## Supplementary material for "Analysis of the ARTIC version 3 and version 4 SARS-CoV-2 primers and their impact on the detection of the G142D amino acid substitution in the spike protein": GISAID Acknowledgements (from another version of this preprint: version 1, 2021-09-28)

We gratefully acknowledge the following Authors from the Originating laboratories responsible for obtaining the specimens, as well as the Submitting laboratories where the genome data were generated and shared via GISAID, on which this research is based.

All Submitters of data may be contacted directly via www.gisaid.org

Authors are sorted alphabetically.

| Accession ID | Originating Laboratory | Submitting Laboratory | Authors |
|---|---|---|---|
| EPI_ISL_1315070 | New South Wales Health Pathology Royal Prince Alfred Hospital | Microbiology RPAH | Foster, C.; Au, J.; Ruiz Silva, M.; Deveson, I.; Bull, R.; Van Hal, S.; Rawlinson, W. |
| EPI_ISL_1337507 | DOHMH Corona | New York City Public Health Laboratory | Jade Wang, et al. |
| EPI_ISL_1409773 | Lighthouse Lab in Cambridge | Wellcome Sanger Institute for the COVID-19 Genomics UK (COG-UK) Consortium | Rob Howes, The Lighthouse Lab in Cambridge and Alex Alderton, Roberto Amato, Jeffrey Barrett, Sonia Goncalves, Ewan Harrison, David K. Jackson, Ian Johnston, Dominic Kwiatkowski, Cordelia Langford, John Sillitoe on behalf of the Wellcome Sanger Institute COVID-19 Surveillance Team |
| EPI_ISL_1419152 | INSACOG-WB | National Institute of Biomedical Genomics – INSACOG | Arindam Maitra, Bhaswati Bandyopadhyay, Nidhan Kumar Biswas, Tamal Ghosh, Sreedhar Chinnaswamy, Ajay Chakraborti, Saumitra Das |
| EPI_ISL_1454606 | Lighthouse Lab in Cambridge | Wellcome Sanger Institute for the COVID-19 Genomics UK (COG-UK) Consortium | Rob Howes, The Lighthouse Lab in Cambridge and Alex Alderton, Roberto Amato, Jeffrey Barrett, Sonia Goncalves, Ewan Harrison, David K. Jackson, Ian Johnston, Dominic Kwiatkowski, Cordelia Langford, John Sillitoe on behalf of the Wellcome Sanger Institute COVID-19 Surveillance Team |
| EPI_ISL_1470937, EPI_ISL_1470959 | Pandemic Response Lab - NYC | Pandemic Response Lab, R&D | Henry Lee, Michael Hammerling, Melissa Hopkins, Cybill del Castillo, Shinyoung Clair Kang, William Ward, Pradeep Bugga, Sol Rey, Dylan Law, Haiping Hao, Jon Laurent |
| EPI_ISL_1473648 | Lighthouse Lab in Cambridge | Wellcome Sanger Institute for the COVID-19 Genomics UK (COG-UK) Consortium | Rob Howes, The Lighthouse Lab in Cambridge and Alex Alderton, Roberto Amato, Jeffrey Barrett, Sonia Goncalves, Ewan Harrison, David K. Jackson, Ian Johnston, Dominic Kwiatkowski, Cordelia Langford, John Sillitoe on behalf of the Wellcome Sanger Institute COVID-19 Surveillance Team |
| EPI_ISL_1482556 | NORTHWELL HEALTH LABORATORIES | Wadsworth Center, New York State Department of Health | Kirsten St. George, Daryl M. Lamson, Alexis Russell, Matthew Shudt, Melissa A Leisner, Jonathan Plitnick, Catharine Prussing, Navjot Singh, John Kelly, Erasmus Schneider, Erica Lasek-Nesselquist |
| EPI_ISL_1494721 | Area of Virology, Serology and Virology Division (SAViD), New South Wales Health Pathology Randwick | Virology Research Laboratory; Area of Virology, Serology and Virology Division (SAViD), New South Wales Health Pathology Randwick | Foster, C.; Au, J.; Ruiz Silva, M.; Deveson, I.; Bull, R.; Van Hal, S.; Rawlinson, W. |
| EPI_ISL_1505036, EPI_ISL_1505038 | Lighthouse Lab in Cambridge | Wellcome Sanger Institute for the COVID-19 Genomics UK (COG-UK) Consortium | Rob Howes, The Lighthouse Lab in Cambridge and Alex Alderton, Roberto Amato, Jeffrey Barrett, Sonia Goncalves, Ewan Harrison, David K. Jackson, Ian Johnston, Dominic Kwiatkowski, Cordelia Langford, John Sillitoe on behalf of the Wellcome Sanger Institute COVID-19 Surveillance Team |
| EPI_ISL_1508996, EPI_ISL_1508997 | PathWest Laboratory Medicine WA | PathWest Laboratory Medicine WA Microbial Surveillance Unit | PathWest Laboratory Medicine WA Microbial Surveillance Unit |
| EPI_ISL_1517918, EPI_ISL_1519271, EPI_ISL_1519290, EPI_ISL_1519300, EPI_ISL_1534758, EPI_ISL_1534917, EPI_ISL_1534938, EPI_ISL_1534945 | Lighthouse Lab in Cambridge | Wellcome Sanger Institute for the COVID-19 Genomics UK (COG-UK) Consortium | Rob Howes, The Lighthouse Lab in Cambridge and Alex Alderton, Roberto Amato, Jeffrey Barrett, Sonia Goncalves, Ewan Harrison, David K. Jackson, Ian Johnston, Dominic Kwiatkowski, Cordelia Langford, John Sillitoe on behalf of the Wellcome Sanger Institute COVID-19 Surveillance Team |
| EPI_ISL_1536300 | Lighthouse Lab in Alderley Park | Wellcome Sanger Institute for the COVID-19 Genomics UK (COG-UK) Consortium | Jacquelyn Wynn, Mairead Hyland, The Lighthouse Lab in Alderley Park and Alex Alderton, Roberto Amato, Jeffrey Barrett, Sonia Goncalves, Ewan Harrison, David K. Jackson, Ian Johnston, Dominic Kwiatkowski, Cordelia Langford, John Sillitoe on behalf of the Wellcome Sanger Institute COVID-19 Surveillance Team |
| EPI_ISL_1537321 | Lighthouse Lab in Cambridge | Wellcome Sanger Institute for the COVID-19 Genomics UK (COG-UK) Consortium | Rob Howes, The Lighthouse Lab in Cambridge and Alex Alderton, Roberto Amato, Jeffrey Barrett, Sonia Goncalves, Ewan Harrison, David K. Jackson, Ian Johnston, Dominic Kwiatkowski, Cordelia Langford, John Sillitoe on behalf of the Wellcome Sanger Institute COVID-19 Surveillance Team |
| EPI_ISL_1543981 | National Public Health Laboratory, National Centre for Infectious Diseases | National Public Health Laboratory, National Centre for Infectious Diseases | Tze Minn Mak, Zhenyang Zhou, Grace Jie Yin Ngan, Royce Ang, Lin Cui, Raymond Tzer Pin Lin |
| EPI_ISL_1544014, EPI_ISL_1544070 | National Centre For Cell Science | National Centre For Cell Science – INSACOG | Dhiraj Paul, Mohak P Gujare, Shivang P. Bhanushali, Mitali Inamdar, Sonal Manik Chavan, Manoj Kumar Bhat, Ajay Pillai, INSACOG Consortium team, Yogesh Shouche |
| EPI_ISL_1544317, EPI_ISL_1544487 | Randox Laboratories | Wellcome Sanger Institute for the COVID-19 Genomics UK (COG-UK) Consortium | Randox Laboratories and Alex Alderton, Roberto Amato, Jeffrey Barrett, Sonia Goncalves, Ewan Harrison, David K. Jackson, Ian Johnston, Dominic Kwiatkowski, Cordelia Langford, John Sillitoe on behalf of the Wellcome Sanger Institute COVID-19 Surveillance Team |
| EPI_ISL_1562503 | Aegis Sciences Corporation | Centers for Disease Control and Prevention Division of Viral Diseases, Pathogen Discovery | Dakota Howard, Dhwani Batra, Peter W. Cook, Kara Moser, Adrian Paskey, Jason Caravas, Benjamin Rambo-Martin, Shatavia Morrison, Christopher Gulvick, Scott Sammons, Yvette Unoarumhi, Darlene Wagner, Matthew Schmerer, Cyndi Clark, Patrick Campbell, Rob Case, Vikramsinha Ghorpade, Holly Houdeshell, Ola Kvalvaag, Dillon Nall, Ethan Sanders, Alec Vest, Shaun Westlund, Matthew Hardison, Clinton R. Paden, Duncan MacCannell |
| EPI_ISL_1564086, EPI_ISL_1564102, EPI_ISL_1564139, EPI_ISL_1564187, EPI_ISL_1564328 | Lighthouse Lab in Cambridge | Wellcome Sanger Institute for the COVID-19 Genomics UK (COG-UK) Consortium | Rob Howes, The Lighthouse Lab in Cambridge and Alex Alderton, Roberto Amato, Jeffrey Barrett, Sonia Goncalves, Ewan Harrison, David K. Jackson, Ian Johnston, Dominic Kwiatkowski, Cordelia Langford, John Sillitoe on behalf of the Wellcome Sanger Institute COVID-19 Surveillance Team |
| EPI_ISL_1564356 | Randox Laboratories | Wellcome Sanger Institute for the COVID-19 Genomics UK (COG-UK) Consortium | Randox Laboratories and Alex Alderton, Roberto Amato, Jeffrey Barrett, Sonia Goncalves, Ewan Harrison, David K. Jackson, Ian Johnston, Dominic Kwiatkowski, Cordelia Langford, John Sillitoe on behalf of the Wellcome Sanger Institute COVID-19 Surveillance Team |
| EPI_ISL_1564772 | Lighthouse Lab in Cambridge | Wellcome Sanger Institute for the COVID-19 Genomics UK (COG-UK) Consortium | Rob Howes, The Lighthouse Lab in Cambridge and Alex Alderton, Roberto Amato, Jeffrey Barrett, Sonia Goncalves, Ewan Harrison, David K. Jackson, Ian Johnston, Dominic Kwiatkowski, Cordelia Langford, John Sillitoe on behalf of the Wellcome Sanger Institute COVID-19 Surveillance Team |
| EPI_ISL_1566762 | Labor Dr. Heidrich & Kollegen MVZ GmbH Hamburg | Robert Koch Institute | unknown |
| EPI_ISL_1583915 | Lighthouse Lab in Glasgow | Wellcome Sanger Institute for the COVID-19 Genomics UK (COG-UK) Consortium | Harper VanSteenhouse, Yumi Kasai, David Gray, Carol Clugston, Anna Dominiczak and Alex Alderton, Roberto Amato, Jeffrey Barrett, Sonia Goncalves, Ewan Harrison, David K. Jackson, Ian Johnston, Dominic Kwiatkowski, Cordelia Langford, John Sillitoe on behalf of the Wellcome Sanger Institute COVID-19 Surveillance Team |
| EPI_ISL_1583942, EPI_ISL_1583979 | Lighthouse Lab in Cambridge | Wellcome Sanger Institute for the COVID-19 Genomics UK (COG-UK) Consortium | Rob Howes, The Lighthouse Lab in Cambridge and Alex Alderton, Roberto Amato, Jeffrey Barrett, Sonia Goncalves, Ewan Harrison, David K. Jackson, Ian Johnston, Dominic Kwiatkowski, Cordelia Langford, John Sillitoe on behalf of the Wellcome Sanger Institute COVID-19 Surveillance Team |
| EPI_ISL_1584529 | Lighthouse Lab in Alderley Park | Wellcome Sanger Institute for the COVID-19 Genomics UK (COG-UK) Consortium | Jacquelyn Wynn, Mairead Hyland, The Lighthouse Lab in Alderley Park and Alex Alderton, Roberto Amato, Jeffrey Barrett, Sonia Goncalves, Ewan Harrison, David K. Jackson, Ian Johnston, Dominic Kwiatkowski, Cordelia Langford, John Sillitoe on behalf of the Wellcome Sanger Institute COVID-19 Surveillance Team |
| EPI_ISL_1589660, EPI_ISL_1589667, EPI_ISL_1589674, EPI_ISL_1589690, EPI_ISL_1589718, EPI_ISL_1589824, EPI_ISL_1589829, EPI_ISL_1589834, EPI_ISL_1589835, EPI_ISL_1589867, EPI_ISL_1589868, EPI_ISL_1589869, EPI_ISL_1589870, EPI_ISL_1589871, EPI_ISL_1589872 | | | |
| see above | INSACOG-WB | National Institute of Biomedical Genomics – INSACOG | Arindam Maitra, Bhaswati Bandyopadhyay, Nidhan Kumar Biswas, Tamal Ghosh, Sreedhar Chinnaswamy, Ajay Chakraborti, Saumitra Das |
| EPI_ISL_1592421, EPI_ISL_1592485 | Helix/Illumina | Centers for Disease Control and Prevention Division of Viral Diseases, Pathogen Discovery | Dakota Howard, Dhwani Batra, Peter W. Cook, Kara Moser, Adrian Paskey, Jason Caravas, Benjamin Rambo-Martin, Shatavia Morrison, Christopher Gulvick, Scott Sammons, Yvette Unoarumhi, Darlene Wagner, Matthew Schmerer, Eileen de Feo, Jan Antico, Christine Tran, Matthew Tolentino, Shannon Wickline, Kim Gietzen, Brad Sickler, Jingtao Liu, Eric Allen, Phil Febbo, Nicole L. Washington, Simon White, Geraint Levan, Kelly Schiabor Barrett, Elizabeth Cirulli, Alexandre Bolze, Ary Ascencio, Charlotte Rivera-Garcia, Ryan Cho, Jason Nguyen, Sherry Wang, Jimmy Ramirez, Tyler Cassens, Efren Sandoval, Magnus Isaksson, William Lee, David Becker, Marc Laurent, James Lu, Clinton R. Paden, Duncan MacCannell |
| EPI_ISL_1594669 | Lighthouse Lab in Milton Keynes | Wellcome Sanger Institute for the COVID-19 Genomics UK (COG-UK) Consortium | The Lighthouse Lab in Milton Keynes and Alex Alderton, Roberto Amato, Jeffrey Barrett, Sonia Goncalves, Ewan Harrison, David K. Jackson, Ian Johnston, Dominic Kwiatkowski, Cordelia Langford, John Sillitoe on behalf of the Wellcome Sanger Institute COVID-19 Surveillance Team |
| EPI_ISL_1594728, EPI_ISL_1594918, EPI_ISL_1595457 | Lighthouse Lab in Cambridge | Wellcome Sanger Institute for the COVID-19 Genomics UK (COG-UK) Consortium | Rob Howes, The Lighthouse Lab in Cambridge and Alex Alderton, Roberto Amato, Jeffrey Barrett, Sonia Goncalves, Ewan Harrison, David K. Jackson, Ian Johnston, Dominic Kwiatkowski, Cordelia Langford, John Sillitoe on behalf of the Wellcome Sanger Institute COVID-19 Surveillance Team |
| EPI_ISL_1611370 | Laboratory Corporation of America | Centers for Disease Control and Prevention Division of Viral Diseases, Pathogen Discovery | Dakota Howard, Dhwani Batra, Peter W. Cook, Kara Moser, Adrian Paskey, Jason Caravas, Benjamin Rambo-Martin, Shatavia Morrison, Christopher Gulvick, Scott Sammons, Yvette Unoarumhi, Darlene Wagner, Matthew Schmerer, Minoo Agarwal, Eyad Almasri, Debbie Boles, Ayla Burns, Nuthawin Charoensri, Oren Cohen, Susan Countryman, Mary Ann Cristobal, Bobbi Croy, Suzanne Dale, Hrushikesh Deshmukh, Amanda Douglas, Vincent Drouillon, Marcia Eisenberg, Howard Engler, Rama Ghatti, Prashant Gupta, Susan Hicks, Jake Humphrey, Lax Iyer, Manoj Jain, Mohan Kolli, Brian Krueger, Tim Kuphal, Stanley Letovsky, Michael Levandoski, Craig Lukasik, Jonathan Meltzer, Brian Norvell, Mindy Nye, Scott Parker, Christos Petropoulos, John Pruitt, Steven Ragan, Scott Ryan, Mike Sapeta, Jana Schroth, Suresh Babu Selvaraju, Goran Stevovic, Amanda Suchanek, Andrea Throop, Lyndon Tilson, Thomas Urban, Joe Voshell, Kimberly Wagner, Jonathan Williams, Mary Williamson, Qian Zeng, Tricia Zwiefelhofer, Clinton R. Paden, Duncan MacCannell |
| EPI_ISL_1614854 | Fulgent Genetics | Centers for Disease Control and Prevention Division of Viral Diseases, Pathogen Discovery | Dakota Howard, Dhwani Batra, Peter W. Cook, Kara Moser, Adrian Paskey, Jason Caravas, Benjamin Rambo-Martin, Shatavia Morrison, Christopher Gulvick, Scott Sammons, Yvette Unoarumhi, Darlene Wagner, Matthew Schmerer, Harry Gao, Mickey Li, John Gao, Joseph Fierro, Benafsh Sapra, Becky Tsai, Yan Meng, Doreen Ng, James Xie, Clinton R. Paden, Duncan MacCannell |
| EPI_ISL_1615349 | Helix/Illumina | Centers for Disease Control and Prevention Division of Viral Diseases, Pathogen Discovery | Dakota Howard, Dhwani Batra, Peter W. Cook, Kara Moser, Adrian Paskey, Jason Caravas, Benjamin Rambo-Martin, Shatavia Morrison, Christopher Gulvick, Scott Sammons, Yvette Unoarumhi, Darlene Wagner, Matthew Schmerer, Eileen de Feo, Jan Antico, Christine Tran, Matthew Tolentino, Shannon Wickline, Kim Gietzen, Brad Sickler, Jingtao Liu, Eric Allen, Phil Febbo, Nicole L. Washington, Simon White, Geraint Levan, Kelly Schiabor Barrett, Elizabeth Cirulli, Alexandre Bolze, Ary Ascencio, Charlotte Rivera-Garcia, Ryan Cho, Jason Nguyen, Sherry Wang, Jimmy Ramirez, Tyler Cassens, Efren Sandoval, Magnus Isaksson, William Lee, David Becker, Marc Laurent, James Lu, Clinton R. Paden, Duncan MacCannell |
| EPI_ISL_1615595, EPI_ISL_1615596, EPI_ISL_1615597, EPI_ISL_1615598 | Area of Virology, Serology and Virology Division (SAViD), New South Wales Health Pathology Randwick | Virology Research Laboratory; Area of Virology, Serology and Virology Division (SAViD), New South Wales Health Pathology Randwick | Foster, C.; Au, J.; Ruiz Silva, M.; Deveson, I.; Bull, R.; Van Hal, S.; Rawlinson, W. |
| EPI_ISL_1615751, EPI_ISL_1615877 | Lighthouse Lab in Cambridge | Wellcome Sanger Institute for the COVID-19 Genomics UK (COG-UK) Consortium | Rob Howes, The Lighthouse Lab in Cambridge and Alex Alderton, Roberto Amato, Jeffrey Barrett, Sonia Goncalves, Ewan Harrison, David K. Jackson, Ian Johnston, Dominic Kwiatkowski, Cordelia Langford, John Sillitoe on behalf of the Wellcome Sanger Institute COVID-19 Surveillance Team |
| EPI_ISL_1620131, EPI_ISL_1620148, | National Public Health Laboratory, National Centre for Infectious Diseases | National Public Health Laboratory, National Centre for Infectious Diseases | Tze Minn Mak, Zhenyang Zhou, Grace Jie Yin Ngan, Royce Ang, Lin Cui, Raymond Tzer Pin Lin |

| | | | |
|---|---|---|---|
| EPI_ISL_1620158, EPI_ISL_1620159 | | | |
| EPI_ISL_1620980 | UW Virology Lab | UW Virology Lab | Pavitra Roychoudhury, Hong Xie, Lasata Shrestha, Shah Mohamed Bakhash, Michelle Lin, Noah R. Baker, Sean Ellis, Saraswathi Sathees, Meei-Li Huang, Keith R Jerome, Alexander Greninger |
| EPI_ISL_1621293 | Middlemore Hospital | Institute of Environmental Science and Research (ESR) | Rachel Boyle, SallyAnn Harbison, Olivia Stroeven, Xiaoyun Ren, Matt Storey, Nikki Freed, Muhammad Faisal, Jing Wang, Hermes Perez, Anja Werno, Antje van der Linden, Arlo Upton, Chris Mansell, David Hammer, Dragana Drinkovic, Gary McAuliffe, Hana Sofia Andersson, James Ussher, Jill Sherwood, Josh Freeman, Julia Howard, Juliet Elvy, Mary DeAlmeida, Matt Blakiston, Matthew Rogers, Max Bloomfield, Michael Addidle, Michelle Balm, Sally Roberts, Sarah Jefferies, Sharmini Muttaiyah, Susan Morpeth, Susan Taylor, Timothy Blackmore, Vani Sathyendran, Veronica Playle, Virginia Hope, Erasmus Smit, Lauren Jelly, Olin Silander, Joep de Ligt |
| EPI_ISL_1621294, EPI_ISL_1621312, EPI_ISL_1621314, EPI_ISL_1621315, EPI_ISL_1621324, EPI_ISL_1621325 | LabPLUS | Institute of Environmental Science and Research (ESR) | Rachel Boyle, SallyAnn Harbison, Olivia Stroeven, Xiaoyun Ren, Matt Storey, Nikki Freed, Muhammad Faisal, Jing Wang, Hermes Perez, Anja Werno, Antje van der Linden, Arlo Upton, Chris Mansell, David Hammer, Dragana Drinkovic, Gary McAuliffe, Hana Sofia Andersson, James Ussher, Jill Sherwood, Josh Freeman, Julia Howard, Juliet Elvy, Mary DeAlmeida, Matt Blakiston, Matthew Rogers, Max Bloomfield, Michael Addidle, Michelle Balm, Sally Roberts, Sarah Jefferies, Sharmini Muttaiyah, Susan Morpeth, Susan Taylor, Timothy Blackmore, Vani Sathyendran, Veronica Playle, Virginia Hope, Erasmus Smit, Lauren Jelly, Olin Silander, Joep de Ligt |
| EPI_ISL_1628185 | UW Virology Lab | UW Virology Lab | Pavitra Roychoudhury, Hong Xie, Lasata Shrestha, Shah Mohamed Bakhash, Michelle Lin, Noah R. Baker, Sean Ellis, Meei-Li Huang, Keith R Jerome, Alexander Greninger |
| EPI_ISL_1631042 | Lighthouse Lab in Alderley Park | Wellcome Sanger Institute for the COVID-19 Genomics UK (COG-UK) Consortium | Jacquelyn Wynn, Mairead Hyland, The Lighthouse Lab in Alderley Park and Alex Alderton, Roberto Amato, Jeffrey Barrett, Sonia Goncalves, Ewan Harrison, David K. Jackson, Ian Johnston, Dominic Kwiatkowski, Cordelia Langford, John Sillitoe on behalf of the Wellcome Sanger Institute COVID-19 Surveillance Team |
| EPI_ISL_1631162 | Lighthouse Lab in Cambridge | Wellcome Sanger Institute for the COVID-19 Genomics UK (COG-UK) Consortium | Rob Howes, The Lighthouse Lab in Cambridge and Alex Alderton, Roberto Amato, Jeffrey Barrett, Sonia Goncalves, Ewan Harrison, David K. Jackson, Ian Johnston, Dominic Kwiatkowski, Cordelia Langford, John Sillitoe on behalf of the Wellcome Sanger Institute COVID-19 Surveillance Team |
| EPI_ISL_1631164, EPI_ISL_1631170 | Randox Laboratories | Wellcome Sanger Institute for the COVID-19 Genomics UK (COG-UK) Consortium | Randox Laboratories and Alex Alderton, Roberto Amato, Jeffrey Barrett, Sonia Goncalves, Ewan Harrison, David K. Jackson, Ian Johnston, Dominic Kwiatkowski, Cordelia Langford, John Sillitoe on behalf of the Wellcome Sanger Institute COVID-19 Surveillance Team |
| EPI_ISL_1631219 | Lighthouse Lab in Cambridge | Wellcome Sanger Institute for the COVID-19 Genomics UK (COG-UK) Consortium | Rob Howes, The Lighthouse Lab in Cambridge and Alex Alderton, Roberto Amato, Jeffrey Barrett, Sonia Goncalves, Ewan Harrison, David K. Jackson, Ian Johnston, Dominic Kwiatkowski, Cordelia Langford, John Sillitoe on behalf of the Wellcome Sanger Institute COVID-19 Surveillance Team |
| EPI_ISL_1631314 | Randox Laboratories | Wellcome Sanger Institute for the COVID-19 Genomics UK (COG-UK) Consortium | Randox Laboratories and Alex Alderton, Roberto Amato, Jeffrey Barrett, Sonia Goncalves, Ewan Harrison, David K. Jackson, Ian Johnston, Dominic Kwiatkowski, Cordelia Langford, John Sillitoe on behalf of the Wellcome Sanger Institute COVID-19 Surveillance Team |
| EPI_ISL_1631645 | Lighthouse Lab in Milton Keynes | Wellcome Sanger Institute for the COVID-19 Genomics UK (COG-UK) Consortium | The Lighthouse Lab in Milton Keynes and Alex Alderton, Roberto Amato, Jeffrey Barrett, Sonia Goncalves, Ewan Harrison, David K. Jackson, Ian Johnston, Dominic Kwiatkowski, Cordelia Langford, John Sillitoe on behalf of the Wellcome Sanger Institute COVID-19 Surveillance Team |
| EPI_ISL_1631836, EPI_ISL_1631914 | Randox Laboratories | Wellcome Sanger Institute for the COVID-19 Genomics UK (COG-UK) Consortium | Randox Laboratories and Alex Alderton, Roberto Amato, Jeffrey Barrett, Sonia Goncalves, Ewan Harrison, David K. Jackson, Ian Johnston, Dominic Kwiatkowski, Cordelia Langford, John Sillitoe on behalf of the Wellcome Sanger Institute COVID-19 Surveillance Team |
| EPI_ISL_1631923, EPI_ISL_1631945 | Lighthouse Lab in Milton Keynes | Wellcome Sanger Institute for the COVID-19 Genomics UK (COG-UK) Consortium | The Lighthouse Lab in Milton Keynes and Alex Alderton, Roberto Amato, Jeffrey Barrett, Sonia Goncalves, Ewan Harrison, David K. Jackson, Ian Johnston, Dominic Kwiatkowski, Cordelia Langford, John Sillitoe on behalf of the Wellcome Sanger Institute COVID-19 Surveillance Team |
| EPI_ISL_1632008 | Lighthouse Lab in Cambridge | Wellcome Sanger Institute for the COVID-19 Genomics UK (COG-UK) Consortium | Rob Howes, The Lighthouse Lab in Cambridge and Alex Alderton, Roberto Amato, Jeffrey Barrett, Sonia Goncalves, Ewan Harrison, David K. Jackson, Ian Johnston, Dominic Kwiatkowski, Cordelia Langford, John Sillitoe on behalf of the Wellcome Sanger Institute COVID-19 Surveillance Team |
| EPI_ISL_1632122, EPI_ISL_1632256 | Randox Laboratories | Wellcome Sanger Institute for the COVID-19 Genomics UK (COG-UK) Consortium | Randox Laboratories and Alex Alderton, Roberto Amato, Jeffrey Barrett, Sonia Goncalves, Ewan Harrison, David K. Jackson, Ian Johnston, Dominic Kwiatkowski, Cordelia Langford, John Sillitoe on behalf of the Wellcome Sanger Institute COVID-19 Surveillance Team |
| EPI_ISL_1632307 | Lighthouse Lab in Glasgow | Wellcome Sanger Institute for the COVID-19 Genomics UK (COG-UK) Consortium | Harper VanSteenhouse, Yumi Kasai, David Gray, Carol Clugston, Anna Dominiczak and Alex Alderton, Roberto Amato, Jeffrey Barrett, Sonia Goncalves, Ewan Harrison, David K. Jackson, Ian Johnston, Dominic Kwiatkowski, Cordelia Langford, John Sillitoe on behalf of the Wellcome Sanger Institute COVID-19 Surveillance Team |
| EPI_ISL_1632316, EPI_ISL_1632335, EPI_ISL_1632384, EPI_ISL_1632412, EPI_ISL_1632426 | Lighthouse Lab in Cambridge | Wellcome Sanger Institute for the COVID-19 Genomics UK (COG-UK) Consortium | Rob Howes, The Lighthouse Lab in Cambridge and Alex Alderton, Roberto Amato, Jeffrey Barrett, Sonia Goncalves, Ewan Harrison, David K. Jackson, Ian Johnston, Dominic Kwiatkowski, Cordelia Langford, John Sillitoe on behalf of the Wellcome Sanger Institute COVID-19 Surveillance Team |
| EPI_ISL_1633348, EPI_ISL_1633353 | Area of Virology, Serology and Virology Division (SAViD), New South Wales Health Pathology Randwick | Virology Research Laboratory; Area of Virology, Serology and Virology Division (SAViD), New South Wales Health Pathology Randwick | Foster, C.; Au, J.; Ruiz Silva, M.; Deveson, I.; Bull, R.; Van Hal, S.; Rawlinson, W. |
| EPI_ISL_1633676, EPI_ISL_1633680 | PathWest Laboratory Medicine WA | PathWest Laboratory Medicine WA Microbial Surveillance Unit | PathWest Laboratory Medicine WA Microbial Surveillance Unit |
| EPI_ISL_1634428, EPI_ISL_1634433 | National Public Health Laboratory, National Centre for Infectious Diseases | National Public Health Laboratory, National Centre for Infectious Diseases | Tze Minn Mak, Zhenyang Zhou, Grace Jie Yin Ngan, Royce Ang, Lin Cui, Raymond Tzer Pin Lin |
| EPI_ISL_1634507, EPI_ISL_1634517 | Lighthouse Lab in Milton Keynes | Wellcome Sanger Institute for the COVID-19 Genomics UK (COG-UK) Consortium | The Lighthouse Lab in Milton Keynes and Alex Alderton, Roberto Amato, Jeffrey Barrett, Sonia Goncalves, Ewan Harrison, David K. Jackson, Ian Johnston, Dominic Kwiatkowski, Cordelia Langford, John Sillitoe on behalf of the Wellcome Sanger Institute COVID-19 Surveillance Team |
| EPI_ISL_1634839, EPI_ISL_1634899, EPI_ISL_1634920, EPI_ISL_1634973, EPI_ISL_1635330 | Lighthouse Lab in Glasgow | Wellcome Sanger Institute for the COVID-19 Genomics UK (COG-UK) Consortium | Harper VanSteenhouse, Yumi Kasai, David Gray, Carol Clugston, Anna Dominiczak and Alex Alderton, Roberto Amato, Jeffrey Barrett, Sonia Goncalves, Ewan Harrison, David K. Jackson, Ian Johnston, Dominic Kwiatkowski, Cordelia Langford, John Sillitoe on behalf of the Wellcome Sanger Institute COVID-19 Surveillance Team |
| EPI_ISL_1635371, EPI_ISL_1635379, EPI_ISL_1635400 | Lighthouse Lab in Cambridge | Wellcome Sanger Institute for the COVID-19 Genomics UK (COG-UK) Consortium | Rob Howes, The Lighthouse Lab in Cambridge and Alex Alderton, Roberto Amato, Jeffrey Barrett, Sonia Goncalves, Ewan Harrison, David K. Jackson, Ian Johnston, Dominic Kwiatkowski, Cordelia Langford, John Sillitoe on behalf of the Wellcome Sanger Institute COVID-19 Surveillance Team |
| EPI_ISL_1635519 | Pandemic Response Lab - NYC | Pandemic Response Lab, R&D | Henry Lee, Michael Hammerling, Melissa Hopkins, Cybill del Castillo, Shinyoung Clair Kang, William Ward, Pradeep Bugga, Sol Rey, Dylan Law, Katharine Nelson, Haiping Hao, Jon Laurent |
| EPI_ISL_1636862 | Liverpool Clinical Laboratories | COVID-19 Genomics UK (COG-UK) Consortium | Sam Haldenby, Alistair Darby, Steve Paterson, Anita Lucaci, Julian Hiscox, M Almsaud, A Alrezaihi, Muhannad Alruwaili, Stuart D Armstrong, Jones Benjamin, Eleanor G Bentley, Anu Chawla, Jordan J Clark, Angela Cowell, Richard Eccles, Isabel García-Dorival, Matthew Gemmell, Alessandro Gerada, PKF Gilmore, Richard Gregory, Ximeng Han, Catherine Hartley, Margaret Hughes, Miren Iturriza-Gomara, James Johnson, L Luu, Jenifer Manson, Charlotte Nelson, Elaine O'Toole, Cassie Olateju, Rebekah Penrice-Randal , Lucille Rainbow, N.P Randle, Trevor Ian Robinson, Parul Sharma, Ghada T Shawli, James P Stewart, Neil Swainston, Ecaterina Vamos, Joanne Watts, Mark Whitehead, Hermione Webster |
| EPI_ISL_1637187 | Northumbria University / South Tees Hospitals NHS Foundation Trust / North Cumbria Integrated Care NHS Foundation Trust / North Tees and Hartlepool NHS Foundation Trust / Newcastle Hospitals NHS Foundation Trust | COVID-19 Genomics UK (COG-UK) Consortium | Darren L Smith,Andrew Nelson,Matthew Bashton,Greg R Young,Joshua Loh,John Allan,Mohammad A Tariq,Giles S Holt,Gary Black,Wen C Yew,Lynn Dover,Paul Baker,Steve Liggett,Sarah Essex,Jane Greenaway,Debra Padgett,Clive Graham,Garren Scott,Edward Barton,Emma Swindells,Brendan Payne,Jennifer Collins,Yusri Taha,Gary Eltringham |
| EPI_ISL_1641213 | MVZ Labor Krone GbR | Robert Koch Institute | unknown |
| EPI_ISL_1647029 | SC (UCO) Igiene e Sanità Pubblica (funzione integrata con SC Microbiologia e Virologia) | ARGO Laboratorio Genomica ed Epigenomica | Licastro D, Dal Monego S, Degasperi M, D'Agaro P, Lombardo F |
| EPI_ISL_1648208, EPI_ISL_1648249 | Quest Diagnostics Incorporated | Centers for Disease Control and Prevention Division of Viral Diseases, Pathogen Discovery | Dakota Howard, Dhwani Batra, Peter W. Cook, Kara Moser, Adrian Paskey, Jason Caravas, Benjamin Rambo-Martin, Shatavia Morrison, Christopher Gulvick, Scott Sammons, Yvette Unoarumhi, Darlene Wagner, Matthew Schmerer, S. H. Rosenthal, A. Gerasimova, R. M. Kagan, B. Anderson, M. Hua, Y. Liu, L.E. Bernstein, K.E. Livingston, A. Perez, I. A. Shlyakhter, R. V. Rolando, R. Owen, P. Tanpaiboon, F. Lacbawan, Clinton R. Paden, Duncan MacCannell |
| EPI_ISL_1652117, EPI_ISL_1652118, EPI_ISL_1652119, EPI_ISL_1652120, EPI_ISL_1652122, EPI_ISL_1652123, EPI_ISL_1652532 | National Public Health Laboratory, National Centre for Infectious Diseases | National Public Health Laboratory, National Centre for Infectious Diseases | Tze Minn Mak, Zhenyang Zhou, Grace Jie Yin Ngan, Royce Ang, Lin Cui, Raymond Tzer Pin Lin |
| EPI_ISL_1652717 | Randox Laboratories | Wellcome Sanger Institute for the COVID-19 Genomics UK (COG-UK) Consortium | Randox Laboratories and Alex Alderton, Roberto Amato, Jeffrey Barrett, Sonia Goncalves, Ewan Harrison, David K. Jackson, Ian Johnston, Dominic Kwiatkowski, Cordelia Langford, John Sillitoe on behalf of the Wellcome Sanger Institute COVID-19 Surveillance Team |
| EPI_ISL_1652724, EPI_ISL_1652738, EPI_ISL_1652761, EPI_ISL_1652763, EPI_ISL_1652850, EPI_ISL_1652890 | Lighthouse Lab in Alderley Park | Wellcome Sanger Institute for the COVID-19 Genomics UK (COG-UK) Consortium | Jacquelyn Wynn, Mairead Hyland, The Lighthouse Lab in Alderley Park and Alex Alderton, Roberto Amato, Jeffrey Barrett, Sonia Goncalves, Ewan Harrison, David K. Jackson, Ian Johnston, Dominic Kwiatkowski, Cordelia Langford, John Sillitoe on behalf of the Wellcome Sanger Institute COVID-19 Surveillance Team |
| EPI_ISL_1652897 | Randox Laboratories | Wellcome Sanger Institute for the COVID-19 Genomics UK (COG-UK) Consortium | Randox Laboratories and Alex Alderton, Roberto Amato, Jeffrey Barrett, Sonia Goncalves, Ewan Harrison, David K. Jackson, Ian Johnston, Dominic Kwiatkowski, Cordelia Langford, John Sillitoe on behalf of the Wellcome Sanger Institute COVID-19 Surveillance Team |
| EPI_ISL_1652964, EPI_ISL_1652981, EPI_ISL_1652993, EPI_ISL_1653090, EPI_ISL_1653104 | Lighthouse Lab in Cambridge | Wellcome Sanger Institute for the COVID-19 Genomics UK (COG-UK) Consortium | Rob Howes, The Lighthouse Lab in Cambridge and Alex Alderton, Roberto Amato, Jeffrey Barrett, Sonia Goncalves, Ewan Harrison, David K. Jackson, Ian Johnston, Dominic Kwiatkowski, Cordelia Langford, John Sillitoe on behalf of the Wellcome Sanger Institute COVID-19 Surveillance Team |
| EPI_ISL_1653187 | Lighthouse Lab in Alderley Park | Wellcome Sanger Institute for the COVID-19 Genomics UK (COG-UK) Consortium | Jacquelyn Wynn, Mairead Hyland, The Lighthouse Lab in Alderley Park and Alex Alderton, Roberto Amato, Jeffrey Barrett, Sonia Goncalves, Ewan Harrison, David K. Jackson, Ian Johnston, Dominic Kwiatkowski, Cordelia Langford, John Sillitoe on behalf of the Wellcome Sanger Institute COVID-19 Surveillance Team |
| EPI_ISL_1653197, | Lighthouse Lab in Cambridge | Wellcome Sanger Institute for the COVID-19 Genomics UK (COG-UK) | Rob Howes, The Lighthouse Lab in Cambridge and Alex Alderton, Roberto Amato, Jeffrey Barrett, Sonia Goncalves, Ewan Harrison, David K. Jackson, Ian Johnston, Dominic Kwiatkowski, Cordelia Langford, John Sillitoe on behalf of the Wellcome |

| | | | |
|---|---|---|---|
| EPI_ISL_1653403, EPI_ISL_1653755 | | Consortium | Sanger Institute COVID-19 Surveillance Team |
| EPI_ISL_1658609, EPI_ISL_1658683 | Viollier AG | Department of Biosystems Science and Engineering, ETH Zurich | Chaoran Chen, Sarah Nadeau, Catharine Aquino, Ivan Topolsky, Philipp Jablonski, Lara Fuhrmann, David Dreifuss, Katharina Jahn, Andreia Cabral de Gouvea, Maria Domenica Moccia, Simon Gruter, Timothy Sykes, Lennart Opitz, Griffin White, Laura Neff, Doris Popovic, Andrea Patrignani, Jay Tracy, Ralph Schlapbach, Christiane Beckmann, Maurice Redondo, Olivier Kobel, Christoph Noppen, Sophie Seidel, Noemie Santamaria de Souza, Niko Beerenwinkel, Tanja Stadler |
| EPI_ISL_1660421, EPI_ISL_1660425, EPI_ISL_1660426, EPI_ISL_1660427, EPI_ISL_1660429, EPI_ISL_1660430, EPI_ISL_1660432, EPI_ISL_1660442, EPI_ISL_1660445, EPI_ISL_1660447, EPI_ISL_1660448, EPI_ISL_1660450, EPI_ISL_1660456, EPI_ISL_1660458, EPI_ISL_1660459 | | | |
| see above | Communicable Disease Laboratory, Public Health Directorate | Communicable Disease Laboratory, Public Health Directorate | Alwasti,H., AlHujairi,Z., AlAbbas,Z., Marhoon,A., Almoamen,G. |
| EPI_ISL_1662284 | Veer Surendra Sai Institute of Medical Sciences and Research, Burla, Sambalpur | Institute of Life Sciences - INSACOG | Sunil K. Raghav, Safal Walia, Arup Ghosh, Atimukta Jha, Amol M. Kanampalliwar, Shifu Aggarwal, Rupesh Dash, Rajeeb Swain, Punit Prasad, INSACOG Consortium, Ajay Parida |
| EPI_ISL_1662291 | Dept. Of Microbiology, Lt.. Baliram Kashyap Memorial Govt. Medical college, Dimrapal, Jagdalpur | Institute of Life Sciences - INSACOG | Sunil K. Raghav, Safal Walia, Arup Ghosh, Atimukta Jha, Amol M. Kanampalliwar, Shifu Aggarwal, Rupesh Dash, Rajeeb Swain, Punit Prasad, INSACOG Consortium, Ajay Parida |
| EPI_ISL_1662451 | Immunogenomics lab, Institute of Life Sciences, Bhubaneswar | Institute of Life Sciences - INSACOG | Sunil K. Raghav, Safal Walia, Arup Ghosh, Atimukta Jha, Amol M. Kanampalliwar, Shifu Aggarwal, Rupesh Dash, Rajeeb Swain, Punit Prasad, INSACOG Consortium, Ajay Parida |
| EPI_ISL_1662783 | Johns Hopkins Hospital Department of Pathology | Johns Hopkins Hospital Department of Pathology | C. Paul Morris, Chun Huai Luo, Adannaya Amadi, Matthew Schwartz, Heba H. Mostafa |
| EPI_ISL_1663247 | BRLSABVM Government Medical College, Rajnandgaon, Chhattisgarh | Institute of Life Sciences - INSACOG | Sunil K. Raghav, Safal Walia, Arup Ghosh, Atimukta Jha, Amol M. Kanampalliwar, Shifu Aggarwal, Rupesh Dash, Rajeeb Swain, Punit Prasad, INSACOG Consortium, Ajay Parida |
| EPI_ISL_1663304, EPI_ISL_1663307, EPI_ISL_1663308, EPI_ISL_1663312 | CIIMS, Bilaspur, Chhattisgarh | Institute of Life Sciences - INSACOG | Sunil K. Raghav, Safal Walia, Arup Ghosh, Atimukta Jha, Amol M. Kanampalliwar, Shifu Aggarwal, Rupesh Dash, Rajeeb Swain, Punit Prasad, INSACOG Consortium, Ajay Parida |
| EPI_ISL_1663366, EPI_ISL_1663367, EPI_ISL_1663375, EPI_ISL_1663376 | NCCS, Pune | Institute of Life Sciences - INSACOG | Sunil K. Raghav, Safal Walia, Arup Ghosh, Atimukta Jha, Amol M. Kanampalliwar, Shifu Aggarwal, Rupesh Dash, Rajeeb Swain, Punit Prasad, INSACOG Consortium, Ajay Parida |
| EPI_ISL_1663491 | Capital hospital, Bhubaneswar | Institute of Life Sciences - INSACOG | Sunil K. Raghav, Safal Walia, Arup Ghosh, Atimukta Jha, Amol M. Kanampalliwar, Shifu Aggarwal, Rupesh Dash, Rajeeb Swain, Punit Prasad, INSACOG Consortium, Ajay Parida |
| EPI_ISL_1663498, EPI_ISL_1663501, EPI_ISL_1663502 | Veer Surendra Sai Institute of Medical Sciences and Research, Burla, Sambalpur | Institute of Life Sciences - INSACOG | Sunil K. Raghav, Safal Walia, Arup Ghosh, Atimukta Jha, Amol M. Kanampalliwar, Shifu Aggarwal, Rupesh Dash, Rajeeb Swain, Punit Prasad, INSACOG Consortium, Ajay Parida |
| EPI_ISL_1663507, EPI_ISL_1663516, EPI_ISL_1663522, EPI_ISL_1663523 | AIIMS, Patna | Institute of Life Sciences - INSACOG | Sunil K. Raghav, Safal Walia, Arup Ghosh, Atimukta Jha, Amol M. Kanampalliwar, Shifu Aggarwal, Rupesh Dash, Rajeeb Swain, Punit Prasad, INSACOG Consortium, Ajay Parida |
| EPI_ISL_1663529, EPI_ISL_1663534 | REGIONAL VRDL, ICMR-RMRC BBSR | Institute of Life Sciences - INSACOG | Sunil K. Raghav, Safal Walia, Arup Ghosh, Atimukta Jha, Amol M. Kanampalliwar, Shifu Aggarwal, Rupesh Dash, Rajeeb Swain, Punit Prasad, INSACOG Consortium, Ajay Parida |
| EPI_ISL_1663541, EPI_ISL_1663543, EPI_ISL_1663548, EPI_ISL_1663549 | MGM Medical College, Jamshedpur | Institute of Life Sciences - INSACOG | Sunil K. Raghav, Safal Walia, Arup Ghosh, Atimukta Jha, Amol M. Kanampalliwar, Shifu Aggarwal, Rupesh Dash, Rajeeb Swain, Punit Prasad, INSACOG Consortium, Ajay Parida |
| EPI_ISL_1663553, EPI_ISL_1663554, EPI_ISL_1663555, EPI_ISL_1663557, EPI_ISL_1663561, EPI_ISL_1663562, EPI_ISL_1663563, EPI_ISL_1663564 | Immunogenomics lab, Institute of Life Sciences, Bhubaneswar | Institute of Life Sciences - INSACOG | Sunil K. Raghav, Safal Walia, Arup Ghosh, Atimukta Jha, Amol M. Kanampalliwar, Shifu Aggarwal, Rupesh Dash, Rajeeb Swain, Punit Prasad, INSACOG Consortium, Ajay Parida |
| EPI_ISL_1666166, EPI_ISL_1666372, EPI_ISL_1666373, EPI_ISL_1666377, EPI_ISL_1666379, EPI_ISL_1666384, EPI_ISL_1666437, EPI_ISL_1666438, EPI_ISL_1666447, EPI_ISL_1666495, EPI_ISL_1667140, EPI_ISL_1667317, EPI_ISL_1667332, EPI_ISL_1667390 | | | |
| see above | Fulgent Genetics | Centers for Disease Control and Prevention Division of Viral Diseases, Pathogen Discovery | Dakota Howard, Dhwani Batra, Peter W. Cook, Kara Moser, Adrian Paskey, Jason Caravas, Benjamin Rambo-Martin, Shatavia Morrison, Christopher Gulvick, Scott Sammons, Yvette Unoarumhi, Darlene Wagner, Matthew Schmerer, Harry Gao, Mickey Li, John Gao, Joseph Fierro, Benafsh Sapra, Becky Tsai, Yan Meng, Doreen Ng, James Xie, Clinton R. Paden, Duncan MacCannell |
| EPI_ISL_1669118, EPI_ISL_1669119, EPI_ISL_1669121 | PathWest Laboratory Medicine WA | PathWest Laboratory Medicine WA Microbial Surveillance Unit | PathWest Laboratory Medicine WA Microbial Surveillance Unit |
| EPI_ISL_1670711, EPI_ISL_1670827 | UW Virology Lab | UW Virology Lab | Pavitra Roychoudhury, Hong Xie, Lasata Shrestha, Shah Mohamed Bakhash, Michelle Lin, Noah R. Baker, Sean Ellis, Meei-Li Huang, Keith R Jerome, Alexander Greninger |
| EPI_ISL_1671604 | NORTHWELL HEALTH LABORATORIES | Wadsworth Center, New York State Department of Health | Kirsten St. George, Daryl M. Lamson, Alexis Russell, Matthew Shudt, Melissa A Leisner, Jonathan Plitnick, Catharine Prussing, Navjot Singh, John Kelly, Erasmus Schneider, Erica Lasek-Nesselquist |
| EPI_ISL_1671957 | SYNLAB | GIGA Medical Genomics | Keith Durkin, Maria Artesi, Sébastien Bontems, Raphaël Boreux, Bouchra Boujemla, Nathalie Renotte, Cécile Meex, Pierrette Melin, Marie-Pierre Hayette, Vincent Bours |
| EPI_ISL_1672867 | Area of Virology, Serology and Virology Division (SAViD), New South Wales Health Pathology Randwick | Virology Research Laboratory; Area of Virology, Serology and Virology Division (SAViD), New South Wales Health Pathology Randwick | Foster, C.; Au, J.; Ruiz Silva, M.; Deveson, I.; Bull, R.; Van Hal, S.; Rawlinson, W. |
| EPI_ISL_1673061, EPI_ISL_1673100, EPI_ISL_1673142 | Lighthouse Lab in Cambridge | Wellcome Sanger Institute for the COVID-19 Genomics UK (COG-UK) Consortium | Rob Howes, The Lighthouse Lab in Cambridge and Alex Alderton, Roberto Amato, Jeffrey Barrett, Sonia Goncalves, Ewan Harrison, David K. Jackson, Ian Johnston, Dominic Kwiatkowski, Cordelia Langford, John Sillitoe on behalf of the Wellcome Sanger Institute COVID-19 Surveillance Team |
| EPI_ISL_1678079 | NYC Department of Health and Mental Hygiene | Centers for Disease Control and Prevention Division of Viral Diseases, Pathogen Discovery | Mili Sheth, Sarah Nobles, Jasmine Padilla, Mark Burroughs, Shoshona Le, Katie Dillon, Peter Cook, Clinton R. Paden, Dhwani Batra, Krista Queen, Kristen Knipe, Dakota Howard, Yvette Unoarumhi, Darlene Wagner, Matthew Schmerer, Ben L. Rambo-Martin, Kristine Lacek, Sam Shepard, Alison Laufer Halpin, Dave Wentworth, Vivien Dugan, Suxiang Tong, Justin Lee |
| EPI_ISL_1679851 | UW Virology Lab | UW Virology Lab | Pavitra Roychoudhury, Hong Xie, Lasata Shrestha, Shah Mohamed Bakhash, Michelle Lin, Noah R. Baker, Sean Ellis, Meei-Li Huang, Keith R Jerome, Alexander Greninger |
| EPI_ISL_1679967, EPI_ISL_1680048 | Helix/Illumina | Centers for Disease Control and Prevention Division of Viral Diseases, Pathogen Discovery | Dakota Howard, Dhwani Batra, Peter W. Cook, Kara Moser, Adrian Paskey, Jason Caravas, Benjamin Rambo-Martin, Shatavia Morrison, Christopher Gulvick, Scott Sammons, Yvette Unoarumhi, Darlene Wagner, Matthew Schmerer, Eileen de Feo, Jan Antico, Christine Tran, Matthew Tolentino, Shannon Wickline, Kim Gietzen, Brad Sickler, Jingtao Liu, Eric Allen, Phil Febbo, Nicole L. Washington, Simon White, Geraint Levan, Kelly Schiabor Barrett, Elizabeth Cirulli, Alexandre Bolze, Ary Ascencio, Charlotte Rivera-Garcia, Ryan Cho, Jason Nguyen, Sherry Wang, Jimmy Ramirez, Tyler Cassens, Efren Sandoval, Magnus Isaksson, William Lee, David Becker, Marc Laurent, James Lu, Clinton R. Paden, Duncan MacCannell |
| EPI_ISL_1681478, EPI_ISL_1681600, EPI_ISL_1682964, EPI_ISL_1683153, EPI_ISL_1683867, EPI_ISL_1683986, EPI_ISL_1684717, EPI_ISL_1685045 | Laboratory Corporation of America | Centers for Disease Control and Prevention Division of Viral Diseases, Pathogen Discovery | Dakota Howard, Dhwani Batra, Peter W. Cook, Kara Moser, Adrian Paskey, Jason Caravas, Benjamin Rambo-Martin, Shatavia Morrison, Christopher Gulvick, Scott Sammons, Yvette Unoarumhi, Darlene Wagner, Matthew Schmerer, Minoo Agarwal, Eyad Almasri, Debbie Boles, Ayla Burns, Nuthawin Charoensri, Oren Cohen, Susan Countryman, Mary Ann Cristobal, Bobbi Croy, Suzanne Dale, Hrushikesh Deshmukh, Amanda Douglas, Vincent Drouillon, Marcia Eisenberg, Howard Engler, Rama Ghatti, Prashant Gupta, Susan Hicks, Jake Humphrey, Lax Iyer, Manoj Jain, Mohan Kolli, Brian Krueger, Tim Kuphal, Stanley Letovsky, Michael Levandoski, Craig Lukasik, Jonathan Meltzer, Brian Norvell, Mindy Nye, Scott Parker, Christos Petropoulos, John Pruitt, Steven Ragan, Scott Ryan, Mike Sapeta, Jana Schroth, Suresh Babu Selvaraju, Goran Stevovic, Amanda Suchanek, Andrea Throop, Lyndon Tilson, Thomas Urban, Joe Voshell, Kimberly Wagner, Jonathan Williams, Mary Williamson, Qian Zeng, Tricia Zwiefelhofer, Clinton R. Paden, Duncan MacCannell |
| EPI_ISL_1685730, EPI_ISL_1690245, EPI_ISL_1690269 | Aegis Sciences Corporation | Centers for Disease Control and Prevention Division of Viral Diseases, Pathogen Discovery | Dakota Howard, Dhwani Batra, Peter W. Cook, Kara Moser, Adrian Paskey, Jason Caravas, Benjamin Rambo-Martin, Shatavia Morrison, Christopher Gulvick, Scott Sammons, Yvette Unoarumhi, Darlene Wagner, Matthew Schmerer, Cyndi Clark, Patrick Campbell, Rob Case, Vikramsinha Ghorpade, Holly Houdeshell, Ola Kvalvaag, Dillon Nall, Ethan Sanders, Alec Vest, Shaun Westlund, Matthew Hardison, Clinton R. Paden, Duncan MacCannell |
| EPI_ISL_1690467 | Helix/Illumina | Centers for Disease Control and Prevention Division of Viral Diseases, Pathogen Discovery | Dakota Howard, Dhwani Batra, Peter W. Cook, Kara Moser, Adrian Paskey, Jason Caravas, Benjamin Rambo-Martin, Shatavia Morrison, Christopher Gulvick, Scott Sammons, Yvette Unoarumhi, Darlene Wagner, Matthew Schmerer, Eileen de Feo, Jan Antico, Christine Tran, Matthew Tolentino, Shannon Wickline, Kim Gietzen, Brad Sickler, Jingtao Liu, Eric Allen, Phil Febbo, Nicole L. Washington, Simon White, Geraint Levan, Kelly Schiabor Barrett, Elizabeth Cirulli, Alexandre Bolze, Ary Ascencio, Charlotte Rivera-Garcia, Ryan Cho, Jason Nguyen, Sherry Wang, Jimmy Ramirez, Tyler Cassens, Efren Sandoval, Magnus Isaksson, William Lee, David Becker, Marc Laurent, James Lu, Clinton R. Paden, Duncan MacCannell |
| EPI_ISL_1697491, EPI_ISL_1697908, EPI_ISL_1697919, EPI_ISL_1697963, EPI_ISL_1697964 | Lighthouse Lab in Cambridge | Wellcome Sanger Institute for the COVID-19 Genomics UK (COG-UK) Consortium | Rob Howes, The Lighthouse Lab in Cambridge and Alex Alderton, Roberto Amato, Jeffrey Barrett, Sonia Goncalves, Ewan Harrison, David K. Jackson, Ian Johnston, Dominic Kwiatkowski, Cordelia Langford, John Sillitoe on behalf of the Wellcome Sanger Institute COVID-19 Surveillance Team |
| EPI_ISL_1697977 | Lighthouse Lab in Milton Keynes | Wellcome Sanger Institute for the COVID-19 Genomics UK (COG-UK) Consortium | The Lighthouse Lab in Milton Keynes and Alex Alderton, Roberto Amato, Jeffrey Barrett, Sonia Goncalves, Ewan Harrison, David K. Jackson, Ian Johnston, Dominic Kwiatkowski, Cordelia Langford, John Sillitoe on behalf of the Wellcome Sanger Institute COVID-19 Surveillance Team |
| EPI_ISL_1697989 | Randox Laboratories | Wellcome Sanger Institute for the COVID-19 Genomics UK (COG-UK) Consortium | Randox Laboratories and Alex Alderton, Roberto Amato, Jeffrey Barrett, Sonia Goncalves, Ewan Harrison, David K. Jackson, Ian Johnston, Dominic Kwiatkowski, Cordelia Langford, John Sillitoe on behalf of the Wellcome Sanger Institute COVID-19 Surveillance Team |
| EPI_ISL_1698079, EPI_ISL_1698093, EPI_ISL_1698106, EPI_ISL_1698144 | Lighthouse Lab in Milton Keynes | Wellcome Sanger Institute for the COVID-19 Genomics UK (COG-UK) Consortium | The Lighthouse Lab in Milton Keynes and Alex Alderton, Roberto Amato, Jeffrey Barrett, Sonia Goncalves, Ewan Harrison, David K. Jackson, Ian Johnston, Dominic Kwiatkowski, Cordelia Langford, John Sillitoe on behalf of the Wellcome Sanger Institute COVID-19 Surveillance Team |
| EPI_ISL_1698227 | Lighthouse Lab in Alderley Park | Wellcome Sanger Institute for the COVID-19 Genomics UK (COG-UK) | Jacquelyn Wynn, Mairead Hyland, The Lighthouse Lab in Alderley Park and Alex Alderton, Roberto Amato, Jeffrey Barrett, Sonia Goncalves, Ewan Harrison, David K. Jackson, Ian Johnston, Dominic Kwiatkowski, Cordelia Langford, John Sillitoe on |

| | | | |
|---|---|---|---|
| | | Consortium | behalf of the Wellcome Sanger Institute COVID-19 Surveillance Team |
| EPI_ISL_1698273, EPI_ISL_1698274, EPI_ISL_1698283 | Lighthouse Lab in Cambridge | Wellcome Sanger Institute for the COVID-19 Genomics UK (COG-UK) Consortium | Rob Howes, The Lighthouse Lab in Cambridge and Alex Alderton, Roberto Amato, Jeffrey Barrett, Sonia Goncalves, Ewan Harrison, David K. Jackson, Ian Johnston, Dominic Kwiatkowski, Cordelia Langford, John Sillitoe on behalf of the Wellcome Sanger Institute COVID-19 Surveillance Team |
| EPI_ISL_1698319 | Lighthouse Lab in Alderley Park | Wellcome Sanger Institute for the COVID-19 Genomics UK (COG-UK) Consortium | Jacquelyn Wynn, Mairead Hyland, The Lighthouse Lab in Alderley Park and Alex Alderton, Roberto Amato, Jeffrey Barrett, Sonia Goncalves, Ewan Harrison, David K. Jackson, Ian Johnston, Dominic Kwiatkowski, Cordelia Langford, John Sillitoe on behalf of the Wellcome Sanger Institute COVID-19 Surveillance Team |
| EPI_ISL_1698322 | Lighthouse Lab in Milton Keynes | Wellcome Sanger Institute for the COVID-19 Genomics UK (COG-UK) Consortium | The Lighthouse Lab in Milton Keynes and Alex Alderton, Roberto Amato, Jeffrey Barrett, Sonia Goncalves, Ewan Harrison, David K. Jackson, Ian Johnston, Dominic Kwiatkowski, Cordelia Langford, John Sillitoe on behalf of the Wellcome Sanger Institute COVID-19 Surveillance Team |
| EPI_ISL_1698325 | Lighthouse Lab in Alderley Park | Wellcome Sanger Institute for the COVID-19 Genomics UK (COG-UK) Consortium | Jacquelyn Wynn, Mairead Hyland, The Lighthouse Lab in Alderley Park and Alex Alderton, Roberto Amato, Jeffrey Barrett, Sonia Goncalves, Ewan Harrison, David K. Jackson, Ian Johnston, Dominic Kwiatkowski, Cordelia Langford, John Sillitoe on behalf of the Wellcome Sanger Institute COVID-19 Surveillance Team |
| EPI_ISL_1698332, EPI_ISL_1698361, EPI_ISL_1698410 | Lighthouse Lab in Cambridge | Wellcome Sanger Institute for the COVID-19 Genomics UK (COG-UK) Consortium | Rob Howes, The Lighthouse Lab in Cambridge and Alex Alderton, Roberto Amato, Jeffrey Barrett, Sonia Goncalves, Ewan Harrison, David K. Jackson, Ian Johnston, Dominic Kwiatkowski, Cordelia Langford, John Sillitoe on behalf of the Wellcome Sanger Institute COVID-19 Surveillance Team |
| EPI_ISL_1698416 | Lighthouse Lab in Alderley Park | Wellcome Sanger Institute for the COVID-19 Genomics UK (COG-UK) Consortium | Jacquelyn Wynn, Mairead Hyland, The Lighthouse Lab in Alderley Park and Alex Alderton, Roberto Amato, Jeffrey Barrett, Sonia Goncalves, Ewan Harrison, David K. Jackson, Ian Johnston, Dominic Kwiatkowski, Cordelia Langford, John Sillitoe on behalf of the Wellcome Sanger Institute COVID-19 Surveillance Team |
| EPI_ISL_1698429, EPI_ISL_1698436 | Lighthouse Lab in Cambridge | Wellcome Sanger Institute for the COVID-19 Genomics UK (COG-UK) Consortium | Rob Howes, The Lighthouse Lab in Cambridge and Alex Alderton, Roberto Amato, Jeffrey Barrett, Sonia Goncalves, Ewan Harrison, David K. Jackson, Ian Johnston, Dominic Kwiatkowski, Cordelia Langford, John Sillitoe on behalf of the Wellcome Sanger Institute COVID-19 Surveillance Team |
| EPI_ISL_1698440 | Lighthouse Lab in Alderley Park | Wellcome Sanger Institute for the COVID-19 Genomics UK (COG-UK) Consortium | Jacquelyn Wynn, Mairead Hyland, The Lighthouse Lab in Alderley Park and Alex Alderton, Roberto Amato, Jeffrey Barrett, Sonia Goncalves, Ewan Harrison, David K. Jackson, Ian Johnston, Dominic Kwiatkowski, Cordelia Langford, John Sillitoe on behalf of the Wellcome Sanger Institute COVID-19 Surveillance Team |
| EPI_ISL_1698521 | Lighthouse Lab in Cambridge | Wellcome Sanger Institute for the COVID-19 Genomics UK (COG-UK) Consortium | Rob Howes, The Lighthouse Lab in Cambridge and Alex Alderton, Roberto Amato, Jeffrey Barrett, Sonia Goncalves, Ewan Harrison, David K. Jackson, Ian Johnston, Dominic Kwiatkowski, Cordelia Langford, John Sillitoe on behalf of the Wellcome Sanger Institute COVID-19 Surveillance Team |
| EPI_ISL_1698543, EPI_ISL_1699101 | Lighthouse Lab in Alderley Park | Wellcome Sanger Institute for the COVID-19 Genomics UK (COG-UK) Consortium | Jacquelyn Wynn, Mairead Hyland, The Lighthouse Lab in Alderley Park and Alex Alderton, Roberto Amato, Jeffrey Barrett, Sonia Goncalves, Ewan Harrison, David K. Jackson, Ian Johnston, Dominic Kwiatkowski, Cordelia Langford, John Sillitoe on behalf of the Wellcome Sanger Institute COVID-19 Surveillance Team |
| EPI_ISL_1699414, EPI_ISL_1699676, EPI_ISL_1699753, EPI_ISL_1699770, EPI_ISL_1699979 | Lighthouse Lab in Milton Keynes | Wellcome Sanger Institute for the COVID-19 Genomics UK (COG-UK) Consortium | The Lighthouse Lab in Milton Keynes and Alex Alderton, Roberto Amato, Jeffrey Barrett, Sonia Goncalves, Ewan Harrison, David K. Jackson, Ian Johnston, Dominic Kwiatkowski, Cordelia Langford, John Sillitoe on behalf of the Wellcome Sanger Institute COVID-19 Surveillance Team |
| EPI_ISL_1700038, EPI_ISL_1700236 | Lighthouse Lab in Alderley Park | Wellcome Sanger Institute for the COVID-19 Genomics UK (COG-UK) Consortium | Jacquelyn Wynn, Mairead Hyland, The Lighthouse Lab in Alderley Park and Alex Alderton, Roberto Amato, Jeffrey Barrett, Sonia Goncalves, Ewan Harrison, David K. Jackson, Ian Johnston, Dominic Kwiatkowski, Cordelia Langford, John Sillitoe on behalf of the Wellcome Sanger Institute COVID-19 Surveillance Team |
| EPI_ISL_1700313, EPI_ISL_1700494 | Randox Laboratories | Wellcome Sanger Institute for the COVID-19 Genomics UK (COG-UK) Consortium | Randox Laboratories and Alex Alderton, Roberto Amato, Jeffrey Barrett, Sonia Goncalves, Ewan Harrison, David K. Jackson, Ian Johnston, Dominic Kwiatkowski, Cordelia Langford, John Sillitoe on behalf of the Wellcome Sanger Institute COVID-19 Surveillance Team |
| EPI_ISL_1701966 | Aegis Sciences Corporation | Centers for Disease Control and Prevention Division of Viral Diseases, Pathogen Discovery | Dakota Howard, Dhwani Batra, Peter W. Cook, Kara Moser, Adrian Paskey, Jason Caravas, Benjamin Rambo-Martin, Shatavia Morrison, Christopher Gulvick, Scott Sammons, Yvette Unoarumhi, Darlene Wagner, Matthew Schmerer, Cyndi Clark, Patrick Campbell, Rob Case, Vikramsinha Ghorpade, Holly Houdeshell, Ola Kvalvaag, Dillon Nall, Ethan Sanders, Alec Vest, Shaun Westlund, Matthew Hardison, Clinton R. Paden, Duncan MacCannell |
| EPI_ISL_1703662, EPI_ISL_1703668, EPI_ISL_1703740, EPI_ISL_1703878, EPI_ISL_1704234, EPI_ISL_1704561, EPI_ISL_1704605, EPI_ISL_1704617, EPI_ISL_1704618, EPI_ISL_1704620, EPI_ISL_1704625, EPI_ISL_1704626, EPI_ISL_1704627, EPI_ISL_1704628, EPI_ISL_1704629, EPI_ISL_1704630, EPI_ISL_1704632, EPI_ISL_1704634, EPI_ISL_1704635, EPI_ISL_1704636, EPI_ISL_1704637, EPI_ISL_1704638, EPI_ISL_1704639 | | | |
| see above | ICMR-National Institute of Virology - INSACOG | NIV Influenza | Dr. Varsha Potdar |
| EPI_ISL_1704830, EPI_ISL_1704831, EPI_ISL_1704832, EPI_ISL_1704833, EPI_ISL_1704834, EPI_ISL_1704835, EPI_ISL_1704836, EPI_ISL_1704837, EPI_ISL_1704838, EPI_ISL_1704839, EPI_ISL_1704840, EPI_ISL_1704841, EPI_ISL_1704842 | | | |
| see above | National Public Health Laboratory, National Centre for Infectious Diseases | National Public Health Laboratory, National Centre for Infectious Diseases | Tze Minn Mak, Zhenyang Zhou, Grace Jie Yin Ngan, Royce Ang, Lin Cui, Raymond Tzer Pin Lin |
| EPI_ISL_1711803 | "OK Public Health Laboratory, Oklahoma State DOH" | Centers for Disease Control and Prevention Division of Viral Diseases, Pathogen Discovery | Mili Sheth, Sarah Nobles, Jasmine Padilla, Mark Burroughs, Shoshona Le, Katie Dillon, Peter Cook, Clinton R. Paden, Dhwani Batra, Krista Queen, Kristen Knipe, Dakota Howard, Yvette Unoarumhi, Darlene Wagner, Matthew Schmerer, Ben L. Rambo-Martin, Kristine Lacek, Sam Shepard, Alison Laufer Halpin, Dave Wentworth, Vivien Dugan, Suxiang Tong, Justin Lee |
| EPI_ISL_1715449, EPI_ISL_1715512, EPI_ISL_1715530, EPI_ISL_1715565 | UW Virology Lab | UW Virology Lab | Pavitra Roychoudhury, Hong Xie, Lasata Shrestha, Tien V. Nguyen, Shah Mohamed Bakhash, Michelle Lin, Noah R. Baker, Sean Ellis, Meei-Li Huang, Keith R Jerome, Alexander Greninger |
| EPI_ISL_1716736 | Laboratory of Immunohematology, Division of Hematology | Greek Genome Center, Biomedical Research Foundation of the Academy of Athens (BRFAA) | Emmanouil Athanasiadis, Ioannis Vatsellas, Theodoros Loupis, Katerina Zoi, Athanasia Mouzaki, Dimitrios Thanos |
| EPI_ISL_1718344, EPI_ISL_1718359 | Lighthouse Lab in Milton Keynes | Wellcome Sanger Institute for the COVID-19 Genomics UK (COG-UK) Consortium | The Lighthouse Lab in Milton Keynes and Alex Alderton, Roberto Amato, Jeffrey Barrett, Sonia Goncalves, Ewan Harrison, David K. Jackson, Ian Johnston, Dominic Kwiatkowski, Cordelia Langford, John Sillitoe on behalf of the Wellcome Sanger Institute COVID-19 Surveillance Team |
| EPI_ISL_1718381, EPI_ISL_1718501 | Lighthouse Lab in Alderley Park | Wellcome Sanger Institute for the COVID-19 Genomics UK (COG-UK) Consortium | Jacquelyn Wynn, Mairead Hyland, The Lighthouse Lab in Alderley Park and Alex Alderton, Roberto Amato, Jeffrey Barrett, Sonia Goncalves, Ewan Harrison, David K. Jackson, Ian Johnston, Dominic Kwiatkowski, Cordelia Langford, John Sillitoe on behalf of the Wellcome Sanger Institute COVID-19 Surveillance Team |
| EPI_ISL_1718509 | Lighthouse Lab in Milton Keynes | Wellcome Sanger Institute for the COVID-19 Genomics UK (COG-UK) Consortium | The Lighthouse Lab in Milton Keynes and Alex Alderton, Roberto Amato, Jeffrey Barrett, Sonia Goncalves, Ewan Harrison, David K. Jackson, Ian Johnston, Dominic Kwiatkowski, Cordelia Langford, John Sillitoe on behalf of the Wellcome Sanger Institute COVID-19 Surveillance Team |
| EPI_ISL_1718585, EPI_ISL_1718588, EPI_ISL_1718597, EPI_ISL_1718598, EPI_ISL_1718630 | Lighthouse Lab in Alderley Park | Wellcome Sanger Institute for the COVID-19 Genomics UK (COG-UK) Consortium | Jacquelyn Wynn, Mairead Hyland, The Lighthouse Lab in Alderley Park and Alex Alderton, Roberto Amato, Jeffrey Barrett, Sonia Goncalves, Ewan Harrison, David K. Jackson, Ian Johnston, Dominic Kwiatkowski, Cordelia Langford, John Sillitoe on behalf of the Wellcome Sanger Institute COVID-19 Surveillance Team |
| EPI_ISL_1718668 | Lighthouse Lab in Milton Keynes | Wellcome Sanger Institute for the COVID-19 Genomics UK (COG-UK) Consortium | The Lighthouse Lab in Milton Keynes and Alex Alderton, Roberto Amato, Jeffrey Barrett, Sonia Goncalves, Ewan Harrison, David K. Jackson, Ian Johnston, Dominic Kwiatkowski, Cordelia Langford, John Sillitoe on behalf of the Wellcome Sanger Institute COVID-19 Surveillance Team |
| EPI_ISL_1718683 | Lighthouse Lab in Alderley Park | Wellcome Sanger Institute for the COVID-19 Genomics UK (COG-UK) Consortium | Jacquelyn Wynn, Mairead Hyland, The Lighthouse Lab in Alderley Park and Alex Alderton, Roberto Amato, Jeffrey Barrett, Sonia Goncalves, Ewan Harrison, David K. Jackson, Ian Johnston, Dominic Kwiatkowski, Cordelia Langford, John Sillitoe on behalf of the Wellcome Sanger Institute COVID-19 Surveillance Team |
| EPI_ISL_1718691 | Lighthouse Lab in Milton Keynes | Wellcome Sanger Institute for the COVID-19 Genomics UK (COG-UK) Consortium | The Lighthouse Lab in Milton Keynes and Alex Alderton, Roberto Amato, Jeffrey Barrett, Sonia Goncalves, Ewan Harrison, David K. Jackson, Ian Johnston, Dominic Kwiatkowski, Cordelia Langford, John Sillitoe on behalf of the Wellcome Sanger Institute COVID-19 Surveillance Team |
| EPI_ISL_1718700 | Randox Laboratories | Wellcome Sanger Institute for the COVID-19 Genomics UK (COG-UK) Consortium | Randox Laboratories and Alex Alderton, Roberto Amato, Jeffrey Barrett, Sonia Goncalves, Ewan Harrison, David K. Jackson, Ian Johnston, Dominic Kwiatkowski, Cordelia Langford, John Sillitoe on behalf of the Wellcome Sanger Institute COVID-19 Surveillance Team |
| EPI_ISL_1718721, EPI_ISL_1718772, EPI_ISL_1718882 | Lighthouse Lab in Alderley Park | Wellcome Sanger Institute for the COVID-19 Genomics UK (COG-UK) Consortium | Jacquelyn Wynn, Mairead Hyland, The Lighthouse Lab in Alderley Park and Alex Alderton, Roberto Amato, Jeffrey Barrett, Sonia Goncalves, Ewan Harrison, David K. Jackson, Ian Johnston, Dominic Kwiatkowski, Cordelia Langford, John Sillitoe on behalf of the Wellcome Sanger Institute COVID-19 Surveillance Team |
| EPI_ISL_1718934, EPI_ISL_1718935 | Lighthouse Lab in Milton Keynes | Wellcome Sanger Institute for the COVID-19 Genomics UK (COG-UK) Consortium | The Lighthouse Lab in Milton Keynes and Alex Alderton, Roberto Amato, Jeffrey Barrett, Sonia Goncalves, Ewan Harrison, David K. Jackson, Ian Johnston, Dominic Kwiatkowski, Cordelia Langford, John Sillitoe on behalf of the Wellcome Sanger Institute COVID-19 Surveillance Team |
| EPI_ISL_1718942 | Lighthouse Lab in Cambridge | Wellcome Sanger Institute for the COVID-19 Genomics UK (COG-UK) Consortium | Rob Howes, The Lighthouse Lab in Cambridge and Alex Alderton, Roberto Amato, Jeffrey Barrett, Sonia Goncalves, Ewan Harrison, David K. Jackson, Ian Johnston, Dominic Kwiatkowski, Cordelia Langford, John Sillitoe on behalf of the Wellcome Sanger Institute COVID-19 Surveillance Team |
| EPI_ISL_1718951 | Lighthouse Lab in Milton Keynes | Wellcome Sanger Institute for the COVID-19 Genomics UK (COG-UK) Consortium | The Lighthouse Lab in Milton Keynes and Alex Alderton, Roberto Amato, Jeffrey Barrett, Sonia Goncalves, Ewan Harrison, David K. Jackson, Ian Johnston, Dominic Kwiatkowski, Cordelia Langford, John Sillitoe on behalf of the Wellcome Sanger Institute COVID-19 Surveillance Team |
| EPI_ISL_1718959 | Lighthouse Lab in Cambridge | Wellcome Sanger Institute for the COVID-19 Genomics UK (COG-UK) Consortium | Rob Howes, The Lighthouse Lab in Cambridge and Alex Alderton, Roberto Amato, Jeffrey Barrett, Sonia Goncalves, Ewan Harrison, David K. Jackson, Ian Johnston, Dominic Kwiatkowski, Cordelia Langford, John Sillitoe on behalf of the Wellcome Sanger Institute COVID-19 Surveillance Team |
| EPI_ISL_1718987 | Lighthouse Lab in Milton Keynes | Wellcome Sanger Institute for the COVID-19 Genomics UK (COG-UK) Consortium | The Lighthouse Lab in Milton Keynes and Alex Alderton, Roberto Amato, Jeffrey Barrett, Sonia Goncalves, Ewan Harrison, David K. Jackson, Ian Johnston, Dominic Kwiatkowski, Cordelia Langford, John Sillitoe on behalf of the Wellcome Sanger Institute COVID-19 Surveillance Team |
| EPI_ISL_1718989, EPI_ISL_1718997 | Lighthouse Lab in Cambridge | Wellcome Sanger Institute for the COVID-19 Genomics UK (COG-UK) Consortium | Rob Howes, The Lighthouse Lab in Cambridge and Alex Alderton, Roberto Amato, Jeffrey Barrett, Sonia Goncalves, Ewan Harrison, David K. Jackson, Ian Johnston, Dominic Kwiatkowski, Cordelia Langford, John Sillitoe on behalf of the Wellcome Sanger Institute COVID-19 Surveillance Team |
| EPI_ISL_1719015, EPI_ISL_1719018 | Lighthouse Lab in Milton Keynes | Wellcome Sanger Institute for the COVID-19 Genomics UK (COG-UK) Consortium | The Lighthouse Lab in Milton Keynes and Alex Alderton, Roberto Amato, Jeffrey Barrett, Sonia Goncalves, Ewan Harrison, David K. Jackson, Ian Johnston, Dominic Kwiatkowski, Cordelia Langford, John Sillitoe on behalf of the Wellcome Sanger Institute COVID-19 Surveillance Team |
| EPI_ISL_1719027, EPI_ISL_1719042 | Lighthouse Lab in Cambridge | Wellcome Sanger Institute for the COVID-19 Genomics UK (COG-UK) Consortium | Rob Howes, The Lighthouse Lab in Cambridge and Alex Alderton, Roberto Amato, Jeffrey Barrett, Sonia Goncalves, Ewan Harrison, David K. Jackson, Ian Johnston, Dominic Kwiatkowski, Cordelia Langford, John Sillitoe on behalf of the Wellcome Sanger Institute COVID-19 Surveillance Team |
| EPI_ISL_1719056 | Lighthouse Lab in Milton Keynes | Wellcome Sanger Institute for the COVID-19 Genomics UK (COG-UK) Consortium | The Lighthouse Lab in Milton Keynes and Alex Alderton, Roberto Amato, Jeffrey Barrett, Sonia Goncalves, Ewan Harrison, David K. Jackson, Ian Johnston, Dominic Kwiatkowski, Cordelia Langford, John Sillitoe on behalf of the Wellcome Sanger Institute COVID-19 Surveillance Team |
| EPI_ISL_1719062 | Lighthouse Lab in Cambridge | Wellcome Sanger Institute for the COVID-19 Genomics UK (COG-UK) Consortium | Rob Howes, The Lighthouse Lab in Cambridge and Alex Alderton, Roberto Amato, Jeffrey Barrett, Sonia Goncalves, Ewan Harrison, David K. Jackson, Ian Johnston, Dominic Kwiatkowski, Cordelia Langford, John Sillitoe on behalf of the Wellcome Sanger Institute COVID-19 Surveillance Team |
| EPI_ISL_1719079 | Lighthouse Lab in Milton Keynes | Wellcome Sanger Institute for the COVID-19 Genomics UK (COG-UK) Consortium | The Lighthouse Lab in Milton Keynes and Alex Alderton, Roberto Amato, Jeffrey Barrett, Sonia Goncalves, Ewan Harrison, David K. Jackson, Ian Johnston, Dominic Kwiatkowski, Cordelia Langford, John Sillitoe on behalf of the Wellcome Sanger Institute COVID-19 Surveillance Team |

| | | | |
|---|---|---|---|
| EPI_ISL_1719099, EPI_ISL_1719134, EPI_ISL_1719142 | Lighthouse Lab in Cambridge | Wellcome Sanger Institute for the COVID-19 Genomics UK (COG-UK) Consortium | Rob Howes, The Lighthouse Lab in Cambridge and Alex Alderton, Roberto Amato, Jeffrey Barrett, Sonia Goncalves, Ewan Harrison, David K. Jackson, Ian Johnston, Dominic Kwiatkowski, Cordelia Langford, John Sillitoe on behalf of the Wellcome Sanger Institute COVID-19 Surveillance Team |
| EPI_ISL_1719162, EPI_ISL_1719212 | Lighthouse Lab in Milton Keynes | Wellcome Sanger Institute for the COVID-19 Genomics UK (COG-UK) Consortium | The Lighthouse Lab in Milton Keynes and Alex Alderton, Roberto Amato, Jeffrey Barrett, Sonia Goncalves, Ewan Harrison, David K. Jackson, Ian Johnston, Dominic Kwiatkowski, Cordelia Langford, John Sillitoe on behalf of the Wellcome Sanger Institute COVID-19 Surveillance Team |
| EPI_ISL_1719573 | Lighthouse Lab in Glasgow | Wellcome Sanger Institute for the COVID-19 Genomics UK (COG-UK) Consortium | Harper VanSteenhouse, Yumi Kasai, David Gray, Carol Clugston, Anna Dominiczak and Alex Alderton, Roberto Amato, Jeffrey Barrett, Sonia Goncalves, Ewan Harrison, David K. Jackson, Ian Johnston, Dominic Kwiatkowski, Cordelia Langford, John Sillitoe on behalf of the Wellcome Sanger Institute COVID-19 Surveillance Team |
| EPI_ISL_1719584 | Lighthouse Lab in Cambridge | Wellcome Sanger Institute for the COVID-19 Genomics UK (COG-UK) Consortium | Rob Howes, The Lighthouse Lab in Cambridge and Alex Alderton, Roberto Amato, Jeffrey Barrett, Sonia Goncalves, Ewan Harrison, David K. Jackson, Ian Johnston, Dominic Kwiatkowski, Cordelia Langford, John Sillitoe on behalf of the Wellcome Sanger Institute COVID-19 Surveillance Team |
| EPI_ISL_1719616 | Lighthouse Lab in Alderley Park | Wellcome Sanger Institute for the COVID-19 Genomics UK (COG-UK) Consortium | Jacquelyn Wynn, Mairead Hyland, The Lighthouse Lab in Alderley Park and Alex Alderton, Roberto Amato, Jeffrey Barrett, Sonia Goncalves, Ewan Harrison, David K. Jackson, Ian Johnston, Dominic Kwiatkowski, Cordelia Langford, John Sillitoe on behalf of the Wellcome Sanger Institute COVID-19 Surveillance Team |
| EPI_ISL_1719618 | Lighthouse Lab in Cambridge | Wellcome Sanger Institute for the COVID-19 Genomics UK (COG-UK) Consortium | Rob Howes, The Lighthouse Lab in Cambridge and Alex Alderton, Roberto Amato, Jeffrey Barrett, Sonia Goncalves, Ewan Harrison, David K. Jackson, Ian Johnston, Dominic Kwiatkowski, Cordelia Langford, John Sillitoe on behalf of the Wellcome Sanger Institute COVID-19 Surveillance Team |
| EPI_ISL_1719728 | Lighthouse Lab in Glasgow | Wellcome Sanger Institute for the COVID-19 Genomics UK (COG-UK) Consortium | Harper VanSteenhouse, Yumi Kasai, David Gray, Carol Clugston, Anna Dominiczak and Alex Alderton, Roberto Amato, Jeffrey Barrett, Sonia Goncalves, Ewan Harrison, David K. Jackson, Ian Johnston, Dominic Kwiatkowski, Cordelia Langford, John Sillitoe on behalf of the Wellcome Sanger Institute COVID-19 Surveillance Team |
| EPI_ISL_1719736 | Lighthouse Lab in Cambridge | Wellcome Sanger Institute for the COVID-19 Genomics UK (COG-UK) Consortium | Rob Howes, The Lighthouse Lab in Cambridge and Alex Alderton, Roberto Amato, Jeffrey Barrett, Sonia Goncalves, Ewan Harrison, David K. Jackson, Ian Johnston, Dominic Kwiatkowski, Cordelia Langford, John Sillitoe on behalf of the Wellcome Sanger Institute COVID-19 Surveillance Team |
| EPI_ISL_1719885, EPI_ISL_1719886, EPI_ISL_1719887, EPI_ISL_1719888, EPI_ISL_1719889, EPI_ISL_1719890, EPI_ISL_1719891, EPI_ISL_1719892, EPI_ISL_1719893, EPI_ISL_1719894, EPI_ISL_1719895, EPI_ISL_1719896, EPI_ISL_1719897, EPI_ISL_1719898, EPI_ISL_1719899, EPI_ISL_1719900 | | | |
| see above | National Public Health Laboratory, National Centre for Infectious Diseases | National Public Health Laboratory, National Centre for Infectious Diseases | Tze Minn Mak, Zhenyang Zhou, Grace Jie Yin Ngan, Royce Ang, Lin Cui, Raymond Tzer Pin Lin |
| EPI_ISL_1722241 | IMD - MVZ Labor Martinsried | Robert Koch Institute | unknown |
| EPI_ISL_1727471 | Limbach - MVZ Humangenetik Ulm | Robert Koch Institute | unknown |
| EPI_ISL_1728331 | SYNLAB MVZ Leverkusen | Robert Koch Institute | unknown |
| EPI_ISL_1730064, EPI_ISL_1730071, EPI_ISL_1730072 | Department of Pathology, University of Cambridge | COVID-19 Genomics UK (COG-UK) Consortium | Aminu S. Jahun, Yasmin Chaudhry, Iliana Georgana, Myra Hosmillo, Rhys Izuagbe, William L. Hamilton, Martin D. Curran, Surendra Parmar, Ian Goodfellow |
| EPI_ISL_1730346 | University College London, Great Ormond Street Hospital for Children NHS Foundation Trust, Imperial College Healthcare NHS Trust | COVID-19 Genomics UK (COG-UK) Consortium | Sergi Castellano, Rachel Williams, Mark Kristiansen, Paola Resende Silva, Sunando Roy, Tony Brooks, Helena Tutill, Paola Niola, Patricia Dyal, Charlotte Williams, Leysa Forrest, Yasmin Panchbhaya, Jacqueline Findlay, Samuel Weeks, Julianne Brown, Kathryn Harris, Paul Randell, James Price, Alison Holmes, Judith Breuer |
| EPI_ISL_1730407 | Northumbria University / South Tees Hospitals NHS Foundation Trust / North Cumbria Integrated Care NHS Foundation Trust / North Tees and Hartlepool NHS Foundation Trust / Newcastle Hospitals NHS Foundation Trust | COVID-19 Genomics UK (COG-UK) Consortium | Darren L Smith,Andrew Nelson,Matthew Bashton,Greg R Young,Joshua Loh,John Allan,Mohammad A Tariq,Giles S Holt,Gary Black,Wen C Yew,Lynn Dover,Paul Baker,Steve Liggett,Sarah Essex,Jane Greenaway,Debra Padgett,Clive Graham,Garren Scott,Edward Barton,Emma Swindells,Brendan Payne,Jennifer Collins,Yusri Taha,Gary Eltringham |
| EPI_ISL_1730635, EPI_ISL_1730640, EPI_ISL_1730648, EPI_ISL_1730650, EPI_ISL_1730652, EPI_ISL_1730654, EPI_ISL_1730662, EPI_ISL_1730804 | Originating lab: Wales Specialist Virology Centre Sequencing lab: Pathogen Genomics Unit | Public Health Wales Microbiology Cardiff Wales Specialist Virology Centre | Catherine Moore, Johnathan Evans, Laura Gifford, Malorie Perry, Simon Cottrell, Angela Marchbank, Alec Birchley, Alexander Adams, Amy Gaskin, Bree Gatica-Wilcox, Jason Coombes, Joel Southgate, Lauren Gilbert, Lee Graham, Nicole Pacchiarini, Sara Kumziene-Summerhayes, Sarah Taylor, Sophie Jones, Sara Rey, Matthew Bull, Joanne Watkins, Sally Corden, Tom Connor |
| EPI_ISL_1730910, EPI_ISL_1730922 | Centre for Enzyme Innovation, University of Portsmouth / Translational Research Laboratory, Portsmouth Hospitals NHS Trust | COVID-19 Genomics UK (COG-UK) Consortium | Angela Beckett,Salman Goudarzi,Christopher Fearn,Kate Cook,Katie Loveson,Sharon Glaysher,Scott Elliott,Samuel Robson |
| EPI_ISL_1731019 | Virology Department, Sheffield Teaching Hospitals NHS Foundation Trust/Department of Infection, Immunity and Cardiovascular Disease, The Medical School, University of Sheffield | COVID-19 Genomics UK (COG-UK) Consortium | Thushan de Silva, Matthew Parker, Nikki Smith, Adri Angyal, Rebecca Brown, Luke Green, Rachel Tucker, Paul Parsons, Danielle Groves, Katie Johnson, Laura Carrilero, Alex Keeley, Dave Partridge, Matthew Wyles, Benjamin Lindsey, Mehmet Yavuz, Mohammad Raza, Cariad Evans |
| EPI_ISL_1731106, EPI_ISL_1731198, EPI_ISL_1731252, EPI_ISL_1731330 | National Virus Reference Laboratory | National Virus Reference Laboratory | Zoe Yandle, Charlene Bennett, Gabriel Gonzalez, Michael Carr, Jonathan Dean, Cillian F De Gascun |
| EPI_ISL_1731752, EPI_ISL_1731753, EPI_ISL_1731754, EPI_ISL_1731755 | State Virus Research and Diagnostic Laboratory (VRDL), AIIMS Raipur | State Virus Research and Diagnostic Laboratory (VRDL), AIIMS Raipur | Pushpendra Singh, Kuldeep Sharma, Priyanka Singh, Somya Sharma, Sanjay Singh Negi, Anudita Bhargava |
| EPI_ISL_1735500 | Helix/Illumina | Centers for Disease Control and Prevention Division of Viral Diseases, Pathogen Discovery | Dakota Howard, Dhwani Batra, Peter W. Cook, Kara Moser, Adrian Paskey, Jason Caravas, Benjamin Rambo-Martin, Shatavia Morrison, Christopher Gulvick, Scott Sammons, Yvette Unoarumhi, Darlene Wagner, Matthew Schmerer, Eileen de Feo, Jan Antico, Christine Tran, Matthew Tolentino, Shannon Wickline, Kim Gietzen, Brad Sickler, Jingtao Liu, Eric Allen, Phil Febbo, Nicole L. Washington, Simon White, Geraint Levan, Kelly Schiabor Barrett, Elizabeth Cirulli, Alexandre Bolze, Ary Ascencio, Charlotte Rivera-Garcia, Ryan Cho, Jason Nguyen, Sherry Wang, Jimmy Ramirez, Tyler Cassens, Efren Sandoval, Magnus Isaksson, William Lee, David Becker, Marc Laurent, James Lu, Clinton R. Paden, Duncan MacCannell |
| EPI_ISL_1736936, EPI_ISL_1737880 | Aegis Sciences Corporation | Centers for Disease Control and Prevention Division of Viral Diseases, Pathogen Discovery | Dakota Howard, Dhwani Batra, Peter W. Cook, Kara Moser, Adrian Paskey, Jason Caravas, Benjamin Rambo-Martin, Shatavia Morrison, Christopher Gulvick, Scott Sammons, Yvette Unoarumhi, Darlene Wagner, Matthew Schmerer, Cyndi Clark, Patrick Campbell, Rob Case, Vikramsinha Ghorpade, Holly Houdeshell, Ola Kvalvaag, Dillon Nall, Ethan Sanders, Alec Vest, Shaun Westlund, Matthew Hardison, Clinton R. Paden, Duncan MacCannell |
| EPI_ISL_1738201 | Quest Diagnostics Incorporated | Centers for Disease Control and Prevention Division of Viral Diseases, Pathogen Discovery | Dakota Howard, Dhwani Batra, Peter W. Cook, Kara Moser, Adrian Paskey, Jason Caravas, Benjamin Rambo-Martin, Shatavia Morrison, Christopher Gulvick, Scott Sammons, Yvette Unoarumhi, Darlene Wagner, Matthew Schmerer, S. H. Rosenthal, A. Gerasimova, R. M. Kagan, B. Anderson, M. Hua, Y. Liu, L.E. Bernstein, K.E. Livingston, A. Perez, I. A. Shlyakhter, R. V. Rolando, R. Owen, P. Tanpaiboon, F. Lacbawan, Clinton R. Paden, Duncan MacCannell |
| EPI_ISL_1740551 | Randox Laboratories | Wellcome Sanger Institute for the COVID-19 Genomics UK (COG-UK) Consortium | Randox Laboratories and Alex Alderton, Roberto Amato, Jeffrey Barrett, Sonia Goncalves, Ewan Harrison, David K. Jackson, Ian Johnston, Dominic Kwiatkowski, Cordelia Langford, John Sillitoe on behalf of the Wellcome Sanger Institute COVID-19 Surveillance Team |
| EPI_ISL_1740580 | Lighthouse Lab in Cambridge | Wellcome Sanger Institute for the COVID-19 Genomics UK (COG-UK) Consortium | Rob Howes, The Lighthouse Lab in Cambridge and Alex Alderton, Roberto Amato, Jeffrey Barrett, Sonia Goncalves, Ewan Harrison, David K. Jackson, Ian Johnston, Dominic Kwiatkowski, Cordelia Langford, John Sillitoe on behalf of the Wellcome Sanger Institute COVID-19 Surveillance Team |
| EPI_ISL_1740600 | Lighthouse Lab in Alderley Park | Wellcome Sanger Institute for the COVID-19 Genomics UK (COG-UK) Consortium | Jacquelyn Wynn, Mairead Hyland, The Lighthouse Lab in Alderley Park and Alex Alderton, Roberto Amato, Jeffrey Barrett, Sonia Goncalves, Ewan Harrison, David K. Jackson, Ian Johnston, Dominic Kwiatkowski, Cordelia Langford, John Sillitoe on behalf of the Wellcome Sanger Institute COVID-19 Surveillance Team |
| EPI_ISL_1740611 | Randox Laboratories | Wellcome Sanger Institute for the COVID-19 Genomics UK (COG-UK) Consortium | Randox Laboratories and Alex Alderton, Roberto Amato, Jeffrey Barrett, Sonia Goncalves, Ewan Harrison, David K. Jackson, Ian Johnston, Dominic Kwiatkowski, Cordelia Langford, John Sillitoe on behalf of the Wellcome Sanger Institute COVID-19 Surveillance Team |
| EPI_ISL_1740616 | Lighthouse Lab in Alderley Park | Wellcome Sanger Institute for the COVID-19 Genomics UK (COG-UK) Consortium | Jacquelyn Wynn, Mairead Hyland, The Lighthouse Lab in Alderley Park and Alex Alderton, Roberto Amato, Jeffrey Barrett, Sonia Goncalves, Ewan Harrison, David K. Jackson, Ian Johnston, Dominic Kwiatkowski, Cordelia Langford, John Sillitoe on behalf of the Wellcome Sanger Institute COVID-19 Surveillance Team |
| EPI_ISL_1740633 | Randox Laboratories | Wellcome Sanger Institute for the COVID-19 Genomics UK (COG-UK) Consortium | Randox Laboratories and Alex Alderton, Roberto Amato, Jeffrey Barrett, Sonia Goncalves, Ewan Harrison, David K. Jackson, Ian Johnston, Dominic Kwiatkowski, Cordelia Langford, John Sillitoe on behalf of the Wellcome Sanger Institute COVID-19 Surveillance Team |
| EPI_ISL_1740654 | Lighthouse Lab in Alderley Park | Wellcome Sanger Institute for the COVID-19 Genomics UK (COG-UK) Consortium | Jacquelyn Wynn, Mairead Hyland, The Lighthouse Lab in Alderley Park and Alex Alderton, Roberto Amato, Jeffrey Barrett, Sonia Goncalves, Ewan Harrison, David K. Jackson, Ian Johnston, Dominic Kwiatkowski, Cordelia Langford, John Sillitoe on behalf of the Wellcome Sanger Institute COVID-19 Surveillance Team |
| EPI_ISL_1740691 | Randox Laboratories | Wellcome Sanger Institute for the COVID-19 Genomics UK (COG-UK) Consortium | Randox Laboratories and Alex Alderton, Roberto Amato, Jeffrey Barrett, Sonia Goncalves, Ewan Harrison, David K. Jackson, Ian Johnston, Dominic Kwiatkowski, Cordelia Langford, John Sillitoe on behalf of the Wellcome Sanger Institute COVID-19 Surveillance Team |
| EPI_ISL_1740707 | Lighthouse Lab in Alderley Park | Wellcome Sanger Institute for the COVID-19 Genomics UK (COG-UK) Consortium | Jacquelyn Wynn, Mairead Hyland, The Lighthouse Lab in Alderley Park and Alex Alderton, Roberto Amato, Jeffrey Barrett, Sonia Goncalves, Ewan Harrison, David K. Jackson, Ian Johnston, Dominic Kwiatkowski, Cordelia Langford, John Sillitoe on behalf of the Wellcome Sanger Institute COVID-19 Surveillance Team |
| EPI_ISL_1740738 | Lighthouse Lab in Glasgow | Wellcome Sanger Institute for the COVID-19 Genomics UK (COG-UK) Consortium | Harper VanSteenhouse, Yumi Kasai, David Gray, Carol Clugston, Anna Dominiczak and Alex Alderton, Roberto Amato, Jeffrey Barrett, Sonia Goncalves, Ewan Harrison, David K. Jackson, Ian Johnston, Dominic Kwiatkowski, Cordelia Langford, John Sillitoe on behalf of the Wellcome Sanger Institute COVID-19 Surveillance Team |
| EPI_ISL_1740750 | Lighthouse Lab in Alderley Park | Wellcome Sanger Institute for the COVID-19 Genomics UK (COG-UK) Consortium | Jacquelyn Wynn, Mairead Hyland, The Lighthouse Lab in Alderley Park and Alex Alderton, Roberto Amato, Jeffrey Barrett, Sonia Goncalves, Ewan Harrison, David K. Jackson, Ian Johnston, Dominic Kwiatkowski, Cordelia Langford, John Sillitoe on behalf of the Wellcome Sanger Institute COVID-19 Surveillance Team |
| EPI_ISL_1740754 | Randox Laboratories | Wellcome Sanger Institute for the COVID-19 Genomics UK (COG-UK) Consortium | Randox Laboratories and Alex Alderton, Roberto Amato, Jeffrey Barrett, Sonia Goncalves, Ewan Harrison, David K. Jackson, Ian Johnston, Dominic Kwiatkowski, Cordelia Langford, John Sillitoe on behalf of the Wellcome Sanger Institute COVID-19 Surveillance Team |
| EPI_ISL_1740763 | Lighthouse Lab in Alderley Park | Wellcome Sanger Institute for the COVID-19 Genomics UK (COG-UK) Consortium | Jacquelyn Wynn, Mairead Hyland, The Lighthouse Lab in Alderley Park and Alex Alderton, Roberto Amato, Jeffrey Barrett, Sonia Goncalves, Ewan Harrison, David K. Jackson, Ian Johnston, Dominic Kwiatkowski, Cordelia Langford, John Sillitoe on behalf of the Wellcome Sanger Institute COVID-19 Surveillance Team |
| EPI_ISL_1740766 | Randox Laboratories | Wellcome Sanger Institute for the COVID-19 Genomics UK (COG-UK) Consortium | Randox Laboratories and Alex Alderton, Roberto Amato, Jeffrey Barrett, Sonia Goncalves, Ewan Harrison, David K. Jackson, Ian Johnston, Dominic Kwiatkowski, Cordelia Langford, John Sillitoe on behalf of the Wellcome Sanger Institute COVID-19 Surveillance Team |
| EPI_ISL_1740778 | Lighthouse Lab in Alderley Park | Wellcome Sanger Institute for the COVID-19 Genomics UK (COG-UK) Consortium | Jacquelyn Wynn, Mairead Hyland, The Lighthouse Lab in Alderley Park and Alex Alderton, Roberto Amato, Jeffrey Barrett, Sonia Goncalves, Ewan Harrison, David K. Jackson, Ian Johnston, Dominic Kwiatkowski, Cordelia Langford, John Sillitoe on behalf of the Wellcome Sanger Institute COVID-19 Surveillance Team |

| | | | |
|---|---|---|---|
| EPI_ISL_1740781 | Lighthouse Lab in Cambridge | Wellcome Sanger Institute for the COVID-19 Genomics UK (COG-UK) Consortium | Rob Howes, The Lighthouse Lab in Cambridge and Alex Alderton, Roberto Amato, Jeffrey Barrett, Sonia Goncalves, Ewan Harrison, David K. Jackson, Ian Johnston, Dominic Kwiatkowski, Cordelia Langford, John Sillitoe on behalf of the Wellcome Sanger Institute COVID-19 Surveillance Team |
| EPI_ISL_1740819 | Lighthouse Lab in Alderley Park | Wellcome Sanger Institute for the COVID-19 Genomics UK (COG-UK) Consortium | Jacquelyn Wynn, Mairead Hyland, The Lighthouse Lab in Alderley Park and Alex Alderton, Roberto Amato, Jeffrey Barrett, Sonia Goncalves, Ewan Harrison, David K. Jackson, Ian Johnston, Dominic Kwiatkowski, Cordelia Langford, John Sillitoe on behalf of the Wellcome Sanger Institute COVID-19 Surveillance Team |
| EPI_ISL_1740858, EPI_ISL_1740882 | Lighthouse Lab in Milton Keynes | Wellcome Sanger Institute for the COVID-19 Genomics UK (COG-UK) Consortium | The Lighthouse Lab in Milton Keynes and Alex Alderton, Roberto Amato, Jeffrey Barrett, Sonia Goncalves, Ewan Harrison, David K. Jackson, Ian Johnston, Dominic Kwiatkowski, Cordelia Langford, John Sillitoe on behalf of the Wellcome Sanger Institute COVID-19 Surveillance Team |
| EPI_ISL_1741081, EPI_ISL_1741129 | Lighthouse Lab in Glasgow | Wellcome Sanger Institute for the COVID-19 Genomics UK (COG-UK) Consortium | Harper VanSteenhouse, Yumi Kasai, David Gray, Carol Clugston, Anna Dominiczak and Alex Alderton, Roberto Amato, Jeffrey Barrett, Sonia Goncalves, Ewan Harrison, David K. Jackson, Ian Johnston, Dominic Kwiatkowski, Cordelia Langford, John Sillitoe on behalf of the Wellcome Sanger Institute COVID-19 Surveillance Team |
| EPI_ISL_1741186 | Lighthouse Lab in Alderley Park | Wellcome Sanger Institute for the COVID-19 Genomics UK (COG-UK) Consortium | Jacquelyn Wynn, Mairead Hyland, The Lighthouse Lab in Alderley Park and Alex Alderton, Roberto Amato, Jeffrey Barrett, Sonia Goncalves, Ewan Harrison, David K. Jackson, Ian Johnston, Dominic Kwiatkowski, Cordelia Langford, John Sillitoe on behalf of the Wellcome Sanger Institute COVID-19 Surveillance Team |
| EPI_ISL_1741213, EPI_ISL_1741259 | Lighthouse Lab in Glasgow | Wellcome Sanger Institute for the COVID-19 Genomics UK (COG-UK) Consortium | Harper VanSteenhouse, Yumi Kasai, David Gray, Carol Clugston, Anna Dominiczak and Alex Alderton, Roberto Amato, Jeffrey Barrett, Sonia Goncalves, Ewan Harrison, David K. Jackson, Ian Johnston, Dominic Kwiatkowski, Cordelia Langford, John Sillitoe on behalf of the Wellcome Sanger Institute COVID-19 Surveillance Team |
| EPI_ISL_1741325 | Randox Laboratories | Wellcome Sanger Institute for the COVID-19 Genomics UK (COG-UK) Consortium | Randox Laboratories and Alex Alderton, Roberto Amato, Jeffrey Barrett, Sonia Goncalves, Ewan Harrison, David K. Jackson, Ian Johnston, Dominic Kwiatkowski, Cordelia Langford, John Sillitoe on behalf of the Wellcome Sanger Institute COVID-19 Surveillance Team |
| EPI_ISL_1741398 | Lighthouse Lab in Milton Keynes | Wellcome Sanger Institute for the COVID-19 Genomics UK (COG-UK) Consortium | The Lighthouse Lab in Milton Keynes and Alex Alderton, Roberto Amato, Jeffrey Barrett, Sonia Goncalves, Ewan Harrison, David K. Jackson, Ian Johnston, Dominic Kwiatkowski, Cordelia Langford, John Sillitoe on behalf of the Wellcome Sanger Institute COVID-19 Surveillance Team |
| EPI_ISL_1741623, EPI_ISL_1741651, EPI_ISL_1741677, EPI_ISL_1741762, EPI_ISL_1741838, EPI_ISL_1741909 | Lighthouse Lab in Glasgow | Wellcome Sanger Institute for the COVID-19 Genomics UK (COG-UK) Consortium | Harper VanSteenhouse, Yumi Kasai, David Gray, Carol Clugston, Anna Dominiczak and Alex Alderton, Roberto Amato, Jeffrey Barrett, Sonia Goncalves, Ewan Harrison, David K. Jackson, Ian Johnston, Dominic Kwiatkowski, Cordelia Langford, John Sillitoe on behalf of the Wellcome Sanger Institute COVID-19 Surveillance Team |
| EPI_ISL_1742834 | J.W. Ruby Memorial Hospital | WVU and Marshall University Combined Genomics Core Facilities | James Denvir, Peter Stoilov, Peter Perrotta, Wesley Kimble, Ryan Percifield |
| EPI_ISL_1743907, EPI_ISL_1744188, EPI_ISL_1744257 | Broad Institute Clinical Research Sequencing Platform | Infectious Disease Program, Broad Institute of Harvard and MIT | Siddle,K.J., Adams,G., Pearlman,L., Gladden-Young,A., Vicente,G., Blumenstiel,B., DeFelice,M., Lee,M., McGovern,S., Lagerborg,K., Rudy,M., DeRuff,K., Carter,A., Normandin,E., Bauer,M., Reilly,S., Tomkins-Tinch,C., Loreth,C., Chaluvadi,S., Meldrim,J., Granger,B., Lemieux,J.E., Birren,B.W., Sabeti,P.C., Larkin,K., Dodge,S., Lennon,N., Madoff,L., Brown,C., Gallagher,G., Smole,S., Park,D.J., Gabriel,S., and MacInnis,B.L. |
| EPI_ISL_1745199 | Platform BIS UZA/UAntwerpen | Labo Klinische Biologie, UZA | Marie Le Mercier, Jasmine Coppens, Basil Britto Xavier, Christine Lammens, Veerle Matheeussen, Herman Goossens |
| EPI_ISL_1750690 | Viollier AG | Department of Biosystems Science and Engineering, ETH Zürich | Chaoran Chen, Sarah Nadeau, Ivan Topolsky, Emmanouil Dermitzakis, Keith Harshman, Ioannis Xenarios, Henri Pegeot, Lorenzo Cerutti, Deborah Penet, Philipp Jablonski, Lara Fuhrmann, David Dreifuss, Katharina Jahn, Christiane Beckmann, Maurice Redondo, Olivier Kobel, Christoph Noppen, Sophie Seidel, Noemie Santamaria de Souza, Niko Beerenwinkel, Tanja Stadler |
| EPI_ISL_1750964 | PathWest Laboratory Medicine WA | PathWest Laboratory Medicine WA Microbial Surveillance Unit | PathWest Laboratory Medicine WA Microbial Surveillance Unit |
| EPI_ISL_1752677 | National Public Health Laboratory, National Centre for Infectious Diseases | National Public Health Laboratory, National Centre for Infectious Diseases | Tze Minn Mak, Zhenyang Zhou, Grace Jie Yin Ngan, Royce Ang, Lin Cui, Raymond Tzer Pin Lin |
| EPI_ISL_1753041, EPI_ISL_1753105, EPI_ISL_1753145 | Quest Diagnostics Incorporated | Centers for Disease Control and Prevention Division of Viral Diseases, Pathogen Discovery | Dakota Howard, Dhwani Batra, Peter W. Cook, Kara Moser, Adrian Paskey, Jason Caravas, Benjamin Rambo-Martin, Shatavia Morrison, Christopher Gulvick, Scott Sammons, Yvette Unoarumhi, Darlene Wagner, Matthew Schmerer, S. H. Rosenthal, A. Gerasimova, R. M. Kagan, B. Anderson, M. Hua, Y. Liu, L.E. Bernstein, K.E. Livingston, A. Perez, I. A. Shlyakhter, R. V. Rolando, R. Owen, P. Tanpaiboon, F. Lacbawan, Clinton R. Paden, Duncan MacCannell |
| EPI_ISL_1753676 | Jessa | Jessa | Cruys et al. on behalf of the Jessa_cmdLab |
| EPI_ISL_1756027, EPI_ISL_1756028, EPI_ISL_1756030, EPI_ISL_1756032 | New South Wales Health Pathology Royal Prince Alfred Hospital | Microbiology RPAH | Foster, C.; Au, J.; Ruiz Silva, M.; Deveson, I.; Bull, R.; Van Hal, S.; Rawlinson, W. |
| EPI_ISL_1758205, EPI_ISL_1758283 | Lighthouse Lab in Milton Keynes | Wellcome Sanger Institute for the COVID-19 Genomics UK (COG-UK) Consortium | The Lighthouse Lab in Milton Keynes and Alex Alderton, Roberto Amato, Jeffrey Barrett, Sonia Goncalves, Ewan Harrison, David K. Jackson, Ian Johnston, Dominic Kwiatkowski, Cordelia Langford, John Sillitoe on behalf of the Wellcome Sanger Institute COVID-19 Surveillance Team |
| EPI_ISL_1758301, EPI_ISL_1758376 | Lighthouse Lab in Alderley Park | Wellcome Sanger Institute for the COVID-19 Genomics UK (COG-UK) Consortium | Jacquelyn Wynn, Mairead Hyland, The Lighthouse Lab in Alderley Park and Alex Alderton, Roberto Amato, Jeffrey Barrett, Sonia Goncalves, Ewan Harrison, David K. Jackson, Ian Johnston, Dominic Kwiatkowski, Cordelia Langford, John Sillitoe on behalf of the Wellcome Sanger Institute COVID-19 Surveillance Team |
| EPI_ISL_1758509, EPI_ISL_1758558, EPI_ISL_1758645 | Lighthouse Lab in Cambridge | Wellcome Sanger Institute for the COVID-19 Genomics UK (COG-UK) Consortium | Rob Howes, The Lighthouse Lab in Cambridge and Alex Alderton, Roberto Amato, Jeffrey Barrett, Sonia Goncalves, Ewan Harrison, David K. Jackson, Ian Johnston, Dominic Kwiatkowski, Cordelia Langford, John Sillitoe on behalf of the Wellcome Sanger Institute COVID-19 Surveillance Team |
| EPI_ISL_1758819, EPI_ISL_1758829 | Lighthouse Lab in Glasgow | Wellcome Sanger Institute for the COVID-19 Genomics UK (COG-UK) Consortium | Harper VanSteenhouse, Yumi Kasai, David Gray, Carol Clugston, Anna Dominiczak and Alex Alderton, Roberto Amato, Jeffrey Barrett, Sonia Goncalves, Ewan Harrison, David K. Jackson, Ian Johnston, Dominic Kwiatkowski, Cordelia Langford, John Sillitoe on behalf of the Wellcome Sanger Institute COVID-19 Surveillance Team |
| EPI_ISL_1758890 | Lighthouse Lab in Alderley Park | Wellcome Sanger Institute for the COVID-19 Genomics UK (COG-UK) Consortium | Jacquelyn Wynn, Mairead Hyland, The Lighthouse Lab in Alderley Park and Alex Alderton, Roberto Amato, Jeffrey Barrett, Sonia Goncalves, Ewan Harrison, David K. Jackson, Ian Johnston, Dominic Kwiatkowski, Cordelia Langford, John Sillitoe on behalf of the Wellcome Sanger Institute COVID-19 Surveillance Team |
| EPI_ISL_1758919 | Lighthouse Lab in Milton Keynes | Wellcome Sanger Institute for the COVID-19 Genomics UK (COG-UK) Consortium | The Lighthouse Lab in Milton Keynes and Alex Alderton, Roberto Amato, Jeffrey Barrett, Sonia Goncalves, Ewan Harrison, David K. Jackson, Ian Johnston, Dominic Kwiatkowski, Cordelia Langford, John Sillitoe on behalf of the Wellcome Sanger Institute COVID-19 Surveillance Team |
| EPI_ISL_1758982 | Randox Laboratories | Wellcome Sanger Institute for the COVID-19 Genomics UK (COG-UK) Consortium | Randox Laboratories and Alex Alderton, Roberto Amato, Jeffrey Barrett, Sonia Goncalves, Ewan Harrison, David K. Jackson, Ian Johnston, Dominic Kwiatkowski, Cordelia Langford, John Sillitoe on behalf of the Wellcome Sanger Institute COVID-19 Surveillance Team |
| EPI_ISL_1759178 | Lighthouse Lab in Milton Keynes | Wellcome Sanger Institute for the COVID-19 Genomics UK (COG-UK) Consortium | The Lighthouse Lab in Milton Keynes and Alex Alderton, Roberto Amato, Jeffrey Barrett, Sonia Goncalves, Ewan Harrison, David K. Jackson, Ian Johnston, Dominic Kwiatkowski, Cordelia Langford, John Sillitoe on behalf of the Wellcome Sanger Institute COVID-19 Surveillance Team |
| EPI_ISL_1759190 | Lighthouse Lab in Alderley Park | Wellcome Sanger Institute for the COVID-19 Genomics UK (COG-UK) Consortium | Jacquelyn Wynn, Mairead Hyland, The Lighthouse Lab in Alderley Park and Alex Alderton, Roberto Amato, Jeffrey Barrett, Sonia Goncalves, Ewan Harrison, David K. Jackson, Ian Johnston, Dominic Kwiatkowski, Cordelia Langford, John Sillitoe on behalf of the Wellcome Sanger Institute COVID-19 Surveillance Team |
| EPI_ISL_1759218 | Lighthouse Lab in Milton Keynes | Wellcome Sanger Institute for the COVID-19 Genomics UK (COG-UK) Consortium | The Lighthouse Lab in Milton Keynes and Alex Alderton, Roberto Amato, Jeffrey Barrett, Sonia Goncalves, Ewan Harrison, David K. Jackson, Ian Johnston, Dominic Kwiatkowski, Cordelia Langford, John Sillitoe on behalf of the Wellcome Sanger Institute COVID-19 Surveillance Team |
| EPI_ISL_1759267 | Lighthouse Lab in Cambridge | Wellcome Sanger Institute for the COVID-19 Genomics UK (COG-UK) Consortium | Rob Howes, The Lighthouse Lab in Cambridge and Alex Alderton, Roberto Amato, Jeffrey Barrett, Sonia Goncalves, Ewan Harrison, David K. Jackson, Ian Johnston, Dominic Kwiatkowski, Cordelia Langford, John Sillitoe on behalf of the Wellcome Sanger Institute COVID-19 Surveillance Team |
| EPI_ISL_1759272 | Lighthouse Lab in Milton Keynes | Wellcome Sanger Institute for the COVID-19 Genomics UK (COG-UK) Consortium | The Lighthouse Lab in Milton Keynes and Alex Alderton, Roberto Amato, Jeffrey Barrett, Sonia Goncalves, Ewan Harrison, David K. Jackson, Ian Johnston, Dominic Kwiatkowski, Cordelia Langford, John Sillitoe on behalf of the Wellcome Sanger Institute COVID-19 Surveillance Team |
| EPI_ISL_1759274 | Lighthouse Lab in Alderley Park | Wellcome Sanger Institute for the COVID-19 Genomics UK (COG-UK) Consortium | Jacquelyn Wynn, Mairead Hyland, The Lighthouse Lab in Alderley Park and Alex Alderton, Roberto Amato, Jeffrey Barrett, Sonia Goncalves, Ewan Harrison, David K. Jackson, Ian Johnston, Dominic Kwiatkowski, Cordelia Langford, John Sillitoe on behalf of the Wellcome Sanger Institute COVID-19 Surveillance Team |
| EPI_ISL_1759278, EPI_ISL_1759335, EPI_ISL_1759819, EPI_ISL_1759847 | Lighthouse Lab in Milton Keynes | Wellcome Sanger Institute for the COVID-19 Genomics UK (COG-UK) Consortium | The Lighthouse Lab in Milton Keynes and Alex Alderton, Roberto Amato, Jeffrey Barrett, Sonia Goncalves, Ewan Harrison, David K. Jackson, Ian Johnston, Dominic Kwiatkowski, Cordelia Langford, John Sillitoe on behalf of the Wellcome Sanger Institute COVID-19 Surveillance Team |
| EPI_ISL_1759884 | Lighthouse Lab in Cambridge | Wellcome Sanger Institute for the COVID-19 Genomics UK (COG-UK) Consortium | Rob Howes, The Lighthouse Lab in Cambridge and Alex Alderton, Roberto Amato, Jeffrey Barrett, Sonia Goncalves, Ewan Harrison, David K. Jackson, Ian Johnston, Dominic Kwiatkowski, Cordelia Langford, John Sillitoe on behalf of the Wellcome Sanger Institute COVID-19 Surveillance Team |
| EPI_ISL_1760176, EPI_ISL_1760178, EPI_ISL_1760180 | Wichita State University - Molecular Diagnostics Lab | Kansas Health and Environmental Lab | Mike Grose, Jonathan Barnell, Ben Olsen, and Phil Adam |
| EPI_ISL_1760647 | Institute for Infectious Diseases, University of Bern, Switzerland | Institute for Infectious Diseases, University of Bern, Switzerland | Alban Ramette, Stefan Neuenschwander, Christian Baumann, Miguel A Terrazos Miani, Cora Sägesser, Pascal Bittel, Peter Keller, Franziska Suter-Riniker, Stephen L Leib |
| EPI_ISL_1761938 | Quest Diagnostics Incorporated | Centers for Disease Control and Prevention Division of Viral Diseases, Pathogen Discovery | Dakota Howard, Dhwani Batra, Peter W. Cook, Kara Moser, Adrian Paskey, Jason Caravas, Benjamin Rambo-Martin, Shatavia Morrison, Christopher Gulvick, Scott Sammons, Yvette Unoarumhi, Darlene Wagner, Matthew Schmerer, S. H. Rosenthal, A. Gerasimova, R. M. Kagan, B. Anderson, M. Hua, Y. Liu, L.E. Bernstein, K.E. Livingston, A. Perez, I. A. Shlyakhter, R. V. Rolando, R. Owen, P. Tanpaiboon, F. Lacbawan, Clinton R. Paden, Duncan MacCannell |
| EPI_ISL_1785273 | National Virus Reference Laboratory | National Virus Reference Laboratory | Zoe Yandle, Charlene Bennett, Gabriel Gonzalez, Michael Carr, Jonathan Dean, Cillian F De Gascun |
| EPI_ISL_1785341 | National Virus Reference Laboratory | National Virus Reference Laboratory | Fiona Crispie, Calum Walsh, Matthew McCabe, Zoe Yandle, Charlene Bennet, Gabriel Gonzalez, Michael Carr, Jonathan Dean, Paul Cotter, Cillian F De Gascun |
| EPI_ISL_1785682 | UW Virology Lab | UW Virology Lab | Pavitra Roychoudhury, Hong Xie, Lasata Shrestha, Tien V. Nguyen, Shah Mohamed Bakhash, Michelle Lin, Noah R. Baker, Sean Ellis, Meei-Li Huang, Keith R Jerome, Alexander Greninger |
| EPI_ISL_1786987 | Broad Institute Clinical Research Sequencing Platform | Infectious Disease Program, Broad Institute of Harvard and MIT | Siddle,K.J., Adams,G., Pearlman,L., Gladden-Young,A., Vicente,G., Blumenstiel,B., DeFelice,M., Lee,M., McGovern,S., Lagerborg,K., Rudy,M., DeRuff,K., Carter,A., Normandin,E., Bauer,M., Reilly,S., Tomkins-Tinch,C., Loreth,C., Chaluvadi,S., Meldrim,J., Granger,B., Lemieux,J.E., Birren,B.W., Sabeti,P.C., Larkin,K., Dodge,S., Lennon,N., Madoff,L., Brown,C., Gallagher,G., Smole,S., Park,D.J., Gabriel,S., and MacInnis,B.L. |
| EPI_ISL_1789323, EPI_ISL_1789403, EPI_ISL_1789408, EPI_ISL_1789424, EPI_ISL_1789426 | UW Virology Lab | UW Virology Lab | Pavitra Roychoudhury, Hong Xie, Lasata Shrestha, Tien V. Nguyen, Shah Mohamed Bakhash, Michelle Lin, Noah R. Baker, Sean Ellis, Meei-Li Huang, Keith R Jerome, Alexander Greninger |
| EPI_ISL_1789541, EPI_ISL_1789542 | WHO National Influenza Centre Russian Federation | WHO National Influenza Centre Russian Federation | Andrey Komissarov, Artem Fadeev, Kseniya Komissarova, Oula Mansour, Mikhail Bakaev, Tamila Musaeva, Maria Timofeeva, Veronika Eder, Maria Pisareva, Daria Danilenko, Ksenia Safina, Elena Nabieva, Georgii Bazykin, Dmitry Lioznov |

| | | | |
|---|---|---|---|
| EPI_ISL_1789673 | Pro-Vitam Diagnostics and Research Laboratory | Pro-Vitam Diagnostics and Research Laboratory | Szilard N. Fejer, Istvan Horvath, Monika Korodi, Kinga Rakosi, Zsuzsanna Jenei |
| EPI_ISL_1790019, EPI_ISL_1790023 | NYU Langone Health | Departments of Pathology and Medicine, New York University School of Medicine | Adriana Heguy, Dacia Dimartino, Emily Guzman, Christian Marier, Peter Meyn, Sitharam Ramaswami, Gael Westby, Paul Zappile, Yutong Zhang, Paolo Cotzia, Guiqing Wang |
| EPI_ISL_1790217, EPI_ISL_1790323, EPI_ISL_1790372 | Randox Laboratories | Wellcome Sanger Institute for the COVID-19 Genomics UK (COG-UK) Consortium | Randox Laboratories and Alex Alderton, Roberto Amato, Jeffrey Barrett, Sonia Goncalves, Ewan Harrison, David K. Jackson, Ian Johnston, Dominic Kwiatkowski, Cordelia Langford, John Sillitoe on behalf of the Wellcome Sanger Institute COVID-19 Surveillance Team |
| EPI_ISL_1790373 | Lighthouse Lab in Glasgow | Wellcome Sanger Institute for the COVID-19 Genomics UK (COG-UK) Consortium | Harper VanSteenhouse, Yumi Kasai, David Gray, Carol Clugston, Anna Dominiczak and Alex Alderton, Roberto Amato, Jeffrey Barrett, Sonia Goncalves, Ewan Harrison, David K. Jackson, Ian Johnston, Dominic Kwiatkowski, Cordelia Langford, John Sillitoe on behalf of the Wellcome Sanger Institute COVID-19 Surveillance Team |
| EPI_ISL_1790378, EPI_ISL_1790415 | Randox Laboratories | Wellcome Sanger Institute for the COVID-19 Genomics UK (COG-UK) Consortium | Randox Laboratories and Alex Alderton, Roberto Amato, Jeffrey Barrett, Sonia Goncalves, Ewan Harrison, David K. Jackson, Ian Johnston, Dominic Kwiatkowski, Cordelia Langford, John Sillitoe on behalf of the Wellcome Sanger Institute COVID-19 Surveillance Team |
| EPI_ISL_1790461 | Lighthouse Lab in Alderley Park | Wellcome Sanger Institute for the COVID-19 Genomics UK (COG-UK) Consortium | Jacquelyn Wynn, Mairead Hyland, The Lighthouse Lab in Alderley Park and Alex Alderton, Roberto Amato, Jeffrey Barrett, Sonia Goncalves, Ewan Harrison, David K. Jackson, Ian Johnston, Dominic Kwiatkowski, Cordelia Langford, John Sillitoe on behalf of the Wellcome Sanger Institute COVID-19 Surveillance Team |
| EPI_ISL_1790473, EPI_ISL_1790475, EPI_ISL_1790529, EPI_ISL_1790532 | Lighthouse Lab in Milton Keynes | Wellcome Sanger Institute for the COVID-19 Genomics UK (COG-UK) Consortium | The Lighthouse Lab in Milton Keynes and Alex Alderton, Roberto Amato, Jeffrey Barrett, Sonia Goncalves, Ewan Harrison, David K. Jackson, Ian Johnston, Dominic Kwiatkowski, Cordelia Langford, John Sillitoe on behalf of the Wellcome Sanger Institute COVID-19 Surveillance Team |
| EPI_ISL_1790538, EPI_ISL_1790547 | Lighthouse Lab in Alderley Park | Wellcome Sanger Institute for the COVID-19 Genomics UK (COG-UK) Consortium | Jacquelyn Wynn, Mairead Hyland, The Lighthouse Lab in Alderley Park and Alex Alderton, Roberto Amato, Jeffrey Barrett, Sonia Goncalves, Ewan Harrison, David K. Jackson, Ian Johnston, Dominic Kwiatkowski, Cordelia Langford, John Sillitoe on behalf of the Wellcome Sanger Institute COVID-19 Surveillance Team |
| EPI_ISL_1790560, EPI_ISL_1790590 | Lighthouse Lab in Milton Keynes | Wellcome Sanger Institute for the COVID-19 Genomics UK (COG-UK) Consortium | The Lighthouse Lab in Milton Keynes and Alex Alderton, Roberto Amato, Jeffrey Barrett, Sonia Goncalves, Ewan Harrison, David K. Jackson, Ian Johnston, Dominic Kwiatkowski, Cordelia Langford, John Sillitoe on behalf of the Wellcome Sanger Institute COVID-19 Surveillance Team |
| EPI_ISL_1790596 | Lighthouse Lab in Alderley Park | Wellcome Sanger Institute for the COVID-19 Genomics UK (COG-UK) Consortium | Jacquelyn Wynn, Mairead Hyland, The Lighthouse Lab in Alderley Park and Alex Alderton, Roberto Amato, Jeffrey Barrett, Sonia Goncalves, Ewan Harrison, David K. Jackson, Ian Johnston, Dominic Kwiatkowski, Cordelia Langford, John Sillitoe on behalf of the Wellcome Sanger Institute COVID-19 Surveillance Team |
| EPI_ISL_1790610, EPI_ISL_1790613, EPI_ISL_1790633, EPI_ISL_1790679, EPI_ISL_1790701, EPI_ISL_1790728 | Lighthouse Lab in Milton Keynes | Wellcome Sanger Institute for the COVID-19 Genomics UK (COG-UK) Consortium | The Lighthouse Lab in Milton Keynes and Alex Alderton, Roberto Amato, Jeffrey Barrett, Sonia Goncalves, Ewan Harrison, David K. Jackson, Ian Johnston, Dominic Kwiatkowski, Cordelia Langford, John Sillitoe on behalf of the Wellcome Sanger Institute COVID-19 Surveillance Team |
| EPI_ISL_1790745 | Lighthouse Lab in Alderley Park | Wellcome Sanger Institute for the COVID-19 Genomics UK (COG-UK) Consortium | Jacquelyn Wynn, Mairead Hyland, The Lighthouse Lab in Alderley Park and Alex Alderton, Roberto Amato, Jeffrey Barrett, Sonia Goncalves, Ewan Harrison, David K. Jackson, Ian Johnston, Dominic Kwiatkowski, Cordelia Langford, John Sillitoe on behalf of the Wellcome Sanger Institute COVID-19 Surveillance Team |
| EPI_ISL_1790753 | Lighthouse Lab in Milton Keynes | Wellcome Sanger Institute for the COVID-19 Genomics UK (COG-UK) Consortium | The Lighthouse Lab in Milton Keynes and Alex Alderton, Roberto Amato, Jeffrey Barrett, Sonia Goncalves, Ewan Harrison, David K. Jackson, Ian Johnston, Dominic Kwiatkowski, Cordelia Langford, John Sillitoe on behalf of the Wellcome Sanger Institute COVID-19 Surveillance Team |
| EPI_ISL_1790771 | Lighthouse Lab in Alderley Park | Wellcome Sanger Institute for the COVID-19 Genomics UK (COG-UK) Consortium | Jacquelyn Wynn, Mairead Hyland, The Lighthouse Lab in Alderley Park and Alex Alderton, Roberto Amato, Jeffrey Barrett, Sonia Goncalves, Ewan Harrison, David K. Jackson, Ian Johnston, Dominic Kwiatkowski, Cordelia Langford, John Sillitoe on behalf of the Wellcome Sanger Institute COVID-19 Surveillance Team |
| EPI_ISL_1790774, EPI_ISL_1790783, EPI_ISL_1790799, EPI_ISL_1790800 | Lighthouse Lab in Milton Keynes | Wellcome Sanger Institute for the COVID-19 Genomics UK (COG-UK) Consortium | The Lighthouse Lab in Milton Keynes and Alex Alderton, Roberto Amato, Jeffrey Barrett, Sonia Goncalves, Ewan Harrison, David K. Jackson, Ian Johnston, Dominic Kwiatkowski, Cordelia Langford, John Sillitoe on behalf of the Wellcome Sanger Institute COVID-19 Surveillance Team |
| EPI_ISL_1790809 | Lighthouse Lab in Alderley Park | Wellcome Sanger Institute for the COVID-19 Genomics UK (COG-UK) Consortium | Jacquelyn Wynn, Mairead Hyland, The Lighthouse Lab in Alderley Park and Alex Alderton, Roberto Amato, Jeffrey Barrett, Sonia Goncalves, Ewan Harrison, David K. Jackson, Ian Johnston, Dominic Kwiatkowski, Cordelia Langford, John Sillitoe on behalf of the Wellcome Sanger Institute COVID-19 Surveillance Team |
| EPI_ISL_1790817, EPI_ISL_1790829, EPI_ISL_1790832, EPI_ISL_1790848, EPI_ISL_1790883, EPI_ISL_1790888, EPI_ISL_1790893, EPI_ISL_1790904, EPI_ISL_1790905, EPI_ISL_1790917, EPI_ISL_1790928, EPI_ISL_1790940, EPI_ISL_1790949, EPI_ISL_1790954 | | | |
| see above | Lighthouse Lab in Milton Keynes | Wellcome Sanger Institute for the COVID-19 Genomics UK (COG-UK) Consortium | The Lighthouse Lab in Milton Keynes and Alex Alderton, Roberto Amato, Jeffrey Barrett, Sonia Goncalves, Ewan Harrison, David K. Jackson, Ian Johnston, Dominic Kwiatkowski, Cordelia Langford, John Sillitoe on behalf of the Wellcome Sanger Institute COVID-19 Surveillance Team |
| EPI_ISL_1790990 | Lighthouse Lab in Alderley Park | Wellcome Sanger Institute for the COVID-19 Genomics UK (COG-UK) Consortium | Jacquelyn Wynn, Mairead Hyland, The Lighthouse Lab in Alderley Park and Alex Alderton, Roberto Amato, Jeffrey Barrett, Sonia Goncalves, Ewan Harrison, David K. Jackson, Ian Johnston, Dominic Kwiatkowski, Cordelia Langford, John Sillitoe on behalf of the Wellcome Sanger Institute COVID-19 Surveillance Team |
| EPI_ISL_1790998, EPI_ISL_1791005 | Lighthouse Lab in Milton Keynes | Wellcome Sanger Institute for the COVID-19 Genomics UK (COG-UK) Consortium | The Lighthouse Lab in Milton Keynes and Alex Alderton, Roberto Amato, Jeffrey Barrett, Sonia Goncalves, Ewan Harrison, David K. Jackson, Ian Johnston, Dominic Kwiatkowski, Cordelia Langford, John Sillitoe on behalf of the Wellcome Sanger Institute COVID-19 Surveillance Team |
| EPI_ISL_1791065, EPI_ISL_1791066 | Kantonsspital Baden AG | Institute of Medical Virology, University of Zurich | Verena Kufner, Gabriela Ziltener, Maryam Zaheri, Stefan Schmutz, Annette Audigé, Maria Grünberg, Kevin Steiner, Jon Huder, Cyril Shah, Riccarda Capaul, Guido Bloemberg, Jürg Böni, Michael Huber, Alexandra Trkola |
| EPI_ISL_1792906, EPI_ISL_1792907, EPI_ISL_1792922 | Dutch COVID-19 response team | National Institute for Public Health and the Environment (RIVM) | Adam Meijer, Harry Vennema, Dirk Eggink, Jeroen Cremer, Sharon van den Brink, Bas van der Veer, AnneMarie van den Brandt, Lisa Wijsman, Kim Freriks, Ryanne Jaarsma, Euníce Then, Lynn Aarts, Sanne Bos, Melissa van Tuil, Robert Kohl, Linda van de Nes, Sjoerd Kuiling, James Groot, Florian Zwagemaker, Dennis Schmitz, Annelies Kroneman, Karim Hajji, Chantal Reusken, on behalf of the national COVID-19 response team |
| EPI_ISL_1794552 | Wisconsin State Laboratory of Hygiene Communicable Disease Division | Wisconsin State Laboratory of Hygiene Communicable Disease Division | Kelsey R. Florek, Abigail C. Shockey, Alicia J. Mooney, Sara Wagner |
| EPI_ISL_1794936 | Scripps Medical Laboratory | Andersen lab at Scripps Research | SEARCH Alliance San Diego with Michael Quigley, Ellen Stefanski, Ian Mchardy |
| EPI_ISL_1797026, EPI_ISL_1797206 | Helix/Illumina | Centers for Disease Control and Prevention Division of Viral Diseases, Pathogen Discovery | Dakota Howard, Dhwani Batra, Peter W. Cook, Kara Moser, Adrian Paskey, Jason Caravas, Benjamin Rambo-Martin, Shatavia Morrison, Christopher Gulvick, Scott Sammons, Yvette Unoarumhi, Darlene Wagner, Matthew Schmerer, Eileen de Feo, Jan Antico, Christine Tran, Matthew Tolentino, Shannon Wickline, Kim Gietzen, Brad Sickler, Jingtao Liu, Eric Allen, Phil Febbo, Nicole L. Washington, Simon White, Geraint Levan, Kelly Schiabor Barrett, Elizabeth Cirulli, Alexandre Bolze, Ary Ascencio, Charlotte Rivera-Garcia, Ryan Cho, Jason Nguyen, Sherry Wang, Jimmy Ramirez, Tyler Cassens, Efren Sandoval, Magnus Isaksson, William Lee, David Becker, Marc Laurent, James Lu, Clinton R. Paden, Duncan MacCannell |
| EPI_ISL_1797626 | Department of Health Technology and Informatics, The Hong Kong Polytechnic University | Department of Health Technology and Informatics, The Hong Kong Polytechnic University | Gilman Kit-Hang Siu, Lam-Kwong Lee, Kenneth Siu-Sing Leung, Jake Siu-Lun Leung, Wing-Hei Lo, Timothy Ting-Leung Ng, Chloe Toi-Mei Chan, Hoi-Ching Jim, Kingsley King-Gee Tam, Hiu-Yin Lao, Denise Sze-Hang Wong, Alan Ka-Lun Wu, Miranda Chong-Yee Yau, Yvette Wai-Man Lai, Kitty Sau-Chun Fung, Sandy Ka-Yee Chau, Barry Kin-Chung Wong, Wing-Kin To, Kristine Luk, Alex Yat-Man Ho, Tak-Lun Que, Kam-Tong Yip, Wing Cheong Yam, David Ho-Keung Shum, Shea Ping Yip |
| EPI_ISL_1798071, EPI_ISL_1798096 | Quest Diagnostics Incorporated | Centers for Disease Control and Prevention Division of Viral Diseases, Pathogen Discovery | Dakota Howard, Dhwani Batra, Peter W. Cook, Kara Moser, Adrian Paskey, Jason Caravas, Benjamin Rambo-Martin, Shatavia Morrison, Christopher Gulvick, Scott Sammons, Yvette Unoarumhi, Darlene Wagner, Matthew Schmerer, S. H. Rosenthal, A. Gerasimova, R. M. Kagan, B. Anderson, M. Hua, Y. Liu, L.E. Bernstein, K.E. Livingston, A. Perez, I. A. Shlyakhter, R. V. Rolando, R. Owen, P. Tanpaiboon, F. Lacbawan, Clinton R. Paden, Duncan MacCannell |
| EPI_ISL_1799003, EPI_ISL_1799336, EPI_ISL_1799474, EPI_ISL_1800829, EPI_ISL_1800981, EPI_ISL_1800987, EPI_ISL_1801017, EPI_ISL_1801360, EPI_ISL_1801691, EPI_ISL_1801976, EPI_ISL_1802058, EPI_ISL_1802476, EPI_ISL_1802644 | | | |
| see above | Laboratory Corporation of America | Centers for Disease Control and Prevention Division of Viral Diseases, Pathogen Discovery | Dakota Howard, Dhwani Batra, Peter W. Cook, Kara Moser, Adrian Paskey, Jason Caravas, Benjamin Rambo-Martin, Shatavia Morrison, Christopher Gulvick, Scott Sammons, Yvette Unoarumhi, Darlene Wagner, Matthew Schmerer, Minoo Agarwal, Eyad Almasri, Debbie Boles, Ayla Burns, Nuthawin Charoensri, Oren Cohen, Susan Countryman, Mary Ann Cristobal, Bobbi Croy, Suzanne Dale, Hrushikesh Deshmukh, Amanda Douglas, Vincent Drouillon, Marcia Eisenberg, Howard Engler, Rama Ghatti, Prashant Gupta, Susan Hicks, Jake Humphrey, Lax Iyer, Manoj Jain, Mohan Kolli, Brian Krueger, Tim Kuphal, Stanley Letovsky, Michael Levandoski, Craig Lukasik, Jonathan Meltzer, Brian Norvell, Mindy Nye, Scott Parker, Christos Petropoulos, John Pruitt, Steven Ragan, Scott Ryan, Mike Sapeta, Jana Schroth, Suresh Babu Selvaraju, Goran Stevovic, Amanda Suchanek, Andrea Throop, Lyndon Tilson, Thomas Urban, Joe Voshell, Kimberly Wagner, Jonathan Williams, Mary Williamson, Qian Zeng, Tricia Zwiefelhofer, Clinton R. Paden, Duncan MacCannell |
| EPI_ISL_1803921, EPI_ISL_1804135, EPI_ISL_1804282 | Helix/Illumina | Centers for Disease Control and Prevention Division of Viral Diseases, Pathogen Discovery | Dakota Howard, Dhwani Batra, Peter W. Cook, Kara Moser, Adrian Paskey, Jason Caravas, Benjamin Rambo-Martin, Shatavia Morrison, Christopher Gulvick, Scott Sammons, Yvette Unoarumhi, Darlene Wagner, Matthew Schmerer, Eileen de Feo, Jan Antico, Christine Tran, Matthew Tolentino, Shannon Wickline, Kim Gietzen, Brad Sickler, Jingtao Liu, Eric Allen, Phil Febbo, Nicole L. Washington, Simon White, Geraint Levan, Kelly Schiabor Barrett, Elizabeth Cirulli, Alexandre Bolze, Ary Ascencio, Charlotte Rivera-Garcia, Ryan Cho, Jason Nguyen, Sherry Wang, Jimmy Ramirez, Tyler Cassens, Efren Sandoval, Magnus Isaksson, William Lee, David Becker, Marc Laurent, James Lu, Clinton R. Paden, Duncan MacCannell |
| EPI_ISL_1805024 | Institute of Virology, Medical Center, University of Freiburg, Freiburg, Germany | Institute of Virology, Clinial Virus Genomics, Medical Center, University of Freiburg, Freiburg, Germany | Jonas Fuchs, Lisa Kern, Lena Jaki, Sandra Reuter, Hajo Grundmann, Marcus Panning |
| EPI_ISL_1805456 | Wichita State University - Molecular Diagnostics Lab | Kansas Health and Environmental Lab | Mike Grose, Jonathan Barnell, Ben Olsen, and Phil Adam |
| EPI_ISL_1805463, EPI_ISL_1805464 | Kansas Health and Environmental Lab | Kansas Health and Environmental Lab | Mike Grose, Jonathan Barnell, Ben Olsen, and Phil Adam |
| EPI_ISL_1805678 | Virginia Division of Consolidated Laboratory Services | Virginia Division of Consolidated Laboratory Services | Virginia DCLS |
| EPI_ISL_1805965 | Lighthouse Lab in Alderley Park | Wellcome Sanger Institute for the COVID-19 Genomics UK (COG-UK) Consortium | Jacquelyn Wynn, Mairead Hyland, The Lighthouse Lab in Alderley Park and Alex Alderton, Roberto Amato, Jeffrey Barrett, Sonia Goncalves, Ewan Harrison, David K. Jackson, Ian Johnston, Dominic Kwiatkowski, Cordelia Langford, John Sillitoe on behalf of the Wellcome Sanger Institute COVID-19 Surveillance Team |
| EPI_ISL_1806011 | Lighthouse Lab in Milton Keynes | Wellcome Sanger Institute for the COVID-19 Genomics UK (COG-UK) Consortium | The Lighthouse Lab in Milton Keynes and Alex Alderton, Roberto Amato, Jeffrey Barrett, Sonia Goncalves, Ewan Harrison, David K. Jackson, Ian Johnston, Dominic Kwiatkowski, Cordelia Langford, John Sillitoe on behalf of the Wellcome Sanger Institute COVID-19 Surveillance Team |
| EPI_ISL_1806086, EPI_ISL_1806094, EPI_ISL_1806236 | Lighthouse Lab in Glasgow | Wellcome Sanger Institute for the COVID-19 Genomics UK (COG-UK) Consortium | Harper VanSteenhouse, Yumi Kasai, David Gray, Carol Clugston, Anna Dominiczak and Alex Alderton, Roberto Amato, Jeffrey Barrett, Sonia Goncalves, Ewan Harrison, David K. Jackson, Ian Johnston, Dominic Kwiatkowski, Cordelia Langford, John Sillitoe on behalf of the Wellcome Sanger Institute COVID-19 Surveillance Team |
| EPI_ISL_1806342 | Lighthouse Lab in Milton Keynes | Wellcome Sanger Institute for the COVID-19 Genomics UK (COG-UK) Consortium | The Lighthouse Lab in Milton Keynes and Alex Alderton, Roberto Amato, Jeffrey Barrett, Sonia Goncalves, Ewan Harrison, David K. Jackson, Ian Johnston, Dominic Kwiatkowski, Cordelia Langford, John Sillitoe on behalf of the Wellcome Sanger Institute COVID-19 Surveillance Team |
| EPI_ISL_1806353 | Lighthouse Lab in Glasgow | Wellcome Sanger Institute for the COVID-19 Genomics UK (COG-UK) Consortium | Harper VanSteenhouse, Yumi Kasai, David Gray, Carol Clugston, Anna Dominiczak and Alex Alderton, Roberto Amato, Jeffrey Barrett, Sonia Goncalves, Ewan Harrison, David K. Jackson, Ian Johnston, Dominic Kwiatkowski, Cordelia Langford, John Sillitoe on behalf of the Wellcome Sanger Institute COVID-19 Surveillance Team |
| EPI_ISL_1806406, EPI_ISL_1806433 | Lighthouse Lab in Milton Keynes | Wellcome Sanger Institute for the COVID-19 Genomics UK (COG-UK) Consortium | The Lighthouse Lab in Milton Keynes and Alex Alderton, Roberto Amato, Jeffrey Barrett, Sonia Goncalves, Ewan Harrison, David K. Jackson, Ian Johnston, Dominic Kwiatkowski, Cordelia Langford, John Sillitoe on behalf of the Wellcome Sanger Institute COVID-19 Surveillance Team |
| EPI_ISL_1806511, EPI_ISL_1806517, | Lighthouse Lab in Alderley Park | Wellcome Sanger Institute for the COVID-19 Genomics UK (COG-UK) Consortium | Jacquelyn Wynn, Mairead Hyland, The Lighthouse Lab in Alderley Park and Alex Alderton, Roberto Amato, Jeffrey Barrett, Sonia Goncalves, Ewan Harrison, David K. Jackson, Ian Johnston, Dominic Kwiatkowski, Cordelia Langford, John Sillitoe on behalf of the Wellcome Sanger Institute COVID-19 Surveillance Team |

| | | | |
|---|---|---|---|
| EPI_ISL_1806521, EPI_ISL_1806550, EPI_ISL_1806561, EPI_ISL_1806594 | | | |
| EPI_ISL_1806601 | Lighthouse Lab in Milton Keynes | Wellcome Sanger Institute for the COVID-19 Genomics UK (COG-UK) Consortium | The Lighthouse Lab in Milton Keynes and Alex Alderton, Roberto Amato, Jeffrey Barrett, Sonia Goncalves, Ewan Harrison, David K. Jackson, Ian Johnston, Dominic Kwiatkowski, Cordelia Langford, John Sillitoe on behalf of the Wellcome Sanger Institute COVID-19 Surveillance Team |
| EPI_ISL_1806633, EPI_ISL_1806670, EPI_ISL_1806695 | Lighthouse Lab in Alderley Park | Wellcome Sanger Institute for the COVID-19 Genomics UK (COG-UK) Consortium | Jacquelyn Wynn, Mairead Hyland, The Lighthouse Lab in Alderley Park and Alex Alderton, Roberto Amato, Jeffrey Barrett, Sonia Goncalves, Ewan Harrison, David K. Jackson, Ian Johnston, Dominic Kwiatkowski, Cordelia Langford, John Sillitoe on behalf of the Wellcome Sanger Institute COVID-19 Surveillance Team |
| EPI_ISL_1806709, EPI_ISL_1806761, EPI_ISL_1806767 | Lighthouse Lab in Milton Keynes | Wellcome Sanger Institute for the COVID-19 Genomics UK (COG-UK) Consortium | The Lighthouse Lab in Milton Keynes and Alex Alderton, Roberto Amato, Jeffrey Barrett, Sonia Goncalves, Ewan Harrison, David K. Jackson, Ian Johnston, Dominic Kwiatkowski, Cordelia Langford, John Sillitoe on behalf of the Wellcome Sanger Institute COVID-19 Surveillance Team |
| EPI_ISL_1806837, EPI_ISL_1806846 | Lighthouse Lab in Alderley Park | Wellcome Sanger Institute for the COVID-19 Genomics UK (COG-UK) Consortium | Jacquelyn Wynn, Mairead Hyland, The Lighthouse Lab in Alderley Park and Alex Alderton, Roberto Amato, Jeffrey Barrett, Sonia Goncalves, Ewan Harrison, David K. Jackson, Ian Johnston, Dominic Kwiatkowski, Cordelia Langford, John Sillitoe on behalf of the Wellcome Sanger Institute COVID-19 Surveillance Team |
| EPI_ISL_1806848 | Lighthouse Lab in Milton Keynes | Wellcome Sanger Institute for the COVID-19 Genomics UK (COG-UK) Consortium | The Lighthouse Lab in Milton Keynes and Alex Alderton, Roberto Amato, Jeffrey Barrett, Sonia Goncalves, Ewan Harrison, David K. Jackson, Ian Johnston, Dominic Kwiatkowski, Cordelia Langford, John Sillitoe on behalf of the Wellcome Sanger Institute COVID-19 Surveillance Team |
| EPI_ISL_1806852 | Lighthouse Lab in Alderley Park | Wellcome Sanger Institute for the COVID-19 Genomics UK (COG-UK) Consortium | Jacquelyn Wynn, Mairead Hyland, The Lighthouse Lab in Alderley Park and Alex Alderton, Roberto Amato, Jeffrey Barrett, Sonia Goncalves, Ewan Harrison, David K. Jackson, Ian Johnston, Dominic Kwiatkowski, Cordelia Langford, John Sillitoe on behalf of the Wellcome Sanger Institute COVID-19 Surveillance Team |
| EPI_ISL_1806892 | Lighthouse Lab in Glasgow | Wellcome Sanger Institute for the COVID-19 Genomics UK (COG-UK) Consortium | Harper VanSteenhouse, Yumi Kasai, David Gray, Carol Clugston, Anna Dominiczak and Alex Alderton, Roberto Amato, Jeffrey Barrett, Sonia Goncalves, Ewan Harrison, David K. Jackson, Ian Johnston, Dominic Kwiatkowski, Cordelia Langford, John Sillitoe on behalf of the Wellcome Sanger Institute COVID-19 Surveillance Team |
| EPI_ISL_1806897, EPI_ISL_1806899, EPI_ISL_1806900, EPI_ISL_1806901, EPI_ISL_1806902, EPI_ISL_1806903, EPI_ISL_1806905 | Lighthouse Lab in Milton Keynes | Wellcome Sanger Institute for the COVID-19 Genomics UK (COG-UK) Consortium | The Lighthouse Lab in Milton Keynes and Alex Alderton, Roberto Amato, Jeffrey Barrett, Sonia Goncalves, Ewan Harrison, David K. Jackson, Ian Johnston, Dominic Kwiatkowski, Cordelia Langford, John Sillitoe on behalf of the Wellcome Sanger Institute COVID-19 Surveillance Team |
| EPI_ISL_1806953 | Randox Laboratories | Wellcome Sanger Institute for the COVID-19 Genomics UK (COG-UK) Consortium | Randox Laboratories and Alex Alderton, Roberto Amato, Jeffrey Barrett, Sonia Goncalves, Ewan Harrison, David K. Jackson, Ian Johnston, Dominic Kwiatkowski, Cordelia Langford, John Sillitoe on behalf of the Wellcome Sanger Institute COVID-19 Surveillance Team |
| EPI_ISL_1807318 | ACT Pathology | Schwessinger Lab | Ashley Jones, Benjamin Schwessinger, Robert Lanfear, Megan McDonald, Ming-Dao Chia, Kevin Murray, Robyn N Hall, Craig Kennedy, Karina Kennedy |
| EPI_ISL_1808683, EPI_ISL_1808686 | Swedish national genomic surveillance program of SARS-CoV-2 | The Public Health Agency of Sweden | Maximilian Riess, Maria Lind Karlberg, Alma Brolund, Swedish national genomic surveillance program of SARS-CoV-2 |
| EPI_ISL_1809681 | Illinois Department of Public Health | Illinois Department of Public Health - Chicago Lab | Vineet K. Dhiman, Ira Heimler, Joel Price |
| EPI_ISL_1811202 | University Hospitals of Geneva, Laboratory of Virology | HUG, Laboratory of Virology and the Health2030 Genome Center | Samuel Cordey, Ana Rita Goncalves, Laurent Kaiser, Lorenzo Cerutti, Henri Pegeot, Melyssa Elies, Deborah Penet, Keith Harshman, Ioannis Xenarios, Emmanouil Dermitzakis |
| EPI_ISL_1816917, EPI_ISL_1816920, EPI_ISL_1816921, EPI_ISL_1816922, EPI_ISL_1816923, EPI_ISL_1816924 | PathWest Laboratory Medicine WA | PathWest Laboratory Medicine WA Microbial Surveillance Unit | PathWest Laboratory Medicine WA Microbial Surveillance Unit |
| EPI_ISL_1816935, EPI_ISL_1816944, EPI_ISL_1816945, EPI_ISL_1816946, EPI_ISL_1816947, EPI_ISL_1816948, EPI_ISL_1816949, EPI_ISL_1816950, EPI_ISL_1816951, EPI_ISL_1816952, EPI_ISL_1816953, EPI_ISL_1816954, EPI_ISL_1816955, EPI_ISL_1816956, EPI_ISL_1816957, EPI_ISL_1816958, EPI_ISL_1816959, EPI_ISL_1816960, EPI_ISL_1816961, EPI_ISL_1816962, EPI_ISL_1816963, EPI_ISL_1816964, EPI_ISL_1816965, EPI_ISL_1816966, EPI_ISL_1816967, EPI_ISL_1816968, EPI_ISL_1816969, EPI_ISL_1816970, EPI_ISL_1816971, EPI_ISL_1816972, EPI_ISL_1816973 | | | |
| see above | National Public Health Laboratory, National Centre for Infectious Diseases | National Public Health Laboratory, National Centre for Infectious Diseases | Tze Minn Mak, Zhenyang Zhou, Grace Jie Yin Ngan, Royce Ang, Lin Cui, Raymond Tzer Pin Lin |
| EPI_ISL_1817558, EPI_ISL_1817559, EPI_ISL_1817560 | PathWest Laboratory Medicine WA | PathWest Laboratory Medicine WA Microbial Surveillance Unit | PathWest Laboratory Medicine WA Microbial Surveillance Unit |
| EPI_ISL_1818383 | Quest Diagnostics Incorporated | Centers for Disease Control and Prevention Division of Viral Diseases, Pathogen Discovery | Dakota Howard, Dhwani Batra, Peter W. Cook, Kara Moser, Adrian Paskey, Jason Caravas, Benjamin Rambo-Martin, Shatavia Morrison, Christopher Gulvick, Scott Sammons, Yvette Unoarumhi, Darlene Wagner, Matthew Schmerer, S. H. Rosenthal, A. Gerasimova, R. M. Kagan, B. Anderson, M. Hua, Y. Liu, L.E. Bernstein, K.E. Livingston, A. Perez, I. A. Shlyakhter, R. V. Rolando, R. Owen, P. Tanpaiboon, F. Lacbawan, Clinton R. Paden, Duncan MacCannell |
| EPI_ISL_1818617, EPI_ISL_1818618 | KEMPEGOWDA INTERNATIONAL AIRPORT | INSACOG-KA, NIMHANS | Chitra Pattabiraman, Pramada Prasad, Anson Kunjumon George, Ananthapadmanabha Kotambail, Darshan Sreenivas, Chetan G K, Gautham Arunachal Udupi, Anita S Desai, V Ravi |
| EPI_ISL_1823618 | KY State Public Health Lab | Centers for Disease Control and Prevention Division of Viral Diseases, Pathogen Discovery | Mili Sheth, Sarah Nobles, Jasmine Padilla, Mark Burroughs, Shoshona Le, Katie Dillon, Peter Cook, Clinton R. Paden, Dhwani Batra, Krista Queen, Kristen Knipe, Dakota Howard, Yvette Unoarumhi, Darlene Wagner, Matthew Schmerer, Ben L. Rambo-Martin, Kristine Lacek, Sam Shepard, Alison Laufer Halpin, Dave Wentworth, Vivien Dugan, Suxiang Tong, Justin Lee |
| EPI_ISL_1824117, EPI_ISL_1824118 | El Camino Hospital | Santa Clara County public Health Laboratory | Santa Clara County Public Health Department |
| EPI_ISL_1824285, EPI_ISL_1824286 | Pathology and Laboratory Medicine Institute, Cleveland Clinic, Ohio, USA | Pathology and Laboratory Medicine Institute, Cleveland Clinic, Ohio, USA | Jessica Spildener, Joy Nakitandwe, Kristen McDonnell, David Plunkett, Zheng Jin Tu, Jay Brock, Yu-Wei Cheng, Gary Procop, Daniel Rhoads, Daniel H. Farkas, David Bosler |
| EPI_ISL_1824604 | PRVKP FK UI | National Institute of Health Research and Development | Vivi Setiawaty, Hana Apsari Pawestri, Subangkit, Kartika Dewi Puspa, Arie Ardiansyah Nugraha, Hartanti Dian Ikawati, Krisna Pangesti, Nelly Puspandari |
| EPI_ISL_1824607 | PKC Jagakarsa | National Institute of Health Research and Development | Vivi Setiawaty, Hana Apsari Pawestri, Subangkit, Kartika Dewi Puspa, Arie Ardiansyah Nugraha, Hartanti Dian Ikawati, Krisna Pangesti, Nelly Puspandari |
| EPI_ISL_1824682, EPI_ISL_1824683, EPI_ISL_1824684 | VITHAS Lab | Microbiology Department. Complexo Hospitalario Universitario de Vigo | Camacho AT, Perez S, Regueiro B |
| EPI_ISL_1825316, EPI_ISL_1825321, EPI_ISL_1825322, EPI_ISL_1825553, EPI_ISL_1825586 | Broad Institute Clinical Research Sequencing Platform | Infectious Disease Program, Broad Institute of Harvard and MIT | Siddle,K.J., Adams,G., Pearlman,L., Gladden-Young,A., Vicente,G., Blumenstiel,B., DeFelice,M., Lee,M., McGovern,S., Lagerborg,K., Rudy,M., DeRuff,K., Carter,A., Normandin,E., Bauer,M., Reilly,S., Tomkins-Tinch,C., Loreth,C., Chaluvadi,S., Meldrim,J., Granger,B., Lemieux,J.E., Birren,B.W., Sabeti,P.C., Larkin,K., Dodge,S., Lennon,N., Madoff,L., Brown,C., Gallagher,G., Smole,S., Park,D.J., Gabriel,S., and MacInnis,B.L. |
| EPI_ISL_1825677 | IHU Méditerranée Infection | IHU Méditerranée Infection | Bernard La Scola et al. |
| EPI_ISL_1826129, EPI_ISL_1826132 | UW Virology Lab | UW Virology Lab | Pavitra Roychoudhury, Hong Xie, Lasata Shrestha, Tien V. Nguyen, Shah Mohamed Bakhash, Michelle Lin, Noah R. Baker, Sean Ellis, Meei-Li Huang, Keith R Jerome, Alexander Greninger |
| EPI_ISL_1827216 | Lab voor klinische biologie | Lab voor klinische biologie | Marija Janevska, Hannelore Hamerlinck, Bruno Verhasselt |
| EPI_ISL_1827723 | UW Virology Lab | UW Virology Lab | Pavitra Roychoudhury, Hong Xie, Lasata Shrestha, Tien V. Nguyen, Shah Mohamed Bakhash, Michelle Lin, Noah R. Baker, Ricardo Perez, Sean Ellis, Nathan Breit, Robert J. Livingston, Meei-Li Huang, Keith R Jerome, Patrick Mathias, Alexander Greninger |
| EPI_ISL_1827727 | Centralny Szpital Kliniczny MSWiA w Warszawie | 1. Virogenetics Laboratory of Virology, Małopolska Centre of Biotechnology, Jagiellonian University. 2. Intercollegiate Faculty of Biotechnology University of Gdansk and Medical University of Gdansk | Lukasz Rabalski, Maciej Kosinski, Natalia Mazur-Panasiuk, Sławomir Butkiewicz, Michał Hampel, Anna Meler, Adam Sybilski, Krzysztof Pyrc, Krystyna Bienkowska-Szewczyk |
| EPI_ISL_1828111, EPI_ISL_1828260, EPI_ISL_1828349, EPI_ISL_1828425, EPI_ISL_1828512, EPI_ISL_1828556, EPI_ISL_1828565, EPI_ISL_1828653, EPI_ISL_1828694 | Pandemic Response Lab - NYC | Pandemic Response Lab, R&D | Henry Lee, Michael Hammerling, Melissa Hopkins, Cybill del Castillo, Shinyoung Clair Kang, William Ward, Pradeep Bugga, Sol Rey, Dylan Law, Katharine Nelson, Haiping Hao, Jon Laurent |
| EPI_ISL_1829206, EPI_ISL_1829207 | Lighthouse Lab in Milton Keynes | Wellcome Sanger Institute for the COVID-19 Genomics UK (COG-UK) Consortium | The Lighthouse Lab in Milton Keynes and Alex Alderton, Roberto Amato, Jeffrey Barrett, Sonia Goncalves, Ewan Harrison, David K. Jackson, Ian Johnston, Dominic Kwiatkowski, Cordelia Langford, John Sillitoe on behalf of the Wellcome Sanger Institute COVID-19 Surveillance Team |
| EPI_ISL_1829212 | Lighthouse Lab in Alderley Park | Wellcome Sanger Institute for the COVID-19 Genomics UK (COG-UK) Consortium | Jacquelyn Wynn, Mairead Hyland, The Lighthouse Lab in Alderley Park and Alex Alderton, Roberto Amato, Jeffrey Barrett, Sonia Goncalves, Ewan Harrison, David K. Jackson, Ian Johnston, Dominic Kwiatkowski, Cordelia Langford, John Sillitoe on behalf of the Wellcome Sanger Institute COVID-19 Surveillance Team |
| EPI_ISL_1829218 | Lighthouse Lab in Milton Keynes | Wellcome Sanger Institute for the COVID-19 Genomics UK (COG-UK) Consortium | The Lighthouse Lab in Milton Keynes and Alex Alderton, Roberto Amato, Jeffrey Barrett, Sonia Goncalves, Ewan Harrison, David K. Jackson, Ian Johnston, Dominic Kwiatkowski, Cordelia Langford, John Sillitoe on behalf of the Wellcome Sanger Institute COVID-19 Surveillance Team |
| EPI_ISL_1829220 | Lighthouse Lab in Alderley Park | Wellcome Sanger Institute for the COVID-19 Genomics UK (COG-UK) Consortium | Jacquelyn Wynn, Mairead Hyland, The Lighthouse Lab in Alderley Park and Alex Alderton, Roberto Amato, Jeffrey Barrett, Sonia Goncalves, Ewan Harrison, David K. Jackson, Ian Johnston, Dominic Kwiatkowski, Cordelia Langford, John Sillitoe on behalf of the Wellcome Sanger Institute COVID-19 Surveillance Team |

| | | | |
|---|---|---|---|
| EPI_ISL_1829225, EPI_ISL_1829226, EPI_ISL_1829228, EPI_ISL_1829230 | Lighthouse Lab in Milton Keynes | Wellcome Sanger Institute for the COVID-19 Genomics UK (COG-UK) Consortium | The Lighthouse Lab in Milton Keynes and Alex Alderton, Roberto Amato, Jeffrey Barrett, Sonia Goncalves, Ewan Harrison, David K. Jackson, Ian Johnston, Dominic Kwiatkowski, Cordelia Langford, John Sillitoe on behalf of the Wellcome Sanger Institute COVID-19 Surveillance Team |
| EPI_ISL_1829233 | Lighthouse Lab in Alderley Park | Wellcome Sanger Institute for the COVID-19 Genomics UK (COG-UK) Consortium | Jacquelyn Wynn, Mairead Hyland, The Lighthouse Lab in Alderley Park and Alex Alderton, Roberto Amato, Jeffrey Barrett, Sonia Goncalves, Ewan Harrison, David K. Jackson, Ian Johnston, Dominic Kwiatkowski, Cordelia Langford, John Sillitoe on behalf of the Wellcome Sanger Institute COVID-19 Surveillance Team |
| EPI_ISL_1829236 | Lighthouse Lab in Milton Keynes | Wellcome Sanger Institute for the COVID-19 Genomics UK (COG-UK) Consortium | The Lighthouse Lab in Milton Keynes and Alex Alderton, Roberto Amato, Jeffrey Barrett, Sonia Goncalves, Ewan Harrison, David K. Jackson, Ian Johnston, Dominic Kwiatkowski, Cordelia Langford, John Sillitoe on behalf of the Wellcome Sanger Institute COVID-19 Surveillance Team |
| EPI_ISL_1829237 | Lighthouse Lab in Alderley Park | Wellcome Sanger Institute for the COVID-19 Genomics UK (COG-UK) Consortium | Jacquelyn Wynn, Mairead Hyland, The Lighthouse Lab in Alderley Park and Alex Alderton, Roberto Amato, Jeffrey Barrett, Sonia Goncalves, Ewan Harrison, David K. Jackson, Ian Johnston, Dominic Kwiatkowski, Cordelia Langford, John Sillitoe on behalf of the Wellcome Sanger Institute COVID-19 Surveillance Team |
| EPI_ISL_1829246, EPI_ISL_1829248, EPI_ISL_1829249, EPI_ISL_1829250 | Lighthouse Lab in Milton Keynes | Wellcome Sanger Institute for the COVID-19 Genomics UK (COG-UK) Consortium | The Lighthouse Lab in Milton Keynes and Alex Alderton, Roberto Amato, Jeffrey Barrett, Sonia Goncalves, Ewan Harrison, David K. Jackson, Ian Johnston, Dominic Kwiatkowski, Cordelia Langford, John Sillitoe on behalf of the Wellcome Sanger Institute COVID-19 Surveillance Team |
| EPI_ISL_1829254, EPI_ISL_1829258 | Lighthouse Lab in Alderley Park | Wellcome Sanger Institute for the COVID-19 Genomics UK (COG-UK) Consortium | Jacquelyn Wynn, Mairead Hyland, The Lighthouse Lab in Alderley Park and Alex Alderton, Roberto Amato, Jeffrey Barrett, Sonia Goncalves, Ewan Harrison, David K. Jackson, Ian Johnston, Dominic Kwiatkowski, Cordelia Langford, John Sillitoe on behalf of the Wellcome Sanger Institute COVID-19 Surveillance Team |
| EPI_ISL_1829259, EPI_ISL_1829261, EPI_ISL_1829268, EPI_ISL_1829277 | Lighthouse Lab in Milton Keynes | Wellcome Sanger Institute for the COVID-19 Genomics UK (COG-UK) Consortium | The Lighthouse Lab in Milton Keynes and Alex Alderton, Roberto Amato, Jeffrey Barrett, Sonia Goncalves, Ewan Harrison, David K. Jackson, Ian Johnston, Dominic Kwiatkowski, Cordelia Langford, John Sillitoe on behalf of the Wellcome Sanger Institute COVID-19 Surveillance Team |
| EPI_ISL_1829279 | Lighthouse Lab in Alderley Park | Wellcome Sanger Institute for the COVID-19 Genomics UK (COG-UK) Consortium | Jacquelyn Wynn, Mairead Hyland, The Lighthouse Lab in Alderley Park and Alex Alderton, Roberto Amato, Jeffrey Barrett, Sonia Goncalves, Ewan Harrison, David K. Jackson, Ian Johnston, Dominic Kwiatkowski, Cordelia Langford, John Sillitoe on behalf of the Wellcome Sanger Institute COVID-19 Surveillance Team |
| EPI_ISL_1829280 | Lighthouse Lab in Milton Keynes | Wellcome Sanger Institute for the COVID-19 Genomics UK (COG-UK) Consortium | The Lighthouse Lab in Milton Keynes and Alex Alderton, Roberto Amato, Jeffrey Barrett, Sonia Goncalves, Ewan Harrison, David K. Jackson, Ian Johnston, Dominic Kwiatkowski, Cordelia Langford, John Sillitoe on behalf of the Wellcome Sanger Institute COVID-19 Surveillance Team |
| EPI_ISL_1829281, EPI_ISL_1829282 | Lighthouse Lab in Alderley Park | Wellcome Sanger Institute for the COVID-19 Genomics UK (COG-UK) Consortium | Jacquelyn Wynn, Mairead Hyland, The Lighthouse Lab in Alderley Park and Alex Alderton, Roberto Amato, Jeffrey Barrett, Sonia Goncalves, Ewan Harrison, David K. Jackson, Ian Johnston, Dominic Kwiatkowski, Cordelia Langford, John Sillitoe on behalf of the Wellcome Sanger Institute COVID-19 Surveillance Team |
| EPI_ISL_1829283, EPI_ISL_1829293, EPI_ISL_1829312, EPI_ISL_1829313, EPI_ISL_1829334, EPI_ISL_1829340, EPI_ISL_1829355 | Lighthouse Lab in Milton Keynes | Wellcome Sanger Institute for the COVID-19 Genomics UK (COG-UK) Consortium | The Lighthouse Lab in Milton Keynes and Alex Alderton, Roberto Amato, Jeffrey Barrett, Sonia Goncalves, Ewan Harrison, David K. Jackson, Ian Johnston, Dominic Kwiatkowski, Cordelia Langford, John Sillitoe on behalf of the Wellcome Sanger Institute COVID-19 Surveillance Team |
| EPI_ISL_1829360 | Lighthouse Lab in Alderley Park | Wellcome Sanger Institute for the COVID-19 Genomics UK (COG-UK) Consortium | Jacquelyn Wynn, Mairead Hyland, The Lighthouse Lab in Alderley Park and Alex Alderton, Roberto Amato, Jeffrey Barrett, Sonia Goncalves, Ewan Harrison, David K. Jackson, Ian Johnston, Dominic Kwiatkowski, Cordelia Langford, John Sillitoe on behalf of the Wellcome Sanger Institute COVID-19 Surveillance Team |
| EPI_ISL_1829370, EPI_ISL_1829372 | Lighthouse Lab in Milton Keynes | Wellcome Sanger Institute for the COVID-19 Genomics UK (COG-UK) Consortium | The Lighthouse Lab in Milton Keynes and Alex Alderton, Roberto Amato, Jeffrey Barrett, Sonia Goncalves, Ewan Harrison, David K. Jackson, Ian Johnston, Dominic Kwiatkowski, Cordelia Langford, John Sillitoe on behalf of the Wellcome Sanger Institute COVID-19 Surveillance Team |
| EPI_ISL_1829382 | Lighthouse Lab in Alderley Park | Wellcome Sanger Institute for the COVID-19 Genomics UK (COG-UK) Consortium | Jacquelyn Wynn, Mairead Hyland, The Lighthouse Lab in Alderley Park and Alex Alderton, Roberto Amato, Jeffrey Barrett, Sonia Goncalves, Ewan Harrison, David K. Jackson, Ian Johnston, Dominic Kwiatkowski, Cordelia Langford, John Sillitoe on behalf of the Wellcome Sanger Institute COVID-19 Surveillance Team |
| EPI_ISL_1829386, EPI_ISL_1829392, EPI_ISL_1829394, EPI_ISL_1829396, EPI_ISL_1829398, EPI_ISL_1829399, EPI_ISL_1829402 | Lighthouse Lab in Milton Keynes | Wellcome Sanger Institute for the COVID-19 Genomics UK (COG-UK) Consortium | The Lighthouse Lab in Milton Keynes and Alex Alderton, Roberto Amato, Jeffrey Barrett, Sonia Goncalves, Ewan Harrison, David K. Jackson, Ian Johnston, Dominic Kwiatkowski, Cordelia Langford, John Sillitoe on behalf of the Wellcome Sanger Institute COVID-19 Surveillance Team |
| EPI_ISL_1829413 | Lighthouse Lab in Alderley Park | Wellcome Sanger Institute for the COVID-19 Genomics UK (COG-UK) Consortium | Jacquelyn Wynn, Mairead Hyland, The Lighthouse Lab in Alderley Park and Alex Alderton, Roberto Amato, Jeffrey Barrett, Sonia Goncalves, Ewan Harrison, David K. Jackson, Ian Johnston, Dominic Kwiatkowski, Cordelia Langford, John Sillitoe on behalf of the Wellcome Sanger Institute COVID-19 Surveillance Team |
| EPI_ISL_1829417, EPI_ISL_1829418, EPI_ISL_1829421, EPI_ISL_1829422, EPI_ISL_1829425, EPI_ISL_1829428 | Lighthouse Lab in Milton Keynes | Wellcome Sanger Institute for the COVID-19 Genomics UK (COG-UK) Consortium | The Lighthouse Lab in Milton Keynes and Alex Alderton, Roberto Amato, Jeffrey Barrett, Sonia Goncalves, Ewan Harrison, David K. Jackson, Ian Johnston, Dominic Kwiatkowski, Cordelia Langford, John Sillitoe on behalf of the Wellcome Sanger Institute COVID-19 Surveillance Team |
| EPI_ISL_1829430, EPI_ISL_1829440 | Lighthouse Lab in Alderley Park | Wellcome Sanger Institute for the COVID-19 Genomics UK (COG-UK) Consortium | Jacquelyn Wynn, Mairead Hyland, The Lighthouse Lab in Alderley Park and Alex Alderton, Roberto Amato, Jeffrey Barrett, Sonia Goncalves, Ewan Harrison, David K. Jackson, Ian Johnston, Dominic Kwiatkowski, Cordelia Langford, John Sillitoe on behalf of the Wellcome Sanger Institute COVID-19 Surveillance Team |
| EPI_ISL_1829449 | Lighthouse Lab in Milton Keynes | Wellcome Sanger Institute for the COVID-19 Genomics UK (COG-UK) Consortium | The Lighthouse Lab in Milton Keynes and Alex Alderton, Roberto Amato, Jeffrey Barrett, Sonia Goncalves, Ewan Harrison, David K. Jackson, Ian Johnston, Dominic Kwiatkowski, Cordelia Langford, John Sillitoe on behalf of the Wellcome Sanger Institute COVID-19 Surveillance Team |
| EPI_ISL_1829458 | Lighthouse Lab in Alderley Park | Wellcome Sanger Institute for the COVID-19 Genomics UK (COG-UK) Consortium | Jacquelyn Wynn, Mairead Hyland, The Lighthouse Lab in Alderley Park and Alex Alderton, Roberto Amato, Jeffrey Barrett, Sonia Goncalves, Ewan Harrison, David K. Jackson, Ian Johnston, Dominic Kwiatkowski, Cordelia Langford, John Sillitoe on behalf of the Wellcome Sanger Institute COVID-19 Surveillance Team |
| EPI_ISL_1829466, EPI_ISL_1829472, EPI_ISL_1829481, EPI_ISL_1829483, EPI_ISL_1829536 | Lighthouse Lab in Milton Keynes | Wellcome Sanger Institute for the COVID-19 Genomics UK (COG-UK) Consortium | The Lighthouse Lab in Milton Keynes and Alex Alderton, Roberto Amato, Jeffrey Barrett, Sonia Goncalves, Ewan Harrison, David K. Jackson, Ian Johnston, Dominic Kwiatkowski, Cordelia Langford, John Sillitoe on behalf of the Wellcome Sanger Institute COVID-19 Surveillance Team |
| EPI_ISL_1829541, EPI_ISL_1829569, EPI_ISL_1829604 | Lighthouse Lab in Glasgow | Wellcome Sanger Institute for the COVID-19 Genomics UK (COG-UK) Consortium | Harper VanSteenhouse, Yumi Kasai, David Gray, Carol Clugston, Anna Dominiczak and Alex Alderton, Roberto Amato, Jeffrey Barrett, Sonia Goncalves, Ewan Harrison, David K. Jackson, Ian Johnston, Dominic Kwiatkowski, Cordelia Langford, John Sillitoe on behalf of the Wellcome Sanger Institute COVID-19 Surveillance Team |
| EPI_ISL_1829729, EPI_ISL_1829744, EPI_ISL_1829774 | Lighthouse Lab in Milton Keynes | Wellcome Sanger Institute for the COVID-19 Genomics UK (COG-UK) Consortium | The Lighthouse Lab in Milton Keynes and Alex Alderton, Roberto Amato, Jeffrey Barrett, Sonia Goncalves, Ewan Harrison, David K. Jackson, Ian Johnston, Dominic Kwiatkowski, Cordelia Langford, John Sillitoe on behalf of the Wellcome Sanger Institute COVID-19 Surveillance Team |
| EPI_ISL_1829780 | Lighthouse Lab in Glasgow | Wellcome Sanger Institute for the COVID-19 Genomics UK (COG-UK) Consortium | Harper VanSteenhouse, Yumi Kasai, David Gray, Carol Clugston, Anna Dominiczak and Alex Alderton, Roberto Amato, Jeffrey Barrett, Sonia Goncalves, Ewan Harrison, David K. Jackson, Ian Johnston, Dominic Kwiatkowski, Cordelia Langford, John Sillitoe on behalf of the Wellcome Sanger Institute COVID-19 Surveillance Team |
| EPI_ISL_1829814 | Randox Laboratories | Wellcome Sanger Institute for the COVID-19 Genomics UK (COG-UK) Consortium | Randox Laboratories and Alex Alderton, Roberto Amato, Jeffrey Barrett, Sonia Goncalves, Ewan Harrison, David K. Jackson, Ian Johnston, Dominic Kwiatkowski, Cordelia Langford, John Sillitoe on behalf of the Wellcome Sanger Institute COVID-19 Surveillance Team |
| EPI_ISL_1829818, EPI_ISL_1829838, EPI_ISL_1829839 | Lighthouse Lab in Alderley Park | Wellcome Sanger Institute for the COVID-19 Genomics UK (COG-UK) Consortium | Jacquelyn Wynn, Mairead Hyland, The Lighthouse Lab in Alderley Park and Alex Alderton, Roberto Amato, Jeffrey Barrett, Sonia Goncalves, Ewan Harrison, David K. Jackson, Ian Johnston, Dominic Kwiatkowski, Cordelia Langford, John Sillitoe on behalf of the Wellcome Sanger Institute COVID-19 Surveillance Team |
| EPI_ISL_1829841 | Randox Laboratories | Wellcome Sanger Institute for the COVID-19 Genomics UK (COG-UK) Consortium | Randox Laboratories and Alex Alderton, Roberto Amato, Jeffrey Barrett, Sonia Goncalves, Ewan Harrison, David K. Jackson, Ian Johnston, Dominic Kwiatkowski, Cordelia Langford, John Sillitoe on behalf of the Wellcome Sanger Institute COVID-19 Surveillance Team |
| EPI_ISL_1829843, EPI_ISL_1829854, EPI_ISL_1829860, EPI_ISL_1829878 | Lighthouse Lab in Alderley Park | Wellcome Sanger Institute for the COVID-19 Genomics UK (COG-UK) Consortium | Jacquelyn Wynn, Mairead Hyland, The Lighthouse Lab in Alderley Park and Alex Alderton, Roberto Amato, Jeffrey Barrett, Sonia Goncalves, Ewan Harrison, David K. Jackson, Ian Johnston, Dominic Kwiatkowski, Cordelia Langford, John Sillitoe on behalf of the Wellcome Sanger Institute COVID-19 Surveillance Team |
| EPI_ISL_1829894 | Randox Laboratories | Wellcome Sanger Institute for the COVID-19 Genomics UK (COG-UK) Consortium | Randox Laboratories and Alex Alderton, Roberto Amato, Jeffrey Barrett, Sonia Goncalves, Ewan Harrison, David K. Jackson, Ian Johnston, Dominic Kwiatkowski, Cordelia Langford, John Sillitoe on behalf of the Wellcome Sanger Institute COVID-19 Surveillance Team |
| EPI_ISL_1829914, EPI_ISL_1829930 | Lighthouse Lab in Alderley Park | Wellcome Sanger Institute for the COVID-19 Genomics UK (COG-UK) Consortium | Jacquelyn Wynn, Mairead Hyland, The Lighthouse Lab in Alderley Park and Alex Alderton, Roberto Amato, Jeffrey Barrett, Sonia Goncalves, Ewan Harrison, David K. Jackson, Ian Johnston, Dominic Kwiatkowski, Cordelia Langford, John Sillitoe on behalf of the Wellcome Sanger Institute COVID-19 Surveillance Team |
| EPI_ISL_1829951, EPI_ISL_1829958 | Randox Laboratories | Wellcome Sanger Institute for the COVID-19 Genomics UK (COG-UK) Consortium | Randox Laboratories and Alex Alderton, Roberto Amato, Jeffrey Barrett, Sonia Goncalves, Ewan Harrison, David K. Jackson, Ian Johnston, Dominic Kwiatkowski, Cordelia Langford, John Sillitoe on behalf of the Wellcome Sanger Institute COVID-19 Surveillance Team |
| EPI_ISL_1829964, EPI_ISL_1829966 | Lighthouse Lab in Alderley Park | Wellcome Sanger Institute for the COVID-19 Genomics UK (COG-UK) Consortium | Jacquelyn Wynn, Mairead Hyland, The Lighthouse Lab in Alderley Park and Alex Alderton, Roberto Amato, Jeffrey Barrett, Sonia Goncalves, Ewan Harrison, David K. Jackson, Ian Johnston, Dominic Kwiatkowski, Cordelia Langford, John Sillitoe on behalf of the Wellcome Sanger Institute COVID-19 Surveillance Team |
| EPI_ISL_1829967 | Randox Laboratories | Wellcome Sanger Institute for the COVID-19 Genomics UK (COG-UK) Consortium | Randox Laboratories and Alex Alderton, Roberto Amato, Jeffrey Barrett, Sonia Goncalves, Ewan Harrison, David K. Jackson, Ian Johnston, Dominic Kwiatkowski, Cordelia Langford, John Sillitoe on behalf of the Wellcome Sanger Institute COVID-19 Surveillance Team |

| | | | |
|---|---|---|---|
| EPI_ISL_1829979, EPI_ISL_1829983, EPI_ISL_1829994 | Lighthouse Lab in Alderley Park | Wellcome Sanger Institute for the COVID-19 Genomics UK (COG-UK) Consortium | Jacquelyn Wynn, Mairead Hyland, The Lighthouse Lab in Alderley Park and Alex Alderton, Roberto Amato, Jeffrey Barrett, Sonia Goncalves, Ewan Harrison, David K. Jackson, Ian Johnston, Dominic Kwiatkowski, Cordelia Langford, John Sillitoe on behalf of the Wellcome Sanger Institute COVID-19 Surveillance Team |
| EPI_ISL_1830001, EPI_ISL_1830033 | Randox Laboratories | Wellcome Sanger Institute for the COVID-19 Genomics UK (COG-UK) Consortium | Randox Laboratories and Alex Alderton, Roberto Amato, Jeffrey Barrett, Sonia Goncalves, Ewan Harrison, David K. Jackson, Ian Johnston, Dominic Kwiatkowski, Cordelia Langford, John Sillitoe on behalf of the Wellcome Sanger Institute COVID-19 Surveillance Team |
| EPI_ISL_1830046 | Lighthouse Lab in Alderley Park | Wellcome Sanger Institute for the COVID-19 Genomics UK (COG-UK) Consortium | Jacquelyn Wynn, Mairead Hyland, The Lighthouse Lab in Alderley Park and Alex Alderton, Roberto Amato, Jeffrey Barrett, Sonia Goncalves, Ewan Harrison, David K. Jackson, Ian Johnston, Dominic Kwiatkowski, Cordelia Langford, John Sillitoe on behalf of the Wellcome Sanger Institute COVID-19 Surveillance Team |
| EPI_ISL_1830056 | Lighthouse Lab in Glasgow | Wellcome Sanger Institute for the COVID-19 Genomics UK (COG-UK) Consortium | Harper VanSteenhouse, Yumi Kasai, David Gray, Carol Clugston, Anna Dominiczak and Alex Alderton, Roberto Amato, Jeffrey Barrett, Sonia Goncalves, Ewan Harrison, David K. Jackson, Ian Johnston, Dominic Kwiatkowski, Cordelia Langford, John Sillitoe on behalf of the Wellcome Sanger Institute COVID-19 Surveillance Team |
| EPI_ISL_1830065, EPI_ISL_1830069, EPI_ISL_1830077, EPI_ISL_1830080 | Lighthouse Lab in Alderley Park | Wellcome Sanger Institute for the COVID-19 Genomics UK (COG-UK) Consortium | Jacquelyn Wynn, Mairead Hyland, The Lighthouse Lab in Alderley Park and Alex Alderton, Roberto Amato, Jeffrey Barrett, Sonia Goncalves, Ewan Harrison, David K. Jackson, Ian Johnston, Dominic Kwiatkowski, Cordelia Langford, John Sillitoe on behalf of the Wellcome Sanger Institute COVID-19 Surveillance Team |
| EPI_ISL_1830108 | Randox Laboratories | Wellcome Sanger Institute for the COVID-19 Genomics UK (COG-UK) Consortium | Randox Laboratories and Alex Alderton, Roberto Amato, Jeffrey Barrett, Sonia Goncalves, Ewan Harrison, David K. Jackson, Ian Johnston, Dominic Kwiatkowski, Cordelia Langford, John Sillitoe on behalf of the Wellcome Sanger Institute COVID-19 Surveillance Team |
| EPI_ISL_1830111, EPI_ISL_1830177, EPI_ISL_1830179, EPI_ISL_1830186 | Lighthouse Lab in Milton Keynes | Wellcome Sanger Institute for the COVID-19 Genomics UK (COG-UK) Consortium | The Lighthouse Lab in Milton Keynes and Alex Alderton, Roberto Amato, Jeffrey Barrett, Sonia Goncalves, Ewan Harrison, David K. Jackson, Ian Johnston, Dominic Kwiatkowski, Cordelia Langford, John Sillitoe on behalf of the Wellcome Sanger Institute COVID-19 Surveillance Team |
| EPI_ISL_1830209, EPI_ISL_1830245 | Randox Laboratories | Wellcome Sanger Institute for the COVID-19 Genomics UK (COG-UK) Consortium | Randox Laboratories and Alex Alderton, Roberto Amato, Jeffrey Barrett, Sonia Goncalves, Ewan Harrison, David K. Jackson, Ian Johnston, Dominic Kwiatkowski, Cordelia Langford, John Sillitoe on behalf of the Wellcome Sanger Institute COVID-19 Surveillance Team |
| EPI_ISL_1830269 | Lighthouse Lab in Milton Keynes | Wellcome Sanger Institute for the COVID-19 Genomics UK (COG-UK) Consortium | The Lighthouse Lab in Milton Keynes and Alex Alderton, Roberto Amato, Jeffrey Barrett, Sonia Goncalves, Ewan Harrison, David K. Jackson, Ian Johnston, Dominic Kwiatkowski, Cordelia Langford, John Sillitoe on behalf of the Wellcome Sanger Institute COVID-19 Surveillance Team |
| EPI_ISL_1830295 | Lighthouse Lab in Glasgow | Wellcome Sanger Institute for the COVID-19 Genomics UK (COG-UK) Consortium | Harper VanSteenhouse, Yumi Kasai, David Gray, Carol Clugston, Anna Dominiczak and Alex Alderton, Roberto Amato, Jeffrey Barrett, Sonia Goncalves, Ewan Harrison, David K. Jackson, Ian Johnston, Dominic Kwiatkowski, Cordelia Langford, John Sillitoe on behalf of the Wellcome Sanger Institute COVID-19 Surveillance Team |
| EPI_ISL_1830305, EPI_ISL_1830328, EPI_ISL_1830341 | Lighthouse Lab in Milton Keynes | Wellcome Sanger Institute for the COVID-19 Genomics UK (COG-UK) Consortium | The Lighthouse Lab in Milton Keynes and Alex Alderton, Roberto Amato, Jeffrey Barrett, Sonia Goncalves, Ewan Harrison, David K. Jackson, Ian Johnston, Dominic Kwiatkowski, Cordelia Langford, John Sillitoe on behalf of the Wellcome Sanger Institute COVID-19 Surveillance Team |
| EPI_ISL_1830356 | Lighthouse Lab in Alderley Park | Wellcome Sanger Institute for the COVID-19 Genomics UK (COG-UK) Consortium | Jacquelyn Wynn, Mairead Hyland, The Lighthouse Lab in Alderley Park and Alex Alderton, Roberto Amato, Jeffrey Barrett, Sonia Goncalves, Ewan Harrison, David K. Jackson, Ian Johnston, Dominic Kwiatkowski, Cordelia Langford, John Sillitoe on behalf of the Wellcome Sanger Institute COVID-19 Surveillance Team |
| EPI_ISL_1830372, EPI_ISL_1830374, EPI_ISL_1830403 | Lighthouse Lab in Milton Keynes | Wellcome Sanger Institute for the COVID-19 Genomics UK (COG-UK) Consortium | The Lighthouse Lab in Milton Keynes and Alex Alderton, Roberto Amato, Jeffrey Barrett, Sonia Goncalves, Ewan Harrison, David K. Jackson, Ian Johnston, Dominic Kwiatkowski, Cordelia Langford, John Sillitoe on behalf of the Wellcome Sanger Institute COVID-19 Surveillance Team |
| EPI_ISL_1830480 | Lighthouse Lab in Alderley Park | Wellcome Sanger Institute for the COVID-19 Genomics UK (COG-UK) Consortium | Jacquelyn Wynn, Mairead Hyland, The Lighthouse Lab in Alderley Park and Alex Alderton, Roberto Amato, Jeffrey Barrett, Sonia Goncalves, Ewan Harrison, David K. Jackson, Ian Johnston, Dominic Kwiatkowski, Cordelia Langford, John Sillitoe on behalf of the Wellcome Sanger Institute COVID-19 Surveillance Team |
| EPI_ISL_1830493 | Lighthouse Lab in Glasgow | Wellcome Sanger Institute for the COVID-19 Genomics UK (COG-UK) Consortium | Harper VanSteenhouse, Yumi Kasai, David Gray, Carol Clugston, Anna Dominiczak and Alex Alderton, Roberto Amato, Jeffrey Barrett, Sonia Goncalves, Ewan Harrison, David K. Jackson, Ian Johnston, Dominic Kwiatkowski, Cordelia Langford, John Sillitoe on behalf of the Wellcome Sanger Institute COVID-19 Surveillance Team |
| EPI_ISL_1830531 | Lighthouse Lab in Milton Keynes | Wellcome Sanger Institute for the COVID-19 Genomics UK (COG-UK) Consortium | The Lighthouse Lab in Milton Keynes and Alex Alderton, Roberto Amato, Jeffrey Barrett, Sonia Goncalves, Ewan Harrison, David K. Jackson, Ian Johnston, Dominic Kwiatkowski, Cordelia Langford, John Sillitoe on behalf of the Wellcome Sanger Institute COVID-19 Surveillance Team |
| EPI_ISL_1830543, EPI_ISL_1830558 | Lighthouse Lab in Alderley Park | Wellcome Sanger Institute for the COVID-19 Genomics UK (COG-UK) Consortium | Jacquelyn Wynn, Mairead Hyland, The Lighthouse Lab in Alderley Park and Alex Alderton, Roberto Amato, Jeffrey Barrett, Sonia Goncalves, Ewan Harrison, David K. Jackson, Ian Johnston, Dominic Kwiatkowski, Cordelia Langford, John Sillitoe on behalf of the Wellcome Sanger Institute COVID-19 Surveillance Team |
| EPI_ISL_1830568, EPI_ISL_1830574 | Lighthouse Lab in Milton Keynes | Wellcome Sanger Institute for the COVID-19 Genomics UK (COG-UK) Consortium | The Lighthouse Lab in Milton Keynes and Alex Alderton, Roberto Amato, Jeffrey Barrett, Sonia Goncalves, Ewan Harrison, David K. Jackson, Ian Johnston, Dominic Kwiatkowski, Cordelia Langford, John Sillitoe on behalf of the Wellcome Sanger Institute COVID-19 Surveillance Team |
| EPI_ISL_1830575 | Lighthouse Lab in Alderley Park | Wellcome Sanger Institute for the COVID-19 Genomics UK (COG-UK) Consortium | Jacquelyn Wynn, Mairead Hyland, The Lighthouse Lab in Alderley Park and Alex Alderton, Roberto Amato, Jeffrey Barrett, Sonia Goncalves, Ewan Harrison, David K. Jackson, Ian Johnston, Dominic Kwiatkowski, Cordelia Langford, John Sillitoe on behalf of the Wellcome Sanger Institute COVID-19 Surveillance Team |
| EPI_ISL_1830578 | Lighthouse Lab in Milton Keynes | Wellcome Sanger Institute for the COVID-19 Genomics UK (COG-UK) Consortium | The Lighthouse Lab in Milton Keynes and Alex Alderton, Roberto Amato, Jeffrey Barrett, Sonia Goncalves, Ewan Harrison, David K. Jackson, Ian Johnston, Dominic Kwiatkowski, Cordelia Langford, John Sillitoe on behalf of the Wellcome Sanger Institute COVID-19 Surveillance Team |
| EPI_ISL_1830586 | Lighthouse Lab in Alderley Park | Wellcome Sanger Institute for the COVID-19 Genomics UK (COG-UK) Consortium | Jacquelyn Wynn, Mairead Hyland, The Lighthouse Lab in Alderley Park and Alex Alderton, Roberto Amato, Jeffrey Barrett, Sonia Goncalves, Ewan Harrison, David K. Jackson, Ian Johnston, Dominic Kwiatkowski, Cordelia Langford, John Sillitoe on behalf of the Wellcome Sanger Institute COVID-19 Surveillance Team |
| EPI_ISL_1830589, EPI_ISL_1830614, EPI_ISL_1830623, EPI_ISL_1830637, EPI_ISL_1830649, EPI_ISL_1830650, EPI_ISL_1830667, EPI_ISL_1830692 | Lighthouse Lab in Milton Keynes | Wellcome Sanger Institute for the COVID-19 Genomics UK (COG-UK) Consortium | The Lighthouse Lab in Milton Keynes and Alex Alderton, Roberto Amato, Jeffrey Barrett, Sonia Goncalves, Ewan Harrison, David K. Jackson, Ian Johnston, Dominic Kwiatkowski, Cordelia Langford, John Sillitoe on behalf of the Wellcome Sanger Institute COVID-19 Surveillance Team |
| EPI_ISL_1830711 | Lighthouse Lab in Alderley Park | Wellcome Sanger Institute for the COVID-19 Genomics UK (COG-UK) Consortium | Jacquelyn Wynn, Mairead Hyland, The Lighthouse Lab in Alderley Park and Alex Alderton, Roberto Amato, Jeffrey Barrett, Sonia Goncalves, Ewan Harrison, David K. Jackson, Ian Johnston, Dominic Kwiatkowski, Cordelia Langford, John Sillitoe on behalf of the Wellcome Sanger Institute COVID-19 Surveillance Team |
| EPI_ISL_1830720 | Lighthouse Lab in Milton Keynes | Wellcome Sanger Institute for the COVID-19 Genomics UK (COG-UK) Consortium | The Lighthouse Lab in Milton Keynes and Alex Alderton, Roberto Amato, Jeffrey Barrett, Sonia Goncalves, Ewan Harrison, David K. Jackson, Ian Johnston, Dominic Kwiatkowski, Cordelia Langford, John Sillitoe on behalf of the Wellcome Sanger Institute COVID-19 Surveillance Team |
| EPI_ISL_1830765, EPI_ISL_1830794, EPI_ISL_1830879, EPI_ISL_1830886 | Lighthouse Lab in Glasgow | Wellcome Sanger Institute for the COVID-19 Genomics UK (COG-UK) Consortium | Harper VanSteenhouse, Yumi Kasai, David Gray, Carol Clugston, Anna Dominiczak and Alex Alderton, Roberto Amato, Jeffrey Barrett, Sonia Goncalves, Ewan Harrison, David K. Jackson, Ian Johnston, Dominic Kwiatkowski, Cordelia Langford, John Sillitoe on behalf of the Wellcome Sanger Institute COVID-19 Surveillance Team |
| EPI_ISL_1831002, EPI_ISL_1831023 | Lighthouse Lab in Alderley Park | Wellcome Sanger Institute for the COVID-19 Genomics UK (COG-UK) Consortium | Jacquelyn Wynn, Mairead Hyland, The Lighthouse Lab in Alderley Park and Alex Alderton, Roberto Amato, Jeffrey Barrett, Sonia Goncalves, Ewan Harrison, David K. Jackson, Ian Johnston, Dominic Kwiatkowski, Cordelia Langford, John Sillitoe on behalf of the Wellcome Sanger Institute COVID-19 Surveillance Team |
| EPI_ISL_1831030, EPI_ISL_1831174 | Lighthouse Lab in Glasgow | Wellcome Sanger Institute for the COVID-19 Genomics UK (COG-UK) Consortium | Harper VanSteenhouse, Yumi Kasai, David Gray, Carol Clugston, Anna Dominiczak and Alex Alderton, Roberto Amato, Jeffrey Barrett, Sonia Goncalves, Ewan Harrison, David K. Jackson, Ian Johnston, Dominic Kwiatkowski, Cordelia Langford, John Sillitoe on behalf of the Wellcome Sanger Institute COVID-19 Surveillance Team |
| EPI_ISL_1831269, EPI_ISL_1831271, EPI_ISL_1831273, EPI_ISL_1831275, EPI_ISL_1831277, EPI_ISL_1831278, EPI_ISL_1831283 | Lighthouse Lab in Milton Keynes | Wellcome Sanger Institute for the COVID-19 Genomics UK (COG-UK) Consortium | The Lighthouse Lab in Milton Keynes and Alex Alderton, Roberto Amato, Jeffrey Barrett, Sonia Goncalves, Ewan Harrison, David K. Jackson, Ian Johnston, Dominic Kwiatkowski, Cordelia Langford, John Sillitoe on behalf of the Wellcome Sanger Institute COVID-19 Surveillance Team |
| EPI_ISL_1831284, EPI_ISL_1831287, EPI_ISL_1831292, EPI_ISL_1831294, EPI_ISL_1831295, EPI_ISL_1831297, EPI_ISL_1831310, EPI_ISL_1831312, EPI_ISL_1831331, EPI_ISL_1831333, EPI_ISL_1831369, EPI_ISL_1831372, EPI_ISL_1831377 | | | |
| see above | Health Services Laboratories | Wellcome Sanger Institute for the COVID-19 Genomics UK (COG-UK) Consortium | Health Services Laboratories and Alex Alderton, Roberto Amato, Jeffrey Barrett, Sonia Goncalves, Ewan Harrison, David K. Jackson, Ian Johnston, Dominic Kwiatkowski, Cordelia Langford, John Sillitoe on behalf of the Wellcome Sanger Institute COVID-19 Surveillance Team |
| EPI_ISL_1832153, EPI_ISL_1832157 | University of Birmingham | COVID-19 Genomics UK (COG-UK) Consortium | Institute of Microbiology, University of Birmingham: Claire McMurray, Joanne Stockton, Samuel Nicholls, Radoslaw Poplawski, Will Rowe, Josh Quick, Nicholas Loman. University of Birmingham Testing Laboratory: Celina M Whalley, Andrew Bosworth, Charlotte Poxon, Kasun Wanigasooriya, Oliver Pickles, Mike Kidd, Alex Richter, Andrew D Beggs PHE Heartlands Lab: Husam Osman, Andrew Bosworth. Queen Elizabeth Hospital: Anna Casey |
| EPI_ISL_1832202, EPI_ISL_1832203, EPI_ISL_1832204, EPI_ISL_1832205, EPI_ISL_1832206, EPI_ISL_1832208, EPI_ISL_1832209, EPI_ISL_1832211, EPI_ISL_1832212, EPI_ISL_1832213, EPI_ISL_1832215, EPI_ISL_1832216, EPI_ISL_1832217, EPI_ISL_1832228 | | | |
| see above | Department of Pathology, University of Cambridge | COVID-19 Genomics UK (COG-UK) Consortium | Aminu S. Jahun, Yasmin Chaudhry, Iliana Georgana, Myra Hosmillo, Rhys Izuagbe, William L. Hamilton, Martin D. Curran, Surendra Parmar, Ian Goodfellow |
| EPI_ISL_1832337, EPI_ISL_1832340 | University Hospitals Of Leicester NHS Trust and DeepSeq Nottingham | COVID-19 Genomics UK (COG-UK) Consortium | Christopher Holmes, Paul Bird, Thomas Helmer, Karlie Fallon, Julian Tang, Jonathan Ball, Patrick McClure, Joeseph Chappell, Nadine Holmes, Matthew Carlisle, Christopher Moore, Fei Sang, Johnny Debebe, Victoria Wright, Matthew Loose |
| EPI_ISL_1832582, EPI_ISL_1832584, EPI_ISL_1832590, EPI_ISL_1832591, EPI_ISL_1832599, EPI_ISL_1832608, EPI_ISL_1832614, EPI_ISL_1832643, EPI_ISL_1832646 | University College London, Great Ormond Street Hospital for Children NHS Foundation Trust, Imperial College Healthcare NHS Trust | COVID-19 Genomics UK (COG-UK) Consortium | Sergi Castellano, Rachel Williams, Mark Kristiansen, Paola Resende Silva, Sunando Roy, Tony Brooks, Helena Tutill, Paola Niola, Patricia Dyal, Charlotte Williams, Leysa Forrest, Yasmin Panchbhaya, Jacqueline Findlay, Samuel Weeks, Julianne Brown, Kathryn Harris, Paul Randell, James Price, Alison Holmes, Judith Breuer |
| EPI_ISL_1832826 | Quadram Institute Bioscience | COVID-19 Genomics UK (COG-UK) Consortium | Dave J. Baker, Gemma L. Kay, Alp Aydin, Thanh Le-Viet, Steven Rudder, Ana P. Tedim, Anastasia Kolyva, Maria Diaz, Leonardo de Oliveira Martins, Nabil-Fareed Alikhan, Lizzie Meadows, Rachael Stanley, Ngozi Elumogo, Muhammed Yasir, |

| | | | |
|---|---|---|---|
| | | | Nicholas M. Thomson, Alexander J Trotter, Rachel Gilroy, Samuel Bloomfield, Claire Stuart, Andrew Bell, Reenesh Prakash, Samir Dervisevic, Alison E. Mather, John Wain, Mark Webber, Andrew J. Page, Justin O'Grady |
| EPI_ISL_1833002 | Queens Medical Centre, Clinical Microbiology Department / DeepSeq Nottingham | COVID-19 Genomics UK (COG-UK) Consortium | Gemma Clark, Wendy Smith, Manjinder Khakh, Vicki M Fleming, Michelle M Lister, Hannah Howson-Wells, Jonathan Ball, Timothy Byaruhanga, Jayasree Dey, Emily Park, Patrick McClure, Joseph Chappell, Theocharis Tsoleridis, Nadine Holmes, Matthew Carlisle, Christopher Moore, Fei Sang, Johnny Debebe, Victoria Wright, Matthew Loose |
| EPI_ISL_1833022 | Lincolnshire Hospitals and DeepSeq Nottingham | COVID-19 Genomics UK (COG-UK) Consortium | Nichola Duckworth, Tim Sloan, Sarah Walsh, Jonathan Ball, Patrick McClure, Joeseph Chappell, Nadine Holmes, Matthew Carlisle, Christopher Moore, Fei Sang, Johnny Debebe, Victoria Wright, Matthew Loose |
| EPI_ISL_1833202 | Wales Specialist Virology Centre Sequencing lab: Pathogen Genomics Unit | Public Health Wales Microbiology Cardiff Wales Specialist Virology Centre | Catherine Moore, Johnathan Evans, Laura Gifford, Malorie Perry, Simon Cottrell, Angela Marchbank, Alec Birchley, Alexander Adams, Amy Gaskin, Bree Gatica-Wilcox, Jason Coombes, Joel Southgate, Lauren Gilbert, Lee Graham, Nicole Pacchiarini, Sara Kumziene-Summerhayes, Sarah Taylor, Sophie Jones, Sara Rey, Matthew Bull, Joanne Watkins, Sally Corden, Tom Connor |
| EPI_ISL_1833278 | Centre for Enzyme Innovation, University of Portsmouth / Translational Research Laboratory, Portsmouth Hospitals NHS Trust | COVID-19 Genomics UK (COG-UK) Consortium | Angela Beckett,Salman Goudarzi,Christopher Fearn,Kate Cook,Katie Loveson,Sharon Glaysher,Scott Elliott,Samuel Robson |
| EPI_ISL_1833679 | Clinical Virology | Clinical Bacteriology | Tim Roloff, Madlen Stange, Helena MB Seth-Smith, Alfredo Mari, Karoline Leuzinger, Julia Bielicki, Manuel Battegay, Hans Hirsch, Adrian Egli |
| EPI_ISL_1833923, EPI_ISL_1833946 | Bioinformatics and Biostatistics Lab, Advanced Sequencing Facility | COVID-19 Genomics UK (COG-UK) Consortium | Aengus Stewart,Jerome Nicod,Chelsea Sawyer,Laura Cubitt,Harshil Patel,Margaret Crawford |
| EPI_ISL_1834352 | Quest Diagnostics Incorporated | Centers for Disease Control and Prevention Division of Viral Diseases, Pathogen Discovery | Dakota Howard, Dhwani Batra, Peter W. Cook, Kara Moser, Adrian Paskey, Jason Caravas, Benjamin Rambo-Martin, Shatavia Morrison, Christopher Gulvick, Scott Sammons, Yvette Unoarumhi, Darlene Wagner, Matthew Schmerer, S. H. Rosenthal, A. Gerasimova, R. M. Kagan, B. Anderson, M. Hua, Y. Liu, L.E. Bernstein, K.E. Livingston, A. Perez, I. A. Shlyakhter, R. V. Rolando, R. Owen, P. Tanpaiboon, F. Lacbawan, Clinton R. Paden, Duncan MacCannell |
| EPI_ISL_1836007, EPI_ISL_1836096 | Aegis Sciences Corporation | Centers for Disease Control and Prevention Division of Viral Diseases, Pathogen Discovery | Dakota Howard, Dhwani Batra, Peter W. Cook, Kara Moser, Adrian Paskey, Jason Caravas, Benjamin Rambo-Martin, Shatavia Morrison, Christopher Gulvick, Scott Sammons, Yvette Unoarumhi, Darlene Wagner, Matthew Schmerer, Cyndi Clark, Patrick Campbell, Rob Case, Vikramsinha Ghorpade, Holly Houdeshell, Ola Kvalvaag, Dillon Nall, Ethan Sanders, Alec Vest, Shaun Westlund, Matthew Hardison, Clinton R. Paden, Duncan MacCannell |
| EPI_ISL_1837991, EPI_ISL_1838012, EPI_ISL_1838023, EPI_ISL_1838042, EPI_ISL_1838045, EPI_ISL_1838063, EPI_ISL_1838073, EPI_ISL_1838080, EPI_ISL_1838083, EPI_ISL_1838086, EPI_ISL_1838109, EPI_ISL_1838114, EPI_ISL_1838123, EPI_ISL_1838125, EPI_ISL_1838134, EPI_ISL_1838151, EPI_ISL_1838194, EPI_ISL_1838217, EPI_ISL_1838225, EPI_ISL_1838229, EPI_ISL_1838260, EPI_ISL_1838262, EPI_ISL_1838265, EPI_ISL_1838282, EPI_ISL_1838285, EPI_ISL_1838292, EPI_ISL_1838312, EPI_ISL_1838315, EPI_ISL_1838322, EPI_ISL_1838327, EPI_ISL_1838337 | | | |
| see above | The National Centre for Cell Science | CSIR-Centre for Cellular and Molecular Biology-INSACOG | Dhiraj Paul, Mohak P Gujare, Shivang P. Bhanushali, Mitali Inamdar, Sonal Manik Chavan, Manoj Kumar Bhat, Ajay Pillai, INSACOG Consortium team, Yogesh Shouche,Payel Mukherjee,Lamuk Zaveri,Tulasi Nagabandi, Ara Sreenivas,Valli Nagalakshmi Undamatla,Shreekant Verma, Amareshwar Vodapalli ,Blessy B John,Viswagithe S L,B Himasri,Onkar Kulkarni,Sofia Banu,Archana Bharadwaj Siva,Sharath Chandra Thota,Karthik Bharadwaj Tallapaka,Rakesh K Mishra,Divya Tej Sowpati |
| EPI_ISL_1838398, EPI_ISL_1838463, EPI_ISL_1838549, EPI_ISL_1838603, EPI_ISL_1838604, EPI_ISL_1838606, EPI_ISL_1838607, EPI_ISL_1838608, EPI_ISL_1838609, EPI_ISL_1838611, EPI_ISL_1838612, EPI_ISL_1838614, EPI_ISL_1838642, EPI_ISL_1838645, EPI_ISL_1838646, EPI_ISL_1838647, EPI_ISL_1838648, EPI_ISL_1838649, EPI_ISL_1838650, EPI_ISL_1838651, EPI_ISL_1838656, EPI_ISL_1838659, EPI_ISL_1838660, EPI_ISL_1838671, EPI_ISL_1838672, EPI_ISL_1838673, EPI_ISL_1838676, EPI_ISL_1838677, EPI_ISL_1838693, EPI_ISL_1838694, EPI_ISL_1838787, EPI_ISL_1838805, EPI_ISL_1838806, EPI_ISL_1838809, EPI_ISL_1838811, EPI_ISL_1838812, EPI_ISL_1838814, EPI_ISL_1838828, EPI_ISL_1838843, EPI_ISL_1838867 | | | |
| see above | CSIR-Centre for Cellular and Molecular Biology | CSIR-Centre for Cellular and Molecular Biology-INSACOG | Payel Mukherjee,Lamuk Zaveri,Tulasi Nagabandi, Ara Sreenivas,Shreekant Verma, Amareshwar Vodapalli ,Blessy B John,Viswagithe S L,B Himasri,Valli Nagalakshmi Undamatla,Onkar Kulkarni,Sofia Banu,Archana Bharadwaj Siva,Sharath Chandra Thota,Karthik Bharadwaj Tallapaka,Rakesh K Mishra,Divya Tej Sowpati |
| EPI_ISL_1839105, EPI_ISL_1839106, EPI_ISL_1839341, EPI_ISL_1839348, EPI_ISL_1839441, EPI_ISL_1839578, EPI_ISL_1839605, EPI_ISL_1839969, EPI_ISL_1840029, EPI_ISL_1840145 | Fulgent Genetics | Centers for Disease Control and Prevention Division of Viral Diseases, Pathogen Discovery | Dakota Howard, Dhwani Batra, Peter W. Cook, Kara Moser, Adrian Paskey, Jason Caravas, Benjamin Rambo-Martin, Shatavia Morrison, Christopher Gulvick, Scott Sammons, Yvette Unoarumhi, Darlene Wagner, Matthew Schmerer, Harry Gao, Mickey Li, John Gao, Joseph Fierro, Benafsh Sapra, Becky Tsai, Yan Meng, Doreen Ng, James Xie, Clinton R. Paden, Duncan MacCannell |
| EPI_ISL_1840480, EPI_ISL_1840498 | Quest Diagnostics Incorporated | Centers for Disease Control and Prevention Division of Viral Diseases, Pathogen Discovery | Dakota Howard, Dhwani Batra, Peter W. Cook, Kara Moser, Adrian Paskey, Jason Caravas, Benjamin Rambo-Martin, Shatavia Morrison, Christopher Gulvick, Scott Sammons, Yvette Unoarumhi, Darlene Wagner, Matthew Schmerer, S. H. Rosenthal, A. Gerasimova, R. M. Kagan, B. Anderson, M. Hua, Y. Liu, L.E. Bernstein, K.E. Livingston, A. Perez, I. A. Shlyakhter, R. V. Rolando, R. Owen, P. Tanpaiboon, F. Lacbawan, Clinton R. Paden, Duncan MacCannell |
| EPI_ISL_1840787 | Azienda Sanitaria dell'Alto Adige Laboratorio Aziendale di Microbiologia e Virologia | Istituto di Genomica Applicata | Elisabetta Pagani, Irene Bianconi, Elisabetta Giacobazzi, Elisa Masi, Stefanie Wieser, Irena Jurman, Vera Vendramin, Gabriele Magris, Eleonora Paparelli, Davide Scaglione, Michele Morgante |
| EPI_ISL_1840801, EPI_ISL_1840814, EPI_ISL_1840828 | Labor Berlin Charité Vivantes GmbH / Institut für Virologie | Charité Universitätsmedizin Berlin, Institut für Virologie/Labor Berlin | Peter Menzel, Christine Stephan, Rolf Schwarzer, Victor M Corman, Barbara Mühlemann, Terry Jones, Christian Drosten |
| EPI_ISL_1841240, EPI_ISL_1841241, EPI_ISL_1841242, EPI_ISL_1841243, EPI_ISL_1841244, EPI_ISL_1841245, EPI_ISL_1841246, EPI_ISL_1841247, EPI_ISL_1841248, EPI_ISL_1841249, EPI_ISL_1841250, EPI_ISL_1841251, EPI_ISL_1841252, EPI_ISL_1841253, EPI_ISL_1841254, EPI_ISL_1841255, EPI_ISL_1841256, EPI_ISL_1841257, EPI_ISL_1841258, EPI_ISL_1841259, EPI_ISL_1841260, EPI_ISL_1841261, EPI_ISL_1841262, EPI_ISL_1841263, EPI_ISL_1841264, EPI_ISL_1841265, EPI_ISL_1841266, EPI_ISL_1841267, EPI_ISL_1841268, EPI_ISL_1841269, EPI_ISL_1841270, EPI_ISL_1841271, EPI_ISL_1841272, EPI_ISL_1841273, EPI_ISL_1841274, EPI_ISL_1841275, EPI_ISL_1841276, EPI_ISL_1841279, EPI_ISL_1841280, EPI_ISL_1841281, EPI_ISL_1841282, EPI_ISL_1841283, EPI_ISL_1841284, EPI_ISL_1841285, EPI_ISL_1841293, EPI_ISL_1841317, EPI_ISL_1841318, EPI_ISL_1841355, EPI_ISL_1841356, EPI_ISL_1841357, EPI_ISL_1841365, EPI_ISL_1841366, EPI_ISL_1841367, EPI_ISL_1841370, EPI_ISL_1841371, EPI_ISL_1841372, EPI_ISL_1841373, EPI_ISL_1841374, EPI_ISL_1841375, EPI_ISL_1841382, EPI_ISL_1841383 | | | |
| see above | ICMR-National Institute of Virology - INSACOG | NIV Influenza | Dr. Varsha Potdar |
| EPI_ISL_1843504 | Labor 28 MVZ GmbH | Robert Koch Institute | unknown |
| EPI_ISL_1843777, EPI_ISL_1843874, EPI_ISL_1843894 | Centogene; Dr. Bauer Laboratoriums GmbH | Robert Koch Institute | unknown |
| EPI_ISL_1845854, EPI_ISL_1846077, EPI_ISL_1846153 | Eurofins LifeCodexx GmbH | Robert Koch Institute | unknown |
| EPI_ISL_1846441 | Uniklinikum Carl Gustav Carus an der TU Dresden; Institut fÃ¼r Virologie | Robert Koch Institute | unknown |
| EPI_ISL_1847444 | MVZ Labor Dr. Limbach & Kollegen GbR | Robert Koch Institute | unknown |
| EPI_ISL_1847807 | SYNLAB MVZ Leinfelden-Echterdingen | Robert Koch Institute | unknown |
| EPI_ISL_1847907 | SYNLAB Labor MÃ¼nchen Zentrum LMZ | Robert Koch Institute | unknown |
| EPI_ISL_1848351 | Oregon State Public Health Laboratory | Oregon State Public Health Laboratory | Rafia Razzaque, Eugene Yeboah, Vanda Makris, Laura Tsaknaridis, John Fontana and Shane Sevey |
| EPI_ISL_1848402 | IMD Labor Frankfurt | Robert Koch Institute | unknown |
| EPI_ISL_1848757 | Labor MÃ¶nchengladbach MVZ Dr. Stein + Kollegen GbR | Robert Koch Institute | unknown |
| EPI_ISL_1849851, EPI_ISL_1850287 | UW Virology Lab | UW Virology Lab | Pavitra Roychoudhury, Hong Xie, Lasata Shrestha, Tien V. Nguyen, Shah Mohamed Bakhash, Michelle Lin, Noah R. Baker, Sean Ellis, Meei-Li Huang, Keith R Jerome, Alexander Greninger |
| EPI_ISL_1850317, EPI_ISL_1850326 | MVZ Medizinisches Labor Hannover GmbH | Robert Koch Institute | unknown |
| EPI_ISL_1850498 | amedes MVZ Hannover | Robert Koch Institute | unknown |
| EPI_ISL_1851102 | MVZ Labor Krone GbR | Robert Koch Institute | unknown |
| EPI_ISL_1851395, EPI_ISL_1851414 | Labor Dr. Heidrich & Kollegen MVZ GmbH Hamburg | Robert Koch Institute | unknown |
| EPI_ISL_1851555 | Synlab MVZ Augsburg | Robert Koch Institute | unknown |
| EPI_ISL_1852081, EPI_ISL_1852099, EPI_ISL_1852126 | MVZ Labor Dr. Limbach & Kollegen GbR | Robert Koch Institute | unknown |
| EPI_ISL_1856819 | UVA Clinical Microbiology Lab | Mathers Lab | Hardik I. Parikh, Katie E. Barry, Amy J. Mathers |
| EPI_ISL_1857081 | Kansas Health and Environmental Lab | Kansas Health and Environmental Lab | Mike Grose, Jonathan Barnell, Ben Olsen, and Phil Adam |
| EPI_ISL_1857556 | Minnesota Department of Health, Public Health Laboratory | Minnesota Department of Health, Public Health Laboratory | Alexandra Lorentz, Jacob Garfin, Matt Plumb, and Xiong Wang |
| EPI_ISL_1857631, EPI_ISL_1857673, EPI_ISL_1857699, EPI_ISL_1857728, EPI_ISL_1857758, EPI_ISL_1857770, EPI_ISL_1857779 | Randox Laboratories | Wellcome Sanger Institute for the COVID-19 Genomics UK (COG-UK) Consortium | Randox Laboratories and Alex Alderton, Roberto Amato, Jeffrey Barrett, Sonia Goncalves, Ewan Harrison, David K. Jackson, Ian Johnston, Dominic Kwiatkowski, Cordelia Langford, John Sillitoe on behalf of the Wellcome Sanger Institute COVID-19 Surveillance Team |
| EPI_ISL_1857806, EPI_ISL_1857812, EPI_ISL_1857859 | Lighthouse Lab in Alderley Park | Wellcome Sanger Institute for the COVID-19 Genomics UK (COG-UK) Consortium | Jacquelyn Wynn, Mairead Hyland, The Lighthouse Lab in Alderley Park and Alex Alderton, Roberto Amato, Jeffrey Barrett, Sonia Goncalves, Ewan Harrison, David K. Jackson, Ian Johnston, Dominic Kwiatkowski, Cordelia Langford, John Sillitoe on behalf of the Wellcome Sanger Institute COVID-19 Surveillance Team |
| EPI_ISL_1857870 | Lighthouse Lab in Milton Keynes | Wellcome Sanger Institute for the COVID-19 Genomics UK (COG-UK) Consortium | The Lighthouse Lab in Milton Keynes and Alex Alderton, Roberto Amato, Jeffrey Barrett, Sonia Goncalves, Ewan Harrison, David K. Jackson, Ian Johnston, Dominic Kwiatkowski, Cordelia Langford, John Sillitoe on behalf of the Wellcome Sanger Institute COVID-19 Surveillance Team |
| EPI_ISL_1857877, EPI_ISL_1857887, EPI_ISL_1857942, | Lighthouse Lab in Alderley Park | Wellcome Sanger Institute for the COVID-19 Genomics UK (COG-UK) Consortium | Jacquelyn Wynn, Mairead Hyland, The Lighthouse Lab in Alderley Park and Alex Alderton, Roberto Amato, Jeffrey Barrett, Sonia Goncalves, Ewan Harrison, David K. Jackson, Ian Johnston, Dominic Kwiatkowski, Cordelia Langford, John Sillitoe on behalf of the Wellcome Sanger Institute COVID-19 Surveillance Team |

| | | | |
|---|---|---|---|
| EPI_ISL_1857943 | | | |
| EPI_ISL_1857944 | Lighthouse Lab in Milton Keynes | Wellcome Sanger Institute for the COVID-19 Genomics UK (COG-UK) Consortium | The Lighthouse Lab in Milton Keynes and Alex Alderton, Roberto Amato, Jeffrey Barrett, Sonia Goncalves, Ewan Harrison, David K. Jackson, Ian Johnston, Dominic Kwiatkowski, Cordelia Langford, John Sillitoe on behalf of the Wellcome Sanger Institute COVID-19 Surveillance Team |
| EPI_ISL_1857977, EPI_ISL_1858006 | Lighthouse Lab in Alderley Park | Wellcome Sanger Institute for the COVID-19 Genomics UK (COG-UK) Consortium | Jacquelyn Wynn, Mairead Hyland, The Lighthouse Lab in Alderley Park and Alex Alderton, Roberto Amato, Jeffrey Barrett, Sonia Goncalves, Ewan Harrison, David K. Jackson, Ian Johnston, Dominic Kwiatkowski, Cordelia Langford, John Sillitoe on behalf of the Wellcome Sanger Institute COVID-19 Surveillance Team |
| EPI_ISL_1858024 | Lighthouse Lab in Milton Keynes | Wellcome Sanger Institute for the COVID-19 Genomics UK (COG-UK) Consortium | The Lighthouse Lab in Milton Keynes and Alex Alderton, Roberto Amato, Jeffrey Barrett, Sonia Goncalves, Ewan Harrison, David K. Jackson, Ian Johnston, Dominic Kwiatkowski, Cordelia Langford, John Sillitoe on behalf of the Wellcome Sanger Institute COVID-19 Surveillance Team |
| EPI_ISL_1858037, EPI_ISL_1858086, EPI_ISL_1858108, EPI_ISL_1858134, EPI_ISL_1858139, EPI_ISL_1858156 | Lighthouse Lab in Alderley Park | Wellcome Sanger Institute for the COVID-19 Genomics UK (COG-UK) Consortium | Jacquelyn Wynn, Mairead Hyland, The Lighthouse Lab in Alderley Park and Alex Alderton, Roberto Amato, Jeffrey Barrett, Sonia Goncalves, Ewan Harrison, David K. Jackson, Ian Johnston, Dominic Kwiatkowski, Cordelia Langford, John Sillitoe on behalf of the Wellcome Sanger Institute COVID-19 Surveillance Team |
| EPI_ISL_1858239 | Lighthouse Lab in Milton Keynes | Wellcome Sanger Institute for the COVID-19 Genomics UK (COG-UK) Consortium | The Lighthouse Lab in Milton Keynes and Alex Alderton, Roberto Amato, Jeffrey Barrett, Sonia Goncalves, Ewan Harrison, David K. Jackson, Ian Johnston, Dominic Kwiatkowski, Cordelia Langford, John Sillitoe on behalf of the Wellcome Sanger Institute COVID-19 Surveillance Team |
| EPI_ISL_1860885, EPI_ISL_1861168 | Department of Virus and Microbiological Special Diagnostics, Statens Serum Institut, Copenhagen, Denmark | Aalborg University | Danish Covid-19 Genome Consortium |
| EPI_ISL_1865437 | Molekylær Medicinsk Afdeling, Aarhus University Hospital, Aarhus, Denmark | Aalborg University | Danish Covid-19 Genome Consortium |
| EPI_ISL_1866210, EPI_ISL_1868349, EPI_ISL_1868655, EPI_ISL_1868943, EPI_ISL_1869673, EPI_ISL_1871129, EPI_ISL_1871787, EPI_ISL_1872528, EPI_ISL_1873137, EPI_ISL_1874259, EPI_ISL_1881663, EPI_ISL_1885762, EPI_ISL_1885979 | | | |
| see above | Department of Virus and Microbiological Special Diagnostics, Statens Serum Institut, Copenhagen, Denmark | Aalborg University | Danish Covid-19 Genome Consortium |
| EPI_ISL_1886104 | Labor Berlin Charité Vivantes GmbH / Institut für Virologie | Charité Universitätsmedizin Berlin, Institut für Virologie/Labor Berlin | Peter Menzel, Christine Stephan, Rolf Schwarzer, Victor M Corman, Barbara Mühlemann, Terry Jones, Christian Drosten |
| EPI_ISL_1889539 | CQRC_QUALITY CONTROL CHEMICAL BIOLOGICAL RISK_AOOR Villa Sofia Cervello Palermo | CQRC_QUALITY CONTROL CHEMICAL BIOLOGICAL RISK_AOOR Villa Sofia Cervello Palermo | Di Gaudio, F., Contino, F., Orlando, V., Buffa, V., Bonvissuto, M., Di Chiara, V., Seidita, G.,Todaro, I., Di Sclafani, A., Lungari, A., Brunacci,G. |
| EPI_ISL_1890934 | Department of Virus and Microbiological Special Diagnostics, Statens Serum Institut, Copenhagen, Denmark | Aalborg University | Danish Covid-19 Genome Consortium |
| EPI_ISL_1890977, EPI_ISL_1890990 | National Virus Reference Laboratory | National Virus Reference Laboratory | Zoe Yandle, Charlene Bennett, Gabriel Gonzalez, Michael Carr, Jonathan Dean, Cillian F De Gascun |
| EPI_ISL_1896403 | Department of Virus and Microbiological Special Diagnostics, Statens Serum Institut, Copenhagen, Denmark | Aalborg University | Danish Covid-19 Genome Consortium |
| EPI_ISL_1897172 | Public Health Ontario Laboratory | Public Health Ontario Laboratory | Vanessa G Allen, Philip Banh, Yao Chen, Richard de Borja, Alireza Eshaghi, Nahuel Fittipaldi, Christine Frantz, Jonathan B Gubbay, Jennifer L Guthrie, Lawrence Heisler, Esha Joshi, Michael Laszloffy, Aimin Li, Michael CY Li, Dean Maxwell, Sandeep Nagra, Samir N Patel, Jared Simpson, Karthikeyan Sivaraman, Ashleigh Sullivan, Yogi Sundaravadanam, Sarah Teatero, Andre Villegas, Matthew Watson, Sandra Zittermann |
| EPI_ISL_1897475 | Oregon State Public Health Laboratory | Oregon State Public Health Laboratory | Rafia Razzaque, Eugene Yeboah, Vanda Makris, Laura Tsaknaridis, John Fontana and Shane Sevey |
| EPI_ISL_1897601, EPI_ISL_1897602, EPI_ISL_1897603, EPI_ISL_1897604, EPI_ISL_1897605, EPI_ISL_1897606, EPI_ISL_1897607, EPI_ISL_1897608, EPI_ISL_1897609, EPI_ISL_1897610, EPI_ISL_1897611, EPI_ISL_1897612, EPI_ISL_1897613, EPI_ISL_1897614, EPI_ISL_1897615, EPI_ISL_1897616, EPI_ISL_1897617, EPI_ISL_1897618, EPI_ISL_1897619, EPI_ISL_1897620, EPI_ISL_1897621 | | | |
| see above | National Public Health Laboratory, National Centre for Infectious Diseases | National Public Health Laboratory, National Centre for Infectious Diseases | Tze Minn Mak, Zhenyang Zhou, Grace Jie Yin Ngan, Royce Ang, Lin Cui, Raymond Tzer Pin Lin |
| EPI_ISL_1899297 | Swedish national genomic surveillance program of SARS-CoV-2 | The Public Health Agency of Sweden | Maximilian Riess, Maria Lind Karlberg, Alma Brolund, Swedish national genomic surveillance program of SARS-CoV-2 |
| EPI_ISL_1904299 | Labor Doz DDr Stefan Mustafa | AGES IMED Vienna | Sara Meschini, Theodhora Ziu, Johanna Schmitt, Elisabeth Walter,Alexander Indra, Stefan Mustafa |
| EPI_ISL_1904449, EPI_ISL_1904454, EPI_ISL_1904457, EPI_ISL_1904459, EPI_ISL_1904460, EPI_ISL_1904462, EPI_ISL_1904463, EPI_ISL_1904464, EPI_ISL_1904465, EPI_ISL_1904466, EPI_ISL_1904468, EPI_ISL_1904469, EPI_ISL_1904470 | | | |
| see above | South Eastern Area Laboratory Services (SEALS) | NSW Health Pathology - Institute of Clinical Pathology and Medical Research; Westmead Hospital; University of Sydney | CIDM-PH et al. |
| EPI_ISL_1904652, EPI_ISL_1904670, EPI_ISL_1904703, EPI_ISL_1904829, EPI_ISL_1904838 | National Virus Reference Laboratory | National Virus Reference Laboratory | Zoe Yandle, Charlene Bennett, Gabriel Gonzalez, Michael Carr, Jonathan Dean, Cillian F De Gascun |
| EPI_ISL_1904852, EPI_ISL_1904857 | LabPLUS | Institute of Environmental Science and Research (ESR) | Rachel Boyle, SallyAnn Harbison, Olivia Stroeven, Xiaoyun Ren, Matt Storey, Nikki Freed, Muhammad Faisal, Jing Wang, Hermes Perez, Anja Werno, Antje van der Linden, Arlo Upton, Chris Mansell, David Hammer, Dragana Drinkovic, Gary McAuliffe, Hana Sofia Andersson, James Ussher, Jill Sherwood, Josh Freeman, Julia Howard, Juliet Elvy, Mary DeAlmeida, Matt Blakiston, Matthew Rogers, Max Bloomfield, Michael Addidle, Michelle Balm, Sally Roberts, Sarah Jefferies, Sharmini Muttaiyah, Susan Morpeth, Susan Taylor, Timothy Blackmore, Vani Sathyendran, Veronica Playle, Virginia Hope, Erasmus Smit, Lauren Jelly, Olin Silander, Joep de Ligt |
| EPI_ISL_1904862, EPI_ISL_1904863, EPI_ISL_1904868, EPI_ISL_1904869, EPI_ISL_1904870 | OLVZ Aalst | OLVZ Aalst | Anne Vankeerberghen |
| EPI_ISL_1904967, EPI_ISL_1904969, EPI_ISL_1904971, EPI_ISL_1904972, EPI_ISL_1904973, EPI_ISL_1904974 | LAM ORIADE ABBAYE ST MARTIN D'HERES | CNR Virus des Infections Respiratoires - France SUD | Antonin Bal, Gregory Destras, Gwendolyne Burfin, Hadrien Regue, Quentin Semanas, Martine Valette, Bruno Lina, Laurence Josset |
| EPI_ISL_1904989, EPI_ISL_1904992, EPI_ISL_1904994, EPI_ISL_1904998 | HOPITAL SAINT ANDRE | CNR Virus des Infections Respiratoires - France SUD | Antonin Bal, Gregory Destras, Gwendolyne Burfin, Hadrien Regue, Quentin Semanas, Martine Valette, Bruno Lina, Laurence Josset |
| EPI_ISL_1906045 | Quest Diagnostics Incorporated | Centers for Disease Control and Prevention Division of Viral Diseases, Pathogen Discovery | Dakota Howard, Dhwani Batra, Peter W. Cook, Kara Moser, Adrian Paskey, Jason Caravas, Benjamin Rambo-Martin, Shatavia Morrison, Christopher Gulvick, Scott Sammons, Yvette Unoarumhi, Darlene Wagner, Matthew Schmerer, S. H. Rosenthal, A. Gerasimova, R. M. Kagan, B. Anderson, M. Hua, Y. Liu, L.E. Bernstein, K.E. Livingston, A. Perez, I. A. Shlyakhter, R. V. Rolando, R. Owen, P. Tanpaiboon, F. Lacbawan, Clinton R. Paden, Duncan MacCannell |
| EPI_ISL_1907995 | Helix/Illumina | Centers for Disease Control and Prevention Division of Viral Diseases, Pathogen Discovery | Dakota Howard, Dhwani Batra, Peter W. Cook, Kara Moser, Adrian Paskey, Jason Caravas, Benjamin Rambo-Martin, Shatavia Morrison, Christopher Gulvick, Scott Sammons, Yvette Unoarumhi, Darlene Wagner, Matthew Schmerer, Eileen de Feo, Jan Antico, Christine Tran, Matthew Tolentino, Shannon Wickline, Kim Gietzen, Brad Sickler, Jingtao Liu, Eric Allen, Phil Febbo, Nicole L. Washington, Simon White, Geraint Levan, Kelly Schiabor Barrett, Elizabeth Cirulli, Alexandre Bolze, Ary Ascencio, Charlotte Rivera-Garcia, Ryan Cho, Jason Nguyen, Sherry Wang, Jimmy Ramirez, Tyler Cassens, Efren Sandoval, Magnus Isaksson, William Lee, David Becker, Marc Laurent, James Lu, Clinton R. Paden, Duncan MacCannell |
| EPI_ISL_1908494 | MD PHL | MD PHL | Maryland Department of Health Laboratories Administration |
| EPI_ISL_1909092, EPI_ISL_1909093, EPI_ISL_1909094, EPI_ISL_1909095 | WSSE Katowice | 1. Academic Center for Pathomorphological and Genetic-Molecular Diagnostics ltd, Bialystok, Poland 2. National Institute of Public Health - National Institute of Hygiene, Warsaw, Poland | Radosław Charkiewicz, Jacek Nikliński, Przemysław Biecek. Joanna Reszeć, Piotr Majewski, Anetta Sulewska, Piotr Karabowicz, Joanna Kiśluk, Janusz Dzięcioł, Konrad Raczkowski, Katarzyna Zacharczuk, Magdalena Nowakowska, Małgorzata Sadkowska-Todys, Tomasz Wołkowicz |
| EPI_ISL_1910791 | Arizona State University | Arizona State University | Peter T. Skidmore, LaRinda A. Holland, Matthew F. Smith, Rabia Maqsood, Nicholas J. Mellor, Joy M. Blain, Valerie Harris, Joshua LaBaer, Vel Murugan, Efrem S. Lim |
| EPI_ISL_1911182, EPI_ISL_1911185, EPI_ISL_1911187, EPI_ISL_1911189, EPI_ISL_1911190, EPI_ISL_1911191 | Area of Virology, Serology and Virology Division (SAViD), New South Wales Health Pathology Randwick | Virology Research Laboratory; Area of Virology, Serology and Virology Division (SAViD), New South Wales Health Pathology Randwick | Foster, C.; Au, J.; Ruiz Silva, M.; Deveson, I.; Bull, R.; Van Hal, S.; Rawlinson, W. |
| EPI_ISL_1911192 | PathWest Laboratory Medicine WA | PathWest Laboratory Medicine WA Microbial Surveillance UNit | PathWest Laboratory Medicine WA Microbial Surveillance UNit |
| EPI_ISL_1911195, EPI_ISL_1911196, EPI_ISL_1911197, EPI_ISL_1911250 | Zhejiang Provincial Center for Disease Control and Prevention Zhoushan Center for Disease Prevention and Contol | Zhejiang Province Center of Disease Control and prevention | Yanjun Zhang, Bing WU, Hongling Wang |
| EPI_ISL_1912013 | Lighthouse Lab in Milton Keynes | Wellcome Sanger Institute for the COVID-19 Genomics UK (COG-UK) Consortium | The Lighthouse Lab in Milton Keynes and Alex Alderton, Roberto Amato, Jeffrey Barrett, Sonia Goncalves, Ewan Harrison, David K. Jackson, Ian Johnston, Dominic Kwiatkowski, Cordelia Langford, John Sillitoe on behalf of the Wellcome Sanger Institute COVID-19 Surveillance Team |
| EPI_ISL_1912014 | Lighthouse Lab in Alderley Park | Wellcome Sanger Institute for the COVID-19 Genomics UK (COG-UK) Consortium | Jacquelyn Wynn, Mairead Hyland, The Lighthouse Lab in Alderley Park and Alex Alderton, Roberto Amato, Jeffrey Barrett, Sonia Goncalves, Ewan Harrison, David K. Jackson, Ian Johnston, Dominic Kwiatkowski, Cordelia Langford, John Sillitoe on behalf of the Wellcome Sanger Institute COVID-19 Surveillance Team |
| EPI_ISL_1912020, EPI_ISL_1912030, | Lighthouse Lab in Milton Keynes | Wellcome Sanger Institute for the COVID-19 Genomics UK (COG-UK) Consortium | The Lighthouse Lab in Milton Keynes and Alex Alderton, Roberto Amato, Jeffrey Barrett, Sonia Goncalves, Ewan Harrison, David K. Jackson, Ian Johnston, Dominic Kwiatkowski, Cordelia Langford, John Sillitoe on behalf of the Wellcome Sanger Institute COVID-19 Surveillance Team |

| | | | |
|---|---|---|---|
| EPI_ISL_1912032, EPI_ISL_1912034, EPI_ISL_1912039, EPI_ISL_1912048, EPI_ISL_1912051, EPI_ISL_1912058 | | | |
| EPI_ISL_1912066 | Lighthouse Lab in Cambridge | Wellcome Sanger Institute for the COVID-19 Genomics UK (COG-UK) Consortium | Rob Howes, The Lighthouse Lab in Cambridge and Alex Alderton, Roberto Amato, Jeffrey Barrett, Sonia Goncalves, Ewan Harrison, David K. Jackson, Ian Johnston, Dominic Kwiatkowski, Cordelia Langford, John Sillitoe on behalf of the Wellcome Sanger Institute COVID-19 Surveillance Team |
| EPI_ISL_1912090, EPI_ISL_1912101 | Lighthouse Lab in Alderley Park | Wellcome Sanger Institute for the COVID-19 Genomics UK (COG-UK) Consortium | Jacquelyn Wynn, Mairead Hyland, The Lighthouse Lab in Alderley Park and Alex Alderton, Roberto Amato, Jeffrey Barrett, Sonia Goncalves, Ewan Harrison, David K. Jackson, Ian Johnston, Dominic Kwiatkowski, Cordelia Langford, John Sillitoe on behalf of the Wellcome Sanger Institute COVID-19 Surveillance Team |
| EPI_ISL_1912114 | Lighthouse Lab in Cambridge | Wellcome Sanger Institute for the COVID-19 Genomics UK (COG-UK) Consortium | Rob Howes, The Lighthouse Lab in Cambridge and Alex Alderton, Roberto Amato, Jeffrey Barrett, Sonia Goncalves, Ewan Harrison, David K. Jackson, Ian Johnston, Dominic Kwiatkowski, Cordelia Langford, John Sillitoe on behalf of the Wellcome Sanger Institute COVID-19 Surveillance Team |
| EPI_ISL_1912117 | Lighthouse Lab in Milton Keynes | Wellcome Sanger Institute for the COVID-19 Genomics UK (COG-UK) Consortium | The Lighthouse Lab in Milton Keynes and Alex Alderton, Roberto Amato, Jeffrey Barrett, Sonia Goncalves, Ewan Harrison, David K. Jackson, Ian Johnston, Dominic Kwiatkowski, Cordelia Langford, John Sillitoe on behalf of the Wellcome Sanger Institute COVID-19 Surveillance Team |
| EPI_ISL_1912119 | Lighthouse Lab in Cambridge | Wellcome Sanger Institute for the COVID-19 Genomics UK (COG-UK) Consortium | Rob Howes, The Lighthouse Lab in Cambridge and Alex Alderton, Roberto Amato, Jeffrey Barrett, Sonia Goncalves, Ewan Harrison, David K. Jackson, Ian Johnston, Dominic Kwiatkowski, Cordelia Langford, John Sillitoe on behalf of the Wellcome Sanger Institute COVID-19 Surveillance Team |
| EPI_ISL_1912140 | Lighthouse Lab in Milton Keynes | Wellcome Sanger Institute for the COVID-19 Genomics UK (COG-UK) Consortium | The Lighthouse Lab in Milton Keynes and Alex Alderton, Roberto Amato, Jeffrey Barrett, Sonia Goncalves, Ewan Harrison, David K. Jackson, Ian Johnston, Dominic Kwiatkowski, Cordelia Langford, John Sillitoe on behalf of the Wellcome Sanger Institute COVID-19 Surveillance Team |
| EPI_ISL_1912157 | Lighthouse Lab in Cambridge | Wellcome Sanger Institute for the COVID-19 Genomics UK (COG-UK) Consortium | Rob Howes, The Lighthouse Lab in Cambridge and Alex Alderton, Roberto Amato, Jeffrey Barrett, Sonia Goncalves, Ewan Harrison, David K. Jackson, Ian Johnston, Dominic Kwiatkowski, Cordelia Langford, John Sillitoe on behalf of the Wellcome Sanger Institute COVID-19 Surveillance Team |
| EPI_ISL_1912159 | Lighthouse Lab in Milton Keynes | Wellcome Sanger Institute for the COVID-19 Genomics UK (COG-UK) Consortium | The Lighthouse Lab in Milton Keynes and Alex Alderton, Roberto Amato, Jeffrey Barrett, Sonia Goncalves, Ewan Harrison, David K. Jackson, Ian Johnston, Dominic Kwiatkowski, Cordelia Langford, John Sillitoe on behalf of the Wellcome Sanger Institute COVID-19 Surveillance Team |
| EPI_ISL_1912160, EPI_ISL_1912162, EPI_ISL_1912163, EPI_ISL_1912170 | Lighthouse Lab in Alderley Park | Wellcome Sanger Institute for the COVID-19 Genomics UK (COG-UK) Consortium | Jacquelyn Wynn, Mairead Hyland, The Lighthouse Lab in Alderley Park and Alex Alderton, Roberto Amato, Jeffrey Barrett, Sonia Goncalves, Ewan Harrison, David K. Jackson, Ian Johnston, Dominic Kwiatkowski, Cordelia Langford, John Sillitoe on behalf of the Wellcome Sanger Institute COVID-19 Surveillance Team |
| EPI_ISL_1912173 | Lighthouse Lab in Cambridge | Wellcome Sanger Institute for the COVID-19 Genomics UK (COG-UK) Consortium | Rob Howes, The Lighthouse Lab in Cambridge and Alex Alderton, Roberto Amato, Jeffrey Barrett, Sonia Goncalves, Ewan Harrison, David K. Jackson, Ian Johnston, Dominic Kwiatkowski, Cordelia Langford, John Sillitoe on behalf of the Wellcome Sanger Institute COVID-19 Surveillance Team |
| EPI_ISL_1912184 | Lighthouse Lab in Alderley Park | Wellcome Sanger Institute for the COVID-19 Genomics UK (COG-UK) Consortium | Jacquelyn Wynn, Mairead Hyland, The Lighthouse Lab in Alderley Park and Alex Alderton, Roberto Amato, Jeffrey Barrett, Sonia Goncalves, Ewan Harrison, David K. Jackson, Ian Johnston, Dominic Kwiatkowski, Cordelia Langford, John Sillitoe on behalf of the Wellcome Sanger Institute COVID-19 Surveillance Team |
| EPI_ISL_1912203, EPI_ISL_1912205 | Lighthouse Lab in Cambridge | Wellcome Sanger Institute for the COVID-19 Genomics UK (COG-UK) Consortium | Rob Howes, The Lighthouse Lab in Cambridge and Alex Alderton, Roberto Amato, Jeffrey Barrett, Sonia Goncalves, Ewan Harrison, David K. Jackson, Ian Johnston, Dominic Kwiatkowski, Cordelia Langford, John Sillitoe on behalf of the Wellcome Sanger Institute COVID-19 Surveillance Team |
| EPI_ISL_1912207 | Lighthouse Lab in Milton Keynes | Wellcome Sanger Institute for the COVID-19 Genomics UK (COG-UK) Consortium | The Lighthouse Lab in Milton Keynes and Alex Alderton, Roberto Amato, Jeffrey Barrett, Sonia Goncalves, Ewan Harrison, David K. Jackson, Ian Johnston, Dominic Kwiatkowski, Cordelia Langford, John Sillitoe on behalf of the Wellcome Sanger Institute COVID-19 Surveillance Team |
| EPI_ISL_1912238 | Lighthouse Lab in Glasgow | Wellcome Sanger Institute for the COVID-19 Genomics UK (COG-UK) Consortium | Harper VanSteenhouse, Yumi Kasai, David Gray, Carol Clugston, Anna Dominiczak and Alex Alderton, Roberto Amato, Jeffrey Barrett, Sonia Goncalves, Ewan Harrison, David K. Jackson, Ian Johnston, Dominic Kwiatkowski, Cordelia Langford, John Sillitoe on behalf of the Wellcome Sanger Institute COVID-19 Surveillance Team |
| EPI_ISL_1912319 | Lighthouse Lab in Milton Keynes | Wellcome Sanger Institute for the COVID-19 Genomics UK (COG-UK) Consortium | The Lighthouse Lab in Milton Keynes and Alex Alderton, Roberto Amato, Jeffrey Barrett, Sonia Goncalves, Ewan Harrison, David K. Jackson, Ian Johnston, Dominic Kwiatkowski, Cordelia Langford, John Sillitoe on behalf of the Wellcome Sanger Institute COVID-19 Surveillance Team |
| EPI_ISL_1912330 | Lighthouse Lab in Glasgow | Wellcome Sanger Institute for the COVID-19 Genomics UK (COG-UK) Consortium | Harper VanSteenhouse, Yumi Kasai, David Gray, Carol Clugston, Anna Dominiczak and Alex Alderton, Roberto Amato, Jeffrey Barrett, Sonia Goncalves, Ewan Harrison, David K. Jackson, Ian Johnston, Dominic Kwiatkowski, Cordelia Langford, John Sillitoe on behalf of the Wellcome Sanger Institute COVID-19 Surveillance Team |
| EPI_ISL_1912361, EPI_ISL_1912364, EPI_ISL_1912365, EPI_ISL_1912375, EPI_ISL_1912394, EPI_ISL_1912396, EPI_ISL_1912401, EPI_ISL_1912409, EPI_ISL_1912413, EPI_ISL_1912418, EPI_ISL_1912430, EPI_ISL_1912443, EPI_ISL_1912455, EPI_ISL_1912461 | | | |
| see above | Lighthouse Lab in Alderley Park | Wellcome Sanger Institute for the COVID-19 Genomics UK (COG-UK) Consortium | Jacquelyn Wynn, Mairead Hyland, The Lighthouse Lab in Alderley Park and Alex Alderton, Roberto Amato, Jeffrey Barrett, Sonia Goncalves, Ewan Harrison, David K. Jackson, Ian Johnston, Dominic Kwiatkowski, Cordelia Langford, John Sillitoe on behalf of the Wellcome Sanger Institute COVID-19 Surveillance Team |
| EPI_ISL_1912472 | Lighthouse Lab in Glasgow | Wellcome Sanger Institute for the COVID-19 Genomics UK (COG-UK) Consortium | Harper VanSteenhouse, Yumi Kasai, David Gray, Carol Clugston, Anna Dominiczak and Alex Alderton, Roberto Amato, Jeffrey Barrett, Sonia Goncalves, Ewan Harrison, David K. Jackson, Ian Johnston, Dominic Kwiatkowski, Cordelia Langford, John Sillitoe on behalf of the Wellcome Sanger Institute COVID-19 Surveillance Team |
| EPI_ISL_1912486, EPI_ISL_1912498, EPI_ISL_1912516, EPI_ISL_1912521, EPI_ISL_1912523, EPI_ISL_1912553, EPI_ISL_1912564 | Lighthouse Lab in Alderley Park | Wellcome Sanger Institute for the COVID-19 Genomics UK (COG-UK) Consortium | Jacquelyn Wynn, Mairead Hyland, The Lighthouse Lab in Alderley Park and Alex Alderton, Roberto Amato, Jeffrey Barrett, Sonia Goncalves, Ewan Harrison, David K. Jackson, Ian Johnston, Dominic Kwiatkowski, Cordelia Langford, John Sillitoe on behalf of the Wellcome Sanger Institute COVID-19 Surveillance Team |
| EPI_ISL_1912648, EPI_ISL_1912669, EPI_ISL_1912682, EPI_ISL_1912695 | Lighthouse Lab in Glasgow | Wellcome Sanger Institute for the COVID-19 Genomics UK (COG-UK) Consortium | Harper VanSteenhouse, Yumi Kasai, David Gray, Carol Clugston, Anna Dominiczak and Alex Alderton, Roberto Amato, Jeffrey Barrett, Sonia Goncalves, Ewan Harrison, David K. Jackson, Ian Johnston, Dominic Kwiatkowski, Cordelia Langford, John Sillitoe on behalf of the Wellcome Sanger Institute COVID-19 Surveillance Team |
| EPI_ISL_1912729 | Lighthouse Lab in Alderley Park | Wellcome Sanger Institute for the COVID-19 Genomics UK (COG-UK) Consortium | Jacquelyn Wynn, Mairead Hyland, The Lighthouse Lab in Alderley Park and Alex Alderton, Roberto Amato, Jeffrey Barrett, Sonia Goncalves, Ewan Harrison, David K. Jackson, Ian Johnston, Dominic Kwiatkowski, Cordelia Langford, John Sillitoe on behalf of the Wellcome Sanger Institute COVID-19 Surveillance Team |
| EPI_ISL_1912732 | Lighthouse Lab in Glasgow | Wellcome Sanger Institute for the COVID-19 Genomics UK (COG-UK) Consortium | Harper VanSteenhouse, Yumi Kasai, David Gray, Carol Clugston, Anna Dominiczak and Alex Alderton, Roberto Amato, Jeffrey Barrett, Sonia Goncalves, Ewan Harrison, David K. Jackson, Ian Johnston, Dominic Kwiatkowski, Cordelia Langford, John Sillitoe on behalf of the Wellcome Sanger Institute COVID-19 Surveillance Team |
| EPI_ISL_1912752, EPI_ISL_1912753, EPI_ISL_1912757, EPI_ISL_1912758 | Lighthouse Lab in Milton Keynes | Wellcome Sanger Institute for the COVID-19 Genomics UK (COG-UK) Consortium | The Lighthouse Lab in Milton Keynes and Alex Alderton, Roberto Amato, Jeffrey Barrett, Sonia Goncalves, Ewan Harrison, David K. Jackson, Ian Johnston, Dominic Kwiatkowski, Cordelia Langford, John Sillitoe on behalf of the Wellcome Sanger Institute COVID-19 Surveillance Team |
| EPI_ISL_1912766, EPI_ISL_1912771, EPI_ISL_1912782 | Randox Laboratories | Wellcome Sanger Institute for the COVID-19 Genomics UK (COG-UK) Consortium | Randox Laboratories and Alex Alderton, Roberto Amato, Jeffrey Barrett, Sonia Goncalves, Ewan Harrison, David K. Jackson, Ian Johnston, Dominic Kwiatkowski, Cordelia Langford, John Sillitoe on behalf of the Wellcome Sanger Institute COVID-19 Surveillance Team |
| EPI_ISL_1913085 | Area of Virology, Serology and Virology Division (SAViD), New South Wales Health Pathology Randwick | Virology Research Laboratory; Area of Virology, Serology and Virology Division (SAViD), New South Wales Health Pathology Randwick | Foster, C.; Au, J.; Ruiz Silva, M.; Deveson, I.; Bull, R.; Van Hal, S.; Rawlinson, W. |
| EPI_ISL_1913109, EPI_ISL_1913110 | Microbiological Diagnostic Unit - Public Health Laboratory (MDU-PHL) | MDU-PHL | Seemann T., Sait, M.L., Sherry, N.L. |
| EPI_ISL_1913205, EPI_ISL_1913206, EPI_ISL_1913207, EPI_ISL_1913209, EPI_ISL_1913211 | Victorian Infectious Diseases Reference Laboratory (VIDRL) | VIDRL and MDU-PHL | Caly L., Seemann T., Sait, M.L., Druce J., Sherry, N.L. |
| EPI_ISL_1913220, EPI_ISL_1913224 | EL CAMINO HOSPITAL | Santa Clara County public Health Laboratory | Santa Clara County Public Health Department |
| EPI_ISL_1914236 | Viollier AG | Department of Biosystems Science and Engineering, ETH Zurich | Chaoran Chen, Sarah Nadeau, Ivan Topolsky, Emmanouil Dermitzakis, Keith Harshman, Ioannis Xenarios, Henri Pegeot, Lorenzo Cerutti, Deborah Penet, Philipp Jablonski, Lara Fuhrmann, David Dreifuss, Katharina Jahn, Christiane Beckmann, Maurice Redondo, Olivier Kobel, Christoph Noppen, Sophie Seidel, Noemie Santamaria de Souza, Niko Beerenwinkel, Tanja Stadler |
| EPI_ISL_1914237 | Viollier AG | Department of Biosystems Science and Engineering, ETH Zurich | Christian Beisel, Sarah Nadeau, Chaoran Chen, Ivan Topolsky, Philipp Jablonski, Lara Fuhrmann, David Dreifuss, Katharina Jahn, Rebecca Denes, Mirjam Feldkamp, Ina Nissen, Natascha Santacroce, Elodie Burcklen, Christiane Beckmann, Maurice Redondo, Olivier Kobel, Christoph Noppen, Sophie Seidel, Noemie Santamaria de Souza, Niko Beerenwinkel, Tanja Stadler |
| EPI_ISL_1914238 | Viollier AG | Department of Biosystems Science and Engineering, ETH Zurich | Chaoran Chen, Sarah Nadeau, Ivan Topolsky, Emmanouil Dermitzakis, Keith Harshman, Ioannis Xenarios, Henri Pegeot, Lorenzo Cerutti, Deborah Penet, Philipp Jablonski, Lara Fuhrmann, David Dreifuss, Katharina Jahn, Christiane Beckmann, Maurice Redondo, Olivier Kobel, Christoph Noppen, Sophie Seidel, Noemie Santamaria de Souza, Niko Beerenwinkel, Tanja Stadler |
| EPI_ISL_1914239 | Viollier AG | Department of Biosystems Science and Engineering, ETH Zurich | Chaoran Chen, Sarah Nadeau, Catharine Aquino, Ivan Topolsky, Philipp Jablonski, Lara Fuhrmann, David Dreifuss, Katharina Jahn, Andreia Cabral de Gouvea, Maria Domenica Moccia, Simon Gruter, Timothy Sykes, Lennart Opitz, Griffin White, Laura Neff, Doris Popovic, Andrea Patrignani, Jay Tracy, Ralph Schlapbach, Christiane Beckmann, Maurice Redondo, Olivier Kobel, Christoph Noppen, Sophie Seidel, Noemie Santamaria de Souza, Niko Beerenwinkel, Tanja Stadler |
| EPI_ISL_1914584, EPI_ISL_1914588, EPI_ISL_1914591, EPI_ISL_1914592, EPI_ISL_1914593, EPI_ISL_1914594, EPI_ISL_1914595, | State Virus Research and Diagnostic Laboratory (VRDL), AIIMS Raipur | State Virus Research and Diagnostic Laboratory (VRDL), AIIMS Raipur | Pushpendra Singh, Kuldeep Sharma, Priyanka Singh, Somya Sharma, Sanjay Singh Negi, Anudita Bhargava |

| | | | |
|---|---|---|---|
| EPI_ISL_1914596, EPI_ISL_1914597, EPI_ISL_1914598 | | | |
| EPI_ISL_1914639, EPI_ISL_1914640, EPI_ISL_1914641, EPI_ISL_1914642, EPI_ISL_1914643, EPI_ISL_1914644 | National Public Health Laboratory, National Centre for Infectious Diseases | National Public Health Laboratory, National Centre for Infectious Diseases | Tze Minn Mak, Zhenyang Zhou, Grace Jie Yin Ngan, Royce Ang, Lin Cui, Raymond Tzer Pin Lin |
| EPI_ISL_1914668 | PathWest Laboratory Medicine WA | PathWest Laboratory Medicine WA Microbial Surveillance Unit | PathWest Laboratory Medicine WA Microbial Surveillance Unit |
| EPI_ISL_1915155, EPI_ISL_1915156 | Labo Analyses Med | National Reference Center for Viruses of Respiratory Infections, Institut Pasteur, Paris | Marion Barbet, Sylvie Behillil, Méline Bizard, Angela Brisebarre, Camille Capel, Vincent Enouf, Louise Lefrançois, Frédéric Lemoine, Christophe Malabat, Corinne Maufrais, Emmanuelle Permal, Etienne Simon-Lorière, Maud Vanpeene, Sylvie Van der Werf ,Fabienne Artur |
| EPI_ISL_1915436, EPI_ISL_1915437, EPI_ISL_1915439 | Institute of Epidemiology, Disease Control and Research (IEDCR) | Institute for Developing Science and Health Initiatives (ideSHi) | Hassan Afrad, Sadia Rahman, Fidausi Qadri, Tahmina Shirin |
| EPI_ISL_1916411, EPI_ISL_1916412, EPI_ISL_1916414, EPI_ISL_1916417, EPI_ISL_1916423, EPI_ISL_1916424 | CSIR-National Environmental Engineering Research Institute | CSIR-Centre for Cellular and Molecular Biology - INSACOG | Onkar Kulkarni, Lamuk Zaveri, Ara Sreenivas, Sofia Banu, Shreekant Verma, Amareshwar Vodapalli , Viswagithe S L, B Himasri, Sharath Chandra Thota, Karthik Bharadwaj Tallapaka, Krishna Khairnar, Rakesh K Mishra,Divya Tej Sowpati |
| EPI_ISL_1916433, EPI_ISL_1916434, EPI_ISL_1916442, EPI_ISL_1916445, EPI_ISL_1916446, EPI_ISL_1916454, EPI_ISL_1916456, EPI_ISL_1916464, EPI_ISL_1916465, EPI_ISL_1916468, EPI_ISL_1916475, EPI_ISL_1916476, EPI_ISL_1916486, EPI_ISL_1916489 | | | |
| see above | CSIR-National Environmental Engineering Research Institute | CSIR-Centre for Cellular and Molecular Biology - INSACOG | Ara Sreenivas, Onkar Kulkarni, Lamuk Zaveri, Sofia Banu, Shreekant Verma, Amareshwar Vodapalli , Viswagithe S L, B Himasri, Sharath Chandra Thota, Karthik Bharadwaj Tallapaka, Krishna Khairnar, Rakesh K Mishra,Divya Tej Sowpati |
| EPI_ISL_1916510 | Institute for Infectious Diseases, University of Bern, Switzerland | Institute for Infectious Diseases, University of Bern, Switzerland | Stefan Neuenschwander, Christian Baumann, Miguel A Terrazos Miani, Cora Sägesser, Pascal Bittel, Peter Keller, Franziska Suter-Riniker, Stephen L Leib, Alban Ramette |
| EPI_ISL_1917450 | Hospital Center Luxembourg | Laboratoire national de sante, Microbiology, Microbial Genomics Platform | Anke Wienecke-Baldacchino, Catherine Ragimbeau,Jessica Tapp, Fatu Djabi, Lise Pignon, Raoul Salmon, Michel Kohnen, Jean-Hugues Francois, Tamir Abdelrahman |
| EPI_ISL_1917963 | Laboratoires Reunis | Laboratoire national de sante, Microbiology, Microbial Genomics Platform | Anke Wienecke-Baldacchino, Catherine Ragimbeau,Jessica Tapp, Fatu Djabi, Lise Pignon, Raoul Salmon, Bernard Weber, Tamir Abdelrahman |
| EPI_ISL_1918919 | Broad Institute Clinical Research Sequencing Platform | Infectious Disease Program, Broad Institute of Harvard and MIT | Siddle,K.J., Adams,G., Pearlman,L., Gladden-Young,A., Vicente,G., Blumenstiel,B., DeFelice,M., Lee,M., McGovern,S., Lagerborg,K., Rudy,M., DeRuff,K., Carter,A., Normandin,E., Bauer,M., Reilly,S., Tomkins-Tinch,C., Loreth,C., Chaluvadi,S., Meldrim,J., Granger,B., Lemieux,J.E., Birren,B.W., Sabeti,P.C., Larkin,K., Dodge,S., Lennon,N., Madoff,L., Brown,C., Gallagher,G., Smole,S., Park,D.J., Gabriel,S., and MacInnis,B.L. |
| EPI_ISL_1919364 | Department of Medical Microbiology - section Molde, Molde Hospital | Norwegian Institute of Public Health, Department of Virology | Kathrine Stene-Johansen, Kamilla Heddeland Instefjord, Hilde Elshaug, Garcia Llorente Ignacio, Jon Bråte, Engebretsen Serina Beate,Pedersen Benedikte Nevjen, Debech Nadia, Atiya R Ali,Marie Paulsen Madsen, Rasmus Riis Kopperud, Hilde Vollan, Karoline Bragstad, Olav Hungnes |
| EPI_ISL_1919655 | SYNLAB | GIGA Medical Genomics | Keith Durkin, Maria Artesi, Sébastien Bontems, Raphaël Boreux, Bouchra Boujemla, Nathalie Renotte, Cécile Meex, Pierrette Melin, Marie-Pierre Hayette, Vincent Bours |
| EPI_ISL_1919755, EPI_ISL_1919757 | Ospedale Santa Caterina Novella | Istituto Zooprofilattico Sperimentale della Puglia e della Basilicata | Parisi A., Bianco A., Capozzi L., Del Sambro L., Simone D., Difato L., Bruno A. |
| EPI_ISL_1919930 | University of Michigan Clinical Microbiology Laboratory | Lauring Lab, University of Michigan, Department of Microbiology and Immunology | Valesano |
| EPI_ISL_1921165 | National Public Health Organization | Institute of Applied Biosciences, Centre for Research and Technology Hellas | Anastasia Chatzidimitriou et al. |
| EPI_ISL_1921169, EPI_ISL_1921174, EPI_ISL_1921190, EPI_ISL_1921197, EPI_ISL_1921211, EPI_ISL_1921217, EPI_ISL_1921227, EPI_ISL_1921241, EPI_ISL_1921251, EPI_ISL_1921253, EPI_ISL_1921262, EPI_ISL_1921263, EPI_ISL_1921278, EPI_ISL_1921279 | | | |
| see above | Lighthouse Lab in Alderley Park | Wellcome Sanger Institute for the COVID-19 Genomics UK (COG-UK) Consortium | Jacquelyn Wynn, Mairead Hyland, The Lighthouse Lab in Alderley Park and Alex Alderton, Roberto Amato, Jeffrey Barrett, Sonia Goncalves, Ewan Harrison, David K. Jackson, Ian Johnston, Dominic Kwiatkowski, Cordelia Langford, John Sillitoe on behalf of the Wellcome Sanger Institute COVID-19 Surveillance Team |
| EPI_ISL_1921288 | National Public Health Organization | Institute of Applied Biosciences, Centre for Research and Technology Hellas | Anastasia Chatzidimitriou et al. |
| EPI_ISL_1921290, EPI_ISL_1921297, EPI_ISL_1921313, EPI_ISL_1921333, EPI_ISL_1921335, EPI_ISL_1921343, EPI_ISL_1921353, EPI_ISL_1921360, EPI_ISL_1921362, EPI_ISL_1921370, EPI_ISL_1921376, EPI_ISL_1921377, EPI_ISL_1921379, EPI_ISL_1921390, EPI_ISL_1921396, EPI_ISL_1921400, EPI_ISL_1921403, EPI_ISL_1921404, EPI_ISL_1921411, EPI_ISL_1921429, EPI_ISL_1921438 | | | |
| see above | Lighthouse Lab in Alderley Park | Wellcome Sanger Institute for the COVID-19 Genomics UK (COG-UK) Consortium | Jacquelyn Wynn, Mairead Hyland, The Lighthouse Lab in Alderley Park and Alex Alderton, Roberto Amato, Jeffrey Barrett, Sonia Goncalves, Ewan Harrison, David K. Jackson, Ian Johnston, Dominic Kwiatkowski, Cordelia Langford, John Sillitoe on behalf of the Wellcome Sanger Institute COVID-19 Surveillance Team |
| EPI_ISL_1921439, EPI_ISL_1921445, EPI_ISL_1921455 | Lighthouse Lab in Milton Keynes | Wellcome Sanger Institute for the COVID-19 Genomics UK (COG-UK) Consortium | The Lighthouse Lab in Milton Keynes and Alex Alderton, Roberto Amato, Jeffrey Barrett, Sonia Goncalves, Ewan Harrison, David K. Jackson, Ian Johnston, Dominic Kwiatkowski, Cordelia Langford, John Sillitoe on behalf of the Wellcome Sanger Institute COVID-19 Surveillance Team |
| EPI_ISL_1921459 | Lighthouse Lab in Alderley Park | Wellcome Sanger Institute for the COVID-19 Genomics UK (COG-UK) Consortium | Jacquelyn Wynn, Mairead Hyland, The Lighthouse Lab in Alderley Park and Alex Alderton, Roberto Amato, Jeffrey Barrett, Sonia Goncalves, Ewan Harrison, David K. Jackson, Ian Johnston, Dominic Kwiatkowski, Cordelia Langford, John Sillitoe on behalf of the Wellcome Sanger Institute COVID-19 Surveillance Team |
| EPI_ISL_1921463 | Randox Laboratories | Wellcome Sanger Institute for the COVID-19 Genomics UK (COG-UK) Consortium | Randox Laboratories and Alex Alderton, Roberto Amato, Jeffrey Barrett, Sonia Goncalves, Ewan Harrison, David K. Jackson, Ian Johnston, Dominic Kwiatkowski, Cordelia Langford, John Sillitoe on behalf of the Wellcome Sanger Institute COVID-19 Surveillance Team |
| EPI_ISL_1921469, EPI_ISL_1921470, EPI_ISL_1921476, EPI_ISL_1921492 | Lighthouse Lab in Milton Keynes | Wellcome Sanger Institute for the COVID-19 Genomics UK (COG-UK) Consortium | The Lighthouse Lab in Milton Keynes and Alex Alderton, Roberto Amato, Jeffrey Barrett, Sonia Goncalves, Ewan Harrison, David K. Jackson, Ian Johnston, Dominic Kwiatkowski, Cordelia Langford, John Sillitoe on behalf of the Wellcome Sanger Institute COVID-19 Surveillance Team |
| EPI_ISL_1921496 | Lighthouse Lab in Alderley Park | Wellcome Sanger Institute for the COVID-19 Genomics UK (COG-UK) Consortium | Jacquelyn Wynn, Mairead Hyland, The Lighthouse Lab in Alderley Park and Alex Alderton, Roberto Amato, Jeffrey Barrett, Sonia Goncalves, Ewan Harrison, David K. Jackson, Ian Johnston, Dominic Kwiatkowski, Cordelia Langford, John Sillitoe on behalf of the Wellcome Sanger Institute COVID-19 Surveillance Team |
| EPI_ISL_1921501, EPI_ISL_1921505, EPI_ISL_1921517, EPI_ISL_1921519, EPI_ISL_1921521 | Lighthouse Lab in Milton Keynes | Wellcome Sanger Institute for the COVID-19 Genomics UK (COG-UK) Consortium | The Lighthouse Lab in Milton Keynes and Alex Alderton, Roberto Amato, Jeffrey Barrett, Sonia Goncalves, Ewan Harrison, David K. Jackson, Ian Johnston, Dominic Kwiatkowski, Cordelia Langford, John Sillitoe on behalf of the Wellcome Sanger Institute COVID-19 Surveillance Team |
| EPI_ISL_1921523 | Randox Laboratories | Wellcome Sanger Institute for the COVID-19 Genomics UK (COG-UK) Consortium | Randox Laboratories and Alex Alderton, Roberto Amato, Jeffrey Barrett, Sonia Goncalves, Ewan Harrison, David K. Jackson, Ian Johnston, Dominic Kwiatkowski, Cordelia Langford, John Sillitoe on behalf of the Wellcome Sanger Institute COVID-19 Surveillance Team |
| EPI_ISL_1921531 | Lighthouse Lab in Milton Keynes | Wellcome Sanger Institute for the COVID-19 Genomics UK (COG-UK) Consortium | The Lighthouse Lab in Milton Keynes and Alex Alderton, Roberto Amato, Jeffrey Barrett, Sonia Goncalves, Ewan Harrison, David K. Jackson, Ian Johnston, Dominic Kwiatkowski, Cordelia Langford, John Sillitoe on behalf of the Wellcome Sanger Institute COVID-19 Surveillance Team |
| EPI_ISL_1921538 | Randox Laboratories | Wellcome Sanger Institute for the COVID-19 Genomics UK (COG-UK) Consortium | Randox Laboratories and Alex Alderton, Roberto Amato, Jeffrey Barrett, Sonia Goncalves, Ewan Harrison, David K. Jackson, Ian Johnston, Dominic Kwiatkowski, Cordelia Langford, John Sillitoe on behalf of the Wellcome Sanger Institute COVID-19 Surveillance Team |
| EPI_ISL_1921542, EPI_ISL_1921548, EPI_ISL_1921556, EPI_ISL_1921563 | Lighthouse Lab in Milton Keynes | Wellcome Sanger Institute for the COVID-19 Genomics UK (COG-UK) Consortium | The Lighthouse Lab in Milton Keynes and Alex Alderton, Roberto Amato, Jeffrey Barrett, Sonia Goncalves, Ewan Harrison, David K. Jackson, Ian Johnston, Dominic Kwiatkowski, Cordelia Langford, John Sillitoe on behalf of the Wellcome Sanger Institute COVID-19 Surveillance Team |
| EPI_ISL_1921568 | Randox Laboratories | Wellcome Sanger Institute for the COVID-19 Genomics UK (COG-UK) Consortium | Randox Laboratories and Alex Alderton, Roberto Amato, Jeffrey Barrett, Sonia Goncalves, Ewan Harrison, David K. Jackson, Ian Johnston, Dominic Kwiatkowski, Cordelia Langford, John Sillitoe on behalf of the Wellcome Sanger Institute COVID-19 Surveillance Team |
| EPI_ISL_1921572 | Lighthouse Lab in Alderley Park | Wellcome Sanger Institute for the COVID-19 Genomics UK (COG-UK) Consortium | Jacquelyn Wynn, Mairead Hyland, The Lighthouse Lab in Alderley Park and Alex Alderton, Roberto Amato, Jeffrey Barrett, Sonia Goncalves, Ewan Harrison, David K. Jackson, Ian Johnston, Dominic Kwiatkowski, Cordelia Langford, John Sillitoe on behalf of the Wellcome Sanger Institute COVID-19 Surveillance Team |
| EPI_ISL_1921579, EPI_ISL_1921586 | Lighthouse Lab in Milton Keynes | Wellcome Sanger Institute for the COVID-19 Genomics UK (COG-UK) Consortium | The Lighthouse Lab in Milton Keynes and Alex Alderton, Roberto Amato, Jeffrey Barrett, Sonia Goncalves, Ewan Harrison, David K. Jackson, Ian Johnston, Dominic Kwiatkowski, Cordelia Langford, John Sillitoe on behalf of the Wellcome Sanger Institute COVID-19 Surveillance Team |
| EPI_ISL_1921596 | Randox Laboratories | Wellcome Sanger Institute for the COVID-19 Genomics UK (COG-UK) Consortium | Randox Laboratories and Alex Alderton, Roberto Amato, Jeffrey Barrett, Sonia Goncalves, Ewan Harrison, David K. Jackson, Ian Johnston, Dominic Kwiatkowski, Cordelia Langford, John Sillitoe on behalf of the Wellcome Sanger Institute COVID-19 Surveillance Team |
| EPI_ISL_1921602 | Lighthouse Lab in Milton Keynes | Wellcome Sanger Institute for the COVID-19 Genomics UK (COG-UK) Consortium | The Lighthouse Lab in Milton Keynes and Alex Alderton, Roberto Amato, Jeffrey Barrett, Sonia Goncalves, Ewan Harrison, David K. Jackson, Ian Johnston, Dominic Kwiatkowski, Cordelia Langford, John Sillitoe on behalf of the Wellcome Sanger Institute COVID-19 Surveillance Team |
| EPI_ISL_1921631 | Randox Laboratories | Wellcome Sanger Institute for the COVID-19 Genomics UK (COG-UK) Consortium | Randox Laboratories and Alex Alderton, Roberto Amato, Jeffrey Barrett, Sonia Goncalves, Ewan Harrison, David K. Jackson, Ian Johnston, Dominic Kwiatkowski, Cordelia Langford, John Sillitoe on behalf of the Wellcome Sanger Institute COVID-19 Surveillance Team |
| EPI_ISL_1921638, EPI_ISL_1921640, EPI_ISL_1921649, EPI_ISL_1921653, EPI_ISL_1921661, EPI_ISL_1921665, | Lighthouse Lab in Milton Keynes | Wellcome Sanger Institute for the COVID-19 Genomics UK (COG-UK) Consortium | The Lighthouse Lab in Milton Keynes and Alex Alderton, Roberto Amato, Jeffrey Barrett, Sonia Goncalves, Ewan Harrison, David K. Jackson, Ian Johnston, Dominic Kwiatkowski, Cordelia Langford, John Sillitoe on behalf of the Wellcome Sanger Institute COVID-19 Surveillance Team |

| | | | |
|---|---|---|---|
| EPI_ISL_1921678, EPI_ISL_1921682, EPI_ISL_1921683, EPI_ISL_1921688 | | | |
| EPI_ISL_1921691 | Randox Laboratories | Wellcome Sanger Institute for the COVID-19 Genomics UK (COG-UK) Consortium | Randox Laboratories and Alex Alderton, Roberto Amato, Jeffrey Barrett, Sonia Goncalves, Ewan Harrison, David K. Jackson, Ian Johnston, Dominic Kwiatkowski, Cordelia Langford, John Sillitoe on behalf of the Wellcome Sanger Institute COVID-19 Surveillance Team |
| EPI_ISL_1921704, EPI_ISL_1921705, EPI_ISL_1921707 | Lighthouse Lab in Milton Keynes | Wellcome Sanger Institute for the COVID-19 Genomics UK (COG-UK) Consortium | The Lighthouse Lab in Milton Keynes and Alex Alderton, Roberto Amato, Jeffrey Barrett, Sonia Goncalves, Ewan Harrison, David K. Jackson, Ian Johnston, Dominic Kwiatkowski, Cordelia Langford, John Sillitoe on behalf of the Wellcome Sanger Institute COVID-19 Surveillance Team |
| EPI_ISL_1921713 | Randox Laboratories | Wellcome Sanger Institute for the COVID-19 Genomics UK (COG-UK) Consortium | Randox Laboratories and Alex Alderton, Roberto Amato, Jeffrey Barrett, Sonia Goncalves, Ewan Harrison, David K. Jackson, Ian Johnston, Dominic Kwiatkowski, Cordelia Langford, John Sillitoe on behalf of the Wellcome Sanger Institute COVID-19 Surveillance Team |
| EPI_ISL_1921719 | Lighthouse Lab in Milton Keynes | Wellcome Sanger Institute for the COVID-19 Genomics UK (COG-UK) Consortium | The Lighthouse Lab in Milton Keynes and Alex Alderton, Roberto Amato, Jeffrey Barrett, Sonia Goncalves, Ewan Harrison, David K. Jackson, Ian Johnston, Dominic Kwiatkowski, Cordelia Langford, John Sillitoe on behalf of the Wellcome Sanger Institute COVID-19 Surveillance Team |
| EPI_ISL_1921731 | Randox Laboratories | Wellcome Sanger Institute for the COVID-19 Genomics UK (COG-UK) Consortium | Randox Laboratories and Alex Alderton, Roberto Amato, Jeffrey Barrett, Sonia Goncalves, Ewan Harrison, David K. Jackson, Ian Johnston, Dominic Kwiatkowski, Cordelia Langford, John Sillitoe on behalf of the Wellcome Sanger Institute COVID-19 Surveillance Team |
| EPI_ISL_1921736 | Lighthouse Lab in Milton Keynes | Wellcome Sanger Institute for the COVID-19 Genomics UK (COG-UK) Consortium | The Lighthouse Lab in Milton Keynes and Alex Alderton, Roberto Amato, Jeffrey Barrett, Sonia Goncalves, Ewan Harrison, David K. Jackson, Ian Johnston, Dominic Kwiatkowski, Cordelia Langford, John Sillitoe on behalf of the Wellcome Sanger Institute COVID-19 Surveillance Team |
| EPI_ISL_1921739, EPI_ISL_1921764, EPI_ISL_1921765, EPI_ISL_1921768 | National Public Health Organization | Institute of Applied Biosciences, Centre for Research and Technology Hellas | Anastasia Chatzidimitriou et al. |
| EPI_ISL_1921855 | Labor Berlin Charite Vivantes GmbH / Institut für Virologie | Charité Universitätsmedizin Berlin, Institut für Virologie/Labor Berlin | Peter Menzel, Christine Stephan, Rolf Schwarzer, Victor M Corman, Barbara Mühlemann, Terry Jones, Christian Drosten |
| EPI_ISL_1922125 | University of Bari Biomedical Sciences and Human Oncology | University of Bari Biomedical Sciences and Human Oncology | Chironna M., Sallustio A., Loconsole D., Accogli M. |
| EPI_ISL_1922480 | Jessa | Jessa | Cruys et al. on behalf of the Jessa_cmdLab |
| EPI_ISL_1922708 | Wichita State University - Molecular Diagnostics Lab | Kansas Health and Environmental Lab | Mike Grose, Jonathan Barnell, Ben Olsen, and Phil Adam |
| EPI_ISL_1922839 | NYU Langone Health | Departments of Pathology and Medicine, New York University School of Medicine | Adriana Heguy, Dacia Dimartino, Emily Guzman, Christian Marier, Peter Meyn, Sitharam Ramaswami, Gael Westby, Paul Zappile, Yutong Zhang, Paolo Cotzia, Guiqing Wang |
| EPI_ISL_1923114 | Aegis Sciences Corporation | Centers for Disease Control and Prevention Division of Viral Diseases, Pathogen Discovery | Dakota Howard, Dhwani Batra, Peter W. Cook, Kara Moser, Adrian Paskey, Jason Caravas, Benjamin Rambo-Martin, Shatavia Morrison, Christopher Gulvick, Scott Sammons, Yvette Unoarumhi, Darlene Wagner, Matthew Schmerer, Cyndi Clark, Patrick Campbell, Rob Case, Vikramsinha Ghorpade, Holly Houdeshell, Ola Kvalvaag, Dillon Nall, Ethan Sanders, Alec Vest, Shaun Westlund, Matthew Hardison, Clinton R. Paden, Duncan MacCannell |
| EPI_ISL_1923331, EPI_ISL_1923362, EPI_ISL_1923401, EPI_ISL_1923785, EPI_ISL_1923911, EPI_ISL_1924007, EPI_ISL_1924031 | Helix/Illumina | Centers for Disease Control and Prevention Division of Viral Diseases, Pathogen Discovery | Dakota Howard, Dhwani Batra, Peter W. Cook, Kara Moser, Adrian Paskey, Jason Caravas, Benjamin Rambo-Martin, Shatavia Morrison, Christopher Gulvick, Scott Sammons, Yvette Unoarumhi, Darlene Wagner, Matthew Schmerer, Eileen de Feo, Jan Antico, Christine Tran, Matthew Tolentino, Shannon Wickline, Kim Gietzen, Brad Sickler, Jingtao Liu, Eric Allen, Phil Febbo, Nicole L. Washington, Simon White, Geraint Levan, Kelly Schiabor Barrett, Elizabeth Cirulli, Alexandre Bolze, Ary Ascencio, Charlotte Rivera-Garcia, Ryan Cho, Jason Nguyen, Sherry Wang, Jimmy Ramirez, Tyler Cassens, Efren Sandoval, Magnus Isaksson, William Lee, David Becker, Marc Laurent, James Lu, Clinton R. Paden, Duncan MacCannell |
| EPI_ISL_1924396, EPI_ISL_1924670, EPI_ISL_1924762, EPI_ISL_1924796, EPI_ISL_1924930, EPI_ISL_1925063 | Quest Diagnostics Incorporated | Centers for Disease Control and Prevention Division of Viral Diseases, Pathogen Discovery | Dakota Howard, Dhwani Batra, Peter W. Cook, Kara Moser, Adrian Paskey, Jason Caravas, Benjamin Rambo-Martin, Shatavia Morrison, Christopher Gulvick, Scott Sammons, Yvette Unoarumhi, Darlene Wagner, Matthew Schmerer, S. H. Rosenthal, A. Gerasimova, R. M. Kagan, B. Anderson, M. Hua, Y. Liu, L.E. Bernstein, K.E. Livingston, A. Perez, I. A. Shlyakhter, R. V. Rolando, R. Owen, P. Tanpaiboon, F. Lacbawan, Clinton R. Paden, Duncan MacCannell |
| EPI_ISL_1925936, EPI_ISL_1926073, EPI_ISL_1926250, EPI_ISL_1926283, EPI_ISL_1926372, EPI_ISL_1926482, EPI_ISL_1926580, EPI_ISL_1926717, EPI_ISL_1926968 | Laboratory Corporation of America | Centers for Disease Control and Prevention Division of Viral Diseases, Pathogen Discovery | Dakota Howard, Dhwani Batra, Peter W. Cook, Kara Moser, Adrian Paskey, Jason Caravas, Benjamin Rambo-Martin, Shatavia Morrison, Christopher Gulvick, Scott Sammons, Yvette Unoarumhi, Darlene Wagner, Matthew Schmerer, Minoo Agarwal, Eyad Almasri, Debbie Boles, Ayla Burns, Nuthawin Charoensri, Oren Cohen, Susan Countryman, Mary Ann Cristobal, Bobbi Croy, Suzanne Dale, Hrushikesh Deshmukh, Amanda Douglas, Vincent Drouillon, Marcia Eisenberg, Howard Engler, Rama Ghatti, Prashant Gupta, Susan Hicks, Jake Humphrey, Lax Iyer, Manoj Jain, Mohan Kolli, Brian Krueger, Tim Kuphal, Stanley Letovsky, Michael Levandoski, Craig Lukasik, Jonathan Meltzer, Brian Norvell, Mindy Nye, Scott Parker, Christos Petropoulos, John Pruitt, Steven Ragan, Scott Ryan, Mike Sapeta, Jana Schroth, Suresh Babu Selvaraju, Goran Stevovic, Amanda Suchanek, Andrea Throop, Lyndon Tilson, Thomas Urban, Joe Voshell, Kimberly Wagner, Jonathan Williams, Mary Williamson, Qian Zeng, Tricia Zwiefelhofer, Clinton R. Paden, Duncan MacCannell |
| EPI_ISL_1927305, EPI_ISL_1927400, EPI_ISL_1927401, EPI_ISL_1927403, EPI_ISL_1927404, EPI_ISL_1927405, EPI_ISL_1927407, EPI_ISL_1927408, EPI_ISL_1927410, EPI_ISL_1927411, EPI_ISL_1927413, EPI_ISL_1927414, EPI_ISL_1927416, EPI_ISL_1927417, EPI_ISL_1927419, EPI_ISL_1927420, EPI_ISL_1927422, EPI_ISL_1927423, EPI_ISL_1927424, EPI_ISL_1927426 | | | |
| see above | SARS-CoV-2 testing team, National Institute of Infectious Diseases | Pathogen Genomics Center, National Institute of Infectious Diseases | Tsuyoshi Sekizuka, Kentaro Itokawa, Rina Tanaka, Masanori Hashino, Nozomu Hanaoka, Masumichi Saito, Naomi Nojiri, Hazuka Y Furihata, Sana Uchikoba, Tsuguto Fujimoto, Makoto Kuroda |
| EPI_ISL_1927603, EPI_ISL_1927971 | Laboratory Corporation of America | Centers for Disease Control and Prevention Division of Viral Diseases, Pathogen Discovery | Dakota Howard, Dhwani Batra, Peter W. Cook, Kara Moser, Adrian Paskey, Jason Caravas, Benjamin Rambo-Martin, Shatavia Morrison, Christopher Gulvick, Scott Sammons, Yvette Unoarumhi, Darlene Wagner, Matthew Schmerer, Minoo Agarwal, Eyad Almasri, Debbie Boles, Ayla Burns, Nuthawin Charoensri, Oren Cohen, Susan Countryman, Mary Ann Cristobal, Bobbi Croy, Suzanne Dale, Hrushikesh Deshmukh, Amanda Douglas, Vincent Drouillon, Marcia Eisenberg, Howard Engler, Rama Ghatti, Prashant Gupta, Susan Hicks, Jake Humphrey, Lax Iyer, Manoj Jain, Mohan Kolli, Brian Krueger, Tim Kuphal, Stanley Letovsky, Michael Levandoski, Craig Lukasik, Jonathan Meltzer, Brian Norvell, Mindy Nye, Scott Parker, Christos Petropoulos, John Pruitt, Steven Ragan, Scott Ryan, Mike Sapeta, Jana Schroth, Suresh Babu Selvaraju, Goran Stevovic, Amanda Suchanek, Andrea Throop, Lyndon Tilson, Thomas Urban, Joe Voshell, Kimberly Wagner, Jonathan Williams, Mary Williamson, Qian Zeng, Tricia Zwiefelhofer, Clinton R. Paden, Duncan MacCannell |
| EPI_ISL_1928407, EPI_ISL_1928417, EPI_ISL_1928439, EPI_ISL_1928457, EPI_ISL_1928462, EPI_ISL_1928477, EPI_ISL_1928484, EPI_ISL_1928506, EPI_ISL_1928507, EPI_ISL_1928509, EPI_ISL_1928512, EPI_ISL_1928513, EPI_ISL_1928515, EPI_ISL_1928520, EPI_ISL_1928522, EPI_ISL_1928525, EPI_ISL_1928527, EPI_ISL_1928530, EPI_ISL_1928531, EPI_ISL_1928534, EPI_ISL_1928536, EPI_ISL_1928537, EPI_ISL_1928539, EPI_ISL_1928540, EPI_ISL_1928541, EPI_ISL_1928543, EPI_ISL_1928544, EPI_ISL_1928547, EPI_ISL_1928549, EPI_ISL_1928550, EPI_ISL_1928552, EPI_ISL_1928553, EPI_ISL_1928555, EPI_ISL_1928556, EPI_ISL_1928558, EPI_ISL_1928559, EPI_ISL_1928561, EPI_ISL_1928562, EPI_ISL_1928565, EPI_ISL_1928566, EPI_ISL_1928568, EPI_ISL_1928569, EPI_ISL_1928571, EPI_ISL_1928572, EPI_ISL_1928573, EPI_ISL_1928575, EPI_ISL_1928576, EPI_ISL_1928578, EPI_ISL_1928579, EPI_ISL_1928581, EPI_ISL_1928583, EPI_ISL_1928584, EPI_ISL_1928586, EPI_ISL_1928588, EPI_ISL_1928590, EPI_ISL_1928592, EPI_ISL_1928593, EPI_ISL_1928595, EPI_ISL_1928596, EPI_ISL_1928598, EPI_ISL_1928599 | | | |
| see above | ICMR-National Institute of Virology - INSACOG | NIV Influenza | Dr. Varsha Potdar |
| EPI_ISL_1928600 | Laboratory Corporation of America | Centers for Disease Control and Prevention Division of Viral Diseases, Pathogen Discovery | Dakota Howard, Dhwani Batra, Peter W. Cook, Kara Moser, Adrian Paskey, Jason Caravas, Benjamin Rambo-Martin, Shatavia Morrison, Christopher Gulvick, Scott Sammons, Yvette Unoarumhi, Darlene Wagner, Matthew Schmerer, Minoo Agarwal, Eyad Almasri, Debbie Boles, Ayla Burns, Nuthawin Charoensri, Oren Cohen, Susan Countryman, Mary Ann Cristobal, Bobbi Croy, Suzanne Dale, Hrushikesh Deshmukh, Amanda Douglas, Vincent Drouillon, Marcia Eisenberg, Howard Engler, Rama Ghatti, Prashant Gupta, Susan Hicks, Jake Humphrey, Lax Iyer, Manoj Jain, Mohan Kolli, Brian Krueger, Tim Kuphal, Stanley Letovsky, Michael Levandoski, Craig Lukasik, Jonathan Meltzer, Brian Norvell, Mindy Nye, Scott Parker, Christos Petropoulos, John Pruitt, Steven Ragan, Scott Ryan, Mike Sapeta, Jana Schroth, Suresh Babu Selvaraju, Goran Stevovic, Amanda Suchanek, Andrea Throop, Lyndon Tilson, Thomas Urban, Joe Voshell, Kimberly Wagner, Jonathan Williams, Mary Williamson, Qian Zeng, Tricia Zwiefelhofer, Clinton R. Paden, Duncan MacCannell |
| EPI_ISL_1928601, EPI_ISL_1928602, EPI_ISL_1928603, EPI_ISL_1928605, EPI_ISL_1928607, EPI_ISL_1928608, EPI_ISL_1928610, EPI_ISL_1928611, EPI_ISL_1928613, EPI_ISL_1928614, EPI_ISL_1928615, EPI_ISL_1928617, EPI_ISL_1928618, EPI_ISL_1928620, EPI_ISL_1928622, EPI_ISL_1928623, EPI_ISL_1928624, EPI_ISL_1928626, EPI_ISL_1928627, EPI_ISL_1928629, EPI_ISL_1928631, EPI_ISL_1928632, EPI_ISL_1928633, EPI_ISL_1928635, EPI_ISL_1928636, EPI_ISL_1928638, EPI_ISL_1928639, EPI_ISL_1928641, EPI_ISL_1928642, EPI_ISL_1928645, EPI_ISL_1928647, EPI_ISL_1928648, EPI_ISL_1928650 | | | |
| see above | ICMR-National Institute of Virology - INSACOG | NIV Influenza | Dr. Varsha Potdar |
| EPI_ISL_1928896, EPI_ISL_1929055, EPI_ISL_1929232, EPI_ISL_1929301, EPI_ISL_1929356, EPI_ISL_1929448, EPI_ISL_1930706, EPI_ISL_1930858, EPI_ISL_1931019, EPI_ISL_1931052, EPI_ISL_1931275, EPI_ISL_1931296, EPI_ISL_1931734, EPI_ISL_1932000, EPI_ISL_1932009 | | | |
| see above | Laboratory Corporation of America | Centers for Disease Control and Prevention Division of Viral Diseases, Pathogen Discovery | Dakota Howard, Dhwani Batra, Peter W. Cook, Kara Moser, Adrian Paskey, Jason Caravas, Benjamin Rambo-Martin, Shatavia Morrison, Christopher Gulvick, Scott Sammons, Yvette Unoarumhi, Darlene Wagner, Matthew Schmerer, Minoo Agarwal, Eyad Almasri, Debbie Boles, Ayla Burns, Nuthawin Charoensri, Oren Cohen, Susan Countryman, Mary Ann Cristobal, Bobbi Croy, Suzanne Dale, Hrushikesh Deshmukh, Amanda Douglas, Vincent Drouillon, Marcia Eisenberg, Howard Engler, Rama Ghatti, Prashant Gupta, Susan Hicks, Jake Humphrey, Lax Iyer, Manoj Jain, Mohan Kolli, Brian Krueger, Tim Kuphal, Stanley Letovsky, Michael Levandoski, Craig Lukasik, Jonathan Meltzer, Brian Norvell, Mindy Nye, Scott Parker, Christos Petropoulos, John Pruitt, Steven Ragan, Scott Ryan, Mike Sapeta, Jana Schroth, Suresh Babu Selvaraju, Goran Stevovic, Amanda Suchanek, Andrea Throop, Lyndon Tilson, Thomas Urban, Joe Voshell, Kimberly Wagner, Jonathan Williams, Mary Williamson, Qian Zeng, Tricia Zwiefelhofer, Clinton R. Paden, Duncan MacCannell |
| EPI_ISL_1935543 | Institute of Microbiology and Immunology, Faculty of Medicine, University of Ljubljana | Institute of Microbiology and Immunology, Faculty of Medicine, University of Ljubljana | Alen Suljič, Samo Zakotnik, Tomaž Mark Zorec, Matic Brvar, Doroteja Vlaj, Andraž Celar, Dominika Šturm, Patricija Pozvek, Špela Pleh, Miša Korva, Mario Poljak, Tatjana Avšič – Županc |
| EPI_ISL_1935601 | Cliniques universitaires Saint-Luc | UCLouvain/IREC/MBLG | Jean Ruelle, Eleonore Ngyuvula, Benoit Kabamba Mukadi |
| EPI_ISL_1936188 | MD PHL | MD PHL | Maryland Department of Health Laboratories Administration |
| EPI_ISL_1936604 | Division of Emerging Infectious Diseases, Bureau of Infectious Diseases Diagnosis Control, Korea Disease Control and Prevention Agency | Division of Emerging Infectious Diseases, Bureau of Infectious Diseases Diagnosis Control, Korea Disease Control and Prevention Agency | Ae Kyung Park, Il-Hwan Kim, Heui Man Kim, Jeong-Min Kim, Jeong-Ah Kim, Chae Young Lee, Jin Sun No, Eun-Jin Kim |
| EPI_ISL_1937844 | NH Dept. of Health and Human Services Public Health Labs | Centers for Disease Control and Prevention Division of Viral Diseases, Pathogen Discovery | Mili Sheth, Sarah Nobles, Jasmine Padilla, Mark Burroughs, Shoshona Le, Katie Dillon, Peter Cook, Clinton R. Paden, Dhwani Batra, Krista Queen, Kristen Knipe, Dakota Howard, Yvette Unoarumhi, Darlene Wagner, Matthew Schmerer, Ben L. Rambo-Martin, Kristine Lacek, Sam Shepard, Alison Laufer Halpin, Dave Wentworth, Vivien Dugan, Suxiang Tong, Justin Lee |
| EPI_ISL_1938308 | Queensland Health Forensic and Scientific Services | Queensland Health Forensic and Scientific Services | Son Nguyen |
| EPI_ISL_1938475, EPI_ISL_1938476, EPI_ISL_1938477 | Institute of Epidemiology, Disease Control and Research (IEDCR) | Institute for Developing Science and Health Initiatives (ideSHi) | Hassan Afrad, Sadia Rahman, Fidausi Qadri, Tahmina Shirin |
| EPI_ISL_1939630, EPI_ISL_1939651 | Lighthouse Lab in Glasgow | Wellcome Sanger Institute for the COVID-19 Genomics UK (COG-UK) Consortium | Harper VanSteenhouse, Yumi Kasai, David Gray, Carol Clugston, Anna Dominiczak and Alex Alderton, Roberto Amato, Jeffrey Barrett, Sonia Goncalves, Ewan Harrison, David K. Jackson, Ian Johnston, Dominic Kwiatkowski, Cordelia Langford, John Sillitoe on behalf of the Wellcome Sanger Institute COVID-19 Surveillance Team |
| EPI_ISL_1939711, EPI_ISL_1939714, EPI_ISL_1939718 | Lighthouse Lab in Milton Keynes | Wellcome Sanger Institute for the COVID-19 Genomics UK (COG-UK) Consortium | The Lighthouse Lab in Milton Keynes and Alex Alderton, Roberto Amato, Jeffrey Barrett, Sonia Goncalves, Ewan Harrison, David K. Jackson, Ian Johnston, Dominic Kwiatkowski, Cordelia Langford, John Sillitoe on behalf of the Wellcome Sanger Institute COVID-19 Surveillance Team |

| | | | |
|---|---|---|---|
| EPI_ISL_1939724, EPI_ISL_1939726, EPI_ISL_1939750, EPI_ISL_1939759, EPI_ISL_1939772, EPI_ISL_1939774, EPI_ISL_1939791, EPI_ISL_1939812 | Lighthouse Lab in Glasgow | Wellcome Sanger Institute for the COVID-19 Genomics UK (COG-UK) Consortium | Harper VanSteenhouse, Yumi Kasai, David Gray, Carol Clugston, Anna Dominiczak and Alex Alderton, Roberto Amato, Jeffrey Barrett, Sonia Goncalves, Ewan Harrison, David K. Jackson, Ian Johnston, Dominic Kwiatkowski, Cordelia Langford, John Sillitoe on behalf of the Wellcome Sanger Institute COVID-19 Surveillance Team |
| EPI_ISL_1939929, EPI_ISL_1939930, EPI_ISL_1939931, EPI_ISL_1939932, EPI_ISL_1939934, EPI_ISL_1939936, EPI_ISL_1939939, EPI_ISL_1939940, EPI_ISL_1939947, EPI_ISL_1939950, EPI_ISL_1939951, EPI_ISL_1939952, EPI_ISL_1939955, EPI_ISL_1939956, EPI_ISL_1939957, EPI_ISL_1939958, EPI_ISL_1939959, EPI_ISL_1939961, EPI_ISL_1939962, EPI_ISL_1939963, EPI_ISL_1939964, EPI_ISL_1939966 | | | |
| see above | JIPMER Puducherry | inStem NCBS - INSACOG | Uma Ramakrishnan Dasaradhi Palakodeti Aswin SaiNarain |
| EPI_ISL_1939969 | KAMAKSHIPALYA | inStem NCBS - INSACOG | Uma Ramakrishnan Dasaradhi Palakodeti Aswin SaiNarain |
| EPI_ISL_1939974 | SUBRAMNAYA NAGAR | inStem NCBS - INSACOG | Uma Ramakrishnan Dasaradhi Palakodeti Aswin SaiNarain |
| EPI_ISL_1939986, EPI_ISL_1939987, EPI_ISL_1939999 | NIMHANS | inStem NCBS - INSACOG | Uma Ramakrishnan Dasaradhi Palakodeti Aswin SaiNarain |
| EPI_ISL_1940001, EPI_ISL_1940002, EPI_ISL_1940003, EPI_ISL_1940004, EPI_ISL_1940008, EPI_ISL_1940011, EPI_ISL_1940013, EPI_ISL_1940014, EPI_ISL_1940015 | HOSAHALLI | inStem NCBS - INSACOG | Uma Ramakrishnan Dasaradhi Palakodeti Aswin SaiNarain |
| EPI_ISL_1940019 | GOVIMDRAJ NAGAR | inStem NCBS - INSACOG | Uma Ramakrishnan Dasaradhi Palakodeti Aswin SaiNarain |
| EPI_ISL_1940026, EPI_ISL_1940036 | A D HALLI | inStem NCBS - INSACOG | Uma Ramakrishnan Dasaradhi Palakodeti Aswin SaiNarain |
| EPI_ISL_1940049 | GOVINDRAJNAGAR | inStem NCBS - INSACOG | Uma Ramakrishnan Dasaradhi Palakodeti Aswin SaiNarain |
| EPI_ISL_1940055, EPI_ISL_1940057 | NIMHANS | inStem NCBS - INSACOG | Uma Ramakrishnan Dasaradhi Palakodeti Aswin SaiNarain |
| EPI_ISL_1940061 | RAILWAY H | inStem NCBS - INSACOG | Uma Ramakrishnan Dasaradhi Palakodeti Aswin SaiNarain |
| EPI_ISL_1940062, EPI_ISL_1940064, EPI_ISL_1940067, EPI_ISL_1940068 | HOSAHALLI | inStem NCBS - INSACOG | Uma Ramakrishnan Dasaradhi Palakodeti Aswin SaiNarain |
| EPI_ISL_1940072, EPI_ISL_1940074, EPI_ISL_1940075 | GOVINDRAJ NAGAR | inStem NCBS - INSACOG | Uma Ramakrishnan Dasaradhi Palakodeti Aswin SaiNarain |
| EPI_ISL_1940092 | AD HALLI | inStem NCBS - INSACOG | Uma Ramakrishnan Dasaradhi Palakodeti Aswin SaiNarain |
| EPI_ISL_1940095, EPI_ISL_1940096, EPI_ISL_1940099 | NIMHANS | inStem NCBS - INSACOG | Uma Ramakrishnan Dasaradhi Palakodeti Aswin SaiNarain |
| EPI_ISL_1940102, EPI_ISL_1940106 | RAILWAY | inStem NCBS - INSACOG | Uma Ramakrishnan Dasaradhi Palakodeti Aswin SaiNarain |
| EPI_ISL_1940107 | NIMHANS | inStem NCBS - INSACOG | Uma Ramakrishnan Dasaradhi Palakodeti Aswin SaiNarain |
| EPI_ISL_1940111 | GOVINDRAJNAGAR | inStem NCBS - INSACOG | Uma Ramakrishnan Dasaradhi Palakodeti Aswin SaiNarain |
| EPI_ISL_1940112, EPI_ISL_1940113, EPI_ISL_1940114, EPI_ISL_1940118, EPI_ISL_1940119, EPI_ISL_1940123, EPI_ISL_1940125, EPI_ISL_1940127, EPI_ISL_1940128 | HOSAHALLI | inStem NCBS - INSACOG | Uma Ramakrishnan Dasaradhi Palakodeti Aswin SaiNarain |
| EPI_ISL_1940134 | NIMHANS | inStem NCBS - INSACOG | Uma Ramakrishnan Dasaradhi Palakodeti Aswin SaiNarain |
| EPI_ISL_1940150 | Wichita State University - Molecular Diagnostics Lab | Kansas Health and Environmental Lab | Mike Grose, Jonathan Barnell, Ben Olsen, and Phil Adam |
| EPI_ISL_1941367, EPI_ISL_1941374, EPI_ISL_1941376 | Landesamt für Verbraucherschutz Sachsen Anhalt, Magdeburg | Institute of Medical Microbiology and Hospital Hygiene | Prof. Dr. Achim Kaasch, Aljoscha Tersteegen |
| EPI_ISL_1941681 | Hospital Universitario Cruces | Biocruces Bizkaia | Ana Belen de la Hoz, Mikel Gallego |
| EPI_ISL_1941773 | LA SOURCE | Laboratory of genomics and metagenomics | Trestan Pillonel, Damien Jacot, Sebastien Aeby, Gilbert Greub, Claire Bertelli |
| EPI_ISL_1941875 | Hospital Universitario Cruces | Biocruces Bizkaia | Ana Belen de la Hoz, Mikel Gallego |
| EPI_ISL_1942165 | Helix/Illumina | Centers for Disease Control and Prevention Division of Viral Diseases, Pathogen Discovery | Dakota Howard, Dhwani Batra, Peter W. Cook, Kara Moser, Adrian Paskey, Jason Caravas, Benjamin Rambo-Martin, Shatavia Morrison, Christopher Gulvick, Scott Sammons, Yvette Unoarumhi, Darlene Wagner, Matthew Schmerer, Eileen de Feo, Jan Antico, Christine Tran, Matthew Tolentino, Shannon Wickline, Kim Gietzen, Brad Sickler, Jingtao Liu, Eric Allen, Phil Febbo, Nicole L. Washington, Simon White, Geraint Levan, Kelly Schiabor Barrett, Elizabeth Cirulli, Alexandre Bolze, Ary Ascencio, Charlotte Rivera-Garcia, Ryan Cho, Jason Nguyen, Sherry Wang, Jimmy Ramirez, Tyler Cassens, Efren Sandoval, Magnus Isaksson, William Lee, David Becker, Marc Laurent, James Lu, Clinton R. Paden, Duncan MacCannell |
| EPI_ISL_1942243 | CQRC_QUALITY CONTROL CHEMICAL BIOLOGICAL RISK_AOOR Villa Sofia Cervello Palermo | CQRC_QUALITY CONTROL CHEMICAL BIOLOGICAL RISK_AOOR Villa Sofia Cervello Palermo | Di Gaudio, F., Contino, F., Orlando, V., Buffa, V., Bonvissuto, M., Di Chiara, V., Seidita, G.,Todaro, I., Di Sclafani, A., Lungari, A., Brunacci,G. |
| EPI_ISL_1942249 | Genome Centre | Genome Centre | Md. Shazid Hasan, Shovon Lal Sarkar, Ali Ahsan Setu, Tanay Chakrabarty, A. S. M. Rubayet Ul Alam, M. Shaminur Rahman, Ovinu Kibria Islam, M. Tanvir Islam, Hassan M. Al-Emran, Iqbal Kabir Jahid, M. Anwar Hossain |
| EPI_ISL_1960012 | NJDOH, Public Health and Environmental Laboratories | NJ_PHEL | Lindsey Bodnar, Shiv K. Verma, Jacquelyn Deverell, Matthew Scarnati, Dana Woell, Byeong Jeong |
| EPI_ISL_1960669, EPI_ISL_1960691, EPI_ISL_1960795, EPI_ISL_1960834, EPI_ISL_1960854 | National Virus Reference Laboratory | National Virus Reference Laboratory | Zoe Yandle, Charlene Bennett, Gabriel Gonzalez, Michael Carr, Jonathan Dean, Cillian F De Gascun |
| EPI_ISL_1962940, EPI_ISL_1962944, EPI_ISL_1962964, EPI_ISL_1962965, EPI_ISL_1962966 | Dutch COVID-19 response team | National Institute for Public Health and the Environment (RIVM) | Adam Meijer, Harry Vennema, Dirk Eggink, Jeroen Cremer, Sharon van den Brink, Bas van der Veer, AnneMarie van den Brandt, Lisa Wijsman, Kim Freriks, Ryanne Jaarsma, Euníce Then, Lynn Aarts, Sanne Bos, Melissa van Tuil, Linda van de Nes, Sjoerd Kuiling, James Groot, Florian Zwagemaker, Dennis Schmitz, Annelies Kroneman, Karim Hajji, Chantal Reusken, on behalf of the national COVID-19 response team |
| EPI_ISL_1963016, EPI_ISL_1963298, EPI_ISL_1963299, EPI_ISL_1963300 | University Hospitals of Geneva, Laboratory of Virology | HUG, Laboratory of Virology and the Health2030 Genome Center | Samuel Cordey, Ana Rita Goncalves, Laurent Kaiser, Lorenzo Cerutti, Henri Pegeot, Melyssa Elies, Deborah Penet, Keith Harshman, Ioannis Xenarios, Emmanouil Dermitzakis |
| EPI_ISL_1963634 | Hong Kong Department of Health | School of Public Health, The University of Hong Kong | Dominic N.C. Tsang, Daniel K.W. Chu, Haogao Gu, Tong Zhang, Leo L.M. Poon, Malik Peiris |
| EPI_ISL_1965786 | Clinical Microbiology Laboratory, NewYork Presbyterian Hospital/Columbia University Irving Medical Center | Uhlemann Laboratory, Columbia University Irving Medical Center | Medini K. Annavajhala, Anne Kelley, Maya Tagliavia, Anne-Catrin Uhlemann |
| EPI_ISL_1968000, EPI_ISL_1968007, EPI_ISL_1968369, EPI_ISL_1968371, EPI_ISL_1968372, EPI_ISL_1968435 | Broad Institute Clinical Research Sequencing Platform | Infectious Disease Program, Broad Institute of Harvard and MIT | Siddle,K.J., Adams,G., Pearlman,L., Gladden-Young,A., Vicente,G., Blumenstiel,B., DeFelice,M., Lee,M., McGovern,S., Lagerborg,K., Rudy,M., DeRuff,K., Carter,A., Normandin,E., Bauer,M., Reilly,S., Tomkins-Tinch,C., Loreth,C., Chaluvadi,S., Meldrim,J., Granger,B., Lemieux,J.E., Birren,B.W., Sabeti,P.C., Larkin,K., Dodge,S., Lennon,N., Madoff,L., Brown,C., Gallagher,G., Smole,S., Park,D.J., Gabriel,S., and MacInnis,B.L. |

| | | | |
|---|---|---|---|
| EPI_ISL_1968884 | Clinical Microbiology, Infection Prevention and Control | Section for Molecular Diagnostics | Björn Hallström, Jonas Björkman |
| EPI_ISL_1969044 | NJDOH, Public Health and Environmental Laboratories | NJ_PHEL | Lindsey Bodnar, Shiv K. Verma, Jacquelyn Deverell, Matthew Scarnati, Dana Woell, Byeong Jeong |
| EPI_ISL_1969243 | Swissbel Hotel Airport | National Institute of Health Research and Development | Vivi Setiawaty, Hana Apsari Pawestri, Subangkit, Kartika Dewi Puspa, Arie Ardiansyah Nugraha, Hartanti Dian Ikawati, Krisna Pangesti, Triyani Soekarso, Nelly Puspandari |
| EPI_ISL_1969244 | Lab RSUP DR Mohammad Hoesin Palembang | National Institute of Health Research and Development | Vivi Setiawaty, Hana Apsari Pawestri, Subangkit, Kartika Dewi Puspa, Arie Ardiansyah Nugraha, Hartanti Dian Ikawati, Krisna Pangesti, Triyani Soekarso, Nelly Puspandari |
| EPI_ISL_1969245 | Lab RSUP DR Mohammad Hoesin Palembang | National Institute of Health Research and Development | Subangkit, Hana Apsari Pawestri, Kartika Dewi Puspa, Arie Ardiansyah Nugraha, Hartanti Dian Ikawati, Krisna Pangesti, Triyani Soekarso, Nelly Puspandari, Vivi Setiawaty |
| EPI_ISL_1969246, EPI_ISL_1969247, EPI_ISL_1969248 | Rumah Sakit Umum Daerah Palangkaraya | National Institute of Health Research and Development | Subangkit, Hana Apsari Pawestri, Kartika Dewi Puspa, Arie Ardiansyah Nugraha, Hartanti Dian Ikawati, Krisna Pangesti, Triyani Soekarso, Nelly Puspandari, Vivi Setiawaty |
| EPI_ISL_1969249 | BBLK Palembang | National Institute of Health Research and Development | Subangkit, Hana Apsari Pawestri, Kartika Dewi Puspa, Arie Ardiansyah Nugraha, Hartanti Dian Ikawati, Krisna Pangesti, Triyani Soekarso, Nelly Puspandari, Vivi Setiawaty |
| EPI_ISL_1969690 | Singapore General Hospital | Department of Microbiology | Nurdyana Abdul Rahman, Kenneth Xin Long Chan, Suphavilai Chayaporn, Kun Lee Lim, Sui Sin Goh, Kian Sing Chan, Lynette Oon, Kern Rei Chng, James Sim Heng Chiak, Niranjan Nagarajan, Karrie Ko |
| EPI_ISL_1969753 | State Virus Research and Diagnostic Laboratory (VRDL), AIIMS Raipur | State Virus Research and Diagnostic Laboratory (VRDL), AIIMS Raipur | Pushpendra Singh, Kuldeep Sharma, Priyanka Singh, Somya Sharma, Sanjay Singh Negi, Anudita Bhargava |
| EPI_ISL_1970089, EPI_ISL_1970092, EPI_ISL_1970094, EPI_ISL_1970095, EPI_ISL_1970099 | Biotechnology Division, NCDC Delhi | NCDC Delhi, Biotechnology Division | Mahesh S Dhar, Kalaiarasan Ponnusamy, Meena Datta, Priyanka Singh, Uma Sharma, Manoj K Singh, Radhakrishnan V. S, Robin Marwal, Hemlata Lall, Hema Gogia, Preeti Madan, Sandhya Kabra, Sujeet K Singh, Partha Rakshit |
| EPI_ISL_1970100, EPI_ISL_1970101, EPI_ISL_1970102, EPI_ISL_1970103, EPI_ISL_1970105, EPI_ISL_1970106, EPI_ISL_1970108, EPI_ISL_1970109, EPI_ISL_1970110, EPI_ISL_1970111, EPI_ISL_1970112, EPI_ISL_1970114, EPI_ISL_1970133, EPI_ISL_1970134, EPI_ISL_1970135, EPI_ISL_1970149, EPI_ISL_1970153, EPI_ISL_1970154, EPI_ISL_1970155, EPI_ISL_1970157, EPI_ISL_1970158, EPI_ISL_1970159, EPI_ISL_1970160, EPI_ISL_1970163, EPI_ISL_1970164, EPI_ISL_1970168, EPI_ISL_1970169, EPI_ISL_1970170, EPI_ISL_1970171, EPI_ISL_1970172, EPI_ISL_1970179, EPI_ISL_1970211, EPI_ISL_1970215, EPI_ISL_1970221, EPI_ISL_1970223, EPI_ISL_1970224, EPI_ISL_1970237, EPI_ISL_1970238, EPI_ISL_1970239, EPI_ISL_1970241, EPI_ISL_1970242 | | | |
| see above | Biotechnology Division, NCDC Delhi | NCDC Delhi, Biotechnology Division | Kalaiarasan Ponnusamy, Meena Datta, Priyanka Singh, Uma Sharma, Manoj K Singh, Radhakrishnan V. S, Robin Marwal, Mahesh S Dhar, Hemlata Lall, Hema Gogia, Preeti Madan, Sandhya Kabra, Sujeet K Singh, Partha Rakshit |
| EPI_ISL_1970243, EPI_ISL_1970247, EPI_ISL_1970249, EPI_ISL_1970250, EPI_ISL_1970251, EPI_ISL_1970267, EPI_ISL_1970269, EPI_ISL_1970274, EPI_ISL_1970276, EPI_ISL_1970277, EPI_ISL_1970278, EPI_ISL_1970279, EPI_ISL_1970281, EPI_ISL_1970286, EPI_ISL_1970288, EPI_ISL_1970310, EPI_ISL_1970313, EPI_ISL_1970314, EPI_ISL_1970318, EPI_ISL_1970319, EPI_ISL_1970323, EPI_ISL_1970324, EPI_ISL_1970327, EPI_ISL_1970328, EPI_ISL_1970332, EPI_ISL_1970333, EPI_ISL_1970334, EPI_ISL_1970335, EPI_ISL_1970336, EPI_ISL_1970337, EPI_ISL_1970340 | | | |
| see above | Biotechnology Division, NCDC Delhi | NCDC Delhi, Biotechnology Division | Meena Datta, Priyanka Singh, Uma Sharma, Manoj K Singh, Radhakrishnan V. S, Robin Marwal, Mahesh S Dhar, Kalaiarasan Ponnusamy, Hemlata Lall, Hema Gogia, Preeti Madan, Sandhya Kabra, Sujeet K Singh, Partha Rakshit |
| EPI_ISL_1970347 | Chongqing Yuzhong District Center For Disease Control And Prevention | Chongqing Municipal Center for Disease Control and Prevention | Sheng Ye, Yun Tang, Yuyue Luo , Zhen Yu, Yingbing Zhou , Hua Ling , Jiaqi Li , Zhangping Tan, Mingyue Wang, Shuang Chen, Wenge Tang, Rong Rong |
| EPI_ISL_1970348 | Chongqing International Travel Health Care Center | Chongqing Municipal Center for Disease Control and Prevention | Sheng Ye, Shuang Chen, Haiyan Wen, Zhen Yu , Lan Zhou, Hua Ling, Dong Wang, Yun Tang, Mingyue Wang, Zhangping Tan, Wenge Tang, Rong Rong |
| EPI_ISL_1970349 | Chongqing International Travel Health Care Center | Chongqing Municipal Center for Disease Control and Prevention | Sheng Ye , Zhangping Tan , Zhaohui Deng，Shuang Chen, Xingdan Luo, Zhen Yu, Yanqing Peng , Hua Ling , Yun Tang, Mingyue Wang, Wenge Tang, Rong Rong |
| EPI_ISL_1970418 | Centre for Dengue Research and AICBU, Department of Immunology and Molecular Medicine | Centre for Dengue Research and AICBU, Department of Immunology and Molecular Medicine | Chandima Jeewandara, Deshni Jayathilaka, Dinuka Ariyaratne, Deshan Madhusanka, Diyanath Ranasinghe, Laksiri Gomes, Gathsaurie Neelika Malavige |
| EPI_ISL_1970422, EPI_ISL_1970427, EPI_ISL_1970429, EPI_ISL_1970431, EPI_ISL_1970433, EPI_ISL_1970434, EPI_ISL_1970435, EPI_ISL_1970437, EPI_ISL_1970438, EPI_ISL_1970439, EPI_ISL_1970440, EPI_ISL_1970441, EPI_ISL_1970442, EPI_ISL_1970443, EPI_ISL_1970444, EPI_ISL_1970445, EPI_ISL_1970447, EPI_ISL_1970448, EPI_ISL_1970449, EPI_ISL_1970450, EPI_ISL_1970451, EPI_ISL_1970452, EPI_ISL_1970453, EPI_ISL_1970454, EPI_ISL_1970455, EPI_ISL_1970456, EPI_ISL_1970457, EPI_ISL_1970458, EPI_ISL_1970459, EPI_ISL_1970461, EPI_ISL_1970462, EPI_ISL_1970463, EPI_ISL_1970464, EPI_ISL_1970465, EPI_ISL_1970466, EPI_ISL_1970467, EPI_ISL_1970468, EPI_ISL_1970469, EPI_ISL_1970470, EPI_ISL_1970471, EPI_ISL_1970472, EPI_ISL_1970473, EPI_ISL_1970474, EPI_ISL_1970475, EPI_ISL_1970476, EPI_ISL_1970477, EPI_ISL_1970478, EPI_ISL_1970479, EPI_ISL_1970480, EPI_ISL_1970481, EPI_ISL_1970482, EPI_ISL_1970484, EPI_ISL_1970485, EPI_ISL_1970486, EPI_ISL_1970488, EPI_ISL_1970489, EPI_ISL_1970491, EPI_ISL_1970492, EPI_ISL_1970493, EPI_ISL_1970494, EPI_ISL_1970495, EPI_ISL_1970496, EPI_ISL_1970497, EPI_ISL_1970499, EPI_ISL_1970500, EPI_ISL_1970501, EPI_ISL_1970502, EPI_ISL_1970503, EPI_ISL_1970504, EPI_ISL_1970505, EPI_ISL_1970506, EPI_ISL_1970507, EPI_ISL_1970508, EPI_ISL_1970509, EPI_ISL_1970510, EPI_ISL_1970511, EPI_ISL_1970512, EPI_ISL_1970513, EPI_ISL_1970514, EPI_ISL_1970515, EPI_ISL_1970517, EPI_ISL_1970518, EPI_ISL_1970519, EPI_ISL_1970520, EPI_ISL_1970522, EPI_ISL_1970523, EPI_ISL_1970524, EPI_ISL_1970525, EPI_ISL_1970526, EPI_ISL_1970527, EPI_ISL_1970529, EPI_ISL_1970530, EPI_ISL_1970531, EPI_ISL_1970534, EPI_ISL_1970535, EPI_ISL_1970536, EPI_ISL_1970538, EPI_ISL_1970540, EPI_ISL_1970541, EPI_ISL_1970542, EPI_ISL_1970543, EPI_ISL_1970544, EPI_ISL_1970545, EPI_ISL_1970546 | | | |
| see above | ICMR-National Institute of Virology - INSACOG | NIV Influenza | Dr. Varsha Potdar |
| EPI_ISL_1970565, EPI_ISL_1970566, EPI_ISL_1970567, EPI_ISL_1970568 | MRC/UVRI & LSHTM Uganda Research Unit | MRC/UVRI & LSHTM Uganda Research Unit | Matthew Cotten, Dan Lule Bugembe, My V.T. Phan, Isaac Sseeewanyana, Patrick Semanda, Susan Nabadda, Pontiano Kaleebu |
| EPI_ISL_1970570, EPI_ISL_1970571, EPI_ISL_1970572, EPI_ISL_1970573, EPI_ISL_1970574, EPI_ISL_1970575 | Virology Laboratory, Scientific Department, Army Medical Center | Virology Laboratory, Scientific Department, Army Medical Center | Silvia Fillo, Riccardo De Sanctis, Antonella Fortunato, Anella Monte, Rossella Brandi, Giulia Campoli, Marzia Cavalli, Lucia Nicosia, Anna Anselmo, Vanessa Vera Fain, Francesco Giordani, Giandomenico Cerreto, Filippo Molinari, Giancarlo Petralito, Florigio Lista |
| EPI_ISL_1970729, EPI_ISL_1970731, EPI_ISL_1970732, EPI_ISL_1970733, EPI_ISL_1970734 | Laboratorio analisi - Policlinico San Pietro | Laboratory of Clinical Microbiology, Virology and Bioemergencies, ASST Fatebenefratelli Sacco - Sacco Hospital | Valeria Micheli, Alessandro Mancon, Alberto Rizzo, Fiorenza Bracchitta, Luca Rizzuto, Maria Rita Gismondo |
| EPI_ISL_1970752 | University of Liège COVID-19 testing center | GIGA Medical Genomics | Keith Durkin, Maria Artesi, Sébastien Bontems, Raphaël Boreux, Bouchra Boujemla, Nathalie Renotte, Cécile Meex, Pierrette Melin, Marie-Pierre Hayette, Vincent Bours |
| EPI_ISL_1971653, EPI_ISL_1971724, EPI_ISL_1971811, EPI_ISL_1971854, EPI_ISL_1971855, EPI_ISL_1971900, EPI_ISL_1971917 | Broad Institute Clinical Research Sequencing Platform | Infectious Disease Program, Broad Institute of Harvard and MIT | Siddle,K.J., Adams,G., Pearlman,L., Gladden-Young,A., Vicente,G., Blumenstiel,B., DeFelice,M., Lee,M., McGovern,S., Lagerborg,K., Rudy,M., DeRuff,K., Carter,A., Normandin,E., Bauer,M., Reilly,S., Tomkins-Tinch,C., Loreth,C., Chaluvadi,S., Meldrim,J., Granger,B., Lemieux,J.E., Birren,B.W., Sabeti,P.C., Larkin,K., Dodge,S., Lennon,N., Madoff,L., Brown,C., Gallagher,G., Smole,S., Park,D.J., Gabriel,S., and MacInnis,B.L. |
| EPI_ISL_1972127, EPI_ISL_1972129 | AZ Zeno | AZ SINT-JAN BRUGGE | Marijke Reynders, Patrick Descheemaeker, Laurien Hoornaert, Thomas Van Landschoot, Jorn Hellemans |
| EPI_ISL_1972356 | Sungai Buloh Hospital | Institute for Medical Research, Infectious Disease Research Centre, National Institutes of Health, Ministry of Health Malaysia | Suppiah J, Kamel K, Mohd Zawawi Z, Azizan MA, Ramly N, Robert F, Thayan R |
| EPI_ISL_1972570 | Hospital Sharp | Microbial Genomics Laboratory | Bruno Gomez-Gil, Julissa Enciso-Ibarra, Alejandra Garcia-Gasca, Daniel Fregoso-Rueda |
| EPI_ISL_1972662 | National Virus Reference Laboratory | National Virus Reference Laboratory | Guerrino Macori, Gabriel Gonzalez, Michael Carr, Zoe Yandle, Charlene Bennett, Jonathan Dean, Seamus Fanning, Cillian F De Gascun |
| EPI_ISL_1972667, EPI_ISL_1972679, EPI_ISL_1972682, EPI_ISL_1972731, EPI_ISL_1972735, EPI_ISL_1972737, EPI_ISL_1972739, EPI_ISL_1972740 | National Virus Reference Laboratory | National Virus Reference Laboratory | Zoe Yandle, Charlene Bennett, Gabriel Gonzalez, Michael Carr, Jonathan Dean, Cillian F De Gascun |
| EPI_ISL_1972833 | National Virus Reference Laboratory | National Virus Reference Laboratory | Fiona Crispie, Calum Walsh, Matthew McCabe, Zoe Yandle, Charlene Bennet, Gabriel Gonzalez, Michael Carr, Jonathan Dean, Paul Cotter, Cillian F De Gascun |
| EPI_ISL_1972908, EPI_ISL_1972909, EPI_ISL_1972911, EPI_ISL_1972912 | PathWest Laboratory Medicine WA | PathWest Laboratory Medicine WA Microbial Surveillance Unit | PathWest Laboratory Medicine WA Microbial Surveillance Unit |
| EPI_ISL_1974711 | Quadram Institute Bioscience | COVID-19 Genomics UK (COG-UK) Consortium | Dave J. Baker, Gemma L. Kay, Alp Aydin, Thanh Le-Viet, Steven Rudder, Ana P. Tedim, Anastasia Kolyva, Maria Diaz, Leonardo de Oliveira Martins, Nabil-Fareed Alikhan, Lizzie Meadows, Rachael Stanley, Ngozi Elumogo, Muhammed Yasir, Nicholas M. Thomson, Alexander J Trotter, Rachel Gilroy, Samuel Bloomfield, Claire Stuart, Andrew Bell, Reenesh Prakash, Samir Dervisevic, Alison E. Mather, John Wain, Mark Webber, Andrew J. Page, Justin O'Grady |
| EPI_ISL_1974723 | University of Birmingham | COVID-19 Genomics UK (COG-UK) Consortium | Institute of Microbiology, University of Birmingham: Claire McMurray, Joanne Stockton, Samuel Nicholls, Radoslaw Poplawski, Will Rowe, Josh Quick, Nicholas Loman. University of Birmingham Testing Laboratory: Celina M Whalley, Andrew Bosworth, Charlotte Poxon, Kasun Wanigasooriya, Oliver Pickles, Mike Kidd, Alex Richter, Andrew D Beggs PHE Heartlands Lab: Husam Osman, Andrew Bosworth. Queen Elizabeth Hospital: Anna Casey |
| EPI_ISL_1974779 | Department of Pathology, University of Cambridge | COVID-19 Genomics UK (COG-UK) Consortium | Aminu S. Jahun, Yasmin Chaudhry, Iliana Georgana, Myra Hosmillo, Rhys Izuagbe, William L. Hamilton, Martin D. Curran, Surendra Parmar, Ian Goodfellow |
| EPI_ISL_1974856 | Virology Department, Royal Infirmary of Edinburgh, NHS Lothian / School of Biological Sciences, University of Edinburgh | COVID-19 Genomics UK (COG-UK) Consortium | McHugh M, Dewar R, Cotton S, Gallagher A, Maloney D, Rooke S, O'Toole Á, Scher E, Hill V, McCrone JT, Colquhoun R, Yu X, Jackson B, Rambaut A, Templeton K |
| EPI_ISL_1975018 | Liverpool Clinical Laboratories | COVID-19 Genomics UK (COG-UK) Consortium | Sam Haldenby, Alistair Darby, Steve Paterson, Anita Lucaci, Julian Hiscox, M Almsaud, A Alrezaihi, Muhannad Alruwaili, Stuart D Armstrong, Jones Benjamin, Eleanor G Bentley, Anu Chawla, Jordan J Clark, Angela Cowell, Richard Eccles, Isabel García-Dorival, Matthew Gemmell, Alessandro Gerada, PKF Gilmore, Richard Gregory, Ximeng Han, Catherine Hartley, Margaret Hughes, Miren Iturriza-Gomara, James Johnson, L Luu, Jenifer Manson, Charlotte Nelson, Elaine O'Toole, Cassie Olateju, Rebekah Penrice-Randal , Lucille Rainbow, N.P Randle, Trevor Ian Robinson, Parul Sharma, Ghada T Shawli, James P Stewart, Neil Swainston, Ecaterina Vamos, Joanne Watts, Mark Whitehead, Hermione Webster |
| EPI_ISL_1975115, EPI_ISL_1975118, EPI_ISL_1975127, EPI_ISL_1975131, EPI_ISL_1975133, EPI_ISL_1975163, | University College London, Great Ormond Street Hospital for Children NHS Foundation Trust, Imperial College Healthcare NHS Trust | COVID-19 Genomics UK (COG-UK) Consortium | Sergi Castellano, Rachel Williams, Mark Kristiansen, Paola Resende Silva, Sunando Roy, Tony Brooks, Helena Tutill, Paola Niola, Patricia Dyal, Charlotte Williams, Leysa Forrest, Yasmin Panchbhaya, Jacqueline Findlay, Samuel Weeks, Julianne Brown, Kathryn Harris, Paul Randell, James Price, Alison Holmes, Judith Breuer |

| | | | |
|---|---|---|---|
| EPI_ISL_1975165, EPI_ISL_1975180 | | | |
| EPI_ISL_1978425, EPI_ISL_1978427, EPI_ISL_1978672, EPI_ISL_1978729, EPI_ISL_1978893 | Northumbria University / South Tees Hospitals NHS Foundation Trust / North Cumbria Integrated Care NHS Foundation Trust / North Tees and Hartlepool NHS Foundation Trust / Newcastle Hospitals NHS Foundation Trust | COVID-19 Genomics UK (COG-UK) Consortium | Darren L Smith,Andrew Nelson,Matthew Bashton,Greg R Young,Joshua Loh,John Allan,Mohammad A Tariq,Giles S Holt,Gary Black,Wen C Yew,Lynn Dover,Paul Baker,Steve Liggett,Sarah Essex,Jane Greenaway,Debra Padgett,Clive Graham,Garren Scott,Edward Barton,Emma Swindells,Brendan Payne,Jennifer Collins,Yusri Taha,Gary Eltringham |
| EPI_ISL_1978960 | Queens Medical Centre, Clinical Microbiology Department / DeepSeq Nottingham | COVID-19 Genomics UK (COG-UK) Consortium | Gemma Clark, Wendy Smith, Manjinder Khakh, Vicki M Fleming, Michelle M Lister, Hannah Howson-Wells, Jonathan Ball, Timothy Byaruhanga, Jayasree Dey, Emily Park, Jack Hill, Patrick McClure, Joseph Chappell, Theocharis Tsoleridis, Nadine Holmes, Matthew Carlisle, Christopher Moore, Fei Sang, Johnny Debebe, Victoria Wright, Matthew Loose |
| EPI_ISL_1979258, EPI_ISL_1979296, EPI_ISL_1979338, EPI_ISL_1979362, EPI_ISL_1979377, EPI_ISL_1979393, EPI_ISL_1979483, EPI_ISL_1979485, EPI_ISL_1979620 | Originating lab: Wales Specialist Virology Centre Sequencing lab: Pathogen Genomics Unit | Public Health Wales Microbiology Cardiff Wales Specialist Virology Centre | Catherine Moore, Johnathan Evans, Laura Gifford, Malorie Perry, Simon Cottrell, Angela Marchbank, Alec Birchley, Alexander Adams, Amy Gaskin, Bree Gatica-Wilcox, Jason Coombes, Joel Southgate, Lauren Gilbert, Lee Graham, Nicole Pacchiarini, Sara Kumziene-Summerhayes, Sarah Taylor, Sophie Jones, Sara Rey, Matthew Bull, Joanne Watkins, Sally Corden, Tom Connor |
| EPI_ISL_1980122, EPI_ISL_1980210 | Centre for Enzyme Innovation, University of Portsmouth / Translational Research Laboratory, Portsmouth Hospitals NHS Trust | COVID-19 Genomics UK (COG-UK) Consortium | Angela Beckett,Salman Goudarzi,Christopher Fearn,Kate Cook,Katie Loveson,Sharon Glaysher,Scott Elliott,Samuel Robson |
| EPI_ISL_1980422, EPI_ISL_1980438 | Virology Department, Sheffield Teaching Hospitals NHS Foundation Trust/Department of Infection, Immunity and Cardiovascular Disease, The Medical School, University of Sheffield | COVID-19 Genomics UK (COG-UK) Consortium | Thushan de Silva, Matthew Parker, Nikki Smith, Adri Angyal, Rebecca Brown, Luke Green, Rachel Tucker, Paul Parsons, Danielle Groves, Katie Johnson, Laura Carrilero, Alex Keeley, Dave Partridge, Matthew Wyles, Benjamin Lindsey, Mehmet Yavuz, Mohammad Raza, Cariad Evans |
| EPI_ISL_1985193, EPI_ISL_1985207 | Lighthouse Lab in Milton Keynes | Wellcome Sanger Institute for the COVID-19 Genomics UK (COG-UK) Consortium | The Lighthouse Lab in Milton Keynes and Alex Alderton, Roberto Amato, Jeffrey Barrett, Sonia Goncalves, Ewan Harrison, David K. Jackson, Ian Johnston, Dominic Kwiatkowski, Cordelia Langford, John Sillitoe on behalf of the Wellcome Sanger Institute COVID-19 Surveillance Team |
| EPI_ISL_1985222 | Lighthouse Lab in Alderley Park | Wellcome Sanger Institute for the COVID-19 Genomics UK (COG-UK) Consortium | Jacquelyn Wynn, Mairead Hyland, The Lighthouse Lab in Alderley Park and Alex Alderton, Roberto Amato, Jeffrey Barrett, Sonia Goncalves, Ewan Harrison, David K. Jackson, Ian Johnston, Dominic Kwiatkowski, Cordelia Langford, John Sillitoe on behalf of the Wellcome Sanger Institute COVID-19 Surveillance Team |
| EPI_ISL_1985243, EPI_ISL_1985265, EPI_ISL_1985275, EPI_ISL_1985284, EPI_ISL_1985351, EPI_ISL_1985369, EPI_ISL_1985380, EPI_ISL_1985403 | Lighthouse Lab in Milton Keynes | Wellcome Sanger Institute for the COVID-19 Genomics UK (COG-UK) Consortium | The Lighthouse Lab in Milton Keynes and Alex Alderton, Roberto Amato, Jeffrey Barrett, Sonia Goncalves, Ewan Harrison, David K. Jackson, Ian Johnston, Dominic Kwiatkowski, Cordelia Langford, John Sillitoe on behalf of the Wellcome Sanger Institute COVID-19 Surveillance Team |
| EPI_ISL_1985411 | Lighthouse Lab in Alderley Park | Wellcome Sanger Institute for the COVID-19 Genomics UK (COG-UK) Consortium | Jacquelyn Wynn, Mairead Hyland, The Lighthouse Lab in Alderley Park and Alex Alderton, Roberto Amato, Jeffrey Barrett, Sonia Goncalves, Ewan Harrison, David K. Jackson, Ian Johnston, Dominic Kwiatkowski, Cordelia Langford, John Sillitoe on behalf of the Wellcome Sanger Institute COVID-19 Surveillance Team |
| EPI_ISL_1985416, EPI_ISL_1985418, EPI_ISL_1985423, EPI_ISL_1985453, EPI_ISL_1985464, EPI_ISL_1985465, EPI_ISL_1985481, EPI_ISL_1985500, EPI_ISL_1985533, EPI_ISL_1985539, EPI_ISL_1985540, EPI_ISL_1985543, EPI_ISL_1985575, EPI_ISL_1985623, EPI_ISL_1985624 | | | |
| see above | Lighthouse Lab in Milton Keynes | Wellcome Sanger Institute for the COVID-19 Genomics UK (COG-UK) Consortium | The Lighthouse Lab in Milton Keynes and Alex Alderton, Roberto Amato, Jeffrey Barrett, Sonia Goncalves, Ewan Harrison, David K. Jackson, Ian Johnston, Dominic Kwiatkowski, Cordelia Langford, John Sillitoe on behalf of the Wellcome Sanger Institute COVID-19 Surveillance Team |
| EPI_ISL_1985692 | Lighthouse Lab in Alderley Park | Wellcome Sanger Institute for the COVID-19 Genomics UK (COG-UK) Consortium | Jacquelyn Wynn, Mairead Hyland, The Lighthouse Lab in Alderley Park and Alex Alderton, Roberto Amato, Jeffrey Barrett, Sonia Goncalves, Ewan Harrison, David K. Jackson, Ian Johnston, Dominic Kwiatkowski, Cordelia Langford, John Sillitoe on behalf of the Wellcome Sanger Institute COVID-19 Surveillance Team |
| EPI_ISL_1985701, EPI_ISL_1985708, EPI_ISL_1985713, EPI_ISL_1985749, EPI_ISL_1985750, EPI_ISL_1985754, EPI_ISL_1985756 | Lighthouse Lab in Milton Keynes | Wellcome Sanger Institute for the COVID-19 Genomics UK (COG-UK) Consortium | The Lighthouse Lab in Milton Keynes and Alex Alderton, Roberto Amato, Jeffrey Barrett, Sonia Goncalves, Ewan Harrison, David K. Jackson, Ian Johnston, Dominic Kwiatkowski, Cordelia Langford, John Sillitoe on behalf of the Wellcome Sanger Institute COVID-19 Surveillance Team |
| EPI_ISL_1985765, EPI_ISL_1985803, EPI_ISL_1985811 | Lighthouse Lab in Alderley Park | Wellcome Sanger Institute for the COVID-19 Genomics UK (COG-UK) Consortium | Jacquelyn Wynn, Mairead Hyland, The Lighthouse Lab in Alderley Park and Alex Alderton, Roberto Amato, Jeffrey Barrett, Sonia Goncalves, Ewan Harrison, David K. Jackson, Ian Johnston, Dominic Kwiatkowski, Cordelia Langford, John Sillitoe on behalf of the Wellcome Sanger Institute COVID-19 Surveillance Team |
| EPI_ISL_1985832, EPI_ISL_1985833, EPI_ISL_1985837, EPI_ISL_1985842, EPI_ISL_1985870, EPI_ISL_1985875, EPI_ISL_1985877, EPI_ISL_1985884, EPI_ISL_1985907, EPI_ISL_1985911, EPI_ISL_1985917 | | | |
| see above | Lighthouse Lab in Milton Keynes | Wellcome Sanger Institute for the COVID-19 Genomics UK (COG-UK) Consortium | The Lighthouse Lab in Milton Keynes and Alex Alderton, Roberto Amato, Jeffrey Barrett, Sonia Goncalves, Ewan Harrison, David K. Jackson, Ian Johnston, Dominic Kwiatkowski, Cordelia Langford, John Sillitoe on behalf of the Wellcome Sanger Institute COVID-19 Surveillance Team |
| EPI_ISL_1985921 | Lighthouse Lab in Alderley Park | Wellcome Sanger Institute for the COVID-19 Genomics UK (COG-UK) Consortium | Jacquelyn Wynn, Mairead Hyland, The Lighthouse Lab in Alderley Park and Alex Alderton, Roberto Amato, Jeffrey Barrett, Sonia Goncalves, Ewan Harrison, David K. Jackson, Ian Johnston, Dominic Kwiatkowski, Cordelia Langford, John Sillitoe on behalf of the Wellcome Sanger Institute COVID-19 Surveillance Team |
| EPI_ISL_1985922, EPI_ISL_1985935, EPI_ISL_1985944 | Lighthouse Lab in Milton Keynes | Wellcome Sanger Institute for the COVID-19 Genomics UK (COG-UK) Consortium | The Lighthouse Lab in Milton Keynes and Alex Alderton, Roberto Amato, Jeffrey Barrett, Sonia Goncalves, Ewan Harrison, David K. Jackson, Ian Johnston, Dominic Kwiatkowski, Cordelia Langford, John Sillitoe on behalf of the Wellcome Sanger Institute COVID-19 Surveillance Team |
| EPI_ISL_1985953, EPI_ISL_1985957 | Lighthouse Lab in Alderley Park | Wellcome Sanger Institute for the COVID-19 Genomics UK (COG-UK) Consortium | Jacquelyn Wynn, Mairead Hyland, The Lighthouse Lab in Alderley Park and Alex Alderton, Roberto Amato, Jeffrey Barrett, Sonia Goncalves, Ewan Harrison, David K. Jackson, Ian Johnston, Dominic Kwiatkowski, Cordelia Langford, John Sillitoe on behalf of the Wellcome Sanger Institute COVID-19 Surveillance Team |
| EPI_ISL_1985964, EPI_ISL_1985969, EPI_ISL_1985974 | Lighthouse Lab in Milton Keynes | Wellcome Sanger Institute for the COVID-19 Genomics UK (COG-UK) Consortium | The Lighthouse Lab in Milton Keynes and Alex Alderton, Roberto Amato, Jeffrey Barrett, Sonia Goncalves, Ewan Harrison, David K. Jackson, Ian Johnston, Dominic Kwiatkowski, Cordelia Langford, John Sillitoe on behalf of the Wellcome Sanger Institute COVID-19 Surveillance Team |
| EPI_ISL_1985984 | Lighthouse Lab in Alderley Park | Wellcome Sanger Institute for the COVID-19 Genomics UK (COG-UK) Consortium | Jacquelyn Wynn, Mairead Hyland, The Lighthouse Lab in Alderley Park and Alex Alderton, Roberto Amato, Jeffrey Barrett, Sonia Goncalves, Ewan Harrison, David K. Jackson, Ian Johnston, Dominic Kwiatkowski, Cordelia Langford, John Sillitoe on behalf of the Wellcome Sanger Institute COVID-19 Surveillance Team |
| EPI_ISL_1985987, EPI_ISL_1985996, EPI_ISL_1985998 | Lighthouse Lab in Milton Keynes | Wellcome Sanger Institute for the COVID-19 Genomics UK (COG-UK) Consortium | The Lighthouse Lab in Milton Keynes and Alex Alderton, Roberto Amato, Jeffrey Barrett, Sonia Goncalves, Ewan Harrison, David K. Jackson, Ian Johnston, Dominic Kwiatkowski, Cordelia Langford, John Sillitoe on behalf of the Wellcome Sanger Institute COVID-19 Surveillance Team |
| EPI_ISL_1986009 | Lighthouse Lab in Alderley Park | Wellcome Sanger Institute for the COVID-19 Genomics UK (COG-UK) Consortium | Jacquelyn Wynn, Mairead Hyland, The Lighthouse Lab in Alderley Park and Alex Alderton, Roberto Amato, Jeffrey Barrett, Sonia Goncalves, Ewan Harrison, David K. Jackson, Ian Johnston, Dominic Kwiatkowski, Cordelia Langford, John Sillitoe on behalf of the Wellcome Sanger Institute COVID-19 Surveillance Team |
| EPI_ISL_1986011, EPI_ISL_1986014 | Lighthouse Lab in Milton Keynes | Wellcome Sanger Institute for the COVID-19 Genomics UK (COG-UK) Consortium | The Lighthouse Lab in Milton Keynes and Alex Alderton, Roberto Amato, Jeffrey Barrett, Sonia Goncalves, Ewan Harrison, David K. Jackson, Ian Johnston, Dominic Kwiatkowski, Cordelia Langford, John Sillitoe on behalf of the Wellcome Sanger Institute COVID-19 Surveillance Team |
| EPI_ISL_1986018, EPI_ISL_1986028, EPI_ISL_1986032 | Lighthouse Lab in Alderley Park | Wellcome Sanger Institute for the COVID-19 Genomics UK (COG-UK) Consortium | Jacquelyn Wynn, Mairead Hyland, The Lighthouse Lab in Alderley Park and Alex Alderton, Roberto Amato, Jeffrey Barrett, Sonia Goncalves, Ewan Harrison, David K. Jackson, Ian Johnston, Dominic Kwiatkowski, Cordelia Langford, John Sillitoe on behalf of the Wellcome Sanger Institute COVID-19 Surveillance Team |
| EPI_ISL_1986033, EPI_ISL_1986052, EPI_ISL_1986053, EPI_ISL_1986054 | Lighthouse Lab in Milton Keynes | Wellcome Sanger Institute for the COVID-19 Genomics UK (COG-UK) Consortium | The Lighthouse Lab in Milton Keynes and Alex Alderton, Roberto Amato, Jeffrey Barrett, Sonia Goncalves, Ewan Harrison, David K. Jackson, Ian Johnston, Dominic Kwiatkowski, Cordelia Langford, John Sillitoe on behalf of the Wellcome Sanger Institute COVID-19 Surveillance Team |
| EPI_ISL_1986058 | Lighthouse Lab in Alderley Park | Wellcome Sanger Institute for the COVID-19 Genomics UK (COG-UK) Consortium | Jacquelyn Wynn, Mairead Hyland, The Lighthouse Lab in Alderley Park and Alex Alderton, Roberto Amato, Jeffrey Barrett, Sonia Goncalves, Ewan Harrison, David K. Jackson, Ian Johnston, Dominic Kwiatkowski, Cordelia Langford, John Sillitoe on behalf of the Wellcome Sanger Institute COVID-19 Surveillance Team |
| EPI_ISL_1986059, EPI_ISL_1986077, EPI_ISL_1986080, EPI_ISL_1986090 | Lighthouse Lab in Milton Keynes | Wellcome Sanger Institute for the COVID-19 Genomics UK (COG-UK) Consortium | The Lighthouse Lab in Milton Keynes and Alex Alderton, Roberto Amato, Jeffrey Barrett, Sonia Goncalves, Ewan Harrison, David K. Jackson, Ian Johnston, Dominic Kwiatkowski, Cordelia Langford, John Sillitoe on behalf of the Wellcome Sanger Institute COVID-19 Surveillance Team |
| EPI_ISL_1986097, EPI_ISL_1986099 | Lighthouse Lab in Alderley Park | Wellcome Sanger Institute for the COVID-19 Genomics UK (COG-UK) Consortium | Jacquelyn Wynn, Mairead Hyland, The Lighthouse Lab in Alderley Park and Alex Alderton, Roberto Amato, Jeffrey Barrett, Sonia Goncalves, Ewan Harrison, David K. Jackson, Ian Johnston, Dominic Kwiatkowski, Cordelia Langford, John Sillitoe on behalf of the Wellcome Sanger Institute COVID-19 Surveillance Team |
| EPI_ISL_1986101 | Lighthouse Lab in Milton Keynes | Wellcome Sanger Institute for the COVID-19 Genomics UK (COG-UK) Consortium | The Lighthouse Lab in Milton Keynes and Alex Alderton, Roberto Amato, Jeffrey Barrett, Sonia Goncalves, Ewan Harrison, David K. Jackson, Ian Johnston, Dominic Kwiatkowski, Cordelia Langford, John Sillitoe on behalf of the Wellcome Sanger Institute COVID-19 Surveillance Team |
| EPI_ISL_1986105, EPI_ISL_1986107 | Lighthouse Lab in Alderley Park | Wellcome Sanger Institute for the COVID-19 Genomics UK (COG-UK) Consortium | Jacquelyn Wynn, Mairead Hyland, The Lighthouse Lab in Alderley Park and Alex Alderton, Roberto Amato, Jeffrey Barrett, Sonia Goncalves, Ewan Harrison, David K. Jackson, Ian Johnston, Dominic Kwiatkowski, Cordelia Langford, John Sillitoe on behalf of the Wellcome Sanger Institute COVID-19 Surveillance Team |
| EPI_ISL_1986120 | Lighthouse Lab in Milton Keynes | Wellcome Sanger Institute for the COVID-19 Genomics UK (COG-UK) | The Lighthouse Lab in Milton Keynes and Alex Alderton, Roberto Amato, Jeffrey Barrett, Sonia Goncalves, Ewan Harrison, David K. Jackson, Ian Johnston, Dominic Kwiatkowski, Cordelia Langford, John Sillitoe on behalf of the Wellcome Sanger |

| | | | |
|---|---|---|---|
| | | Consortium | Institute COVID-19 Surveillance Team |
| EPI_ISL_1986129, EPI_ISL_1986136, EPI_ISL_1986139 | Lighthouse Lab in Alderley Park | Wellcome Sanger Institute for the COVID-19 Genomics UK (COG-UK) Consortium | Jacquelyn Wynn, Mairead Hyland, The Lighthouse Lab in Alderley Park and Alex Alderton, Roberto Amato, Jeffrey Barrett, Sonia Goncalves, Ewan Harrison, David K. Jackson, Ian Johnston, Dominic Kwiatkowski, Cordelia Langford, John Sillitoe on behalf of the Wellcome Sanger Institute COVID-19 Surveillance Team |
| EPI_ISL_1986150, EPI_ISL_1986153 | Lighthouse Lab in Milton Keynes | Wellcome Sanger Institute for the COVID-19 Genomics UK (COG-UK) Consortium | The Lighthouse Lab in Milton Keynes and Alex Alderton, Roberto Amato, Jeffrey Barrett, Sonia Goncalves, Ewan Harrison, David K. Jackson, Ian Johnston, Dominic Kwiatkowski, Cordelia Langford, John Sillitoe on behalf of the Wellcome Sanger Institute COVID-19 Surveillance Team |
| EPI_ISL_1986159, EPI_ISL_1986167 | Lighthouse Lab in Alderley Park | Wellcome Sanger Institute for the COVID-19 Genomics UK (COG-UK) Consortium | Jacquelyn Wynn, Mairead Hyland, The Lighthouse Lab in Alderley Park and Alex Alderton, Roberto Amato, Jeffrey Barrett, Sonia Goncalves, Ewan Harrison, David K. Jackson, Ian Johnston, Dominic Kwiatkowski, Cordelia Langford, John Sillitoe on behalf of the Wellcome Sanger Institute COVID-19 Surveillance Team |
| EPI_ISL_1986183 | Lighthouse Lab in Milton Keynes | Wellcome Sanger Institute for the COVID-19 Genomics UK (COG-UK) Consortium | The Lighthouse Lab in Milton Keynes and Alex Alderton, Roberto Amato, Jeffrey Barrett, Sonia Goncalves, Ewan Harrison, David K. Jackson, Ian Johnston, Dominic Kwiatkowski, Cordelia Langford, John Sillitoe on behalf of the Wellcome Sanger Institute COVID-19 Surveillance Team |
| EPI_ISL_1986184, EPI_ISL_1986218, EPI_ISL_1986220 | Lighthouse Lab in Alderley Park | Wellcome Sanger Institute for the COVID-19 Genomics UK (COG-UK) Consortium | Jacquelyn Wynn, Mairead Hyland, The Lighthouse Lab in Alderley Park and Alex Alderton, Roberto Amato, Jeffrey Barrett, Sonia Goncalves, Ewan Harrison, David K. Jackson, Ian Johnston, Dominic Kwiatkowski, Cordelia Langford, John Sillitoe on behalf of the Wellcome Sanger Institute COVID-19 Surveillance Team |
| EPI_ISL_1986222, EPI_ISL_1986225, EPI_ISL_1986226 | Lighthouse Lab in Milton Keynes | Wellcome Sanger Institute for the COVID-19 Genomics UK (COG-UK) Consortium | The Lighthouse Lab in Milton Keynes and Alex Alderton, Roberto Amato, Jeffrey Barrett, Sonia Goncalves, Ewan Harrison, David K. Jackson, Ian Johnston, Dominic Kwiatkowski, Cordelia Langford, John Sillitoe on behalf of the Wellcome Sanger Institute COVID-19 Surveillance Team |
| EPI_ISL_1986233 | Lighthouse Lab in Alderley Park | Wellcome Sanger Institute for the COVID-19 Genomics UK (COG-UK) Consortium | Jacquelyn Wynn, Mairead Hyland, The Lighthouse Lab in Alderley Park and Alex Alderton, Roberto Amato, Jeffrey Barrett, Sonia Goncalves, Ewan Harrison, David K. Jackson, Ian Johnston, Dominic Kwiatkowski, Cordelia Langford, John Sillitoe on behalf of the Wellcome Sanger Institute COVID-19 Surveillance Team |
| EPI_ISL_1986240, EPI_ISL_1986248 | Lighthouse Lab in Milton Keynes | Wellcome Sanger Institute for the COVID-19 Genomics UK (COG-UK) Consortium | The Lighthouse Lab in Milton Keynes and Alex Alderton, Roberto Amato, Jeffrey Barrett, Sonia Goncalves, Ewan Harrison, David K. Jackson, Ian Johnston, Dominic Kwiatkowski, Cordelia Langford, John Sillitoe on behalf of the Wellcome Sanger Institute COVID-19 Surveillance Team |
| EPI_ISL_1986255 | Lighthouse Lab in Alderley Park | Wellcome Sanger Institute for the COVID-19 Genomics UK (COG-UK) Consortium | Jacquelyn Wynn, Mairead Hyland, The Lighthouse Lab in Alderley Park and Alex Alderton, Roberto Amato, Jeffrey Barrett, Sonia Goncalves, Ewan Harrison, David K. Jackson, Ian Johnston, Dominic Kwiatkowski, Cordelia Langford, John Sillitoe on behalf of the Wellcome Sanger Institute COVID-19 Surveillance Team |
| EPI_ISL_1986261, EPI_ISL_1986262 | Lighthouse Lab in Milton Keynes | Wellcome Sanger Institute for the COVID-19 Genomics UK (COG-UK) Consortium | The Lighthouse Lab in Milton Keynes and Alex Alderton, Roberto Amato, Jeffrey Barrett, Sonia Goncalves, Ewan Harrison, David K. Jackson, Ian Johnston, Dominic Kwiatkowski, Cordelia Langford, John Sillitoe on behalf of the Wellcome Sanger Institute COVID-19 Surveillance Team |
| EPI_ISL_1986281 | Lighthouse Lab in Alderley Park | Wellcome Sanger Institute for the COVID-19 Genomics UK (COG-UK) Consortium | Jacquelyn Wynn, Mairead Hyland, The Lighthouse Lab in Alderley Park and Alex Alderton, Roberto Amato, Jeffrey Barrett, Sonia Goncalves, Ewan Harrison, David K. Jackson, Ian Johnston, Dominic Kwiatkowski, Cordelia Langford, John Sillitoe on behalf of the Wellcome Sanger Institute COVID-19 Surveillance Team |
| EPI_ISL_1986287, EPI_ISL_1986291, EPI_ISL_1986293, EPI_ISL_1986303 | Lighthouse Lab in Milton Keynes | Wellcome Sanger Institute for the COVID-19 Genomics UK (COG-UK) Consortium | The Lighthouse Lab in Milton Keynes and Alex Alderton, Roberto Amato, Jeffrey Barrett, Sonia Goncalves, Ewan Harrison, David K. Jackson, Ian Johnston, Dominic Kwiatkowski, Cordelia Langford, John Sillitoe on behalf of the Wellcome Sanger Institute COVID-19 Surveillance Team |
| EPI_ISL_1986304 | Lighthouse Lab in Alderley Park | Wellcome Sanger Institute for the COVID-19 Genomics UK (COG-UK) Consortium | Jacquelyn Wynn, Mairead Hyland, The Lighthouse Lab in Alderley Park and Alex Alderton, Roberto Amato, Jeffrey Barrett, Sonia Goncalves, Ewan Harrison, David K. Jackson, Ian Johnston, Dominic Kwiatkowski, Cordelia Langford, John Sillitoe on behalf of the Wellcome Sanger Institute COVID-19 Surveillance Team |
| EPI_ISL_1986333, EPI_ISL_1986356, EPI_ISL_1986367, EPI_ISL_1986388 | Lighthouse Lab in Milton Keynes | Wellcome Sanger Institute for the COVID-19 Genomics UK (COG-UK) Consortium | The Lighthouse Lab in Milton Keynes and Alex Alderton, Roberto Amato, Jeffrey Barrett, Sonia Goncalves, Ewan Harrison, David K. Jackson, Ian Johnston, Dominic Kwiatkowski, Cordelia Langford, John Sillitoe on behalf of the Wellcome Sanger Institute COVID-19 Surveillance Team |
| EPI_ISL_1986389 | Lighthouse Lab in Glasgow | Wellcome Sanger Institute for the COVID-19 Genomics UK (COG-UK) Consortium | Harper VanSteenhouse, Yumi Kasai, David Gray, Carol Clugston, Anna Dominiczak and Alex Alderton, Roberto Amato, Jeffrey Barrett, Sonia Goncalves, Ewan Harrison, David K. Jackson, Ian Johnston, Dominic Kwiatkowski, Cordelia Langford, John Sillitoe on behalf of the Wellcome Sanger Institute COVID-19 Surveillance Team |
| EPI_ISL_1986390 | Lighthouse Lab in Alderley Park | Wellcome Sanger Institute for the COVID-19 Genomics UK (COG-UK) Consortium | Jacquelyn Wynn, Mairead Hyland, The Lighthouse Lab in Alderley Park and Alex Alderton, Roberto Amato, Jeffrey Barrett, Sonia Goncalves, Ewan Harrison, David K. Jackson, Ian Johnston, Dominic Kwiatkowski, Cordelia Langford, John Sillitoe on behalf of the Wellcome Sanger Institute COVID-19 Surveillance Team |
| EPI_ISL_1986500 | Lighthouse Lab in Milton Keynes | Wellcome Sanger Institute for the COVID-19 Genomics UK (COG-UK) Consortium | The Lighthouse Lab in Milton Keynes and Alex Alderton, Roberto Amato, Jeffrey Barrett, Sonia Goncalves, Ewan Harrison, David K. Jackson, Ian Johnston, Dominic Kwiatkowski, Cordelia Langford, John Sillitoe on behalf of the Wellcome Sanger Institute COVID-19 Surveillance Team |
| EPI_ISL_1986511, EPI_ISL_1986519 | Lighthouse Lab in Alderley Park | Wellcome Sanger Institute for the COVID-19 Genomics UK (COG-UK) Consortium | Jacquelyn Wynn, Mairead Hyland, The Lighthouse Lab in Alderley Park and Alex Alderton, Roberto Amato, Jeffrey Barrett, Sonia Goncalves, Ewan Harrison, David K. Jackson, Ian Johnston, Dominic Kwiatkowski, Cordelia Langford, John Sillitoe on behalf of the Wellcome Sanger Institute COVID-19 Surveillance Team |
| EPI_ISL_1986555 | Lighthouse Lab in Milton Keynes | Wellcome Sanger Institute for the COVID-19 Genomics UK (COG-UK) Consortium | The Lighthouse Lab in Milton Keynes and Alex Alderton, Roberto Amato, Jeffrey Barrett, Sonia Goncalves, Ewan Harrison, David K. Jackson, Ian Johnston, Dominic Kwiatkowski, Cordelia Langford, John Sillitoe on behalf of the Wellcome Sanger Institute COVID-19 Surveillance Team |
| EPI_ISL_1986571 | Lighthouse Lab in Glasgow | Wellcome Sanger Institute for the COVID-19 Genomics UK (COG-UK) Consortium | Harper VanSteenhouse, Yumi Kasai, David Gray, Carol Clugston, Anna Dominiczak and Alex Alderton, Roberto Amato, Jeffrey Barrett, Sonia Goncalves, Ewan Harrison, David K. Jackson, Ian Johnston, Dominic Kwiatkowski, Cordelia Langford, John Sillitoe on behalf of the Wellcome Sanger Institute COVID-19 Surveillance Team |
| EPI_ISL_1986577 | Lighthouse Lab in Alderley Park | Wellcome Sanger Institute for the COVID-19 Genomics UK (COG-UK) Consortium | Jacquelyn Wynn, Mairead Hyland, The Lighthouse Lab in Alderley Park and Alex Alderton, Roberto Amato, Jeffrey Barrett, Sonia Goncalves, Ewan Harrison, David K. Jackson, Ian Johnston, Dominic Kwiatkowski, Cordelia Langford, John Sillitoe on behalf of the Wellcome Sanger Institute COVID-19 Surveillance Team |
| EPI_ISL_1986578, EPI_ISL_1986584 | Lighthouse Lab in Milton Keynes | Wellcome Sanger Institute for the COVID-19 Genomics UK (COG-UK) Consortium | The Lighthouse Lab in Milton Keynes and Alex Alderton, Roberto Amato, Jeffrey Barrett, Sonia Goncalves, Ewan Harrison, David K. Jackson, Ian Johnston, Dominic Kwiatkowski, Cordelia Langford, John Sillitoe on behalf of the Wellcome Sanger Institute COVID-19 Surveillance Team |
| EPI_ISL_1986589, EPI_ISL_1986593, EPI_ISL_1986617, EPI_ISL_1986626, EPI_ISL_1986639 | Lighthouse Lab in Alderley Park | Wellcome Sanger Institute for the COVID-19 Genomics UK (COG-UK) Consortium | Jacquelyn Wynn, Mairead Hyland, The Lighthouse Lab in Alderley Park and Alex Alderton, Roberto Amato, Jeffrey Barrett, Sonia Goncalves, Ewan Harrison, David K. Jackson, Ian Johnston, Dominic Kwiatkowski, Cordelia Langford, John Sillitoe on behalf of the Wellcome Sanger Institute COVID-19 Surveillance Team |
| EPI_ISL_1986688 | Randox Laboratories | Wellcome Sanger Institute for the COVID-19 Genomics UK (COG-UK) Consortium | Randox Laboratories and Alex Alderton, Roberto Amato, Jeffrey Barrett, Sonia Goncalves, Ewan Harrison, David K. Jackson, Ian Johnston, Dominic Kwiatkowski, Cordelia Langford, John Sillitoe on behalf of the Wellcome Sanger Institute COVID-19 Surveillance Team |
| EPI_ISL_1986692 | Lighthouse Lab in Alderley Park | Wellcome Sanger Institute for the COVID-19 Genomics UK (COG-UK) Consortium | Jacquelyn Wynn, Mairead Hyland, The Lighthouse Lab in Alderley Park and Alex Alderton, Roberto Amato, Jeffrey Barrett, Sonia Goncalves, Ewan Harrison, David K. Jackson, Ian Johnston, Dominic Kwiatkowski, Cordelia Langford, John Sillitoe on behalf of the Wellcome Sanger Institute COVID-19 Surveillance Team |
| EPI_ISL_1986695 | Randox Laboratories | Wellcome Sanger Institute for the COVID-19 Genomics UK (COG-UK) Consortium | Randox Laboratories and Alex Alderton, Roberto Amato, Jeffrey Barrett, Sonia Goncalves, Ewan Harrison, David K. Jackson, Ian Johnston, Dominic Kwiatkowski, Cordelia Langford, John Sillitoe on behalf of the Wellcome Sanger Institute COVID-19 Surveillance Team |
| EPI_ISL_1986704 | Lighthouse Lab in Milton Keynes | Wellcome Sanger Institute for the COVID-19 Genomics UK (COG-UK) Consortium | The Lighthouse Lab in Milton Keynes and Alex Alderton, Roberto Amato, Jeffrey Barrett, Sonia Goncalves, Ewan Harrison, David K. Jackson, Ian Johnston, Dominic Kwiatkowski, Cordelia Langford, John Sillitoe on behalf of the Wellcome Sanger Institute COVID-19 Surveillance Team |
| EPI_ISL_1986707, EPI_ISL_1986712 | Lighthouse Lab in Alderley Park | Wellcome Sanger Institute for the COVID-19 Genomics UK (COG-UK) Consortium | Jacquelyn Wynn, Mairead Hyland, The Lighthouse Lab in Alderley Park and Alex Alderton, Roberto Amato, Jeffrey Barrett, Sonia Goncalves, Ewan Harrison, David K. Jackson, Ian Johnston, Dominic Kwiatkowski, Cordelia Langford, John Sillitoe on behalf of the Wellcome Sanger Institute COVID-19 Surveillance Team |
| EPI_ISL_1986729 | Lighthouse Lab in Milton Keynes | Wellcome Sanger Institute for the COVID-19 Genomics UK (COG-UK) Consortium | The Lighthouse Lab in Milton Keynes and Alex Alderton, Roberto Amato, Jeffrey Barrett, Sonia Goncalves, Ewan Harrison, David K. Jackson, Ian Johnston, Dominic Kwiatkowski, Cordelia Langford, John Sillitoe on behalf of the Wellcome Sanger Institute COVID-19 Surveillance Team |
| EPI_ISL_1986733, EPI_ISL_1986741, EPI_ISL_1986746 | Lighthouse Lab in Alderley Park | Wellcome Sanger Institute for the COVID-19 Genomics UK (COG-UK) Consortium | Jacquelyn Wynn, Mairead Hyland, The Lighthouse Lab in Alderley Park and Alex Alderton, Roberto Amato, Jeffrey Barrett, Sonia Goncalves, Ewan Harrison, David K. Jackson, Ian Johnston, Dominic Kwiatkowski, Cordelia Langford, John Sillitoe on behalf of the Wellcome Sanger Institute COVID-19 Surveillance Team |
| EPI_ISL_1986752, EPI_ISL_1986754 | Lighthouse Lab in Milton Keynes | Wellcome Sanger Institute for the COVID-19 Genomics UK (COG-UK) Consortium | The Lighthouse Lab in Milton Keynes and Alex Alderton, Roberto Amato, Jeffrey Barrett, Sonia Goncalves, Ewan Harrison, David K. Jackson, Ian Johnston, Dominic Kwiatkowski, Cordelia Langford, John Sillitoe on behalf of the Wellcome Sanger Institute COVID-19 Surveillance Team |
| EPI_ISL_1986756 | Lighthouse Lab in Alderley Park | Wellcome Sanger Institute for the COVID-19 Genomics UK (COG-UK) Consortium | Jacquelyn Wynn, Mairead Hyland, The Lighthouse Lab in Alderley Park and Alex Alderton, Roberto Amato, Jeffrey Barrett, Sonia Goncalves, Ewan Harrison, David K. Jackson, Ian Johnston, Dominic Kwiatkowski, Cordelia Langford, John Sillitoe on behalf of the Wellcome Sanger Institute COVID-19 Surveillance Team |
| EPI_ISL_1986757 | Randox Laboratories | Wellcome Sanger Institute for the COVID-19 Genomics UK (COG-UK) Consortium | Randox Laboratories and Alex Alderton, Roberto Amato, Jeffrey Barrett, Sonia Goncalves, Ewan Harrison, David K. Jackson, Ian Johnston, Dominic Kwiatkowski, Cordelia Langford, John Sillitoe on behalf of the Wellcome Sanger Institute COVID-19 Surveillance Team |
| EPI_ISL_1986760, EPI_ISL_1986769 | Lighthouse Lab in Milton Keynes | Wellcome Sanger Institute for the COVID-19 Genomics UK (COG-UK) Consortium | The Lighthouse Lab in Milton Keynes and Alex Alderton, Roberto Amato, Jeffrey Barrett, Sonia Goncalves, Ewan Harrison, David K. Jackson, Ian Johnston, Dominic Kwiatkowski, Cordelia Langford, John Sillitoe on behalf of the Wellcome Sanger Institute COVID-19 Surveillance Team |
| EPI_ISL_1986810, EPI_ISL_1986812, EPI_ISL_1986813 | Lighthouse Lab in Alderley Park | Wellcome Sanger Institute for the COVID-19 Genomics UK (COG-UK) Consortium | Jacquelyn Wynn, Mairead Hyland, The Lighthouse Lab in Alderley Park and Alex Alderton, Roberto Amato, Jeffrey Barrett, Sonia Goncalves, Ewan Harrison, David K. Jackson, Ian Johnston, Dominic Kwiatkowski, Cordelia Langford, John Sillitoe on behalf of the Wellcome Sanger Institute COVID-19 Surveillance Team |
| EPI_ISL_1986827, EPI_ISL_1986834, EPI_ISL_1986850, EPI_ISL_1986851, EPI_ISL_1986856 | Lighthouse Lab in Milton Keynes | Wellcome Sanger Institute for the COVID-19 Genomics UK (COG-UK) Consortium | The Lighthouse Lab in Milton Keynes and Alex Alderton, Roberto Amato, Jeffrey Barrett, Sonia Goncalves, Ewan Harrison, David K. Jackson, Ian Johnston, Dominic Kwiatkowski, Cordelia Langford, John Sillitoe on behalf of the Wellcome Sanger Institute COVID-19 Surveillance Team |
| EPI_ISL_1986858 | Lighthouse Lab in Alderley Park | Wellcome Sanger Institute for the COVID-19 Genomics UK (COG-UK) Consortium | Jacquelyn Wynn, Mairead Hyland, The Lighthouse Lab in Alderley Park and Alex Alderton, Roberto Amato, Jeffrey Barrett, Sonia Goncalves, Ewan Harrison, David K. Jackson, Ian Johnston, Dominic Kwiatkowski, Cordelia Langford, John Sillitoe on behalf of the Wellcome Sanger Institute COVID-19 Surveillance Team |
| EPI_ISL_1986870 | Randox Laboratories | Wellcome Sanger Institute for the COVID-19 Genomics UK (COG-UK) Consortium | Randox Laboratories and Alex Alderton, Roberto Amato, Jeffrey Barrett, Sonia Goncalves, Ewan Harrison, David K. Jackson, Ian Johnston, Dominic Kwiatkowski, Cordelia Langford, John Sillitoe on behalf of the Wellcome Sanger Institute COVID-19 Surveillance Team |

| | | | |
|---|---|---|---|
| EPI_ISL_1986871 | Lighthouse Lab in Alderley Park | Wellcome Sanger Institute for the COVID-19 Genomics UK (COG-UK) Consortium | Jacquelyn Wynn, Mairead Hyland, The Lighthouse Lab in Alderley Park and Alex Alderton, Roberto Amato, Jeffrey Barrett, Sonia Goncalves, Ewan Harrison, David K. Jackson, Ian Johnston, Dominic Kwiatkowski, Cordelia Langford, John Sillitoe on behalf of the Wellcome Sanger Institute COVID-19 Surveillance Team |
| EPI_ISL_1986897 | Lighthouse Lab in Milton Keynes | Wellcome Sanger Institute for the COVID-19 Genomics UK (COG-UK) Consortium | The Lighthouse Lab in Milton Keynes and Alex Alderton, Roberto Amato, Jeffrey Barrett, Sonia Goncalves, Ewan Harrison, David K. Jackson, Ian Johnston, Dominic Kwiatkowski, Cordelia Langford, John Sillitoe on behalf of the Wellcome Sanger Institute COVID-19 Surveillance Team |
| EPI_ISL_1986900, EPI_ISL_1986933 | Lighthouse Lab in Alderley Park | Wellcome Sanger Institute for the COVID-19 Genomics UK (COG-UK) Consortium | Jacquelyn Wynn, Mairead Hyland, The Lighthouse Lab in Alderley Park and Alex Alderton, Roberto Amato, Jeffrey Barrett, Sonia Goncalves, Ewan Harrison, David K. Jackson, Ian Johnston, Dominic Kwiatkowski, Cordelia Langford, John Sillitoe on behalf of the Wellcome Sanger Institute COVID-19 Surveillance Team |
| EPI_ISL_1986950 | Lighthouse Lab in Milton Keynes | Wellcome Sanger Institute for the COVID-19 Genomics UK (COG-UK) Consortium | The Lighthouse Lab in Milton Keynes and Alex Alderton, Roberto Amato, Jeffrey Barrett, Sonia Goncalves, Ewan Harrison, David K. Jackson, Ian Johnston, Dominic Kwiatkowski, Cordelia Langford, John Sillitoe on behalf of the Wellcome Sanger Institute COVID-19 Surveillance Team |
| EPI_ISL_1986954 | Lighthouse Lab in Glasgow | Wellcome Sanger Institute for the COVID-19 Genomics UK (COG-UK) Consortium | Harper VanSteenhouse, Yumi Kasai, David Gray, Carol Clugston, Anna Dominiczak and Alex Alderton, Roberto Amato, Jeffrey Barrett, Sonia Goncalves, Ewan Harrison, David K. Jackson, Ian Johnston, Dominic Kwiatkowski, Cordelia Langford, John Sillitoe on behalf of the Wellcome Sanger Institute COVID-19 Surveillance Team |
| EPI_ISL_1986956 | Randox Laboratories | Wellcome Sanger Institute for the COVID-19 Genomics UK (COG-UK) Consortium | Randox Laboratories and Alex Alderton, Roberto Amato, Jeffrey Barrett, Sonia Goncalves, Ewan Harrison, David K. Jackson, Ian Johnston, Dominic Kwiatkowski, Cordelia Langford, John Sillitoe on behalf of the Wellcome Sanger Institute COVID-19 Surveillance Team |
| EPI_ISL_1986966, EPI_ISL_1986969 | Lighthouse Lab in Milton Keynes | Wellcome Sanger Institute for the COVID-19 Genomics UK (COG-UK) Consortium | The Lighthouse Lab in Milton Keynes and Alex Alderton, Roberto Amato, Jeffrey Barrett, Sonia Goncalves, Ewan Harrison, David K. Jackson, Ian Johnston, Dominic Kwiatkowski, Cordelia Langford, John Sillitoe on behalf of the Wellcome Sanger Institute COVID-19 Surveillance Team |
| EPI_ISL_1986972, EPI_ISL_1986974 | Lighthouse Lab in Alderley Park | Wellcome Sanger Institute for the COVID-19 Genomics UK (COG-UK) Consortium | Jacquelyn Wynn, Mairead Hyland, The Lighthouse Lab in Alderley Park and Alex Alderton, Roberto Amato, Jeffrey Barrett, Sonia Goncalves, Ewan Harrison, David K. Jackson, Ian Johnston, Dominic Kwiatkowski, Cordelia Langford, John Sillitoe on behalf of the Wellcome Sanger Institute COVID-19 Surveillance Team |
| EPI_ISL_1986979 | Randox Laboratories | Wellcome Sanger Institute for the COVID-19 Genomics UK (COG-UK) Consortium | Randox Laboratories and Alex Alderton, Roberto Amato, Jeffrey Barrett, Sonia Goncalves, Ewan Harrison, David K. Jackson, Ian Johnston, Dominic Kwiatkowski, Cordelia Langford, John Sillitoe on behalf of the Wellcome Sanger Institute COVID-19 Surveillance Team |
| EPI_ISL_1986989 | Lighthouse Lab in Glasgow | Wellcome Sanger Institute for the COVID-19 Genomics UK (COG-UK) Consortium | Harper VanSteenhouse, Yumi Kasai, David Gray, Carol Clugston, Anna Dominiczak and Alex Alderton, Roberto Amato, Jeffrey Barrett, Sonia Goncalves, Ewan Harrison, David K. Jackson, Ian Johnston, Dominic Kwiatkowski, Cordelia Langford, John Sillitoe on behalf of the Wellcome Sanger Institute COVID-19 Surveillance Team |
| EPI_ISL_1986993, EPI_ISL_1987000, EPI_ISL_1987001 | Lighthouse Lab in Alderley Park | Wellcome Sanger Institute for the COVID-19 Genomics UK (COG-UK) Consortium | Jacquelyn Wynn, Mairead Hyland, The Lighthouse Lab in Alderley Park and Alex Alderton, Roberto Amato, Jeffrey Barrett, Sonia Goncalves, Ewan Harrison, David K. Jackson, Ian Johnston, Dominic Kwiatkowski, Cordelia Langford, John Sillitoe on behalf of the Wellcome Sanger Institute COVID-19 Surveillance Team |
| EPI_ISL_1987010 | Lighthouse Lab in Milton Keynes | Wellcome Sanger Institute for the COVID-19 Genomics UK (COG-UK) Consortium | The Lighthouse Lab in Milton Keynes and Alex Alderton, Roberto Amato, Jeffrey Barrett, Sonia Goncalves, Ewan Harrison, David K. Jackson, Ian Johnston, Dominic Kwiatkowski, Cordelia Langford, John Sillitoe on behalf of the Wellcome Sanger Institute COVID-19 Surveillance Team |
| EPI_ISL_1987022 | Randox Laboratories | Wellcome Sanger Institute for the COVID-19 Genomics UK (COG-UK) Consortium | Randox Laboratories and Alex Alderton, Roberto Amato, Jeffrey Barrett, Sonia Goncalves, Ewan Harrison, David K. Jackson, Ian Johnston, Dominic Kwiatkowski, Cordelia Langford, John Sillitoe on behalf of the Wellcome Sanger Institute COVID-19 Surveillance Team |
| EPI_ISL_1987041 | Lighthouse Lab in Alderley Park | Wellcome Sanger Institute for the COVID-19 Genomics UK (COG-UK) Consortium | Jacquelyn Wynn, Mairead Hyland, The Lighthouse Lab in Alderley Park and Alex Alderton, Roberto Amato, Jeffrey Barrett, Sonia Goncalves, Ewan Harrison, David K. Jackson, Ian Johnston, Dominic Kwiatkowski, Cordelia Langford, John Sillitoe on behalf of the Wellcome Sanger Institute COVID-19 Surveillance Team |
| EPI_ISL_1987043 | Lighthouse Lab in Milton Keynes | Wellcome Sanger Institute for the COVID-19 Genomics UK (COG-UK) Consortium | The Lighthouse Lab in Milton Keynes and Alex Alderton, Roberto Amato, Jeffrey Barrett, Sonia Goncalves, Ewan Harrison, David K. Jackson, Ian Johnston, Dominic Kwiatkowski, Cordelia Langford, John Sillitoe on behalf of the Wellcome Sanger Institute COVID-19 Surveillance Team |
| EPI_ISL_1987044 | Randox Laboratories | Wellcome Sanger Institute for the COVID-19 Genomics UK (COG-UK) Consortium | Randox Laboratories and Alex Alderton, Roberto Amato, Jeffrey Barrett, Sonia Goncalves, Ewan Harrison, David K. Jackson, Ian Johnston, Dominic Kwiatkowski, Cordelia Langford, John Sillitoe on behalf of the Wellcome Sanger Institute COVID-19 Surveillance Team |
| EPI_ISL_1987050 | Lighthouse Lab in Alderley Park | Wellcome Sanger Institute for the COVID-19 Genomics UK (COG-UK) Consortium | Jacquelyn Wynn, Mairead Hyland, The Lighthouse Lab in Alderley Park and Alex Alderton, Roberto Amato, Jeffrey Barrett, Sonia Goncalves, Ewan Harrison, David K. Jackson, Ian Johnston, Dominic Kwiatkowski, Cordelia Langford, John Sillitoe on behalf of the Wellcome Sanger Institute COVID-19 Surveillance Team |
| EPI_ISL_1987052 | Lighthouse Lab in Milton Keynes | Wellcome Sanger Institute for the COVID-19 Genomics UK (COG-UK) Consortium | The Lighthouse Lab in Milton Keynes and Alex Alderton, Roberto Amato, Jeffrey Barrett, Sonia Goncalves, Ewan Harrison, David K. Jackson, Ian Johnston, Dominic Kwiatkowski, Cordelia Langford, John Sillitoe on behalf of the Wellcome Sanger Institute COVID-19 Surveillance Team |
| EPI_ISL_1987056, EPI_ISL_1987126, EPI_ISL_1987140, EPI_ISL_1987152, EPI_ISL_1987156 | Lighthouse Lab in Alderley Park | Wellcome Sanger Institute for the COVID-19 Genomics UK (COG-UK) Consortium | Jacquelyn Wynn, Mairead Hyland, The Lighthouse Lab in Alderley Park and Alex Alderton, Roberto Amato, Jeffrey Barrett, Sonia Goncalves, Ewan Harrison, David K. Jackson, Ian Johnston, Dominic Kwiatkowski, Cordelia Langford, John Sillitoe on behalf of the Wellcome Sanger Institute COVID-19 Surveillance Team |
| EPI_ISL_1987170, EPI_ISL_1987178, EPI_ISL_1987183 | Randox Laboratories | Wellcome Sanger Institute for the COVID-19 Genomics UK (COG-UK) Consortium | Randox Laboratories and Alex Alderton, Roberto Amato, Jeffrey Barrett, Sonia Goncalves, Ewan Harrison, David K. Jackson, Ian Johnston, Dominic Kwiatkowski, Cordelia Langford, John Sillitoe on behalf of the Wellcome Sanger Institute COVID-19 Surveillance Team |
| EPI_ISL_1987188 | Lighthouse Lab in Alderley Park | Wellcome Sanger Institute for the COVID-19 Genomics UK (COG-UK) Consortium | Jacquelyn Wynn, Mairead Hyland, The Lighthouse Lab in Alderley Park and Alex Alderton, Roberto Amato, Jeffrey Barrett, Sonia Goncalves, Ewan Harrison, David K. Jackson, Ian Johnston, Dominic Kwiatkowski, Cordelia Langford, John Sillitoe on behalf of the Wellcome Sanger Institute COVID-19 Surveillance Team |
| EPI_ISL_1987194 | Lighthouse Lab in Glasgow | Wellcome Sanger Institute for the COVID-19 Genomics UK (COG-UK) Consortium | Harper VanSteenhouse, Yumi Kasai, David Gray, Carol Clugston, Anna Dominiczak and Alex Alderton, Roberto Amato, Jeffrey Barrett, Sonia Goncalves, Ewan Harrison, David K. Jackson, Ian Johnston, Dominic Kwiatkowski, Cordelia Langford, John Sillitoe on behalf of the Wellcome Sanger Institute COVID-19 Surveillance Team |
| EPI_ISL_1987202 | Lighthouse Lab in Milton Keynes | Wellcome Sanger Institute for the COVID-19 Genomics UK (COG-UK) Consortium | The Lighthouse Lab in Milton Keynes and Alex Alderton, Roberto Amato, Jeffrey Barrett, Sonia Goncalves, Ewan Harrison, David K. Jackson, Ian Johnston, Dominic Kwiatkowski, Cordelia Langford, John Sillitoe on behalf of the Wellcome Sanger Institute COVID-19 Surveillance Team |
| EPI_ISL_1987207 | Lighthouse Lab in Alderley Park | Wellcome Sanger Institute for the COVID-19 Genomics UK (COG-UK) Consortium | Jacquelyn Wynn, Mairead Hyland, The Lighthouse Lab in Alderley Park and Alex Alderton, Roberto Amato, Jeffrey Barrett, Sonia Goncalves, Ewan Harrison, David K. Jackson, Ian Johnston, Dominic Kwiatkowski, Cordelia Langford, John Sillitoe on behalf of the Wellcome Sanger Institute COVID-19 Surveillance Team |
| EPI_ISL_1987210 | Randox Laboratories | Wellcome Sanger Institute for the COVID-19 Genomics UK (COG-UK) Consortium | Randox Laboratories and Alex Alderton, Roberto Amato, Jeffrey Barrett, Sonia Goncalves, Ewan Harrison, David K. Jackson, Ian Johnston, Dominic Kwiatkowski, Cordelia Langford, John Sillitoe on behalf of the Wellcome Sanger Institute COVID-19 Surveillance Team |
| EPI_ISL_1987214 | Lighthouse Lab in Alderley Park | Wellcome Sanger Institute for the COVID-19 Genomics UK (COG-UK) Consortium | Jacquelyn Wynn, Mairead Hyland, The Lighthouse Lab in Alderley Park and Alex Alderton, Roberto Amato, Jeffrey Barrett, Sonia Goncalves, Ewan Harrison, David K. Jackson, Ian Johnston, Dominic Kwiatkowski, Cordelia Langford, John Sillitoe on behalf of the Wellcome Sanger Institute COVID-19 Surveillance Team |
| EPI_ISL_1987217 | Lighthouse Lab in Milton Keynes | Wellcome Sanger Institute for the COVID-19 Genomics UK (COG-UK) Consortium | The Lighthouse Lab in Milton Keynes and Alex Alderton, Roberto Amato, Jeffrey Barrett, Sonia Goncalves, Ewan Harrison, David K. Jackson, Ian Johnston, Dominic Kwiatkowski, Cordelia Langford, John Sillitoe on behalf of the Wellcome Sanger Institute COVID-19 Surveillance Team |
| EPI_ISL_1987239 | Lighthouse Lab in Alderley Park | Wellcome Sanger Institute for the COVID-19 Genomics UK (COG-UK) Consortium | Jacquelyn Wynn, Mairead Hyland, The Lighthouse Lab in Alderley Park and Alex Alderton, Roberto Amato, Jeffrey Barrett, Sonia Goncalves, Ewan Harrison, David K. Jackson, Ian Johnston, Dominic Kwiatkowski, Cordelia Langford, John Sillitoe on behalf of the Wellcome Sanger Institute COVID-19 Surveillance Team |
| EPI_ISL_1987244 | Lighthouse Lab in Glasgow | Wellcome Sanger Institute for the COVID-19 Genomics UK (COG-UK) Consortium | Harper VanSteenhouse, Yumi Kasai, David Gray, Carol Clugston, Anna Dominiczak and Alex Alderton, Roberto Amato, Jeffrey Barrett, Sonia Goncalves, Ewan Harrison, David K. Jackson, Ian Johnston, Dominic Kwiatkowski, Cordelia Langford, John Sillitoe on behalf of the Wellcome Sanger Institute COVID-19 Surveillance Team |
| EPI_ISL_1987256 | Randox Laboratories | Wellcome Sanger Institute for the COVID-19 Genomics UK (COG-UK) Consortium | Randox Laboratories and Alex Alderton, Roberto Amato, Jeffrey Barrett, Sonia Goncalves, Ewan Harrison, David K. Jackson, Ian Johnston, Dominic Kwiatkowski, Cordelia Langford, John Sillitoe on behalf of the Wellcome Sanger Institute COVID-19 Surveillance Team |
| EPI_ISL_1987267, EPI_ISL_1987269, EPI_ISL_1987276 | Lighthouse Lab in Alderley Park | Wellcome Sanger Institute for the COVID-19 Genomics UK (COG-UK) Consortium | Jacquelyn Wynn, Mairead Hyland, The Lighthouse Lab in Alderley Park and Alex Alderton, Roberto Amato, Jeffrey Barrett, Sonia Goncalves, Ewan Harrison, David K. Jackson, Ian Johnston, Dominic Kwiatkowski, Cordelia Langford, John Sillitoe on behalf of the Wellcome Sanger Institute COVID-19 Surveillance Team |
| EPI_ISL_1987288 | Lighthouse Lab in Milton Keynes | Wellcome Sanger Institute for the COVID-19 Genomics UK (COG-UK) Consortium | The Lighthouse Lab in Milton Keynes and Alex Alderton, Roberto Amato, Jeffrey Barrett, Sonia Goncalves, Ewan Harrison, David K. Jackson, Ian Johnston, Dominic Kwiatkowski, Cordelia Langford, John Sillitoe on behalf of the Wellcome Sanger Institute COVID-19 Surveillance Team |
| EPI_ISL_1987292 | Lighthouse Lab in Glasgow | Wellcome Sanger Institute for the COVID-19 Genomics UK (COG-UK) Consortium | Harper VanSteenhouse, Yumi Kasai, David Gray, Carol Clugston, Anna Dominiczak and Alex Alderton, Roberto Amato, Jeffrey Barrett, Sonia Goncalves, Ewan Harrison, David K. Jackson, Ian Johnston, Dominic Kwiatkowski, Cordelia Langford, John Sillitoe on behalf of the Wellcome Sanger Institute COVID-19 Surveillance Team |
| EPI_ISL_1987306, EPI_ISL_1987318 | Lighthouse Lab in Milton Keynes | Wellcome Sanger Institute for the COVID-19 Genomics UK (COG-UK) Consortium | The Lighthouse Lab in Milton Keynes and Alex Alderton, Roberto Amato, Jeffrey Barrett, Sonia Goncalves, Ewan Harrison, David K. Jackson, Ian Johnston, Dominic Kwiatkowski, Cordelia Langford, John Sillitoe on behalf of the Wellcome Sanger Institute COVID-19 Surveillance Team |
| EPI_ISL_1987329, EPI_ISL_1987335, EPI_ISL_1987336 | Randox Laboratories | Wellcome Sanger Institute for the COVID-19 Genomics UK (COG-UK) Consortium | Randox Laboratories and Alex Alderton, Roberto Amato, Jeffrey Barrett, Sonia Goncalves, Ewan Harrison, David K. Jackson, Ian Johnston, Dominic Kwiatkowski, Cordelia Langford, John Sillitoe on behalf of the Wellcome Sanger Institute COVID-19 Surveillance Team |
| EPI_ISL_1987341 | Lighthouse Lab in Alderley Park | Wellcome Sanger Institute for the COVID-19 Genomics UK (COG-UK) Consortium | Jacquelyn Wynn, Mairead Hyland, The Lighthouse Lab in Alderley Park and Alex Alderton, Roberto Amato, Jeffrey Barrett, Sonia Goncalves, Ewan Harrison, David K. Jackson, Ian Johnston, Dominic Kwiatkowski, Cordelia Langford, John Sillitoe on behalf of the Wellcome Sanger Institute COVID-19 Surveillance Team |
| EPI_ISL_1987346 | Lighthouse Lab in Milton Keynes | Wellcome Sanger Institute for the COVID-19 Genomics UK (COG-UK) Consortium | The Lighthouse Lab in Milton Keynes and Alex Alderton, Roberto Amato, Jeffrey Barrett, Sonia Goncalves, Ewan Harrison, David K. Jackson, Ian Johnston, Dominic Kwiatkowski, Cordelia Langford, John Sillitoe on behalf of the Wellcome Sanger Institute COVID-19 Surveillance Team |
| EPI_ISL_1987366 | Lighthouse Lab in Alderley Park | Wellcome Sanger Institute for the COVID-19 Genomics UK (COG-UK) Consortium | Jacquelyn Wynn, Mairead Hyland, The Lighthouse Lab in Alderley Park and Alex Alderton, Roberto Amato, Jeffrey Barrett, Sonia Goncalves, Ewan Harrison, David K. Jackson, Ian Johnston, Dominic Kwiatkowski, Cordelia Langford, John Sillitoe on behalf of the Wellcome Sanger Institute COVID-19 Surveillance Team |
| EPI_ISL_1989753, EPI_ISL_1989865, EPI_ISL_1989866, EPI_ISL_1989891, EPI_ISL_1989941, EPI_ISL_1990160 | Quest Diagnostics Incorporated | Centers for Disease Control and Prevention Division of Viral Diseases, Pathogen Discovery | Dakota Howard, Dhwani Batra, Peter W. Cook, Kara Moser, Adrian Paskey, Jason Caravas, Benjamin Rambo-Martin, Shatavia Morrison, Christopher Gulvick, Scott Sammons, Yvette Unoarumhi, Darlene Wagner, Matthew Schmerer, S. H. Rosenthal, A. Gerasimova, R. M. Kagan, B. Anderson, M. Hua, Y. Liu, L.E. Bernstein, K.E. Livingston, A. Perez, I. A. Shlyakhter, R. V. Rolando, R. Owen, P. Tanpaiboon, F. Lacbawan, Clinton R. Paden, Duncan MacCannell |
| EPI_ISL_1990614, EPI_ISL_1990723, EPI_ISL_1990741, EPI_ISL_1990856, | Helix/Illumina | Centers for Disease Control and Prevention Division of Viral Diseases, Pathogen Discovery | Dakota Howard, Dhwani Batra, Peter W. Cook, Kara Moser, Adrian Paskey, Jason Caravas, Benjamin Rambo-Martin, Shatavia Morrison, Christopher Gulvick, Scott Sammons, Yvette Unoarumhi, Darlene Wagner, Matthew Schmerer, Eileen de Feo, Jan Antico, Christine Tran, Matthew Tolentino, Shannon Wickline, Kim Gietzen, Brad Sickler, Jingtao Liu, Eric Allen, Phil Febbo, Nicole L. Washington, Simon White, Geraint Levan, Kelly Schiabor Barrett, Elizabeth Cirulli, Alexandre Bolze, Ary Ascencio, Charlotte Rivera-Garcia, Ryan Cho, Jason Nguyen, Sherry Wang, Jimmy Ramirez, Tyler Cassens, Efren Sandoval, Magnus Isaksson, William Lee, David Becker, Marc Laurent, James Lu, Clinton R. Paden, Duncan MacCannell |

| | | | |
|---|---|---|---|
| EPI_ISL_1990921, EPI_ISL_1991008, EPI_ISL_1991368, EPI_ISL_1991454, EPI_ISL_1991572, EPI_ISL_1991775 | | | |
| EPI_ISL_1992039, EPI_ISL_1992110, EPI_ISL_1992125, EPI_ISL_1992133, EPI_ISL_1992137, EPI_ISL_1992142, EPI_ISL_1992240, EPI_ISL_1992245, EPI_ISL_1992266, EPI_ISL_1992270, EPI_ISL_1992271, EPI_ISL_1992272, EPI_ISL_1992302, EPI_ISL_1992395, EPI_ISL_1992594, EPI_ISL_1992667, EPI_ISL_1992684, EPI_ISL_1992823, EPI_ISL_1992943, EPI_ISL_1992993, EPI_ISL_1993014, EPI_ISL_1993031, EPI_ISL_1993033, EPI_ISL_1993034, EPI_ISL_1993045, EPI_ISL_1993068, EPI_ISL_1993069, EPI_ISL_1993113, EPI_ISL_1993142, EPI_ISL_1993160, EPI_ISL_1993161, EPI_ISL_1993163, EPI_ISL_1993240, EPI_ISL_1993295, EPI_ISL_1993297 | | | |
| see above | Fulgent Genetics | Centers for Disease Control and Prevention Division of Viral Diseases, Pathogen Discovery | Dakota Howard, Dhwani Batra, Peter W. Cook, Kara Moser, Adrian Paskey, Jason Caravas, Benjamin Rambo-Martin, Shatavia Morrison, Christopher Gulvick, Scott Sammons, Yvette Unoarumhi, Darlene Wagner, Matthew Schmerer, Harry Gao, Mickey Li, John Gao, Joseph Fierro, Benafsh Sapra, Becky Tsai, Yan Meng, Doreen Ng, James Xie, Clinton R. Paden, Duncan MacCannell |
| EPI_ISL_1993840, EPI_ISL_1993870, EPI_ISL_1993871, EPI_ISL_1993872, EPI_ISL_1993906, EPI_ISL_1993918 | Clinical Virology | Clinical Bacteriology | Tim Roloff, Madlen Stange, Helena MB Seth-Smith, Alfredo Mari, Karoline Leuzinger, Julia Bielicki, Manuel Battegay, Hans Hirsch, Adrian Egli |
| EPI_ISL_1993936 | UMC Groningen, Clinical Virology, Department of Medical Microbiology and Infection Prevention | UMC Groningen, Clinical Virology, Department of Medical Microbiology and Infection Prevention | Hubert Niesters, Alexander Friedrich, Erley Lizarazo-Forero, Monika Fliss, Lilli Gard, Sigrid Rosema, Coretta Van Leer-Buter, Xuewei Zhou, Marjolein Knoester |
| EPI_ISL_1994063 | Furst Medical Laboratory | Norwegian Institute of Public Health, Department of Virology | Kathrine Stene-Johansen, Kamilla Heddeland Instefjord, Hilde Elshaug, Garcia Llorente Ignacio, Jon Bråte, Engebretsen Serina Beate,Pedersen Benedikte Nevjen, Debech Nadia, Atiya R Ali,Marie Paulsen Madsen, Rasmus Riis Kopperud, Hilde Vollan, Karoline Bragstad, Olav Hungnes |
| EPI_ISL_1994608 | Ampath - Kingsway | KRISP, KZN Research Innovation and Sequencing Platform | Giandhari J, Pillay S, Naidoo Y, Ramphal U, Lessells R, Mdlalose K, York D, Khan S, Emmanuel SJ, Tegally H, Wilkinson E, de Oliveira T |
| EPI_ISL_1994891, EPI_ISL_1994892 | IN State Department of Health Laboratory Services | IN State Department of Health Laboratory Services | Cassandra Campion, Jamie Yeadon, Brian Pope, Lixia Liu, Kyle Brownlee, Melissa Hindenlang, Mark Glazier |
| EPI_ISL_1994923 | Dept. of Microbiology and Infection Control, Akershus University Hospital HF | Dept. of Microbiology and Infection Control, Akershus University Hospital HF | Hege Vangstein Aamot, Alexander Hesselberg Løvestad |
| EPI_ISL_1995876, EPI_ISL_1997014, EPI_ISL_1997354, EPI_ISL_1997404, EPI_ISL_1997765, EPI_ISL_1997967, EPI_ISL_1998231, EPI_ISL_1998831, EPI_ISL_1999092, EPI_ISL_1999775 | Aegis Sciences Corporation | Centers for Disease Control and Prevention Division of Viral Diseases, Pathogen Discovery | Dakota Howard, Dhwani Batra, Peter W. Cook, Kara Moser, Adrian Paskey, Jason Caravas, Benjamin Rambo-Martin, Shatavia Morrison, Christopher Gulvick, Scott Sammons, Yvette Unoarumhi, Darlene Wagner, Matthew Schmerer, Cyndi Clark, Patrick Campbell, Rob Case, Vikramsinha Ghorpade, Holly Houdeshell, Ola Kvalvaag, Dillon Nall, Ethan Sanders, Alec Vest, Shaun Westlund, Matthew Hardison, Clinton R. Paden, Duncan MacCannell |
| EPI_ISL_1999791 | Microbiology Department, Laboratori Clínic Metropolitana Nord. Hospital Universitari Germans Trias i Pujol. | Can Ruti SARS-CoV-2 Sequencing Hub (HUGTiP/IrsiCaixa/IGTP) | Marc Noguera-Julian, Pilar Armengol, Ignacio Blanco, Antoni E Bordoy, Francesc Catala-Moll, Pere-Joan Cardona, Maria Casadellà, Cristina Casañ, Gemma Clara, Bonaventura Clotet, Cristina Esteban, Montserrat Giménez, Mercedes Guerrero, Anna Not, Roger Paredes, Mariona Parera, Verónica Saludes, Alba Sánchez, and Elisa Martró on behalf of the Can Ruti SARS-CoV-2 Sequencing Hub. |
| EPI_ISL_2000028, EPI_ISL_2000090, EPI_ISL_2000128, EPI_ISL_2000209, EPI_ISL_2000226, EPI_ISL_2000262, EPI_ISL_2000388 | Aegis Sciences Corporation | Centers for Disease Control and Prevention Division of Viral Diseases, Pathogen Discovery | Dakota Howard, Dhwani Batra, Peter W. Cook, Kara Moser, Adrian Paskey, Jason Caravas, Benjamin Rambo-Martin, Shatavia Morrison, Christopher Gulvick, Scott Sammons, Yvette Unoarumhi, Darlene Wagner, Matthew Schmerer, Cyndi Clark, Patrick Campbell, Rob Case, Vikramsinha Ghorpade, Holly Houdeshell, Ola Kvalvaag, Dillon Nall, Ethan Sanders, Alec Vest, Shaun Westlund, Matthew Hardison, Clinton R. Paden, Duncan MacCannell |
| EPI_ISL_2000617 | INMI Lazzaro Spallanzani IRCCS | INMI Lazzaro Spallanzani IRCCS | B Bartolini, E Giombini, F Messina, M Rueca, G Bonfiglio, O Butera, CEM Gruber, F Santini, A Di Caro, MR Capobianchi |
| EPI_ISL_2000619 | INMI Lazzaro Spallanzani IRCCS | INMI Lazzaro Spallanzani IRCCS | G Bonfiglio, O Butera, CEM Gruber, F Santini, B Bartolini, E Giombini, F Messina, M Rueca, A Di Caro, MR Capobianchi |
| EPI_ISL_2000620 | INMI Lazzaro Spallanzani IRCCS | INMI Lazzaro Spallanzani IRCCS | O Butera, CEM Gruber, F Santini, B Bartolini, E Giombini, F Messina, M Rueca, G Bonfiglio, MR Capobianchi, A Di Caro |
| EPI_ISL_2000621 | INMI Lazzaro Spallanzani IRCCS | INMI Lazzaro Spallanzani IRCCS | CEM Gruber, F Santini, B Bartolini, E Giombini, F Messina, M Rueca, G Bonfiglio, O Butera, A Di Caro, MR Capobianchi |
| EPI_ISL_2000622 | INMI Lazzaro Spallanzani IRCCS | INMI Lazzaro Spallanzani IRCCS | F Santini, B Bartolini, E Giombini, F Messina, M Rueca, G Bonfiglio, O Butera, CEM Gruber, MR Capobianchi, A Di Caro |
| EPI_ISL_2000623 | INMI Lazzaro Spallanzani IRCCS | INMI Lazzaro Spallanzani IRCCS | B Bartolini, E Giombini, F Messina, M Rueca, G Bonfiglio, O Butera, CEM Gruber, F Santini, A Di Caro, MR Capobianchi |
| EPI_ISL_2000624 | INMI Lazzaro Spallanzani IRCCS | INMI Lazzaro Spallanzani IRCCS | E Giombini, F Messina, M Rueca, G Bonfiglio, O Butera, CEM Gruber, F Santini, B Bartolini, MR Capobianchi, A Di Caro |
| EPI_ISL_2000625 | INMI Lazzaro Spallanzani IRCCS | INMI Lazzaro Spallanzani IRCCS | F Messina, M Rueca, G Bonfiglio, O Butera, CEM Gruber, F Santini, B Bartolini, E Giombini, A Di Caro, MR Capobianchi |
| EPI_ISL_2000626 | INMI Lazzaro Spallanzani IRCCS | INMI Lazzaro Spallanzani IRCCS | M Rueca, G Bonfiglio, O Butera, CEM Gruber, F Santini, B Bartolini, E Giombini, F Messina, MR Capobianchi, A Di Caro |
| EPI_ISL_2000627 | INMI Lazzaro Spallanzani IRCCS | INMI Lazzaro Spallanzani IRCCS | G Bonfiglio, O Butera, CEM Gruber, F Santini, B Bartolini, E Giombini, F Messina, M Rueca, A Di Caro, MR Capobianchi |
| EPI_ISL_2001050 | Hospital Sharp | Microbial Genomics Laboratory | Bruno Gomez-Gil, Julissa Enciso-Ibarra, Alejandra Garcia-Gasca, Daniel Fregoso-Rueda |
| EPI_ISL_2001060 | Queensland Health Forensic and Scientific Services | Queensland Health Forensic and Scientific Services | Son Nguyen |
| EPI_ISL_2001139 | GMERS, Government Medical College, Gandhinagar | Gujarat Biotechnology Research Centre | Nitin Savaliya,Dinesh Kumar,Twinkle Soni,Sonal Sharma ,Zuber Saiyed,Ramesh Pandit,Janvi Raval,Zarna Patel,Gaurishankar Shrimali,Umang Mishra,Nitesh Shah,Chaitanya Joshi,Madhvi Joshi |
| EPI_ISL_2001143 | GMERS, Government Medical College, Gandhinagar | Gujarat Biotechnology Research Centre | Zuber Saiyed,Ramesh Pandit,Janvi Raval,Zarna Patel,Nitin Savaliya,Dinesh Kumar,Twinkle Soni,Sonal Sharma ,Gaurishankar Shrimali,Umang Mishra,Nitesh Shah,Chaitanya Joshi,Madhvi Joshi |
| EPI_ISL_2001183 | GMERS, Government Medical College (GMC), Gotri | Gujarat Biotechnology Research Centre | Twinkle Soni,Sonal Sharma ,Zuber Saiyed,Ramesh Pandit,Janvi Raval,Zarna Patel,Nitin Savaliya,Dinesh Kumar,Bithika Duttaroy,Umang Mishra,Nitesh Shah,Chaitanya Joshi,Madhvi Joshi |
| EPI_ISL_2001185 | GMERS, Government Medical College (GMC), Gotri | Gujarat Biotechnology Research Centre | Zuber Saiyed,Ramesh Pandit,Janvi Raval,Zarna Patel,Nitin Savaliya,Dinesh Kumar,Twinkle Soni,Sonal Sharma ,Bithika Duttaroy,Umang Mishra,Nitesh Shah,Chaitanya Joshi,Madhvi Joshi |
| EPI_ISL_2001186 | GMERS, Government Medical College (GMC), Gotri | Gujarat Biotechnology Research Centre | Ramesh Pandit,Janvi Raval,Zarna Patel,Nitin Savaliya,Dinesh Kumar,Twinkle Soni,Sonal Sharma ,Zuber Saiyed,Bithika Duttaroy,Umang Mishra,Nitesh Shah,Chaitanya Joshi,Madhvi Joshi |
| EPI_ISL_2001187 | VRDL, Goverment Medical College (GMC), Surat | Gujarat Biotechnology Research Centre | Janvi Raval,Zarna Patel,Nitin Savaliya,Dinesh Kumar,Twinkle Soni,Sonal Sharma ,Zuber Saiyed,Ramesh Pandit,Neeta Khandelwal,Umang Mishra,Nitesh Shah,Chaitanya Joshi,Madhvi Joshi |
| EPI_ISL_2001188 | VRDL, Goverment Medical College (GMC), Surat | Gujarat Biotechnology Research Centre | Zarna Patel,Nitin Savaliya,Dinesh Kumar,Twinkle Soni,Sonal Sharma ,Zuber Saiyed,Ramesh Pandit,Janvi Raval,Neeta Khandelwal,Umang Mishra,Nitesh Shah,Chaitanya Joshi,Madhvi Joshi |
| EPI_ISL_2001189 | VRDL, Goverment Medical College (GMC), Surat | Gujarat Biotechnology Research Centre | Nitin Savaliya,Dinesh Kumar,Twinkle Soni,Sonal Sharma ,Zuber Saiyed,Ramesh Pandit,Janvi Raval,Zarna Patel,Neeta Khandelwal,Umang Mishra,Nitesh Shah,Chaitanya Joshi,Madhvi Joshi |
| EPI_ISL_2001190 | VRDL, Goverment Medical College (GMC), Surat | Gujarat Biotechnology Research Centre | Dinesh Kumar,Twinkle Soni,Sonal Sharma ,Zuber Saiyed,Ramesh Pandit,Janvi Raval,Zarna Patel,Nitin Savaliya,Neeta Khandelwal,Umang Mishra,Nitesh Shah,Chaitanya Joshi,Madhvi Joshi |
| EPI_ISL_2001191 | VRDL, Goverment Medical College (GMC), Surat | Gujarat Biotechnology Research Centre | Twinkle Soni,Sonal Sharma ,Zuber Saiyed,Ramesh Pandit,Janvi Raval,Zarna Patel,Nitin Savaliya,Dinesh Kumar,Neeta Khandelwal,Umang Mishra,Nitesh Shah,Chaitanya Joshi,Madhvi Joshi |
| EPI_ISL_2001192 | VRDL, Goverment Medical College (GMC), Surat | Gujarat Biotechnology Research Centre | Sonal Sharma ,Zuber Saiyed,Ramesh Pandit,Janvi Raval,Zarna Patel,Nitin Savaliya,Dinesh Kumar,Twinkle Soni,Neeta Khandelwal,Umang Mishra,Nitesh Shah,Chaitanya Joshi,Madhvi Joshi |
| EPI_ISL_2001193 | VRDL, Goverment Medical College (GMC), Surat | Gujarat Biotechnology Research Centre | Zuber Saiyed,Ramesh Pandit,Janvi Raval,Zarna Patel,Nitin Savaliya,Dinesh Kumar,Twinkle Soni,Sonal Sharma ,Neeta Khandelwal,Umang Mishra,Nitesh Shah,Chaitanya Joshi,Madhvi Joshi |
| EPI_ISL_2001194 | VRDL, Goverment Medical College (GMC), Surat | Gujarat Biotechnology Research Centre | Ramesh Pandit,Janvi Raval,Zarna Patel,Nitin Savaliya,Dinesh Kumar,Twinkle Soni,Sonal Sharma ,Zuber Saiyed,Neeta Khandelwal,Umang Mishra,Nitesh Shah,Chaitanya Joshi,Madhvi Joshi |
| EPI_ISL_2001195 | VRDL, Goverment Medical College (GMC), Surat | Gujarat Biotechnology Research Centre | Janvi Raval,Zarna Patel,Nitin Savaliya,Dinesh Kumar,Twinkle Soni,Sonal Sharma ,Zuber Saiyed,Ramesh Pandit,Neeta Khandelwal,Umang Mishra,Nitesh Shah,Chaitanya Joshi,Madhvi Joshi |
| EPI_ISL_2001196 | VRDL, Goverment Medical College (GMC), Surat | Gujarat Biotechnology Research Centre | Zarna Patel,Nitin Savaliya,Dinesh Kumar,Twinkle Soni,Sonal Sharma ,Zuber Saiyed,Ramesh Pandit,Janvi Raval,Neeta Khandelwal,Umang Mishra,Nitesh Shah,Chaitanya Joshi,Madhvi Joshi |
| EPI_ISL_2001201 | VRDL, Goverment Medical College (GMC), Surat | Gujarat Biotechnology Research Centre | Zuber Saiyed,Ramesh Pandit,Janvi Raval,Zarna Patel,Nitin Savaliya,Dinesh Kumar,Twinkle Soni,Sonal Sharma ,Neeta Khandelwal,Umang Mishra,Nitesh Shah,Chaitanya Joshi,Madhvi Joshi |
| EPI_ISL_2001203, EPI_ISL_2001211 | VRDL, Goverment Medical College (GMC), Surat | Gujarat Biotechnology Research Centre | Janvi Raval,Zarna Patel,Nitin Savaliya,Dinesh Kumar,Twinkle Soni,Sonal Sharma ,Zuber Saiyed,Ramesh Pandit,Neeta Khandelwal,Umang Mishra,Nitesh Shah,Chaitanya Joshi,Madhvi Joshi |
| EPI_ISL_2001212 | VRDL, Goverment Medical College (GMC), Surat | Gujarat Biotechnology Research Centre | Zarna Patel,Nitin Savaliya,Dinesh Kumar,Twinkle Soni,Sonal Sharma ,Zuber Saiyed,Ramesh Pandit,Janvi Raval,Neeta Khandelwal,Umang Mishra,Nitesh Shah,Chaitanya Joshi,Madhvi Joshi |
| EPI_ISL_2001270, EPI_ISL_2001286 | University of Michigan Clinical Microbiology Laboratory | Lauring Lab, University of Michigan, Department of Microbiology and Immunology | Valesano |
| EPI_ISL_2001651, EPI_ISL_2001653 | Intrepid Nepal | Center for Molecular Dynamics Nepal | Rajindra Napit, Prajwol Manandhar, Ajit Poudel, Roji Raut, Anupama Gurung, Saman Pradhan, Kavya Dhital, Sapana Shakya, Dibesh Karmacharya |
| EPI_ISL_2002190, EPI_ISL_2002289, EPI_ISL_2002395 | UW Virology Lab | UW Virology Lab | Pavitra Roychoudhury, Hong Xie, Lasata Shrestha, Tien V. Nguyen, Shah Mohamed Bakhash, Michelle Lin, Noah R. Baker, Sean Ellis, Meei-Li Huang, Keith R Jerome, Alexander Greninger |
| EPI_ISL_2002644 | Centre Hospitalier Universitaire Clermont-Ferrand | CHU Clermont-Ferrand, service de virologie | Bisseux Maxime, Mirand Audrey, Combes Patricia, Henquell Cécile |
| EPI_ISL_2002689, EPI_ISL_2002743, EPI_ISL_2002774, EPI_ISL_2002825, EPI_ISL_2002836, EPI_ISL_2002857, EPI_ISL_2002865, | UW Virology Lab | UW Virology Lab | Pavitra Roychoudhury, Hong Xie, Lasata Shrestha, Tien V. Nguyen, Shah Mohamed Bakhash, Michelle Lin, Noah R. Baker, Sean Ellis, Meei-Li Huang, Keith R Jerome, Alexander Greninger |

| | | | |
|---|---|---|---|
| EPI_ISL_2002878, EPI_ISL_2002924 | | | |
| EPI_ISL_2003297 | University of Liège COVID-19 testing center | GIGA Medical Genomics | Keith Durkin, Maria Artesi, Sébastien Bontems, Raphaël Boreux, Bouchra Boujemla, Nathalie Renotte, Cécile Meex, Pierrette Melin, Marie-Pierre Hayette, Vincent Bours |
| EPI_ISL_2003343 | Department of Clinical Microbiology | GIGA Medical Genomics | Keith Durkin, Maria Artesi, Sébastien Bontems, Raphaël Boreux, Bouchra Boujemla, Nathalie Renotte, Cécile Meex, Pierrette Melin, Marie-Pierre Hayette, Vincent Bours |
| EPI_ISL_2003438 | University of Liège COVID-19 testing center | GIGA Medical Genomics | Keith Durkin, Maria Artesi, Sébastien Bontems, Raphaël Boreux, Bouchra Boujemla, Nathalie Renotte, Cécile Meex, Pierrette Melin, Marie-Pierre Hayette, Vincent Bours |
| EPI_ISL_2003649 | University of Michigan Clinical Microbiology Laboratory | Lauring Lab, University of Michigan, Department of Microbiology and Immunology | Valesano |
| EPI_ISL_2003700 | Virginia Division of Consolidated Laboratory Services | Virginia Division of Consolidated Laboratory Services | Virginia DCLS |
| EPI_ISL_2004296 | CH Setubal | Instituto Nacional de Saude (INSA) | Borges et al |
| EPI_ISL_2004324 | SEEBMO | Instituto Nacional de Saude (INSA) | Borges et al |
| EPI_ISL_2004344 | Randox Laboratories | Wellcome Sanger Institute for the COVID-19 Genomics UK (COG-UK) Consortium | Randox Laboratories and Alex Alderton, Roberto Amato, Jeffrey Barrett, Sonia Goncalves, Ewan Harrison, David K. Jackson, Ian Johnston, Dominic Kwiatkowski, Cordelia Langford, John Sillitoe on behalf of the Wellcome Sanger Institute COVID-19 Surveillance Team |
| EPI_ISL_2004346, EPI_ISL_2004358, EPI_ISL_2004359, EPI_ISL_2004363, EPI_ISL_2004366, EPI_ISL_2004368, EPI_ISL_2004369, EPI_ISL_2004375, EPI_ISL_2004377, EPI_ISL_2004379, EPI_ISL_2004380, EPI_ISL_2004384, EPI_ISL_2004385, EPI_ISL_2004389, EPI_ISL_2004393, EPI_ISL_2004400, EPI_ISL_2004405, EPI_ISL_2004407 | | | |
| see above | Lighthouse Lab in Alderley Park | Wellcome Sanger Institute for the COVID-19 Genomics UK (COG-UK) Consortium | Jacquelyn Wynn, Mairead Hyland, The Lighthouse Lab in Alderley Park and Alex Alderton, Roberto Amato, Jeffrey Barrett, Sonia Goncalves, Ewan Harrison, David K. Jackson, Ian Johnston, Dominic Kwiatkowski, Cordelia Langford, John Sillitoe on behalf of the Wellcome Sanger Institute COVID-19 Surveillance Team |
| EPI_ISL_2004408 | Lighthouse Lab in Glasgow | Wellcome Sanger Institute for the COVID-19 Genomics UK (COG-UK) Consortium | Harper VanSteenhouse, Yumi Kasai, David Gray, Carol Clugston, Anna Dominiczak and Alex Alderton, Roberto Amato, Jeffrey Barrett, Sonia Goncalves, Ewan Harrison, David K. Jackson, Ian Johnston, Dominic Kwiatkowski, Cordelia Langford, John Sillitoe on behalf of the Wellcome Sanger Institute COVID-19 Surveillance Team |
| EPI_ISL_2004415, EPI_ISL_2004416 | Lighthouse Lab in Alderley Park | Wellcome Sanger Institute for the COVID-19 Genomics UK (COG-UK) Consortium | Jacquelyn Wynn, Mairead Hyland, The Lighthouse Lab in Alderley Park and Alex Alderton, Roberto Amato, Jeffrey Barrett, Sonia Goncalves, Ewan Harrison, David K. Jackson, Ian Johnston, Dominic Kwiatkowski, Cordelia Langford, John Sillitoe on behalf of the Wellcome Sanger Institute COVID-19 Surveillance Team |
| EPI_ISL_2004423 | Randox Laboratories | Wellcome Sanger Institute for the COVID-19 Genomics UK (COG-UK) Consortium | Randox Laboratories and Alex Alderton, Roberto Amato, Jeffrey Barrett, Sonia Goncalves, Ewan Harrison, David K. Jackson, Ian Johnston, Dominic Kwiatkowski, Cordelia Langford, John Sillitoe on behalf of the Wellcome Sanger Institute COVID-19 Surveillance Team |
| EPI_ISL_2004426, EPI_ISL_2004440, EPI_ISL_2004442, EPI_ISL_2004445 | Lighthouse Lab in Alderley Park | Wellcome Sanger Institute for the COVID-19 Genomics UK (COG-UK) Consortium | Jacquelyn Wynn, Mairead Hyland, The Lighthouse Lab in Alderley Park and Alex Alderton, Roberto Amato, Jeffrey Barrett, Sonia Goncalves, Ewan Harrison, David K. Jackson, Ian Johnston, Dominic Kwiatkowski, Cordelia Langford, John Sillitoe on behalf of the Wellcome Sanger Institute COVID-19 Surveillance Team |
| EPI_ISL_2004448 | Randox Laboratories | Wellcome Sanger Institute for the COVID-19 Genomics UK (COG-UK) Consortium | Randox Laboratories and Alex Alderton, Roberto Amato, Jeffrey Barrett, Sonia Goncalves, Ewan Harrison, David K. Jackson, Ian Johnston, Dominic Kwiatkowski, Cordelia Langford, John Sillitoe on behalf of the Wellcome Sanger Institute COVID-19 Surveillance Team |
| EPI_ISL_2004450, EPI_ISL_2004457, EPI_ISL_2004458, EPI_ISL_2004461, EPI_ISL_2004462, EPI_ISL_2004470, EPI_ISL_2004475 | Lighthouse Lab in Alderley Park | Wellcome Sanger Institute for the COVID-19 Genomics UK (COG-UK) Consortium | Jacquelyn Wynn, Mairead Hyland, The Lighthouse Lab in Alderley Park and Alex Alderton, Roberto Amato, Jeffrey Barrett, Sonia Goncalves, Ewan Harrison, David K. Jackson, Ian Johnston, Dominic Kwiatkowski, Cordelia Langford, John Sillitoe on behalf of the Wellcome Sanger Institute COVID-19 Surveillance Team |
| EPI_ISL_2004476 | Randox Laboratories | Wellcome Sanger Institute for the COVID-19 Genomics UK (COG-UK) Consortium | Randox Laboratories and Alex Alderton, Roberto Amato, Jeffrey Barrett, Sonia Goncalves, Ewan Harrison, David K. Jackson, Ian Johnston, Dominic Kwiatkowski, Cordelia Langford, John Sillitoe on behalf of the Wellcome Sanger Institute COVID-19 Surveillance Team |
| EPI_ISL_2004481, EPI_ISL_2004488 | Lighthouse Lab in Alderley Park | Wellcome Sanger Institute for the COVID-19 Genomics UK (COG-UK) Consortium | Jacquelyn Wynn, Mairead Hyland, The Lighthouse Lab in Alderley Park and Alex Alderton, Roberto Amato, Jeffrey Barrett, Sonia Goncalves, Ewan Harrison, David K. Jackson, Ian Johnston, Dominic Kwiatkowski, Cordelia Langford, John Sillitoe on behalf of the Wellcome Sanger Institute COVID-19 Surveillance Team |
| EPI_ISL_2004494 | Randox Laboratories | Wellcome Sanger Institute for the COVID-19 Genomics UK (COG-UK) Consortium | Randox Laboratories and Alex Alderton, Roberto Amato, Jeffrey Barrett, Sonia Goncalves, Ewan Harrison, David K. Jackson, Ian Johnston, Dominic Kwiatkowski, Cordelia Langford, John Sillitoe on behalf of the Wellcome Sanger Institute COVID-19 Surveillance Team |
| EPI_ISL_2004502, EPI_ISL_2004506, EPI_ISL_2004507, EPI_ISL_2004509, EPI_ISL_2004510, EPI_ISL_2004512, EPI_ISL_2004513, EPI_ISL_2004517, EPI_ISL_2004519, EPI_ISL_2004529, EPI_ISL_2004531, EPI_ISL_2004533, EPI_ISL_2004537, EPI_ISL_2004539, EPI_ISL_2004543, EPI_ISL_2004544, EPI_ISL_2004547, EPI_ISL_2004552, EPI_ISL_2004554, EPI_ISL_2004558, EPI_ISL_2004564, EPI_ISL_2004571, EPI_ISL_2004572, EPI_ISL_2004573, EPI_ISL_2004575, EPI_ISL_2004579, EPI_ISL_2004582, EPI_ISL_2004586, EPI_ISL_2004596, EPI_ISL_2004606, EPI_ISL_2004610, EPI_ISL_2004612, EPI_ISL_2004616 | | | |
| see above | Lighthouse Lab in Alderley Park | Wellcome Sanger Institute for the COVID-19 Genomics UK (COG-UK) Consortium | Jacquelyn Wynn, Mairead Hyland, The Lighthouse Lab in Alderley Park and Alex Alderton, Roberto Amato, Jeffrey Barrett, Sonia Goncalves, Ewan Harrison, David K. Jackson, Ian Johnston, Dominic Kwiatkowski, Cordelia Langford, John Sillitoe on behalf of the Wellcome Sanger Institute COVID-19 Surveillance Team |
| EPI_ISL_2004627, EPI_ISL_2004628, EPI_ISL_2004648, EPI_ISL_2004653, EPI_ISL_2004655 | Lighthouse Lab in Milton Keynes | Wellcome Sanger Institute for the COVID-19 Genomics UK (COG-UK) Consortium | The Lighthouse Lab in Milton Keynes and Alex Alderton, Roberto Amato, Jeffrey Barrett, Sonia Goncalves, Ewan Harrison, David K. Jackson, Ian Johnston, Dominic Kwiatkowski, Cordelia Langford, John Sillitoe on behalf of the Wellcome Sanger Institute COVID-19 Surveillance Team |
| EPI_ISL_2004658 | Lighthouse Lab in Alderley Park | Wellcome Sanger Institute for the COVID-19 Genomics UK (COG-UK) Consortium | Jacquelyn Wynn, Mairead Hyland, The Lighthouse Lab in Alderley Park and Alex Alderton, Roberto Amato, Jeffrey Barrett, Sonia Goncalves, Ewan Harrison, David K. Jackson, Ian Johnston, Dominic Kwiatkowski, Cordelia Langford, John Sillitoe on behalf of the Wellcome Sanger Institute COVID-19 Surveillance Team |
| EPI_ISL_2004664, EPI_ISL_2004670 | Lighthouse Lab in Milton Keynes | Wellcome Sanger Institute for the COVID-19 Genomics UK (COG-UK) Consortium | The Lighthouse Lab in Milton Keynes and Alex Alderton, Roberto Amato, Jeffrey Barrett, Sonia Goncalves, Ewan Harrison, David K. Jackson, Ian Johnston, Dominic Kwiatkowski, Cordelia Langford, John Sillitoe on behalf of the Wellcome Sanger Institute COVID-19 Surveillance Team |
| EPI_ISL_2004671 | Lighthouse Lab in Alderley Park | Wellcome Sanger Institute for the COVID-19 Genomics UK (COG-UK) Consortium | Jacquelyn Wynn, Mairead Hyland, The Lighthouse Lab in Alderley Park and Alex Alderton, Roberto Amato, Jeffrey Barrett, Sonia Goncalves, Ewan Harrison, David K. Jackson, Ian Johnston, Dominic Kwiatkowski, Cordelia Langford, John Sillitoe on behalf of the Wellcome Sanger Institute COVID-19 Surveillance Team |
| EPI_ISL_2004681, EPI_ISL_2004684, EPI_ISL_2004686, EPI_ISL_2004690, EPI_ISL_2004694, EPI_ISL_2004702, EPI_ISL_2004712, EPI_ISL_2004730 | Lighthouse Lab in Milton Keynes | Wellcome Sanger Institute for the COVID-19 Genomics UK (COG-UK) Consortium | The Lighthouse Lab in Milton Keynes and Alex Alderton, Roberto Amato, Jeffrey Barrett, Sonia Goncalves, Ewan Harrison, David K. Jackson, Ian Johnston, Dominic Kwiatkowski, Cordelia Langford, John Sillitoe on behalf of the Wellcome Sanger Institute COVID-19 Surveillance Team |
| EPI_ISL_2004732 | Lighthouse Lab in Alderley Park | Wellcome Sanger Institute for the COVID-19 Genomics UK (COG-UK) Consortium | Jacquelyn Wynn, Mairead Hyland, The Lighthouse Lab in Alderley Park and Alex Alderton, Roberto Amato, Jeffrey Barrett, Sonia Goncalves, Ewan Harrison, David K. Jackson, Ian Johnston, Dominic Kwiatkowski, Cordelia Langford, John Sillitoe on behalf of the Wellcome Sanger Institute COVID-19 Surveillance Team |
| EPI_ISL_2004740, EPI_ISL_2004742, EPI_ISL_2004745 | Lighthouse Lab in Milton Keynes | Wellcome Sanger Institute for the COVID-19 Genomics UK (COG-UK) Consortium | The Lighthouse Lab in Milton Keynes and Alex Alderton, Roberto Amato, Jeffrey Barrett, Sonia Goncalves, Ewan Harrison, David K. Jackson, Ian Johnston, Dominic Kwiatkowski, Cordelia Langford, John Sillitoe on behalf of the Wellcome Sanger Institute COVID-19 Surveillance Team |
| EPI_ISL_2004747 | Lighthouse Lab in Alderley Park | Wellcome Sanger Institute for the COVID-19 Genomics UK (COG-UK) Consortium | Jacquelyn Wynn, Mairead Hyland, The Lighthouse Lab in Alderley Park and Alex Alderton, Roberto Amato, Jeffrey Barrett, Sonia Goncalves, Ewan Harrison, David K. Jackson, Ian Johnston, Dominic Kwiatkowski, Cordelia Langford, John Sillitoe on behalf of the Wellcome Sanger Institute COVID-19 Surveillance Team |
| EPI_ISL_2004753 | Lighthouse Lab in Milton Keynes | Wellcome Sanger Institute for the COVID-19 Genomics UK (COG-UK) Consortium | The Lighthouse Lab in Milton Keynes and Alex Alderton, Roberto Amato, Jeffrey Barrett, Sonia Goncalves, Ewan Harrison, David K. Jackson, Ian Johnston, Dominic Kwiatkowski, Cordelia Langford, John Sillitoe on behalf of the Wellcome Sanger Institute COVID-19 Surveillance Team |
| EPI_ISL_2004754 | Lighthouse Lab in Alderley Park | Wellcome Sanger Institute for the COVID-19 Genomics UK (COG-UK) Consortium | Jacquelyn Wynn, Mairead Hyland, The Lighthouse Lab in Alderley Park and Alex Alderton, Roberto Amato, Jeffrey Barrett, Sonia Goncalves, Ewan Harrison, David K. Jackson, Ian Johnston, Dominic Kwiatkowski, Cordelia Langford, John Sillitoe on behalf of the Wellcome Sanger Institute COVID-19 Surveillance Team |
| EPI_ISL_2004782, EPI_ISL_2004784, EPI_ISL_2004787, EPI_ISL_2004788, EPI_ISL_2004790, EPI_ISL_2004791, EPI_ISL_2004799, EPI_ISL_2004801, EPI_ISL_2004809, EPI_ISL_2004810 | Lighthouse Lab in Milton Keynes | Wellcome Sanger Institute for the COVID-19 Genomics UK (COG-UK) Consortium | The Lighthouse Lab in Milton Keynes and Alex Alderton, Roberto Amato, Jeffrey Barrett, Sonia Goncalves, Ewan Harrison, David K. Jackson, Ian Johnston, Dominic Kwiatkowski, Cordelia Langford, John Sillitoe on behalf of the Wellcome Sanger Institute COVID-19 Surveillance Team |
| EPI_ISL_2004811 | Lighthouse Lab in Alderley Park | Wellcome Sanger Institute for the COVID-19 Genomics UK (COG-UK) Consortium | Jacquelyn Wynn, Mairead Hyland, The Lighthouse Lab in Alderley Park and Alex Alderton, Roberto Amato, Jeffrey Barrett, Sonia Goncalves, Ewan Harrison, David K. Jackson, Ian Johnston, Dominic Kwiatkowski, Cordelia Langford, John Sillitoe on behalf of the Wellcome Sanger Institute COVID-19 Surveillance Team |
| EPI_ISL_2004819 | Lighthouse Lab in Milton Keynes | Wellcome Sanger Institute for the COVID-19 Genomics UK (COG-UK) Consortium | The Lighthouse Lab in Milton Keynes and Alex Alderton, Roberto Amato, Jeffrey Barrett, Sonia Goncalves, Ewan Harrison, David K. Jackson, Ian Johnston, Dominic Kwiatkowski, Cordelia Langford, John Sillitoe on behalf of the Wellcome Sanger Institute COVID-19 Surveillance Team |
| EPI_ISL_2004821 | Lighthouse Lab in Alderley Park | Wellcome Sanger Institute for the COVID-19 Genomics UK (COG-UK) Consortium | Jacquelyn Wynn, Mairead Hyland, The Lighthouse Lab in Alderley Park and Alex Alderton, Roberto Amato, Jeffrey Barrett, Sonia Goncalves, Ewan Harrison, David K. Jackson, Ian Johnston, Dominic Kwiatkowski, Cordelia Langford, John Sillitoe on behalf of the Wellcome Sanger Institute COVID-19 Surveillance Team |
| EPI_ISL_2004822, EPI_ISL_2004823, EPI_ISL_2004826, EPI_ISL_2004828, EPI_ISL_2004830, EPI_ISL_2004837, EPI_ISL_2004839, EPI_ISL_2004842, EPI_ISL_2004850, EPI_ISL_2004852, EPI_ISL_2004857, EPI_ISL_2004863, EPI_ISL_2004870, EPI_ISL_2004875 | | | |
| see above | Lighthouse Lab in Milton Keynes | Wellcome Sanger Institute for the COVID-19 Genomics UK (COG-UK) Consortium | The Lighthouse Lab in Milton Keynes and Alex Alderton, Roberto Amato, Jeffrey Barrett, Sonia Goncalves, Ewan Harrison, David K. Jackson, Ian Johnston, Dominic Kwiatkowski, Cordelia Langford, John Sillitoe on behalf of the Wellcome Sanger Institute COVID-19 Surveillance Team |

| | | | |
|---|---|---|---|
| EPI_ISL_2004912, EPI_ISL_2004970, EPI_ISL_2005010, EPI_ISL_2005025, EPI_ISL_2005065 | Lighthouse Lab in Alderley Park | Wellcome Sanger Institute for the COVID-19 Genomics UK (COG-UK) Consortium | Jacquelyn Wynn, Mairead Hyland, The Lighthouse Lab in Alderley Park and Alex Alderton, Roberto Amato, Jeffrey Barrett, Sonia Goncalves, Ewan Harrison, David K. Jackson, Ian Johnston, Dominic Kwiatkowski, Cordelia Langford, John Sillitoe on behalf of the Wellcome Sanger Institute COVID-19 Surveillance Team |
| EPI_ISL_2005071 | Lighthouse Lab in Glasgow | Wellcome Sanger Institute for the COVID-19 Genomics UK (COG-UK) Consortium | Harper VanSteenhouse, Yumi Kasai, David Gray, Carol Clugston, Anna Dominiczak and Alex Alderton, Roberto Amato, Jeffrey Barrett, Sonia Goncalves, Ewan Harrison, David K. Jackson, Ian Johnston, Dominic Kwiatkowski, Cordelia Langford, John Sillitoe on behalf of the Wellcome Sanger Institute COVID-19 Surveillance Team |
| EPI_ISL_2005093, EPI_ISL_2005110, EPI_ISL_2005114, EPI_ISL_2005117, EPI_ISL_2005120, EPI_ISL_2005129, EPI_ISL_2005146 | Lighthouse Lab in Alderley Park | Wellcome Sanger Institute for the COVID-19 Genomics UK (COG-UK) Consortium | Jacquelyn Wynn, Mairead Hyland, The Lighthouse Lab in Alderley Park and Alex Alderton, Roberto Amato, Jeffrey Barrett, Sonia Goncalves, Ewan Harrison, David K. Jackson, Ian Johnston, Dominic Kwiatkowski, Cordelia Langford, John Sillitoe on behalf of the Wellcome Sanger Institute COVID-19 Surveillance Team |
| EPI_ISL_2005162 | Randox Laboratories | Wellcome Sanger Institute for the COVID-19 Genomics UK (COG-UK) Consortium | Randox Laboratories and Alex Alderton, Roberto Amato, Jeffrey Barrett, Sonia Goncalves, Ewan Harrison, David K. Jackson, Ian Johnston, Dominic Kwiatkowski, Cordelia Langford, John Sillitoe on behalf of the Wellcome Sanger Institute COVID-19 Surveillance Team |
| EPI_ISL_2005164 | Lighthouse Lab in Glasgow | Wellcome Sanger Institute for the COVID-19 Genomics UK (COG-UK) Consortium | Harper VanSteenhouse, Yumi Kasai, David Gray, Carol Clugston, Anna Dominiczak and Alex Alderton, Roberto Amato, Jeffrey Barrett, Sonia Goncalves, Ewan Harrison, David K. Jackson, Ian Johnston, Dominic Kwiatkowski, Cordelia Langford, John Sillitoe on behalf of the Wellcome Sanger Institute COVID-19 Surveillance Team |
| EPI_ISL_2005192 | Randox Laboratories | Wellcome Sanger Institute for the COVID-19 Genomics UK (COG-UK) Consortium | Randox Laboratories and Alex Alderton, Roberto Amato, Jeffrey Barrett, Sonia Goncalves, Ewan Harrison, David K. Jackson, Ian Johnston, Dominic Kwiatkowski, Cordelia Langford, John Sillitoe on behalf of the Wellcome Sanger Institute COVID-19 Surveillance Team |
| EPI_ISL_2005195, EPI_ISL_2005222 | Lighthouse Lab in Glasgow | Wellcome Sanger Institute for the COVID-19 Genomics UK (COG-UK) Consortium | Harper VanSteenhouse, Yumi Kasai, David Gray, Carol Clugston, Anna Dominiczak and Alex Alderton, Roberto Amato, Jeffrey Barrett, Sonia Goncalves, Ewan Harrison, David K. Jackson, Ian Johnston, Dominic Kwiatkowski, Cordelia Langford, John Sillitoe on behalf of the Wellcome Sanger Institute COVID-19 Surveillance Team |
| EPI_ISL_2005239, EPI_ISL_2005241, EPI_ISL_2005285, EPI_ISL_2005288, EPI_ISL_2005291, EPI_ISL_2005295 | Lighthouse Lab in Alderley Park | Wellcome Sanger Institute for the COVID-19 Genomics UK (COG-UK) Consortium | Jacquelyn Wynn, Mairead Hyland, The Lighthouse Lab in Alderley Park and Alex Alderton, Roberto Amato, Jeffrey Barrett, Sonia Goncalves, Ewan Harrison, David K. Jackson, Ian Johnston, Dominic Kwiatkowski, Cordelia Langford, John Sillitoe on behalf of the Wellcome Sanger Institute COVID-19 Surveillance Team |
| EPI_ISL_2005332 | Lighthouse Lab in Glasgow | Wellcome Sanger Institute for the COVID-19 Genomics UK (COG-UK) Consortium | Harper VanSteenhouse, Yumi Kasai, David Gray, Carol Clugston, Anna Dominiczak and Alex Alderton, Roberto Amato, Jeffrey Barrett, Sonia Goncalves, Ewan Harrison, David K. Jackson, Ian Johnston, Dominic Kwiatkowski, Cordelia Langford, John Sillitoe on behalf of the Wellcome Sanger Institute COVID-19 Surveillance Team |
| EPI_ISL_2005341 | Lighthouse Lab in Milton Keynes | Wellcome Sanger Institute for the COVID-19 Genomics UK (COG-UK) Consortium | The Lighthouse Lab in Milton Keynes and Alex Alderton, Roberto Amato, Jeffrey Barrett, Sonia Goncalves, Ewan Harrison, David K. Jackson, Ian Johnston, Dominic Kwiatkowski, Cordelia Langford, John Sillitoe on behalf of the Wellcome Sanger Institute COVID-19 Surveillance Team |
| EPI_ISL_2005347 | Lighthouse Lab in Alderley Park | Wellcome Sanger Institute for the COVID-19 Genomics UK (COG-UK) Consortium | Jacquelyn Wynn, Mairead Hyland, The Lighthouse Lab in Alderley Park and Alex Alderton, Roberto Amato, Jeffrey Barrett, Sonia Goncalves, Ewan Harrison, David K. Jackson, Ian Johnston, Dominic Kwiatkowski, Cordelia Langford, John Sillitoe on behalf of the Wellcome Sanger Institute COVID-19 Surveillance Team |
| EPI_ISL_2005350 | Health Services Laboratories | Wellcome Sanger Institute for the COVID-19 Genomics UK (COG-UK) Consortium | Health Services Laboratories and Alex Alderton, Roberto Amato, Jeffrey Barrett, Sonia Goncalves, Ewan Harrison, David K. Jackson, Ian Johnston, Dominic Kwiatkowski, Cordelia Langford, John Sillitoe on behalf of the Wellcome Sanger Institute COVID-19 Surveillance Team |
| EPI_ISL_2005355, EPI_ISL_2005359 | Lighthouse Lab in Milton Keynes | Wellcome Sanger Institute for the COVID-19 Genomics UK (COG-UK) Consortium | The Lighthouse Lab in Milton Keynes and Alex Alderton, Roberto Amato, Jeffrey Barrett, Sonia Goncalves, Ewan Harrison, David K. Jackson, Ian Johnston, Dominic Kwiatkowski, Cordelia Langford, John Sillitoe on behalf of the Wellcome Sanger Institute COVID-19 Surveillance Team |
| EPI_ISL_2005413 | Lighthouse Lab in Alderley Park | Wellcome Sanger Institute for the COVID-19 Genomics UK (COG-UK) Consortium | Jacquelyn Wynn, Mairead Hyland, The Lighthouse Lab in Alderley Park and Alex Alderton, Roberto Amato, Jeffrey Barrett, Sonia Goncalves, Ewan Harrison, David K. Jackson, Ian Johnston, Dominic Kwiatkowski, Cordelia Langford, John Sillitoe on behalf of the Wellcome Sanger Institute COVID-19 Surveillance Team |
| EPI_ISL_2005415 | Health Services Laboratories | Wellcome Sanger Institute for the COVID-19 Genomics UK (COG-UK) Consortium | Health Services Laboratories and Alex Alderton, Roberto Amato, Jeffrey Barrett, Sonia Goncalves, Ewan Harrison, David K. Jackson, Ian Johnston, Dominic Kwiatkowski, Cordelia Langford, John Sillitoe on behalf of the Wellcome Sanger Institute COVID-19 Surveillance Team |
| EPI_ISL_2005421 | Lighthouse Lab in Milton Keynes | Wellcome Sanger Institute for the COVID-19 Genomics UK (COG-UK) Consortium | The Lighthouse Lab in Milton Keynes and Alex Alderton, Roberto Amato, Jeffrey Barrett, Sonia Goncalves, Ewan Harrison, David K. Jackson, Ian Johnston, Dominic Kwiatkowski, Cordelia Langford, John Sillitoe on behalf of the Wellcome Sanger Institute COVID-19 Surveillance Team |
| EPI_ISL_2005426, EPI_ISL_2005438 | Lighthouse Lab in Alderley Park | Wellcome Sanger Institute for the COVID-19 Genomics UK (COG-UK) Consortium | Jacquelyn Wynn, Mairead Hyland, The Lighthouse Lab in Alderley Park and Alex Alderton, Roberto Amato, Jeffrey Barrett, Sonia Goncalves, Ewan Harrison, David K. Jackson, Ian Johnston, Dominic Kwiatkowski, Cordelia Langford, John Sillitoe on behalf of the Wellcome Sanger Institute COVID-19 Surveillance Team |
| EPI_ISL_2005441 | Lighthouse Lab in Milton Keynes | Wellcome Sanger Institute for the COVID-19 Genomics UK (COG-UK) Consortium | The Lighthouse Lab in Milton Keynes and Alex Alderton, Roberto Amato, Jeffrey Barrett, Sonia Goncalves, Ewan Harrison, David K. Jackson, Ian Johnston, Dominic Kwiatkowski, Cordelia Langford, John Sillitoe on behalf of the Wellcome Sanger Institute COVID-19 Surveillance Team |
| EPI_ISL_2005455 | Health Services Laboratories | Wellcome Sanger Institute for the COVID-19 Genomics UK (COG-UK) Consortium | Health Services Laboratories and Alex Alderton, Roberto Amato, Jeffrey Barrett, Sonia Goncalves, Ewan Harrison, David K. Jackson, Ian Johnston, Dominic Kwiatkowski, Cordelia Langford, John Sillitoe on behalf of the Wellcome Sanger Institute COVID-19 Surveillance Team |
| EPI_ISL_2005461 | Lighthouse Lab in Milton Keynes | Wellcome Sanger Institute for the COVID-19 Genomics UK (COG-UK) Consortium | The Lighthouse Lab in Milton Keynes and Alex Alderton, Roberto Amato, Jeffrey Barrett, Sonia Goncalves, Ewan Harrison, David K. Jackson, Ian Johnston, Dominic Kwiatkowski, Cordelia Langford, John Sillitoe on behalf of the Wellcome Sanger Institute COVID-19 Surveillance Team |
| EPI_ISL_2005476 | Lighthouse Lab in Alderley Park | Wellcome Sanger Institute for the COVID-19 Genomics UK (COG-UK) Consortium | Jacquelyn Wynn, Mairead Hyland, The Lighthouse Lab in Alderley Park and Alex Alderton, Roberto Amato, Jeffrey Barrett, Sonia Goncalves, Ewan Harrison, David K. Jackson, Ian Johnston, Dominic Kwiatkowski, Cordelia Langford, John Sillitoe on behalf of the Wellcome Sanger Institute COVID-19 Surveillance Team |
| EPI_ISL_2005484 | Lighthouse Lab in Milton Keynes | Wellcome Sanger Institute for the COVID-19 Genomics UK (COG-UK) Consortium | The Lighthouse Lab in Milton Keynes and Alex Alderton, Roberto Amato, Jeffrey Barrett, Sonia Goncalves, Ewan Harrison, David K. Jackson, Ian Johnston, Dominic Kwiatkowski, Cordelia Langford, John Sillitoe on behalf of the Wellcome Sanger Institute COVID-19 Surveillance Team |
| EPI_ISL_2005490, EPI_ISL_2005494, EPI_ISL_2005497 | Health Services Laboratories | Wellcome Sanger Institute for the COVID-19 Genomics UK (COG-UK) Consortium | Health Services Laboratories and Alex Alderton, Roberto Amato, Jeffrey Barrett, Sonia Goncalves, Ewan Harrison, David K. Jackson, Ian Johnston, Dominic Kwiatkowski, Cordelia Langford, John Sillitoe on behalf of the Wellcome Sanger Institute COVID-19 Surveillance Team |
| EPI_ISL_2005510 | Lighthouse Lab in Milton Keynes | Wellcome Sanger Institute for the COVID-19 Genomics UK (COG-UK) Consortium | The Lighthouse Lab in Milton Keynes and Alex Alderton, Roberto Amato, Jeffrey Barrett, Sonia Goncalves, Ewan Harrison, David K. Jackson, Ian Johnston, Dominic Kwiatkowski, Cordelia Langford, John Sillitoe on behalf of the Wellcome Sanger Institute COVID-19 Surveillance Team |
| EPI_ISL_2005519 | Lighthouse Lab in Alderley Park | Wellcome Sanger Institute for the COVID-19 Genomics UK (COG-UK) Consortium | Jacquelyn Wynn, Mairead Hyland, The Lighthouse Lab in Alderley Park and Alex Alderton, Roberto Amato, Jeffrey Barrett, Sonia Goncalves, Ewan Harrison, David K. Jackson, Ian Johnston, Dominic Kwiatkowski, Cordelia Langford, John Sillitoe on behalf of the Wellcome Sanger Institute COVID-19 Surveillance Team |
| EPI_ISL_2005525 | Lighthouse Lab in Milton Keynes | Wellcome Sanger Institute for the COVID-19 Genomics UK (COG-UK) Consortium | The Lighthouse Lab in Milton Keynes and Alex Alderton, Roberto Amato, Jeffrey Barrett, Sonia Goncalves, Ewan Harrison, David K. Jackson, Ian Johnston, Dominic Kwiatkowski, Cordelia Langford, John Sillitoe on behalf of the Wellcome Sanger Institute COVID-19 Surveillance Team |
| EPI_ISL_2005527 | Health Services Laboratories | Wellcome Sanger Institute for the COVID-19 Genomics UK (COG-UK) Consortium | Health Services Laboratories and Alex Alderton, Roberto Amato, Jeffrey Barrett, Sonia Goncalves, Ewan Harrison, David K. Jackson, Ian Johnston, Dominic Kwiatkowski, Cordelia Langford, John Sillitoe on behalf of the Wellcome Sanger Institute COVID-19 Surveillance Team |
| EPI_ISL_2005537 | Lighthouse Lab in Milton Keynes | Wellcome Sanger Institute for the COVID-19 Genomics UK (COG-UK) Consortium | The Lighthouse Lab in Milton Keynes and Alex Alderton, Roberto Amato, Jeffrey Barrett, Sonia Goncalves, Ewan Harrison, David K. Jackson, Ian Johnston, Dominic Kwiatkowski, Cordelia Langford, John Sillitoe on behalf of the Wellcome Sanger Institute COVID-19 Surveillance Team |
| EPI_ISL_2005548 | Lighthouse Lab in Alderley Park | Wellcome Sanger Institute for the COVID-19 Genomics UK (COG-UK) Consortium | Jacquelyn Wynn, Mairead Hyland, The Lighthouse Lab in Alderley Park and Alex Alderton, Roberto Amato, Jeffrey Barrett, Sonia Goncalves, Ewan Harrison, David K. Jackson, Ian Johnston, Dominic Kwiatkowski, Cordelia Langford, John Sillitoe on behalf of the Wellcome Sanger Institute COVID-19 Surveillance Team |
| EPI_ISL_2005569 | Health Services Laboratories | Wellcome Sanger Institute for the COVID-19 Genomics UK (COG-UK) Consortium | Health Services Laboratories and Alex Alderton, Roberto Amato, Jeffrey Barrett, Sonia Goncalves, Ewan Harrison, David K. Jackson, Ian Johnston, Dominic Kwiatkowski, Cordelia Langford, John Sillitoe on behalf of the Wellcome Sanger Institute COVID-19 Surveillance Team |
| EPI_ISL_2005573 | Lighthouse Lab in Alderley Park | Wellcome Sanger Institute for the COVID-19 Genomics UK (COG-UK) Consortium | Jacquelyn Wynn, Mairead Hyland, The Lighthouse Lab in Alderley Park and Alex Alderton, Roberto Amato, Jeffrey Barrett, Sonia Goncalves, Ewan Harrison, David K. Jackson, Ian Johnston, Dominic Kwiatkowski, Cordelia Langford, John Sillitoe on behalf of the Wellcome Sanger Institute COVID-19 Surveillance Team |
| EPI_ISL_2005578, EPI_ISL_2005581 | Lighthouse Lab in Milton Keynes | Wellcome Sanger Institute for the COVID-19 Genomics UK (COG-UK) Consortium | The Lighthouse Lab in Milton Keynes and Alex Alderton, Roberto Amato, Jeffrey Barrett, Sonia Goncalves, Ewan Harrison, David K. Jackson, Ian Johnston, Dominic Kwiatkowski, Cordelia Langford, John Sillitoe on behalf of the Wellcome Sanger Institute COVID-19 Surveillance Team |
| EPI_ISL_2005587 | Health Services Laboratories | Wellcome Sanger Institute for the COVID-19 Genomics UK (COG-UK) Consortium | Health Services Laboratories and Alex Alderton, Roberto Amato, Jeffrey Barrett, Sonia Goncalves, Ewan Harrison, David K. Jackson, Ian Johnston, Dominic Kwiatkowski, Cordelia Langford, John Sillitoe on behalf of the Wellcome Sanger Institute COVID-19 Surveillance Team |
| EPI_ISL_2005588 | Lighthouse Lab in Alderley Park | Wellcome Sanger Institute for the COVID-19 Genomics UK (COG-UK) Consortium | Jacquelyn Wynn, Mairead Hyland, The Lighthouse Lab in Alderley Park and Alex Alderton, Roberto Amato, Jeffrey Barrett, Sonia Goncalves, Ewan Harrison, David K. Jackson, Ian Johnston, Dominic Kwiatkowski, Cordelia Langford, John Sillitoe on behalf of the Wellcome Sanger Institute COVID-19 Surveillance Team |
| EPI_ISL_2005597 | Lighthouse Lab in Milton Keynes | Wellcome Sanger Institute for the COVID-19 Genomics UK (COG-UK) Consortium | The Lighthouse Lab in Milton Keynes and Alex Alderton, Roberto Amato, Jeffrey Barrett, Sonia Goncalves, Ewan Harrison, David K. Jackson, Ian Johnston, Dominic Kwiatkowski, Cordelia Langford, John Sillitoe on behalf of the Wellcome Sanger Institute COVID-19 Surveillance Team |
| EPI_ISL_2005617 | Randox Laboratories | Wellcome Sanger Institute for the COVID-19 Genomics UK (COG-UK) Consortium | Randox Laboratories and Alex Alderton, Roberto Amato, Jeffrey Barrett, Sonia Goncalves, Ewan Harrison, David K. Jackson, Ian Johnston, Dominic Kwiatkowski, Cordelia Langford, John Sillitoe on behalf of the Wellcome Sanger Institute COVID-19 Surveillance Team |
| EPI_ISL_2005637 | Lighthouse Lab in Alderley Park | Wellcome Sanger Institute for the COVID-19 Genomics UK (COG-UK) Consortium | Jacquelyn Wynn, Mairead Hyland, The Lighthouse Lab in Alderley Park and Alex Alderton, Roberto Amato, Jeffrey Barrett, Sonia Goncalves, Ewan Harrison, David K. Jackson, Ian Johnston, Dominic Kwiatkowski, Cordelia Langford, John Sillitoe on behalf of the Wellcome Sanger Institute COVID-19 Surveillance Team |
| EPI_ISL_2005649, EPI_ISL_2005664, EPI_ISL_2005671, EPI_ISL_2005705, EPI_ISL_2005715, EPI_ISL_2005716 | Randox Laboratories | Wellcome Sanger Institute for the COVID-19 Genomics UK (COG-UK) Consortium | Randox Laboratories and Alex Alderton, Roberto Amato, Jeffrey Barrett, Sonia Goncalves, Ewan Harrison, David K. Jackson, Ian Johnston, Dominic Kwiatkowski, Cordelia Langford, John Sillitoe on behalf of the Wellcome Sanger Institute COVID-19 Surveillance Team |

| | | | |
|---|---|---|---|
| EPI_ISL_2005725, EPI_ISL_2005754 | Lighthouse Lab in Milton Keynes | Wellcome Sanger Institute for the COVID-19 Genomics UK (COG-UK) Consortium | The Lighthouse Lab in Milton Keynes and Alex Alderton, Roberto Amato, Jeffrey Barrett, Sonia Goncalves, Ewan Harrison, David K. Jackson, Ian Johnston, Dominic Kwiatkowski, Cordelia Langford, John Sillitoe on behalf of the Wellcome Sanger Institute COVID-19 Surveillance Team |
| EPI_ISL_2005784, EPI_ISL_2005792, EPI_ISL_2005804, EPI_ISL_2005808 | Randox Laboratories | Wellcome Sanger Institute for the COVID-19 Genomics UK (COG-UK) Consortium | Randox Laboratories and Alex Alderton, Roberto Amato, Jeffrey Barrett, Sonia Goncalves, Ewan Harrison, David K. Jackson, Ian Johnston, Dominic Kwiatkowski, Cordelia Langford, John Sillitoe on behalf of the Wellcome Sanger Institute COVID-19 Surveillance Team |
| EPI_ISL_2005819 | Lighthouse Lab in Milton Keynes | Wellcome Sanger Institute for the COVID-19 Genomics UK (COG-UK) Consortium | The Lighthouse Lab in Milton Keynes and Alex Alderton, Roberto Amato, Jeffrey Barrett, Sonia Goncalves, Ewan Harrison, David K. Jackson, Ian Johnston, Dominic Kwiatkowski, Cordelia Langford, John Sillitoe on behalf of the Wellcome Sanger Institute COVID-19 Surveillance Team |
| EPI_ISL_2005820 | Lighthouse Lab in Glasgow | Wellcome Sanger Institute for the COVID-19 Genomics UK (COG-UK) Consortium | Harper VanSteenhouse, Yumi Kasai, David Gray, Carol Clugston, Anna Dominiczak and Alex Alderton, Roberto Amato, Jeffrey Barrett, Sonia Goncalves, Ewan Harrison, David K. Jackson, Ian Johnston, Dominic Kwiatkowski, Cordelia Langford, John Sillitoe on behalf of the Wellcome Sanger Institute COVID-19 Surveillance Team |
| EPI_ISL_2005822, EPI_ISL_2005826, EPI_ISL_2005827, EPI_ISL_2005830, EPI_ISL_2005831, EPI_ISL_2005832, EPI_ISL_2005836 | Lighthouse Lab in Milton Keynes | Wellcome Sanger Institute for the COVID-19 Genomics UK (COG-UK) Consortium | The Lighthouse Lab in Milton Keynes and Alex Alderton, Roberto Amato, Jeffrey Barrett, Sonia Goncalves, Ewan Harrison, David K. Jackson, Ian Johnston, Dominic Kwiatkowski, Cordelia Langford, John Sillitoe on behalf of the Wellcome Sanger Institute COVID-19 Surveillance Team |
| EPI_ISL_2005837, EPI_ISL_2005838, EPI_ISL_2005839 | Randox Laboratories | Wellcome Sanger Institute for the COVID-19 Genomics UK (COG-UK) Consortium | Randox Laboratories and Alex Alderton, Roberto Amato, Jeffrey Barrett, Sonia Goncalves, Ewan Harrison, David K. Jackson, Ian Johnston, Dominic Kwiatkowski, Cordelia Langford, John Sillitoe on behalf of the Wellcome Sanger Institute COVID-19 Surveillance Team |
| EPI_ISL_2005843 | Lighthouse Lab in Milton Keynes | Wellcome Sanger Institute for the COVID-19 Genomics UK (COG-UK) Consortium | The Lighthouse Lab in Milton Keynes and Alex Alderton, Roberto Amato, Jeffrey Barrett, Sonia Goncalves, Ewan Harrison, David K. Jackson, Ian Johnston, Dominic Kwiatkowski, Cordelia Langford, John Sillitoe on behalf of the Wellcome Sanger Institute COVID-19 Surveillance Team |
| EPI_ISL_2005845 | Randox Laboratories | Wellcome Sanger Institute for the COVID-19 Genomics UK (COG-UK) Consortium | Randox Laboratories and Alex Alderton, Roberto Amato, Jeffrey Barrett, Sonia Goncalves, Ewan Harrison, David K. Jackson, Ian Johnston, Dominic Kwiatkowski, Cordelia Langford, John Sillitoe on behalf of the Wellcome Sanger Institute COVID-19 Surveillance Team |
| EPI_ISL_2005849 | Lighthouse Lab in Milton Keynes | Wellcome Sanger Institute for the COVID-19 Genomics UK (COG-UK) Consortium | The Lighthouse Lab in Milton Keynes and Alex Alderton, Roberto Amato, Jeffrey Barrett, Sonia Goncalves, Ewan Harrison, David K. Jackson, Ian Johnston, Dominic Kwiatkowski, Cordelia Langford, John Sillitoe on behalf of the Wellcome Sanger Institute COVID-19 Surveillance Team |
| EPI_ISL_2005851 | Lighthouse Lab in Alderley Park | Wellcome Sanger Institute for the COVID-19 Genomics UK (COG-UK) Consortium | Jacquelyn Wynn, Mairead Hyland, The Lighthouse Lab in Alderley Park and Alex Alderton, Roberto Amato, Jeffrey Barrett, Sonia Goncalves, Ewan Harrison, David K. Jackson, Ian Johnston, Dominic Kwiatkowski, Cordelia Langford, John Sillitoe on behalf of the Wellcome Sanger Institute COVID-19 Surveillance Team |
| EPI_ISL_2005853, EPI_ISL_2005854, EPI_ISL_2005855 | Lighthouse Lab in Milton Keynes | Wellcome Sanger Institute for the COVID-19 Genomics UK (COG-UK) Consortium | The Lighthouse Lab in Milton Keynes and Alex Alderton, Roberto Amato, Jeffrey Barrett, Sonia Goncalves, Ewan Harrison, David K. Jackson, Ian Johnston, Dominic Kwiatkowski, Cordelia Langford, John Sillitoe on behalf of the Wellcome Sanger Institute COVID-19 Surveillance Team |
| EPI_ISL_2005858 | Randox Laboratories | Wellcome Sanger Institute for the COVID-19 Genomics UK (COG-UK) Consortium | Randox Laboratories and Alex Alderton, Roberto Amato, Jeffrey Barrett, Sonia Goncalves, Ewan Harrison, David K. Jackson, Ian Johnston, Dominic Kwiatkowski, Cordelia Langford, John Sillitoe on behalf of the Wellcome Sanger Institute COVID-19 Surveillance Team |
| EPI_ISL_2005860, EPI_ISL_2005861, EPI_ISL_2005862 | Lighthouse Lab in Milton Keynes | Wellcome Sanger Institute for the COVID-19 Genomics UK (COG-UK) Consortium | The Lighthouse Lab in Milton Keynes and Alex Alderton, Roberto Amato, Jeffrey Barrett, Sonia Goncalves, Ewan Harrison, David K. Jackson, Ian Johnston, Dominic Kwiatkowski, Cordelia Langford, John Sillitoe on behalf of the Wellcome Sanger Institute COVID-19 Surveillance Team |
| EPI_ISL_2005866, EPI_ISL_2005868 | Randox Laboratories | Wellcome Sanger Institute for the COVID-19 Genomics UK (COG-UK) Consortium | Randox Laboratories and Alex Alderton, Roberto Amato, Jeffrey Barrett, Sonia Goncalves, Ewan Harrison, David K. Jackson, Ian Johnston, Dominic Kwiatkowski, Cordelia Langford, John Sillitoe on behalf of the Wellcome Sanger Institute COVID-19 Surveillance Team |
| EPI_ISL_2005873, EPI_ISL_2005874, EPI_ISL_2005875 | Lighthouse Lab in Milton Keynes | Wellcome Sanger Institute for the COVID-19 Genomics UK (COG-UK) Consortium | The Lighthouse Lab in Milton Keynes and Alex Alderton, Roberto Amato, Jeffrey Barrett, Sonia Goncalves, Ewan Harrison, David K. Jackson, Ian Johnston, Dominic Kwiatkowski, Cordelia Langford, John Sillitoe on behalf of the Wellcome Sanger Institute COVID-19 Surveillance Team |
| EPI_ISL_2005886 | Randox Laboratories | Wellcome Sanger Institute for the COVID-19 Genomics UK (COG-UK) Consortium | Randox Laboratories and Alex Alderton, Roberto Amato, Jeffrey Barrett, Sonia Goncalves, Ewan Harrison, David K. Jackson, Ian Johnston, Dominic Kwiatkowski, Cordelia Langford, John Sillitoe on behalf of the Wellcome Sanger Institute COVID-19 Surveillance Team |
| EPI_ISL_2005893 | Lighthouse Lab in Glasgow | Wellcome Sanger Institute for the COVID-19 Genomics UK (COG-UK) Consortium | Harper VanSteenhouse, Yumi Kasai, David Gray, Carol Clugston, Anna Dominiczak and Alex Alderton, Roberto Amato, Jeffrey Barrett, Sonia Goncalves, Ewan Harrison, David K. Jackson, Ian Johnston, Dominic Kwiatkowski, Cordelia Langford, John Sillitoe on behalf of the Wellcome Sanger Institute COVID-19 Surveillance Team |
| EPI_ISL_2005898, EPI_ISL_2005905, EPI_ISL_2005906 | Randox Laboratories | Wellcome Sanger Institute for the COVID-19 Genomics UK (COG-UK) Consortium | Randox Laboratories and Alex Alderton, Roberto Amato, Jeffrey Barrett, Sonia Goncalves, Ewan Harrison, David K. Jackson, Ian Johnston, Dominic Kwiatkowski, Cordelia Langford, John Sillitoe on behalf of the Wellcome Sanger Institute COVID-19 Surveillance Team |
| EPI_ISL_2005908, EPI_ISL_2005909 | Lighthouse Lab in Milton Keynes | Wellcome Sanger Institute for the COVID-19 Genomics UK (COG-UK) Consortium | The Lighthouse Lab in Milton Keynes and Alex Alderton, Roberto Amato, Jeffrey Barrett, Sonia Goncalves, Ewan Harrison, David K. Jackson, Ian Johnston, Dominic Kwiatkowski, Cordelia Langford, John Sillitoe on behalf of the Wellcome Sanger Institute COVID-19 Surveillance Team |
| EPI_ISL_2005927 | Lighthouse Lab in Glasgow | Wellcome Sanger Institute for the COVID-19 Genomics UK (COG-UK) Consortium | Harper VanSteenhouse, Yumi Kasai, David Gray, Carol Clugston, Anna Dominiczak and Alex Alderton, Roberto Amato, Jeffrey Barrett, Sonia Goncalves, Ewan Harrison, David K. Jackson, Ian Johnston, Dominic Kwiatkowski, Cordelia Langford, John Sillitoe on behalf of the Wellcome Sanger Institute COVID-19 Surveillance Team |
| EPI_ISL_2005928 | Randox Laboratories | Wellcome Sanger Institute for the COVID-19 Genomics UK (COG-UK) Consortium | Randox Laboratories and Alex Alderton, Roberto Amato, Jeffrey Barrett, Sonia Goncalves, Ewan Harrison, David K. Jackson, Ian Johnston, Dominic Kwiatkowski, Cordelia Langford, John Sillitoe on behalf of the Wellcome Sanger Institute COVID-19 Surveillance Team |
| EPI_ISL_2005942, EPI_ISL_2005944 | Lighthouse Lab in Milton Keynes | Wellcome Sanger Institute for the COVID-19 Genomics UK (COG-UK) Consortium | The Lighthouse Lab in Milton Keynes and Alex Alderton, Roberto Amato, Jeffrey Barrett, Sonia Goncalves, Ewan Harrison, David K. Jackson, Ian Johnston, Dominic Kwiatkowski, Cordelia Langford, John Sillitoe on behalf of the Wellcome Sanger Institute COVID-19 Surveillance Team |
| EPI_ISL_2005947 | Randox Laboratories | Wellcome Sanger Institute for the COVID-19 Genomics UK (COG-UK) Consortium | Randox Laboratories and Alex Alderton, Roberto Amato, Jeffrey Barrett, Sonia Goncalves, Ewan Harrison, David K. Jackson, Ian Johnston, Dominic Kwiatkowski, Cordelia Langford, John Sillitoe on behalf of the Wellcome Sanger Institute COVID-19 Surveillance Team |
| EPI_ISL_2005951, EPI_ISL_2005952, EPI_ISL_2005963, EPI_ISL_2005969, EPI_ISL_2005974, EPI_ISL_2005981, EPI_ISL_2005983, EPI_ISL_2005987 | Lighthouse Lab in Milton Keynes | Wellcome Sanger Institute for the COVID-19 Genomics UK (COG-UK) Consortium | The Lighthouse Lab in Milton Keynes and Alex Alderton, Roberto Amato, Jeffrey Barrett, Sonia Goncalves, Ewan Harrison, David K. Jackson, Ian Johnston, Dominic Kwiatkowski, Cordelia Langford, John Sillitoe on behalf of the Wellcome Sanger Institute COVID-19 Surveillance Team |
| EPI_ISL_2005993 | Randox Laboratories | Wellcome Sanger Institute for the COVID-19 Genomics UK (COG-UK) Consortium | Randox Laboratories and Alex Alderton, Roberto Amato, Jeffrey Barrett, Sonia Goncalves, Ewan Harrison, David K. Jackson, Ian Johnston, Dominic Kwiatkowski, Cordelia Langford, John Sillitoe on behalf of the Wellcome Sanger Institute COVID-19 Surveillance Team |
| EPI_ISL_2005999, EPI_ISL_2006004 | Lighthouse Lab in Milton Keynes | Wellcome Sanger Institute for the COVID-19 Genomics UK (COG-UK) Consortium | The Lighthouse Lab in Milton Keynes and Alex Alderton, Roberto Amato, Jeffrey Barrett, Sonia Goncalves, Ewan Harrison, David K. Jackson, Ian Johnston, Dominic Kwiatkowski, Cordelia Langford, John Sillitoe on behalf of the Wellcome Sanger Institute COVID-19 Surveillance Team |
| EPI_ISL_2006007 | Randox Laboratories | Wellcome Sanger Institute for the COVID-19 Genomics UK (COG-UK) Consortium | Randox Laboratories and Alex Alderton, Roberto Amato, Jeffrey Barrett, Sonia Goncalves, Ewan Harrison, David K. Jackson, Ian Johnston, Dominic Kwiatkowski, Cordelia Langford, John Sillitoe on behalf of the Wellcome Sanger Institute COVID-19 Surveillance Team |
| EPI_ISL_2006009 | Lighthouse Lab in Milton Keynes | Wellcome Sanger Institute for the COVID-19 Genomics UK (COG-UK) Consortium | The Lighthouse Lab in Milton Keynes and Alex Alderton, Roberto Amato, Jeffrey Barrett, Sonia Goncalves, Ewan Harrison, David K. Jackson, Ian Johnston, Dominic Kwiatkowski, Cordelia Langford, John Sillitoe on behalf of the Wellcome Sanger Institute COVID-19 Surveillance Team |
| EPI_ISL_2006010, EPI_ISL_2006012 | Randox Laboratories | Wellcome Sanger Institute for the COVID-19 Genomics UK (COG-UK) Consortium | Randox Laboratories and Alex Alderton, Roberto Amato, Jeffrey Barrett, Sonia Goncalves, Ewan Harrison, David K. Jackson, Ian Johnston, Dominic Kwiatkowski, Cordelia Langford, John Sillitoe on behalf of the Wellcome Sanger Institute COVID-19 Surveillance Team |
| EPI_ISL_2006014, EPI_ISL_2006017, EPI_ISL_2006025, EPI_ISL_2006030, EPI_ISL_2006036, EPI_ISL_2006038, EPI_ISL_2006042 | Lighthouse Lab in Milton Keynes | Wellcome Sanger Institute for the COVID-19 Genomics UK (COG-UK) Consortium | The Lighthouse Lab in Milton Keynes and Alex Alderton, Roberto Amato, Jeffrey Barrett, Sonia Goncalves, Ewan Harrison, David K. Jackson, Ian Johnston, Dominic Kwiatkowski, Cordelia Langford, John Sillitoe on behalf of the Wellcome Sanger Institute COVID-19 Surveillance Team |
| EPI_ISL_2006047 | Lighthouse Lab in Glasgow | Wellcome Sanger Institute for the COVID-19 Genomics UK (COG-UK) Consortium | Harper VanSteenhouse, Yumi Kasai, David Gray, Carol Clugston, Anna Dominiczak and Alex Alderton, Roberto Amato, Jeffrey Barrett, Sonia Goncalves, Ewan Harrison, David K. Jackson, Ian Johnston, Dominic Kwiatkowski, Cordelia Langford, John Sillitoe on behalf of the Wellcome Sanger Institute COVID-19 Surveillance Team |
| EPI_ISL_2006048, EPI_ISL_2006051, EPI_ISL_2006052, EPI_ISL_2006055, EPI_ISL_2006056, EPI_ISL_2006057, EPI_ISL_2006061 | Lighthouse Lab in Milton Keynes | Wellcome Sanger Institute for the COVID-19 Genomics UK (COG-UK) Consortium | The Lighthouse Lab in Milton Keynes and Alex Alderton, Roberto Amato, Jeffrey Barrett, Sonia Goncalves, Ewan Harrison, David K. Jackson, Ian Johnston, Dominic Kwiatkowski, Cordelia Langford, John Sillitoe on behalf of the Wellcome Sanger Institute COVID-19 Surveillance Team |
| EPI_ISL_2006062 | Randox Laboratories | Wellcome Sanger Institute for the COVID-19 Genomics UK (COG-UK) Consortium | Randox Laboratories and Alex Alderton, Roberto Amato, Jeffrey Barrett, Sonia Goncalves, Ewan Harrison, David K. Jackson, Ian Johnston, Dominic Kwiatkowski, Cordelia Langford, John Sillitoe on behalf of the Wellcome Sanger Institute COVID-19 Surveillance Team |
| EPI_ISL_2006064, | Lighthouse Lab in Milton Keynes | Wellcome Sanger Institute for the COVID-19 Genomics UK (COG-UK) Consortium | The Lighthouse Lab in Milton Keynes and Alex Alderton, Roberto Amato, Jeffrey Barrett, Sonia Goncalves, Ewan Harrison, David K. Jackson, Ian Johnston, Dominic Kwiatkowski, Cordelia Langford, John Sillitoe on behalf of the Wellcome Sanger |

| | | | |
|---|---|---|---|
| EPI_ISL_2006067, EPI_ISL_2006070 | | Consortium | Institute COVID-19 Surveillance Team |
| EPI_ISL_2006080, EPI_ISL_2006197, EPI_ISL_2006210, EPI_ISL_2006228, EPI_ISL_2006274 | Lighthouse Lab in Alderley Park | Wellcome Sanger Institute for the COVID-19 Genomics UK (COG-UK) Consortium | Jacquelyn Wynn, Mairead Hyland, The Lighthouse Lab in Alderley Park and Alex Alderton, Roberto Amato, Jeffrey Barrett, Sonia Goncalves, Ewan Harrison, David K. Jackson, Ian Johnston, Dominic Kwiatkowski, Cordelia Langford, John Sillitoe on behalf of the Wellcome Sanger Institute COVID-19 Surveillance Team |
| EPI_ISL_2006317, EPI_ISL_2006366, EPI_ISL_2006372 | Lighthouse Lab in Glasgow | Wellcome Sanger Institute for the COVID-19 Genomics UK (COG-UK) Consortium | Harper VanSteenhouse, Yumi Kasai, David Gray, Carol Clugston, Anna Dominiczak and Alex Alderton, Roberto Amato, Jeffrey Barrett, Sonia Goncalves, Ewan Harrison, David K. Jackson, Ian Johnston, Dominic Kwiatkowski, Cordelia Langford, John Sillitoe on behalf of the Wellcome Sanger Institute COVID-19 Surveillance Team |
| EPI_ISL_2006528, EPI_ISL_2006530 | Lighthouse Lab in Milton Keynes | Wellcome Sanger Institute for the COVID-19 Genomics UK (COG-UK) Consortium | The Lighthouse Lab in Milton Keynes and Alex Alderton, Roberto Amato, Jeffrey Barrett, Sonia Goncalves, Ewan Harrison, David K. Jackson, Ian Johnston, Dominic Kwiatkowski, Cordelia Langford, John Sillitoe on behalf of the Wellcome Sanger Institute COVID-19 Surveillance Team |
| EPI_ISL_2006541, EPI_ISL_2006543, EPI_ISL_2006547, EPI_ISL_2006550, EPI_ISL_2006553, EPI_ISL_2006556, EPI_ISL_2006559, EPI_ISL_2006572, EPI_ISL_2006577, EPI_ISL_2006582, EPI_ISL_2006586, EPI_ISL_2006590, EPI_ISL_2006592, EPI_ISL_2006594, EPI_ISL_2006600, EPI_ISL_2006604, EPI_ISL_2006606, EPI_ISL_2006607, EPI_ISL_2006609, EPI_ISL_2006612, EPI_ISL_2006613, EPI_ISL_2006614, EPI_ISL_2006616, EPI_ISL_2006624, EPI_ISL_2006625, EPI_ISL_2006640, EPI_ISL_2006642, EPI_ISL_2006648, EPI_ISL_2006659, EPI_ISL_2006661 | | | |
| see above | Health Services Laboratories | Wellcome Sanger Institute for the COVID-19 Genomics UK (COG-UK) Consortium | Health Services Laboratories and Alex Alderton, Roberto Amato, Jeffrey Barrett, Sonia Goncalves, Ewan Harrison, David K. Jackson, Ian Johnston, Dominic Kwiatkowski, Cordelia Langford, John Sillitoe on behalf of the Wellcome Sanger Institute COVID-19 Surveillance Team |
| EPI_ISL_2006668 | Lighthouse Lab in Milton Keynes | Wellcome Sanger Institute for the COVID-19 Genomics UK (COG-UK) Consortium | The Lighthouse Lab in Milton Keynes and Alex Alderton, Roberto Amato, Jeffrey Barrett, Sonia Goncalves, Ewan Harrison, David K. Jackson, Ian Johnston, Dominic Kwiatkowski, Cordelia Langford, John Sillitoe on behalf of the Wellcome Sanger Institute COVID-19 Surveillance Team |
| EPI_ISL_2006677 | Max von Pettenkofer Institute, Virology, National Reference Center for Retroviruses, LMU Munich | Laboratory for Functional Genome Analysis; Dept. Genomics; Gene Center of the LMU Munich | Max Muenchhoff; Stefan Krebs; Alexander Graf; Oliver Keppler; Helmut Blum |
| EPI_ISL_2007204 | Jessa | Jessa | Berden et al. on behalf of the Jessa_cmdLab |
| EPI_ISL_2007358 | Akershus University Hospital, Department for Microbiology and Infectious Disease Control | Norwegian Institute of Public Health, Department of Virology | Kathrine Stene-Johansen, Kamilla Heddeland Instefjord, Hilde Elshaug, Garcia Llorente Ignacio, Jon Bråte, Engebretsen Serina Beate,Pedersen Benedikte Nevjen, Debech Nadia, Atiya R Ali,Marie Paulsen Madsen, Rasmus Riis Kopperud, Hilde Vollan, Karoline Bragstad, Olav Hungnes |
| EPI_ISL_2008146, EPI_ISL_2008162, EPI_ISL_2008164, EPI_ISL_2008369 | Massachusetts State Public Health Laboratory | Massachusetts State Public Health Laboratory | Andrew Lang, Timelia Fink, Glen Gallagher, Sandra Smole |
| EPI_ISL_2009735, EPI_ISL_2010071, EPI_ISL_2010188, EPI_ISL_2010380, EPI_ISL_2010430, EPI_ISL_2010478, EPI_ISL_2010812, EPI_ISL_2010872 | Helix/Illumina | Centers for Disease Control and Prevention Division of Viral Diseases, Pathogen Discovery | Dakota Howard, Dhwani Batra, Peter W. Cook, Kara Moser, Adrian Paskey, Jason Caravas, Benjamin Rambo-Martin, Shatavia Morrison, Christopher Gulvick, Scott Sammons, Yvette Unoarumhi, Darlene Wagner, Matthew Schmerer, Eileen de Feo, Jan Antico, Christine Tran, Matthew Tolentino, Shannon Wickline, Kim Gietzen, Brad Sickler, Jingtao Liu, Eric Allen, Phil Febbo, Nicole L. Washington, Simon White, Geraint Levan, Kelly Schiabor Barrett, Elizabeth Cirulli, Alexandre Bolze, Ary Ascencio, Charlotte Rivera-Garcia, Ryan Cho, Jason Nguyen, Sherry Wang, Jimmy Ramirez, Tyler Cassens, Efren Sandoval, Magnus Isaksson, William Lee, David Becker, Marc Laurent, James Lu, Clinton R. Paden, Duncan MacCannell |
| EPI_ISL_2011131 | Quest Diagnostics Incorporated | Centers for Disease Control and Prevention Division of Viral Diseases, Pathogen Discovery | Dakota Howard, Dhwani Batra, Peter W. Cook, Kara Moser, Adrian Paskey, Jason Caravas, Benjamin Rambo-Martin, Shatavia Morrison, Christopher Gulvick, Scott Sammons, Yvette Unoarumhi, Darlene Wagner, Matthew Schmerer, S. H. Rosenthal, A. Gerasimova, R. M. Kagan, B. Anderson, M. Hua, Y. Liu, L.E. Bernstein, K.E. Livingston, A. Perez, I. A. Shlyakhter, R. V. Rolando, R. Owen, P. Tanpaiboon, F. Lacbawan, Clinton R. Paden, Duncan MacCannell |
| EPI_ISL_2011443, EPI_ISL_2011445 | Seattle Flu Study | Seattle Flu Study | Deborah A. Nickerson, Chris D. Frazar, Jover Lee, Benjamin Pelle, Erica Ryke, Matthew Richardson, Amanda Adler, Elisabeth Brandstetter, Peter D. Han, Kairsten Fay, Misja Ilcisin, Kirsten Lacombe, Thomas R. Sibley, Melissa Truong, Caitlin R. Wolf, Karen Cowgill, Stephanie Schrag, Jeff Duchin, Michael Boeckh, Janet A. Englund, Michael Famulare, Barry R. Lutz, Mark J. Rieder, Lea M. Starita, Matthew Thompson, Helen Y. Chu, Trevor Bedford, Jay Shendure |
| EPI_ISL_2011886, EPI_ISL_2012488, EPI_ISL_2012492, EPI_ISL_2012499, EPI_ISL_2012507, EPI_ISL_2012513, EPI_ISL_2012676 | Public Health Ontario Laboratory | Public Health Ontario Laboratory | Vanessa G Allen, Philip Banh, Yao Chen, Richard de Borja, Alireza Eshaghi, Nahuel Fittipaldi, Christine Frantz, Jonathan B Gubbay, Jennifer L Guthrie, Lawrence Heisler, Esha Joshi, Michael Laszloffy, Aimin Li, Michael CY Li, Dean Maxwell, Sandeep Nagra, Samir N Patel, Jared Simpson, Karthikeyan Sivaraman, Ashleigh Sullivan, Yogi Sundaravadanam, Sarah Teatero, Andre Villegas, Matthew Watson, Sandra Zittermann |
| EPI_ISL_2013035 | Plateforme Covid IDF, site Broussais | HEGP - Laboratoire de Virologie | David Veyer, Sandrine Imbeaud, Julien Puech, Nabil Gastli, Hélène Péré, Maxime Wack, Nicolas Robillard |
| EPI_ISL_2013036 | Pathcare-Vermaak Centurion | National Institute for Communicable Diseases of the National Health Laboratory Service | Amoako DG, Tegally H, Scheepers C, Mohale T, Ntuli N, Mahlangu B, Viana R, Glass A,Oliveira TD, Gottberg A, Bhiman JN |
| EPI_ISL_2013037 | Lancet Laboratories | National Institute for Communicable Diseases of the National Health Laboratory Service | Amoako DG, Tegally H, Scheepers C, Mohale T, Ntuli N, Mahlangu B, Viana R, Glass A,Oliveira TD, Gottberg A, Bhiman JN |
| EPI_ISL_2013038 | Pathcare-Vermaak Centurion | National Institute for Communicable Diseases of the National Health Laboratory Service | Amoako DG, Tegally H, Scheepers C, Mohale T, Ntuli N, Mahlangu B, Viana R, Glass A,Oliveira TD, Gottberg A, Bhiman JN |
| EPI_ISL_2013040 | Lancet Laboratories | National Institute for Communicable Diseases of the National Health Laboratory Service | Amoako DG, Tegally H, Scheepers C, Mohale T, Ntuli N, Mahlangu B, Viana R, Glass A,Oliveira TD, Gottberg A, Bhiman JN |
| EPI_ISL_2013703 | Infinity Biologix | Centers for Disease Control and Prevention Division of Viral Diseases, Pathogen Discovery | Dakota Howard, Dhwani Batra, Peter W. Cook, Kara Moser, Adrian Paskey, Jason Caravas, Benjamin Rambo-Martin, Shatavia Morrison, Christopher Gulvick, Scott Sammons, Yvette Unoarumhi, Darlene Wagner, Matthew Schmerer, Christian Bixby, Yihe Wang, Jonathan Schultz, Chirayu Goswami, Russ Hager, Robin Grimwood, Clinton R. Paden, Duncan MacCannell |
| EPI_ISL_2015470 | GOVINDRAJNAGAR | inStem NCBS - INSACOG | Uma Ramakrishnan Dasaradhi Palakodeti Aswin SaiNarain |
| EPI_ISL_2015473, EPI_ISL_2015474, EPI_ISL_2015476, EPI_ISL_2015477, EPI_ISL_2015484, EPI_ISL_2015488 | AGRAHARA PHC | inStem NCBS - INSACOG | Uma Ramakrishnan Dasaradhi Palakodeti Aswin SaiNarain |
| EPI_ISL_2015490 | RAILWAY H | inStem NCBS - INSACOG | Uma Ramakrishnan Dasaradhi Palakodeti Aswin SaiNarain |
| EPI_ISL_2015491 | AGRAHARA PHC | inStem NCBS - INSACOG | Uma Ramakrishnan Dasaradhi Palakodeti Aswin SaiNarain |
| EPI_ISL_2015492 | NIMHANS | inStem NCBS - INSACOG | Uma Ramakrishnan Dasaradhi Palakodeti Aswin SaiNarain |
| EPI_ISL_2015496 | AGRAHARA PHC | inStem NCBS - INSACOG | Uma Ramakrishnan Dasaradhi Palakodeti Aswin SaiNarain |
| EPI_ISL_2015497 | NIMHANS | inStem NCBS - INSACOG | Uma Ramakrishnan Dasaradhi Palakodeti Aswin SaiNarain |
| EPI_ISL_2015502, EPI_ISL_2015504, EPI_ISL_2015505 | CMC Vellore | inStem NCBS - INSACOG | Uma Ramakrishnan Dasaradhi Palakodeti Aswin SaiNarain |
| EPI_ISL_2015506 | HOSA HALLI | inStem NCBS - INSACOG | Uma Ramakrishnan Dasaradhi Palakodeti Aswin SaiNarain |
| EPI_ISL_2015508 | CMC Vellore | inStem NCBS - INSACOG | Uma Ramakrishnan Dasaradhi Palakodeti Aswin SaiNarain |
| EPI_ISL_2015512 | RAILWAY H | inStem NCBS - INSACOG | Uma Ramakrishnan Dasaradhi Palakodeti Aswin SaiNarain |
| EPI_ISL_2015514 | GOVINDRAJNAGAR | inStem NCBS - INSACOG | Uma Ramakrishnan Dasaradhi Palakodeti Aswin SaiNarain |
| EPI_ISL_2015515 | RAILWAY HOSPITAL | inStem NCBS - INSACOG | Uma Ramakrishnan Dasaradhi Palakodeti Aswin SaiNarain |
| EPI_ISL_2015516 | HOSAHALLI | inStem NCBS - INSACOG | Uma Ramakrishnan Dasaradhi Palakodeti Aswin SaiNarain |
| EPI_ISL_2015517 | CMC Vellore | inStem NCBS - INSACOG | Uma Ramakrishnan Dasaradhi Palakodeti Aswin SaiNarain |
| EPI_ISL_2015522 | RAILWAY H | inStem NCBS - INSACOG | Uma Ramakrishnan Dasaradhi Palakodeti Aswin SaiNarain |
| EPI_ISL_2015523 | HOSA HALLI | inStem NCBS - INSACOG | Uma Ramakrishnan Dasaradhi Palakodeti Aswin SaiNarain |
| EPI_ISL_2015526 | CMC Vellore | inStem NCBS - INSACOG | Uma Ramakrishnan Dasaradhi Palakodeti Aswin SaiNarain |
| EPI_ISL_2015527 | HOSAHALLI | inStem NCBS - INSACOG | Uma Ramakrishnan Dasaradhi Palakodeti Aswin SaiNarain |
| EPI_ISL_2015529, EPI_ISL_2015530 | RAILWAY H | inStem NCBS - INSACOG | Uma Ramakrishnan Dasaradhi Palakodeti Aswin SaiNarain |
| EPI_ISL_2015531 | CMC Vellore | inStem NCBS - INSACOG | Uma Ramakrishnan Dasaradhi Palakodeti Aswin SaiNarain |
| EPI_ISL_2015534 | KMC MANIPAL | inStem NCBS - INSACOG | Uma Ramakrishnan Dasaradhi Palakodeti Aswin SaiNarain |
| EPI_ISL_2015535 | AGRAHARA PHC | inStem NCBS - INSACOG | Uma Ramakrishnan Dasaradhi Palakodeti Aswin SaiNarain |
| EPI_ISL_2015536 | CMC Vellore | inStem NCBS - INSACOG | Uma Ramakrishnan Dasaradhi Palakodeti Aswin SaiNarain |
| EPI_ISL_2015538 | GOVINDRAJNAGAR | inStem NCBS - INSACOG | Uma Ramakrishnan Dasaradhi Palakodeti Aswin SaiNarain |

| | | | |
|---|---|---|---|
| EPI_ISL_2015539 | CMC Vellore | inStem NCBS - INSACOG | Uma Ramakrishnan Dasaradhi Palakodeti Aswin SaiNarain |
| EPI_ISL_2015542, EPI_ISL_2015544 | NIMHANS | inStem NCBS - INSACOG | Uma Ramakrishnan Dasaradhi Palakodeti Aswin SaiNarain |
| EPI_ISL_2015560, EPI_ISL_2015561, EPI_ISL_2015563 | CMC Vellore | inStem NCBS - INSACOG | Uma Ramakrishnan Dasaradhi Palakodeti Aswin SaiNarain |
| EPI_ISL_2015565 | HOSAHALLI | inStem NCBS - INSACOG | Uma Ramakrishnan Dasaradhi Palakodeti Aswin SaiNarain |
| EPI_ISL_2015566, EPI_ISL_2015570, EPI_ISL_2015572 | CMC Vellore | inStem NCBS - INSACOG | Uma Ramakrishnan Dasaradhi Palakodeti Aswin SaiNarain |
| EPI_ISL_2015575 | HOSAHALLI | inStem NCBS - INSACOG | Uma Ramakrishnan Dasaradhi Palakodeti Aswin SaiNarain |
| EPI_ISL_2015586 | CMC Vellore | inStem NCBS - INSACOG | Uma Ramakrishnan Dasaradhi Palakodeti Aswin SaiNarain |
| EPI_ISL_2015587 | RAILWAY H | inStem NCBS - INSACOG | Uma Ramakrishnan Dasaradhi Palakodeti Aswin SaiNarain |
| EPI_ISL_2015590, EPI_ISL_2015592 | CMC Vellore | inStem NCBS - INSACOG | Uma Ramakrishnan Dasaradhi Palakodeti Aswin SaiNarain |
| EPI_ISL_2015594 | NIMHANS | inStem NCBS - INSACOG | Uma Ramakrishnan Dasaradhi Palakodeti Aswin SaiNarain |
| EPI_ISL_2015595, EPI_ISL_2015597 | CMC Vellore | inStem NCBS - INSACOG | Uma Ramakrishnan Dasaradhi Palakodeti Aswin SaiNarain |
| EPI_ISL_2015598 | A D HALLI | inStem NCBS - INSACOG | Uma Ramakrishnan Dasaradhi Palakodeti Aswin SaiNarain |
| EPI_ISL_2015602 | CMC Vellore | inStem NCBS - INSACOG | Uma Ramakrishnan Dasaradhi Palakodeti Aswin SaiNarain |
| EPI_ISL_2015604 | NIMHANS | inStem NCBS - INSACOG | Uma Ramakrishnan Dasaradhi Palakodeti Aswin SaiNarain |
| EPI_ISL_2015606, EPI_ISL_2015608, EPI_ISL_2015612 | CMC Vellore | inStem NCBS - INSACOG | Uma Ramakrishnan Dasaradhi Palakodeti Aswin SaiNarain |
| EPI_ISL_2015616 | HOSAHALLI | inStem NCBS - INSACOG | Uma Ramakrishnan Dasaradhi Palakodeti Aswin SaiNarain |
| EPI_ISL_2015619 | GOVINDRAJNAGAR | inStem NCBS - INSACOG | Uma Ramakrishnan Dasaradhi Palakodeti Aswin SaiNarain |
| EPI_ISL_2015620 | RAILWAY | inStem NCBS - INSACOG | Uma Ramakrishnan Dasaradhi Palakodeti Aswin SaiNarain |
| EPI_ISL_2015621 | NIMHANS | inStem NCBS - INSACOG | Uma Ramakrishnan Dasaradhi Palakodeti Aswin SaiNarain |
| EPI_ISL_2015622, EPI_ISL_2015623, EPI_ISL_2015624, EPI_ISL_2015626 | RAILWAY | inStem NCBS - INSACOG | Uma Ramakrishnan Dasaradhi Palakodeti Aswin SaiNarain |
| EPI_ISL_2015627 | Railway | inStem NCBS - INSACOG | Uma Ramakrishnan Dasaradhi Palakodeti Aswin SaiNarain |
| EPI_ISL_2015628 | NIMHANS | inStem NCBS - INSACOG | Uma Ramakrishnan Dasaradhi Palakodeti Aswin SaiNarain |
| EPI_ISL_2015629, EPI_ISL_2015630 | RAILWAY | inStem NCBS - INSACOG | Uma Ramakrishnan Dasaradhi Palakodeti Aswin SaiNarain |
| EPI_ISL_2015631 | NIMHANS | inStem NCBS - INSACOG | Uma Ramakrishnan Dasaradhi Palakodeti Aswin SaiNarain |
| EPI_ISL_2015632 | RAILWAY | inStem NCBS - INSACOG | Uma Ramakrishnan Dasaradhi Palakodeti Aswin SaiNarain |
| EPI_ISL_2015633 | KEMPEGOWDA INTERNATIONAL AIRPORT | inStem NCBS - INSACOG | Uma Ramakrishnan Dasaradhi Palakodeti Aswin SaiNarain |
| EPI_ISL_2015635 | RAILWAY | inStem NCBS - INSACOG | Uma Ramakrishnan Dasaradhi Palakodeti Aswin SaiNarain |
| EPI_ISL_2015636, EPI_ISL_2015637 | NIMHANS | inStem NCBS - INSACOG | Uma Ramakrishnan Dasaradhi Palakodeti Aswin SaiNarain |
| EPI_ISL_2015638 | RAILWAY | inStem NCBS - INSACOG | Uma Ramakrishnan Dasaradhi Palakodeti Aswin SaiNarain |
| EPI_ISL_2015639 | GOVIDRAJNAGAR | inStem NCBS - INSACOG | Uma Ramakrishnan Dasaradhi Palakodeti Aswin SaiNarain |
| EPI_ISL_2015640 | SANJAY GANDHI | inStem NCBS - INSACOG | Uma Ramakrishnan Dasaradhi Palakodeti Aswin SaiNarain |
| EPI_ISL_2015641 | NIMHANS | inStem NCBS - INSACOG | Uma Ramakrishnan Dasaradhi Palakodeti Aswin SaiNarain |
| EPI_ISL_2015692 | National Institute of Public Health - National Institute of Hygiene | 1. National Institute of Public Health - National Institute of Hygiene, Warsaw, Poland 2. Biobank Lab, University of Lodz 3. Laboratory of Respiratory Viruses, Teaching and Clinical Center of the Medical University of Lodz | Dominik Strapagiel, Marta Sobalska-Kwapis, Klaudyna Królikowska, Jakub Lach, Marcin Słomka, Tomasz Płoszaj, Magdalena Traczyk-Borszyńska, Izabela Dróżdż, Maciej Borowiec, Katarzyna Zacharczuk, Magdalena Nowakowska, Małgorzata Sadkowska-Todys, Tomasz Wołkowicz |
| EPI_ISL_2015901, EPI_ISL_2015915 | SELARL MIRIALIS CLUSES | CNR Virus des Infections Respiratoires - France SUD | Antonin Bal, Gregory Destras, Gwendolyne Burfin, Hadrien Regue, Quentin Semanas, Martine Valette, Bruno Lina, Laurence Josset |
| EPI_ISL_2016024 | MIRIALIS CLUSES BECHET | CNR Virus des Infections Respiratoires - France SUD | Antonin Bal, Gregory Destras, Gwendolyne Burfin, Hadrien Regue, Quentin Semanas, Martine Valette, Bruno Lina, Laurence Josset |
| EPI_ISL_2016398, EPI_ISL_2016438 | Broad Institute Clinical Research Sequencing Platform | Infectious Disease Program, Broad Institute of Harvard and MIT | Siddle,K.J., Adams,G., Pearlman,L., Gladden-Young,A., Vicente,G., Blumenstiel,B., DeFelice,M., Lee,M., McGovern,S., Lagerborg,K., Rudy,M., DeRuff,K., Carter,A., Normandin,E., Bauer,M., Reilly,S., Tomkins-Tinch,C., Loreth,C., Chaluvadi,S., Meldrim,J., Granger,B., Lemieux,J.E., Birren,B.W., Sabeti,P.C., Larkin,K., Dodge,S., Lennon,N., Madoff,L., Brown,C., Gallagher,G., Smole,S., Park,D.J., Gabriel,S., and MacInnis,B.L. |
| EPI_ISL_2016711, EPI_ISL_2016712, EPI_ISL_2016714, EPI_ISL_2016715 | Platform BIS UZA/UAntwerpen | Labo Klinische Biologie, UZA | Marie Le Mercier, Jasmine Coppens, Basil Britto Xavier, Christine Lammens, Veerle Matheeussen, Herman Goossens |
| EPI_ISL_2016831 | LHUB-ULB | Labo Klinische Biologie, UZA | Marie Le Mercier, Jasmine Coppens, Basil Britto Xavier, Christine Lammens, Veerle Matheeussen, Herman Goossens |
| EPI_ISL_2017208, EPI_ISL_2017223, EPI_ISL_2017224, EPI_ISL_2017225 | Viollier AG | Department of Biosystems Science and Engineering, ETH Zürich | Christian Beisel, Sarah Nadeau, Chaoran Chen, Ivan Topolsky, Philipp Jablonski, Lara Fuhrmann, David Dreifuss, Katharina Jahn, Rebecca Denes, Mirjam Feldkamp, Ina Nissen, Natascha Santacroce, Elodie Burcklen, Christiane Beckmann, Maurice Redondo, Olivier Kobel, Christoph Noppen, Sophie Seidel, Noemie Santamaria de Souza, Niko Beerenwinkel, Tanja Stadler |
| EPI_ISL_2017485, EPI_ISL_2017486, EPI_ISL_2017487, EPI_ISL_2017488, EPI_ISL_2017489, EPI_ISL_2017490 | OLVZ Aalst | OLVZ Aalst | Anne Vankeerberghen |
| EPI_ISL_2017662, EPI_ISL_2017663 | CLILAB | Microbiology Department | Sara Marti, Aida Gonzalez-Diaz, Laura Calatayud, Jordi Niubó, Miguel Fernandez-Huerta, Carmen Ardanuy, Jordi Camara, M Angeles Domínguez |
| EPI_ISL_2017728 | VRDL, Goverment Medical College (GMC), Surat | Gujarat Biotechnology Research Centre | Nitin Savaliya,Dinesh Kumar,Twinkle Soni,Sonal Sharma ,Zuber Saiyed,Ramesh Pandit,Janvi Raval,Zarna Patel,Neeta Khandelwal,Umang Mishra,Nitesh Shah,Chaitanya Joshi,Madhvi Joshi |
| EPI_ISL_2017731 | VRDL, Goverment Medical College (GMC), Surat | Gujarat Biotechnology Research Centre | Sonal Sharma ,Zuber Saiyed,Ramesh Pandit,Janvi Raval,Zarna Patel,Nitin Savaliya,Dinesh Kumar,Twinkle Soni,Neeta Khandelwal,Umang Mishra,Nitesh Shah,Chaitanya Joshi,Madhvi Joshi |
| EPI_ISL_2017734 | VRDL, Goverment Medical College (GMC), Surat | Gujarat Biotechnology Research Centre | Janvi Raval,Zarna Patel,Nitin Savaliya,Dinesh Kumar,Twinkle Soni,Sonal Sharma ,Zuber Saiyed,Ramesh Pandit,Neeta Khandelwal,Umang Mishra,Nitesh Shah,Chaitanya Joshi,Madhvi Joshi |
| EPI_ISL_2017735 | VRDL, Goverment Medical College (GMC), Surat | Gujarat Biotechnology Research Centre | Zarna Patel,Nitin Savaliya,Dinesh Kumar,Twinkle Soni,Sonal Sharma ,Zuber Saiyed,Ramesh Pandit,Janvi Raval,Neeta Khandelwal,Umang Mishra,Nitesh Shah,Chaitanya Joshi,Madhvi Joshi |
| EPI_ISL_2017737 | VRDL, Goverment Medical College (GMC), Surat | Gujarat Biotechnology Research Centre | Dinesh Kumar,Twinkle Soni,Sonal Sharma ,Zuber Saiyed,Ramesh Pandit,Janvi Raval,Zarna Patel,Nitin Savaliya,Neeta Khandelwal,Umang Mishra,Nitesh Shah,Chaitanya Joshi,Madhvi Joshi |
| EPI_ISL_2017738 | VRDL, Goverment Medical College (GMC), Surat | Gujarat Biotechnology Research Centre | Twinkle Soni,Sonal Sharma ,Zuber Saiyed,Ramesh Pandit,Janvi Raval,Zarna Patel,Nitin Savaliya,Dinesh Kumar,Neeta Khandelwal,Umang Mishra,Nitesh Shah,Chaitanya Joshi,Madhvi Joshi |
| EPI_ISL_2017740 | VRDL, Goverment Medical College (GMC), Surat | Gujarat Biotechnology Research Centre | Zuber Saiyed,Ramesh Pandit,Janvi Raval,Zarna Patel,Nitin Savaliya,Dinesh Kumar,Twinkle Soni,Sonal Sharma ,Neeta Khandelwal,Umang Mishra,Nitesh Shah,Chaitanya Joshi,Madhvi Joshi |
| EPI_ISL_2017744 | VRDL, Goverment Medical College (GMC), Surat | Gujarat Biotechnology Research Centre | Nitin Savaliya,Dinesh Kumar,Twinkle Soni,Sonal Sharma ,Zuber Saiyed,Ramesh Pandit,Janvi Raval,Zarna Patel,Neeta Khandelwal,Umang Mishra,Nitesh Shah,Chaitanya Joshi,Madhvi Joshi |
| EPI_ISL_2017748 | VRDL, Goverment Medical College (GMC) and Hospital, Valsad | Gujarat Biotechnology Research Centre | Zuber Saiyed,Ramesh Pandit,Janvi Raval,Zarna Patel,Nitin Savaliya,Dinesh Kumar,Twinkle Soni,Sonal Sharma ,Vicky Gandhi,Umang Mishra,Nitesh Shah,Chaitanya Joshi,Madhvi Joshi |
| EPI_ISL_2017757 | GMERS, Government Medical College (GMC), Gotri | Gujarat Biotechnology Research Centre | Ramesh Pandit,Janvi Raval,Zarna Patel,Nitin Savaliya,Dinesh Kumar,Twinkle Soni,Sonal Sharma ,Zuber Saiyed,Bithika Duttaroy,Umang Mishra,Nitesh Shah,Chaitanya Joshi,Madhvi Joshi |
| EPI_ISL_2017761 | N.H.L Municipal Medical College, Ahmedabad | Gujarat Biotechnology Research Centre | Dinesh Kumar,Twinkle Soni,Sonal Sharma ,Zuber Saiyed,Ramesh Pandit,Janvi Raval,Zarna Patel,Nitin Savaliya,Jayshri Pethani,Umang Mishra,Nitesh Shah,Chaitanya Joshi,Madhvi Joshi |
| EPI_ISL_2017762 | N.H.L Municipal Medical College, Ahmedabad | Gujarat Biotechnology Research Centre | Twinkle Soni,Sonal Sharma ,Zuber Saiyed,Ramesh Pandit,Janvi Raval,Zarna Patel,Nitin Savaliya,Dinesh Kumar,Jayshri Pethani,Umang Mishra,Nitesh Shah,Chaitanya Joshi,Madhvi Joshi |
| EPI_ISL_2017763 | VRDL, Goverment Medical College (GMC), Surat | Gujarat Biotechnology Research Centre | Sonal Sharma ,Zuber Saiyed,Ramesh Pandit,Janvi Raval,Zarna Patel,Nitin Savaliya,Dinesh Kumar,Twinkle Soni,Neeta Khandelwal,Umang Mishra,Nitesh Shah,Chaitanya Joshi,Madhvi Joshi |
| EPI_ISL_2017764 | VRDL, Goverment Medical College (GMC), Surat | Gujarat Biotechnology Research Centre | Zuber Saiyed,Ramesh Pandit,Janvi Raval,Zarna Patel,Nitin Savaliya,Dinesh Kumar,Twinkle Soni,Sonal Sharma ,Neeta Khandelwal,Umang Mishra,Nitesh Shah,Chaitanya Joshi,Madhvi Joshi |

| | | | |
|---|---|---|---|
| EPI_ISL_2017765 | BJ Medical College and Civil Hospital, Ahmedabad | Gujarat Biotechnology Research Centre | Ramesh Pandit,Janvi Raval,Zarna Patel,Nitin Savaliya,Dinesh Kumar,Twinkle Soni,Sonal Sharma ,Zuber Saiyed,Pranay Shah,Umang Mishra,Nitesh Shah,Chaitanya Joshi,Madhvi Joshi |
| EPI_ISL_2017766 | BJ Medical College and Civil Hospital, Ahmedabad | Gujarat Biotechnology Research Centre | Janvi Raval,Zarna Patel,Nitin Savaliya,Dinesh Kumar,Twinkle Soni,Sonal Sharma ,Zuber Saiyed,Ramesh Pandit,Pranay Shah,Umang Mishra,Nitesh Shah,Chaitanya Joshi,Madhvi Joshi |
| EPI_ISL_2017767 | BJ Medical College and Civil Hospital, Ahmedabad | Gujarat Biotechnology Research Centre | Zarna Patel,Nitin Savaliya,Dinesh Kumar,Twinkle Soni,Sonal Sharma ,Zuber Saiyed,Ramesh Pandit,Janvi Raval,Pranay Shah,Umang Mishra,Nitesh Shah,Chaitanya Joshi,Madhvi Joshi |
| EPI_ISL_2017768 | BJ Medical College and Civil Hospital, Ahmedabad | Gujarat Biotechnology Research Centre | Nitin Savaliya,Dinesh Kumar,Twinkle Soni,Sonal Sharma ,Zuber Saiyed,Ramesh Pandit,Janvi Raval,Zarna Patel,Pranay Shah,Umang Mishra,Nitesh Shah,Chaitanya Joshi,Madhvi Joshi |
| EPI_ISL_2017769 | BJ Medical College and Civil Hospital, Ahmedabad | Gujarat Biotechnology Research Centre | Dinesh Kumar,Twinkle Soni,Sonal Sharma ,Zuber Saiyed,Ramesh Pandit,Janvi Raval,Zarna Patel,Nitin Savaliya,Pranay Shah,Umang Mishra,Nitesh Shah,Chaitanya Joshi,Madhvi Joshi |
| EPI_ISL_2017770 | BJ Medical College and Civil Hospital, Ahmedabad | Gujarat Biotechnology Research Centre | Twinkle Soni,Sonal Sharma ,Zuber Saiyed,Ramesh Pandit,Janvi Raval,Zarna Patel,Nitin Savaliya,Dinesh Kumar,Pranay Shah,Umang Mishra,Nitesh Shah,Chaitanya Joshi,Madhvi Joshi |
| EPI_ISL_2017771 | BJ Medical College and Civil Hospital, Ahmedabad | Gujarat Biotechnology Research Centre | Sonal Sharma ,Zuber Saiyed,Ramesh Pandit,Janvi Raval,Zarna Patel,Nitin Savaliya,Dinesh Kumar,Twinkle Soni,Pranay Shah,Umang Mishra,Nitesh Shah,Chaitanya Joshi,Madhvi Joshi |
| EPI_ISL_2017772 | BJ Medical College and Civil Hospital, Ahmedabad | Gujarat Biotechnology Research Centre | Zuber Saiyed,Ramesh Pandit,Janvi Raval,Zarna Patel,Nitin Savaliya,Dinesh Kumar,Twinkle Soni,Sonal Sharma ,Pranay Shah,Umang Mishra,Nitesh Shah,Chaitanya Joshi,Madhvi Joshi |
| EPI_ISL_2017773 | BJ Medical College and Civil Hospital, Ahmedabad | Gujarat Biotechnology Research Centre | Ramesh Pandit,Janvi Raval,Zarna Patel,Nitin Savaliya,Dinesh Kumar,Twinkle Soni,Sonal Sharma ,Zuber Saiyed,Pranay Shah,Umang Mishra,Nitesh Shah,Chaitanya Joshi,Madhvi Joshi |
| EPI_ISL_2017774 | BJ Medical College and Civil Hospital, Ahmedabad | Gujarat Biotechnology Research Centre | Janvi Raval,Zarna Patel,Nitin Savaliya,Dinesh Kumar,Twinkle Soni,Sonal Sharma ,Zuber Saiyed,Ramesh Pandit,Pranay Shah,Umang Mishra,Nitesh Shah,Chaitanya Joshi,Madhvi Joshi |
| EPI_ISL_2017775 | BJ Medical College and Civil Hospital, Ahmedabad | Gujarat Biotechnology Research Centre | Zarna Patel,Nitin Savaliya,Dinesh Kumar,Twinkle Soni,Sonal Sharma ,Zuber Saiyed,Ramesh Pandit,Janvi Raval,Pranay Shah,Umang Mishra,Nitesh Shah,Chaitanya Joshi,Madhvi Joshi |
| EPI_ISL_2017776 | BJ Medical College and Civil Hospital, Ahmedabad | Gujarat Biotechnology Research Centre | Nitin Savaliya,Dinesh Kumar,Twinkle Soni,Sonal Sharma ,Zuber Saiyed,Ramesh Pandit,Janvi Raval,Zarna Patel,Pranay Shah,Umang Mishra,Nitesh Shah,Chaitanya Joshi,Madhvi Joshi |
| EPI_ISL_2017777 | BJ Medical College and Civil Hospital, Ahmedabad | Gujarat Biotechnology Research Centre | Dinesh Kumar,Twinkle Soni,Sonal Sharma ,Zuber Saiyed,Ramesh Pandit,Janvi Raval,Zarna Patel,Nitin Savaliya,Pranay Shah,Umang Mishra,Nitesh Shah,Chaitanya Joshi,Madhvi Joshi |
| EPI_ISL_2017778 | BJ Medical College and Civil Hospital, Ahmedabad | Gujarat Biotechnology Research Centre | Twinkle Soni,Sonal Sharma ,Zuber Saiyed,Ramesh Pandit,Janvi Raval,Zarna Patel,Nitin Savaliya,Dinesh Kumar,Pranay Shah,Umang Mishra,Nitesh Shah,Chaitanya Joshi,Madhvi Joshi |
| EPI_ISL_2017779 | BJ Medical College and Civil Hospital, Ahmedabad | Gujarat Biotechnology Research Centre | Sonal Sharma ,Zuber Saiyed,Ramesh Pandit,Janvi Raval,Zarna Patel,Nitin Savaliya,Dinesh Kumar,Twinkle Soni,Pranay Shah,Umang Mishra,Nitesh Shah,Chaitanya Joshi,Madhvi Joshi |
| EPI_ISL_2017780 | BJ Medical College and Civil Hospital, Ahmedabad | Gujarat Biotechnology Research Centre | Zuber Saiyed,Ramesh Pandit,Janvi Raval,Zarna Patel,Nitin Savaliya,Dinesh Kumar,Twinkle Soni,Sonal Sharma ,Pranay Shah,Umang Mishra,Nitesh Shah,Chaitanya Joshi,Madhvi Joshi |
| EPI_ISL_2017781 | BJ Medical College and Civil Hospital, Ahmedabad | Gujarat Biotechnology Research Centre | Ramesh Pandit,Janvi Raval,Zarna Patel,Nitin Savaliya,Dinesh Kumar,Twinkle Soni,Sonal Sharma ,Zuber Saiyed,Pranay Shah,Umang Mishra,Nitesh Shah,Chaitanya Joshi,Madhvi Joshi |
| EPI_ISL_2017783 | BJ Medical College and Civil Hospital, Ahmedabad | Gujarat Biotechnology Research Centre | Zarna Patel,Nitin Savaliya,Dinesh Kumar,Twinkle Soni,Sonal Sharma ,Zuber Saiyed,Ramesh Pandit,Janvi Raval,Pranay Shah,Umang Mishra,Nitesh Shah,Chaitanya Joshi,Madhvi Joshi |
| EPI_ISL_2017784 | BJ Medical College and Civil Hospital, Ahmedabad | Gujarat Biotechnology Research Centre | Nitin Savaliya,Dinesh Kumar,Twinkle Soni,Sonal Sharma ,Zuber Saiyed,Ramesh Pandit,Janvi Raval,Zarna Patel,Pranay Shah,Umang Mishra,Nitesh Shah,Chaitanya Joshi,Madhvi Joshi |
| EPI_ISL_2017785 | BJ Medical College and Civil Hospital, Ahmedabad | Gujarat Biotechnology Research Centre | Dinesh Kumar,Twinkle Soni,Sonal Sharma ,Zuber Saiyed,Ramesh Pandit,Janvi Raval,Zarna Patel,Nitin Savaliya,Pranay Shah,Umang Mishra,Nitesh Shah,Chaitanya Joshi,Madhvi Joshi |
| EPI_ISL_2017786 | BJ Medical College and Civil Hospital, Ahmedabad | Gujarat Biotechnology Research Centre | Twinkle Soni,Sonal Sharma ,Zuber Saiyed,Ramesh Pandit,Janvi Raval,Zarna Patel,Nitin Savaliya,Dinesh Kumar,Pranay Shah,Umang Mishra,Nitesh Shah,Chaitanya Joshi,Madhvi Joshi |
| EPI_ISL_2017787 | BJ Medical College and Civil Hospital, Ahmedabad | Gujarat Biotechnology Research Centre | Sonal Sharma ,Zuber Saiyed,Ramesh Pandit,Janvi Raval,Zarna Patel,Nitin Savaliya,Dinesh Kumar,Twinkle Soni,Pranay Shah,Umang Mishra,Nitesh Shah,Chaitanya Joshi,Madhvi Joshi |
| EPI_ISL_2017788 | BJ Medical College and Civil Hospital, Ahmedabad | Gujarat Biotechnology Research Centre | Zuber Saiyed,Ramesh Pandit,Janvi Raval,Zarna Patel,Nitin Savaliya,Dinesh Kumar,Twinkle Soni,Sonal Sharma ,Pranay Shah,Umang Mishra,Nitesh Shah,Chaitanya Joshi,Madhvi Joshi |
| EPI_ISL_2017789 | Gujarat Biotechnology Research Centre | Gujarat Biotechnology Research Centre | Ramesh Pandit,Janvi Raval,Zarna Patel,Nitin Savaliya,Dinesh Kumar,Twinkle Soni,Sonal Sharma ,Zuber Saiyed,Umang Mishra,Nitesh Shah,Chaitanya Joshi,Madhvi Joshi |
| EPI_ISL_2018180 | NM Dept. Health, Scientific Laboratory Division | Centers for Disease Control and Prevention Division of Viral Diseases, Pathogen Discovery | Mili Sheth, Sarah Nobles, Jasmine Padilla, Mark Burroughs, Shoshona Le, Katie Dillon, Peter Cook, Clinton R. Paden, Dhwani Batra, Krista Queen, Kristen Knipe, Dakota Howard, Yvette Unoarumhi, Darlene Wagner, Matthew Schmerer, Ben L. Rambo-Martin, Kristine Lacek, Sam Shepard, Alison Laufer Halpin, Dave Wentworth, Vivien Dugan, Suxiang Tong, Justin Lee |
| EPI_ISL_2018388 | UT-Unified State Labs: Public Health Utah DOH | Centers for Disease Control and Prevention Division of Viral Diseases, Pathogen Discovery | Mili Sheth, Sarah Nobles, Jasmine Padilla, Mark Burroughs, Shoshona Le, Katie Dillon, Peter Cook, Clinton R. Paden, Dhwani Batra, Krista Queen, Kristen Knipe, Dakota Howard, Yvette Unoarumhi, Darlene Wagner, Matthew Schmerer, Ben L. Rambo-Martin, Kristine Lacek, Sam Shepard, Alison Laufer Halpin, Dave Wentworth, Vivien Dugan, Suxiang Tong, Justin Lee |
| EPI_ISL_2018995, EPI_ISL_2019465, EPI_ISL_2019535 | Viollier AG | Department of Biosystems Science and Engineering, ETH Zürich | Chaoran Chen, Sarah Nadeau, Catharine Aquino, Ivan Topolsky, Philipp Jablonski, Lara Fuhrmann, David Dreifuss, Katharina Jahn, Andreia Cabral de Gouvea, Maria Domenica Moccia, Simon Grüter, Timothy Sykes, Lennart Opitz, Griffin White, Laura Neff, Doris Popovic, Andrea Patrignani, Jay Tracy, Ralph Schlapbach, Christiane Beckmann, Maurice Redondo, Olivier Kobel, Christoph Noppen, Sophie Seidel, Noemie Santamaria de Souza, Niko Beerenwinkel, Tanja Stadler |
| EPI_ISL_2020930 | Urban Public Health Centre (UPHC), SECTOR-2, Gandhinagar | Gujarat Biotechnology Research Centre | Amit Kanani,Twinkle Soni,Sonal Sharma ,Zuber Saiyed,Ramesh Pandit,Janvi Raval,Zarna Patel,Nitin Savaliya,Dinesh Kumar,Bhargav Waghela,Bhumika Prajapati,Tejendra Purohit ,Nilen Patel ,Kalpesh Goswami,Amrutlal Patel,Umang Mishra,Nitesh Shah,Chaitanya Joshi,Madhvi Joshi |
| EPI_ISL_2020931 | Urban Public Health Centre (UPHC), SECTOR-2, Gandhinagar | Gujarat Biotechnology Research Centre | Amit Kanani,Sonal Sharma ,Zuber Saiyed,Ramesh Pandit,Janvi Raval,Zarna Patel,Nitin Savaliya,Dinesh Kumar,Twinkle Soni,Bhargav Waghela,Bhumika Prajapati,Tejendra Purohit ,Nilen Patel ,Kalpesh Goswami,Amrutlal Patel,Umang Mishra,Nitesh Shah,Chaitanya Joshi,Madhvi Joshi |
| EPI_ISL_2020932 | Urban Public Health Centre (UPHC), SECTOR-2, Gandhinagar | Gujarat Biotechnology Research Centre | Zuber Saiyed,Ramesh Pandit,Janvi Raval,Zarna Patel,Nitin Savaliya,Dinesh Kumar,Twinkle Soni,Sonal Sharma ,Bhargav Waghela,Bhumika Prajapati,Tejendra Purohit ,Nilen Patel ,Amit Kanani,Kalpesh Goswami,Amrutlal Patel,Umang Mishra,Nitesh Shah,Chaitanya Joshi,Madhvi Joshi |
| EPI_ISL_2020933 | Urban Public Health Centre (UPHC), SECTOR-2, Gandhinagar | Gujarat Biotechnology Research Centre | Ramesh Pandit,Janvi Raval,Zarna Patel,Nitin Savaliya,Dinesh Kumar,Twinkle Soni,Sonal Sharma ,Zuber Saiyed,Bhargav Waghela,Bhumika Prajapati,Tejendra Purohit ,Nilen Patel ,Amit Kanani,Kalpesh Goswami,Amrutlal Patel,Umang Mishra,Nitesh Shah,Chaitanya Joshi,Madhvi Joshi |
| EPI_ISL_2020934 | Urban Public Health Centre (UPHC), SECTOR-2, Gandhinagar | Gujarat Biotechnology Research Centre | Janvi Raval,Zarna Patel,Nitin Savaliya,Dinesh Kumar,Twinkle Soni,Sonal Sharma ,Zuber Saiyed,Ramesh Pandit,Bhargav Waghela,Bhumika Prajapati,Tejendra Purohit ,Nilen Patel ,Amit Kanani,Kalpesh Goswami,Amrutlal Patel,Umang Mishra,Nitesh Shah,Chaitanya Joshi,Madhvi Joshi |
| EPI_ISL_2020935 | Urban Public Health Centre (UPHC), SECTOR-2, Gandhinagar | Gujarat Biotechnology Research Centre | Zarna Patel,Nitin Savaliya,Dinesh Kumar,Twinkle Soni,Sonal Sharma ,Zuber Saiyed,Ramesh Pandit,Janvi Raval,Bhargav Waghela,Bhumika Prajapati,Tejendra Purohit ,Nilen Patel ,Amit Kanani,Kalpesh Goswami,Amrutlal Patel,Umang Mishra,Nitesh Shah,Chaitanya Joshi,Madhvi Joshi |
| EPI_ISL_2020936 | Urban Public Health Centre (UPHC), SECTOR-2, Gandhinagar | Gujarat Biotechnology Research Centre | Nitin Savaliya,Dinesh Kumar,Twinkle Soni,Sonal Sharma ,Zuber Saiyed,Ramesh Pandit,Janvi Raval,Zarna Patel,Bhargav Waghela,Bhumika Prajapati,Tejendra Purohit ,Nilen Patel ,Amit Kanani,Kalpesh Goswami,Amrutlal Patel,Umang Mishra,Nitesh Shah,Chaitanya Joshi,Madhvi Joshi |
| EPI_ISL_2020937 | Urban Public Health Centre (UPHC), SECTOR-2, Gandhinagar | Gujarat Biotechnology Research Centre | Dinesh Kumar,Twinkle Soni,Sonal Sharma ,Zuber Saiyed,Ramesh Pandit,Janvi Raval,Zarna Patel,Nitin Savaliya,Bhargav Waghela,Bhumika Prajapati,Tejendra Purohit ,Nilen Patel ,Amit Kanani,Kalpesh Goswami,Amrutlal Patel,Umang Mishra,Nitesh Shah,Chaitanya Joshi,Madhvi Joshi |
| EPI_ISL_2020938 | Urban Public Health Centre (UPHC), SECTOR-2, Gandhinagar | Gujarat Biotechnology Research Centre | Twinkle Soni,Sonal Sharma ,Zuber Saiyed,Ramesh Pandit,Janvi Raval,Zarna Patel,Nitin Savaliya,Dinesh Kumar,Bhargav Waghela,Bhumika Prajapati,Tejendra Purohit ,Nilen Patel ,Amit Kanani,Kalpesh Goswami,Amrutlal Patel,Umang Mishra,Nitesh Shah,Chaitanya Joshi,Madhvi Joshi |
| EPI_ISL_2020939 | Urban Public Health Centre (UPHC), SECTOR-2, Gandhinagar | Gujarat Biotechnology Research Centre | Bhargav Waghela,Sonal Sharma ,Zuber Saiyed,Ramesh Pandit,Janvi Raval,Zarna Patel,Nitin Savaliya,Dinesh Kumar,Twinkle Soni,Bhumika Prajapati,Tejendra Purohit ,Nilen Patel ,Amit Kanani,Kalpesh Goswami,Amrutlal Patel,Umang Mishra,Nitesh Shah,Chaitanya Joshi,Madhvi Joshi |
| EPI_ISL_2020940 | Urban Public Health Centre (UPHC), SECTOR-2, Gandhinagar | Gujarat Biotechnology Research Centre | Bhargav Waghela,Zuber Saiyed,Ramesh Pandit,Janvi Raval,Zarna Patel,Nitin Savaliya,Dinesh Kumar,Twinkle Soni,Sonal Sharma ,Bhumika Prajapati,Tejendra Purohit ,Nilen Patel ,Amit Kanani,Kalpesh Goswami,Amrutlal Patel,Umang Mishra,Nitesh Shah,Chaitanya Joshi,Madhvi Joshi |
| EPI_ISL_2020941 | Urban Public Health Centre (UPHC), SECTOR-2, Gandhinagar | Gujarat Biotechnology Research Centre | Ramesh Pandit,Janvi Raval,Zarna Patel,Nitin Savaliya,Dinesh Kumar,Twinkle Soni,Sonal Sharma ,Zuber Saiyed,Bhargav Waghela,Bhumika Prajapati,Tejendra Purohit ,Nilen Patel ,Amit Kanani,Kalpesh Goswami,Amrutlal Patel,Umang Mishra,Nitesh Shah,Chaitanya Joshi,Madhvi Joshi |
| EPI_ISL_2020942 | Urban Public Health Centre (UPHC), SECTOR-2, Gandhinagar | Gujarat Biotechnology Research Centre | Janvi Raval,Zarna Patel,Nitin Savaliya,Dinesh Kumar,Twinkle Soni,Sonal Sharma ,Zuber Saiyed,Ramesh Pandit,Bhargav Waghela,Bhumika Prajapati,Tejendra Purohit ,Nilen Patel ,Amit Kanani,Kalpesh Goswami,Amrutlal Patel,Umang Mishra,Nitesh Shah,Chaitanya Joshi,Madhvi Joshi |
| EPI_ISL_2020943 | Urban Public Health Centre (UPHC), SECTOR-2, Gandhinagar | Gujarat Biotechnology Research Centre | Zarna Patel,Nitin Savaliya,Dinesh Kumar,Twinkle Soni,Sonal Sharma ,Zuber Saiyed,Ramesh Pandit,Janvi Raval,Bhargav Waghela,Bhumika Prajapati,Tejendra Purohit ,Nilen Patel ,Amit Kanani,Kalpesh Goswami,Amrutlal Patel,Umang Mishra,Nitesh Shah,Chaitanya Joshi,Madhvi Joshi |
| EPI_ISL_2020944 | Urban Public Health Centre (UPHC), SECTOR-2, Gandhinagar | Gujarat Biotechnology Research Centre | Nitin Savaliya,Dinesh Kumar,Twinkle Soni,Sonal Sharma ,Zuber Saiyed,Ramesh Pandit,Janvi Raval,Zarna Patel,Bhargav Waghela,Bhumika Prajapati,Tejendra Purohit ,Nilen Patel ,Amit Kanani,Kalpesh Goswami,Amrutlal Patel,Umang Mishra,Nitesh Shah,Chaitanya Joshi,Madhvi Joshi |
| EPI_ISL_2020945 | Urban Public Health Centre (UPHC), SECTOR-2, Gandhinagar | Gujarat Biotechnology Research Centre | Bhumika Prajapati,Dinesh Kumar,Twinkle Soni,Sonal Sharma ,Zuber Saiyed,Ramesh Pandit,Janvi Raval,Zarna Patel,Nitin Savaliya,Bhargav Waghela,Tejendra Purohit ,Nilen Patel ,Amit Kanani,Kalpesh Goswami,Amrutlal Patel,Umang Mishra,Nitesh Shah,Chaitanya Joshi,Madhvi Joshi |
| EPI_ISL_2020946 | Urban Public Health Centre (UPHC), SECTOR-2, Gandhinagar | Gujarat Biotechnology Research Centre | Bhumika Prajapati,Twinkle Soni,Sonal Sharma ,Zuber Saiyed,Ramesh Pandit,Janvi Raval,Zarna Patel,Nitin Savaliya,Dinesh Kumar,Bhargav Waghela,Tejendra Purohit ,Nilen Patel ,Amit Kanani,Kalpesh Goswami,Amrutlal Patel,Umang Mishra,Nitesh Shah,Chaitanya Joshi,Madhvi Joshi |
| EPI_ISL_2020947 | Urban Public Health Centre (UPHC), SECTOR-2, Gandhinagar | Gujarat Biotechnology Research Centre | Sonal Sharma ,Zuber Saiyed,Ramesh Pandit,Janvi Raval,Zarna Patel,Nitin Savaliya,Dinesh Kumar,Twinkle Soni,Bhargav Waghela,Bhumika Prajapati,Tejendra Purohit ,Nilen Patel ,Amit Kanani,Kalpesh Goswami,Amrutlal Patel,Umang Mishra,Nitesh Shah,Chaitanya Joshi,Madhvi Joshi |
| EPI_ISL_2020948 | Urban Public Health Centre (UPHC), SECTOR-2, Gandhinagar | Gujarat Biotechnology Research Centre | Tejendra Purohit ,Zuber Saiyed,Ramesh Pandit,Janvi Raval,Zarna Patel,Nitin Savaliya,Dinesh Kumar,Twinkle Soni,Sonal Sharma ,Bhargav Waghela,Bhumika Prajapati,Nilen Patel ,Amit Kanani,Kalpesh Goswami,Amrutlal Patel,Umang Mishra,Nitesh Shah,Chaitanya Joshi,Madhvi Joshi |
| EPI_ISL_2020949 | Urban Public Health Centre (UPHC), SECTOR-2, Gandhinagar | Gujarat Biotechnology Research Centre | Tejendra Purohit ,Ramesh Pandit,Janvi Raval,Zarna Patel,Nitin Savaliya,Dinesh Kumar,Twinkle Soni,Sonal Sharma ,Zuber Saiyed,Bhargav Waghela,Bhumika Prajapati,Nilen Patel ,Amit Kanani,Kalpesh Goswami,Amrutlal Patel,Umang Mishra,Nitesh Shah,Chaitanya Joshi,Madhvi Joshi |
| EPI_ISL_2020950 | Urban Public Health Centre (UPHC), SECTOR-2, Gandhinagar | Gujarat Biotechnology Research Centre | Janvi Raval,Zarna Patel,Nitin Savaliya,Dinesh Kumar,Twinkle Soni,Sonal Sharma ,Zuber Saiyed,Ramesh Pandit,Bhargav Waghela,Bhumika Prajapati,Tejendra Purohit ,Nilen Patel ,Amit Kanani,Kalpesh Goswami,Amrutlal Patel,Umang Mishra,Nitesh Shah,Chaitanya Joshi,Madhvi Joshi |
| EPI_ISL_2020951 | Urban Public Health Centre (UPHC), SECTOR-2, Gandhinagar | Gujarat Biotechnology Research Centre | Zarna Patel,Nitin Savaliya,Dinesh Kumar,Twinkle Soni,Sonal Sharma ,Zuber Saiyed,Ramesh Pandit,Janvi Raval,Bhargav Waghela,Bhumika Prajapati,Tejendra Purohit ,Nilen Patel ,Amit Kanani,Kalpesh Goswami,Amrutlal Patel,Umang Mishra,Nitesh Shah,Chaitanya Joshi,Madhvi Joshi |
| EPI_ISL_2020952 | Urban Public Health Centre (UPHC), SECTOR-2, Gandhinagar | Gujarat Biotechnology Research Centre | Nitin Savaliya,Dinesh Kumar,Twinkle Soni,Sonal Sharma ,Zuber Saiyed,Ramesh Pandit,Janvi Raval,Zarna Patel,Bhargav Waghela,Bhumika Prajapati,Tejendra Purohit ,Nilen Patel ,Amit Kanani,Kalpesh Goswami,Amrutlal Patel,Umang Mishra,Nitesh Shah,Chaitanya Joshi,Madhvi Joshi |
| EPI_ISL_2020953 | Urban Public Health Centre (UPHC), SECTOR-2, Gandhinagar | Gujarat Biotechnology Research Centre | Dinesh Kumar,Twinkle Soni,Sonal Sharma ,Zuber Saiyed,Ramesh Pandit,Janvi Raval,Zarna Patel,Nitin Savaliya,Bhargav Waghela,Bhumika Prajapati,Tejendra Purohit ,Nilen Patel ,Amit Kanani,Kalpesh Goswami,Amrutlal Patel,Umang Mishra,Nitesh Shah,Chaitanya Joshi,Madhvi Joshi |
| EPI_ISL_2020954 | Urban Public Health Centre (UPHC), SECTOR-2, Gandhinagar | Gujarat Biotechnology Research Centre | Twinkle Soni,Sonal Sharma ,Zuber Saiyed,Ramesh Pandit,Janvi Raval,Zarna Patel,Nitin Savaliya,Dinesh Kumar,Bhargav Waghela,Bhumika Prajapati,Tejendra Purohit ,Nilen Patel ,Amit Kanani,Kalpesh Goswami,Amrutlal Patel,Umang Mishra,Nitesh Shah,Chaitanya Joshi,Madhvi Joshi |
| EPI_ISL_2020955 | Urban Public Health Centre (UPHC), SECTOR-2, Gandhinagar | Gujarat Biotechnology Research Centre | Nilen Patel ,Sonal Sharma ,Zuber Saiyed,Ramesh Pandit,Janvi Raval,Zarna Patel,Nitin Savaliya,Dinesh Kumar,Twinkle Soni,Bhargav Waghela,Bhumika Prajapati,Tejendra Purohit ,Amit Kanani,Kalpesh Goswami,Amrutlal Patel,Umang Mishra,Nitesh Shah,Chaitanya Joshi,Madhvi Joshi |
| EPI_ISL_2020956 | Urban Public Health Centre (UPHC), SECTOR-2, Gandhinagar | Gujarat Biotechnology Research Centre | Nilen Patel ,Zuber Saiyed,Ramesh Pandit,Janvi Raval,Zarna Patel,Nitin Savaliya,Dinesh Kumar,Twinkle Soni,Sonal Sharma ,Bhargav Waghela,Bhumika Prajapati,Tejendra Purohit ,Amit Kanani,Kalpesh Goswami,Amrutlal Patel,Umang Mishra,Nitesh Shah,Chaitanya Joshi,Madhvi Joshi |
| EPI_ISL_2020957 | Urban Public Health Centre (UPHC), SECTOR-2, Gandhinagar | Gujarat Biotechnology Research Centre | Kalpesh Goswami,Nitin Savaliya,Dinesh Kumar,Twinkle Soni,Sonal Sharma ,Zuber Saiyed,Ramesh Pandit,Janvi Raval,Zarna Patel,Bhargav Waghela,Bhumika Prajapati,Tejendra Purohit ,Nilen Patel ,Amit Kanani,Amrutlal Patel,Umang |

| | | | |
|---|---|---|---|
| | | | Mishra,Nitesh Shah,Chaitanya Joshi,Madhvi Joshi |
| EPI_ISL_2020958 | Urban Public Health Centre (UPHC), SECTOR-2, Gandhinagar | Gujarat Biotechnology Research Centre | Kalpesh Goswami,Dinesh Kumar,Twinkle Soni,Sonal Sharma ,Zuber Saiyed,Ramesh Pandit,Janvi Raval,Zarna Patel,Nitin Savaliya,Bhargav Waghela,Bhumika Prajapati,Tejendra Purohit ,Nilen Patel ,Amit Kanani,Amrutlal Patel,Umang Mishra,Nitesh Shah,Chaitanya Joshi,Madhvi Joshi |
| EPI_ISL_2020959 | Urban Public Health Centre (UPHC), SECTOR-2, Gandhinagar | Gujarat Biotechnology Research Centre | Amrutlal Patel,Twinkle Soni,Sonal Sharma ,Zuber Saiyed,Ramesh Pandit,Janvi Raval,Zarna Patel,Nitin Savaliya,Dinesh Kumar,Bhargav Waghela,Bhumika Prajapati,Tejendra Purohit ,Nilen Patel ,Amit Kanani,Kalpesh Goswami,Umang Mishra,Nitesh Shah,Chaitanya Joshi,Madhvi Joshi |
| EPI_ISL_2020960 | Urban Public Health Centre (UPHC), SECTOR-2, Gandhinagar | Gujarat Biotechnology Research Centre | Amrutlal Patel,Sonal Sharma ,Zuber Saiyed,Ramesh Pandit,Janvi Raval,Zarna Patel,Nitin Savaliya,Dinesh Kumar,Twinkle Soni,Bhargav Waghela,Bhumika Prajapati,Tejendra Purohit ,Nilen Patel ,Amit Kanani,Kalpesh Goswami,Umang Mishra,Nitesh Shah,Chaitanya Joshi,Madhvi Joshi |
| EPI_ISL_2021047 | Lighthouse Lab in Milton Keynes | Wellcome Sanger Institute for the COVID-19 Genomics UK (COG-UK) Consortium | The Lighthouse Lab in Milton Keynes and Alex Alderton, Roberto Amato, Jeffrey Barrett, Sonia Goncalves, Ewan Harrison, David K. Jackson, Ian Johnston, Dominic Kwiatkowski, Cordelia Langford, John Sillitoe on behalf of the Wellcome Sanger Institute COVID-19 Surveillance Team |
| EPI_ISL_2021051 | Lighthouse Lab in Glasgow | Wellcome Sanger Institute for the COVID-19 Genomics UK (COG-UK) Consortium | Harper VanSteenhouse, Yumi Kasai, David Gray, Carol Clugston, Anna Dominiczak and Alex Alderton, Roberto Amato, Jeffrey Barrett, Sonia Goncalves, Ewan Harrison, David K. Jackson, Ian Johnston, Dominic Kwiatkowski, Cordelia Langford, John Sillitoe on behalf of the Wellcome Sanger Institute COVID-19 Surveillance Team |
| EPI_ISL_2021053 | Lighthouse Lab in Alderley Park | Wellcome Sanger Institute for the COVID-19 Genomics UK (COG-UK) Consortium | Jacquelyn Wynn, Mairead Hyland, The Lighthouse Lab in Alderley Park and Alex Alderton, Roberto Amato, Jeffrey Barrett, Sonia Goncalves, Ewan Harrison, David K. Jackson, Ian Johnston, Dominic Kwiatkowski, Cordelia Langford, John Sillitoe on behalf of the Wellcome Sanger Institute COVID-19 Surveillance Team |
| EPI_ISL_2021055 | Lighthouse Lab in Milton Keynes | Wellcome Sanger Institute for the COVID-19 Genomics UK (COG-UK) Consortium | The Lighthouse Lab in Milton Keynes and Alex Alderton, Roberto Amato, Jeffrey Barrett, Sonia Goncalves, Ewan Harrison, David K. Jackson, Ian Johnston, Dominic Kwiatkowski, Cordelia Langford, John Sillitoe on behalf of the Wellcome Sanger Institute COVID-19 Surveillance Team |
| EPI_ISL_2021061 | Lighthouse Lab in Alderley Park | Wellcome Sanger Institute for the COVID-19 Genomics UK (COG-UK) Consortium | Jacquelyn Wynn, Mairead Hyland, The Lighthouse Lab in Alderley Park and Alex Alderton, Roberto Amato, Jeffrey Barrett, Sonia Goncalves, Ewan Harrison, David K. Jackson, Ian Johnston, Dominic Kwiatkowski, Cordelia Langford, John Sillitoe on behalf of the Wellcome Sanger Institute COVID-19 Surveillance Team |
| EPI_ISL_2021071 | Lighthouse Lab in Milton Keynes | Wellcome Sanger Institute for the COVID-19 Genomics UK (COG-UK) Consortium | The Lighthouse Lab in Milton Keynes and Alex Alderton, Roberto Amato, Jeffrey Barrett, Sonia Goncalves, Ewan Harrison, David K. Jackson, Ian Johnston, Dominic Kwiatkowski, Cordelia Langford, John Sillitoe on behalf of the Wellcome Sanger Institute COVID-19 Surveillance Team |
| EPI_ISL_2021078, EPI_ISL_2021079 | Lighthouse Lab in Glasgow | Wellcome Sanger Institute for the COVID-19 Genomics UK (COG-UK) Consortium | Harper VanSteenhouse, Yumi Kasai, David Gray, Carol Clugston, Anna Dominiczak and Alex Alderton, Roberto Amato, Jeffrey Barrett, Sonia Goncalves, Ewan Harrison, David K. Jackson, Ian Johnston, Dominic Kwiatkowski, Cordelia Langford, John Sillitoe on behalf of the Wellcome Sanger Institute COVID-19 Surveillance Team |
| EPI_ISL_2021083 | Lighthouse Lab in Alderley Park | Wellcome Sanger Institute for the COVID-19 Genomics UK (COG-UK) Consortium | Jacquelyn Wynn, Mairead Hyland, The Lighthouse Lab in Alderley Park and Alex Alderton, Roberto Amato, Jeffrey Barrett, Sonia Goncalves, Ewan Harrison, David K. Jackson, Ian Johnston, Dominic Kwiatkowski, Cordelia Langford, John Sillitoe on behalf of the Wellcome Sanger Institute COVID-19 Surveillance Team |
| EPI_ISL_2021098 | Lighthouse Lab in Glasgow | Wellcome Sanger Institute for the COVID-19 Genomics UK (COG-UK) Consortium | Harper VanSteenhouse, Yumi Kasai, David Gray, Carol Clugston, Anna Dominiczak and Alex Alderton, Roberto Amato, Jeffrey Barrett, Sonia Goncalves, Ewan Harrison, David K. Jackson, Ian Johnston, Dominic Kwiatkowski, Cordelia Langford, John Sillitoe on behalf of the Wellcome Sanger Institute COVID-19 Surveillance Team |
| EPI_ISL_2021104 | Lighthouse Lab in Milton Keynes | Wellcome Sanger Institute for the COVID-19 Genomics UK (COG-UK) Consortium | The Lighthouse Lab in Milton Keynes and Alex Alderton, Roberto Amato, Jeffrey Barrett, Sonia Goncalves, Ewan Harrison, David K. Jackson, Ian Johnston, Dominic Kwiatkowski, Cordelia Langford, John Sillitoe on behalf of the Wellcome Sanger Institute COVID-19 Surveillance Team |
| EPI_ISL_2021108 | Lighthouse Lab in Glasgow | Wellcome Sanger Institute for the COVID-19 Genomics UK (COG-UK) Consortium | Harper VanSteenhouse, Yumi Kasai, David Gray, Carol Clugston, Anna Dominiczak and Alex Alderton, Roberto Amato, Jeffrey Barrett, Sonia Goncalves, Ewan Harrison, David K. Jackson, Ian Johnston, Dominic Kwiatkowski, Cordelia Langford, John Sillitoe on behalf of the Wellcome Sanger Institute COVID-19 Surveillance Team |
| EPI_ISL_2021109 | Lighthouse Lab in Milton Keynes | Wellcome Sanger Institute for the COVID-19 Genomics UK (COG-UK) Consortium | The Lighthouse Lab in Milton Keynes and Alex Alderton, Roberto Amato, Jeffrey Barrett, Sonia Goncalves, Ewan Harrison, David K. Jackson, Ian Johnston, Dominic Kwiatkowski, Cordelia Langford, John Sillitoe on behalf of the Wellcome Sanger Institute COVID-19 Surveillance Team |
| EPI_ISL_2021110, EPI_ISL_2021111, EPI_ISL_2021119, EPI_ISL_2021124 | Lighthouse Lab in Alderley Park | Wellcome Sanger Institute for the COVID-19 Genomics UK (COG-UK) Consortium | Jacquelyn Wynn, Mairead Hyland, The Lighthouse Lab in Alderley Park and Alex Alderton, Roberto Amato, Jeffrey Barrett, Sonia Goncalves, Ewan Harrison, David K. Jackson, Ian Johnston, Dominic Kwiatkowski, Cordelia Langford, John Sillitoe on behalf of the Wellcome Sanger Institute COVID-19 Surveillance Team |
| EPI_ISL_2021132 | Lighthouse Lab in Milton Keynes | Wellcome Sanger Institute for the COVID-19 Genomics UK (COG-UK) Consortium | The Lighthouse Lab in Milton Keynes and Alex Alderton, Roberto Amato, Jeffrey Barrett, Sonia Goncalves, Ewan Harrison, David K. Jackson, Ian Johnston, Dominic Kwiatkowski, Cordelia Langford, John Sillitoe on behalf of the Wellcome Sanger Institute COVID-19 Surveillance Team |
| EPI_ISL_2021144 | Lighthouse Lab in Alderley Park | Wellcome Sanger Institute for the COVID-19 Genomics UK (COG-UK) Consortium | Jacquelyn Wynn, Mairead Hyland, The Lighthouse Lab in Alderley Park and Alex Alderton, Roberto Amato, Jeffrey Barrett, Sonia Goncalves, Ewan Harrison, David K. Jackson, Ian Johnston, Dominic Kwiatkowski, Cordelia Langford, John Sillitoe on behalf of the Wellcome Sanger Institute COVID-19 Surveillance Team |
| EPI_ISL_2021150 | Lighthouse Lab in Glasgow | Wellcome Sanger Institute for the COVID-19 Genomics UK (COG-UK) Consortium | Harper VanSteenhouse, Yumi Kasai, David Gray, Carol Clugston, Anna Dominiczak and Alex Alderton, Roberto Amato, Jeffrey Barrett, Sonia Goncalves, Ewan Harrison, David K. Jackson, Ian Johnston, Dominic Kwiatkowski, Cordelia Langford, John Sillitoe on behalf of the Wellcome Sanger Institute COVID-19 Surveillance Team |
| EPI_ISL_2021152, EPI_ISL_2021154 | Lighthouse Lab in Alderley Park | Wellcome Sanger Institute for the COVID-19 Genomics UK (COG-UK) Consortium | Jacquelyn Wynn, Mairead Hyland, The Lighthouse Lab in Alderley Park and Alex Alderton, Roberto Amato, Jeffrey Barrett, Sonia Goncalves, Ewan Harrison, David K. Jackson, Ian Johnston, Dominic Kwiatkowski, Cordelia Langford, John Sillitoe on behalf of the Wellcome Sanger Institute COVID-19 Surveillance Team |
| EPI_ISL_2021167, EPI_ISL_2021171 | Lighthouse Lab in Milton Keynes | Wellcome Sanger Institute for the COVID-19 Genomics UK (COG-UK) Consortium | The Lighthouse Lab in Milton Keynes and Alex Alderton, Roberto Amato, Jeffrey Barrett, Sonia Goncalves, Ewan Harrison, David K. Jackson, Ian Johnston, Dominic Kwiatkowski, Cordelia Langford, John Sillitoe on behalf of the Wellcome Sanger Institute COVID-19 Surveillance Team |
| EPI_ISL_2021183 | Lighthouse Lab in Alderley Park | Wellcome Sanger Institute for the COVID-19 Genomics UK (COG-UK) Consortium | Jacquelyn Wynn, Mairead Hyland, The Lighthouse Lab in Alderley Park and Alex Alderton, Roberto Amato, Jeffrey Barrett, Sonia Goncalves, Ewan Harrison, David K. Jackson, Ian Johnston, Dominic Kwiatkowski, Cordelia Langford, John Sillitoe on behalf of the Wellcome Sanger Institute COVID-19 Surveillance Team |
| EPI_ISL_2021184 | Lighthouse Lab in Glasgow | Wellcome Sanger Institute for the COVID-19 Genomics UK (COG-UK) Consortium | Harper VanSteenhouse, Yumi Kasai, David Gray, Carol Clugston, Anna Dominiczak and Alex Alderton, Roberto Amato, Jeffrey Barrett, Sonia Goncalves, Ewan Harrison, David K. Jackson, Ian Johnston, Dominic Kwiatkowski, Cordelia Langford, John Sillitoe on behalf of the Wellcome Sanger Institute COVID-19 Surveillance Team |
| EPI_ISL_2021189 | Lighthouse Lab in Milton Keynes | Wellcome Sanger Institute for the COVID-19 Genomics UK (COG-UK) Consortium | The Lighthouse Lab in Milton Keynes and Alex Alderton, Roberto Amato, Jeffrey Barrett, Sonia Goncalves, Ewan Harrison, David K. Jackson, Ian Johnston, Dominic Kwiatkowski, Cordelia Langford, John Sillitoe on behalf of the Wellcome Sanger Institute COVID-19 Surveillance Team |
| EPI_ISL_2021193, EPI_ISL_2021195 | Lighthouse Lab in Alderley Park | Wellcome Sanger Institute for the COVID-19 Genomics UK (COG-UK) Consortium | Jacquelyn Wynn, Mairead Hyland, The Lighthouse Lab in Alderley Park and Alex Alderton, Roberto Amato, Jeffrey Barrett, Sonia Goncalves, Ewan Harrison, David K. Jackson, Ian Johnston, Dominic Kwiatkowski, Cordelia Langford, John Sillitoe on behalf of the Wellcome Sanger Institute COVID-19 Surveillance Team |
| EPI_ISL_2021211 | Lighthouse Lab in Glasgow | Wellcome Sanger Institute for the COVID-19 Genomics UK (COG-UK) Consortium | Harper VanSteenhouse, Yumi Kasai, David Gray, Carol Clugston, Anna Dominiczak and Alex Alderton, Roberto Amato, Jeffrey Barrett, Sonia Goncalves, Ewan Harrison, David K. Jackson, Ian Johnston, Dominic Kwiatkowski, Cordelia Langford, John Sillitoe on behalf of the Wellcome Sanger Institute COVID-19 Surveillance Team |
| EPI_ISL_2021215, EPI_ISL_2021218 | Lighthouse Lab in Alderley Park | Wellcome Sanger Institute for the COVID-19 Genomics UK (COG-UK) Consortium | Jacquelyn Wynn, Mairead Hyland, The Lighthouse Lab in Alderley Park and Alex Alderton, Roberto Amato, Jeffrey Barrett, Sonia Goncalves, Ewan Harrison, David K. Jackson, Ian Johnston, Dominic Kwiatkowski, Cordelia Langford, John Sillitoe on behalf of the Wellcome Sanger Institute COVID-19 Surveillance Team |
| EPI_ISL_2021219, EPI_ISL_2021224 | Lighthouse Lab in Milton Keynes | Wellcome Sanger Institute for the COVID-19 Genomics UK (COG-UK) Consortium | The Lighthouse Lab in Milton Keynes and Alex Alderton, Roberto Amato, Jeffrey Barrett, Sonia Goncalves, Ewan Harrison, David K. Jackson, Ian Johnston, Dominic Kwiatkowski, Cordelia Langford, John Sillitoe on behalf of the Wellcome Sanger Institute COVID-19 Surveillance Team |
| EPI_ISL_2021231 | Lighthouse Lab in Alderley Park | Wellcome Sanger Institute for the COVID-19 Genomics UK (COG-UK) Consortium | Jacquelyn Wynn, Mairead Hyland, The Lighthouse Lab in Alderley Park and Alex Alderton, Roberto Amato, Jeffrey Barrett, Sonia Goncalves, Ewan Harrison, David K. Jackson, Ian Johnston, Dominic Kwiatkowski, Cordelia Langford, John Sillitoe on behalf of the Wellcome Sanger Institute COVID-19 Surveillance Team |
| EPI_ISL_2021232, EPI_ISL_2021238 | Lighthouse Lab in Glasgow | Wellcome Sanger Institute for the COVID-19 Genomics UK (COG-UK) Consortium | Harper VanSteenhouse, Yumi Kasai, David Gray, Carol Clugston, Anna Dominiczak and Alex Alderton, Roberto Amato, Jeffrey Barrett, Sonia Goncalves, Ewan Harrison, David K. Jackson, Ian Johnston, Dominic Kwiatkowski, Cordelia Langford, John Sillitoe on behalf of the Wellcome Sanger Institute COVID-19 Surveillance Team |
| EPI_ISL_2021248, EPI_ISL_2021249 | Lighthouse Lab in Alderley Park | Wellcome Sanger Institute for the COVID-19 Genomics UK (COG-UK) Consortium | Jacquelyn Wynn, Mairead Hyland, The Lighthouse Lab in Alderley Park and Alex Alderton, Roberto Amato, Jeffrey Barrett, Sonia Goncalves, Ewan Harrison, David K. Jackson, Ian Johnston, Dominic Kwiatkowski, Cordelia Langford, John Sillitoe on behalf of the Wellcome Sanger Institute COVID-19 Surveillance Team |
| EPI_ISL_2021253 | Lighthouse Lab in Milton Keynes | Wellcome Sanger Institute for the COVID-19 Genomics UK (COG-UK) Consortium | The Lighthouse Lab in Milton Keynes and Alex Alderton, Roberto Amato, Jeffrey Barrett, Sonia Goncalves, Ewan Harrison, David K. Jackson, Ian Johnston, Dominic Kwiatkowski, Cordelia Langford, John Sillitoe on behalf of the Wellcome Sanger Institute COVID-19 Surveillance Team |
| EPI_ISL_2021255 | Lighthouse Lab in Alderley Park | Wellcome Sanger Institute for the COVID-19 Genomics UK (COG-UK) Consortium | Jacquelyn Wynn, Mairead Hyland, The Lighthouse Lab in Alderley Park and Alex Alderton, Roberto Amato, Jeffrey Barrett, Sonia Goncalves, Ewan Harrison, David K. Jackson, Ian Johnston, Dominic Kwiatkowski, Cordelia Langford, John Sillitoe on behalf of the Wellcome Sanger Institute COVID-19 Surveillance Team |
| EPI_ISL_2021270 | Lighthouse Lab in Milton Keynes | Wellcome Sanger Institute for the COVID-19 Genomics UK (COG-UK) Consortium | The Lighthouse Lab in Milton Keynes and Alex Alderton, Roberto Amato, Jeffrey Barrett, Sonia Goncalves, Ewan Harrison, David K. Jackson, Ian Johnston, Dominic Kwiatkowski, Cordelia Langford, John Sillitoe on behalf of the Wellcome Sanger Institute COVID-19 Surveillance Team |
| EPI_ISL_2021271, EPI_ISL_2021275, EPI_ISL_2021280, EPI_ISL_2021281 | Lighthouse Lab in Alderley Park | Wellcome Sanger Institute for the COVID-19 Genomics UK (COG-UK) Consortium | Jacquelyn Wynn, Mairead Hyland, The Lighthouse Lab in Alderley Park and Alex Alderton, Roberto Amato, Jeffrey Barrett, Sonia Goncalves, Ewan Harrison, David K. Jackson, Ian Johnston, Dominic Kwiatkowski, Cordelia Langford, John Sillitoe on behalf of the Wellcome Sanger Institute COVID-19 Surveillance Team |
| EPI_ISL_2021285 | Lighthouse Lab in Milton Keynes | Wellcome Sanger Institute for the COVID-19 Genomics UK (COG-UK) Consortium | The Lighthouse Lab in Milton Keynes and Alex Alderton, Roberto Amato, Jeffrey Barrett, Sonia Goncalves, Ewan Harrison, David K. Jackson, Ian Johnston, Dominic Kwiatkowski, Cordelia Langford, John Sillitoe on behalf of the Wellcome Sanger Institute COVID-19 Surveillance Team |
| EPI_ISL_2021287 | Lighthouse Lab in Alderley Park | Wellcome Sanger Institute for the COVID-19 Genomics UK (COG-UK) Consortium | Jacquelyn Wynn, Mairead Hyland, The Lighthouse Lab in Alderley Park and Alex Alderton, Roberto Amato, Jeffrey Barrett, Sonia Goncalves, Ewan Harrison, David K. Jackson, Ian Johnston, Dominic Kwiatkowski, Cordelia Langford, John Sillitoe on behalf of the Wellcome Sanger Institute COVID-19 Surveillance Team |
| EPI_ISL_2021289 | Lighthouse Lab in Glasgow | Wellcome Sanger Institute for the COVID-19 Genomics UK (COG-UK) Consortium | Harper VanSteenhouse, Yumi Kasai, David Gray, Carol Clugston, Anna Dominiczak and Alex Alderton, Roberto Amato, Jeffrey Barrett, Sonia Goncalves, Ewan Harrison, David K. Jackson, Ian Johnston, Dominic Kwiatkowski, Cordelia Langford, John Sillitoe on behalf of the Wellcome Sanger Institute COVID-19 Surveillance Team |
| EPI_ISL_2021292, EPI_ISL_2021296 | Lighthouse Lab in Alderley Park | Wellcome Sanger Institute for the COVID-19 Genomics UK (COG-UK) Consortium | Jacquelyn Wynn, Mairead Hyland, The Lighthouse Lab in Alderley Park and Alex Alderton, Roberto Amato, Jeffrey Barrett, Sonia Goncalves, Ewan Harrison, David K. Jackson, Ian Johnston, Dominic Kwiatkowski, Cordelia Langford, John Sillitoe on behalf of the Wellcome Sanger Institute COVID-19 Surveillance Team |
| EPI_ISL_2021312 | Lighthouse Lab in Glasgow | Wellcome Sanger Institute for the COVID-19 Genomics UK (COG-UK) Consortium | Harper VanSteenhouse, Yumi Kasai, David Gray, Carol Clugston, Anna Dominiczak and Alex Alderton, Roberto Amato, Jeffrey Barrett, Sonia Goncalves, Ewan Harrison, David K. Jackson, Ian Johnston, Dominic Kwiatkowski, Cordelia Langford, John Sillitoe on behalf of the Wellcome Sanger Institute COVID-19 Surveillance Team |
| EPI_ISL_2021315, EPI_ISL_2021326, EPI_ISL_2021327, EPI_ISL_2021329 | Lighthouse Lab in Alderley Park | Wellcome Sanger Institute for the COVID-19 Genomics UK (COG-UK) Consortium | Jacquelyn Wynn, Mairead Hyland, The Lighthouse Lab in Alderley Park and Alex Alderton, Roberto Amato, Jeffrey Barrett, Sonia Goncalves, Ewan Harrison, David K. Jackson, Ian Johnston, Dominic Kwiatkowski, Cordelia Langford, John Sillitoe on behalf of the Wellcome Sanger Institute COVID-19 Surveillance Team |
| EPI_ISL_2021344, EPI_ISL_2021347 | Lighthouse Lab in Glasgow | Wellcome Sanger Institute for the COVID-19 Genomics UK (COG-UK) Consortium | Harper VanSteenhouse, Yumi Kasai, David Gray, Carol Clugston, Anna Dominiczak and Alex Alderton, Roberto Amato, Jeffrey Barrett, Sonia Goncalves, Ewan Harrison, David K. Jackson, Ian Johnston, Dominic Kwiatkowski, Cordelia Langford, John Sillitoe on behalf of the Wellcome Sanger Institute COVID-19 Surveillance Team |

| | | | |
|---|---|---|---|
| EPI_ISL_2021349, EPI_ISL_2021374, EPI_ISL_2021376, EPI_ISL_2021377 | Lighthouse Lab in Alderley Park | Wellcome Sanger Institute for the COVID-19 Genomics UK (COG-UK) Consortium | Jacquelyn Wynn, Mairead Hyland, The Lighthouse Lab in Alderley Park and Alex Alderton, Roberto Amato, Jeffrey Barrett, Sonia Goncalves, Ewan Harrison, David K. Jackson, Ian Johnston, Dominic Kwiatkowski, Cordelia Langford, John Sillitoe on behalf of the Wellcome Sanger Institute COVID-19 Surveillance Team |
| EPI_ISL_2021382, EPI_ISL_2021383, EPI_ISL_2021384 | Lighthouse Lab in Milton Keynes | Wellcome Sanger Institute for the COVID-19 Genomics UK (COG-UK) Consortium | The Lighthouse Lab in Milton Keynes and Alex Alderton, Roberto Amato, Jeffrey Barrett, Sonia Goncalves, Ewan Harrison, David K. Jackson, Ian Johnston, Dominic Kwiatkowski, Cordelia Langford, John Sillitoe on behalf of the Wellcome Sanger Institute COVID-19 Surveillance Team |
| EPI_ISL_2021396 | Lighthouse Lab in Alderley Park | Wellcome Sanger Institute for the COVID-19 Genomics UK (COG-UK) Consortium | Jacquelyn Wynn, Mairead Hyland, The Lighthouse Lab in Alderley Park and Alex Alderton, Roberto Amato, Jeffrey Barrett, Sonia Goncalves, Ewan Harrison, David K. Jackson, Ian Johnston, Dominic Kwiatkowski, Cordelia Langford, John Sillitoe on behalf of the Wellcome Sanger Institute COVID-19 Surveillance Team |
| EPI_ISL_2021401 | Lighthouse Lab in Milton Keynes | Wellcome Sanger Institute for the COVID-19 Genomics UK (COG-UK) Consortium | The Lighthouse Lab in Milton Keynes and Alex Alderton, Roberto Amato, Jeffrey Barrett, Sonia Goncalves, Ewan Harrison, David K. Jackson, Ian Johnston, Dominic Kwiatkowski, Cordelia Langford, John Sillitoe on behalf of the Wellcome Sanger Institute COVID-19 Surveillance Team |
| EPI_ISL_2021410, EPI_ISL_2021418 | Lighthouse Lab in Alderley Park | Wellcome Sanger Institute for the COVID-19 Genomics UK (COG-UK) Consortium | Jacquelyn Wynn, Mairead Hyland, The Lighthouse Lab in Alderley Park and Alex Alderton, Roberto Amato, Jeffrey Barrett, Sonia Goncalves, Ewan Harrison, David K. Jackson, Ian Johnston, Dominic Kwiatkowski, Cordelia Langford, John Sillitoe on behalf of the Wellcome Sanger Institute COVID-19 Surveillance Team |
| EPI_ISL_2021419, EPI_ISL_2021422, EPI_ISL_2021425 | Lighthouse Lab in Milton Keynes | Wellcome Sanger Institute for the COVID-19 Genomics UK (COG-UK) Consortium | The Lighthouse Lab in Milton Keynes and Alex Alderton, Roberto Amato, Jeffrey Barrett, Sonia Goncalves, Ewan Harrison, David K. Jackson, Ian Johnston, Dominic Kwiatkowski, Cordelia Langford, John Sillitoe on behalf of the Wellcome Sanger Institute COVID-19 Surveillance Team |
| EPI_ISL_2021432 | Lighthouse Lab in Alderley Park | Wellcome Sanger Institute for the COVID-19 Genomics UK (COG-UK) Consortium | Jacquelyn Wynn, Mairead Hyland, The Lighthouse Lab in Alderley Park and Alex Alderton, Roberto Amato, Jeffrey Barrett, Sonia Goncalves, Ewan Harrison, David K. Jackson, Ian Johnston, Dominic Kwiatkowski, Cordelia Langford, John Sillitoe on behalf of the Wellcome Sanger Institute COVID-19 Surveillance Team |
| EPI_ISL_2021433, EPI_ISL_2021434, EPI_ISL_2021444 | Lighthouse Lab in Milton Keynes | Wellcome Sanger Institute for the COVID-19 Genomics UK (COG-UK) Consortium | The Lighthouse Lab in Milton Keynes and Alex Alderton, Roberto Amato, Jeffrey Barrett, Sonia Goncalves, Ewan Harrison, David K. Jackson, Ian Johnston, Dominic Kwiatkowski, Cordelia Langford, John Sillitoe on behalf of the Wellcome Sanger Institute COVID-19 Surveillance Team |
| EPI_ISL_2021446, EPI_ISL_2021455, EPI_ISL_2021456 | Lighthouse Lab in Alderley Park | Wellcome Sanger Institute for the COVID-19 Genomics UK (COG-UK) Consortium | Jacquelyn Wynn, Mairead Hyland, The Lighthouse Lab in Alderley Park and Alex Alderton, Roberto Amato, Jeffrey Barrett, Sonia Goncalves, Ewan Harrison, David K. Jackson, Ian Johnston, Dominic Kwiatkowski, Cordelia Langford, John Sillitoe on behalf of the Wellcome Sanger Institute COVID-19 Surveillance Team |
| EPI_ISL_2021458, EPI_ISL_2021461 | Lighthouse Lab in Milton Keynes | Wellcome Sanger Institute for the COVID-19 Genomics UK (COG-UK) Consortium | The Lighthouse Lab in Milton Keynes and Alex Alderton, Roberto Amato, Jeffrey Barrett, Sonia Goncalves, Ewan Harrison, David K. Jackson, Ian Johnston, Dominic Kwiatkowski, Cordelia Langford, John Sillitoe on behalf of the Wellcome Sanger Institute COVID-19 Surveillance Team |
| EPI_ISL_2021464 | Lighthouse Lab in Alderley Park | Wellcome Sanger Institute for the COVID-19 Genomics UK (COG-UK) Consortium | Jacquelyn Wynn, Mairead Hyland, The Lighthouse Lab in Alderley Park and Alex Alderton, Roberto Amato, Jeffrey Barrett, Sonia Goncalves, Ewan Harrison, David K. Jackson, Ian Johnston, Dominic Kwiatkowski, Cordelia Langford, John Sillitoe on behalf of the Wellcome Sanger Institute COVID-19 Surveillance Team |
| EPI_ISL_2021467 | Lighthouse Lab in Milton Keynes | Wellcome Sanger Institute for the COVID-19 Genomics UK (COG-UK) Consortium | The Lighthouse Lab in Milton Keynes and Alex Alderton, Roberto Amato, Jeffrey Barrett, Sonia Goncalves, Ewan Harrison, David K. Jackson, Ian Johnston, Dominic Kwiatkowski, Cordelia Langford, John Sillitoe on behalf of the Wellcome Sanger Institute COVID-19 Surveillance Team |
| EPI_ISL_2021468, EPI_ISL_2021473 | Lighthouse Lab in Alderley Park | Wellcome Sanger Institute for the COVID-19 Genomics UK (COG-UK) Consortium | Jacquelyn Wynn, Mairead Hyland, The Lighthouse Lab in Alderley Park and Alex Alderton, Roberto Amato, Jeffrey Barrett, Sonia Goncalves, Ewan Harrison, David K. Jackson, Ian Johnston, Dominic Kwiatkowski, Cordelia Langford, John Sillitoe on behalf of the Wellcome Sanger Institute COVID-19 Surveillance Team |
| EPI_ISL_2021477, EPI_ISL_2021480, EPI_ISL_2021483 | Lighthouse Lab in Milton Keynes | Wellcome Sanger Institute for the COVID-19 Genomics UK (COG-UK) Consortium | The Lighthouse Lab in Milton Keynes and Alex Alderton, Roberto Amato, Jeffrey Barrett, Sonia Goncalves, Ewan Harrison, David K. Jackson, Ian Johnston, Dominic Kwiatkowski, Cordelia Langford, John Sillitoe on behalf of the Wellcome Sanger Institute COVID-19 Surveillance Team |
| EPI_ISL_2021490 | Lighthouse Lab in Alderley Park | Wellcome Sanger Institute for the COVID-19 Genomics UK (COG-UK) Consortium | Jacquelyn Wynn, Mairead Hyland, The Lighthouse Lab in Alderley Park and Alex Alderton, Roberto Amato, Jeffrey Barrett, Sonia Goncalves, Ewan Harrison, David K. Jackson, Ian Johnston, Dominic Kwiatkowski, Cordelia Langford, John Sillitoe on behalf of the Wellcome Sanger Institute COVID-19 Surveillance Team |
| EPI_ISL_2021492 | Lighthouse Lab in Milton Keynes | Wellcome Sanger Institute for the COVID-19 Genomics UK (COG-UK) Consortium | The Lighthouse Lab in Milton Keynes and Alex Alderton, Roberto Amato, Jeffrey Barrett, Sonia Goncalves, Ewan Harrison, David K. Jackson, Ian Johnston, Dominic Kwiatkowski, Cordelia Langford, John Sillitoe on behalf of the Wellcome Sanger Institute COVID-19 Surveillance Team |
| EPI_ISL_2021493, EPI_ISL_2021497, EPI_ISL_2021500, EPI_ISL_2021503 | Lighthouse Lab in Alderley Park | Wellcome Sanger Institute for the COVID-19 Genomics UK (COG-UK) Consortium | Jacquelyn Wynn, Mairead Hyland, The Lighthouse Lab in Alderley Park and Alex Alderton, Roberto Amato, Jeffrey Barrett, Sonia Goncalves, Ewan Harrison, David K. Jackson, Ian Johnston, Dominic Kwiatkowski, Cordelia Langford, John Sillitoe on behalf of the Wellcome Sanger Institute COVID-19 Surveillance Team |
| EPI_ISL_2021508 | Lighthouse Lab in Milton Keynes | Wellcome Sanger Institute for the COVID-19 Genomics UK (COG-UK) Consortium | The Lighthouse Lab in Milton Keynes and Alex Alderton, Roberto Amato, Jeffrey Barrett, Sonia Goncalves, Ewan Harrison, David K. Jackson, Ian Johnston, Dominic Kwiatkowski, Cordelia Langford, John Sillitoe on behalf of the Wellcome Sanger Institute COVID-19 Surveillance Team |
| EPI_ISL_2021519 | Lighthouse Lab in Alderley Park | Wellcome Sanger Institute for the COVID-19 Genomics UK (COG-UK) Consortium | Jacquelyn Wynn, Mairead Hyland, The Lighthouse Lab in Alderley Park and Alex Alderton, Roberto Amato, Jeffrey Barrett, Sonia Goncalves, Ewan Harrison, David K. Jackson, Ian Johnston, Dominic Kwiatkowski, Cordelia Langford, John Sillitoe on behalf of the Wellcome Sanger Institute COVID-19 Surveillance Team |
| EPI_ISL_2021522 | Lighthouse Lab in Milton Keynes | Wellcome Sanger Institute for the COVID-19 Genomics UK (COG-UK) Consortium | The Lighthouse Lab in Milton Keynes and Alex Alderton, Roberto Amato, Jeffrey Barrett, Sonia Goncalves, Ewan Harrison, David K. Jackson, Ian Johnston, Dominic Kwiatkowski, Cordelia Langford, John Sillitoe on behalf of the Wellcome Sanger Institute COVID-19 Surveillance Team |
| EPI_ISL_2021525, EPI_ISL_2021526, EPI_ISL_2021529 | Lighthouse Lab in Alderley Park | Wellcome Sanger Institute for the COVID-19 Genomics UK (COG-UK) Consortium | Jacquelyn Wynn, Mairead Hyland, The Lighthouse Lab in Alderley Park and Alex Alderton, Roberto Amato, Jeffrey Barrett, Sonia Goncalves, Ewan Harrison, David K. Jackson, Ian Johnston, Dominic Kwiatkowski, Cordelia Langford, John Sillitoe on behalf of the Wellcome Sanger Institute COVID-19 Surveillance Team |
| EPI_ISL_2021546, EPI_ISL_2021549 | Lighthouse Lab in Milton Keynes | Wellcome Sanger Institute for the COVID-19 Genomics UK (COG-UK) Consortium | The Lighthouse Lab in Milton Keynes and Alex Alderton, Roberto Amato, Jeffrey Barrett, Sonia Goncalves, Ewan Harrison, David K. Jackson, Ian Johnston, Dominic Kwiatkowski, Cordelia Langford, John Sillitoe on behalf of the Wellcome Sanger Institute COVID-19 Surveillance Team |
| EPI_ISL_2021550, EPI_ISL_2021551, EPI_ISL_2021552 | Lighthouse Lab in Alderley Park | Wellcome Sanger Institute for the COVID-19 Genomics UK (COG-UK) Consortium | Jacquelyn Wynn, Mairead Hyland, The Lighthouse Lab in Alderley Park and Alex Alderton, Roberto Amato, Jeffrey Barrett, Sonia Goncalves, Ewan Harrison, David K. Jackson, Ian Johnston, Dominic Kwiatkowski, Cordelia Langford, John Sillitoe on behalf of the Wellcome Sanger Institute COVID-19 Surveillance Team |
| EPI_ISL_2021558 | Lighthouse Lab in Milton Keynes | Wellcome Sanger Institute for the COVID-19 Genomics UK (COG-UK) Consortium | The Lighthouse Lab in Milton Keynes and Alex Alderton, Roberto Amato, Jeffrey Barrett, Sonia Goncalves, Ewan Harrison, David K. Jackson, Ian Johnston, Dominic Kwiatkowski, Cordelia Langford, John Sillitoe on behalf of the Wellcome Sanger Institute COVID-19 Surveillance Team |
| EPI_ISL_2021561, EPI_ISL_2021568 | Lighthouse Lab in Alderley Park | Wellcome Sanger Institute for the COVID-19 Genomics UK (COG-UK) Consortium | Jacquelyn Wynn, Mairead Hyland, The Lighthouse Lab in Alderley Park and Alex Alderton, Roberto Amato, Jeffrey Barrett, Sonia Goncalves, Ewan Harrison, David K. Jackson, Ian Johnston, Dominic Kwiatkowski, Cordelia Langford, John Sillitoe on behalf of the Wellcome Sanger Institute COVID-19 Surveillance Team |
| EPI_ISL_2021575 | Lighthouse Lab in Milton Keynes | Wellcome Sanger Institute for the COVID-19 Genomics UK (COG-UK) Consortium | The Lighthouse Lab in Milton Keynes and Alex Alderton, Roberto Amato, Jeffrey Barrett, Sonia Goncalves, Ewan Harrison, David K. Jackson, Ian Johnston, Dominic Kwiatkowski, Cordelia Langford, John Sillitoe on behalf of the Wellcome Sanger Institute COVID-19 Surveillance Team |
| EPI_ISL_2021579, EPI_ISL_2021585, EPI_ISL_2021592 | Lighthouse Lab in Alderley Park | Wellcome Sanger Institute for the COVID-19 Genomics UK (COG-UK) Consortium | Jacquelyn Wynn, Mairead Hyland, The Lighthouse Lab in Alderley Park and Alex Alderton, Roberto Amato, Jeffrey Barrett, Sonia Goncalves, Ewan Harrison, David K. Jackson, Ian Johnston, Dominic Kwiatkowski, Cordelia Langford, John Sillitoe on behalf of the Wellcome Sanger Institute COVID-19 Surveillance Team |
| EPI_ISL_2021597 | Lighthouse Lab in Milton Keynes | Wellcome Sanger Institute for the COVID-19 Genomics UK (COG-UK) Consortium | The Lighthouse Lab in Milton Keynes and Alex Alderton, Roberto Amato, Jeffrey Barrett, Sonia Goncalves, Ewan Harrison, David K. Jackson, Ian Johnston, Dominic Kwiatkowski, Cordelia Langford, John Sillitoe on behalf of the Wellcome Sanger Institute COVID-19 Surveillance Team |
| EPI_ISL_2021602 | Lighthouse Lab in Alderley Park | Wellcome Sanger Institute for the COVID-19 Genomics UK (COG-UK) Consortium | Jacquelyn Wynn, Mairead Hyland, The Lighthouse Lab in Alderley Park and Alex Alderton, Roberto Amato, Jeffrey Barrett, Sonia Goncalves, Ewan Harrison, David K. Jackson, Ian Johnston, Dominic Kwiatkowski, Cordelia Langford, John Sillitoe on behalf of the Wellcome Sanger Institute COVID-19 Surveillance Team |
| EPI_ISL_2021603 | Lighthouse Lab in Milton Keynes | Wellcome Sanger Institute for the COVID-19 Genomics UK (COG-UK) Consortium | The Lighthouse Lab in Milton Keynes and Alex Alderton, Roberto Amato, Jeffrey Barrett, Sonia Goncalves, Ewan Harrison, David K. Jackson, Ian Johnston, Dominic Kwiatkowski, Cordelia Langford, John Sillitoe on behalf of the Wellcome Sanger Institute COVID-19 Surveillance Team |
| EPI_ISL_2021609 | Lighthouse Lab in Alderley Park | Wellcome Sanger Institute for the COVID-19 Genomics UK (COG-UK) Consortium | Jacquelyn Wynn, Mairead Hyland, The Lighthouse Lab in Alderley Park and Alex Alderton, Roberto Amato, Jeffrey Barrett, Sonia Goncalves, Ewan Harrison, David K. Jackson, Ian Johnston, Dominic Kwiatkowski, Cordelia Langford, John Sillitoe on behalf of the Wellcome Sanger Institute COVID-19 Surveillance Team |
| EPI_ISL_2021613, EPI_ISL_2021614 | Lighthouse Lab in Milton Keynes | Wellcome Sanger Institute for the COVID-19 Genomics UK (COG-UK) Consortium | The Lighthouse Lab in Milton Keynes and Alex Alderton, Roberto Amato, Jeffrey Barrett, Sonia Goncalves, Ewan Harrison, David K. Jackson, Ian Johnston, Dominic Kwiatkowski, Cordelia Langford, John Sillitoe on behalf of the Wellcome Sanger Institute COVID-19 Surveillance Team |
| EPI_ISL_2021617, EPI_ISL_2021619 | Lighthouse Lab in Alderley Park | Wellcome Sanger Institute for the COVID-19 Genomics UK (COG-UK) Consortium | Jacquelyn Wynn, Mairead Hyland, The Lighthouse Lab in Alderley Park and Alex Alderton, Roberto Amato, Jeffrey Barrett, Sonia Goncalves, Ewan Harrison, David K. Jackson, Ian Johnston, Dominic Kwiatkowski, Cordelia Langford, John Sillitoe on behalf of the Wellcome Sanger Institute COVID-19 Surveillance Team |
| EPI_ISL_2021626 | Lighthouse Lab in Milton Keynes | Wellcome Sanger Institute for the COVID-19 Genomics UK (COG-UK) Consortium | The Lighthouse Lab in Milton Keynes and Alex Alderton, Roberto Amato, Jeffrey Barrett, Sonia Goncalves, Ewan Harrison, David K. Jackson, Ian Johnston, Dominic Kwiatkowski, Cordelia Langford, John Sillitoe on behalf of the Wellcome Sanger Institute COVID-19 Surveillance Team |
| EPI_ISL_2021627 | Lighthouse Lab in Alderley Park | Wellcome Sanger Institute for the COVID-19 Genomics UK (COG-UK) Consortium | Jacquelyn Wynn, Mairead Hyland, The Lighthouse Lab in Alderley Park and Alex Alderton, Roberto Amato, Jeffrey Barrett, Sonia Goncalves, Ewan Harrison, David K. Jackson, Ian Johnston, Dominic Kwiatkowski, Cordelia Langford, John Sillitoe on behalf of the Wellcome Sanger Institute COVID-19 Surveillance Team |
| EPI_ISL_2021632 | Lighthouse Lab in Milton Keynes | Wellcome Sanger Institute for the COVID-19 Genomics UK (COG-UK) Consortium | The Lighthouse Lab in Milton Keynes and Alex Alderton, Roberto Amato, Jeffrey Barrett, Sonia Goncalves, Ewan Harrison, David K. Jackson, Ian Johnston, Dominic Kwiatkowski, Cordelia Langford, John Sillitoe on behalf of the Wellcome Sanger Institute COVID-19 Surveillance Team |
| EPI_ISL_2021633, EPI_ISL_2021640, EPI_ISL_2021647, EPI_ISL_2021657 | Lighthouse Lab in Alderley Park | Wellcome Sanger Institute for the COVID-19 Genomics UK (COG-UK) Consortium | Jacquelyn Wynn, Mairead Hyland, The Lighthouse Lab in Alderley Park and Alex Alderton, Roberto Amato, Jeffrey Barrett, Sonia Goncalves, Ewan Harrison, David K. Jackson, Ian Johnston, Dominic Kwiatkowski, Cordelia Langford, John Sillitoe on behalf of the Wellcome Sanger Institute COVID-19 Surveillance Team |
| EPI_ISL_2021661, EPI_ISL_2021663, EPI_ISL_2021665, EPI_ISL_2021671, EPI_ISL_2021677 | Lighthouse Lab in Milton Keynes | Wellcome Sanger Institute for the COVID-19 Genomics UK (COG-UK) Consortium | The Lighthouse Lab in Milton Keynes and Alex Alderton, Roberto Amato, Jeffrey Barrett, Sonia Goncalves, Ewan Harrison, David K. Jackson, Ian Johnston, Dominic Kwiatkowski, Cordelia Langford, John Sillitoe on behalf of the Wellcome Sanger Institute COVID-19 Surveillance Team |

| | | | |
|---|---|---|---|
| EPI_ISL_2021680 | Lighthouse Lab in Alderley Park | Wellcome Sanger Institute for the COVID-19 Genomics UK (COG-UK) Consortium | Jacquelyn Wynn, Mairead Hyland, The Lighthouse Lab in Alderley Park and Alex Alderton, Roberto Amato, Jeffrey Barrett, Sonia Goncalves, Ewan Harrison, David K. Jackson, Ian Johnston, Dominic Kwiatkowski, Cordelia Langford, John Sillitoe on behalf of the Wellcome Sanger Institute COVID-19 Surveillance Team |
| EPI_ISL_2021685, EPI_ISL_2021688, EPI_ISL_2021690 | Lighthouse Lab in Milton Keynes | Wellcome Sanger Institute for the COVID-19 Genomics UK (COG-UK) Consortium | The Lighthouse Lab in Milton Keynes and Alex Alderton, Roberto Amato, Jeffrey Barrett, Sonia Goncalves, Ewan Harrison, David K. Jackson, Ian Johnston, Dominic Kwiatkowski, Cordelia Langford, John Sillitoe on behalf of the Wellcome Sanger Institute COVID-19 Surveillance Team |
| EPI_ISL_2021692 | Lighthouse Lab in Alderley Park | Wellcome Sanger Institute for the COVID-19 Genomics UK (COG-UK) Consortium | Jacquelyn Wynn, Mairead Hyland, The Lighthouse Lab in Alderley Park and Alex Alderton, Roberto Amato, Jeffrey Barrett, Sonia Goncalves, Ewan Harrison, David K. Jackson, Ian Johnston, Dominic Kwiatkowski, Cordelia Langford, John Sillitoe on behalf of the Wellcome Sanger Institute COVID-19 Surveillance Team |
| EPI_ISL_2021698 | Lighthouse Lab in Glasgow | Wellcome Sanger Institute for the COVID-19 Genomics UK (COG-UK) Consortium | Harper VanSteenhouse, Yumi Kasai, David Gray, Carol Clugston, Anna Dominiczak and Alex Alderton, Roberto Amato, Jeffrey Barrett, Sonia Goncalves, Ewan Harrison, David K. Jackson, Ian Johnston, Dominic Kwiatkowski, Cordelia Langford, John Sillitoe on behalf of the Wellcome Sanger Institute COVID-19 Surveillance Team |
| EPI_ISL_2021711 | Lighthouse Lab in Alderley Park | Wellcome Sanger Institute for the COVID-19 Genomics UK (COG-UK) Consortium | Jacquelyn Wynn, Mairead Hyland, The Lighthouse Lab in Alderley Park and Alex Alderton, Roberto Amato, Jeffrey Barrett, Sonia Goncalves, Ewan Harrison, David K. Jackson, Ian Johnston, Dominic Kwiatkowski, Cordelia Langford, John Sillitoe on behalf of the Wellcome Sanger Institute COVID-19 Surveillance Team |
| EPI_ISL_2021714, EPI_ISL_2021722 | Randox Laboratories | Wellcome Sanger Institute for the COVID-19 Genomics UK (COG-UK) Consortium | Randox Laboratories and Alex Alderton, Roberto Amato, Jeffrey Barrett, Sonia Goncalves, Ewan Harrison, David K. Jackson, Ian Johnston, Dominic Kwiatkowski, Cordelia Langford, John Sillitoe on behalf of the Wellcome Sanger Institute COVID-19 Surveillance Team |
| EPI_ISL_2021728 | Lighthouse Lab in Alderley Park | Wellcome Sanger Institute for the COVID-19 Genomics UK (COG-UK) Consortium | Jacquelyn Wynn, Mairead Hyland, The Lighthouse Lab in Alderley Park and Alex Alderton, Roberto Amato, Jeffrey Barrett, Sonia Goncalves, Ewan Harrison, David K. Jackson, Ian Johnston, Dominic Kwiatkowski, Cordelia Langford, John Sillitoe on behalf of the Wellcome Sanger Institute COVID-19 Surveillance Team |
| EPI_ISL_2021741, EPI_ISL_2021744 | Lighthouse Lab in Milton Keynes | Wellcome Sanger Institute for the COVID-19 Genomics UK (COG-UK) Consortium | The Lighthouse Lab in Milton Keynes and Alex Alderton, Roberto Amato, Jeffrey Barrett, Sonia Goncalves, Ewan Harrison, David K. Jackson, Ian Johnston, Dominic Kwiatkowski, Cordelia Langford, John Sillitoe on behalf of the Wellcome Sanger Institute COVID-19 Surveillance Team |
| EPI_ISL_2021745 | Randox Laboratories | Wellcome Sanger Institute for the COVID-19 Genomics UK (COG-UK) Consortium | Randox Laboratories and Alex Alderton, Roberto Amato, Jeffrey Barrett, Sonia Goncalves, Ewan Harrison, David K. Jackson, Ian Johnston, Dominic Kwiatkowski, Cordelia Langford, John Sillitoe on behalf of the Wellcome Sanger Institute COVID-19 Surveillance Team |
| EPI_ISL_2021747 | Lighthouse Lab in Alderley Park | Wellcome Sanger Institute for the COVID-19 Genomics UK (COG-UK) Consortium | Jacquelyn Wynn, Mairead Hyland, The Lighthouse Lab in Alderley Park and Alex Alderton, Roberto Amato, Jeffrey Barrett, Sonia Goncalves, Ewan Harrison, David K. Jackson, Ian Johnston, Dominic Kwiatkowski, Cordelia Langford, John Sillitoe on behalf of the Wellcome Sanger Institute COVID-19 Surveillance Team |
| EPI_ISL_2021750 | Randox Laboratories | Wellcome Sanger Institute for the COVID-19 Genomics UK (COG-UK) Consortium | Randox Laboratories and Alex Alderton, Roberto Amato, Jeffrey Barrett, Sonia Goncalves, Ewan Harrison, David K. Jackson, Ian Johnston, Dominic Kwiatkowski, Cordelia Langford, John Sillitoe on behalf of the Wellcome Sanger Institute COVID-19 Surveillance Team |
| EPI_ISL_2021753 | Lighthouse Lab in Milton Keynes | Wellcome Sanger Institute for the COVID-19 Genomics UK (COG-UK) Consortium | The Lighthouse Lab in Milton Keynes and Alex Alderton, Roberto Amato, Jeffrey Barrett, Sonia Goncalves, Ewan Harrison, David K. Jackson, Ian Johnston, Dominic Kwiatkowski, Cordelia Langford, John Sillitoe on behalf of the Wellcome Sanger Institute COVID-19 Surveillance Team |
| EPI_ISL_2021759, EPI_ISL_2021763 | Randox Laboratories | Wellcome Sanger Institute for the COVID-19 Genomics UK (COG-UK) Consortium | Randox Laboratories and Alex Alderton, Roberto Amato, Jeffrey Barrett, Sonia Goncalves, Ewan Harrison, David K. Jackson, Ian Johnston, Dominic Kwiatkowski, Cordelia Langford, John Sillitoe on behalf of the Wellcome Sanger Institute COVID-19 Surveillance Team |
| EPI_ISL_2021784 | Lighthouse Lab in Alderley Park | Wellcome Sanger Institute for the COVID-19 Genomics UK (COG-UK) Consortium | Jacquelyn Wynn, Mairead Hyland, The Lighthouse Lab in Alderley Park and Alex Alderton, Roberto Amato, Jeffrey Barrett, Sonia Goncalves, Ewan Harrison, David K. Jackson, Ian Johnston, Dominic Kwiatkowski, Cordelia Langford, John Sillitoe on behalf of the Wellcome Sanger Institute COVID-19 Surveillance Team |
| EPI_ISL_2021801 | Randox Laboratories | Wellcome Sanger Institute for the COVID-19 Genomics UK (COG-UK) Consortium | Randox Laboratories and Alex Alderton, Roberto Amato, Jeffrey Barrett, Sonia Goncalves, Ewan Harrison, David K. Jackson, Ian Johnston, Dominic Kwiatkowski, Cordelia Langford, John Sillitoe on behalf of the Wellcome Sanger Institute COVID-19 Surveillance Team |
| EPI_ISL_2021813, EPI_ISL_2021828 | Lighthouse Lab in Alderley Park | Wellcome Sanger Institute for the COVID-19 Genomics UK (COG-UK) Consortium | Jacquelyn Wynn, Mairead Hyland, The Lighthouse Lab in Alderley Park and Alex Alderton, Roberto Amato, Jeffrey Barrett, Sonia Goncalves, Ewan Harrison, David K. Jackson, Ian Johnston, Dominic Kwiatkowski, Cordelia Langford, John Sillitoe on behalf of the Wellcome Sanger Institute COVID-19 Surveillance Team |
| EPI_ISL_2021832, EPI_ISL_2021834, EPI_ISL_2021839, EPI_ISL_2021841 | Lighthouse Lab in Milton Keynes | Wellcome Sanger Institute for the COVID-19 Genomics UK (COG-UK) Consortium | The Lighthouse Lab in Milton Keynes and Alex Alderton, Roberto Amato, Jeffrey Barrett, Sonia Goncalves, Ewan Harrison, David K. Jackson, Ian Johnston, Dominic Kwiatkowski, Cordelia Langford, John Sillitoe on behalf of the Wellcome Sanger Institute COVID-19 Surveillance Team |
| EPI_ISL_2021842 | Lighthouse Lab in Glasgow | Wellcome Sanger Institute for the COVID-19 Genomics UK (COG-UK) Consortium | Harper VanSteenhouse, Yumi Kasai, David Gray, Carol Clugston, Anna Dominiczak and Alex Alderton, Roberto Amato, Jeffrey Barrett, Sonia Goncalves, Ewan Harrison, David K. Jackson, Ian Johnston, Dominic Kwiatkowski, Cordelia Langford, John Sillitoe on behalf of the Wellcome Sanger Institute COVID-19 Surveillance Team |
| EPI_ISL_2021843 | Lighthouse Lab in Alderley Park | Wellcome Sanger Institute for the COVID-19 Genomics UK (COG-UK) Consortium | Jacquelyn Wynn, Mairead Hyland, The Lighthouse Lab in Alderley Park and Alex Alderton, Roberto Amato, Jeffrey Barrett, Sonia Goncalves, Ewan Harrison, David K. Jackson, Ian Johnston, Dominic Kwiatkowski, Cordelia Langford, John Sillitoe on behalf of the Wellcome Sanger Institute COVID-19 Surveillance Team |
| EPI_ISL_2021844, EPI_ISL_2021846, EPI_ISL_2021848 | Lighthouse Lab in Milton Keynes | Wellcome Sanger Institute for the COVID-19 Genomics UK (COG-UK) Consortium | The Lighthouse Lab in Milton Keynes and Alex Alderton, Roberto Amato, Jeffrey Barrett, Sonia Goncalves, Ewan Harrison, David K. Jackson, Ian Johnston, Dominic Kwiatkowski, Cordelia Langford, John Sillitoe on behalf of the Wellcome Sanger Institute COVID-19 Surveillance Team |
| EPI_ISL_2021852 | Lighthouse Lab in Alderley Park | Wellcome Sanger Institute for the COVID-19 Genomics UK (COG-UK) Consortium | Jacquelyn Wynn, Mairead Hyland, The Lighthouse Lab in Alderley Park and Alex Alderton, Roberto Amato, Jeffrey Barrett, Sonia Goncalves, Ewan Harrison, David K. Jackson, Ian Johnston, Dominic Kwiatkowski, Cordelia Langford, John Sillitoe on behalf of the Wellcome Sanger Institute COVID-19 Surveillance Team |
| EPI_ISL_2021854, EPI_ISL_2021855, EPI_ISL_2021863 | Lighthouse Lab in Milton Keynes | Wellcome Sanger Institute for the COVID-19 Genomics UK (COG-UK) Consortium | The Lighthouse Lab in Milton Keynes and Alex Alderton, Roberto Amato, Jeffrey Barrett, Sonia Goncalves, Ewan Harrison, David K. Jackson, Ian Johnston, Dominic Kwiatkowski, Cordelia Langford, John Sillitoe on behalf of the Wellcome Sanger Institute COVID-19 Surveillance Team |
| EPI_ISL_2021865 | Lighthouse Lab in Alderley Park | Wellcome Sanger Institute for the COVID-19 Genomics UK (COG-UK) Consortium | Jacquelyn Wynn, Mairead Hyland, The Lighthouse Lab in Alderley Park and Alex Alderton, Roberto Amato, Jeffrey Barrett, Sonia Goncalves, Ewan Harrison, David K. Jackson, Ian Johnston, Dominic Kwiatkowski, Cordelia Langford, John Sillitoe on behalf of the Wellcome Sanger Institute COVID-19 Surveillance Team |
| EPI_ISL_2021868 | Lighthouse Lab in Milton Keynes | Wellcome Sanger Institute for the COVID-19 Genomics UK (COG-UK) Consortium | The Lighthouse Lab in Milton Keynes and Alex Alderton, Roberto Amato, Jeffrey Barrett, Sonia Goncalves, Ewan Harrison, David K. Jackson, Ian Johnston, Dominic Kwiatkowski, Cordelia Langford, John Sillitoe on behalf of the Wellcome Sanger Institute COVID-19 Surveillance Team |
| EPI_ISL_2021870 | Lighthouse Lab in Alderley Park | Wellcome Sanger Institute for the COVID-19 Genomics UK (COG-UK) Consortium | Jacquelyn Wynn, Mairead Hyland, The Lighthouse Lab in Alderley Park and Alex Alderton, Roberto Amato, Jeffrey Barrett, Sonia Goncalves, Ewan Harrison, David K. Jackson, Ian Johnston, Dominic Kwiatkowski, Cordelia Langford, John Sillitoe on behalf of the Wellcome Sanger Institute COVID-19 Surveillance Team |
| EPI_ISL_2021871, EPI_ISL_2021872, EPI_ISL_2021874, EPI_ISL_2021878, EPI_ISL_2021880 | Lighthouse Lab in Milton Keynes | Wellcome Sanger Institute for the COVID-19 Genomics UK (COG-UK) Consortium | The Lighthouse Lab in Milton Keynes and Alex Alderton, Roberto Amato, Jeffrey Barrett, Sonia Goncalves, Ewan Harrison, David K. Jackson, Ian Johnston, Dominic Kwiatkowski, Cordelia Langford, John Sillitoe on behalf of the Wellcome Sanger Institute COVID-19 Surveillance Team |
| EPI_ISL_2021882, EPI_ISL_2021883, EPI_ISL_2021890 | Lighthouse Lab in Alderley Park | Wellcome Sanger Institute for the COVID-19 Genomics UK (COG-UK) Consortium | Jacquelyn Wynn, Mairead Hyland, The Lighthouse Lab in Alderley Park and Alex Alderton, Roberto Amato, Jeffrey Barrett, Sonia Goncalves, Ewan Harrison, David K. Jackson, Ian Johnston, Dominic Kwiatkowski, Cordelia Langford, John Sillitoe on behalf of the Wellcome Sanger Institute COVID-19 Surveillance Team |
| EPI_ISL_2021894, EPI_ISL_2021897, EPI_ISL_2021902 | Lighthouse Lab in Milton Keynes | Wellcome Sanger Institute for the COVID-19 Genomics UK (COG-UK) Consortium | The Lighthouse Lab in Milton Keynes and Alex Alderton, Roberto Amato, Jeffrey Barrett, Sonia Goncalves, Ewan Harrison, David K. Jackson, Ian Johnston, Dominic Kwiatkowski, Cordelia Langford, John Sillitoe on behalf of the Wellcome Sanger Institute COVID-19 Surveillance Team |
| EPI_ISL_2021903, EPI_ISL_2021904 | Lighthouse Lab in Alderley Park | Wellcome Sanger Institute for the COVID-19 Genomics UK (COG-UK) Consortium | Jacquelyn Wynn, Mairead Hyland, The Lighthouse Lab in Alderley Park and Alex Alderton, Roberto Amato, Jeffrey Barrett, Sonia Goncalves, Ewan Harrison, David K. Jackson, Ian Johnston, Dominic Kwiatkowski, Cordelia Langford, John Sillitoe on behalf of the Wellcome Sanger Institute COVID-19 Surveillance Team |
| EPI_ISL_2021909, EPI_ISL_2021910 | Lighthouse Lab in Milton Keynes | Wellcome Sanger Institute for the COVID-19 Genomics UK (COG-UK) Consortium | The Lighthouse Lab in Milton Keynes and Alex Alderton, Roberto Amato, Jeffrey Barrett, Sonia Goncalves, Ewan Harrison, David K. Jackson, Ian Johnston, Dominic Kwiatkowski, Cordelia Langford, John Sillitoe on behalf of the Wellcome Sanger Institute COVID-19 Surveillance Team |
| EPI_ISL_2021913, EPI_ISL_2021916 | Lighthouse Lab in Alderley Park | Wellcome Sanger Institute for the COVID-19 Genomics UK (COG-UK) Consortium | Jacquelyn Wynn, Mairead Hyland, The Lighthouse Lab in Alderley Park and Alex Alderton, Roberto Amato, Jeffrey Barrett, Sonia Goncalves, Ewan Harrison, David K. Jackson, Ian Johnston, Dominic Kwiatkowski, Cordelia Langford, John Sillitoe on behalf of the Wellcome Sanger Institute COVID-19 Surveillance Team |
| EPI_ISL_2021924, EPI_ISL_2021930 | Lighthouse Lab in Milton Keynes | Wellcome Sanger Institute for the COVID-19 Genomics UK (COG-UK) Consortium | The Lighthouse Lab in Milton Keynes and Alex Alderton, Roberto Amato, Jeffrey Barrett, Sonia Goncalves, Ewan Harrison, David K. Jackson, Ian Johnston, Dominic Kwiatkowski, Cordelia Langford, John Sillitoe on behalf of the Wellcome Sanger Institute COVID-19 Surveillance Team |
| EPI_ISL_2021933, EPI_ISL_2021934 | Lighthouse Lab in Alderley Park | Wellcome Sanger Institute for the COVID-19 Genomics UK (COG-UK) Consortium | Jacquelyn Wynn, Mairead Hyland, The Lighthouse Lab in Alderley Park and Alex Alderton, Roberto Amato, Jeffrey Barrett, Sonia Goncalves, Ewan Harrison, David K. Jackson, Ian Johnston, Dominic Kwiatkowski, Cordelia Langford, John Sillitoe on behalf of the Wellcome Sanger Institute COVID-19 Surveillance Team |
| EPI_ISL_2021946, EPI_ISL_2021947, EPI_ISL_2021950, EPI_ISL_2021951 | Lighthouse Lab in Milton Keynes | Wellcome Sanger Institute for the COVID-19 Genomics UK (COG-UK) Consortium | The Lighthouse Lab in Milton Keynes and Alex Alderton, Roberto Amato, Jeffrey Barrett, Sonia Goncalves, Ewan Harrison, David K. Jackson, Ian Johnston, Dominic Kwiatkowski, Cordelia Langford, John Sillitoe on behalf of the Wellcome Sanger Institute COVID-19 Surveillance Team |
| EPI_ISL_2021952 | Lighthouse Lab in Alderley Park | Wellcome Sanger Institute for the COVID-19 Genomics UK (COG-UK) Consortium | Jacquelyn Wynn, Mairead Hyland, The Lighthouse Lab in Alderley Park and Alex Alderton, Roberto Amato, Jeffrey Barrett, Sonia Goncalves, Ewan Harrison, David K. Jackson, Ian Johnston, Dominic Kwiatkowski, Cordelia Langford, John Sillitoe on behalf of the Wellcome Sanger Institute COVID-19 Surveillance Team |
| EPI_ISL_2021953, EPI_ISL_2021955 | Lighthouse Lab in Milton Keynes | Wellcome Sanger Institute for the COVID-19 Genomics UK (COG-UK) Consortium | The Lighthouse Lab in Milton Keynes and Alex Alderton, Roberto Amato, Jeffrey Barrett, Sonia Goncalves, Ewan Harrison, David K. Jackson, Ian Johnston, Dominic Kwiatkowski, Cordelia Langford, John Sillitoe on behalf of the Wellcome Sanger Institute COVID-19 Surveillance Team |
| EPI_ISL_2021956 | Lighthouse Lab in Alderley Park | Wellcome Sanger Institute for the COVID-19 Genomics UK (COG-UK) Consortium | Jacquelyn Wynn, Mairead Hyland, The Lighthouse Lab in Alderley Park and Alex Alderton, Roberto Amato, Jeffrey Barrett, Sonia Goncalves, Ewan Harrison, David K. Jackson, Ian Johnston, Dominic Kwiatkowski, Cordelia Langford, John Sillitoe on behalf of the Wellcome Sanger Institute COVID-19 Surveillance Team |
| EPI_ISL_2021958, EPI_ISL_2021960 | Lighthouse Lab in Milton Keynes | Wellcome Sanger Institute for the COVID-19 Genomics UK (COG-UK) Consortium | The Lighthouse Lab in Milton Keynes and Alex Alderton, Roberto Amato, Jeffrey Barrett, Sonia Goncalves, Ewan Harrison, David K. Jackson, Ian Johnston, Dominic Kwiatkowski, Cordelia Langford, John Sillitoe on behalf of the Wellcome Sanger Institute COVID-19 Surveillance Team |
| EPI_ISL_2021963, EPI_ISL_2021971 | Lighthouse Lab in Alderley Park | Wellcome Sanger Institute for the COVID-19 Genomics UK (COG-UK) Consortium | Jacquelyn Wynn, Mairead Hyland, The Lighthouse Lab in Alderley Park and Alex Alderton, Roberto Amato, Jeffrey Barrett, Sonia Goncalves, Ewan Harrison, David K. Jackson, Ian Johnston, Dominic Kwiatkowski, Cordelia Langford, John Sillitoe on behalf of the Wellcome Sanger Institute COVID-19 Surveillance Team |
| EPI_ISL_2021981, EPI_ISL_2021982 | Lighthouse Lab in Milton Keynes | Wellcome Sanger Institute for the COVID-19 Genomics UK (COG-UK) Consortium | The Lighthouse Lab in Milton Keynes and Alex Alderton, Roberto Amato, Jeffrey Barrett, Sonia Goncalves, Ewan Harrison, David K. Jackson, Ian Johnston, Dominic Kwiatkowski, Cordelia Langford, John Sillitoe on behalf of the Wellcome Sanger Institute COVID-19 Surveillance Team |

| | | | |
|---|---|---|---|
| EPI_ISL_2021984 | Lighthouse Lab in Alderley Park | Wellcome Sanger Institute for the COVID-19 Genomics UK (COG-UK) Consortium | Jacquelyn Wynn, Mairead Hyland, The Lighthouse Lab in Alderley Park and Alex Alderton, Roberto Amato, Jeffrey Barrett, Sonia Goncalves, Ewan Harrison, David K. Jackson, Ian Johnston, Dominic Kwiatkowski, Cordelia Langford, John Sillitoe on behalf of the Wellcome Sanger Institute COVID-19 Surveillance Team |
| EPI_ISL_2021985 | Lighthouse Lab in Milton Keynes | Wellcome Sanger Institute for the COVID-19 Genomics UK (COG-UK) Consortium | The Lighthouse Lab in Milton Keynes and Alex Alderton, Roberto Amato, Jeffrey Barrett, Sonia Goncalves, Ewan Harrison, David K. Jackson, Ian Johnston, Dominic Kwiatkowski, Cordelia Langford, John Sillitoe on behalf of the Wellcome Sanger Institute COVID-19 Surveillance Team |
| EPI_ISL_2021989 | Lighthouse Lab in Alderley Park | Wellcome Sanger Institute for the COVID-19 Genomics UK (COG-UK) Consortium | Jacquelyn Wynn, Mairead Hyland, The Lighthouse Lab in Alderley Park and Alex Alderton, Roberto Amato, Jeffrey Barrett, Sonia Goncalves, Ewan Harrison, David K. Jackson, Ian Johnston, Dominic Kwiatkowski, Cordelia Langford, John Sillitoe on behalf of the Wellcome Sanger Institute COVID-19 Surveillance Team |
| EPI_ISL_2021991, EPI_ISL_2022012 | Lighthouse Lab in Milton Keynes | Wellcome Sanger Institute for the COVID-19 Genomics UK (COG-UK) Consortium | The Lighthouse Lab in Milton Keynes and Alex Alderton, Roberto Amato, Jeffrey Barrett, Sonia Goncalves, Ewan Harrison, David K. Jackson, Ian Johnston, Dominic Kwiatkowski, Cordelia Langford, John Sillitoe on behalf of the Wellcome Sanger Institute COVID-19 Surveillance Team |
| EPI_ISL_2022020, EPI_ISL_2022021, EPI_ISL_2022023, EPI_ISL_2022027 | Lighthouse Lab in Alderley Park | Wellcome Sanger Institute for the COVID-19 Genomics UK (COG-UK) Consortium | Jacquelyn Wynn, Mairead Hyland, The Lighthouse Lab in Alderley Park and Alex Alderton, Roberto Amato, Jeffrey Barrett, Sonia Goncalves, Ewan Harrison, David K. Jackson, Ian Johnston, Dominic Kwiatkowski, Cordelia Langford, John Sillitoe on behalf of the Wellcome Sanger Institute COVID-19 Surveillance Team |
| EPI_ISL_2022030, EPI_ISL_2022039, EPI_ISL_2022042, EPI_ISL_2022047, EPI_ISL_2022049, EPI_ISL_2022050, EPI_ISL_2022052 | Lighthouse Lab in Milton Keynes | Wellcome Sanger Institute for the COVID-19 Genomics UK (COG-UK) Consortium | The Lighthouse Lab in Milton Keynes and Alex Alderton, Roberto Amato, Jeffrey Barrett, Sonia Goncalves, Ewan Harrison, David K. Jackson, Ian Johnston, Dominic Kwiatkowski, Cordelia Langford, John Sillitoe on behalf of the Wellcome Sanger Institute COVID-19 Surveillance Team |
| EPI_ISL_2022053 | Lighthouse Lab in Alderley Park | Wellcome Sanger Institute for the COVID-19 Genomics UK (COG-UK) Consortium | Jacquelyn Wynn, Mairead Hyland, The Lighthouse Lab in Alderley Park and Alex Alderton, Roberto Amato, Jeffrey Barrett, Sonia Goncalves, Ewan Harrison, David K. Jackson, Ian Johnston, Dominic Kwiatkowski, Cordelia Langford, John Sillitoe on behalf of the Wellcome Sanger Institute COVID-19 Surveillance Team |
| EPI_ISL_2022059 | Lighthouse Lab in Milton Keynes | Wellcome Sanger Institute for the COVID-19 Genomics UK (COG-UK) Consortium | The Lighthouse Lab in Milton Keynes and Alex Alderton, Roberto Amato, Jeffrey Barrett, Sonia Goncalves, Ewan Harrison, David K. Jackson, Ian Johnston, Dominic Kwiatkowski, Cordelia Langford, John Sillitoe on behalf of the Wellcome Sanger Institute COVID-19 Surveillance Team |
| EPI_ISL_2022066 | Lighthouse Lab in Alderley Park | Wellcome Sanger Institute for the COVID-19 Genomics UK (COG-UK) Consortium | Jacquelyn Wynn, Mairead Hyland, The Lighthouse Lab in Alderley Park and Alex Alderton, Roberto Amato, Jeffrey Barrett, Sonia Goncalves, Ewan Harrison, David K. Jackson, Ian Johnston, Dominic Kwiatkowski, Cordelia Langford, John Sillitoe on behalf of the Wellcome Sanger Institute COVID-19 Surveillance Team |
| EPI_ISL_2022069, EPI_ISL_2022070, EPI_ISL_2022071 | Lighthouse Lab in Milton Keynes | Wellcome Sanger Institute for the COVID-19 Genomics UK (COG-UK) Consortium | The Lighthouse Lab in Milton Keynes and Alex Alderton, Roberto Amato, Jeffrey Barrett, Sonia Goncalves, Ewan Harrison, David K. Jackson, Ian Johnston, Dominic Kwiatkowski, Cordelia Langford, John Sillitoe on behalf of the Wellcome Sanger Institute COVID-19 Surveillance Team |
| EPI_ISL_2022074 | Lighthouse Lab in Alderley Park | Wellcome Sanger Institute for the COVID-19 Genomics UK (COG-UK) Consortium | Jacquelyn Wynn, Mairead Hyland, The Lighthouse Lab in Alderley Park and Alex Alderton, Roberto Amato, Jeffrey Barrett, Sonia Goncalves, Ewan Harrison, David K. Jackson, Ian Johnston, Dominic Kwiatkowski, Cordelia Langford, John Sillitoe on behalf of the Wellcome Sanger Institute COVID-19 Surveillance Team |
| EPI_ISL_2022076 | Lighthouse Lab in Milton Keynes | Wellcome Sanger Institute for the COVID-19 Genomics UK (COG-UK) Consortium | The Lighthouse Lab in Milton Keynes and Alex Alderton, Roberto Amato, Jeffrey Barrett, Sonia Goncalves, Ewan Harrison, David K. Jackson, Ian Johnston, Dominic Kwiatkowski, Cordelia Langford, John Sillitoe on behalf of the Wellcome Sanger Institute COVID-19 Surveillance Team |
| EPI_ISL_2022080 | Lighthouse Lab in Alderley Park | Wellcome Sanger Institute for the COVID-19 Genomics UK (COG-UK) Consortium | Jacquelyn Wynn, Mairead Hyland, The Lighthouse Lab in Alderley Park and Alex Alderton, Roberto Amato, Jeffrey Barrett, Sonia Goncalves, Ewan Harrison, David K. Jackson, Ian Johnston, Dominic Kwiatkowski, Cordelia Langford, John Sillitoe on behalf of the Wellcome Sanger Institute COVID-19 Surveillance Team |
| EPI_ISL_2022085 | Lighthouse Lab in Milton Keynes | Wellcome Sanger Institute for the COVID-19 Genomics UK (COG-UK) Consortium | The Lighthouse Lab in Milton Keynes and Alex Alderton, Roberto Amato, Jeffrey Barrett, Sonia Goncalves, Ewan Harrison, David K. Jackson, Ian Johnston, Dominic Kwiatkowski, Cordelia Langford, John Sillitoe on behalf of the Wellcome Sanger Institute COVID-19 Surveillance Team |
| EPI_ISL_2022086, EPI_ISL_2022087, EPI_ISL_2022097 | Lighthouse Lab in Alderley Park | Wellcome Sanger Institute for the COVID-19 Genomics UK (COG-UK) Consortium | Jacquelyn Wynn, Mairead Hyland, The Lighthouse Lab in Alderley Park and Alex Alderton, Roberto Amato, Jeffrey Barrett, Sonia Goncalves, Ewan Harrison, David K. Jackson, Ian Johnston, Dominic Kwiatkowski, Cordelia Langford, John Sillitoe on behalf of the Wellcome Sanger Institute COVID-19 Surveillance Team |
| EPI_ISL_2022100, EPI_ISL_2022106, EPI_ISL_2022107 | Lighthouse Lab in Milton Keynes | Wellcome Sanger Institute for the COVID-19 Genomics UK (COG-UK) Consortium | The Lighthouse Lab in Milton Keynes and Alex Alderton, Roberto Amato, Jeffrey Barrett, Sonia Goncalves, Ewan Harrison, David K. Jackson, Ian Johnston, Dominic Kwiatkowski, Cordelia Langford, John Sillitoe on behalf of the Wellcome Sanger Institute COVID-19 Surveillance Team |
| EPI_ISL_2022109 | Lighthouse Lab in Alderley Park | Wellcome Sanger Institute for the COVID-19 Genomics UK (COG-UK) Consortium | Jacquelyn Wynn, Mairead Hyland, The Lighthouse Lab in Alderley Park and Alex Alderton, Roberto Amato, Jeffrey Barrett, Sonia Goncalves, Ewan Harrison, David K. Jackson, Ian Johnston, Dominic Kwiatkowski, Cordelia Langford, John Sillitoe on behalf of the Wellcome Sanger Institute COVID-19 Surveillance Team |
| EPI_ISL_2022118, EPI_ISL_2022122 | Lighthouse Lab in Milton Keynes | Wellcome Sanger Institute for the COVID-19 Genomics UK (COG-UK) Consortium | The Lighthouse Lab in Milton Keynes and Alex Alderton, Roberto Amato, Jeffrey Barrett, Sonia Goncalves, Ewan Harrison, David K. Jackson, Ian Johnston, Dominic Kwiatkowski, Cordelia Langford, John Sillitoe on behalf of the Wellcome Sanger Institute COVID-19 Surveillance Team |
| EPI_ISL_2022124 | Lighthouse Lab in Alderley Park | Wellcome Sanger Institute for the COVID-19 Genomics UK (COG-UK) Consortium | Jacquelyn Wynn, Mairead Hyland, The Lighthouse Lab in Alderley Park and Alex Alderton, Roberto Amato, Jeffrey Barrett, Sonia Goncalves, Ewan Harrison, David K. Jackson, Ian Johnston, Dominic Kwiatkowski, Cordelia Langford, John Sillitoe on behalf of the Wellcome Sanger Institute COVID-19 Surveillance Team |
| EPI_ISL_2022131, EPI_ISL_2022146 | Lighthouse Lab in Milton Keynes | Wellcome Sanger Institute for the COVID-19 Genomics UK (COG-UK) Consortium | The Lighthouse Lab in Milton Keynes and Alex Alderton, Roberto Amato, Jeffrey Barrett, Sonia Goncalves, Ewan Harrison, David K. Jackson, Ian Johnston, Dominic Kwiatkowski, Cordelia Langford, John Sillitoe on behalf of the Wellcome Sanger Institute COVID-19 Surveillance Team |
| EPI_ISL_2022149 | Lighthouse Lab in Alderley Park | Wellcome Sanger Institute for the COVID-19 Genomics UK (COG-UK) Consortium | Jacquelyn Wynn, Mairead Hyland, The Lighthouse Lab in Alderley Park and Alex Alderton, Roberto Amato, Jeffrey Barrett, Sonia Goncalves, Ewan Harrison, David K. Jackson, Ian Johnston, Dominic Kwiatkowski, Cordelia Langford, John Sillitoe on behalf of the Wellcome Sanger Institute COVID-19 Surveillance Team |
| EPI_ISL_2022153, EPI_ISL_2022159 | Lighthouse Lab in Milton Keynes | Wellcome Sanger Institute for the COVID-19 Genomics UK (COG-UK) Consortium | The Lighthouse Lab in Milton Keynes and Alex Alderton, Roberto Amato, Jeffrey Barrett, Sonia Goncalves, Ewan Harrison, David K. Jackson, Ian Johnston, Dominic Kwiatkowski, Cordelia Langford, John Sillitoe on behalf of the Wellcome Sanger Institute COVID-19 Surveillance Team |
| EPI_ISL_2022160 | Lighthouse Lab in Alderley Park | Wellcome Sanger Institute for the COVID-19 Genomics UK (COG-UK) Consortium | Jacquelyn Wynn, Mairead Hyland, The Lighthouse Lab in Alderley Park and Alex Alderton, Roberto Amato, Jeffrey Barrett, Sonia Goncalves, Ewan Harrison, David K. Jackson, Ian Johnston, Dominic Kwiatkowski, Cordelia Langford, John Sillitoe on behalf of the Wellcome Sanger Institute COVID-19 Surveillance Team |
| EPI_ISL_2022161 | Lighthouse Lab in Milton Keynes | Wellcome Sanger Institute for the COVID-19 Genomics UK (COG-UK) Consortium | The Lighthouse Lab in Milton Keynes and Alex Alderton, Roberto Amato, Jeffrey Barrett, Sonia Goncalves, Ewan Harrison, David K. Jackson, Ian Johnston, Dominic Kwiatkowski, Cordelia Langford, John Sillitoe on behalf of the Wellcome Sanger Institute COVID-19 Surveillance Team |
| EPI_ISL_2022162, EPI_ISL_2022163 | Lighthouse Lab in Alderley Park | Wellcome Sanger Institute for the COVID-19 Genomics UK (COG-UK) Consortium | Jacquelyn Wynn, Mairead Hyland, The Lighthouse Lab in Alderley Park and Alex Alderton, Roberto Amato, Jeffrey Barrett, Sonia Goncalves, Ewan Harrison, David K. Jackson, Ian Johnston, Dominic Kwiatkowski, Cordelia Langford, John Sillitoe on behalf of the Wellcome Sanger Institute COVID-19 Surveillance Team |
| EPI_ISL_2022167 | Lighthouse Lab in Milton Keynes | Wellcome Sanger Institute for the COVID-19 Genomics UK (COG-UK) Consortium | The Lighthouse Lab in Milton Keynes and Alex Alderton, Roberto Amato, Jeffrey Barrett, Sonia Goncalves, Ewan Harrison, David K. Jackson, Ian Johnston, Dominic Kwiatkowski, Cordelia Langford, John Sillitoe on behalf of the Wellcome Sanger Institute COVID-19 Surveillance Team |
| EPI_ISL_2022171, EPI_ISL_2022172, EPI_ISL_2022181 | Lighthouse Lab in Alderley Park | Wellcome Sanger Institute for the COVID-19 Genomics UK (COG-UK) Consortium | Jacquelyn Wynn, Mairead Hyland, The Lighthouse Lab in Alderley Park and Alex Alderton, Roberto Amato, Jeffrey Barrett, Sonia Goncalves, Ewan Harrison, David K. Jackson, Ian Johnston, Dominic Kwiatkowski, Cordelia Langford, John Sillitoe on behalf of the Wellcome Sanger Institute COVID-19 Surveillance Team |
| EPI_ISL_2022186 | Lighthouse Lab in Milton Keynes | Wellcome Sanger Institute for the COVID-19 Genomics UK (COG-UK) Consortium | The Lighthouse Lab in Milton Keynes and Alex Alderton, Roberto Amato, Jeffrey Barrett, Sonia Goncalves, Ewan Harrison, David K. Jackson, Ian Johnston, Dominic Kwiatkowski, Cordelia Langford, John Sillitoe on behalf of the Wellcome Sanger Institute COVID-19 Surveillance Team |
| EPI_ISL_2022188, EPI_ISL_2022189, EPI_ISL_2022193, EPI_ISL_2022196, EPI_ISL_2022198 | Lighthouse Lab in Alderley Park | Wellcome Sanger Institute for the COVID-19 Genomics UK (COG-UK) Consortium | Jacquelyn Wynn, Mairead Hyland, The Lighthouse Lab in Alderley Park and Alex Alderton, Roberto Amato, Jeffrey Barrett, Sonia Goncalves, Ewan Harrison, David K. Jackson, Ian Johnston, Dominic Kwiatkowski, Cordelia Langford, John Sillitoe on behalf of the Wellcome Sanger Institute COVID-19 Surveillance Team |
| EPI_ISL_2022200 | Lighthouse Lab in Milton Keynes | Wellcome Sanger Institute for the COVID-19 Genomics UK (COG-UK) Consortium | The Lighthouse Lab in Milton Keynes and Alex Alderton, Roberto Amato, Jeffrey Barrett, Sonia Goncalves, Ewan Harrison, David K. Jackson, Ian Johnston, Dominic Kwiatkowski, Cordelia Langford, John Sillitoe on behalf of the Wellcome Sanger Institute COVID-19 Surveillance Team |
| EPI_ISL_2022205, EPI_ISL_2022209, EPI_ISL_2022216, EPI_ISL_2022221 | Lighthouse Lab in Alderley Park | Wellcome Sanger Institute for the COVID-19 Genomics UK (COG-UK) Consortium | Jacquelyn Wynn, Mairead Hyland, The Lighthouse Lab in Alderley Park and Alex Alderton, Roberto Amato, Jeffrey Barrett, Sonia Goncalves, Ewan Harrison, David K. Jackson, Ian Johnston, Dominic Kwiatkowski, Cordelia Langford, John Sillitoe on behalf of the Wellcome Sanger Institute COVID-19 Surveillance Team |
| EPI_ISL_2022224 | Lighthouse Lab in Milton Keynes | Wellcome Sanger Institute for the COVID-19 Genomics UK (COG-UK) Consortium | The Lighthouse Lab in Milton Keynes and Alex Alderton, Roberto Amato, Jeffrey Barrett, Sonia Goncalves, Ewan Harrison, David K. Jackson, Ian Johnston, Dominic Kwiatkowski, Cordelia Langford, John Sillitoe on behalf of the Wellcome Sanger Institute COVID-19 Surveillance Team |
| EPI_ISL_2022226 | Lighthouse Lab in Alderley Park | Wellcome Sanger Institute for the COVID-19 Genomics UK (COG-UK) Consortium | Jacquelyn Wynn, Mairead Hyland, The Lighthouse Lab in Alderley Park and Alex Alderton, Roberto Amato, Jeffrey Barrett, Sonia Goncalves, Ewan Harrison, David K. Jackson, Ian Johnston, Dominic Kwiatkowski, Cordelia Langford, John Sillitoe on behalf of the Wellcome Sanger Institute COVID-19 Surveillance Team |
| EPI_ISL_2022239 | Lighthouse Lab in Milton Keynes | Wellcome Sanger Institute for the COVID-19 Genomics UK (COG-UK) Consortium | The Lighthouse Lab in Milton Keynes and Alex Alderton, Roberto Amato, Jeffrey Barrett, Sonia Goncalves, Ewan Harrison, David K. Jackson, Ian Johnston, Dominic Kwiatkowski, Cordelia Langford, John Sillitoe on behalf of the Wellcome Sanger Institute COVID-19 Surveillance Team |
| EPI_ISL_2022249 | Lighthouse Lab in Alderley Park | Wellcome Sanger Institute for the COVID-19 Genomics UK (COG-UK) Consortium | Jacquelyn Wynn, Mairead Hyland, The Lighthouse Lab in Alderley Park and Alex Alderton, Roberto Amato, Jeffrey Barrett, Sonia Goncalves, Ewan Harrison, David K. Jackson, Ian Johnston, Dominic Kwiatkowski, Cordelia Langford, John Sillitoe on behalf of the Wellcome Sanger Institute COVID-19 Surveillance Team |
| EPI_ISL_2022250, EPI_ISL_2022264 | Lighthouse Lab in Milton Keynes | Wellcome Sanger Institute for the COVID-19 Genomics UK (COG-UK) Consortium | The Lighthouse Lab in Milton Keynes and Alex Alderton, Roberto Amato, Jeffrey Barrett, Sonia Goncalves, Ewan Harrison, David K. Jackson, Ian Johnston, Dominic Kwiatkowski, Cordelia Langford, John Sillitoe on behalf of the Wellcome Sanger Institute COVID-19 Surveillance Team |
| EPI_ISL_2022267, EPI_ISL_2022275 | Lighthouse Lab in Alderley Park | Wellcome Sanger Institute for the COVID-19 Genomics UK (COG-UK) Consortium | Jacquelyn Wynn, Mairead Hyland, The Lighthouse Lab in Alderley Park and Alex Alderton, Roberto Amato, Jeffrey Barrett, Sonia Goncalves, Ewan Harrison, David K. Jackson, Ian Johnston, Dominic Kwiatkowski, Cordelia Langford, John Sillitoe on behalf of the Wellcome Sanger Institute COVID-19 Surveillance Team |
| EPI_ISL_2022280, EPI_ISL_2022290, EPI_ISL_2022295 | Lighthouse Lab in Milton Keynes | Wellcome Sanger Institute for the COVID-19 Genomics UK (COG-UK) Consortium | The Lighthouse Lab in Milton Keynes and Alex Alderton, Roberto Amato, Jeffrey Barrett, Sonia Goncalves, Ewan Harrison, David K. Jackson, Ian Johnston, Dominic Kwiatkowski, Cordelia Langford, John Sillitoe on behalf of the Wellcome Sanger Institute COVID-19 Surveillance Team |

| | | | |
|---|---|---|---|
| EPI_ISL_2022300 | Lighthouse Lab in Alderley Park | Wellcome Sanger Institute for the COVID-19 Genomics UK (COG-UK) Consortium | Jacquelyn Wynn, Mairead Hyland, The Lighthouse Lab in Alderley Park and Alex Alderton, Roberto Amato, Jeffrey Barrett, Sonia Goncalves, Ewan Harrison, David K. Jackson, Ian Johnston, Dominic Kwiatkowski, Cordelia Langford, John Sillitoe on behalf of the Wellcome Sanger Institute COVID-19 Surveillance Team |
| EPI_ISL_2022306 | Lighthouse Lab in Milton Keynes | Wellcome Sanger Institute for the COVID-19 Genomics UK (COG-UK) Consortium | The Lighthouse Lab in Milton Keynes and Alex Alderton, Roberto Amato, Jeffrey Barrett, Sonia Goncalves, Ewan Harrison, David K. Jackson, Ian Johnston, Dominic Kwiatkowski, Cordelia Langford, John Sillitoe on behalf of the Wellcome Sanger Institute COVID-19 Surveillance Team |
| EPI_ISL_2022307 | Lighthouse Lab in Alderley Park | Wellcome Sanger Institute for the COVID-19 Genomics UK (COG-UK) Consortium | Jacquelyn Wynn, Mairead Hyland, The Lighthouse Lab in Alderley Park and Alex Alderton, Roberto Amato, Jeffrey Barrett, Sonia Goncalves, Ewan Harrison, David K. Jackson, Ian Johnston, Dominic Kwiatkowski, Cordelia Langford, John Sillitoe on behalf of the Wellcome Sanger Institute COVID-19 Surveillance Team |
| EPI_ISL_2022308 | Lighthouse Lab in Milton Keynes | Wellcome Sanger Institute for the COVID-19 Genomics UK (COG-UK) Consortium | The Lighthouse Lab in Milton Keynes and Alex Alderton, Roberto Amato, Jeffrey Barrett, Sonia Goncalves, Ewan Harrison, David K. Jackson, Ian Johnston, Dominic Kwiatkowski, Cordelia Langford, John Sillitoe on behalf of the Wellcome Sanger Institute COVID-19 Surveillance Team |
| EPI_ISL_2022315, EPI_ISL_2022318, EPI_ISL_2022325, EPI_ISL_2022329, EPI_ISL_2022335, EPI_ISL_2022336 | Lighthouse Lab in Alderley Park | Wellcome Sanger Institute for the COVID-19 Genomics UK (COG-UK) Consortium | Jacquelyn Wynn, Mairead Hyland, The Lighthouse Lab in Alderley Park and Alex Alderton, Roberto Amato, Jeffrey Barrett, Sonia Goncalves, Ewan Harrison, David K. Jackson, Ian Johnston, Dominic Kwiatkowski, Cordelia Langford, John Sillitoe on behalf of the Wellcome Sanger Institute COVID-19 Surveillance Team |
| EPI_ISL_2022337 | Lighthouse Lab in Milton Keynes | Wellcome Sanger Institute for the COVID-19 Genomics UK (COG-UK) Consortium | The Lighthouse Lab in Milton Keynes and Alex Alderton, Roberto Amato, Jeffrey Barrett, Sonia Goncalves, Ewan Harrison, David K. Jackson, Ian Johnston, Dominic Kwiatkowski, Cordelia Langford, John Sillitoe on behalf of the Wellcome Sanger Institute COVID-19 Surveillance Team |
| EPI_ISL_2022338, EPI_ISL_2022347, EPI_ISL_2022353 | Lighthouse Lab in Alderley Park | Wellcome Sanger Institute for the COVID-19 Genomics UK (COG-UK) Consortium | Jacquelyn Wynn, Mairead Hyland, The Lighthouse Lab in Alderley Park and Alex Alderton, Roberto Amato, Jeffrey Barrett, Sonia Goncalves, Ewan Harrison, David K. Jackson, Ian Johnston, Dominic Kwiatkowski, Cordelia Langford, John Sillitoe on behalf of the Wellcome Sanger Institute COVID-19 Surveillance Team |
| EPI_ISL_2022356 | Lighthouse Lab in Milton Keynes | Wellcome Sanger Institute for the COVID-19 Genomics UK (COG-UK) Consortium | The Lighthouse Lab in Milton Keynes and Alex Alderton, Roberto Amato, Jeffrey Barrett, Sonia Goncalves, Ewan Harrison, David K. Jackson, Ian Johnston, Dominic Kwiatkowski, Cordelia Langford, John Sillitoe on behalf of the Wellcome Sanger Institute COVID-19 Surveillance Team |
| EPI_ISL_2022368, EPI_ISL_2022371 | Lighthouse Lab in Alderley Park | Wellcome Sanger Institute for the COVID-19 Genomics UK (COG-UK) Consortium | Jacquelyn Wynn, Mairead Hyland, The Lighthouse Lab in Alderley Park and Alex Alderton, Roberto Amato, Jeffrey Barrett, Sonia Goncalves, Ewan Harrison, David K. Jackson, Ian Johnston, Dominic Kwiatkowski, Cordelia Langford, John Sillitoe on behalf of the Wellcome Sanger Institute COVID-19 Surveillance Team |
| EPI_ISL_2022375 | Lighthouse Lab in Milton Keynes | Wellcome Sanger Institute for the COVID-19 Genomics UK (COG-UK) Consortium | The Lighthouse Lab in Milton Keynes and Alex Alderton, Roberto Amato, Jeffrey Barrett, Sonia Goncalves, Ewan Harrison, David K. Jackson, Ian Johnston, Dominic Kwiatkowski, Cordelia Langford, John Sillitoe on behalf of the Wellcome Sanger Institute COVID-19 Surveillance Team |
| EPI_ISL_2022377, EPI_ISL_2022379, EPI_ISL_2022382 | Lighthouse Lab in Alderley Park | Wellcome Sanger Institute for the COVID-19 Genomics UK (COG-UK) Consortium | Jacquelyn Wynn, Mairead Hyland, The Lighthouse Lab in Alderley Park and Alex Alderton, Roberto Amato, Jeffrey Barrett, Sonia Goncalves, Ewan Harrison, David K. Jackson, Ian Johnston, Dominic Kwiatkowski, Cordelia Langford, John Sillitoe on behalf of the Wellcome Sanger Institute COVID-19 Surveillance Team |
| EPI_ISL_2022383 | Lighthouse Lab in Milton Keynes | Wellcome Sanger Institute for the COVID-19 Genomics UK (COG-UK) Consortium | The Lighthouse Lab in Milton Keynes and Alex Alderton, Roberto Amato, Jeffrey Barrett, Sonia Goncalves, Ewan Harrison, David K. Jackson, Ian Johnston, Dominic Kwiatkowski, Cordelia Langford, John Sillitoe on behalf of the Wellcome Sanger Institute COVID-19 Surveillance Team |
| EPI_ISL_2022386, EPI_ISL_2022390, EPI_ISL_2022403, EPI_ISL_2022404, EPI_ISL_2022426 | Lighthouse Lab in Alderley Park | Wellcome Sanger Institute for the COVID-19 Genomics UK (COG-UK) Consortium | Jacquelyn Wynn, Mairead Hyland, The Lighthouse Lab in Alderley Park and Alex Alderton, Roberto Amato, Jeffrey Barrett, Sonia Goncalves, Ewan Harrison, David K. Jackson, Ian Johnston, Dominic Kwiatkowski, Cordelia Langford, John Sillitoe on behalf of the Wellcome Sanger Institute COVID-19 Surveillance Team |
| EPI_ISL_2022429 | Lighthouse Lab in Milton Keynes | Wellcome Sanger Institute for the COVID-19 Genomics UK (COG-UK) Consortium | The Lighthouse Lab in Milton Keynes and Alex Alderton, Roberto Amato, Jeffrey Barrett, Sonia Goncalves, Ewan Harrison, David K. Jackson, Ian Johnston, Dominic Kwiatkowski, Cordelia Langford, John Sillitoe on behalf of the Wellcome Sanger Institute COVID-19 Surveillance Team |
| EPI_ISL_2022431 | Lighthouse Lab in Alderley Park | Wellcome Sanger Institute for the COVID-19 Genomics UK (COG-UK) Consortium | Jacquelyn Wynn, Mairead Hyland, The Lighthouse Lab in Alderley Park and Alex Alderton, Roberto Amato, Jeffrey Barrett, Sonia Goncalves, Ewan Harrison, David K. Jackson, Ian Johnston, Dominic Kwiatkowski, Cordelia Langford, John Sillitoe on behalf of the Wellcome Sanger Institute COVID-19 Surveillance Team |
| EPI_ISL_2022494, EPI_ISL_2022501 | Lighthouse Lab in Milton Keynes | Wellcome Sanger Institute for the COVID-19 Genomics UK (COG-UK) Consortium | The Lighthouse Lab in Milton Keynes and Alex Alderton, Roberto Amato, Jeffrey Barrett, Sonia Goncalves, Ewan Harrison, David K. Jackson, Ian Johnston, Dominic Kwiatkowski, Cordelia Langford, John Sillitoe on behalf of the Wellcome Sanger Institute COVID-19 Surveillance Team |
| EPI_ISL_2022507 | Randox Laboratories | Wellcome Sanger Institute for the COVID-19 Genomics UK (COG-UK) Consortium | Randox Laboratories and Alex Alderton, Roberto Amato, Jeffrey Barrett, Sonia Goncalves, Ewan Harrison, David K. Jackson, Ian Johnston, Dominic Kwiatkowski, Cordelia Langford, John Sillitoe on behalf of the Wellcome Sanger Institute COVID-19 Surveillance Team |
| EPI_ISL_2022513 | Lighthouse Lab in Milton Keynes | Wellcome Sanger Institute for the COVID-19 Genomics UK (COG-UK) Consortium | The Lighthouse Lab in Milton Keynes and Alex Alderton, Roberto Amato, Jeffrey Barrett, Sonia Goncalves, Ewan Harrison, David K. Jackson, Ian Johnston, Dominic Kwiatkowski, Cordelia Langford, John Sillitoe on behalf of the Wellcome Sanger Institute COVID-19 Surveillance Team |
| EPI_ISL_2022525 | Randox Laboratories | Wellcome Sanger Institute for the COVID-19 Genomics UK (COG-UK) Consortium | Randox Laboratories and Alex Alderton, Roberto Amato, Jeffrey Barrett, Sonia Goncalves, Ewan Harrison, David K. Jackson, Ian Johnston, Dominic Kwiatkowski, Cordelia Langford, John Sillitoe on behalf of the Wellcome Sanger Institute COVID-19 Surveillance Team |
| EPI_ISL_2022526 | Lighthouse Lab in Milton Keynes | Wellcome Sanger Institute for the COVID-19 Genomics UK (COG-UK) Consortium | The Lighthouse Lab in Milton Keynes and Alex Alderton, Roberto Amato, Jeffrey Barrett, Sonia Goncalves, Ewan Harrison, David K. Jackson, Ian Johnston, Dominic Kwiatkowski, Cordelia Langford, John Sillitoe on behalf of the Wellcome Sanger Institute COVID-19 Surveillance Team |
| EPI_ISL_2022532, EPI_ISL_2022536, EPI_ISL_2022537, EPI_ISL_2022550, EPI_ISL_2022552 | Health Services Laboratories | Wellcome Sanger Institute for the COVID-19 Genomics UK (COG-UK) Consortium | Health Services Laboratories and Alex Alderton, Roberto Amato, Jeffrey Barrett, Sonia Goncalves, Ewan Harrison, David K. Jackson, Ian Johnston, Dominic Kwiatkowski, Cordelia Langford, John Sillitoe on behalf of the Wellcome Sanger Institute COVID-19 Surveillance Team |
| EPI_ISL_2022555 | Lighthouse Lab in Glasgow | Wellcome Sanger Institute for the COVID-19 Genomics UK (COG-UK) Consortium | Harper VanSteenhouse, Yumi Kasai, David Gray, Carol Clugston, Anna Dominiczak and Alex Alderton, Roberto Amato, Jeffrey Barrett, Sonia Goncalves, Ewan Harrison, David K. Jackson, Ian Johnston, Dominic Kwiatkowski, Cordelia Langford, John Sillitoe on behalf of the Wellcome Sanger Institute COVID-19 Surveillance Team |
| EPI_ISL_2022557, EPI_ISL_2022558, EPI_ISL_2022559, EPI_ISL_2022561, EPI_ISL_2022562, EPI_ISL_2022570, EPI_ISL_2022575 | Health Services Laboratories | Wellcome Sanger Institute for the COVID-19 Genomics UK (COG-UK) Consortium | Health Services Laboratories and Alex Alderton, Roberto Amato, Jeffrey Barrett, Sonia Goncalves, Ewan Harrison, David K. Jackson, Ian Johnston, Dominic Kwiatkowski, Cordelia Langford, John Sillitoe on behalf of the Wellcome Sanger Institute COVID-19 Surveillance Team |
| EPI_ISL_2022577 | Lighthouse Lab in Milton Keynes | Wellcome Sanger Institute for the COVID-19 Genomics UK (COG-UK) Consortium | The Lighthouse Lab in Milton Keynes and Alex Alderton, Roberto Amato, Jeffrey Barrett, Sonia Goncalves, Ewan Harrison, David K. Jackson, Ian Johnston, Dominic Kwiatkowski, Cordelia Langford, John Sillitoe on behalf of the Wellcome Sanger Institute COVID-19 Surveillance Team |
| EPI_ISL_2022578 | Randox Laboratories | Wellcome Sanger Institute for the COVID-19 Genomics UK (COG-UK) Consortium | Randox Laboratories and Alex Alderton, Roberto Amato, Jeffrey Barrett, Sonia Goncalves, Ewan Harrison, David K. Jackson, Ian Johnston, Dominic Kwiatkowski, Cordelia Langford, John Sillitoe on behalf of the Wellcome Sanger Institute COVID-19 Surveillance Team |
| EPI_ISL_2022579, EPI_ISL_2022584 | Lighthouse Lab in Milton Keynes | Wellcome Sanger Institute for the COVID-19 Genomics UK (COG-UK) Consortium | The Lighthouse Lab in Milton Keynes and Alex Alderton, Roberto Amato, Jeffrey Barrett, Sonia Goncalves, Ewan Harrison, David K. Jackson, Ian Johnston, Dominic Kwiatkowski, Cordelia Langford, John Sillitoe on behalf of the Wellcome Sanger Institute COVID-19 Surveillance Team |
| EPI_ISL_2022593, EPI_ISL_2022634, EPI_ISL_2022661, EPI_ISL_2022726, EPI_ISL_2022753, EPI_ISL_2022792, EPI_ISL_2022865 | Lighthouse Lab in Glasgow | Wellcome Sanger Institute for the COVID-19 Genomics UK (COG-UK) Consortium | Harper VanSteenhouse, Yumi Kasai, David Gray, Carol Clugston, Anna Dominiczak and Alex Alderton, Roberto Amato, Jeffrey Barrett, Sonia Goncalves, Ewan Harrison, David K. Jackson, Ian Johnston, Dominic Kwiatkowski, Cordelia Langford, John Sillitoe on behalf of the Wellcome Sanger Institute COVID-19 Surveillance Team |
| EPI_ISL_2022902, EPI_ISL_2022913 | Lighthouse Lab in Milton Keynes | Wellcome Sanger Institute for the COVID-19 Genomics UK (COG-UK) Consortium | The Lighthouse Lab in Milton Keynes and Alex Alderton, Roberto Amato, Jeffrey Barrett, Sonia Goncalves, Ewan Harrison, David K. Jackson, Ian Johnston, Dominic Kwiatkowski, Cordelia Langford, John Sillitoe on behalf of the Wellcome Sanger Institute COVID-19 Surveillance Team |
| EPI_ISL_2022978 | Lighthouse Lab in Glasgow | Wellcome Sanger Institute for the COVID-19 Genomics UK (COG-UK) Consortium | Harper VanSteenhouse, Yumi Kasai, David Gray, Carol Clugston, Anna Dominiczak and Alex Alderton, Roberto Amato, Jeffrey Barrett, Sonia Goncalves, Ewan Harrison, David K. Jackson, Ian Johnston, Dominic Kwiatkowski, Cordelia Langford, John Sillitoe on behalf of the Wellcome Sanger Institute COVID-19 Surveillance Team |
| EPI_ISL_2022979 | Lighthouse Lab in Alderley Park | Wellcome Sanger Institute for the COVID-19 Genomics UK (COG-UK) Consortium | Jacquelyn Wynn, Mairead Hyland, The Lighthouse Lab in Alderley Park and Alex Alderton, Roberto Amato, Jeffrey Barrett, Sonia Goncalves, Ewan Harrison, David K. Jackson, Ian Johnston, Dominic Kwiatkowski, Cordelia Langford, John Sillitoe on behalf of the Wellcome Sanger Institute COVID-19 Surveillance Team |
| EPI_ISL_2023030, EPI_ISL_2023031 | Lighthouse Lab in Milton Keynes | Wellcome Sanger Institute for the COVID-19 Genomics UK (COG-UK) Consortium | The Lighthouse Lab in Milton Keynes and Alex Alderton, Roberto Amato, Jeffrey Barrett, Sonia Goncalves, Ewan Harrison, David K. Jackson, Ian Johnston, Dominic Kwiatkowski, Cordelia Langford, John Sillitoe on behalf of the Wellcome Sanger Institute COVID-19 Surveillance Team |
| EPI_ISL_2023939, EPI_ISL_2024372, EPI_ISL_2024800, EPI_ISL_2024848, EPI_ISL_2024988, EPI_ISL_2024994, EPI_ISL_2025050, EPI_ISL_2025253 | Department of Virus and Microbiological Special Diagnostics, Statens Serum Institut, Copenhagen, Denmark | Aalborg University | Danish Covid-19 Genome Consortium |
| EPI_ISL_2025798 | Department of Clinical Microbiology and Center for Genomic Medicine, Rigshospitalet, Copenhagen, Denmark | Aalborg University | Danish Covid-19 Genome Consortium |

EPI_ISL_2026084, EPI_ISL_2026230, EPI_ISL_2026527, EPI_ISL_2026729, EPI_ISL_2026828, EPI_ISL_2027124, EPI_ISL_2027238, EPI_ISL_2027359, EPI_ISL_2027537, EPI_ISL_2027667, EPI_ISL_2027848, EPI_ISL_2028708, EPI_ISL_2028728, EPI_ISL_2028832

| | | | |
|---|---|---|---|
| see above | Department of Virus and Microbiological Special Diagnostics, Statens Serum Institut, Copenhagen, Denmark | Aalborg University | Danish Covid-19 Genome Consortium |
| EPI_ISL_2029113, EPI_ISL_2029114 | HEGP - Laboratoire de Virologie | HEGP - Laboratoire de Virologie | David Veyer, Sandrine Imbeaud, Julien Puech, Hélène Péré, Maxime Wack, Nicolas Robillard |
| EPI_ISL_2029129, EPI_ISL_2029134, EPI_ISL_2029136, EPI_ISL_2029137, EPI_ISL_2029171, EPI_ISL_2029172, EPI_ISL_2029173, EPI_ISL_2029174, EPI_ISL_2029353, EPI_ISL_2029354 | National Virus Reference Laboratory | National Virus Reference Laboratory | Zoe Yandle, Charlene Bennett, Gabriel Gonzalez, Michael Carr, Jonathan Dean, Cillian F De Gascun |
| EPI_ISL_2029687, EPI_ISL_2029713 | Labo Analyses Med | National Reference Center for Viruses of Respiratory Infections, Institut Pasteur, Paris | Marion Barbet, Sylvie Behillil, Méline Bizard, Angela Brisebarre, Camille Capel, Vincent Enouf, Louise Lefrançois, Frédéric Lemoine, Christophe Malabat, Corinne Maufrais, Etienne Simon-Lorière, Maud Vanpeene, Sylvie Van der Werf ,Pierre Netzer |
| EPI_ISL_2029727, EPI_ISL_2029740 | Labo Analyses Med | National Reference Center for Viruses of Respiratory Infections, Institut Pasteur, Paris | Marion Barbet, Sylvie Behillil, Méline Bizard, Angela Brisebarre, Camille Capel, Vincent Enouf, Louise Lefrançois, Frédéric Lemoine, Christophe Malabat, Corinne Maufrais, Pierre Lechat, Etienne Simon-Lorière, Maud Vanpeene, Sylvie Van der Werf ,Jean Boyer |
| EPI_ISL_2029751 | Labo Analyses Med | National Reference Center for Viruses of Respiratory Infections, Institut Pasteur, Paris | Marion Barbet, Sylvie Behillil, Méline Bizard, Angela Brisebarre, Camille Capel, Vincent Enouf, Louise Lefrançois, Frédéric Lemoine, Christophe Malabat, Corinne Maufrais, Etienne Simon-Lorière, Maud Vanpeene, Sylvie Van der Werf ,Judith Zerah |
| EPI_ISL_2029877 | Labo Analyses Med | National Reference Center for Viruses of Respiratory Infections, Institut Pasteur, Paris | Marion Barbet, Sylvie Behillil, Méline Bizard, Angela Brisebarre, Camille Capel, Vincent Enouf, Louise Lefrançois, Frédéric Lemoine, Christophe Malabat, Corinne Maufrais, Victoire Baillet, Etienne Simon-Lorière, Maud Vanpeene, Sylvie Van der Werf ,Judith Zerah |
| EPI_ISL_2029916 | Labo Analyses Med | National Reference Center for Viruses of Respiratory Infections, Institut Pasteur, Paris | Marion Barbet, Sylvie Behillil, Méline Bizard, Angela Brisebarre, Camille Capel, Vincent Enouf, Louise Lefrançois, Frédéric Lemoine, Christophe Malabat, Corinne Maufrais, Etienne Simon-Lorière, Maud Vanpeene, Sylvie Van der Werf ,Elodie Etienne |
| EPI_ISL_2029931 | Hospital | National Reference Center for Viruses of Respiratory Infections, Institut Pasteur, Paris | Marion Barbet, Sylvie Behillil, Méline Bizard, Angela Brisebarre, Camille Capel, Vincent Enouf, Louise Lefrançois, Frédéric Lemoine, Christophe Malabat, Corinne Maufrais, Etienne Simon-Lorière, Maud Vanpeene, Sylvie Van der Werf ,Nabil Gastli |
| EPI_ISL_2031271, EPI_ISL_2031294, EPI_ISL_2031501, EPI_ISL_2031502, EPI_ISL_2031503, EPI_ISL_2031512, EPI_ISL_2031536 | UW Virology Lab | UW Virology Lab | Pavitra Roychoudhury, Hong Xie, Lasata Shrestha, Tien V. Nguyen, Shah Mohamed Bakhash, Michelle Lin, Noah R. Baker, Sean Ellis, Meei-Li Huang, Keith R Jerome, Alexander Greninger |
| EPI_ISL_2032019 | Laboratory of Molecular Biology, Diagnostyka sp. z o.o. | genXone SA, Research & Development Laboratory | Maciej Sykulski, Grzegorz Nowicki, Jakub Grabowski, Natalia Drweska-Matelska, Anna Brylak-Blaszkow, Aleksandra Gidlewicz, Karol Szeszko, Lukasz Krych, Michal Kaszuba |
| EPI_ISL_2032644 | Federal Budget Health Care Institution "Center of Hygiene and Epidemiology in Tver region" | Group of Genomics and Postgenomic Technologies of Central Research Institute of Epidemiology | Samoilov AE, Kaptelova VV, Korneenko EV, Saenko SS, Smirnova YS, Nadtoka MI, Melushenkova TG, Speranskaya AS, Tivanova EV, Kondrasheva LY, Akimkin VG |
| EPI_ISL_2032727, EPI_ISL_2033405 | Swedish national genomic surveillance program of SARS-CoV-2 | The Public Health Agency of Sweden | Maximilian Riess, Maria Lind Karlberg, Alma Brolund, Swedish national genomic surveillance program of SARS-CoV-2 |
| EPI_ISL_2033745 | Translational Health Science and Technology Institute -ESIC medical college and hospital, Faridabad | THSTI Bioassay laboratory | Naseem Ahmed Khan, Heena Shaman, Jigme Wangchuk, Anil Pandey, Akshay Kanakan, Ranjeet Maurya, Janani Srinivasa Vasudevan, Neha Jha, Priyanka Mehta, Rajesh Pandey, Guruprasad R Medigeshi |
| EPI_ISL_2033751, EPI_ISL_2034582, EPI_ISL_2034589 | Swedish national genomic surveillance program of SARS-CoV-2 | The Public Health Agency of Sweden | Maximilian Riess, Maria Lind Karlberg, Alma Brolund, Swedish national genomic surveillance program of SARS-CoV-2 |
| EPI_ISL_2034693 | Laboratory of Molecular Biology, Diagnostyka sp. z o.o. | genXone SA, Research & Development Laboratory | Maciej Sykulski, Grzegorz Nowicki, Jakub Grabowski, Natalia Drweska-Matelska, Anna Brylak-Blaszkow, Aleksandra Gidlewicz, Karol Szeszko, Lukasz Krych, Michal Kaszuba |
| EPI_ISL_2035068 | Yale Clinical Virology Lab | Yale Center for Genomic Analysis | Shrikant Mane, Kaya Bilguvar, Curt Scharfe, Irina Tikhonova, Brooke Sullivan |
| EPI_ISL_2035283 | Laboratory of Molecular Biology, Diagnostyka sp. z o.o. | genXone SA, Research & Development Laboratory | Maciej Sykulski, Grzegorz Nowicki, Jakub Grabowski, Natalia Drweska-Matelska, Anna Brylak-Blaszkow, Aleksandra Gidlewicz, Karol Szeszko, Lukasz Krych, Michal Kaszuba |
| EPI_ISL_2035597 | Missouri State Public Health Laboratory | Missouri State Public Health Laboratory | Matthew Sinn, Joshua Barry, Ashley New |
| EPI_ISL_2035736 | San Gallicano Dermatological Institute I.F.O. | INMI Lazzaro Spallanzani IRCCS | B Bartolini, E Giombini, F Messina, M Rueca, G Bonfiglio, O Butera, CEM Gruber, F Santini, F Pimpinelli, F Ensoli, I Cavallo, A Di Caro, MR Capobianchi |
| EPI_ISL_2035737 | San Gallicano Dermatological Institute I.F.O. | INMI Lazzaro Spallanzani IRCCS | E Giombini, F Messina, M Rueca, G Bonfiglio, O Butera, CEM Gruber, F Santini, B Bartolini, F Pimpinelli, F Ensoli, G Prignano, MR Capobianchi, A Di Caro |
| EPI_ISL_2035738 | San Gallicano Dermatological Institute I.F.O. | INMI Lazzaro Spallanzani IRCCS | F Messina, M Rueca, G Bonfiglio, O Butera, CEM Gruber, F Santini, B Bartolini, E Giombini, F Pimpinelli, F Ensoli, A Mastrofrancesco, A Di Caro, MR Capobianchi |
| EPI_ISL_2035739 | San Gallicano Dermatological Institute I.F.O. | INMI Lazzaro Spallanzani IRCCS | M Rueca, G Bonfiglio, O Butera, CEM Gruber, F Santini, B Bartolini, E Giombini, F Messina, F Pimpinelli, F Ensoli, EG Di Domenico, MR Capobianchi, A Di Caro |
| EPI_ISL_2035740 | San Gallicano Dermatological Institute I.F.O. | INMI Lazzaro Spallanzani IRCCS | G Bonfiglio, O Butera, CEM Gruber, F Santini, B Bartolini, E Giombini, F Messina, M Rueca, F Pimpinelli, F Ensoli, M Pontone, A Di Caro, MR Capobianchi |
| EPI_ISL_2035741 | San Gallicano Dermatological Institute I.F.O. | INMI Lazzaro Spallanzani IRCCS | O Butera, CEM Gruber, F Santini, B Bartolini, E Giombini, F Messina, M Rueca, G Bonfiglio, F Pimpinelli, F Ensoli, G D'Agosto, MR Capobianchi, A Di Caro |
| EPI_ISL_2035742 | San Gallicano Dermatological Institute I.F.O. | INMI Lazzaro Spallanzani IRCCS | CEM Gruber, F Santini, B Bartolini, E Giombini, F Messina, M Rueca, G Bonfiglio, O Butera, F Pimpinelli, F Ensoli, E Trento, A Di Caro, MR Capobianchi |
| EPI_ISL_2035743 | San Gallicano Dermatological Institute I.F.O. | INMI Lazzaro Spallanzani IRCCS | F Santini, B Bartolini, E Giombini, F Messina, M Rueca, G Bonfiglio, O Butera, CEM Gruber, F Pimpinelli, F Ensoli, I Cavallo, MR Capobianchi, A Di Caro |
| EPI_ISL_2035744 | San Gallicano Dermatological Institute I.F.O. | INMI Lazzaro Spallanzani IRCCS | B Bartolini, E Giombini, F Messina, M Rueca, G Bonfiglio, O Butera, CEM Gruber, F Santini, F Pimpinelli, F Ensoli, G Prignano, A Di Caro, MR Capobianchi |
| EPI_ISL_2035745 | San Gallicano Dermatological Institute I.F.O. | INMI Lazzaro Spallanzani IRCCS | E Giombini, F Messina, M Rueca, G Bonfiglio, O Butera, CEM Gruber, F Santini, B Bartolini, F Pimpinelli, F Ensoli, A Mastrofrancesco, MR Capobianchi, A Di Caro |
| EPI_ISL_2035746 | San Gallicano Dermatological Institute I.F.O. | INMI Lazzaro Spallanzani IRCCS | F Messina, M Rueca, G Bonfiglio, O Butera, CEM Gruber, F Santini, B Bartolini, E Giombini, F Pimpinelli, F Ensoli, EG Di Domenico, A Di Caro, MR Capobianchi |
| EPI_ISL_2035747 | San Gallicano Dermatological Institute I.F.O. | INMI Lazzaro Spallanzani IRCCS | M Rueca, G Bonfiglio, O Butera, CEM Gruber, F Santini, B Bartolini, E Giombini, F Messina, F Pimpinelli, F Ensoli, M Pontone, MR Capobianchi, A Di Caro |
| EPI_ISL_2035748 | San Gallicano Dermatological Institute I.F.O. | INMI Lazzaro Spallanzani IRCCS | G Bonfiglio, O Butera, CEM Gruber, F Santini, B Bartolini, E Giombini, F Messina, M Rueca, F Pimpinelli, F Ensoli, G D'Agosto, A Di Caro, MR Capobianchi |
| EPI_ISL_2035749 | San Gallicano Dermatological Institute I.F.O. | INMI Lazzaro Spallanzani IRCCS | O Butera, CEM Gruber, F Santini, B Bartolini, E Giombini, F Messina, M Rueca, G Bonfiglio, F Pimpinelli, F Ensoli, E Trento, MR Capobianchi, A Di Caro |
| EPI_ISL_2035750 | San Gallicano Dermatological Institute I.F.O. | INMI Lazzaro Spallanzani IRCCS | CEM Gruber, F Santini, B Bartolini, E Giombini, F Messina, M Rueca, G Bonfiglio, O Butera, F Pimpinelli, F Ensoli, I Cavallo, A Di Caro, MR Capobianchi |
| EPI_ISL_2035751 | San Gallicano Dermatological Institute I.F.O. | INMI Lazzaro Spallanzani IRCCS | F Santini, B Bartolini, E Giombini, F Messina, M Rueca, G Bonfiglio, O Butera, CEM Gruber, F Pimpinelli, F Ensoli, G Prignano, MR Capobianchi, A Di Caro |
| EPI_ISL_2036262, EPI_ISL_2036263, EPI_ISL_2036264, EPI_ISL_2036265, EPI_ISL_2036266, EPI_ISL_2036267, EPI_ISL_2036268, EPI_ISL_2036269 | Laboratori de Referencia de Catalunya | Laboratori de Referencia de Catalunya | Ramirez A., Bellosillo B., Padilla E., Vilas A., Hernandez JJ., Canal M. |
| EPI_ISL_2036272 | Genome Centre | Genome Centre | Shovon Lal Sarkar, Md. Shazid Hasan, Ali Ahsan Setu, Prosanto Kumar Das, A. S. M. Rubayet Ul Alam, Ovinu Kibria Islam, Md. Tanvir Islam, Hassan Md. Al-Emran, Iqbal Kabir Jahid, M. Anwar Hossain |
| EPI_ISL_2036277, EPI_ISL_2036278, EPI_ISL_2036279, EPI_ISL_2036282, EPI_ISL_2036285, EPI_ISL_2036292, EPI_ISL_2036298, EPI_ISL_2036299, EPI_ISL_2036300, EPI_ISL_2036301, EPI_ISL_2036302, EPI_ISL_2036303, EPI_ISL_2036304 | | | |
| see above | Indian Council of Medical Research-National Institute of Virology, Microbial Containment Complex | Indian Council of Medical Research-National Institute of Virology, Microbial Containment Complex | Pragya D Yadav |
| EPI_ISL_2037012, EPI_ISL_2037044, EPI_ISL_2037052, EPI_ISL_2037056, EPI_ISL_2037057, EPI_ISL_2037069, EPI_ISL_2037173, EPI_ISL_2037251, EPI_ISL_2037276, EPI_ISL_2037293, EPI_ISL_2037360, EPI_ISL_2037365, EPI_ISL_2037367 | | | |
| see above | Pandemic Response Lab - NYC | Pandemic Response Lab, R&D | Henry Lee, Michael Hammerling, Melissa Hopkins, Cybill del Castillo, Shinyoung Clair Kang, William Ward, Pradeep Bugga, Sol Rey, Dylan Law, Katharine Nelson, Haiping Hao, Jon Laurent |
| EPI_ISL_2037457, EPI_ISL_2037490, EPI_ISL_2037527, EPI_ISL_2037537, EPI_ISL_2037573, EPI_ISL_2037581, EPI_ISL_2037634, EPI_ISL_2037663 | UW Virology Lab | UW Virology Lab | Pavitra Roychoudhury, Hong Xie, Lasata Shrestha, Tien V. Nguyen, Shah Mohamed Bakhash, Michelle Lin, Noah R. Baker, Sean Ellis, Meei-Li Huang, Keith R Jerome, Alexander Greninger |
| EPI_ISL_2038558, EPI_ISL_2038729, EPI_ISL_2038753 | Fulgent Genetics | Centers for Disease Control and Prevention Division of Viral Diseases, Pathogen Discovery | Dakota Howard, Dhwani Batra, Peter W. Cook, Kara Moser, Adrian Paskey, Jason Caravas, Benjamin Rambo-Martin, Shatavia Morrison, Christopher Gulvick, Scott Sammons, Yvette Unoarumhi, Darlene Wagner, Matthew Schmerer, Harry Gao, Mickey Li, John Gao, Joseph Fierro, Benafsh Sapra, Becky Tsai, Yan Meng, Doreen Ng, James Xie, Clinton R. Paden, Duncan MacCannell |
| EPI_ISL_2038893 | Federal Budget Health Care Institution "Center of Hygiene and Epidemiology in Tver region" | Group of Genomics and Postgenomic Technologies of Central Research Institute of Epidemiology | Samoilov AE, Kaptelova VV, Korneenko EV, Saenko SS, Smirnova YS, Nadtoka MI, Melushenkova TG, Speranskaya AS, Tivanova EV, Kondrasheva LY, Akimkin VG |

| | | | |
|---|---|---|---|
| EPI_ISL_2038980, EPI_ISL_2038982, EPI_ISL_2038983, EPI_ISL_2039008, EPI_ISL_2039154, EPI_ISL_2039226, EPI_ISL_2039266, EPI_ISL_2039336, EPI_ISL_2039389, EPI_ISL_2039637, EPI_ISL_2039776, EPI_ISL_2039864, EPI_ISL_2039878, EPI_ISL_2040187, EPI_ISL_2040317, EPI_ISL_2040338, EPI_ISL_2040357, EPI_ISL_2040384, EPI_ISL_2040461, EPI_ISL_2040473, EPI_ISL_2040517, EPI_ISL_2040778, EPI_ISL_2040892, EPI_ISL_2040936, EPI_ISL_2040938, EPI_ISL_2040976, EPI_ISL_2041437, EPI_ISL_2041621, EPI_ISL_2041655, EPI_ISL_2041656, EPI_ISL_2041779, EPI_ISL_2041833, EPI_ISL_2041929, EPI_ISL_2041972, EPI_ISL_2042053, EPI_ISL_2042272, EPI_ISL_2042585, EPI_ISL_2042588, EPI_ISL_2042756, EPI_ISL_2042772, EPI_ISL_2042773, EPI_ISL_2042805, EPI_ISL_2042920, EPI_ISL_2042937, EPI_ISL_2042958, EPI_ISL_2042959, EPI_ISL_2043088, EPI_ISL_2043219, EPI_ISL_2043292, EPI_ISL_2043293, EPI_ISL_2043302, EPI_ISL_2043303, EPI_ISL_2043405, EPI_ISL_2043426, EPI_ISL_2043454, EPI_ISL_2043494 | | | |
| see above | Aegis Sciences Corporation | Centers for Disease Control and Prevention Division of Viral Diseases, Pathogen Discovery | Dakota Howard, Dhwani Batra, Peter W. Cook, Kara Moser, Adrian Paskey, Jason Caravas, Benjamin Rambo-Martin, Shatavia Morrison, Christopher Gulvick, Scott Sammons, Yvette Unoarumhi, Darlene Wagner, Matthew Schmerer, Cyndi Clark, Patrick Campbell, Rob Case, Vikramsinha Ghorpade, Holly Houdeshell, Ola Kvalvaag, Dillon Nall, Ethan Sanders, Alec Vest, Shaun Westlund, Matthew Hardison, Clinton R. Paden, Duncan MacCannell |
| EPI_ISL_2043610, EPI_ISL_2043635, EPI_ISL_2043736, EPI_ISL_2043746, EPI_ISL_2043905, EPI_ISL_2043918 | Fulgent Genetics | Centers for Disease Control and Prevention Division of Viral Diseases, Pathogen Discovery | Dakota Howard, Dhwani Batra, Peter W. Cook, Kara Moser, Adrian Paskey, Jason Caravas, Benjamin Rambo-Martin, Shatavia Morrison, Christopher Gulvick, Scott Sammons, Yvette Unoarumhi, Darlene Wagner, Matthew Schmerer, Harry Gao, Mickey Li, John Gao, Joseph Fierro, Benafsh Sapra, Becky Tsai, Yan Meng, Doreen Ng, James Xie, Clinton R. Paden, Duncan MacCannell |
| EPI_ISL_2044007, EPI_ISL_2044013, EPI_ISL_2044036, EPI_ISL_2044077, EPI_ISL_2044210, EPI_ISL_2044211, EPI_ISL_2044343, EPI_ISL_2044454, EPI_ISL_2044588, EPI_ISL_2044622, EPI_ISL_2044694, EPI_ISL_2044695, EPI_ISL_2044706, EPI_ISL_2044786, EPI_ISL_2044941, EPI_ISL_2044942, EPI_ISL_2045014, EPI_ISL_2045112, EPI_ISL_2045385, EPI_ISL_2045440, EPI_ISL_2045668, EPI_ISL_2045783, EPI_ISL_2045852, EPI_ISL_2045856, EPI_ISL_2045932, EPI_ISL_2045959, EPI_ISL_2045974, EPI_ISL_2046278, EPI_ISL_2046354, EPI_ISL_2046357, EPI_ISL_2046388, EPI_ISL_2046570, EPI_ISL_2046793, EPI_ISL_2046823 | | | |
| see above | Laboratory Corporation of America | Centers for Disease Control and Prevention Division of Viral Diseases, Pathogen Discovery | Dakota Howard, Dhwani Batra, Peter W. Cook, Kara Moser, Adrian Paskey, Jason Caravas, Benjamin Rambo-Martin, Shatavia Morrison, Christopher Gulvick, Scott Sammons, Yvette Unoarumhi, Darlene Wagner, Matthew Schmerer, Minoo Agarwal, Eyad Almasri, Debbie Boles, Ayla Burns, Nuthawin Charoensri, Oren Cohen, Susan Countryman, Mary Ann Cristobal, Bobbi Croy, Suzanne Dale, Hrushikesh Deshmukh, Amanda Douglas, Vincent Drouillon, Marcia Eisenberg, Howard Engler, Rama Ghatti, Prashant Gupta, Susan Hicks, Jake Humphrey, Lax Iyer, Manoj Jain, Mohan Kolli, Brian Krueger, Tim Kuphal, Stanley Letovsky, Michael Levandoski, Craig Lukasik, Jonathan Meltzer, Brian Norvell, Mindy Nye, Scott Parker, Christos Petropoulos, John Pruitt, Steven Ragan, Scott Ryan, Mike Sapeta, Jana Schroth, Suresh Babu Selvaraju, Goran Stevovic, Amanda Suchanek, Andrea Throop, Lyndon Tilson, Thomas Urban, Joe Voshell, Kimberly Wagner, Jonathan Williams, Mary Williamson, Qian Zeng, Tricia Zwiefelhofer, Clinton R. Paden, Duncan MacCannell |
| EPI_ISL_2046851, EPI_ISL_2046943, EPI_ISL_2047029, EPI_ISL_2047254, EPI_ISL_2047258 | Quest Diagnostics Incorporated | Centers for Disease Control and Prevention Division of Viral Diseases, Pathogen Discovery | Dakota Howard, Dhwani Batra, Peter W. Cook, Kara Moser, Adrian Paskey, Jason Caravas, Benjamin Rambo-Martin, Shatavia Morrison, Christopher Gulvick, Scott Sammons, Yvette Unoarumhi, Darlene Wagner, Matthew Schmerer, S. H. Rosenthal, A. Gerasimova, R. M. Kagan, B. Anderson, M. Hua, Y. Liu, L.E. Bernstein, K.E. Livingston, A. Perez, I. A. Shlyakhter, R. V. Rolando, R. Owen, P. Tanpaiboon, F. Lacbawan, Clinton R. Paden, Duncan MacCannell |
| EPI_ISL_2047646, EPI_ISL_2047647, EPI_ISL_2047648, EPI_ISL_2047649, EPI_ISL_2047650, EPI_ISL_2047651, EPI_ISL_2047652, EPI_ISL_2047653, EPI_ISL_2047654, EPI_ISL_2047655, EPI_ISL_2047656, EPI_ISL_2047657, EPI_ISL_2047658, EPI_ISL_2047659, EPI_ISL_2047662 | | | |
| see above | Platform BIS UZA/UAntwerpen | Labo Klinische Biologie, UZA | Jasmine Coppens, Marie Le Mercier, Basil Britto Xavier, Christine Lammens, Veerle Matheeussen, Herman Goossens |
| EPI_ISL_2047772, EPI_ISL_2047773 | Hospital Universitari Vall d'Hebron - Vall d'Hebron Institut de Recerca | Hospital Universitari Vall d'Hebron - Vall d'Hebron Institut de Recerca | Cristina Andrés, Maria Piñana, Damir Garcia-Cehic, Ariadna Rando, Juliana Esperalba, Maria Gema Codina, Carla Castillo, Maria Carmen Martin, Tomàs Pumarola, Josep Quer, Andrés Antón |
| EPI_ISL_2080266, EPI_ISL_2080267, EPI_ISL_2080268, EPI_ISL_2080270, EPI_ISL_2080325, EPI_ISL_2080326 | Azienda Sanitaria dell'Alto Adige Laboratorio Aziendale di Microbiologia e Virologia | Istituto di Genomica Applicata | Elisabetta Pagani, Irene Bianconi, Elisabetta Giacobazzi, Elisa Masi, Stefanie Wieser, Irena Jurman, Vera Vendramin, Gabriele Magris, Eleonora Paparelli, Davide Scaglione, Michele Morgante |
| EPI_ISL_2080363, EPI_ISL_2080368, EPI_ISL_2080376, EPI_ISL_2080378, EPI_ISL_2080380, EPI_ISL_2080381, EPI_ISL_2080384, EPI_ISL_2080390, EPI_ISL_2080396, EPI_ISL_2080397, EPI_ISL_2080401, EPI_ISL_2080404, EPI_ISL_2080406, EPI_ISL_2080408, EPI_ISL_2080410, EPI_ISL_2080419 | | | |
| see above | Health Services Laboratories | Wellcome Sanger Institute for the COVID-19 Genomics UK (COG-UK) Consortium | Health Services Laboratories and Alex Alderton, Roberto Amato, Jeffrey Barrett, Sonia Goncalves, Ewan Harrison, David K. Jackson, Ian Johnston, Dominic Kwiatkowski, Cordelia Langford, John Sillitoe on behalf of the Wellcome Sanger Institute COVID-19 Surveillance Team |
| EPI_ISL_2080421 | Lighthouse Lab in Alderley Park | Wellcome Sanger Institute for the COVID-19 Genomics UK (COG-UK) Consortium | Jacquelyn Wynn, Mairead Hyland, The Lighthouse Lab in Alderley Park and Alex Alderton, Roberto Amato, Jeffrey Barrett, Sonia Goncalves, Ewan Harrison, David K. Jackson, Ian Johnston, Dominic Kwiatkowski, Cordelia Langford, John Sillitoe on behalf of the Wellcome Sanger Institute COVID-19 Surveillance Team |
| EPI_ISL_2080423, EPI_ISL_2080424, EPI_ISL_2080426, EPI_ISL_2080428, EPI_ISL_2080433, EPI_ISL_2080435, EPI_ISL_2080437, EPI_ISL_2080440, EPI_ISL_2080441, EPI_ISL_2080443, EPI_ISL_2080445, EPI_ISL_2080446, EPI_ISL_2080447, EPI_ISL_2080449, EPI_ISL_2080451, EPI_ISL_2080459, EPI_ISL_2080462, EPI_ISL_2080465, EPI_ISL_2080470, EPI_ISL_2080471 | | | |
| see above | Health Services Laboratories | Wellcome Sanger Institute for the COVID-19 Genomics UK (COG-UK) Consortium | Health Services Laboratories and Alex Alderton, Roberto Amato, Jeffrey Barrett, Sonia Goncalves, Ewan Harrison, David K. Jackson, Ian Johnston, Dominic Kwiatkowski, Cordelia Langford, John Sillitoe on behalf of the Wellcome Sanger Institute COVID-19 Surveillance Team |
| EPI_ISL_2080476, EPI_ISL_2080477, EPI_ISL_2080478 | Lighthouse Lab in Milton Keynes | Wellcome Sanger Institute for the COVID-19 Genomics UK (COG-UK) Consortium | The Lighthouse Lab in Milton Keynes and Alex Alderton, Roberto Amato, Jeffrey Barrett, Sonia Goncalves, Ewan Harrison, David K. Jackson, Ian Johnston, Dominic Kwiatkowski, Cordelia Langford, John Sillitoe on behalf of the Wellcome Sanger Institute COVID-19 Surveillance Team |
| EPI_ISL_2080488 | Diagnofirm Medical Laboratories | Botswana Harvard HIV Reference Laboratory | Sikhulile Moyo, Wonderful T. Choga, Pamela Smith-Lawrence,Kgomotso Moruisi, Dorcas Maruapula, Keoratile Ntshambiwa, Thongbotho Mphoyakgosi, Boitumelo Zuze, Botshelo Radibe, Legodile Kooepile, Roger Shapiro, Shahin Lockman, Mosepele Mosepele, Joseph Makhema, Madisa Mine, Simani Gaseitsiwe |
| EPI_ISL_2080609 | Tokyo Metropolitan Institute of Public Health | Tokyo Metropolitan Institute of Public Health | Masaki Hayashi, Takako Yamazaki, Akane Negishi, Takeo Sasamoto, Kayo Hagino, Rio Suzuki,Takushi Fujiwara, Hirofumi Miyake, Mami Nagashima, Jun Suzuki, Kenji Sadamasu |
| EPI_ISL_2080768 | OHSU Lab Services Molecular Microbiology Lab | Oregon SARS-CoV-2 Genome Sequencing Center | Brendan L. O'Connell, Sally Grindstaff, Sonia Acharya, Cierra LaBlanc, Kayla Carter, Ruth V. Nichols, Alec J. Hirsch, Donna Hansel, Guang Fan, Xuan Qin, Daniel N. Streblow, William B. Messer, Andrew C. Adey, Benjamin N. Bimber, Brian J. O'Roak |
| EPI_ISL_2081386 | Tampa General Hospital Esoteric Lab | Tampa General Hospital Esoteric Research & Development Lab | Grant Vestal, Deanna Becker, Dominic Uy, Vicki Healer, Amorce Lima, Suzane Silbert |
| EPI_ISL_2081529 | PathWest Laboratory Medicine WA | PathWest Laboratory Medicine WA Microbial Surveillance Unit | PathWest Laboratory Medicine WA Microbial Surveillance Unit |
| EPI_ISL_2082516 | UW Virology Lab | UW Virology Lab | Pavitra Roychoudhury, Hong Xie, Lasata Shrestha, Tien V. Nguyen, Shah Mohamed Bakhash, Michelle Lin, Noah R. Baker, Sean Ellis, Meei-Li Huang, Keith R Jerome, Alexander Greninger |
| EPI_ISL_2083033 | Baylor Scott & White-Grapevine | Baylor Scott & White-Temple | Ari Rao, Linden Morales, Kimberly Walker, Marcus Volz, Shelby Hendrickson |
| EPI_ISL_2086160 | Microbiology Department. Complexo Hospitalario Universitario de Vigo | Microbiology Department. Complexo Hospitalario Universitario de Vigo | Alfaya N, Alonso I, Alvarez M, Cabrera JJ, Carballo R, Cores O, Cortizo S, del-Campo V, Martinez L, Mediero G, Perez S, Potel C, Regueiro B, Rey S, Vasallo FJ |
| EPI_ISL_2086220, EPI_ISL_2086221, EPI_ISL_2086222, EPI_ISL_2086223, EPI_ISL_2086224, EPI_ISL_2086225, EPI_ISL_2086226, EPI_ISL_2086227 | ShipMed Inc | KRISP, KZN Research Innovation and Sequencing Platform | Giandhari J, Pillay S, Naidoo Y, Ramphal U, Lessells R, Emmanuel SJ, Tegally H, Wilkinson E, de Oliveira T |
| EPI_ISL_2086392, EPI_ISL_2086411 | UW Virology Lab | UW Virology Lab | Pavitra Roychoudhury, Hong Xie, Lasata Shrestha, Tien V. Nguyen, Shah Mohamed Bakhash, Michelle Lin, Noah R. Baker, Ricardo Perez, Sean Ellis, Nathan Breit, Robert J. Livingston, Meei-Li Huang, Keith R Jerome, Patrick Mathias, Alexander Greninger |
| EPI_ISL_2087219, EPI_ISL_2087232, EPI_ISL_2087519, EPI_ISL_2087590, EPI_ISL_2087703, EPI_ISL_2087762 | Infinity Biologix | Centers for Disease Control and Prevention Division of Viral Diseases, Pathogen Discovery | Dakota Howard, Dhwani Batra, Peter W. Cook, Kara Moser, Adrian Paskey, Jason Caravas, Benjamin Rambo-Martin, Shatavia Morrison, Christopher Gulvick, Scott Sammons, Yvette Unoarumhi, Darlene Wagner, Matthew Schmerer, Christian Bixby, Yihe Wang, Jonathan Schultz, Chirayu Goswami, Russ Hager, Robin Grimwood, Clinton R. Paden, Duncan MacCannell |
| EPI_ISL_2087953, EPI_ISL_2088054, EPI_ISL_2088086 | National Virus Reference Laboratory | National Virus Reference Laboratory | Zoe Yandle, Charlene Bennett, Gabriel Gonzalez, Michael Carr, Jonathan Dean, Cillian F De Gascun |
| EPI_ISL_2088485, EPI_ISL_2089042, EPI_ISL_2089192, EPI_ISL_2089221, EPI_ISL_2089293 | Infinity Biologix | Centers for Disease Control and Prevention Division of Viral Diseases, Pathogen Discovery | Dakota Howard, Dhwani Batra, Peter W. Cook, Kara Moser, Adrian Paskey, Jason Caravas, Benjamin Rambo-Martin, Shatavia Morrison, Christopher Gulvick, Scott Sammons, Yvette Unoarumhi, Darlene Wagner, Matthew Schmerer, Christian Bixby, Yihe Wang, Jonathan Schultz, Chirayu Goswami, Russ Hager, Robin Grimwood, Clinton R. Paden, Duncan MacCannell |
| EPI_ISL_2089545, EPI_ISL_2089601, EPI_ISL_2089684, EPI_ISL_2089750, EPI_ISL_2089944, EPI_ISL_2090006, EPI_ISL_2090114, EPI_ISL_2090122, EPI_ISL_2090169, EPI_ISL_2090200, EPI_ISL_2090227, EPI_ISL_2090228, EPI_ISL_2090280, EPI_ISL_2090420, EPI_ISL_2090429 | | | |
| see above | Aegis Sciences Corporation | Centers for Disease Control and Prevention Division of Viral Diseases, Pathogen Discovery | Dakota Howard, Dhwani Batra, Peter W. Cook, Kara Moser, Adrian Paskey, Jason Caravas, Benjamin Rambo-Martin, Shatavia Morrison, Christopher Gulvick, Scott Sammons, Yvette Unoarumhi, Darlene Wagner, Matthew Schmerer, Cyndi Clark, Patrick Campbell, Rob Case, Vikramsinha Ghorpade, Holly Houdeshell, Ola Kvalvaag, Dillon Nall, Ethan Sanders, Alec Vest, Shaun Westlund, Matthew Hardison, Clinton R. Paden, Duncan MacCannell |
| EPI_ISL_2090484, EPI_ISL_2090489, EPI_ISL_2090542 | Quest Diagnostics Incorporated | Centers for Disease Control and Prevention Division of Viral Diseases, Pathogen Discovery | Dakota Howard, Dhwani Batra, Peter W. Cook, Kara Moser, Adrian Paskey, Jason Caravas, Benjamin Rambo-Martin, Shatavia Morrison, Christopher Gulvick, Scott Sammons, Yvette Unoarumhi, Darlene Wagner, Matthew Schmerer, S. H. Rosenthal, A. Gerasimova, R. M. Kagan, B. Anderson, M. Hua, Y. Liu, L.E. Bernstein, K.E. Livingston, A. Perez, I. A. Shlyakhter, R. V. Rolando, R. Owen, P. Tanpaiboon, F. Lacbawan, Clinton R. Paden, Duncan MacCannell |
| EPI_ISL_2090620, EPI_ISL_2090621, EPI_ISL_2090622, EPI_ISL_2090623 | WSSE w Warszawie | National Institute of Public Health - National Institute of Hygiene | Wołkowicz Tomasz, Zacharczuk Katarzyna, Sadkowska-Todys Małgorzata, Gierczyńki Rafał |
| EPI_ISL_2090646 | Idaho Bureau of Laboratories | Idaho Bureau of Laboratories | R. Beukelman, Matthew Charles Burns, Aimee Ceniseros, Robert L. Voermans, Christopher Ball |
| EPI_ISL_2090725 | Nebraska Public Health Laboratory | NPHL COVID-19 Response Team | NPHL COVID-19 Response Team |
| EPI_ISL_2090827 | CHOR | UMR PIMIT | Dr David A Wilkinson, Dr Patrick Mavingui, Dr Camille Lebarbenchon, Magali Turpin |
| EPI_ISL_2091014 | Hospital Universitario Cruces | Biocruces Bizkaia | Mikel Gallego, Ana Belen de la Hoz |

| | | | |
|---|---|---|---|
| EPI_ISL_2091020 | Sungai Buloh Hospital | Institute for Medical Research, Infectious Disease Research Centre, National Institutes of Health, Ministry of Health Malaysia | Suppiah J, Kamel K, Mohd Zawawi Z, Azizan MA, Ramly N, Robert F, Thayan R |
| EPI_ISL_2091024 | Sultanah Aminah Hospital, Johor Bahru | Institute for Medical Research, Infectious Disease Research Centre, National Institutes of Health, Ministry of Health Malaysia | Suppiah J, Kamel K, Mohd Zawawi Z, Azizan MA, Ramly N, Robert F, Thayan R |
| EPI_ISL_2091029 | Kansas Health and Environmental Lab | Wichita State University - Molecular Diagnostics Lab | Mike Grose, Jonathan Barnell, Ben Olsen, and Phil Adam |
| EPI_ISL_2091086 | University Hospitals Translational Laboratory (UHTL), University Hospitals | University Hospitals Translational Laboratory (UHTL), University Hospitals | Sadri,N., Alouani,D., Song,X. |
| EPI_ISL_2091462, EPI_ISL_2091464, EPI_ISL_2091474, EPI_ISL_2091476, EPI_ISL_2091485 | Lighthouse Lab in Alderley Park | Wellcome Sanger Institute for the COVID-19 Genomics UK (COG-UK) Consortium | Jacquelyn Wynn, Mairead Hyland, The Lighthouse Lab in Alderley Park and Alex Alderton, Roberto Amato, Jeffrey Barrett, Sonia Goncalves, Ewan Harrison, David K. Jackson, Ian Johnston, Dominic Kwiatkowski, Cordelia Langford, John Sillitoe on behalf of the Wellcome Sanger Institute COVID-19 Surveillance Team |
| EPI_ISL_2091487 | Lighthouse Lab in Milton Keynes | Wellcome Sanger Institute for the COVID-19 Genomics UK (COG-UK) Consortium | The Lighthouse Lab in Milton Keynes and Alex Alderton, Roberto Amato, Jeffrey Barrett, Sonia Goncalves, Ewan Harrison, David K. Jackson, Ian Johnston, Dominic Kwiatkowski, Cordelia Langford, John Sillitoe on behalf of the Wellcome Sanger Institute COVID-19 Surveillance Team |
| EPI_ISL_2091490, EPI_ISL_2091499, EPI_ISL_2091507, EPI_ISL_2091509, EPI_ISL_2091516, EPI_ISL_2091519 | Lighthouse Lab in Alderley Park | Wellcome Sanger Institute for the COVID-19 Genomics UK (COG-UK) Consortium | Jacquelyn Wynn, Mairead Hyland, The Lighthouse Lab in Alderley Park and Alex Alderton, Roberto Amato, Jeffrey Barrett, Sonia Goncalves, Ewan Harrison, David K. Jackson, Ian Johnston, Dominic Kwiatkowski, Cordelia Langford, John Sillitoe on behalf of the Wellcome Sanger Institute COVID-19 Surveillance Team |
| EPI_ISL_2091528 | Lighthouse Lab in Milton Keynes | Wellcome Sanger Institute for the COVID-19 Genomics UK (COG-UK) Consortium | The Lighthouse Lab in Milton Keynes and Alex Alderton, Roberto Amato, Jeffrey Barrett, Sonia Goncalves, Ewan Harrison, David K. Jackson, Ian Johnston, Dominic Kwiatkowski, Cordelia Langford, John Sillitoe on behalf of the Wellcome Sanger Institute COVID-19 Surveillance Team |
| EPI_ISL_2091530, EPI_ISL_2091538, EPI_ISL_2091541 | Lighthouse Lab in Alderley Park | Wellcome Sanger Institute for the COVID-19 Genomics UK (COG-UK) Consortium | Jacquelyn Wynn, Mairead Hyland, The Lighthouse Lab in Alderley Park and Alex Alderton, Roberto Amato, Jeffrey Barrett, Sonia Goncalves, Ewan Harrison, David K. Jackson, Ian Johnston, Dominic Kwiatkowski, Cordelia Langford, John Sillitoe on behalf of the Wellcome Sanger Institute COVID-19 Surveillance Team |
| EPI_ISL_2091542 | Lighthouse Lab in Milton Keynes | Wellcome Sanger Institute for the COVID-19 Genomics UK (COG-UK) Consortium | The Lighthouse Lab in Milton Keynes and Alex Alderton, Roberto Amato, Jeffrey Barrett, Sonia Goncalves, Ewan Harrison, David K. Jackson, Ian Johnston, Dominic Kwiatkowski, Cordelia Langford, John Sillitoe on behalf of the Wellcome Sanger Institute COVID-19 Surveillance Team |
| EPI_ISL_2091548, EPI_ISL_2091550, EPI_ISL_2091552, EPI_ISL_2091560, EPI_ISL_2091568, EPI_ISL_2091570, EPI_ISL_2091572, EPI_ISL_2091574, EPI_ISL_2091575, EPI_ISL_2091577, EPI_ISL_2091579 | | | |
| see above | Lighthouse Lab in Alderley Park | Wellcome Sanger Institute for the COVID-19 Genomics UK (COG-UK) Consortium | Jacquelyn Wynn, Mairead Hyland, The Lighthouse Lab in Alderley Park and Alex Alderton, Roberto Amato, Jeffrey Barrett, Sonia Goncalves, Ewan Harrison, David K. Jackson, Ian Johnston, Dominic Kwiatkowski, Cordelia Langford, John Sillitoe on behalf of the Wellcome Sanger Institute COVID-19 Surveillance Team |
| EPI_ISL_2091586 | Lighthouse Lab in Milton Keynes | Wellcome Sanger Institute for the COVID-19 Genomics UK (COG-UK) Consortium | The Lighthouse Lab in Milton Keynes and Alex Alderton, Roberto Amato, Jeffrey Barrett, Sonia Goncalves, Ewan Harrison, David K. Jackson, Ian Johnston, Dominic Kwiatkowski, Cordelia Langford, John Sillitoe on behalf of the Wellcome Sanger Institute COVID-19 Surveillance Team |
| EPI_ISL_2091595, EPI_ISL_2091601, EPI_ISL_2091604, EPI_ISL_2091609, EPI_ISL_2091612, EPI_ISL_2091618, EPI_ISL_2091620, EPI_ISL_2091621, EPI_ISL_2091623, EPI_ISL_2091626, EPI_ISL_2091631, EPI_ISL_2091634 | | | |
| see above | Lighthouse Lab in Alderley Park | Wellcome Sanger Institute for the COVID-19 Genomics UK (COG-UK) Consortium | Jacquelyn Wynn, Mairead Hyland, The Lighthouse Lab in Alderley Park and Alex Alderton, Roberto Amato, Jeffrey Barrett, Sonia Goncalves, Ewan Harrison, David K. Jackson, Ian Johnston, Dominic Kwiatkowski, Cordelia Langford, John Sillitoe on behalf of the Wellcome Sanger Institute COVID-19 Surveillance Team |
| EPI_ISL_2091636 | Lighthouse Lab in Milton Keynes | Wellcome Sanger Institute for the COVID-19 Genomics UK (COG-UK) Consortium | The Lighthouse Lab in Milton Keynes and Alex Alderton, Roberto Amato, Jeffrey Barrett, Sonia Goncalves, Ewan Harrison, David K. Jackson, Ian Johnston, Dominic Kwiatkowski, Cordelia Langford, John Sillitoe on behalf of the Wellcome Sanger Institute COVID-19 Surveillance Team |
| EPI_ISL_2091637, EPI_ISL_2091638 | Lighthouse Lab in Alderley Park | Wellcome Sanger Institute for the COVID-19 Genomics UK (COG-UK) Consortium | Jacquelyn Wynn, Mairead Hyland, The Lighthouse Lab in Alderley Park and Alex Alderton, Roberto Amato, Jeffrey Barrett, Sonia Goncalves, Ewan Harrison, David K. Jackson, Ian Johnston, Dominic Kwiatkowski, Cordelia Langford, John Sillitoe on behalf of the Wellcome Sanger Institute COVID-19 Surveillance Team |
| EPI_ISL_2091639 | Lighthouse Lab in Milton Keynes | Wellcome Sanger Institute for the COVID-19 Genomics UK (COG-UK) Consortium | The Lighthouse Lab in Milton Keynes and Alex Alderton, Roberto Amato, Jeffrey Barrett, Sonia Goncalves, Ewan Harrison, David K. Jackson, Ian Johnston, Dominic Kwiatkowski, Cordelia Langford, John Sillitoe on behalf of the Wellcome Sanger Institute COVID-19 Surveillance Team |
| EPI_ISL_2091641 | Lighthouse Lab in Alderley Park | Wellcome Sanger Institute for the COVID-19 Genomics UK (COG-UK) Consortium | Jacquelyn Wynn, Mairead Hyland, The Lighthouse Lab in Alderley Park and Alex Alderton, Roberto Amato, Jeffrey Barrett, Sonia Goncalves, Ewan Harrison, David K. Jackson, Ian Johnston, Dominic Kwiatkowski, Cordelia Langford, John Sillitoe on behalf of the Wellcome Sanger Institute COVID-19 Surveillance Team |
| EPI_ISL_2091642 | Lighthouse Lab in Milton Keynes | Wellcome Sanger Institute for the COVID-19 Genomics UK (COG-UK) Consortium | The Lighthouse Lab in Milton Keynes and Alex Alderton, Roberto Amato, Jeffrey Barrett, Sonia Goncalves, Ewan Harrison, David K. Jackson, Ian Johnston, Dominic Kwiatkowski, Cordelia Langford, John Sillitoe on behalf of the Wellcome Sanger Institute COVID-19 Surveillance Team |
| EPI_ISL_2091644, EPI_ISL_2091647, EPI_ISL_2091649, EPI_ISL_2091653, EPI_ISL_2091656, EPI_ISL_2091657 | Lighthouse Lab in Alderley Park | Wellcome Sanger Institute for the COVID-19 Genomics UK (COG-UK) Consortium | Jacquelyn Wynn, Mairead Hyland, The Lighthouse Lab in Alderley Park and Alex Alderton, Roberto Amato, Jeffrey Barrett, Sonia Goncalves, Ewan Harrison, David K. Jackson, Ian Johnston, Dominic Kwiatkowski, Cordelia Langford, John Sillitoe on behalf of the Wellcome Sanger Institute COVID-19 Surveillance Team |
| EPI_ISL_2091665, EPI_ISL_2091666 | Lighthouse Lab in Milton Keynes | Wellcome Sanger Institute for the COVID-19 Genomics UK (COG-UK) Consortium | The Lighthouse Lab in Milton Keynes and Alex Alderton, Roberto Amato, Jeffrey Barrett, Sonia Goncalves, Ewan Harrison, David K. Jackson, Ian Johnston, Dominic Kwiatkowski, Cordelia Langford, John Sillitoe on behalf of the Wellcome Sanger Institute COVID-19 Surveillance Team |
| EPI_ISL_2091672, EPI_ISL_2091680, EPI_ISL_2091684, EPI_ISL_2091685, EPI_ISL_2091686, EPI_ISL_2091690, EPI_ISL_2091692, EPI_ISL_2091694, EPI_ISL_2091696 | Lighthouse Lab in Alderley Park | Wellcome Sanger Institute for the COVID-19 Genomics UK (COG-UK) Consortium | Jacquelyn Wynn, Mairead Hyland, The Lighthouse Lab in Alderley Park and Alex Alderton, Roberto Amato, Jeffrey Barrett, Sonia Goncalves, Ewan Harrison, David K. Jackson, Ian Johnston, Dominic Kwiatkowski, Cordelia Langford, John Sillitoe on behalf of the Wellcome Sanger Institute COVID-19 Surveillance Team |
| EPI_ISL_2091699 | Lighthouse Lab in Milton Keynes | Wellcome Sanger Institute for the COVID-19 Genomics UK (COG-UK) Consortium | The Lighthouse Lab in Milton Keynes and Alex Alderton, Roberto Amato, Jeffrey Barrett, Sonia Goncalves, Ewan Harrison, David K. Jackson, Ian Johnston, Dominic Kwiatkowski, Cordelia Langford, John Sillitoe on behalf of the Wellcome Sanger Institute COVID-19 Surveillance Team |
| EPI_ISL_2091700, EPI_ISL_2091701, EPI_ISL_2091709, EPI_ISL_2091716, EPI_ISL_2091718, EPI_ISL_2091721, EPI_ISL_2091722, EPI_ISL_2091725, EPI_ISL_2091726 | Lighthouse Lab in Alderley Park | Wellcome Sanger Institute for the COVID-19 Genomics UK (COG-UK) Consortium | Jacquelyn Wynn, Mairead Hyland, The Lighthouse Lab in Alderley Park and Alex Alderton, Roberto Amato, Jeffrey Barrett, Sonia Goncalves, Ewan Harrison, David K. Jackson, Ian Johnston, Dominic Kwiatkowski, Cordelia Langford, John Sillitoe on behalf of the Wellcome Sanger Institute COVID-19 Surveillance Team |
| EPI_ISL_2091728 | Lighthouse Lab in Milton Keynes | Wellcome Sanger Institute for the COVID-19 Genomics UK (COG-UK) Consortium | The Lighthouse Lab in Milton Keynes and Alex Alderton, Roberto Amato, Jeffrey Barrett, Sonia Goncalves, Ewan Harrison, David K. Jackson, Ian Johnston, Dominic Kwiatkowski, Cordelia Langford, John Sillitoe on behalf of the Wellcome Sanger Institute COVID-19 Surveillance Team |
| EPI_ISL_2091734 | Lighthouse Lab in Alderley Park | Wellcome Sanger Institute for the COVID-19 Genomics UK (COG-UK) Consortium | Jacquelyn Wynn, Mairead Hyland, The Lighthouse Lab in Alderley Park and Alex Alderton, Roberto Amato, Jeffrey Barrett, Sonia Goncalves, Ewan Harrison, David K. Jackson, Ian Johnston, Dominic Kwiatkowski, Cordelia Langford, John Sillitoe on behalf of the Wellcome Sanger Institute COVID-19 Surveillance Team |
| EPI_ISL_2091746 | Lighthouse Lab in Milton Keynes | Wellcome Sanger Institute for the COVID-19 Genomics UK (COG-UK) Consortium | The Lighthouse Lab in Milton Keynes and Alex Alderton, Roberto Amato, Jeffrey Barrett, Sonia Goncalves, Ewan Harrison, David K. Jackson, Ian Johnston, Dominic Kwiatkowski, Cordelia Langford, John Sillitoe on behalf of the Wellcome Sanger Institute COVID-19 Surveillance Team |
| EPI_ISL_2091753, EPI_ISL_2091754 | Lighthouse Lab in Alderley Park | Wellcome Sanger Institute for the COVID-19 Genomics UK (COG-UK) Consortium | Jacquelyn Wynn, Mairead Hyland, The Lighthouse Lab in Alderley Park and Alex Alderton, Roberto Amato, Jeffrey Barrett, Sonia Goncalves, Ewan Harrison, David K. Jackson, Ian Johnston, Dominic Kwiatkowski, Cordelia Langford, John Sillitoe on behalf of the Wellcome Sanger Institute COVID-19 Surveillance Team |
| EPI_ISL_2091757, EPI_ISL_2091761 | Lighthouse Lab in Milton Keynes | Wellcome Sanger Institute for the COVID-19 Genomics UK (COG-UK) Consortium | The Lighthouse Lab in Milton Keynes and Alex Alderton, Roberto Amato, Jeffrey Barrett, Sonia Goncalves, Ewan Harrison, David K. Jackson, Ian Johnston, Dominic Kwiatkowski, Cordelia Langford, John Sillitoe on behalf of the Wellcome Sanger Institute COVID-19 Surveillance Team |
| EPI_ISL_2091762, EPI_ISL_2091771, EPI_ISL_2091772, EPI_ISL_2091790 | Lighthouse Lab in Alderley Park | Wellcome Sanger Institute for the COVID-19 Genomics UK (COG-UK) Consortium | Jacquelyn Wynn, Mairead Hyland, The Lighthouse Lab in Alderley Park and Alex Alderton, Roberto Amato, Jeffrey Barrett, Sonia Goncalves, Ewan Harrison, David K. Jackson, Ian Johnston, Dominic Kwiatkowski, Cordelia Langford, John Sillitoe on behalf of the Wellcome Sanger Institute COVID-19 Surveillance Team |
| EPI_ISL_2091797, EPI_ISL_2091798, EPI_ISL_2091799, EPI_ISL_2091803, EPI_ISL_2091806, EPI_ISL_2091809 | Lighthouse Lab in Milton Keynes | Wellcome Sanger Institute for the COVID-19 Genomics UK (COG-UK) Consortium | The Lighthouse Lab in Milton Keynes and Alex Alderton, Roberto Amato, Jeffrey Barrett, Sonia Goncalves, Ewan Harrison, David K. Jackson, Ian Johnston, Dominic Kwiatkowski, Cordelia Langford, John Sillitoe on behalf of the Wellcome Sanger Institute COVID-19 Surveillance Team |
| EPI_ISL_2091814 | Lighthouse Lab in Alderley Park | Wellcome Sanger Institute for the COVID-19 Genomics UK (COG-UK) Consortium | Jacquelyn Wynn, Mairead Hyland, The Lighthouse Lab in Alderley Park and Alex Alderton, Roberto Amato, Jeffrey Barrett, Sonia Goncalves, Ewan Harrison, David K. Jackson, Ian Johnston, Dominic Kwiatkowski, Cordelia Langford, John Sillitoe on behalf of the Wellcome Sanger Institute COVID-19 Surveillance Team |
| EPI_ISL_2091815 | Lighthouse Lab in Milton Keynes | Wellcome Sanger Institute for the COVID-19 Genomics UK (COG-UK) | The Lighthouse Lab in Milton Keynes and Alex Alderton, Roberto Amato, Jeffrey Barrett, Sonia Goncalves, Ewan Harrison, David K. Jackson, Ian Johnston, Dominic Kwiatkowski, Cordelia Langford, John Sillitoe on behalf of the Wellcome Sanger |

| | | | |
|---|---|---|---|
| | | Consortium | Institute COVID-19 Surveillance Team |
| EPI_ISL_2091816 | Lighthouse Lab in Alderley Park | Wellcome Sanger Institute for the COVID-19 Genomics UK (COG-UK) Consortium | Jacquelyn Wynn, Mairead Hyland, The Lighthouse Lab in Alderley Park and Alex Alderton, Roberto Amato, Jeffrey Barrett, Sonia Goncalves, Ewan Harrison, David K. Jackson, Ian Johnston, Dominic Kwiatkowski, Cordelia Langford, John Sillitoe on behalf of the Wellcome Sanger Institute COVID-19 Surveillance Team |
| EPI_ISL_2091818, EPI_ISL_2091822 | Lighthouse Lab in Milton Keynes | Wellcome Sanger Institute for the COVID-19 Genomics UK (COG-UK) Consortium | The Lighthouse Lab in Milton Keynes and Alex Alderton, Roberto Amato, Jeffrey Barrett, Sonia Goncalves, Ewan Harrison, David K. Jackson, Ian Johnston, Dominic Kwiatkowski, Cordelia Langford, John Sillitoe on behalf of the Wellcome Sanger Institute COVID-19 Surveillance Team |
| EPI_ISL_2091827 | Lighthouse Lab in Alderley Park | Wellcome Sanger Institute for the COVID-19 Genomics UK (COG-UK) Consortium | Jacquelyn Wynn, Mairead Hyland, The Lighthouse Lab in Alderley Park and Alex Alderton, Roberto Amato, Jeffrey Barrett, Sonia Goncalves, Ewan Harrison, David K. Jackson, Ian Johnston, Dominic Kwiatkowski, Cordelia Langford, John Sillitoe on behalf of the Wellcome Sanger Institute COVID-19 Surveillance Team |
| EPI_ISL_2091839, EPI_ISL_2091840 | Lighthouse Lab in Milton Keynes | Wellcome Sanger Institute for the COVID-19 Genomics UK (COG-UK) Consortium | The Lighthouse Lab in Milton Keynes and Alex Alderton, Roberto Amato, Jeffrey Barrett, Sonia Goncalves, Ewan Harrison, David K. Jackson, Ian Johnston, Dominic Kwiatkowski, Cordelia Langford, John Sillitoe on behalf of the Wellcome Sanger Institute COVID-19 Surveillance Team |
| EPI_ISL_2091841 | Lighthouse Lab in Alderley Park | Wellcome Sanger Institute for the COVID-19 Genomics UK (COG-UK) Consortium | Jacquelyn Wynn, Mairead Hyland, The Lighthouse Lab in Alderley Park and Alex Alderton, Roberto Amato, Jeffrey Barrett, Sonia Goncalves, Ewan Harrison, David K. Jackson, Ian Johnston, Dominic Kwiatkowski, Cordelia Langford, John Sillitoe on behalf of the Wellcome Sanger Institute COVID-19 Surveillance Team |
| EPI_ISL_2091849 | Lighthouse Lab in Milton Keynes | Wellcome Sanger Institute for the COVID-19 Genomics UK (COG-UK) Consortium | The Lighthouse Lab in Milton Keynes and Alex Alderton, Roberto Amato, Jeffrey Barrett, Sonia Goncalves, Ewan Harrison, David K. Jackson, Ian Johnston, Dominic Kwiatkowski, Cordelia Langford, John Sillitoe on behalf of the Wellcome Sanger Institute COVID-19 Surveillance Team |
| EPI_ISL_2091852, EPI_ISL_2091854 | Lighthouse Lab in Alderley Park | Wellcome Sanger Institute for the COVID-19 Genomics UK (COG-UK) Consortium | Jacquelyn Wynn, Mairead Hyland, The Lighthouse Lab in Alderley Park and Alex Alderton, Roberto Amato, Jeffrey Barrett, Sonia Goncalves, Ewan Harrison, David K. Jackson, Ian Johnston, Dominic Kwiatkowski, Cordelia Langford, John Sillitoe on behalf of the Wellcome Sanger Institute COVID-19 Surveillance Team |
| EPI_ISL_2091855 | Lighthouse Lab in Milton Keynes | Wellcome Sanger Institute for the COVID-19 Genomics UK (COG-UK) Consortium | The Lighthouse Lab in Milton Keynes and Alex Alderton, Roberto Amato, Jeffrey Barrett, Sonia Goncalves, Ewan Harrison, David K. Jackson, Ian Johnston, Dominic Kwiatkowski, Cordelia Langford, John Sillitoe on behalf of the Wellcome Sanger Institute COVID-19 Surveillance Team |
| EPI_ISL_2091858 | Lighthouse Lab in Alderley Park | Wellcome Sanger Institute for the COVID-19 Genomics UK (COG-UK) Consortium | Jacquelyn Wynn, Mairead Hyland, The Lighthouse Lab in Alderley Park and Alex Alderton, Roberto Amato, Jeffrey Barrett, Sonia Goncalves, Ewan Harrison, David K. Jackson, Ian Johnston, Dominic Kwiatkowski, Cordelia Langford, John Sillitoe on behalf of the Wellcome Sanger Institute COVID-19 Surveillance Team |
| EPI_ISL_2091867, EPI_ISL_2091883 | Lighthouse Lab in Milton Keynes | Wellcome Sanger Institute for the COVID-19 Genomics UK (COG-UK) Consortium | The Lighthouse Lab in Milton Keynes and Alex Alderton, Roberto Amato, Jeffrey Barrett, Sonia Goncalves, Ewan Harrison, David K. Jackson, Ian Johnston, Dominic Kwiatkowski, Cordelia Langford, John Sillitoe on behalf of the Wellcome Sanger Institute COVID-19 Surveillance Team |
| EPI_ISL_2091884, EPI_ISL_2091885, EPI_ISL_2091893, EPI_ISL_2091896 | Lighthouse Lab in Alderley Park | Wellcome Sanger Institute for the COVID-19 Genomics UK (COG-UK) Consortium | Jacquelyn Wynn, Mairead Hyland, The Lighthouse Lab in Alderley Park and Alex Alderton, Roberto Amato, Jeffrey Barrett, Sonia Goncalves, Ewan Harrison, David K. Jackson, Ian Johnston, Dominic Kwiatkowski, Cordelia Langford, John Sillitoe on behalf of the Wellcome Sanger Institute COVID-19 Surveillance Team |
| EPI_ISL_2091901, EPI_ISL_2091902, EPI_ISL_2091908, EPI_ISL_2091912, EPI_ISL_2091914, EPI_ISL_2091918, EPI_ISL_2091920 | Lighthouse Lab in Milton Keynes | Wellcome Sanger Institute for the COVID-19 Genomics UK (COG-UK) Consortium | The Lighthouse Lab in Milton Keynes and Alex Alderton, Roberto Amato, Jeffrey Barrett, Sonia Goncalves, Ewan Harrison, David K. Jackson, Ian Johnston, Dominic Kwiatkowski, Cordelia Langford, John Sillitoe on behalf of the Wellcome Sanger Institute COVID-19 Surveillance Team |
| EPI_ISL_2091922 | Lighthouse Lab in Alderley Park | Wellcome Sanger Institute for the COVID-19 Genomics UK (COG-UK) Consortium | Jacquelyn Wynn, Mairead Hyland, The Lighthouse Lab in Alderley Park and Alex Alderton, Roberto Amato, Jeffrey Barrett, Sonia Goncalves, Ewan Harrison, David K. Jackson, Ian Johnston, Dominic Kwiatkowski, Cordelia Langford, John Sillitoe on behalf of the Wellcome Sanger Institute COVID-19 Surveillance Team |
| EPI_ISL_2091923, EPI_ISL_2091928 | Lighthouse Lab in Milton Keynes | Wellcome Sanger Institute for the COVID-19 Genomics UK (COG-UK) Consortium | The Lighthouse Lab in Milton Keynes and Alex Alderton, Roberto Amato, Jeffrey Barrett, Sonia Goncalves, Ewan Harrison, David K. Jackson, Ian Johnston, Dominic Kwiatkowski, Cordelia Langford, John Sillitoe on behalf of the Wellcome Sanger Institute COVID-19 Surveillance Team |
| EPI_ISL_2091930 | Lighthouse Lab in Alderley Park | Wellcome Sanger Institute for the COVID-19 Genomics UK (COG-UK) Consortium | Jacquelyn Wynn, Mairead Hyland, The Lighthouse Lab in Alderley Park and Alex Alderton, Roberto Amato, Jeffrey Barrett, Sonia Goncalves, Ewan Harrison, David K. Jackson, Ian Johnston, Dominic Kwiatkowski, Cordelia Langford, John Sillitoe on behalf of the Wellcome Sanger Institute COVID-19 Surveillance Team |
| EPI_ISL_2091935 | Lighthouse Lab in Milton Keynes | Wellcome Sanger Institute for the COVID-19 Genomics UK (COG-UK) Consortium | The Lighthouse Lab in Milton Keynes and Alex Alderton, Roberto Amato, Jeffrey Barrett, Sonia Goncalves, Ewan Harrison, David K. Jackson, Ian Johnston, Dominic Kwiatkowski, Cordelia Langford, John Sillitoe on behalf of the Wellcome Sanger Institute COVID-19 Surveillance Team |
| EPI_ISL_2091943 | Lighthouse Lab in Alderley Park | Wellcome Sanger Institute for the COVID-19 Genomics UK (COG-UK) Consortium | Jacquelyn Wynn, Mairead Hyland, The Lighthouse Lab in Alderley Park and Alex Alderton, Roberto Amato, Jeffrey Barrett, Sonia Goncalves, Ewan Harrison, David K. Jackson, Ian Johnston, Dominic Kwiatkowski, Cordelia Langford, John Sillitoe on behalf of the Wellcome Sanger Institute COVID-19 Surveillance Team |
| EPI_ISL_2091944 | Lighthouse Lab in Milton Keynes | Wellcome Sanger Institute for the COVID-19 Genomics UK (COG-UK) Consortium | The Lighthouse Lab in Milton Keynes and Alex Alderton, Roberto Amato, Jeffrey Barrett, Sonia Goncalves, Ewan Harrison, David K. Jackson, Ian Johnston, Dominic Kwiatkowski, Cordelia Langford, John Sillitoe on behalf of the Wellcome Sanger Institute COVID-19 Surveillance Team |
| EPI_ISL_2091945, EPI_ISL_2091947 | Lighthouse Lab in Alderley Park | Wellcome Sanger Institute for the COVID-19 Genomics UK (COG-UK) Consortium | Jacquelyn Wynn, Mairead Hyland, The Lighthouse Lab in Alderley Park and Alex Alderton, Roberto Amato, Jeffrey Barrett, Sonia Goncalves, Ewan Harrison, David K. Jackson, Ian Johnston, Dominic Kwiatkowski, Cordelia Langford, John Sillitoe on behalf of the Wellcome Sanger Institute COVID-19 Surveillance Team |
| EPI_ISL_2091950 | Lighthouse Lab in Milton Keynes | Wellcome Sanger Institute for the COVID-19 Genomics UK (COG-UK) Consortium | The Lighthouse Lab in Milton Keynes and Alex Alderton, Roberto Amato, Jeffrey Barrett, Sonia Goncalves, Ewan Harrison, David K. Jackson, Ian Johnston, Dominic Kwiatkowski, Cordelia Langford, John Sillitoe on behalf of the Wellcome Sanger Institute COVID-19 Surveillance Team |
| EPI_ISL_2091955, EPI_ISL_2091958, EPI_ISL_2091962 | Lighthouse Lab in Alderley Park | Wellcome Sanger Institute for the COVID-19 Genomics UK (COG-UK) Consortium | Jacquelyn Wynn, Mairead Hyland, The Lighthouse Lab in Alderley Park and Alex Alderton, Roberto Amato, Jeffrey Barrett, Sonia Goncalves, Ewan Harrison, David K. Jackson, Ian Johnston, Dominic Kwiatkowski, Cordelia Langford, John Sillitoe on behalf of the Wellcome Sanger Institute COVID-19 Surveillance Team |
| EPI_ISL_2091971 | Lighthouse Lab in Milton Keynes | Wellcome Sanger Institute for the COVID-19 Genomics UK (COG-UK) Consortium | The Lighthouse Lab in Milton Keynes and Alex Alderton, Roberto Amato, Jeffrey Barrett, Sonia Goncalves, Ewan Harrison, David K. Jackson, Ian Johnston, Dominic Kwiatkowski, Cordelia Langford, John Sillitoe on behalf of the Wellcome Sanger Institute COVID-19 Surveillance Team |
| EPI_ISL_2091974 | Lighthouse Lab in Alderley Park | Wellcome Sanger Institute for the COVID-19 Genomics UK (COG-UK) Consortium | Jacquelyn Wynn, Mairead Hyland, The Lighthouse Lab in Alderley Park and Alex Alderton, Roberto Amato, Jeffrey Barrett, Sonia Goncalves, Ewan Harrison, David K. Jackson, Ian Johnston, Dominic Kwiatkowski, Cordelia Langford, John Sillitoe on behalf of the Wellcome Sanger Institute COVID-19 Surveillance Team |
| EPI_ISL_2091976, EPI_ISL_2091981, EPI_ISL_2091986 | Lighthouse Lab in Milton Keynes | Wellcome Sanger Institute for the COVID-19 Genomics UK (COG-UK) Consortium | The Lighthouse Lab in Milton Keynes and Alex Alderton, Roberto Amato, Jeffrey Barrett, Sonia Goncalves, Ewan Harrison, David K. Jackson, Ian Johnston, Dominic Kwiatkowski, Cordelia Langford, John Sillitoe on behalf of the Wellcome Sanger Institute COVID-19 Surveillance Team |
| EPI_ISL_2091987 | Lighthouse Lab in Alderley Park | Wellcome Sanger Institute for the COVID-19 Genomics UK (COG-UK) Consortium | Jacquelyn Wynn, Mairead Hyland, The Lighthouse Lab in Alderley Park and Alex Alderton, Roberto Amato, Jeffrey Barrett, Sonia Goncalves, Ewan Harrison, David K. Jackson, Ian Johnston, Dominic Kwiatkowski, Cordelia Langford, John Sillitoe on behalf of the Wellcome Sanger Institute COVID-19 Surveillance Team |
| EPI_ISL_2091989, EPI_ISL_2091994, EPI_ISL_2091997 | Lighthouse Lab in Milton Keynes | Wellcome Sanger Institute for the COVID-19 Genomics UK (COG-UK) Consortium | The Lighthouse Lab in Milton Keynes and Alex Alderton, Roberto Amato, Jeffrey Barrett, Sonia Goncalves, Ewan Harrison, David K. Jackson, Ian Johnston, Dominic Kwiatkowski, Cordelia Langford, John Sillitoe on behalf of the Wellcome Sanger Institute COVID-19 Surveillance Team |
| EPI_ISL_2092002, EPI_ISL_2092003, EPI_ISL_2092005 | Lighthouse Lab in Alderley Park | Wellcome Sanger Institute for the COVID-19 Genomics UK (COG-UK) Consortium | Jacquelyn Wynn, Mairead Hyland, The Lighthouse Lab in Alderley Park and Alex Alderton, Roberto Amato, Jeffrey Barrett, Sonia Goncalves, Ewan Harrison, David K. Jackson, Ian Johnston, Dominic Kwiatkowski, Cordelia Langford, John Sillitoe on behalf of the Wellcome Sanger Institute COVID-19 Surveillance Team |
| EPI_ISL_2092007 | Lighthouse Lab in Milton Keynes | Wellcome Sanger Institute for the COVID-19 Genomics UK (COG-UK) Consortium | The Lighthouse Lab in Milton Keynes and Alex Alderton, Roberto Amato, Jeffrey Barrett, Sonia Goncalves, Ewan Harrison, David K. Jackson, Ian Johnston, Dominic Kwiatkowski, Cordelia Langford, John Sillitoe on behalf of the Wellcome Sanger Institute COVID-19 Surveillance Team |
| EPI_ISL_2092008, EPI_ISL_2092010, EPI_ISL_2092013 | Lighthouse Lab in Alderley Park | Wellcome Sanger Institute for the COVID-19 Genomics UK (COG-UK) Consortium | Jacquelyn Wynn, Mairead Hyland, The Lighthouse Lab in Alderley Park and Alex Alderton, Roberto Amato, Jeffrey Barrett, Sonia Goncalves, Ewan Harrison, David K. Jackson, Ian Johnston, Dominic Kwiatkowski, Cordelia Langford, John Sillitoe on behalf of the Wellcome Sanger Institute COVID-19 Surveillance Team |
| EPI_ISL_2092014 | Lighthouse Lab in Milton Keynes | Wellcome Sanger Institute for the COVID-19 Genomics UK (COG-UK) Consortium | The Lighthouse Lab in Milton Keynes and Alex Alderton, Roberto Amato, Jeffrey Barrett, Sonia Goncalves, Ewan Harrison, David K. Jackson, Ian Johnston, Dominic Kwiatkowski, Cordelia Langford, John Sillitoe on behalf of the Wellcome Sanger Institute COVID-19 Surveillance Team |
| EPI_ISL_2092016 | Randox Laboratories | Wellcome Sanger Institute for the COVID-19 Genomics UK (COG-UK) Consortium | Randox Laboratories and Alex Alderton, Roberto Amato, Jeffrey Barrett, Sonia Goncalves, Ewan Harrison, David K. Jackson, Ian Johnston, Dominic Kwiatkowski, Cordelia Langford, John Sillitoe on behalf of the Wellcome Sanger Institute COVID-19 Surveillance Team |
| EPI_ISL_2092018 | Lighthouse Lab in Milton Keynes | Wellcome Sanger Institute for the COVID-19 Genomics UK (COG-UK) Consortium | The Lighthouse Lab in Milton Keynes and Alex Alderton, Roberto Amato, Jeffrey Barrett, Sonia Goncalves, Ewan Harrison, David K. Jackson, Ian Johnston, Dominic Kwiatkowski, Cordelia Langford, John Sillitoe on behalf of the Wellcome Sanger Institute COVID-19 Surveillance Team |
| EPI_ISL_2092019 | Randox Laboratories | Wellcome Sanger Institute for the COVID-19 Genomics UK (COG-UK) Consortium | Randox Laboratories and Alex Alderton, Roberto Amato, Jeffrey Barrett, Sonia Goncalves, Ewan Harrison, David K. Jackson, Ian Johnston, Dominic Kwiatkowski, Cordelia Langford, John Sillitoe on behalf of the Wellcome Sanger Institute COVID-19 Surveillance Team |
| EPI_ISL_2092022, EPI_ISL_2092026 | Lighthouse Lab in Alderley Park | Wellcome Sanger Institute for the COVID-19 Genomics UK (COG-UK) Consortium | Jacquelyn Wynn, Mairead Hyland, The Lighthouse Lab in Alderley Park and Alex Alderton, Roberto Amato, Jeffrey Barrett, Sonia Goncalves, Ewan Harrison, David K. Jackson, Ian Johnston, Dominic Kwiatkowski, Cordelia Langford, John Sillitoe on behalf of the Wellcome Sanger Institute COVID-19 Surveillance Team |
| EPI_ISL_2092027, EPI_ISL_2092033 | Lighthouse Lab in Milton Keynes | Wellcome Sanger Institute for the COVID-19 Genomics UK (COG-UK) Consortium | The Lighthouse Lab in Milton Keynes and Alex Alderton, Roberto Amato, Jeffrey Barrett, Sonia Goncalves, Ewan Harrison, David K. Jackson, Ian Johnston, Dominic Kwiatkowski, Cordelia Langford, John Sillitoe on behalf of the Wellcome Sanger Institute COVID-19 Surveillance Team |
| EPI_ISL_2092034 | Lighthouse Lab in Alderley Park | Wellcome Sanger Institute for the COVID-19 Genomics UK (COG-UK) Consortium | Jacquelyn Wynn, Mairead Hyland, The Lighthouse Lab in Alderley Park and Alex Alderton, Roberto Amato, Jeffrey Barrett, Sonia Goncalves, Ewan Harrison, David K. Jackson, Ian Johnston, Dominic Kwiatkowski, Cordelia Langford, John Sillitoe on behalf of the Wellcome Sanger Institute COVID-19 Surveillance Team |
| EPI_ISL_2092036, EPI_ISL_2092043 | Lighthouse Lab in Milton Keynes | Wellcome Sanger Institute for the COVID-19 Genomics UK (COG-UK) Consortium | The Lighthouse Lab in Milton Keynes and Alex Alderton, Roberto Amato, Jeffrey Barrett, Sonia Goncalves, Ewan Harrison, David K. Jackson, Ian Johnston, Dominic Kwiatkowski, Cordelia Langford, John Sillitoe on behalf of the Wellcome Sanger Institute COVID-19 Surveillance Team |
| EPI_ISL_2092049 | Lighthouse Lab in Alderley Park | Wellcome Sanger Institute for the COVID-19 Genomics UK (COG-UK) Consortium | Jacquelyn Wynn, Mairead Hyland, The Lighthouse Lab in Alderley Park and Alex Alderton, Roberto Amato, Jeffrey Barrett, Sonia Goncalves, Ewan Harrison, David K. Jackson, Ian Johnston, Dominic Kwiatkowski, Cordelia Langford, John Sillitoe on behalf of the Wellcome Sanger Institute COVID-19 Surveillance Team |
| EPI_ISL_2092056 | Lighthouse Lab in Milton Keynes | Wellcome Sanger Institute for the COVID-19 Genomics UK (COG-UK) Consortium | The Lighthouse Lab in Milton Keynes and Alex Alderton, Roberto Amato, Jeffrey Barrett, Sonia Goncalves, Ewan Harrison, David K. Jackson, Ian Johnston, Dominic Kwiatkowski, Cordelia Langford, John Sillitoe on behalf of the Wellcome Sanger Institute COVID-19 Surveillance Team |
| EPI_ISL_2092057, | Lighthouse Lab in Alderley Park | Wellcome Sanger Institute for the COVID-19 Genomics UK (COG-UK) | Jacquelyn Wynn, Mairead Hyland, The Lighthouse Lab in Alderley Park and Alex Alderton, Roberto Amato, Jeffrey Barrett, Sonia Goncalves, Ewan Harrison, David K. Jackson, Ian Johnston, Dominic Kwiatkowski, Cordelia Langford, John Sillitoe on |

| | | | |
|---|---|---|---|
| EPI_ISL_2092059 | | Consortium | behalf of the Wellcome Sanger Institute COVID-19 Surveillance Team |
| EPI_ISL_2092063, EPI_ISL_2092073, EPI_ISL_2092074 | Lighthouse Lab in Milton Keynes | Wellcome Sanger Institute for the COVID-19 Genomics UK (COG-UK) Consortium | The Lighthouse Lab in Milton Keynes and Alex Alderton, Roberto Amato, Jeffrey Barrett, Sonia Goncalves, Ewan Harrison, David K. Jackson, Ian Johnston, Dominic Kwiatkowski, Cordelia Langford, John Sillitoe on behalf of the Wellcome Sanger Institute COVID-19 Surveillance Team |
| EPI_ISL_2092082, EPI_ISL_2092085 | Lighthouse Lab in Alderley Park | Wellcome Sanger Institute for the COVID-19 Genomics UK (COG-UK) Consortium | Jacquelyn Wynn, Mairead Hyland, The Lighthouse Lab in Alderley Park and Alex Alderton, Roberto Amato, Jeffrey Barrett, Sonia Goncalves, Ewan Harrison, David K. Jackson, Ian Johnston, Dominic Kwiatkowski, Cordelia Langford, John Sillitoe on behalf of the Wellcome Sanger Institute COVID-19 Surveillance Team |
| EPI_ISL_2092087, EPI_ISL_2092088, EPI_ISL_2092090 | Lighthouse Lab in Milton Keynes | Wellcome Sanger Institute for the COVID-19 Genomics UK (COG-UK) Consortium | The Lighthouse Lab in Milton Keynes and Alex Alderton, Roberto Amato, Jeffrey Barrett, Sonia Goncalves, Ewan Harrison, David K. Jackson, Ian Johnston, Dominic Kwiatkowski, Cordelia Langford, John Sillitoe on behalf of the Wellcome Sanger Institute COVID-19 Surveillance Team |
| EPI_ISL_2092092 | Lighthouse Lab in Alderley Park | Wellcome Sanger Institute for the COVID-19 Genomics UK (COG-UK) Consortium | Jacquelyn Wynn, Mairead Hyland, The Lighthouse Lab in Alderley Park and Alex Alderton, Roberto Amato, Jeffrey Barrett, Sonia Goncalves, Ewan Harrison, David K. Jackson, Ian Johnston, Dominic Kwiatkowski, Cordelia Langford, John Sillitoe on behalf of the Wellcome Sanger Institute COVID-19 Surveillance Team |
| EPI_ISL_2092094 | Lighthouse Lab in Milton Keynes | Wellcome Sanger Institute for the COVID-19 Genomics UK (COG-UK) Consortium | The Lighthouse Lab in Milton Keynes and Alex Alderton, Roberto Amato, Jeffrey Barrett, Sonia Goncalves, Ewan Harrison, David K. Jackson, Ian Johnston, Dominic Kwiatkowski, Cordelia Langford, John Sillitoe on behalf of the Wellcome Sanger Institute COVID-19 Surveillance Team |
| EPI_ISL_2092096 | Lighthouse Lab in Alderley Park | Wellcome Sanger Institute for the COVID-19 Genomics UK (COG-UK) Consortium | Jacquelyn Wynn, Mairead Hyland, The Lighthouse Lab in Alderley Park and Alex Alderton, Roberto Amato, Jeffrey Barrett, Sonia Goncalves, Ewan Harrison, David K. Jackson, Ian Johnston, Dominic Kwiatkowski, Cordelia Langford, John Sillitoe on behalf of the Wellcome Sanger Institute COVID-19 Surveillance Team |
| EPI_ISL_2092098, EPI_ISL_2092101 | Lighthouse Lab in Milton Keynes | Wellcome Sanger Institute for the COVID-19 Genomics UK (COG-UK) Consortium | The Lighthouse Lab in Milton Keynes and Alex Alderton, Roberto Amato, Jeffrey Barrett, Sonia Goncalves, Ewan Harrison, David K. Jackson, Ian Johnston, Dominic Kwiatkowski, Cordelia Langford, John Sillitoe on behalf of the Wellcome Sanger Institute COVID-19 Surveillance Team |
| EPI_ISL_2092107, EPI_ISL_2092113, EPI_ISL_2092115, EPI_ISL_2092117 | Lighthouse Lab in Alderley Park | Wellcome Sanger Institute for the COVID-19 Genomics UK (COG-UK) Consortium | Jacquelyn Wynn, Mairead Hyland, The Lighthouse Lab in Alderley Park and Alex Alderton, Roberto Amato, Jeffrey Barrett, Sonia Goncalves, Ewan Harrison, David K. Jackson, Ian Johnston, Dominic Kwiatkowski, Cordelia Langford, John Sillitoe on behalf of the Wellcome Sanger Institute COVID-19 Surveillance Team |
| EPI_ISL_2092120 | Randox Laboratories | Wellcome Sanger Institute for the COVID-19 Genomics UK (COG-UK) Consortium | Randox Laboratories and Alex Alderton, Roberto Amato, Jeffrey Barrett, Sonia Goncalves, Ewan Harrison, David K. Jackson, Ian Johnston, Dominic Kwiatkowski, Cordelia Langford, John Sillitoe on behalf of the Wellcome Sanger Institute COVID-19 Surveillance Team |
| EPI_ISL_2092124, EPI_ISL_2092126, EPI_ISL_2092129 | Lighthouse Lab in Alderley Park | Wellcome Sanger Institute for the COVID-19 Genomics UK (COG-UK) Consortium | Jacquelyn Wynn, Mairead Hyland, The Lighthouse Lab in Alderley Park and Alex Alderton, Roberto Amato, Jeffrey Barrett, Sonia Goncalves, Ewan Harrison, David K. Jackson, Ian Johnston, Dominic Kwiatkowski, Cordelia Langford, John Sillitoe on behalf of the Wellcome Sanger Institute COVID-19 Surveillance Team |
| EPI_ISL_2092140 | Randox Laboratories | Wellcome Sanger Institute for the COVID-19 Genomics UK (COG-UK) Consortium | Randox Laboratories and Alex Alderton, Roberto Amato, Jeffrey Barrett, Sonia Goncalves, Ewan Harrison, David K. Jackson, Ian Johnston, Dominic Kwiatkowski, Cordelia Langford, John Sillitoe on behalf of the Wellcome Sanger Institute COVID-19 Surveillance Team |
| EPI_ISL_2092142, EPI_ISL_2092144, EPI_ISL_2092145, EPI_ISL_2092149, EPI_ISL_2092153, EPI_ISL_2092155, EPI_ISL_2092160 | Lighthouse Lab in Alderley Park | Wellcome Sanger Institute for the COVID-19 Genomics UK (COG-UK) Consortium | Jacquelyn Wynn, Mairead Hyland, The Lighthouse Lab in Alderley Park and Alex Alderton, Roberto Amato, Jeffrey Barrett, Sonia Goncalves, Ewan Harrison, David K. Jackson, Ian Johnston, Dominic Kwiatkowski, Cordelia Langford, John Sillitoe on behalf of the Wellcome Sanger Institute COVID-19 Surveillance Team |
| EPI_ISL_2092163 | Randox Laboratories | Wellcome Sanger Institute for the COVID-19 Genomics UK (COG-UK) Consortium | Randox Laboratories and Alex Alderton, Roberto Amato, Jeffrey Barrett, Sonia Goncalves, Ewan Harrison, David K. Jackson, Ian Johnston, Dominic Kwiatkowski, Cordelia Langford, John Sillitoe on behalf of the Wellcome Sanger Institute COVID-19 Surveillance Team |
| EPI_ISL_2092164 | Lighthouse Lab in Alderley Park | Wellcome Sanger Institute for the COVID-19 Genomics UK (COG-UK) Consortium | Jacquelyn Wynn, Mairead Hyland, The Lighthouse Lab in Alderley Park and Alex Alderton, Roberto Amato, Jeffrey Barrett, Sonia Goncalves, Ewan Harrison, David K. Jackson, Ian Johnston, Dominic Kwiatkowski, Cordelia Langford, John Sillitoe on behalf of the Wellcome Sanger Institute COVID-19 Surveillance Team |
| EPI_ISL_2092167 | Randox Laboratories | Wellcome Sanger Institute for the COVID-19 Genomics UK (COG-UK) Consortium | Randox Laboratories and Alex Alderton, Roberto Amato, Jeffrey Barrett, Sonia Goncalves, Ewan Harrison, David K. Jackson, Ian Johnston, Dominic Kwiatkowski, Cordelia Langford, John Sillitoe on behalf of the Wellcome Sanger Institute COVID-19 Surveillance Team |
| EPI_ISL_2092180, EPI_ISL_2092184, EPI_ISL_2092196 | Lighthouse Lab in Alderley Park | Wellcome Sanger Institute for the COVID-19 Genomics UK (COG-UK) Consortium | Jacquelyn Wynn, Mairead Hyland, The Lighthouse Lab in Alderley Park and Alex Alderton, Roberto Amato, Jeffrey Barrett, Sonia Goncalves, Ewan Harrison, David K. Jackson, Ian Johnston, Dominic Kwiatkowski, Cordelia Langford, John Sillitoe on behalf of the Wellcome Sanger Institute COVID-19 Surveillance Team |
| EPI_ISL_2092203, EPI_ISL_2092214 | Randox Laboratories | Wellcome Sanger Institute for the COVID-19 Genomics UK (COG-UK) Consortium | Randox Laboratories and Alex Alderton, Roberto Amato, Jeffrey Barrett, Sonia Goncalves, Ewan Harrison, David K. Jackson, Ian Johnston, Dominic Kwiatkowski, Cordelia Langford, John Sillitoe on behalf of the Wellcome Sanger Institute COVID-19 Surveillance Team |
| EPI_ISL_2092221 | Lighthouse Lab in Alderley Park | Wellcome Sanger Institute for the COVID-19 Genomics UK (COG-UK) Consortium | Jacquelyn Wynn, Mairead Hyland, The Lighthouse Lab in Alderley Park and Alex Alderton, Roberto Amato, Jeffrey Barrett, Sonia Goncalves, Ewan Harrison, David K. Jackson, Ian Johnston, Dominic Kwiatkowski, Cordelia Langford, John Sillitoe on behalf of the Wellcome Sanger Institute COVID-19 Surveillance Team |
| EPI_ISL_2092222 | Randox Laboratories | Wellcome Sanger Institute for the COVID-19 Genomics UK (COG-UK) Consortium | Randox Laboratories and Alex Alderton, Roberto Amato, Jeffrey Barrett, Sonia Goncalves, Ewan Harrison, David K. Jackson, Ian Johnston, Dominic Kwiatkowski, Cordelia Langford, John Sillitoe on behalf of the Wellcome Sanger Institute COVID-19 Surveillance Team |
| EPI_ISL_2092234 | Lighthouse Lab in Alderley Park | Wellcome Sanger Institute for the COVID-19 Genomics UK (COG-UK) Consortium | Jacquelyn Wynn, Mairead Hyland, The Lighthouse Lab in Alderley Park and Alex Alderton, Roberto Amato, Jeffrey Barrett, Sonia Goncalves, Ewan Harrison, David K. Jackson, Ian Johnston, Dominic Kwiatkowski, Cordelia Langford, John Sillitoe on behalf of the Wellcome Sanger Institute COVID-19 Surveillance Team |
| EPI_ISL_2092240 | Randox Laboratories | Wellcome Sanger Institute for the COVID-19 Genomics UK (COG-UK) Consortium | Randox Laboratories and Alex Alderton, Roberto Amato, Jeffrey Barrett, Sonia Goncalves, Ewan Harrison, David K. Jackson, Ian Johnston, Dominic Kwiatkowski, Cordelia Langford, John Sillitoe on behalf of the Wellcome Sanger Institute COVID-19 Surveillance Team |
| EPI_ISL_2092244, EPI_ISL_2092246, EPI_ISL_2092249 | Lighthouse Lab in Alderley Park | Wellcome Sanger Institute for the COVID-19 Genomics UK (COG-UK) Consortium | Jacquelyn Wynn, Mairead Hyland, The Lighthouse Lab in Alderley Park and Alex Alderton, Roberto Amato, Jeffrey Barrett, Sonia Goncalves, Ewan Harrison, David K. Jackson, Ian Johnston, Dominic Kwiatkowski, Cordelia Langford, John Sillitoe on behalf of the Wellcome Sanger Institute COVID-19 Surveillance Team |
| EPI_ISL_2092256 | Randox Laboratories | Wellcome Sanger Institute for the COVID-19 Genomics UK (COG-UK) Consortium | Randox Laboratories and Alex Alderton, Roberto Amato, Jeffrey Barrett, Sonia Goncalves, Ewan Harrison, David K. Jackson, Ian Johnston, Dominic Kwiatkowski, Cordelia Langford, John Sillitoe on behalf of the Wellcome Sanger Institute COVID-19 Surveillance Team |
| EPI_ISL_2092258 | Lighthouse Lab in Alderley Park | Wellcome Sanger Institute for the COVID-19 Genomics UK (COG-UK) Consortium | Jacquelyn Wynn, Mairead Hyland, The Lighthouse Lab in Alderley Park and Alex Alderton, Roberto Amato, Jeffrey Barrett, Sonia Goncalves, Ewan Harrison, David K. Jackson, Ian Johnston, Dominic Kwiatkowski, Cordelia Langford, John Sillitoe on behalf of the Wellcome Sanger Institute COVID-19 Surveillance Team |
| EPI_ISL_2092259 | Randox Laboratories | Wellcome Sanger Institute for the COVID-19 Genomics UK (COG-UK) Consortium | Randox Laboratories and Alex Alderton, Roberto Amato, Jeffrey Barrett, Sonia Goncalves, Ewan Harrison, David K. Jackson, Ian Johnston, Dominic Kwiatkowski, Cordelia Langford, John Sillitoe on behalf of the Wellcome Sanger Institute COVID-19 Surveillance Team |
| EPI_ISL_2092260, EPI_ISL_2092261 | Lighthouse Lab in Alderley Park | Wellcome Sanger Institute for the COVID-19 Genomics UK (COG-UK) Consortium | Jacquelyn Wynn, Mairead Hyland, The Lighthouse Lab in Alderley Park and Alex Alderton, Roberto Amato, Jeffrey Barrett, Sonia Goncalves, Ewan Harrison, David K. Jackson, Ian Johnston, Dominic Kwiatkowski, Cordelia Langford, John Sillitoe on behalf of the Wellcome Sanger Institute COVID-19 Surveillance Team |
| EPI_ISL_2092264, EPI_ISL_2092265, EPI_ISL_2092266 | Randox Laboratories | Wellcome Sanger Institute for the COVID-19 Genomics UK (COG-UK) Consortium | Randox Laboratories and Alex Alderton, Roberto Amato, Jeffrey Barrett, Sonia Goncalves, Ewan Harrison, David K. Jackson, Ian Johnston, Dominic Kwiatkowski, Cordelia Langford, John Sillitoe on behalf of the Wellcome Sanger Institute COVID-19 Surveillance Team |
| EPI_ISL_2092269 | Lighthouse Lab in Alderley Park | Wellcome Sanger Institute for the COVID-19 Genomics UK (COG-UK) Consortium | Jacquelyn Wynn, Mairead Hyland, The Lighthouse Lab in Alderley Park and Alex Alderton, Roberto Amato, Jeffrey Barrett, Sonia Goncalves, Ewan Harrison, David K. Jackson, Ian Johnston, Dominic Kwiatkowski, Cordelia Langford, John Sillitoe on behalf of the Wellcome Sanger Institute COVID-19 Surveillance Team |
| EPI_ISL_2092270 | Randox Laboratories | Wellcome Sanger Institute for the COVID-19 Genomics UK (COG-UK) Consortium | Randox Laboratories and Alex Alderton, Roberto Amato, Jeffrey Barrett, Sonia Goncalves, Ewan Harrison, David K. Jackson, Ian Johnston, Dominic Kwiatkowski, Cordelia Langford, John Sillitoe on behalf of the Wellcome Sanger Institute COVID-19 Surveillance Team |
| EPI_ISL_2092275, EPI_ISL_2092276, EPI_ISL_2092277 | Lighthouse Lab in Alderley Park | Wellcome Sanger Institute for the COVID-19 Genomics UK (COG-UK) Consortium | Jacquelyn Wynn, Mairead Hyland, The Lighthouse Lab in Alderley Park and Alex Alderton, Roberto Amato, Jeffrey Barrett, Sonia Goncalves, Ewan Harrison, David K. Jackson, Ian Johnston, Dominic Kwiatkowski, Cordelia Langford, John Sillitoe on behalf of the Wellcome Sanger Institute COVID-19 Surveillance Team |
| EPI_ISL_2092278 | Randox Laboratories | Wellcome Sanger Institute for the COVID-19 Genomics UK (COG-UK) Consortium | Randox Laboratories and Alex Alderton, Roberto Amato, Jeffrey Barrett, Sonia Goncalves, Ewan Harrison, David K. Jackson, Ian Johnston, Dominic Kwiatkowski, Cordelia Langford, John Sillitoe on behalf of the Wellcome Sanger Institute COVID-19 Surveillance Team |
| EPI_ISL_2092279, EPI_ISL_2092281, EPI_ISL_2092282, EPI_ISL_2092292, EPI_ISL_2092293, EPI_ISL_2092295, EPI_ISL_2092297, EPI_ISL_2092301, EPI_ISL_2092305, EPI_ISL_2092310, EPI_ISL_2092311, EPI_ISL_2092313, EPI_ISL_2092317, EPI_ISL_2092325, EPI_ISL_2092326, EPI_ISL_2092330, EPI_ISL_2092334 | | | |
| see above | Lighthouse Lab in Alderley Park | Wellcome Sanger Institute for the COVID-19 Genomics UK (COG-UK) Consortium | Jacquelyn Wynn, Mairead Hyland, The Lighthouse Lab in Alderley Park and Alex Alderton, Roberto Amato, Jeffrey Barrett, Sonia Goncalves, Ewan Harrison, David K. Jackson, Ian Johnston, Dominic Kwiatkowski, Cordelia Langford, John Sillitoe on behalf of the Wellcome Sanger Institute COVID-19 Surveillance Team |
| EPI_ISL_2092342 | Randox Laboratories | Wellcome Sanger Institute for the COVID-19 Genomics UK (COG-UK) Consortium | Randox Laboratories and Alex Alderton, Roberto Amato, Jeffrey Barrett, Sonia Goncalves, Ewan Harrison, David K. Jackson, Ian Johnston, Dominic Kwiatkowski, Cordelia Langford, John Sillitoe on behalf of the Wellcome Sanger Institute COVID-19 Surveillance Team |
| EPI_ISL_2092350, EPI_ISL_2092352, EPI_ISL_2092355, EPI_ISL_2092357 | Lighthouse Lab in Alderley Park | Wellcome Sanger Institute for the COVID-19 Genomics UK (COG-UK) Consortium | Jacquelyn Wynn, Mairead Hyland, The Lighthouse Lab in Alderley Park and Alex Alderton, Roberto Amato, Jeffrey Barrett, Sonia Goncalves, Ewan Harrison, David K. Jackson, Ian Johnston, Dominic Kwiatkowski, Cordelia Langford, John Sillitoe on behalf of the Wellcome Sanger Institute COVID-19 Surveillance Team |
| EPI_ISL_2092365 | Randox Laboratories | Wellcome Sanger Institute for the COVID-19 Genomics UK (COG-UK) Consortium | Randox Laboratories and Alex Alderton, Roberto Amato, Jeffrey Barrett, Sonia Goncalves, Ewan Harrison, David K. Jackson, Ian Johnston, Dominic Kwiatkowski, Cordelia Langford, John Sillitoe on behalf of the Wellcome Sanger Institute COVID-19 Surveillance Team |
| EPI_ISL_2092368, EPI_ISL_2092371, EPI_ISL_2092372 | Lighthouse Lab in Alderley Park | Wellcome Sanger Institute for the COVID-19 Genomics UK (COG-UK) Consortium | Jacquelyn Wynn, Mairead Hyland, The Lighthouse Lab in Alderley Park and Alex Alderton, Roberto Amato, Jeffrey Barrett, Sonia Goncalves, Ewan Harrison, David K. Jackson, Ian Johnston, Dominic Kwiatkowski, Cordelia Langford, John Sillitoe on behalf of the Wellcome Sanger Institute COVID-19 Surveillance Team |
| EPI_ISL_2092373 | Randox Laboratories | Wellcome Sanger Institute for the COVID-19 Genomics UK (COG-UK) Consortium | Randox Laboratories and Alex Alderton, Roberto Amato, Jeffrey Barrett, Sonia Goncalves, Ewan Harrison, David K. Jackson, Ian Johnston, Dominic Kwiatkowski, Cordelia Langford, John Sillitoe on behalf of the Wellcome Sanger Institute COVID-19 Surveillance Team |

| | | | |
|---|---|---|---|
| EPI_ISL_2092374, EPI_ISL_2092379, EPI_ISL_2092388, EPI_ISL_2092389, EPI_ISL_2092393, EPI_ISL_2092395, EPI_ISL_2092405, EPI_ISL_2092407, EPI_ISL_2092408, EPI_ISL_2092409, EPI_ISL_2092415, EPI_ISL_2092422, EPI_ISL_2092426, EPI_ISL_2092429, EPI_ISL_2092432, EPI_ISL_2092434, EPI_ISL_2092435, EPI_ISL_2092436, EPI_ISL_2092437 | | | |
| see above | Lighthouse Lab in Alderley Park | Wellcome Sanger Institute for the COVID-19 Genomics UK (COG-UK) Consortium | Jacquelyn Wynn, Mairead Hyland, The Lighthouse Lab in Alderley Park and Alex Alderton, Roberto Amato, Jeffrey Barrett, Sonia Goncalves, Ewan Harrison, David K. Jackson, Ian Johnston, Dominic Kwiatkowski, Cordelia Langford, John Sillitoe on behalf of the Wellcome Sanger Institute COVID-19 Surveillance Team |
| EPI_ISL_2092438 | Randox Laboratories | Wellcome Sanger Institute for the COVID-19 Genomics UK (COG-UK) Consortium | Randox Laboratories and Alex Alderton, Roberto Amato, Jeffrey Barrett, Sonia Goncalves, Ewan Harrison, David K. Jackson, Ian Johnston, Dominic Kwiatkowski, Cordelia Langford, John Sillitoe on behalf of the Wellcome Sanger Institute COVID-19 Surveillance Team |
| EPI_ISL_2092456, EPI_ISL_2092464, EPI_ISL_2092465, EPI_ISL_2092475, EPI_ISL_2092484, EPI_ISL_2092496, EPI_ISL_2092516, EPI_ISL_2092526 | Lighthouse Lab in Alderley Park | Wellcome Sanger Institute for the COVID-19 Genomics UK (COG-UK) Consortium | Jacquelyn Wynn, Mairead Hyland, The Lighthouse Lab in Alderley Park and Alex Alderton, Roberto Amato, Jeffrey Barrett, Sonia Goncalves, Ewan Harrison, David K. Jackson, Ian Johnston, Dominic Kwiatkowski, Cordelia Langford, John Sillitoe on behalf of the Wellcome Sanger Institute COVID-19 Surveillance Team |
| EPI_ISL_2092532, EPI_ISL_2092536 | Lighthouse Lab in Glasgow | Wellcome Sanger Institute for the COVID-19 Genomics UK (COG-UK) Consortium | Harper VanSteenhouse, Yumi Kasai, David Gray, Carol Clugston, Anna Dominiczak and Alex Alderton, Roberto Amato, Jeffrey Barrett, Sonia Goncalves, Ewan Harrison, David K. Jackson, Ian Johnston, Dominic Kwiatkowski, Cordelia Langford, John Sillitoe on behalf of the Wellcome Sanger Institute COVID-19 Surveillance Team |
| EPI_ISL_2092548, EPI_ISL_2092553, EPI_ISL_2092565, EPI_ISL_2092586, EPI_ISL_2092595, EPI_ISL_2092599, EPI_ISL_2092612, EPI_ISL_2092613, EPI_ISL_2092645 | Lighthouse Lab in Alderley Park | Wellcome Sanger Institute for the COVID-19 Genomics UK (COG-UK) Consortium | Jacquelyn Wynn, Mairead Hyland, The Lighthouse Lab in Alderley Park and Alex Alderton, Roberto Amato, Jeffrey Barrett, Sonia Goncalves, Ewan Harrison, David K. Jackson, Ian Johnston, Dominic Kwiatkowski, Cordelia Langford, John Sillitoe on behalf of the Wellcome Sanger Institute COVID-19 Surveillance Team |
| EPI_ISL_2092675 | Lighthouse Lab in Milton Keynes | Wellcome Sanger Institute for the COVID-19 Genomics UK (COG-UK) Consortium | The Lighthouse Lab in Milton Keynes and Alex Alderton, Roberto Amato, Jeffrey Barrett, Sonia Goncalves, Ewan Harrison, David K. Jackson, Ian Johnston, Dominic Kwiatkowski, Cordelia Langford, John Sillitoe on behalf of the Wellcome Sanger Institute COVID-19 Surveillance Team |
| EPI_ISL_2092684, EPI_ISL_2092698, EPI_ISL_2092725, EPI_ISL_2092754 | Lighthouse Lab in Glasgow | Wellcome Sanger Institute for the COVID-19 Genomics UK (COG-UK) Consortium | Harper VanSteenhouse, Yumi Kasai, David Gray, Carol Clugston, Anna Dominiczak and Alex Alderton, Roberto Amato, Jeffrey Barrett, Sonia Goncalves, Ewan Harrison, David K. Jackson, Ian Johnston, Dominic Kwiatkowski, Cordelia Langford, John Sillitoe on behalf of the Wellcome Sanger Institute COVID-19 Surveillance Team |
| EPI_ISL_2094352, EPI_ISL_2094353, EPI_ISL_2094354, EPI_ISL_2094386, EPI_ISL_2094387, EPI_ISL_2094388, EPI_ISL_2094389, EPI_ISL_2094390, EPI_ISL_2094391, EPI_ISL_2094392, EPI_ISL_2094393, EPI_ISL_2094394 | | | |
| see above | Dutch COVID-19 response team | National Institute for Public Health and the Environment (RIVM) | Adam Meijer, Harry Vennema, Dirk Eggink, Jeroen Cremer, Sharon van den Brink, Bas van der Veer, AnneMarie van den Brandt, Lisa Wijsman, Kim Freriks, Ryanne Jaarsma, Euníce Then, Lynn Aarts, Sanne Bos, Melissa van Tuil, Linda van de Nes, Sjoerd Kuiling, James Groot, Florian Zwagemaker, Dennis Schmitz, Annelies Kroneman, Karim Hajji, Chantal Reusken, on behalf of the national COVID-19 response team |
| EPI_ISL_2094439, EPI_ISL_2094440, EPI_ISL_2094441 | WSSE w Warszawie | National Institute of Public Health - National Institute of Hygiene | Wołkowicz Tomasz, Zacharczuk Katarzyna, Sadkowska-Todys Małgorzata, Gierczyńki Rafał |
| EPI_ISL_2094505 | LESP Jalisco | Instituto de Diagnostico y Referencia Epidemiologicos (INDRE) | Claudia Wong-Arambula, Abril Rodriguez-Maldonado, Vanessa Rivero-Arredondo, Ariadna Medina-Benitez, Joaquin Quiroz-Mercado, Sergio Rangel-Guerrero, Natividad Cruz-Ortiz, Tatiana Nunez-Garcia, Gisela Barrera-Badillo, Lucia Hernandez-Rivas, Irma Lopez-Martinez, Ernesto Ramirez-Gonzalez. |
| EPI_ISL_2095076, EPI_ISL_2095274, EPI_ISL_2095275, EPI_ISL_2095276, EPI_ISL_2095278, EPI_ISL_2095279, EPI_ISL_2095280, EPI_ISL_2095281, EPI_ISL_2095282 | Max von Pettenkofer Institute, Virology, National Reference Center for Retroviruses, LMU Munich | Laboratory for Functional Genome Analysis; Dept. Genomics; Gene Center of the LMU Munich | Max Muenchhoff; Stefan Krebs; Alexander Graf; Oliver Keppler; Helmut Blum |
| EPI_ISL_2096118 | Washington State Department of Health Public Health Laboratories | Washington State Department of Health Public Health Laboratories | Drew MacKellar, Philip Dykema, Denny Russell, Joenice Gonzalez, Hannah Gray, Geoff Melly, Vanessa De Los Santos, Darren Lucas, JohnAric Peterson, Avi Singh, Rebecca Cao |
| EPI_ISL_2096213 | Maryland Genomics, Institute for Genome Sciences, University of Maryland School of Medicine | Maryland Genomics, Institute for Genome Sciences, University of Maryland School of Medicine | Tallon, Luke J; Sadzewicz, Lisa D; Humphrys, Mike; Ott, Sandra; Roussey, Holly; Mehta, Aditya; Vavikolanu, Kranthi; Fraser, Claire M; Ravel, Jacques |
| EPI_ISL_2096232, EPI_ISL_2096270, EPI_ISL_2096352, EPI_ISL_2096420, EPI_ISL_2096449, EPI_ISL_2096493, EPI_ISL_2096494, EPI_ISL_2096530, EPI_ISL_2096584 | Broad Institute Clinical Research Sequencing Platform | Infectious Disease Program, Broad Institute of Harvard and MIT | Siddle,K.J., Adams,G., Pearlman,L., Gladden-Young,A., Vicente,G., Blumenstiel,B., DeFelice,M., Lee,M., McGovern,S., Lagerborg,K., Rudy,M., DeRuff,K., Carter,A., Normandin,E., Bauer,M., Reilly,S., Tomkins-Tinch,C., Loreth,C., Chaluvadi,S., Meldrim,J., Granger,B., Lemieux,J.E., Birren,B.W., Sabeti,P.C., Larkin,K., Dodge,S., Lennon,N., Madoff,L., Brown,C., Gallagher,G., Smole,S., Park,D.J., Gabriel,S., and MacInnis,B.L. |
| EPI_ISL_2097235 | ALBANY MEDICAL CENTER | Wadsworth Center, New York State Department of Health | Kirsten St. George, Daryl M. Lamson, Alexis Russell, Matthew Shudt, Melissa A Leisner, Jonathan Plitnick, Catharine Prussing, Navjot Singh, John Kelly, Erasmus Schneider, Erica Lasek-Nesselquist |
| EPI_ISL_2097653, EPI_ISL_2097683, EPI_ISL_2097858, EPI_ISL_2098013 | Helix/Illumina | Centers for Disease Control and Prevention Division of Viral Diseases, Pathogen Discovery | Dakota Howard, Dhwani Batra, Peter W. Cook, Kara Moser, Adrian Paskey, Jason Caravas, Benjamin Rambo-Martin, Shatavia Morrison, Christopher Gulvick, Scott Sammons, Yvette Unoarumhi, Darlene Wagner, Matthew Schmerer, Eileen de Feo, Jan Antico, Christine Tran, Matthew Tolentino, Shannon Wickline, Kim Gietzen, Brad Sickler, Jingtao Liu, Eric Allen, Phil Febbo, Nicole L. Washington, Simon White, Geraint Levan, Kelly Schiabor Barrett, Elizabeth Cirulli, Alexandre Bolze, Ary Ascencio, Charlotte Rivera-Garcia, Ryan Cho, Jason Nguyen, Sherry Wang, Jimmy Ramirez, Tyler Cassens, Efren Sandoval, Magnus Isaksson, William Lee, David Becker, Marc Laurent, James Lu, Clinton R. Paden, Duncan MacCannell |
| EPI_ISL_2098125 | ALBANY MEDICAL CENTER | Wadsworth Center, New York State Department of Health | Kirsten St. George, Daryl M. Lamson, Alexis Russell, Matthew Shudt, Melissa A Leisner, Jonathan Plitnick, Catharine Prussing, Navjot Singh, John Kelly, Erasmus Schneider, Erica Lasek-Nesselquist |
| EPI_ISL_2098710, EPI_ISL_2098714, EPI_ISL_2098715, EPI_ISL_2098716, EPI_ISL_2098717, EPI_ISL_2098718, EPI_ISL_2098719, EPI_ISL_2098720, EPI_ISL_2098721, EPI_ISL_2098722 | State Virus Research and Diagnostic Laboratory (VRDL), AIIMS Raipur | State Virus Research and Diagnostic Laboratory (VRDL), AIIMS Raipur | Pushpendra Singh, Kuldeep Sharma, Priyanka Singh, Somya Sharma, MD Rafiiqullah Khan, Jiyan Chandrawanshi, Sanjay Singh Negi, Anudita Bhargava |
| EPI_ISL_2098723 | PathWest Laboratory Medicine WA | PathWest Laboratory Medicine WA Microbial Surveillance Unit | PathWest Laboratory Medicine WA Microbial Surveillance Unit |
| EPI_ISL_2099526 | UMC Groningen, Clinical Virology, Department of Medical Microbiology and Infection Prevention | UMC Groningen, Clinical Virology, Department of Medical Microbiology and Infection Prevention | Hubert Niesters, Alexander Friedrich, Erley Lizarazo-Forero, Monika Fliss, Lilli Gard, Sigrid Rosema, Coretta Van Leer-Buter, Xuewei Zhou, Marjolein Knoester |
| EPI_ISL_2099644 | National Centre for Disease Control | CDFD -INSACOG | To be addded later |
| EPI_ISL_2099853 | National Institute of Infectious Diseases-Prof. Dr. Matei Bals Molecular Diagnostics Laboratory | National Institute of Infectious Diseases-Prof. Dr. Matei Bals Molecular Diagnostics Laboratory | Corina Casangiu, Leontina Banica, Marius Surleac, Ovidiu Vlaicu, Andreea Tudor, Simona Paraschiv, Dan Otelea |
| EPI_ISL_2099869 | Department of Public Health Bucharest | National Institute of Infectious Diseases-Prof. Dr. Matei Bals Molecular Diagnostics Laboratory | Corina Casangiu, Leontina Banica, Marius Surleac, Ovidiu Vlaicu, Andreea Tudor, Simona Paraschiv, Dan Otelea |
| EPI_ISL_2100187, EPI_ISL_2100188, EPI_ISL_2100190, EPI_ISL_2100195, EPI_ISL_2100216 | ICMR-National Institute for Research In Tuberculosis | NIV Influenza | Dr Padmapriyadarsini.C, Dr Luke Elizabeth Hanna, Dr S Siva Kumar, Dr Radha Gopalaswamy, Mr Ramesh K |
| EPI_ISL_2100241, EPI_ISL_2100242 | Viral Respiratory Infections Laboratory, Cantacuzino National Military-Medical Institute | Cantacuzino Institute Virology | Sorin Dinu,Carmen Cherciu, Mihaela Oprea, Mihaela Lazar |
| EPI_ISL_2100243, EPI_ISL_2100244, EPI_ISL_2100245, EPI_ISL_2100246, EPI_ISL_2100247 | Clinical Hospital of Infectious and Tropical Diseases “Dr. Victor Babes” | Cantacuzino Institute Virology | Sorin Dinu,Carmen Cherciu, Mihaela Oprea, Mihaela Lazar |
| EPI_ISL_2100385, | University of Michigan Clinical Microbiology Laboratory | Lauring Lab, University of Michigan, Department of Microbiology | Valesano |

| | | | |
|---|---|---|---|
| EPI_ISL_2100391 | | and Immunology | |
| EPI_ISL_2100635, EPI_ISL_2100636, EPI_ISL_2100637, EPI_ISL_2100638, EPI_ISL_2100639, EPI_ISL_2100640, EPI_ISL_2100641, EPI_ISL_2100642, EPI_ISL_2100643, EPI_ISL_2100644, EPI_ISL_2100645, EPI_ISL_2100646, EPI_ISL_2100647, EPI_ISL_2100648, EPI_ISL_2100649, EPI_ISL_2100650, EPI_ISL_2100651, EPI_ISL_2100652, EPI_ISL_2100653, EPI_ISL_2100654, EPI_ISL_2100655, EPI_ISL_2100656, EPI_ISL_2100657, EPI_ISL_2100658, EPI_ISL_2100659, EPI_ISL_2100660, EPI_ISL_2100661, EPI_ISL_2100662, EPI_ISL_2100663, EPI_ISL_2100664, EPI_ISL_2100665, EPI_ISL_2100666, EPI_ISL_2100667, EPI_ISL_2100668, EPI_ISL_2100669, EPI_ISL_2100670, EPI_ISL_2100671, EPI_ISL_2100672 | | | |
| see above | Delhi North District | CSIR-Institute of Genomics and Integrative Biology | Pooja Sharma*, Bharathram Uppilli*, Animesh Ray, Sushma Rajpoot, Saruchi Wadhwa, Asangala Kamai, Devasish Desai, Manish Kumar, Satish Swain, Sarafaraz Alam, Umang Arora, Sheeba Saifi, Mohammed Ahmed, Ayush Goel, Ashwin Varadarajan, Naveet Wig, Rajesh Pandey, Mohammed Faruq |
| EPI_ISL_2100679, EPI_ISL_2100680, EPI_ISL_2100681, EPI_ISL_2100682, EPI_ISL_2100687, EPI_ISL_2100689, EPI_ISL_2100691, EPI_ISL_2100692, EPI_ISL_2100694, EPI_ISL_2100696, EPI_ISL_2100698, EPI_ISL_2100699 | | | |
| see above | All India Institute of Medical Sciences, Ansari Nagar Delhi | CSIR-Institute of Genomics and Integrative Biology | Pooja Sharma*, Bharathram Uppilli*, Animesh Ray, Sushma Rajpoot, Saruchi Wadhwa, Asangala Kamai, Devasish Desai, Manish Kumar, Satish Swain, Sarafaraz Alam, Umang Arora, Sheeba Saifi, Mohammed Ahmed, Ayush Goel, Ashwin Varadarajan, Naveet Wig, Rajesh Pandey, Mohammed Faruq |
| EPI_ISL_2101018 | Department of Microbiology, Tochigi Prefectural Institute of Public Health and Enviromental Science | Genome Analysis Center, Kamma Memorial Hospital | Fuminori Mizukoshi, Yoshiko Hitomi, Hanako Yazawa, Jun Ishii, Satoko Soma, Kaori Watanabe, Takuya Yazawa, Michiaki Masuda, Hiroshi Kamma |
| EPI_ISL_2101019 | Department of Microbiology, Tochigi Prefectural Institute of Public Health and Environmental Science | Genome Analysis Center, Kamma Memorial Hospital | Fuminori Mizukoshi, Yoshiko Hitomi, Hanako Yazawa, Jun Ishii, Satoko Soma, Kaori Watanabe, Takuya Yazawa, Michiaki Masuda, Hiroshi Kamma |
| EPI_ISL_2101020 | Department of Microbiology, Tochigi Prefectural Institute of Public Health and Enviromental Science | Genome Analysis Center, Kamma Memorial Hospital | Fuminori Mizukoshi, Yoshiko Hitomi, Hanako Yazawa, Jun Ishii, Satoko Soma, Kaori Watanabe, Takuya Yazawa, Michiaki Masuda, Hiroshi Kamma |
| EPI_ISL_2102114 | Medics Labor AG | Clinical Bacteriology | Tim Roloff, Madlen Stange, Helena MB Seth-Smith, Alfredo Mari, Karoline Leuzinger, Julia Bielicki, Manuel Battegay, Hans Hirsch, Adrian Egli |
| EPI_ISL_2102154 | CHUV | Laboratory of genomics and metagenomics | Trestan Pillonel, Damien Jacot, Sébastien Aeby, Gilbert Greub, Claire Bertelli |
| EPI_ISL_2102347 | Maryland Genomics, Institute for Genome Sciences, University of Maryland School of Medicine | Maryland Genomics, Institute for Genome Sciences, University of Maryland School of Medicine | Tallon, Luke J; Sadzewicz, Lisa D; Humphrys, Mike; Ott, Sandra; Roussey, Holly; Mehta, Aditya; Vavikolanu, Kranthi; Fraser, Claire M; Ravel, Jacques |
| EPI_ISL_2103201, EPI_ISL_2103202 | LabPLUS | Institute of Environmental Science and Research (ESR) | Rachel Boyle, SallyAnn Harbison, Olivia Stroeven, Xiaoyun Ren, Matt Storey, Nikki Freed, Muhammad Faisal, Jing Wang, Hermes Perez, Anja Werno, Antje van der Linden, Arlo Upton, Chris Mansell, David Hammer, Dragana Drinkovic, Gary McAuliffe, Hana Sofia Andersson, James Ussher, Jill Sherwood, Josh Freeman, Julia Howard, Juliet Elvy, Mary DeAlmeida, Matt Blakiston, Matthew Rogers, Max Bloomfield, Michael Addidle, Michelle Balm, Sally Roberts, Sarah Jefferies, Sharmini Muttaiyah, Susan Morpeth, Susan Taylor, Timothy Blackmore, Vani Sathyendran, Veronica Playle, Virginia Hope, Erasmus Smit, Lauren Jelly, Olin Silander, Joep de Ligt |
| EPI_ISL_2103264 | Johns Hopkins Hospital Department of Pathology | Johns Hopkins Hospital Department of Pathology | C. Paul Morris, Chun Huai Luo, Adannaya Amadi, Matthew Schwartz, Nicholas Gallagher, Heba H. Mostafa |
| EPI_ISL_2103917, EPI_ISL_2103954, EPI_ISL_2103977, EPI_ISL_2103987, EPI_ISL_2104095, EPI_ISL_2104145, EPI_ISL_2104234, EPI_ISL_2104240, EPI_ISL_2104281, EPI_ISL_2104380 | Broad Institute Clinical Research Sequencing Platform | Infectious Disease Program, Broad Institute of Harvard and MIT | Siddle,K.J., Adams,G., Pearlman,L., Gladden-Young,A., Vicente,G., Blumenstiel,B., DeFelice,M., Lee,M., McGovern,S., Lagerborg,K., Rudy,M., DeRuff,K., Carter,A., Normandin,E., Bauer,M., Reilly,S., Tomkins-Tinch,C., Loreth,C., Chaluvadi,S., Meldrim,J., Granger,B., Lemieux,J.E., Birren,B.W., Sabeti,P.C., Larkin,K., Dodge,S., Lennon,N., Madoff,L., Brown,C., Gallagher,G., Smole,S., Park,D.J., Gabriel,S., and MacInnis,B.L. |
| EPI_ISL_2104479, EPI_ISL_2104494 | University Hospitals of Geneva, Laboratory of Virology | HUG, Laboratory of Virology and the Health2030 Genome Center | Samuel Cordey, Ana Rita Goncalves, Laurent Kaiser, Lorenzo Cerutti, Henri Pegeot, Melyssa Elies, Deborah Penet, Keith Harshman, Ioannis Xenarios, Emmanouil Dermitzakis |
| EPI_ISL_2104743 | Ramathibodi Chakri Naruebodindra Hospital | COVID-19 Network Investigations (CONI) Alliance | Elizabeth Batty, Wasun Chantratita, Thanat Chookajorn, Stefan Fernandez, Angkana Huang, Anthony R. Jones, Khajohn Joonlasak, Chonticha Klungtong, Theerarat Kochakarn, Namfon Kotanan, Krittikorn Kumpornsin, Duangkamon Loesbanluechai, Wudtichai Manasatienkij, Bhakbhoom Panthan, Ekawat Pasomsub, Kingkan Rakmanee, Insee Sensorn, Janjira Thaipadungpanit, Arporn Wangwiwatsin, Treewat Watthanachockchai |
| EPI_ISL_2104866, EPI_ISL_2104900, EPI_ISL_2104962, EPI_ISL_2104997, EPI_ISL_2105091, EPI_ISL_2105092 | UW Virology Lab | UW Virology Lab | Pavitra Roychoudhury, Hong Xie, Lasata Shrestha, Tien V. Nguyen, Shah Mohamed Bakhash, Michelle Lin, Noah R. Baker, Ricardo Perez, Sean Ellis, Nathan Breit, Robert J. Livingston, Meei-Li Huang, Keith R Jerome, Patrick Mathias, Alexander Greninger |
| EPI_ISL_2105565 | Institute of Epidemiology, Disease Control and Research (IEDCR) | ideSHi-IEDCR-icddr,b | Hassan Afrad, Sadia Rahman, Manjur Hossain Khan, Firdausi Qadri, Tahmina Shirin |
| EPI_ISL_2105576, EPI_ISL_2105577, EPI_ISL_2105578, EPI_ISL_2105579, EPI_ISL_2105580, EPI_ISL_2105581 | Delhi North District | CSIR-Institute of Genomics and Integrative Biology | Pooja Sharma*, Bharathram Uppilli*, Animesh Ray, Sushma Rajpoot, Saruchi Wadhwa, Asangala Kamai, Devasish Desai, Manish Kumar, Satish Swain, Sarafaraz Alam, Umang Arora, Sheeba Saifi, Mohammed Ahmed, Ayush Goel, Ashwin Varadarajan, Naveet Wig, Rajesh Pandey, Mohammed Faruq |
| EPI_ISL_2105584, EPI_ISL_2105585 | All India Institute of Medical Sciences, Ansari Nagar Delhi | CSIR-Institute of Genomics and Integrative Biology | Pooja Sharma*, Bharathram Uppilli*, Animesh Ray, Sushma Rajpoot, Saruchi Wadhwa, Asangala Kamai, Devasish Desai, Manish Kumar, Satish Swain, Sarafaraz Alam, Umang Arora, Sheeba Saifi, Mohammed Ahmed, Ayush Goel, Ashwin Varadarajan, Naveet Wig, Rajesh Pandey, Mohammed Faruq |
| EPI_ISL_2105608 | WSSE w Gdansku | 1. Academic Center for Pathomorphological and Genetic-Molecular Diagnostics ltd, Bialystok, Poland 2. National Institute of Public Health - National Institute of Hygiene, Warsaw, Poland | Radosław Charkiewicz, Jacek Nikliński, Przemysław Biecek. Joanna Reszeć, Piotr Majewski, Anetta Sulewska, Piotr Karabowicz, Joanna Kiśluk, Janusz Dzięcioł, Konrad Raczkowski, Katarzyna Zacharczuk, Magdalena Nowakowska, Małgorzata Sadkowska-Todys, Tomasz Wołkowicz |
| EPI_ISL_2105673 | Biolab Diagnostic Laboratories | Biolab Diagnostic Laboratories | Issa Abu-Dayyeh, Ahmad Tibi, Lama Hussein, Shayma Ali, Badia Saddedin, Eiad Atwa, Amid Abdelnour |
| EPI_ISL_2105884 | CSL, Maryland Department of Health | CSL, Maryland Department of Health | Keller,E., Loomis,J., Low,K. |
| EPI_ISL_2106001, EPI_ISL_2106013, EPI_ISL_2106069 | Respiratory Viruses Branch, Centers for Disease Control and Prevention | Respiratory Viruses Branch, Centers for Disease Control and Prevention | Howard,D., Batra,D., Cook,P.W., Moser,K., Paskey,A., Caravas,J., Rambo-Martin,B., Morrison,S., Gulvick,C., Sammons,S., Unoarumhi,Y., Wagner,D., Roychoudhury,M.S., Xie,H., Shrestha,L., Nguyen,T.V., Bakhash,S.M., Lin,M., Baker,N.R., Perez,R., Ellis,S., Breit,N., Livingston,R.J., Huang,M.-L., Jerome,K.R., Mathias,P., Greninger,A., Paden,C.R., MacCannell,D. |
| EPI_ISL_2106198 | Delhi North District | CSIR-Institute of Genomics and Integrative Biology | Pooja Sharma*, Bharathram Uppilli*, Animesh Ray, Sushma Rajpoot, Saruchi Wadhwa, Asangala Kamai, Devasish Desai, Manish Kumar, Satish Swain, Sarafaraz Alam, Umang Arora, Sheeba Saifi, Mohammed Ahmed, Ayush Goel, Ashwin Varadarajan, Naveet Wig, Rajesh Pandey, Mohammed Faruq |
| EPI_ISL_2106292, EPI_ISL_2106299, EPI_ISL_2106311, EPI_ISL_2106323, EPI_ISL_2106331, EPI_ISL_2106344, EPI_ISL_2106347, EPI_ISL_2106363, EPI_ISL_2106388, EPI_ISL_2106401, EPI_ISL_2106415, EPI_ISL_2106449, EPI_ISL_2106469, EPI_ISL_2106526, EPI_ISL_2106566 | | | |
| see above | Pandemic Response Lab - NYC | Pandemic Response Lab, R&D | Henry Lee, Michael Hammerling, Melissa Hopkins, Cybill del Castillo, Shinyoung Clair Kang, William Ward, Pradeep Bugga, Sol Rey, Dylan Law, Katharine Nelson, Haiping Hao, Jon Laurent |
| EPI_ISL_2106967 | Centre for DNA Fingerprinting and Diagnostics | CDFD -INSACOG | Murali Bashyam, Asmita Gupta, Pratyusha Bala, Vinay Donipadi, Divya Vashisht, Ashwin Dalal |
| EPI_ISL_2106975 | BPOM Gorontalo | National Institute of Health Research and Development | Hartanti Dian Ikawati, Hana Apsari Pawestri, Subangkit, Kartika Dewi Puspa, Arie Ardiansyah Nugraha, Triyani Soekarso, Krisna Pangesti, Nelly Puspandari, Vivi Setiawaty |
| EPI_ISL_2107009, EPI_ISL_2107010, EPI_ISL_2107017, EPI_ISL_2107020, EPI_ISL_2107022, EPI_ISL_2107023, EPI_ISL_2107025, EPI_ISL_2107027, EPI_ISL_2107033, EPI_ISL_2107034, EPI_ISL_2107060, EPI_ISL_2107062, EPI_ISL_2107063, EPI_ISL_2107066, EPI_ISL_2107070, EPI_ISL_2107071, EPI_ISL_2107072, EPI_ISL_2107075, EPI_ISL_2107077, EPI_ISL_2107080, EPI_ISL_2107081, EPI_ISL_2107083, EPI_ISL_2107085, EPI_ISL_2107087, EPI_ISL_2107088, EPI_ISL_2107089, EPI_ISL_2107090 | | | |
| see above | National Centre For Cell Science – INSACOG | National Centre For Cell Science | Dhiraj Paul, Shivang P. Bhanushali, Mohak P Gujare, Mitali Inamdar, Manoj Kumar Bhat, Ajay Pillai, INSACOG Consortium team, Yogesh Shouche |
| EPI_ISL_2107443, EPI_ISL_2107444, EPI_ISL_2107446, EPI_ISL_2107447, EPI_ISL_2107448, EPI_ISL_2107451, EPI_ISL_2107454, EPI_ISL_2107455, EPI_ISL_2107456, EPI_ISL_2107458, EPI_ISL_2107499, EPI_ISL_2107500, EPI_ISL_2107501, EPI_ISL_2107506, EPI_ISL_2107529, EPI_ISL_2107535 | | | |
| see above | New South Wales Health Pathology Royal Prince Alfred Hospital | Microbiology RPAH | Foster, C.; Au, J.; Ruiz Silva, M.; Deveson, I.; Bull, R.; Van Hal, S.; Rawlinson, W. |
| EPI_ISL_2107828, EPI_ISL_2107830, EPI_ISL_2107835, EPI_ISL_2107839 | Labor Berlin Charité Vivantes GmbH / Institut für Virologie | Charité Universitätsmedizin Berlin, Institut für Virologie/Labor Berlin | Peter Menzel, Christine Stephan, Rolf Schwarzer, Victor M Corman, Barbara Mühlemann, Terry Jones, Christian Drosten |
| EPI_ISL_2108737, EPI_ISL_2108817 | Bioscientia Labor Wermsdorf | Robert Koch Institute | unknown |
| EPI_ISL_2109111 | Labor 28 MVZ GmbH | Robert Koch Institute | unknown |
| EPI_ISL_2109742 | Bioscientia MVZ Labor Karlsruhe GmbH | Robert Koch Institute | unknown |
| EPI_ISL_2110448, EPI_ISL_2110453, EPI_ISL_2110454, EPI_ISL_2110460, EPI_ISL_2110488, EPI_ISL_2110489, EPI_ISL_2110499 | Eurofins LifeCodexx GmbH | Robert Koch Institute | unknown |
| EPI_ISL_2110643 | Institut Pasteur du Maroc | Functional Genomics platform/CNRST | Melloul Marouane, Chouati Taha, Hemlali Mouhsine, Alaoui Sanaa-amine, Touil Nadia, Elannaz Hicham, Ennibi Khalid, Youbi Mohammed, MERABET MOUAD,Abdelkrim Meziane Bellefquih, NOURLIL Jalal, Maaroufi Abderrahmane, EL FAHIME Elmostafa |
| EPI_ISL_2110715, EPI_ISL_2110737 | Eurofins LifeCodexx GmbH | Robert Koch Institute | unknown |
| EPI_ISL_2111026 | Labor Dr. Wisplinghoff - Kaln | Robert Koch Institute | unknown |
| EPI_ISL_2111142 | Laborarztpraxis Dres. med. Walther Weindel & Kollegen | Robert Koch Institute | unknown |
| EPI_ISL_2111185 | MVZ Labor Krone GbR | Robert Koch Institute | unknown |

| | | | |
|---|---|---|---|
| EPI_ISL_2111238 | MVZ Labor Dr. Fenner und Kollegen (Standort Hamburg) | Robert Koch Institute | unknown |
| EPI_ISL_2111454 | Labor ZOTZ\|KLIMAS; MVZ Dusseldorf-Centrum | Robert Koch Institute | unknown |
| EPI_ISL_2112265 | SYNLAB MVZ Leinfelden-Echterdingen | Robert Koch Institute | unknown |
| EPI_ISL_2112762 | MVZ Medizinisches Labor Hannover GmbH | Robert Koch Institute | unknown |
| EPI_ISL_2112834 | IMD Labor Frankfurt | Robert Koch Institute | unknown |
| EPI_ISL_2114976, EPI_ISL_2115085, EPI_ISL_2115100 | SYNLAB MVZ Heidelberg | Robert Koch Institute | unknown |
| EPI_ISL_2115134, EPI_ISL_2115136, EPI_ISL_2115142, EPI_ISL_2115145 | Dianovis GmbH Greiz | Robert Koch Institute | unknown |
| EPI_ISL_2115162 | MVZ Medizinisches Labor Hannover GmbH | Robert Koch Institute | unknown |
| EPI_ISL_2115263, EPI_ISL_2115271 | MVZ Labor Krone GbR | Robert Koch Institute | unknown |
| EPI_ISL_2115511 | Virologisches Institut des Universitatsklinikums Erlangen | Robert Koch Institute | unknown |
| EPI_ISL_2115877 | amedes MVZ Zentrum fÃ¼r Pathologie und Zytodiagnostik GmbH (KÃ¶ln) | Robert Koch Institute | unknown |
| EPI_ISL_2116115 | Medizinische Laboratorien Dusseldorf | Robert Koch Institute | unknown |
| EPI_ISL_2117300 | SYNLAB Jena Oncoscreen | Robert Koch Institute | unknown |
| EPI_ISL_2117512 | Lighthouse Lab in Alderley Park | Wellcome Sanger Institute for the COVID-19 Genomics UK (COG-UK) Consortium | Jacquelyn Wynn, Mairead Hyland, The Lighthouse Lab in Alderley Park and Alex Alderton, Roberto Amato, Jeffrey Barrett, Sonia Goncalves, Ewan Harrison, David K. Jackson, Ian Johnston, Dominic Kwiatkowski, Cordelia Langford, John Sillitoe on behalf of the Wellcome Sanger Institute COVID-19 Surveillance Team |
| EPI_ISL_2117519, EPI_ISL_2117521 | Lighthouse Lab in Glasgow | Wellcome Sanger Institute for the COVID-19 Genomics UK (COG-UK) Consortium | Harper VanSteenhouse, Yumi Kasai, David Gray, Carol Clugston, Anna Dominiczak and Alex Alderton, Roberto Amato, Jeffrey Barrett, Sonia Goncalves, Ewan Harrison, David K. Jackson, Ian Johnston, Dominic Kwiatkowski, Cordelia Langford, John Sillitoe on behalf of the Wellcome Sanger Institute COVID-19 Surveillance Team |
| EPI_ISL_2117529 | Lighthouse Lab in Milton Keynes | Wellcome Sanger Institute for the COVID-19 Genomics UK (COG-UK) Consortium | The Lighthouse Lab in Milton Keynes and Alex Alderton, Roberto Amato, Jeffrey Barrett, Sonia Goncalves, Ewan Harrison, David K. Jackson, Ian Johnston, Dominic Kwiatkowski, Cordelia Langford, John Sillitoe on behalf of the Wellcome Sanger Institute COVID-19 Surveillance Team |
| EPI_ISL_2117532, EPI_ISL_2117536 | Lighthouse Lab in Glasgow | Wellcome Sanger Institute for the COVID-19 Genomics UK (COG-UK) Consortium | Harper VanSteenhouse, Yumi Kasai, David Gray, Carol Clugston, Anna Dominiczak and Alex Alderton, Roberto Amato, Jeffrey Barrett, Sonia Goncalves, Ewan Harrison, David K. Jackson, Ian Johnston, Dominic Kwiatkowski, Cordelia Langford, John Sillitoe on behalf of the Wellcome Sanger Institute COVID-19 Surveillance Team |
| EPI_ISL_2117538 | Lighthouse Lab in Milton Keynes | Wellcome Sanger Institute for the COVID-19 Genomics UK (COG-UK) Consortium | The Lighthouse Lab in Milton Keynes and Alex Alderton, Roberto Amato, Jeffrey Barrett, Sonia Goncalves, Ewan Harrison, David K. Jackson, Ian Johnston, Dominic Kwiatkowski, Cordelia Langford, John Sillitoe on behalf of the Wellcome Sanger Institute COVID-19 Surveillance Team |
| EPI_ISL_2117539 | Lighthouse Lab in Alderley Park | Wellcome Sanger Institute for the COVID-19 Genomics UK (COG-UK) Consortium | Jacquelyn Wynn, Mairead Hyland, The Lighthouse Lab in Alderley Park and Alex Alderton, Roberto Amato, Jeffrey Barrett, Sonia Goncalves, Ewan Harrison, David K. Jackson, Ian Johnston, Dominic Kwiatkowski, Cordelia Langford, John Sillitoe on behalf of the Wellcome Sanger Institute COVID-19 Surveillance Team |
| EPI_ISL_2117542, EPI_ISL_2117549 | Lighthouse Lab in Milton Keynes | Wellcome Sanger Institute for the COVID-19 Genomics UK (COG-UK) Consortium | The Lighthouse Lab in Milton Keynes and Alex Alderton, Roberto Amato, Jeffrey Barrett, Sonia Goncalves, Ewan Harrison, David K. Jackson, Ian Johnston, Dominic Kwiatkowski, Cordelia Langford, John Sillitoe on behalf of the Wellcome Sanger Institute COVID-19 Surveillance Team |
| EPI_ISL_2117552 | Lighthouse Lab in Alderley Park | Wellcome Sanger Institute for the COVID-19 Genomics UK (COG-UK) Consortium | Jacquelyn Wynn, Mairead Hyland, The Lighthouse Lab in Alderley Park and Alex Alderton, Roberto Amato, Jeffrey Barrett, Sonia Goncalves, Ewan Harrison, David K. Jackson, Ian Johnston, Dominic Kwiatkowski, Cordelia Langford, John Sillitoe on behalf of the Wellcome Sanger Institute COVID-19 Surveillance Team |
| EPI_ISL_2117554 | Lighthouse Lab in Glasgow | Wellcome Sanger Institute for the COVID-19 Genomics UK (COG-UK) Consortium | Harper VanSteenhouse, Yumi Kasai, David Gray, Carol Clugston, Anna Dominiczak and Alex Alderton, Roberto Amato, Jeffrey Barrett, Sonia Goncalves, Ewan Harrison, David K. Jackson, Ian Johnston, Dominic Kwiatkowski, Cordelia Langford, John Sillitoe on behalf of the Wellcome Sanger Institute COVID-19 Surveillance Team |
| EPI_ISL_2117556, EPI_ISL_2117559 | Lighthouse Lab in Milton Keynes | Wellcome Sanger Institute for the COVID-19 Genomics UK (COG-UK) Consortium | The Lighthouse Lab in Milton Keynes and Alex Alderton, Roberto Amato, Jeffrey Barrett, Sonia Goncalves, Ewan Harrison, David K. Jackson, Ian Johnston, Dominic Kwiatkowski, Cordelia Langford, John Sillitoe on behalf of the Wellcome Sanger Institute COVID-19 Surveillance Team |
| EPI_ISL_2117560 | Lighthouse Lab in Alderley Park | Wellcome Sanger Institute for the COVID-19 Genomics UK (COG-UK) Consortium | Jacquelyn Wynn, Mairead Hyland, The Lighthouse Lab in Alderley Park and Alex Alderton, Roberto Amato, Jeffrey Barrett, Sonia Goncalves, Ewan Harrison, David K. Jackson, Ian Johnston, Dominic Kwiatkowski, Cordelia Langford, John Sillitoe on behalf of the Wellcome Sanger Institute COVID-19 Surveillance Team |
| EPI_ISL_2117563 | Lighthouse Lab in Glasgow | Wellcome Sanger Institute for the COVID-19 Genomics UK (COG-UK) Consortium | Harper VanSteenhouse, Yumi Kasai, David Gray, Carol Clugston, Anna Dominiczak and Alex Alderton, Roberto Amato, Jeffrey Barrett, Sonia Goncalves, Ewan Harrison, David K. Jackson, Ian Johnston, Dominic Kwiatkowski, Cordelia Langford, John Sillitoe on behalf of the Wellcome Sanger Institute COVID-19 Surveillance Team |
| EPI_ISL_2117566 | Lighthouse Lab in Milton Keynes | Wellcome Sanger Institute for the COVID-19 Genomics UK (COG-UK) Consortium | The Lighthouse Lab in Milton Keynes and Alex Alderton, Roberto Amato, Jeffrey Barrett, Sonia Goncalves, Ewan Harrison, David K. Jackson, Ian Johnston, Dominic Kwiatkowski, Cordelia Langford, John Sillitoe on behalf of the Wellcome Sanger Institute COVID-19 Surveillance Team |
| EPI_ISL_2117569 | Lighthouse Lab in Alderley Park | Wellcome Sanger Institute for the COVID-19 Genomics UK (COG-UK) Consortium | Jacquelyn Wynn, Mairead Hyland, The Lighthouse Lab in Alderley Park and Alex Alderton, Roberto Amato, Jeffrey Barrett, Sonia Goncalves, Ewan Harrison, David K. Jackson, Ian Johnston, Dominic Kwiatkowski, Cordelia Langford, John Sillitoe on behalf of the Wellcome Sanger Institute COVID-19 Surveillance Team |
| EPI_ISL_2117571, EPI_ISL_2117578, EPI_ISL_2117579 | Lighthouse Lab in Glasgow | Wellcome Sanger Institute for the COVID-19 Genomics UK (COG-UK) Consortium | Harper VanSteenhouse, Yumi Kasai, David Gray, Carol Clugston, Anna Dominiczak and Alex Alderton, Roberto Amato, Jeffrey Barrett, Sonia Goncalves, Ewan Harrison, David K. Jackson, Ian Johnston, Dominic Kwiatkowski, Cordelia Langford, John Sillitoe on behalf of the Wellcome Sanger Institute COVID-19 Surveillance Team |
| EPI_ISL_2117580 | Lighthouse Lab in Alderley Park | Wellcome Sanger Institute for the COVID-19 Genomics UK (COG-UK) Consortium | Jacquelyn Wynn, Mairead Hyland, The Lighthouse Lab in Alderley Park and Alex Alderton, Roberto Amato, Jeffrey Barrett, Sonia Goncalves, Ewan Harrison, David K. Jackson, Ian Johnston, Dominic Kwiatkowski, Cordelia Langford, John Sillitoe on behalf of the Wellcome Sanger Institute COVID-19 Surveillance Team |
| EPI_ISL_2117581 | Lighthouse Lab in Glasgow | Wellcome Sanger Institute for the COVID-19 Genomics UK (COG-UK) Consortium | Harper VanSteenhouse, Yumi Kasai, David Gray, Carol Clugston, Anna Dominiczak and Alex Alderton, Roberto Amato, Jeffrey Barrett, Sonia Goncalves, Ewan Harrison, David K. Jackson, Ian Johnston, Dominic Kwiatkowski, Cordelia Langford, John Sillitoe on behalf of the Wellcome Sanger Institute COVID-19 Surveillance Team |
| EPI_ISL_2117584, EPI_ISL_2117585 | Lighthouse Lab in Milton Keynes | Wellcome Sanger Institute for the COVID-19 Genomics UK (COG-UK) Consortium | The Lighthouse Lab in Milton Keynes and Alex Alderton, Roberto Amato, Jeffrey Barrett, Sonia Goncalves, Ewan Harrison, David K. Jackson, Ian Johnston, Dominic Kwiatkowski, Cordelia Langford, John Sillitoe on behalf of the Wellcome Sanger Institute COVID-19 Surveillance Team |
| EPI_ISL_2117588, EPI_ISL_2117590 | Lighthouse Lab in Alderley Park | Wellcome Sanger Institute for the COVID-19 Genomics UK (COG-UK) Consortium | Jacquelyn Wynn, Mairead Hyland, The Lighthouse Lab in Alderley Park and Alex Alderton, Roberto Amato, Jeffrey Barrett, Sonia Goncalves, Ewan Harrison, David K. Jackson, Ian Johnston, Dominic Kwiatkowski, Cordelia Langford, John Sillitoe on behalf of the Wellcome Sanger Institute COVID-19 Surveillance Team |
| EPI_ISL_2117597, EPI_ISL_2117600 | Lighthouse Lab in Glasgow | Wellcome Sanger Institute for the COVID-19 Genomics UK (COG-UK) Consortium | Harper VanSteenhouse, Yumi Kasai, David Gray, Carol Clugston, Anna Dominiczak and Alex Alderton, Roberto Amato, Jeffrey Barrett, Sonia Goncalves, Ewan Harrison, David K. Jackson, Ian Johnston, Dominic Kwiatkowski, Cordelia Langford, John Sillitoe on behalf of the Wellcome Sanger Institute COVID-19 Surveillance Team |
| EPI_ISL_2117601 | Lighthouse Lab in Milton Keynes | Wellcome Sanger Institute for the COVID-19 Genomics UK (COG-UK) Consortium | The Lighthouse Lab in Milton Keynes and Alex Alderton, Roberto Amato, Jeffrey Barrett, Sonia Goncalves, Ewan Harrison, David K. Jackson, Ian Johnston, Dominic Kwiatkowski, Cordelia Langford, John Sillitoe on behalf of the Wellcome Sanger Institute COVID-19 Surveillance Team |
| EPI_ISL_2117604 | Lighthouse Lab in Alderley Park | Wellcome Sanger Institute for the COVID-19 Genomics UK (COG-UK) Consortium | Jacquelyn Wynn, Mairead Hyland, The Lighthouse Lab in Alderley Park and Alex Alderton, Roberto Amato, Jeffrey Barrett, Sonia Goncalves, Ewan Harrison, David K. Jackson, Ian Johnston, Dominic Kwiatkowski, Cordelia Langford, John Sillitoe on behalf of the Wellcome Sanger Institute COVID-19 Surveillance Team |
| EPI_ISL_2117613, EPI_ISL_2117617 | Lighthouse Lab in Milton Keynes | Wellcome Sanger Institute for the COVID-19 Genomics UK (COG-UK) Consortium | The Lighthouse Lab in Milton Keynes and Alex Alderton, Roberto Amato, Jeffrey Barrett, Sonia Goncalves, Ewan Harrison, David K. Jackson, Ian Johnston, Dominic Kwiatkowski, Cordelia Langford, John Sillitoe on behalf of the Wellcome Sanger Institute COVID-19 Surveillance Team |
| EPI_ISL_2117621 | Lighthouse Lab in Glasgow | Wellcome Sanger Institute for the COVID-19 Genomics UK (COG-UK) Consortium | Harper VanSteenhouse, Yumi Kasai, David Gray, Carol Clugston, Anna Dominiczak and Alex Alderton, Roberto Amato, Jeffrey Barrett, Sonia Goncalves, Ewan Harrison, David K. Jackson, Ian Johnston, Dominic Kwiatkowski, Cordelia Langford, John Sillitoe on behalf of the Wellcome Sanger Institute COVID-19 Surveillance Team |
| EPI_ISL_2117623 | Lighthouse Lab in Alderley Park | Wellcome Sanger Institute for the COVID-19 Genomics UK (COG-UK) Consortium | Jacquelyn Wynn, Mairead Hyland, The Lighthouse Lab in Alderley Park and Alex Alderton, Roberto Amato, Jeffrey Barrett, Sonia Goncalves, Ewan Harrison, David K. Jackson, Ian Johnston, Dominic Kwiatkowski, Cordelia Langford, John Sillitoe on behalf of the Wellcome Sanger Institute COVID-19 Surveillance Team |
| EPI_ISL_2117627 | Lighthouse Lab in Glasgow | Wellcome Sanger Institute for the COVID-19 Genomics UK (COG-UK) Consortium | Harper VanSteenhouse, Yumi Kasai, David Gray, Carol Clugston, Anna Dominiczak and Alex Alderton, Roberto Amato, Jeffrey Barrett, Sonia Goncalves, Ewan Harrison, David K. Jackson, Ian Johnston, Dominic Kwiatkowski, Cordelia Langford, John Sillitoe on behalf of the Wellcome Sanger Institute COVID-19 Surveillance Team |
| EPI_ISL_2117629 | Lighthouse Lab in Milton Keynes | Wellcome Sanger Institute for the COVID-19 Genomics UK (COG-UK) Consortium | The Lighthouse Lab in Milton Keynes and Alex Alderton, Roberto Amato, Jeffrey Barrett, Sonia Goncalves, Ewan Harrison, David K. Jackson, Ian Johnston, Dominic Kwiatkowski, Cordelia Langford, John Sillitoe on behalf of the Wellcome Sanger Institute COVID-19 Surveillance Team |
| EPI_ISL_2117632 | Lighthouse Lab in Glasgow | Wellcome Sanger Institute for the COVID-19 Genomics UK (COG-UK) Consortium | Harper VanSteenhouse, Yumi Kasai, David Gray, Carol Clugston, Anna Dominiczak and Alex Alderton, Roberto Amato, Jeffrey Barrett, Sonia Goncalves, Ewan Harrison, David K. Jackson, Ian Johnston, Dominic Kwiatkowski, Cordelia Langford, John Sillitoe on behalf of the Wellcome Sanger Institute COVID-19 Surveillance Team |
| EPI_ISL_2117635, EPI_ISL_2117638 | Lighthouse Lab in Milton Keynes | Wellcome Sanger Institute for the COVID-19 Genomics UK (COG-UK) Consortium | The Lighthouse Lab in Milton Keynes and Alex Alderton, Roberto Amato, Jeffrey Barrett, Sonia Goncalves, Ewan Harrison, David K. Jackson, Ian Johnston, Dominic Kwiatkowski, Cordelia Langford, John Sillitoe on behalf of the Wellcome Sanger Institute COVID-19 Surveillance Team |
| EPI_ISL_2117639, EPI_ISL_2117643, EPI_ISL_2117644, EPI_ISL_2117645, EPI_ISL_2117648, EPI_ISL_2117649, EPI_ISL_2117651 | Lighthouse Lab in Glasgow | Wellcome Sanger Institute for the COVID-19 Genomics UK (COG-UK) Consortium | Harper VanSteenhouse, Yumi Kasai, David Gray, Carol Clugston, Anna Dominiczak and Alex Alderton, Roberto Amato, Jeffrey Barrett, Sonia Goncalves, Ewan Harrison, David K. Jackson, Ian Johnston, Dominic Kwiatkowski, Cordelia Langford, John Sillitoe on behalf of the Wellcome Sanger Institute COVID-19 Surveillance Team |
| EPI_ISL_2117652, EPI_ISL_2117654 | Lighthouse Lab in Alderley Park | Wellcome Sanger Institute for the COVID-19 Genomics UK (COG-UK) Consortium | Jacquelyn Wynn, Mairead Hyland, The Lighthouse Lab in Alderley Park and Alex Alderton, Roberto Amato, Jeffrey Barrett, Sonia Goncalves, Ewan Harrison, David K. Jackson, Ian Johnston, Dominic Kwiatkowski, Cordelia Langford, John Sillitoe on behalf of the Wellcome Sanger Institute COVID-19 Surveillance Team |
| EPI_ISL_2117656 | Lighthouse Lab in Milton Keynes | Wellcome Sanger Institute for the COVID-19 Genomics UK (COG-UK) | The Lighthouse Lab in Milton Keynes and Alex Alderton, Roberto Amato, Jeffrey Barrett, Sonia Goncalves, Ewan Harrison, David K. Jackson, Ian Johnston, Dominic Kwiatkowski, Cordelia Langford, John Sillitoe on behalf of the Wellcome Sanger |

| | | | |
|---|---|---|---|
| | | Consortium | Institute COVID-19 Surveillance Team |
| EPI_ISL_2117657, EPI_ISL_2117660 | Lighthouse Lab in Glasgow | Wellcome Sanger Institute for the COVID-19 Genomics UK (COG-UK) Consortium | Harper VanSteenhouse, Yumi Kasai, David Gray, Carol Clugston, Anna Dominiczak and Alex Alderton, Roberto Amato, Jeffrey Barrett, Sonia Goncalves, Ewan Harrison, David K. Jackson, Ian Johnston, Dominic Kwiatkowski, Cordelia Langford, John Sillitoe on behalf of the Wellcome Sanger Institute COVID-19 Surveillance Team |
| EPI_ISL_2117665 | Lighthouse Lab in Alderley Park | Wellcome Sanger Institute for the COVID-19 Genomics UK (COG-UK) Consortium | Jacquelyn Wynn, Mairead Hyland, The Lighthouse Lab in Alderley Park and Alex Alderton, Roberto Amato, Jeffrey Barrett, Sonia Goncalves, Ewan Harrison, David K. Jackson, Ian Johnston, Dominic Kwiatkowski, Cordelia Langford, John Sillitoe on behalf of the Wellcome Sanger Institute COVID-19 Surveillance Team |
| EPI_ISL_2117667, EPI_ISL_2117668 | Lighthouse Lab in Glasgow | Wellcome Sanger Institute for the COVID-19 Genomics UK (COG-UK) Consortium | Harper VanSteenhouse, Yumi Kasai, David Gray, Carol Clugston, Anna Dominiczak and Alex Alderton, Roberto Amato, Jeffrey Barrett, Sonia Goncalves, Ewan Harrison, David K. Jackson, Ian Johnston, Dominic Kwiatkowski, Cordelia Langford, John Sillitoe on behalf of the Wellcome Sanger Institute COVID-19 Surveillance Team |
| EPI_ISL_2117672 | Lighthouse Lab in Alderley Park | Wellcome Sanger Institute for the COVID-19 Genomics UK (COG-UK) Consortium | Jacquelyn Wynn, Mairead Hyland, The Lighthouse Lab in Alderley Park and Alex Alderton, Roberto Amato, Jeffrey Barrett, Sonia Goncalves, Ewan Harrison, David K. Jackson, Ian Johnston, Dominic Kwiatkowski, Cordelia Langford, John Sillitoe on behalf of the Wellcome Sanger Institute COVID-19 Surveillance Team |
| EPI_ISL_2117676 | Lighthouse Lab in Milton Keynes | Wellcome Sanger Institute for the COVID-19 Genomics UK (COG-UK) Consortium | The Lighthouse Lab in Milton Keynes and Alex Alderton, Roberto Amato, Jeffrey Barrett, Sonia Goncalves, Ewan Harrison, David K. Jackson, Ian Johnston, Dominic Kwiatkowski, Cordelia Langford, John Sillitoe on behalf of the Wellcome Sanger Institute COVID-19 Surveillance Team |
| EPI_ISL_2117679, EPI_ISL_2117681, EPI_ISL_2117682 | Lighthouse Lab in Glasgow | Wellcome Sanger Institute for the COVID-19 Genomics UK (COG-UK) Consortium | Harper VanSteenhouse, Yumi Kasai, David Gray, Carol Clugston, Anna Dominiczak and Alex Alderton, Roberto Amato, Jeffrey Barrett, Sonia Goncalves, Ewan Harrison, David K. Jackson, Ian Johnston, Dominic Kwiatkowski, Cordelia Langford, John Sillitoe on behalf of the Wellcome Sanger Institute COVID-19 Surveillance Team |
| EPI_ISL_2117684, EPI_ISL_2117687, EPI_ISL_2117688 | Lighthouse Lab in Alderley Park | Wellcome Sanger Institute for the COVID-19 Genomics UK (COG-UK) Consortium | Jacquelyn Wynn, Mairead Hyland, The Lighthouse Lab in Alderley Park and Alex Alderton, Roberto Amato, Jeffrey Barrett, Sonia Goncalves, Ewan Harrison, David K. Jackson, Ian Johnston, Dominic Kwiatkowski, Cordelia Langford, John Sillitoe on behalf of the Wellcome Sanger Institute COVID-19 Surveillance Team |
| EPI_ISL_2117689 | Lighthouse Lab in Milton Keynes | Wellcome Sanger Institute for the COVID-19 Genomics UK (COG-UK) Consortium | The Lighthouse Lab in Milton Keynes and Alex Alderton, Roberto Amato, Jeffrey Barrett, Sonia Goncalves, Ewan Harrison, David K. Jackson, Ian Johnston, Dominic Kwiatkowski, Cordelia Langford, John Sillitoe on behalf of the Wellcome Sanger Institute COVID-19 Surveillance Team |
| EPI_ISL_2117690, EPI_ISL_2117691, EPI_ISL_2117693, EPI_ISL_2117694 | Lighthouse Lab in Alderley Park | Wellcome Sanger Institute for the COVID-19 Genomics UK (COG-UK) Consortium | Jacquelyn Wynn, Mairead Hyland, The Lighthouse Lab in Alderley Park and Alex Alderton, Roberto Amato, Jeffrey Barrett, Sonia Goncalves, Ewan Harrison, David K. Jackson, Ian Johnston, Dominic Kwiatkowski, Cordelia Langford, John Sillitoe on behalf of the Wellcome Sanger Institute COVID-19 Surveillance Team |
| EPI_ISL_2117703, EPI_ISL_2117713, EPI_ISL_2117716 | Lighthouse Lab in Glasgow | Wellcome Sanger Institute for the COVID-19 Genomics UK (COG-UK) Consortium | Harper VanSteenhouse, Yumi Kasai, David Gray, Carol Clugston, Anna Dominiczak and Alex Alderton, Roberto Amato, Jeffrey Barrett, Sonia Goncalves, Ewan Harrison, David K. Jackson, Ian Johnston, Dominic Kwiatkowski, Cordelia Langford, John Sillitoe on behalf of the Wellcome Sanger Institute COVID-19 Surveillance Team |
| EPI_ISL_2117719, EPI_ISL_2117723 | Lighthouse Lab in Alderley Park | Wellcome Sanger Institute for the COVID-19 Genomics UK (COG-UK) Consortium | Jacquelyn Wynn, Mairead Hyland, The Lighthouse Lab in Alderley Park and Alex Alderton, Roberto Amato, Jeffrey Barrett, Sonia Goncalves, Ewan Harrison, David K. Jackson, Ian Johnston, Dominic Kwiatkowski, Cordelia Langford, John Sillitoe on behalf of the Wellcome Sanger Institute COVID-19 Surveillance Team |
| EPI_ISL_2117726, EPI_ISL_2117727 | Lighthouse Lab in Glasgow | Wellcome Sanger Institute for the COVID-19 Genomics UK (COG-UK) Consortium | Harper VanSteenhouse, Yumi Kasai, David Gray, Carol Clugston, Anna Dominiczak and Alex Alderton, Roberto Amato, Jeffrey Barrett, Sonia Goncalves, Ewan Harrison, David K. Jackson, Ian Johnston, Dominic Kwiatkowski, Cordelia Langford, John Sillitoe on behalf of the Wellcome Sanger Institute COVID-19 Surveillance Team |
| EPI_ISL_2117728 | Lighthouse Lab in Milton Keynes | Wellcome Sanger Institute for the COVID-19 Genomics UK (COG-UK) Consortium | The Lighthouse Lab in Milton Keynes and Alex Alderton, Roberto Amato, Jeffrey Barrett, Sonia Goncalves, Ewan Harrison, David K. Jackson, Ian Johnston, Dominic Kwiatkowski, Cordelia Langford, John Sillitoe on behalf of the Wellcome Sanger Institute COVID-19 Surveillance Team |
| EPI_ISL_2117731, EPI_ISL_2117734, EPI_ISL_2117736 | Lighthouse Lab in Glasgow | Wellcome Sanger Institute for the COVID-19 Genomics UK (COG-UK) Consortium | Harper VanSteenhouse, Yumi Kasai, David Gray, Carol Clugston, Anna Dominiczak and Alex Alderton, Roberto Amato, Jeffrey Barrett, Sonia Goncalves, Ewan Harrison, David K. Jackson, Ian Johnston, Dominic Kwiatkowski, Cordelia Langford, John Sillitoe on behalf of the Wellcome Sanger Institute COVID-19 Surveillance Team |
| EPI_ISL_2117740, EPI_ISL_2117743 | Lighthouse Lab in Milton Keynes | Wellcome Sanger Institute for the COVID-19 Genomics UK (COG-UK) Consortium | The Lighthouse Lab in Milton Keynes and Alex Alderton, Roberto Amato, Jeffrey Barrett, Sonia Goncalves, Ewan Harrison, David K. Jackson, Ian Johnston, Dominic Kwiatkowski, Cordelia Langford, John Sillitoe on behalf of the Wellcome Sanger Institute COVID-19 Surveillance Team |
| EPI_ISL_2117745 | Lighthouse Lab in Glasgow | Wellcome Sanger Institute for the COVID-19 Genomics UK (COG-UK) Consortium | Harper VanSteenhouse, Yumi Kasai, David Gray, Carol Clugston, Anna Dominiczak and Alex Alderton, Roberto Amato, Jeffrey Barrett, Sonia Goncalves, Ewan Harrison, David K. Jackson, Ian Johnston, Dominic Kwiatkowski, Cordelia Langford, John Sillitoe on behalf of the Wellcome Sanger Institute COVID-19 Surveillance Team |
| EPI_ISL_2117747 | Lighthouse Lab in Milton Keynes | Wellcome Sanger Institute for the COVID-19 Genomics UK (COG-UK) Consortium | The Lighthouse Lab in Milton Keynes and Alex Alderton, Roberto Amato, Jeffrey Barrett, Sonia Goncalves, Ewan Harrison, David K. Jackson, Ian Johnston, Dominic Kwiatkowski, Cordelia Langford, John Sillitoe on behalf of the Wellcome Sanger Institute COVID-19 Surveillance Team |
| EPI_ISL_2117748 | Lighthouse Lab in Alderley Park | Wellcome Sanger Institute for the COVID-19 Genomics UK (COG-UK) Consortium | Jacquelyn Wynn, Mairead Hyland, The Lighthouse Lab in Alderley Park and Alex Alderton, Roberto Amato, Jeffrey Barrett, Sonia Goncalves, Ewan Harrison, David K. Jackson, Ian Johnston, Dominic Kwiatkowski, Cordelia Langford, John Sillitoe on behalf of the Wellcome Sanger Institute COVID-19 Surveillance Team |
| EPI_ISL_2117749, EPI_ISL_2117751 | Lighthouse Lab in Glasgow | Wellcome Sanger Institute for the COVID-19 Genomics UK (COG-UK) Consortium | Harper VanSteenhouse, Yumi Kasai, David Gray, Carol Clugston, Anna Dominiczak and Alex Alderton, Roberto Amato, Jeffrey Barrett, Sonia Goncalves, Ewan Harrison, David K. Jackson, Ian Johnston, Dominic Kwiatkowski, Cordelia Langford, John Sillitoe on behalf of the Wellcome Sanger Institute COVID-19 Surveillance Team |
| EPI_ISL_2117754 | Lighthouse Lab in Alderley Park | Wellcome Sanger Institute for the COVID-19 Genomics UK (COG-UK) Consortium | Jacquelyn Wynn, Mairead Hyland, The Lighthouse Lab in Alderley Park and Alex Alderton, Roberto Amato, Jeffrey Barrett, Sonia Goncalves, Ewan Harrison, David K. Jackson, Ian Johnston, Dominic Kwiatkowski, Cordelia Langford, John Sillitoe on behalf of the Wellcome Sanger Institute COVID-19 Surveillance Team |
| EPI_ISL_2117755 | Lighthouse Lab in Milton Keynes | Wellcome Sanger Institute for the COVID-19 Genomics UK (COG-UK) Consortium | The Lighthouse Lab in Milton Keynes and Alex Alderton, Roberto Amato, Jeffrey Barrett, Sonia Goncalves, Ewan Harrison, David K. Jackson, Ian Johnston, Dominic Kwiatkowski, Cordelia Langford, John Sillitoe on behalf of the Wellcome Sanger Institute COVID-19 Surveillance Team |
| EPI_ISL_2117760, EPI_ISL_2117761 | Lighthouse Lab in Glasgow | Wellcome Sanger Institute for the COVID-19 Genomics UK (COG-UK) Consortium | Harper VanSteenhouse, Yumi Kasai, David Gray, Carol Clugston, Anna Dominiczak and Alex Alderton, Roberto Amato, Jeffrey Barrett, Sonia Goncalves, Ewan Harrison, David K. Jackson, Ian Johnston, Dominic Kwiatkowski, Cordelia Langford, John Sillitoe on behalf of the Wellcome Sanger Institute COVID-19 Surveillance Team |
| EPI_ISL_2117762 | Lighthouse Lab in Alderley Park | Wellcome Sanger Institute for the COVID-19 Genomics UK (COG-UK) Consortium | Jacquelyn Wynn, Mairead Hyland, The Lighthouse Lab in Alderley Park and Alex Alderton, Roberto Amato, Jeffrey Barrett, Sonia Goncalves, Ewan Harrison, David K. Jackson, Ian Johnston, Dominic Kwiatkowski, Cordelia Langford, John Sillitoe on behalf of the Wellcome Sanger Institute COVID-19 Surveillance Team |
| EPI_ISL_2117767 | Lighthouse Lab in Glasgow | Wellcome Sanger Institute for the COVID-19 Genomics UK (COG-UK) Consortium | Harper VanSteenhouse, Yumi Kasai, David Gray, Carol Clugston, Anna Dominiczak and Alex Alderton, Roberto Amato, Jeffrey Barrett, Sonia Goncalves, Ewan Harrison, David K. Jackson, Ian Johnston, Dominic Kwiatkowski, Cordelia Langford, John Sillitoe on behalf of the Wellcome Sanger Institute COVID-19 Surveillance Team |
| EPI_ISL_2117770, EPI_ISL_2117772 | Lighthouse Lab in Alderley Park | Wellcome Sanger Institute for the COVID-19 Genomics UK (COG-UK) Consortium | Jacquelyn Wynn, Mairead Hyland, The Lighthouse Lab in Alderley Park and Alex Alderton, Roberto Amato, Jeffrey Barrett, Sonia Goncalves, Ewan Harrison, David K. Jackson, Ian Johnston, Dominic Kwiatkowski, Cordelia Langford, John Sillitoe on behalf of the Wellcome Sanger Institute COVID-19 Surveillance Team |
| EPI_ISL_2117773, EPI_ISL_2117779, EPI_ISL_2117781 | Lighthouse Lab in Glasgow | Wellcome Sanger Institute for the COVID-19 Genomics UK (COG-UK) Consortium | Harper VanSteenhouse, Yumi Kasai, David Gray, Carol Clugston, Anna Dominiczak and Alex Alderton, Roberto Amato, Jeffrey Barrett, Sonia Goncalves, Ewan Harrison, David K. Jackson, Ian Johnston, Dominic Kwiatkowski, Cordelia Langford, John Sillitoe on behalf of the Wellcome Sanger Institute COVID-19 Surveillance Team |
| EPI_ISL_2117783 | Lighthouse Lab in Milton Keynes | Wellcome Sanger Institute for the COVID-19 Genomics UK (COG-UK) Consortium | The Lighthouse Lab in Milton Keynes and Alex Alderton, Roberto Amato, Jeffrey Barrett, Sonia Goncalves, Ewan Harrison, David K. Jackson, Ian Johnston, Dominic Kwiatkowski, Cordelia Langford, John Sillitoe on behalf of the Wellcome Sanger Institute COVID-19 Surveillance Team |
| EPI_ISL_2117784, EPI_ISL_2117787, EPI_ISL_2117794 | Lighthouse Lab in Alderley Park | Wellcome Sanger Institute for the COVID-19 Genomics UK (COG-UK) Consortium | Jacquelyn Wynn, Mairead Hyland, The Lighthouse Lab in Alderley Park and Alex Alderton, Roberto Amato, Jeffrey Barrett, Sonia Goncalves, Ewan Harrison, David K. Jackson, Ian Johnston, Dominic Kwiatkowski, Cordelia Langford, John Sillitoe on behalf of the Wellcome Sanger Institute COVID-19 Surveillance Team |
| EPI_ISL_2117799 | Lighthouse Lab in Glasgow | Wellcome Sanger Institute for the COVID-19 Genomics UK (COG-UK) Consortium | Harper VanSteenhouse, Yumi Kasai, David Gray, Carol Clugston, Anna Dominiczak and Alex Alderton, Roberto Amato, Jeffrey Barrett, Sonia Goncalves, Ewan Harrison, David K. Jackson, Ian Johnston, Dominic Kwiatkowski, Cordelia Langford, John Sillitoe on behalf of the Wellcome Sanger Institute COVID-19 Surveillance Team |
| EPI_ISL_2117801, EPI_ISL_2117803 | Lighthouse Lab in Milton Keynes | Wellcome Sanger Institute for the COVID-19 Genomics UK (COG-UK) Consortium | The Lighthouse Lab in Milton Keynes and Alex Alderton, Roberto Amato, Jeffrey Barrett, Sonia Goncalves, Ewan Harrison, David K. Jackson, Ian Johnston, Dominic Kwiatkowski, Cordelia Langford, John Sillitoe on behalf of the Wellcome Sanger Institute COVID-19 Surveillance Team |
| EPI_ISL_2117804, EPI_ISL_2117806, EPI_ISL_2117808 | Lighthouse Lab in Glasgow | Wellcome Sanger Institute for the COVID-19 Genomics UK (COG-UK) Consortium | Harper VanSteenhouse, Yumi Kasai, David Gray, Carol Clugston, Anna Dominiczak and Alex Alderton, Roberto Amato, Jeffrey Barrett, Sonia Goncalves, Ewan Harrison, David K. Jackson, Ian Johnston, Dominic Kwiatkowski, Cordelia Langford, John Sillitoe on behalf of the Wellcome Sanger Institute COVID-19 Surveillance Team |
| EPI_ISL_2117809 | Lighthouse Lab in Alderley Park | Wellcome Sanger Institute for the COVID-19 Genomics UK (COG-UK) Consortium | Jacquelyn Wynn, Mairead Hyland, The Lighthouse Lab in Alderley Park and Alex Alderton, Roberto Amato, Jeffrey Barrett, Sonia Goncalves, Ewan Harrison, David K. Jackson, Ian Johnston, Dominic Kwiatkowski, Cordelia Langford, John Sillitoe on behalf of the Wellcome Sanger Institute COVID-19 Surveillance Team |
| EPI_ISL_2117811, EPI_ISL_2117814 | Lighthouse Lab in Glasgow | Wellcome Sanger Institute for the COVID-19 Genomics UK (COG-UK) Consortium | Harper VanSteenhouse, Yumi Kasai, David Gray, Carol Clugston, Anna Dominiczak and Alex Alderton, Roberto Amato, Jeffrey Barrett, Sonia Goncalves, Ewan Harrison, David K. Jackson, Ian Johnston, Dominic Kwiatkowski, Cordelia Langford, John Sillitoe on behalf of the Wellcome Sanger Institute COVID-19 Surveillance Team |
| EPI_ISL_2117816 | Lighthouse Lab in Milton Keynes | Wellcome Sanger Institute for the COVID-19 Genomics UK (COG-UK) Consortium | The Lighthouse Lab in Milton Keynes and Alex Alderton, Roberto Amato, Jeffrey Barrett, Sonia Goncalves, Ewan Harrison, David K. Jackson, Ian Johnston, Dominic Kwiatkowski, Cordelia Langford, John Sillitoe on behalf of the Wellcome Sanger Institute COVID-19 Surveillance Team |
| EPI_ISL_2117817 | Lighthouse Lab in Glasgow | Wellcome Sanger Institute for the COVID-19 Genomics UK (COG-UK) Consortium | Harper VanSteenhouse, Yumi Kasai, David Gray, Carol Clugston, Anna Dominiczak and Alex Alderton, Roberto Amato, Jeffrey Barrett, Sonia Goncalves, Ewan Harrison, David K. Jackson, Ian Johnston, Dominic Kwiatkowski, Cordelia Langford, John Sillitoe on behalf of the Wellcome Sanger Institute COVID-19 Surveillance Team |
| EPI_ISL_2117819 | Lighthouse Lab in Alderley Park | Wellcome Sanger Institute for the COVID-19 Genomics UK (COG-UK) Consortium | Jacquelyn Wynn, Mairead Hyland, The Lighthouse Lab in Alderley Park and Alex Alderton, Roberto Amato, Jeffrey Barrett, Sonia Goncalves, Ewan Harrison, David K. Jackson, Ian Johnston, Dominic Kwiatkowski, Cordelia Langford, John Sillitoe on behalf of the Wellcome Sanger Institute COVID-19 Surveillance Team |
| EPI_ISL_2117820 | Lighthouse Lab in Glasgow | Wellcome Sanger Institute for the COVID-19 Genomics UK (COG-UK) Consortium | Harper VanSteenhouse, Yumi Kasai, David Gray, Carol Clugston, Anna Dominiczak and Alex Alderton, Roberto Amato, Jeffrey Barrett, Sonia Goncalves, Ewan Harrison, David K. Jackson, Ian Johnston, Dominic Kwiatkowski, Cordelia Langford, John Sillitoe on behalf of the Wellcome Sanger Institute COVID-19 Surveillance Team |
| EPI_ISL_2117822 | Lighthouse Lab in Milton Keynes | Wellcome Sanger Institute for the COVID-19 Genomics UK (COG-UK) Consortium | The Lighthouse Lab in Milton Keynes and Alex Alderton, Roberto Amato, Jeffrey Barrett, Sonia Goncalves, Ewan Harrison, David K. Jackson, Ian Johnston, Dominic Kwiatkowski, Cordelia Langford, John Sillitoe on behalf of the Wellcome Sanger Institute COVID-19 Surveillance Team |
| EPI_ISL_2117823, EPI_ISL_2117828 | Lighthouse Lab in Glasgow | Wellcome Sanger Institute for the COVID-19 Genomics UK (COG-UK) Consortium | Harper VanSteenhouse, Yumi Kasai, David Gray, Carol Clugston, Anna Dominiczak and Alex Alderton, Roberto Amato, Jeffrey Barrett, Sonia Goncalves, Ewan Harrison, David K. Jackson, Ian Johnston, Dominic Kwiatkowski, Cordelia Langford, John Sillitoe on behalf of the Wellcome Sanger Institute COVID-19 Surveillance Team |
| EPI_ISL_2117831 | Lighthouse Lab in Alderley Park | Wellcome Sanger Institute for the COVID-19 Genomics UK (COG-UK) Consortium | Jacquelyn Wynn, Mairead Hyland, The Lighthouse Lab in Alderley Park and Alex Alderton, Roberto Amato, Jeffrey Barrett, Sonia Goncalves, Ewan Harrison, David K. Jackson, Ian Johnston, Dominic Kwiatkowski, Cordelia Langford, John Sillitoe on behalf of the Wellcome Sanger Institute COVID-19 Surveillance Team |
| EPI_ISL_2117837, | Lighthouse Lab in Glasgow | Wellcome Sanger Institute for the COVID-19 Genomics UK (COG-UK) | Harper VanSteenhouse, Yumi Kasai, David Gray, Carol Clugston, Anna Dominiczak and Alex Alderton, Roberto Amato, Jeffrey Barrett, Sonia Goncalves, Ewan Harrison, David K. Jackson, Ian Johnston, Dominic Kwiatkowski, Cordelia Langford, |

| | | | |
|---|---|---|---|
| EPI_ISL_2117842 | | Consortium | John Sillitoe on behalf of the Wellcome Sanger Institute COVID-19 Surveillance Team |
| EPI_ISL_2117843 | Lighthouse Lab in Alderley Park | Wellcome Sanger Institute for the COVID-19 Genomics UK (COG-UK) Consortium | Jacquelyn Wynn, Mairead Hyland, The Lighthouse Lab in Alderley Park and Alex Alderton, Roberto Amato, Jeffrey Barrett, Sonia Goncalves, Ewan Harrison, David K. Jackson, Ian Johnston, Dominic Kwiatkowski, Cordelia Langford, John Sillitoe on behalf of the Wellcome Sanger Institute COVID-19 Surveillance Team |
| EPI_ISL_2117846, EPI_ISL_2117850 | Lighthouse Lab in Milton Keynes | Wellcome Sanger Institute for the COVID-19 Genomics UK (COG-UK) Consortium | The Lighthouse Lab in Milton Keynes and Alex Alderton, Roberto Amato, Jeffrey Barrett, Sonia Goncalves, Ewan Harrison, David K. Jackson, Ian Johnston, Dominic Kwiatkowski, Cordelia Langford, John Sillitoe on behalf of the Wellcome Sanger Institute COVID-19 Surveillance Team |
| EPI_ISL_2117851 | Lighthouse Lab in Alderley Park | Wellcome Sanger Institute for the COVID-19 Genomics UK (COG-UK) Consortium | Jacquelyn Wynn, Mairead Hyland, The Lighthouse Lab in Alderley Park and Alex Alderton, Roberto Amato, Jeffrey Barrett, Sonia Goncalves, Ewan Harrison, David K. Jackson, Ian Johnston, Dominic Kwiatkowski, Cordelia Langford, John Sillitoe on behalf of the Wellcome Sanger Institute COVID-19 Surveillance Team |
| EPI_ISL_2117854, EPI_ISL_2117855, EPI_ISL_2117857 | Lighthouse Lab in Glasgow | Wellcome Sanger Institute for the COVID-19 Genomics UK (COG-UK) Consortium | Harper VanSteenhouse, Yumi Kasai, David Gray, Carol Clugston, Anna Dominiczak and Alex Alderton, Roberto Amato, Jeffrey Barrett, Sonia Goncalves, Ewan Harrison, David K. Jackson, Ian Johnston, Dominic Kwiatkowski, Cordelia Langford, John Sillitoe on behalf of the Wellcome Sanger Institute COVID-19 Surveillance Team |
| EPI_ISL_2117860 | Lighthouse Lab in Alderley Park | Wellcome Sanger Institute for the COVID-19 Genomics UK (COG-UK) Consortium | Jacquelyn Wynn, Mairead Hyland, The Lighthouse Lab in Alderley Park and Alex Alderton, Roberto Amato, Jeffrey Barrett, Sonia Goncalves, Ewan Harrison, David K. Jackson, Ian Johnston, Dominic Kwiatkowski, Cordelia Langford, John Sillitoe on behalf of the Wellcome Sanger Institute COVID-19 Surveillance Team |
| EPI_ISL_2117862 | Lighthouse Lab in Milton Keynes | Wellcome Sanger Institute for the COVID-19 Genomics UK (COG-UK) Consortium | The Lighthouse Lab in Milton Keynes and Alex Alderton, Roberto Amato, Jeffrey Barrett, Sonia Goncalves, Ewan Harrison, David K. Jackson, Ian Johnston, Dominic Kwiatkowski, Cordelia Langford, John Sillitoe on behalf of the Wellcome Sanger Institute COVID-19 Surveillance Team |
| EPI_ISL_2117864, EPI_ISL_2117870 | Lighthouse Lab in Glasgow | Wellcome Sanger Institute for the COVID-19 Genomics UK (COG-UK) Consortium | Harper VanSteenhouse, Yumi Kasai, David Gray, Carol Clugston, Anna Dominiczak and Alex Alderton, Roberto Amato, Jeffrey Barrett, Sonia Goncalves, Ewan Harrison, David K. Jackson, Ian Johnston, Dominic Kwiatkowski, Cordelia Langford, John Sillitoe on behalf of the Wellcome Sanger Institute COVID-19 Surveillance Team |
| EPI_ISL_2117871 | Lighthouse Lab in Alderley Park | Wellcome Sanger Institute for the COVID-19 Genomics UK (COG-UK) Consortium | Jacquelyn Wynn, Mairead Hyland, The Lighthouse Lab in Alderley Park and Alex Alderton, Roberto Amato, Jeffrey Barrett, Sonia Goncalves, Ewan Harrison, David K. Jackson, Ian Johnston, Dominic Kwiatkowski, Cordelia Langford, John Sillitoe on behalf of the Wellcome Sanger Institute COVID-19 Surveillance Team |
| EPI_ISL_2117873 | Lighthouse Lab in Glasgow | Wellcome Sanger Institute for the COVID-19 Genomics UK (COG-UK) Consortium | Harper VanSteenhouse, Yumi Kasai, David Gray, Carol Clugston, Anna Dominiczak and Alex Alderton, Roberto Amato, Jeffrey Barrett, Sonia Goncalves, Ewan Harrison, David K. Jackson, Ian Johnston, Dominic Kwiatkowski, Cordelia Langford, John Sillitoe on behalf of the Wellcome Sanger Institute COVID-19 Surveillance Team |
| EPI_ISL_2117879, EPI_ISL_2117880 | Lighthouse Lab in Alderley Park | Wellcome Sanger Institute for the COVID-19 Genomics UK (COG-UK) Consortium | Jacquelyn Wynn, Mairead Hyland, The Lighthouse Lab in Alderley Park and Alex Alderton, Roberto Amato, Jeffrey Barrett, Sonia Goncalves, Ewan Harrison, David K. Jackson, Ian Johnston, Dominic Kwiatkowski, Cordelia Langford, John Sillitoe on behalf of the Wellcome Sanger Institute COVID-19 Surveillance Team |
| EPI_ISL_2117882 | Lighthouse Lab in Milton Keynes | Wellcome Sanger Institute for the COVID-19 Genomics UK (COG-UK) Consortium | The Lighthouse Lab in Milton Keynes and Alex Alderton, Roberto Amato, Jeffrey Barrett, Sonia Goncalves, Ewan Harrison, David K. Jackson, Ian Johnston, Dominic Kwiatkowski, Cordelia Langford, John Sillitoe on behalf of the Wellcome Sanger Institute COVID-19 Surveillance Team |
| EPI_ISL_2117883, EPI_ISL_2117885 | Lighthouse Lab in Glasgow | Wellcome Sanger Institute for the COVID-19 Genomics UK (COG-UK) Consortium | Harper VanSteenhouse, Yumi Kasai, David Gray, Carol Clugston, Anna Dominiczak and Alex Alderton, Roberto Amato, Jeffrey Barrett, Sonia Goncalves, Ewan Harrison, David K. Jackson, Ian Johnston, Dominic Kwiatkowski, Cordelia Langford, John Sillitoe on behalf of the Wellcome Sanger Institute COVID-19 Surveillance Team |
| EPI_ISL_2117888 | Lighthouse Lab in Milton Keynes | Wellcome Sanger Institute for the COVID-19 Genomics UK (COG-UK) Consortium | The Lighthouse Lab in Milton Keynes and Alex Alderton, Roberto Amato, Jeffrey Barrett, Sonia Goncalves, Ewan Harrison, David K. Jackson, Ian Johnston, Dominic Kwiatkowski, Cordelia Langford, John Sillitoe on behalf of the Wellcome Sanger Institute COVID-19 Surveillance Team |
| EPI_ISL_2117893, EPI_ISL_2117900, EPI_ISL_2117904, EPI_ISL_2117906 | Lighthouse Lab in Glasgow | Wellcome Sanger Institute for the COVID-19 Genomics UK (COG-UK) Consortium | Harper VanSteenhouse, Yumi Kasai, David Gray, Carol Clugston, Anna Dominiczak and Alex Alderton, Roberto Amato, Jeffrey Barrett, Sonia Goncalves, Ewan Harrison, David K. Jackson, Ian Johnston, Dominic Kwiatkowski, Cordelia Langford, John Sillitoe on behalf of the Wellcome Sanger Institute COVID-19 Surveillance Team |
| EPI_ISL_2117908 | Lighthouse Lab in Alderley Park | Wellcome Sanger Institute for the COVID-19 Genomics UK (COG-UK) Consortium | Jacquelyn Wynn, Mairead Hyland, The Lighthouse Lab in Alderley Park and Alex Alderton, Roberto Amato, Jeffrey Barrett, Sonia Goncalves, Ewan Harrison, David K. Jackson, Ian Johnston, Dominic Kwiatkowski, Cordelia Langford, John Sillitoe on behalf of the Wellcome Sanger Institute COVID-19 Surveillance Team |
| EPI_ISL_2117915, EPI_ISL_2117921, EPI_ISL_2117922, EPI_ISL_2117925 | Lighthouse Lab in Glasgow | Wellcome Sanger Institute for the COVID-19 Genomics UK (COG-UK) Consortium | Harper VanSteenhouse, Yumi Kasai, David Gray, Carol Clugston, Anna Dominiczak and Alex Alderton, Roberto Amato, Jeffrey Barrett, Sonia Goncalves, Ewan Harrison, David K. Jackson, Ian Johnston, Dominic Kwiatkowski, Cordelia Langford, John Sillitoe on behalf of the Wellcome Sanger Institute COVID-19 Surveillance Team |
| EPI_ISL_2117928 | Lighthouse Lab in Milton Keynes | Wellcome Sanger Institute for the COVID-19 Genomics UK (COG-UK) Consortium | The Lighthouse Lab in Milton Keynes and Alex Alderton, Roberto Amato, Jeffrey Barrett, Sonia Goncalves, Ewan Harrison, David K. Jackson, Ian Johnston, Dominic Kwiatkowski, Cordelia Langford, John Sillitoe on behalf of the Wellcome Sanger Institute COVID-19 Surveillance Team |
| EPI_ISL_2117931 | Lighthouse Lab in Alderley Park | Wellcome Sanger Institute for the COVID-19 Genomics UK (COG-UK) Consortium | Jacquelyn Wynn, Mairead Hyland, The Lighthouse Lab in Alderley Park and Alex Alderton, Roberto Amato, Jeffrey Barrett, Sonia Goncalves, Ewan Harrison, David K. Jackson, Ian Johnston, Dominic Kwiatkowski, Cordelia Langford, John Sillitoe on behalf of the Wellcome Sanger Institute COVID-19 Surveillance Team |
| EPI_ISL_2117936 | Lighthouse Lab in Milton Keynes | Wellcome Sanger Institute for the COVID-19 Genomics UK (COG-UK) Consortium | The Lighthouse Lab in Milton Keynes and Alex Alderton, Roberto Amato, Jeffrey Barrett, Sonia Goncalves, Ewan Harrison, David K. Jackson, Ian Johnston, Dominic Kwiatkowski, Cordelia Langford, John Sillitoe on behalf of the Wellcome Sanger Institute COVID-19 Surveillance Team |
| EPI_ISL_2117939 | Lighthouse Lab in Alderley Park | Wellcome Sanger Institute for the COVID-19 Genomics UK (COG-UK) Consortium | Jacquelyn Wynn, Mairead Hyland, The Lighthouse Lab in Alderley Park and Alex Alderton, Roberto Amato, Jeffrey Barrett, Sonia Goncalves, Ewan Harrison, David K. Jackson, Ian Johnston, Dominic Kwiatkowski, Cordelia Langford, John Sillitoe on behalf of the Wellcome Sanger Institute COVID-19 Surveillance Team |
| EPI_ISL_2117942, EPI_ISL_2117943 | Lighthouse Lab in Milton Keynes | Wellcome Sanger Institute for the COVID-19 Genomics UK (COG-UK) Consortium | The Lighthouse Lab in Milton Keynes and Alex Alderton, Roberto Amato, Jeffrey Barrett, Sonia Goncalves, Ewan Harrison, David K. Jackson, Ian Johnston, Dominic Kwiatkowski, Cordelia Langford, John Sillitoe on behalf of the Wellcome Sanger Institute COVID-19 Surveillance Team |
| EPI_ISL_2117945, EPI_ISL_2117946, EPI_ISL_2117948, EPI_ISL_2117951 | Lighthouse Lab in Glasgow | Wellcome Sanger Institute for the COVID-19 Genomics UK (COG-UK) Consortium | Harper VanSteenhouse, Yumi Kasai, David Gray, Carol Clugston, Anna Dominiczak and Alex Alderton, Roberto Amato, Jeffrey Barrett, Sonia Goncalves, Ewan Harrison, David K. Jackson, Ian Johnston, Dominic Kwiatkowski, Cordelia Langford, John Sillitoe on behalf of the Wellcome Sanger Institute COVID-19 Surveillance Team |
| EPI_ISL_2117952 | Lighthouse Lab in Milton Keynes | Wellcome Sanger Institute for the COVID-19 Genomics UK (COG-UK) Consortium | The Lighthouse Lab in Milton Keynes and Alex Alderton, Roberto Amato, Jeffrey Barrett, Sonia Goncalves, Ewan Harrison, David K. Jackson, Ian Johnston, Dominic Kwiatkowski, Cordelia Langford, John Sillitoe on behalf of the Wellcome Sanger Institute COVID-19 Surveillance Team |
| EPI_ISL_2117954 | Lighthouse Lab in Alderley Park | Wellcome Sanger Institute for the COVID-19 Genomics UK (COG-UK) Consortium | Jacquelyn Wynn, Mairead Hyland, The Lighthouse Lab in Alderley Park and Alex Alderton, Roberto Amato, Jeffrey Barrett, Sonia Goncalves, Ewan Harrison, David K. Jackson, Ian Johnston, Dominic Kwiatkowski, Cordelia Langford, John Sillitoe on behalf of the Wellcome Sanger Institute COVID-19 Surveillance Team |
| EPI_ISL_2117955 | Lighthouse Lab in Milton Keynes | Wellcome Sanger Institute for the COVID-19 Genomics UK (COG-UK) Consortium | The Lighthouse Lab in Milton Keynes and Alex Alderton, Roberto Amato, Jeffrey Barrett, Sonia Goncalves, Ewan Harrison, David K. Jackson, Ian Johnston, Dominic Kwiatkowski, Cordelia Langford, John Sillitoe on behalf of the Wellcome Sanger Institute COVID-19 Surveillance Team |
| EPI_ISL_2117958 | Lighthouse Lab in Glasgow | Wellcome Sanger Institute for the COVID-19 Genomics UK (COG-UK) Consortium | Harper VanSteenhouse, Yumi Kasai, David Gray, Carol Clugston, Anna Dominiczak and Alex Alderton, Roberto Amato, Jeffrey Barrett, Sonia Goncalves, Ewan Harrison, David K. Jackson, Ian Johnston, Dominic Kwiatkowski, Cordelia Langford, John Sillitoe on behalf of the Wellcome Sanger Institute COVID-19 Surveillance Team |
| EPI_ISL_2117960 | Lighthouse Lab in Milton Keynes | Wellcome Sanger Institute for the COVID-19 Genomics UK (COG-UK) Consortium | The Lighthouse Lab in Milton Keynes and Alex Alderton, Roberto Amato, Jeffrey Barrett, Sonia Goncalves, Ewan Harrison, David K. Jackson, Ian Johnston, Dominic Kwiatkowski, Cordelia Langford, John Sillitoe on behalf of the Wellcome Sanger Institute COVID-19 Surveillance Team |
| EPI_ISL_2117963 | Lighthouse Lab in Glasgow | Wellcome Sanger Institute for the COVID-19 Genomics UK (COG-UK) Consortium | Harper VanSteenhouse, Yumi Kasai, David Gray, Carol Clugston, Anna Dominiczak and Alex Alderton, Roberto Amato, Jeffrey Barrett, Sonia Goncalves, Ewan Harrison, David K. Jackson, Ian Johnston, Dominic Kwiatkowski, Cordelia Langford, John Sillitoe on behalf of the Wellcome Sanger Institute COVID-19 Surveillance Team |
| EPI_ISL_2117964 | Lighthouse Lab in Milton Keynes | Wellcome Sanger Institute for the COVID-19 Genomics UK (COG-UK) Consortium | The Lighthouse Lab in Milton Keynes and Alex Alderton, Roberto Amato, Jeffrey Barrett, Sonia Goncalves, Ewan Harrison, David K. Jackson, Ian Johnston, Dominic Kwiatkowski, Cordelia Langford, John Sillitoe on behalf of the Wellcome Sanger Institute COVID-19 Surveillance Team |
| EPI_ISL_2117966 | Lighthouse Lab in Alderley Park | Wellcome Sanger Institute for the COVID-19 Genomics UK (COG-UK) Consortium | Jacquelyn Wynn, Mairead Hyland, The Lighthouse Lab in Alderley Park and Alex Alderton, Roberto Amato, Jeffrey Barrett, Sonia Goncalves, Ewan Harrison, David K. Jackson, Ian Johnston, Dominic Kwiatkowski, Cordelia Langford, John Sillitoe on behalf of the Wellcome Sanger Institute COVID-19 Surveillance Team |
| EPI_ISL_2117970 | Lighthouse Lab in Glasgow | Wellcome Sanger Institute for the COVID-19 Genomics UK (COG-UK) Consortium | Harper VanSteenhouse, Yumi Kasai, David Gray, Carol Clugston, Anna Dominiczak and Alex Alderton, Roberto Amato, Jeffrey Barrett, Sonia Goncalves, Ewan Harrison, David K. Jackson, Ian Johnston, Dominic Kwiatkowski, Cordelia Langford, John Sillitoe on behalf of the Wellcome Sanger Institute COVID-19 Surveillance Team |
| EPI_ISL_2117972 | Lighthouse Lab in Milton Keynes | Wellcome Sanger Institute for the COVID-19 Genomics UK (COG-UK) Consortium | The Lighthouse Lab in Milton Keynes and Alex Alderton, Roberto Amato, Jeffrey Barrett, Sonia Goncalves, Ewan Harrison, David K. Jackson, Ian Johnston, Dominic Kwiatkowski, Cordelia Langford, John Sillitoe on behalf of the Wellcome Sanger Institute COVID-19 Surveillance Team |
| EPI_ISL_2117977 | Lighthouse Lab in Glasgow | Wellcome Sanger Institute for the COVID-19 Genomics UK (COG-UK) Consortium | Harper VanSteenhouse, Yumi Kasai, David Gray, Carol Clugston, Anna Dominiczak and Alex Alderton, Roberto Amato, Jeffrey Barrett, Sonia Goncalves, Ewan Harrison, David K. Jackson, Ian Johnston, Dominic Kwiatkowski, Cordelia Langford, John Sillitoe on behalf of the Wellcome Sanger Institute COVID-19 Surveillance Team |
| EPI_ISL_2117983 | Lighthouse Lab in Milton Keynes | Wellcome Sanger Institute for the COVID-19 Genomics UK (COG-UK) Consortium | The Lighthouse Lab in Milton Keynes and Alex Alderton, Roberto Amato, Jeffrey Barrett, Sonia Goncalves, Ewan Harrison, David K. Jackson, Ian Johnston, Dominic Kwiatkowski, Cordelia Langford, John Sillitoe on behalf of the Wellcome Sanger Institute COVID-19 Surveillance Team |
| EPI_ISL_2117984 | Lighthouse Lab in Alderley Park | Wellcome Sanger Institute for the COVID-19 Genomics UK (COG-UK) Consortium | Jacquelyn Wynn, Mairead Hyland, The Lighthouse Lab in Alderley Park and Alex Alderton, Roberto Amato, Jeffrey Barrett, Sonia Goncalves, Ewan Harrison, David K. Jackson, Ian Johnston, Dominic Kwiatkowski, Cordelia Langford, John Sillitoe on behalf of the Wellcome Sanger Institute COVID-19 Surveillance Team |
| EPI_ISL_2117985, EPI_ISL_2117988, EPI_ISL_2117990 | Lighthouse Lab in Glasgow | Wellcome Sanger Institute for the COVID-19 Genomics UK (COG-UK) Consortium | Harper VanSteenhouse, Yumi Kasai, David Gray, Carol Clugston, Anna Dominiczak and Alex Alderton, Roberto Amato, Jeffrey Barrett, Sonia Goncalves, Ewan Harrison, David K. Jackson, Ian Johnston, Dominic Kwiatkowski, Cordelia Langford, John Sillitoe on behalf of the Wellcome Sanger Institute COVID-19 Surveillance Team |
| EPI_ISL_2117991 | Lighthouse Lab in Milton Keynes | Wellcome Sanger Institute for the COVID-19 Genomics UK (COG-UK) Consortium | The Lighthouse Lab in Milton Keynes and Alex Alderton, Roberto Amato, Jeffrey Barrett, Sonia Goncalves, Ewan Harrison, David K. Jackson, Ian Johnston, Dominic Kwiatkowski, Cordelia Langford, John Sillitoe on behalf of the Wellcome Sanger Institute COVID-19 Surveillance Team |
| EPI_ISL_2117994, EPI_ISL_2117996 | Lighthouse Lab in Alderley Park | Wellcome Sanger Institute for the COVID-19 Genomics UK (COG-UK) Consortium | Jacquelyn Wynn, Mairead Hyland, The Lighthouse Lab in Alderley Park and Alex Alderton, Roberto Amato, Jeffrey Barrett, Sonia Goncalves, Ewan Harrison, David K. Jackson, Ian Johnston, Dominic Kwiatkowski, Cordelia Langford, John Sillitoe on behalf of the Wellcome Sanger Institute COVID-19 Surveillance Team |
| EPI_ISL_2117998 | Lighthouse Lab in Glasgow | Wellcome Sanger Institute for the COVID-19 Genomics UK (COG-UK) Consortium | Harper VanSteenhouse, Yumi Kasai, David Gray, Carol Clugston, Anna Dominiczak and Alex Alderton, Roberto Amato, Jeffrey Barrett, Sonia Goncalves, Ewan Harrison, David K. Jackson, Ian Johnston, Dominic Kwiatkowski, Cordelia Langford, John Sillitoe on behalf of the Wellcome Sanger Institute COVID-19 Surveillance Team |
| EPI_ISL_2118002 | Lighthouse Lab in Alderley Park | Wellcome Sanger Institute for the COVID-19 Genomics UK (COG-UK) Consortium | Jacquelyn Wynn, Mairead Hyland, The Lighthouse Lab in Alderley Park and Alex Alderton, Roberto Amato, Jeffrey Barrett, Sonia Goncalves, Ewan Harrison, David K. Jackson, Ian Johnston, Dominic Kwiatkowski, Cordelia Langford, John Sillitoe on behalf of the Wellcome Sanger Institute COVID-19 Surveillance Team |
| EPI_ISL_2118007 | Lighthouse Lab in Milton Keynes | Wellcome Sanger Institute for the COVID-19 Genomics UK (COG-UK) Consortium | The Lighthouse Lab in Milton Keynes and Alex Alderton, Roberto Amato, Jeffrey Barrett, Sonia Goncalves, Ewan Harrison, David K. Jackson, Ian Johnston, Dominic Kwiatkowski, Cordelia Langford, John Sillitoe on behalf of the Wellcome Sanger Institute COVID-19 Surveillance Team |

| | | | |
|---|---|---|---|
| EPI_ISL_2118008 | Lighthouse Lab in Glasgow | Wellcome Sanger Institute for the COVID-19 Genomics UK (COG-UK) Consortium | Harper VanSteenhouse, Yumi Kasai, David Gray, Carol Clugston, Anna Dominiczak and Alex Alderton, Roberto Amato, Jeffrey Barrett, Sonia Goncalves, Ewan Harrison, David K. Jackson, Ian Johnston, Dominic Kwiatkowski, Cordelia Langford, John Sillitoe on behalf of the Wellcome Sanger Institute COVID-19 Surveillance Team |
| EPI_ISL_2118011 | Lighthouse Lab in Alderley Park | Wellcome Sanger Institute for the COVID-19 Genomics UK (COG-UK) Consortium | Jacquelyn Wynn, Mairead Hyland, The Lighthouse Lab in Alderley Park and Alex Alderton, Roberto Amato, Jeffrey Barrett, Sonia Goncalves, Ewan Harrison, David K. Jackson, Ian Johnston, Dominic Kwiatkowski, Cordelia Langford, John Sillitoe on behalf of the Wellcome Sanger Institute COVID-19 Surveillance Team |
| EPI_ISL_2118013 | Lighthouse Lab in Glasgow | Wellcome Sanger Institute for the COVID-19 Genomics UK (COG-UK) Consortium | Harper VanSteenhouse, Yumi Kasai, David Gray, Carol Clugston, Anna Dominiczak and Alex Alderton, Roberto Amato, Jeffrey Barrett, Sonia Goncalves, Ewan Harrison, David K. Jackson, Ian Johnston, Dominic Kwiatkowski, Cordelia Langford, John Sillitoe on behalf of the Wellcome Sanger Institute COVID-19 Surveillance Team |
| EPI_ISL_2118016 | Lighthouse Lab in Alderley Park | Wellcome Sanger Institute for the COVID-19 Genomics UK (COG-UK) Consortium | Jacquelyn Wynn, Mairead Hyland, The Lighthouse Lab in Alderley Park and Alex Alderton, Roberto Amato, Jeffrey Barrett, Sonia Goncalves, Ewan Harrison, David K. Jackson, Ian Johnston, Dominic Kwiatkowski, Cordelia Langford, John Sillitoe on behalf of the Wellcome Sanger Institute COVID-19 Surveillance Team |
| EPI_ISL_2118020, EPI_ISL_2118022, EPI_ISL_2118025 | Lighthouse Lab in Glasgow | Wellcome Sanger Institute for the COVID-19 Genomics UK (COG-UK) Consortium | Harper VanSteenhouse, Yumi Kasai, David Gray, Carol Clugston, Anna Dominiczak and Alex Alderton, Roberto Amato, Jeffrey Barrett, Sonia Goncalves, Ewan Harrison, David K. Jackson, Ian Johnston, Dominic Kwiatkowski, Cordelia Langford, John Sillitoe on behalf of the Wellcome Sanger Institute COVID-19 Surveillance Team |
| EPI_ISL_2118027 | Lighthouse Lab in Alderley Park | Wellcome Sanger Institute for the COVID-19 Genomics UK (COG-UK) Consortium | Jacquelyn Wynn, Mairead Hyland, The Lighthouse Lab in Alderley Park and Alex Alderton, Roberto Amato, Jeffrey Barrett, Sonia Goncalves, Ewan Harrison, David K. Jackson, Ian Johnston, Dominic Kwiatkowski, Cordelia Langford, John Sillitoe on behalf of the Wellcome Sanger Institute COVID-19 Surveillance Team |
| EPI_ISL_2118030 | Lighthouse Lab in Glasgow | Wellcome Sanger Institute for the COVID-19 Genomics UK (COG-UK) Consortium | Harper VanSteenhouse, Yumi Kasai, David Gray, Carol Clugston, Anna Dominiczak and Alex Alderton, Roberto Amato, Jeffrey Barrett, Sonia Goncalves, Ewan Harrison, David K. Jackson, Ian Johnston, Dominic Kwiatkowski, Cordelia Langford, John Sillitoe on behalf of the Wellcome Sanger Institute COVID-19 Surveillance Team |
| EPI_ISL_2118032, EPI_ISL_2118036 | Lighthouse Lab in Milton Keynes | Wellcome Sanger Institute for the COVID-19 Genomics UK (COG-UK) Consortium | The Lighthouse Lab in Milton Keynes and Alex Alderton, Roberto Amato, Jeffrey Barrett, Sonia Goncalves, Ewan Harrison, David K. Jackson, Ian Johnston, Dominic Kwiatkowski, Cordelia Langford, John Sillitoe on behalf of the Wellcome Sanger Institute COVID-19 Surveillance Team |
| EPI_ISL_2118038, EPI_ISL_2118042 | Lighthouse Lab in Glasgow | Wellcome Sanger Institute for the COVID-19 Genomics UK (COG-UK) Consortium | Harper VanSteenhouse, Yumi Kasai, David Gray, Carol Clugston, Anna Dominiczak and Alex Alderton, Roberto Amato, Jeffrey Barrett, Sonia Goncalves, Ewan Harrison, David K. Jackson, Ian Johnston, Dominic Kwiatkowski, Cordelia Langford, John Sillitoe on behalf of the Wellcome Sanger Institute COVID-19 Surveillance Team |
| EPI_ISL_2118045 | Lighthouse Lab in Milton Keynes | Wellcome Sanger Institute for the COVID-19 Genomics UK (COG-UK) Consortium | The Lighthouse Lab in Milton Keynes and Alex Alderton, Roberto Amato, Jeffrey Barrett, Sonia Goncalves, Ewan Harrison, David K. Jackson, Ian Johnston, Dominic Kwiatkowski, Cordelia Langford, John Sillitoe on behalf of the Wellcome Sanger Institute COVID-19 Surveillance Team |
| EPI_ISL_2118046, EPI_ISL_2118054 | Lighthouse Lab in Glasgow | Wellcome Sanger Institute for the COVID-19 Genomics UK (COG-UK) Consortium | Harper VanSteenhouse, Yumi Kasai, David Gray, Carol Clugston, Anna Dominiczak and Alex Alderton, Roberto Amato, Jeffrey Barrett, Sonia Goncalves, Ewan Harrison, David K. Jackson, Ian Johnston, Dominic Kwiatkowski, Cordelia Langford, John Sillitoe on behalf of the Wellcome Sanger Institute COVID-19 Surveillance Team |
| EPI_ISL_2118059 | Lighthouse Lab in Alderley Park | Wellcome Sanger Institute for the COVID-19 Genomics UK (COG-UK) Consortium | Jacquelyn Wynn, Mairead Hyland, The Lighthouse Lab in Alderley Park and Alex Alderton, Roberto Amato, Jeffrey Barrett, Sonia Goncalves, Ewan Harrison, David K. Jackson, Ian Johnston, Dominic Kwiatkowski, Cordelia Langford, John Sillitoe on behalf of the Wellcome Sanger Institute COVID-19 Surveillance Team |
| EPI_ISL_2118062 | Lighthouse Lab in Glasgow | Wellcome Sanger Institute for the COVID-19 Genomics UK (COG-UK) Consortium | Harper VanSteenhouse, Yumi Kasai, David Gray, Carol Clugston, Anna Dominiczak and Alex Alderton, Roberto Amato, Jeffrey Barrett, Sonia Goncalves, Ewan Harrison, David K. Jackson, Ian Johnston, Dominic Kwiatkowski, Cordelia Langford, John Sillitoe on behalf of the Wellcome Sanger Institute COVID-19 Surveillance Team |
| EPI_ISL_2118064 | Lighthouse Lab in Alderley Park | Wellcome Sanger Institute for the COVID-19 Genomics UK (COG-UK) Consortium | Jacquelyn Wynn, Mairead Hyland, The Lighthouse Lab in Alderley Park and Alex Alderton, Roberto Amato, Jeffrey Barrett, Sonia Goncalves, Ewan Harrison, David K. Jackson, Ian Johnston, Dominic Kwiatkowski, Cordelia Langford, John Sillitoe on behalf of the Wellcome Sanger Institute COVID-19 Surveillance Team |
| EPI_ISL_2118067 | Lighthouse Lab in Glasgow | Wellcome Sanger Institute for the COVID-19 Genomics UK (COG-UK) Consortium | Harper VanSteenhouse, Yumi Kasai, David Gray, Carol Clugston, Anna Dominiczak and Alex Alderton, Roberto Amato, Jeffrey Barrett, Sonia Goncalves, Ewan Harrison, David K. Jackson, Ian Johnston, Dominic Kwiatkowski, Cordelia Langford, John Sillitoe on behalf of the Wellcome Sanger Institute COVID-19 Surveillance Team |
| EPI_ISL_2118072 | Lighthouse Lab in Alderley Park | Wellcome Sanger Institute for the COVID-19 Genomics UK (COG-UK) Consortium | Jacquelyn Wynn, Mairead Hyland, The Lighthouse Lab in Alderley Park and Alex Alderton, Roberto Amato, Jeffrey Barrett, Sonia Goncalves, Ewan Harrison, David K. Jackson, Ian Johnston, Dominic Kwiatkowski, Cordelia Langford, John Sillitoe on behalf of the Wellcome Sanger Institute COVID-19 Surveillance Team |
| EPI_ISL_2118073 | Lighthouse Lab in Glasgow | Wellcome Sanger Institute for the COVID-19 Genomics UK (COG-UK) Consortium | Harper VanSteenhouse, Yumi Kasai, David Gray, Carol Clugston, Anna Dominiczak and Alex Alderton, Roberto Amato, Jeffrey Barrett, Sonia Goncalves, Ewan Harrison, David K. Jackson, Ian Johnston, Dominic Kwiatkowski, Cordelia Langford, John Sillitoe on behalf of the Wellcome Sanger Institute COVID-19 Surveillance Team |
| EPI_ISL_2118076 | Lighthouse Lab in Alderley Park | Wellcome Sanger Institute for the COVID-19 Genomics UK (COG-UK) Consortium | Jacquelyn Wynn, Mairead Hyland, The Lighthouse Lab in Alderley Park and Alex Alderton, Roberto Amato, Jeffrey Barrett, Sonia Goncalves, Ewan Harrison, David K. Jackson, Ian Johnston, Dominic Kwiatkowski, Cordelia Langford, John Sillitoe on behalf of the Wellcome Sanger Institute COVID-19 Surveillance Team |
| EPI_ISL_2118078, EPI_ISL_2118081 | Lighthouse Lab in Glasgow | Wellcome Sanger Institute for the COVID-19 Genomics UK (COG-UK) Consortium | Harper VanSteenhouse, Yumi Kasai, David Gray, Carol Clugston, Anna Dominiczak and Alex Alderton, Roberto Amato, Jeffrey Barrett, Sonia Goncalves, Ewan Harrison, David K. Jackson, Ian Johnston, Dominic Kwiatkowski, Cordelia Langford, John Sillitoe on behalf of the Wellcome Sanger Institute COVID-19 Surveillance Team |
| EPI_ISL_2118084 | Lighthouse Lab in Milton Keynes | Wellcome Sanger Institute for the COVID-19 Genomics UK (COG-UK) Consortium | The Lighthouse Lab in Milton Keynes and Alex Alderton, Roberto Amato, Jeffrey Barrett, Sonia Goncalves, Ewan Harrison, David K. Jackson, Ian Johnston, Dominic Kwiatkowski, Cordelia Langford, John Sillitoe on behalf of the Wellcome Sanger Institute COVID-19 Surveillance Team |
| EPI_ISL_2118088, EPI_ISL_2118091 | Lighthouse Lab in Glasgow | Wellcome Sanger Institute for the COVID-19 Genomics UK (COG-UK) Consortium | Harper VanSteenhouse, Yumi Kasai, David Gray, Carol Clugston, Anna Dominiczak and Alex Alderton, Roberto Amato, Jeffrey Barrett, Sonia Goncalves, Ewan Harrison, David K. Jackson, Ian Johnston, Dominic Kwiatkowski, Cordelia Langford, John Sillitoe on behalf of the Wellcome Sanger Institute COVID-19 Surveillance Team |
| EPI_ISL_2118093 | Lighthouse Lab in Milton Keynes | Wellcome Sanger Institute for the COVID-19 Genomics UK (COG-UK) Consortium | The Lighthouse Lab in Milton Keynes and Alex Alderton, Roberto Amato, Jeffrey Barrett, Sonia Goncalves, Ewan Harrison, David K. Jackson, Ian Johnston, Dominic Kwiatkowski, Cordelia Langford, John Sillitoe on behalf of the Wellcome Sanger Institute COVID-19 Surveillance Team |
| EPI_ISL_2118094 | Lighthouse Lab in Glasgow | Wellcome Sanger Institute for the COVID-19 Genomics UK (COG-UK) Consortium | Harper VanSteenhouse, Yumi Kasai, David Gray, Carol Clugston, Anna Dominiczak and Alex Alderton, Roberto Amato, Jeffrey Barrett, Sonia Goncalves, Ewan Harrison, David K. Jackson, Ian Johnston, Dominic Kwiatkowski, Cordelia Langford, John Sillitoe on behalf of the Wellcome Sanger Institute COVID-19 Surveillance Team |
| EPI_ISL_2118099 | Lighthouse Lab in Alderley Park | Wellcome Sanger Institute for the COVID-19 Genomics UK (COG-UK) Consortium | Jacquelyn Wynn, Mairead Hyland, The Lighthouse Lab in Alderley Park and Alex Alderton, Roberto Amato, Jeffrey Barrett, Sonia Goncalves, Ewan Harrison, David K. Jackson, Ian Johnston, Dominic Kwiatkowski, Cordelia Langford, John Sillitoe on behalf of the Wellcome Sanger Institute COVID-19 Surveillance Team |
| EPI_ISL_2118105 | Lighthouse Lab in Glasgow | Wellcome Sanger Institute for the COVID-19 Genomics UK (COG-UK) Consortium | Harper VanSteenhouse, Yumi Kasai, David Gray, Carol Clugston, Anna Dominiczak and Alex Alderton, Roberto Amato, Jeffrey Barrett, Sonia Goncalves, Ewan Harrison, David K. Jackson, Ian Johnston, Dominic Kwiatkowski, Cordelia Langford, John Sillitoe on behalf of the Wellcome Sanger Institute COVID-19 Surveillance Team |
| EPI_ISL_2118111 | Lighthouse Lab in Alderley Park | Wellcome Sanger Institute for the COVID-19 Genomics UK (COG-UK) Consortium | Jacquelyn Wynn, Mairead Hyland, The Lighthouse Lab in Alderley Park and Alex Alderton, Roberto Amato, Jeffrey Barrett, Sonia Goncalves, Ewan Harrison, David K. Jackson, Ian Johnston, Dominic Kwiatkowski, Cordelia Langford, John Sillitoe on behalf of the Wellcome Sanger Institute COVID-19 Surveillance Team |
| EPI_ISL_2118118, EPI_ISL_2118119 | Lighthouse Lab in Milton Keynes | Wellcome Sanger Institute for the COVID-19 Genomics UK (COG-UK) Consortium | The Lighthouse Lab in Milton Keynes and Alex Alderton, Roberto Amato, Jeffrey Barrett, Sonia Goncalves, Ewan Harrison, David K. Jackson, Ian Johnston, Dominic Kwiatkowski, Cordelia Langford, John Sillitoe on behalf of the Wellcome Sanger Institute COVID-19 Surveillance Team |
| EPI_ISL_2118120 | Lighthouse Lab in Alderley Park | Wellcome Sanger Institute for the COVID-19 Genomics UK (COG-UK) Consortium | Jacquelyn Wynn, Mairead Hyland, The Lighthouse Lab in Alderley Park and Alex Alderton, Roberto Amato, Jeffrey Barrett, Sonia Goncalves, Ewan Harrison, David K. Jackson, Ian Johnston, Dominic Kwiatkowski, Cordelia Langford, John Sillitoe on behalf of the Wellcome Sanger Institute COVID-19 Surveillance Team |
| EPI_ISL_2118127 | Lighthouse Lab in Glasgow | Wellcome Sanger Institute for the COVID-19 Genomics UK (COG-UK) Consortium | Harper VanSteenhouse, Yumi Kasai, David Gray, Carol Clugston, Anna Dominiczak and Alex Alderton, Roberto Amato, Jeffrey Barrett, Sonia Goncalves, Ewan Harrison, David K. Jackson, Ian Johnston, Dominic Kwiatkowski, Cordelia Langford, John Sillitoe on behalf of the Wellcome Sanger Institute COVID-19 Surveillance Team |
| EPI_ISL_2118129 | Lighthouse Lab in Alderley Park | Wellcome Sanger Institute for the COVID-19 Genomics UK (COG-UK) Consortium | Jacquelyn Wynn, Mairead Hyland, The Lighthouse Lab in Alderley Park and Alex Alderton, Roberto Amato, Jeffrey Barrett, Sonia Goncalves, Ewan Harrison, David K. Jackson, Ian Johnston, Dominic Kwiatkowski, Cordelia Langford, John Sillitoe on behalf of the Wellcome Sanger Institute COVID-19 Surveillance Team |
| EPI_ISL_2118130 | Lighthouse Lab in Glasgow | Wellcome Sanger Institute for the COVID-19 Genomics UK (COG-UK) Consortium | Harper VanSteenhouse, Yumi Kasai, David Gray, Carol Clugston, Anna Dominiczak and Alex Alderton, Roberto Amato, Jeffrey Barrett, Sonia Goncalves, Ewan Harrison, David K. Jackson, Ian Johnston, Dominic Kwiatkowski, Cordelia Langford, John Sillitoe on behalf of the Wellcome Sanger Institute COVID-19 Surveillance Team |
| EPI_ISL_2118132 | Lighthouse Lab in Alderley Park | Wellcome Sanger Institute for the COVID-19 Genomics UK (COG-UK) Consortium | Jacquelyn Wynn, Mairead Hyland, The Lighthouse Lab in Alderley Park and Alex Alderton, Roberto Amato, Jeffrey Barrett, Sonia Goncalves, Ewan Harrison, David K. Jackson, Ian Johnston, Dominic Kwiatkowski, Cordelia Langford, John Sillitoe on behalf of the Wellcome Sanger Institute COVID-19 Surveillance Team |
| EPI_ISL_2118133, EPI_ISL_2118135, EPI_ISL_2118139, EPI_ISL_2118143, EPI_ISL_2118145, EPI_ISL_2118146 | Lighthouse Lab in Glasgow | Wellcome Sanger Institute for the COVID-19 Genomics UK (COG-UK) Consortium | Harper VanSteenhouse, Yumi Kasai, David Gray, Carol Clugston, Anna Dominiczak and Alex Alderton, Roberto Amato, Jeffrey Barrett, Sonia Goncalves, Ewan Harrison, David K. Jackson, Ian Johnston, Dominic Kwiatkowski, Cordelia Langford, John Sillitoe on behalf of the Wellcome Sanger Institute COVID-19 Surveillance Team |
| EPI_ISL_2118148 | Lighthouse Lab in Alderley Park | Wellcome Sanger Institute for the COVID-19 Genomics UK (COG-UK) Consortium | Jacquelyn Wynn, Mairead Hyland, The Lighthouse Lab in Alderley Park and Alex Alderton, Roberto Amato, Jeffrey Barrett, Sonia Goncalves, Ewan Harrison, David K. Jackson, Ian Johnston, Dominic Kwiatkowski, Cordelia Langford, John Sillitoe on behalf of the Wellcome Sanger Institute COVID-19 Surveillance Team |
| EPI_ISL_2118149, EPI_ISL_2118150, EPI_ISL_2118159 | Lighthouse Lab in Glasgow | Wellcome Sanger Institute for the COVID-19 Genomics UK (COG-UK) Consortium | Harper VanSteenhouse, Yumi Kasai, David Gray, Carol Clugston, Anna Dominiczak and Alex Alderton, Roberto Amato, Jeffrey Barrett, Sonia Goncalves, Ewan Harrison, David K. Jackson, Ian Johnston, Dominic Kwiatkowski, Cordelia Langford, John Sillitoe on behalf of the Wellcome Sanger Institute COVID-19 Surveillance Team |
| EPI_ISL_2118164 | Lighthouse Lab in Milton Keynes | Wellcome Sanger Institute for the COVID-19 Genomics UK (COG-UK) Consortium | The Lighthouse Lab in Milton Keynes and Alex Alderton, Roberto Amato, Jeffrey Barrett, Sonia Goncalves, Ewan Harrison, David K. Jackson, Ian Johnston, Dominic Kwiatkowski, Cordelia Langford, John Sillitoe on behalf of the Wellcome Sanger Institute COVID-19 Surveillance Team |
| EPI_ISL_2118167 | Lighthouse Lab in Glasgow | Wellcome Sanger Institute for the COVID-19 Genomics UK (COG-UK) Consortium | Harper VanSteenhouse, Yumi Kasai, David Gray, Carol Clugston, Anna Dominiczak and Alex Alderton, Roberto Amato, Jeffrey Barrett, Sonia Goncalves, Ewan Harrison, David K. Jackson, Ian Johnston, Dominic Kwiatkowski, Cordelia Langford, John Sillitoe on behalf of the Wellcome Sanger Institute COVID-19 Surveillance Team |
| EPI_ISL_2118168 | Lighthouse Lab in Milton Keynes | Wellcome Sanger Institute for the COVID-19 Genomics UK (COG-UK) Consortium | The Lighthouse Lab in Milton Keynes and Alex Alderton, Roberto Amato, Jeffrey Barrett, Sonia Goncalves, Ewan Harrison, David K. Jackson, Ian Johnston, Dominic Kwiatkowski, Cordelia Langford, John Sillitoe on behalf of the Wellcome Sanger Institute COVID-19 Surveillance Team |
| EPI_ISL_2118177 | Lighthouse Lab in Glasgow | Wellcome Sanger Institute for the COVID-19 Genomics UK (COG-UK) Consortium | Harper VanSteenhouse, Yumi Kasai, David Gray, Carol Clugston, Anna Dominiczak and Alex Alderton, Roberto Amato, Jeffrey Barrett, Sonia Goncalves, Ewan Harrison, David K. Jackson, Ian Johnston, Dominic Kwiatkowski, Cordelia Langford, John Sillitoe on behalf of the Wellcome Sanger Institute COVID-19 Surveillance Team |
| EPI_ISL_2118179 | Lighthouse Lab in Milton Keynes | Wellcome Sanger Institute for the COVID-19 Genomics UK (COG-UK) Consortium | The Lighthouse Lab in Milton Keynes and Alex Alderton, Roberto Amato, Jeffrey Barrett, Sonia Goncalves, Ewan Harrison, David K. Jackson, Ian Johnston, Dominic Kwiatkowski, Cordelia Langford, John Sillitoe on behalf of the Wellcome Sanger Institute COVID-19 Surveillance Team |
| EPI_ISL_2118180 | Lighthouse Lab in Glasgow | Wellcome Sanger Institute for the COVID-19 Genomics UK (COG-UK) Consortium | Harper VanSteenhouse, Yumi Kasai, David Gray, Carol Clugston, Anna Dominiczak and Alex Alderton, Roberto Amato, Jeffrey Barrett, Sonia Goncalves, Ewan Harrison, David K. Jackson, Ian Johnston, Dominic Kwiatkowski, Cordelia Langford, John Sillitoe on behalf of the Wellcome Sanger Institute COVID-19 Surveillance Team |
| EPI_ISL_2118182 | Lighthouse Lab in Milton Keynes | Wellcome Sanger Institute for the COVID-19 Genomics UK (COG-UK) Consortium | The Lighthouse Lab in Milton Keynes and Alex Alderton, Roberto Amato, Jeffrey Barrett, Sonia Goncalves, Ewan Harrison, David K. Jackson, Ian Johnston, Dominic Kwiatkowski, Cordelia Langford, John Sillitoe on behalf of the Wellcome Sanger Institute COVID-19 Surveillance Team |
| EPI_ISL_2118185 | Lighthouse Lab in Alderley Park | Wellcome Sanger Institute for the COVID-19 Genomics UK (COG-UK) Consortium | Jacquelyn Wynn, Mairead Hyland, The Lighthouse Lab in Alderley Park and Alex Alderton, Roberto Amato, Jeffrey Barrett, Sonia Goncalves, Ewan Harrison, David K. Jackson, Ian Johnston, Dominic Kwiatkowski, Cordelia Langford, John Sillitoe on behalf of the Wellcome Sanger Institute COVID-19 Surveillance Team |

| | | | |
|---|---|---|---|
| EPI_ISL_2118189 | Lighthouse Lab in Milton Keynes | Wellcome Sanger Institute for the COVID-19 Genomics UK (COG-UK) Consortium | The Lighthouse Lab in Milton Keynes and Alex Alderton, Roberto Amato, Jeffrey Barrett, Sonia Goncalves, Ewan Harrison, David K. Jackson, Ian Johnston, Dominic Kwiatkowski, Cordelia Langford, John Sillitoe on behalf of the Wellcome Sanger Institute COVID-19 Surveillance Team |
| EPI_ISL_2118192 | Lighthouse Lab in Alderley Park | Wellcome Sanger Institute for the COVID-19 Genomics UK (COG-UK) Consortium | Jacquelyn Wynn, Mairead Hyland, The Lighthouse Lab in Alderley Park and Alex Alderton, Roberto Amato, Jeffrey Barrett, Sonia Goncalves, Ewan Harrison, David K. Jackson, Ian Johnston, Dominic Kwiatkowski, Cordelia Langford, John Sillitoe on behalf of the Wellcome Sanger Institute COVID-19 Surveillance Team |
| EPI_ISL_2118193 | Lighthouse Lab in Glasgow | Wellcome Sanger Institute for the COVID-19 Genomics UK (COG-UK) Consortium | Harper VanSteenhouse, Yumi Kasai, David Gray, Carol Clugston, Anna Dominiczak and Alex Alderton, Roberto Amato, Jeffrey Barrett, Sonia Goncalves, Ewan Harrison, David K. Jackson, Ian Johnston, Dominic Kwiatkowski, Cordelia Langford, John Sillitoe on behalf of the Wellcome Sanger Institute COVID-19 Surveillance Team |
| EPI_ISL_2118197 | Lighthouse Lab in Milton Keynes | Wellcome Sanger Institute for the COVID-19 Genomics UK (COG-UK) Consortium | The Lighthouse Lab in Milton Keynes and Alex Alderton, Roberto Amato, Jeffrey Barrett, Sonia Goncalves, Ewan Harrison, David K. Jackson, Ian Johnston, Dominic Kwiatkowski, Cordelia Langford, John Sillitoe on behalf of the Wellcome Sanger Institute COVID-19 Surveillance Team |
| EPI_ISL_2118200 | Lighthouse Lab in Glasgow | Wellcome Sanger Institute for the COVID-19 Genomics UK (COG-UK) Consortium | Harper VanSteenhouse, Yumi Kasai, David Gray, Carol Clugston, Anna Dominiczak and Alex Alderton, Roberto Amato, Jeffrey Barrett, Sonia Goncalves, Ewan Harrison, David K. Jackson, Ian Johnston, Dominic Kwiatkowski, Cordelia Langford, John Sillitoe on behalf of the Wellcome Sanger Institute COVID-19 Surveillance Team |
| EPI_ISL_2118205, EPI_ISL_2118208, EPI_ISL_2118210 | Lighthouse Lab in Milton Keynes | Wellcome Sanger Institute for the COVID-19 Genomics UK (COG-UK) Consortium | The Lighthouse Lab in Milton Keynes and Alex Alderton, Roberto Amato, Jeffrey Barrett, Sonia Goncalves, Ewan Harrison, David K. Jackson, Ian Johnston, Dominic Kwiatkowski, Cordelia Langford, John Sillitoe on behalf of the Wellcome Sanger Institute COVID-19 Surveillance Team |
| EPI_ISL_2118211 | Lighthouse Lab in Alderley Park | Wellcome Sanger Institute for the COVID-19 Genomics UK (COG-UK) Consortium | Jacquelyn Wynn, Mairead Hyland, The Lighthouse Lab in Alderley Park and Alex Alderton, Roberto Amato, Jeffrey Barrett, Sonia Goncalves, Ewan Harrison, David K. Jackson, Ian Johnston, Dominic Kwiatkowski, Cordelia Langford, John Sillitoe on behalf of the Wellcome Sanger Institute COVID-19 Surveillance Team |
| EPI_ISL_2118212, EPI_ISL_2118222 | Lighthouse Lab in Glasgow | Wellcome Sanger Institute for the COVID-19 Genomics UK (COG-UK) Consortium | Harper VanSteenhouse, Yumi Kasai, David Gray, Carol Clugston, Anna Dominiczak and Alex Alderton, Roberto Amato, Jeffrey Barrett, Sonia Goncalves, Ewan Harrison, David K. Jackson, Ian Johnston, Dominic Kwiatkowski, Cordelia Langford, John Sillitoe on behalf of the Wellcome Sanger Institute COVID-19 Surveillance Team |
| EPI_ISL_2118223, EPI_ISL_2118231 | Lighthouse Lab in Milton Keynes | Wellcome Sanger Institute for the COVID-19 Genomics UK (COG-UK) Consortium | The Lighthouse Lab in Milton Keynes and Alex Alderton, Roberto Amato, Jeffrey Barrett, Sonia Goncalves, Ewan Harrison, David K. Jackson, Ian Johnston, Dominic Kwiatkowski, Cordelia Langford, John Sillitoe on behalf of the Wellcome Sanger Institute COVID-19 Surveillance Team |
| EPI_ISL_2118232 | Lighthouse Lab in Glasgow | Wellcome Sanger Institute for the COVID-19 Genomics UK (COG-UK) Consortium | Harper VanSteenhouse, Yumi Kasai, David Gray, Carol Clugston, Anna Dominiczak and Alex Alderton, Roberto Amato, Jeffrey Barrett, Sonia Goncalves, Ewan Harrison, David K. Jackson, Ian Johnston, Dominic Kwiatkowski, Cordelia Langford, John Sillitoe on behalf of the Wellcome Sanger Institute COVID-19 Surveillance Team |
| EPI_ISL_2118234 | Lighthouse Lab in Milton Keynes | Wellcome Sanger Institute for the COVID-19 Genomics UK (COG-UK) Consortium | The Lighthouse Lab in Milton Keynes and Alex Alderton, Roberto Amato, Jeffrey Barrett, Sonia Goncalves, Ewan Harrison, David K. Jackson, Ian Johnston, Dominic Kwiatkowski, Cordelia Langford, John Sillitoe on behalf of the Wellcome Sanger Institute COVID-19 Surveillance Team |
| EPI_ISL_2118235, EPI_ISL_2118238 | Lighthouse Lab in Alderley Park | Wellcome Sanger Institute for the COVID-19 Genomics UK (COG-UK) Consortium | Jacquelyn Wynn, Mairead Hyland, The Lighthouse Lab in Alderley Park and Alex Alderton, Roberto Amato, Jeffrey Barrett, Sonia Goncalves, Ewan Harrison, David K. Jackson, Ian Johnston, Dominic Kwiatkowski, Cordelia Langford, John Sillitoe on behalf of the Wellcome Sanger Institute COVID-19 Surveillance Team |
| EPI_ISL_2118241, EPI_ISL_2118243, EPI_ISL_2118247 | Lighthouse Lab in Glasgow | Wellcome Sanger Institute for the COVID-19 Genomics UK (COG-UK) Consortium | Harper VanSteenhouse, Yumi Kasai, David Gray, Carol Clugston, Anna Dominiczak and Alex Alderton, Roberto Amato, Jeffrey Barrett, Sonia Goncalves, Ewan Harrison, David K. Jackson, Ian Johnston, Dominic Kwiatkowski, Cordelia Langford, John Sillitoe on behalf of the Wellcome Sanger Institute COVID-19 Surveillance Team |
| EPI_ISL_2118249, EPI_ISL_2118256, EPI_ISL_2118261, EPI_ISL_2118262, EPI_ISL_2118270, EPI_ISL_2118272, EPI_ISL_2118310, EPI_ISL_2118318, EPI_ISL_2118320, EPI_ISL_2118322 | Lighthouse Lab in Alderley Park | Wellcome Sanger Institute for the COVID-19 Genomics UK (COG-UK) Consortium | Jacquelyn Wynn, Mairead Hyland, The Lighthouse Lab in Alderley Park and Alex Alderton, Roberto Amato, Jeffrey Barrett, Sonia Goncalves, Ewan Harrison, David K. Jackson, Ian Johnston, Dominic Kwiatkowski, Cordelia Langford, John Sillitoe on behalf of the Wellcome Sanger Institute COVID-19 Surveillance Team |
| EPI_ISL_2118325 | Lighthouse Lab in Milton Keynes | Wellcome Sanger Institute for the COVID-19 Genomics UK (COG-UK) Consortium | The Lighthouse Lab in Milton Keynes and Alex Alderton, Roberto Amato, Jeffrey Barrett, Sonia Goncalves, Ewan Harrison, David K. Jackson, Ian Johnston, Dominic Kwiatkowski, Cordelia Langford, John Sillitoe on behalf of the Wellcome Sanger Institute COVID-19 Surveillance Team |
| EPI_ISL_2118333 | Lighthouse Lab in Alderley Park | Wellcome Sanger Institute for the COVID-19 Genomics UK (COG-UK) Consortium | Jacquelyn Wynn, Mairead Hyland, The Lighthouse Lab in Alderley Park and Alex Alderton, Roberto Amato, Jeffrey Barrett, Sonia Goncalves, Ewan Harrison, David K. Jackson, Ian Johnston, Dominic Kwiatkowski, Cordelia Langford, John Sillitoe on behalf of the Wellcome Sanger Institute COVID-19 Surveillance Team |
| EPI_ISL_2118343 | Lighthouse Lab in Milton Keynes | Wellcome Sanger Institute for the COVID-19 Genomics UK (COG-UK) Consortium | The Lighthouse Lab in Milton Keynes and Alex Alderton, Roberto Amato, Jeffrey Barrett, Sonia Goncalves, Ewan Harrison, David K. Jackson, Ian Johnston, Dominic Kwiatkowski, Cordelia Langford, John Sillitoe on behalf of the Wellcome Sanger Institute COVID-19 Surveillance Team |
| EPI_ISL_2118348, EPI_ISL_2118350, EPI_ISL_2118354, EPI_ISL_2118357 | Lighthouse Lab in Alderley Park | Wellcome Sanger Institute for the COVID-19 Genomics UK (COG-UK) Consortium | Jacquelyn Wynn, Mairead Hyland, The Lighthouse Lab in Alderley Park and Alex Alderton, Roberto Amato, Jeffrey Barrett, Sonia Goncalves, Ewan Harrison, David K. Jackson, Ian Johnston, Dominic Kwiatkowski, Cordelia Langford, John Sillitoe on behalf of the Wellcome Sanger Institute COVID-19 Surveillance Team |
| EPI_ISL_2118359 | Lighthouse Lab in Milton Keynes | Wellcome Sanger Institute for the COVID-19 Genomics UK (COG-UK) Consortium | The Lighthouse Lab in Milton Keynes and Alex Alderton, Roberto Amato, Jeffrey Barrett, Sonia Goncalves, Ewan Harrison, David K. Jackson, Ian Johnston, Dominic Kwiatkowski, Cordelia Langford, John Sillitoe on behalf of the Wellcome Sanger Institute COVID-19 Surveillance Team |
| EPI_ISL_2118360 | Lighthouse Lab in Alderley Park | Wellcome Sanger Institute for the COVID-19 Genomics UK (COG-UK) Consortium | Jacquelyn Wynn, Mairead Hyland, The Lighthouse Lab in Alderley Park and Alex Alderton, Roberto Amato, Jeffrey Barrett, Sonia Goncalves, Ewan Harrison, David K. Jackson, Ian Johnston, Dominic Kwiatkowski, Cordelia Langford, John Sillitoe on behalf of the Wellcome Sanger Institute COVID-19 Surveillance Team |
| EPI_ISL_2118363 | Lighthouse Lab in Milton Keynes | Wellcome Sanger Institute for the COVID-19 Genomics UK (COG-UK) Consortium | The Lighthouse Lab in Milton Keynes and Alex Alderton, Roberto Amato, Jeffrey Barrett, Sonia Goncalves, Ewan Harrison, David K. Jackson, Ian Johnston, Dominic Kwiatkowski, Cordelia Langford, John Sillitoe on behalf of the Wellcome Sanger Institute COVID-19 Surveillance Team |
| EPI_ISL_2118366 | Lighthouse Lab in Alderley Park | Wellcome Sanger Institute for the COVID-19 Genomics UK (COG-UK) Consortium | Jacquelyn Wynn, Mairead Hyland, The Lighthouse Lab in Alderley Park and Alex Alderton, Roberto Amato, Jeffrey Barrett, Sonia Goncalves, Ewan Harrison, David K. Jackson, Ian Johnston, Dominic Kwiatkowski, Cordelia Langford, John Sillitoe on behalf of the Wellcome Sanger Institute COVID-19 Surveillance Team |
| EPI_ISL_2118368 | Lighthouse Lab in Milton Keynes | Wellcome Sanger Institute for the COVID-19 Genomics UK (COG-UK) Consortium | The Lighthouse Lab in Milton Keynes and Alex Alderton, Roberto Amato, Jeffrey Barrett, Sonia Goncalves, Ewan Harrison, David K. Jackson, Ian Johnston, Dominic Kwiatkowski, Cordelia Langford, John Sillitoe on behalf of the Wellcome Sanger Institute COVID-19 Surveillance Team |
| EPI_ISL_2118372, EPI_ISL_2118388 | Lighthouse Lab in Alderley Park | Wellcome Sanger Institute for the COVID-19 Genomics UK (COG-UK) Consortium | Jacquelyn Wynn, Mairead Hyland, The Lighthouse Lab in Alderley Park and Alex Alderton, Roberto Amato, Jeffrey Barrett, Sonia Goncalves, Ewan Harrison, David K. Jackson, Ian Johnston, Dominic Kwiatkowski, Cordelia Langford, John Sillitoe on behalf of the Wellcome Sanger Institute COVID-19 Surveillance Team |
| EPI_ISL_2118406 | Lighthouse Lab in Milton Keynes | Wellcome Sanger Institute for the COVID-19 Genomics UK (COG-UK) Consortium | The Lighthouse Lab in Milton Keynes and Alex Alderton, Roberto Amato, Jeffrey Barrett, Sonia Goncalves, Ewan Harrison, David K. Jackson, Ian Johnston, Dominic Kwiatkowski, Cordelia Langford, John Sillitoe on behalf of the Wellcome Sanger Institute COVID-19 Surveillance Team |
| EPI_ISL_2118416, EPI_ISL_2118421, EPI_ISL_2118422, EPI_ISL_2118427, EPI_ISL_2118428, EPI_ISL_2118430, EPI_ISL_2118433, EPI_ISL_2118434 | Lighthouse Lab in Alderley Park | Wellcome Sanger Institute for the COVID-19 Genomics UK (COG-UK) Consortium | Jacquelyn Wynn, Mairead Hyland, The Lighthouse Lab in Alderley Park and Alex Alderton, Roberto Amato, Jeffrey Barrett, Sonia Goncalves, Ewan Harrison, David K. Jackson, Ian Johnston, Dominic Kwiatkowski, Cordelia Langford, John Sillitoe on behalf of the Wellcome Sanger Institute COVID-19 Surveillance Team |
| EPI_ISL_2118438, EPI_ISL_2118440 | Lighthouse Lab in Milton Keynes | Wellcome Sanger Institute for the COVID-19 Genomics UK (COG-UK) Consortium | The Lighthouse Lab in Milton Keynes and Alex Alderton, Roberto Amato, Jeffrey Barrett, Sonia Goncalves, Ewan Harrison, David K. Jackson, Ian Johnston, Dominic Kwiatkowski, Cordelia Langford, John Sillitoe on behalf of the Wellcome Sanger Institute COVID-19 Surveillance Team |
| EPI_ISL_2118443, EPI_ISL_2118444, EPI_ISL_2118447, EPI_ISL_2118457 | Lighthouse Lab in Alderley Park | Wellcome Sanger Institute for the COVID-19 Genomics UK (COG-UK) Consortium | Jacquelyn Wynn, Mairead Hyland, The Lighthouse Lab in Alderley Park and Alex Alderton, Roberto Amato, Jeffrey Barrett, Sonia Goncalves, Ewan Harrison, David K. Jackson, Ian Johnston, Dominic Kwiatkowski, Cordelia Langford, John Sillitoe on behalf of the Wellcome Sanger Institute COVID-19 Surveillance Team |
| EPI_ISL_2118463 | Lighthouse Lab in Milton Keynes | Wellcome Sanger Institute for the COVID-19 Genomics UK (COG-UK) Consortium | The Lighthouse Lab in Milton Keynes and Alex Alderton, Roberto Amato, Jeffrey Barrett, Sonia Goncalves, Ewan Harrison, David K. Jackson, Ian Johnston, Dominic Kwiatkowski, Cordelia Langford, John Sillitoe on behalf of the Wellcome Sanger Institute COVID-19 Surveillance Team |
| EPI_ISL_2118464 | SYNLAB MVZ Leverkusen | Robert Koch Institute | unknown |
| EPI_ISL_2118465 | Lighthouse Lab in Alderley Park | Wellcome Sanger Institute for the COVID-19 Genomics UK (COG-UK) Consortium | Jacquelyn Wynn, Mairead Hyland, The Lighthouse Lab in Alderley Park and Alex Alderton, Roberto Amato, Jeffrey Barrett, Sonia Goncalves, Ewan Harrison, David K. Jackson, Ian Johnston, Dominic Kwiatkowski, Cordelia Langford, John Sillitoe on behalf of the Wellcome Sanger Institute COVID-19 Surveillance Team |
| EPI_ISL_2118468 | Lighthouse Lab in Milton Keynes | Wellcome Sanger Institute for the COVID-19 Genomics UK (COG-UK) Consortium | The Lighthouse Lab in Milton Keynes and Alex Alderton, Roberto Amato, Jeffrey Barrett, Sonia Goncalves, Ewan Harrison, David K. Jackson, Ian Johnston, Dominic Kwiatkowski, Cordelia Langford, John Sillitoe on behalf of the Wellcome Sanger Institute COVID-19 Surveillance Team |
| EPI_ISL_2118479, EPI_ISL_2118482, EPI_ISL_2118484, EPI_ISL_2118487, EPI_ISL_2118488, EPI_ISL_2118489, EPI_ISL_2118492, EPI_ISL_2118496 | Lighthouse Lab in Alderley Park | Wellcome Sanger Institute for the COVID-19 Genomics UK (COG-UK) Consortium | Jacquelyn Wynn, Mairead Hyland, The Lighthouse Lab in Alderley Park and Alex Alderton, Roberto Amato, Jeffrey Barrett, Sonia Goncalves, Ewan Harrison, David K. Jackson, Ian Johnston, Dominic Kwiatkowski, Cordelia Langford, John Sillitoe on behalf of the Wellcome Sanger Institute COVID-19 Surveillance Team |
| EPI_ISL_2118503 | Lighthouse Lab in Milton Keynes | Wellcome Sanger Institute for the COVID-19 Genomics UK (COG-UK) Consortium | The Lighthouse Lab in Milton Keynes and Alex Alderton, Roberto Amato, Jeffrey Barrett, Sonia Goncalves, Ewan Harrison, David K. Jackson, Ian Johnston, Dominic Kwiatkowski, Cordelia Langford, John Sillitoe on behalf of the Wellcome Sanger Institute COVID-19 Surveillance Team |
| EPI_ISL_2118506 | Lighthouse Lab in Alderley Park | Wellcome Sanger Institute for the COVID-19 Genomics UK (COG-UK) | Jacquelyn Wynn, Mairead Hyland, The Lighthouse Lab in Alderley Park and Alex Alderton, Roberto Amato, Jeffrey Barrett, Sonia Goncalves, Ewan Harrison, David K. Jackson, Ian Johnston, Dominic Kwiatkowski, Cordelia Langford, John Sillitoe on |

| | | | |
|---|---|---|---|
| | | Consortium | behalf of the Wellcome Sanger Institute COVID-19 Surveillance Team |
| EPI_ISL_2118507 | Lighthouse Lab in Milton Keynes | Wellcome Sanger Institute for the COVID-19 Genomics UK (COG-UK) Consortium | The Lighthouse Lab in Milton Keynes and Alex Alderton, Roberto Amato, Jeffrey Barrett, Sonia Goncalves, Ewan Harrison, David K. Jackson, Ian Johnston, Dominic Kwiatkowski, Cordelia Langford, John Sillitoe on behalf of the Wellcome Sanger Institute COVID-19 Surveillance Team |
| EPI_ISL_2118512, EPI_ISL_2118513, EPI_ISL_2118515, EPI_ISL_2118516 | Lighthouse Lab in Alderley Park | Wellcome Sanger Institute for the COVID-19 Genomics UK (COG-UK) Consortium | Jacquelyn Wynn, Mairead Hyland, The Lighthouse Lab in Alderley Park and Alex Alderton, Roberto Amato, Jeffrey Barrett, Sonia Goncalves, Ewan Harrison, David K. Jackson, Ian Johnston, Dominic Kwiatkowski, Cordelia Langford, John Sillitoe on behalf of the Wellcome Sanger Institute COVID-19 Surveillance Team |
| EPI_ISL_2118526 | Lighthouse Lab in Milton Keynes | Wellcome Sanger Institute for the COVID-19 Genomics UK (COG-UK) Consortium | The Lighthouse Lab in Milton Keynes and Alex Alderton, Roberto Amato, Jeffrey Barrett, Sonia Goncalves, Ewan Harrison, David K. Jackson, Ian Johnston, Dominic Kwiatkowski, Cordelia Langford, John Sillitoe on behalf of the Wellcome Sanger Institute COVID-19 Surveillance Team |
| EPI_ISL_2118531, EPI_ISL_2118535, EPI_ISL_2118538, EPI_ISL_2118541, EPI_ISL_2118544, EPI_ISL_2118546 | Lighthouse Lab in Alderley Park | Wellcome Sanger Institute for the COVID-19 Genomics UK (COG-UK) Consortium | Jacquelyn Wynn, Mairead Hyland, The Lighthouse Lab in Alderley Park and Alex Alderton, Roberto Amato, Jeffrey Barrett, Sonia Goncalves, Ewan Harrison, David K. Jackson, Ian Johnston, Dominic Kwiatkowski, Cordelia Langford, John Sillitoe on behalf of the Wellcome Sanger Institute COVID-19 Surveillance Team |
| EPI_ISL_2118549 | Lighthouse Lab in Milton Keynes | Wellcome Sanger Institute for the COVID-19 Genomics UK (COG-UK) Consortium | The Lighthouse Lab in Milton Keynes and Alex Alderton, Roberto Amato, Jeffrey Barrett, Sonia Goncalves, Ewan Harrison, David K. Jackson, Ian Johnston, Dominic Kwiatkowski, Cordelia Langford, John Sillitoe on behalf of the Wellcome Sanger Institute COVID-19 Surveillance Team |
| EPI_ISL_2118552, EPI_ISL_2118556 | Lighthouse Lab in Alderley Park | Wellcome Sanger Institute for the COVID-19 Genomics UK (COG-UK) Consortium | Jacquelyn Wynn, Mairead Hyland, The Lighthouse Lab in Alderley Park and Alex Alderton, Roberto Amato, Jeffrey Barrett, Sonia Goncalves, Ewan Harrison, David K. Jackson, Ian Johnston, Dominic Kwiatkowski, Cordelia Langford, John Sillitoe on behalf of the Wellcome Sanger Institute COVID-19 Surveillance Team |
| EPI_ISL_2118557 | Lighthouse Lab in Milton Keynes | Wellcome Sanger Institute for the COVID-19 Genomics UK (COG-UK) Consortium | The Lighthouse Lab in Milton Keynes and Alex Alderton, Roberto Amato, Jeffrey Barrett, Sonia Goncalves, Ewan Harrison, David K. Jackson, Ian Johnston, Dominic Kwiatkowski, Cordelia Langford, John Sillitoe on behalf of the Wellcome Sanger Institute COVID-19 Surveillance Team |
| EPI_ISL_2118558 | Lighthouse Lab in Alderley Park | Wellcome Sanger Institute for the COVID-19 Genomics UK (COG-UK) Consortium | Jacquelyn Wynn, Mairead Hyland, The Lighthouse Lab in Alderley Park and Alex Alderton, Roberto Amato, Jeffrey Barrett, Sonia Goncalves, Ewan Harrison, David K. Jackson, Ian Johnston, Dominic Kwiatkowski, Cordelia Langford, John Sillitoe on behalf of the Wellcome Sanger Institute COVID-19 Surveillance Team |
| EPI_ISL_2118562 | Lighthouse Lab in Milton Keynes | Wellcome Sanger Institute for the COVID-19 Genomics UK (COG-UK) Consortium | The Lighthouse Lab in Milton Keynes and Alex Alderton, Roberto Amato, Jeffrey Barrett, Sonia Goncalves, Ewan Harrison, David K. Jackson, Ian Johnston, Dominic Kwiatkowski, Cordelia Langford, John Sillitoe on behalf of the Wellcome Sanger Institute COVID-19 Surveillance Team |
| EPI_ISL_2118567, EPI_ISL_2118569 | Lighthouse Lab in Alderley Park | Wellcome Sanger Institute for the COVID-19 Genomics UK (COG-UK) Consortium | Jacquelyn Wynn, Mairead Hyland, The Lighthouse Lab in Alderley Park and Alex Alderton, Roberto Amato, Jeffrey Barrett, Sonia Goncalves, Ewan Harrison, David K. Jackson, Ian Johnston, Dominic Kwiatkowski, Cordelia Langford, John Sillitoe on behalf of the Wellcome Sanger Institute COVID-19 Surveillance Team |
| EPI_ISL_2118570 | Lighthouse Lab in Milton Keynes | Wellcome Sanger Institute for the COVID-19 Genomics UK (COG-UK) Consortium | The Lighthouse Lab in Milton Keynes and Alex Alderton, Roberto Amato, Jeffrey Barrett, Sonia Goncalves, Ewan Harrison, David K. Jackson, Ian Johnston, Dominic Kwiatkowski, Cordelia Langford, John Sillitoe on behalf of the Wellcome Sanger Institute COVID-19 Surveillance Team |
| EPI_ISL_2118573, EPI_ISL_2118575 | Lighthouse Lab in Alderley Park | Wellcome Sanger Institute for the COVID-19 Genomics UK (COG-UK) Consortium | Jacquelyn Wynn, Mairead Hyland, The Lighthouse Lab in Alderley Park and Alex Alderton, Roberto Amato, Jeffrey Barrett, Sonia Goncalves, Ewan Harrison, David K. Jackson, Ian Johnston, Dominic Kwiatkowski, Cordelia Langford, John Sillitoe on behalf of the Wellcome Sanger Institute COVID-19 Surveillance Team |
| EPI_ISL_2118576 | Lighthouse Lab in Milton Keynes | Wellcome Sanger Institute for the COVID-19 Genomics UK (COG-UK) Consortium | The Lighthouse Lab in Milton Keynes and Alex Alderton, Roberto Amato, Jeffrey Barrett, Sonia Goncalves, Ewan Harrison, David K. Jackson, Ian Johnston, Dominic Kwiatkowski, Cordelia Langford, John Sillitoe on behalf of the Wellcome Sanger Institute COVID-19 Surveillance Team |
| EPI_ISL_2118581, EPI_ISL_2118582, EPI_ISL_2118586, EPI_ISL_2118589, EPI_ISL_2118590, EPI_ISL_2118596, EPI_ISL_2118599, EPI_ISL_2118610, EPI_ISL_2118612, EPI_ISL_2118613, EPI_ISL_2118615 | | | |
| see above | Lighthouse Lab in Alderley Park | Wellcome Sanger Institute for the COVID-19 Genomics UK (COG-UK) Consortium | Jacquelyn Wynn, Mairead Hyland, The Lighthouse Lab in Alderley Park and Alex Alderton, Roberto Amato, Jeffrey Barrett, Sonia Goncalves, Ewan Harrison, David K. Jackson, Ian Johnston, Dominic Kwiatkowski, Cordelia Langford, John Sillitoe on behalf of the Wellcome Sanger Institute COVID-19 Surveillance Team |
| EPI_ISL_2118619 | Lighthouse Lab in Milton Keynes | Wellcome Sanger Institute for the COVID-19 Genomics UK (COG-UK) Consortium | The Lighthouse Lab in Milton Keynes and Alex Alderton, Roberto Amato, Jeffrey Barrett, Sonia Goncalves, Ewan Harrison, David K. Jackson, Ian Johnston, Dominic Kwiatkowski, Cordelia Langford, John Sillitoe on behalf of the Wellcome Sanger Institute COVID-19 Surveillance Team |
| EPI_ISL_2118622, EPI_ISL_2118624, EPI_ISL_2118630 | Lighthouse Lab in Alderley Park | Wellcome Sanger Institute for the COVID-19 Genomics UK (COG-UK) Consortium | Jacquelyn Wynn, Mairead Hyland, The Lighthouse Lab in Alderley Park and Alex Alderton, Roberto Amato, Jeffrey Barrett, Sonia Goncalves, Ewan Harrison, David K. Jackson, Ian Johnston, Dominic Kwiatkowski, Cordelia Langford, John Sillitoe on behalf of the Wellcome Sanger Institute COVID-19 Surveillance Team |
| EPI_ISL_2118633 | Lighthouse Lab in Milton Keynes | Wellcome Sanger Institute for the COVID-19 Genomics UK (COG-UK) Consortium | The Lighthouse Lab in Milton Keynes and Alex Alderton, Roberto Amato, Jeffrey Barrett, Sonia Goncalves, Ewan Harrison, David K. Jackson, Ian Johnston, Dominic Kwiatkowski, Cordelia Langford, John Sillitoe on behalf of the Wellcome Sanger Institute COVID-19 Surveillance Team |
| EPI_ISL_2118638, EPI_ISL_2118642, EPI_ISL_2118645, EPI_ISL_2118648, EPI_ISL_2118649, EPI_ISL_2118667, EPI_ISL_2118670, EPI_ISL_2118676, EPI_ISL_2118678 | Lighthouse Lab in Alderley Park | Wellcome Sanger Institute for the COVID-19 Genomics UK (COG-UK) Consortium | Jacquelyn Wynn, Mairead Hyland, The Lighthouse Lab in Alderley Park and Alex Alderton, Roberto Amato, Jeffrey Barrett, Sonia Goncalves, Ewan Harrison, David K. Jackson, Ian Johnston, Dominic Kwiatkowski, Cordelia Langford, John Sillitoe on behalf of the Wellcome Sanger Institute COVID-19 Surveillance Team |
| EPI_ISL_2118681 | Lighthouse Lab in Milton Keynes | Wellcome Sanger Institute for the COVID-19 Genomics UK (COG-UK) Consortium | The Lighthouse Lab in Milton Keynes and Alex Alderton, Roberto Amato, Jeffrey Barrett, Sonia Goncalves, Ewan Harrison, David K. Jackson, Ian Johnston, Dominic Kwiatkowski, Cordelia Langford, John Sillitoe on behalf of the Wellcome Sanger Institute COVID-19 Surveillance Team |
| EPI_ISL_2118682, EPI_ISL_2118684, EPI_ISL_2118685, EPI_ISL_2118687, EPI_ISL_2118689, EPI_ISL_2118694, EPI_ISL_2118696, EPI_ISL_2118704, EPI_ISL_2118705, EPI_ISL_2118708, EPI_ISL_2118709 | | | |
| see above | Lighthouse Lab in Alderley Park | Wellcome Sanger Institute for the COVID-19 Genomics UK (COG-UK) Consortium | Jacquelyn Wynn, Mairead Hyland, The Lighthouse Lab in Alderley Park and Alex Alderton, Roberto Amato, Jeffrey Barrett, Sonia Goncalves, Ewan Harrison, David K. Jackson, Ian Johnston, Dominic Kwiatkowski, Cordelia Langford, John Sillitoe on behalf of the Wellcome Sanger Institute COVID-19 Surveillance Team |
| EPI_ISL_2118715 | Lighthouse Lab in Milton Keynes | Wellcome Sanger Institute for the COVID-19 Genomics UK (COG-UK) Consortium | The Lighthouse Lab in Milton Keynes and Alex Alderton, Roberto Amato, Jeffrey Barrett, Sonia Goncalves, Ewan Harrison, David K. Jackson, Ian Johnston, Dominic Kwiatkowski, Cordelia Langford, John Sillitoe on behalf of the Wellcome Sanger Institute COVID-19 Surveillance Team |
| EPI_ISL_2118721, EPI_ISL_2118722, EPI_ISL_2118724, EPI_ISL_2118728, EPI_ISL_2118732, EPI_ISL_2118738, EPI_ISL_2118742, EPI_ISL_2118745, EPI_ISL_2118748, EPI_ISL_2118751 | Lighthouse Lab in Alderley Park | Wellcome Sanger Institute for the COVID-19 Genomics UK (COG-UK) Consortium | Jacquelyn Wynn, Mairead Hyland, The Lighthouse Lab in Alderley Park and Alex Alderton, Roberto Amato, Jeffrey Barrett, Sonia Goncalves, Ewan Harrison, David K. Jackson, Ian Johnston, Dominic Kwiatkowski, Cordelia Langford, John Sillitoe on behalf of the Wellcome Sanger Institute COVID-19 Surveillance Team |
| EPI_ISL_2118754, EPI_ISL_2118759 | Lighthouse Lab in Milton Keynes | Wellcome Sanger Institute for the COVID-19 Genomics UK (COG-UK) Consortium | The Lighthouse Lab in Milton Keynes and Alex Alderton, Roberto Amato, Jeffrey Barrett, Sonia Goncalves, Ewan Harrison, David K. Jackson, Ian Johnston, Dominic Kwiatkowski, Cordelia Langford, John Sillitoe on behalf of the Wellcome Sanger Institute COVID-19 Surveillance Team |
| EPI_ISL_2118762, EPI_ISL_2118765, EPI_ISL_2118769, EPI_ISL_2118776, EPI_ISL_2118785 | Lighthouse Lab in Alderley Park | Wellcome Sanger Institute for the COVID-19 Genomics UK (COG-UK) Consortium | Jacquelyn Wynn, Mairead Hyland, The Lighthouse Lab in Alderley Park and Alex Alderton, Roberto Amato, Jeffrey Barrett, Sonia Goncalves, Ewan Harrison, David K. Jackson, Ian Johnston, Dominic Kwiatkowski, Cordelia Langford, John Sillitoe on behalf of the Wellcome Sanger Institute COVID-19 Surveillance Team |
| EPI_ISL_2118789 | Lighthouse Lab in Milton Keynes | Wellcome Sanger Institute for the COVID-19 Genomics UK (COG-UK) Consortium | The Lighthouse Lab in Milton Keynes and Alex Alderton, Roberto Amato, Jeffrey Barrett, Sonia Goncalves, Ewan Harrison, David K. Jackson, Ian Johnston, Dominic Kwiatkowski, Cordelia Langford, John Sillitoe on behalf of the Wellcome Sanger Institute COVID-19 Surveillance Team |
| EPI_ISL_2118793, EPI_ISL_2118796 | Lighthouse Lab in Alderley Park | Wellcome Sanger Institute for the COVID-19 Genomics UK (COG-UK) Consortium | Jacquelyn Wynn, Mairead Hyland, The Lighthouse Lab in Alderley Park and Alex Alderton, Roberto Amato, Jeffrey Barrett, Sonia Goncalves, Ewan Harrison, David K. Jackson, Ian Johnston, Dominic Kwiatkowski, Cordelia Langford, John Sillitoe on behalf of the Wellcome Sanger Institute COVID-19 Surveillance Team |
| EPI_ISL_2118800, EPI_ISL_2118801, EPI_ISL_2118804 | Lighthouse Lab in Milton Keynes | Wellcome Sanger Institute for the COVID-19 Genomics UK (COG-UK) Consortium | The Lighthouse Lab in Milton Keynes and Alex Alderton, Roberto Amato, Jeffrey Barrett, Sonia Goncalves, Ewan Harrison, David K. Jackson, Ian Johnston, Dominic Kwiatkowski, Cordelia Langford, John Sillitoe on behalf of the Wellcome Sanger Institute COVID-19 Surveillance Team |
| EPI_ISL_2118807, EPI_ISL_2118811, EPI_ISL_2118816, EPI_ISL_2118817, EPI_ISL_2118819, EPI_ISL_2118829, EPI_ISL_2118830, EPI_ISL_2118831, EPI_ISL_2118832 | Lighthouse Lab in Alderley Park | Wellcome Sanger Institute for the COVID-19 Genomics UK (COG-UK) Consortium | Jacquelyn Wynn, Mairead Hyland, The Lighthouse Lab in Alderley Park and Alex Alderton, Roberto Amato, Jeffrey Barrett, Sonia Goncalves, Ewan Harrison, David K. Jackson, Ian Johnston, Dominic Kwiatkowski, Cordelia Langford, John Sillitoe on behalf of the Wellcome Sanger Institute COVID-19 Surveillance Team |
| EPI_ISL_2118834 | Lighthouse Lab in Milton Keynes | Wellcome Sanger Institute for the COVID-19 Genomics UK (COG-UK) Consortium | The Lighthouse Lab in Milton Keynes and Alex Alderton, Roberto Amato, Jeffrey Barrett, Sonia Goncalves, Ewan Harrison, David K. Jackson, Ian Johnston, Dominic Kwiatkowski, Cordelia Langford, John Sillitoe on behalf of the Wellcome Sanger Institute COVID-19 Surveillance Team |
| EPI_ISL_2118835, EPI_ISL_2118852 | Lighthouse Lab in Alderley Park | Wellcome Sanger Institute for the COVID-19 Genomics UK (COG-UK) Consortium | Jacquelyn Wynn, Mairead Hyland, The Lighthouse Lab in Alderley Park and Alex Alderton, Roberto Amato, Jeffrey Barrett, Sonia Goncalves, Ewan Harrison, David K. Jackson, Ian Johnston, Dominic Kwiatkowski, Cordelia Langford, John Sillitoe on behalf of the Wellcome Sanger Institute COVID-19 Surveillance Team |

| | | | |
|---|---|---|---|
| EPI_ISL_2118858 | Lighthouse Lab in Milton Keynes | Wellcome Sanger Institute for the COVID-19 Genomics UK (COG-UK) Consortium | The Lighthouse Lab in Milton Keynes and Alex Alderton, Roberto Amato, Jeffrey Barrett, Sonia Goncalves, Ewan Harrison, David K. Jackson, Ian Johnston, Dominic Kwiatkowski, Cordelia Langford, John Sillitoe on behalf of the Wellcome Sanger Institute COVID-19 Surveillance Team |
| EPI_ISL_2118882 | Lighthouse Lab in Alderley Park | Wellcome Sanger Institute for the COVID-19 Genomics UK (COG-UK) Consortium | Jacquelyn Wynn, Mairead Hyland, The Lighthouse Lab in Alderley Park and Alex Alderton, Roberto Amato, Jeffrey Barrett, Sonia Goncalves, Ewan Harrison, David K. Jackson, Ian Johnston, Dominic Kwiatkowski, Cordelia Langford, John Sillitoe on behalf of the Wellcome Sanger Institute COVID-19 Surveillance Team |
| EPI_ISL_2118883 | Lighthouse Lab in Milton Keynes | Wellcome Sanger Institute for the COVID-19 Genomics UK (COG-UK) Consortium | The Lighthouse Lab in Milton Keynes and Alex Alderton, Roberto Amato, Jeffrey Barrett, Sonia Goncalves, Ewan Harrison, David K. Jackson, Ian Johnston, Dominic Kwiatkowski, Cordelia Langford, John Sillitoe on behalf of the Wellcome Sanger Institute COVID-19 Surveillance Team |
| EPI_ISL_2118886 | Lighthouse Lab in Alderley Park | Wellcome Sanger Institute for the COVID-19 Genomics UK (COG-UK) Consortium | Jacquelyn Wynn, Mairead Hyland, The Lighthouse Lab in Alderley Park and Alex Alderton, Roberto Amato, Jeffrey Barrett, Sonia Goncalves, Ewan Harrison, David K. Jackson, Ian Johnston, Dominic Kwiatkowski, Cordelia Langford, John Sillitoe on behalf of the Wellcome Sanger Institute COVID-19 Surveillance Team |
| EPI_ISL_2118888 | Lighthouse Lab in Milton Keynes | Wellcome Sanger Institute for the COVID-19 Genomics UK (COG-UK) Consortium | The Lighthouse Lab in Milton Keynes and Alex Alderton, Roberto Amato, Jeffrey Barrett, Sonia Goncalves, Ewan Harrison, David K. Jackson, Ian Johnston, Dominic Kwiatkowski, Cordelia Langford, John Sillitoe on behalf of the Wellcome Sanger Institute COVID-19 Surveillance Team |
| EPI_ISL_2118891 | Lighthouse Lab in Alderley Park | Wellcome Sanger Institute for the COVID-19 Genomics UK (COG-UK) Consortium | Jacquelyn Wynn, Mairead Hyland, The Lighthouse Lab in Alderley Park and Alex Alderton, Roberto Amato, Jeffrey Barrett, Sonia Goncalves, Ewan Harrison, David K. Jackson, Ian Johnston, Dominic Kwiatkowski, Cordelia Langford, John Sillitoe on behalf of the Wellcome Sanger Institute COVID-19 Surveillance Team |
| EPI_ISL_2118896, EPI_ISL_2118899, EPI_ISL_2118909 | Lighthouse Lab in Milton Keynes | Wellcome Sanger Institute for the COVID-19 Genomics UK (COG-UK) Consortium | The Lighthouse Lab in Milton Keynes and Alex Alderton, Roberto Amato, Jeffrey Barrett, Sonia Goncalves, Ewan Harrison, David K. Jackson, Ian Johnston, Dominic Kwiatkowski, Cordelia Langford, John Sillitoe on behalf of the Wellcome Sanger Institute COVID-19 Surveillance Team |
| EPI_ISL_2118915 | Lighthouse Lab in Alderley Park | Wellcome Sanger Institute for the COVID-19 Genomics UK (COG-UK) Consortium | Jacquelyn Wynn, Mairead Hyland, The Lighthouse Lab in Alderley Park and Alex Alderton, Roberto Amato, Jeffrey Barrett, Sonia Goncalves, Ewan Harrison, David K. Jackson, Ian Johnston, Dominic Kwiatkowski, Cordelia Langford, John Sillitoe on behalf of the Wellcome Sanger Institute COVID-19 Surveillance Team |
| EPI_ISL_2118919 | Lighthouse Lab in Milton Keynes | Wellcome Sanger Institute for the COVID-19 Genomics UK (COG-UK) Consortium | The Lighthouse Lab in Milton Keynes and Alex Alderton, Roberto Amato, Jeffrey Barrett, Sonia Goncalves, Ewan Harrison, David K. Jackson, Ian Johnston, Dominic Kwiatkowski, Cordelia Langford, John Sillitoe on behalf of the Wellcome Sanger Institute COVID-19 Surveillance Team |
| EPI_ISL_2118923, EPI_ISL_2118924, EPI_ISL_2118928, EPI_ISL_2118930, EPI_ISL_2118933, EPI_ISL_2118939, EPI_ISL_2118945 | Lighthouse Lab in Alderley Park | Wellcome Sanger Institute for the COVID-19 Genomics UK (COG-UK) Consortium | Jacquelyn Wynn, Mairead Hyland, The Lighthouse Lab in Alderley Park and Alex Alderton, Roberto Amato, Jeffrey Barrett, Sonia Goncalves, Ewan Harrison, David K. Jackson, Ian Johnston, Dominic Kwiatkowski, Cordelia Langford, John Sillitoe on behalf of the Wellcome Sanger Institute COVID-19 Surveillance Team |
| EPI_ISL_2118947, EPI_ISL_2118966, EPI_ISL_2118974, EPI_ISL_2118977, EPI_ISL_2118982 | Lighthouse Lab in Milton Keynes | Wellcome Sanger Institute for the COVID-19 Genomics UK (COG-UK) Consortium | The Lighthouse Lab in Milton Keynes and Alex Alderton, Roberto Amato, Jeffrey Barrett, Sonia Goncalves, Ewan Harrison, David K. Jackson, Ian Johnston, Dominic Kwiatkowski, Cordelia Langford, John Sillitoe on behalf of the Wellcome Sanger Institute COVID-19 Surveillance Team |
| EPI_ISL_2118988, EPI_ISL_2118992, EPI_ISL_2118998 | Lighthouse Lab in Alderley Park | Wellcome Sanger Institute for the COVID-19 Genomics UK (COG-UK) Consortium | Jacquelyn Wynn, Mairead Hyland, The Lighthouse Lab in Alderley Park and Alex Alderton, Roberto Amato, Jeffrey Barrett, Sonia Goncalves, Ewan Harrison, David K. Jackson, Ian Johnston, Dominic Kwiatkowski, Cordelia Langford, John Sillitoe on behalf of the Wellcome Sanger Institute COVID-19 Surveillance Team |
| EPI_ISL_2119001 | Lighthouse Lab in Milton Keynes | Wellcome Sanger Institute for the COVID-19 Genomics UK (COG-UK) Consortium | The Lighthouse Lab in Milton Keynes and Alex Alderton, Roberto Amato, Jeffrey Barrett, Sonia Goncalves, Ewan Harrison, David K. Jackson, Ian Johnston, Dominic Kwiatkowski, Cordelia Langford, John Sillitoe on behalf of the Wellcome Sanger Institute COVID-19 Surveillance Team |
| EPI_ISL_2119006 | Lighthouse Lab in Alderley Park | Wellcome Sanger Institute for the COVID-19 Genomics UK (COG-UK) Consortium | Jacquelyn Wynn, Mairead Hyland, The Lighthouse Lab in Alderley Park and Alex Alderton, Roberto Amato, Jeffrey Barrett, Sonia Goncalves, Ewan Harrison, David K. Jackson, Ian Johnston, Dominic Kwiatkowski, Cordelia Langford, John Sillitoe on behalf of the Wellcome Sanger Institute COVID-19 Surveillance Team |
| EPI_ISL_2119012 | Lighthouse Lab in Milton Keynes | Wellcome Sanger Institute for the COVID-19 Genomics UK (COG-UK) Consortium | The Lighthouse Lab in Milton Keynes and Alex Alderton, Roberto Amato, Jeffrey Barrett, Sonia Goncalves, Ewan Harrison, David K. Jackson, Ian Johnston, Dominic Kwiatkowski, Cordelia Langford, John Sillitoe on behalf of the Wellcome Sanger Institute COVID-19 Surveillance Team |
| EPI_ISL_2119017 | Lighthouse Lab in Alderley Park | Wellcome Sanger Institute for the COVID-19 Genomics UK (COG-UK) Consortium | Jacquelyn Wynn, Mairead Hyland, The Lighthouse Lab in Alderley Park and Alex Alderton, Roberto Amato, Jeffrey Barrett, Sonia Goncalves, Ewan Harrison, David K. Jackson, Ian Johnston, Dominic Kwiatkowski, Cordelia Langford, John Sillitoe on behalf of the Wellcome Sanger Institute COVID-19 Surveillance Team |
| EPI_ISL_2119021 | Lighthouse Lab in Milton Keynes | Wellcome Sanger Institute for the COVID-19 Genomics UK (COG-UK) Consortium | The Lighthouse Lab in Milton Keynes and Alex Alderton, Roberto Amato, Jeffrey Barrett, Sonia Goncalves, Ewan Harrison, David K. Jackson, Ian Johnston, Dominic Kwiatkowski, Cordelia Langford, John Sillitoe on behalf of the Wellcome Sanger Institute COVID-19 Surveillance Team |
| EPI_ISL_2119024, EPI_ISL_2119041, EPI_ISL_2119048 | Lighthouse Lab in Alderley Park | Wellcome Sanger Institute for the COVID-19 Genomics UK (COG-UK) Consortium | Jacquelyn Wynn, Mairead Hyland, The Lighthouse Lab in Alderley Park and Alex Alderton, Roberto Amato, Jeffrey Barrett, Sonia Goncalves, Ewan Harrison, David K. Jackson, Ian Johnston, Dominic Kwiatkowski, Cordelia Langford, John Sillitoe on behalf of the Wellcome Sanger Institute COVID-19 Surveillance Team |
| EPI_ISL_2119085 | Lighthouse Lab in Milton Keynes | Wellcome Sanger Institute for the COVID-19 Genomics UK (COG-UK) Consortium | The Lighthouse Lab in Milton Keynes and Alex Alderton, Roberto Amato, Jeffrey Barrett, Sonia Goncalves, Ewan Harrison, David K. Jackson, Ian Johnston, Dominic Kwiatkowski, Cordelia Langford, John Sillitoe on behalf of the Wellcome Sanger Institute COVID-19 Surveillance Team |
| EPI_ISL_2119088, EPI_ISL_2119099, EPI_ISL_2119100 | Lighthouse Lab in Alderley Park | Wellcome Sanger Institute for the COVID-19 Genomics UK (COG-UK) Consortium | Jacquelyn Wynn, Mairead Hyland, The Lighthouse Lab in Alderley Park and Alex Alderton, Roberto Amato, Jeffrey Barrett, Sonia Goncalves, Ewan Harrison, David K. Jackson, Ian Johnston, Dominic Kwiatkowski, Cordelia Langford, John Sillitoe on behalf of the Wellcome Sanger Institute COVID-19 Surveillance Team |
| EPI_ISL_2119107 | Lighthouse Lab in Milton Keynes | Wellcome Sanger Institute for the COVID-19 Genomics UK (COG-UK) Consortium | The Lighthouse Lab in Milton Keynes and Alex Alderton, Roberto Amato, Jeffrey Barrett, Sonia Goncalves, Ewan Harrison, David K. Jackson, Ian Johnston, Dominic Kwiatkowski, Cordelia Langford, John Sillitoe on behalf of the Wellcome Sanger Institute COVID-19 Surveillance Team |
| EPI_ISL_2119130 | Lighthouse Lab in Alderley Park | Wellcome Sanger Institute for the COVID-19 Genomics UK (COG-UK) Consortium | Jacquelyn Wynn, Mairead Hyland, The Lighthouse Lab in Alderley Park and Alex Alderton, Roberto Amato, Jeffrey Barrett, Sonia Goncalves, Ewan Harrison, David K. Jackson, Ian Johnston, Dominic Kwiatkowski, Cordelia Langford, John Sillitoe on behalf of the Wellcome Sanger Institute COVID-19 Surveillance Team |
| EPI_ISL_2119131, EPI_ISL_2119139 | Lighthouse Lab in Milton Keynes | Wellcome Sanger Institute for the COVID-19 Genomics UK (COG-UK) Consortium | The Lighthouse Lab in Milton Keynes and Alex Alderton, Roberto Amato, Jeffrey Barrett, Sonia Goncalves, Ewan Harrison, David K. Jackson, Ian Johnston, Dominic Kwiatkowski, Cordelia Langford, John Sillitoe on behalf of the Wellcome Sanger Institute COVID-19 Surveillance Team |
| EPI_ISL_2119141, EPI_ISL_2119146, EPI_ISL_2119152, EPI_ISL_2119163, EPI_ISL_2119166 | Lighthouse Lab in Alderley Park | Wellcome Sanger Institute for the COVID-19 Genomics UK (COG-UK) Consortium | Jacquelyn Wynn, Mairead Hyland, The Lighthouse Lab in Alderley Park and Alex Alderton, Roberto Amato, Jeffrey Barrett, Sonia Goncalves, Ewan Harrison, David K. Jackson, Ian Johnston, Dominic Kwiatkowski, Cordelia Langford, John Sillitoe on behalf of the Wellcome Sanger Institute COVID-19 Surveillance Team |
| EPI_ISL_2119174 | Lighthouse Lab in Milton Keynes | Wellcome Sanger Institute for the COVID-19 Genomics UK (COG-UK) Consortium | The Lighthouse Lab in Milton Keynes and Alex Alderton, Roberto Amato, Jeffrey Barrett, Sonia Goncalves, Ewan Harrison, David K. Jackson, Ian Johnston, Dominic Kwiatkowski, Cordelia Langford, John Sillitoe on behalf of the Wellcome Sanger Institute COVID-19 Surveillance Team |
| EPI_ISL_2119183 | Lighthouse Lab in Alderley Park | Wellcome Sanger Institute for the COVID-19 Genomics UK (COG-UK) Consortium | Jacquelyn Wynn, Mairead Hyland, The Lighthouse Lab in Alderley Park and Alex Alderton, Roberto Amato, Jeffrey Barrett, Sonia Goncalves, Ewan Harrison, David K. Jackson, Ian Johnston, Dominic Kwiatkowski, Cordelia Langford, John Sillitoe on behalf of the Wellcome Sanger Institute COVID-19 Surveillance Team |
| EPI_ISL_2119188 | Lighthouse Lab in Milton Keynes | Wellcome Sanger Institute for the COVID-19 Genomics UK (COG-UK) Consortium | The Lighthouse Lab in Milton Keynes and Alex Alderton, Roberto Amato, Jeffrey Barrett, Sonia Goncalves, Ewan Harrison, David K. Jackson, Ian Johnston, Dominic Kwiatkowski, Cordelia Langford, John Sillitoe on behalf of the Wellcome Sanger Institute COVID-19 Surveillance Team |
| EPI_ISL_2119192, EPI_ISL_2119216, EPI_ISL_2119222 | Lighthouse Lab in Alderley Park | Wellcome Sanger Institute for the COVID-19 Genomics UK (COG-UK) Consortium | Jacquelyn Wynn, Mairead Hyland, The Lighthouse Lab in Alderley Park and Alex Alderton, Roberto Amato, Jeffrey Barrett, Sonia Goncalves, Ewan Harrison, David K. Jackson, Ian Johnston, Dominic Kwiatkowski, Cordelia Langford, John Sillitoe on behalf of the Wellcome Sanger Institute COVID-19 Surveillance Team |
| EPI_ISL_2119231, EPI_ISL_2119235 | Lighthouse Lab in Milton Keynes | Wellcome Sanger Institute for the COVID-19 Genomics UK (COG-UK) Consortium | The Lighthouse Lab in Milton Keynes and Alex Alderton, Roberto Amato, Jeffrey Barrett, Sonia Goncalves, Ewan Harrison, David K. Jackson, Ian Johnston, Dominic Kwiatkowski, Cordelia Langford, John Sillitoe on behalf of the Wellcome Sanger Institute COVID-19 Surveillance Team |
| EPI_ISL_2119247, EPI_ISL_2119256, EPI_ISL_2119260 | Lighthouse Lab in Alderley Park | Wellcome Sanger Institute for the COVID-19 Genomics UK (COG-UK) Consortium | Jacquelyn Wynn, Mairead Hyland, The Lighthouse Lab in Alderley Park and Alex Alderton, Roberto Amato, Jeffrey Barrett, Sonia Goncalves, Ewan Harrison, David K. Jackson, Ian Johnston, Dominic Kwiatkowski, Cordelia Langford, John Sillitoe on behalf of the Wellcome Sanger Institute COVID-19 Surveillance Team |
| EPI_ISL_2119261 | Lighthouse Lab in Milton Keynes | Wellcome Sanger Institute for the COVID-19 Genomics UK (COG-UK) Consortium | The Lighthouse Lab in Milton Keynes and Alex Alderton, Roberto Amato, Jeffrey Barrett, Sonia Goncalves, Ewan Harrison, David K. Jackson, Ian Johnston, Dominic Kwiatkowski, Cordelia Langford, John Sillitoe on behalf of the Wellcome Sanger Institute COVID-19 Surveillance Team |
| EPI_ISL_2119265, EPI_ISL_2119271, EPI_ISL_2119275, EPI_ISL_2119280, EPI_ISL_2119289, EPI_ISL_2119301, EPI_ISL_2119311 | Lighthouse Lab in Alderley Park | Wellcome Sanger Institute for the COVID-19 Genomics UK (COG-UK) Consortium | Jacquelyn Wynn, Mairead Hyland, The Lighthouse Lab in Alderley Park and Alex Alderton, Roberto Amato, Jeffrey Barrett, Sonia Goncalves, Ewan Harrison, David K. Jackson, Ian Johnston, Dominic Kwiatkowski, Cordelia Langford, John Sillitoe on behalf of the Wellcome Sanger Institute COVID-19 Surveillance Team |
| EPI_ISL_2119312 | Lighthouse Lab in Milton Keynes | Wellcome Sanger Institute for the COVID-19 Genomics UK (COG-UK) Consortium | The Lighthouse Lab in Milton Keynes and Alex Alderton, Roberto Amato, Jeffrey Barrett, Sonia Goncalves, Ewan Harrison, David K. Jackson, Ian Johnston, Dominic Kwiatkowski, Cordelia Langford, John Sillitoe on behalf of the Wellcome Sanger Institute COVID-19 Surveillance Team |
| EPI_ISL_2119327, EPI_ISL_2119330, EPI_ISL_2119338, EPI_ISL_2119343 | Lighthouse Lab in Alderley Park | Wellcome Sanger Institute for the COVID-19 Genomics UK (COG-UK) Consortium | Jacquelyn Wynn, Mairead Hyland, The Lighthouse Lab in Alderley Park and Alex Alderton, Roberto Amato, Jeffrey Barrett, Sonia Goncalves, Ewan Harrison, David K. Jackson, Ian Johnston, Dominic Kwiatkowski, Cordelia Langford, John Sillitoe on behalf of the Wellcome Sanger Institute COVID-19 Surveillance Team |
| EPI_ISL_2119344 | Lighthouse Lab in Milton Keynes | Wellcome Sanger Institute for the COVID-19 Genomics UK (COG-UK) Consortium | The Lighthouse Lab in Milton Keynes and Alex Alderton, Roberto Amato, Jeffrey Barrett, Sonia Goncalves, Ewan Harrison, David K. Jackson, Ian Johnston, Dominic Kwiatkowski, Cordelia Langford, John Sillitoe on behalf of the Wellcome Sanger Institute COVID-19 Surveillance Team |

| | | | |
|---|---|---|---|
| EPI_ISL_2119346, EPI_ISL_2119349 | Lighthouse Lab in Alderley Park | Wellcome Sanger Institute for the COVID-19 Genomics UK (COG-UK) Consortium | Jacquelyn Wynn, Mairead Hyland, The Lighthouse Lab in Alderley Park and Alex Alderton, Roberto Amato, Jeffrey Barrett, Sonia Goncalves, Ewan Harrison, David K. Jackson, Ian Johnston, Dominic Kwiatkowski, Cordelia Langford, John Sillitoe on behalf of the Wellcome Sanger Institute COVID-19 Surveillance Team |
| EPI_ISL_2119351, EPI_ISL_2119354 | Lighthouse Lab in Glasgow | Wellcome Sanger Institute for the COVID-19 Genomics UK (COG-UK) Consortium | Harper VanSteenhouse, Yumi Kasai, David Gray, Carol Clugston, Anna Dominiczak and Alex Alderton, Roberto Amato, Jeffrey Barrett, Sonia Goncalves, Ewan Harrison, David K. Jackson, Ian Johnston, Dominic Kwiatkowski, Cordelia Langford, John Sillitoe on behalf of the Wellcome Sanger Institute COVID-19 Surveillance Team |
| EPI_ISL_2119356, EPI_ISL_2119371, EPI_ISL_2119376 | Lighthouse Lab in Milton Keynes | Wellcome Sanger Institute for the COVID-19 Genomics UK (COG-UK) Consortium | The Lighthouse Lab in Milton Keynes and Alex Alderton, Roberto Amato, Jeffrey Barrett, Sonia Goncalves, Ewan Harrison, David K. Jackson, Ian Johnston, Dominic Kwiatkowski, Cordelia Langford, John Sillitoe on behalf of the Wellcome Sanger Institute COVID-19 Surveillance Team |
| EPI_ISL_2119389 | Lighthouse Lab in Glasgow | Wellcome Sanger Institute for the COVID-19 Genomics UK (COG-UK) Consortium | Harper VanSteenhouse, Yumi Kasai, David Gray, Carol Clugston, Anna Dominiczak and Alex Alderton, Roberto Amato, Jeffrey Barrett, Sonia Goncalves, Ewan Harrison, David K. Jackson, Ian Johnston, Dominic Kwiatkowski, Cordelia Langford, John Sillitoe on behalf of the Wellcome Sanger Institute COVID-19 Surveillance Team |
| EPI_ISL_2119395 | Lighthouse Lab in Alderley Park | Wellcome Sanger Institute for the COVID-19 Genomics UK (COG-UK) Consortium | Jacquelyn Wynn, Mairead Hyland, The Lighthouse Lab in Alderley Park and Alex Alderton, Roberto Amato, Jeffrey Barrett, Sonia Goncalves, Ewan Harrison, David K. Jackson, Ian Johnston, Dominic Kwiatkowski, Cordelia Langford, John Sillitoe on behalf of the Wellcome Sanger Institute COVID-19 Surveillance Team |
| EPI_ISL_2119400, EPI_ISL_2119405, EPI_ISL_2119412, EPI_ISL_2119414, EPI_ISL_2119419, EPI_ISL_2119427 | Lighthouse Lab in Milton Keynes | Wellcome Sanger Institute for the COVID-19 Genomics UK (COG-UK) Consortium | The Lighthouse Lab in Milton Keynes and Alex Alderton, Roberto Amato, Jeffrey Barrett, Sonia Goncalves, Ewan Harrison, David K. Jackson, Ian Johnston, Dominic Kwiatkowski, Cordelia Langford, John Sillitoe on behalf of the Wellcome Sanger Institute COVID-19 Surveillance Team |
| EPI_ISL_2119430 | Lighthouse Lab in Glasgow | Wellcome Sanger Institute for the COVID-19 Genomics UK (COG-UK) Consortium | Harper VanSteenhouse, Yumi Kasai, David Gray, Carol Clugston, Anna Dominiczak and Alex Alderton, Roberto Amato, Jeffrey Barrett, Sonia Goncalves, Ewan Harrison, David K. Jackson, Ian Johnston, Dominic Kwiatkowski, Cordelia Langford, John Sillitoe on behalf of the Wellcome Sanger Institute COVID-19 Surveillance Team |
| EPI_ISL_2119437, EPI_ISL_2119440 | Lighthouse Lab in Milton Keynes | Wellcome Sanger Institute for the COVID-19 Genomics UK (COG-UK) Consortium | The Lighthouse Lab in Milton Keynes and Alex Alderton, Roberto Amato, Jeffrey Barrett, Sonia Goncalves, Ewan Harrison, David K. Jackson, Ian Johnston, Dominic Kwiatkowski, Cordelia Langford, John Sillitoe on behalf of the Wellcome Sanger Institute COVID-19 Surveillance Team |
| EPI_ISL_2119448 | Lighthouse Lab in Alderley Park | Wellcome Sanger Institute for the COVID-19 Genomics UK (COG-UK) Consortium | Jacquelyn Wynn, Mairead Hyland, The Lighthouse Lab in Alderley Park and Alex Alderton, Roberto Amato, Jeffrey Barrett, Sonia Goncalves, Ewan Harrison, David K. Jackson, Ian Johnston, Dominic Kwiatkowski, Cordelia Langford, John Sillitoe on behalf of the Wellcome Sanger Institute COVID-19 Surveillance Team |
| EPI_ISL_2119460, EPI_ISL_2119462 | Lighthouse Lab in Glasgow | Wellcome Sanger Institute for the COVID-19 Genomics UK (COG-UK) Consortium | Harper VanSteenhouse, Yumi Kasai, David Gray, Carol Clugston, Anna Dominiczak and Alex Alderton, Roberto Amato, Jeffrey Barrett, Sonia Goncalves, Ewan Harrison, David K. Jackson, Ian Johnston, Dominic Kwiatkowski, Cordelia Langford, John Sillitoe on behalf of the Wellcome Sanger Institute COVID-19 Surveillance Team |
| EPI_ISL_2119473 | Lighthouse Lab in Alderley Park | Wellcome Sanger Institute for the COVID-19 Genomics UK (COG-UK) Consortium | Jacquelyn Wynn, Mairead Hyland, The Lighthouse Lab in Alderley Park and Alex Alderton, Roberto Amato, Jeffrey Barrett, Sonia Goncalves, Ewan Harrison, David K. Jackson, Ian Johnston, Dominic Kwiatkowski, Cordelia Langford, John Sillitoe on behalf of the Wellcome Sanger Institute COVID-19 Surveillance Team |
| EPI_ISL_2119482 | Lighthouse Lab in Glasgow | Wellcome Sanger Institute for the COVID-19 Genomics UK (COG-UK) Consortium | Harper VanSteenhouse, Yumi Kasai, David Gray, Carol Clugston, Anna Dominiczak and Alex Alderton, Roberto Amato, Jeffrey Barrett, Sonia Goncalves, Ewan Harrison, David K. Jackson, Ian Johnston, Dominic Kwiatkowski, Cordelia Langford, John Sillitoe on behalf of the Wellcome Sanger Institute COVID-19 Surveillance Team |
| EPI_ISL_2119492 | Lighthouse Lab in Milton Keynes | Wellcome Sanger Institute for the COVID-19 Genomics UK (COG-UK) Consortium | The Lighthouse Lab in Milton Keynes and Alex Alderton, Roberto Amato, Jeffrey Barrett, Sonia Goncalves, Ewan Harrison, David K. Jackson, Ian Johnston, Dominic Kwiatkowski, Cordelia Langford, John Sillitoe on behalf of the Wellcome Sanger Institute COVID-19 Surveillance Team |
| EPI_ISL_2119504 | Lighthouse Lab in Alderley Park | Wellcome Sanger Institute for the COVID-19 Genomics UK (COG-UK) Consortium | Jacquelyn Wynn, Mairead Hyland, The Lighthouse Lab in Alderley Park and Alex Alderton, Roberto Amato, Jeffrey Barrett, Sonia Goncalves, Ewan Harrison, David K. Jackson, Ian Johnston, Dominic Kwiatkowski, Cordelia Langford, John Sillitoe on behalf of the Wellcome Sanger Institute COVID-19 Surveillance Team |
| EPI_ISL_2119510 | Lighthouse Lab in Milton Keynes | Wellcome Sanger Institute for the COVID-19 Genomics UK (COG-UK) Consortium | The Lighthouse Lab in Milton Keynes and Alex Alderton, Roberto Amato, Jeffrey Barrett, Sonia Goncalves, Ewan Harrison, David K. Jackson, Ian Johnston, Dominic Kwiatkowski, Cordelia Langford, John Sillitoe on behalf of the Wellcome Sanger Institute COVID-19 Surveillance Team |
| EPI_ISL_2119513 | Lighthouse Lab in Glasgow | Wellcome Sanger Institute for the COVID-19 Genomics UK (COG-UK) Consortium | Harper VanSteenhouse, Yumi Kasai, David Gray, Carol Clugston, Anna Dominiczak and Alex Alderton, Roberto Amato, Jeffrey Barrett, Sonia Goncalves, Ewan Harrison, David K. Jackson, Ian Johnston, Dominic Kwiatkowski, Cordelia Langford, John Sillitoe on behalf of the Wellcome Sanger Institute COVID-19 Surveillance Team |
| EPI_ISL_2119535, EPI_ISL_2119545, EPI_ISL_2119549 | Lighthouse Lab in Milton Keynes | Wellcome Sanger Institute for the COVID-19 Genomics UK (COG-UK) Consortium | The Lighthouse Lab in Milton Keynes and Alex Alderton, Roberto Amato, Jeffrey Barrett, Sonia Goncalves, Ewan Harrison, David K. Jackson, Ian Johnston, Dominic Kwiatkowski, Cordelia Langford, John Sillitoe on behalf of the Wellcome Sanger Institute COVID-19 Surveillance Team |
| EPI_ISL_2119551 | Lighthouse Lab in Alderley Park | Wellcome Sanger Institute for the COVID-19 Genomics UK (COG-UK) Consortium | Jacquelyn Wynn, Mairead Hyland, The Lighthouse Lab in Alderley Park and Alex Alderton, Roberto Amato, Jeffrey Barrett, Sonia Goncalves, Ewan Harrison, David K. Jackson, Ian Johnston, Dominic Kwiatkowski, Cordelia Langford, John Sillitoe on behalf of the Wellcome Sanger Institute COVID-19 Surveillance Team |
| EPI_ISL_2119565 | Lighthouse Lab in Milton Keynes | Wellcome Sanger Institute for the COVID-19 Genomics UK (COG-UK) Consortium | The Lighthouse Lab in Milton Keynes and Alex Alderton, Roberto Amato, Jeffrey Barrett, Sonia Goncalves, Ewan Harrison, David K. Jackson, Ian Johnston, Dominic Kwiatkowski, Cordelia Langford, John Sillitoe on behalf of the Wellcome Sanger Institute COVID-19 Surveillance Team |
| EPI_ISL_2119569 | Lighthouse Lab in Alderley Park | Wellcome Sanger Institute for the COVID-19 Genomics UK (COG-UK) Consortium | Jacquelyn Wynn, Mairead Hyland, The Lighthouse Lab in Alderley Park and Alex Alderton, Roberto Amato, Jeffrey Barrett, Sonia Goncalves, Ewan Harrison, David K. Jackson, Ian Johnston, Dominic Kwiatkowski, Cordelia Langford, John Sillitoe on behalf of the Wellcome Sanger Institute COVID-19 Surveillance Team |
| EPI_ISL_2119576 | Lighthouse Lab in Glasgow | Wellcome Sanger Institute for the COVID-19 Genomics UK (COG-UK) Consortium | Harper VanSteenhouse, Yumi Kasai, David Gray, Carol Clugston, Anna Dominiczak and Alex Alderton, Roberto Amato, Jeffrey Barrett, Sonia Goncalves, Ewan Harrison, David K. Jackson, Ian Johnston, Dominic Kwiatkowski, Cordelia Langford, John Sillitoe on behalf of the Wellcome Sanger Institute COVID-19 Surveillance Team |
| EPI_ISL_2119580 | Lighthouse Lab in Milton Keynes | Wellcome Sanger Institute for the COVID-19 Genomics UK (COG-UK) Consortium | The Lighthouse Lab in Milton Keynes and Alex Alderton, Roberto Amato, Jeffrey Barrett, Sonia Goncalves, Ewan Harrison, David K. Jackson, Ian Johnston, Dominic Kwiatkowski, Cordelia Langford, John Sillitoe on behalf of the Wellcome Sanger Institute COVID-19 Surveillance Team |
| EPI_ISL_2119582 | Lighthouse Lab in Alderley Park | Wellcome Sanger Institute for the COVID-19 Genomics UK (COG-UK) Consortium | Jacquelyn Wynn, Mairead Hyland, The Lighthouse Lab in Alderley Park and Alex Alderton, Roberto Amato, Jeffrey Barrett, Sonia Goncalves, Ewan Harrison, David K. Jackson, Ian Johnston, Dominic Kwiatkowski, Cordelia Langford, John Sillitoe on behalf of the Wellcome Sanger Institute COVID-19 Surveillance Team |
| EPI_ISL_2119590 | Lighthouse Lab in Milton Keynes | Wellcome Sanger Institute for the COVID-19 Genomics UK (COG-UK) Consortium | The Lighthouse Lab in Milton Keynes and Alex Alderton, Roberto Amato, Jeffrey Barrett, Sonia Goncalves, Ewan Harrison, David K. Jackson, Ian Johnston, Dominic Kwiatkowski, Cordelia Langford, John Sillitoe on behalf of the Wellcome Sanger Institute COVID-19 Surveillance Team |
| EPI_ISL_2119591 | Lighthouse Lab in Alderley Park | Wellcome Sanger Institute for the COVID-19 Genomics UK (COG-UK) Consortium | Jacquelyn Wynn, Mairead Hyland, The Lighthouse Lab in Alderley Park and Alex Alderton, Roberto Amato, Jeffrey Barrett, Sonia Goncalves, Ewan Harrison, David K. Jackson, Ian Johnston, Dominic Kwiatkowski, Cordelia Langford, John Sillitoe on behalf of the Wellcome Sanger Institute COVID-19 Surveillance Team |
| EPI_ISL_2119593 | Lighthouse Lab in Glasgow | Wellcome Sanger Institute for the COVID-19 Genomics UK (COG-UK) Consortium | Harper VanSteenhouse, Yumi Kasai, David Gray, Carol Clugston, Anna Dominiczak and Alex Alderton, Roberto Amato, Jeffrey Barrett, Sonia Goncalves, Ewan Harrison, David K. Jackson, Ian Johnston, Dominic Kwiatkowski, Cordelia Langford, John Sillitoe on behalf of the Wellcome Sanger Institute COVID-19 Surveillance Team |
| EPI_ISL_2119595, EPI_ISL_2119599 | Lighthouse Lab in Alderley Park | Wellcome Sanger Institute for the COVID-19 Genomics UK (COG-UK) Consortium | Jacquelyn Wynn, Mairead Hyland, The Lighthouse Lab in Alderley Park and Alex Alderton, Roberto Amato, Jeffrey Barrett, Sonia Goncalves, Ewan Harrison, David K. Jackson, Ian Johnston, Dominic Kwiatkowski, Cordelia Langford, John Sillitoe on behalf of the Wellcome Sanger Institute COVID-19 Surveillance Team |
| EPI_ISL_2119601, EPI_ISL_2119602, EPI_ISL_2119612, EPI_ISL_2119617, EPI_ISL_2119630 | Lighthouse Lab in Milton Keynes | Wellcome Sanger Institute for the COVID-19 Genomics UK (COG-UK) Consortium | The Lighthouse Lab in Milton Keynes and Alex Alderton, Roberto Amato, Jeffrey Barrett, Sonia Goncalves, Ewan Harrison, David K. Jackson, Ian Johnston, Dominic Kwiatkowski, Cordelia Langford, John Sillitoe on behalf of the Wellcome Sanger Institute COVID-19 Surveillance Team |
| EPI_ISL_2119633 | Lighthouse Lab in Alderley Park | Wellcome Sanger Institute for the COVID-19 Genomics UK (COG-UK) Consortium | Jacquelyn Wynn, Mairead Hyland, The Lighthouse Lab in Alderley Park and Alex Alderton, Roberto Amato, Jeffrey Barrett, Sonia Goncalves, Ewan Harrison, David K. Jackson, Ian Johnston, Dominic Kwiatkowski, Cordelia Langford, John Sillitoe on behalf of the Wellcome Sanger Institute COVID-19 Surveillance Team |
| EPI_ISL_2119635 | Lighthouse Lab in Glasgow | Wellcome Sanger Institute for the COVID-19 Genomics UK (COG-UK) Consortium | Harper VanSteenhouse, Yumi Kasai, David Gray, Carol Clugston, Anna Dominiczak and Alex Alderton, Roberto Amato, Jeffrey Barrett, Sonia Goncalves, Ewan Harrison, David K. Jackson, Ian Johnston, Dominic Kwiatkowski, Cordelia Langford, John Sillitoe on behalf of the Wellcome Sanger Institute COVID-19 Surveillance Team |
| EPI_ISL_2119646 | Lighthouse Lab in Alderley Park | Wellcome Sanger Institute for the COVID-19 Genomics UK (COG-UK) Consortium | Jacquelyn Wynn, Mairead Hyland, The Lighthouse Lab in Alderley Park and Alex Alderton, Roberto Amato, Jeffrey Barrett, Sonia Goncalves, Ewan Harrison, David K. Jackson, Ian Johnston, Dominic Kwiatkowski, Cordelia Langford, John Sillitoe on behalf of the Wellcome Sanger Institute COVID-19 Surveillance Team |
| EPI_ISL_2119649 | Lighthouse Lab in Milton Keynes | Wellcome Sanger Institute for the COVID-19 Genomics UK (COG-UK) Consortium | The Lighthouse Lab in Milton Keynes and Alex Alderton, Roberto Amato, Jeffrey Barrett, Sonia Goncalves, Ewan Harrison, David K. Jackson, Ian Johnston, Dominic Kwiatkowski, Cordelia Langford, John Sillitoe on behalf of the Wellcome Sanger Institute COVID-19 Surveillance Team |
| EPI_ISL_2119654 | Lighthouse Lab in Glasgow | Wellcome Sanger Institute for the COVID-19 Genomics UK (COG-UK) Consortium | Harper VanSteenhouse, Yumi Kasai, David Gray, Carol Clugston, Anna Dominiczak and Alex Alderton, Roberto Amato, Jeffrey Barrett, Sonia Goncalves, Ewan Harrison, David K. Jackson, Ian Johnston, Dominic Kwiatkowski, Cordelia Langford, John Sillitoe on behalf of the Wellcome Sanger Institute COVID-19 Surveillance Team |
| EPI_ISL_2119657, EPI_ISL_2119672, EPI_ISL_2119676, EPI_ISL_2119677 | Lighthouse Lab in Milton Keynes | Wellcome Sanger Institute for the COVID-19 Genomics UK (COG-UK) Consortium | The Lighthouse Lab in Milton Keynes and Alex Alderton, Roberto Amato, Jeffrey Barrett, Sonia Goncalves, Ewan Harrison, David K. Jackson, Ian Johnston, Dominic Kwiatkowski, Cordelia Langford, John Sillitoe on behalf of the Wellcome Sanger Institute COVID-19 Surveillance Team |
| EPI_ISL_2119683 | Lighthouse Lab in Glasgow | Wellcome Sanger Institute for the COVID-19 Genomics UK (COG-UK) Consortium | Harper VanSteenhouse, Yumi Kasai, David Gray, Carol Clugston, Anna Dominiczak and Alex Alderton, Roberto Amato, Jeffrey Barrett, Sonia Goncalves, Ewan Harrison, David K. Jackson, Ian Johnston, Dominic Kwiatkowski, Cordelia Langford, John Sillitoe on behalf of the Wellcome Sanger Institute COVID-19 Surveillance Team |
| EPI_ISL_2119688, EPI_ISL_2119696 | Lighthouse Lab in Milton Keynes | Wellcome Sanger Institute for the COVID-19 Genomics UK (COG-UK) Consortium | The Lighthouse Lab in Milton Keynes and Alex Alderton, Roberto Amato, Jeffrey Barrett, Sonia Goncalves, Ewan Harrison, David K. Jackson, Ian Johnston, Dominic Kwiatkowski, Cordelia Langford, John Sillitoe on behalf of the Wellcome Sanger Institute COVID-19 Surveillance Team |
| EPI_ISL_2119709 | Lighthouse Lab in Alderley Park | Wellcome Sanger Institute for the COVID-19 Genomics UK (COG-UK) Consortium | Jacquelyn Wynn, Mairead Hyland, The Lighthouse Lab in Alderley Park and Alex Alderton, Roberto Amato, Jeffrey Barrett, Sonia Goncalves, Ewan Harrison, David K. Jackson, Ian Johnston, Dominic Kwiatkowski, Cordelia Langford, John Sillitoe on behalf of the Wellcome Sanger Institute COVID-19 Surveillance Team |
| EPI_ISL_2119717 | Lighthouse Lab in Glasgow | Wellcome Sanger Institute for the COVID-19 Genomics UK (COG-UK) Consortium | Harper VanSteenhouse, Yumi Kasai, David Gray, Carol Clugston, Anna Dominiczak and Alex Alderton, Roberto Amato, Jeffrey Barrett, Sonia Goncalves, Ewan Harrison, David K. Jackson, Ian Johnston, Dominic Kwiatkowski, Cordelia Langford, John Sillitoe on behalf of the Wellcome Sanger Institute COVID-19 Surveillance Team |
| EPI_ISL_2119723 | Lighthouse Lab in Milton Keynes | Wellcome Sanger Institute for the COVID-19 Genomics UK (COG-UK) Consortium | The Lighthouse Lab in Milton Keynes and Alex Alderton, Roberto Amato, Jeffrey Barrett, Sonia Goncalves, Ewan Harrison, David K. Jackson, Ian Johnston, Dominic Kwiatkowski, Cordelia Langford, John Sillitoe on behalf of the Wellcome Sanger Institute COVID-19 Surveillance Team |
| EPI_ISL_2119726 | Lighthouse Lab in Alderley Park | Wellcome Sanger Institute for the COVID-19 Genomics UK (COG-UK) Consortium | Jacquelyn Wynn, Mairead Hyland, The Lighthouse Lab in Alderley Park and Alex Alderton, Roberto Amato, Jeffrey Barrett, Sonia Goncalves, Ewan Harrison, David K. Jackson, Ian Johnston, Dominic Kwiatkowski, Cordelia Langford, John Sillitoe on behalf of the Wellcome Sanger Institute COVID-19 Surveillance Team |
| EPI_ISL_2119727, | Lighthouse Lab in Milton Keynes | Wellcome Sanger Institute for the COVID-19 Genomics UK (COG-UK) | The Lighthouse Lab in Milton Keynes and Alex Alderton, Roberto Amato, Jeffrey Barrett, Sonia Goncalves, Ewan Harrison, David K. Jackson, Ian Johnston, Dominic Kwiatkowski, Cordelia Langford, John Sillitoe on behalf of the Wellcome Sanger |

| | | | |
|---|---|---|---|
| EPI_ISL_2119731, EPI_ISL_2119735, EPI_ISL_2119740, EPI_ISL_2119743 | | Consortium | Institute COVID-19 Surveillance Team |
| EPI_ISL_2119748, EPI_ISL_2119749, EPI_ISL_2119754 | Lighthouse Lab in Glasgow | Wellcome Sanger Institute for the COVID-19 Genomics UK (COG-UK) Consortium | Harper VanSteenhouse, Yumi Kasai, David Gray, Carol Clugston, Anna Dominiczak and Alex Alderton, Roberto Amato, Jeffrey Barrett, Sonia Goncalves, Ewan Harrison, David K. Jackson, Ian Johnston, Dominic Kwiatkowski, Cordelia Langford, John Sillitoe on behalf of the Wellcome Sanger Institute COVID-19 Surveillance Team |
| EPI_ISL_2119757, EPI_ISL_2119764 | Lighthouse Lab in Milton Keynes | Wellcome Sanger Institute for the COVID-19 Genomics UK (COG-UK) Consortium | The Lighthouse Lab in Milton Keynes and Alex Alderton, Roberto Amato, Jeffrey Barrett, Sonia Goncalves, Ewan Harrison, David K. Jackson, Ian Johnston, Dominic Kwiatkowski, Cordelia Langford, John Sillitoe on behalf of the Wellcome Sanger Institute COVID-19 Surveillance Team |
| EPI_ISL_2119765 | Lighthouse Lab in Alderley Park | Wellcome Sanger Institute for the COVID-19 Genomics UK (COG-UK) Consortium | Jacquelyn Wynn, Mairead Hyland, The Lighthouse Lab in Alderley Park and Alex Alderton, Roberto Amato, Jeffrey Barrett, Sonia Goncalves, Ewan Harrison, David K. Jackson, Ian Johnston, Dominic Kwiatkowski, Cordelia Langford, John Sillitoe on behalf of the Wellcome Sanger Institute COVID-19 Surveillance Team |
| EPI_ISL_2119767 | Lighthouse Lab in Milton Keynes | Wellcome Sanger Institute for the COVID-19 Genomics UK (COG-UK) Consortium | The Lighthouse Lab in Milton Keynes and Alex Alderton, Roberto Amato, Jeffrey Barrett, Sonia Goncalves, Ewan Harrison, David K. Jackson, Ian Johnston, Dominic Kwiatkowski, Cordelia Langford, John Sillitoe on behalf of the Wellcome Sanger Institute COVID-19 Surveillance Team |
| EPI_ISL_2119772, EPI_ISL_2119775 | Lighthouse Lab in Alderley Park | Wellcome Sanger Institute for the COVID-19 Genomics UK (COG-UK) Consortium | Jacquelyn Wynn, Mairead Hyland, The Lighthouse Lab in Alderley Park and Alex Alderton, Roberto Amato, Jeffrey Barrett, Sonia Goncalves, Ewan Harrison, David K. Jackson, Ian Johnston, Dominic Kwiatkowski, Cordelia Langford, John Sillitoe on behalf of the Wellcome Sanger Institute COVID-19 Surveillance Team |
| EPI_ISL_2119800, EPI_ISL_2119809 | Lighthouse Lab in Milton Keynes | Wellcome Sanger Institute for the COVID-19 Genomics UK (COG-UK) Consortium | The Lighthouse Lab in Milton Keynes and Alex Alderton, Roberto Amato, Jeffrey Barrett, Sonia Goncalves, Ewan Harrison, David K. Jackson, Ian Johnston, Dominic Kwiatkowski, Cordelia Langford, John Sillitoe on behalf of the Wellcome Sanger Institute COVID-19 Surveillance Team |
| EPI_ISL_2119822 | Lighthouse Lab in Alderley Park | Wellcome Sanger Institute for the COVID-19 Genomics UK (COG-UK) Consortium | Jacquelyn Wynn, Mairead Hyland, The Lighthouse Lab in Alderley Park and Alex Alderton, Roberto Amato, Jeffrey Barrett, Sonia Goncalves, Ewan Harrison, David K. Jackson, Ian Johnston, Dominic Kwiatkowski, Cordelia Langford, John Sillitoe on behalf of the Wellcome Sanger Institute COVID-19 Surveillance Team |
| EPI_ISL_2119828 | Lighthouse Lab in Glasgow | Wellcome Sanger Institute for the COVID-19 Genomics UK (COG-UK) Consortium | Harper VanSteenhouse, Yumi Kasai, David Gray, Carol Clugston, Anna Dominiczak and Alex Alderton, Roberto Amato, Jeffrey Barrett, Sonia Goncalves, Ewan Harrison, David K. Jackson, Ian Johnston, Dominic Kwiatkowski, Cordelia Langford, John Sillitoe on behalf of the Wellcome Sanger Institute COVID-19 Surveillance Team |
| EPI_ISL_2119838 | Lighthouse Lab in Alderley Park | Wellcome Sanger Institute for the COVID-19 Genomics UK (COG-UK) Consortium | Jacquelyn Wynn, Mairead Hyland, The Lighthouse Lab in Alderley Park and Alex Alderton, Roberto Amato, Jeffrey Barrett, Sonia Goncalves, Ewan Harrison, David K. Jackson, Ian Johnston, Dominic Kwiatkowski, Cordelia Langford, John Sillitoe on behalf of the Wellcome Sanger Institute COVID-19 Surveillance Team |
| EPI_ISL_2119839, EPI_ISL_2119854, EPI_ISL_2119857 | Lighthouse Lab in Milton Keynes | Wellcome Sanger Institute for the COVID-19 Genomics UK (COG-UK) Consortium | The Lighthouse Lab in Milton Keynes and Alex Alderton, Roberto Amato, Jeffrey Barrett, Sonia Goncalves, Ewan Harrison, David K. Jackson, Ian Johnston, Dominic Kwiatkowski, Cordelia Langford, John Sillitoe on behalf of the Wellcome Sanger Institute COVID-19 Surveillance Team |
| EPI_ISL_2119858 | Lighthouse Lab in Glasgow | Wellcome Sanger Institute for the COVID-19 Genomics UK (COG-UK) Consortium | Harper VanSteenhouse, Yumi Kasai, David Gray, Carol Clugston, Anna Dominiczak and Alex Alderton, Roberto Amato, Jeffrey Barrett, Sonia Goncalves, Ewan Harrison, David K. Jackson, Ian Johnston, Dominic Kwiatkowski, Cordelia Langford, John Sillitoe on behalf of the Wellcome Sanger Institute COVID-19 Surveillance Team |
| EPI_ISL_2119861 | Lighthouse Lab in Milton Keynes | Wellcome Sanger Institute for the COVID-19 Genomics UK (COG-UK) Consortium | The Lighthouse Lab in Milton Keynes and Alex Alderton, Roberto Amato, Jeffrey Barrett, Sonia Goncalves, Ewan Harrison, David K. Jackson, Ian Johnston, Dominic Kwiatkowski, Cordelia Langford, John Sillitoe on behalf of the Wellcome Sanger Institute COVID-19 Surveillance Team |
| EPI_ISL_2119863 | Lighthouse Lab in Glasgow | Wellcome Sanger Institute for the COVID-19 Genomics UK (COG-UK) Consortium | Harper VanSteenhouse, Yumi Kasai, David Gray, Carol Clugston, Anna Dominiczak and Alex Alderton, Roberto Amato, Jeffrey Barrett, Sonia Goncalves, Ewan Harrison, David K. Jackson, Ian Johnston, Dominic Kwiatkowski, Cordelia Langford, John Sillitoe on behalf of the Wellcome Sanger Institute COVID-19 Surveillance Team |
| EPI_ISL_2119865 | Lighthouse Lab in Milton Keynes | Wellcome Sanger Institute for the COVID-19 Genomics UK (COG-UK) Consortium | The Lighthouse Lab in Milton Keynes and Alex Alderton, Roberto Amato, Jeffrey Barrett, Sonia Goncalves, Ewan Harrison, David K. Jackson, Ian Johnston, Dominic Kwiatkowski, Cordelia Langford, John Sillitoe on behalf of the Wellcome Sanger Institute COVID-19 Surveillance Team |
| EPI_ISL_2119870 | Lighthouse Lab in Glasgow | Wellcome Sanger Institute for the COVID-19 Genomics UK (COG-UK) Consortium | Harper VanSteenhouse, Yumi Kasai, David Gray, Carol Clugston, Anna Dominiczak and Alex Alderton, Roberto Amato, Jeffrey Barrett, Sonia Goncalves, Ewan Harrison, David K. Jackson, Ian Johnston, Dominic Kwiatkowski, Cordelia Langford, John Sillitoe on behalf of the Wellcome Sanger Institute COVID-19 Surveillance Team |
| EPI_ISL_2119873, EPI_ISL_2119902 | Lighthouse Lab in Alderley Park | Wellcome Sanger Institute for the COVID-19 Genomics UK (COG-UK) Consortium | Jacquelyn Wynn, Mairead Hyland, The Lighthouse Lab in Alderley Park and Alex Alderton, Roberto Amato, Jeffrey Barrett, Sonia Goncalves, Ewan Harrison, David K. Jackson, Ian Johnston, Dominic Kwiatkowski, Cordelia Langford, John Sillitoe on behalf of the Wellcome Sanger Institute COVID-19 Surveillance Team |
| EPI_ISL_2119906, EPI_ISL_2119907, EPI_ISL_2119910 | Lighthouse Lab in Milton Keynes | Wellcome Sanger Institute for the COVID-19 Genomics UK (COG-UK) Consortium | The Lighthouse Lab in Milton Keynes and Alex Alderton, Roberto Amato, Jeffrey Barrett, Sonia Goncalves, Ewan Harrison, David K. Jackson, Ian Johnston, Dominic Kwiatkowski, Cordelia Langford, John Sillitoe on behalf of the Wellcome Sanger Institute COVID-19 Surveillance Team |
| EPI_ISL_2119915, EPI_ISL_2119919 | Lighthouse Lab in Alderley Park | Wellcome Sanger Institute for the COVID-19 Genomics UK (COG-UK) Consortium | Jacquelyn Wynn, Mairead Hyland, The Lighthouse Lab in Alderley Park and Alex Alderton, Roberto Amato, Jeffrey Barrett, Sonia Goncalves, Ewan Harrison, David K. Jackson, Ian Johnston, Dominic Kwiatkowski, Cordelia Langford, John Sillitoe on behalf of the Wellcome Sanger Institute COVID-19 Surveillance Team |
| EPI_ISL_2119942, EPI_ISL_2119957, EPI_ISL_2119965 | Lighthouse Lab in Milton Keynes | Wellcome Sanger Institute for the COVID-19 Genomics UK (COG-UK) Consortium | The Lighthouse Lab in Milton Keynes and Alex Alderton, Roberto Amato, Jeffrey Barrett, Sonia Goncalves, Ewan Harrison, David K. Jackson, Ian Johnston, Dominic Kwiatkowski, Cordelia Langford, John Sillitoe on behalf of the Wellcome Sanger Institute COVID-19 Surveillance Team |
| EPI_ISL_2119967 | Lighthouse Lab in Glasgow | Wellcome Sanger Institute for the COVID-19 Genomics UK (COG-UK) Consortium | Harper VanSteenhouse, Yumi Kasai, David Gray, Carol Clugston, Anna Dominiczak and Alex Alderton, Roberto Amato, Jeffrey Barrett, Sonia Goncalves, Ewan Harrison, David K. Jackson, Ian Johnston, Dominic Kwiatkowski, Cordelia Langford, John Sillitoe on behalf of the Wellcome Sanger Institute COVID-19 Surveillance Team |
| EPI_ISL_2119979, EPI_ISL_2119982, EPI_ISL_2119985, EPI_ISL_2119990, EPI_ISL_2120000, EPI_ISL_2120006, EPI_ISL_2120008, EPI_ISL_2120013, EPI_ISL_2120027 | Lighthouse Lab in Milton Keynes | Wellcome Sanger Institute for the COVID-19 Genomics UK (COG-UK) Consortium | The Lighthouse Lab in Milton Keynes and Alex Alderton, Roberto Amato, Jeffrey Barrett, Sonia Goncalves, Ewan Harrison, David K. Jackson, Ian Johnston, Dominic Kwiatkowski, Cordelia Langford, John Sillitoe on behalf of the Wellcome Sanger Institute COVID-19 Surveillance Team |
| EPI_ISL_2120035 | Lighthouse Lab in Glasgow | Wellcome Sanger Institute for the COVID-19 Genomics UK (COG-UK) Consortium | Harper VanSteenhouse, Yumi Kasai, David Gray, Carol Clugston, Anna Dominiczak and Alex Alderton, Roberto Amato, Jeffrey Barrett, Sonia Goncalves, Ewan Harrison, David K. Jackson, Ian Johnston, Dominic Kwiatkowski, Cordelia Langford, John Sillitoe on behalf of the Wellcome Sanger Institute COVID-19 Surveillance Team |
| EPI_ISL_2120036, EPI_ISL_2120037, EPI_ISL_2120057, EPI_ISL_2120059, EPI_ISL_2120061, EPI_ISL_2120068, EPI_ISL_2120077, EPI_ISL_2120080, EPI_ISL_2120082, EPI_ISL_2120085, EPI_ISL_2120089, EPI_ISL_2120095, EPI_ISL_2120103, EPI_ISL_2120106, EPI_ISL_2120112, EPI_ISL_2120113, EPI_ISL_2120114, EPI_ISL_2120115 | | | |
| see above | Lighthouse Lab in Milton Keynes | Wellcome Sanger Institute for the COVID-19 Genomics UK (COG-UK) Consortium | The Lighthouse Lab in Milton Keynes and Alex Alderton, Roberto Amato, Jeffrey Barrett, Sonia Goncalves, Ewan Harrison, David K. Jackson, Ian Johnston, Dominic Kwiatkowski, Cordelia Langford, John Sillitoe on behalf of the Wellcome Sanger Institute COVID-19 Surveillance Team |
| EPI_ISL_2120127, EPI_ISL_2120130 | Lighthouse Lab in Alderley Park | Wellcome Sanger Institute for the COVID-19 Genomics UK (COG-UK) Consortium | Jacquelyn Wynn, Mairead Hyland, The Lighthouse Lab in Alderley Park and Alex Alderton, Roberto Amato, Jeffrey Barrett, Sonia Goncalves, Ewan Harrison, David K. Jackson, Ian Johnston, Dominic Kwiatkowski, Cordelia Langford, John Sillitoe on behalf of the Wellcome Sanger Institute COVID-19 Surveillance Team |
| EPI_ISL_2120132 | Lighthouse Lab in Milton Keynes | Wellcome Sanger Institute for the COVID-19 Genomics UK (COG-UK) Consortium | The Lighthouse Lab in Milton Keynes and Alex Alderton, Roberto Amato, Jeffrey Barrett, Sonia Goncalves, Ewan Harrison, David K. Jackson, Ian Johnston, Dominic Kwiatkowski, Cordelia Langford, John Sillitoe on behalf of the Wellcome Sanger Institute COVID-19 Surveillance Team |
| EPI_ISL_2120136 | Lighthouse Lab in Alderley Park | Wellcome Sanger Institute for the COVID-19 Genomics UK (COG-UK) Consortium | Jacquelyn Wynn, Mairead Hyland, The Lighthouse Lab in Alderley Park and Alex Alderton, Roberto Amato, Jeffrey Barrett, Sonia Goncalves, Ewan Harrison, David K. Jackson, Ian Johnston, Dominic Kwiatkowski, Cordelia Langford, John Sillitoe on behalf of the Wellcome Sanger Institute COVID-19 Surveillance Team |
| EPI_ISL_2120150, EPI_ISL_2120152, EPI_ISL_2120157, EPI_ISL_2120168, EPI_ISL_2120173, EPI_ISL_2120176 | Lighthouse Lab in Milton Keynes | Wellcome Sanger Institute for the COVID-19 Genomics UK (COG-UK) Consortium | The Lighthouse Lab in Milton Keynes and Alex Alderton, Roberto Amato, Jeffrey Barrett, Sonia Goncalves, Ewan Harrison, David K. Jackson, Ian Johnston, Dominic Kwiatkowski, Cordelia Langford, John Sillitoe on behalf of the Wellcome Sanger Institute COVID-19 Surveillance Team |
| EPI_ISL_2120178 | Lighthouse Lab in Alderley Park | Wellcome Sanger Institute for the COVID-19 Genomics UK (COG-UK) Consortium | Jacquelyn Wynn, Mairead Hyland, The Lighthouse Lab in Alderley Park and Alex Alderton, Roberto Amato, Jeffrey Barrett, Sonia Goncalves, Ewan Harrison, David K. Jackson, Ian Johnston, Dominic Kwiatkowski, Cordelia Langford, John Sillitoe on behalf of the Wellcome Sanger Institute COVID-19 Surveillance Team |
| EPI_ISL_2120180, EPI_ISL_2120190 | Labormedizin Darmstadt | Robert Koch Institute | unknown |
| EPI_ISL_2120196, EPI_ISL_2120198 | Lighthouse Lab in Milton Keynes | Wellcome Sanger Institute for the COVID-19 Genomics UK (COG-UK) Consortium | The Lighthouse Lab in Milton Keynes and Alex Alderton, Roberto Amato, Jeffrey Barrett, Sonia Goncalves, Ewan Harrison, David K. Jackson, Ian Johnston, Dominic Kwiatkowski, Cordelia Langford, John Sillitoe on behalf of the Wellcome Sanger Institute COVID-19 Surveillance Team |
| EPI_ISL_2120199, EPI_ISL_2120201, EPI_ISL_2120204 | Lighthouse Lab in Alderley Park | Wellcome Sanger Institute for the COVID-19 Genomics UK (COG-UK) Consortium | Jacquelyn Wynn, Mairead Hyland, The Lighthouse Lab in Alderley Park and Alex Alderton, Roberto Amato, Jeffrey Barrett, Sonia Goncalves, Ewan Harrison, David K. Jackson, Ian Johnston, Dominic Kwiatkowski, Cordelia Langford, John Sillitoe on behalf of the Wellcome Sanger Institute COVID-19 Surveillance Team |
| EPI_ISL_2120211, EPI_ISL_2120214, EPI_ISL_2120227 | Lighthouse Lab in Milton Keynes | Wellcome Sanger Institute for the COVID-19 Genomics UK (COG-UK) Consortium | The Lighthouse Lab in Milton Keynes and Alex Alderton, Roberto Amato, Jeffrey Barrett, Sonia Goncalves, Ewan Harrison, David K. Jackson, Ian Johnston, Dominic Kwiatkowski, Cordelia Langford, John Sillitoe on behalf of the Wellcome Sanger Institute COVID-19 Surveillance Team |
| EPI_ISL_2120229 | Lighthouse Lab in Glasgow | Wellcome Sanger Institute for the COVID-19 Genomics UK (COG-UK) Consortium | Harper VanSteenhouse, Yumi Kasai, David Gray, Carol Clugston, Anna Dominiczak and Alex Alderton, Roberto Amato, Jeffrey Barrett, Sonia Goncalves, Ewan Harrison, David K. Jackson, Ian Johnston, Dominic Kwiatkowski, Cordelia Langford, John Sillitoe on behalf of the Wellcome Sanger Institute COVID-19 Surveillance Team |
| EPI_ISL_2120236, EPI_ISL_2120239, EPI_ISL_2120244 | Lighthouse Lab in Milton Keynes | Wellcome Sanger Institute for the COVID-19 Genomics UK (COG-UK) Consortium | The Lighthouse Lab in Milton Keynes and Alex Alderton, Roberto Amato, Jeffrey Barrett, Sonia Goncalves, Ewan Harrison, David K. Jackson, Ian Johnston, Dominic Kwiatkowski, Cordelia Langford, John Sillitoe on behalf of the Wellcome Sanger Institute COVID-19 Surveillance Team |
| EPI_ISL_2120248 | Lighthouse Lab in Alderley Park | Wellcome Sanger Institute for the COVID-19 Genomics UK (COG-UK) Consortium | Jacquelyn Wynn, Mairead Hyland, The Lighthouse Lab in Alderley Park and Alex Alderton, Roberto Amato, Jeffrey Barrett, Sonia Goncalves, Ewan Harrison, David K. Jackson, Ian Johnston, Dominic Kwiatkowski, Cordelia Langford, John Sillitoe on behalf of the Wellcome Sanger Institute COVID-19 Surveillance Team |
| EPI_ISL_2120254, | Lighthouse Lab in Milton Keynes | Wellcome Sanger Institute for the COVID-19 Genomics UK (COG-UK) | The Lighthouse Lab in Milton Keynes and Alex Alderton, Roberto Amato, Jeffrey Barrett, Sonia Goncalves, Ewan Harrison, David K. Jackson, Ian Johnston, Dominic Kwiatkowski, Cordelia Langford, John Sillitoe on behalf of the Wellcome Sanger |

| | | | |
|---|---|---|---|
| EPI_ISL_2120256 | | Consortium | Institute COVID-19 Surveillance Team |
| EPI_ISL_2120258 | Labormedizin Darmstadt | Robert Koch Institute | unknown |
| EPI_ISL_2120264 | Lighthouse Lab in Milton Keynes | Wellcome Sanger Institute for the COVID-19 Genomics UK (COG-UK) Consortium | The Lighthouse Lab in Milton Keynes and Alex Alderton, Roberto Amato, Jeffrey Barrett, Sonia Goncalves, Ewan Harrison, David K. Jackson, Ian Johnston, Dominic Kwiatkowski, Cordelia Langford, John Sillitoe on behalf of the Wellcome Sanger Institute COVID-19 Surveillance Team |
| EPI_ISL_2120265 | Lighthouse Lab in Alderley Park | Wellcome Sanger Institute for the COVID-19 Genomics UK (COG-UK) Consortium | Jacquelyn Wynn, Mairead Hyland, The Lighthouse Lab in Alderley Park and Alex Alderton, Roberto Amato, Jeffrey Barrett, Sonia Goncalves, Ewan Harrison, David K. Jackson, Ian Johnston, Dominic Kwiatkowski, Cordelia Langford, John Sillitoe on behalf of the Wellcome Sanger Institute COVID-19 Surveillance Team |
| EPI_ISL_2120274, EPI_ISL_2120285, EPI_ISL_2120289, EPI_ISL_2120305, EPI_ISL_2120311, EPI_ISL_2120314, EPI_ISL_2120316, EPI_ISL_2120337, EPI_ISL_2120350, EPI_ISL_2120352, EPI_ISL_2120379 | | | |
| see above | Lighthouse Lab in Milton Keynes | Wellcome Sanger Institute for the COVID-19 Genomics UK (COG-UK) Consortium | The Lighthouse Lab in Milton Keynes and Alex Alderton, Roberto Amato, Jeffrey Barrett, Sonia Goncalves, Ewan Harrison, David K. Jackson, Ian Johnston, Dominic Kwiatkowski, Cordelia Langford, John Sillitoe on behalf of the Wellcome Sanger Institute COVID-19 Surveillance Team |
| EPI_ISL_2120382, EPI_ISL_2120385, EPI_ISL_2120387 | Lighthouse Lab in Alderley Park | Wellcome Sanger Institute for the COVID-19 Genomics UK (COG-UK) Consortium | Jacquelyn Wynn, Mairead Hyland, The Lighthouse Lab in Alderley Park and Alex Alderton, Roberto Amato, Jeffrey Barrett, Sonia Goncalves, Ewan Harrison, David K. Jackson, Ian Johnston, Dominic Kwiatkowski, Cordelia Langford, John Sillitoe on behalf of the Wellcome Sanger Institute COVID-19 Surveillance Team |
| EPI_ISL_2120388, EPI_ISL_2120397, EPI_ISL_2120403, EPI_ISL_2120413, EPI_ISL_2120423, EPI_ISL_2120428, EPI_ISL_2120430, EPI_ISL_2120437, EPI_ISL_2120442, EPI_ISL_2120449 | Lighthouse Lab in Milton Keynes | Wellcome Sanger Institute for the COVID-19 Genomics UK (COG-UK) Consortium | The Lighthouse Lab in Milton Keynes and Alex Alderton, Roberto Amato, Jeffrey Barrett, Sonia Goncalves, Ewan Harrison, David K. Jackson, Ian Johnston, Dominic Kwiatkowski, Cordelia Langford, John Sillitoe on behalf of the Wellcome Sanger Institute COVID-19 Surveillance Team |
| EPI_ISL_2120453, EPI_ISL_2120461, EPI_ISL_2120464, EPI_ISL_2120468, EPI_ISL_2120480, EPI_ISL_2120482, EPI_ISL_2120483 | Lighthouse Lab in Alderley Park | Wellcome Sanger Institute for the COVID-19 Genomics UK (COG-UK) Consortium | Jacquelyn Wynn, Mairead Hyland, The Lighthouse Lab in Alderley Park and Alex Alderton, Roberto Amato, Jeffrey Barrett, Sonia Goncalves, Ewan Harrison, David K. Jackson, Ian Johnston, Dominic Kwiatkowski, Cordelia Langford, John Sillitoe on behalf of the Wellcome Sanger Institute COVID-19 Surveillance Team |
| EPI_ISL_2120487 | Lighthouse Lab in Milton Keynes | Wellcome Sanger Institute for the COVID-19 Genomics UK (COG-UK) Consortium | The Lighthouse Lab in Milton Keynes and Alex Alderton, Roberto Amato, Jeffrey Barrett, Sonia Goncalves, Ewan Harrison, David K. Jackson, Ian Johnston, Dominic Kwiatkowski, Cordelia Langford, John Sillitoe on behalf of the Wellcome Sanger Institute COVID-19 Surveillance Team |
| EPI_ISL_2120497 | Lighthouse Lab in Glasgow | Wellcome Sanger Institute for the COVID-19 Genomics UK (COG-UK) Consortium | Harper VanSteenhouse, Yumi Kasai, David Gray, Carol Clugston, Anna Dominiczak and Alex Alderton, Roberto Amato, Jeffrey Barrett, Sonia Goncalves, Ewan Harrison, David K. Jackson, Ian Johnston, Dominic Kwiatkowski, Cordelia Langford, John Sillitoe on behalf of the Wellcome Sanger Institute COVID-19 Surveillance Team |
| EPI_ISL_2120498, EPI_ISL_2120505, EPI_ISL_2120517, EPI_ISL_2120518 | Lighthouse Lab in Alderley Park | Wellcome Sanger Institute for the COVID-19 Genomics UK (COG-UK) Consortium | Jacquelyn Wynn, Mairead Hyland, The Lighthouse Lab in Alderley Park and Alex Alderton, Roberto Amato, Jeffrey Barrett, Sonia Goncalves, Ewan Harrison, David K. Jackson, Ian Johnston, Dominic Kwiatkowski, Cordelia Langford, John Sillitoe on behalf of the Wellcome Sanger Institute COVID-19 Surveillance Team |
| EPI_ISL_2120522, EPI_ISL_2120526 | Lighthouse Lab in Milton Keynes | Wellcome Sanger Institute for the COVID-19 Genomics UK (COG-UK) Consortium | The Lighthouse Lab in Milton Keynes and Alex Alderton, Roberto Amato, Jeffrey Barrett, Sonia Goncalves, Ewan Harrison, David K. Jackson, Ian Johnston, Dominic Kwiatkowski, Cordelia Langford, John Sillitoe on behalf of the Wellcome Sanger Institute COVID-19 Surveillance Team |
| EPI_ISL_2120551, EPI_ISL_2120552, EPI_ISL_2120559, EPI_ISL_2120562, EPI_ISL_2120569, EPI_ISL_2120572 | Lighthouse Lab in Alderley Park | Wellcome Sanger Institute for the COVID-19 Genomics UK (COG-UK) Consortium | Jacquelyn Wynn, Mairead Hyland, The Lighthouse Lab in Alderley Park and Alex Alderton, Roberto Amato, Jeffrey Barrett, Sonia Goncalves, Ewan Harrison, David K. Jackson, Ian Johnston, Dominic Kwiatkowski, Cordelia Langford, John Sillitoe on behalf of the Wellcome Sanger Institute COVID-19 Surveillance Team |
| EPI_ISL_2120578 | Lighthouse Lab in Milton Keynes | Wellcome Sanger Institute for the COVID-19 Genomics UK (COG-UK) Consortium | The Lighthouse Lab in Milton Keynes and Alex Alderton, Roberto Amato, Jeffrey Barrett, Sonia Goncalves, Ewan Harrison, David K. Jackson, Ian Johnston, Dominic Kwiatkowski, Cordelia Langford, John Sillitoe on behalf of the Wellcome Sanger Institute COVID-19 Surveillance Team |
| EPI_ISL_2120586 | MVZ Labor Dr. Limbach & Kollegen GbR | Robert Koch Institute | unknown |
| EPI_ISL_2120588, EPI_ISL_2120591, EPI_ISL_2120604 | Lighthouse Lab in Alderley Park | Wellcome Sanger Institute for the COVID-19 Genomics UK (COG-UK) Consortium | Jacquelyn Wynn, Mairead Hyland, The Lighthouse Lab in Alderley Park and Alex Alderton, Roberto Amato, Jeffrey Barrett, Sonia Goncalves, Ewan Harrison, David K. Jackson, Ian Johnston, Dominic Kwiatkowski, Cordelia Langford, John Sillitoe on behalf of the Wellcome Sanger Institute COVID-19 Surveillance Team |
| EPI_ISL_2120608, EPI_ISL_2120614 | Lighthouse Lab in Milton Keynes | Wellcome Sanger Institute for the COVID-19 Genomics UK (COG-UK) Consortium | The Lighthouse Lab in Milton Keynes and Alex Alderton, Roberto Amato, Jeffrey Barrett, Sonia Goncalves, Ewan Harrison, David K. Jackson, Ian Johnston, Dominic Kwiatkowski, Cordelia Langford, John Sillitoe on behalf of the Wellcome Sanger Institute COVID-19 Surveillance Team |
| EPI_ISL_2120624, EPI_ISL_2120625 | Lighthouse Lab in Alderley Park | Wellcome Sanger Institute for the COVID-19 Genomics UK (COG-UK) Consortium | Jacquelyn Wynn, Mairead Hyland, The Lighthouse Lab in Alderley Park and Alex Alderton, Roberto Amato, Jeffrey Barrett, Sonia Goncalves, Ewan Harrison, David K. Jackson, Ian Johnston, Dominic Kwiatkowski, Cordelia Langford, John Sillitoe on behalf of the Wellcome Sanger Institute COVID-19 Surveillance Team |
| EPI_ISL_2120630 | Lighthouse Lab in Milton Keynes | Wellcome Sanger Institute for the COVID-19 Genomics UK (COG-UK) Consortium | The Lighthouse Lab in Milton Keynes and Alex Alderton, Roberto Amato, Jeffrey Barrett, Sonia Goncalves, Ewan Harrison, David K. Jackson, Ian Johnston, Dominic Kwiatkowski, Cordelia Langford, John Sillitoe on behalf of the Wellcome Sanger Institute COVID-19 Surveillance Team |
| EPI_ISL_2120633 | Lighthouse Lab in Alderley Park | Wellcome Sanger Institute for the COVID-19 Genomics UK (COG-UK) Consortium | Jacquelyn Wynn, Mairead Hyland, The Lighthouse Lab in Alderley Park and Alex Alderton, Roberto Amato, Jeffrey Barrett, Sonia Goncalves, Ewan Harrison, David K. Jackson, Ian Johnston, Dominic Kwiatkowski, Cordelia Langford, John Sillitoe on behalf of the Wellcome Sanger Institute COVID-19 Surveillance Team |
| EPI_ISL_2120636 | Lighthouse Lab in Glasgow | Wellcome Sanger Institute for the COVID-19 Genomics UK (COG-UK) Consortium | Harper VanSteenhouse, Yumi Kasai, David Gray, Carol Clugston, Anna Dominiczak and Alex Alderton, Roberto Amato, Jeffrey Barrett, Sonia Goncalves, Ewan Harrison, David K. Jackson, Ian Johnston, Dominic Kwiatkowski, Cordelia Langford, John Sillitoe on behalf of the Wellcome Sanger Institute COVID-19 Surveillance Team |
| EPI_ISL_2120670 | Bioscientia Labor Wermsdorf | Robert Koch Institute | unknown |
| EPI_ISL_2120710, EPI_ISL_2120712 | Lighthouse Lab in Alderley Park | Wellcome Sanger Institute for the COVID-19 Genomics UK (COG-UK) Consortium | Jacquelyn Wynn, Mairead Hyland, The Lighthouse Lab in Alderley Park and Alex Alderton, Roberto Amato, Jeffrey Barrett, Sonia Goncalves, Ewan Harrison, David K. Jackson, Ian Johnston, Dominic Kwiatkowski, Cordelia Langford, John Sillitoe on behalf of the Wellcome Sanger Institute COVID-19 Surveillance Team |
| EPI_ISL_2120720 | Lighthouse Lab in Glasgow | Wellcome Sanger Institute for the COVID-19 Genomics UK (COG-UK) Consortium | Harper VanSteenhouse, Yumi Kasai, David Gray, Carol Clugston, Anna Dominiczak and Alex Alderton, Roberto Amato, Jeffrey Barrett, Sonia Goncalves, Ewan Harrison, David K. Jackson, Ian Johnston, Dominic Kwiatkowski, Cordelia Langford, John Sillitoe on behalf of the Wellcome Sanger Institute COVID-19 Surveillance Team |
| EPI_ISL_2120727, EPI_ISL_2120732, EPI_ISL_2120742, EPI_ISL_2120757, EPI_ISL_2120763, EPI_ISL_2120764, EPI_ISL_2120766, EPI_ISL_2120768, EPI_ISL_2120769 | Lighthouse Lab in Alderley Park | Wellcome Sanger Institute for the COVID-19 Genomics UK (COG-UK) Consortium | Jacquelyn Wynn, Mairead Hyland, The Lighthouse Lab in Alderley Park and Alex Alderton, Roberto Amato, Jeffrey Barrett, Sonia Goncalves, Ewan Harrison, David K. Jackson, Ian Johnston, Dominic Kwiatkowski, Cordelia Langford, John Sillitoe on behalf of the Wellcome Sanger Institute COVID-19 Surveillance Team |
| EPI_ISL_2120779 | Bioscientia Labor Wermsdorf | Robert Koch Institute | unknown |
| EPI_ISL_2120784 | Lighthouse Lab in Alderley Park | Wellcome Sanger Institute for the COVID-19 Genomics UK (COG-UK) Consortium | Jacquelyn Wynn, Mairead Hyland, The Lighthouse Lab in Alderley Park and Alex Alderton, Roberto Amato, Jeffrey Barrett, Sonia Goncalves, Ewan Harrison, David K. Jackson, Ian Johnston, Dominic Kwiatkowski, Cordelia Langford, John Sillitoe on behalf of the Wellcome Sanger Institute COVID-19 Surveillance Team |
| EPI_ISL_2120790, EPI_ISL_2120798 | Lighthouse Lab in Glasgow | Wellcome Sanger Institute for the COVID-19 Genomics UK (COG-UK) Consortium | Harper VanSteenhouse, Yumi Kasai, David Gray, Carol Clugston, Anna Dominiczak and Alex Alderton, Roberto Amato, Jeffrey Barrett, Sonia Goncalves, Ewan Harrison, David K. Jackson, Ian Johnston, Dominic Kwiatkowski, Cordelia Langford, John Sillitoe on behalf of the Wellcome Sanger Institute COVID-19 Surveillance Team |
| EPI_ISL_2120803, EPI_ISL_2120822, EPI_ISL_2120824, EPI_ISL_2120831, EPI_ISL_2120834 | Lighthouse Lab in Alderley Park | Wellcome Sanger Institute for the COVID-19 Genomics UK (COG-UK) Consortium | Jacquelyn Wynn, Mairead Hyland, The Lighthouse Lab in Alderley Park and Alex Alderton, Roberto Amato, Jeffrey Barrett, Sonia Goncalves, Ewan Harrison, David K. Jackson, Ian Johnston, Dominic Kwiatkowski, Cordelia Langford, John Sillitoe on behalf of the Wellcome Sanger Institute COVID-19 Surveillance Team |
| EPI_ISL_2120846 | Lighthouse Lab in Glasgow | Wellcome Sanger Institute for the COVID-19 Genomics UK (COG-UK) Consortium | Harper VanSteenhouse, Yumi Kasai, David Gray, Carol Clugston, Anna Dominiczak and Alex Alderton, Roberto Amato, Jeffrey Barrett, Sonia Goncalves, Ewan Harrison, David K. Jackson, Ian Johnston, Dominic Kwiatkowski, Cordelia Langford, John Sillitoe on behalf of the Wellcome Sanger Institute COVID-19 Surveillance Team |
| EPI_ISL_2120851, EPI_ISL_2120854 | Lighthouse Lab in Alderley Park | Wellcome Sanger Institute for the COVID-19 Genomics UK (COG-UK) Consortium | Jacquelyn Wynn, Mairead Hyland, The Lighthouse Lab in Alderley Park and Alex Alderton, Roberto Amato, Jeffrey Barrett, Sonia Goncalves, Ewan Harrison, David K. Jackson, Ian Johnston, Dominic Kwiatkowski, Cordelia Langford, John Sillitoe on behalf of the Wellcome Sanger Institute COVID-19 Surveillance Team |
| EPI_ISL_2120861 | Lighthouse Lab in Milton Keynes | Wellcome Sanger Institute for the COVID-19 Genomics UK (COG-UK) Consortium | The Lighthouse Lab in Milton Keynes and Alex Alderton, Roberto Amato, Jeffrey Barrett, Sonia Goncalves, Ewan Harrison, David K. Jackson, Ian Johnston, Dominic Kwiatkowski, Cordelia Langford, John Sillitoe on behalf of the Wellcome Sanger Institute COVID-19 Surveillance Team |
| EPI_ISL_2120872 | Lighthouse Lab in Alderley Park | Wellcome Sanger Institute for the COVID-19 Genomics UK (COG-UK) Consortium | Jacquelyn Wynn, Mairead Hyland, The Lighthouse Lab in Alderley Park and Alex Alderton, Roberto Amato, Jeffrey Barrett, Sonia Goncalves, Ewan Harrison, David K. Jackson, Ian Johnston, Dominic Kwiatkowski, Cordelia Langford, John Sillitoe on behalf of the Wellcome Sanger Institute COVID-19 Surveillance Team |
| EPI_ISL_2120882 | Lighthouse Lab in Milton Keynes | Wellcome Sanger Institute for the COVID-19 Genomics UK (COG-UK) | The Lighthouse Lab in Milton Keynes and Alex Alderton, Roberto Amato, Jeffrey Barrett, Sonia Goncalves, Ewan Harrison, David K. Jackson, Ian Johnston, Dominic Kwiatkowski, Cordelia Langford, John Sillitoe on behalf of the Wellcome Sanger |

| | | | |
|---|---|---|---|
| | | Consortium | Institute COVID-19 Surveillance Team |
| EPI_ISL_2120888, EPI_ISL_2120889 | Lighthouse Lab in Alderley Park | Wellcome Sanger Institute for the COVID-19 Genomics UK (COG-UK) Consortium | Jacquelyn Wynn, Mairead Hyland, The Lighthouse Lab in Alderley Park and Alex Alderton, Roberto Amato, Jeffrey Barrett, Sonia Goncalves, Ewan Harrison, David K. Jackson, Ian Johnston, Dominic Kwiatkowski, Cordelia Langford, John Sillitoe on behalf of the Wellcome Sanger Institute COVID-19 Surveillance Team |
| EPI_ISL_2120891, EPI_ISL_2120902 | Lighthouse Lab in Milton Keynes | Wellcome Sanger Institute for the COVID-19 Genomics UK (COG-UK) Consortium | The Lighthouse Lab in Milton Keynes and Alex Alderton, Roberto Amato, Jeffrey Barrett, Sonia Goncalves, Ewan Harrison, David K. Jackson, Ian Johnston, Dominic Kwiatkowski, Cordelia Langford, John Sillitoe on behalf of the Wellcome Sanger Institute COVID-19 Surveillance Team |
| EPI_ISL_2120911, EPI_ISL_2120913 | Lighthouse Lab in Alderley Park | Wellcome Sanger Institute for the COVID-19 Genomics UK (COG-UK) Consortium | Jacquelyn Wynn, Mairead Hyland, The Lighthouse Lab in Alderley Park and Alex Alderton, Roberto Amato, Jeffrey Barrett, Sonia Goncalves, Ewan Harrison, David K. Jackson, Ian Johnston, Dominic Kwiatkowski, Cordelia Langford, John Sillitoe on behalf of the Wellcome Sanger Institute COVID-19 Surveillance Team |
| EPI_ISL_2120918 | Lighthouse Lab in Milton Keynes | Wellcome Sanger Institute for the COVID-19 Genomics UK (COG-UK) Consortium | The Lighthouse Lab in Milton Keynes and Alex Alderton, Roberto Amato, Jeffrey Barrett, Sonia Goncalves, Ewan Harrison, David K. Jackson, Ian Johnston, Dominic Kwiatkowski, Cordelia Langford, John Sillitoe on behalf of the Wellcome Sanger Institute COVID-19 Surveillance Team |
| EPI_ISL_2120925 | Lighthouse Lab in Alderley Park | Wellcome Sanger Institute for the COVID-19 Genomics UK (COG-UK) Consortium | Jacquelyn Wynn, Mairead Hyland, The Lighthouse Lab in Alderley Park and Alex Alderton, Roberto Amato, Jeffrey Barrett, Sonia Goncalves, Ewan Harrison, David K. Jackson, Ian Johnston, Dominic Kwiatkowski, Cordelia Langford, John Sillitoe on behalf of the Wellcome Sanger Institute COVID-19 Surveillance Team |
| EPI_ISL_2120932 | Lighthouse Lab in Milton Keynes | Wellcome Sanger Institute for the COVID-19 Genomics UK (COG-UK) Consortium | The Lighthouse Lab in Milton Keynes and Alex Alderton, Roberto Amato, Jeffrey Barrett, Sonia Goncalves, Ewan Harrison, David K. Jackson, Ian Johnston, Dominic Kwiatkowski, Cordelia Langford, John Sillitoe on behalf of the Wellcome Sanger Institute COVID-19 Surveillance Team |
| EPI_ISL_2120936, EPI_ISL_2120939, EPI_ISL_2120945, EPI_ISL_2120976 | Lighthouse Lab in Alderley Park | Wellcome Sanger Institute for the COVID-19 Genomics UK (COG-UK) Consortium | Jacquelyn Wynn, Mairead Hyland, The Lighthouse Lab in Alderley Park and Alex Alderton, Roberto Amato, Jeffrey Barrett, Sonia Goncalves, Ewan Harrison, David K. Jackson, Ian Johnston, Dominic Kwiatkowski, Cordelia Langford, John Sillitoe on behalf of the Wellcome Sanger Institute COVID-19 Surveillance Team |
| EPI_ISL_2120979 | Lighthouse Lab in Milton Keynes | Wellcome Sanger Institute for the COVID-19 Genomics UK (COG-UK) Consortium | The Lighthouse Lab in Milton Keynes and Alex Alderton, Roberto Amato, Jeffrey Barrett, Sonia Goncalves, Ewan Harrison, David K. Jackson, Ian Johnston, Dominic Kwiatkowski, Cordelia Langford, John Sillitoe on behalf of the Wellcome Sanger Institute COVID-19 Surveillance Team |
| EPI_ISL_2120987 | Lighthouse Lab in Glasgow | Wellcome Sanger Institute for the COVID-19 Genomics UK (COG-UK) Consortium | Harper VanSteenhouse, Yumi Kasai, David Gray, Carol Clugston, Anna Dominiczak and Alex Alderton, Roberto Amato, Jeffrey Barrett, Sonia Goncalves, Ewan Harrison, David K. Jackson, Ian Johnston, Dominic Kwiatkowski, Cordelia Langford, John Sillitoe on behalf of the Wellcome Sanger Institute COVID-19 Surveillance Team |
| EPI_ISL_2120989, EPI_ISL_2120995, EPI_ISL_2121001 | Lighthouse Lab in Alderley Park | Wellcome Sanger Institute for the COVID-19 Genomics UK (COG-UK) Consortium | Jacquelyn Wynn, Mairead Hyland, The Lighthouse Lab in Alderley Park and Alex Alderton, Roberto Amato, Jeffrey Barrett, Sonia Goncalves, Ewan Harrison, David K. Jackson, Ian Johnston, Dominic Kwiatkowski, Cordelia Langford, John Sillitoe on behalf of the Wellcome Sanger Institute COVID-19 Surveillance Team |
| EPI_ISL_2121007, EPI_ISL_2121011, EPI_ISL_2121021, EPI_ISL_2121025, EPI_ISL_2121032 | Lighthouse Lab in Milton Keynes | Wellcome Sanger Institute for the COVID-19 Genomics UK (COG-UK) Consortium | The Lighthouse Lab in Milton Keynes and Alex Alderton, Roberto Amato, Jeffrey Barrett, Sonia Goncalves, Ewan Harrison, David K. Jackson, Ian Johnston, Dominic Kwiatkowski, Cordelia Langford, John Sillitoe on behalf of the Wellcome Sanger Institute COVID-19 Surveillance Team |
| EPI_ISL_2121042 | Lighthouse Lab in Glasgow | Wellcome Sanger Institute for the COVID-19 Genomics UK (COG-UK) Consortium | Harper VanSteenhouse, Yumi Kasai, David Gray, Carol Clugston, Anna Dominiczak and Alex Alderton, Roberto Amato, Jeffrey Barrett, Sonia Goncalves, Ewan Harrison, David K. Jackson, Ian Johnston, Dominic Kwiatkowski, Cordelia Langford, John Sillitoe on behalf of the Wellcome Sanger Institute COVID-19 Surveillance Team |
| EPI_ISL_2121056, EPI_ISL_2121066, EPI_ISL_2121076, EPI_ISL_2121078 | Lighthouse Lab in Milton Keynes | Wellcome Sanger Institute for the COVID-19 Genomics UK (COG-UK) Consortium | The Lighthouse Lab in Milton Keynes and Alex Alderton, Roberto Amato, Jeffrey Barrett, Sonia Goncalves, Ewan Harrison, David K. Jackson, Ian Johnston, Dominic Kwiatkowski, Cordelia Langford, John Sillitoe on behalf of the Wellcome Sanger Institute COVID-19 Surveillance Team |
| EPI_ISL_2121095 | Lighthouse Lab in Alderley Park | Wellcome Sanger Institute for the COVID-19 Genomics UK (COG-UK) Consortium | Jacquelyn Wynn, Mairead Hyland, The Lighthouse Lab in Alderley Park and Alex Alderton, Roberto Amato, Jeffrey Barrett, Sonia Goncalves, Ewan Harrison, David K. Jackson, Ian Johnston, Dominic Kwiatkowski, Cordelia Langford, John Sillitoe on behalf of the Wellcome Sanger Institute COVID-19 Surveillance Team |
| EPI_ISL_2121111 | Lighthouse Lab in Milton Keynes | Wellcome Sanger Institute for the COVID-19 Genomics UK (COG-UK) Consortium | The Lighthouse Lab in Milton Keynes and Alex Alderton, Roberto Amato, Jeffrey Barrett, Sonia Goncalves, Ewan Harrison, David K. Jackson, Ian Johnston, Dominic Kwiatkowski, Cordelia Langford, John Sillitoe on behalf of the Wellcome Sanger Institute COVID-19 Surveillance Team |
| EPI_ISL_2121120 | Lighthouse Lab in Glasgow | Wellcome Sanger Institute for the COVID-19 Genomics UK (COG-UK) Consortium | Harper VanSteenhouse, Yumi Kasai, David Gray, Carol Clugston, Anna Dominiczak and Alex Alderton, Roberto Amato, Jeffrey Barrett, Sonia Goncalves, Ewan Harrison, David K. Jackson, Ian Johnston, Dominic Kwiatkowski, Cordelia Langford, John Sillitoe on behalf of the Wellcome Sanger Institute COVID-19 Surveillance Team |
| EPI_ISL_2121121 | LabKom - Labor Augsburg MVZ GmbH | Robert Koch Institute | unknown |
| EPI_ISL_2121129 | Lighthouse Lab in Glasgow | Wellcome Sanger Institute for the COVID-19 Genomics UK (COG-UK) Consortium | Harper VanSteenhouse, Yumi Kasai, David Gray, Carol Clugston, Anna Dominiczak and Alex Alderton, Roberto Amato, Jeffrey Barrett, Sonia Goncalves, Ewan Harrison, David K. Jackson, Ian Johnston, Dominic Kwiatkowski, Cordelia Langford, John Sillitoe on behalf of the Wellcome Sanger Institute COVID-19 Surveillance Team |
| EPI_ISL_2121157, EPI_ISL_2121165, EPI_ISL_2121166 | Lighthouse Lab in Milton Keynes | Wellcome Sanger Institute for the COVID-19 Genomics UK (COG-UK) Consortium | The Lighthouse Lab in Milton Keynes and Alex Alderton, Roberto Amato, Jeffrey Barrett, Sonia Goncalves, Ewan Harrison, David K. Jackson, Ian Johnston, Dominic Kwiatkowski, Cordelia Langford, John Sillitoe on behalf of the Wellcome Sanger Institute COVID-19 Surveillance Team |
| EPI_ISL_2121188 | Lighthouse Lab in Alderley Park | Wellcome Sanger Institute for the COVID-19 Genomics UK (COG-UK) Consortium | Jacquelyn Wynn, Mairead Hyland, The Lighthouse Lab in Alderley Park and Alex Alderton, Roberto Amato, Jeffrey Barrett, Sonia Goncalves, Ewan Harrison, David K. Jackson, Ian Johnston, Dominic Kwiatkowski, Cordelia Langford, John Sillitoe on behalf of the Wellcome Sanger Institute COVID-19 Surveillance Team |
| EPI_ISL_2121198, EPI_ISL_2121210 | Lighthouse Lab in Milton Keynes | Wellcome Sanger Institute for the COVID-19 Genomics UK (COG-UK) Consortium | The Lighthouse Lab in Milton Keynes and Alex Alderton, Roberto Amato, Jeffrey Barrett, Sonia Goncalves, Ewan Harrison, David K. Jackson, Ian Johnston, Dominic Kwiatkowski, Cordelia Langford, John Sillitoe on behalf of the Wellcome Sanger Institute COVID-19 Surveillance Team |
| EPI_ISL_2121218, EPI_ISL_2121225 | Lighthouse Lab in Glasgow | Wellcome Sanger Institute for the COVID-19 Genomics UK (COG-UK) Consortium | Harper VanSteenhouse, Yumi Kasai, David Gray, Carol Clugston, Anna Dominiczak and Alex Alderton, Roberto Amato, Jeffrey Barrett, Sonia Goncalves, Ewan Harrison, David K. Jackson, Ian Johnston, Dominic Kwiatkowski, Cordelia Langford, John Sillitoe on behalf of the Wellcome Sanger Institute COVID-19 Surveillance Team |
| EPI_ISL_2121230, EPI_ISL_2121237, EPI_ISL_2121242, EPI_ISL_2121246, EPI_ISL_2121266, EPI_ISL_2121273, EPI_ISL_2121277, EPI_ISL_2121286, EPI_ISL_2121291 | Lighthouse Lab in Milton Keynes | Wellcome Sanger Institute for the COVID-19 Genomics UK (COG-UK) Consortium | The Lighthouse Lab in Milton Keynes and Alex Alderton, Roberto Amato, Jeffrey Barrett, Sonia Goncalves, Ewan Harrison, David K. Jackson, Ian Johnston, Dominic Kwiatkowski, Cordelia Langford, John Sillitoe on behalf of the Wellcome Sanger Institute COVID-19 Surveillance Team |
| EPI_ISL_2121307 | LabKom - Labor an der Salzbrucke MVZ GmbH | Robert Koch Institute | unknown |
| EPI_ISL_2121309, EPI_ISL_2121315, EPI_ISL_2121318 | Lighthouse Lab in Milton Keynes | Wellcome Sanger Institute for the COVID-19 Genomics UK (COG-UK) Consortium | The Lighthouse Lab in Milton Keynes and Alex Alderton, Roberto Amato, Jeffrey Barrett, Sonia Goncalves, Ewan Harrison, David K. Jackson, Ian Johnston, Dominic Kwiatkowski, Cordelia Langford, John Sillitoe on behalf of the Wellcome Sanger Institute COVID-19 Surveillance Team |
| EPI_ISL_2121330 | Lighthouse Lab in Glasgow | Wellcome Sanger Institute for the COVID-19 Genomics UK (COG-UK) Consortium | Harper VanSteenhouse, Yumi Kasai, David Gray, Carol Clugston, Anna Dominiczak and Alex Alderton, Roberto Amato, Jeffrey Barrett, Sonia Goncalves, Ewan Harrison, David K. Jackson, Ian Johnston, Dominic Kwiatkowski, Cordelia Langford, John Sillitoe on behalf of the Wellcome Sanger Institute COVID-19 Surveillance Team |
| EPI_ISL_2121338 | Lighthouse Lab in Milton Keynes | Wellcome Sanger Institute for the COVID-19 Genomics UK (COG-UK) Consortium | The Lighthouse Lab in Milton Keynes and Alex Alderton, Roberto Amato, Jeffrey Barrett, Sonia Goncalves, Ewan Harrison, David K. Jackson, Ian Johnston, Dominic Kwiatkowski, Cordelia Langford, John Sillitoe on behalf of the Wellcome Sanger Institute COVID-19 Surveillance Team |
| EPI_ISL_2121343 | Lighthouse Lab in Glasgow | Wellcome Sanger Institute for the COVID-19 Genomics UK (COG-UK) Consortium | Harper VanSteenhouse, Yumi Kasai, David Gray, Carol Clugston, Anna Dominiczak and Alex Alderton, Roberto Amato, Jeffrey Barrett, Sonia Goncalves, Ewan Harrison, David K. Jackson, Ian Johnston, Dominic Kwiatkowski, Cordelia Langford, John Sillitoe on behalf of the Wellcome Sanger Institute COVID-19 Surveillance Team |
| EPI_ISL_2121347, EPI_ISL_2121355 | Lighthouse Lab in Milton Keynes | Wellcome Sanger Institute for the COVID-19 Genomics UK (COG-UK) Consortium | The Lighthouse Lab in Milton Keynes and Alex Alderton, Roberto Amato, Jeffrey Barrett, Sonia Goncalves, Ewan Harrison, David K. Jackson, Ian Johnston, Dominic Kwiatkowski, Cordelia Langford, John Sillitoe on behalf of the Wellcome Sanger Institute COVID-19 Surveillance Team |
| EPI_ISL_2121370 | Lighthouse Lab in Alderley Park | Wellcome Sanger Institute for the COVID-19 Genomics UK (COG-UK) Consortium | Jacquelyn Wynn, Mairead Hyland, The Lighthouse Lab in Alderley Park and Alex Alderton, Roberto Amato, Jeffrey Barrett, Sonia Goncalves, Ewan Harrison, David K. Jackson, Ian Johnston, Dominic Kwiatkowski, Cordelia Langford, John Sillitoe on behalf of the Wellcome Sanger Institute COVID-19 Surveillance Team |
| EPI_ISL_2121373, EPI_ISL_2121374, EPI_ISL_2121376, EPI_ISL_2121397 | Lighthouse Lab in Milton Keynes | Wellcome Sanger Institute for the COVID-19 Genomics UK (COG-UK) Consortium | The Lighthouse Lab in Milton Keynes and Alex Alderton, Roberto Amato, Jeffrey Barrett, Sonia Goncalves, Ewan Harrison, David K. Jackson, Ian Johnston, Dominic Kwiatkowski, Cordelia Langford, John Sillitoe on behalf of the Wellcome Sanger Institute COVID-19 Surveillance Team |
| EPI_ISL_2121408 | Lighthouse Lab in Glasgow | Wellcome Sanger Institute for the COVID-19 Genomics UK (COG-UK) Consortium | Harper VanSteenhouse, Yumi Kasai, David Gray, Carol Clugston, Anna Dominiczak and Alex Alderton, Roberto Amato, Jeffrey Barrett, Sonia Goncalves, Ewan Harrison, David K. Jackson, Ian Johnston, Dominic Kwiatkowski, Cordelia Langford, John Sillitoe on behalf of the Wellcome Sanger Institute COVID-19 Surveillance Team |
| EPI_ISL_2121413, EPI_ISL_2121420, EPI_ISL_2121425, EPI_ISL_2121429, EPI_ISL_2121436, EPI_ISL_2121437, EPI_ISL_2121473, EPI_ISL_2121479, EPI_ISL_2121483, EPI_ISL_2121484, EPI_ISL_2121498, EPI_ISL_2121500, EPI_ISL_2121509, EPI_ISL_2121513, EPI_ISL_2121514, EPI_ISL_2121525, EPI_ISL_2121527, EPI_ISL_2121533, EPI_ISL_2121536 | | | |
| see above | Lighthouse Lab in Milton Keynes | Wellcome Sanger Institute for the COVID-19 Genomics UK (COG-UK) Consortium | The Lighthouse Lab in Milton Keynes and Alex Alderton, Roberto Amato, Jeffrey Barrett, Sonia Goncalves, Ewan Harrison, David K. Jackson, Ian Johnston, Dominic Kwiatkowski, Cordelia Langford, John Sillitoe on behalf of the Wellcome Sanger Institute COVID-19 Surveillance Team |
| EPI_ISL_2121546 | Lighthouse Lab in Alderley Park | Wellcome Sanger Institute for the COVID-19 Genomics UK (COG-UK) Consortium | Jacquelyn Wynn, Mairead Hyland, The Lighthouse Lab in Alderley Park and Alex Alderton, Roberto Amato, Jeffrey Barrett, Sonia Goncalves, Ewan Harrison, David K. Jackson, Ian Johnston, Dominic Kwiatkowski, Cordelia Langford, John Sillitoe on behalf of the Wellcome Sanger Institute COVID-19 Surveillance Team |
| EPI_ISL_2121551 | Lighthouse Lab in Glasgow | Wellcome Sanger Institute for the COVID-19 Genomics UK (COG-UK) Consortium | Harper VanSteenhouse, Yumi Kasai, David Gray, Carol Clugston, Anna Dominiczak and Alex Alderton, Roberto Amato, Jeffrey Barrett, Sonia Goncalves, Ewan Harrison, David K. Jackson, Ian Johnston, Dominic Kwiatkowski, Cordelia Langford, John Sillitoe on behalf of the Wellcome Sanger Institute COVID-19 Surveillance Team |
| EPI_ISL_2121556 | Labor 28 MVZ GmbH | Robert Koch Institute | unknown |
| EPI_ISL_2121557, EPI_ISL_2121576, EPI_ISL_2121577 | Lighthouse Lab in Milton Keynes | Wellcome Sanger Institute for the COVID-19 Genomics UK (COG-UK) Consortium | The Lighthouse Lab in Milton Keynes and Alex Alderton, Roberto Amato, Jeffrey Barrett, Sonia Goncalves, Ewan Harrison, David K. Jackson, Ian Johnston, Dominic Kwiatkowski, Cordelia Langford, John Sillitoe on behalf of the Wellcome Sanger Institute COVID-19 Surveillance Team |

| | | | |
|---|---|---|---|
| EPI_ISL_2121579 | Lighthouse Lab in Glasgow | Wellcome Sanger Institute for the COVID-19 Genomics UK (COG-UK) Consortium | Harper VanSteenhouse, Yumi Kasai, David Gray, Carol Clugston, Anna Dominiczak and Alex Alderton, Roberto Amato, Jeffrey Barrett, Sonia Goncalves, Ewan Harrison, David K. Jackson, Ian Johnston, Dominic Kwiatkowski, Cordelia Langford, John Sillitoe on behalf of the Wellcome Sanger Institute COVID-19 Surveillance Team |
| EPI_ISL_2121580, EPI_ISL_2121591, EPI_ISL_2121593, EPI_ISL_2121598, EPI_ISL_2121618 | Lighthouse Lab in Milton Keynes | Wellcome Sanger Institute for the COVID-19 Genomics UK (COG-UK) Consortium | The Lighthouse Lab in Milton Keynes and Alex Alderton, Roberto Amato, Jeffrey Barrett, Sonia Goncalves, Ewan Harrison, David K. Jackson, Ian Johnston, Dominic Kwiatkowski, Cordelia Langford, John Sillitoe on behalf of the Wellcome Sanger Institute COVID-19 Surveillance Team |
| EPI_ISL_2121620, EPI_ISL_2121637 | Lighthouse Lab in Glasgow | Wellcome Sanger Institute for the COVID-19 Genomics UK (COG-UK) Consortium | Harper VanSteenhouse, Yumi Kasai, David Gray, Carol Clugston, Anna Dominiczak and Alex Alderton, Roberto Amato, Jeffrey Barrett, Sonia Goncalves, Ewan Harrison, David K. Jackson, Ian Johnston, Dominic Kwiatkowski, Cordelia Langford, John Sillitoe on behalf of the Wellcome Sanger Institute COVID-19 Surveillance Team |
| EPI_ISL_2121639, EPI_ISL_2121650, EPI_ISL_2121662 | Lighthouse Lab in Milton Keynes | Wellcome Sanger Institute for the COVID-19 Genomics UK (COG-UK) Consortium | The Lighthouse Lab in Milton Keynes and Alex Alderton, Roberto Amato, Jeffrey Barrett, Sonia Goncalves, Ewan Harrison, David K. Jackson, Ian Johnston, Dominic Kwiatkowski, Cordelia Langford, John Sillitoe on behalf of the Wellcome Sanger Institute COVID-19 Surveillance Team |
| EPI_ISL_2121663 | Lighthouse Lab in Glasgow | Wellcome Sanger Institute for the COVID-19 Genomics UK (COG-UK) Consortium | Harper VanSteenhouse, Yumi Kasai, David Gray, Carol Clugston, Anna Dominiczak and Alex Alderton, Roberto Amato, Jeffrey Barrett, Sonia Goncalves, Ewan Harrison, David K. Jackson, Ian Johnston, Dominic Kwiatkowski, Cordelia Langford, John Sillitoe on behalf of the Wellcome Sanger Institute COVID-19 Surveillance Team |
| EPI_ISL_2121668, EPI_ISL_2121671, EPI_ISL_2121680, EPI_ISL_2121684, EPI_ISL_2121688, EPI_ISL_2121691 | Lighthouse Lab in Milton Keynes | Wellcome Sanger Institute for the COVID-19 Genomics UK (COG-UK) Consortium | The Lighthouse Lab in Milton Keynes and Alex Alderton, Roberto Amato, Jeffrey Barrett, Sonia Goncalves, Ewan Harrison, David K. Jackson, Ian Johnston, Dominic Kwiatkowski, Cordelia Langford, John Sillitoe on behalf of the Wellcome Sanger Institute COVID-19 Surveillance Team |
| EPI_ISL_2121699, EPI_ISL_2121702, EPI_ISL_2121705 | Lighthouse Lab in Glasgow | Wellcome Sanger Institute for the COVID-19 Genomics UK (COG-UK) Consortium | Harper VanSteenhouse, Yumi Kasai, David Gray, Carol Clugston, Anna Dominiczak and Alex Alderton, Roberto Amato, Jeffrey Barrett, Sonia Goncalves, Ewan Harrison, David K. Jackson, Ian Johnston, Dominic Kwiatkowski, Cordelia Langford, John Sillitoe on behalf of the Wellcome Sanger Institute COVID-19 Surveillance Team |
| EPI_ISL_2121710, EPI_ISL_2121719 | Lighthouse Lab in Milton Keynes | Wellcome Sanger Institute for the COVID-19 Genomics UK (COG-UK) Consortium | The Lighthouse Lab in Milton Keynes and Alex Alderton, Roberto Amato, Jeffrey Barrett, Sonia Goncalves, Ewan Harrison, David K. Jackson, Ian Johnston, Dominic Kwiatkowski, Cordelia Langford, John Sillitoe on behalf of the Wellcome Sanger Institute COVID-19 Surveillance Team |
| EPI_ISL_2121721, EPI_ISL_2121724 | Lighthouse Lab in Glasgow | Wellcome Sanger Institute for the COVID-19 Genomics UK (COG-UK) Consortium | Harper VanSteenhouse, Yumi Kasai, David Gray, Carol Clugston, Anna Dominiczak and Alex Alderton, Roberto Amato, Jeffrey Barrett, Sonia Goncalves, Ewan Harrison, David K. Jackson, Ian Johnston, Dominic Kwiatkowski, Cordelia Langford, John Sillitoe on behalf of the Wellcome Sanger Institute COVID-19 Surveillance Team |
| EPI_ISL_2121725 | Lighthouse Lab in Milton Keynes | Wellcome Sanger Institute for the COVID-19 Genomics UK (COG-UK) Consortium | The Lighthouse Lab in Milton Keynes and Alex Alderton, Roberto Amato, Jeffrey Barrett, Sonia Goncalves, Ewan Harrison, David K. Jackson, Ian Johnston, Dominic Kwiatkowski, Cordelia Langford, John Sillitoe on behalf of the Wellcome Sanger Institute COVID-19 Surveillance Team |
| EPI_ISL_2121727, EPI_ISL_2121729, EPI_ISL_2121732, EPI_ISL_2121738 | Lighthouse Lab in Glasgow | Wellcome Sanger Institute for the COVID-19 Genomics UK (COG-UK) Consortium | Harper VanSteenhouse, Yumi Kasai, David Gray, Carol Clugston, Anna Dominiczak and Alex Alderton, Roberto Amato, Jeffrey Barrett, Sonia Goncalves, Ewan Harrison, David K. Jackson, Ian Johnston, Dominic Kwiatkowski, Cordelia Langford, John Sillitoe on behalf of the Wellcome Sanger Institute COVID-19 Surveillance Team |
| EPI_ISL_2121743, EPI_ISL_2121748, EPI_ISL_2121750, EPI_ISL_2121756, EPI_ISL_2121761, EPI_ISL_2121762, EPI_ISL_2121764, EPI_ISL_2121767, EPI_ISL_2121769, EPI_ISL_2121773 | Lighthouse Lab in Milton Keynes | Wellcome Sanger Institute for the COVID-19 Genomics UK (COG-UK) Consortium | The Lighthouse Lab in Milton Keynes and Alex Alderton, Roberto Amato, Jeffrey Barrett, Sonia Goncalves, Ewan Harrison, David K. Jackson, Ian Johnston, Dominic Kwiatkowski, Cordelia Langford, John Sillitoe on behalf of the Wellcome Sanger Institute COVID-19 Surveillance Team |
| EPI_ISL_2121777 | Lighthouse Lab in Glasgow | Wellcome Sanger Institute for the COVID-19 Genomics UK (COG-UK) Consortium | Harper VanSteenhouse, Yumi Kasai, David Gray, Carol Clugston, Anna Dominiczak and Alex Alderton, Roberto Amato, Jeffrey Barrett, Sonia Goncalves, Ewan Harrison, David K. Jackson, Ian Johnston, Dominic Kwiatkowski, Cordelia Langford, John Sillitoe on behalf of the Wellcome Sanger Institute COVID-19 Surveillance Team |
| EPI_ISL_2121778, EPI_ISL_2121781, EPI_ISL_2121782 | Lighthouse Lab in Milton Keynes | Wellcome Sanger Institute for the COVID-19 Genomics UK (COG-UK) Consortium | The Lighthouse Lab in Milton Keynes and Alex Alderton, Roberto Amato, Jeffrey Barrett, Sonia Goncalves, Ewan Harrison, David K. Jackson, Ian Johnston, Dominic Kwiatkowski, Cordelia Langford, John Sillitoe on behalf of the Wellcome Sanger Institute COVID-19 Surveillance Team |
| EPI_ISL_2121787, EPI_ISL_2121788, EPI_ISL_2121791, EPI_ISL_2121795 | Lighthouse Lab in Glasgow | Wellcome Sanger Institute for the COVID-19 Genomics UK (COG-UK) Consortium | Harper VanSteenhouse, Yumi Kasai, David Gray, Carol Clugston, Anna Dominiczak and Alex Alderton, Roberto Amato, Jeffrey Barrett, Sonia Goncalves, Ewan Harrison, David K. Jackson, Ian Johnston, Dominic Kwiatkowski, Cordelia Langford, John Sillitoe on behalf of the Wellcome Sanger Institute COVID-19 Surveillance Team |
| EPI_ISL_2121806 | Lighthouse Lab in Milton Keynes | Wellcome Sanger Institute for the COVID-19 Genomics UK (COG-UK) Consortium | The Lighthouse Lab in Milton Keynes and Alex Alderton, Roberto Amato, Jeffrey Barrett, Sonia Goncalves, Ewan Harrison, David K. Jackson, Ian Johnston, Dominic Kwiatkowski, Cordelia Langford, John Sillitoe on behalf of the Wellcome Sanger Institute COVID-19 Surveillance Team |
| EPI_ISL_2121809, EPI_ISL_2121813, EPI_ISL_2121817 | Lighthouse Lab in Glasgow | Wellcome Sanger Institute for the COVID-19 Genomics UK (COG-UK) Consortium | Harper VanSteenhouse, Yumi Kasai, David Gray, Carol Clugston, Anna Dominiczak and Alex Alderton, Roberto Amato, Jeffrey Barrett, Sonia Goncalves, Ewan Harrison, David K. Jackson, Ian Johnston, Dominic Kwiatkowski, Cordelia Langford, John Sillitoe on behalf of the Wellcome Sanger Institute COVID-19 Surveillance Team |
| EPI_ISL_2121818 | Bioscientia Labor Wermsdorf | Robert Koch Institute | unknown |
| EPI_ISL_2121826 | Lighthouse Lab in Milton Keynes | Wellcome Sanger Institute for the COVID-19 Genomics UK (COG-UK) Consortium | The Lighthouse Lab in Milton Keynes and Alex Alderton, Roberto Amato, Jeffrey Barrett, Sonia Goncalves, Ewan Harrison, David K. Jackson, Ian Johnston, Dominic Kwiatkowski, Cordelia Langford, John Sillitoe on behalf of the Wellcome Sanger Institute COVID-19 Surveillance Team |
| EPI_ISL_2121830, EPI_ISL_2121831 | Lighthouse Lab in Glasgow | Wellcome Sanger Institute for the COVID-19 Genomics UK (COG-UK) Consortium | Harper VanSteenhouse, Yumi Kasai, David Gray, Carol Clugston, Anna Dominiczak and Alex Alderton, Roberto Amato, Jeffrey Barrett, Sonia Goncalves, Ewan Harrison, David K. Jackson, Ian Johnston, Dominic Kwiatkowski, Cordelia Langford, John Sillitoe on behalf of the Wellcome Sanger Institute COVID-19 Surveillance Team |
| EPI_ISL_2121838 | Lighthouse Lab in Milton Keynes | Wellcome Sanger Institute for the COVID-19 Genomics UK (COG-UK) Consortium | The Lighthouse Lab in Milton Keynes and Alex Alderton, Roberto Amato, Jeffrey Barrett, Sonia Goncalves, Ewan Harrison, David K. Jackson, Ian Johnston, Dominic Kwiatkowski, Cordelia Langford, John Sillitoe on behalf of the Wellcome Sanger Institute COVID-19 Surveillance Team |
| EPI_ISL_2121840, EPI_ISL_2121844 | Lighthouse Lab in Glasgow | Wellcome Sanger Institute for the COVID-19 Genomics UK (COG-UK) Consortium | Harper VanSteenhouse, Yumi Kasai, David Gray, Carol Clugston, Anna Dominiczak and Alex Alderton, Roberto Amato, Jeffrey Barrett, Sonia Goncalves, Ewan Harrison, David K. Jackson, Ian Johnston, Dominic Kwiatkowski, Cordelia Langford, John Sillitoe on behalf of the Wellcome Sanger Institute COVID-19 Surveillance Team |
| EPI_ISL_2121847 | Lighthouse Lab in Milton Keynes | Wellcome Sanger Institute for the COVID-19 Genomics UK (COG-UK) Consortium | The Lighthouse Lab in Milton Keynes and Alex Alderton, Roberto Amato, Jeffrey Barrett, Sonia Goncalves, Ewan Harrison, David K. Jackson, Ian Johnston, Dominic Kwiatkowski, Cordelia Langford, John Sillitoe on behalf of the Wellcome Sanger Institute COVID-19 Surveillance Team |
| EPI_ISL_2121849 | Lighthouse Lab in Glasgow | Wellcome Sanger Institute for the COVID-19 Genomics UK (COG-UK) Consortium | Harper VanSteenhouse, Yumi Kasai, David Gray, Carol Clugston, Anna Dominiczak and Alex Alderton, Roberto Amato, Jeffrey Barrett, Sonia Goncalves, Ewan Harrison, David K. Jackson, Ian Johnston, Dominic Kwiatkowski, Cordelia Langford, John Sillitoe on behalf of the Wellcome Sanger Institute COVID-19 Surveillance Team |
| EPI_ISL_2121855, EPI_ISL_2121856, EPI_ISL_2121863, EPI_ISL_2121867 | Lighthouse Lab in Milton Keynes | Wellcome Sanger Institute for the COVID-19 Genomics UK (COG-UK) Consortium | The Lighthouse Lab in Milton Keynes and Alex Alderton, Roberto Amato, Jeffrey Barrett, Sonia Goncalves, Ewan Harrison, David K. Jackson, Ian Johnston, Dominic Kwiatkowski, Cordelia Langford, John Sillitoe on behalf of the Wellcome Sanger Institute COVID-19 Surveillance Team |
| EPI_ISL_2121869 | Lighthouse Lab in Glasgow | Wellcome Sanger Institute for the COVID-19 Genomics UK (COG-UK) Consortium | Harper VanSteenhouse, Yumi Kasai, David Gray, Carol Clugston, Anna Dominiczak and Alex Alderton, Roberto Amato, Jeffrey Barrett, Sonia Goncalves, Ewan Harrison, David K. Jackson, Ian Johnston, Dominic Kwiatkowski, Cordelia Langford, John Sillitoe on behalf of the Wellcome Sanger Institute COVID-19 Surveillance Team |
| EPI_ISL_2121870, EPI_ISL_2121872, EPI_ISL_2121877 | Lighthouse Lab in Milton Keynes | Wellcome Sanger Institute for the COVID-19 Genomics UK (COG-UK) Consortium | The Lighthouse Lab in Milton Keynes and Alex Alderton, Roberto Amato, Jeffrey Barrett, Sonia Goncalves, Ewan Harrison, David K. Jackson, Ian Johnston, Dominic Kwiatkowski, Cordelia Langford, John Sillitoe on behalf of the Wellcome Sanger Institute COVID-19 Surveillance Team |
| EPI_ISL_2121881 | Lighthouse Lab in Glasgow | Wellcome Sanger Institute for the COVID-19 Genomics UK (COG-UK) Consortium | Harper VanSteenhouse, Yumi Kasai, David Gray, Carol Clugston, Anna Dominiczak and Alex Alderton, Roberto Amato, Jeffrey Barrett, Sonia Goncalves, Ewan Harrison, David K. Jackson, Ian Johnston, Dominic Kwiatkowski, Cordelia Langford, John Sillitoe on behalf of the Wellcome Sanger Institute COVID-19 Surveillance Team |
| EPI_ISL_2121887 | Labor 28 MVZ GmbH | Robert Koch Institute | unknown |
| EPI_ISL_2121893 | Lighthouse Lab in Milton Keynes | Wellcome Sanger Institute for the COVID-19 Genomics UK (COG-UK) Consortium | The Lighthouse Lab in Milton Keynes and Alex Alderton, Roberto Amato, Jeffrey Barrett, Sonia Goncalves, Ewan Harrison, David K. Jackson, Ian Johnston, Dominic Kwiatkowski, Cordelia Langford, John Sillitoe on behalf of the Wellcome Sanger Institute COVID-19 Surveillance Team |
| EPI_ISL_2121894, EPI_ISL_2121896 | Lighthouse Lab in Glasgow | Wellcome Sanger Institute for the COVID-19 Genomics UK (COG-UK) Consortium | Harper VanSteenhouse, Yumi Kasai, David Gray, Carol Clugston, Anna Dominiczak and Alex Alderton, Roberto Amato, Jeffrey Barrett, Sonia Goncalves, Ewan Harrison, David K. Jackson, Ian Johnston, Dominic Kwiatkowski, Cordelia Langford, John Sillitoe on behalf of the Wellcome Sanger Institute COVID-19 Surveillance Team |
| EPI_ISL_2121903 | Labor 28 MVZ GmbH | Robert Koch Institute | unknown |
| EPI_ISL_2121906 | Lighthouse Lab in Milton Keynes | Wellcome Sanger Institute for the COVID-19 Genomics UK (COG-UK) Consortium | The Lighthouse Lab in Milton Keynes and Alex Alderton, Roberto Amato, Jeffrey Barrett, Sonia Goncalves, Ewan Harrison, David K. Jackson, Ian Johnston, Dominic Kwiatkowski, Cordelia Langford, John Sillitoe on behalf of the Wellcome Sanger Institute COVID-19 Surveillance Team |
| EPI_ISL_2121917 | Lighthouse Lab in Glasgow | Wellcome Sanger Institute for the COVID-19 Genomics UK (COG-UK) Consortium | Harper VanSteenhouse, Yumi Kasai, David Gray, Carol Clugston, Anna Dominiczak and Alex Alderton, Roberto Amato, Jeffrey Barrett, Sonia Goncalves, Ewan Harrison, David K. Jackson, Ian Johnston, Dominic Kwiatkowski, Cordelia Langford, John Sillitoe on behalf of the Wellcome Sanger Institute COVID-19 Surveillance Team |
| EPI_ISL_2121922 | Lighthouse Lab in Milton Keynes | Wellcome Sanger Institute for the COVID-19 Genomics UK (COG-UK) Consortium | The Lighthouse Lab in Milton Keynes and Alex Alderton, Roberto Amato, Jeffrey Barrett, Sonia Goncalves, Ewan Harrison, David K. Jackson, Ian Johnston, Dominic Kwiatkowski, Cordelia Langford, John Sillitoe on behalf of the Wellcome Sanger Institute COVID-19 Surveillance Team |
| EPI_ISL_2121923, | Lighthouse Lab in Glasgow | Wellcome Sanger Institute for the COVID-19 Genomics UK (COG-UK) | Harper VanSteenhouse, Yumi Kasai, David Gray, Carol Clugston, Anna Dominiczak and Alex Alderton, Roberto Amato, Jeffrey Barrett, Sonia Goncalves, Ewan Harrison, David K. Jackson, Ian Johnston, Dominic Kwiatkowski, Cordelia Langford, |

| | | | |
|---|---|---|---|
| EPI_ISL_2121928 | | Consortium | John Sillitoe on behalf of the Wellcome Sanger Institute COVID-19 Surveillance Team |
| EPI_ISL_2121933 | Lighthouse Lab in Milton Keynes | Wellcome Sanger Institute for the COVID-19 Genomics UK (COG-UK) Consortium | The Lighthouse Lab in Milton Keynes and Alex Alderton, Roberto Amato, Jeffrey Barrett, Sonia Goncalves, Ewan Harrison, David K. Jackson, Ian Johnston, Dominic Kwiatkowski, Cordelia Langford, John Sillitoe on behalf of the Wellcome Sanger Institute COVID-19 Surveillance Team |
| EPI_ISL_2121950 | Lighthouse Lab in Glasgow | Wellcome Sanger Institute for the COVID-19 Genomics UK (COG-UK) Consortium | Harper VanSteenhouse, Yumi Kasai, David Gray, Carol Clugston, Anna Dominiczak and Alex Alderton, Roberto Amato, Jeffrey Barrett, Sonia Goncalves, Ewan Harrison, David K. Jackson, Ian Johnston, Dominic Kwiatkowski, Cordelia Langford, John Sillitoe on behalf of the Wellcome Sanger Institute COVID-19 Surveillance Team |
| EPI_ISL_2121952, EPI_ISL_2121966 | Lighthouse Lab in Milton Keynes | Wellcome Sanger Institute for the COVID-19 Genomics UK (COG-UK) Consortium | The Lighthouse Lab in Milton Keynes and Alex Alderton, Roberto Amato, Jeffrey Barrett, Sonia Goncalves, Ewan Harrison, David K. Jackson, Ian Johnston, Dominic Kwiatkowski, Cordelia Langford, John Sillitoe on behalf of the Wellcome Sanger Institute COVID-19 Surveillance Team |
| EPI_ISL_2121968, EPI_ISL_2121970, EPI_ISL_2121971 | Lighthouse Lab in Glasgow | Wellcome Sanger Institute for the COVID-19 Genomics UK (COG-UK) Consortium | Harper VanSteenhouse, Yumi Kasai, David Gray, Carol Clugston, Anna Dominiczak and Alex Alderton, Roberto Amato, Jeffrey Barrett, Sonia Goncalves, Ewan Harrison, David K. Jackson, Ian Johnston, Dominic Kwiatkowski, Cordelia Langford, John Sillitoe on behalf of the Wellcome Sanger Institute COVID-19 Surveillance Team |
| EPI_ISL_2121974, EPI_ISL_2121976, EPI_ISL_2121977 | Lighthouse Lab in Milton Keynes | Wellcome Sanger Institute for the COVID-19 Genomics UK (COG-UK) Consortium | The Lighthouse Lab in Milton Keynes and Alex Alderton, Roberto Amato, Jeffrey Barrett, Sonia Goncalves, Ewan Harrison, David K. Jackson, Ian Johnston, Dominic Kwiatkowski, Cordelia Langford, John Sillitoe on behalf of the Wellcome Sanger Institute COVID-19 Surveillance Team |
| EPI_ISL_2121982 | Lighthouse Lab in Glasgow | Wellcome Sanger Institute for the COVID-19 Genomics UK (COG-UK) Consortium | Harper VanSteenhouse, Yumi Kasai, David Gray, Carol Clugston, Anna Dominiczak and Alex Alderton, Roberto Amato, Jeffrey Barrett, Sonia Goncalves, Ewan Harrison, David K. Jackson, Ian Johnston, Dominic Kwiatkowski, Cordelia Langford, John Sillitoe on behalf of the Wellcome Sanger Institute COVID-19 Surveillance Team |
| EPI_ISL_2121989 | Lighthouse Lab in Milton Keynes | Wellcome Sanger Institute for the COVID-19 Genomics UK (COG-UK) Consortium | The Lighthouse Lab in Milton Keynes and Alex Alderton, Roberto Amato, Jeffrey Barrett, Sonia Goncalves, Ewan Harrison, David K. Jackson, Ian Johnston, Dominic Kwiatkowski, Cordelia Langford, John Sillitoe on behalf of the Wellcome Sanger Institute COVID-19 Surveillance Team |
| EPI_ISL_2121994 | Lighthouse Lab in Glasgow | Wellcome Sanger Institute for the COVID-19 Genomics UK (COG-UK) Consortium | Harper VanSteenhouse, Yumi Kasai, David Gray, Carol Clugston, Anna Dominiczak and Alex Alderton, Roberto Amato, Jeffrey Barrett, Sonia Goncalves, Ewan Harrison, David K. Jackson, Ian Johnston, Dominic Kwiatkowski, Cordelia Langford, John Sillitoe on behalf of the Wellcome Sanger Institute COVID-19 Surveillance Team |
| EPI_ISL_2122007, EPI_ISL_2122009 | Lighthouse Lab in Milton Keynes | Wellcome Sanger Institute for the COVID-19 Genomics UK (COG-UK) Consortium | The Lighthouse Lab in Milton Keynes and Alex Alderton, Roberto Amato, Jeffrey Barrett, Sonia Goncalves, Ewan Harrison, David K. Jackson, Ian Johnston, Dominic Kwiatkowski, Cordelia Langford, John Sillitoe on behalf of the Wellcome Sanger Institute COVID-19 Surveillance Team |
| EPI_ISL_2122013, EPI_ISL_2122016 | Lighthouse Lab in Glasgow | Wellcome Sanger Institute for the COVID-19 Genomics UK (COG-UK) Consortium | Harper VanSteenhouse, Yumi Kasai, David Gray, Carol Clugston, Anna Dominiczak and Alex Alderton, Roberto Amato, Jeffrey Barrett, Sonia Goncalves, Ewan Harrison, David K. Jackson, Ian Johnston, Dominic Kwiatkowski, Cordelia Langford, John Sillitoe on behalf of the Wellcome Sanger Institute COVID-19 Surveillance Team |
| EPI_ISL_2122018 | Lighthouse Lab in Milton Keynes | Wellcome Sanger Institute for the COVID-19 Genomics UK (COG-UK) Consortium | The Lighthouse Lab in Milton Keynes and Alex Alderton, Roberto Amato, Jeffrey Barrett, Sonia Goncalves, Ewan Harrison, David K. Jackson, Ian Johnston, Dominic Kwiatkowski, Cordelia Langford, John Sillitoe on behalf of the Wellcome Sanger Institute COVID-19 Surveillance Team |
| EPI_ISL_2122020 | Bioscientia Labor Wermsdorf | Robert Koch Institute | unknown |
| EPI_ISL_2122021 | Lighthouse Lab in Glasgow | Wellcome Sanger Institute for the COVID-19 Genomics UK (COG-UK) Consortium | Harper VanSteenhouse, Yumi Kasai, David Gray, Carol Clugston, Anna Dominiczak and Alex Alderton, Roberto Amato, Jeffrey Barrett, Sonia Goncalves, Ewan Harrison, David K. Jackson, Ian Johnston, Dominic Kwiatkowski, Cordelia Langford, John Sillitoe on behalf of the Wellcome Sanger Institute COVID-19 Surveillance Team |
| EPI_ISL_2122024, EPI_ISL_2122050, EPI_ISL_2122051 | Lighthouse Lab in Milton Keynes | Wellcome Sanger Institute for the COVID-19 Genomics UK (COG-UK) Consortium | The Lighthouse Lab in Milton Keynes and Alex Alderton, Roberto Amato, Jeffrey Barrett, Sonia Goncalves, Ewan Harrison, David K. Jackson, Ian Johnston, Dominic Kwiatkowski, Cordelia Langford, John Sillitoe on behalf of the Wellcome Sanger Institute COVID-19 Surveillance Team |
| EPI_ISL_2122055 | Lighthouse Lab in Glasgow | Wellcome Sanger Institute for the COVID-19 Genomics UK (COG-UK) Consortium | Harper VanSteenhouse, Yumi Kasai, David Gray, Carol Clugston, Anna Dominiczak and Alex Alderton, Roberto Amato, Jeffrey Barrett, Sonia Goncalves, Ewan Harrison, David K. Jackson, Ian Johnston, Dominic Kwiatkowski, Cordelia Langford, John Sillitoe on behalf of the Wellcome Sanger Institute COVID-19 Surveillance Team |
| EPI_ISL_2122056, EPI_ISL_2122058, EPI_ISL_2122063, EPI_ISL_2122071 | Lighthouse Lab in Milton Keynes | Wellcome Sanger Institute for the COVID-19 Genomics UK (COG-UK) Consortium | The Lighthouse Lab in Milton Keynes and Alex Alderton, Roberto Amato, Jeffrey Barrett, Sonia Goncalves, Ewan Harrison, David K. Jackson, Ian Johnston, Dominic Kwiatkowski, Cordelia Langford, John Sillitoe on behalf of the Wellcome Sanger Institute COVID-19 Surveillance Team |
| EPI_ISL_2122073 | Lighthouse Lab in Glasgow | Wellcome Sanger Institute for the COVID-19 Genomics UK (COG-UK) Consortium | Harper VanSteenhouse, Yumi Kasai, David Gray, Carol Clugston, Anna Dominiczak and Alex Alderton, Roberto Amato, Jeffrey Barrett, Sonia Goncalves, Ewan Harrison, David K. Jackson, Ian Johnston, Dominic Kwiatkowski, Cordelia Langford, John Sillitoe on behalf of the Wellcome Sanger Institute COVID-19 Surveillance Team |
| EPI_ISL_2122076, EPI_ISL_2122089, EPI_ISL_2122091 | Lighthouse Lab in Milton Keynes | Wellcome Sanger Institute for the COVID-19 Genomics UK (COG-UK) Consortium | The Lighthouse Lab in Milton Keynes and Alex Alderton, Roberto Amato, Jeffrey Barrett, Sonia Goncalves, Ewan Harrison, David K. Jackson, Ian Johnston, Dominic Kwiatkowski, Cordelia Langford, John Sillitoe on behalf of the Wellcome Sanger Institute COVID-19 Surveillance Team |
| EPI_ISL_2122098 | Lighthouse Lab in Glasgow | Wellcome Sanger Institute for the COVID-19 Genomics UK (COG-UK) Consortium | Harper VanSteenhouse, Yumi Kasai, David Gray, Carol Clugston, Anna Dominiczak and Alex Alderton, Roberto Amato, Jeffrey Barrett, Sonia Goncalves, Ewan Harrison, David K. Jackson, Ian Johnston, Dominic Kwiatkowski, Cordelia Langford, John Sillitoe on behalf of the Wellcome Sanger Institute COVID-19 Surveillance Team |
| EPI_ISL_2122100, EPI_ISL_2122102, EPI_ISL_2122108, EPI_ISL_2122116, EPI_ISL_2122119 | Lighthouse Lab in Milton Keynes | Wellcome Sanger Institute for the COVID-19 Genomics UK (COG-UK) Consortium | The Lighthouse Lab in Milton Keynes and Alex Alderton, Roberto Amato, Jeffrey Barrett, Sonia Goncalves, Ewan Harrison, David K. Jackson, Ian Johnston, Dominic Kwiatkowski, Cordelia Langford, John Sillitoe on behalf of the Wellcome Sanger Institute COVID-19 Surveillance Team |
| EPI_ISL_2122125 | Randox Laboratories | Wellcome Sanger Institute for the COVID-19 Genomics UK (COG-UK) Consortium | Randox Laboratories and Alex Alderton, Roberto Amato, Jeffrey Barrett, Sonia Goncalves, Ewan Harrison, David K. Jackson, Ian Johnston, Dominic Kwiatkowski, Cordelia Langford, John Sillitoe on behalf of the Wellcome Sanger Institute COVID-19 Surveillance Team |
| EPI_ISL_2122130 | Lighthouse Lab in Alderley Park | Wellcome Sanger Institute for the COVID-19 Genomics UK (COG-UK) Consortium | Jacquelyn Wynn, Mairead Hyland, The Lighthouse Lab in Alderley Park and Alex Alderton, Roberto Amato, Jeffrey Barrett, Sonia Goncalves, Ewan Harrison, David K. Jackson, Ian Johnston, Dominic Kwiatkowski, Cordelia Langford, John Sillitoe on behalf of the Wellcome Sanger Institute COVID-19 Surveillance Team |
| EPI_ISL_2122148, EPI_ISL_2122150 | Lighthouse Lab in Milton Keynes | Wellcome Sanger Institute for the COVID-19 Genomics UK (COG-UK) Consortium | The Lighthouse Lab in Milton Keynes and Alex Alderton, Roberto Amato, Jeffrey Barrett, Sonia Goncalves, Ewan Harrison, David K. Jackson, Ian Johnston, Dominic Kwiatkowski, Cordelia Langford, John Sillitoe on behalf of the Wellcome Sanger Institute COVID-19 Surveillance Team |
| EPI_ISL_2122153, EPI_ISL_2122169, EPI_ISL_2122174 | Lighthouse Lab in Glasgow | Wellcome Sanger Institute for the COVID-19 Genomics UK (COG-UK) Consortium | Harper VanSteenhouse, Yumi Kasai, David Gray, Carol Clugston, Anna Dominiczak and Alex Alderton, Roberto Amato, Jeffrey Barrett, Sonia Goncalves, Ewan Harrison, David K. Jackson, Ian Johnston, Dominic Kwiatkowski, Cordelia Langford, John Sillitoe on behalf of the Wellcome Sanger Institute COVID-19 Surveillance Team |
| EPI_ISL_2122180 | Lighthouse Lab in Milton Keynes | Wellcome Sanger Institute for the COVID-19 Genomics UK (COG-UK) Consortium | The Lighthouse Lab in Milton Keynes and Alex Alderton, Roberto Amato, Jeffrey Barrett, Sonia Goncalves, Ewan Harrison, David K. Jackson, Ian Johnston, Dominic Kwiatkowski, Cordelia Langford, John Sillitoe on behalf of the Wellcome Sanger Institute COVID-19 Surveillance Team |
| EPI_ISL_2122186 | Lighthouse Lab in Glasgow | Wellcome Sanger Institute for the COVID-19 Genomics UK (COG-UK) Consortium | Harper VanSteenhouse, Yumi Kasai, David Gray, Carol Clugston, Anna Dominiczak and Alex Alderton, Roberto Amato, Jeffrey Barrett, Sonia Goncalves, Ewan Harrison, David K. Jackson, Ian Johnston, Dominic Kwiatkowski, Cordelia Langford, John Sillitoe on behalf of the Wellcome Sanger Institute COVID-19 Surveillance Team |
| EPI_ISL_2122202, EPI_ISL_2122205 | Lighthouse Lab in Milton Keynes | Wellcome Sanger Institute for the COVID-19 Genomics UK (COG-UK) Consortium | The Lighthouse Lab in Milton Keynes and Alex Alderton, Roberto Amato, Jeffrey Barrett, Sonia Goncalves, Ewan Harrison, David K. Jackson, Ian Johnston, Dominic Kwiatkowski, Cordelia Langford, John Sillitoe on behalf of the Wellcome Sanger Institute COVID-19 Surveillance Team |
| EPI_ISL_2122206, EPI_ISL_2122217, EPI_ISL_2122218 | Lighthouse Lab in Alderley Park | Wellcome Sanger Institute for the COVID-19 Genomics UK (COG-UK) Consortium | Jacquelyn Wynn, Mairead Hyland, The Lighthouse Lab in Alderley Park and Alex Alderton, Roberto Amato, Jeffrey Barrett, Sonia Goncalves, Ewan Harrison, David K. Jackson, Ian Johnston, Dominic Kwiatkowski, Cordelia Langford, John Sillitoe on behalf of the Wellcome Sanger Institute COVID-19 Surveillance Team |
| EPI_ISL_2122220 | Lighthouse Lab in Glasgow | Wellcome Sanger Institute for the COVID-19 Genomics UK (COG-UK) Consortium | Harper VanSteenhouse, Yumi Kasai, David Gray, Carol Clugston, Anna Dominiczak and Alex Alderton, Roberto Amato, Jeffrey Barrett, Sonia Goncalves, Ewan Harrison, David K. Jackson, Ian Johnston, Dominic Kwiatkowski, Cordelia Langford, John Sillitoe on behalf of the Wellcome Sanger Institute COVID-19 Surveillance Team |
| EPI_ISL_2122229, EPI_ISL_2122235, EPI_ISL_2122245 | Lighthouse Lab in Alderley Park | Wellcome Sanger Institute for the COVID-19 Genomics UK (COG-UK) Consortium | Jacquelyn Wynn, Mairead Hyland, The Lighthouse Lab in Alderley Park and Alex Alderton, Roberto Amato, Jeffrey Barrett, Sonia Goncalves, Ewan Harrison, David K. Jackson, Ian Johnston, Dominic Kwiatkowski, Cordelia Langford, John Sillitoe on behalf of the Wellcome Sanger Institute COVID-19 Surveillance Team |
| EPI_ISL_2122250 | Lighthouse Lab in Glasgow | Wellcome Sanger Institute for the COVID-19 Genomics UK (COG-UK) Consortium | Harper VanSteenhouse, Yumi Kasai, David Gray, Carol Clugston, Anna Dominiczak and Alex Alderton, Roberto Amato, Jeffrey Barrett, Sonia Goncalves, Ewan Harrison, David K. Jackson, Ian Johnston, Dominic Kwiatkowski, Cordelia Langford, John Sillitoe on behalf of the Wellcome Sanger Institute COVID-19 Surveillance Team |
| EPI_ISL_2122276 | Lighthouse Lab in Alderley Park | Wellcome Sanger Institute for the COVID-19 Genomics UK (COG-UK) Consortium | Jacquelyn Wynn, Mairead Hyland, The Lighthouse Lab in Alderley Park and Alex Alderton, Roberto Amato, Jeffrey Barrett, Sonia Goncalves, Ewan Harrison, David K. Jackson, Ian Johnston, Dominic Kwiatkowski, Cordelia Langford, John Sillitoe on behalf of the Wellcome Sanger Institute COVID-19 Surveillance Team |
| EPI_ISL_2122279 | Lighthouse Lab in Milton Keynes | Wellcome Sanger Institute for the COVID-19 Genomics UK (COG-UK) Consortium | The Lighthouse Lab in Milton Keynes and Alex Alderton, Roberto Amato, Jeffrey Barrett, Sonia Goncalves, Ewan Harrison, David K. Jackson, Ian Johnston, Dominic Kwiatkowski, Cordelia Langford, John Sillitoe on behalf of the Wellcome Sanger Institute COVID-19 Surveillance Team |
| EPI_ISL_2122286 | Lighthouse Lab in Alderley Park | Wellcome Sanger Institute for the COVID-19 Genomics UK (COG-UK) Consortium | Jacquelyn Wynn, Mairead Hyland, The Lighthouse Lab in Alderley Park and Alex Alderton, Roberto Amato, Jeffrey Barrett, Sonia Goncalves, Ewan Harrison, David K. Jackson, Ian Johnston, Dominic Kwiatkowski, Cordelia Langford, John Sillitoe on behalf of the Wellcome Sanger Institute COVID-19 Surveillance Team |
| EPI_ISL_2122308 | Lighthouse Lab in Milton Keynes | Wellcome Sanger Institute for the COVID-19 Genomics UK (COG-UK) Consortium | The Lighthouse Lab in Milton Keynes and Alex Alderton, Roberto Amato, Jeffrey Barrett, Sonia Goncalves, Ewan Harrison, David K. Jackson, Ian Johnston, Dominic Kwiatkowski, Cordelia Langford, John Sillitoe on behalf of the Wellcome Sanger Institute COVID-19 Surveillance Team |
| EPI_ISL_2122316 | Lighthouse Lab in Glasgow | Wellcome Sanger Institute for the COVID-19 Genomics UK (COG-UK) Consortium | Harper VanSteenhouse, Yumi Kasai, David Gray, Carol Clugston, Anna Dominiczak and Alex Alderton, Roberto Amato, Jeffrey Barrett, Sonia Goncalves, Ewan Harrison, David K. Jackson, Ian Johnston, Dominic Kwiatkowski, Cordelia Langford, John Sillitoe on behalf of the Wellcome Sanger Institute COVID-19 Surveillance Team |
| EPI_ISL_2122319 | Lighthouse Lab in Milton Keynes | Wellcome Sanger Institute for the COVID-19 Genomics UK (COG-UK) Consortium | The Lighthouse Lab in Milton Keynes and Alex Alderton, Roberto Amato, Jeffrey Barrett, Sonia Goncalves, Ewan Harrison, David K. Jackson, Ian Johnston, Dominic Kwiatkowski, Cordelia Langford, John Sillitoe on behalf of the Wellcome Sanger Institute COVID-19 Surveillance Team |
| EPI_ISL_2122320, EPI_ISL_2122338 | Lighthouse Lab in Glasgow | Wellcome Sanger Institute for the COVID-19 Genomics UK (COG-UK) Consortium | Harper VanSteenhouse, Yumi Kasai, David Gray, Carol Clugston, Anna Dominiczak and Alex Alderton, Roberto Amato, Jeffrey Barrett, Sonia Goncalves, Ewan Harrison, David K. Jackson, Ian Johnston, Dominic Kwiatkowski, Cordelia Langford, John Sillitoe on behalf of the Wellcome Sanger Institute COVID-19 Surveillance Team |
| EPI_ISL_2122343 | Lighthouse Lab in Alderley Park | Wellcome Sanger Institute for the COVID-19 Genomics UK (COG-UK) Consortium | Jacquelyn Wynn, Mairead Hyland, The Lighthouse Lab in Alderley Park and Alex Alderton, Roberto Amato, Jeffrey Barrett, Sonia Goncalves, Ewan Harrison, David K. Jackson, Ian Johnston, Dominic Kwiatkowski, Cordelia Langford, John Sillitoe on behalf of the Wellcome Sanger Institute COVID-19 Surveillance Team |
| EPI_ISL_2122346 | Lighthouse Lab in Glasgow | Wellcome Sanger Institute for the COVID-19 Genomics UK (COG-UK) Consortium | Harper VanSteenhouse, Yumi Kasai, David Gray, Carol Clugston, Anna Dominiczak and Alex Alderton, Roberto Amato, Jeffrey Barrett, Sonia Goncalves, Ewan Harrison, David K. Jackson, Ian Johnston, Dominic Kwiatkowski, Cordelia Langford, John Sillitoe on behalf of the Wellcome Sanger Institute COVID-19 Surveillance Team |

| | | | |
|---|---|---|---|
| EPI_ISL_2122354 | Lighthouse Lab in Alderley Park | Wellcome Sanger Institute for the COVID-19 Genomics UK (COG-UK) Consortium | Jacquelyn Wynn, Mairead Hyland, The Lighthouse Lab in Alderley Park and Alex Alderton, Roberto Amato, Jeffrey Barrett, Sonia Goncalves, Ewan Harrison, David K. Jackson, Ian Johnston, Dominic Kwiatkowski, Cordelia Langford, John Sillitoe on behalf of the Wellcome Sanger Institute COVID-19 Surveillance Team |
| EPI_ISL_2122371 | Lighthouse Lab in Glasgow | Wellcome Sanger Institute for the COVID-19 Genomics UK (COG-UK) Consortium | Harper VanSteenhouse, Yumi Kasai, David Gray, Carol Clugston, Anna Dominiczak and Alex Alderton, Roberto Amato, Jeffrey Barrett, Sonia Goncalves, Ewan Harrison, David K. Jackson, Ian Johnston, Dominic Kwiatkowski, Cordelia Langford, John Sillitoe on behalf of the Wellcome Sanger Institute COVID-19 Surveillance Team |
| EPI_ISL_2122383, EPI_ISL_2122398 | Lighthouse Lab in Alderley Park | Wellcome Sanger Institute for the COVID-19 Genomics UK (COG-UK) Consortium | Jacquelyn Wynn, Mairead Hyland, The Lighthouse Lab in Alderley Park and Alex Alderton, Roberto Amato, Jeffrey Barrett, Sonia Goncalves, Ewan Harrison, David K. Jackson, Ian Johnston, Dominic Kwiatkowski, Cordelia Langford, John Sillitoe on behalf of the Wellcome Sanger Institute COVID-19 Surveillance Team |
| EPI_ISL_2122402 | Lighthouse Lab in Milton Keynes | Wellcome Sanger Institute for the COVID-19 Genomics UK (COG-UK) Consortium | The Lighthouse Lab in Milton Keynes and Alex Alderton, Roberto Amato, Jeffrey Barrett, Sonia Goncalves, Ewan Harrison, David K. Jackson, Ian Johnston, Dominic Kwiatkowski, Cordelia Langford, John Sillitoe on behalf of the Wellcome Sanger Institute COVID-19 Surveillance Team |
| EPI_ISL_2122406 | Lighthouse Lab in Glasgow | Wellcome Sanger Institute for the COVID-19 Genomics UK (COG-UK) Consortium | Harper VanSteenhouse, Yumi Kasai, David Gray, Carol Clugston, Anna Dominiczak and Alex Alderton, Roberto Amato, Jeffrey Barrett, Sonia Goncalves, Ewan Harrison, David K. Jackson, Ian Johnston, Dominic Kwiatkowski, Cordelia Langford, John Sillitoe on behalf of the Wellcome Sanger Institute COVID-19 Surveillance Team |
| EPI_ISL_2122414 | Lighthouse Lab in Alderley Park | Wellcome Sanger Institute for the COVID-19 Genomics UK (COG-UK) Consortium | Jacquelyn Wynn, Mairead Hyland, The Lighthouse Lab in Alderley Park and Alex Alderton, Roberto Amato, Jeffrey Barrett, Sonia Goncalves, Ewan Harrison, David K. Jackson, Ian Johnston, Dominic Kwiatkowski, Cordelia Langford, John Sillitoe on behalf of the Wellcome Sanger Institute COVID-19 Surveillance Team |
| EPI_ISL_2122418 | Lighthouse Lab in Glasgow | Wellcome Sanger Institute for the COVID-19 Genomics UK (COG-UK) Consortium | Harper VanSteenhouse, Yumi Kasai, David Gray, Carol Clugston, Anna Dominiczak and Alex Alderton, Roberto Amato, Jeffrey Barrett, Sonia Goncalves, Ewan Harrison, David K. Jackson, Ian Johnston, Dominic Kwiatkowski, Cordelia Langford, John Sillitoe on behalf of the Wellcome Sanger Institute COVID-19 Surveillance Team |
| EPI_ISL_2122421, EPI_ISL_2122438, EPI_ISL_2122461 | Lighthouse Lab in Alderley Park | Wellcome Sanger Institute for the COVID-19 Genomics UK (COG-UK) Consortium | Jacquelyn Wynn, Mairead Hyland, The Lighthouse Lab in Alderley Park and Alex Alderton, Roberto Amato, Jeffrey Barrett, Sonia Goncalves, Ewan Harrison, David K. Jackson, Ian Johnston, Dominic Kwiatkowski, Cordelia Langford, John Sillitoe on behalf of the Wellcome Sanger Institute COVID-19 Surveillance Team |
| EPI_ISL_2122468, EPI_ISL_2122470, EPI_ISL_2122473 | Lighthouse Lab in Milton Keynes | Wellcome Sanger Institute for the COVID-19 Genomics UK (COG-UK) Consortium | The Lighthouse Lab in Milton Keynes and Alex Alderton, Roberto Amato, Jeffrey Barrett, Sonia Goncalves, Ewan Harrison, David K. Jackson, Ian Johnston, Dominic Kwiatkowski, Cordelia Langford, John Sillitoe on behalf of the Wellcome Sanger Institute COVID-19 Surveillance Team |
| EPI_ISL_2122479, EPI_ISL_2122480 | Lighthouse Lab in Glasgow | Wellcome Sanger Institute for the COVID-19 Genomics UK (COG-UK) Consortium | Harper VanSteenhouse, Yumi Kasai, David Gray, Carol Clugston, Anna Dominiczak and Alex Alderton, Roberto Amato, Jeffrey Barrett, Sonia Goncalves, Ewan Harrison, David K. Jackson, Ian Johnston, Dominic Kwiatkowski, Cordelia Langford, John Sillitoe on behalf of the Wellcome Sanger Institute COVID-19 Surveillance Team |
| EPI_ISL_2122484 | Lighthouse Lab in Milton Keynes | Wellcome Sanger Institute for the COVID-19 Genomics UK (COG-UK) Consortium | The Lighthouse Lab in Milton Keynes and Alex Alderton, Roberto Amato, Jeffrey Barrett, Sonia Goncalves, Ewan Harrison, David K. Jackson, Ian Johnston, Dominic Kwiatkowski, Cordelia Langford, John Sillitoe on behalf of the Wellcome Sanger Institute COVID-19 Surveillance Team |
| EPI_ISL_2122502, EPI_ISL_2122503 | Randox Laboratories | Wellcome Sanger Institute for the COVID-19 Genomics UK (COG-UK) Consortium | Randox Laboratories and Alex Alderton, Roberto Amato, Jeffrey Barrett, Sonia Goncalves, Ewan Harrison, David K. Jackson, Ian Johnston, Dominic Kwiatkowski, Cordelia Langford, John Sillitoe on behalf of the Wellcome Sanger Institute COVID-19 Surveillance Team |
| EPI_ISL_2122510 | Lighthouse Lab in Alderley Park | Wellcome Sanger Institute for the COVID-19 Genomics UK (COG-UK) Consortium | Jacquelyn Wynn, Mairead Hyland, The Lighthouse Lab in Alderley Park and Alex Alderton, Roberto Amato, Jeffrey Barrett, Sonia Goncalves, Ewan Harrison, David K. Jackson, Ian Johnston, Dominic Kwiatkowski, Cordelia Langford, John Sillitoe on behalf of the Wellcome Sanger Institute COVID-19 Surveillance Team |
| EPI_ISL_2122512 | Lighthouse Lab in Glasgow | Wellcome Sanger Institute for the COVID-19 Genomics UK (COG-UK) Consortium | Harper VanSteenhouse, Yumi Kasai, David Gray, Carol Clugston, Anna Dominiczak and Alex Alderton, Roberto Amato, Jeffrey Barrett, Sonia Goncalves, Ewan Harrison, David K. Jackson, Ian Johnston, Dominic Kwiatkowski, Cordelia Langford, John Sillitoe on behalf of the Wellcome Sanger Institute COVID-19 Surveillance Team |
| EPI_ISL_2122523, EPI_ISL_2122529, EPI_ISL_2122530 | Lighthouse Lab in Alderley Park | Wellcome Sanger Institute for the COVID-19 Genomics UK (COG-UK) Consortium | Jacquelyn Wynn, Mairead Hyland, The Lighthouse Lab in Alderley Park and Alex Alderton, Roberto Amato, Jeffrey Barrett, Sonia Goncalves, Ewan Harrison, David K. Jackson, Ian Johnston, Dominic Kwiatkowski, Cordelia Langford, John Sillitoe on behalf of the Wellcome Sanger Institute COVID-19 Surveillance Team |
| EPI_ISL_2122543 | Lighthouse Lab in Glasgow | Wellcome Sanger Institute for the COVID-19 Genomics UK (COG-UK) Consortium | Harper VanSteenhouse, Yumi Kasai, David Gray, Carol Clugston, Anna Dominiczak and Alex Alderton, Roberto Amato, Jeffrey Barrett, Sonia Goncalves, Ewan Harrison, David K. Jackson, Ian Johnston, Dominic Kwiatkowski, Cordelia Langford, John Sillitoe on behalf of the Wellcome Sanger Institute COVID-19 Surveillance Team |
| EPI_ISL_2122547 | Lighthouse Lab in Alderley Park | Wellcome Sanger Institute for the COVID-19 Genomics UK (COG-UK) Consortium | Jacquelyn Wynn, Mairead Hyland, The Lighthouse Lab in Alderley Park and Alex Alderton, Roberto Amato, Jeffrey Barrett, Sonia Goncalves, Ewan Harrison, David K. Jackson, Ian Johnston, Dominic Kwiatkowski, Cordelia Langford, John Sillitoe on behalf of the Wellcome Sanger Institute COVID-19 Surveillance Team |
| EPI_ISL_2122554 | Lighthouse Lab in Glasgow | Wellcome Sanger Institute for the COVID-19 Genomics UK (COG-UK) Consortium | Harper VanSteenhouse, Yumi Kasai, David Gray, Carol Clugston, Anna Dominiczak and Alex Alderton, Roberto Amato, Jeffrey Barrett, Sonia Goncalves, Ewan Harrison, David K. Jackson, Ian Johnston, Dominic Kwiatkowski, Cordelia Langford, John Sillitoe on behalf of the Wellcome Sanger Institute COVID-19 Surveillance Team |
| EPI_ISL_2122562 | Lighthouse Lab in Milton Keynes | Wellcome Sanger Institute for the COVID-19 Genomics UK (COG-UK) Consortium | The Lighthouse Lab in Milton Keynes and Alex Alderton, Roberto Amato, Jeffrey Barrett, Sonia Goncalves, Ewan Harrison, David K. Jackson, Ian Johnston, Dominic Kwiatkowski, Cordelia Langford, John Sillitoe on behalf of the Wellcome Sanger Institute COVID-19 Surveillance Team |
| EPI_ISL_2122571, EPI_ISL_2122579 | Lighthouse Lab in Glasgow | Wellcome Sanger Institute for the COVID-19 Genomics UK (COG-UK) Consortium | Harper VanSteenhouse, Yumi Kasai, David Gray, Carol Clugston, Anna Dominiczak and Alex Alderton, Roberto Amato, Jeffrey Barrett, Sonia Goncalves, Ewan Harrison, David K. Jackson, Ian Johnston, Dominic Kwiatkowski, Cordelia Langford, John Sillitoe on behalf of the Wellcome Sanger Institute COVID-19 Surveillance Team |
| EPI_ISL_2122586 | Lighthouse Lab in Milton Keynes | Wellcome Sanger Institute for the COVID-19 Genomics UK (COG-UK) Consortium | The Lighthouse Lab in Milton Keynes and Alex Alderton, Roberto Amato, Jeffrey Barrett, Sonia Goncalves, Ewan Harrison, David K. Jackson, Ian Johnston, Dominic Kwiatkowski, Cordelia Langford, John Sillitoe on behalf of the Wellcome Sanger Institute COVID-19 Surveillance Team |
| EPI_ISL_2122587 | Lighthouse Lab in Alderley Park | Wellcome Sanger Institute for the COVID-19 Genomics UK (COG-UK) Consortium | Jacquelyn Wynn, Mairead Hyland, The Lighthouse Lab in Alderley Park and Alex Alderton, Roberto Amato, Jeffrey Barrett, Sonia Goncalves, Ewan Harrison, David K. Jackson, Ian Johnston, Dominic Kwiatkowski, Cordelia Langford, John Sillitoe on behalf of the Wellcome Sanger Institute COVID-19 Surveillance Team |
| EPI_ISL_2122596 | Lighthouse Lab in Glasgow | Wellcome Sanger Institute for the COVID-19 Genomics UK (COG-UK) Consortium | Harper VanSteenhouse, Yumi Kasai, David Gray, Carol Clugston, Anna Dominiczak and Alex Alderton, Roberto Amato, Jeffrey Barrett, Sonia Goncalves, Ewan Harrison, David K. Jackson, Ian Johnston, Dominic Kwiatkowski, Cordelia Langford, John Sillitoe on behalf of the Wellcome Sanger Institute COVID-19 Surveillance Team |
| EPI_ISL_2122599 | MVZ Labor Dr. Limbach & Kollegen GbR | Robert Koch Institute | unknown |
| EPI_ISL_2122602, EPI_ISL_2122629 | Lighthouse Lab in Alderley Park | Wellcome Sanger Institute for the COVID-19 Genomics UK (COG-UK) Consortium | Jacquelyn Wynn, Mairead Hyland, The Lighthouse Lab in Alderley Park and Alex Alderton, Roberto Amato, Jeffrey Barrett, Sonia Goncalves, Ewan Harrison, David K. Jackson, Ian Johnston, Dominic Kwiatkowski, Cordelia Langford, John Sillitoe on behalf of the Wellcome Sanger Institute COVID-19 Surveillance Team |
| EPI_ISL_2122636 | Randox Laboratories | Wellcome Sanger Institute for the COVID-19 Genomics UK (COG-UK) Consortium | Randox Laboratories and Alex Alderton, Roberto Amato, Jeffrey Barrett, Sonia Goncalves, Ewan Harrison, David K. Jackson, Ian Johnston, Dominic Kwiatkowski, Cordelia Langford, John Sillitoe on behalf of the Wellcome Sanger Institute COVID-19 Surveillance Team |
| EPI_ISL_2122639 | Lighthouse Lab in Milton Keynes | Wellcome Sanger Institute for the COVID-19 Genomics UK (COG-UK) Consortium | The Lighthouse Lab in Milton Keynes and Alex Alderton, Roberto Amato, Jeffrey Barrett, Sonia Goncalves, Ewan Harrison, David K. Jackson, Ian Johnston, Dominic Kwiatkowski, Cordelia Langford, John Sillitoe on behalf of the Wellcome Sanger Institute COVID-19 Surveillance Team |
| EPI_ISL_2122654, EPI_ISL_2122655 | Lighthouse Lab in Alderley Park | Wellcome Sanger Institute for the COVID-19 Genomics UK (COG-UK) Consortium | Jacquelyn Wynn, Mairead Hyland, The Lighthouse Lab in Alderley Park and Alex Alderton, Roberto Amato, Jeffrey Barrett, Sonia Goncalves, Ewan Harrison, David K. Jackson, Ian Johnston, Dominic Kwiatkowski, Cordelia Langford, John Sillitoe on behalf of the Wellcome Sanger Institute COVID-19 Surveillance Team |
| EPI_ISL_2122659 | Lighthouse Lab in Glasgow | Wellcome Sanger Institute for the COVID-19 Genomics UK (COG-UK) Consortium | Harper VanSteenhouse, Yumi Kasai, David Gray, Carol Clugston, Anna Dominiczak and Alex Alderton, Roberto Amato, Jeffrey Barrett, Sonia Goncalves, Ewan Harrison, David K. Jackson, Ian Johnston, Dominic Kwiatkowski, Cordelia Langford, John Sillitoe on behalf of the Wellcome Sanger Institute COVID-19 Surveillance Team |
| EPI_ISL_2122662 | Lighthouse Lab in Alderley Park | Wellcome Sanger Institute for the COVID-19 Genomics UK (COG-UK) Consortium | Jacquelyn Wynn, Mairead Hyland, The Lighthouse Lab in Alderley Park and Alex Alderton, Roberto Amato, Jeffrey Barrett, Sonia Goncalves, Ewan Harrison, David K. Jackson, Ian Johnston, Dominic Kwiatkowski, Cordelia Langford, John Sillitoe on behalf of the Wellcome Sanger Institute COVID-19 Surveillance Team |
| EPI_ISL_2122665 | Lighthouse Lab in Milton Keynes | Wellcome Sanger Institute for the COVID-19 Genomics UK (COG-UK) Consortium | The Lighthouse Lab in Milton Keynes and Alex Alderton, Roberto Amato, Jeffrey Barrett, Sonia Goncalves, Ewan Harrison, David K. Jackson, Ian Johnston, Dominic Kwiatkowski, Cordelia Langford, John Sillitoe on behalf of the Wellcome Sanger Institute COVID-19 Surveillance Team |
| EPI_ISL_2122669 | Lighthouse Lab in Glasgow | Wellcome Sanger Institute for the COVID-19 Genomics UK (COG-UK) Consortium | Harper VanSteenhouse, Yumi Kasai, David Gray, Carol Clugston, Anna Dominiczak and Alex Alderton, Roberto Amato, Jeffrey Barrett, Sonia Goncalves, Ewan Harrison, David K. Jackson, Ian Johnston, Dominic Kwiatkowski, Cordelia Langford, John Sillitoe on behalf of the Wellcome Sanger Institute COVID-19 Surveillance Team |
| EPI_ISL_2122677, EPI_ISL_2122680 | Lighthouse Lab in Alderley Park | Wellcome Sanger Institute for the COVID-19 Genomics UK (COG-UK) Consortium | Jacquelyn Wynn, Mairead Hyland, The Lighthouse Lab in Alderley Park and Alex Alderton, Roberto Amato, Jeffrey Barrett, Sonia Goncalves, Ewan Harrison, David K. Jackson, Ian Johnston, Dominic Kwiatkowski, Cordelia Langford, John Sillitoe on behalf of the Wellcome Sanger Institute COVID-19 Surveillance Team |
| EPI_ISL_2122681 | Lighthouse Lab in Milton Keynes | Wellcome Sanger Institute for the COVID-19 Genomics UK (COG-UK) Consortium | The Lighthouse Lab in Milton Keynes and Alex Alderton, Roberto Amato, Jeffrey Barrett, Sonia Goncalves, Ewan Harrison, David K. Jackson, Ian Johnston, Dominic Kwiatkowski, Cordelia Langford, John Sillitoe on behalf of the Wellcome Sanger Institute COVID-19 Surveillance Team |
| EPI_ISL_2122687, EPI_ISL_2122689 | Lighthouse Lab in Alderley Park | Wellcome Sanger Institute for the COVID-19 Genomics UK (COG-UK) Consortium | Jacquelyn Wynn, Mairead Hyland, The Lighthouse Lab in Alderley Park and Alex Alderton, Roberto Amato, Jeffrey Barrett, Sonia Goncalves, Ewan Harrison, David K. Jackson, Ian Johnston, Dominic Kwiatkowski, Cordelia Langford, John Sillitoe on behalf of the Wellcome Sanger Institute COVID-19 Surveillance Team |
| EPI_ISL_2122701, EPI_ISL_2122702, EPI_ISL_2122704 | Lighthouse Lab in Milton Keynes | Wellcome Sanger Institute for the COVID-19 Genomics UK (COG-UK) Consortium | The Lighthouse Lab in Milton Keynes and Alex Alderton, Roberto Amato, Jeffrey Barrett, Sonia Goncalves, Ewan Harrison, David K. Jackson, Ian Johnston, Dominic Kwiatkowski, Cordelia Langford, John Sillitoe on behalf of the Wellcome Sanger Institute COVID-19 Surveillance Team |
| EPI_ISL_2122710, EPI_ISL_2122724 | Lighthouse Lab in Glasgow | Wellcome Sanger Institute for the COVID-19 Genomics UK (COG-UK) Consortium | Harper VanSteenhouse, Yumi Kasai, David Gray, Carol Clugston, Anna Dominiczak and Alex Alderton, Roberto Amato, Jeffrey Barrett, Sonia Goncalves, Ewan Harrison, David K. Jackson, Ian Johnston, Dominic Kwiatkowski, Cordelia Langford, John Sillitoe on behalf of the Wellcome Sanger Institute COVID-19 Surveillance Team |
| EPI_ISL_2122726 | MVZ Labor Dr. Limbach & Kollegen GbR | Robert Koch Institute | unknown |
| EPI_ISL_2122728 | Lighthouse Lab in Glasgow | Wellcome Sanger Institute for the COVID-19 Genomics UK (COG-UK) Consortium | Harper VanSteenhouse, Yumi Kasai, David Gray, Carol Clugston, Anna Dominiczak and Alex Alderton, Roberto Amato, Jeffrey Barrett, Sonia Goncalves, Ewan Harrison, David K. Jackson, Ian Johnston, Dominic Kwiatkowski, Cordelia Langford, John Sillitoe on behalf of the Wellcome Sanger Institute COVID-19 Surveillance Team |
| EPI_ISL_2122746 | MDI Limbach Berlin GmbH; MVZ Labor Berlin | Robert Koch Institute | unknown |
| EPI_ISL_2122807, EPI_ISL_2122809, EPI_ISL_2122813 | Lighthouse Lab in Glasgow | Wellcome Sanger Institute for the COVID-19 Genomics UK (COG-UK) Consortium | Harper VanSteenhouse, Yumi Kasai, David Gray, Carol Clugston, Anna Dominiczak and Alex Alderton, Roberto Amato, Jeffrey Barrett, Sonia Goncalves, Ewan Harrison, David K. Jackson, Ian Johnston, Dominic Kwiatkowski, Cordelia Langford, John Sillitoe on behalf of the Wellcome Sanger Institute COVID-19 Surveillance Team |
| EPI_ISL_2122852 | Landesuntersuchungsanstalt fur das Gesundheits- und VeterinÃ¤rwesen (LUA) Sachsen | Robert Koch Institute | unknown |
| EPI_ISL_2122856, EPI_ISL_2122872, EPI_ISL_2122903, EPI_ISL_2122913 | Lighthouse Lab in Glasgow | Wellcome Sanger Institute for the COVID-19 Genomics UK (COG-UK) Consortium | Harper VanSteenhouse, Yumi Kasai, David Gray, Carol Clugston, Anna Dominiczak and Alex Alderton, Roberto Amato, Jeffrey Barrett, Sonia Goncalves, Ewan Harrison, David K. Jackson, Ian Johnston, Dominic Kwiatkowski, Cordelia Langford, John Sillitoe on behalf of the Wellcome Sanger Institute COVID-19 Surveillance Team |

| | | | |
|---|---|---|---|
| EPI_ISL_2123120, EPI_ISL_2123158, EPI_ISL_2123187, EPI_ISL_2123202, EPI_ISL_2123224, EPI_ISL_2123237, EPI_ISL_2123266, EPI_ISL_2123293, EPI_ISL_2123341, EPI_ISL_2123353 | Centogene; Dr. Bauer Laboratoriums GmbH | Robert Koch Institute | unknown |
| EPI_ISL_2123430, EPI_ISL_2123440, EPI_ISL_2123503, EPI_ISL_2123616, EPI_ISL_2123713 | SYNLAB Labor MÃ¼nchen Zentrum LMZ | Robert Koch Institute | unknown |
| EPI_ISL_2123724, EPI_ISL_2123736, EPI_ISL_2123791, EPI_ISL_2123794, EPI_ISL_2123795, EPI_ISL_2123806 | Eurofins LifeCodexx GmbH | Robert Koch Institute | unknown |
| EPI_ISL_2124051 | SYNLAB Labor MÃ¼nchen Zentrum LMZ | Robert Koch Institute | unknown |
| EPI_ISL_2124152, EPI_ISL_2124170 | SYNLAB Labor Munchen Zentrum LMZ | Robert Koch Institute | unknown |
| EPI_ISL_2124215 | SYNLAB MVZ Berlin | Robert Koch Institute | unknown |
| EPI_ISL_2124226, EPI_ISL_2124236 | SYNLAB Labor Munchen Zentrum LMZ | Robert Koch Institute | unknown |
| EPI_ISL_2124368 | MVZ Labor Dr. Reising-Ackermann und Kollegen | Robert Koch Institute | unknown |
| EPI_ISL_2125017 | Labor Dr. Wisplinghoff - Koln | Robert Koch Institute | unknown |
| EPI_ISL_2125018, EPI_ISL_2125024 | Fraunhofer-Institut fur Zelltherapie und Immunologie IZI AG Next-Generation Diagnostics | Robert Koch Institute | unknown |
| EPI_ISL_2125129 | Labor Dr. Spranger | Robert Koch Institute | unknown |
| EPI_ISL_2125446, EPI_ISL_2125447, EPI_ISL_2125449 | University of Birmingham | COVID-19 Genomics UK (COG-UK) Consortium | Institute of Microbiology, University of Birmingham: Claire McMurray, Joanne Stockton, Samuel Nicholls, Radoslaw Poplawski, Will Rowe, Josh Quick, Nicholas Loman. University of Birmingham Testing Laboratory: Celina M Whalley, Andrew Bosworth, Charlotte Poxon, Kasun Wanigasooriya, Oliver Pickles, Mike Kidd, Alex Richter, Andrew D Beggs PHE Heartlands Lab: Husam Osman, Andrew Bosworth. Queen Elizabeth Hospital: Anna Casey |
| EPI_ISL_2125456, EPI_ISL_2125458, EPI_ISL_2125459, EPI_ISL_2125461, EPI_ISL_2125463, EPI_ISL_2125465 | Department of Pathology, University of Cambridge | COVID-19 Genomics UK (COG-UK) Consortium | Aminu S. Jahun, Yasmin Chaudhry, Iliana Georgana, Myra Hosmillo, Rhys Izuagbe, William L. Hamilton, Martin D. Curran, Surendra Parmar, Ian Goodfellow |
| EPI_ISL_2125466 | Labor Dr. Schumacher MVZ | Robert Koch Institute | unknown |
| EPI_ISL_2125484 | Department of Pathology, University of Cambridge | COVID-19 Genomics UK (COG-UK) Consortium | Aminu S. Jahun, Yasmin Chaudhry, Iliana Georgana, Myra Hosmillo, Rhys Izuagbe, William L. Hamilton, Martin D. Curran, Surendra Parmar, Ian Goodfellow |
| EPI_ISL_2125642, EPI_ISL_2125659, EPI_ISL_2125661, EPI_ISL_2125662, EPI_ISL_2125711, EPI_ISL_2125713, EPI_ISL_2125715, EPI_ISL_2125717, EPI_ISL_2125744, EPI_ISL_2125745, EPI_ISL_2125748, EPI_ISL_2125749 | | | |
| see above | Virology Department, Royal Infirmary of Edinburgh, NHS Lothian / School of Biological Sciences, University of Edinburgh | COVID-19 Genomics UK (COG-UK) Consortium | McHugh M, Dewar R, Cotton S, Gallagher A, Maloney D, Rooke S, O'Toole Á, Scher E, Hill V, McCrone JT, Colquhoun R, Yu X, Jackson B, Rambaut A, Templeton K |
| EPI_ISL_2125827 | Bioscientia Labor Wermsdorf | Robert Koch Institute | unknown |
| EPI_ISL_2125917 | Labor 28 MVZ GmbH | Robert Koch Institute | unknown |
| EPI_ISL_2125921 | University Hospitals Of Leicester NHS Trust and DeepSeq Nottingham | COVID-19 Genomics UK (COG-UK) Consortium | Christopher Holmes, Paul Bird, Thomas Helmer, Karlie Fallon, Julian Tang, Jonathan Ball, Patrick McClure, Joeseph Chappell, Nadine Holmes, Matthew Carlisle, Christopher Moore, Fei Sang, Johnny Debebe, Victoria Wright, Matthew Loose |
| EPI_ISL_2125945 | LabKom - MVZ Labor Bochum MLB GmbH | Robert Koch Institute | unknown |
| EPI_ISL_2126019, EPI_ISL_2126024 | Barts Health NHS Trust | COVID-19 Genomics UK (COG-UK) Consortium | CUTINO-MOGUEL, Maria-Teresa; HARRINGTON, David; OWOYEMI, Dola; KULASEGARAN-SHYLINI, Raghavendran; BROAD, Claire; KELE, Beatrix |
| EPI_ISL_2126133 | Labor Dr. Schumacher MVZ | Robert Koch Institute | unknown |
| EPI_ISL_2126142, EPI_ISL_2126147, EPI_ISL_2126149 | University College London, Great Ormond Street Hospital for Children NHS Foundation Trust, Imperial College Healthcare NHS Trust | COVID-19 Genomics UK (COG-UK) Consortium | Sergi Castellano, Rachel Williams, Mark Kristiansen, Paola Resende Silva, Sunando Roy, Tony Brooks, Helena Tutill, Paola Niola, Patricia Dyal, Charlotte Williams, Leysa Forrest, Yasmin Panchbhaya, Jacqueline Findlay, Samuel Weeks, Julianne Brown, Kathryn Harris, Paul Randell, James Price, Alison Holmes, Judith Breuer |
| EPI_ISL_2126150 | Bioscientia MVZ Labor Karlsruhe GmbH | Robert Koch Institute | unknown |
| EPI_ISL_2126151, EPI_ISL_2126152, EPI_ISL_2126154, EPI_ISL_2126157, EPI_ISL_2126159, EPI_ISL_2126168, EPI_ISL_2126170, EPI_ISL_2126175 | University College London, Great Ormond Street Hospital for Children NHS Foundation Trust, Imperial College Healthcare NHS Trust | COVID-19 Genomics UK (COG-UK) Consortium | Sergi Castellano, Rachel Williams, Mark Kristiansen, Paola Resende Silva, Sunando Roy, Tony Brooks, Helena Tutill, Paola Niola, Patricia Dyal, Charlotte Williams, Leysa Forrest, Yasmin Panchbhaya, Jacqueline Findlay, Samuel Weeks, Julianne Brown, Kathryn Harris, Paul Randell, James Price, Alison Holmes, Judith Breuer |
| EPI_ISL_2126212, EPI_ISL_2126253, EPI_ISL_2126321, EPI_ISL_2126374, EPI_ISL_2126425 | Northumbria University / South Tees Hospitals NHS Foundation Trust / North Cumbria Integrated Care NHS Foundation Trust / North Tees and Hartlepool NHS Foundation Trust / Newcastle Hospitals NHS Foundation Trust | COVID-19 Genomics UK (COG-UK) Consortium | Darren L Smith,Andrew Nelson,Matthew Bashton,Greg R Young,Joshua Loh,John Allan,Mohammad A Tariq,Giles S Holt,Gary Black,Wen C Yew,Lynn Dover,Paul Baker,Steve Liggett,Sarah Essex,Jane Greenaway,Debra Padgett,Clive Graham,Garren Scott,Edward Barton,Emma Swindells,Brendan Payne,Jennifer Collins,Yusri Taha,Gary Eltringham |
| EPI_ISL_2126453, EPI_ISL_2126465 | SYNLAB MVZ Weiden | Robert Koch Institute | unknown |
| EPI_ISL_2126817, EPI_ISL_2126819 | Northumbria University / South Tees Hospitals NHS Foundation Trust / North Cumbria Integrated Care NHS Foundation Trust / North Tees and Hartlepool NHS Foundation Trust / Newcastle Hospitals NHS Foundation Trust | COVID-19 Genomics UK (COG-UK) Consortium | Darren L Smith,Andrew Nelson,Matthew Bashton,Greg R Young,Joshua Loh,John Allan,Mohammad A Tariq,Giles S Holt,Gary Black,Wen C Yew,Lynn Dover,Paul Baker,Steve Liggett,Sarah Essex,Jane Greenaway,Debra Padgett,Clive Graham,Garren Scott,Edward Barton,Emma Swindells,Brendan Payne,Jennifer Collins,Yusri Taha,Gary Eltringham |
| EPI_ISL_2126843 | SYNLAB MVZ Trier | Robert Koch Institute | unknown |
| EPI_ISL_2126933, EPI_ISL_2127073 | Northumbria University / South Tees Hospitals NHS Foundation Trust / North Cumbria Integrated Care NHS Foundation Trust / North Tees and Hartlepool NHS Foundation Trust / Newcastle Hospitals NHS Foundation Trust | COVID-19 Genomics UK (COG-UK) Consortium | Darren L Smith,Andrew Nelson,Matthew Bashton,Greg R Young,Joshua Loh,John Allan,Mohammad A Tariq,Giles S Holt,Gary Black,Wen C Yew,Lynn Dover,Paul Baker,Steve Liggett,Sarah Essex,Jane Greenaway,Debra Padgett,Clive Graham,Garren Scott,Edward Barton,Emma Swindells,Brendan Payne,Jennifer Collins,Yusri Taha,Gary Eltringham |
| EPI_ISL_2127123 | MVZ Labor Dr. Fenner und Kollegen (Standort Hamburg) | Robert Koch Institute | unknown |
| EPI_ISL_2127147 | Northumbria University / South Tees Hospitals NHS Foundation Trust / North Cumbria Integrated Care NHS Foundation Trust / North Tees and Hartlepool NHS Foundation Trust / Newcastle Hospitals NHS Foundation Trust | COVID-19 Genomics UK (COG-UK) Consortium | Darren L Smith,Andrew Nelson,Matthew Bashton,Greg R Young,Joshua Loh,John Allan,Mohammad A Tariq,Giles S Holt,Gary Black,Wen C Yew,Lynn Dover,Paul Baker,Steve Liggett,Sarah Essex,Jane Greenaway,Debra Padgett,Clive Graham,Garren Scott,Edward Barton,Emma Swindells,Brendan Payne,Jennifer Collins,Yusri Taha,Gary Eltringham |
| EPI_ISL_2127172 | MVZ Labor Dr. Fenner und Kollegen (Standort Hamburg) | Robert Koch Institute | unknown |
| EPI_ISL_2127187, EPI_ISL_2127189, EPI_ISL_2127195, EPI_ISL_2127214, EPI_ISL_2127289, EPI_ISL_2127292, EPI_ISL_2127303, EPI_ISL_2127336, EPI_ISL_2127340, EPI_ISL_2127346, EPI_ISL_2127449, EPI_ISL_2127467, EPI_ISL_2127480, EPI_ISL_2127491, EPI_ISL_2127503 | | | |
| see above | Northumbria University / South Tees Hospitals NHS Foundation Trust / North Cumbria Integrated Care NHS Foundation Trust / North Tees and Hartlepool NHS Foundation Trust / Newcastle Hospitals NHS Foundation Trust | COVID-19 Genomics UK (COG-UK) Consortium | Darren L Smith,Andrew Nelson,Matthew Bashton,Greg R Young,Joshua Loh,John Allan,Mohammad A Tariq,Giles S Holt,Gary Black,Wen C Yew,Lynn Dover,Paul Baker,Steve Liggett,Sarah Essex,Jane Greenaway,Debra Padgett,Clive Graham,Garren Scott,Edward Barton,Emma Swindells,Brendan Payne,Jennifer Collins,Yusri Taha,Gary Eltringham |
| EPI_ISL_2127606, | Queens Medical Centre, Clinical Microbiology Department / | COVID-19 Genomics UK (COG-UK) Consortium | Gemma Clark, Wendy Smith, Manjinder Khakh, Vicki M Fleming, Michelle M Lister, Hannah Howson-Wells, Jonathan Ball, Timothy Byaruhanga, Jayasree Dey, Emily Park, Jack Hill, Patrick McClure, Joseph Chappell, Theocharis Tsoleridis, Nadine |

| | | | |
|---|---|---|---|
| EPI_ISL_2127610, EPI_ISL_2127621, EPI_ISL_2127623, EPI_ISL_2127624 | DeepSeq Nottingham | | Holmes, Matthew Carlisle, Christopher Moore, Fei Sang, Johnny Debebe, Victoria Wright, Matthew Loose |
| EPI_ISL_2127734, EPI_ISL_2127759, EPI_ISL_2127792, EPI_ISL_2127798, EPI_ISL_2127804, EPI_ISL_2127805, EPI_ISL_2127811, EPI_ISL_2127831, EPI_ISL_2127871 | Wales Specialist Virology Centre Sequencing lab: Pathogen Genomics Unit | Public Health Wales Microbiology Cardiff Wales Specialist Virology Centre | Catherine Moore, Johnathan Evans, Laura Gifford, Malorie Perry, Simon Cottrell, Angela Marchbank, Alec Birchley, Alexander Adams, Amy Gaskin, Bree Gatica-Wilcox, Jason Coombes, Joel Southgate, Lauren Gilbert, Lee Graham, Nicole Pacchiarini, Sara Kumziene-Summerhayes, Sarah Taylor, Sophie Jones, Sara Rey, Matthew Bull, Joanne Watkins, Sally Corden, Tom Connor |
| EPI_ISL_2128467, EPI_ISL_2128473, EPI_ISL_2128478, EPI_ISL_2128479 | Virology Department, Sheffield Teaching Hospitals NHS Foundation Trust/Department of Infection, Immunity and Cardiovascular Disease, The Medical School, University of Sheffield | COVID-19 Genomics UK (COG-UK) Consortium | Thushan de Silva, Matthew Parker, Nikki Smith, Adri Angyal, Rebecca Brown, Luke Green, Rachel Tucker, Paul Parsons, Danielle Groves, Katie Johnson, Laura Carrilero, Alex Keeley, Dave Partridge, Matthew Wyles, Benjamin Lindsey, Mehmet Yavuz, Mohammad Raza, Cariad Evans |
| EPI_ISL_2128517 | Bioinformatics and Biostatistics Lab, Advanced Sequencing Facility | COVID-19 Genomics UK (COG-UK) Consortium | Aengus Stewart,Jerome Nicod,Chelsea Sawyer,Laura Cubitt,Harshil Patel,Margaret Crawford |
| EPI_ISL_2129020, EPI_ISL_2129024, EPI_ISL_2129029, EPI_ISL_2129038, EPI_ISL_2129039, EPI_ISL_2129047, EPI_ISL_2129050, EPI_ISL_2129064, EPI_ISL_2129066, EPI_ISL_2129067, EPI_ISL_2129069 | | | |
| see above | Medizinische Laboratorien Dusseldorf | Robert Koch Institute | unknown |
| EPI_ISL_2129595 | MVZ Dr. Eberhard & Partner Dortmund | Robert Koch Institute | unknown |
| EPI_ISL_2129645 | Limbach - MVZ Labor Eveld & Kollegen Essen | Robert Koch Institute | unknown |
| EPI_ISL_2130246 | SYNLAB MVZ Trier | Robert Koch Institute | unknown |
| EPI_ISL_2130723 | amedes MVZ fÃ¼r Labordiagnostik Rhein-Main | Robert Koch Institute | unknown |
| EPI_ISL_2130761 | amedes MVZ fÃ¼r Laboratoriumsmedizin Hamburg | Robert Koch Institute | unknown |
| EPI_ISL_2131156 | Bundeswehrkrankenhaus Berlin | Bundeswehr Institute of Microbiology | Markus Antwerpen, Alexandra Rehn, Mathias Walter, Malena Bestehorn-Willmann, Mike Pillukat, Sabine Zange, Enrico Georgi, Roman Wölfel |
| EPI_ISL_2131214 | WSSE Warszawa | 1. National Institute of Public Health - National Institute of Hygiene, Warsaw, Poland 2. Biobank Lab, University of Lodz 3. Laboratory of Respiratory Viruses, Teaching and Clinical Center of the Medical University of Lodz | Dominik Strapagiel, Marta Sobalska-Kwapis, Klaudyna Królikowska, Jakub Lach, Marcin Słomka, Tomasz Płoszaj, Magdalena Traczyk-Borszyńska, Izabela Dróżdż, Maciej Borowiec, Katarzyna Zacharczuk, Magdalena Nowakowska, Małgorzata Sadkowska-Todys, Tomasz Wołkowicz |
| EPI_ISL_2131296, EPI_ISL_2131297, EPI_ISL_2131298, EPI_ISL_2131303, EPI_ISL_2131310, EPI_ISL_2131345 | Platform BIS UZA/UAntwerpen | Labo Klinische Biologie, UZA | Jasmine Coppens, Marie Le Mercier, Basil Britto Xavier, Christine Lammens, Veerle Matheeussen, Herman Goossens |
| EPI_ISL_2131509, EPI_ISL_2131510, EPI_ISL_2131514, EPI_ISL_2131516, EPI_ISL_2131517, EPI_ISL_2131518, EPI_ISL_2131519, EPI_ISL_2131520, EPI_ISL_2131521, EPI_ISL_2131522, EPI_ISL_2131523, EPI_ISL_2131524, EPI_ISL_2131525, EPI_ISL_2131526, EPI_ISL_2131527, EPI_ISL_2131528, EPI_ISL_2131529, EPI_ISL_2131530, EPI_ISL_2131531, EPI_ISL_2131532, EPI_ISL_2131533, EPI_ISL_2131534, EPI_ISL_2131538, EPI_ISL_2131540, EPI_ISL_2131541, EPI_ISL_2131542, EPI_ISL_2131543, EPI_ISL_2131544, EPI_ISL_2131546, EPI_ISL_2131547, EPI_ISL_2131548, EPI_ISL_2131549, EPI_ISL_2131550, EPI_ISL_2131551, EPI_ISL_2131552, EPI_ISL_2131553, EPI_ISL_2131554, EPI_ISL_2131555, EPI_ISL_2131556, EPI_ISL_2131557, EPI_ISL_2131558, EPI_ISL_2131559, EPI_ISL_2131560, EPI_ISL_2131561, EPI_ISL_2131562, EPI_ISL_2131563, EPI_ISL_2131564, EPI_ISL_2131565, EPI_ISL_2131566, EPI_ISL_2131567, EPI_ISL_2131568, EPI_ISL_2131569, EPI_ISL_2131570, EPI_ISL_2131571, EPI_ISL_2131572 | | | |
| see above | ICMR-National Institute of Virology - INSACOG | NIV Influenza | Dr. Varsha Potdar |
| EPI_ISL_2131667, EPI_ISL_2131668, EPI_ISL_2131671, EPI_ISL_2131672, EPI_ISL_2131673, EPI_ISL_2131676, EPI_ISL_2131677, EPI_ISL_2131690, EPI_ISL_2131691, EPI_ISL_2131692, EPI_ISL_2131693, EPI_ISL_2131694, EPI_ISL_2131695, EPI_ISL_2131696, EPI_ISL_2131697, EPI_ISL_2131698, EPI_ISL_2131699, EPI_ISL_2131700, EPI_ISL_2131701, EPI_ISL_2131702, EPI_ISL_2131703, EPI_ISL_2131704, EPI_ISL_2131705, EPI_ISL_2131706, EPI_ISL_2131707, EPI_ISL_2131708, EPI_ISL_2131709, EPI_ISL_2131710, EPI_ISL_2131711, EPI_ISL_2131712, EPI_ISL_2131713, EPI_ISL_2131714, EPI_ISL_2131715, EPI_ISL_2131716, EPI_ISL_2131717, EPI_ISL_2131718, EPI_ISL_2131719, EPI_ISL_2131720, EPI_ISL_2131721, EPI_ISL_2131722, EPI_ISL_2131723, EPI_ISL_2131724, EPI_ISL_2131725, EPI_ISL_2131726, EPI_ISL_2131727, EPI_ISL_2131728, EPI_ISL_2131729, EPI_ISL_2131730, EPI_ISL_2131731, EPI_ISL_2131732, EPI_ISL_2131733, EPI_ISL_2131734, EPI_ISL_2131735, EPI_ISL_2131736, EPI_ISL_2131737, EPI_ISL_2131738, EPI_ISL_2131739, EPI_ISL_2131740, EPI_ISL_2131741, EPI_ISL_2131742, EPI_ISL_2131743, EPI_ISL_2131744, EPI_ISL_2131745, EPI_ISL_2131746, EPI_ISL_2131747, EPI_ISL_2131748, EPI_ISL_2131749, EPI_ISL_2131750, EPI_ISL_2131751, EPI_ISL_2131752, EPI_ISL_2131753, EPI_ISL_2131754, EPI_ISL_2131755, EPI_ISL_2131756, EPI_ISL_2131757, EPI_ISL_2131758, EPI_ISL_2131759, EPI_ISL_2131760, EPI_ISL_2131761, EPI_ISL_2131762, EPI_ISL_2131763, EPI_ISL_2131764, EPI_ISL_2131765, EPI_ISL_2131766, EPI_ISL_2131767, EPI_ISL_2131768, EPI_ISL_2131769, EPI_ISL_2131770, EPI_ISL_2131771, EPI_ISL_2131772, EPI_ISL_2131773, EPI_ISL_2131774, EPI_ISL_2131775, EPI_ISL_2131776, EPI_ISL_2131777, EPI_ISL_2131778, EPI_ISL_2131779, EPI_ISL_2131780, EPI_ISL_2131781, EPI_ISL_2131782, EPI_ISL_2131783, EPI_ISL_2131784, EPI_ISL_2131785, EPI_ISL_2131786, EPI_ISL_2131787, EPI_ISL_2131788, EPI_ISL_2131795, EPI_ISL_2131796 | | | |
| see above | SARS-CoV-2 testing team, National Institute of Infectious Diseases | Pathogen Genomics Center, National Institute of Infectious Diseases | Tsuyoshi Sekizuka, Kentaro Itokawa, Rina Tanaka, Masanori Hashino, Nozomu Hanaoka, Masumichi Saito, Naomi Nojiri, Hazuka Y Furihata, Sana Uchikoba, Tsuguto Fujimoto, Makoto Kuroda |
| EPI_ISL_2131856, EPI_ISL_2131879, EPI_ISL_2131885, EPI_ISL_2131897, EPI_ISL_2131943, EPI_ISL_2131950, EPI_ISL_2131965, EPI_ISL_2131977, EPI_ISL_2132053, EPI_ISL_2132109, EPI_ISL_2132111, EPI_ISL_2132152, EPI_ISL_2132155, EPI_ISL_2132165, EPI_ISL_2132169, EPI_ISL_2132188, EPI_ISL_2132191, EPI_ISL_2132215 | | | |
| see above | National Virus Reference Laboratory | National Virus Reference Laboratory | Zoe Yandle, Charlene Bennett, Gabriel Gonzalez, Michael Carr, Jonathan Dean, Cillian F De Gascun |
| EPI_ISL_2132653 | UMC Groningen, Clinical Virology, Department of Medical Microbiology and Infection Prevention | UMC Groningen, Clinical Virology, Department of Medical Microbiology and Infection Prevention | Hubert Niesters, Alexander Friedrich, Erley Lizarazo-Forero, Monika Fliss, Lilli Gard, Sigrid Rosema, Coretta Van Leer-Buter, Xuewei Zhou, Marjolein Knoester |
| EPI_ISL_2133258, EPI_ISL_2133308, EPI_ISL_2133309, EPI_ISL_2133312, EPI_ISL_2133434, EPI_ISL_2133436, EPI_ISL_2133456, EPI_ISL_2133475 | Quest Diagnostics Incorporated | Centers for Disease Control and Prevention Division of Viral Diseases, Pathogen Discovery | Dakota Howard, Dhwani Batra, Peter W. Cook, Kara Moser, Adrian Paskey, Jason Caravas, Benjamin Rambo-Martin, Shatavia Morrison, Christopher Gulvick, Scott Sammons, Yvette Unoarumhi, Darlene Wagner, Matthew Schmerer, S. H. Rosenthal, A. Gerasimova, R. M. Kagan, B. Anderson, M. Hua, Y. Liu, L.E. Bernstein, K.E. Livingston, A. Perez, I. A. Shlyakhter, R. V. Rolando, R. Owen, P. Tanpaiboon, F. Lacbawan, Clinton R. Paden, Duncan MacCannell |
| EPI_ISL_2133939, EPI_ISL_2133949, EPI_ISL_2133987, EPI_ISL_2133992, EPI_ISL_2134010, EPI_ISL_2134011, EPI_ISL_2134012, EPI_ISL_2134013, EPI_ISL_2134014, EPI_ISL_2134065, EPI_ISL_2134067, EPI_ISL_2134078, EPI_ISL_2134105, EPI_ISL_2134126, EPI_ISL_2134268, EPI_ISL_2134296, EPI_ISL_2134343, EPI_ISL_2134346, EPI_ISL_2134347, EPI_ISL_2134421, EPI_ISL_2134475, EPI_ISL_2134533, EPI_ISL_2134535 | | | |
| see above | Fulgent Genetics | Centers for Disease Control and Prevention Division of Viral Diseases, Pathogen Discovery | Dakota Howard, Dhwani Batra, Peter W. Cook, Kara Moser, Adrian Paskey, Jason Caravas, Benjamin Rambo-Martin, Shatavia Morrison, Christopher Gulvick, Scott Sammons, Yvette Unoarumhi, Darlene Wagner, Matthew Schmerer, Harry Gao, Mickey Li, John Gao, Joseph Fierro, Benafsh Sapra, Becky Tsai, Yan Meng, Doreen Ng, James Xie, Clinton R. Paden, Duncan MacCannell |
| EPI_ISL_2134586, EPI_ISL_2134610 | IN State Department of Health Laboratory Services | IN State Department of Health Laboratory Services | Cassandra Campion, Jamie Yeadon, Brian Pope, Lixia Liu, Kyle Brownlee, Melissa Hindenlang, Mark Glazier |
| EPI_ISL_2134689 | Platform BIS UZA/UAntwerpen | Labo Klinische Biologie, UZA | Marie Le Mercier, Jasmine Coppens, Basil Britto Xavier, Christine Lammens, Veerle Matheeussen, Herman Goossens |
| EPI_ISL_2134882, EPI_ISL_2134892, EPI_ISL_2134901, EPI_ISL_2134906, EPI_ISL_2134911, EPI_ISL_2134921, EPI_ISL_2134928, EPI_ISL_2134933 | HOSPITAL UNIVERSITARIO SON ESPASES | HOSPITAL UNIVERSITARIO SON ESPASES | Carla López-Causapé, Pablo Fraile-Ribot, Antonio Oliver, SeqCovid |
| EPI_ISL_2135087 | Illinois Department of Public Health - Springfield Lab | Illinois Department of Public Health - Springfield Lab | Bryan Sim, Gordon McCall |
| EPI_ISL_2135481, EPI_ISL_2135482, EPI_ISL_2135484, EPI_ISL_2135485, EPI_ISL_2135487, EPI_ISL_2135488, EPI_ISL_2135490 | Epimol – Molecular Epidemiology Group (UV-FISABIO) | SeqCOVID-SPAIN consortium/IBV(CSIC) | María Alma Bracho and SeqCOVID-SPAIN consortium |
| EPI_ISL_2135836, EPI_ISL_2135842, EPI_ISL_2135843, EPI_ISL_2135844, EPI_ISL_2135845, EPI_ISL_2135846 | Viral Respiratory Lab, National Institute for Biomedical Research (INRB) | Pathogen Sequencing Lab, National Institute for Biomedical Research (INRB) | Placide Mbala-Kingebeni, Edith Nkwembe, Eddy Kinganda-Lusamaki, Amuri Aziza, Francisca Muyembe Mawete, Emmanuel Lokilo Lofiko, Jean Claude Makangara, Catherine Pratt, Matthias Pauthner, Josh Quick, Allison Black, James Hadfield, Trevor Bedford, Ian Goodfellow, Andrew Rambaut, Nick Loman, Kristian Andersen, Michael Wiley, Steve Ahuka-Mundeke, Jean-Jacques Muyembe Tamfum |
| EPI_ISL_2136247 | University of Michigan Clinical Microbiology Laboratory | Lauring Lab, University of Michigan, Department of Microbiology and Immunology | Valesano |
| EPI_ISL_2137225 | Austrian Agency for Health and Food Safety (AGES) | Bergthaler laboratory, CeMM Research Center for Molecular Medicine of the Austrian Academy of Sciences | Lukas Endler, Anna Schedl, Fabian Amman, Petr Triska, Thomas Penz, Benedikt Agerer, Maelle Le Moing, Michael Schuster, Bekir Erguner, Jan Laine, Martin Senekowitsch, Christoph Bock, Andreas Bergthaler |
| EPI_ISL_2137845, EPI_ISL_2137846, | Lighthouse Lab in Alderley Park | Wellcome Sanger Institute for the COVID-19 Genomics UK (COG-UK) Consortium | Jacquelyn Wynn, Mairead Hyland, The Lighthouse Lab in Alderley Park and Alex Alderton, Roberto Amato, Jeffrey Barrett, Sonia Goncalves, Ewan Harrison, David K. Jackson, Ian Johnston, Dominic Kwiatkowski, Cordelia Langford, John Sillitoe on behalf of the Wellcome Sanger Institute COVID-19 Surveillance Team |

| | | | |
|---|---|---|---|
| EPI_ISL_2137850 | | | |
| EPI_ISL_2137851, EPI_ISL_2137855, EPI_ISL_2137856, EPI_ISL_2137859, EPI_ISL_2137861, EPI_ISL_2137862, EPI_ISL_2137864, EPI_ISL_2137865, EPI_ISL_2137866, EPI_ISL_2137867, EPI_ISL_2137870, EPI_ISL_2137871, EPI_ISL_2137874 | | | |
| see above | Lighthouse Lab in Milton Keynes | Wellcome Sanger Institute for the COVID-19 Genomics UK (COG-UK) Consortium | The Lighthouse Lab in Milton Keynes and Alex Alderton, Roberto Amato, Jeffrey Barrett, Sonia Goncalves, Ewan Harrison, David K. Jackson, Ian Johnston, Dominic Kwiatkowski, Cordelia Langford, John Sillitoe on behalf of the Wellcome Sanger Institute COVID-19 Surveillance Team |
| EPI_ISL_2137875, EPI_ISL_2137877 | Lighthouse Lab in Alderley Park | Wellcome Sanger Institute for the COVID-19 Genomics UK (COG-UK) Consortium | Jacquelyn Wynn, Mairead Hyland, The Lighthouse Lab in Alderley Park and Alex Alderton, Roberto Amato, Jeffrey Barrett, Sonia Goncalves, Ewan Harrison, David K. Jackson, Ian Johnston, Dominic Kwiatkowski, Cordelia Langford, John Sillitoe on behalf of the Wellcome Sanger Institute COVID-19 Surveillance Team |
| EPI_ISL_2137878 | Lighthouse Lab in Milton Keynes | Wellcome Sanger Institute for the COVID-19 Genomics UK (COG-UK) Consortium | The Lighthouse Lab in Milton Keynes and Alex Alderton, Roberto Amato, Jeffrey Barrett, Sonia Goncalves, Ewan Harrison, David K. Jackson, Ian Johnston, Dominic Kwiatkowski, Cordelia Langford, John Sillitoe on behalf of the Wellcome Sanger Institute COVID-19 Surveillance Team |
| EPI_ISL_2137879 | Lighthouse Lab in Alderley Park | Wellcome Sanger Institute for the COVID-19 Genomics UK (COG-UK) Consortium | Jacquelyn Wynn, Mairead Hyland, The Lighthouse Lab in Alderley Park and Alex Alderton, Roberto Amato, Jeffrey Barrett, Sonia Goncalves, Ewan Harrison, David K. Jackson, Ian Johnston, Dominic Kwiatkowski, Cordelia Langford, John Sillitoe on behalf of the Wellcome Sanger Institute COVID-19 Surveillance Team |
| EPI_ISL_2137880, EPI_ISL_2137881 | Lighthouse Lab in Milton Keynes | Wellcome Sanger Institute for the COVID-19 Genomics UK (COG-UK) Consortium | The Lighthouse Lab in Milton Keynes and Alex Alderton, Roberto Amato, Jeffrey Barrett, Sonia Goncalves, Ewan Harrison, David K. Jackson, Ian Johnston, Dominic Kwiatkowski, Cordelia Langford, John Sillitoe on behalf of the Wellcome Sanger Institute COVID-19 Surveillance Team |
| EPI_ISL_2137882, EPI_ISL_2137887 | Lighthouse Lab in Alderley Park | Wellcome Sanger Institute for the COVID-19 Genomics UK (COG-UK) Consortium | Jacquelyn Wynn, Mairead Hyland, The Lighthouse Lab in Alderley Park and Alex Alderton, Roberto Amato, Jeffrey Barrett, Sonia Goncalves, Ewan Harrison, David K. Jackson, Ian Johnston, Dominic Kwiatkowski, Cordelia Langford, John Sillitoe on behalf of the Wellcome Sanger Institute COVID-19 Surveillance Team |
| EPI_ISL_2137889, EPI_ISL_2137892 | Lighthouse Lab in Milton Keynes | Wellcome Sanger Institute for the COVID-19 Genomics UK (COG-UK) Consortium | The Lighthouse Lab in Milton Keynes and Alex Alderton, Roberto Amato, Jeffrey Barrett, Sonia Goncalves, Ewan Harrison, David K. Jackson, Ian Johnston, Dominic Kwiatkowski, Cordelia Langford, John Sillitoe on behalf of the Wellcome Sanger Institute COVID-19 Surveillance Team |
| EPI_ISL_2137893, EPI_ISL_2137894 | Lighthouse Lab in Alderley Park | Wellcome Sanger Institute for the COVID-19 Genomics UK (COG-UK) Consortium | Jacquelyn Wynn, Mairead Hyland, The Lighthouse Lab in Alderley Park and Alex Alderton, Roberto Amato, Jeffrey Barrett, Sonia Goncalves, Ewan Harrison, David K. Jackson, Ian Johnston, Dominic Kwiatkowski, Cordelia Langford, John Sillitoe on behalf of the Wellcome Sanger Institute COVID-19 Surveillance Team |
| EPI_ISL_2137895 | Lighthouse Lab in Milton Keynes | Wellcome Sanger Institute for the COVID-19 Genomics UK (COG-UK) Consortium | The Lighthouse Lab in Milton Keynes and Alex Alderton, Roberto Amato, Jeffrey Barrett, Sonia Goncalves, Ewan Harrison, David K. Jackson, Ian Johnston, Dominic Kwiatkowski, Cordelia Langford, John Sillitoe on behalf of the Wellcome Sanger Institute COVID-19 Surveillance Team |
| EPI_ISL_2137898, EPI_ISL_2137900 | Lighthouse Lab in Alderley Park | Wellcome Sanger Institute for the COVID-19 Genomics UK (COG-UK) Consortium | Jacquelyn Wynn, Mairead Hyland, The Lighthouse Lab in Alderley Park and Alex Alderton, Roberto Amato, Jeffrey Barrett, Sonia Goncalves, Ewan Harrison, David K. Jackson, Ian Johnston, Dominic Kwiatkowski, Cordelia Langford, John Sillitoe on behalf of the Wellcome Sanger Institute COVID-19 Surveillance Team |
| EPI_ISL_2137901 | Lighthouse Lab in Milton Keynes | Wellcome Sanger Institute for the COVID-19 Genomics UK (COG-UK) Consortium | The Lighthouse Lab in Milton Keynes and Alex Alderton, Roberto Amato, Jeffrey Barrett, Sonia Goncalves, Ewan Harrison, David K. Jackson, Ian Johnston, Dominic Kwiatkowski, Cordelia Langford, John Sillitoe on behalf of the Wellcome Sanger Institute COVID-19 Surveillance Team |
| EPI_ISL_2137903 | Lighthouse Lab in Alderley Park | Wellcome Sanger Institute for the COVID-19 Genomics UK (COG-UK) Consortium | Jacquelyn Wynn, Mairead Hyland, The Lighthouse Lab in Alderley Park and Alex Alderton, Roberto Amato, Jeffrey Barrett, Sonia Goncalves, Ewan Harrison, David K. Jackson, Ian Johnston, Dominic Kwiatkowski, Cordelia Langford, John Sillitoe on behalf of the Wellcome Sanger Institute COVID-19 Surveillance Team |
| EPI_ISL_2137905, EPI_ISL_2137908, EPI_ISL_2137909 | Lighthouse Lab in Milton Keynes | Wellcome Sanger Institute for the COVID-19 Genomics UK (COG-UK) Consortium | The Lighthouse Lab in Milton Keynes and Alex Alderton, Roberto Amato, Jeffrey Barrett, Sonia Goncalves, Ewan Harrison, David K. Jackson, Ian Johnston, Dominic Kwiatkowski, Cordelia Langford, John Sillitoe on behalf of the Wellcome Sanger Institute COVID-19 Surveillance Team |
| EPI_ISL_2137910, EPI_ISL_2137911 | Lighthouse Lab in Alderley Park | Wellcome Sanger Institute for the COVID-19 Genomics UK (COG-UK) Consortium | Jacquelyn Wynn, Mairead Hyland, The Lighthouse Lab in Alderley Park and Alex Alderton, Roberto Amato, Jeffrey Barrett, Sonia Goncalves, Ewan Harrison, David K. Jackson, Ian Johnston, Dominic Kwiatkowski, Cordelia Langford, John Sillitoe on behalf of the Wellcome Sanger Institute COVID-19 Surveillance Team |
| EPI_ISL_2137914, EPI_ISL_2137915, EPI_ISL_2137916 | Lighthouse Lab in Milton Keynes | Wellcome Sanger Institute for the COVID-19 Genomics UK (COG-UK) Consortium | The Lighthouse Lab in Milton Keynes and Alex Alderton, Roberto Amato, Jeffrey Barrett, Sonia Goncalves, Ewan Harrison, David K. Jackson, Ian Johnston, Dominic Kwiatkowski, Cordelia Langford, John Sillitoe on behalf of the Wellcome Sanger Institute COVID-19 Surveillance Team |
| EPI_ISL_2137919, EPI_ISL_2137920 | Lighthouse Lab in Alderley Park | Wellcome Sanger Institute for the COVID-19 Genomics UK (COG-UK) Consortium | Jacquelyn Wynn, Mairead Hyland, The Lighthouse Lab in Alderley Park and Alex Alderton, Roberto Amato, Jeffrey Barrett, Sonia Goncalves, Ewan Harrison, David K. Jackson, Ian Johnston, Dominic Kwiatkowski, Cordelia Langford, John Sillitoe on behalf of the Wellcome Sanger Institute COVID-19 Surveillance Team |
| EPI_ISL_2137924 | Lighthouse Lab in Milton Keynes | Wellcome Sanger Institute for the COVID-19 Genomics UK (COG-UK) Consortium | The Lighthouse Lab in Milton Keynes and Alex Alderton, Roberto Amato, Jeffrey Barrett, Sonia Goncalves, Ewan Harrison, David K. Jackson, Ian Johnston, Dominic Kwiatkowski, Cordelia Langford, John Sillitoe on behalf of the Wellcome Sanger Institute COVID-19 Surveillance Team |
| EPI_ISL_2137925 | Lighthouse Lab in Alderley Park | Wellcome Sanger Institute for the COVID-19 Genomics UK (COG-UK) Consortium | Jacquelyn Wynn, Mairead Hyland, The Lighthouse Lab in Alderley Park and Alex Alderton, Roberto Amato, Jeffrey Barrett, Sonia Goncalves, Ewan Harrison, David K. Jackson, Ian Johnston, Dominic Kwiatkowski, Cordelia Langford, John Sillitoe on behalf of the Wellcome Sanger Institute COVID-19 Surveillance Team |
| EPI_ISL_2137927 | Lighthouse Lab in Milton Keynes | Wellcome Sanger Institute for the COVID-19 Genomics UK (COG-UK) Consortium | The Lighthouse Lab in Milton Keynes and Alex Alderton, Roberto Amato, Jeffrey Barrett, Sonia Goncalves, Ewan Harrison, David K. Jackson, Ian Johnston, Dominic Kwiatkowski, Cordelia Langford, John Sillitoe on behalf of the Wellcome Sanger Institute COVID-19 Surveillance Team |
| EPI_ISL_2137928 | Lighthouse Lab in Alderley Park | Wellcome Sanger Institute for the COVID-19 Genomics UK (COG-UK) Consortium | Jacquelyn Wynn, Mairead Hyland, The Lighthouse Lab in Alderley Park and Alex Alderton, Roberto Amato, Jeffrey Barrett, Sonia Goncalves, Ewan Harrison, David K. Jackson, Ian Johnston, Dominic Kwiatkowski, Cordelia Langford, John Sillitoe on behalf of the Wellcome Sanger Institute COVID-19 Surveillance Team |
| EPI_ISL_2137931, EPI_ISL_2137932 | Lighthouse Lab in Milton Keynes | Wellcome Sanger Institute for the COVID-19 Genomics UK (COG-UK) Consortium | The Lighthouse Lab in Milton Keynes and Alex Alderton, Roberto Amato, Jeffrey Barrett, Sonia Goncalves, Ewan Harrison, David K. Jackson, Ian Johnston, Dominic Kwiatkowski, Cordelia Langford, John Sillitoe on behalf of the Wellcome Sanger Institute COVID-19 Surveillance Team |
| EPI_ISL_2137933 | Lighthouse Lab in Alderley Park | Wellcome Sanger Institute for the COVID-19 Genomics UK (COG-UK) Consortium | Jacquelyn Wynn, Mairead Hyland, The Lighthouse Lab in Alderley Park and Alex Alderton, Roberto Amato, Jeffrey Barrett, Sonia Goncalves, Ewan Harrison, David K. Jackson, Ian Johnston, Dominic Kwiatkowski, Cordelia Langford, John Sillitoe on behalf of the Wellcome Sanger Institute COVID-19 Surveillance Team |
| EPI_ISL_2137939, EPI_ISL_2137940, EPI_ISL_2137944, EPI_ISL_2137946, EPI_ISL_2137948 | Lighthouse Lab in Milton Keynes | Wellcome Sanger Institute for the COVID-19 Genomics UK (COG-UK) Consortium | The Lighthouse Lab in Milton Keynes and Alex Alderton, Roberto Amato, Jeffrey Barrett, Sonia Goncalves, Ewan Harrison, David K. Jackson, Ian Johnston, Dominic Kwiatkowski, Cordelia Langford, John Sillitoe on behalf of the Wellcome Sanger Institute COVID-19 Surveillance Team |
| EPI_ISL_2137950 | Lighthouse Lab in Alderley Park | Wellcome Sanger Institute for the COVID-19 Genomics UK (COG-UK) Consortium | Jacquelyn Wynn, Mairead Hyland, The Lighthouse Lab in Alderley Park and Alex Alderton, Roberto Amato, Jeffrey Barrett, Sonia Goncalves, Ewan Harrison, David K. Jackson, Ian Johnston, Dominic Kwiatkowski, Cordelia Langford, John Sillitoe on behalf of the Wellcome Sanger Institute COVID-19 Surveillance Team |
| EPI_ISL_2137951 | Lighthouse Lab in Milton Keynes | Wellcome Sanger Institute for the COVID-19 Genomics UK (COG-UK) Consortium | The Lighthouse Lab in Milton Keynes and Alex Alderton, Roberto Amato, Jeffrey Barrett, Sonia Goncalves, Ewan Harrison, David K. Jackson, Ian Johnston, Dominic Kwiatkowski, Cordelia Langford, John Sillitoe on behalf of the Wellcome Sanger Institute COVID-19 Surveillance Team |
| EPI_ISL_2137952 | Lighthouse Lab in Alderley Park | Wellcome Sanger Institute for the COVID-19 Genomics UK (COG-UK) Consortium | Jacquelyn Wynn, Mairead Hyland, The Lighthouse Lab in Alderley Park and Alex Alderton, Roberto Amato, Jeffrey Barrett, Sonia Goncalves, Ewan Harrison, David K. Jackson, Ian Johnston, Dominic Kwiatkowski, Cordelia Langford, John Sillitoe on behalf of the Wellcome Sanger Institute COVID-19 Surveillance Team |
| EPI_ISL_2137955, EPI_ISL_2137958, EPI_ISL_2137959, EPI_ISL_2137961 | Lighthouse Lab in Milton Keynes | Wellcome Sanger Institute for the COVID-19 Genomics UK (COG-UK) Consortium | The Lighthouse Lab in Milton Keynes and Alex Alderton, Roberto Amato, Jeffrey Barrett, Sonia Goncalves, Ewan Harrison, David K. Jackson, Ian Johnston, Dominic Kwiatkowski, Cordelia Langford, John Sillitoe on behalf of the Wellcome Sanger Institute COVID-19 Surveillance Team |
| EPI_ISL_2137963 | Lighthouse Lab in Alderley Park | Wellcome Sanger Institute for the COVID-19 Genomics UK (COG-UK) Consortium | Jacquelyn Wynn, Mairead Hyland, The Lighthouse Lab in Alderley Park and Alex Alderton, Roberto Amato, Jeffrey Barrett, Sonia Goncalves, Ewan Harrison, David K. Jackson, Ian Johnston, Dominic Kwiatkowski, Cordelia Langford, John Sillitoe on behalf of the Wellcome Sanger Institute COVID-19 Surveillance Team |
| EPI_ISL_2137964, EPI_ISL_2137966 | Lighthouse Lab in Milton Keynes | Wellcome Sanger Institute for the COVID-19 Genomics UK (COG-UK) Consortium | The Lighthouse Lab in Milton Keynes and Alex Alderton, Roberto Amato, Jeffrey Barrett, Sonia Goncalves, Ewan Harrison, David K. Jackson, Ian Johnston, Dominic Kwiatkowski, Cordelia Langford, John Sillitoe on behalf of the Wellcome Sanger Institute COVID-19 Surveillance Team |
| EPI_ISL_2137967, EPI_ISL_2137968 | Lighthouse Lab in Alderley Park | Wellcome Sanger Institute for the COVID-19 Genomics UK (COG-UK) Consortium | Jacquelyn Wynn, Mairead Hyland, The Lighthouse Lab in Alderley Park and Alex Alderton, Roberto Amato, Jeffrey Barrett, Sonia Goncalves, Ewan Harrison, David K. Jackson, Ian Johnston, Dominic Kwiatkowski, Cordelia Langford, John Sillitoe on behalf of the Wellcome Sanger Institute COVID-19 Surveillance Team |
| EPI_ISL_2137971, EPI_ISL_2137973 | Lighthouse Lab in Milton Keynes | Wellcome Sanger Institute for the COVID-19 Genomics UK (COG-UK) Consortium | The Lighthouse Lab in Milton Keynes and Alex Alderton, Roberto Amato, Jeffrey Barrett, Sonia Goncalves, Ewan Harrison, David K. Jackson, Ian Johnston, Dominic Kwiatkowski, Cordelia Langford, John Sillitoe on behalf of the Wellcome Sanger Institute COVID-19 Surveillance Team |
| EPI_ISL_2137978 | Lighthouse Lab in Alderley Park | Wellcome Sanger Institute for the COVID-19 Genomics UK (COG-UK) Consortium | Jacquelyn Wynn, Mairead Hyland, The Lighthouse Lab in Alderley Park and Alex Alderton, Roberto Amato, Jeffrey Barrett, Sonia Goncalves, Ewan Harrison, David K. Jackson, Ian Johnston, Dominic Kwiatkowski, Cordelia Langford, John Sillitoe on behalf of the Wellcome Sanger Institute COVID-19 Surveillance Team |
| EPI_ISL_2137981, EPI_ISL_2137986, EPI_ISL_2137987, EPI_ISL_2137988, EPI_ISL_2137989, EPI_ISL_2137994 | Lighthouse Lab in Milton Keynes | Wellcome Sanger Institute for the COVID-19 Genomics UK (COG-UK) Consortium | The Lighthouse Lab in Milton Keynes and Alex Alderton, Roberto Amato, Jeffrey Barrett, Sonia Goncalves, Ewan Harrison, David K. Jackson, Ian Johnston, Dominic Kwiatkowski, Cordelia Langford, John Sillitoe on behalf of the Wellcome Sanger Institute COVID-19 Surveillance Team |
| EPI_ISL_2137997, EPI_ISL_2137998 | Lighthouse Lab in Alderley Park | Wellcome Sanger Institute for the COVID-19 Genomics UK (COG-UK) Consortium | Jacquelyn Wynn, Mairead Hyland, The Lighthouse Lab in Alderley Park and Alex Alderton, Roberto Amato, Jeffrey Barrett, Sonia Goncalves, Ewan Harrison, David K. Jackson, Ian Johnston, Dominic Kwiatkowski, Cordelia Langford, John Sillitoe on behalf of the Wellcome Sanger Institute COVID-19 Surveillance Team |
| EPI_ISL_2137999 | Lighthouse Lab in Milton Keynes | Wellcome Sanger Institute for the COVID-19 Genomics UK (COG-UK) Consortium | The Lighthouse Lab in Milton Keynes and Alex Alderton, Roberto Amato, Jeffrey Barrett, Sonia Goncalves, Ewan Harrison, David K. Jackson, Ian Johnston, Dominic Kwiatkowski, Cordelia Langford, John Sillitoe on behalf of the Wellcome Sanger Institute COVID-19 Surveillance Team |
| EPI_ISL_2138000 | Lighthouse Lab in Alderley Park | Wellcome Sanger Institute for the COVID-19 Genomics UK (COG-UK) Consortium | Jacquelyn Wynn, Mairead Hyland, The Lighthouse Lab in Alderley Park and Alex Alderton, Roberto Amato, Jeffrey Barrett, Sonia Goncalves, Ewan Harrison, David K. Jackson, Ian Johnston, Dominic Kwiatkowski, Cordelia Langford, John Sillitoe on behalf of the Wellcome Sanger Institute COVID-19 Surveillance Team |
| EPI_ISL_2138007, EPI_ISL_2138008, EPI_ISL_2138009, EPI_ISL_2138013, EPI_ISL_2138014, EPI_ISL_2138015, EPI_ISL_2138021, EPI_ISL_2138025 | Lighthouse Lab in Milton Keynes | Wellcome Sanger Institute for the COVID-19 Genomics UK (COG-UK) Consortium | The Lighthouse Lab in Milton Keynes and Alex Alderton, Roberto Amato, Jeffrey Barrett, Sonia Goncalves, Ewan Harrison, David K. Jackson, Ian Johnston, Dominic Kwiatkowski, Cordelia Langford, John Sillitoe on behalf of the Wellcome Sanger Institute COVID-19 Surveillance Team |

| | | | |
|---|---|---|---|
| EPI_ISL_2138028 | Lighthouse Lab in Alderley Park | Wellcome Sanger Institute for the COVID-19 Genomics UK (COG-UK) Consortium | Jacquelyn Wynn, Mairead Hyland, The Lighthouse Lab in Alderley Park and Alex Alderton, Roberto Amato, Jeffrey Barrett, Sonia Goncalves, Ewan Harrison, David K. Jackson, Ian Johnston, Dominic Kwiatkowski, Cordelia Langford, John Sillitoe on behalf of the Wellcome Sanger Institute COVID-19 Surveillance Team |
| EPI_ISL_2138029, EPI_ISL_2138030 | Lighthouse Lab in Milton Keynes | Wellcome Sanger Institute for the COVID-19 Genomics UK (COG-UK) Consortium | The Lighthouse Lab in Milton Keynes and Alex Alderton, Roberto Amato, Jeffrey Barrett, Sonia Goncalves, Ewan Harrison, David K. Jackson, Ian Johnston, Dominic Kwiatkowski, Cordelia Langford, John Sillitoe on behalf of the Wellcome Sanger Institute COVID-19 Surveillance Team |
| EPI_ISL_2138031, EPI_ISL_2138034 | Lighthouse Lab in Alderley Park | Wellcome Sanger Institute for the COVID-19 Genomics UK (COG-UK) Consortium | Jacquelyn Wynn, Mairead Hyland, The Lighthouse Lab in Alderley Park and Alex Alderton, Roberto Amato, Jeffrey Barrett, Sonia Goncalves, Ewan Harrison, David K. Jackson, Ian Johnston, Dominic Kwiatkowski, Cordelia Langford, John Sillitoe on behalf of the Wellcome Sanger Institute COVID-19 Surveillance Team |
| EPI_ISL_2138037, EPI_ISL_2138040 | Lighthouse Lab in Milton Keynes | Wellcome Sanger Institute for the COVID-19 Genomics UK (COG-UK) Consortium | The Lighthouse Lab in Milton Keynes and Alex Alderton, Roberto Amato, Jeffrey Barrett, Sonia Goncalves, Ewan Harrison, David K. Jackson, Ian Johnston, Dominic Kwiatkowski, Cordelia Langford, John Sillitoe on behalf of the Wellcome Sanger Institute COVID-19 Surveillance Team |
| EPI_ISL_2138043 | Lighthouse Lab in Alderley Park | Wellcome Sanger Institute for the COVID-19 Genomics UK (COG-UK) Consortium | Jacquelyn Wynn, Mairead Hyland, The Lighthouse Lab in Alderley Park and Alex Alderton, Roberto Amato, Jeffrey Barrett, Sonia Goncalves, Ewan Harrison, David K. Jackson, Ian Johnston, Dominic Kwiatkowski, Cordelia Langford, John Sillitoe on behalf of the Wellcome Sanger Institute COVID-19 Surveillance Team |
| EPI_ISL_2138045 | Lighthouse Lab in Milton Keynes | Wellcome Sanger Institute for the COVID-19 Genomics UK (COG-UK) Consortium | The Lighthouse Lab in Milton Keynes and Alex Alderton, Roberto Amato, Jeffrey Barrett, Sonia Goncalves, Ewan Harrison, David K. Jackson, Ian Johnston, Dominic Kwiatkowski, Cordelia Langford, John Sillitoe on behalf of the Wellcome Sanger Institute COVID-19 Surveillance Team |
| EPI_ISL_2138046, EPI_ISL_2138049 | Lighthouse Lab in Alderley Park | Wellcome Sanger Institute for the COVID-19 Genomics UK (COG-UK) Consortium | Jacquelyn Wynn, Mairead Hyland, The Lighthouse Lab in Alderley Park and Alex Alderton, Roberto Amato, Jeffrey Barrett, Sonia Goncalves, Ewan Harrison, David K. Jackson, Ian Johnston, Dominic Kwiatkowski, Cordelia Langford, John Sillitoe on behalf of the Wellcome Sanger Institute COVID-19 Surveillance Team |
| EPI_ISL_2138050 | Lighthouse Lab in Glasgow | Wellcome Sanger Institute for the COVID-19 Genomics UK (COG-UK) Consortium | Harper VanSteenhouse, Yumi Kasai, David Gray, Carol Clugston, Anna Dominiczak and Alex Alderton, Roberto Amato, Jeffrey Barrett, Sonia Goncalves, Ewan Harrison, David K. Jackson, Ian Johnston, Dominic Kwiatkowski, Cordelia Langford, John Sillitoe on behalf of the Wellcome Sanger Institute COVID-19 Surveillance Team |
| EPI_ISL_2138051, EPI_ISL_2138055, EPI_ISL_2138056, EPI_ISL_2138059 | Lighthouse Lab in Milton Keynes | Wellcome Sanger Institute for the COVID-19 Genomics UK (COG-UK) Consortium | The Lighthouse Lab in Milton Keynes and Alex Alderton, Roberto Amato, Jeffrey Barrett, Sonia Goncalves, Ewan Harrison, David K. Jackson, Ian Johnston, Dominic Kwiatkowski, Cordelia Langford, John Sillitoe on behalf of the Wellcome Sanger Institute COVID-19 Surveillance Team |
| EPI_ISL_2138063 | Lighthouse Lab in Alderley Park | Wellcome Sanger Institute for the COVID-19 Genomics UK (COG-UK) Consortium | Jacquelyn Wynn, Mairead Hyland, The Lighthouse Lab in Alderley Park and Alex Alderton, Roberto Amato, Jeffrey Barrett, Sonia Goncalves, Ewan Harrison, David K. Jackson, Ian Johnston, Dominic Kwiatkowski, Cordelia Langford, John Sillitoe on behalf of the Wellcome Sanger Institute COVID-19 Surveillance Team |
| EPI_ISL_2138067, EPI_ISL_2138068, EPI_ISL_2138070, EPI_ISL_2138072 | Lighthouse Lab in Milton Keynes | Wellcome Sanger Institute for the COVID-19 Genomics UK (COG-UK) Consortium | The Lighthouse Lab in Milton Keynes and Alex Alderton, Roberto Amato, Jeffrey Barrett, Sonia Goncalves, Ewan Harrison, David K. Jackson, Ian Johnston, Dominic Kwiatkowski, Cordelia Langford, John Sillitoe on behalf of the Wellcome Sanger Institute COVID-19 Surveillance Team |
| EPI_ISL_2138073 | Lighthouse Lab in Alderley Park | Wellcome Sanger Institute for the COVID-19 Genomics UK (COG-UK) Consortium | Jacquelyn Wynn, Mairead Hyland, The Lighthouse Lab in Alderley Park and Alex Alderton, Roberto Amato, Jeffrey Barrett, Sonia Goncalves, Ewan Harrison, David K. Jackson, Ian Johnston, Dominic Kwiatkowski, Cordelia Langford, John Sillitoe on behalf of the Wellcome Sanger Institute COVID-19 Surveillance Team |
| EPI_ISL_2138077, EPI_ISL_2138079 | Lighthouse Lab in Milton Keynes | Wellcome Sanger Institute for the COVID-19 Genomics UK (COG-UK) Consortium | The Lighthouse Lab in Milton Keynes and Alex Alderton, Roberto Amato, Jeffrey Barrett, Sonia Goncalves, Ewan Harrison, David K. Jackson, Ian Johnston, Dominic Kwiatkowski, Cordelia Langford, John Sillitoe on behalf of the Wellcome Sanger Institute COVID-19 Surveillance Team |
| EPI_ISL_2138082 | Lighthouse Lab in Alderley Park | Wellcome Sanger Institute for the COVID-19 Genomics UK (COG-UK) Consortium | Jacquelyn Wynn, Mairead Hyland, The Lighthouse Lab in Alderley Park and Alex Alderton, Roberto Amato, Jeffrey Barrett, Sonia Goncalves, Ewan Harrison, David K. Jackson, Ian Johnston, Dominic Kwiatkowski, Cordelia Langford, John Sillitoe on behalf of the Wellcome Sanger Institute COVID-19 Surveillance Team |
| EPI_ISL_2138089, EPI_ISL_2138091, EPI_ISL_2138097 | Lighthouse Lab in Milton Keynes | Wellcome Sanger Institute for the COVID-19 Genomics UK (COG-UK) Consortium | The Lighthouse Lab in Milton Keynes and Alex Alderton, Roberto Amato, Jeffrey Barrett, Sonia Goncalves, Ewan Harrison, David K. Jackson, Ian Johnston, Dominic Kwiatkowski, Cordelia Langford, John Sillitoe on behalf of the Wellcome Sanger Institute COVID-19 Surveillance Team |
| EPI_ISL_2138098 | Lighthouse Lab in Alderley Park | Wellcome Sanger Institute for the COVID-19 Genomics UK (COG-UK) Consortium | Jacquelyn Wynn, Mairead Hyland, The Lighthouse Lab in Alderley Park and Alex Alderton, Roberto Amato, Jeffrey Barrett, Sonia Goncalves, Ewan Harrison, David K. Jackson, Ian Johnston, Dominic Kwiatkowski, Cordelia Langford, John Sillitoe on behalf of the Wellcome Sanger Institute COVID-19 Surveillance Team |
| EPI_ISL_2138103 | Lighthouse Lab in Milton Keynes | Wellcome Sanger Institute for the COVID-19 Genomics UK (COG-UK) Consortium | The Lighthouse Lab in Milton Keynes and Alex Alderton, Roberto Amato, Jeffrey Barrett, Sonia Goncalves, Ewan Harrison, David K. Jackson, Ian Johnston, Dominic Kwiatkowski, Cordelia Langford, John Sillitoe on behalf of the Wellcome Sanger Institute COVID-19 Surveillance Team |
| EPI_ISL_2138107 | Lighthouse Lab in Alderley Park | Wellcome Sanger Institute for the COVID-19 Genomics UK (COG-UK) Consortium | Jacquelyn Wynn, Mairead Hyland, The Lighthouse Lab in Alderley Park and Alex Alderton, Roberto Amato, Jeffrey Barrett, Sonia Goncalves, Ewan Harrison, David K. Jackson, Ian Johnston, Dominic Kwiatkowski, Cordelia Langford, John Sillitoe on behalf of the Wellcome Sanger Institute COVID-19 Surveillance Team |
| EPI_ISL_2138108 | Lighthouse Lab in Milton Keynes | Wellcome Sanger Institute for the COVID-19 Genomics UK (COG-UK) Consortium | The Lighthouse Lab in Milton Keynes and Alex Alderton, Roberto Amato, Jeffrey Barrett, Sonia Goncalves, Ewan Harrison, David K. Jackson, Ian Johnston, Dominic Kwiatkowski, Cordelia Langford, John Sillitoe on behalf of the Wellcome Sanger Institute COVID-19 Surveillance Team |
| EPI_ISL_2138109, EPI_ISL_2138112 | Lighthouse Lab in Alderley Park | Wellcome Sanger Institute for the COVID-19 Genomics UK (COG-UK) Consortium | Jacquelyn Wynn, Mairead Hyland, The Lighthouse Lab in Alderley Park and Alex Alderton, Roberto Amato, Jeffrey Barrett, Sonia Goncalves, Ewan Harrison, David K. Jackson, Ian Johnston, Dominic Kwiatkowski, Cordelia Langford, John Sillitoe on behalf of the Wellcome Sanger Institute COVID-19 Surveillance Team |
| EPI_ISL_2138115, EPI_ISL_2138118, EPI_ISL_2138121, EPI_ISL_2138124, EPI_ISL_2138126, EPI_ISL_2138127, EPI_ISL_2138135 | Lighthouse Lab in Milton Keynes | Wellcome Sanger Institute for the COVID-19 Genomics UK (COG-UK) Consortium | The Lighthouse Lab in Milton Keynes and Alex Alderton, Roberto Amato, Jeffrey Barrett, Sonia Goncalves, Ewan Harrison, David K. Jackson, Ian Johnston, Dominic Kwiatkowski, Cordelia Langford, John Sillitoe on behalf of the Wellcome Sanger Institute COVID-19 Surveillance Team |
| EPI_ISL_2138137, EPI_ISL_2138139 | Lighthouse Lab in Alderley Park | Wellcome Sanger Institute for the COVID-19 Genomics UK (COG-UK) Consortium | Jacquelyn Wynn, Mairead Hyland, The Lighthouse Lab in Alderley Park and Alex Alderton, Roberto Amato, Jeffrey Barrett, Sonia Goncalves, Ewan Harrison, David K. Jackson, Ian Johnston, Dominic Kwiatkowski, Cordelia Langford, John Sillitoe on behalf of the Wellcome Sanger Institute COVID-19 Surveillance Team |
| EPI_ISL_2138141 | Lighthouse Lab in Milton Keynes | Wellcome Sanger Institute for the COVID-19 Genomics UK (COG-UK) Consortium | The Lighthouse Lab in Milton Keynes and Alex Alderton, Roberto Amato, Jeffrey Barrett, Sonia Goncalves, Ewan Harrison, David K. Jackson, Ian Johnston, Dominic Kwiatkowski, Cordelia Langford, John Sillitoe on behalf of the Wellcome Sanger Institute COVID-19 Surveillance Team |
| EPI_ISL_2138143 | Lighthouse Lab in Alderley Park | Wellcome Sanger Institute for the COVID-19 Genomics UK (COG-UK) Consortium | Jacquelyn Wynn, Mairead Hyland, The Lighthouse Lab in Alderley Park and Alex Alderton, Roberto Amato, Jeffrey Barrett, Sonia Goncalves, Ewan Harrison, David K. Jackson, Ian Johnston, Dominic Kwiatkowski, Cordelia Langford, John Sillitoe on behalf of the Wellcome Sanger Institute COVID-19 Surveillance Team |
| EPI_ISL_2138145 | Lighthouse Lab in Milton Keynes | Wellcome Sanger Institute for the COVID-19 Genomics UK (COG-UK) Consortium | The Lighthouse Lab in Milton Keynes and Alex Alderton, Roberto Amato, Jeffrey Barrett, Sonia Goncalves, Ewan Harrison, David K. Jackson, Ian Johnston, Dominic Kwiatkowski, Cordelia Langford, John Sillitoe on behalf of the Wellcome Sanger Institute COVID-19 Surveillance Team |
| EPI_ISL_2138147 | Lighthouse Lab in Alderley Park | Wellcome Sanger Institute for the COVID-19 Genomics UK (COG-UK) Consortium | Jacquelyn Wynn, Mairead Hyland, The Lighthouse Lab in Alderley Park and Alex Alderton, Roberto Amato, Jeffrey Barrett, Sonia Goncalves, Ewan Harrison, David K. Jackson, Ian Johnston, Dominic Kwiatkowski, Cordelia Langford, John Sillitoe on behalf of the Wellcome Sanger Institute COVID-19 Surveillance Team |
| EPI_ISL_2138153, EPI_ISL_2138154, EPI_ISL_2138156, EPI_ISL_2138160, EPI_ISL_2138162 | Lighthouse Lab in Milton Keynes | Wellcome Sanger Institute for the COVID-19 Genomics UK (COG-UK) Consortium | The Lighthouse Lab in Milton Keynes and Alex Alderton, Roberto Amato, Jeffrey Barrett, Sonia Goncalves, Ewan Harrison, David K. Jackson, Ian Johnston, Dominic Kwiatkowski, Cordelia Langford, John Sillitoe on behalf of the Wellcome Sanger Institute COVID-19 Surveillance Team |
| EPI_ISL_2138163, EPI_ISL_2138164 | Lighthouse Lab in Alderley Park | Wellcome Sanger Institute for the COVID-19 Genomics UK (COG-UK) Consortium | Jacquelyn Wynn, Mairead Hyland, The Lighthouse Lab in Alderley Park and Alex Alderton, Roberto Amato, Jeffrey Barrett, Sonia Goncalves, Ewan Harrison, David K. Jackson, Ian Johnston, Dominic Kwiatkowski, Cordelia Langford, John Sillitoe on behalf of the Wellcome Sanger Institute COVID-19 Surveillance Team |
| EPI_ISL_2138165 | Lighthouse Lab in Glasgow | Wellcome Sanger Institute for the COVID-19 Genomics UK (COG-UK) Consortium | Harper VanSteenhouse, Yumi Kasai, David Gray, Carol Clugston, Anna Dominiczak and Alex Alderton, Roberto Amato, Jeffrey Barrett, Sonia Goncalves, Ewan Harrison, David K. Jackson, Ian Johnston, Dominic Kwiatkowski, Cordelia Langford, John Sillitoe on behalf of the Wellcome Sanger Institute COVID-19 Surveillance Team |
| EPI_ISL_2138167 | Lighthouse Lab in Alderley Park | Wellcome Sanger Institute for the COVID-19 Genomics UK (COG-UK) Consortium | Jacquelyn Wynn, Mairead Hyland, The Lighthouse Lab in Alderley Park and Alex Alderton, Roberto Amato, Jeffrey Barrett, Sonia Goncalves, Ewan Harrison, David K. Jackson, Ian Johnston, Dominic Kwiatkowski, Cordelia Langford, John Sillitoe on behalf of the Wellcome Sanger Institute COVID-19 Surveillance Team |
| EPI_ISL_2138168 | Lighthouse Lab in Milton Keynes | Wellcome Sanger Institute for the COVID-19 Genomics UK (COG-UK) Consortium | The Lighthouse Lab in Milton Keynes and Alex Alderton, Roberto Amato, Jeffrey Barrett, Sonia Goncalves, Ewan Harrison, David K. Jackson, Ian Johnston, Dominic Kwiatkowski, Cordelia Langford, John Sillitoe on behalf of the Wellcome Sanger Institute COVID-19 Surveillance Team |
| EPI_ISL_2138173 | Lighthouse Lab in Alderley Park | Wellcome Sanger Institute for the COVID-19 Genomics UK (COG-UK) Consortium | Jacquelyn Wynn, Mairead Hyland, The Lighthouse Lab in Alderley Park and Alex Alderton, Roberto Amato, Jeffrey Barrett, Sonia Goncalves, Ewan Harrison, David K. Jackson, Ian Johnston, Dominic Kwiatkowski, Cordelia Langford, John Sillitoe on behalf of the Wellcome Sanger Institute COVID-19 Surveillance Team |
| EPI_ISL_2138177 | Lighthouse Lab in Milton Keynes | Wellcome Sanger Institute for the COVID-19 Genomics UK (COG-UK) Consortium | The Lighthouse Lab in Milton Keynes and Alex Alderton, Roberto Amato, Jeffrey Barrett, Sonia Goncalves, Ewan Harrison, David K. Jackson, Ian Johnston, Dominic Kwiatkowski, Cordelia Langford, John Sillitoe on behalf of the Wellcome Sanger Institute COVID-19 Surveillance Team |
| EPI_ISL_2138178 | Lighthouse Lab in Alderley Park | Wellcome Sanger Institute for the COVID-19 Genomics UK (COG-UK) Consortium | Jacquelyn Wynn, Mairead Hyland, The Lighthouse Lab in Alderley Park and Alex Alderton, Roberto Amato, Jeffrey Barrett, Sonia Goncalves, Ewan Harrison, David K. Jackson, Ian Johnston, Dominic Kwiatkowski, Cordelia Langford, John Sillitoe on behalf of the Wellcome Sanger Institute COVID-19 Surveillance Team |
| EPI_ISL_2138179 | Lighthouse Lab in Milton Keynes | Wellcome Sanger Institute for the COVID-19 Genomics UK (COG-UK) Consortium | The Lighthouse Lab in Milton Keynes and Alex Alderton, Roberto Amato, Jeffrey Barrett, Sonia Goncalves, Ewan Harrison, David K. Jackson, Ian Johnston, Dominic Kwiatkowski, Cordelia Langford, John Sillitoe on behalf of the Wellcome Sanger Institute COVID-19 Surveillance Team |
| EPI_ISL_2138181 | Lighthouse Lab in Alderley Park | Wellcome Sanger Institute for the COVID-19 Genomics UK (COG-UK) Consortium | Jacquelyn Wynn, Mairead Hyland, The Lighthouse Lab in Alderley Park and Alex Alderton, Roberto Amato, Jeffrey Barrett, Sonia Goncalves, Ewan Harrison, David K. Jackson, Ian Johnston, Dominic Kwiatkowski, Cordelia Langford, John Sillitoe on behalf of the Wellcome Sanger Institute COVID-19 Surveillance Team |
| EPI_ISL_2138186, EPI_ISL_2138188, EPI_ISL_2138189, EPI_ISL_2138195 | Lighthouse Lab in Milton Keynes | Wellcome Sanger Institute for the COVID-19 Genomics UK (COG-UK) Consortium | The Lighthouse Lab in Milton Keynes and Alex Alderton, Roberto Amato, Jeffrey Barrett, Sonia Goncalves, Ewan Harrison, David K. Jackson, Ian Johnston, Dominic Kwiatkowski, Cordelia Langford, John Sillitoe on behalf of the Wellcome Sanger Institute COVID-19 Surveillance Team |
| EPI_ISL_2138198 | Lighthouse Lab in Alderley Park | Wellcome Sanger Institute for the COVID-19 Genomics UK (COG-UK) Consortium | Jacquelyn Wynn, Mairead Hyland, The Lighthouse Lab in Alderley Park and Alex Alderton, Roberto Amato, Jeffrey Barrett, Sonia Goncalves, Ewan Harrison, David K. Jackson, Ian Johnston, Dominic Kwiatkowski, Cordelia Langford, John Sillitoe on behalf of the Wellcome Sanger Institute COVID-19 Surveillance Team |
| EPI_ISL_2138199, EPI_ISL_2138200, | Lighthouse Lab in Milton Keynes | Wellcome Sanger Institute for the COVID-19 Genomics UK (COG-UK) Consortium | The Lighthouse Lab in Milton Keynes and Alex Alderton, Roberto Amato, Jeffrey Barrett, Sonia Goncalves, Ewan Harrison, David K. Jackson, Ian Johnston, Dominic Kwiatkowski, Cordelia Langford, John Sillitoe on behalf of the Wellcome Sanger Institute COVID-19 Surveillance Team |

| | | | |
|---|---|---|---|
| EPI_ISL_2138203, EPI_ISL_2138204 | | | |
| EPI_ISL_2138205, EPI_ISL_2138208 | Lighthouse Lab in Alderley Park | Wellcome Sanger Institute for the COVID-19 Genomics UK (COG-UK) Consortium | Jacquelyn Wynn, Mairead Hyland, The Lighthouse Lab in Alderley Park and Alex Alderton, Roberto Amato, Jeffrey Barrett, Sonia Goncalves, Ewan Harrison, David K. Jackson, Ian Johnston, Dominic Kwiatkowski, Cordelia Langford, John Sillitoe on behalf of the Wellcome Sanger Institute COVID-19 Surveillance Team |
| EPI_ISL_2138213 | Lighthouse Lab in Milton Keynes | Wellcome Sanger Institute for the COVID-19 Genomics UK (COG-UK) Consortium | The Lighthouse Lab in Milton Keynes and Alex Alderton, Roberto Amato, Jeffrey Barrett, Sonia Goncalves, Ewan Harrison, David K. Jackson, Ian Johnston, Dominic Kwiatkowski, Cordelia Langford, John Sillitoe on behalf of the Wellcome Sanger Institute COVID-19 Surveillance Team |
| EPI_ISL_2138214 | Lighthouse Lab in Alderley Park | Wellcome Sanger Institute for the COVID-19 Genomics UK (COG-UK) Consortium | Jacquelyn Wynn, Mairead Hyland, The Lighthouse Lab in Alderley Park and Alex Alderton, Roberto Amato, Jeffrey Barrett, Sonia Goncalves, Ewan Harrison, David K. Jackson, Ian Johnston, Dominic Kwiatkowski, Cordelia Langford, John Sillitoe on behalf of the Wellcome Sanger Institute COVID-19 Surveillance Team |
| EPI_ISL_2138216 | Lighthouse Lab in Milton Keynes | Wellcome Sanger Institute for the COVID-19 Genomics UK (COG-UK) Consortium | The Lighthouse Lab in Milton Keynes and Alex Alderton, Roberto Amato, Jeffrey Barrett, Sonia Goncalves, Ewan Harrison, David K. Jackson, Ian Johnston, Dominic Kwiatkowski, Cordelia Langford, John Sillitoe on behalf of the Wellcome Sanger Institute COVID-19 Surveillance Team |
| EPI_ISL_2138219 | Lighthouse Lab in Alderley Park | Wellcome Sanger Institute for the COVID-19 Genomics UK (COG-UK) Consortium | Jacquelyn Wynn, Mairead Hyland, The Lighthouse Lab in Alderley Park and Alex Alderton, Roberto Amato, Jeffrey Barrett, Sonia Goncalves, Ewan Harrison, David K. Jackson, Ian Johnston, Dominic Kwiatkowski, Cordelia Langford, John Sillitoe on behalf of the Wellcome Sanger Institute COVID-19 Surveillance Team |
| EPI_ISL_2138220 | Lighthouse Lab in Milton Keynes | Wellcome Sanger Institute for the COVID-19 Genomics UK (COG-UK) Consortium | The Lighthouse Lab in Milton Keynes and Alex Alderton, Roberto Amato, Jeffrey Barrett, Sonia Goncalves, Ewan Harrison, David K. Jackson, Ian Johnston, Dominic Kwiatkowski, Cordelia Langford, John Sillitoe on behalf of the Wellcome Sanger Institute COVID-19 Surveillance Team |
| EPI_ISL_2138224 | Lighthouse Lab in Alderley Park | Wellcome Sanger Institute for the COVID-19 Genomics UK (COG-UK) Consortium | Jacquelyn Wynn, Mairead Hyland, The Lighthouse Lab in Alderley Park and Alex Alderton, Roberto Amato, Jeffrey Barrett, Sonia Goncalves, Ewan Harrison, David K. Jackson, Ian Johnston, Dominic Kwiatkowski, Cordelia Langford, John Sillitoe on behalf of the Wellcome Sanger Institute COVID-19 Surveillance Team |
| EPI_ISL_2138225, EPI_ISL_2138227, EPI_ISL_2138230, EPI_ISL_2138233 | Lighthouse Lab in Milton Keynes | Wellcome Sanger Institute for the COVID-19 Genomics UK (COG-UK) Consortium | The Lighthouse Lab in Milton Keynes and Alex Alderton, Roberto Amato, Jeffrey Barrett, Sonia Goncalves, Ewan Harrison, David K. Jackson, Ian Johnston, Dominic Kwiatkowski, Cordelia Langford, John Sillitoe on behalf of the Wellcome Sanger Institute COVID-19 Surveillance Team |
| EPI_ISL_2138238 | Lighthouse Lab in Alderley Park | Wellcome Sanger Institute for the COVID-19 Genomics UK (COG-UK) Consortium | Jacquelyn Wynn, Mairead Hyland, The Lighthouse Lab in Alderley Park and Alex Alderton, Roberto Amato, Jeffrey Barrett, Sonia Goncalves, Ewan Harrison, David K. Jackson, Ian Johnston, Dominic Kwiatkowski, Cordelia Langford, John Sillitoe on behalf of the Wellcome Sanger Institute COVID-19 Surveillance Team |
| EPI_ISL_2138239 | Lighthouse Lab in Milton Keynes | Wellcome Sanger Institute for the COVID-19 Genomics UK (COG-UK) Consortium | The Lighthouse Lab in Milton Keynes and Alex Alderton, Roberto Amato, Jeffrey Barrett, Sonia Goncalves, Ewan Harrison, David K. Jackson, Ian Johnston, Dominic Kwiatkowski, Cordelia Langford, John Sillitoe on behalf of the Wellcome Sanger Institute COVID-19 Surveillance Team |
| EPI_ISL_2138242 | Lighthouse Lab in Alderley Park | Wellcome Sanger Institute for the COVID-19 Genomics UK (COG-UK) Consortium | Jacquelyn Wynn, Mairead Hyland, The Lighthouse Lab in Alderley Park and Alex Alderton, Roberto Amato, Jeffrey Barrett, Sonia Goncalves, Ewan Harrison, David K. Jackson, Ian Johnston, Dominic Kwiatkowski, Cordelia Langford, John Sillitoe on behalf of the Wellcome Sanger Institute COVID-19 Surveillance Team |
| EPI_ISL_2138243, EPI_ISL_2138245, EPI_ISL_2138250, EPI_ISL_2138253, EPI_ISL_2138254 | Lighthouse Lab in Milton Keynes | Wellcome Sanger Institute for the COVID-19 Genomics UK (COG-UK) Consortium | The Lighthouse Lab in Milton Keynes and Alex Alderton, Roberto Amato, Jeffrey Barrett, Sonia Goncalves, Ewan Harrison, David K. Jackson, Ian Johnston, Dominic Kwiatkowski, Cordelia Langford, John Sillitoe on behalf of the Wellcome Sanger Institute COVID-19 Surveillance Team |
| EPI_ISL_2138255, EPI_ISL_2138260 | Lighthouse Lab in Alderley Park | Wellcome Sanger Institute for the COVID-19 Genomics UK (COG-UK) Consortium | Jacquelyn Wynn, Mairead Hyland, The Lighthouse Lab in Alderley Park and Alex Alderton, Roberto Amato, Jeffrey Barrett, Sonia Goncalves, Ewan Harrison, David K. Jackson, Ian Johnston, Dominic Kwiatkowski, Cordelia Langford, John Sillitoe on behalf of the Wellcome Sanger Institute COVID-19 Surveillance Team |
| EPI_ISL_2138261, EPI_ISL_2138267 | Lighthouse Lab in Milton Keynes | Wellcome Sanger Institute for the COVID-19 Genomics UK (COG-UK) Consortium | The Lighthouse Lab in Milton Keynes and Alex Alderton, Roberto Amato, Jeffrey Barrett, Sonia Goncalves, Ewan Harrison, David K. Jackson, Ian Johnston, Dominic Kwiatkowski, Cordelia Langford, John Sillitoe on behalf of the Wellcome Sanger Institute COVID-19 Surveillance Team |
| EPI_ISL_2138268, EPI_ISL_2138272 | Lighthouse Lab in Alderley Park | Wellcome Sanger Institute for the COVID-19 Genomics UK (COG-UK) Consortium | Jacquelyn Wynn, Mairead Hyland, The Lighthouse Lab in Alderley Park and Alex Alderton, Roberto Amato, Jeffrey Barrett, Sonia Goncalves, Ewan Harrison, David K. Jackson, Ian Johnston, Dominic Kwiatkowski, Cordelia Langford, John Sillitoe on behalf of the Wellcome Sanger Institute COVID-19 Surveillance Team |
| EPI_ISL_2138275 | Lighthouse Lab in Milton Keynes | Wellcome Sanger Institute for the COVID-19 Genomics UK (COG-UK) Consortium | The Lighthouse Lab in Milton Keynes and Alex Alderton, Roberto Amato, Jeffrey Barrett, Sonia Goncalves, Ewan Harrison, David K. Jackson, Ian Johnston, Dominic Kwiatkowski, Cordelia Langford, John Sillitoe on behalf of the Wellcome Sanger Institute COVID-19 Surveillance Team |
| EPI_ISL_2138276 | Lighthouse Lab in Alderley Park | Wellcome Sanger Institute for the COVID-19 Genomics UK (COG-UK) Consortium | Jacquelyn Wynn, Mairead Hyland, The Lighthouse Lab in Alderley Park and Alex Alderton, Roberto Amato, Jeffrey Barrett, Sonia Goncalves, Ewan Harrison, David K. Jackson, Ian Johnston, Dominic Kwiatkowski, Cordelia Langford, John Sillitoe on behalf of the Wellcome Sanger Institute COVID-19 Surveillance Team |
| EPI_ISL_2138277, EPI_ISL_2138278, EPI_ISL_2138279, EPI_ISL_2138281, EPI_ISL_2138282, EPI_ISL_2138284 | Lighthouse Lab in Milton Keynes | Wellcome Sanger Institute for the COVID-19 Genomics UK (COG-UK) Consortium | The Lighthouse Lab in Milton Keynes and Alex Alderton, Roberto Amato, Jeffrey Barrett, Sonia Goncalves, Ewan Harrison, David K. Jackson, Ian Johnston, Dominic Kwiatkowski, Cordelia Langford, John Sillitoe on behalf of the Wellcome Sanger Institute COVID-19 Surveillance Team |
| EPI_ISL_2138289, EPI_ISL_2138291, EPI_ISL_2138293, EPI_ISL_2138294 | Lighthouse Lab in Alderley Park | Wellcome Sanger Institute for the COVID-19 Genomics UK (COG-UK) Consortium | Jacquelyn Wynn, Mairead Hyland, The Lighthouse Lab in Alderley Park and Alex Alderton, Roberto Amato, Jeffrey Barrett, Sonia Goncalves, Ewan Harrison, David K. Jackson, Ian Johnston, Dominic Kwiatkowski, Cordelia Langford, John Sillitoe on behalf of the Wellcome Sanger Institute COVID-19 Surveillance Team |
| EPI_ISL_2138295 | Lighthouse Lab in Glasgow | Wellcome Sanger Institute for the COVID-19 Genomics UK (COG-UK) Consortium | Harper VanSteenhouse, Yumi Kasai, David Gray, Carol Clugston, Anna Dominiczak and Alex Alderton, Roberto Amato, Jeffrey Barrett, Sonia Goncalves, Ewan Harrison, David K. Jackson, Ian Johnston, Dominic Kwiatkowski, Cordelia Langford, John Sillitoe on behalf of the Wellcome Sanger Institute COVID-19 Surveillance Team |
| EPI_ISL_2138296 | Lighthouse Lab in Alderley Park | Wellcome Sanger Institute for the COVID-19 Genomics UK (COG-UK) Consortium | Jacquelyn Wynn, Mairead Hyland, The Lighthouse Lab in Alderley Park and Alex Alderton, Roberto Amato, Jeffrey Barrett, Sonia Goncalves, Ewan Harrison, David K. Jackson, Ian Johnston, Dominic Kwiatkowski, Cordelia Langford, John Sillitoe on behalf of the Wellcome Sanger Institute COVID-19 Surveillance Team |
| EPI_ISL_2138302 | Lighthouse Lab in Milton Keynes | Wellcome Sanger Institute for the COVID-19 Genomics UK (COG-UK) Consortium | The Lighthouse Lab in Milton Keynes and Alex Alderton, Roberto Amato, Jeffrey Barrett, Sonia Goncalves, Ewan Harrison, David K. Jackson, Ian Johnston, Dominic Kwiatkowski, Cordelia Langford, John Sillitoe on behalf of the Wellcome Sanger Institute COVID-19 Surveillance Team |
| EPI_ISL_2138306, EPI_ISL_2138307 | Lighthouse Lab in Alderley Park | Wellcome Sanger Institute for the COVID-19 Genomics UK (COG-UK) Consortium | Jacquelyn Wynn, Mairead Hyland, The Lighthouse Lab in Alderley Park and Alex Alderton, Roberto Amato, Jeffrey Barrett, Sonia Goncalves, Ewan Harrison, David K. Jackson, Ian Johnston, Dominic Kwiatkowski, Cordelia Langford, John Sillitoe on behalf of the Wellcome Sanger Institute COVID-19 Surveillance Team |
| EPI_ISL_2138310 | Lighthouse Lab in Milton Keynes | Wellcome Sanger Institute for the COVID-19 Genomics UK (COG-UK) Consortium | The Lighthouse Lab in Milton Keynes and Alex Alderton, Roberto Amato, Jeffrey Barrett, Sonia Goncalves, Ewan Harrison, David K. Jackson, Ian Johnston, Dominic Kwiatkowski, Cordelia Langford, John Sillitoe on behalf of the Wellcome Sanger Institute COVID-19 Surveillance Team |
| EPI_ISL_2138311 | Lighthouse Lab in Alderley Park | Wellcome Sanger Institute for the COVID-19 Genomics UK (COG-UK) Consortium | Jacquelyn Wynn, Mairead Hyland, The Lighthouse Lab in Alderley Park and Alex Alderton, Roberto Amato, Jeffrey Barrett, Sonia Goncalves, Ewan Harrison, David K. Jackson, Ian Johnston, Dominic Kwiatkowski, Cordelia Langford, John Sillitoe on behalf of the Wellcome Sanger Institute COVID-19 Surveillance Team |
| EPI_ISL_2138313, EPI_ISL_2138314 | Lighthouse Lab in Milton Keynes | Wellcome Sanger Institute for the COVID-19 Genomics UK (COG-UK) Consortium | The Lighthouse Lab in Milton Keynes and Alex Alderton, Roberto Amato, Jeffrey Barrett, Sonia Goncalves, Ewan Harrison, David K. Jackson, Ian Johnston, Dominic Kwiatkowski, Cordelia Langford, John Sillitoe on behalf of the Wellcome Sanger Institute COVID-19 Surveillance Team |
| EPI_ISL_2138315, EPI_ISL_2138318, EPI_ISL_2138319, EPI_ISL_2138321, EPI_ISL_2138322 | Lighthouse Lab in Alderley Park | Wellcome Sanger Institute for the COVID-19 Genomics UK (COG-UK) Consortium | Jacquelyn Wynn, Mairead Hyland, The Lighthouse Lab in Alderley Park and Alex Alderton, Roberto Amato, Jeffrey Barrett, Sonia Goncalves, Ewan Harrison, David K. Jackson, Ian Johnston, Dominic Kwiatkowski, Cordelia Langford, John Sillitoe on behalf of the Wellcome Sanger Institute COVID-19 Surveillance Team |
| EPI_ISL_2138323, EPI_ISL_2138324, EPI_ISL_2138326, EPI_ISL_2138327 | Lighthouse Lab in Milton Keynes | Wellcome Sanger Institute for the COVID-19 Genomics UK (COG-UK) Consortium | The Lighthouse Lab in Milton Keynes and Alex Alderton, Roberto Amato, Jeffrey Barrett, Sonia Goncalves, Ewan Harrison, David K. Jackson, Ian Johnston, Dominic Kwiatkowski, Cordelia Langford, John Sillitoe on behalf of the Wellcome Sanger Institute COVID-19 Surveillance Team |
| EPI_ISL_2138331 | Lighthouse Lab in Alderley Park | Wellcome Sanger Institute for the COVID-19 Genomics UK (COG-UK) Consortium | Jacquelyn Wynn, Mairead Hyland, The Lighthouse Lab in Alderley Park and Alex Alderton, Roberto Amato, Jeffrey Barrett, Sonia Goncalves, Ewan Harrison, David K. Jackson, Ian Johnston, Dominic Kwiatkowski, Cordelia Langford, John Sillitoe on behalf of the Wellcome Sanger Institute COVID-19 Surveillance Team |
| EPI_ISL_2138332, EPI_ISL_2138335 | Lighthouse Lab in Milton Keynes | Wellcome Sanger Institute for the COVID-19 Genomics UK (COG-UK) Consortium | The Lighthouse Lab in Milton Keynes and Alex Alderton, Roberto Amato, Jeffrey Barrett, Sonia Goncalves, Ewan Harrison, David K. Jackson, Ian Johnston, Dominic Kwiatkowski, Cordelia Langford, John Sillitoe on behalf of the Wellcome Sanger Institute COVID-19 Surveillance Team |
| EPI_ISL_2138339 | Lighthouse Lab in Alderley Park | Wellcome Sanger Institute for the COVID-19 Genomics UK (COG-UK) Consortium | Jacquelyn Wynn, Mairead Hyland, The Lighthouse Lab in Alderley Park and Alex Alderton, Roberto Amato, Jeffrey Barrett, Sonia Goncalves, Ewan Harrison, David K. Jackson, Ian Johnston, Dominic Kwiatkowski, Cordelia Langford, John Sillitoe on behalf of the Wellcome Sanger Institute COVID-19 Surveillance Team |
| EPI_ISL_2138340, EPI_ISL_2138342, EPI_ISL_2138345 | Lighthouse Lab in Milton Keynes | Wellcome Sanger Institute for the COVID-19 Genomics UK (COG-UK) Consortium | The Lighthouse Lab in Milton Keynes and Alex Alderton, Roberto Amato, Jeffrey Barrett, Sonia Goncalves, Ewan Harrison, David K. Jackson, Ian Johnston, Dominic Kwiatkowski, Cordelia Langford, John Sillitoe on behalf of the Wellcome Sanger Institute COVID-19 Surveillance Team |
| EPI_ISL_2138353, EPI_ISL_2138354, EPI_ISL_2138355, EPI_ISL_2138356, EPI_ISL_2138363 | Lighthouse Lab in Alderley Park | Wellcome Sanger Institute for the COVID-19 Genomics UK (COG-UK) Consortium | Jacquelyn Wynn, Mairead Hyland, The Lighthouse Lab in Alderley Park and Alex Alderton, Roberto Amato, Jeffrey Barrett, Sonia Goncalves, Ewan Harrison, David K. Jackson, Ian Johnston, Dominic Kwiatkowski, Cordelia Langford, John Sillitoe on behalf of the Wellcome Sanger Institute COVID-19 Surveillance Team |
| EPI_ISL_2138364, EPI_ISL_2138366, EPI_ISL_2138368, EPI_ISL_2138371 | Lighthouse Lab in Milton Keynes | Wellcome Sanger Institute for the COVID-19 Genomics UK (COG-UK) Consortium | The Lighthouse Lab in Milton Keynes and Alex Alderton, Roberto Amato, Jeffrey Barrett, Sonia Goncalves, Ewan Harrison, David K. Jackson, Ian Johnston, Dominic Kwiatkowski, Cordelia Langford, John Sillitoe on behalf of the Wellcome Sanger Institute COVID-19 Surveillance Team |
| EPI_ISL_2138372 | Lighthouse Lab in Alderley Park | Wellcome Sanger Institute for the COVID-19 Genomics UK (COG-UK) Consortium | Jacquelyn Wynn, Mairead Hyland, The Lighthouse Lab in Alderley Park and Alex Alderton, Roberto Amato, Jeffrey Barrett, Sonia Goncalves, Ewan Harrison, David K. Jackson, Ian Johnston, Dominic Kwiatkowski, Cordelia Langford, John Sillitoe on behalf of the Wellcome Sanger Institute COVID-19 Surveillance Team |

| | | | |
|---|---|---|---|
| EPI_ISL_2138374, EPI_ISL_2138375, EPI_ISL_2138378, EPI_ISL_2138381 | Lighthouse Lab in Milton Keynes | Wellcome Sanger Institute for the COVID-19 Genomics UK (COG-UK) Consortium | The Lighthouse Lab in Milton Keynes and Alex Alderton, Roberto Amato, Jeffrey Barrett, Sonia Goncalves, Ewan Harrison, David K. Jackson, Ian Johnston, Dominic Kwiatkowski, Cordelia Langford, John Sillitoe on behalf of the Wellcome Sanger Institute COVID-19 Surveillance Team |
| EPI_ISL_2138385 | Health Services Laboratories | Wellcome Sanger Institute for the COVID-19 Genomics UK (COG-UK) Consortium | Health Services Laboratories and Alex Alderton, Roberto Amato, Jeffrey Barrett, Sonia Goncalves, Ewan Harrison, David K. Jackson, Ian Johnston, Dominic Kwiatkowski, Cordelia Langford, John Sillitoe on behalf of the Wellcome Sanger Institute COVID-19 Surveillance Team |
| EPI_ISL_2138386 | Lighthouse Lab in Alderley Park | Wellcome Sanger Institute for the COVID-19 Genomics UK (COG-UK) Consortium | Jacquelyn Wynn, Mairead Hyland, The Lighthouse Lab in Alderley Park and Alex Alderton, Roberto Amato, Jeffrey Barrett, Sonia Goncalves, Ewan Harrison, David K. Jackson, Ian Johnston, Dominic Kwiatkowski, Cordelia Langford, John Sillitoe on behalf of the Wellcome Sanger Institute COVID-19 Surveillance Team |
| EPI_ISL_2138392 | Lighthouse Lab in Milton Keynes | Wellcome Sanger Institute for the COVID-19 Genomics UK (COG-UK) Consortium | The Lighthouse Lab in Milton Keynes and Alex Alderton, Roberto Amato, Jeffrey Barrett, Sonia Goncalves, Ewan Harrison, David K. Jackson, Ian Johnston, Dominic Kwiatkowski, Cordelia Langford, John Sillitoe on behalf of the Wellcome Sanger Institute COVID-19 Surveillance Team |
| EPI_ISL_2138398 | Lighthouse Lab in Alderley Park | Wellcome Sanger Institute for the COVID-19 Genomics UK (COG-UK) Consortium | Jacquelyn Wynn, Mairead Hyland, The Lighthouse Lab in Alderley Park and Alex Alderton, Roberto Amato, Jeffrey Barrett, Sonia Goncalves, Ewan Harrison, David K. Jackson, Ian Johnston, Dominic Kwiatkowski, Cordelia Langford, John Sillitoe on behalf of the Wellcome Sanger Institute COVID-19 Surveillance Team |
| EPI_ISL_2138400, EPI_ISL_2138401 | Lighthouse Lab in Milton Keynes | Wellcome Sanger Institute for the COVID-19 Genomics UK (COG-UK) Consortium | The Lighthouse Lab in Milton Keynes and Alex Alderton, Roberto Amato, Jeffrey Barrett, Sonia Goncalves, Ewan Harrison, David K. Jackson, Ian Johnston, Dominic Kwiatkowski, Cordelia Langford, John Sillitoe on behalf of the Wellcome Sanger Institute COVID-19 Surveillance Team |
| EPI_ISL_2138406, EPI_ISL_2138407, EPI_ISL_2138409 | Lighthouse Lab in Alderley Park | Wellcome Sanger Institute for the COVID-19 Genomics UK (COG-UK) Consortium | Jacquelyn Wynn, Mairead Hyland, The Lighthouse Lab in Alderley Park and Alex Alderton, Roberto Amato, Jeffrey Barrett, Sonia Goncalves, Ewan Harrison, David K. Jackson, Ian Johnston, Dominic Kwiatkowski, Cordelia Langford, John Sillitoe on behalf of the Wellcome Sanger Institute COVID-19 Surveillance Team |
| EPI_ISL_2138412, EPI_ISL_2138413, EPI_ISL_2138417 | Lighthouse Lab in Milton Keynes | Wellcome Sanger Institute for the COVID-19 Genomics UK (COG-UK) Consortium | The Lighthouse Lab in Milton Keynes and Alex Alderton, Roberto Amato, Jeffrey Barrett, Sonia Goncalves, Ewan Harrison, David K. Jackson, Ian Johnston, Dominic Kwiatkowski, Cordelia Langford, John Sillitoe on behalf of the Wellcome Sanger Institute COVID-19 Surveillance Team |
| EPI_ISL_2138420 | Lighthouse Lab in Alderley Park | Wellcome Sanger Institute for the COVID-19 Genomics UK (COG-UK) Consortium | Jacquelyn Wynn, Mairead Hyland, The Lighthouse Lab in Alderley Park and Alex Alderton, Roberto Amato, Jeffrey Barrett, Sonia Goncalves, Ewan Harrison, David K. Jackson, Ian Johnston, Dominic Kwiatkowski, Cordelia Langford, John Sillitoe on behalf of the Wellcome Sanger Institute COVID-19 Surveillance Team |
| EPI_ISL_2138424, EPI_ISL_2138427 | Lighthouse Lab in Milton Keynes | Wellcome Sanger Institute for the COVID-19 Genomics UK (COG-UK) Consortium | The Lighthouse Lab in Milton Keynes and Alex Alderton, Roberto Amato, Jeffrey Barrett, Sonia Goncalves, Ewan Harrison, David K. Jackson, Ian Johnston, Dominic Kwiatkowski, Cordelia Langford, John Sillitoe on behalf of the Wellcome Sanger Institute COVID-19 Surveillance Team |
| EPI_ISL_2138428, EPI_ISL_2138430 | Lighthouse Lab in Alderley Park | Wellcome Sanger Institute for the COVID-19 Genomics UK (COG-UK) Consortium | Jacquelyn Wynn, Mairead Hyland, The Lighthouse Lab in Alderley Park and Alex Alderton, Roberto Amato, Jeffrey Barrett, Sonia Goncalves, Ewan Harrison, David K. Jackson, Ian Johnston, Dominic Kwiatkowski, Cordelia Langford, John Sillitoe on behalf of the Wellcome Sanger Institute COVID-19 Surveillance Team |
| EPI_ISL_2138433 | Lighthouse Lab in Milton Keynes | Wellcome Sanger Institute for the COVID-19 Genomics UK (COG-UK) Consortium | The Lighthouse Lab in Milton Keynes and Alex Alderton, Roberto Amato, Jeffrey Barrett, Sonia Goncalves, Ewan Harrison, David K. Jackson, Ian Johnston, Dominic Kwiatkowski, Cordelia Langford, John Sillitoe on behalf of the Wellcome Sanger Institute COVID-19 Surveillance Team |
| EPI_ISL_2138434 | Health Services Laboratories | Wellcome Sanger Institute for the COVID-19 Genomics UK (COG-UK) Consortium | Health Services Laboratories and Alex Alderton, Roberto Amato, Jeffrey Barrett, Sonia Goncalves, Ewan Harrison, David K. Jackson, Ian Johnston, Dominic Kwiatkowski, Cordelia Langford, John Sillitoe on behalf of the Wellcome Sanger Institute COVID-19 Surveillance Team |
| EPI_ISL_2138438, EPI_ISL_2138439 | Lighthouse Lab in Milton Keynes | Wellcome Sanger Institute for the COVID-19 Genomics UK (COG-UK) Consortium | The Lighthouse Lab in Milton Keynes and Alex Alderton, Roberto Amato, Jeffrey Barrett, Sonia Goncalves, Ewan Harrison, David K. Jackson, Ian Johnston, Dominic Kwiatkowski, Cordelia Langford, John Sillitoe on behalf of the Wellcome Sanger Institute COVID-19 Surveillance Team |
| EPI_ISL_2138440 | Lighthouse Lab in Alderley Park | Wellcome Sanger Institute for the COVID-19 Genomics UK (COG-UK) Consortium | Jacquelyn Wynn, Mairead Hyland, The Lighthouse Lab in Alderley Park and Alex Alderton, Roberto Amato, Jeffrey Barrett, Sonia Goncalves, Ewan Harrison, David K. Jackson, Ian Johnston, Dominic Kwiatkowski, Cordelia Langford, John Sillitoe on behalf of the Wellcome Sanger Institute COVID-19 Surveillance Team |
| EPI_ISL_2138445 | Lighthouse Lab in Milton Keynes | Wellcome Sanger Institute for the COVID-19 Genomics UK (COG-UK) Consortium | The Lighthouse Lab in Milton Keynes and Alex Alderton, Roberto Amato, Jeffrey Barrett, Sonia Goncalves, Ewan Harrison, David K. Jackson, Ian Johnston, Dominic Kwiatkowski, Cordelia Langford, John Sillitoe on behalf of the Wellcome Sanger Institute COVID-19 Surveillance Team |
| EPI_ISL_2138446 | Lighthouse Lab in Alderley Park | Wellcome Sanger Institute for the COVID-19 Genomics UK (COG-UK) Consortium | Jacquelyn Wynn, Mairead Hyland, The Lighthouse Lab in Alderley Park and Alex Alderton, Roberto Amato, Jeffrey Barrett, Sonia Goncalves, Ewan Harrison, David K. Jackson, Ian Johnston, Dominic Kwiatkowski, Cordelia Langford, John Sillitoe on behalf of the Wellcome Sanger Institute COVID-19 Surveillance Team |
| EPI_ISL_2138447, EPI_ISL_2138449, EPI_ISL_2138452 | Lighthouse Lab in Milton Keynes | Wellcome Sanger Institute for the COVID-19 Genomics UK (COG-UK) Consortium | The Lighthouse Lab in Milton Keynes and Alex Alderton, Roberto Amato, Jeffrey Barrett, Sonia Goncalves, Ewan Harrison, David K. Jackson, Ian Johnston, Dominic Kwiatkowski, Cordelia Langford, John Sillitoe on behalf of the Wellcome Sanger Institute COVID-19 Surveillance Team |
| EPI_ISL_2138456, EPI_ISL_2138460 | Lighthouse Lab in Alderley Park | Wellcome Sanger Institute for the COVID-19 Genomics UK (COG-UK) Consortium | Jacquelyn Wynn, Mairead Hyland, The Lighthouse Lab in Alderley Park and Alex Alderton, Roberto Amato, Jeffrey Barrett, Sonia Goncalves, Ewan Harrison, David K. Jackson, Ian Johnston, Dominic Kwiatkowski, Cordelia Langford, John Sillitoe on behalf of the Wellcome Sanger Institute COVID-19 Surveillance Team |
| EPI_ISL_2138463, EPI_ISL_2138465, EPI_ISL_2138466 | Lighthouse Lab in Milton Keynes | Wellcome Sanger Institute for the COVID-19 Genomics UK (COG-UK) Consortium | The Lighthouse Lab in Milton Keynes and Alex Alderton, Roberto Amato, Jeffrey Barrett, Sonia Goncalves, Ewan Harrison, David K. Jackson, Ian Johnston, Dominic Kwiatkowski, Cordelia Langford, John Sillitoe on behalf of the Wellcome Sanger Institute COVID-19 Surveillance Team |
| EPI_ISL_2138471 | Health Services Laboratories | Wellcome Sanger Institute for the COVID-19 Genomics UK (COG-UK) Consortium | Health Services Laboratories and Alex Alderton, Roberto Amato, Jeffrey Barrett, Sonia Goncalves, Ewan Harrison, David K. Jackson, Ian Johnston, Dominic Kwiatkowski, Cordelia Langford, John Sillitoe on behalf of the Wellcome Sanger Institute COVID-19 Surveillance Team |
| EPI_ISL_2138476, EPI_ISL_2138477, EPI_ISL_2138480, EPI_ISL_2138481 | Lighthouse Lab in Alderley Park | Wellcome Sanger Institute for the COVID-19 Genomics UK (COG-UK) Consortium | Jacquelyn Wynn, Mairead Hyland, The Lighthouse Lab in Alderley Park and Alex Alderton, Roberto Amato, Jeffrey Barrett, Sonia Goncalves, Ewan Harrison, David K. Jackson, Ian Johnston, Dominic Kwiatkowski, Cordelia Langford, John Sillitoe on behalf of the Wellcome Sanger Institute COVID-19 Surveillance Team |
| EPI_ISL_2138485 | Health Services Laboratories | Wellcome Sanger Institute for the COVID-19 Genomics UK (COG-UK) Consortium | Health Services Laboratories and Alex Alderton, Roberto Amato, Jeffrey Barrett, Sonia Goncalves, Ewan Harrison, David K. Jackson, Ian Johnston, Dominic Kwiatkowski, Cordelia Langford, John Sillitoe on behalf of the Wellcome Sanger Institute COVID-19 Surveillance Team |
| EPI_ISL_2138490 | Lighthouse Lab in Milton Keynes | Wellcome Sanger Institute for the COVID-19 Genomics UK (COG-UK) Consortium | The Lighthouse Lab in Milton Keynes and Alex Alderton, Roberto Amato, Jeffrey Barrett, Sonia Goncalves, Ewan Harrison, David K. Jackson, Ian Johnston, Dominic Kwiatkowski, Cordelia Langford, John Sillitoe on behalf of the Wellcome Sanger Institute COVID-19 Surveillance Team |
| EPI_ISL_2138491, EPI_ISL_2138494 | Lighthouse Lab in Alderley Park | Wellcome Sanger Institute for the COVID-19 Genomics UK (COG-UK) Consortium | Jacquelyn Wynn, Mairead Hyland, The Lighthouse Lab in Alderley Park and Alex Alderton, Roberto Amato, Jeffrey Barrett, Sonia Goncalves, Ewan Harrison, David K. Jackson, Ian Johnston, Dominic Kwiatkowski, Cordelia Langford, John Sillitoe on behalf of the Wellcome Sanger Institute COVID-19 Surveillance Team |
| EPI_ISL_2138495, EPI_ISL_2138497, EPI_ISL_2138500 | Lighthouse Lab in Milton Keynes | Wellcome Sanger Institute for the COVID-19 Genomics UK (COG-UK) Consortium | The Lighthouse Lab in Milton Keynes and Alex Alderton, Roberto Amato, Jeffrey Barrett, Sonia Goncalves, Ewan Harrison, David K. Jackson, Ian Johnston, Dominic Kwiatkowski, Cordelia Langford, John Sillitoe on behalf of the Wellcome Sanger Institute COVID-19 Surveillance Team |
| EPI_ISL_2138503 | Health Services Laboratories | Wellcome Sanger Institute for the COVID-19 Genomics UK (COG-UK) Consortium | Health Services Laboratories and Alex Alderton, Roberto Amato, Jeffrey Barrett, Sonia Goncalves, Ewan Harrison, David K. Jackson, Ian Johnston, Dominic Kwiatkowski, Cordelia Langford, John Sillitoe on behalf of the Wellcome Sanger Institute COVID-19 Surveillance Team |
| EPI_ISL_2138504 | Lighthouse Lab in Milton Keynes | Wellcome Sanger Institute for the COVID-19 Genomics UK (COG-UK) Consortium | The Lighthouse Lab in Milton Keynes and Alex Alderton, Roberto Amato, Jeffrey Barrett, Sonia Goncalves, Ewan Harrison, David K. Jackson, Ian Johnston, Dominic Kwiatkowski, Cordelia Langford, John Sillitoe on behalf of the Wellcome Sanger Institute COVID-19 Surveillance Team |
| EPI_ISL_2138508 | Lighthouse Lab in Alderley Park | Wellcome Sanger Institute for the COVID-19 Genomics UK (COG-UK) Consortium | Jacquelyn Wynn, Mairead Hyland, The Lighthouse Lab in Alderley Park and Alex Alderton, Roberto Amato, Jeffrey Barrett, Sonia Goncalves, Ewan Harrison, David K. Jackson, Ian Johnston, Dominic Kwiatkowski, Cordelia Langford, John Sillitoe on behalf of the Wellcome Sanger Institute COVID-19 Surveillance Team |
| EPI_ISL_2138510 | Lighthouse Lab in Milton Keynes | Wellcome Sanger Institute for the COVID-19 Genomics UK (COG-UK) Consortium | The Lighthouse Lab in Milton Keynes and Alex Alderton, Roberto Amato, Jeffrey Barrett, Sonia Goncalves, Ewan Harrison, David K. Jackson, Ian Johnston, Dominic Kwiatkowski, Cordelia Langford, John Sillitoe on behalf of the Wellcome Sanger Institute COVID-19 Surveillance Team |
| EPI_ISL_2138511 | Lighthouse Lab in Alderley Park | Wellcome Sanger Institute for the COVID-19 Genomics UK (COG-UK) Consortium | Jacquelyn Wynn, Mairead Hyland, The Lighthouse Lab in Alderley Park and Alex Alderton, Roberto Amato, Jeffrey Barrett, Sonia Goncalves, Ewan Harrison, David K. Jackson, Ian Johnston, Dominic Kwiatkowski, Cordelia Langford, John Sillitoe on behalf of the Wellcome Sanger Institute COVID-19 Surveillance Team |
| EPI_ISL_2138516 | Health Services Laboratories | Wellcome Sanger Institute for the COVID-19 Genomics UK (COG-UK) Consortium | Health Services Laboratories and Alex Alderton, Roberto Amato, Jeffrey Barrett, Sonia Goncalves, Ewan Harrison, David K. Jackson, Ian Johnston, Dominic Kwiatkowski, Cordelia Langford, John Sillitoe on behalf of the Wellcome Sanger Institute COVID-19 Surveillance Team |
| EPI_ISL_2138518, EPI_ISL_2138519 | Lighthouse Lab in Milton Keynes | Wellcome Sanger Institute for the COVID-19 Genomics UK (COG-UK) Consortium | The Lighthouse Lab in Milton Keynes and Alex Alderton, Roberto Amato, Jeffrey Barrett, Sonia Goncalves, Ewan Harrison, David K. Jackson, Ian Johnston, Dominic Kwiatkowski, Cordelia Langford, John Sillitoe on behalf of the Wellcome Sanger Institute COVID-19 Surveillance Team |
| EPI_ISL_2138520, EPI_ISL_2138528 | Lighthouse Lab in Alderley Park | Wellcome Sanger Institute for the COVID-19 Genomics UK (COG-UK) Consortium | Jacquelyn Wynn, Mairead Hyland, The Lighthouse Lab in Alderley Park and Alex Alderton, Roberto Amato, Jeffrey Barrett, Sonia Goncalves, Ewan Harrison, David K. Jackson, Ian Johnston, Dominic Kwiatkowski, Cordelia Langford, John Sillitoe on behalf of the Wellcome Sanger Institute COVID-19 Surveillance Team |
| EPI_ISL_2138529 | Lighthouse Lab in Milton Keynes | Wellcome Sanger Institute for the COVID-19 Genomics UK (COG-UK) Consortium | The Lighthouse Lab in Milton Keynes and Alex Alderton, Roberto Amato, Jeffrey Barrett, Sonia Goncalves, Ewan Harrison, David K. Jackson, Ian Johnston, Dominic Kwiatkowski, Cordelia Langford, John Sillitoe on behalf of the Wellcome Sanger Institute COVID-19 Surveillance Team |
| EPI_ISL_2138532, EPI_ISL_2138533 | Lighthouse Lab in Alderley Park | Wellcome Sanger Institute for the COVID-19 Genomics UK (COG-UK) Consortium | Jacquelyn Wynn, Mairead Hyland, The Lighthouse Lab in Alderley Park and Alex Alderton, Roberto Amato, Jeffrey Barrett, Sonia Goncalves, Ewan Harrison, David K. Jackson, Ian Johnston, Dominic Kwiatkowski, Cordelia Langford, John Sillitoe on behalf of the Wellcome Sanger Institute COVID-19 Surveillance Team |
| EPI_ISL_2138534, EPI_ISL_2138535, EPI_ISL_2138536, EPI_ISL_2138537 | Lighthouse Lab in Milton Keynes | Wellcome Sanger Institute for the COVID-19 Genomics UK (COG-UK) Consortium | The Lighthouse Lab in Milton Keynes and Alex Alderton, Roberto Amato, Jeffrey Barrett, Sonia Goncalves, Ewan Harrison, David K. Jackson, Ian Johnston, Dominic Kwiatkowski, Cordelia Langford, John Sillitoe on behalf of the Wellcome Sanger Institute COVID-19 Surveillance Team |
| EPI_ISL_2138538 | Lighthouse Lab in Alderley Park | Wellcome Sanger Institute for the COVID-19 Genomics UK (COG-UK) Consortium | Jacquelyn Wynn, Mairead Hyland, The Lighthouse Lab in Alderley Park and Alex Alderton, Roberto Amato, Jeffrey Barrett, Sonia Goncalves, Ewan Harrison, David K. Jackson, Ian Johnston, Dominic Kwiatkowski, Cordelia Langford, John Sillitoe on behalf of the Wellcome Sanger Institute COVID-19 Surveillance Team |
| EPI_ISL_2138539, EPI_ISL_2138540 | Lighthouse Lab in Milton Keynes | Wellcome Sanger Institute for the COVID-19 Genomics UK (COG-UK) Consortium | The Lighthouse Lab in Milton Keynes and Alex Alderton, Roberto Amato, Jeffrey Barrett, Sonia Goncalves, Ewan Harrison, David K. Jackson, Ian Johnston, Dominic Kwiatkowski, Cordelia Langford, John Sillitoe on behalf of the Wellcome Sanger Institute COVID-19 Surveillance Team |
| EPI_ISL_2138544 | Lighthouse Lab in Alderley Park | Wellcome Sanger Institute for the COVID-19 Genomics UK (COG-UK) Consortium | Jacquelyn Wynn, Mairead Hyland, The Lighthouse Lab in Alderley Park and Alex Alderton, Roberto Amato, Jeffrey Barrett, Sonia Goncalves, Ewan Harrison, David K. Jackson, Ian Johnston, Dominic Kwiatkowski, Cordelia Langford, John Sillitoe on behalf of the Wellcome Sanger Institute COVID-19 Surveillance Team |
| EPI_ISL_2138545, EPI_ISL_2138547, | Lighthouse Lab in Milton Keynes | Wellcome Sanger Institute for the COVID-19 Genomics UK (COG-UK) Consortium | The Lighthouse Lab in Milton Keynes and Alex Alderton, Roberto Amato, Jeffrey Barrett, Sonia Goncalves, Ewan Harrison, David K. Jackson, Ian Johnston, Dominic Kwiatkowski, Cordelia Langford, John Sillitoe on behalf of the Wellcome Sanger Institute COVID-19 Surveillance Team |

| | | | |
|---|---|---|---|
| EPI_ISL_2138548 | | | |
| EPI_ISL_2138549 | Health Services Laboratories | Wellcome Sanger Institute for the COVID-19 Genomics UK (COG-UK) Consortium | Health Services Laboratories and Alex Alderton, Roberto Amato, Jeffrey Barrett, Sonia Goncalves, Ewan Harrison, David K. Jackson, Ian Johnston, Dominic Kwiatkowski, Cordelia Langford, John Sillitoe on behalf of the Wellcome Sanger Institute COVID-19 Surveillance Team |
| EPI_ISL_2138550 | Lighthouse Lab in Milton Keynes | Wellcome Sanger Institute for the COVID-19 Genomics UK (COG-UK) Consortium | The Lighthouse Lab in Milton Keynes and Alex Alderton, Roberto Amato, Jeffrey Barrett, Sonia Goncalves, Ewan Harrison, David K. Jackson, Ian Johnston, Dominic Kwiatkowski, Cordelia Langford, John Sillitoe on behalf of the Wellcome Sanger Institute COVID-19 Surveillance Team |
| EPI_ISL_2138553 | Lighthouse Lab in Alderley Park | Wellcome Sanger Institute for the COVID-19 Genomics UK (COG-UK) Consortium | Jacquelyn Wynn, Mairead Hyland, The Lighthouse Lab in Alderley Park and Alex Alderton, Roberto Amato, Jeffrey Barrett, Sonia Goncalves, Ewan Harrison, David K. Jackson, Ian Johnston, Dominic Kwiatkowski, Cordelia Langford, John Sillitoe on behalf of the Wellcome Sanger Institute COVID-19 Surveillance Team |
| EPI_ISL_2138555 | Lighthouse Lab in Milton Keynes | Wellcome Sanger Institute for the COVID-19 Genomics UK (COG-UK) Consortium | The Lighthouse Lab in Milton Keynes and Alex Alderton, Roberto Amato, Jeffrey Barrett, Sonia Goncalves, Ewan Harrison, David K. Jackson, Ian Johnston, Dominic Kwiatkowski, Cordelia Langford, John Sillitoe on behalf of the Wellcome Sanger Institute COVID-19 Surveillance Team |
| EPI_ISL_2138560 | Lighthouse Lab in Alderley Park | Wellcome Sanger Institute for the COVID-19 Genomics UK (COG-UK) Consortium | Jacquelyn Wynn, Mairead Hyland, The Lighthouse Lab in Alderley Park and Alex Alderton, Roberto Amato, Jeffrey Barrett, Sonia Goncalves, Ewan Harrison, David K. Jackson, Ian Johnston, Dominic Kwiatkowski, Cordelia Langford, John Sillitoe on behalf of the Wellcome Sanger Institute COVID-19 Surveillance Team |
| EPI_ISL_2138561 | Health Services Laboratories | Wellcome Sanger Institute for the COVID-19 Genomics UK (COG-UK) Consortium | Health Services Laboratories and Alex Alderton, Roberto Amato, Jeffrey Barrett, Sonia Goncalves, Ewan Harrison, David K. Jackson, Ian Johnston, Dominic Kwiatkowski, Cordelia Langford, John Sillitoe on behalf of the Wellcome Sanger Institute COVID-19 Surveillance Team |
| EPI_ISL_2138562 | Lighthouse Lab in Alderley Park | Wellcome Sanger Institute for the COVID-19 Genomics UK (COG-UK) Consortium | Jacquelyn Wynn, Mairead Hyland, The Lighthouse Lab in Alderley Park and Alex Alderton, Roberto Amato, Jeffrey Barrett, Sonia Goncalves, Ewan Harrison, David K. Jackson, Ian Johnston, Dominic Kwiatkowski, Cordelia Langford, John Sillitoe on behalf of the Wellcome Sanger Institute COVID-19 Surveillance Team |
| EPI_ISL_2138567 | Lighthouse Lab in Milton Keynes | Wellcome Sanger Institute for the COVID-19 Genomics UK (COG-UK) Consortium | The Lighthouse Lab in Milton Keynes and Alex Alderton, Roberto Amato, Jeffrey Barrett, Sonia Goncalves, Ewan Harrison, David K. Jackson, Ian Johnston, Dominic Kwiatkowski, Cordelia Langford, John Sillitoe on behalf of the Wellcome Sanger Institute COVID-19 Surveillance Team |
| EPI_ISL_2138571 | Lighthouse Lab in Alderley Park | Wellcome Sanger Institute for the COVID-19 Genomics UK (COG-UK) Consortium | Jacquelyn Wynn, Mairead Hyland, The Lighthouse Lab in Alderley Park and Alex Alderton, Roberto Amato, Jeffrey Barrett, Sonia Goncalves, Ewan Harrison, David K. Jackson, Ian Johnston, Dominic Kwiatkowski, Cordelia Langford, John Sillitoe on behalf of the Wellcome Sanger Institute COVID-19 Surveillance Team |
| EPI_ISL_2138576 | Lighthouse Lab in Milton Keynes | Wellcome Sanger Institute for the COVID-19 Genomics UK (COG-UK) Consortium | The Lighthouse Lab in Milton Keynes and Alex Alderton, Roberto Amato, Jeffrey Barrett, Sonia Goncalves, Ewan Harrison, David K. Jackson, Ian Johnston, Dominic Kwiatkowski, Cordelia Langford, John Sillitoe on behalf of the Wellcome Sanger Institute COVID-19 Surveillance Team |
| EPI_ISL_2138578, EPI_ISL_2138587 | Lighthouse Lab in Alderley Park | Wellcome Sanger Institute for the COVID-19 Genomics UK (COG-UK) Consortium | Jacquelyn Wynn, Mairead Hyland, The Lighthouse Lab in Alderley Park and Alex Alderton, Roberto Amato, Jeffrey Barrett, Sonia Goncalves, Ewan Harrison, David K. Jackson, Ian Johnston, Dominic Kwiatkowski, Cordelia Langford, John Sillitoe on behalf of the Wellcome Sanger Institute COVID-19 Surveillance Team |
| EPI_ISL_2138588 | Lighthouse Lab in Milton Keynes | Wellcome Sanger Institute for the COVID-19 Genomics UK (COG-UK) Consortium | The Lighthouse Lab in Milton Keynes and Alex Alderton, Roberto Amato, Jeffrey Barrett, Sonia Goncalves, Ewan Harrison, David K. Jackson, Ian Johnston, Dominic Kwiatkowski, Cordelia Langford, John Sillitoe on behalf of the Wellcome Sanger Institute COVID-19 Surveillance Team |
| EPI_ISL_2138589 | Lighthouse Lab in Alderley Park | Wellcome Sanger Institute for the COVID-19 Genomics UK (COG-UK) Consortium | Jacquelyn Wynn, Mairead Hyland, The Lighthouse Lab in Alderley Park and Alex Alderton, Roberto Amato, Jeffrey Barrett, Sonia Goncalves, Ewan Harrison, David K. Jackson, Ian Johnston, Dominic Kwiatkowski, Cordelia Langford, John Sillitoe on behalf of the Wellcome Sanger Institute COVID-19 Surveillance Team |
| EPI_ISL_2138593, EPI_ISL_2138595 | Lighthouse Lab in Milton Keynes | Wellcome Sanger Institute for the COVID-19 Genomics UK (COG-UK) Consortium | The Lighthouse Lab in Milton Keynes and Alex Alderton, Roberto Amato, Jeffrey Barrett, Sonia Goncalves, Ewan Harrison, David K. Jackson, Ian Johnston, Dominic Kwiatkowski, Cordelia Langford, John Sillitoe on behalf of the Wellcome Sanger Institute COVID-19 Surveillance Team |
| EPI_ISL_2138600 | Health Services Laboratories | Wellcome Sanger Institute for the COVID-19 Genomics UK (COG-UK) Consortium | Health Services Laboratories and Alex Alderton, Roberto Amato, Jeffrey Barrett, Sonia Goncalves, Ewan Harrison, David K. Jackson, Ian Johnston, Dominic Kwiatkowski, Cordelia Langford, John Sillitoe on behalf of the Wellcome Sanger Institute COVID-19 Surveillance Team |
| EPI_ISL_2138602 | Lighthouse Lab in Alderley Park | Wellcome Sanger Institute for the COVID-19 Genomics UK (COG-UK) Consortium | Jacquelyn Wynn, Mairead Hyland, The Lighthouse Lab in Alderley Park and Alex Alderton, Roberto Amato, Jeffrey Barrett, Sonia Goncalves, Ewan Harrison, David K. Jackson, Ian Johnston, Dominic Kwiatkowski, Cordelia Langford, John Sillitoe on behalf of the Wellcome Sanger Institute COVID-19 Surveillance Team |
| EPI_ISL_2138605 | Lighthouse Lab in Milton Keynes | Wellcome Sanger Institute for the COVID-19 Genomics UK (COG-UK) Consortium | The Lighthouse Lab in Milton Keynes and Alex Alderton, Roberto Amato, Jeffrey Barrett, Sonia Goncalves, Ewan Harrison, David K. Jackson, Ian Johnston, Dominic Kwiatkowski, Cordelia Langford, John Sillitoe on behalf of the Wellcome Sanger Institute COVID-19 Surveillance Team |
| EPI_ISL_2138608, EPI_ISL_2138609 | Lighthouse Lab in Alderley Park | Wellcome Sanger Institute for the COVID-19 Genomics UK (COG-UK) Consortium | Jacquelyn Wynn, Mairead Hyland, The Lighthouse Lab in Alderley Park and Alex Alderton, Roberto Amato, Jeffrey Barrett, Sonia Goncalves, Ewan Harrison, David K. Jackson, Ian Johnston, Dominic Kwiatkowski, Cordelia Langford, John Sillitoe on behalf of the Wellcome Sanger Institute COVID-19 Surveillance Team |
| EPI_ISL_2138615 | Lighthouse Lab in Milton Keynes | Wellcome Sanger Institute for the COVID-19 Genomics UK (COG-UK) Consortium | The Lighthouse Lab in Milton Keynes and Alex Alderton, Roberto Amato, Jeffrey Barrett, Sonia Goncalves, Ewan Harrison, David K. Jackson, Ian Johnston, Dominic Kwiatkowski, Cordelia Langford, John Sillitoe on behalf of the Wellcome Sanger Institute COVID-19 Surveillance Team |
| EPI_ISL_2138616 | Lighthouse Lab in Alderley Park | Wellcome Sanger Institute for the COVID-19 Genomics UK (COG-UK) Consortium | Jacquelyn Wynn, Mairead Hyland, The Lighthouse Lab in Alderley Park and Alex Alderton, Roberto Amato, Jeffrey Barrett, Sonia Goncalves, Ewan Harrison, David K. Jackson, Ian Johnston, Dominic Kwiatkowski, Cordelia Langford, John Sillitoe on behalf of the Wellcome Sanger Institute COVID-19 Surveillance Team |
| EPI_ISL_2138617 | Health Services Laboratories | Wellcome Sanger Institute for the COVID-19 Genomics UK (COG-UK) Consortium | Health Services Laboratories and Alex Alderton, Roberto Amato, Jeffrey Barrett, Sonia Goncalves, Ewan Harrison, David K. Jackson, Ian Johnston, Dominic Kwiatkowski, Cordelia Langford, John Sillitoe on behalf of the Wellcome Sanger Institute COVID-19 Surveillance Team |
| EPI_ISL_2138618 | Lighthouse Lab in Milton Keynes | Wellcome Sanger Institute for the COVID-19 Genomics UK (COG-UK) Consortium | The Lighthouse Lab in Milton Keynes and Alex Alderton, Roberto Amato, Jeffrey Barrett, Sonia Goncalves, Ewan Harrison, David K. Jackson, Ian Johnston, Dominic Kwiatkowski, Cordelia Langford, John Sillitoe on behalf of the Wellcome Sanger Institute COVID-19 Surveillance Team |
| EPI_ISL_2138620 | Lighthouse Lab in Alderley Park | Wellcome Sanger Institute for the COVID-19 Genomics UK (COG-UK) Consortium | Jacquelyn Wynn, Mairead Hyland, The Lighthouse Lab in Alderley Park and Alex Alderton, Roberto Amato, Jeffrey Barrett, Sonia Goncalves, Ewan Harrison, David K. Jackson, Ian Johnston, Dominic Kwiatkowski, Cordelia Langford, John Sillitoe on behalf of the Wellcome Sanger Institute COVID-19 Surveillance Team |
| EPI_ISL_2138621 | Lighthouse Lab in Milton Keynes | Wellcome Sanger Institute for the COVID-19 Genomics UK (COG-UK) Consortium | The Lighthouse Lab in Milton Keynes and Alex Alderton, Roberto Amato, Jeffrey Barrett, Sonia Goncalves, Ewan Harrison, David K. Jackson, Ian Johnston, Dominic Kwiatkowski, Cordelia Langford, John Sillitoe on behalf of the Wellcome Sanger Institute COVID-19 Surveillance Team |
| EPI_ISL_2138622 | Health Services Laboratories | Wellcome Sanger Institute for the COVID-19 Genomics UK (COG-UK) Consortium | Health Services Laboratories and Alex Alderton, Roberto Amato, Jeffrey Barrett, Sonia Goncalves, Ewan Harrison, David K. Jackson, Ian Johnston, Dominic Kwiatkowski, Cordelia Langford, John Sillitoe on behalf of the Wellcome Sanger Institute COVID-19 Surveillance Team |
| EPI_ISL_2138628, EPI_ISL_2138631, EPI_ISL_2138634 | Lighthouse Lab in Milton Keynes | Wellcome Sanger Institute for the COVID-19 Genomics UK (COG-UK) Consortium | The Lighthouse Lab in Milton Keynes and Alex Alderton, Roberto Amato, Jeffrey Barrett, Sonia Goncalves, Ewan Harrison, David K. Jackson, Ian Johnston, Dominic Kwiatkowski, Cordelia Langford, John Sillitoe on behalf of the Wellcome Sanger Institute COVID-19 Surveillance Team |
| EPI_ISL_2138635 | Lighthouse Lab in Alderley Park | Wellcome Sanger Institute for the COVID-19 Genomics UK (COG-UK) Consortium | Jacquelyn Wynn, Mairead Hyland, The Lighthouse Lab in Alderley Park and Alex Alderton, Roberto Amato, Jeffrey Barrett, Sonia Goncalves, Ewan Harrison, David K. Jackson, Ian Johnston, Dominic Kwiatkowski, Cordelia Langford, John Sillitoe on behalf of the Wellcome Sanger Institute COVID-19 Surveillance Team |
| EPI_ISL_2138642 | Lighthouse Lab in Milton Keynes | Wellcome Sanger Institute for the COVID-19 Genomics UK (COG-UK) Consortium | The Lighthouse Lab in Milton Keynes and Alex Alderton, Roberto Amato, Jeffrey Barrett, Sonia Goncalves, Ewan Harrison, David K. Jackson, Ian Johnston, Dominic Kwiatkowski, Cordelia Langford, John Sillitoe on behalf of the Wellcome Sanger Institute COVID-19 Surveillance Team |
| EPI_ISL_2138644 | Lighthouse Lab in Alderley Park | Wellcome Sanger Institute for the COVID-19 Genomics UK (COG-UK) Consortium | Jacquelyn Wynn, Mairead Hyland, The Lighthouse Lab in Alderley Park and Alex Alderton, Roberto Amato, Jeffrey Barrett, Sonia Goncalves, Ewan Harrison, David K. Jackson, Ian Johnston, Dominic Kwiatkowski, Cordelia Langford, John Sillitoe on behalf of the Wellcome Sanger Institute COVID-19 Surveillance Team |
| EPI_ISL_2138646, EPI_ISL_2138648 | Lighthouse Lab in Milton Keynes | Wellcome Sanger Institute for the COVID-19 Genomics UK (COG-UK) Consortium | The Lighthouse Lab in Milton Keynes and Alex Alderton, Roberto Amato, Jeffrey Barrett, Sonia Goncalves, Ewan Harrison, David K. Jackson, Ian Johnston, Dominic Kwiatkowski, Cordelia Langford, John Sillitoe on behalf of the Wellcome Sanger Institute COVID-19 Surveillance Team |
| EPI_ISL_2138650 | Lighthouse Lab in Alderley Park | Wellcome Sanger Institute for the COVID-19 Genomics UK (COG-UK) Consortium | Jacquelyn Wynn, Mairead Hyland, The Lighthouse Lab in Alderley Park and Alex Alderton, Roberto Amato, Jeffrey Barrett, Sonia Goncalves, Ewan Harrison, David K. Jackson, Ian Johnston, Dominic Kwiatkowski, Cordelia Langford, John Sillitoe on behalf of the Wellcome Sanger Institute COVID-19 Surveillance Team |
| EPI_ISL_2138651 | Lighthouse Lab in Milton Keynes | Wellcome Sanger Institute for the COVID-19 Genomics UK (COG-UK) Consortium | The Lighthouse Lab in Milton Keynes and Alex Alderton, Roberto Amato, Jeffrey Barrett, Sonia Goncalves, Ewan Harrison, David K. Jackson, Ian Johnston, Dominic Kwiatkowski, Cordelia Langford, John Sillitoe on behalf of the Wellcome Sanger Institute COVID-19 Surveillance Team |
| EPI_ISL_2138653 | Lighthouse Lab in Alderley Park | Wellcome Sanger Institute for the COVID-19 Genomics UK (COG-UK) Consortium | Jacquelyn Wynn, Mairead Hyland, The Lighthouse Lab in Alderley Park and Alex Alderton, Roberto Amato, Jeffrey Barrett, Sonia Goncalves, Ewan Harrison, David K. Jackson, Ian Johnston, Dominic Kwiatkowski, Cordelia Langford, John Sillitoe on behalf of the Wellcome Sanger Institute COVID-19 Surveillance Team |
| EPI_ISL_2138658, EPI_ISL_2138661, EPI_ISL_2138664, EPI_ISL_2138667, EPI_ISL_2138672, EPI_ISL_2138673 | Lighthouse Lab in Milton Keynes | Wellcome Sanger Institute for the COVID-19 Genomics UK (COG-UK) Consortium | The Lighthouse Lab in Milton Keynes and Alex Alderton, Roberto Amato, Jeffrey Barrett, Sonia Goncalves, Ewan Harrison, David K. Jackson, Ian Johnston, Dominic Kwiatkowski, Cordelia Langford, John Sillitoe on behalf of the Wellcome Sanger Institute COVID-19 Surveillance Team |
| EPI_ISL_2138675 | Lighthouse Lab in Alderley Park | Wellcome Sanger Institute for the COVID-19 Genomics UK (COG-UK) Consortium | Jacquelyn Wynn, Mairead Hyland, The Lighthouse Lab in Alderley Park and Alex Alderton, Roberto Amato, Jeffrey Barrett, Sonia Goncalves, Ewan Harrison, David K. Jackson, Ian Johnston, Dominic Kwiatkowski, Cordelia Langford, John Sillitoe on behalf of the Wellcome Sanger Institute COVID-19 Surveillance Team |
| EPI_ISL_2138677, EPI_ISL_2138679 | Lighthouse Lab in Milton Keynes | Wellcome Sanger Institute for the COVID-19 Genomics UK (COG-UK) Consortium | The Lighthouse Lab in Milton Keynes and Alex Alderton, Roberto Amato, Jeffrey Barrett, Sonia Goncalves, Ewan Harrison, David K. Jackson, Ian Johnston, Dominic Kwiatkowski, Cordelia Langford, John Sillitoe on behalf of the Wellcome Sanger Institute COVID-19 Surveillance Team |
| EPI_ISL_2138681 | Health Services Laboratories | Wellcome Sanger Institute for the COVID-19 Genomics UK (COG-UK) Consortium | Health Services Laboratories and Alex Alderton, Roberto Amato, Jeffrey Barrett, Sonia Goncalves, Ewan Harrison, David K. Jackson, Ian Johnston, Dominic Kwiatkowski, Cordelia Langford, John Sillitoe on behalf of the Wellcome Sanger Institute COVID-19 Surveillance Team |
| EPI_ISL_2138682 | Lighthouse Lab in Milton Keynes | Wellcome Sanger Institute for the COVID-19 Genomics UK (COG-UK) Consortium | The Lighthouse Lab in Milton Keynes and Alex Alderton, Roberto Amato, Jeffrey Barrett, Sonia Goncalves, Ewan Harrison, David K. Jackson, Ian Johnston, Dominic Kwiatkowski, Cordelia Langford, John Sillitoe on behalf of the Wellcome Sanger Institute COVID-19 Surveillance Team |
| EPI_ISL_2138683 | Lighthouse Lab in Alderley Park | Wellcome Sanger Institute for the COVID-19 Genomics UK (COG-UK) Consortium | Jacquelyn Wynn, Mairead Hyland, The Lighthouse Lab in Alderley Park and Alex Alderton, Roberto Amato, Jeffrey Barrett, Sonia Goncalves, Ewan Harrison, David K. Jackson, Ian Johnston, Dominic Kwiatkowski, Cordelia Langford, John Sillitoe on behalf of the Wellcome Sanger Institute COVID-19 Surveillance Team |
| EPI_ISL_2138684 | Health Services Laboratories | Wellcome Sanger Institute for the COVID-19 Genomics UK (COG-UK) Consortium | Health Services Laboratories and Alex Alderton, Roberto Amato, Jeffrey Barrett, Sonia Goncalves, Ewan Harrison, David K. Jackson, Ian Johnston, Dominic Kwiatkowski, Cordelia Langford, John Sillitoe on behalf of the Wellcome Sanger Institute COVID-19 Surveillance Team |
| EPI_ISL_2138685 | Lighthouse Lab in Milton Keynes | Wellcome Sanger Institute for the COVID-19 Genomics UK (COG-UK) Consortium | The Lighthouse Lab in Milton Keynes and Alex Alderton, Roberto Amato, Jeffrey Barrett, Sonia Goncalves, Ewan Harrison, David K. Jackson, Ian Johnston, Dominic Kwiatkowski, Cordelia Langford, John Sillitoe on behalf of the Wellcome Sanger Institute COVID-19 Surveillance Team |
| EPI_ISL_2138686 | Lighthouse Lab in Alderley Park | Wellcome Sanger Institute for the COVID-19 Genomics UK (COG-UK) Consortium | Jacquelyn Wynn, Mairead Hyland, The Lighthouse Lab in Alderley Park and Alex Alderton, Roberto Amato, Jeffrey Barrett, Sonia Goncalves, Ewan Harrison, David K. Jackson, Ian Johnston, Dominic Kwiatkowski, Cordelia Langford, John Sillitoe on behalf of the Wellcome Sanger Institute COVID-19 Surveillance Team |

| | | | |
|---|---|---|---|
| EPI_ISL_2138687 | Lighthouse Lab in Milton Keynes | Wellcome Sanger Institute for the COVID-19 Genomics UK (COG-UK) Consortium | The Lighthouse Lab in Milton Keynes and Alex Alderton, Roberto Amato, Jeffrey Barrett, Sonia Goncalves, Ewan Harrison, David K. Jackson, Ian Johnston, Dominic Kwiatkowski, Cordelia Langford, John Sillitoe on behalf of the Wellcome Sanger Institute COVID-19 Surveillance Team |
| EPI_ISL_2138692, EPI_ISL_2138693 | Lighthouse Lab in Alderley Park | Wellcome Sanger Institute for the COVID-19 Genomics UK (COG-UK) Consortium | Jacquelyn Wynn, Mairead Hyland, The Lighthouse Lab in Alderley Park and Alex Alderton, Roberto Amato, Jeffrey Barrett, Sonia Goncalves, Ewan Harrison, David K. Jackson, Ian Johnston, Dominic Kwiatkowski, Cordelia Langford, John Sillitoe on behalf of the Wellcome Sanger Institute COVID-19 Surveillance Team |
| EPI_ISL_2138694 | Health Services Laboratories | Wellcome Sanger Institute for the COVID-19 Genomics UK (COG-UK) Consortium | Health Services Laboratories and Alex Alderton, Roberto Amato, Jeffrey Barrett, Sonia Goncalves, Ewan Harrison, David K. Jackson, Ian Johnston, Dominic Kwiatkowski, Cordelia Langford, John Sillitoe on behalf of the Wellcome Sanger Institute COVID-19 Surveillance Team |
| EPI_ISL_2138695 | Lighthouse Lab in Milton Keynes | Wellcome Sanger Institute for the COVID-19 Genomics UK (COG-UK) Consortium | The Lighthouse Lab in Milton Keynes and Alex Alderton, Roberto Amato, Jeffrey Barrett, Sonia Goncalves, Ewan Harrison, David K. Jackson, Ian Johnston, Dominic Kwiatkowski, Cordelia Langford, John Sillitoe on behalf of the Wellcome Sanger Institute COVID-19 Surveillance Team |
| EPI_ISL_2138697, EPI_ISL_2138698 | Lighthouse Lab in Alderley Park | Wellcome Sanger Institute for the COVID-19 Genomics UK (COG-UK) Consortium | Jacquelyn Wynn, Mairead Hyland, The Lighthouse Lab in Alderley Park and Alex Alderton, Roberto Amato, Jeffrey Barrett, Sonia Goncalves, Ewan Harrison, David K. Jackson, Ian Johnston, Dominic Kwiatkowski, Cordelia Langford, John Sillitoe on behalf of the Wellcome Sanger Institute COVID-19 Surveillance Team |
| EPI_ISL_2138702, EPI_ISL_2138703, EPI_ISL_2138704, EPI_ISL_2138706, EPI_ISL_2138707, EPI_ISL_2138711, EPI_ISL_2138712, EPI_ISL_2138714, EPI_ISL_2138716 | Lighthouse Lab in Milton Keynes | Wellcome Sanger Institute for the COVID-19 Genomics UK (COG-UK) Consortium | The Lighthouse Lab in Milton Keynes and Alex Alderton, Roberto Amato, Jeffrey Barrett, Sonia Goncalves, Ewan Harrison, David K. Jackson, Ian Johnston, Dominic Kwiatkowski, Cordelia Langford, John Sillitoe on behalf of the Wellcome Sanger Institute COVID-19 Surveillance Team |
| EPI_ISL_2139194, EPI_ISL_2139219, EPI_ISL_2139411, EPI_ISL_2139416 | Public Health Ontario Laboratory | Public Health Ontario Laboratory | Vanessa G Allen, Philip Banh, Yao Chen, Richard de Borja, Alireza Eshaghi, Nahuel Fittipaldi, Christine Frantz, Jonathan B Gubbay, Jennifer L Guthrie, Lawrence Heisler, Esha Joshi, Michael Laszloffy, Aimin Li, Michael CY Li, Dean Maxwell, Sandeep Nagra, Samir N Patel, Jared Simpson, Karthikeyan Sivaraman, Ashleigh Sullivan, Yogi Sundaravadanam, Sarah Teatero, Andre Villegas, Matthew Watson, Sandra Zittermann |
| EPI_ISL_2139737 | University of Liège COVID-19 testing center | GIGA Medical Genomics | Keith Durkin, Maria Artesi, Bouchra Boujemla, Nathalie Renotte, Cécile Meex, Sébastien Bontems, Fabrice Bureau, Laurent Gillet, Wouter Coppieters, Marie-Pierre Hayette, Vincent Bours |
| EPI_ISL_2140020, EPI_ISL_2140036 | Illinois Department of Public Health | Illinois Department of Public Health - Chicago Lab | Vineet K. Dhiman, Ira Heimler, Joel Price |
| EPI_ISL_2140071 | Servicio Virosis Respiratorias-Departamento Virología-INEI | Instituto Nacional Enfermedades Infecciosas C.G.Malbran | Baumeister E., Avaro M., Benedetti E., Russo M., Dattero ME, Pontoriero A., Cisterna D., Molina V., Perandones C., Tuduri E., Lorenzo F., Poklepovich T., Campos J. |
| EPI_ISL_2140237 | Zakład Diagnostyki Laboratoryjnej SPWSZ | 1. Tricity SARS-CoV-2 sequencing consortium: University of Gdansk, Medical University of Gdansk, Vaxican Ltd., Invicta Ltd. 2. National Institute of Public Health - National Institute of Hygiene, Warsaw, Poland | Maciej Kosinski, Celina Cybulska, Krystyna Bienkowska Szewczyk, Maciej Grzybek, Karolina Gackowska, Katarzyna Groth, Lukasz Rabalski, Katarzyna Zacharczuk, Magdalena Nowakowska, Małgorzata Sadkowska-Todys, Tomasz Wołkowicz |
| EPI_ISL_2140238, EPI_ISL_2140239, EPI_ISL_2140240, EPI_ISL_2140241, EPI_ISL_2140242, EPI_ISL_2140243, EPI_ISL_2140244, EPI_ISL_2140245 | Laboratorium Analiz Lekarskich ALAB | 1. Tricity SARS-CoV-2 sequencing consortium: University of Gdansk, Medical University of Gdansk, Vaxican Ltd., Invicta Ltd. 2. National Institute of Public Health - National Institute of Hygiene, Warsaw, Poland | Maciej Kosinski, Celina Cybulska, Krystyna Bienkowska Szewczyk, Maciej Grzybek, Karolina Gackowska, Katarzyna Groth, Lukasz Rabalski, Katarzyna Zacharczuk, Magdalena Nowakowska, Małgorzata Sadkowska-Todys, Tomasz Wołkowicz |
| EPI_ISL_2140365, EPI_ISL_2140366, EPI_ISL_2140367 | Diagnostyka Sp. z o.o. | 1. Tricity SARS-CoV-2 sequencing consortium: University of Gdansk, Medical University of Gdansk, Vaxican Ltd., Invicta Ltd. 2. National Institute of Public Health - National Institute of Hygiene, Warsaw, Poland | Maciej Kosinski, Celina Cybulska, Krystyna Bienkowska Szewczyk, Maciej Grzybek, Karolina Gackowska, Katarzyna Groth, Lukasz Rabalski, Katarzyna Zacharczuk, Magdalena Nowakowska, Małgorzata Sadkowska-Todys, Tomasz Wołkowicz |
| EPI_ISL_2140524 | UCK | 1. Tricity SARS-CoV-2 sequencing consortium: University of Gdansk, Medical University of Gdansk, Vaxican Ltd., Invicta Ltd. 2. National Institute of Public Health - National Institute of Hygiene, Warsaw, Poland | Maciej Kosinski, Celina Cybulska, Krystyna Bienkowska Szewczyk, Maciej Grzybek, Karolina Gackowska, Katarzyna Groth, Lukasz Rabalski, Katarzyna Zacharczuk, Magdalena Nowakowska, Małgorzata Sadkowska-Todys, Tomasz Wołkowicz |
| EPI_ISL_2140611, EPI_ISL_2140641 | Microbiology Department, Laboratori Clínic Metropolitana Nord. Hospital Universitari Germans Trias i Pujol | Can Ruti SARS-CoV-2 Sequencing Hub (HUGTiP/IrsiCaixa/IGTP) | Marc Noguera-Julian, Pilar Armengol, Ignacio Blanco, Antoni E Bordoy, Francesc Catala-Moll, Pere-Joan Cardona, Maria Casadellà, Cristina Casañ, Gemma Clara, Bonaventura Clotet, Cristina Esteban, Montserrat Giménez, Mercedes Guerrero, Anna Not, Roger Paredes, Mariona Parera, Verónica Saludes, Alba Sánchez, and Elisa Martró on behalf of the Can Ruti SARS-CoV-2 Sequencing Hub. |
| EPI_ISL_2140878 | The Ohio State University Applied Microbiology Services Laboratory | The Ohio State University Applied Microbiology Services Laboratory | Seth A. Faith PhD |
| EPI_ISL_2140890, EPI_ISL_2140894, EPI_ISL_2140904, EPI_ISL_2140908, EPI_ISL_2140909, EPI_ISL_2140941, EPI_ISL_2140944, EPI_ISL_2140948, EPI_ISL_2141245, EPI_ISL_2141270, EPI_ISL_2141271, EPI_ISL_2141272 | | | |
| see above | Public Health Ontario Laboratory | Public Health Ontario Laboratory | Vanessa G Allen, Philip Banh, Yao Chen, Richard de Borja, Alireza Eshaghi, Nahuel Fittipaldi, Christine Frantz, Jonathan B Gubbay, Jennifer L Guthrie, Lawrence Heisler, Esha Joshi, Michael Laszloffy, Aimin Li, Michael CY Li, Dean Maxwell, Sandeep Nagra, Samir N Patel, Jared Simpson, Karthikeyan Sivaraman, Ashleigh Sullivan, Yogi Sundaravadanam, Sarah Teatero, Andre Villegas, Matthew Watson, Sandra Zittermann |
| EPI_ISL_2141442 | TXDSHS | TXDSHS | Rashmi Tuladhar, Bonnie Oh, Jenny Zhang, Maliha Rahman, Mayela Pedrueza, Anita Pokharel, Karen Bobier, Lorraine Rodriguez, Myong Koag, Chun Wang, Rachel Lee, Grace Kubin |
| EPI_ISL_2142418, EPI_ISL_2142435 | HOPITAL SAINT ANDRE | CNR Virus des Infections Respiratoires - France SUD | Antonin Bal, Gregory Destras, Gwendolyne Burfin, Hadrien Regue, Quentin Semanas, Martine Valette, Bruno Lina, Laurence Josset |
| EPI_ISL_2142939, EPI_ISL_2142947, EPI_ISL_2143008, EPI_ISL_2143023, EPI_ISL_2143064, EPI_ISL_2143096, EPI_ISL_2143229, EPI_ISL_2143359, EPI_ISL_2143363, EPI_ISL_2143449 | Quest Diagnostics Incorporated | Centers for Disease Control and Prevention Division of Viral Diseases, Pathogen Discovery | Dakota Howard, Dhwani Batra, Peter W. Cook, Kara Moser, Adrian Paskey, Jason Caravas, Benjamin Rambo-Martin, Shatavia Morrison, Christopher Gulvick, Scott Sammons, Yvette Unoarumhi, Darlene Wagner, Matthew Schmerer, S. H. Rosenthal, A. Gerasimova, R. M. Kagan, B. Anderson, M. Hua, Y. Liu, L.E. Bernstein, K.E. Livingston, A. Perez, I. A. Shlyakhter, R. V. Rolando, R. Owen, P. Tanpaiboon, F. Lacbawan, Clinton R. Paden, Duncan MacCannell |
| EPI_ISL_2143486, EPI_ISL_2143633 | CHU LILLE | CHU Lille - Laboratoire de Virologie | AIT YAHYA Emilie, ALIDJINOU Enagnon Kazali, BOCKET Laurence, CREPIN Michel, DEMAY Christophe, ENGELMANN Ilka, GEFFROY Sandrine, GUIGON Aurélie, LAMBERT Valérie, LAZREK Mouna, NOBILLIAUX Florian, PREVOST Brigitte, THUILLIER Caroline, TINEZ Claire |
| EPI_ISL_2143679, EPI_ISL_2143750, EPI_ISL_2143921 | Quest Diagnostics Incorporated | Centers for Disease Control and Prevention Division of Viral Diseases, Pathogen Discovery | Dakota Howard, Dhwani Batra, Peter W. Cook, Kara Moser, Adrian Paskey, Jason Caravas, Benjamin Rambo-Martin, Shatavia Morrison, Christopher Gulvick, Scott Sammons, Yvette Unoarumhi, Darlene Wagner, Matthew Schmerer, S. H. Rosenthal, A. Gerasimova, R. M. Kagan, B. Anderson, M. Hua, Y. Liu, L.E. Bernstein, K.E. Livingston, A. Perez, I. A. Shlyakhter, R. V. Rolando, R. Owen, P. Tanpaiboon, F. Lacbawan, Clinton R. Paden, Duncan MacCannell |
| EPI_ISL_2144255, EPI_ISL_2144340, EPI_ISL_2144363, EPI_ISL_2144594, EPI_ISL_2145008, EPI_ISL_2145113, EPI_ISL_2145142 | Helix/Illumina | Centers for Disease Control and Prevention Division of Viral Diseases, Pathogen Discovery | Dakota Howard, Dhwani Batra, Peter W. Cook, Kara Moser, Adrian Paskey, Jason Caravas, Benjamin Rambo-Martin, Shatavia Morrison, Christopher Gulvick, Scott Sammons, Yvette Unoarumhi, Darlene Wagner, Matthew Schmerer, Eileen de Feo, Jan Antico, Christine Tran, Matthew Tolentino, Shannon Wickline, Kim Gietzen, Brad Sickler, Jingtao Liu, Eric Allen, Phil Febbo, Nicole L. Washington, Simon White, Geraint Levan, Kelly Schiabor Barrett, Elizabeth Cirulli, Alexandre Bolze, Ary Ascencio, Charlotte Rivera-Garcia, Ryan Cho, Jason Nguyen, Sherry Wang, Jimmy Ramirez, Tyler Cassens, Efren Sandoval, Magnus Isaksson, William Lee, David Becker, Marc Laurent, James Lu, Clinton R. Paden, Duncan MacCannell |
| EPI_ISL_2145143 | Dutch COVID-19 response team | Erasmus Medical Center | Bas Oude Munnink, Reina Sikkema, David Nieuwenhuijse, Irina Chestakova, Anne van der Linden, Marjan Boter, Emmanuelle Munger, Corine GeurtsvanKessel, Annemiek van der Eijk, Richard Molenkamp, Marion Koopmans, on behalf of the Dutch national COVID-19 respo |
| EPI_ISL_2145272 | Helix/Illumina | Centers for Disease Control and Prevention Division of Viral Diseases, Pathogen Discovery | Dakota Howard, Dhwani Batra, Peter W. Cook, Kara Moser, Adrian Paskey, Jason Caravas, Benjamin Rambo-Martin, Shatavia Morrison, Christopher Gulvick, Scott Sammons, Yvette Unoarumhi, Darlene Wagner, Matthew Schmerer, Eileen de Feo, Jan Antico, Christine Tran, Matthew Tolentino, Shannon Wickline, Kim Gietzen, Brad Sickler, Jingtao Liu, Eric Allen, Phil Febbo, Nicole L. Washington, Simon White, Geraint Levan, Kelly Schiabor Barrett, Elizabeth Cirulli, Alexandre Bolze, Ary Ascencio, Charlotte Rivera-Garcia, Ryan Cho, Jason Nguyen, Sherry Wang, Jimmy Ramirez, Tyler Cassens, Efren Sandoval, Magnus Isaksson, William Lee, David Becker, Marc Laurent, James Lu, Clinton R. Paden, Duncan MacCannell |
| EPI_ISL_2145682, EPI_ISL_2145703 | Dutch COVID-19 response team | Erasmus Medical Center | Bas Oude Munnink, Reina Sikkema, David Nieuwenhuijse, Irina Chestakova, Anne van der Linden, Marjan Boter, Emmanuelle Munger, Corine GeurtsvanKessel, Annemiek van der Eijk, Richard Molenkamp, Marion Koopmans, on behalf of the Dutch national COVID-19 respo |
| EPI_ISL_2145734 | Helix/Illumina | Centers for Disease Control and Prevention Division of Viral Diseases, Pathogen Discovery | Dakota Howard, Dhwani Batra, Peter W. Cook, Kara Moser, Adrian Paskey, Jason Caravas, Benjamin Rambo-Martin, Shatavia Morrison, Christopher Gulvick, Scott Sammons, Yvette Unoarumhi, Darlene Wagner, Matthew Schmerer, Eileen de Feo, Jan Antico, Christine Tran, Matthew Tolentino, Shannon Wickline, Kim Gietzen, Brad Sickler, Jingtao Liu, Eric Allen, Phil Febbo, Nicole L. Washington, Simon White, Geraint Levan, Kelly Schiabor Barrett, Elizabeth Cirulli, Alexandre Bolze, Ary Ascencio, Charlotte Rivera-Garcia, Ryan Cho, Jason Nguyen, Sherry Wang, Jimmy Ramirez, Tyler Cassens, Efren Sandoval, Magnus Isaksson, William Lee, David Becker, Marc Laurent, James Lu, Clinton R. Paden, Duncan MacCannell |
| EPI_ISL_2145782, EPI_ISL_2145783 | Dutch COVID-19 response team | Erasmus Medical Center | Bas Oude Munnink, Reina Sikkema, David Nieuwenhuijse, Irina Chestakova, Anne van der Linden, Marjan Boter, Emmanuelle Munger, Corine GeurtsvanKessel, Annemiek van der Eijk, Richard Molenkamp, Marion Koopmans, on behalf of the Dutch national COVID-19 respo |
| EPI_ISL_2145844, EPI_ISL_2145852, EPI_ISL_2145882, EPI_ISL_2145895, EPI_ISL_2145921, EPI_ISL_2146150, EPI_ISL_2146175, EPI_ISL_2146201, EPI_ISL_2146219, EPI_ISL_2146328, EPI_ISL_2146490, EPI_ISL_2146496, EPI_ISL_2146559, EPI_ISL_2146561, EPI_ISL_2146606, EPI_ISL_2146607, EPI_ISL_2146619, EPI_ISL_2146645, EPI_ISL_2146687, EPI_ISL_2146693, EPI_ISL_2146755, EPI_ISL_2146764, EPI_ISL_2146810, EPI_ISL_2146890, EPI_ISL_2146898, EPI_ISL_2146944, EPI_ISL_2146965, EPI_ISL_2147072, EPI_ISL_2147105, EPI_ISL_2147217, EPI_ISL_2147408, EPI_ISL_2147487, EPI_ISL_2147489, EPI_ISL_2147511, EPI_ISL_2147592, EPI_ISL_2147593, EPI_ISL_2147680, EPI_ISL_2147707, EPI_ISL_2147709, EPI_ISL_2147821, EPI_ISL_2147874, EPI_ISL_2147930, EPI_ISL_2147983, EPI_ISL_2147992, EPI_ISL_2148058, EPI_ISL_2148082, EPI_ISL_2148120, EPI_ISL_2148121, EPI_ISL_2148496, EPI_ISL_2148548, EPI_ISL_2148604, EPI_ISL_2148677, EPI_ISL_2148801, EPI_ISL_2148833, EPI_ISL_2148878, EPI_ISL_2149165, EPI_ISL_2149183, EPI_ISL_2149224, EPI_ISL_2149225, EPI_ISL_2149227, EPI_ISL_2149329, EPI_ISL_2149359, EPI_ISL_2149395, EPI_ISL_2149523, EPI_ISL_2149524, EPI_ISL_2149545, EPI_ISL_2149546, EPI_ISL_2149579, EPI_ISL_2149646, EPI_ISL_2149654, EPI_ISL_2149672, EPI_ISL_2149677, EPI_ISL_2149713, EPI_ISL_2149715, EPI_ISL_2149796, EPI_ISL_2149802, EPI_ISL_2149931, EPI_ISL_2150060, EPI_ISL_2150095, EPI_ISL_2150121, EPI_ISL_2150164, EPI_ISL_2150210, EPI_ISL_2150223, EPI_ISL_2150247, EPI_ISL_2150411, EPI_ISL_2150417, EPI_ISL_2150516, EPI_ISL_2150584, EPI_ISL_2150585, EPI_ISL_2150587, EPI_ISL_2150880 | | | |
| see above | Aegis Sciences Corporation | Centers for Disease Control and Prevention Division of Viral | Dakota Howard, Dhwani Batra, Peter W. Cook, Kara Moser, Adrian Paskey, Jason Caravas, Benjamin Rambo-Martin, Shatavia Morrison, Christopher Gulvick, Scott Sammons, Yvette Unoarumhi, Darlene Wagner, Matthew Schmerer, Cyndi Clark, |

| | | | |
|---|---|---|---|
| | | Diseases, Pathogen Discovery | Patrick Campbell, Rob Case, Vikramsinha Ghorpade, Holly Houdeshell, Ola Kvalvaag, Dillon Nall, Ethan Sanders, Alec Vest, Shaun Westlund, Matthew Hardison, Clinton R. Paden, Duncan MacCannell |
| EPI_ISL_2151096, EPI_ISL_2151097, EPI_ISL_2151098, EPI_ISL_2151099, EPI_ISL_2151100, EPI_ISL_2151101, EPI_ISL_2151102 | INSACOG-Sikkim | National Institute of Biomedical Genomics – INSACOG | Arindam Maitra, Tara D Sharma, Nidhan Kumar Biswas, Kaden Zangmu Bhutia, Sreedhar Chinnaswamy Tshering Ongmu Bhutia, Saumitra Das |
| EPI_ISL_2151164 | MS Palya PHC | inStem NCBS - INSACOG | Uma Ramakrishnan Dasaradhi Palakodeti Aswin SaiNarain |
| EPI_ISL_2151166, EPI_ISL_2151169, EPI_ISL_2151171, EPI_ISL_2151172, EPI_ISL_2151174, EPI_ISL_2151177, EPI_ISL_2151179, EPI_ISL_2151180 | Kodigehalli PHC | inStem NCBS - INSACOG | Uma Ramakrishnan Dasaradhi Palakodeti Aswin SaiNarain |
| EPI_ISL_2151181 | Sahakara Nagar PHC | inStem NCBS - INSACOG | Uma Ramakrishnan Dasaradhi Palakodeti Aswin SaiNarain |
| EPI_ISL_2151183, EPI_ISL_2151184, EPI_ISL_2151185, EPI_ISL_2151190, EPI_ISL_2151191, EPI_ISL_2151194, EPI_ISL_2151197, EPI_ISL_2151198, EPI_ISL_2151199, EPI_ISL_2151200, EPI_ISL_2151201, EPI_ISL_2151202, EPI_ISL_2151203, EPI_ISL_2151204, EPI_ISL_2151211, EPI_ISL_2151213, EPI_ISL_2151215, EPI_ISL_2151216, EPI_ISL_2151219, EPI_ISL_2151223, EPI_ISL_2151225, EPI_ISL_2151227, EPI_ISL_2151229, EPI_ISL_2151236, EPI_ISL_2151237, EPI_ISL_2151238, EPI_ISL_2151239, EPI_ISL_2151240, EPI_ISL_2151241, EPI_ISL_2151242, EPI_ISL_2151244 | | | |
| see above | Kodigehalli PHC | inStem NCBS - INSACOG | Uma Ramakrishnan Dasaradhi Palakodeti Aswin SaiNarain |
| EPI_ISL_2151245 | RAILWAY HOSPITAL | inStem NCBS - INSACOG | Uma Ramakrishnan Dasaradhi Palakodeti Aswin SaiNarain |
| EPI_ISL_2151246 | SANJAY GANDHI INSTITUTE OF TRAUMA AND ORTHOPAEDICS | inStem NCBS - INSACOG | Uma Ramakrishnan Dasaradhi Palakodeti Aswin SaiNarain |
| EPI_ISL_2151247, EPI_ISL_2151250 | NIMHANS | inStem NCBS - INSACOG | Uma Ramakrishnan Dasaradhi Palakodeti Aswin SaiNarain |
| EPI_ISL_2151251, EPI_ISL_2151252 | RAILWAY HOSPITAL | inStem NCBS - INSACOG | Uma Ramakrishnan Dasaradhi Palakodeti Aswin SaiNarain |
| EPI_ISL_2151253, EPI_ISL_2151255, EPI_ISL_2151256 | NIMHANS | inStem NCBS - INSACOG | Uma Ramakrishnan Dasaradhi Palakodeti Aswin SaiNarain |
| EPI_ISL_2151258 | SANJAY GANDHI INSTITUTE OF TRAUMA AND ORTHOPAEDICS | inStem NCBS - INSACOG | Uma Ramakrishnan Dasaradhi Palakodeti Aswin SaiNarain |
| EPI_ISL_2151259, EPI_ISL_2151260, EPI_ISL_2151261, EPI_ISL_2151262, EPI_ISL_2151263 | RAILWAY HOSPITAL | inStem NCBS - INSACOG | Uma Ramakrishnan Dasaradhi Palakodeti Aswin SaiNarain |
| EPI_ISL_2151264, EPI_ISL_2151266 | RH HOSAHALLI | inStem NCBS - INSACOG | Uma Ramakrishnan Dasaradhi Palakodeti Aswin SaiNarain |
| EPI_ISL_2151267 | SRI JAYADEVA INSTITUTE OF CARDIOLOGY | inStem NCBS - INSACOG | Uma Ramakrishnan Dasaradhi Palakodeti Aswin SaiNarain |
| EPI_ISL_2151269, EPI_ISL_2151270, EPI_ISL_2151271 | NIMHANS | inStem NCBS - INSACOG | Uma Ramakrishnan Dasaradhi Palakodeti Aswin SaiNarain |
| EPI_ISL_2151273 | RAILWAY HOSPITAL | inStem NCBS - INSACOG | Uma Ramakrishnan Dasaradhi Palakodeti Aswin SaiNarain |
| EPI_ISL_2151274, EPI_ISL_2151275 | JJR NAGAR UPHC | inStem NCBS - INSACOG | Uma Ramakrishnan Dasaradhi Palakodeti Aswin SaiNarain |
| EPI_ISL_2151279 | NIMHANS | inStem NCBS - INSACOG | Uma Ramakrishnan Dasaradhi Palakodeti Aswin SaiNarain |
| EPI_ISL_2151280 | SANJAY GANDHI INSTITUTE OF TRAUMA AND ORTHOPAEDICS | inStem NCBS - INSACOG | Uma Ramakrishnan Dasaradhi Palakodeti Aswin SaiNarain |
| EPI_ISL_2151281, EPI_ISL_2151282 | RAILWAY HOSPITAL | inStem NCBS - INSACOG | Uma Ramakrishnan Dasaradhi Palakodeti Aswin SaiNarain |
| EPI_ISL_2151284 | NIMHANS | inStem NCBS - INSACOG | Uma Ramakrishnan Dasaradhi Palakodeti Aswin SaiNarain |
| EPI_ISL_2151285, EPI_ISL_2151286 | RAILWAY HOSPITAL | inStem NCBS - INSACOG | Uma Ramakrishnan Dasaradhi Palakodeti Aswin SaiNarain |
| EPI_ISL_2151289, EPI_ISL_2151290, EPI_ISL_2151291, EPI_ISL_2151292, EPI_ISL_2151294 | NIMHANS | inStem NCBS - INSACOG | Uma Ramakrishnan Dasaradhi Palakodeti Aswin SaiNarain |
| EPI_ISL_2151295 | PHC AGARA | inStem NCBS - INSACOG | Uma Ramakrishnan Dasaradhi Palakodeti Aswin SaiNarain |
| EPI_ISL_2151296 | RAILWAY HOSPITAL | inStem NCBS - INSACOG | Uma Ramakrishnan Dasaradhi Palakodeti Aswin SaiNarain |
| EPI_ISL_2151297, EPI_ISL_2151298 | RAILWAY | inStem NCBS - INSACOG | Uma Ramakrishnan Dasaradhi Palakodeti Aswin SaiNarain |
| EPI_ISL_2151299, EPI_ISL_2151301 | RAILWAY HOSPITAL | inStem NCBS - INSACOG | Uma Ramakrishnan Dasaradhi Palakodeti Aswin SaiNarain |
| EPI_ISL_2151302, EPI_ISL_2151304, EPI_ISL_2151305, EPI_ISL_2151306, EPI_ISL_2151307, EPI_ISL_2151309, EPI_ISL_2151311, EPI_ISL_2151312 | NIMHANS | inStem NCBS - INSACOG | Uma Ramakrishnan Dasaradhi Palakodeti Aswin SaiNarain |
| EPI_ISL_2151313 | GOVINDRAJ NAGAR UPHC | inStem NCBS - INSACOG | Uma Ramakrishnan Dasaradhi Palakodeti Aswin SaiNarain |
| EPI_ISL_2151315, EPI_ISL_2151316, EPI_ISL_2151317, EPI_ISL_2151318, EPI_ISL_2151319 | RAILWAY HOSPITAL | inStem NCBS - INSACOG | Uma Ramakrishnan Dasaradhi Palakodeti Aswin SaiNarain |
| EPI_ISL_2151320, EPI_ISL_2151321, EPI_ISL_2151322 | RAILWAY | inStem NCBS - INSACOG | Uma Ramakrishnan Dasaradhi Palakodeti Aswin SaiNarain |
| EPI_ISL_2151323 | NIMHANS | inStem NCBS - INSACOG | Uma Ramakrishnan Dasaradhi Palakodeti Aswin SaiNarain |
| EPI_ISL_2151324 | UPHC PANTHARPALYA | inStem NCBS - INSACOG | Uma Ramakrishnan Dasaradhi Palakodeti Aswin SaiNarain |
| EPI_ISL_2151326 | RAILWAY HOSPITAL | inStem NCBS - INSACOG | Uma Ramakrishnan Dasaradhi Palakodeti Aswin SaiNarain |
| EPI_ISL_2151328 | RAILWAY | inStem NCBS - INSACOG | Uma Ramakrishnan Dasaradhi Palakodeti Aswin SaiNarain |
| EPI_ISL_2151329, EPI_ISL_2151330, EPI_ISL_2151331 | NIMHANS | inStem NCBS - INSACOG | Uma Ramakrishnan Dasaradhi Palakodeti Aswin SaiNarain |
| EPI_ISL_2151333 | RAILWAY | inStem NCBS - INSACOG | Uma Ramakrishnan Dasaradhi Palakodeti Aswin SaiNarain |
| EPI_ISL_2151871 | Hôpital Avicenne | Department of Virology, Henri Mondor University Hospital, Assistance Publique Hôpitaux de Paris, Université Paris-Est Créteil, INSERM U955 | Christophe Rodriguez, Slim Fourati, Vanessa Demontant, Guillaume Gricourt, Melissa N'Debi, Alexandre Soulier, Elisabeth Trawinski, Jean-Michel Pawlotsky |
| EPI_ISL_2152567, | BROUSSAIS | Department of Virology, Henri Mondor University Hospital, | Christophe Rodriguez, Slim Fourati, Vanessa Demontant, Guillaume Gricourt, Melissa N'Debi, Alexandre Soulier, Elisabeth Trawinski, Jean-Michel Pawlotsky |

| | | | |
|---|---|---|---|
| EPI_ISL_2152568, EPI_ISL_2152569, EPI_ISL_2152570, EPI_ISL_2152571 | | Assistance Publique Hôpitaux de Paris, Université Paris-Est Créteil, INSERM U955 | |
| EPI_ISL_2152656 | Lighthouse Lab in Alderley Park | Wellcome Sanger Institute for the COVID-19 Genomics UK (COG-UK) Consortium | Jacquelyn Wynn, Mairead Hyland, The Lighthouse Lab in Alderley Park and Alex Alderton, Roberto Amato, Jeffrey Barrett, Sonia Goncalves, Ewan Harrison, David K. Jackson, Ian Johnston, Dominic Kwiatkowski, Cordelia Langford, John Sillitoe on behalf of the Wellcome Sanger Institute COVID-19 Surveillance Team |
| EPI_ISL_2152658, EPI_ISL_2152660, EPI_ISL_2152661 | Lighthouse Lab in Milton Keynes | Wellcome Sanger Institute for the COVID-19 Genomics UK (COG-UK) Consortium | The Lighthouse Lab in Milton Keynes and Alex Alderton, Roberto Amato, Jeffrey Barrett, Sonia Goncalves, Ewan Harrison, David K. Jackson, Ian Johnston, Dominic Kwiatkowski, Cordelia Langford, John Sillitoe on behalf of the Wellcome Sanger Institute COVID-19 Surveillance Team |
| EPI_ISL_2152670, EPI_ISL_2152672, EPI_ISL_2152676, EPI_ISL_2152684, EPI_ISL_2152686, EPI_ISL_2152697, EPI_ISL_2152700, EPI_ISL_2152703, EPI_ISL_2152707, EPI_ISL_2152708, EPI_ISL_2152710, EPI_ISL_2152720, EPI_ISL_2152721, EPI_ISL_2152730, EPI_ISL_2152731, EPI_ISL_2152739, EPI_ISL_2152744, EPI_ISL_2152746, EPI_ISL_2152755, EPI_ISL_2152761, EPI_ISL_2152764, EPI_ISL_2152765, EPI_ISL_2152769, EPI_ISL_2152770, EPI_ISL_2152773, EPI_ISL_2152784, EPI_ISL_2152786, EPI_ISL_2152789, EPI_ISL_2152790, EPI_ISL_2152794, EPI_ISL_2152800, EPI_ISL_2152801, EPI_ISL_2152807, EPI_ISL_2152811, EPI_ISL_2152812, EPI_ISL_2152817, EPI_ISL_2152821, EPI_ISL_2152826, EPI_ISL_2152828, EPI_ISL_2152830, EPI_ISL_2152837, EPI_ISL_2152849, EPI_ISL_2152853, EPI_ISL_2152856, EPI_ISL_2152857, EPI_ISL_2152858, EPI_ISL_2152860, EPI_ISL_2152862, EPI_ISL_2152863, EPI_ISL_2152871, EPI_ISL_2152872, EPI_ISL_2152875, EPI_ISL_2152877, EPI_ISL_2152882, EPI_ISL_2152885, EPI_ISL_2152886, EPI_ISL_2152887 | | | |
| see above | Lighthouse Lab in Glasgow | Wellcome Sanger Institute for the COVID-19 Genomics UK (COG-UK) Consortium | Harper VanSteenhouse, Yumi Kasai, David Gray, Carol Clugston, Anna Dominiczak and Alex Alderton, Roberto Amato, Jeffrey Barrett, Sonia Goncalves, Ewan Harrison, David K. Jackson, Ian Johnston, Dominic Kwiatkowski, Cordelia Langford, John Sillitoe on behalf of the Wellcome Sanger Institute COVID-19 Surveillance Team |
| EPI_ISL_2152889 | Lighthouse Lab in Milton Keynes | Wellcome Sanger Institute for the COVID-19 Genomics UK (COG-UK) Consortium | The Lighthouse Lab in Milton Keynes and Alex Alderton, Roberto Amato, Jeffrey Barrett, Sonia Goncalves, Ewan Harrison, David K. Jackson, Ian Johnston, Dominic Kwiatkowski, Cordelia Langford, John Sillitoe on behalf of the Wellcome Sanger Institute COVID-19 Surveillance Team |
| EPI_ISL_2152890 | Lighthouse Lab in Glasgow | Wellcome Sanger Institute for the COVID-19 Genomics UK (COG-UK) Consortium | Harper VanSteenhouse, Yumi Kasai, David Gray, Carol Clugston, Anna Dominiczak and Alex Alderton, Roberto Amato, Jeffrey Barrett, Sonia Goncalves, Ewan Harrison, David K. Jackson, Ian Johnston, Dominic Kwiatkowski, Cordelia Langford, John Sillitoe on behalf of the Wellcome Sanger Institute COVID-19 Surveillance Team |
| EPI_ISL_2152896 | Lighthouse Lab in Milton Keynes | Wellcome Sanger Institute for the COVID-19 Genomics UK (COG-UK) Consortium | The Lighthouse Lab in Milton Keynes and Alex Alderton, Roberto Amato, Jeffrey Barrett, Sonia Goncalves, Ewan Harrison, David K. Jackson, Ian Johnston, Dominic Kwiatkowski, Cordelia Langford, John Sillitoe on behalf of the Wellcome Sanger Institute COVID-19 Surveillance Team |
| EPI_ISL_2152899, EPI_ISL_2152901, EPI_ISL_2152908, EPI_ISL_2152917, EPI_ISL_2152920, EPI_ISL_2152924, EPI_ISL_2152926, EPI_ISL_2152930, EPI_ISL_2152931, EPI_ISL_2152932, EPI_ISL_2152934, EPI_ISL_2152936, EPI_ISL_2152937, EPI_ISL_2152938, EPI_ISL_2152939, EPI_ISL_2152942, EPI_ISL_2152945, EPI_ISL_2152952, EPI_ISL_2152954, EPI_ISL_2152956, EPI_ISL_2152962, EPI_ISL_2152967, EPI_ISL_2152970, EPI_ISL_2152974, EPI_ISL_2152982, EPI_ISL_2152998, EPI_ISL_2153004, EPI_ISL_2153005, EPI_ISL_2153009, EPI_ISL_2153012 | | | |
| see above | Lighthouse Lab in Glasgow | Wellcome Sanger Institute for the COVID-19 Genomics UK (COG-UK) Consortium | Harper VanSteenhouse, Yumi Kasai, David Gray, Carol Clugston, Anna Dominiczak and Alex Alderton, Roberto Amato, Jeffrey Barrett, Sonia Goncalves, Ewan Harrison, David K. Jackson, Ian Johnston, Dominic Kwiatkowski, Cordelia Langford, John Sillitoe on behalf of the Wellcome Sanger Institute COVID-19 Surveillance Team |
| EPI_ISL_2153016 | Lighthouse Lab in Milton Keynes | Wellcome Sanger Institute for the COVID-19 Genomics UK (COG-UK) Consortium | The Lighthouse Lab in Milton Keynes and Alex Alderton, Roberto Amato, Jeffrey Barrett, Sonia Goncalves, Ewan Harrison, David K. Jackson, Ian Johnston, Dominic Kwiatkowski, Cordelia Langford, John Sillitoe on behalf of the Wellcome Sanger Institute COVID-19 Surveillance Team |
| EPI_ISL_2153025, EPI_ISL_2153026, EPI_ISL_2153031 | Lighthouse Lab in Glasgow | Wellcome Sanger Institute for the COVID-19 Genomics UK (COG-UK) Consortium | Harper VanSteenhouse, Yumi Kasai, David Gray, Carol Clugston, Anna Dominiczak and Alex Alderton, Roberto Amato, Jeffrey Barrett, Sonia Goncalves, Ewan Harrison, David K. Jackson, Ian Johnston, Dominic Kwiatkowski, Cordelia Langford, John Sillitoe on behalf of the Wellcome Sanger Institute COVID-19 Surveillance Team |
| EPI_ISL_2153044 | Lighthouse Lab in Milton Keynes | Wellcome Sanger Institute for the COVID-19 Genomics UK (COG-UK) Consortium | The Lighthouse Lab in Milton Keynes and Alex Alderton, Roberto Amato, Jeffrey Barrett, Sonia Goncalves, Ewan Harrison, David K. Jackson, Ian Johnston, Dominic Kwiatkowski, Cordelia Langford, John Sillitoe on behalf of the Wellcome Sanger Institute COVID-19 Surveillance Team |
| EPI_ISL_2153065, EPI_ISL_2153066 | Israel Central Virology laboratory | Israel Central Virology laboratory | Neta Zuckerman, Efrat Dahan Bucris, Michal Mandelboim, Yaniv Lustig, Dana Bar-Ilan, Miranda Geva, Oran Erster, Danit Sofer, Ella Mendelson, Orna Mor |
| EPI_ISL_2153476 | Botswana Harvard HIV Reference Laboratory | Botswana Harvard HIV Reference Laboratory | Sikhulile Moyo, Wonderful T. Choga, Pamela Smith-Lawrence, Kgomotso Moruisi, Dorcas Maruapula, Keoratile Ntshambiwa, Thongbotho Mphoyakgosi, Boitumelo Zuze, Botshelo Radibe, Legodile Kooepile, Ontlametse T. Bareng, Roger Shapiro, Shahin Lockman, Joseph Makhema, Madisa Mine, Mosepele Mosepele, Simani Gaseitsiwe |
| EPI_ISL_2155359 | Laboratorio HUB -Azienda Ospedaliero Universitaria - AOU – Cagliari | Laboratorio SPOKE Biologia Molecolare -Azienda Ospedaliero Universitaria - AOU – Cagliari | Germano Orrù, Sara Fais, Valentina Medda, Alessandra Scano, Miriam Loddo, Riccardo Cappai, Ferdinando Coghe |
| EPI_ISL_2155360, EPI_ISL_2155361 | Laboratorio HUB -Azienda Ospedaliero Universitaria - AOU – Cagliari | Laboratorio SPOKE Biologia Molecolare -Azienda Ospedaliero Universitaria - AOU – Cagliari | Sara Fais, Valentina Medda, Alessandra Scano, Germano Orrù, Miriam Loddo, Riccardo Cappai, Ferdinando Coghe |
| EPI_ISL_2155362, EPI_ISL_2155363, EPI_ISL_2155364, EPI_ISL_2155365, EPI_ISL_2155366, EPI_ISL_2155367, EPI_ISL_2155368, EPI_ISL_2155369 | Laboratorio HUB -Azienda Ospedaliero Universitaria - AOU – Cagliari | Laboratorio SPOKE Biologia Molecolare -Azienda Ospedaliero Universitaria - AOU – Cagliari | Germano Orrù, Sara Fais, Valentina Medda, Alessandra Scano, Miriam Loddo, Riccardo Cappai, Ferdinando Coghe |
| EPI_ISL_2156833, EPI_ISL_2156836, EPI_ISL_2156837, EPI_ISL_2156840, EPI_ISL_2156843 | CSIR-Centre for Cellular and Molecular Biology | CSIR-Centre for Cellular and Molecular Biology-INSACOG | Tulasi Nagabandi, Lamuk Zaveri,Ara Sreenivas,Shreekant Verma, Amareshwar Vodapalli ,Blessy B John,Viswagithe S L,B Himasri,Valli Nagalakshmi Undamatla,Payel Mukherjee,Sofia Banu,Archana Bharadwaj Siva,Sharath Chandra Thota,Karthik Bharadwaj Tallapaka,Rakesh K Mishra,Divya Tej Sowpati |
| EPI_ISL_2156844, EPI_ISL_2156845, EPI_ISL_2156846, EPI_ISL_2156849, EPI_ISL_2156850, EPI_ISL_2156851, EPI_ISL_2156852, EPI_ISL_2156856, EPI_ISL_2156857, EPI_ISL_2156858, EPI_ISL_2156859, EPI_ISL_2156860, EPI_ISL_2156862, EPI_ISL_2156863, EPI_ISL_2156864, EPI_ISL_2156865, EPI_ISL_2156866, EPI_ISL_2156867, EPI_ISL_2156868, EPI_ISL_2156869, EPI_ISL_2156870 | | | |
| see above | CSIR-Centre for Cellular and Molecular Biology | CSIR-Centre for Cellular and Molecular Biology-INSACOG | Blessy B John,Viswagithe S L,Tulasi Nagabandi, Lamuk Zaveri,Ara Sreenivas,Shreekant Verma, Amareshwar Vodapalli ,B Himasri,Valli Nagalakshmi Undamatla,Payel Mukherjee,Sofia Banu,Archana Bharadwaj Siva,Sharath Chandra Thota,Karthik Bharadwaj Tallapaka,Rakesh K Mishra,Divya Tej Sowpati |
| EPI_ISL_2156874, EPI_ISL_2156875, EPI_ISL_2156876 | CSIR-Centre for Cellular and Molecular Biology | CSIR-Centre for Cellular and Molecular Biology-INSACOG | Tulasi Nagabandi, Lamuk Zaveri,Ara Sreenivas,Shreekant Verma, Amareshwar Vodapalli ,Blessy B John,Viswagithe S L,B Himasri,Valli Nagalakshmi Undamatla,Payel Mukherjee,Sofia Banu,Archana Bharadwaj Siva,Sharath Chandra Thota,Karthik Bharadwaj Tallapaka,Rakesh K Mishra,Divya Tej Sowpati |
| EPI_ISL_2156877, EPI_ISL_2156878, EPI_ISL_2156879, EPI_ISL_2156880, EPI_ISL_2156882 | CSIR-Centre for Cellular and Molecular Biology | CSIR-Centre for Cellular and Molecular Biology-INSACOG | Blessy B John,Viswagithe S L,Tulasi Nagabandi, Lamuk Zaveri,Ara Sreenivas,Shreekant Verma, Amareshwar Vodapalli ,B Himasri,Valli Nagalakshmi Undamatla,Payel Mukherjee,Sofia Banu,Archana Bharadwaj Siva,Sharath Chandra Thota,Karthik Bharadwaj Tallapaka,Rakesh K Mishra,Divya Tej Sowpati |
| EPI_ISL_2156884, EPI_ISL_2156885, EPI_ISL_2156886, EPI_ISL_2156887, EPI_ISL_2156888, EPI_ISL_2156891, EPI_ISL_2156892, EPI_ISL_2156893, EPI_ISL_2156894, EPI_ISL_2156895, EPI_ISL_2156898, EPI_ISL_2156899, EPI_ISL_2156900, EPI_ISL_2156901, EPI_ISL_2156902, EPI_ISL_2156905, EPI_ISL_2156906, EPI_ISL_2156907, EPI_ISL_2156908, EPI_ISL_2156913 | | | |
| see above | CSIR-Centre for Cellular and Molecular Biology | CSIR-Centre for Cellular and Molecular Biology-INSACOG | Ara Sreenivas,Shreekant Verma,Tulasi Nagabandi, Lamuk Zaveri, Amareshwar Vodapalli ,Blessy B John,Viswagithe S L,B Himasri,Valli Nagalakshmi Undamatla,Payel Mukherjee,Sofia Banu,Archana Bharadwaj Siva,Sharath Chandra Thota,Karthik Bharadwaj Tallapaka,Rakesh K Mishra,Divya Tej Sowpati |
| EPI_ISL_2156920, EPI_ISL_2156932 | Mapmygenome | CSIR-Centre for Cellular and Molecular Biology-INSACOG | Lamuk Zaveri,Tulasi Nagabandi, Ara Sreenivas,Shreekant Verma, Amareshwar Vodapalli ,Blessy B John,Viswagithe S L,B Himasri,Valli Nagalakshmi Undamatla,Payel Mukherjee,Sofia Banu,Archana Bharadwaj Siva,Sharath Chandra Thota,Karthik Bharadwaj Tallapaka,Rakesh K Mishra,Divya Tej Sowpati |
| EPI_ISL_2156939, EPI_ISL_2156940, EPI_ISL_2156941, EPI_ISL_2156942, EPI_ISL_2156943, EPI_ISL_2156944, EPI_ISL_2156947, EPI_ISL_2156948, EPI_ISL_2156949, EPI_ISL_2156950 | CSIR-Centre for Cellular and Molecular Biology | CSIR-Centre for Cellular and Molecular Biology-INSACOG | Blessy B John,Viswagithe S L,Tulasi Nagabandi, Lamuk Zaveri,Ara Sreenivas,Shreekant Verma, Amareshwar Vodapalli ,B Himasri,Valli Nagalakshmi Undamatla,Payel Mukherjee,Sofia Banu,Archana Bharadwaj Siva,Sharath Chandra Thota,Karthik Bharadwaj Tallapaka,Rakesh K Mishra,Divya Tej Sowpati |
| EPI_ISL_2156958, EPI_ISL_2156964, EPI_ISL_2156968, EPI_ISL_2156969 | CSIR-Centre for Cellular and Molecular Biology | CSIR-Centre for Cellular and Molecular Biology-INSACOG | Ara Sreenivas,Shreekant Verma,Tulasi Nagabandi, Lamuk Zaveri, Amareshwar Vodapalli ,Blessy B John,Viswagithe S L,B Himasri,Valli Nagalakshmi Undamatla,Payel Mukherjee,Sofia Banu,Archana Bharadwaj Siva,Sharath Chandra Thota,Karthik Bharadwaj Tallapaka,Rakesh K Mishra,Divya Tej Sowpati |
| EPI_ISL_2156984, EPI_ISL_2156990 | CSIR-Centre for Cellular and Molecular Biology | CSIR-Centre for Cellular and Molecular Biology-INSACOG | Tulasi Nagabandi, Lamuk Zaveri,Ara Sreenivas,Shreekant Verma, Amareshwar Vodapalli ,Blessy B John,Viswagithe S L,B Himasri,Valli Nagalakshmi Undamatla,Payel Mukherjee,Sofia Banu,Archana Bharadwaj Siva,Sharath Chandra Thota,Karthik Bharadwaj Tallapaka,Rakesh K Mishra,Divya Tej Sowpati |
| EPI_ISL_2156991, EPI_ISL_2156994, EPI_ISL_2156995 | CSIR-Centre for Cellular and Molecular Biology | CSIR-Centre for Cellular and Molecular Biology-INSACOG | Payel Mukherjee,Lamuk Zaveri,Tulasi Nagabandi, Ara Sreenivas,Shreekant Verma, Amareshwar Vodapalli ,Blessy B John,Viswagithe S L,B Himasri,Valli Nagalakshmi Undamatla,Sofia Banu,Archana Bharadwaj Siva,Sharath Chandra Thota,Karthik Bharadwaj Tallapaka,Rakesh K Mishra,Divya Tej Sowpati |
| EPI_ISL_2156996, EPI_ISL_2156997, EPI_ISL_2156999, EPI_ISL_2157000, EPI_ISL_2157001 | CSIR-Centre for Cellular and Molecular Biology | CSIR-Centre for Cellular and Molecular Biology-INSACOG | B Himasri,Valli Nagalakshmi Undamatla,Lamuk Zaveri,Tulasi Nagabandi, Ara Sreenivas,Shreekant Verma, Amareshwar Vodapalli ,Blessy B John,Viswagithe S L,Payel Mukherjee,Sofia Banu,Archana Bharadwaj Siva,Sharath Chandra Thota,Karthik Bharadwaj Tallapaka,Rakesh K Mishra,Divya Tej Sowpati |
| EPI_ISL_2157010, EPI_ISL_2157017, EPI_ISL_2157018 | CSIR-Centre for Cellular and Molecular Biology | CSIR-Centre for Cellular and Molecular Biology-INSACOG | Blessy B John,Viswagithe S L,Tulasi Nagabandi, Lamuk Zaveri,Ara Sreenivas,Shreekant Verma, Amareshwar Vodapalli ,B Himasri,Valli Nagalakshmi Undamatla,Payel Mukherjee,Sofia Banu,Archana Bharadwaj Siva,Sharath Chandra Thota,Karthik Bharadwaj Tallapaka,Rakesh K Mishra,Divya Tej Sowpati |
| EPI_ISL_2157019, | Mapmygenome | CSIR-Centre for Cellular and Molecular Biology-INSACOG | Blessy B John,Viswagithe S L,Tulasi Nagabandi, Lamuk Zaveri,Ara Sreenivas,Shreekant Verma, Amareshwar Vodapalli ,B Himasri,Valli Nagalakshmi Undamatla,Payel Mukherjee,Sofia Banu,Archana Bharadwaj Siva,Sharath Chandra Thota,Karthik |

| | | | |
|---|---|---|---|
| EPI_ISL_2157021 | | | Bharadwaj Tallapaka,Rakesh K Mishra,Divya Tej Sowpati |
| EPI_ISL_2157036, EPI_ISL_2157037, EPI_ISL_2157038, EPI_ISL_2157039 | CSIR-Centre for Cellular and Molecular Biology | CSIR-Centre for Cellular and Molecular Biology-INSACOG | Ara Sreenivas,Shreekant Verma,Tulasi Nagabandi, Lamuk Zaveri, Amareshwar Vodapalli ,Blessy B John,Viswagithe S L,B Himasri,Valli Nagalakshmi Undamatla,Payel Mukherjee,Sofia Banu,Archana Bharadwaj Siva,Sharath Chandra Thota,Karthik Bharadwaj Tallapaka,Rakesh K Mishra,Divya Tej Sowpati |
| EPI_ISL_2157044, EPI_ISL_2157046, EPI_ISL_2157047, EPI_ISL_2157050 | CSIR-Centre for Cellular and Molecular Biology | CSIR-Centre for Cellular and Molecular Biology-INSACOG | Lamuk Zaveri,Tulasi Nagabandi, Ara Sreenivas,Shreekant Verma, Amareshwar Vodapalli ,Blessy B John,Viswagithe S L,B Himasri,Valli Nagalakshmi Undamatla,Payel Mukherjee,Sofia Banu,Archana Bharadwaj Siva,Sharath Chandra Thota,Karthik Bharadwaj Tallapaka,Rakesh K Mishra,Divya Tej Sowpati |
| EPI_ISL_2157051, EPI_ISL_2157053, EPI_ISL_2157054, EPI_ISL_2157057, EPI_ISL_2157058, EPI_ISL_2157060, EPI_ISL_2157063 | CSIR-Centre for Cellular and Molecular Biology | CSIR-Centre for Cellular and Molecular Biology-INSACOG | Tulasi Nagabandi, Lamuk Zaveri,Ara Sreenivas,Shreekant Verma, Amareshwar Vodapalli ,Blessy B John,Viswagithe S L,B Himasri,Valli Nagalakshmi Undamatla,Payel Mukherjee,Sofia Banu,Archana Bharadwaj Siva,Sharath Chandra Thota,Karthik Bharadwaj Tallapaka,Rakesh K Mishra,Divya Tej Sowpati |
| EPI_ISL_2157065, EPI_ISL_2157066, EPI_ISL_2157067, EPI_ISL_2157068 | CSIR-Centre for Cellular and Molecular Biology | CSIR-Centre for Cellular and Molecular Biology-INSACOG | Payel Mukherjee,Lamuk Zaveri,Tulasi Nagabandi, Ara Sreenivas,Shreekant Verma, Amareshwar Vodapalli ,Blessy B John,Viswagithe S L,B Himasri,Valli Nagalakshmi Undamatla,Sofia Banu,Archana Bharadwaj Siva,Sharath Chandra Thota,Karthik Bharadwaj Tallapaka,Rakesh K Mishra,Divya Tej Sowpati |
| EPI_ISL_2157076 | CSIR-Centre for Cellular and Molecular Biology | CSIR-Centre for Cellular and Molecular Biology-INSACOG | B Himasri,Valli Nagalakshmi Undamatla,Lamuk Zaveri,Tulasi Nagabandi, Ara Sreenivas,Shreekant Verma, Amareshwar Vodapalli ,Blessy B John,Viswagithe S L,Payel Mukherjee,Sofia Banu,Archana Bharadwaj Siva,Sharath Chandra Thota,Karthik Bharadwaj Tallapaka,Rakesh K Mishra,Divya Tej Sowpati |
| EPI_ISL_2157077, EPI_ISL_2157078, EPI_ISL_2157079, EPI_ISL_2157080, EPI_ISL_2157081, EPI_ISL_2157082, EPI_ISL_2157085 | CSIR-Centre for Cellular and Molecular Biology | CSIR-Centre for Cellular and Molecular Biology-INSACOG | Payel Mukherjee,Lamuk Zaveri,Tulasi Nagabandi, Ara Sreenivas,Shreekant Verma, Amareshwar Vodapalli ,Blessy B John,Viswagithe S L,B Himasri,Valli Nagalakshmi Undamatla,Sofia Banu,Archana Bharadwaj Siva,Sharath Chandra Thota,Karthik Bharadwaj Tallapaka,Rakesh K Mishra,Divya Tej Sowpati |
| EPI_ISL_2157093 | CSIR-Centre for Cellular and Molecular Biology | CSIR-Centre for Cellular and Molecular Biology-INSACOG | Lamuk Zaveri,Tulasi Nagabandi, Ara Sreenivas,Shreekant Verma, Amareshwar Vodapalli ,Blessy B John,Viswagithe S L,B Himasri,Valli Nagalakshmi Undamatla,Payel Mukherjee,Sofia Banu,Archana Bharadwaj Siva,Sharath Chandra Thota,Karthik Bharadwaj Tallapaka,Rakesh K Mishra,Divya Tej Sowpati |
| EPI_ISL_2157099, EPI_ISL_2157102, EPI_ISL_2157103, EPI_ISL_2157104, EPI_ISL_2157105, EPI_ISL_2157106, EPI_ISL_2157107 | CSIR-Centre for Cellular and Molecular Biology | CSIR-Centre for Cellular and Molecular Biology-INSACOG | Payel Mukherjee,Lamuk Zaveri,Tulasi Nagabandi, Ara Sreenivas,Shreekant Verma, Amareshwar Vodapalli ,Blessy B John,Viswagithe S L,B Himasri,Valli Nagalakshmi Undamatla,Sofia Banu,Archana Bharadwaj Siva,Sharath Chandra Thota,Karthik Bharadwaj Tallapaka,Rakesh K Mishra,Divya Tej Sowpati |
| EPI_ISL_2157111, EPI_ISL_2157112, EPI_ISL_2157113, EPI_ISL_2157114 | CSIR-Centre for Cellular and Molecular Biology | CSIR-Centre for Cellular and Molecular Biology-INSACOG | Tulasi Nagabandi, Lamuk Zaveri,Ara Sreenivas,Shreekant Verma, Amareshwar Vodapalli ,Blessy B John,Viswagithe S L,B Himasri,Valli Nagalakshmi Undamatla,Payel Mukherjee,Sofia Banu,Archana Bharadwaj Siva,Sharath Chandra Thota,Karthik Bharadwaj Tallapaka,Rakesh K Mishra,Divya Tej Sowpati |
| EPI_ISL_2157118 | CSIR-Centre for Cellular and Molecular Biology | CSIR-Centre for Cellular and Molecular Biology-INSACOG | Payel Mukherjee,Lamuk Zaveri,Tulasi Nagabandi, Ara Sreenivas,Shreekant Verma, Amareshwar Vodapalli ,Blessy B John,Viswagithe S L,B Himasri,Valli Nagalakshmi Undamatla,Sofia Banu,Archana Bharadwaj Siva,Sharath Chandra Thota,Karthik Bharadwaj Tallapaka,Rakesh K Mishra,Divya Tej Sowpati |
| EPI_ISL_2157121, EPI_ISL_2157123, EPI_ISL_2157124, EPI_ISL_2157125, EPI_ISL_2157129, EPI_ISL_2157130 | CSIR-Centre for Cellular and Molecular Biology | CSIR-Centre for Cellular and Molecular Biology-INSACOG | Lamuk Zaveri,Tulasi Nagabandi, Ara Sreenivas,Shreekant Verma, Amareshwar Vodapalli ,Blessy B John,Viswagithe S L,B Himasri,Valli Nagalakshmi Undamatla,Payel Mukherjee,Sofia Banu,Archana Bharadwaj Siva,Sharath Chandra Thota,Karthik Bharadwaj Tallapaka,Rakesh K Mishra,Divya Tej Sowpati |
| EPI_ISL_2157154, EPI_ISL_2157155 | CSIR-Centre for Cellular and Molecular Biology | CSIR-Centre for Cellular and Molecular Biology-INSACOG | Payel Mukherjee,Lamuk Zaveri,Tulasi Nagabandi, Ara Sreenivas,Shreekant Verma, Amareshwar Vodapalli ,Blessy B John,Viswagithe S L,B Himasri,Valli Nagalakshmi Undamatla,Sofia Banu,Archana Bharadwaj Siva,Sharath Chandra Thota,Karthik Bharadwaj Tallapaka,Rakesh K Mishra,Divya Tej Sowpati |
| EPI_ISL_2157168 | Mapmygenome | CSIR-Centre for Cellular and Molecular Biology - INSACOG | Ara Sreenivas, Onkar Kulkarni, Lamuk Zaveri, Sofia Banu, Shreekant Verma, Amareshwar Vodapalli , Viswagithe S L, B Himasri, Sharath Chandra Thota, Karthik Bharadwaj Tallapaka, Rakesh K Mishra,Divya Tej Sowpati |
| EPI_ISL_2157177 | Mapmygenome | CSIR-Centre for Cellular and Molecular Biology - INSACOG | Onkar Kulkarni, Ara Sreenivas, Lamuk Zaveri, Sofia Banu, Shreekant Verma, Amareshwar Vodapalli , Viswagithe S L, B Himasri, Sharath Chandra Thota, Karthik Bharadwaj Tallapaka, Rakesh K Mishra,Divya Tej Sowpati |
| EPI_ISL_2157184 | Mapmygenome | CSIR-Centre for Cellular and Molecular Biology - INSACOG | Lamuk Zaveri, Ara Sreenivas, Onkar Kulkarni, Sofia Banu, Shreekant Verma, Amareshwar Vodapalli , Viswagithe S L, B Himasri, Sharath Chandra Thota, Karthik Bharadwaj Tallapaka, Rakesh K Mishra,Divya Tej Sowpati |
| EPI_ISL_2157190, EPI_ISL_2157194 | Mapmygenome | CSIR-Centre for Cellular and Molecular Biology - INSACOG | Sofia Banu, Ara Sreenivas, Onkar Kulkarni, Lamuk Zaveri, Shreekant Verma, Amareshwar Vodapalli , Viswagithe S L, B Himasri, Sharath Chandra Thota, Karthik Bharadwaj Tallapaka, Rakesh K Mishra,Divya Tej Sowpati |
| EPI_ISL_2157216 | Kansas Health and Environmental Lab | Kansas Health and Environmental Lab | Mike Grose, Katherine Wiggins, Jonathan Barnell, Ben Olsen, and Phil Adam |
| EPI_ISL_2158402, EPI_ISL_2158539 | National Virus Reference Laboratory | National Virus Reference Laboratory | Zoe Yandle, Charlene Bennett, Gabriel Gonzalez, Michael Carr, Jonathan Dean, Cillian F De Gascun |
| EPI_ISL_2158614, EPI_ISL_2158616, EPI_ISL_2158617 | Department of Veterinary Science, National Institute of Infectious Diseases | Department of Veterinary Science, National Institute of Infectious Diseases | Yudai Kuroda, Tsukasa Yamamoto, Keita Ishijima, Souichi Yamada, Shuetsu Fukushi, Ken Maeda |
| EPI_ISL_2158913, EPI_ISL_2158914, EPI_ISL_2158920, EPI_ISL_2159166, EPI_ISL_2159199, EPI_ISL_2159217, EPI_ISL_2159264, EPI_ISL_2159352 | Aegis Sciences Corporation | Centers for Disease Control and Prevention Division of Viral Diseases, Pathogen Discovery | Dakota Howard, Dhwani Batra, Peter W. Cook, Kara Moser, Adrian Paskey, Jason Caravas, Benjamin Rambo-Martin, Shatavia Morrison, Christopher Gulvick, Scott Sammons, Yvette Unoarumhi, Darlene Wagner, Matthew Schmerer, Cyndi Clark, Patrick Campbell, Rob Case, Vikramsinha Ghorpade, Holly Houdeshell, Ola Kvalvaag, Dillon Nall, Ethan Sanders, Alec Vest, Shaun Westlund, Matthew Hardison, Clinton R. Paden, Duncan MacCannell |
| EPI_ISL_2159577, EPI_ISL_2159751, EPI_ISL_2159811, EPI_ISL_2159870, EPI_ISL_2160115, EPI_ISL_2160284, EPI_ISL_2160446, EPI_ISL_2160619, EPI_ISL_2160637 | Helix/Illumina | Centers for Disease Control and Prevention Division of Viral Diseases, Pathogen Discovery | Dakota Howard, Dhwani Batra, Peter W. Cook, Kara Moser, Adrian Paskey, Jason Caravas, Benjamin Rambo-Martin, Shatavia Morrison, Christopher Gulvick, Scott Sammons, Yvette Unoarumhi, Darlene Wagner, Matthew Schmerer, Eileen de Feo, Jan Antico, Christine Tran, Matthew Tolentino, Shannon Wickline, Kim Gietzen, Brad Sickler, Jingtao Liu, Eric Allen, Phil Febbo, Nicole L. Washington, Simon White, Geraint Levan, Kelly Schiabor Barrett, Elizabeth Cirulli, Alexandre Bolze, Ary Ascencio, Charlotte Rivera-Garcia, Ryan Cho, Jason Nguyen, Sherry Wang, Jimmy Ramirez, Tyler Cassens, Efren Sandoval, Magnus Isaksson, William Lee, David Becker, Marc Laurent, James Lu, Clinton R. Paden, Duncan MacCannell |
| EPI_ISL_2160766, EPI_ISL_2160771, EPI_ISL_2160774, EPI_ISL_2160792, EPI_ISL_2160804, EPI_ISL_2160840, EPI_ISL_2160848, EPI_ISL_2160849, EPI_ISL_2160889, EPI_ISL_2160900, EPI_ISL_2160908, EPI_ISL_2160946, EPI_ISL_2161002 | | | |
| see above | Fulgent Genetics | Centers for Disease Control and Prevention Division of Viral Diseases, Pathogen Discovery | Dakota Howard, Dhwani Batra, Peter W. Cook, Kara Moser, Adrian Paskey, Jason Caravas, Benjamin Rambo-Martin, Shatavia Morrison, Christopher Gulvick, Scott Sammons, Yvette Unoarumhi, Darlene Wagner, Matthew Schmerer, Harry Gao, Mickey Li, John Gao, Joseph Fierro, Benafsh Sapra, Becky Tsai, Yan Meng, Doreen Ng, James Xie, Clinton R. Paden, Duncan MacCannell |
| EPI_ISL_2161520, EPI_ISL_2161528 | Maryland Genomics, Institute for Genome Sciences, University of Maryland School of Medicine | Maryland Genomics, Institute for Genome Sciences, University of Maryland School of Medicine | Tallon, Luke J; Sadzewicz, Lisa D; Humphrys, Mike; Ott, Sandra; Roussey, Holly; Mehta, Aditya; Vavikolanu, Kranthi; Fraser, Claire M; Ravel, Jacques |
| EPI_ISL_2161784 | NL-Dr. Leonard A. Miller Centre for Health Services | National Microbiology Laboratory (NML) | Anna Majer, Shari Tyson, Grace Seo, Philip Mabon, Elsie Grudeski, Rhiannon Huzarewich, Russell Mandes, Anneliese Landgraff, Jennifer Tanner, Natalie Knox, Morag Graham, Gary Van Domselaar, Robert Needle, Yang Yu, Adel Malek, Laura Gilbert, George Zahariadis, Nathalie Bastien, Yan Li, Timothy Booth, Darian Hole, Madison Chapel, Kirsten Biggar, Kerri Smith, CanCOGeN's metadata curation team, Public Health Agency of Canada CanCOGeN team |
| EPI_ISL_2161818, EPI_ISL_2161848, EPI_ISL_2161871, EPI_ISL_2161913, EPI_ISL_2161915, EPI_ISL_2161925, EPI_ISL_2161928, EPI_ISL_2161967, EPI_ISL_2161978 | UW Virology Lab | UW Virology Lab | Pavitra Roychoudhury, Hong Xie, Lasata Shrestha, Tien V. Nguyen, Shah Mohamed Bakhash, Michelle Lin, Noah R. Baker, Ricardo Perez, Sean Ellis, Nathan Breit, Robert J. Livingston, Meei-Li Huang, Keith R Jerome, Patrick Mathias, Alexander Greninger |
| EPI_ISL_2162072, EPI_ISL_2162120 | NS-QEII Health Sciences Centre | National Microbiology Laboratory (NML) | Anna Majer, Shari Tyson, Grace Seo, Philip Mabon, Elsie Grudeski, Rhiannon Huzarewich, Russell Mandes, Anneliese Landgraff, Jennifer Tanner, Natalie Knox, Morag Graham, Gary Van Domselaar, Todd Hatchette, Jason LeBlanc, Janice Pettipas, Dan Gaston, Nathalie Bastien, Yan Li, Timothy Booth, Darian Hole, Madison Chapel, Kirsten Biggar, CanCOGeN's metadata curation team, Public Health Agency of Canada CanCOGeN team |

| | | | |
|---|---|---|---|
| EPI_ISL_2162255, EPI_ISL_2162266, EPI_ISL_2162276 | Hospital Universitari Vall d'Hebron - Vall d'Hebron Institut de Recerca | Hospital Universitari Vall d'Hebron - Vall d'Hebron Institut de Recerca | Cristina Andrés, Maria Piñana, Alejandra González-Sánchez, Damir Garcia-Cehic, Ariadna Rando, Juliana Esperalba, Maria Gema Codina, Carla Castillo, Maria Carmen Martin, Tomàs Pumarola, Josep Quer, Andrés Antón |
| EPI_ISL_2162392, EPI_ISL_2162393, EPI_ISL_2162395, EPI_ISL_2162411, EPI_ISL_2162417, EPI_ISL_2162423, EPI_ISL_2162425, EPI_ISL_2162427, EPI_ISL_2162429, EPI_ISL_2162435, EPI_ISL_2162437, EPI_ISL_2162438, EPI_ISL_2162441, EPI_ISL_2162444, EPI_ISL_2162445, EPI_ISL_2162455, EPI_ISL_2162461, EPI_ISL_2162466 | | | |
| see above | National Centre For Cell Science | National Centre For Cell Science – INSACOG | Dhiraj Paul, Mohak P Gujare, Mitali Inamdar, Shivang P. Bhanushali, Manoj Kumar Bhat, Ajay Pillai, INSACOG Consortium team, Yogesh Shouche |
| EPI_ISL_2170576 | Alberta Precision Labs (APL) | Public Health Agency of Canada (PHAC) National Microbiology Laboratory | Buss, E, Croxen M, Deo A, Dieu P, Gill K, Ferrato C, Khan F, Koleva P, Li V, Lloyd C, Lynch T, Ma R, Murphy S, Pabbaraju K, Shokoples S, Tipples G, Thayer J, Whitehouse M, Wong A, Yu C, Zelyas N |
| EPI_ISL_2170808, EPI_ISL_2170809, EPI_ISL_2170810 | CAP SARRIÀ | Banc de Sang i Teixits | Francisco Vidal, Irene Corrales, Nina Borràs, Carlos Hobeich, Natália Comes, Lorena Ramírez, Noemí Gonzalez, Silvia Sauleda, Maria Glòria Soria |
| EPI_ISL_2170814 | CAP PRATS DE LLUÇANÈS | Banc de Sang i Teixits | Francisco Vidal, Irene Corrales, Nina Borràs, Carlos Hobeich, Natália Comes, Lorena Ramírez, Noemí Gonzalez, Silvia Sauleda, Maria Glòria Soria |
| EPI_ISL_2171242, EPI_ISL_2171265, EPI_ISL_2171266 | Janamaitri Hospital | Molecular and Genomics Research Lab, Dhulikhel Hospital, Kathmandu University Hospital | Rajeev Shrestha, Nishan Katuwal, Navin Adhikari, Dipesh Tamrakar, Meghnath Dhimal, Pradip Gyanwali, Saroj Bhattarai, Surendra Kumar Madhup |
| EPI_ISL_2171267, EPI_ISL_2171281 | Dhulikhel Hospital, Kathmandu University Hospital | Molecular and Genomics Research Lab, Dhulikhel Hospital, Kathmandu University Hospital | Rajeev Shrestha, Nishan Katuwal, Navin Adhikari, Dipesh Tamrakar, Meghnath Dhimal, Pradip Gyanwali, Saroj Bhattarai, Surendra Kumar Madhup |
| EPI_ISL_2171675, EPI_ISL_2171677 | Lighthouse Lab in Alderley Park | Wellcome Sanger Institute for the COVID-19 Genomics UK (COG-UK) Consortium | Jacquelyn Wynn, Mairead Hyland, The Lighthouse Lab in Alderley Park and Alex Alderton, Roberto Amato, Jeffrey Barrett, Sonia Goncalves, Ewan Harrison, David K. Jackson, Ian Johnston, Dominic Kwiatkowski, Cordelia Langford, John Sillitoe on behalf of the Wellcome Sanger Institute COVID-19 Surveillance Team |
| EPI_ISL_2171681 | Lighthouse Lab in Glasgow | Wellcome Sanger Institute for the COVID-19 Genomics UK (COG-UK) Consortium | Harper VanSteenhouse, Yumi Kasai, David Gray, Carol Clugston, Anna Dominiczak and Alex Alderton, Roberto Amato, Jeffrey Barrett, Sonia Goncalves, Ewan Harrison, David K. Jackson, Ian Johnston, Dominic Kwiatkowski, Cordelia Langford, John Sillitoe on behalf of the Wellcome Sanger Institute COVID-19 Surveillance Team |
| EPI_ISL_2171683, EPI_ISL_2171684, EPI_ISL_2171685, EPI_ISL_2171686, EPI_ISL_2171688, EPI_ISL_2171689, EPI_ISL_2171690, EPI_ISL_2171692, EPI_ISL_2171698, EPI_ISL_2171699, EPI_ISL_2171701, EPI_ISL_2171702, EPI_ISL_2171704, EPI_ISL_2171705 | | | |
| see above | Lighthouse Lab in Alderley Park | Wellcome Sanger Institute for the COVID-19 Genomics UK (COG-UK) Consortium | Jacquelyn Wynn, Mairead Hyland, The Lighthouse Lab in Alderley Park and Alex Alderton, Roberto Amato, Jeffrey Barrett, Sonia Goncalves, Ewan Harrison, David K. Jackson, Ian Johnston, Dominic Kwiatkowski, Cordelia Langford, John Sillitoe on behalf of the Wellcome Sanger Institute COVID-19 Surveillance Team |
| EPI_ISL_2171706, EPI_ISL_2171707 | Lighthouse Lab in Glasgow | Wellcome Sanger Institute for the COVID-19 Genomics UK (COG-UK) Consortium | Harper VanSteenhouse, Yumi Kasai, David Gray, Carol Clugston, Anna Dominiczak and Alex Alderton, Roberto Amato, Jeffrey Barrett, Sonia Goncalves, Ewan Harrison, David K. Jackson, Ian Johnston, Dominic Kwiatkowski, Cordelia Langford, John Sillitoe on behalf of the Wellcome Sanger Institute COVID-19 Surveillance Team |
| EPI_ISL_2171711, EPI_ISL_2171713, EPI_ISL_2171714, EPI_ISL_2171716, EPI_ISL_2171717, EPI_ISL_2171718, EPI_ISL_2171719 | Lighthouse Lab in Alderley Park | Wellcome Sanger Institute for the COVID-19 Genomics UK (COG-UK) Consortium | Jacquelyn Wynn, Mairead Hyland, The Lighthouse Lab in Alderley Park and Alex Alderton, Roberto Amato, Jeffrey Barrett, Sonia Goncalves, Ewan Harrison, David K. Jackson, Ian Johnston, Dominic Kwiatkowski, Cordelia Langford, John Sillitoe on behalf of the Wellcome Sanger Institute COVID-19 Surveillance Team |
| EPI_ISL_2171721 | Lighthouse Lab in Glasgow | Wellcome Sanger Institute for the COVID-19 Genomics UK (COG-UK) Consortium | Harper VanSteenhouse, Yumi Kasai, David Gray, Carol Clugston, Anna Dominiczak and Alex Alderton, Roberto Amato, Jeffrey Barrett, Sonia Goncalves, Ewan Harrison, David K. Jackson, Ian Johnston, Dominic Kwiatkowski, Cordelia Langford, John Sillitoe on behalf of the Wellcome Sanger Institute COVID-19 Surveillance Team |
| EPI_ISL_2171724, EPI_ISL_2171727, EPI_ISL_2171728, EPI_ISL_2171729, EPI_ISL_2171730, EPI_ISL_2171733, EPI_ISL_2171735, EPI_ISL_2171741, EPI_ISL_2171743, EPI_ISL_2171744 | Lighthouse Lab in Alderley Park | Wellcome Sanger Institute for the COVID-19 Genomics UK (COG-UK) Consortium | Jacquelyn Wynn, Mairead Hyland, The Lighthouse Lab in Alderley Park and Alex Alderton, Roberto Amato, Jeffrey Barrett, Sonia Goncalves, Ewan Harrison, David K. Jackson, Ian Johnston, Dominic Kwiatkowski, Cordelia Langford, John Sillitoe on behalf of the Wellcome Sanger Institute COVID-19 Surveillance Team |
| EPI_ISL_2171748 | Lighthouse Lab in Glasgow | Wellcome Sanger Institute for the COVID-19 Genomics UK (COG-UK) Consortium | Harper VanSteenhouse, Yumi Kasai, David Gray, Carol Clugston, Anna Dominiczak and Alex Alderton, Roberto Amato, Jeffrey Barrett, Sonia Goncalves, Ewan Harrison, David K. Jackson, Ian Johnston, Dominic Kwiatkowski, Cordelia Langford, John Sillitoe on behalf of the Wellcome Sanger Institute COVID-19 Surveillance Team |
| EPI_ISL_2171750, EPI_ISL_2171753, EPI_ISL_2171761 | Lighthouse Lab in Alderley Park | Wellcome Sanger Institute for the COVID-19 Genomics UK (COG-UK) Consortium | Jacquelyn Wynn, Mairead Hyland, The Lighthouse Lab in Alderley Park and Alex Alderton, Roberto Amato, Jeffrey Barrett, Sonia Goncalves, Ewan Harrison, David K. Jackson, Ian Johnston, Dominic Kwiatkowski, Cordelia Langford, John Sillitoe on behalf of the Wellcome Sanger Institute COVID-19 Surveillance Team |
| EPI_ISL_2171762, EPI_ISL_2171763, EPI_ISL_2171764 | Lighthouse Lab in Glasgow | Wellcome Sanger Institute for the COVID-19 Genomics UK (COG-UK) Consortium | Harper VanSteenhouse, Yumi Kasai, David Gray, Carol Clugston, Anna Dominiczak and Alex Alderton, Roberto Amato, Jeffrey Barrett, Sonia Goncalves, Ewan Harrison, David K. Jackson, Ian Johnston, Dominic Kwiatkowski, Cordelia Langford, John Sillitoe on behalf of the Wellcome Sanger Institute COVID-19 Surveillance Team |
| EPI_ISL_2171765, EPI_ISL_2171767, EPI_ISL_2171772, EPI_ISL_2171775, EPI_ISL_2171779, EPI_ISL_2171780, EPI_ISL_2171781 | Lighthouse Lab in Alderley Park | Wellcome Sanger Institute for the COVID-19 Genomics UK (COG-UK) Consortium | Jacquelyn Wynn, Mairead Hyland, The Lighthouse Lab in Alderley Park and Alex Alderton, Roberto Amato, Jeffrey Barrett, Sonia Goncalves, Ewan Harrison, David K. Jackson, Ian Johnston, Dominic Kwiatkowski, Cordelia Langford, John Sillitoe on behalf of the Wellcome Sanger Institute COVID-19 Surveillance Team |
| EPI_ISL_2171784 | Lighthouse Lab in Glasgow | Wellcome Sanger Institute for the COVID-19 Genomics UK (COG-UK) Consortium | Harper VanSteenhouse, Yumi Kasai, David Gray, Carol Clugston, Anna Dominiczak and Alex Alderton, Roberto Amato, Jeffrey Barrett, Sonia Goncalves, Ewan Harrison, David K. Jackson, Ian Johnston, Dominic Kwiatkowski, Cordelia Langford, John Sillitoe on behalf of the Wellcome Sanger Institute COVID-19 Surveillance Team |
| EPI_ISL_2171785, EPI_ISL_2171787, EPI_ISL_2171788, EPI_ISL_2171789, EPI_ISL_2171790, EPI_ISL_2171791, EPI_ISL_2171792, EPI_ISL_2171793 | Lighthouse Lab in Alderley Park | Wellcome Sanger Institute for the COVID-19 Genomics UK (COG-UK) Consortium | Jacquelyn Wynn, Mairead Hyland, The Lighthouse Lab in Alderley Park and Alex Alderton, Roberto Amato, Jeffrey Barrett, Sonia Goncalves, Ewan Harrison, David K. Jackson, Ian Johnston, Dominic Kwiatkowski, Cordelia Langford, John Sillitoe on behalf of the Wellcome Sanger Institute COVID-19 Surveillance Team |
| EPI_ISL_2171804 | Lighthouse Lab in Glasgow | Wellcome Sanger Institute for the COVID-19 Genomics UK (COG-UK) Consortium | Harper VanSteenhouse, Yumi Kasai, David Gray, Carol Clugston, Anna Dominiczak and Alex Alderton, Roberto Amato, Jeffrey Barrett, Sonia Goncalves, Ewan Harrison, David K. Jackson, Ian Johnston, Dominic Kwiatkowski, Cordelia Langford, John Sillitoe on behalf of the Wellcome Sanger Institute COVID-19 Surveillance Team |
| EPI_ISL_2171810, EPI_ISL_2171814 | Lighthouse Lab in Alderley Park | Wellcome Sanger Institute for the COVID-19 Genomics UK (COG-UK) Consortium | Jacquelyn Wynn, Mairead Hyland, The Lighthouse Lab in Alderley Park and Alex Alderton, Roberto Amato, Jeffrey Barrett, Sonia Goncalves, Ewan Harrison, David K. Jackson, Ian Johnston, Dominic Kwiatkowski, Cordelia Langford, John Sillitoe on behalf of the Wellcome Sanger Institute COVID-19 Surveillance Team |
| EPI_ISL_2171815 | Lighthouse Lab in Glasgow | Wellcome Sanger Institute for the COVID-19 Genomics UK (COG-UK) Consortium | Harper VanSteenhouse, Yumi Kasai, David Gray, Carol Clugston, Anna Dominiczak and Alex Alderton, Roberto Amato, Jeffrey Barrett, Sonia Goncalves, Ewan Harrison, David K. Jackson, Ian Johnston, Dominic Kwiatkowski, Cordelia Langford, John Sillitoe on behalf of the Wellcome Sanger Institute COVID-19 Surveillance Team |
| EPI_ISL_2171816 | Lighthouse Lab in Alderley Park | Wellcome Sanger Institute for the COVID-19 Genomics UK (COG-UK) Consortium | Jacquelyn Wynn, Mairead Hyland, The Lighthouse Lab in Alderley Park and Alex Alderton, Roberto Amato, Jeffrey Barrett, Sonia Goncalves, Ewan Harrison, David K. Jackson, Ian Johnston, Dominic Kwiatkowski, Cordelia Langford, John Sillitoe on behalf of the Wellcome Sanger Institute COVID-19 Surveillance Team |
| EPI_ISL_2171820, EPI_ISL_2171822 | Lighthouse Lab in Glasgow | Wellcome Sanger Institute for the COVID-19 Genomics UK (COG-UK) Consortium | Harper VanSteenhouse, Yumi Kasai, David Gray, Carol Clugston, Anna Dominiczak and Alex Alderton, Roberto Amato, Jeffrey Barrett, Sonia Goncalves, Ewan Harrison, David K. Jackson, Ian Johnston, Dominic Kwiatkowski, Cordelia Langford, John Sillitoe on behalf of the Wellcome Sanger Institute COVID-19 Surveillance Team |
| EPI_ISL_2171823, EPI_ISL_2171825, EPI_ISL_2171826 | Lighthouse Lab in Alderley Park | Wellcome Sanger Institute for the COVID-19 Genomics UK (COG-UK) Consortium | Jacquelyn Wynn, Mairead Hyland, The Lighthouse Lab in Alderley Park and Alex Alderton, Roberto Amato, Jeffrey Barrett, Sonia Goncalves, Ewan Harrison, David K. Jackson, Ian Johnston, Dominic Kwiatkowski, Cordelia Langford, John Sillitoe on behalf of the Wellcome Sanger Institute COVID-19 Surveillance Team |
| EPI_ISL_2171827 | Lighthouse Lab in Glasgow | Wellcome Sanger Institute for the COVID-19 Genomics UK (COG-UK) Consortium | Harper VanSteenhouse, Yumi Kasai, David Gray, Carol Clugston, Anna Dominiczak and Alex Alderton, Roberto Amato, Jeffrey Barrett, Sonia Goncalves, Ewan Harrison, David K. Jackson, Ian Johnston, Dominic Kwiatkowski, Cordelia Langford, John Sillitoe on behalf of the Wellcome Sanger Institute COVID-19 Surveillance Team |
| EPI_ISL_2171828, EPI_ISL_2171830, EPI_ISL_2171831, EPI_ISL_2171834, EPI_ISL_2171838 | Lighthouse Lab in Alderley Park | Wellcome Sanger Institute for the COVID-19 Genomics UK (COG-UK) Consortium | Jacquelyn Wynn, Mairead Hyland, The Lighthouse Lab in Alderley Park and Alex Alderton, Roberto Amato, Jeffrey Barrett, Sonia Goncalves, Ewan Harrison, David K. Jackson, Ian Johnston, Dominic Kwiatkowski, Cordelia Langford, John Sillitoe on behalf of the Wellcome Sanger Institute COVID-19 Surveillance Team |
| EPI_ISL_2171840 | Lighthouse Lab in Glasgow | Wellcome Sanger Institute for the COVID-19 Genomics UK (COG-UK) Consortium | Harper VanSteenhouse, Yumi Kasai, David Gray, Carol Clugston, Anna Dominiczak and Alex Alderton, Roberto Amato, Jeffrey Barrett, Sonia Goncalves, Ewan Harrison, David K. Jackson, Ian Johnston, Dominic Kwiatkowski, Cordelia Langford, John Sillitoe on behalf of the Wellcome Sanger Institute COVID-19 Surveillance Team |
| EPI_ISL_2171844, EPI_ISL_2171845, EPI_ISL_2171846, EPI_ISL_2171852 | Lighthouse Lab in Alderley Park | Wellcome Sanger Institute for the COVID-19 Genomics UK (COG-UK) Consortium | Jacquelyn Wynn, Mairead Hyland, The Lighthouse Lab in Alderley Park and Alex Alderton, Roberto Amato, Jeffrey Barrett, Sonia Goncalves, Ewan Harrison, David K. Jackson, Ian Johnston, Dominic Kwiatkowski, Cordelia Langford, John Sillitoe on behalf of the Wellcome Sanger Institute COVID-19 Surveillance Team |

| | | | |
|---|---|---|---|
| EPI_ISL_2171853, EPI_ISL_2171854 | Lighthouse Lab in Glasgow | Wellcome Sanger Institute for the COVID-19 Genomics UK (COG-UK) Consortium | Harper VanSteenhouse, Yumi Kasai, David Gray, Carol Clugston, Anna Dominiczak and Alex Alderton, Roberto Amato, Jeffrey Barrett, Sonia Goncalves, Ewan Harrison, David K. Jackson, Ian Johnston, Dominic Kwiatkowski, Cordelia Langford, John Sillitoe on behalf of the Wellcome Sanger Institute COVID-19 Surveillance Team |
| EPI_ISL_2171858, EPI_ISL_2171860 | Lighthouse Lab in Alderley Park | Wellcome Sanger Institute for the COVID-19 Genomics UK (COG-UK) Consortium | Jacquelyn Wynn, Mairead Hyland, The Lighthouse Lab in Alderley Park and Alex Alderton, Roberto Amato, Jeffrey Barrett, Sonia Goncalves, Ewan Harrison, David K. Jackson, Ian Johnston, Dominic Kwiatkowski, Cordelia Langford, John Sillitoe on behalf of the Wellcome Sanger Institute COVID-19 Surveillance Team |
| EPI_ISL_2171861 | Lighthouse Lab in Glasgow | Wellcome Sanger Institute for the COVID-19 Genomics UK (COG-UK) Consortium | Harper VanSteenhouse, Yumi Kasai, David Gray, Carol Clugston, Anna Dominiczak and Alex Alderton, Roberto Amato, Jeffrey Barrett, Sonia Goncalves, Ewan Harrison, David K. Jackson, Ian Johnston, Dominic Kwiatkowski, Cordelia Langford, John Sillitoe on behalf of the Wellcome Sanger Institute COVID-19 Surveillance Team |
| EPI_ISL_2171862 | Lighthouse Lab in Alderley Park | Wellcome Sanger Institute for the COVID-19 Genomics UK (COG-UK) Consortium | Jacquelyn Wynn, Mairead Hyland, The Lighthouse Lab in Alderley Park and Alex Alderton, Roberto Amato, Jeffrey Barrett, Sonia Goncalves, Ewan Harrison, David K. Jackson, Ian Johnston, Dominic Kwiatkowski, Cordelia Langford, John Sillitoe on behalf of the Wellcome Sanger Institute COVID-19 Surveillance Team |
| EPI_ISL_2171863 | Lighthouse Lab in Milton Keynes | Wellcome Sanger Institute for the COVID-19 Genomics UK (COG-UK) Consortium | The Lighthouse Lab in Milton Keynes and Alex Alderton, Roberto Amato, Jeffrey Barrett, Sonia Goncalves, Ewan Harrison, David K. Jackson, Ian Johnston, Dominic Kwiatkowski, Cordelia Langford, John Sillitoe on behalf of the Wellcome Sanger Institute COVID-19 Surveillance Team |
| EPI_ISL_2171864, EPI_ISL_2171865, EPI_ISL_2171866 | Lighthouse Lab in Alderley Park | Wellcome Sanger Institute for the COVID-19 Genomics UK (COG-UK) Consortium | Jacquelyn Wynn, Mairead Hyland, The Lighthouse Lab in Alderley Park and Alex Alderton, Roberto Amato, Jeffrey Barrett, Sonia Goncalves, Ewan Harrison, David K. Jackson, Ian Johnston, Dominic Kwiatkowski, Cordelia Langford, John Sillitoe on behalf of the Wellcome Sanger Institute COVID-19 Surveillance Team |
| EPI_ISL_2171867 | Lighthouse Lab in Glasgow | Wellcome Sanger Institute for the COVID-19 Genomics UK (COG-UK) Consortium | Harper VanSteenhouse, Yumi Kasai, David Gray, Carol Clugston, Anna Dominiczak and Alex Alderton, Roberto Amato, Jeffrey Barrett, Sonia Goncalves, Ewan Harrison, David K. Jackson, Ian Johnston, Dominic Kwiatkowski, Cordelia Langford, John Sillitoe on behalf of the Wellcome Sanger Institute COVID-19 Surveillance Team |
| EPI_ISL_2171868, EPI_ISL_2171869, EPI_ISL_2171870 | Lighthouse Lab in Alderley Park | Wellcome Sanger Institute for the COVID-19 Genomics UK (COG-UK) Consortium | Jacquelyn Wynn, Mairead Hyland, The Lighthouse Lab in Alderley Park and Alex Alderton, Roberto Amato, Jeffrey Barrett, Sonia Goncalves, Ewan Harrison, David K. Jackson, Ian Johnston, Dominic Kwiatkowski, Cordelia Langford, John Sillitoe on behalf of the Wellcome Sanger Institute COVID-19 Surveillance Team |
| EPI_ISL_2171871, EPI_ISL_2171873, EPI_ISL_2171875, EPI_ISL_2171876 | Lighthouse Lab in Glasgow | Wellcome Sanger Institute for the COVID-19 Genomics UK (COG-UK) Consortium | Harper VanSteenhouse, Yumi Kasai, David Gray, Carol Clugston, Anna Dominiczak and Alex Alderton, Roberto Amato, Jeffrey Barrett, Sonia Goncalves, Ewan Harrison, David K. Jackson, Ian Johnston, Dominic Kwiatkowski, Cordelia Langford, John Sillitoe on behalf of the Wellcome Sanger Institute COVID-19 Surveillance Team |
| EPI_ISL_2171877, EPI_ISL_2171879 | Lighthouse Lab in Alderley Park | Wellcome Sanger Institute for the COVID-19 Genomics UK (COG-UK) Consortium | Jacquelyn Wynn, Mairead Hyland, The Lighthouse Lab in Alderley Park and Alex Alderton, Roberto Amato, Jeffrey Barrett, Sonia Goncalves, Ewan Harrison, David K. Jackson, Ian Johnston, Dominic Kwiatkowski, Cordelia Langford, John Sillitoe on behalf of the Wellcome Sanger Institute COVID-19 Surveillance Team |
| EPI_ISL_2171881 | Lighthouse Lab in Glasgow | Wellcome Sanger Institute for the COVID-19 Genomics UK (COG-UK) Consortium | Harper VanSteenhouse, Yumi Kasai, David Gray, Carol Clugston, Anna Dominiczak and Alex Alderton, Roberto Amato, Jeffrey Barrett, Sonia Goncalves, Ewan Harrison, David K. Jackson, Ian Johnston, Dominic Kwiatkowski, Cordelia Langford, John Sillitoe on behalf of the Wellcome Sanger Institute COVID-19 Surveillance Team |
| EPI_ISL_2171882 | Lighthouse Lab in Alderley Park | Wellcome Sanger Institute for the COVID-19 Genomics UK (COG-UK) Consortium | Jacquelyn Wynn, Mairead Hyland, The Lighthouse Lab in Alderley Park and Alex Alderton, Roberto Amato, Jeffrey Barrett, Sonia Goncalves, Ewan Harrison, David K. Jackson, Ian Johnston, Dominic Kwiatkowski, Cordelia Langford, John Sillitoe on behalf of the Wellcome Sanger Institute COVID-19 Surveillance Team |
| EPI_ISL_2171883, EPI_ISL_2171885, EPI_ISL_2171886 | Lighthouse Lab in Glasgow | Wellcome Sanger Institute for the COVID-19 Genomics UK (COG-UK) Consortium | Harper VanSteenhouse, Yumi Kasai, David Gray, Carol Clugston, Anna Dominiczak and Alex Alderton, Roberto Amato, Jeffrey Barrett, Sonia Goncalves, Ewan Harrison, David K. Jackson, Ian Johnston, Dominic Kwiatkowski, Cordelia Langford, John Sillitoe on behalf of the Wellcome Sanger Institute COVID-19 Surveillance Team |
| EPI_ISL_2171887 | Lighthouse Lab in Milton Keynes | Wellcome Sanger Institute for the COVID-19 Genomics UK (COG-UK) Consortium | The Lighthouse Lab in Milton Keynes and Alex Alderton, Roberto Amato, Jeffrey Barrett, Sonia Goncalves, Ewan Harrison, David K. Jackson, Ian Johnston, Dominic Kwiatkowski, Cordelia Langford, John Sillitoe on behalf of the Wellcome Sanger Institute COVID-19 Surveillance Team |
| EPI_ISL_2171888 | Lighthouse Lab in Alderley Park | Wellcome Sanger Institute for the COVID-19 Genomics UK (COG-UK) Consortium | Jacquelyn Wynn, Mairead Hyland, The Lighthouse Lab in Alderley Park and Alex Alderton, Roberto Amato, Jeffrey Barrett, Sonia Goncalves, Ewan Harrison, David K. Jackson, Ian Johnston, Dominic Kwiatkowski, Cordelia Langford, John Sillitoe on behalf of the Wellcome Sanger Institute COVID-19 Surveillance Team |
| EPI_ISL_2171889, EPI_ISL_2171890 | Lighthouse Lab in Glasgow | Wellcome Sanger Institute for the COVID-19 Genomics UK (COG-UK) Consortium | Harper VanSteenhouse, Yumi Kasai, David Gray, Carol Clugston, Anna Dominiczak and Alex Alderton, Roberto Amato, Jeffrey Barrett, Sonia Goncalves, Ewan Harrison, David K. Jackson, Ian Johnston, Dominic Kwiatkowski, Cordelia Langford, John Sillitoe on behalf of the Wellcome Sanger Institute COVID-19 Surveillance Team |
| EPI_ISL_2171891 | Lighthouse Lab in Alderley Park | Wellcome Sanger Institute for the COVID-19 Genomics UK (COG-UK) Consortium | Jacquelyn Wynn, Mairead Hyland, The Lighthouse Lab in Alderley Park and Alex Alderton, Roberto Amato, Jeffrey Barrett, Sonia Goncalves, Ewan Harrison, David K. Jackson, Ian Johnston, Dominic Kwiatkowski, Cordelia Langford, John Sillitoe on behalf of the Wellcome Sanger Institute COVID-19 Surveillance Team |
| EPI_ISL_2171892 | Lighthouse Lab in Milton Keynes | Wellcome Sanger Institute for the COVID-19 Genomics UK (COG-UK) Consortium | The Lighthouse Lab in Milton Keynes and Alex Alderton, Roberto Amato, Jeffrey Barrett, Sonia Goncalves, Ewan Harrison, David K. Jackson, Ian Johnston, Dominic Kwiatkowski, Cordelia Langford, John Sillitoe on behalf of the Wellcome Sanger Institute COVID-19 Surveillance Team |
| EPI_ISL_2171893 | Lighthouse Lab in Glasgow | Wellcome Sanger Institute for the COVID-19 Genomics UK (COG-UK) Consortium | Harper VanSteenhouse, Yumi Kasai, David Gray, Carol Clugston, Anna Dominiczak and Alex Alderton, Roberto Amato, Jeffrey Barrett, Sonia Goncalves, Ewan Harrison, David K. Jackson, Ian Johnston, Dominic Kwiatkowski, Cordelia Langford, John Sillitoe on behalf of the Wellcome Sanger Institute COVID-19 Surveillance Team |
| EPI_ISL_2171894, EPI_ISL_2171898, EPI_ISL_2171901, EPI_ISL_2171903, EPI_ISL_2171904, EPI_ISL_2171907 | Lighthouse Lab in Alderley Park | Wellcome Sanger Institute for the COVID-19 Genomics UK (COG-UK) Consortium | Jacquelyn Wynn, Mairead Hyland, The Lighthouse Lab in Alderley Park and Alex Alderton, Roberto Amato, Jeffrey Barrett, Sonia Goncalves, Ewan Harrison, David K. Jackson, Ian Johnston, Dominic Kwiatkowski, Cordelia Langford, John Sillitoe on behalf of the Wellcome Sanger Institute COVID-19 Surveillance Team |
| EPI_ISL_2171911, EPI_ISL_2171912 | Lighthouse Lab in Glasgow | Wellcome Sanger Institute for the COVID-19 Genomics UK (COG-UK) Consortium | Harper VanSteenhouse, Yumi Kasai, David Gray, Carol Clugston, Anna Dominiczak and Alex Alderton, Roberto Amato, Jeffrey Barrett, Sonia Goncalves, Ewan Harrison, David K. Jackson, Ian Johnston, Dominic Kwiatkowski, Cordelia Langford, John Sillitoe on behalf of the Wellcome Sanger Institute COVID-19 Surveillance Team |
| EPI_ISL_2171913 | Lighthouse Lab in Alderley Park | Wellcome Sanger Institute for the COVID-19 Genomics UK (COG-UK) Consortium | Jacquelyn Wynn, Mairead Hyland, The Lighthouse Lab in Alderley Park and Alex Alderton, Roberto Amato, Jeffrey Barrett, Sonia Goncalves, Ewan Harrison, David K. Jackson, Ian Johnston, Dominic Kwiatkowski, Cordelia Langford, John Sillitoe on behalf of the Wellcome Sanger Institute COVID-19 Surveillance Team |
| EPI_ISL_2171914 | Lighthouse Lab in Glasgow | Wellcome Sanger Institute for the COVID-19 Genomics UK (COG-UK) Consortium | Harper VanSteenhouse, Yumi Kasai, David Gray, Carol Clugston, Anna Dominiczak and Alex Alderton, Roberto Amato, Jeffrey Barrett, Sonia Goncalves, Ewan Harrison, David K. Jackson, Ian Johnston, Dominic Kwiatkowski, Cordelia Langford, John Sillitoe on behalf of the Wellcome Sanger Institute COVID-19 Surveillance Team |
| EPI_ISL_2171915 | Lighthouse Lab in Alderley Park | Wellcome Sanger Institute for the COVID-19 Genomics UK (COG-UK) Consortium | Jacquelyn Wynn, Mairead Hyland, The Lighthouse Lab in Alderley Park and Alex Alderton, Roberto Amato, Jeffrey Barrett, Sonia Goncalves, Ewan Harrison, David K. Jackson, Ian Johnston, Dominic Kwiatkowski, Cordelia Langford, John Sillitoe on behalf of the Wellcome Sanger Institute COVID-19 Surveillance Team |
| EPI_ISL_2171918, EPI_ISL_2171920 | Lighthouse Lab in Glasgow | Wellcome Sanger Institute for the COVID-19 Genomics UK (COG-UK) Consortium | Harper VanSteenhouse, Yumi Kasai, David Gray, Carol Clugston, Anna Dominiczak and Alex Alderton, Roberto Amato, Jeffrey Barrett, Sonia Goncalves, Ewan Harrison, David K. Jackson, Ian Johnston, Dominic Kwiatkowski, Cordelia Langford, John Sillitoe on behalf of the Wellcome Sanger Institute COVID-19 Surveillance Team |
| EPI_ISL_2171922, EPI_ISL_2171923, EPI_ISL_2171925 | Lighthouse Lab in Alderley Park | Wellcome Sanger Institute for the COVID-19 Genomics UK (COG-UK) Consortium | Jacquelyn Wynn, Mairead Hyland, The Lighthouse Lab in Alderley Park and Alex Alderton, Roberto Amato, Jeffrey Barrett, Sonia Goncalves, Ewan Harrison, David K. Jackson, Ian Johnston, Dominic Kwiatkowski, Cordelia Langford, John Sillitoe on behalf of the Wellcome Sanger Institute COVID-19 Surveillance Team |
| EPI_ISL_2171928 | Lighthouse Lab in Glasgow | Wellcome Sanger Institute for the COVID-19 Genomics UK (COG-UK) Consortium | Harper VanSteenhouse, Yumi Kasai, David Gray, Carol Clugston, Anna Dominiczak and Alex Alderton, Roberto Amato, Jeffrey Barrett, Sonia Goncalves, Ewan Harrison, David K. Jackson, Ian Johnston, Dominic Kwiatkowski, Cordelia Langford, John Sillitoe on behalf of the Wellcome Sanger Institute COVID-19 Surveillance Team |
| EPI_ISL_2171931, EPI_ISL_2171932, EPI_ISL_2171934 | Lighthouse Lab in Alderley Park | Wellcome Sanger Institute for the COVID-19 Genomics UK (COG-UK) Consortium | Jacquelyn Wynn, Mairead Hyland, The Lighthouse Lab in Alderley Park and Alex Alderton, Roberto Amato, Jeffrey Barrett, Sonia Goncalves, Ewan Harrison, David K. Jackson, Ian Johnston, Dominic Kwiatkowski, Cordelia Langford, John Sillitoe on behalf of the Wellcome Sanger Institute COVID-19 Surveillance Team |
| EPI_ISL_2171936, EPI_ISL_2171937, EPI_ISL_2171939, EPI_ISL_2171940 | Lighthouse Lab in Glasgow | Wellcome Sanger Institute for the COVID-19 Genomics UK (COG-UK) Consortium | Harper VanSteenhouse, Yumi Kasai, David Gray, Carol Clugston, Anna Dominiczak and Alex Alderton, Roberto Amato, Jeffrey Barrett, Sonia Goncalves, Ewan Harrison, David K. Jackson, Ian Johnston, Dominic Kwiatkowski, Cordelia Langford, John Sillitoe on behalf of the Wellcome Sanger Institute COVID-19 Surveillance Team |
| EPI_ISL_2171941 | Lighthouse Lab in Milton Keynes | Wellcome Sanger Institute for the COVID-19 Genomics UK (COG-UK) Consortium | The Lighthouse Lab in Milton Keynes and Alex Alderton, Roberto Amato, Jeffrey Barrett, Sonia Goncalves, Ewan Harrison, David K. Jackson, Ian Johnston, Dominic Kwiatkowski, Cordelia Langford, John Sillitoe on behalf of the Wellcome Sanger Institute COVID-19 Surveillance Team |
| EPI_ISL_2171942, EPI_ISL_2171945, EPI_ISL_2171946 | Lighthouse Lab in Alderley Park | Wellcome Sanger Institute for the COVID-19 Genomics UK (COG-UK) Consortium | Jacquelyn Wynn, Mairead Hyland, The Lighthouse Lab in Alderley Park and Alex Alderton, Roberto Amato, Jeffrey Barrett, Sonia Goncalves, Ewan Harrison, David K. Jackson, Ian Johnston, Dominic Kwiatkowski, Cordelia Langford, John Sillitoe on behalf of the Wellcome Sanger Institute COVID-19 Surveillance Team |
| EPI_ISL_2171951 | Lighthouse Lab in Glasgow | Wellcome Sanger Institute for the COVID-19 Genomics UK (COG-UK) Consortium | Harper VanSteenhouse, Yumi Kasai, David Gray, Carol Clugston, Anna Dominiczak and Alex Alderton, Roberto Amato, Jeffrey Barrett, Sonia Goncalves, Ewan Harrison, David K. Jackson, Ian Johnston, Dominic Kwiatkowski, Cordelia Langford, John Sillitoe on behalf of the Wellcome Sanger Institute COVID-19 Surveillance Team |
| EPI_ISL_2171952, EPI_ISL_2171953, EPI_ISL_2171954, EPI_ISL_2171955, EPI_ISL_2171956 | Lighthouse Lab in Alderley Park | Wellcome Sanger Institute for the COVID-19 Genomics UK (COG-UK) Consortium | Jacquelyn Wynn, Mairead Hyland, The Lighthouse Lab in Alderley Park and Alex Alderton, Roberto Amato, Jeffrey Barrett, Sonia Goncalves, Ewan Harrison, David K. Jackson, Ian Johnston, Dominic Kwiatkowski, Cordelia Langford, John Sillitoe on behalf of the Wellcome Sanger Institute COVID-19 Surveillance Team |
| EPI_ISL_2171957 | Lighthouse Lab in Glasgow | Wellcome Sanger Institute for the COVID-19 Genomics UK (COG-UK) Consortium | Harper VanSteenhouse, Yumi Kasai, David Gray, Carol Clugston, Anna Dominiczak and Alex Alderton, Roberto Amato, Jeffrey Barrett, Sonia Goncalves, Ewan Harrison, David K. Jackson, Ian Johnston, Dominic Kwiatkowski, Cordelia Langford, John Sillitoe on behalf of the Wellcome Sanger Institute COVID-19 Surveillance Team |
| EPI_ISL_2171958 | Lighthouse Lab in Alderley Park | Wellcome Sanger Institute for the COVID-19 Genomics UK (COG-UK) Consortium | Jacquelyn Wynn, Mairead Hyland, The Lighthouse Lab in Alderley Park and Alex Alderton, Roberto Amato, Jeffrey Barrett, Sonia Goncalves, Ewan Harrison, David K. Jackson, Ian Johnston, Dominic Kwiatkowski, Cordelia Langford, John Sillitoe on behalf of the Wellcome Sanger Institute COVID-19 Surveillance Team |
| EPI_ISL_2171959, EPI_ISL_2171961 | Lighthouse Lab in Glasgow | Wellcome Sanger Institute for the COVID-19 Genomics UK (COG-UK) Consortium | Harper VanSteenhouse, Yumi Kasai, David Gray, Carol Clugston, Anna Dominiczak and Alex Alderton, Roberto Amato, Jeffrey Barrett, Sonia Goncalves, Ewan Harrison, David K. Jackson, Ian Johnston, Dominic Kwiatkowski, Cordelia Langford, John Sillitoe on behalf of the Wellcome Sanger Institute COVID-19 Surveillance Team |
| EPI_ISL_2171962 | Lighthouse Lab in Alderley Park | Wellcome Sanger Institute for the COVID-19 Genomics UK (COG-UK) Consortium | Jacquelyn Wynn, Mairead Hyland, The Lighthouse Lab in Alderley Park and Alex Alderton, Roberto Amato, Jeffrey Barrett, Sonia Goncalves, Ewan Harrison, David K. Jackson, Ian Johnston, Dominic Kwiatkowski, Cordelia Langford, John Sillitoe on behalf of the Wellcome Sanger Institute COVID-19 Surveillance Team |
| EPI_ISL_2171963 | Lighthouse Lab in Glasgow | Wellcome Sanger Institute for the COVID-19 Genomics UK (COG-UK) Consortium | Harper VanSteenhouse, Yumi Kasai, David Gray, Carol Clugston, Anna Dominiczak and Alex Alderton, Roberto Amato, Jeffrey Barrett, Sonia Goncalves, Ewan Harrison, David K. Jackson, Ian Johnston, Dominic Kwiatkowski, Cordelia Langford, John Sillitoe on behalf of the Wellcome Sanger Institute COVID-19 Surveillance Team |

| | | | |
|---|---|---|---|
| EPI_ISL_2171965 | Lighthouse Lab in Alderley Park | Wellcome Sanger Institute for the COVID-19 Genomics UK (COG-UK) Consortium | Jacquelyn Wynn, Mairead Hyland, The Lighthouse Lab in Alderley Park and Alex Alderton, Roberto Amato, Jeffrey Barrett, Sonia Goncalves, Ewan Harrison, David K. Jackson, Ian Johnston, Dominic Kwiatkowski, Cordelia Langford, John Sillitoe on behalf of the Wellcome Sanger Institute COVID-19 Surveillance Team |
| EPI_ISL_2171966 | Lighthouse Lab in Glasgow | Wellcome Sanger Institute for the COVID-19 Genomics UK (COG-UK) Consortium | Harper VanSteenhouse, Yumi Kasai, David Gray, Carol Clugston, Anna Dominiczak and Alex Alderton, Roberto Amato, Jeffrey Barrett, Sonia Goncalves, Ewan Harrison, David K. Jackson, Ian Johnston, Dominic Kwiatkowski, Cordelia Langford, John Sillitoe on behalf of the Wellcome Sanger Institute COVID-19 Surveillance Team |
| EPI_ISL_2171968, EPI_ISL_2171969 | Lighthouse Lab in Alderley Park | Wellcome Sanger Institute for the COVID-19 Genomics UK (COG-UK) Consortium | Jacquelyn Wynn, Mairead Hyland, The Lighthouse Lab in Alderley Park and Alex Alderton, Roberto Amato, Jeffrey Barrett, Sonia Goncalves, Ewan Harrison, David K. Jackson, Ian Johnston, Dominic Kwiatkowski, Cordelia Langford, John Sillitoe on behalf of the Wellcome Sanger Institute COVID-19 Surveillance Team |
| EPI_ISL_2171970, EPI_ISL_2171971 | Lighthouse Lab in Glasgow | Wellcome Sanger Institute for the COVID-19 Genomics UK (COG-UK) Consortium | Harper VanSteenhouse, Yumi Kasai, David Gray, Carol Clugston, Anna Dominiczak and Alex Alderton, Roberto Amato, Jeffrey Barrett, Sonia Goncalves, Ewan Harrison, David K. Jackson, Ian Johnston, Dominic Kwiatkowski, Cordelia Langford, John Sillitoe on behalf of the Wellcome Sanger Institute COVID-19 Surveillance Team |
| EPI_ISL_2171972, EPI_ISL_2171974, EPI_ISL_2171975 | Lighthouse Lab in Alderley Park | Wellcome Sanger Institute for the COVID-19 Genomics UK (COG-UK) Consortium | Jacquelyn Wynn, Mairead Hyland, The Lighthouse Lab in Alderley Park and Alex Alderton, Roberto Amato, Jeffrey Barrett, Sonia Goncalves, Ewan Harrison, David K. Jackson, Ian Johnston, Dominic Kwiatkowski, Cordelia Langford, John Sillitoe on behalf of the Wellcome Sanger Institute COVID-19 Surveillance Team |
| EPI_ISL_2171976 | Lighthouse Lab in Glasgow | Wellcome Sanger Institute for the COVID-19 Genomics UK (COG-UK) Consortium | Harper VanSteenhouse, Yumi Kasai, David Gray, Carol Clugston, Anna Dominiczak and Alex Alderton, Roberto Amato, Jeffrey Barrett, Sonia Goncalves, Ewan Harrison, David K. Jackson, Ian Johnston, Dominic Kwiatkowski, Cordelia Langford, John Sillitoe on behalf of the Wellcome Sanger Institute COVID-19 Surveillance Team |
| EPI_ISL_2171979, EPI_ISL_2171985 | Lighthouse Lab in Alderley Park | Wellcome Sanger Institute for the COVID-19 Genomics UK (COG-UK) Consortium | Jacquelyn Wynn, Mairead Hyland, The Lighthouse Lab in Alderley Park and Alex Alderton, Roberto Amato, Jeffrey Barrett, Sonia Goncalves, Ewan Harrison, David K. Jackson, Ian Johnston, Dominic Kwiatkowski, Cordelia Langford, John Sillitoe on behalf of the Wellcome Sanger Institute COVID-19 Surveillance Team |
| EPI_ISL_2171986 | Lighthouse Lab in Milton Keynes | Wellcome Sanger Institute for the COVID-19 Genomics UK (COG-UK) Consortium | The Lighthouse Lab in Milton Keynes and Alex Alderton, Roberto Amato, Jeffrey Barrett, Sonia Goncalves, Ewan Harrison, David K. Jackson, Ian Johnston, Dominic Kwiatkowski, Cordelia Langford, John Sillitoe on behalf of the Wellcome Sanger Institute COVID-19 Surveillance Team |
| EPI_ISL_2171987, EPI_ISL_2171989, EPI_ISL_2171990, EPI_ISL_2171991 | Lighthouse Lab in Alderley Park | Wellcome Sanger Institute for the COVID-19 Genomics UK (COG-UK) Consortium | Jacquelyn Wynn, Mairead Hyland, The Lighthouse Lab in Alderley Park and Alex Alderton, Roberto Amato, Jeffrey Barrett, Sonia Goncalves, Ewan Harrison, David K. Jackson, Ian Johnston, Dominic Kwiatkowski, Cordelia Langford, John Sillitoe on behalf of the Wellcome Sanger Institute COVID-19 Surveillance Team |
| EPI_ISL_2171994, EPI_ISL_2171995 | Lighthouse Lab in Glasgow | Wellcome Sanger Institute for the COVID-19 Genomics UK (COG-UK) Consortium | Harper VanSteenhouse, Yumi Kasai, David Gray, Carol Clugston, Anna Dominiczak and Alex Alderton, Roberto Amato, Jeffrey Barrett, Sonia Goncalves, Ewan Harrison, David K. Jackson, Ian Johnston, Dominic Kwiatkowski, Cordelia Langford, John Sillitoe on behalf of the Wellcome Sanger Institute COVID-19 Surveillance Team |
| EPI_ISL_2171996, EPI_ISL_2171998, EPI_ISL_2171999, EPI_ISL_2172000 | Lighthouse Lab in Alderley Park | Wellcome Sanger Institute for the COVID-19 Genomics UK (COG-UK) Consortium | Jacquelyn Wynn, Mairead Hyland, The Lighthouse Lab in Alderley Park and Alex Alderton, Roberto Amato, Jeffrey Barrett, Sonia Goncalves, Ewan Harrison, David K. Jackson, Ian Johnston, Dominic Kwiatkowski, Cordelia Langford, John Sillitoe on behalf of the Wellcome Sanger Institute COVID-19 Surveillance Team |
| EPI_ISL_2172001 | Lighthouse Lab in Glasgow | Wellcome Sanger Institute for the COVID-19 Genomics UK (COG-UK) Consortium | Harper VanSteenhouse, Yumi Kasai, David Gray, Carol Clugston, Anna Dominiczak and Alex Alderton, Roberto Amato, Jeffrey Barrett, Sonia Goncalves, Ewan Harrison, David K. Jackson, Ian Johnston, Dominic Kwiatkowski, Cordelia Langford, John Sillitoe on behalf of the Wellcome Sanger Institute COVID-19 Surveillance Team |
| EPI_ISL_2172002, EPI_ISL_2172003, EPI_ISL_2172004 | Lighthouse Lab in Alderley Park | Wellcome Sanger Institute for the COVID-19 Genomics UK (COG-UK) Consortium | Jacquelyn Wynn, Mairead Hyland, The Lighthouse Lab in Alderley Park and Alex Alderton, Roberto Amato, Jeffrey Barrett, Sonia Goncalves, Ewan Harrison, David K. Jackson, Ian Johnston, Dominic Kwiatkowski, Cordelia Langford, John Sillitoe on behalf of the Wellcome Sanger Institute COVID-19 Surveillance Team |
| EPI_ISL_2172006, EPI_ISL_2172008, EPI_ISL_2172009, EPI_ISL_2172011, EPI_ISL_2172012, EPI_ISL_2172015 | Lighthouse Lab in Glasgow | Wellcome Sanger Institute for the COVID-19 Genomics UK (COG-UK) Consortium | Harper VanSteenhouse, Yumi Kasai, David Gray, Carol Clugston, Anna Dominiczak and Alex Alderton, Roberto Amato, Jeffrey Barrett, Sonia Goncalves, Ewan Harrison, David K. Jackson, Ian Johnston, Dominic Kwiatkowski, Cordelia Langford, John Sillitoe on behalf of the Wellcome Sanger Institute COVID-19 Surveillance Team |
| EPI_ISL_2172016, EPI_ISL_2172017, EPI_ISL_2172018, EPI_ISL_2172020, EPI_ISL_2172024, EPI_ISL_2172026 | Lighthouse Lab in Alderley Park | Wellcome Sanger Institute for the COVID-19 Genomics UK (COG-UK) Consortium | Jacquelyn Wynn, Mairead Hyland, The Lighthouse Lab in Alderley Park and Alex Alderton, Roberto Amato, Jeffrey Barrett, Sonia Goncalves, Ewan Harrison, David K. Jackson, Ian Johnston, Dominic Kwiatkowski, Cordelia Langford, John Sillitoe on behalf of the Wellcome Sanger Institute COVID-19 Surveillance Team |
| EPI_ISL_2172027 | Lighthouse Lab in Glasgow | Wellcome Sanger Institute for the COVID-19 Genomics UK (COG-UK) Consortium | Harper VanSteenhouse, Yumi Kasai, David Gray, Carol Clugston, Anna Dominiczak and Alex Alderton, Roberto Amato, Jeffrey Barrett, Sonia Goncalves, Ewan Harrison, David K. Jackson, Ian Johnston, Dominic Kwiatkowski, Cordelia Langford, John Sillitoe on behalf of the Wellcome Sanger Institute COVID-19 Surveillance Team |
| EPI_ISL_2172029, EPI_ISL_2172031, EPI_ISL_2172032 | Lighthouse Lab in Alderley Park | Wellcome Sanger Institute for the COVID-19 Genomics UK (COG-UK) Consortium | Jacquelyn Wynn, Mairead Hyland, The Lighthouse Lab in Alderley Park and Alex Alderton, Roberto Amato, Jeffrey Barrett, Sonia Goncalves, Ewan Harrison, David K. Jackson, Ian Johnston, Dominic Kwiatkowski, Cordelia Langford, John Sillitoe on behalf of the Wellcome Sanger Institute COVID-19 Surveillance Team |
| EPI_ISL_2172037 | Lighthouse Lab in Glasgow | Wellcome Sanger Institute for the COVID-19 Genomics UK (COG-UK) Consortium | Harper VanSteenhouse, Yumi Kasai, David Gray, Carol Clugston, Anna Dominiczak and Alex Alderton, Roberto Amato, Jeffrey Barrett, Sonia Goncalves, Ewan Harrison, David K. Jackson, Ian Johnston, Dominic Kwiatkowski, Cordelia Langford, John Sillitoe on behalf of the Wellcome Sanger Institute COVID-19 Surveillance Team |
| EPI_ISL_2172039, EPI_ISL_2172040, EPI_ISL_2172042, EPI_ISL_2172043, EPI_ISL_2172044 | Lighthouse Lab in Alderley Park | Wellcome Sanger Institute for the COVID-19 Genomics UK (COG-UK) Consortium | Jacquelyn Wynn, Mairead Hyland, The Lighthouse Lab in Alderley Park and Alex Alderton, Roberto Amato, Jeffrey Barrett, Sonia Goncalves, Ewan Harrison, David K. Jackson, Ian Johnston, Dominic Kwiatkowski, Cordelia Langford, John Sillitoe on behalf of the Wellcome Sanger Institute COVID-19 Surveillance Team |
| EPI_ISL_2172045 | Lighthouse Lab in Glasgow | Wellcome Sanger Institute for the COVID-19 Genomics UK (COG-UK) Consortium | Harper VanSteenhouse, Yumi Kasai, David Gray, Carol Clugston, Anna Dominiczak and Alex Alderton, Roberto Amato, Jeffrey Barrett, Sonia Goncalves, Ewan Harrison, David K. Jackson, Ian Johnston, Dominic Kwiatkowski, Cordelia Langford, John Sillitoe on behalf of the Wellcome Sanger Institute COVID-19 Surveillance Team |
| EPI_ISL_2172046, EPI_ISL_2172047 | Lighthouse Lab in Alderley Park | Wellcome Sanger Institute for the COVID-19 Genomics UK (COG-UK) Consortium | Jacquelyn Wynn, Mairead Hyland, The Lighthouse Lab in Alderley Park and Alex Alderton, Roberto Amato, Jeffrey Barrett, Sonia Goncalves, Ewan Harrison, David K. Jackson, Ian Johnston, Dominic Kwiatkowski, Cordelia Langford, John Sillitoe on behalf of the Wellcome Sanger Institute COVID-19 Surveillance Team |
| EPI_ISL_2172048 | Lighthouse Lab in Glasgow | Wellcome Sanger Institute for the COVID-19 Genomics UK (COG-UK) Consortium | Harper VanSteenhouse, Yumi Kasai, David Gray, Carol Clugston, Anna Dominiczak and Alex Alderton, Roberto Amato, Jeffrey Barrett, Sonia Goncalves, Ewan Harrison, David K. Jackson, Ian Johnston, Dominic Kwiatkowski, Cordelia Langford, John Sillitoe on behalf of the Wellcome Sanger Institute COVID-19 Surveillance Team |
| EPI_ISL_2172049, EPI_ISL_2172050, EPI_ISL_2172051, EPI_ISL_2172052, EPI_ISL_2172055 | Lighthouse Lab in Alderley Park | Wellcome Sanger Institute for the COVID-19 Genomics UK (COG-UK) Consortium | Jacquelyn Wynn, Mairead Hyland, The Lighthouse Lab in Alderley Park and Alex Alderton, Roberto Amato, Jeffrey Barrett, Sonia Goncalves, Ewan Harrison, David K. Jackson, Ian Johnston, Dominic Kwiatkowski, Cordelia Langford, John Sillitoe on behalf of the Wellcome Sanger Institute COVID-19 Surveillance Team |
| EPI_ISL_2172056, EPI_ISL_2172060, EPI_ISL_2172061 | Lighthouse Lab in Glasgow | Wellcome Sanger Institute for the COVID-19 Genomics UK (COG-UK) Consortium | Harper VanSteenhouse, Yumi Kasai, David Gray, Carol Clugston, Anna Dominiczak and Alex Alderton, Roberto Amato, Jeffrey Barrett, Sonia Goncalves, Ewan Harrison, David K. Jackson, Ian Johnston, Dominic Kwiatkowski, Cordelia Langford, John Sillitoe on behalf of the Wellcome Sanger Institute COVID-19 Surveillance Team |
| EPI_ISL_2172062 | Lighthouse Lab in Alderley Park | Wellcome Sanger Institute for the COVID-19 Genomics UK (COG-UK) Consortium | Jacquelyn Wynn, Mairead Hyland, The Lighthouse Lab in Alderley Park and Alex Alderton, Roberto Amato, Jeffrey Barrett, Sonia Goncalves, Ewan Harrison, David K. Jackson, Ian Johnston, Dominic Kwiatkowski, Cordelia Langford, John Sillitoe on behalf of the Wellcome Sanger Institute COVID-19 Surveillance Team |
| EPI_ISL_2172064 | Lighthouse Lab in Glasgow | Wellcome Sanger Institute for the COVID-19 Genomics UK (COG-UK) Consortium | Harper VanSteenhouse, Yumi Kasai, David Gray, Carol Clugston, Anna Dominiczak and Alex Alderton, Roberto Amato, Jeffrey Barrett, Sonia Goncalves, Ewan Harrison, David K. Jackson, Ian Johnston, Dominic Kwiatkowski, Cordelia Langford, John Sillitoe on behalf of the Wellcome Sanger Institute COVID-19 Surveillance Team |
| EPI_ISL_2172065, EPI_ISL_2172067, EPI_ISL_2172068 | Lighthouse Lab in Alderley Park | Wellcome Sanger Institute for the COVID-19 Genomics UK (COG-UK) Consortium | Jacquelyn Wynn, Mairead Hyland, The Lighthouse Lab in Alderley Park and Alex Alderton, Roberto Amato, Jeffrey Barrett, Sonia Goncalves, Ewan Harrison, David K. Jackson, Ian Johnston, Dominic Kwiatkowski, Cordelia Langford, John Sillitoe on behalf of the Wellcome Sanger Institute COVID-19 Surveillance Team |
| EPI_ISL_2172069, EPI_ISL_2172070, EPI_ISL_2172071 | Lighthouse Lab in Glasgow | Wellcome Sanger Institute for the COVID-19 Genomics UK (COG-UK) Consortium | Harper VanSteenhouse, Yumi Kasai, David Gray, Carol Clugston, Anna Dominiczak and Alex Alderton, Roberto Amato, Jeffrey Barrett, Sonia Goncalves, Ewan Harrison, David K. Jackson, Ian Johnston, Dominic Kwiatkowski, Cordelia Langford, John Sillitoe on behalf of the Wellcome Sanger Institute COVID-19 Surveillance Team |
| EPI_ISL_2172073 | Lighthouse Lab in Alderley Park | Wellcome Sanger Institute for the COVID-19 Genomics UK (COG-UK) Consortium | Jacquelyn Wynn, Mairead Hyland, The Lighthouse Lab in Alderley Park and Alex Alderton, Roberto Amato, Jeffrey Barrett, Sonia Goncalves, Ewan Harrison, David K. Jackson, Ian Johnston, Dominic Kwiatkowski, Cordelia Langford, John Sillitoe on behalf of the Wellcome Sanger Institute COVID-19 Surveillance Team |
| EPI_ISL_2172076 | Lighthouse Lab in Glasgow | Wellcome Sanger Institute for the COVID-19 Genomics UK (COG-UK) Consortium | Harper VanSteenhouse, Yumi Kasai, David Gray, Carol Clugston, Anna Dominiczak and Alex Alderton, Roberto Amato, Jeffrey Barrett, Sonia Goncalves, Ewan Harrison, David K. Jackson, Ian Johnston, Dominic Kwiatkowski, Cordelia Langford, John Sillitoe on behalf of the Wellcome Sanger Institute COVID-19 Surveillance Team |
| EPI_ISL_2172077, EPI_ISL_2172079, EPI_ISL_2172080, EPI_ISL_2172083, EPI_ISL_2172084 | Lighthouse Lab in Alderley Park | Wellcome Sanger Institute for the COVID-19 Genomics UK (COG-UK) Consortium | Jacquelyn Wynn, Mairead Hyland, The Lighthouse Lab in Alderley Park and Alex Alderton, Roberto Amato, Jeffrey Barrett, Sonia Goncalves, Ewan Harrison, David K. Jackson, Ian Johnston, Dominic Kwiatkowski, Cordelia Langford, John Sillitoe on behalf of the Wellcome Sanger Institute COVID-19 Surveillance Team |
| EPI_ISL_2172085 | Lighthouse Lab in Glasgow | Wellcome Sanger Institute for the COVID-19 Genomics UK (COG-UK) Consortium | Harper VanSteenhouse, Yumi Kasai, David Gray, Carol Clugston, Anna Dominiczak and Alex Alderton, Roberto Amato, Jeffrey Barrett, Sonia Goncalves, Ewan Harrison, David K. Jackson, Ian Johnston, Dominic Kwiatkowski, Cordelia Langford, John Sillitoe on behalf of the Wellcome Sanger Institute COVID-19 Surveillance Team |
| EPI_ISL_2172087, EPI_ISL_2172091 | Lighthouse Lab in Alderley Park | Wellcome Sanger Institute for the COVID-19 Genomics UK (COG-UK) Consortium | Jacquelyn Wynn, Mairead Hyland, The Lighthouse Lab in Alderley Park and Alex Alderton, Roberto Amato, Jeffrey Barrett, Sonia Goncalves, Ewan Harrison, David K. Jackson, Ian Johnston, Dominic Kwiatkowski, Cordelia Langford, John Sillitoe on behalf of the Wellcome Sanger Institute COVID-19 Surveillance Team |
| EPI_ISL_2172099, EPI_ISL_2172104, | Lighthouse Lab in Milton Keynes | Wellcome Sanger Institute for the COVID-19 Genomics UK (COG-UK) Consortium | The Lighthouse Lab in Milton Keynes and Alex Alderton, Roberto Amato, Jeffrey Barrett, Sonia Goncalves, Ewan Harrison, David K. Jackson, Ian Johnston, Dominic Kwiatkowski, Cordelia Langford, John Sillitoe on behalf of the Wellcome Sanger Institute COVID-19 Surveillance Team |

| | | | |
|---|---|---|---|
| EPI_ISL_2172107 | | | |
| EPI_ISL_2172116, EPI_ISL_2172121, EPI_ISL_2172139 | Lighthouse Lab in Alderley Park | Wellcome Sanger Institute for the COVID-19 Genomics UK (COG-UK) Consortium | Jacquelyn Wynn, Mairead Hyland, The Lighthouse Lab in Alderley Park and Alex Alderton, Roberto Amato, Jeffrey Barrett, Sonia Goncalves, Ewan Harrison, David K. Jackson, Ian Johnston, Dominic Kwiatkowski, Cordelia Langford, John Sillitoe on behalf of the Wellcome Sanger Institute COVID-19 Surveillance Team |
| EPI_ISL_2172143 | Lighthouse Lab in Milton Keynes | Wellcome Sanger Institute for the COVID-19 Genomics UK (COG-UK) Consortium | The Lighthouse Lab in Milton Keynes and Alex Alderton, Roberto Amato, Jeffrey Barrett, Sonia Goncalves, Ewan Harrison, David K. Jackson, Ian Johnston, Dominic Kwiatkowski, Cordelia Langford, John Sillitoe on behalf of the Wellcome Sanger Institute COVID-19 Surveillance Team |
| EPI_ISL_2172153 | Lighthouse Lab in Alderley Park | Wellcome Sanger Institute for the COVID-19 Genomics UK (COG-UK) Consortium | Jacquelyn Wynn, Mairead Hyland, The Lighthouse Lab in Alderley Park and Alex Alderton, Roberto Amato, Jeffrey Barrett, Sonia Goncalves, Ewan Harrison, David K. Jackson, Ian Johnston, Dominic Kwiatkowski, Cordelia Langford, John Sillitoe on behalf of the Wellcome Sanger Institute COVID-19 Surveillance Team |
| EPI_ISL_2172174 | Lighthouse Lab in Milton Keynes | Wellcome Sanger Institute for the COVID-19 Genomics UK (COG-UK) Consortium | The Lighthouse Lab in Milton Keynes and Alex Alderton, Roberto Amato, Jeffrey Barrett, Sonia Goncalves, Ewan Harrison, David K. Jackson, Ian Johnston, Dominic Kwiatkowski, Cordelia Langford, John Sillitoe on behalf of the Wellcome Sanger Institute COVID-19 Surveillance Team |
| EPI_ISL_2172182 | Lighthouse Lab in Alderley Park | Wellcome Sanger Institute for the COVID-19 Genomics UK (COG-UK) Consortium | Jacquelyn Wynn, Mairead Hyland, The Lighthouse Lab in Alderley Park and Alex Alderton, Roberto Amato, Jeffrey Barrett, Sonia Goncalves, Ewan Harrison, David K. Jackson, Ian Johnston, Dominic Kwiatkowski, Cordelia Langford, John Sillitoe on behalf of the Wellcome Sanger Institute COVID-19 Surveillance Team |
| EPI_ISL_2172196, EPI_ISL_2172210 | Lighthouse Lab in Milton Keynes | Wellcome Sanger Institute for the COVID-19 Genomics UK (COG-UK) Consortium | The Lighthouse Lab in Milton Keynes and Alex Alderton, Roberto Amato, Jeffrey Barrett, Sonia Goncalves, Ewan Harrison, David K. Jackson, Ian Johnston, Dominic Kwiatkowski, Cordelia Langford, John Sillitoe on behalf of the Wellcome Sanger Institute COVID-19 Surveillance Team |
| EPI_ISL_2172223, EPI_ISL_2172229 | Lighthouse Lab in Alderley Park | Wellcome Sanger Institute for the COVID-19 Genomics UK (COG-UK) Consortium | Jacquelyn Wynn, Mairead Hyland, The Lighthouse Lab in Alderley Park and Alex Alderton, Roberto Amato, Jeffrey Barrett, Sonia Goncalves, Ewan Harrison, David K. Jackson, Ian Johnston, Dominic Kwiatkowski, Cordelia Langford, John Sillitoe on behalf of the Wellcome Sanger Institute COVID-19 Surveillance Team |
| EPI_ISL_2172230, EPI_ISL_2172249, EPI_ISL_2172253 | Lighthouse Lab in Milton Keynes | Wellcome Sanger Institute for the COVID-19 Genomics UK (COG-UK) Consortium | The Lighthouse Lab in Milton Keynes and Alex Alderton, Roberto Amato, Jeffrey Barrett, Sonia Goncalves, Ewan Harrison, David K. Jackson, Ian Johnston, Dominic Kwiatkowski, Cordelia Langford, John Sillitoe on behalf of the Wellcome Sanger Institute COVID-19 Surveillance Team |
| EPI_ISL_2172255 | Lighthouse Lab in Alderley Park | Wellcome Sanger Institute for the COVID-19 Genomics UK (COG-UK) Consortium | Jacquelyn Wynn, Mairead Hyland, The Lighthouse Lab in Alderley Park and Alex Alderton, Roberto Amato, Jeffrey Barrett, Sonia Goncalves, Ewan Harrison, David K. Jackson, Ian Johnston, Dominic Kwiatkowski, Cordelia Langford, John Sillitoe on behalf of the Wellcome Sanger Institute COVID-19 Surveillance Team |
| EPI_ISL_2172263 | Lighthouse Lab in Milton Keynes | Wellcome Sanger Institute for the COVID-19 Genomics UK (COG-UK) Consortium | The Lighthouse Lab in Milton Keynes and Alex Alderton, Roberto Amato, Jeffrey Barrett, Sonia Goncalves, Ewan Harrison, David K. Jackson, Ian Johnston, Dominic Kwiatkowski, Cordelia Langford, John Sillitoe on behalf of the Wellcome Sanger Institute COVID-19 Surveillance Team |
| EPI_ISL_2172265, EPI_ISL_2172279 | Lighthouse Lab in Alderley Park | Wellcome Sanger Institute for the COVID-19 Genomics UK (COG-UK) Consortium | Jacquelyn Wynn, Mairead Hyland, The Lighthouse Lab in Alderley Park and Alex Alderton, Roberto Amato, Jeffrey Barrett, Sonia Goncalves, Ewan Harrison, David K. Jackson, Ian Johnston, Dominic Kwiatkowski, Cordelia Langford, John Sillitoe on behalf of the Wellcome Sanger Institute COVID-19 Surveillance Team |
| EPI_ISL_2172318 | Lighthouse Lab in Milton Keynes | Wellcome Sanger Institute for the COVID-19 Genomics UK (COG-UK) Consortium | The Lighthouse Lab in Milton Keynes and Alex Alderton, Roberto Amato, Jeffrey Barrett, Sonia Goncalves, Ewan Harrison, David K. Jackson, Ian Johnston, Dominic Kwiatkowski, Cordelia Langford, John Sillitoe on behalf of the Wellcome Sanger Institute COVID-19 Surveillance Team |
| EPI_ISL_2172324 | Lighthouse Lab in Alderley Park | Wellcome Sanger Institute for the COVID-19 Genomics UK (COG-UK) Consortium | Jacquelyn Wynn, Mairead Hyland, The Lighthouse Lab in Alderley Park and Alex Alderton, Roberto Amato, Jeffrey Barrett, Sonia Goncalves, Ewan Harrison, David K. Jackson, Ian Johnston, Dominic Kwiatkowski, Cordelia Langford, John Sillitoe on behalf of the Wellcome Sanger Institute COVID-19 Surveillance Team |
| EPI_ISL_2172325 | Lighthouse Lab in Milton Keynes | Wellcome Sanger Institute for the COVID-19 Genomics UK (COG-UK) Consortium | The Lighthouse Lab in Milton Keynes and Alex Alderton, Roberto Amato, Jeffrey Barrett, Sonia Goncalves, Ewan Harrison, David K. Jackson, Ian Johnston, Dominic Kwiatkowski, Cordelia Langford, John Sillitoe on behalf of the Wellcome Sanger Institute COVID-19 Surveillance Team |
| EPI_ISL_2172328 | Lighthouse Lab in Alderley Park | Wellcome Sanger Institute for the COVID-19 Genomics UK (COG-UK) Consortium | Jacquelyn Wynn, Mairead Hyland, The Lighthouse Lab in Alderley Park and Alex Alderton, Roberto Amato, Jeffrey Barrett, Sonia Goncalves, Ewan Harrison, David K. Jackson, Ian Johnston, Dominic Kwiatkowski, Cordelia Langford, John Sillitoe on behalf of the Wellcome Sanger Institute COVID-19 Surveillance Team |
| EPI_ISL_2172348, EPI_ISL_2172360 | Lighthouse Lab in Milton Keynes | Wellcome Sanger Institute for the COVID-19 Genomics UK (COG-UK) Consortium | The Lighthouse Lab in Milton Keynes and Alex Alderton, Roberto Amato, Jeffrey Barrett, Sonia Goncalves, Ewan Harrison, David K. Jackson, Ian Johnston, Dominic Kwiatkowski, Cordelia Langford, John Sillitoe on behalf of the Wellcome Sanger Institute COVID-19 Surveillance Team |
| EPI_ISL_2172366 | Lighthouse Lab in Alderley Park | Wellcome Sanger Institute for the COVID-19 Genomics UK (COG-UK) Consortium | Jacquelyn Wynn, Mairead Hyland, The Lighthouse Lab in Alderley Park and Alex Alderton, Roberto Amato, Jeffrey Barrett, Sonia Goncalves, Ewan Harrison, David K. Jackson, Ian Johnston, Dominic Kwiatkowski, Cordelia Langford, John Sillitoe on behalf of the Wellcome Sanger Institute COVID-19 Surveillance Team |
| EPI_ISL_2172372 | Lighthouse Lab in Milton Keynes | Wellcome Sanger Institute for the COVID-19 Genomics UK (COG-UK) Consortium | The Lighthouse Lab in Milton Keynes and Alex Alderton, Roberto Amato, Jeffrey Barrett, Sonia Goncalves, Ewan Harrison, David K. Jackson, Ian Johnston, Dominic Kwiatkowski, Cordelia Langford, John Sillitoe on behalf of the Wellcome Sanger Institute COVID-19 Surveillance Team |
| EPI_ISL_2172374, EPI_ISL_2172380, EPI_ISL_2172381, EPI_ISL_2172386, EPI_ISL_2172401 | Lighthouse Lab in Alderley Park | Wellcome Sanger Institute for the COVID-19 Genomics UK (COG-UK) Consortium | Jacquelyn Wynn, Mairead Hyland, The Lighthouse Lab in Alderley Park and Alex Alderton, Roberto Amato, Jeffrey Barrett, Sonia Goncalves, Ewan Harrison, David K. Jackson, Ian Johnston, Dominic Kwiatkowski, Cordelia Langford, John Sillitoe on behalf of the Wellcome Sanger Institute COVID-19 Surveillance Team |
| EPI_ISL_2172542, EPI_ISL_2172586, EPI_ISL_2172683, EPI_ISL_2172791, EPI_ISL_2173209, EPI_ISL_2173553, EPI_ISL_2174330, EPI_ISL_2174379, EPI_ISL_2174463, EPI_ISL_2174512, EPI_ISL_2174862, EPI_ISL_2175337, EPI_ISL_2175485, EPI_ISL_2175486, EPI_ISL_2175543, EPI_ISL_2175701, EPI_ISL_2175863, EPI_ISL_2176057, EPI_ISL_2176964, EPI_ISL_2177177, EPI_ISL_2177191, EPI_ISL_2177275 | | | |
| see above | Department of Virus and Microbiological Special Diagnostics, Statens Serum Institut, Copenhagen, Denmark | Aalborg University | Danish Covid-19 Genome Consortium |
| EPI_ISL_2177412 | TXDSHS | TXDSHS | Rashmi Tuladhar, Bonnie Oh, Jenny Zhang, Maliha Rahman, Mayela Pedrueza, Anita Pokharel, Karen Bobier, Lorraine Rodriguez, Myong Koag, Chun Wang, Rachel Lee, Grace Kubin |
| EPI_ISL_2177545, EPI_ISL_2178114 | Department of Virus and Microbiological Special Diagnostics, Statens Serum Institut, Copenhagen, Denmark | Aalborg University | Danish Covid-19 Genome Consortium |
| EPI_ISL_2178978 | OLVZ Aalst | OLVZ Aalst | Astrid Holderbeke |
| EPI_ISL_2179170 | DOHMH PHL | New York City Public Health Laboratory | Jade Wang, et al. |
| EPI_ISL_2179186 | DOHMH Jamaica | New York City Public Health Laboratory | Jade Wang, et al. |
| EPI_ISL_2179878, EPI_ISL_2179901 | Fulgent Genetics | Centers for Disease Control and Prevention Division of Viral Diseases, Pathogen Discovery | Dakota Howard, Dhwani Batra, Peter W. Cook, Kara Moser, Adrian Paskey, Jason Caravas, Benjamin Rambo-Martin, Shatavia Morrison, Christopher Gulvick, Scott Sammons, Yvette Unoarumhi, Darlene Wagner, Matthew Schmerer, Harry Gao, Mickey Li, John Gao, Joseph Fierro, Benafsh Sapra, Becky Tsai, Yan Meng, Doreen Ng, James Xie, Clinton R. Paden, Duncan MacCannell |
| EPI_ISL_2180454, EPI_ISL_2180455, EPI_ISL_2180456, EPI_ISL_2180457 | Institute of Epidemiology, Disease Control and Research (IEDCR) | IEDCR-ideSHi-icddr,b | Hassan Afrad, Sadia Rahman, Manjur Hossain Khan, Firdausi Qadri, Tahmina Shirin |
| EPI_ISL_2180461, EPI_ISL_2180479, EPI_ISL_2180501, EPI_ISL_2180516 | Quest Diagnostics Incorporated | Centers for Disease Control and Prevention Division of Viral Diseases, Pathogen Discovery | Dakota Howard, Dhwani Batra, Peter W. Cook, Kara Moser, Adrian Paskey, Jason Caravas, Benjamin Rambo-Martin, Shatavia Morrison, Christopher Gulvick, Scott Sammons, Yvette Unoarumhi, Darlene Wagner, Matthew Schmerer, S. H. Rosenthal, A. Gerasimova, R. M. Kagan, B. Anderson, M. Hua, Y. Liu, L.E. Bernstein, K.E. Livingston, A. Perez, I. A. Shlyakhter, R. V. Rolando, R. Owen, P. Tanpaiboon, F. Lacbawan, Clinton R. Paden, Duncan MacCannell |
| EPI_ISL_2180629, EPI_ISL_2180642, EPI_ISL_2180715, EPI_ISL_2180786, EPI_ISL_2180899, EPI_ISL_2181104, EPI_ISL_2181113, EPI_ISL_2181136, EPI_ISL_2181219, EPI_ISL_2181262, EPI_ISL_2181306, EPI_ISL_2181311, EPI_ISL_2181323, EPI_ISL_2181324, EPI_ISL_2181391, EPI_ISL_2181435, EPI_ISL_2181534 | | | |
| see above | Aegis Sciences Corporation | Centers for Disease Control and Prevention Division of Viral Diseases, Pathogen Discovery | Dakota Howard, Dhwani Batra, Peter W. Cook, Kara Moser, Adrian Paskey, Jason Caravas, Benjamin Rambo-Martin, Shatavia Morrison, Christopher Gulvick, Scott Sammons, Yvette Unoarumhi, Darlene Wagner, Matthew Schmerer, Cyndi Clark, Patrick Campbell, Rob Case, Vikramsinha Ghorpade, Holly Houdeshell, Ola Kvalvaag, Dillon Nall, Ethan Sanders, Alec Vest, Shaun Westlund, Matthew Hardison, Clinton R. Paden, Duncan MacCannell |
| EPI_ISL_2181699, EPI_ISL_2181727, EPI_ISL_2181917, EPI_ISL_2182032, EPI_ISL_2182035, EPI_ISL_2182078, EPI_ISL_2182225, EPI_ISL_2182270, EPI_ISL_2182444, EPI_ISL_2182485, EPI_ISL_2182642, EPI_ISL_2182878, EPI_ISL_2182881 | | | |
| see above | Laboratory Corporation of America | Centers for Disease Control and Prevention Division of Viral Diseases, Pathogen Discovery | Dakota Howard, Dhwani Batra, Peter W. Cook, Kara Moser, Adrian Paskey, Jason Caravas, Benjamin Rambo-Martin, Shatavia Morrison, Christopher Gulvick, Scott Sammons, Yvette Unoarumhi, Darlene Wagner, Matthew Schmerer, Minoo Agarwal, Eyad Almasri, Debbie Boles, Ayla Burns, Nuthawin Charoensri, Oren Cohen, Susan Countryman, Mary Ann Cristobal, Bobbi Croy, Suzanne Dale, Hrushikesh Deshmukh, Amanda Douglas, Vincent Drouillon, Marcia Eisenberg, Howard Engler, Rama Ghatti, Prashant Gupta, Susan Hicks, Jake Humphrey, Lax Iyer, Manoj Jain, Mohan Kolli, Brian Krueger, Tim Kuphal, Stanley Letovsky, Michael Levandoski, Craig Lukasik, Jonathan Meltzer, Brian Norvell, Mindy Nye, Scott Parker, Christos Petropoulos, John Pruitt, Steven Ragan, Scott Ryan, Mike Sapeta, Jana Schroth, Suresh Babu Selvaraju, Goran Stevovic, Amanda Suchanek, Andrea Throop, Lyndon Tilson, Thomas Urban, Joe Voshell, Kimberly Wagner, Jonathan Williams, Mary Williamson, Qian Zeng, Tricia Zwiefelhofer, Clinton R. Paden, Duncan MacCannell |
| EPI_ISL_2182923 | Israel Central Virology laboratory | Israel National Consortium for SARS-CoV-2 sequencing | Neta Zuckerman, Efrat Dahan Bucris, Michal Mandelboim, Dana Bar-Ilan, Miranda Geva, Netanel Abu, Oran Erster, Efrat Glick-Saar, Omri Nayshool, Gideon Rechavi, Ella Mendelson, Orna Mor |
| EPI_ISL_2182993 | Laboratory Corporation of America | Centers for Disease Control and Prevention Division of Viral Diseases, Pathogen Discovery | Dakota Howard, Dhwani Batra, Peter W. Cook, Kara Moser, Adrian Paskey, Jason Caravas, Benjamin Rambo-Martin, Shatavia Morrison, Christopher Gulvick, Scott Sammons, Yvette Unoarumhi, Darlene Wagner, Matthew Schmerer, Minoo Agarwal, Eyad Almasri, Debbie Boles, Ayla Burns, Nuthawin Charoensri, Oren Cohen, Susan Countryman, Mary Ann Cristobal, Bobbi Croy, Suzanne Dale, Hrushikesh Deshmukh, Amanda Douglas, Vincent Drouillon, Marcia Eisenberg, Howard Engler, Rama Ghatti, Prashant Gupta, Susan Hicks, Jake Humphrey, Lax Iyer, Manoj Jain, Mohan Kolli, Brian Krueger, Tim Kuphal, Stanley Letovsky, Michael Levandoski, Craig Lukasik, Jonathan Meltzer, Brian Norvell, Mindy Nye, Scott Parker, Christos Petropoulos, John Pruitt, Steven Ragan, Scott Ryan, Mike Sapeta, Jana Schroth, Suresh Babu Selvaraju, Goran Stevovic, Amanda Suchanek, Andrea Throop, Lyndon Tilson, Thomas Urban, Joe Voshell, Kimberly Wagner, Jonathan Williams, Mary Williamson, Qian Zeng, Tricia Zwiefelhofer, Clinton R. Paden, Duncan MacCannell |
| EPI_ISL_2183026, EPI_ISL_2183029, EPI_ISL_2183057, EPI_ISL_2183059, EPI_ISL_2183060, EPI_ISL_2183062 | Israel Central Virology laboratory | Israel National Consortium for SARS-CoV-2 sequencing | Neta Zuckerman, Efrat Dahan Bucris, Michal Mandelboim, Dana Bar-Ilan, Miranda Geva, Netanel Abu, Oran Erster, Efrat Glick-Saar, Omri Nayshool, Gideon Rechavi, Ella Mendelson, Orna Mor |
| EPI_ISL_2183100 | Laboratory Corporation of America | Centers for Disease Control and Prevention Division of Viral Diseases, Pathogen Discovery | Dakota Howard, Dhwani Batra, Peter W. Cook, Kara Moser, Adrian Paskey, Jason Caravas, Benjamin Rambo-Martin, Shatavia Morrison, Christopher Gulvick, Scott Sammons, Yvette Unoarumhi, Darlene Wagner, Matthew Schmerer, Minoo Agarwal, Eyad Almasri, Debbie Boles, Ayla Burns, Nuthawin Charoensri, Oren Cohen, Susan Countryman, Mary Ann Cristobal, Bobbi Croy, Suzanne Dale, Hrushikesh Deshmukh, Amanda Douglas, Vincent Drouillon, Marcia Eisenberg, Howard Engler, Rama Ghatti, Prashant Gupta, Susan Hicks, Jake Humphrey, Lax Iyer, Manoj Jain, Mohan Kolli, Brian Krueger, Tim Kuphal, Stanley Letovsky, Michael Levandoski, Craig Lukasik, Jonathan Meltzer, Brian Norvell, Mindy Nye, Scott Parker, Christos Petropoulos, John Pruitt, Steven Ragan, Scott Ryan, Mike Sapeta, Jana Schroth, Suresh Babu Selvaraju, Goran Stevovic, Amanda Suchanek, Andrea Throop, Lyndon Tilson, Thomas Urban, Joe Voshell, Kimberly Wagner, Jonathan Williams, Mary Williamson, Qian Zeng, Tricia Zwiefelhofer, Clinton R. Paden, Duncan MacCannell |
| EPI_ISL_2183112, EPI_ISL_2183114, EPI_ISL_2183116, EPI_ISL_2183117 | Israel Central Virology laboratory | Israel National Consortium for SARS-CoV-2 sequencing | Neta Zuckerman, Efrat Dahan Bucris, Michal Mandelboim, Dana Bar-Ilan, Miranda Geva, Netanel Abu, Oran Erster, Efrat Glick-Saar, Omri Nayshool, Gideon Rechavi, Ella Mendelson, Orna Mor |

| | | | |
|---|---|---|---|
| EPI_ISL_2183147 | Laboratory Corporation of America | Centers for Disease Control and Prevention Division of Viral Diseases, Pathogen Discovery | Dakota Howard, Dhwani Batra, Peter W. Cook, Kara Moser, Adrian Paskey, Jason Caravas, Benjamin Rambo-Martin, Shatavia Morrison, Christopher Gulvick, Scott Sammons, Yvette Unoarumhi, Darlene Wagner, Matthew Schmerer, Minoo Agarwal, Eyad Almasri, Debbie Boles, Ayla Burns, Nuthawin Charoensri, Oren Cohen, Susan Countryman, Mary Ann Cristobal, Bobbi Croy, Suzanne Dale, Hrushikesh Deshmukh, Amanda Douglas, Vincent Drouillon, Marcia Eisenberg, Howard Engler, Rama Ghatti, Prashant Gupta, Susan Hicks, Jake Humphrey, Lax Iyer, Manoj Jain, Mohan Kolli, Brian Krueger, Tim Kuphal, Stanley Letovsky, Michael Levandoski, Craig Lukasik, Jonathan Meltzer, Brian Norvell, Mindy Nye, Scott Parker, Christos Petropoulos, John Pruitt, Steven Ragan, Scott Ryan, Mike Sapeta, Jana Schroth, Suresh Babu Selvaraju, Goran Stevovic, Amanda Suchanek, Andrea Throop, Lyndon Tilson, Thomas Urban, Joe Voshell, Kimberly Wagner, Jonathan Williams, Mary Williamson, Qian Zeng, Tricia Zwiefelhofer, Clinton R. Paden, Duncan MacCannell |
| EPI_ISL_2183341, EPI_ISL_2183420, EPI_ISL_2183422, EPI_ISL_2183424, EPI_ISL_2183427, EPI_ISL_2183429, EPI_ISL_2183431, EPI_ISL_2183433 | Israel Central Virology laboratory | Israel National Consortium for SARS-CoV-2 sequencing | Neta Zuckerman, Efrat Dahan Bucris, Michal Mandelboim, Dana Bar-Ilan, Miranda Geva, Netanel Abu, Oran Erster, Efrat Glick-Saar, Omri Nayshool, Gideon Rechavi, Ella Mendelson, Orna Mor |
| EPI_ISL_2183543, EPI_ISL_2183545 | Laboratory Corporation of America | Centers for Disease Control and Prevention Division of Viral Diseases, Pathogen Discovery | Dakota Howard, Dhwani Batra, Peter W. Cook, Kara Moser, Adrian Paskey, Jason Caravas, Benjamin Rambo-Martin, Shatavia Morrison, Christopher Gulvick, Scott Sammons, Yvette Unoarumhi, Darlene Wagner, Matthew Schmerer, Minoo Agarwal, Eyad Almasri, Debbie Boles, Ayla Burns, Nuthawin Charoensri, Oren Cohen, Susan Countryman, Mary Ann Cristobal, Bobbi Croy, Suzanne Dale, Hrushikesh Deshmukh, Amanda Douglas, Vincent Drouillon, Marcia Eisenberg, Howard Engler, Rama Ghatti, Prashant Gupta, Susan Hicks, Jake Humphrey, Lax Iyer, Manoj Jain, Mohan Kolli, Brian Krueger, Tim Kuphal, Stanley Letovsky, Michael Levandoski, Craig Lukasik, Jonathan Meltzer, Brian Norvell, Mindy Nye, Scott Parker, Christos Petropoulos, John Pruitt, Steven Ragan, Scott Ryan, Mike Sapeta, Jana Schroth, Suresh Babu Selvaraju, Goran Stevovic, Amanda Suchanek, Andrea Throop, Lyndon Tilson, Thomas Urban, Joe Voshell, Kimberly Wagner, Jonathan Williams, Mary Williamson, Qian Zeng, Tricia Zwiefelhofer, Clinton R. Paden, Duncan MacCannell |
| EPI_ISL_2183613, EPI_ISL_2183615, EPI_ISL_2183679, EPI_ISL_2183691, EPI_ISL_2183692, EPI_ISL_2183694, EPI_ISL_2183702, EPI_ISL_2183704, EPI_ISL_2183706 | Israel Central Virology laboratory | Israel National Consortium for SARS-CoV-2 sequencing | Neta Zuckerman, Efrat Dahan Bucris, Michal Mandelboim, Dana Bar-Ilan, Miranda Geva, Netanel Abu, Oran Erster, Efrat Glick-Saar, Omri Nayshool, Gideon Rechavi, Ella Mendelson, Orna Mor |
| EPI_ISL_2183710 | Laboratory Corporation of America | Centers for Disease Control and Prevention Division of Viral Diseases, Pathogen Discovery | Dakota Howard, Dhwani Batra, Peter W. Cook, Kara Moser, Adrian Paskey, Jason Caravas, Benjamin Rambo-Martin, Shatavia Morrison, Christopher Gulvick, Scott Sammons, Yvette Unoarumhi, Darlene Wagner, Matthew Schmerer, Minoo Agarwal, Eyad Almasri, Debbie Boles, Ayla Burns, Nuthawin Charoensri, Oren Cohen, Susan Countryman, Mary Ann Cristobal, Bobbi Croy, Suzanne Dale, Hrushikesh Deshmukh, Amanda Douglas, Vincent Drouillon, Marcia Eisenberg, Howard Engler, Rama Ghatti, Prashant Gupta, Susan Hicks, Jake Humphrey, Lax Iyer, Manoj Jain, Mohan Kolli, Brian Krueger, Tim Kuphal, Stanley Letovsky, Michael Levandoski, Craig Lukasik, Jonathan Meltzer, Brian Norvell, Mindy Nye, Scott Parker, Christos Petropoulos, John Pruitt, Steven Ragan, Scott Ryan, Mike Sapeta, Jana Schroth, Suresh Babu Selvaraju, Goran Stevovic, Amanda Suchanek, Andrea Throop, Lyndon Tilson, Thomas Urban, Joe Voshell, Kimberly Wagner, Jonathan Williams, Mary Williamson, Qian Zeng, Tricia Zwiefelhofer, Clinton R. Paden, Duncan MacCannell |
| EPI_ISL_2183721, EPI_ISL_2183727, EPI_ISL_2183736, EPI_ISL_2183749 | Israel Central Virology laboratory | Israel National Consortium for SARS-CoV-2 sequencing | Neta Zuckerman, Efrat Dahan Bucris, Michal Mandelboim, Dana Bar-Ilan, Miranda Geva, Netanel Abu, Oran Erster, Efrat Glick-Saar, Omri Nayshool, Gideon Rechavi, Ella Mendelson, Orna Mor |
| EPI_ISL_2183751 | Laboratory Corporation of America | Centers for Disease Control and Prevention Division of Viral Diseases, Pathogen Discovery | Dakota Howard, Dhwani Batra, Peter W. Cook, Kara Moser, Adrian Paskey, Jason Caravas, Benjamin Rambo-Martin, Shatavia Morrison, Christopher Gulvick, Scott Sammons, Yvette Unoarumhi, Darlene Wagner, Matthew Schmerer, Minoo Agarwal, Eyad Almasri, Debbie Boles, Ayla Burns, Nuthawin Charoensri, Oren Cohen, Susan Countryman, Mary Ann Cristobal, Bobbi Croy, Suzanne Dale, Hrushikesh Deshmukh, Amanda Douglas, Vincent Drouillon, Marcia Eisenberg, Howard Engler, Rama Ghatti, Prashant Gupta, Susan Hicks, Jake Humphrey, Lax Iyer, Manoj Jain, Mohan Kolli, Brian Krueger, Tim Kuphal, Stanley Letovsky, Michael Levandoski, Craig Lukasik, Jonathan Meltzer, Brian Norvell, Mindy Nye, Scott Parker, Christos Petropoulos, John Pruitt, Steven Ragan, Scott Ryan, Mike Sapeta, Jana Schroth, Suresh Babu Selvaraju, Goran Stevovic, Amanda Suchanek, Andrea Throop, Lyndon Tilson, Thomas Urban, Joe Voshell, Kimberly Wagner, Jonathan Williams, Mary Williamson, Qian Zeng, Tricia Zwiefelhofer, Clinton R. Paden, Duncan MacCannell |
| EPI_ISL_2183777, EPI_ISL_2183779, EPI_ISL_2183802, EPI_ISL_2183805 | Israel Central Virology laboratory | Israel National Consortium for SARS-CoV-2 sequencing | Neta Zuckerman, Efrat Dahan Bucris, Michal Mandelboim, Dana Bar-Ilan, Miranda Geva, Netanel Abu, Oran Erster, Efrat Glick-Saar, Omri Nayshool, Gideon Rechavi, Ella Mendelson, Orna Mor |
| EPI_ISL_2183985, EPI_ISL_2184012, EPI_ISL_2184064, EPI_ISL_2184123, EPI_ISL_2184258, EPI_ISL_2184285, EPI_ISL_2184378, EPI_ISL_2184555, EPI_ISL_2184611, EPI_ISL_2184696, EPI_ISL_2184842, EPI_ISL_2184984, EPI_ISL_2185013, EPI_ISL_2185018, EPI_ISL_2185032, EPI_ISL_2185173, EPI_ISL_2185215, EPI_ISL_2185277, EPI_ISL_2185452 | | | |
| see above | Laboratory Corporation of America | Centers for Disease Control and Prevention Division of Viral Diseases, Pathogen Discovery | Dakota Howard, Dhwani Batra, Peter W. Cook, Kara Moser, Adrian Paskey, Jason Caravas, Benjamin Rambo-Martin, Shatavia Morrison, Christopher Gulvick, Scott Sammons, Yvette Unoarumhi, Darlene Wagner, Matthew Schmerer, Minoo Agarwal, Eyad Almasri, Debbie Boles, Ayla Burns, Nuthawin Charoensri, Oren Cohen, Susan Countryman, Mary Ann Cristobal, Bobbi Croy, Suzanne Dale, Hrushikesh Deshmukh, Amanda Douglas, Vincent Drouillon, Marcia Eisenberg, Howard Engler, Rama Ghatti, Prashant Gupta, Susan Hicks, Jake Humphrey, Lax Iyer, Manoj Jain, Mohan Kolli, Brian Krueger, Tim Kuphal, Stanley Letovsky, Michael Levandoski, Craig Lukasik, Jonathan Meltzer, Brian Norvell, Mindy Nye, Scott Parker, Christos Petropoulos, John Pruitt, Steven Ragan, Scott Ryan, Mike Sapeta, Jana Schroth, Suresh Babu Selvaraju, Goran Stevovic, Amanda Suchanek, Andrea Throop, Lyndon Tilson, Thomas Urban, Joe Voshell, Kimberly Wagner, Jonathan Williams, Mary Williamson, Qian Zeng, Tricia Zwiefelhofer, Clinton R. Paden, Duncan MacCannell |
| EPI_ISL_2185596, EPI_ISL_2185612, EPI_ISL_2185623 | NJDOH, Public Health and Environmental Laboratories | NJ_PHEL | Lindsey Bodnar, Shiv K. Verma, Jacquelyn Deverell, Matthew Scarnati, Dana Woell, Byeong Jeong |
| EPI_ISL_2186510, EPI_ISL_2186522, EPI_ISL_2186603, EPI_ISL_2186947, EPI_ISL_2187138 | Infinity Biologix | Centers for Disease Control and Prevention Division of Viral Diseases, Pathogen Discovery | Dakota Howard, Dhwani Batra, Peter W. Cook, Kara Moser, Adrian Paskey, Jason Caravas, Benjamin Rambo-Martin, Shatavia Morrison, Christopher Gulvick, Scott Sammons, Yvette Unoarumhi, Darlene Wagner, Matthew Schmerer, Christian Bixby, Yihe Wang, Jonathan Schultz, Chirayu Goswami, Russ Hager, Robin Grimwood, Clinton R. Paden, Duncan MacCannell |
| EPI_ISL_2187341 | Helix/Illumina | Centers for Disease Control and Prevention Division of Viral Diseases, Pathogen Discovery | Dakota Howard, Dhwani Batra, Peter W. Cook, Kara Moser, Adrian Paskey, Jason Caravas, Benjamin Rambo-Martin, Shatavia Morrison, Christopher Gulvick, Scott Sammons, Yvette Unoarumhi, Darlene Wagner, Matthew Schmerer, Eileen de Feo, Jan Antico, Christine Tran, Matthew Tolentino, Shannon Wickline, Kim Gietzen, Brad Sickler, Jingtao Liu, Eric Allen, Phil Febbo, Nicole L. Washington, Simon White, Geraint Levan, Kelly Schiabor Barrett, Elizabeth Cirulli, Alexandre Bolze, Ary Ascencio, Charlotte Rivera-Garcia, Ryan Cho, Jason Nguyen, Sherry Wang, Jimmy Ramirez, Tyler Cassens, Efren Sandoval, Magnus Isaksson, William Lee, David Becker, Marc Laurent, James Lu, Clinton R. Paden, Duncan MacCannell |
| EPI_ISL_2187476, EPI_ISL_2187606 | Aegis Sciences Corporation | Centers for Disease Control and Prevention Division of Viral Diseases, Pathogen Discovery | Dakota Howard, Dhwani Batra, Peter W. Cook, Kara Moser, Adrian Paskey, Jason Caravas, Benjamin Rambo-Martin, Shatavia Morrison, Christopher Gulvick, Scott Sammons, Yvette Unoarumhi, Darlene Wagner, Matthew Schmerer, Cyndi Clark, Patrick Campbell, Rob Case, Vikramsinha Ghorpade, Holly Houdeshell, Ola Kvalvaag, Dillon Nall, Ethan Sanders, Alec Vest, Shaun Westlund, Matthew Hardison, Clinton R. Paden, Duncan MacCannell |
| EPI_ISL_2188352 | Hospital | National Reference Center for Viruses of Respiratory Infections, Institut Pasteur, Paris | Marion Barbet, Sylvie Behillil, Méline Bizard, Angela Brisebarre, Camille Capel, Vincent Enouf, Louise Lefrançois, Frédéric Lemoine, Christophe Malabat, Corinne Maufrais, Etienne Simon-Lorière, Maud Vanpeene, Sylvie Van der Werf ,Laura Djamdjian |
| EPI_ISL_2188418 | Hospital | National Reference Center for Viruses of Respiratory Infections, Institut Pasteur, Paris | Marion Barbet, Sylvie Behillil, Méline Bizard, Angela Brisebarre, Camille Capel, Vincent Enouf, Louise Lefrançois, Frédéric Lemoine, Christophe Malabat, Corinne Maufrais, Amaury Vaysse, Etienne Simon-Lorière, Maud Vanpeene, Sylvie Van der Werf ,Raphael Guiheneuf |
| EPI_ISL_2188419 | Hospital | National Reference Center for Viruses of Respiratory Infections, Institut Pasteur, Paris | Marion Barbet, Sylvie Behillil, Méline Bizard, Angela Brisebarre, Camille Capel, Vincent Enouf, Louise Lefrançois, Frédéric Lemoine, Christophe Malabat, Corinne Maufrais, Etienne Simon-Lorière, Maud Vanpeene, Sylvie Van der Werf ,Raphael Guiheneuf |
| EPI_ISL_2188450, EPI_ISL_2188451, EPI_ISL_2188452 | Hospital | National Reference Center for Viruses of Respiratory Infections, Institut Pasteur, Paris | Marion Barbet, Sylvie Behillil, Méline Bizard, Angela Brisebarre, Camille Capel, Vincent Enouf, Louise Lefrançois, Frédéric Lemoine, Christophe Malabat, Corinne Maufrais, Etienne Simon-Lorière, Maud Vanpeene, Sylvie Van der Werf ,Nabil Gastli |
| EPI_ISL_2189572, EPI_ISL_2189596, EPI_ISL_2189638, EPI_ISL_2189639, EPI_ISL_2189641, EPI_ISL_2189642, EPI_ISL_2189643, EPI_ISL_2189644, EPI_ISL_2189646, EPI_ISL_2189649, EPI_ISL_2189650, EPI_ISL_2189652, EPI_ISL_2189654, EPI_ISL_2189656, EPI_ISL_2189657, EPI_ISL_2189658, EPI_ISL_2189662, EPI_ISL_2189663, EPI_ISL_2189669, EPI_ISL_2189670, EPI_ISL_2189676, EPI_ISL_2189679, EPI_ISL_2189686, EPI_ISL_2189691, EPI_ISL_2189697, EPI_ISL_2189698, EPI_ISL_2189699, EPI_ISL_2189704, EPI_ISL_2189705, EPI_ISL_2189708, EPI_ISL_2189710, EPI_ISL_2189711, EPI_ISL_2189712, EPI_ISL_2189713 | | | |
| see above | ICMR-National Institute of Virology | NCDC Delhi, Biotechnology Division | V.Vipat,Meena Datta,S.Jadhav,Priyanka Singh,U.Saha ,M.Das,Uma Sharma,G.Divekar,T.Sanjeevi,Manoj K Singh,S Jadhav,Radhakrishnan V. S,A Walimbe ,Robin Marwal,V Malik,Mahesh S Dhar,K.Iyengar,Kalaiarasan Ponnusamy,K P.Shinde,Hemlata Lall,K.Korabu,Z. Sayyed,Hema Gogia, S.Shelkande,R.Verma,Preeti Madan,S.Bhorekar,B.Apoorva,Sandhya Kabra,A.Titkare,H.Kengale,Sujeet K Singh,V.Autude,A.Awhale,Partha Rakshit,M L Choudhary |
| EPI_ISL_2189716, EPI_ISL_2189717, EPI_ISL_2189718, EPI_ISL_2189723, EPI_ISL_2189724, EPI_ISL_2189726, EPI_ISL_2189728, EPI_ISL_2189731, EPI_ISL_2189738, EPI_ISL_2189744, EPI_ISL_2189748, EPI_ISL_2189749, EPI_ISL_2189755, EPI_ISL_2189761, EPI_ISL_2189765, EPI_ISL_2189766, EPI_ISL_2189768, EPI_ISL_2189770, EPI_ISL_2189772, EPI_ISL_2189773, EPI_ISL_2189778, EPI_ISL_2189780, EPI_ISL_2189786, EPI_ISL_2189787, EPI_ISL_2189788, EPI_ISL_2189791, EPI_ISL_2189792, EPI_ISL_2189795, EPI_ISL_2189796, EPI_ISL_2189800, EPI_ISL_2189803, EPI_ISL_2189806, EPI_ISL_2189809, EPI_ISL_2189810, EPI_ISL_2189811, EPI_ISL_2189813, EPI_ISL_2189814, EPI_ISL_2189817, EPI_ISL_2189818, EPI_ISL_2189819, EPI_ISL_2189823, EPI_ISL_2189825, EPI_ISL_2189827, EPI_ISL_2189828, EPI_ISL_2189829, EPI_ISL_2189830, EPI_ISL_2189831, EPI_ISL_2189834, EPI_ISL_2189836, EPI_ISL_2189837, EPI_ISL_2189840, EPI_ISL_2189841, EPI_ISL_2189843, EPI_ISL_2189845, EPI_ISL_2189849, EPI_ISL_2189850, EPI_ISL_2189852, EPI_ISL_2189857, EPI_ISL_2189859, EPI_ISL_2189860, EPI_ISL_2189863 | | | |
| see above | ICMR-National Institute of Virology | NCDC Delhi, Biotechnology Division | Meena Datta,V.Vipat,S.Jadhav,Priyanka Singh,U.Saha ,M.Das,Uma Sharma,G.Divekar,T.Sanjeevi,Manoj K Singh,S Jadhav,Radhakrishnan V. S,A Walimbe ,Robin Marwal,V Malik,Mahesh S Dhar,K.Iyengar,Kalaiarasan Ponnusamy,K P.Shinde,Hemlata Lall,K.Korabu,Z. Sayyed,Hema Gogia, S.Shelkande,R.Verma,Preeti Madan,S.Bhorekar,B.Apoorva,Sandhya Kabra,A.Titkare,H.Kengale,Sujeet K Singh,V.Autude,A.Awhale,M L Choudhary,Partha Rakshit |
| EPI_ISL_2190101 | Department of Public Health Bucharest | National Institute of Infectious Diseases-Prof. Dr. Matei Bals Molecular Diagnostics Laboratory | Corina Casangiu, Leontina Banica, Marius Surleac, Ovidiu Vlaicu, Andreea Tudor, Simona Paraschiv, Dan Otelea |
| EPI_ISL_2190106 | National Institute of Infectious Diseases-Prof. Dr. Matei Bals Molecular Diagnostics Laboratory | National Institute of Infectious Diseases-Prof. Dr. Matei Bals Molecular Diagnostics Laboratory | Corina Casangiu, Leontina Banica, Marius Surleac, Ovidiu Vlaicu, Andreea Tudor, Simona Paraschiv, Dan Otelea |
| EPI_ISL_2192265, EPI_ISL_2192404, EPI_ISL_2192405, EPI_ISL_2192420, EPI_ISL_2192463 | Lab voor klinische biologie | Lab voor klinische biologie | Marija Janevska, Hannelore Hamerlinck, Bruno Verhasselt |
| EPI_ISL_2192655, EPI_ISL_2192706, EPI_ISL_2192713, EPI_ISL_2192746, EPI_ISL_2192788, EPI_ISL_2192796, EPI_ISL_2192812, EPI_ISL_2192912, EPI_ISL_2192922, EPI_ISL_2192925, EPI_ISL_2192955, EPI_ISL_2192973, EPI_ISL_2193007, EPI_ISL_2193047, EPI_ISL_2193085, EPI_ISL_2193090 | | | |
| see above | Broad Institute Clinical Research Sequencing Platform | Infectious Disease Program, Broad Institute of Harvard and MIT | Siddle,K.J., Adams,G., Pearlman,L., Gladden-Young,A., Vicente,G., Blumenstiel,B., DeFelice,M., Lee,M., McGovern,S., Lagerborg,K., Rudy,M., DeRuff,K., Carter,A., Normandin,E., Bauer,M., Reilly,S., Tomkins-Tinch,C., Loreth,C., Chaluvadi,S., Meldrim,J., Granger,B., Lemieux,J.E., Birren,B.W., Sabeti,P.C., Larkin,K., Dodge,S., Lennon,N., Madoff,L., Brown,C., Gallagher,G., Smole,S., Park,D.J., Gabriel,S., and MacInnis,B.L. |
| EPI_ISL_2193354, EPI_ISL_2193355, EPI_ISL_2193356, EPI_ISL_2193357, | Ampath Laboratories | National Institute for Communicable Diseases of the National Health Laboratory Service | Amoako DG, Scheepers C, Mohale T, Ntuli N, Mahlangu B, Viana R, Glass A, Marshall T, Gottberg A, Bhiman JN |

| | | | |
|---|---|---|---|
| EPI_ISL_2193358, EPI_ISL_2193359, EPI_ISL_2193360, EPI_ISL_2193361 | | | |
| EPI_ISL_2193386 | University Hospitals Translational Laboratory (UHTL), University Hospitals | University Hospitals Translational Laboratory (UHTL), University Hospitals | Sadri,N., Alouani,D., Song,X. |
| EPI_ISL_2193825, EPI_ISL_2193835, EPI_ISL_2193895, EPI_ISL_2193904, EPI_ISL_2193909, EPI_ISL_2193922, EPI_ISL_2193937 | National Virus Reference Laboratory | National Virus Reference Laboratory | Zoe Yandle, Charlene Bennett, Gabriel Gonzalez, Michael Carr, Jonathan Dean, Cillian F De Gascun |
| EPI_ISL_2193979 | Wyoming Public Health Laboratory | Wyoming Public Health Laboratory | Jim Mildenberger, Wanda Manley, Noah Hull, Taylor Fearing, Lynette Gumbleton, Channing Weber, Ashley Norberg, Chayse Rowley, Marley Goetz, Brian Dominguez, Elliot Thomasson, Sam Britz, Cari Sloma, and Rob Christensen |
| EPI_ISL_2194081, EPI_ISL_2194082, EPI_ISL_2194173 | UW Virology Lab | UW Virology Lab | Pavitra Roychoudhury, Hong Xie, Lasata Shrestha, Tien V. Nguyen, Shah Mohamed Bakhash, Michelle Lin, Noah R. Baker, Ricardo Perez, Sean Ellis, Nathan Breit, Robert J. Livingston, Meei-Li Huang, Keith R Jerome, Patrick Mathias, Alexander Greninger |
| EPI_ISL_2194237 | Istituto Zooprofilattico Sperimentale del Mezzogiorno | TIGEM | Antonio Grimaldi Patrizia Annunziata Francesco Panariello Biancamaria Pierri Claudia Tiberio Teresa Giuliano Valentina Bouche Chiara Colantuono Maria Concetta Cuomo Denise Di Concilio Lucio Di Filippo Anna Manfredi Marcello Salvi Antonio Limone Luigi Atripaldi Pellegrino Cerino Andrea Ballabio Davide Cacchiarelli |
| EPI_ISL_2195974, EPI_ISL_2195975 | J.W. Ruby Memorial Hospital | WVU and Marshall University Combined Genomics Core Facilities | James Denvir, Peter Stoilov, Peter Perrotta, Wesley Kimble, Ryan Percifield |
| EPI_ISL_2198344, EPI_ISL_2198345, EPI_ISL_2198347, EPI_ISL_2198349 | Lighthouse Lab in Milton Keynes | Wellcome Sanger Institute for the COVID-19 Genomics UK (COG-UK) Consortium | The Lighthouse Lab in Milton Keynes and Alex Alderton, Roberto Amato, Jeffrey Barrett, Sonia Goncalves, Ewan Harrison, David K. Jackson, Ian Johnston, Dominic Kwiatkowski, Cordelia Langford, John Sillitoe on behalf of the Wellcome Sanger Institute COVID-19 Surveillance Team |
| EPI_ISL_2198350 | Lighthouse Lab in Alderley Park | Wellcome Sanger Institute for the COVID-19 Genomics UK (COG-UK) Consortium | Jacquelyn Wynn, Mairead Hyland, The Lighthouse Lab in Alderley Park and Alex Alderton, Roberto Amato, Jeffrey Barrett, Sonia Goncalves, Ewan Harrison, David K. Jackson, Ian Johnston, Dominic Kwiatkowski, Cordelia Langford, John Sillitoe on behalf of the Wellcome Sanger Institute COVID-19 Surveillance Team |
| EPI_ISL_2198352, EPI_ISL_2198354, EPI_ISL_2198355, EPI_ISL_2198356, EPI_ISL_2198358 | Lighthouse Lab in Milton Keynes | Wellcome Sanger Institute for the COVID-19 Genomics UK (COG-UK) Consortium | The Lighthouse Lab in Milton Keynes and Alex Alderton, Roberto Amato, Jeffrey Barrett, Sonia Goncalves, Ewan Harrison, David K. Jackson, Ian Johnston, Dominic Kwiatkowski, Cordelia Langford, John Sillitoe on behalf of the Wellcome Sanger Institute COVID-19 Surveillance Team |
| EPI_ISL_2198362 | Lighthouse Lab in Alderley Park | Wellcome Sanger Institute for the COVID-19 Genomics UK (COG-UK) Consortium | Jacquelyn Wynn, Mairead Hyland, The Lighthouse Lab in Alderley Park and Alex Alderton, Roberto Amato, Jeffrey Barrett, Sonia Goncalves, Ewan Harrison, David K. Jackson, Ian Johnston, Dominic Kwiatkowski, Cordelia Langford, John Sillitoe on behalf of the Wellcome Sanger Institute COVID-19 Surveillance Team |
| EPI_ISL_2198364 | Lighthouse Lab in Milton Keynes | Wellcome Sanger Institute for the COVID-19 Genomics UK (COG-UK) Consortium | The Lighthouse Lab in Milton Keynes and Alex Alderton, Roberto Amato, Jeffrey Barrett, Sonia Goncalves, Ewan Harrison, David K. Jackson, Ian Johnston, Dominic Kwiatkowski, Cordelia Langford, John Sillitoe on behalf of the Wellcome Sanger Institute COVID-19 Surveillance Team |
| EPI_ISL_2198366 | Lighthouse Lab in Alderley Park | Wellcome Sanger Institute for the COVID-19 Genomics UK (COG-UK) Consortium | Jacquelyn Wynn, Mairead Hyland, The Lighthouse Lab in Alderley Park and Alex Alderton, Roberto Amato, Jeffrey Barrett, Sonia Goncalves, Ewan Harrison, David K. Jackson, Ian Johnston, Dominic Kwiatkowski, Cordelia Langford, John Sillitoe on behalf of the Wellcome Sanger Institute COVID-19 Surveillance Team |
| EPI_ISL_2198369, EPI_ISL_2198375, EPI_ISL_2198376 | Lighthouse Lab in Milton Keynes | Wellcome Sanger Institute for the COVID-19 Genomics UK (COG-UK) Consortium | The Lighthouse Lab in Milton Keynes and Alex Alderton, Roberto Amato, Jeffrey Barrett, Sonia Goncalves, Ewan Harrison, David K. Jackson, Ian Johnston, Dominic Kwiatkowski, Cordelia Langford, John Sillitoe on behalf of the Wellcome Sanger Institute COVID-19 Surveillance Team |
| EPI_ISL_2198380, EPI_ISL_2198381 | Lighthouse Lab in Alderley Park | Wellcome Sanger Institute for the COVID-19 Genomics UK (COG-UK) Consortium | Jacquelyn Wynn, Mairead Hyland, The Lighthouse Lab in Alderley Park and Alex Alderton, Roberto Amato, Jeffrey Barrett, Sonia Goncalves, Ewan Harrison, David K. Jackson, Ian Johnston, Dominic Kwiatkowski, Cordelia Langford, John Sillitoe on behalf of the Wellcome Sanger Institute COVID-19 Surveillance Team |
| EPI_ISL_2198383, EPI_ISL_2198384, EPI_ISL_2198385 | Lighthouse Lab in Milton Keynes | Wellcome Sanger Institute for the COVID-19 Genomics UK (COG-UK) Consortium | The Lighthouse Lab in Milton Keynes and Alex Alderton, Roberto Amato, Jeffrey Barrett, Sonia Goncalves, Ewan Harrison, David K. Jackson, Ian Johnston, Dominic Kwiatkowski, Cordelia Langford, John Sillitoe on behalf of the Wellcome Sanger Institute COVID-19 Surveillance Team |
| EPI_ISL_2198386, EPI_ISL_2198387, EPI_ISL_2198388 | Lighthouse Lab in Alderley Park | Wellcome Sanger Institute for the COVID-19 Genomics UK (COG-UK) Consortium | Jacquelyn Wynn, Mairead Hyland, The Lighthouse Lab in Alderley Park and Alex Alderton, Roberto Amato, Jeffrey Barrett, Sonia Goncalves, Ewan Harrison, David K. Jackson, Ian Johnston, Dominic Kwiatkowski, Cordelia Langford, John Sillitoe on behalf of the Wellcome Sanger Institute COVID-19 Surveillance Team |
| EPI_ISL_2198391, EPI_ISL_2198394, EPI_ISL_2198395 | Lighthouse Lab in Milton Keynes | Wellcome Sanger Institute for the COVID-19 Genomics UK (COG-UK) Consortium | The Lighthouse Lab in Milton Keynes and Alex Alderton, Roberto Amato, Jeffrey Barrett, Sonia Goncalves, Ewan Harrison, David K. Jackson, Ian Johnston, Dominic Kwiatkowski, Cordelia Langford, John Sillitoe on behalf of the Wellcome Sanger Institute COVID-19 Surveillance Team |
| EPI_ISL_2198396, EPI_ISL_2198397 | Lighthouse Lab in Alderley Park | Wellcome Sanger Institute for the COVID-19 Genomics UK (COG-UK) Consortium | Jacquelyn Wynn, Mairead Hyland, The Lighthouse Lab in Alderley Park and Alex Alderton, Roberto Amato, Jeffrey Barrett, Sonia Goncalves, Ewan Harrison, David K. Jackson, Ian Johnston, Dominic Kwiatkowski, Cordelia Langford, John Sillitoe on behalf of the Wellcome Sanger Institute COVID-19 Surveillance Team |
| EPI_ISL_2198398 | Lighthouse Lab in Milton Keynes | Wellcome Sanger Institute for the COVID-19 Genomics UK (COG-UK) Consortium | The Lighthouse Lab in Milton Keynes and Alex Alderton, Roberto Amato, Jeffrey Barrett, Sonia Goncalves, Ewan Harrison, David K. Jackson, Ian Johnston, Dominic Kwiatkowski, Cordelia Langford, John Sillitoe on behalf of the Wellcome Sanger Institute COVID-19 Surveillance Team |
| EPI_ISL_2198400 | Lighthouse Lab in Alderley Park | Wellcome Sanger Institute for the COVID-19 Genomics UK (COG-UK) Consortium | Jacquelyn Wynn, Mairead Hyland, The Lighthouse Lab in Alderley Park and Alex Alderton, Roberto Amato, Jeffrey Barrett, Sonia Goncalves, Ewan Harrison, David K. Jackson, Ian Johnston, Dominic Kwiatkowski, Cordelia Langford, John Sillitoe on behalf of the Wellcome Sanger Institute COVID-19 Surveillance Team |
| EPI_ISL_2198403 | Lighthouse Lab in Milton Keynes | Wellcome Sanger Institute for the COVID-19 Genomics UK (COG-UK) Consortium | The Lighthouse Lab in Milton Keynes and Alex Alderton, Roberto Amato, Jeffrey Barrett, Sonia Goncalves, Ewan Harrison, David K. Jackson, Ian Johnston, Dominic Kwiatkowski, Cordelia Langford, John Sillitoe on behalf of the Wellcome Sanger Institute COVID-19 Surveillance Team |
| EPI_ISL_2198406 | Lighthouse Lab in Alderley Park | Wellcome Sanger Institute for the COVID-19 Genomics UK (COG-UK) Consortium | Jacquelyn Wynn, Mairead Hyland, The Lighthouse Lab in Alderley Park and Alex Alderton, Roberto Amato, Jeffrey Barrett, Sonia Goncalves, Ewan Harrison, David K. Jackson, Ian Johnston, Dominic Kwiatkowski, Cordelia Langford, John Sillitoe on behalf of the Wellcome Sanger Institute COVID-19 Surveillance Team |
| EPI_ISL_2198408, EPI_ISL_2198411 | Lighthouse Lab in Milton Keynes | Wellcome Sanger Institute for the COVID-19 Genomics UK (COG-UK) Consortium | The Lighthouse Lab in Milton Keynes and Alex Alderton, Roberto Amato, Jeffrey Barrett, Sonia Goncalves, Ewan Harrison, David K. Jackson, Ian Johnston, Dominic Kwiatkowski, Cordelia Langford, John Sillitoe on behalf of the Wellcome Sanger Institute COVID-19 Surveillance Team |
| EPI_ISL_2198413 | Lighthouse Lab in Alderley Park | Wellcome Sanger Institute for the COVID-19 Genomics UK (COG-UK) Consortium | Jacquelyn Wynn, Mairead Hyland, The Lighthouse Lab in Alderley Park and Alex Alderton, Roberto Amato, Jeffrey Barrett, Sonia Goncalves, Ewan Harrison, David K. Jackson, Ian Johnston, Dominic Kwiatkowski, Cordelia Langford, John Sillitoe on behalf of the Wellcome Sanger Institute COVID-19 Surveillance Team |
| EPI_ISL_2198415 | Lighthouse Lab in Milton Keynes | Wellcome Sanger Institute for the COVID-19 Genomics UK (COG-UK) Consortium | The Lighthouse Lab in Milton Keynes and Alex Alderton, Roberto Amato, Jeffrey Barrett, Sonia Goncalves, Ewan Harrison, David K. Jackson, Ian Johnston, Dominic Kwiatkowski, Cordelia Langford, John Sillitoe on behalf of the Wellcome Sanger Institute COVID-19 Surveillance Team |
| EPI_ISL_2198417 | Lighthouse Lab in Alderley Park | Wellcome Sanger Institute for the COVID-19 Genomics UK (COG-UK) Consortium | Jacquelyn Wynn, Mairead Hyland, The Lighthouse Lab in Alderley Park and Alex Alderton, Roberto Amato, Jeffrey Barrett, Sonia Goncalves, Ewan Harrison, David K. Jackson, Ian Johnston, Dominic Kwiatkowski, Cordelia Langford, John Sillitoe on behalf of the Wellcome Sanger Institute COVID-19 Surveillance Team |
| EPI_ISL_2198418 | Lighthouse Lab in Milton Keynes | Wellcome Sanger Institute for the COVID-19 Genomics UK (COG-UK) Consortium | The Lighthouse Lab in Milton Keynes and Alex Alderton, Roberto Amato, Jeffrey Barrett, Sonia Goncalves, Ewan Harrison, David K. Jackson, Ian Johnston, Dominic Kwiatkowski, Cordelia Langford, John Sillitoe on behalf of the Wellcome Sanger Institute COVID-19 Surveillance Team |
| EPI_ISL_2198420 | Lighthouse Lab in Alderley Park | Wellcome Sanger Institute for the COVID-19 Genomics UK (COG-UK) Consortium | Jacquelyn Wynn, Mairead Hyland, The Lighthouse Lab in Alderley Park and Alex Alderton, Roberto Amato, Jeffrey Barrett, Sonia Goncalves, Ewan Harrison, David K. Jackson, Ian Johnston, Dominic Kwiatkowski, Cordelia Langford, John Sillitoe on behalf of the Wellcome Sanger Institute COVID-19 Surveillance Team |
| EPI_ISL_2198422, EPI_ISL_2198427 | Lighthouse Lab in Milton Keynes | Wellcome Sanger Institute for the COVID-19 Genomics UK (COG-UK) Consortium | The Lighthouse Lab in Milton Keynes and Alex Alderton, Roberto Amato, Jeffrey Barrett, Sonia Goncalves, Ewan Harrison, David K. Jackson, Ian Johnston, Dominic Kwiatkowski, Cordelia Langford, John Sillitoe on behalf of the Wellcome Sanger Institute COVID-19 Surveillance Team |
| EPI_ISL_2198430 | Lighthouse Lab in Alderley Park | Wellcome Sanger Institute for the COVID-19 Genomics UK (COG-UK) Consortium | Jacquelyn Wynn, Mairead Hyland, The Lighthouse Lab in Alderley Park and Alex Alderton, Roberto Amato, Jeffrey Barrett, Sonia Goncalves, Ewan Harrison, David K. Jackson, Ian Johnston, Dominic Kwiatkowski, Cordelia Langford, John Sillitoe on behalf of the Wellcome Sanger Institute COVID-19 Surveillance Team |
| EPI_ISL_2198432, EPI_ISL_2198434, EPI_ISL_2198436 | Lighthouse Lab in Milton Keynes | Wellcome Sanger Institute for the COVID-19 Genomics UK (COG-UK) Consortium | The Lighthouse Lab in Milton Keynes and Alex Alderton, Roberto Amato, Jeffrey Barrett, Sonia Goncalves, Ewan Harrison, David K. Jackson, Ian Johnston, Dominic Kwiatkowski, Cordelia Langford, John Sillitoe on behalf of the Wellcome Sanger Institute COVID-19 Surveillance Team |
| EPI_ISL_2198439 | Lighthouse Lab in Alderley Park | Wellcome Sanger Institute for the COVID-19 Genomics UK (COG-UK) Consortium | Jacquelyn Wynn, Mairead Hyland, The Lighthouse Lab in Alderley Park and Alex Alderton, Roberto Amato, Jeffrey Barrett, Sonia Goncalves, Ewan Harrison, David K. Jackson, Ian Johnston, Dominic Kwiatkowski, Cordelia Langford, John Sillitoe on behalf of the Wellcome Sanger Institute COVID-19 Surveillance Team |
| EPI_ISL_2198441, EPI_ISL_2198443, EPI_ISL_2198447, EPI_ISL_2198448, EPI_ISL_2198449, EPI_ISL_2198452, EPI_ISL_2198453 | Lighthouse Lab in Milton Keynes | Wellcome Sanger Institute for the COVID-19 Genomics UK (COG-UK) Consortium | The Lighthouse Lab in Milton Keynes and Alex Alderton, Roberto Amato, Jeffrey Barrett, Sonia Goncalves, Ewan Harrison, David K. Jackson, Ian Johnston, Dominic Kwiatkowski, Cordelia Langford, John Sillitoe on behalf of the Wellcome Sanger Institute COVID-19 Surveillance Team |
| EPI_ISL_2198454 | Lighthouse Lab in Alderley Park | Wellcome Sanger Institute for the COVID-19 Genomics UK (COG-UK) Consortium | Jacquelyn Wynn, Mairead Hyland, The Lighthouse Lab in Alderley Park and Alex Alderton, Roberto Amato, Jeffrey Barrett, Sonia Goncalves, Ewan Harrison, David K. Jackson, Ian Johnston, Dominic Kwiatkowski, Cordelia Langford, John Sillitoe on behalf of the Wellcome Sanger Institute COVID-19 Surveillance Team |
| EPI_ISL_2198455, EPI_ISL_2198456, EPI_ISL_2198465, | Lighthouse Lab in Milton Keynes | Wellcome Sanger Institute for the COVID-19 Genomics UK (COG-UK) Consortium | The Lighthouse Lab in Milton Keynes and Alex Alderton, Roberto Amato, Jeffrey Barrett, Sonia Goncalves, Ewan Harrison, David K. Jackson, Ian Johnston, Dominic Kwiatkowski, Cordelia Langford, John Sillitoe on behalf of the Wellcome Sanger Institute COVID-19 Surveillance Team |

| | | | |
|---|---|---|---|
| EPI_ISL_2198467, EPI_ISL_2198470, EPI_ISL_2198471, EPI_ISL_2198472 | | | |
| EPI_ISL_2198476 | Lighthouse Lab in Alderley Park | Wellcome Sanger Institute for the COVID-19 Genomics UK (COG-UK) Consortium | Jacquelyn Wynn, Mairead Hyland, The Lighthouse Lab in Alderley Park and Alex Alderton, Roberto Amato, Jeffrey Barrett, Sonia Goncalves, Ewan Harrison, David K. Jackson, Ian Johnston, Dominic Kwiatkowski, Cordelia Langford, John Sillitoe on behalf of the Wellcome Sanger Institute COVID-19 Surveillance Team |
| EPI_ISL_2198477, EPI_ISL_2198478 | Lighthouse Lab in Milton Keynes | Wellcome Sanger Institute for the COVID-19 Genomics UK (COG-UK) Consortium | The Lighthouse Lab in Milton Keynes and Alex Alderton, Roberto Amato, Jeffrey Barrett, Sonia Goncalves, Ewan Harrison, David K. Jackson, Ian Johnston, Dominic Kwiatkowski, Cordelia Langford, John Sillitoe on behalf of the Wellcome Sanger Institute COVID-19 Surveillance Team |
| EPI_ISL_2198482 | Lighthouse Lab in Alderley Park | Wellcome Sanger Institute for the COVID-19 Genomics UK (COG-UK) Consortium | Jacquelyn Wynn, Mairead Hyland, The Lighthouse Lab in Alderley Park and Alex Alderton, Roberto Amato, Jeffrey Barrett, Sonia Goncalves, Ewan Harrison, David K. Jackson, Ian Johnston, Dominic Kwiatkowski, Cordelia Langford, John Sillitoe on behalf of the Wellcome Sanger Institute COVID-19 Surveillance Team |
| EPI_ISL_2198483, EPI_ISL_2198486, EPI_ISL_2198487 | Lighthouse Lab in Milton Keynes | Wellcome Sanger Institute for the COVID-19 Genomics UK (COG-UK) Consortium | The Lighthouse Lab in Milton Keynes and Alex Alderton, Roberto Amato, Jeffrey Barrett, Sonia Goncalves, Ewan Harrison, David K. Jackson, Ian Johnston, Dominic Kwiatkowski, Cordelia Langford, John Sillitoe on behalf of the Wellcome Sanger Institute COVID-19 Surveillance Team |
| EPI_ISL_2198489 | Lighthouse Lab in Alderley Park | Wellcome Sanger Institute for the COVID-19 Genomics UK (COG-UK) Consortium | Jacquelyn Wynn, Mairead Hyland, The Lighthouse Lab in Alderley Park and Alex Alderton, Roberto Amato, Jeffrey Barrett, Sonia Goncalves, Ewan Harrison, David K. Jackson, Ian Johnston, Dominic Kwiatkowski, Cordelia Langford, John Sillitoe on behalf of the Wellcome Sanger Institute COVID-19 Surveillance Team |
| EPI_ISL_2198492, EPI_ISL_2198493 | Lighthouse Lab in Milton Keynes | Wellcome Sanger Institute for the COVID-19 Genomics UK (COG-UK) Consortium | The Lighthouse Lab in Milton Keynes and Alex Alderton, Roberto Amato, Jeffrey Barrett, Sonia Goncalves, Ewan Harrison, David K. Jackson, Ian Johnston, Dominic Kwiatkowski, Cordelia Langford, John Sillitoe on behalf of the Wellcome Sanger Institute COVID-19 Surveillance Team |
| EPI_ISL_2198496 | Lighthouse Lab in Alderley Park | Wellcome Sanger Institute for the COVID-19 Genomics UK (COG-UK) Consortium | Jacquelyn Wynn, Mairead Hyland, The Lighthouse Lab in Alderley Park and Alex Alderton, Roberto Amato, Jeffrey Barrett, Sonia Goncalves, Ewan Harrison, David K. Jackson, Ian Johnston, Dominic Kwiatkowski, Cordelia Langford, John Sillitoe on behalf of the Wellcome Sanger Institute COVID-19 Surveillance Team |
| EPI_ISL_2198499, EPI_ISL_2198502 | Lighthouse Lab in Milton Keynes | Wellcome Sanger Institute for the COVID-19 Genomics UK (COG-UK) Consortium | The Lighthouse Lab in Milton Keynes and Alex Alderton, Roberto Amato, Jeffrey Barrett, Sonia Goncalves, Ewan Harrison, David K. Jackson, Ian Johnston, Dominic Kwiatkowski, Cordelia Langford, John Sillitoe on behalf of the Wellcome Sanger Institute COVID-19 Surveillance Team |
| EPI_ISL_2198504 | Lighthouse Lab in Alderley Park | Wellcome Sanger Institute for the COVID-19 Genomics UK (COG-UK) Consortium | Jacquelyn Wynn, Mairead Hyland, The Lighthouse Lab in Alderley Park and Alex Alderton, Roberto Amato, Jeffrey Barrett, Sonia Goncalves, Ewan Harrison, David K. Jackson, Ian Johnston, Dominic Kwiatkowski, Cordelia Langford, John Sillitoe on behalf of the Wellcome Sanger Institute COVID-19 Surveillance Team |
| EPI_ISL_2198506, EPI_ISL_2198514 | Lighthouse Lab in Milton Keynes | Wellcome Sanger Institute for the COVID-19 Genomics UK (COG-UK) Consortium | The Lighthouse Lab in Milton Keynes and Alex Alderton, Roberto Amato, Jeffrey Barrett, Sonia Goncalves, Ewan Harrison, David K. Jackson, Ian Johnston, Dominic Kwiatkowski, Cordelia Langford, John Sillitoe on behalf of the Wellcome Sanger Institute COVID-19 Surveillance Team |
| EPI_ISL_2198515 | Lighthouse Lab in Alderley Park | Wellcome Sanger Institute for the COVID-19 Genomics UK (COG-UK) Consortium | Jacquelyn Wynn, Mairead Hyland, The Lighthouse Lab in Alderley Park and Alex Alderton, Roberto Amato, Jeffrey Barrett, Sonia Goncalves, Ewan Harrison, David K. Jackson, Ian Johnston, Dominic Kwiatkowski, Cordelia Langford, John Sillitoe on behalf of the Wellcome Sanger Institute COVID-19 Surveillance Team |
| EPI_ISL_2198516 | Lighthouse Lab in Milton Keynes | Wellcome Sanger Institute for the COVID-19 Genomics UK (COG-UK) Consortium | The Lighthouse Lab in Milton Keynes and Alex Alderton, Roberto Amato, Jeffrey Barrett, Sonia Goncalves, Ewan Harrison, David K. Jackson, Ian Johnston, Dominic Kwiatkowski, Cordelia Langford, John Sillitoe on behalf of the Wellcome Sanger Institute COVID-19 Surveillance Team |
| EPI_ISL_2198518 | Lighthouse Lab in Alderley Park | Wellcome Sanger Institute for the COVID-19 Genomics UK (COG-UK) Consortium | Jacquelyn Wynn, Mairead Hyland, The Lighthouse Lab in Alderley Park and Alex Alderton, Roberto Amato, Jeffrey Barrett, Sonia Goncalves, Ewan Harrison, David K. Jackson, Ian Johnston, Dominic Kwiatkowski, Cordelia Langford, John Sillitoe on behalf of the Wellcome Sanger Institute COVID-19 Surveillance Team |
| EPI_ISL_2198519 | Lighthouse Lab in Milton Keynes | Wellcome Sanger Institute for the COVID-19 Genomics UK (COG-UK) Consortium | The Lighthouse Lab in Milton Keynes and Alex Alderton, Roberto Amato, Jeffrey Barrett, Sonia Goncalves, Ewan Harrison, David K. Jackson, Ian Johnston, Dominic Kwiatkowski, Cordelia Langford, John Sillitoe on behalf of the Wellcome Sanger Institute COVID-19 Surveillance Team |
| EPI_ISL_2198520 | Lighthouse Lab in Alderley Park | Wellcome Sanger Institute for the COVID-19 Genomics UK (COG-UK) Consortium | Jacquelyn Wynn, Mairead Hyland, The Lighthouse Lab in Alderley Park and Alex Alderton, Roberto Amato, Jeffrey Barrett, Sonia Goncalves, Ewan Harrison, David K. Jackson, Ian Johnston, Dominic Kwiatkowski, Cordelia Langford, John Sillitoe on behalf of the Wellcome Sanger Institute COVID-19 Surveillance Team |
| EPI_ISL_2198521, EPI_ISL_2198522, EPI_ISL_2198524, EPI_ISL_2198526, EPI_ISL_2198535 | Lighthouse Lab in Milton Keynes | Wellcome Sanger Institute for the COVID-19 Genomics UK (COG-UK) Consortium | The Lighthouse Lab in Milton Keynes and Alex Alderton, Roberto Amato, Jeffrey Barrett, Sonia Goncalves, Ewan Harrison, David K. Jackson, Ian Johnston, Dominic Kwiatkowski, Cordelia Langford, John Sillitoe on behalf of the Wellcome Sanger Institute COVID-19 Surveillance Team |
| EPI_ISL_2198538 | Lighthouse Lab in Alderley Park | Wellcome Sanger Institute for the COVID-19 Genomics UK (COG-UK) Consortium | Jacquelyn Wynn, Mairead Hyland, The Lighthouse Lab in Alderley Park and Alex Alderton, Roberto Amato, Jeffrey Barrett, Sonia Goncalves, Ewan Harrison, David K. Jackson, Ian Johnston, Dominic Kwiatkowski, Cordelia Langford, John Sillitoe on behalf of the Wellcome Sanger Institute COVID-19 Surveillance Team |
| EPI_ISL_2198540, EPI_ISL_2198541, EPI_ISL_2198542, EPI_ISL_2198545, EPI_ISL_2198546 | Lighthouse Lab in Milton Keynes | Wellcome Sanger Institute for the COVID-19 Genomics UK (COG-UK) Consortium | The Lighthouse Lab in Milton Keynes and Alex Alderton, Roberto Amato, Jeffrey Barrett, Sonia Goncalves, Ewan Harrison, David K. Jackson, Ian Johnston, Dominic Kwiatkowski, Cordelia Langford, John Sillitoe on behalf of the Wellcome Sanger Institute COVID-19 Surveillance Team |
| EPI_ISL_2198547 | Lighthouse Lab in Alderley Park | Wellcome Sanger Institute for the COVID-19 Genomics UK (COG-UK) Consortium | Jacquelyn Wynn, Mairead Hyland, The Lighthouse Lab in Alderley Park and Alex Alderton, Roberto Amato, Jeffrey Barrett, Sonia Goncalves, Ewan Harrison, David K. Jackson, Ian Johnston, Dominic Kwiatkowski, Cordelia Langford, John Sillitoe on behalf of the Wellcome Sanger Institute COVID-19 Surveillance Team |
| EPI_ISL_2198548, EPI_ISL_2198553, EPI_ISL_2198555, EPI_ISL_2198556 | Lighthouse Lab in Milton Keynes | Wellcome Sanger Institute for the COVID-19 Genomics UK (COG-UK) Consortium | The Lighthouse Lab in Milton Keynes and Alex Alderton, Roberto Amato, Jeffrey Barrett, Sonia Goncalves, Ewan Harrison, David K. Jackson, Ian Johnston, Dominic Kwiatkowski, Cordelia Langford, John Sillitoe on behalf of the Wellcome Sanger Institute COVID-19 Surveillance Team |
| EPI_ISL_2198557, EPI_ISL_2198558, EPI_ISL_2198559 | Lighthouse Lab in Alderley Park | Wellcome Sanger Institute for the COVID-19 Genomics UK (COG-UK) Consortium | Jacquelyn Wynn, Mairead Hyland, The Lighthouse Lab in Alderley Park and Alex Alderton, Roberto Amato, Jeffrey Barrett, Sonia Goncalves, Ewan Harrison, David K. Jackson, Ian Johnston, Dominic Kwiatkowski, Cordelia Langford, John Sillitoe on behalf of the Wellcome Sanger Institute COVID-19 Surveillance Team |
| EPI_ISL_2198566 | Lighthouse Lab in Milton Keynes | Wellcome Sanger Institute for the COVID-19 Genomics UK (COG-UK) Consortium | The Lighthouse Lab in Milton Keynes and Alex Alderton, Roberto Amato, Jeffrey Barrett, Sonia Goncalves, Ewan Harrison, David K. Jackson, Ian Johnston, Dominic Kwiatkowski, Cordelia Langford, John Sillitoe on behalf of the Wellcome Sanger Institute COVID-19 Surveillance Team |
| EPI_ISL_2198569 | Lighthouse Lab in Alderley Park | Wellcome Sanger Institute for the COVID-19 Genomics UK (COG-UK) Consortium | Jacquelyn Wynn, Mairead Hyland, The Lighthouse Lab in Alderley Park and Alex Alderton, Roberto Amato, Jeffrey Barrett, Sonia Goncalves, Ewan Harrison, David K. Jackson, Ian Johnston, Dominic Kwiatkowski, Cordelia Langford, John Sillitoe on behalf of the Wellcome Sanger Institute COVID-19 Surveillance Team |
| EPI_ISL_2198570 | Lighthouse Lab in Milton Keynes | Wellcome Sanger Institute for the COVID-19 Genomics UK (COG-UK) Consortium | The Lighthouse Lab in Milton Keynes and Alex Alderton, Roberto Amato, Jeffrey Barrett, Sonia Goncalves, Ewan Harrison, David K. Jackson, Ian Johnston, Dominic Kwiatkowski, Cordelia Langford, John Sillitoe on behalf of the Wellcome Sanger Institute COVID-19 Surveillance Team |
| EPI_ISL_2198571 | Lighthouse Lab in Alderley Park | Wellcome Sanger Institute for the COVID-19 Genomics UK (COG-UK) Consortium | Jacquelyn Wynn, Mairead Hyland, The Lighthouse Lab in Alderley Park and Alex Alderton, Roberto Amato, Jeffrey Barrett, Sonia Goncalves, Ewan Harrison, David K. Jackson, Ian Johnston, Dominic Kwiatkowski, Cordelia Langford, John Sillitoe on behalf of the Wellcome Sanger Institute COVID-19 Surveillance Team |
| EPI_ISL_2198572, EPI_ISL_2198576, EPI_ISL_2198577, EPI_ISL_2198581, EPI_ISL_2198583 | Lighthouse Lab in Milton Keynes | Wellcome Sanger Institute for the COVID-19 Genomics UK (COG-UK) Consortium | The Lighthouse Lab in Milton Keynes and Alex Alderton, Roberto Amato, Jeffrey Barrett, Sonia Goncalves, Ewan Harrison, David K. Jackson, Ian Johnston, Dominic Kwiatkowski, Cordelia Langford, John Sillitoe on behalf of the Wellcome Sanger Institute COVID-19 Surveillance Team |
| EPI_ISL_2198584 | Lighthouse Lab in Alderley Park | Wellcome Sanger Institute for the COVID-19 Genomics UK (COG-UK) Consortium | Jacquelyn Wynn, Mairead Hyland, The Lighthouse Lab in Alderley Park and Alex Alderton, Roberto Amato, Jeffrey Barrett, Sonia Goncalves, Ewan Harrison, David K. Jackson, Ian Johnston, Dominic Kwiatkowski, Cordelia Langford, John Sillitoe on behalf of the Wellcome Sanger Institute COVID-19 Surveillance Team |
| EPI_ISL_2198587, EPI_ISL_2198589 | Lighthouse Lab in Milton Keynes | Wellcome Sanger Institute for the COVID-19 Genomics UK (COG-UK) Consortium | The Lighthouse Lab in Milton Keynes and Alex Alderton, Roberto Amato, Jeffrey Barrett, Sonia Goncalves, Ewan Harrison, David K. Jackson, Ian Johnston, Dominic Kwiatkowski, Cordelia Langford, John Sillitoe on behalf of the Wellcome Sanger Institute COVID-19 Surveillance Team |
| EPI_ISL_2198590 | Lighthouse Lab in Alderley Park | Wellcome Sanger Institute for the COVID-19 Genomics UK (COG-UK) Consortium | Jacquelyn Wynn, Mairead Hyland, The Lighthouse Lab in Alderley Park and Alex Alderton, Roberto Amato, Jeffrey Barrett, Sonia Goncalves, Ewan Harrison, David K. Jackson, Ian Johnston, Dominic Kwiatkowski, Cordelia Langford, John Sillitoe on behalf of the Wellcome Sanger Institute COVID-19 Surveillance Team |
| EPI_ISL_2198591 | Lighthouse Lab in Milton Keynes | Wellcome Sanger Institute for the COVID-19 Genomics UK (COG-UK) Consortium | The Lighthouse Lab in Milton Keynes and Alex Alderton, Roberto Amato, Jeffrey Barrett, Sonia Goncalves, Ewan Harrison, David K. Jackson, Ian Johnston, Dominic Kwiatkowski, Cordelia Langford, John Sillitoe on behalf of the Wellcome Sanger Institute COVID-19 Surveillance Team |
| EPI_ISL_2198593 | Lighthouse Lab in Alderley Park | Wellcome Sanger Institute for the COVID-19 Genomics UK (COG-UK) Consortium | Jacquelyn Wynn, Mairead Hyland, The Lighthouse Lab in Alderley Park and Alex Alderton, Roberto Amato, Jeffrey Barrett, Sonia Goncalves, Ewan Harrison, David K. Jackson, Ian Johnston, Dominic Kwiatkowski, Cordelia Langford, John Sillitoe on behalf of the Wellcome Sanger Institute COVID-19 Surveillance Team |
| EPI_ISL_2198595, EPI_ISL_2198600, EPI_ISL_2198601 | Lighthouse Lab in Milton Keynes | Wellcome Sanger Institute for the COVID-19 Genomics UK (COG-UK) Consortium | The Lighthouse Lab in Milton Keynes and Alex Alderton, Roberto Amato, Jeffrey Barrett, Sonia Goncalves, Ewan Harrison, David K. Jackson, Ian Johnston, Dominic Kwiatkowski, Cordelia Langford, John Sillitoe on behalf of the Wellcome Sanger Institute COVID-19 Surveillance Team |
| EPI_ISL_2198603 | Lighthouse Lab in Alderley Park | Wellcome Sanger Institute for the COVID-19 Genomics UK (COG-UK) Consortium | Jacquelyn Wynn, Mairead Hyland, The Lighthouse Lab in Alderley Park and Alex Alderton, Roberto Amato, Jeffrey Barrett, Sonia Goncalves, Ewan Harrison, David K. Jackson, Ian Johnston, Dominic Kwiatkowski, Cordelia Langford, John Sillitoe on behalf of the Wellcome Sanger Institute COVID-19 Surveillance Team |
| EPI_ISL_2198610, EPI_ISL_2198612, EPI_ISL_2198615, EPI_ISL_2198618, EPI_ISL_2198621, EPI_ISL_2198625, EPI_ISL_2198629 | Lighthouse Lab in Milton Keynes | Wellcome Sanger Institute for the COVID-19 Genomics UK (COG-UK) Consortium | The Lighthouse Lab in Milton Keynes and Alex Alderton, Roberto Amato, Jeffrey Barrett, Sonia Goncalves, Ewan Harrison, David K. Jackson, Ian Johnston, Dominic Kwiatkowski, Cordelia Langford, John Sillitoe on behalf of the Wellcome Sanger Institute COVID-19 Surveillance Team |
| EPI_ISL_2198630 | Lighthouse Lab in Alderley Park | Wellcome Sanger Institute for the COVID-19 Genomics UK (COG-UK) Consortium | Jacquelyn Wynn, Mairead Hyland, The Lighthouse Lab in Alderley Park and Alex Alderton, Roberto Amato, Jeffrey Barrett, Sonia Goncalves, Ewan Harrison, David K. Jackson, Ian Johnston, Dominic Kwiatkowski, Cordelia Langford, John Sillitoe on behalf of the Wellcome Sanger Institute COVID-19 Surveillance Team |
| EPI_ISL_2198633, | Lighthouse Lab in Milton Keynes | Wellcome Sanger Institute for the COVID-19 Genomics UK (COG-UK) | The Lighthouse Lab in Milton Keynes and Alex Alderton, Roberto Amato, Jeffrey Barrett, Sonia Goncalves, Ewan Harrison, David K. Jackson, Ian Johnston, Dominic Kwiatkowski, Cordelia Langford, John Sillitoe on behalf of the Wellcome Sanger |

| | | | |
|---|---|---|---|
| EPI_ISL_2198634, EPI_ISL_2198636, EPI_ISL_2198639, EPI_ISL_2198642, EPI_ISL_2198643, EPI_ISL_2198644 | | Consortium | Institute COVID-19 Surveillance Team |
| EPI_ISL_2198645 | Lighthouse Lab in Alderley Park | Wellcome Sanger Institute for the COVID-19 Genomics UK (COG-UK) Consortium | Jacquelyn Wynn, Mairead Hyland, The Lighthouse Lab in Alderley Park and Alex Alderton, Roberto Amato, Jeffrey Barrett, Sonia Goncalves, Ewan Harrison, David K. Jackson, Ian Johnston, Dominic Kwiatkowski, Cordelia Langford, John Sillitoe on behalf of the Wellcome Sanger Institute COVID-19 Surveillance Team |
| EPI_ISL_2198647, EPI_ISL_2198649 | Lighthouse Lab in Milton Keynes | Wellcome Sanger Institute for the COVID-19 Genomics UK (COG-UK) Consortium | The Lighthouse Lab in Milton Keynes and Alex Alderton, Roberto Amato, Jeffrey Barrett, Sonia Goncalves, Ewan Harrison, David K. Jackson, Ian Johnston, Dominic Kwiatkowski, Cordelia Langford, John Sillitoe on behalf of the Wellcome Sanger Institute COVID-19 Surveillance Team |
| EPI_ISL_2198650, EPI_ISL_2198651 | Lighthouse Lab in Alderley Park | Wellcome Sanger Institute for the COVID-19 Genomics UK (COG-UK) Consortium | Jacquelyn Wynn, Mairead Hyland, The Lighthouse Lab in Alderley Park and Alex Alderton, Roberto Amato, Jeffrey Barrett, Sonia Goncalves, Ewan Harrison, David K. Jackson, Ian Johnston, Dominic Kwiatkowski, Cordelia Langford, John Sillitoe on behalf of the Wellcome Sanger Institute COVID-19 Surveillance Team |
| EPI_ISL_2198654, EPI_ISL_2198655 | Lighthouse Lab in Milton Keynes | Wellcome Sanger Institute for the COVID-19 Genomics UK (COG-UK) Consortium | The Lighthouse Lab in Milton Keynes and Alex Alderton, Roberto Amato, Jeffrey Barrett, Sonia Goncalves, Ewan Harrison, David K. Jackson, Ian Johnston, Dominic Kwiatkowski, Cordelia Langford, John Sillitoe on behalf of the Wellcome Sanger Institute COVID-19 Surveillance Team |
| EPI_ISL_2198656 | Lighthouse Lab in Alderley Park | Wellcome Sanger Institute for the COVID-19 Genomics UK (COG-UK) Consortium | Jacquelyn Wynn, Mairead Hyland, The Lighthouse Lab in Alderley Park and Alex Alderton, Roberto Amato, Jeffrey Barrett, Sonia Goncalves, Ewan Harrison, David K. Jackson, Ian Johnston, Dominic Kwiatkowski, Cordelia Langford, John Sillitoe on behalf of the Wellcome Sanger Institute COVID-19 Surveillance Team |
| EPI_ISL_2198657 | Lighthouse Lab in Milton Keynes | Wellcome Sanger Institute for the COVID-19 Genomics UK (COG-UK) Consortium | The Lighthouse Lab in Milton Keynes and Alex Alderton, Roberto Amato, Jeffrey Barrett, Sonia Goncalves, Ewan Harrison, David K. Jackson, Ian Johnston, Dominic Kwiatkowski, Cordelia Langford, John Sillitoe on behalf of the Wellcome Sanger Institute COVID-19 Surveillance Team |
| EPI_ISL_2198658 | Lighthouse Lab in Alderley Park | Wellcome Sanger Institute for the COVID-19 Genomics UK (COG-UK) Consortium | Jacquelyn Wynn, Mairead Hyland, The Lighthouse Lab in Alderley Park and Alex Alderton, Roberto Amato, Jeffrey Barrett, Sonia Goncalves, Ewan Harrison, David K. Jackson, Ian Johnston, Dominic Kwiatkowski, Cordelia Langford, John Sillitoe on behalf of the Wellcome Sanger Institute COVID-19 Surveillance Team |
| EPI_ISL_2198661, EPI_ISL_2198663 | Lighthouse Lab in Milton Keynes | Wellcome Sanger Institute for the COVID-19 Genomics UK (COG-UK) Consortium | The Lighthouse Lab in Milton Keynes and Alex Alderton, Roberto Amato, Jeffrey Barrett, Sonia Goncalves, Ewan Harrison, David K. Jackson, Ian Johnston, Dominic Kwiatkowski, Cordelia Langford, John Sillitoe on behalf of the Wellcome Sanger Institute COVID-19 Surveillance Team |
| EPI_ISL_2198664 | Lighthouse Lab in Alderley Park | Wellcome Sanger Institute for the COVID-19 Genomics UK (COG-UK) Consortium | Jacquelyn Wynn, Mairead Hyland, The Lighthouse Lab in Alderley Park and Alex Alderton, Roberto Amato, Jeffrey Barrett, Sonia Goncalves, Ewan Harrison, David K. Jackson, Ian Johnston, Dominic Kwiatkowski, Cordelia Langford, John Sillitoe on behalf of the Wellcome Sanger Institute COVID-19 Surveillance Team |
| EPI_ISL_2198667, EPI_ISL_2198669 | Lighthouse Lab in Milton Keynes | Wellcome Sanger Institute for the COVID-19 Genomics UK (COG-UK) Consortium | The Lighthouse Lab in Milton Keynes and Alex Alderton, Roberto Amato, Jeffrey Barrett, Sonia Goncalves, Ewan Harrison, David K. Jackson, Ian Johnston, Dominic Kwiatkowski, Cordelia Langford, John Sillitoe on behalf of the Wellcome Sanger Institute COVID-19 Surveillance Team |
| EPI_ISL_2198671 | Lighthouse Lab in Glasgow | Wellcome Sanger Institute for the COVID-19 Genomics UK (COG-UK) Consortium | Harper VanSteenhouse, Yumi Kasai, David Gray, Carol Clugston, Anna Dominiczak and Alex Alderton, Roberto Amato, Jeffrey Barrett, Sonia Goncalves, Ewan Harrison, David K. Jackson, Ian Johnston, Dominic Kwiatkowski, Cordelia Langford, John Sillitoe on behalf of the Wellcome Sanger Institute COVID-19 Surveillance Team |
| EPI_ISL_2198673 | Lighthouse Lab in Alderley Park | Wellcome Sanger Institute for the COVID-19 Genomics UK (COG-UK) Consortium | Jacquelyn Wynn, Mairead Hyland, The Lighthouse Lab in Alderley Park and Alex Alderton, Roberto Amato, Jeffrey Barrett, Sonia Goncalves, Ewan Harrison, David K. Jackson, Ian Johnston, Dominic Kwiatkowski, Cordelia Langford, John Sillitoe on behalf of the Wellcome Sanger Institute COVID-19 Surveillance Team |
| EPI_ISL_2198675 | Lighthouse Lab in Milton Keynes | Wellcome Sanger Institute for the COVID-19 Genomics UK (COG-UK) Consortium | The Lighthouse Lab in Milton Keynes and Alex Alderton, Roberto Amato, Jeffrey Barrett, Sonia Goncalves, Ewan Harrison, David K. Jackson, Ian Johnston, Dominic Kwiatkowski, Cordelia Langford, John Sillitoe on behalf of the Wellcome Sanger Institute COVID-19 Surveillance Team |
| EPI_ISL_2198677, EPI_ISL_2198678 | Lighthouse Lab in Alderley Park | Wellcome Sanger Institute for the COVID-19 Genomics UK (COG-UK) Consortium | Jacquelyn Wynn, Mairead Hyland, The Lighthouse Lab in Alderley Park and Alex Alderton, Roberto Amato, Jeffrey Barrett, Sonia Goncalves, Ewan Harrison, David K. Jackson, Ian Johnston, Dominic Kwiatkowski, Cordelia Langford, John Sillitoe on behalf of the Wellcome Sanger Institute COVID-19 Surveillance Team |
| EPI_ISL_2198679 | Lighthouse Lab in Milton Keynes | Wellcome Sanger Institute for the COVID-19 Genomics UK (COG-UK) Consortium | The Lighthouse Lab in Milton Keynes and Alex Alderton, Roberto Amato, Jeffrey Barrett, Sonia Goncalves, Ewan Harrison, David K. Jackson, Ian Johnston, Dominic Kwiatkowski, Cordelia Langford, John Sillitoe on behalf of the Wellcome Sanger Institute COVID-19 Surveillance Team |
| EPI_ISL_2198680, EPI_ISL_2198681 | Lighthouse Lab in Alderley Park | Wellcome Sanger Institute for the COVID-19 Genomics UK (COG-UK) Consortium | Jacquelyn Wynn, Mairead Hyland, The Lighthouse Lab in Alderley Park and Alex Alderton, Roberto Amato, Jeffrey Barrett, Sonia Goncalves, Ewan Harrison, David K. Jackson, Ian Johnston, Dominic Kwiatkowski, Cordelia Langford, John Sillitoe on behalf of the Wellcome Sanger Institute COVID-19 Surveillance Team |
| EPI_ISL_2198682 | Lighthouse Lab in Milton Keynes | Wellcome Sanger Institute for the COVID-19 Genomics UK (COG-UK) Consortium | The Lighthouse Lab in Milton Keynes and Alex Alderton, Roberto Amato, Jeffrey Barrett, Sonia Goncalves, Ewan Harrison, David K. Jackson, Ian Johnston, Dominic Kwiatkowski, Cordelia Langford, John Sillitoe on behalf of the Wellcome Sanger Institute COVID-19 Surveillance Team |
| EPI_ISL_2198686 | Lighthouse Lab in Alderley Park | Wellcome Sanger Institute for the COVID-19 Genomics UK (COG-UK) Consortium | Jacquelyn Wynn, Mairead Hyland, The Lighthouse Lab in Alderley Park and Alex Alderton, Roberto Amato, Jeffrey Barrett, Sonia Goncalves, Ewan Harrison, David K. Jackson, Ian Johnston, Dominic Kwiatkowski, Cordelia Langford, John Sillitoe on behalf of the Wellcome Sanger Institute COVID-19 Surveillance Team |
| EPI_ISL_2198687, EPI_ISL_2198688 | Lighthouse Lab in Milton Keynes | Wellcome Sanger Institute for the COVID-19 Genomics UK (COG-UK) Consortium | The Lighthouse Lab in Milton Keynes and Alex Alderton, Roberto Amato, Jeffrey Barrett, Sonia Goncalves, Ewan Harrison, David K. Jackson, Ian Johnston, Dominic Kwiatkowski, Cordelia Langford, John Sillitoe on behalf of the Wellcome Sanger Institute COVID-19 Surveillance Team |
| EPI_ISL_2198689 | Lighthouse Lab in Alderley Park | Wellcome Sanger Institute for the COVID-19 Genomics UK (COG-UK) Consortium | Jacquelyn Wynn, Mairead Hyland, The Lighthouse Lab in Alderley Park and Alex Alderton, Roberto Amato, Jeffrey Barrett, Sonia Goncalves, Ewan Harrison, David K. Jackson, Ian Johnston, Dominic Kwiatkowski, Cordelia Langford, John Sillitoe on behalf of the Wellcome Sanger Institute COVID-19 Surveillance Team |
| EPI_ISL_2198691 | Lighthouse Lab in Milton Keynes | Wellcome Sanger Institute for the COVID-19 Genomics UK (COG-UK) Consortium | The Lighthouse Lab in Milton Keynes and Alex Alderton, Roberto Amato, Jeffrey Barrett, Sonia Goncalves, Ewan Harrison, David K. Jackson, Ian Johnston, Dominic Kwiatkowski, Cordelia Langford, John Sillitoe on behalf of the Wellcome Sanger Institute COVID-19 Surveillance Team |
| EPI_ISL_2198692 | Lighthouse Lab in Alderley Park | Wellcome Sanger Institute for the COVID-19 Genomics UK (COG-UK) Consortium | Jacquelyn Wynn, Mairead Hyland, The Lighthouse Lab in Alderley Park and Alex Alderton, Roberto Amato, Jeffrey Barrett, Sonia Goncalves, Ewan Harrison, David K. Jackson, Ian Johnston, Dominic Kwiatkowski, Cordelia Langford, John Sillitoe on behalf of the Wellcome Sanger Institute COVID-19 Surveillance Team |
| EPI_ISL_2198700 | Lighthouse Lab in Milton Keynes | Wellcome Sanger Institute for the COVID-19 Genomics UK (COG-UK) Consortium | The Lighthouse Lab in Milton Keynes and Alex Alderton, Roberto Amato, Jeffrey Barrett, Sonia Goncalves, Ewan Harrison, David K. Jackson, Ian Johnston, Dominic Kwiatkowski, Cordelia Langford, John Sillitoe on behalf of the Wellcome Sanger Institute COVID-19 Surveillance Team |
| EPI_ISL_2198701, EPI_ISL_2198702 | Lighthouse Lab in Alderley Park | Wellcome Sanger Institute for the COVID-19 Genomics UK (COG-UK) Consortium | Jacquelyn Wynn, Mairead Hyland, The Lighthouse Lab in Alderley Park and Alex Alderton, Roberto Amato, Jeffrey Barrett, Sonia Goncalves, Ewan Harrison, David K. Jackson, Ian Johnston, Dominic Kwiatkowski, Cordelia Langford, John Sillitoe on behalf of the Wellcome Sanger Institute COVID-19 Surveillance Team |
| EPI_ISL_2198704, EPI_ISL_2198706, EPI_ISL_2198707 | Lighthouse Lab in Milton Keynes | Wellcome Sanger Institute for the COVID-19 Genomics UK (COG-UK) Consortium | The Lighthouse Lab in Milton Keynes and Alex Alderton, Roberto Amato, Jeffrey Barrett, Sonia Goncalves, Ewan Harrison, David K. Jackson, Ian Johnston, Dominic Kwiatkowski, Cordelia Langford, John Sillitoe on behalf of the Wellcome Sanger Institute COVID-19 Surveillance Team |
| EPI_ISL_2198714, EPI_ISL_2198715 | Lighthouse Lab in Alderley Park | Wellcome Sanger Institute for the COVID-19 Genomics UK (COG-UK) Consortium | Jacquelyn Wynn, Mairead Hyland, The Lighthouse Lab in Alderley Park and Alex Alderton, Roberto Amato, Jeffrey Barrett, Sonia Goncalves, Ewan Harrison, David K. Jackson, Ian Johnston, Dominic Kwiatkowski, Cordelia Langford, John Sillitoe on behalf of the Wellcome Sanger Institute COVID-19 Surveillance Team |
| EPI_ISL_2198716, EPI_ISL_2198717, EPI_ISL_2198718 | Lighthouse Lab in Milton Keynes | Wellcome Sanger Institute for the COVID-19 Genomics UK (COG-UK) Consortium | The Lighthouse Lab in Milton Keynes and Alex Alderton, Roberto Amato, Jeffrey Barrett, Sonia Goncalves, Ewan Harrison, David K. Jackson, Ian Johnston, Dominic Kwiatkowski, Cordelia Langford, John Sillitoe on behalf of the Wellcome Sanger Institute COVID-19 Surveillance Team |
| EPI_ISL_2198719 | Lighthouse Lab in Alderley Park | Wellcome Sanger Institute for the COVID-19 Genomics UK (COG-UK) Consortium | Jacquelyn Wynn, Mairead Hyland, The Lighthouse Lab in Alderley Park and Alex Alderton, Roberto Amato, Jeffrey Barrett, Sonia Goncalves, Ewan Harrison, David K. Jackson, Ian Johnston, Dominic Kwiatkowski, Cordelia Langford, John Sillitoe on behalf of the Wellcome Sanger Institute COVID-19 Surveillance Team |
| EPI_ISL_2198723 | Lighthouse Lab in Milton Keynes | Wellcome Sanger Institute for the COVID-19 Genomics UK (COG-UK) Consortium | The Lighthouse Lab in Milton Keynes and Alex Alderton, Roberto Amato, Jeffrey Barrett, Sonia Goncalves, Ewan Harrison, David K. Jackson, Ian Johnston, Dominic Kwiatkowski, Cordelia Langford, John Sillitoe on behalf of the Wellcome Sanger Institute COVID-19 Surveillance Team |
| EPI_ISL_2198727 | Lighthouse Lab in Alderley Park | Wellcome Sanger Institute for the COVID-19 Genomics UK (COG-UK) Consortium | Jacquelyn Wynn, Mairead Hyland, The Lighthouse Lab in Alderley Park and Alex Alderton, Roberto Amato, Jeffrey Barrett, Sonia Goncalves, Ewan Harrison, David K. Jackson, Ian Johnston, Dominic Kwiatkowski, Cordelia Langford, John Sillitoe on behalf of the Wellcome Sanger Institute COVID-19 Surveillance Team |
| EPI_ISL_2198728 | Lighthouse Lab in Milton Keynes | Wellcome Sanger Institute for the COVID-19 Genomics UK (COG-UK) Consortium | The Lighthouse Lab in Milton Keynes and Alex Alderton, Roberto Amato, Jeffrey Barrett, Sonia Goncalves, Ewan Harrison, David K. Jackson, Ian Johnston, Dominic Kwiatkowski, Cordelia Langford, John Sillitoe on behalf of the Wellcome Sanger Institute COVID-19 Surveillance Team |
| EPI_ISL_2198729 | Lighthouse Lab in Alderley Park | Wellcome Sanger Institute for the COVID-19 Genomics UK (COG-UK) Consortium | Jacquelyn Wynn, Mairead Hyland, The Lighthouse Lab in Alderley Park and Alex Alderton, Roberto Amato, Jeffrey Barrett, Sonia Goncalves, Ewan Harrison, David K. Jackson, Ian Johnston, Dominic Kwiatkowski, Cordelia Langford, John Sillitoe on behalf of the Wellcome Sanger Institute COVID-19 Surveillance Team |
| EPI_ISL_2198730 | Lighthouse Lab in Milton Keynes | Wellcome Sanger Institute for the COVID-19 Genomics UK (COG-UK) Consortium | The Lighthouse Lab in Milton Keynes and Alex Alderton, Roberto Amato, Jeffrey Barrett, Sonia Goncalves, Ewan Harrison, David K. Jackson, Ian Johnston, Dominic Kwiatkowski, Cordelia Langford, John Sillitoe on behalf of the Wellcome Sanger Institute COVID-19 Surveillance Team |
| EPI_ISL_2198732, EPI_ISL_2198733 | Lighthouse Lab in Alderley Park | Wellcome Sanger Institute for the COVID-19 Genomics UK (COG-UK) Consortium | Jacquelyn Wynn, Mairead Hyland, The Lighthouse Lab in Alderley Park and Alex Alderton, Roberto Amato, Jeffrey Barrett, Sonia Goncalves, Ewan Harrison, David K. Jackson, Ian Johnston, Dominic Kwiatkowski, Cordelia Langford, John Sillitoe on behalf of the Wellcome Sanger Institute COVID-19 Surveillance Team |
| EPI_ISL_2198737 | Lighthouse Lab in Milton Keynes | Wellcome Sanger Institute for the COVID-19 Genomics UK (COG-UK) Consortium | The Lighthouse Lab in Milton Keynes and Alex Alderton, Roberto Amato, Jeffrey Barrett, Sonia Goncalves, Ewan Harrison, David K. Jackson, Ian Johnston, Dominic Kwiatkowski, Cordelia Langford, John Sillitoe on behalf of the Wellcome Sanger Institute COVID-19 Surveillance Team |
| EPI_ISL_2198738 | Lighthouse Lab in Glasgow | Wellcome Sanger Institute for the COVID-19 Genomics UK (COG-UK) Consortium | Harper VanSteenhouse, Yumi Kasai, David Gray, Carol Clugston, Anna Dominiczak and Alex Alderton, Roberto Amato, Jeffrey Barrett, Sonia Goncalves, Ewan Harrison, David K. Jackson, Ian Johnston, Dominic Kwiatkowski, Cordelia Langford, John Sillitoe on behalf of the Wellcome Sanger Institute COVID-19 Surveillance Team |
| EPI_ISL_2198739, EPI_ISL_2198740 | Lighthouse Lab in Alderley Park | Wellcome Sanger Institute for the COVID-19 Genomics UK (COG-UK) Consortium | Jacquelyn Wynn, Mairead Hyland, The Lighthouse Lab in Alderley Park and Alex Alderton, Roberto Amato, Jeffrey Barrett, Sonia Goncalves, Ewan Harrison, David K. Jackson, Ian Johnston, Dominic Kwiatkowski, Cordelia Langford, John Sillitoe on behalf of the Wellcome Sanger Institute COVID-19 Surveillance Team |
| EPI_ISL_2198742, EPI_ISL_2198743, EPI_ISL_2198747, EPI_ISL_2198748 | Lighthouse Lab in Milton Keynes | Wellcome Sanger Institute for the COVID-19 Genomics UK (COG-UK) Consortium | The Lighthouse Lab in Milton Keynes and Alex Alderton, Roberto Amato, Jeffrey Barrett, Sonia Goncalves, Ewan Harrison, David K. Jackson, Ian Johnston, Dominic Kwiatkowski, Cordelia Langford, John Sillitoe on behalf of the Wellcome Sanger Institute COVID-19 Surveillance Team |
| EPI_ISL_2198749 | Lighthouse Lab in Alderley Park | Wellcome Sanger Institute for the COVID-19 Genomics UK (COG-UK) Consortium | Jacquelyn Wynn, Mairead Hyland, The Lighthouse Lab in Alderley Park and Alex Alderton, Roberto Amato, Jeffrey Barrett, Sonia Goncalves, Ewan Harrison, David K. Jackson, Ian Johnston, Dominic Kwiatkowski, Cordelia Langford, John Sillitoe on behalf of the Wellcome Sanger Institute COVID-19 Surveillance Team |
| EPI_ISL_2198750 | Lighthouse Lab in Milton Keynes | Wellcome Sanger Institute for the COVID-19 Genomics UK (COG-UK) Consortium | The Lighthouse Lab in Milton Keynes and Alex Alderton, Roberto Amato, Jeffrey Barrett, Sonia Goncalves, Ewan Harrison, David K. Jackson, Ian Johnston, Dominic Kwiatkowski, Cordelia Langford, John Sillitoe on behalf of the Wellcome Sanger Institute COVID-19 Surveillance Team |
| EPI_ISL_2198751 | Lighthouse Lab in Alderley Park | Wellcome Sanger Institute for the COVID-19 Genomics UK (COG-UK) Consortium | Jacquelyn Wynn, Mairead Hyland, The Lighthouse Lab in Alderley Park and Alex Alderton, Roberto Amato, Jeffrey Barrett, Sonia Goncalves, Ewan Harrison, David K. Jackson, Ian Johnston, Dominic Kwiatkowski, Cordelia Langford, John Sillitoe on behalf of the Wellcome Sanger Institute COVID-19 Surveillance Team |

| | | | |
|---|---|---|---|
| EPI_ISL_2198753 | Lighthouse Lab in Glasgow | Wellcome Sanger Institute for the COVID-19 Genomics UK (COG-UK) Consortium | Harper VanSteenhouse, Yumi Kasai, David Gray, Carol Clugston, Anna Dominiczak and Alex Alderton, Roberto Amato, Jeffrey Barrett, Sonia Goncalves, Ewan Harrison, David K. Jackson, Ian Johnston, Dominic Kwiatkowski, Cordelia Langford, John Sillitoe on behalf of the Wellcome Sanger Institute COVID-19 Surveillance Team |
| EPI_ISL_2198754 | Lighthouse Lab in Alderley Park | Wellcome Sanger Institute for the COVID-19 Genomics UK (COG-UK) Consortium | Jacquelyn Wynn, Mairead Hyland, The Lighthouse Lab in Alderley Park and Alex Alderton, Roberto Amato, Jeffrey Barrett, Sonia Goncalves, Ewan Harrison, David K. Jackson, Ian Johnston, Dominic Kwiatkowski, Cordelia Langford, John Sillitoe on behalf of the Wellcome Sanger Institute COVID-19 Surveillance Team |
| EPI_ISL_2198755 | Lighthouse Lab in Glasgow | Wellcome Sanger Institute for the COVID-19 Genomics UK (COG-UK) Consortium | Harper VanSteenhouse, Yumi Kasai, David Gray, Carol Clugston, Anna Dominiczak and Alex Alderton, Roberto Amato, Jeffrey Barrett, Sonia Goncalves, Ewan Harrison, David K. Jackson, Ian Johnston, Dominic Kwiatkowski, Cordelia Langford, John Sillitoe on behalf of the Wellcome Sanger Institute COVID-19 Surveillance Team |
| EPI_ISL_2198756, EPI_ISL_2198760 | Lighthouse Lab in Alderley Park | Wellcome Sanger Institute for the COVID-19 Genomics UK (COG-UK) Consortium | Jacquelyn Wynn, Mairead Hyland, The Lighthouse Lab in Alderley Park and Alex Alderton, Roberto Amato, Jeffrey Barrett, Sonia Goncalves, Ewan Harrison, David K. Jackson, Ian Johnston, Dominic Kwiatkowski, Cordelia Langford, John Sillitoe on behalf of the Wellcome Sanger Institute COVID-19 Surveillance Team |
| EPI_ISL_2198761 | Lighthouse Lab in Milton Keynes | Wellcome Sanger Institute for the COVID-19 Genomics UK (COG-UK) Consortium | The Lighthouse Lab in Milton Keynes and Alex Alderton, Roberto Amato, Jeffrey Barrett, Sonia Goncalves, Ewan Harrison, David K. Jackson, Ian Johnston, Dominic Kwiatkowski, Cordelia Langford, John Sillitoe on behalf of the Wellcome Sanger Institute COVID-19 Surveillance Team |
| EPI_ISL_2198763 | Lighthouse Lab in Alderley Park | Wellcome Sanger Institute for the COVID-19 Genomics UK (COG-UK) Consortium | Jacquelyn Wynn, Mairead Hyland, The Lighthouse Lab in Alderley Park and Alex Alderton, Roberto Amato, Jeffrey Barrett, Sonia Goncalves, Ewan Harrison, David K. Jackson, Ian Johnston, Dominic Kwiatkowski, Cordelia Langford, John Sillitoe on behalf of the Wellcome Sanger Institute COVID-19 Surveillance Team |
| EPI_ISL_2198764, EPI_ISL_2198766 | Lighthouse Lab in Milton Keynes | Wellcome Sanger Institute for the COVID-19 Genomics UK (COG-UK) Consortium | The Lighthouse Lab in Milton Keynes and Alex Alderton, Roberto Amato, Jeffrey Barrett, Sonia Goncalves, Ewan Harrison, David K. Jackson, Ian Johnston, Dominic Kwiatkowski, Cordelia Langford, John Sillitoe on behalf of the Wellcome Sanger Institute COVID-19 Surveillance Team |
| EPI_ISL_2198767 | Lighthouse Lab in Alderley Park | Wellcome Sanger Institute for the COVID-19 Genomics UK (COG-UK) Consortium | Jacquelyn Wynn, Mairead Hyland, The Lighthouse Lab in Alderley Park and Alex Alderton, Roberto Amato, Jeffrey Barrett, Sonia Goncalves, Ewan Harrison, David K. Jackson, Ian Johnston, Dominic Kwiatkowski, Cordelia Langford, John Sillitoe on behalf of the Wellcome Sanger Institute COVID-19 Surveillance Team |
| EPI_ISL_2198768, EPI_ISL_2198770, EPI_ISL_2198772 | Lighthouse Lab in Milton Keynes | Wellcome Sanger Institute for the COVID-19 Genomics UK (COG-UK) Consortium | The Lighthouse Lab in Milton Keynes and Alex Alderton, Roberto Amato, Jeffrey Barrett, Sonia Goncalves, Ewan Harrison, David K. Jackson, Ian Johnston, Dominic Kwiatkowski, Cordelia Langford, John Sillitoe on behalf of the Wellcome Sanger Institute COVID-19 Surveillance Team |
| EPI_ISL_2198775 | Lighthouse Lab in Alderley Park | Wellcome Sanger Institute for the COVID-19 Genomics UK (COG-UK) Consortium | Jacquelyn Wynn, Mairead Hyland, The Lighthouse Lab in Alderley Park and Alex Alderton, Roberto Amato, Jeffrey Barrett, Sonia Goncalves, Ewan Harrison, David K. Jackson, Ian Johnston, Dominic Kwiatkowski, Cordelia Langford, John Sillitoe on behalf of the Wellcome Sanger Institute COVID-19 Surveillance Team |
| EPI_ISL_2198778 | Lighthouse Lab in Milton Keynes | Wellcome Sanger Institute for the COVID-19 Genomics UK (COG-UK) Consortium | The Lighthouse Lab in Milton Keynes and Alex Alderton, Roberto Amato, Jeffrey Barrett, Sonia Goncalves, Ewan Harrison, David K. Jackson, Ian Johnston, Dominic Kwiatkowski, Cordelia Langford, John Sillitoe on behalf of the Wellcome Sanger Institute COVID-19 Surveillance Team |
| EPI_ISL_2198781, EPI_ISL_2198785 | Lighthouse Lab in Alderley Park | Wellcome Sanger Institute for the COVID-19 Genomics UK (COG-UK) Consortium | Jacquelyn Wynn, Mairead Hyland, The Lighthouse Lab in Alderley Park and Alex Alderton, Roberto Amato, Jeffrey Barrett, Sonia Goncalves, Ewan Harrison, David K. Jackson, Ian Johnston, Dominic Kwiatkowski, Cordelia Langford, John Sillitoe on behalf of the Wellcome Sanger Institute COVID-19 Surveillance Team |
| EPI_ISL_2198789 | Lighthouse Lab in Glasgow | Wellcome Sanger Institute for the COVID-19 Genomics UK (COG-UK) Consortium | Harper VanSteenhouse, Yumi Kasai, David Gray, Carol Clugston, Anna Dominiczak and Alex Alderton, Roberto Amato, Jeffrey Barrett, Sonia Goncalves, Ewan Harrison, David K. Jackson, Ian Johnston, Dominic Kwiatkowski, Cordelia Langford, John Sillitoe on behalf of the Wellcome Sanger Institute COVID-19 Surveillance Team |
| EPI_ISL_2198792, EPI_ISL_2198794 | Lighthouse Lab in Milton Keynes | Wellcome Sanger Institute for the COVID-19 Genomics UK (COG-UK) Consortium | The Lighthouse Lab in Milton Keynes and Alex Alderton, Roberto Amato, Jeffrey Barrett, Sonia Goncalves, Ewan Harrison, David K. Jackson, Ian Johnston, Dominic Kwiatkowski, Cordelia Langford, John Sillitoe on behalf of the Wellcome Sanger Institute COVID-19 Surveillance Team |
| EPI_ISL_2198796 | Lighthouse Lab in Alderley Park | Wellcome Sanger Institute for the COVID-19 Genomics UK (COG-UK) Consortium | Jacquelyn Wynn, Mairead Hyland, The Lighthouse Lab in Alderley Park and Alex Alderton, Roberto Amato, Jeffrey Barrett, Sonia Goncalves, Ewan Harrison, David K. Jackson, Ian Johnston, Dominic Kwiatkowski, Cordelia Langford, John Sillitoe on behalf of the Wellcome Sanger Institute COVID-19 Surveillance Team |
| EPI_ISL_2198799, EPI_ISL_2198800 | Lighthouse Lab in Milton Keynes | Wellcome Sanger Institute for the COVID-19 Genomics UK (COG-UK) Consortium | The Lighthouse Lab in Milton Keynes and Alex Alderton, Roberto Amato, Jeffrey Barrett, Sonia Goncalves, Ewan Harrison, David K. Jackson, Ian Johnston, Dominic Kwiatkowski, Cordelia Langford, John Sillitoe on behalf of the Wellcome Sanger Institute COVID-19 Surveillance Team |
| EPI_ISL_2198804 | Lighthouse Lab in Glasgow | Wellcome Sanger Institute for the COVID-19 Genomics UK (COG-UK) Consortium | Harper VanSteenhouse, Yumi Kasai, David Gray, Carol Clugston, Anna Dominiczak and Alex Alderton, Roberto Amato, Jeffrey Barrett, Sonia Goncalves, Ewan Harrison, David K. Jackson, Ian Johnston, Dominic Kwiatkowski, Cordelia Langford, John Sillitoe on behalf of the Wellcome Sanger Institute COVID-19 Surveillance Team |
| EPI_ISL_2198806, EPI_ISL_2198808 | Lighthouse Lab in Alderley Park | Wellcome Sanger Institute for the COVID-19 Genomics UK (COG-UK) Consortium | Jacquelyn Wynn, Mairead Hyland, The Lighthouse Lab in Alderley Park and Alex Alderton, Roberto Amato, Jeffrey Barrett, Sonia Goncalves, Ewan Harrison, David K. Jackson, Ian Johnston, Dominic Kwiatkowski, Cordelia Langford, John Sillitoe on behalf of the Wellcome Sanger Institute COVID-19 Surveillance Team |
| EPI_ISL_2198810 | Lighthouse Lab in Milton Keynes | Wellcome Sanger Institute for the COVID-19 Genomics UK (COG-UK) Consortium | The Lighthouse Lab in Milton Keynes and Alex Alderton, Roberto Amato, Jeffrey Barrett, Sonia Goncalves, Ewan Harrison, David K. Jackson, Ian Johnston, Dominic Kwiatkowski, Cordelia Langford, John Sillitoe on behalf of the Wellcome Sanger Institute COVID-19 Surveillance Team |
| EPI_ISL_2198811, EPI_ISL_2198812 | Lighthouse Lab in Alderley Park | Wellcome Sanger Institute for the COVID-19 Genomics UK (COG-UK) Consortium | Jacquelyn Wynn, Mairead Hyland, The Lighthouse Lab in Alderley Park and Alex Alderton, Roberto Amato, Jeffrey Barrett, Sonia Goncalves, Ewan Harrison, David K. Jackson, Ian Johnston, Dominic Kwiatkowski, Cordelia Langford, John Sillitoe on behalf of the Wellcome Sanger Institute COVID-19 Surveillance Team |
| EPI_ISL_2198816 | Lighthouse Lab in Milton Keynes | Wellcome Sanger Institute for the COVID-19 Genomics UK (COG-UK) Consortium | The Lighthouse Lab in Milton Keynes and Alex Alderton, Roberto Amato, Jeffrey Barrett, Sonia Goncalves, Ewan Harrison, David K. Jackson, Ian Johnston, Dominic Kwiatkowski, Cordelia Langford, John Sillitoe on behalf of the Wellcome Sanger Institute COVID-19 Surveillance Team |
| EPI_ISL_2198817 | Lighthouse Lab in Glasgow | Wellcome Sanger Institute for the COVID-19 Genomics UK (COG-UK) Consortium | Harper VanSteenhouse, Yumi Kasai, David Gray, Carol Clugston, Anna Dominiczak and Alex Alderton, Roberto Amato, Jeffrey Barrett, Sonia Goncalves, Ewan Harrison, David K. Jackson, Ian Johnston, Dominic Kwiatkowski, Cordelia Langford, John Sillitoe on behalf of the Wellcome Sanger Institute COVID-19 Surveillance Team |
| EPI_ISL_2198818 | Lighthouse Lab in Milton Keynes | Wellcome Sanger Institute for the COVID-19 Genomics UK (COG-UK) Consortium | The Lighthouse Lab in Milton Keynes and Alex Alderton, Roberto Amato, Jeffrey Barrett, Sonia Goncalves, Ewan Harrison, David K. Jackson, Ian Johnston, Dominic Kwiatkowski, Cordelia Langford, John Sillitoe on behalf of the Wellcome Sanger Institute COVID-19 Surveillance Team |
| EPI_ISL_2198820 | Lighthouse Lab in Alderley Park | Wellcome Sanger Institute for the COVID-19 Genomics UK (COG-UK) Consortium | Jacquelyn Wynn, Mairead Hyland, The Lighthouse Lab in Alderley Park and Alex Alderton, Roberto Amato, Jeffrey Barrett, Sonia Goncalves, Ewan Harrison, David K. Jackson, Ian Johnston, Dominic Kwiatkowski, Cordelia Langford, John Sillitoe on behalf of the Wellcome Sanger Institute COVID-19 Surveillance Team |
| EPI_ISL_2198823, EPI_ISL_2198830, EPI_ISL_2198834 | Lighthouse Lab in Milton Keynes | Wellcome Sanger Institute for the COVID-19 Genomics UK (COG-UK) Consortium | The Lighthouse Lab in Milton Keynes and Alex Alderton, Roberto Amato, Jeffrey Barrett, Sonia Goncalves, Ewan Harrison, David K. Jackson, Ian Johnston, Dominic Kwiatkowski, Cordelia Langford, John Sillitoe on behalf of the Wellcome Sanger Institute COVID-19 Surveillance Team |
| EPI_ISL_2198835, EPI_ISL_2198845 | Lighthouse Lab in Alderley Park | Wellcome Sanger Institute for the COVID-19 Genomics UK (COG-UK) Consortium | Jacquelyn Wynn, Mairead Hyland, The Lighthouse Lab in Alderley Park and Alex Alderton, Roberto Amato, Jeffrey Barrett, Sonia Goncalves, Ewan Harrison, David K. Jackson, Ian Johnston, Dominic Kwiatkowski, Cordelia Langford, John Sillitoe on behalf of the Wellcome Sanger Institute COVID-19 Surveillance Team |
| EPI_ISL_2198847, EPI_ISL_2198849, EPI_ISL_2198851, EPI_ISL_2198854 | Lighthouse Lab in Milton Keynes | Wellcome Sanger Institute for the COVID-19 Genomics UK (COG-UK) Consortium | The Lighthouse Lab in Milton Keynes and Alex Alderton, Roberto Amato, Jeffrey Barrett, Sonia Goncalves, Ewan Harrison, David K. Jackson, Ian Johnston, Dominic Kwiatkowski, Cordelia Langford, John Sillitoe on behalf of the Wellcome Sanger Institute COVID-19 Surveillance Team |
| EPI_ISL_2198855 | Lighthouse Lab in Glasgow | Wellcome Sanger Institute for the COVID-19 Genomics UK (COG-UK) Consortium | Harper VanSteenhouse, Yumi Kasai, David Gray, Carol Clugston, Anna Dominiczak and Alex Alderton, Roberto Amato, Jeffrey Barrett, Sonia Goncalves, Ewan Harrison, David K. Jackson, Ian Johnston, Dominic Kwiatkowski, Cordelia Langford, John Sillitoe on behalf of the Wellcome Sanger Institute COVID-19 Surveillance Team |
| EPI_ISL_2198856 | Lighthouse Lab in Alderley Park | Wellcome Sanger Institute for the COVID-19 Genomics UK (COG-UK) Consortium | Jacquelyn Wynn, Mairead Hyland, The Lighthouse Lab in Alderley Park and Alex Alderton, Roberto Amato, Jeffrey Barrett, Sonia Goncalves, Ewan Harrison, David K. Jackson, Ian Johnston, Dominic Kwiatkowski, Cordelia Langford, John Sillitoe on behalf of the Wellcome Sanger Institute COVID-19 Surveillance Team |
| EPI_ISL_2198857 | Lighthouse Lab in Milton Keynes | Wellcome Sanger Institute for the COVID-19 Genomics UK (COG-UK) Consortium | The Lighthouse Lab in Milton Keynes and Alex Alderton, Roberto Amato, Jeffrey Barrett, Sonia Goncalves, Ewan Harrison, David K. Jackson, Ian Johnston, Dominic Kwiatkowski, Cordelia Langford, John Sillitoe on behalf of the Wellcome Sanger Institute COVID-19 Surveillance Team |
| EPI_ISL_2198860 | Lighthouse Lab in Alderley Park | Wellcome Sanger Institute for the COVID-19 Genomics UK (COG-UK) Consortium | Jacquelyn Wynn, Mairead Hyland, The Lighthouse Lab in Alderley Park and Alex Alderton, Roberto Amato, Jeffrey Barrett, Sonia Goncalves, Ewan Harrison, David K. Jackson, Ian Johnston, Dominic Kwiatkowski, Cordelia Langford, John Sillitoe on behalf of the Wellcome Sanger Institute COVID-19 Surveillance Team |
| EPI_ISL_2198861, EPI_ISL_2198866, EPI_ISL_2198869 | Lighthouse Lab in Milton Keynes | Wellcome Sanger Institute for the COVID-19 Genomics UK (COG-UK) Consortium | The Lighthouse Lab in Milton Keynes and Alex Alderton, Roberto Amato, Jeffrey Barrett, Sonia Goncalves, Ewan Harrison, David K. Jackson, Ian Johnston, Dominic Kwiatkowski, Cordelia Langford, John Sillitoe on behalf of the Wellcome Sanger Institute COVID-19 Surveillance Team |
| EPI_ISL_2198870 | Lighthouse Lab in Alderley Park | Wellcome Sanger Institute for the COVID-19 Genomics UK (COG-UK) Consortium | Jacquelyn Wynn, Mairead Hyland, The Lighthouse Lab in Alderley Park and Alex Alderton, Roberto Amato, Jeffrey Barrett, Sonia Goncalves, Ewan Harrison, David K. Jackson, Ian Johnston, Dominic Kwiatkowski, Cordelia Langford, John Sillitoe on behalf of the Wellcome Sanger Institute COVID-19 Surveillance Team |
| EPI_ISL_2198874, EPI_ISL_2198875 | Lighthouse Lab in Milton Keynes | Wellcome Sanger Institute for the COVID-19 Genomics UK (COG-UK) Consortium | The Lighthouse Lab in Milton Keynes and Alex Alderton, Roberto Amato, Jeffrey Barrett, Sonia Goncalves, Ewan Harrison, David K. Jackson, Ian Johnston, Dominic Kwiatkowski, Cordelia Langford, John Sillitoe on behalf of the Wellcome Sanger Institute COVID-19 Surveillance Team |
| EPI_ISL_2198876, EPI_ISL_2198879 | Lighthouse Lab in Alderley Park | Wellcome Sanger Institute for the COVID-19 Genomics UK (COG-UK) Consortium | Jacquelyn Wynn, Mairead Hyland, The Lighthouse Lab in Alderley Park and Alex Alderton, Roberto Amato, Jeffrey Barrett, Sonia Goncalves, Ewan Harrison, David K. Jackson, Ian Johnston, Dominic Kwiatkowski, Cordelia Langford, John Sillitoe on behalf of the Wellcome Sanger Institute COVID-19 Surveillance Team |
| EPI_ISL_2198880 | Lighthouse Lab in Glasgow | Wellcome Sanger Institute for the COVID-19 Genomics UK (COG-UK) Consortium | Harper VanSteenhouse, Yumi Kasai, David Gray, Carol Clugston, Anna Dominiczak and Alex Alderton, Roberto Amato, Jeffrey Barrett, Sonia Goncalves, Ewan Harrison, David K. Jackson, Ian Johnston, Dominic Kwiatkowski, Cordelia Langford, John Sillitoe on behalf of the Wellcome Sanger Institute COVID-19 Surveillance Team |
| EPI_ISL_2198881, EPI_ISL_2198885, EPI_ISL_2198887, EPI_ISL_2198889 | Lighthouse Lab in Milton Keynes | Wellcome Sanger Institute for the COVID-19 Genomics UK (COG-UK) Consortium | The Lighthouse Lab in Milton Keynes and Alex Alderton, Roberto Amato, Jeffrey Barrett, Sonia Goncalves, Ewan Harrison, David K. Jackson, Ian Johnston, Dominic Kwiatkowski, Cordelia Langford, John Sillitoe on behalf of the Wellcome Sanger Institute COVID-19 Surveillance Team |
| EPI_ISL_2198892 | Lighthouse Lab in Alderley Park | Wellcome Sanger Institute for the COVID-19 Genomics UK (COG-UK) Consortium | Jacquelyn Wynn, Mairead Hyland, The Lighthouse Lab in Alderley Park and Alex Alderton, Roberto Amato, Jeffrey Barrett, Sonia Goncalves, Ewan Harrison, David K. Jackson, Ian Johnston, Dominic Kwiatkowski, Cordelia Langford, John Sillitoe on behalf of the Wellcome Sanger Institute COVID-19 Surveillance Team |
| EPI_ISL_2198893 | Lighthouse Lab in Milton Keynes | Wellcome Sanger Institute for the COVID-19 Genomics UK (COG-UK) Consortium | The Lighthouse Lab in Milton Keynes and Alex Alderton, Roberto Amato, Jeffrey Barrett, Sonia Goncalves, Ewan Harrison, David K. Jackson, Ian Johnston, Dominic Kwiatkowski, Cordelia Langford, John Sillitoe on behalf of the Wellcome Sanger Institute COVID-19 Surveillance Team |
| EPI_ISL_2198894, EPI_ISL_2198900, EPI_ISL_2198901 | Lighthouse Lab in Alderley Park | Wellcome Sanger Institute for the COVID-19 Genomics UK (COG-UK) Consortium | Jacquelyn Wynn, Mairead Hyland, The Lighthouse Lab in Alderley Park and Alex Alderton, Roberto Amato, Jeffrey Barrett, Sonia Goncalves, Ewan Harrison, David K. Jackson, Ian Johnston, Dominic Kwiatkowski, Cordelia Langford, John Sillitoe on behalf of the Wellcome Sanger Institute COVID-19 Surveillance Team |
| EPI_ISL_2198903 | Lighthouse Lab in Milton Keynes | Wellcome Sanger Institute for the COVID-19 Genomics UK (COG-UK) Consortium | The Lighthouse Lab in Milton Keynes and Alex Alderton, Roberto Amato, Jeffrey Barrett, Sonia Goncalves, Ewan Harrison, David K. Jackson, Ian Johnston, Dominic Kwiatkowski, Cordelia Langford, John Sillitoe on behalf of the Wellcome Sanger Institute COVID-19 Surveillance Team |
| EPI_ISL_2198908, EPI_ISL_2198909 | Lighthouse Lab in Alderley Park | Wellcome Sanger Institute for the COVID-19 Genomics UK (COG-UK) Consortium | Jacquelyn Wynn, Mairead Hyland, The Lighthouse Lab in Alderley Park and Alex Alderton, Roberto Amato, Jeffrey Barrett, Sonia Goncalves, Ewan Harrison, David K. Jackson, Ian Johnston, Dominic Kwiatkowski, Cordelia Langford, John Sillitoe on behalf of the Wellcome Sanger Institute COVID-19 Surveillance Team |

| | | | |
|---|---|---|---|
| EPI_ISL_2198911, EPI_ISL_2198913, EPI_ISL_2198914 | Lighthouse Lab in Milton Keynes | Wellcome Sanger Institute for the COVID-19 Genomics UK (COG-UK) Consortium | The Lighthouse Lab in Milton Keynes and Alex Alderton, Roberto Amato, Jeffrey Barrett, Sonia Goncalves, Ewan Harrison, David K. Jackson, Ian Johnston, Dominic Kwiatkowski, Cordelia Langford, John Sillitoe on behalf of the Wellcome Sanger Institute COVID-19 Surveillance Team |
| EPI_ISL_2198917, EPI_ISL_2198919, EPI_ISL_2198920, EPI_ISL_2198921, EPI_ISL_2198922, EPI_ISL_2198926 | Lighthouse Lab in Alderley Park | Wellcome Sanger Institute for the COVID-19 Genomics UK (COG-UK) Consortium | Jacquelyn Wynn, Mairead Hyland, The Lighthouse Lab in Alderley Park and Alex Alderton, Roberto Amato, Jeffrey Barrett, Sonia Goncalves, Ewan Harrison, David K. Jackson, Ian Johnston, Dominic Kwiatkowski, Cordelia Langford, John Sillitoe on behalf of the Wellcome Sanger Institute COVID-19 Surveillance Team |
| EPI_ISL_2198929, EPI_ISL_2198933, EPI_ISL_2198935 | Lighthouse Lab in Milton Keynes | Wellcome Sanger Institute for the COVID-19 Genomics UK (COG-UK) Consortium | The Lighthouse Lab in Milton Keynes and Alex Alderton, Roberto Amato, Jeffrey Barrett, Sonia Goncalves, Ewan Harrison, David K. Jackson, Ian Johnston, Dominic Kwiatkowski, Cordelia Langford, John Sillitoe on behalf of the Wellcome Sanger Institute COVID-19 Surveillance Team |
| EPI_ISL_2198936, EPI_ISL_2198938, EPI_ISL_2198939 | Lighthouse Lab in Alderley Park | Wellcome Sanger Institute for the COVID-19 Genomics UK (COG-UK) Consortium | Jacquelyn Wynn, Mairead Hyland, The Lighthouse Lab in Alderley Park and Alex Alderton, Roberto Amato, Jeffrey Barrett, Sonia Goncalves, Ewan Harrison, David K. Jackson, Ian Johnston, Dominic Kwiatkowski, Cordelia Langford, John Sillitoe on behalf of the Wellcome Sanger Institute COVID-19 Surveillance Team |
| EPI_ISL_2198941, EPI_ISL_2198942, EPI_ISL_2198944 | Lighthouse Lab in Milton Keynes | Wellcome Sanger Institute for the COVID-19 Genomics UK (COG-UK) Consortium | The Lighthouse Lab in Milton Keynes and Alex Alderton, Roberto Amato, Jeffrey Barrett, Sonia Goncalves, Ewan Harrison, David K. Jackson, Ian Johnston, Dominic Kwiatkowski, Cordelia Langford, John Sillitoe on behalf of the Wellcome Sanger Institute COVID-19 Surveillance Team |
| EPI_ISL_2198947 | Lighthouse Lab in Glasgow | Wellcome Sanger Institute for the COVID-19 Genomics UK (COG-UK) Consortium | Harper VanSteenhouse, Yumi Kasai, David Gray, Carol Clugston, Anna Dominiczak and Alex Alderton, Roberto Amato, Jeffrey Barrett, Sonia Goncalves, Ewan Harrison, David K. Jackson, Ian Johnston, Dominic Kwiatkowski, Cordelia Langford, John Sillitoe on behalf of the Wellcome Sanger Institute COVID-19 Surveillance Team |
| EPI_ISL_2198948, EPI_ISL_2198952, EPI_ISL_2198953 | Lighthouse Lab in Alderley Park | Wellcome Sanger Institute for the COVID-19 Genomics UK (COG-UK) Consortium | Jacquelyn Wynn, Mairead Hyland, The Lighthouse Lab in Alderley Park and Alex Alderton, Roberto Amato, Jeffrey Barrett, Sonia Goncalves, Ewan Harrison, David K. Jackson, Ian Johnston, Dominic Kwiatkowski, Cordelia Langford, John Sillitoe on behalf of the Wellcome Sanger Institute COVID-19 Surveillance Team |
| EPI_ISL_2198960 | Lighthouse Lab in Milton Keynes | Wellcome Sanger Institute for the COVID-19 Genomics UK (COG-UK) Consortium | The Lighthouse Lab in Milton Keynes and Alex Alderton, Roberto Amato, Jeffrey Barrett, Sonia Goncalves, Ewan Harrison, David K. Jackson, Ian Johnston, Dominic Kwiatkowski, Cordelia Langford, John Sillitoe on behalf of the Wellcome Sanger Institute COVID-19 Surveillance Team |
| EPI_ISL_2198964, EPI_ISL_2198965 | Lighthouse Lab in Alderley Park | Wellcome Sanger Institute for the COVID-19 Genomics UK (COG-UK) Consortium | Jacquelyn Wynn, Mairead Hyland, The Lighthouse Lab in Alderley Park and Alex Alderton, Roberto Amato, Jeffrey Barrett, Sonia Goncalves, Ewan Harrison, David K. Jackson, Ian Johnston, Dominic Kwiatkowski, Cordelia Langford, John Sillitoe on behalf of the Wellcome Sanger Institute COVID-19 Surveillance Team |
| EPI_ISL_2198966 | Lighthouse Lab in Glasgow | Wellcome Sanger Institute for the COVID-19 Genomics UK (COG-UK) Consortium | Harper VanSteenhouse, Yumi Kasai, David Gray, Carol Clugston, Anna Dominiczak and Alex Alderton, Roberto Amato, Jeffrey Barrett, Sonia Goncalves, Ewan Harrison, David K. Jackson, Ian Johnston, Dominic Kwiatkowski, Cordelia Langford, John Sillitoe on behalf of the Wellcome Sanger Institute COVID-19 Surveillance Team |
| EPI_ISL_2198970 | Lighthouse Lab in Alderley Park | Wellcome Sanger Institute for the COVID-19 Genomics UK (COG-UK) Consortium | Jacquelyn Wynn, Mairead Hyland, The Lighthouse Lab in Alderley Park and Alex Alderton, Roberto Amato, Jeffrey Barrett, Sonia Goncalves, Ewan Harrison, David K. Jackson, Ian Johnston, Dominic Kwiatkowski, Cordelia Langford, John Sillitoe on behalf of the Wellcome Sanger Institute COVID-19 Surveillance Team |
| EPI_ISL_2198971, EPI_ISL_2198973, EPI_ISL_2198974 | Lighthouse Lab in Milton Keynes | Wellcome Sanger Institute for the COVID-19 Genomics UK (COG-UK) Consortium | The Lighthouse Lab in Milton Keynes and Alex Alderton, Roberto Amato, Jeffrey Barrett, Sonia Goncalves, Ewan Harrison, David K. Jackson, Ian Johnston, Dominic Kwiatkowski, Cordelia Langford, John Sillitoe on behalf of the Wellcome Sanger Institute COVID-19 Surveillance Team |
| EPI_ISL_2198975 | Lighthouse Lab in Alderley Park | Wellcome Sanger Institute for the COVID-19 Genomics UK (COG-UK) Consortium | Jacquelyn Wynn, Mairead Hyland, The Lighthouse Lab in Alderley Park and Alex Alderton, Roberto Amato, Jeffrey Barrett, Sonia Goncalves, Ewan Harrison, David K. Jackson, Ian Johnston, Dominic Kwiatkowski, Cordelia Langford, John Sillitoe on behalf of the Wellcome Sanger Institute COVID-19 Surveillance Team |
| EPI_ISL_2198978 | Lighthouse Lab in Milton Keynes | Wellcome Sanger Institute for the COVID-19 Genomics UK (COG-UK) Consortium | The Lighthouse Lab in Milton Keynes and Alex Alderton, Roberto Amato, Jeffrey Barrett, Sonia Goncalves, Ewan Harrison, David K. Jackson, Ian Johnston, Dominic Kwiatkowski, Cordelia Langford, John Sillitoe on behalf of the Wellcome Sanger Institute COVID-19 Surveillance Team |
| EPI_ISL_2198980 | Lighthouse Lab in Alderley Park | Wellcome Sanger Institute for the COVID-19 Genomics UK (COG-UK) Consortium | Jacquelyn Wynn, Mairead Hyland, The Lighthouse Lab in Alderley Park and Alex Alderton, Roberto Amato, Jeffrey Barrett, Sonia Goncalves, Ewan Harrison, David K. Jackson, Ian Johnston, Dominic Kwiatkowski, Cordelia Langford, John Sillitoe on behalf of the Wellcome Sanger Institute COVID-19 Surveillance Team |
| EPI_ISL_2198981 | Lighthouse Lab in Milton Keynes | Wellcome Sanger Institute for the COVID-19 Genomics UK (COG-UK) Consortium | The Lighthouse Lab in Milton Keynes and Alex Alderton, Roberto Amato, Jeffrey Barrett, Sonia Goncalves, Ewan Harrison, David K. Jackson, Ian Johnston, Dominic Kwiatkowski, Cordelia Langford, John Sillitoe on behalf of the Wellcome Sanger Institute COVID-19 Surveillance Team |
| EPI_ISL_2198986, EPI_ISL_2198987, EPI_ISL_2198988 | Lighthouse Lab in Alderley Park | Wellcome Sanger Institute for the COVID-19 Genomics UK (COG-UK) Consortium | Jacquelyn Wynn, Mairead Hyland, The Lighthouse Lab in Alderley Park and Alex Alderton, Roberto Amato, Jeffrey Barrett, Sonia Goncalves, Ewan Harrison, David K. Jackson, Ian Johnston, Dominic Kwiatkowski, Cordelia Langford, John Sillitoe on behalf of the Wellcome Sanger Institute COVID-19 Surveillance Team |
| EPI_ISL_2198993, EPI_ISL_2198994, EPI_ISL_2198997, EPI_ISL_2199001 | Lighthouse Lab in Milton Keynes | Wellcome Sanger Institute for the COVID-19 Genomics UK (COG-UK) Consortium | The Lighthouse Lab in Milton Keynes and Alex Alderton, Roberto Amato, Jeffrey Barrett, Sonia Goncalves, Ewan Harrison, David K. Jackson, Ian Johnston, Dominic Kwiatkowski, Cordelia Langford, John Sillitoe on behalf of the Wellcome Sanger Institute COVID-19 Surveillance Team |
| EPI_ISL_2199003 | Lighthouse Lab in Alderley Park | Wellcome Sanger Institute for the COVID-19 Genomics UK (COG-UK) Consortium | Jacquelyn Wynn, Mairead Hyland, The Lighthouse Lab in Alderley Park and Alex Alderton, Roberto Amato, Jeffrey Barrett, Sonia Goncalves, Ewan Harrison, David K. Jackson, Ian Johnston, Dominic Kwiatkowski, Cordelia Langford, John Sillitoe on behalf of the Wellcome Sanger Institute COVID-19 Surveillance Team |
| EPI_ISL_2199005, EPI_ISL_2199007, EPI_ISL_2199009 | Lighthouse Lab in Glasgow | Wellcome Sanger Institute for the COVID-19 Genomics UK (COG-UK) Consortium | Harper VanSteenhouse, Yumi Kasai, David Gray, Carol Clugston, Anna Dominiczak and Alex Alderton, Roberto Amato, Jeffrey Barrett, Sonia Goncalves, Ewan Harrison, David K. Jackson, Ian Johnston, Dominic Kwiatkowski, Cordelia Langford, John Sillitoe on behalf of the Wellcome Sanger Institute COVID-19 Surveillance Team |
| EPI_ISL_2199013 | Lighthouse Lab in Milton Keynes | Wellcome Sanger Institute for the COVID-19 Genomics UK (COG-UK) Consortium | The Lighthouse Lab in Milton Keynes and Alex Alderton, Roberto Amato, Jeffrey Barrett, Sonia Goncalves, Ewan Harrison, David K. Jackson, Ian Johnston, Dominic Kwiatkowski, Cordelia Langford, John Sillitoe on behalf of the Wellcome Sanger Institute COVID-19 Surveillance Team |
| EPI_ISL_2199014, EPI_ISL_2199017 | Lighthouse Lab in Glasgow | Wellcome Sanger Institute for the COVID-19 Genomics UK (COG-UK) Consortium | Harper VanSteenhouse, Yumi Kasai, David Gray, Carol Clugston, Anna Dominiczak and Alex Alderton, Roberto Amato, Jeffrey Barrett, Sonia Goncalves, Ewan Harrison, David K. Jackson, Ian Johnston, Dominic Kwiatkowski, Cordelia Langford, John Sillitoe on behalf of the Wellcome Sanger Institute COVID-19 Surveillance Team |
| EPI_ISL_2199018 | Lighthouse Lab in Milton Keynes | Wellcome Sanger Institute for the COVID-19 Genomics UK (COG-UK) Consortium | The Lighthouse Lab in Milton Keynes and Alex Alderton, Roberto Amato, Jeffrey Barrett, Sonia Goncalves, Ewan Harrison, David K. Jackson, Ian Johnston, Dominic Kwiatkowski, Cordelia Langford, John Sillitoe on behalf of the Wellcome Sanger Institute COVID-19 Surveillance Team |
| EPI_ISL_2199019, EPI_ISL_2199023 | Lighthouse Lab in Glasgow | Wellcome Sanger Institute for the COVID-19 Genomics UK (COG-UK) Consortium | Harper VanSteenhouse, Yumi Kasai, David Gray, Carol Clugston, Anna Dominiczak and Alex Alderton, Roberto Amato, Jeffrey Barrett, Sonia Goncalves, Ewan Harrison, David K. Jackson, Ian Johnston, Dominic Kwiatkowski, Cordelia Langford, John Sillitoe on behalf of the Wellcome Sanger Institute COVID-19 Surveillance Team |
| EPI_ISL_2199024 | Lighthouse Lab in Alderley Park | Wellcome Sanger Institute for the COVID-19 Genomics UK (COG-UK) Consortium | Jacquelyn Wynn, Mairead Hyland, The Lighthouse Lab in Alderley Park and Alex Alderton, Roberto Amato, Jeffrey Barrett, Sonia Goncalves, Ewan Harrison, David K. Jackson, Ian Johnston, Dominic Kwiatkowski, Cordelia Langford, John Sillitoe on behalf of the Wellcome Sanger Institute COVID-19 Surveillance Team |
| EPI_ISL_2199027 | Lighthouse Lab in Milton Keynes | Wellcome Sanger Institute for the COVID-19 Genomics UK (COG-UK) Consortium | The Lighthouse Lab in Milton Keynes and Alex Alderton, Roberto Amato, Jeffrey Barrett, Sonia Goncalves, Ewan Harrison, David K. Jackson, Ian Johnston, Dominic Kwiatkowski, Cordelia Langford, John Sillitoe on behalf of the Wellcome Sanger Institute COVID-19 Surveillance Team |
| EPI_ISL_2199032 | Lighthouse Lab in Alderley Park | Wellcome Sanger Institute for the COVID-19 Genomics UK (COG-UK) Consortium | Jacquelyn Wynn, Mairead Hyland, The Lighthouse Lab in Alderley Park and Alex Alderton, Roberto Amato, Jeffrey Barrett, Sonia Goncalves, Ewan Harrison, David K. Jackson, Ian Johnston, Dominic Kwiatkowski, Cordelia Langford, John Sillitoe on behalf of the Wellcome Sanger Institute COVID-19 Surveillance Team |
| EPI_ISL_2199033 | Lighthouse Lab in Glasgow | Wellcome Sanger Institute for the COVID-19 Genomics UK (COG-UK) Consortium | Harper VanSteenhouse, Yumi Kasai, David Gray, Carol Clugston, Anna Dominiczak and Alex Alderton, Roberto Amato, Jeffrey Barrett, Sonia Goncalves, Ewan Harrison, David K. Jackson, Ian Johnston, Dominic Kwiatkowski, Cordelia Langford, John Sillitoe on behalf of the Wellcome Sanger Institute COVID-19 Surveillance Team |
| EPI_ISL_2199034, EPI_ISL_2199035 | Lighthouse Lab in Alderley Park | Wellcome Sanger Institute for the COVID-19 Genomics UK (COG-UK) Consortium | Jacquelyn Wynn, Mairead Hyland, The Lighthouse Lab in Alderley Park and Alex Alderton, Roberto Amato, Jeffrey Barrett, Sonia Goncalves, Ewan Harrison, David K. Jackson, Ian Johnston, Dominic Kwiatkowski, Cordelia Langford, John Sillitoe on behalf of the Wellcome Sanger Institute COVID-19 Surveillance Team |
| EPI_ISL_2199036 | Lighthouse Lab in Milton Keynes | Wellcome Sanger Institute for the COVID-19 Genomics UK (COG-UK) Consortium | The Lighthouse Lab in Milton Keynes and Alex Alderton, Roberto Amato, Jeffrey Barrett, Sonia Goncalves, Ewan Harrison, David K. Jackson, Ian Johnston, Dominic Kwiatkowski, Cordelia Langford, John Sillitoe on behalf of the Wellcome Sanger Institute COVID-19 Surveillance Team |
| EPI_ISL_2199040, EPI_ISL_2199041, EPI_ISL_2199042, EPI_ISL_2199046, EPI_ISL_2199050 | Lighthouse Lab in Glasgow | Wellcome Sanger Institute for the COVID-19 Genomics UK (COG-UK) Consortium | Harper VanSteenhouse, Yumi Kasai, David Gray, Carol Clugston, Anna Dominiczak and Alex Alderton, Roberto Amato, Jeffrey Barrett, Sonia Goncalves, Ewan Harrison, David K. Jackson, Ian Johnston, Dominic Kwiatkowski, Cordelia Langford, John Sillitoe on behalf of the Wellcome Sanger Institute COVID-19 Surveillance Team |
| EPI_ISL_2199051, EPI_ISL_2199055, EPI_ISL_2199061 | Lighthouse Lab in Milton Keynes | Wellcome Sanger Institute for the COVID-19 Genomics UK (COG-UK) Consortium | The Lighthouse Lab in Milton Keynes and Alex Alderton, Roberto Amato, Jeffrey Barrett, Sonia Goncalves, Ewan Harrison, David K. Jackson, Ian Johnston, Dominic Kwiatkowski, Cordelia Langford, John Sillitoe on behalf of the Wellcome Sanger Institute COVID-19 Surveillance Team |
| EPI_ISL_2199063, EPI_ISL_2199065, EPI_ISL_2199067, EPI_ISL_2199070 | Lighthouse Lab in Glasgow | Wellcome Sanger Institute for the COVID-19 Genomics UK (COG-UK) Consortium | Harper VanSteenhouse, Yumi Kasai, David Gray, Carol Clugston, Anna Dominiczak and Alex Alderton, Roberto Amato, Jeffrey Barrett, Sonia Goncalves, Ewan Harrison, David K. Jackson, Ian Johnston, Dominic Kwiatkowski, Cordelia Langford, John Sillitoe on behalf of the Wellcome Sanger Institute COVID-19 Surveillance Team |
| EPI_ISL_2199072 | Lighthouse Lab in Alderley Park | Wellcome Sanger Institute for the COVID-19 Genomics UK (COG-UK) Consortium | Jacquelyn Wynn, Mairead Hyland, The Lighthouse Lab in Alderley Park and Alex Alderton, Roberto Amato, Jeffrey Barrett, Sonia Goncalves, Ewan Harrison, David K. Jackson, Ian Johnston, Dominic Kwiatkowski, Cordelia Langford, John Sillitoe on behalf of the Wellcome Sanger Institute COVID-19 Surveillance Team |
| EPI_ISL_2199075 | Lighthouse Lab in Milton Keynes | Wellcome Sanger Institute for the COVID-19 Genomics UK (COG-UK) Consortium | The Lighthouse Lab in Milton Keynes and Alex Alderton, Roberto Amato, Jeffrey Barrett, Sonia Goncalves, Ewan Harrison, David K. Jackson, Ian Johnston, Dominic Kwiatkowski, Cordelia Langford, John Sillitoe on behalf of the Wellcome Sanger Institute COVID-19 Surveillance Team |
| EPI_ISL_2199077 | Lighthouse Lab in Alderley Park | Wellcome Sanger Institute for the COVID-19 Genomics UK (COG-UK) Consortium | Jacquelyn Wynn, Mairead Hyland, The Lighthouse Lab in Alderley Park and Alex Alderton, Roberto Amato, Jeffrey Barrett, Sonia Goncalves, Ewan Harrison, David K. Jackson, Ian Johnston, Dominic Kwiatkowski, Cordelia Langford, John Sillitoe on behalf of the Wellcome Sanger Institute COVID-19 Surveillance Team |
| EPI_ISL_2199078 | Lighthouse Lab in Glasgow | Wellcome Sanger Institute for the COVID-19 Genomics UK (COG-UK) Consortium | Harper VanSteenhouse, Yumi Kasai, David Gray, Carol Clugston, Anna Dominiczak and Alex Alderton, Roberto Amato, Jeffrey Barrett, Sonia Goncalves, Ewan Harrison, David K. Jackson, Ian Johnston, Dominic Kwiatkowski, Cordelia Langford, John Sillitoe on behalf of the Wellcome Sanger Institute COVID-19 Surveillance Team |

| | | | |
|---|---|---|---|
| EPI_ISL_2199079 | Lighthouse Lab in Alderley Park | Wellcome Sanger Institute for the COVID-19 Genomics UK (COG-UK) Consortium | Jacquelyn Wynn, Mairead Hyland, The Lighthouse Lab in Alderley Park and Alex Alderton, Roberto Amato, Jeffrey Barrett, Sonia Goncalves, Ewan Harrison, David K. Jackson, Ian Johnston, Dominic Kwiatkowski, Cordelia Langford, John Sillitoe on behalf of the Wellcome Sanger Institute COVID-19 Surveillance Team |
| EPI_ISL_2199081, EPI_ISL_2199082 | Lighthouse Lab in Glasgow | Wellcome Sanger Institute for the COVID-19 Genomics UK (COG-UK) Consortium | Harper VanSteenhouse, Yumi Kasai, David Gray, Carol Clugston, Anna Dominiczak and Alex Alderton, Roberto Amato, Jeffrey Barrett, Sonia Goncalves, Ewan Harrison, David K. Jackson, Ian Johnston, Dominic Kwiatkowski, Cordelia Langford, John Sillitoe on behalf of the Wellcome Sanger Institute COVID-19 Surveillance Team |
| EPI_ISL_2199083, EPI_ISL_2199085 | Lighthouse Lab in Alderley Park | Wellcome Sanger Institute for the COVID-19 Genomics UK (COG-UK) Consortium | Jacquelyn Wynn, Mairead Hyland, The Lighthouse Lab in Alderley Park and Alex Alderton, Roberto Amato, Jeffrey Barrett, Sonia Goncalves, Ewan Harrison, David K. Jackson, Ian Johnston, Dominic Kwiatkowski, Cordelia Langford, John Sillitoe on behalf of the Wellcome Sanger Institute COVID-19 Surveillance Team |
| EPI_ISL_2199087 | Lighthouse Lab in Glasgow | Wellcome Sanger Institute for the COVID-19 Genomics UK (COG-UK) Consortium | Harper VanSteenhouse, Yumi Kasai, David Gray, Carol Clugston, Anna Dominiczak and Alex Alderton, Roberto Amato, Jeffrey Barrett, Sonia Goncalves, Ewan Harrison, David K. Jackson, Ian Johnston, Dominic Kwiatkowski, Cordelia Langford, John Sillitoe on behalf of the Wellcome Sanger Institute COVID-19 Surveillance Team |
| EPI_ISL_2199088 | Lighthouse Lab in Alderley Park | Wellcome Sanger Institute for the COVID-19 Genomics UK (COG-UK) Consortium | Jacquelyn Wynn, Mairead Hyland, The Lighthouse Lab in Alderley Park and Alex Alderton, Roberto Amato, Jeffrey Barrett, Sonia Goncalves, Ewan Harrison, David K. Jackson, Ian Johnston, Dominic Kwiatkowski, Cordelia Langford, John Sillitoe on behalf of the Wellcome Sanger Institute COVID-19 Surveillance Team |
| EPI_ISL_2199089 | Lighthouse Lab in Glasgow | Wellcome Sanger Institute for the COVID-19 Genomics UK (COG-UK) Consortium | Harper VanSteenhouse, Yumi Kasai, David Gray, Carol Clugston, Anna Dominiczak and Alex Alderton, Roberto Amato, Jeffrey Barrett, Sonia Goncalves, Ewan Harrison, David K. Jackson, Ian Johnston, Dominic Kwiatkowski, Cordelia Langford, John Sillitoe on behalf of the Wellcome Sanger Institute COVID-19 Surveillance Team |
| EPI_ISL_2199090 | Lighthouse Lab in Alderley Park | Wellcome Sanger Institute for the COVID-19 Genomics UK (COG-UK) Consortium | Jacquelyn Wynn, Mairead Hyland, The Lighthouse Lab in Alderley Park and Alex Alderton, Roberto Amato, Jeffrey Barrett, Sonia Goncalves, Ewan Harrison, David K. Jackson, Ian Johnston, Dominic Kwiatkowski, Cordelia Langford, John Sillitoe on behalf of the Wellcome Sanger Institute COVID-19 Surveillance Team |
| EPI_ISL_2199094, EPI_ISL_2199095 | Lighthouse Lab in Glasgow | Wellcome Sanger Institute for the COVID-19 Genomics UK (COG-UK) Consortium | Harper VanSteenhouse, Yumi Kasai, David Gray, Carol Clugston, Anna Dominiczak and Alex Alderton, Roberto Amato, Jeffrey Barrett, Sonia Goncalves, Ewan Harrison, David K. Jackson, Ian Johnston, Dominic Kwiatkowski, Cordelia Langford, John Sillitoe on behalf of the Wellcome Sanger Institute COVID-19 Surveillance Team |
| EPI_ISL_2199098 | Lighthouse Lab in Milton Keynes | Wellcome Sanger Institute for the COVID-19 Genomics UK (COG-UK) Consortium | The Lighthouse Lab in Milton Keynes and Alex Alderton, Roberto Amato, Jeffrey Barrett, Sonia Goncalves, Ewan Harrison, David K. Jackson, Ian Johnston, Dominic Kwiatkowski, Cordelia Langford, John Sillitoe on behalf of the Wellcome Sanger Institute COVID-19 Surveillance Team |
| EPI_ISL_2199100 | Lighthouse Lab in Alderley Park | Wellcome Sanger Institute for the COVID-19 Genomics UK (COG-UK) Consortium | Jacquelyn Wynn, Mairead Hyland, The Lighthouse Lab in Alderley Park and Alex Alderton, Roberto Amato, Jeffrey Barrett, Sonia Goncalves, Ewan Harrison, David K. Jackson, Ian Johnston, Dominic Kwiatkowski, Cordelia Langford, John Sillitoe on behalf of the Wellcome Sanger Institute COVID-19 Surveillance Team |
| EPI_ISL_2199101 | Lighthouse Lab in Milton Keynes | Wellcome Sanger Institute for the COVID-19 Genomics UK (COG-UK) Consortium | The Lighthouse Lab in Milton Keynes and Alex Alderton, Roberto Amato, Jeffrey Barrett, Sonia Goncalves, Ewan Harrison, David K. Jackson, Ian Johnston, Dominic Kwiatkowski, Cordelia Langford, John Sillitoe on behalf of the Wellcome Sanger Institute COVID-19 Surveillance Team |
| EPI_ISL_2199104, EPI_ISL_2199107 | Lighthouse Lab in Glasgow | Wellcome Sanger Institute for the COVID-19 Genomics UK (COG-UK) Consortium | Harper VanSteenhouse, Yumi Kasai, David Gray, Carol Clugston, Anna Dominiczak and Alex Alderton, Roberto Amato, Jeffrey Barrett, Sonia Goncalves, Ewan Harrison, David K. Jackson, Ian Johnston, Dominic Kwiatkowski, Cordelia Langford, John Sillitoe on behalf of the Wellcome Sanger Institute COVID-19 Surveillance Team |
| EPI_ISL_2199108 | Lighthouse Lab in Alderley Park | Wellcome Sanger Institute for the COVID-19 Genomics UK (COG-UK) Consortium | Jacquelyn Wynn, Mairead Hyland, The Lighthouse Lab in Alderley Park and Alex Alderton, Roberto Amato, Jeffrey Barrett, Sonia Goncalves, Ewan Harrison, David K. Jackson, Ian Johnston, Dominic Kwiatkowski, Cordelia Langford, John Sillitoe on behalf of the Wellcome Sanger Institute COVID-19 Surveillance Team |
| EPI_ISL_2199110 | Lighthouse Lab in Milton Keynes | Wellcome Sanger Institute for the COVID-19 Genomics UK (COG-UK) Consortium | The Lighthouse Lab in Milton Keynes and Alex Alderton, Roberto Amato, Jeffrey Barrett, Sonia Goncalves, Ewan Harrison, David K. Jackson, Ian Johnston, Dominic Kwiatkowski, Cordelia Langford, John Sillitoe on behalf of the Wellcome Sanger Institute COVID-19 Surveillance Team |
| EPI_ISL_2199111, EPI_ISL_2199114 | Lighthouse Lab in Alderley Park | Wellcome Sanger Institute for the COVID-19 Genomics UK (COG-UK) Consortium | Jacquelyn Wynn, Mairead Hyland, The Lighthouse Lab in Alderley Park and Alex Alderton, Roberto Amato, Jeffrey Barrett, Sonia Goncalves, Ewan Harrison, David K. Jackson, Ian Johnston, Dominic Kwiatkowski, Cordelia Langford, John Sillitoe on behalf of the Wellcome Sanger Institute COVID-19 Surveillance Team |
| EPI_ISL_2199118 | Lighthouse Lab in Glasgow | Wellcome Sanger Institute for the COVID-19 Genomics UK (COG-UK) Consortium | Harper VanSteenhouse, Yumi Kasai, David Gray, Carol Clugston, Anna Dominiczak and Alex Alderton, Roberto Amato, Jeffrey Barrett, Sonia Goncalves, Ewan Harrison, David K. Jackson, Ian Johnston, Dominic Kwiatkowski, Cordelia Langford, John Sillitoe on behalf of the Wellcome Sanger Institute COVID-19 Surveillance Team |
| EPI_ISL_2199119 | Lighthouse Lab in Alderley Park | Wellcome Sanger Institute for the COVID-19 Genomics UK (COG-UK) Consortium | Jacquelyn Wynn, Mairead Hyland, The Lighthouse Lab in Alderley Park and Alex Alderton, Roberto Amato, Jeffrey Barrett, Sonia Goncalves, Ewan Harrison, David K. Jackson, Ian Johnston, Dominic Kwiatkowski, Cordelia Langford, John Sillitoe on behalf of the Wellcome Sanger Institute COVID-19 Surveillance Team |
| EPI_ISL_2199120 | Lighthouse Lab in Glasgow | Wellcome Sanger Institute for the COVID-19 Genomics UK (COG-UK) Consortium | Harper VanSteenhouse, Yumi Kasai, David Gray, Carol Clugston, Anna Dominiczak and Alex Alderton, Roberto Amato, Jeffrey Barrett, Sonia Goncalves, Ewan Harrison, David K. Jackson, Ian Johnston, Dominic Kwiatkowski, Cordelia Langford, John Sillitoe on behalf of the Wellcome Sanger Institute COVID-19 Surveillance Team |
| EPI_ISL_2199121, EPI_ISL_2199122, EPI_ISL_2199123 | Lighthouse Lab in Alderley Park | Wellcome Sanger Institute for the COVID-19 Genomics UK (COG-UK) Consortium | Jacquelyn Wynn, Mairead Hyland, The Lighthouse Lab in Alderley Park and Alex Alderton, Roberto Amato, Jeffrey Barrett, Sonia Goncalves, Ewan Harrison, David K. Jackson, Ian Johnston, Dominic Kwiatkowski, Cordelia Langford, John Sillitoe on behalf of the Wellcome Sanger Institute COVID-19 Surveillance Team |
| EPI_ISL_2199124, EPI_ISL_2199126 | Lighthouse Lab in Milton Keynes | Wellcome Sanger Institute for the COVID-19 Genomics UK (COG-UK) Consortium | The Lighthouse Lab in Milton Keynes and Alex Alderton, Roberto Amato, Jeffrey Barrett, Sonia Goncalves, Ewan Harrison, David K. Jackson, Ian Johnston, Dominic Kwiatkowski, Cordelia Langford, John Sillitoe on behalf of the Wellcome Sanger Institute COVID-19 Surveillance Team |
| EPI_ISL_2199129 | Lighthouse Lab in Glasgow | Wellcome Sanger Institute for the COVID-19 Genomics UK (COG-UK) Consortium | Harper VanSteenhouse, Yumi Kasai, David Gray, Carol Clugston, Anna Dominiczak and Alex Alderton, Roberto Amato, Jeffrey Barrett, Sonia Goncalves, Ewan Harrison, David K. Jackson, Ian Johnston, Dominic Kwiatkowski, Cordelia Langford, John Sillitoe on behalf of the Wellcome Sanger Institute COVID-19 Surveillance Team |
| EPI_ISL_2199130 | Lighthouse Lab in Alderley Park | Wellcome Sanger Institute for the COVID-19 Genomics UK (COG-UK) Consortium | Jacquelyn Wynn, Mairead Hyland, The Lighthouse Lab in Alderley Park and Alex Alderton, Roberto Amato, Jeffrey Barrett, Sonia Goncalves, Ewan Harrison, David K. Jackson, Ian Johnston, Dominic Kwiatkowski, Cordelia Langford, John Sillitoe on behalf of the Wellcome Sanger Institute COVID-19 Surveillance Team |
| EPI_ISL_2199131 | Lighthouse Lab in Milton Keynes | Wellcome Sanger Institute for the COVID-19 Genomics UK (COG-UK) Consortium | The Lighthouse Lab in Milton Keynes and Alex Alderton, Roberto Amato, Jeffrey Barrett, Sonia Goncalves, Ewan Harrison, David K. Jackson, Ian Johnston, Dominic Kwiatkowski, Cordelia Langford, John Sillitoe on behalf of the Wellcome Sanger Institute COVID-19 Surveillance Team |
| EPI_ISL_2199132, EPI_ISL_2199133, EPI_ISL_2199138, EPI_ISL_2199143, EPI_ISL_2199151, EPI_ISL_2199155 | Lighthouse Lab in Glasgow | Wellcome Sanger Institute for the COVID-19 Genomics UK (COG-UK) Consortium | Harper VanSteenhouse, Yumi Kasai, David Gray, Carol Clugston, Anna Dominiczak and Alex Alderton, Roberto Amato, Jeffrey Barrett, Sonia Goncalves, Ewan Harrison, David K. Jackson, Ian Johnston, Dominic Kwiatkowski, Cordelia Langford, John Sillitoe on behalf of the Wellcome Sanger Institute COVID-19 Surveillance Team |
| EPI_ISL_2199156 | Lighthouse Lab in Alderley Park | Wellcome Sanger Institute for the COVID-19 Genomics UK (COG-UK) Consortium | Jacquelyn Wynn, Mairead Hyland, The Lighthouse Lab in Alderley Park and Alex Alderton, Roberto Amato, Jeffrey Barrett, Sonia Goncalves, Ewan Harrison, David K. Jackson, Ian Johnston, Dominic Kwiatkowski, Cordelia Langford, John Sillitoe on behalf of the Wellcome Sanger Institute COVID-19 Surveillance Team |
| EPI_ISL_2199160 | Lighthouse Lab in Milton Keynes | Wellcome Sanger Institute for the COVID-19 Genomics UK (COG-UK) Consortium | The Lighthouse Lab in Milton Keynes and Alex Alderton, Roberto Amato, Jeffrey Barrett, Sonia Goncalves, Ewan Harrison, David K. Jackson, Ian Johnston, Dominic Kwiatkowski, Cordelia Langford, John Sillitoe on behalf of the Wellcome Sanger Institute COVID-19 Surveillance Team |
| EPI_ISL_2199161, EPI_ISL_2199164 | Lighthouse Lab in Alderley Park | Wellcome Sanger Institute for the COVID-19 Genomics UK (COG-UK) Consortium | Jacquelyn Wynn, Mairead Hyland, The Lighthouse Lab in Alderley Park and Alex Alderton, Roberto Amato, Jeffrey Barrett, Sonia Goncalves, Ewan Harrison, David K. Jackson, Ian Johnston, Dominic Kwiatkowski, Cordelia Langford, John Sillitoe on behalf of the Wellcome Sanger Institute COVID-19 Surveillance Team |
| EPI_ISL_2199167 | Lighthouse Lab in Milton Keynes | Wellcome Sanger Institute for the COVID-19 Genomics UK (COG-UK) Consortium | The Lighthouse Lab in Milton Keynes and Alex Alderton, Roberto Amato, Jeffrey Barrett, Sonia Goncalves, Ewan Harrison, David K. Jackson, Ian Johnston, Dominic Kwiatkowski, Cordelia Langford, John Sillitoe on behalf of the Wellcome Sanger Institute COVID-19 Surveillance Team |
| EPI_ISL_2199169, EPI_ISL_2199172, EPI_ISL_2199174 | Lighthouse Lab in Glasgow | Wellcome Sanger Institute for the COVID-19 Genomics UK (COG-UK) Consortium | Harper VanSteenhouse, Yumi Kasai, David Gray, Carol Clugston, Anna Dominiczak and Alex Alderton, Roberto Amato, Jeffrey Barrett, Sonia Goncalves, Ewan Harrison, David K. Jackson, Ian Johnston, Dominic Kwiatkowski, Cordelia Langford, John Sillitoe on behalf of the Wellcome Sanger Institute COVID-19 Surveillance Team |
| EPI_ISL_2199177 | Lighthouse Lab in Milton Keynes | Wellcome Sanger Institute for the COVID-19 Genomics UK (COG-UK) Consortium | The Lighthouse Lab in Milton Keynes and Alex Alderton, Roberto Amato, Jeffrey Barrett, Sonia Goncalves, Ewan Harrison, David K. Jackson, Ian Johnston, Dominic Kwiatkowski, Cordelia Langford, John Sillitoe on behalf of the Wellcome Sanger Institute COVID-19 Surveillance Team |
| EPI_ISL_2199180 | Lighthouse Lab in Alderley Park | Wellcome Sanger Institute for the COVID-19 Genomics UK (COG-UK) Consortium | Jacquelyn Wynn, Mairead Hyland, The Lighthouse Lab in Alderley Park and Alex Alderton, Roberto Amato, Jeffrey Barrett, Sonia Goncalves, Ewan Harrison, David K. Jackson, Ian Johnston, Dominic Kwiatkowski, Cordelia Langford, John Sillitoe on behalf of the Wellcome Sanger Institute COVID-19 Surveillance Team |
| EPI_ISL_2199182 | Lighthouse Lab in Milton Keynes | Wellcome Sanger Institute for the COVID-19 Genomics UK (COG-UK) Consortium | The Lighthouse Lab in Milton Keynes and Alex Alderton, Roberto Amato, Jeffrey Barrett, Sonia Goncalves, Ewan Harrison, David K. Jackson, Ian Johnston, Dominic Kwiatkowski, Cordelia Langford, John Sillitoe on behalf of the Wellcome Sanger Institute COVID-19 Surveillance Team |
| EPI_ISL_2199183, EPI_ISL_2199185, EPI_ISL_2199187 | Lighthouse Lab in Glasgow | Wellcome Sanger Institute for the COVID-19 Genomics UK (COG-UK) Consortium | Harper VanSteenhouse, Yumi Kasai, David Gray, Carol Clugston, Anna Dominiczak and Alex Alderton, Roberto Amato, Jeffrey Barrett, Sonia Goncalves, Ewan Harrison, David K. Jackson, Ian Johnston, Dominic Kwiatkowski, Cordelia Langford, John Sillitoe on behalf of the Wellcome Sanger Institute COVID-19 Surveillance Team |
| EPI_ISL_2199191, EPI_ISL_2199198 | Lighthouse Lab in Alderley Park | Wellcome Sanger Institute for the COVID-19 Genomics UK (COG-UK) Consortium | Jacquelyn Wynn, Mairead Hyland, The Lighthouse Lab in Alderley Park and Alex Alderton, Roberto Amato, Jeffrey Barrett, Sonia Goncalves, Ewan Harrison, David K. Jackson, Ian Johnston, Dominic Kwiatkowski, Cordelia Langford, John Sillitoe on behalf of the Wellcome Sanger Institute COVID-19 Surveillance Team |
| EPI_ISL_2199203 | Lighthouse Lab in Glasgow | Wellcome Sanger Institute for the COVID-19 Genomics UK (COG-UK) Consortium | Harper VanSteenhouse, Yumi Kasai, David Gray, Carol Clugston, Anna Dominiczak and Alex Alderton, Roberto Amato, Jeffrey Barrett, Sonia Goncalves, Ewan Harrison, David K. Jackson, Ian Johnston, Dominic Kwiatkowski, Cordelia Langford, John Sillitoe on behalf of the Wellcome Sanger Institute COVID-19 Surveillance Team |
| EPI_ISL_2199205 | Lighthouse Lab in Alderley Park | Wellcome Sanger Institute for the COVID-19 Genomics UK (COG-UK) Consortium | Jacquelyn Wynn, Mairead Hyland, The Lighthouse Lab in Alderley Park and Alex Alderton, Roberto Amato, Jeffrey Barrett, Sonia Goncalves, Ewan Harrison, David K. Jackson, Ian Johnston, Dominic Kwiatkowski, Cordelia Langford, John Sillitoe on behalf of the Wellcome Sanger Institute COVID-19 Surveillance Team |
| EPI_ISL_2199209 | Lighthouse Lab in Milton Keynes | Wellcome Sanger Institute for the COVID-19 Genomics UK (COG-UK) Consortium | The Lighthouse Lab in Milton Keynes and Alex Alderton, Roberto Amato, Jeffrey Barrett, Sonia Goncalves, Ewan Harrison, David K. Jackson, Ian Johnston, Dominic Kwiatkowski, Cordelia Langford, John Sillitoe on behalf of the Wellcome Sanger Institute COVID-19 Surveillance Team |
| EPI_ISL_2199212 | Lighthouse Lab in Glasgow | Wellcome Sanger Institute for the COVID-19 Genomics UK (COG-UK) Consortium | Harper VanSteenhouse, Yumi Kasai, David Gray, Carol Clugston, Anna Dominiczak and Alex Alderton, Roberto Amato, Jeffrey Barrett, Sonia Goncalves, Ewan Harrison, David K. Jackson, Ian Johnston, Dominic Kwiatkowski, Cordelia Langford, John Sillitoe on behalf of the Wellcome Sanger Institute COVID-19 Surveillance Team |
| EPI_ISL_2199214 | Lighthouse Lab in Milton Keynes | Wellcome Sanger Institute for the COVID-19 Genomics UK (COG-UK) Consortium | The Lighthouse Lab in Milton Keynes and Alex Alderton, Roberto Amato, Jeffrey Barrett, Sonia Goncalves, Ewan Harrison, David K. Jackson, Ian Johnston, Dominic Kwiatkowski, Cordelia Langford, John Sillitoe on behalf of the Wellcome Sanger Institute COVID-19 Surveillance Team |
| EPI_ISL_2199219 | Lighthouse Lab in Alderley Park | Wellcome Sanger Institute for the COVID-19 Genomics UK (COG-UK) Consortium | Jacquelyn Wynn, Mairead Hyland, The Lighthouse Lab in Alderley Park and Alex Alderton, Roberto Amato, Jeffrey Barrett, Sonia Goncalves, Ewan Harrison, David K. Jackson, Ian Johnston, Dominic Kwiatkowski, Cordelia Langford, John Sillitoe on behalf of the Wellcome Sanger Institute COVID-19 Surveillance Team |
| EPI_ISL_2199222 | Lighthouse Lab in Glasgow | Wellcome Sanger Institute for the COVID-19 Genomics UK (COG-UK) Consortium | Harper VanSteenhouse, Yumi Kasai, David Gray, Carol Clugston, Anna Dominiczak and Alex Alderton, Roberto Amato, Jeffrey Barrett, Sonia Goncalves, Ewan Harrison, David K. Jackson, Ian Johnston, Dominic Kwiatkowski, Cordelia Langford, John Sillitoe on behalf of the Wellcome Sanger Institute COVID-19 Surveillance Team |
| EPI_ISL_2199226 | Lighthouse Lab in Milton Keynes | Wellcome Sanger Institute for the COVID-19 Genomics UK (COG-UK) Consortium | The Lighthouse Lab in Milton Keynes and Alex Alderton, Roberto Amato, Jeffrey Barrett, Sonia Goncalves, Ewan Harrison, David K. Jackson, Ian Johnston, Dominic Kwiatkowski, Cordelia Langford, John Sillitoe on behalf of the Wellcome Sanger Institute COVID-19 Surveillance Team |
| EPI_ISL_2199230, | Lighthouse Lab in Glasgow | Wellcome Sanger Institute for the COVID-19 Genomics UK (COG-UK) | Harper VanSteenhouse, Yumi Kasai, David Gray, Carol Clugston, Anna Dominiczak and Alex Alderton, Roberto Amato, Jeffrey Barrett, Sonia Goncalves, Ewan Harrison, David K. Jackson, Ian Johnston, Dominic Kwiatkowski, Cordelia Langford, |

| | | | |
|---|---|---|---|
| EPI_ISL_2199231 | | Consortium | John Sillitoe on behalf of the Wellcome Sanger Institute COVID-19 Surveillance Team |
| EPI_ISL_2199232 | Lighthouse Lab in Milton Keynes | Wellcome Sanger Institute for the COVID-19 Genomics UK (COG-UK) Consortium | The Lighthouse Lab in Milton Keynes and Alex Alderton, Roberto Amato, Jeffrey Barrett, Sonia Goncalves, Ewan Harrison, David K. Jackson, Ian Johnston, Dominic Kwiatkowski, Cordelia Langford, John Sillitoe on behalf of the Wellcome Sanger Institute COVID-19 Surveillance Team |
| EPI_ISL_2199234 | Lighthouse Lab in Glasgow | Wellcome Sanger Institute for the COVID-19 Genomics UK (COG-UK) Consortium | Harper VanSteenhouse, Yumi Kasai, David Gray, Carol Clugston, Anna Dominiczak and Alex Alderton, Roberto Amato, Jeffrey Barrett, Sonia Goncalves, Ewan Harrison, David K. Jackson, Ian Johnston, Dominic Kwiatkowski, Cordelia Langford, John Sillitoe on behalf of the Wellcome Sanger Institute COVID-19 Surveillance Team |
| EPI_ISL_2199236, EPI_ISL_2199242 | Lighthouse Lab in Alderley Park | Wellcome Sanger Institute for the COVID-19 Genomics UK (COG-UK) Consortium | Jacquelyn Wynn, Mairead Hyland, The Lighthouse Lab in Alderley Park and Alex Alderton, Roberto Amato, Jeffrey Barrett, Sonia Goncalves, Ewan Harrison, David K. Jackson, Ian Johnston, Dominic Kwiatkowski, Cordelia Langford, John Sillitoe on behalf of the Wellcome Sanger Institute COVID-19 Surveillance Team |
| EPI_ISL_2199246 | Lighthouse Lab in Glasgow | Wellcome Sanger Institute for the COVID-19 Genomics UK (COG-UK) Consortium | Harper VanSteenhouse, Yumi Kasai, David Gray, Carol Clugston, Anna Dominiczak and Alex Alderton, Roberto Amato, Jeffrey Barrett, Sonia Goncalves, Ewan Harrison, David K. Jackson, Ian Johnston, Dominic Kwiatkowski, Cordelia Langford, John Sillitoe on behalf of the Wellcome Sanger Institute COVID-19 Surveillance Team |
| EPI_ISL_2199247, EPI_ISL_2199250, EPI_ISL_2199251, EPI_ISL_2199253, EPI_ISL_2199254, EPI_ISL_2199257 | Lighthouse Lab in Alderley Park | Wellcome Sanger Institute for the COVID-19 Genomics UK (COG-UK) Consortium | Jacquelyn Wynn, Mairead Hyland, The Lighthouse Lab in Alderley Park and Alex Alderton, Roberto Amato, Jeffrey Barrett, Sonia Goncalves, Ewan Harrison, David K. Jackson, Ian Johnston, Dominic Kwiatkowski, Cordelia Langford, John Sillitoe on behalf of the Wellcome Sanger Institute COVID-19 Surveillance Team |
| EPI_ISL_2199261 | Lighthouse Lab in Glasgow | Wellcome Sanger Institute for the COVID-19 Genomics UK (COG-UK) Consortium | Harper VanSteenhouse, Yumi Kasai, David Gray, Carol Clugston, Anna Dominiczak and Alex Alderton, Roberto Amato, Jeffrey Barrett, Sonia Goncalves, Ewan Harrison, David K. Jackson, Ian Johnston, Dominic Kwiatkowski, Cordelia Langford, John Sillitoe on behalf of the Wellcome Sanger Institute COVID-19 Surveillance Team |
| EPI_ISL_2199263, EPI_ISL_2199268 | Lighthouse Lab in Milton Keynes | Wellcome Sanger Institute for the COVID-19 Genomics UK (COG-UK) Consortium | The Lighthouse Lab in Milton Keynes and Alex Alderton, Roberto Amato, Jeffrey Barrett, Sonia Goncalves, Ewan Harrison, David K. Jackson, Ian Johnston, Dominic Kwiatkowski, Cordelia Langford, John Sillitoe on behalf of the Wellcome Sanger Institute COVID-19 Surveillance Team |
| EPI_ISL_2199271 | Lighthouse Lab in Alderley Park | Wellcome Sanger Institute for the COVID-19 Genomics UK (COG-UK) Consortium | Jacquelyn Wynn, Mairead Hyland, The Lighthouse Lab in Alderley Park and Alex Alderton, Roberto Amato, Jeffrey Barrett, Sonia Goncalves, Ewan Harrison, David K. Jackson, Ian Johnston, Dominic Kwiatkowski, Cordelia Langford, John Sillitoe on behalf of the Wellcome Sanger Institute COVID-19 Surveillance Team |
| EPI_ISL_2199276 | Lighthouse Lab in Glasgow | Wellcome Sanger Institute for the COVID-19 Genomics UK (COG-UK) Consortium | Harper VanSteenhouse, Yumi Kasai, David Gray, Carol Clugston, Anna Dominiczak and Alex Alderton, Roberto Amato, Jeffrey Barrett, Sonia Goncalves, Ewan Harrison, David K. Jackson, Ian Johnston, Dominic Kwiatkowski, Cordelia Langford, John Sillitoe on behalf of the Wellcome Sanger Institute COVID-19 Surveillance Team |
| EPI_ISL_2199280, EPI_ISL_2199283, EPI_ISL_2199285 | Lighthouse Lab in Alderley Park | Wellcome Sanger Institute for the COVID-19 Genomics UK (COG-UK) Consortium | Jacquelyn Wynn, Mairead Hyland, The Lighthouse Lab in Alderley Park and Alex Alderton, Roberto Amato, Jeffrey Barrett, Sonia Goncalves, Ewan Harrison, David K. Jackson, Ian Johnston, Dominic Kwiatkowski, Cordelia Langford, John Sillitoe on behalf of the Wellcome Sanger Institute COVID-19 Surveillance Team |
| EPI_ISL_2199289, EPI_ISL_2199291 | Lighthouse Lab in Milton Keynes | Wellcome Sanger Institute for the COVID-19 Genomics UK (COG-UK) Consortium | The Lighthouse Lab in Milton Keynes and Alex Alderton, Roberto Amato, Jeffrey Barrett, Sonia Goncalves, Ewan Harrison, David K. Jackson, Ian Johnston, Dominic Kwiatkowski, Cordelia Langford, John Sillitoe on behalf of the Wellcome Sanger Institute COVID-19 Surveillance Team |
| EPI_ISL_2199293, EPI_ISL_2199294 | Lighthouse Lab in Glasgow | Wellcome Sanger Institute for the COVID-19 Genomics UK (COG-UK) Consortium | Harper VanSteenhouse, Yumi Kasai, David Gray, Carol Clugston, Anna Dominiczak and Alex Alderton, Roberto Amato, Jeffrey Barrett, Sonia Goncalves, Ewan Harrison, David K. Jackson, Ian Johnston, Dominic Kwiatkowski, Cordelia Langford, John Sillitoe on behalf of the Wellcome Sanger Institute COVID-19 Surveillance Team |
| EPI_ISL_2199295 | Lighthouse Lab in Alderley Park | Wellcome Sanger Institute for the COVID-19 Genomics UK (COG-UK) Consortium | Jacquelyn Wynn, Mairead Hyland, The Lighthouse Lab in Alderley Park and Alex Alderton, Roberto Amato, Jeffrey Barrett, Sonia Goncalves, Ewan Harrison, David K. Jackson, Ian Johnston, Dominic Kwiatkowski, Cordelia Langford, John Sillitoe on behalf of the Wellcome Sanger Institute COVID-19 Surveillance Team |
| EPI_ISL_2199296 | Lighthouse Lab in Glasgow | Wellcome Sanger Institute for the COVID-19 Genomics UK (COG-UK) Consortium | Harper VanSteenhouse, Yumi Kasai, David Gray, Carol Clugston, Anna Dominiczak and Alex Alderton, Roberto Amato, Jeffrey Barrett, Sonia Goncalves, Ewan Harrison, David K. Jackson, Ian Johnston, Dominic Kwiatkowski, Cordelia Langford, John Sillitoe on behalf of the Wellcome Sanger Institute COVID-19 Surveillance Team |
| EPI_ISL_2199301 | Lighthouse Lab in Milton Keynes | Wellcome Sanger Institute for the COVID-19 Genomics UK (COG-UK) Consortium | The Lighthouse Lab in Milton Keynes and Alex Alderton, Roberto Amato, Jeffrey Barrett, Sonia Goncalves, Ewan Harrison, David K. Jackson, Ian Johnston, Dominic Kwiatkowski, Cordelia Langford, John Sillitoe on behalf of the Wellcome Sanger Institute COVID-19 Surveillance Team |
| EPI_ISL_2199302 | Lighthouse Lab in Alderley Park | Wellcome Sanger Institute for the COVID-19 Genomics UK (COG-UK) Consortium | Jacquelyn Wynn, Mairead Hyland, The Lighthouse Lab in Alderley Park and Alex Alderton, Roberto Amato, Jeffrey Barrett, Sonia Goncalves, Ewan Harrison, David K. Jackson, Ian Johnston, Dominic Kwiatkowski, Cordelia Langford, John Sillitoe on behalf of the Wellcome Sanger Institute COVID-19 Surveillance Team |
| EPI_ISL_2199304, EPI_ISL_2199305 | Lighthouse Lab in Glasgow | Wellcome Sanger Institute for the COVID-19 Genomics UK (COG-UK) Consortium | Harper VanSteenhouse, Yumi Kasai, David Gray, Carol Clugston, Anna Dominiczak and Alex Alderton, Roberto Amato, Jeffrey Barrett, Sonia Goncalves, Ewan Harrison, David K. Jackson, Ian Johnston, Dominic Kwiatkowski, Cordelia Langford, John Sillitoe on behalf of the Wellcome Sanger Institute COVID-19 Surveillance Team |
| EPI_ISL_2199306, EPI_ISL_2199309 | Lighthouse Lab in Milton Keynes | Wellcome Sanger Institute for the COVID-19 Genomics UK (COG-UK) Consortium | The Lighthouse Lab in Milton Keynes and Alex Alderton, Roberto Amato, Jeffrey Barrett, Sonia Goncalves, Ewan Harrison, David K. Jackson, Ian Johnston, Dominic Kwiatkowski, Cordelia Langford, John Sillitoe on behalf of the Wellcome Sanger Institute COVID-19 Surveillance Team |
| EPI_ISL_2199311, EPI_ISL_2199313 | Lighthouse Lab in Alderley Park | Wellcome Sanger Institute for the COVID-19 Genomics UK (COG-UK) Consortium | Jacquelyn Wynn, Mairead Hyland, The Lighthouse Lab in Alderley Park and Alex Alderton, Roberto Amato, Jeffrey Barrett, Sonia Goncalves, Ewan Harrison, David K. Jackson, Ian Johnston, Dominic Kwiatkowski, Cordelia Langford, John Sillitoe on behalf of the Wellcome Sanger Institute COVID-19 Surveillance Team |
| EPI_ISL_2199314, EPI_ISL_2199322, EPI_ISL_2199325 | Lighthouse Lab in Milton Keynes | Wellcome Sanger Institute for the COVID-19 Genomics UK (COG-UK) Consortium | The Lighthouse Lab in Milton Keynes and Alex Alderton, Roberto Amato, Jeffrey Barrett, Sonia Goncalves, Ewan Harrison, David K. Jackson, Ian Johnston, Dominic Kwiatkowski, Cordelia Langford, John Sillitoe on behalf of the Wellcome Sanger Institute COVID-19 Surveillance Team |
| EPI_ISL_2199331 | Lighthouse Lab in Glasgow | Wellcome Sanger Institute for the COVID-19 Genomics UK (COG-UK) Consortium | Harper VanSteenhouse, Yumi Kasai, David Gray, Carol Clugston, Anna Dominiczak and Alex Alderton, Roberto Amato, Jeffrey Barrett, Sonia Goncalves, Ewan Harrison, David K. Jackson, Ian Johnston, Dominic Kwiatkowski, Cordelia Langford, John Sillitoe on behalf of the Wellcome Sanger Institute COVID-19 Surveillance Team |
| EPI_ISL_2199332 | Lighthouse Lab in Alderley Park | Wellcome Sanger Institute for the COVID-19 Genomics UK (COG-UK) Consortium | Jacquelyn Wynn, Mairead Hyland, The Lighthouse Lab in Alderley Park and Alex Alderton, Roberto Amato, Jeffrey Barrett, Sonia Goncalves, Ewan Harrison, David K. Jackson, Ian Johnston, Dominic Kwiatkowski, Cordelia Langford, John Sillitoe on behalf of the Wellcome Sanger Institute COVID-19 Surveillance Team |
| EPI_ISL_2199333 | Lighthouse Lab in Milton Keynes | Wellcome Sanger Institute for the COVID-19 Genomics UK (COG-UK) Consortium | The Lighthouse Lab in Milton Keynes and Alex Alderton, Roberto Amato, Jeffrey Barrett, Sonia Goncalves, Ewan Harrison, David K. Jackson, Ian Johnston, Dominic Kwiatkowski, Cordelia Langford, John Sillitoe on behalf of the Wellcome Sanger Institute COVID-19 Surveillance Team |
| EPI_ISL_2199337 | Lighthouse Lab in Glasgow | Wellcome Sanger Institute for the COVID-19 Genomics UK (COG-UK) Consortium | Harper VanSteenhouse, Yumi Kasai, David Gray, Carol Clugston, Anna Dominiczak and Alex Alderton, Roberto Amato, Jeffrey Barrett, Sonia Goncalves, Ewan Harrison, David K. Jackson, Ian Johnston, Dominic Kwiatkowski, Cordelia Langford, John Sillitoe on behalf of the Wellcome Sanger Institute COVID-19 Surveillance Team |
| EPI_ISL_2199342, EPI_ISL_2199345, EPI_ISL_2199347, EPI_ISL_2199352, EPI_ISL_2199356, EPI_ISL_2199359, EPI_ISL_2199360, EPI_ISL_2199361, EPI_ISL_2199366, EPI_ISL_2199367 | Lighthouse Lab in Alderley Park | Wellcome Sanger Institute for the COVID-19 Genomics UK (COG-UK) Consortium | Jacquelyn Wynn, Mairead Hyland, The Lighthouse Lab in Alderley Park and Alex Alderton, Roberto Amato, Jeffrey Barrett, Sonia Goncalves, Ewan Harrison, David K. Jackson, Ian Johnston, Dominic Kwiatkowski, Cordelia Langford, John Sillitoe on behalf of the Wellcome Sanger Institute COVID-19 Surveillance Team |
| EPI_ISL_2199369 | Lighthouse Lab in Milton Keynes | Wellcome Sanger Institute for the COVID-19 Genomics UK (COG-UK) Consortium | The Lighthouse Lab in Milton Keynes and Alex Alderton, Roberto Amato, Jeffrey Barrett, Sonia Goncalves, Ewan Harrison, David K. Jackson, Ian Johnston, Dominic Kwiatkowski, Cordelia Langford, John Sillitoe on behalf of the Wellcome Sanger Institute COVID-19 Surveillance Team |
| EPI_ISL_2199370, EPI_ISL_2199371, EPI_ISL_2199373, EPI_ISL_2199375, EPI_ISL_2199377, EPI_ISL_2199378, EPI_ISL_2199380 | Lighthouse Lab in Alderley Park | Wellcome Sanger Institute for the COVID-19 Genomics UK (COG-UK) Consortium | Jacquelyn Wynn, Mairead Hyland, The Lighthouse Lab in Alderley Park and Alex Alderton, Roberto Amato, Jeffrey Barrett, Sonia Goncalves, Ewan Harrison, David K. Jackson, Ian Johnston, Dominic Kwiatkowski, Cordelia Langford, John Sillitoe on behalf of the Wellcome Sanger Institute COVID-19 Surveillance Team |
| EPI_ISL_2199383 | Lighthouse Lab in Milton Keynes | Wellcome Sanger Institute for the COVID-19 Genomics UK (COG-UK) Consortium | The Lighthouse Lab in Milton Keynes and Alex Alderton, Roberto Amato, Jeffrey Barrett, Sonia Goncalves, Ewan Harrison, David K. Jackson, Ian Johnston, Dominic Kwiatkowski, Cordelia Langford, John Sillitoe on behalf of the Wellcome Sanger Institute COVID-19 Surveillance Team |
| EPI_ISL_2199386, EPI_ISL_2199388, EPI_ISL_2199390, EPI_ISL_2199392, EPI_ISL_2199395, EPI_ISL_2199398, EPI_ISL_2199399, EPI_ISL_2199401, EPI_ISL_2199404, EPI_ISL_2199407, EPI_ISL_2199408, EPI_ISL_2199410, EPI_ISL_2199416, EPI_ISL_2199417, EPI_ISL_2199421, EPI_ISL_2199424, EPI_ISL_2199425, EPI_ISL_2199428 | | | |
| see above | Lighthouse Lab in Alderley Park | Wellcome Sanger Institute for the COVID-19 Genomics UK (COG-UK) Consortium | Jacquelyn Wynn, Mairead Hyland, The Lighthouse Lab in Alderley Park and Alex Alderton, Roberto Amato, Jeffrey Barrett, Sonia Goncalves, Ewan Harrison, David K. Jackson, Ian Johnston, Dominic Kwiatkowski, Cordelia Langford, John Sillitoe on behalf of the Wellcome Sanger Institute COVID-19 Surveillance Team |
| EPI_ISL_2199430 | Lighthouse Lab in Milton Keynes | Wellcome Sanger Institute for the COVID-19 Genomics UK (COG-UK) Consortium | The Lighthouse Lab in Milton Keynes and Alex Alderton, Roberto Amato, Jeffrey Barrett, Sonia Goncalves, Ewan Harrison, David K. Jackson, Ian Johnston, Dominic Kwiatkowski, Cordelia Langford, John Sillitoe on behalf of the Wellcome Sanger Institute COVID-19 Surveillance Team |
| EPI_ISL_2199432, EPI_ISL_2199434, EPI_ISL_2199435, EPI_ISL_2199436, EPI_ISL_2199440, EPI_ISL_2199441, EPI_ISL_2199442, EPI_ISL_2199445, EPI_ISL_2199447, EPI_ISL_2199452, EPI_ISL_2199454, EPI_ISL_2199457 | | | |
| see above | Lighthouse Lab in Alderley Park | Wellcome Sanger Institute for the COVID-19 Genomics UK (COG-UK) Consortium | Jacquelyn Wynn, Mairead Hyland, The Lighthouse Lab in Alderley Park and Alex Alderton, Roberto Amato, Jeffrey Barrett, Sonia Goncalves, Ewan Harrison, David K. Jackson, Ian Johnston, Dominic Kwiatkowski, Cordelia Langford, John Sillitoe on behalf of the Wellcome Sanger Institute COVID-19 Surveillance Team |
| EPI_ISL_2199463 | Lighthouse Lab in Milton Keynes | Wellcome Sanger Institute for the COVID-19 Genomics UK (COG-UK) Consortium | The Lighthouse Lab in Milton Keynes and Alex Alderton, Roberto Amato, Jeffrey Barrett, Sonia Goncalves, Ewan Harrison, David K. Jackson, Ian Johnston, Dominic Kwiatkowski, Cordelia Langford, John Sillitoe on behalf of the Wellcome Sanger Institute COVID-19 Surveillance Team |
| EPI_ISL_2199467, EPI_ISL_2199469 | Lighthouse Lab in Alderley Park | Wellcome Sanger Institute for the COVID-19 Genomics UK (COG-UK) Consortium | Jacquelyn Wynn, Mairead Hyland, The Lighthouse Lab in Alderley Park and Alex Alderton, Roberto Amato, Jeffrey Barrett, Sonia Goncalves, Ewan Harrison, David K. Jackson, Ian Johnston, Dominic Kwiatkowski, Cordelia Langford, John Sillitoe on behalf of the Wellcome Sanger Institute COVID-19 Surveillance Team |
| EPI_ISL_2199470 | Lighthouse Lab in Milton Keynes | Wellcome Sanger Institute for the COVID-19 Genomics UK (COG-UK) Consortium | The Lighthouse Lab in Milton Keynes and Alex Alderton, Roberto Amato, Jeffrey Barrett, Sonia Goncalves, Ewan Harrison, David K. Jackson, Ian Johnston, Dominic Kwiatkowski, Cordelia Langford, John Sillitoe on behalf of the Wellcome Sanger Institute COVID-19 Surveillance Team |
| EPI_ISL_2199471, EPI_ISL_2199474, EPI_ISL_2199475, EPI_ISL_2199476, EPI_ISL_2199477, EPI_ISL_2199479, EPI_ISL_2199481, EPI_ISL_2199483, EPI_ISL_2199484, EPI_ISL_2199485, EPI_ISL_2199486, EPI_ISL_2199487, EPI_ISL_2199489, EPI_ISL_2199491, EPI_ISL_2199493, EPI_ISL_2199494, EPI_ISL_2199495, EPI_ISL_2199496, EPI_ISL_2199497, EPI_ISL_2199499, EPI_ISL_2199500, EPI_ISL_2199501, EPI_ISL_2199503 | | | |
| see above | Lighthouse Lab in Alderley Park | Wellcome Sanger Institute for the COVID-19 Genomics UK (COG-UK) | Jacquelyn Wynn, Mairead Hyland, The Lighthouse Lab in Alderley Park and Alex Alderton, Roberto Amato, Jeffrey Barrett, Sonia Goncalves, Ewan Harrison, David K. Jackson, Ian Johnston, Dominic Kwiatkowski, Cordelia Langford, John Sillitoe on |

| | | | |
|---|---|---|---|
| | | Consortium | behalf of the Wellcome Sanger Institute COVID-19 Surveillance Team |
| EPI_ISL_2199504 | Lighthouse Lab in Milton Keynes | Wellcome Sanger Institute for the COVID-19 Genomics UK (COG-UK) Consortium | The Lighthouse Lab in Milton Keynes and Alex Alderton, Roberto Amato, Jeffrey Barrett, Sonia Goncalves, Ewan Harrison, David K. Jackson, Ian Johnston, Dominic Kwiatkowski, Cordelia Langford, John Sillitoe on behalf of the Wellcome Sanger Institute COVID-19 Surveillance Team |
| EPI_ISL_2199507, EPI_ISL_2199509, EPI_ISL_2199511, EPI_ISL_2199512, EPI_ISL_2199514, EPI_ISL_2199515, EPI_ISL_2199516 | Lighthouse Lab in Alderley Park | Wellcome Sanger Institute for the COVID-19 Genomics UK (COG-UK) Consortium | Jacquelyn Wynn, Mairead Hyland, The Lighthouse Lab in Alderley Park and Alex Alderton, Roberto Amato, Jeffrey Barrett, Sonia Goncalves, Ewan Harrison, David K. Jackson, Ian Johnston, Dominic Kwiatkowski, Cordelia Langford, John Sillitoe on behalf of the Wellcome Sanger Institute COVID-19 Surveillance Team |
| EPI_ISL_2199519 | Lighthouse Lab in Milton Keynes | Wellcome Sanger Institute for the COVID-19 Genomics UK (COG-UK) Consortium | The Lighthouse Lab in Milton Keynes and Alex Alderton, Roberto Amato, Jeffrey Barrett, Sonia Goncalves, Ewan Harrison, David K. Jackson, Ian Johnston, Dominic Kwiatkowski, Cordelia Langford, John Sillitoe on behalf of the Wellcome Sanger Institute COVID-19 Surveillance Team |
| EPI_ISL_2199521, EPI_ISL_2199523, EPI_ISL_2199524, EPI_ISL_2199525, EPI_ISL_2199526, EPI_ISL_2199527, EPI_ISL_2199528, EPI_ISL_2199529, EPI_ISL_2199533, EPI_ISL_2199536, EPI_ISL_2199538, EPI_ISL_2199540, EPI_ISL_2199541, EPI_ISL_2199543, EPI_ISL_2199544, EPI_ISL_2199546, EPI_ISL_2199547, EPI_ISL_2199548 | | | |
| see above | Lighthouse Lab in Alderley Park | Wellcome Sanger Institute for the COVID-19 Genomics UK (COG-UK) Consortium | Jacquelyn Wynn, Mairead Hyland, The Lighthouse Lab in Alderley Park and Alex Alderton, Roberto Amato, Jeffrey Barrett, Sonia Goncalves, Ewan Harrison, David K. Jackson, Ian Johnston, Dominic Kwiatkowski, Cordelia Langford, John Sillitoe on behalf of the Wellcome Sanger Institute COVID-19 Surveillance Team |
| EPI_ISL_2199549 | Lighthouse Lab in Milton Keynes | Wellcome Sanger Institute for the COVID-19 Genomics UK (COG-UK) Consortium | The Lighthouse Lab in Milton Keynes and Alex Alderton, Roberto Amato, Jeffrey Barrett, Sonia Goncalves, Ewan Harrison, David K. Jackson, Ian Johnston, Dominic Kwiatkowski, Cordelia Langford, John Sillitoe on behalf of the Wellcome Sanger Institute COVID-19 Surveillance Team |
| EPI_ISL_2199550, EPI_ISL_2199551, EPI_ISL_2199552, EPI_ISL_2199555, EPI_ISL_2199557, EPI_ISL_2199561, EPI_ISL_2199562, EPI_ISL_2199564, EPI_ISL_2199565, EPI_ISL_2199567, EPI_ISL_2199573, EPI_ISL_2199574, EPI_ISL_2199575, EPI_ISL_2199576, EPI_ISL_2199577, EPI_ISL_2199578, EPI_ISL_2199579, EPI_ISL_2199580, EPI_ISL_2199588, EPI_ISL_2199590, EPI_ISL_2199593, EPI_ISL_2199594, EPI_ISL_2199597, EPI_ISL_2199598, EPI_ISL_2199599, EPI_ISL_2199600, EPI_ISL_2199601, EPI_ISL_2199607, EPI_ISL_2199611, EPI_ISL_2199612, EPI_ISL_2199613 | | | |
| see above | Lighthouse Lab in Alderley Park | Wellcome Sanger Institute for the COVID-19 Genomics UK (COG-UK) Consortium | Jacquelyn Wynn, Mairead Hyland, The Lighthouse Lab in Alderley Park and Alex Alderton, Roberto Amato, Jeffrey Barrett, Sonia Goncalves, Ewan Harrison, David K. Jackson, Ian Johnston, Dominic Kwiatkowski, Cordelia Langford, John Sillitoe on behalf of the Wellcome Sanger Institute COVID-19 Surveillance Team |
| EPI_ISL_2199614 | Lighthouse Lab in Milton Keynes | Wellcome Sanger Institute for the COVID-19 Genomics UK (COG-UK) Consortium | The Lighthouse Lab in Milton Keynes and Alex Alderton, Roberto Amato, Jeffrey Barrett, Sonia Goncalves, Ewan Harrison, David K. Jackson, Ian Johnston, Dominic Kwiatkowski, Cordelia Langford, John Sillitoe on behalf of the Wellcome Sanger Institute COVID-19 Surveillance Team |
| EPI_ISL_2199615, EPI_ISL_2199617, EPI_ISL_2199618, EPI_ISL_2199621, EPI_ISL_2199626, EPI_ISL_2199628, EPI_ISL_2199629 | Lighthouse Lab in Alderley Park | Wellcome Sanger Institute for the COVID-19 Genomics UK (COG-UK) Consortium | Jacquelyn Wynn, Mairead Hyland, The Lighthouse Lab in Alderley Park and Alex Alderton, Roberto Amato, Jeffrey Barrett, Sonia Goncalves, Ewan Harrison, David K. Jackson, Ian Johnston, Dominic Kwiatkowski, Cordelia Langford, John Sillitoe on behalf of the Wellcome Sanger Institute COVID-19 Surveillance Team |
| EPI_ISL_2199630 | Lighthouse Lab in Milton Keynes | Wellcome Sanger Institute for the COVID-19 Genomics UK (COG-UK) Consortium | The Lighthouse Lab in Milton Keynes and Alex Alderton, Roberto Amato, Jeffrey Barrett, Sonia Goncalves, Ewan Harrison, David K. Jackson, Ian Johnston, Dominic Kwiatkowski, Cordelia Langford, John Sillitoe on behalf of the Wellcome Sanger Institute COVID-19 Surveillance Team |
| EPI_ISL_2199633, EPI_ISL_2199637, EPI_ISL_2199638, EPI_ISL_2199639, EPI_ISL_2199640, EPI_ISL_2199642, EPI_ISL_2199643, EPI_ISL_2199644, EPI_ISL_2199645, EPI_ISL_2199647, EPI_ISL_2199654, EPI_ISL_2199658, EPI_ISL_2199662, EPI_ISL_2199663, EPI_ISL_2199664, EPI_ISL_2199665, EPI_ISL_2199674, EPI_ISL_2199675, EPI_ISL_2199676, EPI_ISL_2199677, EPI_ISL_2199678, EPI_ISL_2199679, EPI_ISL_2199681, EPI_ISL_2199682 | | | |
| see above | Lighthouse Lab in Alderley Park | Wellcome Sanger Institute for the COVID-19 Genomics UK (COG-UK) Consortium | Jacquelyn Wynn, Mairead Hyland, The Lighthouse Lab in Alderley Park and Alex Alderton, Roberto Amato, Jeffrey Barrett, Sonia Goncalves, Ewan Harrison, David K. Jackson, Ian Johnston, Dominic Kwiatkowski, Cordelia Langford, John Sillitoe on behalf of the Wellcome Sanger Institute COVID-19 Surveillance Team |
| EPI_ISL_2199684, EPI_ISL_2199685, EPI_ISL_2199696, EPI_ISL_2199698, EPI_ISL_2199699, EPI_ISL_2199700, EPI_ISL_2199704, EPI_ISL_2199713, EPI_ISL_2199718, EPI_ISL_2199724, EPI_ISL_2199725, EPI_ISL_2199726, EPI_ISL_2199727, EPI_ISL_2199729, EPI_ISL_2199735, EPI_ISL_2199739, EPI_ISL_2199741, EPI_ISL_2199742, EPI_ISL_2199759, EPI_ISL_2199761, EPI_ISL_2199768, EPI_ISL_2199772, EPI_ISL_2199774, EPI_ISL_2199776, EPI_ISL_2199777, EPI_ISL_2199780, EPI_ISL_2199786, EPI_ISL_2199789, EPI_ISL_2199791, EPI_ISL_2199793, EPI_ISL_2199795, EPI_ISL_2199803, EPI_ISL_2199804, EPI_ISL_2199808, EPI_ISL_2199812, EPI_ISL_2199819, EPI_ISL_2199825, EPI_ISL_2199829, EPI_ISL_2199833, EPI_ISL_2199836, EPI_ISL_2199843, EPI_ISL_2199850, EPI_ISL_2199851, EPI_ISL_2199853, EPI_ISL_2199854, EPI_ISL_2199855, EPI_ISL_2199857, EPI_ISL_2199867, EPI_ISL_2199874, EPI_ISL_2199878, EPI_ISL_2199880, EPI_ISL_2199883, EPI_ISL_2199896, EPI_ISL_2199898, EPI_ISL_2199899, EPI_ISL_2199900, EPI_ISL_2199903, EPI_ISL_2199906, EPI_ISL_2199909, EPI_ISL_2199912, EPI_ISL_2199914, EPI_ISL_2199917, EPI_ISL_2199921, EPI_ISL_2199925, EPI_ISL_2199932, EPI_ISL_2199933, EPI_ISL_2199935, EPI_ISL_2199939, EPI_ISL_2199941, EPI_ISL_2199942, EPI_ISL_2199946, EPI_ISL_2199953, EPI_ISL_2199955, EPI_ISL_2199957, EPI_ISL_2199958, EPI_ISL_2199962, EPI_ISL_2199964, EPI_ISL_2199965, EPI_ISL_2199966, EPI_ISL_2199967, EPI_ISL_2199973, EPI_ISL_2199975, EPI_ISL_2199977 | | | |
| see above | Lighthouse Lab in Glasgow | Wellcome Sanger Institute for the COVID-19 Genomics UK (COG-UK) Consortium | Harper VanSteenhouse, Yumi Kasai, David Gray, Carol Clugston, Anna Dominiczak and Alex Alderton, Roberto Amato, Jeffrey Barrett, Sonia Goncalves, Ewan Harrison, David K. Jackson, Ian Johnston, Dominic Kwiatkowski, Cordelia Langford, John Sillitoe on behalf of the Wellcome Sanger Institute COVID-19 Surveillance Team |
| EPI_ISL_2199979, EPI_ISL_2199980, EPI_ISL_2199981, EPI_ISL_2199982 | Lighthouse Lab in Milton Keynes | Wellcome Sanger Institute for the COVID-19 Genomics UK (COG-UK) Consortium | The Lighthouse Lab in Milton Keynes and Alex Alderton, Roberto Amato, Jeffrey Barrett, Sonia Goncalves, Ewan Harrison, David K. Jackson, Ian Johnston, Dominic Kwiatkowski, Cordelia Langford, John Sillitoe on behalf of the Wellcome Sanger Institute COVID-19 Surveillance Team |
| EPI_ISL_2200633 | Schweizer Paraplegiker Zentrum | Institute of Medical Virology | Verena Kufner, Gabriela Ziltener, Maryam Zaheri, Stefan Schmutz, Annette Audigé, Maria Grünberg, Kevin Steiner, Jon Huder, Cyril Shah, Riccarda Capaul, Guido Bloemberg, Jürg Böni, Michael Huber, Alexandra Trkola |
| EPI_ISL_2201543, EPI_ISL_2201546, EPI_ISL_2201789, EPI_ISL_2202123 | Aegis Sciences Corporation | Centers for Disease Control and Prevention Division of Viral Diseases, Pathogen Discovery | Dakota Howard, Dhwani Batra, Peter W. Cook, Kara Moser, Adrian Paskey, Jason Caravas, Benjamin Rambo-Martin, Shatavia Morrison, Christopher Gulvick, Scott Sammons, Yvette Unoarumhi, Darlene Wagner, Matthew Schmerer, Cyndi Clark, Patrick Campbell, Rob Case, Vikramsinha Ghorpade, Holly Houdeshell, Ola Kvalvaag, Dillon Nall, Ethan Sanders, Alec Vest, Shaun Westlund, Matthew Hardison, Clinton R. Paden, Duncan MacCannell |
| EPI_ISL_2202807 | NYU Langone Health | Departments of Pathology and Medicine, New York University School of Medicine | Adriana Heguy, Dacia Dimartino, Emily Guzman, Christian Marier, Peter Meyn, Sitharam Ramaswami, Gael Westby, Paul Zappile, Yutong Zhang, Paolo Cotzia, Guiqing Wang |
| EPI_ISL_2203031, EPI_ISL_2203200, EPI_ISL_2203265, EPI_ISL_2203267 | Aegis Sciences Corporation | Centers for Disease Control and Prevention Division of Viral Diseases, Pathogen Discovery | Dakota Howard, Dhwani Batra, Peter W. Cook, Kara Moser, Adrian Paskey, Jason Caravas, Benjamin Rambo-Martin, Shatavia Morrison, Christopher Gulvick, Scott Sammons, Yvette Unoarumhi, Darlene Wagner, Matthew Schmerer, Cyndi Clark, Patrick Campbell, Rob Case, Vikramsinha Ghorpade, Holly Houdeshell, Ola Kvalvaag, Dillon Nall, Ethan Sanders, Alec Vest, Shaun Westlund, Matthew Hardison, Clinton R. Paden, Duncan MacCannell |
| EPI_ISL_2203739, EPI_ISL_2203742, EPI_ISL_2203788, EPI_ISL_2203953, EPI_ISL_2204104 | Quest Diagnostics Incorporated | Centers for Disease Control and Prevention Division of Viral Diseases, Pathogen Discovery | Dakota Howard, Dhwani Batra, Peter W. Cook, Kara Moser, Adrian Paskey, Jason Caravas, Benjamin Rambo-Martin, Shatavia Morrison, Christopher Gulvick, Scott Sammons, Yvette Unoarumhi, Darlene Wagner, Matthew Schmerer, S. H. Rosenthal, A. Gerasimova, R. M. Kagan, B. Anderson, M. Hua, Y. Liu, L.E. Bernstein, K.E. Livingston, A. Perez, I. A. Shlyakhter, R. V. Rolando, R. Owen, P. Tanpaiboon, F. Lacbawan, Clinton R. Paden, Duncan MacCannell |
| EPI_ISL_2204729, EPI_ISL_2204788, EPI_ISL_2204930 | Aegis Sciences Corporation | Centers for Disease Control and Prevention Division of Viral Diseases, Pathogen Discovery | Dakota Howard, Dhwani Batra, Peter W. Cook, Kara Moser, Adrian Paskey, Jason Caravas, Benjamin Rambo-Martin, Shatavia Morrison, Christopher Gulvick, Scott Sammons, Yvette Unoarumhi, Darlene Wagner, Matthew Schmerer, Cyndi Clark, Patrick Campbell, Rob Case, Vikramsinha Ghorpade, Holly Houdeshell, Ola Kvalvaag, Dillon Nall, Ethan Sanders, Alec Vest, Shaun Westlund, Matthew Hardison, Clinton R. Paden, Duncan MacCannell |
| EPI_ISL_2205005, EPI_ISL_2205007 | Nebraska Public Health Laboratory | NPHL COVID-19 Response Team | NPHL COVID-19 Response Team |
| EPI_ISL_2205865 | Swedish national genomic surveillance program of SARS-CoV-2 | The Public Health Agency of Sweden | Maximilian Riess, Maria Lind Karlberg, Alma Brolund, Swedish national genomic surveillance program of SARS-CoV-2 |
| EPI_ISL_2207016 | Nebraska Public Health Laboratory | NPHL COVID-19 Response Team | NPHL COVID-19 Response Team |
| EPI_ISL_2209223 | Swedish national genomic surveillance program of SARS-CoV-2 | The Public Health Agency of Sweden | Maximilian Riess, Maria Lind Karlberg, Alma Brolund, Swedish national genomic surveillance program of SARS-CoV-2 |
| EPI_ISL_2212778 | Viollier AG | Department of Biosystems Science and Engineering, ETH Zürich | Christian Beisel, Sarah Nadeau, Chaoran Chen, Ivan Topolsky, Philipp Jablonski, Lara Fuhrmann, David Dreifuss, Katharina Jahn, Rebecca Denes, Mirjam Feldkamp, Ina Nissen, Natascha Santacroce, Elodie Burcklen, Christiane Beckmann, Maurice Redondo, Olivier Kobel, Christoph Noppen, Sophie Seidel, Noemie Santamaria de Souza, Niko Beerenwinkel, Tanja Stadler |
| EPI_ISL_2213568, EPI_ISL_2213571, EPI_ISL_2213577, EPI_ISL_2213579, EPI_ISL_2213582, EPI_ISL_2213584 | University Hospitals of Geneva, Laboratory of Virology | HUG, Laboratory of Virology and the Health2030 Genome Center | Samuel Cordey, Ana Rita Goncalves, Laurent Kaiser, Lorenzo Cerutti, Henri Pegeot, Melyssa Elies, Deborah Penet, Keith Harshman, Ioannis Xenarios, Emmanouil Dermitzakis |
| EPI_ISL_2220505, EPI_ISL_2220507, EPI_ISL_2220513, EPI_ISL_2220515, EPI_ISL_2220518, EPI_ISL_2220520 | Dutch COVID-19 response team | National Institute for Public Health and the Environment (RIVM) | Adam Meijer, Harry Vennema, Dirk Eggink, Jeroen Cremer, Sharon van den Brink, Bas van der Veer, AnneMarie van den Brandt, Lisa Wijsman, Kim Freriks, Ryanne Jaarsma, Euníce Then, Lynn Aarts, Sanne Bos, Melissa van Tuil, Linda van de Nes, Sjoerd Kuiling, James Groot, Florian Zwagemaker, Dennis Schmitz, Annelies Kroneman, Karim Hajji, Chantal Reusken, on behalf of the national COVID-19 response team |
| EPI_ISL_2224331 | University of Mississippi Medical Center, Department of Pathology | University of Mississippi Medical Center, Molecular and Genomics Core Facility | Ashley C. Johnson, Ithiel J. Frame, Krishna K. Ayyalasomayajula, Michael R. Garrett, D. Ashley Robinson |
| EPI_ISL_2224471, EPI_ISL_2224473 | MS Public Health Laboratory | University of Mississippi Medical Center, Molecular and Genomics Core Facility | Ashley C. Johnson, Ithiel J. Frame, Krishna K. Ayyalasomayajula, Michael R. Garrett, D. Ashley Robinson |
| EPI_ISL_2225235, EPI_ISL_2225238 | Houston Methodist Hospital | Houston Methodist Hospital | Randall J. Olsen, Paul A. Christensen, S. Wesley Long, Sishir Subedi, Robert Olson, Marcus Nguyen, James J. Davis, Matthew Ojeda Saavedra, Prasanti Yerramilli, Layne Pruitt, Kristina Reppond, Madison N. Shyer, Jessica Cambric, Ryan Gadd, Ilya J. Finkelstein, Jimmy Gollihar, and James M. Musser |
| EPI_ISL_2225532 | Wyoming Public Health Laboratory | Wyoming Public Health Laboratory | Jim Mildenberger, Taylor Fearing, Channing Weber, Ashley Norberg, Chayse Rowley, Marley Goetz, Brian Dominguez, Elliot Thomasson, Brittany Oher, Sam Britz, Cari Sloma, and Rob Christensen |
| EPI_ISL_2225632, EPI_ISL_2225633, EPI_ISL_2225637, EPI_ISL_2225642, EPI_ISL_2225644, EPI_ISL_2225645, EPI_ISL_2225647, EPI_ISL_2225649, EPI_ISL_2225654, EPI_ISL_2225658, EPI_ISL_2225660, EPI_ISL_2225661, EPI_ISL_2225662, EPI_ISL_2225664, EPI_ISL_2225666, EPI_ISL_2225669, EPI_ISL_2225671, EPI_ISL_2225672, EPI_ISL_2225673, EPI_ISL_2225675, EPI_ISL_2225676, EPI_ISL_2225677, EPI_ISL_2225678, EPI_ISL_2225679, EPI_ISL_2225681, EPI_ISL_2225682, EPI_ISL_2225684, EPI_ISL_2225685, EPI_ISL_2225686, EPI_ISL_2225688, EPI_ISL_2225689, EPI_ISL_2225690, EPI_ISL_2225691, EPI_ISL_2225692, EPI_ISL_2225693 | | | |
| see above | National Centre For Cell Science | National Centre For Cell Science – INSACOG | Dhiraj Paul, Shivang P. Bhanushali, Mitali Inamdar, Mohak P Gujare, Manoj Kumar Bhat, Ajay Pillai, INSACOG Consortium team, Yogesh Shouche |

| | | | |
|---|---|---|---|
| EPI_ISL_2225843 | Akershus University Hospital, Department for Microbiology and Infectious Disease Control | Norwegian Institute of Public Health, Department of Virology | Kathrine Stene-Johansen, Kamilla Heddeland Instefjord, Hilde Elshaug, Garcia Llorente Ignacio, Jon Bråte, Engebretsen Serina Beate,Pedersen Benedikte Nevjen, Debech Nadia, Atiya R Ali,Marie Paulsen Madsen, Rasmus Riis Kopperud, Hilde Vollan, Karoline Bragstad, Olav Hungnes |
| EPI_ISL_2226262 | Mapmygenome | CSIR-Centre for Cellular and Molecular Biology-INSACOG | Lamuk Zaveri, Ara Sreenivas, Onkar Kulkarni, Sofia Banu, Shreekant Verma, Amareshwar Vodapalli , Viswagithe S L, B Himasri, Sharath Chandra Thota, Karthik Bharadwaj Tallapaka, Rakesh K Mishra,Divya Tej Sowpati |
| EPI_ISL_2226348 | CHU POITIERS | CHU Poitiers | Caroline MICHAUD, Maxime PICHON, Manon PRAT, Valentin BON-BARET, Birama N'DIAYE, Magali GARCIA, Agnes BEBY-DEFAUX, Nicolas LEVEQUE |
| EPI_ISL_2226818, EPI_ISL_2226819, EPI_ISL_2226821 | Wyoming Public Health Laboratory | Wyoming Public Health Laboratory | Jim Mildenberger, Wanda Manley, Noah Hull, Taylor Fearing, Lynette Gumbleton, Channing Weber, Ashley Norberg, Chayse Rowley, Marley Goetz, Brian Dominguez, Elliot Thomasson, Sam Britz, Cari Sloma, and Rob Christensen |
| EPI_ISL_2227204, EPI_ISL_2227268, EPI_ISL_2227269 | COVID-19 National Reference Laboratoty, Pasteur Institute of Iran | Genetics Research Center, University of Social Welfare and Rehabilitation Sciences | Zohreh Fattahi, Marzieh Mohseni, Kimia Kahrizi, Mahsa Tavakoli,Tahmineh Jalali, Mohammad Hassan Pouriayevali, Mostafa Salehi-Vaziri, Hossein Najmabadi. |
| EPI_ISL_2227270 | COVID-19 National Reference Laboratoty, Pasteur Institute of Iran. | Genetics Research Center, University of Social Welfare and Rehabilitation Sciences | Zohreh Fattahi, Marzieh Mohseni, Kimia Kahrizi, Mahsa Tavakoli,Tahmineh Jalali, Mohammad Hassan Pouriayevali, Mostafa Salehi-Vaziri, Hossein Najmabadi. |
| EPI_ISL_2227271 | COVID-19 National Reference Laboratory, Pasteur Institute of Iran. | Genetics Research Center, University of Social Welfare and Rehabilitation Sciences | Zohreh Fattahi, Marzieh Mohseni, Kimia Kahrizi, Mahsa Tavakoli,Tahmineh Jalali, Mohammad Hassan Pouriayevali, Mostafa Salehi-Vaziri, Hossein Najmabadi. |
| EPI_ISL_2227272 | COVID-19 National Reference Laboratory, Pasteur Institute of Iran | Genetics Research Center, University of Social Welfare and Rehabilitation Sciences | Zohreh Fattahi, Marzieh Mohseni, Kimia Kahrizi, Mahsa Tavakoli,Tahmineh Jalali, Mohammad Hassan Pouriayevali, Mostafa Salehi-Vaziri, Hossein Najmabadi. |
| EPI_ISL_2227358, EPI_ISL_2227359 | Labor Berlin Charité Vivantes GmbH / Institut für Virologie | Charité Universitätsmedizin Berlin, Institut für Virologie/Labor Berlin | Peter Menzel, Christine Stephan, Rolf Schwarzer, Victor M Corman, Barbara Mühlemann, Terry Jones, Christian Drosten |
| EPI_ISL_2227398, EPI_ISL_2227400 | Wichita State University - Molecular Diagnostics Lab | Kansas Health and Environmental Lab | Mike Grose, Jonathan Barnell, Ben Olsen, and Phil Adam |
| EPI_ISL_2227435 | MD PHL | MD PHL | Maryland Department of Health Laboratories Administration |
| EPI_ISL_2227694, EPI_ISL_2227695 | OHSU Lab Services Molecular Microbiology Lab | Oregon SARS-CoV-2 Genome Sequencing Center | Brendan L. O'Connell, Sally Grindstaff, Sonia Acharya, Cierra LaBlanc, Kayla Carter, Ruth V. Nichols, Alec J. Hirsch, Donna Hansel, Guang Fan, Xuan Qin, Daniel N. Streblow, William B. Messer, Andrew C. Adey, Benjamin N. Bimber, Brian J. O'Roak |
| EPI_ISL_2228090 | CT Department of Public Health | CT Department of Public Health | Claire_Pearson,Tu_N_Nguyen |
| EPI_ISL_2228674, EPI_ISL_2228675 | LABM Unilabs EYLAU Corbeil | Department of Virology, Henri Mondor University Hospital, Assistance Publique Hôpitaux de Paris, Université Paris-Est Créteil, INSERM U955 | Christophe Rodriguez, Slim Fourati, Vanessa Demontant, Guillaume Gricourt, Melissa N'Debi, Alexandre Soulier, Elisabeth Trawinski, Jean-Michel Pawlotsky |
| EPI_ISL_2228707 | CH. ROBERT BALLANGER | Department of Virology, Henri Mondor University Hospital, Assistance Publique Hôpitaux de Paris, Université Paris-Est Créteil, INSERM U955 | Christophe Rodriguez, Slim Fourati, Vanessa Demontant, Guillaume Gricourt, Melissa N'Debi, Alexandre Soulier, Elisabeth Trawinski, Jean-Michel Pawlotsky |
| EPI_ISL_2228723, EPI_ISL_2228724 | Hôpital Avicenne | Department of Virology, Henri Mondor University Hospital, Assistance Publique Hôpitaux de Paris, Université Paris-Est Créteil, INSERM U955 | Christophe Rodriguez, Slim Fourati, Vanessa Demontant, Guillaume Gricourt, Melissa N'Debi, Alexandre Soulier, Elisabeth Trawinski, Jean-Michel Pawlotsky |
| EPI_ISL_2228856, EPI_ISL_2228858, EPI_ISL_2228859 | Cerballiance Paris IDF est Saint-Denis | Department of Virology, Henri Mondor University Hospital, Assistance Publique Hôpitaux de Paris, Université Paris-Est Créteil, INSERM U955 | Christophe Rodriguez, Slim Fourati, Vanessa Demontant, Guillaume Gricourt, Melissa N'Debi, Alexandre Soulier, Elisabeth Trawinski, Jean-Michel Pawlotsky |
| EPI_ISL_2228954 | SC Dept of Health and Env. Control-Bureau of Laboratories | Centers for Disease Control and Prevention Division of Viral Diseases, Pathogen Discovery | Mili Sheth, Sarah Nobles, Jasmine Padilla, Mark Burroughs, Shoshona Le, Katie Dillon, Peter Cook, Clinton R. Paden, Dhwani Batra, Krista Queen, Kristen Knipe, Dakota Howard, Yvette Unoarumhi, Darlene Wagner, Matthew Schmerer, Ben L. Rambo-Martin, Kristine Lacek, Sam Shepard, Alison Laufer Halpin, Dave Wentworth, Vivien Dugan, Suxiang Tong, Justin Lee |
| EPI_ISL_2228962 | DE Public Health Laboratory | Centers for Disease Control and Prevention Division of Viral Diseases, Pathogen Discovery | Mili Sheth, Sarah Nobles, Jasmine Padilla, Mark Burroughs, Shoshona Le, Katie Dillon, Peter Cook, Clinton R. Paden, Dhwani Batra, Krista Queen, Kristen Knipe, Dakota Howard, Yvette Unoarumhi, Darlene Wagner, Matthew Schmerer, Ben L. Rambo-Martin, Kristine Lacek, Sam Shepard, Alison Laufer Halpin, Dave Wentworth, Vivien Dugan, Suxiang Tong, Justin Lee |
| EPI_ISL_2229053 | BIOMNIS EUROFINS IVRY | Department of Virology, Henri Mondor University Hospital, Assistance Publique Hôpitaux de Paris, Université Paris-Est Créteil, INSERM U955 | Christophe Rodriguez, Slim Fourati, Vanessa Demontant, Guillaume Gricourt, Melissa N'Debi, Alexandre Soulier, Elisabeth Trawinski, Jean-Michel Pawlotsky |
| EPI_ISL_2229105 | Guam Public Health Laboratory | Centers for Disease Control and Prevention Division of Viral Diseases, Pathogen Discovery | Mili Sheth, Sarah Nobles, Jasmine Padilla, Mark Burroughs, Shoshona Le, Katie Dillon, Peter Cook, Clinton R. Paden, Dhwani Batra, Krista Queen, Kristen Knipe, Dakota Howard, Yvette Unoarumhi, Darlene Wagner, Matthew Schmerer, Ben L. Rambo-Martin, Kristine Lacek, Sam Shepard, Alison Laufer Halpin, Dave Wentworth, Vivien Dugan, Suxiang Tong, Justin Lee |
| EPI_ISL_2229118 | "AR Dept. of Health-PHL, Molecular Diagnostics" | Centers for Disease Control and Prevention Division of Viral Diseases, Pathogen Discovery | Mili Sheth, Sarah Nobles, Jasmine Padilla, Mark Burroughs, Shoshona Le, Katie Dillon, Peter Cook, Clinton R. Paden, Dhwani Batra, Krista Queen, Kristen Knipe, Dakota Howard, Yvette Unoarumhi, Darlene Wagner, Matthew Schmerer, Ben L. Rambo-Martin, Kristine Lacek, Sam Shepard, Alison Laufer Halpin, Dave Wentworth, Vivien Dugan, Suxiang Tong, Justin Lee |
| EPI_ISL_2229145, EPI_ISL_2229146 | "CDPH, Viral and Rickettsial Disease Laboratory" | Centers for Disease Control and Prevention Division of Viral Diseases, Pathogen Discovery | Mili Sheth, Sarah Nobles, Jasmine Padilla, Mark Burroughs, Shoshona Le, Katie Dillon, Peter Cook, Clinton R. Paden, Dhwani Batra, Krista Queen, Kristen Knipe, Dakota Howard, Yvette Unoarumhi, Darlene Wagner, Matthew Schmerer, Ben L. Rambo-Martin, Kristine Lacek, Sam Shepard, Alison Laufer Halpin, Dave Wentworth, Vivien Dugan, Suxiang Tong, Justin Lee |
| EPI_ISL_2229243 | UT-Unified State Labs: Public Health Utah DOH | Centers for Disease Control and Prevention Division of Viral Diseases, Pathogen Discovery | Mili Sheth, Sarah Nobles, Jasmine Padilla, Mark Burroughs, Shoshona Le, Katie Dillon, Peter Cook, Clinton R. Paden, Dhwani Batra, Krista Queen, Kristen Knipe, Dakota Howard, Yvette Unoarumhi, Darlene Wagner, Matthew Schmerer, Ben L. Rambo-Martin, Kristine Lacek, Sam Shepard, Alison Laufer Halpin, Dave Wentworth, Vivien Dugan, Suxiang Tong, Justin Lee |
| EPI_ISL_2229272 | LA Office of Public Health Laboratories | Centers for Disease Control and Prevention Division of Viral Diseases, Pathogen Discovery | Mili Sheth, Sarah Nobles, Jasmine Padilla, Mark Burroughs, Shoshona Le, Katie Dillon, Peter Cook, Clinton R. Paden, Dhwani Batra, Krista Queen, Kristen Knipe, Dakota Howard, Yvette Unoarumhi, Darlene Wagner, Matthew Schmerer, Ben L. Rambo-Martin, Kristine Lacek, Sam Shepard, Alison Laufer Halpin, Dave Wentworth, Vivien Dugan, Suxiang Tong, Justin Lee |
| EPI_ISL_2229350 | LSUHS Emerging Viral Threat Laboratory | LSUHS Emerging Viral Threat Laboratory | Jeremy P. Kamil, Jennifer L. Carroll, Maarten Van Diest, Rona S. Scott,Gregory L. Ware, Alexander Mijalis, Andrew D. Yurochko, Christopher G. Kevil, John A. Vanchiere |
| EPI_ISL_2229383 | Willis-Knighton Medical Center Hospital Laboratory | LSUHS Emerging Viral Threat Laboratory | Rona S. Scott, Gregory L. Ware, Alexander Mijalis, Jeremy P. Kamil, Jennifer L. Carroll, Maarten Van Diest, Andrew D. Yurochko, Christopher G. Kevil, John A. Vanchiere |
| EPI_ISL_2229435 | Virginia Division of Consolidated Laboratory Services | Virginia Division of Consolidated Laboratory Services | Virginia DCLS |
| EPI_ISL_2229531, EPI_ISL_2229775 | Massachusetts State Public Health Laboratory | Massachusetts State Public Health Laboratory | Andrew Lang, Timelia Fink, Glen Gallagher, Sandra Smole |
| EPI_ISL_2230013 | Florida Bureau of Public Health Laboratories | Florida Bureau of Public Health Laboratories | Sarah Schmedes, Jason Blanton |
| EPI_ISL_2230038 | The Jackson Laboratory | The Jackson Laboratory | Long J, Bergeron D, Renzette N, Adams M, Omerza G, Kelly K, Li L |
| EPI_ISL_2230722 | Southern Nevada Public Health Laboratory | Southern Nevada Public Health Laboratory | Michael Picker |
| EPI_ISL_2231198 | Sharp HealthCare Laboratory | Andersen lab at Scripps Research | SEARCH Alliance San Diego with Aaron Harding, Jacquelyn Berumen, Cathy Woerle, Liam McGinnis, Art Mendoza, Omid Bakhtar |
| EPI_ISL_2231590 | AP SSO | CSIR-Centre for Cellular and Molecular Biology-INSACOG | Lamuk Zaveri,Tulasi Nagabandi, Ara Sreenivas,Shreekant Verma, Amareshwar Vodapalli ,Blessy B John,Viswagithe S L,B Himasri,Valli Nagalakshmi Undamatla,Payel Mukherjee,Sofia Banu,Archana Bharadwaj Siva,Sharath Chandra Thota,Karthik Bharadwaj Tallapaka,Rakesh K Mishra,Divya Tej Sowpati |
| EPI_ISL_2231597 | AP SSO | CSIR-Centre for Cellular and Molecular Biology-INSACOG | Payel Mukherjee,Lamuk Zaveri,Tulasi Nagabandi, Ara Sreenivas,Shreekant Verma, Amareshwar Vodapalli ,Blessy B John,Viswagithe S L,B Himasri,Valli Nagalakshmi Undamatla,Sofia Banu,Archana Bharadwaj Siva,Sharath Chandra Thota,Karthik Bharadwaj Tallapaka,Rakesh K Mishra,Divya Tej Sowpati |
| EPI_ISL_2231598 | AP SSO | CSIR-Centre for Cellular and Molecular Biology-INSACOG | B Himasri,Valli Nagalakshmi Undamatla,Lamuk Zaveri,Tulasi Nagabandi, Ara Sreenivas,Shreekant Verma, Amareshwar Vodapalli ,Blessy B John,Viswagithe S L,Payel Mukherjee,Sofia Banu,Archana Bharadwaj Siva,Sharath Chandra Thota,Karthik Bharadwaj Tallapaka,Rakesh K Mishra,Divya Tej Sowpati |
| EPI_ISL_2231599 | AP SSO | CSIR-Centre for Cellular and Molecular Biology-INSACOG | Lamuk Zaveri,Tulasi Nagabandi, Ara Sreenivas,Shreekant Verma, Amareshwar Vodapalli ,Blessy B John,Viswagithe S L,B Himasri,Valli Nagalakshmi Undamatla,Payel Mukherjee,Sofia Banu,Archana Bharadwaj Siva,Sharath Chandra Thota,Karthik Bharadwaj Tallapaka,Rakesh K Mishra,Divya Tej Sowpati |
| EPI_ISL_2231601 | AP SSO | CSIR-Centre for Cellular and Molecular Biology-INSACOG | Payel Mukherjee,Lamuk Zaveri,Tulasi Nagabandi, Ara Sreenivas,Shreekant Verma, Amareshwar Vodapalli ,Blessy B John,Viswagithe S L,B Himasri,Valli Nagalakshmi Undamatla,Sofia Banu,Archana Bharadwaj Siva,Sharath Chandra Thota,Karthik Bharadwaj Tallapaka,Rakesh K Mishra,Divya Tej Sowpati |
| EPI_ISL_2231605 | AP SSO | CSIR-Centre for Cellular and Molecular Biology-INSACOG | B Himasri,Valli Nagalakshmi Undamatla,Lamuk Zaveri,Tulasi Nagabandi, Ara Sreenivas,Shreekant Verma, Amareshwar Vodapalli ,Blessy B John,Viswagithe S L,Payel Mukherjee,Sofia Banu,Archana Bharadwaj Siva,Sharath Chandra Thota,Karthik Bharadwaj Tallapaka,Rakesh K Mishra,Divya Tej Sowpati |
| EPI_ISL_2231607, EPI_ISL_2231608, EPI_ISL_2231609, EPI_ISL_2231611 | AP SSO | CSIR-Centre for Cellular and Molecular Biology-INSACOG | Lamuk Zaveri,Tulasi Nagabandi, Ara Sreenivas,Shreekant Verma, Amareshwar Vodapalli ,Blessy B John,Viswagithe S L,B Himasri,Valli Nagalakshmi Undamatla,Payel Mukherjee,Sofia Banu,Archana Bharadwaj Siva,Sharath Chandra Thota,Karthik Bharadwaj Tallapaka,Rakesh K Mishra,Divya Tej Sowpati |
| EPI_ISL_2231612, EPI_ISL_2231614 | AP SSO | CSIR-Centre for Cellular and Molecular Biology-INSACOG | Payel Mukherjee,Lamuk Zaveri,Tulasi Nagabandi, Ara Sreenivas,Shreekant Verma, Amareshwar Vodapalli ,Blessy B John,Viswagithe S L,B Himasri,Valli Nagalakshmi Undamatla,Sofia Banu,Archana Bharadwaj Siva,Sharath Chandra Thota,Karthik Bharadwaj Tallapaka,Rakesh K Mishra,Divya Tej Sowpati |
| EPI_ISL_2231615, EPI_ISL_2231616 | AP SSO | CSIR-Centre for Cellular and Molecular Biology-INSACOG | Lamuk Zaveri,Tulasi Nagabandi, Ara Sreenivas,Shreekant Verma, Amareshwar Vodapalli ,Blessy B John,Viswagithe S L,B Himasri,Valli Nagalakshmi Undamatla,Payel Mukherjee,Sofia Banu,Archana Bharadwaj Siva,Sharath Chandra Thota,Karthik Bharadwaj Tallapaka,Rakesh K Mishra,Divya Tej Sowpati |
| EPI_ISL_2231627 | AP SSO | CSIR-Centre for Cellular and Molecular Biology-INSACOG | B Himasri,Valli Nagalakshmi Undamatla,Lamuk Zaveri,Tulasi Nagabandi, Ara Sreenivas,Shreekant Verma, Amareshwar Vodapalli ,Blessy B John,Viswagithe S L,Payel Mukherjee,Sofia Banu,Archana Bharadwaj Siva,Sharath Chandra Thota,Karthik Bharadwaj Tallapaka,Rakesh K Mishra,Divya Tej Sowpati |
| EPI_ISL_2231628 | AP SSO | CSIR-Centre for Cellular and Molecular Biology-INSACOG | Payel Mukherjee,Lamuk Zaveri,Tulasi Nagabandi, Ara Sreenivas,Shreekant Verma, Amareshwar Vodapalli ,Blessy B John,Viswagithe S L,B Himasri,Valli Nagalakshmi Undamatla,Sofia Banu,Archana Bharadwaj Siva,Sharath Chandra Thota,Karthik Bharadwaj Tallapaka,Rakesh K Mishra,Divya Tej Sowpati |
| EPI_ISL_2231632 | AP SSO | CSIR-Centre for Cellular and Molecular Biology-INSACOG | Lamuk Zaveri,Tulasi Nagabandi, Ara Sreenivas,Shreekant Verma, Amareshwar Vodapalli ,Blessy B John,Viswagithe S L,B Himasri,Valli Nagalakshmi Undamatla,Payel Mukherjee,Sofia Banu,Archana Bharadwaj Siva,Sharath Chandra Thota,Karthik Bharadwaj Tallapaka,Rakesh K Mishra,Divya Tej Sowpati |
| EPI_ISL_2231633 | AP SSO | CSIR-Centre for Cellular and Molecular Biology-INSACOG | Tulasi Nagabandi, Lamuk Zaveri,Ara Sreenivas,Shreekant Verma, Amareshwar Vodapalli ,Blessy B John,Viswagithe S L,B Himasri,Valli Nagalakshmi Undamatla,Payel Mukherjee,Sofia Banu,Archana Bharadwaj Siva,Sharath Chandra Thota,Karthik Bharadwaj Tallapaka,Rakesh K Mishra,Divya Tej Sowpati |

| | | | |
|---|---|---|---|
| EPI_ISL_2231634, EPI_ISL_2231635 | AP SSO | CSIR-Centre for Cellular and Molecular Biology-INSACOG | Lamuk Zaveri,Tulasi Nagabandi, Ara Sreenivas,Shreekant Verma, Amareshwar Vodapalli ,Blessy B John,Viswagithe S L,B Himasri,Valli Nagalakshmi Undamatla,Payel Mukherjee,Sofia Banu,Archana Bharadwaj Siva,Sharath Chandra Thota,Karthik Bharadwaj Tallapaka,Rakesh K Mishra,Divya Tej Sowpati |
| EPI_ISL_2231636 | AP SSO | CSIR-Centre for Cellular and Molecular Biology-INSACOG | Tulasi Nagabandi, Lamuk Zaveri,Ara Sreenivas,Shreekant Verma, Amareshwar Vodapalli ,Blessy B John,Viswagithe S L,B Himasri,Valli Nagalakshmi Undamatla,Payel Mukherjee,Sofia Banu,Archana Bharadwaj Siva,Sharath Chandra Thota,Karthik Bharadwaj Tallapaka,Rakesh K Mishra,Divya Tej Sowpati |
| EPI_ISL_2231639 | AP SSO | CSIR-Centre for Cellular and Molecular Biology-INSACOG | Lamuk Zaveri,Tulasi Nagabandi, Ara Sreenivas,Shreekant Verma, Amareshwar Vodapalli ,Blessy B John,Viswagithe S L,B Himasri,Valli Nagalakshmi Undamatla,Payel Mukherjee,Sofia Banu,Archana Bharadwaj Siva,Sharath Chandra Thota,Karthik Bharadwaj Tallapaka,Rakesh K Mishra,Divya Tej Sowpati |
| EPI_ISL_2231640 | AP SSO | CSIR-Centre for Cellular and Molecular Biology-INSACOG | Tulasi Nagabandi, Lamuk Zaveri,Ara Sreenivas,Shreekant Verma, Amareshwar Vodapalli ,Blessy B John,Viswagithe S L,B Himasri,Valli Nagalakshmi Undamatla,Payel Mukherjee,Sofia Banu,Archana Bharadwaj Siva,Sharath Chandra Thota,Karthik Bharadwaj Tallapaka,Rakesh K Mishra,Divya Tej Sowpati |
| EPI_ISL_2231641 | AP SSO | CSIR-Centre for Cellular and Molecular Biology-INSACOG | Payel Mukherjee,Lamuk Zaveri,Tulasi Nagabandi, Ara Sreenivas,Shreekant Verma, Amareshwar Vodapalli ,Blessy B John,Viswagithe S L,B Himasri,Valli Nagalakshmi Undamatla,Sofia Banu,Archana Bharadwaj Siva,Sharath Chandra Thota,Karthik Bharadwaj Tallapaka,Rakesh K Mishra,Divya Tej Sowpati |
| EPI_ISL_2231642 | AP SSO | CSIR-Centre for Cellular and Molecular Biology-INSACOG | Lamuk Zaveri,Tulasi Nagabandi, Ara Sreenivas,Shreekant Verma, Amareshwar Vodapalli ,Blessy B John,Viswagithe S L,B Himasri,Valli Nagalakshmi Undamatla,Payel Mukherjee,Sofia Banu,Archana Bharadwaj Siva,Sharath Chandra Thota,Karthik Bharadwaj Tallapaka,Rakesh K Mishra,Divya Tej Sowpati |
| EPI_ISL_2231645 | AP SSO | CSIR-Centre for Cellular and Molecular Biology-INSACOG | Payel Mukherjee,Lamuk Zaveri,Tulasi Nagabandi, Ara Sreenivas,Shreekant Verma, Amareshwar Vodapalli ,Blessy B John,Viswagithe S L,B Himasri,Valli Nagalakshmi Undamatla,Sofia Banu,Archana Bharadwaj Siva,Sharath Chandra Thota,Karthik Bharadwaj Tallapaka,Rakesh K Mishra,Divya Tej Sowpati |
| EPI_ISL_2231650, EPI_ISL_2231651, EPI_ISL_2231652, EPI_ISL_2231654 | AP SSO | CSIR-Centre for Cellular and Molecular Biology-INSACOG | Tulasi Nagabandi, Lamuk Zaveri,Ara Sreenivas,Shreekant Verma, Amareshwar Vodapalli ,Blessy B John,Viswagithe S L,B Himasri,Valli Nagalakshmi Undamatla,Payel Mukherjee,Sofia Banu,Archana Bharadwaj Siva,Sharath Chandra Thota,Karthik Bharadwaj Tallapaka,Rakesh K Mishra,Divya Tej Sowpati |
| EPI_ISL_2231656 | AP SSO | CSIR-Centre for Cellular and Molecular Biology-INSACOG | Payel Mukherjee,Lamuk Zaveri,Tulasi Nagabandi, Ara Sreenivas,Shreekant Verma, Amareshwar Vodapalli ,Blessy B John,Viswagithe S L,B Himasri,Valli Nagalakshmi Undamatla,Sofia Banu,Archana Bharadwaj Siva,Sharath Chandra Thota,Karthik Bharadwaj Tallapaka,Rakesh K Mishra,Divya Tej Sowpati |
| EPI_ISL_2231659, EPI_ISL_2231660, EPI_ISL_2231665, EPI_ISL_2231666, EPI_ISL_2231667, EPI_ISL_2231669 | AP SSO | CSIR-Centre for Cellular and Molecular Biology-INSACOG | Tulasi Nagabandi, Lamuk Zaveri,Ara Sreenivas,Shreekant Verma, Amareshwar Vodapalli ,Blessy B John,Viswagithe S L,B Himasri,Valli Nagalakshmi Undamatla,Payel Mukherjee,Sofia Banu,Archana Bharadwaj Siva,Sharath Chandra Thota,Karthik Bharadwaj Tallapaka,Rakesh K Mishra,Divya Tej Sowpati |
| EPI_ISL_2231672 | AP SSO | CSIR-Centre for Cellular and Molecular Biology-INSACOG | Payel Mukherjee,Lamuk Zaveri,Tulasi Nagabandi, Ara Sreenivas,Shreekant Verma, Amareshwar Vodapalli ,Blessy B John,Viswagithe S L,B Himasri,Valli Nagalakshmi Undamatla,Sofia Banu,Archana Bharadwaj Siva,Sharath Chandra Thota,Karthik Bharadwaj Tallapaka,Rakesh K Mishra,Divya Tej Sowpati |
| EPI_ISL_2231677, EPI_ISL_2231678 | AP SSO | CSIR-Centre for Cellular and Molecular Biology-INSACOG | Lamuk Zaveri,Tulasi Nagabandi, Ara Sreenivas,Shreekant Verma, Amareshwar Vodapalli ,Blessy B John,Viswagithe S L,B Himasri,Valli Nagalakshmi Undamatla,Payel Mukherjee,Sofia Banu,Archana Bharadwaj Siva,Sharath Chandra Thota,Karthik Bharadwaj Tallapaka,Rakesh K Mishra,Divya Tej Sowpati |
| EPI_ISL_2231680 | AP SSO | CSIR-Centre for Cellular and Molecular Biology-INSACOG | Tulasi Nagabandi, Lamuk Zaveri,Ara Sreenivas,Shreekant Verma, Amareshwar Vodapalli ,Blessy B John,Viswagithe S L,B Himasri,Valli Nagalakshmi Undamatla,Payel Mukherjee,Sofia Banu,Archana Bharadwaj Siva,Sharath Chandra Thota,Karthik Bharadwaj Tallapaka,Rakesh K Mishra,Divya Tej Sowpati |
| EPI_ISL_2231681 | AP SSO | CSIR-Centre for Cellular and Molecular Biology-INSACOG | Lamuk Zaveri,Tulasi Nagabandi, Ara Sreenivas,Shreekant Verma, Amareshwar Vodapalli ,Blessy B John,Viswagithe S L,B Himasri,Valli Nagalakshmi Undamatla,Payel Mukherjee,Sofia Banu,Archana Bharadwaj Siva,Sharath Chandra Thota,Karthik Bharadwaj Tallapaka,Rakesh K Mishra,Divya Tej Sowpati |
| EPI_ISL_2231684 | AP SSO | CSIR-Centre for Cellular and Molecular Biology-INSACOG | Payel Mukherjee,Lamuk Zaveri,Tulasi Nagabandi, Ara Sreenivas,Shreekant Verma, Amareshwar Vodapalli ,Blessy B John,Viswagithe S L,B Himasri,Valli Nagalakshmi Undamatla,Sofia Banu,Archana Bharadwaj Siva,Sharath Chandra Thota,Karthik Bharadwaj Tallapaka,Rakesh K Mishra,Divya Tej Sowpati |
| EPI_ISL_2231685 | AP SSO | CSIR-Centre for Cellular and Molecular Biology-INSACOG | Lamuk Zaveri,Tulasi Nagabandi, Ara Sreenivas,Shreekant Verma, Amareshwar Vodapalli ,Blessy B John,Viswagithe S L,B Himasri,Valli Nagalakshmi Undamatla,Payel Mukherjee,Sofia Banu,Archana Bharadwaj Siva,Sharath Chandra Thota,Karthik Bharadwaj Tallapaka,Rakesh K Mishra,Divya Tej Sowpati |
| EPI_ISL_2231687, EPI_ISL_2231690 | AP SSO | CSIR-Centre for Cellular and Molecular Biology-INSACOG | Payel Mukherjee,Lamuk Zaveri,Tulasi Nagabandi, Ara Sreenivas,Shreekant Verma, Amareshwar Vodapalli ,Blessy B John,Viswagithe S L,B Himasri,Valli Nagalakshmi Undamatla,Sofia Banu,Archana Bharadwaj Siva,Sharath Chandra Thota,Karthik Bharadwaj Tallapaka,Rakesh K Mishra,Divya Tej Sowpati |
| EPI_ISL_2231691 | AP SSO | CSIR-Centre for Cellular and Molecular Biology-INSACOG | Lamuk Zaveri,Tulasi Nagabandi, Ara Sreenivas,Shreekant Verma, Amareshwar Vodapalli ,Blessy B John,Viswagithe S L,B Himasri,Valli Nagalakshmi Undamatla,Payel Mukherjee,Sofia Banu,Archana Bharadwaj Siva,Sharath Chandra Thota,Karthik Bharadwaj Tallapaka,Rakesh K Mishra,Divya Tej Sowpati |
| EPI_ISL_2231695 | AP SSO | CSIR-Centre for Cellular and Molecular Biology-INSACOG | Tulasi Nagabandi, Lamuk Zaveri,Ara Sreenivas,Shreekant Verma, Amareshwar Vodapalli ,Blessy B John,Viswagithe S L,B Himasri,Valli Nagalakshmi Undamatla,Payel Mukherjee,Sofia Banu,Archana Bharadwaj Siva,Sharath Chandra Thota,Karthik Bharadwaj Tallapaka,Rakesh K Mishra,Divya Tej Sowpati |
| EPI_ISL_2231696 | AP SSO | CSIR-Centre for Cellular and Molecular Biology-INSACOG | Payel Mukherjee,Lamuk Zaveri,Tulasi Nagabandi, Ara Sreenivas,Shreekant Verma, Amareshwar Vodapalli ,Blessy B John,Viswagithe S L,B Himasri,Valli Nagalakshmi Undamatla,Sofia Banu,Archana Bharadwaj Siva,Sharath Chandra Thota,Karthik Bharadwaj Tallapaka,Rakesh K Mishra,Divya Tej Sowpati |
| EPI_ISL_2231697 | AP SSO | CSIR-Centre for Cellular and Molecular Biology-INSACOG | Lamuk Zaveri,Tulasi Nagabandi, Ara Sreenivas,Shreekant Verma, Amareshwar Vodapalli ,Blessy B John,Viswagithe S L,B Himasri,Valli Nagalakshmi Undamatla,Payel Mukherjee,Sofia Banu,Archana Bharadwaj Siva,Sharath Chandra Thota,Karthik Bharadwaj Tallapaka,Rakesh K Mishra,Divya Tej Sowpati |
| EPI_ISL_2231701 | AP SSO | CSIR-Centre for Cellular and Molecular Biology-INSACOG | Payel Mukherjee,Lamuk Zaveri,Tulasi Nagabandi, Ara Sreenivas,Shreekant Verma, Amareshwar Vodapalli ,Blessy B John,Viswagithe S L,B Himasri,Valli Nagalakshmi Undamatla,Sofia Banu,Archana Bharadwaj Siva,Sharath Chandra Thota,Karthik Bharadwaj Tallapaka,Rakesh K Mishra,Divya Tej Sowpati |
| EPI_ISL_2231702 | AP SSO | CSIR-Centre for Cellular and Molecular Biology-INSACOG | Lamuk Zaveri,Tulasi Nagabandi, Ara Sreenivas,Shreekant Verma, Amareshwar Vodapalli ,Blessy B John,Viswagithe S L,B Himasri,Valli Nagalakshmi Undamatla,Payel Mukherjee,Sofia Banu,Archana Bharadwaj Siva,Sharath Chandra Thota,Karthik Bharadwaj Tallapaka,Rakesh K Mishra,Divya Tej Sowpati |
| EPI_ISL_2231703, EPI_ISL_2231706, EPI_ISL_2231707 | AP SSO | CSIR-Centre for Cellular and Molecular Biology-INSACOG | Payel Mukherjee,Lamuk Zaveri,Tulasi Nagabandi, Ara Sreenivas,Shreekant Verma, Amareshwar Vodapalli ,Blessy B John,Viswagithe S L,B Himasri,Valli Nagalakshmi Undamatla,Sofia Banu,Archana Bharadwaj Siva,Sharath Chandra Thota,Karthik Bharadwaj Tallapaka,Rakesh K Mishra,Divya Tej Sowpati |
| EPI_ISL_2231711 | AP SSO | CSIR-Centre for Cellular and Molecular Biology-INSACOG | B Himasri,Valli Nagalakshmi Undamatla,Lamuk Zaveri,Tulasi Nagabandi, Ara Sreenivas,Shreekant Verma, Amareshwar Vodapalli ,Blessy B John,Viswagithe S L,Payel Mukherjee,Sofia Banu,Archana Bharadwaj Siva,Sharath Chandra Thota,Karthik Bharadwaj Tallapaka,Rakesh K Mishra,Divya Tej Sowpati |
| EPI_ISL_2231712 | AP SSO | CSIR-Centre for Cellular and Molecular Biology-INSACOG | Tulasi Nagabandi, Lamuk Zaveri,Ara Sreenivas,Shreekant Verma, Amareshwar Vodapalli ,Blessy B John,Viswagithe S L,B Himasri,Valli Nagalakshmi Undamatla,Payel Mukherjee,Sofia Banu,Archana Bharadwaj Siva,Sharath Chandra Thota,Karthik Bharadwaj Tallapaka,Rakesh K Mishra,Divya Tej Sowpati |
| EPI_ISL_2231713 | AP SSO | CSIR-Centre for Cellular and Molecular Biology-INSACOG | Payel Mukherjee,Lamuk Zaveri,Tulasi Nagabandi, Ara Sreenivas,Shreekant Verma, Amareshwar Vodapalli ,Blessy B John,Viswagithe S L,B Himasri,Valli Nagalakshmi Undamatla,Sofia Banu,Archana Bharadwaj Siva,Sharath Chandra Thota,Karthik Bharadwaj Tallapaka,Rakesh K Mishra,Divya Tej Sowpati |
| EPI_ISL_2231716 | AP SSO | CSIR-Centre for Cellular and Molecular Biology-INSACOG | Lamuk Zaveri,Tulasi Nagabandi, Ara Sreenivas,Shreekant Verma, Amareshwar Vodapalli ,Blessy B John,Viswagithe S L,B Himasri,Valli Nagalakshmi Undamatla,Payel Mukherjee,Sofia Banu,Archana Bharadwaj Siva,Sharath Chandra Thota,Karthik Bharadwaj Tallapaka,Rakesh K Mishra,Divya Tej Sowpati |
| EPI_ISL_2231721, EPI_ISL_2231726 | AP SSO | CSIR-Centre for Cellular and Molecular Biology-INSACOG | Payel Mukherjee,Lamuk Zaveri,Tulasi Nagabandi, Ara Sreenivas,Shreekant Verma, Amareshwar Vodapalli ,Blessy B John,Viswagithe S L,B Himasri,Valli Nagalakshmi Undamatla,Sofia Banu,Archana Bharadwaj Siva,Sharath Chandra Thota,Karthik Bharadwaj Tallapaka,Rakesh K Mishra,Divya Tej Sowpati |
| EPI_ISL_2231728 | AP SSO | CSIR-Centre for Cellular and Molecular Biology-INSACOG | Tulasi Nagabandi, Lamuk Zaveri,Ara Sreenivas,Shreekant Verma, Amareshwar Vodapalli ,Blessy B John,Viswagithe S L,B Himasri,Valli Nagalakshmi Undamatla,Payel Mukherjee,Sofia Banu,Archana Bharadwaj Siva,Sharath Chandra Thota,Karthik Bharadwaj Tallapaka,Rakesh K Mishra,Divya Tej Sowpati |
| EPI_ISL_2231730, EPI_ISL_2231732 | AP SSO | CSIR-Centre for Cellular and Molecular Biology-INSACOG | Payel Mukherjee,Lamuk Zaveri,Tulasi Nagabandi, Ara Sreenivas,Shreekant Verma, Amareshwar Vodapalli ,Blessy B John,Viswagithe S L,B Himasri,Valli Nagalakshmi Undamatla,Sofia Banu,Archana Bharadwaj Siva,Sharath Chandra Thota,Karthik Bharadwaj Tallapaka,Rakesh K Mishra,Divya Tej Sowpati |
| EPI_ISL_2231738 | AP SSO | CSIR-Centre for Cellular and Molecular Biology-INSACOG | Lamuk Zaveri,Tulasi Nagabandi, Ara Sreenivas,Shreekant Verma, Amareshwar Vodapalli ,Blessy B John,Viswagithe S L,B Himasri,Valli Nagalakshmi Undamatla,Payel Mukherjee,Sofia Banu,Archana Bharadwaj Siva,Sharath Chandra Thota,Karthik Bharadwaj Tallapaka,Rakesh K Mishra,Divya Tej Sowpati |
| EPI_ISL_2231739 | AP SSO | CSIR-Centre for Cellular and Molecular Biology-INSACOG | B Himasri,Valli Nagalakshmi Undamatla,Lamuk Zaveri,Tulasi Nagabandi, Ara Sreenivas,Shreekant Verma, Amareshwar Vodapalli ,Blessy B John,Viswagithe S L,Payel Mukherjee,Sofia Banu,Archana Bharadwaj Siva,Sharath Chandra Thota,Karthik Bharadwaj Tallapaka,Rakesh K Mishra,Divya Tej Sowpati |
| EPI_ISL_2231746, EPI_ISL_2231747, EPI_ISL_2231748, EPI_ISL_2231752 | AP SSO | CSIR-Centre for Cellular and Molecular Biology-INSACOG | Payel Mukherjee,Lamuk Zaveri,Tulasi Nagabandi, Ara Sreenivas,Shreekant Verma, Amareshwar Vodapalli ,Blessy B John,Viswagithe S L,B Himasri,Valli Nagalakshmi Undamatla,Sofia Banu,Archana Bharadwaj Siva,Sharath Chandra Thota,Karthik Bharadwaj Tallapaka,Rakesh K Mishra,Divya Tej Sowpati |
| EPI_ISL_2231754 | AP SSO | CSIR-Centre for Cellular and Molecular Biology-INSACOG | B Himasri,Valli Nagalakshmi Undamatla,Lamuk Zaveri,Tulasi Nagabandi, Ara Sreenivas,Shreekant Verma, Amareshwar Vodapalli ,Blessy B John,Viswagithe S L,Payel Mukherjee,Sofia Banu,Archana Bharadwaj Siva,Sharath Chandra Thota,Karthik Bharadwaj Tallapaka,Rakesh K Mishra,Divya Tej Sowpati |
| EPI_ISL_2231756, EPI_ISL_2231757, EPI_ISL_2231759, EPI_ISL_2231761 | AP SSO | CSIR-Centre for Cellular and Molecular Biology-INSACOG | Tulasi Nagabandi, Lamuk Zaveri,Ara Sreenivas,Shreekant Verma, Amareshwar Vodapalli ,Blessy B John,Viswagithe S L,B Himasri,Valli Nagalakshmi Undamatla,Payel Mukherjee,Sofia Banu,Archana Bharadwaj Siva,Sharath Chandra Thota,Karthik Bharadwaj Tallapaka,Rakesh K Mishra,Divya Tej Sowpati |
| EPI_ISL_2231767 | AP SSO | CSIR-Centre for Cellular and Molecular Biology-INSACOG | Payel Mukherjee,Lamuk Zaveri,Tulasi Nagabandi, Ara Sreenivas,Shreekant Verma, Amareshwar Vodapalli ,Blessy B John,Viswagithe S L,B Himasri,Valli Nagalakshmi Undamatla,Sofia Banu,Archana Bharadwaj Siva,Sharath Chandra Thota,Karthik Bharadwaj Tallapaka,Rakesh K Mishra,Divya Tej Sowpati |
| EPI_ISL_2231769, EPI_ISL_2231772, EPI_ISL_2231773, EPI_ISL_2231777, EPI_ISL_2231779, EPI_ISL_2231782, EPI_ISL_2231784, EPI_ISL_2231794 | AP SSO | CSIR-Centre for Cellular and Molecular Biology-INSACOG | Tulasi Nagabandi, Lamuk Zaveri,Ara Sreenivas,Shreekant Verma, Amareshwar Vodapalli ,Blessy B John,Viswagithe S L,B Himasri,Valli Nagalakshmi Undamatla,Payel Mukherjee,Sofia Banu,Archana Bharadwaj Siva,Sharath Chandra Thota,Karthik Bharadwaj Tallapaka,Rakesh K Mishra,Divya Tej Sowpati |

| | | | |
|---|---|---|---|
| EPI_ISL_2231798, EPI_ISL_2231799 | AP SSO | CSIR-Centre for Cellular and Molecular Biology-INSACOG | Payel Mukherjee,Lamuk Zaveri,Tulasi Nagabandi, Ara Sreenivas,Shreekant Verma, Amareshwar Vodapalli ,Blessy B John,Viswagithe S L,B Himasri,Valli Nagalakshmi Undamatla,Sofia Banu,Archana Bharadwaj Siva,Sharath Chandra Thota,Karthik Bharadwaj Tallapaka,Rakesh K Mishra,Divya Tej Sowpati |
| EPI_ISL_2231808, EPI_ISL_2231809, EPI_ISL_2231812, EPI_ISL_2231821, EPI_ISL_2231822, EPI_ISL_2231823, EPI_ISL_2231830, EPI_ISL_2231831, EPI_ISL_2231832, EPI_ISL_2231841, EPI_ISL_2231848, EPI_ISL_2231849, EPI_ISL_2231850, EPI_ISL_2231853, EPI_ISL_2231854, EPI_ISL_2231855, EPI_ISL_2231864, EPI_ISL_2231866, EPI_ISL_2231867, EPI_ISL_2231869 | | | |
| see above | AP SSO | CSIR-Centre for Cellular and Molecular Biology-INSACOG | Tulasi Nagabandi, Lamuk Zaveri,Ara Sreenivas,Shreekant Verma, Amareshwar Vodapalli ,Blessy B John,Viswagithe S L,B Himasri,Valli Nagalakshmi Undamatla,Payel Mukherjee,Sofia Banu,Archana Bharadwaj Siva,Sharath Chandra Thota,Karthik Bharadwaj Tallapaka,Rakesh K Mishra,Divya Tej Sowpati |
| EPI_ISL_2232671 | Ministry of Health Turkey | Ministry of Health Turkey | Fatma Bayrakdar, Yasemin Cosgun, Suleyman Yalcin, Gulay Korukluoglu |
| EPI_ISL_2232832 | Austrian Agency for Health and Food Safety (AGES) | Bergthaler laboratory, CeMM Research Center for Molecular Medicine of the Austrian Academy of Sciences | Lukas Endler, Anna Schedl, Fabian Amman, Petr Triska, Thomas Penz, Benedikt Agerer, Maelle Le Moing, Michael Schuster, Bekir Erguner, Jan Laine, Martin Senekowitsch, Christoph Bock, Andreas Bergthaler |
| EPI_ISL_2233082 | Nebraska Public Health Laboratory | NPHL COVID-19 Response Team | NPHL COVID-19 Response Team |
| EPI_ISL_2233089 | RSUD Abdul Moeis | National Institute of Health Research and Development | Vivi Setiawaty, Hartanti Dian Ikawati, Hana Apsari Pawestri, Subangkit, Kartika Dewi Puspa, Arie Ardiansyah Nugraha, Triyani Soekarso, Krisna Pangesti, Nelly Puspandari |
| EPI_ISL_2233090 | Dinkes Samarinda | National Institute of Health Research and Development | Vivi Setiawaty, Hartanti Dian Ikawati, Hana Apsari Pawestri, Subangkit, Kartika Dewi Puspa, Arie Ardiansyah Nugraha, Triyani Soekarso, Krisna Pangesti, Nelly Puspandari |
| EPI_ISL_2233091, EPI_ISL_2233092, EPI_ISL_2233093 | Kantor Kesehatan Pelabuhan Kelas II Cilacap | National Institute of Health Research and Development | Kartika Dewi Puspa, Subangkit, Arie Ardiansyah Nugraha, Hartanti Dian Ikawati, Hana Apsari Pawestri, Triyani Soekarso, Krisna Pangesti, Nelly Puspandari, Vivi Setiawaty |
| EPI_ISL_2233094 | Labkesda Prov Kal Tim | National Institute of Health Research and Development | Kartika Dewi Puspa, Subangkit, Arie Ardiansyah Nugraha, Hartanti Dian Ikawati, Hana Apsari Pawestri, Triyani Soekarso, Krisna Pangesti, Nelly Puspandari, Vivi Setiawaty |
| EPI_ISL_2233095 | Hotel Horison | National Institute of Health Research and Development | Kartika Dewi Puspa, Subangkit, Arie Ardiansyah Nugraha, Hartanti Dian Ikawati, Hana Apsari Pawestri, Triyani Soekarso, Krisna Pangesti, Nelly Puspandari, Vivi Setiawaty |
| EPI_ISL_2233096, EPI_ISL_2233097, EPI_ISL_2233098, EPI_ISL_2233099, EPI_ISL_2233100, EPI_ISL_2233101, EPI_ISL_2233102, EPI_ISL_2233103, EPI_ISL_2233104, EPI_ISL_2233105 | Kantor Kesehatan Pelabuhan Kelas II Cilacap | National Institute of Health Research and Development | Subangkit, Hana Apsari Pawestri, Hartanti Dian Ikawati, Kartika Dewi Puspa, Arie Ardiansyah Nugraha, Triyani Soekarso, Krisna Pangesti, Nelly Puspandari, Vivi Setiawaty |
| EPI_ISL_2233145, EPI_ISL_2233146 | National Centre For Cell Science | National Centre For Cell Science – INSACOG | Dhiraj Paul, Shivang P. Bhanushali, Mitali Inamdar, Mohak P Gujare, Manoj Kumar Bhat, Ajay Pillai, INSACOG Consortium team, Yogesh Shouche |
| EPI_ISL_2233209, EPI_ISL_2233280 | UW Virology Lab | UW Virology Lab | Pavitra Roychoudhury, Hong Xie, Lasata Shrestha, Tien V. Nguyen, Shah Mohamed Bakhash, Michelle Lin, Noah R. Baker, Sean Ellis, Meei-Li Huang, Keith R Jerome, Alexander Greninger |
| EPI_ISL_2233382 | Hospital Labuan | Institute for Medical Research, Infectious Disease Research Centre, National Institutes of Health, Ministry of Health Malaysia | Suppiah J, Kamel K, Mohd Zawawi Z, Azizan MA, Ramly N, Robert F, Thayan R |
| EPI_ISL_2233495, EPI_ISL_2233544, EPI_ISL_2233550, EPI_ISL_2233595, EPI_ISL_2233667 | National Centre for Disease Control | CDFD -INSACOG | Murali Bashyam, Asmita Gupta, Pratyusha Bala, Vinay Donipadi, Divya Vashisht, Ashwin Dalal |
| EPI_ISL_2234965, EPI_ISL_2234986, EPI_ISL_2234993, EPI_ISL_2235011, EPI_ISL_2235025, EPI_ISL_2235027, EPI_ISL_2235032, EPI_ISL_2235043, EPI_ISL_2235047, EPI_ISL_2235061, EPI_ISL_2235077, EPI_ISL_2235113, EPI_ISL_2235124, EPI_ISL_2235135, EPI_ISL_2235207, EPI_ISL_2235209, EPI_ISL_2235216 | | | |
| see above | Pandemic Response Lab - NYC | Pandemic Response Lab, R&D | Henry Lee, Michael Hammerling, Melissa Hopkins, Cybill del Castillo, Shinyoung Clair Kang, William Ward, Pradeep Bugga, Sol Rey, Dylan Law, Katharine Nelson, Haiping Hao, Jon Laurent |
| EPI_ISL_2235305 | AZ Zeno Knokke-Blankenberge | AZ SINT-JAN BRUGGE | Marijke Reynders, Patrick Descheemaeker, Laurien Hoornaert, Thomas Van Landschoot, Jorn Hellemans, Marie Madeleine Chabert-Consen |
| EPI_ISL_2235328 | SIESP CHIETI - DRIVE IN CHIETI | Istituto Zooprofilattico Sperimentale dell'Abruzzo e Molise "G. Caporale" | Lorusso A, Marcacci M, Di Domenico M, Ancora M, Curini V, Di Lollo Valeria, Mangone I, Rinaldi A, Delli Compagni E, Scialabba S, Caporale M, Di Pasquale A, Cammà C, Puglia I, Calistri P, Savini G |
| EPI_ISL_2235528, EPI_ISL_2235530 | Lighthouse Lab in Alderley Park | Wellcome Sanger Institute for the COVID-19 Genomics UK (COG-UK) Consortium | Jacquelyn Wynn, Mairead Hyland, The Lighthouse Lab in Alderley Park and Alex Alderton, Roberto Amato, Jeffrey Barrett, Sonia Goncalves, Ewan Harrison, David K. Jackson, Ian Johnston, Dominic Kwiatkowski, Cordelia Langford, John Sillitoe on behalf of the Wellcome Sanger Institute COVID-19 Surveillance Team |
| EPI_ISL_2235531 | Lighthouse Lab in Milton Keynes | Wellcome Sanger Institute for the COVID-19 Genomics UK (COG-UK) Consortium | The Lighthouse Lab in Milton Keynes and Alex Alderton, Roberto Amato, Jeffrey Barrett, Sonia Goncalves, Ewan Harrison, David K. Jackson, Ian Johnston, Dominic Kwiatkowski, Cordelia Langford, John Sillitoe on behalf of the Wellcome Sanger Institute COVID-19 Surveillance Team |
| EPI_ISL_2235532, EPI_ISL_2235533, EPI_ISL_2235534, EPI_ISL_2235536 | Lighthouse Lab in Glasgow | Wellcome Sanger Institute for the COVID-19 Genomics UK (COG-UK) Consortium | Harper VanSteenhouse, Yumi Kasai, David Gray, Carol Clugston, Anna Dominiczak and Alex Alderton, Roberto Amato, Jeffrey Barrett, Sonia Goncalves, Ewan Harrison, David K. Jackson, Ian Johnston, Dominic Kwiatkowski, Cordelia Langford, John Sillitoe on behalf of the Wellcome Sanger Institute COVID-19 Surveillance Team |
| EPI_ISL_2235537 | Lighthouse Lab in Alderley Park | Wellcome Sanger Institute for the COVID-19 Genomics UK (COG-UK) Consortium | Jacquelyn Wynn, Mairead Hyland, The Lighthouse Lab in Alderley Park and Alex Alderton, Roberto Amato, Jeffrey Barrett, Sonia Goncalves, Ewan Harrison, David K. Jackson, Ian Johnston, Dominic Kwiatkowski, Cordelia Langford, John Sillitoe on behalf of the Wellcome Sanger Institute COVID-19 Surveillance Team |
| EPI_ISL_2235538, EPI_ISL_2235543 | Lighthouse Lab in Glasgow | Wellcome Sanger Institute for the COVID-19 Genomics UK (COG-UK) Consortium | Harper VanSteenhouse, Yumi Kasai, David Gray, Carol Clugston, Anna Dominiczak and Alex Alderton, Roberto Amato, Jeffrey Barrett, Sonia Goncalves, Ewan Harrison, David K. Jackson, Ian Johnston, Dominic Kwiatkowski, Cordelia Langford, John Sillitoe on behalf of the Wellcome Sanger Institute COVID-19 Surveillance Team |
| EPI_ISL_2235545 | Health Services Laboratories | Wellcome Sanger Institute for the COVID-19 Genomics UK (COG-UK) Consortium | Health Services Laboratories and Alex Alderton, Roberto Amato, Jeffrey Barrett, Sonia Goncalves, Ewan Harrison, David K. Jackson, Ian Johnston, Dominic Kwiatkowski, Cordelia Langford, John Sillitoe on behalf of the Wellcome Sanger Institute COVID-19 Surveillance Team |
| EPI_ISL_2235546 | Lighthouse Lab in Glasgow | Wellcome Sanger Institute for the COVID-19 Genomics UK (COG-UK) Consortium | Harper VanSteenhouse, Yumi Kasai, David Gray, Carol Clugston, Anna Dominiczak and Alex Alderton, Roberto Amato, Jeffrey Barrett, Sonia Goncalves, Ewan Harrison, David K. Jackson, Ian Johnston, Dominic Kwiatkowski, Cordelia Langford, John Sillitoe on behalf of the Wellcome Sanger Institute COVID-19 Surveillance Team |
| EPI_ISL_2235549, EPI_ISL_2235551 | Lighthouse Lab in Milton Keynes | Wellcome Sanger Institute for the COVID-19 Genomics UK (COG-UK) Consortium | The Lighthouse Lab in Milton Keynes and Alex Alderton, Roberto Amato, Jeffrey Barrett, Sonia Goncalves, Ewan Harrison, David K. Jackson, Ian Johnston, Dominic Kwiatkowski, Cordelia Langford, John Sillitoe on behalf of the Wellcome Sanger Institute COVID-19 Surveillance Team |
| EPI_ISL_2235554, EPI_ISL_2235555, EPI_ISL_2235556 | Lighthouse Lab in Glasgow | Wellcome Sanger Institute for the COVID-19 Genomics UK (COG-UK) Consortium | Harper VanSteenhouse, Yumi Kasai, David Gray, Carol Clugston, Anna Dominiczak and Alex Alderton, Roberto Amato, Jeffrey Barrett, Sonia Goncalves, Ewan Harrison, David K. Jackson, Ian Johnston, Dominic Kwiatkowski, Cordelia Langford, John Sillitoe on behalf of the Wellcome Sanger Institute COVID-19 Surveillance Team |
| EPI_ISL_2235557 | Lighthouse Lab in Milton Keynes | Wellcome Sanger Institute for the COVID-19 Genomics UK (COG-UK) Consortium | The Lighthouse Lab in Milton Keynes and Alex Alderton, Roberto Amato, Jeffrey Barrett, Sonia Goncalves, Ewan Harrison, David K. Jackson, Ian Johnston, Dominic Kwiatkowski, Cordelia Langford, John Sillitoe on behalf of the Wellcome Sanger Institute COVID-19 Surveillance Team |
| EPI_ISL_2235558 | Lighthouse Lab in Glasgow | Wellcome Sanger Institute for the COVID-19 Genomics UK (COG-UK) Consortium | Harper VanSteenhouse, Yumi Kasai, David Gray, Carol Clugston, Anna Dominiczak and Alex Alderton, Roberto Amato, Jeffrey Barrett, Sonia Goncalves, Ewan Harrison, David K. Jackson, Ian Johnston, Dominic Kwiatkowski, Cordelia Langford, John Sillitoe on behalf of the Wellcome Sanger Institute COVID-19 Surveillance Team |
| EPI_ISL_2235559 | Lighthouse Lab in Milton Keynes | Wellcome Sanger Institute for the COVID-19 Genomics UK (COG-UK) Consortium | The Lighthouse Lab in Milton Keynes and Alex Alderton, Roberto Amato, Jeffrey Barrett, Sonia Goncalves, Ewan Harrison, David K. Jackson, Ian Johnston, Dominic Kwiatkowski, Cordelia Langford, John Sillitoe on behalf of the Wellcome Sanger Institute COVID-19 Surveillance Team |
| EPI_ISL_2235560, EPI_ISL_2235562 | Lighthouse Lab in Alderley Park | Wellcome Sanger Institute for the COVID-19 Genomics UK (COG-UK) Consortium | Jacquelyn Wynn, Mairead Hyland, The Lighthouse Lab in Alderley Park and Alex Alderton, Roberto Amato, Jeffrey Barrett, Sonia Goncalves, Ewan Harrison, David K. Jackson, Ian Johnston, Dominic Kwiatkowski, Cordelia Langford, John Sillitoe on behalf of the Wellcome Sanger Institute COVID-19 Surveillance Team |
| EPI_ISL_2235563 | Lighthouse Lab in Glasgow | Wellcome Sanger Institute for the COVID-19 Genomics UK (COG-UK) Consortium | Harper VanSteenhouse, Yumi Kasai, David Gray, Carol Clugston, Anna Dominiczak and Alex Alderton, Roberto Amato, Jeffrey Barrett, Sonia Goncalves, Ewan Harrison, David K. Jackson, Ian Johnston, Dominic Kwiatkowski, Cordelia Langford, John Sillitoe on behalf of the Wellcome Sanger Institute COVID-19 Surveillance Team |
| EPI_ISL_2235564, EPI_ISL_2235567 | Lighthouse Lab in Milton Keynes | Wellcome Sanger Institute for the COVID-19 Genomics UK (COG-UK) Consortium | The Lighthouse Lab in Milton Keynes and Alex Alderton, Roberto Amato, Jeffrey Barrett, Sonia Goncalves, Ewan Harrison, David K. Jackson, Ian Johnston, Dominic Kwiatkowski, Cordelia Langford, John Sillitoe on behalf of the Wellcome Sanger Institute COVID-19 Surveillance Team |
| EPI_ISL_2235568 | Lighthouse Lab in Alderley Park | Wellcome Sanger Institute for the COVID-19 Genomics UK (COG-UK) Consortium | Jacquelyn Wynn, Mairead Hyland, The Lighthouse Lab in Alderley Park and Alex Alderton, Roberto Amato, Jeffrey Barrett, Sonia Goncalves, Ewan Harrison, David K. Jackson, Ian Johnston, Dominic Kwiatkowski, Cordelia Langford, John Sillitoe on behalf of the Wellcome Sanger Institute COVID-19 Surveillance Team |
| EPI_ISL_2235569 | Lighthouse Lab in Milton Keynes | Wellcome Sanger Institute for the COVID-19 Genomics UK (COG-UK) Consortium | The Lighthouse Lab in Milton Keynes and Alex Alderton, Roberto Amato, Jeffrey Barrett, Sonia Goncalves, Ewan Harrison, David K. Jackson, Ian Johnston, Dominic Kwiatkowski, Cordelia Langford, John Sillitoe on behalf of the Wellcome Sanger Institute COVID-19 Surveillance Team |
| EPI_ISL_2235576 | Lighthouse Lab in Glasgow | Wellcome Sanger Institute for the COVID-19 Genomics UK (COG-UK) Consortium | Harper VanSteenhouse, Yumi Kasai, David Gray, Carol Clugston, Anna Dominiczak and Alex Alderton, Roberto Amato, Jeffrey Barrett, Sonia Goncalves, Ewan Harrison, David K. Jackson, Ian Johnston, Dominic Kwiatkowski, Cordelia Langford, John Sillitoe on behalf of the Wellcome Sanger Institute COVID-19 Surveillance Team |
| EPI_ISL_2235577 | Lighthouse Lab in Alderley Park | Wellcome Sanger Institute for the COVID-19 Genomics UK (COG-UK) Consortium | Jacquelyn Wynn, Mairead Hyland, The Lighthouse Lab in Alderley Park and Alex Alderton, Roberto Amato, Jeffrey Barrett, Sonia Goncalves, Ewan Harrison, David K. Jackson, Ian Johnston, Dominic Kwiatkowski, Cordelia Langford, John Sillitoe on behalf of the Wellcome Sanger Institute COVID-19 Surveillance Team |
| EPI_ISL_2235579 | Lighthouse Lab in Glasgow | Wellcome Sanger Institute for the COVID-19 Genomics UK (COG-UK) Consortium | Harper VanSteenhouse, Yumi Kasai, David Gray, Carol Clugston, Anna Dominiczak and Alex Alderton, Roberto Amato, Jeffrey Barrett, Sonia Goncalves, Ewan Harrison, David K. Jackson, Ian Johnston, Dominic Kwiatkowski, Cordelia Langford, John Sillitoe on behalf of the Wellcome Sanger Institute COVID-19 Surveillance Team |
| EPI_ISL_2235580 | Health Services Laboratories | Wellcome Sanger Institute for the COVID-19 Genomics UK (COG-UK) Consortium | Health Services Laboratories and Alex Alderton, Roberto Amato, Jeffrey Barrett, Sonia Goncalves, Ewan Harrison, David K. Jackson, Ian Johnston, Dominic Kwiatkowski, Cordelia Langford, John Sillitoe on behalf of the Wellcome Sanger Institute COVID-19 Surveillance Team |
| EPI_ISL_2235585 | Lighthouse Lab in Glasgow | Wellcome Sanger Institute for the COVID-19 Genomics UK (COG-UK) Consortium | Harper VanSteenhouse, Yumi Kasai, David Gray, Carol Clugston, Anna Dominiczak and Alex Alderton, Roberto Amato, Jeffrey Barrett, Sonia Goncalves, Ewan Harrison, David K. Jackson, Ian Johnston, Dominic Kwiatkowski, Cordelia Langford, John Sillitoe on behalf of the Wellcome Sanger Institute COVID-19 Surveillance Team |
| EPI_ISL_2235586 | Lighthouse Lab in Alderley Park | Wellcome Sanger Institute for the COVID-19 Genomics UK (COG-UK) Consortium | Jacquelyn Wynn, Mairead Hyland, The Lighthouse Lab in Alderley Park and Alex Alderton, Roberto Amato, Jeffrey Barrett, Sonia Goncalves, Ewan Harrison, David K. Jackson, Ian Johnston, Dominic Kwiatkowski, Cordelia Langford, John Sillitoe on behalf of the Wellcome Sanger Institute COVID-19 Surveillance Team |

| | | | |
|---|---|---|---|
| EPI_ISL_2235587, EPI_ISL_2235589, EPI_ISL_2235590, EPI_ISL_2235593, EPI_ISL_2235595 | Lighthouse Lab in Glasgow | Wellcome Sanger Institute for the COVID-19 Genomics UK (COG-UK) Consortium | Harper VanSteenhouse, Yumi Kasai, David Gray, Carol Clugston, Anna Dominiczak and Alex Alderton, Roberto Amato, Jeffrey Barrett, Sonia Goncalves, Ewan Harrison, David K. Jackson, Ian Johnston, Dominic Kwiatkowski, Cordelia Langford, John Sillitoe on behalf of the Wellcome Sanger Institute COVID-19 Surveillance Team |
| EPI_ISL_2235596 | Lighthouse Lab in Alderley Park | Wellcome Sanger Institute for the COVID-19 Genomics UK (COG-UK) Consortium | Jacquelyn Wynn, Mairead Hyland, The Lighthouse Lab in Alderley Park and Alex Alderton, Roberto Amato, Jeffrey Barrett, Sonia Goncalves, Ewan Harrison, David K. Jackson, Ian Johnston, Dominic Kwiatkowski, Cordelia Langford, John Sillitoe on behalf of the Wellcome Sanger Institute COVID-19 Surveillance Team |
| EPI_ISL_2235597, EPI_ISL_2235598 | Lighthouse Lab in Milton Keynes | Wellcome Sanger Institute for the COVID-19 Genomics UK (COG-UK) Consortium | The Lighthouse Lab in Milton Keynes and Alex Alderton, Roberto Amato, Jeffrey Barrett, Sonia Goncalves, Ewan Harrison, David K. Jackson, Ian Johnston, Dominic Kwiatkowski, Cordelia Langford, John Sillitoe on behalf of the Wellcome Sanger Institute COVID-19 Surveillance Team |
| EPI_ISL_2235599 | Health Services Laboratories | Wellcome Sanger Institute for the COVID-19 Genomics UK (COG-UK) Consortium | Health Services Laboratories and Alex Alderton, Roberto Amato, Jeffrey Barrett, Sonia Goncalves, Ewan Harrison, David K. Jackson, Ian Johnston, Dominic Kwiatkowski, Cordelia Langford, John Sillitoe on behalf of the Wellcome Sanger Institute COVID-19 Surveillance Team |
| EPI_ISL_2235600 | Lighthouse Lab in Alderley Park | Wellcome Sanger Institute for the COVID-19 Genomics UK (COG-UK) Consortium | Jacquelyn Wynn, Mairead Hyland, The Lighthouse Lab in Alderley Park and Alex Alderton, Roberto Amato, Jeffrey Barrett, Sonia Goncalves, Ewan Harrison, David K. Jackson, Ian Johnston, Dominic Kwiatkowski, Cordelia Langford, John Sillitoe on behalf of the Wellcome Sanger Institute COVID-19 Surveillance Team |
| EPI_ISL_2235601, EPI_ISL_2235602 | Lighthouse Lab in Glasgow | Wellcome Sanger Institute for the COVID-19 Genomics UK (COG-UK) Consortium | Harper VanSteenhouse, Yumi Kasai, David Gray, Carol Clugston, Anna Dominiczak and Alex Alderton, Roberto Amato, Jeffrey Barrett, Sonia Goncalves, Ewan Harrison, David K. Jackson, Ian Johnston, Dominic Kwiatkowski, Cordelia Langford, John Sillitoe on behalf of the Wellcome Sanger Institute COVID-19 Surveillance Team |
| EPI_ISL_2235603, EPI_ISL_2235604 | Health Services Laboratories | Wellcome Sanger Institute for the COVID-19 Genomics UK (COG-UK) Consortium | Health Services Laboratories and Alex Alderton, Roberto Amato, Jeffrey Barrett, Sonia Goncalves, Ewan Harrison, David K. Jackson, Ian Johnston, Dominic Kwiatkowski, Cordelia Langford, John Sillitoe on behalf of the Wellcome Sanger Institute COVID-19 Surveillance Team |
| EPI_ISL_2235607, EPI_ISL_2235608, EPI_ISL_2235614 | Lighthouse Lab in Glasgow | Wellcome Sanger Institute for the COVID-19 Genomics UK (COG-UK) Consortium | Harper VanSteenhouse, Yumi Kasai, David Gray, Carol Clugston, Anna Dominiczak and Alex Alderton, Roberto Amato, Jeffrey Barrett, Sonia Goncalves, Ewan Harrison, David K. Jackson, Ian Johnston, Dominic Kwiatkowski, Cordelia Langford, John Sillitoe on behalf of the Wellcome Sanger Institute COVID-19 Surveillance Team |
| EPI_ISL_2235616 | Health Services Laboratories | Wellcome Sanger Institute for the COVID-19 Genomics UK (COG-UK) Consortium | Health Services Laboratories and Alex Alderton, Roberto Amato, Jeffrey Barrett, Sonia Goncalves, Ewan Harrison, David K. Jackson, Ian Johnston, Dominic Kwiatkowski, Cordelia Langford, John Sillitoe on behalf of the Wellcome Sanger Institute COVID-19 Surveillance Team |
| EPI_ISL_2235617 | Lighthouse Lab in Glasgow | Wellcome Sanger Institute for the COVID-19 Genomics UK (COG-UK) Consortium | Harper VanSteenhouse, Yumi Kasai, David Gray, Carol Clugston, Anna Dominiczak and Alex Alderton, Roberto Amato, Jeffrey Barrett, Sonia Goncalves, Ewan Harrison, David K. Jackson, Ian Johnston, Dominic Kwiatkowski, Cordelia Langford, John Sillitoe on behalf of the Wellcome Sanger Institute COVID-19 Surveillance Team |
| EPI_ISL_2235621 | Lighthouse Lab in Milton Keynes | Wellcome Sanger Institute for the COVID-19 Genomics UK (COG-UK) Consortium | The Lighthouse Lab in Milton Keynes and Alex Alderton, Roberto Amato, Jeffrey Barrett, Sonia Goncalves, Ewan Harrison, David K. Jackson, Ian Johnston, Dominic Kwiatkowski, Cordelia Langford, John Sillitoe on behalf of the Wellcome Sanger Institute COVID-19 Surveillance Team |
| EPI_ISL_2235622 | Health Services Laboratories | Wellcome Sanger Institute for the COVID-19 Genomics UK (COG-UK) Consortium | Health Services Laboratories and Alex Alderton, Roberto Amato, Jeffrey Barrett, Sonia Goncalves, Ewan Harrison, David K. Jackson, Ian Johnston, Dominic Kwiatkowski, Cordelia Langford, John Sillitoe on behalf of the Wellcome Sanger Institute COVID-19 Surveillance Team |
| EPI_ISL_2235627 | Lighthouse Lab in Milton Keynes | Wellcome Sanger Institute for the COVID-19 Genomics UK (COG-UK) Consortium | The Lighthouse Lab in Milton Keynes and Alex Alderton, Roberto Amato, Jeffrey Barrett, Sonia Goncalves, Ewan Harrison, David K. Jackson, Ian Johnston, Dominic Kwiatkowski, Cordelia Langford, John Sillitoe on behalf of the Wellcome Sanger Institute COVID-19 Surveillance Team |
| EPI_ISL_2235628, EPI_ISL_2235629 | Lighthouse Lab in Alderley Park | Wellcome Sanger Institute for the COVID-19 Genomics UK (COG-UK) Consortium | Jacquelyn Wynn, Mairead Hyland, The Lighthouse Lab in Alderley Park and Alex Alderton, Roberto Amato, Jeffrey Barrett, Sonia Goncalves, Ewan Harrison, David K. Jackson, Ian Johnston, Dominic Kwiatkowski, Cordelia Langford, John Sillitoe on behalf of the Wellcome Sanger Institute COVID-19 Surveillance Team |
| EPI_ISL_2235631, EPI_ISL_2235632 | Lighthouse Lab in Glasgow | Wellcome Sanger Institute for the COVID-19 Genomics UK (COG-UK) Consortium | Harper VanSteenhouse, Yumi Kasai, David Gray, Carol Clugston, Anna Dominiczak and Alex Alderton, Roberto Amato, Jeffrey Barrett, Sonia Goncalves, Ewan Harrison, David K. Jackson, Ian Johnston, Dominic Kwiatkowski, Cordelia Langford, John Sillitoe on behalf of the Wellcome Sanger Institute COVID-19 Surveillance Team |
| EPI_ISL_2235633 | Health Services Laboratories | Wellcome Sanger Institute for the COVID-19 Genomics UK (COG-UK) Consortium | Health Services Laboratories and Alex Alderton, Roberto Amato, Jeffrey Barrett, Sonia Goncalves, Ewan Harrison, David K. Jackson, Ian Johnston, Dominic Kwiatkowski, Cordelia Langford, John Sillitoe on behalf of the Wellcome Sanger Institute COVID-19 Surveillance Team |
| EPI_ISL_2235635, EPI_ISL_2235636 | Lighthouse Lab in Alderley Park | Wellcome Sanger Institute for the COVID-19 Genomics UK (COG-UK) Consortium | Jacquelyn Wynn, Mairead Hyland, The Lighthouse Lab in Alderley Park and Alex Alderton, Roberto Amato, Jeffrey Barrett, Sonia Goncalves, Ewan Harrison, David K. Jackson, Ian Johnston, Dominic Kwiatkowski, Cordelia Langford, John Sillitoe on behalf of the Wellcome Sanger Institute COVID-19 Surveillance Team |
| EPI_ISL_2235638, EPI_ISL_2235639, EPI_ISL_2235644 | Health Services Laboratories | Wellcome Sanger Institute for the COVID-19 Genomics UK (COG-UK) Consortium | Health Services Laboratories and Alex Alderton, Roberto Amato, Jeffrey Barrett, Sonia Goncalves, Ewan Harrison, David K. Jackson, Ian Johnston, Dominic Kwiatkowski, Cordelia Langford, John Sillitoe on behalf of the Wellcome Sanger Institute COVID-19 Surveillance Team |
| EPI_ISL_2235646, EPI_ISL_2235648 | Lighthouse Lab in Glasgow | Wellcome Sanger Institute for the COVID-19 Genomics UK (COG-UK) Consortium | Harper VanSteenhouse, Yumi Kasai, David Gray, Carol Clugston, Anna Dominiczak and Alex Alderton, Roberto Amato, Jeffrey Barrett, Sonia Goncalves, Ewan Harrison, David K. Jackson, Ian Johnston, Dominic Kwiatkowski, Cordelia Langford, John Sillitoe on behalf of the Wellcome Sanger Institute COVID-19 Surveillance Team |
| EPI_ISL_2235650 | Lighthouse Lab in Milton Keynes | Wellcome Sanger Institute for the COVID-19 Genomics UK (COG-UK) Consortium | The Lighthouse Lab in Milton Keynes and Alex Alderton, Roberto Amato, Jeffrey Barrett, Sonia Goncalves, Ewan Harrison, David K. Jackson, Ian Johnston, Dominic Kwiatkowski, Cordelia Langford, John Sillitoe on behalf of the Wellcome Sanger Institute COVID-19 Surveillance Team |
| EPI_ISL_2235652 | Health Services Laboratories | Wellcome Sanger Institute for the COVID-19 Genomics UK (COG-UK) Consortium | Health Services Laboratories and Alex Alderton, Roberto Amato, Jeffrey Barrett, Sonia Goncalves, Ewan Harrison, David K. Jackson, Ian Johnston, Dominic Kwiatkowski, Cordelia Langford, John Sillitoe on behalf of the Wellcome Sanger Institute COVID-19 Surveillance Team |
| EPI_ISL_2235659 | Lighthouse Lab in Alderley Park | Wellcome Sanger Institute for the COVID-19 Genomics UK (COG-UK) Consortium | Jacquelyn Wynn, Mairead Hyland, The Lighthouse Lab in Alderley Park and Alex Alderton, Roberto Amato, Jeffrey Barrett, Sonia Goncalves, Ewan Harrison, David K. Jackson, Ian Johnston, Dominic Kwiatkowski, Cordelia Langford, John Sillitoe on behalf of the Wellcome Sanger Institute COVID-19 Surveillance Team |
| EPI_ISL_2235660 | Lighthouse Lab in Milton Keynes | Wellcome Sanger Institute for the COVID-19 Genomics UK (COG-UK) Consortium | The Lighthouse Lab in Milton Keynes and Alex Alderton, Roberto Amato, Jeffrey Barrett, Sonia Goncalves, Ewan Harrison, David K. Jackson, Ian Johnston, Dominic Kwiatkowski, Cordelia Langford, John Sillitoe on behalf of the Wellcome Sanger Institute COVID-19 Surveillance Team |
| EPI_ISL_2235661, EPI_ISL_2235665 | Lighthouse Lab in Glasgow | Wellcome Sanger Institute for the COVID-19 Genomics UK (COG-UK) Consortium | Harper VanSteenhouse, Yumi Kasai, David Gray, Carol Clugston, Anna Dominiczak and Alex Alderton, Roberto Amato, Jeffrey Barrett, Sonia Goncalves, Ewan Harrison, David K. Jackson, Ian Johnston, Dominic Kwiatkowski, Cordelia Langford, John Sillitoe on behalf of the Wellcome Sanger Institute COVID-19 Surveillance Team |
| EPI_ISL_2235666 | Lighthouse Lab in Milton Keynes | Wellcome Sanger Institute for the COVID-19 Genomics UK (COG-UK) Consortium | The Lighthouse Lab in Milton Keynes and Alex Alderton, Roberto Amato, Jeffrey Barrett, Sonia Goncalves, Ewan Harrison, David K. Jackson, Ian Johnston, Dominic Kwiatkowski, Cordelia Langford, John Sillitoe on behalf of the Wellcome Sanger Institute COVID-19 Surveillance Team |
| EPI_ISL_2235667, EPI_ISL_2235670 | Lighthouse Lab in Glasgow | Wellcome Sanger Institute for the COVID-19 Genomics UK (COG-UK) Consortium | Harper VanSteenhouse, Yumi Kasai, David Gray, Carol Clugston, Anna Dominiczak and Alex Alderton, Roberto Amato, Jeffrey Barrett, Sonia Goncalves, Ewan Harrison, David K. Jackson, Ian Johnston, Dominic Kwiatkowski, Cordelia Langford, John Sillitoe on behalf of the Wellcome Sanger Institute COVID-19 Surveillance Team |
| EPI_ISL_2235671 | Lighthouse Lab in Alderley Park | Wellcome Sanger Institute for the COVID-19 Genomics UK (COG-UK) Consortium | Jacquelyn Wynn, Mairead Hyland, The Lighthouse Lab in Alderley Park and Alex Alderton, Roberto Amato, Jeffrey Barrett, Sonia Goncalves, Ewan Harrison, David K. Jackson, Ian Johnston, Dominic Kwiatkowski, Cordelia Langford, John Sillitoe on behalf of the Wellcome Sanger Institute COVID-19 Surveillance Team |
| EPI_ISL_2235672 | Lighthouse Lab in Glasgow | Wellcome Sanger Institute for the COVID-19 Genomics UK (COG-UK) Consortium | Harper VanSteenhouse, Yumi Kasai, David Gray, Carol Clugston, Anna Dominiczak and Alex Alderton, Roberto Amato, Jeffrey Barrett, Sonia Goncalves, Ewan Harrison, David K. Jackson, Ian Johnston, Dominic Kwiatkowski, Cordelia Langford, John Sillitoe on behalf of the Wellcome Sanger Institute COVID-19 Surveillance Team |
| EPI_ISL_2235673 | Lighthouse Lab in Milton Keynes | Wellcome Sanger Institute for the COVID-19 Genomics UK (COG-UK) Consortium | The Lighthouse Lab in Milton Keynes and Alex Alderton, Roberto Amato, Jeffrey Barrett, Sonia Goncalves, Ewan Harrison, David K. Jackson, Ian Johnston, Dominic Kwiatkowski, Cordelia Langford, John Sillitoe on behalf of the Wellcome Sanger Institute COVID-19 Surveillance Team |
| EPI_ISL_2235674, EPI_ISL_2235677 | Lighthouse Lab in Alderley Park | Wellcome Sanger Institute for the COVID-19 Genomics UK (COG-UK) Consortium | Jacquelyn Wynn, Mairead Hyland, The Lighthouse Lab in Alderley Park and Alex Alderton, Roberto Amato, Jeffrey Barrett, Sonia Goncalves, Ewan Harrison, David K. Jackson, Ian Johnston, Dominic Kwiatkowski, Cordelia Langford, John Sillitoe on behalf of the Wellcome Sanger Institute COVID-19 Surveillance Team |
| EPI_ISL_2235678, EPI_ISL_2235680, EPI_ISL_2235682, EPI_ISL_2235683 | Lighthouse Lab in Glasgow | Wellcome Sanger Institute for the COVID-19 Genomics UK (COG-UK) Consortium | Harper VanSteenhouse, Yumi Kasai, David Gray, Carol Clugston, Anna Dominiczak and Alex Alderton, Roberto Amato, Jeffrey Barrett, Sonia Goncalves, Ewan Harrison, David K. Jackson, Ian Johnston, Dominic Kwiatkowski, Cordelia Langford, John Sillitoe on behalf of the Wellcome Sanger Institute COVID-19 Surveillance Team |
| EPI_ISL_2235685 | Lighthouse Lab in Alderley Park | Wellcome Sanger Institute for the COVID-19 Genomics UK (COG-UK) Consortium | Jacquelyn Wynn, Mairead Hyland, The Lighthouse Lab in Alderley Park and Alex Alderton, Roberto Amato, Jeffrey Barrett, Sonia Goncalves, Ewan Harrison, David K. Jackson, Ian Johnston, Dominic Kwiatkowski, Cordelia Langford, John Sillitoe on behalf of the Wellcome Sanger Institute COVID-19 Surveillance Team |
| EPI_ISL_2235687, EPI_ISL_2235688, EPI_ISL_2235689, EPI_ISL_2235690, EPI_ISL_2235693 | Lighthouse Lab in Glasgow | Wellcome Sanger Institute for the COVID-19 Genomics UK (COG-UK) Consortium | Harper VanSteenhouse, Yumi Kasai, David Gray, Carol Clugston, Anna Dominiczak and Alex Alderton, Roberto Amato, Jeffrey Barrett, Sonia Goncalves, Ewan Harrison, David K. Jackson, Ian Johnston, Dominic Kwiatkowski, Cordelia Langford, John Sillitoe on behalf of the Wellcome Sanger Institute COVID-19 Surveillance Team |
| EPI_ISL_2235700, EPI_ISL_2235702 | Lighthouse Lab in Alderley Park | Wellcome Sanger Institute for the COVID-19 Genomics UK (COG-UK) Consortium | Jacquelyn Wynn, Mairead Hyland, The Lighthouse Lab in Alderley Park and Alex Alderton, Roberto Amato, Jeffrey Barrett, Sonia Goncalves, Ewan Harrison, David K. Jackson, Ian Johnston, Dominic Kwiatkowski, Cordelia Langford, John Sillitoe on behalf of the Wellcome Sanger Institute COVID-19 Surveillance Team |
| EPI_ISL_2235706 | Lighthouse Lab in Milton Keynes | Wellcome Sanger Institute for the COVID-19 Genomics UK (COG-UK) Consortium | The Lighthouse Lab in Milton Keynes and Alex Alderton, Roberto Amato, Jeffrey Barrett, Sonia Goncalves, Ewan Harrison, David K. Jackson, Ian Johnston, Dominic Kwiatkowski, Cordelia Langford, John Sillitoe on behalf of the Wellcome Sanger Institute COVID-19 Surveillance Team |
| EPI_ISL_2235707 | Health Services Laboratories | Wellcome Sanger Institute for the COVID-19 Genomics UK (COG-UK) Consortium | Health Services Laboratories and Alex Alderton, Roberto Amato, Jeffrey Barrett, Sonia Goncalves, Ewan Harrison, David K. Jackson, Ian Johnston, Dominic Kwiatkowski, Cordelia Langford, John Sillitoe on behalf of the Wellcome Sanger Institute COVID-19 Surveillance Team |
| EPI_ISL_2235708 | Lighthouse Lab in Glasgow | Wellcome Sanger Institute for the COVID-19 Genomics UK (COG-UK) Consortium | Harper VanSteenhouse, Yumi Kasai, David Gray, Carol Clugston, Anna Dominiczak and Alex Alderton, Roberto Amato, Jeffrey Barrett, Sonia Goncalves, Ewan Harrison, David K. Jackson, Ian Johnston, Dominic Kwiatkowski, Cordelia Langford, John Sillitoe on behalf of the Wellcome Sanger Institute COVID-19 Surveillance Team |
| EPI_ISL_2235710 | Health Services Laboratories | Wellcome Sanger Institute for the COVID-19 Genomics UK (COG-UK) Consortium | Health Services Laboratories and Alex Alderton, Roberto Amato, Jeffrey Barrett, Sonia Goncalves, Ewan Harrison, David K. Jackson, Ian Johnston, Dominic Kwiatkowski, Cordelia Langford, John Sillitoe on behalf of the Wellcome Sanger Institute COVID-19 Surveillance Team |
| EPI_ISL_2235712 | Lighthouse Lab in Alderley Park | Wellcome Sanger Institute for the COVID-19 Genomics UK (COG-UK) Consortium | Jacquelyn Wynn, Mairead Hyland, The Lighthouse Lab in Alderley Park and Alex Alderton, Roberto Amato, Jeffrey Barrett, Sonia Goncalves, Ewan Harrison, David K. Jackson, Ian Johnston, Dominic Kwiatkowski, Cordelia Langford, John Sillitoe on behalf of the Wellcome Sanger Institute COVID-19 Surveillance Team |
| EPI_ISL_2235715, EPI_ISL_2235716, EPI_ISL_2235719 | Lighthouse Lab in Glasgow | Wellcome Sanger Institute for the COVID-19 Genomics UK (COG-UK) Consortium | Harper VanSteenhouse, Yumi Kasai, David Gray, Carol Clugston, Anna Dominiczak and Alex Alderton, Roberto Amato, Jeffrey Barrett, Sonia Goncalves, Ewan Harrison, David K. Jackson, Ian Johnston, Dominic Kwiatkowski, Cordelia Langford, John Sillitoe on behalf of the Wellcome Sanger Institute COVID-19 Surveillance Team |
| EPI_ISL_2235720 | Lighthouse Lab in Milton Keynes | Wellcome Sanger Institute for the COVID-19 Genomics UK (COG-UK) | The Lighthouse Lab in Milton Keynes and Alex Alderton, Roberto Amato, Jeffrey Barrett, Sonia Goncalves, Ewan Harrison, David K. Jackson, Ian Johnston, Dominic Kwiatkowski, Cordelia Langford, John Sillitoe on behalf of the Wellcome Sanger |

| | | | |
|---|---|---|---|
| | | Consortium | Institute COVID-19 Surveillance Team |
| EPI_ISL_2235721 | Lighthouse Lab in Alderley Park | Wellcome Sanger Institute for the COVID-19 Genomics UK (COG-UK) Consortium | Jacquelyn Wynn, Mairead Hyland, The Lighthouse Lab in Alderley Park and Alex Alderton, Roberto Amato, Jeffrey Barrett, Sonia Goncalves, Ewan Harrison, David K. Jackson, Ian Johnston, Dominic Kwiatkowski, Cordelia Langford, John Sillitoe on behalf of the Wellcome Sanger Institute COVID-19 Surveillance Team |
| EPI_ISL_2235722, EPI_ISL_2235723 | Health Services Laboratories | Wellcome Sanger Institute for the COVID-19 Genomics UK (COG-UK) Consortium | Health Services Laboratories and Alex Alderton, Roberto Amato, Jeffrey Barrett, Sonia Goncalves, Ewan Harrison, David K. Jackson, Ian Johnston, Dominic Kwiatkowski, Cordelia Langford, John Sillitoe on behalf of the Wellcome Sanger Institute COVID-19 Surveillance Team |
| EPI_ISL_2235724 | Lighthouse Lab in Glasgow | Wellcome Sanger Institute for the COVID-19 Genomics UK (COG-UK) Consortium | Harper VanSteenhouse, Yumi Kasai, David Gray, Carol Clugston, Anna Dominiczak and Alex Alderton, Roberto Amato, Jeffrey Barrett, Sonia Goncalves, Ewan Harrison, David K. Jackson, Ian Johnston, Dominic Kwiatkowski, Cordelia Langford, John Sillitoe on behalf of the Wellcome Sanger Institute COVID-19 Surveillance Team |
| EPI_ISL_2235726 | Lighthouse Lab in Milton Keynes | Wellcome Sanger Institute for the COVID-19 Genomics UK (COG-UK) Consortium | The Lighthouse Lab in Milton Keynes and Alex Alderton, Roberto Amato, Jeffrey Barrett, Sonia Goncalves, Ewan Harrison, David K. Jackson, Ian Johnston, Dominic Kwiatkowski, Cordelia Langford, John Sillitoe on behalf of the Wellcome Sanger Institute COVID-19 Surveillance Team |
| EPI_ISL_2235728 | Health Services Laboratories | Wellcome Sanger Institute for the COVID-19 Genomics UK (COG-UK) Consortium | Health Services Laboratories and Alex Alderton, Roberto Amato, Jeffrey Barrett, Sonia Goncalves, Ewan Harrison, David K. Jackson, Ian Johnston, Dominic Kwiatkowski, Cordelia Langford, John Sillitoe on behalf of the Wellcome Sanger Institute COVID-19 Surveillance Team |
| EPI_ISL_2235729 | Lighthouse Lab in Milton Keynes | Wellcome Sanger Institute for the COVID-19 Genomics UK (COG-UK) Consortium | The Lighthouse Lab in Milton Keynes and Alex Alderton, Roberto Amato, Jeffrey Barrett, Sonia Goncalves, Ewan Harrison, David K. Jackson, Ian Johnston, Dominic Kwiatkowski, Cordelia Langford, John Sillitoe on behalf of the Wellcome Sanger Institute COVID-19 Surveillance Team |
| EPI_ISL_2235730 | Health Services Laboratories | Wellcome Sanger Institute for the COVID-19 Genomics UK (COG-UK) Consortium | Health Services Laboratories and Alex Alderton, Roberto Amato, Jeffrey Barrett, Sonia Goncalves, Ewan Harrison, David K. Jackson, Ian Johnston, Dominic Kwiatkowski, Cordelia Langford, John Sillitoe on behalf of the Wellcome Sanger Institute COVID-19 Surveillance Team |
| EPI_ISL_2235731 | Lighthouse Lab in Milton Keynes | Wellcome Sanger Institute for the COVID-19 Genomics UK (COG-UK) Consortium | The Lighthouse Lab in Milton Keynes and Alex Alderton, Roberto Amato, Jeffrey Barrett, Sonia Goncalves, Ewan Harrison, David K. Jackson, Ian Johnston, Dominic Kwiatkowski, Cordelia Langford, John Sillitoe on behalf of the Wellcome Sanger Institute COVID-19 Surveillance Team |
| EPI_ISL_2235733 | Lighthouse Lab in Glasgow | Wellcome Sanger Institute for the COVID-19 Genomics UK (COG-UK) Consortium | Harper VanSteenhouse, Yumi Kasai, David Gray, Carol Clugston, Anna Dominiczak and Alex Alderton, Roberto Amato, Jeffrey Barrett, Sonia Goncalves, Ewan Harrison, David K. Jackson, Ian Johnston, Dominic Kwiatkowski, Cordelia Langford, John Sillitoe on behalf of the Wellcome Sanger Institute COVID-19 Surveillance Team |
| EPI_ISL_2235734 | Lighthouse Lab in Milton Keynes | Wellcome Sanger Institute for the COVID-19 Genomics UK (COG-UK) Consortium | The Lighthouse Lab in Milton Keynes and Alex Alderton, Roberto Amato, Jeffrey Barrett, Sonia Goncalves, Ewan Harrison, David K. Jackson, Ian Johnston, Dominic Kwiatkowski, Cordelia Langford, John Sillitoe on behalf of the Wellcome Sanger Institute COVID-19 Surveillance Team |
| EPI_ISL_2235738, EPI_ISL_2235740 | Lighthouse Lab in Glasgow | Wellcome Sanger Institute for the COVID-19 Genomics UK (COG-UK) Consortium | Harper VanSteenhouse, Yumi Kasai, David Gray, Carol Clugston, Anna Dominiczak and Alex Alderton, Roberto Amato, Jeffrey Barrett, Sonia Goncalves, Ewan Harrison, David K. Jackson, Ian Johnston, Dominic Kwiatkowski, Cordelia Langford, John Sillitoe on behalf of the Wellcome Sanger Institute COVID-19 Surveillance Team |
| EPI_ISL_2235742 | Lighthouse Lab in Alderley Park | Wellcome Sanger Institute for the COVID-19 Genomics UK (COG-UK) Consortium | Jacquelyn Wynn, Mairead Hyland, The Lighthouse Lab in Alderley Park and Alex Alderton, Roberto Amato, Jeffrey Barrett, Sonia Goncalves, Ewan Harrison, David K. Jackson, Ian Johnston, Dominic Kwiatkowski, Cordelia Langford, John Sillitoe on behalf of the Wellcome Sanger Institute COVID-19 Surveillance Team |
| EPI_ISL_2235743 | Health Services Laboratories | Wellcome Sanger Institute for the COVID-19 Genomics UK (COG-UK) Consortium | Health Services Laboratories and Alex Alderton, Roberto Amato, Jeffrey Barrett, Sonia Goncalves, Ewan Harrison, David K. Jackson, Ian Johnston, Dominic Kwiatkowski, Cordelia Langford, John Sillitoe on behalf of the Wellcome Sanger Institute COVID-19 Surveillance Team |
| EPI_ISL_2235746 | Lighthouse Lab in Alderley Park | Wellcome Sanger Institute for the COVID-19 Genomics UK (COG-UK) Consortium | Jacquelyn Wynn, Mairead Hyland, The Lighthouse Lab in Alderley Park and Alex Alderton, Roberto Amato, Jeffrey Barrett, Sonia Goncalves, Ewan Harrison, David K. Jackson, Ian Johnston, Dominic Kwiatkowski, Cordelia Langford, John Sillitoe on behalf of the Wellcome Sanger Institute COVID-19 Surveillance Team |
| EPI_ISL_2235747, EPI_ISL_2235750 | Lighthouse Lab in Glasgow | Wellcome Sanger Institute for the COVID-19 Genomics UK (COG-UK) Consortium | Harper VanSteenhouse, Yumi Kasai, David Gray, Carol Clugston, Anna Dominiczak and Alex Alderton, Roberto Amato, Jeffrey Barrett, Sonia Goncalves, Ewan Harrison, David K. Jackson, Ian Johnston, Dominic Kwiatkowski, Cordelia Langford, John Sillitoe on behalf of the Wellcome Sanger Institute COVID-19 Surveillance Team |
| EPI_ISL_2235751 | Lighthouse Lab in Milton Keynes | Wellcome Sanger Institute for the COVID-19 Genomics UK (COG-UK) Consortium | The Lighthouse Lab in Milton Keynes and Alex Alderton, Roberto Amato, Jeffrey Barrett, Sonia Goncalves, Ewan Harrison, David K. Jackson, Ian Johnston, Dominic Kwiatkowski, Cordelia Langford, John Sillitoe on behalf of the Wellcome Sanger Institute COVID-19 Surveillance Team |
| EPI_ISL_2235754 | Health Services Laboratories | Wellcome Sanger Institute for the COVID-19 Genomics UK (COG-UK) Consortium | Health Services Laboratories and Alex Alderton, Roberto Amato, Jeffrey Barrett, Sonia Goncalves, Ewan Harrison, David K. Jackson, Ian Johnston, Dominic Kwiatkowski, Cordelia Langford, John Sillitoe on behalf of the Wellcome Sanger Institute COVID-19 Surveillance Team |
| EPI_ISL_2235757, EPI_ISL_2235762, EPI_ISL_2235763 | Lighthouse Lab in Alderley Park | Wellcome Sanger Institute for the COVID-19 Genomics UK (COG-UK) Consortium | Jacquelyn Wynn, Mairead Hyland, The Lighthouse Lab in Alderley Park and Alex Alderton, Roberto Amato, Jeffrey Barrett, Sonia Goncalves, Ewan Harrison, David K. Jackson, Ian Johnston, Dominic Kwiatkowski, Cordelia Langford, John Sillitoe on behalf of the Wellcome Sanger Institute COVID-19 Surveillance Team |
| EPI_ISL_2235765 | Lighthouse Lab in Milton Keynes | Wellcome Sanger Institute for the COVID-19 Genomics UK (COG-UK) Consortium | The Lighthouse Lab in Milton Keynes and Alex Alderton, Roberto Amato, Jeffrey Barrett, Sonia Goncalves, Ewan Harrison, David K. Jackson, Ian Johnston, Dominic Kwiatkowski, Cordelia Langford, John Sillitoe on behalf of the Wellcome Sanger Institute COVID-19 Surveillance Team |
| EPI_ISL_2235766, EPI_ISL_2235767 | Lighthouse Lab in Glasgow | Wellcome Sanger Institute for the COVID-19 Genomics UK (COG-UK) Consortium | Harper VanSteenhouse, Yumi Kasai, David Gray, Carol Clugston, Anna Dominiczak and Alex Alderton, Roberto Amato, Jeffrey Barrett, Sonia Goncalves, Ewan Harrison, David K. Jackson, Ian Johnston, Dominic Kwiatkowski, Cordelia Langford, John Sillitoe on behalf of the Wellcome Sanger Institute COVID-19 Surveillance Team |
| EPI_ISL_2235768, EPI_ISL_2235770 | Lighthouse Lab in Milton Keynes | Wellcome Sanger Institute for the COVID-19 Genomics UK (COG-UK) Consortium | The Lighthouse Lab in Milton Keynes and Alex Alderton, Roberto Amato, Jeffrey Barrett, Sonia Goncalves, Ewan Harrison, David K. Jackson, Ian Johnston, Dominic Kwiatkowski, Cordelia Langford, John Sillitoe on behalf of the Wellcome Sanger Institute COVID-19 Surveillance Team |
| EPI_ISL_2235771 | Lighthouse Lab in Glasgow | Wellcome Sanger Institute for the COVID-19 Genomics UK (COG-UK) Consortium | Harper VanSteenhouse, Yumi Kasai, David Gray, Carol Clugston, Anna Dominiczak and Alex Alderton, Roberto Amato, Jeffrey Barrett, Sonia Goncalves, Ewan Harrison, David K. Jackson, Ian Johnston, Dominic Kwiatkowski, Cordelia Langford, John Sillitoe on behalf of the Wellcome Sanger Institute COVID-19 Surveillance Team |
| EPI_ISL_2235773, EPI_ISL_2235775 | Lighthouse Lab in Alderley Park | Wellcome Sanger Institute for the COVID-19 Genomics UK (COG-UK) Consortium | Jacquelyn Wynn, Mairead Hyland, The Lighthouse Lab in Alderley Park and Alex Alderton, Roberto Amato, Jeffrey Barrett, Sonia Goncalves, Ewan Harrison, David K. Jackson, Ian Johnston, Dominic Kwiatkowski, Cordelia Langford, John Sillitoe on behalf of the Wellcome Sanger Institute COVID-19 Surveillance Team |
| EPI_ISL_2235776 | Lighthouse Lab in Glasgow | Wellcome Sanger Institute for the COVID-19 Genomics UK (COG-UK) Consortium | Harper VanSteenhouse, Yumi Kasai, David Gray, Carol Clugston, Anna Dominiczak and Alex Alderton, Roberto Amato, Jeffrey Barrett, Sonia Goncalves, Ewan Harrison, David K. Jackson, Ian Johnston, Dominic Kwiatkowski, Cordelia Langford, John Sillitoe on behalf of the Wellcome Sanger Institute COVID-19 Surveillance Team |
| EPI_ISL_2235777 | Lighthouse Lab in Alderley Park | Wellcome Sanger Institute for the COVID-19 Genomics UK (COG-UK) Consortium | Jacquelyn Wynn, Mairead Hyland, The Lighthouse Lab in Alderley Park and Alex Alderton, Roberto Amato, Jeffrey Barrett, Sonia Goncalves, Ewan Harrison, David K. Jackson, Ian Johnston, Dominic Kwiatkowski, Cordelia Langford, John Sillitoe on behalf of the Wellcome Sanger Institute COVID-19 Surveillance Team |
| EPI_ISL_2235779 | Lighthouse Lab in Glasgow | Wellcome Sanger Institute for the COVID-19 Genomics UK (COG-UK) Consortium | Harper VanSteenhouse, Yumi Kasai, David Gray, Carol Clugston, Anna Dominiczak and Alex Alderton, Roberto Amato, Jeffrey Barrett, Sonia Goncalves, Ewan Harrison, David K. Jackson, Ian Johnston, Dominic Kwiatkowski, Cordelia Langford, John Sillitoe on behalf of the Wellcome Sanger Institute COVID-19 Surveillance Team |
| EPI_ISL_2235780 | Lighthouse Lab in Milton Keynes | Wellcome Sanger Institute for the COVID-19 Genomics UK (COG-UK) Consortium | The Lighthouse Lab in Milton Keynes and Alex Alderton, Roberto Amato, Jeffrey Barrett, Sonia Goncalves, Ewan Harrison, David K. Jackson, Ian Johnston, Dominic Kwiatkowski, Cordelia Langford, John Sillitoe on behalf of the Wellcome Sanger Institute COVID-19 Surveillance Team |
| EPI_ISL_2235783 | Lighthouse Lab in Glasgow | Wellcome Sanger Institute for the COVID-19 Genomics UK (COG-UK) Consortium | Harper VanSteenhouse, Yumi Kasai, David Gray, Carol Clugston, Anna Dominiczak and Alex Alderton, Roberto Amato, Jeffrey Barrett, Sonia Goncalves, Ewan Harrison, David K. Jackson, Ian Johnston, Dominic Kwiatkowski, Cordelia Langford, John Sillitoe on behalf of the Wellcome Sanger Institute COVID-19 Surveillance Team |
| EPI_ISL_2235784, EPI_ISL_2235786, EPI_ISL_2235787 | Health Services Laboratories | Wellcome Sanger Institute for the COVID-19 Genomics UK (COG-UK) Consortium | Health Services Laboratories and Alex Alderton, Roberto Amato, Jeffrey Barrett, Sonia Goncalves, Ewan Harrison, David K. Jackson, Ian Johnston, Dominic Kwiatkowski, Cordelia Langford, John Sillitoe on behalf of the Wellcome Sanger Institute COVID-19 Surveillance Team |
| EPI_ISL_2235788 | Lighthouse Lab in Glasgow | Wellcome Sanger Institute for the COVID-19 Genomics UK (COG-UK) Consortium | Harper VanSteenhouse, Yumi Kasai, David Gray, Carol Clugston, Anna Dominiczak and Alex Alderton, Roberto Amato, Jeffrey Barrett, Sonia Goncalves, Ewan Harrison, David K. Jackson, Ian Johnston, Dominic Kwiatkowski, Cordelia Langford, John Sillitoe on behalf of the Wellcome Sanger Institute COVID-19 Surveillance Team |
| EPI_ISL_2235794, EPI_ISL_2235795 | Lighthouse Lab in Alderley Park | Wellcome Sanger Institute for the COVID-19 Genomics UK (COG-UK) Consortium | Jacquelyn Wynn, Mairead Hyland, The Lighthouse Lab in Alderley Park and Alex Alderton, Roberto Amato, Jeffrey Barrett, Sonia Goncalves, Ewan Harrison, David K. Jackson, Ian Johnston, Dominic Kwiatkowski, Cordelia Langford, John Sillitoe on behalf of the Wellcome Sanger Institute COVID-19 Surveillance Team |
| EPI_ISL_2235797 | Lighthouse Lab in Glasgow | Wellcome Sanger Institute for the COVID-19 Genomics UK (COG-UK) Consortium | Harper VanSteenhouse, Yumi Kasai, David Gray, Carol Clugston, Anna Dominiczak and Alex Alderton, Roberto Amato, Jeffrey Barrett, Sonia Goncalves, Ewan Harrison, David K. Jackson, Ian Johnston, Dominic Kwiatkowski, Cordelia Langford, John Sillitoe on behalf of the Wellcome Sanger Institute COVID-19 Surveillance Team |
| EPI_ISL_2235798 | Lighthouse Lab in Milton Keynes | Wellcome Sanger Institute for the COVID-19 Genomics UK (COG-UK) Consortium | The Lighthouse Lab in Milton Keynes and Alex Alderton, Roberto Amato, Jeffrey Barrett, Sonia Goncalves, Ewan Harrison, David K. Jackson, Ian Johnston, Dominic Kwiatkowski, Cordelia Langford, John Sillitoe on behalf of the Wellcome Sanger Institute COVID-19 Surveillance Team |
| EPI_ISL_2235799 | Lighthouse Lab in Glasgow | Wellcome Sanger Institute for the COVID-19 Genomics UK (COG-UK) Consortium | Harper VanSteenhouse, Yumi Kasai, David Gray, Carol Clugston, Anna Dominiczak and Alex Alderton, Roberto Amato, Jeffrey Barrett, Sonia Goncalves, Ewan Harrison, David K. Jackson, Ian Johnston, Dominic Kwiatkowski, Cordelia Langford, John Sillitoe on behalf of the Wellcome Sanger Institute COVID-19 Surveillance Team |
| EPI_ISL_2235800, EPI_ISL_2235801 | Lighthouse Lab in Milton Keynes | Wellcome Sanger Institute for the COVID-19 Genomics UK (COG-UK) Consortium | The Lighthouse Lab in Milton Keynes and Alex Alderton, Roberto Amato, Jeffrey Barrett, Sonia Goncalves, Ewan Harrison, David K. Jackson, Ian Johnston, Dominic Kwiatkowski, Cordelia Langford, John Sillitoe on behalf of the Wellcome Sanger Institute COVID-19 Surveillance Team |
| EPI_ISL_2235804 | Lighthouse Lab in Alderley Park | Wellcome Sanger Institute for the COVID-19 Genomics UK (COG-UK) Consortium | Jacquelyn Wynn, Mairead Hyland, The Lighthouse Lab in Alderley Park and Alex Alderton, Roberto Amato, Jeffrey Barrett, Sonia Goncalves, Ewan Harrison, David K. Jackson, Ian Johnston, Dominic Kwiatkowski, Cordelia Langford, John Sillitoe on behalf of the Wellcome Sanger Institute COVID-19 Surveillance Team |
| EPI_ISL_2235805 | Lighthouse Lab in Milton Keynes | Wellcome Sanger Institute for the COVID-19 Genomics UK (COG-UK) Consortium | The Lighthouse Lab in Milton Keynes and Alex Alderton, Roberto Amato, Jeffrey Barrett, Sonia Goncalves, Ewan Harrison, David K. Jackson, Ian Johnston, Dominic Kwiatkowski, Cordelia Langford, John Sillitoe on behalf of the Wellcome Sanger Institute COVID-19 Surveillance Team |
| EPI_ISL_2235807 | Lighthouse Lab in Glasgow | Wellcome Sanger Institute for the COVID-19 Genomics UK (COG-UK) Consortium | Harper VanSteenhouse, Yumi Kasai, David Gray, Carol Clugston, Anna Dominiczak and Alex Alderton, Roberto Amato, Jeffrey Barrett, Sonia Goncalves, Ewan Harrison, David K. Jackson, Ian Johnston, Dominic Kwiatkowski, Cordelia Langford, John Sillitoe on behalf of the Wellcome Sanger Institute COVID-19 Surveillance Team |
| EPI_ISL_2235810 | Lighthouse Lab in Milton Keynes | Wellcome Sanger Institute for the COVID-19 Genomics UK (COG-UK) Consortium | The Lighthouse Lab in Milton Keynes and Alex Alderton, Roberto Amato, Jeffrey Barrett, Sonia Goncalves, Ewan Harrison, David K. Jackson, Ian Johnston, Dominic Kwiatkowski, Cordelia Langford, John Sillitoe on behalf of the Wellcome Sanger Institute COVID-19 Surveillance Team |
| EPI_ISL_2235813, EPI_ISL_2235815 | Lighthouse Lab in Glasgow | Wellcome Sanger Institute for the COVID-19 Genomics UK (COG-UK) Consortium | Harper VanSteenhouse, Yumi Kasai, David Gray, Carol Clugston, Anna Dominiczak and Alex Alderton, Roberto Amato, Jeffrey Barrett, Sonia Goncalves, Ewan Harrison, David K. Jackson, Ian Johnston, Dominic Kwiatkowski, Cordelia Langford, John Sillitoe on behalf of the Wellcome Sanger Institute COVID-19 Surveillance Team |
| EPI_ISL_2235816 | Lighthouse Lab in Alderley Park | Wellcome Sanger Institute for the COVID-19 Genomics UK (COG-UK) Consortium | Jacquelyn Wynn, Mairead Hyland, The Lighthouse Lab in Alderley Park and Alex Alderton, Roberto Amato, Jeffrey Barrett, Sonia Goncalves, Ewan Harrison, David K. Jackson, Ian Johnston, Dominic Kwiatkowski, Cordelia Langford, John Sillitoe on behalf of the Wellcome Sanger Institute COVID-19 Surveillance Team |
| EPI_ISL_2235821 | Health Services Laboratories | Wellcome Sanger Institute for the COVID-19 Genomics UK (COG-UK) Consortium | Health Services Laboratories and Alex Alderton, Roberto Amato, Jeffrey Barrett, Sonia Goncalves, Ewan Harrison, David K. Jackson, Ian Johnston, Dominic Kwiatkowski, Cordelia Langford, John Sillitoe on behalf of the Wellcome Sanger Institute COVID-19 Surveillance Team |
| EPI_ISL_2235823 | Lighthouse Lab in Alderley Park | Wellcome Sanger Institute for the COVID-19 Genomics UK (COG-UK) Consortium | Jacquelyn Wynn, Mairead Hyland, The Lighthouse Lab in Alderley Park and Alex Alderton, Roberto Amato, Jeffrey Barrett, Sonia Goncalves, Ewan Harrison, David K. Jackson, Ian Johnston, Dominic Kwiatkowski, Cordelia Langford, John Sillitoe on behalf of the Wellcome Sanger Institute COVID-19 Surveillance Team |
| EPI_ISL_2235824, | Lighthouse Lab in Glasgow | Wellcome Sanger Institute for the COVID-19 Genomics UK (COG-UK) | Harper VanSteenhouse, Yumi Kasai, David Gray, Carol Clugston, Anna Dominiczak and Alex Alderton, Roberto Amato, Jeffrey Barrett, Sonia Goncalves, Ewan Harrison, David K. Jackson, Ian Johnston, Dominic Kwiatkowski, Cordelia Langford, |

| | | | |
|---|---|---|---|
| EPI_ISL_2235826 | | Consortium | John Sillitoe on behalf of the Wellcome Sanger Institute COVID-19 Surveillance Team |
| EPI_ISL_2235827 | Lighthouse Lab in Alderley Park | Wellcome Sanger Institute for the COVID-19 Genomics UK (COG-UK) Consortium | Jacquelyn Wynn, Mairead Hyland, The Lighthouse Lab in Alderley Park and Alex Alderton, Roberto Amato, Jeffrey Barrett, Sonia Goncalves, Ewan Harrison, David K. Jackson, Ian Johnston, Dominic Kwiatkowski, Cordelia Langford, John Sillitoe on behalf of the Wellcome Sanger Institute COVID-19 Surveillance Team |
| EPI_ISL_2235828 | Lighthouse Lab in Milton Keynes | Wellcome Sanger Institute for the COVID-19 Genomics UK (COG-UK) Consortium | The Lighthouse Lab in Milton Keynes and Alex Alderton, Roberto Amato, Jeffrey Barrett, Sonia Goncalves, Ewan Harrison, David K. Jackson, Ian Johnston, Dominic Kwiatkowski, Cordelia Langford, John Sillitoe on behalf of the Wellcome Sanger Institute COVID-19 Surveillance Team |
| EPI_ISL_2235829, EPI_ISL_2235830 | Lighthouse Lab in Glasgow | Wellcome Sanger Institute for the COVID-19 Genomics UK (COG-UK) Consortium | Harper VanSteenhouse, Yumi Kasai, David Gray, Carol Clugston, Anna Dominiczak and Alex Alderton, Roberto Amato, Jeffrey Barrett, Sonia Goncalves, Ewan Harrison, David K. Jackson, Ian Johnston, Dominic Kwiatkowski, Cordelia Langford, John Sillitoe on behalf of the Wellcome Sanger Institute COVID-19 Surveillance Team |
| EPI_ISL_2235832 | Lighthouse Lab in Alderley Park | Wellcome Sanger Institute for the COVID-19 Genomics UK (COG-UK) Consortium | Jacquelyn Wynn, Mairead Hyland, The Lighthouse Lab in Alderley Park and Alex Alderton, Roberto Amato, Jeffrey Barrett, Sonia Goncalves, Ewan Harrison, David K. Jackson, Ian Johnston, Dominic Kwiatkowski, Cordelia Langford, John Sillitoe on behalf of the Wellcome Sanger Institute COVID-19 Surveillance Team |
| EPI_ISL_2235833, EPI_ISL_2235834 | Lighthouse Lab in Milton Keynes | Wellcome Sanger Institute for the COVID-19 Genomics UK (COG-UK) Consortium | The Lighthouse Lab in Milton Keynes and Alex Alderton, Roberto Amato, Jeffrey Barrett, Sonia Goncalves, Ewan Harrison, David K. Jackson, Ian Johnston, Dominic Kwiatkowski, Cordelia Langford, John Sillitoe on behalf of the Wellcome Sanger Institute COVID-19 Surveillance Team |
| EPI_ISL_2235836, EPI_ISL_2235838, EPI_ISL_2235841, EPI_ISL_2235844 | Lighthouse Lab in Glasgow | Wellcome Sanger Institute for the COVID-19 Genomics UK (COG-UK) Consortium | Harper VanSteenhouse, Yumi Kasai, David Gray, Carol Clugston, Anna Dominiczak and Alex Alderton, Roberto Amato, Jeffrey Barrett, Sonia Goncalves, Ewan Harrison, David K. Jackson, Ian Johnston, Dominic Kwiatkowski, Cordelia Langford, John Sillitoe on behalf of the Wellcome Sanger Institute COVID-19 Surveillance Team |
| EPI_ISL_2235845, EPI_ISL_2235846 | Lighthouse Lab in Milton Keynes | Wellcome Sanger Institute for the COVID-19 Genomics UK (COG-UK) Consortium | The Lighthouse Lab in Milton Keynes and Alex Alderton, Roberto Amato, Jeffrey Barrett, Sonia Goncalves, Ewan Harrison, David K. Jackson, Ian Johnston, Dominic Kwiatkowski, Cordelia Langford, John Sillitoe on behalf of the Wellcome Sanger Institute COVID-19 Surveillance Team |
| EPI_ISL_2235849 | Lighthouse Lab in Alderley Park | Wellcome Sanger Institute for the COVID-19 Genomics UK (COG-UK) Consortium | Jacquelyn Wynn, Mairead Hyland, The Lighthouse Lab in Alderley Park and Alex Alderton, Roberto Amato, Jeffrey Barrett, Sonia Goncalves, Ewan Harrison, David K. Jackson, Ian Johnston, Dominic Kwiatkowski, Cordelia Langford, John Sillitoe on behalf of the Wellcome Sanger Institute COVID-19 Surveillance Team |
| EPI_ISL_2235850 | Lighthouse Lab in Glasgow | Wellcome Sanger Institute for the COVID-19 Genomics UK (COG-UK) Consortium | Harper VanSteenhouse, Yumi Kasai, David Gray, Carol Clugston, Anna Dominiczak and Alex Alderton, Roberto Amato, Jeffrey Barrett, Sonia Goncalves, Ewan Harrison, David K. Jackson, Ian Johnston, Dominic Kwiatkowski, Cordelia Langford, |
| EPI_ISL_2235851 | Health Services Laboratories | Wellcome Sanger Institute for the COVID-19 Genomics UK (COG-UK) Consortium | Health Services Laboratories and Alex Alderton, Roberto Amato, Jeffrey Barrett, Sonia Goncalves, Ewan Harrison, David K. Jackson, Ian Johnston, Dominic Kwiatkowski, Cordelia Langford, John Sillitoe on behalf of the Wellcome Sanger Institute COVID-19 Surveillance Team |
| EPI_ISL_2235852, EPI_ISL_2235855 | Lighthouse Lab in Glasgow | Wellcome Sanger Institute for the COVID-19 Genomics UK (COG-UK) Consortium | Harper VanSteenhouse, Yumi Kasai, David Gray, Carol Clugston, Anna Dominiczak and Alex Alderton, Roberto Amato, Jeffrey Barrett, Sonia Goncalves, Ewan Harrison, David K. Jackson, Ian Johnston, Dominic Kwiatkowski, Cordelia Langford, John Sillitoe on behalf of the Wellcome Sanger Institute COVID-19 Surveillance Team |
| EPI_ISL_2235856 | Lighthouse Lab in Milton Keynes | Wellcome Sanger Institute for the COVID-19 Genomics UK (COG-UK) Consortium | The Lighthouse Lab in Milton Keynes and Alex Alderton, Roberto Amato, Jeffrey Barrett, Sonia Goncalves, Ewan Harrison, David K. Jackson, Ian Johnston, Dominic Kwiatkowski, Cordelia Langford, John Sillitoe on behalf of the Wellcome Sanger Institute COVID-19 Surveillance Team |
| EPI_ISL_2235858 | Lighthouse Lab in Alderley Park | Wellcome Sanger Institute for the COVID-19 Genomics UK (COG-UK) Consortium | Jacquelyn Wynn, Mairead Hyland, The Lighthouse Lab in Alderley Park and Alex Alderton, Roberto Amato, Jeffrey Barrett, Sonia Goncalves, Ewan Harrison, David K. Jackson, Ian Johnston, Dominic Kwiatkowski, Cordelia Langford, John Sillitoe on behalf of the Wellcome Sanger Institute COVID-19 Surveillance Team |
| EPI_ISL_2235862 | Lighthouse Lab in Glasgow | Wellcome Sanger Institute for the COVID-19 Genomics UK (COG-UK) Consortium | Harper VanSteenhouse, Yumi Kasai, David Gray, Carol Clugston, Anna Dominiczak and Alex Alderton, Roberto Amato, Jeffrey Barrett, Sonia Goncalves, Ewan Harrison, David K. Jackson, Ian Johnston, Dominic Kwiatkowski, Cordelia Langford, John Sillitoe on behalf of the Wellcome Sanger Institute COVID-19 Surveillance Team |
| EPI_ISL_2235863, EPI_ISL_2235865, EPI_ISL_2235866 | Lighthouse Lab in Alderley Park | Wellcome Sanger Institute for the COVID-19 Genomics UK (COG-UK) Consortium | Jacquelyn Wynn, Mairead Hyland, The Lighthouse Lab in Alderley Park and Alex Alderton, Roberto Amato, Jeffrey Barrett, Sonia Goncalves, Ewan Harrison, David K. Jackson, Ian Johnston, Dominic Kwiatkowski, Cordelia Langford, John Sillitoe on behalf of the Wellcome Sanger Institute COVID-19 Surveillance Team |
| EPI_ISL_2235869 | Lighthouse Lab in Glasgow | Wellcome Sanger Institute for the COVID-19 Genomics UK (COG-UK) Consortium | Harper VanSteenhouse, Yumi Kasai, David Gray, Carol Clugston, Anna Dominiczak and Alex Alderton, Roberto Amato, Jeffrey Barrett, Sonia Goncalves, Ewan Harrison, David K. Jackson, Ian Johnston, Dominic Kwiatkowski, Cordelia Langford, John Sillitoe on behalf of the Wellcome Sanger Institute COVID-19 Surveillance Team |
| EPI_ISL_2235871 | Lighthouse Lab in Alderley Park | Wellcome Sanger Institute for the COVID-19 Genomics UK (COG-UK) Consortium | Jacquelyn Wynn, Mairead Hyland, The Lighthouse Lab in Alderley Park and Alex Alderton, Roberto Amato, Jeffrey Barrett, Sonia Goncalves, Ewan Harrison, David K. Jackson, Ian Johnston, Dominic Kwiatkowski, Cordelia Langford, John Sillitoe on behalf of the Wellcome Sanger Institute COVID-19 Surveillance Team |
| EPI_ISL_2235873 | Lighthouse Lab in Milton Keynes | Wellcome Sanger Institute for the COVID-19 Genomics UK (COG-UK) Consortium | The Lighthouse Lab in Milton Keynes and Alex Alderton, Roberto Amato, Jeffrey Barrett, Sonia Goncalves, Ewan Harrison, David K. Jackson, Ian Johnston, Dominic Kwiatkowski, Cordelia Langford, John Sillitoe on behalf of the Wellcome Sanger Institute COVID-19 Surveillance Team |
| EPI_ISL_2235874 | Lighthouse Lab in Glasgow | Wellcome Sanger Institute for the COVID-19 Genomics UK (COG-UK) Consortium | Harper VanSteenhouse, Yumi Kasai, David Gray, Carol Clugston, Anna Dominiczak and Alex Alderton, Roberto Amato, Jeffrey Barrett, Sonia Goncalves, Ewan Harrison, David K. Jackson, Ian Johnston, Dominic Kwiatkowski, Cordelia Langford, John Sillitoe on behalf of the Wellcome Sanger Institute COVID-19 Surveillance Team |
| EPI_ISL_2235876, EPI_ISL_2235877, EPI_ISL_2235880, EPI_ISL_2235881, EPI_ISL_2235883, EPI_ISL_2235885, EPI_ISL_2235888, EPI_ISL_2235889, EPI_ISL_2235890, EPI_ISL_2235891 | Lighthouse Lab in Alderley Park | Wellcome Sanger Institute for the COVID-19 Genomics UK (COG-UK) Consortium | Jacquelyn Wynn, Mairead Hyland, The Lighthouse Lab in Alderley Park and Alex Alderton, Roberto Amato, Jeffrey Barrett, Sonia Goncalves, Ewan Harrison, David K. Jackson, Ian Johnston, Dominic Kwiatkowski, Cordelia Langford, John Sillitoe on behalf of the Wellcome Sanger Institute COVID-19 Surveillance Team |
| EPI_ISL_2235894, EPI_ISL_2235896, EPI_ISL_2235897 | Lighthouse Lab in Milton Keynes | Wellcome Sanger Institute for the COVID-19 Genomics UK (COG-UK) Consortium | The Lighthouse Lab in Milton Keynes and Alex Alderton, Roberto Amato, Jeffrey Barrett, Sonia Goncalves, Ewan Harrison, David K. Jackson, Ian Johnston, Dominic Kwiatkowski, Cordelia Langford, John Sillitoe on behalf of the Wellcome Sanger Institute COVID-19 Surveillance Team |
| EPI_ISL_2235898, EPI_ISL_2235900 | Lighthouse Lab in Glasgow | Wellcome Sanger Institute for the COVID-19 Genomics UK (COG-UK) Consortium | Harper VanSteenhouse, Yumi Kasai, David Gray, Carol Clugston, Anna Dominiczak and Alex Alderton, Roberto Amato, Jeffrey Barrett, Sonia Goncalves, Ewan Harrison, David K. Jackson, Ian Johnston, Dominic Kwiatkowski, Cordelia Langford, John Sillitoe on behalf of the Wellcome Sanger Institute COVID-19 Surveillance Team |
| EPI_ISL_2235903, EPI_ISL_2235905 | Lighthouse Lab in Alderley Park | Wellcome Sanger Institute for the COVID-19 Genomics UK (COG-UK) Consortium | Jacquelyn Wynn, Mairead Hyland, The Lighthouse Lab in Alderley Park and Alex Alderton, Roberto Amato, Jeffrey Barrett, Sonia Goncalves, Ewan Harrison, David K. Jackson, Ian Johnston, Dominic Kwiatkowski, Cordelia Langford, John Sillitoe on behalf of the Wellcome Sanger Institute COVID-19 Surveillance Team |
| EPI_ISL_2235906 | Lighthouse Lab in Milton Keynes | Wellcome Sanger Institute for the COVID-19 Genomics UK (COG-UK) Consortium | The Lighthouse Lab in Milton Keynes and Alex Alderton, Roberto Amato, Jeffrey Barrett, Sonia Goncalves, Ewan Harrison, David K. Jackson, Ian Johnston, Dominic Kwiatkowski, Cordelia Langford, John Sillitoe on behalf of the Wellcome Sanger Institute COVID-19 Surveillance Team |
| EPI_ISL_2235909 | Lighthouse Lab in Alderley Park | Wellcome Sanger Institute for the COVID-19 Genomics UK (COG-UK) Consortium | Jacquelyn Wynn, Mairead Hyland, The Lighthouse Lab in Alderley Park and Alex Alderton, Roberto Amato, Jeffrey Barrett, Sonia Goncalves, Ewan Harrison, David K. Jackson, Ian Johnston, Dominic Kwiatkowski, Cordelia Langford, John Sillitoe on behalf of the Wellcome Sanger Institute COVID-19 Surveillance Team |
| EPI_ISL_2235910 | Lighthouse Lab in Milton Keynes | Wellcome Sanger Institute for the COVID-19 Genomics UK (COG-UK) Consortium | The Lighthouse Lab in Milton Keynes and Alex Alderton, Roberto Amato, Jeffrey Barrett, Sonia Goncalves, Ewan Harrison, David K. Jackson, Ian Johnston, Dominic Kwiatkowski, Cordelia Langford, John Sillitoe on behalf of the Wellcome Sanger Institute COVID-19 Surveillance Team |
| EPI_ISL_2235911, EPI_ISL_2235914 | Lighthouse Lab in Alderley Park | Wellcome Sanger Institute for the COVID-19 Genomics UK (COG-UK) Consortium | Jacquelyn Wynn, Mairead Hyland, The Lighthouse Lab in Alderley Park and Alex Alderton, Roberto Amato, Jeffrey Barrett, Sonia Goncalves, Ewan Harrison, David K. Jackson, Ian Johnston, Dominic Kwiatkowski, Cordelia Langford, John Sillitoe on behalf of the Wellcome Sanger Institute COVID-19 Surveillance Team |
| EPI_ISL_2235921 | Lighthouse Lab in Milton Keynes | Wellcome Sanger Institute for the COVID-19 Genomics UK (COG-UK) Consortium | The Lighthouse Lab in Milton Keynes and Alex Alderton, Roberto Amato, Jeffrey Barrett, Sonia Goncalves, Ewan Harrison, David K. Jackson, Ian Johnston, Dominic Kwiatkowski, Cordelia Langford, John Sillitoe on behalf of the Wellcome Sanger Institute COVID-19 Surveillance Team |
| EPI_ISL_2235922 | Lighthouse Lab in Glasgow | Wellcome Sanger Institute for the COVID-19 Genomics UK (COG-UK) Consortium | Harper VanSteenhouse, Yumi Kasai, David Gray, Carol Clugston, Anna Dominiczak and Alex Alderton, Roberto Amato, Jeffrey Barrett, Sonia Goncalves, Ewan Harrison, David K. Jackson, Ian Johnston, Dominic Kwiatkowski, Cordelia Langford, John Sillitoe on behalf of the Wellcome Sanger Institute COVID-19 Surveillance Team |
| EPI_ISL_2235924, EPI_ISL_2235926, EPI_ISL_2235927, EPI_ISL_2235929 | Lighthouse Lab in Alderley Park | Wellcome Sanger Institute for the COVID-19 Genomics UK (COG-UK) Consortium | Jacquelyn Wynn, Mairead Hyland, The Lighthouse Lab in Alderley Park and Alex Alderton, Roberto Amato, Jeffrey Barrett, Sonia Goncalves, Ewan Harrison, David K. Jackson, Ian Johnston, Dominic Kwiatkowski, Cordelia Langford, John Sillitoe on behalf of the Wellcome Sanger Institute COVID-19 Surveillance Team |
| EPI_ISL_2235931 | Lighthouse Lab in Milton Keynes | Wellcome Sanger Institute for the COVID-19 Genomics UK (COG-UK) Consortium | The Lighthouse Lab in Milton Keynes and Alex Alderton, Roberto Amato, Jeffrey Barrett, Sonia Goncalves, Ewan Harrison, David K. Jackson, Ian Johnston, Dominic Kwiatkowski, Cordelia Langford, John Sillitoe on behalf of the Wellcome Sanger Institute COVID-19 Surveillance Team |
| EPI_ISL_2235937, EPI_ISL_2235941 | Lighthouse Lab in Alderley Park | Wellcome Sanger Institute for the COVID-19 Genomics UK (COG-UK) Consortium | Jacquelyn Wynn, Mairead Hyland, The Lighthouse Lab in Alderley Park and Alex Alderton, Roberto Amato, Jeffrey Barrett, Sonia Goncalves, Ewan Harrison, David K. Jackson, Ian Johnston, Dominic Kwiatkowski, Cordelia Langford, John Sillitoe on behalf of the Wellcome Sanger Institute COVID-19 Surveillance Team |
| EPI_ISL_2235942 | Lighthouse Lab in Milton Keynes | Wellcome Sanger Institute for the COVID-19 Genomics UK (COG-UK) Consortium | The Lighthouse Lab in Milton Keynes and Alex Alderton, Roberto Amato, Jeffrey Barrett, Sonia Goncalves, Ewan Harrison, David K. Jackson, Ian Johnston, Dominic Kwiatkowski, Cordelia Langford, John Sillitoe on behalf of the Wellcome Sanger Institute COVID-19 Surveillance Team |
| EPI_ISL_2235943, EPI_ISL_2235944, EPI_ISL_2235945, EPI_ISL_2235946, EPI_ISL_2235947, EPI_ISL_2235948, EPI_ISL_2235949 | Lighthouse Lab in Alderley Park | Wellcome Sanger Institute for the COVID-19 Genomics UK (COG-UK) Consortium | Jacquelyn Wynn, Mairead Hyland, The Lighthouse Lab in Alderley Park and Alex Alderton, Roberto Amato, Jeffrey Barrett, Sonia Goncalves, Ewan Harrison, David K. Jackson, Ian Johnston, Dominic Kwiatkowski, Cordelia Langford, John Sillitoe on behalf of the Wellcome Sanger Institute COVID-19 Surveillance Team |
| EPI_ISL_2235950 | Lighthouse Lab in Milton Keynes | Wellcome Sanger Institute for the COVID-19 Genomics UK (COG-UK) Consortium | The Lighthouse Lab in Milton Keynes and Alex Alderton, Roberto Amato, Jeffrey Barrett, Sonia Goncalves, Ewan Harrison, David K. Jackson, Ian Johnston, Dominic Kwiatkowski, Cordelia Langford, John Sillitoe on behalf of the Wellcome Sanger Institute COVID-19 Surveillance Team |
| EPI_ISL_2235951 | Lighthouse Lab in Alderley Park | Wellcome Sanger Institute for the COVID-19 Genomics UK (COG-UK) Consortium | Jacquelyn Wynn, Mairead Hyland, The Lighthouse Lab in Alderley Park and Alex Alderton, Roberto Amato, Jeffrey Barrett, Sonia Goncalves, Ewan Harrison, David K. Jackson, Ian Johnston, Dominic Kwiatkowski, Cordelia Langford, John Sillitoe on behalf of the Wellcome Sanger Institute COVID-19 Surveillance Team |
| EPI_ISL_2235952 | Lighthouse Lab in Glasgow | Wellcome Sanger Institute for the COVID-19 Genomics UK (COG-UK) Consortium | Harper VanSteenhouse, Yumi Kasai, David Gray, Carol Clugston, Anna Dominiczak and Alex Alderton, Roberto Amato, Jeffrey Barrett, Sonia Goncalves, Ewan Harrison, David K. Jackson, Ian Johnston, Dominic Kwiatkowski, Cordelia Langford, John Sillitoe on behalf of the Wellcome Sanger Institute COVID-19 Surveillance Team |

| | | | |
|---|---|---|---|
| EPI_ISL_2235956 | Lighthouse Lab in Alderley Park | Wellcome Sanger Institute for the COVID-19 Genomics UK (COG-UK) Consortium | Jacquelyn Wynn, Mairead Hyland, The Lighthouse Lab in Alderley Park and Alex Alderton, Roberto Amato, Jeffrey Barrett, Sonia Goncalves, Ewan Harrison, David K. Jackson, Ian Johnston, Dominic Kwiatkowski, Cordelia Langford, John Sillitoe on behalf of the Wellcome Sanger Institute COVID-19 Surveillance Team |
| EPI_ISL_2235957, EPI_ISL_2235958, EPI_ISL_2235959 | Lighthouse Lab in Glasgow | Wellcome Sanger Institute for the COVID-19 Genomics UK (COG-UK) Consortium | Harper VanSteenhouse, Yumi Kasai, David Gray, Carol Clugston, Anna Dominiczak and Alex Alderton, Roberto Amato, Jeffrey Barrett, Sonia Goncalves, Ewan Harrison, David K. Jackson, Ian Johnston, Dominic Kwiatkowski, Cordelia Langford, John Sillitoe on behalf of the Wellcome Sanger Institute COVID-19 Surveillance Team |
| EPI_ISL_2235960 | Lighthouse Lab in Alderley Park | Wellcome Sanger Institute for the COVID-19 Genomics UK (COG-UK) Consortium | Jacquelyn Wynn, Mairead Hyland, The Lighthouse Lab in Alderley Park and Alex Alderton, Roberto Amato, Jeffrey Barrett, Sonia Goncalves, Ewan Harrison, David K. Jackson, Ian Johnston, Dominic Kwiatkowski, Cordelia Langford, John Sillitoe on behalf of the Wellcome Sanger Institute COVID-19 Surveillance Team |
| EPI_ISL_2235962 | Lighthouse Lab in Glasgow | Wellcome Sanger Institute for the COVID-19 Genomics UK (COG-UK) Consortium | Harper VanSteenhouse, Yumi Kasai, David Gray, Carol Clugston, Anna Dominiczak and Alex Alderton, Roberto Amato, Jeffrey Barrett, Sonia Goncalves, Ewan Harrison, David K. Jackson, Ian Johnston, Dominic Kwiatkowski, Cordelia Langford, John Sillitoe on behalf of the Wellcome Sanger Institute COVID-19 Surveillance Team |
| EPI_ISL_2235963, EPI_ISL_2235966, EPI_ISL_2235967, EPI_ISL_2235969 | Lighthouse Lab in Alderley Park | Wellcome Sanger Institute for the COVID-19 Genomics UK (COG-UK) Consortium | Jacquelyn Wynn, Mairead Hyland, The Lighthouse Lab in Alderley Park and Alex Alderton, Roberto Amato, Jeffrey Barrett, Sonia Goncalves, Ewan Harrison, David K. Jackson, Ian Johnston, Dominic Kwiatkowski, Cordelia Langford, John Sillitoe on behalf of the Wellcome Sanger Institute COVID-19 Surveillance Team |
| EPI_ISL_2235970, EPI_ISL_2235971, EPI_ISL_2235972 | Lighthouse Lab in Milton Keynes | Wellcome Sanger Institute for the COVID-19 Genomics UK (COG-UK) Consortium | The Lighthouse Lab in Milton Keynes and Alex Alderton, Roberto Amato, Jeffrey Barrett, Sonia Goncalves, Ewan Harrison, David K. Jackson, Ian Johnston, Dominic Kwiatkowski, Cordelia Langford, John Sillitoe on behalf of the Wellcome Sanger Institute COVID-19 Surveillance Team |
| EPI_ISL_2235973, EPI_ISL_2235974 | Lighthouse Lab in Alderley Park | Wellcome Sanger Institute for the COVID-19 Genomics UK (COG-UK) Consortium | Jacquelyn Wynn, Mairead Hyland, The Lighthouse Lab in Alderley Park and Alex Alderton, Roberto Amato, Jeffrey Barrett, Sonia Goncalves, Ewan Harrison, David K. Jackson, Ian Johnston, Dominic Kwiatkowski, Cordelia Langford, John Sillitoe on behalf of the Wellcome Sanger Institute COVID-19 Surveillance Team |
| EPI_ISL_2235975 | Lighthouse Lab in Glasgow | Wellcome Sanger Institute for the COVID-19 Genomics UK (COG-UK) Consortium | Harper VanSteenhouse, Yumi Kasai, David Gray, Carol Clugston, Anna Dominiczak and Alex Alderton, Roberto Amato, Jeffrey Barrett, Sonia Goncalves, Ewan Harrison, David K. Jackson, Ian Johnston, Dominic Kwiatkowski, Cordelia Langford, John Sillitoe on behalf of the Wellcome Sanger Institute COVID-19 Surveillance Team |
| EPI_ISL_2235977, EPI_ISL_2235978, EPI_ISL_2235979 | Lighthouse Lab in Alderley Park | Wellcome Sanger Institute for the COVID-19 Genomics UK (COG-UK) Consortium | Jacquelyn Wynn, Mairead Hyland, The Lighthouse Lab in Alderley Park and Alex Alderton, Roberto Amato, Jeffrey Barrett, Sonia Goncalves, Ewan Harrison, David K. Jackson, Ian Johnston, Dominic Kwiatkowski, Cordelia Langford, John Sillitoe on behalf of the Wellcome Sanger Institute COVID-19 Surveillance Team |
| EPI_ISL_2235980, EPI_ISL_2235982 | Lighthouse Lab in Glasgow | Wellcome Sanger Institute for the COVID-19 Genomics UK (COG-UK) Consortium | Harper VanSteenhouse, Yumi Kasai, David Gray, Carol Clugston, Anna Dominiczak and Alex Alderton, Roberto Amato, Jeffrey Barrett, Sonia Goncalves, Ewan Harrison, David K. Jackson, Ian Johnston, Dominic Kwiatkowski, Cordelia Langford, John Sillitoe on behalf of the Wellcome Sanger Institute COVID-19 Surveillance Team |
| EPI_ISL_2235985 | Lighthouse Lab in Alderley Park | Wellcome Sanger Institute for the COVID-19 Genomics UK (COG-UK) Consortium | Jacquelyn Wynn, Mairead Hyland, The Lighthouse Lab in Alderley Park and Alex Alderton, Roberto Amato, Jeffrey Barrett, Sonia Goncalves, Ewan Harrison, David K. Jackson, Ian Johnston, Dominic Kwiatkowski, Cordelia Langford, John Sillitoe on behalf of the Wellcome Sanger Institute COVID-19 Surveillance Team |
| EPI_ISL_2235986, EPI_ISL_2235987 | Lighthouse Lab in Glasgow | Wellcome Sanger Institute for the COVID-19 Genomics UK (COG-UK) Consortium | Harper VanSteenhouse, Yumi Kasai, David Gray, Carol Clugston, Anna Dominiczak and Alex Alderton, Roberto Amato, Jeffrey Barrett, Sonia Goncalves, Ewan Harrison, David K. Jackson, Ian Johnston, Dominic Kwiatkowski, Cordelia Langford, John Sillitoe on behalf of the Wellcome Sanger Institute COVID-19 Surveillance Team |
| EPI_ISL_2235989 | Lighthouse Lab in Alderley Park | Wellcome Sanger Institute for the COVID-19 Genomics UK (COG-UK) Consortium | Jacquelyn Wynn, Mairead Hyland, The Lighthouse Lab in Alderley Park and Alex Alderton, Roberto Amato, Jeffrey Barrett, Sonia Goncalves, Ewan Harrison, David K. Jackson, Ian Johnston, Dominic Kwiatkowski, Cordelia Langford, John Sillitoe on behalf of the Wellcome Sanger Institute COVID-19 Surveillance Team |
| EPI_ISL_2235993 | Lighthouse Lab in Glasgow | Wellcome Sanger Institute for the COVID-19 Genomics UK (COG-UK) Consortium | Harper VanSteenhouse, Yumi Kasai, David Gray, Carol Clugston, Anna Dominiczak and Alex Alderton, Roberto Amato, Jeffrey Barrett, Sonia Goncalves, Ewan Harrison, David K. Jackson, Ian Johnston, Dominic Kwiatkowski, Cordelia Langford, John Sillitoe on behalf of the Wellcome Sanger Institute COVID-19 Surveillance Team |
| EPI_ISL_2235994 | Lighthouse Lab in Milton Keynes | Wellcome Sanger Institute for the COVID-19 Genomics UK (COG-UK) Consortium | The Lighthouse Lab in Milton Keynes and Alex Alderton, Roberto Amato, Jeffrey Barrett, Sonia Goncalves, Ewan Harrison, David K. Jackson, Ian Johnston, Dominic Kwiatkowski, Cordelia Langford, John Sillitoe on behalf of the Wellcome Sanger Institute COVID-19 Surveillance Team |
| EPI_ISL_2235995, EPI_ISL_2235997 | Lighthouse Lab in Glasgow | Wellcome Sanger Institute for the COVID-19 Genomics UK (COG-UK) Consortium | Harper VanSteenhouse, Yumi Kasai, David Gray, Carol Clugston, Anna Dominiczak and Alex Alderton, Roberto Amato, Jeffrey Barrett, Sonia Goncalves, Ewan Harrison, David K. Jackson, Ian Johnston, Dominic Kwiatkowski, Cordelia Langford, John Sillitoe on behalf of the Wellcome Sanger Institute COVID-19 Surveillance Team |
| EPI_ISL_2235998, EPI_ISL_2236000, EPI_ISL_2236001 | Lighthouse Lab in Alderley Park | Wellcome Sanger Institute for the COVID-19 Genomics UK (COG-UK) Consortium | Jacquelyn Wynn, Mairead Hyland, The Lighthouse Lab in Alderley Park and Alex Alderton, Roberto Amato, Jeffrey Barrett, Sonia Goncalves, Ewan Harrison, David K. Jackson, Ian Johnston, Dominic Kwiatkowski, Cordelia Langford, John Sillitoe on behalf of the Wellcome Sanger Institute COVID-19 Surveillance Team |
| EPI_ISL_2236002 | Lighthouse Lab in Glasgow | Wellcome Sanger Institute for the COVID-19 Genomics UK (COG-UK) Consortium | Harper VanSteenhouse, Yumi Kasai, David Gray, Carol Clugston, Anna Dominiczak and Alex Alderton, Roberto Amato, Jeffrey Barrett, Sonia Goncalves, Ewan Harrison, David K. Jackson, Ian Johnston, Dominic Kwiatkowski, Cordelia Langford, John Sillitoe on behalf of the Wellcome Sanger Institute COVID-19 Surveillance Team |
| EPI_ISL_2236004 | Lighthouse Lab in Alderley Park | Wellcome Sanger Institute for the COVID-19 Genomics UK (COG-UK) Consortium | Jacquelyn Wynn, Mairead Hyland, The Lighthouse Lab in Alderley Park and Alex Alderton, Roberto Amato, Jeffrey Barrett, Sonia Goncalves, Ewan Harrison, David K. Jackson, Ian Johnston, Dominic Kwiatkowski, Cordelia Langford, John Sillitoe on behalf of the Wellcome Sanger Institute COVID-19 Surveillance Team |
| EPI_ISL_2236010 | Lighthouse Lab in Milton Keynes | Wellcome Sanger Institute for the COVID-19 Genomics UK (COG-UK) Consortium | The Lighthouse Lab in Milton Keynes and Alex Alderton, Roberto Amato, Jeffrey Barrett, Sonia Goncalves, Ewan Harrison, David K. Jackson, Ian Johnston, Dominic Kwiatkowski, Cordelia Langford, John Sillitoe on behalf of the Wellcome Sanger Institute COVID-19 Surveillance Team |
| EPI_ISL_2236012 | Lighthouse Lab in Alderley Park | Wellcome Sanger Institute for the COVID-19 Genomics UK (COG-UK) Consortium | Jacquelyn Wynn, Mairead Hyland, The Lighthouse Lab in Alderley Park and Alex Alderton, Roberto Amato, Jeffrey Barrett, Sonia Goncalves, Ewan Harrison, David K. Jackson, Ian Johnston, Dominic Kwiatkowski, Cordelia Langford, John Sillitoe on behalf of the Wellcome Sanger Institute COVID-19 Surveillance Team |
| EPI_ISL_2236015 | Lighthouse Lab in Glasgow | Wellcome Sanger Institute for the COVID-19 Genomics UK (COG-UK) Consortium | Harper VanSteenhouse, Yumi Kasai, David Gray, Carol Clugston, Anna Dominiczak and Alex Alderton, Roberto Amato, Jeffrey Barrett, Sonia Goncalves, Ewan Harrison, David K. Jackson, Ian Johnston, Dominic Kwiatkowski, Cordelia Langford, John Sillitoe on behalf of the Wellcome Sanger Institute COVID-19 Surveillance Team |
| EPI_ISL_2236018, EPI_ISL_2236019 | Lighthouse Lab in Alderley Park | Wellcome Sanger Institute for the COVID-19 Genomics UK (COG-UK) Consortium | Jacquelyn Wynn, Mairead Hyland, The Lighthouse Lab in Alderley Park and Alex Alderton, Roberto Amato, Jeffrey Barrett, Sonia Goncalves, Ewan Harrison, David K. Jackson, Ian Johnston, Dominic Kwiatkowski, Cordelia Langford, John Sillitoe on behalf of the Wellcome Sanger Institute COVID-19 Surveillance Team |
| EPI_ISL_2236020 | Lighthouse Lab in Glasgow | Wellcome Sanger Institute for the COVID-19 Genomics UK (COG-UK) Consortium | Harper VanSteenhouse, Yumi Kasai, David Gray, Carol Clugston, Anna Dominiczak and Alex Alderton, Roberto Amato, Jeffrey Barrett, Sonia Goncalves, Ewan Harrison, David K. Jackson, Ian Johnston, Dominic Kwiatkowski, Cordelia Langford, John Sillitoe on behalf of the Wellcome Sanger Institute COVID-19 Surveillance Team |
| EPI_ISL_2236021, EPI_ISL_2236022, EPI_ISL_2236027, EPI_ISL_2236028 | Lighthouse Lab in Alderley Park | Wellcome Sanger Institute for the COVID-19 Genomics UK (COG-UK) Consortium | Jacquelyn Wynn, Mairead Hyland, The Lighthouse Lab in Alderley Park and Alex Alderton, Roberto Amato, Jeffrey Barrett, Sonia Goncalves, Ewan Harrison, David K. Jackson, Ian Johnston, Dominic Kwiatkowski, Cordelia Langford, John Sillitoe on behalf of the Wellcome Sanger Institute COVID-19 Surveillance Team |
| EPI_ISL_2236034 | Lighthouse Lab in Milton Keynes | Wellcome Sanger Institute for the COVID-19 Genomics UK (COG-UK) Consortium | The Lighthouse Lab in Milton Keynes and Alex Alderton, Roberto Amato, Jeffrey Barrett, Sonia Goncalves, Ewan Harrison, David K. Jackson, Ian Johnston, Dominic Kwiatkowski, Cordelia Langford, John Sillitoe on behalf of the Wellcome Sanger Institute COVID-19 Surveillance Team |
| EPI_ISL_2236038 | Lighthouse Lab in Glasgow | Wellcome Sanger Institute for the COVID-19 Genomics UK (COG-UK) Consortium | Harper VanSteenhouse, Yumi Kasai, David Gray, Carol Clugston, Anna Dominiczak and Alex Alderton, Roberto Amato, Jeffrey Barrett, Sonia Goncalves, Ewan Harrison, David K. Jackson, Ian Johnston, Dominic Kwiatkowski, Cordelia Langford, John Sillitoe on behalf of the Wellcome Sanger Institute COVID-19 Surveillance Team |
| EPI_ISL_2236039 | Lighthouse Lab in Alderley Park | Wellcome Sanger Institute for the COVID-19 Genomics UK (COG-UK) Consortium | Jacquelyn Wynn, Mairead Hyland, The Lighthouse Lab in Alderley Park and Alex Alderton, Roberto Amato, Jeffrey Barrett, Sonia Goncalves, Ewan Harrison, David K. Jackson, Ian Johnston, Dominic Kwiatkowski, Cordelia Langford, John Sillitoe on behalf of the Wellcome Sanger Institute COVID-19 Surveillance Team |
| EPI_ISL_2236040, EPI_ISL_2236041 | Lighthouse Lab in Glasgow | Wellcome Sanger Institute for the COVID-19 Genomics UK (COG-UK) Consortium | Harper VanSteenhouse, Yumi Kasai, David Gray, Carol Clugston, Anna Dominiczak and Alex Alderton, Roberto Amato, Jeffrey Barrett, Sonia Goncalves, Ewan Harrison, David K. Jackson, Ian Johnston, Dominic Kwiatkowski, Cordelia Langford, John Sillitoe on behalf of the Wellcome Sanger Institute COVID-19 Surveillance Team |
| EPI_ISL_2236042, EPI_ISL_2236043, EPI_ISL_2236044, EPI_ISL_2236045 | Lighthouse Lab in Alderley Park | Wellcome Sanger Institute for the COVID-19 Genomics UK (COG-UK) Consortium | Jacquelyn Wynn, Mairead Hyland, The Lighthouse Lab in Alderley Park and Alex Alderton, Roberto Amato, Jeffrey Barrett, Sonia Goncalves, Ewan Harrison, David K. Jackson, Ian Johnston, Dominic Kwiatkowski, Cordelia Langford, John Sillitoe on behalf of the Wellcome Sanger Institute COVID-19 Surveillance Team |
| EPI_ISL_2236046 | Lighthouse Lab in Glasgow | Wellcome Sanger Institute for the COVID-19 Genomics UK (COG-UK) Consortium | Harper VanSteenhouse, Yumi Kasai, David Gray, Carol Clugston, Anna Dominiczak and Alex Alderton, Roberto Amato, Jeffrey Barrett, Sonia Goncalves, Ewan Harrison, David K. Jackson, Ian Johnston, Dominic Kwiatkowski, Cordelia Langford, John Sillitoe on behalf of the Wellcome Sanger Institute COVID-19 Surveillance Team |
| EPI_ISL_2236047, EPI_ISL_2236048, EPI_ISL_2236050, EPI_ISL_2236051 | Lighthouse Lab in Alderley Park | Wellcome Sanger Institute for the COVID-19 Genomics UK (COG-UK) Consortium | Jacquelyn Wynn, Mairead Hyland, The Lighthouse Lab in Alderley Park and Alex Alderton, Roberto Amato, Jeffrey Barrett, Sonia Goncalves, Ewan Harrison, David K. Jackson, Ian Johnston, Dominic Kwiatkowski, Cordelia Langford, John Sillitoe on behalf of the Wellcome Sanger Institute COVID-19 Surveillance Team |
| EPI_ISL_2236052, EPI_ISL_2236053 | Lighthouse Lab in Milton Keynes | Wellcome Sanger Institute for the COVID-19 Genomics UK (COG-UK) Consortium | The Lighthouse Lab in Milton Keynes and Alex Alderton, Roberto Amato, Jeffrey Barrett, Sonia Goncalves, Ewan Harrison, David K. Jackson, Ian Johnston, Dominic Kwiatkowski, Cordelia Langford, John Sillitoe on behalf of the Wellcome Sanger Institute COVID-19 Surveillance Team |
| EPI_ISL_2236056 | Lighthouse Lab in Alderley Park | Wellcome Sanger Institute for the COVID-19 Genomics UK (COG-UK) Consortium | Jacquelyn Wynn, Mairead Hyland, The Lighthouse Lab in Alderley Park and Alex Alderton, Roberto Amato, Jeffrey Barrett, Sonia Goncalves, Ewan Harrison, David K. Jackson, Ian Johnston, Dominic Kwiatkowski, Cordelia Langford, John Sillitoe on behalf of the Wellcome Sanger Institute COVID-19 Surveillance Team |
| EPI_ISL_2236057, EPI_ISL_2236058, EPI_ISL_2236060 | Lighthouse Lab in Glasgow | Wellcome Sanger Institute for the COVID-19 Genomics UK (COG-UK) Consortium | Harper VanSteenhouse, Yumi Kasai, David Gray, Carol Clugston, Anna Dominiczak and Alex Alderton, Roberto Amato, Jeffrey Barrett, Sonia Goncalves, Ewan Harrison, David K. Jackson, Ian Johnston, Dominic Kwiatkowski, Cordelia Langford, John Sillitoe on behalf of the Wellcome Sanger Institute COVID-19 Surveillance Team |
| EPI_ISL_2236061, EPI_ISL_2236062 | Lighthouse Lab in Alderley Park | Wellcome Sanger Institute for the COVID-19 Genomics UK (COG-UK) Consortium | Jacquelyn Wynn, Mairead Hyland, The Lighthouse Lab in Alderley Park and Alex Alderton, Roberto Amato, Jeffrey Barrett, Sonia Goncalves, Ewan Harrison, David K. Jackson, Ian Johnston, Dominic Kwiatkowski, Cordelia Langford, John Sillitoe on behalf of the Wellcome Sanger Institute COVID-19 Surveillance Team |
| EPI_ISL_2236064 | Lighthouse Lab in Glasgow | Wellcome Sanger Institute for the COVID-19 Genomics UK (COG-UK) Consortium | Harper VanSteenhouse, Yumi Kasai, David Gray, Carol Clugston, Anna Dominiczak and Alex Alderton, Roberto Amato, Jeffrey Barrett, Sonia Goncalves, Ewan Harrison, David K. Jackson, Ian Johnston, Dominic Kwiatkowski, Cordelia Langford, John Sillitoe on behalf of the Wellcome Sanger Institute COVID-19 Surveillance Team |
| EPI_ISL_2236066 | Lighthouse Lab in Alderley Park | Wellcome Sanger Institute for the COVID-19 Genomics UK (COG-UK) Consortium | Jacquelyn Wynn, Mairead Hyland, The Lighthouse Lab in Alderley Park and Alex Alderton, Roberto Amato, Jeffrey Barrett, Sonia Goncalves, Ewan Harrison, David K. Jackson, Ian Johnston, Dominic Kwiatkowski, Cordelia Langford, John Sillitoe on behalf of the Wellcome Sanger Institute COVID-19 Surveillance Team |
| EPI_ISL_2236067 | Lighthouse Lab in Milton Keynes | Wellcome Sanger Institute for the COVID-19 Genomics UK (COG-UK) Consortium | The Lighthouse Lab in Milton Keynes and Alex Alderton, Roberto Amato, Jeffrey Barrett, Sonia Goncalves, Ewan Harrison, David K. Jackson, Ian Johnston, Dominic Kwiatkowski, Cordelia Langford, John Sillitoe on behalf of the Wellcome Sanger Institute COVID-19 Surveillance Team |
| EPI_ISL_2236068, | Lighthouse Lab in Alderley Park | Wellcome Sanger Institute for the COVID-19 Genomics UK (COG-UK) | Jacquelyn Wynn, Mairead Hyland, The Lighthouse Lab in Alderley Park and Alex Alderton, Roberto Amato, Jeffrey Barrett, Sonia Goncalves, Ewan Harrison, David K. Jackson, Ian Johnston, Dominic Kwiatkowski, Cordelia Langford, John Sillitoe on |

| | | | |
|---|---|---|---|
| EPI_ISL_2236069, EPI_ISL_2236072, EPI_ISL_2236076, EPI_ISL_2236077 | | Consortium | behalf of the Wellcome Sanger Institute COVID-19 Surveillance Team |
| EPI_ISL_2236078, EPI_ISL_2236079, EPI_ISL_2236080 | Lighthouse Lab in Glasgow | Wellcome Sanger Institute for the COVID-19 Genomics UK (COG-UK) Consortium | Harper VanSteenhouse, Yumi Kasai, David Gray, Carol Clugston, Anna Dominiczak and Alex Alderton, Roberto Amato, Jeffrey Barrett, Sonia Goncalves, Ewan Harrison, David K. Jackson, Ian Johnston, Dominic Kwiatkowski, Cordelia Langford, John Sillitoe on behalf of the Wellcome Sanger Institute COVID-19 Surveillance Team |
| EPI_ISL_2236082 | Lighthouse Lab in Alderley Park | Wellcome Sanger Institute for the COVID-19 Genomics UK (COG-UK) Consortium | Jacquelyn Wynn, Mairead Hyland, The Lighthouse Lab in Alderley Park and Alex Alderton, Roberto Amato, Jeffrey Barrett, Sonia Goncalves, Ewan Harrison, David K. Jackson, Ian Johnston, Dominic Kwiatkowski, Cordelia Langford, John Sillitoe on behalf of the Wellcome Sanger Institute COVID-19 Surveillance Team |
| EPI_ISL_2236083, EPI_ISL_2236084 | Lighthouse Lab in Milton Keynes | Wellcome Sanger Institute for the COVID-19 Genomics UK (COG-UK) Consortium | The Lighthouse Lab in Milton Keynes and Alex Alderton, Roberto Amato, Jeffrey Barrett, Sonia Goncalves, Ewan Harrison, David K. Jackson, Ian Johnston, Dominic Kwiatkowski, Cordelia Langford, John Sillitoe on behalf of the Wellcome Sanger Institute COVID-19 Surveillance Team |
| EPI_ISL_2236085, EPI_ISL_2236087 | Lighthouse Lab in Alderley Park | Wellcome Sanger Institute for the COVID-19 Genomics UK (COG-UK) Consortium | Jacquelyn Wynn, Mairead Hyland, The Lighthouse Lab in Alderley Park and Alex Alderton, Roberto Amato, Jeffrey Barrett, Sonia Goncalves, Ewan Harrison, David K. Jackson, Ian Johnston, Dominic Kwiatkowski, Cordelia Langford, John Sillitoe on behalf of the Wellcome Sanger Institute COVID-19 Surveillance Team |
| EPI_ISL_2236088 | Lighthouse Lab in Milton Keynes | Wellcome Sanger Institute for the COVID-19 Genomics UK (COG-UK) Consortium | The Lighthouse Lab in Milton Keynes and Alex Alderton, Roberto Amato, Jeffrey Barrett, Sonia Goncalves, Ewan Harrison, David K. Jackson, Ian Johnston, Dominic Kwiatkowski, Cordelia Langford, John Sillitoe on behalf of the Wellcome Sanger Institute COVID-19 Surveillance Team |
| EPI_ISL_2236089, EPI_ISL_2236090 | Lighthouse Lab in Alderley Park | Wellcome Sanger Institute for the COVID-19 Genomics UK (COG-UK) Consortium | Jacquelyn Wynn, Mairead Hyland, The Lighthouse Lab in Alderley Park and Alex Alderton, Roberto Amato, Jeffrey Barrett, Sonia Goncalves, Ewan Harrison, David K. Jackson, Ian Johnston, Dominic Kwiatkowski, Cordelia Langford, John Sillitoe on behalf of the Wellcome Sanger Institute COVID-19 Surveillance Team |
| EPI_ISL_2236092 | Lighthouse Lab in Glasgow | Wellcome Sanger Institute for the COVID-19 Genomics UK (COG-UK) Consortium | Harper VanSteenhouse, Yumi Kasai, David Gray, Carol Clugston, Anna Dominiczak and Alex Alderton, Roberto Amato, Jeffrey Barrett, Sonia Goncalves, Ewan Harrison, David K. Jackson, Ian Johnston, Dominic Kwiatkowski, Cordelia Langford, John Sillitoe on behalf of the Wellcome Sanger Institute COVID-19 Surveillance Team |
| EPI_ISL_2236093, EPI_ISL_2236094, EPI_ISL_2236096, EPI_ISL_2236098, EPI_ISL_2236099 | Lighthouse Lab in Alderley Park | Wellcome Sanger Institute for the COVID-19 Genomics UK (COG-UK) Consortium | Jacquelyn Wynn, Mairead Hyland, The Lighthouse Lab in Alderley Park and Alex Alderton, Roberto Amato, Jeffrey Barrett, Sonia Goncalves, Ewan Harrison, David K. Jackson, Ian Johnston, Dominic Kwiatkowski, Cordelia Langford, John Sillitoe on behalf of the Wellcome Sanger Institute COVID-19 Surveillance Team |
| EPI_ISL_2236101 | Lighthouse Lab in Milton Keynes | Wellcome Sanger Institute for the COVID-19 Genomics UK (COG-UK) Consortium | The Lighthouse Lab in Milton Keynes and Alex Alderton, Roberto Amato, Jeffrey Barrett, Sonia Goncalves, Ewan Harrison, David K. Jackson, Ian Johnston, Dominic Kwiatkowski, Cordelia Langford, John Sillitoe on behalf of the Wellcome Sanger Institute COVID-19 Surveillance Team |
| EPI_ISL_2236102 | Lighthouse Lab in Alderley Park | Wellcome Sanger Institute for the COVID-19 Genomics UK (COG-UK) Consortium | Jacquelyn Wynn, Mairead Hyland, The Lighthouse Lab in Alderley Park and Alex Alderton, Roberto Amato, Jeffrey Barrett, Sonia Goncalves, Ewan Harrison, David K. Jackson, Ian Johnston, Dominic Kwiatkowski, Cordelia Langford, John Sillitoe on behalf of the Wellcome Sanger Institute COVID-19 Surveillance Team |
| EPI_ISL_2236104 | Lighthouse Lab in Milton Keynes | Wellcome Sanger Institute for the COVID-19 Genomics UK (COG-UK) Consortium | The Lighthouse Lab in Milton Keynes and Alex Alderton, Roberto Amato, Jeffrey Barrett, Sonia Goncalves, Ewan Harrison, David K. Jackson, Ian Johnston, Dominic Kwiatkowski, Cordelia Langford, John Sillitoe on behalf of the Wellcome Sanger Institute COVID-19 Surveillance Team |
| EPI_ISL_2236105 | Lighthouse Lab in Glasgow | Wellcome Sanger Institute for the COVID-19 Genomics UK (COG-UK) Consortium | Harper VanSteenhouse, Yumi Kasai, David Gray, Carol Clugston, Anna Dominiczak and Alex Alderton, Roberto Amato, Jeffrey Barrett, Sonia Goncalves, Ewan Harrison, David K. Jackson, Ian Johnston, Dominic Kwiatkowski, Cordelia Langford, John Sillitoe on behalf of the Wellcome Sanger Institute COVID-19 Surveillance Team |
| EPI_ISL_2236108 | Lighthouse Lab in Milton Keynes | Wellcome Sanger Institute for the COVID-19 Genomics UK (COG-UK) Consortium | The Lighthouse Lab in Milton Keynes and Alex Alderton, Roberto Amato, Jeffrey Barrett, Sonia Goncalves, Ewan Harrison, David K. Jackson, Ian Johnston, Dominic Kwiatkowski, Cordelia Langford, John Sillitoe on behalf of the Wellcome Sanger Institute COVID-19 Surveillance Team |
| EPI_ISL_2236110, EPI_ISL_2236111 | Lighthouse Lab in Glasgow | Wellcome Sanger Institute for the COVID-19 Genomics UK (COG-UK) Consortium | Harper VanSteenhouse, Yumi Kasai, David Gray, Carol Clugston, Anna Dominiczak and Alex Alderton, Roberto Amato, Jeffrey Barrett, Sonia Goncalves, Ewan Harrison, David K. Jackson, Ian Johnston, Dominic Kwiatkowski, Cordelia Langford, John Sillitoe on behalf of the Wellcome Sanger Institute COVID-19 Surveillance Team |
| EPI_ISL_2236114 | Lighthouse Lab in Alderley Park | Wellcome Sanger Institute for the COVID-19 Genomics UK (COG-UK) Consortium | Jacquelyn Wynn, Mairead Hyland, The Lighthouse Lab in Alderley Park and Alex Alderton, Roberto Amato, Jeffrey Barrett, Sonia Goncalves, Ewan Harrison, David K. Jackson, Ian Johnston, Dominic Kwiatkowski, Cordelia Langford, John Sillitoe on behalf of the Wellcome Sanger Institute COVID-19 Surveillance Team |
| EPI_ISL_2236115 | Lighthouse Lab in Glasgow | Wellcome Sanger Institute for the COVID-19 Genomics UK (COG-UK) Consortium | Harper VanSteenhouse, Yumi Kasai, David Gray, Carol Clugston, Anna Dominiczak and Alex Alderton, Roberto Amato, Jeffrey Barrett, Sonia Goncalves, Ewan Harrison, David K. Jackson, Ian Johnston, Dominic Kwiatkowski, Cordelia Langford, John Sillitoe on behalf of the Wellcome Sanger Institute COVID-19 Surveillance Team |
| EPI_ISL_2236116 | Lighthouse Lab in Alderley Park | Wellcome Sanger Institute for the COVID-19 Genomics UK (COG-UK) Consortium | Jacquelyn Wynn, Mairead Hyland, The Lighthouse Lab in Alderley Park and Alex Alderton, Roberto Amato, Jeffrey Barrett, Sonia Goncalves, Ewan Harrison, David K. Jackson, Ian Johnston, Dominic Kwiatkowski, Cordelia Langford, John Sillitoe on behalf of the Wellcome Sanger Institute COVID-19 Surveillance Team |
| EPI_ISL_2236118 | Lighthouse Lab in Milton Keynes | Wellcome Sanger Institute for the COVID-19 Genomics UK (COG-UK) Consortium | The Lighthouse Lab in Milton Keynes and Alex Alderton, Roberto Amato, Jeffrey Barrett, Sonia Goncalves, Ewan Harrison, David K. Jackson, Ian Johnston, Dominic Kwiatkowski, Cordelia Langford, John Sillitoe on behalf of the Wellcome Sanger Institute COVID-19 Surveillance Team |
| EPI_ISL_2236119, EPI_ISL_2236122 | Lighthouse Lab in Alderley Park | Wellcome Sanger Institute for the COVID-19 Genomics UK (COG-UK) Consortium | Jacquelyn Wynn, Mairead Hyland, The Lighthouse Lab in Alderley Park and Alex Alderton, Roberto Amato, Jeffrey Barrett, Sonia Goncalves, Ewan Harrison, David K. Jackson, Ian Johnston, Dominic Kwiatkowski, Cordelia Langford, John Sillitoe on behalf of the Wellcome Sanger Institute COVID-19 Surveillance Team |
| EPI_ISL_2236123 | Lighthouse Lab in Glasgow | Wellcome Sanger Institute for the COVID-19 Genomics UK (COG-UK) Consortium | Harper VanSteenhouse, Yumi Kasai, David Gray, Carol Clugston, Anna Dominiczak and Alex Alderton, Roberto Amato, Jeffrey Barrett, Sonia Goncalves, Ewan Harrison, David K. Jackson, Ian Johnston, Dominic Kwiatkowski, Cordelia Langford, John Sillitoe on behalf of the Wellcome Sanger Institute COVID-19 Surveillance Team |
| EPI_ISL_2236124, EPI_ISL_2236126 | Lighthouse Lab in Alderley Park | Wellcome Sanger Institute for the COVID-19 Genomics UK (COG-UK) Consortium | Jacquelyn Wynn, Mairead Hyland, The Lighthouse Lab in Alderley Park and Alex Alderton, Roberto Amato, Jeffrey Barrett, Sonia Goncalves, Ewan Harrison, David K. Jackson, Ian Johnston, Dominic Kwiatkowski, Cordelia Langford, John Sillitoe on behalf of the Wellcome Sanger Institute COVID-19 Surveillance Team |
| EPI_ISL_2236127 | Lighthouse Lab in Glasgow | Wellcome Sanger Institute for the COVID-19 Genomics UK (COG-UK) Consortium | Harper VanSteenhouse, Yumi Kasai, David Gray, Carol Clugston, Anna Dominiczak and Alex Alderton, Roberto Amato, Jeffrey Barrett, Sonia Goncalves, Ewan Harrison, David K. Jackson, Ian Johnston, Dominic Kwiatkowski, Cordelia Langford, John Sillitoe on behalf of the Wellcome Sanger Institute COVID-19 Surveillance Team |
| EPI_ISL_2236129, EPI_ISL_2236131, EPI_ISL_2236132 | Lighthouse Lab in Alderley Park | Wellcome Sanger Institute for the COVID-19 Genomics UK (COG-UK) Consortium | Jacquelyn Wynn, Mairead Hyland, The Lighthouse Lab in Alderley Park and Alex Alderton, Roberto Amato, Jeffrey Barrett, Sonia Goncalves, Ewan Harrison, David K. Jackson, Ian Johnston, Dominic Kwiatkowski, Cordelia Langford, John Sillitoe on behalf of the Wellcome Sanger Institute COVID-19 Surveillance Team |
| EPI_ISL_2236134 | Lighthouse Lab in Milton Keynes | Wellcome Sanger Institute for the COVID-19 Genomics UK (COG-UK) Consortium | The Lighthouse Lab in Milton Keynes and Alex Alderton, Roberto Amato, Jeffrey Barrett, Sonia Goncalves, Ewan Harrison, David K. Jackson, Ian Johnston, Dominic Kwiatkowski, Cordelia Langford, John Sillitoe on behalf of the Wellcome Sanger Institute COVID-19 Surveillance Team |
| EPI_ISL_2236135, EPI_ISL_2236136 | Lighthouse Lab in Alderley Park | Wellcome Sanger Institute for the COVID-19 Genomics UK (COG-UK) Consortium | Jacquelyn Wynn, Mairead Hyland, The Lighthouse Lab in Alderley Park and Alex Alderton, Roberto Amato, Jeffrey Barrett, Sonia Goncalves, Ewan Harrison, David K. Jackson, Ian Johnston, Dominic Kwiatkowski, Cordelia Langford, John Sillitoe on behalf of the Wellcome Sanger Institute COVID-19 Surveillance Team |
| EPI_ISL_2236137 | Lighthouse Lab in Milton Keynes | Wellcome Sanger Institute for the COVID-19 Genomics UK (COG-UK) Consortium | The Lighthouse Lab in Milton Keynes and Alex Alderton, Roberto Amato, Jeffrey Barrett, Sonia Goncalves, Ewan Harrison, David K. Jackson, Ian Johnston, Dominic Kwiatkowski, Cordelia Langford, John Sillitoe on behalf of the Wellcome Sanger Institute COVID-19 Surveillance Team |
| EPI_ISL_2236140 | Lighthouse Lab in Alderley Park | Wellcome Sanger Institute for the COVID-19 Genomics UK (COG-UK) Consortium | Jacquelyn Wynn, Mairead Hyland, The Lighthouse Lab in Alderley Park and Alex Alderton, Roberto Amato, Jeffrey Barrett, Sonia Goncalves, Ewan Harrison, David K. Jackson, Ian Johnston, Dominic Kwiatkowski, Cordelia Langford, John Sillitoe on behalf of the Wellcome Sanger Institute COVID-19 Surveillance Team |
| EPI_ISL_2236141 | Lighthouse Lab in Milton Keynes | Wellcome Sanger Institute for the COVID-19 Genomics UK (COG-UK) Consortium | The Lighthouse Lab in Milton Keynes and Alex Alderton, Roberto Amato, Jeffrey Barrett, Sonia Goncalves, Ewan Harrison, David K. Jackson, Ian Johnston, Dominic Kwiatkowski, Cordelia Langford, John Sillitoe on behalf of the Wellcome Sanger Institute COVID-19 Surveillance Team |
| EPI_ISL_2236142, EPI_ISL_2236143, EPI_ISL_2236144 | Lighthouse Lab in Alderley Park | Wellcome Sanger Institute for the COVID-19 Genomics UK (COG-UK) Consortium | Jacquelyn Wynn, Mairead Hyland, The Lighthouse Lab in Alderley Park and Alex Alderton, Roberto Amato, Jeffrey Barrett, Sonia Goncalves, Ewan Harrison, David K. Jackson, Ian Johnston, Dominic Kwiatkowski, Cordelia Langford, John Sillitoe on behalf of the Wellcome Sanger Institute COVID-19 Surveillance Team |
| EPI_ISL_2236146 | Lighthouse Lab in Glasgow | Wellcome Sanger Institute for the COVID-19 Genomics UK (COG-UK) Consortium | Harper VanSteenhouse, Yumi Kasai, David Gray, Carol Clugston, Anna Dominiczak and Alex Alderton, Roberto Amato, Jeffrey Barrett, Sonia Goncalves, Ewan Harrison, David K. Jackson, Ian Johnston, Dominic Kwiatkowski, Cordelia Langford, John Sillitoe on behalf of the Wellcome Sanger Institute COVID-19 Surveillance Team |
| EPI_ISL_2236147 | Lighthouse Lab in Alderley Park | Wellcome Sanger Institute for the COVID-19 Genomics UK (COG-UK) Consortium | Jacquelyn Wynn, Mairead Hyland, The Lighthouse Lab in Alderley Park and Alex Alderton, Roberto Amato, Jeffrey Barrett, Sonia Goncalves, Ewan Harrison, David K. Jackson, Ian Johnston, Dominic Kwiatkowski, Cordelia Langford, John Sillitoe on behalf of the Wellcome Sanger Institute COVID-19 Surveillance Team |
| EPI_ISL_2236149 | Lighthouse Lab in Milton Keynes | Wellcome Sanger Institute for the COVID-19 Genomics UK (COG-UK) Consortium | The Lighthouse Lab in Milton Keynes and Alex Alderton, Roberto Amato, Jeffrey Barrett, Sonia Goncalves, Ewan Harrison, David K. Jackson, Ian Johnston, Dominic Kwiatkowski, Cordelia Langford, John Sillitoe on behalf of the Wellcome Sanger Institute COVID-19 Surveillance Team |
| EPI_ISL_2236150, EPI_ISL_2236152, EPI_ISL_2236153 | Lighthouse Lab in Glasgow | Wellcome Sanger Institute for the COVID-19 Genomics UK (COG-UK) Consortium | Harper VanSteenhouse, Yumi Kasai, David Gray, Carol Clugston, Anna Dominiczak and Alex Alderton, Roberto Amato, Jeffrey Barrett, Sonia Goncalves, Ewan Harrison, David K. Jackson, Ian Johnston, Dominic Kwiatkowski, Cordelia Langford, John Sillitoe on behalf of the Wellcome Sanger Institute COVID-19 Surveillance Team |
| EPI_ISL_2236154 | Lighthouse Lab in Alderley Park | Wellcome Sanger Institute for the COVID-19 Genomics UK (COG-UK) Consortium | Jacquelyn Wynn, Mairead Hyland, The Lighthouse Lab in Alderley Park and Alex Alderton, Roberto Amato, Jeffrey Barrett, Sonia Goncalves, Ewan Harrison, David K. Jackson, Ian Johnston, Dominic Kwiatkowski, Cordelia Langford, John Sillitoe on behalf of the Wellcome Sanger Institute COVID-19 Surveillance Team |
| EPI_ISL_2236159 | Lighthouse Lab in Milton Keynes | Wellcome Sanger Institute for the COVID-19 Genomics UK (COG-UK) Consortium | The Lighthouse Lab in Milton Keynes and Alex Alderton, Roberto Amato, Jeffrey Barrett, Sonia Goncalves, Ewan Harrison, David K. Jackson, Ian Johnston, Dominic Kwiatkowski, Cordelia Langford, John Sillitoe on behalf of the Wellcome Sanger Institute COVID-19 Surveillance Team |
| EPI_ISL_2236160 | Lighthouse Lab in Glasgow | Wellcome Sanger Institute for the COVID-19 Genomics UK (COG-UK) Consortium | Harper VanSteenhouse, Yumi Kasai, David Gray, Carol Clugston, Anna Dominiczak and Alex Alderton, Roberto Amato, Jeffrey Barrett, Sonia Goncalves, Ewan Harrison, David K. Jackson, Ian Johnston, Dominic Kwiatkowski, Cordelia Langford, John Sillitoe on behalf of the Wellcome Sanger Institute COVID-19 Surveillance Team |
| EPI_ISL_2236161 | Lighthouse Lab in Alderley Park | Wellcome Sanger Institute for the COVID-19 Genomics UK (COG-UK) Consortium | Jacquelyn Wynn, Mairead Hyland, The Lighthouse Lab in Alderley Park and Alex Alderton, Roberto Amato, Jeffrey Barrett, Sonia Goncalves, Ewan Harrison, David K. Jackson, Ian Johnston, Dominic Kwiatkowski, Cordelia Langford, John Sillitoe on behalf of the Wellcome Sanger Institute COVID-19 Surveillance Team |
| EPI_ISL_2236164, EPI_ISL_2236165 | Lighthouse Lab in Glasgow | Wellcome Sanger Institute for the COVID-19 Genomics UK (COG-UK) Consortium | Harper VanSteenhouse, Yumi Kasai, David Gray, Carol Clugston, Anna Dominiczak and Alex Alderton, Roberto Amato, Jeffrey Barrett, Sonia Goncalves, Ewan Harrison, David K. Jackson, Ian Johnston, Dominic Kwiatkowski, Cordelia Langford, John Sillitoe on behalf of the Wellcome Sanger Institute COVID-19 Surveillance Team |
| EPI_ISL_2236166, EPI_ISL_2236171 | Lighthouse Lab in Alderley Park | Wellcome Sanger Institute for the COVID-19 Genomics UK (COG-UK) Consortium | Jacquelyn Wynn, Mairead Hyland, The Lighthouse Lab in Alderley Park and Alex Alderton, Roberto Amato, Jeffrey Barrett, Sonia Goncalves, Ewan Harrison, David K. Jackson, Ian Johnston, Dominic Kwiatkowski, Cordelia Langford, John Sillitoe on behalf of the Wellcome Sanger Institute COVID-19 Surveillance Team |
| EPI_ISL_2236172, EPI_ISL_2236173 | Lighthouse Lab in Milton Keynes | Wellcome Sanger Institute for the COVID-19 Genomics UK (COG-UK) Consortium | The Lighthouse Lab in Milton Keynes and Alex Alderton, Roberto Amato, Jeffrey Barrett, Sonia Goncalves, Ewan Harrison, David K. Jackson, Ian Johnston, Dominic Kwiatkowski, Cordelia Langford, John Sillitoe on behalf of the Wellcome Sanger Institute COVID-19 Surveillance Team |
| EPI_ISL_2236174, | Lighthouse Lab in Alderley Park | Wellcome Sanger Institute for the COVID-19 Genomics UK (COG-UK) | Jacquelyn Wynn, Mairead Hyland, The Lighthouse Lab in Alderley Park and Alex Alderton, Roberto Amato, Jeffrey Barrett, Sonia Goncalves, Ewan Harrison, David K. Jackson, Ian Johnston, Dominic Kwiatkowski, Cordelia Langford, John Sillitoe on |

| | | | |
|---|---|---|---|
| EPI_ISL_2236176, EPI_ISL_2236177 | | Consortium | behalf of the Wellcome Sanger Institute COVID-19 Surveillance Team |
| EPI_ISL_2236179 | Lighthouse Lab in Milton Keynes | Wellcome Sanger Institute for the COVID-19 Genomics UK (COG-UK) Consortium | The Lighthouse Lab in Milton Keynes and Alex Alderton, Roberto Amato, Jeffrey Barrett, Sonia Goncalves, Ewan Harrison, David K. Jackson, Ian Johnston, Dominic Kwiatkowski, Cordelia Langford, John Sillitoe on behalf of the Wellcome Sanger Institute COVID-19 Surveillance Team |
| EPI_ISL_2236180, EPI_ISL_2236181 | Lighthouse Lab in Alderley Park | Wellcome Sanger Institute for the COVID-19 Genomics UK (COG-UK) Consortium | Jacquelyn Wynn, Mairead Hyland, The Lighthouse Lab in Alderley Park and Alex Alderton, Roberto Amato, Jeffrey Barrett, Sonia Goncalves, Ewan Harrison, David K. Jackson, Ian Johnston, Dominic Kwiatkowski, Cordelia Langford, John Sillitoe on behalf of the Wellcome Sanger Institute COVID-19 Surveillance Team |
| EPI_ISL_2236182 | Lighthouse Lab in Glasgow | Wellcome Sanger Institute for the COVID-19 Genomics UK (COG-UK) Consortium | Harper VanSteenhouse, Yumi Kasai, David Gray, Carol Clugston, Anna Dominiczak and Alex Alderton, Roberto Amato, Jeffrey Barrett, Sonia Goncalves, Ewan Harrison, David K. Jackson, Ian Johnston, Dominic Kwiatkowski, Cordelia Langford, John Sillitoe on behalf of the Wellcome Sanger Institute COVID-19 Surveillance Team |
| EPI_ISL_2236184 | Lighthouse Lab in Alderley Park | Wellcome Sanger Institute for the COVID-19 Genomics UK (COG-UK) Consortium | Jacquelyn Wynn, Mairead Hyland, The Lighthouse Lab in Alderley Park and Alex Alderton, Roberto Amato, Jeffrey Barrett, Sonia Goncalves, Ewan Harrison, David K. Jackson, Ian Johnston, Dominic Kwiatkowski, Cordelia Langford, John Sillitoe on behalf of the Wellcome Sanger Institute COVID-19 Surveillance Team |
| EPI_ISL_2236186 | Lighthouse Lab in Glasgow | Wellcome Sanger Institute for the COVID-19 Genomics UK (COG-UK) Consortium | Harper VanSteenhouse, Yumi Kasai, David Gray, Carol Clugston, Anna Dominiczak and Alex Alderton, Roberto Amato, Jeffrey Barrett, Sonia Goncalves, Ewan Harrison, David K. Jackson, Ian Johnston, Dominic Kwiatkowski, Cordelia Langford, John Sillitoe on behalf of the Wellcome Sanger Institute COVID-19 Surveillance Team |
| EPI_ISL_2236187, EPI_ISL_2236189, EPI_ISL_2236190 | Lighthouse Lab in Alderley Park | Wellcome Sanger Institute for the COVID-19 Genomics UK (COG-UK) Consortium | Jacquelyn Wynn, Mairead Hyland, The Lighthouse Lab in Alderley Park and Alex Alderton, Roberto Amato, Jeffrey Barrett, Sonia Goncalves, Ewan Harrison, David K. Jackson, Ian Johnston, Dominic Kwiatkowski, Cordelia Langford, John Sillitoe on behalf of the Wellcome Sanger Institute COVID-19 Surveillance Team |
| EPI_ISL_2236191 | Lighthouse Lab in Glasgow | Wellcome Sanger Institute for the COVID-19 Genomics UK (COG-UK) Consortium | Harper VanSteenhouse, Yumi Kasai, David Gray, Carol Clugston, Anna Dominiczak and Alex Alderton, Roberto Amato, Jeffrey Barrett, Sonia Goncalves, Ewan Harrison, David K. Jackson, Ian Johnston, Dominic Kwiatkowski, Cordelia Langford, John Sillitoe on behalf of the Wellcome Sanger Institute COVID-19 Surveillance Team |
| EPI_ISL_2236192 | Lighthouse Lab in Alderley Park | Wellcome Sanger Institute for the COVID-19 Genomics UK (COG-UK) Consortium | Jacquelyn Wynn, Mairead Hyland, The Lighthouse Lab in Alderley Park and Alex Alderton, Roberto Amato, Jeffrey Barrett, Sonia Goncalves, Ewan Harrison, David K. Jackson, Ian Johnston, Dominic Kwiatkowski, Cordelia Langford, John Sillitoe on behalf of the Wellcome Sanger Institute COVID-19 Surveillance Team |
| EPI_ISL_2236194 | Lighthouse Lab in Milton Keynes | Wellcome Sanger Institute for the COVID-19 Genomics UK (COG-UK) Consortium | The Lighthouse Lab in Milton Keynes and Alex Alderton, Roberto Amato, Jeffrey Barrett, Sonia Goncalves, Ewan Harrison, David K. Jackson, Ian Johnston, Dominic Kwiatkowski, Cordelia Langford, John Sillitoe on behalf of the Wellcome Sanger Institute COVID-19 Surveillance Team |
| EPI_ISL_2236198, EPI_ISL_2236200, EPI_ISL_2236201, EPI_ISL_2236205 | Lighthouse Lab in Alderley Park | Wellcome Sanger Institute for the COVID-19 Genomics UK (COG-UK) Consortium | Jacquelyn Wynn, Mairead Hyland, The Lighthouse Lab in Alderley Park and Alex Alderton, Roberto Amato, Jeffrey Barrett, Sonia Goncalves, Ewan Harrison, David K. Jackson, Ian Johnston, Dominic Kwiatkowski, Cordelia Langford, John Sillitoe on behalf of the Wellcome Sanger Institute COVID-19 Surveillance Team |
| EPI_ISL_2236206 | Lighthouse Lab in Milton Keynes | Wellcome Sanger Institute for the COVID-19 Genomics UK (COG-UK) Consortium | The Lighthouse Lab in Milton Keynes and Alex Alderton, Roberto Amato, Jeffrey Barrett, Sonia Goncalves, Ewan Harrison, David K. Jackson, Ian Johnston, Dominic Kwiatkowski, Cordelia Langford, John Sillitoe on behalf of the Wellcome Sanger Institute COVID-19 Surveillance Team |
| EPI_ISL_2236207 | Lighthouse Lab in Alderley Park | Wellcome Sanger Institute for the COVID-19 Genomics UK (COG-UK) Consortium | Jacquelyn Wynn, Mairead Hyland, The Lighthouse Lab in Alderley Park and Alex Alderton, Roberto Amato, Jeffrey Barrett, Sonia Goncalves, Ewan Harrison, David K. Jackson, Ian Johnston, Dominic Kwiatkowski, Cordelia Langford, John Sillitoe on behalf of the Wellcome Sanger Institute COVID-19 Surveillance Team |
| EPI_ISL_2236208 | Lighthouse Lab in Milton Keynes | Wellcome Sanger Institute for the COVID-19 Genomics UK (COG-UK) Consortium | The Lighthouse Lab in Milton Keynes and Alex Alderton, Roberto Amato, Jeffrey Barrett, Sonia Goncalves, Ewan Harrison, David K. Jackson, Ian Johnston, Dominic Kwiatkowski, Cordelia Langford, John Sillitoe on behalf of the Wellcome Sanger Institute COVID-19 Surveillance Team |
| EPI_ISL_2236209, EPI_ISL_2236212, EPI_ISL_2236215, EPI_ISL_2236217, EPI_ISL_2236218, EPI_ISL_2236220, EPI_ISL_2236222, EPI_ISL_2236223, EPI_ISL_2236225, EPI_ISL_2236226, EPI_ISL_2236227, EPI_ISL_2236228, EPI_ISL_2236229, EPI_ISL_2236230, EPI_ISL_2236231, EPI_ISL_2236232 | | | |
| see above | Lighthouse Lab in Alderley Park | Wellcome Sanger Institute for the COVID-19 Genomics UK (COG-UK) Consortium | Jacquelyn Wynn, Mairead Hyland, The Lighthouse Lab in Alderley Park and Alex Alderton, Roberto Amato, Jeffrey Barrett, Sonia Goncalves, Ewan Harrison, David K. Jackson, Ian Johnston, Dominic Kwiatkowski, Cordelia Langford, John Sillitoe on behalf of the Wellcome Sanger Institute COVID-19 Surveillance Team |
| EPI_ISL_2236236 | Lighthouse Lab in Milton Keynes | Wellcome Sanger Institute for the COVID-19 Genomics UK (COG-UK) Consortium | The Lighthouse Lab in Milton Keynes and Alex Alderton, Roberto Amato, Jeffrey Barrett, Sonia Goncalves, Ewan Harrison, David K. Jackson, Ian Johnston, Dominic Kwiatkowski, Cordelia Langford, John Sillitoe on behalf of the Wellcome Sanger Institute COVID-19 Surveillance Team |
| EPI_ISL_2236238, EPI_ISL_2236239, EPI_ISL_2236240, EPI_ISL_2236243, EPI_ISL_2236247, EPI_ISL_2236249, EPI_ISL_2236250, EPI_ISL_2236251, EPI_ISL_2236254, EPI_ISL_2236255 | Lighthouse Lab in Alderley Park | Wellcome Sanger Institute for the COVID-19 Genomics UK (COG-UK) Consortium | Jacquelyn Wynn, Mairead Hyland, The Lighthouse Lab in Alderley Park and Alex Alderton, Roberto Amato, Jeffrey Barrett, Sonia Goncalves, Ewan Harrison, David K. Jackson, Ian Johnston, Dominic Kwiatkowski, Cordelia Langford, John Sillitoe on behalf of the Wellcome Sanger Institute COVID-19 Surveillance Team |
| EPI_ISL_2236256 | Lighthouse Lab in Glasgow | Wellcome Sanger Institute for the COVID-19 Genomics UK (COG-UK) Consortium | Harper VanSteenhouse, Yumi Kasai, David Gray, Carol Clugston, Anna Dominiczak and Alex Alderton, Roberto Amato, Jeffrey Barrett, Sonia Goncalves, Ewan Harrison, David K. Jackson, Ian Johnston, Dominic Kwiatkowski, Cordelia Langford, John Sillitoe on behalf of the Wellcome Sanger Institute COVID-19 Surveillance Team |
| EPI_ISL_2236257 | Lighthouse Lab in Milton Keynes | Wellcome Sanger Institute for the COVID-19 Genomics UK (COG-UK) Consortium | The Lighthouse Lab in Milton Keynes and Alex Alderton, Roberto Amato, Jeffrey Barrett, Sonia Goncalves, Ewan Harrison, David K. Jackson, Ian Johnston, Dominic Kwiatkowski, Cordelia Langford, John Sillitoe on behalf of the Wellcome Sanger Institute COVID-19 Surveillance Team |
| EPI_ISL_2236261, EPI_ISL_2236262, EPI_ISL_2236264, EPI_ISL_2236265, EPI_ISL_2236266, EPI_ISL_2236267, EPI_ISL_2236268, EPI_ISL_2236270, EPI_ISL_2236271, EPI_ISL_2236273, EPI_ISL_2236275, EPI_ISL_2236277 | | | |
| see above | Lighthouse Lab in Alderley Park | Wellcome Sanger Institute for the COVID-19 Genomics UK (COG-UK) Consortium | Jacquelyn Wynn, Mairead Hyland, The Lighthouse Lab in Alderley Park and Alex Alderton, Roberto Amato, Jeffrey Barrett, Sonia Goncalves, Ewan Harrison, David K. Jackson, Ian Johnston, Dominic Kwiatkowski, Cordelia Langford, John Sillitoe on behalf of the Wellcome Sanger Institute COVID-19 Surveillance Team |
| EPI_ISL_2236278 | Lighthouse Lab in Glasgow | Wellcome Sanger Institute for the COVID-19 Genomics UK (COG-UK) Consortium | Harper VanSteenhouse, Yumi Kasai, David Gray, Carol Clugston, Anna Dominiczak and Alex Alderton, Roberto Amato, Jeffrey Barrett, Sonia Goncalves, Ewan Harrison, David K. Jackson, Ian Johnston, Dominic Kwiatkowski, Cordelia Langford, John Sillitoe on behalf of the Wellcome Sanger Institute COVID-19 Surveillance Team |
| EPI_ISL_2236279, EPI_ISL_2236280, EPI_ISL_2236283, EPI_ISL_2236284, EPI_ISL_2236286, EPI_ISL_2236290, EPI_ISL_2236291, EPI_ISL_2236293, EPI_ISL_2236294, EPI_ISL_2236296, EPI_ISL_2236297, EPI_ISL_2236298, EPI_ISL_2236299, EPI_ISL_2236300, EPI_ISL_2236301, EPI_ISL_2236302 | | | |
| see above | Lighthouse Lab in Alderley Park | Wellcome Sanger Institute for the COVID-19 Genomics UK (COG-UK) Consortium | Jacquelyn Wynn, Mairead Hyland, The Lighthouse Lab in Alderley Park and Alex Alderton, Roberto Amato, Jeffrey Barrett, Sonia Goncalves, Ewan Harrison, David K. Jackson, Ian Johnston, Dominic Kwiatkowski, Cordelia Langford, John Sillitoe on behalf of the Wellcome Sanger Institute COVID-19 Surveillance Team |
| EPI_ISL_2236303 | Lighthouse Lab in Milton Keynes | Wellcome Sanger Institute for the COVID-19 Genomics UK (COG-UK) Consortium | The Lighthouse Lab in Milton Keynes and Alex Alderton, Roberto Amato, Jeffrey Barrett, Sonia Goncalves, Ewan Harrison, David K. Jackson, Ian Johnston, Dominic Kwiatkowski, Cordelia Langford, John Sillitoe on behalf of the Wellcome Sanger Institute COVID-19 Surveillance Team |
| EPI_ISL_2236304 | Lighthouse Lab in Glasgow | Wellcome Sanger Institute for the COVID-19 Genomics UK (COG-UK) Consortium | Harper VanSteenhouse, Yumi Kasai, David Gray, Carol Clugston, Anna Dominiczak and Alex Alderton, Roberto Amato, Jeffrey Barrett, Sonia Goncalves, Ewan Harrison, David K. Jackson, Ian Johnston, Dominic Kwiatkowski, Cordelia Langford, John Sillitoe on behalf of the Wellcome Sanger Institute COVID-19 Surveillance Team |
| EPI_ISL_2236306, EPI_ISL_2236307, EPI_ISL_2236308, EPI_ISL_2236309, EPI_ISL_2236313, EPI_ISL_2236314, EPI_ISL_2236315, EPI_ISL_2236317, EPI_ISL_2236318, EPI_ISL_2236319, EPI_ISL_2236320, EPI_ISL_2236321, EPI_ISL_2236323, EPI_ISL_2236324, EPI_ISL_2236325, EPI_ISL_2236326, EPI_ISL_2236327, EPI_ISL_2236328, EPI_ISL_2236329, EPI_ISL_2236331, EPI_ISL_2236334 | | | |
| see above | Lighthouse Lab in Alderley Park | Wellcome Sanger Institute for the COVID-19 Genomics UK (COG-UK) Consortium | Jacquelyn Wynn, Mairead Hyland, The Lighthouse Lab in Alderley Park and Alex Alderton, Roberto Amato, Jeffrey Barrett, Sonia Goncalves, Ewan Harrison, David K. Jackson, Ian Johnston, Dominic Kwiatkowski, Cordelia Langford, John Sillitoe on behalf of the Wellcome Sanger Institute COVID-19 Surveillance Team |
| EPI_ISL_2236336 | Lighthouse Lab in Milton Keynes | Wellcome Sanger Institute for the COVID-19 Genomics UK (COG-UK) Consortium | The Lighthouse Lab in Milton Keynes and Alex Alderton, Roberto Amato, Jeffrey Barrett, Sonia Goncalves, Ewan Harrison, David K. Jackson, Ian Johnston, Dominic Kwiatkowski, Cordelia Langford, John Sillitoe on behalf of the Wellcome Sanger Institute COVID-19 Surveillance Team |
| EPI_ISL_2236337, EPI_ISL_2236339 | Lighthouse Lab in Alderley Park | Wellcome Sanger Institute for the COVID-19 Genomics UK (COG-UK) Consortium | Jacquelyn Wynn, Mairead Hyland, The Lighthouse Lab in Alderley Park and Alex Alderton, Roberto Amato, Jeffrey Barrett, Sonia Goncalves, Ewan Harrison, David K. Jackson, Ian Johnston, Dominic Kwiatkowski, Cordelia Langford, John Sillitoe, behalf of the Wellcome Sanger Institute COVID-19 Surveillance Team |
| EPI_ISL_2236340, EPI_ISL_2236342 | Lighthouse Lab in Glasgow | Wellcome Sanger Institute for the COVID-19 Genomics UK (COG-UK) Consortium | Harper VanSteenhouse, Yumi Kasai, David Gray, Carol Clugston, Anna Dominiczak and Alex Alderton, Roberto Amato, Jeffrey Barrett, Sonia Goncalves, Ewan Harrison, David K. Jackson, Ian Johnston, Dominic Kwiatkowski, Cordelia Langford, John Sillitoe on behalf of the Wellcome Sanger Institute COVID-19 Surveillance Team |
| EPI_ISL_2236344 | Lighthouse Lab in Milton Keynes | Wellcome Sanger Institute for the COVID-19 Genomics UK (COG-UK) Consortium | The Lighthouse Lab in Milton Keynes and Alex Alderton, Roberto Amato, Jeffrey Barrett, Sonia Goncalves, Ewan Harrison, David K. Jackson, Ian Johnston, Dominic Kwiatkowski, Cordelia Langford, John Sillitoe on behalf of the Wellcome Sanger Institute COVID-19 Surveillance Team |
| EPI_ISL_2236345 | Lighthouse Lab in Alderley Park | Wellcome Sanger Institute for the COVID-19 Genomics UK (COG-UK) Consortium | Jacquelyn Wynn, Mairead Hyland, The Lighthouse Lab in Alderley Park and Alex Alderton, Roberto Amato, Jeffrey Barrett, Sonia Goncalves, Ewan Harrison, David K. Jackson, Ian Johnston, Dominic Kwiatkowski, Cordelia Langford, John Sillitoe on behalf of the Wellcome Sanger Institute COVID-19 Surveillance Team |
| EPI_ISL_2236346 | Lighthouse Lab in Glasgow | Wellcome Sanger Institute for the COVID-19 Genomics UK (COG-UK) Consortium | Harper VanSteenhouse, Yumi Kasai, David Gray, Carol Clugston, Anna Dominiczak and Alex Alderton, Roberto Amato, Jeffrey Barrett, Sonia Goncalves, Ewan Harrison, David K. Jackson, Ian Johnston, Dominic Kwiatkowski, Cordelia Langford, John Sillitoe on behalf of the Wellcome Sanger Institute COVID-19 Surveillance Team |
| EPI_ISL_2236347, EPI_ISL_2236348, EPI_ISL_2236349, EPI_ISL_2236351, EPI_ISL_2236352, EPI_ISL_2236353, EPI_ISL_2236354, EPI_ISL_2236355 | Lighthouse Lab in Alderley Park | Wellcome Sanger Institute for the COVID-19 Genomics UK (COG-UK) Consortium | Jacquelyn Wynn, Mairead Hyland, The Lighthouse Lab in Alderley Park and Alex Alderton, Roberto Amato, Jeffrey Barrett, Sonia Goncalves, Ewan Harrison, David K. Jackson, Ian Johnston, Dominic Kwiatkowski, Cordelia Langford, John Sillitoe on behalf of the Wellcome Sanger Institute COVID-19 Surveillance Team |
| EPI_ISL_2236357 | Lighthouse Lab in Milton Keynes | Wellcome Sanger Institute for the COVID-19 Genomics UK (COG-UK) Consortium | The Lighthouse Lab in Milton Keynes and Alex Alderton, Roberto Amato, Jeffrey Barrett, Sonia Goncalves, Ewan Harrison, David K. Jackson, Ian Johnston, Dominic Kwiatkowski, Cordelia Langford, John Sillitoe on behalf of the Wellcome Sanger Institute COVID-19 Surveillance Team |
| EPI_ISL_2236359, EPI_ISL_2236362, EPI_ISL_2236364, EPI_ISL_2236367, EPI_ISL_2236369, EPI_ISL_2236370, EPI_ISL_2236371, EPI_ISL_2236372, EPI_ISL_2236374, EPI_ISL_2236375, EPI_ISL_2236379, EPI_ISL_2236382, EPI_ISL_2236384, EPI_ISL_2236385, EPI_ISL_2236387, EPI_ISL_2236389, EPI_ISL_2236391 | | | |
| see above | Lighthouse Lab in Alderley Park | Wellcome Sanger Institute for the COVID-19 Genomics UK (COG-UK) Consortium | Jacquelyn Wynn, Mairead Hyland, The Lighthouse Lab in Alderley Park and Alex Alderton, Roberto Amato, Jeffrey Barrett, Sonia Goncalves, Ewan Harrison, David K. Jackson, Ian Johnston, Dominic Kwiatkowski, Cordelia Langford, John Sillitoe on behalf of the Wellcome Sanger Institute COVID-19 Surveillance Team |
| EPI_ISL_2236392, EPI_ISL_2236394 | Lighthouse Lab in Milton Keynes | Wellcome Sanger Institute for the COVID-19 Genomics UK (COG-UK) Consortium | The Lighthouse Lab in Milton Keynes and Alex Alderton, Roberto Amato, Jeffrey Barrett, Sonia Goncalves, Ewan Harrison, David K. Jackson, Ian Johnston, Dominic Kwiatkowski, Cordelia Langford, John Sillitoe on behalf of the Wellcome Sanger Institute COVID-19 Surveillance Team |
| EPI_ISL_2236396, EPI_ISL_2236397, EPI_ISL_2236398, EPI_ISL_2236399, EPI_ISL_2236400, EPI_ISL_2236401, EPI_ISL_2236402, EPI_ISL_2236404, EPI_ISL_2236405, EPI_ISL_2236407, EPI_ISL_2236410, EPI_ISL_2236412, EPI_ISL_2236414, EPI_ISL_2236415, EPI_ISL_2236416, EPI_ISL_2236417, EPI_ISL_2236421, EPI_ISL_2236422, EPI_ISL_2236426, EPI_ISL_2236427, EPI_ISL_2236428, EPI_ISL_2236430, | | | |

| | | | |
|---|---|---|---|
| EPI_ISL_2236432 | | | |
| see above | Lighthouse Lab in Alderley Park | Wellcome Sanger Institute for the COVID-19 Genomics UK (COG-UK) Consortium | Jacquelyn Wynn, Mairead Hyland, The Lighthouse Lab in Alderley Park and Alex Alderton, Roberto Amato, Jeffrey Barrett, Sonia Goncalves, Ewan Harrison, David K. Jackson, Ian Johnston, Dominic Kwiatkowski, Cordelia Langford, John Sillitoe on behalf of the Wellcome Sanger Institute COVID-19 Surveillance Team |
| EPI_ISL_2236433 | Lighthouse Lab in Milton Keynes | Wellcome Sanger Institute for the COVID-19 Genomics UK (COG-UK) Consortium | The Lighthouse Lab in Milton Keynes and Alex Alderton, Roberto Amato, Jeffrey Barrett, Sonia Goncalves, Ewan Harrison, David K. Jackson, Ian Johnston, Dominic Kwiatkowski, Cordelia Langford, John Sillitoe on behalf of the Wellcome Sanger Institute COVID-19 Surveillance Team |
| EPI_ISL_2236434, EPI_ISL_2236435, EPI_ISL_2236436, EPI_ISL_2236437, EPI_ISL_2236439, EPI_ISL_2236440, EPI_ISL_2236441, EPI_ISL_2236442, EPI_ISL_2236447, EPI_ISL_2236449 | Lighthouse Lab in Alderley Park | Wellcome Sanger Institute for the COVID-19 Genomics UK (COG-UK) Consortium | Jacquelyn Wynn, Mairead Hyland, The Lighthouse Lab in Alderley Park and Alex Alderton, Roberto Amato, Jeffrey Barrett, Sonia Goncalves, Ewan Harrison, David K. Jackson, Ian Johnston, Dominic Kwiatkowski, Cordelia Langford, John Sillitoe on behalf of the Wellcome Sanger Institute COVID-19 Surveillance Team |
| EPI_ISL_2236450 | Lighthouse Lab in Glasgow | Wellcome Sanger Institute for the COVID-19 Genomics UK (COG-UK) Consortium | Harper VanSteenhouse, Yumi Kasai, David Gray, Carol Clugston, Anna Dominiczak and Alex Alderton, Roberto Amato, Jeffrey Barrett, Sonia Goncalves, Ewan Harrison, David K. Jackson, Ian Johnston, Dominic Kwiatkowski, Cordelia Langford, John Sillitoe on behalf of the Wellcome Sanger Institute COVID-19 Surveillance Team |
| EPI_ISL_2236451, EPI_ISL_2236452, EPI_ISL_2236453, EPI_ISL_2236454, EPI_ISL_2236457, EPI_ISL_2236458, EPI_ISL_2236461 | Lighthouse Lab in Alderley Park | Wellcome Sanger Institute for the COVID-19 Genomics UK (COG-UK) Consortium | Jacquelyn Wynn, Mairead Hyland, The Lighthouse Lab in Alderley Park and Alex Alderton, Roberto Amato, Jeffrey Barrett, Sonia Goncalves, Ewan Harrison, David K. Jackson, Ian Johnston, Dominic Kwiatkowski, Cordelia Langford, John Sillitoe on behalf of the Wellcome Sanger Institute COVID-19 Surveillance Team |
| EPI_ISL_2236463 | Lighthouse Lab in Milton Keynes | Wellcome Sanger Institute for the COVID-19 Genomics UK (COG-UK) Consortium | The Lighthouse Lab in Milton Keynes and Alex Alderton, Roberto Amato, Jeffrey Barrett, Sonia Goncalves, Ewan Harrison, David K. Jackson, Ian Johnston, Dominic Kwiatkowski, Cordelia Langford, John Sillitoe on behalf of the Wellcome Sanger Institute COVID-19 Surveillance Team |
| EPI_ISL_2236464, EPI_ISL_2236465, EPI_ISL_2236466, EPI_ISL_2236468, EPI_ISL_2236469, EPI_ISL_2236471, EPI_ISL_2236472, EPI_ISL_2236473 | Lighthouse Lab in Alderley Park | Wellcome Sanger Institute for the COVID-19 Genomics UK (COG-UK) Consortium | Jacquelyn Wynn, Mairead Hyland, The Lighthouse Lab in Alderley Park and Alex Alderton, Roberto Amato, Jeffrey Barrett, Sonia Goncalves, Ewan Harrison, David K. Jackson, Ian Johnston, Dominic Kwiatkowski, Cordelia Langford, John Sillitoe on behalf of the Wellcome Sanger Institute COVID-19 Surveillance Team |
| EPI_ISL_2236475, EPI_ISL_2236478, EPI_ISL_2236480, EPI_ISL_2236483, EPI_ISL_2236484, EPI_ISL_2236485, EPI_ISL_2236486, EPI_ISL_2236487, EPI_ISL_2236488, EPI_ISL_2236491, EPI_ISL_2236495, EPI_ISL_2236497, EPI_ISL_2236501, EPI_ISL_2236505, EPI_ISL_2236506 | | | |
| see above | Lighthouse Lab in Milton Keynes | Wellcome Sanger Institute for the COVID-19 Genomics UK (COG-UK) Consortium | The Lighthouse Lab in Milton Keynes and Alex Alderton, Roberto Amato, Jeffrey Barrett, Sonia Goncalves, Ewan Harrison, David K. Jackson, Ian Johnston, Dominic Kwiatkowski, Cordelia Langford, John Sillitoe on behalf of the Wellcome Sanger Institute COVID-19 Surveillance Team |
| EPI_ISL_2236507 | Lighthouse Lab in Glasgow | Wellcome Sanger Institute for the COVID-19 Genomics UK (COG-UK) Consortium | Harper VanSteenhouse, Yumi Kasai, David Gray, Carol Clugston, Anna Dominiczak and Alex Alderton, Roberto Amato, Jeffrey Barrett, Sonia Goncalves, Ewan Harrison, David K. Jackson, Ian Johnston, Dominic Kwiatkowski, Cordelia Langford, John Sillitoe on behalf of the Wellcome Sanger Institute COVID-19 Surveillance Team |
| EPI_ISL_2236508, EPI_ISL_2236509, EPI_ISL_2236511, EPI_ISL_2236512, EPI_ISL_2236513, EPI_ISL_2236514 | Lighthouse Lab in Milton Keynes | Wellcome Sanger Institute for the COVID-19 Genomics UK (COG-UK) Consortium | The Lighthouse Lab in Milton Keynes and Alex Alderton, Roberto Amato, Jeffrey Barrett, Sonia Goncalves, Ewan Harrison, David K. Jackson, Ian Johnston, Dominic Kwiatkowski, Cordelia Langford, John Sillitoe on behalf of the Wellcome Sanger Institute COVID-19 Surveillance Team |
| EPI_ISL_2236515, EPI_ISL_2236518 | Lighthouse Lab in Glasgow | Wellcome Sanger Institute for the COVID-19 Genomics UK (COG-UK) Consortium | Harper VanSteenhouse, Yumi Kasai, David Gray, Carol Clugston, Anna Dominiczak and Alex Alderton, Roberto Amato, Jeffrey Barrett, Sonia Goncalves, Ewan Harrison, David K. Jackson, Ian Johnston, Dominic Kwiatkowski, Cordelia Langford, John Sillitoe on behalf of the Wellcome Sanger Institute COVID-19 Surveillance Team |
| EPI_ISL_2236520 | Lighthouse Lab in Milton Keynes | Wellcome Sanger Institute for the COVID-19 Genomics UK (COG-UK) Consortium | The Lighthouse Lab in Milton Keynes and Alex Alderton, Roberto Amato, Jeffrey Barrett, Sonia Goncalves, Ewan Harrison, David K. Jackson, Ian Johnston, Dominic Kwiatkowski, Cordelia Langford, John Sillitoe on behalf of the Wellcome Sanger Institute COVID-19 Surveillance Team |
| EPI_ISL_2236523 | Lighthouse Lab in Glasgow | Wellcome Sanger Institute for the COVID-19 Genomics UK (COG-UK) Consortium | Harper VanSteenhouse, Yumi Kasai, David Gray, Carol Clugston, Anna Dominiczak and Alex Alderton, Roberto Amato, Jeffrey Barrett, Sonia Goncalves, Ewan Harrison, David K. Jackson, Ian Johnston, Dominic Kwiatkowski, Cordelia Langford, John Sillitoe on behalf of the Wellcome Sanger Institute COVID-19 Surveillance Team |
| EPI_ISL_2236524 | Lighthouse Lab in Milton Keynes | Wellcome Sanger Institute for the COVID-19 Genomics UK (COG-UK) Consortium | The Lighthouse Lab in Milton Keynes and Alex Alderton, Roberto Amato, Jeffrey Barrett, Sonia Goncalves, Ewan Harrison, David K. Jackson, Ian Johnston, Dominic Kwiatkowski, Cordelia Langford, John Sillitoe on behalf of the Wellcome Sanger Institute COVID-19 Surveillance Team |
| EPI_ISL_2236525 | Lighthouse Lab in Glasgow | Wellcome Sanger Institute for the COVID-19 Genomics UK (COG-UK) Consortium | Harper VanSteenhouse, Yumi Kasai, David Gray, Carol Clugston, Anna Dominiczak and Alex Alderton, Roberto Amato, Jeffrey Barrett, Sonia Goncalves, Ewan Harrison, David K. Jackson, Ian Johnston, Dominic Kwiatkowski, Cordelia Langford, John Sillitoe on behalf of the Wellcome Sanger Institute COVID-19 Surveillance Team |
| EPI_ISL_2236526, EPI_ISL_2236529, EPI_ISL_2236530, EPI_ISL_2236534, EPI_ISL_2236535, EPI_ISL_2236536, EPI_ISL_2236537 | Lighthouse Lab in Milton Keynes | Wellcome Sanger Institute for the COVID-19 Genomics UK (COG-UK) Consortium | The Lighthouse Lab in Milton Keynes and Alex Alderton, Roberto Amato, Jeffrey Barrett, Sonia Goncalves, Ewan Harrison, David K. Jackson, Ian Johnston, Dominic Kwiatkowski, Cordelia Langford, John Sillitoe on behalf of the Wellcome Sanger Institute COVID-19 Surveillance Team |
| EPI_ISL_2236539, EPI_ISL_2236541 | Lighthouse Lab in Glasgow | Wellcome Sanger Institute for the COVID-19 Genomics UK (COG-UK) Consortium | Harper VanSteenhouse, Yumi Kasai, David Gray, Carol Clugston, Anna Dominiczak and Alex Alderton, Roberto Amato, Jeffrey Barrett, Sonia Goncalves, Ewan Harrison, David K. Jackson, Ian Johnston, Dominic Kwiatkowski, Cordelia Langford, John Sillitoe on behalf of the Wellcome Sanger Institute COVID-19 Surveillance Team |
| EPI_ISL_2236543, EPI_ISL_2236544, EPI_ISL_2236546, EPI_ISL_2236547 | Lighthouse Lab in Milton Keynes | Wellcome Sanger Institute for the COVID-19 Genomics UK (COG-UK) Consortium | The Lighthouse Lab in Milton Keynes and Alex Alderton, Roberto Amato, Jeffrey Barrett, Sonia Goncalves, Ewan Harrison, David K. Jackson, Ian Johnston, Dominic Kwiatkowski, Cordelia Langford, John Sillitoe on behalf of the Wellcome Sanger Institute COVID-19 Surveillance Team |
| EPI_ISL_2236549 | Lighthouse Lab in Glasgow | Wellcome Sanger Institute for the COVID-19 Genomics UK (COG-UK) Consortium | Harper VanSteenhouse, Yumi Kasai, David Gray, Carol Clugston, Anna Dominiczak and Alex Alderton, Roberto Amato, Jeffrey Barrett, Sonia Goncalves, Ewan Harrison, David K. Jackson, Ian Johnston, Dominic Kwiatkowski, Cordelia Langford, John Sillitoe on behalf of the Wellcome Sanger Institute COVID-19 Surveillance Team |
| EPI_ISL_2236550, EPI_ISL_2236552, EPI_ISL_2236557 | Lighthouse Lab in Milton Keynes | Wellcome Sanger Institute for the COVID-19 Genomics UK (COG-UK) Consortium | The Lighthouse Lab in Milton Keynes and Alex Alderton, Roberto Amato, Jeffrey Barrett, Sonia Goncalves, Ewan Harrison, David K. Jackson, Ian Johnston, Dominic Kwiatkowski, Cordelia Langford, John Sillitoe on behalf of the Wellcome Sanger Institute COVID-19 Surveillance Team |
| EPI_ISL_2236558, EPI_ISL_2236559 | Lighthouse Lab in Glasgow | Wellcome Sanger Institute for the COVID-19 Genomics UK (COG-UK) Consortium | Harper VanSteenhouse, Yumi Kasai, David Gray, Carol Clugston, Anna Dominiczak and Alex Alderton, Roberto Amato, Jeffrey Barrett, Sonia Goncalves, Ewan Harrison, David K. Jackson, Ian Johnston, Dominic Kwiatkowski, Cordelia Langford, John Sillitoe on behalf of the Wellcome Sanger Institute COVID-19 Surveillance Team |
| EPI_ISL_2236560, EPI_ISL_2236561, EPI_ISL_2236562, EPI_ISL_2236563 | Lighthouse Lab in Milton Keynes | Wellcome Sanger Institute for the COVID-19 Genomics UK (COG-UK) Consortium | The Lighthouse Lab in Milton Keynes and Alex Alderton, Roberto Amato, Jeffrey Barrett, Sonia Goncalves, Ewan Harrison, David K. Jackson, Ian Johnston, Dominic Kwiatkowski, Cordelia Langford, John Sillitoe on behalf of the Wellcome Sanger Institute COVID-19 Surveillance Team |
| EPI_ISL_2236564 | Lighthouse Lab in Glasgow | Wellcome Sanger Institute for the COVID-19 Genomics UK (COG-UK) Consortium | Harper VanSteenhouse, Yumi Kasai, David Gray, Carol Clugston, Anna Dominiczak and Alex Alderton, Roberto Amato, Jeffrey Barrett, Sonia Goncalves, Ewan Harrison, David K. Jackson, Ian Johnston, Dominic Kwiatkowski, Cordelia Langford, John Sillitoe on behalf of the Wellcome Sanger Institute COVID-19 Surveillance Team |
| EPI_ISL_2236565, EPI_ISL_2236567, EPI_ISL_2236568 | Lighthouse Lab in Milton Keynes | Wellcome Sanger Institute for the COVID-19 Genomics UK (COG-UK) Consortium | The Lighthouse Lab in Milton Keynes and Alex Alderton, Roberto Amato, Jeffrey Barrett, Sonia Goncalves, Ewan Harrison, David K. Jackson, Ian Johnston, Dominic Kwiatkowski, Cordelia Langford, John Sillitoe on behalf of the Wellcome Sanger Institute COVID-19 Surveillance Team |
| EPI_ISL_2236569 | Lighthouse Lab in Glasgow | Wellcome Sanger Institute for the COVID-19 Genomics UK (COG-UK) Consortium | Harper VanSteenhouse, Yumi Kasai, David Gray, Carol Clugston, Anna Dominiczak and Alex Alderton, Roberto Amato, Jeffrey Barrett, Sonia Goncalves, Ewan Harrison, David K. Jackson, Ian Johnston, Dominic Kwiatkowski, Cordelia Langford, John Sillitoe on behalf of the Wellcome Sanger Institute COVID-19 Surveillance Team |
| EPI_ISL_2236570, EPI_ISL_2236571 | Lighthouse Lab in Milton Keynes | Wellcome Sanger Institute for the COVID-19 Genomics UK (COG-UK) Consortium | The Lighthouse Lab in Milton Keynes and Alex Alderton, Roberto Amato, Jeffrey Barrett, Sonia Goncalves, Ewan Harrison, David K. Jackson, Ian Johnston, Dominic Kwiatkowski, Cordelia Langford, John Sillitoe on behalf of the Wellcome Sanger Institute COVID-19 Surveillance Team |
| EPI_ISL_2236572, EPI_ISL_2236573, EPI_ISL_2236574, EPI_ISL_2236577 | Lighthouse Lab in Glasgow | Wellcome Sanger Institute for the COVID-19 Genomics UK (COG-UK) Consortium | Harper VanSteenhouse, Yumi Kasai, David Gray, Carol Clugston, Anna Dominiczak and Alex Alderton, Roberto Amato, Jeffrey Barrett, Sonia Goncalves, Ewan Harrison, David K. Jackson, Ian Johnston, Dominic Kwiatkowski, Cordelia Langford, John Sillitoe on behalf of the Wellcome Sanger Institute COVID-19 Surveillance Team |
| EPI_ISL_2236579, EPI_ISL_2236580, EPI_ISL_2236581, EPI_ISL_2236582 | Lighthouse Lab in Milton Keynes | Wellcome Sanger Institute for the COVID-19 Genomics UK (COG-UK) Consortium | The Lighthouse Lab in Milton Keynes and Alex Alderton, Roberto Amato, Jeffrey Barrett, Sonia Goncalves, Ewan Harrison, David K. Jackson, Ian Johnston, Dominic Kwiatkowski, Cordelia Langford, John Sillitoe on behalf of the Wellcome Sanger Institute COVID-19 Surveillance Team |
| EPI_ISL_2236584 | Lighthouse Lab in Glasgow | Wellcome Sanger Institute for the COVID-19 Genomics UK (COG-UK) | Harper VanSteenhouse, Yumi Kasai, David Gray, Carol Clugston, Anna Dominiczak and Alex Alderton, Roberto Amato, Jeffrey Barrett, Sonia Goncalves, Ewan Harrison, David K. Jackson, Ian Johnston, Dominic Kwiatkowski, Cordelia Langford, |

| | | | |
|---|---|---|---|
| | | Consortium | John Sillitoe on behalf of the Wellcome Sanger Institute COVID-19 Surveillance Team |
| EPI_ISL_2236587, EPI_ISL_2236588, EPI_ISL_2236589, EPI_ISL_2236590, EPI_ISL_2236592 | Lighthouse Lab in Milton Keynes | Wellcome Sanger Institute for the COVID-19 Genomics UK (COG-UK) Consortium | The Lighthouse Lab in Milton Keynes and Alex Alderton, Roberto Amato, Jeffrey Barrett, Sonia Goncalves, Ewan Harrison, David K. Jackson, Ian Johnston, Dominic Kwiatkowski, Cordelia Langford, John Sillitoe on behalf of the Wellcome Sanger Institute COVID-19 Surveillance Team |
| EPI_ISL_2236595 | Lighthouse Lab in Glasgow | Wellcome Sanger Institute for the COVID-19 Genomics UK (COG-UK) Consortium | Harper VanSteenhouse, Yumi Kasai, David Gray, Carol Clugston, Anna Dominiczak and Alex Alderton, Roberto Amato, Jeffrey Barrett, Sonia Goncalves, Ewan Harrison, David K. Jackson, Ian Johnston, Dominic Kwiatkowski, Cordelia Langford, John Sillitoe on behalf of the Wellcome Sanger Institute COVID-19 Surveillance Team |
| EPI_ISL_2236596, EPI_ISL_2236599, EPI_ISL_2236600, EPI_ISL_2236601, EPI_ISL_2236602, EPI_ISL_2236603, EPI_ISL_2236604, EPI_ISL_2236606, EPI_ISL_2236608, EPI_ISL_2236609, EPI_ISL_2236611, EPI_ISL_2236612, EPI_ISL_2236613, EPI_ISL_2236614, EPI_ISL_2236616 | | | |
| see above | Lighthouse Lab in Milton Keynes | Wellcome Sanger Institute for the COVID-19 Genomics UK (COG-UK) Consortium | The Lighthouse Lab in Milton Keynes and Alex Alderton, Roberto Amato, Jeffrey Barrett, Sonia Goncalves, Ewan Harrison, David K. Jackson, Ian Johnston, Dominic Kwiatkowski, Cordelia Langford, John Sillitoe on behalf of the Wellcome Sanger Institute COVID-19 Surveillance Team |
| EPI_ISL_2236617 | Lighthouse Lab in Glasgow | Wellcome Sanger Institute for the COVID-19 Genomics UK (COG-UK) Consortium | Harper VanSteenhouse, Yumi Kasai, David Gray, Carol Clugston, Anna Dominiczak and Alex Alderton, Roberto Amato, Jeffrey Barrett, Sonia Goncalves, Ewan Harrison, David K. Jackson, Ian Johnston, Dominic Kwiatkowski, Cordelia Langford, John Sillitoe on behalf of the Wellcome Sanger Institute COVID-19 Surveillance Team |
| EPI_ISL_2236618, EPI_ISL_2236620, EPI_ISL_2236623, EPI_ISL_2236625 | Lighthouse Lab in Milton Keynes | Wellcome Sanger Institute for the COVID-19 Genomics UK (COG-UK) Consortium | The Lighthouse Lab in Milton Keynes and Alex Alderton, Roberto Amato, Jeffrey Barrett, Sonia Goncalves, Ewan Harrison, David K. Jackson, Ian Johnston, Dominic Kwiatkowski, Cordelia Langford, John Sillitoe on behalf of the Wellcome Sanger Institute COVID-19 Surveillance Team |
| EPI_ISL_2236626 | Lighthouse Lab in Glasgow | Wellcome Sanger Institute for the COVID-19 Genomics UK (COG-UK) Consortium | Harper VanSteenhouse, Yumi Kasai, David Gray, Carol Clugston, Anna Dominiczak and Alex Alderton, Roberto Amato, Jeffrey Barrett, Sonia Goncalves, Ewan Harrison, David K. Jackson, Ian Johnston, Dominic Kwiatkowski, Cordelia Langford, John Sillitoe on behalf of the Wellcome Sanger Institute COVID-19 Surveillance Team |
| EPI_ISL_2236627, EPI_ISL_2236629, EPI_ISL_2236631, EPI_ISL_2236632 | Lighthouse Lab in Milton Keynes | Wellcome Sanger Institute for the COVID-19 Genomics UK (COG-UK) Consortium | The Lighthouse Lab in Milton Keynes and Alex Alderton, Roberto Amato, Jeffrey Barrett, Sonia Goncalves, Ewan Harrison, David K. Jackson, Ian Johnston, Dominic Kwiatkowski, Cordelia Langford, John Sillitoe on behalf of the Wellcome Sanger Institute COVID-19 Surveillance Team |
| EPI_ISL_2236633, EPI_ISL_2236634 | Lighthouse Lab in Glasgow | Wellcome Sanger Institute for the COVID-19 Genomics UK (COG-UK) Consortium | Harper VanSteenhouse, Yumi Kasai, David Gray, Carol Clugston, Anna Dominiczak and Alex Alderton, Roberto Amato, Jeffrey Barrett, Sonia Goncalves, Ewan Harrison, David K. Jackson, Ian Johnston, Dominic Kwiatkowski, Cordelia Langford, John Sillitoe on behalf of the Wellcome Sanger Institute COVID-19 Surveillance Team |
| EPI_ISL_2236636, EPI_ISL_2236640, EPI_ISL_2236641, EPI_ISL_2236642, EPI_ISL_2236644, EPI_ISL_2236645, EPI_ISL_2236646, EPI_ISL_2236648, EPI_ISL_2236649, EPI_ISL_2236653, EPI_ISL_2236655, EPI_ISL_2236657, EPI_ISL_2236658, EPI_ISL_2236662, EPI_ISL_2236663, EPI_ISL_2236665, EPI_ISL_2236668, EPI_ISL_2236670 | | | |
| see above | Lighthouse Lab in Milton Keynes | Wellcome Sanger Institute for the COVID-19 Genomics UK (COG-UK) Consortium | The Lighthouse Lab in Milton Keynes and Alex Alderton, Roberto Amato, Jeffrey Barrett, Sonia Goncalves, Ewan Harrison, David K. Jackson, Ian Johnston, Dominic Kwiatkowski, Cordelia Langford, John Sillitoe on behalf of the Wellcome Sanger Institute COVID-19 Surveillance Team |
| EPI_ISL_2236674 | Lighthouse Lab in Glasgow | Wellcome Sanger Institute for the COVID-19 Genomics UK (COG-UK) Consortium | Harper VanSteenhouse, Yumi Kasai, David Gray, Carol Clugston, Anna Dominiczak and Alex Alderton, Roberto Amato, Jeffrey Barrett, Sonia Goncalves, Ewan Harrison, David K. Jackson, Ian Johnston, Dominic Kwiatkowski, Cordelia Langford, John Sillitoe on behalf of the Wellcome Sanger Institute COVID-19 Surveillance Team |
| EPI_ISL_2236675, EPI_ISL_2236676 | Lighthouse Lab in Milton Keynes | Wellcome Sanger Institute for the COVID-19 Genomics UK (COG-UK) Consortium | The Lighthouse Lab in Milton Keynes and Alex Alderton, Roberto Amato, Jeffrey Barrett, Sonia Goncalves, Ewan Harrison, David K. Jackson, Ian Johnston, Dominic Kwiatkowski, Cordelia Langford, John Sillitoe on behalf of the Wellcome Sanger Institute COVID-19 Surveillance Team |
| EPI_ISL_2236683 | Lighthouse Lab in Glasgow | Wellcome Sanger Institute for the COVID-19 Genomics UK (COG-UK) Consortium | Harper VanSteenhouse, Yumi Kasai, David Gray, Carol Clugston, Anna Dominiczak and Alex Alderton, Roberto Amato, Jeffrey Barrett, Sonia Goncalves, Ewan Harrison, David K. Jackson, Ian Johnston, Dominic Kwiatkowski, Cordelia Langford, John Sillitoe on behalf of the Wellcome Sanger Institute COVID-19 Surveillance Team |
| EPI_ISL_2236684, EPI_ISL_2236685 | Lighthouse Lab in Milton Keynes | Wellcome Sanger Institute for the COVID-19 Genomics UK (COG-UK) Consortium | The Lighthouse Lab in Milton Keynes and Alex Alderton, Roberto Amato, Jeffrey Barrett, Sonia Goncalves, Ewan Harrison, David K. Jackson, Ian Johnston, Dominic Kwiatkowski, Cordelia Langford, John Sillitoe on behalf of the Wellcome Sanger Institute COVID-19 Surveillance Team |
| EPI_ISL_2236687 | Lighthouse Lab in Glasgow | Wellcome Sanger Institute for the COVID-19 Genomics UK (COG-UK) Consortium | Harper VanSteenhouse, Yumi Kasai, David Gray, Carol Clugston, Anna Dominiczak and Alex Alderton, Roberto Amato, Jeffrey Barrett, Sonia Goncalves, Ewan Harrison, David K. Jackson, Ian Johnston, Dominic Kwiatkowski, Cordelia Langford, John Sillitoe on behalf of the Wellcome Sanger Institute COVID-19 Surveillance Team |
| EPI_ISL_2236692, EPI_ISL_2236695, EPI_ISL_2236696, EPI_ISL_2236699, EPI_ISL_2236700, EPI_ISL_2236701 | Lighthouse Lab in Milton Keynes | Wellcome Sanger Institute for the COVID-19 Genomics UK (COG-UK) Consortium | The Lighthouse Lab in Milton Keynes and Alex Alderton, Roberto Amato, Jeffrey Barrett, Sonia Goncalves, Ewan Harrison, David K. Jackson, Ian Johnston, Dominic Kwiatkowski, Cordelia Langford, John Sillitoe on behalf of the Wellcome Sanger Institute COVID-19 Surveillance Team |
| EPI_ISL_2236702 | Lighthouse Lab in Glasgow | Wellcome Sanger Institute for the COVID-19 Genomics UK (COG-UK) Consortium | Harper VanSteenhouse, Yumi Kasai, David Gray, Carol Clugston, Anna Dominiczak and Alex Alderton, Roberto Amato, Jeffrey Barrett, Sonia Goncalves, Ewan Harrison, David K. Jackson, Ian Johnston, Dominic Kwiatkowski, Cordelia Langford, John Sillitoe on behalf of the Wellcome Sanger Institute COVID-19 Surveillance Team |
| EPI_ISL_2236703, EPI_ISL_2236708 | Lighthouse Lab in Milton Keynes | Wellcome Sanger Institute for the COVID-19 Genomics UK (COG-UK) Consortium | The Lighthouse Lab in Milton Keynes and Alex Alderton, Roberto Amato, Jeffrey Barrett, Sonia Goncalves, Ewan Harrison, David K. Jackson, Ian Johnston, Dominic Kwiatkowski, Cordelia Langford, John Sillitoe on behalf of the Wellcome Sanger Institute COVID-19 Surveillance Team |
| EPI_ISL_2236709, EPI_ISL_2236710 | Lighthouse Lab in Glasgow | Wellcome Sanger Institute for the COVID-19 Genomics UK (COG-UK) Consortium | Harper VanSteenhouse, Yumi Kasai, David Gray, Carol Clugston, Anna Dominiczak and Alex Alderton, Roberto Amato, Jeffrey Barrett, Sonia Goncalves, Ewan Harrison, David K. Jackson, Ian Johnston, Dominic Kwiatkowski, Cordelia Langford, John Sillitoe on behalf of the Wellcome Sanger Institute COVID-19 Surveillance Team |
| EPI_ISL_2236711, EPI_ISL_2236712, EPI_ISL_2236713 | Lighthouse Lab in Milton Keynes | Wellcome Sanger Institute for the COVID-19 Genomics UK (COG-UK) Consortium | The Lighthouse Lab in Milton Keynes and Alex Alderton, Roberto Amato, Jeffrey Barrett, Sonia Goncalves, Ewan Harrison, David K. Jackson, Ian Johnston, Dominic Kwiatkowski, Cordelia Langford, John Sillitoe on behalf of the Wellcome Sanger Institute COVID-19 Surveillance Team |
| EPI_ISL_2236714 | Lighthouse Lab in Glasgow | Wellcome Sanger Institute for the COVID-19 Genomics UK (COG-UK) Consortium | Harper VanSteenhouse, Yumi Kasai, David Gray, Carol Clugston, Anna Dominiczak and Alex Alderton, Roberto Amato, Jeffrey Barrett, Sonia Goncalves, Ewan Harrison, David K. Jackson, Ian Johnston, Dominic Kwiatkowski, Cordelia Langford, John Sillitoe on behalf of the Wellcome Sanger Institute COVID-19 Surveillance Team |
| EPI_ISL_2236715, EPI_ISL_2236717, EPI_ISL_2236718, EPI_ISL_2236722, EPI_ISL_2236723, EPI_ISL_2236724, EPI_ISL_2236725, EPI_ISL_2236726, EPI_ISL_2236730, EPI_ISL_2236731, EPI_ISL_2236733, EPI_ISL_2236735 | | | |
| see above | Lighthouse Lab in Milton Keynes | Wellcome Sanger Institute for the COVID-19 Genomics UK (COG-UK) Consortium | The Lighthouse Lab in Milton Keynes and Alex Alderton, Roberto Amato, Jeffrey Barrett, Sonia Goncalves, Ewan Harrison, David K. Jackson, Ian Johnston, Dominic Kwiatkowski, Cordelia Langford, John Sillitoe on behalf of the Wellcome Sanger Institute COVID-19 Surveillance Team |
| EPI_ISL_2236737 | Lighthouse Lab in Glasgow | Wellcome Sanger Institute for the COVID-19 Genomics UK (COG-UK) Consortium | Harper VanSteenhouse, Yumi Kasai, David Gray, Carol Clugston, Anna Dominiczak and Alex Alderton, Roberto Amato, Jeffrey Barrett, Sonia Goncalves, Ewan Harrison, David K. Jackson, Ian Johnston, Dominic Kwiatkowski, Cordelia Langford, John Sillitoe on behalf of the Wellcome Sanger Institute COVID-19 Surveillance Team |
| EPI_ISL_2236739, EPI_ISL_2236742, EPI_ISL_2236743 | Lighthouse Lab in Milton Keynes | Wellcome Sanger Institute for the COVID-19 Genomics UK (COG-UK) Consortium | The Lighthouse Lab in Milton Keynes and Alex Alderton, Roberto Amato, Jeffrey Barrett, Sonia Goncalves, Ewan Harrison, David K. Jackson, Ian Johnston, Dominic Kwiatkowski, Cordelia Langford, John Sillitoe on behalf of the Wellcome Sanger Institute COVID-19 Surveillance Team |
| EPI_ISL_2236744 | Lighthouse Lab in Glasgow | Wellcome Sanger Institute for the COVID-19 Genomics UK (COG-UK) Consortium | Harper VanSteenhouse, Yumi Kasai, David Gray, Carol Clugston, Anna Dominiczak and Alex Alderton, Roberto Amato, Jeffrey Barrett, Sonia Goncalves, Ewan Harrison, David K. Jackson, Ian Johnston, Dominic Kwiatkowski, Cordelia Langford, John Sillitoe on behalf of the Wellcome Sanger Institute COVID-19 Surveillance Team |
| EPI_ISL_2236745, EPI_ISL_2236747, EPI_ISL_2236753 | Lighthouse Lab in Milton Keynes | Wellcome Sanger Institute for the COVID-19 Genomics UK (COG-UK) Consortium | The Lighthouse Lab in Milton Keynes and Alex Alderton, Roberto Amato, Jeffrey Barrett, Sonia Goncalves, Ewan Harrison, David K. Jackson, Ian Johnston, Dominic Kwiatkowski, Cordelia Langford, John Sillitoe on behalf of the Wellcome Sanger Institute COVID-19 Surveillance Team |
| EPI_ISL_2236756 | Lighthouse Lab in Glasgow | Wellcome Sanger Institute for the COVID-19 Genomics UK (COG-UK) Consortium | Harper VanSteenhouse, Yumi Kasai, David Gray, Carol Clugston, Anna Dominiczak and Alex Alderton, Roberto Amato, Jeffrey Barrett, Sonia Goncalves, Ewan Harrison, David K. Jackson, Ian Johnston, Dominic Kwiatkowski, Cordelia Langford, John Sillitoe on behalf of the Wellcome Sanger Institute COVID-19 Surveillance Team |
| EPI_ISL_2236758 | Lighthouse Lab in Milton Keynes | Wellcome Sanger Institute for the COVID-19 Genomics UK (COG-UK) Consortium | The Lighthouse Lab in Milton Keynes and Alex Alderton, Roberto Amato, Jeffrey Barrett, Sonia Goncalves, Ewan Harrison, David K. Jackson, Ian Johnston, Dominic Kwiatkowski, Cordelia Langford, John Sillitoe on behalf of the Wellcome Sanger Institute COVID-19 Surveillance Team |
| EPI_ISL_2236759 | Lighthouse Lab in Glasgow | Wellcome Sanger Institute for the COVID-19 Genomics UK (COG-UK) Consortium | Harper VanSteenhouse, Yumi Kasai, David Gray, Carol Clugston, Anna Dominiczak and Alex Alderton, Roberto Amato, Jeffrey Barrett, Sonia Goncalves, Ewan Harrison, David K. Jackson, Ian Johnston, Dominic Kwiatkowski, Cordelia Langford, John Sillitoe on behalf of the Wellcome Sanger Institute COVID-19 Surveillance Team |
| EPI_ISL_2236761, EPI_ISL_2236762, EPI_ISL_2236763 | Lighthouse Lab in Milton Keynes | Wellcome Sanger Institute for the COVID-19 Genomics UK (COG-UK) Consortium | The Lighthouse Lab in Milton Keynes and Alex Alderton, Roberto Amato, Jeffrey Barrett, Sonia Goncalves, Ewan Harrison, David K. Jackson, Ian Johnston, Dominic Kwiatkowski, Cordelia Langford, John Sillitoe on behalf of the Wellcome Sanger Institute COVID-19 Surveillance Team |
| EPI_ISL_2236765, EPI_ISL_2236770, EPI_ISL_2236771, EPI_ISL_2236773 | Lighthouse Lab in Glasgow | Wellcome Sanger Institute for the COVID-19 Genomics UK (COG-UK) Consortium | Harper VanSteenhouse, Yumi Kasai, David Gray, Carol Clugston, Anna Dominiczak and Alex Alderton, Roberto Amato, Jeffrey Barrett, Sonia Goncalves, Ewan Harrison, David K. Jackson, Ian Johnston, Dominic Kwiatkowski, Cordelia Langford, John Sillitoe on behalf of the Wellcome Sanger Institute COVID-19 Surveillance Team |
| EPI_ISL_2236774 | Lighthouse Lab in Milton Keynes | Wellcome Sanger Institute for the COVID-19 Genomics UK (COG-UK) Consortium | The Lighthouse Lab in Milton Keynes and Alex Alderton, Roberto Amato, Jeffrey Barrett, Sonia Goncalves, Ewan Harrison, David K. Jackson, Ian Johnston, Dominic Kwiatkowski, Cordelia Langford, John Sillitoe on behalf of the Wellcome Sanger Institute COVID-19 Surveillance Team |
| EPI_ISL_2236776 | Lighthouse Lab in Glasgow | Wellcome Sanger Institute for the COVID-19 Genomics UK (COG-UK) Consortium | Harper VanSteenhouse, Yumi Kasai, David Gray, Carol Clugston, Anna Dominiczak and Alex Alderton, Roberto Amato, Jeffrey Barrett, Sonia Goncalves, Ewan Harrison, David K. Jackson, Ian Johnston, Dominic Kwiatkowski, Cordelia Langford, John Sillitoe on behalf of the Wellcome Sanger Institute COVID-19 Surveillance Team |
| EPI_ISL_2236777, EPI_ISL_2236778, EPI_ISL_2236779, EPI_ISL_2236786, EPI_ISL_2236787, EPI_ISL_2236788, EPI_ISL_2236789, EPI_ISL_2236790, EPI_ISL_2236791, | Lighthouse Lab in Milton Keynes | Wellcome Sanger Institute for the COVID-19 Genomics UK (COG-UK) Consortium | The Lighthouse Lab in Milton Keynes and Alex Alderton, Roberto Amato, Jeffrey Barrett, Sonia Goncalves, Ewan Harrison, David K. Jackson, Ian Johnston, Dominic Kwiatkowski, Cordelia Langford, John Sillitoe on behalf of the Wellcome Sanger Institute COVID-19 Surveillance Team |

| | | | |
|---|---|---|---|
| EPI_ISL_2236792 | | | |
| EPI_ISL_2236796 | Lighthouse Lab in Glasgow | Wellcome Sanger Institute for the COVID-19 Genomics UK (COG-UK) Consortium | Harper VanSteenhouse, Yumi Kasai, David Gray, Carol Clugston, Anna Dominiczak and Alex Alderton, Roberto Amato, Jeffrey Barrett, Sonia Goncalves, Ewan Harrison, David K. Jackson, Ian Johnston, Dominic Kwiatkowski, Cordelia Langford, John Sillitoe on behalf of the Wellcome Sanger Institute COVID-19 Surveillance Team |
| EPI_ISL_2236797, EPI_ISL_2236798 | Lighthouse Lab in Milton Keynes | Wellcome Sanger Institute for the COVID-19 Genomics UK (COG-UK) Consortium | The Lighthouse Lab in Milton Keynes and Alex Alderton, Roberto Amato, Jeffrey Barrett, Sonia Goncalves, Ewan Harrison, David K. Jackson, Ian Johnston, Dominic Kwiatkowski, Cordelia Langford, John Sillitoe on behalf of the Wellcome Sanger Institute COVID-19 Surveillance Team |
| EPI_ISL_2236799, EPI_ISL_2236800, EPI_ISL_2236801 | Lighthouse Lab in Glasgow | Wellcome Sanger Institute for the COVID-19 Genomics UK (COG-UK) Consortium | Harper VanSteenhouse, Yumi Kasai, David Gray, Carol Clugston, Anna Dominiczak and Alex Alderton, Roberto Amato, Jeffrey Barrett, Sonia Goncalves, Ewan Harrison, David K. Jackson, Ian Johnston, Dominic Kwiatkowski, Cordelia Langford, John Sillitoe on behalf of the Wellcome Sanger Institute COVID-19 Surveillance Team |
| EPI_ISL_2236802, EPI_ISL_2236803, EPI_ISL_2236805, EPI_ISL_2236807 | Lighthouse Lab in Milton Keynes | Wellcome Sanger Institute for the COVID-19 Genomics UK (COG-UK) Consortium | The Lighthouse Lab in Milton Keynes and Alex Alderton, Roberto Amato, Jeffrey Barrett, Sonia Goncalves, Ewan Harrison, David K. Jackson, Ian Johnston, Dominic Kwiatkowski, Cordelia Langford, John Sillitoe on behalf of the Wellcome Sanger Institute COVID-19 Surveillance Team |
| EPI_ISL_2236808 | Lighthouse Lab in Glasgow | Wellcome Sanger Institute for the COVID-19 Genomics UK (COG-UK) Consortium | Harper VanSteenhouse, Yumi Kasai, David Gray, Carol Clugston, Anna Dominiczak and Alex Alderton, Roberto Amato, Jeffrey Barrett, Sonia Goncalves, Ewan Harrison, David K. Jackson, Ian Johnston, Dominic Kwiatkowski, Cordelia Langford, John Sillitoe on behalf of the Wellcome Sanger Institute COVID-19 Surveillance Team |
| EPI_ISL_2236810, EPI_ISL_2236811, EPI_ISL_2236814, EPI_ISL_2236816 | Lighthouse Lab in Milton Keynes | Wellcome Sanger Institute for the COVID-19 Genomics UK (COG-UK) Consortium | The Lighthouse Lab in Milton Keynes and Alex Alderton, Roberto Amato, Jeffrey Barrett, Sonia Goncalves, Ewan Harrison, David K. Jackson, Ian Johnston, Dominic Kwiatkowski, Cordelia Langford, John Sillitoe on behalf of the Wellcome Sanger Institute COVID-19 Surveillance Team |
| EPI_ISL_2236817 | Lighthouse Lab in Glasgow | Wellcome Sanger Institute for the COVID-19 Genomics UK (COG-UK) Consortium | Harper VanSteenhouse, Yumi Kasai, David Gray, Carol Clugston, Anna Dominiczak and Alex Alderton, Roberto Amato, Jeffrey Barrett, Sonia Goncalves, Ewan Harrison, David K. Jackson, Ian Johnston, Dominic Kwiatkowski, Cordelia Langford, John Sillitoe on behalf of the Wellcome Sanger Institute COVID-19 Surveillance Team |
| EPI_ISL_2236819 | Lighthouse Lab in Milton Keynes | Wellcome Sanger Institute for the COVID-19 Genomics UK (COG-UK) Consortium | The Lighthouse Lab in Milton Keynes and Alex Alderton, Roberto Amato, Jeffrey Barrett, Sonia Goncalves, Ewan Harrison, David K. Jackson, Ian Johnston, Dominic Kwiatkowski, Cordelia Langford, John Sillitoe on behalf of the Wellcome Sanger Institute COVID-19 Surveillance Team |
| EPI_ISL_2236825, EPI_ISL_2236826 | Lighthouse Lab in Glasgow | Wellcome Sanger Institute for the COVID-19 Genomics UK (COG-UK) Consortium | Harper VanSteenhouse, Yumi Kasai, David Gray, Carol Clugston, Anna Dominiczak and Alex Alderton, Roberto Amato, Jeffrey Barrett, Sonia Goncalves, Ewan Harrison, David K. Jackson, Ian Johnston, Dominic Kwiatkowski, Cordelia Langford, John Sillitoe on behalf of the Wellcome Sanger Institute COVID-19 Surveillance Team |
| EPI_ISL_2236829 | Lighthouse Lab in Alderley Park | Wellcome Sanger Institute for the COVID-19 Genomics UK (COG-UK) Consortium | Jacquelyn Wynn, Mairead Hyland, The Lighthouse Lab in Alderley Park and Alex Alderton, Roberto Amato, Jeffrey Barrett, Sonia Goncalves, Ewan Harrison, David K. Jackson, Ian Johnston, Dominic Kwiatkowski, Cordelia Langford, John Sillitoe on behalf of the Wellcome Sanger Institute COVID-19 Surveillance Team |
| EPI_ISL_2236831, EPI_ISL_2236833 | Lighthouse Lab in Glasgow | Wellcome Sanger Institute for the COVID-19 Genomics UK (COG-UK) Consortium | Harper VanSteenhouse, Yumi Kasai, David Gray, Carol Clugston, Anna Dominiczak and Alex Alderton, Roberto Amato, Jeffrey Barrett, Sonia Goncalves, Ewan Harrison, David K. Jackson, Ian Johnston, Dominic Kwiatkowski, Cordelia Langford, John Sillitoe on behalf of the Wellcome Sanger Institute COVID-19 Surveillance Team |
| EPI_ISL_2236836, EPI_ISL_2236837 | Lighthouse Lab in Alderley Park | Wellcome Sanger Institute for the COVID-19 Genomics UK (COG-UK) Consortium | Jacquelyn Wynn, Mairead Hyland, The Lighthouse Lab in Alderley Park and Alex Alderton, Roberto Amato, Jeffrey Barrett, Sonia Goncalves, Ewan Harrison, David K. Jackson, Ian Johnston, Dominic Kwiatkowski, Cordelia Langford, John Sillitoe on behalf of the Wellcome Sanger Institute COVID-19 Surveillance Team |
| EPI_ISL_2236839 | Lighthouse Lab in Glasgow | Wellcome Sanger Institute for the COVID-19 Genomics UK (COG-UK) Consortium | Harper VanSteenhouse, Yumi Kasai, David Gray, Carol Clugston, Anna Dominiczak and Alex Alderton, Roberto Amato, Jeffrey Barrett, Sonia Goncalves, Ewan Harrison, David K. Jackson, Ian Johnston, Dominic Kwiatkowski, Cordelia Langford, John Sillitoe on behalf of the Wellcome Sanger Institute COVID-19 Surveillance Team |
| EPI_ISL_2236840, EPI_ISL_2236841, EPI_ISL_2236843, EPI_ISL_2236844, EPI_ISL_2236845, EPI_ISL_2236849 | Lighthouse Lab in Alderley Park | Wellcome Sanger Institute for the COVID-19 Genomics UK (COG-UK) Consortium | Jacquelyn Wynn, Mairead Hyland, The Lighthouse Lab in Alderley Park and Alex Alderton, Roberto Amato, Jeffrey Barrett, Sonia Goncalves, Ewan Harrison, David K. Jackson, Ian Johnston, Dominic Kwiatkowski, Cordelia Langford, John Sillitoe on behalf of the Wellcome Sanger Institute COVID-19 Surveillance Team |
| EPI_ISL_2236853, EPI_ISL_2236854, EPI_ISL_2236855, EPI_ISL_2236856 | Lighthouse Lab in Glasgow | Wellcome Sanger Institute for the COVID-19 Genomics UK (COG-UK) Consortium | Harper VanSteenhouse, Yumi Kasai, David Gray, Carol Clugston, Anna Dominiczak and Alex Alderton, Roberto Amato, Jeffrey Barrett, Sonia Goncalves, Ewan Harrison, David K. Jackson, Ian Johnston, Dominic Kwiatkowski, Cordelia Langford, John Sillitoe on behalf of the Wellcome Sanger Institute COVID-19 Surveillance Team |
| EPI_ISL_2236857 | Lighthouse Lab in Alderley Park | Wellcome Sanger Institute for the COVID-19 Genomics UK (COG-UK) Consortium | Jacquelyn Wynn, Mairead Hyland, The Lighthouse Lab in Alderley Park and Alex Alderton, Roberto Amato, Jeffrey Barrett, Sonia Goncalves, Ewan Harrison, David K. Jackson, Ian Johnston, Dominic Kwiatkowski, Cordelia Langford, John Sillitoe on behalf of the Wellcome Sanger Institute COVID-19 Surveillance Team |
| EPI_ISL_2236858, EPI_ISL_2236863 | Lighthouse Lab in Milton Keynes | Wellcome Sanger Institute for the COVID-19 Genomics UK (COG-UK) Consortium | The Lighthouse Lab in Milton Keynes and Alex Alderton, Roberto Amato, Jeffrey Barrett, Sonia Goncalves, Ewan Harrison, David K. Jackson, Ian Johnston, Dominic Kwiatkowski, Cordelia Langford, John Sillitoe on behalf of the Wellcome Sanger Institute COVID-19 Surveillance Team |
| EPI_ISL_2236866 | Lighthouse Lab in Alderley Park | Wellcome Sanger Institute for the COVID-19 Genomics UK (COG-UK) Consortium | Jacquelyn Wynn, Mairead Hyland, The Lighthouse Lab in Alderley Park and Alex Alderton, Roberto Amato, Jeffrey Barrett, Sonia Goncalves, Ewan Harrison, David K. Jackson, Ian Johnston, Dominic Kwiatkowski, Cordelia Langford, John Sillitoe on behalf of the Wellcome Sanger Institute COVID-19 Surveillance Team |
| EPI_ISL_2236876 | Lighthouse Lab in Glasgow | Wellcome Sanger Institute for the COVID-19 Genomics UK (COG-UK) Consortium | Harper VanSteenhouse, Yumi Kasai, David Gray, Carol Clugston, Anna Dominiczak and Alex Alderton, Roberto Amato, Jeffrey Barrett, Sonia Goncalves, Ewan Harrison, David K. Jackson, Ian Johnston, Dominic Kwiatkowski, Cordelia Langford, John Sillitoe on behalf of the Wellcome Sanger Institute COVID-19 Surveillance Team |
| EPI_ISL_2236877 | Lighthouse Lab in Milton Keynes | Wellcome Sanger Institute for the COVID-19 Genomics UK (COG-UK) Consortium | The Lighthouse Lab in Milton Keynes and Alex Alderton, Roberto Amato, Jeffrey Barrett, Sonia Goncalves, Ewan Harrison, David K. Jackson, Ian Johnston, Dominic Kwiatkowski, Cordelia Langford, John Sillitoe on behalf of the Wellcome Sanger Institute COVID-19 Surveillance Team |
| EPI_ISL_2236880, EPI_ISL_2236882, EPI_ISL_2236883 | Lighthouse Lab in Glasgow | Wellcome Sanger Institute for the COVID-19 Genomics UK (COG-UK) Consortium | Harper VanSteenhouse, Yumi Kasai, David Gray, Carol Clugston, Anna Dominiczak and Alex Alderton, Roberto Amato, Jeffrey Barrett, Sonia Goncalves, Ewan Harrison, David K. Jackson, Ian Johnston, Dominic Kwiatkowski, Cordelia Langford, John Sillitoe on behalf of the Wellcome Sanger Institute COVID-19 Surveillance Team |
| EPI_ISL_2236886, EPI_ISL_2236889 | Lighthouse Lab in Milton Keynes | Wellcome Sanger Institute for the COVID-19 Genomics UK (COG-UK) Consortium | The Lighthouse Lab in Milton Keynes and Alex Alderton, Roberto Amato, Jeffrey Barrett, Sonia Goncalves, Ewan Harrison, David K. Jackson, Ian Johnston, Dominic Kwiatkowski, Cordelia Langford, John Sillitoe on behalf of the Wellcome Sanger Institute COVID-19 Surveillance Team |
| EPI_ISL_2236893 | Lighthouse Lab in Glasgow | Wellcome Sanger Institute for the COVID-19 Genomics UK (COG-UK) Consortium | Harper VanSteenhouse, Yumi Kasai, David Gray, Carol Clugston, Anna Dominiczak and Alex Alderton, Roberto Amato, Jeffrey Barrett, Sonia Goncalves, Ewan Harrison, David K. Jackson, Ian Johnston, Dominic Kwiatkowski, Cordelia Langford, John Sillitoe on behalf of the Wellcome Sanger Institute COVID-19 Surveillance Team |
| EPI_ISL_2236894, EPI_ISL_2236898, EPI_ISL_2236899, EPI_ISL_2236900, EPI_ISL_2236902 | Lighthouse Lab in Alderley Park | Wellcome Sanger Institute for the COVID-19 Genomics UK (COG-UK) Consortium | Jacquelyn Wynn, Mairead Hyland, The Lighthouse Lab in Alderley Park and Alex Alderton, Roberto Amato, Jeffrey Barrett, Sonia Goncalves, Ewan Harrison, David K. Jackson, Ian Johnston, Dominic Kwiatkowski, Cordelia Langford, John Sillitoe on behalf of the Wellcome Sanger Institute COVID-19 Surveillance Team |
| EPI_ISL_2236904 | Lighthouse Lab in Milton Keynes | Wellcome Sanger Institute for the COVID-19 Genomics UK (COG-UK) Consortium | The Lighthouse Lab in Milton Keynes and Alex Alderton, Roberto Amato, Jeffrey Barrett, Sonia Goncalves, Ewan Harrison, David K. Jackson, Ian Johnston, Dominic Kwiatkowski, Cordelia Langford, John Sillitoe on behalf of the Wellcome Sanger Institute COVID-19 Surveillance Team |
| EPI_ISL_2236905 | Lighthouse Lab in Alderley Park | Wellcome Sanger Institute for the COVID-19 Genomics UK (COG-UK) Consortium | Jacquelyn Wynn, Mairead Hyland, The Lighthouse Lab in Alderley Park and Alex Alderton, Roberto Amato, Jeffrey Barrett, Sonia Goncalves, Ewan Harrison, David K. Jackson, Ian Johnston, Dominic Kwiatkowski, Cordelia Langford, John Sillitoe on behalf of the Wellcome Sanger Institute COVID-19 Surveillance Team |
| EPI_ISL_2236906 | Lighthouse Lab in Glasgow | Wellcome Sanger Institute for the COVID-19 Genomics UK (COG-UK) Consortium | Harper VanSteenhouse, Yumi Kasai, David Gray, Carol Clugston, Anna Dominiczak and Alex Alderton, Roberto Amato, Jeffrey Barrett, Sonia Goncalves, Ewan Harrison, David K. Jackson, Ian Johnston, Dominic Kwiatkowski, Cordelia Langford, John Sillitoe on behalf of the Wellcome Sanger Institute COVID-19 Surveillance Team |
| EPI_ISL_2236907, EPI_ISL_2236908 | Lighthouse Lab in Alderley Park | Wellcome Sanger Institute for the COVID-19 Genomics UK (COG-UK) Consortium | Jacquelyn Wynn, Mairead Hyland, The Lighthouse Lab in Alderley Park and Alex Alderton, Roberto Amato, Jeffrey Barrett, Sonia Goncalves, Ewan Harrison, David K. Jackson, Ian Johnston, Dominic Kwiatkowski, Cordelia Langford, John Sillitoe on behalf of the Wellcome Sanger Institute COVID-19 Surveillance Team |
| EPI_ISL_2236909, EPI_ISL_2236912 | Lighthouse Lab in Glasgow | Wellcome Sanger Institute for the COVID-19 Genomics UK (COG-UK) Consortium | Harper VanSteenhouse, Yumi Kasai, David Gray, Carol Clugston, Anna Dominiczak and Alex Alderton, Roberto Amato, Jeffrey Barrett, Sonia Goncalves, Ewan Harrison, David K. Jackson, Ian Johnston, Dominic Kwiatkowski, Cordelia Langford, John Sillitoe on behalf of the Wellcome Sanger Institute COVID-19 Surveillance Team |
| EPI_ISL_2236915 | Lighthouse Lab in Alderley Park | Wellcome Sanger Institute for the COVID-19 Genomics UK (COG-UK) Consortium | Jacquelyn Wynn, Mairead Hyland, The Lighthouse Lab in Alderley Park and Alex Alderton, Roberto Amato, Jeffrey Barrett, Sonia Goncalves, Ewan Harrison, David K. Jackson, Ian Johnston, Dominic Kwiatkowski, Cordelia Langford, John Sillitoe on behalf of the Wellcome Sanger Institute COVID-19 Surveillance Team |
| EPI_ISL_2236916, EPI_ISL_2236918 | Lighthouse Lab in Glasgow | Wellcome Sanger Institute for the COVID-19 Genomics UK (COG-UK) Consortium | Harper VanSteenhouse, Yumi Kasai, David Gray, Carol Clugston, Anna Dominiczak and Alex Alderton, Roberto Amato, Jeffrey Barrett, Sonia Goncalves, Ewan Harrison, David K. Jackson, Ian Johnston, Dominic Kwiatkowski, Cordelia Langford, John Sillitoe on behalf of the Wellcome Sanger Institute COVID-19 Surveillance Team |
| EPI_ISL_2236925 | Lighthouse Lab in Milton Keynes | Wellcome Sanger Institute for the COVID-19 Genomics UK (COG-UK) Consortium | The Lighthouse Lab in Milton Keynes and Alex Alderton, Roberto Amato, Jeffrey Barrett, Sonia Goncalves, Ewan Harrison, David K. Jackson, Ian Johnston, Dominic Kwiatkowski, Cordelia Langford, John Sillitoe on behalf of the Wellcome Sanger Institute COVID-19 Surveillance Team |
| EPI_ISL_2236930 | Lighthouse Lab in Alderley Park | Wellcome Sanger Institute for the COVID-19 Genomics UK (COG-UK) Consortium | Jacquelyn Wynn, Mairead Hyland, The Lighthouse Lab in Alderley Park and Alex Alderton, Roberto Amato, Jeffrey Barrett, Sonia Goncalves, Ewan Harrison, David K. Jackson, Ian Johnston, Dominic Kwiatkowski, Cordelia Langford, John Sillitoe on behalf of the Wellcome Sanger Institute COVID-19 Surveillance Team |
| EPI_ISL_2236931, EPI_ISL_2236932 | Lighthouse Lab in Milton Keynes | Wellcome Sanger Institute for the COVID-19 Genomics UK (COG-UK) Consortium | The Lighthouse Lab in Milton Keynes and Alex Alderton, Roberto Amato, Jeffrey Barrett, Sonia Goncalves, Ewan Harrison, David K. Jackson, Ian Johnston, Dominic Kwiatkowski, Cordelia Langford, John Sillitoe on behalf of the Wellcome Sanger Institute COVID-19 Surveillance Team |
| EPI_ISL_2236933, EPI_ISL_2236935 | Lighthouse Lab in Glasgow | Wellcome Sanger Institute for the COVID-19 Genomics UK (COG-UK) Consortium | Harper VanSteenhouse, Yumi Kasai, David Gray, Carol Clugston, Anna Dominiczak and Alex Alderton, Roberto Amato, Jeffrey Barrett, Sonia Goncalves, Ewan Harrison, David K. Jackson, Ian Johnston, Dominic Kwiatkowski, Cordelia Langford, John Sillitoe on behalf of the Wellcome Sanger Institute COVID-19 Surveillance Team |
| EPI_ISL_2236938 | Lighthouse Lab in Milton Keynes | Wellcome Sanger Institute for the COVID-19 Genomics UK (COG-UK) Consortium | The Lighthouse Lab in Milton Keynes and Alex Alderton, Roberto Amato, Jeffrey Barrett, Sonia Goncalves, Ewan Harrison, David K. Jackson, Ian Johnston, Dominic Kwiatkowski, Cordelia Langford, John Sillitoe on behalf of the Wellcome Sanger Institute COVID-19 Surveillance Team |
| EPI_ISL_2236940 | Lighthouse Lab in Glasgow | Wellcome Sanger Institute for the COVID-19 Genomics UK (COG-UK) Consortium | Harper VanSteenhouse, Yumi Kasai, David Gray, Carol Clugston, Anna Dominiczak and Alex Alderton, Roberto Amato, Jeffrey Barrett, Sonia Goncalves, Ewan Harrison, David K. Jackson, Ian Johnston, Dominic Kwiatkowski, Cordelia Langford, John Sillitoe on behalf of the Wellcome Sanger Institute COVID-19 Surveillance Team |
| EPI_ISL_2236942 | Lighthouse Lab in Alderley Park | Wellcome Sanger Institute for the COVID-19 Genomics UK (COG-UK) Consortium | Jacquelyn Wynn, Mairead Hyland, The Lighthouse Lab in Alderley Park and Alex Alderton, Roberto Amato, Jeffrey Barrett, Sonia Goncalves, Ewan Harrison, David K. Jackson, Ian Johnston, Dominic Kwiatkowski, Cordelia Langford, John Sillitoe on behalf of the Wellcome Sanger Institute COVID-19 Surveillance Team |
| EPI_ISL_2236944 | Lighthouse Lab in Glasgow | Wellcome Sanger Institute for the COVID-19 Genomics UK (COG-UK) | Harper VanSteenhouse, Yumi Kasai, David Gray, Carol Clugston, Anna Dominiczak and Alex Alderton, Roberto Amato, Jeffrey Barrett, Sonia Goncalves, Ewan Harrison, David K. Jackson, Ian Johnston, Dominic Kwiatkowski, Cordelia Langford, |

| | | | |
|---|---|---|---|
| | | Consortium | John Sillitoe on behalf of the Wellcome Sanger Institute COVID-19 Surveillance Team |
| EPI_ISL_2236946 | Lighthouse Lab in Alderley Park | Wellcome Sanger Institute for the COVID-19 Genomics UK (COG-UK) Consortium | Jacquelyn Wynn, Mairead Hyland, The Lighthouse Lab in Alderley Park and Alex Alderton, Roberto Amato, Jeffrey Barrett, Sonia Goncalves, Ewan Harrison, David K. Jackson, Ian Johnston, Dominic Kwiatkowski, Cordelia Langford, John Sillitoe on behalf of the Wellcome Sanger Institute COVID-19 Surveillance Team |
| EPI_ISL_2236949 | Lighthouse Lab in Glasgow | Wellcome Sanger Institute for the COVID-19 Genomics UK (COG-UK) Consortium | Harper VanSteenhouse, Yumi Kasai, David Gray, Carol Clugston, Anna Dominiczak and Alex Alderton, Roberto Amato, Jeffrey Barrett, Sonia Goncalves, Ewan Harrison, David K. Jackson, Ian Johnston, Dominic Kwiatkowski, Cordelia Langford, John Sillitoe on behalf of the Wellcome Sanger Institute COVID-19 Surveillance Team |
| EPI_ISL_2236952, EPI_ISL_2236953, EPI_ISL_2236958, EPI_ISL_2236960 | Lighthouse Lab in Alderley Park | Wellcome Sanger Institute for the COVID-19 Genomics UK (COG-UK) Consortium | Jacquelyn Wynn, Mairead Hyland, The Lighthouse Lab in Alderley Park and Alex Alderton, Roberto Amato, Jeffrey Barrett, Sonia Goncalves, Ewan Harrison, David K. Jackson, Ian Johnston, Dominic Kwiatkowski, Cordelia Langford, John Sillitoe on behalf of the Wellcome Sanger Institute COVID-19 Surveillance Team |
| EPI_ISL_2236961, EPI_ISL_2236965 | Lighthouse Lab in Glasgow | Wellcome Sanger Institute for the COVID-19 Genomics UK (COG-UK) Consortium | Harper VanSteenhouse, Yumi Kasai, David Gray, Carol Clugston, Anna Dominiczak and Alex Alderton, Roberto Amato, Jeffrey Barrett, Sonia Goncalves, Ewan Harrison, David K. Jackson, Ian Johnston, Dominic Kwiatkowski, Cordelia Langford, John Sillitoe on behalf of the Wellcome Sanger Institute COVID-19 Surveillance Team |
| EPI_ISL_2236969 | Lighthouse Lab in Alderley Park | Wellcome Sanger Institute for the COVID-19 Genomics UK (COG-UK) Consortium | Jacquelyn Wynn, Mairead Hyland, The Lighthouse Lab in Alderley Park and Alex Alderton, Roberto Amato, Jeffrey Barrett, Sonia Goncalves, Ewan Harrison, David K. Jackson, Ian Johnston, Dominic Kwiatkowski, Cordelia Langford, John Sillitoe on behalf of the Wellcome Sanger Institute COVID-19 Surveillance Team |
| EPI_ISL_2236970 | Lighthouse Lab in Milton Keynes | Wellcome Sanger Institute for the COVID-19 Genomics UK (COG-UK) Consortium | The Lighthouse Lab in Milton Keynes and Alex Alderton, Roberto Amato, Jeffrey Barrett, Sonia Goncalves, Ewan Harrison, David K. Jackson, Ian Johnston, Dominic Kwiatkowski, Cordelia Langford, John Sillitoe on behalf of the Wellcome Sanger Institute COVID-19 Surveillance Team |
| EPI_ISL_2236971 | Lighthouse Lab in Alderley Park | Wellcome Sanger Institute for the COVID-19 Genomics UK (COG-UK) Consortium | Jacquelyn Wynn, Mairead Hyland, The Lighthouse Lab in Alderley Park and Alex Alderton, Roberto Amato, Jeffrey Barrett, Sonia Goncalves, Ewan Harrison, David K. Jackson, Ian Johnston, Dominic Kwiatkowski, Cordelia Langford, John Sillitoe on behalf of the Wellcome Sanger Institute COVID-19 Surveillance Team |
| EPI_ISL_2236972 | Lighthouse Lab in Glasgow | Wellcome Sanger Institute for the COVID-19 Genomics UK (COG-UK) Consortium | Harper VanSteenhouse, Yumi Kasai, David Gray, Carol Clugston, Anna Dominiczak and Alex Alderton, Roberto Amato, Jeffrey Barrett, Sonia Goncalves, Ewan Harrison, David K. Jackson, Ian Johnston, Dominic Kwiatkowski, Cordelia Langford, John Sillitoe on behalf of the Wellcome Sanger Institute COVID-19 Surveillance Team |
| EPI_ISL_2236973 | Lighthouse Lab in Milton Keynes | Wellcome Sanger Institute for the COVID-19 Genomics UK (COG-UK) Consortium | The Lighthouse Lab in Milton Keynes and Alex Alderton, Roberto Amato, Jeffrey Barrett, Sonia Goncalves, Ewan Harrison, David K. Jackson, Ian Johnston, Dominic Kwiatkowski, Cordelia Langford, John Sillitoe on behalf of the Wellcome Sanger Institute COVID-19 Surveillance Team |
| EPI_ISL_2236975, EPI_ISL_2236976, EPI_ISL_2236977, EPI_ISL_2236978 | Lighthouse Lab in Glasgow | Wellcome Sanger Institute for the COVID-19 Genomics UK (COG-UK) Consortium | Harper VanSteenhouse, Yumi Kasai, David Gray, Carol Clugston, Anna Dominiczak and Alex Alderton, Roberto Amato, Jeffrey Barrett, Sonia Goncalves, Ewan Harrison, David K. Jackson, Ian Johnston, Dominic Kwiatkowski, Cordelia Langford, John Sillitoe on behalf of the Wellcome Sanger Institute COVID-19 Surveillance Team |
| EPI_ISL_2236980 | Lighthouse Lab in Milton Keynes | Wellcome Sanger Institute for the COVID-19 Genomics UK (COG-UK) Consortium | The Lighthouse Lab in Milton Keynes and Alex Alderton, Roberto Amato, Jeffrey Barrett, Sonia Goncalves, Ewan Harrison, David K. Jackson, Ian Johnston, Dominic Kwiatkowski, Cordelia Langford, John Sillitoe on behalf of the Wellcome Sanger Institute COVID-19 Surveillance Team |
| EPI_ISL_2236984 | Lighthouse Lab in Alderley Park | Wellcome Sanger Institute for the COVID-19 Genomics UK (COG-UK) Consortium | Jacquelyn Wynn, Mairead Hyland, The Lighthouse Lab in Alderley Park and Alex Alderton, Roberto Amato, Jeffrey Barrett, Sonia Goncalves, Ewan Harrison, David K. Jackson, Ian Johnston, Dominic Kwiatkowski, Cordelia Langford, John Sillitoe on behalf of the Wellcome Sanger Institute COVID-19 Surveillance Team |
| EPI_ISL_2236985 | Lighthouse Lab in Glasgow | Wellcome Sanger Institute for the COVID-19 Genomics UK (COG-UK) Consortium | Harper VanSteenhouse, Yumi Kasai, David Gray, Carol Clugston, Anna Dominiczak and Alex Alderton, Roberto Amato, Jeffrey Barrett, Sonia Goncalves, Ewan Harrison, David K. Jackson, Ian Johnston, Dominic Kwiatkowski, Cordelia Langford, John Sillitoe on behalf of the Wellcome Sanger Institute COVID-19 Surveillance Team |
| EPI_ISL_2236987, EPI_ISL_2236990 | Lighthouse Lab in Alderley Park | Wellcome Sanger Institute for the COVID-19 Genomics UK (COG-UK) Consortium | Jacquelyn Wynn, Mairead Hyland, The Lighthouse Lab in Alderley Park and Alex Alderton, Roberto Amato, Jeffrey Barrett, Sonia Goncalves, Ewan Harrison, David K. Jackson, Ian Johnston, Dominic Kwiatkowski, Cordelia Langford, John Sillitoe on behalf of the Wellcome Sanger Institute COVID-19 Surveillance Team |
| EPI_ISL_2236991 | Lighthouse Lab in Glasgow | Wellcome Sanger Institute for the COVID-19 Genomics UK (COG-UK) Consortium | Harper VanSteenhouse, Yumi Kasai, David Gray, Carol Clugston, Anna Dominiczak and Alex Alderton, Roberto Amato, Jeffrey Barrett, Sonia Goncalves, Ewan Harrison, David K. Jackson, Ian Johnston, Dominic Kwiatkowski, Cordelia Langford, John Sillitoe on behalf of the Wellcome Sanger Institute COVID-19 Surveillance Team |
| EPI_ISL_2236992 | Lighthouse Lab in Alderley Park | Wellcome Sanger Institute for the COVID-19 Genomics UK (COG-UK) Consortium | Jacquelyn Wynn, Mairead Hyland, The Lighthouse Lab in Alderley Park and Alex Alderton, Roberto Amato, Jeffrey Barrett, Sonia Goncalves, Ewan Harrison, David K. Jackson, Ian Johnston, Dominic Kwiatkowski, Cordelia Langford, John Sillitoe on behalf of the Wellcome Sanger Institute COVID-19 Surveillance Team |
| EPI_ISL_2236994 | Lighthouse Lab in Milton Keynes | Wellcome Sanger Institute for the COVID-19 Genomics UK (COG-UK) Consortium | The Lighthouse Lab in Milton Keynes and Alex Alderton, Roberto Amato, Jeffrey Barrett, Sonia Goncalves, Ewan Harrison, David K. Jackson, Ian Johnston, Dominic Kwiatkowski, Cordelia Langford, John Sillitoe on behalf of the Wellcome Sanger Institute COVID-19 Surveillance Team |
| EPI_ISL_2237000 | Lighthouse Lab in Alderley Park | Wellcome Sanger Institute for the COVID-19 Genomics UK (COG-UK) Consortium | Jacquelyn Wynn, Mairead Hyland, The Lighthouse Lab in Alderley Park and Alex Alderton, Roberto Amato, Jeffrey Barrett, Sonia Goncalves, Ewan Harrison, David K. Jackson, Ian Johnston, Dominic Kwiatkowski, Cordelia Langford, John Sillitoe on behalf of the Wellcome Sanger Institute COVID-19 Surveillance Team |
| EPI_ISL_2237001, EPI_ISL_2237010 | Lighthouse Lab in Glasgow | Wellcome Sanger Institute for the COVID-19 Genomics UK (COG-UK) Consortium | Harper VanSteenhouse, Yumi Kasai, David Gray, Carol Clugston, Anna Dominiczak and Alex Alderton, Roberto Amato, Jeffrey Barrett, Sonia Goncalves, Ewan Harrison, David K. Jackson, Ian Johnston, Dominic Kwiatkowski, Cordelia Langford, John Sillitoe on behalf of the Wellcome Sanger Institute COVID-19 Surveillance Team |
| EPI_ISL_2237016, EPI_ISL_2237017, EPI_ISL_2237019 | Lighthouse Lab in Milton Keynes | Wellcome Sanger Institute for the COVID-19 Genomics UK (COG-UK) Consortium | The Lighthouse Lab in Milton Keynes and Alex Alderton, Roberto Amato, Jeffrey Barrett, Sonia Goncalves, Ewan Harrison, David K. Jackson, Ian Johnston, Dominic Kwiatkowski, Cordelia Langford, John Sillitoe on behalf of the Wellcome Sanger Institute COVID-19 Surveillance Team |
| EPI_ISL_2237026, EPI_ISL_2237027 | Lighthouse Lab in Alderley Park | Wellcome Sanger Institute for the COVID-19 Genomics UK (COG-UK) Consortium | Jacquelyn Wynn, Mairead Hyland, The Lighthouse Lab in Alderley Park and Alex Alderton, Roberto Amato, Jeffrey Barrett, Sonia Goncalves, Ewan Harrison, David K. Jackson, Ian Johnston, Dominic Kwiatkowski, Cordelia Langford, John Sillitoe on behalf of the Wellcome Sanger Institute COVID-19 Surveillance Team |
| EPI_ISL_2237029, EPI_ISL_2237031, EPI_ISL_2237033 | Lighthouse Lab in Glasgow | Wellcome Sanger Institute for the COVID-19 Genomics UK (COG-UK) Consortium | Harper VanSteenhouse, Yumi Kasai, David Gray, Carol Clugston, Anna Dominiczak and Alex Alderton, Roberto Amato, Jeffrey Barrett, Sonia Goncalves, Ewan Harrison, David K. Jackson, Ian Johnston, Dominic Kwiatkowski, Cordelia Langford, John Sillitoe on behalf of the Wellcome Sanger Institute COVID-19 Surveillance Team |
| EPI_ISL_2237035, EPI_ISL_2237036, EPI_ISL_2237038 | Lighthouse Lab in Alderley Park | Wellcome Sanger Institute for the COVID-19 Genomics UK (COG-UK) Consortium | Jacquelyn Wynn, Mairead Hyland, The Lighthouse Lab in Alderley Park and Alex Alderton, Roberto Amato, Jeffrey Barrett, Sonia Goncalves, Ewan Harrison, David K. Jackson, Ian Johnston, Dominic Kwiatkowski, Cordelia Langford, John Sillitoe on behalf of the Wellcome Sanger Institute COVID-19 Surveillance Team |
| EPI_ISL_2237039 | Lighthouse Lab in Glasgow | Wellcome Sanger Institute for the COVID-19 Genomics UK (COG-UK) Consortium | Harper VanSteenhouse, Yumi Kasai, David Gray, Carol Clugston, Anna Dominiczak and Alex Alderton, Roberto Amato, Jeffrey Barrett, Sonia Goncalves, Ewan Harrison, David K. Jackson, Ian Johnston, Dominic Kwiatkowski, Cordelia Langford, John Sillitoe on behalf of the Wellcome Sanger Institute COVID-19 Surveillance Team |
| EPI_ISL_2237040 | Lighthouse Lab in Alderley Park | Wellcome Sanger Institute for the COVID-19 Genomics UK (COG-UK) Consortium | Jacquelyn Wynn, Mairead Hyland, The Lighthouse Lab in Alderley Park and Alex Alderton, Roberto Amato, Jeffrey Barrett, Sonia Goncalves, Ewan Harrison, David K. Jackson, Ian Johnston, Dominic Kwiatkowski, Cordelia Langford, John Sillitoe on behalf of the Wellcome Sanger Institute COVID-19 Surveillance Team |
| EPI_ISL_2237043 | Lighthouse Lab in Glasgow | Wellcome Sanger Institute for the COVID-19 Genomics UK (COG-UK) Consortium | Harper VanSteenhouse, Yumi Kasai, David Gray, Carol Clugston, Anna Dominiczak and Alex Alderton, Roberto Amato, Jeffrey Barrett, Sonia Goncalves, Ewan Harrison, David K. Jackson, Ian Johnston, Dominic Kwiatkowski, Cordelia Langford, John Sillitoe on behalf of the Wellcome Sanger Institute COVID-19 Surveillance Team |
| EPI_ISL_2237044 | Lighthouse Lab in Milton Keynes | Wellcome Sanger Institute for the COVID-19 Genomics UK (COG-UK) Consortium | The Lighthouse Lab in Milton Keynes and Alex Alderton, Roberto Amato, Jeffrey Barrett, Sonia Goncalves, Ewan Harrison, David K. Jackson, Ian Johnston, Dominic Kwiatkowski, Cordelia Langford, John Sillitoe on behalf of the Wellcome Sanger Institute COVID-19 Surveillance Team |
| EPI_ISL_2237045 | Lighthouse Lab in Alderley Park | Wellcome Sanger Institute for the COVID-19 Genomics UK (COG-UK) Consortium | Jacquelyn Wynn, Mairead Hyland, The Lighthouse Lab in Alderley Park and Alex Alderton, Roberto Amato, Jeffrey Barrett, Sonia Goncalves, Ewan Harrison, David K. Jackson, Ian Johnston, Dominic Kwiatkowski, Cordelia Langford, John Sillitoe on behalf of the Wellcome Sanger Institute COVID-19 Surveillance Team |
| EPI_ISL_2237046 | Lighthouse Lab in Milton Keynes | Wellcome Sanger Institute for the COVID-19 Genomics UK (COG-UK) Consortium | The Lighthouse Lab in Milton Keynes and Alex Alderton, Roberto Amato, Jeffrey Barrett, Sonia Goncalves, Ewan Harrison, David K. Jackson, Ian Johnston, Dominic Kwiatkowski, Cordelia Langford, John Sillitoe on behalf of the Wellcome Sanger Institute COVID-19 Surveillance Team |
| EPI_ISL_2237049, EPI_ISL_2237052 | Lighthouse Lab in Glasgow | Wellcome Sanger Institute for the COVID-19 Genomics UK (COG-UK) Consortium | Harper VanSteenhouse, Yumi Kasai, David Gray, Carol Clugston, Anna Dominiczak and Alex Alderton, Roberto Amato, Jeffrey Barrett, Sonia Goncalves, Ewan Harrison, David K. Jackson, Ian Johnston, Dominic Kwiatkowski, Cordelia Langford, John Sillitoe on behalf of the Wellcome Sanger Institute COVID-19 Surveillance Team |
| EPI_ISL_2237055 | Lighthouse Lab in Milton Keynes | Wellcome Sanger Institute for the COVID-19 Genomics UK (COG-UK) Consortium | The Lighthouse Lab in Milton Keynes and Alex Alderton, Roberto Amato, Jeffrey Barrett, Sonia Goncalves, Ewan Harrison, David K. Jackson, Ian Johnston, Dominic Kwiatkowski, Cordelia Langford, John Sillitoe on behalf of the Wellcome Sanger Institute COVID-19 Surveillance Team |
| EPI_ISL_2237056 | Lighthouse Lab in Glasgow | Wellcome Sanger Institute for the COVID-19 Genomics UK (COG-UK) Consortium | Harper VanSteenhouse, Yumi Kasai, David Gray, Carol Clugston, Anna Dominiczak and Alex Alderton, Roberto Amato, Jeffrey Barrett, Sonia Goncalves, Ewan Harrison, David K. Jackson, Ian Johnston, Dominic Kwiatkowski, Cordelia Langford, John Sillitoe on behalf of the Wellcome Sanger Institute COVID-19 Surveillance Team |
| EPI_ISL_2237057, EPI_ISL_2237059 | Lighthouse Lab in Alderley Park | Wellcome Sanger Institute for the COVID-19 Genomics UK (COG-UK) Consortium | Jacquelyn Wynn, Mairead Hyland, The Lighthouse Lab in Alderley Park and Alex Alderton, Roberto Amato, Jeffrey Barrett, Sonia Goncalves, Ewan Harrison, David K. Jackson, Ian Johnston, Dominic Kwiatkowski, Cordelia Langford, John Sillitoe on behalf of the Wellcome Sanger Institute COVID-19 Surveillance Team |
| EPI_ISL_2237060 | Lighthouse Lab in Glasgow | Wellcome Sanger Institute for the COVID-19 Genomics UK (COG-UK) Consortium | Harper VanSteenhouse, Yumi Kasai, David Gray, Carol Clugston, Anna Dominiczak and Alex Alderton, Roberto Amato, Jeffrey Barrett, Sonia Goncalves, Ewan Harrison, David K. Jackson, Ian Johnston, Dominic Kwiatkowski, Cordelia Langford, John Sillitoe on behalf of the Wellcome Sanger Institute COVID-19 Surveillance Team |
| EPI_ISL_2237062 | Lighthouse Lab in Alderley Park | Wellcome Sanger Institute for the COVID-19 Genomics UK (COG-UK) Consortium | Jacquelyn Wynn, Mairead Hyland, The Lighthouse Lab in Alderley Park and Alex Alderton, Roberto Amato, Jeffrey Barrett, Sonia Goncalves, Ewan Harrison, David K. Jackson, Ian Johnston, Dominic Kwiatkowski, Cordelia Langford, John Sillitoe on behalf of the Wellcome Sanger Institute COVID-19 Surveillance Team |
| EPI_ISL_2237066, EPI_ISL_2237067, EPI_ISL_2237073 | Lighthouse Lab in Milton Keynes | Wellcome Sanger Institute for the COVID-19 Genomics UK (COG-UK) Consortium | The Lighthouse Lab in Milton Keynes and Alex Alderton, Roberto Amato, Jeffrey Barrett, Sonia Goncalves, Ewan Harrison, David K. Jackson, Ian Johnston, Dominic Kwiatkowski, Cordelia Langford, John Sillitoe on behalf of the Wellcome Sanger Institute COVID-19 Surveillance Team |
| EPI_ISL_2237077, EPI_ISL_2237078 | Lighthouse Lab in Glasgow | Wellcome Sanger Institute for the COVID-19 Genomics UK (COG-UK) Consortium | Harper VanSteenhouse, Yumi Kasai, David Gray, Carol Clugston, Anna Dominiczak and Alex Alderton, Roberto Amato, Jeffrey Barrett, Sonia Goncalves, Ewan Harrison, David K. Jackson, Ian Johnston, Dominic Kwiatkowski, Cordelia Langford, John Sillitoe on behalf of the Wellcome Sanger Institute COVID-19 Surveillance Team |
| EPI_ISL_2237081, EPI_ISL_2237083, EPI_ISL_2237088 | Lighthouse Lab in Alderley Park | Wellcome Sanger Institute for the COVID-19 Genomics UK (COG-UK) Consortium | Jacquelyn Wynn, Mairead Hyland, The Lighthouse Lab in Alderley Park and Alex Alderton, Roberto Amato, Jeffrey Barrett, Sonia Goncalves, Ewan Harrison, David K. Jackson, Ian Johnston, Dominic Kwiatkowski, Cordelia Langford, John Sillitoe on behalf of the Wellcome Sanger Institute COVID-19 Surveillance Team |
| EPI_ISL_2237089 | Lighthouse Lab in Glasgow | Wellcome Sanger Institute for the COVID-19 Genomics UK (COG-UK) Consortium | Harper VanSteenhouse, Yumi Kasai, David Gray, Carol Clugston, Anna Dominiczak and Alex Alderton, Roberto Amato, Jeffrey Barrett, Sonia Goncalves, Ewan Harrison, David K. Jackson, Ian Johnston, Dominic Kwiatkowski, Cordelia Langford, John Sillitoe on behalf of the Wellcome Sanger Institute COVID-19 Surveillance Team |
| EPI_ISL_2237090, EPI_ISL_2237091 | Lighthouse Lab in Alderley Park | Wellcome Sanger Institute for the COVID-19 Genomics UK (COG-UK) Consortium | Jacquelyn Wynn, Mairead Hyland, The Lighthouse Lab in Alderley Park and Alex Alderton, Roberto Amato, Jeffrey Barrett, Sonia Goncalves, Ewan Harrison, David K. Jackson, Ian Johnston, Dominic Kwiatkowski, Cordelia Langford, John Sillitoe on behalf of the Wellcome Sanger Institute COVID-19 Surveillance Team |
| EPI_ISL_2237092 | Lighthouse Lab in Glasgow | Wellcome Sanger Institute for the COVID-19 Genomics UK (COG-UK) Consortium | Harper VanSteenhouse, Yumi Kasai, David Gray, Carol Clugston, Anna Dominiczak and Alex Alderton, Roberto Amato, Jeffrey Barrett, Sonia Goncalves, Ewan Harrison, David K. Jackson, Ian Johnston, Dominic Kwiatkowski, Cordelia Langford, John Sillitoe on behalf of the Wellcome Sanger Institute COVID-19 Surveillance Team |

| | | | |
|---|---|---|---|
| EPI_ISL_2237095 | Lighthouse Lab in Milton Keynes | Wellcome Sanger Institute for the COVID-19 Genomics UK (COG-UK) Consortium | The Lighthouse Lab in Milton Keynes and Alex Alderton, Roberto Amato, Jeffrey Barrett, Sonia Goncalves, Ewan Harrison, David K. Jackson, Ian Johnston, Dominic Kwiatkowski, Cordelia Langford, John Sillitoe on behalf of the Wellcome Sanger Institute COVID-19 Surveillance Team |
| EPI_ISL_2237098 | Lighthouse Lab in Alderley Park | Wellcome Sanger Institute for the COVID-19 Genomics UK (COG-UK) Consortium | Jacquelyn Wynn, Mairead Hyland, The Lighthouse Lab in Alderley Park and Alex Alderton, Roberto Amato, Jeffrey Barrett, Sonia Goncalves, Ewan Harrison, David K. Jackson, Ian Johnston, Dominic Kwiatkowski, Cordelia Langford, John Sillitoe on behalf of the Wellcome Sanger Institute COVID-19 Surveillance Team |
| EPI_ISL_2237101 | Lighthouse Lab in Glasgow | Wellcome Sanger Institute for the COVID-19 Genomics UK (COG-UK) Consortium | Harper VanSteenhouse, Yumi Kasai, David Gray, Carol Clugston, Anna Dominiczak and Alex Alderton, Roberto Amato, Jeffrey Barrett, Sonia Goncalves, Ewan Harrison, David K. Jackson, Ian Johnston, Dominic Kwiatkowski, Cordelia Langford, John Sillitoe on behalf of the Wellcome Sanger Institute COVID-19 Surveillance Team |
| EPI_ISL_2237103 | Lighthouse Lab in Alderley Park | Wellcome Sanger Institute for the COVID-19 Genomics UK (COG-UK) Consortium | Jacquelyn Wynn, Mairead Hyland, The Lighthouse Lab in Alderley Park and Alex Alderton, Roberto Amato, Jeffrey Barrett, Sonia Goncalves, Ewan Harrison, David K. Jackson, Ian Johnston, Dominic Kwiatkowski, Cordelia Langford, John Sillitoe on behalf of the Wellcome Sanger Institute COVID-19 Surveillance Team |
| EPI_ISL_2237105 | Lighthouse Lab in Glasgow | Wellcome Sanger Institute for the COVID-19 Genomics UK (COG-UK) Consortium | Harper VanSteenhouse, Yumi Kasai, David Gray, Carol Clugston, Anna Dominiczak and Alex Alderton, Roberto Amato, Jeffrey Barrett, Sonia Goncalves, Ewan Harrison, David K. Jackson, Ian Johnston, Dominic Kwiatkowski, Cordelia Langford, John Sillitoe on behalf of the Wellcome Sanger Institute COVID-19 Surveillance Team |
| EPI_ISL_2237107, EPI_ISL_2237110 | Lighthouse Lab in Alderley Park | Wellcome Sanger Institute for the COVID-19 Genomics UK (COG-UK) Consortium | Jacquelyn Wynn, Mairead Hyland, The Lighthouse Lab in Alderley Park and Alex Alderton, Roberto Amato, Jeffrey Barrett, Sonia Goncalves, Ewan Harrison, David K. Jackson, Ian Johnston, Dominic Kwiatkowski, Cordelia Langford, John Sillitoe on behalf of the Wellcome Sanger Institute COVID-19 Surveillance Team |
| EPI_ISL_2237115 | Lighthouse Lab in Milton Keynes | Wellcome Sanger Institute for the COVID-19 Genomics UK (COG-UK) Consortium | The Lighthouse Lab in Milton Keynes and Alex Alderton, Roberto Amato, Jeffrey Barrett, Sonia Goncalves, Ewan Harrison, David K. Jackson, Ian Johnston, Dominic Kwiatkowski, Cordelia Langford, John Sillitoe on behalf of the Wellcome Sanger Institute COVID-19 Surveillance Team |
| EPI_ISL_2237116 | Lighthouse Lab in Glasgow | Wellcome Sanger Institute for the COVID-19 Genomics UK (COG-UK) Consortium | Harper VanSteenhouse, Yumi Kasai, David Gray, Carol Clugston, Anna Dominiczak and Alex Alderton, Roberto Amato, Jeffrey Barrett, Sonia Goncalves, Ewan Harrison, David K. Jackson, Ian Johnston, Dominic Kwiatkowski, Cordelia Langford, John Sillitoe on behalf of the Wellcome Sanger Institute COVID-19 Surveillance Team |
| EPI_ISL_2237117, EPI_ISL_2237118, EPI_ISL_2237119 | Lighthouse Lab in Alderley Park | Wellcome Sanger Institute for the COVID-19 Genomics UK (COG-UK) Consortium | Jacquelyn Wynn, Mairead Hyland, The Lighthouse Lab in Alderley Park and Alex Alderton, Roberto Amato, Jeffrey Barrett, Sonia Goncalves, Ewan Harrison, David K. Jackson, Ian Johnston, Dominic Kwiatkowski, Cordelia Langford, John Sillitoe on behalf of the Wellcome Sanger Institute COVID-19 Surveillance Team |
| EPI_ISL_2237122 | Lighthouse Lab in Glasgow | Wellcome Sanger Institute for the COVID-19 Genomics UK (COG-UK) Consortium | Harper VanSteenhouse, Yumi Kasai, David Gray, Carol Clugston, Anna Dominiczak and Alex Alderton, Roberto Amato, Jeffrey Barrett, Sonia Goncalves, Ewan Harrison, David K. Jackson, Ian Johnston, Dominic Kwiatkowski, Cordelia Langford, John Sillitoe on behalf of the Wellcome Sanger Institute COVID-19 Surveillance Team |
| EPI_ISL_2237125 | Lighthouse Lab in Alderley Park | Wellcome Sanger Institute for the COVID-19 Genomics UK (COG-UK) Consortium | Jacquelyn Wynn, Mairead Hyland, The Lighthouse Lab in Alderley Park and Alex Alderton, Roberto Amato, Jeffrey Barrett, Sonia Goncalves, Ewan Harrison, David K. Jackson, Ian Johnston, Dominic Kwiatkowski, Cordelia Langford, John Sillitoe on behalf of the Wellcome Sanger Institute COVID-19 Surveillance Team |
| EPI_ISL_2237126 | Lighthouse Lab in Glasgow | Wellcome Sanger Institute for the COVID-19 Genomics UK (COG-UK) Consortium | Harper VanSteenhouse, Yumi Kasai, David Gray, Carol Clugston, Anna Dominiczak and Alex Alderton, Roberto Amato, Jeffrey Barrett, Sonia Goncalves, Ewan Harrison, David K. Jackson, Ian Johnston, Dominic Kwiatkowski, Cordelia Langford, John Sillitoe on behalf of the Wellcome Sanger Institute COVID-19 Surveillance Team |
| EPI_ISL_2237128 | Lighthouse Lab in Milton Keynes | Wellcome Sanger Institute for the COVID-19 Genomics UK (COG-UK) Consortium | The Lighthouse Lab in Milton Keynes and Alex Alderton, Roberto Amato, Jeffrey Barrett, Sonia Goncalves, Ewan Harrison, David K. Jackson, Ian Johnston, Dominic Kwiatkowski, Cordelia Langford, John Sillitoe on behalf of the Wellcome Sanger Institute COVID-19 Surveillance Team |
| EPI_ISL_2237129, EPI_ISL_2237130, EPI_ISL_2237133 | Lighthouse Lab in Alderley Park | Wellcome Sanger Institute for the COVID-19 Genomics UK (COG-UK) Consortium | Jacquelyn Wynn, Mairead Hyland, The Lighthouse Lab in Alderley Park and Alex Alderton, Roberto Amato, Jeffrey Barrett, Sonia Goncalves, Ewan Harrison, David K. Jackson, Ian Johnston, Dominic Kwiatkowski, Cordelia Langford, John Sillitoe on behalf of the Wellcome Sanger Institute COVID-19 Surveillance Team |
| EPI_ISL_2237135 | Lighthouse Lab in Glasgow | Wellcome Sanger Institute for the COVID-19 Genomics UK (COG-UK) Consortium | Harper VanSteenhouse, Yumi Kasai, David Gray, Carol Clugston, Anna Dominiczak and Alex Alderton, Roberto Amato, Jeffrey Barrett, Sonia Goncalves, Ewan Harrison, David K. Jackson, Ian Johnston, Dominic Kwiatkowski, Cordelia Langford, John Sillitoe on behalf of the Wellcome Sanger Institute COVID-19 Surveillance Team |
| EPI_ISL_2237136 | Lighthouse Lab in Milton Keynes | Wellcome Sanger Institute for the COVID-19 Genomics UK (COG-UK) Consortium | The Lighthouse Lab in Milton Keynes and Alex Alderton, Roberto Amato, Jeffrey Barrett, Sonia Goncalves, Ewan Harrison, David K. Jackson, Ian Johnston, Dominic Kwiatkowski, Cordelia Langford, John Sillitoe on behalf of the Wellcome Sanger Institute COVID-19 Surveillance Team |
| EPI_ISL_2237137 | Lighthouse Lab in Glasgow | Wellcome Sanger Institute for the COVID-19 Genomics UK (COG-UK) Consortium | Harper VanSteenhouse, Yumi Kasai, David Gray, Carol Clugston, Anna Dominiczak and Alex Alderton, Roberto Amato, Jeffrey Barrett, Sonia Goncalves, Ewan Harrison, David K. Jackson, Ian Johnston, Dominic Kwiatkowski, Cordelia Langford, John Sillitoe on behalf of the Wellcome Sanger Institute COVID-19 Surveillance Team |
| EPI_ISL_2237142, EPI_ISL_2237143 | Lighthouse Lab in Milton Keynes | Wellcome Sanger Institute for the COVID-19 Genomics UK (COG-UK) Consortium | The Lighthouse Lab in Milton Keynes and Alex Alderton, Roberto Amato, Jeffrey Barrett, Sonia Goncalves, Ewan Harrison, David K. Jackson, Ian Johnston, Dominic Kwiatkowski, Cordelia Langford, John Sillitoe on behalf of the Wellcome Sanger Institute COVID-19 Surveillance Team |
| EPI_ISL_2237145, EPI_ISL_2237148, EPI_ISL_2237152, EPI_ISL_2237154, EPI_ISL_2237156 | Lighthouse Lab in Alderley Park | Wellcome Sanger Institute for the COVID-19 Genomics UK (COG-UK) Consortium | Jacquelyn Wynn, Mairead Hyland, The Lighthouse Lab in Alderley Park and Alex Alderton, Roberto Amato, Jeffrey Barrett, Sonia Goncalves, Ewan Harrison, David K. Jackson, Ian Johnston, Dominic Kwiatkowski, Cordelia Langford, John Sillitoe on behalf of the Wellcome Sanger Institute COVID-19 Surveillance Team |
| EPI_ISL_2237158 | Lighthouse Lab in Glasgow | Wellcome Sanger Institute for the COVID-19 Genomics UK (COG-UK) Consortium | Harper VanSteenhouse, Yumi Kasai, David Gray, Carol Clugston, Anna Dominiczak and Alex Alderton, Roberto Amato, Jeffrey Barrett, Sonia Goncalves, Ewan Harrison, David K. Jackson, Ian Johnston, Dominic Kwiatkowski, Cordelia Langford, John Sillitoe on behalf of the Wellcome Sanger Institute COVID-19 Surveillance Team |
| EPI_ISL_2237159, EPI_ISL_2237163, EPI_ISL_2237166, EPI_ISL_2237170 | Lighthouse Lab in Alderley Park | Wellcome Sanger Institute for the COVID-19 Genomics UK (COG-UK) Consortium | Jacquelyn Wynn, Mairead Hyland, The Lighthouse Lab in Alderley Park and Alex Alderton, Roberto Amato, Jeffrey Barrett, Sonia Goncalves, Ewan Harrison, David K. Jackson, Ian Johnston, Dominic Kwiatkowski, Cordelia Langford, John Sillitoe on behalf of the Wellcome Sanger Institute COVID-19 Surveillance Team |
| EPI_ISL_2237174 | Lighthouse Lab in Glasgow | Wellcome Sanger Institute for the COVID-19 Genomics UK (COG-UK) Consortium | Harper VanSteenhouse, Yumi Kasai, David Gray, Carol Clugston, Anna Dominiczak and Alex Alderton, Roberto Amato, Jeffrey Barrett, Sonia Goncalves, Ewan Harrison, David K. Jackson, Ian Johnston, Dominic Kwiatkowski, Cordelia Langford, John Sillitoe on behalf of the Wellcome Sanger Institute COVID-19 Surveillance Team |
| EPI_ISL_2237176 | Lighthouse Lab in Alderley Park | Wellcome Sanger Institute for the COVID-19 Genomics UK (COG-UK) Consortium | Jacquelyn Wynn, Mairead Hyland, The Lighthouse Lab in Alderley Park and Alex Alderton, Roberto Amato, Jeffrey Barrett, Sonia Goncalves, Ewan Harrison, David K. Jackson, Ian Johnston, Dominic Kwiatkowski, Cordelia Langford, John Sillitoe on behalf of the Wellcome Sanger Institute COVID-19 Surveillance Team |
| EPI_ISL_2237177 | Lighthouse Lab in Glasgow | Wellcome Sanger Institute for the COVID-19 Genomics UK (COG-UK) Consortium | Harper VanSteenhouse, Yumi Kasai, David Gray, Carol Clugston, Anna Dominiczak and Alex Alderton, Roberto Amato, Jeffrey Barrett, Sonia Goncalves, Ewan Harrison, David K. Jackson, Ian Johnston, Dominic Kwiatkowski, Cordelia Langford, John Sillitoe on behalf of the Wellcome Sanger Institute COVID-19 Surveillance Team |
| EPI_ISL_2237178, EPI_ISL_2237179 | Lighthouse Lab in Alderley Park | Wellcome Sanger Institute for the COVID-19 Genomics UK (COG-UK) Consortium | Jacquelyn Wynn, Mairead Hyland, The Lighthouse Lab in Alderley Park and Alex Alderton, Roberto Amato, Jeffrey Barrett, Sonia Goncalves, Ewan Harrison, David K. Jackson, Ian Johnston, Dominic Kwiatkowski, Cordelia Langford, John Sillitoe on behalf of the Wellcome Sanger Institute COVID-19 Surveillance Team |
| EPI_ISL_2237180 | Lighthouse Lab in Glasgow | Wellcome Sanger Institute for the COVID-19 Genomics UK (COG-UK) Consortium | Harper VanSteenhouse, Yumi Kasai, David Gray, Carol Clugston, Anna Dominiczak and Alex Alderton, Roberto Amato, Jeffrey Barrett, Sonia Goncalves, Ewan Harrison, David K. Jackson, Ian Johnston, Dominic Kwiatkowski, Cordelia Langford, John Sillitoe on behalf of the Wellcome Sanger Institute COVID-19 Surveillance Team |
| EPI_ISL_2237181 | Lighthouse Lab in Alderley Park | Wellcome Sanger Institute for the COVID-19 Genomics UK (COG-UK) Consortium | Jacquelyn Wynn, Mairead Hyland, The Lighthouse Lab in Alderley Park and Alex Alderton, Roberto Amato, Jeffrey Barrett, Sonia Goncalves, Ewan Harrison, David K. Jackson, Ian Johnston, Dominic Kwiatkowski, Cordelia Langford, John Sillitoe on behalf of the Wellcome Sanger Institute COVID-19 Surveillance Team |
| EPI_ISL_2237182 | Lighthouse Lab in Glasgow | Wellcome Sanger Institute for the COVID-19 Genomics UK (COG-UK) Consortium | Harper VanSteenhouse, Yumi Kasai, David Gray, Carol Clugston, Anna Dominiczak and Alex Alderton, Roberto Amato, Jeffrey Barrett, Sonia Goncalves, Ewan Harrison, David K. Jackson, Ian Johnston, Dominic Kwiatkowski, Cordelia Langford, John Sillitoe on behalf of the Wellcome Sanger Institute COVID-19 Surveillance Team |
| EPI_ISL_2237184, EPI_ISL_2237185, EPI_ISL_2237186, EPI_ISL_2237187 | Lighthouse Lab in Alderley Park | Wellcome Sanger Institute for the COVID-19 Genomics UK (COG-UK) Consortium | Jacquelyn Wynn, Mairead Hyland, The Lighthouse Lab in Alderley Park and Alex Alderton, Roberto Amato, Jeffrey Barrett, Sonia Goncalves, Ewan Harrison, David K. Jackson, Ian Johnston, Dominic Kwiatkowski, Cordelia Langford, John Sillitoe on behalf of the Wellcome Sanger Institute COVID-19 Surveillance Team |
| EPI_ISL_2237189 | Lighthouse Lab in Glasgow | Wellcome Sanger Institute for the COVID-19 Genomics UK (COG-UK) Consortium | Harper VanSteenhouse, Yumi Kasai, David Gray, Carol Clugston, Anna Dominiczak and Alex Alderton, Roberto Amato, Jeffrey Barrett, Sonia Goncalves, Ewan Harrison, David K. Jackson, Ian Johnston, Dominic Kwiatkowski, Cordelia Langford, John Sillitoe on behalf of the Wellcome Sanger Institute COVID-19 Surveillance Team |
| EPI_ISL_2237190, EPI_ISL_2237191 | Lighthouse Lab in Alderley Park | Wellcome Sanger Institute for the COVID-19 Genomics UK (COG-UK) Consortium | Jacquelyn Wynn, Mairead Hyland, The Lighthouse Lab in Alderley Park and Alex Alderton, Roberto Amato, Jeffrey Barrett, Sonia Goncalves, Ewan Harrison, David K. Jackson, Ian Johnston, Dominic Kwiatkowski, Cordelia Langford, John Sillitoe on behalf of the Wellcome Sanger Institute COVID-19 Surveillance Team |
| EPI_ISL_2237192, EPI_ISL_2237196 | Lighthouse Lab in Glasgow | Wellcome Sanger Institute for the COVID-19 Genomics UK (COG-UK) Consortium | Harper VanSteenhouse, Yumi Kasai, David Gray, Carol Clugston, Anna Dominiczak and Alex Alderton, Roberto Amato, Jeffrey Barrett, Sonia Goncalves, Ewan Harrison, David K. Jackson, Ian Johnston, Dominic Kwiatkowski, Cordelia Langford, John Sillitoe on behalf of the Wellcome Sanger Institute COVID-19 Surveillance Team |
| EPI_ISL_2237197 | Lighthouse Lab in Alderley Park | Wellcome Sanger Institute for the COVID-19 Genomics UK (COG-UK) Consortium | Jacquelyn Wynn, Mairead Hyland, The Lighthouse Lab in Alderley Park and Alex Alderton, Roberto Amato, Jeffrey Barrett, Sonia Goncalves, Ewan Harrison, David K. Jackson, Ian Johnston, Dominic Kwiatkowski, Cordelia Langford, John Sillitoe on behalf of the Wellcome Sanger Institute COVID-19 Surveillance Team |
| EPI_ISL_2237198, EPI_ISL_2237199 | Lighthouse Lab in Glasgow | Wellcome Sanger Institute for the COVID-19 Genomics UK (COG-UK) Consortium | Harper VanSteenhouse, Yumi Kasai, David Gray, Carol Clugston, Anna Dominiczak and Alex Alderton, Roberto Amato, Jeffrey Barrett, Sonia Goncalves, Ewan Harrison, David K. Jackson, Ian Johnston, Dominic Kwiatkowski, Cordelia Langford, John Sillitoe on behalf of the Wellcome Sanger Institute COVID-19 Surveillance Team |
| EPI_ISL_2237200, EPI_ISL_2237201, EPI_ISL_2237202, EPI_ISL_2237204, EPI_ISL_2237205, EPI_ISL_2237207, EPI_ISL_2237210, EPI_ISL_2237211, EPI_ISL_2237212 | Lighthouse Lab in Alderley Park | Wellcome Sanger Institute for the COVID-19 Genomics UK (COG-UK) Consortium | Jacquelyn Wynn, Mairead Hyland, The Lighthouse Lab in Alderley Park and Alex Alderton, Roberto Amato, Jeffrey Barrett, Sonia Goncalves, Ewan Harrison, David K. Jackson, Ian Johnston, Dominic Kwiatkowski, Cordelia Langford, John Sillitoe on behalf of the Wellcome Sanger Institute COVID-19 Surveillance Team |
| EPI_ISL_2237213 | Lighthouse Lab in Glasgow | Wellcome Sanger Institute for the COVID-19 Genomics UK (COG-UK) Consortium | Harper VanSteenhouse, Yumi Kasai, David Gray, Carol Clugston, Anna Dominiczak and Alex Alderton, Roberto Amato, Jeffrey Barrett, Sonia Goncalves, Ewan Harrison, David K. Jackson, Ian Johnston, Dominic Kwiatkowski, Cordelia Langford, John Sillitoe on behalf of the Wellcome Sanger Institute COVID-19 Surveillance Team |
| EPI_ISL_2237214, EPI_ISL_2237215, EPI_ISL_2237216, EPI_ISL_2237217, EPI_ISL_2237218, EPI_ISL_2237219, | Lighthouse Lab in Alderley Park | Wellcome Sanger Institute for the COVID-19 Genomics UK (COG-UK) Consortium | Jacquelyn Wynn, Mairead Hyland, The Lighthouse Lab in Alderley Park and Alex Alderton, Roberto Amato, Jeffrey Barrett, Sonia Goncalves, Ewan Harrison, David K. Jackson, Ian Johnston, Dominic Kwiatkowski, Cordelia Langford, John Sillitoe on behalf of the Wellcome Sanger Institute COVID-19 Surveillance Team |

| | | | |
|---|---|---|---|
| EPI_ISL_2237220, EPI_ISL_2237227, EPI_ISL_2237228 | | | |
| EPI_ISL_2237229 | Lighthouse Lab in Glasgow | Wellcome Sanger Institute for the COVID-19 Genomics UK (COG-UK) Consortium | Harper VanSteenhouse, Yumi Kasai, David Gray, Carol Clugston, Anna Dominiczak and Alex Alderton, Roberto Amato, Jeffrey Barrett, Sonia Goncalves, Ewan Harrison, David K. Jackson, Ian Johnston, Dominic Kwiatkowski, Cordelia Langford, John Sillitoe on behalf of the Wellcome Sanger Institute COVID-19 Surveillance Team |
| EPI_ISL_2237232, EPI_ISL_2237233, EPI_ISL_2237236 | Lighthouse Lab in Alderley Park | Wellcome Sanger Institute for the COVID-19 Genomics UK (COG-UK) Consortium | Jacquelyn Wynn, Mairead Hyland, The Lighthouse Lab in Alderley Park and Alex Alderton, Roberto Amato, Jeffrey Barrett, Sonia Goncalves, Ewan Harrison, David K. Jackson, Ian Johnston, Dominic Kwiatkowski, Cordelia Langford, John Sillitoe on behalf of the Wellcome Sanger Institute COVID-19 Surveillance Team |
| EPI_ISL_2237237 | Lighthouse Lab in Glasgow | Wellcome Sanger Institute for the COVID-19 Genomics UK (COG-UK) Consortium | Harper VanSteenhouse, Yumi Kasai, David Gray, Carol Clugston, Anna Dominiczak and Alex Alderton, Roberto Amato, Jeffrey Barrett, Sonia Goncalves, Ewan Harrison, David K. Jackson, Ian Johnston, Dominic Kwiatkowski, Cordelia Langford, John Sillitoe on behalf of the Wellcome Sanger Institute COVID-19 Surveillance Team |
| EPI_ISL_2237238, EPI_ISL_2237239 | Lighthouse Lab in Alderley Park | Wellcome Sanger Institute for the COVID-19 Genomics UK (COG-UK) Consortium | Jacquelyn Wynn, Mairead Hyland, The Lighthouse Lab in Alderley Park and Alex Alderton, Roberto Amato, Jeffrey Barrett, Sonia Goncalves, Ewan Harrison, David K. Jackson, Ian Johnston, Dominic Kwiatkowski, Cordelia Langford, John Sillitoe on behalf of the Wellcome Sanger Institute COVID-19 Surveillance Team |
| EPI_ISL_2237242 | Lighthouse Lab in Glasgow | Wellcome Sanger Institute for the COVID-19 Genomics UK (COG-UK) Consortium | Harper VanSteenhouse, Yumi Kasai, David Gray, Carol Clugston, Anna Dominiczak and Alex Alderton, Roberto Amato, Jeffrey Barrett, Sonia Goncalves, Ewan Harrison, David K. Jackson, Ian Johnston, Dominic Kwiatkowski, Cordelia Langford, John Sillitoe on behalf of the Wellcome Sanger Institute COVID-19 Surveillance Team |
| EPI_ISL_2237243 | Lighthouse Lab in Alderley Park | Wellcome Sanger Institute for the COVID-19 Genomics UK (COG-UK) Consortium | Jacquelyn Wynn, Mairead Hyland, The Lighthouse Lab in Alderley Park and Alex Alderton, Roberto Amato, Jeffrey Barrett, Sonia Goncalves, Ewan Harrison, David K. Jackson, Ian Johnston, Dominic Kwiatkowski, Cordelia Langford, John Sillitoe on behalf of the Wellcome Sanger Institute COVID-19 Surveillance Team |
| EPI_ISL_2237245 | Lighthouse Lab in Glasgow | Wellcome Sanger Institute for the COVID-19 Genomics UK (COG-UK) Consortium | Harper VanSteenhouse, Yumi Kasai, David Gray, Carol Clugston, Anna Dominiczak and Alex Alderton, Roberto Amato, Jeffrey Barrett, Sonia Goncalves, Ewan Harrison, David K. Jackson, Ian Johnston, Dominic Kwiatkowski, Cordelia Langford, John Sillitoe on behalf of the Wellcome Sanger Institute COVID-19 Surveillance Team |
| EPI_ISL_2237246, EPI_ISL_2237248 | Lighthouse Lab in Alderley Park | Wellcome Sanger Institute for the COVID-19 Genomics UK (COG-UK) Consortium | Jacquelyn Wynn, Mairead Hyland, The Lighthouse Lab in Alderley Park and Alex Alderton, Roberto Amato, Jeffrey Barrett, Sonia Goncalves, Ewan Harrison, David K. Jackson, Ian Johnston, Dominic Kwiatkowski, Cordelia Langford, John Sillitoe on behalf of the Wellcome Sanger Institute COVID-19 Surveillance Team |
| EPI_ISL_2237249 | Lighthouse Lab in Glasgow | Wellcome Sanger Institute for the COVID-19 Genomics UK (COG-UK) Consortium | Harper VanSteenhouse, Yumi Kasai, David Gray, Carol Clugston, Anna Dominiczak and Alex Alderton, Roberto Amato, Jeffrey Barrett, Sonia Goncalves, Ewan Harrison, David K. Jackson, Ian Johnston, Dominic Kwiatkowski, Cordelia Langford, John Sillitoe on behalf of the Wellcome Sanger Institute COVID-19 Surveillance Team |
| EPI_ISL_2237250, EPI_ISL_2237251 | Lighthouse Lab in Alderley Park | Wellcome Sanger Institute for the COVID-19 Genomics UK (COG-UK) Consortium | Jacquelyn Wynn, Mairead Hyland, The Lighthouse Lab in Alderley Park and Alex Alderton, Roberto Amato, Jeffrey Barrett, Sonia Goncalves, Ewan Harrison, David K. Jackson, Ian Johnston, Dominic Kwiatkowski, Cordelia Langford, John Sillitoe on behalf of the Wellcome Sanger Institute COVID-19 Surveillance Team |
| EPI_ISL_2237253 | Lighthouse Lab in Glasgow | Wellcome Sanger Institute for the COVID-19 Genomics UK (COG-UK) Consortium | Harper VanSteenhouse, Yumi Kasai, David Gray, Carol Clugston, Anna Dominiczak and Alex Alderton, Roberto Amato, Jeffrey Barrett, Sonia Goncalves, Ewan Harrison, David K. Jackson, Ian Johnston, Dominic Kwiatkowski, Cordelia Langford, John Sillitoe on behalf of the Wellcome Sanger Institute COVID-19 Surveillance Team |
| EPI_ISL_2237255, EPI_ISL_2237258, EPI_ISL_2237259, EPI_ISL_2237260, EPI_ISL_2237261 | Lighthouse Lab in Alderley Park | Wellcome Sanger Institute for the COVID-19 Genomics UK (COG-UK) Consortium | Jacquelyn Wynn, Mairead Hyland, The Lighthouse Lab in Alderley Park and Alex Alderton, Roberto Amato, Jeffrey Barrett, Sonia Goncalves, Ewan Harrison, David K. Jackson, Ian Johnston, Dominic Kwiatkowski, Cordelia Langford, John Sillitoe on behalf of the Wellcome Sanger Institute COVID-19 Surveillance Team |
| EPI_ISL_2237262 | Lighthouse Lab in Glasgow | Wellcome Sanger Institute for the COVID-19 Genomics UK (COG-UK) Consortium | Harper VanSteenhouse, Yumi Kasai, David Gray, Carol Clugston, Anna Dominiczak and Alex Alderton, Roberto Amato, Jeffrey Barrett, Sonia Goncalves, Ewan Harrison, David K. Jackson, Ian Johnston, Dominic Kwiatkowski, Cordelia Langford, John Sillitoe on behalf of the Wellcome Sanger Institute COVID-19 Surveillance Team |
| EPI_ISL_2237263, EPI_ISL_2237264, EPI_ISL_2237267, EPI_ISL_2237268, EPI_ISL_2237275, EPI_ISL_2237277, EPI_ISL_2237278, EPI_ISL_2237280 | Lighthouse Lab in Alderley Park | Wellcome Sanger Institute for the COVID-19 Genomics UK (COG-UK) Consortium | Jacquelyn Wynn, Mairead Hyland, The Lighthouse Lab in Alderley Park and Alex Alderton, Roberto Amato, Jeffrey Barrett, Sonia Goncalves, Ewan Harrison, David K. Jackson, Ian Johnston, Dominic Kwiatkowski, Cordelia Langford, John Sillitoe on behalf of the Wellcome Sanger Institute COVID-19 Surveillance Team |
| EPI_ISL_2237282, EPI_ISL_2237283, EPI_ISL_2237284, EPI_ISL_2237286 | Lighthouse Lab in Glasgow | Wellcome Sanger Institute for the COVID-19 Genomics UK (COG-UK) Consortium | Harper VanSteenhouse, Yumi Kasai, David Gray, Carol Clugston, Anna Dominiczak and Alex Alderton, Roberto Amato, Jeffrey Barrett, Sonia Goncalves, Ewan Harrison, David K. Jackson, Ian Johnston, Dominic Kwiatkowski, Cordelia Langford, John Sillitoe on behalf of the Wellcome Sanger Institute COVID-19 Surveillance Team |
| EPI_ISL_2237287, EPI_ISL_2237292, EPI_ISL_2237294, EPI_ISL_2237295, EPI_ISL_2237296, EPI_ISL_2237298, EPI_ISL_2237299, EPI_ISL_2237300 | Lighthouse Lab in Alderley Park | Wellcome Sanger Institute for the COVID-19 Genomics UK (COG-UK) Consortium | Jacquelyn Wynn, Mairead Hyland, The Lighthouse Lab in Alderley Park and Alex Alderton, Roberto Amato, Jeffrey Barrett, Sonia Goncalves, Ewan Harrison, David K. Jackson, Ian Johnston, Dominic Kwiatkowski, Cordelia Langford, John Sillitoe on behalf of the Wellcome Sanger Institute COVID-19 Surveillance Team |
| EPI_ISL_2237302 | Lighthouse Lab in Glasgow | Wellcome Sanger Institute for the COVID-19 Genomics UK (COG-UK) Consortium | Harper VanSteenhouse, Yumi Kasai, David Gray, Carol Clugston, Anna Dominiczak and Alex Alderton, Roberto Amato, Jeffrey Barrett, Sonia Goncalves, Ewan Harrison, David K. Jackson, Ian Johnston, Dominic Kwiatkowski, Cordelia Langford, John Sillitoe on behalf of the Wellcome Sanger Institute COVID-19 Surveillance Team |
| EPI_ISL_2237303, EPI_ISL_2237305, EPI_ISL_2237306, EPI_ISL_2237307, EPI_ISL_2237308 | Lighthouse Lab in Alderley Park | Wellcome Sanger Institute for the COVID-19 Genomics UK (COG-UK) Consortium | Jacquelyn Wynn, Mairead Hyland, The Lighthouse Lab in Alderley Park and Alex Alderton, Roberto Amato, Jeffrey Barrett, Sonia Goncalves, Ewan Harrison, David K. Jackson, Ian Johnston, Dominic Kwiatkowski, Cordelia Langford, John Sillitoe on behalf of the Wellcome Sanger Institute COVID-19 Surveillance Team |
| EPI_ISL_2237309, EPI_ISL_2237311 | Lighthouse Lab in Glasgow | Wellcome Sanger Institute for the COVID-19 Genomics UK (COG-UK) Consortium | Harper VanSteenhouse, Yumi Kasai, David Gray, Carol Clugston, Anna Dominiczak and Alex Alderton, Roberto Amato, Jeffrey Barrett, Sonia Goncalves, Ewan Harrison, David K. Jackson, Ian Johnston, Dominic Kwiatkowski, Cordelia Langford, John Sillitoe on behalf of the Wellcome Sanger Institute COVID-19 Surveillance Team |
| EPI_ISL_2237313 | Lighthouse Lab in Alderley Park | Wellcome Sanger Institute for the COVID-19 Genomics UK (COG-UK) Consortium | Jacquelyn Wynn, Mairead Hyland, The Lighthouse Lab in Alderley Park and Alex Alderton, Roberto Amato, Jeffrey Barrett, Sonia Goncalves, Ewan Harrison, David K. Jackson, Ian Johnston, Dominic Kwiatkowski, Cordelia Langford, John Sillitoe on behalf of the Wellcome Sanger Institute COVID-19 Surveillance Team |
| EPI_ISL_2237315 | Lighthouse Lab in Glasgow | Wellcome Sanger Institute for the COVID-19 Genomics UK (COG-UK) Consortium | Harper VanSteenhouse, Yumi Kasai, David Gray, Carol Clugston, Anna Dominiczak and Alex Alderton, Roberto Amato, Jeffrey Barrett, Sonia Goncalves, Ewan Harrison, David K. Jackson, Ian Johnston, Dominic Kwiatkowski, Cordelia Langford, John Sillitoe on behalf of the Wellcome Sanger Institute COVID-19 Surveillance Team |
| EPI_ISL_2237317, EPI_ISL_2237320 | Lighthouse Lab in Alderley Park | Wellcome Sanger Institute for the COVID-19 Genomics UK (COG-UK) Consortium | Jacquelyn Wynn, Mairead Hyland, The Lighthouse Lab in Alderley Park and Alex Alderton, Roberto Amato, Jeffrey Barrett, Sonia Goncalves, Ewan Harrison, David K. Jackson, Ian Johnston, Dominic Kwiatkowski, Cordelia Langford, John Sillitoe on behalf of the Wellcome Sanger Institute COVID-19 Surveillance Team |
| EPI_ISL_2237321 | Lighthouse Lab in Glasgow | Wellcome Sanger Institute for the COVID-19 Genomics UK (COG-UK) Consortium | Harper VanSteenhouse, Yumi Kasai, David Gray, Carol Clugston, Anna Dominiczak and Alex Alderton, Roberto Amato, Jeffrey Barrett, Sonia Goncalves, Ewan Harrison, David K. Jackson, Ian Johnston, Dominic Kwiatkowski, Cordelia Langford, John Sillitoe on behalf of the Wellcome Sanger Institute COVID-19 Surveillance Team |
| EPI_ISL_2237327, EPI_ISL_2237331, EPI_ISL_2237337, EPI_ISL_2237339, EPI_ISL_2237342 | Lighthouse Lab in Alderley Park | Wellcome Sanger Institute for the COVID-19 Genomics UK (COG-UK) Consortium | Jacquelyn Wynn, Mairead Hyland, The Lighthouse Lab in Alderley Park and Alex Alderton, Roberto Amato, Jeffrey Barrett, Sonia Goncalves, Ewan Harrison, David K. Jackson, Ian Johnston, Dominic Kwiatkowski, Cordelia Langford, John Sillitoe on behalf of the Wellcome Sanger Institute COVID-19 Surveillance Team |
| EPI_ISL_2237344, EPI_ISL_2237345, EPI_ISL_2237346 | Lighthouse Lab in Glasgow | Wellcome Sanger Institute for the COVID-19 Genomics UK (COG-UK) Consortium | Harper VanSteenhouse, Yumi Kasai, David Gray, Carol Clugston, Anna Dominiczak and Alex Alderton, Roberto Amato, Jeffrey Barrett, Sonia Goncalves, Ewan Harrison, David K. Jackson, Ian Johnston, Dominic Kwiatkowski, Cordelia Langford, John Sillitoe on behalf of the Wellcome Sanger Institute COVID-19 Surveillance Team |
| EPI_ISL_2237347 | Lighthouse Lab in Alderley Park | Wellcome Sanger Institute for the COVID-19 Genomics UK (COG-UK) Consortium | Jacquelyn Wynn, Mairead Hyland, The Lighthouse Lab in Alderley Park and Alex Alderton, Roberto Amato, Jeffrey Barrett, Sonia Goncalves, Ewan Harrison, David K. Jackson, Ian Johnston, Dominic Kwiatkowski, Cordelia Langford, John Sillitoe on behalf of the Wellcome Sanger Institute COVID-19 Surveillance Team |
| EPI_ISL_2237349, EPI_ISL_2237352 | Lighthouse Lab in Glasgow | Wellcome Sanger Institute for the COVID-19 Genomics UK (COG-UK) Consortium | Harper VanSteenhouse, Yumi Kasai, David Gray, Carol Clugston, Anna Dominiczak and Alex Alderton, Roberto Amato, Jeffrey Barrett, Sonia Goncalves, Ewan Harrison, David K. Jackson, Ian Johnston, Dominic Kwiatkowski, Cordelia Langford, John Sillitoe on behalf of the Wellcome Sanger Institute COVID-19 Surveillance Team |
| EPI_ISL_2237353, EPI_ISL_2237354, EPI_ISL_2237357, EPI_ISL_2237358, EPI_ISL_2237360, EPI_ISL_2237361, EPI_ISL_2237362, EPI_ISL_2237368 | Lighthouse Lab in Alderley Park | Wellcome Sanger Institute for the COVID-19 Genomics UK (COG-UK) Consortium | Jacquelyn Wynn, Mairead Hyland, The Lighthouse Lab in Alderley Park and Alex Alderton, Roberto Amato, Jeffrey Barrett, Sonia Goncalves, Ewan Harrison, David K. Jackson, Ian Johnston, Dominic Kwiatkowski, Cordelia Langford, John Sillitoe on behalf of the Wellcome Sanger Institute COVID-19 Surveillance Team |
| EPI_ISL_2237369, EPI_ISL_2237370 | Lighthouse Lab in Glasgow | Wellcome Sanger Institute for the COVID-19 Genomics UK (COG-UK) Consortium | Harper VanSteenhouse, Yumi Kasai, David Gray, Carol Clugston, Anna Dominiczak and Alex Alderton, Roberto Amato, Jeffrey Barrett, Sonia Goncalves, Ewan Harrison, David K. Jackson, Ian Johnston, Dominic Kwiatkowski, Cordelia Langford, John Sillitoe on behalf of the Wellcome Sanger Institute COVID-19 Surveillance Team |
| EPI_ISL_2237371, EPI_ISL_2237373 | Lighthouse Lab in Alderley Park | Wellcome Sanger Institute for the COVID-19 Genomics UK (COG-UK) Consortium | Jacquelyn Wynn, Mairead Hyland, The Lighthouse Lab in Alderley Park and Alex Alderton, Roberto Amato, Jeffrey Barrett, Sonia Goncalves, Ewan Harrison, David K. Jackson, Ian Johnston, Dominic Kwiatkowski, Cordelia Langford, John Sillitoe on behalf of the Wellcome Sanger Institute COVID-19 Surveillance Team |
| EPI_ISL_2237376 | Lighthouse Lab in Glasgow | Wellcome Sanger Institute for the COVID-19 Genomics UK (COG-UK) Consortium | Harper VanSteenhouse, Yumi Kasai, David Gray, Carol Clugston, Anna Dominiczak and Alex Alderton, Roberto Amato, Jeffrey Barrett, Sonia Goncalves, Ewan Harrison, David K. Jackson, Ian Johnston, Dominic Kwiatkowski, Cordelia Langford, John Sillitoe on behalf of the Wellcome Sanger Institute COVID-19 Surveillance Team |

| | | | |
|---|---|---|---|
| EPI_ISL_2237377, EPI_ISL_2237379 | Lighthouse Lab in Alderley Park | Wellcome Sanger Institute for the COVID-19 Genomics UK (COG-UK) Consortium | Jacquelyn Wynn, Mairead Hyland, The Lighthouse Lab in Alderley Park and Alex Alderton, Roberto Amato, Jeffrey Barrett, Sonia Goncalves, Ewan Harrison, David K. Jackson, Ian Johnston, Dominic Kwiatkowski, Cordelia Langford, John Sillitoe on behalf of the Wellcome Sanger Institute COVID-19 Surveillance Team |
| EPI_ISL_2237380, EPI_ISL_2237382, EPI_ISL_2237385 | Lighthouse Lab in Glasgow | Wellcome Sanger Institute for the COVID-19 Genomics UK (COG-UK) Consortium | Harper VanSteenhouse, Yumi Kasai, David Gray, Carol Clugston, Anna Dominiczak and Alex Alderton, Roberto Amato, Jeffrey Barrett, Sonia Goncalves, Ewan Harrison, David K. Jackson, Ian Johnston, Dominic Kwiatkowski, Cordelia Langford, John Sillitoe on behalf of the Wellcome Sanger Institute COVID-19 Surveillance Team |
| EPI_ISL_2237388, EPI_ISL_2237390 | Lighthouse Lab in Alderley Park | Wellcome Sanger Institute for the COVID-19 Genomics UK (COG-UK) Consortium | Jacquelyn Wynn, Mairead Hyland, The Lighthouse Lab in Alderley Park and Alex Alderton, Roberto Amato, Jeffrey Barrett, Sonia Goncalves, Ewan Harrison, David K. Jackson, Ian Johnston, Dominic Kwiatkowski, Cordelia Langford, John Sillitoe on behalf of the Wellcome Sanger Institute COVID-19 Surveillance Team |
| EPI_ISL_2237391 | Lighthouse Lab in Glasgow | Wellcome Sanger Institute for the COVID-19 Genomics UK (COG-UK) Consortium | Harper VanSteenhouse, Yumi Kasai, David Gray, Carol Clugston, Anna Dominiczak and Alex Alderton, Roberto Amato, Jeffrey Barrett, Sonia Goncalves, Ewan Harrison, David K. Jackson, Ian Johnston, Dominic Kwiatkowski, Cordelia Langford, John Sillitoe on behalf of the Wellcome Sanger Institute COVID-19 Surveillance Team |
| EPI_ISL_2237393, EPI_ISL_2237394, EPI_ISL_2237395, EPI_ISL_2237396, EPI_ISL_2237398 | Lighthouse Lab in Alderley Park | Wellcome Sanger Institute for the COVID-19 Genomics UK (COG-UK) Consortium | Jacquelyn Wynn, Mairead Hyland, The Lighthouse Lab in Alderley Park and Alex Alderton, Roberto Amato, Jeffrey Barrett, Sonia Goncalves, Ewan Harrison, David K. Jackson, Ian Johnston, Dominic Kwiatkowski, Cordelia Langford, John Sillitoe on behalf of the Wellcome Sanger Institute COVID-19 Surveillance Team |
| EPI_ISL_2237399, EPI_ISL_2237400 | Lighthouse Lab in Glasgow | Wellcome Sanger Institute for the COVID-19 Genomics UK (COG-UK) Consortium | Harper VanSteenhouse, Yumi Kasai, David Gray, Carol Clugston, Anna Dominiczak and Alex Alderton, Roberto Amato, Jeffrey Barrett, Sonia Goncalves, Ewan Harrison, David K. Jackson, Ian Johnston, Dominic Kwiatkowski, Cordelia Langford, John Sillitoe on behalf of the Wellcome Sanger Institute COVID-19 Surveillance Team |
| EPI_ISL_2237404 | Lighthouse Lab in Alderley Park | Wellcome Sanger Institute for the COVID-19 Genomics UK (COG-UK) Consortium | Jacquelyn Wynn, Mairead Hyland, The Lighthouse Lab in Alderley Park and Alex Alderton, Roberto Amato, Jeffrey Barrett, Sonia Goncalves, Ewan Harrison, David K. Jackson, Ian Johnston, Dominic Kwiatkowski, Cordelia Langford, John Sillitoe on behalf of the Wellcome Sanger Institute COVID-19 Surveillance Team |
| EPI_ISL_2237405, EPI_ISL_2237406 | Lighthouse Lab in Glasgow | Wellcome Sanger Institute for the COVID-19 Genomics UK (COG-UK) Consortium | Harper VanSteenhouse, Yumi Kasai, David Gray, Carol Clugston, Anna Dominiczak and Alex Alderton, Roberto Amato, Jeffrey Barrett, Sonia Goncalves, Ewan Harrison, David K. Jackson, Ian Johnston, Dominic Kwiatkowski, Cordelia Langford, John Sillitoe on behalf of the Wellcome Sanger Institute COVID-19 Surveillance Team |
| EPI_ISL_2237407, EPI_ISL_2237408, EPI_ISL_2237409 | Lighthouse Lab in Alderley Park | Wellcome Sanger Institute for the COVID-19 Genomics UK (COG-UK) Consortium | Jacquelyn Wynn, Mairead Hyland, The Lighthouse Lab in Alderley Park and Alex Alderton, Roberto Amato, Jeffrey Barrett, Sonia Goncalves, Ewan Harrison, David K. Jackson, Ian Johnston, Dominic Kwiatkowski, Cordelia Langford, John Sillitoe on behalf of the Wellcome Sanger Institute COVID-19 Surveillance Team |
| EPI_ISL_2237410, EPI_ISL_2237413 | Lighthouse Lab in Glasgow | Wellcome Sanger Institute for the COVID-19 Genomics UK (COG-UK) Consortium | Harper VanSteenhouse, Yumi Kasai, David Gray, Carol Clugston, Anna Dominiczak and Alex Alderton, Roberto Amato, Jeffrey Barrett, Sonia Goncalves, Ewan Harrison, David K. Jackson, Ian Johnston, Dominic Kwiatkowski, Cordelia Langford, John Sillitoe on behalf of the Wellcome Sanger Institute COVID-19 Surveillance Team |
| EPI_ISL_2237417 | Lighthouse Lab in Milton Keynes | Wellcome Sanger Institute for the COVID-19 Genomics UK (COG-UK) Consortium | The Lighthouse Lab in Milton Keynes and Alex Alderton, Roberto Amato, Jeffrey Barrett, Sonia Goncalves, Ewan Harrison, David K. Jackson, Ian Johnston, Dominic Kwiatkowski, Cordelia Langford, John Sillitoe on behalf of the Wellcome Sanger Institute COVID-19 Surveillance Team |
| EPI_ISL_2237418 | Lighthouse Lab in Alderley Park | Wellcome Sanger Institute for the COVID-19 Genomics UK (COG-UK) Consortium | Jacquelyn Wynn, Mairead Hyland, The Lighthouse Lab in Alderley Park and Alex Alderton, Roberto Amato, Jeffrey Barrett, Sonia Goncalves, Ewan Harrison, David K. Jackson, Ian Johnston, Dominic Kwiatkowski, Cordelia Langford, John Sillitoe on behalf of the Wellcome Sanger Institute COVID-19 Surveillance Team |
| EPI_ISL_2237419, EPI_ISL_2237425, EPI_ISL_2237426 | Lighthouse Lab in Milton Keynes | Wellcome Sanger Institute for the COVID-19 Genomics UK (COG-UK) Consortium | The Lighthouse Lab in Milton Keynes and Alex Alderton, Roberto Amato, Jeffrey Barrett, Sonia Goncalves, Ewan Harrison, David K. Jackson, Ian Johnston, Dominic Kwiatkowski, Cordelia Langford, John Sillitoe on behalf of the Wellcome Sanger Institute COVID-19 Surveillance Team |
| EPI_ISL_2237435 | Lighthouse Lab in Alderley Park | Wellcome Sanger Institute for the COVID-19 Genomics UK (COG-UK) Consortium | Jacquelyn Wynn, Mairead Hyland, The Lighthouse Lab in Alderley Park and Alex Alderton, Roberto Amato, Jeffrey Barrett, Sonia Goncalves, Ewan Harrison, David K. Jackson, Ian Johnston, Dominic Kwiatkowski, Cordelia Langford, John Sillitoe on behalf of the Wellcome Sanger Institute COVID-19 Surveillance Team |
| EPI_ISL_2237437 | Lighthouse Lab in Glasgow | Wellcome Sanger Institute for the COVID-19 Genomics UK (COG-UK) Consortium | Harper VanSteenhouse, Yumi Kasai, David Gray, Carol Clugston, Anna Dominiczak and Alex Alderton, Roberto Amato, Jeffrey Barrett, Sonia Goncalves, Ewan Harrison, David K. Jackson, Ian Johnston, Dominic Kwiatkowski, Cordelia Langford, John Sillitoe on behalf of the Wellcome Sanger Institute COVID-19 Surveillance Team |
| EPI_ISL_2237438 | Lighthouse Lab in Alderley Park | Wellcome Sanger Institute for the COVID-19 Genomics UK (COG-UK) Consortium | Jacquelyn Wynn, Mairead Hyland, The Lighthouse Lab in Alderley Park and Alex Alderton, Roberto Amato, Jeffrey Barrett, Sonia Goncalves, Ewan Harrison, David K. Jackson, Ian Johnston, Dominic Kwiatkowski, Cordelia Langford, John Sillitoe on behalf of the Wellcome Sanger Institute COVID-19 Surveillance Team |
| EPI_ISL_2237440 | Lighthouse Lab in Glasgow | Wellcome Sanger Institute for the COVID-19 Genomics UK (COG-UK) Consortium | Harper VanSteenhouse, Yumi Kasai, David Gray, Carol Clugston, Anna Dominiczak and Alex Alderton, Roberto Amato, Jeffrey Barrett, Sonia Goncalves, Ewan Harrison, David K. Jackson, Ian Johnston, Dominic Kwiatkowski, Cordelia Langford, John Sillitoe on behalf of the Wellcome Sanger Institute COVID-19 Surveillance Team |
| EPI_ISL_2237442 | Lighthouse Lab in Milton Keynes | Wellcome Sanger Institute for the COVID-19 Genomics UK (COG-UK) Consortium | The Lighthouse Lab in Milton Keynes and Alex Alderton, Roberto Amato, Jeffrey Barrett, Sonia Goncalves, Ewan Harrison, David K. Jackson, Ian Johnston, Dominic Kwiatkowski, Cordelia Langford, John Sillitoe on behalf of the Wellcome Sanger Institute COVID-19 Surveillance Team |
| EPI_ISL_2237443 | Lighthouse Lab in Alderley Park | Wellcome Sanger Institute for the COVID-19 Genomics UK (COG-UK) Consortium | Jacquelyn Wynn, Mairead Hyland, The Lighthouse Lab in Alderley Park and Alex Alderton, Roberto Amato, Jeffrey Barrett, Sonia Goncalves, Ewan Harrison, David K. Jackson, Ian Johnston, Dominic Kwiatkowski, Cordelia Langford, John Sillitoe on behalf of the Wellcome Sanger Institute COVID-19 Surveillance Team |
| EPI_ISL_2237444 | Lighthouse Lab in Milton Keynes | Wellcome Sanger Institute for the COVID-19 Genomics UK (COG-UK) Consortium | The Lighthouse Lab in Milton Keynes and Alex Alderton, Roberto Amato, Jeffrey Barrett, Sonia Goncalves, Ewan Harrison, David K. Jackson, Ian Johnston, Dominic Kwiatkowski, Cordelia Langford, John Sillitoe on behalf of the Wellcome Sanger Institute COVID-19 Surveillance Team |
| EPI_ISL_2237445 | Lighthouse Lab in Glasgow | Wellcome Sanger Institute for the COVID-19 Genomics UK (COG-UK) Consortium | Harper VanSteenhouse, Yumi Kasai, David Gray, Carol Clugston, Anna Dominiczak and Alex Alderton, Roberto Amato, Jeffrey Barrett, Sonia Goncalves, Ewan Harrison, David K. Jackson, Ian Johnston, Dominic Kwiatkowski, Cordelia Langford, John Sillitoe on behalf of the Wellcome Sanger Institute COVID-19 Surveillance Team |
| EPI_ISL_2237446 | Lighthouse Lab in Alderley Park | Wellcome Sanger Institute for the COVID-19 Genomics UK (COG-UK) Consortium | Jacquelyn Wynn, Mairead Hyland, The Lighthouse Lab in Alderley Park and Alex Alderton, Roberto Amato, Jeffrey Barrett, Sonia Goncalves, Ewan Harrison, David K. Jackson, Ian Johnston, Dominic Kwiatkowski, Cordelia Langford, John Sillitoe on behalf of the Wellcome Sanger Institute COVID-19 Surveillance Team |
| EPI_ISL_2237447 | Lighthouse Lab in Glasgow | Wellcome Sanger Institute for the COVID-19 Genomics UK (COG-UK) Consortium | Harper VanSteenhouse, Yumi Kasai, David Gray, Carol Clugston, Anna Dominiczak and Alex Alderton, Roberto Amato, Jeffrey Barrett, Sonia Goncalves, Ewan Harrison, David K. Jackson, Ian Johnston, Dominic Kwiatkowski, Cordelia Langford, John Sillitoe on behalf of the Wellcome Sanger Institute COVID-19 Surveillance Team |
| EPI_ISL_2237448 | Lighthouse Lab in Alderley Park | Wellcome Sanger Institute for the COVID-19 Genomics UK (COG-UK) Consortium | Jacquelyn Wynn, Mairead Hyland, The Lighthouse Lab in Alderley Park and Alex Alderton, Roberto Amato, Jeffrey Barrett, Sonia Goncalves, Ewan Harrison, David K. Jackson, Ian Johnston, Dominic Kwiatkowski, Cordelia Langford, John Sillitoe on behalf of the Wellcome Sanger Institute COVID-19 Surveillance Team |
| EPI_ISL_2237450 | Lighthouse Lab in Milton Keynes | Wellcome Sanger Institute for the COVID-19 Genomics UK (COG-UK) Consortium | The Lighthouse Lab in Milton Keynes and Alex Alderton, Roberto Amato, Jeffrey Barrett, Sonia Goncalves, Ewan Harrison, David K. Jackson, Ian Johnston, Dominic Kwiatkowski, Cordelia Langford, John Sillitoe on behalf of the Wellcome Sanger Institute COVID-19 Surveillance Team |
| EPI_ISL_2237454 | Lighthouse Lab in Alderley Park | Wellcome Sanger Institute for the COVID-19 Genomics UK (COG-UK) Consortium | Jacquelyn Wynn, Mairead Hyland, The Lighthouse Lab in Alderley Park and Alex Alderton, Roberto Amato, Jeffrey Barrett, Sonia Goncalves, Ewan Harrison, David K. Jackson, Ian Johnston, Dominic Kwiatkowski, Cordelia Langford, John Sillitoe on behalf of the Wellcome Sanger Institute COVID-19 Surveillance Team |
| EPI_ISL_2237455 | Lighthouse Lab in Glasgow | Wellcome Sanger Institute for the COVID-19 Genomics UK (COG-UK) Consortium | Harper VanSteenhouse, Yumi Kasai, David Gray, Carol Clugston, Anna Dominiczak and Alex Alderton, Roberto Amato, Jeffrey Barrett, Sonia Goncalves, Ewan Harrison, David K. Jackson, Ian Johnston, Dominic Kwiatkowski, Cordelia Langford, John Sillitoe on behalf of the Wellcome Sanger Institute COVID-19 Surveillance Team |
| EPI_ISL_2237456 | Lighthouse Lab in Milton Keynes | Wellcome Sanger Institute for the COVID-19 Genomics UK (COG-UK) Consortium | The Lighthouse Lab in Milton Keynes and Alex Alderton, Roberto Amato, Jeffrey Barrett, Sonia Goncalves, Ewan Harrison, David K. Jackson, Ian Johnston, Dominic Kwiatkowski, Cordelia Langford, John Sillitoe on behalf of the Wellcome Sanger Institute COVID-19 Surveillance Team |
| EPI_ISL_2237461, EPI_ISL_2237465 | Lighthouse Lab in Alderley Park | Wellcome Sanger Institute for the COVID-19 Genomics UK (COG-UK) Consortium | Jacquelyn Wynn, Mairead Hyland, The Lighthouse Lab in Alderley Park and Alex Alderton, Roberto Amato, Jeffrey Barrett, Sonia Goncalves, Ewan Harrison, David K. Jackson, Ian Johnston, Dominic Kwiatkowski, Cordelia Langford, John Sillitoe on behalf of the Wellcome Sanger Institute COVID-19 Surveillance Team |
| EPI_ISL_2237466, EPI_ISL_2237468 | Lighthouse Lab in Glasgow | Wellcome Sanger Institute for the COVID-19 Genomics UK (COG-UK) Consortium | Harper VanSteenhouse, Yumi Kasai, David Gray, Carol Clugston, Anna Dominiczak and Alex Alderton, Roberto Amato, Jeffrey Barrett, Sonia Goncalves, Ewan Harrison, David K. Jackson, Ian Johnston, Dominic Kwiatkowski, Cordelia Langford, John Sillitoe on behalf of the Wellcome Sanger Institute COVID-19 Surveillance Team |
| EPI_ISL_2237469 | Lighthouse Lab in Alderley Park | Wellcome Sanger Institute for the COVID-19 Genomics UK (COG-UK) Consortium | Jacquelyn Wynn, Mairead Hyland, The Lighthouse Lab in Alderley Park and Alex Alderton, Roberto Amato, Jeffrey Barrett, Sonia Goncalves, Ewan Harrison, David K. Jackson, Ian Johnston, Dominic Kwiatkowski, Cordelia Langford, John Sillitoe on behalf of the Wellcome Sanger Institute COVID-19 Surveillance Team |
| EPI_ISL_2237470, EPI_ISL_2237474, EPI_ISL_2237478 | Lighthouse Lab in Milton Keynes | Wellcome Sanger Institute for the COVID-19 Genomics UK (COG-UK) Consortium | The Lighthouse Lab in Milton Keynes and Alex Alderton, Roberto Amato, Jeffrey Barrett, Sonia Goncalves, Ewan Harrison, David K. Jackson, Ian Johnston, Dominic Kwiatkowski, Cordelia Langford, John Sillitoe on behalf of the Wellcome Sanger Institute COVID-19 Surveillance Team |
| EPI_ISL_2237483 | Lighthouse Lab in Glasgow | Wellcome Sanger Institute for the COVID-19 Genomics UK (COG-UK) Consortium | Harper VanSteenhouse, Yumi Kasai, David Gray, Carol Clugston, Anna Dominiczak and Alex Alderton, Roberto Amato, Jeffrey Barrett, Sonia Goncalves, Ewan Harrison, David K. Jackson, Ian Johnston, Dominic Kwiatkowski, Cordelia Langford, John Sillitoe on behalf of the Wellcome Sanger Institute COVID-19 Surveillance Team |
| EPI_ISL_2237484 | Lighthouse Lab in Milton Keynes | Wellcome Sanger Institute for the COVID-19 Genomics UK (COG-UK) Consortium | The Lighthouse Lab in Milton Keynes and Alex Alderton, Roberto Amato, Jeffrey Barrett, Sonia Goncalves, Ewan Harrison, David K. Jackson, Ian Johnston, Dominic Kwiatkowski, Cordelia Langford, John Sillitoe on behalf of the Wellcome Sanger Institute COVID-19 Surveillance Team |
| EPI_ISL_2237486, EPI_ISL_2237487 | Lighthouse Lab in Alderley Park | Wellcome Sanger Institute for the COVID-19 Genomics UK (COG-UK) Consortium | Jacquelyn Wynn, Mairead Hyland, The Lighthouse Lab in Alderley Park and Alex Alderton, Roberto Amato, Jeffrey Barrett, Sonia Goncalves, Ewan Harrison, David K. Jackson, Ian Johnston, Dominic Kwiatkowski, Cordelia Langford, John Sillitoe on behalf of the Wellcome Sanger Institute COVID-19 Surveillance Team |
| EPI_ISL_2237490, EPI_ISL_2237492 | Lighthouse Lab in Milton Keynes | Wellcome Sanger Institute for the COVID-19 Genomics UK (COG-UK) Consortium | The Lighthouse Lab in Milton Keynes and Alex Alderton, Roberto Amato, Jeffrey Barrett, Sonia Goncalves, Ewan Harrison, David K. Jackson, Ian Johnston, Dominic Kwiatkowski, Cordelia Langford, John Sillitoe on behalf of the Wellcome Sanger Institute COVID-19 Surveillance Team |
| EPI_ISL_2237494, EPI_ISL_2237500 | Lighthouse Lab in Glasgow | Wellcome Sanger Institute for the COVID-19 Genomics UK (COG-UK) Consortium | Harper VanSteenhouse, Yumi Kasai, David Gray, Carol Clugston, Anna Dominiczak and Alex Alderton, Roberto Amato, Jeffrey Barrett, Sonia Goncalves, Ewan Harrison, David K. Jackson, Ian Johnston, Dominic Kwiatkowski, Cordelia Langford, John Sillitoe on behalf of the Wellcome Sanger Institute COVID-19 Surveillance Team |
| EPI_ISL_2237502 | Lighthouse Lab in Milton Keynes | Wellcome Sanger Institute for the COVID-19 Genomics UK (COG-UK) Consortium | The Lighthouse Lab in Milton Keynes and Alex Alderton, Roberto Amato, Jeffrey Barrett, Sonia Goncalves, Ewan Harrison, David K. Jackson, Ian Johnston, Dominic Kwiatkowski, Cordelia Langford, John Sillitoe on behalf of the Wellcome Sanger Institute COVID-19 Surveillance Team |
| EPI_ISL_2237503, EPI_ISL_2237504 | Lighthouse Lab in Alderley Park | Wellcome Sanger Institute for the COVID-19 Genomics UK (COG-UK) Consortium | Jacquelyn Wynn, Mairead Hyland, The Lighthouse Lab in Alderley Park and Alex Alderton, Roberto Amato, Jeffrey Barrett, Sonia Goncalves, Ewan Harrison, David K. Jackson, Ian Johnston, Dominic Kwiatkowski, Cordelia Langford, John Sillitoe on behalf of the Wellcome Sanger Institute COVID-19 Surveillance Team |
| EPI_ISL_2237508 | Lighthouse Lab in Milton Keynes | Wellcome Sanger Institute for the COVID-19 Genomics UK (COG-UK) Consortium | The Lighthouse Lab in Milton Keynes and Alex Alderton, Roberto Amato, Jeffrey Barrett, Sonia Goncalves, Ewan Harrison, David K. Jackson, Ian Johnston, Dominic Kwiatkowski, Cordelia Langford, John Sillitoe on behalf of the Wellcome Sanger Institute COVID-19 Surveillance Team |
| EPI_ISL_2237509 | Lighthouse Lab in Glasgow | Wellcome Sanger Institute for the COVID-19 Genomics UK (COG-UK) Consortium | Harper VanSteenhouse, Yumi Kasai, David Gray, Carol Clugston, Anna Dominiczak and Alex Alderton, Roberto Amato, Jeffrey Barrett, Sonia Goncalves, Ewan Harrison, David K. Jackson, Ian Johnston, Dominic Kwiatkowski, Cordelia Langford, John Sillitoe on behalf of the Wellcome Sanger Institute COVID-19 Surveillance Team |
| EPI_ISL_2237511 | Lighthouse Lab in Milton Keynes | Wellcome Sanger Institute for the COVID-19 Genomics UK (COG-UK) Consortium | The Lighthouse Lab in Milton Keynes and Alex Alderton, Roberto Amato, Jeffrey Barrett, Sonia Goncalves, Ewan Harrison, David K. Jackson, Ian Johnston, Dominic Kwiatkowski, Cordelia Langford, John Sillitoe on behalf of the Wellcome Sanger Institute COVID-19 Surveillance Team |
| EPI_ISL_2237515 | Lighthouse Lab in Alderley Park | Wellcome Sanger Institute for the COVID-19 Genomics UK (COG-UK) | Jacquelyn Wynn, Mairead Hyland, The Lighthouse Lab in Alderley Park and Alex Alderton, Roberto Amato, Jeffrey Barrett, Sonia Goncalves, Ewan Harrison, David K. Jackson, Ian Johnston, Dominic Kwiatkowski, Cordelia Langford, John Sillitoe on |

| | | | |
|---|---|---|---|
| | | Consortium | behalf of the Wellcome Sanger Institute COVID-19 Surveillance Team |
| EPI_ISL_2237516, EPI_ISL_2237517 | Lighthouse Lab in Milton Keynes | Wellcome Sanger Institute for the COVID-19 Genomics UK (COG-UK) Consortium | The Lighthouse Lab in Milton Keynes and Alex Alderton, Roberto Amato, Jeffrey Barrett, Sonia Goncalves, Ewan Harrison, David K. Jackson, Ian Johnston, Dominic Kwiatkowski, Cordelia Langford, John Sillitoe on behalf of the Wellcome Sanger Institute COVID-19 Surveillance Team |
| EPI_ISL_2237518, EPI_ISL_2237519 | Lighthouse Lab in Alderley Park | Wellcome Sanger Institute for the COVID-19 Genomics UK (COG-UK) Consortium | Jacquelyn Wynn, Mairead Hyland, The Lighthouse Lab in Alderley Park and Alex Alderton, Roberto Amato, Jeffrey Barrett, Sonia Goncalves, Ewan Harrison, David K. Jackson, Ian Johnston, Dominic Kwiatkowski, Cordelia Langford, John Sillitoe on behalf of the Wellcome Sanger Institute COVID-19 Surveillance Team |
| EPI_ISL_2237523, EPI_ISL_2237524, EPI_ISL_2237527 | Lighthouse Lab in Milton Keynes | Wellcome Sanger Institute for the COVID-19 Genomics UK (COG-UK) Consortium | The Lighthouse Lab in Milton Keynes and Alex Alderton, Roberto Amato, Jeffrey Barrett, Sonia Goncalves, Ewan Harrison, David K. Jackson, Ian Johnston, Dominic Kwiatkowski, Cordelia Langford, John Sillitoe on behalf of the Wellcome Sanger Institute COVID-19 Surveillance Team |
| EPI_ISL_2237529 | Lighthouse Lab in Glasgow | Wellcome Sanger Institute for the COVID-19 Genomics UK (COG-UK) Consortium | Harper VanSteenhouse, Yumi Kasai, David Gray, Carol Clugston, Anna Dominiczak and Alex Alderton, Roberto Amato, Jeffrey Barrett, Sonia Goncalves, Ewan Harrison, David K. Jackson, Ian Johnston, Dominic Kwiatkowski, Cordelia Langford, John Sillitoe on behalf of the Wellcome Sanger Institute COVID-19 Surveillance Team |
| EPI_ISL_2237530 | Lighthouse Lab in Alderley Park | Wellcome Sanger Institute for the COVID-19 Genomics UK (COG-UK) Consortium | Jacquelyn Wynn, Mairead Hyland, The Lighthouse Lab in Alderley Park and Alex Alderton, Roberto Amato, Jeffrey Barrett, Sonia Goncalves, Ewan Harrison, David K. Jackson, Ian Johnston, Dominic Kwiatkowski, Cordelia Langford, John Sillitoe on behalf of the Wellcome Sanger Institute COVID-19 Surveillance Team |
| EPI_ISL_2237531 | Lighthouse Lab in Milton Keynes | Wellcome Sanger Institute for the COVID-19 Genomics UK (COG-UK) Consortium | The Lighthouse Lab in Milton Keynes and Alex Alderton, Roberto Amato, Jeffrey Barrett, Sonia Goncalves, Ewan Harrison, David K. Jackson, Ian Johnston, Dominic Kwiatkowski, Cordelia Langford, John Sillitoe on behalf of the Wellcome Sanger Institute COVID-19 Surveillance Team |
| EPI_ISL_2237532 | Lighthouse Lab in Glasgow | Wellcome Sanger Institute for the COVID-19 Genomics UK (COG-UK) Consortium | Harper VanSteenhouse, Yumi Kasai, David Gray, Carol Clugston, Anna Dominiczak and Alex Alderton, Roberto Amato, Jeffrey Barrett, Sonia Goncalves, Ewan Harrison, David K. Jackson, Ian Johnston, Dominic Kwiatkowski, Cordelia Langford, John Sillitoe on behalf of the Wellcome Sanger Institute COVID-19 Surveillance Team |
| EPI_ISL_2237534, EPI_ISL_2237535, EPI_ISL_2237536 | Lighthouse Lab in Milton Keynes | Wellcome Sanger Institute for the COVID-19 Genomics UK (COG-UK) Consortium | The Lighthouse Lab in Milton Keynes and Alex Alderton, Roberto Amato, Jeffrey Barrett, Sonia Goncalves, Ewan Harrison, David K. Jackson, Ian Johnston, Dominic Kwiatkowski, Cordelia Langford, John Sillitoe on behalf of the Wellcome Sanger Institute COVID-19 Surveillance Team |
| EPI_ISL_2237537 | Lighthouse Lab in Alderley Park | Wellcome Sanger Institute for the COVID-19 Genomics UK (COG-UK) Consortium | Jacquelyn Wynn, Mairead Hyland, The Lighthouse Lab in Alderley Park and Alex Alderton, Roberto Amato, Jeffrey Barrett, Sonia Goncalves, Ewan Harrison, David K. Jackson, Ian Johnston, Dominic Kwiatkowski, Cordelia Langford, John Sillitoe on behalf of the Wellcome Sanger Institute COVID-19 Surveillance Team |
| EPI_ISL_2237539 | Lighthouse Lab in Glasgow | Wellcome Sanger Institute for the COVID-19 Genomics UK (COG-UK) Consortium | Harper VanSteenhouse, Yumi Kasai, David Gray, Carol Clugston, Anna Dominiczak and Alex Alderton, Roberto Amato, Jeffrey Barrett, Sonia Goncalves, Ewan Harrison, David K. Jackson, Ian Johnston, Dominic Kwiatkowski, Cordelia Langford, John Sillitoe on behalf of the Wellcome Sanger Institute COVID-19 Surveillance Team |
| EPI_ISL_2237540, EPI_ISL_2237542, EPI_ISL_2237543 | Lighthouse Lab in Milton Keynes | Wellcome Sanger Institute for the COVID-19 Genomics UK (COG-UK) Consortium | The Lighthouse Lab in Milton Keynes and Alex Alderton, Roberto Amato, Jeffrey Barrett, Sonia Goncalves, Ewan Harrison, David K. Jackson, Ian Johnston, Dominic Kwiatkowski, Cordelia Langford, John Sillitoe on behalf of the Wellcome Sanger Institute COVID-19 Surveillance Team |
| EPI_ISL_2237546 | Lighthouse Lab in Glasgow | Wellcome Sanger Institute for the COVID-19 Genomics UK (COG-UK) Consortium | Harper VanSteenhouse, Yumi Kasai, David Gray, Carol Clugston, Anna Dominiczak and Alex Alderton, Roberto Amato, Jeffrey Barrett, Sonia Goncalves, Ewan Harrison, David K. Jackson, Ian Johnston, Dominic Kwiatkowski, Cordelia Langford, John Sillitoe on behalf of the Wellcome Sanger Institute COVID-19 Surveillance Team |
| EPI_ISL_2237547, EPI_ISL_2237548, EPI_ISL_2237556 | Lighthouse Lab in Alderley Park | Wellcome Sanger Institute for the COVID-19 Genomics UK (COG-UK) Consortium | Jacquelyn Wynn, Mairead Hyland, The Lighthouse Lab in Alderley Park and Alex Alderton, Roberto Amato, Jeffrey Barrett, Sonia Goncalves, Ewan Harrison, David K. Jackson, Ian Johnston, Dominic Kwiatkowski, Cordelia Langford, John Sillitoe on behalf of the Wellcome Sanger Institute COVID-19 Surveillance Team |
| EPI_ISL_2237558 | Lighthouse Lab in Glasgow | Wellcome Sanger Institute for the COVID-19 Genomics UK (COG-UK) Consortium | Harper VanSteenhouse, Yumi Kasai, David Gray, Carol Clugston, Anna Dominiczak and Alex Alderton, Roberto Amato, Jeffrey Barrett, Sonia Goncalves, Ewan Harrison, David K. Jackson, Ian Johnston, Dominic Kwiatkowski, Cordelia Langford, John Sillitoe on behalf of the Wellcome Sanger Institute COVID-19 Surveillance Team |
| EPI_ISL_2237560 | Lighthouse Lab in Alderley Park | Wellcome Sanger Institute for the COVID-19 Genomics UK (COG-UK) Consortium | Jacquelyn Wynn, Mairead Hyland, The Lighthouse Lab in Alderley Park and Alex Alderton, Roberto Amato, Jeffrey Barrett, Sonia Goncalves, Ewan Harrison, David K. Jackson, Ian Johnston, Dominic Kwiatkowski, Cordelia Langford, John Sillitoe on behalf of the Wellcome Sanger Institute COVID-19 Surveillance Team |
| EPI_ISL_2237562, EPI_ISL_2237564 | Lighthouse Lab in Milton Keynes | Wellcome Sanger Institute for the COVID-19 Genomics UK (COG-UK) Consortium | The Lighthouse Lab in Milton Keynes and Alex Alderton, Roberto Amato, Jeffrey Barrett, Sonia Goncalves, Ewan Harrison, David K. Jackson, Ian Johnston, Dominic Kwiatkowski, Cordelia Langford, John Sillitoe on behalf of the Wellcome Sanger Institute COVID-19 Surveillance Team |
| EPI_ISL_2237567 | Lighthouse Lab in Alderley Park | Wellcome Sanger Institute for the COVID-19 Genomics UK (COG-UK) Consortium | Jacquelyn Wynn, Mairead Hyland, The Lighthouse Lab in Alderley Park and Alex Alderton, Roberto Amato, Jeffrey Barrett, Sonia Goncalves, Ewan Harrison, David K. Jackson, Ian Johnston, Dominic Kwiatkowski, Cordelia Langford, John Sillitoe on behalf of the Wellcome Sanger Institute COVID-19 Surveillance Team |
| EPI_ISL_2237568, EPI_ISL_2237569 | Lighthouse Lab in Milton Keynes | Wellcome Sanger Institute for the COVID-19 Genomics UK (COG-UK) Consortium | The Lighthouse Lab in Milton Keynes and Alex Alderton, Roberto Amato, Jeffrey Barrett, Sonia Goncalves, Ewan Harrison, David K. Jackson, Ian Johnston, Dominic Kwiatkowski, Cordelia Langford, John Sillitoe on behalf of the Wellcome Sanger Institute COVID-19 Surveillance Team |
| EPI_ISL_2237572, EPI_ISL_2237574 | Lighthouse Lab in Alderley Park | Wellcome Sanger Institute for the COVID-19 Genomics UK (COG-UK) Consortium | Jacquelyn Wynn, Mairead Hyland, The Lighthouse Lab in Alderley Park and Alex Alderton, Roberto Amato, Jeffrey Barrett, Sonia Goncalves, Ewan Harrison, David K. Jackson, Ian Johnston, Dominic Kwiatkowski, Cordelia Langford, John Sillitoe on behalf of the Wellcome Sanger Institute COVID-19 Surveillance Team |
| EPI_ISL_2237575 | Lighthouse Lab in Glasgow | Wellcome Sanger Institute for the COVID-19 Genomics UK (COG-UK) Consortium | Harper VanSteenhouse, Yumi Kasai, David Gray, Carol Clugston, Anna Dominiczak and Alex Alderton, Roberto Amato, Jeffrey Barrett, Sonia Goncalves, Ewan Harrison, David K. Jackson, Ian Johnston, Dominic Kwiatkowski, Cordelia Langford, John Sillitoe on behalf of the Wellcome Sanger Institute COVID-19 Surveillance Team |
| EPI_ISL_2237581 | Lighthouse Lab in Milton Keynes | Wellcome Sanger Institute for the COVID-19 Genomics UK (COG-UK) Consortium | The Lighthouse Lab in Milton Keynes and Alex Alderton, Roberto Amato, Jeffrey Barrett, Sonia Goncalves, Ewan Harrison, David K. Jackson, Ian Johnston, Dominic Kwiatkowski, Cordelia Langford, John Sillitoe on behalf of the Wellcome Sanger Institute COVID-19 Surveillance Team |
| EPI_ISL_2237584 | Lighthouse Lab in Alderley Park | Wellcome Sanger Institute for the COVID-19 Genomics UK (COG-UK) Consortium | Jacquelyn Wynn, Mairead Hyland, The Lighthouse Lab in Alderley Park and Alex Alderton, Roberto Amato, Jeffrey Barrett, Sonia Goncalves, Ewan Harrison, David K. Jackson, Ian Johnston, Dominic Kwiatkowski, Cordelia Langford, John Sillitoe on behalf of the Wellcome Sanger Institute COVID-19 Surveillance Team |
| EPI_ISL_2237588, EPI_ISL_2237590, EPI_ISL_2237591, EPI_ISL_2237596 | Lighthouse Lab in Milton Keynes | Wellcome Sanger Institute for the COVID-19 Genomics UK (COG-UK) Consortium | The Lighthouse Lab in Milton Keynes and Alex Alderton, Roberto Amato, Jeffrey Barrett, Sonia Goncalves, Ewan Harrison, David K. Jackson, Ian Johnston, Dominic Kwiatkowski, Cordelia Langford, John Sillitoe on behalf of the Wellcome Sanger Institute COVID-19 Surveillance Team |
| EPI_ISL_2237600, EPI_ISL_2237601, EPI_ISL_2237605, EPI_ISL_2237606 | Lighthouse Lab in Alderley Park | Wellcome Sanger Institute for the COVID-19 Genomics UK (COG-UK) Consortium | Jacquelyn Wynn, Mairead Hyland, The Lighthouse Lab in Alderley Park and Alex Alderton, Roberto Amato, Jeffrey Barrett, Sonia Goncalves, Ewan Harrison, David K. Jackson, Ian Johnston, Dominic Kwiatkowski, Cordelia Langford, John Sillitoe on behalf of the Wellcome Sanger Institute COVID-19 Surveillance Team |
| EPI_ISL_2237607, EPI_ISL_2237608 | Lighthouse Lab in Milton Keynes | Wellcome Sanger Institute for the COVID-19 Genomics UK (COG-UK) Consortium | The Lighthouse Lab in Milton Keynes and Alex Alderton, Roberto Amato, Jeffrey Barrett, Sonia Goncalves, Ewan Harrison, David K. Jackson, Ian Johnston, Dominic Kwiatkowski, Cordelia Langford, John Sillitoe on behalf of the Wellcome Sanger Institute COVID-19 Surveillance Team |
| EPI_ISL_2237610 | Lighthouse Lab in Glasgow | Wellcome Sanger Institute for the COVID-19 Genomics UK (COG-UK) Consortium | Harper VanSteenhouse, Yumi Kasai, David Gray, Carol Clugston, Anna Dominiczak and Alex Alderton, Roberto Amato, Jeffrey Barrett, Sonia Goncalves, Ewan Harrison, David K. Jackson, Ian Johnston, Dominic Kwiatkowski, Cordelia Langford, John Sillitoe on behalf of the Wellcome Sanger Institute COVID-19 Surveillance Team |
| EPI_ISL_2237611 | Lighthouse Lab in Alderley Park | Wellcome Sanger Institute for the COVID-19 Genomics UK (COG-UK) Consortium | Jacquelyn Wynn, Mairead Hyland, The Lighthouse Lab in Alderley Park and Alex Alderton, Roberto Amato, Jeffrey Barrett, Sonia Goncalves, Ewan Harrison, David K. Jackson, Ian Johnston, Dominic Kwiatkowski, Cordelia Langford, John Sillitoe on behalf of the Wellcome Sanger Institute COVID-19 Surveillance Team |
| EPI_ISL_2237613 | Lighthouse Lab in Milton Keynes | Wellcome Sanger Institute for the COVID-19 Genomics UK (COG-UK) Consortium | The Lighthouse Lab in Milton Keynes and Alex Alderton, Roberto Amato, Jeffrey Barrett, Sonia Goncalves, Ewan Harrison, David K. Jackson, Ian Johnston, Dominic Kwiatkowski, Cordelia Langford, John Sillitoe on behalf of the Wellcome Sanger Institute COVID-19 Surveillance Team |
| EPI_ISL_2237614 | Lighthouse Lab in Glasgow | Wellcome Sanger Institute for the COVID-19 Genomics UK (COG-UK) Consortium | Harper VanSteenhouse, Yumi Kasai, David Gray, Carol Clugston, Anna Dominiczak and Alex Alderton, Roberto Amato, Jeffrey Barrett, Sonia Goncalves, Ewan Harrison, David K. Jackson, Ian Johnston, Dominic Kwiatkowski, Cordelia Langford, John Sillitoe on behalf of the Wellcome Sanger Institute COVID-19 Surveillance Team |
| EPI_ISL_2237615 | Lighthouse Lab in Milton Keynes | Wellcome Sanger Institute for the COVID-19 Genomics UK (COG-UK) Consortium | The Lighthouse Lab in Milton Keynes and Alex Alderton, Roberto Amato, Jeffrey Barrett, Sonia Goncalves, Ewan Harrison, David K. Jackson, Ian Johnston, Dominic Kwiatkowski, Cordelia Langford, John Sillitoe on behalf of the Wellcome Sanger Institute COVID-19 Surveillance Team |
| EPI_ISL_2237618 | Lighthouse Lab in Alderley Park | Wellcome Sanger Institute for the COVID-19 Genomics UK (COG-UK) Consortium | Jacquelyn Wynn, Mairead Hyland, The Lighthouse Lab in Alderley Park and Alex Alderton, Roberto Amato, Jeffrey Barrett, Sonia Goncalves, Ewan Harrison, David K. Jackson, Ian Johnston, Dominic Kwiatkowski, Cordelia Langford, John Sillitoe on behalf of the Wellcome Sanger Institute COVID-19 Surveillance Team |
| EPI_ISL_2237622, EPI_ISL_2237623 | Lighthouse Lab in Glasgow | Wellcome Sanger Institute for the COVID-19 Genomics UK (COG-UK) Consortium | Harper VanSteenhouse, Yumi Kasai, David Gray, Carol Clugston, Anna Dominiczak and Alex Alderton, Roberto Amato, Jeffrey Barrett, Sonia Goncalves, Ewan Harrison, David K. Jackson, Ian Johnston, Dominic Kwiatkowski, Cordelia Langford, John Sillitoe on behalf of the Wellcome Sanger Institute COVID-19 Surveillance Team |
| EPI_ISL_2237625, EPI_ISL_2237627, EPI_ISL_2237628 | Lighthouse Lab in Alderley Park | Wellcome Sanger Institute for the COVID-19 Genomics UK (COG-UK) Consortium | Jacquelyn Wynn, Mairead Hyland, The Lighthouse Lab in Alderley Park and Alex Alderton, Roberto Amato, Jeffrey Barrett, Sonia Goncalves, Ewan Harrison, David K. Jackson, Ian Johnston, Dominic Kwiatkowski, Cordelia Langford, John Sillitoe on behalf of the Wellcome Sanger Institute COVID-19 Surveillance Team |
| EPI_ISL_2237631, EPI_ISL_2237633 | Lighthouse Lab in Milton Keynes | Wellcome Sanger Institute for the COVID-19 Genomics UK (COG-UK) Consortium | The Lighthouse Lab in Milton Keynes and Alex Alderton, Roberto Amato, Jeffrey Barrett, Sonia Goncalves, Ewan Harrison, David K. Jackson, Ian Johnston, Dominic Kwiatkowski, Cordelia Langford, John Sillitoe on behalf of the Wellcome Sanger Institute COVID-19 Surveillance Team |
| EPI_ISL_2237635 | Lighthouse Lab in Glasgow | Wellcome Sanger Institute for the COVID-19 Genomics UK (COG-UK) Consortium | Harper VanSteenhouse, Yumi Kasai, David Gray, Carol Clugston, Anna Dominiczak and Alex Alderton, Roberto Amato, Jeffrey Barrett, Sonia Goncalves, Ewan Harrison, David K. Jackson, Ian Johnston, Dominic Kwiatkowski, Cordelia Langford, John Sillitoe on behalf of the Wellcome Sanger Institute COVID-19 Surveillance Team |
| EPI_ISL_2237636, EPI_ISL_2237638, EPI_ISL_2237639, EPI_ISL_2237640 | Lighthouse Lab in Milton Keynes | Wellcome Sanger Institute for the COVID-19 Genomics UK (COG-UK) Consortium | The Lighthouse Lab in Milton Keynes and Alex Alderton, Roberto Amato, Jeffrey Barrett, Sonia Goncalves, Ewan Harrison, David K. Jackson, Ian Johnston, Dominic Kwiatkowski, Cordelia Langford, John Sillitoe on behalf of the Wellcome Sanger Institute COVID-19 Surveillance Team |
| EPI_ISL_2237641 | Lighthouse Lab in Glasgow | Wellcome Sanger Institute for the COVID-19 Genomics UK (COG-UK) Consortium | Harper VanSteenhouse, Yumi Kasai, David Gray, Carol Clugston, Anna Dominiczak and Alex Alderton, Roberto Amato, Jeffrey Barrett, Sonia Goncalves, Ewan Harrison, David K. Jackson, Ian Johnston, Dominic Kwiatkowski, Cordelia Langford, John Sillitoe on behalf of the Wellcome Sanger Institute COVID-19 Surveillance Team |
| EPI_ISL_2237642, EPI_ISL_2237643 | Lighthouse Lab in Milton Keynes | Wellcome Sanger Institute for the COVID-19 Genomics UK (COG-UK) Consortium | The Lighthouse Lab in Milton Keynes and Alex Alderton, Roberto Amato, Jeffrey Barrett, Sonia Goncalves, Ewan Harrison, David K. Jackson, Ian Johnston, Dominic Kwiatkowski, Cordelia Langford, John Sillitoe on behalf of the Wellcome Sanger Institute COVID-19 Surveillance Team |
| EPI_ISL_2237647 | Lighthouse Lab in Glasgow | Wellcome Sanger Institute for the COVID-19 Genomics UK (COG-UK) Consortium | Harper VanSteenhouse, Yumi Kasai, David Gray, Carol Clugston, Anna Dominiczak and Alex Alderton, Roberto Amato, Jeffrey Barrett, Sonia Goncalves, Ewan Harrison, David K. Jackson, Ian Johnston, Dominic Kwiatkowski, Cordelia Langford, John Sillitoe on behalf of the Wellcome Sanger Institute COVID-19 Surveillance Team |
| EPI_ISL_2237648, EPI_ISL_2237651 | Lighthouse Lab in Milton Keynes | Wellcome Sanger Institute for the COVID-19 Genomics UK (COG-UK) Consortium | The Lighthouse Lab in Milton Keynes and Alex Alderton, Roberto Amato, Jeffrey Barrett, Sonia Goncalves, Ewan Harrison, David K. Jackson, Ian Johnston, Dominic Kwiatkowski, Cordelia Langford, John Sillitoe on behalf of the Wellcome Sanger Institute COVID-19 Surveillance Team |

| | | | |
|---|---|---|---|
| EPI_ISL_2237655 | Lighthouse Lab in Glasgow | Wellcome Sanger Institute for the COVID-19 Genomics UK (COG-UK) Consortium | Harper VanSteenhouse, Yumi Kasai, David Gray, Carol Clugston, Anna Dominiczak and Alex Alderton, Roberto Amato, Jeffrey Barrett, Sonia Goncalves, Ewan Harrison, David K. Jackson, Ian Johnston, Dominic Kwiatkowski, Cordelia Langford, John Sillitoe on behalf of the Wellcome Sanger Institute COVID-19 Surveillance Team |
| EPI_ISL_2237656, EPI_ISL_2237657 | Lighthouse Lab in Alderley Park | Wellcome Sanger Institute for the COVID-19 Genomics UK (COG-UK) Consortium | Jacquelyn Wynn, Mairead Hyland, The Lighthouse Lab in Alderley Park and Alex Alderton, Roberto Amato, Jeffrey Barrett, Sonia Goncalves, Ewan Harrison, David K. Jackson, Ian Johnston, Dominic Kwiatkowski, Cordelia Langford, John Sillitoe on behalf of the Wellcome Sanger Institute COVID-19 Surveillance Team |
| EPI_ISL_2237658, EPI_ISL_2237659 | Lighthouse Lab in Milton Keynes | Wellcome Sanger Institute for the COVID-19 Genomics UK (COG-UK) Consortium | The Lighthouse Lab in Milton Keynes and Alex Alderton, Roberto Amato, Jeffrey Barrett, Sonia Goncalves, Ewan Harrison, David K. Jackson, Ian Johnston, Dominic Kwiatkowski, Cordelia Langford, John Sillitoe on behalf of the Wellcome Sanger Institute COVID-19 Surveillance Team |
| EPI_ISL_2237662, EPI_ISL_2237663 | Lighthouse Lab in Alderley Park | Wellcome Sanger Institute for the COVID-19 Genomics UK (COG-UK) Consortium | Jacquelyn Wynn, Mairead Hyland, The Lighthouse Lab in Alderley Park and Alex Alderton, Roberto Amato, Jeffrey Barrett, Sonia Goncalves, Ewan Harrison, David K. Jackson, Ian Johnston, Dominic Kwiatkowski, Cordelia Langford, John Sillitoe on behalf of the Wellcome Sanger Institute COVID-19 Surveillance Team |
| EPI_ISL_2237664 | Lighthouse Lab in Glasgow | Wellcome Sanger Institute for the COVID-19 Genomics UK (COG-UK) Consortium | Harper VanSteenhouse, Yumi Kasai, David Gray, Carol Clugston, Anna Dominiczak and Alex Alderton, Roberto Amato, Jeffrey Barrett, Sonia Goncalves, Ewan Harrison, David K. Jackson, Ian Johnston, Dominic Kwiatkowski, Cordelia Langford, John Sillitoe on behalf of the Wellcome Sanger Institute COVID-19 Surveillance Team |
| EPI_ISL_2237665 | Lighthouse Lab in Alderley Park | Wellcome Sanger Institute for the COVID-19 Genomics UK (COG-UK) Consortium | Jacquelyn Wynn, Mairead Hyland, The Lighthouse Lab in Alderley Park and Alex Alderton, Roberto Amato, Jeffrey Barrett, Sonia Goncalves, Ewan Harrison, David K. Jackson, Ian Johnston, Dominic Kwiatkowski, Cordelia Langford, John Sillitoe on behalf of the Wellcome Sanger Institute COVID-19 Surveillance Team |
| EPI_ISL_2237667 | Lighthouse Lab in Glasgow | Wellcome Sanger Institute for the COVID-19 Genomics UK (COG-UK) Consortium | Harper VanSteenhouse, Yumi Kasai, David Gray, Carol Clugston, Anna Dominiczak and Alex Alderton, Roberto Amato, Jeffrey Barrett, Sonia Goncalves, Ewan Harrison, David K. Jackson, Ian Johnston, Dominic Kwiatkowski, Cordelia Langford, John Sillitoe on behalf of the Wellcome Sanger Institute COVID-19 Surveillance Team |
| EPI_ISL_2237669, EPI_ISL_2237671, EPI_ISL_2237672, EPI_ISL_2237673, EPI_ISL_2237674 | Lighthouse Lab in Milton Keynes | Wellcome Sanger Institute for the COVID-19 Genomics UK (COG-UK) Consortium | The Lighthouse Lab in Milton Keynes and Alex Alderton, Roberto Amato, Jeffrey Barrett, Sonia Goncalves, Ewan Harrison, David K. Jackson, Ian Johnston, Dominic Kwiatkowski, Cordelia Langford, John Sillitoe on behalf of the Wellcome Sanger Institute COVID-19 Surveillance Team |
| EPI_ISL_2237675 | Lighthouse Lab in Glasgow | Wellcome Sanger Institute for the COVID-19 Genomics UK (COG-UK) Consortium | Harper VanSteenhouse, Yumi Kasai, David Gray, Carol Clugston, Anna Dominiczak and Alex Alderton, Roberto Amato, Jeffrey Barrett, Sonia Goncalves, Ewan Harrison, David K. Jackson, Ian Johnston, Dominic Kwiatkowski, Cordelia Langford, John Sillitoe on behalf of the Wellcome Sanger Institute COVID-19 Surveillance Team |
| EPI_ISL_2237676 | Lighthouse Lab in Milton Keynes | Wellcome Sanger Institute for the COVID-19 Genomics UK (COG-UK) Consortium | The Lighthouse Lab in Milton Keynes and Alex Alderton, Roberto Amato, Jeffrey Barrett, Sonia Goncalves, Ewan Harrison, David K. Jackson, Ian Johnston, Dominic Kwiatkowski, Cordelia Langford, John Sillitoe on behalf of the Wellcome Sanger Institute COVID-19 Surveillance Team |
| EPI_ISL_2237678 | Lighthouse Lab in Alderley Park | Wellcome Sanger Institute for the COVID-19 Genomics UK (COG-UK) Consortium | Jacquelyn Wynn, Mairead Hyland, The Lighthouse Lab in Alderley Park and Alex Alderton, Roberto Amato, Jeffrey Barrett, Sonia Goncalves, Ewan Harrison, David K. Jackson, Ian Johnston, Dominic Kwiatkowski, Cordelia Langford, John Sillitoe on behalf of the Wellcome Sanger Institute COVID-19 Surveillance Team |
| EPI_ISL_2237679, EPI_ISL_2237680, EPI_ISL_2237682, EPI_ISL_2237683 | Lighthouse Lab in Milton Keynes | Wellcome Sanger Institute for the COVID-19 Genomics UK (COG-UK) Consortium | The Lighthouse Lab in Milton Keynes and Alex Alderton, Roberto Amato, Jeffrey Barrett, Sonia Goncalves, Ewan Harrison, David K. Jackson, Ian Johnston, Dominic Kwiatkowski, Cordelia Langford, John Sillitoe on behalf of the Wellcome Sanger Institute COVID-19 Surveillance Team |
| EPI_ISL_2237686 | Lighthouse Lab in Glasgow | Wellcome Sanger Institute for the COVID-19 Genomics UK (COG-UK) Consortium | Harper VanSteenhouse, Yumi Kasai, David Gray, Carol Clugston, Anna Dominiczak and Alex Alderton, Roberto Amato, Jeffrey Barrett, Sonia Goncalves, Ewan Harrison, David K. Jackson, Ian Johnston, Dominic Kwiatkowski, Cordelia Langford, John Sillitoe on behalf of the Wellcome Sanger Institute COVID-19 Surveillance Team |
| EPI_ISL_2237688, EPI_ISL_2237689, EPI_ISL_2237690 | Lighthouse Lab in Milton Keynes | Wellcome Sanger Institute for the COVID-19 Genomics UK (COG-UK) Consortium | The Lighthouse Lab in Milton Keynes and Alex Alderton, Roberto Amato, Jeffrey Barrett, Sonia Goncalves, Ewan Harrison, David K. Jackson, Ian Johnston, Dominic Kwiatkowski, Cordelia Langford, John Sillitoe on behalf of the Wellcome Sanger Institute COVID-19 Surveillance Team |
| EPI_ISL_2237692, EPI_ISL_2237694 | Lighthouse Lab in Alderley Park | Wellcome Sanger Institute for the COVID-19 Genomics UK (COG-UK) Consortium | Jacquelyn Wynn, Mairead Hyland, The Lighthouse Lab in Alderley Park and Alex Alderton, Roberto Amato, Jeffrey Barrett, Sonia Goncalves, Ewan Harrison, David K. Jackson, Ian Johnston, Dominic Kwiatkowski, Cordelia Langford, John Sillitoe on behalf of the Wellcome Sanger Institute COVID-19 Surveillance Team |
| EPI_ISL_2237695 | Lighthouse Lab in Milton Keynes | Wellcome Sanger Institute for the COVID-19 Genomics UK (COG-UK) Consortium | The Lighthouse Lab in Milton Keynes and Alex Alderton, Roberto Amato, Jeffrey Barrett, Sonia Goncalves, Ewan Harrison, David K. Jackson, Ian Johnston, Dominic Kwiatkowski, Cordelia Langford, John Sillitoe on behalf of the Wellcome Sanger Institute COVID-19 Surveillance Team |
| EPI_ISL_2237698, EPI_ISL_2237699 | Lighthouse Lab in Glasgow | Wellcome Sanger Institute for the COVID-19 Genomics UK (COG-UK) Consortium | Harper VanSteenhouse, Yumi Kasai, David Gray, Carol Clugston, Anna Dominiczak and Alex Alderton, Roberto Amato, Jeffrey Barrett, Sonia Goncalves, Ewan Harrison, David K. Jackson, Ian Johnston, Dominic Kwiatkowski, Cordelia Langford, John Sillitoe on behalf of the Wellcome Sanger Institute COVID-19 Surveillance Team |
| EPI_ISL_2237700 | Lighthouse Lab in Milton Keynes | Wellcome Sanger Institute for the COVID-19 Genomics UK (COG-UK) Consortium | The Lighthouse Lab in Milton Keynes and Alex Alderton, Roberto Amato, Jeffrey Barrett, Sonia Goncalves, Ewan Harrison, David K. Jackson, Ian Johnston, Dominic Kwiatkowski, Cordelia Langford, John Sillitoe on behalf of the Wellcome Sanger Institute COVID-19 Surveillance Team |
| EPI_ISL_2237703 | Lighthouse Lab in Alderley Park | Wellcome Sanger Institute for the COVID-19 Genomics UK (COG-UK) Consortium | Jacquelyn Wynn, Mairead Hyland, The Lighthouse Lab in Alderley Park and Alex Alderton, Roberto Amato, Jeffrey Barrett, Sonia Goncalves, Ewan Harrison, David K. Jackson, Ian Johnston, Dominic Kwiatkowski, Cordelia Langford, John Sillitoe on behalf of the Wellcome Sanger Institute COVID-19 Surveillance Team |
| EPI_ISL_2237704 | Lighthouse Lab in Glasgow | Wellcome Sanger Institute for the COVID-19 Genomics UK (COG-UK) Consortium | Harper VanSteenhouse, Yumi Kasai, David Gray, Carol Clugston, Anna Dominiczak and Alex Alderton, Roberto Amato, Jeffrey Barrett, Sonia Goncalves, Ewan Harrison, David K. Jackson, Ian Johnston, Dominic Kwiatkowski, Cordelia Langford, John Sillitoe on behalf of the Wellcome Sanger Institute COVID-19 Surveillance Team |
| EPI_ISL_2237705, EPI_ISL_2237706, EPI_ISL_2237707, EPI_ISL_2237711, EPI_ISL_2237712 | Lighthouse Lab in Milton Keynes | Wellcome Sanger Institute for the COVID-19 Genomics UK (COG-UK) Consortium | The Lighthouse Lab in Milton Keynes and Alex Alderton, Roberto Amato, Jeffrey Barrett, Sonia Goncalves, Ewan Harrison, David K. Jackson, Ian Johnston, Dominic Kwiatkowski, Cordelia Langford, John Sillitoe on behalf of the Wellcome Sanger Institute COVID-19 Surveillance Team |
| EPI_ISL_2237713 | Lighthouse Lab in Glasgow | Wellcome Sanger Institute for the COVID-19 Genomics UK (COG-UK) Consortium | Harper VanSteenhouse, Yumi Kasai, David Gray, Carol Clugston, Anna Dominiczak and Alex Alderton, Roberto Amato, Jeffrey Barrett, Sonia Goncalves, Ewan Harrison, David K. Jackson, Ian Johnston, Dominic Kwiatkowski, Cordelia Langford, John Sillitoe on behalf of the Wellcome Sanger Institute COVID-19 Surveillance Team |
| EPI_ISL_2237715 | Lighthouse Lab in Milton Keynes | Wellcome Sanger Institute for the COVID-19 Genomics UK (COG-UK) Consortium | The Lighthouse Lab in Milton Keynes and Alex Alderton, Roberto Amato, Jeffrey Barrett, Sonia Goncalves, Ewan Harrison, David K. Jackson, Ian Johnston, Dominic Kwiatkowski, Cordelia Langford, John Sillitoe on behalf of the Wellcome Sanger Institute COVID-19 Surveillance Team |
| EPI_ISL_2237716 | Lighthouse Lab in Glasgow | Wellcome Sanger Institute for the COVID-19 Genomics UK (COG-UK) Consortium | Harper VanSteenhouse, Yumi Kasai, David Gray, Carol Clugston, Anna Dominiczak and Alex Alderton, Roberto Amato, Jeffrey Barrett, Sonia Goncalves, Ewan Harrison, David K. Jackson, Ian Johnston, Dominic Kwiatkowski, Cordelia Langford, John Sillitoe on behalf of the Wellcome Sanger Institute COVID-19 Surveillance Team |
| EPI_ISL_2237719, EPI_ISL_2237720, EPI_ISL_2237721, EPI_ISL_2237723 | Lighthouse Lab in Alderley Park | Wellcome Sanger Institute for the COVID-19 Genomics UK (COG-UK) Consortium | Jacquelyn Wynn, Mairead Hyland, The Lighthouse Lab in Alderley Park and Alex Alderton, Roberto Amato, Jeffrey Barrett, Sonia Goncalves, Ewan Harrison, David K. Jackson, Ian Johnston, Dominic Kwiatkowski, Cordelia Langford, John Sillitoe on behalf of the Wellcome Sanger Institute COVID-19 Surveillance Team |
| EPI_ISL_2237725, EPI_ISL_2237727, EPI_ISL_2237729, EPI_ISL_2237737 | Lighthouse Lab in Milton Keynes | Wellcome Sanger Institute for the COVID-19 Genomics UK (COG-UK) Consortium | The Lighthouse Lab in Milton Keynes and Alex Alderton, Roberto Amato, Jeffrey Barrett, Sonia Goncalves, Ewan Harrison, David K. Jackson, Ian Johnston, Dominic Kwiatkowski, Cordelia Langford, John Sillitoe on behalf of the Wellcome Sanger Institute COVID-19 Surveillance Team |
| EPI_ISL_2237739 | Lighthouse Lab in Alderley Park | Wellcome Sanger Institute for the COVID-19 Genomics UK (COG-UK) Consortium | Jacquelyn Wynn, Mairead Hyland, The Lighthouse Lab in Alderley Park and Alex Alderton, Roberto Amato, Jeffrey Barrett, Sonia Goncalves, Ewan Harrison, David K. Jackson, Ian Johnston, Dominic Kwiatkowski, Cordelia Langford, John Sillitoe on behalf of the Wellcome Sanger Institute COVID-19 Surveillance Team |
| EPI_ISL_2237744 | Lighthouse Lab in Glasgow | Wellcome Sanger Institute for the COVID-19 Genomics UK (COG-UK) Consortium | Harper VanSteenhouse, Yumi Kasai, David Gray, Carol Clugston, Anna Dominiczak and Alex Alderton, Roberto Amato, Jeffrey Barrett, Sonia Goncalves, Ewan Harrison, David K. Jackson, Ian Johnston, Dominic Kwiatkowski, Cordelia Langford, John Sillitoe on behalf of the Wellcome Sanger Institute COVID-19 Surveillance Team |
| EPI_ISL_2237747, EPI_ISL_2237748 | Lighthouse Lab in Milton Keynes | Wellcome Sanger Institute for the COVID-19 Genomics UK (COG-UK) Consortium | The Lighthouse Lab in Milton Keynes and Alex Alderton, Roberto Amato, Jeffrey Barrett, Sonia Goncalves, Ewan Harrison, David K. Jackson, Ian Johnston, Dominic Kwiatkowski, Cordelia Langford, John Sillitoe on behalf of the Wellcome Sanger Institute COVID-19 Surveillance Team |
| EPI_ISL_2237758 | Lighthouse Lab in Glasgow | Wellcome Sanger Institute for the COVID-19 Genomics UK (COG-UK) Consortium | Harper VanSteenhouse, Yumi Kasai, David Gray, Carol Clugston, Anna Dominiczak and Alex Alderton, Roberto Amato, Jeffrey Barrett, Sonia Goncalves, Ewan Harrison, David K. Jackson, Ian Johnston, Dominic Kwiatkowski, Cordelia Langford, John Sillitoe on behalf of the Wellcome Sanger Institute COVID-19 Surveillance Team |
| EPI_ISL_2237760 | Lighthouse Lab in Milton Keynes | Wellcome Sanger Institute for the COVID-19 Genomics UK (COG-UK) Consortium | The Lighthouse Lab in Milton Keynes and Alex Alderton, Roberto Amato, Jeffrey Barrett, Sonia Goncalves, Ewan Harrison, David K. Jackson, Ian Johnston, Dominic Kwiatkowski, Cordelia Langford, John Sillitoe on behalf of the Wellcome Sanger Institute COVID-19 Surveillance Team |
| EPI_ISL_2237761 | Lighthouse Lab in Glasgow | Wellcome Sanger Institute for the COVID-19 Genomics UK (COG-UK) Consortium | Harper VanSteenhouse, Yumi Kasai, David Gray, Carol Clugston, Anna Dominiczak and Alex Alderton, Roberto Amato, Jeffrey Barrett, Sonia Goncalves, Ewan Harrison, David K. Jackson, Ian Johnston, Dominic Kwiatkowski, Cordelia Langford, John Sillitoe on behalf of the Wellcome Sanger Institute COVID-19 Surveillance Team |
| EPI_ISL_2237764 | Lighthouse Lab in Alderley Park | Wellcome Sanger Institute for the COVID-19 Genomics UK (COG-UK) Consortium | Jacquelyn Wynn, Mairead Hyland, The Lighthouse Lab in Alderley Park and Alex Alderton, Roberto Amato, Jeffrey Barrett, Sonia Goncalves, Ewan Harrison, David K. Jackson, Ian Johnston, Dominic Kwiatkowski, Cordelia Langford, John Sillitoe on behalf of the Wellcome Sanger Institute COVID-19 Surveillance Team |
| EPI_ISL_2237765 | Lighthouse Lab in Glasgow | Wellcome Sanger Institute for the COVID-19 Genomics UK (COG-UK) Consortium | Harper VanSteenhouse, Yumi Kasai, David Gray, Carol Clugston, Anna Dominiczak and Alex Alderton, Roberto Amato, Jeffrey Barrett, Sonia Goncalves, Ewan Harrison, David K. Jackson, Ian Johnston, Dominic Kwiatkowski, Cordelia Langford, John Sillitoe on behalf of the Wellcome Sanger Institute COVID-19 Surveillance Team |
| EPI_ISL_2237767, EPI_ISL_2237768 | Lighthouse Lab in Milton Keynes | Wellcome Sanger Institute for the COVID-19 Genomics UK (COG-UK) Consortium | The Lighthouse Lab in Milton Keynes and Alex Alderton, Roberto Amato, Jeffrey Barrett, Sonia Goncalves, Ewan Harrison, David K. Jackson, Ian Johnston, Dominic Kwiatkowski, Cordelia Langford, John Sillitoe on behalf of the Wellcome Sanger Institute COVID-19 Surveillance Team |
| EPI_ISL_2237769, EPI_ISL_2237772 | Lighthouse Lab in Glasgow | Wellcome Sanger Institute for the COVID-19 Genomics UK (COG-UK) Consortium | Harper VanSteenhouse, Yumi Kasai, David Gray, Carol Clugston, Anna Dominiczak and Alex Alderton, Roberto Amato, Jeffrey Barrett, Sonia Goncalves, Ewan Harrison, David K. Jackson, Ian Johnston, Dominic Kwiatkowski, Cordelia Langford, John Sillitoe on behalf of the Wellcome Sanger Institute COVID-19 Surveillance Team |
| EPI_ISL_2237777, EPI_ISL_2237779 | Lighthouse Lab in Alderley Park | Wellcome Sanger Institute for the COVID-19 Genomics UK (COG-UK) Consortium | Jacquelyn Wynn, Mairead Hyland, The Lighthouse Lab in Alderley Park and Alex Alderton, Roberto Amato, Jeffrey Barrett, Sonia Goncalves, Ewan Harrison, David K. Jackson, Ian Johnston, Dominic Kwiatkowski, Cordelia Langford, John Sillitoe on behalf of the Wellcome Sanger Institute COVID-19 Surveillance Team |
| EPI_ISL_2237780, EPI_ISL_2237785, EPI_ISL_2237791, EPI_ISL_2237792 | Lighthouse Lab in Milton Keynes | Wellcome Sanger Institute for the COVID-19 Genomics UK (COG-UK) Consortium | The Lighthouse Lab in Milton Keynes and Alex Alderton, Roberto Amato, Jeffrey Barrett, Sonia Goncalves, Ewan Harrison, David K. Jackson, Ian Johnston, Dominic Kwiatkowski, Cordelia Langford, John Sillitoe on behalf of the Wellcome Sanger Institute COVID-19 Surveillance Team |
| EPI_ISL_2237793, EPI_ISL_2237796, | Lighthouse Lab in Glasgow | Wellcome Sanger Institute for the COVID-19 Genomics UK (COG-UK) Consortium | Harper VanSteenhouse, Yumi Kasai, David Gray, Carol Clugston, Anna Dominiczak and Alex Alderton, Roberto Amato, Jeffrey Barrett, Sonia Goncalves, Ewan Harrison, David K. Jackson, Ian Johnston, Dominic Kwiatkowski, Cordelia Langford, John Sillitoe on behalf of the Wellcome Sanger Institute COVID-19 Surveillance Team |

| | | | |
|---|---|---|---|
| EPI_ISL_2237797, EPI_ISL_2237798 | | | |
| EPI_ISL_2237801 | Lighthouse Lab in Milton Keynes | Wellcome Sanger Institute for the COVID-19 Genomics UK (COG-UK) Consortium | The Lighthouse Lab in Milton Keynes and Alex Alderton, Roberto Amato, Jeffrey Barrett, Sonia Goncalves, Ewan Harrison, David K. Jackson, Ian Johnston, Dominic Kwiatkowski, Cordelia Langford, John Sillitoe on behalf of the Wellcome Sanger Institute COVID-19 Surveillance Team |
| EPI_ISL_2237802, EPI_ISL_2237803 | Lighthouse Lab in Glasgow | Wellcome Sanger Institute for the COVID-19 Genomics UK (COG-UK) Consortium | Harper VanSteenhouse, Yumi Kasai, David Gray, Carol Clugston, Anna Dominiczak and Alex Alderton, Roberto Amato, Jeffrey Barrett, Sonia Goncalves, Ewan Harrison, David K. Jackson, Ian Johnston, Dominic Kwiatkowski, Cordelia Langford, John Sillitoe on behalf of the Wellcome Sanger Institute COVID-19 Surveillance Team |
| EPI_ISL_2237804 | Lighthouse Lab in Alderley Park | Wellcome Sanger Institute for the COVID-19 Genomics UK (COG-UK) Consortium | Jacquelyn Wynn, Mairead Hyland, The Lighthouse Lab in Alderley Park and Alex Alderton, Roberto Amato, Jeffrey Barrett, Sonia Goncalves, Ewan Harrison, David K. Jackson, Ian Johnston, Dominic Kwiatkowski, Cordelia Langford, John Sillitoe on behalf of the Wellcome Sanger Institute COVID-19 Surveillance Team |
| EPI_ISL_2237805, EPI_ISL_2237807 | Lighthouse Lab in Milton Keynes | Wellcome Sanger Institute for the COVID-19 Genomics UK (COG-UK) Consortium | The Lighthouse Lab in Milton Keynes and Alex Alderton, Roberto Amato, Jeffrey Barrett, Sonia Goncalves, Ewan Harrison, David K. Jackson, Ian Johnston, Dominic Kwiatkowski, Cordelia Langford, John Sillitoe on behalf of the Wellcome Sanger Institute COVID-19 Surveillance Team |
| EPI_ISL_2237814, EPI_ISL_2237816, EPI_ISL_2237820 | Lighthouse Lab in Glasgow | Wellcome Sanger Institute for the COVID-19 Genomics UK (COG-UK) Consortium | Harper VanSteenhouse, Yumi Kasai, David Gray, Carol Clugston, Anna Dominiczak and Alex Alderton, Roberto Amato, Jeffrey Barrett, Sonia Goncalves, Ewan Harrison, David K. Jackson, Ian Johnston, Dominic Kwiatkowski, Cordelia Langford, John Sillitoe on behalf of the Wellcome Sanger Institute COVID-19 Surveillance Team |
| EPI_ISL_2237822 | Lighthouse Lab in Milton Keynes | Wellcome Sanger Institute for the COVID-19 Genomics UK (COG-UK) Consortium | The Lighthouse Lab in Milton Keynes and Alex Alderton, Roberto Amato, Jeffrey Barrett, Sonia Goncalves, Ewan Harrison, David K. Jackson, Ian Johnston, Dominic Kwiatkowski, Cordelia Langford, John Sillitoe on behalf of the Wellcome Sanger Institute COVID-19 Surveillance Team |
| EPI_ISL_2237823 | Lighthouse Lab in Alderley Park | Wellcome Sanger Institute for the COVID-19 Genomics UK (COG-UK) Consortium | Jacquelyn Wynn, Mairead Hyland, The Lighthouse Lab in Alderley Park and Alex Alderton, Roberto Amato, Jeffrey Barrett, Sonia Goncalves, Ewan Harrison, David K. Jackson, Ian Johnston, Dominic Kwiatkowski, Cordelia Langford, John Sillitoe on behalf of the Wellcome Sanger Institute COVID-19 Surveillance Team |
| EPI_ISL_2237824, EPI_ISL_2237825 | Lighthouse Lab in Milton Keynes | Wellcome Sanger Institute for the COVID-19 Genomics UK (COG-UK) Consortium | The Lighthouse Lab in Milton Keynes and Alex Alderton, Roberto Amato, Jeffrey Barrett, Sonia Goncalves, Ewan Harrison, David K. Jackson, Ian Johnston, Dominic Kwiatkowski, Cordelia Langford, John Sillitoe on behalf of the Wellcome Sanger Institute COVID-19 Surveillance Team |
| EPI_ISL_2237827, EPI_ISL_2237828 | Lighthouse Lab in Alderley Park | Wellcome Sanger Institute for the COVID-19 Genomics UK (COG-UK) Consortium | Jacquelyn Wynn, Mairead Hyland, The Lighthouse Lab in Alderley Park and Alex Alderton, Roberto Amato, Jeffrey Barrett, Sonia Goncalves, Ewan Harrison, David K. Jackson, Ian Johnston, Dominic Kwiatkowski, Cordelia Langford, John Sillitoe on behalf of the Wellcome Sanger Institute COVID-19 Surveillance Team |
| EPI_ISL_2237830 | Lighthouse Lab in Glasgow | Wellcome Sanger Institute for the COVID-19 Genomics UK (COG-UK) Consortium | Harper VanSteenhouse, Yumi Kasai, David Gray, Carol Clugston, Anna Dominiczak and Alex Alderton, Roberto Amato, Jeffrey Barrett, Sonia Goncalves, Ewan Harrison, David K. Jackson, Ian Johnston, Dominic Kwiatkowski, Cordelia Langford, John Sillitoe on behalf of the Wellcome Sanger Institute COVID-19 Surveillance Team |
| EPI_ISL_2237833 | Lighthouse Lab in Alderley Park | Wellcome Sanger Institute for the COVID-19 Genomics UK (COG-UK) Consortium | Jacquelyn Wynn, Mairead Hyland, The Lighthouse Lab in Alderley Park and Alex Alderton, Roberto Amato, Jeffrey Barrett, Sonia Goncalves, Ewan Harrison, David K. Jackson, Ian Johnston, Dominic Kwiatkowski, Cordelia Langford, John Sillitoe on behalf of the Wellcome Sanger Institute COVID-19 Surveillance Team |
| EPI_ISL_2237834 | Lighthouse Lab in Glasgow | Wellcome Sanger Institute for the COVID-19 Genomics UK (COG-UK) Consortium | Harper VanSteenhouse, Yumi Kasai, David Gray, Carol Clugston, Anna Dominiczak and Alex Alderton, Roberto Amato, Jeffrey Barrett, Sonia Goncalves, Ewan Harrison, David K. Jackson, Ian Johnston, Dominic Kwiatkowski, Cordelia Langford, John Sillitoe on behalf of the Wellcome Sanger Institute COVID-19 Surveillance Team |
| EPI_ISL_2237836, EPI_ISL_2237837 | Lighthouse Lab in Milton Keynes | Wellcome Sanger Institute for the COVID-19 Genomics UK (COG-UK) Consortium | The Lighthouse Lab in Milton Keynes and Alex Alderton, Roberto Amato, Jeffrey Barrett, Sonia Goncalves, Ewan Harrison, David K. Jackson, Ian Johnston, Dominic Kwiatkowski, Cordelia Langford, John Sillitoe on behalf of the Wellcome Sanger Institute COVID-19 Surveillance Team |
| EPI_ISL_2237840, EPI_ISL_2237845 | Lighthouse Lab in Glasgow | Wellcome Sanger Institute for the COVID-19 Genomics UK (COG-UK) Consortium | Harper VanSteenhouse, Yumi Kasai, David Gray, Carol Clugston, Anna Dominiczak and Alex Alderton, Roberto Amato, Jeffrey Barrett, Sonia Goncalves, Ewan Harrison, David K. Jackson, Ian Johnston, Dominic Kwiatkowski, Cordelia Langford, John Sillitoe on behalf of the Wellcome Sanger Institute COVID-19 Surveillance Team |
| EPI_ISL_2237846, EPI_ISL_2237847 | Lighthouse Lab in Milton Keynes | Wellcome Sanger Institute for the COVID-19 Genomics UK (COG-UK) Consortium | The Lighthouse Lab in Milton Keynes and Alex Alderton, Roberto Amato, Jeffrey Barrett, Sonia Goncalves, Ewan Harrison, David K. Jackson, Ian Johnston, Dominic Kwiatkowski, Cordelia Langford, John Sillitoe on behalf of the Wellcome Sanger Institute COVID-19 Surveillance Team |
| EPI_ISL_2237849 | Lighthouse Lab in Alderley Park | Wellcome Sanger Institute for the COVID-19 Genomics UK (COG-UK) Consortium | Jacquelyn Wynn, Mairead Hyland, The Lighthouse Lab in Alderley Park and Alex Alderton, Roberto Amato, Jeffrey Barrett, Sonia Goncalves, Ewan Harrison, David K. Jackson, Ian Johnston, Dominic Kwiatkowski, Cordelia Langford, John Sillitoe on behalf of the Wellcome Sanger Institute COVID-19 Surveillance Team |
| EPI_ISL_2237850 | Lighthouse Lab in Milton Keynes | Wellcome Sanger Institute for the COVID-19 Genomics UK (COG-UK) Consortium | The Lighthouse Lab in Milton Keynes and Alex Alderton, Roberto Amato, Jeffrey Barrett, Sonia Goncalves, Ewan Harrison, David K. Jackson, Ian Johnston, Dominic Kwiatkowski, Cordelia Langford, John Sillitoe on behalf of the Wellcome Sanger Institute COVID-19 Surveillance Team |
| EPI_ISL_2237851 | Lighthouse Lab in Alderley Park | Wellcome Sanger Institute for the COVID-19 Genomics UK (COG-UK) Consortium | Jacquelyn Wynn, Mairead Hyland, The Lighthouse Lab in Alderley Park and Alex Alderton, Roberto Amato, Jeffrey Barrett, Sonia Goncalves, Ewan Harrison, David K. Jackson, Ian Johnston, Dominic Kwiatkowski, Cordelia Langford, John Sillitoe on behalf of the Wellcome Sanger Institute COVID-19 Surveillance Team |
| EPI_ISL_2237853 | Lighthouse Lab in Milton Keynes | Wellcome Sanger Institute for the COVID-19 Genomics UK (COG-UK) Consortium | The Lighthouse Lab in Milton Keynes and Alex Alderton, Roberto Amato, Jeffrey Barrett, Sonia Goncalves, Ewan Harrison, David K. Jackson, Ian Johnston, Dominic Kwiatkowski, Cordelia Langford, John Sillitoe on behalf of the Wellcome Sanger Institute COVID-19 Surveillance Team |
| EPI_ISL_2237854 | Lighthouse Lab in Glasgow | Wellcome Sanger Institute for the COVID-19 Genomics UK (COG-UK) Consortium | Harper VanSteenhouse, Yumi Kasai, David Gray, Carol Clugston, Anna Dominiczak and Alex Alderton, Roberto Amato, Jeffrey Barrett, Sonia Goncalves, Ewan Harrison, David K. Jackson, Ian Johnston, Dominic Kwiatkowski, Cordelia Langford, John Sillitoe on behalf of the Wellcome Sanger Institute COVID-19 Surveillance Team |
| EPI_ISL_2237855 | Lighthouse Lab in Milton Keynes | Wellcome Sanger Institute for the COVID-19 Genomics UK (COG-UK) Consortium | The Lighthouse Lab in Milton Keynes and Alex Alderton, Roberto Amato, Jeffrey Barrett, Sonia Goncalves, Ewan Harrison, David K. Jackson, Ian Johnston, Dominic Kwiatkowski, Cordelia Langford, John Sillitoe on behalf of the Wellcome Sanger Institute COVID-19 Surveillance Team |
| EPI_ISL_2237861, EPI_ISL_2237863 | Lighthouse Lab in Glasgow | Wellcome Sanger Institute for the COVID-19 Genomics UK (COG-UK) Consortium | Harper VanSteenhouse, Yumi Kasai, David Gray, Carol Clugston, Anna Dominiczak and Alex Alderton, Roberto Amato, Jeffrey Barrett, Sonia Goncalves, Ewan Harrison, David K. Jackson, Ian Johnston, Dominic Kwiatkowski, Cordelia Langford, John Sillitoe on behalf of the Wellcome Sanger Institute COVID-19 Surveillance Team |
| EPI_ISL_2237866 | Lighthouse Lab in Milton Keynes | Wellcome Sanger Institute for the COVID-19 Genomics UK (COG-UK) Consortium | The Lighthouse Lab in Milton Keynes and Alex Alderton, Roberto Amato, Jeffrey Barrett, Sonia Goncalves, Ewan Harrison, David K. Jackson, Ian Johnston, Dominic Kwiatkowski, Cordelia Langford, John Sillitoe on behalf of the Wellcome Sanger Institute COVID-19 Surveillance Team |
| EPI_ISL_2237867 | Lighthouse Lab in Glasgow | Wellcome Sanger Institute for the COVID-19 Genomics UK (COG-UK) Consortium | Harper VanSteenhouse, Yumi Kasai, David Gray, Carol Clugston, Anna Dominiczak and Alex Alderton, Roberto Amato, Jeffrey Barrett, Sonia Goncalves, Ewan Harrison, David K. Jackson, Ian Johnston, Dominic Kwiatkowski, Cordelia Langford, John Sillitoe on behalf of the Wellcome Sanger Institute COVID-19 Surveillance Team |
| EPI_ISL_2237868, EPI_ISL_2237870 | Lighthouse Lab in Alderley Park | Wellcome Sanger Institute for the COVID-19 Genomics UK (COG-UK) Consortium | Jacquelyn Wynn, Mairead Hyland, The Lighthouse Lab in Alderley Park and Alex Alderton, Roberto Amato, Jeffrey Barrett, Sonia Goncalves, Ewan Harrison, David K. Jackson, Ian Johnston, Dominic Kwiatkowski, Cordelia Langford, John Sillitoe on behalf of the Wellcome Sanger Institute COVID-19 Surveillance Team |
| EPI_ISL_2237871, EPI_ISL_2237876 | Lighthouse Lab in Milton Keynes | Wellcome Sanger Institute for the COVID-19 Genomics UK (COG-UK) Consortium | The Lighthouse Lab in Milton Keynes and Alex Alderton, Roberto Amato, Jeffrey Barrett, Sonia Goncalves, Ewan Harrison, David K. Jackson, Ian Johnston, Dominic Kwiatkowski, Cordelia Langford, John Sillitoe on behalf of the Wellcome Sanger Institute COVID-19 Surveillance Team |
| EPI_ISL_2237878 | Lighthouse Lab in Alderley Park | Wellcome Sanger Institute for the COVID-19 Genomics UK (COG-UK) Consortium | Jacquelyn Wynn, Mairead Hyland, The Lighthouse Lab in Alderley Park and Alex Alderton, Roberto Amato, Jeffrey Barrett, Sonia Goncalves, Ewan Harrison, David K. Jackson, Ian Johnston, Dominic Kwiatkowski, Cordelia Langford, John Sillitoe on behalf of the Wellcome Sanger Institute COVID-19 Surveillance Team |
| EPI_ISL_2237879 | Lighthouse Lab in Glasgow | Wellcome Sanger Institute for the COVID-19 Genomics UK (COG-UK) Consortium | Harper VanSteenhouse, Yumi Kasai, David Gray, Carol Clugston, Anna Dominiczak and Alex Alderton, Roberto Amato, Jeffrey Barrett, Sonia Goncalves, Ewan Harrison, David K. Jackson, Ian Johnston, Dominic Kwiatkowski, Cordelia Langford, John Sillitoe on behalf of the Wellcome Sanger Institute COVID-19 Surveillance Team |
| EPI_ISL_2237882, EPI_ISL_2237883, EPI_ISL_2237885 | Lighthouse Lab in Alderley Park | Wellcome Sanger Institute for the COVID-19 Genomics UK (COG-UK) Consortium | Jacquelyn Wynn, Mairead Hyland, The Lighthouse Lab in Alderley Park and Alex Alderton, Roberto Amato, Jeffrey Barrett, Sonia Goncalves, Ewan Harrison, David K. Jackson, Ian Johnston, Dominic Kwiatkowski, Cordelia Langford, John Sillitoe on behalf of the Wellcome Sanger Institute COVID-19 Surveillance Team |
| EPI_ISL_2237887 | Lighthouse Lab in Milton Keynes | Wellcome Sanger Institute for the COVID-19 Genomics UK (COG-UK) Consortium | The Lighthouse Lab in Milton Keynes and Alex Alderton, Roberto Amato, Jeffrey Barrett, Sonia Goncalves, Ewan Harrison, David K. Jackson, Ian Johnston, Dominic Kwiatkowski, Cordelia Langford, John Sillitoe on behalf of the Wellcome Sanger Institute COVID-19 Surveillance Team |
| EPI_ISL_2237889 | Lighthouse Lab in Glasgow | Wellcome Sanger Institute for the COVID-19 Genomics UK (COG-UK) Consortium | Harper VanSteenhouse, Yumi Kasai, David Gray, Carol Clugston, Anna Dominiczak and Alex Alderton, Roberto Amato, Jeffrey Barrett, Sonia Goncalves, Ewan Harrison, David K. Jackson, Ian Johnston, Dominic Kwiatkowski, Cordelia Langford, John Sillitoe on behalf of the Wellcome Sanger Institute COVID-19 Surveillance Team |
| EPI_ISL_2237890 | Lighthouse Lab in Milton Keynes | Wellcome Sanger Institute for the COVID-19 Genomics UK (COG-UK) Consortium | The Lighthouse Lab in Milton Keynes and Alex Alderton, Roberto Amato, Jeffrey Barrett, Sonia Goncalves, Ewan Harrison, David K. Jackson, Ian Johnston, Dominic Kwiatkowski, Cordelia Langford, John Sillitoe on behalf of the Wellcome Sanger Institute COVID-19 Surveillance Team |
| EPI_ISL_2237891, EPI_ISL_2237893 | Lighthouse Lab in Glasgow | Wellcome Sanger Institute for the COVID-19 Genomics UK (COG-UK) Consortium | Harper VanSteenhouse, Yumi Kasai, David Gray, Carol Clugston, Anna Dominiczak and Alex Alderton, Roberto Amato, Jeffrey Barrett, Sonia Goncalves, Ewan Harrison, David K. Jackson, Ian Johnston, Dominic Kwiatkowski, Cordelia Langford, John Sillitoe on behalf of the Wellcome Sanger Institute COVID-19 Surveillance Team |
| EPI_ISL_2237894 | Lighthouse Lab in Alderley Park | Wellcome Sanger Institute for the COVID-19 Genomics UK (COG-UK) Consortium | Jacquelyn Wynn, Mairead Hyland, The Lighthouse Lab in Alderley Park and Alex Alderton, Roberto Amato, Jeffrey Barrett, Sonia Goncalves, Ewan Harrison, David K. Jackson, Ian Johnston, Dominic Kwiatkowski, Cordelia Langford, John Sillitoe on behalf of the Wellcome Sanger Institute COVID-19 Surveillance Team |
| EPI_ISL_2237895, EPI_ISL_2237896 | Lighthouse Lab in Milton Keynes | Wellcome Sanger Institute for the COVID-19 Genomics UK (COG-UK) Consortium | The Lighthouse Lab in Milton Keynes and Alex Alderton, Roberto Amato, Jeffrey Barrett, Sonia Goncalves, Ewan Harrison, David K. Jackson, Ian Johnston, Dominic Kwiatkowski, Cordelia Langford, John Sillitoe on behalf of the Wellcome Sanger Institute COVID-19 Surveillance Team |
| EPI_ISL_2237898 | Lighthouse Lab in Glasgow | Wellcome Sanger Institute for the COVID-19 Genomics UK (COG-UK) Consortium | Harper VanSteenhouse, Yumi Kasai, David Gray, Carol Clugston, Anna Dominiczak and Alex Alderton, Roberto Amato, Jeffrey Barrett, Sonia Goncalves, Ewan Harrison, David K. Jackson, Ian Johnston, Dominic Kwiatkowski, Cordelia Langford, John Sillitoe on behalf of the Wellcome Sanger Institute COVID-19 Surveillance Team |
| EPI_ISL_2237905 | Lighthouse Lab in Milton Keynes | Wellcome Sanger Institute for the COVID-19 Genomics UK (COG-UK) Consortium | The Lighthouse Lab in Milton Keynes and Alex Alderton, Roberto Amato, Jeffrey Barrett, Sonia Goncalves, Ewan Harrison, David K. Jackson, Ian Johnston, Dominic Kwiatkowski, Cordelia Langford, John Sillitoe on behalf of the Wellcome Sanger Institute COVID-19 Surveillance Team |
| EPI_ISL_2237909, EPI_ISL_2237910, EPI_ISL_2237913, EPI_ISL_2237914 | Lighthouse Lab in Glasgow | Wellcome Sanger Institute for the COVID-19 Genomics UK (COG-UK) Consortium | Harper VanSteenhouse, Yumi Kasai, David Gray, Carol Clugston, Anna Dominiczak and Alex Alderton, Roberto Amato, Jeffrey Barrett, Sonia Goncalves, Ewan Harrison, David K. Jackson, Ian Johnston, Dominic Kwiatkowski, Cordelia Langford, John Sillitoe on behalf of the Wellcome Sanger Institute COVID-19 Surveillance Team |
| EPI_ISL_2237919 | Lighthouse Lab in Milton Keynes | Wellcome Sanger Institute for the COVID-19 Genomics UK (COG-UK) Consortium | The Lighthouse Lab in Milton Keynes and Alex Alderton, Roberto Amato, Jeffrey Barrett, Sonia Goncalves, Ewan Harrison, David K. Jackson, Ian Johnston, Dominic Kwiatkowski, Cordelia Langford, John Sillitoe on behalf of the Wellcome Sanger Institute COVID-19 Surveillance Team |
| EPI_ISL_2237920 | Lighthouse Lab in Alderley Park | Wellcome Sanger Institute for the COVID-19 Genomics UK (COG-UK) Consortium | Jacquelyn Wynn, Mairead Hyland, The Lighthouse Lab in Alderley Park and Alex Alderton, Roberto Amato, Jeffrey Barrett, Sonia Goncalves, Ewan Harrison, David K. Jackson, Ian Johnston, Dominic Kwiatkowski, Cordelia Langford, John Sillitoe on behalf of the Wellcome Sanger Institute COVID-19 Surveillance Team |
| EPI_ISL_2237926 | Lighthouse Lab in Glasgow | Wellcome Sanger Institute for the COVID-19 Genomics UK (COG-UK) Consortium | Harper VanSteenhouse, Yumi Kasai, David Gray, Carol Clugston, Anna Dominiczak and Alex Alderton, Roberto Amato, Jeffrey Barrett, Sonia Goncalves, Ewan Harrison, David K. Jackson, Ian Johnston, Dominic Kwiatkowski, Cordelia Langford, John Sillitoe on behalf of the Wellcome Sanger Institute COVID-19 Surveillance Team |
| EPI_ISL_2237929 | Lighthouse Lab in Milton Keynes | Wellcome Sanger Institute for the COVID-19 Genomics UK (COG-UK) Consortium | The Lighthouse Lab in Milton Keynes and Alex Alderton, Roberto Amato, Jeffrey Barrett, Sonia Goncalves, Ewan Harrison, David K. Jackson, Ian Johnston, Dominic Kwiatkowski, Cordelia Langford, John Sillitoe on behalf of the Wellcome Sanger Institute COVID-19 Surveillance Team |

| | | | |
|---|---|---|---|
| EPI_ISL_2237932 | Lighthouse Lab in Glasgow | Wellcome Sanger Institute for the COVID-19 Genomics UK (COG-UK) Consortium | Harper VanSteenhouse, Yumi Kasai, David Gray, Carol Clugston, Anna Dominiczak and Alex Alderton, Roberto Amato, Jeffrey Barrett, Sonia Goncalves, Ewan Harrison, David K. Jackson, Ian Johnston, Dominic Kwiatkowski, Cordelia Langford, John Sillitoe on behalf of the Wellcome Sanger Institute COVID-19 Surveillance Team |
| EPI_ISL_2237933 | Lighthouse Lab in Milton Keynes | Wellcome Sanger Institute for the COVID-19 Genomics UK (COG-UK) Consortium | The Lighthouse Lab in Milton Keynes and Alex Alderton, Roberto Amato, Jeffrey Barrett, Sonia Goncalves, Ewan Harrison, David K. Jackson, Ian Johnston, Dominic Kwiatkowski, Cordelia Langford, John Sillitoe on behalf of the Wellcome Sanger Institute COVID-19 Surveillance Team |
| EPI_ISL_2237934 | Lighthouse Lab in Alderley Park | Wellcome Sanger Institute for the COVID-19 Genomics UK (COG-UK) Consortium | Jacquelyn Wynn, Mairead Hyland, The Lighthouse Lab in Alderley Park and Alex Alderton, Roberto Amato, Jeffrey Barrett, Sonia Goncalves, Ewan Harrison, David K. Jackson, Ian Johnston, Dominic Kwiatkowski, Cordelia Langford, John Sillitoe on behalf of the Wellcome Sanger Institute COVID-19 Surveillance Team |
| EPI_ISL_2237935 | Lighthouse Lab in Milton Keynes | Wellcome Sanger Institute for the COVID-19 Genomics UK (COG-UK) Consortium | The Lighthouse Lab in Milton Keynes and Alex Alderton, Roberto Amato, Jeffrey Barrett, Sonia Goncalves, Ewan Harrison, David K. Jackson, Ian Johnston, Dominic Kwiatkowski, Cordelia Langford, John Sillitoe on behalf of the Wellcome Sanger Institute COVID-19 Surveillance Team |
| EPI_ISL_2237939, EPI_ISL_2237941 | Lighthouse Lab in Glasgow | Wellcome Sanger Institute for the COVID-19 Genomics UK (COG-UK) Consortium | Harper VanSteenhouse, Yumi Kasai, David Gray, Carol Clugston, Anna Dominiczak and Alex Alderton, Roberto Amato, Jeffrey Barrett, Sonia Goncalves, Ewan Harrison, David K. Jackson, Ian Johnston, Dominic Kwiatkowski, Cordelia Langford, John Sillitoe on behalf of the Wellcome Sanger Institute COVID-19 Surveillance Team |
| EPI_ISL_2237942, EPI_ISL_2237943, EPI_ISL_2237944 | Lighthouse Lab in Milton Keynes | Wellcome Sanger Institute for the COVID-19 Genomics UK (COG-UK) Consortium | The Lighthouse Lab in Milton Keynes and Alex Alderton, Roberto Amato, Jeffrey Barrett, Sonia Goncalves, Ewan Harrison, David K. Jackson, Ian Johnston, Dominic Kwiatkowski, Cordelia Langford, John Sillitoe on behalf of the Wellcome Sanger Institute COVID-19 Surveillance Team |
| EPI_ISL_2237945 | Lighthouse Lab in Alderley Park | Wellcome Sanger Institute for the COVID-19 Genomics UK (COG-UK) Consortium | Jacquelyn Wynn, Mairead Hyland, The Lighthouse Lab in Alderley Park and Alex Alderton, Roberto Amato, Jeffrey Barrett, Sonia Goncalves, Ewan Harrison, David K. Jackson, Ian Johnston, Dominic Kwiatkowski, Cordelia Langford, John Sillitoe on behalf of the Wellcome Sanger Institute COVID-19 Surveillance Team |
| EPI_ISL_2237947, EPI_ISL_2237949, EPI_ISL_2237950, EPI_ISL_2237951 | Lighthouse Lab in Milton Keynes | Wellcome Sanger Institute for the COVID-19 Genomics UK (COG-UK) Consortium | The Lighthouse Lab in Milton Keynes and Alex Alderton, Roberto Amato, Jeffrey Barrett, Sonia Goncalves, Ewan Harrison, David K. Jackson, Ian Johnston, Dominic Kwiatkowski, Cordelia Langford, John Sillitoe on behalf of the Wellcome Sanger Institute COVID-19 Surveillance Team |
| EPI_ISL_2237952 | Lighthouse Lab in Glasgow | Wellcome Sanger Institute for the COVID-19 Genomics UK (COG-UK) Consortium | Harper VanSteenhouse, Yumi Kasai, David Gray, Carol Clugston, Anna Dominiczak and Alex Alderton, Roberto Amato, Jeffrey Barrett, Sonia Goncalves, Ewan Harrison, David K. Jackson, Ian Johnston, Dominic Kwiatkowski, Cordelia Langford, John Sillitoe on behalf of the Wellcome Sanger Institute COVID-19 Surveillance Team |
| EPI_ISL_2237955 | Lighthouse Lab in Alderley Park | Wellcome Sanger Institute for the COVID-19 Genomics UK (COG-UK) Consortium | Jacquelyn Wynn, Mairead Hyland, The Lighthouse Lab in Alderley Park and Alex Alderton, Roberto Amato, Jeffrey Barrett, Sonia Goncalves, Ewan Harrison, David K. Jackson, Ian Johnston, Dominic Kwiatkowski, Cordelia Langford, John Sillitoe on behalf of the Wellcome Sanger Institute COVID-19 Surveillance Team |
| EPI_ISL_2237956 | Lighthouse Lab in Glasgow | Wellcome Sanger Institute for the COVID-19 Genomics UK (COG-UK) Consortium | Harper VanSteenhouse, Yumi Kasai, David Gray, Carol Clugston, Anna Dominiczak and Alex Alderton, Roberto Amato, Jeffrey Barrett, Sonia Goncalves, Ewan Harrison, David K. Jackson, Ian Johnston, Dominic Kwiatkowski, Cordelia Langford, John Sillitoe on behalf of the Wellcome Sanger Institute COVID-19 Surveillance Team |
| EPI_ISL_2237959 | Lighthouse Lab in Milton Keynes | Wellcome Sanger Institute for the COVID-19 Genomics UK (COG-UK) Consortium | The Lighthouse Lab in Milton Keynes and Alex Alderton, Roberto Amato, Jeffrey Barrett, Sonia Goncalves, Ewan Harrison, David K. Jackson, Ian Johnston, Dominic Kwiatkowski, Cordelia Langford, John Sillitoe on behalf of the Wellcome Sanger Institute COVID-19 Surveillance Team |
| EPI_ISL_2237960 | Lighthouse Lab in Glasgow | Wellcome Sanger Institute for the COVID-19 Genomics UK (COG-UK) Consortium | Harper VanSteenhouse, Yumi Kasai, David Gray, Carol Clugston, Anna Dominiczak and Alex Alderton, Roberto Amato, Jeffrey Barrett, Sonia Goncalves, Ewan Harrison, David K. Jackson, Ian Johnston, Dominic Kwiatkowski, Cordelia Langford, John Sillitoe on behalf of the Wellcome Sanger Institute COVID-19 Surveillance Team |
| EPI_ISL_2237962 | Lighthouse Lab in Milton Keynes | Wellcome Sanger Institute for the COVID-19 Genomics UK (COG-UK) Consortium | The Lighthouse Lab in Milton Keynes and Alex Alderton, Roberto Amato, Jeffrey Barrett, Sonia Goncalves, Ewan Harrison, David K. Jackson, Ian Johnston, Dominic Kwiatkowski, Cordelia Langford, John Sillitoe on behalf of the Wellcome Sanger Institute COVID-19 Surveillance Team |
| EPI_ISL_2237966 | Lighthouse Lab in Glasgow | Wellcome Sanger Institute for the COVID-19 Genomics UK (COG-UK) Consortium | Harper VanSteenhouse, Yumi Kasai, David Gray, Carol Clugston, Anna Dominiczak and Alex Alderton, Roberto Amato, Jeffrey Barrett, Sonia Goncalves, Ewan Harrison, David K. Jackson, Ian Johnston, Dominic Kwiatkowski, Cordelia Langford, John Sillitoe on behalf of the Wellcome Sanger Institute COVID-19 Surveillance Team |
| EPI_ISL_2237969 | Lighthouse Lab in Milton Keynes | Wellcome Sanger Institute for the COVID-19 Genomics UK (COG-UK) Consortium | The Lighthouse Lab in Milton Keynes and Alex Alderton, Roberto Amato, Jeffrey Barrett, Sonia Goncalves, Ewan Harrison, David K. Jackson, Ian Johnston, Dominic Kwiatkowski, Cordelia Langford, John Sillitoe on behalf of the Wellcome Sanger Institute COVID-19 Surveillance Team |
| EPI_ISL_2237970, EPI_ISL_2237971 | Lighthouse Lab in Glasgow | Wellcome Sanger Institute for the COVID-19 Genomics UK (COG-UK) Consortium | Harper VanSteenhouse, Yumi Kasai, David Gray, Carol Clugston, Anna Dominiczak and Alex Alderton, Roberto Amato, Jeffrey Barrett, Sonia Goncalves, Ewan Harrison, David K. Jackson, Ian Johnston, Dominic Kwiatkowski, Cordelia Langford, John Sillitoe on behalf of the Wellcome Sanger Institute COVID-19 Surveillance Team |
| EPI_ISL_2237975, EPI_ISL_2237976, EPI_ISL_2237977 | Lighthouse Lab in Milton Keynes | Wellcome Sanger Institute for the COVID-19 Genomics UK (COG-UK) Consortium | The Lighthouse Lab in Milton Keynes and Alex Alderton, Roberto Amato, Jeffrey Barrett, Sonia Goncalves, Ewan Harrison, David K. Jackson, Ian Johnston, Dominic Kwiatkowski, Cordelia Langford, John Sillitoe on behalf of the Wellcome Sanger Institute COVID-19 Surveillance Team |
| EPI_ISL_2237979, EPI_ISL_2237981 | Lighthouse Lab in Glasgow | Wellcome Sanger Institute for the COVID-19 Genomics UK (COG-UK) Consortium | Harper VanSteenhouse, Yumi Kasai, David Gray, Carol Clugston, Anna Dominiczak and Alex Alderton, Roberto Amato, Jeffrey Barrett, Sonia Goncalves, Ewan Harrison, David K. Jackson, Ian Johnston, Dominic Kwiatkowski, Cordelia Langford, John Sillitoe on behalf of the Wellcome Sanger Institute COVID-19 Surveillance Team |
| EPI_ISL_2237982, EPI_ISL_2237983 | Lighthouse Lab in Milton Keynes | Wellcome Sanger Institute for the COVID-19 Genomics UK (COG-UK) Consortium | The Lighthouse Lab in Milton Keynes and Alex Alderton, Roberto Amato, Jeffrey Barrett, Sonia Goncalves, Ewan Harrison, David K. Jackson, Ian Johnston, Dominic Kwiatkowski, Cordelia Langford, John Sillitoe on behalf of the Wellcome Sanger Institute COVID-19 Surveillance Team |
| EPI_ISL_2237985, EPI_ISL_2237986, EPI_ISL_2237987 | Lighthouse Lab in Glasgow | Wellcome Sanger Institute for the COVID-19 Genomics UK (COG-UK) Consortium | Harper VanSteenhouse, Yumi Kasai, David Gray, Carol Clugston, Anna Dominiczak and Alex Alderton, Roberto Amato, Jeffrey Barrett, Sonia Goncalves, Ewan Harrison, David K. Jackson, Ian Johnston, Dominic Kwiatkowski, Cordelia Langford, John Sillitoe on behalf of the Wellcome Sanger Institute COVID-19 Surveillance Team |
| EPI_ISL_2237991 | Lighthouse Lab in Milton Keynes | Wellcome Sanger Institute for the COVID-19 Genomics UK (COG-UK) Consortium | The Lighthouse Lab in Milton Keynes and Alex Alderton, Roberto Amato, Jeffrey Barrett, Sonia Goncalves, Ewan Harrison, David K. Jackson, Ian Johnston, Dominic Kwiatkowski, Cordelia Langford, John Sillitoe on behalf of the Wellcome Sanger Institute COVID-19 Surveillance Team |
| EPI_ISL_2237992 | Lighthouse Lab in Glasgow | Wellcome Sanger Institute for the COVID-19 Genomics UK (COG-UK) Consortium | Harper VanSteenhouse, Yumi Kasai, David Gray, Carol Clugston, Anna Dominiczak and Alex Alderton, Roberto Amato, Jeffrey Barrett, Sonia Goncalves, Ewan Harrison, David K. Jackson, Ian Johnston, Dominic Kwiatkowski, Cordelia Langford, John Sillitoe on behalf of the Wellcome Sanger Institute COVID-19 Surveillance Team |
| EPI_ISL_2237993 | Lighthouse Lab in Alderley Park | Wellcome Sanger Institute for the COVID-19 Genomics UK (COG-UK) Consortium | Jacquelyn Wynn, Mairead Hyland, The Lighthouse Lab in Alderley Park and Alex Alderton, Roberto Amato, Jeffrey Barrett, Sonia Goncalves, Ewan Harrison, David K. Jackson, Ian Johnston, Dominic Kwiatkowski, Cordelia Langford, John Sillitoe on behalf of the Wellcome Sanger Institute COVID-19 Surveillance Team |
| EPI_ISL_2237994 | Lighthouse Lab in Milton Keynes | Wellcome Sanger Institute for the COVID-19 Genomics UK (COG-UK) Consortium | The Lighthouse Lab in Milton Keynes and Alex Alderton, Roberto Amato, Jeffrey Barrett, Sonia Goncalves, Ewan Harrison, David K. Jackson, Ian Johnston, Dominic Kwiatkowski, Cordelia Langford, John Sillitoe on behalf of the Wellcome Sanger Institute COVID-19 Surveillance Team |
| EPI_ISL_2237996 | Lighthouse Lab in Alderley Park | Wellcome Sanger Institute for the COVID-19 Genomics UK (COG-UK) Consortium | Jacquelyn Wynn, Mairead Hyland, The Lighthouse Lab in Alderley Park and Alex Alderton, Roberto Amato, Jeffrey Barrett, Sonia Goncalves, Ewan Harrison, David K. Jackson, Ian Johnston, Dominic Kwiatkowski, Cordelia Langford, John Sillitoe on behalf of the Wellcome Sanger Institute COVID-19 Surveillance Team |
| EPI_ISL_2237999 | Lighthouse Lab in Glasgow | Wellcome Sanger Institute for the COVID-19 Genomics UK (COG-UK) Consortium | Harper VanSteenhouse, Yumi Kasai, David Gray, Carol Clugston, Anna Dominiczak and Alex Alderton, Roberto Amato, Jeffrey Barrett, Sonia Goncalves, Ewan Harrison, David K. Jackson, Ian Johnston, Dominic Kwiatkowski, Cordelia Langford, John Sillitoe on behalf of the Wellcome Sanger Institute COVID-19 Surveillance Team |
| EPI_ISL_2238001, EPI_ISL_2238003 | Lighthouse Lab in Milton Keynes | Wellcome Sanger Institute for the COVID-19 Genomics UK (COG-UK) Consortium | The Lighthouse Lab in Milton Keynes and Alex Alderton, Roberto Amato, Jeffrey Barrett, Sonia Goncalves, Ewan Harrison, David K. Jackson, Ian Johnston, Dominic Kwiatkowski, Cordelia Langford, John Sillitoe on behalf of the Wellcome Sanger Institute COVID-19 Surveillance Team |
| EPI_ISL_2238004 | Lighthouse Lab in Glasgow | Wellcome Sanger Institute for the COVID-19 Genomics UK (COG-UK) Consortium | Harper VanSteenhouse, Yumi Kasai, David Gray, Carol Clugston, Anna Dominiczak and Alex Alderton, Roberto Amato, Jeffrey Barrett, Sonia Goncalves, Ewan Harrison, David K. Jackson, Ian Johnston, Dominic Kwiatkowski, Cordelia Langford, John Sillitoe on behalf of the Wellcome Sanger Institute COVID-19 Surveillance Team |
| EPI_ISL_2238007, EPI_ISL_2238008 | Lighthouse Lab in Milton Keynes | Wellcome Sanger Institute for the COVID-19 Genomics UK (COG-UK) Consortium | The Lighthouse Lab in Milton Keynes and Alex Alderton, Roberto Amato, Jeffrey Barrett, Sonia Goncalves, Ewan Harrison, David K. Jackson, Ian Johnston, Dominic Kwiatkowski, Cordelia Langford, John Sillitoe on behalf of the Wellcome Sanger Institute COVID-19 Surveillance Team |
| EPI_ISL_2238010 | Lighthouse Lab in Glasgow | Wellcome Sanger Institute for the COVID-19 Genomics UK (COG-UK) Consortium | Harper VanSteenhouse, Yumi Kasai, David Gray, Carol Clugston, Anna Dominiczak and Alex Alderton, Roberto Amato, Jeffrey Barrett, Sonia Goncalves, Ewan Harrison, David K. Jackson, Ian Johnston, Dominic Kwiatkowski, Cordelia Langford, John Sillitoe on behalf of the Wellcome Sanger Institute COVID-19 Surveillance Team |
| EPI_ISL_2238011 | Lighthouse Lab in Alderley Park | Wellcome Sanger Institute for the COVID-19 Genomics UK (COG-UK) Consortium | Jacquelyn Wynn, Mairead Hyland, The Lighthouse Lab in Alderley Park and Alex Alderton, Roberto Amato, Jeffrey Barrett, Sonia Goncalves, Ewan Harrison, David K. Jackson, Ian Johnston, Dominic Kwiatkowski, Cordelia Langford, John Sillitoe on behalf of the Wellcome Sanger Institute COVID-19 Surveillance Team |
| EPI_ISL_2238013, EPI_ISL_2238015 | Lighthouse Lab in Milton Keynes | Wellcome Sanger Institute for the COVID-19 Genomics UK (COG-UK) Consortium | The Lighthouse Lab in Milton Keynes and Alex Alderton, Roberto Amato, Jeffrey Barrett, Sonia Goncalves, Ewan Harrison, David K. Jackson, Ian Johnston, Dominic Kwiatkowski, Cordelia Langford, John Sillitoe on behalf of the Wellcome Sanger Institute COVID-19 Surveillance Team |
| EPI_ISL_2238021 | Lighthouse Lab in Alderley Park | Wellcome Sanger Institute for the COVID-19 Genomics UK (COG-UK) Consortium | Jacquelyn Wynn, Mairead Hyland, The Lighthouse Lab in Alderley Park and Alex Alderton, Roberto Amato, Jeffrey Barrett, Sonia Goncalves, Ewan Harrison, David K. Jackson, Ian Johnston, Dominic Kwiatkowski, Cordelia Langford, John Sillitoe on behalf of the Wellcome Sanger Institute COVID-19 Surveillance Team |
| EPI_ISL_2238024, EPI_ISL_2238025 | Lighthouse Lab in Glasgow | Wellcome Sanger Institute for the COVID-19 Genomics UK (COG-UK) Consortium | Harper VanSteenhouse, Yumi Kasai, David Gray, Carol Clugston, Anna Dominiczak and Alex Alderton, Roberto Amato, Jeffrey Barrett, Sonia Goncalves, Ewan Harrison, David K. Jackson, Ian Johnston, Dominic Kwiatkowski, Cordelia Langford, John Sillitoe on behalf of the Wellcome Sanger Institute COVID-19 Surveillance Team |
| EPI_ISL_2238032, EPI_ISL_2238034 | Lighthouse Lab in Alderley Park | Wellcome Sanger Institute for the COVID-19 Genomics UK (COG-UK) Consortium | Jacquelyn Wynn, Mairead Hyland, The Lighthouse Lab in Alderley Park and Alex Alderton, Roberto Amato, Jeffrey Barrett, Sonia Goncalves, Ewan Harrison, David K. Jackson, Ian Johnston, Dominic Kwiatkowski, Cordelia Langford, John Sillitoe on behalf of the Wellcome Sanger Institute COVID-19 Surveillance Team |
| EPI_ISL_2238036 | Lighthouse Lab in Milton Keynes | Wellcome Sanger Institute for the COVID-19 Genomics UK (COG-UK) Consortium | The Lighthouse Lab in Milton Keynes and Alex Alderton, Roberto Amato, Jeffrey Barrett, Sonia Goncalves, Ewan Harrison, David K. Jackson, Ian Johnston, Dominic Kwiatkowski, Cordelia Langford, John Sillitoe on behalf of the Wellcome Sanger Institute COVID-19 Surveillance Team |
| EPI_ISL_2238037 | Lighthouse Lab in Glasgow | Wellcome Sanger Institute for the COVID-19 Genomics UK (COG-UK) Consortium | Harper VanSteenhouse, Yumi Kasai, David Gray, Carol Clugston, Anna Dominiczak and Alex Alderton, Roberto Amato, Jeffrey Barrett, Sonia Goncalves, Ewan Harrison, David K. Jackson, Ian Johnston, Dominic Kwiatkowski, Cordelia Langford, John Sillitoe on behalf of the Wellcome Sanger Institute COVID-19 Surveillance Team |
| EPI_ISL_2238038 | Lighthouse Lab in Alderley Park | Wellcome Sanger Institute for the COVID-19 Genomics UK (COG-UK) Consortium | Jacquelyn Wynn, Mairead Hyland, The Lighthouse Lab in Alderley Park and Alex Alderton, Roberto Amato, Jeffrey Barrett, Sonia Goncalves, Ewan Harrison, David K. Jackson, Ian Johnston, Dominic Kwiatkowski, Cordelia Langford, John Sillitoe on behalf of the Wellcome Sanger Institute COVID-19 Surveillance Team |
| EPI_ISL_2238039, EPI_ISL_2238041 | Lighthouse Lab in Glasgow | Wellcome Sanger Institute for the COVID-19 Genomics UK (COG-UK) Consortium | Harper VanSteenhouse, Yumi Kasai, David Gray, Carol Clugston, Anna Dominiczak and Alex Alderton, Roberto Amato, Jeffrey Barrett, Sonia Goncalves, Ewan Harrison, David K. Jackson, Ian Johnston, Dominic Kwiatkowski, Cordelia Langford, John Sillitoe on behalf of the Wellcome Sanger Institute COVID-19 Surveillance Team |
| EPI_ISL_2238042 | Lighthouse Lab in Alderley Park | Wellcome Sanger Institute for the COVID-19 Genomics UK (COG-UK) Consortium | Jacquelyn Wynn, Mairead Hyland, The Lighthouse Lab in Alderley Park and Alex Alderton, Roberto Amato, Jeffrey Barrett, Sonia Goncalves, Ewan Harrison, David K. Jackson, Ian Johnston, Dominic Kwiatkowski, Cordelia Langford, John Sillitoe on behalf of the Wellcome Sanger Institute COVID-19 Surveillance Team |
| EPI_ISL_2238045 | Lighthouse Lab in Glasgow | Wellcome Sanger Institute for the COVID-19 Genomics UK (COG-UK) Consortium | Harper VanSteenhouse, Yumi Kasai, David Gray, Carol Clugston, Anna Dominiczak and Alex Alderton, Roberto Amato, Jeffrey Barrett, Sonia Goncalves, Ewan Harrison, David K. Jackson, Ian Johnston, Dominic Kwiatkowski, Cordelia Langford, John Sillitoe on behalf of the Wellcome Sanger Institute COVID-19 Surveillance Team |
| EPI_ISL_2238047 | Lighthouse Lab in Milton Keynes | Wellcome Sanger Institute for the COVID-19 Genomics UK (COG-UK) Consortium | The Lighthouse Lab in Milton Keynes and Alex Alderton, Roberto Amato, Jeffrey Barrett, Sonia Goncalves, Ewan Harrison, David K. Jackson, Ian Johnston, Dominic Kwiatkowski, Cordelia Langford, John Sillitoe on behalf of the Wellcome Sanger Institute COVID-19 Surveillance Team |

| | | | |
|---|---|---|---|
| EPI_ISL_2238049 | Lighthouse Lab in Glasgow | Wellcome Sanger Institute for the COVID-19 Genomics UK (COG-UK) Consortium | Harper VanSteenhouse, Yumi Kasai, David Gray, Carol Clugston, Anna Dominiczak and Alex Alderton, Roberto Amato, Jeffrey Barrett, Sonia Goncalves, Ewan Harrison, David K. Jackson, Ian Johnston, Dominic Kwiatkowski, Cordelia Langford, John Sillitoe on behalf of the Wellcome Sanger Institute COVID-19 Surveillance Team |
| EPI_ISL_2238050 | Lighthouse Lab in Milton Keynes | Wellcome Sanger Institute for the COVID-19 Genomics UK (COG-UK) Consortium | The Lighthouse Lab in Milton Keynes and Alex Alderton, Roberto Amato, Jeffrey Barrett, Sonia Goncalves, Ewan Harrison, David K. Jackson, Ian Johnston, Dominic Kwiatkowski, Cordelia Langford, John Sillitoe on behalf of the Wellcome Sanger Institute COVID-19 Surveillance Team |
| EPI_ISL_2238051 | Lighthouse Lab in Alderley Park | Wellcome Sanger Institute for the COVID-19 Genomics UK (COG-UK) Consortium | Jacquelyn Wynn, Mairead Hyland, The Lighthouse Lab in Alderley Park and Alex Alderton, Roberto Amato, Jeffrey Barrett, Sonia Goncalves, Ewan Harrison, David K. Jackson, Ian Johnston, Dominic Kwiatkowski, Cordelia Langford, John Sillitoe on behalf of the Wellcome Sanger Institute COVID-19 Surveillance Team |
| EPI_ISL_2238052 | Lighthouse Lab in Milton Keynes | Wellcome Sanger Institute for the COVID-19 Genomics UK (COG-UK) Consortium | The Lighthouse Lab in Milton Keynes and Alex Alderton, Roberto Amato, Jeffrey Barrett, Sonia Goncalves, Ewan Harrison, David K. Jackson, Ian Johnston, Dominic Kwiatkowski, Cordelia Langford, John Sillitoe on behalf of the Wellcome Sanger Institute COVID-19 Surveillance Team |
| EPI_ISL_2238054 | Lighthouse Lab in Glasgow | Wellcome Sanger Institute for the COVID-19 Genomics UK (COG-UK) Consortium | Harper VanSteenhouse, Yumi Kasai, David Gray, Carol Clugston, Anna Dominiczak and Alex Alderton, Roberto Amato, Jeffrey Barrett, Sonia Goncalves, Ewan Harrison, David K. Jackson, Ian Johnston, Dominic Kwiatkowski, Cordelia Langford, John Sillitoe on behalf of the Wellcome Sanger Institute COVID-19 Surveillance Team |
| EPI_ISL_2238055 | Lighthouse Lab in Milton Keynes | Wellcome Sanger Institute for the COVID-19 Genomics UK (COG-UK) Consortium | The Lighthouse Lab in Milton Keynes and Alex Alderton, Roberto Amato, Jeffrey Barrett, Sonia Goncalves, Ewan Harrison, David K. Jackson, Ian Johnston, Dominic Kwiatkowski, Cordelia Langford, John Sillitoe on behalf of the Wellcome Sanger Institute COVID-19 Surveillance Team |
| EPI_ISL_2238060 | Lighthouse Lab in Alderley Park | Wellcome Sanger Institute for the COVID-19 Genomics UK (COG-UK) Consortium | Jacquelyn Wynn, Mairead Hyland, The Lighthouse Lab in Alderley Park and Alex Alderton, Roberto Amato, Jeffrey Barrett, Sonia Goncalves, Ewan Harrison, David K. Jackson, Ian Johnston, Dominic Kwiatkowski, Cordelia Langford, John Sillitoe on behalf of the Wellcome Sanger Institute COVID-19 Surveillance Team |
| EPI_ISL_2238061 | Lighthouse Lab in Glasgow | Wellcome Sanger Institute for the COVID-19 Genomics UK (COG-UK) Consortium | Harper VanSteenhouse, Yumi Kasai, David Gray, Carol Clugston, Anna Dominiczak and Alex Alderton, Roberto Amato, Jeffrey Barrett, Sonia Goncalves, Ewan Harrison, David K. Jackson, Ian Johnston, Dominic Kwiatkowski, Cordelia Langford, John Sillitoe on behalf of the Wellcome Sanger Institute COVID-19 Surveillance Team |
| EPI_ISL_2238063, EPI_ISL_2238066 | Lighthouse Lab in Milton Keynes | Wellcome Sanger Institute for the COVID-19 Genomics UK (COG-UK) Consortium | The Lighthouse Lab in Milton Keynes and Alex Alderton, Roberto Amato, Jeffrey Barrett, Sonia Goncalves, Ewan Harrison, David K. Jackson, Ian Johnston, Dominic Kwiatkowski, Cordelia Langford, John Sillitoe on behalf of the Wellcome Sanger Institute COVID-19 Surveillance Team |
| EPI_ISL_2238071 | Lighthouse Lab in Glasgow | Wellcome Sanger Institute for the COVID-19 Genomics UK (COG-UK) Consortium | Harper VanSteenhouse, Yumi Kasai, David Gray, Carol Clugston, Anna Dominiczak and Alex Alderton, Roberto Amato, Jeffrey Barrett, Sonia Goncalves, Ewan Harrison, David K. Jackson, Ian Johnston, Dominic Kwiatkowski, Cordelia Langford, John Sillitoe on behalf of the Wellcome Sanger Institute COVID-19 Surveillance Team |
| EPI_ISL_2238073, EPI_ISL_2238074 | Lighthouse Lab in Milton Keynes | Wellcome Sanger Institute for the COVID-19 Genomics UK (COG-UK) Consortium | The Lighthouse Lab in Milton Keynes and Alex Alderton, Roberto Amato, Jeffrey Barrett, Sonia Goncalves, Ewan Harrison, David K. Jackson, Ian Johnston, Dominic Kwiatkowski, Cordelia Langford, John Sillitoe on behalf of the Wellcome Sanger Institute COVID-19 Surveillance Team |
| EPI_ISL_2238076 | Lighthouse Lab in Glasgow | Wellcome Sanger Institute for the COVID-19 Genomics UK (COG-UK) Consortium | Harper VanSteenhouse, Yumi Kasai, David Gray, Carol Clugston, Anna Dominiczak and Alex Alderton, Roberto Amato, Jeffrey Barrett, Sonia Goncalves, Ewan Harrison, David K. Jackson, Ian Johnston, Dominic Kwiatkowski, Cordelia Langford, John Sillitoe on behalf of the Wellcome Sanger Institute COVID-19 Surveillance Team |
| EPI_ISL_2238079, EPI_ISL_2238082, EPI_ISL_2238085 | Lighthouse Lab in Milton Keynes | Wellcome Sanger Institute for the COVID-19 Genomics UK (COG-UK) Consortium | The Lighthouse Lab in Milton Keynes and Alex Alderton, Roberto Amato, Jeffrey Barrett, Sonia Goncalves, Ewan Harrison, David K. Jackson, Ian Johnston, Dominic Kwiatkowski, Cordelia Langford, John Sillitoe on behalf of the Wellcome Sanger Institute COVID-19 Surveillance Team |
| EPI_ISL_2238086 | Lighthouse Lab in Glasgow | Wellcome Sanger Institute for the COVID-19 Genomics UK (COG-UK) Consortium | Harper VanSteenhouse, Yumi Kasai, David Gray, Carol Clugston, Anna Dominiczak and Alex Alderton, Roberto Amato, Jeffrey Barrett, Sonia Goncalves, Ewan Harrison, David K. Jackson, Ian Johnston, Dominic Kwiatkowski, Cordelia Langford, John Sillitoe on behalf of the Wellcome Sanger Institute COVID-19 Surveillance Team |
| EPI_ISL_2238089, EPI_ISL_2238091 | Lighthouse Lab in Milton Keynes | Wellcome Sanger Institute for the COVID-19 Genomics UK (COG-UK) Consortium | The Lighthouse Lab in Milton Keynes and Alex Alderton, Roberto Amato, Jeffrey Barrett, Sonia Goncalves, Ewan Harrison, David K. Jackson, Ian Johnston, Dominic Kwiatkowski, Cordelia Langford, John Sillitoe on behalf of the Wellcome Sanger Institute COVID-19 Surveillance Team |
| EPI_ISL_2238092, EPI_ISL_2238094 | Lighthouse Lab in Glasgow | Wellcome Sanger Institute for the COVID-19 Genomics UK (COG-UK) Consortium | Harper VanSteenhouse, Yumi Kasai, David Gray, Carol Clugston, Anna Dominiczak and Alex Alderton, Roberto Amato, Jeffrey Barrett, Sonia Goncalves, Ewan Harrison, David K. Jackson, Ian Johnston, Dominic Kwiatkowski, Cordelia Langford, John Sillitoe on behalf of the Wellcome Sanger Institute COVID-19 Surveillance Team |
| EPI_ISL_2238095, EPI_ISL_2238096, EPI_ISL_2238099 | Lighthouse Lab in Milton Keynes | Wellcome Sanger Institute for the COVID-19 Genomics UK (COG-UK) Consortium | The Lighthouse Lab in Milton Keynes and Alex Alderton, Roberto Amato, Jeffrey Barrett, Sonia Goncalves, Ewan Harrison, David K. Jackson, Ian Johnston, Dominic Kwiatkowski, Cordelia Langford, John Sillitoe on behalf of the Wellcome Sanger Institute COVID-19 Surveillance Team |
| EPI_ISL_2238100, EPI_ISL_2238102 | Lighthouse Lab in Glasgow | Wellcome Sanger Institute for the COVID-19 Genomics UK (COG-UK) Consortium | Harper VanSteenhouse, Yumi Kasai, David Gray, Carol Clugston, Anna Dominiczak and Alex Alderton, Roberto Amato, Jeffrey Barrett, Sonia Goncalves, Ewan Harrison, David K. Jackson, Ian Johnston, Dominic Kwiatkowski, Cordelia Langford, John Sillitoe on behalf of the Wellcome Sanger Institute COVID-19 Surveillance Team |
| EPI_ISL_2238104, EPI_ISL_2238105, EPI_ISL_2238114, EPI_ISL_2238116, EPI_ISL_2238118 | Lighthouse Lab in Milton Keynes | Wellcome Sanger Institute for the COVID-19 Genomics UK (COG-UK) Consortium | The Lighthouse Lab in Milton Keynes and Alex Alderton, Roberto Amato, Jeffrey Barrett, Sonia Goncalves, Ewan Harrison, David K. Jackson, Ian Johnston, Dominic Kwiatkowski, Cordelia Langford, John Sillitoe on behalf of the Wellcome Sanger Institute COVID-19 Surveillance Team |
| EPI_ISL_2238123 | Lighthouse Lab in Glasgow | Wellcome Sanger Institute for the COVID-19 Genomics UK (COG-UK) Consortium | Harper VanSteenhouse, Yumi Kasai, David Gray, Carol Clugston, Anna Dominiczak and Alex Alderton, Roberto Amato, Jeffrey Barrett, Sonia Goncalves, Ewan Harrison, David K. Jackson, Ian Johnston, Dominic Kwiatkowski, Cordelia Langford, John Sillitoe on behalf of the Wellcome Sanger Institute COVID-19 Surveillance Team |
| EPI_ISL_2238124 | Lighthouse Lab in Milton Keynes | Wellcome Sanger Institute for the COVID-19 Genomics UK (COG-UK) Consortium | The Lighthouse Lab in Milton Keynes and Alex Alderton, Roberto Amato, Jeffrey Barrett, Sonia Goncalves, Ewan Harrison, David K. Jackson, Ian Johnston, Dominic Kwiatkowski, Cordelia Langford, John Sillitoe on behalf of the Wellcome Sanger Institute COVID-19 Surveillance Team |
| EPI_ISL_2238125 | Lighthouse Lab in Glasgow | Wellcome Sanger Institute for the COVID-19 Genomics UK (COG-UK) Consortium | Harper VanSteenhouse, Yumi Kasai, David Gray, Carol Clugston, Anna Dominiczak and Alex Alderton, Roberto Amato, Jeffrey Barrett, Sonia Goncalves, Ewan Harrison, David K. Jackson, Ian Johnston, Dominic Kwiatkowski, Cordelia Langford, John Sillitoe on behalf of the Wellcome Sanger Institute COVID-19 Surveillance Team |
| EPI_ISL_2238127 | Lighthouse Lab in Milton Keynes | Wellcome Sanger Institute for the COVID-19 Genomics UK (COG-UK) Consortium | The Lighthouse Lab in Milton Keynes and Alex Alderton, Roberto Amato, Jeffrey Barrett, Sonia Goncalves, Ewan Harrison, David K. Jackson, Ian Johnston, Dominic Kwiatkowski, Cordelia Langford, John Sillitoe on behalf of the Wellcome Sanger Institute COVID-19 Surveillance Team |
| EPI_ISL_2238128, EPI_ISL_2238130 | Lighthouse Lab in Glasgow | Wellcome Sanger Institute for the COVID-19 Genomics UK (COG-UK) Consortium | Harper VanSteenhouse, Yumi Kasai, David Gray, Carol Clugston, Anna Dominiczak and Alex Alderton, Roberto Amato, Jeffrey Barrett, Sonia Goncalves, Ewan Harrison, David K. Jackson, Ian Johnston, Dominic Kwiatkowski, Cordelia Langford, John Sillitoe on behalf of the Wellcome Sanger Institute COVID-19 Surveillance Team |
| EPI_ISL_2238131 | Lighthouse Lab in Milton Keynes | Wellcome Sanger Institute for the COVID-19 Genomics UK (COG-UK) Consortium | The Lighthouse Lab in Milton Keynes and Alex Alderton, Roberto Amato, Jeffrey Barrett, Sonia Goncalves, Ewan Harrison, David K. Jackson, Ian Johnston, Dominic Kwiatkowski, Cordelia Langford, John Sillitoe on behalf of the Wellcome Sanger Institute COVID-19 Surveillance Team |
| EPI_ISL_2238135, EPI_ISL_2238136 | Lighthouse Lab in Glasgow | Wellcome Sanger Institute for the COVID-19 Genomics UK (COG-UK) Consortium | Harper VanSteenhouse, Yumi Kasai, David Gray, Carol Clugston, Anna Dominiczak and Alex Alderton, Roberto Amato, Jeffrey Barrett, Sonia Goncalves, Ewan Harrison, David K. Jackson, Ian Johnston, Dominic Kwiatkowski, Cordelia Langford, John Sillitoe on behalf of the Wellcome Sanger Institute COVID-19 Surveillance Team |
| EPI_ISL_2238137, EPI_ISL_2238138 | Lighthouse Lab in Milton Keynes | Wellcome Sanger Institute for the COVID-19 Genomics UK (COG-UK) Consortium | The Lighthouse Lab in Milton Keynes and Alex Alderton, Roberto Amato, Jeffrey Barrett, Sonia Goncalves, Ewan Harrison, David K. Jackson, Ian Johnston, Dominic Kwiatkowski, Cordelia Langford, John Sillitoe on behalf of the Wellcome Sanger Institute COVID-19 Surveillance Team |
| EPI_ISL_2238139 | Lighthouse Lab in Glasgow | Wellcome Sanger Institute for the COVID-19 Genomics UK (COG-UK) Consortium | Harper VanSteenhouse, Yumi Kasai, David Gray, Carol Clugston, Anna Dominiczak and Alex Alderton, Roberto Amato, Jeffrey Barrett, Sonia Goncalves, Ewan Harrison, David K. Jackson, Ian Johnston, Dominic Kwiatkowski, Cordelia Langford, John Sillitoe on behalf of the Wellcome Sanger Institute COVID-19 Surveillance Team |
| EPI_ISL_2238140 | Lighthouse Lab in Milton Keynes | Wellcome Sanger Institute for the COVID-19 Genomics UK (COG-UK) Consortium | The Lighthouse Lab in Milton Keynes and Alex Alderton, Roberto Amato, Jeffrey Barrett, Sonia Goncalves, Ewan Harrison, David K. Jackson, Ian Johnston, Dominic Kwiatkowski, Cordelia Langford, John Sillitoe on behalf of the Wellcome Sanger Institute COVID-19 Surveillance Team |
| EPI_ISL_2238143, EPI_ISL_2238145, EPI_ISL_2238149, EPI_ISL_2238150 | Lighthouse Lab in Glasgow | Wellcome Sanger Institute for the COVID-19 Genomics UK (COG-UK) Consortium | Harper VanSteenhouse, Yumi Kasai, David Gray, Carol Clugston, Anna Dominiczak and Alex Alderton, Roberto Amato, Jeffrey Barrett, Sonia Goncalves, Ewan Harrison, David K. Jackson, Ian Johnston, Dominic Kwiatkowski, Cordelia Langford, John Sillitoe on behalf of the Wellcome Sanger Institute COVID-19 Surveillance Team |
| EPI_ISL_2238151 | Lighthouse Lab in Milton Keynes | Wellcome Sanger Institute for the COVID-19 Genomics UK (COG-UK) Consortium | The Lighthouse Lab in Milton Keynes and Alex Alderton, Roberto Amato, Jeffrey Barrett, Sonia Goncalves, Ewan Harrison, David K. Jackson, Ian Johnston, Dominic Kwiatkowski, Cordelia Langford, John Sillitoe on behalf of the Wellcome Sanger Institute COVID-19 Surveillance Team |
| EPI_ISL_2238153 | Lighthouse Lab in Glasgow | Wellcome Sanger Institute for the COVID-19 Genomics UK (COG-UK) Consortium | Harper VanSteenhouse, Yumi Kasai, David Gray, Carol Clugston, Anna Dominiczak and Alex Alderton, Roberto Amato, Jeffrey Barrett, Sonia Goncalves, Ewan Harrison, David K. Jackson, Ian Johnston, Dominic Kwiatkowski, Cordelia Langford, John Sillitoe on behalf of the Wellcome Sanger Institute COVID-19 Surveillance Team |
| EPI_ISL_2238154 | Lighthouse Lab in Milton Keynes | Wellcome Sanger Institute for the COVID-19 Genomics UK (COG-UK) Consortium | The Lighthouse Lab in Milton Keynes and Alex Alderton, Roberto Amato, Jeffrey Barrett, Sonia Goncalves, Ewan Harrison, David K. Jackson, Ian Johnston, Dominic Kwiatkowski, Cordelia Langford, John Sillitoe on behalf of the Wellcome Sanger Institute COVID-19 Surveillance Team |
| EPI_ISL_2238156, EPI_ISL_2238162 | Lighthouse Lab in Glasgow | Wellcome Sanger Institute for the COVID-19 Genomics UK (COG-UK) Consortium | Harper VanSteenhouse, Yumi Kasai, David Gray, Carol Clugston, Anna Dominiczak and Alex Alderton, Roberto Amato, Jeffrey Barrett, Sonia Goncalves, Ewan Harrison, David K. Jackson, Ian Johnston, Dominic Kwiatkowski, Cordelia Langford, John Sillitoe on behalf of the Wellcome Sanger Institute COVID-19 Surveillance Team |
| EPI_ISL_2238163 | Lighthouse Lab in Milton Keynes | Wellcome Sanger Institute for the COVID-19 Genomics UK (COG-UK) Consortium | The Lighthouse Lab in Milton Keynes and Alex Alderton, Roberto Amato, Jeffrey Barrett, Sonia Goncalves, Ewan Harrison, David K. Jackson, Ian Johnston, Dominic Kwiatkowski, Cordelia Langford, John Sillitoe on behalf of the Wellcome Sanger Institute COVID-19 Surveillance Team |
| EPI_ISL_2238164 | Lighthouse Lab in Glasgow | Wellcome Sanger Institute for the COVID-19 Genomics UK (COG-UK) Consortium | Harper VanSteenhouse, Yumi Kasai, David Gray, Carol Clugston, Anna Dominiczak and Alex Alderton, Roberto Amato, Jeffrey Barrett, Sonia Goncalves, Ewan Harrison, David K. Jackson, Ian Johnston, Dominic Kwiatkowski, Cordelia Langford, John Sillitoe on behalf of the Wellcome Sanger Institute COVID-19 Surveillance Team |
| EPI_ISL_2238165, EPI_ISL_2238167 | Lighthouse Lab in Milton Keynes | Wellcome Sanger Institute for the COVID-19 Genomics UK (COG-UK) Consortium | The Lighthouse Lab in Milton Keynes and Alex Alderton, Roberto Amato, Jeffrey Barrett, Sonia Goncalves, Ewan Harrison, David K. Jackson, Ian Johnston, Dominic Kwiatkowski, Cordelia Langford, John Sillitoe on behalf of the Wellcome Sanger Institute COVID-19 Surveillance Team |
| EPI_ISL_2238168 | Lighthouse Lab in Glasgow | Wellcome Sanger Institute for the COVID-19 Genomics UK (COG-UK) Consortium | Harper VanSteenhouse, Yumi Kasai, David Gray, Carol Clugston, Anna Dominiczak and Alex Alderton, Roberto Amato, Jeffrey Barrett, Sonia Goncalves, Ewan Harrison, David K. Jackson, Ian Johnston, Dominic Kwiatkowski, Cordelia Langford, John Sillitoe on behalf of the Wellcome Sanger Institute COVID-19 Surveillance Team |
| EPI_ISL_2238169 | Lighthouse Lab in Milton Keynes | Wellcome Sanger Institute for the COVID-19 Genomics UK (COG-UK) Consortium | The Lighthouse Lab in Milton Keynes and Alex Alderton, Roberto Amato, Jeffrey Barrett, Sonia Goncalves, Ewan Harrison, David K. Jackson, Ian Johnston, Dominic Kwiatkowski, Cordelia Langford, John Sillitoe on behalf of the Wellcome Sanger Institute COVID-19 Surveillance Team |
| EPI_ISL_2238172, EPI_ISL_2238173 | Lighthouse Lab in Glasgow | Wellcome Sanger Institute for the COVID-19 Genomics UK (COG-UK) Consortium | Harper VanSteenhouse, Yumi Kasai, David Gray, Carol Clugston, Anna Dominiczak and Alex Alderton, Roberto Amato, Jeffrey Barrett, Sonia Goncalves, Ewan Harrison, David K. Jackson, Ian Johnston, Dominic Kwiatkowski, Cordelia Langford, John Sillitoe on behalf of the Wellcome Sanger Institute COVID-19 Surveillance Team |
| EPI_ISL_2238174 | Lighthouse Lab in Milton Keynes | Wellcome Sanger Institute for the COVID-19 Genomics UK (COG-UK) Consortium | The Lighthouse Lab in Milton Keynes and Alex Alderton, Roberto Amato, Jeffrey Barrett, Sonia Goncalves, Ewan Harrison, David K. Jackson, Ian Johnston, Dominic Kwiatkowski, Cordelia Langford, John Sillitoe on behalf of the Wellcome Sanger Institute COVID-19 Surveillance Team |
| EPI_ISL_2238176 | Lighthouse Lab in Glasgow | Wellcome Sanger Institute for the COVID-19 Genomics UK (COG-UK) Consortium | Harper VanSteenhouse, Yumi Kasai, David Gray, Carol Clugston, Anna Dominiczak and Alex Alderton, Roberto Amato, Jeffrey Barrett, Sonia Goncalves, Ewan Harrison, David K. Jackson, Ian Johnston, Dominic Kwiatkowski, Cordelia Langford, John Sillitoe on behalf of the Wellcome Sanger Institute COVID-19 Surveillance Team |

| | | | |
|---|---|---|---|
| EPI_ISL_2238178, EPI_ISL_2238183 | Lighthouse Lab in Milton Keynes | Wellcome Sanger Institute for the COVID-19 Genomics UK (COG-UK) Consortium | The Lighthouse Lab in Milton Keynes and Alex Alderton, Roberto Amato, Jeffrey Barrett, Sonia Goncalves, Ewan Harrison, David K. Jackson, Ian Johnston, Dominic Kwiatkowski, Cordelia Langford, John Sillitoe on behalf of the Wellcome Sanger Institute COVID-19 Surveillance Team |
| EPI_ISL_2238184, EPI_ISL_2238186, EPI_ISL_2238188, EPI_ISL_2238199, EPI_ISL_2238203 | Lighthouse Lab in Glasgow | Wellcome Sanger Institute for the COVID-19 Genomics UK (COG-UK) Consortium | Harper VanSteenhouse, Yumi Kasai, David Gray, Carol Clugston, Anna Dominiczak and Alex Alderton, Roberto Amato, Jeffrey Barrett, Sonia Goncalves, Ewan Harrison, David K. Jackson, Ian Johnston, Dominic Kwiatkowski, Cordelia Langford, John Sillitoe on behalf of the Wellcome Sanger Institute COVID-19 Surveillance Team |
| EPI_ISL_2238205 | Lighthouse Lab in Milton Keynes | Wellcome Sanger Institute for the COVID-19 Genomics UK (COG-UK) Consortium | The Lighthouse Lab in Milton Keynes and Alex Alderton, Roberto Amato, Jeffrey Barrett, Sonia Goncalves, Ewan Harrison, David K. Jackson, Ian Johnston, Dominic Kwiatkowski, Cordelia Langford, John Sillitoe on behalf of the Wellcome Sanger Institute COVID-19 Surveillance Team |
| EPI_ISL_2238206, EPI_ISL_2238212, EPI_ISL_2238217 | Lighthouse Lab in Glasgow | Wellcome Sanger Institute for the COVID-19 Genomics UK (COG-UK) Consortium | Harper VanSteenhouse, Yumi Kasai, David Gray, Carol Clugston, Anna Dominiczak and Alex Alderton, Roberto Amato, Jeffrey Barrett, Sonia Goncalves, Ewan Harrison, David K. Jackson, Ian Johnston, Dominic Kwiatkowski, Cordelia Langford, John Sillitoe on behalf of the Wellcome Sanger Institute COVID-19 Surveillance Team |
| EPI_ISL_2238218 | Lighthouse Lab in Milton Keynes | Wellcome Sanger Institute for the COVID-19 Genomics UK (COG-UK) Consortium | The Lighthouse Lab in Milton Keynes and Alex Alderton, Roberto Amato, Jeffrey Barrett, Sonia Goncalves, Ewan Harrison, David K. Jackson, Ian Johnston, Dominic Kwiatkowski, Cordelia Langford, John Sillitoe on behalf of the Wellcome Sanger Institute COVID-19 Surveillance Team |
| EPI_ISL_2238219, EPI_ISL_2238221, EPI_ISL_2238224, EPI_ISL_2238225 | Lighthouse Lab in Glasgow | Wellcome Sanger Institute for the COVID-19 Genomics UK (COG-UK) Consortium | Harper VanSteenhouse, Yumi Kasai, David Gray, Carol Clugston, Anna Dominiczak and Alex Alderton, Roberto Amato, Jeffrey Barrett, Sonia Goncalves, Ewan Harrison, David K. Jackson, Ian Johnston, Dominic Kwiatkowski, Cordelia Langford, John Sillitoe on behalf of the Wellcome Sanger Institute COVID-19 Surveillance Team |
| EPI_ISL_2238230 | Lighthouse Lab in Milton Keynes | Wellcome Sanger Institute for the COVID-19 Genomics UK (COG-UK) Consortium | The Lighthouse Lab in Milton Keynes and Alex Alderton, Roberto Amato, Jeffrey Barrett, Sonia Goncalves, Ewan Harrison, David K. Jackson, Ian Johnston, Dominic Kwiatkowski, Cordelia Langford, John Sillitoe on behalf of the Wellcome Sanger Institute COVID-19 Surveillance Team |
| EPI_ISL_2238234 | Lighthouse Lab in Glasgow | Wellcome Sanger Institute for the COVID-19 Genomics UK (COG-UK) Consortium | Harper VanSteenhouse, Yumi Kasai, David Gray, Carol Clugston, Anna Dominiczak and Alex Alderton, Roberto Amato, Jeffrey Barrett, Sonia Goncalves, Ewan Harrison, David K. Jackson, Ian Johnston, Dominic Kwiatkowski, Cordelia Langford, John Sillitoe on behalf of the Wellcome Sanger Institute COVID-19 Surveillance Team |
| EPI_ISL_2238235, EPI_ISL_2238236, EPI_ISL_2238238 | Lighthouse Lab in Milton Keynes | Wellcome Sanger Institute for the COVID-19 Genomics UK (COG-UK) Consortium | The Lighthouse Lab in Milton Keynes and Alex Alderton, Roberto Amato, Jeffrey Barrett, Sonia Goncalves, Ewan Harrison, David K. Jackson, Ian Johnston, Dominic Kwiatkowski, Cordelia Langford, John Sillitoe on behalf of the Wellcome Sanger Institute COVID-19 Surveillance Team |
| EPI_ISL_2238242, EPI_ISL_2238244, EPI_ISL_2238245, EPI_ISL_2238247, EPI_ISL_2238250 | Lighthouse Lab in Glasgow | Wellcome Sanger Institute for the COVID-19 Genomics UK (COG-UK) Consortium | Harper VanSteenhouse, Yumi Kasai, David Gray, Carol Clugston, Anna Dominiczak and Alex Alderton, Roberto Amato, Jeffrey Barrett, Sonia Goncalves, Ewan Harrison, David K. Jackson, Ian Johnston, Dominic Kwiatkowski, Cordelia Langford, John Sillitoe on behalf of the Wellcome Sanger Institute COVID-19 Surveillance Team |
| EPI_ISL_2238251 | Lighthouse Lab in Alderley Park | Wellcome Sanger Institute for the COVID-19 Genomics UK (COG-UK) Consortium | Jacquelyn Wynn, Mairead Hyland, The Lighthouse Lab in Alderley Park and Alex Alderton, Roberto Amato, Jeffrey Barrett, Sonia Goncalves, Ewan Harrison, David K. Jackson, Ian Johnston, Dominic Kwiatkowski, Cordelia Langford, John Sillitoe on behalf of the Wellcome Sanger Institute COVID-19 Surveillance Team |
| EPI_ISL_2238253 | Lighthouse Lab in Milton Keynes | Wellcome Sanger Institute for the COVID-19 Genomics UK (COG-UK) Consortium | The Lighthouse Lab in Milton Keynes and Alex Alderton, Roberto Amato, Jeffrey Barrett, Sonia Goncalves, Ewan Harrison, David K. Jackson, Ian Johnston, Dominic Kwiatkowski, Cordelia Langford, John Sillitoe on behalf of the Wellcome Sanger Institute COVID-19 Surveillance Team |
| EPI_ISL_2238257 | Lighthouse Lab in Glasgow | Wellcome Sanger Institute for the COVID-19 Genomics UK (COG-UK) Consortium | Harper VanSteenhouse, Yumi Kasai, David Gray, Carol Clugston, Anna Dominiczak and Alex Alderton, Roberto Amato, Jeffrey Barrett, Sonia Goncalves, Ewan Harrison, David K. Jackson, Ian Johnston, Dominic Kwiatkowski, Cordelia Langford, John Sillitoe on behalf of the Wellcome Sanger Institute COVID-19 Surveillance Team |
| EPI_ISL_2238259 | Lighthouse Lab in Milton Keynes | Wellcome Sanger Institute for the COVID-19 Genomics UK (COG-UK) Consortium | The Lighthouse Lab in Milton Keynes and Alex Alderton, Roberto Amato, Jeffrey Barrett, Sonia Goncalves, Ewan Harrison, David K. Jackson, Ian Johnston, Dominic Kwiatkowski, Cordelia Langford, John Sillitoe on behalf of the Wellcome Sanger Institute COVID-19 Surveillance Team |
| EPI_ISL_2238261 | Lighthouse Lab in Alderley Park | Wellcome Sanger Institute for the COVID-19 Genomics UK (COG-UK) Consortium | Jacquelyn Wynn, Mairead Hyland, The Lighthouse Lab in Alderley Park and Alex Alderton, Roberto Amato, Jeffrey Barrett, Sonia Goncalves, Ewan Harrison, David K. Jackson, Ian Johnston, Dominic Kwiatkowski, Cordelia Langford, John Sillitoe on behalf of the Wellcome Sanger Institute COVID-19 Surveillance Team |
| EPI_ISL_2238263, EPI_ISL_2238266 | Lighthouse Lab in Glasgow | Wellcome Sanger Institute for the COVID-19 Genomics UK (COG-UK) Consortium | Harper VanSteenhouse, Yumi Kasai, David Gray, Carol Clugston, Anna Dominiczak and Alex Alderton, Roberto Amato, Jeffrey Barrett, Sonia Goncalves, Ewan Harrison, David K. Jackson, Ian Johnston, Dominic Kwiatkowski, Cordelia Langford, John Sillitoe on behalf of the Wellcome Sanger Institute COVID-19 Surveillance Team |
| EPI_ISL_2238267 | Lighthouse Lab in Milton Keynes | Wellcome Sanger Institute for the COVID-19 Genomics UK (COG-UK) Consortium | The Lighthouse Lab in Milton Keynes and Alex Alderton, Roberto Amato, Jeffrey Barrett, Sonia Goncalves, Ewan Harrison, David K. Jackson, Ian Johnston, Dominic Kwiatkowski, Cordelia Langford, John Sillitoe on behalf of the Wellcome Sanger Institute COVID-19 Surveillance Team |
| EPI_ISL_2238268, EPI_ISL_2238269 | Lighthouse Lab in Glasgow | Wellcome Sanger Institute for the COVID-19 Genomics UK (COG-UK) Consortium | Harper VanSteenhouse, Yumi Kasai, David Gray, Carol Clugston, Anna Dominiczak and Alex Alderton, Roberto Amato, Jeffrey Barrett, Sonia Goncalves, Ewan Harrison, David K. Jackson, Ian Johnston, Dominic Kwiatkowski, Cordelia Langford, John Sillitoe on behalf of the Wellcome Sanger Institute COVID-19 Surveillance Team |
| EPI_ISL_2238272 | Lighthouse Lab in Milton Keynes | Wellcome Sanger Institute for the COVID-19 Genomics UK (COG-UK) Consortium | The Lighthouse Lab in Milton Keynes and Alex Alderton, Roberto Amato, Jeffrey Barrett, Sonia Goncalves, Ewan Harrison, David K. Jackson, Ian Johnston, Dominic Kwiatkowski, Cordelia Langford, John Sillitoe on behalf of the Wellcome Sanger Institute COVID-19 Surveillance Team |
| EPI_ISL_2238274 | Lighthouse Lab in Glasgow | Wellcome Sanger Institute for the COVID-19 Genomics UK (COG-UK) Consortium | Harper VanSteenhouse, Yumi Kasai, David Gray, Carol Clugston, Anna Dominiczak and Alex Alderton, Roberto Amato, Jeffrey Barrett, Sonia Goncalves, Ewan Harrison, David K. Jackson, Ian Johnston, Dominic Kwiatkowski, Cordelia Langford, John Sillitoe on behalf of the Wellcome Sanger Institute COVID-19 Surveillance Team |
| EPI_ISL_2238276 | Lighthouse Lab in Milton Keynes | Wellcome Sanger Institute for the COVID-19 Genomics UK (COG-UK) Consortium | The Lighthouse Lab in Milton Keynes and Alex Alderton, Roberto Amato, Jeffrey Barrett, Sonia Goncalves, Ewan Harrison, David K. Jackson, Ian Johnston, Dominic Kwiatkowski, Cordelia Langford, John Sillitoe on behalf of the Wellcome Sanger Institute COVID-19 Surveillance Team |
| EPI_ISL_2238278 | Lighthouse Lab in Glasgow | Wellcome Sanger Institute for the COVID-19 Genomics UK (COG-UK) Consortium | Harper VanSteenhouse, Yumi Kasai, David Gray, Carol Clugston, Anna Dominiczak and Alex Alderton, Roberto Amato, Jeffrey Barrett, Sonia Goncalves, Ewan Harrison, David K. Jackson, Ian Johnston, Dominic Kwiatkowski, Cordelia Langford, John Sillitoe on behalf of the Wellcome Sanger Institute COVID-19 Surveillance Team |
| EPI_ISL_2238280, EPI_ISL_2238283, EPI_ISL_2238284 | Lighthouse Lab in Milton Keynes | Wellcome Sanger Institute for the COVID-19 Genomics UK (COG-UK) Consortium | The Lighthouse Lab in Milton Keynes and Alex Alderton, Roberto Amato, Jeffrey Barrett, Sonia Goncalves, Ewan Harrison, David K. Jackson, Ian Johnston, Dominic Kwiatkowski, Cordelia Langford, John Sillitoe on behalf of the Wellcome Sanger Institute COVID-19 Surveillance Team |
| EPI_ISL_2238285, EPI_ISL_2238286 | Lighthouse Lab in Glasgow | Wellcome Sanger Institute for the COVID-19 Genomics UK (COG-UK) Consortium | Harper VanSteenhouse, Yumi Kasai, David Gray, Carol Clugston, Anna Dominiczak and Alex Alderton, Roberto Amato, Jeffrey Barrett, Sonia Goncalves, Ewan Harrison, David K. Jackson, Ian Johnston, Dominic Kwiatkowski, Cordelia Langford, John Sillitoe on behalf of the Wellcome Sanger Institute COVID-19 Surveillance Team |
| EPI_ISL_2238287, EPI_ISL_2238288 | Lighthouse Lab in Milton Keynes | Wellcome Sanger Institute for the COVID-19 Genomics UK (COG-UK) Consortium | The Lighthouse Lab in Milton Keynes and Alex Alderton, Roberto Amato, Jeffrey Barrett, Sonia Goncalves, Ewan Harrison, David K. Jackson, Ian Johnston, Dominic Kwiatkowski, Cordelia Langford, John Sillitoe on behalf of the Wellcome Sanger Institute COVID-19 Surveillance Team |
| EPI_ISL_2238289, EPI_ISL_2238291, EPI_ISL_2238298, EPI_ISL_2238301 | Lighthouse Lab in Glasgow | Wellcome Sanger Institute for the COVID-19 Genomics UK (COG-UK) Consortium | Harper VanSteenhouse, Yumi Kasai, David Gray, Carol Clugston, Anna Dominiczak and Alex Alderton, Roberto Amato, Jeffrey Barrett, Sonia Goncalves, Ewan Harrison, David K. Jackson, Ian Johnston, Dominic Kwiatkowski, Cordelia Langford, John Sillitoe on behalf of the Wellcome Sanger Institute COVID-19 Surveillance Team |
| EPI_ISL_2238304, EPI_ISL_2238305 | Lighthouse Lab in Milton Keynes | Wellcome Sanger Institute for the COVID-19 Genomics UK (COG-UK) Consortium | The Lighthouse Lab in Milton Keynes and Alex Alderton, Roberto Amato, Jeffrey Barrett, Sonia Goncalves, Ewan Harrison, David K. Jackson, Ian Johnston, Dominic Kwiatkowski, Cordelia Langford, John Sillitoe on behalf of the Wellcome Sanger Institute COVID-19 Surveillance Team |
| EPI_ISL_2238308 | Lighthouse Lab in Glasgow | Wellcome Sanger Institute for the COVID-19 Genomics UK (COG-UK) Consortium | Harper VanSteenhouse, Yumi Kasai, David Gray, Carol Clugston, Anna Dominiczak and Alex Alderton, Roberto Amato, Jeffrey Barrett, Sonia Goncalves, Ewan Harrison, David K. Jackson, Ian Johnston, Dominic Kwiatkowski, Cordelia Langford, John Sillitoe on behalf of the Wellcome Sanger Institute COVID-19 Surveillance Team |
| EPI_ISL_2238310 | Lighthouse Lab in Milton Keynes | Wellcome Sanger Institute for the COVID-19 Genomics UK (COG-UK) Consortium | The Lighthouse Lab in Milton Keynes and Alex Alderton, Roberto Amato, Jeffrey Barrett, Sonia Goncalves, Ewan Harrison, David K. Jackson, Ian Johnston, Dominic Kwiatkowski, Cordelia Langford, John Sillitoe on behalf of the Wellcome Sanger Institute COVID-19 Surveillance Team |
| EPI_ISL_2238314 | Lighthouse Lab in Glasgow | Wellcome Sanger Institute for the COVID-19 Genomics UK (COG-UK) Consortium | Harper VanSteenhouse, Yumi Kasai, David Gray, Carol Clugston, Anna Dominiczak and Alex Alderton, Roberto Amato, Jeffrey Barrett, Sonia Goncalves, Ewan Harrison, David K. Jackson, Ian Johnston, Dominic Kwiatkowski, Cordelia Langford, John Sillitoe on behalf of the Wellcome Sanger Institute COVID-19 Surveillance Team |
| EPI_ISL_2238316, EPI_ISL_2238317, EPI_ISL_2238319, EPI_ISL_2238325, EPI_ISL_2238327, EPI_ISL_2238330 | Lighthouse Lab in Milton Keynes | Wellcome Sanger Institute for the COVID-19 Genomics UK (COG-UK) Consortium | The Lighthouse Lab in Milton Keynes and Alex Alderton, Roberto Amato, Jeffrey Barrett, Sonia Goncalves, Ewan Harrison, David K. Jackson, Ian Johnston, Dominic Kwiatkowski, Cordelia Langford, John Sillitoe on behalf of the Wellcome Sanger Institute COVID-19 Surveillance Team |
| EPI_ISL_2238334 | Lighthouse Lab in Alderley Park | Wellcome Sanger Institute for the COVID-19 Genomics UK (COG-UK) Consortium | Jacquelyn Wynn, Mairead Hyland, The Lighthouse Lab in Alderley Park and Alex Alderton, Roberto Amato, Jeffrey Barrett, Sonia Goncalves, Ewan Harrison, David K. Jackson, Ian Johnston, Dominic Kwiatkowski, Cordelia Langford, John Sillitoe on behalf of the Wellcome Sanger Institute COVID-19 Surveillance Team |
| EPI_ISL_2238336 | Lighthouse Lab in Glasgow | Wellcome Sanger Institute for the COVID-19 Genomics UK (COG-UK) Consortium | Harper VanSteenhouse, Yumi Kasai, David Gray, Carol Clugston, Anna Dominiczak and Alex Alderton, Roberto Amato, Jeffrey Barrett, Sonia Goncalves, Ewan Harrison, David K. Jackson, Ian Johnston, Dominic Kwiatkowski, Cordelia Langford, John Sillitoe on behalf of the Wellcome Sanger Institute COVID-19 Surveillance Team |
| EPI_ISL_2238338 | Lighthouse Lab in Milton Keynes | Wellcome Sanger Institute for the COVID-19 Genomics UK (COG-UK) Consortium | The Lighthouse Lab in Milton Keynes and Alex Alderton, Roberto Amato, Jeffrey Barrett, Sonia Goncalves, Ewan Harrison, David K. Jackson, Ian Johnston, Dominic Kwiatkowski, Cordelia Langford, John Sillitoe on behalf of the Wellcome Sanger Institute COVID-19 Surveillance Team |
| EPI_ISL_2238339 | Lighthouse Lab in Glasgow | Wellcome Sanger Institute for the COVID-19 Genomics UK (COG-UK) Consortium | Harper VanSteenhouse, Yumi Kasai, David Gray, Carol Clugston, Anna Dominiczak and Alex Alderton, Roberto Amato, Jeffrey Barrett, Sonia Goncalves, Ewan Harrison, David K. Jackson, Ian Johnston, Dominic Kwiatkowski, Cordelia Langford, John Sillitoe on behalf of the Wellcome Sanger Institute COVID-19 Surveillance Team |
| EPI_ISL_2238342, EPI_ISL_2238345, EPI_ISL_2238346 | Lighthouse Lab in Milton Keynes | Wellcome Sanger Institute for the COVID-19 Genomics UK (COG-UK) Consortium | The Lighthouse Lab in Milton Keynes and Alex Alderton, Roberto Amato, Jeffrey Barrett, Sonia Goncalves, Ewan Harrison, David K. Jackson, Ian Johnston, Dominic Kwiatkowski, Cordelia Langford, John Sillitoe on behalf of the Wellcome Sanger Institute COVID-19 Surveillance Team |
| EPI_ISL_2238353, EPI_ISL_2238355 | Lighthouse Lab in Glasgow | Wellcome Sanger Institute for the COVID-19 Genomics UK (COG-UK) Consortium | Harper VanSteenhouse, Yumi Kasai, David Gray, Carol Clugston, Anna Dominiczak and Alex Alderton, Roberto Amato, Jeffrey Barrett, Sonia Goncalves, Ewan Harrison, David K. Jackson, Ian Johnston, Dominic Kwiatkowski, Cordelia Langford, John Sillitoe on behalf of the Wellcome Sanger Institute COVID-19 Surveillance Team |
| EPI_ISL_2238363, | Lighthouse Lab in Alderley Park | Wellcome Sanger Institute for the COVID-19 Genomics UK (COG-UK) | Jacquelyn Wynn, Mairead Hyland, The Lighthouse Lab in Alderley Park and Alex Alderton, Roberto Amato, Jeffrey Barrett, Sonia Goncalves, Ewan Harrison, David K. Jackson, Ian Johnston, Dominic Kwiatkowski, Cordelia Langford, John Sillitoe on |

| | | | |
|---|---|---|---|
| EPI_ISL_2238364, EPI_ISL_2238366, EPI_ISL_2238381, EPI_ISL_2238384 | | Consortium | behalf of the Wellcome Sanger Institute COVID-19 Surveillance Team |
| EPI_ISL_2238385, EPI_ISL_2238389, EPI_ISL_2238406 | Lighthouse Lab in Glasgow | Wellcome Sanger Institute for the COVID-19 Genomics UK (COG-UK) Consortium | Harper VanSteenhouse, Yumi Kasai, David Gray, Carol Clugston, Anna Dominiczak and Alex Alderton, Roberto Amato, Jeffrey Barrett, Sonia Goncalves, Ewan Harrison, David K. Jackson, Ian Johnston, Dominic Kwiatkowski, Cordelia Langford, John Sillitoe on behalf of the Wellcome Sanger Institute COVID-19 Surveillance Team |
| EPI_ISL_2238411, EPI_ISL_2238415, EPI_ISL_2238421, EPI_ISL_2238422, EPI_ISL_2238423 | Lighthouse Lab in Alderley Park | Wellcome Sanger Institute for the COVID-19 Genomics UK (COG-UK) Consortium | Jacquelyn Wynn, Mairead Hyland, The Lighthouse Lab in Alderley Park and Alex Alderton, Roberto Amato, Jeffrey Barrett, Sonia Goncalves, Ewan Harrison, David K. Jackson, Ian Johnston, Dominic Kwiatkowski, Cordelia Langford, John Sillitoe on behalf of the Wellcome Sanger Institute COVID-19 Surveillance Team |
| EPI_ISL_2238425 | Lighthouse Lab in Glasgow | Wellcome Sanger Institute for the COVID-19 Genomics UK (COG-UK) Consortium | Harper VanSteenhouse, Yumi Kasai, David Gray, Carol Clugston, Anna Dominiczak and Alex Alderton, Roberto Amato, Jeffrey Barrett, Sonia Goncalves, Ewan Harrison, David K. Jackson, Ian Johnston, Dominic Kwiatkowski, Cordelia Langford, John Sillitoe on behalf of the Wellcome Sanger Institute COVID-19 Surveillance Team |
| EPI_ISL_2238426, EPI_ISL_2238431, EPI_ISL_2238433 | Lighthouse Lab in Alderley Park | Wellcome Sanger Institute for the COVID-19 Genomics UK (COG-UK) Consortium | Jacquelyn Wynn, Mairead Hyland, The Lighthouse Lab in Alderley Park and Alex Alderton, Roberto Amato, Jeffrey Barrett, Sonia Goncalves, Ewan Harrison, David K. Jackson, Ian Johnston, Dominic Kwiatkowski, Cordelia Langford, John Sillitoe on behalf of the Wellcome Sanger Institute COVID-19 Surveillance Team |
| EPI_ISL_2238436, EPI_ISL_2238440 | Lighthouse Lab in Glasgow | Wellcome Sanger Institute for the COVID-19 Genomics UK (COG-UK) Consortium | Harper VanSteenhouse, Yumi Kasai, David Gray, Carol Clugston, Anna Dominiczak and Alex Alderton, Roberto Amato, Jeffrey Barrett, Sonia Goncalves, Ewan Harrison, David K. Jackson, Ian Johnston, Dominic Kwiatkowski, Cordelia Langford, John Sillitoe on behalf of the Wellcome Sanger Institute COVID-19 Surveillance Team |
| EPI_ISL_2238441 | Lighthouse Lab in Alderley Park | Wellcome Sanger Institute for the COVID-19 Genomics UK (COG-UK) Consortium | Jacquelyn Wynn, Mairead Hyland, The Lighthouse Lab in Alderley Park and Alex Alderton, Roberto Amato, Jeffrey Barrett, Sonia Goncalves, Ewan Harrison, David K. Jackson, Ian Johnston, Dominic Kwiatkowski, Cordelia Langford, John Sillitoe on behalf of the Wellcome Sanger Institute COVID-19 Surveillance Team |
| EPI_ISL_2238451, EPI_ISL_2238455, EPI_ISL_2238460, EPI_ISL_2238464, EPI_ISL_2238465 | Lighthouse Lab in Glasgow | Wellcome Sanger Institute for the COVID-19 Genomics UK (COG-UK) Consortium | Harper VanSteenhouse, Yumi Kasai, David Gray, Carol Clugston, Anna Dominiczak and Alex Alderton, Roberto Amato, Jeffrey Barrett, Sonia Goncalves, Ewan Harrison, David K. Jackson, Ian Johnston, Dominic Kwiatkowski, Cordelia Langford, John Sillitoe on behalf of the Wellcome Sanger Institute COVID-19 Surveillance Team |
| EPI_ISL_2238466 | Lighthouse Lab in Alderley Park | Wellcome Sanger Institute for the COVID-19 Genomics UK (COG-UK) Consortium | Jacquelyn Wynn, Mairead Hyland, The Lighthouse Lab in Alderley Park and Alex Alderton, Roberto Amato, Jeffrey Barrett, Sonia Goncalves, Ewan Harrison, David K. Jackson, Ian Johnston, Dominic Kwiatkowski, Cordelia Langford, John Sillitoe on behalf of the Wellcome Sanger Institute COVID-19 Surveillance Team |
| EPI_ISL_2238473, EPI_ISL_2238476 | Lighthouse Lab in Glasgow | Wellcome Sanger Institute for the COVID-19 Genomics UK (COG-UK) Consortium | Harper VanSteenhouse, Yumi Kasai, David Gray, Carol Clugston, Anna Dominiczak and Alex Alderton, Roberto Amato, Jeffrey Barrett, Sonia Goncalves, Ewan Harrison, David K. Jackson, Ian Johnston, Dominic Kwiatkowski, Cordelia Langford, John Sillitoe on behalf of the Wellcome Sanger Institute COVID-19 Surveillance Team |
| EPI_ISL_2238480 | Lighthouse Lab in Alderley Park | Wellcome Sanger Institute for the COVID-19 Genomics UK (COG-UK) Consortium | Jacquelyn Wynn, Mairead Hyland, The Lighthouse Lab in Alderley Park and Alex Alderton, Roberto Amato, Jeffrey Barrett, Sonia Goncalves, Ewan Harrison, David K. Jackson, Ian Johnston, Dominic Kwiatkowski, Cordelia Langford, John Sillitoe on behalf of the Wellcome Sanger Institute COVID-19 Surveillance Team |
| EPI_ISL_2238481 | Lighthouse Lab in Glasgow | Wellcome Sanger Institute for the COVID-19 Genomics UK (COG-UK) Consortium | Harper VanSteenhouse, Yumi Kasai, David Gray, Carol Clugston, Anna Dominiczak and Alex Alderton, Roberto Amato, Jeffrey Barrett, Sonia Goncalves, Ewan Harrison, David K. Jackson, Ian Johnston, Dominic Kwiatkowski, Cordelia Langford, John Sillitoe on behalf of the Wellcome Sanger Institute COVID-19 Surveillance Team |
| EPI_ISL_2238486 | Lighthouse Lab in Alderley Park | Wellcome Sanger Institute for the COVID-19 Genomics UK (COG-UK) Consortium | Jacquelyn Wynn, Mairead Hyland, The Lighthouse Lab in Alderley Park and Alex Alderton, Roberto Amato, Jeffrey Barrett, Sonia Goncalves, Ewan Harrison, David K. Jackson, Ian Johnston, Dominic Kwiatkowski, Cordelia Langford, John Sillitoe on behalf of the Wellcome Sanger Institute COVID-19 Surveillance Team |
| EPI_ISL_2238499, EPI_ISL_2238504, EPI_ISL_2238519 | Lighthouse Lab in Glasgow | Wellcome Sanger Institute for the COVID-19 Genomics UK (COG-UK) Consortium | Harper VanSteenhouse, Yumi Kasai, David Gray, Carol Clugston, Anna Dominiczak and Alex Alderton, Roberto Amato, Jeffrey Barrett, Sonia Goncalves, Ewan Harrison, David K. Jackson, Ian Johnston, Dominic Kwiatkowski, Cordelia Langford, John Sillitoe on behalf of the Wellcome Sanger Institute COVID-19 Surveillance Team |
| EPI_ISL_2238526 | Lighthouse Lab in Alderley Park | Wellcome Sanger Institute for the COVID-19 Genomics UK (COG-UK) Consortium | Jacquelyn Wynn, Mairead Hyland, The Lighthouse Lab in Alderley Park and Alex Alderton, Roberto Amato, Jeffrey Barrett, Sonia Goncalves, Ewan Harrison, David K. Jackson, Ian Johnston, Dominic Kwiatkowski, Cordelia Langford, John Sillitoe on behalf of the Wellcome Sanger Institute COVID-19 Surveillance Team |
| EPI_ISL_2238527, EPI_ISL_2238528, EPI_ISL_2238529 | Lighthouse Lab in Glasgow | Wellcome Sanger Institute for the COVID-19 Genomics UK (COG-UK) Consortium | Harper VanSteenhouse, Yumi Kasai, David Gray, Carol Clugston, Anna Dominiczak and Alex Alderton, Roberto Amato, Jeffrey Barrett, Sonia Goncalves, Ewan Harrison, David K. Jackson, Ian Johnston, Dominic Kwiatkowski, Cordelia Langford, John Sillitoe on behalf of the Wellcome Sanger Institute COVID-19 Surveillance Team |
| EPI_ISL_2238537, EPI_ISL_2238545, EPI_ISL_2238548, EPI_ISL_2238553, EPI_ISL_2238556 | Lighthouse Lab in Alderley Park | Wellcome Sanger Institute for the COVID-19 Genomics UK (COG-UK) Consortium | Jacquelyn Wynn, Mairead Hyland, The Lighthouse Lab in Alderley Park and Alex Alderton, Roberto Amato, Jeffrey Barrett, Sonia Goncalves, Ewan Harrison, David K. Jackson, Ian Johnston, Dominic Kwiatkowski, Cordelia Langford, John Sillitoe on behalf of the Wellcome Sanger Institute COVID-19 Surveillance Team |
| EPI_ISL_2238557, EPI_ISL_2238560 | Lighthouse Lab in Glasgow | Wellcome Sanger Institute for the COVID-19 Genomics UK (COG-UK) Consortium | Harper VanSteenhouse, Yumi Kasai, David Gray, Carol Clugston, Anna Dominiczak and Alex Alderton, Roberto Amato, Jeffrey Barrett, Sonia Goncalves, Ewan Harrison, David K. Jackson, Ian Johnston, Dominic Kwiatkowski, Cordelia Langford, John Sillitoe on behalf of the Wellcome Sanger Institute COVID-19 Surveillance Team |
| EPI_ISL_2238561 | Lighthouse Lab in Alderley Park | Wellcome Sanger Institute for the COVID-19 Genomics UK (COG-UK) Consortium | Jacquelyn Wynn, Mairead Hyland, The Lighthouse Lab in Alderley Park and Alex Alderton, Roberto Amato, Jeffrey Barrett, Sonia Goncalves, Ewan Harrison, David K. Jackson, Ian Johnston, Dominic Kwiatkowski, Cordelia Langford, John Sillitoe on behalf of the Wellcome Sanger Institute COVID-19 Surveillance Team |
| EPI_ISL_2238568, EPI_ISL_2238577, EPI_ISL_2238589 | Lighthouse Lab in Glasgow | Wellcome Sanger Institute for the COVID-19 Genomics UK (COG-UK) Consortium | Harper VanSteenhouse, Yumi Kasai, David Gray, Carol Clugston, Anna Dominiczak and Alex Alderton, Roberto Amato, Jeffrey Barrett, Sonia Goncalves, Ewan Harrison, David K. Jackson, Ian Johnston, Dominic Kwiatkowski, Cordelia Langford, John Sillitoe on behalf of the Wellcome Sanger Institute COVID-19 Surveillance Team |
| EPI_ISL_2238594 | Lighthouse Lab in Alderley Park | Wellcome Sanger Institute for the COVID-19 Genomics UK (COG-UK) Consortium | Jacquelyn Wynn, Mairead Hyland, The Lighthouse Lab in Alderley Park and Alex Alderton, Roberto Amato, Jeffrey Barrett, Sonia Goncalves, Ewan Harrison, David K. Jackson, Ian Johnston, Dominic Kwiatkowski, Cordelia Langford, John Sillitoe on behalf of the Wellcome Sanger Institute COVID-19 Surveillance Team |
| EPI_ISL_2238598 | Lighthouse Lab in Glasgow | Wellcome Sanger Institute for the COVID-19 Genomics UK (COG-UK) Consortium | Harper VanSteenhouse, Yumi Kasai, David Gray, Carol Clugston, Anna Dominiczak and Alex Alderton, Roberto Amato, Jeffrey Barrett, Sonia Goncalves, Ewan Harrison, David K. Jackson, Ian Johnston, Dominic Kwiatkowski, Cordelia Langford, John Sillitoe on behalf of the Wellcome Sanger Institute COVID-19 Surveillance Team |
| EPI_ISL_2238618, EPI_ISL_2238619, EPI_ISL_2238620 | Lighthouse Lab in Alderley Park | Wellcome Sanger Institute for the COVID-19 Genomics UK (COG-UK) Consortium | Jacquelyn Wynn, Mairead Hyland, The Lighthouse Lab in Alderley Park and Alex Alderton, Roberto Amato, Jeffrey Barrett, Sonia Goncalves, Ewan Harrison, David K. Jackson, Ian Johnston, Dominic Kwiatkowski, Cordelia Langford, John Sillitoe on behalf of the Wellcome Sanger Institute COVID-19 Surveillance Team |
| EPI_ISL_2238622, EPI_ISL_2238630 | Lighthouse Lab in Glasgow | Wellcome Sanger Institute for the COVID-19 Genomics UK (COG-UK) Consortium | Harper VanSteenhouse, Yumi Kasai, David Gray, Carol Clugston, Anna Dominiczak and Alex Alderton, Roberto Amato, Jeffrey Barrett, Sonia Goncalves, Ewan Harrison, David K. Jackson, Ian Johnston, Dominic Kwiatkowski, Cordelia Langford, John Sillitoe on behalf of the Wellcome Sanger Institute COVID-19 Surveillance Team |
| EPI_ISL_2238633 | Lighthouse Lab in Alderley Park | Wellcome Sanger Institute for the COVID-19 Genomics UK (COG-UK) Consortium | Jacquelyn Wynn, Mairead Hyland, The Lighthouse Lab in Alderley Park and Alex Alderton, Roberto Amato, Jeffrey Barrett, Sonia Goncalves, Ewan Harrison, David K. Jackson, Ian Johnston, Dominic Kwiatkowski, Cordelia Langford, John Sillitoe on behalf of the Wellcome Sanger Institute COVID-19 Surveillance Team |
| EPI_ISL_2238637 | Lighthouse Lab in Glasgow | Wellcome Sanger Institute for the COVID-19 Genomics UK (COG-UK) Consortium | Harper VanSteenhouse, Yumi Kasai, David Gray, Carol Clugston, Anna Dominiczak and Alex Alderton, Roberto Amato, Jeffrey Barrett, Sonia Goncalves, Ewan Harrison, David K. Jackson, Ian Johnston, Dominic Kwiatkowski, Cordelia Langford, John Sillitoe on behalf of the Wellcome Sanger Institute COVID-19 Surveillance Team |
| EPI_ISL_2238638, EPI_ISL_2238640, EPI_ISL_2238641, EPI_ISL_2238649, EPI_ISL_2238659, EPI_ISL_2238669 | Lighthouse Lab in Alderley Park | Wellcome Sanger Institute for the COVID-19 Genomics UK (COG-UK) Consortium | Jacquelyn Wynn, Mairead Hyland, The Lighthouse Lab in Alderley Park and Alex Alderton, Roberto Amato, Jeffrey Barrett, Sonia Goncalves, Ewan Harrison, David K. Jackson, Ian Johnston, Dominic Kwiatkowski, Cordelia Langford, John Sillitoe on behalf of the Wellcome Sanger Institute COVID-19 Surveillance Team |
| EPI_ISL_2238671 | Lighthouse Lab in Glasgow | Wellcome Sanger Institute for the COVID-19 Genomics UK (COG-UK) Consortium | Harper VanSteenhouse, Yumi Kasai, David Gray, Carol Clugston, Anna Dominiczak and Alex Alderton, Roberto Amato, Jeffrey Barrett, Sonia Goncalves, Ewan Harrison, David K. Jackson, Ian Johnston, Dominic Kwiatkowski, Cordelia Langford, John Sillitoe on behalf of the Wellcome Sanger Institute COVID-19 Surveillance Team |
| EPI_ISL_2238676, EPI_ISL_2238678 | Lighthouse Lab in Alderley Park | Wellcome Sanger Institute for the COVID-19 Genomics UK (COG-UK) Consortium | Jacquelyn Wynn, Mairead Hyland, The Lighthouse Lab in Alderley Park and Alex Alderton, Roberto Amato, Jeffrey Barrett, Sonia Goncalves, Ewan Harrison, David K. Jackson, Ian Johnston, Dominic Kwiatkowski, Cordelia Langford, John Sillitoe on behalf of the Wellcome Sanger Institute COVID-19 Surveillance Team |
| EPI_ISL_2238682, EPI_ISL_2238685, EPI_ISL_2238693, EPI_ISL_2238695 | Lighthouse Lab in Milton Keynes | Wellcome Sanger Institute for the COVID-19 Genomics UK (COG-UK) Consortium | The Lighthouse Lab in Milton Keynes and Alex Alderton, Roberto Amato, Jeffrey Barrett, Sonia Goncalves, Ewan Harrison, David K. Jackson, Ian Johnston, Dominic Kwiatkowski, Cordelia Langford, John Sillitoe on behalf of the Wellcome Sanger Institute COVID-19 Surveillance Team |
| EPI_ISL_2238730, EPI_ISL_2238733, EPI_ISL_2238736 | University of Birmingham | COVID-19 Genomics UK (COG-UK) Consortium | Institute of Microbiology, University of Birmingham: Claire McMurray, Joanne Stockton, Samuel Nicholls, Radoslaw Poplawski, Will Rowe, Josh Quick, Nicholas Loman. University of Birmingham Testing Laboratory: Celina M Whalley, Andrew Bosworth, Charlotte Poxon, Kasun Wanigasooriya, Oliver Pickles, Mike Kidd, Alex Richter, Andrew D Beggs PHE Heartlands Lab: Husam Osman, Andrew Bosworth. Queen Elizabeth Hospital: Anna Casey |
| EPI_ISL_2238739, EPI_ISL_2238742, EPI_ISL_2238743, EPI_ISL_2238744, EPI_ISL_2238745, EPI_ISL_2238746, EPI_ISL_2238749 | Department of Pathology, University of Cambridge | COVID-19 Genomics UK (COG-UK) Consortium | Aminu S. Jahun, Yasmin Chaudhry, Iliana Georgana, Myra Hosmillo, Rhys Izuagbe, William L. Hamilton, Martin D. Curran, Surendra Parmar, Ian Goodfellow |
| EPI_ISL_2238812, EPI_ISL_2238813, EPI_ISL_2238814, | Virology Department, Royal Infirmary of Edinburgh, NHS Lothian / School of Biological Sciences, University of Edinburgh | COVID-19 Genomics UK (COG-UK) Consortium | McHugh M, Dewar R, Cotton S, Gallagher A, Maloney D, Rooke S, O'Toole Á, Scher E, Hill V, McCrone JT, Colquhoun R, Yu X, Jackson B, Rambaut A, Templeton K |

| | | | |
|---|---|---|---|
| EPI_ISL_2238815, EPI_ISL_2238816, EPI_ISL_2238818 | | | |
| EPI_ISL_2238840 | University of Exeter | COVID-19 Genomics UK (COG-UK) Consortium | Ben Temperton,Aaron Jeffries,Michelle Michelsen,Joanna Warwick-Dugdale,Audrey Farbos,Robyn Manley,Stephen Michell,Jane Masoli |
| EPI_ISL_2238862, EPI_ISL_2238864, EPI_ISL_2238865, EPI_ISL_2238866, EPI_ISL_2238869, EPI_ISL_2238870 | University Hospitals Of Leicester NHS Trust and DeepSeq Nottingham | COVID-19 Genomics UK (COG-UK) Consortium | Christopher Holmes, Paul Bird, Thomas Helmer, Karlie Fallon, Julian Tang, Jonathan Ball, Patrick McClure, Joeseph Chappell, Nadine Holmes, Matthew Carlisle, Christopher Moore, Fei Sang, Johnny Debebe, Victoria Wright, Matthew Loose |
| EPI_ISL_2238871, EPI_ISL_2238872, EPI_ISL_2238875, EPI_ISL_2238876, EPI_ISL_2238877, EPI_ISL_2238881, EPI_ISL_2238886, EPI_ISL_2238889, EPI_ISL_2238890, EPI_ISL_2238891 | Liverpool Clinical Laboratories | COVID-19 Genomics UK (COG-UK) Consortium | Sam Haldenby, Alistair Darby, Steve Paterson, Anita Lucaci, Julian Hiscox, M Almsaud, A Alrezaihi, Muhannad Alruwaili, Stuart D Armstrong, Jones Benjamin, Eleanor G Bentley, Anu Chawla, Jordan J Clark, Angela Cowell, Richard Eccles, Isabel García-Dorival, Matthew Gemmell, Alessandro Gerada, PKF Gilmore, Richard Gregory, Ximeng Han, Catherine Hartley, Margaret Hughes, Miren Iturriza-Gomara, James Johnson, L Luu, Jenifer Manson, Charlotte Nelson, Elaine O'Toole, Cassie Olateju, Rebekah Penrice-Randal , Lucille Rainbow, N.P Randle, Trevor Ian Robinson, Parul Sharma, Ghada T Shawli, James P Stewart, Neil Swainston, Ecaterina Vamos, Joanne Watts, Mark Whitehead, Hermione Webster |
| EPI_ISL_2238945 | Barts Health NHS Trust | COVID-19 Genomics UK (COG-UK) Consortium | CUTINO-MOGUEL, Maria-Teresa; HARRINGTON, David; OWOYEMI, Dola; KULASEGARAN-SHYLINI, Raghavendran; BROAD, Claire; KELE, Beatrix |
| EPI_ISL_2238949, EPI_ISL_2238951, EPI_ISL_2238954, EPI_ISL_2238955, EPI_ISL_2238956, EPI_ISL_2238957, EPI_ISL_2238959, EPI_ISL_2238960, EPI_ISL_2238961, EPI_ISL_2238963, EPI_ISL_2238964, EPI_ISL_2238965, EPI_ISL_2238966, EPI_ISL_2238968, EPI_ISL_2238969, EPI_ISL_2238971, EPI_ISL_2238972, EPI_ISL_2238973, EPI_ISL_2238979, EPI_ISL_2238981, EPI_ISL_2238982 | | | |
| see above | University College London, Great Ormond Street Hospital for Children NHS Foundation Trust, Imperial College Healthcare NHS Trust | COVID-19 Genomics UK (COG-UK) Consortium | Sergi Castellano, Rachel Williams, Mark Kristiansen, Paola Resende Silva, Sunando Roy, Tony Brooks, Helena Tutill, Paola Niola, Patricia Dyal, Charlotte Williams, Leysa Forrest, Yasmin Panchbhaya, Jacqueline Findlay, Samuel Weeks, Julianne Brown, Kathryn Harris, Paul Randell, James Price, Alison Holmes, Judith Breuer |
| EPI_ISL_2239321, EPI_ISL_2239324, EPI_ISL_2239341, EPI_ISL_2239374, EPI_ISL_2239384, EPI_ISL_2239561, EPI_ISL_2239568, EPI_ISL_2239605, EPI_ISL_2239606, EPI_ISL_2239650, EPI_ISL_2239651, EPI_ISL_2239660, EPI_ISL_2239667, EPI_ISL_2239683, EPI_ISL_2239702, EPI_ISL_2239708, EPI_ISL_2239724, EPI_ISL_2239725, EPI_ISL_2239734, EPI_ISL_2239746, EPI_ISL_2239748, EPI_ISL_2239752, EPI_ISL_2239763, EPI_ISL_2239765, EPI_ISL_2239769, EPI_ISL_2239773, EPI_ISL_2239774, EPI_ISL_2239788, EPI_ISL_2239789, EPI_ISL_2239795, EPI_ISL_2239802, EPI_ISL_2239813, EPI_ISL_2239820, EPI_ISL_2239859, EPI_ISL_2239866, EPI_ISL_2239931, EPI_ISL_2239933, EPI_ISL_2239935, EPI_ISL_2239940, EPI_ISL_2239944, EPI_ISL_2239946, EPI_ISL_2239948, EPI_ISL_2239950, EPI_ISL_2239955, EPI_ISL_2239973, EPI_ISL_2240016, EPI_ISL_2240095, EPI_ISL_2240130, EPI_ISL_2240136, EPI_ISL_2240141, EPI_ISL_2240145, EPI_ISL_2240147, EPI_ISL_2240165, EPI_ISL_2240168, EPI_ISL_2240186, EPI_ISL_2240187, EPI_ISL_2240190, EPI_ISL_2240192, EPI_ISL_2240195, EPI_ISL_2240197, EPI_ISL_2240209, EPI_ISL_2240213, EPI_ISL_2240218, EPI_ISL_2240267, EPI_ISL_2240306, EPI_ISL_2240354, EPI_ISL_2240357, EPI_ISL_2240362, EPI_ISL_2240408, EPI_ISL_2240410 | | | |
| see above | Northumbria University / South Tees Hospitals NHS Foundation Trust / North Cumbria Integrated Care NHS Foundation Trust / North Tees and Hartlepool NHS Foundation Trust / Newcastle Hospitals NHS Foundation Trust | COVID-19 Genomics UK (COG-UK) Consortium | Darren L Smith,Andrew Nelson,Matthew Bashton,Greg R Young,Joshua Loh,John Allan,Mohammad A Tariq,Giles S Holt,Gary Black,Wen C Yew,Lynn Dover,Paul Baker,Steve Liggett,Sarah Essex,Jane Greenaway,Debra Padgett,Clive Graham,Garren Scott,Edward Barton,Emma Swindells,Brendan Payne,Jennifer Collins,Yusri Taha,Gary Eltringham |
| EPI_ISL_2240447 | Lincolnshire Hospitals and DeepSeq Nottingham | COVID-19 Genomics UK (COG-UK) Consortium | Nichola Duckworth, Tim Sloan, Sarah Walsh, Jonathan Ball, Patrick McClure, Joeseph Chappell, Nadine Holmes, Matthew Carlisle, Christopher Moore, Fei Sang, Johnny Debebe, Victoria Wright, Matthew Loose |
| EPI_ISL_2240453, EPI_ISL_2240454, EPI_ISL_2240458, EPI_ISL_2240459, EPI_ISL_2240460 | Queens Medical Centre, Clinical Microbiology Department / DeepSeq Nottingham | COVID-19 Genomics UK (COG-UK) Consortium | Gemma Clark, Wendy Smith, Manjinder Khakh, Vicki M Fleming, Michelle M Lister, Hannah Howson-Wells, Jonathan Ball, Timothy Byaruhanga, Jayasree Dey, Emily Park, Jack Hill, Patrick McClure, Joseph Chappell, Theocharis Tsoleridis, Nadine Holmes, Matthew Carlisle, Christopher Moore, Fei Sang, Johnny Debebe, Victoria Wright, Matthew Loose |
| EPI_ISL_2240501 | Oxford Viromics, NDM, University of Oxford; Oxford University Hospitals; Basingstoke and North Hampshire Hospital | COVID-19 Genomics UK (COG-UK) Consortium | Tanya Golubchik, David Bonsall, George Macintyre, Amy Trebes, Mariateresa de Cesare, Catrin Moore, Alex Mobbs, Anita Justice, Robert Shaw, Monique Andersson, Timothy Peto, Emma Wise, Nathan Moore, Jessica Lynch, Nick Cortes, Matilde Mori, Stephen Kidd, David Buck, John Todd, Christophe Fraser |
| EPI_ISL_2240511, EPI_ISL_2240518, EPI_ISL_2240541, EPI_ISL_2240547, EPI_ISL_2240553, EPI_ISL_2240564, EPI_ISL_2240568, EPI_ISL_2240579, EPI_ISL_2240580, EPI_ISL_2240582, EPI_ISL_2240589, EPI_ISL_2240593, EPI_ISL_2240594, EPI_ISL_2240595, EPI_ISL_2240597, EPI_ISL_2240598, EPI_ISL_2240599, EPI_ISL_2240600, EPI_ISL_2240601, EPI_ISL_2240604, EPI_ISL_2240612, EPI_ISL_2240618, EPI_ISL_2240624, EPI_ISL_2240628, EPI_ISL_2240630, EPI_ISL_2240631, EPI_ISL_2240638, EPI_ISL_2240639, EPI_ISL_2240641, EPI_ISL_2240642, EPI_ISL_2240651, EPI_ISL_2240658 | | | |
| see above | Originating lab: Wales Specialist Virology Centre Sequencing lab: Pathogen Genomics Unit | Public Health Wales Microbiology Cardiff Wales Specialist Virology Centre | Catherine Moore, Johnathan Evans, Laura Gifford, Malorie Perry, Simon Cottrell, Angela Marchbank, Alec Birchley, Alexander Adams, Amy Gaskin, Bree Gatica-Wilcox, Jason Coombes, Joel Southgate, Lauren Gilbert, Lee Graham, Nicole Pacchiarini, Sara Kumziene-Summerhayes, Sarah Taylor, Sophie Jones, Sara Rey, Matthew Bull, Joanne Watkins, Sally Corden, Tom Connor |
| EPI_ISL_2240664 | Centre for Enzyme Innovation, University of Portsmouth / Translational Research Laboratory, Portsmouth Hospitals NHS Trust / Hampshire Hospitals NHS Foundation Trust / Bournemouth University / University Hospitals Dorset NHS Foundation Trust / Brighton and Sussex University Hospitals NHS Trust / Dartford and Gravesham NHS Trust / Isle of Wight NHS Trust / Maidstone and Tunbridge Wells NHS Trust / University Hospital Southampton NHS Foundation Trust | COVID-19 Genomics UK (COG-UK) Consortium | Samuel Robson, Angela Beckett, Salman Goudarzi, Christopher Fearn, Kate Cook, Katie Loveson, Sharon Glaysher, Scott Elliott, Kelly Bicknell, Sarah Wyllie, Allyson Lloyd, Robert Impey, Anoop Chauhan, Stephen Kidd, Nathan Moore, Nick Cortes, Claire Thomas, Anna Mantzaouratou, Sarah Buchan, Magdalena Barrow, Andrew Butt, Liz Sheridan, Jonnie Seymour, Dorian Crudgington, Ben Macklin, Mohammed Hassan-Ibrahim, Cassandra Malone, Benjamin Cogger, Kevin Tucker, Samirakhon Raupova, Rachael Jeremiah, Anibolina Castigador, Emily Macnaughton, Karen Withell, Kordo Saeed, Jacqui Prieto, Adhyana Mahanama, Buddhini Samaraweera, Siona Silvieri, Emanuela Pelosi, Eleri Wilson-Davies, Sarah Jeremiah, Helen Wheeler, Matthew Harvey, Thea Sass, Helen Umpleby, Stephen Aplin |
| EPI_ISL_2240693, EPI_ISL_2240700, EPI_ISL_2240726 | Virology Department, Sheffield Teaching Hospitals NHS Foundation Trust/Department of Infection, Immunity and Cardiovascular Disease, The Medical School, University of Sheffield | COVID-19 Genomics UK (COG-UK) Consortium | Thushan de Silva, Matthew Parker, Nikki Smith, Adri Angyal, Rebecca Brown, Luke Green, Rachel Tucker, Paul Parsons, Danielle Groves, Katie Johnson, Laura Carrilero, Alex Keeley, Dave Partridge, Matthew Wyles, Benjamin Lindsey, Mehmet Yavuz, Mohammad Raza, Cariad Evans |
| EPI_ISL_2240824 | Laboratorio HUB -Azienda Ospedaliero Universitaria - AOU – Cagliari | Laboratorio SPOKE Biologia Molecolare -Azienda Ospedaliero Universitaria - AOU – Cagliari | Alessandra Scano, Sara Fais, Valentina Medda, Germano Orrù, Miriam Loddo, Riccardo Cappai, Ferdinando Coghe |
| EPI_ISL_2240828, EPI_ISL_2240834, EPI_ISL_2240843, EPI_ISL_2240844, EPI_ISL_2240882, EPI_ISL_2240899, EPI_ISL_2240912, EPI_ISL_2240923, EPI_ISL_2240935, EPI_ISL_2241055, EPI_ISL_2241056, EPI_ISL_2241058, EPI_ISL_2241078, EPI_ISL_2241082, EPI_ISL_2241118, EPI_ISL_2241134, EPI_ISL_2241192, EPI_ISL_2241253, EPI_ISL_2241268, EPI_ISL_2241284 | | | |
| see above | National Virus Reference Laboratory | National Virus Reference Laboratory | Zoe Yandle, Charlene Bennett, Gabriel Gonzalez, Michael Carr, Jonathan Dean, Cillian F De Gascun |
| EPI_ISL_2241351 | Wojewodzka Stacja Sanitarno-Epidemiologiczna w Olsztynie, Laboratorium Badan Epidemiologiczno-Klinicznych | Wojewodzka Stacja Sanitarno-Epidemiologiczna w Olsztynie, Laboratorium Badan Epidemiologiczno-Klinicznych | Sylwia Krzetowska, Monika Czerminska, Ewa Liszewska, Tomasz Jakubczak, Marta Lukian, Patryk Bielecki, Aleksandra Kobiatko, Emilia Tarabasz, Paulina Rozycka, Barbara Dolinska |
| EPI_ISL_2245238 | University of Liège COVID-19 testing center | GIGA Medical Genomics | Keith Durkin, Maria Artesi, Bouchra Boujemla, Nathalie Renotte, Cécile Meex, Sébastien Bontems, Fabrice Bureau, Laurent Gillet, Wouter Coppieters, Marie-Pierre Hayette, Vincent Bours |
| EPI_ISL_2246063, EPI_ISL_2246064, EPI_ISL_2246065, EPI_ISL_2246066, EPI_ISL_2246115, EPI_ISL_2246121 | Platform BIS UZA/UAntwerpen | Labo Klinische Biologie, UZA | Marie Le Mercier, Jasmine Coppens, Basil Britto Xavier, Christine Lammens, Veerle Matheeussen, Herman Goossens |
| EPI_ISL_2246823 | National Virus Reference Laboratory | National Virus Reference Laboratory | Zoe Yandle, Charlene Bennett, Gabriel Gonzalez, Michael Carr, Jonathan Dean, Cillian F De Gascun |
| EPI_ISL_2246827, EPI_ISL_2246828 | LESP Ciudad de Mexico /LANS | Instituto de Diagnostico y Referencia Epidemiologicos (INDRE) | Claudia Wong-Arambula, Abril Rodriguez-Maldonado, Vanessa Rivero-Arredondo, Ariadna Medina-Benitez, Joaquin Quiroz-Mercado, Sergio Rangel-Guerrero, Natividad Cruz-Ortiz, Tatiana Nunez-Garcia, Gisela Barrera-Badillo, Lucia Hernandez-Rivas, Irma Lopez-Martinez, Ernesto Ramirez-Gonzalez. |
| EPI_ISL_2246829, EPI_ISL_2246830, EPI_ISL_2246831, EPI_ISL_2246832, EPI_ISL_2246833 | LESP Michoacan | Instituto de Diagnostico y Referencia Epidemiologicos (INDRE) | Claudia Wong-Arambula, Abril Rodriguez-Maldonado, Vanessa Rivero-Arredondo, Ariadna Medina-Benitez, Joaquin Quiroz-Mercado, Sergio Rangel-Guerrero, Natividad Cruz-Ortiz, Tatiana Nunez-Garcia, Gisela Barrera-Badillo, Lucia Hernandez-Rivas, Irma Lopez-Martinez, Ernesto Ramirez-Gonzalez. |
| EPI_ISL_2246834, EPI_ISL_2246835, EPI_ISL_2246836, EPI_ISL_2246837 | LESP Sinaloa | Instituto de Diagnostico y Referencia Epidemiologicos (INDRE) | Claudia Wong-Arambula, Abril Rodriguez-Maldonado, Vanessa Rivero-Arredondo, Ariadna Medina-Benitez, Joaquin Quiroz-Mercado, Sergio Rangel-Guerrero, Natividad Cruz-Ortiz, Tatiana Nunez-Garcia, Gisela Barrera-Badillo, Lucia Hernandez-Rivas, Irma Lopez-Martinez, Ernesto Ramirez-Gonzalez. |
| EPI_ISL_2246838 | LESP Ciudad de Mexico | Instituto de Diagnostico y Referencia Epidemiologicos (INDRE) | Claudia Wong-Arambula, Abril Rodriguez-Maldonado, Vanessa Rivero-Arredondo, Ariadna Medina-Benitez, Joaquin Quiroz-Mercado, Sergio Rangel-Guerrero, Natividad Cruz-Ortiz, Tatiana Nunez-Garcia, Gisela Barrera-Badillo, Lucia Hernandez-Rivas, Irma Lopez-Martinez, Ernesto Ramirez-Gonzalez. |
| EPI_ISL_2246887, EPI_ISL_2246888, EPI_ISL_2246889 | Florida Bureau of Public Health Laboratories | Florida Bureau of Public Health Laboratories | Sarah Schmedes, Jason Blanton |
| EPI_ISL_2247027 | WSSE w Krakowie | 1. Academic Center for Pathomorphological and Genetic-Molecular Diagnostics ltd, Bialystok, Poland 2. National Institute of Public Health - National Institute of Hygiene, Warsaw, Poland | Radosław Charkiewicz, Jacek Nikliński, Przemysław Biecek. Joanna Reszeć, Piotr Majewski, Anetta Sulewska, Piotr Karabowicz, Joanna Kiśluk, Janusz Dzięcioł, Konrad Raczkowski, Katarzyna Zacharczuk, Magdalena Nowakowska, Małgorzata Sadkowska-Todys, Tomasz Wołkowicz |
| EPI_ISL_2247090, EPI_ISL_2247111, EPI_ISL_2247115, EPI_ISL_2247178, EPI_ISL_2247186, EPI_ISL_2247198, EPI_ISL_2247242, | Quest Diagnostics Incorporated | Centers for Disease Control and Prevention Division of Viral Diseases, Pathogen Discovery | Dakota Howard, Dhwani Batra, Peter W. Cook, Kara Moser, Adrian Paskey, Jason Caravas, Benjamin Rambo-Martin, Shatavia Morrison, Christopher Gulvick, Scott Sammons, Yvette Unoarumhi, Darlene Wagner, Matthew Schmerer, S. H. Rosenthal, A. Gerasimova, R. M. Kagan, B. Anderson, M. Hua, Y. Liu, L.E. Bernstein, K.E. Livingston, A. Perez, I. A. Shlyakhter, R. V. Rolando, R. Owen, P. Tanpaiboon, F. Lacbawan, Clinton R. Paden, Duncan MacCannell |

| | | | |
|---|---|---|---|
| EPI_ISL_2247275, EPI_ISL_2247281 | | | |
| EPI_ISL_2247378, EPI_ISL_2247397, EPI_ISL_2247537, EPI_ISL_2247575, EPI_ISL_2247606, EPI_ISL_2247615, EPI_ISL_2247625, EPI_ISL_2247647, EPI_ISL_2247704, EPI_ISL_2248038, EPI_ISL_2248071, EPI_ISL_2248220, EPI_ISL_2248308, EPI_ISL_2248458 | | | |
| see above | Helix/Illumina | Centers for Disease Control and Prevention Division of Viral Diseases, Pathogen Discovery | Dakota Howard, Dhwani Batra, Peter W. Cook, Kara Moser, Adrian Paskey, Jason Caravas, Benjamin Rambo-Martin, Shatavia Morrison, Christopher Gulvick, Scott Sammons, Yvette Unoarumhi, Darlene Wagner, Matthew Schmerer, Eileen de Feo, Jan Antico, Christine Tran, Matthew Tolentino, Shannon Wickline, Kim Gietzen, Brad Sickler, Jingtao Liu, Eric Allen, Phil Febbo, Nicole L. Washington, Simon White, Geraint Levan, Kelly Schiabor Barrett, Elizabeth Cirulli, Alexandre Bolze, Ary Ascencio, Charlotte Rivera-Garcia, Ryan Cho, Jason Nguyen, Sherry Wang, Jimmy Ramirez, Tyler Cassens, Efren Sandoval, Magnus Isaksson, William Lee, David Becker, Marc Laurent, James Lu, Clinton R. Paden, Duncan MacCannell |
| EPI_ISL_2248483, EPI_ISL_2248541, EPI_ISL_2248619, EPI_ISL_2248631, EPI_ISL_2248662 | Quest Diagnostics Incorporated | Centers for Disease Control and Prevention Division of Viral Diseases, Pathogen Discovery | Dakota Howard, Dhwani Batra, Peter W. Cook, Kara Moser, Adrian Paskey, Jason Caravas, Benjamin Rambo-Martin, Shatavia Morrison, Christopher Gulvick, Scott Sammons, Yvette Unoarumhi, Darlene Wagner, Matthew Schmerer, S. H. Rosenthal, A. Gerasimova, R. M. Kagan, B. Anderson, M. Hua, Y. Liu, L.E. Bernstein, K.E. Livingston, A. Perez, I. A. Shlyakhter, R. V. Rolando, R. Owen, P. Tanpaiboon, F. Lacbawan, Clinton R. Paden, Duncan MacCannell |
| EPI_ISL_2248749, EPI_ISL_2248750 | Kansas Health and Environmental Lab | Kansas Health and Environmental Lab | Mike Grose, Jonathan Barnell, Ben Olsen, and Phil Adam |
| EPI_ISL_2248796, EPI_ISL_2248799, EPI_ISL_2248815, EPI_ISL_2248821, EPI_ISL_2248826, EPI_ISL_2248831, EPI_ISL_2248832, EPI_ISL_2248833, EPI_ISL_2248834 | CHLC | Instituto Nacional de Saude (INSA) | Borges et al |
| EPI_ISL_2248851 | Cintramedica | Instituto Nacional de Saude (INSA) | Borges et al |
| EPI_ISL_2248870, EPI_ISL_2248871, EPI_ISL_2248874 | Germano de Sousa | Instituto Nacional de Saude (INSA) | Borges et al |
| EPI_ISL_2248930, EPI_ISL_2248935 | Joaquim Chaves Saude | Instituto Nacional de Saude (INSA) | Borges et al |
| EPI_ISL_2248966 | ARS Algarve - Laboratorio Laura Ayres | Instituto Nacional de Saude (INSA) | Borges et al |
| EPI_ISL_2248985, EPI_ISL_2249005, EPI_ISL_2249015 | Synlab | Instituto Nacional de Saude (INSA) | Borges et al |
| EPI_ISL_2249024, EPI_ISL_2249037, EPI_ISL_2249038 | UNILABS | Instituto Nacional de Saude (INSA) | Borges et al |
| EPI_ISL_2249064 | ARS Algarve - Laboratorio Laura Ayres | Instituto Nacional de Saude (INSA) and Institute of Biomedicine (iBiMed), Universidade de Aveiro | Borges et al |
| EPI_ISL_2249115 | ULS Litoral Alentejano | Instituto Nacional de Saude (INSA) and Institute of Biomedicine (iBiMed), Universidade de Aveiro | Borges et al |
| EPI_ISL_2249132 | ABC Algarve | Instituto Nacional de Saude (INSA) and Centro de Investigaçao em Biodiversidade e Recursos Geneticos (CIBIO), Universidade do Porto | Borges et al |
| EPI_ISL_2249174 | CHULN - H Santa Maria | Instituto Nacional de Saude (INSA) and Centro de Investigaçao em Biodiversidade e Recursos Geneticos (CIBIO), Universidade do Porto | Borges et al |
| EPI_ISL_2249202, EPI_ISL_2249204 | ULS Litoral Alentejano | Instituto Nacional de Saude (INSA) and Centro de Investigaçao em Biodiversidade e Recursos Geneticos (CIBIO), Universidade do Porto | Borges et al |
| EPI_ISL_2249246, EPI_ISL_2249247, EPI_ISL_2249248, EPI_ISL_2249249, EPI_ISL_2249252, EPI_ISL_2249253, EPI_ISL_2249255 | Hospital Sharp | Microbial Genomics Laboratory | Bruno Gomez-Gil, Julissa Enciso-Ibarra, Alejandra Garcia-Gasca, Daniel Fregoso-Rueda |
| EPI_ISL_2249257 | New South Wales Health Pathology Royal Prince Alfred Hospital | Microbiology RPAH | Foster, C.; Au, J.; Ruiz Silva, M.; Deveson, I.; Bull, R.; Van Hal, S.; Rawlinson, W. |
| EPI_ISL_2249523, EPI_ISL_2249544 | Minnesota Department of Health, Public Health Laboratory | Minnesota Department of Health, Public Health Laboratory | Alexandra Lorentz, Jacob Garfin, Matt Plumb, and Xiong Wang |
| EPI_ISL_2249559 | M Health Fairview | Minnesota Department of Health, Public Health Laboratory | Alexandra Lorentz, Jacob Garfin, Matt Plumb, and Xiong Wang |
| EPI_ISL_2250038, EPI_ISL_2250073 | Arizona State University | Arizona State University | Peter T. Skidmore, LaRinda A. Holland, Matthew F. Smith, Rabia Maqsood, Nicholas J. Mellor, Joy M. Blain, Valerie Harris, Joshua LaBaer, Vel Murugan, Efrem S. Lim |
| EPI_ISL_2250189, EPI_ISL_2250190, EPI_ISL_2250191, EPI_ISL_2250193, EPI_ISL_2250196, EPI_ISL_2250197, EPI_ISL_2250198, EPI_ISL_2250199, EPI_ISL_2250200, EPI_ISL_2250202, EPI_ISL_2250204, EPI_ISL_2250206, EPI_ISL_2250207, EPI_ISL_2250208, EPI_ISL_2250209, EPI_ISL_2250211, EPI_ISL_2250216, EPI_ISL_2250217, EPI_ISL_2250218, EPI_ISL_2250219, EPI_ISL_2250220 | | | |
| see above | Royal Darwin Hospital Pathology | Microbiological Diagnostic Unit Public Health Laboratory (MDU-PHL) | Meumann, E., Caly L., Seemann T., Sait, M.L., Druce J., Sherry, N.L. |
| EPI_ISL_2250230 | Microbiological Diagnostic Unit - Public Health Laboratory (MDU-PHL) | Microbiological Diagnostic Unit Public Health Laboratory (MDU-PHL) | Seemann T., Sait, M.L., Sherry, N.L. |
| EPI_ISL_2250723, EPI_ISL_2250746, EPI_ISL_2250748, EPI_ISL_2251194, EPI_ISL_2251195, EPI_ISL_2251508, EPI_ISL_2251529, EPI_ISL_2251546, EPI_ISL_2251555, EPI_ISL_2251569, EPI_ISL_2251598, EPI_ISL_2251719, EPI_ISL_2251786, EPI_ISL_2251794, EPI_ISL_2251797, EPI_ISL_2251798, EPI_ISL_2251803, EPI_ISL_2251805, EPI_ISL_2251813, EPI_ISL_2251814, EPI_ISL_2251815, EPI_ISL_2251817, EPI_ISL_2251819, EPI_ISL_2251827, EPI_ISL_2251836, EPI_ISL_2251841, EPI_ISL_2251843, EPI_ISL_2251846, EPI_ISL_2251849, EPI_ISL_2251850, EPI_ISL_2251852, EPI_ISL_2251854, EPI_ISL_2251855, EPI_ISL_2251857, EPI_ISL_2251858, EPI_ISL_2251859, EPI_ISL_2251956, EPI_ISL_2251958, EPI_ISL_2251960, EPI_ISL_2251961, EPI_ISL_2251966, EPI_ISL_2251968, EPI_ISL_2251970, EPI_ISL_2251971, EPI_ISL_2252024, EPI_ISL_2252249, EPI_ISL_2252253, EPI_ISL_2252276, EPI_ISL_2252278, EPI_ISL_2252301, EPI_ISL_2252329, EPI_ISL_2252332, EPI_ISL_2252354, EPI_ISL_2252372, EPI_ISL_2252401, EPI_ISL_2252402, EPI_ISL_2252407, EPI_ISL_2252412, EPI_ISL_2252424, EPI_ISL_2252436, EPI_ISL_2252488, EPI_ISL_2252500, EPI_ISL_2252557, EPI_ISL_2252569, EPI_ISL_2252598, EPI_ISL_2252611, EPI_ISL_2252614, EPI_ISL_2252618, EPI_ISL_2252668, EPI_ISL_2252678, EPI_ISL_2252688, EPI_ISL_2252694, EPI_ISL_2252719, EPI_ISL_2252720, EPI_ISL_2252722, EPI_ISL_2252737, EPI_ISL_2252742, EPI_ISL_2252754, EPI_ISL_2252764, EPI_ISL_2252768, EPI_ISL_2252776, EPI_ISL_2252783, EPI_ISL_2252813, EPI_ISL_2252830, EPI_ISL_2252866, EPI_ISL_2252867, EPI_ISL_2252868, EPI_ISL_2252870, EPI_ISL_2252900, EPI_ISL_2252933, EPI_ISL_2253032, EPI_ISL_2253043, EPI_ISL_2253051, EPI_ISL_2253054, EPI_ISL_2253066, EPI_ISL_2253071, EPI_ISL_2253077, EPI_ISL_2253098 | | | |
| see above | Public Health Ontario Laboratory | Public Health Ontario Laboratory | Vanessa G Allen, Philip Banh, Yao Chen, Richard de Borja, Alireza Eshaghi, Nahuel Fittipaldi, Christine Frantz, Jonathan B Gubbay, Jennifer L Guthrie, Lawrence Heisler, Esha Joshi, Michael Laszloffy, Aimin Li, Michael CY Li, Dean Maxwell, Sandeep Nagra, Samir N Patel, Jared Simpson, Karthikeyan Sivaraman, Ashleigh Sullivan, Yogi Sundaravadanam, Sarah Teatero, Andre Villegas, Matthew Watson, Sandra Zittermann |
| EPI_ISL_2253111, EPI_ISL_2253112, EPI_ISL_2253113, EPI_ISL_2253114, EPI_ISL_2253115, EPI_ISL_2253118, EPI_ISL_2253121, EPI_ISL_2253122, EPI_ISL_2253123 | SARS-CoV-2 testing team, National Institute of Infectious Diseases | Pathogen Genomics Center, National Institute of Infectious Diseases | Tsuyoshi Sekizuka, Kentaro Itokawa, Rina Tanaka, Masanori Hashino, Nozomu Hanaoka, Masumichi Saito, Naomi Nojiri, Hazuka Y Furihata, Sana Uchikoba, Tsuguto Fujimoto, Makoto Kuroda |
| EPI_ISL_2253222, EPI_ISL_2253225 | Health Services Laboratories | Wellcome Sanger Institute for the COVID-19 Genomics UK (COG-UK) Consortium | Health Services Laboratories and Alex Alderton, Roberto Amato, Jeffrey Barrett, Sonia Goncalves, Ewan Harrison, David K. Jackson, Ian Johnston, Dominic Kwiatkowski, Cordelia Langford, John Sillitoe on behalf of the Wellcome Sanger Institute COVID-19 Surveillance Team |
| EPI_ISL_2253226, EPI_ISL_2253227 | Lighthouse Lab in Milton Keynes | Wellcome Sanger Institute for the COVID-19 Genomics UK (COG-UK) Consortium | The Lighthouse Lab in Milton Keynes and Alex Alderton, Roberto Amato, Jeffrey Barrett, Sonia Goncalves, Ewan Harrison, David K. Jackson, Ian Johnston, Dominic Kwiatkowski, Cordelia Langford, John Sillitoe on behalf of the Wellcome Sanger Institute COVID-19 Surveillance Team |
| EPI_ISL_2253228 | Health Services Laboratories | Wellcome Sanger Institute for the COVID-19 Genomics UK (COG-UK) Consortium | Health Services Laboratories and Alex Alderton, Roberto Amato, Jeffrey Barrett, Sonia Goncalves, Ewan Harrison, David K. Jackson, Ian Johnston, Dominic Kwiatkowski, Cordelia Langford, John Sillitoe on behalf of the Wellcome Sanger Institute COVID-19 Surveillance Team |
| EPI_ISL_2253229, EPI_ISL_2253233, EPI_ISL_2253235 | Lighthouse Lab in Milton Keynes | Wellcome Sanger Institute for the COVID-19 Genomics UK (COG-UK) Consortium | The Lighthouse Lab in Milton Keynes and Alex Alderton, Roberto Amato, Jeffrey Barrett, Sonia Goncalves, Ewan Harrison, David K. Jackson, Ian Johnston, Dominic Kwiatkowski, Cordelia Langford, John Sillitoe on behalf of the Wellcome Sanger Institute COVID-19 Surveillance Team |
| EPI_ISL_2253236 | Health Services Laboratories | Wellcome Sanger Institute for the COVID-19 Genomics UK (COG-UK) Consortium | Health Services Laboratories and Alex Alderton, Roberto Amato, Jeffrey Barrett, Sonia Goncalves, Ewan Harrison, David K. Jackson, Ian Johnston, Dominic Kwiatkowski, Cordelia Langford, John Sillitoe on behalf of the Wellcome Sanger Institute COVID-19 Surveillance Team |
| EPI_ISL_2253238 | Lighthouse Lab in Milton Keynes | Wellcome Sanger Institute for the COVID-19 Genomics UK (COG-UK) Consortium | The Lighthouse Lab in Milton Keynes and Alex Alderton, Roberto Amato, Jeffrey Barrett, Sonia Goncalves, Ewan Harrison, David K. Jackson, Ian Johnston, Dominic Kwiatkowski, Cordelia Langford, John Sillitoe on behalf of the Wellcome Sanger Institute COVID-19 Surveillance Team |
| EPI_ISL_2253240 | Health Services Laboratories | Wellcome Sanger Institute for the COVID-19 Genomics UK (COG-UK) Consortium | Health Services Laboratories and Alex Alderton, Roberto Amato, Jeffrey Barrett, Sonia Goncalves, Ewan Harrison, David K. Jackson, Ian Johnston, Dominic Kwiatkowski, Cordelia Langford, John Sillitoe on behalf of the Wellcome Sanger Institute COVID-19 Surveillance Team |
| EPI_ISL_2253241, EPI_ISL_2253242, | Lighthouse Lab in Milton Keynes | Wellcome Sanger Institute for the COVID-19 Genomics UK (COG-UK) Consortium | The Lighthouse Lab in Milton Keynes and Alex Alderton, Roberto Amato, Jeffrey Barrett, Sonia Goncalves, Ewan Harrison, David K. Jackson, Ian Johnston, Dominic Kwiatkowski, Cordelia Langford, John Sillitoe on behalf of the Wellcome Sanger Institute COVID-19 Surveillance Team |

| | | | |
|---|---|---|---|
| EPI_ISL_2253243 | | | |
| EPI_ISL_2253244, EPI_ISL_2253246 | Health Services Laboratories | Wellcome Sanger Institute for the COVID-19 Genomics UK (COG-UK) Consortium | Health Services Laboratories and Alex Alderton, Roberto Amato, Jeffrey Barrett, Sonia Goncalves, Ewan Harrison, David K. Jackson, Ian Johnston, Dominic Kwiatkowski, Cordelia Langford, John Sillitoe on behalf of the Wellcome Sanger Institute COVID-19 Surveillance Team |
| EPI_ISL_2253247, EPI_ISL_2253249, EPI_ISL_2253250, EPI_ISL_2253253, EPI_ISL_2253256 | Lighthouse Lab in Milton Keynes | Wellcome Sanger Institute for the COVID-19 Genomics UK (COG-UK) Consortium | The Lighthouse Lab in Milton Keynes and Alex Alderton, Roberto Amato, Jeffrey Barrett, Sonia Goncalves, Ewan Harrison, David K. Jackson, Ian Johnston, Dominic Kwiatkowski, Cordelia Langford, John Sillitoe on behalf of the Wellcome Sanger Institute COVID-19 Surveillance Team |
| EPI_ISL_2253257 | Health Services Laboratories | Wellcome Sanger Institute for the COVID-19 Genomics UK (COG-UK) Consortium | Health Services Laboratories and Alex Alderton, Roberto Amato, Jeffrey Barrett, Sonia Goncalves, Ewan Harrison, David K. Jackson, Ian Johnston, Dominic Kwiatkowski, Cordelia Langford, John Sillitoe on behalf of the Wellcome Sanger Institute COVID-19 Surveillance Team |
| EPI_ISL_2253258, EPI_ISL_2253259 | Lighthouse Lab in Milton Keynes | Wellcome Sanger Institute for the COVID-19 Genomics UK (COG-UK) Consortium | The Lighthouse Lab in Milton Keynes and Alex Alderton, Roberto Amato, Jeffrey Barrett, Sonia Goncalves, Ewan Harrison, David K. Jackson, Ian Johnston, Dominic Kwiatkowski, Cordelia Langford, John Sillitoe on behalf of the Wellcome Sanger Institute COVID-19 Surveillance Team |
| EPI_ISL_2253261 | Health Services Laboratories | Wellcome Sanger Institute for the COVID-19 Genomics UK (COG-UK) Consortium | Health Services Laboratories and Alex Alderton, Roberto Amato, Jeffrey Barrett, Sonia Goncalves, Ewan Harrison, David K. Jackson, Ian Johnston, Dominic Kwiatkowski, Cordelia Langford, John Sillitoe on behalf of the Wellcome Sanger Institute COVID-19 Surveillance Team |
| EPI_ISL_2253264 | Lighthouse Lab in Glasgow | Wellcome Sanger Institute for the COVID-19 Genomics UK (COG-UK) Consortium | Harper VanSteenhouse, Yumi Kasai, David Gray, Carol Clugston, Anna Dominiczak and Alex Alderton, Roberto Amato, Jeffrey Barrett, Sonia Goncalves, Ewan Harrison, David K. Jackson, Ian Johnston, Dominic Kwiatkowski, Cordelia Langford, John Sillitoe on behalf of the Wellcome Sanger Institute COVID-19 Surveillance Team |
| EPI_ISL_2253265, EPI_ISL_2253267 | Health Services Laboratories | Wellcome Sanger Institute for the COVID-19 Genomics UK (COG-UK) Consortium | Health Services Laboratories and Alex Alderton, Roberto Amato, Jeffrey Barrett, Sonia Goncalves, Ewan Harrison, David K. Jackson, Ian Johnston, Dominic Kwiatkowski, Cordelia Langford, John Sillitoe on behalf of the Wellcome Sanger Institute COVID-19 Surveillance Team |
| EPI_ISL_2253270, EPI_ISL_2253271, EPI_ISL_2253273, EPI_ISL_2253274 | Lighthouse Lab in Milton Keynes | Wellcome Sanger Institute for the COVID-19 Genomics UK (COG-UK) Consortium | The Lighthouse Lab in Milton Keynes and Alex Alderton, Roberto Amato, Jeffrey Barrett, Sonia Goncalves, Ewan Harrison, David K. Jackson, Ian Johnston, Dominic Kwiatkowski, Cordelia Langford, John Sillitoe on behalf of the Wellcome Sanger Institute COVID-19 Surveillance Team |
| EPI_ISL_2253275, EPI_ISL_2253276 | Health Services Laboratories | Wellcome Sanger Institute for the COVID-19 Genomics UK (COG-UK) Consortium | Health Services Laboratories and Alex Alderton, Roberto Amato, Jeffrey Barrett, Sonia Goncalves, Ewan Harrison, David K. Jackson, Ian Johnston, Dominic Kwiatkowski, Cordelia Langford, John Sillitoe on behalf of the Wellcome Sanger Institute COVID-19 Surveillance Team |
| EPI_ISL_2253277 | Lighthouse Lab in Milton Keynes | Wellcome Sanger Institute for the COVID-19 Genomics UK (COG-UK) Consortium | The Lighthouse Lab in Milton Keynes and Alex Alderton, Roberto Amato, Jeffrey Barrett, Sonia Goncalves, Ewan Harrison, David K. Jackson, Ian Johnston, Dominic Kwiatkowski, Cordelia Langford, John Sillitoe on behalf of the Wellcome Sanger Institute COVID-19 Surveillance Team |
| EPI_ISL_2253278 | Health Services Laboratories | Wellcome Sanger Institute for the COVID-19 Genomics UK (COG-UK) Consortium | Health Services Laboratories and Alex Alderton, Roberto Amato, Jeffrey Barrett, Sonia Goncalves, Ewan Harrison, David K. Jackson, Ian Johnston, Dominic Kwiatkowski, Cordelia Langford, John Sillitoe on behalf of the Wellcome Sanger Institute COVID-19 Surveillance Team |
| EPI_ISL_2253279, EPI_ISL_2253280, EPI_ISL_2253282 | Lighthouse Lab in Milton Keynes | Wellcome Sanger Institute for the COVID-19 Genomics UK (COG-UK) Consortium | The Lighthouse Lab in Milton Keynes and Alex Alderton, Roberto Amato, Jeffrey Barrett, Sonia Goncalves, Ewan Harrison, David K. Jackson, Ian Johnston, Dominic Kwiatkowski, Cordelia Langford, John Sillitoe on behalf of the Wellcome Sanger Institute COVID-19 Surveillance Team |
| EPI_ISL_2253283 | Health Services Laboratories | Wellcome Sanger Institute for the COVID-19 Genomics UK (COG-UK) Consortium | Health Services Laboratories and Alex Alderton, Roberto Amato, Jeffrey Barrett, Sonia Goncalves, Ewan Harrison, David K. Jackson, Ian Johnston, Dominic Kwiatkowski, Cordelia Langford, John Sillitoe on behalf of the Wellcome Sanger Institute COVID-19 Surveillance Team |
| EPI_ISL_2253284 | Lighthouse Lab in Milton Keynes | Wellcome Sanger Institute for the COVID-19 Genomics UK (COG-UK) Consortium | The Lighthouse Lab in Milton Keynes and Alex Alderton, Roberto Amato, Jeffrey Barrett, Sonia Goncalves, Ewan Harrison, David K. Jackson, Ian Johnston, Dominic Kwiatkowski, Cordelia Langford, John Sillitoe on behalf of the Wellcome Sanger Institute COVID-19 Surveillance Team |
| EPI_ISL_2253285, EPI_ISL_2253286 | Health Services Laboratories | Wellcome Sanger Institute for the COVID-19 Genomics UK (COG-UK) Consortium | Health Services Laboratories and Alex Alderton, Roberto Amato, Jeffrey Barrett, Sonia Goncalves, Ewan Harrison, David K. Jackson, Ian Johnston, Dominic Kwiatkowski, Cordelia Langford, John Sillitoe on behalf of the Wellcome Sanger Institute COVID-19 Surveillance Team |
| EPI_ISL_2253287 | Lighthouse Lab in Milton Keynes | Wellcome Sanger Institute for the COVID-19 Genomics UK (COG-UK) Consortium | The Lighthouse Lab in Milton Keynes and Alex Alderton, Roberto Amato, Jeffrey Barrett, Sonia Goncalves, Ewan Harrison, David K. Jackson, Ian Johnston, Dominic Kwiatkowski, Cordelia Langford, John Sillitoe on behalf of the Wellcome Sanger Institute COVID-19 Surveillance Team |
| EPI_ISL_2253289, EPI_ISL_2253292, EPI_ISL_2253293 | Health Services Laboratories | Wellcome Sanger Institute for the COVID-19 Genomics UK (COG-UK) Consortium | Health Services Laboratories and Alex Alderton, Roberto Amato, Jeffrey Barrett, Sonia Goncalves, Ewan Harrison, David K. Jackson, Ian Johnston, Dominic Kwiatkowski, Cordelia Langford, John Sillitoe on behalf of the Wellcome Sanger Institute COVID-19 Surveillance Team |
| EPI_ISL_2253294, EPI_ISL_2253295, EPI_ISL_2253296 | Lighthouse Lab in Milton Keynes | Wellcome Sanger Institute for the COVID-19 Genomics UK (COG-UK) Consortium | The Lighthouse Lab in Milton Keynes and Alex Alderton, Roberto Amato, Jeffrey Barrett, Sonia Goncalves, Ewan Harrison, David K. Jackson, Ian Johnston, Dominic Kwiatkowski, Cordelia Langford, John Sillitoe on behalf of the Wellcome Sanger Institute COVID-19 Surveillance Team |
| EPI_ISL_2253297, EPI_ISL_2253298 | Health Services Laboratories | Wellcome Sanger Institute for the COVID-19 Genomics UK (COG-UK) Consortium | Health Services Laboratories and Alex Alderton, Roberto Amato, Jeffrey Barrett, Sonia Goncalves, Ewan Harrison, David K. Jackson, Ian Johnston, Dominic Kwiatkowski, Cordelia Langford, John Sillitoe on behalf of the Wellcome Sanger Institute COVID-19 Surveillance Team |
| EPI_ISL_2253300 | Lighthouse Lab in Milton Keynes | Wellcome Sanger Institute for the COVID-19 Genomics UK (COG-UK) Consortium | The Lighthouse Lab in Milton Keynes and Alex Alderton, Roberto Amato, Jeffrey Barrett, Sonia Goncalves, Ewan Harrison, David K. Jackson, Ian Johnston, Dominic Kwiatkowski, Cordelia Langford, John Sillitoe on behalf of the Wellcome Sanger Institute COVID-19 Surveillance Team |
| EPI_ISL_2253302, EPI_ISL_2253303, EPI_ISL_2253304 | Health Services Laboratories | Wellcome Sanger Institute for the COVID-19 Genomics UK (COG-UK) Consortium | Health Services Laboratories and Alex Alderton, Roberto Amato, Jeffrey Barrett, Sonia Goncalves, Ewan Harrison, David K. Jackson, Ian Johnston, Dominic Kwiatkowski, Cordelia Langford, John Sillitoe on behalf of the Wellcome Sanger Institute COVID-19 Surveillance Team |
| EPI_ISL_2253305 | Lighthouse Lab in Milton Keynes | Wellcome Sanger Institute for the COVID-19 Genomics UK (COG-UK) Consortium | The Lighthouse Lab in Milton Keynes and Alex Alderton, Roberto Amato, Jeffrey Barrett, Sonia Goncalves, Ewan Harrison, David K. Jackson, Ian Johnston, Dominic Kwiatkowski, Cordelia Langford, John Sillitoe on behalf of the Wellcome Sanger Institute COVID-19 Surveillance Team |
| EPI_ISL_2253306 | Health Services Laboratories | Wellcome Sanger Institute for the COVID-19 Genomics UK (COG-UK) Consortium | Health Services Laboratories and Alex Alderton, Roberto Amato, Jeffrey Barrett, Sonia Goncalves, Ewan Harrison, David K. Jackson, Ian Johnston, Dominic Kwiatkowski, Cordelia Langford, John Sillitoe on behalf of the Wellcome Sanger Institute COVID-19 Surveillance Team |
| EPI_ISL_2253307 | Lighthouse Lab in Milton Keynes | Wellcome Sanger Institute for the COVID-19 Genomics UK (COG-UK) Consortium | The Lighthouse Lab in Milton Keynes and Alex Alderton, Roberto Amato, Jeffrey Barrett, Sonia Goncalves, Ewan Harrison, David K. Jackson, Ian Johnston, Dominic Kwiatkowski, Cordelia Langford, John Sillitoe on behalf of the Wellcome Sanger Institute COVID-19 Surveillance Team |
| EPI_ISL_2253308 | Health Services Laboratories | Wellcome Sanger Institute for the COVID-19 Genomics UK (COG-UK) Consortium | Health Services Laboratories and Alex Alderton, Roberto Amato, Jeffrey Barrett, Sonia Goncalves, Ewan Harrison, David K. Jackson, Ian Johnston, Dominic Kwiatkowski, Cordelia Langford, John Sillitoe on behalf of the Wellcome Sanger Institute COVID-19 Surveillance Team |
| EPI_ISL_2253309 | Lighthouse Lab in Milton Keynes | Wellcome Sanger Institute for the COVID-19 Genomics UK (COG-UK) Consortium | The Lighthouse Lab in Milton Keynes and Alex Alderton, Roberto Amato, Jeffrey Barrett, Sonia Goncalves, Ewan Harrison, David K. Jackson, Ian Johnston, Dominic Kwiatkowski, Cordelia Langford, John Sillitoe on behalf of the Wellcome Sanger Institute COVID-19 Surveillance Team |
| EPI_ISL_2253310, EPI_ISL_2253311 | Health Services Laboratories | Wellcome Sanger Institute for the COVID-19 Genomics UK (COG-UK) Consortium | Health Services Laboratories and Alex Alderton, Roberto Amato, Jeffrey Barrett, Sonia Goncalves, Ewan Harrison, David K. Jackson, Ian Johnston, Dominic Kwiatkowski, Cordelia Langford, John Sillitoe on behalf of the Wellcome Sanger Institute COVID-19 Surveillance Team |
| EPI_ISL_2253313, EPI_ISL_2253314 | Lighthouse Lab in Milton Keynes | Wellcome Sanger Institute for the COVID-19 Genomics UK (COG-UK) Consortium | The Lighthouse Lab in Milton Keynes and Alex Alderton, Roberto Amato, Jeffrey Barrett, Sonia Goncalves, Ewan Harrison, David K. Jackson, Ian Johnston, Dominic Kwiatkowski, Cordelia Langford, John Sillitoe on behalf of the Wellcome Sanger Institute COVID-19 Surveillance Team |
| EPI_ISL_2253316, EPI_ISL_2253319, EPI_ISL_2253320 | Health Services Laboratories | Wellcome Sanger Institute for the COVID-19 Genomics UK (COG-UK) Consortium | Health Services Laboratories and Alex Alderton, Roberto Amato, Jeffrey Barrett, Sonia Goncalves, Ewan Harrison, David K. Jackson, Ian Johnston, Dominic Kwiatkowski, Cordelia Langford, John Sillitoe on behalf of the Wellcome Sanger Institute COVID-19 Surveillance Team |
| EPI_ISL_2253322 | Lighthouse Lab in Milton Keynes | Wellcome Sanger Institute for the COVID-19 Genomics UK (COG-UK) Consortium | The Lighthouse Lab in Milton Keynes and Alex Alderton, Roberto Amato, Jeffrey Barrett, Sonia Goncalves, Ewan Harrison, David K. Jackson, Ian Johnston, Dominic Kwiatkowski, Cordelia Langford, John Sillitoe on behalf of the Wellcome Sanger Institute COVID-19 Surveillance Team |
| EPI_ISL_2253326 | Health Services Laboratories | Wellcome Sanger Institute for the COVID-19 Genomics UK (COG-UK) Consortium | Health Services Laboratories and Alex Alderton, Roberto Amato, Jeffrey Barrett, Sonia Goncalves, Ewan Harrison, David K. Jackson, Ian Johnston, Dominic Kwiatkowski, Cordelia Langford, John Sillitoe on behalf of the Wellcome Sanger Institute COVID-19 Surveillance Team |
| EPI_ISL_2253328, EPI_ISL_2253329 | Lighthouse Lab in Milton Keynes | Wellcome Sanger Institute for the COVID-19 Genomics UK (COG-UK) Consortium | The Lighthouse Lab in Milton Keynes and Alex Alderton, Roberto Amato, Jeffrey Barrett, Sonia Goncalves, Ewan Harrison, David K. Jackson, Ian Johnston, Dominic Kwiatkowski, Cordelia Langford, John Sillitoe on behalf of the Wellcome Sanger Institute COVID-19 Surveillance Team |
| EPI_ISL_2253330, EPI_ISL_2253332, EPI_ISL_2253334, EPI_ISL_2253335, EPI_ISL_2253340 | Health Services Laboratories | Wellcome Sanger Institute for the COVID-19 Genomics UK (COG-UK) Consortium | Health Services Laboratories and Alex Alderton, Roberto Amato, Jeffrey Barrett, Sonia Goncalves, Ewan Harrison, David K. Jackson, Ian Johnston, Dominic Kwiatkowski, Cordelia Langford, John Sillitoe on behalf of the Wellcome Sanger Institute COVID-19 Surveillance Team |
| EPI_ISL_2253342 | Lighthouse Lab in Milton Keynes | Wellcome Sanger Institute for the COVID-19 Genomics UK (COG-UK) Consortium | The Lighthouse Lab in Milton Keynes and Alex Alderton, Roberto Amato, Jeffrey Barrett, Sonia Goncalves, Ewan Harrison, David K. Jackson, Ian Johnston, Dominic Kwiatkowski, Cordelia Langford, John Sillitoe on behalf of the Wellcome Sanger Institute COVID-19 Surveillance Team |
| EPI_ISL_2253343, EPI_ISL_2253346, EPI_ISL_2253349, EPI_ISL_2253350 | Health Services Laboratories | Wellcome Sanger Institute for the COVID-19 Genomics UK (COG-UK) Consortium | Health Services Laboratories and Alex Alderton, Roberto Amato, Jeffrey Barrett, Sonia Goncalves, Ewan Harrison, David K. Jackson, Ian Johnston, Dominic Kwiatkowski, Cordelia Langford, John Sillitoe on behalf of the Wellcome Sanger Institute COVID-19 Surveillance Team |
| EPI_ISL_2253351, EPI_ISL_2253353, EPI_ISL_2253357, EPI_ISL_2253358 | Lighthouse Lab in Milton Keynes | Wellcome Sanger Institute for the COVID-19 Genomics UK (COG-UK) Consortium | The Lighthouse Lab in Milton Keynes and Alex Alderton, Roberto Amato, Jeffrey Barrett, Sonia Goncalves, Ewan Harrison, David K. Jackson, Ian Johnston, Dominic Kwiatkowski, Cordelia Langford, John Sillitoe on behalf of the Wellcome Sanger Institute COVID-19 Surveillance Team |
| EPI_ISL_2253360 | Health Services Laboratories | Wellcome Sanger Institute for the COVID-19 Genomics UK (COG-UK) Consortium | Health Services Laboratories and Alex Alderton, Roberto Amato, Jeffrey Barrett, Sonia Goncalves, Ewan Harrison, David K. Jackson, Ian Johnston, Dominic Kwiatkowski, Cordelia Langford, John Sillitoe on behalf of the Wellcome Sanger Institute COVID-19 Surveillance Team |
| EPI_ISL_2253361, | Lighthouse Lab in Milton Keynes | Wellcome Sanger Institute for the COVID-19 Genomics UK (COG-UK) | The Lighthouse Lab in Milton Keynes and Alex Alderton, Roberto Amato, Jeffrey Barrett, Sonia Goncalves, Ewan Harrison, David K. Jackson, Ian Johnston, Dominic Kwiatkowski, Cordelia Langford, John Sillitoe on behalf of the Wellcome Sanger |

| | | | |
|---|---|---|---|
| EPI_ISL_2253363 | | Consortium | Institute COVID-19 Surveillance Team |
| EPI_ISL_2253364, EPI_ISL_2253365, EPI_ISL_2253366 | Health Services Laboratories | Wellcome Sanger Institute for the COVID-19 Genomics UK (COG-UK) Consortium | Health Services Laboratories and Alex Alderton, Roberto Amato, Jeffrey Barrett, Sonia Goncalves, Ewan Harrison, David K. Jackson, Ian Johnston, Dominic Kwiatkowski, Cordelia Langford, John Sillitoe on behalf of the Wellcome Sanger Institute COVID-19 Surveillance Team |
| EPI_ISL_2253368, EPI_ISL_2253370, EPI_ISL_2253373, EPI_ISL_2253377, EPI_ISL_2253378 | Lighthouse Lab in Milton Keynes | Wellcome Sanger Institute for the COVID-19 Genomics UK (COG-UK) Consortium | The Lighthouse Lab in Milton Keynes and Alex Alderton, Roberto Amato, Jeffrey Barrett, Sonia Goncalves, Ewan Harrison, David K. Jackson, Ian Johnston, Dominic Kwiatkowski, Cordelia Langford, John Sillitoe on behalf of the Wellcome Sanger Institute COVID-19 Surveillance Team |
| EPI_ISL_2253383, EPI_ISL_2253385 | Health Services Laboratories | Wellcome Sanger Institute for the COVID-19 Genomics UK (COG-UK) Consortium | Health Services Laboratories and Alex Alderton, Roberto Amato, Jeffrey Barrett, Sonia Goncalves, Ewan Harrison, David K. Jackson, Ian Johnston, Dominic Kwiatkowski, Cordelia Langford, John Sillitoe on behalf of the Wellcome Sanger Institute COVID-19 Surveillance Team |
| EPI_ISL_2253387 | Lighthouse Lab in Milton Keynes | Wellcome Sanger Institute for the COVID-19 Genomics UK (COG-UK) Consortium | The Lighthouse Lab in Milton Keynes and Alex Alderton, Roberto Amato, Jeffrey Barrett, Sonia Goncalves, Ewan Harrison, David K. Jackson, Ian Johnston, Dominic Kwiatkowski, Cordelia Langford, John Sillitoe on behalf of the Wellcome Sanger Institute COVID-19 Surveillance Team |
| EPI_ISL_2253389 | Lighthouse Lab in Alderley Park | Wellcome Sanger Institute for the COVID-19 Genomics UK (COG-UK) Consortium | Jacquelyn Wynn, Mairead Hyland, The Lighthouse Lab in Alderley Park and Alex Alderton, Roberto Amato, Jeffrey Barrett, Sonia Goncalves, Ewan Harrison, David K. Jackson, Ian Johnston, Dominic Kwiatkowski, Cordelia Langford, John Sillitoe on behalf of the Wellcome Sanger Institute COVID-19 Surveillance Team |
| EPI_ISL_2253391 | Lighthouse Lab in Milton Keynes | Wellcome Sanger Institute for the COVID-19 Genomics UK (COG-UK) Consortium | The Lighthouse Lab in Milton Keynes and Alex Alderton, Roberto Amato, Jeffrey Barrett, Sonia Goncalves, Ewan Harrison, David K. Jackson, Ian Johnston, Dominic Kwiatkowski, Cordelia Langford, John Sillitoe on behalf of the Wellcome Sanger Institute COVID-19 Surveillance Team |
| EPI_ISL_2253392, EPI_ISL_2253393, EPI_ISL_2253394 | Lighthouse Lab in Alderley Park | Wellcome Sanger Institute for the COVID-19 Genomics UK (COG-UK) Consortium | Jacquelyn Wynn, Mairead Hyland, The Lighthouse Lab in Alderley Park and Alex Alderton, Roberto Amato, Jeffrey Barrett, Sonia Goncalves, Ewan Harrison, David K. Jackson, Ian Johnston, Dominic Kwiatkowski, Cordelia Langford, John Sillitoe on behalf of the Wellcome Sanger Institute COVID-19 Surveillance Team |
| EPI_ISL_2253396 | Health Services Laboratories | Wellcome Sanger Institute for the COVID-19 Genomics UK (COG-UK) Consortium | Health Services Laboratories and Alex Alderton, Roberto Amato, Jeffrey Barrett, Sonia Goncalves, Ewan Harrison, David K. Jackson, Ian Johnston, Dominic Kwiatkowski, Cordelia Langford, John Sillitoe on behalf of the Wellcome Sanger Institute COVID-19 Surveillance Team |
| EPI_ISL_2253397 | Lighthouse Lab in Milton Keynes | Wellcome Sanger Institute for the COVID-19 Genomics UK (COG-UK) Consortium | The Lighthouse Lab in Milton Keynes and Alex Alderton, Roberto Amato, Jeffrey Barrett, Sonia Goncalves, Ewan Harrison, David K. Jackson, Ian Johnston, Dominic Kwiatkowski, Cordelia Langford, John Sillitoe on behalf of the Wellcome Sanger Institute COVID-19 Surveillance Team |
| EPI_ISL_2253398, EPI_ISL_2253399 | Lighthouse Lab in Alderley Park | Wellcome Sanger Institute for the COVID-19 Genomics UK (COG-UK) Consortium | Jacquelyn Wynn, Mairead Hyland, The Lighthouse Lab in Alderley Park and Alex Alderton, Roberto Amato, Jeffrey Barrett, Sonia Goncalves, Ewan Harrison, David K. Jackson, Ian Johnston, Dominic Kwiatkowski, Cordelia Langford, John Sillitoe on behalf of the Wellcome Sanger Institute COVID-19 Surveillance Team |
| EPI_ISL_2253400 | Lighthouse Lab in Milton Keynes | Wellcome Sanger Institute for the COVID-19 Genomics UK (COG-UK) Consortium | The Lighthouse Lab in Milton Keynes and Alex Alderton, Roberto Amato, Jeffrey Barrett, Sonia Goncalves, Ewan Harrison, David K. Jackson, Ian Johnston, Dominic Kwiatkowski, Cordelia Langford, John Sillitoe on behalf of the Wellcome Sanger Institute COVID-19 Surveillance Team |
| EPI_ISL_2253404 | Lighthouse Lab in Alderley Park | Wellcome Sanger Institute for the COVID-19 Genomics UK (COG-UK) Consortium | Jacquelyn Wynn, Mairead Hyland, The Lighthouse Lab in Alderley Park and Alex Alderton, Roberto Amato, Jeffrey Barrett, Sonia Goncalves, Ewan Harrison, David K. Jackson, Ian Johnston, Dominic Kwiatkowski, Cordelia Langford, John Sillitoe on behalf of the Wellcome Sanger Institute COVID-19 Surveillance Team |
| EPI_ISL_2253405 | Lighthouse Lab in Milton Keynes | Wellcome Sanger Institute for the COVID-19 Genomics UK (COG-UK) Consortium | The Lighthouse Lab in Milton Keynes and Alex Alderton, Roberto Amato, Jeffrey Barrett, Sonia Goncalves, Ewan Harrison, David K. Jackson, Ian Johnston, Dominic Kwiatkowski, Cordelia Langford, John Sillitoe on behalf of the Wellcome Sanger Institute COVID-19 Surveillance Team |
| EPI_ISL_2253407, EPI_ISL_2253408 | Lighthouse Lab in Alderley Park | Wellcome Sanger Institute for the COVID-19 Genomics UK (COG-UK) Consortium | Jacquelyn Wynn, Mairead Hyland, The Lighthouse Lab in Alderley Park and Alex Alderton, Roberto Amato, Jeffrey Barrett, Sonia Goncalves, Ewan Harrison, David K. Jackson, Ian Johnston, Dominic Kwiatkowski, Cordelia Langford, John Sillitoe on behalf of the Wellcome Sanger Institute COVID-19 Surveillance Team |
| EPI_ISL_2253409 | Health Services Laboratories | Wellcome Sanger Institute for the COVID-19 Genomics UK (COG-UK) Consortium | Health Services Laboratories and Alex Alderton, Roberto Amato, Jeffrey Barrett, Sonia Goncalves, Ewan Harrison, David K. Jackson, Ian Johnston, Dominic Kwiatkowski, Cordelia Langford, John Sillitoe on behalf of the Wellcome Sanger Institute COVID-19 Surveillance Team |
| EPI_ISL_2253410 | Lighthouse Lab in Alderley Park | Wellcome Sanger Institute for the COVID-19 Genomics UK (COG-UK) Consortium | Jacquelyn Wynn, Mairead Hyland, The Lighthouse Lab in Alderley Park and Alex Alderton, Roberto Amato, Jeffrey Barrett, Sonia Goncalves, Ewan Harrison, David K. Jackson, Ian Johnston, Dominic Kwiatkowski, Cordelia Langford, John Sillitoe on behalf of the Wellcome Sanger Institute COVID-19 Surveillance Team |
| EPI_ISL_2253411, EPI_ISL_2253413 | Lighthouse Lab in Milton Keynes | Wellcome Sanger Institute for the COVID-19 Genomics UK (COG-UK) Consortium | The Lighthouse Lab in Milton Keynes and Alex Alderton, Roberto Amato, Jeffrey Barrett, Sonia Goncalves, Ewan Harrison, David K. Jackson, Ian Johnston, Dominic Kwiatkowski, Cordelia Langford, John Sillitoe on behalf of the Wellcome Sanger Institute COVID-19 Surveillance Team |
| EPI_ISL_2253414, EPI_ISL_2253415 | Lighthouse Lab in Alderley Park | Wellcome Sanger Institute for the COVID-19 Genomics UK (COG-UK) Consortium | Jacquelyn Wynn, Mairead Hyland, The Lighthouse Lab in Alderley Park and Alex Alderton, Roberto Amato, Jeffrey Barrett, Sonia Goncalves, Ewan Harrison, David K. Jackson, Ian Johnston, Dominic Kwiatkowski, Cordelia Langford, John Sillitoe on behalf of the Wellcome Sanger Institute COVID-19 Surveillance Team |
| EPI_ISL_2253416 | Lighthouse Lab in Milton Keynes | Wellcome Sanger Institute for the COVID-19 Genomics UK (COG-UK) Consortium | The Lighthouse Lab in Milton Keynes and Alex Alderton, Roberto Amato, Jeffrey Barrett, Sonia Goncalves, Ewan Harrison, David K. Jackson, Ian Johnston, Dominic Kwiatkowski, Cordelia Langford, John Sillitoe on behalf of the Wellcome Sanger Institute COVID-19 Surveillance Team |
| EPI_ISL_2253417, EPI_ISL_2253418 | Lighthouse Lab in Alderley Park | Wellcome Sanger Institute for the COVID-19 Genomics UK (COG-UK) Consortium | Jacquelyn Wynn, Mairead Hyland, The Lighthouse Lab in Alderley Park and Alex Alderton, Roberto Amato, Jeffrey Barrett, Sonia Goncalves, Ewan Harrison, David K. Jackson, Ian Johnston, Dominic Kwiatkowski, Cordelia Langford, John Sillitoe on behalf of the Wellcome Sanger Institute COVID-19 Surveillance Team |
| EPI_ISL_2253419 | Lighthouse Lab in Milton Keynes | Wellcome Sanger Institute for the COVID-19 Genomics UK (COG-UK) Consortium | The Lighthouse Lab in Milton Keynes and Alex Alderton, Roberto Amato, Jeffrey Barrett, Sonia Goncalves, Ewan Harrison, David K. Jackson, Ian Johnston, Dominic Kwiatkowski, Cordelia Langford, John Sillitoe on behalf of the Wellcome Sanger Institute COVID-19 Surveillance Team |
| EPI_ISL_2253420, EPI_ISL_2253421, EPI_ISL_2253422, EPI_ISL_2253425 | Lighthouse Lab in Alderley Park | Wellcome Sanger Institute for the COVID-19 Genomics UK (COG-UK) Consortium | Jacquelyn Wynn, Mairead Hyland, The Lighthouse Lab in Alderley Park and Alex Alderton, Roberto Amato, Jeffrey Barrett, Sonia Goncalves, Ewan Harrison, David K. Jackson, Ian Johnston, Dominic Kwiatkowski, Cordelia Langford, John Sillitoe on behalf of the Wellcome Sanger Institute COVID-19 Surveillance Team |
| EPI_ISL_2253426 | Lighthouse Lab in Milton Keynes | Wellcome Sanger Institute for the COVID-19 Genomics UK (COG-UK) Consortium | The Lighthouse Lab in Milton Keynes and Alex Alderton, Roberto Amato, Jeffrey Barrett, Sonia Goncalves, Ewan Harrison, David K. Jackson, Ian Johnston, Dominic Kwiatkowski, Cordelia Langford, John Sillitoe on behalf of the Wellcome Sanger Institute COVID-19 Surveillance Team |
| EPI_ISL_2253427 | Lighthouse Lab in Alderley Park | Wellcome Sanger Institute for the COVID-19 Genomics UK (COG-UK) Consortium | Jacquelyn Wynn, Mairead Hyland, The Lighthouse Lab in Alderley Park and Alex Alderton, Roberto Amato, Jeffrey Barrett, Sonia Goncalves, Ewan Harrison, David K. Jackson, Ian Johnston, Dominic Kwiatkowski, Cordelia Langford, John Sillitoe on behalf of the Wellcome Sanger Institute COVID-19 Surveillance Team |
| EPI_ISL_2253428, EPI_ISL_2253429 | Lighthouse Lab in Milton Keynes | Wellcome Sanger Institute for the COVID-19 Genomics UK (COG-UK) Consortium | The Lighthouse Lab in Milton Keynes and Alex Alderton, Roberto Amato, Jeffrey Barrett, Sonia Goncalves, Ewan Harrison, David K. Jackson, Ian Johnston, Dominic Kwiatkowski, Cordelia Langford, John Sillitoe on behalf of the Wellcome Sanger Institute COVID-19 Surveillance Team |
| EPI_ISL_2253430 | Lighthouse Lab in Glasgow | Wellcome Sanger Institute for the COVID-19 Genomics UK (COG-UK) Consortium | Harper VanSteenhouse, Yumi Kasai, David Gray, Carol Clugston, Anna Dominiczak and Alex Alderton, Roberto Amato, Jeffrey Barrett, Sonia Goncalves, Ewan Harrison, David K. Jackson, Ian Johnston, Dominic Kwiatkowski, Cordelia Langford, John Sillitoe on behalf of the Wellcome Sanger Institute COVID-19 Surveillance Team |
| EPI_ISL_2253431 | Health Services Laboratories | Wellcome Sanger Institute for the COVID-19 Genomics UK (COG-UK) Consortium | Health Services Laboratories and Alex Alderton, Roberto Amato, Jeffrey Barrett, Sonia Goncalves, Ewan Harrison, David K. Jackson, Ian Johnston, Dominic Kwiatkowski, Cordelia Langford, John Sillitoe on behalf of the Wellcome Sanger Institute COVID-19 Surveillance Team |
| EPI_ISL_2253432 | Lighthouse Lab in Milton Keynes | Wellcome Sanger Institute for the COVID-19 Genomics UK (COG-UK) Consortium | The Lighthouse Lab in Milton Keynes and Alex Alderton, Roberto Amato, Jeffrey Barrett, Sonia Goncalves, Ewan Harrison, David K. Jackson, Ian Johnston, Dominic Kwiatkowski, Cordelia Langford, John Sillitoe on behalf of the Wellcome Sanger Institute COVID-19 Surveillance Team |
| EPI_ISL_2253433, EPI_ISL_2253435, EPI_ISL_2253436, EPI_ISL_2253437, EPI_ISL_2253438 | Lighthouse Lab in Alderley Park | Wellcome Sanger Institute for the COVID-19 Genomics UK (COG-UK) Consortium | Jacquelyn Wynn, Mairead Hyland, The Lighthouse Lab in Alderley Park and Alex Alderton, Roberto Amato, Jeffrey Barrett, Sonia Goncalves, Ewan Harrison, David K. Jackson, Ian Johnston, Dominic Kwiatkowski, Cordelia Langford, John Sillitoe on behalf of the Wellcome Sanger Institute COVID-19 Surveillance Team |
| EPI_ISL_2253439 | Lighthouse Lab in Milton Keynes | Wellcome Sanger Institute for the COVID-19 Genomics UK (COG-UK) Consortium | The Lighthouse Lab in Milton Keynes and Alex Alderton, Roberto Amato, Jeffrey Barrett, Sonia Goncalves, Ewan Harrison, David K. Jackson, Ian Johnston, Dominic Kwiatkowski, Cordelia Langford, John Sillitoe on behalf of the Wellcome Sanger Institute COVID-19 Surveillance Team |
| EPI_ISL_2253440 | Lighthouse Lab in Alderley Park | Wellcome Sanger Institute for the COVID-19 Genomics UK (COG-UK) Consortium | Jacquelyn Wynn, Mairead Hyland, The Lighthouse Lab in Alderley Park and Alex Alderton, Roberto Amato, Jeffrey Barrett, Sonia Goncalves, Ewan Harrison, David K. Jackson, Ian Johnston, Dominic Kwiatkowski, Cordelia Langford, John Sillitoe on behalf of the Wellcome Sanger Institute COVID-19 Surveillance Team |
| EPI_ISL_2253443 | Lighthouse Lab in Milton Keynes | Wellcome Sanger Institute for the COVID-19 Genomics UK (COG-UK) Consortium | The Lighthouse Lab in Milton Keynes and Alex Alderton, Roberto Amato, Jeffrey Barrett, Sonia Goncalves, Ewan Harrison, David K. Jackson, Ian Johnston, Dominic Kwiatkowski, Cordelia Langford, John Sillitoe on behalf of the Wellcome Sanger Institute COVID-19 Surveillance Team |
| EPI_ISL_2253444, EPI_ISL_2253446, EPI_ISL_2253447 | Lighthouse Lab in Alderley Park | Wellcome Sanger Institute for the COVID-19 Genomics UK (COG-UK) Consortium | Jacquelyn Wynn, Mairead Hyland, The Lighthouse Lab in Alderley Park and Alex Alderton, Roberto Amato, Jeffrey Barrett, Sonia Goncalves, Ewan Harrison, David K. Jackson, Ian Johnston, Dominic Kwiatkowski, Cordelia Langford, John Sillitoe on behalf of the Wellcome Sanger Institute COVID-19 Surveillance Team |
| EPI_ISL_2253448 | Lighthouse Lab in Milton Keynes | Wellcome Sanger Institute for the COVID-19 Genomics UK (COG-UK) Consortium | The Lighthouse Lab in Milton Keynes and Alex Alderton, Roberto Amato, Jeffrey Barrett, Sonia Goncalves, Ewan Harrison, David K. Jackson, Ian Johnston, Dominic Kwiatkowski, Cordelia Langford, John Sillitoe on behalf of the Wellcome Sanger Institute COVID-19 Surveillance Team |
| EPI_ISL_2253449 | Lighthouse Lab in Alderley Park | Wellcome Sanger Institute for the COVID-19 Genomics UK (COG-UK) Consortium | Jacquelyn Wynn, Mairead Hyland, The Lighthouse Lab in Alderley Park and Alex Alderton, Roberto Amato, Jeffrey Barrett, Sonia Goncalves, Ewan Harrison, David K. Jackson, Ian Johnston, Dominic Kwiatkowski, Cordelia Langford, John Sillitoe on behalf of the Wellcome Sanger Institute COVID-19 Surveillance Team |
| EPI_ISL_2253451 | Lighthouse Lab in Glasgow | Wellcome Sanger Institute for the COVID-19 Genomics UK (COG-UK) Consortium | Harper VanSteenhouse, Yumi Kasai, David Gray, Carol Clugston, Anna Dominiczak and Alex Alderton, Roberto Amato, Jeffrey Barrett, Sonia Goncalves, Ewan Harrison, David K. Jackson, Ian Johnston, Dominic Kwiatkowski, Cordelia Langford, John Sillitoe on behalf of the Wellcome Sanger Institute COVID-19 Surveillance Team |
| EPI_ISL_2253452, EPI_ISL_2253453, EPI_ISL_2253454 | Lighthouse Lab in Alderley Park | Wellcome Sanger Institute for the COVID-19 Genomics UK (COG-UK) Consortium | Jacquelyn Wynn, Mairead Hyland, The Lighthouse Lab in Alderley Park and Alex Alderton, Roberto Amato, Jeffrey Barrett, Sonia Goncalves, Ewan Harrison, David K. Jackson, Ian Johnston, Dominic Kwiatkowski, Cordelia Langford, John Sillitoe on behalf of the Wellcome Sanger Institute COVID-19 Surveillance Team |
| EPI_ISL_2253456 | Lighthouse Lab in Milton Keynes | Wellcome Sanger Institute for the COVID-19 Genomics UK (COG-UK) Consortium | The Lighthouse Lab in Milton Keynes and Alex Alderton, Roberto Amato, Jeffrey Barrett, Sonia Goncalves, Ewan Harrison, David K. Jackson, Ian Johnston, Dominic Kwiatkowski, Cordelia Langford, John Sillitoe on behalf of the Wellcome Sanger Institute COVID-19 Surveillance Team |
| EPI_ISL_2253457 | Health Services Laboratories | Wellcome Sanger Institute for the COVID-19 Genomics UK (COG-UK) Consortium | Health Services Laboratories and Alex Alderton, Roberto Amato, Jeffrey Barrett, Sonia Goncalves, Ewan Harrison, David K. Jackson, Ian Johnston, Dominic Kwiatkowski, Cordelia Langford, John Sillitoe on behalf of the Wellcome Sanger Institute COVID-19 Surveillance Team |
| EPI_ISL_2253458 | Lighthouse Lab in Milton Keynes | Wellcome Sanger Institute for the COVID-19 Genomics UK (COG-UK) | The Lighthouse Lab in Milton Keynes and Alex Alderton, Roberto Amato, Jeffrey Barrett, Sonia Goncalves, Ewan Harrison, David K. Jackson, Ian Johnston, Dominic Kwiatkowski, Cordelia Langford, John Sillitoe on behalf of the Wellcome Sanger |

| | | | |
|---|---|---|---|
| | | Consortium | Institute COVID-19 Surveillance Team |
| EPI_ISL_2253459 | Lighthouse Lab in Alderley Park | Wellcome Sanger Institute for the COVID-19 Genomics UK (COG-UK) Consortium | Jacquelyn Wynn, Mairead Hyland, The Lighthouse Lab in Alderley Park and Alex Alderton, Roberto Amato, Jeffrey Barrett, Sonia Goncalves, Ewan Harrison, David K. Jackson, Ian Johnston, Dominic Kwiatkowski, Cordelia Langford, John Sillitoe on behalf of the Wellcome Sanger Institute COVID-19 Surveillance Team |
| EPI_ISL_2253460 | Health Services Laboratories | Wellcome Sanger Institute for the COVID-19 Genomics UK (COG-UK) Consortium | Health Services Laboratories and Alex Alderton, Roberto Amato, Jeffrey Barrett, Sonia Goncalves, Ewan Harrison, David K. Jackson, Ian Johnston, Dominic Kwiatkowski, Cordelia Langford, John Sillitoe on behalf of the Wellcome Sanger Institute COVID-19 Surveillance Team |
| EPI_ISL_2253463, EPI_ISL_2253464 | Lighthouse Lab in Alderley Park | Wellcome Sanger Institute for the COVID-19 Genomics UK (COG-UK) Consortium | Jacquelyn Wynn, Mairead Hyland, The Lighthouse Lab in Alderley Park and Alex Alderton, Roberto Amato, Jeffrey Barrett, Sonia Goncalves, Ewan Harrison, David K. Jackson, Ian Johnston, Dominic Kwiatkowski, Cordelia Langford, John Sillitoe on behalf of the Wellcome Sanger Institute COVID-19 Surveillance Team |
| EPI_ISL_2253465 | Lighthouse Lab in Glasgow | Wellcome Sanger Institute for the COVID-19 Genomics UK (COG-UK) Consortium | Harper VanSteenhouse, Yumi Kasai, David Gray, Carol Clugston, Anna Dominiczak and Alex Alderton, Roberto Amato, Jeffrey Barrett, Sonia Goncalves, Ewan Harrison, David K. Jackson, Ian Johnston, Dominic Kwiatkowski, Cordelia Langford, John Sillitoe on behalf of the Wellcome Sanger Institute COVID-19 Surveillance Team |
| EPI_ISL_2253468 | Lighthouse Lab in Milton Keynes | Wellcome Sanger Institute for the COVID-19 Genomics UK (COG-UK) Consortium | The Lighthouse Lab in Milton Keynes and Alex Alderton, Roberto Amato, Jeffrey Barrett, Sonia Goncalves, Ewan Harrison, David K. Jackson, Ian Johnston, Dominic Kwiatkowski, Cordelia Langford, John Sillitoe on behalf of the Wellcome Sanger Institute COVID-19 Surveillance Team |
| EPI_ISL_2253469, EPI_ISL_2253472, EPI_ISL_2253474, EPI_ISL_2253475 | Lighthouse Lab in Alderley Park | Wellcome Sanger Institute for the COVID-19 Genomics UK (COG-UK) Consortium | Jacquelyn Wynn, Mairead Hyland, The Lighthouse Lab in Alderley Park and Alex Alderton, Roberto Amato, Jeffrey Barrett, Sonia Goncalves, Ewan Harrison, David K. Jackson, Ian Johnston, Dominic Kwiatkowski, Cordelia Langford, John Sillitoe on behalf of the Wellcome Sanger Institute COVID-19 Surveillance Team |
| EPI_ISL_2253477 | Lighthouse Lab in Milton Keynes | Wellcome Sanger Institute for the COVID-19 Genomics UK (COG-UK) Consortium | The Lighthouse Lab in Milton Keynes and Alex Alderton, Roberto Amato, Jeffrey Barrett, Sonia Goncalves, Ewan Harrison, David K. Jackson, Ian Johnston, Dominic Kwiatkowski, Cordelia Langford, John Sillitoe on behalf of the Wellcome Sanger Institute COVID-19 Surveillance Team |
| EPI_ISL_2253479 | Lighthouse Lab in Alderley Park | Wellcome Sanger Institute for the COVID-19 Genomics UK (COG-UK) Consortium | Jacquelyn Wynn, Mairead Hyland, The Lighthouse Lab in Alderley Park and Alex Alderton, Roberto Amato, Jeffrey Barrett, Sonia Goncalves, Ewan Harrison, David K. Jackson, Ian Johnston, Dominic Kwiatkowski, Cordelia Langford, John Sillitoe on behalf of the Wellcome Sanger Institute COVID-19 Surveillance Team |
| EPI_ISL_2253480, EPI_ISL_2253481, EPI_ISL_2253482 | Lighthouse Lab in Milton Keynes | Wellcome Sanger Institute for the COVID-19 Genomics UK (COG-UK) Consortium | The Lighthouse Lab in Milton Keynes and Alex Alderton, Roberto Amato, Jeffrey Barrett, Sonia Goncalves, Ewan Harrison, David K. Jackson, Ian Johnston, Dominic Kwiatkowski, Cordelia Langford, John Sillitoe on behalf of the Wellcome Sanger Institute COVID-19 Surveillance Team |
| EPI_ISL_2253483, EPI_ISL_2253484 | Lighthouse Lab in Alderley Park | Wellcome Sanger Institute for the COVID-19 Genomics UK (COG-UK) Consortium | Jacquelyn Wynn, Mairead Hyland, The Lighthouse Lab in Alderley Park and Alex Alderton, Roberto Amato, Jeffrey Barrett, Sonia Goncalves, Ewan Harrison, David K. Jackson, Ian Johnston, Dominic Kwiatkowski, Cordelia Langford, John Sillitoe on behalf of the Wellcome Sanger Institute COVID-19 Surveillance Team |
| EPI_ISL_2253486 | Health Services Laboratories | Wellcome Sanger Institute for the COVID-19 Genomics UK (COG-UK) Consortium | Health Services Laboratories and Alex Alderton, Roberto Amato, Jeffrey Barrett, Sonia Goncalves, Ewan Harrison, David K. Jackson, Ian Johnston, Dominic Kwiatkowski, Cordelia Langford, John Sillitoe on behalf of the Wellcome Sanger Institute COVID-19 Surveillance Team |
| EPI_ISL_2253487 | Lighthouse Lab in Milton Keynes | Wellcome Sanger Institute for the COVID-19 Genomics UK (COG-UK) Consortium | The Lighthouse Lab in Milton Keynes and Alex Alderton, Roberto Amato, Jeffrey Barrett, Sonia Goncalves, Ewan Harrison, David K. Jackson, Ian Johnston, Dominic Kwiatkowski, Cordelia Langford, John Sillitoe on behalf of the Wellcome Sanger Institute COVID-19 Surveillance Team |
| EPI_ISL_2253488 | Lighthouse Lab in Alderley Park | Wellcome Sanger Institute for the COVID-19 Genomics UK (COG-UK) Consortium | Jacquelyn Wynn, Mairead Hyland, The Lighthouse Lab in Alderley Park and Alex Alderton, Roberto Amato, Jeffrey Barrett, Sonia Goncalves, Ewan Harrison, David K. Jackson, Ian Johnston, Dominic Kwiatkowski, Cordelia Langford, John Sillitoe on behalf of the Wellcome Sanger Institute COVID-19 Surveillance Team |
| EPI_ISL_2253490 | Lighthouse Lab in Milton Keynes | Wellcome Sanger Institute for the COVID-19 Genomics UK (COG-UK) Consortium | The Lighthouse Lab in Milton Keynes and Alex Alderton, Roberto Amato, Jeffrey Barrett, Sonia Goncalves, Ewan Harrison, David K. Jackson, Ian Johnston, Dominic Kwiatkowski, Cordelia Langford, John Sillitoe on behalf of the Wellcome Sanger Institute COVID-19 Surveillance Team |
| EPI_ISL_2253491 | Health Services Laboratories | Wellcome Sanger Institute for the COVID-19 Genomics UK (COG-UK) Consortium | Health Services Laboratories and Alex Alderton, Roberto Amato, Jeffrey Barrett, Sonia Goncalves, Ewan Harrison, David K. Jackson, Ian Johnston, Dominic Kwiatkowski, Cordelia Langford, John Sillitoe on behalf of the Wellcome Sanger Institute COVID-19 Surveillance Team |
| EPI_ISL_2253492 | Lighthouse Lab in Alderley Park | Wellcome Sanger Institute for the COVID-19 Genomics UK (COG-UK) Consortium | Jacquelyn Wynn, Mairead Hyland, The Lighthouse Lab in Alderley Park and Alex Alderton, Roberto Amato, Jeffrey Barrett, Sonia Goncalves, Ewan Harrison, David K. Jackson, Ian Johnston, Dominic Kwiatkowski, Cordelia Langford, John Sillitoe on behalf of the Wellcome Sanger Institute COVID-19 Surveillance Team |
| EPI_ISL_2253493 | Lighthouse Lab in Milton Keynes | Wellcome Sanger Institute for the COVID-19 Genomics UK (COG-UK) Consortium | The Lighthouse Lab in Milton Keynes and Alex Alderton, Roberto Amato, Jeffrey Barrett, Sonia Goncalves, Ewan Harrison, David K. Jackson, Ian Johnston, Dominic Kwiatkowski, Cordelia Langford, John Sillitoe on behalf of the Wellcome Sanger Institute COVID-19 Surveillance Team |
| EPI_ISL_2253494 | Lighthouse Lab in Glasgow | Wellcome Sanger Institute for the COVID-19 Genomics UK (COG-UK) Consortium | Harper VanSteenhouse, Yumi Kasai, David Gray, Carol Clugston, Anna Dominiczak and Alex Alderton, Roberto Amato, Jeffrey Barrett, Sonia Goncalves, Ewan Harrison, David K. Jackson, Ian Johnston, Dominic Kwiatkowski, Cordelia Langford, John Sillitoe on behalf of the Wellcome Sanger Institute COVID-19 Surveillance Team |
| EPI_ISL_2253495 | Lighthouse Lab in Alderley Park | Wellcome Sanger Institute for the COVID-19 Genomics UK (COG-UK) Consortium | Jacquelyn Wynn, Mairead Hyland, The Lighthouse Lab in Alderley Park and Alex Alderton, Roberto Amato, Jeffrey Barrett, Sonia Goncalves, Ewan Harrison, David K. Jackson, Ian Johnston, Dominic Kwiatkowski, Cordelia Langford, John Sillitoe on behalf of the Wellcome Sanger Institute COVID-19 Surveillance Team |
| EPI_ISL_2253496 | Lighthouse Lab in Milton Keynes | Wellcome Sanger Institute for the COVID-19 Genomics UK (COG-UK) Consortium | The Lighthouse Lab in Milton Keynes and Alex Alderton, Roberto Amato, Jeffrey Barrett, Sonia Goncalves, Ewan Harrison, David K. Jackson, Ian Johnston, Dominic Kwiatkowski, Cordelia Langford, John Sillitoe on behalf of the Wellcome Sanger Institute COVID-19 Surveillance Team |
| EPI_ISL_2253497, EPI_ISL_2253498 | Lighthouse Lab in Alderley Park | Wellcome Sanger Institute for the COVID-19 Genomics UK (COG-UK) Consortium | Jacquelyn Wynn, Mairead Hyland, The Lighthouse Lab in Alderley Park and Alex Alderton, Roberto Amato, Jeffrey Barrett, Sonia Goncalves, Ewan Harrison, David K. Jackson, Ian Johnston, Dominic Kwiatkowski, Cordelia Langford, John Sillitoe on behalf of the Wellcome Sanger Institute COVID-19 Surveillance Team |
| EPI_ISL_2253499 | Health Services Laboratories | Wellcome Sanger Institute for the COVID-19 Genomics UK (COG-UK) Consortium | Health Services Laboratories and Alex Alderton, Roberto Amato, Jeffrey Barrett, Sonia Goncalves, Ewan Harrison, David K. Jackson, Ian Johnston, Dominic Kwiatkowski, Cordelia Langford, John Sillitoe on behalf of the Wellcome Sanger Institute COVID-19 Surveillance Team |
| EPI_ISL_2253500, EPI_ISL_2253501, EPI_ISL_2253502, EPI_ISL_2253503, EPI_ISL_2253505 | Lighthouse Lab in Alderley Park | Wellcome Sanger Institute for the COVID-19 Genomics UK (COG-UK) Consortium | Jacquelyn Wynn, Mairead Hyland, The Lighthouse Lab in Alderley Park and Alex Alderton, Roberto Amato, Jeffrey Barrett, Sonia Goncalves, Ewan Harrison, David K. Jackson, Ian Johnston, Dominic Kwiatkowski, Cordelia Langford, John Sillitoe on behalf of the Wellcome Sanger Institute COVID-19 Surveillance Team |
| EPI_ISL_2253507, EPI_ISL_2253508 | Lighthouse Lab in Milton Keynes | Wellcome Sanger Institute for the COVID-19 Genomics UK (COG-UK) Consortium | The Lighthouse Lab in Milton Keynes and Alex Alderton, Roberto Amato, Jeffrey Barrett, Sonia Goncalves, Ewan Harrison, David K. Jackson, Ian Johnston, Dominic Kwiatkowski, Cordelia Langford, John Sillitoe on behalf of the Wellcome Sanger Institute COVID-19 Surveillance Team |
| EPI_ISL_2253509, EPI_ISL_2253510 | Lighthouse Lab in Alderley Park | Wellcome Sanger Institute for the COVID-19 Genomics UK (COG-UK) Consortium | Jacquelyn Wynn, Mairead Hyland, The Lighthouse Lab in Alderley Park and Alex Alderton, Roberto Amato, Jeffrey Barrett, Sonia Goncalves, Ewan Harrison, David K. Jackson, Ian Johnston, Dominic Kwiatkowski, Cordelia Langford, John Sillitoe on behalf of the Wellcome Sanger Institute COVID-19 Surveillance Team |
| EPI_ISL_2253511, EPI_ISL_2253512 | Lighthouse Lab in Milton Keynes | Wellcome Sanger Institute for the COVID-19 Genomics UK (COG-UK) Consortium | The Lighthouse Lab in Milton Keynes and Alex Alderton, Roberto Amato, Jeffrey Barrett, Sonia Goncalves, Ewan Harrison, David K. Jackson, Ian Johnston, Dominic Kwiatkowski, Cordelia Langford, John Sillitoe on behalf of the Wellcome Sanger Institute COVID-19 Surveillance Team |
| EPI_ISL_2253514, EPI_ISL_2253515, EPI_ISL_2253516, EPI_ISL_2253517 | Lighthouse Lab in Alderley Park | Wellcome Sanger Institute for the COVID-19 Genomics UK (COG-UK) Consortium | Jacquelyn Wynn, Mairead Hyland, The Lighthouse Lab in Alderley Park and Alex Alderton, Roberto Amato, Jeffrey Barrett, Sonia Goncalves, Ewan Harrison, David K. Jackson, Ian Johnston, Dominic Kwiatkowski, Cordelia Langford, John Sillitoe on behalf of the Wellcome Sanger Institute COVID-19 Surveillance Team |
| EPI_ISL_2253519 | Health Services Laboratories | Wellcome Sanger Institute for the COVID-19 Genomics UK (COG-UK) Consortium | Health Services Laboratories and Alex Alderton, Roberto Amato, Jeffrey Barrett, Sonia Goncalves, Ewan Harrison, David K. Jackson, Ian Johnston, Dominic Kwiatkowski, Cordelia Langford, John Sillitoe on behalf of the Wellcome Sanger Institute COVID-19 Surveillance Team |
| EPI_ISL_2253521 | Lighthouse Lab in Alderley Park | Wellcome Sanger Institute for the COVID-19 Genomics UK (COG-UK) Consortium | Jacquelyn Wynn, Mairead Hyland, The Lighthouse Lab in Alderley Park and Alex Alderton, Roberto Amato, Jeffrey Barrett, Sonia Goncalves, Ewan Harrison, David K. Jackson, Ian Johnston, Dominic Kwiatkowski, Cordelia Langford, John Sillitoe on behalf of the Wellcome Sanger Institute COVID-19 Surveillance Team |
| EPI_ISL_2253524 | Lighthouse Lab in Milton Keynes | Wellcome Sanger Institute for the COVID-19 Genomics UK (COG-UK) Consortium | The Lighthouse Lab in Milton Keynes and Alex Alderton, Roberto Amato, Jeffrey Barrett, Sonia Goncalves, Ewan Harrison, David K. Jackson, Ian Johnston, Dominic Kwiatkowski, Cordelia Langford, John Sillitoe on behalf of the Wellcome Sanger Institute COVID-19 Surveillance Team |
| EPI_ISL_2253531, EPI_ISL_2253532 | Lighthouse Lab in Alderley Park | Wellcome Sanger Institute for the COVID-19 Genomics UK (COG-UK) Consortium | Jacquelyn Wynn, Mairead Hyland, The Lighthouse Lab in Alderley Park and Alex Alderton, Roberto Amato, Jeffrey Barrett, Sonia Goncalves, Ewan Harrison, David K. Jackson, Ian Johnston, Dominic Kwiatkowski, Cordelia Langford, John Sillitoe on behalf of the Wellcome Sanger Institute COVID-19 Surveillance Team |
| EPI_ISL_2253533 | Lighthouse Lab in Milton Keynes | Wellcome Sanger Institute for the COVID-19 Genomics UK (COG-UK) Consortium | The Lighthouse Lab in Milton Keynes and Alex Alderton, Roberto Amato, Jeffrey Barrett, Sonia Goncalves, Ewan Harrison, David K. Jackson, Ian Johnston, Dominic Kwiatkowski, Cordelia Langford, John Sillitoe on behalf of the Wellcome Sanger Institute COVID-19 Surveillance Team |
| EPI_ISL_2253534, EPI_ISL_2253536 | Lighthouse Lab in Alderley Park | Wellcome Sanger Institute for the COVID-19 Genomics UK (COG-UK) Consortium | Jacquelyn Wynn, Mairead Hyland, The Lighthouse Lab in Alderley Park and Alex Alderton, Roberto Amato, Jeffrey Barrett, Sonia Goncalves, Ewan Harrison, David K. Jackson, Ian Johnston, Dominic Kwiatkowski, Cordelia Langford, John Sillitoe on behalf of the Wellcome Sanger Institute COVID-19 Surveillance Team |
| EPI_ISL_2253537 | Lighthouse Lab in Milton Keynes | Wellcome Sanger Institute for the COVID-19 Genomics UK (COG-UK) Consortium | The Lighthouse Lab in Milton Keynes and Alex Alderton, Roberto Amato, Jeffrey Barrett, Sonia Goncalves, Ewan Harrison, David K. Jackson, Ian Johnston, Dominic Kwiatkowski, Cordelia Langford, John Sillitoe on behalf of the Wellcome Sanger Institute COVID-19 Surveillance Team |
| EPI_ISL_2253539 | Lighthouse Lab in Glasgow | Wellcome Sanger Institute for the COVID-19 Genomics UK (COG-UK) Consortium | Harper VanSteenhouse, Yumi Kasai, David Gray, Carol Clugston, Anna Dominiczak and Alex Alderton, Roberto Amato, Jeffrey Barrett, Sonia Goncalves, Ewan Harrison, David K. Jackson, Ian Johnston, Dominic Kwiatkowski, Cordelia Langford, John Sillitoe on behalf of the Wellcome Sanger Institute COVID-19 Surveillance Team |
| EPI_ISL_2253540 | Lighthouse Lab in Milton Keynes | Wellcome Sanger Institute for the COVID-19 Genomics UK (COG-UK) Consortium | The Lighthouse Lab in Milton Keynes and Alex Alderton, Roberto Amato, Jeffrey Barrett, Sonia Goncalves, Ewan Harrison, David K. Jackson, Ian Johnston, Dominic Kwiatkowski, Cordelia Langford, John Sillitoe on behalf of the Wellcome Sanger Institute COVID-19 Surveillance Team |
| EPI_ISL_2253541 | Lighthouse Lab in Alderley Park | Wellcome Sanger Institute for the COVID-19 Genomics UK (COG-UK) Consortium | Jacquelyn Wynn, Mairead Hyland, The Lighthouse Lab in Alderley Park and Alex Alderton, Roberto Amato, Jeffrey Barrett, Sonia Goncalves, Ewan Harrison, David K. Jackson, Ian Johnston, Dominic Kwiatkowski, Cordelia Langford, John Sillitoe on behalf of the Wellcome Sanger Institute COVID-19 Surveillance Team |
| EPI_ISL_2253543 | Lighthouse Lab in Milton Keynes | Wellcome Sanger Institute for the COVID-19 Genomics UK (COG-UK) Consortium | The Lighthouse Lab in Milton Keynes and Alex Alderton, Roberto Amato, Jeffrey Barrett, Sonia Goncalves, Ewan Harrison, David K. Jackson, Ian Johnston, Dominic Kwiatkowski, Cordelia Langford, John Sillitoe on behalf of the Wellcome Sanger Institute COVID-19 Surveillance Team |
| EPI_ISL_2253544, EPI_ISL_2253546, EPI_ISL_2253547 | Lighthouse Lab in Alderley Park | Wellcome Sanger Institute for the COVID-19 Genomics UK (COG-UK) Consortium | Jacquelyn Wynn, Mairead Hyland, The Lighthouse Lab in Alderley Park and Alex Alderton, Roberto Amato, Jeffrey Barrett, Sonia Goncalves, Ewan Harrison, David K. Jackson, Ian Johnston, Dominic Kwiatkowski, Cordelia Langford, John Sillitoe on behalf of the Wellcome Sanger Institute COVID-19 Surveillance Team |
| EPI_ISL_2253550 | Lighthouse Lab in Milton Keynes | Wellcome Sanger Institute for the COVID-19 Genomics UK (COG-UK) Consortium | The Lighthouse Lab in Milton Keynes and Alex Alderton, Roberto Amato, Jeffrey Barrett, Sonia Goncalves, Ewan Harrison, David K. Jackson, Ian Johnston, Dominic Kwiatkowski, Cordelia Langford, John Sillitoe on behalf of the Wellcome Sanger Institute COVID-19 Surveillance Team |
| EPI_ISL_2253552, | Lighthouse Lab in Alderley Park | Wellcome Sanger Institute for the COVID-19 Genomics UK (COG-UK) | Jacquelyn Wynn, Mairead Hyland, The Lighthouse Lab in Alderley Park and Alex Alderton, Roberto Amato, Jeffrey Barrett, Sonia Goncalves, Ewan Harrison, David K. Jackson, Ian Johnston, Dominic Kwiatkowski, Cordelia Langford, John Sillitoe on |

| | | | |
|---|---|---|---|
| EPI_ISL_2253554, EPI_ISL_2253555, EPI_ISL_2253558, EPI_ISL_2253561, EPI_ISL_2253562 | | Consortium | behalf of the Wellcome Sanger Institute COVID-19 Surveillance Team |
| EPI_ISL_2253563, EPI_ISL_2253564 | Lighthouse Lab in Milton Keynes | Wellcome Sanger Institute for the COVID-19 Genomics UK (COG-UK) Consortium | The Lighthouse Lab in Milton Keynes and Alex Alderton, Roberto Amato, Jeffrey Barrett, Sonia Goncalves, Ewan Harrison, David K. Jackson, Ian Johnston, Dominic Kwiatkowski, Cordelia Langford, John Sillitoe on behalf of the Wellcome Sanger Institute COVID-19 Surveillance Team |
| EPI_ISL_2253568, EPI_ISL_2253569, EPI_ISL_2253570, EPI_ISL_2253571 | Lighthouse Lab in Alderley Park | Wellcome Sanger Institute for the COVID-19 Genomics UK (COG-UK) Consortium | Jacquelyn Wynn, Mairead Hyland, The Lighthouse Lab in Alderley Park and Alex Alderton, Roberto Amato, Jeffrey Barrett, Sonia Goncalves, Ewan Harrison, David K. Jackson, Ian Johnston, Dominic Kwiatkowski, Cordelia Langford, John Sillitoe on behalf of the Wellcome Sanger Institute COVID-19 Surveillance Team |
| EPI_ISL_2253572, EPI_ISL_2253573 | Lighthouse Lab in Milton Keynes | Wellcome Sanger Institute for the COVID-19 Genomics UK (COG-UK) Consortium | The Lighthouse Lab in Milton Keynes and Alex Alderton, Roberto Amato, Jeffrey Barrett, Sonia Goncalves, Ewan Harrison, David K. Jackson, Ian Johnston, Dominic Kwiatkowski, Cordelia Langford, John Sillitoe on behalf of the Wellcome Sanger Institute COVID-19 Surveillance Team |
| EPI_ISL_2253575 | Lighthouse Lab in Alderley Park | Wellcome Sanger Institute for the COVID-19 Genomics UK (COG-UK) Consortium | Jacquelyn Wynn, Mairead Hyland, The Lighthouse Lab in Alderley Park and Alex Alderton, Roberto Amato, Jeffrey Barrett, Sonia Goncalves, Ewan Harrison, David K. Jackson, Ian Johnston, Dominic Kwiatkowski, Cordelia Langford, John Sillitoe on behalf of the Wellcome Sanger Institute COVID-19 Surveillance Team |
| EPI_ISL_2253576 | Lighthouse Lab in Milton Keynes | Wellcome Sanger Institute for the COVID-19 Genomics UK (COG-UK) Consortium | The Lighthouse Lab in Milton Keynes and Alex Alderton, Roberto Amato, Jeffrey Barrett, Sonia Goncalves, Ewan Harrison, David K. Jackson, Ian Johnston, Dominic Kwiatkowski, Cordelia Langford, John Sillitoe on behalf of the Wellcome Sanger Institute COVID-19 Surveillance Team |
| EPI_ISL_2253578 | Health Services Laboratories | Wellcome Sanger Institute for the COVID-19 Genomics UK (COG-UK) Consortium | Health Services Laboratories and Alex Alderton, Roberto Amato, Jeffrey Barrett, Sonia Goncalves, Ewan Harrison, David K. Jackson, Ian Johnston, Dominic Kwiatkowski, Cordelia Langford, John Sillitoe on behalf of the Wellcome Sanger Institute COVID-19 Surveillance Team |
| EPI_ISL_2253580 | Lighthouse Lab in Milton Keynes | Wellcome Sanger Institute for the COVID-19 Genomics UK (COG-UK) Consortium | The Lighthouse Lab in Milton Keynes and Alex Alderton, Roberto Amato, Jeffrey Barrett, Sonia Goncalves, Ewan Harrison, David K. Jackson, Ian Johnston, Dominic Kwiatkowski, Cordelia Langford, John Sillitoe on behalf of the Wellcome Sanger Institute COVID-19 Surveillance Team |
| EPI_ISL_2253581, EPI_ISL_2253582 | Lighthouse Lab in Alderley Park | Wellcome Sanger Institute for the COVID-19 Genomics UK (COG-UK) Consortium | Jacquelyn Wynn, Mairead Hyland, The Lighthouse Lab in Alderley Park and Alex Alderton, Roberto Amato, Jeffrey Barrett, Sonia Goncalves, Ewan Harrison, David K. Jackson, Ian Johnston, Dominic Kwiatkowski, Cordelia Langford, John Sillitoe on behalf of the Wellcome Sanger Institute COVID-19 Surveillance Team |
| EPI_ISL_2253583 | Lighthouse Lab in Milton Keynes | Wellcome Sanger Institute for the COVID-19 Genomics UK (COG-UK) Consortium | The Lighthouse Lab in Milton Keynes and Alex Alderton, Roberto Amato, Jeffrey Barrett, Sonia Goncalves, Ewan Harrison, David K. Jackson, Ian Johnston, Dominic Kwiatkowski, Cordelia Langford, John Sillitoe on behalf of the Wellcome Sanger Institute COVID-19 Surveillance Team |
| EPI_ISL_2253584, EPI_ISL_2253585, EPI_ISL_2253586, EPI_ISL_2253587, EPI_ISL_2253588, EPI_ISL_2253589, EPI_ISL_2253590, EPI_ISL_2253591, EPI_ISL_2253592, EPI_ISL_2253593, EPI_ISL_2253594, EPI_ISL_2253595, EPI_ISL_2253596, EPI_ISL_2253597, EPI_ISL_2253598 | | | |
| see above | Lighthouse Lab in Alderley Park | Wellcome Sanger Institute for the COVID-19 Genomics UK (COG-UK) Consortium | Jacquelyn Wynn, Mairead Hyland, The Lighthouse Lab in Alderley Park and Alex Alderton, Roberto Amato, Jeffrey Barrett, Sonia Goncalves, Ewan Harrison, David K. Jackson, Ian Johnston, Dominic Kwiatkowski, Cordelia Langford, John Sillitoe on behalf of the Wellcome Sanger Institute COVID-19 Surveillance Team |
| EPI_ISL_2253599, EPI_ISL_2253600 | Lighthouse Lab in Milton Keynes | Wellcome Sanger Institute for the COVID-19 Genomics UK (COG-UK) Consortium | The Lighthouse Lab in Milton Keynes and Alex Alderton, Roberto Amato, Jeffrey Barrett, Sonia Goncalves, Ewan Harrison, David K. Jackson, Ian Johnston, Dominic Kwiatkowski, Cordelia Langford, John Sillitoe on behalf of the Wellcome Sanger Institute COVID-19 Surveillance Team |
| EPI_ISL_2253601, EPI_ISL_2253603 | Lighthouse Lab in Alderley Park | Wellcome Sanger Institute for the COVID-19 Genomics UK (COG-UK) Consortium | Jacquelyn Wynn, Mairead Hyland, The Lighthouse Lab in Alderley Park and Alex Alderton, Roberto Amato, Jeffrey Barrett, Sonia Goncalves, Ewan Harrison, David K. Jackson, Ian Johnston, Dominic Kwiatkowski, Cordelia Langford, John Sillitoe on behalf of the Wellcome Sanger Institute COVID-19 Surveillance Team |
| EPI_ISL_2253604 | Lighthouse Lab in Milton Keynes | Wellcome Sanger Institute for the COVID-19 Genomics UK (COG-UK) Consortium | The Lighthouse Lab in Milton Keynes and Alex Alderton, Roberto Amato, Jeffrey Barrett, Sonia Goncalves, Ewan Harrison, David K. Jackson, Ian Johnston, Dominic Kwiatkowski, Cordelia Langford, John Sillitoe on behalf of the Wellcome Sanger Institute COVID-19 Surveillance Team |
| EPI_ISL_2253605 | Lighthouse Lab in Alderley Park | Wellcome Sanger Institute for the COVID-19 Genomics UK (COG-UK) Consortium | Jacquelyn Wynn, Mairead Hyland, The Lighthouse Lab in Alderley Park and Alex Alderton, Roberto Amato, Jeffrey Barrett, Sonia Goncalves, Ewan Harrison, David K. Jackson, Ian Johnston, Dominic Kwiatkowski, Cordelia Langford, John Sillitoe on behalf of the Wellcome Sanger Institute COVID-19 Surveillance Team |
| EPI_ISL_2253606 | Lighthouse Lab in Milton Keynes | Wellcome Sanger Institute for the COVID-19 Genomics UK (COG-UK) Consortium | The Lighthouse Lab in Milton Keynes and Alex Alderton, Roberto Amato, Jeffrey Barrett, Sonia Goncalves, Ewan Harrison, David K. Jackson, Ian Johnston, Dominic Kwiatkowski, Cordelia Langford, John Sillitoe on behalf of the Wellcome Sanger Institute COVID-19 Surveillance Team |
| EPI_ISL_2253607 | Lighthouse Lab in Alderley Park | Wellcome Sanger Institute for the COVID-19 Genomics UK (COG-UK) Consortium | Jacquelyn Wynn, Mairead Hyland, The Lighthouse Lab in Alderley Park and Alex Alderton, Roberto Amato, Jeffrey Barrett, Sonia Goncalves, Ewan Harrison, David K. Jackson, Ian Johnston, Dominic Kwiatkowski, Cordelia Langford, John Sillitoe on behalf of the Wellcome Sanger Institute COVID-19 Surveillance Team |
| EPI_ISL_2253609 | Lighthouse Lab in Glasgow | Wellcome Sanger Institute for the COVID-19 Genomics UK (COG-UK) Consortium | Harper VanSteenhouse, Yumi Kasai, David Gray, Carol Clugston, Anna Dominiczak and Alex Alderton, Roberto Amato, Jeffrey Barrett, Sonia Goncalves, Ewan Harrison, David K. Jackson, Ian Johnston, Dominic Kwiatkowski, Cordelia Langford, John Sillitoe on behalf of the Wellcome Sanger Institute COVID-19 Surveillance Team |
| EPI_ISL_2253610 | Lighthouse Lab in Alderley Park | Wellcome Sanger Institute for the COVID-19 Genomics UK (COG-UK) Consortium | Jacquelyn Wynn, Mairead Hyland, The Lighthouse Lab in Alderley Park and Alex Alderton, Roberto Amato, Jeffrey Barrett, Sonia Goncalves, Ewan Harrison, David K. Jackson, Ian Johnston, Dominic Kwiatkowski, Cordelia Langford, John Sillitoe on behalf of the Wellcome Sanger Institute COVID-19 Surveillance Team |
| EPI_ISL_2253611 | Lighthouse Lab in Glasgow | Wellcome Sanger Institute for the COVID-19 Genomics UK (COG-UK) Consortium | Harper VanSteenhouse, Yumi Kasai, David Gray, Carol Clugston, Anna Dominiczak and Alex Alderton, Roberto Amato, Jeffrey Barrett, Sonia Goncalves, Ewan Harrison, David K. Jackson, Ian Johnston, Dominic Kwiatkowski, Cordelia Langford, John Sillitoe on behalf of the Wellcome Sanger Institute COVID-19 Surveillance Team |
| EPI_ISL_2253613 | Lighthouse Lab in Alderley Park | Wellcome Sanger Institute for the COVID-19 Genomics UK (COG-UK) Consortium | Jacquelyn Wynn, Mairead Hyland, The Lighthouse Lab in Alderley Park and Alex Alderton, Roberto Amato, Jeffrey Barrett, Sonia Goncalves, Ewan Harrison, David K. Jackson, Ian Johnston, Dominic Kwiatkowski, Cordelia Langford, John Sillitoe on behalf of the Wellcome Sanger Institute COVID-19 Surveillance Team |
| EPI_ISL_2253614 | Health Services Laboratories | Wellcome Sanger Institute for the COVID-19 Genomics UK (COG-UK) Consortium | Health Services Laboratories and Alex Alderton, Roberto Amato, Jeffrey Barrett, Sonia Goncalves, Ewan Harrison, David K. Jackson, Ian Johnston, Dominic Kwiatkowski, Cordelia Langford, John Sillitoe on behalf of the Wellcome Sanger Institute COVID-19 Surveillance Team |
| EPI_ISL_2253615 | Lighthouse Lab in Alderley Park | Wellcome Sanger Institute for the COVID-19 Genomics UK (COG-UK) Consortium | Jacquelyn Wynn, Mairead Hyland, The Lighthouse Lab in Alderley Park and Alex Alderton, Roberto Amato, Jeffrey Barrett, Sonia Goncalves, Ewan Harrison, David K. Jackson, Ian Johnston, Dominic Kwiatkowski, Cordelia Langford, John Sillitoe on behalf of the Wellcome Sanger Institute COVID-19 Surveillance Team |
| EPI_ISL_2253618 | Lighthouse Lab in Milton Keynes | Wellcome Sanger Institute for the COVID-19 Genomics UK (COG-UK) Consortium | The Lighthouse Lab in Milton Keynes and Alex Alderton, Roberto Amato, Jeffrey Barrett, Sonia Goncalves, Ewan Harrison, David K. Jackson, Ian Johnston, Dominic Kwiatkowski, Cordelia Langford, John Sillitoe on behalf of the Wellcome Sanger Institute COVID-19 Surveillance Team |
| EPI_ISL_2253619, EPI_ISL_2253620 | Lighthouse Lab in Alderley Park | Wellcome Sanger Institute for the COVID-19 Genomics UK (COG-UK) Consortium | Jacquelyn Wynn, Mairead Hyland, The Lighthouse Lab in Alderley Park and Alex Alderton, Roberto Amato, Jeffrey Barrett, Sonia Goncalves, Ewan Harrison, David K. Jackson, Ian Johnston, Dominic Kwiatkowski, Cordelia Langford, John Sillitoe on behalf of the Wellcome Sanger Institute COVID-19 Surveillance Team |
| EPI_ISL_2253621 | Lighthouse Lab in Glasgow | Wellcome Sanger Institute for the COVID-19 Genomics UK (COG-UK) Consortium | Harper VanSteenhouse, Yumi Kasai, David Gray, Carol Clugston, Anna Dominiczak and Alex Alderton, Roberto Amato, Jeffrey Barrett, Sonia Goncalves, Ewan Harrison, David K. Jackson, Ian Johnston, Dominic Kwiatkowski, Cordelia Langford, John Sillitoe on behalf of the Wellcome Sanger Institute COVID-19 Surveillance Team |
| EPI_ISL_2253623, EPI_ISL_2253624, EPI_ISL_2253625, EPI_ISL_2253626 | Lighthouse Lab in Alderley Park | Wellcome Sanger Institute for the COVID-19 Genomics UK (COG-UK) Consortium | Jacquelyn Wynn, Mairead Hyland, The Lighthouse Lab in Alderley Park and Alex Alderton, Roberto Amato, Jeffrey Barrett, Sonia Goncalves, Ewan Harrison, David K. Jackson, Ian Johnston, Dominic Kwiatkowski, Cordelia Langford, John Sillitoe on behalf of the Wellcome Sanger Institute COVID-19 Surveillance Team |
| EPI_ISL_2253627 | Lighthouse Lab in Glasgow | Wellcome Sanger Institute for the COVID-19 Genomics UK (COG-UK) Consortium | Harper VanSteenhouse, Yumi Kasai, David Gray, Carol Clugston, Anna Dominiczak and Alex Alderton, Roberto Amato, Jeffrey Barrett, Sonia Goncalves, Ewan Harrison, David K. Jackson, Ian Johnston, Dominic Kwiatkowski, Cordelia Langford, John Sillitoe on behalf of the Wellcome Sanger Institute COVID-19 Surveillance Team |
| EPI_ISL_2253628 | Lighthouse Lab in Alderley Park | Wellcome Sanger Institute for the COVID-19 Genomics UK (COG-UK) Consortium | Jacquelyn Wynn, Mairead Hyland, The Lighthouse Lab in Alderley Park and Alex Alderton, Roberto Amato, Jeffrey Barrett, Sonia Goncalves, Ewan Harrison, David K. Jackson, Ian Johnston, Dominic Kwiatkowski, Cordelia Langford, John Sillitoe on behalf of the Wellcome Sanger Institute COVID-19 Surveillance Team |
| EPI_ISL_2253629, EPI_ISL_2253635 | Lighthouse Lab in Milton Keynes | Wellcome Sanger Institute for the COVID-19 Genomics UK (COG-UK) Consortium | The Lighthouse Lab in Milton Keynes and Alex Alderton, Roberto Amato, Jeffrey Barrett, Sonia Goncalves, Ewan Harrison, David K. Jackson, Ian Johnston, Dominic Kwiatkowski, Cordelia Langford, John Sillitoe on behalf of the Wellcome Sanger Institute COVID-19 Surveillance Team |
| EPI_ISL_2253636 | Lighthouse Lab in Alderley Park | Wellcome Sanger Institute for the COVID-19 Genomics UK (COG-UK) Consortium | Jacquelyn Wynn, Mairead Hyland, The Lighthouse Lab in Alderley Park and Alex Alderton, Roberto Amato, Jeffrey Barrett, Sonia Goncalves, Ewan Harrison, David K. Jackson, Ian Johnston, Dominic Kwiatkowski, Cordelia Langford, John Sillitoe on behalf of the Wellcome Sanger Institute COVID-19 Surveillance Team |
| EPI_ISL_2253638, EPI_ISL_2253640 | Lighthouse Lab in Milton Keynes | Wellcome Sanger Institute for the COVID-19 Genomics UK (COG-UK) Consortium | The Lighthouse Lab in Milton Keynes and Alex Alderton, Roberto Amato, Jeffrey Barrett, Sonia Goncalves, Ewan Harrison, David K. Jackson, Ian Johnston, Dominic Kwiatkowski, Cordelia Langford, John Sillitoe on behalf of the Wellcome Sanger Institute COVID-19 Surveillance Team |
| EPI_ISL_2253645, EPI_ISL_2253646, EPI_ISL_2253647, EPI_ISL_2253648, EPI_ISL_2253649, EPI_ISL_2253651, EPI_ISL_2253653, EPI_ISL_2253654 | Lighthouse Lab in Alderley Park | Wellcome Sanger Institute for the COVID-19 Genomics UK (COG-UK) Consortium | Jacquelyn Wynn, Mairead Hyland, The Lighthouse Lab in Alderley Park and Alex Alderton, Roberto Amato, Jeffrey Barrett, Sonia Goncalves, Ewan Harrison, David K. Jackson, Ian Johnston, Dominic Kwiatkowski, Cordelia Langford, John Sillitoe on behalf of the Wellcome Sanger Institute COVID-19 Surveillance Team |
| EPI_ISL_2253656 | Lighthouse Lab in Milton Keynes | Wellcome Sanger Institute for the COVID-19 Genomics UK (COG-UK) Consortium | The Lighthouse Lab in Milton Keynes and Alex Alderton, Roberto Amato, Jeffrey Barrett, Sonia Goncalves, Ewan Harrison, David K. Jackson, Ian Johnston, Dominic Kwiatkowski, Cordelia Langford, John Sillitoe on behalf of the Wellcome Sanger Institute COVID-19 Surveillance Team |
| EPI_ISL_2253658, EPI_ISL_2253659 | Lighthouse Lab in Alderley Park | Wellcome Sanger Institute for the COVID-19 Genomics UK (COG-UK) Consortium | Jacquelyn Wynn, Mairead Hyland, The Lighthouse Lab in Alderley Park and Alex Alderton, Roberto Amato, Jeffrey Barrett, Sonia Goncalves, Ewan Harrison, David K. Jackson, Ian Johnston, Dominic Kwiatkowski, Cordelia Langford, John Sillitoe on behalf of the Wellcome Sanger Institute COVID-19 Surveillance Team |
| EPI_ISL_2253660 | Health Services Laboratories | Wellcome Sanger Institute for the COVID-19 Genomics UK (COG-UK) Consortium | Health Services Laboratories and Alex Alderton, Roberto Amato, Jeffrey Barrett, Sonia Goncalves, Ewan Harrison, David K. Jackson, Ian Johnston, Dominic Kwiatkowski, Cordelia Langford, John Sillitoe on behalf of the Wellcome Sanger Institute COVID-19 Surveillance Team |
| EPI_ISL_2253664, EPI_ISL_2253665, EPI_ISL_2253666, EPI_ISL_2253667, EPI_ISL_2253669, EPI_ISL_2253672 | Lighthouse Lab in Alderley Park | Wellcome Sanger Institute for the COVID-19 Genomics UK (COG-UK) Consortium | Jacquelyn Wynn, Mairead Hyland, The Lighthouse Lab in Alderley Park and Alex Alderton, Roberto Amato, Jeffrey Barrett, Sonia Goncalves, Ewan Harrison, David K. Jackson, Ian Johnston, Dominic Kwiatkowski, Cordelia Langford, John Sillitoe on behalf of the Wellcome Sanger Institute COVID-19 Surveillance Team |
| EPI_ISL_2253673 | Lighthouse Lab in Milton Keynes | Wellcome Sanger Institute for the COVID-19 Genomics UK (COG-UK) | The Lighthouse Lab in Milton Keynes and Alex Alderton, Roberto Amato, Jeffrey Barrett, Sonia Goncalves, Ewan Harrison, David K. Jackson, Ian Johnston, Dominic Kwiatkowski, Cordelia Langford, John Sillitoe on behalf of the Wellcome Sanger |

| | | | |
|---|---|---|---|
| | | Consortium | Institute COVID-19 Surveillance Team |
| EPI_ISL_2253674, EPI_ISL_2253676 | Lighthouse Lab in Alderley Park | Wellcome Sanger Institute for the COVID-19 Genomics UK (COG-UK) Consortium | Jacquelyn Wynn, Mairead Hyland, The Lighthouse Lab in Alderley Park and Alex Alderton, Roberto Amato, Jeffrey Barrett, Sonia Goncalves, Ewan Harrison, David K. Jackson, Ian Johnston, Dominic Kwiatkowski, Cordelia Langford, John Sillitoe on behalf of the Wellcome Sanger Institute COVID-19 Surveillance Team |
| EPI_ISL_2253681, EPI_ISL_2253682, EPI_ISL_2253685, EPI_ISL_2253687, EPI_ISL_2253691, EPI_ISL_2253692, EPI_ISL_2253693, EPI_ISL_2253694, EPI_ISL_2253700, EPI_ISL_2253702, EPI_ISL_2253703, EPI_ISL_2253704, EPI_ISL_2253709, EPI_ISL_2253710, EPI_ISL_2253713, EPI_ISL_2253714, EPI_ISL_2253715, EPI_ISL_2253718, EPI_ISL_2253719, EPI_ISL_2253721, EPI_ISL_2253725, EPI_ISL_2253732, EPI_ISL_2253733, EPI_ISL_2253737, EPI_ISL_2253739, EPI_ISL_2253742, EPI_ISL_2253745, EPI_ISL_2253747, EPI_ISL_2253748, EPI_ISL_2253750, EPI_ISL_2253753, EPI_ISL_2253754, EPI_ISL_2253760, EPI_ISL_2253761, EPI_ISL_2253765, EPI_ISL_2253767, EPI_ISL_2253771, EPI_ISL_2253775, EPI_ISL_2253779, EPI_ISL_2253780, EPI_ISL_2253785, EPI_ISL_2253787, EPI_ISL_2253788, EPI_ISL_2253790, EPI_ISL_2253791, EPI_ISL_2253792, EPI_ISL_2253793, EPI_ISL_2253796, EPI_ISL_2253797, EPI_ISL_2253799, EPI_ISL_2253800, EPI_ISL_2253801, EPI_ISL_2253803, EPI_ISL_2253804, EPI_ISL_2253805, EPI_ISL_2253806, EPI_ISL_2253807, EPI_ISL_2253808, EPI_ISL_2253809, EPI_ISL_2253813, EPI_ISL_2253815, EPI_ISL_2253819, EPI_ISL_2253823, EPI_ISL_2253827, EPI_ISL_2253829, EPI_ISL_2253830, EPI_ISL_2253833, EPI_ISL_2253834, EPI_ISL_2253836, EPI_ISL_2253837, EPI_ISL_2253843, EPI_ISL_2253845, EPI_ISL_2253846, EPI_ISL_2253854, EPI_ISL_2253858, EPI_ISL_2253860, EPI_ISL_2253863, EPI_ISL_2253864, EPI_ISL_2253867, EPI_ISL_2253869, EPI_ISL_2253871, EPI_ISL_2253874, EPI_ISL_2253879, EPI_ISL_2253884, EPI_ISL_2253887, EPI_ISL_2253888, EPI_ISL_2253889, EPI_ISL_2253891, EPI_ISL_2253892, EPI_ISL_2253897, EPI_ISL_2253902, EPI_ISL_2253906, EPI_ISL_2253910, EPI_ISL_2253911, EPI_ISL_2253913, EPI_ISL_2253915, EPI_ISL_2253923, EPI_ISL_2253928, EPI_ISL_2253929, EPI_ISL_2253932, EPI_ISL_2253937 | | | |
| see above | Lighthouse Lab in Milton Keynes | Wellcome Sanger Institute for the COVID-19 Genomics UK (COG-UK) Consortium | The Lighthouse Lab in Milton Keynes and Alex Alderton, Roberto Amato, Jeffrey Barrett, Sonia Goncalves, Ewan Harrison, David K. Jackson, Ian Johnston, Dominic Kwiatkowski, Cordelia Langford, John Sillitoe on behalf of the Wellcome Sanger Institute COVID-19 Surveillance Team |
| EPI_ISL_2253939, EPI_ISL_2253940, EPI_ISL_2253942, EPI_ISL_2253944, EPI_ISL_2253945, EPI_ISL_2253946, EPI_ISL_2253948, EPI_ISL_2253950 | Lighthouse Lab in Alderley Park | Wellcome Sanger Institute for the COVID-19 Genomics UK (COG-UK) Consortium | Jacquelyn Wynn, Mairead Hyland, The Lighthouse Lab in Alderley Park and Alex Alderton, Roberto Amato, Jeffrey Barrett, Sonia Goncalves, Ewan Harrison, David K. Jackson, Ian Johnston, Dominic Kwiatkowski, Cordelia Langford, John Sillitoe on behalf of the Wellcome Sanger Institute COVID-19 Surveillance Team |
| EPI_ISL_2253952 | Lighthouse Lab in Glasgow | Wellcome Sanger Institute for the COVID-19 Genomics UK (COG-UK) Consortium | Harper VanSteenhouse, Yumi Kasai, David Gray, Carol Clugston, Anna Dominiczak and Alex Alderton, Roberto Amato, Jeffrey Barrett, Sonia Goncalves, Ewan Harrison, David K. Jackson, Ian Johnston, Dominic Kwiatkowski, Cordelia Langford, John Sillitoe on behalf of the Wellcome Sanger Institute COVID-19 Surveillance Team |
| EPI_ISL_2253953 | Lighthouse Lab in Milton Keynes | Wellcome Sanger Institute for the COVID-19 Genomics UK (COG-UK) Consortium | The Lighthouse Lab in Milton Keynes and Alex Alderton, Roberto Amato, Jeffrey Barrett, Sonia Goncalves, Ewan Harrison, David K. Jackson, Ian Johnston, Dominic Kwiatkowski, Cordelia Langford, John Sillitoe on behalf of the Wellcome Sanger Institute COVID-19 Surveillance Team |
| EPI_ISL_2253955, EPI_ISL_2253959, EPI_ISL_2253960 | Lighthouse Lab in Alderley Park | Wellcome Sanger Institute for the COVID-19 Genomics UK (COG-UK) Consortium | Jacquelyn Wynn, Mairead Hyland, The Lighthouse Lab in Alderley Park and Alex Alderton, Roberto Amato, Jeffrey Barrett, Sonia Goncalves, Ewan Harrison, David K. Jackson, Ian Johnston, Dominic Kwiatkowski, Cordelia Langford, John Sillitoe on behalf of the Wellcome Sanger Institute COVID-19 Surveillance Team |
| EPI_ISL_2253962 | Lighthouse Lab in Milton Keynes | Wellcome Sanger Institute for the COVID-19 Genomics UK (COG-UK) Consortium | The Lighthouse Lab in Milton Keynes and Alex Alderton, Roberto Amato, Jeffrey Barrett, Sonia Goncalves, Ewan Harrison, David K. Jackson, Ian Johnston, Dominic Kwiatkowski, Cordelia Langford, John Sillitoe on behalf of the Wellcome Sanger Institute COVID-19 Surveillance Team |
| EPI_ISL_2253964, EPI_ISL_2253965, EPI_ISL_2253966, EPI_ISL_2253967, EPI_ISL_2253968, EPI_ISL_2253969, EPI_ISL_2253971, EPI_ISL_2253972, EPI_ISL_2253973 | Lighthouse Lab in Alderley Park | Wellcome Sanger Institute for the COVID-19 Genomics UK (COG-UK) Consortium | Jacquelyn Wynn, Mairead Hyland, The Lighthouse Lab in Alderley Park and Alex Alderton, Roberto Amato, Jeffrey Barrett, Sonia Goncalves, Ewan Harrison, David K. Jackson, Ian Johnston, Dominic Kwiatkowski, Cordelia Langford, John Sillitoe on behalf of the Wellcome Sanger Institute COVID-19 Surveillance Team |
| EPI_ISL_2253977 | Lighthouse Lab in Milton Keynes | Wellcome Sanger Institute for the COVID-19 Genomics UK (COG-UK) Consortium | The Lighthouse Lab in Milton Keynes and Alex Alderton, Roberto Amato, Jeffrey Barrett, Sonia Goncalves, Ewan Harrison, David K. Jackson, Ian Johnston, Dominic Kwiatkowski, Cordelia Langford, John Sillitoe on behalf of the Wellcome Sanger Institute COVID-19 Surveillance Team |
| EPI_ISL_2253978, EPI_ISL_2253979 | Lighthouse Lab in Alderley Park | Wellcome Sanger Institute for the COVID-19 Genomics UK (COG-UK) Consortium | Jacquelyn Wynn, Mairead Hyland, The Lighthouse Lab in Alderley Park and Alex Alderton, Roberto Amato, Jeffrey Barrett, Sonia Goncalves, Ewan Harrison, David K. Jackson, Ian Johnston, Dominic Kwiatkowski, Cordelia Langford, John Sillitoe on behalf of the Wellcome Sanger Institute COVID-19 Surveillance Team |
| EPI_ISL_2253981 | Lighthouse Lab in Glasgow | Wellcome Sanger Institute for the COVID-19 Genomics UK (COG-UK) Consortium | Harper VanSteenhouse, Yumi Kasai, David Gray, Carol Clugston, Anna Dominiczak and Alex Alderton, Roberto Amato, Jeffrey Barrett, Sonia Goncalves, Ewan Harrison, David K. Jackson, Ian Johnston, Dominic Kwiatkowski, Cordelia Langford, John Sillitoe on behalf of the Wellcome Sanger Institute COVID-19 Surveillance Team |
| EPI_ISL_2253984 | Lighthouse Lab in Milton Keynes | Wellcome Sanger Institute for the COVID-19 Genomics UK (COG-UK) Consortium | The Lighthouse Lab in Milton Keynes and Alex Alderton, Roberto Amato, Jeffrey Barrett, Sonia Goncalves, Ewan Harrison, David K. Jackson, Ian Johnston, Dominic Kwiatkowski, Cordelia Langford, John Sillitoe on behalf of the Wellcome Sanger Institute COVID-19 Surveillance Team |
| EPI_ISL_2253985, EPI_ISL_2253986, EPI_ISL_2253989, EPI_ISL_2253991 | Lighthouse Lab in Alderley Park | Wellcome Sanger Institute for the COVID-19 Genomics UK (COG-UK) Consortium | Jacquelyn Wynn, Mairead Hyland, The Lighthouse Lab in Alderley Park and Alex Alderton, Roberto Amato, Jeffrey Barrett, Sonia Goncalves, Ewan Harrison, David K. Jackson, Ian Johnston, Dominic Kwiatkowski, Cordelia Langford, John Sillitoe on behalf of the Wellcome Sanger Institute COVID-19 Surveillance Team |
| EPI_ISL_2253992 | Lighthouse Lab in Glasgow | Wellcome Sanger Institute for the COVID-19 Genomics UK (COG-UK) Consortium | Harper VanSteenhouse, Yumi Kasai, David Gray, Carol Clugston, Anna Dominiczak and Alex Alderton, Roberto Amato, Jeffrey Barrett, Sonia Goncalves, Ewan Harrison, David K. Jackson, Ian Johnston, Dominic Kwiatkowski, Cordelia Langford, John Sillitoe on behalf of the Wellcome Sanger Institute COVID-19 Surveillance Team |
| EPI_ISL_2253993, EPI_ISL_2253994 | Lighthouse Lab in Alderley Park | Wellcome Sanger Institute for the COVID-19 Genomics UK (COG-UK) Consortium | Jacquelyn Wynn, Mairead Hyland, The Lighthouse Lab in Alderley Park and Alex Alderton, Roberto Amato, Jeffrey Barrett, Sonia Goncalves, Ewan Harrison, David K. Jackson, Ian Johnston, Dominic Kwiatkowski, Cordelia Langford, John Sillitoe on behalf of the Wellcome Sanger Institute COVID-19 Surveillance Team |
| EPI_ISL_2253995 | Lighthouse Lab in Milton Keynes | Wellcome Sanger Institute for the COVID-19 Genomics UK (COG-UK) Consortium | The Lighthouse Lab in Milton Keynes and Alex Alderton, Roberto Amato, Jeffrey Barrett, Sonia Goncalves, Ewan Harrison, David K. Jackson, Ian Johnston, Dominic Kwiatkowski, Cordelia Langford, John Sillitoe on behalf of the Wellcome Sanger Institute COVID-19 Surveillance Team |
| EPI_ISL_2253998 | Lighthouse Lab in Alderley Park | Wellcome Sanger Institute for the COVID-19 Genomics UK (COG-UK) Consortium | Jacquelyn Wynn, Mairead Hyland, The Lighthouse Lab in Alderley Park and Alex Alderton, Roberto Amato, Jeffrey Barrett, Sonia Goncalves, Ewan Harrison, David K. Jackson, Ian Johnston, Dominic Kwiatkowski, Cordelia Langford, John Sillitoe on behalf of the Wellcome Sanger Institute COVID-19 Surveillance Team |
| EPI_ISL_2254000 | Lighthouse Lab in Milton Keynes | Wellcome Sanger Institute for the COVID-19 Genomics UK (COG-UK) Consortium | The Lighthouse Lab in Milton Keynes and Alex Alderton, Roberto Amato, Jeffrey Barrett, Sonia Goncalves, Ewan Harrison, David K. Jackson, Ian Johnston, Dominic Kwiatkowski, Cordelia Langford, John Sillitoe on behalf of the Wellcome Sanger Institute COVID-19 Surveillance Team |
| EPI_ISL_2254002, EPI_ISL_2254003, EPI_ISL_2254008 | Lighthouse Lab in Alderley Park | Wellcome Sanger Institute for the COVID-19 Genomics UK (COG-UK) Consortium | Jacquelyn Wynn, Mairead Hyland, The Lighthouse Lab in Alderley Park and Alex Alderton, Roberto Amato, Jeffrey Barrett, Sonia Goncalves, Ewan Harrison, David K. Jackson, Ian Johnston, Dominic Kwiatkowski, Cordelia Langford, John Sillitoe on behalf of the Wellcome Sanger Institute COVID-19 Surveillance Team |
| EPI_ISL_2254010 | Lighthouse Lab in Milton Keynes | Wellcome Sanger Institute for the COVID-19 Genomics UK (COG-UK) Consortium | The Lighthouse Lab in Milton Keynes and Alex Alderton, Roberto Amato, Jeffrey Barrett, Sonia Goncalves, Ewan Harrison, David K. Jackson, Ian Johnston, Dominic Kwiatkowski, Cordelia Langford, John Sillitoe on behalf of the Wellcome Sanger Institute COVID-19 Surveillance Team |
| EPI_ISL_2254011 | Lighthouse Lab in Alderley Park | Wellcome Sanger Institute for the COVID-19 Genomics UK (COG-UK) Consortium | Jacquelyn Wynn, Mairead Hyland, The Lighthouse Lab in Alderley Park and Alex Alderton, Roberto Amato, Jeffrey Barrett, Sonia Goncalves, Ewan Harrison, David K. Jackson, Ian Johnston, Dominic Kwiatkowski, Cordelia Langford, John Sillitoe on behalf of the Wellcome Sanger Institute COVID-19 Surveillance Team |
| EPI_ISL_2254012 | Lighthouse Lab in Milton Keynes | Wellcome Sanger Institute for the COVID-19 Genomics UK (COG-UK) Consortium | The Lighthouse Lab in Milton Keynes and Alex Alderton, Roberto Amato, Jeffrey Barrett, Sonia Goncalves, Ewan Harrison, David K. Jackson, Ian Johnston, Dominic Kwiatkowski, Cordelia Langford, John Sillitoe on behalf of the Wellcome Sanger Institute COVID-19 Surveillance Team |
| EPI_ISL_2254013 | Lighthouse Lab in Alderley Park | Wellcome Sanger Institute for the COVID-19 Genomics UK (COG-UK) Consortium | Jacquelyn Wynn, Mairead Hyland, The Lighthouse Lab in Alderley Park and Alex Alderton, Roberto Amato, Jeffrey Barrett, Sonia Goncalves, Ewan Harrison, David K. Jackson, Ian Johnston, Dominic Kwiatkowski, Cordelia Langford, John Sillitoe on behalf of the Wellcome Sanger Institute COVID-19 Surveillance Team |
| EPI_ISL_2254014, EPI_ISL_2254015 | Lighthouse Lab in Milton Keynes | Wellcome Sanger Institute for the COVID-19 Genomics UK (COG-UK) Consortium | The Lighthouse Lab in Milton Keynes and Alex Alderton, Roberto Amato, Jeffrey Barrett, Sonia Goncalves, Ewan Harrison, David K. Jackson, Ian Johnston, Dominic Kwiatkowski, Cordelia Langford, John Sillitoe on behalf of the Wellcome Sanger Institute COVID-19 Surveillance Team |
| EPI_ISL_2254018, EPI_ISL_2254022, EPI_ISL_2254023, EPI_ISL_2254024, EPI_ISL_2254027 | Lighthouse Lab in Alderley Park | Wellcome Sanger Institute for the COVID-19 Genomics UK (COG-UK) Consortium | Jacquelyn Wynn, Mairead Hyland, The Lighthouse Lab in Alderley Park and Alex Alderton, Roberto Amato, Jeffrey Barrett, Sonia Goncalves, Ewan Harrison, David K. Jackson, Ian Johnston, Dominic Kwiatkowski, Cordelia Langford, John Sillitoe on behalf of the Wellcome Sanger Institute COVID-19 Surveillance Team |
| EPI_ISL_2254028 | Lighthouse Lab in Milton Keynes | Wellcome Sanger Institute for the COVID-19 Genomics UK (COG-UK) Consortium | The Lighthouse Lab in Milton Keynes and Alex Alderton, Roberto Amato, Jeffrey Barrett, Sonia Goncalves, Ewan Harrison, David K. Jackson, Ian Johnston, Dominic Kwiatkowski, Cordelia Langford, John Sillitoe on behalf of the Wellcome Sanger Institute COVID-19 Surveillance Team |
| EPI_ISL_2254030, EPI_ISL_2254031, EPI_ISL_2254034, EPI_ISL_2254035 | Lighthouse Lab in Alderley Park | Wellcome Sanger Institute for the COVID-19 Genomics UK (COG-UK) Consortium | Jacquelyn Wynn, Mairead Hyland, The Lighthouse Lab in Alderley Park and Alex Alderton, Roberto Amato, Jeffrey Barrett, Sonia Goncalves, Ewan Harrison, David K. Jackson, Ian Johnston, Dominic Kwiatkowski, Cordelia Langford, John Sillitoe on behalf of the Wellcome Sanger Institute COVID-19 Surveillance Team |
| EPI_ISL_2254036 | Lighthouse Lab in Glasgow | Wellcome Sanger Institute for the COVID-19 Genomics UK (COG-UK) Consortium | Harper VanSteenhouse, Yumi Kasai, David Gray, Carol Clugston, Anna Dominiczak and Alex Alderton, Roberto Amato, Jeffrey Barrett, Sonia Goncalves, Ewan Harrison, David K. Jackson, Ian Johnston, Dominic Kwiatkowski, Cordelia Langford, John Sillitoe on behalf of the Wellcome Sanger Institute COVID-19 Surveillance Team |
| EPI_ISL_2254037, EPI_ISL_2254041, EPI_ISL_2254042, EPI_ISL_2254043, EPI_ISL_2254047, EPI_ISL_2254050 | Lighthouse Lab in Alderley Park | Wellcome Sanger Institute for the COVID-19 Genomics UK (COG-UK) Consortium | Jacquelyn Wynn, Mairead Hyland, The Lighthouse Lab in Alderley Park and Alex Alderton, Roberto Amato, Jeffrey Barrett, Sonia Goncalves, Ewan Harrison, David K. Jackson, Ian Johnston, Dominic Kwiatkowski, Cordelia Langford, John Sillitoe on behalf of the Wellcome Sanger Institute COVID-19 Surveillance Team |
| EPI_ISL_2254053 | Lighthouse Lab in Milton Keynes | Wellcome Sanger Institute for the COVID-19 Genomics UK (COG-UK) Consortium | The Lighthouse Lab in Milton Keynes and Alex Alderton, Roberto Amato, Jeffrey Barrett, Sonia Goncalves, Ewan Harrison, David K. Jackson, Ian Johnston, Dominic Kwiatkowski, Cordelia Langford, John Sillitoe on behalf of the Wellcome Sanger Institute COVID-19 Surveillance Team |
| EPI_ISL_2254054 | Lighthouse Lab in Glasgow | Wellcome Sanger Institute for the COVID-19 Genomics UK (COG-UK) Consortium | Harper VanSteenhouse, Yumi Kasai, David Gray, Carol Clugston, Anna Dominiczak and Alex Alderton, Roberto Amato, Jeffrey Barrett, Sonia Goncalves, Ewan Harrison, David K. Jackson, Ian Johnston, Dominic Kwiatkowski, Cordelia Langford, John Sillitoe on behalf of the Wellcome Sanger Institute COVID-19 Surveillance Team |
| EPI_ISL_2254056, | Lighthouse Lab in Alderley Park | Wellcome Sanger Institute for the COVID-19 Genomics UK (COG-UK) | Jacquelyn Wynn, Mairead Hyland, The Lighthouse Lab in Alderley Park and Alex Alderton, Roberto Amato, Jeffrey Barrett, Sonia Goncalves, Ewan Harrison, David K. Jackson, Ian Johnston, Dominic Kwiatkowski, Cordelia Langford, John Sillitoe on |

| | | | |
|---|---|---|---|
| EPI_ISL_2254058 | | Consortium | behalf of the Wellcome Sanger Institute COVID-19 Surveillance Team |
| EPI_ISL_2254060 | Lighthouse Lab in Milton Keynes | Wellcome Sanger Institute for the COVID-19 Genomics UK (COG-UK) Consortium | The Lighthouse Lab in Milton Keynes and Alex Alderton, Roberto Amato, Jeffrey Barrett, Sonia Goncalves, Ewan Harrison, David K. Jackson, Ian Johnston, Dominic Kwiatkowski, Cordelia Langford, John Sillitoe on behalf of the Wellcome Sanger Institute COVID-19 Surveillance Team |
| EPI_ISL_2254061 | Lighthouse Lab in Glasgow | Wellcome Sanger Institute for the COVID-19 Genomics UK (COG-UK) Consortium | Harper VanSteenhouse, Yumi Kasai, David Gray, Carol Clugston, Anna Dominiczak and Alex Alderton, Roberto Amato, Jeffrey Barrett, Sonia Goncalves, Ewan Harrison, David K. Jackson, Ian Johnston, Dominic Kwiatkowski, Cordelia Langford, John Sillitoe on behalf of the Wellcome Sanger Institute COVID-19 Surveillance Team |
| EPI_ISL_2254062, EPI_ISL_2254063, EPI_ISL_2254065, EPI_ISL_2254066, EPI_ISL_2254068, EPI_ISL_2254069 | Lighthouse Lab in Alderley Park | Wellcome Sanger Institute for the COVID-19 Genomics UK (COG-UK) Consortium | Jacquelyn Wynn, Mairead Hyland, The Lighthouse Lab in Alderley Park and Alex Alderton, Roberto Amato, Jeffrey Barrett, Sonia Goncalves, Ewan Harrison, David K. Jackson, Ian Johnston, Dominic Kwiatkowski, Cordelia Langford, John Sillitoe on behalf of the Wellcome Sanger Institute COVID-19 Surveillance Team |
| EPI_ISL_2254071 | Lighthouse Lab in Milton Keynes | Wellcome Sanger Institute for the COVID-19 Genomics UK (COG-UK) Consortium | The Lighthouse Lab in Milton Keynes and Alex Alderton, Roberto Amato, Jeffrey Barrett, Sonia Goncalves, Ewan Harrison, David K. Jackson, Ian Johnston, Dominic Kwiatkowski, Cordelia Langford, John Sillitoe on behalf of the Wellcome Sanger Institute COVID-19 Surveillance Team |
| EPI_ISL_2254083 | Randox Laboratories | Wellcome Sanger Institute for the COVID-19 Genomics UK (COG-UK) Consortium | Randox Laboratories and Alex Alderton, Roberto Amato, Jeffrey Barrett, Sonia Goncalves, Ewan Harrison, David K. Jackson, Ian Johnston, Dominic Kwiatkowski, Cordelia Langford, John Sillitoe on behalf of the Wellcome Sanger Institute COVID-19 Surveillance Team |
| EPI_ISL_2254086 | Lighthouse Lab in Glasgow | Wellcome Sanger Institute for the COVID-19 Genomics UK (COG-UK) Consortium | Harper VanSteenhouse, Yumi Kasai, David Gray, Carol Clugston, Anna Dominiczak and Alex Alderton, Roberto Amato, Jeffrey Barrett, Sonia Goncalves, Ewan Harrison, David K. Jackson, Ian Johnston, Dominic Kwiatkowski, Cordelia Langford, John Sillitoe on behalf of the Wellcome Sanger Institute COVID-19 Surveillance Team |
| EPI_ISL_2254094 | Randox Laboratories | Wellcome Sanger Institute for the COVID-19 Genomics UK (COG-UK) Consortium | Randox Laboratories and Alex Alderton, Roberto Amato, Jeffrey Barrett, Sonia Goncalves, Ewan Harrison, David K. Jackson, Ian Johnston, Dominic Kwiatkowski, Cordelia Langford, John Sillitoe on behalf of the Wellcome Sanger Institute COVID-19 Surveillance Team |
| EPI_ISL_2254106, EPI_ISL_2254121, EPI_ISL_2254122 | Lighthouse Lab in Milton Keynes | Wellcome Sanger Institute for the COVID-19 Genomics UK (COG-UK) Consortium | The Lighthouse Lab in Milton Keynes and Alex Alderton, Roberto Amato, Jeffrey Barrett, Sonia Goncalves, Ewan Harrison, David K. Jackson, Ian Johnston, Dominic Kwiatkowski, Cordelia Langford, John Sillitoe on behalf of the Wellcome Sanger Institute COVID-19 Surveillance Team |
| EPI_ISL_2254135 | Microbiology Department, Laboratori Clínic Metropolitana Nord. Hospital Universitari Germans Trias i Pujol. | Can Ruti SARS-CoV-2 Sequencing Hub (HUGTiP/IrsiCaixa/IGTP) | Marc Noguera-Julian, Pilar Armengol, Ignacio Blanco, Antoni E Bordoy, Francesc Catala-Moll, Pere-Joan Cardona, Maria Casadellà, Cristina Casañ, Gemma Clara, Bonaventura Clotet, Cristina Esteban, Montserrat Giménez, Mercedes Guerrero, Anna Not, Roger Paredes, Mariona Parera, Verónica Saludes, Alba Sánchez, and Elisa Martró on behalf of the Can Ruti SARS-CoV-2 Sequencing Hub. |
| EPI_ISL_2254398 | Respiratory Viruses Branch, Centers for Disease Control and Prevention | Respiratory Viruses Branch, Centers for Disease Control and Prevention | Howard,D., Batra,D., Cook,P.W., Moser,K., Paskey,A., Caravas,J., Rambo-Martin,B., Morrison,S., Gulvick,C., Sammons,S., Unoarumhi,Y., Wagner,D., Schmerer,M., Clark,C., Campbell,P., Case,R., Ghorpade,V., Houdeshell,H., Kvalvaag,O., Nall,D., Sanders,E., Vest,A., Westlund,S., Hardison,M., Paden,C.R., MacCannell,D. |
| EPI_ISL_2257706 | Swedish national genomic surveillance program of SARS-CoV-2 | The Public Health Agency of Sweden | Maximilian Riess, Maria Lind Karlberg, Alma Brolund, Swedish national genomic surveillance program of SARS-CoV-2 |
| EPI_ISL_2259251 | Quest Diagnostics | Los Angeles County Public Health Laboratories | P. Hemarajata et al. |
| EPI_ISL_2259439 | Providence Little Company of Mary Medical Center | Los Angeles County Public Health Laboratories | P. Hemarajata et al. |
| EPI_ISL_2259546, EPI_ISL_2259576, EPI_ISL_2259579, EPI_ISL_2259582 | Eurofins LifeCodexx GmbH | Robert Koch Institute | unknown |
| EPI_ISL_2259641, EPI_ISL_2259642, EPI_ISL_2259653 | MVZ Labor Dr. Limbach & Kollegen GbR | Robert Koch Institute | unknown |
| EPI_ISL_2259713 | Niedersächsisches Landesgesundheitsamt (NLGA) | Robert Koch Institute | unknown |
| EPI_ISL_2259771 | MVZ Labor Dr. Reising-Ackermann und Kollegen | Robert Koch Institute | unknown |
| EPI_ISL_2259850, EPI_ISL_2259953, EPI_ISL_2259962, EPI_ISL_2259990, EPI_ISL_2260028, EPI_ISL_2260038 | CENTOGENE Frankfurt Laboratory: Niederlassung Industriepark Höchst | Robert Koch Institute | unknown |
| EPI_ISL_2260059 | MDI Limbach Berlin GmbH; MVZ Labor Berlin | Robert Koch Institute | unknown |
| EPI_ISL_2260092, EPI_ISL_2260146, EPI_ISL_2260299, EPI_ISL_2260301, EPI_ISL_2260332 | Labor 28 MVZ GmbH | Robert Koch Institute | unknown |
| EPI_ISL_2260379 | USC Clinical Lab | Los Angeles County Public Health Laboratories | P. Hemarajata et al. |
| EPI_ISL_2260430, EPI_ISL_2260533, EPI_ISL_2260540, EPI_ISL_2260543 | Bayerisches Landesamt für Gesundheit und Lebensmittelsicherheit (LGL) | Robert Koch Institute | unknown |
| EPI_ISL_2260662 | Sonic - MVZ Medizinisches Labor Bremen GmbH | Robert Koch Institute | unknown |
| EPI_ISL_2260688 | LabKom - Labor Mainz MVZ GmbH | Robert Koch Institute | unknown |
| EPI_ISL_2260689 | Sonic - MVZ Medizinisches Labor Bremen GmbH | Robert Koch Institute | unknown |
| EPI_ISL_2260703 | LabKom - Labor Mainz MVZ GmbH | Robert Koch Institute | unknown |
| EPI_ISL_2260735, EPI_ISL_2260774 | Sonic - MVZ Medizinisches Labor Bremen GmbH | Robert Koch Institute | unknown |
| EPI_ISL_2260789 | Bioscientia Labor Wermsdorf | Robert Koch Institute | unknown |
| EPI_ISL_2260807, EPI_ISL_2260829, EPI_ISL_2260879 | Sonic - MVZ Medizinisches Labor Bremen GmbH | Robert Koch Institute | unknown |
| EPI_ISL_2260917 | MVZ Labor Dr. Limbach & Kollegen GbR | Robert Koch Institute | unknown |
| EPI_ISL_2260987 | Procomcure Biotech Germany GmbH | Robert Koch Institute | unknown |
| EPI_ISL_2261019 | MVZ Dr. Eberhard & Partner Dortmund | Robert Koch Institute | unknown |
| EPI_ISL_2261471 | MVZ Labor Dr. Quade & Kollegen GmbH | Robert Koch Institute | unknown |
| EPI_ISL_2262247, EPI_ISL_2262249 | MVZ Labor Dr. Limbach & Kollegen GbR | Robert Koch Institute | unknown |
| EPI_ISL_2262263 | RS Umum Kartika Pulomas | National Institute of Health Research and Development | Vivi Setiawaty, Hana Apsari Pawestri, Hartanti Dian Ikawati, Subangkit, Kartika Dewi Puspa, Arie Ardiansyah Nugraha, Triyani Soekarso, Krisna Pangesti, Nelly Puspandari |
| EPI_ISL_2262283, EPI_ISL_2262285, EPI_ISL_2262288, EPI_ISL_2262289, EPI_ISL_2262291 | RSPI Prof. Dr. Sulianti Saroso | National Institute of Health Research and Development | Hana Apsari Pawestri, Kartika Dewi Puspa, Arie Ardiansyah Nugraha, Subangkit, Hartanti Dian Ikawati, Triyani Soekarso, Krisna Pangesti, Nelly Puspandari, Vivi Setiawaty |
| EPI_ISL_2262310, EPI_ISL_2262315, EPI_ISL_2262318 | Fraunhofer-Institut für Zelltherapie und Immunologie IZI AG Next-Generation Diagnostics | Robert Koch Institute | unknown |
| EPI_ISL_2262371 | Städtisches Klinikum Dresden Institut für Klinische Chemie und Laboratoriumsmedizin | Robert Koch Institute | unknown |
| EPI_ISL_2262379 | Uniklinikum Carl Gustav Carus an der TU Dresden; Institut für Virologie | Robert Koch Institute | unknown |
| EPI_ISL_2262483 | MVZ Labor Dr. Quade & Kollegen GmbH | Robert Koch Institute | unknown |
| EPI_ISL_2262724, EPI_ISL_2262751, EPI_ISL_2262763 | Bayerisches Landesamt für Gesundheit und Lebensmittelsicherheit (LGL) | Robert Koch Institute | unknown |
| EPI_ISL_2263043, EPI_ISL_2263047 | Limbach - MVZ Labor Eveld & Kollegen Essen | Robert Koch Institute | unknown |
| EPI_ISL_2263053 | MVZ Labor Dr. Reising-Ackermann und Kollegen | Robert Koch Institute | unknown |

| | | | |
|---|---|---|---|
| EPI_ISL_2263116 | MDI Limbach Berlin GmbH; MVZ Labor Berlin | Robert Koch Institute | unknown |
| EPI_ISL_2263203 | Clemenshospital - Alexianer Clemenshospital Münster | Robert Koch Institute | unknown |
| EPI_ISL_2263654 | SYNLAB MVZ Leinfelden-Echterdingen | Robert Koch Institute | unknown |
| EPI_ISL_2263790 | SYNLAB MVZ Trier | Robert Koch Institute | unknown |
| EPI_ISL_2264324 | SYNLAB MVZ Leinfelden-Echterdingen | Robert Koch Institute | unknown |
| EPI_ISL_2264489, EPI_ISL_2264511 | Universitätsklinikum Bonn - Institut für Virologie | Robert Koch Institute | unknown |
| EPI_ISL_2264533 | Universitätsklinikum des Saarlandes - Institut für Virologie | Robert Koch Institute | unknown |
| EPI_ISL_2264534 | Universitätsklinikum Bonn - Institut für Virologie | Robert Koch Institute | unknown |
| EPI_ISL_2264556 | Limbach - MVZ Labor Eveld & Kollegen Essen | Robert Koch Institute | unknown |
| EPI_ISL_2264558, EPI_ISL_2264578, EPI_ISL_2264588 | MVZ Labor Dr. Limbach & Kollegen GbR | Robert Koch Institute | unknown |
| EPI_ISL_2264591 | Clemenshospital - Alexianer Clemenshospital Münster | Robert Koch Institute | unknown |
| EPI_ISL_2264719, EPI_ISL_2264740 | MDI Limbach Berlin GmbH; MVZ Labor Berlin | Robert Koch Institute | unknown |
| EPI_ISL_2264766 | Ospedale Santa Caterina Novella | Istituto Zooprofilattico Sperimentale della Puglia e della Basilicata | Parisi A., Bianco A., Capozzi L., Del Sambro L., Simone D., Difato L., Bruno A. R. |
| EPI_ISL_2264831 | Presidio di Brindisi Di Summa - Perrino | Istituto Zooprofilattico Sperimentale della Puglia e della Basilicata | Parisi A., Bianco A., Capozzi L., Del Sambro L., Simone D., Difato L., Santoro A., Ridolfi D., Giannico A. |
| EPI_ISL_2264832 | Labor ZOTZ\|KLIMAS; MVZ Düsseldorf-Centrum | Robert Koch Institute | unknown |
| EPI_ISL_2265025 | MVZ Dr. Eberhard & Partner Dortmund | Robert Koch Institute | unknown |
| EPI_ISL_2265125 | SYNLAB Labor München Zentrum LMZ | Robert Koch Institute | unknown |
| EPI_ISL_2265521 | Labor Prof. Dr. G. Enders MVZ GbR | Robert Koch Institute | unknown |
| EPI_ISL_2265617 | Labor Dr. Schumacher MVZ | Robert Koch Institute | unknown |
| EPI_ISL_2266083 | Department für Labormedizin Abteilung III Bereich Molekulare Diagnostik Universitätsklinikum Halle(Saale) | Robert Koch Institute | unknown |
| EPI_ISL_2266232 | amedes MVZ DIAMEDIS Sennestadt | Robert Koch Institute | unknown |
| EPI_ISL_2266662 | Medizinische Laboratorien Düsseldorf | Robert Koch Institute | unknown |
| EPI_ISL_2266918 | Synlab MVZ Augsburg | Robert Koch Institute | unknown |
| EPI_ISL_2267012 | SYNLAB MVZ Berlin | Robert Koch Institute | unknown |
| EPI_ISL_2267203, EPI_ISL_2267288 | Synlab MVZ Augsburg | Robert Koch Institute | unknown |
| EPI_ISL_2267618, EPI_ISL_2267676, EPI_ISL_2267677, EPI_ISL_2267689 | SYNLAB MVZ Weiden | Robert Koch Institute | unknown |
| EPI_ISL_2267823 | SYNLAB MVZ Berlin | Robert Koch Institute | unknown |
| EPI_ISL_2267834, EPI_ISL_2267842, EPI_ISL_2267850 | SYNLAB MVZ Weiden | Robert Koch Institute | unknown |
| EPI_ISL_2267893 | SYNLAB MVZ Dachau | Robert Koch Institute | unknown |
| EPI_ISL_2267952 | CENTOGENE Frankfurt Laboratory: Niederlassung Industriepark Höchst | Robert Koch Institute | unknown |
| EPI_ISL_2268011 | Aegis Sciences Corporation | Centers for Disease Control and Prevention Division of Viral Diseases, Pathogen Discovery | Dakota Howard, Dhwani Batra, Peter W. Cook, Kara Moser, Adrian Paskey, Jason Caravas, Benjamin Rambo-Martin, Shatavia Morrison, Christopher Gulvick, Scott Sammons, Yvette Unoarumhi, Darlene Wagner, Matthew Schmerer, Cyndi Clark, Patrick Campbell, Rob Case, Vikramsinha Ghorpade, Holly Houdeshell, Ola Kvalvaag, Dillon Nall, Ethan Sanders, Alec Vest, Shaun Westlund, Matthew Hardison, Clinton R. Paden, Duncan MacCannell |
| EPI_ISL_2268083, EPI_ISL_2268136 | CENTOGENE Frankfurt Laboratory: Niederlassung Industriepark Höchst | Robert Koch Institute | unknown |
| EPI_ISL_2268203 | Aegis Sciences Corporation | Centers for Disease Control and Prevention Division of Viral Diseases, Pathogen Discovery | Dakota Howard, Dhwani Batra, Peter W. Cook, Kara Moser, Adrian Paskey, Jason Caravas, Benjamin Rambo-Martin, Shatavia Morrison, Christopher Gulvick, Scott Sammons, Yvette Unoarumhi, Darlene Wagner, Matthew Schmerer, Cyndi Clark, Patrick Campbell, Rob Case, Vikramsinha Ghorpade, Holly Houdeshell, Ola Kvalvaag, Dillon Nall, Ethan Sanders, Alec Vest, Shaun Westlund, Matthew Hardison, Clinton R. Paden, Duncan MacCannell |
| EPI_ISL_2268239 | CENTOGENE Frankfurt Laboratory: Niederlassung Industriepark Höchst | Robert Koch Institute | unknown |
| EPI_ISL_2268256 | Aegis Sciences Corporation | Centers for Disease Control and Prevention Division of Viral Diseases, Pathogen Discovery | Dakota Howard, Dhwani Batra, Peter W. Cook, Kara Moser, Adrian Paskey, Jason Caravas, Benjamin Rambo-Martin, Shatavia Morrison, Christopher Gulvick, Scott Sammons, Yvette Unoarumhi, Darlene Wagner, Matthew Schmerer, Cyndi Clark, Patrick Campbell, Rob Case, Vikramsinha Ghorpade, Holly Houdeshell, Ola Kvalvaag, Dillon Nall, Ethan Sanders, Alec Vest, Shaun Westlund, Matthew Hardison, Clinton R. Paden, Duncan MacCannell |
| EPI_ISL_2268308, EPI_ISL_2268322 | CENTOGENE Frankfurt Laboratory: Niederlassung Industriepark Höchst | Robert Koch Institute | unknown |
| EPI_ISL_2268455, EPI_ISL_2268523, EPI_ISL_2268536, EPI_ISL_2268576, EPI_ISL_2268595, EPI_ISL_2268602, EPI_ISL_2268674, EPI_ISL_2268692, EPI_ISL_2268710, EPI_ISL_2268814, EPI_ISL_2268838, EPI_ISL_2268860, EPI_ISL_2268877, EPI_ISL_2268891, EPI_ISL_2268916, EPI_ISL_2268937, EPI_ISL_2268940, EPI_ISL_2268946, EPI_ISL_2268963, EPI_ISL_2268970, EPI_ISL_2269027, EPI_ISL_2269042, EPI_ISL_2269044, EPI_ISL_2269051, EPI_ISL_2269062 | | | |
| see above | Quest Diagnostics Incorporated | Centers for Disease Control and Prevention Division of Viral Diseases, Pathogen Discovery | Dakota Howard, Dhwani Batra, Peter W. Cook, Kara Moser, Adrian Paskey, Jason Caravas, Benjamin Rambo-Martin, Shatavia Morrison, Christopher Gulvick, Scott Sammons, Yvette Unoarumhi, Darlene Wagner, Matthew Schmerer, S. H. Rosenthal, A. Gerasimova, R. M. Kagan, B. Anderson, M. Hua, Y. Liu, L.E. Bernstein, K.E. Livingston, A. Perez, I. A. Shlyakhter, R. V. Rolando, R. Owen, P. Tanpaiboon, F. Lacbawan, Clinton R. Paden, Duncan MacCannell |
| EPI_ISL_2269218, EPI_ISL_2269230, EPI_ISL_2269231, EPI_ISL_2269237, EPI_ISL_2269238, EPI_ISL_2269239, EPI_ISL_2269243, EPI_ISL_2269249, EPI_ISL_2269317, EPI_ISL_2269326, EPI_ISL_2269337, EPI_ISL_2269391, EPI_ISL_2269392, EPI_ISL_2269393, EPI_ISL_2269394, EPI_ISL_2269396, EPI_ISL_2269397, EPI_ISL_2269449, EPI_ISL_2269450, EPI_ISL_2269451, EPI_ISL_2269454, EPI_ISL_2269456, EPI_ISL_2269457, EPI_ISL_2269485, EPI_ISL_2269488 | | | |
| see above | Fulgent Genetics | Centers for Disease Control and Prevention Division of Viral Diseases, Pathogen Discovery | Dakota Howard, Dhwani Batra, Peter W. Cook, Kara Moser, Adrian Paskey, Jason Caravas, Benjamin Rambo-Martin, Shatavia Morrison, Christopher Gulvick, Scott Sammons, Yvette Unoarumhi, Darlene Wagner, Matthew Schmerer, Harry Gao, Mickey Li, John Gao, Joseph Fierro, Benafsh Sapra, Becky Tsai, Yan Meng, Doreen Ng, James Xie, Clinton R. Paden, Duncan MacCannell |
| EPI_ISL_2269491 | Mobile County Health Department | Synergy Laboratories | Megan Cornwell |
| EPI_ISL_2269513 | Fulgent Genetics | Centers for Disease Control and Prevention Division of Viral Diseases, Pathogen Discovery | Dakota Howard, Dhwani Batra, Peter W. Cook, Kara Moser, Adrian Paskey, Jason Caravas, Benjamin Rambo-Martin, Shatavia Morrison, Christopher Gulvick, Scott Sammons, Yvette Unoarumhi, Darlene Wagner, Matthew Schmerer, Harry Gao, Mickey Li, John Gao, Joseph Fierro, Benafsh Sapra, Becky Tsai, Yan Meng, Doreen Ng, James Xie, Clinton R. Paden, Duncan MacCannell |
| EPI_ISL_2269619, EPI_ISL_2269712, EPI_ISL_2269822, EPI_ISL_2270252 | Helix/Illumina | Centers for Disease Control and Prevention Division of Viral Diseases, Pathogen Discovery | Dakota Howard, Dhwani Batra, Peter W. Cook, Kara Moser, Adrian Paskey, Jason Caravas, Benjamin Rambo-Martin, Shatavia Morrison, Christopher Gulvick, Scott Sammons, Yvette Unoarumhi, Darlene Wagner, Matthew Schmerer, Eileen de Feo, Jan Antico, Christine Tran, Matthew Tolentino, Shannon Wickline, Kim Gietzen, Brad Sickler, Jingtao Liu, Eric Allen, Phil Febbo, Nicole L. Washington, Simon White, Geraint Levan, Kelly Schiabor Barrett, Elizabeth Cirulli, Alexandre Bolze, Ary Ascencio, Charlotte Rivera-Garcia, Ryan Cho, Jason Nguyen, Sherry Wang, Jimmy Ramirez, Tyler Cassens, Efren Sandoval, Magnus Isaksson, William Lee, David Becker, Marc Laurent, James Lu, Clinton R. Paden, Duncan MacCannell |
| EPI_ISL_2270355 | Mobile County Health Department | Synergy Laboratories | Megan Cornwell |
| EPI_ISL_2270430, EPI_ISL_2270494, EPI_ISL_2270532, EPI_ISL_2270535, EPI_ISL_2270570, EPI_ISL_2270601, EPI_ISL_2270707, EPI_ISL_2270722, EPI_ISL_2270737 | Helix/Illumina | Centers for Disease Control and Prevention Division of Viral Diseases, Pathogen Discovery | Dakota Howard, Dhwani Batra, Peter W. Cook, Kara Moser, Adrian Paskey, Jason Caravas, Benjamin Rambo-Martin, Shatavia Morrison, Christopher Gulvick, Scott Sammons, Yvette Unoarumhi, Darlene Wagner, Matthew Schmerer, Eileen de Feo, Jan Antico, Christine Tran, Matthew Tolentino, Shannon Wickline, Kim Gietzen, Brad Sickler, Jingtao Liu, Eric Allen, Phil Febbo, Nicole L. Washington, Simon White, Geraint Levan, Kelly Schiabor Barrett, Elizabeth Cirulli, Alexandre Bolze, Ary Ascencio, Charlotte Rivera-Garcia, Ryan Cho, Jason Nguyen, Sherry Wang, Jimmy Ramirez, Tyler Cassens, Efren Sandoval, Magnus Isaksson, William Lee, David Becker, Marc Laurent, James Lu, Clinton R. Paden, Duncan MacCannell |
| EPI_ISL_2270762 | Mobile County Health Department | Synergy Laboratories | Megan Cornwell |
| EPI_ISL_2271720, EPI_ISL_2271721, EPI_ISL_2271722 | Kansas Health and Environmental Lab | Kansas Health and Environmental Lab | Mike Grose, Katherine Wiggins, Jonathan Barnell, Ben Olsen, and Phil Adam |
| EPI_ISL_2271876, EPI_ISL_2271877 | MD PHL | MD PHL | Maryland Department of Health Laboratories Administration |
| EPI_ISL_2272477, EPI_ISL_2272479, EPI_ISL_2272480, EPI_ISL_2272481, EPI_ISL_2272483, EPI_ISL_2272484, EPI_ISL_2272486, EPI_ISL_2272487, EPI_ISL_2272489, EPI_ISL_2272494, EPI_ISL_2272495, EPI_ISL_2272497 | | | |
| see above | Indian Council of Medical Research-National Institute of Virology, Microbial Containment Complex | Indian Council of Medical Research-National Institute of Virology, Microbial Containment Complex | Pragya D Yadav |

| | | | |
|---|---|---|---|
| EPI_ISL_2272555 | Alaska State Virology Laboratory | Alaska State Virology Laboratory | Stephanie DeRonde, Elva House, Jacob Zidek, Lisa Smith, Ph.D., Jack Chen, Ph.D. |
| EPI_ISL_2272597, EPI_ISL_2272598, EPI_ISL_2272599, EPI_ISL_2272600, EPI_ISL_2272601, EPI_ISL_2272602 | MEPHI, Aix Marseille University | MEPHI, Aix Marseille University | Anthony LEVASSEUR |
| EPI_ISL_2272661, EPI_ISL_2272669 | UW Virology Lab | UW Virology Lab | Pavitra Roychoudhury, Hong Xie, Lasata Shrestha, Tien V. Nguyen, Shah Mohamed Bakhash, Michelle Lin, Noah R. Baker, Ricardo Perez, Sean Ellis, Nathan Breit, Robert J. Livingston, Meei-Li Huang, Keith R Jerome, Patrick Mathias, Alexander Greninger |
| EPI_ISL_2272670, EPI_ISL_2272671, EPI_ISL_2272672, EPI_ISL_2272678, EPI_ISL_2272679, EPI_ISL_2272687, EPI_ISL_2272690, EPI_ISL_2272760, EPI_ISL_2272768, EPI_ISL_2272769, EPI_ISL_2272775 | | | |
| see above | Platform BIS UZA/UAntwerpen | Labo Klinische Biologie, UZA | Marie Le Mercier, Jasmine Coppens, Basil Britto Xavier, Christine Lammens, Veerle Matheeussen, Herman Goossens |
| EPI_ISL_2272778, EPI_ISL_2272779 | Algemeen Medisch Labo | Labo Klinische Biologie, UZA | Marie Le Mercier, Jasmine Coppens, Basil Britto Xavier, Christine Lammens, Veerle Matheeussen, Herman Goossens |
| EPI_ISL_2272780, EPI_ISL_2272781, EPI_ISL_2272783, EPI_ISL_2272784, EPI_ISL_2272785, EPI_ISL_2272786 | Clinical Hospital of Infectious and Tropical Diseases “Dr. Victor Babes” | Cantacuzino Institute Virology | Sorin Dinu, Usein Codruta, Carmen Cherciu, Mihaela Oprea, Mihaela Lazar |
| EPI_ISL_2272789, EPI_ISL_2272790, EPI_ISL_2272791, EPI_ISL_2272794, EPI_ISL_2272795, EPI_ISL_2272796, EPI_ISL_2272798, EPI_ISL_2272799, EPI_ISL_2272800, EPI_ISL_2272801, EPI_ISL_2272802, EPI_ISL_2272803, EPI_ISL_2272804, EPI_ISL_2272805, EPI_ISL_2272807, EPI_ISL_2272808, EPI_ISL_2272809, EPI_ISL_2272810, EPI_ISL_2272811, EPI_ISL_2272812, EPI_ISL_2272814, EPI_ISL_2272815, EPI_ISL_2272817, EPI_ISL_2272819, EPI_ISL_2272820, EPI_ISL_2272821, EPI_ISL_2272824, EPI_ISL_2272827, EPI_ISL_2272828, EPI_ISL_2272829, EPI_ISL_2272830, EPI_ISL_2272831, EPI_ISL_2272832, EPI_ISL_2272833, EPI_ISL_2272834, EPI_ISL_2272835, EPI_ISL_2272836, EPI_ISL_2272838, EPI_ISL_2272839, EPI_ISL_2272840, EPI_ISL_2272841, EPI_ISL_2272843, EPI_ISL_2272844, EPI_ISL_2272845, EPI_ISL_2272846, EPI_ISL_2272847, EPI_ISL_2272848, EPI_ISL_2272849, EPI_ISL_2272850, EPI_ISL_2272851, EPI_ISL_2272852, EPI_ISL_2272853, EPI_ISL_2272854, EPI_ISL_2272855, EPI_ISL_2272856, EPI_ISL_2272857, EPI_ISL_2272858, EPI_ISL_2272859, EPI_ISL_2272860, EPI_ISL_2272861, EPI_ISL_2272862, EPI_ISL_2272863, EPI_ISL_2272864, EPI_ISL_2272865, EPI_ISL_2272866, EPI_ISL_2272867, EPI_ISL_2272868, EPI_ISL_2272869, EPI_ISL_2272870, EPI_ISL_2272871, EPI_ISL_2272872, EPI_ISL_2272873, EPI_ISL_2272874, EPI_ISL_2272876, EPI_ISL_2272877, EPI_ISL_2272878, EPI_ISL_2272879, EPI_ISL_2272881, EPI_ISL_2272882, EPI_ISL_2272883, EPI_ISL_2272884, EPI_ISL_2272885, EPI_ISL_2272886, EPI_ISL_2272887, EPI_ISL_2272888, EPI_ISL_2272890, EPI_ISL_2272893, EPI_ISL_2272895, EPI_ISL_2272897, EPI_ISL_2272898, EPI_ISL_2272899, EPI_ISL_2272900, EPI_ISL_2272901, EPI_ISL_2272903, EPI_ISL_2272904, EPI_ISL_2272905, EPI_ISL_2272908, EPI_ISL_2272910, EPI_ISL_2272911, EPI_ISL_2272912, EPI_ISL_2272913, EPI_ISL_2272914, EPI_ISL_2272916, EPI_ISL_2272917, EPI_ISL_2272918, EPI_ISL_2272919, EPI_ISL_2272920, EPI_ISL_2272922, EPI_ISL_2272923, EPI_ISL_2272924, EPI_ISL_2272926, EPI_ISL_2272928, EPI_ISL_2272932, EPI_ISL_2272933, EPI_ISL_2272936, EPI_ISL_2272937 | | | |
| see above | JIPMER Puducherry | inStem NCBS - INSACOG | Uma Ramakrishnan Dasaradhi Palakodeti Aswin SaiNarain |
| EPI_ISL_2272938, EPI_ISL_2272939, EPI_ISL_2272940, EPI_ISL_2272943, EPI_ISL_2272947 | Wichita State University - Molecular Diagnostics Lab | Kansas Health and Environmental Lab | Mike Grose, Katherine Wiggins, Jonathan Barnell, Ben Olsen, and Phil Adam |
| EPI_ISL_2273505 | National Laboratory for Health, Environment and Food, OMM, Maribor | NLZOH (National Laboratory for Health, Environment and Food) / CISLD (Clinical Institute of Special Laboratory Diagnostics), University Children's Hospital, University Medical Center Ljubljana | Sandra Janezic, Aleksander Mahnic, Maja Rupnik, Tjasa Žohar Čretnik, Alenka Štorman, Nika Gobec, Aleksander Kocuvan, Kaja Tominc, Maša Jarčič, Mojca Cimerman, Nika Volmajer, Andrej Golle / Jernej Kovač, Barbara Jenko Bizjan, Tine Tesovnik, Robert Šket, Katarina Kozmos, Ana Grom, Maruša Debeljak, Marko Pokorn, Tadej Battelino |
| EPI_ISL_2273760, EPI_ISL_2273792 | UW Virology Lab | UW Virology Lab | Pavitra Roychoudhury, Hong Xie, Lasata Shrestha, Tien V. Nguyen, Shah Mohamed Bakhash, Michelle Lin, Noah R. Baker, Ricardo Perez, Sean Ellis, Nathan Breit, Robert J. Livingston, Meei-Li Huang, Keith R Jerome, Patrick Mathias, Alexander Greninger |
| EPI_ISL_2273847 | City of Milwaukee Health Department Laboratory | City of Milwaukee Health Department Laboratory | Sanjib Bhattacharyya, Manjeet Khubbar, Amy Bauer, Jennifer Lentz, Samantha Scott |
| EPI_ISL_2274244, EPI_ISL_2274246, EPI_ISL_2274249 | Queensland Health Forensic and Scientific Services | Queensland Health Forensic and Scientific Services | Son Nguyen |
| EPI_ISL_2274250 | New South Wales Health Pathology Royal Prince Alfred Hospital | Microbiology RPAH | Foster, C.; Au, J.; Ruiz Silva, M.; Deveson, I.; Bull, R.; Van Hal, S.; Rawlinson, W. |
| EPI_ISL_2274379, EPI_ISL_2274381, EPI_ISL_2274383, EPI_ISL_2274397, EPI_ISL_2274399, EPI_ISL_2274404 | Ministry of Public Health / Hamad Medical Corporation | Biomedical Research Center (BRC), Qatar University / Qatar Genome Project (QGP) | BRC: Fatiha M. Benslimane, Heba A. Al-Khatib, Oal Al-Jamal, Dana Al-Batesh, Hadi M. Yassine, Asmaa A. Al-Thani. MOPH and HMC: Abdullatif Al-Khal, Muna A. S. Al-Maslamani, Mashael A. Al-Bader, Hamda Alromaihi, Roberto Bertollini, Peter V. Coyle, Einas A. E. Al-Kuwari, Hamad E. Al-Romaihi, Salih Al-Marri, Mohammed Al-Thani, Reham A. El-Kahlout. QBB: Tasneem Al-Hamad, Dina Elgakhlab QGP: Fatima H. Al-Kuwari, Chadi Saad |
| EPI_ISL_2274445 | Labormedizinisches Zentrum Dr Risch | Clinical Bacteriology | Tim Roloff, Madlen Stange, Helena MB Seth-Smith, Alfredo Mari, Karoline Leuzinger, Julia Bielicki, Nadia Wohlwend,Martin Risch, Lorenz Risch, Manuel Battegay, Hans Hirsch, Adrian Egli |
| EPI_ISL_2274706 | NYC Pandemic Response Lab | Wadsworth Center, New York State Department of Health | Kirsten St. George, Daryl M. Lamson, Alexis Russell, Matthew Shudt, Melissa A Leisner, Jonathan Plitnick, Catharine Prussing, Navjot Singh, John Kelly, Erasmus Schneider, Erica Lasek-Nesselquist |
| EPI_ISL_2274979 | Aeroporto Internacional de Guarulhos | Instituto Adolfo Lutz, Interdiciplinary Procedures Center, Strategic Laboratory | Claudio Tavares Sacchi, Claudia Regina Gonçalves, Erica Valessa Ramos Gomes, Karoline Rodrigues Campos, Caio Vinicius Dias Lopes, Leonardo Jose Tadeu de Araujo |
| EPI_ISL_2274980, EPI_ISL_2274982, EPI_ISL_2274989, EPI_ISL_2274991, EPI_ISL_2274992, EPI_ISL_2274994 | Lighthouse Lab in Milton Keynes | Wellcome Sanger Institute for the COVID-19 Genomics UK (COG-UK) Consortium | The Lighthouse Lab in Milton Keynes and Alex Alderton, Roberto Amato, Jeffrey Barrett, Sonia Goncalves, Ewan Harrison, David K. Jackson, Ian Johnston, Dominic Kwiatkowski, Cordelia Langford, John Sillitoe on behalf of the Wellcome Sanger Institute COVID-19 Surveillance Team |
| EPI_ISL_2274995 | Lighthouse Lab in Alderley Park | Wellcome Sanger Institute for the COVID-19 Genomics UK (COG-UK) Consortium | Jacquelyn Wynn, Mairead Hyland, The Lighthouse Lab in Alderley Park and Alex Alderton, Roberto Amato, Jeffrey Barrett, Sonia Goncalves, Ewan Harrison, David K. Jackson, Ian Johnston, Dominic Kwiatkowski, Cordelia Langford, John Sillitoe on behalf of the Wellcome Sanger Institute COVID-19 Surveillance Team |
| EPI_ISL_2274997 | Lighthouse Lab in Milton Keynes | Wellcome Sanger Institute for the COVID-19 Genomics UK (COG-UK) Consortium | The Lighthouse Lab in Milton Keynes and Alex Alderton, Roberto Amato, Jeffrey Barrett, Sonia Goncalves, Ewan Harrison, David K. Jackson, Ian Johnston, Dominic Kwiatkowski, Cordelia Langford, John Sillitoe on behalf of the Wellcome Sanger Institute COVID-19 Surveillance Team |
| EPI_ISL_2275002 | Lighthouse Lab in Alderley Park | Wellcome Sanger Institute for the COVID-19 Genomics UK (COG-UK) Consortium | Jacquelyn Wynn, Mairead Hyland, The Lighthouse Lab in Alderley Park and Alex Alderton, Roberto Amato, Jeffrey Barrett, Sonia Goncalves, Ewan Harrison, David K. Jackson, Ian Johnston, Dominic Kwiatkowski, Cordelia Langford, John Sillitoe on behalf of the Wellcome Sanger Institute COVID-19 Surveillance Team |
| EPI_ISL_2275004, EPI_ISL_2275007, EPI_ISL_2275011 | Lighthouse Lab in Milton Keynes | Wellcome Sanger Institute for the COVID-19 Genomics UK (COG-UK) Consortium | The Lighthouse Lab in Milton Keynes and Alex Alderton, Roberto Amato, Jeffrey Barrett, Sonia Goncalves, Ewan Harrison, David K. Jackson, Ian Johnston, Dominic Kwiatkowski, Cordelia Langford, John Sillitoe on behalf of the Wellcome Sanger Institute COVID-19 Surveillance Team |
| EPI_ISL_2275012 | Lighthouse Lab in Alderley Park | Wellcome Sanger Institute for the COVID-19 Genomics UK (COG-UK) Consortium | Jacquelyn Wynn, Mairead Hyland, The Lighthouse Lab in Alderley Park and Alex Alderton, Roberto Amato, Jeffrey Barrett, Sonia Goncalves, Ewan Harrison, David K. Jackson, Ian Johnston, Dominic Kwiatkowski, Cordelia Langford, John Sillitoe on behalf of the Wellcome Sanger Institute COVID-19 Surveillance Team |
| EPI_ISL_2275015, EPI_ISL_2275016, EPI_ISL_2275017, EPI_ISL_2275018, EPI_ISL_2275021, EPI_ISL_2275022, EPI_ISL_2275023, EPI_ISL_2275025, EPI_ISL_2275026, EPI_ISL_2275027 | Lighthouse Lab in Milton Keynes | Wellcome Sanger Institute for the COVID-19 Genomics UK (COG-UK) Consortium | The Lighthouse Lab in Milton Keynes and Alex Alderton, Roberto Amato, Jeffrey Barrett, Sonia Goncalves, Ewan Harrison, David K. Jackson, Ian Johnston, Dominic Kwiatkowski, Cordelia Langford, John Sillitoe on behalf of the Wellcome Sanger Institute COVID-19 Surveillance Team |
| EPI_ISL_2275030 | Lighthouse Lab in Glasgow | Wellcome Sanger Institute for the COVID-19 Genomics UK (COG-UK) Consortium | Harper VanSteenhouse, Yumi Kasai, David Gray, Carol Clugston, Anna Dominiczak and Alex Alderton, Roberto Amato, Jeffrey Barrett, Sonia Goncalves, Ewan Harrison, David K. Jackson, Ian Johnston, Dominic Kwiatkowski, Cordelia Langford, John Sillitoe on behalf of the Wellcome Sanger Institute COVID-19 Surveillance Team |
| EPI_ISL_2275031, EPI_ISL_2275036, EPI_ISL_2275039, EPI_ISL_2275040, EPI_ISL_2275042, EPI_ISL_2275043, EPI_ISL_2275044, EPI_ISL_2275046, EPI_ISL_2275048, EPI_ISL_2275049, EPI_ISL_2275050, EPI_ISL_2275051 | | | |
| see above | Lighthouse Lab in Milton Keynes | Wellcome Sanger Institute for the COVID-19 Genomics UK (COG-UK) Consortium | The Lighthouse Lab in Milton Keynes and Alex Alderton, Roberto Amato, Jeffrey Barrett, Sonia Goncalves, Ewan Harrison, David K. Jackson, Ian Johnston, Dominic Kwiatkowski, Cordelia Langford, John Sillitoe on behalf of the Wellcome Sanger Institute COVID-19 Surveillance Team |
| EPI_ISL_2275052 | Lighthouse Lab in Alderley Park | Wellcome Sanger Institute for the COVID-19 Genomics UK (COG-UK) Consortium | Jacquelyn Wynn, Mairead Hyland, The Lighthouse Lab in Alderley Park and Alex Alderton, Roberto Amato, Jeffrey Barrett, Sonia Goncalves, Ewan Harrison, David K. Jackson, Ian Johnston, Dominic Kwiatkowski, Cordelia Langford, John Sillitoe on behalf of the Wellcome Sanger Institute COVID-19 Surveillance Team |
| EPI_ISL_2275054 | Lighthouse Lab in Milton Keynes | Wellcome Sanger Institute for the COVID-19 Genomics UK (COG-UK) Consortium | The Lighthouse Lab in Milton Keynes and Alex Alderton, Roberto Amato, Jeffrey Barrett, Sonia Goncalves, Ewan Harrison, David K. Jackson, Ian Johnston, Dominic Kwiatkowski, Cordelia Langford, John Sillitoe on behalf of the Wellcome Sanger Institute COVID-19 Surveillance Team |
| EPI_ISL_2275055 | Lighthouse Lab in Alderley Park | Wellcome Sanger Institute for the COVID-19 Genomics UK (COG-UK) Consortium | Jacquelyn Wynn, Mairead Hyland, The Lighthouse Lab in Alderley Park and Alex Alderton, Roberto Amato, Jeffrey Barrett, Sonia Goncalves, Ewan Harrison, David K. Jackson, Ian Johnston, Dominic Kwiatkowski, Cordelia Langford, John Sillitoe on behalf of the Wellcome Sanger Institute COVID-19 Surveillance Team |
| EPI_ISL_2275056 | Lighthouse Lab in Milton Keynes | Wellcome Sanger Institute for the COVID-19 Genomics UK (COG-UK) Consortium | The Lighthouse Lab in Milton Keynes and Alex Alderton, Roberto Amato, Jeffrey Barrett, Sonia Goncalves, Ewan Harrison, David K. Jackson, Ian Johnston, Dominic Kwiatkowski, Cordelia Langford, John Sillitoe on behalf of the Wellcome Sanger Institute COVID-19 Surveillance Team |
| EPI_ISL_2275057 | Lighthouse Lab in Alderley Park | Wellcome Sanger Institute for the COVID-19 Genomics UK (COG-UK) Consortium | Jacquelyn Wynn, Mairead Hyland, The Lighthouse Lab in Alderley Park and Alex Alderton, Roberto Amato, Jeffrey Barrett, Sonia Goncalves, Ewan Harrison, David K. Jackson, Ian Johnston, Dominic Kwiatkowski, Cordelia Langford, John Sillitoe on behalf of the Wellcome Sanger Institute COVID-19 Surveillance Team |
| EPI_ISL_2275059, EPI_ISL_2275060, | Lighthouse Lab in Milton Keynes | Wellcome Sanger Institute for the COVID-19 Genomics UK (COG-UK) Consortium | The Lighthouse Lab in Milton Keynes and Alex Alderton, Roberto Amato, Jeffrey Barrett, Sonia Goncalves, Ewan Harrison, David K. Jackson, Ian Johnston, Dominic Kwiatkowski, Cordelia Langford, John Sillitoe on behalf of the Wellcome Sanger Institute COVID-19 Surveillance Team |

| | | | |
|---|---|---|---|
| EPI_ISL_2275061 | | | |
| EPI_ISL_2275062 | Lighthouse Lab in Alderley Park | Wellcome Sanger Institute for the COVID-19 Genomics UK (COG-UK) Consortium | Jacquelyn Wynn, Mairead Hyland, The Lighthouse Lab in Alderley Park and Alex Alderton, Roberto Amato, Jeffrey Barrett, Sonia Goncalves, Ewan Harrison, David K. Jackson, Ian Johnston, Dominic Kwiatkowski, Cordelia Langford, John Sillitoe on behalf of the Wellcome Sanger Institute COVID-19 Surveillance Team |
| EPI_ISL_2275063, EPI_ISL_2275065, EPI_ISL_2275068, EPI_ISL_2275069, EPI_ISL_2275071, EPI_ISL_2275072, EPI_ISL_2275073, EPI_ISL_2275074, EPI_ISL_2275075, EPI_ISL_2275076, EPI_ISL_2275077, EPI_ISL_2275080, EPI_ISL_2275082, EPI_ISL_2275083, EPI_ISL_2275085, EPI_ISL_2275089, EPI_ISL_2275090, EPI_ISL_2275093, EPI_ISL_2275094, EPI_ISL_2275095, EPI_ISL_2275096, EPI_ISL_2275097, EPI_ISL_2275098, EPI_ISL_2275100 | | | |
| see above | Lighthouse Lab in Milton Keynes | Wellcome Sanger Institute for the COVID-19 Genomics UK (COG-UK) Consortium | The Lighthouse Lab in Milton Keynes and Alex Alderton, Roberto Amato, Jeffrey Barrett, Sonia Goncalves, Ewan Harrison, David K. Jackson, Ian Johnston, Dominic Kwiatkowski, Cordelia Langford, John Sillitoe on behalf of the Wellcome Sanger Institute COVID-19 Surveillance Team |
| EPI_ISL_2275103 | Lighthouse Lab in Alderley Park | Wellcome Sanger Institute for the COVID-19 Genomics UK (COG-UK) Consortium | Jacquelyn Wynn, Mairead Hyland, The Lighthouse Lab in Alderley Park and Alex Alderton, Roberto Amato, Jeffrey Barrett, Sonia Goncalves, Ewan Harrison, David K. Jackson, Ian Johnston, Dominic Kwiatkowski, Cordelia Langford, John Sillitoe on behalf of the Wellcome Sanger Institute COVID-19 Surveillance Team |
| EPI_ISL_2275106, EPI_ISL_2275107, EPI_ISL_2275108 | Lighthouse Lab in Milton Keynes | Wellcome Sanger Institute for the COVID-19 Genomics UK (COG-UK) Consortium | The Lighthouse Lab in Milton Keynes and Alex Alderton, Roberto Amato, Jeffrey Barrett, Sonia Goncalves, Ewan Harrison, David K. Jackson, Ian Johnston, Dominic Kwiatkowski, Cordelia Langford, John Sillitoe on behalf of the Wellcome Sanger Institute COVID-19 Surveillance Team |
| EPI_ISL_2275109, EPI_ISL_2275111 | Lighthouse Lab in Alderley Park | Wellcome Sanger Institute for the COVID-19 Genomics UK (COG-UK) Consortium | Jacquelyn Wynn, Mairead Hyland, The Lighthouse Lab in Alderley Park and Alex Alderton, Roberto Amato, Jeffrey Barrett, Sonia Goncalves, Ewan Harrison, David K. Jackson, Ian Johnston, Dominic Kwiatkowski, Cordelia Langford, John Sillitoe on behalf of the Wellcome Sanger Institute COVID-19 Surveillance Team |
| EPI_ISL_2275112, EPI_ISL_2275114, EPI_ISL_2275116, EPI_ISL_2275117, EPI_ISL_2275118, EPI_ISL_2275120, EPI_ISL_2275121, EPI_ISL_2275123, EPI_ISL_2275124, EPI_ISL_2275125 | Lighthouse Lab in Milton Keynes | Wellcome Sanger Institute for the COVID-19 Genomics UK (COG-UK) Consortium | The Lighthouse Lab in Milton Keynes and Alex Alderton, Roberto Amato, Jeffrey Barrett, Sonia Goncalves, Ewan Harrison, David K. Jackson, Ian Johnston, Dominic Kwiatkowski, Cordelia Langford, John Sillitoe on behalf of the Wellcome Sanger Institute COVID-19 Surveillance Team |
| EPI_ISL_2275128 | Lighthouse Lab in Alderley Park | Wellcome Sanger Institute for the COVID-19 Genomics UK (COG-UK) Consortium | Jacquelyn Wynn, Mairead Hyland, The Lighthouse Lab in Alderley Park and Alex Alderton, Roberto Amato, Jeffrey Barrett, Sonia Goncalves, Ewan Harrison, David K. Jackson, Ian Johnston, Dominic Kwiatkowski, Cordelia Langford, John Sillitoe on behalf of the Wellcome Sanger Institute COVID-19 Surveillance Team |
| EPI_ISL_2275130, EPI_ISL_2275135 | Lighthouse Lab in Milton Keynes | Wellcome Sanger Institute for the COVID-19 Genomics UK (COG-UK) Consortium | The Lighthouse Lab in Milton Keynes and Alex Alderton, Roberto Amato, Jeffrey Barrett, Sonia Goncalves, Ewan Harrison, David K. Jackson, Ian Johnston, Dominic Kwiatkowski, Cordelia Langford, John Sillitoe on behalf of the Wellcome Sanger Institute COVID-19 Surveillance Team |
| EPI_ISL_2275137, EPI_ISL_2275139 | Lighthouse Lab in Alderley Park | Wellcome Sanger Institute for the COVID-19 Genomics UK (COG-UK) Consortium | Jacquelyn Wynn, Mairead Hyland, The Lighthouse Lab in Alderley Park and Alex Alderton, Roberto Amato, Jeffrey Barrett, Sonia Goncalves, Ewan Harrison, David K. Jackson, Ian Johnston, Dominic Kwiatkowski, Cordelia Langford, John Sillitoe on behalf of the Wellcome Sanger Institute COVID-19 Surveillance Team |
| EPI_ISL_2275140, EPI_ISL_2275143, EPI_ISL_2275144, EPI_ISL_2275145, EPI_ISL_2275146, EPI_ISL_2275147 | Lighthouse Lab in Milton Keynes | Wellcome Sanger Institute for the COVID-19 Genomics UK (COG-UK) Consortium | The Lighthouse Lab in Milton Keynes and Alex Alderton, Roberto Amato, Jeffrey Barrett, Sonia Goncalves, Ewan Harrison, David K. Jackson, Ian Johnston, Dominic Kwiatkowski, Cordelia Langford, John Sillitoe on behalf of the Wellcome Sanger Institute COVID-19 Surveillance Team |
| EPI_ISL_2275148 | Lighthouse Lab in Alderley Park | Wellcome Sanger Institute for the COVID-19 Genomics UK (COG-UK) Consortium | Jacquelyn Wynn, Mairead Hyland, The Lighthouse Lab in Alderley Park and Alex Alderton, Roberto Amato, Jeffrey Barrett, Sonia Goncalves, Ewan Harrison, David K. Jackson, Ian Johnston, Dominic Kwiatkowski, Cordelia Langford, John Sillitoe on behalf of the Wellcome Sanger Institute COVID-19 Surveillance Team |
| EPI_ISL_2275149, EPI_ISL_2275150, EPI_ISL_2275151, EPI_ISL_2275152, EPI_ISL_2275154 | Lighthouse Lab in Milton Keynes | Wellcome Sanger Institute for the COVID-19 Genomics UK (COG-UK) Consortium | The Lighthouse Lab in Milton Keynes and Alex Alderton, Roberto Amato, Jeffrey Barrett, Sonia Goncalves, Ewan Harrison, David K. Jackson, Ian Johnston, Dominic Kwiatkowski, Cordelia Langford, John Sillitoe on behalf of the Wellcome Sanger Institute COVID-19 Surveillance Team |
| EPI_ISL_2275155 | Lighthouse Lab in Alderley Park | Wellcome Sanger Institute for the COVID-19 Genomics UK (COG-UK) Consortium | Jacquelyn Wynn, Mairead Hyland, The Lighthouse Lab in Alderley Park and Alex Alderton, Roberto Amato, Jeffrey Barrett, Sonia Goncalves, Ewan Harrison, David K. Jackson, Ian Johnston, Dominic Kwiatkowski, Cordelia Langford, John Sillitoe on behalf of the Wellcome Sanger Institute COVID-19 Surveillance Team |
| EPI_ISL_2275157 | Lighthouse Lab in Milton Keynes | Wellcome Sanger Institute for the COVID-19 Genomics UK (COG-UK) Consortium | The Lighthouse Lab in Milton Keynes and Alex Alderton, Roberto Amato, Jeffrey Barrett, Sonia Goncalves, Ewan Harrison, David K. Jackson, Ian Johnston, Dominic Kwiatkowski, Cordelia Langford, John Sillitoe on behalf of the Wellcome Sanger Institute COVID-19 Surveillance Team |
| EPI_ISL_2275158 | Lighthouse Lab in Glasgow | Wellcome Sanger Institute for the COVID-19 Genomics UK (COG-UK) Consortium | Harper VanSteenhouse, Yumi Kasai, David Gray, Carol Clugston, Anna Dominiczak and Alex Alderton, Roberto Amato, Jeffrey Barrett, Sonia Goncalves, Ewan Harrison, David K. Jackson, Ian Johnston, Dominic Kwiatkowski, Cordelia Langford, John Sillitoe on behalf of the Wellcome Sanger Institute COVID-19 Surveillance Team |
| EPI_ISL_2275159, EPI_ISL_2275162, EPI_ISL_2275164, EPI_ISL_2275165, EPI_ISL_2275167, EPI_ISL_2275168, EPI_ISL_2275169, EPI_ISL_2275170 | Lighthouse Lab in Milton Keynes | Wellcome Sanger Institute for the COVID-19 Genomics UK (COG-UK) Consortium | The Lighthouse Lab in Milton Keynes and Alex Alderton, Roberto Amato, Jeffrey Barrett, Sonia Goncalves, Ewan Harrison, David K. Jackson, Ian Johnston, Dominic Kwiatkowski, Cordelia Langford, John Sillitoe on behalf of the Wellcome Sanger Institute COVID-19 Surveillance Team |
| EPI_ISL_2275171, EPI_ISL_2275172 | Lighthouse Lab in Alderley Park | Wellcome Sanger Institute for the COVID-19 Genomics UK (COG-UK) Consortium | Jacquelyn Wynn, Mairead Hyland, The Lighthouse Lab in Alderley Park and Alex Alderton, Roberto Amato, Jeffrey Barrett, Sonia Goncalves, Ewan Harrison, David K. Jackson, Ian Johnston, Dominic Kwiatkowski, Cordelia Langford, John Sillitoe on behalf of the Wellcome Sanger Institute COVID-19 Surveillance Team |
| EPI_ISL_2275173, EPI_ISL_2275177 | Lighthouse Lab in Milton Keynes | Wellcome Sanger Institute for the COVID-19 Genomics UK (COG-UK) Consortium | The Lighthouse Lab in Milton Keynes and Alex Alderton, Roberto Amato, Jeffrey Barrett, Sonia Goncalves, Ewan Harrison, David K. Jackson, Ian Johnston, Dominic Kwiatkowski, Cordelia Langford, John Sillitoe on behalf of the Wellcome Sanger Institute COVID-19 Surveillance Team |
| EPI_ISL_2275179 | Lighthouse Lab in Alderley Park | Wellcome Sanger Institute for the COVID-19 Genomics UK (COG-UK) Consortium | Jacquelyn Wynn, Mairead Hyland, The Lighthouse Lab in Alderley Park and Alex Alderton, Roberto Amato, Jeffrey Barrett, Sonia Goncalves, Ewan Harrison, David K. Jackson, Ian Johnston, Dominic Kwiatkowski, Cordelia Langford, John Sillitoe on behalf of the Wellcome Sanger Institute COVID-19 Surveillance Team |
| EPI_ISL_2275180, EPI_ISL_2275182, EPI_ISL_2275183, EPI_ISL_2275184 | Lighthouse Lab in Milton Keynes | Wellcome Sanger Institute for the COVID-19 Genomics UK (COG-UK) Consortium | The Lighthouse Lab in Milton Keynes and Alex Alderton, Roberto Amato, Jeffrey Barrett, Sonia Goncalves, Ewan Harrison, David K. Jackson, Ian Johnston, Dominic Kwiatkowski, Cordelia Langford, John Sillitoe on behalf of the Wellcome Sanger Institute COVID-19 Surveillance Team |
| EPI_ISL_2275185 | Lighthouse Lab in Alderley Park | Wellcome Sanger Institute for the COVID-19 Genomics UK (COG-UK) Consortium | Jacquelyn Wynn, Mairead Hyland, The Lighthouse Lab in Alderley Park and Alex Alderton, Roberto Amato, Jeffrey Barrett, Sonia Goncalves, Ewan Harrison, David K. Jackson, Ian Johnston, Dominic Kwiatkowski, Cordelia Langford, John Sillitoe on behalf of the Wellcome Sanger Institute COVID-19 Surveillance Team |
| EPI_ISL_2275187 | Lighthouse Lab in Milton Keynes | Wellcome Sanger Institute for the COVID-19 Genomics UK (COG-UK) Consortium | The Lighthouse Lab in Milton Keynes and Alex Alderton, Roberto Amato, Jeffrey Barrett, Sonia Goncalves, Ewan Harrison, David K. Jackson, Ian Johnston, Dominic Kwiatkowski, Cordelia Langford, John Sillitoe on behalf of the Wellcome Sanger Institute COVID-19 Surveillance Team |
| EPI_ISL_2275188, EPI_ISL_2275189, EPI_ISL_2275190 | Lighthouse Lab in Alderley Park | Wellcome Sanger Institute for the COVID-19 Genomics UK (COG-UK) Consortium | Jacquelyn Wynn, Mairead Hyland, The Lighthouse Lab in Alderley Park and Alex Alderton, Roberto Amato, Jeffrey Barrett, Sonia Goncalves, Ewan Harrison, David K. Jackson, Ian Johnston, Dominic Kwiatkowski, Cordelia Langford, John Sillitoe on behalf of the Wellcome Sanger Institute COVID-19 Surveillance Team |
| EPI_ISL_2275191 | Lighthouse Lab in Milton Keynes | Wellcome Sanger Institute for the COVID-19 Genomics UK (COG-UK) Consortium | The Lighthouse Lab in Milton Keynes and Alex Alderton, Roberto Amato, Jeffrey Barrett, Sonia Goncalves, Ewan Harrison, David K. Jackson, Ian Johnston, Dominic Kwiatkowski, Cordelia Langford, John Sillitoe on behalf of the Wellcome Sanger Institute COVID-19 Surveillance Team |
| EPI_ISL_2275192 | Lighthouse Lab in Glasgow | Wellcome Sanger Institute for the COVID-19 Genomics UK (COG-UK) Consortium | Harper VanSteenhouse, Yumi Kasai, David Gray, Carol Clugston, Anna Dominiczak and Alex Alderton, Roberto Amato, Jeffrey Barrett, Sonia Goncalves, Ewan Harrison, David K. Jackson, Ian Johnston, Dominic Kwiatkowski, Cordelia Langford, John Sillitoe on behalf of the Wellcome Sanger Institute COVID-19 Surveillance Team |
| EPI_ISL_2275193 | Lighthouse Lab in Alderley Park | Wellcome Sanger Institute for the COVID-19 Genomics UK (COG-UK) Consortium | Jacquelyn Wynn, Mairead Hyland, The Lighthouse Lab in Alderley Park and Alex Alderton, Roberto Amato, Jeffrey Barrett, Sonia Goncalves, Ewan Harrison, David K. Jackson, Ian Johnston, Dominic Kwiatkowski, Cordelia Langford, John Sillitoe on behalf of the Wellcome Sanger Institute COVID-19 Surveillance Team |
| EPI_ISL_2275194 | Lighthouse Lab in Glasgow | Wellcome Sanger Institute for the COVID-19 Genomics UK (COG-UK) Consortium | Harper VanSteenhouse, Yumi Kasai, David Gray, Carol Clugston, Anna Dominiczak and Alex Alderton, Roberto Amato, Jeffrey Barrett, Sonia Goncalves, Ewan Harrison, David K. Jackson, Ian Johnston, Dominic Kwiatkowski, Cordelia Langford, John Sillitoe on behalf of the Wellcome Sanger Institute COVID-19 Surveillance Team |
| EPI_ISL_2275195 | Lighthouse Lab in Alderley Park | Wellcome Sanger Institute for the COVID-19 Genomics UK (COG-UK) Consortium | Jacquelyn Wynn, Mairead Hyland, The Lighthouse Lab in Alderley Park and Alex Alderton, Roberto Amato, Jeffrey Barrett, Sonia Goncalves, Ewan Harrison, David K. Jackson, Ian Johnston, Dominic Kwiatkowski, Cordelia Langford, John Sillitoe on behalf of the Wellcome Sanger Institute COVID-19 Surveillance Team |
| EPI_ISL_2275196 | Lighthouse Lab in Milton Keynes | Wellcome Sanger Institute for the COVID-19 Genomics UK (COG-UK) Consortium | The Lighthouse Lab in Milton Keynes and Alex Alderton, Roberto Amato, Jeffrey Barrett, Sonia Goncalves, Ewan Harrison, David K. Jackson, Ian Johnston, Dominic Kwiatkowski, Cordelia Langford, John Sillitoe on behalf of the Wellcome Sanger Institute COVID-19 Surveillance Team |
| EPI_ISL_2275198 | Lighthouse Lab in Alderley Park | Wellcome Sanger Institute for the COVID-19 Genomics UK (COG-UK) Consortium | Jacquelyn Wynn, Mairead Hyland, The Lighthouse Lab in Alderley Park and Alex Alderton, Roberto Amato, Jeffrey Barrett, Sonia Goncalves, Ewan Harrison, David K. Jackson, Ian Johnston, Dominic Kwiatkowski, Cordelia Langford, John Sillitoe on behalf of the Wellcome Sanger Institute COVID-19 Surveillance Team |
| EPI_ISL_2275199, EPI_ISL_2275200 | Lighthouse Lab in Milton Keynes | Wellcome Sanger Institute for the COVID-19 Genomics UK (COG-UK) Consortium | The Lighthouse Lab in Milton Keynes and Alex Alderton, Roberto Amato, Jeffrey Barrett, Sonia Goncalves, Ewan Harrison, David K. Jackson, Ian Johnston, Dominic Kwiatkowski, Cordelia Langford, John Sillitoe on behalf of the Wellcome Sanger Institute COVID-19 Surveillance Team |
| EPI_ISL_2275201, EPI_ISL_2275202, EPI_ISL_2275204, EPI_ISL_2275205, EPI_ISL_2275206 | Lighthouse Lab in Alderley Park | Wellcome Sanger Institute for the COVID-19 Genomics UK (COG-UK) Consortium | Jacquelyn Wynn, Mairead Hyland, The Lighthouse Lab in Alderley Park and Alex Alderton, Roberto Amato, Jeffrey Barrett, Sonia Goncalves, Ewan Harrison, David K. Jackson, Ian Johnston, Dominic Kwiatkowski, Cordelia Langford, John Sillitoe on behalf of the Wellcome Sanger Institute COVID-19 Surveillance Team |

| | | | |
|---|---|---|---|
| EPI_ISL_2275207, EPI_ISL_2275209 | Lighthouse Lab in Glasgow | Wellcome Sanger Institute for the COVID-19 Genomics UK (COG-UK) Consortium | Harper VanSteenhouse, Yumi Kasai, David Gray, Carol Clugston, Anna Dominiczak and Alex Alderton, Roberto Amato, Jeffrey Barrett, Sonia Goncalves, Ewan Harrison, David K. Jackson, Ian Johnston, Dominic Kwiatkowski, Cordelia Langford, John Sillitoe on behalf of the Wellcome Sanger Institute COVID-19 Surveillance Team |
| EPI_ISL_2275210, EPI_ISL_2275211 | Lighthouse Lab in Alderley Park | Wellcome Sanger Institute for the COVID-19 Genomics UK (COG-UK) Consortium | Jacquelyn Wynn, Mairead Hyland, The Lighthouse Lab in Alderley Park and Alex Alderton, Roberto Amato, Jeffrey Barrett, Sonia Goncalves, Ewan Harrison, David K. Jackson, Ian Johnston, Dominic Kwiatkowski, Cordelia Langford, John Sillitoe on behalf of the Wellcome Sanger Institute COVID-19 Surveillance Team |
| EPI_ISL_2275213, EPI_ISL_2275214 | Lighthouse Lab in Glasgow | Wellcome Sanger Institute for the COVID-19 Genomics UK (COG-UK) Consortium | Harper VanSteenhouse, Yumi Kasai, David Gray, Carol Clugston, Anna Dominiczak and Alex Alderton, Roberto Amato, Jeffrey Barrett, Sonia Goncalves, Ewan Harrison, David K. Jackson, Ian Johnston, Dominic Kwiatkowski, Cordelia Langford, John Sillitoe on behalf of the Wellcome Sanger Institute COVID-19 Surveillance Team |
| EPI_ISL_2275216 | Lighthouse Lab in Milton Keynes | Wellcome Sanger Institute for the COVID-19 Genomics UK (COG-UK) Consortium | The Lighthouse Lab in Milton Keynes and Alex Alderton, Roberto Amato, Jeffrey Barrett, Sonia Goncalves, Ewan Harrison, David K. Jackson, Ian Johnston, Dominic Kwiatkowski, Cordelia Langford, John Sillitoe on behalf of the Wellcome Sanger Institute COVID-19 Surveillance Team |
| EPI_ISL_2275218 | Lighthouse Lab in Glasgow | Wellcome Sanger Institute for the COVID-19 Genomics UK (COG-UK) Consortium | Harper VanSteenhouse, Yumi Kasai, David Gray, Carol Clugston, Anna Dominiczak and Alex Alderton, Roberto Amato, Jeffrey Barrett, Sonia Goncalves, Ewan Harrison, David K. Jackson, Ian Johnston, Dominic Kwiatkowski, Cordelia Langford, John Sillitoe on behalf of the Wellcome Sanger Institute COVID-19 Surveillance Team |
| EPI_ISL_2275222, EPI_ISL_2275223, EPI_ISL_2275224 | Lighthouse Lab in Milton Keynes | Wellcome Sanger Institute for the COVID-19 Genomics UK (COG-UK) Consortium | The Lighthouse Lab in Milton Keynes and Alex Alderton, Roberto Amato, Jeffrey Barrett, Sonia Goncalves, Ewan Harrison, David K. Jackson, Ian Johnston, Dominic Kwiatkowski, Cordelia Langford, John Sillitoe on behalf of the Wellcome Sanger Institute COVID-19 Surveillance Team |
| EPI_ISL_2275225, EPI_ISL_2275227 | Lighthouse Lab in Glasgow | Wellcome Sanger Institute for the COVID-19 Genomics UK (COG-UK) Consortium | Harper VanSteenhouse, Yumi Kasai, David Gray, Carol Clugston, Anna Dominiczak and Alex Alderton, Roberto Amato, Jeffrey Barrett, Sonia Goncalves, Ewan Harrison, David K. Jackson, Ian Johnston, Dominic Kwiatkowski, Cordelia Langford, John Sillitoe on behalf of the Wellcome Sanger Institute COVID-19 Surveillance Team |
| EPI_ISL_2275229 | Lighthouse Lab in Alderley Park | Wellcome Sanger Institute for the COVID-19 Genomics UK (COG-UK) Consortium | Jacquelyn Wynn, Mairead Hyland, The Lighthouse Lab in Alderley Park and Alex Alderton, Roberto Amato, Jeffrey Barrett, Sonia Goncalves, Ewan Harrison, David K. Jackson, Ian Johnston, Dominic Kwiatkowski, Cordelia Langford, John Sillitoe on behalf of the Wellcome Sanger Institute COVID-19 Surveillance Team |
| EPI_ISL_2275230 | Lighthouse Lab in Milton Keynes | Wellcome Sanger Institute for the COVID-19 Genomics UK (COG-UK) Consortium | The Lighthouse Lab in Milton Keynes and Alex Alderton, Roberto Amato, Jeffrey Barrett, Sonia Goncalves, Ewan Harrison, David K. Jackson, Ian Johnston, Dominic Kwiatkowski, Cordelia Langford, John Sillitoe on behalf of the Wellcome Sanger Institute COVID-19 Surveillance Team |
| EPI_ISL_2275231, EPI_ISL_2275232 | Lighthouse Lab in Alderley Park | Wellcome Sanger Institute for the COVID-19 Genomics UK (COG-UK) Consortium | Jacquelyn Wynn, Mairead Hyland, The Lighthouse Lab in Alderley Park and Alex Alderton, Roberto Amato, Jeffrey Barrett, Sonia Goncalves, Ewan Harrison, David K. Jackson, Ian Johnston, Dominic Kwiatkowski, Cordelia Langford, John Sillitoe on behalf of the Wellcome Sanger Institute COVID-19 Surveillance Team |
| EPI_ISL_2275233 | Lighthouse Lab in Milton Keynes | Wellcome Sanger Institute for the COVID-19 Genomics UK (COG-UK) Consortium | The Lighthouse Lab in Milton Keynes and Alex Alderton, Roberto Amato, Jeffrey Barrett, Sonia Goncalves, Ewan Harrison, David K. Jackson, Ian Johnston, Dominic Kwiatkowski, Cordelia Langford, John Sillitoe on behalf of the Wellcome Sanger Institute COVID-19 Surveillance Team |
| EPI_ISL_2275234 | Lighthouse Lab in Glasgow | Wellcome Sanger Institute for the COVID-19 Genomics UK (COG-UK) Consortium | Harper VanSteenhouse, Yumi Kasai, David Gray, Carol Clugston, Anna Dominiczak and Alex Alderton, Roberto Amato, Jeffrey Barrett, Sonia Goncalves, Ewan Harrison, David K. Jackson, Ian Johnston, Dominic Kwiatkowski, Cordelia Langford, John Sillitoe on behalf of the Wellcome Sanger Institute COVID-19 Surveillance Team |
| EPI_ISL_2275235 | Lighthouse Lab in Milton Keynes | Wellcome Sanger Institute for the COVID-19 Genomics UK (COG-UK) Consortium | The Lighthouse Lab in Milton Keynes and Alex Alderton, Roberto Amato, Jeffrey Barrett, Sonia Goncalves, Ewan Harrison, David K. Jackson, Ian Johnston, Dominic Kwiatkowski, Cordelia Langford, John Sillitoe on behalf of the Wellcome Sanger Institute COVID-19 Surveillance Team |
| EPI_ISL_2275236, EPI_ISL_2275237, EPI_ISL_2275238, EPI_ISL_2275239 | Lighthouse Lab in Alderley Park | Wellcome Sanger Institute for the COVID-19 Genomics UK (COG-UK) Consortium | Jacquelyn Wynn, Mairead Hyland, The Lighthouse Lab in Alderley Park and Alex Alderton, Roberto Amato, Jeffrey Barrett, Sonia Goncalves, Ewan Harrison, David K. Jackson, Ian Johnston, Dominic Kwiatkowski, Cordelia Langford, John Sillitoe on behalf of the Wellcome Sanger Institute COVID-19 Surveillance Team |
| EPI_ISL_2275240, EPI_ISL_2275241 | Lighthouse Lab in Glasgow | Wellcome Sanger Institute for the COVID-19 Genomics UK (COG-UK) Consortium | Harper VanSteenhouse, Yumi Kasai, David Gray, Carol Clugston, Anna Dominiczak and Alex Alderton, Roberto Amato, Jeffrey Barrett, Sonia Goncalves, Ewan Harrison, David K. Jackson, Ian Johnston, Dominic Kwiatkowski, Cordelia Langford, John Sillitoe on behalf of the Wellcome Sanger Institute COVID-19 Surveillance Team |
| EPI_ISL_2275242, EPI_ISL_2275243, EPI_ISL_2275245 | Lighthouse Lab in Alderley Park | Wellcome Sanger Institute for the COVID-19 Genomics UK (COG-UK) Consortium | Jacquelyn Wynn, Mairead Hyland, The Lighthouse Lab in Alderley Park and Alex Alderton, Roberto Amato, Jeffrey Barrett, Sonia Goncalves, Ewan Harrison, David K. Jackson, Ian Johnston, Dominic Kwiatkowski, Cordelia Langford, John Sillitoe on behalf of the Wellcome Sanger Institute COVID-19 Surveillance Team |
| EPI_ISL_2275246 | Lighthouse Lab in Milton Keynes | Wellcome Sanger Institute for the COVID-19 Genomics UK (COG-UK) Consortium | The Lighthouse Lab in Milton Keynes and Alex Alderton, Roberto Amato, Jeffrey Barrett, Sonia Goncalves, Ewan Harrison, David K. Jackson, Ian Johnston, Dominic Kwiatkowski, Cordelia Langford, John Sillitoe on behalf of the Wellcome Sanger Institute COVID-19 Surveillance Team |
| EPI_ISL_2275247 | Lighthouse Lab in Alderley Park | Wellcome Sanger Institute for the COVID-19 Genomics UK (COG-UK) Consortium | Jacquelyn Wynn, Mairead Hyland, The Lighthouse Lab in Alderley Park and Alex Alderton, Roberto Amato, Jeffrey Barrett, Sonia Goncalves, Ewan Harrison, David K. Jackson, Ian Johnston, Dominic Kwiatkowski, Cordelia Langford, John Sillitoe on behalf of the Wellcome Sanger Institute COVID-19 Surveillance Team |
| EPI_ISL_2275248 | Lighthouse Lab in Glasgow | Wellcome Sanger Institute for the COVID-19 Genomics UK (COG-UK) Consortium | Harper VanSteenhouse, Yumi Kasai, David Gray, Carol Clugston, Anna Dominiczak and Alex Alderton, Roberto Amato, Jeffrey Barrett, Sonia Goncalves, Ewan Harrison, David K. Jackson, Ian Johnston, Dominic Kwiatkowski, Cordelia Langford, John Sillitoe on behalf of the Wellcome Sanger Institute COVID-19 Surveillance Team |
| EPI_ISL_2275249 | Lighthouse Lab in Milton Keynes | Wellcome Sanger Institute for the COVID-19 Genomics UK (COG-UK) Consortium | The Lighthouse Lab in Milton Keynes and Alex Alderton, Roberto Amato, Jeffrey Barrett, Sonia Goncalves, Ewan Harrison, David K. Jackson, Ian Johnston, Dominic Kwiatkowski, Cordelia Langford, John Sillitoe on behalf of the Wellcome Sanger Institute COVID-19 Surveillance Team |
| EPI_ISL_2275250, EPI_ISL_2275251 | Lighthouse Lab in Glasgow | Wellcome Sanger Institute for the COVID-19 Genomics UK (COG-UK) Consortium | Harper VanSteenhouse, Yumi Kasai, David Gray, Carol Clugston, Anna Dominiczak and Alex Alderton, Roberto Amato, Jeffrey Barrett, Sonia Goncalves, Ewan Harrison, David K. Jackson, Ian Johnston, Dominic Kwiatkowski, Cordelia Langford, John Sillitoe on behalf of the Wellcome Sanger Institute COVID-19 Surveillance Team |
| EPI_ISL_2275252 | Lighthouse Lab in Alderley Park | Wellcome Sanger Institute for the COVID-19 Genomics UK (COG-UK) Consortium | Jacquelyn Wynn, Mairead Hyland, The Lighthouse Lab in Alderley Park and Alex Alderton, Roberto Amato, Jeffrey Barrett, Sonia Goncalves, Ewan Harrison, David K. Jackson, Ian Johnston, Dominic Kwiatkowski, Cordelia Langford, John Sillitoe on behalf of the Wellcome Sanger Institute COVID-19 Surveillance Team |
| EPI_ISL_2275253 | Lighthouse Lab in Milton Keynes | Wellcome Sanger Institute for the COVID-19 Genomics UK (COG-UK) Consortium | The Lighthouse Lab in Milton Keynes and Alex Alderton, Roberto Amato, Jeffrey Barrett, Sonia Goncalves, Ewan Harrison, David K. Jackson, Ian Johnston, Dominic Kwiatkowski, Cordelia Langford, John Sillitoe on behalf of the Wellcome Sanger Institute COVID-19 Surveillance Team |
| EPI_ISL_2275254 | Lighthouse Lab in Glasgow | Wellcome Sanger Institute for the COVID-19 Genomics UK (COG-UK) Consortium | Harper VanSteenhouse, Yumi Kasai, David Gray, Carol Clugston, Anna Dominiczak and Alex Alderton, Roberto Amato, Jeffrey Barrett, Sonia Goncalves, Ewan Harrison, David K. Jackson, Ian Johnston, Dominic Kwiatkowski, Cordelia Langford, John Sillitoe on behalf of the Wellcome Sanger Institute COVID-19 Surveillance Team |
| EPI_ISL_2275256 | Lighthouse Lab in Alderley Park | Wellcome Sanger Institute for the COVID-19 Genomics UK (COG-UK) Consortium | Jacquelyn Wynn, Mairead Hyland, The Lighthouse Lab in Alderley Park and Alex Alderton, Roberto Amato, Jeffrey Barrett, Sonia Goncalves, Ewan Harrison, David K. Jackson, Ian Johnston, Dominic Kwiatkowski, Cordelia Langford, John Sillitoe on behalf of the Wellcome Sanger Institute COVID-19 Surveillance Team |
| EPI_ISL_2275257 | Lighthouse Lab in Glasgow | Wellcome Sanger Institute for the COVID-19 Genomics UK (COG-UK) Consortium | Harper VanSteenhouse, Yumi Kasai, David Gray, Carol Clugston, Anna Dominiczak and Alex Alderton, Roberto Amato, Jeffrey Barrett, Sonia Goncalves, Ewan Harrison, David K. Jackson, Ian Johnston, Dominic Kwiatkowski, Cordelia Langford, John Sillitoe on behalf of the Wellcome Sanger Institute COVID-19 Surveillance Team |
| EPI_ISL_2275258, EPI_ISL_2275259, EPI_ISL_2275260 | Lighthouse Lab in Milton Keynes | Wellcome Sanger Institute for the COVID-19 Genomics UK (COG-UK) Consortium | The Lighthouse Lab in Milton Keynes and Alex Alderton, Roberto Amato, Jeffrey Barrett, Sonia Goncalves, Ewan Harrison, David K. Jackson, Ian Johnston, Dominic Kwiatkowski, Cordelia Langford, John Sillitoe on behalf of the Wellcome Sanger Institute COVID-19 Surveillance Team |
| EPI_ISL_2275262, EPI_ISL_2275264 | Lighthouse Lab in Alderley Park | Wellcome Sanger Institute for the COVID-19 Genomics UK (COG-UK) Consortium | Jacquelyn Wynn, Mairead Hyland, The Lighthouse Lab in Alderley Park and Alex Alderton, Roberto Amato, Jeffrey Barrett, Sonia Goncalves, Ewan Harrison, David K. Jackson, Ian Johnston, Dominic Kwiatkowski, Cordelia Langford, John Sillitoe on behalf of the Wellcome Sanger Institute COVID-19 Surveillance Team |
| EPI_ISL_2275265, EPI_ISL_2275266 | Lighthouse Lab in Glasgow | Wellcome Sanger Institute for the COVID-19 Genomics UK (COG-UK) Consortium | Harper VanSteenhouse, Yumi Kasai, David Gray, Carol Clugston, Anna Dominiczak and Alex Alderton, Roberto Amato, Jeffrey Barrett, Sonia Goncalves, Ewan Harrison, David K. Jackson, Ian Johnston, Dominic Kwiatkowski, Cordelia Langford, John Sillitoe on behalf of the Wellcome Sanger Institute COVID-19 Surveillance Team |
| EPI_ISL_2275268, EPI_ISL_2275269 | Lighthouse Lab in Alderley Park | Wellcome Sanger Institute for the COVID-19 Genomics UK (COG-UK) Consortium | Jacquelyn Wynn, Mairead Hyland, The Lighthouse Lab in Alderley Park and Alex Alderton, Roberto Amato, Jeffrey Barrett, Sonia Goncalves, Ewan Harrison, David K. Jackson, Ian Johnston, Dominic Kwiatkowski, Cordelia Langford, John Sillitoe on behalf of the Wellcome Sanger Institute COVID-19 Surveillance Team |
| EPI_ISL_2275270 | Lighthouse Lab in Milton Keynes | Wellcome Sanger Institute for the COVID-19 Genomics UK (COG-UK) Consortium | The Lighthouse Lab in Milton Keynes and Alex Alderton, Roberto Amato, Jeffrey Barrett, Sonia Goncalves, Ewan Harrison, David K. Jackson, Ian Johnston, Dominic Kwiatkowski, Cordelia Langford, John Sillitoe on behalf of the Wellcome Sanger Institute COVID-19 Surveillance Team |
| EPI_ISL_2275271, EPI_ISL_2275272 | Lighthouse Lab in Alderley Park | Wellcome Sanger Institute for the COVID-19 Genomics UK (COG-UK) Consortium | Jacquelyn Wynn, Mairead Hyland, The Lighthouse Lab in Alderley Park and Alex Alderton, Roberto Amato, Jeffrey Barrett, Sonia Goncalves, Ewan Harrison, David K. Jackson, Ian Johnston, Dominic Kwiatkowski, Cordelia Langford, John Sillitoe on behalf of the Wellcome Sanger Institute COVID-19 Surveillance Team |
| EPI_ISL_2275274, EPI_ISL_2275276, EPI_ISL_2275278 | Lighthouse Lab in Glasgow | Wellcome Sanger Institute for the COVID-19 Genomics UK (COG-UK) Consortium | Harper VanSteenhouse, Yumi Kasai, David Gray, Carol Clugston, Anna Dominiczak and Alex Alderton, Roberto Amato, Jeffrey Barrett, Sonia Goncalves, Ewan Harrison, David K. Jackson, Ian Johnston, Dominic Kwiatkowski, Cordelia Langford, John Sillitoe on behalf of the Wellcome Sanger Institute COVID-19 Surveillance Team |
| EPI_ISL_2275279 | Lighthouse Lab in Alderley Park | Wellcome Sanger Institute for the COVID-19 Genomics UK (COG-UK) Consortium | Jacquelyn Wynn, Mairead Hyland, The Lighthouse Lab in Alderley Park and Alex Alderton, Roberto Amato, Jeffrey Barrett, Sonia Goncalves, Ewan Harrison, David K. Jackson, Ian Johnston, Dominic Kwiatkowski, Cordelia Langford, John Sillitoe on behalf of the Wellcome Sanger Institute COVID-19 Surveillance Team |
| EPI_ISL_2275280 | Lighthouse Lab in Milton Keynes | Wellcome Sanger Institute for the COVID-19 Genomics UK (COG-UK) Consortium | The Lighthouse Lab in Milton Keynes and Alex Alderton, Roberto Amato, Jeffrey Barrett, Sonia Goncalves, Ewan Harrison, David K. Jackson, Ian Johnston, Dominic Kwiatkowski, Cordelia Langford, John Sillitoe on behalf of the Wellcome Sanger Institute COVID-19 Surveillance Team |
| EPI_ISL_2275282 | Lighthouse Lab in Glasgow | Wellcome Sanger Institute for the COVID-19 Genomics UK (COG-UK) Consortium | Harper VanSteenhouse, Yumi Kasai, David Gray, Carol Clugston, Anna Dominiczak and Alex Alderton, Roberto Amato, Jeffrey Barrett, Sonia Goncalves, Ewan Harrison, David K. Jackson, Ian Johnston, Dominic Kwiatkowski, Cordelia Langford, John Sillitoe on behalf of the Wellcome Sanger Institute COVID-19 Surveillance Team |
| EPI_ISL_2275284, EPI_ISL_2275285 | Lighthouse Lab in Milton Keynes | Wellcome Sanger Institute for the COVID-19 Genomics UK (COG-UK) Consortium | The Lighthouse Lab in Milton Keynes and Alex Alderton, Roberto Amato, Jeffrey Barrett, Sonia Goncalves, Ewan Harrison, David K. Jackson, Ian Johnston, Dominic Kwiatkowski, Cordelia Langford, John Sillitoe on behalf of the Wellcome Sanger Institute COVID-19 Surveillance Team |
| EPI_ISL_2275286 | Lighthouse Lab in Glasgow | Wellcome Sanger Institute for the COVID-19 Genomics UK (COG-UK) Consortium | Harper VanSteenhouse, Yumi Kasai, David Gray, Carol Clugston, Anna Dominiczak and Alex Alderton, Roberto Amato, Jeffrey Barrett, Sonia Goncalves, Ewan Harrison, David K. Jackson, Ian Johnston, Dominic Kwiatkowski, Cordelia Langford, John Sillitoe on behalf of the Wellcome Sanger Institute COVID-19 Surveillance Team |
| EPI_ISL_2275287 | Lighthouse Lab in Milton Keynes | Wellcome Sanger Institute for the COVID-19 Genomics UK (COG-UK) Consortium | The Lighthouse Lab in Milton Keynes and Alex Alderton, Roberto Amato, Jeffrey Barrett, Sonia Goncalves, Ewan Harrison, David K. Jackson, Ian Johnston, Dominic Kwiatkowski, Cordelia Langford, John Sillitoe on behalf of the Wellcome Sanger Institute COVID-19 Surveillance Team |
| EPI_ISL_2275288 | Lighthouse Lab in Alderley Park | Wellcome Sanger Institute for the COVID-19 Genomics UK (COG-UK) Consortium | Jacquelyn Wynn, Mairead Hyland, The Lighthouse Lab in Alderley Park and Alex Alderton, Roberto Amato, Jeffrey Barrett, Sonia Goncalves, Ewan Harrison, David K. Jackson, Ian Johnston, Dominic Kwiatkowski, Cordelia Langford, John Sillitoe on behalf of the Wellcome Sanger Institute COVID-19 Surveillance Team |
| EPI_ISL_2275289 | Lighthouse Lab in Glasgow | Wellcome Sanger Institute for the COVID-19 Genomics UK (COG-UK) Consortium | Harper VanSteenhouse, Yumi Kasai, David Gray, Carol Clugston, Anna Dominiczak and Alex Alderton, Roberto Amato, Jeffrey Barrett, Sonia Goncalves, Ewan Harrison, David K. Jackson, Ian Johnston, Dominic Kwiatkowski, Cordelia Langford, John Sillitoe on behalf of the Wellcome Sanger Institute COVID-19 Surveillance Team |
| EPI_ISL_2275294 | Lighthouse Lab in Milton Keynes | Wellcome Sanger Institute for the COVID-19 Genomics UK (COG-UK) Consortium | The Lighthouse Lab in Milton Keynes and Alex Alderton, Roberto Amato, Jeffrey Barrett, Sonia Goncalves, Ewan Harrison, David K. Jackson, Ian Johnston, Dominic Kwiatkowski, Cordelia Langford, John Sillitoe on behalf of the Wellcome Sanger Institute COVID-19 Surveillance Team |
| EPI_ISL_2275295, EPI_ISL_2275298 | Lighthouse Lab in Glasgow | Wellcome Sanger Institute for the COVID-19 Genomics UK (COG-UK) Consortium | Harper VanSteenhouse, Yumi Kasai, David Gray, Carol Clugston, Anna Dominiczak and Alex Alderton, Roberto Amato, Jeffrey Barrett, Sonia Goncalves, Ewan Harrison, David K. Jackson, Ian Johnston, Dominic Kwiatkowski, Cordelia Langford, John Sillitoe on behalf of the Wellcome Sanger Institute COVID-19 Surveillance Team |

| | | | |
|---|---|---|---|
| EPI_ISL_2275300 | Lighthouse Lab in Milton Keynes | Wellcome Sanger Institute for the COVID-19 Genomics UK (COG-UK) Consortium | The Lighthouse Lab in Milton Keynes and Alex Alderton, Roberto Amato, Jeffrey Barrett, Sonia Goncalves, Ewan Harrison, David K. Jackson, Ian Johnston, Dominic Kwiatkowski, Cordelia Langford, John Sillitoe on behalf of the Wellcome Sanger Institute COVID-19 Surveillance Team |
| EPI_ISL_2275302 | Lighthouse Lab in Glasgow | Wellcome Sanger Institute for the COVID-19 Genomics UK (COG-UK) Consortium | Harper VanSteenhouse, Yumi Kasai, David Gray, Carol Clugston, Anna Dominiczak and Alex Alderton, Roberto Amato, Jeffrey Barrett, Sonia Goncalves, Ewan Harrison, David K. Jackson, Ian Johnston, Dominic Kwiatkowski, Cordelia Langford, John Sillitoe on behalf of the Wellcome Sanger Institute COVID-19 Surveillance Team |
| EPI_ISL_2275305 | Lighthouse Lab in Alderley Park | Wellcome Sanger Institute for the COVID-19 Genomics UK (COG-UK) Consortium | Jacquelyn Wynn, Mairead Hyland, The Lighthouse Lab in Alderley Park and Alex Alderton, Roberto Amato, Jeffrey Barrett, Sonia Goncalves, Ewan Harrison, David K. Jackson, Ian Johnston, Dominic Kwiatkowski, Cordelia Langford, John Sillitoe on behalf of the Wellcome Sanger Institute COVID-19 Surveillance Team |
| EPI_ISL_2275306 | Lighthouse Lab in Glasgow | Wellcome Sanger Institute for the COVID-19 Genomics UK (COG-UK) Consortium | Harper VanSteenhouse, Yumi Kasai, David Gray, Carol Clugston, Anna Dominiczak and Alex Alderton, Roberto Amato, Jeffrey Barrett, Sonia Goncalves, Ewan Harrison, David K. Jackson, Ian Johnston, Dominic Kwiatkowski, Cordelia Langford, John Sillitoe on behalf of the Wellcome Sanger Institute COVID-19 Surveillance Team |
| EPI_ISL_2275308, EPI_ISL_2275309 | Lighthouse Lab in Alderley Park | Wellcome Sanger Institute for the COVID-19 Genomics UK (COG-UK) Consortium | Jacquelyn Wynn, Mairead Hyland, The Lighthouse Lab in Alderley Park and Alex Alderton, Roberto Amato, Jeffrey Barrett, Sonia Goncalves, Ewan Harrison, David K. Jackson, Ian Johnston, Dominic Kwiatkowski, Cordelia Langford, John Sillitoe on behalf of the Wellcome Sanger Institute COVID-19 Surveillance Team |
| EPI_ISL_2275310, EPI_ISL_2275311 | Lighthouse Lab in Milton Keynes | Wellcome Sanger Institute for the COVID-19 Genomics UK (COG-UK) Consortium | The Lighthouse Lab in Milton Keynes and Alex Alderton, Roberto Amato, Jeffrey Barrett, Sonia Goncalves, Ewan Harrison, David K. Jackson, Ian Johnston, Dominic Kwiatkowski, Cordelia Langford, John Sillitoe on behalf of the Wellcome Sanger Institute COVID-19 Surveillance Team |
| EPI_ISL_2275312 | Lighthouse Lab in Alderley Park | Wellcome Sanger Institute for the COVID-19 Genomics UK (COG-UK) Consortium | Jacquelyn Wynn, Mairead Hyland, The Lighthouse Lab in Alderley Park and Alex Alderton, Roberto Amato, Jeffrey Barrett, Sonia Goncalves, Ewan Harrison, David K. Jackson, Ian Johnston, Dominic Kwiatkowski, Cordelia Langford, John Sillitoe on behalf of the Wellcome Sanger Institute COVID-19 Surveillance Team |
| EPI_ISL_2275316 | Lighthouse Lab in Glasgow | Wellcome Sanger Institute for the COVID-19 Genomics UK (COG-UK) Consortium | Harper VanSteenhouse, Yumi Kasai, David Gray, Carol Clugston, Anna Dominiczak and Alex Alderton, Roberto Amato, Jeffrey Barrett, Sonia Goncalves, Ewan Harrison, David K. Jackson, Ian Johnston, Dominic Kwiatkowski, Cordelia Langford, John Sillitoe on behalf of the Wellcome Sanger Institute COVID-19 Surveillance Team |
| EPI_ISL_2275321 | Lighthouse Lab in Alderley Park | Wellcome Sanger Institute for the COVID-19 Genomics UK (COG-UK) Consortium | Jacquelyn Wynn, Mairead Hyland, The Lighthouse Lab in Alderley Park and Alex Alderton, Roberto Amato, Jeffrey Barrett, Sonia Goncalves, Ewan Harrison, David K. Jackson, Ian Johnston, Dominic Kwiatkowski, Cordelia Langford, John Sillitoe on behalf of the Wellcome Sanger Institute COVID-19 Surveillance Team |
| EPI_ISL_2275322 | Lighthouse Lab in Milton Keynes | Wellcome Sanger Institute for the COVID-19 Genomics UK (COG-UK) Consortium | The Lighthouse Lab in Milton Keynes and Alex Alderton, Roberto Amato, Jeffrey Barrett, Sonia Goncalves, Ewan Harrison, David K. Jackson, Ian Johnston, Dominic Kwiatkowski, Cordelia Langford, John Sillitoe on behalf of the Wellcome Sanger Institute COVID-19 Surveillance Team |
| EPI_ISL_2275323 | Lighthouse Lab in Alderley Park | Wellcome Sanger Institute for the COVID-19 Genomics UK (COG-UK) Consortium | Jacquelyn Wynn, Mairead Hyland, The Lighthouse Lab in Alderley Park and Alex Alderton, Roberto Amato, Jeffrey Barrett, Sonia Goncalves, Ewan Harrison, David K. Jackson, Ian Johnston, Dominic Kwiatkowski, Cordelia Langford, John Sillitoe on behalf of the Wellcome Sanger Institute COVID-19 Surveillance Team |
| EPI_ISL_2275324 | Lighthouse Lab in Glasgow | Wellcome Sanger Institute for the COVID-19 Genomics UK (COG-UK) Consortium | Harper VanSteenhouse, Yumi Kasai, David Gray, Carol Clugston, Anna Dominiczak and Alex Alderton, Roberto Amato, Jeffrey Barrett, Sonia Goncalves, Ewan Harrison, David K. Jackson, Ian Johnston, Dominic Kwiatkowski, Cordelia Langford, John Sillitoe on behalf of the Wellcome Sanger Institute COVID-19 Surveillance Team |
| EPI_ISL_2275326, EPI_ISL_2275327, EPI_ISL_2275328 | Lighthouse Lab in Alderley Park | Wellcome Sanger Institute for the COVID-19 Genomics UK (COG-UK) Consortium | Jacquelyn Wynn, Mairead Hyland, The Lighthouse Lab in Alderley Park and Alex Alderton, Roberto Amato, Jeffrey Barrett, Sonia Goncalves, Ewan Harrison, David K. Jackson, Ian Johnston, Dominic Kwiatkowski, Cordelia Langford, John Sillitoe on behalf of the Wellcome Sanger Institute COVID-19 Surveillance Team |
| EPI_ISL_2275329 | Lighthouse Lab in Glasgow | Wellcome Sanger Institute for the COVID-19 Genomics UK (COG-UK) Consortium | Harper VanSteenhouse, Yumi Kasai, David Gray, Carol Clugston, Anna Dominiczak and Alex Alderton, Roberto Amato, Jeffrey Barrett, Sonia Goncalves, Ewan Harrison, David K. Jackson, Ian Johnston, Dominic Kwiatkowski, Cordelia Langford, John Sillitoe on behalf of the Wellcome Sanger Institute COVID-19 Surveillance Team |
| EPI_ISL_2275330 | Lighthouse Lab in Alderley Park | Wellcome Sanger Institute for the COVID-19 Genomics UK (COG-UK) Consortium | Jacquelyn Wynn, Mairead Hyland, The Lighthouse Lab in Alderley Park and Alex Alderton, Roberto Amato, Jeffrey Barrett, Sonia Goncalves, Ewan Harrison, David K. Jackson, Ian Johnston, Dominic Kwiatkowski, Cordelia Langford, John Sillitoe on behalf of the Wellcome Sanger Institute COVID-19 Surveillance Team |
| EPI_ISL_2275331 | Lighthouse Lab in Milton Keynes | Wellcome Sanger Institute for the COVID-19 Genomics UK (COG-UK) Consortium | The Lighthouse Lab in Milton Keynes and Alex Alderton, Roberto Amato, Jeffrey Barrett, Sonia Goncalves, Ewan Harrison, David K. Jackson, Ian Johnston, Dominic Kwiatkowski, Cordelia Langford, John Sillitoe on behalf of the Wellcome Sanger Institute COVID-19 Surveillance Team |
| EPI_ISL_2275332 | Lighthouse Lab in Glasgow | Wellcome Sanger Institute for the COVID-19 Genomics UK (COG-UK) Consortium | Harper VanSteenhouse, Yumi Kasai, David Gray, Carol Clugston, Anna Dominiczak and Alex Alderton, Roberto Amato, Jeffrey Barrett, Sonia Goncalves, Ewan Harrison, David K. Jackson, Ian Johnston, Dominic Kwiatkowski, Cordelia Langford, John Sillitoe on behalf of the Wellcome Sanger Institute COVID-19 Surveillance Team |
| EPI_ISL_2275333, EPI_ISL_2275334 | Lighthouse Lab in Milton Keynes | Wellcome Sanger Institute for the COVID-19 Genomics UK (COG-UK) Consortium | The Lighthouse Lab in Milton Keynes and Alex Alderton, Roberto Amato, Jeffrey Barrett, Sonia Goncalves, Ewan Harrison, David K. Jackson, Ian Johnston, Dominic Kwiatkowski, Cordelia Langford, John Sillitoe on behalf of the Wellcome Sanger Institute COVID-19 Surveillance Team |
| EPI_ISL_2275335, EPI_ISL_2275336 | Lighthouse Lab in Glasgow | Wellcome Sanger Institute for the COVID-19 Genomics UK (COG-UK) Consortium | Harper VanSteenhouse, Yumi Kasai, David Gray, Carol Clugston, Anna Dominiczak and Alex Alderton, Roberto Amato, Jeffrey Barrett, Sonia Goncalves, Ewan Harrison, David K. Jackson, Ian Johnston, Dominic Kwiatkowski, Cordelia Langford, John Sillitoe on behalf of the Wellcome Sanger Institute COVID-19 Surveillance Team |
| EPI_ISL_2275337 | Lighthouse Lab in Alderley Park | Wellcome Sanger Institute for the COVID-19 Genomics UK (COG-UK) Consortium | Jacquelyn Wynn, Mairead Hyland, The Lighthouse Lab in Alderley Park and Alex Alderton, Roberto Amato, Jeffrey Barrett, Sonia Goncalves, Ewan Harrison, David K. Jackson, Ian Johnston, Dominic Kwiatkowski, Cordelia Langford, John Sillitoe on behalf of the Wellcome Sanger Institute COVID-19 Surveillance Team |
| EPI_ISL_2275338 | Lighthouse Lab in Glasgow | Wellcome Sanger Institute for the COVID-19 Genomics UK (COG-UK) Consortium | Harper VanSteenhouse, Yumi Kasai, David Gray, Carol Clugston, Anna Dominiczak and Alex Alderton, Roberto Amato, Jeffrey Barrett, Sonia Goncalves, Ewan Harrison, David K. Jackson, Ian Johnston, Dominic Kwiatkowski, Cordelia Langford, John Sillitoe on behalf of the Wellcome Sanger Institute COVID-19 Surveillance Team |
| EPI_ISL_2275341 | Lighthouse Lab in Alderley Park | Wellcome Sanger Institute for the COVID-19 Genomics UK (COG-UK) Consortium | Jacquelyn Wynn, Mairead Hyland, The Lighthouse Lab in Alderley Park and Alex Alderton, Roberto Amato, Jeffrey Barrett, Sonia Goncalves, Ewan Harrison, David K. Jackson, Ian Johnston, Dominic Kwiatkowski, Cordelia Langford, John Sillitoe on behalf of the Wellcome Sanger Institute COVID-19 Surveillance Team |
| EPI_ISL_2275342, EPI_ISL_2275344 | Lighthouse Lab in Milton Keynes | Wellcome Sanger Institute for the COVID-19 Genomics UK (COG-UK) Consortium | The Lighthouse Lab in Milton Keynes and Alex Alderton, Roberto Amato, Jeffrey Barrett, Sonia Goncalves, Ewan Harrison, David K. Jackson, Ian Johnston, Dominic Kwiatkowski, Cordelia Langford, John Sillitoe on behalf of the Wellcome Sanger Institute COVID-19 Surveillance Team |
| EPI_ISL_2275345, EPI_ISL_2275346, EPI_ISL_2275347, EPI_ISL_2275348, EPI_ISL_2275349 | Lighthouse Lab in Alderley Park | Wellcome Sanger Institute for the COVID-19 Genomics UK (COG-UK) Consortium | Jacquelyn Wynn, Mairead Hyland, The Lighthouse Lab in Alderley Park and Alex Alderton, Roberto Amato, Jeffrey Barrett, Sonia Goncalves, Ewan Harrison, David K. Jackson, Ian Johnston, Dominic Kwiatkowski, Cordelia Langford, John Sillitoe on behalf of the Wellcome Sanger Institute COVID-19 Surveillance Team |
| EPI_ISL_2275351 | Lighthouse Lab in Glasgow | Wellcome Sanger Institute for the COVID-19 Genomics UK (COG-UK) Consortium | Harper VanSteenhouse, Yumi Kasai, David Gray, Carol Clugston, Anna Dominiczak and Alex Alderton, Roberto Amato, Jeffrey Barrett, Sonia Goncalves, Ewan Harrison, David K. Jackson, Ian Johnston, Dominic Kwiatkowski, Cordelia Langford, John Sillitoe on behalf of the Wellcome Sanger Institute COVID-19 Surveillance Team |
| EPI_ISL_2275354 | Lighthouse Lab in Alderley Park | Wellcome Sanger Institute for the COVID-19 Genomics UK (COG-UK) Consortium | Jacquelyn Wynn, Mairead Hyland, The Lighthouse Lab in Alderley Park and Alex Alderton, Roberto Amato, Jeffrey Barrett, Sonia Goncalves, Ewan Harrison, David K. Jackson, Ian Johnston, Dominic Kwiatkowski, Cordelia Langford, John Sillitoe on behalf of the Wellcome Sanger Institute COVID-19 Surveillance Team |
| EPI_ISL_2275355 | Lighthouse Lab in Milton Keynes | Wellcome Sanger Institute for the COVID-19 Genomics UK (COG-UK) Consortium | The Lighthouse Lab in Milton Keynes and Alex Alderton, Roberto Amato, Jeffrey Barrett, Sonia Goncalves, Ewan Harrison, David K. Jackson, Ian Johnston, Dominic Kwiatkowski, Cordelia Langford, John Sillitoe on behalf of the Wellcome Sanger Institute COVID-19 Surveillance Team |
| EPI_ISL_2275358, EPI_ISL_2275360 | Lighthouse Lab in Alderley Park | Wellcome Sanger Institute for the COVID-19 Genomics UK (COG-UK) Consortium | Jacquelyn Wynn, Mairead Hyland, The Lighthouse Lab in Alderley Park and Alex Alderton, Roberto Amato, Jeffrey Barrett, Sonia Goncalves, Ewan Harrison, David K. Jackson, Ian Johnston, Dominic Kwiatkowski, Cordelia Langford, John Sillitoe on behalf of the Wellcome Sanger Institute COVID-19 Surveillance Team |
| EPI_ISL_2275361 | Lighthouse Lab in Milton Keynes | Wellcome Sanger Institute for the COVID-19 Genomics UK (COG-UK) Consortium | The Lighthouse Lab in Milton Keynes and Alex Alderton, Roberto Amato, Jeffrey Barrett, Sonia Goncalves, Ewan Harrison, David K. Jackson, Ian Johnston, Dominic Kwiatkowski, Cordelia Langford, John Sillitoe on behalf of the Wellcome Sanger Institute COVID-19 Surveillance Team |
| EPI_ISL_2275364, EPI_ISL_2275366 | Lighthouse Lab in Alderley Park | Wellcome Sanger Institute for the COVID-19 Genomics UK (COG-UK) Consortium | Jacquelyn Wynn, Mairead Hyland, The Lighthouse Lab in Alderley Park and Alex Alderton, Roberto Amato, Jeffrey Barrett, Sonia Goncalves, Ewan Harrison, David K. Jackson, Ian Johnston, Dominic Kwiatkowski, Cordelia Langford, John Sillitoe on behalf of the Wellcome Sanger Institute COVID-19 Surveillance Team |
| EPI_ISL_2275367, EPI_ISL_2275368 | Lighthouse Lab in Milton Keynes | Wellcome Sanger Institute for the COVID-19 Genomics UK (COG-UK) Consortium | The Lighthouse Lab in Milton Keynes and Alex Alderton, Roberto Amato, Jeffrey Barrett, Sonia Goncalves, Ewan Harrison, David K. Jackson, Ian Johnston, Dominic Kwiatkowski, Cordelia Langford, John Sillitoe on behalf of the Wellcome Sanger Institute COVID-19 Surveillance Team |
| EPI_ISL_2275370, EPI_ISL_2275371 | Lighthouse Lab in Glasgow | Wellcome Sanger Institute for the COVID-19 Genomics UK (COG-UK) Consortium | Harper VanSteenhouse, Yumi Kasai, David Gray, Carol Clugston, Anna Dominiczak and Alex Alderton, Roberto Amato, Jeffrey Barrett, Sonia Goncalves, Ewan Harrison, David K. Jackson, Ian Johnston, Dominic Kwiatkowski, Cordelia Langford, John Sillitoe on behalf of the Wellcome Sanger Institute COVID-19 Surveillance Team |
| EPI_ISL_2275372, EPI_ISL_2275374 | Lighthouse Lab in Milton Keynes | Wellcome Sanger Institute for the COVID-19 Genomics UK (COG-UK) Consortium | The Lighthouse Lab in Milton Keynes and Alex Alderton, Roberto Amato, Jeffrey Barrett, Sonia Goncalves, Ewan Harrison, David K. Jackson, Ian Johnston, Dominic Kwiatkowski, Cordelia Langford, John Sillitoe on behalf of the Wellcome Sanger Institute COVID-19 Surveillance Team |
| EPI_ISL_2275375, EPI_ISL_2275376, EPI_ISL_2275377 | Lighthouse Lab in Alderley Park | Wellcome Sanger Institute for the COVID-19 Genomics UK (COG-UK) Consortium | Jacquelyn Wynn, Mairead Hyland, The Lighthouse Lab in Alderley Park and Alex Alderton, Roberto Amato, Jeffrey Barrett, Sonia Goncalves, Ewan Harrison, David K. Jackson, Ian Johnston, Dominic Kwiatkowski, Cordelia Langford, John Sillitoe on behalf of the Wellcome Sanger Institute COVID-19 Surveillance Team |
| EPI_ISL_2275378, EPI_ISL_2275379 | Lighthouse Lab in Milton Keynes | Wellcome Sanger Institute for the COVID-19 Genomics UK (COG-UK) Consortium | The Lighthouse Lab in Milton Keynes and Alex Alderton, Roberto Amato, Jeffrey Barrett, Sonia Goncalves, Ewan Harrison, David K. Jackson, Ian Johnston, Dominic Kwiatkowski, Cordelia Langford, John Sillitoe on behalf of the Wellcome Sanger Institute COVID-19 Surveillance Team |
| EPI_ISL_2275380 | Lighthouse Lab in Alderley Park | Wellcome Sanger Institute for the COVID-19 Genomics UK (COG-UK) Consortium | Jacquelyn Wynn, Mairead Hyland, The Lighthouse Lab in Alderley Park and Alex Alderton, Roberto Amato, Jeffrey Barrett, Sonia Goncalves, Ewan Harrison, David K. Jackson, Ian Johnston, Dominic Kwiatkowski, Cordelia Langford, John Sillitoe on behalf of the Wellcome Sanger Institute COVID-19 Surveillance Team |
| EPI_ISL_2275381, EPI_ISL_2275382 | Lighthouse Lab in Glasgow | Wellcome Sanger Institute for the COVID-19 Genomics UK (COG-UK) Consortium | Harper VanSteenhouse, Yumi Kasai, David Gray, Carol Clugston, Anna Dominiczak and Alex Alderton, Roberto Amato, Jeffrey Barrett, Sonia Goncalves, Ewan Harrison, David K. Jackson, Ian Johnston, Dominic Kwiatkowski, Cordelia Langford, John Sillitoe on behalf of the Wellcome Sanger Institute COVID-19 Surveillance Team |
| EPI_ISL_2275385 | Lighthouse Lab in Alderley Park | Wellcome Sanger Institute for the COVID-19 Genomics UK (COG-UK) Consortium | Jacquelyn Wynn, Mairead Hyland, The Lighthouse Lab in Alderley Park and Alex Alderton, Roberto Amato, Jeffrey Barrett, Sonia Goncalves, Ewan Harrison, David K. Jackson, Ian Johnston, Dominic Kwiatkowski, Cordelia Langford, John Sillitoe on behalf of the Wellcome Sanger Institute COVID-19 Surveillance Team |
| EPI_ISL_2275387, EPI_ISL_2275389, EPI_ISL_2275390 | Lighthouse Lab in Glasgow | Wellcome Sanger Institute for the COVID-19 Genomics UK (COG-UK) Consortium | Harper VanSteenhouse, Yumi Kasai, David Gray, Carol Clugston, Anna Dominiczak and Alex Alderton, Roberto Amato, Jeffrey Barrett, Sonia Goncalves, Ewan Harrison, David K. Jackson, Ian Johnston, Dominic Kwiatkowski, Cordelia Langford, John Sillitoe on behalf of the Wellcome Sanger Institute COVID-19 Surveillance Team |
| EPI_ISL_2275391, EPI_ISL_2275392 | Lighthouse Lab in Milton Keynes | Wellcome Sanger Institute for the COVID-19 Genomics UK (COG-UK) Consortium | The Lighthouse Lab in Milton Keynes and Alex Alderton, Roberto Amato, Jeffrey Barrett, Sonia Goncalves, Ewan Harrison, David K. Jackson, Ian Johnston, Dominic Kwiatkowski, Cordelia Langford, John Sillitoe on behalf of the Wellcome Sanger Institute COVID-19 Surveillance Team |
| EPI_ISL_2275393, EPI_ISL_2275396 | Lighthouse Lab in Glasgow | Wellcome Sanger Institute for the COVID-19 Genomics UK (COG-UK) Consortium | Harper VanSteenhouse, Yumi Kasai, David Gray, Carol Clugston, Anna Dominiczak and Alex Alderton, Roberto Amato, Jeffrey Barrett, Sonia Goncalves, Ewan Harrison, David K. Jackson, Ian Johnston, Dominic Kwiatkowski, Cordelia Langford, John Sillitoe on behalf of the Wellcome Sanger Institute COVID-19 Surveillance Team |
| EPI_ISL_2275398 | Lighthouse Lab in Alderley Park | Wellcome Sanger Institute for the COVID-19 Genomics UK (COG-UK) Consortium | Jacquelyn Wynn, Mairead Hyland, The Lighthouse Lab in Alderley Park and Alex Alderton, Roberto Amato, Jeffrey Barrett, Sonia Goncalves, Ewan Harrison, David K. Jackson, Ian Johnston, Dominic Kwiatkowski, Cordelia Langford, John Sillitoe on behalf of the Wellcome Sanger Institute COVID-19 Surveillance Team |
| EPI_ISL_2275399 | Lighthouse Lab in Glasgow | Wellcome Sanger Institute for the COVID-19 Genomics UK (COG-UK) | Harper VanSteenhouse, Yumi Kasai, David Gray, Carol Clugston, Anna Dominiczak and Alex Alderton, Roberto Amato, Jeffrey Barrett, Sonia Goncalves, Ewan Harrison, David K. Jackson, Ian Johnston, Dominic Kwiatkowski, Cordelia Langford, |

| | | | |
|---|---|---|---|
| | | Consortium | John Sillitoe on behalf of the Wellcome Sanger Institute COVID-19 Surveillance Team |
| EPI_ISL_2275400, EPI_ISL_2275401 | Lighthouse Lab in Milton Keynes | Wellcome Sanger Institute for the COVID-19 Genomics UK (COG-UK) Consortium | The Lighthouse Lab in Milton Keynes and Alex Alderton, Roberto Amato, Jeffrey Barrett, Sonia Goncalves, Ewan Harrison, David K. Jackson, Ian Johnston, Dominic Kwiatkowski, Cordelia Langford, John Sillitoe on behalf of the Wellcome Sanger Institute COVID-19 Surveillance Team |
| EPI_ISL_2275402 | Lighthouse Lab in Alderley Park | Wellcome Sanger Institute for the COVID-19 Genomics UK (COG-UK) Consortium | Jacquelyn Wynn, Mairead Hyland, The Lighthouse Lab in Alderley Park and Alex Alderton, Roberto Amato, Jeffrey Barrett, Sonia Goncalves, Ewan Harrison, David K. Jackson, Ian Johnston, Dominic Kwiatkowski, Cordelia Langford, John Sillitoe on behalf of the Wellcome Sanger Institute COVID-19 Surveillance Team |
| EPI_ISL_2275403 | Lighthouse Lab in Milton Keynes | Wellcome Sanger Institute for the COVID-19 Genomics UK (COG-UK) Consortium | The Lighthouse Lab in Milton Keynes and Alex Alderton, Roberto Amato, Jeffrey Barrett, Sonia Goncalves, Ewan Harrison, David K. Jackson, Ian Johnston, Dominic Kwiatkowski, Cordelia Langford, John Sillitoe on behalf of the Wellcome Sanger Institute COVID-19 Surveillance Team |
| EPI_ISL_2275408, EPI_ISL_2275410, EPI_ISL_2275411, EPI_ISL_2275414, EPI_ISL_2275418, EPI_ISL_2275419, EPI_ISL_2275421, EPI_ISL_2275422, EPI_ISL_2275424, EPI_ISL_2275434, EPI_ISL_2275435, EPI_ISL_2275438, EPI_ISL_2275439, EPI_ISL_2275444, EPI_ISL_2275456, EPI_ISL_2275457, EPI_ISL_2275461, EPI_ISL_2275462, EPI_ISL_2275472, EPI_ISL_2275476, EPI_ISL_2275478, EPI_ISL_2275481, EPI_ISL_2275485, EPI_ISL_2275489, EPI_ISL_2275492, EPI_ISL_2275500, EPI_ISL_2275506, EPI_ISL_2275510, EPI_ISL_2275511, EPI_ISL_2275512, EPI_ISL_2275516, EPI_ISL_2275519, EPI_ISL_2275521, EPI_ISL_2275523, EPI_ISL_2275524, EPI_ISL_2275529, EPI_ISL_2275530, EPI_ISL_2275532, EPI_ISL_2275539, EPI_ISL_2275541, EPI_ISL_2275542, EPI_ISL_2275552, EPI_ISL_2275557, EPI_ISL_2275559, EPI_ISL_2275569, EPI_ISL_2275570, EPI_ISL_2275571, EPI_ISL_2275573, EPI_ISL_2275574, EPI_ISL_2275581, EPI_ISL_2275583, EPI_ISL_2275584, EPI_ISL_2275586, EPI_ISL_2275588, EPI_ISL_2275589, EPI_ISL_2275590, EPI_ISL_2275591, EPI_ISL_2275594, EPI_ISL_2275596, EPI_ISL_2275597, EPI_ISL_2275598, EPI_ISL_2275599, EPI_ISL_2275603, EPI_ISL_2275605, EPI_ISL_2275609, EPI_ISL_2275610, EPI_ISL_2275612, EPI_ISL_2275613, EPI_ISL_2275614, EPI_ISL_2275615, EPI_ISL_2275617, EPI_ISL_2275620, EPI_ISL_2275621, EPI_ISL_2275625, EPI_ISL_2275627, EPI_ISL_2275628, EPI_ISL_2275631, EPI_ISL_2275635, EPI_ISL_2275637, EPI_ISL_2275638, EPI_ISL_2275640, EPI_ISL_2275646, EPI_ISL_2275651, EPI_ISL_2275652, EPI_ISL_2275653, EPI_ISL_2275654, EPI_ISL_2275658, EPI_ISL_2275659, EPI_ISL_2275662, EPI_ISL_2275664, EPI_ISL_2275666, EPI_ISL_2275670, EPI_ISL_2275671, EPI_ISL_2275673, EPI_ISL_2275678, EPI_ISL_2275679, EPI_ISL_2275680, EPI_ISL_2275683, EPI_ISL_2275685, EPI_ISL_2275686, EPI_ISL_2275687, EPI_ISL_2275691, EPI_ISL_2275697, EPI_ISL_2275699, EPI_ISL_2275703, EPI_ISL_2275710, EPI_ISL_2275711, EPI_ISL_2275713, EPI_ISL_2275715, EPI_ISL_2275719, EPI_ISL_2275720, EPI_ISL_2275721, EPI_ISL_2275724, EPI_ISL_2275726, EPI_ISL_2275727, EPI_ISL_2275728, EPI_ISL_2275731, EPI_ISL_2275733, EPI_ISL_2275734, EPI_ISL_2275736, EPI_ISL_2275739, EPI_ISL_2275740, EPI_ISL_2275743, EPI_ISL_2275744, EPI_ISL_2275746 | | | |
| see above | Lighthouse Lab in Glasgow | Wellcome Sanger Institute for the COVID-19 Genomics UK (COG-UK) Consortium | Harper VanSteenhouse, Yumi Kasai, David Gray, Carol Clugston, Anna Dominiczak and Alex Alderton, Roberto Amato, Jeffrey Barrett, Sonia Goncalves, Ewan Harrison, David K. Jackson, Ian Johnston, Dominic Kwiatkowski, Cordelia Langford, John Sillitoe on behalf of the Wellcome Sanger Institute COVID-19 Surveillance Team |
| EPI_ISL_2276096 | Kansas Health and Environmental Lab | Kansas Health and Environmental Lab | Mike Grose, Katherine Wiggins, Jonathan Barnell, Ben Olsen, and Phil Adam |
| EPI_ISL_2277663, EPI_ISL_2277672 | Institute for Infectious Diseases | Institute for Infectious Diseases | Stefan Neuenschwander, Christian Baumann, Miguel A Terrazos Miani, Cora Sägesser, Pascal Bittel, Peter Keller, Franziska Suter-Riniker, Stephen L Leib, Alban Ramette |
| EPI_ISL_2278013, EPI_ISL_2278015, EPI_ISL_2278019, EPI_ISL_2278020, EPI_ISL_2278021, EPI_ISL_2278024, EPI_ISL_2278026, EPI_ISL_2278027, EPI_ISL_2278031, EPI_ISL_2278036, EPI_ISL_2278038, EPI_ISL_2278041, EPI_ISL_2278044, EPI_ISL_2278046, EPI_ISL_2278048, EPI_ISL_2278051, EPI_ISL_2278052, EPI_ISL_2278053, EPI_ISL_2278054, EPI_ISL_2278055, EPI_ISL_2278061, EPI_ISL_2278064, EPI_ISL_2278065, EPI_ISL_2278067, EPI_ISL_2278073, EPI_ISL_2278081, EPI_ISL_2278082, EPI_ISL_2278083, EPI_ISL_2278085, EPI_ISL_2278088, EPI_ISL_2278089, EPI_ISL_2278094, EPI_ISL_2278098, EPI_ISL_2278099, EPI_ISL_2278101, EPI_ISL_2278102, EPI_ISL_2278107, EPI_ISL_2278109, EPI_ISL_2278120, EPI_ISL_2278121, EPI_ISL_2278125, EPI_ISL_2278127, EPI_ISL_2278144, EPI_ISL_2278157, EPI_ISL_2278163, EPI_ISL_2278173 | | | |
| see above | Lighthouse Laboratory Plymouth | Wellcome Sanger Institute for the COVID-19 Genomics UK (COG-UK) Consortium | Lighthouse Laboratory Plymouth and Alex Alderton, Roberto Amato, Jeffrey Barrett, Sonia Goncalves, Ewan Harrison, David K. Jackson, Ian Johnston, Dominic Kwiatkowski, Cordelia Langford, John Sillitoe on behalf of the Wellcome Sanger Institute COVID-19 Surveillance Team |
| EPI_ISL_2279192, EPI_ISL_2279263 | Wisconsin State Laboratory of Hygiene Communicable Disease Division | Wisconsin State Laboratory of Hygiene Communicable Disease Division | Abigail C. Shockey, Alicia J. Mooney, Erika M. Hanson, Tonya Danz, Richard Griesser, Sara Wagner, Kelsey R. Florek |
| EPI_ISL_2279524, EPI_ISL_2279552 | URMC LABS | Wadsworth Center, New York State Department of Health | Kirsten St. George, Daryl M. Lamson, Alexis Russell, Matthew Shudt, Melissa A Leisner, Jonathan Plitnick, Catharine Prussing, Navjot Singh, John Kelly, Erasmus Schneider, Erica Lasek-Nesselquist |
| EPI_ISL_2280307, EPI_ISL_2280313, EPI_ISL_2280383, EPI_ISL_2280386, EPI_ISL_2280418, EPI_ISL_2280435, EPI_ISL_2280441, EPI_ISL_2280466, EPI_ISL_2280469, EPI_ISL_2280481, EPI_ISL_2280488, EPI_ISL_2280550, EPI_ISL_2280568, EPI_ISL_2280590, EPI_ISL_2280613, EPI_ISL_2280633, EPI_ISL_2280634, EPI_ISL_2280703, EPI_ISL_2280723, EPI_ISL_2280735, EPI_ISL_2280749, EPI_ISL_2280750, EPI_ISL_2280764, EPI_ISL_2280860, EPI_ISL_2280877, EPI_ISL_2281098, EPI_ISL_2281190 | | | |
| see above | Aegis Sciences Corporation | Centers for Disease Control and Prevention Division of Viral Diseases, Pathogen Discovery | Dakota Howard, Dhwani Batra, Peter W. Cook, Kara Moser, Adrian Paskey, Jason Caravas, Benjamin Rambo-Martin, Shatavia Morrison, Christopher Gulvick, Scott Sammons, Yvette Unoarumhi, Darlene Wagner, Matthew Schmerer, Cyndi Clark, Patrick Campbell, Rob Case, Vikramsinha Ghorpade, Holly Houdeshell, Ola Kvalvaag, Dillon Nall, Ethan Sanders, Alec Vest, Shaun Westlund, Matthew Hardison, Clinton R. Paden, Duncan MacCannell |
| EPI_ISL_2281283, EPI_ISL_2281295, EPI_ISL_2281368, EPI_ISL_2281391, EPI_ISL_2281401 | Microbiology Department, Laboratori Clínic Metropolitana Nord. Hospital Universitari Germans Trias i Pujol | Can Ruti SARS-CoV-2 Sequencing Hub (HUGTiP/IrsiCaixa/IGTP) | Marc Noguera-Julian, Pilar Armengol, Ignacio Blanco, Antoni E Bordoy, Francesc Catala-Moll, Pere-Joan Cardona, Maria Casadellà, Cristina Casañ, Gemma Clara, Bonaventura Clotet, Cristina Esteban, Montserrat Giménez, Mercedes Guerrero, Anna Not, Roger Paredes, Mariona Parera, Verónica Saludes, Alba Sánchez, and Elisa Martró on behalf of the Can Ruti SARS-CoV-2 Sequencing Hub. |
| EPI_ISL_2281845 | NJDOH, Public Health and Environmental Laboratories | NJ_PHEL | Lindsey Bodnar, Shiv K. Verma, Jacquelyn Deverell, Ryan Pachucki, Dana Woell, Byeong Jeong |
| EPI_ISL_2281907 | Istituto Zooprofilattico Sperimentale del Mezzogiorno | TIGEM | Antonio Grimaldi Patrizia Annunziata Francesco Panariello Biancamaria Pierri Claudia Tiberio Teresa Giuliano Valentina Bouche Chiara Colantuono Maria Concetta Cuomo Denise Di Concilio Lucio Di Filippo Anna Manfredi Marcello Salvi Antonio Limone Luigi Atripaldi Pellegrino Cerino Andrea Ballabio Davide Cacchiarelli |
| EPI_ISL_2282038, EPI_ISL_2282040 | NJDOH, Public Health and Environmental Laboratories | NJ_PHEL | Lindsey Bodnar, Shiv K. Verma, Jacquelyn Deverell, Matthew Scarnati, Dana Woell, Byeong Jeong |
| EPI_ISL_2282452, EPI_ISL_2282534, EPI_ISL_2282565, EPI_ISL_2282836 | Infinity Biologix | Centers for Disease Control and Prevention Division of Viral Diseases, Pathogen Discovery | Dakota Howard, Dhwani Batra, Peter W. Cook, Kara Moser, Adrian Paskey, Jason Caravas, Benjamin Rambo-Martin, Shatavia Morrison, Christopher Gulvick, Scott Sammons, Yvette Unoarumhi, Darlene Wagner, Matthew Schmerer, Christian Bixby, Yihe Wang, Jonathan Schultz, Chirayu Goswami, Russ Hager, Robin Grimwood, Clinton R. Paden, Duncan MacCannell |
| EPI_ISL_2283860 | Aegis Sciences Corporation | Centers for Disease Control and Prevention Division of Viral Diseases, Pathogen Discovery | Dakota Howard, Dhwani Batra, Peter W. Cook, Kara Moser, Adrian Paskey, Jason Caravas, Benjamin Rambo-Martin, Shatavia Morrison, Christopher Gulvick, Scott Sammons, Yvette Unoarumhi, Darlene Wagner, Matthew Schmerer, Cyndi Clark, Patrick Campbell, Rob Case, Vikramsinha Ghorpade, Holly Houdeshell, Ola Kvalvaag, Dillon Nall, Ethan Sanders, Alec Vest, Shaun Westlund, Matthew Hardison, Clinton R. Paden, Duncan MacCannell |
| EPI_ISL_2284223, EPI_ISL_2284263, EPI_ISL_2284264, EPI_ISL_2284332, EPI_ISL_2284347, EPI_ISL_2284369, EPI_ISL_2284378, EPI_ISL_2284388 | National Virus Reference Laboratory | National Virus Reference Laboratory | Fiona Crispie, Calum Walsh, Matthew McCabe, Zoe Yandle, Charlene Bennet, Gabriel Gonzalez, Michael Carr, Jonathan Dean, Paul Cotter, Cillian F De Gascun |
| EPI_ISL_2284693, EPI_ISL_2284699 | Utah Public Health Laboratory | Utah Public Health Laboratory | Erin L. Young, Kelly F. Oakeson, Tara Gallagher |
| EPI_ISL_2284863 | Wichita State University - Molecular Diagnostics Lab | Kansas Health and Environmental Lab | Mike Grose, Katherine Wiggins, Jonathan Barnell, Ben Olsen, and Phil Adam |
| EPI_ISL_2284885, EPI_ISL_2284886, EPI_ISL_2284887, EPI_ISL_2284888, EPI_ISL_2284889, EPI_ISL_2284890, EPI_ISL_2284891, EPI_ISL_2284892, EPI_ISL_2284893, EPI_ISL_2284894 | Institute of Epidemiology, Disease Control and Research (IEDCR) | IEDCR-ideSHi-icddr,b | Hassan Afrad, Sadia Rahman, Manjur Hossain Khan, Firdausi Qadri, Tahmina Shirin |
| EPI_ISL_2284911, EPI_ISL_2284917, EPI_ISL_2284918, EPI_ISL_2284919, EPI_ISL_2284972 | Hospital Universitari Vall d'Hebron - Vall d'Hebron Institut de Recerca | Hospital Universitari Vall d'Hebron - Vall d'Hebron Institut de Recerca | Cristina Andrés, Maria Piñana, Alejandra González-Sánchez, Damir Garcia-Cehic, Ariadna Rando, Juliana Esperalba, Maria Gema Codina, Carla Castillo, Maria Carmen Martin, Tomàs Pumarola, Josep Quer, Andrés Antón |
| EPI_ISL_2285004, EPI_ISL_2285023, EPI_ISL_2285039, EPI_ISL_2285071, EPI_ISL_2285073 | Mako Medical | Centers for Disease Control and Prevention Division of Viral Diseases, Pathogen Discovery | Dakota Howard, Dhwani Batra, Peter W. Cook, Kara Moser, Adrian Paskey, Jason Caravas, Benjamin Rambo-Martin, Shatavia Morrison, Christopher Gulvick, Scott Sammons, Yvette Unoarumhi, Darlene Wagner, Matthew Schmerer,Matthew Tugwell, Lauren Moon, Clinton R. Paden, Duncan MacCannell |
| EPI_ISL_2285121 | Microbiology Department. Complexo Hospitalario Universitario de Vigo | Microbiology Department. Complexo Hospitalario Universitario de Vigo | Alfaya N, Alonso I, Alvarez M, Cabrera JJ, Carballo R, Cores O, Cortizo S, del-Campo V, Martinez L, Mediero G, Perez S, Potel C, Regueiro B, Rey S, Vasallo FJ |
| EPI_ISL_2285157, EPI_ISL_2285158, EPI_ISL_2285159, EPI_ISL_2285160, EPI_ISL_2285161 | VITHAS Lab | Microbiology Department. Complexo Hospitalario Universitario de Vigo | Camacho AT, Perez S, Regueiro B |
| EPI_ISL_2285774 | Wyoming Public Health Laboratory | Wyoming Public Health Laboratory | Jim Mildenberger, Taylor Fearing, Channing Weber, Ashley Norberg, Chayse Rowley, Marley Goetz, Brian Dominguez, Elliot Thomasson, Brittany Oher, Sam Britz, Cari Sloma, and Rob Christensen |
| EPI_ISL_2285862, EPI_ISL_2285863 | National Influenza Centre | National Influenza Centre | William K. Ampofo, Michael Marks, Ivy A. Asante, Sharon Hsu, Benjamiin B. Lindsey, Benjamin H. Foulkes, Mildred Adusei-Poku, Linda Boatemaa, Lorreta Kwasah, Joseph Oliver-Commey, Ernest Asiedu, Franklin Asiedu-Bekoe, Gordon Awandare, Joyce Ngoi, Dennis Laryea, Mathew D. Parker, Thushan I de Silva, |
| EPI_ISL_2286253, EPI_ISL_2286257 | MEPHI, Aix Marseille University | MEPHI, Aix Marseille University | Anthony LEVASSEUR |
| EPI_ISL_2287856, EPI_ISL_2287859 | New Mexico Department of Health Scientific Laboratory | New Mexico Department of Health Scientific Laboratory | Ellie Johnson, Anastacia Griego-Fisher, D'eldra Malone, Jennifer Benoit |
| EPI_ISL_2288947, EPI_ISL_2288948, EPI_ISL_2288973 | FUNDACIO ALTHAIA-MANRESA | Banc de Sang i Teixits | Francisco Vidal, Irene Corrales, Nina Borràs, Carlos Hobeich, Natália Comes, Lorena Ramírez, Noemí Gonzalez, Silvia Sauleda, Maria Glòria Soria |

| | | | |
|---|---|---|---|
| EPI_ISL_2288982 | BST | Banc de Sang i Teixits | Francisco Vidal, Irene Corrales, Nina Borràs, Carlos Hobeich, Natália Comes, Lorena Ramírez, Noemí Gonzalez, Silvia Sauleda, Maria Glòria Soria |
| EPI_ISL_2289002 | University of Liège COVID-19 testing center | GIGA Medical Genomics | Keith Durkin, Maria Artesi, Bouchra Boujemla, Nathalie Renotte, Cécile Meex, Sébastien Bontems, Fabrice Bureau, Laurent Gillet, Wouter Coppieters, Marie-Pierre Hayette, Vincent Bours |
| EPI_ISL_2289248, EPI_ISL_2289249, EPI_ISL_2289250 | LBM UNIBIO VALENTIN | CNR Virus des Infections Respiratoires - France SUD | Antonin Bal, Gregory Destras, Gwendolyne Burfin, Hadrien Regue, Quentin Semanas, Martine Valette, Bruno Lina, Laurence Josset |
| EPI_ISL_2289352 | MIRIALIS CLUSES BECHET | CNR Virus des Infections Respiratoires - France SUD | Antonin Bal, Gregory Destras, Gwendolyne Burfin, Hadrien Regue, Quentin Semanas, Martine Valette, Bruno Lina, Laurence Josset |
| EPI_ISL_2289683 | CERBALLIANCE LAURENT DU VAR | CNR Virus des Infections Respiratoires - France SUD | Antonin Bal, Gregory Destras, Gwendolyne Burfin, Hadrien Regue, Quentin Semanas, Martine Valette, Bruno Lina, Laurence Josset |
| EPI_ISL_2290622 | Uniwersyteckie Centrum Kliniczne | 1. National Institute of Public Health - National Institute of Hygiene, Warsaw, Poland 2. Biobank Lab, University of Lodz 3. Laboratory of Respiratory Viruses, Teaching and Clinical Center of the Medical University of Lodz | Dominik Strapagiel, Marta Sobalska-Kwapis, Klaudyna Królikowska, Jakub Lach, Marcin Słomka, Tomasz Płoszaj, Magdalena Traczyk-Borszyńska, Izabela Dróżdż, Maciej Borowiec, Katarzyna Zacharczuk, Magdalena Nowakowska, Małgorzata Sadkowska-Todys, Tomasz Wołkowicz |
| EPI_ISL_2290665, EPI_ISL_2290666 | Invicta | 1. National Institute of Public Health - National Institute of Hygiene, Warsaw, Poland 2. Biobank Lab, University of Lodz 3. Laboratory of Respiratory Viruses, Teaching and Clinical Center of the Medical University of Lodz | Dominik Strapagiel, Marta Sobalska-Kwapis, Klaudyna Królikowska, Jakub Lach, Marcin Słomka, Tomasz Płoszaj, Magdalena Traczyk-Borszyńska, Izabela Dróżdż, Maciej Borowiec, Katarzyna Zacharczuk, Magdalena Nowakowska, Małgorzata Sadkowska-Todys, Tomasz Wołkowicz |
| EPI_ISL_2290768, EPI_ISL_2290769 | MSHS Clinical Microbiology Laboratories | MSHS Pathogen Surveillance Program | Ana S. Gonzalez-Reiche, Hala Alshammary, Mitchell J. Sullivan, Brianne Ciferri, Ajay Obla, Angela Amoako, Mahmoud Awawda, Daniel Floda, Julia Matthews, Ashley Salimbangon, Levy Sominsky, Katherine Beach, Kayla Russo, Charles Gleason, Shelcie Fabre, Giulio Kleiner, Zenab Khan, Bremy Alburquerque, Adriana van de Guchte, Komal Srivastava, Matthew M. Hernandez, Jayeeta Dutta, Denise Jurczyszak, Nancy Francoeur, Betsaida Salom Melo, Irina Oussenko, Gintaras Deikus, Juan Soto, Shwetha Hara Sridhar, Ying-Chih Wang, Kathryn Twyman, Deena R. Altman, Robert Sebra, Adolfo García-Sastre, Marta Luksza, Gopi Patel, Sarah Schaefer, Melissa Gitman, Michael D. Nowak, Alberto Paniz-Mondolfi, Emilia Mia Sordillo, Viviana Simon, Harm van Bakel |
| EPI_ISL_2290872, EPI_ISL_2291743, EPI_ISL_2291745, EPI_ISL_2291749, EPI_ISL_2291802, EPI_ISL_2292130, EPI_ISL_2292131, EPI_ISL_2292137, EPI_ISL_2292170, EPI_ISL_2292173, EPI_ISL_2292394, EPI_ISL_2292414, EPI_ISL_2292490, EPI_ISL_2292558, EPI_ISL_2292612 | | | |
| see above | Utah Public Health Laboratory | Utah Public Health Laboratory | Erin L. Young, Kelly F. Oakeson, Tara Gallagher |
| EPI_ISL_2292722, EPI_ISL_2292727 | Platform BIS UZA/UAntwerpen | Labo Klinische Biologie, UZA | Jasmine Coppens, Marie Le Mercier, Basil Britto Xavier, Christine Lammens, Veerle Matheeussen, Herman Goossens |
| EPI_ISL_2292738, EPI_ISL_2292902, EPI_ISL_2292937, EPI_ISL_2292941, EPI_ISL_2292972 | Utah Public Health Laboratory | Utah Public Health Laboratory | Erin L. Young, Kelly F. Oakeson, Tara Gallagher |
| EPI_ISL_2293245 | Heilig hart Lier | Imelda Ziekenhuis | Johan Frans, Hanne Valgaeren, Dagmar Obbels |
| EPI_ISL_2293281 | Hospital | National Reference Center for Viruses of Respiratory Infections, Institut Pasteur, Paris | Marion Barbet, Sylvie Behillil, Méline Bizard, Angela Brisebarre, Camille Capel, Vincent Enouf, Louise Lefrançois, Frédéric Lemoine, Christophe Malabat, Corinne Maufrais, Pierre Lechat, Etienne Simon-Lorière, Maud Vanpeene, Sylvie Van der Werf ,HéLèNe Roger |
| EPI_ISL_2293284 | Hospital | National Reference Center for Viruses of Respiratory Infections, Institut Pasteur, Paris | Marion Barbet, Sylvie Behillil, Méline Bizard, Angela Brisebarre, Camille Capel, Vincent Enouf, Louise Lefrançois, Frédéric Lemoine, Christophe Malabat, Corinne Maufrais, Pierre Lechat, Etienne Simon-Lorière, Maud Vanpeene, Sylvie Van der Werf ,Laura Djamdjian |
| EPI_ISL_2293285 | Hospital | National Reference Center for Viruses of Respiratory Infections, Institut Pasteur, Paris | Marion Barbet, Sylvie Behillil, Méline Bizard, Angela Brisebarre, Camille Capel, Vincent Enouf, Louise Lefrançois, Frédéric Lemoine, Christophe Malabat, Corinne Maufrais, Etienne Simon-Lorière, Maud Vanpeene, Sylvie Van der Werf ,Laura Djamdjian |
| EPI_ISL_2293303 | Hospital | National Reference Center for Viruses of Respiratory Infections, Institut Pasteur, Paris | Marion Barbet, Sylvie Behillil, Méline Bizard, Angela Brisebarre, Camille Capel, Vincent Enouf, Louise Lefrançois, Frédéric Lemoine, Christophe Malabat, Corinne Maufrais, Pierre Lechat, Etienne Simon-Lorière, Maud Vanpeene, Sylvie Van der Werf ,Elodie Gleize |
| EPI_ISL_2293304, EPI_ISL_2293305, EPI_ISL_2293306 | Hospital | National Reference Center for Viruses of Respiratory Infections, Institut Pasteur, Paris | Marion Barbet, Sylvie Behillil, Méline Bizard, Angela Brisebarre, Camille Capel, Vincent Enouf, Louise Lefrançois, Frédéric Lemoine, Christophe Malabat, Corinne Maufrais, Etienne Simon-Lorière, Maud Vanpeene, Sylvie Van der Werf ,Elodie Gleize |
| EPI_ISL_2293307 | Hospital | National Reference Center for Viruses of Respiratory Infections, Institut Pasteur, Paris | Marion Barbet, Sylvie Behillil, Méline Bizard, Angela Brisebarre, Camille Capel, Vincent Enouf, Louise Lefrançois, Frédéric Lemoine, Christophe Malabat, Corinne Maufrais, Pierre Lechat, Etienne Simon-Lorière, Maud Vanpeene, Sylvie Van der Werf ,Elodie Gleize |
| EPI_ISL_2293308, EPI_ISL_2293309 | Hospital | National Reference Center for Viruses of Respiratory Infections, Institut Pasteur, Paris | Marion Barbet, Sylvie Behillil, Méline Bizard, Angela Brisebarre, Camille Capel, Vincent Enouf, Louise Lefrançois, Frédéric Lemoine, Christophe Malabat, Corinne Maufrais, Etienne Simon-Lorière, Maud Vanpeene, Sylvie Van der Werf ,Elodie Gleize |
| EPI_ISL_2295285 | OK Public Health Laboratory, Oklahoma State DOH | Genomics and Discovery, Respiratory Viruses Branch, Division of Viral Diseases, Centers for Disease Control and Prevention | Jing Zhang, Anna Kelleher, Ying Tao, Yan Li, Brian Lynch, Krista Queen, Anna Uehara, Peter Cook, Han Jia Justin Ng, Rachel Marine, Clinton R. Paden, Dhwani Batra, Haibin Wang, Tara Coalter, Jasmine Padilla, Morgan Davis, Mili Sheth, Sarah Nobles, Mark Burroughs, Justin Lee, Adam Retchless, Suxiang Tong |
| EPI_ISL_2296394, EPI_ISL_2296447 | Yale Clinical Virology Lab | Grubaugh Lab - Yale School of Public Health | Joseph Fauver, Mallery Breban, Isabel Ott, Tara Alpert, Mary Petrone, Anderson Brito, Chantal Vogels, Annie Watkins, Chaney Kalinich, Jessica Rothman, Marie L. Landry, Nathan Grubaugh |
| EPI_ISL_2296521 | Premier Medical Lab | Minnesota Department of Health, Public Health Laboratory | Alexandra Lorentz, Jacob Garfin, Matt Plumb, and Xiong Wang |
| EPI_ISL_2296691 | Lighthouse Lab in Milton Keynes | Wellcome Sanger Institute for the COVID-19 Genomics UK (COG-UK) Consortium | The Lighthouse Lab in Milton Keynes and Alex Alderton, Roberto Amato, Jeffrey Barrett, Sonia Goncalves, Ewan Harrison, David K. Jackson, Ian Johnston, Dominic Kwiatkowski, Cordelia Langford, John Sillitoe on behalf of the Wellcome Sanger Institute COVID-19 Surveillance Team |
| EPI_ISL_2296694 | Lighthouse Lab in Alderley Park | Wellcome Sanger Institute for the COVID-19 Genomics UK (COG-UK) Consortium | Jacquelyn Wynn, Mairead Hyland, The Lighthouse Lab in Alderley Park and Alex Alderton, Roberto Amato, Jeffrey Barrett, Sonia Goncalves, Ewan Harrison, David K. Jackson, Ian Johnston, Dominic Kwiatkowski, Cordelia Langford, John Sillitoe on behalf of the Wellcome Sanger Institute COVID-19 Surveillance Team |
| EPI_ISL_2296695 | Lighthouse Lab in Milton Keynes | Wellcome Sanger Institute for the COVID-19 Genomics UK (COG-UK) Consortium | The Lighthouse Lab in Milton Keynes and Alex Alderton, Roberto Amato, Jeffrey Barrett, Sonia Goncalves, Ewan Harrison, David K. Jackson, Ian Johnston, Dominic Kwiatkowski, Cordelia Langford, John Sillitoe on behalf of the Wellcome Sanger Institute COVID-19 Surveillance Team |
| EPI_ISL_2296700 | Lighthouse Lab in Alderley Park | Wellcome Sanger Institute for the COVID-19 Genomics UK (COG-UK) Consortium | Jacquelyn Wynn, Mairead Hyland, The Lighthouse Lab in Alderley Park and Alex Alderton, Roberto Amato, Jeffrey Barrett, Sonia Goncalves, Ewan Harrison, David K. Jackson, Ian Johnston, Dominic Kwiatkowski, Cordelia Langford, John Sillitoe on behalf of the Wellcome Sanger Institute COVID-19 Surveillance Team |
| EPI_ISL_2296701 | Lighthouse Lab in Milton Keynes | Wellcome Sanger Institute for the COVID-19 Genomics UK (COG-UK) Consortium | The Lighthouse Lab in Milton Keynes and Alex Alderton, Roberto Amato, Jeffrey Barrett, Sonia Goncalves, Ewan Harrison, David K. Jackson, Ian Johnston, Dominic Kwiatkowski, Cordelia Langford, John Sillitoe on behalf of the Wellcome Sanger Institute COVID-19 Surveillance Team |
| EPI_ISL_2296702 | Lighthouse Lab in Alderley Park | Wellcome Sanger Institute for the COVID-19 Genomics UK (COG-UK) Consortium | Jacquelyn Wynn, Mairead Hyland, The Lighthouse Lab in Alderley Park and Alex Alderton, Roberto Amato, Jeffrey Barrett, Sonia Goncalves, Ewan Harrison, David K. Jackson, Ian Johnston, Dominic Kwiatkowski, Cordelia Langford, John Sillitoe on behalf of the Wellcome Sanger Institute COVID-19 Surveillance Team |
| EPI_ISL_2296704 | Lighthouse Lab in Glasgow | Wellcome Sanger Institute for the COVID-19 Genomics UK (COG-UK) Consortium | Harper VanSteenhouse, Yumi Kasai, David Gray, Carol Clugston, Anna Dominiczak and Alex Alderton, Roberto Amato, Jeffrey Barrett, Sonia Goncalves, Ewan Harrison, David K. Jackson, Ian Johnston, Dominic Kwiatkowski, Cordelia Langford, John Sillitoe on behalf of the Wellcome Sanger Institute COVID-19 Surveillance Team |
| EPI_ISL_2296705, EPI_ISL_2296706 | Hospital General Universitario Gregorio Marañón | Hospital General Universitario Gregorio Marañón | Sergio Buenestado Serrano, Pedro Sola Campoy, Laura Pérez-Lago, Cristina Rodriguez-Grande, Pilar Catalán, Patricia Muñoz, Darío García de Viedma |
| EPI_ISL_2296745, EPI_ISL_2296934, EPI_ISL_2297070, EPI_ISL_2297089, EPI_ISL_2297212, EPI_ISL_2298360, EPI_ISL_2298552, EPI_ISL_2298630, EPI_ISL_2298846, EPI_ISL_2299149, EPI_ISL_2299162 | | | |
| see above | Department of Virus and Microbiological Special Diagnostics, Statens Serum Institut, Copenhagen, Denmark | Aalborg University | Danish Covid-19 Genome Consortium |
| EPI_ISL_2299323, EPI_ISL_2299371, EPI_ISL_2299395, EPI_ISL_2299425, EPI_ISL_2299513 | UW Virology Lab | UW Virology Lab | Pavitra Roychoudhury, Hong Xie, Lasata Shrestha, Tien V. Nguyen, Shah Mohamed Bakhash, Michelle Lin, Noah R. Baker, Sean Ellis, Meei-Li Huang, Keith R Jerome, Alexander Greninger |
| EPI_ISL_2299834, EPI_ISL_2299973, EPI_ISL_2299977, EPI_ISL_2299991 | Department of Virus and Microbiological Special Diagnostics, Statens Serum Institut, Copenhagen, Denmark | Aalborg University | Danish Covid-19 Genome Consortium |
| EPI_ISL_2300398, EPI_ISL_2300529, EPI_ISL_2300590, EPI_ISL_2300597 | Illinois Department of Public Health | Illinois Department of Public Health - Chicago Lab | Vineet K. Dhiman, Ira Heimler, Joel Price |
| EPI_ISL_2300683, EPI_ISL_2300806, EPI_ISL_2300839 | Department of Virus and Microbiological Special Diagnostics, Statens Serum Institut, Copenhagen, Denmark | Aalborg University | Danish Covid-19 Genome Consortium |
| EPI_ISL_2301382 | UCK | 1. Tricity SARS-CoV-2 sequencing consortium: University of Gdansk, Medical University of Gdansk, Vaxican Ltd., Invicta Ltd. 2. National Institute of Public Health - National Institute of Hygiene, Warsaw, Poland | Maciej Kosinski, Celina Cybulska, Krystyna Bienkowska Szewczyk, Maciej Grzybek, Karolina Gackowska, Marcin Lubocki, Katarzyna Groth, Lukasz Rabalski, Katarzyna Zacharczuk, Magdalena Nowakowska, Małgorzata Sadkowska-Todys, Tomasz Wołkowicz |
| EPI_ISL_2301430 | National Platform bis UMONS/Jolimont | National Platform bis UMONS/Jolimont | François Dufrasne, Guillaume Bayon-Vicente, Florian Juszczak, Gautier Detry, Ruddy Wattiez |
| EPI_ISL_2303934, EPI_ISL_2303935, EPI_ISL_2303972, EPI_ISL_2303973, EPI_ISL_2303974, EPI_ISL_2303975, EPI_ISL_2303976, EPI_ISL_2303977, EPI_ISL_2303978, EPI_ISL_2303980, EPI_ISL_2303981, EPI_ISL_2303982, EPI_ISL_2303983, EPI_ISL_2303984 | | | |
| see above | Dutch COVID-19 response team | National Institute for Public Health and the Environment (RIVM) | Adam Meijer, Harry Vennema, Dirk Eggink, Jeroen Cremer, Sharon van den Brink, Bas van der Veer, AnneMarie van den Brandt, Lisa Wijsman, Kim Freriks, Ryanne Jaarsma, Euníce Then, Lynn Aarts, Sanne Bos, Melissa van Tuil, Linda van de Nes, Sjoerd Kuiling, James Groot, Florian Zwagemaker, Dennis Schmitz, Annelies Kroneman, Karim Hajji, Chantal Reusken, on behalf of the national COVID-19 response team |

| | | | |
|---|---|---|---|
| EPI_ISL_2304047, EPI_ISL_2304050, EPI_ISL_2304057, EPI_ISL_2304058, EPI_ISL_2304062, EPI_ISL_2304073, EPI_ISL_2304079, EPI_ISL_2304082, EPI_ISL_2304085 | National Centre For Cell Science | National Centre For Cell Science – INSACOG | Dhiraj Paul, Mohak P Gujare, Shivang P. Bhanushali, Mitali Inamdar, Manoj Kumar Bhat, Ajay Pillai, INSACOG Consortium team, Yogesh Shouche |
| EPI_ISL_2304086, EPI_ISL_2304087, EPI_ISL_2304088, EPI_ISL_2304089, EPI_ISL_2304092, EPI_ISL_2304093, EPI_ISL_2304094, EPI_ISL_2304095, EPI_ISL_2304096, EPI_ISL_2304097, EPI_ISL_2304098, EPI_ISL_2304100, EPI_ISL_2304101, EPI_ISL_2304108, EPI_ISL_2304109, EPI_ISL_2304110, EPI_ISL_2304112, EPI_ISL_2304113, EPI_ISL_2304114, EPI_ISL_2304116, EPI_ISL_2304117, EPI_ISL_2304118, EPI_ISL_2304120, EPI_ISL_2304121, EPI_ISL_2304124, EPI_ISL_2304125, EPI_ISL_2304126, EPI_ISL_2304128 | | | |
| see above | Armed Forces Medical College Pune | National Centre For Cell Science – INSACOG | Dhiraj Paul, Mohak P Gujare, Shivang P. Bhanushali, Mitali Inamdar, Manoj Kumar Bhat, Ajay Pillai, INSACOG Consortium team, Yogesh Shouche |
| EPI_ISL_2304141, EPI_ISL_2304142, EPI_ISL_2304143 | National Platform bis UMONS/Jolimont | National Platform bis UMONS/Jolimont | François Dufrasne, Guillaume Bayon-Vicente, Florian Juszczak, Gautier Detry, Ruddy Wattiez |
| EPI_ISL_2304211, EPI_ISL_2304240, EPI_ISL_2304258 | Missouri State Public Health Laboratory | Missouri State Public Health Laboratory | Matthew Sinn, Joshua Barry, Ashley New |
| EPI_ISL_2304324, EPI_ISL_2304327 | Alameda County Public Health Department | Alameda County Public Health Department | Kristina Hsieh, Maria Elma, Kevin Libuit |
| EPI_ISL_2304357, EPI_ISL_2304405, EPI_ISL_2304415, EPI_ISL_2304444, EPI_ISL_2304453, EPI_ISL_2304456 | Indiana Animal Disease Diagnostic Laboratory | Carpi Laboratory - Purdue University | Jack Dorman, Ilinca I Ciubotariu, Nicole M Perry, Jobin J Kattoor, Lev Gorenstein, Abebe A Fola, G Kenitra Hendrix, Rebecca P Wilkes, Giovanna Carpi |
| EPI_ISL_2304461, EPI_ISL_2304462, EPI_ISL_2304463, EPI_ISL_2304465 | Institute of Epidemiology, Disease Control and Research (IEDCR) | IEDCR-ideSHi-icddr,b | Hassan Afrad, Sadia Rahman, Manjur Hossain Khan, Firdausi Qadri, Tahmina Shirin |
| EPI_ISL_2304503, EPI_ISL_2304539, EPI_ISL_2304553, EPI_ISL_2304586, EPI_ISL_2304589, EPI_ISL_2304642, EPI_ISL_2304688 | Quest Diagnostics Incorporated | Centers for Disease Control and Prevention Division of Viral Diseases, Pathogen Discovery | Dakota Howard, Dhwani Batra, Peter W. Cook, Kara Moser, Adrian Paskey, Jason Caravas, Benjamin Rambo-Martin, Shatavia Morrison, Christopher Gulvick, Scott Sammons, Yvette Unoarumhi, Darlene Wagner, Matthew Schmerer, S. H. Rosenthal, A. Gerasimova, R. M. Kagan, B. Anderson, M. Hua, Y. Liu, L.E. Bernstein, K.E. Livingston, A. Perez, I. A. Shlyakhter, R. V. Rolando, R. Owen, P. Tanpaiboon, F. Lacbawan, Clinton R. Paden, Duncan MacCannell |
| EPI_ISL_2304947, EPI_ISL_2304971, EPI_ISL_2305001, EPI_ISL_2305002, EPI_ISL_2305106, EPI_ISL_2305108, EPI_ISL_2305131, EPI_ISL_2305139, EPI_ISL_2305173, EPI_ISL_2305203, EPI_ISL_2305210, EPI_ISL_2305221, EPI_ISL_2305235, EPI_ISL_2305283, EPI_ISL_2305308, EPI_ISL_2305370, EPI_ISL_2305371, EPI_ISL_2305445, EPI_ISL_2305488, EPI_ISL_2305497, EPI_ISL_2305531, EPI_ISL_2305543, EPI_ISL_2305590, EPI_ISL_2305596, EPI_ISL_2305611, EPI_ISL_2305629, EPI_ISL_2305630, EPI_ISL_2305762, EPI_ISL_2306019, EPI_ISL_2306061, EPI_ISL_2306122, EPI_ISL_2306159, EPI_ISL_2306160, EPI_ISL_2306180, EPI_ISL_2306181, EPI_ISL_2306182, EPI_ISL_2306190, EPI_ISL_2306264, EPI_ISL_2306281, EPI_ISL_2306304, EPI_ISL_2306323, EPI_ISL_2306330, EPI_ISL_2306338, EPI_ISL_2306395, EPI_ISL_2306428, EPI_ISL_2306457, EPI_ISL_2306511, EPI_ISL_2306521, EPI_ISL_2306601, EPI_ISL_2306605, EPI_ISL_2306627, EPI_ISL_2306632, EPI_ISL_2306709, EPI_ISL_2306773, EPI_ISL_2306819, EPI_ISL_2306894, EPI_ISL_2306914, EPI_ISL_2306920, EPI_ISL_2307000, EPI_ISL_2307031, EPI_ISL_2307070, EPI_ISL_2307079 | | | |
| see above | Laboratory Corporation of America | Centers for Disease Control and Prevention Division of Viral Diseases, Pathogen Discovery | Dakota Howard, Dhwani Batra, Peter W. Cook, Kara Moser, Adrian Paskey, Jason Caravas, Benjamin Rambo-Martin, Shatavia Morrison, Christopher Gulvick, Scott Sammons, Yvette Unoarumhi, Darlene Wagner, Matthew Schmerer, Minoo Agarwal, Eyad Almasri, Debbie Boles, Ayla Burns, Nuthawin Charoensri, Oren Cohen, Susan Countryman, Mary Ann Cristobal, Bobbi Croy, Suzanne Dale, Hrushikesh Deshmukh, Amanda Douglas, Vincent Drouillon, Marcia Eisenberg, Howard Engler, Rama Ghatti, Prashant Gupta, Susan Hicks, Jake Humphrey, Lax Iyer, Manoj Jain, Mohan Kolli, Brian Krueger, Tim Kuphal, Stanley Letovsky, Michael Levandoski, Craig Lukasik, Jonathan Meltzer, Brian Norvell, Mindy Nye, Scott Parker, Christos Petropoulos, John Pruitt, Steven Ragan, Scott Ryan, Mike Sapeta, Jana Schroth, Suresh Babu Selvaraju, Goran Stevovic, Amanda Suchanek, Andrea Throop, Lyndon Tilson, Thomas Urban, Joe Voshell, Kimberly Wagner, Jonathan Williams, Mary Williamson, Qian Zeng, Tricia Zwiefelhofer, Clinton R. Paden, Duncan MacCannell |
| EPI_ISL_2307102, EPI_ISL_2307104, EPI_ISL_2307105, EPI_ISL_2307107, EPI_ISL_2307108, EPI_ISL_2307109, EPI_ISL_2307110, EPI_ISL_2307111, EPI_ISL_2307112, EPI_ISL_2307113, EPI_ISL_2307114, EPI_ISL_2307115, EPI_ISL_2307116, EPI_ISL_2307117, EPI_ISL_2307118, EPI_ISL_2307119, EPI_ISL_2307120, EPI_ISL_2307121, EPI_ISL_2307122, EPI_ISL_2307123, EPI_ISL_2307124, EPI_ISL_2307125, EPI_ISL_2307126, EPI_ISL_2307127, EPI_ISL_2307128, EPI_ISL_2307129, EPI_ISL_2307130, EPI_ISL_2307131, EPI_ISL_2307132, EPI_ISL_2307133, EPI_ISL_2307134, EPI_ISL_2307135, EPI_ISL_2307136 | | | |
| see above | State Virus Research and Diagnostic Laboratory (VRDL), AIIMS Raipur | State Virus Research and Diagnostic Laboratory (VRDL), AIIMS Raipur | Pushpendra Singh, Kuldeep Sharma, Priyanka Singh, Somya Sharma, MD Rafiiqullah Khan, Jiyan Chandrawanshi, Sanjay Singh Negi, Anudita Bhargava |
| EPI_ISL_2307175, EPI_ISL_2307195, EPI_ISL_2307271 | Mako Medical | Centers for Disease Control and Prevention Division of Viral Diseases, Pathogen Discovery | Dakota Howard, Dhwani Batra, Peter W. Cook, Kara Moser, Adrian Paskey, Jason Caravas, Benjamin Rambo-Martin, Shatavia Morrison, Christopher Gulvick, Scott Sammons, Yvette Unoarumhi, Darlene Wagner, Matthew Schmerer,Matthew Tugwell, Lauren Moon, Clinton R. Paden, Duncan MacCannell |
| EPI_ISL_2307592, EPI_ISL_2307656 | Viollier AG | Department of Biosystems Science and Engineering, ETH Zürich | Chaoran Chen, Sarah Nadeau, Catharine Aquino, Ivan Topolsky, Philipp Jablonski, Lara Fuhrmann, David Dreifuss, Katharina Jahn, Daniel Ehrsam, Isabel Stürmer, Andreia Cabral de Gouvea, Maria Domenica Moccia, Simon Grüter, Timothy Sykes, Lennart Opitz, Griffin White, Laura Neff, Doris Popovic, Andrea Patrignani, Jay Tracy, Ralph Schlapbach, Christiane Beckmann, Maurice Redondo, Olivier Kobel, Christoph Noppen, Sophie Seidel, Noemie Santamaria de Souza, Niko Beerenwinkel, Tanja Stadler |
| EPI_ISL_2307932, EPI_ISL_2308031 | Viollier AG | Department of Biosystems Science and Engineering, ETH Zürich | Christian Beisel, Sarah Nadeau, Chaoran Chen, Ivan Topolsky, Philipp Jablonski, Lara Fuhrmann, David Dreifuss, Katharina Jahn, Rebecca Denes, Mirjam Feldkamp, Ina Nissen, Natascha Santacroce, Elodie Burcklen, Christiane Beckmann, Maurice Redondo, Olivier Kobel, Christoph Noppen, Sophie Seidel, Noemie Santamaria de Souza, Niko Beerenwinkel, Tanja Stadler |
| EPI_ISL_2308061 | Viollier AG | Department of Biosystems Science and Engineering, ETH Zürich | Chaoran Chen, Sarah Nadeau, Catharine Aquino, Ivan Topolsky, Philipp Jablonski, Lara Fuhrmann, David Dreifuss, Katharina Jahn, Daniel Ehrsam, Isabel Stürmer, Andreia Cabral de Gouvea, Maria Domenica Moccia, Simon Grüter, Timothy Sykes, Lennart Opitz, Griffin White, Laura Neff, Doris Popovic, Andrea Patrignani, Jay Tracy, Ralph Schlapbach, Christiane Beckmann, Maurice Redondo, Olivier Kobel, Christoph Noppen, Sophie Seidel, Noemie Santamaria de Souza, Niko Beerenwinkel, Tanja Stadler |
| EPI_ISL_2308249 | National Institute of Infectious Diseases-Prof. Dr. Matei Bals Molecular Diagnostics Laboratory | National Institute of Infectious Diseases-Prof. Dr. Matei Bals Molecular Diagnostics Laboratory | Corina Casangiu, Leontina Banica, Marius Surleac, Ovidiu Vlaicu, Andreea Tudor, Simona Paraschiv, Dan Otelea |
| EPI_ISL_2308273 | Indian Council of Medical Research-National Institute of Virology, Microbial Containment Complex | Indian Council of Medical Research-National Institute of Virology, Microbial Containment Complex | Pragya D Yadav |
| EPI_ISL_2308278 | Laboratori de Referencia de Catalunya | Laboratori de Referencia de Catalunya | Ramirez A., Bellosillo B., Padilla E., Vilas A., Hernandez JJ., Canal M. |
| EPI_ISL_2308534, EPI_ISL_2308539 | Nebraska Public Health Laboratory | NPHL COVID-19 Response Team | NPHL COVID-19 Response Team |
| EPI_ISL_2309138, EPI_ISL_2309139, EPI_ISL_2309413, EPI_ISL_2309414, EPI_ISL_2309914, EPI_ISL_2309986, EPI_ISL_2310076, EPI_ISL_2310077, EPI_ISL_2310169, EPI_ISL_2310174, EPI_ISL_2310175, EPI_ISL_2310292, EPI_ISL_2310423, EPI_ISL_2310438, EPI_ISL_2310440, EPI_ISL_2310485, EPI_ISL_2310490, EPI_ISL_2310491, EPI_ISL_2310524, EPI_ISL_2310526, EPI_ISL_2310645, EPI_ISL_2310748, EPI_ISL_2310749, EPI_ISL_2310750, EPI_ISL_2311063, EPI_ISL_2311064, EPI_ISL_2311065, EPI_ISL_2311066, EPI_ISL_2311067, EPI_ISL_2311123 | | | |
| see above | Colorado Department of Public Health and Environment | Colorado Department of Public Health and Environment | Laura Bankers, Molly C. Hetherington-Rauth, Diana Ir, Alexandria Rossheim, Shannon R. Matzinger, Sarah Elizabeth Totten, Emily A. Travanty |
| EPI_ISL_2311763 | GA Department of Public Health Laboratory | Genomics and Discovery, Respiratory Viruses Branch, Division of Viral Diseases, Centers for Disease Control and Prevention | Jing Zhang, Anna Kelleher, Ying Tao, Yan Li, Brian Lynch, Krista Queen, Anna Uehara, Peter Cook, Han Jia Justin Ng, Rachel Marine, Clinton R. Paden, Dhwani Batra, Haibin Wang, Tara Coalter, Jasmine Padilla, Morgan Davis, Mili Sheth, Sarah Nobles, Mark Burroughs, Justin Lee, Adam Retchless, Suxiang Tong |
| EPI_ISL_2311769 | OK Public Health Laboratory, Oklahoma State DOH | Genomics and Discovery, Respiratory Viruses Branch, Division of Viral Diseases, Centers for Disease Control and Prevention | Jing Zhang, Anna Kelleher, Ying Tao, Yan Li, Brian Lynch, Krista Queen, Anna Uehara, Peter Cook, Han Jia Justin Ng, Rachel Marine, Clinton R. Paden, Dhwani Batra, Haibin Wang, Tara Coalter, Jasmine Padilla, Morgan Davis, Mili Sheth, Sarah Nobles, Mark Burroughs, Justin Lee, Adam Retchless, Suxiang Tong |
| EPI_ISL_2311861 | Fundação Ezequiel Dias | Coordenação Geral de Laboratórios de Saúde Pública (CGLAB/DAEVS/SVS/MS) | Vagner Fonseca, et al. |
| EPI_ISL_2311890, EPI_ISL_2311898, EPI_ISL_2311908, EPI_ISL_2311913, EPI_ISL_2311938, EPI_ISL_2311970, EPI_ISL_2311975, EPI_ISL_2311995, EPI_ISL_2312020, EPI_ISL_2312060, EPI_ISL_2312066, EPI_ISL_2312080, EPI_ISL_2312101, EPI_ISL_2312105, EPI_ISL_2312151, EPI_ISL_2312223, EPI_ISL_2312225, EPI_ISL_2312231, EPI_ISL_2312242, EPI_ISL_2312252, EPI_ISL_2312258, EPI_ISL_2312266, EPI_ISL_2312273, EPI_ISL_2312278, EPI_ISL_2312284, EPI_ISL_2312348, EPI_ISL_2312358, EPI_ISL_2312368, EPI_ISL_2312380, EPI_ISL_2312414, EPI_ISL_2312415, EPI_ISL_2312420 | | | |
| see above | Broad Institute Clinical Research Sequencing Platform | Infectious Disease Program, Broad Institute of Harvard and MIT | Siddle,K.J., Adams,G., Pearlman,L., Gladden-Young,A., Vicente,G., Blumenstiel,B., DeFelice,M., Lee,M., McGovern,S., Lagerborg,K., Rudy,M., DeRuff,K., Carter,A., Normandin,E., Bauer,M., Reilly,S., Tomkins-Tinch,C., Loreth,C., Chaluvadi,S., Meldrim,J., Granger,B., Lemieux,J.E., Birren,B.W., Sabeti,P.C., Larkin,K., Dodge,S., Lennon,N., Madoff,L., Brown,C., Gallagher,G., Smole,S., Park,D.J., Gabriel,S., and MacInnis,B.L. |
| EPI_ISL_2312655 | Laboratorio Unico Provinciale Dipartimento Biotecnologie, Trasfusionale e di Laboratorio | U.O. Microbiologia Laboratorio Unico Centro Servizi AUSL della Romagna | Sara Ghisellini, Michela Boni, Giorgio Dirani, Silvia Zannoli, Francesca Taddei, Giulia Gatti, Ilaria Poggianti, Vittorio Sambri |
| EPI_ISL_2313086 | Department of Virology | Department of Virology | Massab Umair, Aamer Ikram, Muhammad Salman, Muhammad Ammar, Adnan Haider |
| EPI_ISL_2313095, EPI_ISL_2313096 | Houston Health Department, Disease Prevention and Control | Houston Health Department, Disease Prevention and Control | Penn,R., Brown,P., Lara,A., Lai,Y. |
| EPI_ISL_2313142 | Limbach - MVZ Labor Eveld & Kollegen Essen | Robert Koch Institute | unknown |
| EPI_ISL_2313176, EPI_ISL_2313224 | MVZ Labor Dr. Limbach & Kollegen GbR | Robert Koch Institute | unknown |
| EPI_ISL_2313256, EPI_ISL_2313281 | MVZ Dr. Eberhard & Partner Dortmund | Robert Koch Institute | unknown |
| EPI_ISL_2313284 | Laborarztpraxis Osnabrück | Robert Koch Institute | unknown |
| EPI_ISL_2313294, EPI_ISL_2313316, | MVZ Dr. Eberhard & Partner Dortmund | Robert Koch Institute | unknown |

| | | | |
|---|---|---|---|
| EPI_ISL_2313335 | | | |
| EPI_ISL_2313341, EPI_ISL_2313343, EPI_ISL_2313345, EPI_ISL_2313346, EPI_ISL_2313347 | Robert Koch-Institut ZBS1 | Robert Koch Institute | Annika Brinkmann |
| EPI_ISL_2313392 | Sonic - Labor Staber Heilbronn | Robert Koch Institute | unknown |
| EPI_ISL_2313446, EPI_ISL_2313507, EPI_ISL_2313650, EPI_ISL_2313798, EPI_ISL_2313822 | Labor 28 MVZ GmbH | Robert Koch Institute | unknown |
| EPI_ISL_2313845 | Bioscientia Labor Wermsdorf | Robert Koch Institute | unknown |
| EPI_ISL_2313909 | Landesuntersuchungsanstalt für das Gesundheits- und Veterinärwesen (LUA) Sachsen | Robert Koch Institute | unknown |
| EPI_ISL_2313935 | LABOR CHEMNITZ – Zentrum für Diagnostik am Klinikum Chemnitz | Robert Koch Institute | unknown |
| EPI_ISL_2313955 | Städtisches Klinikum Dresden Institut für Klinische Chemie und Laboratoriumsmedizin | Robert Koch Institute | unknown |
| EPI_ISL_2313991, EPI_ISL_2314020, EPI_ISL_2314031 | Labormedizin Darmstadt | Robert Koch Institute | unknown |
| EPI_ISL_2314038 | MVZ Labor Dr. Limbach & Kollegen GbR | Robert Koch Institute | unknown |
| EPI_ISL_2314081 | Universitätsklinikum Frankfurt - Institut für Medizinische Virologie | Robert Koch Institute | unknown |
| EPI_ISL_2314097 | Bioscientia MVZ Labor Karlsruhe GmbH | Robert Koch Institute | unknown |
| EPI_ISL_2314169 | Universitätsklinikum Frankfurt - Institut für Medizinische Virologie | Robert Koch Institute | unknown |
| EPI_ISL_2314250, EPI_ISL_2314283, EPI_ISL_2314284, EPI_ISL_2314288 | Bioscientia MVZ Labor Karlsruhe GmbH | Robert Koch Institute | unknown |
| EPI_ISL_2314308 | Sonic - Labor Dr. von Froreich GmbH | Robert Koch Institute | unknown |
| EPI_ISL_2314310 | Sonic - Labor Lademannbogen MVZ GmbH | Robert Koch Institute | unknown |
| EPI_ISL_2314355, EPI_ISL_2314377 | Eurofins LifeCodexx GmbH | Robert Koch Institute | unknown |
| EPI_ISL_2314589 | SYNLAB MVZ Leinfelden-Echterdingen | Robert Koch Institute | unknown |
| EPI_ISL_2314667, EPI_ISL_2314669, EPI_ISL_2314671 | Labor Mönchengladbach MVZ Dr. Stein + Kollegen GbR | Robert Koch Institute | unknown |
| EPI_ISL_2314690 | MVZ Medizinisches Labor Hannover GmbH | Robert Koch Institute | unknown |
| EPI_ISL_2314761 | MDI Limbach Berlin GmbH; MVZ Labor Berlin | Robert Koch Institute | unknown |
| EPI_ISL_2314768 | MVZ Medizinisches Labor Hannover GmbH | Robert Koch Institute | unknown |
| EPI_ISL_2314851 | Landesgesundheitsamt Baden-Württemberg | Robert Koch Institute | unknown |
| EPI_ISL_2314888, EPI_ISL_2315009 | Diagnosticum - Labor Neukirchen | Robert Koch Institute | unknown |
| EPI_ISL_2315110 | SYNLAB Labor München Zentrum LMZ | Robert Koch Institute | unknown |
| EPI_ISL_2315525 | MVZ Labor Dr. Quade & Kollegen GmbH | Robert Koch Institute | unknown |
| EPI_ISL_2315614, EPI_ISL_2315623, EPI_ISL_2315628, EPI_ISL_2315635, EPI_ISL_2315646 | Eurofins LifeCodexx GmbH | Robert Koch Institute | unknown |
| EPI_ISL_2315667 | Universitätsklinikum Bonn - Institut für Virologie | Robert Koch Institute | unknown |
| EPI_ISL_2315816 | Bayerisches Landesamt für Gesundheit und Lebensmittelsicherheit (LGL) | Robert Koch Institute | unknown |
| EPI_ISL_2316279 | Lighthouse Lab in Alderley Park | Wellcome Sanger Institute for the COVID-19 Genomics UK (COG-UK) Consortium | Jacquelyn Wynn, Mairead Hyland, The Lighthouse Lab in Alderley Park and Alex Alderton, Roberto Amato, Jeffrey Barrett, Sonia Goncalves, Ewan Harrison, David K. Jackson, Ian Johnston, Dominic Kwiatkowski, Cordelia Langford, John Sillitoe on behalf of the Wellcome Sanger Institute COVID-19 Surveillance Team |
| EPI_ISL_2316281 | Health Services Laboratories | Wellcome Sanger Institute for the COVID-19 Genomics UK (COG-UK) Consortium | Health Services Laboratories and Alex Alderton, Roberto Amato, Jeffrey Barrett, Sonia Goncalves, Ewan Harrison, David K. Jackson, Ian Johnston, Dominic Kwiatkowski, Cordelia Langford, John Sillitoe on behalf of the Wellcome Sanger Institute COVID-19 Surveillance Team |
| EPI_ISL_2316284, EPI_ISL_2316285, EPI_ISL_2316287, EPI_ISL_2316288 | Lighthouse Lab in Milton Keynes | Wellcome Sanger Institute for the COVID-19 Genomics UK (COG-UK) Consortium | The Lighthouse Lab in Milton Keynes and Alex Alderton, Roberto Amato, Jeffrey Barrett, Sonia Goncalves, Ewan Harrison, David K. Jackson, Ian Johnston, Dominic Kwiatkowski, Cordelia Langford, John Sillitoe on behalf of the Wellcome Sanger Institute COVID-19 Surveillance Team |
| EPI_ISL_2316290 | Lighthouse Lab in Alderley Park | Wellcome Sanger Institute for the COVID-19 Genomics UK (COG-UK) Consortium | Jacquelyn Wynn, Mairead Hyland, The Lighthouse Lab in Alderley Park and Alex Alderton, Roberto Amato, Jeffrey Barrett, Sonia Goncalves, Ewan Harrison, David K. Jackson, Ian Johnston, Dominic Kwiatkowski, Cordelia Langford, John Sillitoe on behalf of the Wellcome Sanger Institute COVID-19 Surveillance Team |
| EPI_ISL_2316293 | Health Services Laboratories | Wellcome Sanger Institute for the COVID-19 Genomics UK (COG-UK) Consortium | Health Services Laboratories and Alex Alderton, Roberto Amato, Jeffrey Barrett, Sonia Goncalves, Ewan Harrison, David K. Jackson, Ian Johnston, Dominic Kwiatkowski, Cordelia Langford, John Sillitoe on behalf of the Wellcome Sanger Institute COVID-19 Surveillance Team |
| EPI_ISL_2316294, EPI_ISL_2316295 | Lighthouse Lab in Alderley Park | Wellcome Sanger Institute for the COVID-19 Genomics UK (COG-UK) Consortium | Jacquelyn Wynn, Mairead Hyland, The Lighthouse Lab in Alderley Park and Alex Alderton, Roberto Amato, Jeffrey Barrett, Sonia Goncalves, Ewan Harrison, David K. Jackson, Ian Johnston, Dominic Kwiatkowski, Cordelia Langford, John Sillitoe on behalf of the Wellcome Sanger Institute COVID-19 Surveillance Team |
| EPI_ISL_2316297 | Health Services Laboratories | Wellcome Sanger Institute for the COVID-19 Genomics UK (COG-UK) Consortium | Health Services Laboratories and Alex Alderton, Roberto Amato, Jeffrey Barrett, Sonia Goncalves, Ewan Harrison, David K. Jackson, Ian Johnston, Dominic Kwiatkowski, Cordelia Langford, John Sillitoe on behalf of the Wellcome Sanger Institute COVID-19 Surveillance Team |
| EPI_ISL_2316298, EPI_ISL_2316300, EPI_ISL_2316304 | Lighthouse Lab in Alderley Park | Wellcome Sanger Institute for the COVID-19 Genomics UK (COG-UK) Consortium | Jacquelyn Wynn, Mairead Hyland, The Lighthouse Lab in Alderley Park and Alex Alderton, Roberto Amato, Jeffrey Barrett, Sonia Goncalves, Ewan Harrison, David K. Jackson, Ian Johnston, Dominic Kwiatkowski, Cordelia Langford, John Sillitoe on behalf of the Wellcome Sanger Institute COVID-19 Surveillance Team |
| EPI_ISL_2316306 | Health Services Laboratories | Wellcome Sanger Institute for the COVID-19 Genomics UK (COG-UK) Consortium | Health Services Laboratories and Alex Alderton, Roberto Amato, Jeffrey Barrett, Sonia Goncalves, Ewan Harrison, David K. Jackson, Ian Johnston, Dominic Kwiatkowski, Cordelia Langford, John Sillitoe on behalf of the Wellcome Sanger Institute COVID-19 Surveillance Team |
| EPI_ISL_2316307, EPI_ISL_2316309 | Lighthouse Lab in Alderley Park | Wellcome Sanger Institute for the COVID-19 Genomics UK (COG-UK) Consortium | Jacquelyn Wynn, Mairead Hyland, The Lighthouse Lab in Alderley Park and Alex Alderton, Roberto Amato, Jeffrey Barrett, Sonia Goncalves, Ewan Harrison, David K. Jackson, Ian Johnston, Dominic Kwiatkowski, Cordelia Langford, John Sillitoe on behalf of the Wellcome Sanger Institute COVID-19 Surveillance Team |
| EPI_ISL_2316313, EPI_ISL_2316314, EPI_ISL_2316316, EPI_ISL_2316317 | Lighthouse Lab in Milton Keynes | Wellcome Sanger Institute for the COVID-19 Genomics UK (COG-UK) Consortium | The Lighthouse Lab in Milton Keynes and Alex Alderton, Roberto Amato, Jeffrey Barrett, Sonia Goncalves, Ewan Harrison, David K. Jackson, Ian Johnston, Dominic Kwiatkowski, Cordelia Langford, John Sillitoe on behalf of the Wellcome Sanger Institute COVID-19 Surveillance Team |
| EPI_ISL_2316318 | Lighthouse Lab in Alderley Park | Wellcome Sanger Institute for the COVID-19 Genomics UK (COG-UK) Consortium | Jacquelyn Wynn, Mairead Hyland, The Lighthouse Lab in Alderley Park and Alex Alderton, Roberto Amato, Jeffrey Barrett, Sonia Goncalves, Ewan Harrison, David K. Jackson, Ian Johnston, Dominic Kwiatkowski, Cordelia Langford, John Sillitoe on behalf of the Wellcome Sanger Institute COVID-19 Surveillance Team |
| EPI_ISL_2316322 | Lighthouse Lab in Milton Keynes | Wellcome Sanger Institute for the COVID-19 Genomics UK (COG-UK) Consortium | The Lighthouse Lab in Milton Keynes and Alex Alderton, Roberto Amato, Jeffrey Barrett, Sonia Goncalves, Ewan Harrison, David K. Jackson, Ian Johnston, Dominic Kwiatkowski, Cordelia Langford, John Sillitoe on behalf of the Wellcome Sanger Institute COVID-19 Surveillance Team |
| EPI_ISL_2316323 | Lighthouse Lab in Alderley Park | Wellcome Sanger Institute for the COVID-19 Genomics UK (COG-UK) Consortium | Jacquelyn Wynn, Mairead Hyland, The Lighthouse Lab in Alderley Park and Alex Alderton, Roberto Amato, Jeffrey Barrett, Sonia Goncalves, Ewan Harrison, David K. Jackson, Ian Johnston, Dominic Kwiatkowski, Cordelia Langford, John Sillitoe on behalf of the Wellcome Sanger Institute COVID-19 Surveillance Team |
| EPI_ISL_2316325 | Lighthouse Lab in Milton Keynes | Wellcome Sanger Institute for the COVID-19 Genomics UK (COG-UK) Consortium | The Lighthouse Lab in Milton Keynes and Alex Alderton, Roberto Amato, Jeffrey Barrett, Sonia Goncalves, Ewan Harrison, David K. Jackson, Ian Johnston, Dominic Kwiatkowski, Cordelia Langford, John Sillitoe on behalf of the Wellcome Sanger Institute COVID-19 Surveillance Team |

| | | | |
|---|---|---|---|
| EPI_ISL_2316326 | Lighthouse Lab in Alderley Park | Wellcome Sanger Institute for the COVID-19 Genomics UK (COG-UK) Consortium | Jacquelyn Wynn, Mairead Hyland, The Lighthouse Lab in Alderley Park and Alex Alderton, Roberto Amato, Jeffrey Barrett, Sonia Goncalves, Ewan Harrison, David K. Jackson, Ian Johnston, Dominic Kwiatkowski, Cordelia Langford, John Sillitoe on behalf of the Wellcome Sanger Institute COVID-19 Surveillance Team |
| EPI_ISL_2316329 | Lighthouse Lab in Milton Keynes | Wellcome Sanger Institute for the COVID-19 Genomics UK (COG-UK) Consortium | The Lighthouse Lab in Milton Keynes and Alex Alderton, Roberto Amato, Jeffrey Barrett, Sonia Goncalves, Ewan Harrison, David K. Jackson, Ian Johnston, Dominic Kwiatkowski, Cordelia Langford, John Sillitoe on behalf of the Wellcome Sanger Institute COVID-19 Surveillance Team |
| EPI_ISL_2316330 | Lighthouse Lab in Alderley Park | Wellcome Sanger Institute for the COVID-19 Genomics UK (COG-UK) Consortium | Jacquelyn Wynn, Mairead Hyland, The Lighthouse Lab in Alderley Park and Alex Alderton, Roberto Amato, Jeffrey Barrett, Sonia Goncalves, Ewan Harrison, David K. Jackson, Ian Johnston, Dominic Kwiatkowski, Cordelia Langford, John Sillitoe on behalf of the Wellcome Sanger Institute COVID-19 Surveillance Team |
| EPI_ISL_2316332 | Lighthouse Lab in Milton Keynes | Wellcome Sanger Institute for the COVID-19 Genomics UK (COG-UK) Consortium | The Lighthouse Lab in Milton Keynes and Alex Alderton, Roberto Amato, Jeffrey Barrett, Sonia Goncalves, Ewan Harrison, David K. Jackson, Ian Johnston, Dominic Kwiatkowski, Cordelia Langford, John Sillitoe on behalf of the Wellcome Sanger Institute COVID-19 Surveillance Team |
| EPI_ISL_2316333, EPI_ISL_2316335 | Lighthouse Lab in Alderley Park | Wellcome Sanger Institute for the COVID-19 Genomics UK (COG-UK) Consortium | Jacquelyn Wynn, Mairead Hyland, The Lighthouse Lab in Alderley Park and Alex Alderton, Roberto Amato, Jeffrey Barrett, Sonia Goncalves, Ewan Harrison, David K. Jackson, Ian Johnston, Dominic Kwiatkowski, Cordelia Langford, John Sillitoe on behalf of the Wellcome Sanger Institute COVID-19 Surveillance Team |
| EPI_ISL_2316336 | Health Services Laboratories | Wellcome Sanger Institute for the COVID-19 Genomics UK (COG-UK) Consortium | Health Services Laboratories and Alex Alderton, Roberto Amato, Jeffrey Barrett, Sonia Goncalves, Ewan Harrison, David K. Jackson, Ian Johnston, Dominic Kwiatkowski, Cordelia Langford, John Sillitoe on behalf of the Wellcome Sanger Institute COVID-19 Surveillance Team |
| EPI_ISL_2316340 | Lighthouse Lab in Milton Keynes | Wellcome Sanger Institute for the COVID-19 Genomics UK (COG-UK) Consortium | The Lighthouse Lab in Milton Keynes and Alex Alderton, Roberto Amato, Jeffrey Barrett, Sonia Goncalves, Ewan Harrison, David K. Jackson, Ian Johnston, Dominic Kwiatkowski, Cordelia Langford, John Sillitoe on behalf of the Wellcome Sanger Institute COVID-19 Surveillance Team |
| EPI_ISL_2316342, EPI_ISL_2316345 | Lighthouse Lab in Alderley Park | Wellcome Sanger Institute for the COVID-19 Genomics UK (COG-UK) Consortium | Jacquelyn Wynn, Mairead Hyland, The Lighthouse Lab in Alderley Park and Alex Alderton, Roberto Amato, Jeffrey Barrett, Sonia Goncalves, Ewan Harrison, David K. Jackson, Ian Johnston, Dominic Kwiatkowski, Cordelia Langford, John Sillitoe on behalf of the Wellcome Sanger Institute COVID-19 Surveillance Team |
| EPI_ISL_2316346 | Lighthouse Lab in Milton Keynes | Wellcome Sanger Institute for the COVID-19 Genomics UK (COG-UK) Consortium | The Lighthouse Lab in Milton Keynes and Alex Alderton, Roberto Amato, Jeffrey Barrett, Sonia Goncalves, Ewan Harrison, David K. Jackson, Ian Johnston, Dominic Kwiatkowski, Cordelia Langford, John Sillitoe on behalf of the Wellcome Sanger Institute COVID-19 Surveillance Team |
| EPI_ISL_2316348 | Labor für medizinische Mikrobiologie Bakteriologie Mykologie Virologie & Infektionsserologie - Labor Mölbis | Robert Koch Institute | unknown |
| EPI_ISL_2316353 | Lighthouse Lab in Alderley Park | Wellcome Sanger Institute for the COVID-19 Genomics UK (COG-UK) Consortium | Jacquelyn Wynn, Mairead Hyland, The Lighthouse Lab in Alderley Park and Alex Alderton, Roberto Amato, Jeffrey Barrett, Sonia Goncalves, Ewan Harrison, David K. Jackson, Ian Johnston, Dominic Kwiatkowski, Cordelia Langford, John Sillitoe on behalf of the Wellcome Sanger Institute COVID-19 Surveillance Team |
| EPI_ISL_2316354 | Lighthouse Lab in Milton Keynes | Wellcome Sanger Institute for the COVID-19 Genomics UK (COG-UK) Consortium | The Lighthouse Lab in Milton Keynes and Alex Alderton, Roberto Amato, Jeffrey Barrett, Sonia Goncalves, Ewan Harrison, David K. Jackson, Ian Johnston, Dominic Kwiatkowski, Cordelia Langford, John Sillitoe on behalf of the Wellcome Sanger Institute COVID-19 Surveillance Team |
| EPI_ISL_2316359 | Health Services Laboratories | Wellcome Sanger Institute for the COVID-19 Genomics UK (COG-UK) Consortium | Health Services Laboratories and Alex Alderton, Roberto Amato, Jeffrey Barrett, Sonia Goncalves, Ewan Harrison, David K. Jackson, Ian Johnston, Dominic Kwiatkowski, Cordelia Langford, John Sillitoe on behalf of the Wellcome Sanger Institute COVID-19 Surveillance Team |
| EPI_ISL_2316361, EPI_ISL_2316362 | Lighthouse Lab in Alderley Park | Wellcome Sanger Institute for the COVID-19 Genomics UK (COG-UK) Consortium | Jacquelyn Wynn, Mairead Hyland, The Lighthouse Lab in Alderley Park and Alex Alderton, Roberto Amato, Jeffrey Barrett, Sonia Goncalves, Ewan Harrison, David K. Jackson, Ian Johnston, Dominic Kwiatkowski, Cordelia Langford, John Sillitoe on behalf of the Wellcome Sanger Institute COVID-19 Surveillance Team |
| EPI_ISL_2316365 | Health Services Laboratories | Wellcome Sanger Institute for the COVID-19 Genomics UK (COG-UK) Consortium | Health Services Laboratories and Alex Alderton, Roberto Amato, Jeffrey Barrett, Sonia Goncalves, Ewan Harrison, David K. Jackson, Ian Johnston, Dominic Kwiatkowski, Cordelia Langford, John Sillitoe on behalf of the Wellcome Sanger Institute COVID-19 Surveillance Team |
| EPI_ISL_2316366, EPI_ISL_2316369, EPI_ISL_2316371, EPI_ISL_2316372, EPI_ISL_2316373, EPI_ISL_2316375, EPI_ISL_2316376, EPI_ISL_2316378 | Lighthouse Lab in Alderley Park | Wellcome Sanger Institute for the COVID-19 Genomics UK (COG-UK) Consortium | Jacquelyn Wynn, Mairead Hyland, The Lighthouse Lab in Alderley Park and Alex Alderton, Roberto Amato, Jeffrey Barrett, Sonia Goncalves, Ewan Harrison, David K. Jackson, Ian Johnston, Dominic Kwiatkowski, Cordelia Langford, John Sillitoe on behalf of the Wellcome Sanger Institute COVID-19 Surveillance Team |
| EPI_ISL_2316379 | Lighthouse Lab in Milton Keynes | Wellcome Sanger Institute for the COVID-19 Genomics UK (COG-UK) Consortium | The Lighthouse Lab in Milton Keynes and Alex Alderton, Roberto Amato, Jeffrey Barrett, Sonia Goncalves, Ewan Harrison, David K. Jackson, Ian Johnston, Dominic Kwiatkowski, Cordelia Langford, John Sillitoe on behalf of the Wellcome Sanger Institute COVID-19 Surveillance Team |
| EPI_ISL_2316381 | Lighthouse Lab in Alderley Park | Wellcome Sanger Institute for the COVID-19 Genomics UK (COG-UK) Consortium | Jacquelyn Wynn, Mairead Hyland, The Lighthouse Lab in Alderley Park and Alex Alderton, Roberto Amato, Jeffrey Barrett, Sonia Goncalves, Ewan Harrison, David K. Jackson, Ian Johnston, Dominic Kwiatkowski, Cordelia Langford, John Sillitoe on behalf of the Wellcome Sanger Institute COVID-19 Surveillance Team |
| EPI_ISL_2316384 | Health Services Laboratories | Wellcome Sanger Institute for the COVID-19 Genomics UK (COG-UK) Consortium | Health Services Laboratories and Alex Alderton, Roberto Amato, Jeffrey Barrett, Sonia Goncalves, Ewan Harrison, David K. Jackson, Ian Johnston, Dominic Kwiatkowski, Cordelia Langford, John Sillitoe on behalf of the Wellcome Sanger Institute COVID-19 Surveillance Team |
| EPI_ISL_2316387, EPI_ISL_2316389, EPI_ISL_2316392, EPI_ISL_2316393 | Lighthouse Lab in Alderley Park | Wellcome Sanger Institute for the COVID-19 Genomics UK (COG-UK) Consortium | Jacquelyn Wynn, Mairead Hyland, The Lighthouse Lab in Alderley Park and Alex Alderton, Roberto Amato, Jeffrey Barrett, Sonia Goncalves, Ewan Harrison, David K. Jackson, Ian Johnston, Dominic Kwiatkowski, Cordelia Langford, John Sillitoe on behalf of the Wellcome Sanger Institute COVID-19 Surveillance Team |
| EPI_ISL_2316395 | Lighthouse Lab in Milton Keynes | Wellcome Sanger Institute for the COVID-19 Genomics UK (COG-UK) Consortium | The Lighthouse Lab in Milton Keynes and Alex Alderton, Roberto Amato, Jeffrey Barrett, Sonia Goncalves, Ewan Harrison, David K. Jackson, Ian Johnston, Dominic Kwiatkowski, Cordelia Langford, John Sillitoe on behalf of the Wellcome Sanger Institute COVID-19 Surveillance Team |
| EPI_ISL_2316397 | Eurofins MVZ Labor Gelsenkirchen | Robert Koch Institute | unknown |
| EPI_ISL_2316398 | Lighthouse Lab in Milton Keynes | Wellcome Sanger Institute for the COVID-19 Genomics UK (COG-UK) Consortium | The Lighthouse Lab in Milton Keynes and Alex Alderton, Roberto Amato, Jeffrey Barrett, Sonia Goncalves, Ewan Harrison, David K. Jackson, Ian Johnston, Dominic Kwiatkowski, Cordelia Langford, John Sillitoe on behalf of the Wellcome Sanger Institute COVID-19 Surveillance Team |
| EPI_ISL_2316399 | Eurofins MVZ Labor Gelsenkirchen | Robert Koch Institute | unknown |
| EPI_ISL_2316400 | Lighthouse Lab in Milton Keynes | Wellcome Sanger Institute for the COVID-19 Genomics UK (COG-UK) Consortium | The Lighthouse Lab in Milton Keynes and Alex Alderton, Roberto Amato, Jeffrey Barrett, Sonia Goncalves, Ewan Harrison, David K. Jackson, Ian Johnston, Dominic Kwiatkowski, Cordelia Langford, John Sillitoe on behalf of the Wellcome Sanger Institute COVID-19 Surveillance Team |
| EPI_ISL_2316401 | Lighthouse Lab in Alderley Park | Wellcome Sanger Institute for the COVID-19 Genomics UK (COG-UK) Consortium | Jacquelyn Wynn, Mairead Hyland, The Lighthouse Lab in Alderley Park and Alex Alderton, Roberto Amato, Jeffrey Barrett, Sonia Goncalves, Ewan Harrison, David K. Jackson, Ian Johnston, Dominic Kwiatkowski, Cordelia Langford, John Sillitoe on behalf of the Wellcome Sanger Institute COVID-19 Surveillance Team |
| EPI_ISL_2316403 | Lighthouse Lab in Milton Keynes | Wellcome Sanger Institute for the COVID-19 Genomics UK (COG-UK) Consortium | The Lighthouse Lab in Milton Keynes and Alex Alderton, Roberto Amato, Jeffrey Barrett, Sonia Goncalves, Ewan Harrison, David K. Jackson, Ian Johnston, Dominic Kwiatkowski, Cordelia Langford, John Sillitoe on behalf of the Wellcome Sanger Institute COVID-19 Surveillance Team |
| EPI_ISL_2316404, EPI_ISL_2316405 | Lighthouse Lab in Alderley Park | Wellcome Sanger Institute for the COVID-19 Genomics UK (COG-UK) Consortium | Jacquelyn Wynn, Mairead Hyland, The Lighthouse Lab in Alderley Park and Alex Alderton, Roberto Amato, Jeffrey Barrett, Sonia Goncalves, Ewan Harrison, David K. Jackson, Ian Johnston, Dominic Kwiatkowski, Cordelia Langford, John Sillitoe on behalf of the Wellcome Sanger Institute COVID-19 Surveillance Team |
| EPI_ISL_2316406 | Eurofins MVZ Labor Gelsenkirchen | Robert Koch Institute | unknown |
| EPI_ISL_2316407 | Lighthouse Lab in Alderley Park | Wellcome Sanger Institute for the COVID-19 Genomics UK (COG-UK) Consortium | Jacquelyn Wynn, Mairead Hyland, The Lighthouse Lab in Alderley Park and Alex Alderton, Roberto Amato, Jeffrey Barrett, Sonia Goncalves, Ewan Harrison, David K. Jackson, Ian Johnston, Dominic Kwiatkowski, Cordelia Langford, John Sillitoe on behalf of the Wellcome Sanger Institute COVID-19 Surveillance Team |
| EPI_ISL_2316408 | Lighthouse Lab in Milton Keynes | Wellcome Sanger Institute for the COVID-19 Genomics UK (COG-UK) Consortium | The Lighthouse Lab in Milton Keynes and Alex Alderton, Roberto Amato, Jeffrey Barrett, Sonia Goncalves, Ewan Harrison, David K. Jackson, Ian Johnston, Dominic Kwiatkowski, Cordelia Langford, John Sillitoe on behalf of the Wellcome Sanger Institute COVID-19 Surveillance Team |
| EPI_ISL_2316411 | Health Services Laboratories | Wellcome Sanger Institute for the COVID-19 Genomics UK (COG-UK) Consortium | Health Services Laboratories and Alex Alderton, Roberto Amato, Jeffrey Barrett, Sonia Goncalves, Ewan Harrison, David K. Jackson, Ian Johnston, Dominic Kwiatkowski, Cordelia Langford, John Sillitoe on behalf of the Wellcome Sanger Institute COVID-19 Surveillance Team |
| EPI_ISL_2316412 | Eurofins MVZ Labor Gelsenkirchen | Robert Koch Institute | unknown |
| EPI_ISL_2316413 | Health Services Laboratories | Wellcome Sanger Institute for the COVID-19 Genomics UK (COG-UK) Consortium | Health Services Laboratories and Alex Alderton, Roberto Amato, Jeffrey Barrett, Sonia Goncalves, Ewan Harrison, David K. Jackson, Ian Johnston, Dominic Kwiatkowski, Cordelia Langford, John Sillitoe on behalf of the Wellcome Sanger Institute COVID-19 Surveillance Team |
| EPI_ISL_2316414, EPI_ISL_2316416, EPI_ISL_2316419 | Lighthouse Lab in Alderley Park | Wellcome Sanger Institute for the COVID-19 Genomics UK (COG-UK) Consortium | Jacquelyn Wynn, Mairead Hyland, The Lighthouse Lab in Alderley Park and Alex Alderton, Roberto Amato, Jeffrey Barrett, Sonia Goncalves, Ewan Harrison, David K. Jackson, Ian Johnston, Dominic Kwiatkowski, Cordelia Langford, John Sillitoe on behalf of the Wellcome Sanger Institute COVID-19 Surveillance Team |
| EPI_ISL_2316420 | Lighthouse Lab in Milton Keynes | Wellcome Sanger Institute for the COVID-19 Genomics UK (COG-UK) Consortium | The Lighthouse Lab in Milton Keynes and Alex Alderton, Roberto Amato, Jeffrey Barrett, Sonia Goncalves, Ewan Harrison, David K. Jackson, Ian Johnston, Dominic Kwiatkowski, Cordelia Langford, John Sillitoe on behalf of the Wellcome Sanger Institute COVID-19 Surveillance Team |
| EPI_ISL_2316422 | Lighthouse Lab in Alderley Park | Wellcome Sanger Institute for the COVID-19 Genomics UK (COG-UK) Consortium | Jacquelyn Wynn, Mairead Hyland, The Lighthouse Lab in Alderley Park and Alex Alderton, Roberto Amato, Jeffrey Barrett, Sonia Goncalves, Ewan Harrison, David K. Jackson, Ian Johnston, Dominic Kwiatkowski, Cordelia Langford, John Sillitoe on behalf of the Wellcome Sanger Institute COVID-19 Surveillance Team |
| EPI_ISL_2316423 | Lighthouse Lab in Milton Keynes | Wellcome Sanger Institute for the COVID-19 Genomics UK (COG-UK) Consortium | The Lighthouse Lab in Milton Keynes and Alex Alderton, Roberto Amato, Jeffrey Barrett, Sonia Goncalves, Ewan Harrison, David K. Jackson, Ian Johnston, Dominic Kwiatkowski, Cordelia Langford, John Sillitoe on behalf of the Wellcome Sanger Institute COVID-19 Surveillance Team |
| EPI_ISL_2316424 | Lighthouse Lab in Alderley Park | Wellcome Sanger Institute for the COVID-19 Genomics UK (COG-UK) Consortium | Jacquelyn Wynn, Mairead Hyland, The Lighthouse Lab in Alderley Park and Alex Alderton, Roberto Amato, Jeffrey Barrett, Sonia Goncalves, Ewan Harrison, David K. Jackson, Ian Johnston, Dominic Kwiatkowski, Cordelia Langford, John Sillitoe on behalf of the Wellcome Sanger Institute COVID-19 Surveillance Team |
| EPI_ISL_2316426 | Lighthouse Lab in Milton Keynes | Wellcome Sanger Institute for the COVID-19 Genomics UK (COG-UK) Consortium | The Lighthouse Lab in Milton Keynes and Alex Alderton, Roberto Amato, Jeffrey Barrett, Sonia Goncalves, Ewan Harrison, David K. Jackson, Ian Johnston, Dominic Kwiatkowski, Cordelia Langford, John Sillitoe on behalf of the Wellcome Sanger Institute COVID-19 Surveillance Team |
| EPI_ISL_2316427 | Lighthouse Lab in Alderley Park | Wellcome Sanger Institute for the COVID-19 Genomics UK (COG-UK) Consortium | Jacquelyn Wynn, Mairead Hyland, The Lighthouse Lab in Alderley Park and Alex Alderton, Roberto Amato, Jeffrey Barrett, Sonia Goncalves, Ewan Harrison, David K. Jackson, Ian Johnston, Dominic Kwiatkowski, Cordelia Langford, John Sillitoe on behalf of the Wellcome Sanger Institute COVID-19 Surveillance Team |
| EPI_ISL_2316428 | Lighthouse Lab in Milton Keynes | Wellcome Sanger Institute for the COVID-19 Genomics UK (COG-UK) Consortium | The Lighthouse Lab in Milton Keynes and Alex Alderton, Roberto Amato, Jeffrey Barrett, Sonia Goncalves, Ewan Harrison, David K. Jackson, Ian Johnston, Dominic Kwiatkowski, Cordelia Langford, John Sillitoe on behalf of the Wellcome Sanger Institute COVID-19 Surveillance Team |
| EPI_ISL_2316430 | Health Services Laboratories | Wellcome Sanger Institute for the COVID-19 Genomics UK (COG-UK) Consortium | Health Services Laboratories and Alex Alderton, Roberto Amato, Jeffrey Barrett, Sonia Goncalves, Ewan Harrison, David K. Jackson, Ian Johnston, Dominic Kwiatkowski, Cordelia Langford, John Sillitoe on behalf of the Wellcome Sanger Institute COVID-19 Surveillance Team |

| | | | |
|---|---|---|---|
| EPI_ISL_2316431, EPI_ISL_2316432 | Lighthouse Lab in Milton Keynes | Wellcome Sanger Institute for the COVID-19 Genomics UK (COG-UK) Consortium | The Lighthouse Lab in Milton Keynes and Alex Alderton, Roberto Amato, Jeffrey Barrett, Sonia Goncalves, Ewan Harrison, David K. Jackson, Ian Johnston, Dominic Kwiatkowski, Cordelia Langford, John Sillitoe on behalf of the Wellcome Sanger Institute COVID-19 Surveillance Team |
| EPI_ISL_2316433 | MVZ Medizinisches Labor Hannover GmbH | Robert Koch Institute | unknown |
| EPI_ISL_2316434, EPI_ISL_2316437, EPI_ISL_2316441 | Lighthouse Lab in Alderley Park | Wellcome Sanger Institute for the COVID-19 Genomics UK (COG-UK) Consortium | Jacquelyn Wynn, Mairead Hyland, The Lighthouse Lab in Alderley Park and Alex Alderton, Roberto Amato, Jeffrey Barrett, Sonia Goncalves, Ewan Harrison, David K. Jackson, Ian Johnston, Dominic Kwiatkowski, Cordelia Langford, John Sillitoe on behalf of the Wellcome Sanger Institute COVID-19 Surveillance Team |
| EPI_ISL_2316442, EPI_ISL_2316444 | Lighthouse Lab in Milton Keynes | Wellcome Sanger Institute for the COVID-19 Genomics UK (COG-UK) Consortium | The Lighthouse Lab in Milton Keynes and Alex Alderton, Roberto Amato, Jeffrey Barrett, Sonia Goncalves, Ewan Harrison, David K. Jackson, Ian Johnston, Dominic Kwiatkowski, Cordelia Langford, John Sillitoe on behalf of the Wellcome Sanger Institute COVID-19 Surveillance Team |
| EPI_ISL_2316445 | Lighthouse Lab in Alderley Park | Wellcome Sanger Institute for the COVID-19 Genomics UK (COG-UK) Consortium | Jacquelyn Wynn, Mairead Hyland, The Lighthouse Lab in Alderley Park and Alex Alderton, Roberto Amato, Jeffrey Barrett, Sonia Goncalves, Ewan Harrison, David K. Jackson, Ian Johnston, Dominic Kwiatkowski, Cordelia Langford, John Sillitoe on behalf of the Wellcome Sanger Institute COVID-19 Surveillance Team |
| EPI_ISL_2316447 | Lighthouse Lab in Milton Keynes | Wellcome Sanger Institute for the COVID-19 Genomics UK (COG-UK) Consortium | The Lighthouse Lab in Milton Keynes and Alex Alderton, Roberto Amato, Jeffrey Barrett, Sonia Goncalves, Ewan Harrison, David K. Jackson, Ian Johnston, Dominic Kwiatkowski, Cordelia Langford, John Sillitoe on behalf of the Wellcome Sanger Institute COVID-19 Surveillance Team |
| EPI_ISL_2316448 | Lighthouse Lab in Alderley Park | Wellcome Sanger Institute for the COVID-19 Genomics UK (COG-UK) Consortium | Jacquelyn Wynn, Mairead Hyland, The Lighthouse Lab in Alderley Park and Alex Alderton, Roberto Amato, Jeffrey Barrett, Sonia Goncalves, Ewan Harrison, David K. Jackson, Ian Johnston, Dominic Kwiatkowski, Cordelia Langford, John Sillitoe on behalf of the Wellcome Sanger Institute COVID-19 Surveillance Team |
| EPI_ISL_2316449 | MVZ Medizinisches Labor Hannover GmbH | Robert Koch Institute | unknown |
| EPI_ISL_2316450, EPI_ISL_2316451 | Lighthouse Lab in Milton Keynes | Wellcome Sanger Institute for the COVID-19 Genomics UK (COG-UK) Consortium | The Lighthouse Lab in Milton Keynes and Alex Alderton, Roberto Amato, Jeffrey Barrett, Sonia Goncalves, Ewan Harrison, David K. Jackson, Ian Johnston, Dominic Kwiatkowski, Cordelia Langford, John Sillitoe on behalf of the Wellcome Sanger Institute COVID-19 Surveillance Team |
| EPI_ISL_2316452, EPI_ISL_2316453 | Lighthouse Lab in Alderley Park | Wellcome Sanger Institute for the COVID-19 Genomics UK (COG-UK) Consortium | Jacquelyn Wynn, Mairead Hyland, The Lighthouse Lab in Alderley Park and Alex Alderton, Roberto Amato, Jeffrey Barrett, Sonia Goncalves, Ewan Harrison, David K. Jackson, Ian Johnston, Dominic Kwiatkowski, Cordelia Langford, John Sillitoe on behalf of the Wellcome Sanger Institute COVID-19 Surveillance Team |
| EPI_ISL_2316457 | Lighthouse Lab in Milton Keynes | Wellcome Sanger Institute for the COVID-19 Genomics UK (COG-UK) Consortium | The Lighthouse Lab in Milton Keynes and Alex Alderton, Roberto Amato, Jeffrey Barrett, Sonia Goncalves, Ewan Harrison, David K. Jackson, Ian Johnston, Dominic Kwiatkowski, Cordelia Langford, John Sillitoe on behalf of the Wellcome Sanger Institute COVID-19 Surveillance Team |
| EPI_ISL_2316458 | Lighthouse Lab in Alderley Park | Wellcome Sanger Institute for the COVID-19 Genomics UK (COG-UK) Consortium | Jacquelyn Wynn, Mairead Hyland, The Lighthouse Lab in Alderley Park and Alex Alderton, Roberto Amato, Jeffrey Barrett, Sonia Goncalves, Ewan Harrison, David K. Jackson, Ian Johnston, Dominic Kwiatkowski, Cordelia Langford, John Sillitoe on behalf of the Wellcome Sanger Institute COVID-19 Surveillance Team |
| EPI_ISL_2316459 | Health Services Laboratories | Wellcome Sanger Institute for the COVID-19 Genomics UK (COG-UK) Consortium | Health Services Laboratories and Alex Alderton, Roberto Amato, Jeffrey Barrett, Sonia Goncalves, Ewan Harrison, David K. Jackson, Ian Johnston, Dominic Kwiatkowski, Cordelia Langford, John Sillitoe on behalf of the Wellcome Sanger Institute COVID-19 Surveillance Team |
| EPI_ISL_2316461 | Lighthouse Lab in Alderley Park | Wellcome Sanger Institute for the COVID-19 Genomics UK (COG-UK) Consortium | Jacquelyn Wynn, Mairead Hyland, The Lighthouse Lab in Alderley Park and Alex Alderton, Roberto Amato, Jeffrey Barrett, Sonia Goncalves, Ewan Harrison, David K. Jackson, Ian Johnston, Dominic Kwiatkowski, Cordelia Langford, John Sillitoe on behalf of the Wellcome Sanger Institute COVID-19 Surveillance Team |
| EPI_ISL_2316462 | Lighthouse Lab in Milton Keynes | Wellcome Sanger Institute for the COVID-19 Genomics UK (COG-UK) Consortium | The Lighthouse Lab in Milton Keynes and Alex Alderton, Roberto Amato, Jeffrey Barrett, Sonia Goncalves, Ewan Harrison, David K. Jackson, Ian Johnston, Dominic Kwiatkowski, Cordelia Langford, John Sillitoe on behalf of the Wellcome Sanger Institute COVID-19 Surveillance Team |
| EPI_ISL_2316463, EPI_ISL_2316464 | Lighthouse Lab in Alderley Park | Wellcome Sanger Institute for the COVID-19 Genomics UK (COG-UK) Consortium | Jacquelyn Wynn, Mairead Hyland, The Lighthouse Lab in Alderley Park and Alex Alderton, Roberto Amato, Jeffrey Barrett, Sonia Goncalves, Ewan Harrison, David K. Jackson, Ian Johnston, Dominic Kwiatkowski, Cordelia Langford, John Sillitoe on behalf of the Wellcome Sanger Institute COVID-19 Surveillance Team |
| EPI_ISL_2316465 | Lighthouse Lab in Milton Keynes | Wellcome Sanger Institute for the COVID-19 Genomics UK (COG-UK) Consortium | The Lighthouse Lab in Milton Keynes and Alex Alderton, Roberto Amato, Jeffrey Barrett, Sonia Goncalves, Ewan Harrison, David K. Jackson, Ian Johnston, Dominic Kwiatkowski, Cordelia Langford, John Sillitoe on behalf of the Wellcome Sanger Institute COVID-19 Surveillance Team |
| EPI_ISL_2316468, EPI_ISL_2316469, EPI_ISL_2316471, EPI_ISL_2316473 | Lighthouse Lab in Alderley Park | Wellcome Sanger Institute for the COVID-19 Genomics UK (COG-UK) Consortium | Jacquelyn Wynn, Mairead Hyland, The Lighthouse Lab in Alderley Park and Alex Alderton, Roberto Amato, Jeffrey Barrett, Sonia Goncalves, Ewan Harrison, David K. Jackson, Ian Johnston, Dominic Kwiatkowski, Cordelia Langford, John Sillitoe on behalf of the Wellcome Sanger Institute COVID-19 Surveillance Team |
| EPI_ISL_2316475 | Lighthouse Lab in Milton Keynes | Wellcome Sanger Institute for the COVID-19 Genomics UK (COG-UK) Consortium | The Lighthouse Lab in Milton Keynes and Alex Alderton, Roberto Amato, Jeffrey Barrett, Sonia Goncalves, Ewan Harrison, David K. Jackson, Ian Johnston, Dominic Kwiatkowski, Cordelia Langford, John Sillitoe on behalf of the Wellcome Sanger Institute COVID-19 Surveillance Team |
| EPI_ISL_2316476 | Lighthouse Lab in Alderley Park | Wellcome Sanger Institute for the COVID-19 Genomics UK (COG-UK) Consortium | Jacquelyn Wynn, Mairead Hyland, The Lighthouse Lab in Alderley Park and Alex Alderton, Roberto Amato, Jeffrey Barrett, Sonia Goncalves, Ewan Harrison, David K. Jackson, Ian Johnston, Dominic Kwiatkowski, Cordelia Langford, John Sillitoe on behalf of the Wellcome Sanger Institute COVID-19 Surveillance Team |
| EPI_ISL_2316479, EPI_ISL_2316480, EPI_ISL_2316482, EPI_ISL_2316483, EPI_ISL_2316486 | Lighthouse Lab in Milton Keynes | Wellcome Sanger Institute for the COVID-19 Genomics UK (COG-UK) Consortium | The Lighthouse Lab in Milton Keynes and Alex Alderton, Roberto Amato, Jeffrey Barrett, Sonia Goncalves, Ewan Harrison, David K. Jackson, Ian Johnston, Dominic Kwiatkowski, Cordelia Langford, John Sillitoe on behalf of the Wellcome Sanger Institute COVID-19 Surveillance Team |
| EPI_ISL_2316487 | Lighthouse Lab in Alderley Park | Wellcome Sanger Institute for the COVID-19 Genomics UK (COG-UK) Consortium | Jacquelyn Wynn, Mairead Hyland, The Lighthouse Lab in Alderley Park and Alex Alderton, Roberto Amato, Jeffrey Barrett, Sonia Goncalves, Ewan Harrison, David K. Jackson, Ian Johnston, Dominic Kwiatkowski, Cordelia Langford, John Sillitoe on behalf of the Wellcome Sanger Institute COVID-19 Surveillance Team |
| EPI_ISL_2316489 | Lighthouse Lab in Milton Keynes | Wellcome Sanger Institute for the COVID-19 Genomics UK (COG-UK) Consortium | The Lighthouse Lab in Milton Keynes and Alex Alderton, Roberto Amato, Jeffrey Barrett, Sonia Goncalves, Ewan Harrison, David K. Jackson, Ian Johnston, Dominic Kwiatkowski, Cordelia Langford, John Sillitoe on behalf of the Wellcome Sanger Institute COVID-19 Surveillance Team |
| EPI_ISL_2316491 | Lighthouse Lab in Alderley Park | Wellcome Sanger Institute for the COVID-19 Genomics UK (COG-UK) Consortium | Jacquelyn Wynn, Mairead Hyland, The Lighthouse Lab in Alderley Park and Alex Alderton, Roberto Amato, Jeffrey Barrett, Sonia Goncalves, Ewan Harrison, David K. Jackson, Ian Johnston, Dominic Kwiatkowski, Cordelia Langford, John Sillitoe on behalf of the Wellcome Sanger Institute COVID-19 Surveillance Team |
| EPI_ISL_2316495 | Lighthouse Lab in Milton Keynes | Wellcome Sanger Institute for the COVID-19 Genomics UK (COG-UK) Consortium | The Lighthouse Lab in Milton Keynes and Alex Alderton, Roberto Amato, Jeffrey Barrett, Sonia Goncalves, Ewan Harrison, David K. Jackson, Ian Johnston, Dominic Kwiatkowski, Cordelia Langford, John Sillitoe on behalf of the Wellcome Sanger Institute COVID-19 Surveillance Team |
| EPI_ISL_2316496, EPI_ISL_2316498, EPI_ISL_2316500, EPI_ISL_2316501 | Lighthouse Lab in Alderley Park | Wellcome Sanger Institute for the COVID-19 Genomics UK (COG-UK) Consortium | Jacquelyn Wynn, Mairead Hyland, The Lighthouse Lab in Alderley Park and Alex Alderton, Roberto Amato, Jeffrey Barrett, Sonia Goncalves, Ewan Harrison, David K. Jackson, Ian Johnston, Dominic Kwiatkowski, Cordelia Langford, John Sillitoe on behalf of the Wellcome Sanger Institute COVID-19 Surveillance Team |
| EPI_ISL_2316502, EPI_ISL_2316505, EPI_ISL_2316507 | Health Services Laboratories | Wellcome Sanger Institute for the COVID-19 Genomics UK (COG-UK) Consortium | Health Services Laboratories and Alex Alderton, Roberto Amato, Jeffrey Barrett, Sonia Goncalves, Ewan Harrison, David K. Jackson, Ian Johnston, Dominic Kwiatkowski, Cordelia Langford, John Sillitoe on behalf of the Wellcome Sanger Institute COVID-19 Surveillance Team |
| EPI_ISL_2316509, EPI_ISL_2316512, EPI_ISL_2316513, EPI_ISL_2316514 | Lighthouse Lab in Alderley Park | Wellcome Sanger Institute for the COVID-19 Genomics UK (COG-UK) Consortium | Jacquelyn Wynn, Mairead Hyland, The Lighthouse Lab in Alderley Park and Alex Alderton, Roberto Amato, Jeffrey Barrett, Sonia Goncalves, Ewan Harrison, David K. Jackson, Ian Johnston, Dominic Kwiatkowski, Cordelia Langford, John Sillitoe on behalf of the Wellcome Sanger Institute COVID-19 Surveillance Team |
| EPI_ISL_2316517 | Health Services Laboratories | Wellcome Sanger Institute for the COVID-19 Genomics UK (COG-UK) Consortium | Health Services Laboratories and Alex Alderton, Roberto Amato, Jeffrey Barrett, Sonia Goncalves, Ewan Harrison, David K. Jackson, Ian Johnston, Dominic Kwiatkowski, Cordelia Langford, John Sillitoe on behalf of the Wellcome Sanger Institute COVID-19 Surveillance Team |
| EPI_ISL_2316519 | Lighthouse Lab in Alderley Park | Wellcome Sanger Institute for the COVID-19 Genomics UK (COG-UK) Consortium | Jacquelyn Wynn, Mairead Hyland, The Lighthouse Lab in Alderley Park and Alex Alderton, Roberto Amato, Jeffrey Barrett, Sonia Goncalves, Ewan Harrison, David K. Jackson, Ian Johnston, Dominic Kwiatkowski, Cordelia Langford, John Sillitoe on behalf of the Wellcome Sanger Institute COVID-19 Surveillance Team |
| EPI_ISL_2316520 | Lighthouse Lab in Milton Keynes | Wellcome Sanger Institute for the COVID-19 Genomics UK (COG-UK) Consortium | The Lighthouse Lab in Milton Keynes and Alex Alderton, Roberto Amato, Jeffrey Barrett, Sonia Goncalves, Ewan Harrison, David K. Jackson, Ian Johnston, Dominic Kwiatkowski, Cordelia Langford, John Sillitoe on behalf of the Wellcome Sanger Institute COVID-19 Surveillance Team |
| EPI_ISL_2316523 | Lighthouse Lab in Glasgow | Wellcome Sanger Institute for the COVID-19 Genomics UK (COG-UK) Consortium | Harper VanSteenhouse, Yumi Kasai, David Gray, Carol Clugston, Anna Dominiczak and Alex Alderton, Roberto Amato, Jeffrey Barrett, Sonia Goncalves, Ewan Harrison, David K. Jackson, Ian Johnston, Dominic Kwiatkowski, Cordelia Langford, John Sillitoe on behalf of the Wellcome Sanger Institute COVID-19 Surveillance Team |
| EPI_ISL_2316524 | Lighthouse Lab in Alderley Park | Wellcome Sanger Institute for the COVID-19 Genomics UK (COG-UK) Consortium | Jacquelyn Wynn, Mairead Hyland, The Lighthouse Lab in Alderley Park and Alex Alderton, Roberto Amato, Jeffrey Barrett, Sonia Goncalves, Ewan Harrison, David K. Jackson, Ian Johnston, Dominic Kwiatkowski, Cordelia Langford, John Sillitoe on behalf of the Wellcome Sanger Institute COVID-19 Surveillance Team |
| EPI_ISL_2316526 | Lighthouse Lab in Milton Keynes | Wellcome Sanger Institute for the COVID-19 Genomics UK (COG-UK) Consortium | The Lighthouse Lab in Milton Keynes and Alex Alderton, Roberto Amato, Jeffrey Barrett, Sonia Goncalves, Ewan Harrison, David K. Jackson, Ian Johnston, Dominic Kwiatkowski, Cordelia Langford, John Sillitoe on behalf of the Wellcome Sanger Institute COVID-19 Surveillance Team |
| EPI_ISL_2316527 | Lighthouse Lab in Alderley Park | Wellcome Sanger Institute for the COVID-19 Genomics UK (COG-UK) Consortium | Jacquelyn Wynn, Mairead Hyland, The Lighthouse Lab in Alderley Park and Alex Alderton, Roberto Amato, Jeffrey Barrett, Sonia Goncalves, Ewan Harrison, David K. Jackson, Ian Johnston, Dominic Kwiatkowski, Cordelia Langford, John Sillitoe on behalf of the Wellcome Sanger Institute COVID-19 Surveillance Team |
| EPI_ISL_2316533 | Health Services Laboratories | Wellcome Sanger Institute for the COVID-19 Genomics UK (COG-UK) Consortium | Health Services Laboratories and Alex Alderton, Roberto Amato, Jeffrey Barrett, Sonia Goncalves, Ewan Harrison, David K. Jackson, Ian Johnston, Dominic Kwiatkowski, Cordelia Langford, John Sillitoe on behalf of the Wellcome Sanger Institute COVID-19 Surveillance Team |
| EPI_ISL_2316536, EPI_ISL_2316538, EPI_ISL_2316539, EPI_ISL_2316541 | Lighthouse Lab in Alderley Park | Wellcome Sanger Institute for the COVID-19 Genomics UK (COG-UK) Consortium | Jacquelyn Wynn, Mairead Hyland, The Lighthouse Lab in Alderley Park and Alex Alderton, Roberto Amato, Jeffrey Barrett, Sonia Goncalves, Ewan Harrison, David K. Jackson, Ian Johnston, Dominic Kwiatkowski, Cordelia Langford, John Sillitoe on behalf of the Wellcome Sanger Institute COVID-19 Surveillance Team |
| EPI_ISL_2316544 | Lighthouse Lab in Milton Keynes | Wellcome Sanger Institute for the COVID-19 Genomics UK (COG-UK) Consortium | The Lighthouse Lab in Milton Keynes and Alex Alderton, Roberto Amato, Jeffrey Barrett, Sonia Goncalves, Ewan Harrison, David K. Jackson, Ian Johnston, Dominic Kwiatkowski, Cordelia Langford, John Sillitoe on behalf of the Wellcome Sanger Institute COVID-19 Surveillance Team |
| EPI_ISL_2316545, EPI_ISL_2316548, EPI_ISL_2316550, EPI_ISL_2316551, EPI_ISL_2316553, EPI_ISL_2316556, | Lighthouse Lab in Alderley Park | Wellcome Sanger Institute for the COVID-19 Genomics UK (COG-UK) Consortium | Jacquelyn Wynn, Mairead Hyland, The Lighthouse Lab in Alderley Park and Alex Alderton, Roberto Amato, Jeffrey Barrett, Sonia Goncalves, Ewan Harrison, David K. Jackson, Ian Johnston, Dominic Kwiatkowski, Cordelia Langford, John Sillitoe on behalf of the Wellcome Sanger Institute COVID-19 Surveillance Team |

| | | | |
|---|---|---|---|
| EPI_ISL_2316562, EPI_ISL_2316563 | | | |
| EPI_ISL_2316564 | Lighthouse Lab in Milton Keynes | Wellcome Sanger Institute for the COVID-19 Genomics UK (COG-UK) Consortium | The Lighthouse Lab in Milton Keynes and Alex Alderton, Roberto Amato, Jeffrey Barrett, Sonia Goncalves, Ewan Harrison, David K. Jackson, Ian Johnston, Dominic Kwiatkowski, Cordelia Langford, John Sillitoe on behalf of the Wellcome Sanger Institute COVID-19 Surveillance Team |
| EPI_ISL_2316566, EPI_ISL_2316568, EPI_ISL_2316569 | Lighthouse Lab in Alderley Park | Wellcome Sanger Institute for the COVID-19 Genomics UK (COG-UK) Consortium | Jacquelyn Wynn, Mairead Hyland, The Lighthouse Lab in Alderley Park and Alex Alderton, Roberto Amato, Jeffrey Barrett, Sonia Goncalves, Ewan Harrison, David K. Jackson, Ian Johnston, Dominic Kwiatkowski, Cordelia Langford, John Sillitoe on behalf of the Wellcome Sanger Institute COVID-19 Surveillance Team |
| EPI_ISL_2316570 | Lighthouse Lab in Milton Keynes | Wellcome Sanger Institute for the COVID-19 Genomics UK (COG-UK) Consortium | The Lighthouse Lab in Milton Keynes and Alex Alderton, Roberto Amato, Jeffrey Barrett, Sonia Goncalves, Ewan Harrison, David K. Jackson, Ian Johnston, Dominic Kwiatkowski, Cordelia Langford, John Sillitoe on behalf of the Wellcome Sanger Institute COVID-19 Surveillance Team |
| EPI_ISL_2316575, EPI_ISL_2316576, EPI_ISL_2316579 | Lighthouse Lab in Alderley Park | Wellcome Sanger Institute for the COVID-19 Genomics UK (COG-UK) Consortium | Jacquelyn Wynn, Mairead Hyland, The Lighthouse Lab in Alderley Park and Alex Alderton, Roberto Amato, Jeffrey Barrett, Sonia Goncalves, Ewan Harrison, David K. Jackson, Ian Johnston, Dominic Kwiatkowski, Cordelia Langford, John Sillitoe on behalf of the Wellcome Sanger Institute COVID-19 Surveillance Team |
| EPI_ISL_2316581 | Lighthouse Lab in Milton Keynes | Wellcome Sanger Institute for the COVID-19 Genomics UK (COG-UK) Consortium | The Lighthouse Lab in Milton Keynes and Alex Alderton, Roberto Amato, Jeffrey Barrett, Sonia Goncalves, Ewan Harrison, David K. Jackson, Ian Johnston, Dominic Kwiatkowski, Cordelia Langford, John Sillitoe on behalf of the Wellcome Sanger Institute COVID-19 Surveillance Team |
| EPI_ISL_2316582, EPI_ISL_2316584, EPI_ISL_2316586 | Lighthouse Lab in Alderley Park | Wellcome Sanger Institute for the COVID-19 Genomics UK (COG-UK) Consortium | Jacquelyn Wynn, Mairead Hyland, The Lighthouse Lab in Alderley Park and Alex Alderton, Roberto Amato, Jeffrey Barrett, Sonia Goncalves, Ewan Harrison, David K. Jackson, Ian Johnston, Dominic Kwiatkowski, Cordelia Langford, John Sillitoe on behalf of the Wellcome Sanger Institute COVID-19 Surveillance Team |
| EPI_ISL_2316587 | Lighthouse Lab in Milton Keynes | Wellcome Sanger Institute for the COVID-19 Genomics UK (COG-UK) Consortium | The Lighthouse Lab in Milton Keynes and Alex Alderton, Roberto Amato, Jeffrey Barrett, Sonia Goncalves, Ewan Harrison, David K. Jackson, Ian Johnston, Dominic Kwiatkowski, Cordelia Langford, John Sillitoe on behalf of the Wellcome Sanger Institute COVID-19 Surveillance Team |
| EPI_ISL_2316596, EPI_ISL_2316598, EPI_ISL_2316601, EPI_ISL_2316603, EPI_ISL_2316604, EPI_ISL_2316606, EPI_ISL_2316611 | Lighthouse Lab in Alderley Park | Wellcome Sanger Institute for the COVID-19 Genomics UK (COG-UK) Consortium | Jacquelyn Wynn, Mairead Hyland, The Lighthouse Lab in Alderley Park and Alex Alderton, Roberto Amato, Jeffrey Barrett, Sonia Goncalves, Ewan Harrison, David K. Jackson, Ian Johnston, Dominic Kwiatkowski, Cordelia Langford, John Sillitoe on behalf of the Wellcome Sanger Institute COVID-19 Surveillance Team |
| EPI_ISL_2316616 | Lighthouse Lab in Milton Keynes | Wellcome Sanger Institute for the COVID-19 Genomics UK (COG-UK) Consortium | The Lighthouse Lab in Milton Keynes and Alex Alderton, Roberto Amato, Jeffrey Barrett, Sonia Goncalves, Ewan Harrison, David K. Jackson, Ian Johnston, Dominic Kwiatkowski, Cordelia Langford, John Sillitoe on behalf of the Wellcome Sanger Institute COVID-19 Surveillance Team |
| EPI_ISL_2316618 | Lighthouse Lab in Alderley Park | Wellcome Sanger Institute for the COVID-19 Genomics UK (COG-UK) Consortium | Jacquelyn Wynn, Mairead Hyland, The Lighthouse Lab in Alderley Park and Alex Alderton, Roberto Amato, Jeffrey Barrett, Sonia Goncalves, Ewan Harrison, David K. Jackson, Ian Johnston, Dominic Kwiatkowski, Cordelia Langford, John Sillitoe on behalf of the Wellcome Sanger Institute COVID-19 Surveillance Team |
| EPI_ISL_2316619, EPI_ISL_2316621, EPI_ISL_2316623, EPI_ISL_2316624, EPI_ISL_2316629, EPI_ISL_2316630 | Lighthouse Lab in Milton Keynes | Wellcome Sanger Institute for the COVID-19 Genomics UK (COG-UK) Consortium | The Lighthouse Lab in Milton Keynes and Alex Alderton, Roberto Amato, Jeffrey Barrett, Sonia Goncalves, Ewan Harrison, David K. Jackson, Ian Johnston, Dominic Kwiatkowski, Cordelia Langford, John Sillitoe on behalf of the Wellcome Sanger Institute COVID-19 Surveillance Team |
| EPI_ISL_2316632, EPI_ISL_2316633, EPI_ISL_2316634, EPI_ISL_2316637 | Lighthouse Lab in Alderley Park | Wellcome Sanger Institute for the COVID-19 Genomics UK (COG-UK) Consortium | Jacquelyn Wynn, Mairead Hyland, The Lighthouse Lab in Alderley Park and Alex Alderton, Roberto Amato, Jeffrey Barrett, Sonia Goncalves, Ewan Harrison, David K. Jackson, Ian Johnston, Dominic Kwiatkowski, Cordelia Langford, John Sillitoe on behalf of the Wellcome Sanger Institute COVID-19 Surveillance Team |
| EPI_ISL_2316640 | Lighthouse Lab in Glasgow | Wellcome Sanger Institute for the COVID-19 Genomics UK (COG-UK) Consortium | Harper VanSteenhouse, Yumi Kasai, David Gray, Carol Clugston, Anna Dominiczak and Alex Alderton, Roberto Amato, Jeffrey Barrett, Sonia Goncalves, Ewan Harrison, David K. Jackson, Ian Johnston, Dominic Kwiatkowski, Cordelia Langford, John Sillitoe on behalf of the Wellcome Sanger Institute COVID-19 Surveillance Team |
| EPI_ISL_2316641 | Lighthouse Lab in Alderley Park | Wellcome Sanger Institute for the COVID-19 Genomics UK (COG-UK) Consortium | Jacquelyn Wynn, Mairead Hyland, The Lighthouse Lab in Alderley Park and Alex Alderton, Roberto Amato, Jeffrey Barrett, Sonia Goncalves, Ewan Harrison, David K. Jackson, Ian Johnston, Dominic Kwiatkowski, Cordelia Langford, John Sillitoe on behalf of the Wellcome Sanger Institute COVID-19 Surveillance Team |
| EPI_ISL_2316646 | Lighthouse Lab in Milton Keynes | Wellcome Sanger Institute for the COVID-19 Genomics UK (COG-UK) Consortium | The Lighthouse Lab in Milton Keynes and Alex Alderton, Roberto Amato, Jeffrey Barrett, Sonia Goncalves, Ewan Harrison, David K. Jackson, Ian Johnston, Dominic Kwiatkowski, Cordelia Langford, John Sillitoe on behalf of the Wellcome Sanger Institute COVID-19 Surveillance Team |
| EPI_ISL_2316651 | Lighthouse Lab in Alderley Park | Wellcome Sanger Institute for the COVID-19 Genomics UK (COG-UK) Consortium | Jacquelyn Wynn, Mairead Hyland, The Lighthouse Lab in Alderley Park and Alex Alderton, Roberto Amato, Jeffrey Barrett, Sonia Goncalves, Ewan Harrison, David K. Jackson, Ian Johnston, Dominic Kwiatkowski, Cordelia Langford, John Sillitoe on behalf of the Wellcome Sanger Institute COVID-19 Surveillance Team |
| EPI_ISL_2316652, EPI_ISL_2316653 | Lighthouse Lab in Milton Keynes | Wellcome Sanger Institute for the COVID-19 Genomics UK (COG-UK) Consortium | The Lighthouse Lab in Milton Keynes and Alex Alderton, Roberto Amato, Jeffrey Barrett, Sonia Goncalves, Ewan Harrison, David K. Jackson, Ian Johnston, Dominic Kwiatkowski, Cordelia Langford, John Sillitoe on behalf of the Wellcome Sanger Institute COVID-19 Surveillance Team |
| EPI_ISL_2316656 | Lighthouse Lab in Alderley Park | Wellcome Sanger Institute for the COVID-19 Genomics UK (COG-UK) Consortium | Jacquelyn Wynn, Mairead Hyland, The Lighthouse Lab in Alderley Park and Alex Alderton, Roberto Amato, Jeffrey Barrett, Sonia Goncalves, Ewan Harrison, David K. Jackson, Ian Johnston, Dominic Kwiatkowski, Cordelia Langford, John Sillitoe on behalf of the Wellcome Sanger Institute COVID-19 Surveillance Team |
| EPI_ISL_2316658, EPI_ISL_2316659 | Lighthouse Lab in Milton Keynes | Wellcome Sanger Institute for the COVID-19 Genomics UK (COG-UK) Consortium | The Lighthouse Lab in Milton Keynes and Alex Alderton, Roberto Amato, Jeffrey Barrett, Sonia Goncalves, Ewan Harrison, David K. Jackson, Ian Johnston, Dominic Kwiatkowski, Cordelia Langford, John Sillitoe on behalf of the Wellcome Sanger Institute COVID-19 Surveillance Team |
| EPI_ISL_2316660 | Lighthouse Lab in Alderley Park | Wellcome Sanger Institute for the COVID-19 Genomics UK (COG-UK) Consortium | Jacquelyn Wynn, Mairead Hyland, The Lighthouse Lab in Alderley Park and Alex Alderton, Roberto Amato, Jeffrey Barrett, Sonia Goncalves, Ewan Harrison, David K. Jackson, Ian Johnston, Dominic Kwiatkowski, Cordelia Langford, John Sillitoe on behalf of the Wellcome Sanger Institute COVID-19 Surveillance Team |
| EPI_ISL_2316662 | Lighthouse Lab in Milton Keynes | Wellcome Sanger Institute for the COVID-19 Genomics UK (COG-UK) Consortium | The Lighthouse Lab in Milton Keynes and Alex Alderton, Roberto Amato, Jeffrey Barrett, Sonia Goncalves, Ewan Harrison, David K. Jackson, Ian Johnston, Dominic Kwiatkowski, Cordelia Langford, John Sillitoe on behalf of the Wellcome Sanger Institute COVID-19 Surveillance Team |
| EPI_ISL_2316663 | Lighthouse Lab in Alderley Park | Wellcome Sanger Institute for the COVID-19 Genomics UK (COG-UK) Consortium | Jacquelyn Wynn, Mairead Hyland, The Lighthouse Lab in Alderley Park and Alex Alderton, Roberto Amato, Jeffrey Barrett, Sonia Goncalves, Ewan Harrison, David K. Jackson, Ian Johnston, Dominic Kwiatkowski, Cordelia Langford, John Sillitoe on behalf of the Wellcome Sanger Institute COVID-19 Surveillance Team |
| EPI_ISL_2316665, EPI_ISL_2316668 | Lighthouse Lab in Glasgow | Wellcome Sanger Institute for the COVID-19 Genomics UK (COG-UK) Consortium | Harper VanSteenhouse, Yumi Kasai, David Gray, Carol Clugston, Anna Dominiczak and Alex Alderton, Roberto Amato, Jeffrey Barrett, Sonia Goncalves, Ewan Harrison, David K. Jackson, Ian Johnston, Dominic Kwiatkowski, Cordelia Langford, John Sillitoe on behalf of the Wellcome Sanger Institute COVID-19 Surveillance Team |
| EPI_ISL_2316669 | Lighthouse Lab in Milton Keynes | Wellcome Sanger Institute for the COVID-19 Genomics UK (COG-UK) Consortium | The Lighthouse Lab in Milton Keynes and Alex Alderton, Roberto Amato, Jeffrey Barrett, Sonia Goncalves, Ewan Harrison, David K. Jackson, Ian Johnston, Dominic Kwiatkowski, Cordelia Langford, John Sillitoe on behalf of the Wellcome Sanger Institute COVID-19 Surveillance Team |
| EPI_ISL_2316674 | Lighthouse Lab in Glasgow | Wellcome Sanger Institute for the COVID-19 Genomics UK (COG-UK) Consortium | Harper VanSteenhouse, Yumi Kasai, David Gray, Carol Clugston, Anna Dominiczak and Alex Alderton, Roberto Amato, Jeffrey Barrett, Sonia Goncalves, Ewan Harrison, David K. Jackson, Ian Johnston, Dominic Kwiatkowski, Cordelia Langford, John Sillitoe on behalf of the Wellcome Sanger Institute COVID-19 Surveillance Team |
| EPI_ISL_2316676 | Lighthouse Lab in Milton Keynes | Wellcome Sanger Institute for the COVID-19 Genomics UK (COG-UK) Consortium | The Lighthouse Lab in Milton Keynes and Alex Alderton, Roberto Amato, Jeffrey Barrett, Sonia Goncalves, Ewan Harrison, David K. Jackson, Ian Johnston, Dominic Kwiatkowski, Cordelia Langford, John Sillitoe on behalf of the Wellcome Sanger Institute COVID-19 Surveillance Team |
| EPI_ISL_2316679, EPI_ISL_2316680, EPI_ISL_2316682, EPI_ISL_2316684, EPI_ISL_2316686, EPI_ISL_2316687, EPI_ISL_2316689, EPI_ISL_2316690 | Lighthouse Lab in Glasgow | Wellcome Sanger Institute for the COVID-19 Genomics UK (COG-UK) Consortium | Harper VanSteenhouse, Yumi Kasai, David Gray, Carol Clugston, Anna Dominiczak and Alex Alderton, Roberto Amato, Jeffrey Barrett, Sonia Goncalves, Ewan Harrison, David K. Jackson, Ian Johnston, Dominic Kwiatkowski, Cordelia Langford, John Sillitoe on behalf of the Wellcome Sanger Institute COVID-19 Surveillance Team |
| EPI_ISL_2316692 | Lighthouse Lab in Milton Keynes | Wellcome Sanger Institute for the COVID-19 Genomics UK (COG-UK) Consortium | The Lighthouse Lab in Milton Keynes and Alex Alderton, Roberto Amato, Jeffrey Barrett, Sonia Goncalves, Ewan Harrison, David K. Jackson, Ian Johnston, Dominic Kwiatkowski, Cordelia Langford, John Sillitoe on behalf of the Wellcome Sanger Institute COVID-19 Surveillance Team |
| EPI_ISL_2316693 | Lighthouse Lab in Glasgow | Wellcome Sanger Institute for the COVID-19 Genomics UK (COG-UK) Consortium | Harper VanSteenhouse, Yumi Kasai, David Gray, Carol Clugston, Anna Dominiczak and Alex Alderton, Roberto Amato, Jeffrey Barrett, Sonia Goncalves, Ewan Harrison, David K. Jackson, Ian Johnston, Dominic Kwiatkowski, Cordelia Langford, John Sillitoe on behalf of the Wellcome Sanger Institute COVID-19 Surveillance Team |
| EPI_ISL_2316695 | Lighthouse Lab in Milton Keynes | Wellcome Sanger Institute for the COVID-19 Genomics UK (COG-UK) Consortium | The Lighthouse Lab in Milton Keynes and Alex Alderton, Roberto Amato, Jeffrey Barrett, Sonia Goncalves, Ewan Harrison, David K. Jackson, Ian Johnston, Dominic Kwiatkowski, Cordelia Langford, John Sillitoe on behalf of the Wellcome Sanger Institute COVID-19 Surveillance Team |
| EPI_ISL_2316696, EPI_ISL_2316699, EPI_ISL_2316701, EPI_ISL_2316704, EPI_ISL_2316707 | Lighthouse Lab in Glasgow | Wellcome Sanger Institute for the COVID-19 Genomics UK (COG-UK) Consortium | Harper VanSteenhouse, Yumi Kasai, David Gray, Carol Clugston, Anna Dominiczak and Alex Alderton, Roberto Amato, Jeffrey Barrett, Sonia Goncalves, Ewan Harrison, David K. Jackson, Ian Johnston, Dominic Kwiatkowski, Cordelia Langford, John Sillitoe on behalf of the Wellcome Sanger Institute COVID-19 Surveillance Team |
| EPI_ISL_2316709, EPI_ISL_2316712 | Lighthouse Lab in Milton Keynes | Wellcome Sanger Institute for the COVID-19 Genomics UK (COG-UK) Consortium | The Lighthouse Lab in Milton Keynes and Alex Alderton, Roberto Amato, Jeffrey Barrett, Sonia Goncalves, Ewan Harrison, David K. Jackson, Ian Johnston, Dominic Kwiatkowski, Cordelia Langford, John Sillitoe on behalf of the Wellcome Sanger Institute COVID-19 Surveillance Team |
| EPI_ISL_2316717, EPI_ISL_2316720 | Lighthouse Lab in Glasgow | Wellcome Sanger Institute for the COVID-19 Genomics UK (COG-UK) Consortium | Harper VanSteenhouse, Yumi Kasai, David Gray, Carol Clugston, Anna Dominiczak and Alex Alderton, Roberto Amato, Jeffrey Barrett, Sonia Goncalves, Ewan Harrison, David K. Jackson, Ian Johnston, Dominic Kwiatkowski, Cordelia Langford, John Sillitoe on behalf of the Wellcome Sanger Institute COVID-19 Surveillance Team |
| EPI_ISL_2316723, EPI_ISL_2316730, EPI_ISL_2316734 | Lighthouse Lab in Milton Keynes | Wellcome Sanger Institute for the COVID-19 Genomics UK (COG-UK) Consortium | The Lighthouse Lab in Milton Keynes and Alex Alderton, Roberto Amato, Jeffrey Barrett, Sonia Goncalves, Ewan Harrison, David K. Jackson, Ian Johnston, Dominic Kwiatkowski, Cordelia Langford, John Sillitoe on behalf of the Wellcome Sanger Institute COVID-19 Surveillance Team |

| | | | |
|---|---|---|---|
| EPI_ISL_2316740 | Lighthouse Lab in Glasgow | Wellcome Sanger Institute for the COVID-19 Genomics UK (COG-UK) Consortium | Harper VanSteenhouse, Yumi Kasai, David Gray, Carol Clugston, Anna Dominiczak and Alex Alderton, Roberto Amato, Jeffrey Barrett, Sonia Goncalves, Ewan Harrison, David K. Jackson, Ian Johnston, Dominic Kwiatkowski, Cordelia Langford, John Sillitoe on behalf of the Wellcome Sanger Institute COVID-19 Surveillance Team |
| EPI_ISL_2316742 | Lighthouse Lab in Milton Keynes | Wellcome Sanger Institute for the COVID-19 Genomics UK (COG-UK) Consortium | The Lighthouse Lab in Milton Keynes and Alex Alderton, Roberto Amato, Jeffrey Barrett, Sonia Goncalves, Ewan Harrison, David K. Jackson, Ian Johnston, Dominic Kwiatkowski, Cordelia Langford, John Sillitoe on behalf of the Wellcome Sanger Institute COVID-19 Surveillance Team |
| EPI_ISL_2316745, EPI_ISL_2316746, EPI_ISL_2316747 | Lighthouse Lab in Glasgow | Wellcome Sanger Institute for the COVID-19 Genomics UK (COG-UK) Consortium | Harper VanSteenhouse, Yumi Kasai, David Gray, Carol Clugston, Anna Dominiczak and Alex Alderton, Roberto Amato, Jeffrey Barrett, Sonia Goncalves, Ewan Harrison, David K. Jackson, Ian Johnston, Dominic Kwiatkowski, Cordelia Langford, John Sillitoe on behalf of the Wellcome Sanger Institute COVID-19 Surveillance Team |
| EPI_ISL_2316749, EPI_ISL_2316753, EPI_ISL_2316755 | Lighthouse Lab in Milton Keynes | Wellcome Sanger Institute for the COVID-19 Genomics UK (COG-UK) Consortium | The Lighthouse Lab in Milton Keynes and Alex Alderton, Roberto Amato, Jeffrey Barrett, Sonia Goncalves, Ewan Harrison, David K. Jackson, Ian Johnston, Dominic Kwiatkowski, Cordelia Langford, John Sillitoe on behalf of the Wellcome Sanger Institute COVID-19 Surveillance Team |
| EPI_ISL_2316756, EPI_ISL_2316758 | Lighthouse Lab in Glasgow | Wellcome Sanger Institute for the COVID-19 Genomics UK (COG-UK) Consortium | Harper VanSteenhouse, Yumi Kasai, David Gray, Carol Clugston, Anna Dominiczak and Alex Alderton, Roberto Amato, Jeffrey Barrett, Sonia Goncalves, Ewan Harrison, David K. Jackson, Ian Johnston, Dominic Kwiatkowski, Cordelia Langford, John Sillitoe on behalf of the Wellcome Sanger Institute COVID-19 Surveillance Team |
| EPI_ISL_2316763 | Lighthouse Lab in Milton Keynes | Wellcome Sanger Institute for the COVID-19 Genomics UK (COG-UK) Consortium | The Lighthouse Lab in Milton Keynes and Alex Alderton, Roberto Amato, Jeffrey Barrett, Sonia Goncalves, Ewan Harrison, David K. Jackson, Ian Johnston, Dominic Kwiatkowski, Cordelia Langford, John Sillitoe on behalf of the Wellcome Sanger Institute COVID-19 Surveillance Team |
| EPI_ISL_2316765 | Lighthouse Lab in Glasgow | Wellcome Sanger Institute for the COVID-19 Genomics UK (COG-UK) Consortium | Harper VanSteenhouse, Yumi Kasai, David Gray, Carol Clugston, Anna Dominiczak and Alex Alderton, Roberto Amato, Jeffrey Barrett, Sonia Goncalves, Ewan Harrison, David K. Jackson, Ian Johnston, Dominic Kwiatkowski, Cordelia Langford, John Sillitoe on behalf of the Wellcome Sanger Institute COVID-19 Surveillance Team |
| EPI_ISL_2316766 | Lighthouse Lab in Milton Keynes | Wellcome Sanger Institute for the COVID-19 Genomics UK (COG-UK) Consortium | The Lighthouse Lab in Milton Keynes and Alex Alderton, Roberto Amato, Jeffrey Barrett, Sonia Goncalves, Ewan Harrison, David K. Jackson, Ian Johnston, Dominic Kwiatkowski, Cordelia Langford, John Sillitoe on behalf of the Wellcome Sanger Institute COVID-19 Surveillance Team |
| EPI_ISL_2316767, EPI_ISL_2316773 | Lighthouse Lab in Glasgow | Wellcome Sanger Institute for the COVID-19 Genomics UK (COG-UK) Consortium | Harper VanSteenhouse, Yumi Kasai, David Gray, Carol Clugston, Anna Dominiczak and Alex Alderton, Roberto Amato, Jeffrey Barrett, Sonia Goncalves, Ewan Harrison, David K. Jackson, Ian Johnston, Dominic Kwiatkowski, Cordelia Langford, John Sillitoe on behalf of the Wellcome Sanger Institute COVID-19 Surveillance Team |
| EPI_ISL_2316774, EPI_ISL_2316776 | Lighthouse Lab in Milton Keynes | Wellcome Sanger Institute for the COVID-19 Genomics UK (COG-UK) Consortium | The Lighthouse Lab in Milton Keynes and Alex Alderton, Roberto Amato, Jeffrey Barrett, Sonia Goncalves, Ewan Harrison, David K. Jackson, Ian Johnston, Dominic Kwiatkowski, Cordelia Langford, John Sillitoe on behalf of the Wellcome Sanger Institute COVID-19 Surveillance Team |
| EPI_ISL_2316777, EPI_ISL_2316781, EPI_ISL_2316787, EPI_ISL_2316790, EPI_ISL_2316791, EPI_ISL_2316793 | Lighthouse Lab in Glasgow | Wellcome Sanger Institute for the COVID-19 Genomics UK (COG-UK) Consortium | Harper VanSteenhouse, Yumi Kasai, David Gray, Carol Clugston, Anna Dominiczak and Alex Alderton, Roberto Amato, Jeffrey Barrett, Sonia Goncalves, Ewan Harrison, David K. Jackson, Ian Johnston, Dominic Kwiatkowski, Cordelia Langford, John Sillitoe on behalf of the Wellcome Sanger Institute COVID-19 Surveillance Team |
| EPI_ISL_2316794 | Lighthouse Lab in Milton Keynes | Wellcome Sanger Institute for the COVID-19 Genomics UK (COG-UK) Consortium | The Lighthouse Lab in Milton Keynes and Alex Alderton, Roberto Amato, Jeffrey Barrett, Sonia Goncalves, Ewan Harrison, David K. Jackson, Ian Johnston, Dominic Kwiatkowski, Cordelia Langford, John Sillitoe on behalf of the Wellcome Sanger Institute COVID-19 Surveillance Team |
| EPI_ISL_2316796, EPI_ISL_2316797, EPI_ISL_2316800 | Lighthouse Lab in Glasgow | Wellcome Sanger Institute for the COVID-19 Genomics UK (COG-UK) Consortium | Harper VanSteenhouse, Yumi Kasai, David Gray, Carol Clugston, Anna Dominiczak and Alex Alderton, Roberto Amato, Jeffrey Barrett, Sonia Goncalves, Ewan Harrison, David K. Jackson, Ian Johnston, Dominic Kwiatkowski, Cordelia Langford, John Sillitoe on behalf of the Wellcome Sanger Institute COVID-19 Surveillance Team |
| EPI_ISL_2316801 | Lighthouse Lab in Milton Keynes | Wellcome Sanger Institute for the COVID-19 Genomics UK (COG-UK) Consortium | The Lighthouse Lab in Milton Keynes and Alex Alderton, Roberto Amato, Jeffrey Barrett, Sonia Goncalves, Ewan Harrison, David K. Jackson, Ian Johnston, Dominic Kwiatkowski, Cordelia Langford, John Sillitoe on behalf of the Wellcome Sanger Institute COVID-19 Surveillance Team |
| EPI_ISL_2316805, EPI_ISL_2316806 | Lighthouse Lab in Glasgow | Wellcome Sanger Institute for the COVID-19 Genomics UK (COG-UK) Consortium | Harper VanSteenhouse, Yumi Kasai, David Gray, Carol Clugston, Anna Dominiczak and Alex Alderton, Roberto Amato, Jeffrey Barrett, Sonia Goncalves, Ewan Harrison, David K. Jackson, Ian Johnston, Dominic Kwiatkowski, Cordelia Langford, John Sillitoe on behalf of the Wellcome Sanger Institute COVID-19 Surveillance Team |
| EPI_ISL_2316809, EPI_ISL_2316812 | Lighthouse Lab in Milton Keynes | Wellcome Sanger Institute for the COVID-19 Genomics UK (COG-UK) Consortium | The Lighthouse Lab in Milton Keynes and Alex Alderton, Roberto Amato, Jeffrey Barrett, Sonia Goncalves, Ewan Harrison, David K. Jackson, Ian Johnston, Dominic Kwiatkowski, Cordelia Langford, John Sillitoe on behalf of the Wellcome Sanger Institute COVID-19 Surveillance Team |
| EPI_ISL_2316813, EPI_ISL_2316818, EPI_ISL_2316819 | Lighthouse Lab in Glasgow | Wellcome Sanger Institute for the COVID-19 Genomics UK (COG-UK) Consortium | Harper VanSteenhouse, Yumi Kasai, David Gray, Carol Clugston, Anna Dominiczak and Alex Alderton, Roberto Amato, Jeffrey Barrett, Sonia Goncalves, Ewan Harrison, David K. Jackson, Ian Johnston, Dominic Kwiatkowski, Cordelia Langford, John Sillitoe on behalf of the Wellcome Sanger Institute COVID-19 Surveillance Team |
| EPI_ISL_2316824, EPI_ISL_2316825, EPI_ISL_2316827 | Lighthouse Lab in Milton Keynes | Wellcome Sanger Institute for the COVID-19 Genomics UK (COG-UK) Consortium | The Lighthouse Lab in Milton Keynes and Alex Alderton, Roberto Amato, Jeffrey Barrett, Sonia Goncalves, Ewan Harrison, David K. Jackson, Ian Johnston, Dominic Kwiatkowski, Cordelia Langford, John Sillitoe on behalf of the Wellcome Sanger Institute COVID-19 Surveillance Team |
| EPI_ISL_2316828, EPI_ISL_2316831, EPI_ISL_2316832, EPI_ISL_2316835 | Lighthouse Lab in Glasgow | Wellcome Sanger Institute for the COVID-19 Genomics UK (COG-UK) Consortium | Harper VanSteenhouse, Yumi Kasai, David Gray, Carol Clugston, Anna Dominiczak and Alex Alderton, Roberto Amato, Jeffrey Barrett, Sonia Goncalves, Ewan Harrison, David K. Jackson, Ian Johnston, Dominic Kwiatkowski, Cordelia Langford, John Sillitoe on behalf of the Wellcome Sanger Institute COVID-19 Surveillance Team |
| EPI_ISL_2316836 | Eurofins LifeCodexx GmbH | Robert Koch Institute | unknown |
| EPI_ISL_2316837 | Lighthouse Lab in Milton Keynes | Wellcome Sanger Institute for the COVID-19 Genomics UK (COG-UK) Consortium | The Lighthouse Lab in Milton Keynes and Alex Alderton, Roberto Amato, Jeffrey Barrett, Sonia Goncalves, Ewan Harrison, David K. Jackson, Ian Johnston, Dominic Kwiatkowski, Cordelia Langford, John Sillitoe on behalf of the Wellcome Sanger Institute COVID-19 Surveillance Team |
| EPI_ISL_2316838, EPI_ISL_2316841, EPI_ISL_2316842 | Lighthouse Lab in Glasgow | Wellcome Sanger Institute for the COVID-19 Genomics UK (COG-UK) Consortium | Harper VanSteenhouse, Yumi Kasai, David Gray, Carol Clugston, Anna Dominiczak and Alex Alderton, Roberto Amato, Jeffrey Barrett, Sonia Goncalves, Ewan Harrison, David K. Jackson, Ian Johnston, Dominic Kwiatkowski, Cordelia Langford, John Sillitoe on behalf of the Wellcome Sanger Institute COVID-19 Surveillance Team |
| EPI_ISL_2316843 | Eurofins LifeCodexx GmbH | Robert Koch Institute | unknown |
| EPI_ISL_2316844 | Lighthouse Lab in Glasgow | Wellcome Sanger Institute for the COVID-19 Genomics UK (COG-UK) Consortium | Harper VanSteenhouse, Yumi Kasai, David Gray, Carol Clugston, Anna Dominiczak and Alex Alderton, Roberto Amato, Jeffrey Barrett, Sonia Goncalves, Ewan Harrison, David K. Jackson, Ian Johnston, Dominic Kwiatkowski, Cordelia Langford, John Sillitoe on behalf of the Wellcome Sanger Institute COVID-19 Surveillance Team |
| EPI_ISL_2316846 | Lighthouse Lab in Milton Keynes | Wellcome Sanger Institute for the COVID-19 Genomics UK (COG-UK) Consortium | The Lighthouse Lab in Milton Keynes and Alex Alderton, Roberto Amato, Jeffrey Barrett, Sonia Goncalves, Ewan Harrison, David K. Jackson, Ian Johnston, Dominic Kwiatkowski, Cordelia Langford, John Sillitoe on behalf of the Wellcome Sanger Institute COVID-19 Surveillance Team |
| EPI_ISL_2316849 | Eurofins LifeCodexx GmbH | Robert Koch Institute | unknown |
| EPI_ISL_2316850, EPI_ISL_2316852 | Lighthouse Lab in Glasgow | Wellcome Sanger Institute for the COVID-19 Genomics UK (COG-UK) Consortium | Harper VanSteenhouse, Yumi Kasai, David Gray, Carol Clugston, Anna Dominiczak and Alex Alderton, Roberto Amato, Jeffrey Barrett, Sonia Goncalves, Ewan Harrison, David K. Jackson, Ian Johnston, Dominic Kwiatkowski, Cordelia Langford, John Sillitoe on behalf of the Wellcome Sanger Institute COVID-19 Surveillance Team |
| EPI_ISL_2316854 | Lighthouse Lab in Milton Keynes | Wellcome Sanger Institute for the COVID-19 Genomics UK (COG-UK) Consortium | The Lighthouse Lab in Milton Keynes and Alex Alderton, Roberto Amato, Jeffrey Barrett, Sonia Goncalves, Ewan Harrison, David K. Jackson, Ian Johnston, Dominic Kwiatkowski, Cordelia Langford, John Sillitoe on behalf of the Wellcome Sanger Institute COVID-19 Surveillance Team |
| EPI_ISL_2316855, EPI_ISL_2316857, EPI_ISL_2316862 | Lighthouse Lab in Glasgow | Wellcome Sanger Institute for the COVID-19 Genomics UK (COG-UK) Consortium | Harper VanSteenhouse, Yumi Kasai, David Gray, Carol Clugston, Anna Dominiczak and Alex Alderton, Roberto Amato, Jeffrey Barrett, Sonia Goncalves, Ewan Harrison, David K. Jackson, Ian Johnston, Dominic Kwiatkowski, Cordelia Langford, John Sillitoe on behalf of the Wellcome Sanger Institute COVID-19 Surveillance Team |
| EPI_ISL_2316864 | Lighthouse Lab in Milton Keynes | Wellcome Sanger Institute for the COVID-19 Genomics UK (COG-UK) Consortium | The Lighthouse Lab in Milton Keynes and Alex Alderton, Roberto Amato, Jeffrey Barrett, Sonia Goncalves, Ewan Harrison, David K. Jackson, Ian Johnston, Dominic Kwiatkowski, Cordelia Langford, John Sillitoe on behalf of the Wellcome Sanger Institute COVID-19 Surveillance Team |
| EPI_ISL_2316865 | Lighthouse Lab in Glasgow | Wellcome Sanger Institute for the COVID-19 Genomics UK (COG-UK) Consortium | Harper VanSteenhouse, Yumi Kasai, David Gray, Carol Clugston, Anna Dominiczak and Alex Alderton, Roberto Amato, Jeffrey Barrett, Sonia Goncalves, Ewan Harrison, David K. Jackson, Ian Johnston, Dominic Kwiatkowski, Cordelia Langford, John Sillitoe on behalf of the Wellcome Sanger Institute COVID-19 Surveillance Team |
| EPI_ISL_2316866 | Lighthouse Lab in Milton Keynes | Wellcome Sanger Institute for the COVID-19 Genomics UK (COG-UK) Consortium | The Lighthouse Lab in Milton Keynes and Alex Alderton, Roberto Amato, Jeffrey Barrett, Sonia Goncalves, Ewan Harrison, David K. Jackson, Ian Johnston, Dominic Kwiatkowski, Cordelia Langford, John Sillitoe on behalf of the Wellcome Sanger Institute COVID-19 Surveillance Team |
| EPI_ISL_2316867, EPI_ISL_2316868, EPI_ISL_2316870, EPI_ISL_2316872, EPI_ISL_2316873 | Lighthouse Lab in Glasgow | Wellcome Sanger Institute for the COVID-19 Genomics UK (COG-UK) Consortium | Harper VanSteenhouse, Yumi Kasai, David Gray, Carol Clugston, Anna Dominiczak and Alex Alderton, Roberto Amato, Jeffrey Barrett, Sonia Goncalves, Ewan Harrison, David K. Jackson, Ian Johnston, Dominic Kwiatkowski, Cordelia Langford, John Sillitoe on behalf of the Wellcome Sanger Institute COVID-19 Surveillance Team |
| EPI_ISL_2316875 | Lighthouse Lab in Milton Keynes | Wellcome Sanger Institute for the COVID-19 Genomics UK (COG-UK) Consortium | The Lighthouse Lab in Milton Keynes and Alex Alderton, Roberto Amato, Jeffrey Barrett, Sonia Goncalves, Ewan Harrison, David K. Jackson, Ian Johnston, Dominic Kwiatkowski, Cordelia Langford, John Sillitoe on behalf of the Wellcome Sanger Institute COVID-19 Surveillance Team |
| EPI_ISL_2316876, EPI_ISL_2316878, EPI_ISL_2316879 | Lighthouse Lab in Glasgow | Wellcome Sanger Institute for the COVID-19 Genomics UK (COG-UK) Consortium | Harper VanSteenhouse, Yumi Kasai, David Gray, Carol Clugston, Anna Dominiczak and Alex Alderton, Roberto Amato, Jeffrey Barrett, Sonia Goncalves, Ewan Harrison, David K. Jackson, Ian Johnston, Dominic Kwiatkowski, Cordelia Langford, John Sillitoe on behalf of the Wellcome Sanger Institute COVID-19 Surveillance Team |
| EPI_ISL_2316881 | Lighthouse Lab in Milton Keynes | Wellcome Sanger Institute for the COVID-19 Genomics UK (COG-UK) Consortium | The Lighthouse Lab in Milton Keynes and Alex Alderton, Roberto Amato, Jeffrey Barrett, Sonia Goncalves, Ewan Harrison, David K. Jackson, Ian Johnston, Dominic Kwiatkowski, Cordelia Langford, John Sillitoe on behalf of the Wellcome Sanger Institute COVID-19 Surveillance Team |
| EPI_ISL_2316882 | Lighthouse Lab in Glasgow | Wellcome Sanger Institute for the COVID-19 Genomics UK (COG-UK) Consortium | Harper VanSteenhouse, Yumi Kasai, David Gray, Carol Clugston, Anna Dominiczak and Alex Alderton, Roberto Amato, Jeffrey Barrett, Sonia Goncalves, Ewan Harrison, David K. Jackson, Ian Johnston, Dominic Kwiatkowski, Cordelia Langford, John Sillitoe on behalf of the Wellcome Sanger Institute COVID-19 Surveillance Team |
| EPI_ISL_2316883, EPI_ISL_2316886, EPI_ISL_2316887 | Lighthouse Lab in Milton Keynes | Wellcome Sanger Institute for the COVID-19 Genomics UK (COG-UK) Consortium | The Lighthouse Lab in Milton Keynes and Alex Alderton, Roberto Amato, Jeffrey Barrett, Sonia Goncalves, Ewan Harrison, David K. Jackson, Ian Johnston, Dominic Kwiatkowski, Cordelia Langford, John Sillitoe on behalf of the Wellcome Sanger Institute COVID-19 Surveillance Team |
| EPI_ISL_2316888 | Eurofins LifeCodexx GmbH | Robert Koch Institute | unknown |
| EPI_ISL_2316889, | Lighthouse Lab in Glasgow | Wellcome Sanger Institute for the COVID-19 Genomics UK (COG-UK) | Harper VanSteenhouse, Yumi Kasai, David Gray, Carol Clugston, Anna Dominiczak and Alex Alderton, Roberto Amato, Jeffrey Barrett, Sonia Goncalves, Ewan Harrison, David K. Jackson, Ian Johnston, Dominic Kwiatkowski, Cordelia Langford, |

| | | | |
|---|---|---|---|
| EPI_ISL_2316892, EPI_ISL_2316893, EPI_ISL_2316895 | | Consortium | John Sillitoe on behalf of the Wellcome Sanger Institute COVID-19 Surveillance Team |
| EPI_ISL_2316897 | Lighthouse Lab in Milton Keynes | Wellcome Sanger Institute for the COVID-19 Genomics UK (COG-UK) Consortium | The Lighthouse Lab in Milton Keynes and Alex Alderton, Roberto Amato, Jeffrey Barrett, Sonia Goncalves, Ewan Harrison, David K. Jackson, Ian Johnston, Dominic Kwiatkowski, Cordelia Langford, John Sillitoe on behalf of the Wellcome Sanger Institute COVID-19 Surveillance Team |
| EPI_ISL_2316898 | Eurofins LifeCodexx GmbH | Robert Koch Institute | unknown |
| EPI_ISL_2316899, EPI_ISL_2316900 | Lighthouse Lab in Glasgow | Wellcome Sanger Institute for the COVID-19 Genomics UK (COG-UK) Consortium | Harper VanSteenhouse, Yumi Kasai, David Gray, Carol Clugston, Anna Dominiczak and Alex Alderton, Roberto Amato, Jeffrey Barrett, Sonia Goncalves, Ewan Harrison, David K. Jackson, Ian Johnston, Dominic Kwiatkowski, Cordelia Langford, John Sillitoe on behalf of the Wellcome Sanger Institute COVID-19 Surveillance Team |
| EPI_ISL_2316901 | Eurofins LifeCodexx GmbH | Robert Koch Institute | unknown |
| EPI_ISL_2316903, EPI_ISL_2316907, EPI_ISL_2316908, EPI_ISL_2316910 | Lighthouse Lab in Glasgow | Wellcome Sanger Institute for the COVID-19 Genomics UK (COG-UK) Consortium | Harper VanSteenhouse, Yumi Kasai, David Gray, Carol Clugston, Anna Dominiczak and Alex Alderton, Roberto Amato, Jeffrey Barrett, Sonia Goncalves, Ewan Harrison, David K. Jackson, Ian Johnston, Dominic Kwiatkowski, Cordelia Langford, John Sillitoe on behalf of the Wellcome Sanger Institute COVID-19 Surveillance Team |
| EPI_ISL_2316912 | Lighthouse Lab in Milton Keynes | Wellcome Sanger Institute for the COVID-19 Genomics UK (COG-UK) Consortium | The Lighthouse Lab in Milton Keynes and Alex Alderton, Roberto Amato, Jeffrey Barrett, Sonia Goncalves, Ewan Harrison, David K. Jackson, Ian Johnston, Dominic Kwiatkowski, Cordelia Langford, John Sillitoe on behalf of the Wellcome Sanger Institute COVID-19 Surveillance Team |
| EPI_ISL_2316913, EPI_ISL_2316919 | Lighthouse Lab in Glasgow | Wellcome Sanger Institute for the COVID-19 Genomics UK (COG-UK) Consortium | Harper VanSteenhouse, Yumi Kasai, David Gray, Carol Clugston, Anna Dominiczak and Alex Alderton, Roberto Amato, Jeffrey Barrett, Sonia Goncalves, Ewan Harrison, David K. Jackson, Ian Johnston, Dominic Kwiatkowski, Cordelia Langford, John Sillitoe on behalf of the Wellcome Sanger Institute COVID-19 Surveillance Team |
| EPI_ISL_2316920 | Lighthouse Lab in Milton Keynes | Wellcome Sanger Institute for the COVID-19 Genomics UK (COG-UK) Consortium | The Lighthouse Lab in Milton Keynes and Alex Alderton, Roberto Amato, Jeffrey Barrett, Sonia Goncalves, Ewan Harrison, David K. Jackson, Ian Johnston, Dominic Kwiatkowski, Cordelia Langford, John Sillitoe on behalf of the Wellcome Sanger Institute COVID-19 Surveillance Team |
| EPI_ISL_2316922 | Lighthouse Lab in Glasgow | Wellcome Sanger Institute for the COVID-19 Genomics UK (COG-UK) Consortium | Harper VanSteenhouse, Yumi Kasai, David Gray, Carol Clugston, Anna Dominiczak and Alex Alderton, Roberto Amato, Jeffrey Barrett, Sonia Goncalves, Ewan Harrison, David K. Jackson, Ian Johnston, Dominic Kwiatkowski, Cordelia Langford, John Sillitoe on behalf of the Wellcome Sanger Institute COVID-19 Surveillance Team |
| EPI_ISL_2316923, EPI_ISL_2316924 | Lighthouse Lab in Milton Keynes | Wellcome Sanger Institute for the COVID-19 Genomics UK (COG-UK) Consortium | The Lighthouse Lab in Milton Keynes and Alex Alderton, Roberto Amato, Jeffrey Barrett, Sonia Goncalves, Ewan Harrison, David K. Jackson, Ian Johnston, Dominic Kwiatkowski, Cordelia Langford, John Sillitoe on behalf of the Wellcome Sanger Institute COVID-19 Surveillance Team |
| EPI_ISL_2316926, EPI_ISL_2316927 | Lighthouse Lab in Glasgow | Wellcome Sanger Institute for the COVID-19 Genomics UK (COG-UK) Consortium | Harper VanSteenhouse, Yumi Kasai, David Gray, Carol Clugston, Anna Dominiczak and Alex Alderton, Roberto Amato, Jeffrey Barrett, Sonia Goncalves, Ewan Harrison, David K. Jackson, Ian Johnston, Dominic Kwiatkowski, Cordelia Langford, John Sillitoe on behalf of the Wellcome Sanger Institute COVID-19 Surveillance Team |
| EPI_ISL_2316928 | Eurofins LifeCodexx GmbH | Robert Koch Institute | unknown |
| EPI_ISL_2316929, EPI_ISL_2316932 | Lighthouse Lab in Glasgow | Wellcome Sanger Institute for the COVID-19 Genomics UK (COG-UK) Consortium | Harper VanSteenhouse, Yumi Kasai, David Gray, Carol Clugston, Anna Dominiczak and Alex Alderton, Roberto Amato, Jeffrey Barrett, Sonia Goncalves, Ewan Harrison, David K. Jackson, Ian Johnston, Dominic Kwiatkowski, Cordelia Langford, John Sillitoe on behalf of the Wellcome Sanger Institute COVID-19 Surveillance Team |
| EPI_ISL_2316934 | Eurofins LifeCodexx GmbH | Robert Koch Institute | unknown |
| EPI_ISL_2316936 | Lighthouse Lab in Milton Keynes | Wellcome Sanger Institute for the COVID-19 Genomics UK (COG-UK) Consortium | The Lighthouse Lab in Milton Keynes and Alex Alderton, Roberto Amato, Jeffrey Barrett, Sonia Goncalves, Ewan Harrison, David K. Jackson, Ian Johnston, Dominic Kwiatkowski, Cordelia Langford, John Sillitoe on behalf of the Wellcome Sanger Institute COVID-19 Surveillance Team |
| EPI_ISL_2316937, EPI_ISL_2316939 | Eurofins LifeCodexx GmbH | Robert Koch Institute | unknown |
| EPI_ISL_2316943 | Lighthouse Lab in Milton Keynes | Wellcome Sanger Institute for the COVID-19 Genomics UK (COG-UK) Consortium | The Lighthouse Lab in Milton Keynes and Alex Alderton, Roberto Amato, Jeffrey Barrett, Sonia Goncalves, Ewan Harrison, David K. Jackson, Ian Johnston, Dominic Kwiatkowski, Cordelia Langford, John Sillitoe on behalf of the Wellcome Sanger Institute COVID-19 Surveillance Team |
| EPI_ISL_2316944, EPI_ISL_2316947, EPI_ISL_2316949, EPI_ISL_2316951, EPI_ISL_2316954 | Lighthouse Lab in Glasgow | Wellcome Sanger Institute for the COVID-19 Genomics UK (COG-UK) Consortium | Harper VanSteenhouse, Yumi Kasai, David Gray, Carol Clugston, Anna Dominiczak and Alex Alderton, Roberto Amato, Jeffrey Barrett, Sonia Goncalves, Ewan Harrison, David K. Jackson, Ian Johnston, Dominic Kwiatkowski, Cordelia Langford, John Sillitoe on behalf of the Wellcome Sanger Institute COVID-19 Surveillance Team |
| EPI_ISL_2316956 | Lighthouse Lab in Milton Keynes | Wellcome Sanger Institute for the COVID-19 Genomics UK (COG-UK) Consortium | The Lighthouse Lab in Milton Keynes and Alex Alderton, Roberto Amato, Jeffrey Barrett, Sonia Goncalves, Ewan Harrison, David K. Jackson, Ian Johnston, Dominic Kwiatkowski, Cordelia Langford, John Sillitoe on behalf of the Wellcome Sanger Institute COVID-19 Surveillance Team |
| EPI_ISL_2316957, EPI_ISL_2316958 | Lighthouse Lab in Glasgow | Wellcome Sanger Institute for the COVID-19 Genomics UK (COG-UK) Consortium | Harper VanSteenhouse, Yumi Kasai, David Gray, Carol Clugston, Anna Dominiczak and Alex Alderton, Roberto Amato, Jeffrey Barrett, Sonia Goncalves, Ewan Harrison, David K. Jackson, Ian Johnston, Dominic Kwiatkowski, Cordelia Langford, John Sillitoe on behalf of the Wellcome Sanger Institute COVID-19 Surveillance Team |
| EPI_ISL_2316960 | Lighthouse Lab in Milton Keynes | Wellcome Sanger Institute for the COVID-19 Genomics UK (COG-UK) Consortium | The Lighthouse Lab in Milton Keynes and Alex Alderton, Roberto Amato, Jeffrey Barrett, Sonia Goncalves, Ewan Harrison, David K. Jackson, Ian Johnston, Dominic Kwiatkowski, Cordelia Langford, John Sillitoe on behalf of the Wellcome Sanger Institute COVID-19 Surveillance Team |
| EPI_ISL_2316961, EPI_ISL_2316963, EPI_ISL_2316964, EPI_ISL_2316965 | Lighthouse Lab in Glasgow | Wellcome Sanger Institute for the COVID-19 Genomics UK (COG-UK) Consortium | Harper VanSteenhouse, Yumi Kasai, David Gray, Carol Clugston, Anna Dominiczak and Alex Alderton, Roberto Amato, Jeffrey Barrett, Sonia Goncalves, Ewan Harrison, David K. Jackson, Ian Johnston, Dominic Kwiatkowski, Cordelia Langford, John Sillitoe on behalf of the Wellcome Sanger Institute COVID-19 Surveillance Team |
| EPI_ISL_2316967 | Lighthouse Lab in Milton Keynes | Wellcome Sanger Institute for the COVID-19 Genomics UK (COG-UK) Consortium | The Lighthouse Lab in Milton Keynes and Alex Alderton, Roberto Amato, Jeffrey Barrett, Sonia Goncalves, Ewan Harrison, David K. Jackson, Ian Johnston, Dominic Kwiatkowski, Cordelia Langford, John Sillitoe on behalf of the Wellcome Sanger Institute COVID-19 Surveillance Team |
| EPI_ISL_2316970, EPI_ISL_2316971 | Lighthouse Lab in Glasgow | Wellcome Sanger Institute for the COVID-19 Genomics UK (COG-UK) Consortium | Harper VanSteenhouse, Yumi Kasai, David Gray, Carol Clugston, Anna Dominiczak and Alex Alderton, Roberto Amato, Jeffrey Barrett, Sonia Goncalves, Ewan Harrison, David K. Jackson, Ian Johnston, Dominic Kwiatkowski, Cordelia Langford, John Sillitoe on behalf of the Wellcome Sanger Institute COVID-19 Surveillance Team |
| EPI_ISL_2316972, EPI_ISL_2316975 | Lighthouse Lab in Milton Keynes | Wellcome Sanger Institute for the COVID-19 Genomics UK (COG-UK) Consortium | The Lighthouse Lab in Milton Keynes and Alex Alderton, Roberto Amato, Jeffrey Barrett, Sonia Goncalves, Ewan Harrison, David K. Jackson, Ian Johnston, Dominic Kwiatkowski, Cordelia Langford, John Sillitoe on behalf of the Wellcome Sanger Institute COVID-19 Surveillance Team |
| EPI_ISL_2316976, EPI_ISL_2316978, EPI_ISL_2316979, EPI_ISL_2316980 | Lighthouse Lab in Glasgow | Wellcome Sanger Institute for the COVID-19 Genomics UK (COG-UK) Consortium | Harper VanSteenhouse, Yumi Kasai, David Gray, Carol Clugston, Anna Dominiczak and Alex Alderton, Roberto Amato, Jeffrey Barrett, Sonia Goncalves, Ewan Harrison, David K. Jackson, Ian Johnston, Dominic Kwiatkowski, Cordelia Langford, John Sillitoe on behalf of the Wellcome Sanger Institute COVID-19 Surveillance Team |
| EPI_ISL_2316982 | Lighthouse Lab in Milton Keynes | Wellcome Sanger Institute for the COVID-19 Genomics UK (COG-UK) Consortium | The Lighthouse Lab in Milton Keynes and Alex Alderton, Roberto Amato, Jeffrey Barrett, Sonia Goncalves, Ewan Harrison, David K. Jackson, Ian Johnston, Dominic Kwiatkowski, Cordelia Langford, John Sillitoe on behalf of the Wellcome Sanger Institute COVID-19 Surveillance Team |
| EPI_ISL_2316983, EPI_ISL_2316985 | Lighthouse Lab in Glasgow | Wellcome Sanger Institute for the COVID-19 Genomics UK (COG-UK) Consortium | Harper VanSteenhouse, Yumi Kasai, David Gray, Carol Clugston, Anna Dominiczak and Alex Alderton, Roberto Amato, Jeffrey Barrett, Sonia Goncalves, Ewan Harrison, David K. Jackson, Ian Johnston, Dominic Kwiatkowski, Cordelia Langford, John Sillitoe on behalf of the Wellcome Sanger Institute COVID-19 Surveillance Team |
| EPI_ISL_2316988 | Eurofins LifeCodexx GmbH | Robert Koch Institute | unknown |
| EPI_ISL_2316991 | Lighthouse Lab in Milton Keynes | Wellcome Sanger Institute for the COVID-19 Genomics UK (COG-UK) Consortium | The Lighthouse Lab in Milton Keynes and Alex Alderton, Roberto Amato, Jeffrey Barrett, Sonia Goncalves, Ewan Harrison, David K. Jackson, Ian Johnston, Dominic Kwiatkowski, Cordelia Langford, John Sillitoe on behalf of the Wellcome Sanger Institute COVID-19 Surveillance Team |
| EPI_ISL_2316992, EPI_ISL_2316993 | Lighthouse Lab in Glasgow | Wellcome Sanger Institute for the COVID-19 Genomics UK (COG-UK) Consortium | Harper VanSteenhouse, Yumi Kasai, David Gray, Carol Clugston, Anna Dominiczak and Alex Alderton, Roberto Amato, Jeffrey Barrett, Sonia Goncalves, Ewan Harrison, David K. Jackson, Ian Johnston, Dominic Kwiatkowski, Cordelia Langford, John Sillitoe on behalf of the Wellcome Sanger Institute COVID-19 Surveillance Team |
| EPI_ISL_2316994 | Eurofins LifeCodexx GmbH | Robert Koch Institute | unknown |
| EPI_ISL_2316995, EPI_ISL_2316999 | Lighthouse Lab in Glasgow | Wellcome Sanger Institute for the COVID-19 Genomics UK (COG-UK) Consortium | Harper VanSteenhouse, Yumi Kasai, David Gray, Carol Clugston, Anna Dominiczak and Alex Alderton, Roberto Amato, Jeffrey Barrett, Sonia Goncalves, Ewan Harrison, David K. Jackson, Ian Johnston, Dominic Kwiatkowski, Cordelia Langford, John Sillitoe on behalf of the Wellcome Sanger Institute COVID-19 Surveillance Team |
| EPI_ISL_2317000 | Lighthouse Lab in Milton Keynes | Wellcome Sanger Institute for the COVID-19 Genomics UK (COG-UK) Consortium | The Lighthouse Lab in Milton Keynes and Alex Alderton, Roberto Amato, Jeffrey Barrett, Sonia Goncalves, Ewan Harrison, David K. Jackson, Ian Johnston, Dominic Kwiatkowski, Cordelia Langford, John Sillitoe on behalf of the Wellcome Sanger Institute COVID-19 Surveillance Team |
| EPI_ISL_2317002 | Lighthouse Lab in Glasgow | Wellcome Sanger Institute for the COVID-19 Genomics UK (COG-UK) Consortium | Harper VanSteenhouse, Yumi Kasai, David Gray, Carol Clugston, Anna Dominiczak and Alex Alderton, Roberto Amato, Jeffrey Barrett, Sonia Goncalves, Ewan Harrison, David K. Jackson, Ian Johnston, Dominic Kwiatkowski, Cordelia Langford, John Sillitoe on behalf of the Wellcome Sanger Institute COVID-19 Surveillance Team |
| EPI_ISL_2317005, EPI_ISL_2317006, EPI_ISL_2317009, EPI_ISL_2317010, EPI_ISL_2317027, EPI_ISL_2317029, EPI_ISL_2317033 | Eurofins LifeCodexx GmbH | Robert Koch Institute | unknown |
| EPI_ISL_2317037 | Lighthouse Lab in Milton Keynes | Wellcome Sanger Institute for the COVID-19 Genomics UK (COG-UK) Consortium | The Lighthouse Lab in Milton Keynes and Alex Alderton, Roberto Amato, Jeffrey Barrett, Sonia Goncalves, Ewan Harrison, David K. Jackson, Ian Johnston, Dominic Kwiatkowski, Cordelia Langford, John Sillitoe on behalf of the Wellcome Sanger Institute COVID-19 Surveillance Team |
| EPI_ISL_2317038, EPI_ISL_2317039 | Lighthouse Lab in Glasgow | Wellcome Sanger Institute for the COVID-19 Genomics UK (COG-UK) Consortium | Harper VanSteenhouse, Yumi Kasai, David Gray, Carol Clugston, Anna Dominiczak and Alex Alderton, Roberto Amato, Jeffrey Barrett, Sonia Goncalves, Ewan Harrison, David K. Jackson, Ian Johnston, Dominic Kwiatkowski, Cordelia Langford, John Sillitoe on behalf of the Wellcome Sanger Institute COVID-19 Surveillance Team |
| EPI_ISL_2317040 | Lighthouse Lab in Milton Keynes | Wellcome Sanger Institute for the COVID-19 Genomics UK (COG-UK) Consortium | The Lighthouse Lab in Milton Keynes and Alex Alderton, Roberto Amato, Jeffrey Barrett, Sonia Goncalves, Ewan Harrison, David K. Jackson, Ian Johnston, Dominic Kwiatkowski, Cordelia Langford, John Sillitoe on behalf of the Wellcome Sanger Institute COVID-19 Surveillance Team |
| EPI_ISL_2317042, EPI_ISL_2317043 | Lighthouse Lab in Glasgow | Wellcome Sanger Institute for the COVID-19 Genomics UK (COG-UK) Consortium | Harper VanSteenhouse, Yumi Kasai, David Gray, Carol Clugston, Anna Dominiczak and Alex Alderton, Roberto Amato, Jeffrey Barrett, Sonia Goncalves, Ewan Harrison, David K. Jackson, Ian Johnston, Dominic Kwiatkowski, Cordelia Langford, John Sillitoe on behalf of the Wellcome Sanger Institute COVID-19 Surveillance Team |
| EPI_ISL_2317044 | Lighthouse Lab in Milton Keynes | Wellcome Sanger Institute for the COVID-19 Genomics UK (COG-UK) | The Lighthouse Lab in Milton Keynes and Alex Alderton, Roberto Amato, Jeffrey Barrett, Sonia Goncalves, Ewan Harrison, David K. Jackson, Ian Johnston, Dominic Kwiatkowski, Cordelia Langford, John Sillitoe on behalf of the Wellcome Sanger |

| | | | |
|---|---|---|---|
| | | Consortium | Institute COVID-19 Surveillance Team |
| EPI_ISL_2317046, EPI_ISL_2317047 | Lighthouse Lab in Glasgow | Wellcome Sanger Institute for the COVID-19 Genomics UK (COG-UK) Consortium | Harper VanSteenhouse, Yumi Kasai, David Gray, Carol Clugston, Anna Dominiczak and Alex Alderton, Roberto Amato, Jeffrey Barrett, Sonia Goncalves, Ewan Harrison, David K. Jackson, Ian Johnston, Dominic Kwiatkowski, Cordelia Langford, John Sillitoe on behalf of the Wellcome Sanger Institute COVID-19 Surveillance Team |
| EPI_ISL_2317048 | Lighthouse Lab in Milton Keynes | Wellcome Sanger Institute for the COVID-19 Genomics UK (COG-UK) Consortium | The Lighthouse Lab in Milton Keynes and Alex Alderton, Roberto Amato, Jeffrey Barrett, Sonia Goncalves, Ewan Harrison, David K. Jackson, Ian Johnston, Dominic Kwiatkowski, Cordelia Langford, John Sillitoe on behalf of the Wellcome Sanger Institute COVID-19 Surveillance Team |
| EPI_ISL_2317050, EPI_ISL_2317051, EPI_ISL_2317053, EPI_ISL_2317054, EPI_ISL_2317056, EPI_ISL_2317057, EPI_ISL_2317058, EPI_ISL_2317061, EPI_ISL_2317062, EPI_ISL_2317066 | Lighthouse Lab in Glasgow | Wellcome Sanger Institute for the COVID-19 Genomics UK (COG-UK) Consortium | Harper VanSteenhouse, Yumi Kasai, David Gray, Carol Clugston, Anna Dominiczak and Alex Alderton, Roberto Amato, Jeffrey Barrett, Sonia Goncalves, Ewan Harrison, David K. Jackson, Ian Johnston, Dominic Kwiatkowski, Cordelia Langford, John Sillitoe on behalf of the Wellcome Sanger Institute COVID-19 Surveillance Team |
| EPI_ISL_2317067 | Lighthouse Lab in Milton Keynes | Wellcome Sanger Institute for the COVID-19 Genomics UK (COG-UK) Consortium | The Lighthouse Lab in Milton Keynes and Alex Alderton, Roberto Amato, Jeffrey Barrett, Sonia Goncalves, Ewan Harrison, David K. Jackson, Ian Johnston, Dominic Kwiatkowski, Cordelia Langford, John Sillitoe on behalf of the Wellcome Sanger Institute COVID-19 Surveillance Team |
| EPI_ISL_2317068, EPI_ISL_2317070 | Lighthouse Lab in Glasgow | Wellcome Sanger Institute for the COVID-19 Genomics UK (COG-UK) Consortium | Harper VanSteenhouse, Yumi Kasai, David Gray, Carol Clugston, Anna Dominiczak and Alex Alderton, Roberto Amato, Jeffrey Barrett, Sonia Goncalves, Ewan Harrison, David K. Jackson, Ian Johnston, Dominic Kwiatkowski, Cordelia Langford, John Sillitoe on behalf of the Wellcome Sanger Institute COVID-19 Surveillance Team |
| EPI_ISL_2317073, EPI_ISL_2317075, EPI_ISL_2317076 | Lighthouse Lab in Milton Keynes | Wellcome Sanger Institute for the COVID-19 Genomics UK (COG-UK) Consortium | The Lighthouse Lab in Milton Keynes and Alex Alderton, Roberto Amato, Jeffrey Barrett, Sonia Goncalves, Ewan Harrison, David K. Jackson, Ian Johnston, Dominic Kwiatkowski, Cordelia Langford, John Sillitoe on behalf of the Wellcome Sanger Institute COVID-19 Surveillance Team |
| EPI_ISL_2317078 | Lighthouse Lab in Glasgow | Wellcome Sanger Institute for the COVID-19 Genomics UK (COG-UK) Consortium | Harper VanSteenhouse, Yumi Kasai, David Gray, Carol Clugston, Anna Dominiczak and Alex Alderton, Roberto Amato, Jeffrey Barrett, Sonia Goncalves, Ewan Harrison, David K. Jackson, Ian Johnston, Dominic Kwiatkowski, Cordelia Langford, John Sillitoe on behalf of the Wellcome Sanger Institute COVID-19 Surveillance Team |
| EPI_ISL_2317080 | Lighthouse Lab in Milton Keynes | Wellcome Sanger Institute for the COVID-19 Genomics UK (COG-UK) Consortium | The Lighthouse Lab in Milton Keynes and Alex Alderton, Roberto Amato, Jeffrey Barrett, Sonia Goncalves, Ewan Harrison, David K. Jackson, Ian Johnston, Dominic Kwiatkowski, Cordelia Langford, John Sillitoe on behalf of the Wellcome Sanger Institute COVID-19 Surveillance Team |
| EPI_ISL_2317083, EPI_ISL_2317084, EPI_ISL_2317086, EPI_ISL_2317088, EPI_ISL_2317089 | Lighthouse Lab in Glasgow | Wellcome Sanger Institute for the COVID-19 Genomics UK (COG-UK) Consortium | Harper VanSteenhouse, Yumi Kasai, David Gray, Carol Clugston, Anna Dominiczak and Alex Alderton, Roberto Amato, Jeffrey Barrett, Sonia Goncalves, Ewan Harrison, David K. Jackson, Ian Johnston, Dominic Kwiatkowski, Cordelia Langford, John Sillitoe on behalf of the Wellcome Sanger Institute COVID-19 Surveillance Team |
| EPI_ISL_2317090 | Lighthouse Lab in Milton Keynes | Wellcome Sanger Institute for the COVID-19 Genomics UK (COG-UK) Consortium | The Lighthouse Lab in Milton Keynes and Alex Alderton, Roberto Amato, Jeffrey Barrett, Sonia Goncalves, Ewan Harrison, David K. Jackson, Ian Johnston, Dominic Kwiatkowski, Cordelia Langford, John Sillitoe on behalf of the Wellcome Sanger Institute COVID-19 Surveillance Team |
| EPI_ISL_2317091 | Lighthouse Lab in Glasgow | Wellcome Sanger Institute for the COVID-19 Genomics UK (COG-UK) Consortium | Harper VanSteenhouse, Yumi Kasai, David Gray, Carol Clugston, Anna Dominiczak and Alex Alderton, Roberto Amato, Jeffrey Barrett, Sonia Goncalves, Ewan Harrison, David K. Jackson, Ian Johnston, Dominic Kwiatkowski, Cordelia Langford, John Sillitoe on behalf of the Wellcome Sanger Institute COVID-19 Surveillance Team |
| EPI_ISL_2317092, EPI_ISL_2317094 | Lighthouse Lab in Milton Keynes | Wellcome Sanger Institute for the COVID-19 Genomics UK (COG-UK) Consortium | The Lighthouse Lab in Milton Keynes and Alex Alderton, Roberto Amato, Jeffrey Barrett, Sonia Goncalves, Ewan Harrison, David K. Jackson, Ian Johnston, Dominic Kwiatkowski, Cordelia Langford, John Sillitoe on behalf of the Wellcome Sanger Institute COVID-19 Surveillance Team |
| EPI_ISL_2317097, EPI_ISL_2317098, EPI_ISL_2317101 | Lighthouse Lab in Glasgow | Wellcome Sanger Institute for the COVID-19 Genomics UK (COG-UK) Consortium | Harper VanSteenhouse, Yumi Kasai, David Gray, Carol Clugston, Anna Dominiczak and Alex Alderton, Roberto Amato, Jeffrey Barrett, Sonia Goncalves, Ewan Harrison, David K. Jackson, Ian Johnston, Dominic Kwiatkowski, Cordelia Langford, John Sillitoe on behalf of the Wellcome Sanger Institute COVID-19 Surveillance Team |
| EPI_ISL_2317102, EPI_ISL_2317105 | Lighthouse Lab in Milton Keynes | Wellcome Sanger Institute for the COVID-19 Genomics UK (COG-UK) Consortium | The Lighthouse Lab in Milton Keynes and Alex Alderton, Roberto Amato, Jeffrey Barrett, Sonia Goncalves, Ewan Harrison, David K. Jackson, Ian Johnston, Dominic Kwiatkowski, Cordelia Langford, John Sillitoe on behalf of the Wellcome Sanger Institute COVID-19 Surveillance Team |
| EPI_ISL_2317107 | Lighthouse Lab in Glasgow | Wellcome Sanger Institute for the COVID-19 Genomics UK (COG-UK) Consortium | Harper VanSteenhouse, Yumi Kasai, David Gray, Carol Clugston, Anna Dominiczak and Alex Alderton, Roberto Amato, Jeffrey Barrett, Sonia Goncalves, Ewan Harrison, David K. Jackson, Ian Johnston, Dominic Kwiatkowski, Cordelia Langford, John Sillitoe on behalf of the Wellcome Sanger Institute COVID-19 Surveillance Team |
| EPI_ISL_2317108 | Lighthouse Lab in Milton Keynes | Wellcome Sanger Institute for the COVID-19 Genomics UK (COG-UK) Consortium | The Lighthouse Lab in Milton Keynes and Alex Alderton, Roberto Amato, Jeffrey Barrett, Sonia Goncalves, Ewan Harrison, David K. Jackson, Ian Johnston, Dominic Kwiatkowski, Cordelia Langford, John Sillitoe on behalf of the Wellcome Sanger Institute COVID-19 Surveillance Team |
| EPI_ISL_2317109 | Lighthouse Lab in Glasgow | Wellcome Sanger Institute for the COVID-19 Genomics UK (COG-UK) Consortium | Harper VanSteenhouse, Yumi Kasai, David Gray, Carol Clugston, Anna Dominiczak and Alex Alderton, Roberto Amato, Jeffrey Barrett, Sonia Goncalves, Ewan Harrison, David K. Jackson, Ian Johnston, Dominic Kwiatkowski, Cordelia Langford, John Sillitoe on behalf of the Wellcome Sanger Institute COVID-19 Surveillance Team |
| EPI_ISL_2317110 | Lighthouse Lab in Milton Keynes | Wellcome Sanger Institute for the COVID-19 Genomics UK (COG-UK) Consortium | The Lighthouse Lab in Milton Keynes and Alex Alderton, Roberto Amato, Jeffrey Barrett, Sonia Goncalves, Ewan Harrison, David K. Jackson, Ian Johnston, Dominic Kwiatkowski, Cordelia Langford, John Sillitoe on behalf of the Wellcome Sanger Institute COVID-19 Surveillance Team |
| EPI_ISL_2317111, EPI_ISL_2317112, EPI_ISL_2317114 | Lighthouse Lab in Glasgow | Wellcome Sanger Institute for the COVID-19 Genomics UK (COG-UK) Consortium | Harper VanSteenhouse, Yumi Kasai, David Gray, Carol Clugston, Anna Dominiczak and Alex Alderton, Roberto Amato, Jeffrey Barrett, Sonia Goncalves, Ewan Harrison, David K. Jackson, Ian Johnston, Dominic Kwiatkowski, Cordelia Langford, John Sillitoe on behalf of the Wellcome Sanger Institute COVID-19 Surveillance Team |
| EPI_ISL_2317115, EPI_ISL_2317117 | Lighthouse Lab in Milton Keynes | Wellcome Sanger Institute for the COVID-19 Genomics UK (COG-UK) Consortium | The Lighthouse Lab in Milton Keynes and Alex Alderton, Roberto Amato, Jeffrey Barrett, Sonia Goncalves, Ewan Harrison, David K. Jackson, Ian Johnston, Dominic Kwiatkowski, Cordelia Langford, John Sillitoe on behalf of the Wellcome Sanger Institute COVID-19 Surveillance Team |
| EPI_ISL_2317119 | Lighthouse Lab in Glasgow | Wellcome Sanger Institute for the COVID-19 Genomics UK (COG-UK) Consortium | Harper VanSteenhouse, Yumi Kasai, David Gray, Carol Clugston, Anna Dominiczak and Alex Alderton, Roberto Amato, Jeffrey Barrett, Sonia Goncalves, Ewan Harrison, David K. Jackson, Ian Johnston, Dominic Kwiatkowski, Cordelia Langford, John Sillitoe on behalf of the Wellcome Sanger Institute COVID-19 Surveillance Team |
| EPI_ISL_2317121, EPI_ISL_2317122 | Lighthouse Lab in Milton Keynes | Wellcome Sanger Institute for the COVID-19 Genomics UK (COG-UK) Consortium | The Lighthouse Lab in Milton Keynes and Alex Alderton, Roberto Amato, Jeffrey Barrett, Sonia Goncalves, Ewan Harrison, David K. Jackson, Ian Johnston, Dominic Kwiatkowski, Cordelia Langford, John Sillitoe on behalf of the Wellcome Sanger Institute COVID-19 Surveillance Team |
| EPI_ISL_2317123, EPI_ISL_2317124, EPI_ISL_2317125 | Lighthouse Lab in Glasgow | Wellcome Sanger Institute for the COVID-19 Genomics UK (COG-UK) Consortium | Harper VanSteenhouse, Yumi Kasai, David Gray, Carol Clugston, Anna Dominiczak and Alex Alderton, Roberto Amato, Jeffrey Barrett, Sonia Goncalves, Ewan Harrison, David K. Jackson, Ian Johnston, Dominic Kwiatkowski, Cordelia Langford, John Sillitoe on behalf of the Wellcome Sanger Institute COVID-19 Surveillance Team |
| EPI_ISL_2317126 | Lighthouse Lab in Milton Keynes | Wellcome Sanger Institute for the COVID-19 Genomics UK (COG-UK) Consortium | The Lighthouse Lab in Milton Keynes and Alex Alderton, Roberto Amato, Jeffrey Barrett, Sonia Goncalves, Ewan Harrison, David K. Jackson, Ian Johnston, Dominic Kwiatkowski, Cordelia Langford, John Sillitoe on behalf of the Wellcome Sanger Institute COVID-19 Surveillance Team |
| EPI_ISL_2317127, EPI_ISL_2317131 | Lighthouse Lab in Glasgow | Wellcome Sanger Institute for the COVID-19 Genomics UK (COG-UK) Consortium | Harper VanSteenhouse, Yumi Kasai, David Gray, Carol Clugston, Anna Dominiczak and Alex Alderton, Roberto Amato, Jeffrey Barrett, Sonia Goncalves, Ewan Harrison, David K. Jackson, Ian Johnston, Dominic Kwiatkowski, Cordelia Langford, John Sillitoe on behalf of the Wellcome Sanger Institute COVID-19 Surveillance Team |
| EPI_ISL_2317133 | Lighthouse Lab in Milton Keynes | Wellcome Sanger Institute for the COVID-19 Genomics UK (COG-UK) Consortium | The Lighthouse Lab in Milton Keynes and Alex Alderton, Roberto Amato, Jeffrey Barrett, Sonia Goncalves, Ewan Harrison, David K. Jackson, Ian Johnston, Dominic Kwiatkowski, Cordelia Langford, John Sillitoe on behalf of the Wellcome Sanger Institute COVID-19 Surveillance Team |
| EPI_ISL_2317136, EPI_ISL_2317137, EPI_ISL_2317138, EPI_ISL_2317139, EPI_ISL_2317141, EPI_ISL_2317144 | Lighthouse Lab in Glasgow | Wellcome Sanger Institute for the COVID-19 Genomics UK (COG-UK) Consortium | Harper VanSteenhouse, Yumi Kasai, David Gray, Carol Clugston, Anna Dominiczak and Alex Alderton, Roberto Amato, Jeffrey Barrett, Sonia Goncalves, Ewan Harrison, David K. Jackson, Ian Johnston, Dominic Kwiatkowski, Cordelia Langford, John Sillitoe on behalf of the Wellcome Sanger Institute COVID-19 Surveillance Team |
| EPI_ISL_2317146 | Lighthouse Lab in Milton Keynes | Wellcome Sanger Institute for the COVID-19 Genomics UK (COG-UK) Consortium | The Lighthouse Lab in Milton Keynes and Alex Alderton, Roberto Amato, Jeffrey Barrett, Sonia Goncalves, Ewan Harrison, David K. Jackson, Ian Johnston, Dominic Kwiatkowski, Cordelia Langford, John Sillitoe on behalf of the Wellcome Sanger Institute COVID-19 Surveillance Team |
| EPI_ISL_2317150 | Lighthouse Lab in Glasgow | Wellcome Sanger Institute for the COVID-19 Genomics UK (COG-UK) Consortium | Harper VanSteenhouse, Yumi Kasai, David Gray, Carol Clugston, Anna Dominiczak and Alex Alderton, Roberto Amato, Jeffrey Barrett, Sonia Goncalves, Ewan Harrison, David K. Jackson, Ian Johnston, Dominic Kwiatkowski, Cordelia Langford, John Sillitoe on behalf of the Wellcome Sanger Institute COVID-19 Surveillance Team |
| EPI_ISL_2317151, EPI_ISL_2317152 | Lighthouse Lab in Milton Keynes | Wellcome Sanger Institute for the COVID-19 Genomics UK (COG-UK) Consortium | The Lighthouse Lab in Milton Keynes and Alex Alderton, Roberto Amato, Jeffrey Barrett, Sonia Goncalves, Ewan Harrison, David K. Jackson, Ian Johnston, Dominic Kwiatkowski, Cordelia Langford, John Sillitoe on behalf of the Wellcome Sanger Institute COVID-19 Surveillance Team |
| EPI_ISL_2317153 | Lighthouse Lab in Glasgow | Wellcome Sanger Institute for the COVID-19 Genomics UK (COG-UK) Consortium | Harper VanSteenhouse, Yumi Kasai, David Gray, Carol Clugston, Anna Dominiczak and Alex Alderton, Roberto Amato, Jeffrey Barrett, Sonia Goncalves, Ewan Harrison, David K. Jackson, Ian Johnston, Dominic Kwiatkowski, Cordelia Langford, John Sillitoe on behalf of the Wellcome Sanger Institute COVID-19 Surveillance Team |
| EPI_ISL_2317154 | Lighthouse Lab in Milton Keynes | Wellcome Sanger Institute for the COVID-19 Genomics UK (COG-UK) Consortium | The Lighthouse Lab in Milton Keynes and Alex Alderton, Roberto Amato, Jeffrey Barrett, Sonia Goncalves, Ewan Harrison, David K. Jackson, Ian Johnston, Dominic Kwiatkowski, Cordelia Langford, John Sillitoe on behalf of the Wellcome Sanger Institute COVID-19 Surveillance Team |
| EPI_ISL_2317155, EPI_ISL_2317157, EPI_ISL_2317158, EPI_ISL_2317159, EPI_ISL_2317160, EPI_ISL_2317161 | Lighthouse Lab in Glasgow | Wellcome Sanger Institute for the COVID-19 Genomics UK (COG-UK) Consortium | Harper VanSteenhouse, Yumi Kasai, David Gray, Carol Clugston, Anna Dominiczak and Alex Alderton, Roberto Amato, Jeffrey Barrett, Sonia Goncalves, Ewan Harrison, David K. Jackson, Ian Johnston, Dominic Kwiatkowski, Cordelia Langford, John Sillitoe on behalf of the Wellcome Sanger Institute COVID-19 Surveillance Team |
| EPI_ISL_2317162 | Lighthouse Laboratory Plymouth | Wellcome Sanger Institute for the COVID-19 Genomics UK (COG-UK) Consortium | Lighthouse Laboratory Plymouth and Alex Alderton, Roberto Amato, Jeffrey Barrett, Sonia Goncalves, Ewan Harrison, David K. Jackson, Ian Johnston, Dominic Kwiatkowski, Cordelia Langford, John Sillitoe on behalf of the Wellcome Sanger Institute COVID-19 Surveillance Team |
| EPI_ISL_2317163, EPI_ISL_2317164, EPI_ISL_2317166 | Lighthouse Lab in Milton Keynes | Wellcome Sanger Institute for the COVID-19 Genomics UK (COG-UK) Consortium | The Lighthouse Lab in Milton Keynes and Alex Alderton, Roberto Amato, Jeffrey Barrett, Sonia Goncalves, Ewan Harrison, David K. Jackson, Ian Johnston, Dominic Kwiatkowski, Cordelia Langford, John Sillitoe on behalf of the Wellcome Sanger Institute COVID-19 Surveillance Team |

| | | | |
|---|---|---|---|
| EPI_ISL_2317171 | Health Services Laboratories | Wellcome Sanger Institute for the COVID-19 Genomics UK (COG-UK) Consortium | Health Services Laboratories and Alex Alderton, Roberto Amato, Jeffrey Barrett, Sonia Goncalves, Ewan Harrison, David K. Jackson, Ian Johnston, Dominic Kwiatkowski, Cordelia Langford, John Sillitoe on behalf of the Wellcome Sanger Institute COVID-19 Surveillance Team |
| EPI_ISL_2317172 | Lighthouse Lab in Milton Keynes | Wellcome Sanger Institute for the COVID-19 Genomics UK (COG-UK) Consortium | The Lighthouse Lab in Milton Keynes and Alex Alderton, Roberto Amato, Jeffrey Barrett, Sonia Goncalves, Ewan Harrison, David K. Jackson, Ian Johnston, Dominic Kwiatkowski, Cordelia Langford, John Sillitoe on behalf of the Wellcome Sanger Institute COVID-19 Surveillance Team |
| EPI_ISL_2317173, EPI_ISL_2317174 | Lighthouse Lab in Alderley Park | Wellcome Sanger Institute for the COVID-19 Genomics UK (COG-UK) Consortium | Jacquelyn Wynn, Mairead Hyland, The Lighthouse Lab in Alderley Park and Alex Alderton, Roberto Amato, Jeffrey Barrett, Sonia Goncalves, Ewan Harrison, David K. Jackson, Ian Johnston, Dominic Kwiatkowski, Cordelia Langford, John Sillitoe on behalf of the Wellcome Sanger Institute COVID-19 Surveillance Team |
| EPI_ISL_2317175 | Lighthouse Lab in Milton Keynes | Wellcome Sanger Institute for the COVID-19 Genomics UK (COG-UK) Consortium | The Lighthouse Lab in Milton Keynes and Alex Alderton, Roberto Amato, Jeffrey Barrett, Sonia Goncalves, Ewan Harrison, David K. Jackson, Ian Johnston, Dominic Kwiatkowski, Cordelia Langford, John Sillitoe on behalf of the Wellcome Sanger Institute COVID-19 Surveillance Team |
| EPI_ISL_2317176 | Lighthouse Lab in Alderley Park | Wellcome Sanger Institute for the COVID-19 Genomics UK (COG-UK) Consortium | Jacquelyn Wynn, Mairead Hyland, The Lighthouse Lab in Alderley Park and Alex Alderton, Roberto Amato, Jeffrey Barrett, Sonia Goncalves, Ewan Harrison, David K. Jackson, Ian Johnston, Dominic Kwiatkowski, Cordelia Langford, John Sillitoe on behalf of the Wellcome Sanger Institute COVID-19 Surveillance Team |
| EPI_ISL_2317177 | Lighthouse Lab in Milton Keynes | Wellcome Sanger Institute for the COVID-19 Genomics UK (COG-UK) Consortium | The Lighthouse Lab in Milton Keynes and Alex Alderton, Roberto Amato, Jeffrey Barrett, Sonia Goncalves, Ewan Harrison, David K. Jackson, Ian Johnston, Dominic Kwiatkowski, Cordelia Langford, John Sillitoe on behalf of the Wellcome Sanger Institute COVID-19 Surveillance Team |
| EPI_ISL_2317178, EPI_ISL_2317179, EPI_ISL_2317181 | Lighthouse Lab in Alderley Park | Wellcome Sanger Institute for the COVID-19 Genomics UK (COG-UK) Consortium | Jacquelyn Wynn, Mairead Hyland, The Lighthouse Lab in Alderley Park and Alex Alderton, Roberto Amato, Jeffrey Barrett, Sonia Goncalves, Ewan Harrison, David K. Jackson, Ian Johnston, Dominic Kwiatkowski, Cordelia Langford, John Sillitoe on behalf of the Wellcome Sanger Institute COVID-19 Surveillance Team |
| EPI_ISL_2317183 | Lighthouse Lab in Milton Keynes | Wellcome Sanger Institute for the COVID-19 Genomics UK (COG-UK) Consortium | The Lighthouse Lab in Milton Keynes and Alex Alderton, Roberto Amato, Jeffrey Barrett, Sonia Goncalves, Ewan Harrison, David K. Jackson, Ian Johnston, Dominic Kwiatkowski, Cordelia Langford, John Sillitoe on behalf of the Wellcome Sanger Institute COVID-19 Surveillance Team |
| EPI_ISL_2317185 | Lighthouse Lab in Alderley Park | Wellcome Sanger Institute for the COVID-19 Genomics UK (COG-UK) Consortium | Jacquelyn Wynn, Mairead Hyland, The Lighthouse Lab in Alderley Park and Alex Alderton, Roberto Amato, Jeffrey Barrett, Sonia Goncalves, Ewan Harrison, David K. Jackson, Ian Johnston, Dominic Kwiatkowski, Cordelia Langford, John Sillitoe on behalf of the Wellcome Sanger Institute COVID-19 Surveillance Team |
| EPI_ISL_2317186 | Lighthouse Lab in Milton Keynes | Wellcome Sanger Institute for the COVID-19 Genomics UK (COG-UK) Consortium | The Lighthouse Lab in Milton Keynes and Alex Alderton, Roberto Amato, Jeffrey Barrett, Sonia Goncalves, Ewan Harrison, David K. Jackson, Ian Johnston, Dominic Kwiatkowski, Cordelia Langford, John Sillitoe on behalf of the Wellcome Sanger Institute COVID-19 Surveillance Team |
| EPI_ISL_2317188 | Health Services Laboratories | Wellcome Sanger Institute for the COVID-19 Genomics UK (COG-UK) Consortium | Health Services Laboratories and Alex Alderton, Roberto Amato, Jeffrey Barrett, Sonia Goncalves, Ewan Harrison, David K. Jackson, Ian Johnston, Dominic Kwiatkowski, Cordelia Langford, John Sillitoe on behalf of the Wellcome Sanger Institute COVID-19 Surveillance Team |
| EPI_ISL_2317190 | Lighthouse Laboratory Plymouth | Wellcome Sanger Institute for the COVID-19 Genomics UK (COG-UK) Consortium | Lighthouse Laboratory Plymouth and Alex Alderton, Roberto Amato, Jeffrey Barrett, Sonia Goncalves, Ewan Harrison, David K. Jackson, Ian Johnston, Dominic Kwiatkowski, Cordelia Langford, John Sillitoe on behalf of the Wellcome Sanger Institute COVID-19 Surveillance Team |
| EPI_ISL_2317194 | Lighthouse Lab in Milton Keynes | Wellcome Sanger Institute for the COVID-19 Genomics UK (COG-UK) Consortium | The Lighthouse Lab in Milton Keynes and Alex Alderton, Roberto Amato, Jeffrey Barrett, Sonia Goncalves, Ewan Harrison, David K. Jackson, Ian Johnston, Dominic Kwiatkowski, Cordelia Langford, John Sillitoe on behalf of the Wellcome Sanger Institute COVID-19 Surveillance Team |
| EPI_ISL_2317195 | Lighthouse Lab in Alderley Park | Wellcome Sanger Institute for the COVID-19 Genomics UK (COG-UK) Consortium | Jacquelyn Wynn, Mairead Hyland, The Lighthouse Lab in Alderley Park and Alex Alderton, Roberto Amato, Jeffrey Barrett, Sonia Goncalves, Ewan Harrison, David K. Jackson, Ian Johnston, Dominic Kwiatkowski, Cordelia Langford, John Sillitoe on behalf of the Wellcome Sanger Institute COVID-19 Surveillance Team |
| EPI_ISL_2317196 | Lighthouse Lab in Milton Keynes | Wellcome Sanger Institute for the COVID-19 Genomics UK (COG-UK) Consortium | The Lighthouse Lab in Milton Keynes and Alex Alderton, Roberto Amato, Jeffrey Barrett, Sonia Goncalves, Ewan Harrison, David K. Jackson, Ian Johnston, Dominic Kwiatkowski, Cordelia Langford, John Sillitoe on behalf of the Wellcome Sanger Institute COVID-19 Surveillance Team |
| EPI_ISL_2317198 | Lighthouse Lab in Alderley Park | Wellcome Sanger Institute for the COVID-19 Genomics UK (COG-UK) Consortium | Jacquelyn Wynn, Mairead Hyland, The Lighthouse Lab in Alderley Park and Alex Alderton, Roberto Amato, Jeffrey Barrett, Sonia Goncalves, Ewan Harrison, David K. Jackson, Ian Johnston, Dominic Kwiatkowski, Cordelia Langford, John Sillitoe on behalf of the Wellcome Sanger Institute COVID-19 Surveillance Team |
| EPI_ISL_2317199 | Health Services Laboratories | Wellcome Sanger Institute for the COVID-19 Genomics UK (COG-UK) Consortium | Health Services Laboratories and Alex Alderton, Roberto Amato, Jeffrey Barrett, Sonia Goncalves, Ewan Harrison, David K. Jackson, Ian Johnston, Dominic Kwiatkowski, Cordelia Langford, John Sillitoe on behalf of the Wellcome Sanger Institute COVID-19 Surveillance Team |
| EPI_ISL_2317202 | Lighthouse Laboratory Plymouth | Wellcome Sanger Institute for the COVID-19 Genomics UK (COG-UK) Consortium | Lighthouse Laboratory Plymouth and Alex Alderton, Roberto Amato, Jeffrey Barrett, Sonia Goncalves, Ewan Harrison, David K. Jackson, Ian Johnston, Dominic Kwiatkowski, Cordelia Langford, John Sillitoe on behalf of the Wellcome Sanger Institute COVID-19 Surveillance Team |
| EPI_ISL_2317203 | Lighthouse Lab in Alderley Park | Wellcome Sanger Institute for the COVID-19 Genomics UK (COG-UK) Consortium | Jacquelyn Wynn, Mairead Hyland, The Lighthouse Lab in Alderley Park and Alex Alderton, Roberto Amato, Jeffrey Barrett, Sonia Goncalves, Ewan Harrison, David K. Jackson, Ian Johnston, Dominic Kwiatkowski, Cordelia Langford, John Sillitoe on behalf of the Wellcome Sanger Institute COVID-19 Surveillance Team |
| EPI_ISL_2317204 | Health Services Laboratories | Wellcome Sanger Institute for the COVID-19 Genomics UK (COG-UK) Consortium | Health Services Laboratories and Alex Alderton, Roberto Amato, Jeffrey Barrett, Sonia Goncalves, Ewan Harrison, David K. Jackson, Ian Johnston, Dominic Kwiatkowski, Cordelia Langford, John Sillitoe on behalf of the Wellcome Sanger Institute COVID-19 Surveillance Team |
| EPI_ISL_2317206 | Lighthouse Lab in Alderley Park | Wellcome Sanger Institute for the COVID-19 Genomics UK (COG-UK) Consortium | Jacquelyn Wynn, Mairead Hyland, The Lighthouse Lab in Alderley Park and Alex Alderton, Roberto Amato, Jeffrey Barrett, Sonia Goncalves, Ewan Harrison, David K. Jackson, Ian Johnston, Dominic Kwiatkowski, Cordelia Langford, John Sillitoe on behalf of the Wellcome Sanger Institute COVID-19 Surveillance Team |
| EPI_ISL_2317209, EPI_ISL_2317212 | Lighthouse Lab in Milton Keynes | Wellcome Sanger Institute for the COVID-19 Genomics UK (COG-UK) Consortium | The Lighthouse Lab in Milton Keynes and Alex Alderton, Roberto Amato, Jeffrey Barrett, Sonia Goncalves, Ewan Harrison, David K. Jackson, Ian Johnston, Dominic Kwiatkowski, Cordelia Langford, John Sillitoe on behalf of the Wellcome Sanger Institute COVID-19 Surveillance Team |
| EPI_ISL_2317213, EPI_ISL_2317214 | Lighthouse Lab in Alderley Park | Wellcome Sanger Institute for the COVID-19 Genomics UK (COG-UK) Consortium | Jacquelyn Wynn, Mairead Hyland, The Lighthouse Lab in Alderley Park and Alex Alderton, Roberto Amato, Jeffrey Barrett, Sonia Goncalves, Ewan Harrison, David K. Jackson, Ian Johnston, Dominic Kwiatkowski, Cordelia Langford, John Sillitoe on behalf of the Wellcome Sanger Institute COVID-19 Surveillance Team |
| EPI_ISL_2317218 | Health Services Laboratories | Wellcome Sanger Institute for the COVID-19 Genomics UK (COG-UK) Consortium | Health Services Laboratories and Alex Alderton, Roberto Amato, Jeffrey Barrett, Sonia Goncalves, Ewan Harrison, David K. Jackson, Ian Johnston, Dominic Kwiatkowski, Cordelia Langford, John Sillitoe on behalf of the Wellcome Sanger Institute COVID-19 Surveillance Team |
| EPI_ISL_2317219 | Lighthouse Lab in Alderley Park | Wellcome Sanger Institute for the COVID-19 Genomics UK (COG-UK) Consortium | Jacquelyn Wynn, Mairead Hyland, The Lighthouse Lab in Alderley Park and Alex Alderton, Roberto Amato, Jeffrey Barrett, Sonia Goncalves, Ewan Harrison, David K. Jackson, Ian Johnston, Dominic Kwiatkowski, Cordelia Langford, John Sillitoe on behalf of the Wellcome Sanger Institute COVID-19 Surveillance Team |
| EPI_ISL_2317220, EPI_ISL_2317222, EPI_ISL_2317225 | Lighthouse Lab in Milton Keynes | Wellcome Sanger Institute for the COVID-19 Genomics UK (COG-UK) Consortium | The Lighthouse Lab in Milton Keynes and Alex Alderton, Roberto Amato, Jeffrey Barrett, Sonia Goncalves, Ewan Harrison, David K. Jackson, Ian Johnston, Dominic Kwiatkowski, Cordelia Langford, John Sillitoe on behalf of the Wellcome Sanger Institute COVID-19 Surveillance Team |
| EPI_ISL_2317227 | Lighthouse Lab in Alderley Park | Wellcome Sanger Institute for the COVID-19 Genomics UK (COG-UK) Consortium | Jacquelyn Wynn, Mairead Hyland, The Lighthouse Lab in Alderley Park and Alex Alderton, Roberto Amato, Jeffrey Barrett, Sonia Goncalves, Ewan Harrison, David K. Jackson, Ian Johnston, Dominic Kwiatkowski, Cordelia Langford, John Sillitoe on behalf of the Wellcome Sanger Institute COVID-19 Surveillance Team |
| EPI_ISL_2317229, EPI_ISL_2317230, EPI_ISL_2317231 | Lighthouse Lab in Milton Keynes | Wellcome Sanger Institute for the COVID-19 Genomics UK (COG-UK) Consortium | The Lighthouse Lab in Milton Keynes and Alex Alderton, Roberto Amato, Jeffrey Barrett, Sonia Goncalves, Ewan Harrison, David K. Jackson, Ian Johnston, Dominic Kwiatkowski, Cordelia Langford, John Sillitoe on behalf of the Wellcome Sanger Institute COVID-19 Surveillance Team |
| EPI_ISL_2317232 | Lighthouse Lab in Alderley Park | Wellcome Sanger Institute for the COVID-19 Genomics UK (COG-UK) Consortium | Jacquelyn Wynn, Mairead Hyland, The Lighthouse Lab in Alderley Park and Alex Alderton, Roberto Amato, Jeffrey Barrett, Sonia Goncalves, Ewan Harrison, David K. Jackson, Ian Johnston, Dominic Kwiatkowski, Cordelia Langford, John Sillitoe on behalf of the Wellcome Sanger Institute COVID-19 Surveillance Team |
| EPI_ISL_2317234 | Lighthouse Lab in Milton Keynes | Wellcome Sanger Institute for the COVID-19 Genomics UK (COG-UK) Consortium | The Lighthouse Lab in Milton Keynes and Alex Alderton, Roberto Amato, Jeffrey Barrett, Sonia Goncalves, Ewan Harrison, David K. Jackson, Ian Johnston, Dominic Kwiatkowski, Cordelia Langford, John Sillitoe on behalf of the Wellcome Sanger Institute COVID-19 Surveillance Team |
| EPI_ISL_2317235 | Lighthouse Lab in Alderley Park | Wellcome Sanger Institute for the COVID-19 Genomics UK (COG-UK) Consortium | Jacquelyn Wynn, Mairead Hyland, The Lighthouse Lab in Alderley Park and Alex Alderton, Roberto Amato, Jeffrey Barrett, Sonia Goncalves, Ewan Harrison, David K. Jackson, Ian Johnston, Dominic Kwiatkowski, Cordelia Langford, John Sillitoe on behalf of the Wellcome Sanger Institute COVID-19 Surveillance Team |
| EPI_ISL_2317236 | Health Services Laboratories | Wellcome Sanger Institute for the COVID-19 Genomics UK (COG-UK) Consortium | Health Services Laboratories and Alex Alderton, Roberto Amato, Jeffrey Barrett, Sonia Goncalves, Ewan Harrison, David K. Jackson, Ian Johnston, Dominic Kwiatkowski, Cordelia Langford, John Sillitoe on behalf of the Wellcome Sanger Institute COVID-19 Surveillance Team |
| EPI_ISL_2317237 | Lighthouse Lab in Alderley Park | Wellcome Sanger Institute for the COVID-19 Genomics UK (COG-UK) Consortium | Jacquelyn Wynn, Mairead Hyland, The Lighthouse Lab in Alderley Park and Alex Alderton, Roberto Amato, Jeffrey Barrett, Sonia Goncalves, Ewan Harrison, David K. Jackson, Ian Johnston, Dominic Kwiatkowski, Cordelia Langford, John Sillitoe on behalf of the Wellcome Sanger Institute COVID-19 Surveillance Team |
| EPI_ISL_2317239 | Lighthouse Lab in Milton Keynes | Wellcome Sanger Institute for the COVID-19 Genomics UK (COG-UK) Consortium | The Lighthouse Lab in Milton Keynes and Alex Alderton, Roberto Amato, Jeffrey Barrett, Sonia Goncalves, Ewan Harrison, David K. Jackson, Ian Johnston, Dominic Kwiatkowski, Cordelia Langford, John Sillitoe on behalf of the Wellcome Sanger Institute COVID-19 Surveillance Team |
| EPI_ISL_2317240 | Lighthouse Lab in Alderley Park | Wellcome Sanger Institute for the COVID-19 Genomics UK (COG-UK) Consortium | Jacquelyn Wynn, Mairead Hyland, The Lighthouse Lab in Alderley Park and Alex Alderton, Roberto Amato, Jeffrey Barrett, Sonia Goncalves, Ewan Harrison, David K. Jackson, Ian Johnston, Dominic Kwiatkowski, Cordelia Langford, John Sillitoe on behalf of the Wellcome Sanger Institute COVID-19 Surveillance Team |
| EPI_ISL_2317241 | Health Services Laboratories | Wellcome Sanger Institute for the COVID-19 Genomics UK (COG-UK) Consortium | Health Services Laboratories and Alex Alderton, Roberto Amato, Jeffrey Barrett, Sonia Goncalves, Ewan Harrison, David K. Jackson, Ian Johnston, Dominic Kwiatkowski, Cordelia Langford, John Sillitoe on behalf of the Wellcome Sanger Institute COVID-19 Surveillance Team |
| EPI_ISL_2317242, EPI_ISL_2317243 | Lighthouse Lab in Milton Keynes | Wellcome Sanger Institute for the COVID-19 Genomics UK (COG-UK) Consortium | The Lighthouse Lab in Milton Keynes and Alex Alderton, Roberto Amato, Jeffrey Barrett, Sonia Goncalves, Ewan Harrison, David K. Jackson, Ian Johnston, Dominic Kwiatkowski, Cordelia Langford, John Sillitoe on behalf of the Wellcome Sanger Institute COVID-19 Surveillance Team |
| EPI_ISL_2317244, EPI_ISL_2317246, EPI_ISL_2317247 | Lighthouse Lab in Alderley Park | Wellcome Sanger Institute for the COVID-19 Genomics UK (COG-UK) Consortium | Jacquelyn Wynn, Mairead Hyland, The Lighthouse Lab in Alderley Park and Alex Alderton, Roberto Amato, Jeffrey Barrett, Sonia Goncalves, Ewan Harrison, David K. Jackson, Ian Johnston, Dominic Kwiatkowski, Cordelia Langford, John Sillitoe on behalf of the Wellcome Sanger Institute COVID-19 Surveillance Team |
| EPI_ISL_2317249 | Lighthouse Lab in Milton Keynes | Wellcome Sanger Institute for the COVID-19 Genomics UK (COG-UK) Consortium | The Lighthouse Lab in Milton Keynes and Alex Alderton, Roberto Amato, Jeffrey Barrett, Sonia Goncalves, Ewan Harrison, David K. Jackson, Ian Johnston, Dominic Kwiatkowski, Cordelia Langford, John Sillitoe on behalf of the Wellcome Sanger Institute COVID-19 Surveillance Team |
| EPI_ISL_2317250, EPI_ISL_2317251 | Lighthouse Lab in Alderley Park | Wellcome Sanger Institute for the COVID-19 Genomics UK (COG-UK) Consortium | Jacquelyn Wynn, Mairead Hyland, The Lighthouse Lab in Alderley Park and Alex Alderton, Roberto Amato, Jeffrey Barrett, Sonia Goncalves, Ewan Harrison, David K. Jackson, Ian Johnston, Dominic Kwiatkowski, Cordelia Langford, John Sillitoe on behalf of the Wellcome Sanger Institute COVID-19 Surveillance Team |
| EPI_ISL_2317253 | Lighthouse Lab in Milton Keynes | Wellcome Sanger Institute for the COVID-19 Genomics UK (COG-UK) Consortium | The Lighthouse Lab in Milton Keynes and Alex Alderton, Roberto Amato, Jeffrey Barrett, Sonia Goncalves, Ewan Harrison, David K. Jackson, Ian Johnston, Dominic Kwiatkowski, Cordelia Langford, John Sillitoe on behalf of the Wellcome Sanger Institute COVID-19 Surveillance Team |
| EPI_ISL_2317256 | Lighthouse Lab in Alderley Park | Wellcome Sanger Institute for the COVID-19 Genomics UK (COG-UK) Consortium | Jacquelyn Wynn, Mairead Hyland, The Lighthouse Lab in Alderley Park and Alex Alderton, Roberto Amato, Jeffrey Barrett, Sonia Goncalves, Ewan Harrison, David K. Jackson, Ian Johnston, Dominic Kwiatkowski, Cordelia Langford, John Sillitoe on behalf of the Wellcome Sanger Institute COVID-19 Surveillance Team |
| EPI_ISL_2317257 | Health Services Laboratories | Wellcome Sanger Institute for the COVID-19 Genomics UK (COG-UK) Consortium | Health Services Laboratories and Alex Alderton, Roberto Amato, Jeffrey Barrett, Sonia Goncalves, Ewan Harrison, David K. Jackson, Ian Johnston, Dominic Kwiatkowski, Cordelia Langford, John Sillitoe on behalf of the Wellcome Sanger Institute COVID-19 Surveillance Team |
| EPI_ISL_2317258 | Lighthouse Laboratory Plymouth | Wellcome Sanger Institute for the COVID-19 Genomics UK (COG-UK) | Lighthouse Laboratory Plymouth and Alex Alderton, Roberto Amato, Jeffrey Barrett, Sonia Goncalves, Ewan Harrison, David K. Jackson, Ian Johnston, Dominic Kwiatkowski, Cordelia Langford, John Sillitoe on behalf of the Wellcome Sanger |

| | | | |
|---|---|---|---|
| | | Consortium | Institute COVID-19 Surveillance Team |
| EPI_ISL_2317260 | Lighthouse Lab in Milton Keynes | Wellcome Sanger Institute for the COVID-19 Genomics UK (COG-UK) Consortium | The Lighthouse Lab in Milton Keynes and Alex Alderton, Roberto Amato, Jeffrey Barrett, Sonia Goncalves, Ewan Harrison, David K. Jackson, Ian Johnston, Dominic Kwiatkowski, Cordelia Langford, John Sillitoe on behalf of the Wellcome Sanger Institute COVID-19 Surveillance Team |
| EPI_ISL_2317263 | Lighthouse Lab in Alderley Park | Wellcome Sanger Institute for the COVID-19 Genomics UK (COG-UK) Consortium | Jacquelyn Wynn, Mairead Hyland, The Lighthouse Lab in Alderley Park and Alex Alderton, Roberto Amato, Jeffrey Barrett, Sonia Goncalves, Ewan Harrison, David K. Jackson, Ian Johnston, Dominic Kwiatkowski, Cordelia Langford, John Sillitoe on behalf of the Wellcome Sanger Institute COVID-19 Surveillance Team |
| EPI_ISL_2317264 | Lighthouse Lab in Milton Keynes | Wellcome Sanger Institute for the COVID-19 Genomics UK (COG-UK) Consortium | The Lighthouse Lab in Milton Keynes and Alex Alderton, Roberto Amato, Jeffrey Barrett, Sonia Goncalves, Ewan Harrison, David K. Jackson, Ian Johnston, Dominic Kwiatkowski, Cordelia Langford, John Sillitoe on behalf of the Wellcome Sanger Institute COVID-19 Surveillance Team |
| EPI_ISL_2317265 | Lighthouse Lab in Alderley Park | Wellcome Sanger Institute for the COVID-19 Genomics UK (COG-UK) Consortium | Jacquelyn Wynn, Mairead Hyland, The Lighthouse Lab in Alderley Park and Alex Alderton, Roberto Amato, Jeffrey Barrett, Sonia Goncalves, Ewan Harrison, David K. Jackson, Ian Johnston, Dominic Kwiatkowski, Cordelia Langford, John Sillitoe on behalf of the Wellcome Sanger Institute COVID-19 Surveillance Team |
| EPI_ISL_2317266 | Lighthouse Laboratory Plymouth | Wellcome Sanger Institute for the COVID-19 Genomics UK (COG-UK) Consortium | Lighthouse Laboratory Plymouth and Alex Alderton, Roberto Amato, Jeffrey Barrett, Sonia Goncalves, Ewan Harrison, David K. Jackson, Ian Johnston, Dominic Kwiatkowski, Cordelia Langford, John Sillitoe on behalf of the Wellcome Sanger Institute COVID-19 Surveillance Team |
| EPI_ISL_2317267 | Lighthouse Lab in Alderley Park | Wellcome Sanger Institute for the COVID-19 Genomics UK (COG-UK) Consortium | Jacquelyn Wynn, Mairead Hyland, The Lighthouse Lab in Alderley Park and Alex Alderton, Roberto Amato, Jeffrey Barrett, Sonia Goncalves, Ewan Harrison, David K. Jackson, Ian Johnston, Dominic Kwiatkowski, Cordelia Langford, John Sillitoe on behalf of the Wellcome Sanger Institute COVID-19 Surveillance Team |
| EPI_ISL_2317268, EPI_ISL_2317269 | Lighthouse Lab in Milton Keynes | Wellcome Sanger Institute for the COVID-19 Genomics UK (COG-UK) Consortium | The Lighthouse Lab in Milton Keynes and Alex Alderton, Roberto Amato, Jeffrey Barrett, Sonia Goncalves, Ewan Harrison, David K. Jackson, Ian Johnston, Dominic Kwiatkowski, Cordelia Langford, John Sillitoe on behalf of the Wellcome Sanger Institute COVID-19 Surveillance Team |
| EPI_ISL_2317270 | Lighthouse Lab in Alderley Park | Wellcome Sanger Institute for the COVID-19 Genomics UK (COG-UK) Consortium | Jacquelyn Wynn, Mairead Hyland, The Lighthouse Lab in Alderley Park and Alex Alderton, Roberto Amato, Jeffrey Barrett, Sonia Goncalves, Ewan Harrison, David K. Jackson, Ian Johnston, Dominic Kwiatkowski, Cordelia Langford, John Sillitoe on behalf of the Wellcome Sanger Institute COVID-19 Surveillance Team |
| EPI_ISL_2317273, EPI_ISL_2317276, EPI_ISL_2317277 | Lighthouse Lab in Milton Keynes | Wellcome Sanger Institute for the COVID-19 Genomics UK (COG-UK) Consortium | The Lighthouse Lab in Milton Keynes and Alex Alderton, Roberto Amato, Jeffrey Barrett, Sonia Goncalves, Ewan Harrison, David K. Jackson, Ian Johnston, Dominic Kwiatkowski, Cordelia Langford, John Sillitoe on behalf of the Wellcome Sanger Institute COVID-19 Surveillance Team |
| EPI_ISL_2317279 | Health Services Laboratories | Wellcome Sanger Institute for the COVID-19 Genomics UK (COG-UK) Consortium | Health Services Laboratories and Alex Alderton, Roberto Amato, Jeffrey Barrett, Sonia Goncalves, Ewan Harrison, David K. Jackson, Ian Johnston, Dominic Kwiatkowski, Cordelia Langford, John Sillitoe on behalf of the Wellcome Sanger Institute COVID-19 Surveillance Team |
| EPI_ISL_2317283 | Lighthouse Laboratory Plymouth | Wellcome Sanger Institute for the COVID-19 Genomics UK (COG-UK) Consortium | Lighthouse Laboratory Plymouth and Alex Alderton, Roberto Amato, Jeffrey Barrett, Sonia Goncalves, Ewan Harrison, David K. Jackson, Ian Johnston, Dominic Kwiatkowski, Cordelia Langford, John Sillitoe on behalf of the Wellcome Sanger Institute COVID-19 Surveillance Team |
| EPI_ISL_2317284 | Lighthouse Lab in Milton Keynes | Wellcome Sanger Institute for the COVID-19 Genomics UK (COG-UK) Consortium | The Lighthouse Lab in Milton Keynes and Alex Alderton, Roberto Amato, Jeffrey Barrett, Sonia Goncalves, Ewan Harrison, David K. Jackson, Ian Johnston, Dominic Kwiatkowski, Cordelia Langford, John Sillitoe on behalf of the Wellcome Sanger Institute COVID-19 Surveillance Team |
| EPI_ISL_2317285 | Lighthouse Lab in Alderley Park | Wellcome Sanger Institute for the COVID-19 Genomics UK (COG-UK) Consortium | Jacquelyn Wynn, Mairead Hyland, The Lighthouse Lab in Alderley Park and Alex Alderton, Roberto Amato, Jeffrey Barrett, Sonia Goncalves, Ewan Harrison, David K. Jackson, Ian Johnston, Dominic Kwiatkowski, Cordelia Langford, John Sillitoe on behalf of the Wellcome Sanger Institute COVID-19 Surveillance Team |
| EPI_ISL_2317286 | Health Services Laboratories | Wellcome Sanger Institute for the COVID-19 Genomics UK (COG-UK) Consortium | Health Services Laboratories and Alex Alderton, Roberto Amato, Jeffrey Barrett, Sonia Goncalves, Ewan Harrison, David K. Jackson, Ian Johnston, Dominic Kwiatkowski, Cordelia Langford, John Sillitoe on behalf of the Wellcome Sanger Institute COVID-19 Surveillance Team |
| EPI_ISL_2317287, EPI_ISL_2317289 | Lighthouse Lab in Alderley Park | Wellcome Sanger Institute for the COVID-19 Genomics UK (COG-UK) Consortium | Jacquelyn Wynn, Mairead Hyland, The Lighthouse Lab in Alderley Park and Alex Alderton, Roberto Amato, Jeffrey Barrett, Sonia Goncalves, Ewan Harrison, David K. Jackson, Ian Johnston, Dominic Kwiatkowski, Cordelia Langford, John Sillitoe on behalf of the Wellcome Sanger Institute COVID-19 Surveillance Team |
| EPI_ISL_2317290 | Lighthouse Lab in Milton Keynes | Wellcome Sanger Institute for the COVID-19 Genomics UK (COG-UK) Consortium | The Lighthouse Lab in Milton Keynes and Alex Alderton, Roberto Amato, Jeffrey Barrett, Sonia Goncalves, Ewan Harrison, David K. Jackson, Ian Johnston, Dominic Kwiatkowski, Cordelia Langford, John Sillitoe on behalf of the Wellcome Sanger Institute COVID-19 Surveillance Team |
| EPI_ISL_2317292, EPI_ISL_2317294, EPI_ISL_2317295 | Lighthouse Lab in Alderley Park | Wellcome Sanger Institute for the COVID-19 Genomics UK (COG-UK) Consortium | Jacquelyn Wynn, Mairead Hyland, The Lighthouse Lab in Alderley Park and Alex Alderton, Roberto Amato, Jeffrey Barrett, Sonia Goncalves, Ewan Harrison, David K. Jackson, Ian Johnston, Dominic Kwiatkowski, Cordelia Langford, John Sillitoe on behalf of the Wellcome Sanger Institute COVID-19 Surveillance Team |
| EPI_ISL_2317296, EPI_ISL_2317297 | Health Services Laboratories | Wellcome Sanger Institute for the COVID-19 Genomics UK (COG-UK) Consortium | Health Services Laboratories and Alex Alderton, Roberto Amato, Jeffrey Barrett, Sonia Goncalves, Ewan Harrison, David K. Jackson, Ian Johnston, Dominic Kwiatkowski, Cordelia Langford, John Sillitoe on behalf of the Wellcome Sanger Institute COVID-19 Surveillance Team |
| EPI_ISL_2317298, EPI_ISL_2317299 | Lighthouse Lab in Milton Keynes | Wellcome Sanger Institute for the COVID-19 Genomics UK (COG-UK) Consortium | The Lighthouse Lab in Milton Keynes and Alex Alderton, Roberto Amato, Jeffrey Barrett, Sonia Goncalves, Ewan Harrison, David K. Jackson, Ian Johnston, Dominic Kwiatkowski, Cordelia Langford, John Sillitoe on behalf of the Wellcome Sanger Institute COVID-19 Surveillance Team |
| EPI_ISL_2317300 | Lighthouse Lab in Alderley Park | Wellcome Sanger Institute for the COVID-19 Genomics UK (COG-UK) Consortium | Jacquelyn Wynn, Mairead Hyland, The Lighthouse Lab in Alderley Park and Alex Alderton, Roberto Amato, Jeffrey Barrett, Sonia Goncalves, Ewan Harrison, David K. Jackson, Ian Johnston, Dominic Kwiatkowski, Cordelia Langford, John Sillitoe on behalf of the Wellcome Sanger Institute COVID-19 Surveillance Team |
| EPI_ISL_2317302 | Lighthouse Laboratory Plymouth | Wellcome Sanger Institute for the COVID-19 Genomics UK (COG-UK) Consortium | Lighthouse Laboratory Plymouth and Alex Alderton, Roberto Amato, Jeffrey Barrett, Sonia Goncalves, Ewan Harrison, David K. Jackson, Ian Johnston, Dominic Kwiatkowski, Cordelia Langford, John Sillitoe on behalf of the Wellcome Sanger Institute COVID-19 Surveillance Team |
| EPI_ISL_2317305 | Lighthouse Lab in Alderley Park | Wellcome Sanger Institute for the COVID-19 Genomics UK (COG-UK) Consortium | Jacquelyn Wynn, Mairead Hyland, The Lighthouse Lab in Alderley Park and Alex Alderton, Roberto Amato, Jeffrey Barrett, Sonia Goncalves, Ewan Harrison, David K. Jackson, Ian Johnston, Dominic Kwiatkowski, Cordelia Langford, John Sillitoe on behalf of the Wellcome Sanger Institute COVID-19 Surveillance Team |
| EPI_ISL_2317309 | Lighthouse Laboratory Plymouth | Wellcome Sanger Institute for the COVID-19 Genomics UK (COG-UK) Consortium | Lighthouse Laboratory Plymouth and Alex Alderton, Roberto Amato, Jeffrey Barrett, Sonia Goncalves, Ewan Harrison, David K. Jackson, Ian Johnston, Dominic Kwiatkowski, Cordelia Langford, John Sillitoe on behalf of the Wellcome Sanger Institute COVID-19 Surveillance Team |
| EPI_ISL_2317310 | Lighthouse Lab in Alderley Park | Wellcome Sanger Institute for the COVID-19 Genomics UK (COG-UK) Consortium | Jacquelyn Wynn, Mairead Hyland, The Lighthouse Lab in Alderley Park and Alex Alderton, Roberto Amato, Jeffrey Barrett, Sonia Goncalves, Ewan Harrison, David K. Jackson, Ian Johnston, Dominic Kwiatkowski, Cordelia Langford, John Sillitoe on behalf of the Wellcome Sanger Institute COVID-19 Surveillance Team |
| EPI_ISL_2317311 | Lighthouse Lab in Milton Keynes | Wellcome Sanger Institute for the COVID-19 Genomics UK (COG-UK) Consortium | The Lighthouse Lab in Milton Keynes and Alex Alderton, Roberto Amato, Jeffrey Barrett, Sonia Goncalves, Ewan Harrison, David K. Jackson, Ian Johnston, Dominic Kwiatkowski, Cordelia Langford, John Sillitoe on behalf of the Wellcome Sanger Institute COVID-19 Surveillance Team |
| EPI_ISL_2317312 | Lighthouse Lab in Alderley Park | Wellcome Sanger Institute for the COVID-19 Genomics UK (COG-UK) Consortium | Jacquelyn Wynn, Mairead Hyland, The Lighthouse Lab in Alderley Park and Alex Alderton, Roberto Amato, Jeffrey Barrett, Sonia Goncalves, Ewan Harrison, David K. Jackson, Ian Johnston, Dominic Kwiatkowski, Cordelia Langford, John Sillitoe on behalf of the Wellcome Sanger Institute COVID-19 Surveillance Team |
| EPI_ISL_2317317 | Lighthouse Laboratory Plymouth | Wellcome Sanger Institute for the COVID-19 Genomics UK (COG-UK) Consortium | Lighthouse Laboratory Plymouth and Alex Alderton, Roberto Amato, Jeffrey Barrett, Sonia Goncalves, Ewan Harrison, David K. Jackson, Ian Johnston, Dominic Kwiatkowski, Cordelia Langford, John Sillitoe on behalf of the Wellcome Sanger Institute COVID-19 Surveillance Team |
| EPI_ISL_2317321, EPI_ISL_2317328 | Lighthouse Lab in Alderley Park | Wellcome Sanger Institute for the COVID-19 Genomics UK (COG-UK) Consortium | Jacquelyn Wynn, Mairead Hyland, The Lighthouse Lab in Alderley Park and Alex Alderton, Roberto Amato, Jeffrey Barrett, Sonia Goncalves, Ewan Harrison, David K. Jackson, Ian Johnston, Dominic Kwiatkowski, Cordelia Langford, John Sillitoe on behalf of the Wellcome Sanger Institute COVID-19 Surveillance Team |
| EPI_ISL_2317329, EPI_ISL_2317330 | Lighthouse Lab in Milton Keynes | Wellcome Sanger Institute for the COVID-19 Genomics UK (COG-UK) Consortium | The Lighthouse Lab in Milton Keynes and Alex Alderton, Roberto Amato, Jeffrey Barrett, Sonia Goncalves, Ewan Harrison, David K. Jackson, Ian Johnston, Dominic Kwiatkowski, Cordelia Langford, John Sillitoe on behalf of the Wellcome Sanger Institute COVID-19 Surveillance Team |
| EPI_ISL_2317331, EPI_ISL_2317332 | Lighthouse Lab in Alderley Park | Wellcome Sanger Institute for the COVID-19 Genomics UK (COG-UK) Consortium | Jacquelyn Wynn, Mairead Hyland, The Lighthouse Lab in Alderley Park and Alex Alderton, Roberto Amato, Jeffrey Barrett, Sonia Goncalves, Ewan Harrison, David K. Jackson, Ian Johnston, Dominic Kwiatkowski, Cordelia Langford, John Sillitoe on behalf of the Wellcome Sanger Institute COVID-19 Surveillance Team |
| EPI_ISL_2317333 | Health Services Laboratories | Wellcome Sanger Institute for the COVID-19 Genomics UK (COG-UK) Consortium | Health Services Laboratories and Alex Alderton, Roberto Amato, Jeffrey Barrett, Sonia Goncalves, Ewan Harrison, David K. Jackson, Ian Johnston, Dominic Kwiatkowski, Cordelia Langford, John Sillitoe on behalf of the Wellcome Sanger Institute COVID-19 Surveillance Team |
| EPI_ISL_2317334 | Lighthouse Lab in Milton Keynes | Wellcome Sanger Institute for the COVID-19 Genomics UK (COG-UK) Consortium | The Lighthouse Lab in Milton Keynes and Alex Alderton, Roberto Amato, Jeffrey Barrett, Sonia Goncalves, Ewan Harrison, David K. Jackson, Ian Johnston, Dominic Kwiatkowski, Cordelia Langford, John Sillitoe on behalf of the Wellcome Sanger Institute COVID-19 Surveillance Team |
| EPI_ISL_2317335 | Randox Laboratories | Wellcome Sanger Institute for the COVID-19 Genomics UK (COG-UK) Consortium | Randox Laboratories and Alex Alderton, Roberto Amato, Jeffrey Barrett, Sonia Goncalves, Ewan Harrison, David K. Jackson, Ian Johnston, Dominic Kwiatkowski, Cordelia Langford, John Sillitoe on behalf of the Wellcome Sanger Institute COVID-19 Surveillance Team |
| EPI_ISL_2317338, EPI_ISL_2317339 | Lighthouse Lab in Alderley Park | Wellcome Sanger Institute for the COVID-19 Genomics UK (COG-UK) Consortium | Jacquelyn Wynn, Mairead Hyland, The Lighthouse Lab in Alderley Park and Alex Alderton, Roberto Amato, Jeffrey Barrett, Sonia Goncalves, Ewan Harrison, David K. Jackson, Ian Johnston, Dominic Kwiatkowski, Cordelia Langford, John Sillitoe on behalf of the Wellcome Sanger Institute COVID-19 Surveillance Team |
| EPI_ISL_2317340, EPI_ISL_2317341 | Lighthouse Lab in Milton Keynes | Wellcome Sanger Institute for the COVID-19 Genomics UK (COG-UK) Consortium | The Lighthouse Lab in Milton Keynes and Alex Alderton, Roberto Amato, Jeffrey Barrett, Sonia Goncalves, Ewan Harrison, David K. Jackson, Ian Johnston, Dominic Kwiatkowski, Cordelia Langford, John Sillitoe on behalf of the Wellcome Sanger Institute COVID-19 Surveillance Team |
| EPI_ISL_2317344 | Lighthouse Laboratory Plymouth | Wellcome Sanger Institute for the COVID-19 Genomics UK (COG-UK) Consortium | Lighthouse Laboratory Plymouth and Alex Alderton, Roberto Amato, Jeffrey Barrett, Sonia Goncalves, Ewan Harrison, David K. Jackson, Ian Johnston, Dominic Kwiatkowski, Cordelia Langford, John Sillitoe on behalf of the Wellcome Sanger Institute COVID-19 Surveillance Team |
| EPI_ISL_2317345, EPI_ISL_2317346, EPI_ISL_2317350 | Lighthouse Lab in Alderley Park | Wellcome Sanger Institute for the COVID-19 Genomics UK (COG-UK) Consortium | Jacquelyn Wynn, Mairead Hyland, The Lighthouse Lab in Alderley Park and Alex Alderton, Roberto Amato, Jeffrey Barrett, Sonia Goncalves, Ewan Harrison, David K. Jackson, Ian Johnston, Dominic Kwiatkowski, Cordelia Langford, John Sillitoe on behalf of the Wellcome Sanger Institute COVID-19 Surveillance Team |
| EPI_ISL_2317351 | Health Services Laboratories | Wellcome Sanger Institute for the COVID-19 Genomics UK (COG-UK) Consortium | Health Services Laboratories and Alex Alderton, Roberto Amato, Jeffrey Barrett, Sonia Goncalves, Ewan Harrison, David K. Jackson, Ian Johnston, Dominic Kwiatkowski, Cordelia Langford, John Sillitoe on behalf of the Wellcome Sanger Institute COVID-19 Surveillance Team |
| EPI_ISL_2317352, EPI_ISL_2317353 | Lighthouse Lab in Milton Keynes | Wellcome Sanger Institute for the COVID-19 Genomics UK (COG-UK) Consortium | The Lighthouse Lab in Milton Keynes and Alex Alderton, Roberto Amato, Jeffrey Barrett, Sonia Goncalves, Ewan Harrison, David K. Jackson, Ian Johnston, Dominic Kwiatkowski, Cordelia Langford, John Sillitoe on behalf of the Wellcome Sanger Institute COVID-19 Surveillance Team |
| EPI_ISL_2317355 | Lighthouse Lab in Alderley Park | Wellcome Sanger Institute for the COVID-19 Genomics UK (COG-UK) Consortium | Jacquelyn Wynn, Mairead Hyland, The Lighthouse Lab in Alderley Park and Alex Alderton, Roberto Amato, Jeffrey Barrett, Sonia Goncalves, Ewan Harrison, David K. Jackson, Ian Johnston, Dominic Kwiatkowski, Cordelia Langford, John Sillitoe on behalf of the Wellcome Sanger Institute COVID-19 Surveillance Team |
| EPI_ISL_2317357 | Health Services Laboratories | Wellcome Sanger Institute for the COVID-19 Genomics UK (COG-UK) Consortium | Health Services Laboratories and Alex Alderton, Roberto Amato, Jeffrey Barrett, Sonia Goncalves, Ewan Harrison, David K. Jackson, Ian Johnston, Dominic Kwiatkowski, Cordelia Langford, John Sillitoe on behalf of the Wellcome Sanger Institute COVID-19 Surveillance Team |
| EPI_ISL_2317367 | Lighthouse Lab in Milton Keynes | Wellcome Sanger Institute for the COVID-19 Genomics UK (COG-UK) Consortium | The Lighthouse Lab in Milton Keynes and Alex Alderton, Roberto Amato, Jeffrey Barrett, Sonia Goncalves, Ewan Harrison, David K. Jackson, Ian Johnston, Dominic Kwiatkowski, Cordelia Langford, John Sillitoe on behalf of the Wellcome Sanger Institute COVID-19 Surveillance Team |
| EPI_ISL_2317368 | Lighthouse Lab in Alderley Park | Wellcome Sanger Institute for the COVID-19 Genomics UK (COG-UK) Consortium | Jacquelyn Wynn, Mairead Hyland, The Lighthouse Lab in Alderley Park and Alex Alderton, Roberto Amato, Jeffrey Barrett, Sonia Goncalves, Ewan Harrison, David K. Jackson, Ian Johnston, Dominic Kwiatkowski, Cordelia Langford, John Sillitoe on behalf of the Wellcome Sanger Institute COVID-19 Surveillance Team |
| EPI_ISL_2317369 | Lighthouse Laboratory Plymouth | Wellcome Sanger Institute for the COVID-19 Genomics UK (COG-UK) | Lighthouse Laboratory Plymouth and Alex Alderton, Roberto Amato, Jeffrey Barrett, Sonia Goncalves, Ewan Harrison, David K. Jackson, Ian Johnston, Dominic Kwiatkowski, Cordelia Langford, John Sillitoe on behalf of the Wellcome Sanger |

| | | | |
|---|---|---|---|
| | | Consortium | Institute COVID-19 Surveillance Team |
| EPI_ISL_2317370, EPI_ISL_2317371 | Lighthouse Lab in Alderley Park | Wellcome Sanger Institute for the COVID-19 Genomics UK (COG-UK) Consortium | Jacquelyn Wynn, Mairead Hyland, The Lighthouse Lab in Alderley Park and Alex Alderton, Roberto Amato, Jeffrey Barrett, Sonia Goncalves, Ewan Harrison, David K. Jackson, Ian Johnston, Dominic Kwiatkowski, Cordelia Langford, John Sillitoe on behalf of the Wellcome Sanger Institute COVID-19 Surveillance Team |
| EPI_ISL_2317377 | Lighthouse Lab in Milton Keynes | Wellcome Sanger Institute for the COVID-19 Genomics UK (COG-UK) Consortium | The Lighthouse Lab in Milton Keynes and Alex Alderton, Roberto Amato, Jeffrey Barrett, Sonia Goncalves, Ewan Harrison, David K. Jackson, Ian Johnston, Dominic Kwiatkowski, Cordelia Langford, John Sillitoe on behalf of the Wellcome Sanger Institute COVID-19 Surveillance Team |
| EPI_ISL_2317378, EPI_ISL_2317380 | Health Services Laboratories | Wellcome Sanger Institute for the COVID-19 Genomics UK (COG-UK) Consortium | Health Services Laboratories and Alex Alderton, Roberto Amato, Jeffrey Barrett, Sonia Goncalves, Ewan Harrison, David K. Jackson, Ian Johnston, Dominic Kwiatkowski, Cordelia Langford, John Sillitoe on behalf of the Wellcome Sanger Institute COVID-19 Surveillance Team |
| EPI_ISL_2317381, EPI_ISL_2317382, EPI_ISL_2317383, EPI_ISL_2317384, EPI_ISL_2317385, EPI_ISL_2317386 | Lighthouse Lab in Alderley Park | Wellcome Sanger Institute for the COVID-19 Genomics UK (COG-UK) Consortium | Jacquelyn Wynn, Mairead Hyland, The Lighthouse Lab in Alderley Park and Alex Alderton, Roberto Amato, Jeffrey Barrett, Sonia Goncalves, Ewan Harrison, David K. Jackson, Ian Johnston, Dominic Kwiatkowski, Cordelia Langford, John Sillitoe on behalf of the Wellcome Sanger Institute COVID-19 Surveillance Team |
| EPI_ISL_2317387 | Health Services Laboratories | Wellcome Sanger Institute for the COVID-19 Genomics UK (COG-UK) Consortium | Health Services Laboratories and Alex Alderton, Roberto Amato, Jeffrey Barrett, Sonia Goncalves, Ewan Harrison, David K. Jackson, Ian Johnston, Dominic Kwiatkowski, Cordelia Langford, John Sillitoe on behalf of the Wellcome Sanger Institute COVID-19 Surveillance Team |
| EPI_ISL_2317389 | Lighthouse Lab in Milton Keynes | Wellcome Sanger Institute for the COVID-19 Genomics UK (COG-UK) Consortium | The Lighthouse Lab in Milton Keynes and Alex Alderton, Roberto Amato, Jeffrey Barrett, Sonia Goncalves, Ewan Harrison, David K. Jackson, Ian Johnston, Dominic Kwiatkowski, Cordelia Langford, John Sillitoe on behalf of the Wellcome Sanger Institute COVID-19 Surveillance Team |
| EPI_ISL_2317390, EPI_ISL_2317393, EPI_ISL_2317394 | Lighthouse Lab in Alderley Park | Wellcome Sanger Institute for the COVID-19 Genomics UK (COG-UK) Consortium | Jacquelyn Wynn, Mairead Hyland, The Lighthouse Lab in Alderley Park and Alex Alderton, Roberto Amato, Jeffrey Barrett, Sonia Goncalves, Ewan Harrison, David K. Jackson, Ian Johnston, Dominic Kwiatkowski, Cordelia Langford, John Sillitoe on behalf of the Wellcome Sanger Institute COVID-19 Surveillance Team |
| EPI_ISL_2317395 | Lighthouse Lab in Milton Keynes | Wellcome Sanger Institute for the COVID-19 Genomics UK (COG-UK) Consortium | The Lighthouse Lab in Milton Keynes and Alex Alderton, Roberto Amato, Jeffrey Barrett, Sonia Goncalves, Ewan Harrison, David K. Jackson, Ian Johnston, Dominic Kwiatkowski, Cordelia Langford, John Sillitoe on behalf of the Wellcome Sanger Institute COVID-19 Surveillance Team |
| EPI_ISL_2317398, EPI_ISL_2317399 | Lighthouse Lab in Alderley Park | Wellcome Sanger Institute for the COVID-19 Genomics UK (COG-UK) Consortium | Jacquelyn Wynn, Mairead Hyland, The Lighthouse Lab in Alderley Park and Alex Alderton, Roberto Amato, Jeffrey Barrett, Sonia Goncalves, Ewan Harrison, David K. Jackson, Ian Johnston, Dominic Kwiatkowski, Cordelia Langford, John Sillitoe on behalf of the Wellcome Sanger Institute COVID-19 Surveillance Team |
| EPI_ISL_2317400, EPI_ISL_2317401 | Lighthouse Lab in Milton Keynes | Wellcome Sanger Institute for the COVID-19 Genomics UK (COG-UK) Consortium | The Lighthouse Lab in Milton Keynes and Alex Alderton, Roberto Amato, Jeffrey Barrett, Sonia Goncalves, Ewan Harrison, David K. Jackson, Ian Johnston, Dominic Kwiatkowski, Cordelia Langford, John Sillitoe on behalf of the Wellcome Sanger Institute COVID-19 Surveillance Team |
| EPI_ISL_2317404 | Lighthouse Lab in Alderley Park | Wellcome Sanger Institute for the COVID-19 Genomics UK (COG-UK) Consortium | Jacquelyn Wynn, Mairead Hyland, The Lighthouse Lab in Alderley Park and Alex Alderton, Roberto Amato, Jeffrey Barrett, Sonia Goncalves, Ewan Harrison, David K. Jackson, Ian Johnston, Dominic Kwiatkowski, Cordelia Langford, John Sillitoe on behalf of the Wellcome Sanger Institute COVID-19 Surveillance Team |
| EPI_ISL_2317406 | Lighthouse Lab in Milton Keynes | Wellcome Sanger Institute for the COVID-19 Genomics UK (COG-UK) Consortium | The Lighthouse Lab in Milton Keynes and Alex Alderton, Roberto Amato, Jeffrey Barrett, Sonia Goncalves, Ewan Harrison, David K. Jackson, Ian Johnston, Dominic Kwiatkowski, Cordelia Langford, John Sillitoe on behalf of the Wellcome Sanger Institute COVID-19 Surveillance Team |
| EPI_ISL_2317408 | Lighthouse Lab in Alderley Park | Wellcome Sanger Institute for the COVID-19 Genomics UK (COG-UK) Consortium | Jacquelyn Wynn, Mairead Hyland, The Lighthouse Lab in Alderley Park and Alex Alderton, Roberto Amato, Jeffrey Barrett, Sonia Goncalves, Ewan Harrison, David K. Jackson, Ian Johnston, Dominic Kwiatkowski, Cordelia Langford, John Sillitoe on behalf of the Wellcome Sanger Institute COVID-19 Surveillance Team |
| EPI_ISL_2317411 | Health Services Laboratories | Wellcome Sanger Institute for the COVID-19 Genomics UK (COG-UK) Consortium | Health Services Laboratories and Alex Alderton, Roberto Amato, Jeffrey Barrett, Sonia Goncalves, Ewan Harrison, David K. Jackson, Ian Johnston, Dominic Kwiatkowski, Cordelia Langford, John Sillitoe on behalf of the Wellcome Sanger Institute COVID-19 Surveillance Team |
| EPI_ISL_2317414, EPI_ISL_2317415, EPI_ISL_2317420, EPI_ISL_2317421, EPI_ISL_2317422, EPI_ISL_2317425 | Lighthouse Lab in Milton Keynes | Wellcome Sanger Institute for the COVID-19 Genomics UK (COG-UK) Consortium | The Lighthouse Lab in Milton Keynes and Alex Alderton, Roberto Amato, Jeffrey Barrett, Sonia Goncalves, Ewan Harrison, David K. Jackson, Ian Johnston, Dominic Kwiatkowski, Cordelia Langford, John Sillitoe on behalf of the Wellcome Sanger Institute COVID-19 Surveillance Team |
| EPI_ISL_2317426, EPI_ISL_2317428 | Lighthouse Lab in Alderley Park | Wellcome Sanger Institute for the COVID-19 Genomics UK (COG-UK) Consortium | Jacquelyn Wynn, Mairead Hyland, The Lighthouse Lab in Alderley Park and Alex Alderton, Roberto Amato, Jeffrey Barrett, Sonia Goncalves, Ewan Harrison, David K. Jackson, Ian Johnston, Dominic Kwiatkowski, Cordelia Langford, John Sillitoe on behalf of the Wellcome Sanger Institute COVID-19 Surveillance Team |
| EPI_ISL_2317429 | Lighthouse Lab in Milton Keynes | Wellcome Sanger Institute for the COVID-19 Genomics UK (COG-UK) Consortium | The Lighthouse Lab in Milton Keynes and Alex Alderton, Roberto Amato, Jeffrey Barrett, Sonia Goncalves, Ewan Harrison, David K. Jackson, Ian Johnston, Dominic Kwiatkowski, Cordelia Langford, John Sillitoe on behalf of the Wellcome Sanger Institute COVID-19 Surveillance Team |
| EPI_ISL_2317430, EPI_ISL_2317431, EPI_ISL_2317432 | Lighthouse Lab in Alderley Park | Wellcome Sanger Institute for the COVID-19 Genomics UK (COG-UK) Consortium | Jacquelyn Wynn, Mairead Hyland, The Lighthouse Lab in Alderley Park and Alex Alderton, Roberto Amato, Jeffrey Barrett, Sonia Goncalves, Ewan Harrison, David K. Jackson, Ian Johnston, Dominic Kwiatkowski, Cordelia Langford, John Sillitoe on behalf of the Wellcome Sanger Institute COVID-19 Surveillance Team |
| EPI_ISL_2317439, EPI_ISL_2317440, EPI_ISL_2317441 | Lighthouse Lab in Milton Keynes | Wellcome Sanger Institute for the COVID-19 Genomics UK (COG-UK) Consortium | The Lighthouse Lab in Milton Keynes and Alex Alderton, Roberto Amato, Jeffrey Barrett, Sonia Goncalves, Ewan Harrison, David K. Jackson, Ian Johnston, Dominic Kwiatkowski, Cordelia Langford, John Sillitoe on behalf of the Wellcome Sanger Institute COVID-19 Surveillance Team |
| EPI_ISL_2317442 | Lighthouse Lab in Alderley Park | Wellcome Sanger Institute for the COVID-19 Genomics UK (COG-UK) Consortium | Jacquelyn Wynn, Mairead Hyland, The Lighthouse Lab in Alderley Park and Alex Alderton, Roberto Amato, Jeffrey Barrett, Sonia Goncalves, Ewan Harrison, David K. Jackson, Ian Johnston, Dominic Kwiatkowski, Cordelia Langford, John Sillitoe on behalf of the Wellcome Sanger Institute COVID-19 Surveillance Team |
| EPI_ISL_2317443 | Lighthouse Lab in Milton Keynes | Wellcome Sanger Institute for the COVID-19 Genomics UK (COG-UK) Consortium | The Lighthouse Lab in Milton Keynes and Alex Alderton, Roberto Amato, Jeffrey Barrett, Sonia Goncalves, Ewan Harrison, David K. Jackson, Ian Johnston, Dominic Kwiatkowski, Cordelia Langford, John Sillitoe on behalf of the Wellcome Sanger Institute COVID-19 Surveillance Team |
| EPI_ISL_2317444 | Lighthouse Lab in Alderley Park | Wellcome Sanger Institute for the COVID-19 Genomics UK (COG-UK) Consortium | Jacquelyn Wynn, Mairead Hyland, The Lighthouse Lab in Alderley Park and Alex Alderton, Roberto Amato, Jeffrey Barrett, Sonia Goncalves, Ewan Harrison, David K. Jackson, Ian Johnston, Dominic Kwiatkowski, Cordelia Langford, John Sillitoe on behalf of the Wellcome Sanger Institute COVID-19 Surveillance Team |
| EPI_ISL_2317445, EPI_ISL_2317446, EPI_ISL_2317447, EPI_ISL_2317449, EPI_ISL_2317450 | Lighthouse Lab in Milton Keynes | Wellcome Sanger Institute for the COVID-19 Genomics UK (COG-UK) Consortium | The Lighthouse Lab in Milton Keynes and Alex Alderton, Roberto Amato, Jeffrey Barrett, Sonia Goncalves, Ewan Harrison, David K. Jackson, Ian Johnston, Dominic Kwiatkowski, Cordelia Langford, John Sillitoe on behalf of the Wellcome Sanger Institute COVID-19 Surveillance Team |
| EPI_ISL_2317451 | Lighthouse Lab in Alderley Park | Wellcome Sanger Institute for the COVID-19 Genomics UK (COG-UK) Consortium | Jacquelyn Wynn, Mairead Hyland, The Lighthouse Lab in Alderley Park and Alex Alderton, Roberto Amato, Jeffrey Barrett, Sonia Goncalves, Ewan Harrison, David K. Jackson, Ian Johnston, Dominic Kwiatkowski, Cordelia Langford, John Sillitoe on behalf of the Wellcome Sanger Institute COVID-19 Surveillance Team |
| EPI_ISL_2317452, EPI_ISL_2317453, EPI_ISL_2317454 | Lighthouse Lab in Milton Keynes | Wellcome Sanger Institute for the COVID-19 Genomics UK (COG-UK) Consortium | The Lighthouse Lab in Milton Keynes and Alex Alderton, Roberto Amato, Jeffrey Barrett, Sonia Goncalves, Ewan Harrison, David K. Jackson, Ian Johnston, Dominic Kwiatkowski, Cordelia Langford, John Sillitoe on behalf of the Wellcome Sanger Institute COVID-19 Surveillance Team |
| EPI_ISL_2317455 | Lighthouse Lab in Alderley Park | Wellcome Sanger Institute for the COVID-19 Genomics UK (COG-UK) Consortium | Jacquelyn Wynn, Mairead Hyland, The Lighthouse Lab in Alderley Park and Alex Alderton, Roberto Amato, Jeffrey Barrett, Sonia Goncalves, Ewan Harrison, David K. Jackson, Ian Johnston, Dominic Kwiatkowski, Cordelia Langford, John Sillitoe on behalf of the Wellcome Sanger Institute COVID-19 Surveillance Team |
| EPI_ISL_2317456, EPI_ISL_2317458, EPI_ISL_2317459, EPI_ISL_2317462, EPI_ISL_2317463, EPI_ISL_2317465, EPI_ISL_2317467, EPI_ISL_2317468, EPI_ISL_2317471, EPI_ISL_2317472 | Lighthouse Lab in Milton Keynes | Wellcome Sanger Institute for the COVID-19 Genomics UK (COG-UK) Consortium | The Lighthouse Lab in Milton Keynes and Alex Alderton, Roberto Amato, Jeffrey Barrett, Sonia Goncalves, Ewan Harrison, David K. Jackson, Ian Johnston, Dominic Kwiatkowski, Cordelia Langford, John Sillitoe on behalf of the Wellcome Sanger Institute COVID-19 Surveillance Team |
| EPI_ISL_2317473 | Lighthouse Lab in Alderley Park | Wellcome Sanger Institute for the COVID-19 Genomics UK (COG-UK) Consortium | Jacquelyn Wynn, Mairead Hyland, The Lighthouse Lab in Alderley Park and Alex Alderton, Roberto Amato, Jeffrey Barrett, Sonia Goncalves, Ewan Harrison, David K. Jackson, Ian Johnston, Dominic Kwiatkowski, Cordelia Langford, John Sillitoe on behalf of the Wellcome Sanger Institute COVID-19 Surveillance Team |
| EPI_ISL_2317474, EPI_ISL_2317475, EPI_ISL_2317476, EPI_ISL_2317477, EPI_ISL_2317478, EPI_ISL_2317482 | Lighthouse Lab in Milton Keynes | Wellcome Sanger Institute for the COVID-19 Genomics UK (COG-UK) Consortium | The Lighthouse Lab in Milton Keynes and Alex Alderton, Roberto Amato, Jeffrey Barrett, Sonia Goncalves, Ewan Harrison, David K. Jackson, Ian Johnston, Dominic Kwiatkowski, Cordelia Langford, John Sillitoe on behalf of the Wellcome Sanger Institute COVID-19 Surveillance Team |
| EPI_ISL_2317483 | Lighthouse Lab in Alderley Park | Wellcome Sanger Institute for the COVID-19 Genomics UK (COG-UK) Consortium | Jacquelyn Wynn, Mairead Hyland, The Lighthouse Lab in Alderley Park and Alex Alderton, Roberto Amato, Jeffrey Barrett, Sonia Goncalves, Ewan Harrison, David K. Jackson, Ian Johnston, Dominic Kwiatkowski, Cordelia Langford, John Sillitoe on behalf of the Wellcome Sanger Institute COVID-19 Surveillance Team |
| EPI_ISL_2317484, EPI_ISL_2317485 | Lighthouse Lab in Milton Keynes | Wellcome Sanger Institute for the COVID-19 Genomics UK (COG-UK) Consortium | The Lighthouse Lab in Milton Keynes and Alex Alderton, Roberto Amato, Jeffrey Barrett, Sonia Goncalves, Ewan Harrison, David K. Jackson, Ian Johnston, Dominic Kwiatkowski, Cordelia Langford, John Sillitoe on behalf of the Wellcome Sanger Institute COVID-19 Surveillance Team |
| EPI_ISL_2317486 | Lighthouse Lab in Alderley Park | Wellcome Sanger Institute for the COVID-19 Genomics UK (COG-UK) Consortium | Jacquelyn Wynn, Mairead Hyland, The Lighthouse Lab in Alderley Park and Alex Alderton, Roberto Amato, Jeffrey Barrett, Sonia Goncalves, Ewan Harrison, David K. Jackson, Ian Johnston, Dominic Kwiatkowski, Cordelia Langford, John Sillitoe on behalf of the Wellcome Sanger Institute COVID-19 Surveillance Team |
| EPI_ISL_2317489 | Lighthouse Lab in Milton Keynes | Wellcome Sanger Institute for the COVID-19 Genomics UK (COG-UK) Consortium | The Lighthouse Lab in Milton Keynes and Alex Alderton, Roberto Amato, Jeffrey Barrett, Sonia Goncalves, Ewan Harrison, David K. Jackson, Ian Johnston, Dominic Kwiatkowski, Cordelia Langford, John Sillitoe on behalf of the Wellcome Sanger Institute COVID-19 Surveillance Team |
| EPI_ISL_2317491 | Lighthouse Lab in Alderley Park | Wellcome Sanger Institute for the COVID-19 Genomics UK (COG-UK) | Jacquelyn Wynn, Mairead Hyland, The Lighthouse Lab in Alderley Park and Alex Alderton, Roberto Amato, Jeffrey Barrett, Sonia Goncalves, Ewan Harrison, David K. Jackson, Ian Johnston, Dominic Kwiatkowski, Cordelia Langford, John Sillitoe on |

| | | | |
|---|---|---|---|
| | | Consortium | behalf of the Wellcome Sanger Institute COVID-19 Surveillance Team |
| EPI_ISL_2317492 | Lighthouse Lab in Milton Keynes | Wellcome Sanger Institute for the COVID-19 Genomics UK (COG-UK) Consortium | The Lighthouse Lab in Milton Keynes and Alex Alderton, Roberto Amato, Jeffrey Barrett, Sonia Goncalves, Ewan Harrison, David K. Jackson, Ian Johnston, Dominic Kwiatkowski, Cordelia Langford, John Sillitoe on behalf of the Wellcome Sanger Institute COVID-19 Surveillance Team |
| EPI_ISL_2317494 | Lighthouse Lab in Alderley Park | Wellcome Sanger Institute for the COVID-19 Genomics UK (COG-UK) Consortium | Jacquelyn Wynn, Mairead Hyland, The Lighthouse Lab in Alderley Park and Alex Alderton, Roberto Amato, Jeffrey Barrett, Sonia Goncalves, Ewan Harrison, David K. Jackson, Ian Johnston, Dominic Kwiatkowski, Cordelia Langford, John Sillitoe on behalf of the Wellcome Sanger Institute COVID-19 Surveillance Team |
| EPI_ISL_2317496, EPI_ISL_2317497, EPI_ISL_2317501, EPI_ISL_2317502, EPI_ISL_2317503, EPI_ISL_2317505, EPI_ISL_2317506 | Lighthouse Lab in Milton Keynes | Wellcome Sanger Institute for the COVID-19 Genomics UK (COG-UK) Consortium | The Lighthouse Lab in Milton Keynes and Alex Alderton, Roberto Amato, Jeffrey Barrett, Sonia Goncalves, Ewan Harrison, David K. Jackson, Ian Johnston, Dominic Kwiatkowski, Cordelia Langford, John Sillitoe on behalf of the Wellcome Sanger Institute COVID-19 Surveillance Team |
| EPI_ISL_2317507 | Lighthouse Lab in Alderley Park | Wellcome Sanger Institute for the COVID-19 Genomics UK (COG-UK) Consortium | Jacquelyn Wynn, Mairead Hyland, The Lighthouse Lab in Alderley Park and Alex Alderton, Roberto Amato, Jeffrey Barrett, Sonia Goncalves, Ewan Harrison, David K. Jackson, Ian Johnston, Dominic Kwiatkowski, Cordelia Langford, John Sillitoe on behalf of the Wellcome Sanger Institute COVID-19 Surveillance Team |
| EPI_ISL_2317508, EPI_ISL_2317509, EPI_ISL_2317510, EPI_ISL_2317512, EPI_ISL_2317513, EPI_ISL_2317515, EPI_ISL_2317518, EPI_ISL_2317521, EPI_ISL_2317523 | Lighthouse Lab in Milton Keynes | Wellcome Sanger Institute for the COVID-19 Genomics UK (COG-UK) Consortium | The Lighthouse Lab in Milton Keynes and Alex Alderton, Roberto Amato, Jeffrey Barrett, Sonia Goncalves, Ewan Harrison, David K. Jackson, Ian Johnston, Dominic Kwiatkowski, Cordelia Langford, John Sillitoe on behalf of the Wellcome Sanger Institute COVID-19 Surveillance Team |
| EPI_ISL_2317524 | Lighthouse Lab in Alderley Park | Wellcome Sanger Institute for the COVID-19 Genomics UK (COG-UK) Consortium | Jacquelyn Wynn, Mairead Hyland, The Lighthouse Lab in Alderley Park and Alex Alderton, Roberto Amato, Jeffrey Barrett, Sonia Goncalves, Ewan Harrison, David K. Jackson, Ian Johnston, Dominic Kwiatkowski, Cordelia Langford, John Sillitoe on behalf of the Wellcome Sanger Institute COVID-19 Surveillance Team |
| EPI_ISL_2317525 | Lighthouse Lab in Milton Keynes | Wellcome Sanger Institute for the COVID-19 Genomics UK (COG-UK) Consortium | The Lighthouse Lab in Milton Keynes and Alex Alderton, Roberto Amato, Jeffrey Barrett, Sonia Goncalves, Ewan Harrison, David K. Jackson, Ian Johnston, Dominic Kwiatkowski, Cordelia Langford, John Sillitoe on behalf of the Wellcome Sanger Institute COVID-19 Surveillance Team |
| EPI_ISL_2317526 | Lighthouse Lab in Alderley Park | Wellcome Sanger Institute for the COVID-19 Genomics UK (COG-UK) Consortium | Jacquelyn Wynn, Mairead Hyland, The Lighthouse Lab in Alderley Park and Alex Alderton, Roberto Amato, Jeffrey Barrett, Sonia Goncalves, Ewan Harrison, David K. Jackson, Ian Johnston, Dominic Kwiatkowski, Cordelia Langford, John Sillitoe on behalf of the Wellcome Sanger Institute COVID-19 Surveillance Team |
| EPI_ISL_2317527, EPI_ISL_2317528, EPI_ISL_2317532, EPI_ISL_2317533, EPI_ISL_2317535, EPI_ISL_2317536 | Lighthouse Lab in Milton Keynes | Wellcome Sanger Institute for the COVID-19 Genomics UK (COG-UK) Consortium | The Lighthouse Lab in Milton Keynes and Alex Alderton, Roberto Amato, Jeffrey Barrett, Sonia Goncalves, Ewan Harrison, David K. Jackson, Ian Johnston, Dominic Kwiatkowski, Cordelia Langford, John Sillitoe on behalf of the Wellcome Sanger Institute COVID-19 Surveillance Team |
| EPI_ISL_2317537, EPI_ISL_2317538, EPI_ISL_2317539 | Lighthouse Lab in Alderley Park | Wellcome Sanger Institute for the COVID-19 Genomics UK (COG-UK) Consortium | Jacquelyn Wynn, Mairead Hyland, The Lighthouse Lab in Alderley Park and Alex Alderton, Roberto Amato, Jeffrey Barrett, Sonia Goncalves, Ewan Harrison, David K. Jackson, Ian Johnston, Dominic Kwiatkowski, Cordelia Langford, John Sillitoe on behalf of the Wellcome Sanger Institute COVID-19 Surveillance Team |
| EPI_ISL_2317541, EPI_ISL_2317542, EPI_ISL_2317543, EPI_ISL_2317545, EPI_ISL_2317548, EPI_ISL_2317549, EPI_ISL_2317552, EPI_ISL_2317553, EPI_ISL_2317554, EPI_ISL_2317555 | Lighthouse Lab in Milton Keynes | Wellcome Sanger Institute for the COVID-19 Genomics UK (COG-UK) Consortium | The Lighthouse Lab in Milton Keynes and Alex Alderton, Roberto Amato, Jeffrey Barrett, Sonia Goncalves, Ewan Harrison, David K. Jackson, Ian Johnston, Dominic Kwiatkowski, Cordelia Langford, John Sillitoe on behalf of the Wellcome Sanger Institute COVID-19 Surveillance Team |
| EPI_ISL_2317556 | Lighthouse Lab in Alderley Park | Wellcome Sanger Institute for the COVID-19 Genomics UK (COG-UK) Consortium | Jacquelyn Wynn, Mairead Hyland, The Lighthouse Lab in Alderley Park and Alex Alderton, Roberto Amato, Jeffrey Barrett, Sonia Goncalves, Ewan Harrison, David K. Jackson, Ian Johnston, Dominic Kwiatkowski, Cordelia Langford, John Sillitoe on behalf of the Wellcome Sanger Institute COVID-19 Surveillance Team |
| EPI_ISL_2317558, EPI_ISL_2317559, EPI_ISL_2317562 | Lighthouse Lab in Milton Keynes | Wellcome Sanger Institute for the COVID-19 Genomics UK (COG-UK) Consortium | The Lighthouse Lab in Milton Keynes and Alex Alderton, Roberto Amato, Jeffrey Barrett, Sonia Goncalves, Ewan Harrison, David K. Jackson, Ian Johnston, Dominic Kwiatkowski, Cordelia Langford, John Sillitoe on behalf of the Wellcome Sanger Institute COVID-19 Surveillance Team |
| EPI_ISL_2317563 | Lighthouse Lab in Alderley Park | Wellcome Sanger Institute for the COVID-19 Genomics UK (COG-UK) Consortium | Jacquelyn Wynn, Mairead Hyland, The Lighthouse Lab in Alderley Park and Alex Alderton, Roberto Amato, Jeffrey Barrett, Sonia Goncalves, Ewan Harrison, David K. Jackson, Ian Johnston, Dominic Kwiatkowski, Cordelia Langford, John Sillitoe on behalf of the Wellcome Sanger Institute COVID-19 Surveillance Team |
| EPI_ISL_2317564, EPI_ISL_2317566, EPI_ISL_2317567, EPI_ISL_2317568, EPI_ISL_2317569, EPI_ISL_2317570, EPI_ISL_2317571, EPI_ISL_2317573 | Lighthouse Lab in Milton Keynes | Wellcome Sanger Institute for the COVID-19 Genomics UK (COG-UK) Consortium | The Lighthouse Lab in Milton Keynes and Alex Alderton, Roberto Amato, Jeffrey Barrett, Sonia Goncalves, Ewan Harrison, David K. Jackson, Ian Johnston, Dominic Kwiatkowski, Cordelia Langford, John Sillitoe on behalf of the Wellcome Sanger Institute COVID-19 Surveillance Team |
| EPI_ISL_2317575, EPI_ISL_2317576 | Lighthouse Lab in Alderley Park | Wellcome Sanger Institute for the COVID-19 Genomics UK (COG-UK) Consortium | Jacquelyn Wynn, Mairead Hyland, The Lighthouse Lab in Alderley Park and Alex Alderton, Roberto Amato, Jeffrey Barrett, Sonia Goncalves, Ewan Harrison, David K. Jackson, Ian Johnston, Dominic Kwiatkowski, Cordelia Langford, John Sillitoe on behalf of the Wellcome Sanger Institute COVID-19 Surveillance Team |
| EPI_ISL_2317577, EPI_ISL_2317578, EPI_ISL_2317579 | Lighthouse Lab in Milton Keynes | Wellcome Sanger Institute for the COVID-19 Genomics UK (COG-UK) Consortium | The Lighthouse Lab in Milton Keynes and Alex Alderton, Roberto Amato, Jeffrey Barrett, Sonia Goncalves, Ewan Harrison, David K. Jackson, Ian Johnston, Dominic Kwiatkowski, Cordelia Langford, John Sillitoe on behalf of the Wellcome Sanger Institute COVID-19 Surveillance Team |
| EPI_ISL_2317581 | Lighthouse Lab in Alderley Park | Wellcome Sanger Institute for the COVID-19 Genomics UK (COG-UK) Consortium | Jacquelyn Wynn, Mairead Hyland, The Lighthouse Lab in Alderley Park and Alex Alderton, Roberto Amato, Jeffrey Barrett, Sonia Goncalves, Ewan Harrison, David K. Jackson, Ian Johnston, Dominic Kwiatkowski, Cordelia Langford, John Sillitoe on behalf of the Wellcome Sanger Institute COVID-19 Surveillance Team |
| EPI_ISL_2317582, EPI_ISL_2317589 | Lighthouse Lab in Milton Keynes | Wellcome Sanger Institute for the COVID-19 Genomics UK (COG-UK) Consortium | The Lighthouse Lab in Milton Keynes and Alex Alderton, Roberto Amato, Jeffrey Barrett, Sonia Goncalves, Ewan Harrison, David K. Jackson, Ian Johnston, Dominic Kwiatkowski, Cordelia Langford, John Sillitoe on behalf of the Wellcome Sanger Institute COVID-19 Surveillance Team |
| EPI_ISL_2317591, EPI_ISL_2317592, EPI_ISL_2317595 | Lighthouse Lab in Alderley Park | Wellcome Sanger Institute for the COVID-19 Genomics UK (COG-UK) Consortium | Jacquelyn Wynn, Mairead Hyland, The Lighthouse Lab in Alderley Park and Alex Alderton, Roberto Amato, Jeffrey Barrett, Sonia Goncalves, Ewan Harrison, David K. Jackson, Ian Johnston, Dominic Kwiatkowski, Cordelia Langford, John Sillitoe on behalf of the Wellcome Sanger Institute COVID-19 Surveillance Team |
| EPI_ISL_2317596, EPI_ISL_2317597 | Lighthouse Lab in Milton Keynes | Wellcome Sanger Institute for the COVID-19 Genomics UK (COG-UK) Consortium | The Lighthouse Lab in Milton Keynes and Alex Alderton, Roberto Amato, Jeffrey Barrett, Sonia Goncalves, Ewan Harrison, David K. Jackson, Ian Johnston, Dominic Kwiatkowski, Cordelia Langford, John Sillitoe on behalf of the Wellcome Sanger Institute COVID-19 Surveillance Team |
| EPI_ISL_2317598 | Lighthouse Lab in Alderley Park | Wellcome Sanger Institute for the COVID-19 Genomics UK (COG-UK) Consortium | Jacquelyn Wynn, Mairead Hyland, The Lighthouse Lab in Alderley Park and Alex Alderton, Roberto Amato, Jeffrey Barrett, Sonia Goncalves, Ewan Harrison, David K. Jackson, Ian Johnston, Dominic Kwiatkowski, Cordelia Langford, John Sillitoe on behalf of the Wellcome Sanger Institute COVID-19 Surveillance Team |
| EPI_ISL_2317600 | Lighthouse Lab in Milton Keynes | Wellcome Sanger Institute for the COVID-19 Genomics UK (COG-UK) Consortium | The Lighthouse Lab in Milton Keynes and Alex Alderton, Roberto Amato, Jeffrey Barrett, Sonia Goncalves, Ewan Harrison, David K. Jackson, Ian Johnston, Dominic Kwiatkowski, Cordelia Langford, John Sillitoe on behalf of the Wellcome Sanger Institute COVID-19 Surveillance Team |
| EPI_ISL_2317601 | Lighthouse Lab in Alderley Park | Wellcome Sanger Institute for the COVID-19 Genomics UK (COG-UK) Consortium | Jacquelyn Wynn, Mairead Hyland, The Lighthouse Lab in Alderley Park and Alex Alderton, Roberto Amato, Jeffrey Barrett, Sonia Goncalves, Ewan Harrison, David K. Jackson, Ian Johnston, Dominic Kwiatkowski, Cordelia Langford, John Sillitoe on behalf of the Wellcome Sanger Institute COVID-19 Surveillance Team |
| EPI_ISL_2317602 | Lighthouse Lab in Milton Keynes | Wellcome Sanger Institute for the COVID-19 Genomics UK (COG-UK) Consortium | The Lighthouse Lab in Milton Keynes and Alex Alderton, Roberto Amato, Jeffrey Barrett, Sonia Goncalves, Ewan Harrison, David K. Jackson, Ian Johnston, Dominic Kwiatkowski, Cordelia Langford, John Sillitoe on behalf of the Wellcome Sanger Institute COVID-19 Surveillance Team |
| EPI_ISL_2317603 | Lighthouse Lab in Alderley Park | Wellcome Sanger Institute for the COVID-19 Genomics UK (COG-UK) Consortium | Jacquelyn Wynn, Mairead Hyland, The Lighthouse Lab in Alderley Park and Alex Alderton, Roberto Amato, Jeffrey Barrett, Sonia Goncalves, Ewan Harrison, David K. Jackson, Ian Johnston, Dominic Kwiatkowski, Cordelia Langford, John Sillitoe on behalf of the Wellcome Sanger Institute COVID-19 Surveillance Team |
| EPI_ISL_2317605, EPI_ISL_2317606, EPI_ISL_2317607, EPI_ISL_2317608, EPI_ISL_2317609, EPI_ISL_2317610, EPI_ISL_2317611, EPI_ISL_2317612 | Lighthouse Lab in Milton Keynes | Wellcome Sanger Institute for the COVID-19 Genomics UK (COG-UK) Consortium | The Lighthouse Lab in Milton Keynes and Alex Alderton, Roberto Amato, Jeffrey Barrett, Sonia Goncalves, Ewan Harrison, David K. Jackson, Ian Johnston, Dominic Kwiatkowski, Cordelia Langford, John Sillitoe on behalf of the Wellcome Sanger Institute COVID-19 Surveillance Team |
| EPI_ISL_2317614 | Lighthouse Lab in Alderley Park | Wellcome Sanger Institute for the COVID-19 Genomics UK (COG-UK) Consortium | Jacquelyn Wynn, Mairead Hyland, The Lighthouse Lab in Alderley Park and Alex Alderton, Roberto Amato, Jeffrey Barrett, Sonia Goncalves, Ewan Harrison, David K. Jackson, Ian Johnston, Dominic Kwiatkowski, Cordelia Langford, John Sillitoe on behalf of the Wellcome Sanger Institute COVID-19 Surveillance Team |

| | | | |
|---|---|---|---|
| EPI_ISL_2317616 | Lighthouse Lab in Milton Keynes | Wellcome Sanger Institute for the COVID-19 Genomics UK (COG-UK) Consortium | The Lighthouse Lab in Milton Keynes and Alex Alderton, Roberto Amato, Jeffrey Barrett, Sonia Goncalves, Ewan Harrison, David K. Jackson, Ian Johnston, Dominic Kwiatkowski, Cordelia Langford, John Sillitoe on behalf of the Wellcome Sanger Institute COVID-19 Surveillance Team |
| EPI_ISL_2317620, EPI_ISL_2317623, EPI_ISL_2317624, EPI_ISL_2317625 | Lighthouse Lab in Alderley Park | Wellcome Sanger Institute for the COVID-19 Genomics UK (COG-UK) Consortium | Jacquelyn Wynn, Mairead Hyland, The Lighthouse Lab in Alderley Park and Alex Alderton, Roberto Amato, Jeffrey Barrett, Sonia Goncalves, Ewan Harrison, David K. Jackson, Ian Johnston, Dominic Kwiatkowski, Cordelia Langford, John Sillitoe on behalf of the Wellcome Sanger Institute COVID-19 Surveillance Team |
| EPI_ISL_2317626, EPI_ISL_2317627, EPI_ISL_2317628 | Lighthouse Lab in Milton Keynes | Wellcome Sanger Institute for the COVID-19 Genomics UK (COG-UK) Consortium | The Lighthouse Lab in Milton Keynes and Alex Alderton, Roberto Amato, Jeffrey Barrett, Sonia Goncalves, Ewan Harrison, David K. Jackson, Ian Johnston, Dominic Kwiatkowski, Cordelia Langford, John Sillitoe on behalf of the Wellcome Sanger Institute COVID-19 Surveillance Team |
| EPI_ISL_2317630 | Lighthouse Lab in Alderley Park | Wellcome Sanger Institute for the COVID-19 Genomics UK (COG-UK) Consortium | Jacquelyn Wynn, Mairead Hyland, The Lighthouse Lab in Alderley Park and Alex Alderton, Roberto Amato, Jeffrey Barrett, Sonia Goncalves, Ewan Harrison, David K. Jackson, Ian Johnston, Dominic Kwiatkowski, Cordelia Langford, John Sillitoe on behalf of the Wellcome Sanger Institute COVID-19 Surveillance Team |
| EPI_ISL_2317631, EPI_ISL_2317632, EPI_ISL_2317636 | Lighthouse Lab in Milton Keynes | Wellcome Sanger Institute for the COVID-19 Genomics UK (COG-UK) Consortium | The Lighthouse Lab in Milton Keynes and Alex Alderton, Roberto Amato, Jeffrey Barrett, Sonia Goncalves, Ewan Harrison, David K. Jackson, Ian Johnston, Dominic Kwiatkowski, Cordelia Langford, John Sillitoe on behalf of the Wellcome Sanger Institute COVID-19 Surveillance Team |
| EPI_ISL_2317637 | Lighthouse Lab in Alderley Park | Wellcome Sanger Institute for the COVID-19 Genomics UK (COG-UK) Consortium | Jacquelyn Wynn, Mairead Hyland, The Lighthouse Lab in Alderley Park and Alex Alderton, Roberto Amato, Jeffrey Barrett, Sonia Goncalves, Ewan Harrison, David K. Jackson, Ian Johnston, Dominic Kwiatkowski, Cordelia Langford, John Sillitoe on behalf of the Wellcome Sanger Institute COVID-19 Surveillance Team |
| EPI_ISL_2317639, EPI_ISL_2317640, EPI_ISL_2317641, EPI_ISL_2317645 | Lighthouse Lab in Milton Keynes | Wellcome Sanger Institute for the COVID-19 Genomics UK (COG-UK) Consortium | The Lighthouse Lab in Milton Keynes and Alex Alderton, Roberto Amato, Jeffrey Barrett, Sonia Goncalves, Ewan Harrison, David K. Jackson, Ian Johnston, Dominic Kwiatkowski, Cordelia Langford, John Sillitoe on behalf of the Wellcome Sanger Institute COVID-19 Surveillance Team |
| EPI_ISL_2317647 | Lighthouse Lab in Alderley Park | Wellcome Sanger Institute for the COVID-19 Genomics UK (COG-UK) Consortium | Jacquelyn Wynn, Mairead Hyland, The Lighthouse Lab in Alderley Park and Alex Alderton, Roberto Amato, Jeffrey Barrett, Sonia Goncalves, Ewan Harrison, David K. Jackson, Ian Johnston, Dominic Kwiatkowski, Cordelia Langford, John Sillitoe on behalf of the Wellcome Sanger Institute COVID-19 Surveillance Team |
| EPI_ISL_2317648 | Lighthouse Lab in Milton Keynes | Wellcome Sanger Institute for the COVID-19 Genomics UK (COG-UK) Consortium | The Lighthouse Lab in Milton Keynes and Alex Alderton, Roberto Amato, Jeffrey Barrett, Sonia Goncalves, Ewan Harrison, David K. Jackson, Ian Johnston, Dominic Kwiatkowski, Cordelia Langford, John Sillitoe on behalf of the Wellcome Sanger Institute COVID-19 Surveillance Team |
| EPI_ISL_2317649 | Lighthouse Lab in Alderley Park | Wellcome Sanger Institute for the COVID-19 Genomics UK (COG-UK) Consortium | Jacquelyn Wynn, Mairead Hyland, The Lighthouse Lab in Alderley Park and Alex Alderton, Roberto Amato, Jeffrey Barrett, Sonia Goncalves, Ewan Harrison, David K. Jackson, Ian Johnston, Dominic Kwiatkowski, Cordelia Langford, John Sillitoe on behalf of the Wellcome Sanger Institute COVID-19 Surveillance Team |
| EPI_ISL_2317651, EPI_ISL_2317653 | Lighthouse Lab in Milton Keynes | Wellcome Sanger Institute for the COVID-19 Genomics UK (COG-UK) Consortium | The Lighthouse Lab in Milton Keynes and Alex Alderton, Roberto Amato, Jeffrey Barrett, Sonia Goncalves, Ewan Harrison, David K. Jackson, Ian Johnston, Dominic Kwiatkowski, Cordelia Langford, John Sillitoe on behalf of the Wellcome Sanger Institute COVID-19 Surveillance Team |
| EPI_ISL_2317655, EPI_ISL_2317656 | Lighthouse Lab in Alderley Park | Wellcome Sanger Institute for the COVID-19 Genomics UK (COG-UK) Consortium | Jacquelyn Wynn, Mairead Hyland, The Lighthouse Lab in Alderley Park and Alex Alderton, Roberto Amato, Jeffrey Barrett, Sonia Goncalves, Ewan Harrison, David K. Jackson, Ian Johnston, Dominic Kwiatkowski, Cordelia Langford, John Sillitoe on behalf of the Wellcome Sanger Institute COVID-19 Surveillance Team |
| EPI_ISL_2317660 | Lighthouse Lab in Milton Keynes | Wellcome Sanger Institute for the COVID-19 Genomics UK (COG-UK) Consortium | The Lighthouse Lab in Milton Keynes and Alex Alderton, Roberto Amato, Jeffrey Barrett, Sonia Goncalves, Ewan Harrison, David K. Jackson, Ian Johnston, Dominic Kwiatkowski, Cordelia Langford, John Sillitoe on behalf of the Wellcome Sanger Institute COVID-19 Surveillance Team |
| EPI_ISL_2317661, EPI_ISL_2317662, EPI_ISL_2317663 | Lighthouse Lab in Alderley Park | Wellcome Sanger Institute for the COVID-19 Genomics UK (COG-UK) Consortium | Jacquelyn Wynn, Mairead Hyland, The Lighthouse Lab in Alderley Park and Alex Alderton, Roberto Amato, Jeffrey Barrett, Sonia Goncalves, Ewan Harrison, David K. Jackson, Ian Johnston, Dominic Kwiatkowski, Cordelia Langford, John Sillitoe on behalf of the Wellcome Sanger Institute COVID-19 Surveillance Team |
| EPI_ISL_2317666, EPI_ISL_2317667, EPI_ISL_2317668 | Lighthouse Lab in Milton Keynes | Wellcome Sanger Institute for the COVID-19 Genomics UK (COG-UK) Consortium | The Lighthouse Lab in Milton Keynes and Alex Alderton, Roberto Amato, Jeffrey Barrett, Sonia Goncalves, Ewan Harrison, David K. Jackson, Ian Johnston, Dominic Kwiatkowski, Cordelia Langford, John Sillitoe on behalf of the Wellcome Sanger Institute COVID-19 Surveillance Team |
| EPI_ISL_2317670 | Lighthouse Lab in Alderley Park | Wellcome Sanger Institute for the COVID-19 Genomics UK (COG-UK) Consortium | Jacquelyn Wynn, Mairead Hyland, The Lighthouse Lab in Alderley Park and Alex Alderton, Roberto Amato, Jeffrey Barrett, Sonia Goncalves, Ewan Harrison, David K. Jackson, Ian Johnston, Dominic Kwiatkowski, Cordelia Langford, John Sillitoe on behalf of the Wellcome Sanger Institute COVID-19 Surveillance Team |
| EPI_ISL_2317671 | Lighthouse Lab in Milton Keynes | Wellcome Sanger Institute for the COVID-19 Genomics UK (COG-UK) Consortium | The Lighthouse Lab in Milton Keynes and Alex Alderton, Roberto Amato, Jeffrey Barrett, Sonia Goncalves, Ewan Harrison, David K. Jackson, Ian Johnston, Dominic Kwiatkowski, Cordelia Langford, John Sillitoe on behalf of the Wellcome Sanger Institute COVID-19 Surveillance Team |
| EPI_ISL_2317672 | Lighthouse Lab in Alderley Park | Wellcome Sanger Institute for the COVID-19 Genomics UK (COG-UK) Consortium | Jacquelyn Wynn, Mairead Hyland, The Lighthouse Lab in Alderley Park and Alex Alderton, Roberto Amato, Jeffrey Barrett, Sonia Goncalves, Ewan Harrison, David K. Jackson, Ian Johnston, Dominic Kwiatkowski, Cordelia Langford, John Sillitoe on behalf of the Wellcome Sanger Institute COVID-19 Surveillance Team |
| EPI_ISL_2317673 | Lighthouse Lab in Milton Keynes | Wellcome Sanger Institute for the COVID-19 Genomics UK (COG-UK) Consortium | The Lighthouse Lab in Milton Keynes and Alex Alderton, Roberto Amato, Jeffrey Barrett, Sonia Goncalves, Ewan Harrison, David K. Jackson, Ian Johnston, Dominic Kwiatkowski, Cordelia Langford, John Sillitoe on behalf of the Wellcome Sanger Institute COVID-19 Surveillance Team |
| EPI_ISL_2317674 | Lighthouse Lab in Alderley Park | Wellcome Sanger Institute for the COVID-19 Genomics UK (COG-UK) Consortium | Jacquelyn Wynn, Mairead Hyland, The Lighthouse Lab in Alderley Park and Alex Alderton, Roberto Amato, Jeffrey Barrett, Sonia Goncalves, Ewan Harrison, David K. Jackson, Ian Johnston, Dominic Kwiatkowski, Cordelia Langford, John Sillitoe on behalf of the Wellcome Sanger Institute COVID-19 Surveillance Team |
| EPI_ISL_2317675 | Lighthouse Lab in Milton Keynes | Wellcome Sanger Institute for the COVID-19 Genomics UK (COG-UK) Consortium | The Lighthouse Lab in Milton Keynes and Alex Alderton, Roberto Amato, Jeffrey Barrett, Sonia Goncalves, Ewan Harrison, David K. Jackson, Ian Johnston, Dominic Kwiatkowski, Cordelia Langford, John Sillitoe on behalf of the Wellcome Sanger Institute COVID-19 Surveillance Team |
| EPI_ISL_2317676, EPI_ISL_2317677, EPI_ISL_2317678, EPI_ISL_2317681 | Lighthouse Lab in Alderley Park | Wellcome Sanger Institute for the COVID-19 Genomics UK (COG-UK) Consortium | Jacquelyn Wynn, Mairead Hyland, The Lighthouse Lab in Alderley Park and Alex Alderton, Roberto Amato, Jeffrey Barrett, Sonia Goncalves, Ewan Harrison, David K. Jackson, Ian Johnston, Dominic Kwiatkowski, Cordelia Langford, John Sillitoe on behalf of the Wellcome Sanger Institute COVID-19 Surveillance Team |
| EPI_ISL_2317682, EPI_ISL_2317683, EPI_ISL_2317685, EPI_ISL_2317686, EPI_ISL_2317687 | Lighthouse Lab in Milton Keynes | Wellcome Sanger Institute for the COVID-19 Genomics UK (COG-UK) Consortium | The Lighthouse Lab in Milton Keynes and Alex Alderton, Roberto Amato, Jeffrey Barrett, Sonia Goncalves, Ewan Harrison, David K. Jackson, Ian Johnston, Dominic Kwiatkowski, Cordelia Langford, John Sillitoe on behalf of the Wellcome Sanger Institute COVID-19 Surveillance Team |
| EPI_ISL_2317690 | Lighthouse Lab in Alderley Park | Wellcome Sanger Institute for the COVID-19 Genomics UK (COG-UK) Consortium | Jacquelyn Wynn, Mairead Hyland, The Lighthouse Lab in Alderley Park and Alex Alderton, Roberto Amato, Jeffrey Barrett, Sonia Goncalves, Ewan Harrison, David K. Jackson, Ian Johnston, Dominic Kwiatkowski, Cordelia Langford, John Sillitoe on behalf of the Wellcome Sanger Institute COVID-19 Surveillance Team |
| EPI_ISL_2317691, EPI_ISL_2317693, EPI_ISL_2317694, EPI_ISL_2317695, EPI_ISL_2317696, EPI_ISL_2317697, EPI_ISL_2317698, EPI_ISL_2317702, EPI_ISL_2317704 | Lighthouse Lab in Milton Keynes | Wellcome Sanger Institute for the COVID-19 Genomics UK (COG-UK) Consortium | The Lighthouse Lab in Milton Keynes and Alex Alderton, Roberto Amato, Jeffrey Barrett, Sonia Goncalves, Ewan Harrison, David K. Jackson, Ian Johnston, Dominic Kwiatkowski, Cordelia Langford, John Sillitoe on behalf of the Wellcome Sanger Institute COVID-19 Surveillance Team |
| EPI_ISL_2317707 | Lighthouse Lab in Alderley Park | Wellcome Sanger Institute for the COVID-19 Genomics UK (COG-UK) Consortium | Jacquelyn Wynn, Mairead Hyland, The Lighthouse Lab in Alderley Park and Alex Alderton, Roberto Amato, Jeffrey Barrett, Sonia Goncalves, Ewan Harrison, David K. Jackson, Ian Johnston, Dominic Kwiatkowski, Cordelia Langford, John Sillitoe on behalf of the Wellcome Sanger Institute COVID-19 Surveillance Team |
| EPI_ISL_2317709 | Lighthouse Lab in Milton Keynes | Wellcome Sanger Institute for the COVID-19 Genomics UK (COG-UK) Consortium | The Lighthouse Lab in Milton Keynes and Alex Alderton, Roberto Amato, Jeffrey Barrett, Sonia Goncalves, Ewan Harrison, David K. Jackson, Ian Johnston, Dominic Kwiatkowski, Cordelia Langford, John Sillitoe on behalf of the Wellcome Sanger Institute COVID-19 Surveillance Team |
| EPI_ISL_2317712 | Lighthouse Lab in Alderley Park | Wellcome Sanger Institute for the COVID-19 Genomics UK (COG-UK) Consortium | Jacquelyn Wynn, Mairead Hyland, The Lighthouse Lab in Alderley Park and Alex Alderton, Roberto Amato, Jeffrey Barrett, Sonia Goncalves, Ewan Harrison, David K. Jackson, Ian Johnston, Dominic Kwiatkowski, Cordelia Langford, John Sillitoe on behalf of the Wellcome Sanger Institute COVID-19 Surveillance Team |
| EPI_ISL_2317713 | Lighthouse Lab in Milton Keynes | Wellcome Sanger Institute for the COVID-19 Genomics UK (COG-UK) Consortium | The Lighthouse Lab in Milton Keynes and Alex Alderton, Roberto Amato, Jeffrey Barrett, Sonia Goncalves, Ewan Harrison, David K. Jackson, Ian Johnston, Dominic Kwiatkowski, Cordelia Langford, John Sillitoe on behalf of the Wellcome Sanger Institute COVID-19 Surveillance Team |
| EPI_ISL_2317714 | Lighthouse Lab in Alderley Park | Wellcome Sanger Institute for the COVID-19 Genomics UK (COG-UK) Consortium | Jacquelyn Wynn, Mairead Hyland, The Lighthouse Lab in Alderley Park and Alex Alderton, Roberto Amato, Jeffrey Barrett, Sonia Goncalves, Ewan Harrison, David K. Jackson, Ian Johnston, Dominic Kwiatkowski, Cordelia Langford, John Sillitoe on behalf of the Wellcome Sanger Institute COVID-19 Surveillance Team |
| EPI_ISL_2317716, EPI_ISL_2317720, EPI_ISL_2317721, EPI_ISL_2317722, EPI_ISL_2317723, EPI_ISL_2317724 | Lighthouse Lab in Milton Keynes | Wellcome Sanger Institute for the COVID-19 Genomics UK (COG-UK) Consortium | The Lighthouse Lab in Milton Keynes and Alex Alderton, Roberto Amato, Jeffrey Barrett, Sonia Goncalves, Ewan Harrison, David K. Jackson, Ian Johnston, Dominic Kwiatkowski, Cordelia Langford, John Sillitoe on behalf of the Wellcome Sanger Institute COVID-19 Surveillance Team |
| EPI_ISL_2317725, EPI_ISL_2317727 | Lighthouse Lab in Alderley Park | Wellcome Sanger Institute for the COVID-19 Genomics UK (COG-UK) Consortium | Jacquelyn Wynn, Mairead Hyland, The Lighthouse Lab in Alderley Park and Alex Alderton, Roberto Amato, Jeffrey Barrett, Sonia Goncalves, Ewan Harrison, David K. Jackson, Ian Johnston, Dominic Kwiatkowski, Cordelia Langford, John Sillitoe on behalf of the Wellcome Sanger Institute COVID-19 Surveillance Team |
| EPI_ISL_2317728, EPI_ISL_2317729 | Lighthouse Lab in Milton Keynes | Wellcome Sanger Institute for the COVID-19 Genomics UK (COG-UK) Consortium | The Lighthouse Lab in Milton Keynes and Alex Alderton, Roberto Amato, Jeffrey Barrett, Sonia Goncalves, Ewan Harrison, David K. Jackson, Ian Johnston, Dominic Kwiatkowski, Cordelia Langford, John Sillitoe on behalf of the Wellcome Sanger Institute COVID-19 Surveillance Team |
| EPI_ISL_2317730, EPI_ISL_2317731 | Lighthouse Lab in Alderley Park | Wellcome Sanger Institute for the COVID-19 Genomics UK (COG-UK) Consortium | Jacquelyn Wynn, Mairead Hyland, The Lighthouse Lab in Alderley Park and Alex Alderton, Roberto Amato, Jeffrey Barrett, Sonia Goncalves, Ewan Harrison, David K. Jackson, Ian Johnston, Dominic Kwiatkowski, Cordelia Langford, John Sillitoe on behalf of the Wellcome Sanger Institute COVID-19 Surveillance Team |
| EPI_ISL_2317732 | Lighthouse Lab in Milton Keynes | Wellcome Sanger Institute for the COVID-19 Genomics UK (COG-UK) Consortium | The Lighthouse Lab in Milton Keynes and Alex Alderton, Roberto Amato, Jeffrey Barrett, Sonia Goncalves, Ewan Harrison, David K. Jackson, Ian Johnston, Dominic Kwiatkowski, Cordelia Langford, John Sillitoe on behalf of the Wellcome Sanger Institute COVID-19 Surveillance Team |
| EPI_ISL_2317733 | Lighthouse Lab in Alderley Park | Wellcome Sanger Institute for the COVID-19 Genomics UK (COG-UK) | Jacquelyn Wynn, Mairead Hyland, The Lighthouse Lab in Alderley Park and Alex Alderton, Roberto Amato, Jeffrey Barrett, Sonia Goncalves, Ewan Harrison, David K. Jackson, Ian Johnston, Dominic Kwiatkowski, Cordelia Langford, John Sillitoe on |

| | | | |
|---|---|---|---|
| | | Consortium | behalf of the Wellcome Sanger Institute COVID-19 Surveillance Team |
| EPI_ISL_2317734 | Lighthouse Lab in Milton Keynes | Wellcome Sanger Institute for the COVID-19 Genomics UK (COG-UK) Consortium | The Lighthouse Lab in Milton Keynes and Alex Alderton, Roberto Amato, Jeffrey Barrett, Sonia Goncalves, Ewan Harrison, David K. Jackson, Ian Johnston, Dominic Kwiatkowski, Cordelia Langford, John Sillitoe on behalf of the Wellcome Sanger Institute COVID-19 Surveillance Team |
| EPI_ISL_2317735, EPI_ISL_2317738 | Lighthouse Lab in Alderley Park | Wellcome Sanger Institute for the COVID-19 Genomics UK (COG-UK) Consortium | Jacquelyn Wynn, Mairead Hyland, The Lighthouse Lab in Alderley Park and Alex Alderton, Roberto Amato, Jeffrey Barrett, Sonia Goncalves, Ewan Harrison, David K. Jackson, Ian Johnston, Dominic Kwiatkowski, Cordelia Langford, John Sillitoe on behalf of the Wellcome Sanger Institute COVID-19 Surveillance Team |
| EPI_ISL_2317739, EPI_ISL_2317741 | Lighthouse Lab in Milton Keynes | Wellcome Sanger Institute for the COVID-19 Genomics UK (COG-UK) Consortium | The Lighthouse Lab in Milton Keynes and Alex Alderton, Roberto Amato, Jeffrey Barrett, Sonia Goncalves, Ewan Harrison, David K. Jackson, Ian Johnston, Dominic Kwiatkowski, Cordelia Langford, John Sillitoe on behalf of the Wellcome Sanger Institute COVID-19 Surveillance Team |
| EPI_ISL_2317743, EPI_ISL_2317744 | Lighthouse Lab in Alderley Park | Wellcome Sanger Institute for the COVID-19 Genomics UK (COG-UK) Consortium | Jacquelyn Wynn, Mairead Hyland, The Lighthouse Lab in Alderley Park and Alex Alderton, Roberto Amato, Jeffrey Barrett, Sonia Goncalves, Ewan Harrison, David K. Jackson, Ian Johnston, Dominic Kwiatkowski, Cordelia Langford, John Sillitoe on behalf of the Wellcome Sanger Institute COVID-19 Surveillance Team |
| EPI_ISL_2317745, EPI_ISL_2317747, EPI_ISL_2317748 | Lighthouse Lab in Milton Keynes | Wellcome Sanger Institute for the COVID-19 Genomics UK (COG-UK) Consortium | The Lighthouse Lab in Milton Keynes and Alex Alderton, Roberto Amato, Jeffrey Barrett, Sonia Goncalves, Ewan Harrison, David K. Jackson, Ian Johnston, Dominic Kwiatkowski, Cordelia Langford, John Sillitoe on behalf of the Wellcome Sanger Institute COVID-19 Surveillance Team |
| EPI_ISL_2317749, EPI_ISL_2317750 | Lighthouse Lab in Alderley Park | Wellcome Sanger Institute for the COVID-19 Genomics UK (COG-UK) Consortium | Jacquelyn Wynn, Mairead Hyland, The Lighthouse Lab in Alderley Park and Alex Alderton, Roberto Amato, Jeffrey Barrett, Sonia Goncalves, Ewan Harrison, David K. Jackson, Ian Johnston, Dominic Kwiatkowski, Cordelia Langford, John Sillitoe on behalf of the Wellcome Sanger Institute COVID-19 Surveillance Team |
| EPI_ISL_2317751 | Lighthouse Lab in Milton Keynes | Wellcome Sanger Institute for the COVID-19 Genomics UK (COG-UK) Consortium | The Lighthouse Lab in Milton Keynes and Alex Alderton, Roberto Amato, Jeffrey Barrett, Sonia Goncalves, Ewan Harrison, David K. Jackson, Ian Johnston, Dominic Kwiatkowski, Cordelia Langford, John Sillitoe on behalf of the Wellcome Sanger Institute COVID-19 Surveillance Team |
| EPI_ISL_2317753, EPI_ISL_2317754 | Lighthouse Lab in Alderley Park | Wellcome Sanger Institute for the COVID-19 Genomics UK (COG-UK) Consortium | Jacquelyn Wynn, Mairead Hyland, The Lighthouse Lab in Alderley Park and Alex Alderton, Roberto Amato, Jeffrey Barrett, Sonia Goncalves, Ewan Harrison, David K. Jackson, Ian Johnston, Dominic Kwiatkowski, Cordelia Langford, John Sillitoe on behalf of the Wellcome Sanger Institute COVID-19 Surveillance Team |
| EPI_ISL_2317755, EPI_ISL_2317757 | Lighthouse Lab in Milton Keynes | Wellcome Sanger Institute for the COVID-19 Genomics UK (COG-UK) Consortium | The Lighthouse Lab in Milton Keynes and Alex Alderton, Roberto Amato, Jeffrey Barrett, Sonia Goncalves, Ewan Harrison, David K. Jackson, Ian Johnston, Dominic Kwiatkowski, Cordelia Langford, John Sillitoe on behalf of the Wellcome Sanger Institute COVID-19 Surveillance Team |
| EPI_ISL_2317758, EPI_ISL_2317759, EPI_ISL_2317760, EPI_ISL_2317763, EPI_ISL_2317764, EPI_ISL_2317766, EPI_ISL_2317768, EPI_ISL_2317772 | Lighthouse Lab in Alderley Park | Wellcome Sanger Institute for the COVID-19 Genomics UK (COG-UK) Consortium | Jacquelyn Wynn, Mairead Hyland, The Lighthouse Lab in Alderley Park and Alex Alderton, Roberto Amato, Jeffrey Barrett, Sonia Goncalves, Ewan Harrison, David K. Jackson, Ian Johnston, Dominic Kwiatkowski, Cordelia Langford, John Sillitoe on behalf of the Wellcome Sanger Institute COVID-19 Surveillance Team |
| EPI_ISL_2317773 | Lighthouse Lab in Milton Keynes | Wellcome Sanger Institute for the COVID-19 Genomics UK (COG-UK) Consortium | The Lighthouse Lab in Milton Keynes and Alex Alderton, Roberto Amato, Jeffrey Barrett, Sonia Goncalves, Ewan Harrison, David K. Jackson, Ian Johnston, Dominic Kwiatkowski, Cordelia Langford, John Sillitoe on behalf of the Wellcome Sanger Institute COVID-19 Surveillance Team |
| EPI_ISL_2317774 | Randox Laboratories | Wellcome Sanger Institute for the COVID-19 Genomics UK (COG-UK) Consortium | Randox Laboratories and Alex Alderton, Roberto Amato, Jeffrey Barrett, Sonia Goncalves, Ewan Harrison, David K. Jackson, Ian Johnston, Dominic Kwiatkowski, Cordelia Langford, John Sillitoe on behalf of the Wellcome Sanger Institute COVID-19 Surveillance Team |
| EPI_ISL_2317775, EPI_ISL_2317777, EPI_ISL_2317778 | Lighthouse Lab in Alderley Park | Wellcome Sanger Institute for the COVID-19 Genomics UK (COG-UK) Consortium | Jacquelyn Wynn, Mairead Hyland, The Lighthouse Lab in Alderley Park and Alex Alderton, Roberto Amato, Jeffrey Barrett, Sonia Goncalves, Ewan Harrison, David K. Jackson, Ian Johnston, Dominic Kwiatkowski, Cordelia Langford, John Sillitoe on behalf of the Wellcome Sanger Institute COVID-19 Surveillance Team |
| EPI_ISL_2317780 | Lighthouse Lab in Milton Keynes | Wellcome Sanger Institute for the COVID-19 Genomics UK (COG-UK) Consortium | The Lighthouse Lab in Milton Keynes and Alex Alderton, Roberto Amato, Jeffrey Barrett, Sonia Goncalves, Ewan Harrison, David K. Jackson, Ian Johnston, Dominic Kwiatkowski, Cordelia Langford, John Sillitoe on behalf of the Wellcome Sanger Institute COVID-19 Surveillance Team |
| EPI_ISL_2317782, EPI_ISL_2317783 | Lighthouse Lab in Alderley Park | Wellcome Sanger Institute for the COVID-19 Genomics UK (COG-UK) Consortium | Jacquelyn Wynn, Mairead Hyland, The Lighthouse Lab in Alderley Park and Alex Alderton, Roberto Amato, Jeffrey Barrett, Sonia Goncalves, Ewan Harrison, David K. Jackson, Ian Johnston, Dominic Kwiatkowski, Cordelia Langford, John Sillitoe on behalf of the Wellcome Sanger Institute COVID-19 Surveillance Team |
| EPI_ISL_2317784 | Randox Laboratories | Wellcome Sanger Institute for the COVID-19 Genomics UK (COG-UK) Consortium | Randox Laboratories and Alex Alderton, Roberto Amato, Jeffrey Barrett, Sonia Goncalves, Ewan Harrison, David K. Jackson, Ian Johnston, Dominic Kwiatkowski, Cordelia Langford, John Sillitoe on behalf of the Wellcome Sanger Institute COVID-19 Surveillance Team |
| EPI_ISL_2317786, EPI_ISL_2317788, EPI_ISL_2317789, EPI_ISL_2317790, EPI_ISL_2317792 | Lighthouse Lab in Alderley Park | Wellcome Sanger Institute for the COVID-19 Genomics UK (COG-UK) Consortium | Jacquelyn Wynn, Mairead Hyland, The Lighthouse Lab in Alderley Park and Alex Alderton, Roberto Amato, Jeffrey Barrett, Sonia Goncalves, Ewan Harrison, David K. Jackson, Ian Johnston, Dominic Kwiatkowski, Cordelia Langford, John Sillitoe on behalf of the Wellcome Sanger Institute COVID-19 Surveillance Team |
| EPI_ISL_2317793 | Lighthouse Lab in Milton Keynes | Wellcome Sanger Institute for the COVID-19 Genomics UK (COG-UK) Consortium | The Lighthouse Lab in Milton Keynes and Alex Alderton, Roberto Amato, Jeffrey Barrett, Sonia Goncalves, Ewan Harrison, David K. Jackson, Ian Johnston, Dominic Kwiatkowski, Cordelia Langford, John Sillitoe on behalf of the Wellcome Sanger Institute COVID-19 Surveillance Team |
| EPI_ISL_2317794 | Lighthouse Lab in Alderley Park | Wellcome Sanger Institute for the COVID-19 Genomics UK (COG-UK) Consortium | Jacquelyn Wynn, Mairead Hyland, The Lighthouse Lab in Alderley Park and Alex Alderton, Roberto Amato, Jeffrey Barrett, Sonia Goncalves, Ewan Harrison, David K. Jackson, Ian Johnston, Dominic Kwiatkowski, Cordelia Langford, John Sillitoe on behalf of the Wellcome Sanger Institute COVID-19 Surveillance Team |
| EPI_ISL_2317797, EPI_ISL_2317798 | Lighthouse Lab in Milton Keynes | Wellcome Sanger Institute for the COVID-19 Genomics UK (COG-UK) Consortium | The Lighthouse Lab in Milton Keynes and Alex Alderton, Roberto Amato, Jeffrey Barrett, Sonia Goncalves, Ewan Harrison, David K. Jackson, Ian Johnston, Dominic Kwiatkowski, Cordelia Langford, John Sillitoe on behalf of the Wellcome Sanger Institute COVID-19 Surveillance Team |
| EPI_ISL_2317800 | Lighthouse Lab in Alderley Park | Wellcome Sanger Institute for the COVID-19 Genomics UK (COG-UK) Consortium | Jacquelyn Wynn, Mairead Hyland, The Lighthouse Lab in Alderley Park and Alex Alderton, Roberto Amato, Jeffrey Barrett, Sonia Goncalves, Ewan Harrison, David K. Jackson, Ian Johnston, Dominic Kwiatkowski, Cordelia Langford, John Sillitoe on behalf of the Wellcome Sanger Institute COVID-19 Surveillance Team |
| EPI_ISL_2317801, EPI_ISL_2317802, EPI_ISL_2317803 | Lighthouse Lab in Milton Keynes | Wellcome Sanger Institute for the COVID-19 Genomics UK (COG-UK) Consortium | The Lighthouse Lab in Milton Keynes and Alex Alderton, Roberto Amato, Jeffrey Barrett, Sonia Goncalves, Ewan Harrison, David K. Jackson, Ian Johnston, Dominic Kwiatkowski, Cordelia Langford, John Sillitoe on behalf of the Wellcome Sanger Institute COVID-19 Surveillance Team |
| EPI_ISL_2317804 | Lighthouse Lab in Alderley Park | Wellcome Sanger Institute for the COVID-19 Genomics UK (COG-UK) Consortium | Jacquelyn Wynn, Mairead Hyland, The Lighthouse Lab in Alderley Park and Alex Alderton, Roberto Amato, Jeffrey Barrett, Sonia Goncalves, Ewan Harrison, David K. Jackson, Ian Johnston, Dominic Kwiatkowski, Cordelia Langford, John Sillitoe on behalf of the Wellcome Sanger Institute COVID-19 Surveillance Team |
| EPI_ISL_2317805 | Lighthouse Lab in Milton Keynes | Wellcome Sanger Institute for the COVID-19 Genomics UK (COG-UK) Consortium | The Lighthouse Lab in Milton Keynes and Alex Alderton, Roberto Amato, Jeffrey Barrett, Sonia Goncalves, Ewan Harrison, David K. Jackson, Ian Johnston, Dominic Kwiatkowski, Cordelia Langford, John Sillitoe on behalf of the Wellcome Sanger Institute COVID-19 Surveillance Team |
| EPI_ISL_2317806, EPI_ISL_2317807 | Lighthouse Lab in Alderley Park | Wellcome Sanger Institute for the COVID-19 Genomics UK (COG-UK) Consortium | Jacquelyn Wynn, Mairead Hyland, The Lighthouse Lab in Alderley Park and Alex Alderton, Roberto Amato, Jeffrey Barrett, Sonia Goncalves, Ewan Harrison, David K. Jackson, Ian Johnston, Dominic Kwiatkowski, Cordelia Langford, John Sillitoe on behalf of the Wellcome Sanger Institute COVID-19 Surveillance Team |
| EPI_ISL_2317809 | Lighthouse Lab in Milton Keynes | Wellcome Sanger Institute for the COVID-19 Genomics UK (COG-UK) Consortium | The Lighthouse Lab in Milton Keynes and Alex Alderton, Roberto Amato, Jeffrey Barrett, Sonia Goncalves, Ewan Harrison, David K. Jackson, Ian Johnston, Dominic Kwiatkowski, Cordelia Langford, John Sillitoe on behalf of the Wellcome Sanger Institute COVID-19 Surveillance Team |
| EPI_ISL_2317811 | Randox Laboratories | Wellcome Sanger Institute for the COVID-19 Genomics UK (COG-UK) Consortium | Randox Laboratories and Alex Alderton, Roberto Amato, Jeffrey Barrett, Sonia Goncalves, Ewan Harrison, David K. Jackson, Ian Johnston, Dominic Kwiatkowski, Cordelia Langford, John Sillitoe on behalf of the Wellcome Sanger Institute COVID-19 Surveillance Team |
| EPI_ISL_2317812, EPI_ISL_2317813, EPI_ISL_2317814 | Lighthouse Lab in Alderley Park | Wellcome Sanger Institute for the COVID-19 Genomics UK (COG-UK) Consortium | Jacquelyn Wynn, Mairead Hyland, The Lighthouse Lab in Alderley Park and Alex Alderton, Roberto Amato, Jeffrey Barrett, Sonia Goncalves, Ewan Harrison, David K. Jackson, Ian Johnston, Dominic Kwiatkowski, Cordelia Langford, John Sillitoe on behalf of the Wellcome Sanger Institute COVID-19 Surveillance Team |
| EPI_ISL_2317815 | Lighthouse Lab in Milton Keynes | Wellcome Sanger Institute for the COVID-19 Genomics UK (COG-UK) Consortium | The Lighthouse Lab in Milton Keynes and Alex Alderton, Roberto Amato, Jeffrey Barrett, Sonia Goncalves, Ewan Harrison, David K. Jackson, Ian Johnston, Dominic Kwiatkowski, Cordelia Langford, John Sillitoe on behalf of the Wellcome Sanger Institute COVID-19 Surveillance Team |
| EPI_ISL_2317816, EPI_ISL_2317817, EPI_ISL_2317819, EPI_ISL_2317820, EPI_ISL_2317822, EPI_ISL_2317823 | Lighthouse Lab in Alderley Park | Wellcome Sanger Institute for the COVID-19 Genomics UK (COG-UK) Consortium | Jacquelyn Wynn, Mairead Hyland, The Lighthouse Lab in Alderley Park and Alex Alderton, Roberto Amato, Jeffrey Barrett, Sonia Goncalves, Ewan Harrison, David K. Jackson, Ian Johnston, Dominic Kwiatkowski, Cordelia Langford, John Sillitoe on behalf of the Wellcome Sanger Institute COVID-19 Surveillance Team |
| EPI_ISL_2317825, EPI_ISL_2317826 | Lighthouse Lab in Milton Keynes | Wellcome Sanger Institute for the COVID-19 Genomics UK (COG-UK) Consortium | The Lighthouse Lab in Milton Keynes and Alex Alderton, Roberto Amato, Jeffrey Barrett, Sonia Goncalves, Ewan Harrison, David K. Jackson, Ian Johnston, Dominic Kwiatkowski, Cordelia Langford, John Sillitoe on behalf of the Wellcome Sanger Institute COVID-19 Surveillance Team |
| EPI_ISL_2317828 | Lighthouse Lab in Alderley Park | Wellcome Sanger Institute for the COVID-19 Genomics UK (COG-UK) Consortium | Jacquelyn Wynn, Mairead Hyland, The Lighthouse Lab in Alderley Park and Alex Alderton, Roberto Amato, Jeffrey Barrett, Sonia Goncalves, Ewan Harrison, David K. Jackson, Ian Johnston, Dominic Kwiatkowski, Cordelia Langford, John Sillitoe on behalf of the Wellcome Sanger Institute COVID-19 Surveillance Team |
| EPI_ISL_2317829, EPI_ISL_2317830 | Lighthouse Lab in Milton Keynes | Wellcome Sanger Institute for the COVID-19 Genomics UK (COG-UK) Consortium | The Lighthouse Lab in Milton Keynes and Alex Alderton, Roberto Amato, Jeffrey Barrett, Sonia Goncalves, Ewan Harrison, David K. Jackson, Ian Johnston, Dominic Kwiatkowski, Cordelia Langford, John Sillitoe on behalf of the Wellcome Sanger Institute COVID-19 Surveillance Team |
| EPI_ISL_2317832, EPI_ISL_2317833 | Lighthouse Lab in Alderley Park | Wellcome Sanger Institute for the COVID-19 Genomics UK (COG-UK) Consortium | Jacquelyn Wynn, Mairead Hyland, The Lighthouse Lab in Alderley Park and Alex Alderton, Roberto Amato, Jeffrey Barrett, Sonia Goncalves, Ewan Harrison, David K. Jackson, Ian Johnston, Dominic Kwiatkowski, Cordelia Langford, John Sillitoe on behalf of the Wellcome Sanger Institute COVID-19 Surveillance Team |
| EPI_ISL_2317834 | Lighthouse Lab in Milton Keynes | Wellcome Sanger Institute for the COVID-19 Genomics UK (COG-UK) Consortium | The Lighthouse Lab in Milton Keynes and Alex Alderton, Roberto Amato, Jeffrey Barrett, Sonia Goncalves, Ewan Harrison, David K. Jackson, Ian Johnston, Dominic Kwiatkowski, Cordelia Langford, John Sillitoe on behalf of the Wellcome Sanger Institute COVID-19 Surveillance Team |
| EPI_ISL_2317836 | Lighthouse Lab in Alderley Park | Wellcome Sanger Institute for the COVID-19 Genomics UK (COG-UK) Consortium | Jacquelyn Wynn, Mairead Hyland, The Lighthouse Lab in Alderley Park and Alex Alderton, Roberto Amato, Jeffrey Barrett, Sonia Goncalves, Ewan Harrison, David K. Jackson, Ian Johnston, Dominic Kwiatkowski, Cordelia Langford, John Sillitoe on behalf of the Wellcome Sanger Institute COVID-19 Surveillance Team |
| EPI_ISL_2317837 | Lighthouse Lab in Milton Keynes | Wellcome Sanger Institute for the COVID-19 Genomics UK (COG-UK) Consortium | The Lighthouse Lab in Milton Keynes and Alex Alderton, Roberto Amato, Jeffrey Barrett, Sonia Goncalves, Ewan Harrison, David K. Jackson, Ian Johnston, Dominic Kwiatkowski, Cordelia Langford, John Sillitoe on behalf of the Wellcome Sanger Institute COVID-19 Surveillance Team |
| EPI_ISL_2317839, | Lighthouse Lab in Alderley Park | Wellcome Sanger Institute for the COVID-19 Genomics UK (COG-UK) | Jacquelyn Wynn, Mairead Hyland, The Lighthouse Lab in Alderley Park and Alex Alderton, Roberto Amato, Jeffrey Barrett, Sonia Goncalves, Ewan Harrison, David K. Jackson, Ian Johnston, Dominic Kwiatkowski, Cordelia Langford, John Sillitoe on |

| | | | |
|---|---|---|---|
| EPI_ISL_2317840, EPI_ISL_2317841, EPI_ISL_2317844, EPI_ISL_2317849 | | Consortium | behalf of the Wellcome Sanger Institute COVID-19 Surveillance Team |
| EPI_ISL_2317851 | Randox Laboratories | Wellcome Sanger Institute for the COVID-19 Genomics UK (COG-UK) Consortium | Randox Laboratories and Alex Alderton, Roberto Amato, Jeffrey Barrett, Sonia Goncalves, Ewan Harrison, David K. Jackson, Ian Johnston, Dominic Kwiatkowski, Cordelia Langford, John Sillitoe on behalf of the Wellcome Sanger Institute COVID-19 Surveillance Team |
| EPI_ISL_2317852 | Lighthouse Lab in Milton Keynes | Wellcome Sanger Institute for the COVID-19 Genomics UK (COG-UK) Consortium | The Lighthouse Lab in Milton Keynes and Alex Alderton, Roberto Amato, Jeffrey Barrett, Sonia Goncalves, Ewan Harrison, David K. Jackson, Ian Johnston, Dominic Kwiatkowski, Cordelia Langford, John Sillitoe on behalf of the Wellcome Sanger Institute COVID-19 Surveillance Team |
| EPI_ISL_2317853, EPI_ISL_2317856, EPI_ISL_2317857, EPI_ISL_2317859, EPI_ISL_2317860, EPI_ISL_2317861, EPI_ISL_2317863 | Lighthouse Lab in Alderley Park | Wellcome Sanger Institute for the COVID-19 Genomics UK (COG-UK) Consortium | Jacquelyn Wynn, Mairead Hyland, The Lighthouse Lab in Alderley Park and Alex Alderton, Roberto Amato, Jeffrey Barrett, Sonia Goncalves, Ewan Harrison, David K. Jackson, Ian Johnston, Dominic Kwiatkowski, Cordelia Langford, John Sillitoe on behalf of the Wellcome Sanger Institute COVID-19 Surveillance Team |
| EPI_ISL_2317864, EPI_ISL_2317865 | Lighthouse Lab in Milton Keynes | Wellcome Sanger Institute for the COVID-19 Genomics UK (COG-UK) Consortium | The Lighthouse Lab in Milton Keynes and Alex Alderton, Roberto Amato, Jeffrey Barrett, Sonia Goncalves, Ewan Harrison, David K. Jackson, Ian Johnston, Dominic Kwiatkowski, Cordelia Langford, John Sillitoe on behalf of the Wellcome Sanger Institute COVID-19 Surveillance Team |
| EPI_ISL_2317866, EPI_ISL_2317867, EPI_ISL_2317868, EPI_ISL_2317871, EPI_ISL_2317876, EPI_ISL_2317877 | Lighthouse Lab in Alderley Park | Wellcome Sanger Institute for the COVID-19 Genomics UK (COG-UK) Consortium | Jacquelyn Wynn, Mairead Hyland, The Lighthouse Lab in Alderley Park and Alex Alderton, Roberto Amato, Jeffrey Barrett, Sonia Goncalves, Ewan Harrison, David K. Jackson, Ian Johnston, Dominic Kwiatkowski, Cordelia Langford, John Sillitoe on behalf of the Wellcome Sanger Institute COVID-19 Surveillance Team |
| EPI_ISL_2317883 | Lighthouse Lab in Milton Keynes | Wellcome Sanger Institute for the COVID-19 Genomics UK (COG-UK) Consortium | The Lighthouse Lab in Milton Keynes and Alex Alderton, Roberto Amato, Jeffrey Barrett, Sonia Goncalves, Ewan Harrison, David K. Jackson, Ian Johnston, Dominic Kwiatkowski, Cordelia Langford, John Sillitoe on behalf of the Wellcome Sanger Institute COVID-19 Surveillance Team |
| EPI_ISL_2317884, EPI_ISL_2317885 | Lighthouse Lab in Alderley Park | Wellcome Sanger Institute for the COVID-19 Genomics UK (COG-UK) Consortium | Jacquelyn Wynn, Mairead Hyland, The Lighthouse Lab in Alderley Park and Alex Alderton, Roberto Amato, Jeffrey Barrett, Sonia Goncalves, Ewan Harrison, David K. Jackson, Ian Johnston, Dominic Kwiatkowski, Cordelia Langford, John Sillitoe on behalf of the Wellcome Sanger Institute COVID-19 Surveillance Team |
| EPI_ISL_2317887, EPI_ISL_2317892 | Lighthouse Lab in Milton Keynes | Wellcome Sanger Institute for the COVID-19 Genomics UK (COG-UK) Consortium | The Lighthouse Lab in Milton Keynes and Alex Alderton, Roberto Amato, Jeffrey Barrett, Sonia Goncalves, Ewan Harrison, David K. Jackson, Ian Johnston, Dominic Kwiatkowski, Cordelia Langford, John Sillitoe on behalf of the Wellcome Sanger Institute COVID-19 Surveillance Team |
| EPI_ISL_2317893, EPI_ISL_2317894, EPI_ISL_2317895, EPI_ISL_2317897, EPI_ISL_2317900, EPI_ISL_2317901, EPI_ISL_2317902, EPI_ISL_2317904, EPI_ISL_2317905, EPI_ISL_2317908, EPI_ISL_2317910, EPI_ISL_2317912, EPI_ISL_2317914, EPI_ISL_2317918, EPI_ISL_2317920, EPI_ISL_2317921 | | | |
| see above | Lighthouse Lab in Alderley Park | Wellcome Sanger Institute for the COVID-19 Genomics UK (COG-UK) Consortium | Jacquelyn Wynn, Mairead Hyland, The Lighthouse Lab in Alderley Park and Alex Alderton, Roberto Amato, Jeffrey Barrett, Sonia Goncalves, Ewan Harrison, David K. Jackson, Ian Johnston, Dominic Kwiatkowski, Cordelia Langford, John Sillitoe on behalf of the Wellcome Sanger Institute COVID-19 Surveillance Team |
| EPI_ISL_2317922 | Lighthouse Lab in Milton Keynes | Wellcome Sanger Institute for the COVID-19 Genomics UK (COG-UK) Consortium | The Lighthouse Lab in Milton Keynes and Alex Alderton, Roberto Amato, Jeffrey Barrett, Sonia Goncalves, Ewan Harrison, David K. Jackson, Ian Johnston, Dominic Kwiatkowski, Cordelia Langford, John Sillitoe on behalf of the Wellcome Sanger Institute COVID-19 Surveillance Team |
| EPI_ISL_2317923, EPI_ISL_2317924, EPI_ISL_2317925, EPI_ISL_2317926, EPI_ISL_2317927, EPI_ISL_2317928 | Lighthouse Lab in Alderley Park | Wellcome Sanger Institute for the COVID-19 Genomics UK (COG-UK) Consortium | Jacquelyn Wynn, Mairead Hyland, The Lighthouse Lab in Alderley Park and Alex Alderton, Roberto Amato, Jeffrey Barrett, Sonia Goncalves, Ewan Harrison, David K. Jackson, Ian Johnston, Dominic Kwiatkowski, Cordelia Langford, John Sillitoe on behalf of the Wellcome Sanger Institute COVID-19 Surveillance Team |
| EPI_ISL_2317929 | Lighthouse Lab in Milton Keynes | Wellcome Sanger Institute for the COVID-19 Genomics UK (COG-UK) Consortium | The Lighthouse Lab in Milton Keynes and Alex Alderton, Roberto Amato, Jeffrey Barrett, Sonia Goncalves, Ewan Harrison, David K. Jackson, Ian Johnston, Dominic Kwiatkowski, Cordelia Langford, John Sillitoe on behalf of the Wellcome Sanger Institute COVID-19 Surveillance Team |
| EPI_ISL_2317930 | Randox Laboratories | Wellcome Sanger Institute for the COVID-19 Genomics UK (COG-UK) Consortium | Randox Laboratories and Alex Alderton, Roberto Amato, Jeffrey Barrett, Sonia Goncalves, Ewan Harrison, David K. Jackson, Ian Johnston, Dominic Kwiatkowski, Cordelia Langford, John Sillitoe on behalf of the Wellcome Sanger Institute COVID-19 Surveillance Team |
| EPI_ISL_2317931, EPI_ISL_2317932, EPI_ISL_2317933, EPI_ISL_2317934 | Lighthouse Lab in Alderley Park | Wellcome Sanger Institute for the COVID-19 Genomics UK (COG-UK) Consortium | Jacquelyn Wynn, Mairead Hyland, The Lighthouse Lab in Alderley Park and Alex Alderton, Roberto Amato, Jeffrey Barrett, Sonia Goncalves, Ewan Harrison, David K. Jackson, Ian Johnston, Dominic Kwiatkowski, Cordelia Langford, John Sillitoe on behalf of the Wellcome Sanger Institute COVID-19 Surveillance Team |
| EPI_ISL_2317935 | Lighthouse Lab in Milton Keynes | Wellcome Sanger Institute for the COVID-19 Genomics UK (COG-UK) Consortium | The Lighthouse Lab in Milton Keynes and Alex Alderton, Roberto Amato, Jeffrey Barrett, Sonia Goncalves, Ewan Harrison, David K. Jackson, Ian Johnston, Dominic Kwiatkowski, Cordelia Langford, John Sillitoe on behalf of the Wellcome Sanger Institute COVID-19 Surveillance Team |
| EPI_ISL_2317936 | Lighthouse Lab in Alderley Park | Wellcome Sanger Institute for the COVID-19 Genomics UK (COG-UK) Consortium | Jacquelyn Wynn, Mairead Hyland, The Lighthouse Lab in Alderley Park and Alex Alderton, Roberto Amato, Jeffrey Barrett, Sonia Goncalves, Ewan Harrison, David K. Jackson, Ian Johnston, Dominic Kwiatkowski, Cordelia Langford, John Sillitoe on behalf of the Wellcome Sanger Institute COVID-19 Surveillance Team |
| EPI_ISL_2317937 | Lighthouse Lab in Milton Keynes | Wellcome Sanger Institute for the COVID-19 Genomics UK (COG-UK) Consortium | The Lighthouse Lab in Milton Keynes and Alex Alderton, Roberto Amato, Jeffrey Barrett, Sonia Goncalves, Ewan Harrison, David K. Jackson, Ian Johnston, Dominic Kwiatkowski, Cordelia Langford, John Sillitoe on behalf of the Wellcome Sanger Institute COVID-19 Surveillance Team |
| EPI_ISL_2317938, EPI_ISL_2317939, EPI_ISL_2317940, EPI_ISL_2317944, EPI_ISL_2317946, EPI_ISL_2317948 | Lighthouse Lab in Alderley Park | Wellcome Sanger Institute for the COVID-19 Genomics UK (COG-UK) Consortium | Jacquelyn Wynn, Mairead Hyland, The Lighthouse Lab in Alderley Park and Alex Alderton, Roberto Amato, Jeffrey Barrett, Sonia Goncalves, Ewan Harrison, David K. Jackson, Ian Johnston, Dominic Kwiatkowski, Cordelia Langford, John Sillitoe on behalf of the Wellcome Sanger Institute COVID-19 Surveillance Team |
| EPI_ISL_2317949 | Lighthouse Lab in Milton Keynes | Wellcome Sanger Institute for the COVID-19 Genomics UK (COG-UK) Consortium | The Lighthouse Lab in Milton Keynes and Alex Alderton, Roberto Amato, Jeffrey Barrett, Sonia Goncalves, Ewan Harrison, David K. Jackson, Ian Johnston, Dominic Kwiatkowski, Cordelia Langford, John Sillitoe on behalf of the Wellcome Sanger Institute COVID-19 Surveillance Team |
| EPI_ISL_2317950, EPI_ISL_2317951, EPI_ISL_2317952, EPI_ISL_2317953 | Lighthouse Lab in Alderley Park | Wellcome Sanger Institute for the COVID-19 Genomics UK (COG-UK) Consortium | Jacquelyn Wynn, Mairead Hyland, The Lighthouse Lab in Alderley Park and Alex Alderton, Roberto Amato, Jeffrey Barrett, Sonia Goncalves, Ewan Harrison, David K. Jackson, Ian Johnston, Dominic Kwiatkowski, Cordelia Langford, John Sillitoe on behalf of the Wellcome Sanger Institute COVID-19 Surveillance Team |
| EPI_ISL_2317954 | Randox Laboratories | Wellcome Sanger Institute for the COVID-19 Genomics UK (COG-UK) Consortium | Randox Laboratories and Alex Alderton, Roberto Amato, Jeffrey Barrett, Sonia Goncalves, Ewan Harrison, David K. Jackson, Ian Johnston, Dominic Kwiatkowski, Cordelia Langford, John Sillitoe on behalf of the Wellcome Sanger Institute COVID-19 Surveillance Team |
| EPI_ISL_2317955 | Lighthouse Lab in Milton Keynes | Wellcome Sanger Institute for the COVID-19 Genomics UK (COG-UK) Consortium | The Lighthouse Lab in Milton Keynes and Alex Alderton, Roberto Amato, Jeffrey Barrett, Sonia Goncalves, Ewan Harrison, David K. Jackson, Ian Johnston, Dominic Kwiatkowski, Cordelia Langford, John Sillitoe on behalf of the Wellcome Sanger Institute COVID-19 Surveillance Team |
| EPI_ISL_2317957, EPI_ISL_2317958, EPI_ISL_2317959 | Lighthouse Lab in Alderley Park | Wellcome Sanger Institute for the COVID-19 Genomics UK (COG-UK) Consortium | Jacquelyn Wynn, Mairead Hyland, The Lighthouse Lab in Alderley Park and Alex Alderton, Roberto Amato, Jeffrey Barrett, Sonia Goncalves, Ewan Harrison, David K. Jackson, Ian Johnston, Dominic Kwiatkowski, Cordelia Langford, John Sillitoe on behalf of the Wellcome Sanger Institute COVID-19 Surveillance Team |
| EPI_ISL_2317960 | Lighthouse Lab in Milton Keynes | Wellcome Sanger Institute for the COVID-19 Genomics UK (COG-UK) Consortium | The Lighthouse Lab in Milton Keynes and Alex Alderton, Roberto Amato, Jeffrey Barrett, Sonia Goncalves, Ewan Harrison, David K. Jackson, Ian Johnston, Dominic Kwiatkowski, Cordelia Langford, John Sillitoe on behalf of the Wellcome Sanger Institute COVID-19 Surveillance Team |
| EPI_ISL_2317961, EPI_ISL_2317962, EPI_ISL_2317963, EPI_ISL_2317964, EPI_ISL_2317965, EPI_ISL_2317967, EPI_ISL_2317968 | Lighthouse Lab in Alderley Park | Wellcome Sanger Institute for the COVID-19 Genomics UK (COG-UK) Consortium | Jacquelyn Wynn, Mairead Hyland, The Lighthouse Lab in Alderley Park and Alex Alderton, Roberto Amato, Jeffrey Barrett, Sonia Goncalves, Ewan Harrison, David K. Jackson, Ian Johnston, Dominic Kwiatkowski, Cordelia Langford, John Sillitoe on behalf of the Wellcome Sanger Institute COVID-19 Surveillance Team |
| EPI_ISL_2317969 | Lighthouse Lab in Milton Keynes | Wellcome Sanger Institute for the COVID-19 Genomics UK (COG-UK) Consortium | The Lighthouse Lab in Milton Keynes and Alex Alderton, Roberto Amato, Jeffrey Barrett, Sonia Goncalves, Ewan Harrison, David K. Jackson, Ian Johnston, Dominic Kwiatkowski, Cordelia Langford, John Sillitoe on behalf of the Wellcome Sanger Institute COVID-19 Surveillance Team |
| EPI_ISL_2317970 | Lighthouse Lab in Alderley Park | Wellcome Sanger Institute for the COVID-19 Genomics UK (COG-UK) Consortium | Jacquelyn Wynn, Mairead Hyland, The Lighthouse Lab in Alderley Park and Alex Alderton, Roberto Amato, Jeffrey Barrett, Sonia Goncalves, Ewan Harrison, David K. Jackson, Ian Johnston, Dominic Kwiatkowski, Cordelia Langford, John Sillitoe on behalf of the Wellcome Sanger Institute COVID-19 Surveillance Team |
| EPI_ISL_2317971 | Lighthouse Lab in Milton Keynes | Wellcome Sanger Institute for the COVID-19 Genomics UK (COG-UK) Consortium | The Lighthouse Lab in Milton Keynes and Alex Alderton, Roberto Amato, Jeffrey Barrett, Sonia Goncalves, Ewan Harrison, David K. Jackson, Ian Johnston, Dominic Kwiatkowski, Cordelia Langford, John Sillitoe on behalf of the Wellcome Sanger Institute COVID-19 Surveillance Team |
| EPI_ISL_2317972 | Lighthouse Lab in Alderley Park | Wellcome Sanger Institute for the COVID-19 Genomics UK (COG-UK) Consortium | Jacquelyn Wynn, Mairead Hyland, The Lighthouse Lab in Alderley Park and Alex Alderton, Roberto Amato, Jeffrey Barrett, Sonia Goncalves, Ewan Harrison, David K. Jackson, Ian Johnston, Dominic Kwiatkowski, Cordelia Langford, John Sillitoe on behalf of the Wellcome Sanger Institute COVID-19 Surveillance Team |
| EPI_ISL_2317973 | Lighthouse Lab in Milton Keynes | Wellcome Sanger Institute for the COVID-19 Genomics UK (COG-UK) Consortium | The Lighthouse Lab in Milton Keynes and Alex Alderton, Roberto Amato, Jeffrey Barrett, Sonia Goncalves, Ewan Harrison, David K. Jackson, Ian Johnston, Dominic Kwiatkowski, Cordelia Langford, John Sillitoe on behalf of the Wellcome Sanger Institute COVID-19 Surveillance Team |
| EPI_ISL_2317974 | Lighthouse Lab in Alderley Park | Wellcome Sanger Institute for the COVID-19 Genomics UK (COG-UK) Consortium | Jacquelyn Wynn, Mairead Hyland, The Lighthouse Lab in Alderley Park and Alex Alderton, Roberto Amato, Jeffrey Barrett, Sonia Goncalves, Ewan Harrison, David K. Jackson, Ian Johnston, Dominic Kwiatkowski, Cordelia Langford, John Sillitoe on behalf of the Wellcome Sanger Institute COVID-19 Surveillance Team |
| EPI_ISL_2317975 | Lighthouse Lab in Milton Keynes | Wellcome Sanger Institute for the COVID-19 Genomics UK (COG-UK) | The Lighthouse Lab in Milton Keynes and Alex Alderton, Roberto Amato, Jeffrey Barrett, Sonia Goncalves, Ewan Harrison, David K. Jackson, Ian Johnston, Dominic Kwiatkowski, Cordelia Langford, John Sillitoe on behalf of the Wellcome Sanger |

| | | | |
|---|---|---|---|
| | | Consortium | Institute COVID-19 Surveillance Team |
| EPI_ISL_2317976, EPI_ISL_2317977 | Lighthouse Lab in Alderley Park | Wellcome Sanger Institute for the COVID-19 Genomics UK (COG-UK) Consortium | Jacquelyn Wynn, Mairead Hyland, The Lighthouse Lab in Alderley Park and Alex Alderton, Roberto Amato, Jeffrey Barrett, Sonia Goncalves, Ewan Harrison, David K. Jackson, Ian Johnston, Dominic Kwiatkowski, Cordelia Langford, John Sillitoe on behalf of the Wellcome Sanger Institute COVID-19 Surveillance Team |
| EPI_ISL_2317980 | Randox Laboratories | Wellcome Sanger Institute for the COVID-19 Genomics UK (COG-UK) Consortium | Randox Laboratories and Alex Alderton, Roberto Amato, Jeffrey Barrett, Sonia Goncalves, Ewan Harrison, David K. Jackson, Ian Johnston, Dominic Kwiatkowski, Cordelia Langford, John Sillitoe on behalf of the Wellcome Sanger Institute COVID-19 Surveillance Team |
| EPI_ISL_2317982 | Lighthouse Lab in Milton Keynes | Wellcome Sanger Institute for the COVID-19 Genomics UK (COG-UK) Consortium | The Lighthouse Lab in Milton Keynes and Alex Alderton, Roberto Amato, Jeffrey Barrett, Sonia Goncalves, Ewan Harrison, David K. Jackson, Ian Johnston, Dominic Kwiatkowski, Cordelia Langford, John Sillitoe on behalf of the Wellcome Sanger Institute COVID-19 Surveillance Team |
| EPI_ISL_2317983 | Lighthouse Lab in Alderley Park | Wellcome Sanger Institute for the COVID-19 Genomics UK (COG-UK) Consortium | Jacquelyn Wynn, Mairead Hyland, The Lighthouse Lab in Alderley Park and Alex Alderton, Roberto Amato, Jeffrey Barrett, Sonia Goncalves, Ewan Harrison, David K. Jackson, Ian Johnston, Dominic Kwiatkowski, Cordelia Langford, John Sillitoe on behalf of the Wellcome Sanger Institute COVID-19 Surveillance Team |
| EPI_ISL_2317984, EPI_ISL_2317985 | Lighthouse Lab in Milton Keynes | Wellcome Sanger Institute for the COVID-19 Genomics UK (COG-UK) Consortium | The Lighthouse Lab in Milton Keynes and Alex Alderton, Roberto Amato, Jeffrey Barrett, Sonia Goncalves, Ewan Harrison, David K. Jackson, Ian Johnston, Dominic Kwiatkowski, Cordelia Langford, John Sillitoe on behalf of the Wellcome Sanger Institute COVID-19 Surveillance Team |
| EPI_ISL_2317986, EPI_ISL_2317988, EPI_ISL_2317989, EPI_ISL_2317990, EPI_ISL_2317992, EPI_ISL_2317993, EPI_ISL_2317996, EPI_ISL_2317999, EPI_ISL_2318002 | Lighthouse Lab in Alderley Park | Wellcome Sanger Institute for the COVID-19 Genomics UK (COG-UK) Consortium | Jacquelyn Wynn, Mairead Hyland, The Lighthouse Lab in Alderley Park and Alex Alderton, Roberto Amato, Jeffrey Barrett, Sonia Goncalves, Ewan Harrison, David K. Jackson, Ian Johnston, Dominic Kwiatkowski, Cordelia Langford, John Sillitoe on behalf of the Wellcome Sanger Institute COVID-19 Surveillance Team |
| EPI_ISL_2318003 | Lighthouse Lab in Milton Keynes | Wellcome Sanger Institute for the COVID-19 Genomics UK (COG-UK) Consortium | The Lighthouse Lab in Milton Keynes and Alex Alderton, Roberto Amato, Jeffrey Barrett, Sonia Goncalves, Ewan Harrison, David K. Jackson, Ian Johnston, Dominic Kwiatkowski, Cordelia Langford, John Sillitoe on behalf of the Wellcome Sanger Institute COVID-19 Surveillance Team |
| EPI_ISL_2318004, EPI_ISL_2318007, EPI_ISL_2318010 | Lighthouse Lab in Alderley Park | Wellcome Sanger Institute for the COVID-19 Genomics UK (COG-UK) Consortium | Jacquelyn Wynn, Mairead Hyland, The Lighthouse Lab in Alderley Park and Alex Alderton, Roberto Amato, Jeffrey Barrett, Sonia Goncalves, Ewan Harrison, David K. Jackson, Ian Johnston, Dominic Kwiatkowski, Cordelia Langford, John Sillitoe on behalf of the Wellcome Sanger Institute COVID-19 Surveillance Team |
| EPI_ISL_2318011 | Lighthouse Lab in Milton Keynes | Wellcome Sanger Institute for the COVID-19 Genomics UK (COG-UK) Consortium | The Lighthouse Lab in Milton Keynes and Alex Alderton, Roberto Amato, Jeffrey Barrett, Sonia Goncalves, Ewan Harrison, David K. Jackson, Ian Johnston, Dominic Kwiatkowski, Cordelia Langford, John Sillitoe on behalf of the Wellcome Sanger Institute COVID-19 Surveillance Team |
| EPI_ISL_2318013 | Lighthouse Lab in Alderley Park | Wellcome Sanger Institute for the COVID-19 Genomics UK (COG-UK) Consortium | Jacquelyn Wynn, Mairead Hyland, The Lighthouse Lab in Alderley Park and Alex Alderton, Roberto Amato, Jeffrey Barrett, Sonia Goncalves, Ewan Harrison, David K. Jackson, Ian Johnston, Dominic Kwiatkowski, Cordelia Langford, John Sillitoe on behalf of the Wellcome Sanger Institute COVID-19 Surveillance Team |
| EPI_ISL_2318016, EPI_ISL_2318017 | Lighthouse Lab in Glasgow | Wellcome Sanger Institute for the COVID-19 Genomics UK (COG-UK) Consortium | Harper VanSteenhouse, Yumi Kasai, David Gray, Carol Clugston, Anna Dominiczak and Alex Alderton, Roberto Amato, Jeffrey Barrett, Sonia Goncalves, Ewan Harrison, David K. Jackson, Ian Johnston, Dominic Kwiatkowski, Cordelia Langford, John Sillitoe on behalf of the Wellcome Sanger Institute COVID-19 Surveillance Team |
| EPI_ISL_2318018, EPI_ISL_2318019 | Lighthouse Lab in Alderley Park | Wellcome Sanger Institute for the COVID-19 Genomics UK (COG-UK) Consortium | Jacquelyn Wynn, Mairead Hyland, The Lighthouse Lab in Alderley Park and Alex Alderton, Roberto Amato, Jeffrey Barrett, Sonia Goncalves, Ewan Harrison, David K. Jackson, Ian Johnston, Dominic Kwiatkowski, Cordelia Langford, John Sillitoe on behalf of the Wellcome Sanger Institute COVID-19 Surveillance Team |
| EPI_ISL_2318020, EPI_ISL_2318021 | Lighthouse Lab in Glasgow | Wellcome Sanger Institute for the COVID-19 Genomics UK (COG-UK) Consortium | Harper VanSteenhouse, Yumi Kasai, David Gray, Carol Clugston, Anna Dominiczak and Alex Alderton, Roberto Amato, Jeffrey Barrett, Sonia Goncalves, Ewan Harrison, David K. Jackson, Ian Johnston, Dominic Kwiatkowski, Cordelia Langford, John Sillitoe on behalf of the Wellcome Sanger Institute COVID-19 Surveillance Team |
| EPI_ISL_2318022, EPI_ISL_2318023 | Lighthouse Lab in Alderley Park | Wellcome Sanger Institute for the COVID-19 Genomics UK (COG-UK) Consortium | Jacquelyn Wynn, Mairead Hyland, The Lighthouse Lab in Alderley Park and Alex Alderton, Roberto Amato, Jeffrey Barrett, Sonia Goncalves, Ewan Harrison, David K. Jackson, Ian Johnston, Dominic Kwiatkowski, Cordelia Langford, John Sillitoe on behalf of the Wellcome Sanger Institute COVID-19 Surveillance Team |
| EPI_ISL_2318024 | Lighthouse Lab in Glasgow | Wellcome Sanger Institute for the COVID-19 Genomics UK (COG-UK) Consortium | Harper VanSteenhouse, Yumi Kasai, David Gray, Carol Clugston, Anna Dominiczak and Alex Alderton, Roberto Amato, Jeffrey Barrett, Sonia Goncalves, Ewan Harrison, David K. Jackson, Ian Johnston, Dominic Kwiatkowski, Cordelia Langford, John Sillitoe on behalf of the Wellcome Sanger Institute COVID-19 Surveillance Team |
| EPI_ISL_2318025, EPI_ISL_2318026, EPI_ISL_2318027, EPI_ISL_2318029, EPI_ISL_2318030, EPI_ISL_2318031 | Lighthouse Lab in Alderley Park | Wellcome Sanger Institute for the COVID-19 Genomics UK (COG-UK) Consortium | Jacquelyn Wynn, Mairead Hyland, The Lighthouse Lab in Alderley Park and Alex Alderton, Roberto Amato, Jeffrey Barrett, Sonia Goncalves, Ewan Harrison, David K. Jackson, Ian Johnston, Dominic Kwiatkowski, Cordelia Langford, John Sillitoe on behalf of the Wellcome Sanger Institute COVID-19 Surveillance Team |
| EPI_ISL_2318033 | Health Services Laboratories | Wellcome Sanger Institute for the COVID-19 Genomics UK (COG-UK) Consortium | Health Services Laboratories and Alex Alderton, Roberto Amato, Jeffrey Barrett, Sonia Goncalves, Ewan Harrison, David K. Jackson, Ian Johnston, Dominic Kwiatkowski, Cordelia Langford, John Sillitoe on behalf of the Wellcome Sanger Institute COVID-19 Surveillance Team |
| EPI_ISL_2318035, EPI_ISL_2318036, EPI_ISL_2318038, EPI_ISL_2318039, EPI_ISL_2318040 | Lighthouse Lab in Alderley Park | Wellcome Sanger Institute for the COVID-19 Genomics UK (COG-UK) Consortium | Jacquelyn Wynn, Mairead Hyland, The Lighthouse Lab in Alderley Park and Alex Alderton, Roberto Amato, Jeffrey Barrett, Sonia Goncalves, Ewan Harrison, David K. Jackson, Ian Johnston, Dominic Kwiatkowski, Cordelia Langford, John Sillitoe on behalf of the Wellcome Sanger Institute COVID-19 Surveillance Team |
| EPI_ISL_2318042 | Health Services Laboratories | Wellcome Sanger Institute for the COVID-19 Genomics UK (COG-UK) Consortium | Health Services Laboratories and Alex Alderton, Roberto Amato, Jeffrey Barrett, Sonia Goncalves, Ewan Harrison, David K. Jackson, Ian Johnston, Dominic Kwiatkowski, Cordelia Langford, John Sillitoe on behalf of the Wellcome Sanger Institute COVID-19 Surveillance Team |
| EPI_ISL_2318045, EPI_ISL_2318046, EPI_ISL_2318047, EPI_ISL_2318049, EPI_ISL_2318050, EPI_ISL_2318051, EPI_ISL_2318053 | Lighthouse Lab in Alderley Park | Wellcome Sanger Institute for the COVID-19 Genomics UK (COG-UK) Consortium | Jacquelyn Wynn, Mairead Hyland, The Lighthouse Lab in Alderley Park and Alex Alderton, Roberto Amato, Jeffrey Barrett, Sonia Goncalves, Ewan Harrison, David K. Jackson, Ian Johnston, Dominic Kwiatkowski, Cordelia Langford, John Sillitoe on behalf of the Wellcome Sanger Institute COVID-19 Surveillance Team |
| EPI_ISL_2318054 | Lighthouse Lab in Glasgow | Wellcome Sanger Institute for the COVID-19 Genomics UK (COG-UK) Consortium | Harper VanSteenhouse, Yumi Kasai, David Gray, Carol Clugston, Anna Dominiczak and Alex Alderton, Roberto Amato, Jeffrey Barrett, Sonia Goncalves, Ewan Harrison, David K. Jackson, Ian Johnston, Dominic Kwiatkowski, Cordelia Langford, John Sillitoe on behalf of the Wellcome Sanger Institute COVID-19 Surveillance Team |
| EPI_ISL_2318056, EPI_ISL_2318058, EPI_ISL_2318062 | Lighthouse Lab in Alderley Park | Wellcome Sanger Institute for the COVID-19 Genomics UK (COG-UK) Consortium | Jacquelyn Wynn, Mairead Hyland, The Lighthouse Lab in Alderley Park and Alex Alderton, Roberto Amato, Jeffrey Barrett, Sonia Goncalves, Ewan Harrison, David K. Jackson, Ian Johnston, Dominic Kwiatkowski, Cordelia Langford, John Sillitoe on behalf of the Wellcome Sanger Institute COVID-19 Surveillance Team |
| EPI_ISL_2318063, EPI_ISL_2318065 | Lighthouse Lab in Glasgow | Wellcome Sanger Institute for the COVID-19 Genomics UK (COG-UK) Consortium | Harper VanSteenhouse, Yumi Kasai, David Gray, Carol Clugston, Anna Dominiczak and Alex Alderton, Roberto Amato, Jeffrey Barrett, Sonia Goncalves, Ewan Harrison, David K. Jackson, Ian Johnston, Dominic Kwiatkowski, Cordelia Langford, John Sillitoe on behalf of the Wellcome Sanger Institute COVID-19 Surveillance Team |
| EPI_ISL_2318066 | Health Services Laboratories | Wellcome Sanger Institute for the COVID-19 Genomics UK (COG-UK) Consortium | Health Services Laboratories and Alex Alderton, Roberto Amato, Jeffrey Barrett, Sonia Goncalves, Ewan Harrison, David K. Jackson, Ian Johnston, Dominic Kwiatkowski, Cordelia Langford, John Sillitoe on behalf of the Wellcome Sanger Institute COVID-19 Surveillance Team |
| EPI_ISL_2318067 | Lighthouse Lab in Glasgow | Wellcome Sanger Institute for the COVID-19 Genomics UK (COG-UK) Consortium | Harper VanSteenhouse, Yumi Kasai, David Gray, Carol Clugston, Anna Dominiczak and Alex Alderton, Roberto Amato, Jeffrey Barrett, Sonia Goncalves, Ewan Harrison, David K. Jackson, Ian Johnston, Dominic Kwiatkowski, Cordelia Langford, John Sillitoe on behalf of the Wellcome Sanger Institute COVID-19 Surveillance Team |
| EPI_ISL_2318068 | Lighthouse Lab in Alderley Park | Wellcome Sanger Institute for the COVID-19 Genomics UK (COG-UK) Consortium | Jacquelyn Wynn, Mairead Hyland, The Lighthouse Lab in Alderley Park and Alex Alderton, Roberto Amato, Jeffrey Barrett, Sonia Goncalves, Ewan Harrison, David K. Jackson, Ian Johnston, Dominic Kwiatkowski, Cordelia Langford, John Sillitoe on behalf of the Wellcome Sanger Institute COVID-19 Surveillance Team |
| EPI_ISL_2318069 | Lighthouse Lab in Glasgow | Wellcome Sanger Institute for the COVID-19 Genomics UK (COG-UK) Consortium | Harper VanSteenhouse, Yumi Kasai, David Gray, Carol Clugston, Anna Dominiczak and Alex Alderton, Roberto Amato, Jeffrey Barrett, Sonia Goncalves, Ewan Harrison, David K. Jackson, Ian Johnston, Dominic Kwiatkowski, Cordelia Langford, John Sillitoe on behalf of the Wellcome Sanger Institute COVID-19 Surveillance Team |
| EPI_ISL_2318071, EPI_ISL_2318072, EPI_ISL_2318075 | Lighthouse Lab in Alderley Park | Wellcome Sanger Institute for the COVID-19 Genomics UK (COG-UK) Consortium | Jacquelyn Wynn, Mairead Hyland, The Lighthouse Lab in Alderley Park and Alex Alderton, Roberto Amato, Jeffrey Barrett, Sonia Goncalves, Ewan Harrison, David K. Jackson, Ian Johnston, Dominic Kwiatkowski, Cordelia Langford, John Sillitoe on behalf of the Wellcome Sanger Institute COVID-19 Surveillance Team |
| EPI_ISL_2318076 | Lighthouse Lab in Glasgow | Wellcome Sanger Institute for the COVID-19 Genomics UK (COG-UK) Consortium | Harper VanSteenhouse, Yumi Kasai, David Gray, Carol Clugston, Anna Dominiczak and Alex Alderton, Roberto Amato, Jeffrey Barrett, Sonia Goncalves, Ewan Harrison, David K. Jackson, Ian Johnston, Dominic Kwiatkowski, Cordelia Langford, John Sillitoe on behalf of the Wellcome Sanger Institute COVID-19 Surveillance Team |
| EPI_ISL_2318077, EPI_ISL_2318078 | Lighthouse Lab in Alderley Park | Wellcome Sanger Institute for the COVID-19 Genomics UK (COG-UK) Consortium | Jacquelyn Wynn, Mairead Hyland, The Lighthouse Lab in Alderley Park and Alex Alderton, Roberto Amato, Jeffrey Barrett, Sonia Goncalves, Ewan Harrison, David K. Jackson, Ian Johnston, Dominic Kwiatkowski, Cordelia Langford, John Sillitoe on behalf of the Wellcome Sanger Institute COVID-19 Surveillance Team |
| EPI_ISL_2318081 | Lighthouse Lab in Glasgow | Wellcome Sanger Institute for the COVID-19 Genomics UK (COG-UK) Consortium | Harper VanSteenhouse, Yumi Kasai, David Gray, Carol Clugston, Anna Dominiczak and Alex Alderton, Roberto Amato, Jeffrey Barrett, Sonia Goncalves, Ewan Harrison, David K. Jackson, Ian Johnston, Dominic Kwiatkowski, Cordelia Langford, John Sillitoe on behalf of the Wellcome Sanger Institute COVID-19 Surveillance Team |
| EPI_ISL_2318082, EPI_ISL_2318084 | Lighthouse Lab in Alderley Park | Wellcome Sanger Institute for the COVID-19 Genomics UK (COG-UK) Consortium | Jacquelyn Wynn, Mairead Hyland, The Lighthouse Lab in Alderley Park and Alex Alderton, Roberto Amato, Jeffrey Barrett, Sonia Goncalves, Ewan Harrison, David K. Jackson, Ian Johnston, Dominic Kwiatkowski, Cordelia Langford, John Sillitoe on behalf of the Wellcome Sanger Institute COVID-19 Surveillance Team |
| EPI_ISL_2318085 | Lighthouse Lab in Glasgow | Wellcome Sanger Institute for the COVID-19 Genomics UK (COG-UK) Consortium | Harper VanSteenhouse, Yumi Kasai, David Gray, Carol Clugston, Anna Dominiczak and Alex Alderton, Roberto Amato, Jeffrey Barrett, Sonia Goncalves, Ewan Harrison, David K. Jackson, Ian Johnston, Dominic Kwiatkowski, Cordelia Langford, John Sillitoe on behalf of the Wellcome Sanger Institute COVID-19 Surveillance Team |
| EPI_ISL_2318086, EPI_ISL_2318089 | Lighthouse Lab in Alderley Park | Wellcome Sanger Institute for the COVID-19 Genomics UK (COG-UK) Consortium | Jacquelyn Wynn, Mairead Hyland, The Lighthouse Lab in Alderley Park and Alex Alderton, Roberto Amato, Jeffrey Barrett, Sonia Goncalves, Ewan Harrison, David K. Jackson, Ian Johnston, Dominic Kwiatkowski, Cordelia Langford, John Sillitoe on behalf of the Wellcome Sanger Institute COVID-19 Surveillance Team |
| EPI_ISL_2318090 | Lighthouse Lab in Glasgow | Wellcome Sanger Institute for the COVID-19 Genomics UK (COG-UK) Consortium | Harper VanSteenhouse, Yumi Kasai, David Gray, Carol Clugston, Anna Dominiczak and Alex Alderton, Roberto Amato, Jeffrey Barrett, Sonia Goncalves, Ewan Harrison, David K. Jackson, Ian Johnston, Dominic Kwiatkowski, Cordelia Langford, John Sillitoe on behalf of the Wellcome Sanger Institute COVID-19 Surveillance Team |
| EPI_ISL_2318091, | Lighthouse Lab in Alderley Park | Wellcome Sanger Institute for the COVID-19 Genomics UK (COG-UK) | Jacquelyn Wynn, Mairead Hyland, The Lighthouse Lab in Alderley Park and Alex Alderton, Roberto Amato, Jeffrey Barrett, Sonia Goncalves, Ewan Harrison, David K. Jackson, Ian Johnston, Dominic Kwiatkowski, Cordelia Langford, John Sillitoe on |

| | | | |
|---|---|---|---|
| EPI_ISL_2318092, EPI_ISL_2318094, EPI_ISL_2318095, EPI_ISL_2318096, EPI_ISL_2318097, EPI_ISL_2318098, EPI_ISL_2318100, EPI_ISL_2318101 | | Consortium | behalf of the Wellcome Sanger Institute COVID-19 Surveillance Team |
| EPI_ISL_2318102 | Lighthouse Lab in Glasgow | Wellcome Sanger Institute for the COVID-19 Genomics UK (COG-UK) Consortium | Harper VanSteenhouse, Yumi Kasai, David Gray, Carol Clugston, Anna Dominiczak and Alex Alderton, Roberto Amato, Jeffrey Barrett, Sonia Goncalves, Ewan Harrison, David K. Jackson, Ian Johnston, Dominic Kwiatkowski, Cordelia Langford, John Sillitoe on behalf of the Wellcome Sanger Institute COVID-19 Surveillance Team |
| EPI_ISL_2318103 | Health Services Laboratories | Wellcome Sanger Institute for the COVID-19 Genomics UK (COG-UK) Consortium | Health Services Laboratories and Alex Alderton, Roberto Amato, Jeffrey Barrett, Sonia Goncalves, Ewan Harrison, David K. Jackson, Ian Johnston, Dominic Kwiatkowski, Cordelia Langford, John Sillitoe on behalf of the Wellcome Sanger Institute COVID-19 Surveillance Team |
| EPI_ISL_2318104, EPI_ISL_2318106, EPI_ISL_2318108 | Lighthouse Lab in Alderley Park | Wellcome Sanger Institute for the COVID-19 Genomics UK (COG-UK) Consortium | Jacquelyn Wynn, Mairead Hyland, The Lighthouse Lab in Alderley Park and Alex Alderton, Roberto Amato, Jeffrey Barrett, Sonia Goncalves, Ewan Harrison, David K. Jackson, Ian Johnston, Dominic Kwiatkowski, Cordelia Langford, John Sillitoe on behalf of the Wellcome Sanger Institute COVID-19 Surveillance Team |
| EPI_ISL_2318111 | Lighthouse Lab in Glasgow | Wellcome Sanger Institute for the COVID-19 Genomics UK (COG-UK) Consortium | Harper VanSteenhouse, Yumi Kasai, David Gray, Carol Clugston, Anna Dominiczak and Alex Alderton, Roberto Amato, Jeffrey Barrett, Sonia Goncalves, Ewan Harrison, David K. Jackson, Ian Johnston, Dominic Kwiatkowski, Cordelia Langford, John Sillitoe on behalf of the Wellcome Sanger Institute COVID-19 Surveillance Team |
| EPI_ISL_2318112 | Health Services Laboratories | Wellcome Sanger Institute for the COVID-19 Genomics UK (COG-UK) Consortium | Health Services Laboratories and Alex Alderton, Roberto Amato, Jeffrey Barrett, Sonia Goncalves, Ewan Harrison, David K. Jackson, Ian Johnston, Dominic Kwiatkowski, Cordelia Langford, John Sillitoe on behalf of the Wellcome Sanger Institute COVID-19 Surveillance Team |
| EPI_ISL_2318113 | Lighthouse Lab in Alderley Park | Wellcome Sanger Institute for the COVID-19 Genomics UK (COG-UK) Consortium | Jacquelyn Wynn, Mairead Hyland, The Lighthouse Lab in Alderley Park and Alex Alderton, Roberto Amato, Jeffrey Barrett, Sonia Goncalves, Ewan Harrison, David K. Jackson, Ian Johnston, Dominic Kwiatkowski, Cordelia Langford, John Sillitoe on behalf of the Wellcome Sanger Institute COVID-19 Surveillance Team |
| EPI_ISL_2318114 | Health Services Laboratories | Wellcome Sanger Institute for the COVID-19 Genomics UK (COG-UK) Consortium | Health Services Laboratories and Alex Alderton, Roberto Amato, Jeffrey Barrett, Sonia Goncalves, Ewan Harrison, David K. Jackson, Ian Johnston, Dominic Kwiatkowski, Cordelia Langford, John Sillitoe on behalf of the Wellcome Sanger Institute COVID-19 Surveillance Team |
| EPI_ISL_2318115, EPI_ISL_2318116 | Lighthouse Lab in Alderley Park | Wellcome Sanger Institute for the COVID-19 Genomics UK (COG-UK) Consortium | Jacquelyn Wynn, Mairead Hyland, The Lighthouse Lab in Alderley Park and Alex Alderton, Roberto Amato, Jeffrey Barrett, Sonia Goncalves, Ewan Harrison, David K. Jackson, Ian Johnston, Dominic Kwiatkowski, Cordelia Langford, John Sillitoe on behalf of the Wellcome Sanger Institute COVID-19 Surveillance Team |
| EPI_ISL_2318118, EPI_ISL_2318120 | Lighthouse Lab in Glasgow | Wellcome Sanger Institute for the COVID-19 Genomics UK (COG-UK) Consortium | Harper VanSteenhouse, Yumi Kasai, David Gray, Carol Clugston, Anna Dominiczak and Alex Alderton, Roberto Amato, Jeffrey Barrett, Sonia Goncalves, Ewan Harrison, David K. Jackson, Ian Johnston, Dominic Kwiatkowski, Cordelia Langford, John Sillitoe on behalf of the Wellcome Sanger Institute COVID-19 Surveillance Team |
| EPI_ISL_2318121, EPI_ISL_2318122 | Lighthouse Lab in Alderley Park | Wellcome Sanger Institute for the COVID-19 Genomics UK (COG-UK) Consortium | Jacquelyn Wynn, Mairead Hyland, The Lighthouse Lab in Alderley Park and Alex Alderton, Roberto Amato, Jeffrey Barrett, Sonia Goncalves, Ewan Harrison, David K. Jackson, Ian Johnston, Dominic Kwiatkowski, Cordelia Langford, John Sillitoe on behalf of the Wellcome Sanger Institute COVID-19 Surveillance Team |
| EPI_ISL_2318123 | Lighthouse Lab in Glasgow | Wellcome Sanger Institute for the COVID-19 Genomics UK (COG-UK) Consortium | Harper VanSteenhouse, Yumi Kasai, David Gray, Carol Clugston, Anna Dominiczak and Alex Alderton, Roberto Amato, Jeffrey Barrett, Sonia Goncalves, Ewan Harrison, David K. Jackson, Ian Johnston, Dominic Kwiatkowski, Cordelia Langford, John Sillitoe on behalf of the Wellcome Sanger Institute COVID-19 Surveillance Team |
| EPI_ISL_2318125, EPI_ISL_2318126, EPI_ISL_2318127, EPI_ISL_2318128, EPI_ISL_2318129 | Lighthouse Lab in Alderley Park | Wellcome Sanger Institute for the COVID-19 Genomics UK (COG-UK) Consortium | Jacquelyn Wynn, Mairead Hyland, The Lighthouse Lab in Alderley Park and Alex Alderton, Roberto Amato, Jeffrey Barrett, Sonia Goncalves, Ewan Harrison, David K. Jackson, Ian Johnston, Dominic Kwiatkowski, Cordelia Langford, John Sillitoe on behalf of the Wellcome Sanger Institute COVID-19 Surveillance Team |
| EPI_ISL_2318130 | Lighthouse Lab in Glasgow | Wellcome Sanger Institute for the COVID-19 Genomics UK (COG-UK) Consortium | Harper VanSteenhouse, Yumi Kasai, David Gray, Carol Clugston, Anna Dominiczak and Alex Alderton, Roberto Amato, Jeffrey Barrett, Sonia Goncalves, Ewan Harrison, David K. Jackson, Ian Johnston, Dominic Kwiatkowski, Cordelia Langford, John Sillitoe on behalf of the Wellcome Sanger Institute COVID-19 Surveillance Team |
| EPI_ISL_2318133, EPI_ISL_2318134, EPI_ISL_2318135, EPI_ISL_2318136, EPI_ISL_2318137, EPI_ISL_2318138, EPI_ISL_2318140 | Lighthouse Lab in Alderley Park | Wellcome Sanger Institute for the COVID-19 Genomics UK (COG-UK) Consortium | Jacquelyn Wynn, Mairead Hyland, The Lighthouse Lab in Alderley Park and Alex Alderton, Roberto Amato, Jeffrey Barrett, Sonia Goncalves, Ewan Harrison, David K. Jackson, Ian Johnston, Dominic Kwiatkowski, Cordelia Langford, John Sillitoe on behalf of the Wellcome Sanger Institute COVID-19 Surveillance Team |
| EPI_ISL_2318141 | Lighthouse Lab in Glasgow | Wellcome Sanger Institute for the COVID-19 Genomics UK (COG-UK) Consortium | Harper VanSteenhouse, Yumi Kasai, David Gray, Carol Clugston, Anna Dominiczak and Alex Alderton, Roberto Amato, Jeffrey Barrett, Sonia Goncalves, Ewan Harrison, David K. Jackson, Ian Johnston, Dominic Kwiatkowski, Cordelia Langford, John Sillitoe on behalf of the Wellcome Sanger Institute COVID-19 Surveillance Team |
| EPI_ISL_2318142, EPI_ISL_2318144 | Lighthouse Lab in Alderley Park | Wellcome Sanger Institute for the COVID-19 Genomics UK (COG-UK) Consortium | Jacquelyn Wynn, Mairead Hyland, The Lighthouse Lab in Alderley Park and Alex Alderton, Roberto Amato, Jeffrey Barrett, Sonia Goncalves, Ewan Harrison, David K. Jackson, Ian Johnston, Dominic Kwiatkowski, Cordelia Langford, John Sillitoe on behalf of the Wellcome Sanger Institute COVID-19 Surveillance Team |
| EPI_ISL_2318145 | Lighthouse Lab in Glasgow | Wellcome Sanger Institute for the COVID-19 Genomics UK (COG-UK) Consortium | Harper VanSteenhouse, Yumi Kasai, David Gray, Carol Clugston, Anna Dominiczak and Alex Alderton, Roberto Amato, Jeffrey Barrett, Sonia Goncalves, Ewan Harrison, David K. Jackson, Ian Johnston, Dominic Kwiatkowski, Cordelia Langford, John Sillitoe on behalf of the Wellcome Sanger Institute COVID-19 Surveillance Team |
| EPI_ISL_2318146, EPI_ISL_2318147, EPI_ISL_2318148, EPI_ISL_2318149 | Lighthouse Lab in Alderley Park | Wellcome Sanger Institute for the COVID-19 Genomics UK (COG-UK) Consortium | Jacquelyn Wynn, Mairead Hyland, The Lighthouse Lab in Alderley Park and Alex Alderton, Roberto Amato, Jeffrey Barrett, Sonia Goncalves, Ewan Harrison, David K. Jackson, Ian Johnston, Dominic Kwiatkowski, Cordelia Langford, John Sillitoe on behalf of the Wellcome Sanger Institute COVID-19 Surveillance Team |
| EPI_ISL_2318150 | Lighthouse Lab in Glasgow | Wellcome Sanger Institute for the COVID-19 Genomics UK (COG-UK) Consortium | Harper VanSteenhouse, Yumi Kasai, David Gray, Carol Clugston, Anna Dominiczak and Alex Alderton, Roberto Amato, Jeffrey Barrett, Sonia Goncalves, Ewan Harrison, David K. Jackson, Ian Johnston, Dominic Kwiatkowski, Cordelia Langford, John Sillitoe on behalf of the Wellcome Sanger Institute COVID-19 Surveillance Team |
| EPI_ISL_2318151 | Lighthouse Lab in Alderley Park | Wellcome Sanger Institute for the COVID-19 Genomics UK (COG-UK) Consortium | Jacquelyn Wynn, Mairead Hyland, The Lighthouse Lab in Alderley Park and Alex Alderton, Roberto Amato, Jeffrey Barrett, Sonia Goncalves, Ewan Harrison, David K. Jackson, Ian Johnston, Dominic Kwiatkowski, Cordelia Langford, John Sillitoe on behalf of the Wellcome Sanger Institute COVID-19 Surveillance Team |
| EPI_ISL_2318153 | Lighthouse Lab in Glasgow | Wellcome Sanger Institute for the COVID-19 Genomics UK (COG-UK) Consortium | Harper VanSteenhouse, Yumi Kasai, David Gray, Carol Clugston, Anna Dominiczak and Alex Alderton, Roberto Amato, Jeffrey Barrett, Sonia Goncalves, Ewan Harrison, David K. Jackson, Ian Johnston, Dominic Kwiatkowski, Cordelia Langford, John Sillitoe on behalf of the Wellcome Sanger Institute COVID-19 Surveillance Team |
| EPI_ISL_2318155, EPI_ISL_2318159, EPI_ISL_2318160, EPI_ISL_2318161, EPI_ISL_2318162, EPI_ISL_2318163 | Lighthouse Lab in Alderley Park | Wellcome Sanger Institute for the COVID-19 Genomics UK (COG-UK) Consortium | Jacquelyn Wynn, Mairead Hyland, The Lighthouse Lab in Alderley Park and Alex Alderton, Roberto Amato, Jeffrey Barrett, Sonia Goncalves, Ewan Harrison, David K. Jackson, Ian Johnston, Dominic Kwiatkowski, Cordelia Langford, John Sillitoe on behalf of the Wellcome Sanger Institute COVID-19 Surveillance Team |
| EPI_ISL_2318164 | Lighthouse Lab in Glasgow | Wellcome Sanger Institute for the COVID-19 Genomics UK (COG-UK) Consortium | Harper VanSteenhouse, Yumi Kasai, David Gray, Carol Clugston, Anna Dominiczak and Alex Alderton, Roberto Amato, Jeffrey Barrett, Sonia Goncalves, Ewan Harrison, David K. Jackson, Ian Johnston, Dominic Kwiatkowski, Cordelia Langford, John Sillitoe on behalf of the Wellcome Sanger Institute COVID-19 Surveillance Team |
| EPI_ISL_2318165, EPI_ISL_2318166, EPI_ISL_2318168, EPI_ISL_2318169, EPI_ISL_2318170, EPI_ISL_2318171, EPI_ISL_2318172 | Lighthouse Lab in Alderley Park | Wellcome Sanger Institute for the COVID-19 Genomics UK (COG-UK) Consortium | Jacquelyn Wynn, Mairead Hyland, The Lighthouse Lab in Alderley Park and Alex Alderton, Roberto Amato, Jeffrey Barrett, Sonia Goncalves, Ewan Harrison, David K. Jackson, Ian Johnston, Dominic Kwiatkowski, Cordelia Langford, John Sillitoe on behalf of the Wellcome Sanger Institute COVID-19 Surveillance Team |
| EPI_ISL_2318173, EPI_ISL_2318174, EPI_ISL_2318175 | Lighthouse Lab in Glasgow | Wellcome Sanger Institute for the COVID-19 Genomics UK (COG-UK) Consortium | Harper VanSteenhouse, Yumi Kasai, David Gray, Carol Clugston, Anna Dominiczak and Alex Alderton, Roberto Amato, Jeffrey Barrett, Sonia Goncalves, Ewan Harrison, David K. Jackson, Ian Johnston, Dominic Kwiatkowski, Cordelia Langford, John Sillitoe on behalf of the Wellcome Sanger Institute COVID-19 Surveillance Team |
| EPI_ISL_2318176, EPI_ISL_2318178, EPI_ISL_2318179, EPI_ISL_2318180 | Lighthouse Lab in Alderley Park | Wellcome Sanger Institute for the COVID-19 Genomics UK (COG-UK) Consortium | Jacquelyn Wynn, Mairead Hyland, The Lighthouse Lab in Alderley Park and Alex Alderton, Roberto Amato, Jeffrey Barrett, Sonia Goncalves, Ewan Harrison, David K. Jackson, Ian Johnston, Dominic Kwiatkowski, Cordelia Langford, John Sillitoe on behalf of the Wellcome Sanger Institute COVID-19 Surveillance Team |
| EPI_ISL_2318181 | Lighthouse Lab in Glasgow | Wellcome Sanger Institute for the COVID-19 Genomics UK (COG-UK) Consortium | Harper VanSteenhouse, Yumi Kasai, David Gray, Carol Clugston, Anna Dominiczak and Alex Alderton, Roberto Amato, Jeffrey Barrett, Sonia Goncalves, Ewan Harrison, David K. Jackson, Ian Johnston, Dominic Kwiatkowski, Cordelia Langford, John Sillitoe on behalf of the Wellcome Sanger Institute COVID-19 Surveillance Team |
| EPI_ISL_2318182, EPI_ISL_2318184, EPI_ISL_2318185, EPI_ISL_2318186, EPI_ISL_2318187, EPI_ISL_2318188, EPI_ISL_2318189, EPI_ISL_2318190, EPI_ISL_2318191, EPI_ISL_2318192, EPI_ISL_2318193, EPI_ISL_2318194, EPI_ISL_2318195, EPI_ISL_2318197, EPI_ISL_2318199 | | | |
| see above | Lighthouse Lab in Alderley Park | Wellcome Sanger Institute for the COVID-19 Genomics UK (COG-UK) Consortium | Jacquelyn Wynn, Mairead Hyland, The Lighthouse Lab in Alderley Park and Alex Alderton, Roberto Amato, Jeffrey Barrett, Sonia Goncalves, Ewan Harrison, David K. Jackson, Ian Johnston, Dominic Kwiatkowski, Cordelia Langford, John Sillitoe on behalf of the Wellcome Sanger Institute COVID-19 Surveillance Team |
| EPI_ISL_2318200 | Lighthouse Lab in Glasgow | Wellcome Sanger Institute for the COVID-19 Genomics UK (COG-UK) Consortium | Harper VanSteenhouse, Yumi Kasai, David Gray, Carol Clugston, Anna Dominiczak and Alex Alderton, Roberto Amato, Jeffrey Barrett, Sonia Goncalves, Ewan Harrison, David K. Jackson, Ian Johnston, Dominic Kwiatkowski, Cordelia Langford, John Sillitoe on behalf of the Wellcome Sanger Institute COVID-19 Surveillance Team |
| EPI_ISL_2318201, EPI_ISL_2318204, EPI_ISL_2318205 | Lighthouse Lab in Alderley Park | Wellcome Sanger Institute for the COVID-19 Genomics UK (COG-UK) Consortium | Jacquelyn Wynn, Mairead Hyland, The Lighthouse Lab in Alderley Park and Alex Alderton, Roberto Amato, Jeffrey Barrett, Sonia Goncalves, Ewan Harrison, David K. Jackson, Ian Johnston, Dominic Kwiatkowski, Cordelia Langford, John Sillitoe on behalf of the Wellcome Sanger Institute COVID-19 Surveillance Team |
| EPI_ISL_2318206 | Health Services Laboratories | Wellcome Sanger Institute for the COVID-19 Genomics UK (COG-UK) Consortium | Health Services Laboratories and Alex Alderton, Roberto Amato, Jeffrey Barrett, Sonia Goncalves, Ewan Harrison, David K. Jackson, Ian Johnston, Dominic Kwiatkowski, Cordelia Langford, John Sillitoe on behalf of the Wellcome Sanger Institute COVID-19 Surveillance Team |
| EPI_ISL_2318208 | Lighthouse Lab in Glasgow | Wellcome Sanger Institute for the COVID-19 Genomics UK (COG-UK) | Harper VanSteenhouse, Yumi Kasai, David Gray, Carol Clugston, Anna Dominiczak and Alex Alderton, Roberto Amato, Jeffrey Barrett, Sonia Goncalves, Ewan Harrison, David K. Jackson, Ian Johnston, Dominic Kwiatkowski, Cordelia Langford, |

| | | | |
|---|---|---|---|
| | | Consortium | John Sillitoe on behalf of the Wellcome Sanger Institute COVID-19 Surveillance Team |
| EPI_ISL_2318209, EPI_ISL_2318210, EPI_ISL_2318211, EPI_ISL_2318212, EPI_ISL_2318214, EPI_ISL_2318215, EPI_ISL_2318219, EPI_ISL_2318220, EPI_ISL_2318222, EPI_ISL_2318223 | Lighthouse Lab in Alderley Park | Wellcome Sanger Institute for the COVID-19 Genomics UK (COG-UK) Consortium | Jacquelyn Wynn, Mairead Hyland, The Lighthouse Lab in Alderley Park and Alex Alderton, Roberto Amato, Jeffrey Barrett, Sonia Goncalves, Ewan Harrison, David K. Jackson, Ian Johnston, Dominic Kwiatkowski, Cordelia Langford, John Sillitoe on behalf of the Wellcome Sanger Institute COVID-19 Surveillance Team |
| EPI_ISL_2318224 | Lighthouse Lab in Glasgow | Wellcome Sanger Institute for the COVID-19 Genomics UK (COG-UK) Consortium | Harper VanSteenhouse, Yumi Kasai, David Gray, Carol Clugston, Anna Dominiczak and Alex Alderton, Roberto Amato, Jeffrey Barrett, Sonia Goncalves, Ewan Harrison, David K. Jackson, Ian Johnston, Dominic Kwiatkowski, Cordelia Langford, John Sillitoe on behalf of the Wellcome Sanger Institute COVID-19 Surveillance Team |
| EPI_ISL_2318225, EPI_ISL_2318227, EPI_ISL_2318229, EPI_ISL_2318230 | Lighthouse Lab in Alderley Park | Wellcome Sanger Institute for the COVID-19 Genomics UK (COG-UK) Consortium | Jacquelyn Wynn, Mairead Hyland, The Lighthouse Lab in Alderley Park and Alex Alderton, Roberto Amato, Jeffrey Barrett, Sonia Goncalves, Ewan Harrison, David K. Jackson, Ian Johnston, Dominic Kwiatkowski, Cordelia Langford, John Sillitoe on behalf of the Wellcome Sanger Institute COVID-19 Surveillance Team |
| EPI_ISL_2318231 | Lighthouse Lab in Glasgow | Wellcome Sanger Institute for the COVID-19 Genomics UK (COG-UK) Consortium | Harper VanSteenhouse, Yumi Kasai, David Gray, Carol Clugston, Anna Dominiczak and Alex Alderton, Roberto Amato, Jeffrey Barrett, Sonia Goncalves, Ewan Harrison, David K. Jackson, Ian Johnston, Dominic Kwiatkowski, Cordelia Langford, John Sillitoe on behalf of the Wellcome Sanger Institute COVID-19 Surveillance Team |
| EPI_ISL_2318232, EPI_ISL_2318235, EPI_ISL_2318236, EPI_ISL_2318238, EPI_ISL_2318239, EPI_ISL_2318240 | Lighthouse Lab in Alderley Park | Wellcome Sanger Institute for the COVID-19 Genomics UK (COG-UK) Consortium | Jacquelyn Wynn, Mairead Hyland, The Lighthouse Lab in Alderley Park and Alex Alderton, Roberto Amato, Jeffrey Barrett, Sonia Goncalves, Ewan Harrison, David K. Jackson, Ian Johnston, Dominic Kwiatkowski, Cordelia Langford, John Sillitoe on behalf of the Wellcome Sanger Institute COVID-19 Surveillance Team |
| EPI_ISL_2318241 | Lighthouse Lab in Glasgow | Wellcome Sanger Institute for the COVID-19 Genomics UK (COG-UK) Consortium | Harper VanSteenhouse, Yumi Kasai, David Gray, Carol Clugston, Anna Dominiczak and Alex Alderton, Roberto Amato, Jeffrey Barrett, Sonia Goncalves, Ewan Harrison, David K. Jackson, Ian Johnston, Dominic Kwiatkowski, Cordelia Langford, John Sillitoe on behalf of the Wellcome Sanger Institute COVID-19 Surveillance Team |
| EPI_ISL_2318242, EPI_ISL_2318246, EPI_ISL_2318247 | Lighthouse Lab in Alderley Park | Wellcome Sanger Institute for the COVID-19 Genomics UK (COG-UK) Consortium | Jacquelyn Wynn, Mairead Hyland, The Lighthouse Lab in Alderley Park and Alex Alderton, Roberto Amato, Jeffrey Barrett, Sonia Goncalves, Ewan Harrison, David K. Jackson, Ian Johnston, Dominic Kwiatkowski, Cordelia Langford, John Sillitoe on behalf of the Wellcome Sanger Institute COVID-19 Surveillance Team |
| EPI_ISL_2318248 | Health Services Laboratories | Wellcome Sanger Institute for the COVID-19 Genomics UK (COG-UK) Consortium | Health Services Laboratories and Alex Alderton, Roberto Amato, Jeffrey Barrett, Sonia Goncalves, Ewan Harrison, David K. Jackson, Ian Johnston, Dominic Kwiatkowski, Cordelia Langford, John Sillitoe on behalf of the Wellcome Sanger Institute COVID-19 Surveillance Team |
| EPI_ISL_2318249 | Lighthouse Lab in Glasgow | Wellcome Sanger Institute for the COVID-19 Genomics UK (COG-UK) Consortium | Harper VanSteenhouse, Yumi Kasai, David Gray, Carol Clugston, Anna Dominiczak and Alex Alderton, Roberto Amato, Jeffrey Barrett, Sonia Goncalves, Ewan Harrison, David K. Jackson, Ian Johnston, Dominic Kwiatkowski, Cordelia Langford, John Sillitoe on behalf of the Wellcome Sanger Institute COVID-19 Surveillance Team |
| EPI_ISL_2318250 | Lighthouse Lab in Alderley Park | Wellcome Sanger Institute for the COVID-19 Genomics UK (COG-UK) Consortium | Jacquelyn Wynn, Mairead Hyland, The Lighthouse Lab in Alderley Park and Alex Alderton, Roberto Amato, Jeffrey Barrett, Sonia Goncalves, Ewan Harrison, David K. Jackson, Ian Johnston, Dominic Kwiatkowski, Cordelia Langford, John Sillitoe on behalf of the Wellcome Sanger Institute COVID-19 Surveillance Team |
| EPI_ISL_2318251 | Lighthouse Lab in Glasgow | Wellcome Sanger Institute for the COVID-19 Genomics UK (COG-UK) Consortium | Harper VanSteenhouse, Yumi Kasai, David Gray, Carol Clugston, Anna Dominiczak and Alex Alderton, Roberto Amato, Jeffrey Barrett, Sonia Goncalves, Ewan Harrison, David K. Jackson, Ian Johnston, Dominic Kwiatkowski, Cordelia Langford, John Sillitoe on behalf of the Wellcome Sanger Institute COVID-19 Surveillance Team |
| EPI_ISL_2318252, EPI_ISL_2318253, EPI_ISL_2318254 | Lighthouse Lab in Alderley Park | Wellcome Sanger Institute for the COVID-19 Genomics UK (COG-UK) Consortium | Jacquelyn Wynn, Mairead Hyland, The Lighthouse Lab in Alderley Park and Alex Alderton, Roberto Amato, Jeffrey Barrett, Sonia Goncalves, Ewan Harrison, David K. Jackson, Ian Johnston, Dominic Kwiatkowski, Cordelia Langford, John Sillitoe on behalf of the Wellcome Sanger Institute COVID-19 Surveillance Team |
| EPI_ISL_2318255 | Lighthouse Lab in Glasgow | Wellcome Sanger Institute for the COVID-19 Genomics UK (COG-UK) Consortium | Harper VanSteenhouse, Yumi Kasai, David Gray, Carol Clugston, Anna Dominiczak and Alex Alderton, Roberto Amato, Jeffrey Barrett, Sonia Goncalves, Ewan Harrison, David K. Jackson, Ian Johnston, Dominic Kwiatkowski, Cordelia Langford, John Sillitoe on behalf of the Wellcome Sanger Institute COVID-19 Surveillance Team |
| EPI_ISL_2318256, EPI_ISL_2318258, EPI_ISL_2318259, EPI_ISL_2318260, EPI_ISL_2318261, EPI_ISL_2318262, EPI_ISL_2318263 | Lighthouse Lab in Alderley Park | Wellcome Sanger Institute for the COVID-19 Genomics UK (COG-UK) Consortium | Jacquelyn Wynn, Mairead Hyland, The Lighthouse Lab in Alderley Park and Alex Alderton, Roberto Amato, Jeffrey Barrett, Sonia Goncalves, Ewan Harrison, David K. Jackson, Ian Johnston, Dominic Kwiatkowski, Cordelia Langford, John Sillitoe on behalf of the Wellcome Sanger Institute COVID-19 Surveillance Team |
| EPI_ISL_2318265 | Health Services Laboratories | Wellcome Sanger Institute for the COVID-19 Genomics UK (COG-UK) Consortium | Health Services Laboratories and Alex Alderton, Roberto Amato, Jeffrey Barrett, Sonia Goncalves, Ewan Harrison, David K. Jackson, Ian Johnston, Dominic Kwiatkowski, Cordelia Langford, John Sillitoe on behalf of the Wellcome Sanger Institute COVID-19 Surveillance Team |
| EPI_ISL_2318266, EPI_ISL_2318267, EPI_ISL_2318268 | Lighthouse Lab in Alderley Park | Wellcome Sanger Institute for the COVID-19 Genomics UK (COG-UK) Consortium | Jacquelyn Wynn, Mairead Hyland, The Lighthouse Lab in Alderley Park and Alex Alderton, Roberto Amato, Jeffrey Barrett, Sonia Goncalves, Ewan Harrison, David K. Jackson, Ian Johnston, Dominic Kwiatkowski, Cordelia Langford, John Sillitoe on behalf of the Wellcome Sanger Institute COVID-19 Surveillance Team |
| EPI_ISL_2318269 | Lighthouse Lab in Glasgow | Wellcome Sanger Institute for the COVID-19 Genomics UK (COG-UK) Consortium | Harper VanSteenhouse, Yumi Kasai, David Gray, Carol Clugston, Anna Dominiczak and Alex Alderton, Roberto Amato, Jeffrey Barrett, Sonia Goncalves, Ewan Harrison, David K. Jackson, Ian Johnston, Dominic Kwiatkowski, Cordelia Langford, John Sillitoe on behalf of the Wellcome Sanger Institute COVID-19 Surveillance Team |
| EPI_ISL_2318270, EPI_ISL_2318271, EPI_ISL_2318272, EPI_ISL_2318273, EPI_ISL_2318274, EPI_ISL_2318275, EPI_ISL_2318276, EPI_ISL_2318277 | Lighthouse Lab in Alderley Park | Wellcome Sanger Institute for the COVID-19 Genomics UK (COG-UK) Consortium | Jacquelyn Wynn, Mairead Hyland, The Lighthouse Lab in Alderley Park and Alex Alderton, Roberto Amato, Jeffrey Barrett, Sonia Goncalves, Ewan Harrison, David K. Jackson, Ian Johnston, Dominic Kwiatkowski, Cordelia Langford, John Sillitoe on behalf of the Wellcome Sanger Institute COVID-19 Surveillance Team |
| EPI_ISL_2318278, EPI_ISL_2318280 | Lighthouse Lab in Glasgow | Wellcome Sanger Institute for the COVID-19 Genomics UK (COG-UK) Consortium | Harper VanSteenhouse, Yumi Kasai, David Gray, Carol Clugston, Anna Dominiczak and Alex Alderton, Roberto Amato, Jeffrey Barrett, Sonia Goncalves, Ewan Harrison, David K. Jackson, Ian Johnston, Dominic Kwiatkowski, Cordelia Langford, John Sillitoe on behalf of the Wellcome Sanger Institute COVID-19 Surveillance Team |
| EPI_ISL_2318281, EPI_ISL_2318282, EPI_ISL_2318284, EPI_ISL_2318285, EPI_ISL_2318286, EPI_ISL_2318288, EPI_ISL_2318289 | Lighthouse Lab in Alderley Park | Wellcome Sanger Institute for the COVID-19 Genomics UK (COG-UK) Consortium | Jacquelyn Wynn, Mairead Hyland, The Lighthouse Lab in Alderley Park and Alex Alderton, Roberto Amato, Jeffrey Barrett, Sonia Goncalves, Ewan Harrison, David K. Jackson, Ian Johnston, Dominic Kwiatkowski, Cordelia Langford, John Sillitoe on behalf of the Wellcome Sanger Institute COVID-19 Surveillance Team |
| EPI_ISL_2318290 | Lighthouse Lab in Glasgow | Wellcome Sanger Institute for the COVID-19 Genomics UK (COG-UK) Consortium | Harper VanSteenhouse, Yumi Kasai, David Gray, Carol Clugston, Anna Dominiczak and Alex Alderton, Roberto Amato, Jeffrey Barrett, Sonia Goncalves, Ewan Harrison, David K. Jackson, Ian Johnston, Dominic Kwiatkowski, Cordelia Langford, John Sillitoe on behalf of the Wellcome Sanger Institute COVID-19 Surveillance Team |
| EPI_ISL_2318291, EPI_ISL_2318293, EPI_ISL_2318294, EPI_ISL_2318295, EPI_ISL_2318296, EPI_ISL_2318297, EPI_ISL_2318298, EPI_ISL_2318300 | Lighthouse Lab in Alderley Park | Wellcome Sanger Institute for the COVID-19 Genomics UK (COG-UK) Consortium | Jacquelyn Wynn, Mairead Hyland, The Lighthouse Lab in Alderley Park and Alex Alderton, Roberto Amato, Jeffrey Barrett, Sonia Goncalves, Ewan Harrison, David K. Jackson, Ian Johnston, Dominic Kwiatkowski, Cordelia Langford, John Sillitoe on behalf of the Wellcome Sanger Institute COVID-19 Surveillance Team |
| EPI_ISL_2318301 | Lighthouse Lab in Glasgow | Wellcome Sanger Institute for the COVID-19 Genomics UK (COG-UK) Consortium | Harper VanSteenhouse, Yumi Kasai, David Gray, Carol Clugston, Anna Dominiczak and Alex Alderton, Roberto Amato, Jeffrey Barrett, Sonia Goncalves, Ewan Harrison, David K. Jackson, Ian Johnston, Dominic Kwiatkowski, Cordelia Langford, John Sillitoe on behalf of the Wellcome Sanger Institute COVID-19 Surveillance Team |
| EPI_ISL_2318302, EPI_ISL_2318303, EPI_ISL_2318304, EPI_ISL_2318305, EPI_ISL_2318306, EPI_ISL_2318307, EPI_ISL_2318308, EPI_ISL_2318311 | Lighthouse Lab in Alderley Park | Wellcome Sanger Institute for the COVID-19 Genomics UK (COG-UK) Consortium | Jacquelyn Wynn, Mairead Hyland, The Lighthouse Lab in Alderley Park and Alex Alderton, Roberto Amato, Jeffrey Barrett, Sonia Goncalves, Ewan Harrison, David K. Jackson, Ian Johnston, Dominic Kwiatkowski, Cordelia Langford, John Sillitoe on behalf of the Wellcome Sanger Institute COVID-19 Surveillance Team |
| EPI_ISL_2318312 | Lighthouse Lab in Glasgow | Wellcome Sanger Institute for the COVID-19 Genomics UK (COG-UK) Consortium | Harper VanSteenhouse, Yumi Kasai, David Gray, Carol Clugston, Anna Dominiczak and Alex Alderton, Roberto Amato, Jeffrey Barrett, Sonia Goncalves, Ewan Harrison, David K. Jackson, Ian Johnston, Dominic Kwiatkowski, Cordelia Langford, John Sillitoe on behalf of the Wellcome Sanger Institute COVID-19 Surveillance Team |
| EPI_ISL_2318313, EPI_ISL_2318315 | Lighthouse Lab in Alderley Park | Wellcome Sanger Institute for the COVID-19 Genomics UK (COG-UK) Consortium | Jacquelyn Wynn, Mairead Hyland, The Lighthouse Lab in Alderley Park and Alex Alderton, Roberto Amato, Jeffrey Barrett, Sonia Goncalves, Ewan Harrison, David K. Jackson, Ian Johnston, Dominic Kwiatkowski, Cordelia Langford, John Sillitoe on behalf of the Wellcome Sanger Institute COVID-19 Surveillance Team |

| | | | |
|---|---|---|---|
| EPI_ISL_2318316, EPI_ISL_2318320 | Lighthouse Lab in Glasgow | Wellcome Sanger Institute for the COVID-19 Genomics UK (COG-UK) Consortium | Harper VanSteenhouse, Yumi Kasai, David Gray, Carol Clugston, Anna Dominiczak and Alex Alderton, Roberto Amato, Jeffrey Barrett, Sonia Goncalves, Ewan Harrison, David K. Jackson, Ian Johnston, Dominic Kwiatkowski, Cordelia Langford, John Sillitoe on behalf of the Wellcome Sanger Institute COVID-19 Surveillance Team |
| EPI_ISL_2318321, EPI_ISL_2318322, EPI_ISL_2318323, EPI_ISL_2318324, EPI_ISL_2318325, EPI_ISL_2318326, EPI_ISL_2318327, EPI_ISL_2318328, EPI_ISL_2318329, EPI_ISL_2318330 | Lighthouse Lab in Alderley Park | Wellcome Sanger Institute for the COVID-19 Genomics UK (COG-UK) Consortium | Jacquelyn Wynn, Mairead Hyland, The Lighthouse Lab in Alderley Park and Alex Alderton, Roberto Amato, Jeffrey Barrett, Sonia Goncalves, Ewan Harrison, David K. Jackson, Ian Johnston, Dominic Kwiatkowski, Cordelia Langford, John Sillitoe on behalf of the Wellcome Sanger Institute COVID-19 Surveillance Team |
| EPI_ISL_2318331 | Lighthouse Lab in Glasgow | Wellcome Sanger Institute for the COVID-19 Genomics UK (COG-UK) Consortium | Harper VanSteenhouse, Yumi Kasai, David Gray, Carol Clugston, Anna Dominiczak and Alex Alderton, Roberto Amato, Jeffrey Barrett, Sonia Goncalves, Ewan Harrison, David K. Jackson, Ian Johnston, Dominic Kwiatkowski, Cordelia Langford, John Sillitoe on behalf of the Wellcome Sanger Institute COVID-19 Surveillance Team |
| EPI_ISL_2318332, EPI_ISL_2318333, EPI_ISL_2318335, EPI_ISL_2318338, EPI_ISL_2318339 | Lighthouse Lab in Alderley Park | Wellcome Sanger Institute for the COVID-19 Genomics UK (COG-UK) Consortium | Jacquelyn Wynn, Mairead Hyland, The Lighthouse Lab in Alderley Park and Alex Alderton, Roberto Amato, Jeffrey Barrett, Sonia Goncalves, Ewan Harrison, David K. Jackson, Ian Johnston, Dominic Kwiatkowski, Cordelia Langford, John Sillitoe on behalf of the Wellcome Sanger Institute COVID-19 Surveillance Team |
| EPI_ISL_2318340 | Lighthouse Lab in Glasgow | Wellcome Sanger Institute for the COVID-19 Genomics UK (COG-UK) Consortium | Harper VanSteenhouse, Yumi Kasai, David Gray, Carol Clugston, Anna Dominiczak and Alex Alderton, Roberto Amato, Jeffrey Barrett, Sonia Goncalves, Ewan Harrison, David K. Jackson, Ian Johnston, Dominic Kwiatkowski, Cordelia Langford, John Sillitoe on behalf of the Wellcome Sanger Institute COVID-19 Surveillance Team |
| EPI_ISL_2318341 | Health Services Laboratories | Wellcome Sanger Institute for the COVID-19 Genomics UK (COG-UK) Consortium | Health Services Laboratories and Alex Alderton, Roberto Amato, Jeffrey Barrett, Sonia Goncalves, Ewan Harrison, David K. Jackson, Ian Johnston, Dominic Kwiatkowski, Cordelia Langford, John Sillitoe on behalf of the Wellcome Sanger Institute COVID-19 Surveillance Team |
| EPI_ISL_2318342 | Lighthouse Lab in Alderley Park | Wellcome Sanger Institute for the COVID-19 Genomics UK (COG-UK) Consortium | Jacquelyn Wynn, Mairead Hyland, The Lighthouse Lab in Alderley Park and Alex Alderton, Roberto Amato, Jeffrey Barrett, Sonia Goncalves, Ewan Harrison, David K. Jackson, Ian Johnston, Dominic Kwiatkowski, Cordelia Langford, John Sillitoe on behalf of the Wellcome Sanger Institute COVID-19 Surveillance Team |
| EPI_ISL_2318343 | Lighthouse Lab in Glasgow | Wellcome Sanger Institute for the COVID-19 Genomics UK (COG-UK) Consortium | Harper VanSteenhouse, Yumi Kasai, David Gray, Carol Clugston, Anna Dominiczak and Alex Alderton, Roberto Amato, Jeffrey Barrett, Sonia Goncalves, Ewan Harrison, David K. Jackson, Ian Johnston, Dominic Kwiatkowski, Cordelia Langford, John Sillitoe on behalf of the Wellcome Sanger Institute COVID-19 Surveillance Team |
| EPI_ISL_2318347, EPI_ISL_2318349, EPI_ISL_2318351, EPI_ISL_2318352, EPI_ISL_2318353, EPI_ISL_2318354 | Lighthouse Lab in Alderley Park | Wellcome Sanger Institute for the COVID-19 Genomics UK (COG-UK) Consortium | Jacquelyn Wynn, Mairead Hyland, The Lighthouse Lab in Alderley Park and Alex Alderton, Roberto Amato, Jeffrey Barrett, Sonia Goncalves, Ewan Harrison, David K. Jackson, Ian Johnston, Dominic Kwiatkowski, Cordelia Langford, John Sillitoe on behalf of the Wellcome Sanger Institute COVID-19 Surveillance Team |
| EPI_ISL_2318358 | Lighthouse Lab in Glasgow | Wellcome Sanger Institute for the COVID-19 Genomics UK (COG-UK) Consortium | Harper VanSteenhouse, Yumi Kasai, David Gray, Carol Clugston, Anna Dominiczak and Alex Alderton, Roberto Amato, Jeffrey Barrett, Sonia Goncalves, Ewan Harrison, David K. Jackson, Ian Johnston, Dominic Kwiatkowski, Cordelia Langford, John Sillitoe on behalf of the Wellcome Sanger Institute COVID-19 Surveillance Team |
| EPI_ISL_2318359, EPI_ISL_2318360, EPI_ISL_2318362, EPI_ISL_2318363, EPI_ISL_2318364, EPI_ISL_2318365 | Lighthouse Lab in Alderley Park | Wellcome Sanger Institute for the COVID-19 Genomics UK (COG-UK) Consortium | Jacquelyn Wynn, Mairead Hyland, The Lighthouse Lab in Alderley Park and Alex Alderton, Roberto Amato, Jeffrey Barrett, Sonia Goncalves, Ewan Harrison, David K. Jackson, Ian Johnston, Dominic Kwiatkowski, Cordelia Langford, John Sillitoe on behalf of the Wellcome Sanger Institute COVID-19 Surveillance Team |
| EPI_ISL_2318366, EPI_ISL_2318367 | Lighthouse Lab in Glasgow | Wellcome Sanger Institute for the COVID-19 Genomics UK (COG-UK) Consortium | Harper VanSteenhouse, Yumi Kasai, David Gray, Carol Clugston, Anna Dominiczak and Alex Alderton, Roberto Amato, Jeffrey Barrett, Sonia Goncalves, Ewan Harrison, David K. Jackson, Ian Johnston, Dominic Kwiatkowski, Cordelia Langford, John Sillitoe on behalf of the Wellcome Sanger Institute COVID-19 Surveillance Team |
| EPI_ISL_2318368, EPI_ISL_2318370 | Lighthouse Lab in Alderley Park | Wellcome Sanger Institute for the COVID-19 Genomics UK (COG-UK) Consortium | Jacquelyn Wynn, Mairead Hyland, The Lighthouse Lab in Alderley Park and Alex Alderton, Roberto Amato, Jeffrey Barrett, Sonia Goncalves, Ewan Harrison, David K. Jackson, Ian Johnston, Dominic Kwiatkowski, Cordelia Langford, John Sillitoe on behalf of the Wellcome Sanger Institute COVID-19 Surveillance Team |
| EPI_ISL_2318373 | Lighthouse Lab in Glasgow | Wellcome Sanger Institute for the COVID-19 Genomics UK (COG-UK) Consortium | Harper VanSteenhouse, Yumi Kasai, David Gray, Carol Clugston, Anna Dominiczak and Alex Alderton, Roberto Amato, Jeffrey Barrett, Sonia Goncalves, Ewan Harrison, David K. Jackson, Ian Johnston, Dominic Kwiatkowski, Cordelia Langford, John Sillitoe on behalf of the Wellcome Sanger Institute COVID-19 Surveillance Team |
| EPI_ISL_2318374, EPI_ISL_2318375, EPI_ISL_2318378, EPI_ISL_2318379 | Lighthouse Lab in Alderley Park | Wellcome Sanger Institute for the COVID-19 Genomics UK (COG-UK) Consortium | Jacquelyn Wynn, Mairead Hyland, The Lighthouse Lab in Alderley Park and Alex Alderton, Roberto Amato, Jeffrey Barrett, Sonia Goncalves, Ewan Harrison, David K. Jackson, Ian Johnston, Dominic Kwiatkowski, Cordelia Langford, John Sillitoe on behalf of the Wellcome Sanger Institute COVID-19 Surveillance Team |
| EPI_ISL_2318380 | Lighthouse Lab in Glasgow | Wellcome Sanger Institute for the COVID-19 Genomics UK (COG-UK) Consortium | Harper VanSteenhouse, Yumi Kasai, David Gray, Carol Clugston, Anna Dominiczak and Alex Alderton, Roberto Amato, Jeffrey Barrett, Sonia Goncalves, Ewan Harrison, David K. Jackson, Ian Johnston, Dominic Kwiatkowski, Cordelia Langford, John Sillitoe on behalf of the Wellcome Sanger Institute COVID-19 Surveillance Team |
| EPI_ISL_2318382, EPI_ISL_2318384, EPI_ISL_2318385 | Lighthouse Lab in Alderley Park | Wellcome Sanger Institute for the COVID-19 Genomics UK (COG-UK) Consortium | Jacquelyn Wynn, Mairead Hyland, The Lighthouse Lab in Alderley Park and Alex Alderton, Roberto Amato, Jeffrey Barrett, Sonia Goncalves, Ewan Harrison, David K. Jackson, Ian Johnston, Dominic Kwiatkowski, Cordelia Langford, John Sillitoe on behalf of the Wellcome Sanger Institute COVID-19 Surveillance Team |
| EPI_ISL_2318386 | Lighthouse Lab in Glasgow | Wellcome Sanger Institute for the COVID-19 Genomics UK (COG-UK) Consortium | Harper VanSteenhouse, Yumi Kasai, David Gray, Carol Clugston, Anna Dominiczak and Alex Alderton, Roberto Amato, Jeffrey Barrett, Sonia Goncalves, Ewan Harrison, David K. Jackson, Ian Johnston, Dominic Kwiatkowski, Cordelia Langford, John Sillitoe on behalf of the Wellcome Sanger Institute COVID-19 Surveillance Team |
| EPI_ISL_2318387 | Lighthouse Lab in Alderley Park | Wellcome Sanger Institute for the COVID-19 Genomics UK (COG-UK) Consortium | Jacquelyn Wynn, Mairead Hyland, The Lighthouse Lab in Alderley Park and Alex Alderton, Roberto Amato, Jeffrey Barrett, Sonia Goncalves, Ewan Harrison, David K. Jackson, Ian Johnston, Dominic Kwiatkowski, Cordelia Langford, John Sillitoe on behalf of the Wellcome Sanger Institute COVID-19 Surveillance Team |
| EPI_ISL_2318388, EPI_ISL_2318390 | Lighthouse Lab in Glasgow | Wellcome Sanger Institute for the COVID-19 Genomics UK (COG-UK) Consortium | Harper VanSteenhouse, Yumi Kasai, David Gray, Carol Clugston, Anna Dominiczak and Alex Alderton, Roberto Amato, Jeffrey Barrett, Sonia Goncalves, Ewan Harrison, David K. Jackson, Ian Johnston, Dominic Kwiatkowski, Cordelia Langford, John Sillitoe on behalf of the Wellcome Sanger Institute COVID-19 Surveillance Team |
| EPI_ISL_2318391 | Lighthouse Lab in Alderley Park | Wellcome Sanger Institute for the COVID-19 Genomics UK (COG-UK) Consortium | Jacquelyn Wynn, Mairead Hyland, The Lighthouse Lab in Alderley Park and Alex Alderton, Roberto Amato, Jeffrey Barrett, Sonia Goncalves, Ewan Harrison, David K. Jackson, Ian Johnston, Dominic Kwiatkowski, Cordelia Langford, John Sillitoe on behalf of the Wellcome Sanger Institute COVID-19 Surveillance Team |
| EPI_ISL_2318392 | Lighthouse Lab in Glasgow | Wellcome Sanger Institute for the COVID-19 Genomics UK (COG-UK) Consortium | Harper VanSteenhouse, Yumi Kasai, David Gray, Carol Clugston, Anna Dominiczak and Alex Alderton, Roberto Amato, Jeffrey Barrett, Sonia Goncalves, Ewan Harrison, David K. Jackson, Ian Johnston, Dominic Kwiatkowski, Cordelia Langford, John Sillitoe on behalf of the Wellcome Sanger Institute COVID-19 Surveillance Team |
| EPI_ISL_2318393, EPI_ISL_2318395, EPI_ISL_2318396, EPI_ISL_2318398 | Lighthouse Lab in Alderley Park | Wellcome Sanger Institute for the COVID-19 Genomics UK (COG-UK) Consortium | Jacquelyn Wynn, Mairead Hyland, The Lighthouse Lab in Alderley Park and Alex Alderton, Roberto Amato, Jeffrey Barrett, Sonia Goncalves, Ewan Harrison, David K. Jackson, Ian Johnston, Dominic Kwiatkowski, Cordelia Langford, John Sillitoe on behalf of the Wellcome Sanger Institute COVID-19 Surveillance Team |
| EPI_ISL_2318399, EPI_ISL_2318400 | Lighthouse Lab in Glasgow | Wellcome Sanger Institute for the COVID-19 Genomics UK (COG-UK) Consortium | Harper VanSteenhouse, Yumi Kasai, David Gray, Carol Clugston, Anna Dominiczak and Alex Alderton, Roberto Amato, Jeffrey Barrett, Sonia Goncalves, Ewan Harrison, David K. Jackson, Ian Johnston, Dominic Kwiatkowski, Cordelia Langford, John Sillitoe on behalf of the Wellcome Sanger Institute COVID-19 Surveillance Team |
| EPI_ISL_2318401, EPI_ISL_2318402, EPI_ISL_2318404, EPI_ISL_2318405, EPI_ISL_2318406, EPI_ISL_2318407, EPI_ISL_2318408, EPI_ISL_2318409, EPI_ISL_2318410, EPI_ISL_2318411 | Lighthouse Lab in Alderley Park | Wellcome Sanger Institute for the COVID-19 Genomics UK (COG-UK) Consortium | Jacquelyn Wynn, Mairead Hyland, The Lighthouse Lab in Alderley Park and Alex Alderton, Roberto Amato, Jeffrey Barrett, Sonia Goncalves, Ewan Harrison, David K. Jackson, Ian Johnston, Dominic Kwiatkowski, Cordelia Langford, John Sillitoe on behalf of the Wellcome Sanger Institute COVID-19 Surveillance Team |
| EPI_ISL_2318412 | Lighthouse Lab in Glasgow | Wellcome Sanger Institute for the COVID-19 Genomics UK (COG-UK) Consortium | Harper VanSteenhouse, Yumi Kasai, David Gray, Carol Clugston, Anna Dominiczak and Alex Alderton, Roberto Amato, Jeffrey Barrett, Sonia Goncalves, Ewan Harrison, David K. Jackson, Ian Johnston, Dominic Kwiatkowski, Cordelia Langford, John Sillitoe on behalf of the Wellcome Sanger Institute COVID-19 Surveillance Team |
| EPI_ISL_2318414 | Lighthouse Lab in Alderley Park | Wellcome Sanger Institute for the COVID-19 Genomics UK (COG-UK) Consortium | Jacquelyn Wynn, Mairead Hyland, The Lighthouse Lab in Alderley Park and Alex Alderton, Roberto Amato, Jeffrey Barrett, Sonia Goncalves, Ewan Harrison, David K. Jackson, Ian Johnston, Dominic Kwiatkowski, Cordelia Langford, John Sillitoe on behalf of the Wellcome Sanger Institute COVID-19 Surveillance Team |
| EPI_ISL_2318415 | Lighthouse Lab in Glasgow | Wellcome Sanger Institute for the COVID-19 Genomics UK (COG-UK) Consortium | Harper VanSteenhouse, Yumi Kasai, David Gray, Carol Clugston, Anna Dominiczak and Alex Alderton, Roberto Amato, Jeffrey Barrett, Sonia Goncalves, Ewan Harrison, David K. Jackson, Ian Johnston, Dominic Kwiatkowski, Cordelia Langford, John Sillitoe on behalf of the Wellcome Sanger Institute COVID-19 Surveillance Team |
| EPI_ISL_2318416, EPI_ISL_2318417, EPI_ISL_2318418 | Lighthouse Lab in Alderley Park | Wellcome Sanger Institute for the COVID-19 Genomics UK (COG-UK) Consortium | Jacquelyn Wynn, Mairead Hyland, The Lighthouse Lab in Alderley Park and Alex Alderton, Roberto Amato, Jeffrey Barrett, Sonia Goncalves, Ewan Harrison, David K. Jackson, Ian Johnston, Dominic Kwiatkowski, Cordelia Langford, John Sillitoe on behalf of the Wellcome Sanger Institute COVID-19 Surveillance Team |
| EPI_ISL_2318419 | Lighthouse Lab in Glasgow | Wellcome Sanger Institute for the COVID-19 Genomics UK (COG-UK) Consortium | Harper VanSteenhouse, Yumi Kasai, David Gray, Carol Clugston, Anna Dominiczak and Alex Alderton, Roberto Amato, Jeffrey Barrett, Sonia Goncalves, Ewan Harrison, David K. Jackson, Ian Johnston, Dominic Kwiatkowski, Cordelia Langford, John Sillitoe on behalf of the Wellcome Sanger Institute COVID-19 Surveillance Team |
| EPI_ISL_2318420, EPI_ISL_2318421, EPI_ISL_2318422, | Lighthouse Lab in Alderley Park | Wellcome Sanger Institute for the COVID-19 Genomics UK (COG-UK) Consortium | Jacquelyn Wynn, Mairead Hyland, The Lighthouse Lab in Alderley Park and Alex Alderton, Roberto Amato, Jeffrey Barrett, Sonia Goncalves, Ewan Harrison, David K. Jackson, Ian Johnston, Dominic Kwiatkowski, Cordelia Langford, John Sillitoe on behalf of the Wellcome Sanger Institute COVID-19 Surveillance Team |

| | | | |
|---|---|---|---|
| EPI_ISL_2318424, EPI_ISL_2318425, EPI_ISL_2318426, EPI_ISL_2318427, EPI_ISL_2318429, EPI_ISL_2318431 | | | |
| EPI_ISL_2318434 | Lighthouse Lab in Glasgow | Wellcome Sanger Institute for the COVID-19 Genomics UK (COG-UK) Consortium | Harper VanSteenhouse, Yumi Kasai, David Gray, Carol Clugston, Anna Dominiczak and Alex Alderton, Roberto Amato, Jeffrey Barrett, Sonia Goncalves, Ewan Harrison, David K. Jackson, Ian Johnston, Dominic Kwiatkowski, Cordelia Langford, John Sillitoe on behalf of the Wellcome Sanger Institute COVID-19 Surveillance Team |
| EPI_ISL_2318435, EPI_ISL_2318436, EPI_ISL_2318438, EPI_ISL_2318439, EPI_ISL_2318440, EPI_ISL_2318441, EPI_ISL_2318442, EPI_ISL_2318444, EPI_ISL_2318446, EPI_ISL_2318447, EPI_ISL_2318448 | | | |
| see above | Lighthouse Lab in Alderley Park | Wellcome Sanger Institute for the COVID-19 Genomics UK (COG-UK) Consortium | Jacquelyn Wynn, Mairead Hyland, The Lighthouse Lab in Alderley Park and Alex Alderton, Roberto Amato, Jeffrey Barrett, Sonia Goncalves, Ewan Harrison, David K. Jackson, Ian Johnston, Dominic Kwiatkowski, Cordelia Langford, John Sillitoe on behalf of the Wellcome Sanger Institute COVID-19 Surveillance Team |
| EPI_ISL_2318450 | Lighthouse Lab in Glasgow | Wellcome Sanger Institute for the COVID-19 Genomics UK (COG-UK) Consortium | Harper VanSteenhouse, Yumi Kasai, David Gray, Carol Clugston, Anna Dominiczak and Alex Alderton, Roberto Amato, Jeffrey Barrett, Sonia Goncalves, Ewan Harrison, David K. Jackson, Ian Johnston, Dominic Kwiatkowski, Cordelia Langford, John Sillitoe on behalf of the Wellcome Sanger Institute COVID-19 Surveillance Team |
| EPI_ISL_2318451, EPI_ISL_2318452, EPI_ISL_2318454, EPI_ISL_2318455, EPI_ISL_2318456, EPI_ISL_2318457, EPI_ISL_2318458, EPI_ISL_2318459, EPI_ISL_2318460, EPI_ISL_2318461, EPI_ISL_2318462, EPI_ISL_2318465, EPI_ISL_2318466, EPI_ISL_2318467 | | | |
| see above | Lighthouse Lab in Alderley Park | Wellcome Sanger Institute for the COVID-19 Genomics UK (COG-UK) Consortium | Jacquelyn Wynn, Mairead Hyland, The Lighthouse Lab in Alderley Park and Alex Alderton, Roberto Amato, Jeffrey Barrett, Sonia Goncalves, Ewan Harrison, David K. Jackson, Ian Johnston, Dominic Kwiatkowski, Cordelia Langford, John Sillitoe on behalf of the Wellcome Sanger Institute COVID-19 Surveillance Team |
| EPI_ISL_2318468, EPI_ISL_2318471, EPI_ISL_2318472, EPI_ISL_2318473, EPI_ISL_2318474 | Lighthouse Lab in Glasgow | Wellcome Sanger Institute for the COVID-19 Genomics UK (COG-UK) Consortium | Harper VanSteenhouse, Yumi Kasai, David Gray, Carol Clugston, Anna Dominiczak and Alex Alderton, Roberto Amato, Jeffrey Barrett, Sonia Goncalves, Ewan Harrison, David K. Jackson, Ian Johnston, Dominic Kwiatkowski, Cordelia Langford, John Sillitoe on behalf of the Wellcome Sanger Institute COVID-19 Surveillance Team |
| EPI_ISL_2318477, EPI_ISL_2318480, EPI_ISL_2318484, EPI_ISL_2318485 | Lighthouse Lab in Alderley Park | Wellcome Sanger Institute for the COVID-19 Genomics UK (COG-UK) Consortium | Jacquelyn Wynn, Mairead Hyland, The Lighthouse Lab in Alderley Park and Alex Alderton, Roberto Amato, Jeffrey Barrett, Sonia Goncalves, Ewan Harrison, David K. Jackson, Ian Johnston, Dominic Kwiatkowski, Cordelia Langford, John Sillitoe on behalf of the Wellcome Sanger Institute COVID-19 Surveillance Team |
| EPI_ISL_2318486, EPI_ISL_2318487, EPI_ISL_2318488, EPI_ISL_2318489 | Lighthouse Lab in Glasgow | Wellcome Sanger Institute for the COVID-19 Genomics UK (COG-UK) Consortium | Harper VanSteenhouse, Yumi Kasai, David Gray, Carol Clugston, Anna Dominiczak and Alex Alderton, Roberto Amato, Jeffrey Barrett, Sonia Goncalves, Ewan Harrison, David K. Jackson, Ian Johnston, Dominic Kwiatkowski, Cordelia Langford, John Sillitoe on behalf of the Wellcome Sanger Institute COVID-19 Surveillance Team |
| EPI_ISL_2318490, EPI_ISL_2318491, EPI_ISL_2318492, EPI_ISL_2318493, EPI_ISL_2318495, EPI_ISL_2318498, EPI_ISL_2318499 | Lighthouse Lab in Alderley Park | Wellcome Sanger Institute for the COVID-19 Genomics UK (COG-UK) Consortium | Jacquelyn Wynn, Mairead Hyland, The Lighthouse Lab in Alderley Park and Alex Alderton, Roberto Amato, Jeffrey Barrett, Sonia Goncalves, Ewan Harrison, David K. Jackson, Ian Johnston, Dominic Kwiatkowski, Cordelia Langford, John Sillitoe on behalf of the Wellcome Sanger Institute COVID-19 Surveillance Team |
| EPI_ISL_2318500, EPI_ISL_2318501 | Lighthouse Lab in Glasgow | Wellcome Sanger Institute for the COVID-19 Genomics UK (COG-UK) Consortium | Harper VanSteenhouse, Yumi Kasai, David Gray, Carol Clugston, Anna Dominiczak and Alex Alderton, Roberto Amato, Jeffrey Barrett, Sonia Goncalves, Ewan Harrison, David K. Jackson, Ian Johnston, Dominic Kwiatkowski, Cordelia Langford, John Sillitoe on behalf of the Wellcome Sanger Institute COVID-19 Surveillance Team |
| EPI_ISL_2318502 | Lighthouse Lab in Alderley Park | Wellcome Sanger Institute for the COVID-19 Genomics UK (COG-UK) Consortium | Jacquelyn Wynn, Mairead Hyland, The Lighthouse Lab in Alderley Park and Alex Alderton, Roberto Amato, Jeffrey Barrett, Sonia Goncalves, Ewan Harrison, David K. Jackson, Ian Johnston, Dominic Kwiatkowski, Cordelia Langford, John Sillitoe on behalf of the Wellcome Sanger Institute COVID-19 Surveillance Team |
| EPI_ISL_2318504 | Lighthouse Lab in Glasgow | Wellcome Sanger Institute for the COVID-19 Genomics UK (COG-UK) Consortium | Harper VanSteenhouse, Yumi Kasai, David Gray, Carol Clugston, Anna Dominiczak and Alex Alderton, Roberto Amato, Jeffrey Barrett, Sonia Goncalves, Ewan Harrison, David K. Jackson, Ian Johnston, Dominic Kwiatkowski, Cordelia Langford, John Sillitoe on behalf of the Wellcome Sanger Institute COVID-19 Surveillance Team |
| EPI_ISL_2318505, EPI_ISL_2318508 | Lighthouse Lab in Alderley Park | Wellcome Sanger Institute for the COVID-19 Genomics UK (COG-UK) Consortium | Jacquelyn Wynn, Mairead Hyland, The Lighthouse Lab in Alderley Park and Alex Alderton, Roberto Amato, Jeffrey Barrett, Sonia Goncalves, Ewan Harrison, David K. Jackson, Ian Johnston, Dominic Kwiatkowski, Cordelia Langford, John Sillitoe on behalf of the Wellcome Sanger Institute COVID-19 Surveillance Team |
| EPI_ISL_2318510 | Lighthouse Lab in Glasgow | Wellcome Sanger Institute for the COVID-19 Genomics UK (COG-UK) Consortium | Harper VanSteenhouse, Yumi Kasai, David Gray, Carol Clugston, Anna Dominiczak and Alex Alderton, Roberto Amato, Jeffrey Barrett, Sonia Goncalves, Ewan Harrison, David K. Jackson, Ian Johnston, Dominic Kwiatkowski, Cordelia Langford, John Sillitoe on behalf of the Wellcome Sanger Institute COVID-19 Surveillance Team |
| EPI_ISL_2318511, EPI_ISL_2318516, EPI_ISL_2318517, EPI_ISL_2318518 | Lighthouse Lab in Alderley Park | Wellcome Sanger Institute for the COVID-19 Genomics UK (COG-UK) Consortium | Jacquelyn Wynn, Mairead Hyland, The Lighthouse Lab in Alderley Park and Alex Alderton, Roberto Amato, Jeffrey Barrett, Sonia Goncalves, Ewan Harrison, David K. Jackson, Ian Johnston, Dominic Kwiatkowski, Cordelia Langford, John Sillitoe on behalf of the Wellcome Sanger Institute COVID-19 Surveillance Team |
| EPI_ISL_2318519 | Lighthouse Lab in Glasgow | Wellcome Sanger Institute for the COVID-19 Genomics UK (COG-UK) Consortium | Harper VanSteenhouse, Yumi Kasai, David Gray, Carol Clugston, Anna Dominiczak and Alex Alderton, Roberto Amato, Jeffrey Barrett, Sonia Goncalves, Ewan Harrison, David K. Jackson, Ian Johnston, Dominic Kwiatkowski, Cordelia Langford, John Sillitoe on behalf of the Wellcome Sanger Institute COVID-19 Surveillance Team |
| EPI_ISL_2318522, EPI_ISL_2318523, EPI_ISL_2318524, EPI_ISL_2318525, EPI_ISL_2318528, EPI_ISL_2318530, EPI_ISL_2318531, EPI_ISL_2318534, EPI_ISL_2318535 | Lighthouse Lab in Alderley Park | Wellcome Sanger Institute for the COVID-19 Genomics UK (COG-UK) Consortium | Jacquelyn Wynn, Mairead Hyland, The Lighthouse Lab in Alderley Park and Alex Alderton, Roberto Amato, Jeffrey Barrett, Sonia Goncalves, Ewan Harrison, David K. Jackson, Ian Johnston, Dominic Kwiatkowski, Cordelia Langford, John Sillitoe on behalf of the Wellcome Sanger Institute COVID-19 Surveillance Team |
| EPI_ISL_2318537 | Lighthouse Lab in Glasgow | Wellcome Sanger Institute for the COVID-19 Genomics UK (COG-UK) Consortium | Harper VanSteenhouse, Yumi Kasai, David Gray, Carol Clugston, Anna Dominiczak and Alex Alderton, Roberto Amato, Jeffrey Barrett, Sonia Goncalves, Ewan Harrison, David K. Jackson, Ian Johnston, Dominic Kwiatkowski, Cordelia Langford, John Sillitoe on behalf of the Wellcome Sanger Institute COVID-19 Surveillance Team |
| EPI_ISL_2318539, EPI_ISL_2318540, EPI_ISL_2318541, EPI_ISL_2318542, EPI_ISL_2318543 | Lighthouse Lab in Alderley Park | Wellcome Sanger Institute for the COVID-19 Genomics UK (COG-UK) Consortium | Jacquelyn Wynn, Mairead Hyland, The Lighthouse Lab in Alderley Park and Alex Alderton, Roberto Amato, Jeffrey Barrett, Sonia Goncalves, Ewan Harrison, David K. Jackson, Ian Johnston, Dominic Kwiatkowski, Cordelia Langford, John Sillitoe on behalf of the Wellcome Sanger Institute COVID-19 Surveillance Team |
| EPI_ISL_2318544 | Lighthouse Lab in Glasgow | Wellcome Sanger Institute for the COVID-19 Genomics UK (COG-UK) Consortium | Harper VanSteenhouse, Yumi Kasai, David Gray, Carol Clugston, Anna Dominiczak and Alex Alderton, Roberto Amato, Jeffrey Barrett, Sonia Goncalves, Ewan Harrison, David K. Jackson, Ian Johnston, Dominic Kwiatkowski, Cordelia Langford, John Sillitoe on behalf of the Wellcome Sanger Institute COVID-19 Surveillance Team |
| EPI_ISL_2318546, EPI_ISL_2318547, EPI_ISL_2318549, EPI_ISL_2318550, EPI_ISL_2318551, EPI_ISL_2318552, EPI_ISL_2318554, EPI_ISL_2318556, EPI_ISL_2318557, EPI_ISL_2318559, EPI_ISL_2318560, EPI_ISL_2318561, EPI_ISL_2318562, EPI_ISL_2318563, EPI_ISL_2318564, EPI_ISL_2318566, EPI_ISL_2318568, EPI_ISL_2318569, EPI_ISL_2318570, EPI_ISL_2318572, EPI_ISL_2318574 | | | |
| see above | Lighthouse Lab in Alderley Park | Wellcome Sanger Institute for the COVID-19 Genomics UK (COG-UK) Consortium | Jacquelyn Wynn, Mairead Hyland, The Lighthouse Lab in Alderley Park and Alex Alderton, Roberto Amato, Jeffrey Barrett, Sonia Goncalves, Ewan Harrison, David K. Jackson, Ian Johnston, Dominic Kwiatkowski, Cordelia Langford, John Sillitoe on behalf of the Wellcome Sanger Institute COVID-19 Surveillance Team |
| EPI_ISL_2318576 | Lighthouse Lab in Glasgow | Wellcome Sanger Institute for the COVID-19 Genomics UK (COG-UK) Consortium | Harper VanSteenhouse, Yumi Kasai, David Gray, Carol Clugston, Anna Dominiczak and Alex Alderton, Roberto Amato, Jeffrey Barrett, Sonia Goncalves, Ewan Harrison, David K. Jackson, Ian Johnston, Dominic Kwiatkowski, Cordelia Langford, John Sillitoe on behalf of the Wellcome Sanger Institute COVID-19 Surveillance Team |
| EPI_ISL_2318578 | Lighthouse Lab in Alderley Park | Wellcome Sanger Institute for the COVID-19 Genomics UK (COG-UK) Consortium | Jacquelyn Wynn, Mairead Hyland, The Lighthouse Lab in Alderley Park and Alex Alderton, Roberto Amato, Jeffrey Barrett, Sonia Goncalves, Ewan Harrison, David K. Jackson, Ian Johnston, Dominic Kwiatkowski, Cordelia Langford, John Sillitoe on behalf of the Wellcome Sanger Institute COVID-19 Surveillance Team |
| EPI_ISL_2318581 | Lighthouse Lab in Glasgow | Wellcome Sanger Institute for the COVID-19 Genomics UK (COG-UK) Consortium | Harper VanSteenhouse, Yumi Kasai, David Gray, Carol Clugston, Anna Dominiczak and Alex Alderton, Roberto Amato, Jeffrey Barrett, Sonia Goncalves, Ewan Harrison, David K. Jackson, Ian Johnston, Dominic Kwiatkowski, Cordelia Langford, John Sillitoe on behalf of the Wellcome Sanger Institute COVID-19 Surveillance Team |
| EPI_ISL_2318582, EPI_ISL_2318583, EPI_ISL_2318584 | Lighthouse Lab in Alderley Park | Wellcome Sanger Institute for the COVID-19 Genomics UK (COG-UK) Consortium | Jacquelyn Wynn, Mairead Hyland, The Lighthouse Lab in Alderley Park and Alex Alderton, Roberto Amato, Jeffrey Barrett, Sonia Goncalves, Ewan Harrison, David K. Jackson, Ian Johnston, Dominic Kwiatkowski, Cordelia Langford, John Sillitoe on behalf of the Wellcome Sanger Institute COVID-19 Surveillance Team |
| EPI_ISL_2318585 | Lighthouse Lab in Glasgow | Wellcome Sanger Institute for the COVID-19 Genomics UK (COG-UK) Consortium | Harper VanSteenhouse, Yumi Kasai, David Gray, Carol Clugston, Anna Dominiczak and Alex Alderton, Roberto Amato, Jeffrey Barrett, Sonia Goncalves, Ewan Harrison, David K. Jackson, Ian Johnston, Dominic Kwiatkowski, Cordelia Langford, John Sillitoe on behalf of the Wellcome Sanger Institute COVID-19 Surveillance Team |
| EPI_ISL_2318588, EPI_ISL_2318589, EPI_ISL_2318590, EPI_ISL_2318591, EPI_ISL_2318593 | Lighthouse Lab in Alderley Park | Wellcome Sanger Institute for the COVID-19 Genomics UK (COG-UK) Consortium | Jacquelyn Wynn, Mairead Hyland, The Lighthouse Lab in Alderley Park and Alex Alderton, Roberto Amato, Jeffrey Barrett, Sonia Goncalves, Ewan Harrison, David K. Jackson, Ian Johnston, Dominic Kwiatkowski, Cordelia Langford, John Sillitoe on behalf of the Wellcome Sanger Institute COVID-19 Surveillance Team |
| EPI_ISL_2318594 | Lighthouse Lab in Glasgow | Wellcome Sanger Institute for the COVID-19 Genomics UK (COG-UK) Consortium | Harper VanSteenhouse, Yumi Kasai, David Gray, Carol Clugston, Anna Dominiczak and Alex Alderton, Roberto Amato, Jeffrey Barrett, Sonia Goncalves, Ewan Harrison, David K. Jackson, Ian Johnston, Dominic Kwiatkowski, Cordelia Langford, John Sillitoe on behalf of the Wellcome Sanger Institute COVID-19 Surveillance Team |
| EPI_ISL_2318595, EPI_ISL_2318596, EPI_ISL_2318598, EPI_ISL_2318599, EPI_ISL_2318601, EPI_ISL_2318602, EPI_ISL_2318605, EPI_ISL_2318606, EPI_ISL_2318608, EPI_ISL_2318609, EPI_ISL_2318610, EPI_ISL_2318611, EPI_ISL_2318612, EPI_ISL_2318613 | | | |
| see above | Lighthouse Lab in Alderley Park | Wellcome Sanger Institute for the COVID-19 Genomics UK (COG-UK) Consortium | Jacquelyn Wynn, Mairead Hyland, The Lighthouse Lab in Alderley Park and Alex Alderton, Roberto Amato, Jeffrey Barrett, Sonia Goncalves, Ewan Harrison, David K. Jackson, Ian Johnston, Dominic Kwiatkowski, Cordelia Langford, John Sillitoe on behalf of the Wellcome Sanger Institute COVID-19 Surveillance Team |
| EPI_ISL_2318615 | Lighthouse Lab in Glasgow | Wellcome Sanger Institute for the COVID-19 Genomics UK (COG-UK) Consortium | Harper VanSteenhouse, Yumi Kasai, David Gray, Carol Clugston, Anna Dominiczak and Alex Alderton, Roberto Amato, Jeffrey Barrett, Sonia Goncalves, Ewan Harrison, David K. Jackson, Ian Johnston, Dominic Kwiatkowski, Cordelia Langford, John Sillitoe on behalf of the Wellcome Sanger Institute COVID-19 Surveillance Team |

| | | | |
|---|---|---|---|
| EPI_ISL_2318616, EPI_ISL_2318618, EPI_ISL_2318619, EPI_ISL_2318620, EPI_ISL_2318621, EPI_ISL_2318623 | Lighthouse Lab in Alderley Park | Wellcome Sanger Institute for the COVID-19 Genomics UK (COG-UK) Consortium | Jacquelyn Wynn, Mairead Hyland, The Lighthouse Lab in Alderley Park and Alex Alderton, Roberto Amato, Jeffrey Barrett, Sonia Goncalves, Ewan Harrison, David K. Jackson, Ian Johnston, Dominic Kwiatkowski, Cordelia Langford, John Sillitoe on behalf of the Wellcome Sanger Institute COVID-19 Surveillance Team |
| EPI_ISL_2318625 | Lighthouse Lab in Glasgow | Wellcome Sanger Institute for the COVID-19 Genomics UK (COG-UK) Consortium | Harper VanSteenhouse, Yumi Kasai, David Gray, Carol Clugston, Anna Dominiczak and Alex Alderton, Roberto Amato, Jeffrey Barrett, Sonia Goncalves, Ewan Harrison, David K. Jackson, Ian Johnston, Dominic Kwiatkowski, Cordelia Langford, John Sillitoe on behalf of the Wellcome Sanger Institute COVID-19 Surveillance Team |
| EPI_ISL_2318627, EPI_ISL_2318630, EPI_ISL_2318632, EPI_ISL_2318634, EPI_ISL_2318635, EPI_ISL_2318636 | Lighthouse Lab in Alderley Park | Wellcome Sanger Institute for the COVID-19 Genomics UK (COG-UK) Consortium | Jacquelyn Wynn, Mairead Hyland, The Lighthouse Lab in Alderley Park and Alex Alderton, Roberto Amato, Jeffrey Barrett, Sonia Goncalves, Ewan Harrison, David K. Jackson, Ian Johnston, Dominic Kwiatkowski, Cordelia Langford, John Sillitoe on behalf of the Wellcome Sanger Institute COVID-19 Surveillance Team |
| EPI_ISL_2318637 | Lighthouse Lab in Glasgow | Wellcome Sanger Institute for the COVID-19 Genomics UK (COG-UK) Consortium | Harper VanSteenhouse, Yumi Kasai, David Gray, Carol Clugston, Anna Dominiczak and Alex Alderton, Roberto Amato, Jeffrey Barrett, Sonia Goncalves, Ewan Harrison, David K. Jackson, Ian Johnston, Dominic Kwiatkowski, Cordelia Langford, John Sillitoe on behalf of the Wellcome Sanger Institute COVID-19 Surveillance Team |
| EPI_ISL_2318638, EPI_ISL_2318639 | Lighthouse Lab in Alderley Park | Wellcome Sanger Institute for the COVID-19 Genomics UK (COG-UK) Consortium | Jacquelyn Wynn, Mairead Hyland, The Lighthouse Lab in Alderley Park and Alex Alderton, Roberto Amato, Jeffrey Barrett, Sonia Goncalves, Ewan Harrison, David K. Jackson, Ian Johnston, Dominic Kwiatkowski, Cordelia Langford, John Sillitoe on behalf of the Wellcome Sanger Institute COVID-19 Surveillance Team |
| EPI_ISL_2318641 | Lighthouse Lab in Glasgow | Wellcome Sanger Institute for the COVID-19 Genomics UK (COG-UK) Consortium | Harper VanSteenhouse, Yumi Kasai, David Gray, Carol Clugston, Anna Dominiczak and Alex Alderton, Roberto Amato, Jeffrey Barrett, Sonia Goncalves, Ewan Harrison, David K. Jackson, Ian Johnston, Dominic Kwiatkowski, Cordelia Langford, John Sillitoe on behalf of the Wellcome Sanger Institute COVID-19 Surveillance Team |
| EPI_ISL_2318642, EPI_ISL_2318643, EPI_ISL_2318646, EPI_ISL_2318647, EPI_ISL_2318648 | Lighthouse Lab in Alderley Park | Wellcome Sanger Institute for the COVID-19 Genomics UK (COG-UK) Consortium | Jacquelyn Wynn, Mairead Hyland, The Lighthouse Lab in Alderley Park and Alex Alderton, Roberto Amato, Jeffrey Barrett, Sonia Goncalves, Ewan Harrison, David K. Jackson, Ian Johnston, Dominic Kwiatkowski, Cordelia Langford, John Sillitoe on behalf of the Wellcome Sanger Institute COVID-19 Surveillance Team |
| EPI_ISL_2318649 | Lighthouse Lab in Glasgow | Wellcome Sanger Institute for the COVID-19 Genomics UK (COG-UK) Consortium | Harper VanSteenhouse, Yumi Kasai, David Gray, Carol Clugston, Anna Dominiczak and Alex Alderton, Roberto Amato, Jeffrey Barrett, Sonia Goncalves, Ewan Harrison, David K. Jackson, Ian Johnston, Dominic Kwiatkowski, Cordelia Langford, John Sillitoe on behalf of the Wellcome Sanger Institute COVID-19 Surveillance Team |
| EPI_ISL_2318651, EPI_ISL_2318652, EPI_ISL_2318653, EPI_ISL_2318655, EPI_ISL_2318656, EPI_ISL_2318657 | Lighthouse Lab in Alderley Park | Wellcome Sanger Institute for the COVID-19 Genomics UK (COG-UK) Consortium | Jacquelyn Wynn, Mairead Hyland, The Lighthouse Lab in Alderley Park and Alex Alderton, Roberto Amato, Jeffrey Barrett, Sonia Goncalves, Ewan Harrison, David K. Jackson, Ian Johnston, Dominic Kwiatkowski, Cordelia Langford, John Sillitoe on behalf of the Wellcome Sanger Institute COVID-19 Surveillance Team |
| EPI_ISL_2318658 | Lighthouse Lab in Glasgow | Wellcome Sanger Institute for the COVID-19 Genomics UK (COG-UK) Consortium | Harper VanSteenhouse, Yumi Kasai, David Gray, Carol Clugston, Anna Dominiczak and Alex Alderton, Roberto Amato, Jeffrey Barrett, Sonia Goncalves, Ewan Harrison, David K. Jackson, Ian Johnston, Dominic Kwiatkowski, Cordelia Langford, John Sillitoe on behalf of the Wellcome Sanger Institute COVID-19 Surveillance Team |
| EPI_ISL_2318660, EPI_ISL_2318664 | Lighthouse Lab in Alderley Park | Wellcome Sanger Institute for the COVID-19 Genomics UK (COG-UK) Consortium | Jacquelyn Wynn, Mairead Hyland, The Lighthouse Lab in Alderley Park and Alex Alderton, Roberto Amato, Jeffrey Barrett, Sonia Goncalves, Ewan Harrison, David K. Jackson, Ian Johnston, Dominic Kwiatkowski, Cordelia Langford, John Sillitoe on behalf of the Wellcome Sanger Institute COVID-19 Surveillance Team |
| EPI_ISL_2318665 | Lighthouse Lab in Glasgow | Wellcome Sanger Institute for the COVID-19 Genomics UK (COG-UK) Consortium | Harper VanSteenhouse, Yumi Kasai, David Gray, Carol Clugston, Anna Dominiczak and Alex Alderton, Roberto Amato, Jeffrey Barrett, Sonia Goncalves, Ewan Harrison, David K. Jackson, Ian Johnston, Dominic Kwiatkowski, Cordelia Langford, John Sillitoe on behalf of the Wellcome Sanger Institute COVID-19 Surveillance Team |
| EPI_ISL_2318667, EPI_ISL_2318671, EPI_ISL_2318673, EPI_ISL_2318674, EPI_ISL_2318675, EPI_ISL_2318679, EPI_ISL_2318680, EPI_ISL_2318682, EPI_ISL_2318683, EPI_ISL_2318684, EPI_ISL_2318685 | | | |
| see above | Lighthouse Lab in Alderley Park | Wellcome Sanger Institute for the COVID-19 Genomics UK (COG-UK) Consortium | Jacquelyn Wynn, Mairead Hyland, The Lighthouse Lab in Alderley Park and Alex Alderton, Roberto Amato, Jeffrey Barrett, Sonia Goncalves, Ewan Harrison, David K. Jackson, Ian Johnston, Dominic Kwiatkowski, Cordelia Langford, John Sillitoe on behalf of the Wellcome Sanger Institute COVID-19 Surveillance Team |
| EPI_ISL_2318686, EPI_ISL_2318688 | Lighthouse Lab in Glasgow | Wellcome Sanger Institute for the COVID-19 Genomics UK (COG-UK) Consortium | Harper VanSteenhouse, Yumi Kasai, David Gray, Carol Clugston, Anna Dominiczak and Alex Alderton, Roberto Amato, Jeffrey Barrett, Sonia Goncalves, Ewan Harrison, David K. Jackson, Ian Johnston, Dominic Kwiatkowski, Cordelia Langford, John Sillitoe on behalf of the Wellcome Sanger Institute COVID-19 Surveillance Team |
| EPI_ISL_2318690, EPI_ISL_2318691, EPI_ISL_2318692, EPI_ISL_2318696, EPI_ISL_2318697 | Lighthouse Lab in Alderley Park | Wellcome Sanger Institute for the COVID-19 Genomics UK (COG-UK) Consortium | Jacquelyn Wynn, Mairead Hyland, The Lighthouse Lab in Alderley Park and Alex Alderton, Roberto Amato, Jeffrey Barrett, Sonia Goncalves, Ewan Harrison, David K. Jackson, Ian Johnston, Dominic Kwiatkowski, Cordelia Langford, John Sillitoe on behalf of the Wellcome Sanger Institute COVID-19 Surveillance Team |
| EPI_ISL_2318700 | Lighthouse Lab in Glasgow | Wellcome Sanger Institute for the COVID-19 Genomics UK (COG-UK) Consortium | Harper VanSteenhouse, Yumi Kasai, David Gray, Carol Clugston, Anna Dominiczak and Alex Alderton, Roberto Amato, Jeffrey Barrett, Sonia Goncalves, Ewan Harrison, David K. Jackson, Ian Johnston, Dominic Kwiatkowski, Cordelia Langford, John Sillitoe on behalf of the Wellcome Sanger Institute COVID-19 Surveillance Team |
| EPI_ISL_2318703 | Lighthouse Lab in Alderley Park | Wellcome Sanger Institute for the COVID-19 Genomics UK (COG-UK) Consortium | Jacquelyn Wynn, Mairead Hyland, The Lighthouse Lab in Alderley Park and Alex Alderton, Roberto Amato, Jeffrey Barrett, Sonia Goncalves, Ewan Harrison, David K. Jackson, Ian Johnston, Dominic Kwiatkowski, Cordelia Langford, John Sillitoe on behalf of the Wellcome Sanger Institute COVID-19 Surveillance Team |
| EPI_ISL_2318704 | Lighthouse Lab in Glasgow | Wellcome Sanger Institute for the COVID-19 Genomics UK (COG-UK) Consortium | Harper VanSteenhouse, Yumi Kasai, David Gray, Carol Clugston, Anna Dominiczak and Alex Alderton, Roberto Amato, Jeffrey Barrett, Sonia Goncalves, Ewan Harrison, David K. Jackson, Ian Johnston, Dominic Kwiatkowski, Cordelia Langford, John Sillitoe on behalf of the Wellcome Sanger Institute COVID-19 Surveillance Team |
| EPI_ISL_2318706, EPI_ISL_2318707, EPI_ISL_2318708, EPI_ISL_2318709, EPI_ISL_2318711 | Lighthouse Lab in Alderley Park | Wellcome Sanger Institute for the COVID-19 Genomics UK (COG-UK) Consortium | Jacquelyn Wynn, Mairead Hyland, The Lighthouse Lab in Alderley Park and Alex Alderton, Roberto Amato, Jeffrey Barrett, Sonia Goncalves, Ewan Harrison, David K. Jackson, Ian Johnston, Dominic Kwiatkowski, Cordelia Langford, John Sillitoe on behalf of the Wellcome Sanger Institute COVID-19 Surveillance Team |
| EPI_ISL_2318712 | Lighthouse Lab in Glasgow | Wellcome Sanger Institute for the COVID-19 Genomics UK (COG-UK) Consortium | Harper VanSteenhouse, Yumi Kasai, David Gray, Carol Clugston, Anna Dominiczak and Alex Alderton, Roberto Amato, Jeffrey Barrett, Sonia Goncalves, Ewan Harrison, David K. Jackson, Ian Johnston, Dominic Kwiatkowski, Cordelia Langford, John Sillitoe on behalf of the Wellcome Sanger Institute COVID-19 Surveillance Team |
| EPI_ISL_2318714 | Lighthouse Lab in Alderley Park | Wellcome Sanger Institute for the COVID-19 Genomics UK (COG-UK) Consortium | Jacquelyn Wynn, Mairead Hyland, The Lighthouse Lab in Alderley Park and Alex Alderton, Roberto Amato, Jeffrey Barrett, Sonia Goncalves, Ewan Harrison, David K. Jackson, Ian Johnston, Dominic Kwiatkowski, Cordelia Langford, John Sillitoe on behalf of the Wellcome Sanger Institute COVID-19 Surveillance Team |
| EPI_ISL_2318715, EPI_ISL_2318717, EPI_ISL_2318718, EPI_ISL_2318719, EPI_ISL_2318720, EPI_ISL_2318722, EPI_ISL_2318723, EPI_ISL_2318725, EPI_ISL_2318728, EPI_ISL_2318731, EPI_ISL_2318736, EPI_ISL_2318737, EPI_ISL_2318740, EPI_ISL_2318741, EPI_ISL_2318743, EPI_ISL_2318744, EPI_ISL_2318746, EPI_ISL_2318748, EPI_ISL_2318753, EPI_ISL_2318756, EPI_ISL_2318758, EPI_ISL_2318759, EPI_ISL_2318760, EPI_ISL_2318762, EPI_ISL_2318764, EPI_ISL_2318766, EPI_ISL_2318767, EPI_ISL_2318768, EPI_ISL_2318769, EPI_ISL_2318770, EPI_ISL_2318773, EPI_ISL_2318774, EPI_ISL_2318775, EPI_ISL_2318777, EPI_ISL_2318780, EPI_ISL_2318783, EPI_ISL_2318784, EPI_ISL_2318787, EPI_ISL_2318788, EPI_ISL_2318789, EPI_ISL_2318793, EPI_ISL_2318795, EPI_ISL_2318796, EPI_ISL_2318797, EPI_ISL_2318798, EPI_ISL_2318801, EPI_ISL_2318802, EPI_ISL_2318805, EPI_ISL_2318806, EPI_ISL_2318808, EPI_ISL_2318809, EPI_ISL_2318811, EPI_ISL_2318812, EPI_ISL_2318816, EPI_ISL_2318817, EPI_ISL_2318818, EPI_ISL_2318819, EPI_ISL_2318821, EPI_ISL_2318822, EPI_ISL_2318824, EPI_ISL_2318827, EPI_ISL_2318829, EPI_ISL_2318830, EPI_ISL_2318832, EPI_ISL_2318833, EPI_ISL_2318835, EPI_ISL_2318836, EPI_ISL_2318838, EPI_ISL_2318840, EPI_ISL_2318841, EPI_ISL_2318843, EPI_ISL_2318847, EPI_ISL_2318852, EPI_ISL_2318853, EPI_ISL_2318854, EPI_ISL_2318856, EPI_ISL_2318857, EPI_ISL_2318858, EPI_ISL_2318859, EPI_ISL_2318861, EPI_ISL_2318863, EPI_ISL_2318864, EPI_ISL_2318865, EPI_ISL_2318866, EPI_ISL_2318867, EPI_ISL_2318868, EPI_ISL_2318869, EPI_ISL_2318870, EPI_ISL_2318872, EPI_ISL_2318875, EPI_ISL_2318876, EPI_ISL_2318877, EPI_ISL_2318878, EPI_ISL_2318880, EPI_ISL_2318881, EPI_ISL_2318882, EPI_ISL_2318883, EPI_ISL_2318885, EPI_ISL_2318886, EPI_ISL_2318887, EPI_ISL_2318888, EPI_ISL_2318890, EPI_ISL_2318891, EPI_ISL_2318892, EPI_ISL_2318894, EPI_ISL_2318895, EPI_ISL_2318896 | | | |
| see above | Lighthouse Lab in Glasgow | Wellcome Sanger Institute for the COVID-19 Genomics UK (COG-UK) Consortium | Harper VanSteenhouse, Yumi Kasai, David Gray, Carol Clugston, Anna Dominiczak and Alex Alderton, Roberto Amato, Jeffrey Barrett, Sonia Goncalves, Ewan Harrison, David K. Jackson, Ian Johnston, Dominic Kwiatkowski, Cordelia Langford, John Sillitoe on behalf of the Wellcome Sanger Institute COVID-19 Surveillance Team |
| EPI_ISL_2318901, EPI_ISL_2318902, EPI_ISL_2318924, EPI_ISL_2318925, EPI_ISL_2318927, EPI_ISL_2318930, EPI_ISL_2318971, EPI_ISL_2318972 | Azienda Sanitaria dell'Alto Adige Laboratorio Aziendale di Microbiologia e Virologia | Istituto di Genomica Applicata | Elisabetta Pagani, Irene Bianconi, Elisabetta Giacobazzi, Elisa Masi, Stefanie Wieser, Irena Jurman, Vera Vendramin, Gabriele Magris, Eleonora Paparelli, Davide Scaglione, Michele Morgante |
| EPI_ISL_2318999 | Megalab, Molecular and Cytogenetics Diagnostics | Department for Virology, Molecular Biology and Genome Research, R. G. Lugar Center for Public Health Research, National Center for Disease Control and Public Health (NCDC) of Georgia. | Giorgi Tomashvili, Gvantsa Brachveli, Meri Pantsulaia, Giorgi Gogoladze, Nino Berishvili, Tata Imnadze, Ana Papkiauri, Gvantsa Chanturia, Ann Machablishvili, Nato Kotaria, Marine Murtskhvaladze, Lela Sabadze, Mari Gavashelidze, Tamar Jashiashvili, Tea Tevdoradze, Ketevan Sidamonidze, Ekaterine Khmaladze, Ekaterine Zhghenti, Roena Sukhiashvili, Mariam Zakalashvili, Lela Urushadze, Magda Dgebuadze, Davit Tsaguria, Nino Chikhovani, Ekaterine Zangaladze, Adam Kotorashvili, Maia Alkhazashvili, Irma Burjanadze, Anna Kasradze, Khatuna Zakhashvili, Paata Imnadze, Amiran Gamkrelidze. |
| EPI_ISL_2319006 | Department for Virology, Molecular Biology and Genome Research, R. G. Lugar Center for Public Health Research, National Center for Disease Control and Public Health (NCDC) of Georgia. | Department for Virology, Molecular Biology and Genome Research, R. G. Lugar Center for Public Health Research, National Center for Disease Control and Public Health (NCDC) of Georgia. | Meri Pantsulaia, Giorgi Tomashvili, Gvantsa Brachveli, Giorgi Gogoladze, Nino Berishvili, Tata Imnadze, Ana Papkiauri, Gvantsa Chanturia, Ann Machablishvili, Nato Kotaria, Marine Murtskhvaladze, Lela Sabadze, Mari Gavashelidze, Tamar Jashiashvili, Tea Tevdoradze, Ketevan Sidamonidze, Ekaterine Khmaladze, Ekaterine Zhghenti, Roena Sukhiashvili, Mariam Zakalashvili, Lela Urushadze, Magda Dgebuadze, Davit Tsaguria, Ekaterine Zangaladze, Adam Kotorashvili, Maia Alkhazashvili, Irma Burjanadze, Anna Kasradze, Khatuna Zakhashvili, Paata Imnadze, Amiran Gamkrelidze. |
| EPI_ISL_2319007 | Department for Virology, Molecular Biology and Genome Research, R. G. Lugar Center for Public Health Research, National Center for Disease Control and Public Health (NCDC) of Georgia. | Department for Virology, Molecular Biology and Genome Research, R. G. Lugar Center for Public Health Research, National Center for Disease Control and Public Health (NCDC) of Georgia. | Giorgi Tomashvili, Meri Pantsulaia, Gvantsa Brachveli, Giorgi Gogoladze, Nino Berishvili, Tata Imnadze, Ana Papkiauri, Gvantsa Chanturia, Ann Machablishvili, Nato Kotaria, Marine Murtskhvaladze, Lela Sabadze, Mari Gavashelidze, Tamar Jashiashvili, Tea Tevdoradze, Ketevan Sidamonidze, Ekaterine Khmaladze, Ekaterine Zhghenti, Roena Sukhiashvili, Mariam Zakalashvili, Lela Urushadze, Magda Dgebuadze, Davit Tsaguria, Ekaterine Zangaladze, Adam Kotorashvili, Maia Alkhazashvili, Irma Burjanadze, Anna Kasradze, Khatuna Zakhashvili, Paata Imnadze, Amiran Gamkrelidze. |
| EPI_ISL_2319009 | Diagnostyka. Laboratoria Medyczne. | 1. ViroGenetics - BSL3 Laboratory of Virology, Małopolska Centre of Biotechnology, Jagiellonian University; 2. genXone SA, Research & Development Laboratory | Mazur-Panasiuk,N., Nowicki G, Gromowski,T., Drweska-Matelska N, Grabowski J, Brylak A, Gidlewicz A, Szeszko K, Sykulski M, Krych L, Kowalski,M., Wydmanski W., Szulc,P., Januszczak S, Labaj,P.P., Kaszuba M, Pyrc,K. |
| EPI_ISL_2319503 | Maryland Genomics, Institute for Genome Sciences, University of Maryland School of Medicine | Maryland Genomics, Institute for Genome Sciences, University of Maryland School of Medicine | Tallon, Luke J; Sadzewicz, Lisa D; Humphrys, Mike; Ott, Sandra; Roussey, Holly; Mehta, Aditya; Vavikolanu, Kranthi; Fraser, Claire M; Ravel, Jacques |
| EPI_ISL_2319973 | Reditus Laboratories | Reditus Laboratories | Joshua J. Geltz, Ph.D., Robert M. Sgambelluri, Ph.D., Rex Dyer, Ph.D., Cassy Philips, M.S., Alexa Eichelberger, M.S. |

| | | | |
|---|---|---|---|
| EPI_ISL_2320021, EPI_ISL_2320131, EPI_ISL_2320147, EPI_ISL_2320156, EPI_ISL_2320176, EPI_ISL_2320200, EPI_ISL_2320383, EPI_ISL_2320446, EPI_ISL_2320572, EPI_ISL_2320699, EPI_ISL_2320767, EPI_ISL_2320883, EPI_ISL_2321076, EPI_ISL_2321081, EPI_ISL_2321132 | | | |
| see above | Helix/Illumina | Centers for Disease Control and Prevention Division of Viral Diseases, Pathogen Discovery | Dakota Howard, Dhwani Batra, Peter W. Cook, Kara Moser, Adrian Paskey, Jason Caravas, Benjamin Rambo-Martin, Shatavia Morrison, Christopher Gulvick, Scott Sammons, Yvette Unoarumhi, Darlene Wagner, Matthew Schmerer, Eileen de Feo, Jan Antico, Christine Tran, Matthew Tolentino, Shannon Wickline, Kim Gietzen, Brad Sickler, Jingtao Liu, Eric Allen, Phil Febbo, Nicole L. Washington, Simon White, Geraint Levan, Kelly Schiabor Barrett, Elizabeth Cirulli, Alexandre Bolze, Ary Ascencio, Charlotte Rivera-Garcia, Ryan Cho, Jason Nguyen, Sherry Wang, Jimmy Ramirez, Tyler Cassens, Efren Sandoval, Magnus Isaksson, William Lee, David Becker, Marc Laurent, James Lu, Clinton R. Paden, Duncan MacCannell |
| EPI_ISL_2321160, EPI_ISL_2321164, EPI_ISL_2321165, EPI_ISL_2321166, EPI_ISL_2321168, EPI_ISL_2321169 | Royal Darwin Hospital Pathology | MDU-PHL | Meumann, E., Caly L., Seemann T., Sait, M.L., Druce J., Sherry, N.L. |
| EPI_ISL_2321171 | Microbiological Diagnostic Unit - Public Health Laboratory (MDU-PHL) | MDU-PHL | Seemann T., Sait, M.L., Sherry, N.L. |
| EPI_ISL_2321313, EPI_ISL_2321328 | Aegis Sciences Corporation | Centers for Disease Control and Prevention Division of Viral Diseases, Pathogen Discovery | Dakota Howard, Dhwani Batra, Peter W. Cook, Kara Moser, Adrian Paskey, Jason Caravas, Benjamin Rambo-Martin, Shatavia Morrison, Christopher Gulvick, Scott Sammons, Yvette Unoarumhi, Darlene Wagner, Matthew Schmerer, Cyndi Clark, Patrick Campbell, Rob Case, Vikramsinha Ghorpade, Holly Houdeshell, Ola Kvalvaag, Dillon Nall, Ethan Sanders, Alec Vest, Shaun Westlund, Matthew Hardison, Clinton R. Paden, Duncan MacCannell |
| EPI_ISL_2322717 | MD PHL | MD PHL | Maryland Department of Health Laboratories Administration |
| EPI_ISL_2322922 | UNILABS - LAUSANNE | Laboratory of genomics and metagenomics | Trestan Pillonel, Damien Jacot, Sébastien Aeby, Gilbert Greub, Claire Bertelli |
| EPI_ISL_2322968 | EHC MORGES | Laboratory of genomics and metagenomics | Trestan Pillonel, Damien Jacot, Sébastien Aeby, Gilbert Greub, Claire Bertelli |
| EPI_ISL_2322989 | Heilig hart Lier | Imeda Hospital | Johan Frans, Dagmar Obbels, Hanne Valgaeren |
| EPI_ISL_2322996 | Imeda Hospital | Imeda Hospital | Johan Frans, Dagmar Obbels, Hanne Valgaeren |
| EPI_ISL_2323023 | Mapmygenome | CSIR-Centre for Cellular and Molecular Biology - INSACOG | Lamuk Zaveri, Ara Sreenivas, Onkar Kulkarni, Sofia Banu, Shreekant Verma, Amareshwar Vodapalli , Viswagithe S L, B Himasri, Sharath Chandra Thota, Karthik Bharadwaj Tallapaka, Rakesh K Mishra,Divya Tej Sowpati |
| EPI_ISL_2323024 | AP SSO | CSIR-Centre for Cellular and Molecular Biology-INSACOG | Tulasi Nagabandi, Lamuk Zaveri,Ara Sreenivas,Shreekant Verma, Amareshwar Vodapalli ,Blessy B John,Viswagithe S L,B Himasri,Valli Nagalakshmi Undamatla,Payel Mukherjee,Sofia Banu,Archana Bharadwaj Siva,Sharath Chandra Thota,Karthik Bharadwaj Tallapaka,Rakesh K Mishra,Divya Tej Sowpati |
| EPI_ISL_2323315 | Furst Medical Laboratory | Norwegian Institute of Public Health, Department of Virology | Kathrine Stene-Johansen, Kamilla Heddeland Instefjord, Hilde Elshaug, Garcia Llorente Ignacio, Jon Bråte, Engebretsen Serina Beate, Line Victoria Moen, Pedersen Benedikte Nevjen, Debech Nadia, Atiya R Ali,Marie Paulsen Madsen, Rasmus Riis Kopperud, Hilde Vollan, Karoline Bragstad, Olav Hungnes |
| EPI_ISL_2323325, EPI_ISL_2323326 | Ostfold Hospital Trust - Kalnes, Centre for Laboratory Medicine, Section for gene technology and infection serology | Norwegian Institute of Public Health, Department of Virology | Kathrine Stene-Johansen, Kamilla Heddeland Instefjord, Hilde Elshaug, Garcia Llorente Ignacio, Jon Bråte, Engebretsen Serina Beate, Line Victoria Moen, Pedersen Benedikte Nevjen, Debech Nadia, Atiya R Ali,Marie Paulsen Madsen, Rasmus Riis Kopperud, Hilde Vollan, Karoline Bragstad, Olav Hungnes |
| EPI_ISL_2323341, EPI_ISL_2323347 | Akershus University Hospital, Department for Microbiology and Infectious Disease Control | Norwegian Institute of Public Health, Department of Virology | Kathrine Stene-Johansen, Kamilla Heddeland Instefjord, Hilde Elshaug, Garcia Llorente Ignacio, Jon Bråte, Engebretsen Serina Beate, Line Victoria Moen, Pedersen Benedikte Nevjen, Debech Nadia, Atiya R Ali,Marie Paulsen Madsen, Rasmus Riis Kopperud, Hilde Vollan, Karoline Bragstad, Olav Hungnes |
| EPI_ISL_2323452 | GA Department of Public Health Laboratory | Genomics and Discovery, Respiratory Viruses Branch, Division of Viral Diseases, Centers for Disease Control and Prevention | Jing Zhang, Anna Kelleher, Ying Tao, Yan Li, Brian Lynch, Krista Queen, Anna Uehara, Peter Cook, Han Jia Justin Ng, Rachel Marine, Clinton R. Paden, Dhwani Batra, Haibin Wang, Tara Coalter, Jasmine Padilla, Morgan Davis, Mili Sheth, Sarah Nobles, Mark Burroughs, Justin Lee, Adam Retchless, Suxiang Tong |
| EPI_ISL_2323453, EPI_ISL_2323454 | AR Dept. of Health-PHL, Molecular Diagnostics | Genomics and Discovery, Respiratory Viruses Branch, Division of Viral Diseases, Centers for Disease Control and Prevention | Jing Zhang, Anna Kelleher, Ying Tao, Yan Li, Brian Lynch, Krista Queen, Anna Uehara, Peter Cook, Han Jia Justin Ng, Rachel Marine, Clinton R. Paden, Dhwani Batra, Haibin Wang, Tara Coalter, Jasmine Padilla, Morgan Davis, Mili Sheth, Sarah Nobles, Mark Burroughs, Justin Lee, Adam Retchless, Suxiang Tong |
| EPI_ISL_2323583, EPI_ISL_2323642, EPI_ISL_2323770, EPI_ISL_2323988, EPI_ISL_2324007, EPI_ISL_2324032, EPI_ISL_2324115 | Infinity Biologix | Centers for Disease Control and Prevention Division of Viral Diseases, Pathogen Discovery | Dakota Howard, Dhwani Batra, Peter W. Cook, Kara Moser, Adrian Paskey, Jason Caravas, Benjamin Rambo-Martin, Shatavia Morrison, Christopher Gulvick, Scott Sammons, Yvette Unoarumhi, Darlene Wagner, Matthew Schmerer, Christian Bixby, Yihe Wang, Jonathan Schultz, Chirayu Goswami, Russ Hager, Robin Grimwood, Clinton R. Paden, Duncan MacCannell |
| EPI_ISL_2324268 | Institute for Medical and Chemical Laboratory Diagnostics, Kepler Universitätsklinikum | Bergthaler laboratory, CeMM Research Center for Molecular Medicine of the Austrian Academy of Sciences | Lukas Endler, Anna Schedl, Fabian Amman, Petr Triska, Thomas Penz, Benedikt Agerer, Maelle Le Moing, Michael Schuster, Bekir Erguner, Jan Laine, Martin Senekowitsch, Christoph Bock, Andreas Bergthaler |
| EPI_ISL_2324450, EPI_ISL_2324484, EPI_ISL_2324505, EPI_ISL_2324507, EPI_ISL_2324547, EPI_ISL_2324551 | Laboratory Corporation of America | Centers for Disease Control and Prevention Division of Viral Diseases, Pathogen Discovery | Dakota Howard, Dhwani Batra, Peter W. Cook, Kara Moser, Adrian Paskey, Jason Caravas, Benjamin Rambo-Martin, Shatavia Morrison, Christopher Gulvick, Scott Sammons, Yvette Unoarumhi, Darlene Wagner, Matthew Schmerer, Minoo Agarwal, Eyad Almasri, Debbie Boles, Ayla Burns, Nuthawin Charoensri, Oren Cohen, Susan Countryman, Mary Ann Cristobal, Bobbi Croy, Suzanne Dale, Hrushikesh Deshmukh, Amanda Douglas, Vincent Drouillon, Marcia Eisenberg, Howard Engler, Rama Ghatti, Prashant Gupta, Susan Hicks, Jake Humphrey, Lax Iyer, Lisa Pfefferle, Manoj Jain, Matthew Robinson, Mohan Kolli, Brian Krueger, Tim Kuphal, Stanley Letovsky, Michael Levandoski, Craig Lukasik, Jonathan Meltzer, Brian Norvell, Mindy Nye, Scott Parker, Christos Petropoulos, John Pruitt, Steven Ragan, Scott Ryan, Mike Sapeta, Jana Schroth, Suresh Babu Selvaraju, Goran Stevovic, Amanda Suchanek, Andrea Throop, Lyndon Tilson, Thomas Urban, Joe Voshell, Kimberly Wagner, Jonathan Williams, Mary Williamson, Qian Zeng, Tricia Zwiefelhofer, Clinton R. Paden, Duncan MacCannell |
| EPI_ISL_2324942, EPI_ISL_2324943, EPI_ISL_2324944, EPI_ISL_2324948 | National Institute of Public Health | National Institute of Public Health | Helena Jirincova, Jaromira Vecerova, Timotej Suri, Dusan Trnka, Alexander Nagy |
| EPI_ISL_2329654 | Pathogen Genomics Center, National Institute of Infectious Diseases | Pathogen Genomics Center, National Institute of Infectious Diseases | Tsuyoshi Sekizuka, Kentaro Itokawa, Rina Tanaka, Masanori Hashino, Makoto Kuroda |
| EPI_ISL_2329929, EPI_ISL_2329930, EPI_ISL_2329931, EPI_ISL_2329932, EPI_ISL_2329933 | SARS-CoV-2 testing team, National Institute of Infectious Diseases | Pathogen Genomics Center, National Institute of Infectious Diseases | Tsuyoshi Sekizuka, Kentaro Itokawa, Rina Tanaka, Masanori Hashino, Nozomu Hanaoka, Masumichi Saito, Naomi Nojiri, Hazuka Y Furihata, Sana Uchikoba, Tsuguto Fujimoto, Makoto Kuroda |
| EPI_ISL_2330490 | Pathogen Genomics Center, National Institute of Infectious Diseases | Pathogen Genomics Center, National Institute of Infectious Diseases | Tsuyoshi Sekizuka, Kentaro Itokawa, Rina Tanaka, Masanori Hashino, Makoto Kuroda |
| EPI_ISL_2331496, EPI_ISL_2331507 | Johns Hopkins Hospital Department of Pathology | Johns Hopkins Hospital Department of Pathology | C. Paul Morris, Chun Huai Luo, Adannaya Amadi, Matthew Schwartz, Nicholas Gallagher, Heba H. Mostafa |
| EPI_ISL_2331830 | Platform BIS UZA/UAntwerpen | Labo Klinische Biologie, UZA | Marie Le Mercier, Jasmine Coppens, Basil Britto Xavier, Christine Lammens, Veerle Matheeussen, Herman Goossens |
| EPI_ISL_2332624 | LBM ALPHABIO, Marseille | LBM ALPHABIO, Marseille | Vincent GARCIA |
| EPI_ISL_2332699 | Maple Grove Hospital | Minnesota Department of Health, Public Health Laboratory | Alexandra Lorentz, Jacob Garfin, Matt Plumb, and Xiong Wang |
| EPI_ISL_2332704, EPI_ISL_2332706, EPI_ISL_2332707, EPI_ISL_2332708, EPI_ISL_2332709, EPI_ISL_2332711, EPI_ISL_2332713, EPI_ISL_2332714, EPI_ISL_2332716, EPI_ISL_2332717, EPI_ISL_2332718, EPI_ISL_2332719, EPI_ISL_2332721, EPI_ISL_2332722, EPI_ISL_2332723, EPI_ISL_2332724, EPI_ISL_2332725, EPI_ISL_2332726, EPI_ISL_2332727, EPI_ISL_2332728, EPI_ISL_2332729, EPI_ISL_2332730, EPI_ISL_2332731, EPI_ISL_2332736, EPI_ISL_2332737, EPI_ISL_2332739, EPI_ISL_2332743, EPI_ISL_2332744, EPI_ISL_2332745, EPI_ISL_2332746, EPI_ISL_2332747, EPI_ISL_2332748, EPI_ISL_2332749, EPI_ISL_2332750, EPI_ISL_2332751, EPI_ISL_2332752, EPI_ISL_2332753, EPI_ISL_2332754, EPI_ISL_2332755, EPI_ISL_2332756, EPI_ISL_2332757, EPI_ISL_2332758, EPI_ISL_2332759, EPI_ISL_2332760, EPI_ISL_2332761, EPI_ISL_2332763, EPI_ISL_2332764, EPI_ISL_2332765, EPI_ISL_2332768, EPI_ISL_2332772, EPI_ISL_2332773, EPI_ISL_2332778, EPI_ISL_2332779, EPI_ISL_2332780, EPI_ISL_2332784, EPI_ISL_2332786, EPI_ISL_2332787, EPI_ISL_2332788 | | | |
| see above | National Centre For Cell Science | National Centre For Cell Science – INSACOG | Dhiraj Paul, Shivang P. Bhanushali, Mitali Inamdar, Mohak P Gujare, Manoj Kumar Bhat, Ajay Pillai, INSACOG Consortium team, Yogesh Shouche |
| EPI_ISL_2332851, EPI_ISL_2332856, EPI_ISL_2332860, EPI_ISL_2332868, EPI_ISL_2332873, EPI_ISL_2332888, EPI_ISL_2332889, EPI_ISL_2332929, EPI_ISL_2332964, EPI_ISL_2333011, EPI_ISL_2333013, EPI_ISL_2333049, EPI_ISL_2333067, EPI_ISL_2333069 | | | |
| see above | UW Virology Lab | UW Virology Lab | Pavitra Roychoudhury, Hong Xie, Lasata Shrestha, Tien V. Nguyen, Shah Mohamed Bakhash, Michelle Lin, Noah R. Baker, Ricardo Perez, Sean Ellis, Nathan Breit, Robert J. Livingston, Meei-Li Huang, Keith R Jerome, Patrick Mathias, Alexander Greninger |
| EPI_ISL_2333503 | Furst Medical Laboratory | Norwegian Institute of Public Health, Department of Virology | 'Kathrine Stene-Johansen, Kamilla Heddeland Instefjord, Hilde Elshaug, Garcia Llorente Ignacio, Jon Bråte, Engebretsen Serina Beate,Pedersen Benedikte Nevjen, Debech Nadia, Line Victoria Moen, Atiya R Ali,Marie Paulsen Madsen, Rasmus Riis Kopperud, Hilde Vollan, Karoline Bragstad, Olav Hungnes |
| EPI_ISL_2333504, EPI_ISL_2333505, EPI_ISL_2333506, EPI_ISL_2333507 | Akershus University Hospital, Department for Microbiology and Infectious Disease Control | Norwegian Institute of Public Health, Department of Virology | 'Kathrine Stene-Johansen, Kamilla Heddeland Instefjord, Hilde Elshaug, Garcia Llorente Ignacio, Jon Bråte, Engebretsen Serina Beate,Pedersen Benedikte Nevjen, Debech Nadia, Line Victoria Moen, Atiya R Ali,Marie Paulsen Madsen, Rasmus Riis Kopperud, Hilde Vollan, Karoline Bragstad, Olav Hungnes |
| EPI_ISL_2333508, EPI_ISL_2333509 | Furst Medical Laboratory | Norwegian Institute of Public Health, Department of Virology | 'Kathrine Stene-Johansen, Kamilla Heddeland Instefjord, Hilde Elshaug, Garcia Llorente Ignacio, Jon Bråte, Engebretsen Serina Beate,Pedersen Benedikte Nevjen, Debech Nadia, Line Victoria Moen, Atiya R Ali,Marie Paulsen Madsen, Rasmus Riis Kopperud, Hilde Vollan, Karoline Bragstad, Olav Hungnes |
| EPI_ISL_2333510 | Akershus University Hospital, Department for Microbiology and Infectious Disease Control | Norwegian Institute of Public Health, Department of Virology | 'Kathrine Stene-Johansen, Kamilla Heddeland Instefjord, Hilde Elshaug, Garcia Llorente Ignacio, Jon Bråte, Engebretsen Serina Beate,Pedersen Benedikte Nevjen, Debech Nadia, Line Victoria Moen, Atiya R Ali,Marie Paulsen Madsen, Rasmus Riis Kopperud, Hilde Vollan, Karoline Bragstad, Olav Hungnes |
| EPI_ISL_2333511, EPI_ISL_2333512 | Furst Medical Laboratory | Norwegian Institute of Public Health, Department of Virology | 'Kathrine Stene-Johansen, Kamilla Heddeland Instefjord, Hilde Elshaug, Garcia Llorente Ignacio, Jon Bråte, Engebretsen Serina Beate,Pedersen Benedikte Nevjen, Debech Nadia, Line Victoria Moen, Atiya R Ali,Marie Paulsen Madsen, Rasmus Riis Kopperud, Hilde Vollan, Karoline Bragstad, Olav Hungnes |
| EPI_ISL_2333747 | Biogroup Bio Lam-LCD Saint-Denis | Department of Virology, Henri Mondor University Hospital, Assistance Publique Hôpitaux de Paris, Université Paris-Est Créteil, INSERM U955 | Christophe Rodriguez, Slim Fourati, Vanessa Demontant, Guillaume Gricourt, Melissa N'Debi, Alexandre Soulier, Elisabeth Trawinski, Jean-Michel Pawlotsky |
| EPI_ISL_2333757 | Hôpital Avicenne | Department of Virology, Henri Mondor University Hospital, Assistance Publique Hôpitaux de Paris, Université Paris-Est Créteil, INSERM U955 | Christophe Rodriguez, Slim Fourati, Vanessa Demontant, Guillaume Gricourt, Melissa N'Debi, Alexandre Soulier, Elisabeth Trawinski, Jean-Michel Pawlotsky |
| EPI_ISL_2333922 | MS Public Health Laboratory | University of Mississippi Medical Center, Molecular and Genomics Core Facility | Ashley C. Johnson, Ithiel J. Frame, Krishna K. Ayyalasomayajula, Michael R. Garrett, D. Ashley Robinson |

| | | | |
|---|---|---|---|
| EPI_ISL_2333982 | GROUPE HOSPITALIER SUD ILE DE FRANCE | Department of Virology, Henri Mondor University Hospital, Assistance Publique Hôpitaux de Paris, Université Paris-Est Créteil, INSERM U955 | Christophe Rodriguez, Slim Fourati, Vanessa Demontant, Guillaume Gricourt, Melissa N'Debi, Alexandre Soulier, Elisabeth Trawinski, Jean-Michel Pawlotsky |
| EPI_ISL_2335415 | Pathogen Genomics Center, National Institute of Infectious Diseases | Pathogen Genomics Center, National Institute of Infectious Diseases | Tsuyoshi Sekizuka, Kentaro Itokawa, Rina Tanaka, Masanori Hashino, Makoto Kuroda |
| EPI_ISL_2336215 | Washington State Department of Health Public Health Laboratories | Washington State Department of Health Public Health Laboratories | Drew MacKellar, Philip Dykema, Denny Russell, Joenice Gonzalez, Hannah Gray, Geoff Melly, Vanessa De Los Santos, Darren Lucas, JohnAric Peterson, Avi Singh, Rebecca Cao |
| EPI_ISL_2336637, EPI_ISL_2337043 | SARS-CoV-2 testing team, National Institute of Infectious Diseases | Pathogen Genomics Center, National Institute of Infectious Diseases | Tsuyoshi Sekizuka, Kentaro Itokawa, Rina Tanaka, Masanori Hashino, Nozomu Hanaoka, Masumichi Saito, Naomi Nojiri, Hazuka Y Furihata, Sana Uchikoba, Tsuguto Fujimoto, Makoto Kuroda |
| EPI_ISL_2337257 | Chiba Prefectural Institute of Public Health | Pathogen Genomics Center, National Institute of Infectious Diseases | Tsuyoshi Sekizuka, Kentaro Itokawa, Rina Tanaka, Masanori Hashino, Makoto Kuroda |
| EPI_ISL_2337324, EPI_ISL_2337429, EPI_ISL_2337471, EPI_ISL_2337524 | Pathogen Genomics Center, National Institute of Infectious Diseases | Pathogen Genomics Center, National Institute of Infectious Diseases | Tsuyoshi Sekizuka, Kentaro Itokawa, Rina Tanaka, Masanori Hashino, Makoto Kuroda |
| EPI_ISL_2339438, EPI_ISL_2339452, EPI_ISL_2339458, EPI_ISL_2339465, EPI_ISL_2339467, EPI_ISL_2339481, EPI_ISL_2339482, EPI_ISL_2339501 | Labor Berlin Charité Vivantes GmbH / Institut für Virologie | Charité Universitätsmedizin Berlin, Institut für Virologie/Labor Berlin | Peter Menzel, Christine Stephan, Rolf Schwarzer, Victor M Corman, Barbara Mühlemann, Terry Jones, Christian Drosten |
| EPI_ISL_2340135 | Laboratorio Aziendale di Microbiologia e Virologia, Azienda Sanitaria dell'Alto Adige | Laboratorio Aziendale di Microbiologia e Virologia, Azienda Sanitaria dell'Alto Adige | Irene Bianconi, Elisabetta Giacobazzi, Elisa Masi, Stefanie Wieser, Anne Picard, Claudia Volpato, Chiara Cantaloni, Elisabetta Pagani |
| EPI_ISL_2340136, EPI_ISL_2340137, EPI_ISL_2340138 | CHMT | Instituto Nacional de Saude (INSA) | Borges et al |
| EPI_ISL_2340146, EPI_ISL_2340147, EPI_ISL_2340152, EPI_ISL_2340153 | Germano de Sousa | Instituto Nacional de Saude (INSA) | Borges et al |
| EPI_ISL_2340227 | H Divino Espirito Santo - Ponta Delgada | Instituto Nacional de Saude (INSA) | Borges et al |
| EPI_ISL_2340230 | ULS Litoral Alentejano | Instituto Nacional de Saude (INSA) | Borges et al |
| EPI_ISL_2340336, EPI_ISL_2340339, EPI_ISL_2340340, EPI_ISL_2340341, EPI_ISL_2340348, EPI_ISL_2340358 | Germano de Sousa | Instituto Nacional de Saude (INSA) and BioSystems & Integrative Sciences Institute (BioISI) Genomics Unit, FCUL | Borges et al |
| EPI_ISL_2340461 | Instituto Nacional de Saude (INSA) | Instituto Nacional de Saude (INSA) and BioSystems & Integrative Sciences Institute (BioISI) Genomics Unit, FCUL | Borges et al |
| EPI_ISL_2340672 | University of Wisconsin-Madison AIDS Vaccine Research Laboratories | University of Wisconsin-Madison AIDS Vaccine Research Laboratories | Gage Moreno, Katarina Braun, et al. AIDS Vaccine Research Laboratories |
| EPI_ISL_2340686 | UW Virology Lab | UW Virology Lab | Pavitra Roychoudhury, Hong Xie, Lasata Shrestha, Tien V. Nguyen, Shah Mohamed Bakhash, Michelle Lin, Noah R. Baker, Sean Ellis, Meei-Li Huang, Keith R Jerome, Alexander Greninger |
| EPI_ISL_2340841 | University of Wisconsin-Madison AIDS Vaccine Research Laboratories | University of Wisconsin-Madison AIDS Vaccine Research Laboratories | Gage Moreno, Katarina Braun, et al. AIDS Vaccine Research Laboratories |
| EPI_ISL_2341495, EPI_ISL_2341560 | Santa Clara County Public Health Laboratory | Chan-Zuckerberg Biohub | CZB Cliahub Consortium |
| EPI_ISL_2341631 | CA DPH Viral and Rickettsial Disease Laboratory | Chan-Zuckerberg Biohub | CZB Cliahub Consortium |
| EPI_ISL_2341641 | Santa Clara County Public Health Laboratory | Chan-Zuckerberg Biohub | CZB Cliahub Consortium |
| EPI_ISL_2341666 | Veer Surendra Sai Institute of Medical Sciences and Research, Burla, Sambalpur | Institute of Life Sciences - INSACOG | Sunil K. Raghav, Safal Walia, Arup Ghosh, Atimukta Jha, Amol M. Kanampalliwar, Omprakash Shiriwas, Sana Fatma, Shifu Aggarwal, Rupesh Dash, Rajeeb Swain, Punit Prasad, INSACOG Consortium, Ajay Parida |
| EPI_ISL_2341679 | Pt. Jawahar Lal Nehru Memorial Medical College, Raipur | Institute of Life Sciences - INSACOG | Sunil K. Raghav, Safal Walia, Arup Ghosh, Atimukta Jha, Amol M. Kanampalliwar, Omprakash Shiriwas, Sana Fatma, Shifu Aggarwal, Rupesh Dash, Rajeeb Swain, Punit Prasad, INSACOG Consortium, Ajay Parida |
| EPI_ISL_2341682 | Institute of Life Sciences, Bhubaneswar | Institute of Life Sciences - INSACOG | Sunil K. Raghav, Safal Walia, Arup Ghosh, Atimukta Jha, Amol M. Kanampalliwar, Omprakash Shiriwas, Sana Fatma, Shifu Aggarwal, Rupesh Dash, Rajeeb Swain, Punit Prasad, INSACOG Consortium, Ajay Parida |
| EPI_ISL_2341685 | S.C.B. Medical College and Hospital, Cuttack | Institute of Life Sciences - INSACOG | Sunil K. Raghav, Safal Walia, Arup Ghosh, Atimukta Jha, Amol M. Kanampalliwar, Omprakash Shiriwas, Sana Fatma, Shifu Aggarwal, Rupesh Dash, Rajeeb Swain, Punit Prasad, INSACOG Consortium, Ajay Parida |
| EPI_ISL_2341690 | FM Medical College, Balasore | Institute of Life Sciences - INSACOG | Sunil K. Raghav, Safal Walia, Arup Ghosh, Atimukta Jha, Amol M. Kanampalliwar, Omprakash Shiriwas, Sana Fatma, Shifu Aggarwal, Rupesh Dash, Rajeeb Swain, Punit Prasad, INSACOG Consortium, Ajay Parida |
| EPI_ISL_2341692, EPI_ISL_2341693 | Pt. Jawahar Lal Nehru Memorial Medical College, Raipur | Institute of Life Sciences - INSACOG | Sunil K. Raghav, Safal Walia, Arup Ghosh, Atimukta Jha, Amol M. Kanampalliwar, Omprakash Shiriwas, Sana Fatma, Shifu Aggarwal, Rupesh Dash, Rajeeb Swain, Punit Prasad, INSACOG Consortium, Ajay Parida |
| EPI_ISL_2341694 | FM Medical College, Balasore | Institute of Life Sciences - INSACOG | Sunil K. Raghav, Safal Walia, Arup Ghosh, Atimukta Jha, Amol M. Kanampalliwar, Omprakash Shiriwas, Sana Fatma, Shifu Aggarwal, Rupesh Dash, Rajeeb Swain, Punit Prasad, INSACOG Consortium, Ajay Parida |
| EPI_ISL_2341695 | Institute of Life Sciences, Bhubaneswar | Institute of Life Sciences - INSACOG | Sunil K. Raghav, Safal Walia, Arup Ghosh, Atimukta Jha, Amol M. Kanampalliwar, Omprakash Shiriwas, Sana Fatma, Shifu Aggarwal, Rupesh Dash, Rajeeb Swain, Punit Prasad, INSACOG Consortium, Ajay Parida |
| EPI_ISL_2341697 | S.C.B. Medical College and Hospital, Cuttack | Institute of Life Sciences - INSACOG | Sunil K. Raghav, Safal Walia, Arup Ghosh, Atimukta Jha, Amol M. Kanampalliwar, Omprakash Shiriwas, Sana Fatma, Shifu Aggarwal, Rupesh Dash, Rajeeb Swain, Punit Prasad, INSACOG Consortium, Ajay Parida |
| EPI_ISL_2341698, EPI_ISL_2341699, EPI_ISL_2341701, EPI_ISL_2341704, EPI_ISL_2341706 | Institute of Life Sciences, Bhubaneswar | Institute of Life Sciences - INSACOG | Sunil K. Raghav, Safal Walia, Arup Ghosh, Atimukta Jha, Amol M. Kanampalliwar, Omprakash Shiriwas, Sana Fatma, Shifu Aggarwal, Rupesh Dash, Rajeeb Swain, Punit Prasad, INSACOG Consortium, Ajay Parida |
| EPI_ISL_2341708, EPI_ISL_2341709 | Pt. Jawahar Lal Nehru Memorial Medical College, Raipur | Institute of Life Sciences - INSACOG | Sunil K. Raghav, Safal Walia, Arup Ghosh, Atimukta Jha, Amol M. Kanampalliwar, Omprakash Shiriwas, Sana Fatma, Shifu Aggarwal, Rupesh Dash, Rajeeb Swain, Punit Prasad, INSACOG Consortium, Ajay Parida |
| EPI_ISL_2341712 | Institute of Life Sciences, Bhubaneswar | Institute of Life Sciences - INSACOG | Sunil K. Raghav, Safal Walia, Arup Ghosh, Atimukta Jha, Amol M. Kanampalliwar, Omprakash Shiriwas, Sana Fatma, Shifu Aggarwal, Rupesh Dash, Rajeeb Swain, Punit Prasad, INSACOG Consortium, Ajay Parida |
| EPI_ISL_2341714, EPI_ISL_2341717 | S.C.B. Medical College and Hospital, Cuttack | Institute of Life Sciences - INSACOG | Sunil K. Raghav, Safal Walia, Arup Ghosh, Atimukta Jha, Amol M. Kanampalliwar, Omprakash Shiriwas, Sana Fatma, Shifu Aggarwal, Rupesh Dash, Rajeeb Swain, Punit Prasad, INSACOG Consortium, Ajay Parida |
| EPI_ISL_2341721, EPI_ISL_2341722, EPI_ISL_2341724 | Pt. Jawahar Lal Nehru Memorial Medical College, Raipur | Institute of Life Sciences - INSACOG | Sunil K. Raghav, Safal Walia, Arup Ghosh, Atimukta Jha, Amol M. Kanampalliwar, Omprakash Shiriwas, Sana Fatma, Shifu Aggarwal, Rupesh Dash, Rajeeb Swain, Punit Prasad, INSACOG Consortium, Ajay Parida |
| EPI_ISL_2341727 | SLN Medical College and Hospital, Koraput | Institute of Life Sciences - INSACOG | Sunil K. Raghav, Safal Walia, Arup Ghosh, Atimukta Jha, Amol M. Kanampalliwar, Omprakash Shiriwas, Sana Fatma, Shifu Aggarwal, Rupesh Dash, Rajeeb Swain, Punit Prasad, INSACOG Consortium, Ajay Parida |
| EPI_ISL_2341728 | S.C.B. Medical College and Hospital, Cuttack | Institute of Life Sciences - INSACOG | Sunil K. Raghav, Safal Walia, Arup Ghosh, Atimukta Jha, Amol M. Kanampalliwar, Omprakash Shiriwas, Sana Fatma, Shifu Aggarwal, Rupesh Dash, Rajeeb Swain, Punit Prasad, INSACOG Consortium, Ajay Parida |
| EPI_ISL_2341783, EPI_ISL_2341785 | Institute of Life Sciences, Bhubaneswar | Institute of Life Sciences - INSACOG | Sunil K. Raghav, Safal Walia, Arup Ghosh, Atimukta Jha, Amol M. Kanampalliwar, Omprakash Shiriwas, Sana Fatma, Shifu Aggarwal, Rupesh Dash, Rajeeb Swain, Punit Prasad, INSACOG Consortium, Ajay Parida |
| EPI_ISL_2341788 | Capital Hospital, Bhubaneswar | Institute of Life Sciences - INSACOG | Sunil K. Raghav, Safal Walia, Arup Ghosh, Atimukta Jha, Amol M. Kanampalliwar, Omprakash Shiriwas, Sana Fatma, Shifu Aggarwal, Rupesh Dash, Rajeeb Swain, Punit Prasad, INSACOG Consortium, Ajay Parida |
| EPI_ISL_2341790 | Pt. Jawahar Lal Nehru Memorial Medical College, Raipur | Institute of Life Sciences - INSACOG | Sunil K. Raghav, Safal Walia, Arup Ghosh, Atimukta Jha, Amol M. Kanampalliwar, Omprakash Shiriwas, Sana Fatma, Shifu Aggarwal, Rupesh Dash, Rajeeb Swain, Punit Prasad, INSACOG Consortium, Ajay Parida |
| EPI_ISL_2341795 | S.C.B. Medical College and Hospital, Cuttack | Institute of Life Sciences - INSACOG | Sunil K. Raghav, Safal Walia, Arup Ghosh, Atimukta Jha, Amol M. Kanampalliwar, Omprakash Shiriwas, Sana Fatma, Shifu Aggarwal, Rupesh Dash, Rajeeb Swain, Punit Prasad, INSACOG Consortium, Ajay Parida |
| EPI_ISL_2341806 | Capital Hospital, Bhubaneswar | Institute of Life Sciences - INSACOG | Sunil K. Raghav, Safal Walia, Arup Ghosh, Atimukta Jha, Amol M. Kanampalliwar, Omprakash Shiriwas, Sana Fatma, Shifu Aggarwal, Rupesh Dash, Rajeeb Swain, Punit Prasad, INSACOG Consortium, Ajay Parida |
| EPI_ISL_2341842 | Institute of Life Sciences, Bhubaneswar | Institute of Life Sciences - INSACOG | Sunil K. Raghav, Safal Walia, Arup Ghosh, Atimukta Jha, Amol M. Kanampalliwar, Omprakash Shiriwas, Sana Fatma, Shifu Aggarwal, Rupesh Dash, Rajeeb Swain, Punit Prasad, INSACOG Consortium, Ajay Parida |
| EPI_ISL_2341850 | FM Medical College, Balasore | Institute of Life Sciences - INSACOG | Sunil K. Raghav, Safal Walia, Arup Ghosh, Atimukta Jha, Amol M. Kanampalliwar, Omprakash Shiriwas, Sana Fatma, Shifu Aggarwal, Rupesh Dash, Rajeeb Swain, Punit Prasad, INSACOG Consortium, Ajay Parida |
| EPI_ISL_2341852, EPI_ISL_2341853, EPI_ISL_2341855, EPI_ISL_2341856, EPI_ISL_2341859 | S.C.B. Medical College and Hospital, Cuttack | Institute of Life Sciences - INSACOG | Sunil K. Raghav, Safal Walia, Arup Ghosh, Atimukta Jha, Amol M. Kanampalliwar, Omprakash Shiriwas, Sana Fatma, Shifu Aggarwal, Rupesh Dash, Rajeeb Swain, Punit Prasad, INSACOG Consortium, Ajay Parida |

EPI_ISL_2341862, EPI_ISL_2341863, EPI_ISL_2341864, EPI_ISL_2341866, EPI_ISL_2341867, EPI_ISL_2341869, EPI_ISL_2341871, EPI_ISL_2341872, EPI_ISL_2341875, EPI_ISL_2341876, EPI_ISL_2341880, EPI_ISL_2341881, EPI_ISL_2341883, EPI_ISL_2341885, EPI_ISL_2341887, EPI_ISL_2341889, EPI_ISL_2341890, EPI_ISL_2341891, EPI_ISL_2341892, EPI_ISL_2341896, EPI_ISL_2341897, EPI_ISL_2341898,

EPI_ISL_2341899, EPI_ISL_2341900, EPI_ISL_2341901, EPI_ISL_2341902, EPI_ISL_2341904, EPI_ISL_2341905, EPI_ISL_2341906, EPI_ISL_2341907, EPI_ISL_2341909, EPI_ISL_2341910, EPI_ISL_2341911, EPI_ISL_2341912, EPI_ISL_2341913, EPI_ISL_2341914, EPI_ISL_2341920, EPI_ISL_2341921

| | | | |
|---|---|---|---|
| see above | Pt. Jawahar Lal Nehru Memorial Medical College, Raipur | Institute of Life Sciences - INSACOG | Sunil K. Raghav, Safal Walia, Arup Ghosh, Atimukta Jha, Amol M. Kanampalliwar, Omprakash Shiriwas, Sana Fatma, Shifu Aggarwal, Rupesh Dash, Rajeeb Swain, Punit Prasad, INSACOG Consortium, Ajay Parida |
| EPI_ISL_2341923 | Capital Hospital, Bhubaneswar | Institute of Life Sciences - INSACOG | Sunil K. Raghav, Safal Walia, Arup Ghosh, Atimukta Jha, Amol M. Kanampalliwar, Omprakash Shiriwas, Sana Fatma, Shifu Aggarwal, Rupesh Dash, Rajeeb Swain, Punit Prasad, INSACOG Consortium, Ajay Parida |
| EPI_ISL_2341924, EPI_ISL_2341925, EPI_ISL_2341926 | Institute of Life Sciences, Bhubaneswar | Institute of Life Sciences - INSACOG | Sunil K. Raghav, Safal Walia, Arup Ghosh, Atimukta Jha, Amol M. Kanampalliwar, Omprakash Shiriwas, Sana Fatma, Shifu Aggarwal, Rupesh Dash, Rajeeb Swain, Punit Prasad, INSACOG Consortium, Ajay Parida |
| EPI_ISL_2341931, EPI_ISL_2341932, EPI_ISL_2341934, EPI_ISL_2341935, EPI_ISL_2341936 | FM Medical College, Balasore | Institute of Life Sciences - INSACOG | Sunil K. Raghav, Safal Walia, Arup Ghosh, Atimukta Jha, Amol M. Kanampalliwar, Omprakash Shiriwas, Sana Fatma, Shifu Aggarwal, Rupesh Dash, Rajeeb Swain, Punit Prasad, INSACOG Consortium, Ajay Parida |
| EPI_ISL_2341938 | S.C.B. Medical College and Hospital, Cuttack | Institute of Life Sciences - INSACOG | Sunil K. Raghav, Safal Walia, Arup Ghosh, Atimukta Jha, Amol M. Kanampalliwar, Omprakash Shiriwas, Sana Fatma, Shifu Aggarwal, Rupesh Dash, Rajeeb Swain, Punit Prasad, INSACOG Consortium, Ajay Parida |
| EPI_ISL_2341941, EPI_ISL_2341942 | Institute of Life Sciences, Bhubaneswar | Institute of Life Sciences - INSACOG | Sunil K. Raghav, Safal Walia, Arup Ghosh, Atimukta Jha, Amol M. Kanampalliwar, Omprakash Shiriwas, Sana Fatma, Shifu Aggarwal, Rupesh Dash, Rajeeb Swain, Punit Prasad, INSACOG Consortium, Ajay Parida |
| EPI_ISL_2341993 | Tata Main Hospital, Jharkhand | Institute of Life Sciences - INSACOG | Sunil K. Raghav, Safal Walia, Arup Ghosh, Atimukta Jha, Amol M. Kanampalliwar, Omprakash Shiriwas, Sana Fatma, Shifu Aggarwal, Rupesh Dash, Rajeeb Swain, Punit Prasad, INSACOG Consortium, Ajay Parida |
| EPI_ISL_2341995 | Itki Aarogyashala, Ranchi | Institute of Life Sciences - INSACOG | Sunil K. Raghav, Safal Walia, Arup Ghosh, Atimukta Jha, Amol M. Kanampalliwar, Omprakash Shiriwas, Sana Fatma, Shifu Aggarwal, Rupesh Dash, Rajeeb Swain, Punit Prasad, INSACOG Consortium, Ajay Parida |
| EPI_ISL_2341996 | District Head Quarters Hospital, Nayagarh | Institute of Life Sciences - INSACOG | Sunil K. Raghav, Safal Walia, Arup Ghosh, Atimukta Jha, Amol M. Kanampalliwar, Omprakash Shiriwas, Sana Fatma, Shifu Aggarwal, Rupesh Dash, Rajeeb Swain, Punit Prasad, INSACOG Consortium, Ajay Parida |
| EPI_ISL_2341997 | Tata Main Hospital, Jharkhand | Institute of Life Sciences - INSACOG | Sunil K. Raghav, Safal Walia, Arup Ghosh, Atimukta Jha, Amol M. Kanampalliwar, Omprakash Shiriwas, Sana Fatma, Shifu Aggarwal, Rupesh Dash, Rajeeb Swain, Punit Prasad, INSACOG Consortium, Ajay Parida |
| EPI_ISL_2341998 | Institute of Life Sciences, Bhubaneswar | Institute of Life Sciences - INSACOG | Sunil K. Raghav, Safal Walia, Arup Ghosh, Atimukta Jha, Amol M. Kanampalliwar, Omprakash Shiriwas, Sana Fatma, Shifu Aggarwal, Rupesh Dash, Rajeeb Swain, Punit Prasad, INSACOG Consortium, Ajay Parida |
| EPI_ISL_2342000 | Tata Main Hospital, Jharkhand | Institute of Life Sciences - INSACOG | Sunil K. Raghav, Safal Walia, Arup Ghosh, Atimukta Jha, Amol M. Kanampalliwar, Omprakash Shiriwas, Sana Fatma, Shifu Aggarwal, Rupesh Dash, Rajeeb Swain, Punit Prasad, INSACOG Consortium, Ajay Parida |
| EPI_ISL_2342002 | PRM Medical College & Hospital, Baripada | Institute of Life Sciences - INSACOG | Sunil K. Raghav, Safal Walia, Arup Ghosh, Atimukta Jha, Amol M. Kanampalliwar, Omprakash Shiriwas, Sana Fatma, Shifu Aggarwal, Rupesh Dash, Rajeeb Swain, Punit Prasad, INSACOG Consortium, Ajay Parida |
| EPI_ISL_2342004 | Rajendra Institute of Medical Sciences, Ranchi | Institute of Life Sciences - INSACOG | Sunil K. Raghav, Safal Walia, Arup Ghosh, Atimukta Jha, Amol M. Kanampalliwar, Omprakash Shiriwas, Sana Fatma, Shifu Aggarwal, Rupesh Dash, Rajeeb Swain, Punit Prasad, INSACOG Consortium, Ajay Parida |
| EPI_ISL_2342006 | Sheikh Bhikhari Medical College, Hazaribag | Institute of Life Sciences - INSACOG | Sunil K. Raghav, Safal Walia, Arup Ghosh, Atimukta Jha, Amol M. Kanampalliwar, Omprakash Shiriwas, Sana Fatma, Shifu Aggarwal, Rupesh Dash, Rajeeb Swain, Punit Prasad, INSACOG Consortium, Ajay Parida |
| EPI_ISL_2342007, EPI_ISL_2342009 | Old District Headquarter Hospital, Jharsuguda | Institute of Life Sciences - INSACOG | Sunil K. Raghav, Safal Walia, Arup Ghosh, Atimukta Jha, Amol M. Kanampalliwar, Omprakash Shiriwas, Sana Fatma, Shifu Aggarwal, Rupesh Dash, Rajeeb Swain, Punit Prasad, INSACOG Consortium, Ajay Parida |
| EPI_ISL_2342010 | Sheikh Bhikhari Medical College, Hazaribag | Institute of Life Sciences - INSACOG | Sunil K. Raghav, Safal Walia, Arup Ghosh, Atimukta Jha, Amol M. Kanampalliwar, Omprakash Shiriwas, Sana Fatma, Shifu Aggarwal, Rupesh Dash, Rajeeb Swain, Punit Prasad, INSACOG Consortium, Ajay Parida |
| EPI_ISL_2342011 | FM Medical College, Balasore | Institute of Life Sciences - INSACOG | Sunil K. Raghav, Safal Walia, Arup Ghosh, Atimukta Jha, Amol M. Kanampalliwar, Omprakash Shiriwas, Sana Fatma, Shifu Aggarwal, Rupesh Dash, Rajeeb Swain, Punit Prasad, INSACOG Consortium, Ajay Parida |
| EPI_ISL_2342012 | Tata Main Hospital, Jharkhand | Institute of Life Sciences - INSACOG | Sunil K. Raghav, Safal Walia, Arup Ghosh, Atimukta Jha, Amol M. Kanampalliwar, Omprakash Shiriwas, Sana Fatma, Shifu Aggarwal, Rupesh Dash, Rajeeb Swain, Punit Prasad, INSACOG Consortium, Ajay Parida |
| EPI_ISL_2342013 | Rajendra Institute of Medical Sciences, Ranchi | Institute of Life Sciences - INSACOG | Sunil K. Raghav, Safal Walia, Arup Ghosh, Atimukta Jha, Amol M. Kanampalliwar, Omprakash Shiriwas, Sana Fatma, Shifu Aggarwal, Rupesh Dash, Rajeeb Swain, Punit Prasad, INSACOG Consortium, Ajay Parida |
| EPI_ISL_2342015 | Tata Main Hospital, Jharkhand | Institute of Life Sciences - INSACOG | Sunil K. Raghav, Safal Walia, Arup Ghosh, Atimukta Jha, Amol M. Kanampalliwar, Omprakash Shiriwas, Sana Fatma, Shifu Aggarwal, Rupesh Dash, Rajeeb Swain, Punit Prasad, INSACOG Consortium, Ajay Parida |
| EPI_ISL_2342016 | Sheikh Bhikhari Medical College, Hazaribag | Institute of Life Sciences - INSACOG | Sunil K. Raghav, Safal Walia, Arup Ghosh, Atimukta Jha, Amol M. Kanampalliwar, Omprakash Shiriwas, Sana Fatma, Shifu Aggarwal, Rupesh Dash, Rajeeb Swain, Punit Prasad, INSACOG Consortium, Ajay Parida |
| EPI_ISL_2342017 | Institute of Life Sciences, Bhubaneswar | Institute of Life Sciences - INSACOG | Sunil K. Raghav, Safal Walia, Arup Ghosh, Atimukta Jha, Amol M. Kanampalliwar, Omprakash Shiriwas, Sana Fatma, Shifu Aggarwal, Rupesh Dash, Rajeeb Swain, Punit Prasad, INSACOG Consortium, Ajay Parida |
| EPI_ISL_2342019 | Chief District Medical Office, Cuttack | Institute of Life Sciences - INSACOG | Sunil K. Raghav, Safal Walia, Arup Ghosh, Atimukta Jha, Amol M. Kanampalliwar, Omprakash Shiriwas, Sana Fatma, Shifu Aggarwal, Rupesh Dash, Rajeeb Swain, Punit Prasad, INSACOG Consortium, Ajay Parida |
| EPI_ISL_2342020 | PRM Medical College & Hospital, Baripada | Institute of Life Sciences - INSACOG | Sunil K. Raghav, Safal Walia, Arup Ghosh, Atimukta Jha, Amol M. Kanampalliwar, Omprakash Shiriwas, Sana Fatma, Shifu Aggarwal, Rupesh Dash, Rajeeb Swain, Punit Prasad, INSACOG Consortium, Ajay Parida |
| EPI_ISL_2342021 | Institute of Life Sciences, Bhubaneswar | Institute of Life Sciences - INSACOG | Sunil K. Raghav, Safal Walia, Arup Ghosh, Atimukta Jha, Amol M. Kanampalliwar, Omprakash Shiriwas, Sana Fatma, Shifu Aggarwal, Rupesh Dash, Rajeeb Swain, Punit Prasad, INSACOG Consortium, Ajay Parida |
| EPI_ISL_2342022 | Sheikh Bhikhari Medical College, Hazaribag | Institute of Life Sciences - INSACOG | Sunil K. Raghav, Safal Walia, Arup Ghosh, Atimukta Jha, Amol M. Kanampalliwar, Omprakash Shiriwas, Sana Fatma, Shifu Aggarwal, Rupesh Dash, Rajeeb Swain, Punit Prasad, INSACOG Consortium, Ajay Parida |
| EPI_ISL_2342023 | Old District Headquarter Hospital, Jharsuguda | Institute of Life Sciences - INSACOG | Sunil K. Raghav, Safal Walia, Arup Ghosh, Atimukta Jha, Amol M. Kanampalliwar, Omprakash Shiriwas, Sana Fatma, Shifu Aggarwal, Rupesh Dash, Rajeeb Swain, Punit Prasad, INSACOG Consortium, Ajay Parida |
| EPI_ISL_2342024 | FM Medical College, Balasore | Institute of Life Sciences - INSACOG | Sunil K. Raghav, Safal Walia, Arup Ghosh, Atimukta Jha, Amol M. Kanampalliwar, Omprakash Shiriwas, Sana Fatma, Shifu Aggarwal, Rupesh Dash, Rajeeb Swain, Punit Prasad, INSACOG Consortium, Ajay Parida |
| EPI_ISL_2342025 | Sheikh Bhikhari Medical College, Hazaribag | Institute of Life Sciences - INSACOG | Sunil K. Raghav, Safal Walia, Arup Ghosh, Atimukta Jha, Amol M. Kanampalliwar, Omprakash Shiriwas, Sana Fatma, Shifu Aggarwal, Rupesh Dash, Rajeeb Swain, Punit Prasad, INSACOG Consortium, Ajay Parida |
| EPI_ISL_2342026 | FM Medical College, Balasore | Institute of Life Sciences - INSACOG | Sunil K. Raghav, Safal Walia, Arup Ghosh, Atimukta Jha, Amol M. Kanampalliwar, Omprakash Shiriwas, Sana Fatma, Shifu Aggarwal, Rupesh Dash, Rajeeb Swain, Punit Prasad, INSACOG Consortium, Ajay Parida |
| EPI_ISL_2342027 | Tata Main Hospital, Jharkhand | Institute of Life Sciences - INSACOG | Sunil K. Raghav, Safal Walia, Arup Ghosh, Atimukta Jha, Amol M. Kanampalliwar, Omprakash Shiriwas, Sana Fatma, Shifu Aggarwal, Rupesh Dash, Rajeeb Swain, Punit Prasad, INSACOG Consortium, Ajay Parida |
| EPI_ISL_2342028 | Institute of Life Sciences, Bhubaneswar | Institute of Life Sciences - INSACOG | Sunil K. Raghav, Safal Walia, Arup Ghosh, Atimukta Jha, Amol M. Kanampalliwar, Omprakash Shiriwas, Sana Fatma, Shifu Aggarwal, Rupesh Dash, Rajeeb Swain, Punit Prasad, INSACOG Consortium, Ajay Parida |
| EPI_ISL_2342029 | Tata Main Hospital, Jharkhand | Institute of Life Sciences - INSACOG | Sunil K. Raghav, Safal Walia, Arup Ghosh, Atimukta Jha, Amol M. Kanampalliwar, Omprakash Shiriwas, Sana Fatma, Shifu Aggarwal, Rupesh Dash, Rajeeb Swain, Punit Prasad, INSACOG Consortium, Ajay Parida |
| EPI_ISL_2342032 | Itki Aarogyashala, Ranchi | Institute of Life Sciences - INSACOG | Sunil K. Raghav, Safal Walia, Arup Ghosh, Atimukta Jha, Amol M. Kanampalliwar, Omprakash Shiriwas, Sana Fatma, Shifu Aggarwal, Rupesh Dash, Rajeeb Swain, Punit Prasad, INSACOG Consortium, Ajay Parida |
| EPI_ISL_2342033 | Tata Main Hospital, Jharkhand | Institute of Life Sciences - INSACOG | Sunil K. Raghav, Safal Walia, Arup Ghosh, Atimukta Jha, Amol M. Kanampalliwar, Omprakash Shiriwas, Sana Fatma, Shifu Aggarwal, Rupesh Dash, Rajeeb Swain, Punit Prasad, INSACOG Consortium, Ajay Parida |
| EPI_ISL_2342034 | Old District Headquarter Hospital, Jharsuguda | Institute of Life Sciences - INSACOG | Sunil K. Raghav, Safal Walia, Arup Ghosh, Atimukta Jha, Amol M. Kanampalliwar, Omprakash Shiriwas, Sana Fatma, Shifu Aggarwal, Rupesh Dash, Rajeeb Swain, Punit Prasad, INSACOG Consortium, Ajay Parida |
| EPI_ISL_2342035 | FM Medical College, Balasore | Institute of Life Sciences - INSACOG | Sunil K. Raghav, Safal Walia, Arup Ghosh, Atimukta Jha, Amol M. Kanampalliwar, Omprakash Shiriwas, Sana Fatma, Shifu Aggarwal, Rupesh Dash, Rajeeb Swain, Punit Prasad, INSACOG Consortium, Ajay Parida |
| EPI_ISL_2342036 | Tata Main Hospital, Jharkhand | Institute of Life Sciences - INSACOG | Sunil K. Raghav, Safal Walia, Arup Ghosh, Atimukta Jha, Amol M. Kanampalliwar, Omprakash Shiriwas, Sana Fatma, Shifu Aggarwal, Rupesh Dash, Rajeeb Swain, Punit Prasad, INSACOG Consortium, Ajay Parida |
| EPI_ISL_2342037 | Itki Aarogyashala, Ranchi | Institute of Life Sciences - INSACOG | Sunil K. Raghav, Safal Walia, Arup Ghosh, Atimukta Jha, Amol M. Kanampalliwar, Omprakash Shiriwas, Sana Fatma, Shifu Aggarwal, Rupesh Dash, Rajeeb Swain, Punit Prasad, INSACOG Consortium, Ajay Parida |
| EPI_ISL_2342038 | District Headquarter Hospital, Puri | Institute of Life Sciences - INSACOG | Sunil K. Raghav, Safal Walia, Arup Ghosh, Atimukta Jha, Amol M. Kanampalliwar, Omprakash Shiriwas, Sana Fatma, Shifu Aggarwal, Rupesh Dash, Rajeeb Swain, Punit Prasad, INSACOG Consortium, Ajay Parida |
| EPI_ISL_2342040 | Tata Main Hospital, Jharkhand | Institute of Life Sciences - INSACOG | Sunil K. Raghav, Safal Walia, Arup Ghosh, Atimukta Jha, Amol M. Kanampalliwar, Omprakash Shiriwas, Sana Fatma, Shifu Aggarwal, Rupesh Dash, Rajeeb Swain, Punit Prasad, INSACOG Consortium, Ajay Parida |
| EPI_ISL_2342043 | Sheikh Bhikhari Medical College, Hazaribag | Institute of Life Sciences - INSACOG | Sunil K. Raghav, Safal Walia, Arup Ghosh, Atimukta Jha, Amol M. Kanampalliwar, Omprakash Shiriwas, Sana Fatma, Shifu Aggarwal, Rupesh Dash, Rajeeb Swain, Punit Prasad, INSACOG Consortium, Ajay Parida |
| EPI_ISL_2342044 | Old District Headquarter Hospital, Jharsuguda | Institute of Life Sciences - INSACOG | Sunil K. Raghav, Safal Walia, Arup Ghosh, Atimukta Jha, Amol M. Kanampalliwar, Omprakash Shiriwas, Sana Fatma, Shifu Aggarwal, Rupesh Dash, Rajeeb Swain, Punit Prasad, INSACOG Consortium, Ajay Parida |
| EPI_ISL_2342045 | Sheikh Bhikhari Medical College, Hazaribag | Institute of Life Sciences - INSACOG | Sunil K. Raghav, Safal Walia, Arup Ghosh, Atimukta Jha, Amol M. Kanampalliwar, Omprakash Shiriwas, Sana Fatma, Shifu Aggarwal, Rupesh Dash, Rajeeb Swain, Punit Prasad, INSACOG Consortium, Ajay Parida |
| EPI_ISL_2342046 | Mahatma Gandhi Memorial Medical College & Hospital Jamshedpur | Institute of Life Sciences - INSACOG | Sunil K. Raghav, Safal Walia, Arup Ghosh, Atimukta Jha, Amol M. Kanampalliwar, Omprakash Shiriwas, Sana Fatma, Shifu Aggarwal, Rupesh Dash, Rajeeb Swain, Punit Prasad, INSACOG Consortium, Ajay Parida |
| EPI_ISL_2342047 | District Head Quarters Hospital, Nayagarh | Institute of Life Sciences - INSACOG | Sunil K. Raghav, Safal Walia, Arup Ghosh, Atimukta Jha, Amol M. Kanampalliwar, Omprakash Shiriwas, Sana Fatma, Shifu Aggarwal, Rupesh Dash, Rajeeb Swain, Punit Prasad, INSACOG Consortium, Ajay Parida |
| EPI_ISL_2342049 | Mahatma Gandhi Memorial Medical College & Hospital Jamshedpur | Institute of Life Sciences - INSACOG | Sunil K. Raghav, Safal Walia, Arup Ghosh, Atimukta Jha, Amol M. Kanampalliwar, Omprakash Shiriwas, Sana Fatma, Shifu Aggarwal, Rupesh Dash, Rajeeb Swain, Punit Prasad, INSACOG Consortium, Ajay Parida |
| EPI_ISL_2342050 | Old District Headquarter Hospital, Jharsuguda | Institute of Life Sciences - INSACOG | Sunil K. Raghav, Safal Walia, Arup Ghosh, Atimukta Jha, Amol M. Kanampalliwar, Omprakash Shiriwas, Sana Fatma, Shifu Aggarwal, Rupesh Dash, Rajeeb Swain, Punit Prasad, INSACOG Consortium, Ajay Parida |
| EPI_ISL_2342051 | Mahatma Gandhi Memorial Medical College & Hospital Jamshedpur | Institute of Life Sciences - INSACOG | Sunil K. Raghav, Safal Walia, Arup Ghosh, Atimukta Jha, Amol M. Kanampalliwar, Omprakash Shiriwas, Sana Fatma, Shifu Aggarwal, Rupesh Dash, Rajeeb Swain, Punit Prasad, INSACOG Consortium, Ajay Parida |
| EPI_ISL_2342052 | FM Medical College, Balasore | Institute of Life Sciences - INSACOG | Sunil K. Raghav, Safal Walia, Arup Ghosh, Atimukta Jha, Amol M. Kanampalliwar, Omprakash Shiriwas, Sana Fatma, Shifu Aggarwal, Rupesh Dash, Rajeeb Swain, Punit Prasad, INSACOG Consortium, Ajay Parida |
| EPI_ISL_2342053 | Itki Aarogyashala, Ranchi | Institute of Life Sciences - INSACOG | Sunil K. Raghav, Safal Walia, Arup Ghosh, Atimukta Jha, Amol M. Kanampalliwar, Omprakash Shiriwas, Sana Fatma, Shifu Aggarwal, Rupesh Dash, Rajeeb Swain, Punit Prasad, INSACOG Consortium, Ajay Parida |
| EPI_ISL_2342054 | Rajendra Institute of Medical Sciences, Ranchi | Institute of Life Sciences - INSACOG | Sunil K. Raghav, Safal Walia, Arup Ghosh, Atimukta Jha, Amol M. Kanampalliwar, Omprakash Shiriwas, Sana Fatma, Shifu Aggarwal, Rupesh Dash, Rajeeb Swain, Punit Prasad, INSACOG Consortium, Ajay Parida |
| EPI_ISL_2342055 | Sheikh Bhikhari Medical College, Hazaribag | Institute of Life Sciences - INSACOG | Sunil K. Raghav, Safal Walia, Arup Ghosh, Atimukta Jha, Amol M. Kanampalliwar, Omprakash Shiriwas, Sana Fatma, Shifu Aggarwal, Rupesh Dash, Rajeeb Swain, Punit Prasad, INSACOG Consortium, Ajay Parida |
| EPI_ISL_2342056 | FM Medical College, Balasore | Institute of Life Sciences - INSACOG | Sunil K. Raghav, Safal Walia, Arup Ghosh, Atimukta Jha, Amol M. Kanampalliwar, Omprakash Shiriwas, Sana Fatma, Shifu Aggarwal, Rupesh Dash, Rajeeb Swain, Punit Prasad, INSACOG Consortium, Ajay Parida |
| EPI_ISL_2342058 | PRM Medical College & Hospital, Baripada | Institute of Life Sciences - INSACOG | Sunil K. Raghav, Safal Walia, Arup Ghosh, Atimukta Jha, Amol M. Kanampalliwar, Omprakash Shiriwas, Sana Fatma, Shifu Aggarwal, Rupesh Dash, Rajeeb Swain, Punit Prasad, INSACOG Consortium, Ajay Parida |
| EPI_ISL_2342061 | Tata Main Hospital, Jharkhand | Institute of Life Sciences - INSACOG | Sunil K. Raghav, Safal Walia, Arup Ghosh, Atimukta Jha, Amol M. Kanampalliwar, Omprakash Shiriwas, Sana Fatma, Shifu Aggarwal, Rupesh Dash, Rajeeb Swain, Punit Prasad, INSACOG Consortium, Ajay Parida |
| EPI_ISL_2342062 | Institute of Life Sciences, Bhubaneswar | Institute of Life Sciences - INSACOG | Sunil K. Raghav, Safal Walia, Arup Ghosh, Atimukta Jha, Amol M. Kanampalliwar, Omprakash Shiriwas, Sana Fatma, Shifu Aggarwal, Rupesh Dash, Rajeeb Swain, Punit Prasad, INSACOG Consortium, Ajay Parida |
| EPI_ISL_2342063 | Mahatma Gandhi Memorial Medical College & Hospital Jamshedpur | Institute of Life Sciences - INSACOG | Sunil K. Raghav, Safal Walia, Arup Ghosh, Atimukta Jha, Amol M. Kanampalliwar, Omprakash Shiriwas, Sana Fatma, Shifu Aggarwal, Rupesh Dash, Rajeeb Swain, Punit Prasad, INSACOG Consortium, Ajay Parida |
| EPI_ISL_2342064 | Rajendra Institute of Medical Sciences, Ranchi | Institute of Life Sciences - INSACOG | Sunil K. Raghav, Safal Walia, Arup Ghosh, Atimukta Jha, Amol M. Kanampalliwar, Omprakash Shiriwas, Sana Fatma, Shifu Aggarwal, Rupesh Dash, Rajeeb Swain, Punit Prasad, INSACOG Consortium, Ajay Parida |
| EPI_ISL_2342065 | District Headquarter Hospital, Puri | Institute of Life Sciences - INSACOG | Sunil K. Raghav, Safal Walia, Arup Ghosh, Atimukta Jha, Amol M. Kanampalliwar, Omprakash Shiriwas, Sana Fatma, Shifu Aggarwal, Rupesh Dash, Rajeeb Swain, Punit Prasad, INSACOG Consortium, Ajay Parida |
| EPI_ISL_2342066, EPI_ISL_2342067 | Rajendra Institute of Medical Sciences, Ranchi | Institute of Life Sciences - INSACOG | Sunil K. Raghav, Safal Walia, Arup Ghosh, Atimukta Jha, Amol M. Kanampalliwar, Omprakash Shiriwas, Sana Fatma, Shifu Aggarwal, Rupesh Dash, Rajeeb Swain, Punit Prasad, INSACOG Consortium, Ajay Parida |
| EPI_ISL_2342068 | Tata Main Hospital, Jharkhand | Institute of Life Sciences - INSACOG | Sunil K. Raghav, Safal Walia, Arup Ghosh, Atimukta Jha, Amol M. Kanampalliwar, Omprakash Shiriwas, Sana Fatma, Shifu Aggarwal, Rupesh Dash, Rajeeb Swain, Punit Prasad, INSACOG Consortium, Ajay Parida |
| EPI_ISL_2342069 | Mahatma Gandhi Memorial Medical College & Hospital Jamshedpur | Institute of Life Sciences - INSACOG | Sunil K. Raghav, Safal Walia, Arup Ghosh, Atimukta Jha, Amol M. Kanampalliwar, Omprakash Shiriwas, Sana Fatma, Shifu Aggarwal, Rupesh Dash, Rajeeb Swain, Punit Prasad, INSACOG Consortium, Ajay Parida |
| EPI_ISL_2342070 | District Headquarter Hospital, Puri | Institute of Life Sciences - INSACOG | Sunil K. Raghav, Safal Walia, Arup Ghosh, Atimukta Jha, Amol M. Kanampalliwar, Omprakash Shiriwas, Sana Fatma, Shifu Aggarwal, Rupesh Dash, Rajeeb Swain, Punit Prasad, INSACOG Consortium, Ajay Parida |
| EPI_ISL_2342071 | Itki Aarogyashala, Ranchi | Institute of Life Sciences - INSACOG | Sunil K. Raghav, Safal Walia, Arup Ghosh, Atimukta Jha, Amol M. Kanampalliwar, Omprakash Shiriwas, Sana Fatma, Shifu Aggarwal, Rupesh Dash, Rajeeb Swain, Punit Prasad, INSACOG Consortium, Ajay Parida |

| | | | |
|---|---|---|---|
| EPI_ISL_2342072 | Institute of Life Sciences, Bhubaneswar | Institute of Life Sciences - INSACOG | Sunil K. Raghav, Safal Walia, Arup Ghosh, Atimukta Jha, Amol M. Kanampalliwar, Omprakash Shiriwas, Sana Fatma, Shifu Aggarwal, Rupesh Dash, Rajeeb Swain, Punit Prasad, INSACOG Consortium, Ajay Parida |
| EPI_ISL_2342073 | District Headquarter Hospital, Puri | Institute of Life Sciences - INSACOG | Sunil K. Raghav, Safal Walia, Arup Ghosh, Atimukta Jha, Amol M. Kanampalliwar, Omprakash Shiriwas, Sana Fatma, Shifu Aggarwal, Rupesh Dash, Rajeeb Swain, Punit Prasad, INSACOG Consortium, Ajay Parida |
| EPI_ISL_2342074 | Institute of Life Sciences, Bhubaneswar | Institute of Life Sciences - INSACOG | Sunil K. Raghav, Safal Walia, Arup Ghosh, Atimukta Jha, Amol M. Kanampalliwar, Omprakash Shiriwas, Sana Fatma, Shifu Aggarwal, Rupesh Dash, Rajeeb Swain, Punit Prasad, INSACOG Consortium, Ajay Parida |
| EPI_ISL_2342075 | Mahatma Gandhi Memorial Medical College & Hospital Jamshedpur | Institute of Life Sciences - INSACOG | Sunil K. Raghav, Safal Walia, Arup Ghosh, Atimukta Jha, Amol M. Kanampalliwar, Omprakash Shiriwas, Sana Fatma, Shifu Aggarwal, Rupesh Dash, Rajeeb Swain, Punit Prasad, INSACOG Consortium, Ajay Parida |
| EPI_ISL_2342077, EPI_ISL_2342078 | Tata Main Hospital, Jharkhand | Institute of Life Sciences - INSACOG | Sunil K. Raghav, Safal Walia, Arup Ghosh, Atimukta Jha, Amol M. Kanampalliwar, Omprakash Shiriwas, Sana Fatma, Shifu Aggarwal, Rupesh Dash, Rajeeb Swain, Punit Prasad, INSACOG Consortium, Ajay Parida |
| EPI_ISL_2342079 | District Headquarter Hospital, Puri | Institute of Life Sciences - INSACOG | Sunil K. Raghav, Safal Walia, Arup Ghosh, Atimukta Jha, Amol M. Kanampalliwar, Omprakash Shiriwas, Sana Fatma, Shifu Aggarwal, Rupesh Dash, Rajeeb Swain, Punit Prasad, INSACOG Consortium, Ajay Parida |
| EPI_ISL_2342080 | FM Medical College, Balasore | Institute of Life Sciences - INSACOG | Sunil K. Raghav, Safal Walia, Arup Ghosh, Atimukta Jha, Amol M. Kanampalliwar, Omprakash Shiriwas, Sana Fatma, Shifu Aggarwal, Rupesh Dash, Rajeeb Swain, Punit Prasad, INSACOG Consortium, Ajay Parida |
| EPI_ISL_2342081 | Itki Aarogyashala, Ranchi | Institute of Life Sciences - INSACOG | Sunil K. Raghav, Safal Walia, Arup Ghosh, Atimukta Jha, Amol M. Kanampalliwar, Omprakash Shiriwas, Sana Fatma, Shifu Aggarwal, Rupesh Dash, Rajeeb Swain, Punit Prasad, INSACOG Consortium, Ajay Parida |
| EPI_ISL_2342083 | PRM Medical College & Hospital, Baripada | Institute of Life Sciences - INSACOG | Sunil K. Raghav, Safal Walia, Arup Ghosh, Atimukta Jha, Amol M. Kanampalliwar, Omprakash Shiriwas, Sana Fatma, Shifu Aggarwal, Rupesh Dash, Rajeeb Swain, Punit Prasad, INSACOG Consortium, Ajay Parida |
| EPI_ISL_2342084 | Itki Aarogyashala, Ranchi | Institute of Life Sciences - INSACOG | Sunil K. Raghav, Safal Walia, Arup Ghosh, Atimukta Jha, Amol M. Kanampalliwar, Omprakash Shiriwas, Sana Fatma, Shifu Aggarwal, Rupesh Dash, Rajeeb Swain, Punit Prasad, INSACOG Consortium, Ajay Parida |
| EPI_ISL_2342086 | District Headquarter Hospital, Puri | Institute of Life Sciences - INSACOG | Sunil K. Raghav, Safal Walia, Arup Ghosh, Atimukta Jha, Amol M. Kanampalliwar, Omprakash Shiriwas, Sana Fatma, Shifu Aggarwal, Rupesh Dash, Rajeeb Swain, Punit Prasad, INSACOG Consortium, Ajay Parida |
| EPI_ISL_2342089 | Tata Main Hospital, Jharkhand | Institute of Life Sciences - INSACOG | Sunil K. Raghav, Safal Walia, Arup Ghosh, Atimukta Jha, Amol M. Kanampalliwar, Omprakash Shiriwas, Sana Fatma, Shifu Aggarwal, Rupesh Dash, Rajeeb Swain, Punit Prasad, INSACOG Consortium, Ajay Parida |
| EPI_ISL_2342090 | Itki Aarogyashala, Ranchi | Institute of Life Sciences - INSACOG | Sunil K. Raghav, Safal Walia, Arup Ghosh, Atimukta Jha, Amol M. Kanampalliwar, Omprakash Shiriwas, Sana Fatma, Shifu Aggarwal, Rupesh Dash, Rajeeb Swain, Punit Prasad, INSACOG Consortium, Ajay Parida |
| EPI_ISL_2342091, EPI_ISL_2342092 | Sheikh Bhikhari Medical College, Hazaribag | Institute of Life Sciences - INSACOG | Sunil K. Raghav, Safal Walia, Arup Ghosh, Atimukta Jha, Amol M. Kanampalliwar, Omprakash Shiriwas, Sana Fatma, Shifu Aggarwal, Rupesh Dash, Rajeeb Swain, Punit Prasad, INSACOG Consortium, Ajay Parida |
| EPI_ISL_2342093 | Tata Main Hospital, Jharkhand | Institute of Life Sciences - INSACOG | Sunil K. Raghav, Safal Walia, Arup Ghosh, Atimukta Jha, Amol M. Kanampalliwar, Omprakash Shiriwas, Sana Fatma, Shifu Aggarwal, Rupesh Dash, Rajeeb Swain, Punit Prasad, INSACOG Consortium, Ajay Parida |
| EPI_ISL_2342094 | Itki Aarogyashala, Ranchi | Institute of Life Sciences - INSACOG | Sunil K. Raghav, Safal Walia, Arup Ghosh, Atimukta Jha, Amol M. Kanampalliwar, Omprakash Shiriwas, Sana Fatma, Shifu Aggarwal, Rupesh Dash, Rajeeb Swain, Punit Prasad, INSACOG Consortium, Ajay Parida |
| EPI_ISL_2342100 | District Headquarters Hospital, Deogarh | Institute of Life Sciences - INSACOG | Sunil K. Raghav, Safal Walia, Arup Ghosh, Atimukta Jha, Amol M. Kanampalliwar, Omprakash Shiriwas, Sana Fatma, Shifu Aggarwal, Rupesh Dash, Rajeeb Swain, Punit Prasad, INSACOG Consortium, Ajay Parida |
| EPI_ISL_2342101 | Itki Aarogyashala, Ranchi | Institute of Life Sciences - INSACOG | Sunil K. Raghav, Safal Walia, Arup Ghosh, Atimukta Jha, Amol M. Kanampalliwar, Omprakash Shiriwas, Sana Fatma, Shifu Aggarwal, Rupesh Dash, Rajeeb Swain, Punit Prasad, INSACOG Consortium, Ajay Parida |
| EPI_ISL_2342107 | Tata Main Hospital, Jharkhand | Institute of Life Sciences - INSACOG | Sunil K. Raghav, Safal Walia, Arup Ghosh, Atimukta Jha, Amol M. Kanampalliwar, Omprakash Shiriwas, Sana Fatma, Shifu Aggarwal, Rupesh Dash, Rajeeb Swain, Punit Prasad, INSACOG Consortium, Ajay Parida |
| EPI_ISL_2342111 | Old District Headquarter Hospital, Jharsuguda | Institute of Life Sciences - INSACOG | Sunil K. Raghav, Safal Walia, Arup Ghosh, Atimukta Jha, Amol M. Kanampalliwar, Omprakash Shiriwas, Sana Fatma, Shifu Aggarwal, Rupesh Dash, Rajeeb Swain, Punit Prasad, INSACOG Consortium, Ajay Parida |
| EPI_ISL_2342115 | Institute of Life Sciences, Bhubaneswar | Institute of Life Sciences - INSACOG | Sunil K. Raghav, Safal Walia, Arup Ghosh, Atimukta Jha, Amol M. Kanampalliwar, Omprakash Shiriwas, Sana Fatma, Shifu Aggarwal, Rupesh Dash, Rajeeb Swain, Punit Prasad, INSACOG Consortium, Ajay Parida |
| EPI_ISL_2342116 | Itki Aarogyashala, Ranchi | Institute of Life Sciences - INSACOG | Sunil K. Raghav, Safal Walia, Arup Ghosh, Atimukta Jha, Amol M. Kanampalliwar, Omprakash Shiriwas, Sana Fatma, Shifu Aggarwal, Rupesh Dash, Rajeeb Swain, Punit Prasad, INSACOG Consortium, Ajay Parida |
| EPI_ISL_2342117, EPI_ISL_2342118 | Tata Main Hospital, Jharkhand | Institute of Life Sciences - INSACOG | Sunil K. Raghav, Safal Walia, Arup Ghosh, Atimukta Jha, Amol M. Kanampalliwar, Omprakash Shiriwas, Sana Fatma, Shifu Aggarwal, Rupesh Dash, Rajeeb Swain, Punit Prasad, INSACOG Consortium, Ajay Parida |
| EPI_ISL_2342119 | Itki Aarogyashala, Ranchi | Institute of Life Sciences - INSACOG | Sunil K. Raghav, Safal Walia, Arup Ghosh, Atimukta Jha, Amol M. Kanampalliwar, Omprakash Shiriwas, Sana Fatma, Shifu Aggarwal, Rupesh Dash, Rajeeb Swain, Punit Prasad, INSACOG Consortium, Ajay Parida |
| EPI_ISL_2342120 | Institute of Life Sciences, Bhubaneswar | Institute of Life Sciences - INSACOG | Sunil K. Raghav, Safal Walia, Arup Ghosh, Atimukta Jha, Amol M. Kanampalliwar, Omprakash Shiriwas, Sana Fatma, Shifu Aggarwal, Rupesh Dash, Rajeeb Swain, Punit Prasad, INSACOG Consortium, Ajay Parida |
| EPI_ISL_2342121 | Sheikh Bhikhari Medical College, Hazaribag | Institute of Life Sciences - INSACOG | Sunil K. Raghav, Safal Walia, Arup Ghosh, Atimukta Jha, Amol M. Kanampalliwar, Omprakash Shiriwas, Sana Fatma, Shifu Aggarwal, Rupesh Dash, Rajeeb Swain, Punit Prasad, INSACOG Consortium, Ajay Parida |
| EPI_ISL_2342123 | Tata Main Hospital, Jharkhand | Institute of Life Sciences - INSACOG | Sunil K. Raghav, Safal Walia, Arup Ghosh, Atimukta Jha, Amol M. Kanampalliwar, Omprakash Shiriwas, Sana Fatma, Shifu Aggarwal, Rupesh Dash, Rajeeb Swain, Punit Prasad, INSACOG Consortium, Ajay Parida |
| EPI_ISL_2342125 | Sheikh Bhikhari Medical College, Hazaribag | Institute of Life Sciences - INSACOG | Sunil K. Raghav, Safal Walia, Arup Ghosh, Atimukta Jha, Amol M. Kanampalliwar, Omprakash Shiriwas, Sana Fatma, Shifu Aggarwal, Rupesh Dash, Rajeeb Swain, Punit Prasad, INSACOG Consortium, Ajay Parida |
| EPI_ISL_2342126 | Itki Aarogyashala, Ranchi | Institute of Life Sciences - INSACOG | Sunil K. Raghav, Safal Walia, Arup Ghosh, Atimukta Jha, Amol M. Kanampalliwar, Omprakash Shiriwas, Sana Fatma, Shifu Aggarwal, Rupesh Dash, Rajeeb Swain, Punit Prasad, INSACOG Consortium, Ajay Parida |
| EPI_ISL_2342127 | Tata Main Hospital, Jharkhand | Institute of Life Sciences - INSACOG | Sunil K. Raghav, Safal Walia, Arup Ghosh, Atimukta Jha, Amol M. Kanampalliwar, Omprakash Shiriwas, Sana Fatma, Shifu Aggarwal, Rupesh Dash, Rajeeb Swain, Punit Prasad, INSACOG Consortium, Ajay Parida |
| EPI_ISL_2342128 | Chief District Medical Office, Cuttack | Institute of Life Sciences - INSACOG | Sunil K. Raghav, Safal Walia, Arup Ghosh, Atimukta Jha, Amol M. Kanampalliwar, Omprakash Shiriwas, Sana Fatma, Shifu Aggarwal, Rupesh Dash, Rajeeb Swain, Punit Prasad, INSACOG Consortium, Ajay Parida |
| EPI_ISL_2342242, EPI_ISL_2342249 | Tata Main Hospital, Jharkhand | Institute of Life Sciences - INSACOG | Sunil K. Raghav, Safal Walia, Arup Ghosh, Atimukta Jha, Amol M. Kanampalliwar, Omprakash Shiriwas, Sana Fatma, Shifu Aggarwal, Rupesh Dash, Rajeeb Swain, Punit Prasad, INSACOG Consortium, Ajay Parida |
| EPI_ISL_2342284 | Institute of Life Sciences, Bhubaneswar | Institute of Life Sciences - INSACOG | Sunil K. Raghav, Safal Walia, Arup Ghosh, Atimukta Jha, Amol M. Kanampalliwar, Omprakash Shiriwas, Sana Fatma, Shifu Aggarwal, Rupesh Dash, Rajeeb Swain, Punit Prasad, INSACOG Consortium, Ajay Parida |
| EPI_ISL_2342286 | District Headquarters Hospital, Deogarh | Institute of Life Sciences - INSACOG | Sunil K. Raghav, Safal Walia, Arup Ghosh, Atimukta Jha, Amol M. Kanampalliwar, Omprakash Shiriwas, Sana Fatma, Shifu Aggarwal, Rupesh Dash, Rajeeb Swain, Punit Prasad, INSACOG Consortium, Ajay Parida |
| EPI_ISL_2342293, EPI_ISL_2342294, EPI_ISL_2342295, EPI_ISL_2342296, EPI_ISL_2342297 | Mahatma Gandhi Memorial Medical College & Hospital Jamshedpur | Institute of Life Sciences - INSACOG | Sunil K. Raghav, Safal Walia, Arup Ghosh, Atimukta Jha, Amol M. Kanampalliwar, Omprakash Shiriwas, Sana Fatma, Shifu Aggarwal, Rupesh Dash, Rajeeb Swain, Punit Prasad, INSACOG Consortium, Ajay Parida |
| EPI_ISL_2342298 | Rajendra Institute of Medical Sciences, Ranchi | Institute of Life Sciences - INSACOG | Sunil K. Raghav, Safal Walia, Arup Ghosh, Atimukta Jha, Amol M. Kanampalliwar, Omprakash Shiriwas, Sana Fatma, Shifu Aggarwal, Rupesh Dash, Rajeeb Swain, Punit Prasad, INSACOG Consortium, Ajay Parida |
| EPI_ISL_2342299, EPI_ISL_2342300, EPI_ISL_2342301, EPI_ISL_2342302, EPI_ISL_2342303, EPI_ISL_2342304, EPI_ISL_2342305, EPI_ISL_2342306 | District Headquarter Hospital, Puri | Institute of Life Sciences - INSACOG | Sunil K. Raghav, Safal Walia, Arup Ghosh, Atimukta Jha, Amol M. Kanampalliwar, Omprakash Shiriwas, Sana Fatma, Shifu Aggarwal, Rupesh Dash, Rajeeb Swain, Punit Prasad, INSACOG Consortium, Ajay Parida |
| EPI_ISL_2342307 | PRM Medical College & Hospital, Baripada | Institute of Life Sciences - INSACOG | Sunil K. Raghav, Safal Walia, Arup Ghosh, Atimukta Jha, Amol M. Kanampalliwar, Omprakash Shiriwas, Sana Fatma, Shifu Aggarwal, Rupesh Dash, Rajeeb Swain, Punit Prasad, INSACOG Consortium, Ajay Parida |
| EPI_ISL_2342308, EPI_ISL_2342309, EPI_ISL_2342310, EPI_ISL_2342311, EPI_ISL_2342312, EPI_ISL_2342313, EPI_ISL_2342314, EPI_ISL_2342315, EPI_ISL_2342316, EPI_ISL_2342317, EPI_ISL_2342318, EPI_ISL_2342319, EPI_ISL_2342320, EPI_ISL_2342321, EPI_ISL_2342324, EPI_ISL_2342325, EPI_ISL_2342326, EPI_ISL_2342327 | | | |
| see above | Sheikh Bhikhari Medical College, Hazaribag | Institute of Life Sciences - INSACOG | Sunil K. Raghav, Safal Walia, Arup Ghosh, Atimukta Jha, Amol M. Kanampalliwar, Omprakash Shiriwas, Sana Fatma, Shifu Aggarwal, Rupesh Dash, Rajeeb Swain, Punit Prasad, INSACOG Consortium, Ajay Parida |
| EPI_ISL_2342329, EPI_ISL_2342331, EPI_ISL_2342332, EPI_ISL_2342333, EPI_ISL_2342334, EPI_ISL_2342335, EPI_ISL_2342336, EPI_ISL_2342337, EPI_ISL_2342338, EPI_ISL_2342340, EPI_ISL_2342343, EPI_ISL_2342344, EPI_ISL_2342345, EPI_ISL_2342346, EPI_ISL_2342347, EPI_ISL_2342348, EPI_ISL_2342349, EPI_ISL_2342350, EPI_ISL_2342351, EPI_ISL_2342352, EPI_ISL_2342353 | | | |
| see above | Tata Main Hospital, Jharkhand | Institute of Life Sciences - INSACOG | Sunil K. Raghav, Safal Walia, Arup Ghosh, Atimukta Jha, Amol M. Kanampalliwar, Omprakash Shiriwas, Sana Fatma, Shifu Aggarwal, Rupesh Dash, Rajeeb Swain, Punit Prasad, INSACOG Consortium, Ajay Parida |
| EPI_ISL_2342354 | Chief District Medical Office, Cuttack | Institute of Life Sciences - INSACOG | Sunil K. Raghav, Safal Walia, Arup Ghosh, Atimukta Jha, Amol M. Kanampalliwar, Omprakash Shiriwas, Sana Fatma, Shifu Aggarwal, Rupesh Dash, Rajeeb Swain, Punit Prasad, INSACOG Consortium, Ajay Parida |
| EPI_ISL_2342355 | Itki Aarogyashala, Ranchi | Institute of Life Sciences - INSACOG | Sunil K. Raghav, Safal Walia, Arup Ghosh, Atimukta Jha, Amol M. Kanampalliwar, Omprakash Shiriwas, Sana Fatma, Shifu Aggarwal, Rupesh Dash, Rajeeb Swain, Punit Prasad, INSACOG Consortium, Ajay Parida |
| EPI_ISL_2342357 | District Headquarters Hospital, Deogarh | Institute of Life Sciences - INSACOG | Sunil K. Raghav, Safal Walia, Arup Ghosh, Atimukta Jha, Amol M. Kanampalliwar, Omprakash Shiriwas, Sana Fatma, Shifu Aggarwal, Rupesh Dash, Rajeeb Swain, Punit Prasad, INSACOG Consortium, Ajay Parida |
| EPI_ISL_2342358 | Old District Headquarter Hospital, Jharsuguda | Institute of Life Sciences - INSACOG | Sunil K. Raghav, Safal Walia, Arup Ghosh, Atimukta Jha, Amol M. Kanampalliwar, Omprakash Shiriwas, Sana Fatma, Shifu Aggarwal, Rupesh Dash, Rajeeb Swain, Punit Prasad, INSACOG Consortium, Ajay Parida |
| EPI_ISL_2342359, EPI_ISL_2342360, EPI_ISL_2342361, EPI_ISL_2342362, EPI_ISL_2342363 | Institute of Life Sciences, Bhubaneswar | Institute of Life Sciences - INSACOG | Sunil K. Raghav, Safal Walia, Arup Ghosh, Atimukta Jha, Amol M. Kanampalliwar, Omprakash Shiriwas, Sana Fatma, Shifu Aggarwal, Rupesh Dash, Rajeeb Swain, Punit Prasad, INSACOG Consortium, Ajay Parida |
| EPI_ISL_2342514 | Institute for Infectious Diseases, University of Bern, Switzerland | Institute for Infectious Diseases, University of Bern, Switzerland | Stefan Neuenschwander, Christian Baumann, Miguel A Terrazos Miani, Cora Sägesser, Pascal Bittel, Peter Keller, Franziska Suter-Riniker, Stephen L Leib, Alban Ramette |
| EPI_ISL_2342604 | Institute of Life Sciences, Bhubaneswar | Institute of Life Sciences - INSACOG | Sunil K. Raghav, Safal Walia, Arup Ghosh, Atimukta Jha, Amol M. Kanampalliwar, Omprakash Shiriwas, Sana Fatma, Shifu Aggarwal, Rupesh Dash, Rajeeb Swain, Punit Prasad, INSACOG Consortium, Ajay Parida |
| EPI_ISL_2342611 | Rajendra Institute of Medical Sciences, Ranchi | Institute of Life Sciences - INSACOG | Sunil K. Raghav, Safal Walia, Arup Ghosh, Atimukta Jha, Amol M. Kanampalliwar, Omprakash Shiriwas, Sana Fatma, Shifu Aggarwal, Rupesh Dash, Rajeeb Swain, Punit Prasad, INSACOG Consortium, Ajay Parida |
| EPI_ISL_2342613, EPI_ISL_2342616 | MGM Medical College & Hospital, Jamshedpur | Institute of Life Sciences - INSACOG | Sunil K. Raghav, Safal Walia, Arup Ghosh, Atimukta Jha, Amol M. Kanampalliwar, Omprakash Shiriwas, Sana Fatma, Shifu Aggarwal, Rupesh Dash, Rajeeb Swain, Punit Prasad, INSACOG Consortium, Ajay Parida |
| EPI_ISL_2342617 | BRLSABVM Government Medical College, Rajnandgaon, Chhattisgarh | Institute of Life Sciences - INSACOG | Sunil K. Raghav, Safal Walia, Arup Ghosh, Atimukta Jha, Amol M. Kanampalliwar, Omprakash Shiriwas, Sana Fatma, Shifu Aggarwal, Rupesh Dash, Rajeeb Swain, Punit Prasad, INSACOG Consortium, Ajay Parida |
| EPI_ISL_2342619 | MGM Medical College & Hospital, Jamshedpur | Institute of Life Sciences - INSACOG | Sunil K. Raghav, Safal Walia, Arup Ghosh, Atimukta Jha, Amol M. Kanampalliwar, Omprakash Shiriwas, Sana Fatma, Shifu Aggarwal, Rupesh Dash, Rajeeb Swain, Punit Prasad, INSACOG Consortium, Ajay Parida |
| EPI_ISL_2342621 | BRLSABVM Government Medical College, Rajnandgaon, Chhattisgarh | Institute of Life Sciences - INSACOG | Sunil K. Raghav, Safal Walia, Arup Ghosh, Atimukta Jha, Amol M. Kanampalliwar, Omprakash Shiriwas, Sana Fatma, Shifu Aggarwal, Rupesh Dash, Rajeeb Swain, Punit Prasad, INSACOG Consortium, Ajay Parida |
| EPI_ISL_2342624 | MGM Medical College & Hospital, Jamshedpur | Institute of Life Sciences - INSACOG | Sunil K. Raghav, Safal Walia, Arup Ghosh, Atimukta Jha, Amol M. Kanampalliwar, Omprakash Shiriwas, Sana Fatma, Shifu Aggarwal, Rupesh Dash, Rajeeb Swain, Punit Prasad, INSACOG Consortium, Ajay Parida |
| EPI_ISL_2342625 | BRLSABVM Government Medical College, Rajnandgaon, Chhattisgarh | Institute of Life Sciences - INSACOG | Sunil K. Raghav, Safal Walia, Arup Ghosh, Atimukta Jha, Amol M. Kanampalliwar, Omprakash Shiriwas, Sana Fatma, Shifu Aggarwal, Rupesh Dash, Rajeeb Swain, Punit Prasad, INSACOG Consortium, Ajay Parida |
| EPI_ISL_2342626, EPI_ISL_2342628, EPI_ISL_2342629, EPI_ISL_2342631 | MGM Medical College & Hospital, Jamshedpur | Institute of Life Sciences - INSACOG | Sunil K. Raghav, Safal Walia, Arup Ghosh, Atimukta Jha, Amol M. Kanampalliwar, Omprakash Shiriwas, Sana Fatma, Shifu Aggarwal, Rupesh Dash, Rajeeb Swain, Punit Prasad, INSACOG Consortium, Ajay Parida |
| EPI_ISL_2342633 | PRM Medical College & Hospital, Baripada | Institute of Life Sciences - INSACOG | Sunil K. Raghav, Safal Walia, Arup Ghosh, Atimukta Jha, Amol M. Kanampalliwar, Omprakash Shiriwas, Sana Fatma, Shifu Aggarwal, Rupesh Dash, Rajeeb Swain, Punit Prasad, INSACOG Consortium, Ajay Parida |

| | | | |
|---|---|---|---|
| EPI_ISL_2342636 | BRLSABVM Government Medical College, Rajnandgaon, Chhattisgarh | Institute of Life Sciences - INSACOG | Sunil K. Raghav, Safal Walia, Arup Ghosh, Atimukta Jha, Amol M. Kanampalliwar, Omprakash Shiriwas, Sana Fatma, Shifu Aggarwal, Rupesh Dash, Rajeeb Swain, Punit Prasad, INSACOG Consortium, Ajay Parida |
| EPI_ISL_2342637 | MGM Medical College & Hospital, Jamshedpur | Institute of Life Sciences - INSACOG | Sunil K. Raghav, Safal Walia, Arup Ghosh, Atimukta Jha, Amol M. Kanampalliwar, Omprakash Shiriwas, Sana Fatma, Shifu Aggarwal, Rupesh Dash, Rajeeb Swain, Punit Prasad, INSACOG Consortium, Ajay Parida |
| EPI_ISL_2342639 | Institute of Life Sciences, Bhubaneswar | Institute of Life Sciences - INSACOG | Sunil K. Raghav, Safal Walia, Arup Ghosh, Atimukta Jha, Amol M. Kanampalliwar, Omprakash Shiriwas, Sana Fatma, Shifu Aggarwal, Rupesh Dash, Rajeeb Swain, Punit Prasad, INSACOG Consortium, Ajay Parida |
| EPI_ISL_2342642, EPI_ISL_2342644 | NCCS, Pune | Institute of Life Sciences - INSACOG | Sunil K. Raghav, Safal Walia, Arup Ghosh, Atimukta Jha, Amol M. Kanampalliwar, Omprakash Shiriwas, Sana Fatma, Shifu Aggarwal, Rupesh Dash, Rajeeb Swain, Punit Prasad, INSACOG Consortium, Ajay Parida |
| EPI_ISL_2342645 | Institute of Life Sciences, Bhubaneswar | Institute of Life Sciences - INSACOG | Sunil K. Raghav, Safal Walia, Arup Ghosh, Atimukta Jha, Amol M. Kanampalliwar, Omprakash Shiriwas, Sana Fatma, Shifu Aggarwal, Rupesh Dash, Rajeeb Swain, Punit Prasad, INSACOG Consortium, Ajay Parida |
| EPI_ISL_2342646 | MGM Medical College & Hospital, Jamshedpur | Institute of Life Sciences - INSACOG | Sunil K. Raghav, Safal Walia, Arup Ghosh, Atimukta Jha, Amol M. Kanampalliwar, Omprakash Shiriwas, Sana Fatma, Shifu Aggarwal, Rupesh Dash, Rajeeb Swain, Punit Prasad, INSACOG Consortium, Ajay Parida |
| EPI_ISL_2342647 | BRLSABVM Government Medical College, Rajnandgaon, Chhattisgarh | Institute of Life Sciences - INSACOG | Sunil K. Raghav, Safal Walia, Arup Ghosh, Atimukta Jha, Amol M. Kanampalliwar, Omprakash Shiriwas, Sana Fatma, Shifu Aggarwal, Rupesh Dash, Rajeeb Swain, Punit Prasad, INSACOG Consortium, Ajay Parida |
| EPI_ISL_2342648 | MGM Medical College & Hospital, Jamshedpur | Institute of Life Sciences - INSACOG | Sunil K. Raghav, Safal Walia, Arup Ghosh, Atimukta Jha, Amol M. Kanampalliwar, Omprakash Shiriwas, Sana Fatma, Shifu Aggarwal, Rupesh Dash, Rajeeb Swain, Punit Prasad, INSACOG Consortium, Ajay Parida |
| EPI_ISL_2342651 | BRLSABVM Government Medical College, Rajnandgaon, Chhattisgarh | Institute of Life Sciences - INSACOG | Sunil K. Raghav, Safal Walia, Arup Ghosh, Atimukta Jha, Amol M. Kanampalliwar, Omprakash Shiriwas, Sana Fatma, Shifu Aggarwal, Rupesh Dash, Rajeeb Swain, Punit Prasad, INSACOG Consortium, Ajay Parida |
| EPI_ISL_2342652 | Institute of Life Sciences, Bhubaneswar | Institute of Life Sciences - INSACOG | Sunil K. Raghav, Safal Walia, Arup Ghosh, Atimukta Jha, Amol M. Kanampalliwar, Omprakash Shiriwas, Sana Fatma, Shifu Aggarwal, Rupesh Dash, Rajeeb Swain, Punit Prasad, INSACOG Consortium, Ajay Parida |
| EPI_ISL_2342653 | MGM Medical College & Hospital, Jamshedpur | Institute of Life Sciences - INSACOG | Sunil K. Raghav, Safal Walia, Arup Ghosh, Atimukta Jha, Amol M. Kanampalliwar, Omprakash Shiriwas, Sana Fatma, Shifu Aggarwal, Rupesh Dash, Rajeeb Swain, Punit Prasad, INSACOG Consortium, Ajay Parida |
| EPI_ISL_2342656 | NCCS, Pune | Institute of Life Sciences - INSACOG | Sunil K. Raghav, Safal Walia, Arup Ghosh, Atimukta Jha, Amol M. Kanampalliwar, Omprakash Shiriwas, Sana Fatma, Shifu Aggarwal, Rupesh Dash, Rajeeb Swain, Punit Prasad, INSACOG Consortium, Ajay Parida |
| EPI_ISL_2342658 | MGM Medical College & Hospital, Jamshedpur | Institute of Life Sciences - INSACOG | Sunil K. Raghav, Safal Walia, Arup Ghosh, Atimukta Jha, Amol M. Kanampalliwar, Omprakash Shiriwas, Sana Fatma, Shifu Aggarwal, Rupesh Dash, Rajeeb Swain, Punit Prasad, INSACOG Consortium, Ajay Parida |
| EPI_ISL_2342659, EPI_ISL_2342661 | BRLSABVM Government Medical College, Rajnandgaon, Chhattisgarh | Institute of Life Sciences - INSACOG | Sunil K. Raghav, Safal Walia, Arup Ghosh, Atimukta Jha, Amol M. Kanampalliwar, Omprakash Shiriwas, Sana Fatma, Shifu Aggarwal, Rupesh Dash, Rajeeb Swain, Punit Prasad, INSACOG Consortium, Ajay Parida |
| EPI_ISL_2342662, EPI_ISL_2342663 | MGM Medical College & Hospital, Jamshedpur | Institute of Life Sciences - INSACOG | Sunil K. Raghav, Safal Walia, Arup Ghosh, Atimukta Jha, Amol M. Kanampalliwar, Omprakash Shiriwas, Sana Fatma, Shifu Aggarwal, Rupesh Dash, Rajeeb Swain, Punit Prasad, INSACOG Consortium, Ajay Parida |
| EPI_ISL_2342664 | Rajendra Institute of Medical Sciences, Ranchi | Institute of Life Sciences - INSACOG | Sunil K. Raghav, Safal Walia, Arup Ghosh, Atimukta Jha, Amol M. Kanampalliwar, Omprakash Shiriwas, Sana Fatma, Shifu Aggarwal, Rupesh Dash, Rajeeb Swain, Punit Prasad, INSACOG Consortium, Ajay Parida |
| EPI_ISL_2342666 | Institute of Life Sciences, Bhubaneswar | Institute of Life Sciences - INSACOG | Sunil K. Raghav, Safal Walia, Arup Ghosh, Atimukta Jha, Amol M. Kanampalliwar, Omprakash Shiriwas, Sana Fatma, Shifu Aggarwal, Rupesh Dash, Rajeeb Swain, Punit Prasad, INSACOG Consortium, Ajay Parida |
| EPI_ISL_2342668 | MGM Medical College & Hospital, Jamshedpur | Institute of Life Sciences - INSACOG | Sunil K. Raghav, Safal Walia, Arup Ghosh, Atimukta Jha, Amol M. Kanampalliwar, Omprakash Shiriwas, Sana Fatma, Shifu Aggarwal, Rupesh Dash, Rajeeb Swain, Punit Prasad, INSACOG Consortium, Ajay Parida |
| EPI_ISL_2342669, EPI_ISL_2342670 | BRLSABVM Government Medical College, Rajnandgaon, Chhattisgarh | Institute of Life Sciences - INSACOG | Sunil K. Raghav, Safal Walia, Arup Ghosh, Atimukta Jha, Amol M. Kanampalliwar, Omprakash Shiriwas, Sana Fatma, Shifu Aggarwal, Rupesh Dash, Rajeeb Swain, Punit Prasad, INSACOG Consortium, Ajay Parida |
| EPI_ISL_2342671 | Institute of Life Sciences, Bhubaneswar | Institute of Life Sciences - INSACOG | Sunil K. Raghav, Safal Walia, Arup Ghosh, Atimukta Jha, Amol M. Kanampalliwar, Omprakash Shiriwas, Sana Fatma, Shifu Aggarwal, Rupesh Dash, Rajeeb Swain, Punit Prasad, INSACOG Consortium, Ajay Parida |
| EPI_ISL_2342677 | MGM Medical College & Hospital, Jamshedpur | Institute of Life Sciences - INSACOG | Sunil K. Raghav, Safal Walia, Arup Ghosh, Atimukta Jha, Amol M. Kanampalliwar, Omprakash Shiriwas, Sana Fatma, Shifu Aggarwal, Rupesh Dash, Rajeeb Swain, Punit Prasad, INSACOG Consortium, Ajay Parida |
| EPI_ISL_2342682 | Rajendra Institute of Medical Sciences, Ranchi | Institute of Life Sciences - INSACOG | Sunil K. Raghav, Safal Walia, Arup Ghosh, Atimukta Jha, Amol M. Kanampalliwar, Omprakash Shiriwas, Sana Fatma, Shifu Aggarwal, Rupesh Dash, Rajeeb Swain, Punit Prasad, INSACOG Consortium, Ajay Parida |
| EPI_ISL_2342688 | MGM Medical College & Hospital, Jamshedpur | Institute of Life Sciences - INSACOG | Sunil K. Raghav, Safal Walia, Arup Ghosh, Atimukta Jha, Amol M. Kanampalliwar, Omprakash Shiriwas, Sana Fatma, Shifu Aggarwal, Rupesh Dash, Rajeeb Swain, Punit Prasad, INSACOG Consortium, Ajay Parida |
| EPI_ISL_2342695 | Institute of Life Sciences, Bhubaneswar | Institute of Life Sciences - INSACOG | Sunil K. Raghav, Safal Walia, Arup Ghosh, Atimukta Jha, Amol M. Kanampalliwar, Omprakash Shiriwas, Sana Fatma, Shifu Aggarwal, Rupesh Dash, Rajeeb Swain, Punit Prasad, INSACOG Consortium, Ajay Parida |
| EPI_ISL_2342701, EPI_ISL_2342704, EPI_ISL_2342705 | MGM Medical College & Hospital, Jamshedpur | Institute of Life Sciences - INSACOG | Sunil K. Raghav, Safal Walia, Arup Ghosh, Atimukta Jha, Amol M. Kanampalliwar, Omprakash Shiriwas, Sana Fatma, Shifu Aggarwal, Rupesh Dash, Rajeeb Swain, Punit Prasad, INSACOG Consortium, Ajay Parida |
| EPI_ISL_2342708 | BRLSABVM Government Medical College, Rajnandgaon, Chhattisgarh | Institute of Life Sciences - INSACOG | Sunil K. Raghav, Safal Walia, Arup Ghosh, Atimukta Jha, Amol M. Kanampalliwar, Omprakash Shiriwas, Sana Fatma, Shifu Aggarwal, Rupesh Dash, Rajeeb Swain, Punit Prasad, INSACOG Consortium, Ajay Parida |
| EPI_ISL_2342716 | PRM Medical College & Hospital, Baripada | Institute of Life Sciences - INSACOG | Sunil K. Raghav, Safal Walia, Arup Ghosh, Atimukta Jha, Amol M. Kanampalliwar, Omprakash Shiriwas, Sana Fatma, Shifu Aggarwal, Rupesh Dash, Rajeeb Swain, Punit Prasad, INSACOG Consortium, Ajay Parida |
| EPI_ISL_2342720 | MGM Medical College & Hospital, Jamshedpur | Institute of Life Sciences - INSACOG | Sunil K. Raghav, Safal Walia, Arup Ghosh, Atimukta Jha, Amol M. Kanampalliwar, Omprakash Shiriwas, Sana Fatma, Shifu Aggarwal, Rupesh Dash, Rajeeb Swain, Punit Prasad, INSACOG Consortium, Ajay Parida |
| EPI_ISL_2342722 | NCCS, Pune | Institute of Life Sciences - INSACOG | Sunil K. Raghav, Safal Walia, Arup Ghosh, Atimukta Jha, Amol M. Kanampalliwar, Omprakash Shiriwas, Sana Fatma, Shifu Aggarwal, Rupesh Dash, Rajeeb Swain, Punit Prasad, INSACOG Consortium, Ajay Parida |
| EPI_ISL_2342834, EPI_ISL_2342857, EPI_ISL_2342863, EPI_ISL_2342864, EPI_ISL_2342866, EPI_ISL_2342867, EPI_ISL_2342868, EPI_ISL_2342869, EPI_ISL_2342870, EPI_ISL_2342871, EPI_ISL_2342873, EPI_ISL_2342875, EPI_ISL_2342876, EPI_ISL_2342877, EPI_ISL_2342878, EPI_ISL_2342879, EPI_ISL_2342880, EPI_ISL_2342881 | | | |
| see above | BRLSABVM Government Medical College, Rajnandgaon, Chhattisgarh | Institute of Life Sciences - INSACOG | Sunil K. Raghav, Safal Walia, Arup Ghosh, Atimukta Jha, Amol M. Kanampalliwar, Omprakash Shiriwas, Sana Fatma, Shifu Aggarwal, Rupesh Dash, Rajeeb Swain, Punit Prasad, INSACOG Consortium, Ajay Parida |
| EPI_ISL_2342919, EPI_ISL_2342930, EPI_ISL_2342931, EPI_ISL_2342932, EPI_ISL_2342937, EPI_ISL_2342938, EPI_ISL_2342946, EPI_ISL_2342947, EPI_ISL_2342953, EPI_ISL_2342956, EPI_ISL_2342964 | | | |
| see above | NCCS, Pune | Institute of Life Sciences - INSACOG | Sunil K. Raghav, Safal Walia, Arup Ghosh, Atimukta Jha, Amol M. Kanampalliwar, Omprakash Shiriwas, Sana Fatma, Shifu Aggarwal, Rupesh Dash, Rajeeb Swain, Punit Prasad, INSACOG Consortium, Ajay Parida |
| EPI_ISL_2342971, EPI_ISL_2342972, EPI_ISL_2342973, EPI_ISL_2342974, EPI_ISL_2342975, EPI_ISL_2342976, EPI_ISL_2342977, EPI_ISL_2342978, EPI_ISL_2342979 | MGM Medical College & Hospital, Jamshedpur | Institute of Life Sciences - INSACOG | Sunil K. Raghav, Safal Walia, Arup Ghosh, Atimukta Jha, Amol M. Kanampalliwar, Omprakash Shiriwas, Sana Fatma, Shifu Aggarwal, Rupesh Dash, Rajeeb Swain, Punit Prasad, INSACOG Consortium, Ajay Parida |
| EPI_ISL_2342980 | Institute of Life Sciences, Bhubaneswar | Institute of Life Sciences - INSACOG | Sunil K. Raghav, Safal Walia, Arup Ghosh, Atimukta Jha, Amol M. Kanampalliwar, Omprakash Shiriwas, Sana Fatma, Shifu Aggarwal, Rupesh Dash, Rajeeb Swain, Punit Prasad, INSACOG Consortium, Ajay Parida |
| EPI_ISL_2342999 | LESP Michoacan/LANS | Instituto de Diagnostico y Referencia Epidemiologicos (INDRE) | Claudia Wong-Arambula, Abril Rodriguez-Maldonado, Vanessa Rivero-Arredondo, Ariadna Medina-Benitez, Joaquin Quiroz-Mercado, Sergio Rangel-Guerrero, Natividad Cruz-Ortiz, Tatiana Nunez-Garcia, Gisela Barrera-Badillo, Lucia Hernandez-Rivas, Irma Lopez-Martinez, Ernesto Ramirez-Gonzalez. |
| EPI_ISL_2343005 | LESP Sinaloa | Instituto de Diagnostico y Referencia Epidemiologicos (INDRE) | Claudia Wong-Arambula, Abril Rodriguez-Maldonado, Vanessa Rivero-Arredondo, Ariadna Medina-Benitez, Joaquin Quiroz-Mercado, Sergio Rangel-Guerrero, Natividad Cruz-Ortiz, Tatiana Nunez-Garcia, Gisela Barrera-Badillo, Lucia Hernandez-Rivas, Irma Lopez-Martinez, Ernesto Ramirez-Gonzalez. |
| EPI_ISL_2343033, EPI_ISL_2343068, EPI_ISL_2343125 | University Hospitals of Geneva, Laboratory of Virology | HUG, Laboratory of Virology and the Health2030 Genome Center | Samuel Cordey, Ana Rita Goncalves, Laurent Kaiser, Lorenzo Cerutti, Henri Pegeot, Melyssa Elies, Deborah Penet, Keith Harshman, Ioannis Xenarios, Emmanouil Dermitzakis |
| EPI_ISL_2346118 | Oregon State Public Health Laboratory | Oregon State Public Health Laboratory | Rafia Razzaque, Eugene Yeboah, Vanda Makris, Laura Tsaknaridis, John Fontana and Shane Sevey |
| EPI_ISL_2346160 | Emory Molecular Diagnostics Laboratory, Emory Healthcare | Piantadosi Lab, Emory Department of Pathology | Ahmed Babiker, Anne Piantadosi |
| EPI_ISL_2346305, EPI_ISL_2346306, EPI_ISL_2346307 | Plateforme de testing Namuroise | Plateforme de testing Namuroise | Lesly Nyinkeu Kemamen; Otto Gaetan ; Denis Olivier ; Degosserie Jonathan ; Mullier François |
| EPI_ISL_2346472, EPI_ISL_2346473 | Lighthouse Lab in Milton Keynes | Wellcome Sanger Institute for the COVID-19 Genomics UK (COG-UK) Consortium | The Lighthouse Lab in Milton Keynes and Alex Alderton, Roberto Amato, Jeffrey Barrett, Sonia Goncalves, Ewan Harrison, David K. Jackson, Ian Johnston, Dominic Kwiatkowski, Cordelia Langford, John Sillitoe on behalf of the Wellcome Sanger Institute COVID-19 Surveillance Team |
| EPI_ISL_2346474, EPI_ISL_2346475 | Lighthouse Lab in Alderley Park | Wellcome Sanger Institute for the COVID-19 Genomics UK (COG-UK) Consortium | Jacquelyn Wynn, Mairead Hyland, The Lighthouse Lab in Alderley Park and Alex Alderton, Roberto Amato, Jeffrey Barrett, Sonia Goncalves, Ewan Harrison, David K. Jackson, Ian Johnston, Dominic Kwiatkowski, Cordelia Langford, John Sillitoe on behalf of the Wellcome Sanger Institute COVID-19 Surveillance Team |
| EPI_ISL_2346479, EPI_ISL_2346480, EPI_ISL_2346481 | Lighthouse Lab in Milton Keynes | Wellcome Sanger Institute for the COVID-19 Genomics UK (COG-UK) Consortium | The Lighthouse Lab in Milton Keynes and Alex Alderton, Roberto Amato, Jeffrey Barrett, Sonia Goncalves, Ewan Harrison, David K. Jackson, Ian Johnston, Dominic Kwiatkowski, Cordelia Langford, John Sillitoe on behalf of the Wellcome Sanger Institute COVID-19 Surveillance Team |
| EPI_ISL_2346482 | Lighthouse Lab in Alderley Park | Wellcome Sanger Institute for the COVID-19 Genomics UK (COG-UK) Consortium | Jacquelyn Wynn, Mairead Hyland, The Lighthouse Lab in Alderley Park and Alex Alderton, Roberto Amato, Jeffrey Barrett, Sonia Goncalves, Ewan Harrison, David K. Jackson, Ian Johnston, Dominic Kwiatkowski, Cordelia Langford, John Sillitoe on behalf of the Wellcome Sanger Institute COVID-19 Surveillance Team |
| EPI_ISL_2346483 | Lighthouse Lab in Glasgow | Wellcome Sanger Institute for the COVID-19 Genomics UK (COG-UK) Consortium | Harper VanSteenhouse, Yumi Kasai, David Gray, Carol Clugston, Anna Dominiczak and Alex Alderton, Roberto Amato, Jeffrey Barrett, Sonia Goncalves, Ewan Harrison, David K. Jackson, Ian Johnston, Dominic Kwiatkowski, Cordelia Langford, John Sillitoe on behalf of the Wellcome Sanger Institute COVID-19 Surveillance Team |
| EPI_ISL_2346484 | Lighthouse Lab in Milton Keynes | Wellcome Sanger Institute for the COVID-19 Genomics UK (COG-UK) Consortium | The Lighthouse Lab in Milton Keynes and Alex Alderton, Roberto Amato, Jeffrey Barrett, Sonia Goncalves, Ewan Harrison, David K. Jackson, Ian Johnston, Dominic Kwiatkowski, Cordelia Langford, John Sillitoe on behalf of the Wellcome Sanger Institute COVID-19 Surveillance Team |
| EPI_ISL_2346485 | Lighthouse Lab in Alderley Park | Wellcome Sanger Institute for the COVID-19 Genomics UK (COG-UK) Consortium | Jacquelyn Wynn, Mairead Hyland, The Lighthouse Lab in Alderley Park and Alex Alderton, Roberto Amato, Jeffrey Barrett, Sonia Goncalves, Ewan Harrison, David K. Jackson, Ian Johnston, Dominic Kwiatkowski, Cordelia Langford, John Sillitoe on behalf of the Wellcome Sanger Institute COVID-19 Surveillance Team |
| EPI_ISL_2346486 | Lighthouse Lab in Milton Keynes | Wellcome Sanger Institute for the COVID-19 Genomics UK (COG-UK) Consortium | The Lighthouse Lab in Milton Keynes and Alex Alderton, Roberto Amato, Jeffrey Barrett, Sonia Goncalves, Ewan Harrison, David K. Jackson, Ian Johnston, Dominic Kwiatkowski, Cordelia Langford, John Sillitoe on behalf of the Wellcome Sanger Institute COVID-19 Surveillance Team |
| EPI_ISL_2346487 | Lighthouse Lab in Alderley Park | Wellcome Sanger Institute for the COVID-19 Genomics UK (COG-UK) Consortium | Jacquelyn Wynn, Mairead Hyland, The Lighthouse Lab in Alderley Park and Alex Alderton, Roberto Amato, Jeffrey Barrett, Sonia Goncalves, Ewan Harrison, David K. Jackson, Ian Johnston, Dominic Kwiatkowski, Cordelia Langford, John Sillitoe on behalf of the Wellcome Sanger Institute COVID-19 Surveillance Team |
| EPI_ISL_2346489, EPI_ISL_2346491 | Lighthouse Lab in Milton Keynes | Wellcome Sanger Institute for the COVID-19 Genomics UK (COG-UK) Consortium | The Lighthouse Lab in Milton Keynes and Alex Alderton, Roberto Amato, Jeffrey Barrett, Sonia Goncalves, Ewan Harrison, David K. Jackson, Ian Johnston, Dominic Kwiatkowski, Cordelia Langford, John Sillitoe on behalf of the Wellcome Sanger Institute COVID-19 Surveillance Team |

| | | | |
|---|---|---|---|
| EPI_ISL_2346492, EPI_ISL_2346495 | Lighthouse Lab in Alderley Park | Wellcome Sanger Institute for the COVID-19 Genomics UK (COG-UK) Consortium | Jacquelyn Wynn, Mairead Hyland, The Lighthouse Lab in Alderley Park and Alex Alderton, Roberto Amato, Jeffrey Barrett, Sonia Goncalves, Ewan Harrison, David K. Jackson, Ian Johnston, Dominic Kwiatkowski, Cordelia Langford, John Sillitoe on behalf of the Wellcome Sanger Institute COVID-19 Surveillance Team |
| EPI_ISL_2346496 | Lighthouse Lab in Milton Keynes | Wellcome Sanger Institute for the COVID-19 Genomics UK (COG-UK) Consortium | The Lighthouse Lab in Milton Keynes and Alex Alderton, Roberto Amato, Jeffrey Barrett, Sonia Goncalves, Ewan Harrison, David K. Jackson, Ian Johnston, Dominic Kwiatkowski, Cordelia Langford, John Sillitoe on behalf of the Wellcome Sanger Institute COVID-19 Surveillance Team |
| EPI_ISL_2346497, EPI_ISL_2346499 | Lighthouse Lab in Alderley Park | Wellcome Sanger Institute for the COVID-19 Genomics UK (COG-UK) Consortium | Jacquelyn Wynn, Mairead Hyland, The Lighthouse Lab in Alderley Park and Alex Alderton, Roberto Amato, Jeffrey Barrett, Sonia Goncalves, Ewan Harrison, David K. Jackson, Ian Johnston, Dominic Kwiatkowski, Cordelia Langford, John Sillitoe on behalf of the Wellcome Sanger Institute COVID-19 Surveillance Team |
| EPI_ISL_2346500 | Lighthouse Lab in Milton Keynes | Wellcome Sanger Institute for the COVID-19 Genomics UK (COG-UK) Consortium | The Lighthouse Lab in Milton Keynes and Alex Alderton, Roberto Amato, Jeffrey Barrett, Sonia Goncalves, Ewan Harrison, David K. Jackson, Ian Johnston, Dominic Kwiatkowski, Cordelia Langford, John Sillitoe on behalf of the Wellcome Sanger Institute COVID-19 Surveillance Team |
| EPI_ISL_2346502 | Lighthouse Lab in Alderley Park | Wellcome Sanger Institute for the COVID-19 Genomics UK (COG-UK) Consortium | Jacquelyn Wynn, Mairead Hyland, The Lighthouse Lab in Alderley Park and Alex Alderton, Roberto Amato, Jeffrey Barrett, Sonia Goncalves, Ewan Harrison, David K. Jackson, Ian Johnston, Dominic Kwiatkowski, Cordelia Langford, John Sillitoe on behalf of the Wellcome Sanger Institute COVID-19 Surveillance Team |
| EPI_ISL_2346503 | Lighthouse Lab in Milton Keynes | Wellcome Sanger Institute for the COVID-19 Genomics UK (COG-UK) Consortium | The Lighthouse Lab in Milton Keynes and Alex Alderton, Roberto Amato, Jeffrey Barrett, Sonia Goncalves, Ewan Harrison, David K. Jackson, Ian Johnston, Dominic Kwiatkowski, Cordelia Langford, John Sillitoe on behalf of the Wellcome Sanger Institute COVID-19 Surveillance Team |
| EPI_ISL_2346504 | Lighthouse Lab in Alderley Park | Wellcome Sanger Institute for the COVID-19 Genomics UK (COG-UK) Consortium | Jacquelyn Wynn, Mairead Hyland, The Lighthouse Lab in Alderley Park and Alex Alderton, Roberto Amato, Jeffrey Barrett, Sonia Goncalves, Ewan Harrison, David K. Jackson, Ian Johnston, Dominic Kwiatkowski, Cordelia Langford, John Sillitoe on behalf of the Wellcome Sanger Institute COVID-19 Surveillance Team |
| EPI_ISL_2346505, EPI_ISL_2346507 | Lighthouse Lab in Milton Keynes | Wellcome Sanger Institute for the COVID-19 Genomics UK (COG-UK) Consortium | The Lighthouse Lab in Milton Keynes and Alex Alderton, Roberto Amato, Jeffrey Barrett, Sonia Goncalves, Ewan Harrison, David K. Jackson, Ian Johnston, Dominic Kwiatkowski, Cordelia Langford, John Sillitoe on behalf of the Wellcome Sanger Institute COVID-19 Surveillance Team |
| EPI_ISL_2346508, EPI_ISL_2346509 | Lighthouse Lab in Alderley Park | Wellcome Sanger Institute for the COVID-19 Genomics UK (COG-UK) Consortium | Jacquelyn Wynn, Mairead Hyland, The Lighthouse Lab in Alderley Park and Alex Alderton, Roberto Amato, Jeffrey Barrett, Sonia Goncalves, Ewan Harrison, David K. Jackson, Ian Johnston, Dominic Kwiatkowski, Cordelia Langford, John Sillitoe on behalf of the Wellcome Sanger Institute COVID-19 Surveillance Team |
| EPI_ISL_2346510, EPI_ISL_2346511 | Lighthouse Lab in Milton Keynes | Wellcome Sanger Institute for the COVID-19 Genomics UK (COG-UK) Consortium | The Lighthouse Lab in Milton Keynes and Alex Alderton, Roberto Amato, Jeffrey Barrett, Sonia Goncalves, Ewan Harrison, David K. Jackson, Ian Johnston, Dominic Kwiatkowski, Cordelia Langford, John Sillitoe on behalf of the Wellcome Sanger Institute COVID-19 Surveillance Team |
| EPI_ISL_2346514, EPI_ISL_2346516, EPI_ISL_2346517, EPI_ISL_2346518, EPI_ISL_2346522 | Lighthouse Lab in Alderley Park | Wellcome Sanger Institute for the COVID-19 Genomics UK (COG-UK) Consortium | Jacquelyn Wynn, Mairead Hyland, The Lighthouse Lab in Alderley Park and Alex Alderton, Roberto Amato, Jeffrey Barrett, Sonia Goncalves, Ewan Harrison, David K. Jackson, Ian Johnston, Dominic Kwiatkowski, Cordelia Langford, John Sillitoe on behalf of the Wellcome Sanger Institute COVID-19 Surveillance Team |
| EPI_ISL_2346523, EPI_ISL_2346524 | Lighthouse Lab in Glasgow | Wellcome Sanger Institute for the COVID-19 Genomics UK (COG-UK) Consortium | Harper VanSteenhouse, Yumi Kasai, David Gray, Carol Clugston, Anna Dominiczak and Alex Alderton, Roberto Amato, Jeffrey Barrett, Sonia Goncalves, Ewan Harrison, David K. Jackson, Ian Johnston, Dominic Kwiatkowski, Cordelia Langford, John Sillitoe on behalf of the Wellcome Sanger Institute COVID-19 Surveillance Team |
| EPI_ISL_2346525 | Lighthouse Lab in Milton Keynes | Wellcome Sanger Institute for the COVID-19 Genomics UK (COG-UK) Consortium | The Lighthouse Lab in Milton Keynes and Alex Alderton, Roberto Amato, Jeffrey Barrett, Sonia Goncalves, Ewan Harrison, David K. Jackson, Ian Johnston, Dominic Kwiatkowski, Cordelia Langford, John Sillitoe on behalf of the Wellcome Sanger Institute COVID-19 Surveillance Team |
| EPI_ISL_2346526 | Lighthouse Lab in Alderley Park | Wellcome Sanger Institute for the COVID-19 Genomics UK (COG-UK) Consortium | Jacquelyn Wynn, Mairead Hyland, The Lighthouse Lab in Alderley Park and Alex Alderton, Roberto Amato, Jeffrey Barrett, Sonia Goncalves, Ewan Harrison, David K. Jackson, Ian Johnston, Dominic Kwiatkowski, Cordelia Langford, John Sillitoe on behalf of the Wellcome Sanger Institute COVID-19 Surveillance Team |
| EPI_ISL_2346527, EPI_ISL_2346530, EPI_ISL_2346531, EPI_ISL_2346534, EPI_ISL_2346538, EPI_ISL_2346539, EPI_ISL_2346540 | Lighthouse Lab in Milton Keynes | Wellcome Sanger Institute for the COVID-19 Genomics UK (COG-UK) Consortium | The Lighthouse Lab in Milton Keynes and Alex Alderton, Roberto Amato, Jeffrey Barrett, Sonia Goncalves, Ewan Harrison, David K. Jackson, Ian Johnston, Dominic Kwiatkowski, Cordelia Langford, John Sillitoe on behalf of the Wellcome Sanger Institute COVID-19 Surveillance Team |
| EPI_ISL_2346541, EPI_ISL_2346543, EPI_ISL_2346544, EPI_ISL_2346545 | Lighthouse Lab in Alderley Park | Wellcome Sanger Institute for the COVID-19 Genomics UK (COG-UK) Consortium | Jacquelyn Wynn, Mairead Hyland, The Lighthouse Lab in Alderley Park and Alex Alderton, Roberto Amato, Jeffrey Barrett, Sonia Goncalves, Ewan Harrison, David K. Jackson, Ian Johnston, Dominic Kwiatkowski, Cordelia Langford, John Sillitoe on behalf of the Wellcome Sanger Institute COVID-19 Surveillance Team |
| EPI_ISL_2346546, EPI_ISL_2346547, EPI_ISL_2346548 | Lighthouse Lab in Milton Keynes | Wellcome Sanger Institute for the COVID-19 Genomics UK (COG-UK) Consortium | The Lighthouse Lab in Milton Keynes and Alex Alderton, Roberto Amato, Jeffrey Barrett, Sonia Goncalves, Ewan Harrison, David K. Jackson, Ian Johnston, Dominic Kwiatkowski, Cordelia Langford, John Sillitoe on behalf of the Wellcome Sanger Institute COVID-19 Surveillance Team |
| EPI_ISL_2346554, EPI_ISL_2346555, EPI_ISL_2346556, EPI_ISL_2346557 | Lighthouse Lab in Alderley Park | Wellcome Sanger Institute for the COVID-19 Genomics UK (COG-UK) Consortium | Jacquelyn Wynn, Mairead Hyland, The Lighthouse Lab in Alderley Park and Alex Alderton, Roberto Amato, Jeffrey Barrett, Sonia Goncalves, Ewan Harrison, David K. Jackson, Ian Johnston, Dominic Kwiatkowski, Cordelia Langford, John Sillitoe on behalf of the Wellcome Sanger Institute COVID-19 Surveillance Team |
| EPI_ISL_2346560, EPI_ISL_2346561 | Lighthouse Lab in Milton Keynes | Wellcome Sanger Institute for the COVID-19 Genomics UK (COG-UK) Consortium | The Lighthouse Lab in Milton Keynes and Alex Alderton, Roberto Amato, Jeffrey Barrett, Sonia Goncalves, Ewan Harrison, David K. Jackson, Ian Johnston, Dominic Kwiatkowski, Cordelia Langford, John Sillitoe on behalf of the Wellcome Sanger Institute COVID-19 Surveillance Team |
| EPI_ISL_2346564 | Lighthouse Lab in Alderley Park | Wellcome Sanger Institute for the COVID-19 Genomics UK (COG-UK) Consortium | Jacquelyn Wynn, Mairead Hyland, The Lighthouse Lab in Alderley Park and Alex Alderton, Roberto Amato, Jeffrey Barrett, Sonia Goncalves, Ewan Harrison, David K. Jackson, Ian Johnston, Dominic Kwiatkowski, Cordelia Langford, John Sillitoe on behalf of the Wellcome Sanger Institute COVID-19 Surveillance Team |
| EPI_ISL_2346565, EPI_ISL_2346566 | Lighthouse Lab in Milton Keynes | Wellcome Sanger Institute for the COVID-19 Genomics UK (COG-UK) Consortium | The Lighthouse Lab in Milton Keynes and Alex Alderton, Roberto Amato, Jeffrey Barrett, Sonia Goncalves, Ewan Harrison, David K. Jackson, Ian Johnston, Dominic Kwiatkowski, Cordelia Langford, John Sillitoe on behalf of the Wellcome Sanger Institute COVID-19 Surveillance Team |
| EPI_ISL_2346567, EPI_ISL_2346569, EPI_ISL_2346572 | Lighthouse Lab in Alderley Park | Wellcome Sanger Institute for the COVID-19 Genomics UK (COG-UK) Consortium | Jacquelyn Wynn, Mairead Hyland, The Lighthouse Lab in Alderley Park and Alex Alderton, Roberto Amato, Jeffrey Barrett, Sonia Goncalves, Ewan Harrison, David K. Jackson, Ian Johnston, Dominic Kwiatkowski, Cordelia Langford, John Sillitoe on behalf of the Wellcome Sanger Institute COVID-19 Surveillance Team |
| EPI_ISL_2346573, EPI_ISL_2346574, EPI_ISL_2346577, EPI_ISL_2346578, EPI_ISL_2346579, EPI_ISL_2346580, EPI_ISL_2346581, EPI_ISL_2346582, EPI_ISL_2346586 | Lighthouse Lab in Milton Keynes | Wellcome Sanger Institute for the COVID-19 Genomics UK (COG-UK) Consortium | The Lighthouse Lab in Milton Keynes and Alex Alderton, Roberto Amato, Jeffrey Barrett, Sonia Goncalves, Ewan Harrison, David K. Jackson, Ian Johnston, Dominic Kwiatkowski, Cordelia Langford, John Sillitoe on behalf of the Wellcome Sanger Institute COVID-19 Surveillance Team |
| EPI_ISL_2346588 | Lighthouse Lab in Alderley Park | Wellcome Sanger Institute for the COVID-19 Genomics UK (COG-UK) Consortium | Jacquelyn Wynn, Mairead Hyland, The Lighthouse Lab in Alderley Park and Alex Alderton, Roberto Amato, Jeffrey Barrett, Sonia Goncalves, Ewan Harrison, David K. Jackson, Ian Johnston, Dominic Kwiatkowski, Cordelia Langford, John Sillitoe on behalf of the Wellcome Sanger Institute COVID-19 Surveillance Team |
| EPI_ISL_2346589, EPI_ISL_2346590 | Lighthouse Lab in Milton Keynes | Wellcome Sanger Institute for the COVID-19 Genomics UK (COG-UK) Consortium | The Lighthouse Lab in Milton Keynes and Alex Alderton, Roberto Amato, Jeffrey Barrett, Sonia Goncalves, Ewan Harrison, David K. Jackson, Ian Johnston, Dominic Kwiatkowski, Cordelia Langford, John Sillitoe on behalf of the Wellcome Sanger Institute COVID-19 Surveillance Team |
| EPI_ISL_2346591, EPI_ISL_2346592, EPI_ISL_2346593 | Lighthouse Lab in Alderley Park | Wellcome Sanger Institute for the COVID-19 Genomics UK (COG-UK) Consortium | Jacquelyn Wynn, Mairead Hyland, The Lighthouse Lab in Alderley Park and Alex Alderton, Roberto Amato, Jeffrey Barrett, Sonia Goncalves, Ewan Harrison, David K. Jackson, Ian Johnston, Dominic Kwiatkowski, Cordelia Langford, John Sillitoe on behalf of the Wellcome Sanger Institute COVID-19 Surveillance Team |
| EPI_ISL_2346595, EPI_ISL_2346596 | Lighthouse Lab in Milton Keynes | Wellcome Sanger Institute for the COVID-19 Genomics UK (COG-UK) Consortium | The Lighthouse Lab in Milton Keynes and Alex Alderton, Roberto Amato, Jeffrey Barrett, Sonia Goncalves, Ewan Harrison, David K. Jackson, Ian Johnston, Dominic Kwiatkowski, Cordelia Langford, John Sillitoe on behalf of the Wellcome Sanger Institute COVID-19 Surveillance Team |
| EPI_ISL_2346597 | Lighthouse Lab in Alderley Park | Wellcome Sanger Institute for the COVID-19 Genomics UK (COG-UK) Consortium | Jacquelyn Wynn, Mairead Hyland, The Lighthouse Lab in Alderley Park and Alex Alderton, Roberto Amato, Jeffrey Barrett, Sonia Goncalves, Ewan Harrison, David K. Jackson, Ian Johnston, Dominic Kwiatkowski, Cordelia Langford, John Sillitoe on behalf of the Wellcome Sanger Institute COVID-19 Surveillance Team |
| EPI_ISL_2346598, EPI_ISL_2346601 | Lighthouse Lab in Milton Keynes | Wellcome Sanger Institute for the COVID-19 Genomics UK (COG-UK) Consortium | The Lighthouse Lab in Milton Keynes and Alex Alderton, Roberto Amato, Jeffrey Barrett, Sonia Goncalves, Ewan Harrison, David K. Jackson, Ian Johnston, Dominic Kwiatkowski, Cordelia Langford, John Sillitoe on behalf of the Wellcome Sanger Institute COVID-19 Surveillance Team |
| EPI_ISL_2346602 | Lighthouse Lab in Alderley Park | Wellcome Sanger Institute for the COVID-19 Genomics UK (COG-UK) Consortium | Jacquelyn Wynn, Mairead Hyland, The Lighthouse Lab in Alderley Park and Alex Alderton, Roberto Amato, Jeffrey Barrett, Sonia Goncalves, Ewan Harrison, David K. Jackson, Ian Johnston, Dominic Kwiatkowski, Cordelia Langford, John Sillitoe on behalf of the Wellcome Sanger Institute COVID-19 Surveillance Team |
| EPI_ISL_2346604 | Lighthouse Lab in Milton Keynes | Wellcome Sanger Institute for the COVID-19 Genomics UK (COG-UK) Consortium | The Lighthouse Lab in Milton Keynes and Alex Alderton, Roberto Amato, Jeffrey Barrett, Sonia Goncalves, Ewan Harrison, David K. Jackson, Ian Johnston, Dominic Kwiatkowski, Cordelia Langford, John Sillitoe on behalf of the Wellcome Sanger Institute COVID-19 Surveillance Team |
| EPI_ISL_2346605, EPI_ISL_2346606 | Lighthouse Lab in Alderley Park | Wellcome Sanger Institute for the COVID-19 Genomics UK (COG-UK) Consortium | Jacquelyn Wynn, Mairead Hyland, The Lighthouse Lab in Alderley Park and Alex Alderton, Roberto Amato, Jeffrey Barrett, Sonia Goncalves, Ewan Harrison, David K. Jackson, Ian Johnston, Dominic Kwiatkowski, Cordelia Langford, John Sillitoe on behalf of the Wellcome Sanger Institute COVID-19 Surveillance Team |
| EPI_ISL_2346608, EPI_ISL_2346610, EPI_ISL_2346611, EPI_ISL_2346612 | Lighthouse Lab in Milton Keynes | Wellcome Sanger Institute for the COVID-19 Genomics UK (COG-UK) Consortium | The Lighthouse Lab in Milton Keynes and Alex Alderton, Roberto Amato, Jeffrey Barrett, Sonia Goncalves, Ewan Harrison, David K. Jackson, Ian Johnston, Dominic Kwiatkowski, Cordelia Langford, John Sillitoe on behalf of the Wellcome Sanger Institute COVID-19 Surveillance Team |
| EPI_ISL_2346613 | Lighthouse Lab in Alderley Park | Wellcome Sanger Institute for the COVID-19 Genomics UK (COG-UK) Consortium | Jacquelyn Wynn, Mairead Hyland, The Lighthouse Lab in Alderley Park and Alex Alderton, Roberto Amato, Jeffrey Barrett, Sonia Goncalves, Ewan Harrison, David K. Jackson, Ian Johnston, Dominic Kwiatkowski, Cordelia Langford, John Sillitoe on behalf of the Wellcome Sanger Institute COVID-19 Surveillance Team |
| EPI_ISL_2346615, EPI_ISL_2346616, EPI_ISL_2346617 | Lighthouse Lab in Milton Keynes | Wellcome Sanger Institute for the COVID-19 Genomics UK (COG-UK) Consortium | The Lighthouse Lab in Milton Keynes and Alex Alderton, Roberto Amato, Jeffrey Barrett, Sonia Goncalves, Ewan Harrison, David K. Jackson, Ian Johnston, Dominic Kwiatkowski, Cordelia Langford, John Sillitoe on behalf of the Wellcome Sanger Institute COVID-19 Surveillance Team |

| | | | |
|---|---|---|---|
| EPI_ISL_2346618 | Lighthouse Lab in Glasgow | Wellcome Sanger Institute for the COVID-19 Genomics UK (COG-UK) Consortium | Harper VanSteenhouse, Yumi Kasai, David Gray, Carol Clugston, Anna Dominiczak and Alex Alderton, Roberto Amato, Jeffrey Barrett, Sonia Goncalves, Ewan Harrison, David K. Jackson, Ian Johnston, Dominic Kwiatkowski, Cordelia Langford, John Sillitoe on behalf of the Wellcome Sanger Institute COVID-19 Surveillance Team |
| EPI_ISL_2346619, EPI_ISL_2346620 | Lighthouse Lab in Alderley Park | Wellcome Sanger Institute for the COVID-19 Genomics UK (COG-UK) Consortium | Jacquelyn Wynn, Mairead Hyland, The Lighthouse Lab in Alderley Park and Alex Alderton, Roberto Amato, Jeffrey Barrett, Sonia Goncalves, Ewan Harrison, David K. Jackson, Ian Johnston, Dominic Kwiatkowski, Cordelia Langford, John Sillitoe on behalf of the Wellcome Sanger Institute COVID-19 Surveillance Team |
| EPI_ISL_2346623 | Lighthouse Lab in Milton Keynes | Wellcome Sanger Institute for the COVID-19 Genomics UK (COG-UK) Consortium | The Lighthouse Lab in Milton Keynes and Alex Alderton, Roberto Amato, Jeffrey Barrett, Sonia Goncalves, Ewan Harrison, David K. Jackson, Ian Johnston, Dominic Kwiatkowski, Cordelia Langford, John Sillitoe on behalf of the Wellcome Sanger Institute COVID-19 Surveillance Team |
| EPI_ISL_2346624, EPI_ISL_2346627 | Lighthouse Lab in Alderley Park | Wellcome Sanger Institute for the COVID-19 Genomics UK (COG-UK) Consortium | Jacquelyn Wynn, Mairead Hyland, The Lighthouse Lab in Alderley Park and Alex Alderton, Roberto Amato, Jeffrey Barrett, Sonia Goncalves, Ewan Harrison, David K. Jackson, Ian Johnston, Dominic Kwiatkowski, Cordelia Langford, John Sillitoe on behalf of the Wellcome Sanger Institute COVID-19 Surveillance Team |
| EPI_ISL_2346628 | Lighthouse Lab in Milton Keynes | Wellcome Sanger Institute for the COVID-19 Genomics UK (COG-UK) Consortium | The Lighthouse Lab in Milton Keynes and Alex Alderton, Roberto Amato, Jeffrey Barrett, Sonia Goncalves, Ewan Harrison, David K. Jackson, Ian Johnston, Dominic Kwiatkowski, Cordelia Langford, John Sillitoe on behalf of the Wellcome Sanger Institute COVID-19 Surveillance Team |
| EPI_ISL_2346629 | Lighthouse Lab in Alderley Park | Wellcome Sanger Institute for the COVID-19 Genomics UK (COG-UK) Consortium | Jacquelyn Wynn, Mairead Hyland, The Lighthouse Lab in Alderley Park and Alex Alderton, Roberto Amato, Jeffrey Barrett, Sonia Goncalves, Ewan Harrison, David K. Jackson, Ian Johnston, Dominic Kwiatkowski, Cordelia Langford, John Sillitoe on behalf of the Wellcome Sanger Institute COVID-19 Surveillance Team |
| EPI_ISL_2346630, EPI_ISL_2346631, EPI_ISL_2346632 | Lighthouse Lab in Milton Keynes | Wellcome Sanger Institute for the COVID-19 Genomics UK (COG-UK) Consortium | The Lighthouse Lab in Milton Keynes and Alex Alderton, Roberto Amato, Jeffrey Barrett, Sonia Goncalves, Ewan Harrison, David K. Jackson, Ian Johnston, Dominic Kwiatkowski, Cordelia Langford, John Sillitoe on behalf of the Wellcome Sanger Institute COVID-19 Surveillance Team |
| EPI_ISL_2346633, EPI_ISL_2346634, EPI_ISL_2346635, EPI_ISL_2346636, EPI_ISL_2346637 | Lighthouse Lab in Alderley Park | Wellcome Sanger Institute for the COVID-19 Genomics UK (COG-UK) Consortium | Jacquelyn Wynn, Mairead Hyland, The Lighthouse Lab in Alderley Park and Alex Alderton, Roberto Amato, Jeffrey Barrett, Sonia Goncalves, Ewan Harrison, David K. Jackson, Ian Johnston, Dominic Kwiatkowski, Cordelia Langford, John Sillitoe on behalf of the Wellcome Sanger Institute COVID-19 Surveillance Team |
| EPI_ISL_2346638 | Lighthouse Lab in Milton Keynes | Wellcome Sanger Institute for the COVID-19 Genomics UK (COG-UK) Consortium | The Lighthouse Lab in Milton Keynes and Alex Alderton, Roberto Amato, Jeffrey Barrett, Sonia Goncalves, Ewan Harrison, David K. Jackson, Ian Johnston, Dominic Kwiatkowski, Cordelia Langford, John Sillitoe on behalf of the Wellcome Sanger Institute COVID-19 Surveillance Team |
| EPI_ISL_2346639, EPI_ISL_2346644 | Lighthouse Lab in Alderley Park | Wellcome Sanger Institute for the COVID-19 Genomics UK (COG-UK) Consortium | Jacquelyn Wynn, Mairead Hyland, The Lighthouse Lab in Alderley Park and Alex Alderton, Roberto Amato, Jeffrey Barrett, Sonia Goncalves, Ewan Harrison, David K. Jackson, Ian Johnston, Dominic Kwiatkowski, Cordelia Langford, John Sillitoe on behalf of the Wellcome Sanger Institute COVID-19 Surveillance Team |
| EPI_ISL_2346646 | Lighthouse Lab in Glasgow | Wellcome Sanger Institute for the COVID-19 Genomics UK (COG-UK) Consortium | Harper VanSteenhouse, Yumi Kasai, David Gray, Carol Clugston, Anna Dominiczak and Alex Alderton, Roberto Amato, Jeffrey Barrett, Sonia Goncalves, Ewan Harrison, David K. Jackson, Ian Johnston, Dominic Kwiatkowski, Cordelia Langford, John Sillitoe on behalf of the Wellcome Sanger Institute COVID-19 Surveillance Team |
| EPI_ISL_2346647, EPI_ISL_2346648 | Lighthouse Lab in Milton Keynes | Wellcome Sanger Institute for the COVID-19 Genomics UK (COG-UK) Consortium | The Lighthouse Lab in Milton Keynes and Alex Alderton, Roberto Amato, Jeffrey Barrett, Sonia Goncalves, Ewan Harrison, David K. Jackson, Ian Johnston, Dominic Kwiatkowski, Cordelia Langford, John Sillitoe on behalf of the Wellcome Sanger Institute COVID-19 Surveillance Team |
| EPI_ISL_2346650, EPI_ISL_2346651, EPI_ISL_2346652 | Lighthouse Lab in Alderley Park | Wellcome Sanger Institute for the COVID-19 Genomics UK (COG-UK) Consortium | Jacquelyn Wynn, Mairead Hyland, The Lighthouse Lab in Alderley Park and Alex Alderton, Roberto Amato, Jeffrey Barrett, Sonia Goncalves, Ewan Harrison, David K. Jackson, Ian Johnston, Dominic Kwiatkowski, Cordelia Langford, John Sillitoe on behalf of the Wellcome Sanger Institute COVID-19 Surveillance Team |
| EPI_ISL_2346653, EPI_ISL_2346654 | Lighthouse Lab in Milton Keynes | Wellcome Sanger Institute for the COVID-19 Genomics UK (COG-UK) Consortium | The Lighthouse Lab in Milton Keynes and Alex Alderton, Roberto Amato, Jeffrey Barrett, Sonia Goncalves, Ewan Harrison, David K. Jackson, Ian Johnston, Dominic Kwiatkowski, Cordelia Langford, John Sillitoe on behalf of the Wellcome Sanger Institute COVID-19 Surveillance Team |
| EPI_ISL_2346655, EPI_ISL_2346656 | Lighthouse Lab in Alderley Park | Wellcome Sanger Institute for the COVID-19 Genomics UK (COG-UK) Consortium | Jacquelyn Wynn, Mairead Hyland, The Lighthouse Lab in Alderley Park and Alex Alderton, Roberto Amato, Jeffrey Barrett, Sonia Goncalves, Ewan Harrison, David K. Jackson, Ian Johnston, Dominic Kwiatkowski, Cordelia Langford, John Sillitoe on behalf of the Wellcome Sanger Institute COVID-19 Surveillance Team |
| EPI_ISL_2346657, EPI_ISL_2346658 | Lighthouse Lab in Milton Keynes | Wellcome Sanger Institute for the COVID-19 Genomics UK (COG-UK) Consortium | The Lighthouse Lab in Milton Keynes and Alex Alderton, Roberto Amato, Jeffrey Barrett, Sonia Goncalves, Ewan Harrison, David K. Jackson, Ian Johnston, Dominic Kwiatkowski, Cordelia Langford, John Sillitoe on behalf of the Wellcome Sanger Institute COVID-19 Surveillance Team |
| EPI_ISL_2346659, EPI_ISL_2346660, EPI_ISL_2346661 | Lighthouse Lab in Alderley Park | Wellcome Sanger Institute for the COVID-19 Genomics UK (COG-UK) Consortium | Jacquelyn Wynn, Mairead Hyland, The Lighthouse Lab in Alderley Park and Alex Alderton, Roberto Amato, Jeffrey Barrett, Sonia Goncalves, Ewan Harrison, David K. Jackson, Ian Johnston, Dominic Kwiatkowski, Cordelia Langford, John Sillitoe on behalf of the Wellcome Sanger Institute COVID-19 Surveillance Team |
| EPI_ISL_2346662 | Lighthouse Lab in Milton Keynes | Wellcome Sanger Institute for the COVID-19 Genomics UK (COG-UK) Consortium | The Lighthouse Lab in Milton Keynes and Alex Alderton, Roberto Amato, Jeffrey Barrett, Sonia Goncalves, Ewan Harrison, David K. Jackson, Ian Johnston, Dominic Kwiatkowski, Cordelia Langford, John Sillitoe on behalf of the Wellcome Sanger Institute COVID-19 Surveillance Team |
| EPI_ISL_2346663 | Lighthouse Lab in Alderley Park | Wellcome Sanger Institute for the COVID-19 Genomics UK (COG-UK) Consortium | Jacquelyn Wynn, Mairead Hyland, The Lighthouse Lab in Alderley Park and Alex Alderton, Roberto Amato, Jeffrey Barrett, Sonia Goncalves, Ewan Harrison, David K. Jackson, Ian Johnston, Dominic Kwiatkowski, Cordelia Langford, John Sillitoe on behalf of the Wellcome Sanger Institute COVID-19 Surveillance Team |
| EPI_ISL_2346665 | Lighthouse Lab in Milton Keynes | Wellcome Sanger Institute for the COVID-19 Genomics UK (COG-UK) Consortium | The Lighthouse Lab in Milton Keynes and Alex Alderton, Roberto Amato, Jeffrey Barrett, Sonia Goncalves, Ewan Harrison, David K. Jackson, Ian Johnston, Dominic Kwiatkowski, Cordelia Langford, John Sillitoe on behalf of the Wellcome Sanger Institute COVID-19 Surveillance Team |
| EPI_ISL_2346668, EPI_ISL_2346669 | Lighthouse Lab in Alderley Park | Wellcome Sanger Institute for the COVID-19 Genomics UK (COG-UK) Consortium | Jacquelyn Wynn, Mairead Hyland, The Lighthouse Lab in Alderley Park and Alex Alderton, Roberto Amato, Jeffrey Barrett, Sonia Goncalves, Ewan Harrison, David K. Jackson, Ian Johnston, Dominic Kwiatkowski, Cordelia Langford, John Sillitoe on behalf of the Wellcome Sanger Institute COVID-19 Surveillance Team |
| EPI_ISL_2346670, EPI_ISL_2346671 | Lighthouse Lab in Milton Keynes | Wellcome Sanger Institute for the COVID-19 Genomics UK (COG-UK) Consortium | The Lighthouse Lab in Milton Keynes and Alex Alderton, Roberto Amato, Jeffrey Barrett, Sonia Goncalves, Ewan Harrison, David K. Jackson, Ian Johnston, Dominic Kwiatkowski, Cordelia Langford, John Sillitoe on behalf of the Wellcome Sanger Institute COVID-19 Surveillance Team |
| EPI_ISL_2346672 | Lighthouse Lab in Alderley Park | Wellcome Sanger Institute for the COVID-19 Genomics UK (COG-UK) Consortium | Jacquelyn Wynn, Mairead Hyland, The Lighthouse Lab in Alderley Park and Alex Alderton, Roberto Amato, Jeffrey Barrett, Sonia Goncalves, Ewan Harrison, David K. Jackson, Ian Johnston, Dominic Kwiatkowski, Cordelia Langford, John Sillitoe on behalf of the Wellcome Sanger Institute COVID-19 Surveillance Team |
| EPI_ISL_2346673 | Lighthouse Lab in Milton Keynes | Wellcome Sanger Institute for the COVID-19 Genomics UK (COG-UK) Consortium | The Lighthouse Lab in Milton Keynes and Alex Alderton, Roberto Amato, Jeffrey Barrett, Sonia Goncalves, Ewan Harrison, David K. Jackson, Ian Johnston, Dominic Kwiatkowski, Cordelia Langford, John Sillitoe on behalf of the Wellcome Sanger Institute COVID-19 Surveillance Team |
| EPI_ISL_2346675, EPI_ISL_2346676 | Lighthouse Lab in Alderley Park | Wellcome Sanger Institute for the COVID-19 Genomics UK (COG-UK) Consortium | Jacquelyn Wynn, Mairead Hyland, The Lighthouse Lab in Alderley Park and Alex Alderton, Roberto Amato, Jeffrey Barrett, Sonia Goncalves, Ewan Harrison, David K. Jackson, Ian Johnston, Dominic Kwiatkowski, Cordelia Langford, John Sillitoe on behalf of the Wellcome Sanger Institute COVID-19 Surveillance Team |
| EPI_ISL_2346678, EPI_ISL_2346679, EPI_ISL_2346680, EPI_ISL_2346681, EPI_ISL_2346682, EPI_ISL_2346683 | Lighthouse Lab in Milton Keynes | Wellcome Sanger Institute for the COVID-19 Genomics UK (COG-UK) Consortium | The Lighthouse Lab in Milton Keynes and Alex Alderton, Roberto Amato, Jeffrey Barrett, Sonia Goncalves, Ewan Harrison, David K. Jackson, Ian Johnston, Dominic Kwiatkowski, Cordelia Langford, John Sillitoe on behalf of the Wellcome Sanger Institute COVID-19 Surveillance Team |
| EPI_ISL_2346684 | Lighthouse Lab in Alderley Park | Wellcome Sanger Institute for the COVID-19 Genomics UK (COG-UK) Consortium | Jacquelyn Wynn, Mairead Hyland, The Lighthouse Lab in Alderley Park and Alex Alderton, Roberto Amato, Jeffrey Barrett, Sonia Goncalves, Ewan Harrison, David K. Jackson, Ian Johnston, Dominic Kwiatkowski, Cordelia Langford, John Sillitoe on behalf of the Wellcome Sanger Institute COVID-19 Surveillance Team |
| EPI_ISL_2346685 | Lighthouse Lab in Milton Keynes | Wellcome Sanger Institute for the COVID-19 Genomics UK (COG-UK) Consortium | The Lighthouse Lab in Milton Keynes and Alex Alderton, Roberto Amato, Jeffrey Barrett, Sonia Goncalves, Ewan Harrison, David K. Jackson, Ian Johnston, Dominic Kwiatkowski, Cordelia Langford, John Sillitoe on behalf of the Wellcome Sanger Institute COVID-19 Surveillance Team |
| EPI_ISL_2346687 | Lighthouse Lab in Glasgow | Wellcome Sanger Institute for the COVID-19 Genomics UK (COG-UK) Consortium | Harper VanSteenhouse, Yumi Kasai, David Gray, Carol Clugston, Anna Dominiczak and Alex Alderton, Roberto Amato, Jeffrey Barrett, Sonia Goncalves, Ewan Harrison, David K. Jackson, Ian Johnston, Dominic Kwiatkowski, Cordelia Langford, John Sillitoe on behalf of the Wellcome Sanger Institute COVID-19 Surveillance Team |
| EPI_ISL_2346689 | Lighthouse Lab in Milton Keynes | Wellcome Sanger Institute for the COVID-19 Genomics UK (COG-UK) Consortium | The Lighthouse Lab in Milton Keynes and Alex Alderton, Roberto Amato, Jeffrey Barrett, Sonia Goncalves, Ewan Harrison, David K. Jackson, Ian Johnston, Dominic Kwiatkowski, Cordelia Langford, John Sillitoe on behalf of the Wellcome Sanger Institute COVID-19 Surveillance Team |
| EPI_ISL_2346690 | Lighthouse Lab in Alderley Park | Wellcome Sanger Institute for the COVID-19 Genomics UK (COG-UK) Consortium | Jacquelyn Wynn, Mairead Hyland, The Lighthouse Lab in Alderley Park and Alex Alderton, Roberto Amato, Jeffrey Barrett, Sonia Goncalves, Ewan Harrison, David K. Jackson, Ian Johnston, Dominic Kwiatkowski, Cordelia Langford, John Sillitoe on behalf of the Wellcome Sanger Institute COVID-19 Surveillance Team |
| EPI_ISL_2346691 | Lighthouse Lab in Milton Keynes | Wellcome Sanger Institute for the COVID-19 Genomics UK (COG-UK) Consortium | The Lighthouse Lab in Milton Keynes and Alex Alderton, Roberto Amato, Jeffrey Barrett, Sonia Goncalves, Ewan Harrison, David K. Jackson, Ian Johnston, Dominic Kwiatkowski, Cordelia Langford, John Sillitoe on behalf of the Wellcome Sanger Institute COVID-19 Surveillance Team |
| EPI_ISL_2346692 | Lighthouse Lab in Alderley Park | Wellcome Sanger Institute for the COVID-19 Genomics UK (COG-UK) Consortium | Jacquelyn Wynn, Mairead Hyland, The Lighthouse Lab in Alderley Park and Alex Alderton, Roberto Amato, Jeffrey Barrett, Sonia Goncalves, Ewan Harrison, David K. Jackson, Ian Johnston, Dominic Kwiatkowski, Cordelia Langford, John Sillitoe on behalf of the Wellcome Sanger Institute COVID-19 Surveillance Team |
| EPI_ISL_2346694, EPI_ISL_2346695, EPI_ISL_2346696, EPI_ISL_2346697 | Lighthouse Lab in Milton Keynes | Wellcome Sanger Institute for the COVID-19 Genomics UK (COG-UK) Consortium | The Lighthouse Lab in Milton Keynes and Alex Alderton, Roberto Amato, Jeffrey Barrett, Sonia Goncalves, Ewan Harrison, David K. Jackson, Ian Johnston, Dominic Kwiatkowski, Cordelia Langford, John Sillitoe on behalf of the Wellcome Sanger Institute COVID-19 Surveillance Team |
| EPI_ISL_2346698 | Lighthouse Lab in Alderley Park | Wellcome Sanger Institute for the COVID-19 Genomics UK (COG-UK) Consortium | Jacquelyn Wynn, Mairead Hyland, The Lighthouse Lab in Alderley Park and Alex Alderton, Roberto Amato, Jeffrey Barrett, Sonia Goncalves, Ewan Harrison, David K. Jackson, Ian Johnston, Dominic Kwiatkowski, Cordelia Langford, John Sillitoe on behalf of the Wellcome Sanger Institute COVID-19 Surveillance Team |
| EPI_ISL_2346702 | Lighthouse Lab in Milton Keynes | Wellcome Sanger Institute for the COVID-19 Genomics UK (COG-UK) Consortium | The Lighthouse Lab in Milton Keynes and Alex Alderton, Roberto Amato, Jeffrey Barrett, Sonia Goncalves, Ewan Harrison, David K. Jackson, Ian Johnston, Dominic Kwiatkowski, Cordelia Langford, John Sillitoe on behalf of the Wellcome Sanger Institute COVID-19 Surveillance Team |
| EPI_ISL_2346703 | Lighthouse Lab in Alderley Park | Wellcome Sanger Institute for the COVID-19 Genomics UK (COG-UK) Consortium | Jacquelyn Wynn, Mairead Hyland, The Lighthouse Lab in Alderley Park and Alex Alderton, Roberto Amato, Jeffrey Barrett, Sonia Goncalves, Ewan Harrison, David K. Jackson, Ian Johnston, Dominic Kwiatkowski, Cordelia Langford, John Sillitoe on behalf of the Wellcome Sanger Institute COVID-19 Surveillance Team |
| EPI_ISL_2346704, EPI_ISL_2346706, EPI_ISL_2346707, EPI_ISL_2346708, EPI_ISL_2346712 | Lighthouse Lab in Milton Keynes | Wellcome Sanger Institute for the COVID-19 Genomics UK (COG-UK) Consortium | The Lighthouse Lab in Milton Keynes and Alex Alderton, Roberto Amato, Jeffrey Barrett, Sonia Goncalves, Ewan Harrison, David K. Jackson, Ian Johnston, Dominic Kwiatkowski, Cordelia Langford, John Sillitoe on behalf of the Wellcome Sanger Institute COVID-19 Surveillance Team |
| EPI_ISL_2346715 | Lighthouse Lab in Alderley Park | Wellcome Sanger Institute for the COVID-19 Genomics UK (COG-UK) Consortium | Jacquelyn Wynn, Mairead Hyland, The Lighthouse Lab in Alderley Park and Alex Alderton, Roberto Amato, Jeffrey Barrett, Sonia Goncalves, Ewan Harrison, David K. Jackson, Ian Johnston, Dominic Kwiatkowski, Cordelia Langford, John Sillitoe on behalf of the Wellcome Sanger Institute COVID-19 Surveillance Team |

| | | | |
|---|---|---|---|
| EPI_ISL_2346718, EPI_ISL_2346720 | Lighthouse Lab in Milton Keynes | Wellcome Sanger Institute for the COVID-19 Genomics UK (COG-UK) Consortium | The Lighthouse Lab in Milton Keynes and Alex Alderton, Roberto Amato, Jeffrey Barrett, Sonia Goncalves, Ewan Harrison, David K. Jackson, Ian Johnston, Dominic Kwiatkowski, Cordelia Langford, John Sillitoe on behalf of the Wellcome Sanger Institute COVID-19 Surveillance Team |
| EPI_ISL_2346721, EPI_ISL_2346723 | Lighthouse Lab in Alderley Park | Wellcome Sanger Institute for the COVID-19 Genomics UK (COG-UK) Consortium | Jacquelyn Wynn, Mairead Hyland, The Lighthouse Lab in Alderley Park and Alex Alderton, Roberto Amato, Jeffrey Barrett, Sonia Goncalves, Ewan Harrison, David K. Jackson, Ian Johnston, Dominic Kwiatkowski, Cordelia Langford, John Sillitoe on behalf of the Wellcome Sanger Institute COVID-19 Surveillance Team |
| EPI_ISL_2346724 | Lighthouse Lab in Milton Keynes | Wellcome Sanger Institute for the COVID-19 Genomics UK (COG-UK) Consortium | The Lighthouse Lab in Milton Keynes and Alex Alderton, Roberto Amato, Jeffrey Barrett, Sonia Goncalves, Ewan Harrison, David K. Jackson, Ian Johnston, Dominic Kwiatkowski, Cordelia Langford, John Sillitoe on behalf of the Wellcome Sanger Institute COVID-19 Surveillance Team |
| EPI_ISL_2346725 | Lighthouse Lab in Alderley Park | Wellcome Sanger Institute for the COVID-19 Genomics UK (COG-UK) Consortium | Jacquelyn Wynn, Mairead Hyland, The Lighthouse Lab in Alderley Park and Alex Alderton, Roberto Amato, Jeffrey Barrett, Sonia Goncalves, Ewan Harrison, David K. Jackson, Ian Johnston, Dominic Kwiatkowski, Cordelia Langford, John Sillitoe on behalf of the Wellcome Sanger Institute COVID-19 Surveillance Team |
| EPI_ISL_2346726 | Lighthouse Lab in Milton Keynes | Wellcome Sanger Institute for the COVID-19 Genomics UK (COG-UK) Consortium | The Lighthouse Lab in Milton Keynes and Alex Alderton, Roberto Amato, Jeffrey Barrett, Sonia Goncalves, Ewan Harrison, David K. Jackson, Ian Johnston, Dominic Kwiatkowski, Cordelia Langford, John Sillitoe on behalf of the Wellcome Sanger Institute COVID-19 Surveillance Team |
| EPI_ISL_2346728, EPI_ISL_2346730 | Lighthouse Lab in Alderley Park | Wellcome Sanger Institute for the COVID-19 Genomics UK (COG-UK) Consortium | Jacquelyn Wynn, Mairead Hyland, The Lighthouse Lab in Alderley Park and Alex Alderton, Roberto Amato, Jeffrey Barrett, Sonia Goncalves, Ewan Harrison, David K. Jackson, Ian Johnston, Dominic Kwiatkowski, Cordelia Langford, John Sillitoe on behalf of the Wellcome Sanger Institute COVID-19 Surveillance Team |
| EPI_ISL_2346731 | Lighthouse Lab in Milton Keynes | Wellcome Sanger Institute for the COVID-19 Genomics UK (COG-UK) Consortium | The Lighthouse Lab in Milton Keynes and Alex Alderton, Roberto Amato, Jeffrey Barrett, Sonia Goncalves, Ewan Harrison, David K. Jackson, Ian Johnston, Dominic Kwiatkowski, Cordelia Langford, John Sillitoe on behalf of the Wellcome Sanger Institute COVID-19 Surveillance Team |
| EPI_ISL_2346733, EPI_ISL_2346734, EPI_ISL_2346735 | Lighthouse Lab in Alderley Park | Wellcome Sanger Institute for the COVID-19 Genomics UK (COG-UK) Consortium | Jacquelyn Wynn, Mairead Hyland, The Lighthouse Lab in Alderley Park and Alex Alderton, Roberto Amato, Jeffrey Barrett, Sonia Goncalves, Ewan Harrison, David K. Jackson, Ian Johnston, Dominic Kwiatkowski, Cordelia Langford, John Sillitoe on behalf of the Wellcome Sanger Institute COVID-19 Surveillance Team |
| EPI_ISL_2346736 | Lighthouse Lab in Milton Keynes | Wellcome Sanger Institute for the COVID-19 Genomics UK (COG-UK) Consortium | The Lighthouse Lab in Milton Keynes and Alex Alderton, Roberto Amato, Jeffrey Barrett, Sonia Goncalves, Ewan Harrison, David K. Jackson, Ian Johnston, Dominic Kwiatkowski, Cordelia Langford, John Sillitoe on behalf of the Wellcome Sanger Institute COVID-19 Surveillance Team |
| EPI_ISL_2346738 | Lighthouse Lab in Alderley Park | Wellcome Sanger Institute for the COVID-19 Genomics UK (COG-UK) Consortium | Jacquelyn Wynn, Mairead Hyland, The Lighthouse Lab in Alderley Park and Alex Alderton, Roberto Amato, Jeffrey Barrett, Sonia Goncalves, Ewan Harrison, David K. Jackson, Ian Johnston, Dominic Kwiatkowski, Cordelia Langford, John Sillitoe on behalf of the Wellcome Sanger Institute COVID-19 Surveillance Team |
| EPI_ISL_2346739 | Lighthouse Lab in Milton Keynes | Wellcome Sanger Institute for the COVID-19 Genomics UK (COG-UK) Consortium | The Lighthouse Lab in Milton Keynes and Alex Alderton, Roberto Amato, Jeffrey Barrett, Sonia Goncalves, Ewan Harrison, David K. Jackson, Ian Johnston, Dominic Kwiatkowski, Cordelia Langford, John Sillitoe on behalf of the Wellcome Sanger Institute COVID-19 Surveillance Team |
| EPI_ISL_2346740, EPI_ISL_2346741, EPI_ISL_2346742, EPI_ISL_2346744, EPI_ISL_2346747, EPI_ISL_2346748, EPI_ISL_2346749, EPI_ISL_2346750, EPI_ISL_2346751, EPI_ISL_2346753, EPI_ISL_2346754, EPI_ISL_2346755, EPI_ISL_2346756, EPI_ISL_2346759 | | | |
| see above | Lighthouse Lab in Alderley Park | Wellcome Sanger Institute for the COVID-19 Genomics UK (COG-UK) Consortium | Jacquelyn Wynn, Mairead Hyland, The Lighthouse Lab in Alderley Park and Alex Alderton, Roberto Amato, Jeffrey Barrett, Sonia Goncalves, Ewan Harrison, David K. Jackson, Ian Johnston, Dominic Kwiatkowski, Cordelia Langford, John Sillitoe on behalf of the Wellcome Sanger Institute COVID-19 Surveillance Team |
| EPI_ISL_2346760 | Lighthouse Lab in Milton Keynes | Wellcome Sanger Institute for the COVID-19 Genomics UK (COG-UK) Consortium | The Lighthouse Lab in Milton Keynes and Alex Alderton, Roberto Amato, Jeffrey Barrett, Sonia Goncalves, Ewan Harrison, David K. Jackson, Ian Johnston, Dominic Kwiatkowski, Cordelia Langford, John Sillitoe on behalf of the Wellcome Sanger Institute COVID-19 Surveillance Team |
| EPI_ISL_2346762, EPI_ISL_2346763, EPI_ISL_2346764, EPI_ISL_2346766, EPI_ISL_2346767, EPI_ISL_2346768 | Lighthouse Lab in Alderley Park | Wellcome Sanger Institute for the COVID-19 Genomics UK (COG-UK) Consortium | Jacquelyn Wynn, Mairead Hyland, The Lighthouse Lab in Alderley Park and Alex Alderton, Roberto Amato, Jeffrey Barrett, Sonia Goncalves, Ewan Harrison, David K. Jackson, Ian Johnston, Dominic Kwiatkowski, Cordelia Langford, John Sillitoe on behalf of the Wellcome Sanger Institute COVID-19 Surveillance Team |
| EPI_ISL_2346769, EPI_ISL_2346770 | Lighthouse Lab in Milton Keynes | Wellcome Sanger Institute for the COVID-19 Genomics UK (COG-UK) Consortium | The Lighthouse Lab in Milton Keynes and Alex Alderton, Roberto Amato, Jeffrey Barrett, Sonia Goncalves, Ewan Harrison, David K. Jackson, Ian Johnston, Dominic Kwiatkowski, Cordelia Langford, John Sillitoe on behalf of the Wellcome Sanger Institute COVID-19 Surveillance Team |
| EPI_ISL_2346772 | Lighthouse Lab in Alderley Park | Wellcome Sanger Institute for the COVID-19 Genomics UK (COG-UK) Consortium | Jacquelyn Wynn, Mairead Hyland, The Lighthouse Lab in Alderley Park and Alex Alderton, Roberto Amato, Jeffrey Barrett, Sonia Goncalves, Ewan Harrison, David K. Jackson, Ian Johnston, Dominic Kwiatkowski, Cordelia Langford, John Sillitoe on behalf of the Wellcome Sanger Institute COVID-19 Surveillance Team |
| EPI_ISL_2346774 | Lighthouse Lab in Milton Keynes | Wellcome Sanger Institute for the COVID-19 Genomics UK (COG-UK) Consortium | The Lighthouse Lab in Milton Keynes and Alex Alderton, Roberto Amato, Jeffrey Barrett, Sonia Goncalves, Ewan Harrison, David K. Jackson, Ian Johnston, Dominic Kwiatkowski, Cordelia Langford, John Sillitoe on behalf of the Wellcome Sanger Institute COVID-19 Surveillance Team |
| EPI_ISL_2346775, EPI_ISL_2346776, EPI_ISL_2346777 | Lighthouse Lab in Alderley Park | Wellcome Sanger Institute for the COVID-19 Genomics UK (COG-UK) Consortium | Jacquelyn Wynn, Mairead Hyland, The Lighthouse Lab in Alderley Park and Alex Alderton, Roberto Amato, Jeffrey Barrett, Sonia Goncalves, Ewan Harrison, David K. Jackson, Ian Johnston, Dominic Kwiatkowski, Cordelia Langford, John Sillitoe on behalf of the Wellcome Sanger Institute COVID-19 Surveillance Team |
| EPI_ISL_2346779 | Lighthouse Lab in Milton Keynes | Wellcome Sanger Institute for the COVID-19 Genomics UK (COG-UK) Consortium | The Lighthouse Lab in Milton Keynes and Alex Alderton, Roberto Amato, Jeffrey Barrett, Sonia Goncalves, Ewan Harrison, David K. Jackson, Ian Johnston, Dominic Kwiatkowski, Cordelia Langford, John Sillitoe on behalf of the Wellcome Sanger Institute COVID-19 Surveillance Team |
| EPI_ISL_2346782, EPI_ISL_2346783, EPI_ISL_2346784, EPI_ISL_2346787, EPI_ISL_2346788, EPI_ISL_2346789, EPI_ISL_2346790 | Lighthouse Lab in Alderley Park | Wellcome Sanger Institute for the COVID-19 Genomics UK (COG-UK) Consortium | Jacquelyn Wynn, Mairead Hyland, The Lighthouse Lab in Alderley Park and Alex Alderton, Roberto Amato, Jeffrey Barrett, Sonia Goncalves, Ewan Harrison, David K. Jackson, Ian Johnston, Dominic Kwiatkowski, Cordelia Langford, John Sillitoe on behalf of the Wellcome Sanger Institute COVID-19 Surveillance Team |
| EPI_ISL_2346791 | Lighthouse Lab in Milton Keynes | Wellcome Sanger Institute for the COVID-19 Genomics UK (COG-UK) Consortium | The Lighthouse Lab in Milton Keynes and Alex Alderton, Roberto Amato, Jeffrey Barrett, Sonia Goncalves, Ewan Harrison, David K. Jackson, Ian Johnston, Dominic Kwiatkowski, Cordelia Langford, John Sillitoe on behalf of the Wellcome Sanger Institute COVID-19 Surveillance Team |
| EPI_ISL_2346793 | Lighthouse Lab in Alderley Park | Wellcome Sanger Institute for the COVID-19 Genomics UK (COG-UK) Consortium | Jacquelyn Wynn, Mairead Hyland, The Lighthouse Lab in Alderley Park and Alex Alderton, Roberto Amato, Jeffrey Barrett, Sonia Goncalves, Ewan Harrison, David K. Jackson, Ian Johnston, Dominic Kwiatkowski, Cordelia Langford, John Sillitoe on behalf of the Wellcome Sanger Institute COVID-19 Surveillance Team |
| EPI_ISL_2346794 | Lighthouse Lab in Milton Keynes | Wellcome Sanger Institute for the COVID-19 Genomics UK (COG-UK) Consortium | The Lighthouse Lab in Milton Keynes and Alex Alderton, Roberto Amato, Jeffrey Barrett, Sonia Goncalves, Ewan Harrison, David K. Jackson, Ian Johnston, Dominic Kwiatkowski, Cordelia Langford, John Sillitoe on behalf of the Wellcome Sanger Institute COVID-19 Surveillance Team |
| EPI_ISL_2346795 | Lighthouse Lab in Alderley Park | Wellcome Sanger Institute for the COVID-19 Genomics UK (COG-UK) Consortium | Jacquelyn Wynn, Mairead Hyland, The Lighthouse Lab in Alderley Park and Alex Alderton, Roberto Amato, Jeffrey Barrett, Sonia Goncalves, Ewan Harrison, David K. Jackson, Ian Johnston, Dominic Kwiatkowski, Cordelia Langford, John Sillitoe on behalf of the Wellcome Sanger Institute COVID-19 Surveillance Team |
| EPI_ISL_2346797 | Lighthouse Lab in Milton Keynes | Wellcome Sanger Institute for the COVID-19 Genomics UK (COG-UK) Consortium | The Lighthouse Lab in Milton Keynes and Alex Alderton, Roberto Amato, Jeffrey Barrett, Sonia Goncalves, Ewan Harrison, David K. Jackson, Ian Johnston, Dominic Kwiatkowski, Cordelia Langford, John Sillitoe on behalf of the Wellcome Sanger Institute COVID-19 Surveillance Team |
| EPI_ISL_2346800, EPI_ISL_2346802, EPI_ISL_2346803, EPI_ISL_2346804 | Lighthouse Lab in Alderley Park | Wellcome Sanger Institute for the COVID-19 Genomics UK (COG-UK) Consortium | Jacquelyn Wynn, Mairead Hyland, The Lighthouse Lab in Alderley Park and Alex Alderton, Roberto Amato, Jeffrey Barrett, Sonia Goncalves, Ewan Harrison, David K. Jackson, Ian Johnston, Dominic Kwiatkowski, Cordelia Langford, John Sillitoe on behalf of the Wellcome Sanger Institute COVID-19 Surveillance Team |
| EPI_ISL_2346806 | Lighthouse Lab in Milton Keynes | Wellcome Sanger Institute for the COVID-19 Genomics UK (COG-UK) Consortium | The Lighthouse Lab in Milton Keynes and Alex Alderton, Roberto Amato, Jeffrey Barrett, Sonia Goncalves, Ewan Harrison, David K. Jackson, Ian Johnston, Dominic Kwiatkowski, Cordelia Langford, John Sillitoe on behalf of the Wellcome Sanger Institute COVID-19 Surveillance Team |
| EPI_ISL_2346808 | Lighthouse Lab in Alderley Park | Wellcome Sanger Institute for the COVID-19 Genomics UK (COG-UK) Consortium | Jacquelyn Wynn, Mairead Hyland, The Lighthouse Lab in Alderley Park and Alex Alderton, Roberto Amato, Jeffrey Barrett, Sonia Goncalves, Ewan Harrison, David K. Jackson, Ian Johnston, Dominic Kwiatkowski, Cordelia Langford, John Sillitoe on behalf of the Wellcome Sanger Institute COVID-19 Surveillance Team |
| EPI_ISL_2346809, EPI_ISL_2346810 | Lighthouse Lab in Milton Keynes | Wellcome Sanger Institute for the COVID-19 Genomics UK (COG-UK) Consortium | The Lighthouse Lab in Milton Keynes and Alex Alderton, Roberto Amato, Jeffrey Barrett, Sonia Goncalves, Ewan Harrison, David K. Jackson, Ian Johnston, Dominic Kwiatkowski, Cordelia Langford, John Sillitoe on behalf of the Wellcome Sanger Institute COVID-19 Surveillance Team |
| EPI_ISL_2346811, EPI_ISL_2346812, EPI_ISL_2346814, EPI_ISL_2346815, EPI_ISL_2346816, EPI_ISL_2346817, EPI_ISL_2346818 | Lighthouse Lab in Alderley Park | Wellcome Sanger Institute for the COVID-19 Genomics UK (COG-UK) Consortium | Jacquelyn Wynn, Mairead Hyland, The Lighthouse Lab in Alderley Park and Alex Alderton, Roberto Amato, Jeffrey Barrett, Sonia Goncalves, Ewan Harrison, David K. Jackson, Ian Johnston, Dominic Kwiatkowski, Cordelia Langford, John Sillitoe on behalf of the Wellcome Sanger Institute COVID-19 Surveillance Team |
| EPI_ISL_2346819, EPI_ISL_2346820 | Lighthouse Lab in Milton Keynes | Wellcome Sanger Institute for the COVID-19 Genomics UK (COG-UK) Consortium | The Lighthouse Lab in Milton Keynes and Alex Alderton, Roberto Amato, Jeffrey Barrett, Sonia Goncalves, Ewan Harrison, David K. Jackson, Ian Johnston, Dominic Kwiatkowski, Cordelia Langford, John Sillitoe on behalf of the Wellcome Sanger Institute COVID-19 Surveillance Team |
| EPI_ISL_2346821 | Lighthouse Lab in Alderley Park | Wellcome Sanger Institute for the COVID-19 Genomics UK (COG-UK) Consortium | Jacquelyn Wynn, Mairead Hyland, The Lighthouse Lab in Alderley Park and Alex Alderton, Roberto Amato, Jeffrey Barrett, Sonia Goncalves, Ewan Harrison, David K. Jackson, Ian Johnston, Dominic Kwiatkowski, Cordelia Langford, John Sillitoe on behalf of the Wellcome Sanger Institute COVID-19 Surveillance Team |
| EPI_ISL_2346822 | Lighthouse Lab in Milton Keynes | Wellcome Sanger Institute for the COVID-19 Genomics UK (COG-UK) Consortium | The Lighthouse Lab in Milton Keynes and Alex Alderton, Roberto Amato, Jeffrey Barrett, Sonia Goncalves, Ewan Harrison, David K. Jackson, Ian Johnston, Dominic Kwiatkowski, Cordelia Langford, John Sillitoe on behalf of the Wellcome Sanger Institute COVID-19 Surveillance Team |
| EPI_ISL_2346823, EPI_ISL_2346824, EPI_ISL_2346825, EPI_ISL_2346826, EPI_ISL_2346827, EPI_ISL_2346829 | Lighthouse Lab in Alderley Park | Wellcome Sanger Institute for the COVID-19 Genomics UK (COG-UK) Consortium | Jacquelyn Wynn, Mairead Hyland, The Lighthouse Lab in Alderley Park and Alex Alderton, Roberto Amato, Jeffrey Barrett, Sonia Goncalves, Ewan Harrison, David K. Jackson, Ian Johnston, Dominic Kwiatkowski, Cordelia Langford, John Sillitoe on behalf of the Wellcome Sanger Institute COVID-19 Surveillance Team |
| EPI_ISL_2346830 | Lighthouse Lab in Milton Keynes | Wellcome Sanger Institute for the COVID-19 Genomics UK (COG-UK) Consortium | The Lighthouse Lab in Milton Keynes and Alex Alderton, Roberto Amato, Jeffrey Barrett, Sonia Goncalves, Ewan Harrison, David K. Jackson, Ian Johnston, Dominic Kwiatkowski, Cordelia Langford, John Sillitoe on behalf of the Wellcome Sanger Institute COVID-19 Surveillance Team |
| EPI_ISL_2346832 | Lighthouse Lab in Alderley Park | Wellcome Sanger Institute for the COVID-19 Genomics UK (COG-UK) | Jacquelyn Wynn, Mairead Hyland, The Lighthouse Lab in Alderley Park and Alex Alderton, Roberto Amato, Jeffrey Barrett, Sonia Goncalves, Ewan Harrison, David K. Jackson, Ian Johnston, Dominic Kwiatkowski, Cordelia Langford, John Sillitoe on |

| | | | |
|---|---|---|---|
| | | Consortium | behalf of the Wellcome Sanger Institute COVID-19 Surveillance Team |
| EPI_ISL_2346833 | Lighthouse Lab in Milton Keynes | Wellcome Sanger Institute for the COVID-19 Genomics UK (COG-UK) Consortium | The Lighthouse Lab in Milton Keynes and Alex Alderton, Roberto Amato, Jeffrey Barrett, Sonia Goncalves, Ewan Harrison, David K. Jackson, Ian Johnston, Dominic Kwiatkowski, Cordelia Langford, John Sillitoe on behalf of the Wellcome Sanger Institute COVID-19 Surveillance Team |
| EPI_ISL_2346834, EPI_ISL_2346835, EPI_ISL_2346836 | Lighthouse Lab in Alderley Park | Wellcome Sanger Institute for the COVID-19 Genomics UK (COG-UK) Consortium | Jacquelyn Wynn, Mairead Hyland, The Lighthouse Lab in Alderley Park and Alex Alderton, Roberto Amato, Jeffrey Barrett, Sonia Goncalves, Ewan Harrison, David K. Jackson, Ian Johnston, Dominic Kwiatkowski, Cordelia Langford, John Sillitoe on behalf of the Wellcome Sanger Institute COVID-19 Surveillance Team |
| EPI_ISL_2346837, EPI_ISL_2346838 | Lighthouse Lab in Milton Keynes | Wellcome Sanger Institute for the COVID-19 Genomics UK (COG-UK) Consortium | The Lighthouse Lab in Milton Keynes and Alex Alderton, Roberto Amato, Jeffrey Barrett, Sonia Goncalves, Ewan Harrison, David K. Jackson, Ian Johnston, Dominic Kwiatkowski, Cordelia Langford, John Sillitoe on behalf of the Wellcome Sanger Institute COVID-19 Surveillance Team |
| EPI_ISL_2346839, EPI_ISL_2346842, EPI_ISL_2346843, EPI_ISL_2346844, EPI_ISL_2346845, EPI_ISL_2346846 | Lighthouse Lab in Alderley Park | Wellcome Sanger Institute for the COVID-19 Genomics UK (COG-UK) Consortium | Jacquelyn Wynn, Mairead Hyland, The Lighthouse Lab in Alderley Park and Alex Alderton, Roberto Amato, Jeffrey Barrett, Sonia Goncalves, Ewan Harrison, David K. Jackson, Ian Johnston, Dominic Kwiatkowski, Cordelia Langford, John Sillitoe on behalf of the Wellcome Sanger Institute COVID-19 Surveillance Team |
| EPI_ISL_2346847 | Lighthouse Lab in Milton Keynes | Wellcome Sanger Institute for the COVID-19 Genomics UK (COG-UK) Consortium | The Lighthouse Lab in Milton Keynes and Alex Alderton, Roberto Amato, Jeffrey Barrett, Sonia Goncalves, Ewan Harrison, David K. Jackson, Ian Johnston, Dominic Kwiatkowski, Cordelia Langford, John Sillitoe on behalf of the Wellcome Sanger Institute COVID-19 Surveillance Team |
| EPI_ISL_2346848 | Lighthouse Lab in Alderley Park | Wellcome Sanger Institute for the COVID-19 Genomics UK (COG-UK) Consortium | Jacquelyn Wynn, Mairead Hyland, The Lighthouse Lab in Alderley Park and Alex Alderton, Roberto Amato, Jeffrey Barrett, Sonia Goncalves, Ewan Harrison, David K. Jackson, Ian Johnston, Dominic Kwiatkowski, Cordelia Langford, John Sillitoe on behalf of the Wellcome Sanger Institute COVID-19 Surveillance Team |
| EPI_ISL_2346849 | Lighthouse Lab in Milton Keynes | Wellcome Sanger Institute for the COVID-19 Genomics UK (COG-UK) Consortium | The Lighthouse Lab in Milton Keynes and Alex Alderton, Roberto Amato, Jeffrey Barrett, Sonia Goncalves, Ewan Harrison, David K. Jackson, Ian Johnston, Dominic Kwiatkowski, Cordelia Langford, John Sillitoe on behalf of the Wellcome Sanger Institute COVID-19 Surveillance Team |
| EPI_ISL_2346850, EPI_ISL_2346851, EPI_ISL_2346853, EPI_ISL_2346854, EPI_ISL_2346855, EPI_ISL_2346856, EPI_ISL_2346857 | Lighthouse Lab in Alderley Park | Wellcome Sanger Institute for the COVID-19 Genomics UK (COG-UK) Consortium | Jacquelyn Wynn, Mairead Hyland, The Lighthouse Lab in Alderley Park and Alex Alderton, Roberto Amato, Jeffrey Barrett, Sonia Goncalves, Ewan Harrison, David K. Jackson, Ian Johnston, Dominic Kwiatkowski, Cordelia Langford, John Sillitoe on behalf of the Wellcome Sanger Institute COVID-19 Surveillance Team |
| EPI_ISL_2346858 | Lighthouse Lab in Milton Keynes | Wellcome Sanger Institute for the COVID-19 Genomics UK (COG-UK) Consortium | The Lighthouse Lab in Milton Keynes and Alex Alderton, Roberto Amato, Jeffrey Barrett, Sonia Goncalves, Ewan Harrison, David K. Jackson, Ian Johnston, Dominic Kwiatkowski, Cordelia Langford, John Sillitoe on behalf of the Wellcome Sanger Institute COVID-19 Surveillance Team |
| EPI_ISL_2346859, EPI_ISL_2346860, EPI_ISL_2346861 | Lighthouse Lab in Alderley Park | Wellcome Sanger Institute for the COVID-19 Genomics UK (COG-UK) Consortium | Jacquelyn Wynn, Mairead Hyland, The Lighthouse Lab in Alderley Park and Alex Alderton, Roberto Amato, Jeffrey Barrett, Sonia Goncalves, Ewan Harrison, David K. Jackson, Ian Johnston, Dominic Kwiatkowski, Cordelia Langford, John Sillitoe on behalf of the Wellcome Sanger Institute COVID-19 Surveillance Team |
| EPI_ISL_2346862 | Lighthouse Lab in Milton Keynes | Wellcome Sanger Institute for the COVID-19 Genomics UK (COG-UK) Consortium | The Lighthouse Lab in Milton Keynes and Alex Alderton, Roberto Amato, Jeffrey Barrett, Sonia Goncalves, Ewan Harrison, David K. Jackson, Ian Johnston, Dominic Kwiatkowski, Cordelia Langford, John Sillitoe on behalf of the Wellcome Sanger Institute COVID-19 Surveillance Team |
| EPI_ISL_2346863, EPI_ISL_2346866, EPI_ISL_2346868, EPI_ISL_2346869, EPI_ISL_2346870, EPI_ISL_2346871 | Lighthouse Lab in Alderley Park | Wellcome Sanger Institute for the COVID-19 Genomics UK (COG-UK) Consortium | Jacquelyn Wynn, Mairead Hyland, The Lighthouse Lab in Alderley Park and Alex Alderton, Roberto Amato, Jeffrey Barrett, Sonia Goncalves, Ewan Harrison, David K. Jackson, Ian Johnston, Dominic Kwiatkowski, Cordelia Langford, John Sillitoe on behalf of the Wellcome Sanger Institute COVID-19 Surveillance Team |
| EPI_ISL_2346872 | Lighthouse Lab in Milton Keynes | Wellcome Sanger Institute for the COVID-19 Genomics UK (COG-UK) Consortium | The Lighthouse Lab in Milton Keynes and Alex Alderton, Roberto Amato, Jeffrey Barrett, Sonia Goncalves, Ewan Harrison, David K. Jackson, Ian Johnston, Dominic Kwiatkowski, Cordelia Langford, John Sillitoe on behalf of the Wellcome Sanger Institute COVID-19 Surveillance Team |
| EPI_ISL_2346873, EPI_ISL_2346874, EPI_ISL_2346875, EPI_ISL_2346877, EPI_ISL_2346878, EPI_ISL_2346879, EPI_ISL_2346880 | Lighthouse Lab in Alderley Park | Wellcome Sanger Institute for the COVID-19 Genomics UK (COG-UK) Consortium | Jacquelyn Wynn, Mairead Hyland, The Lighthouse Lab in Alderley Park and Alex Alderton, Roberto Amato, Jeffrey Barrett, Sonia Goncalves, Ewan Harrison, David K. Jackson, Ian Johnston, Dominic Kwiatkowski, Cordelia Langford, John Sillitoe on behalf of the Wellcome Sanger Institute COVID-19 Surveillance Team |
| EPI_ISL_2346882 | Lighthouse Lab in Milton Keynes | Wellcome Sanger Institute for the COVID-19 Genomics UK (COG-UK) Consortium | The Lighthouse Lab in Milton Keynes and Alex Alderton, Roberto Amato, Jeffrey Barrett, Sonia Goncalves, Ewan Harrison, David K. Jackson, Ian Johnston, Dominic Kwiatkowski, Cordelia Langford, John Sillitoe on behalf of the Wellcome Sanger Institute COVID-19 Surveillance Team |
| EPI_ISL_2346883, EPI_ISL_2346884, EPI_ISL_2346885, EPI_ISL_2346887 | Lighthouse Lab in Alderley Park | Wellcome Sanger Institute for the COVID-19 Genomics UK (COG-UK) Consortium | Jacquelyn Wynn, Mairead Hyland, The Lighthouse Lab in Alderley Park and Alex Alderton, Roberto Amato, Jeffrey Barrett, Sonia Goncalves, Ewan Harrison, David K. Jackson, Ian Johnston, Dominic Kwiatkowski, Cordelia Langford, John Sillitoe on behalf of the Wellcome Sanger Institute COVID-19 Surveillance Team |
| EPI_ISL_2346888 | Lighthouse Lab in Milton Keynes | Wellcome Sanger Institute for the COVID-19 Genomics UK (COG-UK) Consortium | The Lighthouse Lab in Milton Keynes and Alex Alderton, Roberto Amato, Jeffrey Barrett, Sonia Goncalves, Ewan Harrison, David K. Jackson, Ian Johnston, Dominic Kwiatkowski, Cordelia Langford, John Sillitoe on behalf of the Wellcome Sanger Institute COVID-19 Surveillance Team |
| EPI_ISL_2346889, EPI_ISL_2346891, EPI_ISL_2346892 | Lighthouse Lab in Alderley Park | Wellcome Sanger Institute for the COVID-19 Genomics UK (COG-UK) Consortium | Jacquelyn Wynn, Mairead Hyland, The Lighthouse Lab in Alderley Park and Alex Alderton, Roberto Amato, Jeffrey Barrett, Sonia Goncalves, Ewan Harrison, David K. Jackson, Ian Johnston, Dominic Kwiatkowski, Cordelia Langford, John Sillitoe on behalf of the Wellcome Sanger Institute COVID-19 Surveillance Team |
| EPI_ISL_2346893 | Lighthouse Lab in Milton Keynes | Wellcome Sanger Institute for the COVID-19 Genomics UK (COG-UK) Consortium | The Lighthouse Lab in Milton Keynes and Alex Alderton, Roberto Amato, Jeffrey Barrett, Sonia Goncalves, Ewan Harrison, David K. Jackson, Ian Johnston, Dominic Kwiatkowski, Cordelia Langford, John Sillitoe on behalf of the Wellcome Sanger Institute COVID-19 Surveillance Team |
| EPI_ISL_2346894, EPI_ISL_2346895, EPI_ISL_2346896, EPI_ISL_2346897, EPI_ISL_2346898, EPI_ISL_2346899, EPI_ISL_2346900, EPI_ISL_2346901, EPI_ISL_2346902, EPI_ISL_2346903 | Lighthouse Lab in Alderley Park | Wellcome Sanger Institute for the COVID-19 Genomics UK (COG-UK) Consortium | Jacquelyn Wynn, Mairead Hyland, The Lighthouse Lab in Alderley Park and Alex Alderton, Roberto Amato, Jeffrey Barrett, Sonia Goncalves, Ewan Harrison, David K. Jackson, Ian Johnston, Dominic Kwiatkowski, Cordelia Langford, John Sillitoe on behalf of the Wellcome Sanger Institute COVID-19 Surveillance Team |
| EPI_ISL_2346904 | Lighthouse Lab in Milton Keynes | Wellcome Sanger Institute for the COVID-19 Genomics UK (COG-UK) Consortium | The Lighthouse Lab in Milton Keynes and Alex Alderton, Roberto Amato, Jeffrey Barrett, Sonia Goncalves, Ewan Harrison, David K. Jackson, Ian Johnston, Dominic Kwiatkowski, Cordelia Langford, John Sillitoe on behalf of the Wellcome Sanger Institute COVID-19 Surveillance Team |
| EPI_ISL_2346906 | Lighthouse Lab in Alderley Park | Wellcome Sanger Institute for the COVID-19 Genomics UK (COG-UK) Consortium | Jacquelyn Wynn, Mairead Hyland, The Lighthouse Lab in Alderley Park and Alex Alderton, Roberto Amato, Jeffrey Barrett, Sonia Goncalves, Ewan Harrison, David K. Jackson, Ian Johnston, Dominic Kwiatkowski, Cordelia Langford, John Sillitoe on behalf of the Wellcome Sanger Institute COVID-19 Surveillance Team |
| EPI_ISL_2346907 | Lighthouse Lab in Milton Keynes | Wellcome Sanger Institute for the COVID-19 Genomics UK (COG-UK) Consortium | The Lighthouse Lab in Milton Keynes and Alex Alderton, Roberto Amato, Jeffrey Barrett, Sonia Goncalves, Ewan Harrison, David K. Jackson, Ian Johnston, Dominic Kwiatkowski, Cordelia Langford, John Sillitoe on behalf of the Wellcome Sanger Institute COVID-19 Surveillance Team |
| EPI_ISL_2346908, EPI_ISL_2346909 | Lighthouse Lab in Alderley Park | Wellcome Sanger Institute for the COVID-19 Genomics UK (COG-UK) Consortium | Jacquelyn Wynn, Mairead Hyland, The Lighthouse Lab in Alderley Park and Alex Alderton, Roberto Amato, Jeffrey Barrett, Sonia Goncalves, Ewan Harrison, David K. Jackson, Ian Johnston, Dominic Kwiatkowski, Cordelia Langford, John Sillitoe on behalf of the Wellcome Sanger Institute COVID-19 Surveillance Team |
| EPI_ISL_2346910 | Lighthouse Lab in Milton Keynes | Wellcome Sanger Institute for the COVID-19 Genomics UK (COG-UK) Consortium | The Lighthouse Lab in Milton Keynes and Alex Alderton, Roberto Amato, Jeffrey Barrett, Sonia Goncalves, Ewan Harrison, David K. Jackson, Ian Johnston, Dominic Kwiatkowski, Cordelia Langford, John Sillitoe on behalf of the Wellcome Sanger Institute COVID-19 Surveillance Team |
| EPI_ISL_2346912, EPI_ISL_2346913, EPI_ISL_2346914, EPI_ISL_2346915, EPI_ISL_2346916, EPI_ISL_2346917, EPI_ISL_2346918 | Lighthouse Lab in Alderley Park | Wellcome Sanger Institute for the COVID-19 Genomics UK (COG-UK) Consortium | Jacquelyn Wynn, Mairead Hyland, The Lighthouse Lab in Alderley Park and Alex Alderton, Roberto Amato, Jeffrey Barrett, Sonia Goncalves, Ewan Harrison, David K. Jackson, Ian Johnston, Dominic Kwiatkowski, Cordelia Langford, John Sillitoe on behalf of the Wellcome Sanger Institute COVID-19 Surveillance Team |
| EPI_ISL_2346921, EPI_ISL_2346922 | Lighthouse Lab in Milton Keynes | Wellcome Sanger Institute for the COVID-19 Genomics UK (COG-UK) Consortium | The Lighthouse Lab in Milton Keynes and Alex Alderton, Roberto Amato, Jeffrey Barrett, Sonia Goncalves, Ewan Harrison, David K. Jackson, Ian Johnston, Dominic Kwiatkowski, Cordelia Langford, John Sillitoe on behalf of the Wellcome Sanger Institute COVID-19 Surveillance Team |
| EPI_ISL_2346923, EPI_ISL_2346924, EPI_ISL_2346925, EPI_ISL_2346926, EPI_ISL_2346927, EPI_ISL_2346929, EPI_ISL_2346930, EPI_ISL_2346931, EPI_ISL_2346932, EPI_ISL_2346933, EPI_ISL_2346934, EPI_ISL_2346937, EPI_ISL_2346938, EPI_ISL_2346940, EPI_ISL_2346942, EPI_ISL_2346943, EPI_ISL_2346945 | | | |
| see above | Lighthouse Lab in Alderley Park | Wellcome Sanger Institute for the COVID-19 Genomics UK (COG-UK) Consortium | Jacquelyn Wynn, Mairead Hyland, The Lighthouse Lab in Alderley Park and Alex Alderton, Roberto Amato, Jeffrey Barrett, Sonia Goncalves, Ewan Harrison, David K. Jackson, Ian Johnston, Dominic Kwiatkowski, Cordelia Langford, John Sillitoe on behalf of the Wellcome Sanger Institute COVID-19 Surveillance Team |
| EPI_ISL_2346946 | Lighthouse Lab in Milton Keynes | Wellcome Sanger Institute for the COVID-19 Genomics UK (COG-UK) Consortium | The Lighthouse Lab in Milton Keynes and Alex Alderton, Roberto Amato, Jeffrey Barrett, Sonia Goncalves, Ewan Harrison, David K. Jackson, Ian Johnston, Dominic Kwiatkowski, Cordelia Langford, John Sillitoe on behalf of the Wellcome Sanger Institute COVID-19 Surveillance Team |

EPI_ISL_2346947, EPI_ISL_2346949, EPI_ISL_2346950, EPI_ISL_2346951, EPI_ISL_2346952, EPI_ISL_2346954, EPI_ISL_2346956, EPI_ISL_2346958, EPI_ISL_2346959, EPI_ISL_2346961, EPI_ISL_2346962

| | | | |
|---|---|---|---|
| see above | Lighthouse Lab in Alderley Park | Wellcome Sanger Institute for the COVID-19 Genomics UK (COG-UK) Consortium | Jacquelyn Wynn, Mairead Hyland, The Lighthouse Lab in Alderley Park and Alex Alderton, Roberto Amato, Jeffrey Barrett, Sonia Goncalves, Ewan Harrison, David K. Jackson, Ian Johnston, Dominic Kwiatkowski, Cordelia Langford, John Sillitoe on behalf of the Wellcome Sanger Institute COVID-19 Surveillance Team |
| EPI_ISL_2346964 | Lighthouse Lab in Milton Keynes | Wellcome Sanger Institute for the COVID-19 Genomics UK (COG-UK) Consortium | The Lighthouse Lab in Milton Keynes and Alex Alderton, Roberto Amato, Jeffrey Barrett, Sonia Goncalves, Ewan Harrison, David K. Jackson, Ian Johnston, Dominic Kwiatkowski, Cordelia Langford, John Sillitoe on behalf of the Wellcome Sanger Institute COVID-19 Surveillance Team |
| EPI_ISL_2346965, EPI_ISL_2346966, EPI_ISL_2346967 | Lighthouse Lab in Alderley Park | Wellcome Sanger Institute for the COVID-19 Genomics UK (COG-UK) Consortium | Jacquelyn Wynn, Mairead Hyland, The Lighthouse Lab in Alderley Park and Alex Alderton, Roberto Amato, Jeffrey Barrett, Sonia Goncalves, Ewan Harrison, David K. Jackson, Ian Johnston, Dominic Kwiatkowski, Cordelia Langford, John Sillitoe on behalf of the Wellcome Sanger Institute COVID-19 Surveillance Team |
| EPI_ISL_2346969 | Lighthouse Lab in Milton Keynes | Wellcome Sanger Institute for the COVID-19 Genomics UK (COG-UK) Consortium | The Lighthouse Lab in Milton Keynes and Alex Alderton, Roberto Amato, Jeffrey Barrett, Sonia Goncalves, Ewan Harrison, David K. Jackson, Ian Johnston, Dominic Kwiatkowski, Cordelia Langford, John Sillitoe on behalf of the Wellcome Sanger Institute COVID-19 Surveillance Team |
| EPI_ISL_2346970, EPI_ISL_2346972 | Lighthouse Lab in Alderley Park | Wellcome Sanger Institute for the COVID-19 Genomics UK (COG-UK) Consortium | Jacquelyn Wynn, Mairead Hyland, The Lighthouse Lab in Alderley Park and Alex Alderton, Roberto Amato, Jeffrey Barrett, Sonia Goncalves, Ewan Harrison, David K. Jackson, Ian Johnston, Dominic Kwiatkowski, Cordelia Langford, John Sillitoe on behalf of the Wellcome Sanger Institute COVID-19 Surveillance Team |
| EPI_ISL_2346973 | Lighthouse Lab in Milton Keynes | Wellcome Sanger Institute for the COVID-19 Genomics UK (COG-UK) Consortium | The Lighthouse Lab in Milton Keynes and Alex Alderton, Roberto Amato, Jeffrey Barrett, Sonia Goncalves, Ewan Harrison, David K. Jackson, Ian Johnston, Dominic Kwiatkowski, Cordelia Langford, John Sillitoe on behalf of the Wellcome Sanger Institute COVID-19 Surveillance Team |
| EPI_ISL_2346974, EPI_ISL_2346975, EPI_ISL_2346976, EPI_ISL_2346977 | Lighthouse Lab in Alderley Park | Wellcome Sanger Institute for the COVID-19 Genomics UK (COG-UK) Consortium | Jacquelyn Wynn, Mairead Hyland, The Lighthouse Lab in Alderley Park and Alex Alderton, Roberto Amato, Jeffrey Barrett, Sonia Goncalves, Ewan Harrison, David K. Jackson, Ian Johnston, Dominic Kwiatkowski, Cordelia Langford, John Sillitoe on behalf of the Wellcome Sanger Institute COVID-19 Surveillance Team |
| EPI_ISL_2346978 | Lighthouse Lab in Milton Keynes | Wellcome Sanger Institute for the COVID-19 Genomics UK (COG-UK) Consortium | The Lighthouse Lab in Milton Keynes and Alex Alderton, Roberto Amato, Jeffrey Barrett, Sonia Goncalves, Ewan Harrison, David K. Jackson, Ian Johnston, Dominic Kwiatkowski, Cordelia Langford, John Sillitoe on behalf of the Wellcome Sanger Institute COVID-19 Surveillance Team |
| EPI_ISL_2346979, EPI_ISL_2346981, EPI_ISL_2346982, EPI_ISL_2346983, EPI_ISL_2346986 | Lighthouse Lab in Alderley Park | Wellcome Sanger Institute for the COVID-19 Genomics UK (COG-UK) Consortium | Jacquelyn Wynn, Mairead Hyland, The Lighthouse Lab in Alderley Park and Alex Alderton, Roberto Amato, Jeffrey Barrett, Sonia Goncalves, Ewan Harrison, David K. Jackson, Ian Johnston, Dominic Kwiatkowski, Cordelia Langford, John Sillitoe on behalf of the Wellcome Sanger Institute COVID-19 Surveillance Team |
| EPI_ISL_2346987 | Lighthouse Lab in Milton Keynes | Wellcome Sanger Institute for the COVID-19 Genomics UK (COG-UK) Consortium | The Lighthouse Lab in Milton Keynes and Alex Alderton, Roberto Amato, Jeffrey Barrett, Sonia Goncalves, Ewan Harrison, David K. Jackson, Ian Johnston, Dominic Kwiatkowski, Cordelia Langford, John Sillitoe on behalf of the Wellcome Sanger Institute COVID-19 Surveillance Team |
| EPI_ISL_2346988, EPI_ISL_2346989, EPI_ISL_2346990, EPI_ISL_2346991, EPI_ISL_2346992, EPI_ISL_2346993, EPI_ISL_2346994, EPI_ISL_2346995, EPI_ISL_2346996, EPI_ISL_2346997 | Lighthouse Lab in Alderley Park | Wellcome Sanger Institute for the COVID-19 Genomics UK (COG-UK) Consortium | Jacquelyn Wynn, Mairead Hyland, The Lighthouse Lab in Alderley Park and Alex Alderton, Roberto Amato, Jeffrey Barrett, Sonia Goncalves, Ewan Harrison, David K. Jackson, Ian Johnston, Dominic Kwiatkowski, Cordelia Langford, John Sillitoe on behalf of the Wellcome Sanger Institute COVID-19 Surveillance Team |
| EPI_ISL_2346998 | Lighthouse Lab in Milton Keynes | Wellcome Sanger Institute for the COVID-19 Genomics UK (COG-UK) Consortium | The Lighthouse Lab in Milton Keynes and Alex Alderton, Roberto Amato, Jeffrey Barrett, Sonia Goncalves, Ewan Harrison, David K. Jackson, Ian Johnston, Dominic Kwiatkowski, Cordelia Langford, John Sillitoe on behalf of the Wellcome Sanger Institute COVID-19 Surveillance Team |
| EPI_ISL_2347002, EPI_ISL_2347004, EPI_ISL_2347005, EPI_ISL_2347006, EPI_ISL_2347008 | Lighthouse Lab in Alderley Park | Wellcome Sanger Institute for the COVID-19 Genomics UK (COG-UK) Consortium | Jacquelyn Wynn, Mairead Hyland, The Lighthouse Lab in Alderley Park and Alex Alderton, Roberto Amato, Jeffrey Barrett, Sonia Goncalves, Ewan Harrison, David K. Jackson, Ian Johnston, Dominic Kwiatkowski, Cordelia Langford, John Sillitoe on behalf of the Wellcome Sanger Institute COVID-19 Surveillance Team |
| EPI_ISL_2347010, EPI_ISL_2347012, EPI_ISL_2347014 | Lighthouse Lab in Milton Keynes | Wellcome Sanger Institute for the COVID-19 Genomics UK (COG-UK) Consortium | The Lighthouse Lab in Milton Keynes and Alex Alderton, Roberto Amato, Jeffrey Barrett, Sonia Goncalves, Ewan Harrison, David K. Jackson, Ian Johnston, Dominic Kwiatkowski, Cordelia Langford, John Sillitoe on behalf of the Wellcome Sanger Institute COVID-19 Surveillance Team |
| EPI_ISL_2347015 | Lighthouse Lab in Alderley Park | Wellcome Sanger Institute for the COVID-19 Genomics UK (COG-UK) Consortium | Jacquelyn Wynn, Mairead Hyland, The Lighthouse Lab in Alderley Park and Alex Alderton, Roberto Amato, Jeffrey Barrett, Sonia Goncalves, Ewan Harrison, David K. Jackson, Ian Johnston, Dominic Kwiatkowski, Cordelia Langford, John Sillitoe on behalf of the Wellcome Sanger Institute COVID-19 Surveillance Team |
| EPI_ISL_2347016, EPI_ISL_2347017 | Lighthouse Lab in Milton Keynes | Wellcome Sanger Institute for the COVID-19 Genomics UK (COG-UK) Consortium | The Lighthouse Lab in Milton Keynes and Alex Alderton, Roberto Amato, Jeffrey Barrett, Sonia Goncalves, Ewan Harrison, David K. Jackson, Ian Johnston, Dominic Kwiatkowski, Cordelia Langford, John Sillitoe on behalf of the Wellcome Sanger Institute COVID-19 Surveillance Team |
| EPI_ISL_2347018, EPI_ISL_2347019, EPI_ISL_2347020, EPI_ISL_2347021 | Lighthouse Lab in Alderley Park | Wellcome Sanger Institute for the COVID-19 Genomics UK (COG-UK) Consortium | Jacquelyn Wynn, Mairead Hyland, The Lighthouse Lab in Alderley Park and Alex Alderton, Roberto Amato, Jeffrey Barrett, Sonia Goncalves, Ewan Harrison, David K. Jackson, Ian Johnston, Dominic Kwiatkowski, Cordelia Langford, John Sillitoe on behalf of the Wellcome Sanger Institute COVID-19 Surveillance Team |
| EPI_ISL_2347022 | Lighthouse Lab in Milton Keynes | Wellcome Sanger Institute for the COVID-19 Genomics UK (COG-UK) Consortium | The Lighthouse Lab in Milton Keynes and Alex Alderton, Roberto Amato, Jeffrey Barrett, Sonia Goncalves, Ewan Harrison, David K. Jackson, Ian Johnston, Dominic Kwiatkowski, Cordelia Langford, John Sillitoe on behalf of the Wellcome Sanger Institute COVID-19 Surveillance Team |
| EPI_ISL_2347024, EPI_ISL_2347026, EPI_ISL_2347027 | Lighthouse Lab in Alderley Park | Wellcome Sanger Institute for the COVID-19 Genomics UK (COG-UK) Consortium | Jacquelyn Wynn, Mairead Hyland, The Lighthouse Lab in Alderley Park and Alex Alderton, Roberto Amato, Jeffrey Barrett, Sonia Goncalves, Ewan Harrison, David K. Jackson, Ian Johnston, Dominic Kwiatkowski, Cordelia Langford, John Sillitoe on behalf of the Wellcome Sanger Institute COVID-19 Surveillance Team |
| EPI_ISL_2347028 | Lighthouse Lab in Milton Keynes | Wellcome Sanger Institute for the COVID-19 Genomics UK (COG-UK) Consortium | The Lighthouse Lab in Milton Keynes and Alex Alderton, Roberto Amato, Jeffrey Barrett, Sonia Goncalves, Ewan Harrison, David K. Jackson, Ian Johnston, Dominic Kwiatkowski, Cordelia Langford, John Sillitoe on behalf of the Wellcome Sanger Institute COVID-19 Surveillance Team |
| EPI_ISL_2347029, EPI_ISL_2347030, EPI_ISL_2347031, EPI_ISL_2347032, EPI_ISL_2347033 | Lighthouse Lab in Alderley Park | Wellcome Sanger Institute for the COVID-19 Genomics UK (COG-UK) Consortium | Jacquelyn Wynn, Mairead Hyland, The Lighthouse Lab in Alderley Park and Alex Alderton, Roberto Amato, Jeffrey Barrett, Sonia Goncalves, Ewan Harrison, David K. Jackson, Ian Johnston, Dominic Kwiatkowski, Cordelia Langford, John Sillitoe on behalf of the Wellcome Sanger Institute COVID-19 Surveillance Team |
| EPI_ISL_2347034 | Lighthouse Lab in Milton Keynes | Wellcome Sanger Institute for the COVID-19 Genomics UK (COG-UK) Consortium | The Lighthouse Lab in Milton Keynes and Alex Alderton, Roberto Amato, Jeffrey Barrett, Sonia Goncalves, Ewan Harrison, David K. Jackson, Ian Johnston, Dominic Kwiatkowski, Cordelia Langford, John Sillitoe on behalf of the Wellcome Sanger Institute COVID-19 Surveillance Team |
| EPI_ISL_2347035, EPI_ISL_2347037, EPI_ISL_2347038, EPI_ISL_2347041 | Lighthouse Lab in Alderley Park | Wellcome Sanger Institute for the COVID-19 Genomics UK (COG-UK) Consortium | Jacquelyn Wynn, Mairead Hyland, The Lighthouse Lab in Alderley Park and Alex Alderton, Roberto Amato, Jeffrey Barrett, Sonia Goncalves, Ewan Harrison, David K. Jackson, Ian Johnston, Dominic Kwiatkowski, Cordelia Langford, John Sillitoe on behalf of the Wellcome Sanger Institute COVID-19 Surveillance Team |
| EPI_ISL_2347043 | Lighthouse Lab in Milton Keynes | Wellcome Sanger Institute for the COVID-19 Genomics UK (COG-UK) Consortium | The Lighthouse Lab in Milton Keynes and Alex Alderton, Roberto Amato, Jeffrey Barrett, Sonia Goncalves, Ewan Harrison, David K. Jackson, Ian Johnston, Dominic Kwiatkowski, Cordelia Langford, John Sillitoe on behalf of the Wellcome Sanger Institute COVID-19 Surveillance Team |

EPI_ISL_2347046, EPI_ISL_2347048, EPI_ISL_2347050, EPI_ISL_2347052, EPI_ISL_2347054, EPI_ISL_2347055, EPI_ISL_2347056, EPI_ISL_2347057, EPI_ISL_2347058, EPI_ISL_2347060, EPI_ISL_2347061, EPI_ISL_2347062, EPI_ISL_2347063, EPI_ISL_2347064, EPI_ISL_2347066

| | | | |
|---|---|---|---|
| see above | Lighthouse Lab in Alderley Park | Wellcome Sanger Institute for the COVID-19 Genomics UK (COG-UK) Consortium | Jacquelyn Wynn, Mairead Hyland, The Lighthouse Lab in Alderley Park and Alex Alderton, Roberto Amato, Jeffrey Barrett, Sonia Goncalves, Ewan Harrison, David K. Jackson, Ian Johnston, Dominic Kwiatkowski, Cordelia Langford, John Sillitoe on behalf of the Wellcome Sanger Institute COVID-19 Surveillance Team |
| EPI_ISL_2347067 | Lighthouse Lab in Milton Keynes | Wellcome Sanger Institute for the COVID-19 Genomics UK (COG-UK) Consortium | The Lighthouse Lab in Milton Keynes and Alex Alderton, Roberto Amato, Jeffrey Barrett, Sonia Goncalves, Ewan Harrison, David K. Jackson, Ian Johnston, Dominic Kwiatkowski, Cordelia Langford, John Sillitoe on behalf of the Wellcome Sanger Institute COVID-19 Surveillance Team |
| EPI_ISL_2347068 | Lighthouse Lab in Alderley Park | Wellcome Sanger Institute for the COVID-19 Genomics UK (COG-UK) Consortium | Jacquelyn Wynn, Mairead Hyland, The Lighthouse Lab in Alderley Park and Alex Alderton, Roberto Amato, Jeffrey Barrett, Sonia Goncalves, Ewan Harrison, David K. Jackson, Ian Johnston, Dominic Kwiatkowski, Cordelia Langford, John Sillitoe on behalf of the Wellcome Sanger Institute COVID-19 Surveillance Team |
| EPI_ISL_2347070 | Lighthouse Lab in Milton Keynes | Wellcome Sanger Institute for the COVID-19 Genomics UK (COG-UK) Consortium | The Lighthouse Lab in Milton Keynes and Alex Alderton, Roberto Amato, Jeffrey Barrett, Sonia Goncalves, Ewan Harrison, David K. Jackson, Ian Johnston, Dominic Kwiatkowski, Cordelia Langford, John Sillitoe on behalf of the Wellcome Sanger Institute COVID-19 Surveillance Team |
| EPI_ISL_2347071, EPI_ISL_2347072, EPI_ISL_2347074, EPI_ISL_2347075, EPI_ISL_2347077, EPI_ISL_2347078, EPI_ISL_2347079, EPI_ISL_2347080 | Lighthouse Lab in Alderley Park | Wellcome Sanger Institute for the COVID-19 Genomics UK (COG-UK) Consortium | Jacquelyn Wynn, Mairead Hyland, The Lighthouse Lab in Alderley Park and Alex Alderton, Roberto Amato, Jeffrey Barrett, Sonia Goncalves, Ewan Harrison, David K. Jackson, Ian Johnston, Dominic Kwiatkowski, Cordelia Langford, John Sillitoe on behalf of the Wellcome Sanger Institute COVID-19 Surveillance Team |
| EPI_ISL_2347082, EPI_ISL_2347084 | Lighthouse Lab in Milton Keynes | Wellcome Sanger Institute for the COVID-19 Genomics UK (COG-UK) Consortium | The Lighthouse Lab in Milton Keynes and Alex Alderton, Roberto Amato, Jeffrey Barrett, Sonia Goncalves, Ewan Harrison, David K. Jackson, Ian Johnston, Dominic Kwiatkowski, Cordelia Langford, John Sillitoe on behalf of the Wellcome Sanger Institute COVID-19 Surveillance Team |
| EPI_ISL_2347085, | Lighthouse Lab in Alderley Park | Wellcome Sanger Institute for the COVID-19 Genomics UK (COG-UK) | Jacquelyn Wynn, Mairead Hyland, The Lighthouse Lab in Alderley Park and Alex Alderton, Roberto Amato, Jeffrey Barrett, Sonia Goncalves, Ewan Harrison, David K. Jackson, Ian Johnston, Dominic Kwiatkowski, Cordelia Langford, John Sillitoe on |

| | | | |
|---|---|---|---|
| EPI_ISL_2347086 | | Consortium | behalf of the Wellcome Sanger Institute COVID-19 Surveillance Team |
| EPI_ISL_2347088, EPI_ISL_2347089, EPI_ISL_2347090, EPI_ISL_2347092, EPI_ISL_2347093, EPI_ISL_2347094, EPI_ISL_2347096, EPI_ISL_2347097, EPI_ISL_2347098, EPI_ISL_2347099, EPI_ISL_2347100, EPI_ISL_2347101, EPI_ISL_2347102, EPI_ISL_2347103, EPI_ISL_2347104, EPI_ISL_2347105, EPI_ISL_2347107, EPI_ISL_2347108, EPI_ISL_2347109, EPI_ISL_2347110, EPI_ISL_2347112, EPI_ISL_2347113, EPI_ISL_2347114, EPI_ISL_2347115, EPI_ISL_2347117, EPI_ISL_2347118, EPI_ISL_2347119, EPI_ISL_2347120, EPI_ISL_2347121, EPI_ISL_2347122, EPI_ISL_2347123, EPI_ISL_2347128, EPI_ISL_2347130, EPI_ISL_2347131, EPI_ISL_2347132, EPI_ISL_2347133, EPI_ISL_2347134, EPI_ISL_2347135, EPI_ISL_2347136, EPI_ISL_2347137, EPI_ISL_2347139, EPI_ISL_2347142, EPI_ISL_2347144, EPI_ISL_2347145, EPI_ISL_2347146, EPI_ISL_2347147, EPI_ISL_2347151, EPI_ISL_2347152, EPI_ISL_2347153, EPI_ISL_2347154, EPI_ISL_2347155, EPI_ISL_2347157, EPI_ISL_2347158, EPI_ISL_2347160, EPI_ISL_2347162, EPI_ISL_2347163, EPI_ISL_2347164, EPI_ISL_2347165, EPI_ISL_2347166, EPI_ISL_2347167, EPI_ISL_2347168, EPI_ISL_2347169, EPI_ISL_2347170, EPI_ISL_2347171, EPI_ISL_2347172, EPI_ISL_2347173, EPI_ISL_2347174, EPI_ISL_2347176, EPI_ISL_2347177, EPI_ISL_2347178 | | | |
| see above | Lighthouse Lab in Milton Keynes | Wellcome Sanger Institute for the COVID-19 Genomics UK (COG-UK) Consortium | The Lighthouse Lab in Milton Keynes and Alex Alderton, Roberto Amato, Jeffrey Barrett, Sonia Goncalves, Ewan Harrison, David K. Jackson, Ian Johnston, Dominic Kwiatkowski, Cordelia Langford, John Sillitoe on behalf of the Wellcome Sanger Institute COVID-19 Surveillance Team |
| EPI_ISL_2347179 | Lighthouse Lab in Glasgow | Wellcome Sanger Institute for the COVID-19 Genomics UK (COG-UK) Consortium | Harper VanSteenhouse, Yumi Kasai, David Gray, Carol Clugston, Anna Dominiczak and Alex Alderton, Roberto Amato, Jeffrey Barrett, Sonia Goncalves, Ewan Harrison, David K. Jackson, Ian Johnston, Dominic Kwiatkowski, Cordelia Langford, John Sillitoe on behalf of the Wellcome Sanger Institute COVID-19 Surveillance Team |
| EPI_ISL_2347180, EPI_ISL_2347181, EPI_ISL_2347182, EPI_ISL_2347183, EPI_ISL_2347184, EPI_ISL_2347186, EPI_ISL_2347187, EPI_ISL_2347188, EPI_ISL_2347189, EPI_ISL_2347190, EPI_ISL_2347191, EPI_ISL_2347194, EPI_ISL_2347195, EPI_ISL_2347196, EPI_ISL_2347198, EPI_ISL_2347199, EPI_ISL_2347201, EPI_ISL_2347202, EPI_ISL_2347203, EPI_ISL_2347204, EPI_ISL_2347205, EPI_ISL_2347207, EPI_ISL_2347209, EPI_ISL_2347211, EPI_ISL_2347212, EPI_ISL_2347215, EPI_ISL_2347218, EPI_ISL_2347219, EPI_ISL_2347220, EPI_ISL_2347221, EPI_ISL_2347222, EPI_ISL_2347223, EPI_ISL_2347224, EPI_ISL_2347226, EPI_ISL_2347227, EPI_ISL_2347228, EPI_ISL_2347229, EPI_ISL_2347230, EPI_ISL_2347231, EPI_ISL_2347232, EPI_ISL_2347233, EPI_ISL_2347235, EPI_ISL_2347236, EPI_ISL_2347238, EPI_ISL_2347239, EPI_ISL_2347241, EPI_ISL_2347242, EPI_ISL_2347243, EPI_ISL_2347244, EPI_ISL_2347246, EPI_ISL_2347248, EPI_ISL_2347249, EPI_ISL_2347250, EPI_ISL_2347251, EPI_ISL_2347252, EPI_ISL_2347254, EPI_ISL_2347256, EPI_ISL_2347261, EPI_ISL_2347263, EPI_ISL_2347265, EPI_ISL_2347266 | | | |
| see above | Lighthouse Lab in Milton Keynes | Wellcome Sanger Institute for the COVID-19 Genomics UK (COG-UK) Consortium | The Lighthouse Lab in Milton Keynes and Alex Alderton, Roberto Amato, Jeffrey Barrett, Sonia Goncalves, Ewan Harrison, David K. Jackson, Ian Johnston, Dominic Kwiatkowski, Cordelia Langford, John Sillitoe on behalf of the Wellcome Sanger Institute COVID-19 Surveillance Team |
| EPI_ISL_2347268 | Lighthouse Lab in Glasgow | Wellcome Sanger Institute for the COVID-19 Genomics UK (COG-UK) Consortium | Harper VanSteenhouse, Yumi Kasai, David Gray, Carol Clugston, Anna Dominiczak and Alex Alderton, Roberto Amato, Jeffrey Barrett, Sonia Goncalves, Ewan Harrison, David K. Jackson, Ian Johnston, Dominic Kwiatkowski, Cordelia Langford, John Sillitoe on behalf of the Wellcome Sanger Institute COVID-19 Surveillance Team |
| EPI_ISL_2347269, EPI_ISL_2347271, EPI_ISL_2347272, EPI_ISL_2347273, EPI_ISL_2347274, EPI_ISL_2347275, EPI_ISL_2347277, EPI_ISL_2347278, EPI_ISL_2347279, EPI_ISL_2347280, EPI_ISL_2347281, EPI_ISL_2347284, EPI_ISL_2347285, EPI_ISL_2347288, EPI_ISL_2347289, EPI_ISL_2347290, EPI_ISL_2347291, EPI_ISL_2347295, EPI_ISL_2347297, EPI_ISL_2347298, EPI_ISL_2347300, EPI_ISL_2347301, EPI_ISL_2347302, EPI_ISL_2347303, EPI_ISL_2347306, EPI_ISL_2347307, EPI_ISL_2347313, EPI_ISL_2347316, EPI_ISL_2347317, EPI_ISL_2347318, EPI_ISL_2347319, EPI_ISL_2347320, EPI_ISL_2347322, EPI_ISL_2347323, EPI_ISL_2347324, EPI_ISL_2347325, EPI_ISL_2347326, EPI_ISL_2347327, EPI_ISL_2347329, EPI_ISL_2347330, EPI_ISL_2347332 | | | |
| see above | Lighthouse Lab in Milton Keynes | Wellcome Sanger Institute for the COVID-19 Genomics UK (COG-UK) Consortium | The Lighthouse Lab in Milton Keynes and Alex Alderton, Roberto Amato, Jeffrey Barrett, Sonia Goncalves, Ewan Harrison, David K. Jackson, Ian Johnston, Dominic Kwiatkowski, Cordelia Langford, John Sillitoe on behalf of the Wellcome Sanger Institute COVID-19 Surveillance Team |
| EPI_ISL_2347335, EPI_ISL_2347336, EPI_ISL_2347337, EPI_ISL_2347339, EPI_ISL_2347340, EPI_ISL_2347341, EPI_ISL_2347343, EPI_ISL_2347345 | Lighthouse Lab in Alderley Park | Wellcome Sanger Institute for the COVID-19 Genomics UK (COG-UK) Consortium | Jacquelyn Wynn, Mairead Hyland, The Lighthouse Lab in Alderley Park and Alex Alderton, Roberto Amato, Jeffrey Barrett, Sonia Goncalves, Ewan Harrison, David K. Jackson, Ian Johnston, Dominic Kwiatkowski, Cordelia Langford, John Sillitoe on behalf of the Wellcome Sanger Institute COVID-19 Surveillance Team |
| EPI_ISL_2347347 | Lighthouse Lab in Glasgow | Wellcome Sanger Institute for the COVID-19 Genomics UK (COG-UK) Consortium | Harper VanSteenhouse, Yumi Kasai, David Gray, Carol Clugston, Anna Dominiczak and Alex Alderton, Roberto Amato, Jeffrey Barrett, Sonia Goncalves, Ewan Harrison, David K. Jackson, Ian Johnston, Dominic Kwiatkowski, Cordelia Langford, John Sillitoe on behalf of the Wellcome Sanger Institute COVID-19 Surveillance Team |
| EPI_ISL_2347348, EPI_ISL_2347351, EPI_ISL_2347352, EPI_ISL_2347354, EPI_ISL_2347355, EPI_ISL_2347356, EPI_ISL_2347357, EPI_ISL_2347358, EPI_ISL_2347359, EPI_ISL_2347360, EPI_ISL_2347361, EPI_ISL_2347362, EPI_ISL_2347363, EPI_ISL_2347364, EPI_ISL_2347365, EPI_ISL_2347366, EPI_ISL_2347367, EPI_ISL_2347368, EPI_ISL_2347369, EPI_ISL_2347370 | | | |
| see above | Lighthouse Lab in Alderley Park | Wellcome Sanger Institute for the COVID-19 Genomics UK (COG-UK) Consortium | Jacquelyn Wynn, Mairead Hyland, The Lighthouse Lab in Alderley Park and Alex Alderton, Roberto Amato, Jeffrey Barrett, Sonia Goncalves, Ewan Harrison, David K. Jackson, Ian Johnston, Dominic Kwiatkowski, Cordelia Langford, John Sillitoe on behalf of the Wellcome Sanger Institute COVID-19 Surveillance Team |
| EPI_ISL_2347373 | Health Services Laboratories | Wellcome Sanger Institute for the COVID-19 Genomics UK (COG-UK) Consortium | Health Services Laboratories and Alex Alderton, Roberto Amato, Jeffrey Barrett, Sonia Goncalves, Ewan Harrison, David K. Jackson, Ian Johnston, Dominic Kwiatkowski, Cordelia Langford, John Sillitoe on behalf of the Wellcome Sanger Institute COVID-19 Surveillance Team |
| EPI_ISL_2347375, EPI_ISL_2347376, EPI_ISL_2347377, EPI_ISL_2347378, EPI_ISL_2347379, EPI_ISL_2347380, EPI_ISL_2347381 | Lighthouse Lab in Alderley Park | Wellcome Sanger Institute for the COVID-19 Genomics UK (COG-UK) Consortium | Jacquelyn Wynn, Mairead Hyland, The Lighthouse Lab in Alderley Park and Alex Alderton, Roberto Amato, Jeffrey Barrett, Sonia Goncalves, Ewan Harrison, David K. Jackson, Ian Johnston, Dominic Kwiatkowski, Cordelia Langford, John Sillitoe on behalf of the Wellcome Sanger Institute COVID-19 Surveillance Team |
| EPI_ISL_2347382 | Health Services Laboratories | Wellcome Sanger Institute for the COVID-19 Genomics UK (COG-UK) Consortium | Health Services Laboratories and Alex Alderton, Roberto Amato, Jeffrey Barrett, Sonia Goncalves, Ewan Harrison, David K. Jackson, Ian Johnston, Dominic Kwiatkowski, Cordelia Langford, John Sillitoe on behalf of the Wellcome Sanger Institute COVID-19 Surveillance Team |
| EPI_ISL_2347383, EPI_ISL_2347384, EPI_ISL_2347385 | Lighthouse Lab in Alderley Park | Wellcome Sanger Institute for the COVID-19 Genomics UK (COG-UK) Consortium | Jacquelyn Wynn, Mairead Hyland, The Lighthouse Lab in Alderley Park and Alex Alderton, Roberto Amato, Jeffrey Barrett, Sonia Goncalves, Ewan Harrison, David K. Jackson, Ian Johnston, Dominic Kwiatkowski, Cordelia Langford, John Sillitoe on behalf of the Wellcome Sanger Institute COVID-19 Surveillance Team |
| EPI_ISL_2347386 | Health Services Laboratories | Wellcome Sanger Institute for the COVID-19 Genomics UK (COG-UK) Consortium | Health Services Laboratories and Alex Alderton, Roberto Amato, Jeffrey Barrett, Sonia Goncalves, Ewan Harrison, David K. Jackson, Ian Johnston, Dominic Kwiatkowski, Cordelia Langford, John Sillitoe on behalf of the Wellcome Sanger Institute COVID-19 Surveillance Team |
| EPI_ISL_2347387 | Lighthouse Lab in Alderley Park | Wellcome Sanger Institute for the COVID-19 Genomics UK (COG-UK) Consortium | Jacquelyn Wynn, Mairead Hyland, The Lighthouse Lab in Alderley Park and Alex Alderton, Roberto Amato, Jeffrey Barrett, Sonia Goncalves, Ewan Harrison, David K. Jackson, Ian Johnston, Dominic Kwiatkowski, Cordelia Langford, John Sillitoe on behalf of the Wellcome Sanger Institute COVID-19 Surveillance Team |
| EPI_ISL_2347388 | Lighthouse Lab in Milton Keynes | Wellcome Sanger Institute for the COVID-19 Genomics UK (COG-UK) Consortium | The Lighthouse Lab in Milton Keynes and Alex Alderton, Roberto Amato, Jeffrey Barrett, Sonia Goncalves, Ewan Harrison, David K. Jackson, Ian Johnston, Dominic Kwiatkowski, Cordelia Langford, John Sillitoe on behalf of the Wellcome Sanger Institute COVID-19 Surveillance Team |
| EPI_ISL_2347389, EPI_ISL_2347390, EPI_ISL_2347392, EPI_ISL_2347393, EPI_ISL_2347394, EPI_ISL_2347395, EPI_ISL_2347396, EPI_ISL_2347398, EPI_ISL_2347402 | Lighthouse Lab in Alderley Park | Wellcome Sanger Institute for the COVID-19 Genomics UK (COG-UK) Consortium | Jacquelyn Wynn, Mairead Hyland, The Lighthouse Lab in Alderley Park and Alex Alderton, Roberto Amato, Jeffrey Barrett, Sonia Goncalves, Ewan Harrison, David K. Jackson, Ian Johnston, Dominic Kwiatkowski, Cordelia Langford, John Sillitoe on behalf of the Wellcome Sanger Institute COVID-19 Surveillance Team |
| EPI_ISL_2347403 | Health Services Laboratories | Wellcome Sanger Institute for the COVID-19 Genomics UK (COG-UK) Consortium | Health Services Laboratories and Alex Alderton, Roberto Amato, Jeffrey Barrett, Sonia Goncalves, Ewan Harrison, David K. Jackson, Ian Johnston, Dominic Kwiatkowski, Cordelia Langford, John Sillitoe on behalf of the Wellcome Sanger Institute COVID-19 Surveillance Team |
| EPI_ISL_2347404, EPI_ISL_2347405, EPI_ISL_2347406, EPI_ISL_2347407 | Lighthouse Lab in Alderley Park | Wellcome Sanger Institute for the COVID-19 Genomics UK (COG-UK) Consortium | Jacquelyn Wynn, Mairead Hyland, The Lighthouse Lab in Alderley Park and Alex Alderton, Roberto Amato, Jeffrey Barrett, Sonia Goncalves, Ewan Harrison, David K. Jackson, Ian Johnston, Dominic Kwiatkowski, Cordelia Langford, John Sillitoe on behalf of the Wellcome Sanger Institute COVID-19 Surveillance Team |
| EPI_ISL_2347408 | Health Services Laboratories | Wellcome Sanger Institute for the COVID-19 Genomics UK (COG-UK) Consortium | Health Services Laboratories and Alex Alderton, Roberto Amato, Jeffrey Barrett, Sonia Goncalves, Ewan Harrison, David K. Jackson, Ian Johnston, Dominic Kwiatkowski, Cordelia Langford, John Sillitoe on behalf of the Wellcome Sanger Institute COVID-19 Surveillance Team |
| EPI_ISL_2347409 | Lighthouse Lab in Alderley Park | Wellcome Sanger Institute for the COVID-19 Genomics UK (COG-UK) Consortium | Jacquelyn Wynn, Mairead Hyland, The Lighthouse Lab in Alderley Park and Alex Alderton, Roberto Amato, Jeffrey Barrett, Sonia Goncalves, Ewan Harrison, David K. Jackson, Ian Johnston, Dominic Kwiatkowski, Cordelia Langford, John Sillitoe on behalf of the Wellcome Sanger Institute COVID-19 Surveillance Team |
| EPI_ISL_2347410 | Health Services Laboratories | Wellcome Sanger Institute for the COVID-19 Genomics UK (COG-UK) Consortium | Health Services Laboratories and Alex Alderton, Roberto Amato, Jeffrey Barrett, Sonia Goncalves, Ewan Harrison, David K. Jackson, Ian Johnston, Dominic Kwiatkowski, Cordelia Langford, John Sillitoe on behalf of the Wellcome Sanger Institute COVID-19 Surveillance Team |
| EPI_ISL_2347411, EPI_ISL_2347412 | Lighthouse Lab in Milton Keynes | Wellcome Sanger Institute for the COVID-19 Genomics UK (COG-UK) Consortium | The Lighthouse Lab in Milton Keynes and Alex Alderton, Roberto Amato, Jeffrey Barrett, Sonia Goncalves, Ewan Harrison, David K. Jackson, Ian Johnston, Dominic Kwiatkowski, Cordelia Langford, John Sillitoe on behalf of the Wellcome Sanger Institute COVID-19 Surveillance Team |
| EPI_ISL_2347414, EPI_ISL_2347415, EPI_ISL_2347417 | Lighthouse Lab in Alderley Park | Wellcome Sanger Institute for the COVID-19 Genomics UK (COG-UK) Consortium | Jacquelyn Wynn, Mairead Hyland, The Lighthouse Lab in Alderley Park and Alex Alderton, Roberto Amato, Jeffrey Barrett, Sonia Goncalves, Ewan Harrison, David K. Jackson, Ian Johnston, Dominic Kwiatkowski, Cordelia Langford, John Sillitoe on behalf of the Wellcome Sanger Institute COVID-19 Surveillance Team |
| EPI_ISL_2347418 | Lighthouse Lab in Glasgow | Wellcome Sanger Institute for the COVID-19 Genomics UK (COG-UK) Consortium | Harper VanSteenhouse, Yumi Kasai, David Gray, Carol Clugston, Anna Dominiczak and Alex Alderton, Roberto Amato, Jeffrey Barrett, Sonia Goncalves, Ewan Harrison, David K. Jackson, Ian Johnston, Dominic Kwiatkowski, Cordelia Langford, John Sillitoe on behalf of the Wellcome Sanger Institute COVID-19 Surveillance Team |
| EPI_ISL_2347419, EPI_ISL_2347420, EPI_ISL_2347422, EPI_ISL_2347423, EPI_ISL_2347425 | Lighthouse Lab in Alderley Park | Wellcome Sanger Institute for the COVID-19 Genomics UK (COG-UK) Consortium | Jacquelyn Wynn, Mairead Hyland, The Lighthouse Lab in Alderley Park and Alex Alderton, Roberto Amato, Jeffrey Barrett, Sonia Goncalves, Ewan Harrison, David K. Jackson, Ian Johnston, Dominic Kwiatkowski, Cordelia Langford, John Sillitoe on behalf of the Wellcome Sanger Institute COVID-19 Surveillance Team |
| EPI_ISL_2347426 | Lighthouse Lab in Glasgow | Wellcome Sanger Institute for the COVID-19 Genomics UK (COG-UK) Consortium | Harper VanSteenhouse, Yumi Kasai, David Gray, Carol Clugston, Anna Dominiczak and Alex Alderton, Roberto Amato, Jeffrey Barrett, Sonia Goncalves, Ewan Harrison, David K. Jackson, Ian Johnston, Dominic Kwiatkowski, Cordelia Langford, John Sillitoe on behalf of the Wellcome Sanger Institute COVID-19 Surveillance Team |
| EPI_ISL_2347428, EPI_ISL_2347429, EPI_ISL_2347430, EPI_ISL_2347431, EPI_ISL_2347432, EPI_ISL_2347433, EPI_ISL_2347434, EPI_ISL_2347435 | Lighthouse Lab in Alderley Park | Wellcome Sanger Institute for the COVID-19 Genomics UK (COG-UK) Consortium | Jacquelyn Wynn, Mairead Hyland, The Lighthouse Lab in Alderley Park and Alex Alderton, Roberto Amato, Jeffrey Barrett, Sonia Goncalves, Ewan Harrison, David K. Jackson, Ian Johnston, Dominic Kwiatkowski, Cordelia Langford, John Sillitoe on behalf of the Wellcome Sanger Institute COVID-19 Surveillance Team |

| | | | |
|---|---|---|---|
| EPI_ISL_2347436 | Health Services Laboratories | Wellcome Sanger Institute for the COVID-19 Genomics UK (COG-UK) Consortium | Health Services Laboratories and Alex Alderton, Roberto Amato, Jeffrey Barrett, Sonia Goncalves, Ewan Harrison, David K. Jackson, Ian Johnston, Dominic Kwiatkowski, Cordelia Langford, John Sillitoe on behalf of the Wellcome Sanger Institute COVID-19 Surveillance Team |
| EPI_ISL_2347438, EPI_ISL_2347439 | Lighthouse Lab in Alderley Park | Wellcome Sanger Institute for the COVID-19 Genomics UK (COG-UK) Consortium | Jacquelyn Wynn, Mairead Hyland, The Lighthouse Lab in Alderley Park and Alex Alderton, Roberto Amato, Jeffrey Barrett, Sonia Goncalves, Ewan Harrison, David K. Jackson, Ian Johnston, Dominic Kwiatkowski, Cordelia Langford, John Sillitoe on behalf of the Wellcome Sanger Institute COVID-19 Surveillance Team |
| EPI_ISL_2347441 | Lighthouse Lab in Glasgow | Wellcome Sanger Institute for the COVID-19 Genomics UK (COG-UK) Consortium | Harper VanSteenhouse, Yumi Kasai, David Gray, Carol Clugston, Anna Dominiczak and Alex Alderton, Roberto Amato, Jeffrey Barrett, Sonia Goncalves, Ewan Harrison, David K. Jackson, Ian Johnston, Dominic Kwiatkowski, Cordelia Langford, John Sillitoe on behalf of the Wellcome Sanger Institute COVID-19 Surveillance Team |
| EPI_ISL_2347442, EPI_ISL_2347443 | Lighthouse Lab in Alderley Park | Wellcome Sanger Institute for the COVID-19 Genomics UK (COG-UK) Consortium | Jacquelyn Wynn, Mairead Hyland, The Lighthouse Lab in Alderley Park and Alex Alderton, Roberto Amato, Jeffrey Barrett, Sonia Goncalves, Ewan Harrison, David K. Jackson, Ian Johnston, Dominic Kwiatkowski, Cordelia Langford, John Sillitoe on behalf of the Wellcome Sanger Institute COVID-19 Surveillance Team |
| EPI_ISL_2347445 | Health Services Laboratories | Wellcome Sanger Institute for the COVID-19 Genomics UK (COG-UK) Consortium | Health Services Laboratories and Alex Alderton, Roberto Amato, Jeffrey Barrett, Sonia Goncalves, Ewan Harrison, David K. Jackson, Ian Johnston, Dominic Kwiatkowski, Cordelia Langford, John Sillitoe on behalf of the Wellcome Sanger Institute COVID-19 Surveillance Team |
| EPI_ISL_2347446, EPI_ISL_2347447, EPI_ISL_2347448, EPI_ISL_2347449, EPI_ISL_2347450, EPI_ISL_2347451 | Lighthouse Lab in Alderley Park | Wellcome Sanger Institute for the COVID-19 Genomics UK (COG-UK) Consortium | Jacquelyn Wynn, Mairead Hyland, The Lighthouse Lab in Alderley Park and Alex Alderton, Roberto Amato, Jeffrey Barrett, Sonia Goncalves, Ewan Harrison, David K. Jackson, Ian Johnston, Dominic Kwiatkowski, Cordelia Langford, John Sillitoe on behalf of the Wellcome Sanger Institute COVID-19 Surveillance Team |
| EPI_ISL_2347452 | Health Services Laboratories | Wellcome Sanger Institute for the COVID-19 Genomics UK (COG-UK) Consortium | Health Services Laboratories and Alex Alderton, Roberto Amato, Jeffrey Barrett, Sonia Goncalves, Ewan Harrison, David K. Jackson, Ian Johnston, Dominic Kwiatkowski, Cordelia Langford, John Sillitoe on behalf of the Wellcome Sanger Institute COVID-19 Surveillance Team |
| EPI_ISL_2347454, EPI_ISL_2347455, EPI_ISL_2347456, EPI_ISL_2347457, EPI_ISL_2347458, EPI_ISL_2347459, EPI_ISL_2347460, EPI_ISL_2347461, EPI_ISL_2347463, EPI_ISL_2347465, EPI_ISL_2347466, EPI_ISL_2347467, EPI_ISL_2347468, EPI_ISL_2347470, EPI_ISL_2347472 | | | |
| see above | Lighthouse Lab in Alderley Park | Wellcome Sanger Institute for the COVID-19 Genomics UK (COG-UK) Consortium | Jacquelyn Wynn, Mairead Hyland, The Lighthouse Lab in Alderley Park and Alex Alderton, Roberto Amato, Jeffrey Barrett, Sonia Goncalves, Ewan Harrison, David K. Jackson, Ian Johnston, Dominic Kwiatkowski, Cordelia Langford, John Sillitoe on behalf of the Wellcome Sanger Institute COVID-19 Surveillance Team |
| EPI_ISL_2347473 | Health Services Laboratories | Wellcome Sanger Institute for the COVID-19 Genomics UK (COG-UK) Consortium | Health Services Laboratories and Alex Alderton, Roberto Amato, Jeffrey Barrett, Sonia Goncalves, Ewan Harrison, David K. Jackson, Ian Johnston, Dominic Kwiatkowski, Cordelia Langford, John Sillitoe on behalf of the Wellcome Sanger Institute COVID-19 Surveillance Team |
| EPI_ISL_2347474, EPI_ISL_2347475, EPI_ISL_2347476, EPI_ISL_2347477, EPI_ISL_2347479 | Lighthouse Lab in Alderley Park | Wellcome Sanger Institute for the COVID-19 Genomics UK (COG-UK) Consortium | Jacquelyn Wynn, Mairead Hyland, The Lighthouse Lab in Alderley Park and Alex Alderton, Roberto Amato, Jeffrey Barrett, Sonia Goncalves, Ewan Harrison, David K. Jackson, Ian Johnston, Dominic Kwiatkowski, Cordelia Langford, John Sillitoe on behalf of the Wellcome Sanger Institute COVID-19 Surveillance Team |
| EPI_ISL_2347480 | Health Services Laboratories | Wellcome Sanger Institute for the COVID-19 Genomics UK (COG-UK) Consortium | Health Services Laboratories and Alex Alderton, Roberto Amato, Jeffrey Barrett, Sonia Goncalves, Ewan Harrison, David K. Jackson, Ian Johnston, Dominic Kwiatkowski, Cordelia Langford, John Sillitoe on behalf of the Wellcome Sanger Institute COVID-19 Surveillance Team |
| EPI_ISL_2347481, EPI_ISL_2347482, EPI_ISL_2347483, EPI_ISL_2347484, EPI_ISL_2347485, EPI_ISL_2347486, EPI_ISL_2347487 | Lighthouse Lab in Alderley Park | Wellcome Sanger Institute for the COVID-19 Genomics UK (COG-UK) Consortium | Jacquelyn Wynn, Mairead Hyland, The Lighthouse Lab in Alderley Park and Alex Alderton, Roberto Amato, Jeffrey Barrett, Sonia Goncalves, Ewan Harrison, David K. Jackson, Ian Johnston, Dominic Kwiatkowski, Cordelia Langford, John Sillitoe on behalf of the Wellcome Sanger Institute COVID-19 Surveillance Team |
| EPI_ISL_2347488 | Health Services Laboratories | Wellcome Sanger Institute for the COVID-19 Genomics UK (COG-UK) Consortium | Health Services Laboratories and Alex Alderton, Roberto Amato, Jeffrey Barrett, Sonia Goncalves, Ewan Harrison, David K. Jackson, Ian Johnston, Dominic Kwiatkowski, Cordelia Langford, John Sillitoe on behalf of the Wellcome Sanger Institute COVID-19 Surveillance Team |
| EPI_ISL_2347489, EPI_ISL_2347492, EPI_ISL_2347493, EPI_ISL_2347494, EPI_ISL_2347495, EPI_ISL_2347496, EPI_ISL_2347497, EPI_ISL_2347498, EPI_ISL_2347501, EPI_ISL_2347502, EPI_ISL_2347503, EPI_ISL_2347504, EPI_ISL_2347505 | | | |
| see above | Lighthouse Lab in Alderley Park | Wellcome Sanger Institute for the COVID-19 Genomics UK (COG-UK) Consortium | Jacquelyn Wynn, Mairead Hyland, The Lighthouse Lab in Alderley Park and Alex Alderton, Roberto Amato, Jeffrey Barrett, Sonia Goncalves, Ewan Harrison, David K. Jackson, Ian Johnston, Dominic Kwiatkowski, Cordelia Langford, John Sillitoe on behalf of the Wellcome Sanger Institute COVID-19 Surveillance Team |
| EPI_ISL_2347506 | Health Services Laboratories | Wellcome Sanger Institute for the COVID-19 Genomics UK (COG-UK) Consortium | Health Services Laboratories and Alex Alderton, Roberto Amato, Jeffrey Barrett, Sonia Goncalves, Ewan Harrison, David K. Jackson, Ian Johnston, Dominic Kwiatkowski, Cordelia Langford, John Sillitoe on behalf of the Wellcome Sanger Institute COVID-19 Surveillance Team |
| EPI_ISL_2347508, EPI_ISL_2347509, EPI_ISL_2347510, EPI_ISL_2347511 | Lighthouse Lab in Alderley Park | Wellcome Sanger Institute for the COVID-19 Genomics UK (COG-UK) Consortium | Jacquelyn Wynn, Mairead Hyland, The Lighthouse Lab in Alderley Park and Alex Alderton, Roberto Amato, Jeffrey Barrett, Sonia Goncalves, Ewan Harrison, David K. Jackson, Ian Johnston, Dominic Kwiatkowski, Cordelia Langford, John Sillitoe on behalf of the Wellcome Sanger Institute COVID-19 Surveillance Team |
| EPI_ISL_2347512 | Health Services Laboratories | Wellcome Sanger Institute for the COVID-19 Genomics UK (COG-UK) Consortium | Health Services Laboratories and Alex Alderton, Roberto Amato, Jeffrey Barrett, Sonia Goncalves, Ewan Harrison, David K. Jackson, Ian Johnston, Dominic Kwiatkowski, Cordelia Langford, John Sillitoe on behalf of the Wellcome Sanger Institute COVID-19 Surveillance Team |
| EPI_ISL_2347513, EPI_ISL_2347514, EPI_ISL_2347516, EPI_ISL_2347517, EPI_ISL_2347518, EPI_ISL_2347520, EPI_ISL_2347521, EPI_ISL_2347522 | Lighthouse Lab in Alderley Park | Wellcome Sanger Institute for the COVID-19 Genomics UK (COG-UK) Consortium | Jacquelyn Wynn, Mairead Hyland, The Lighthouse Lab in Alderley Park and Alex Alderton, Roberto Amato, Jeffrey Barrett, Sonia Goncalves, Ewan Harrison, David K. Jackson, Ian Johnston, Dominic Kwiatkowski, Cordelia Langford, John Sillitoe on behalf of the Wellcome Sanger Institute COVID-19 Surveillance Team |
| EPI_ISL_2347523 | Health Services Laboratories | Wellcome Sanger Institute for the COVID-19 Genomics UK (COG-UK) Consortium | Health Services Laboratories and Alex Alderton, Roberto Amato, Jeffrey Barrett, Sonia Goncalves, Ewan Harrison, David K. Jackson, Ian Johnston, Dominic Kwiatkowski, Cordelia Langford, John Sillitoe on behalf of the Wellcome Sanger Institute COVID-19 Surveillance Team |
| EPI_ISL_2347524, EPI_ISL_2347525, EPI_ISL_2347527, EPI_ISL_2347528, EPI_ISL_2347530 | Lighthouse Lab in Alderley Park | Wellcome Sanger Institute for the COVID-19 Genomics UK (COG-UK) Consortium | Jacquelyn Wynn, Mairead Hyland, The Lighthouse Lab in Alderley Park and Alex Alderton, Roberto Amato, Jeffrey Barrett, Sonia Goncalves, Ewan Harrison, David K. Jackson, Ian Johnston, Dominic Kwiatkowski, Cordelia Langford, John Sillitoe on behalf of the Wellcome Sanger Institute COVID-19 Surveillance Team |
| EPI_ISL_2347531 | Health Services Laboratories | Wellcome Sanger Institute for the COVID-19 Genomics UK (COG-UK) Consortium | Health Services Laboratories and Alex Alderton, Roberto Amato, Jeffrey Barrett, Sonia Goncalves, Ewan Harrison, David K. Jackson, Ian Johnston, Dominic Kwiatkowski, Cordelia Langford, John Sillitoe on behalf of the Wellcome Sanger Institute COVID-19 Surveillance Team |
| EPI_ISL_2347532, EPI_ISL_2347533, EPI_ISL_2347534, EPI_ISL_2347535, EPI_ISL_2347537, EPI_ISL_2347538, EPI_ISL_2347539, EPI_ISL_2347540 | Lighthouse Lab in Alderley Park | Wellcome Sanger Institute for the COVID-19 Genomics UK (COG-UK) Consortium | Jacquelyn Wynn, Mairead Hyland, The Lighthouse Lab in Alderley Park and Alex Alderton, Roberto Amato, Jeffrey Barrett, Sonia Goncalves, Ewan Harrison, David K. Jackson, Ian Johnston, Dominic Kwiatkowski, Cordelia Langford, John Sillitoe on behalf of the Wellcome Sanger Institute COVID-19 Surveillance Team |
| EPI_ISL_2347541 | Health Services Laboratories | Wellcome Sanger Institute for the COVID-19 Genomics UK (COG-UK) Consortium | Health Services Laboratories and Alex Alderton, Roberto Amato, Jeffrey Barrett, Sonia Goncalves, Ewan Harrison, David K. Jackson, Ian Johnston, Dominic Kwiatkowski, Cordelia Langford, John Sillitoe on behalf of the Wellcome Sanger Institute COVID-19 Surveillance Team |
| EPI_ISL_2347543, EPI_ISL_2347544, EPI_ISL_2347545, EPI_ISL_2347546, EPI_ISL_2347547, EPI_ISL_2347549, EPI_ISL_2347550, EPI_ISL_2347551, EPI_ISL_2347552, EPI_ISL_2347553, EPI_ISL_2347554, EPI_ISL_2347555, EPI_ISL_2347556 | | | |
| see above | Lighthouse Lab in Alderley Park | Wellcome Sanger Institute for the COVID-19 Genomics UK (COG-UK) Consortium | Jacquelyn Wynn, Mairead Hyland, The Lighthouse Lab in Alderley Park and Alex Alderton, Roberto Amato, Jeffrey Barrett, Sonia Goncalves, Ewan Harrison, David K. Jackson, Ian Johnston, Dominic Kwiatkowski, Cordelia Langford, John Sillitoe on behalf of the Wellcome Sanger Institute COVID-19 Surveillance Team |
| EPI_ISL_2347558 | Health Services Laboratories | Wellcome Sanger Institute for the COVID-19 Genomics UK (COG-UK) Consortium | Health Services Laboratories and Alex Alderton, Roberto Amato, Jeffrey Barrett, Sonia Goncalves, Ewan Harrison, David K. Jackson, Ian Johnston, Dominic Kwiatkowski, Cordelia Langford, John Sillitoe on behalf of the Wellcome Sanger Institute COVID-19 Surveillance Team |
| EPI_ISL_2347559, EPI_ISL_2347560, EPI_ISL_2347561, EPI_ISL_2347563, EPI_ISL_2347567 | Lighthouse Lab in Alderley Park | Wellcome Sanger Institute for the COVID-19 Genomics UK (COG-UK) Consortium | Jacquelyn Wynn, Mairead Hyland, The Lighthouse Lab in Alderley Park and Alex Alderton, Roberto Amato, Jeffrey Barrett, Sonia Goncalves, Ewan Harrison, David K. Jackson, Ian Johnston, Dominic Kwiatkowski, Cordelia Langford, John Sillitoe on behalf of the Wellcome Sanger Institute COVID-19 Surveillance Team |
| EPI_ISL_2347568, EPI_ISL_2347569 | Health Services Laboratories | Wellcome Sanger Institute for the COVID-19 Genomics UK (COG-UK) Consortium | Health Services Laboratories and Alex Alderton, Roberto Amato, Jeffrey Barrett, Sonia Goncalves, Ewan Harrison, David K. Jackson, Ian Johnston, Dominic Kwiatkowski, Cordelia Langford, John Sillitoe on behalf of the Wellcome Sanger Institute COVID-19 Surveillance Team |
| EPI_ISL_2347570 | Lighthouse Lab in Alderley Park | Wellcome Sanger Institute for the COVID-19 Genomics UK (COG-UK) Consortium | Jacquelyn Wynn, Mairead Hyland, The Lighthouse Lab in Alderley Park and Alex Alderton, Roberto Amato, Jeffrey Barrett, Sonia Goncalves, Ewan Harrison, David K. Jackson, Ian Johnston, Dominic Kwiatkowski, Cordelia Langford, John Sillitoe on behalf of the Wellcome Sanger Institute COVID-19 Surveillance Team |
| EPI_ISL_2347571 | Lighthouse Lab in Milton Keynes | Wellcome Sanger Institute for the COVID-19 Genomics UK (COG-UK) Consortium | The Lighthouse Lab in Milton Keynes and Alex Alderton, Roberto Amato, Jeffrey Barrett, Sonia Goncalves, Ewan Harrison, David K. Jackson, Ian Johnston, Dominic Kwiatkowski, Cordelia Langford, John Sillitoe on behalf of the Wellcome Sanger Institute COVID-19 Surveillance Team |
| EPI_ISL_2347572, EPI_ISL_2347573, EPI_ISL_2347574, | Lighthouse Lab in Alderley Park | Wellcome Sanger Institute for the COVID-19 Genomics UK (COG-UK) Consortium | Jacquelyn Wynn, Mairead Hyland, The Lighthouse Lab in Alderley Park and Alex Alderton, Roberto Amato, Jeffrey Barrett, Sonia Goncalves, Ewan Harrison, David K. Jackson, Ian Johnston, Dominic Kwiatkowski, Cordelia Langford, John Sillitoe on behalf of the Wellcome Sanger Institute COVID-19 Surveillance Team |

| | | | |
|---|---|---|---|
| EPI_ISL_2347575, EPI_ISL_2347578, EPI_ISL_2347579 | | | |
| EPI_ISL_2347580 | Health Services Laboratories | Wellcome Sanger Institute for the COVID-19 Genomics UK (COG-UK) Consortium | Health Services Laboratories and Alex Alderton, Roberto Amato, Jeffrey Barrett, Sonia Goncalves, Ewan Harrison, David K. Jackson, Ian Johnston, Dominic Kwiatkowski, Cordelia Langford, John Sillitoe on behalf of the Wellcome Sanger Institute COVID-19 Surveillance Team |
| EPI_ISL_2347582, EPI_ISL_2347583, EPI_ISL_2347584, EPI_ISL_2347585, EPI_ISL_2347586, EPI_ISL_2347587, EPI_ISL_2347588, EPI_ISL_2347589, EPI_ISL_2347590, EPI_ISL_2347591, EPI_ISL_2347592, EPI_ISL_2347593, EPI_ISL_2347594, EPI_ISL_2347595, EPI_ISL_2347597, EPI_ISL_2347599, EPI_ISL_2347600, EPI_ISL_2347601, EPI_ISL_2347603, EPI_ISL_2347604, EPI_ISL_2347606, EPI_ISL_2347607, EPI_ISL_2347608, EPI_ISL_2347609, EPI_ISL_2347610, EPI_ISL_2347611, EPI_ISL_2347613, EPI_ISL_2347614, EPI_ISL_2347615, EPI_ISL_2347616 | | | |
| see above | Lighthouse Lab in Alderley Park | Wellcome Sanger Institute for the COVID-19 Genomics UK (COG-UK) Consortium | Jacquelyn Wynn, Mairead Hyland, The Lighthouse Lab in Alderley Park and Alex Alderton, Roberto Amato, Jeffrey Barrett, Sonia Goncalves, Ewan Harrison, David K. Jackson, Ian Johnston, Dominic Kwiatkowski, Cordelia Langford, John Sillitoe on behalf of the Wellcome Sanger Institute COVID-19 Surveillance Team |
| EPI_ISL_2347618 | Lighthouse Laboratory Plymouth | Wellcome Sanger Institute for the COVID-19 Genomics UK (COG-UK) Consortium | Lighthouse Laboratory Plymouth and Alex Alderton, Roberto Amato, Jeffrey Barrett, Sonia Goncalves, Ewan Harrison, David K. Jackson, Ian Johnston, Dominic Kwiatkowski, Cordelia Langford, John Sillitoe on behalf of the Wellcome Sanger Institute COVID-19 Surveillance Team |
| EPI_ISL_2347619, EPI_ISL_2347622, EPI_ISL_2347624, EPI_ISL_2347625, EPI_ISL_2347627, EPI_ISL_2347629, EPI_ISL_2347630, EPI_ISL_2347632, EPI_ISL_2347633, EPI_ISL_2347637, EPI_ISL_2347639, EPI_ISL_2347640, EPI_ISL_2347641, EPI_ISL_2347642 | | | |
| see above | Lighthouse Lab in Alderley Park | Wellcome Sanger Institute for the COVID-19 Genomics UK (COG-UK) Consortium | Jacquelyn Wynn, Mairead Hyland, The Lighthouse Lab in Alderley Park and Alex Alderton, Roberto Amato, Jeffrey Barrett, Sonia Goncalves, Ewan Harrison, David K. Jackson, Ian Johnston, Dominic Kwiatkowski, Cordelia Langford, John Sillitoe on behalf of the Wellcome Sanger Institute COVID-19 Surveillance Team |
| EPI_ISL_2347643 | Lighthouse Laboratory Plymouth | Wellcome Sanger Institute for the COVID-19 Genomics UK (COG-UK) Consortium | Lighthouse Laboratory Plymouth and Alex Alderton, Roberto Amato, Jeffrey Barrett, Sonia Goncalves, Ewan Harrison, David K. Jackson, Ian Johnston, Dominic Kwiatkowski, Cordelia Langford, John Sillitoe on behalf of the Wellcome Sanger Institute COVID-19 Surveillance Team |
| EPI_ISL_2347644, EPI_ISL_2347645, EPI_ISL_2347646, EPI_ISL_2347647, EPI_ISL_2347649, EPI_ISL_2347650, EPI_ISL_2347651 | Lighthouse Lab in Alderley Park | Wellcome Sanger Institute for the COVID-19 Genomics UK (COG-UK) Consortium | Jacquelyn Wynn, Mairead Hyland, The Lighthouse Lab in Alderley Park and Alex Alderton, Roberto Amato, Jeffrey Barrett, Sonia Goncalves, Ewan Harrison, David K. Jackson, Ian Johnston, Dominic Kwiatkowski, Cordelia Langford, John Sillitoe on behalf of the Wellcome Sanger Institute COVID-19 Surveillance Team |
| EPI_ISL_2347655 | Lighthouse Lab in Glasgow | Wellcome Sanger Institute for the COVID-19 Genomics UK (COG-UK) Consortium | Harper VanSteenhouse, Yumi Kasai, David Gray, Carol Clugston, Anna Dominiczak and Alex Alderton, Roberto Amato, Jeffrey Barrett, Sonia Goncalves, Ewan Harrison, David K. Jackson, Ian Johnston, Dominic Kwiatkowski, Cordelia Langford, John Sillitoe on behalf of the Wellcome Sanger Institute COVID-19 Surveillance Team |
| EPI_ISL_2347656, EPI_ISL_2347659, EPI_ISL_2347661, EPI_ISL_2347662 | Lighthouse Lab in Alderley Park | Wellcome Sanger Institute for the COVID-19 Genomics UK (COG-UK) Consortium | Jacquelyn Wynn, Mairead Hyland, The Lighthouse Lab in Alderley Park and Alex Alderton, Roberto Amato, Jeffrey Barrett, Sonia Goncalves, Ewan Harrison, David K. Jackson, Ian Johnston, Dominic Kwiatkowski, Cordelia Langford, John Sillitoe on behalf of the Wellcome Sanger Institute COVID-19 Surveillance Team |
| EPI_ISL_2347663 | Lighthouse Lab in Glasgow | Wellcome Sanger Institute for the COVID-19 Genomics UK (COG-UK) Consortium | Harper VanSteenhouse, Yumi Kasai, David Gray, Carol Clugston, Anna Dominiczak and Alex Alderton, Roberto Amato, Jeffrey Barrett, Sonia Goncalves, Ewan Harrison, David K. Jackson, Ian Johnston, Dominic Kwiatkowski, Cordelia Langford, John Sillitoe on behalf of the Wellcome Sanger Institute COVID-19 Surveillance Team |
| EPI_ISL_2347665, EPI_ISL_2347666, EPI_ISL_2347667, EPI_ISL_2347668 | Lighthouse Lab in Alderley Park | Wellcome Sanger Institute for the COVID-19 Genomics UK (COG-UK) Consortium | Jacquelyn Wynn, Mairead Hyland, The Lighthouse Lab in Alderley Park and Alex Alderton, Roberto Amato, Jeffrey Barrett, Sonia Goncalves, Ewan Harrison, David K. Jackson, Ian Johnston, Dominic Kwiatkowski, Cordelia Langford, John Sillitoe on behalf of the Wellcome Sanger Institute COVID-19 Surveillance Team |
| EPI_ISL_2347669 | Lighthouse Laboratory Plymouth | Wellcome Sanger Institute for the COVID-19 Genomics UK (COG-UK) Consortium | Lighthouse Laboratory Plymouth and Alex Alderton, Roberto Amato, Jeffrey Barrett, Sonia Goncalves, Ewan Harrison, David K. Jackson, Ian Johnston, Dominic Kwiatkowski, Cordelia Langford, John Sillitoe on behalf of the Wellcome Sanger Institute COVID-19 Surveillance Team |
| EPI_ISL_2347670, EPI_ISL_2347671, EPI_ISL_2347672 | Lighthouse Lab in Alderley Park | Wellcome Sanger Institute for the COVID-19 Genomics UK (COG-UK) Consortium | Jacquelyn Wynn, Mairead Hyland, The Lighthouse Lab in Alderley Park and Alex Alderton, Roberto Amato, Jeffrey Barrett, Sonia Goncalves, Ewan Harrison, David K. Jackson, Ian Johnston, Dominic Kwiatkowski, Cordelia Langford, John Sillitoe on behalf of the Wellcome Sanger Institute COVID-19 Surveillance Team |
| EPI_ISL_2347673 | Lighthouse Lab in Glasgow | Wellcome Sanger Institute for the COVID-19 Genomics UK (COG-UK) Consortium | Harper VanSteenhouse, Yumi Kasai, David Gray, Carol Clugston, Anna Dominiczak and Alex Alderton, Roberto Amato, Jeffrey Barrett, Sonia Goncalves, Ewan Harrison, David K. Jackson, Ian Johnston, Dominic Kwiatkowski, Cordelia Langford, John Sillitoe on behalf of the Wellcome Sanger Institute COVID-19 Surveillance Team |
| EPI_ISL_2347674, EPI_ISL_2347678, EPI_ISL_2347680 | Lighthouse Lab in Alderley Park | Wellcome Sanger Institute for the COVID-19 Genomics UK (COG-UK) Consortium | Jacquelyn Wynn, Mairead Hyland, The Lighthouse Lab in Alderley Park and Alex Alderton, Roberto Amato, Jeffrey Barrett, Sonia Goncalves, Ewan Harrison, David K. Jackson, Ian Johnston, Dominic Kwiatkowski, Cordelia Langford, John Sillitoe on behalf of the Wellcome Sanger Institute COVID-19 Surveillance Team |
| EPI_ISL_2347683 | Health Services Laboratories | Wellcome Sanger Institute for the COVID-19 Genomics UK (COG-UK) Consortium | Health Services Laboratories and Alex Alderton, Roberto Amato, Jeffrey Barrett, Sonia Goncalves, Ewan Harrison, David K. Jackson, Ian Johnston, Dominic Kwiatkowski, Cordelia Langford, John Sillitoe on behalf of the Wellcome Sanger Institute COVID-19 Surveillance Team |
| EPI_ISL_2347685, EPI_ISL_2347686, EPI_ISL_2347687, EPI_ISL_2347689, EPI_ISL_2347691, EPI_ISL_2347692, EPI_ISL_2347693, EPI_ISL_2347695, EPI_ISL_2347696, EPI_ISL_2347699, EPI_ISL_2347701, EPI_ISL_2347702 | | | |
| see above | Lighthouse Lab in Alderley Park | Wellcome Sanger Institute for the COVID-19 Genomics UK (COG-UK) Consortium | Jacquelyn Wynn, Mairead Hyland, The Lighthouse Lab in Alderley Park and Alex Alderton, Roberto Amato, Jeffrey Barrett, Sonia Goncalves, Ewan Harrison, David K. Jackson, Ian Johnston, Dominic Kwiatkowski, Cordelia Langford, John Sillitoe on behalf of the Wellcome Sanger Institute COVID-19 Surveillance Team |
| EPI_ISL_2347704 | Lighthouse Laboratory Plymouth | Wellcome Sanger Institute for the COVID-19 Genomics UK (COG-UK) Consortium | Lighthouse Laboratory Plymouth and Alex Alderton, Roberto Amato, Jeffrey Barrett, Sonia Goncalves, Ewan Harrison, David K. Jackson, Ian Johnston, Dominic Kwiatkowski, Cordelia Langford, John Sillitoe on behalf of the Wellcome Sanger Institute COVID-19 Surveillance Team |
| EPI_ISL_2347705, EPI_ISL_2347706, EPI_ISL_2347713, EPI_ISL_2347714, EPI_ISL_2347716, EPI_ISL_2347717, EPI_ISL_2347718, EPI_ISL_2347722, EPI_ISL_2347723, EPI_ISL_2347724, EPI_ISL_2347725, EPI_ISL_2347727 | | | |
| see above | Lighthouse Lab in Alderley Park | Wellcome Sanger Institute for the COVID-19 Genomics UK (COG-UK) Consortium | Jacquelyn Wynn, Mairead Hyland, The Lighthouse Lab in Alderley Park and Alex Alderton, Roberto Amato, Jeffrey Barrett, Sonia Goncalves, Ewan Harrison, David K. Jackson, Ian Johnston, Dominic Kwiatkowski, Cordelia Langford, John Sillitoe on behalf of the Wellcome Sanger Institute COVID-19 Surveillance Team |
| EPI_ISL_2347728 | Lighthouse Laboratory Plymouth | Wellcome Sanger Institute for the COVID-19 Genomics UK (COG-UK) Consortium | Lighthouse Laboratory Plymouth and Alex Alderton, Roberto Amato, Jeffrey Barrett, Sonia Goncalves, Ewan Harrison, David K. Jackson, Ian Johnston, Dominic Kwiatkowski, Cordelia Langford, John Sillitoe on behalf of the Wellcome Sanger Institute COVID-19 Surveillance Team |
| EPI_ISL_2347729 | Lighthouse Lab in Alderley Park | Wellcome Sanger Institute for the COVID-19 Genomics UK (COG-UK) Consortium | Jacquelyn Wynn, Mairead Hyland, The Lighthouse Lab in Alderley Park and Alex Alderton, Roberto Amato, Jeffrey Barrett, Sonia Goncalves, Ewan Harrison, David K. Jackson, Ian Johnston, Dominic Kwiatkowski, Cordelia Langford, John Sillitoe on behalf of the Wellcome Sanger Institute COVID-19 Surveillance Team |
| EPI_ISL_2347730 | Lighthouse Laboratory Plymouth | Wellcome Sanger Institute for the COVID-19 Genomics UK (COG-UK) Consortium | Lighthouse Laboratory Plymouth and Alex Alderton, Roberto Amato, Jeffrey Barrett, Sonia Goncalves, Ewan Harrison, David K. Jackson, Ian Johnston, Dominic Kwiatkowski, Cordelia Langford, John Sillitoe on behalf of the Wellcome Sanger Institute COVID-19 Surveillance Team |
| EPI_ISL_2347731, EPI_ISL_2347734, EPI_ISL_2347735, EPI_ISL_2347737, EPI_ISL_2347738, EPI_ISL_2347739, EPI_ISL_2347740, EPI_ISL_2347742, EPI_ISL_2347743, EPI_ISL_2347746, EPI_ISL_2347747, EPI_ISL_2347748 | | | |
| see above | Lighthouse Lab in Alderley Park | Wellcome Sanger Institute for the COVID-19 Genomics UK (COG-UK) Consortium | Jacquelyn Wynn, Mairead Hyland, The Lighthouse Lab in Alderley Park and Alex Alderton, Roberto Amato, Jeffrey Barrett, Sonia Goncalves, Ewan Harrison, David K. Jackson, Ian Johnston, Dominic Kwiatkowski, Cordelia Langford, John Sillitoe on behalf of the Wellcome Sanger Institute COVID-19 Surveillance Team |
| EPI_ISL_2347749 | Lighthouse Lab in Glasgow | Wellcome Sanger Institute for the COVID-19 Genomics UK (COG-UK) Consortium | Harper VanSteenhouse, Yumi Kasai, David Gray, Carol Clugston, Anna Dominiczak and Alex Alderton, Roberto Amato, Jeffrey Barrett, Sonia Goncalves, Ewan Harrison, David K. Jackson, Ian Johnston, Dominic Kwiatkowski, Cordelia Langford, John Sillitoe on behalf of the Wellcome Sanger Institute COVID-19 Surveillance Team |
| EPI_ISL_2347750, EPI_ISL_2347752, EPI_ISL_2347754, EPI_ISL_2347755, EPI_ISL_2347757, EPI_ISL_2347758, EPI_ISL_2347759, EPI_ISL_2347760, EPI_ISL_2347761, EPI_ISL_2347762, EPI_ISL_2347763, EPI_ISL_2347764, EPI_ISL_2347765, EPI_ISL_2347767, EPI_ISL_2347768 | | | |
| see above | Lighthouse Lab in Alderley Park | Wellcome Sanger Institute for the COVID-19 Genomics UK (COG-UK) Consortium | Jacquelyn Wynn, Mairead Hyland, The Lighthouse Lab in Alderley Park and Alex Alderton, Roberto Amato, Jeffrey Barrett, Sonia Goncalves, Ewan Harrison, David K. Jackson, Ian Johnston, Dominic Kwiatkowski, Cordelia Langford, John Sillitoe on behalf of the Wellcome Sanger Institute COVID-19 Surveillance Team |
| EPI_ISL_2347770 | Health Services Laboratories | Wellcome Sanger Institute for the COVID-19 Genomics UK (COG-UK) Consortium | Health Services Laboratories and Alex Alderton, Roberto Amato, Jeffrey Barrett, Sonia Goncalves, Ewan Harrison, David K. Jackson, Ian Johnston, Dominic Kwiatkowski, Cordelia Langford, John Sillitoe on behalf of the Wellcome Sanger Institute COVID-19 Surveillance Team |
| EPI_ISL_2347771, EPI_ISL_2347772, EPI_ISL_2347775 | Lighthouse Lab in Alderley Park | Wellcome Sanger Institute for the COVID-19 Genomics UK (COG-UK) Consortium | Jacquelyn Wynn, Mairead Hyland, The Lighthouse Lab in Alderley Park and Alex Alderton, Roberto Amato, Jeffrey Barrett, Sonia Goncalves, Ewan Harrison, David K. Jackson, Ian Johnston, Dominic Kwiatkowski, Cordelia Langford, John Sillitoe on behalf of the Wellcome Sanger Institute COVID-19 Surveillance Team |
| EPI_ISL_2347776 | Health Services Laboratories | Wellcome Sanger Institute for the COVID-19 Genomics UK (COG-UK) Consortium | Health Services Laboratories and Alex Alderton, Roberto Amato, Jeffrey Barrett, Sonia Goncalves, Ewan Harrison, David K. Jackson, Ian Johnston, Dominic Kwiatkowski, Cordelia Langford, John Sillitoe on behalf of the Wellcome Sanger Institute COVID-19 Surveillance Team |
| EPI_ISL_2347777, EPI_ISL_2347778, EPI_ISL_2347781, EPI_ISL_2347782, EPI_ISL_2347783 | Lighthouse Lab in Alderley Park | Wellcome Sanger Institute for the COVID-19 Genomics UK (COG-UK) Consortium | Jacquelyn Wynn, Mairead Hyland, The Lighthouse Lab in Alderley Park and Alex Alderton, Roberto Amato, Jeffrey Barrett, Sonia Goncalves, Ewan Harrison, David K. Jackson, Ian Johnston, Dominic Kwiatkowski, Cordelia Langford, John Sillitoe on behalf of the Wellcome Sanger Institute COVID-19 Surveillance Team |
| EPI_ISL_2347784 | Lighthouse Lab in Glasgow | Wellcome Sanger Institute for the COVID-19 Genomics UK (COG-UK) Consortium | Harper VanSteenhouse, Yumi Kasai, David Gray, Carol Clugston, Anna Dominiczak and Alex Alderton, Roberto Amato, Jeffrey Barrett, Sonia Goncalves, Ewan Harrison, David K. Jackson, Ian Johnston, Dominic Kwiatkowski, Cordelia Langford, John Sillitoe on behalf of the Wellcome Sanger Institute COVID-19 Surveillance Team |
| EPI_ISL_2347786, EPI_ISL_2347787, EPI_ISL_2347788, EPI_ISL_2347789, EPI_ISL_2347790 | Lighthouse Lab in Alderley Park | Wellcome Sanger Institute for the COVID-19 Genomics UK (COG-UK) Consortium | Jacquelyn Wynn, Mairead Hyland, The Lighthouse Lab in Alderley Park and Alex Alderton, Roberto Amato, Jeffrey Barrett, Sonia Goncalves, Ewan Harrison, David K. Jackson, Ian Johnston, Dominic Kwiatkowski, Cordelia Langford, John Sillitoe on behalf of the Wellcome Sanger Institute COVID-19 Surveillance Team |
| EPI_ISL_2347791, EPI_ISL_2347792 | Lighthouse Laboratory Plymouth | Wellcome Sanger Institute for the COVID-19 Genomics UK (COG-UK) Consortium | Lighthouse Laboratory Plymouth and Alex Alderton, Roberto Amato, Jeffrey Barrett, Sonia Goncalves, Ewan Harrison, David K. Jackson, Ian Johnston, Dominic Kwiatkowski, Cordelia Langford, John Sillitoe on behalf of the Wellcome Sanger Institute COVID-19 Surveillance Team |
| EPI_ISL_2347795, EPI_ISL_2347796, EPI_ISL_2347798, | Lighthouse Lab in Alderley Park | Wellcome Sanger Institute for the COVID-19 Genomics UK (COG-UK) Consortium | Jacquelyn Wynn, Mairead Hyland, The Lighthouse Lab in Alderley Park and Alex Alderton, Roberto Amato, Jeffrey Barrett, Sonia Goncalves, Ewan Harrison, David K. Jackson, Ian Johnston, Dominic Kwiatkowski, Cordelia Langford, John Sillitoe on behalf of the Wellcome Sanger Institute COVID-19 Surveillance Team |

| | | | |
|---|---|---|---|
| EPI_ISL_2347799, EPI_ISL_2347801, EPI_ISL_2347802, EPI_ISL_2347803 | | | |
| EPI_ISL_2347804 | Health Services Laboratories | Wellcome Sanger Institute for the COVID-19 Genomics UK (COG-UK) Consortium | Health Services Laboratories and Alex Alderton, Roberto Amato, Jeffrey Barrett, Sonia Goncalves, Ewan Harrison, David K. Jackson, Ian Johnston, Dominic Kwiatkowski, Cordelia Langford, John Sillitoe on behalf of the Wellcome Sanger Institute COVID-19 Surveillance Team |
| EPI_ISL_2347806, EPI_ISL_2347809, EPI_ISL_2347810, EPI_ISL_2347812, EPI_ISL_2347814, EPI_ISL_2347815, EPI_ISL_2347816, EPI_ISL_2347817, EPI_ISL_2347818, EPI_ISL_2347820, EPI_ISL_2347822, EPI_ISL_2347824 | | | |
| see above | Lighthouse Lab in Alderley Park | Wellcome Sanger Institute for the COVID-19 Genomics UK (COG-UK) Consortium | Jacquelyn Wynn, Mairead Hyland, The Lighthouse Lab in Alderley Park and Alex Alderton, Roberto Amato, Jeffrey Barrett, Sonia Goncalves, Ewan Harrison, David K. Jackson, Ian Johnston, Dominic Kwiatkowski, Cordelia Langford, John Sillitoe on behalf of the Wellcome Sanger Institute COVID-19 Surveillance Team |
| EPI_ISL_2347825 | Lighthouse Lab in Glasgow | Wellcome Sanger Institute for the COVID-19 Genomics UK (COG-UK) Consortium | Harper VanSteenhouse, Yumi Kasai, David Gray, Carol Clugston, Anna Dominiczak and Alex Alderton, Roberto Amato, Jeffrey Barrett, Sonia Goncalves, Ewan Harrison, David K. Jackson, Ian Johnston, Dominic Kwiatkowski, Cordelia Langford, John Sillitoe on behalf of the Wellcome Sanger Institute COVID-19 Surveillance Team |
| EPI_ISL_2347826, EPI_ISL_2347827, EPI_ISL_2347829, EPI_ISL_2347830, EPI_ISL_2347831 | Lighthouse Lab in Alderley Park | Wellcome Sanger Institute for the COVID-19 Genomics UK (COG-UK) Consortium | Jacquelyn Wynn, Mairead Hyland, The Lighthouse Lab in Alderley Park and Alex Alderton, Roberto Amato, Jeffrey Barrett, Sonia Goncalves, Ewan Harrison, David K. Jackson, Ian Johnston, Dominic Kwiatkowski, Cordelia Langford, John Sillitoe on behalf of the Wellcome Sanger Institute COVID-19 Surveillance Team |
| EPI_ISL_2347832 | Lighthouse Lab in Glasgow | Wellcome Sanger Institute for the COVID-19 Genomics UK (COG-UK) Consortium | Harper VanSteenhouse, Yumi Kasai, David Gray, Carol Clugston, Anna Dominiczak and Alex Alderton, Roberto Amato, Jeffrey Barrett, Sonia Goncalves, Ewan Harrison, David K. Jackson, Ian Johnston, Dominic Kwiatkowski, Cordelia Langford, John Sillitoe on behalf of the Wellcome Sanger Institute COVID-19 Surveillance Team |
| EPI_ISL_2347833, EPI_ISL_2347834, EPI_ISL_2347835, EPI_ISL_2347836, EPI_ISL_2347837, EPI_ISL_2347838, EPI_ISL_2347839, EPI_ISL_2347840, EPI_ISL_2347841, EPI_ISL_2347842 | Lighthouse Lab in Alderley Park | Wellcome Sanger Institute for the COVID-19 Genomics UK (COG-UK) Consortium | Jacquelyn Wynn, Mairead Hyland, The Lighthouse Lab in Alderley Park and Alex Alderton, Roberto Amato, Jeffrey Barrett, Sonia Goncalves, Ewan Harrison, David K. Jackson, Ian Johnston, Dominic Kwiatkowski, Cordelia Langford, John Sillitoe on behalf of the Wellcome Sanger Institute COVID-19 Surveillance Team |
| EPI_ISL_2347843 | Lighthouse Laboratory Plymouth | Wellcome Sanger Institute for the COVID-19 Genomics UK (COG-UK) Consortium | Lighthouse Laboratory Plymouth and Alex Alderton, Roberto Amato, Jeffrey Barrett, Sonia Goncalves, Ewan Harrison, David K. Jackson, Ian Johnston, Dominic Kwiatkowski, Cordelia Langford, John Sillitoe on behalf of the Wellcome Sanger Institute COVID-19 Surveillance Team |
| EPI_ISL_2347844, EPI_ISL_2347845, EPI_ISL_2347846, EPI_ISL_2347850, EPI_ISL_2347853, EPI_ISL_2347854, EPI_ISL_2347855, EPI_ISL_2347856, EPI_ISL_2347858, EPI_ISL_2347859, EPI_ISL_2347860 | | | |
| see above | Lighthouse Lab in Alderley Park | Wellcome Sanger Institute for the COVID-19 Genomics UK (COG-UK) Consortium | Jacquelyn Wynn, Mairead Hyland, The Lighthouse Lab in Alderley Park and Alex Alderton, Roberto Amato, Jeffrey Barrett, Sonia Goncalves, Ewan Harrison, David K. Jackson, Ian Johnston, Dominic Kwiatkowski, Cordelia Langford, John Sillitoe on behalf of the Wellcome Sanger Institute COVID-19 Surveillance Team |
| EPI_ISL_2347861 | Health Services Laboratories | Wellcome Sanger Institute for the COVID-19 Genomics UK (COG-UK) Consortium | Health Services Laboratories and Alex Alderton, Roberto Amato, Jeffrey Barrett, Sonia Goncalves, Ewan Harrison, David K. Jackson, Ian Johnston, Dominic Kwiatkowski, Cordelia Langford, John Sillitoe on behalf of the Wellcome Sanger Institute COVID-19 Surveillance Team |
| EPI_ISL_2347862 | Lighthouse Lab in Alderley Park | Wellcome Sanger Institute for the COVID-19 Genomics UK (COG-UK) Consortium | Jacquelyn Wynn, Mairead Hyland, The Lighthouse Lab in Alderley Park and Alex Alderton, Roberto Amato, Jeffrey Barrett, Sonia Goncalves, Ewan Harrison, David K. Jackson, Ian Johnston, Dominic Kwiatkowski, Cordelia Langford, John Sillitoe on behalf of the Wellcome Sanger Institute COVID-19 Surveillance Team |
| EPI_ISL_2347865 | Lighthouse Lab in Milton Keynes | Wellcome Sanger Institute for the COVID-19 Genomics UK (COG-UK) Consortium | The Lighthouse Lab in Milton Keynes and Alex Alderton, Roberto Amato, Jeffrey Barrett, Sonia Goncalves, Ewan Harrison, David K. Jackson, Ian Johnston, Dominic Kwiatkowski, Cordelia Langford, John Sillitoe on behalf of the Wellcome Sanger Institute COVID-19 Surveillance Team |
| EPI_ISL_2347866 | Lighthouse Laboratory Plymouth | Wellcome Sanger Institute for the COVID-19 Genomics UK (COG-UK) Consortium | Lighthouse Laboratory Plymouth and Alex Alderton, Roberto Amato, Jeffrey Barrett, Sonia Goncalves, Ewan Harrison, David K. Jackson, Ian Johnston, Dominic Kwiatkowski, Cordelia Langford, John Sillitoe on behalf of the Wellcome Sanger Institute COVID-19 Surveillance Team |
| EPI_ISL_2347867, EPI_ISL_2347870 | Lighthouse Lab in Alderley Park | Wellcome Sanger Institute for the COVID-19 Genomics UK (COG-UK) Consortium | Jacquelyn Wynn, Mairead Hyland, The Lighthouse Lab in Alderley Park and Alex Alderton, Roberto Amato, Jeffrey Barrett, Sonia Goncalves, Ewan Harrison, David K. Jackson, Ian Johnston, Dominic Kwiatkowski, Cordelia Langford, John Sillitoe on behalf of the Wellcome Sanger Institute COVID-19 Surveillance Team |
| EPI_ISL_2347871 | Lighthouse Laboratory Plymouth | Wellcome Sanger Institute for the COVID-19 Genomics UK (COG-UK) Consortium | Lighthouse Laboratory Plymouth and Alex Alderton, Roberto Amato, Jeffrey Barrett, Sonia Goncalves, Ewan Harrison, David K. Jackson, Ian Johnston, Dominic Kwiatkowski, Cordelia Langford, John Sillitoe on behalf of the Wellcome Sanger Institute COVID-19 Surveillance Team |
| EPI_ISL_2347872 | Health Services Laboratories | Wellcome Sanger Institute for the COVID-19 Genomics UK (COG-UK) Consortium | Health Services Laboratories and Alex Alderton, Roberto Amato, Jeffrey Barrett, Sonia Goncalves, Ewan Harrison, David K. Jackson, Ian Johnston, Dominic Kwiatkowski, Cordelia Langford, John Sillitoe on behalf of the Wellcome Sanger Institute COVID-19 Surveillance Team |
| EPI_ISL_2347874 | Lighthouse Lab in Milton Keynes | Wellcome Sanger Institute for the COVID-19 Genomics UK (COG-UK) Consortium | The Lighthouse Lab in Milton Keynes and Alex Alderton, Roberto Amato, Jeffrey Barrett, Sonia Goncalves, Ewan Harrison, David K. Jackson, Ian Johnston, Dominic Kwiatkowski, Cordelia Langford, John Sillitoe on behalf of the Wellcome Sanger Institute COVID-19 Surveillance Team |
| EPI_ISL_2347875, EPI_ISL_2347876, EPI_ISL_2347877 | Lighthouse Lab in Alderley Park | Wellcome Sanger Institute for the COVID-19 Genomics UK (COG-UK) Consortium | Jacquelyn Wynn, Mairead Hyland, The Lighthouse Lab in Alderley Park and Alex Alderton, Roberto Amato, Jeffrey Barrett, Sonia Goncalves, Ewan Harrison, David K. Jackson, Ian Johnston, Dominic Kwiatkowski, Cordelia Langford, John Sillitoe on behalf of the Wellcome Sanger Institute COVID-19 Surveillance Team |
| EPI_ISL_2347879 | Lighthouse Lab in Milton Keynes | Wellcome Sanger Institute for the COVID-19 Genomics UK (COG-UK) Consortium | The Lighthouse Lab in Milton Keynes and Alex Alderton, Roberto Amato, Jeffrey Barrett, Sonia Goncalves, Ewan Harrison, David K. Jackson, Ian Johnston, Dominic Kwiatkowski, Cordelia Langford, John Sillitoe on behalf of the Wellcome Sanger Institute COVID-19 Surveillance Team |
| EPI_ISL_2347881 | Lighthouse Lab in Alderley Park | Wellcome Sanger Institute for the COVID-19 Genomics UK (COG-UK) Consortium | Jacquelyn Wynn, Mairead Hyland, The Lighthouse Lab in Alderley Park and Alex Alderton, Roberto Amato, Jeffrey Barrett, Sonia Goncalves, Ewan Harrison, David K. Jackson, Ian Johnston, Dominic Kwiatkowski, Cordelia Langford, John Sillitoe on behalf of the Wellcome Sanger Institute COVID-19 Surveillance Team |
| EPI_ISL_2347882, EPI_ISL_2347883 | Lighthouse Lab in Milton Keynes | Wellcome Sanger Institute for the COVID-19 Genomics UK (COG-UK) Consortium | The Lighthouse Lab in Milton Keynes and Alex Alderton, Roberto Amato, Jeffrey Barrett, Sonia Goncalves, Ewan Harrison, David K. Jackson, Ian Johnston, Dominic Kwiatkowski, Cordelia Langford, John Sillitoe on behalf of the Wellcome Sanger Institute COVID-19 Surveillance Team |
| EPI_ISL_2347884, EPI_ISL_2347885 | Lighthouse Lab in Alderley Park | Wellcome Sanger Institute for the COVID-19 Genomics UK (COG-UK) Consortium | Jacquelyn Wynn, Mairead Hyland, The Lighthouse Lab in Alderley Park and Alex Alderton, Roberto Amato, Jeffrey Barrett, Sonia Goncalves, Ewan Harrison, David K. Jackson, Ian Johnston, Dominic Kwiatkowski, Cordelia Langford, John Sillitoe on behalf of the Wellcome Sanger Institute COVID-19 Surveillance Team |
| EPI_ISL_2347893 | Health Services Laboratories | Wellcome Sanger Institute for the COVID-19 Genomics UK (COG-UK) Consortium | Health Services Laboratories and Alex Alderton, Roberto Amato, Jeffrey Barrett, Sonia Goncalves, Ewan Harrison, David K. Jackson, Ian Johnston, Dominic Kwiatkowski, Cordelia Langford, John Sillitoe on behalf of the Wellcome Sanger Institute COVID-19 Surveillance Team |
| EPI_ISL_2347894 | Lighthouse Laboratory Plymouth | Wellcome Sanger Institute for the COVID-19 Genomics UK (COG-UK) Consortium | Lighthouse Laboratory Plymouth and Alex Alderton, Roberto Amato, Jeffrey Barrett, Sonia Goncalves, Ewan Harrison, David K. Jackson, Ian Johnston, Dominic Kwiatkowski, Cordelia Langford, John Sillitoe on behalf of the Wellcome Sanger Institute COVID-19 Surveillance Team |
| EPI_ISL_2347895 | Lighthouse Lab in Milton Keynes | Wellcome Sanger Institute for the COVID-19 Genomics UK (COG-UK) Consortium | The Lighthouse Lab in Milton Keynes and Alex Alderton, Roberto Amato, Jeffrey Barrett, Sonia Goncalves, Ewan Harrison, David K. Jackson, Ian Johnston, Dominic Kwiatkowski, Cordelia Langford, John Sillitoe on behalf of the Wellcome Sanger Institute COVID-19 Surveillance Team |
| EPI_ISL_2347896 | Lighthouse Lab in Alderley Park | Wellcome Sanger Institute for the COVID-19 Genomics UK (COG-UK) Consortium | Jacquelyn Wynn, Mairead Hyland, The Lighthouse Lab in Alderley Park and Alex Alderton, Roberto Amato, Jeffrey Barrett, Sonia Goncalves, Ewan Harrison, David K. Jackson, Ian Johnston, Dominic Kwiatkowski, Cordelia Langford, John Sillitoe on behalf of the Wellcome Sanger Institute COVID-19 Surveillance Team |
| EPI_ISL_2347898 | Lighthouse Lab in Milton Keynes | Wellcome Sanger Institute for the COVID-19 Genomics UK (COG-UK) Consortium | The Lighthouse Lab in Milton Keynes and Alex Alderton, Roberto Amato, Jeffrey Barrett, Sonia Goncalves, Ewan Harrison, David K. Jackson, Ian Johnston, Dominic Kwiatkowski, Cordelia Langford, John Sillitoe on behalf of the Wellcome Sanger Institute COVID-19 Surveillance Team |
| EPI_ISL_2347899, EPI_ISL_2347900 | Lighthouse Lab in Alderley Park | Wellcome Sanger Institute for the COVID-19 Genomics UK (COG-UK) Consortium | Jacquelyn Wynn, Mairead Hyland, The Lighthouse Lab in Alderley Park and Alex Alderton, Roberto Amato, Jeffrey Barrett, Sonia Goncalves, Ewan Harrison, David K. Jackson, Ian Johnston, Dominic Kwiatkowski, Cordelia Langford, John Sillitoe on behalf of the Wellcome Sanger Institute COVID-19 Surveillance Team |
| EPI_ISL_2347901 | Health Services Laboratories | Wellcome Sanger Institute for the COVID-19 Genomics UK (COG-UK) Consortium | Health Services Laboratories and Alex Alderton, Roberto Amato, Jeffrey Barrett, Sonia Goncalves, Ewan Harrison, David K. Jackson, Ian Johnston, Dominic Kwiatkowski, Cordelia Langford, John Sillitoe on behalf of the Wellcome Sanger Institute COVID-19 Surveillance Team |
| EPI_ISL_2347902 | Lighthouse Lab in Alderley Park | Wellcome Sanger Institute for the COVID-19 Genomics UK (COG-UK) Consortium | Jacquelyn Wynn, Mairead Hyland, The Lighthouse Lab in Alderley Park and Alex Alderton, Roberto Amato, Jeffrey Barrett, Sonia Goncalves, Ewan Harrison, David K. Jackson, Ian Johnston, Dominic Kwiatkowski, Cordelia Langford, John Sillitoe on behalf of the Wellcome Sanger Institute COVID-19 Surveillance Team |
| EPI_ISL_2347903 | Lighthouse Lab in Milton Keynes | Wellcome Sanger Institute for the COVID-19 Genomics UK (COG-UK) Consortium | The Lighthouse Lab in Milton Keynes and Alex Alderton, Roberto Amato, Jeffrey Barrett, Sonia Goncalves, Ewan Harrison, David K. Jackson, Ian Johnston, Dominic Kwiatkowski, Cordelia Langford, John Sillitoe on behalf of the Wellcome Sanger Institute COVID-19 Surveillance Team |
| EPI_ISL_2347905 | Lighthouse Laboratory Plymouth | Wellcome Sanger Institute for the COVID-19 Genomics UK (COG-UK) Consortium | Lighthouse Laboratory Plymouth and Alex Alderton, Roberto Amato, Jeffrey Barrett, Sonia Goncalves, Ewan Harrison, David K. Jackson, Ian Johnston, Dominic Kwiatkowski, Cordelia Langford, John Sillitoe on behalf of the Wellcome Sanger Institute COVID-19 Surveillance Team |
| EPI_ISL_2347906, EPI_ISL_2347907, EPI_ISL_2347908 | Lighthouse Lab in Alderley Park | Wellcome Sanger Institute for the COVID-19 Genomics UK (COG-UK) Consortium | Jacquelyn Wynn, Mairead Hyland, The Lighthouse Lab in Alderley Park and Alex Alderton, Roberto Amato, Jeffrey Barrett, Sonia Goncalves, Ewan Harrison, David K. Jackson, Ian Johnston, Dominic Kwiatkowski, Cordelia Langford, John Sillitoe on behalf of the Wellcome Sanger Institute COVID-19 Surveillance Team |
| EPI_ISL_2347909 | Lighthouse Lab in Milton Keynes | Wellcome Sanger Institute for the COVID-19 Genomics UK (COG-UK) Consortium | The Lighthouse Lab in Milton Keynes and Alex Alderton, Roberto Amato, Jeffrey Barrett, Sonia Goncalves, Ewan Harrison, David K. Jackson, Ian Johnston, Dominic Kwiatkowski, Cordelia Langford, John Sillitoe on behalf of the Wellcome Sanger Institute COVID-19 Surveillance Team |
| EPI_ISL_2347910 | Lighthouse Lab in Alderley Park | Wellcome Sanger Institute for the COVID-19 Genomics UK (COG-UK) Consortium | Jacquelyn Wynn, Mairead Hyland, The Lighthouse Lab in Alderley Park and Alex Alderton, Roberto Amato, Jeffrey Barrett, Sonia Goncalves, Ewan Harrison, David K. Jackson, Ian Johnston, Dominic Kwiatkowski, Cordelia Langford, John Sillitoe on behalf of the Wellcome Sanger Institute COVID-19 Surveillance Team |
| EPI_ISL_2347911 | Lighthouse Laboratory Plymouth | Wellcome Sanger Institute for the COVID-19 Genomics UK (COG-UK) Consortium | Lighthouse Laboratory Plymouth and Alex Alderton, Roberto Amato, Jeffrey Barrett, Sonia Goncalves, Ewan Harrison, David K. Jackson, Ian Johnston, Dominic Kwiatkowski, Cordelia Langford, John Sillitoe on behalf of the Wellcome Sanger Institute COVID-19 Surveillance Team |
| EPI_ISL_2347912, EPI_ISL_2347913 | Health Services Laboratories | Wellcome Sanger Institute for the COVID-19 Genomics UK (COG-UK) Consortium | Health Services Laboratories and Alex Alderton, Roberto Amato, Jeffrey Barrett, Sonia Goncalves, Ewan Harrison, David K. Jackson, Ian Johnston, Dominic Kwiatkowski, Cordelia Langford, John Sillitoe on behalf of the Wellcome Sanger Institute COVID-19 Surveillance Team |
| EPI_ISL_2347914, EPI_ISL_2347916, | Lighthouse Lab in Alderley Park | Wellcome Sanger Institute for the COVID-19 Genomics UK (COG-UK) Consortium | Jacquelyn Wynn, Mairead Hyland, The Lighthouse Lab in Alderley Park and Alex Alderton, Roberto Amato, Jeffrey Barrett, Sonia Goncalves, Ewan Harrison, David K. Jackson, Ian Johnston, Dominic Kwiatkowski, Cordelia Langford, John Sillitoe on behalf of the Wellcome Sanger Institute COVID-19 Surveillance Team |

| | | | |
|---|---|---|---|
| EPI_ISL_2347918 | | | |
| EPI_ISL_2347919 | Health Services Laboratories | Wellcome Sanger Institute for the COVID-19 Genomics UK (COG-UK) Consortium | Health Services Laboratories and Alex Alderton, Roberto Amato, Jeffrey Barrett, Sonia Goncalves, Ewan Harrison, David K. Jackson, Ian Johnston, Dominic Kwiatkowski, Cordelia Langford, John Sillitoe on behalf of the Wellcome Sanger Institute COVID-19 Surveillance Team |
| EPI_ISL_2347920 | Lighthouse Lab in Milton Keynes | Wellcome Sanger Institute for the COVID-19 Genomics UK (COG-UK) Consortium | The Lighthouse Lab in Milton Keynes and Alex Alderton, Roberto Amato, Jeffrey Barrett, Sonia Goncalves, Ewan Harrison, David K. Jackson, Ian Johnston, Dominic Kwiatkowski, Cordelia Langford, John Sillitoe on behalf of the Wellcome Sanger Institute COVID-19 Surveillance Team |
| EPI_ISL_2347922 | Health Services Laboratories | Wellcome Sanger Institute for the COVID-19 Genomics UK (COG-UK) Consortium | Health Services Laboratories and Alex Alderton, Roberto Amato, Jeffrey Barrett, Sonia Goncalves, Ewan Harrison, David K. Jackson, Ian Johnston, Dominic Kwiatkowski, Cordelia Langford, John Sillitoe on behalf of the Wellcome Sanger Institute COVID-19 Surveillance Team |
| EPI_ISL_2347923 | Lighthouse Lab in Milton Keynes | Wellcome Sanger Institute for the COVID-19 Genomics UK (COG-UK) Consortium | The Lighthouse Lab in Milton Keynes and Alex Alderton, Roberto Amato, Jeffrey Barrett, Sonia Goncalves, Ewan Harrison, David K. Jackson, Ian Johnston, Dominic Kwiatkowski, Cordelia Langford, John Sillitoe on behalf of the Wellcome Sanger Institute COVID-19 Surveillance Team |
| EPI_ISL_2347924 | Lighthouse Laboratory Plymouth | Wellcome Sanger Institute for the COVID-19 Genomics UK (COG-UK) Consortium | Lighthouse Laboratory Plymouth and Alex Alderton, Roberto Amato, Jeffrey Barrett, Sonia Goncalves, Ewan Harrison, David K. Jackson, Ian Johnston, Dominic Kwiatkowski, Cordelia Langford, John Sillitoe on behalf of the Wellcome Sanger Institute COVID-19 Surveillance Team |
| EPI_ISL_2347925, EPI_ISL_2347926, EPI_ISL_2347927, EPI_ISL_2347929 | Lighthouse Lab in Alderley Park | Wellcome Sanger Institute for the COVID-19 Genomics UK (COG-UK) Consortium | Jacquelyn Wynn, Mairead Hyland, The Lighthouse Lab in Alderley Park and Alex Alderton, Roberto Amato, Jeffrey Barrett, Sonia Goncalves, Ewan Harrison, David K. Jackson, Ian Johnston, Dominic Kwiatkowski, Cordelia Langford, John Sillitoe on behalf of the Wellcome Sanger Institute COVID-19 Surveillance Team |
| EPI_ISL_2347930 | Health Services Laboratories | Wellcome Sanger Institute for the COVID-19 Genomics UK (COG-UK) Consortium | Health Services Laboratories and Alex Alderton, Roberto Amato, Jeffrey Barrett, Sonia Goncalves, Ewan Harrison, David K. Jackson, Ian Johnston, Dominic Kwiatkowski, Cordelia Langford, John Sillitoe on behalf of the Wellcome Sanger Institute COVID-19 Surveillance Team |
| EPI_ISL_2347931 | Lighthouse Lab in Alderley Park | Wellcome Sanger Institute for the COVID-19 Genomics UK (COG-UK) Consortium | Jacquelyn Wynn, Mairead Hyland, The Lighthouse Lab in Alderley Park and Alex Alderton, Roberto Amato, Jeffrey Barrett, Sonia Goncalves, Ewan Harrison, David K. Jackson, Ian Johnston, Dominic Kwiatkowski, Cordelia Langford, John Sillitoe on behalf of the Wellcome Sanger Institute COVID-19 Surveillance Team |
| EPI_ISL_2347933, EPI_ISL_2347934, EPI_ISL_2347935, EPI_ISL_2347936 | Lighthouse Lab in Milton Keynes | Wellcome Sanger Institute for the COVID-19 Genomics UK (COG-UK) Consortium | The Lighthouse Lab in Milton Keynes and Alex Alderton, Roberto Amato, Jeffrey Barrett, Sonia Goncalves, Ewan Harrison, David K. Jackson, Ian Johnston, Dominic Kwiatkowski, Cordelia Langford, John Sillitoe on behalf of the Wellcome Sanger Institute COVID-19 Surveillance Team |
| EPI_ISL_2347938 | Lighthouse Lab in Alderley Park | Wellcome Sanger Institute for the COVID-19 Genomics UK (COG-UK) Consortium | Jacquelyn Wynn, Mairead Hyland, The Lighthouse Lab in Alderley Park and Alex Alderton, Roberto Amato, Jeffrey Barrett, Sonia Goncalves, Ewan Harrison, David K. Jackson, Ian Johnston, Dominic Kwiatkowski, Cordelia Langford, John Sillitoe on behalf of the Wellcome Sanger Institute COVID-19 Surveillance Team |
| EPI_ISL_2347939 | Lighthouse Lab in Milton Keynes | Wellcome Sanger Institute for the COVID-19 Genomics UK (COG-UK) Consortium | The Lighthouse Lab in Milton Keynes and Alex Alderton, Roberto Amato, Jeffrey Barrett, Sonia Goncalves, Ewan Harrison, David K. Jackson, Ian Johnston, Dominic Kwiatkowski, Cordelia Langford, John Sillitoe on behalf of the Wellcome Sanger Institute COVID-19 Surveillance Team |
| EPI_ISL_2347940 | Lighthouse Lab in Alderley Park | Wellcome Sanger Institute for the COVID-19 Genomics UK (COG-UK) Consortium | Jacquelyn Wynn, Mairead Hyland, The Lighthouse Lab in Alderley Park and Alex Alderton, Roberto Amato, Jeffrey Barrett, Sonia Goncalves, Ewan Harrison, David K. Jackson, Ian Johnston, Dominic Kwiatkowski, Cordelia Langford, John Sillitoe on behalf of the Wellcome Sanger Institute COVID-19 Surveillance Team |
| EPI_ISL_2347941 | Health Services Laboratories | Wellcome Sanger Institute for the COVID-19 Genomics UK (COG-UK) Consortium | Health Services Laboratories and Alex Alderton, Roberto Amato, Jeffrey Barrett, Sonia Goncalves, Ewan Harrison, David K. Jackson, Ian Johnston, Dominic Kwiatkowski, Cordelia Langford, John Sillitoe on behalf of the Wellcome Sanger Institute COVID-19 Surveillance Team |
| EPI_ISL_2347943, EPI_ISL_2347944 | Lighthouse Lab in Milton Keynes | Wellcome Sanger Institute for the COVID-19 Genomics UK (COG-UK) Consortium | The Lighthouse Lab in Milton Keynes and Alex Alderton, Roberto Amato, Jeffrey Barrett, Sonia Goncalves, Ewan Harrison, David K. Jackson, Ian Johnston, Dominic Kwiatkowski, Cordelia Langford, John Sillitoe on behalf of the Wellcome Sanger Institute COVID-19 Surveillance Team |
| EPI_ISL_2347945 | Lighthouse Lab in Alderley Park | Wellcome Sanger Institute for the COVID-19 Genomics UK (COG-UK) Consortium | Jacquelyn Wynn, Mairead Hyland, The Lighthouse Lab in Alderley Park and Alex Alderton, Roberto Amato, Jeffrey Barrett, Sonia Goncalves, Ewan Harrison, David K. Jackson, Ian Johnston, Dominic Kwiatkowski, Cordelia Langford, John Sillitoe on behalf of the Wellcome Sanger Institute COVID-19 Surveillance Team |
| EPI_ISL_2347946 | Lighthouse Lab in Milton Keynes | Wellcome Sanger Institute for the COVID-19 Genomics UK (COG-UK) Consortium | The Lighthouse Lab in Milton Keynes and Alex Alderton, Roberto Amato, Jeffrey Barrett, Sonia Goncalves, Ewan Harrison, David K. Jackson, Ian Johnston, Dominic Kwiatkowski, Cordelia Langford, John Sillitoe on behalf of the Wellcome Sanger Institute COVID-19 Surveillance Team |
| EPI_ISL_2347947 | Lighthouse Lab in Alderley Park | Wellcome Sanger Institute for the COVID-19 Genomics UK (COG-UK) Consortium | Jacquelyn Wynn, Mairead Hyland, The Lighthouse Lab in Alderley Park and Alex Alderton, Roberto Amato, Jeffrey Barrett, Sonia Goncalves, Ewan Harrison, David K. Jackson, Ian Johnston, Dominic Kwiatkowski, Cordelia Langford, John Sillitoe on behalf of the Wellcome Sanger Institute COVID-19 Surveillance Team |
| EPI_ISL_2347948 | Lighthouse Lab in Milton Keynes | Wellcome Sanger Institute for the COVID-19 Genomics UK (COG-UK) Consortium | The Lighthouse Lab in Milton Keynes and Alex Alderton, Roberto Amato, Jeffrey Barrett, Sonia Goncalves, Ewan Harrison, David K. Jackson, Ian Johnston, Dominic Kwiatkowski, Cordelia Langford, John Sillitoe on behalf of the Wellcome Sanger Institute COVID-19 Surveillance Team |
| EPI_ISL_2347950, EPI_ISL_2347952 | Lighthouse Lab in Alderley Park | Wellcome Sanger Institute for the COVID-19 Genomics UK (COG-UK) Consortium | Jacquelyn Wynn, Mairead Hyland, The Lighthouse Lab in Alderley Park and Alex Alderton, Roberto Amato, Jeffrey Barrett, Sonia Goncalves, Ewan Harrison, David K. Jackson, Ian Johnston, Dominic Kwiatkowski, Cordelia Langford, John Sillitoe on behalf of the Wellcome Sanger Institute COVID-19 Surveillance Team |
| EPI_ISL_2347955 | Lighthouse Lab in Milton Keynes | Wellcome Sanger Institute for the COVID-19 Genomics UK (COG-UK) Consortium | The Lighthouse Lab in Milton Keynes and Alex Alderton, Roberto Amato, Jeffrey Barrett, Sonia Goncalves, Ewan Harrison, David K. Jackson, Ian Johnston, Dominic Kwiatkowski, Cordelia Langford, John Sillitoe on behalf of the Wellcome Sanger Institute COVID-19 Surveillance Team |
| EPI_ISL_2347956 | Health Services Laboratories | Wellcome Sanger Institute for the COVID-19 Genomics UK (COG-UK) Consortium | Health Services Laboratories and Alex Alderton, Roberto Amato, Jeffrey Barrett, Sonia Goncalves, Ewan Harrison, David K. Jackson, Ian Johnston, Dominic Kwiatkowski, Cordelia Langford, John Sillitoe on behalf of the Wellcome Sanger Institute COVID-19 Surveillance Team |
| EPI_ISL_2347957, EPI_ISL_2347958, EPI_ISL_2347959, EPI_ISL_2347960, EPI_ISL_2347963, EPI_ISL_2347964, EPI_ISL_2347965 | Lighthouse Lab in Alderley Park | Wellcome Sanger Institute for the COVID-19 Genomics UK (COG-UK) Consortium | Jacquelyn Wynn, Mairead Hyland, The Lighthouse Lab in Alderley Park and Alex Alderton, Roberto Amato, Jeffrey Barrett, Sonia Goncalves, Ewan Harrison, David K. Jackson, Ian Johnston, Dominic Kwiatkowski, Cordelia Langford, John Sillitoe on behalf of the Wellcome Sanger Institute COVID-19 Surveillance Team |
| EPI_ISL_2347967 | Health Services Laboratories | Wellcome Sanger Institute for the COVID-19 Genomics UK (COG-UK) Consortium | Health Services Laboratories and Alex Alderton, Roberto Amato, Jeffrey Barrett, Sonia Goncalves, Ewan Harrison, David K. Jackson, Ian Johnston, Dominic Kwiatkowski, Cordelia Langford, John Sillitoe on behalf of the Wellcome Sanger Institute COVID-19 Surveillance Team |
| EPI_ISL_2347969, EPI_ISL_2347970 | Lighthouse Lab in Alderley Park | Wellcome Sanger Institute for the COVID-19 Genomics UK (COG-UK) Consortium | Jacquelyn Wynn, Mairead Hyland, The Lighthouse Lab in Alderley Park and Alex Alderton, Roberto Amato, Jeffrey Barrett, Sonia Goncalves, Ewan Harrison, David K. Jackson, Ian Johnston, Dominic Kwiatkowski, Cordelia Langford, John Sillitoe on behalf of the Wellcome Sanger Institute COVID-19 Surveillance Team |
| EPI_ISL_2347971 | Lighthouse Lab in Milton Keynes | Wellcome Sanger Institute for the COVID-19 Genomics UK (COG-UK) Consortium | The Lighthouse Lab in Milton Keynes and Alex Alderton, Roberto Amato, Jeffrey Barrett, Sonia Goncalves, Ewan Harrison, David K. Jackson, Ian Johnston, Dominic Kwiatkowski, Cordelia Langford, John Sillitoe on behalf of the Wellcome Sanger Institute COVID-19 Surveillance Team |
| EPI_ISL_2347972, EPI_ISL_2347973, EPI_ISL_2347974 | Lighthouse Lab in Alderley Park | Wellcome Sanger Institute for the COVID-19 Genomics UK (COG-UK) Consortium | Jacquelyn Wynn, Mairead Hyland, The Lighthouse Lab in Alderley Park and Alex Alderton, Roberto Amato, Jeffrey Barrett, Sonia Goncalves, Ewan Harrison, David K. Jackson, Ian Johnston, Dominic Kwiatkowski, Cordelia Langford, John Sillitoe on behalf of the Wellcome Sanger Institute COVID-19 Surveillance Team |
| EPI_ISL_2347976 | Lighthouse Lab in Milton Keynes | Wellcome Sanger Institute for the COVID-19 Genomics UK (COG-UK) Consortium | The Lighthouse Lab in Milton Keynes and Alex Alderton, Roberto Amato, Jeffrey Barrett, Sonia Goncalves, Ewan Harrison, David K. Jackson, Ian Johnston, Dominic Kwiatkowski, Cordelia Langford, John Sillitoe on behalf of the Wellcome Sanger Institute COVID-19 Surveillance Team |
| EPI_ISL_2347977, EPI_ISL_2347978, EPI_ISL_2347979, EPI_ISL_2347982, EPI_ISL_2347986 | Lighthouse Lab in Alderley Park | Wellcome Sanger Institute for the COVID-19 Genomics UK (COG-UK) Consortium | Jacquelyn Wynn, Mairead Hyland, The Lighthouse Lab in Alderley Park and Alex Alderton, Roberto Amato, Jeffrey Barrett, Sonia Goncalves, Ewan Harrison, David K. Jackson, Ian Johnston, Dominic Kwiatkowski, Cordelia Langford, John Sillitoe on behalf of the Wellcome Sanger Institute COVID-19 Surveillance Team |
| EPI_ISL_2347987 | Lighthouse Lab in Milton Keynes | Wellcome Sanger Institute for the COVID-19 Genomics UK (COG-UK) Consortium | The Lighthouse Lab in Milton Keynes and Alex Alderton, Roberto Amato, Jeffrey Barrett, Sonia Goncalves, Ewan Harrison, David K. Jackson, Ian Johnston, Dominic Kwiatkowski, Cordelia Langford, John Sillitoe on behalf of the Wellcome Sanger Institute COVID-19 Surveillance Team |
| EPI_ISL_2347990, EPI_ISL_2347991, EPI_ISL_2347994, EPI_ISL_2347995 | Lighthouse Lab in Alderley Park | Wellcome Sanger Institute for the COVID-19 Genomics UK (COG-UK) Consortium | Jacquelyn Wynn, Mairead Hyland, The Lighthouse Lab in Alderley Park and Alex Alderton, Roberto Amato, Jeffrey Barrett, Sonia Goncalves, Ewan Harrison, David K. Jackson, Ian Johnston, Dominic Kwiatkowski, Cordelia Langford, John Sillitoe on behalf of the Wellcome Sanger Institute COVID-19 Surveillance Team |
| EPI_ISL_2347996 | Lighthouse Lab in Milton Keynes | Wellcome Sanger Institute for the COVID-19 Genomics UK (COG-UK) Consortium | The Lighthouse Lab in Milton Keynes and Alex Alderton, Roberto Amato, Jeffrey Barrett, Sonia Goncalves, Ewan Harrison, David K. Jackson, Ian Johnston, Dominic Kwiatkowski, Cordelia Langford, John Sillitoe on behalf of the Wellcome Sanger Institute COVID-19 Surveillance Team |
| EPI_ISL_2347998 | Lighthouse Laboratory Plymouth | Wellcome Sanger Institute for the COVID-19 Genomics UK (COG-UK) Consortium | Lighthouse Laboratory Plymouth and Alex Alderton, Roberto Amato, Jeffrey Barrett, Sonia Goncalves, Ewan Harrison, David K. Jackson, Ian Johnston, Dominic Kwiatkowski, Cordelia Langford, John Sillitoe on behalf of the Wellcome Sanger Institute COVID-19 Surveillance Team |
| EPI_ISL_2348000, EPI_ISL_2348001, EPI_ISL_2348002, EPI_ISL_2348003 | Lighthouse Lab in Alderley Park | Wellcome Sanger Institute for the COVID-19 Genomics UK (COG-UK) Consortium | Jacquelyn Wynn, Mairead Hyland, The Lighthouse Lab in Alderley Park and Alex Alderton, Roberto Amato, Jeffrey Barrett, Sonia Goncalves, Ewan Harrison, David K. Jackson, Ian Johnston, Dominic Kwiatkowski, Cordelia Langford, John Sillitoe on behalf of the Wellcome Sanger Institute COVID-19 Surveillance Team |
| EPI_ISL_2348004 | Health Services Laboratories | Wellcome Sanger Institute for the COVID-19 Genomics UK (COG-UK) Consortium | Health Services Laboratories and Alex Alderton, Roberto Amato, Jeffrey Barrett, Sonia Goncalves, Ewan Harrison, David K. Jackson, Ian Johnston, Dominic Kwiatkowski, Cordelia Langford, John Sillitoe on behalf of the Wellcome Sanger Institute COVID-19 Surveillance Team |
| EPI_ISL_2348005 | Lighthouse Lab in Alderley Park | Wellcome Sanger Institute for the COVID-19 Genomics UK (COG-UK) Consortium | Jacquelyn Wynn, Mairead Hyland, The Lighthouse Lab in Alderley Park and Alex Alderton, Roberto Amato, Jeffrey Barrett, Sonia Goncalves, Ewan Harrison, David K. Jackson, Ian Johnston, Dominic Kwiatkowski, Cordelia Langford, John Sillitoe on behalf of the Wellcome Sanger Institute COVID-19 Surveillance Team |
| EPI_ISL_2348007 | Health Services Laboratories | Wellcome Sanger Institute for the COVID-19 Genomics UK (COG-UK) Consortium | Health Services Laboratories and Alex Alderton, Roberto Amato, Jeffrey Barrett, Sonia Goncalves, Ewan Harrison, David K. Jackson, Ian Johnston, Dominic Kwiatkowski, Cordelia Langford, John Sillitoe on behalf of the Wellcome Sanger Institute COVID-19 Surveillance Team |
| EPI_ISL_2348008, EPI_ISL_2348009 | Lighthouse Lab in Alderley Park | Wellcome Sanger Institute for the COVID-19 Genomics UK (COG-UK) Consortium | Jacquelyn Wynn, Mairead Hyland, The Lighthouse Lab in Alderley Park and Alex Alderton, Roberto Amato, Jeffrey Barrett, Sonia Goncalves, Ewan Harrison, David K. Jackson, Ian Johnston, Dominic Kwiatkowski, Cordelia Langford, John Sillitoe on behalf of the Wellcome Sanger Institute COVID-19 Surveillance Team |
| EPI_ISL_2348015 | Lighthouse Lab in Milton Keynes | Wellcome Sanger Institute for the COVID-19 Genomics UK (COG-UK) | The Lighthouse Lab in Milton Keynes and Alex Alderton, Roberto Amato, Jeffrey Barrett, Sonia Goncalves, Ewan Harrison, David K. Jackson, Ian Johnston, Dominic Kwiatkowski, Cordelia Langford, John Sillitoe on behalf of the Wellcome Sanger |

| | | | |
|---|---|---|---|
| | | Consortium | Institute COVID-19 Surveillance Team |
| EPI_ISL_2348016 | Health Services Laboratories | Wellcome Sanger Institute for the COVID-19 Genomics UK (COG-UK) Consortium | Health Services Laboratories and Alex Alderton, Roberto Amato, Jeffrey Barrett, Sonia Goncalves, Ewan Harrison, David K. Jackson, Ian Johnston, Dominic Kwiatkowski, Cordelia Langford, John Sillitoe on behalf of the Wellcome Sanger Institute COVID-19 Surveillance Team |
| EPI_ISL_2348018 | Lighthouse Lab in Alderley Park | Wellcome Sanger Institute for the COVID-19 Genomics UK (COG-UK) Consortium | Jacquelyn Wynn, Mairead Hyland, The Lighthouse Lab in Alderley Park and Alex Alderton, Roberto Amato, Jeffrey Barrett, Sonia Goncalves, Ewan Harrison, David K. Jackson, Ian Johnston, Dominic Kwiatkowski, Cordelia Langford, John Sillitoe on behalf of the Wellcome Sanger Institute COVID-19 Surveillance Team |
| EPI_ISL_2348019, EPI_ISL_2348020 | Lighthouse Lab in Milton Keynes | Wellcome Sanger Institute for the COVID-19 Genomics UK (COG-UK) Consortium | The Lighthouse Lab in Milton Keynes and Alex Alderton, Roberto Amato, Jeffrey Barrett, Sonia Goncalves, Ewan Harrison, David K. Jackson, Ian Johnston, Dominic Kwiatkowski, Cordelia Langford, John Sillitoe on behalf of the Wellcome Sanger Institute COVID-19 Surveillance Team |
| EPI_ISL_2348021, EPI_ISL_2348023, EPI_ISL_2348024, EPI_ISL_2348025, EPI_ISL_2348026, EPI_ISL_2348027 | Lighthouse Lab in Alderley Park | Wellcome Sanger Institute for the COVID-19 Genomics UK (COG-UK) Consortium | Jacquelyn Wynn, Mairead Hyland, The Lighthouse Lab in Alderley Park and Alex Alderton, Roberto Amato, Jeffrey Barrett, Sonia Goncalves, Ewan Harrison, David K. Jackson, Ian Johnston, Dominic Kwiatkowski, Cordelia Langford, John Sillitoe on behalf of the Wellcome Sanger Institute COVID-19 Surveillance Team |
| EPI_ISL_2348028 | Lighthouse Lab in Milton Keynes | Wellcome Sanger Institute for the COVID-19 Genomics UK (COG-UK) Consortium | The Lighthouse Lab in Milton Keynes and Alex Alderton, Roberto Amato, Jeffrey Barrett, Sonia Goncalves, Ewan Harrison, David K. Jackson, Ian Johnston, Dominic Kwiatkowski, Cordelia Langford, John Sillitoe on behalf of the Wellcome Sanger Institute COVID-19 Surveillance Team |
| EPI_ISL_2348029 | Health Services Laboratories | Wellcome Sanger Institute for the COVID-19 Genomics UK (COG-UK) Consortium | Health Services Laboratories and Alex Alderton, Roberto Amato, Jeffrey Barrett, Sonia Goncalves, Ewan Harrison, David K. Jackson, Ian Johnston, Dominic Kwiatkowski, Cordelia Langford, John Sillitoe on behalf of the Wellcome Sanger Institute COVID-19 Surveillance Team |
| EPI_ISL_2348031, EPI_ISL_2348032 | Lighthouse Lab in Alderley Park | Wellcome Sanger Institute for the COVID-19 Genomics UK (COG-UK) Consortium | Jacquelyn Wynn, Mairead Hyland, The Lighthouse Lab in Alderley Park and Alex Alderton, Roberto Amato, Jeffrey Barrett, Sonia Goncalves, Ewan Harrison, David K. Jackson, Ian Johnston, Dominic Kwiatkowski, Cordelia Langford, John Sillitoe on behalf of the Wellcome Sanger Institute COVID-19 Surveillance Team |
| EPI_ISL_2348033 | Lighthouse Lab in Milton Keynes | Wellcome Sanger Institute for the COVID-19 Genomics UK (COG-UK) Consortium | The Lighthouse Lab in Milton Keynes and Alex Alderton, Roberto Amato, Jeffrey Barrett, Sonia Goncalves, Ewan Harrison, David K. Jackson, Ian Johnston, Dominic Kwiatkowski, Cordelia Langford, John Sillitoe on behalf of the Wellcome Sanger Institute COVID-19 Surveillance Team |
| EPI_ISL_2348034 | Lighthouse Lab in Alderley Park | Wellcome Sanger Institute for the COVID-19 Genomics UK (COG-UK) Consortium | Jacquelyn Wynn, Mairead Hyland, The Lighthouse Lab in Alderley Park and Alex Alderton, Roberto Amato, Jeffrey Barrett, Sonia Goncalves, Ewan Harrison, David K. Jackson, Ian Johnston, Dominic Kwiatkowski, Cordelia Langford, John Sillitoe on behalf of the Wellcome Sanger Institute COVID-19 Surveillance Team |
| EPI_ISL_2348035 | Lighthouse Lab in Milton Keynes | Wellcome Sanger Institute for the COVID-19 Genomics UK (COG-UK) Consortium | The Lighthouse Lab in Milton Keynes and Alex Alderton, Roberto Amato, Jeffrey Barrett, Sonia Goncalves, Ewan Harrison, David K. Jackson, Ian Johnston, Dominic Kwiatkowski, Cordelia Langford, John Sillitoe on behalf of the Wellcome Sanger Institute COVID-19 Surveillance Team |
| EPI_ISL_2348036 | Lighthouse Lab in Alderley Park | Wellcome Sanger Institute for the COVID-19 Genomics UK (COG-UK) Consortium | Jacquelyn Wynn, Mairead Hyland, The Lighthouse Lab in Alderley Park and Alex Alderton, Roberto Amato, Jeffrey Barrett, Sonia Goncalves, Ewan Harrison, David K. Jackson, Ian Johnston, Dominic Kwiatkowski, Cordelia Langford, John Sillitoe on behalf of the Wellcome Sanger Institute COVID-19 Surveillance Team |
| EPI_ISL_2348037 | Lighthouse Laboratory Plymouth | Wellcome Sanger Institute for the COVID-19 Genomics UK (COG-UK) Consortium | Lighthouse Laboratory Plymouth and Alex Alderton, Roberto Amato, Jeffrey Barrett, Sonia Goncalves, Ewan Harrison, David K. Jackson, Ian Johnston, Dominic Kwiatkowski, Cordelia Langford, John Sillitoe on behalf of the Wellcome Sanger Institute COVID-19 Surveillance Team |
| EPI_ISL_2348040, EPI_ISL_2348041, EPI_ISL_2348042 | Lighthouse Lab in Milton Keynes | Wellcome Sanger Institute for the COVID-19 Genomics UK (COG-UK) Consortium | The Lighthouse Lab in Milton Keynes and Alex Alderton, Roberto Amato, Jeffrey Barrett, Sonia Goncalves, Ewan Harrison, David K. Jackson, Ian Johnston, Dominic Kwiatkowski, Cordelia Langford, John Sillitoe on behalf of the Wellcome Sanger Institute COVID-19 Surveillance Team |
| EPI_ISL_2348043, EPI_ISL_2348044, EPI_ISL_2348045 | Lighthouse Lab in Alderley Park | Wellcome Sanger Institute for the COVID-19 Genomics UK (COG-UK) Consortium | Jacquelyn Wynn, Mairead Hyland, The Lighthouse Lab in Alderley Park and Alex Alderton, Roberto Amato, Jeffrey Barrett, Sonia Goncalves, Ewan Harrison, David K. Jackson, Ian Johnston, Dominic Kwiatkowski, Cordelia Langford, John Sillitoe on behalf of the Wellcome Sanger Institute COVID-19 Surveillance Team |
| EPI_ISL_2348046 | Health Services Laboratories | Wellcome Sanger Institute for the COVID-19 Genomics UK (COG-UK) Consortium | Health Services Laboratories and Alex Alderton, Roberto Amato, Jeffrey Barrett, Sonia Goncalves, Ewan Harrison, David K. Jackson, Ian Johnston, Dominic Kwiatkowski, Cordelia Langford, John Sillitoe on behalf of the Wellcome Sanger Institute COVID-19 Surveillance Team |
| EPI_ISL_2348047, EPI_ISL_2348048, EPI_ISL_2348049 | Lighthouse Lab in Alderley Park | Wellcome Sanger Institute for the COVID-19 Genomics UK (COG-UK) Consortium | Jacquelyn Wynn, Mairead Hyland, The Lighthouse Lab in Alderley Park and Alex Alderton, Roberto Amato, Jeffrey Barrett, Sonia Goncalves, Ewan Harrison, David K. Jackson, Ian Johnston, Dominic Kwiatkowski, Cordelia Langford, John Sillitoe on behalf of the Wellcome Sanger Institute COVID-19 Surveillance Team |
| EPI_ISL_2348050, EPI_ISL_2348051 | Lighthouse Lab in Milton Keynes | Wellcome Sanger Institute for the COVID-19 Genomics UK (COG-UK) Consortium | The Lighthouse Lab in Milton Keynes and Alex Alderton, Roberto Amato, Jeffrey Barrett, Sonia Goncalves, Ewan Harrison, David K. Jackson, Ian Johnston, Dominic Kwiatkowski, Cordelia Langford, John Sillitoe on behalf of the Wellcome Sanger Institute COVID-19 Surveillance Team |
| EPI_ISL_2348052 | Lighthouse Lab in Alderley Park | Wellcome Sanger Institute for the COVID-19 Genomics UK (COG-UK) Consortium | Jacquelyn Wynn, Mairead Hyland, The Lighthouse Lab in Alderley Park and Alex Alderton, Roberto Amato, Jeffrey Barrett, Sonia Goncalves, Ewan Harrison, David K. Jackson, Ian Johnston, Dominic Kwiatkowski, Cordelia Langford, John Sillitoe on behalf of the Wellcome Sanger Institute COVID-19 Surveillance Team |
| EPI_ISL_2348053 | Lighthouse Laboratory Plymouth | Wellcome Sanger Institute for the COVID-19 Genomics UK (COG-UK) Consortium | Lighthouse Laboratory Plymouth and Alex Alderton, Roberto Amato, Jeffrey Barrett, Sonia Goncalves, Ewan Harrison, David K. Jackson, Ian Johnston, Dominic Kwiatkowski, Cordelia Langford, John Sillitoe on behalf of the Wellcome Sanger Institute COVID-19 Surveillance Team |
| EPI_ISL_2348054, EPI_ISL_2348055 | Lighthouse Lab in Alderley Park | Wellcome Sanger Institute for the COVID-19 Genomics UK (COG-UK) Consortium | Jacquelyn Wynn, Mairead Hyland, The Lighthouse Lab in Alderley Park and Alex Alderton, Roberto Amato, Jeffrey Barrett, Sonia Goncalves, Ewan Harrison, David K. Jackson, Ian Johnston, Dominic Kwiatkowski, Cordelia Langford, John Sillitoe on behalf of the Wellcome Sanger Institute COVID-19 Surveillance Team |
| EPI_ISL_2348056 | Lighthouse Laboratory Plymouth | Wellcome Sanger Institute for the COVID-19 Genomics UK (COG-UK) Consortium | Lighthouse Laboratory Plymouth and Alex Alderton, Roberto Amato, Jeffrey Barrett, Sonia Goncalves, Ewan Harrison, David K. Jackson, Ian Johnston, Dominic Kwiatkowski, Cordelia Langford, John Sillitoe on behalf of the Wellcome Sanger Institute COVID-19 Surveillance Team |
| EPI_ISL_2348057 | Health Services Laboratories | Wellcome Sanger Institute for the COVID-19 Genomics UK (COG-UK) Consortium | Health Services Laboratories and Alex Alderton, Roberto Amato, Jeffrey Barrett, Sonia Goncalves, Ewan Harrison, David K. Jackson, Ian Johnston, Dominic Kwiatkowski, Cordelia Langford, John Sillitoe on behalf of the Wellcome Sanger Institute COVID-19 Surveillance Team |
| EPI_ISL_2348059, EPI_ISL_2348060 | Lighthouse Lab in Alderley Park | Wellcome Sanger Institute for the COVID-19 Genomics UK (COG-UK) Consortium | Jacquelyn Wynn, Mairead Hyland, The Lighthouse Lab in Alderley Park and Alex Alderton, Roberto Amato, Jeffrey Barrett, Sonia Goncalves, Ewan Harrison, David K. Jackson, Ian Johnston, Dominic Kwiatkowski, Cordelia Langford, John Sillitoe on behalf of the Wellcome Sanger Institute COVID-19 Surveillance Team |
| EPI_ISL_2348061 | Lighthouse Laboratory Plymouth | Wellcome Sanger Institute for the COVID-19 Genomics UK (COG-UK) Consortium | Lighthouse Laboratory Plymouth and Alex Alderton, Roberto Amato, Jeffrey Barrett, Sonia Goncalves, Ewan Harrison, David K. Jackson, Ian Johnston, Dominic Kwiatkowski, Cordelia Langford, John Sillitoe on behalf of the Wellcome Sanger Institute COVID-19 Surveillance Team |
| EPI_ISL_2348062, EPI_ISL_2348063 | Lighthouse Lab in Milton Keynes | Wellcome Sanger Institute for the COVID-19 Genomics UK (COG-UK) Consortium | The Lighthouse Lab in Milton Keynes and Alex Alderton, Roberto Amato, Jeffrey Barrett, Sonia Goncalves, Ewan Harrison, David K. Jackson, Ian Johnston, Dominic Kwiatkowski, Cordelia Langford, John Sillitoe on behalf of the Wellcome Sanger Institute COVID-19 Surveillance Team |
| EPI_ISL_2348064 | Lighthouse Lab in Alderley Park | Wellcome Sanger Institute for the COVID-19 Genomics UK (COG-UK) Consortium | Jacquelyn Wynn, Mairead Hyland, The Lighthouse Lab in Alderley Park and Alex Alderton, Roberto Amato, Jeffrey Barrett, Sonia Goncalves, Ewan Harrison, David K. Jackson, Ian Johnston, Dominic Kwiatkowski, Cordelia Langford, John Sillitoe on behalf of the Wellcome Sanger Institute COVID-19 Surveillance Team |
| EPI_ISL_2348065 | Health Services Laboratories | Wellcome Sanger Institute for the COVID-19 Genomics UK (COG-UK) Consortium | Health Services Laboratories and Alex Alderton, Roberto Amato, Jeffrey Barrett, Sonia Goncalves, Ewan Harrison, David K. Jackson, Ian Johnston, Dominic Kwiatkowski, Cordelia Langford, John Sillitoe on behalf of the Wellcome Sanger Institute COVID-19 Surveillance Team |
| EPI_ISL_2348066 | Lighthouse Lab in Alderley Park | Wellcome Sanger Institute for the COVID-19 Genomics UK (COG-UK) Consortium | Jacquelyn Wynn, Mairead Hyland, The Lighthouse Lab in Alderley Park and Alex Alderton, Roberto Amato, Jeffrey Barrett, Sonia Goncalves, Ewan Harrison, David K. Jackson, Ian Johnston, Dominic Kwiatkowski, Cordelia Langford, John Sillitoe on behalf of the Wellcome Sanger Institute COVID-19 Surveillance Team |
| EPI_ISL_2348068 | Lighthouse Lab in Milton Keynes | Wellcome Sanger Institute for the COVID-19 Genomics UK (COG-UK) Consortium | The Lighthouse Lab in Milton Keynes and Alex Alderton, Roberto Amato, Jeffrey Barrett, Sonia Goncalves, Ewan Harrison, David K. Jackson, Ian Johnston, Dominic Kwiatkowski, Cordelia Langford, John Sillitoe on behalf of the Wellcome Sanger Institute COVID-19 Surveillance Team |
| EPI_ISL_2348069, EPI_ISL_2348070 | Lighthouse Lab in Alderley Park | Wellcome Sanger Institute for the COVID-19 Genomics UK (COG-UK) Consortium | Jacquelyn Wynn, Mairead Hyland, The Lighthouse Lab in Alderley Park and Alex Alderton, Roberto Amato, Jeffrey Barrett, Sonia Goncalves, Ewan Harrison, David K. Jackson, Ian Johnston, Dominic Kwiatkowski, Cordelia Langford, John Sillitoe on behalf of the Wellcome Sanger Institute COVID-19 Surveillance Team |
| EPI_ISL_2348071 | Lighthouse Lab in Milton Keynes | Wellcome Sanger Institute for the COVID-19 Genomics UK (COG-UK) Consortium | The Lighthouse Lab in Milton Keynes and Alex Alderton, Roberto Amato, Jeffrey Barrett, Sonia Goncalves, Ewan Harrison, David K. Jackson, Ian Johnston, Dominic Kwiatkowski, Cordelia Langford, John Sillitoe on behalf of the Wellcome Sanger Institute COVID-19 Surveillance Team |
| EPI_ISL_2348072 | Lighthouse Lab in Alderley Park | Wellcome Sanger Institute for the COVID-19 Genomics UK (COG-UK) Consortium | Jacquelyn Wynn, Mairead Hyland, The Lighthouse Lab in Alderley Park and Alex Alderton, Roberto Amato, Jeffrey Barrett, Sonia Goncalves, Ewan Harrison, David K. Jackson, Ian Johnston, Dominic Kwiatkowski, Cordelia Langford, John Sillitoe on behalf of the Wellcome Sanger Institute COVID-19 Surveillance Team |
| EPI_ISL_2348073 | Lighthouse Laboratory Plymouth | Wellcome Sanger Institute for the COVID-19 Genomics UK (COG-UK) Consortium | Lighthouse Laboratory Plymouth and Alex Alderton, Roberto Amato, Jeffrey Barrett, Sonia Goncalves, Ewan Harrison, David K. Jackson, Ian Johnston, Dominic Kwiatkowski, Cordelia Langford, John Sillitoe on behalf of the Wellcome Sanger Institute COVID-19 Surveillance Team |
| EPI_ISL_2348075, EPI_ISL_2348076, EPI_ISL_2348077, EPI_ISL_2348078, EPI_ISL_2348079, EPI_ISL_2348080, EPI_ISL_2348083, EPI_ISL_2348085 | Lighthouse Lab in Alderley Park | Wellcome Sanger Institute for the COVID-19 Genomics UK (COG-UK) Consortium | Jacquelyn Wynn, Mairead Hyland, The Lighthouse Lab in Alderley Park and Alex Alderton, Roberto Amato, Jeffrey Barrett, Sonia Goncalves, Ewan Harrison, David K. Jackson, Ian Johnston, Dominic Kwiatkowski, Cordelia Langford, John Sillitoe on behalf of the Wellcome Sanger Institute COVID-19 Surveillance Team |
| EPI_ISL_2348088 | Lighthouse Lab in Milton Keynes | Wellcome Sanger Institute for the COVID-19 Genomics UK (COG-UK) Consortium | The Lighthouse Lab in Milton Keynes and Alex Alderton, Roberto Amato, Jeffrey Barrett, Sonia Goncalves, Ewan Harrison, David K. Jackson, Ian Johnston, Dominic Kwiatkowski, Cordelia Langford, John Sillitoe on behalf of the Wellcome Sanger Institute COVID-19 Surveillance Team |
| EPI_ISL_2348089 | Lighthouse Lab in Alderley Park | Wellcome Sanger Institute for the COVID-19 Genomics UK (COG-UK) Consortium | Jacquelyn Wynn, Mairead Hyland, The Lighthouse Lab in Alderley Park and Alex Alderton, Roberto Amato, Jeffrey Barrett, Sonia Goncalves, Ewan Harrison, David K. Jackson, Ian Johnston, Dominic Kwiatkowski, Cordelia Langford, John Sillitoe on behalf of the Wellcome Sanger Institute COVID-19 Surveillance Team |
| EPI_ISL_2348090 | Health Services Laboratories | Wellcome Sanger Institute for the COVID-19 Genomics UK (COG-UK) Consortium | Health Services Laboratories and Alex Alderton, Roberto Amato, Jeffrey Barrett, Sonia Goncalves, Ewan Harrison, David K. Jackson, Ian Johnston, Dominic Kwiatkowski, Cordelia Langford, John Sillitoe on behalf of the Wellcome Sanger Institute COVID-19 Surveillance Team |
| EPI_ISL_2348091 | Lighthouse Lab in Alderley Park | Wellcome Sanger Institute for the COVID-19 Genomics UK (COG-UK) Consortium | Jacquelyn Wynn, Mairead Hyland, The Lighthouse Lab in Alderley Park and Alex Alderton, Roberto Amato, Jeffrey Barrett, Sonia Goncalves, Ewan Harrison, David K. Jackson, Ian Johnston, Dominic Kwiatkowski, Cordelia Langford, John Sillitoe on behalf of the Wellcome Sanger Institute COVID-19 Surveillance Team |
| EPI_ISL_2348094 | Health Services Laboratories | Wellcome Sanger Institute for the COVID-19 Genomics UK (COG-UK) Consortium | Health Services Laboratories and Alex Alderton, Roberto Amato, Jeffrey Barrett, Sonia Goncalves, Ewan Harrison, David K. Jackson, Ian Johnston, Dominic Kwiatkowski, Cordelia Langford, John Sillitoe on behalf of the Wellcome Sanger Institute COVID-19 Surveillance Team |

| | | | |
|---|---|---|---|
| EPI_ISL_2348095, EPI_ISL_2348097 | Lighthouse Lab in Alderley Park | Wellcome Sanger Institute for the COVID-19 Genomics UK (COG-UK) Consortium | Jacquelyn Wynn, Mairead Hyland, The Lighthouse Lab in Alderley Park and Alex Alderton, Roberto Amato, Jeffrey Barrett, Sonia Goncalves, Ewan Harrison, David K. Jackson, Ian Johnston, Dominic Kwiatkowski, Cordelia Langford, John Sillitoe on behalf of the Wellcome Sanger Institute COVID-19 Surveillance Team |
| EPI_ISL_2348098 | Health Services Laboratories | Wellcome Sanger Institute for the COVID-19 Genomics UK (COG-UK) Consortium | Health Services Laboratories and Alex Alderton, Roberto Amato, Jeffrey Barrett, Sonia Goncalves, Ewan Harrison, David K. Jackson, Ian Johnston, Dominic Kwiatkowski, Cordelia Langford, John Sillitoe on behalf of the Wellcome Sanger Institute COVID-19 Surveillance Team |
| EPI_ISL_2348099, EPI_ISL_2348100, EPI_ISL_2348102 | Lighthouse Lab in Alderley Park | Wellcome Sanger Institute for the COVID-19 Genomics UK (COG-UK) Consortium | Jacquelyn Wynn, Mairead Hyland, The Lighthouse Lab in Alderley Park and Alex Alderton, Roberto Amato, Jeffrey Barrett, Sonia Goncalves, Ewan Harrison, David K. Jackson, Ian Johnston, Dominic Kwiatkowski, Cordelia Langford, John Sillitoe on behalf of the Wellcome Sanger Institute COVID-19 Surveillance Team |
| EPI_ISL_2348103 | Lighthouse Lab in Milton Keynes | Wellcome Sanger Institute for the COVID-19 Genomics UK (COG-UK) Consortium | The Lighthouse Lab in Milton Keynes and Alex Alderton, Roberto Amato, Jeffrey Barrett, Sonia Goncalves, Ewan Harrison, David K. Jackson, Ian Johnston, Dominic Kwiatkowski, Cordelia Langford, John Sillitoe on behalf of the Wellcome Sanger Institute COVID-19 Surveillance Team |
| EPI_ISL_2348104 | Lighthouse Lab in Alderley Park | Wellcome Sanger Institute for the COVID-19 Genomics UK (COG-UK) Consortium | Jacquelyn Wynn, Mairead Hyland, The Lighthouse Lab in Alderley Park and Alex Alderton, Roberto Amato, Jeffrey Barrett, Sonia Goncalves, Ewan Harrison, David K. Jackson, Ian Johnston, Dominic Kwiatkowski, Cordelia Langford, John Sillitoe on behalf of the Wellcome Sanger Institute COVID-19 Surveillance Team |
| EPI_ISL_2348105 | Lighthouse Lab in Milton Keynes | Wellcome Sanger Institute for the COVID-19 Genomics UK (COG-UK) Consortium | The Lighthouse Lab in Milton Keynes and Alex Alderton, Roberto Amato, Jeffrey Barrett, Sonia Goncalves, Ewan Harrison, David K. Jackson, Ian Johnston, Dominic Kwiatkowski, Cordelia Langford, John Sillitoe on behalf of the Wellcome Sanger Institute COVID-19 Surveillance Team |
| EPI_ISL_2348106, EPI_ISL_2348107, EPI_ISL_2348108, EPI_ISL_2348110 | Lighthouse Lab in Alderley Park | Wellcome Sanger Institute for the COVID-19 Genomics UK (COG-UK) Consortium | Jacquelyn Wynn, Mairead Hyland, The Lighthouse Lab in Alderley Park and Alex Alderton, Roberto Amato, Jeffrey Barrett, Sonia Goncalves, Ewan Harrison, David K. Jackson, Ian Johnston, Dominic Kwiatkowski, Cordelia Langford, John Sillitoe on behalf of the Wellcome Sanger Institute COVID-19 Surveillance Team |
| EPI_ISL_2348111 | Health Services Laboratories | Wellcome Sanger Institute for the COVID-19 Genomics UK (COG-UK) Consortium | Health Services Laboratories and Alex Alderton, Roberto Amato, Jeffrey Barrett, Sonia Goncalves, Ewan Harrison, David K. Jackson, Ian Johnston, Dominic Kwiatkowski, Cordelia Langford, John Sillitoe on behalf of the Wellcome Sanger Institute COVID-19 Surveillance Team |
| EPI_ISL_2348112 | Lighthouse Lab in Milton Keynes | Wellcome Sanger Institute for the COVID-19 Genomics UK (COG-UK) Consortium | The Lighthouse Lab in Milton Keynes and Alex Alderton, Roberto Amato, Jeffrey Barrett, Sonia Goncalves, Ewan Harrison, David K. Jackson, Ian Johnston, Dominic Kwiatkowski, Cordelia Langford, John Sillitoe on behalf of the Wellcome Sanger Institute COVID-19 Surveillance Team |
| EPI_ISL_2348113, EPI_ISL_2348114, EPI_ISL_2348116, EPI_ISL_2348117 | Lighthouse Lab in Alderley Park | Wellcome Sanger Institute for the COVID-19 Genomics UK (COG-UK) Consortium | Jacquelyn Wynn, Mairead Hyland, The Lighthouse Lab in Alderley Park and Alex Alderton, Roberto Amato, Jeffrey Barrett, Sonia Goncalves, Ewan Harrison, David K. Jackson, Ian Johnston, Dominic Kwiatkowski, Cordelia Langford, John Sillitoe on behalf of the Wellcome Sanger Institute COVID-19 Surveillance Team |
| EPI_ISL_2348118 | Lighthouse Laboratory Plymouth | Wellcome Sanger Institute for the COVID-19 Genomics UK (COG-UK) Consortium | Lighthouse Laboratory Plymouth and Alex Alderton, Roberto Amato, Jeffrey Barrett, Sonia Goncalves, Ewan Harrison, David K. Jackson, Ian Johnston, Dominic Kwiatkowski, Cordelia Langford, John Sillitoe on behalf of the Wellcome Sanger Institute COVID-19 Surveillance Team |
| EPI_ISL_2348119, EPI_ISL_2348120 | Lighthouse Lab in Alderley Park | Wellcome Sanger Institute for the COVID-19 Genomics UK (COG-UK) Consortium | Jacquelyn Wynn, Mairead Hyland, The Lighthouse Lab in Alderley Park and Alex Alderton, Roberto Amato, Jeffrey Barrett, Sonia Goncalves, Ewan Harrison, David K. Jackson, Ian Johnston, Dominic Kwiatkowski, Cordelia Langford, John Sillitoe on behalf of the Wellcome Sanger Institute COVID-19 Surveillance Team |
| EPI_ISL_2348122 | Lighthouse Lab in Milton Keynes | Wellcome Sanger Institute for the COVID-19 Genomics UK (COG-UK) Consortium | The Lighthouse Lab in Milton Keynes and Alex Alderton, Roberto Amato, Jeffrey Barrett, Sonia Goncalves, Ewan Harrison, David K. Jackson, Ian Johnston, Dominic Kwiatkowski, Cordelia Langford, John Sillitoe on behalf of the Wellcome Sanger Institute COVID-19 Surveillance Team |
| EPI_ISL_2348123, EPI_ISL_2348124, EPI_ISL_2348125 | Lighthouse Lab in Alderley Park | Wellcome Sanger Institute for the COVID-19 Genomics UK (COG-UK) Consortium | Jacquelyn Wynn, Mairead Hyland, The Lighthouse Lab in Alderley Park and Alex Alderton, Roberto Amato, Jeffrey Barrett, Sonia Goncalves, Ewan Harrison, David K. Jackson, Ian Johnston, Dominic Kwiatkowski, Cordelia Langford, John Sillitoe on behalf of the Wellcome Sanger Institute COVID-19 Surveillance Team |
| EPI_ISL_2348127 | Lighthouse Lab in Milton Keynes | Wellcome Sanger Institute for the COVID-19 Genomics UK (COG-UK) Consortium | The Lighthouse Lab in Milton Keynes and Alex Alderton, Roberto Amato, Jeffrey Barrett, Sonia Goncalves, Ewan Harrison, David K. Jackson, Ian Johnston, Dominic Kwiatkowski, Cordelia Langford, John Sillitoe on behalf of the Wellcome Sanger Institute COVID-19 Surveillance Team |
| EPI_ISL_2348128 | Lighthouse Lab in Alderley Park | Wellcome Sanger Institute for the COVID-19 Genomics UK (COG-UK) Consortium | Jacquelyn Wynn, Mairead Hyland, The Lighthouse Lab in Alderley Park and Alex Alderton, Roberto Amato, Jeffrey Barrett, Sonia Goncalves, Ewan Harrison, David K. Jackson, Ian Johnston, Dominic Kwiatkowski, Cordelia Langford, John Sillitoe on behalf of the Wellcome Sanger Institute COVID-19 Surveillance Team |
| EPI_ISL_2348130 | Lighthouse Lab in Milton Keynes | Wellcome Sanger Institute for the COVID-19 Genomics UK (COG-UK) Consortium | The Lighthouse Lab in Milton Keynes and Alex Alderton, Roberto Amato, Jeffrey Barrett, Sonia Goncalves, Ewan Harrison, David K. Jackson, Ian Johnston, Dominic Kwiatkowski, Cordelia Langford, John Sillitoe on behalf of the Wellcome Sanger Institute COVID-19 Surveillance Team |
| EPI_ISL_2348131 | Lighthouse Lab in Alderley Park | Wellcome Sanger Institute for the COVID-19 Genomics UK (COG-UK) Consortium | Jacquelyn Wynn, Mairead Hyland, The Lighthouse Lab in Alderley Park and Alex Alderton, Roberto Amato, Jeffrey Barrett, Sonia Goncalves, Ewan Harrison, David K. Jackson, Ian Johnston, Dominic Kwiatkowski, Cordelia Langford, John Sillitoe on behalf of the Wellcome Sanger Institute COVID-19 Surveillance Team |
| EPI_ISL_2348132, EPI_ISL_2348134 | Lighthouse Lab in Milton Keynes | Wellcome Sanger Institute for the COVID-19 Genomics UK (COG-UK) Consortium | The Lighthouse Lab in Milton Keynes and Alex Alderton, Roberto Amato, Jeffrey Barrett, Sonia Goncalves, Ewan Harrison, David K. Jackson, Ian Johnston, Dominic Kwiatkowski, Cordelia Langford, John Sillitoe on behalf of the Wellcome Sanger Institute COVID-19 Surveillance Team |
| EPI_ISL_2348135, EPI_ISL_2348137, EPI_ISL_2348138, EPI_ISL_2348139, EPI_ISL_2348141, EPI_ISL_2348142, EPI_ISL_2348143, EPI_ISL_2348145, EPI_ISL_2348146 | Lighthouse Lab in Alderley Park | Wellcome Sanger Institute for the COVID-19 Genomics UK (COG-UK) Consortium | Jacquelyn Wynn, Mairead Hyland, The Lighthouse Lab in Alderley Park and Alex Alderton, Roberto Amato, Jeffrey Barrett, Sonia Goncalves, Ewan Harrison, David K. Jackson, Ian Johnston, Dominic Kwiatkowski, Cordelia Langford, John Sillitoe on behalf of the Wellcome Sanger Institute COVID-19 Surveillance Team |
| EPI_ISL_2348150 | Lighthouse Laboratory Plymouth | Wellcome Sanger Institute for the COVID-19 Genomics UK (COG-UK) Consortium | Lighthouse Laboratory Plymouth and Alex Alderton, Roberto Amato, Jeffrey Barrett, Sonia Goncalves, Ewan Harrison, David K. Jackson, Ian Johnston, Dominic Kwiatkowski, Cordelia Langford, John Sillitoe on behalf of the Wellcome Sanger Institute COVID-19 Surveillance Team |
| EPI_ISL_2348152 | Lighthouse Lab in Milton Keynes | Wellcome Sanger Institute for the COVID-19 Genomics UK (COG-UK) Consortium | The Lighthouse Lab in Milton Keynes and Alex Alderton, Roberto Amato, Jeffrey Barrett, Sonia Goncalves, Ewan Harrison, David K. Jackson, Ian Johnston, Dominic Kwiatkowski, Cordelia Langford, John Sillitoe on behalf of the Wellcome Sanger Institute COVID-19 Surveillance Team |
| EPI_ISL_2348155, EPI_ISL_2348156 | Lighthouse Lab in Alderley Park | Wellcome Sanger Institute for the COVID-19 Genomics UK (COG-UK) Consortium | Jacquelyn Wynn, Mairead Hyland, The Lighthouse Lab in Alderley Park and Alex Alderton, Roberto Amato, Jeffrey Barrett, Sonia Goncalves, Ewan Harrison, David K. Jackson, Ian Johnston, Dominic Kwiatkowski, Cordelia Langford, John Sillitoe on behalf of the Wellcome Sanger Institute COVID-19 Surveillance Team |
| EPI_ISL_2348159 | Lighthouse Laboratory Plymouth | Wellcome Sanger Institute for the COVID-19 Genomics UK (COG-UK) Consortium | Lighthouse Laboratory Plymouth and Alex Alderton, Roberto Amato, Jeffrey Barrett, Sonia Goncalves, Ewan Harrison, David K. Jackson, Ian Johnston, Dominic Kwiatkowski, Cordelia Langford, John Sillitoe on behalf of the Wellcome Sanger Institute COVID-19 Surveillance Team |
| EPI_ISL_2348161 | Lighthouse Lab in Alderley Park | Wellcome Sanger Institute for the COVID-19 Genomics UK (COG-UK) Consortium | Jacquelyn Wynn, Mairead Hyland, The Lighthouse Lab in Alderley Park and Alex Alderton, Roberto Amato, Jeffrey Barrett, Sonia Goncalves, Ewan Harrison, David K. Jackson, Ian Johnston, Dominic Kwiatkowski, Cordelia Langford, John Sillitoe on behalf of the Wellcome Sanger Institute COVID-19 Surveillance Team |
| EPI_ISL_2348162 | Health Services Laboratories | Wellcome Sanger Institute for the COVID-19 Genomics UK (COG-UK) Consortium | Health Services Laboratories and Alex Alderton, Roberto Amato, Jeffrey Barrett, Sonia Goncalves, Ewan Harrison, David K. Jackson, Ian Johnston, Dominic Kwiatkowski, Cordelia Langford, John Sillitoe on behalf of the Wellcome Sanger Institute COVID-19 Surveillance Team |
| EPI_ISL_2348163 | Lighthouse Lab in Milton Keynes | Wellcome Sanger Institute for the COVID-19 Genomics UK (COG-UK) Consortium | The Lighthouse Lab in Milton Keynes and Alex Alderton, Roberto Amato, Jeffrey Barrett, Sonia Goncalves, Ewan Harrison, David K. Jackson, Ian Johnston, Dominic Kwiatkowski, Cordelia Langford, John Sillitoe on behalf of the Wellcome Sanger Institute COVID-19 Surveillance Team |
| EPI_ISL_2348164, EPI_ISL_2348166, EPI_ISL_2348167, EPI_ISL_2348169, EPI_ISL_2348170 | Lighthouse Lab in Alderley Park | Wellcome Sanger Institute for the COVID-19 Genomics UK (COG-UK) Consortium | Jacquelyn Wynn, Mairead Hyland, The Lighthouse Lab in Alderley Park and Alex Alderton, Roberto Amato, Jeffrey Barrett, Sonia Goncalves, Ewan Harrison, David K. Jackson, Ian Johnston, Dominic Kwiatkowski, Cordelia Langford, John Sillitoe on behalf of the Wellcome Sanger Institute COVID-19 Surveillance Team |
| EPI_ISL_2348171 | Lighthouse Lab in Milton Keynes | Wellcome Sanger Institute for the COVID-19 Genomics UK (COG-UK) Consortium | The Lighthouse Lab in Milton Keynes and Alex Alderton, Roberto Amato, Jeffrey Barrett, Sonia Goncalves, Ewan Harrison, David K. Jackson, Ian Johnston, Dominic Kwiatkowski, Cordelia Langford, John Sillitoe on behalf of the Wellcome Sanger Institute COVID-19 Surveillance Team |
| EPI_ISL_2348172 | Lighthouse Lab in Alderley Park | Wellcome Sanger Institute for the COVID-19 Genomics UK (COG-UK) Consortium | Jacquelyn Wynn, Mairead Hyland, The Lighthouse Lab in Alderley Park and Alex Alderton, Roberto Amato, Jeffrey Barrett, Sonia Goncalves, Ewan Harrison, David K. Jackson, Ian Johnston, Dominic Kwiatkowski, Cordelia Langford, John Sillitoe on behalf of the Wellcome Sanger Institute COVID-19 Surveillance Team |
| EPI_ISL_2348173 | Lighthouse Lab in Milton Keynes | Wellcome Sanger Institute for the COVID-19 Genomics UK (COG-UK) Consortium | The Lighthouse Lab in Milton Keynes and Alex Alderton, Roberto Amato, Jeffrey Barrett, Sonia Goncalves, Ewan Harrison, David K. Jackson, Ian Johnston, Dominic Kwiatkowski, Cordelia Langford, John Sillitoe on behalf of the Wellcome Sanger Institute COVID-19 Surveillance Team |
| EPI_ISL_2348174, EPI_ISL_2348175, EPI_ISL_2348177, EPI_ISL_2348178, EPI_ISL_2348179, EPI_ISL_2348182, EPI_ISL_2348183, EPI_ISL_2348184, EPI_ISL_2348185, EPI_ISL_2348186, EPI_ISL_2348188 | | | |
| see above | Lighthouse Lab in Alderley Park | Wellcome Sanger Institute for the COVID-19 Genomics UK (COG-UK) Consortium | Jacquelyn Wynn, Mairead Hyland, The Lighthouse Lab in Alderley Park and Alex Alderton, Roberto Amato, Jeffrey Barrett, Sonia Goncalves, Ewan Harrison, David K. Jackson, Ian Johnston, Dominic Kwiatkowski, Cordelia Langford, John Sillitoe on behalf of the Wellcome Sanger Institute COVID-19 Surveillance Team |
| EPI_ISL_2348190, EPI_ISL_2348192 | Lighthouse Lab in Milton Keynes | Wellcome Sanger Institute for the COVID-19 Genomics UK (COG-UK) Consortium | The Lighthouse Lab in Milton Keynes and Alex Alderton, Roberto Amato, Jeffrey Barrett, Sonia Goncalves, Ewan Harrison, David K. Jackson, Ian Johnston, Dominic Kwiatkowski, Cordelia Langford, John Sillitoe on behalf of the Wellcome Sanger Institute COVID-19 Surveillance Team |
| EPI_ISL_2348195 | Lighthouse Lab in Alderley Park | Wellcome Sanger Institute for the COVID-19 Genomics UK (COG-UK) Consortium | Jacquelyn Wynn, Mairead Hyland, The Lighthouse Lab in Alderley Park and Alex Alderton, Roberto Amato, Jeffrey Barrett, Sonia Goncalves, Ewan Harrison, David K. Jackson, Ian Johnston, Dominic Kwiatkowski, Cordelia Langford, John Sillitoe on behalf of the Wellcome Sanger Institute COVID-19 Surveillance Team |
| EPI_ISL_2348198 | Lighthouse Lab in Milton Keynes | Wellcome Sanger Institute for the COVID-19 Genomics UK (COG-UK) Consortium | The Lighthouse Lab in Milton Keynes and Alex Alderton, Roberto Amato, Jeffrey Barrett, Sonia Goncalves, Ewan Harrison, David K. Jackson, Ian Johnston, Dominic Kwiatkowski, Cordelia Langford, John Sillitoe on behalf of the Wellcome Sanger Institute COVID-19 Surveillance Team |
| EPI_ISL_2348200, EPI_ISL_2348201, EPI_ISL_2348202, EPI_ISL_2348203 | Lighthouse Lab in Alderley Park | Wellcome Sanger Institute for the COVID-19 Genomics UK (COG-UK) Consortium | Jacquelyn Wynn, Mairead Hyland, The Lighthouse Lab in Alderley Park and Alex Alderton, Roberto Amato, Jeffrey Barrett, Sonia Goncalves, Ewan Harrison, David K. Jackson, Ian Johnston, Dominic Kwiatkowski, Cordelia Langford, John Sillitoe on behalf of the Wellcome Sanger Institute COVID-19 Surveillance Team |
| EPI_ISL_2348205, EPI_ISL_2348207, EPI_ISL_2348208, | Lighthouse Lab in Milton Keynes | Wellcome Sanger Institute for the COVID-19 Genomics UK (COG-UK) Consortium | The Lighthouse Lab in Milton Keynes and Alex Alderton, Roberto Amato, Jeffrey Barrett, Sonia Goncalves, Ewan Harrison, David K. Jackson, Ian Johnston, Dominic Kwiatkowski, Cordelia Langford, John Sillitoe on behalf of the Wellcome Sanger Institute COVID-19 Surveillance Team |

| | | | |
|---|---|---|---|
| EPI_ISL_2348209, EPI_ISL_2348214 | | | |
| EPI_ISL_2348216 | Lighthouse Lab in Alderley Park | Wellcome Sanger Institute for the COVID-19 Genomics UK (COG-UK) Consortium | Jacquelyn Wynn, Mairead Hyland, The Lighthouse Lab in Alderley Park and Alex Alderton, Roberto Amato, Jeffrey Barrett, Sonia Goncalves, Ewan Harrison, David K. Jackson, Ian Johnston, Dominic Kwiatkowski, Cordelia Langford, John Sillitoe on behalf of the Wellcome Sanger Institute COVID-19 Surveillance Team |
| EPI_ISL_2348217 | Lighthouse Lab in Milton Keynes | Wellcome Sanger Institute for the COVID-19 Genomics UK (COG-UK) Consortium | The Lighthouse Lab in Milton Keynes and Alex Alderton, Roberto Amato, Jeffrey Barrett, Sonia Goncalves, Ewan Harrison, David K. Jackson, Ian Johnston, Dominic Kwiatkowski, Cordelia Langford, John Sillitoe on behalf of the Wellcome Sanger Institute COVID-19 Surveillance Team |
| EPI_ISL_2348218, EPI_ISL_2348219 | Lighthouse Lab in Alderley Park | Wellcome Sanger Institute for the COVID-19 Genomics UK (COG-UK) Consortium | Jacquelyn Wynn, Mairead Hyland, The Lighthouse Lab in Alderley Park and Alex Alderton, Roberto Amato, Jeffrey Barrett, Sonia Goncalves, Ewan Harrison, David K. Jackson, Ian Johnston, Dominic Kwiatkowski, Cordelia Langford, John Sillitoe on behalf of the Wellcome Sanger Institute COVID-19 Surveillance Team |
| EPI_ISL_2348221 | Lighthouse Lab in Milton Keynes | Wellcome Sanger Institute for the COVID-19 Genomics UK (COG-UK) Consortium | The Lighthouse Lab in Milton Keynes and Alex Alderton, Roberto Amato, Jeffrey Barrett, Sonia Goncalves, Ewan Harrison, David K. Jackson, Ian Johnston, Dominic Kwiatkowski, Cordelia Langford, John Sillitoe on behalf of the Wellcome Sanger Institute COVID-19 Surveillance Team |
| EPI_ISL_2348222 | Lighthouse Lab in Alderley Park | Wellcome Sanger Institute for the COVID-19 Genomics UK (COG-UK) Consortium | Jacquelyn Wynn, Mairead Hyland, The Lighthouse Lab in Alderley Park and Alex Alderton, Roberto Amato, Jeffrey Barrett, Sonia Goncalves, Ewan Harrison, David K. Jackson, Ian Johnston, Dominic Kwiatkowski, Cordelia Langford, John Sillitoe on behalf of the Wellcome Sanger Institute COVID-19 Surveillance Team |
| EPI_ISL_2348223 | Lighthouse Lab in Milton Keynes | Wellcome Sanger Institute for the COVID-19 Genomics UK (COG-UK) Consortium | The Lighthouse Lab in Milton Keynes and Alex Alderton, Roberto Amato, Jeffrey Barrett, Sonia Goncalves, Ewan Harrison, David K. Jackson, Ian Johnston, Dominic Kwiatkowski, Cordelia Langford, John Sillitoe on behalf of the Wellcome Sanger Institute COVID-19 Surveillance Team |
| EPI_ISL_2348224 | Lighthouse Lab in Alderley Park | Wellcome Sanger Institute for the COVID-19 Genomics UK (COG-UK) Consortium | Jacquelyn Wynn, Mairead Hyland, The Lighthouse Lab in Alderley Park and Alex Alderton, Roberto Amato, Jeffrey Barrett, Sonia Goncalves, Ewan Harrison, David K. Jackson, Ian Johnston, Dominic Kwiatkowski, Cordelia Langford, John Sillitoe on behalf of the Wellcome Sanger Institute COVID-19 Surveillance Team |
| EPI_ISL_2348225 | Lighthouse Lab in Milton Keynes | Wellcome Sanger Institute for the COVID-19 Genomics UK (COG-UK) Consortium | The Lighthouse Lab in Milton Keynes and Alex Alderton, Roberto Amato, Jeffrey Barrett, Sonia Goncalves, Ewan Harrison, David K. Jackson, Ian Johnston, Dominic Kwiatkowski, Cordelia Langford, John Sillitoe on behalf of the Wellcome Sanger Institute COVID-19 Surveillance Team |
| EPI_ISL_2348226 | Lighthouse Lab in Glasgow | Wellcome Sanger Institute for the COVID-19 Genomics UK (COG-UK) Consortium | Harper VanSteenhouse, Yumi Kasai, David Gray, Carol Clugston, Anna Dominiczak and Alex Alderton, Roberto Amato, Jeffrey Barrett, Sonia Goncalves, Ewan Harrison, David K. Jackson, Ian Johnston, Dominic Kwiatkowski, Cordelia Langford, John Sillitoe on behalf of the Wellcome Sanger Institute COVID-19 Surveillance Team |
| EPI_ISL_2348227, EPI_ISL_2348228 | Lighthouse Lab in Milton Keynes | Wellcome Sanger Institute for the COVID-19 Genomics UK (COG-UK) Consortium | The Lighthouse Lab in Milton Keynes and Alex Alderton, Roberto Amato, Jeffrey Barrett, Sonia Goncalves, Ewan Harrison, David K. Jackson, Ian Johnston, Dominic Kwiatkowski, Cordelia Langford, John Sillitoe on behalf of the Wellcome Sanger Institute COVID-19 Surveillance Team |
| EPI_ISL_2348229 | Lighthouse Lab in Alderley Park | Wellcome Sanger Institute for the COVID-19 Genomics UK (COG-UK) Consortium | Jacquelyn Wynn, Mairead Hyland, The Lighthouse Lab in Alderley Park and Alex Alderton, Roberto Amato, Jeffrey Barrett, Sonia Goncalves, Ewan Harrison, David K. Jackson, Ian Johnston, Dominic Kwiatkowski, Cordelia Langford, John Sillitoe on behalf of the Wellcome Sanger Institute COVID-19 Surveillance Team |
| EPI_ISL_2348230 | Lighthouse Lab in Milton Keynes | Wellcome Sanger Institute for the COVID-19 Genomics UK (COG-UK) Consortium | The Lighthouse Lab in Milton Keynes and Alex Alderton, Roberto Amato, Jeffrey Barrett, Sonia Goncalves, Ewan Harrison, David K. Jackson, Ian Johnston, Dominic Kwiatkowski, Cordelia Langford, John Sillitoe on behalf of the Wellcome Sanger Institute COVID-19 Surveillance Team |
| EPI_ISL_2348231, EPI_ISL_2348232, EPI_ISL_2348233 | Lighthouse Lab in Alderley Park | Wellcome Sanger Institute for the COVID-19 Genomics UK (COG-UK) Consortium | Jacquelyn Wynn, Mairead Hyland, The Lighthouse Lab in Alderley Park and Alex Alderton, Roberto Amato, Jeffrey Barrett, Sonia Goncalves, Ewan Harrison, David K. Jackson, Ian Johnston, Dominic Kwiatkowski, Cordelia Langford, John Sillitoe on behalf of the Wellcome Sanger Institute COVID-19 Surveillance Team |
| EPI_ISL_2348237 | Lighthouse Lab in Milton Keynes | Wellcome Sanger Institute for the COVID-19 Genomics UK (COG-UK) Consortium | The Lighthouse Lab in Milton Keynes and Alex Alderton, Roberto Amato, Jeffrey Barrett, Sonia Goncalves, Ewan Harrison, David K. Jackson, Ian Johnston, Dominic Kwiatkowski, Cordelia Langford, John Sillitoe on behalf of the Wellcome Sanger Institute COVID-19 Surveillance Team |
| EPI_ISL_2348240 | Lighthouse Lab in Alderley Park | Wellcome Sanger Institute for the COVID-19 Genomics UK (COG-UK) Consortium | Jacquelyn Wynn, Mairead Hyland, The Lighthouse Lab in Alderley Park and Alex Alderton, Roberto Amato, Jeffrey Barrett, Sonia Goncalves, Ewan Harrison, David K. Jackson, Ian Johnston, Dominic Kwiatkowski, Cordelia Langford, John Sillitoe on behalf of the Wellcome Sanger Institute COVID-19 Surveillance Team |
| EPI_ISL_2348242, EPI_ISL_2348243, EPI_ISL_2348246, EPI_ISL_2348247, EPI_ISL_2348248 | Lighthouse Lab in Milton Keynes | Wellcome Sanger Institute for the COVID-19 Genomics UK (COG-UK) Consortium | The Lighthouse Lab in Milton Keynes and Alex Alderton, Roberto Amato, Jeffrey Barrett, Sonia Goncalves, Ewan Harrison, David K. Jackson, Ian Johnston, Dominic Kwiatkowski, Cordelia Langford, John Sillitoe on behalf of the Wellcome Sanger Institute COVID-19 Surveillance Team |
| EPI_ISL_2348251 | Lighthouse Lab in Glasgow | Wellcome Sanger Institute for the COVID-19 Genomics UK (COG-UK) Consortium | Harper VanSteenhouse, Yumi Kasai, David Gray, Carol Clugston, Anna Dominiczak and Alex Alderton, Roberto Amato, Jeffrey Barrett, Sonia Goncalves, Ewan Harrison, David K. Jackson, Ian Johnston, Dominic Kwiatkowski, Cordelia Langford, John Sillitoe on behalf of the Wellcome Sanger Institute COVID-19 Surveillance Team |
| EPI_ISL_2348254 | Lighthouse Lab in Alderley Park | Wellcome Sanger Institute for the COVID-19 Genomics UK (COG-UK) Consortium | Jacquelyn Wynn, Mairead Hyland, The Lighthouse Lab in Alderley Park and Alex Alderton, Roberto Amato, Jeffrey Barrett, Sonia Goncalves, Ewan Harrison, David K. Jackson, Ian Johnston, Dominic Kwiatkowski, Cordelia Langford, John Sillitoe on behalf of the Wellcome Sanger Institute COVID-19 Surveillance Team |
| EPI_ISL_2348255 | Lighthouse Lab in Milton Keynes | Wellcome Sanger Institute for the COVID-19 Genomics UK (COG-UK) Consortium | The Lighthouse Lab in Milton Keynes and Alex Alderton, Roberto Amato, Jeffrey Barrett, Sonia Goncalves, Ewan Harrison, David K. Jackson, Ian Johnston, Dominic Kwiatkowski, Cordelia Langford, John Sillitoe on behalf of the Wellcome Sanger Institute COVID-19 Surveillance Team |
| EPI_ISL_2348258 | Lighthouse Lab in Alderley Park | Wellcome Sanger Institute for the COVID-19 Genomics UK (COG-UK) Consortium | Jacquelyn Wynn, Mairead Hyland, The Lighthouse Lab in Alderley Park and Alex Alderton, Roberto Amato, Jeffrey Barrett, Sonia Goncalves, Ewan Harrison, David K. Jackson, Ian Johnston, Dominic Kwiatkowski, Cordelia Langford, John Sillitoe on behalf of the Wellcome Sanger Institute COVID-19 Surveillance Team |
| EPI_ISL_2348259, EPI_ISL_2348260 | Lighthouse Lab in Milton Keynes | Wellcome Sanger Institute for the COVID-19 Genomics UK (COG-UK) Consortium | The Lighthouse Lab in Milton Keynes and Alex Alderton, Roberto Amato, Jeffrey Barrett, Sonia Goncalves, Ewan Harrison, David K. Jackson, Ian Johnston, Dominic Kwiatkowski, Cordelia Langford, John Sillitoe on behalf of the Wellcome Sanger Institute COVID-19 Surveillance Team |
| EPI_ISL_2348262 | Lighthouse Lab in Alderley Park | Wellcome Sanger Institute for the COVID-19 Genomics UK (COG-UK) Consortium | Jacquelyn Wynn, Mairead Hyland, The Lighthouse Lab in Alderley Park and Alex Alderton, Roberto Amato, Jeffrey Barrett, Sonia Goncalves, Ewan Harrison, David K. Jackson, Ian Johnston, Dominic Kwiatkowski, Cordelia Langford, John Sillitoe on behalf of the Wellcome Sanger Institute COVID-19 Surveillance Team |
| EPI_ISL_2348263 | Lighthouse Lab in Milton Keynes | Wellcome Sanger Institute for the COVID-19 Genomics UK (COG-UK) Consortium | The Lighthouse Lab in Milton Keynes and Alex Alderton, Roberto Amato, Jeffrey Barrett, Sonia Goncalves, Ewan Harrison, David K. Jackson, Ian Johnston, Dominic Kwiatkowski, Cordelia Langford, John Sillitoe on behalf of the Wellcome Sanger Institute COVID-19 Surveillance Team |
| EPI_ISL_2348264, EPI_ISL_2348265 | Lighthouse Lab in Alderley Park | Wellcome Sanger Institute for the COVID-19 Genomics UK (COG-UK) Consortium | Jacquelyn Wynn, Mairead Hyland, The Lighthouse Lab in Alderley Park and Alex Alderton, Roberto Amato, Jeffrey Barrett, Sonia Goncalves, Ewan Harrison, David K. Jackson, Ian Johnston, Dominic Kwiatkowski, Cordelia Langford, John Sillitoe on behalf of the Wellcome Sanger Institute COVID-19 Surveillance Team |
| EPI_ISL_2348266, EPI_ISL_2348267 | Lighthouse Lab in Milton Keynes | Wellcome Sanger Institute for the COVID-19 Genomics UK (COG-UK) Consortium | The Lighthouse Lab in Milton Keynes and Alex Alderton, Roberto Amato, Jeffrey Barrett, Sonia Goncalves, Ewan Harrison, David K. Jackson, Ian Johnston, Dominic Kwiatkowski, Cordelia Langford, John Sillitoe on behalf of the Wellcome Sanger Institute COVID-19 Surveillance Team |
| EPI_ISL_2348271, EPI_ISL_2348272, EPI_ISL_2348274, EPI_ISL_2348275 | Lighthouse Lab in Alderley Park | Wellcome Sanger Institute for the COVID-19 Genomics UK (COG-UK) Consortium | Jacquelyn Wynn, Mairead Hyland, The Lighthouse Lab in Alderley Park and Alex Alderton, Roberto Amato, Jeffrey Barrett, Sonia Goncalves, Ewan Harrison, David K. Jackson, Ian Johnston, Dominic Kwiatkowski, Cordelia Langford, John Sillitoe on behalf of the Wellcome Sanger Institute COVID-19 Surveillance Team |
| EPI_ISL_2348276 | Lighthouse Lab in Glasgow | Wellcome Sanger Institute for the COVID-19 Genomics UK (COG-UK) Consortium | Harper VanSteenhouse, Yumi Kasai, David Gray, Carol Clugston, Anna Dominiczak and Alex Alderton, Roberto Amato, Jeffrey Barrett, Sonia Goncalves, Ewan Harrison, David K. Jackson, Ian Johnston, Dominic Kwiatkowski, Cordelia Langford, John Sillitoe on behalf of the Wellcome Sanger Institute COVID-19 Surveillance Team |
| EPI_ISL_2348277 | Lighthouse Lab in Alderley Park | Wellcome Sanger Institute for the COVID-19 Genomics UK (COG-UK) Consortium | Jacquelyn Wynn, Mairead Hyland, The Lighthouse Lab in Alderley Park and Alex Alderton, Roberto Amato, Jeffrey Barrett, Sonia Goncalves, Ewan Harrison, David K. Jackson, Ian Johnston, Dominic Kwiatkowski, Cordelia Langford, John Sillitoe on behalf of the Wellcome Sanger Institute COVID-19 Surveillance Team |
| EPI_ISL_2348278 | Lighthouse Lab in Milton Keynes | Wellcome Sanger Institute for the COVID-19 Genomics UK (COG-UK) Consortium | The Lighthouse Lab in Milton Keynes and Alex Alderton, Roberto Amato, Jeffrey Barrett, Sonia Goncalves, Ewan Harrison, David K. Jackson, Ian Johnston, Dominic Kwiatkowski, Cordelia Langford, John Sillitoe on behalf of the Wellcome Sanger Institute COVID-19 Surveillance Team |
| EPI_ISL_2348280 | Lighthouse Lab in Alderley Park | Wellcome Sanger Institute for the COVID-19 Genomics UK (COG-UK) Consortium | Jacquelyn Wynn, Mairead Hyland, The Lighthouse Lab in Alderley Park and Alex Alderton, Roberto Amato, Jeffrey Barrett, Sonia Goncalves, Ewan Harrison, David K. Jackson, Ian Johnston, Dominic Kwiatkowski, Cordelia Langford, John Sillitoe on behalf of the Wellcome Sanger Institute COVID-19 Surveillance Team |
| EPI_ISL_2348282 | Lighthouse Lab in Glasgow | Wellcome Sanger Institute for the COVID-19 Genomics UK (COG-UK) Consortium | Harper VanSteenhouse, Yumi Kasai, David Gray, Carol Clugston, Anna Dominiczak and Alex Alderton, Roberto Amato, Jeffrey Barrett, Sonia Goncalves, Ewan Harrison, David K. Jackson, Ian Johnston, Dominic Kwiatkowski, Cordelia Langford, John Sillitoe on behalf of the Wellcome Sanger Institute COVID-19 Surveillance Team |
| EPI_ISL_2348285, EPI_ISL_2348286, EPI_ISL_2348287, EPI_ISL_2348288 | Lighthouse Lab in Alderley Park | Wellcome Sanger Institute for the COVID-19 Genomics UK (COG-UK) Consortium | Jacquelyn Wynn, Mairead Hyland, The Lighthouse Lab in Alderley Park and Alex Alderton, Roberto Amato, Jeffrey Barrett, Sonia Goncalves, Ewan Harrison, David K. Jackson, Ian Johnston, Dominic Kwiatkowski, Cordelia Langford, John Sillitoe on behalf of the Wellcome Sanger Institute COVID-19 Surveillance Team |
| EPI_ISL_2348289 | Lighthouse Lab in Milton Keynes | Wellcome Sanger Institute for the COVID-19 Genomics UK (COG-UK) Consortium | The Lighthouse Lab in Milton Keynes and Alex Alderton, Roberto Amato, Jeffrey Barrett, Sonia Goncalves, Ewan Harrison, David K. Jackson, Ian Johnston, Dominic Kwiatkowski, Cordelia Langford, John Sillitoe on behalf of the Wellcome Sanger Institute COVID-19 Surveillance Team |
| EPI_ISL_2348290, EPI_ISL_2348292 | Lighthouse Lab in Alderley Park | Wellcome Sanger Institute for the COVID-19 Genomics UK (COG-UK) Consortium | Jacquelyn Wynn, Mairead Hyland, The Lighthouse Lab in Alderley Park and Alex Alderton, Roberto Amato, Jeffrey Barrett, Sonia Goncalves, Ewan Harrison, David K. Jackson, Ian Johnston, Dominic Kwiatkowski, Cordelia Langford, John Sillitoe on behalf of the Wellcome Sanger Institute COVID-19 Surveillance Team |
| EPI_ISL_2348293, EPI_ISL_2348294, EPI_ISL_2348295 | Lighthouse Lab in Milton Keynes | Wellcome Sanger Institute for the COVID-19 Genomics UK (COG-UK) Consortium | The Lighthouse Lab in Milton Keynes and Alex Alderton, Roberto Amato, Jeffrey Barrett, Sonia Goncalves, Ewan Harrison, David K. Jackson, Ian Johnston, Dominic Kwiatkowski, Cordelia Langford, John Sillitoe on behalf of the Wellcome Sanger Institute COVID-19 Surveillance Team |
| EPI_ISL_2348297 | Lighthouse Lab in Alderley Park | Wellcome Sanger Institute for the COVID-19 Genomics UK (COG-UK) Consortium | Jacquelyn Wynn, Mairead Hyland, The Lighthouse Lab in Alderley Park and Alex Alderton, Roberto Amato, Jeffrey Barrett, Sonia Goncalves, Ewan Harrison, David K. Jackson, Ian Johnston, Dominic Kwiatkowski, Cordelia Langford, John Sillitoe on behalf of the Wellcome Sanger Institute COVID-19 Surveillance Team |
| EPI_ISL_2348298, EPI_ISL_2348299, EPI_ISL_2348300 | Lighthouse Lab in Milton Keynes | Wellcome Sanger Institute for the COVID-19 Genomics UK (COG-UK) Consortium | The Lighthouse Lab in Milton Keynes and Alex Alderton, Roberto Amato, Jeffrey Barrett, Sonia Goncalves, Ewan Harrison, David K. Jackson, Ian Johnston, Dominic Kwiatkowski, Cordelia Langford, John Sillitoe on behalf of the Wellcome Sanger Institute COVID-19 Surveillance Team |
| EPI_ISL_2348301, EPI_ISL_2348304 | Lighthouse Lab in Alderley Park | Wellcome Sanger Institute for the COVID-19 Genomics UK (COG-UK) Consortium | Jacquelyn Wynn, Mairead Hyland, The Lighthouse Lab in Alderley Park and Alex Alderton, Roberto Amato, Jeffrey Barrett, Sonia Goncalves, Ewan Harrison, David K. Jackson, Ian Johnston, Dominic Kwiatkowski, Cordelia Langford, John Sillitoe on behalf of the Wellcome Sanger Institute COVID-19 Surveillance Team |
| EPI_ISL_2348305 | Lighthouse Lab in Milton Keynes | Wellcome Sanger Institute for the COVID-19 Genomics UK (COG-UK) Consortium | The Lighthouse Lab in Milton Keynes and Alex Alderton, Roberto Amato, Jeffrey Barrett, Sonia Goncalves, Ewan Harrison, David K. Jackson, Ian Johnston, Dominic Kwiatkowski, Cordelia Langford, John Sillitoe on behalf of the Wellcome Sanger Institute COVID-19 Surveillance Team |
| EPI_ISL_2348306, EPI_ISL_2348307, | Lighthouse Lab in Alderley Park | Wellcome Sanger Institute for the COVID-19 Genomics UK (COG-UK) Consortium | Jacquelyn Wynn, Mairead Hyland, The Lighthouse Lab in Alderley Park and Alex Alderton, Roberto Amato, Jeffrey Barrett, Sonia Goncalves, Ewan Harrison, David K. Jackson, Ian Johnston, Dominic Kwiatkowski, Cordelia Langford, John Sillitoe on behalf of the Wellcome Sanger Institute COVID-19 Surveillance Team |

| | | | |
|---|---|---|---|
| EPI_ISL_2348308, EPI_ISL_2348311, EPI_ISL_2348315, EPI_ISL_2348316, EPI_ISL_2348317 | | | |
| EPI_ISL_2348318, EPI_ISL_2348319, EPI_ISL_2348320, EPI_ISL_2348321 | Lighthouse Lab in Milton Keynes | Wellcome Sanger Institute for the COVID-19 Genomics UK (COG-UK) Consortium | The Lighthouse Lab in Milton Keynes and Alex Alderton, Roberto Amato, Jeffrey Barrett, Sonia Goncalves, Ewan Harrison, David K. Jackson, Ian Johnston, Dominic Kwiatkowski, Cordelia Langford, John Sillitoe on behalf of the Wellcome Sanger Institute COVID-19 Surveillance Team |
| EPI_ISL_2348323 | Lighthouse Lab in Alderley Park | Wellcome Sanger Institute for the COVID-19 Genomics UK (COG-UK) Consortium | Jacquelyn Wynn, Mairead Hyland, The Lighthouse Lab in Alderley Park and Alex Alderton, Roberto Amato, Jeffrey Barrett, Sonia Goncalves, Ewan Harrison, David K. Jackson, Ian Johnston, Dominic Kwiatkowski, Cordelia Langford, John Sillitoe on behalf of the Wellcome Sanger Institute COVID-19 Surveillance Team |
| EPI_ISL_2348324, EPI_ISL_2348325 | Lighthouse Lab in Milton Keynes | Wellcome Sanger Institute for the COVID-19 Genomics UK (COG-UK) Consortium | The Lighthouse Lab in Milton Keynes and Alex Alderton, Roberto Amato, Jeffrey Barrett, Sonia Goncalves, Ewan Harrison, David K. Jackson, Ian Johnston, Dominic Kwiatkowski, Cordelia Langford, John Sillitoe on behalf of the Wellcome Sanger Institute COVID-19 Surveillance Team |
| EPI_ISL_2348327, EPI_ISL_2348328 | Lighthouse Lab in Alderley Park | Wellcome Sanger Institute for the COVID-19 Genomics UK (COG-UK) Consortium | Jacquelyn Wynn, Mairead Hyland, The Lighthouse Lab in Alderley Park and Alex Alderton, Roberto Amato, Jeffrey Barrett, Sonia Goncalves, Ewan Harrison, David K. Jackson, Ian Johnston, Dominic Kwiatkowski, Cordelia Langford, John Sillitoe on behalf of the Wellcome Sanger Institute COVID-19 Surveillance Team |
| EPI_ISL_2348330 | Lighthouse Lab in Milton Keynes | Wellcome Sanger Institute for the COVID-19 Genomics UK (COG-UK) Consortium | The Lighthouse Lab in Milton Keynes and Alex Alderton, Roberto Amato, Jeffrey Barrett, Sonia Goncalves, Ewan Harrison, David K. Jackson, Ian Johnston, Dominic Kwiatkowski, Cordelia Langford, John Sillitoe on behalf of the Wellcome Sanger Institute COVID-19 Surveillance Team |
| EPI_ISL_2348331 | Lighthouse Lab in Alderley Park | Wellcome Sanger Institute for the COVID-19 Genomics UK (COG-UK) Consortium | Jacquelyn Wynn, Mairead Hyland, The Lighthouse Lab in Alderley Park and Alex Alderton, Roberto Amato, Jeffrey Barrett, Sonia Goncalves, Ewan Harrison, David K. Jackson, Ian Johnston, Dominic Kwiatkowski, Cordelia Langford, John Sillitoe on behalf of the Wellcome Sanger Institute COVID-19 Surveillance Team |
| EPI_ISL_2348332, EPI_ISL_2348333, EPI_ISL_2348334, EPI_ISL_2348335 | Lighthouse Lab in Milton Keynes | Wellcome Sanger Institute for the COVID-19 Genomics UK (COG-UK) Consortium | The Lighthouse Lab in Milton Keynes and Alex Alderton, Roberto Amato, Jeffrey Barrett, Sonia Goncalves, Ewan Harrison, David K. Jackson, Ian Johnston, Dominic Kwiatkowski, Cordelia Langford, John Sillitoe on behalf of the Wellcome Sanger Institute COVID-19 Surveillance Team |
| EPI_ISL_2348336, EPI_ISL_2348338 | Lighthouse Lab in Alderley Park | Wellcome Sanger Institute for the COVID-19 Genomics UK (COG-UK) Consortium | Jacquelyn Wynn, Mairead Hyland, The Lighthouse Lab in Alderley Park and Alex Alderton, Roberto Amato, Jeffrey Barrett, Sonia Goncalves, Ewan Harrison, David K. Jackson, Ian Johnston, Dominic Kwiatkowski, Cordelia Langford, John Sillitoe on behalf of the Wellcome Sanger Institute COVID-19 Surveillance Team |
| EPI_ISL_2348339, EPI_ISL_2348341 | Lighthouse Lab in Milton Keynes | Wellcome Sanger Institute for the COVID-19 Genomics UK (COG-UK) Consortium | The Lighthouse Lab in Milton Keynes and Alex Alderton, Roberto Amato, Jeffrey Barrett, Sonia Goncalves, Ewan Harrison, David K. Jackson, Ian Johnston, Dominic Kwiatkowski, Cordelia Langford, John Sillitoe on behalf of the Wellcome Sanger Institute COVID-19 Surveillance Team |
| EPI_ISL_2348342, EPI_ISL_2348344, EPI_ISL_2348345, EPI_ISL_2348346, EPI_ISL_2348348, EPI_ISL_2348349 | Lighthouse Lab in Alderley Park | Wellcome Sanger Institute for the COVID-19 Genomics UK (COG-UK) Consortium | Jacquelyn Wynn, Mairead Hyland, The Lighthouse Lab in Alderley Park and Alex Alderton, Roberto Amato, Jeffrey Barrett, Sonia Goncalves, Ewan Harrison, David K. Jackson, Ian Johnston, Dominic Kwiatkowski, Cordelia Langford, John Sillitoe on behalf of the Wellcome Sanger Institute COVID-19 Surveillance Team |
| EPI_ISL_2348351, EPI_ISL_2348353 | Lighthouse Lab in Milton Keynes | Wellcome Sanger Institute for the COVID-19 Genomics UK (COG-UK) Consortium | The Lighthouse Lab in Milton Keynes and Alex Alderton, Roberto Amato, Jeffrey Barrett, Sonia Goncalves, Ewan Harrison, David K. Jackson, Ian Johnston, Dominic Kwiatkowski, Cordelia Langford, John Sillitoe on behalf of the Wellcome Sanger Institute COVID-19 Surveillance Team |
| EPI_ISL_2348354, EPI_ISL_2348356 | Lighthouse Lab in Alderley Park | Wellcome Sanger Institute for the COVID-19 Genomics UK (COG-UK) Consortium | Jacquelyn Wynn, Mairead Hyland, The Lighthouse Lab in Alderley Park and Alex Alderton, Roberto Amato, Jeffrey Barrett, Sonia Goncalves, Ewan Harrison, David K. Jackson, Ian Johnston, Dominic Kwiatkowski, Cordelia Langford, John Sillitoe on behalf of the Wellcome Sanger Institute COVID-19 Surveillance Team |
| EPI_ISL_2348357, EPI_ISL_2348358, EPI_ISL_2348360 | Lighthouse Lab in Milton Keynes | Wellcome Sanger Institute for the COVID-19 Genomics UK (COG-UK) Consortium | The Lighthouse Lab in Milton Keynes and Alex Alderton, Roberto Amato, Jeffrey Barrett, Sonia Goncalves, Ewan Harrison, David K. Jackson, Ian Johnston, Dominic Kwiatkowski, Cordelia Langford, John Sillitoe on behalf of the Wellcome Sanger Institute COVID-19 Surveillance Team |
| EPI_ISL_2348362, EPI_ISL_2348363, EPI_ISL_2348364, EPI_ISL_2348365 | Lighthouse Lab in Alderley Park | Wellcome Sanger Institute for the COVID-19 Genomics UK (COG-UK) Consortium | Jacquelyn Wynn, Mairead Hyland, The Lighthouse Lab in Alderley Park and Alex Alderton, Roberto Amato, Jeffrey Barrett, Sonia Goncalves, Ewan Harrison, David K. Jackson, Ian Johnston, Dominic Kwiatkowski, Cordelia Langford, John Sillitoe on behalf of the Wellcome Sanger Institute COVID-19 Surveillance Team |
| EPI_ISL_2348366 | Lighthouse Lab in Milton Keynes | Wellcome Sanger Institute for the COVID-19 Genomics UK (COG-UK) Consortium | The Lighthouse Lab in Milton Keynes and Alex Alderton, Roberto Amato, Jeffrey Barrett, Sonia Goncalves, Ewan Harrison, David K. Jackson, Ian Johnston, Dominic Kwiatkowski, Cordelia Langford, John Sillitoe on behalf of the Wellcome Sanger Institute COVID-19 Surveillance Team |
| EPI_ISL_2348367 | Lighthouse Lab in Alderley Park | Wellcome Sanger Institute for the COVID-19 Genomics UK (COG-UK) Consortium | Jacquelyn Wynn, Mairead Hyland, The Lighthouse Lab in Alderley Park and Alex Alderton, Roberto Amato, Jeffrey Barrett, Sonia Goncalves, Ewan Harrison, David K. Jackson, Ian Johnston, Dominic Kwiatkowski, Cordelia Langford, John Sillitoe on behalf of the Wellcome Sanger Institute COVID-19 Surveillance Team |
| EPI_ISL_2348368 | Lighthouse Lab in Milton Keynes | Wellcome Sanger Institute for the COVID-19 Genomics UK (COG-UK) Consortium | The Lighthouse Lab in Milton Keynes and Alex Alderton, Roberto Amato, Jeffrey Barrett, Sonia Goncalves, Ewan Harrison, David K. Jackson, Ian Johnston, Dominic Kwiatkowski, Cordelia Langford, John Sillitoe on behalf of the Wellcome Sanger Institute COVID-19 Surveillance Team |
| EPI_ISL_2348369, EPI_ISL_2348371, EPI_ISL_2348372 | Lighthouse Lab in Alderley Park | Wellcome Sanger Institute for the COVID-19 Genomics UK (COG-UK) Consortium | Jacquelyn Wynn, Mairead Hyland, The Lighthouse Lab in Alderley Park and Alex Alderton, Roberto Amato, Jeffrey Barrett, Sonia Goncalves, Ewan Harrison, David K. Jackson, Ian Johnston, Dominic Kwiatkowski, Cordelia Langford, John Sillitoe on behalf of the Wellcome Sanger Institute COVID-19 Surveillance Team |
| EPI_ISL_2348374 | Lighthouse Lab in Glasgow | Wellcome Sanger Institute for the COVID-19 Genomics UK (COG-UK) Consortium | Harper VanSteenhouse, Yumi Kasai, David Gray, Carol Clugston, Anna Dominiczak and Alex Alderton, Roberto Amato, Jeffrey Barrett, Sonia Goncalves, Ewan Harrison, David K. Jackson, Ian Johnston, Dominic Kwiatkowski, Cordelia Langford, John Sillitoe on behalf of the Wellcome Sanger Institute COVID-19 Surveillance Team |
| EPI_ISL_2348375 | Lighthouse Lab in Milton Keynes | Wellcome Sanger Institute for the COVID-19 Genomics UK (COG-UK) Consortium | The Lighthouse Lab in Milton Keynes and Alex Alderton, Roberto Amato, Jeffrey Barrett, Sonia Goncalves, Ewan Harrison, David K. Jackson, Ian Johnston, Dominic Kwiatkowski, Cordelia Langford, John Sillitoe on behalf of the Wellcome Sanger Institute COVID-19 Surveillance Team |
| EPI_ISL_2348376, EPI_ISL_2348377, EPI_ISL_2348379, EPI_ISL_2348382, EPI_ISL_2348383 | Lighthouse Lab in Alderley Park | Wellcome Sanger Institute for the COVID-19 Genomics UK (COG-UK) Consortium | Jacquelyn Wynn, Mairead Hyland, The Lighthouse Lab in Alderley Park and Alex Alderton, Roberto Amato, Jeffrey Barrett, Sonia Goncalves, Ewan Harrison, David K. Jackson, Ian Johnston, Dominic Kwiatkowski, Cordelia Langford, John Sillitoe on behalf of the Wellcome Sanger Institute COVID-19 Surveillance Team |
| EPI_ISL_2348385 | Lighthouse Lab in Milton Keynes | Wellcome Sanger Institute for the COVID-19 Genomics UK (COG-UK) Consortium | The Lighthouse Lab in Milton Keynes and Alex Alderton, Roberto Amato, Jeffrey Barrett, Sonia Goncalves, Ewan Harrison, David K. Jackson, Ian Johnston, Dominic Kwiatkowski, Cordelia Langford, John Sillitoe on behalf of the Wellcome Sanger Institute COVID-19 Surveillance Team |
| EPI_ISL_2348388 | Lighthouse Lab in Glasgow | Wellcome Sanger Institute for the COVID-19 Genomics UK (COG-UK) Consortium | Harper VanSteenhouse, Yumi Kasai, David Gray, Carol Clugston, Anna Dominiczak and Alex Alderton, Roberto Amato, Jeffrey Barrett, Sonia Goncalves, Ewan Harrison, David K. Jackson, Ian Johnston, Dominic Kwiatkowski, Cordelia Langford, John Sillitoe on behalf of the Wellcome Sanger Institute COVID-19 Surveillance Team |
| EPI_ISL_2348389 | Lighthouse Lab in Milton Keynes | Wellcome Sanger Institute for the COVID-19 Genomics UK (COG-UK) Consortium | The Lighthouse Lab in Milton Keynes and Alex Alderton, Roberto Amato, Jeffrey Barrett, Sonia Goncalves, Ewan Harrison, David K. Jackson, Ian Johnston, Dominic Kwiatkowski, Cordelia Langford, John Sillitoe on behalf of the Wellcome Sanger Institute COVID-19 Surveillance Team |
| EPI_ISL_2348391, EPI_ISL_2348392 | Lighthouse Lab in Alderley Park | Wellcome Sanger Institute for the COVID-19 Genomics UK (COG-UK) Consortium | Jacquelyn Wynn, Mairead Hyland, The Lighthouse Lab in Alderley Park and Alex Alderton, Roberto Amato, Jeffrey Barrett, Sonia Goncalves, Ewan Harrison, David K. Jackson, Ian Johnston, Dominic Kwiatkowski, Cordelia Langford, John Sillitoe on behalf of the Wellcome Sanger Institute COVID-19 Surveillance Team |
| EPI_ISL_2348393, EPI_ISL_2348395, EPI_ISL_2348397, EPI_ISL_2348399 | Lighthouse Lab in Milton Keynes | Wellcome Sanger Institute for the COVID-19 Genomics UK (COG-UK) Consortium | The Lighthouse Lab in Milton Keynes and Alex Alderton, Roberto Amato, Jeffrey Barrett, Sonia Goncalves, Ewan Harrison, David K. Jackson, Ian Johnston, Dominic Kwiatkowski, Cordelia Langford, John Sillitoe on behalf of the Wellcome Sanger Institute COVID-19 Surveillance Team |
| EPI_ISL_2348400, EPI_ISL_2348401 | Lighthouse Lab in Alderley Park | Wellcome Sanger Institute for the COVID-19 Genomics UK (COG-UK) Consortium | Jacquelyn Wynn, Mairead Hyland, The Lighthouse Lab in Alderley Park and Alex Alderton, Roberto Amato, Jeffrey Barrett, Sonia Goncalves, Ewan Harrison, David K. Jackson, Ian Johnston, Dominic Kwiatkowski, Cordelia Langford, John Sillitoe on behalf of the Wellcome Sanger Institute COVID-19 Surveillance Team |
| EPI_ISL_2348402, EPI_ISL_2348404, EPI_ISL_2348405 | Lighthouse Lab in Milton Keynes | Wellcome Sanger Institute for the COVID-19 Genomics UK (COG-UK) Consortium | The Lighthouse Lab in Milton Keynes and Alex Alderton, Roberto Amato, Jeffrey Barrett, Sonia Goncalves, Ewan Harrison, David K. Jackson, Ian Johnston, Dominic Kwiatkowski, Cordelia Langford, John Sillitoe on behalf of the Wellcome Sanger Institute COVID-19 Surveillance Team |
| EPI_ISL_2348407 | Lighthouse Lab in Alderley Park | Wellcome Sanger Institute for the COVID-19 Genomics UK (COG-UK) Consortium | Jacquelyn Wynn, Mairead Hyland, The Lighthouse Lab in Alderley Park and Alex Alderton, Roberto Amato, Jeffrey Barrett, Sonia Goncalves, Ewan Harrison, David K. Jackson, Ian Johnston, Dominic Kwiatkowski, Cordelia Langford, John Sillitoe on behalf of the Wellcome Sanger Institute COVID-19 Surveillance Team |
| EPI_ISL_2348408 | Lighthouse Lab in Milton Keynes | Wellcome Sanger Institute for the COVID-19 Genomics UK (COG-UK) Consortium | The Lighthouse Lab in Milton Keynes and Alex Alderton, Roberto Amato, Jeffrey Barrett, Sonia Goncalves, Ewan Harrison, David K. Jackson, Ian Johnston, Dominic Kwiatkowski, Cordelia Langford, John Sillitoe on behalf of the Wellcome Sanger Institute COVID-19 Surveillance Team |
| EPI_ISL_2348409 | Lighthouse Lab in Alderley Park | Wellcome Sanger Institute for the COVID-19 Genomics UK (COG-UK) Consortium | Jacquelyn Wynn, Mairead Hyland, The Lighthouse Lab in Alderley Park and Alex Alderton, Roberto Amato, Jeffrey Barrett, Sonia Goncalves, Ewan Harrison, David K. Jackson, Ian Johnston, Dominic Kwiatkowski, Cordelia Langford, John Sillitoe on behalf of the Wellcome Sanger Institute COVID-19 Surveillance Team |
| EPI_ISL_2348410, EPI_ISL_2348412 | Lighthouse Lab in Milton Keynes | Wellcome Sanger Institute for the COVID-19 Genomics UK (COG-UK) Consortium | The Lighthouse Lab in Milton Keynes and Alex Alderton, Roberto Amato, Jeffrey Barrett, Sonia Goncalves, Ewan Harrison, David K. Jackson, Ian Johnston, Dominic Kwiatkowski, Cordelia Langford, John Sillitoe on behalf of the Wellcome Sanger Institute COVID-19 Surveillance Team |
| EPI_ISL_2348414 | Lighthouse Lab in Alderley Park | Wellcome Sanger Institute for the COVID-19 Genomics UK (COG-UK) Consortium | Jacquelyn Wynn, Mairead Hyland, The Lighthouse Lab in Alderley Park and Alex Alderton, Roberto Amato, Jeffrey Barrett, Sonia Goncalves, Ewan Harrison, David K. Jackson, Ian Johnston, Dominic Kwiatkowski, Cordelia Langford, John Sillitoe on behalf of the Wellcome Sanger Institute COVID-19 Surveillance Team |
| EPI_ISL_2348416, EPI_ISL_2348417, EPI_ISL_2348419 | Lighthouse Lab in Milton Keynes | Wellcome Sanger Institute for the COVID-19 Genomics UK (COG-UK) Consortium | The Lighthouse Lab in Milton Keynes and Alex Alderton, Roberto Amato, Jeffrey Barrett, Sonia Goncalves, Ewan Harrison, David K. Jackson, Ian Johnston, Dominic Kwiatkowski, Cordelia Langford, John Sillitoe on behalf of the Wellcome Sanger Institute COVID-19 Surveillance Team |
| EPI_ISL_2348422, EPI_ISL_2348424, EPI_ISL_2348425, | Lighthouse Lab in Alderley Park | Wellcome Sanger Institute for the COVID-19 Genomics UK (COG-UK) Consortium | Jacquelyn Wynn, Mairead Hyland, The Lighthouse Lab in Alderley Park and Alex Alderton, Roberto Amato, Jeffrey Barrett, Sonia Goncalves, Ewan Harrison, David K. Jackson, Ian Johnston, Dominic Kwiatkowski, Cordelia Langford, John Sillitoe on behalf of the Wellcome Sanger Institute COVID-19 Surveillance Team |

| | | | |
|---|---|---|---|
| EPI_ISL_2348426, EPI_ISL_2348428, EPI_ISL_2348429 | | | |
| EPI_ISL_2348430, EPI_ISL_2348431, EPI_ISL_2348432 | Lighthouse Lab in Milton Keynes | Wellcome Sanger Institute for the COVID-19 Genomics UK (COG-UK) Consortium | The Lighthouse Lab in Milton Keynes and Alex Alderton, Roberto Amato, Jeffrey Barrett, Sonia Goncalves, Ewan Harrison, David K. Jackson, Ian Johnston, Dominic Kwiatkowski, Cordelia Langford, John Sillitoe on behalf of the Wellcome Sanger Institute COVID-19 Surveillance Team |
| EPI_ISL_2348433 | Lighthouse Lab in Alderley Park | Wellcome Sanger Institute for the COVID-19 Genomics UK (COG-UK) Consortium | Jacquelyn Wynn, Mairead Hyland, The Lighthouse Lab in Alderley Park and Alex Alderton, Roberto Amato, Jeffrey Barrett, Sonia Goncalves, Ewan Harrison, David K. Jackson, Ian Johnston, Dominic Kwiatkowski, Cordelia Langford, John Sillitoe on behalf of the Wellcome Sanger Institute COVID-19 Surveillance Team |
| EPI_ISL_2348434 | Lighthouse Lab in Milton Keynes | Wellcome Sanger Institute for the COVID-19 Genomics UK (COG-UK) Consortium | The Lighthouse Lab in Milton Keynes and Alex Alderton, Roberto Amato, Jeffrey Barrett, Sonia Goncalves, Ewan Harrison, David K. Jackson, Ian Johnston, Dominic Kwiatkowski, Cordelia Langford, John Sillitoe on behalf of the Wellcome Sanger Institute COVID-19 Surveillance Team |
| EPI_ISL_2348435, EPI_ISL_2348436, EPI_ISL_2348437 | Lighthouse Lab in Alderley Park | Wellcome Sanger Institute for the COVID-19 Genomics UK (COG-UK) Consortium | Jacquelyn Wynn, Mairead Hyland, The Lighthouse Lab in Alderley Park and Alex Alderton, Roberto Amato, Jeffrey Barrett, Sonia Goncalves, Ewan Harrison, David K. Jackson, Ian Johnston, Dominic Kwiatkowski, Cordelia Langford, John Sillitoe on behalf of the Wellcome Sanger Institute COVID-19 Surveillance Team |
| EPI_ISL_2348439, EPI_ISL_2348443, EPI_ISL_2348444, EPI_ISL_2348445 | Lighthouse Lab in Milton Keynes | Wellcome Sanger Institute for the COVID-19 Genomics UK (COG-UK) Consortium | The Lighthouse Lab in Milton Keynes and Alex Alderton, Roberto Amato, Jeffrey Barrett, Sonia Goncalves, Ewan Harrison, David K. Jackson, Ian Johnston, Dominic Kwiatkowski, Cordelia Langford, John Sillitoe on behalf of the Wellcome Sanger Institute COVID-19 Surveillance Team |
| EPI_ISL_2348446, EPI_ISL_2348447, EPI_ISL_2348448 | Lighthouse Lab in Alderley Park | Wellcome Sanger Institute for the COVID-19 Genomics UK (COG-UK) Consortium | Jacquelyn Wynn, Mairead Hyland, The Lighthouse Lab in Alderley Park and Alex Alderton, Roberto Amato, Jeffrey Barrett, Sonia Goncalves, Ewan Harrison, David K. Jackson, Ian Johnston, Dominic Kwiatkowski, Cordelia Langford, John Sillitoe on behalf of the Wellcome Sanger Institute COVID-19 Surveillance Team |
| EPI_ISL_2348449 | Lighthouse Lab in Milton Keynes | Wellcome Sanger Institute for the COVID-19 Genomics UK (COG-UK) Consortium | The Lighthouse Lab in Milton Keynes and Alex Alderton, Roberto Amato, Jeffrey Barrett, Sonia Goncalves, Ewan Harrison, David K. Jackson, Ian Johnston, Dominic Kwiatkowski, Cordelia Langford, John Sillitoe on behalf of the Wellcome Sanger Institute COVID-19 Surveillance Team |
| EPI_ISL_2348451, EPI_ISL_2348452, EPI_ISL_2348454, EPI_ISL_2348457 | Lighthouse Lab in Alderley Park | Wellcome Sanger Institute for the COVID-19 Genomics UK (COG-UK) Consortium | Jacquelyn Wynn, Mairead Hyland, The Lighthouse Lab in Alderley Park and Alex Alderton, Roberto Amato, Jeffrey Barrett, Sonia Goncalves, Ewan Harrison, David K. Jackson, Ian Johnston, Dominic Kwiatkowski, Cordelia Langford, John Sillitoe on behalf of the Wellcome Sanger Institute COVID-19 Surveillance Team |
| EPI_ISL_2348458 | Lighthouse Lab in Milton Keynes | Wellcome Sanger Institute for the COVID-19 Genomics UK (COG-UK) Consortium | The Lighthouse Lab in Milton Keynes and Alex Alderton, Roberto Amato, Jeffrey Barrett, Sonia Goncalves, Ewan Harrison, David K. Jackson, Ian Johnston, Dominic Kwiatkowski, Cordelia Langford, John Sillitoe on behalf of the Wellcome Sanger Institute COVID-19 Surveillance Team |
| EPI_ISL_2348459 | Lighthouse Lab in Alderley Park | Wellcome Sanger Institute for the COVID-19 Genomics UK (COG-UK) Consortium | Jacquelyn Wynn, Mairead Hyland, The Lighthouse Lab in Alderley Park and Alex Alderton, Roberto Amato, Jeffrey Barrett, Sonia Goncalves, Ewan Harrison, David K. Jackson, Ian Johnston, Dominic Kwiatkowski, Cordelia Langford, John Sillitoe on behalf of the Wellcome Sanger Institute COVID-19 Surveillance Team |
| EPI_ISL_2348460 | Lighthouse Lab in Milton Keynes | Wellcome Sanger Institute for the COVID-19 Genomics UK (COG-UK) Consortium | The Lighthouse Lab in Milton Keynes and Alex Alderton, Roberto Amato, Jeffrey Barrett, Sonia Goncalves, Ewan Harrison, David K. Jackson, Ian Johnston, Dominic Kwiatkowski, Cordelia Langford, John Sillitoe on behalf of the Wellcome Sanger Institute COVID-19 Surveillance Team |
| EPI_ISL_2348461, EPI_ISL_2348462, EPI_ISL_2348465 | Lighthouse Lab in Alderley Park | Wellcome Sanger Institute for the COVID-19 Genomics UK (COG-UK) Consortium | Jacquelyn Wynn, Mairead Hyland, The Lighthouse Lab in Alderley Park and Alex Alderton, Roberto Amato, Jeffrey Barrett, Sonia Goncalves, Ewan Harrison, David K. Jackson, Ian Johnston, Dominic Kwiatkowski, Cordelia Langford, John Sillitoe on behalf of the Wellcome Sanger Institute COVID-19 Surveillance Team |
| EPI_ISL_2348466 | Lighthouse Lab in Milton Keynes | Wellcome Sanger Institute for the COVID-19 Genomics UK (COG-UK) Consortium | The Lighthouse Lab in Milton Keynes and Alex Alderton, Roberto Amato, Jeffrey Barrett, Sonia Goncalves, Ewan Harrison, David K. Jackson, Ian Johnston, Dominic Kwiatkowski, Cordelia Langford, John Sillitoe on behalf of the Wellcome Sanger Institute COVID-19 Surveillance Team |
| EPI_ISL_2348467 | Lighthouse Lab in Alderley Park | Wellcome Sanger Institute for the COVID-19 Genomics UK (COG-UK) Consortium | Jacquelyn Wynn, Mairead Hyland, The Lighthouse Lab in Alderley Park and Alex Alderton, Roberto Amato, Jeffrey Barrett, Sonia Goncalves, Ewan Harrison, David K. Jackson, Ian Johnston, Dominic Kwiatkowski, Cordelia Langford, John Sillitoe on behalf of the Wellcome Sanger Institute COVID-19 Surveillance Team |
| EPI_ISL_2348468, EPI_ISL_2348471, EPI_ISL_2348472, EPI_ISL_2348473 | Lighthouse Lab in Milton Keynes | Wellcome Sanger Institute for the COVID-19 Genomics UK (COG-UK) Consortium | The Lighthouse Lab in Milton Keynes and Alex Alderton, Roberto Amato, Jeffrey Barrett, Sonia Goncalves, Ewan Harrison, David K. Jackson, Ian Johnston, Dominic Kwiatkowski, Cordelia Langford, John Sillitoe on behalf of the Wellcome Sanger Institute COVID-19 Surveillance Team |
| EPI_ISL_2348474 | Lighthouse Lab in Alderley Park | Wellcome Sanger Institute for the COVID-19 Genomics UK (COG-UK) Consortium | Jacquelyn Wynn, Mairead Hyland, The Lighthouse Lab in Alderley Park and Alex Alderton, Roberto Amato, Jeffrey Barrett, Sonia Goncalves, Ewan Harrison, David K. Jackson, Ian Johnston, Dominic Kwiatkowski, Cordelia Langford, John Sillitoe on behalf of the Wellcome Sanger Institute COVID-19 Surveillance Team |
| EPI_ISL_2348477, EPI_ISL_2348480, EPI_ISL_2348484, EPI_ISL_2348485 | Lighthouse Lab in Milton Keynes | Wellcome Sanger Institute for the COVID-19 Genomics UK (COG-UK) Consortium | The Lighthouse Lab in Milton Keynes and Alex Alderton, Roberto Amato, Jeffrey Barrett, Sonia Goncalves, Ewan Harrison, David K. Jackson, Ian Johnston, Dominic Kwiatkowski, Cordelia Langford, John Sillitoe on behalf of the Wellcome Sanger Institute COVID-19 Surveillance Team |
| EPI_ISL_2348499 | Megalab, Molecular and Cytogenetics Diagnostics | Department for Virology, Molecular Biology and Genome Research, R. G. Lugar Center for Public Health Research, National Center for Disease Control and Public Health (NCDC) of Georgia. | Giorgi Tomashvili, Giorgi Gogoladze, Gvantsa Brachveli, Meri Pantsulaia, Nino Berishvili, Tata Imnadze, Ana Papkiauri, Gvantsa Chanturia, Ann Machablishvili, Nato Kotaria, Marine Murtskhvaladze, Lela Sabadze, Mari Gavashelidze, Tamar Jashiashvili, Tea Tevdoradze, Ketevan Sidamonidze, Ekaterine Khmaladze, Ekaterine Zhghenti, Roena Sukhiashvili, Mariam Zakalashvili, Lela Urushadze, Magda Dgebuadze, Davit Tsaguria, Nino Chikhovani, Ekaterine Zangaladze, Adam Kotorashvili, Maia Alkhazashvili, Irma Burjanadze, Anna Kasradze, Khatuna Zakhashvili, Paata Imnadze, Amiran Gamkrelidze. |
| EPI_ISL_2348573 | SYNLAB | GIGA Medical Genomics | Keith Durkin, Maria Artesi, Sébastien Bontems, Raphaël Boreux, Bouchra Boujemla, Nathalie Renotte, Cécile Meex, Pierrette Melin, Marie-Pierre Hayette, Vincent Bours |
| EPI_ISL_2348574, EPI_ISL_2348575, EPI_ISL_2348576, EPI_ISL_2348577, EPI_ISL_2348578, EPI_ISL_2348579, EPI_ISL_2348580, EPI_ISL_2348581, EPI_ISL_2348582, EPI_ISL_2348583, EPI_ISL_2348584, EPI_ISL_2348585, EPI_ISL_2348587, EPI_ISL_2348588, EPI_ISL_2348589, EPI_ISL_2348590, EPI_ISL_2348591, EPI_ISL_2348592 | | | |
| see above | Labo Luc Olivier | GIGA Medical Genomics | Keith Durkin, Maria Artesi, Sébastien Bontems, Raphaël Boreux, Bouchra Boujemla, Nathalie Renotte, Cécile Meex, Pierrette Melin, Marie-Pierre Hayette, Vincent Bours |
| EPI_ISL_2348811, EPI_ISL_2348812, EPI_ISL_2348814, EPI_ISL_2348816, EPI_ISL_2348817, EPI_ISL_2348818, EPI_ISL_2348828, EPI_ISL_2348829 | Institute for Urban Disease Control and Prevention | COVID-19 Network Investigations (CONI) Alliance | Elizabeth Batty, Wasun Chantratita, Thanat Chookajorn, Stefan Fernandez, Angkana Huang, Anthony R. Jones, Khajohn Joonlasak, Chonticha Klungtong, Theerarat Kochakarn, Namfon Kotanan, Krittikorn Kumpornsin, Duangkamon Loesbanluechai, Wudtichai Manasatienkij, Bhakbhoom Panthan, Ekawat Pasomsub, Kingkan Rakmanee, Insee Sensorn, Janjira Thaipadungpanit, Arporn Wangwiwatsin, Treewat Watthanachockchai, Kamolthip Atsawawaranunt, Prayuth Kaewmalang, Amornmas Kongklieng, Pukkaporn Parnwijitkul, Vichan Pawun |
| EPI_ISL_2348848, EPI_ISL_2348862 | Cliniques universitaires Saint-Luc | UCLouvain/IREC/MBLG | Jean Ruelle, Ophélie Simon, Nicolas Pinte, Benoit Kabamba Mukadi |
| EPI_ISL_2348923, EPI_ISL_2348924, EPI_ISL_2348925 | Hôpitaux Iris Sud Etterbeek-Ixelles | UCLouvain/IREC/MBLG | Jean Ruelle, Ophélie Simon, Nicolas Pinte, Benoit Kabamba Mukadi |
| EPI_ISL_2349615, EPI_ISL_2349619, EPI_ISL_2349632, EPI_ISL_2349636, EPI_ISL_2349641, EPI_ISL_2349669, EPI_ISL_2349673, EPI_ISL_2349679, EPI_ISL_2349685 | Pandemic Response Lab - NYC | Pandemic Response Lab, R&D | Henry Lee, Michael Hammerling, Melissa Hopkins, Cybill del Castillo, Shinyoung Clair Kang, William Ward, Pradeep Bugga, Sol Rey, Dylan Law, Katharine Nelson, Haiping Hao, Jon Laurent |
| EPI_ISL_2349711, EPI_ISL_2349712, EPI_ISL_2349713, EPI_ISL_2349714, EPI_ISL_2349715, EPI_ISL_2349716, EPI_ISL_2349717, EPI_ISL_2349718, EPI_ISL_2349719, EPI_ISL_2349720, EPI_ISL_2349721, EPI_ISL_2349722, EPI_ISL_2349723, EPI_ISL_2349724, EPI_ISL_2349725, EPI_ISL_2349726, EPI_ISL_2349727, EPI_ISL_2349728, EPI_ISL_2349729, EPI_ISL_2349730, EPI_ISL_2349731, EPI_ISL_2349732, EPI_ISL_2349733, EPI_ISL_2349734, EPI_ISL_2349735, EPI_ISL_2349736, EPI_ISL_2349737, EPI_ISL_2349738, EPI_ISL_2349739, EPI_ISL_2349740, EPI_ISL_2349741, EPI_ISL_2349742, EPI_ISL_2349743, EPI_ISL_2349744, EPI_ISL_2349745, EPI_ISL_2349746, EPI_ISL_2349747, EPI_ISL_2349748, EPI_ISL_2349749, EPI_ISL_2349750, EPI_ISL_2349751, EPI_ISL_2349752, EPI_ISL_2349753, EPI_ISL_2349754, EPI_ISL_2349755, EPI_ISL_2349756, EPI_ISL_2349757, EPI_ISL_2349758, EPI_ISL_2349759, EPI_ISL_2349760, EPI_ISL_2349761, EPI_ISL_2349762, EPI_ISL_2349763, EPI_ISL_2349764, EPI_ISL_2349765, EPI_ISL_2349766, EPI_ISL_2349767, EPI_ISL_2349782, EPI_ISL_2349783, EPI_ISL_2349784, EPI_ISL_2349785, EPI_ISL_2349786, EPI_ISL_2349787, EPI_ISL_2349788, EPI_ISL_2349789, EPI_ISL_2349791 | | | |
| see above | National Public Health Laboratory, National Centre for Infectious Diseases | National Public Health Laboratory, National Centre for Infectious Diseases | Tze Minn Mak, Zhenyang Zhou, Grace Jie Yin Ngan, Royce Ang, Lin Cui, Raymond Tzer Pin Lin |
| EPI_ISL_2349806, EPI_ISL_2349807, EPI_ISL_2349808, EPI_ISL_2349809, EPI_ISL_2349810, EPI_ISL_2349813, EPI_ISL_2349814, EPI_ISL_2349815, EPI_ISL_2349816, EPI_ISL_2349817, EPI_ISL_2349818, EPI_ISL_2349819, EPI_ISL_2349820, EPI_ISL_2349821, EPI_ISL_2349822, EPI_ISL_2349823, EPI_ISL_2349824, EPI_ISL_2349825, EPI_ISL_2349826, EPI_ISL_2349827, EPI_ISL_2349828, EPI_ISL_2349829, EPI_ISL_2349830, EPI_ISL_2349831, EPI_ISL_2349835, EPI_ISL_2349836, EPI_ISL_2349837, EPI_ISL_2349838, EPI_ISL_2349839, EPI_ISL_2349840, EPI_ISL_2349841, EPI_ISL_2349842, EPI_ISL_2349843, EPI_ISL_2349844, EPI_ISL_2349845, EPI_ISL_2349846, EPI_ISL_2349847, EPI_ISL_2349848, EPI_ISL_2349849, EPI_ISL_2349850, EPI_ISL_2349851, EPI_ISL_2349852, EPI_ISL_2349853, EPI_ISL_2349854, EPI_ISL_2349855, EPI_ISL_2349856, EPI_ISL_2349857, EPI_ISL_2349858, EPI_ISL_2349859, EPI_ISL_2349860, EPI_ISL_2349861, EPI_ISL_2349862, EPI_ISL_2349863, EPI_ISL_2349864, EPI_ISL_2349865, EPI_ISL_2349866, EPI_ISL_2349867, EPI_ISL_2349868, EPI_ISL_2349869, EPI_ISL_2349870, EPI_ISL_2349871, EPI_ISL_2349872, EPI_ISL_2349873, EPI_ISL_2349874, EPI_ISL_2349875, EPI_ISL_2349876, EPI_ISL_2349877, EPI_ISL_2349878 | | | |
| see above | National Public Health Laboratory, National Centre for Infectious Diseases | National Public Health Laboratory, National Centre for Infectious Diseases | Tze Minn Mak, Zhenyang Zhou, Royce Ang, Lin Cui, Raymond Tzer Pin Lin |
| EPI_ISL_2349933 | Salud Digna | Instituto Nacional de Medicina Genomica | Hidalgo-Miranda A, Cedro-Tanda A, Mendoza-Vargas A, Reyes-Grajeda JP, Abraham Campos-Romero, Moreno-Camacho José Luis, Rodriguez-Gallegos Jorge, Luna-Ruiz Marco, Gonzalez-Barrera D, Rangel-DeLeon D, Munguia-Garza P, Ramirez-Vega O, Escobar-Arrazola, M, Herrera-Montalvo LA. |
| EPI_ISL_2350787, EPI_ISL_2350789, EPI_ISL_2350790 | Translational Health Science and Technology Institute -ESIC medical college and hospital, Faridabad | THSTI Bioassay laboratory | Naseem Ahmed Khan, Heena Shaman, Janmejay Singh, Kamal Pargai, Jigme Wangchuk, Anil Pandey, Akshay Kanakan, Ranjeet Maurya, Janani Srinivasa Vasudevan, Neha Jha, Priyanka Mehta, Rajesh Pandey, Guruprasad R Medigeshi |
| EPI_ISL_2350905, EPI_ISL_2350908, | Wojewodzka Stacja Sanitarno-Epidemiologiczna w Olsztynie, Laboratorium Badan Epidemiologiczno-Klinicznych | Wojewodzka Stacja Sanitarno-Epidemiologiczna w Olsztynie, Laboratorium Badan Epidemiologiczno-Klinicznych | Sylwia Krzetowska, Monika Czerminska, Ewa Liszewska, Tomasz Jakubczak, Marta Lukian, Patryk Bielecki, Aleksandra Kobiatko, Emilia Tarabasz, Paulina Rozycka, Barbara Dolinska |

| | | | |
|---|---|---|---|
| EPI_ISL_2350914 | | | |
| EPI_ISL_2350915 | Orebro University Hospital | Orebro University Hospital | Sundqvist M et al |
| EPI_ISL_2351289, EPI_ISL_2351330, EPI_ISL_2351357 | Microbiology Department, Laboratori Clínic Metropolitana Nord. Hospital Universitari Germans Trias i Pujol. | Can Ruti SARS-CoV-2 Sequencing Hub (HUGTiP/IrsiCaixa/IGTP) | Marc Noguera-Julian, Pilar Armengol, Ignacio Blanco, Antoni E Bordoy, Francesc Catala-Moll, Pere-Joan Cardona, Maria Casadellà, Cristina Casañ, Gemma Clara, Bonaventura Clotet, Cristina Esteban, Montserrat Giménez, Mercedes Guerrero, Anna Not, Roger Paredes, Mariona Parera, Verónica Saludes, Alba Sánchez, and Elisa Martró on behalf of the Can Ruti SARS-CoV-2 Sequencing Hub. |
| EPI_ISL_2351383 | Althaia. Xarxa Assistencial Universitària de Manresa | IrsiCaixa | Marc Noguera-Julian, Mariona Parera, Maria Casadellà, Pilar Armengol, Francesc Catala-Moll, Roger Paredes, Bonaventura Clotet Gloria Trujillo, Rafel Perez Vidal, Jaume Trapé, Carolina Gonzalez-Fernandez, Miquel Micó, Antonia Flor, Roger Paredes, Eulalia Grau, Bonaventura Clotet |
| EPI_ISL_2352009, EPI_ISL_2352011 | Hospital Universitari Bellvitge | Microbiology Department | Sara Marti, Aida Gonzalez-Diaz, Laura Calatayud, Jordi Niubó, Miguel Fernandez-Huerta, Carmen Ardanuy, Jordi Camara, M Angeles Domínguez |
| EPI_ISL_2352689, EPI_ISL_2352690, EPI_ISL_2352691, EPI_ISL_2352692, EPI_ISL_2352693, EPI_ISL_2352694 | Lighthouse Lab in Milton Keynes | Wellcome Sanger Institute for the COVID-19 Genomics UK (COG-UK) Consortium | The Lighthouse Lab in Milton Keynes and Alex Alderton, Roberto Amato, Jeffrey Barrett, Sonia Goncalves, Ewan Harrison, David K. Jackson, Ian Johnston, Dominic Kwiatkowski, Cordelia Langford, John Sillitoe on behalf of the Wellcome Sanger Institute COVID-19 Surveillance Team |
| EPI_ISL_2352695 | Lighthouse Lab in Alderley Park | Wellcome Sanger Institute for the COVID-19 Genomics UK (COG-UK) Consortium | Jacquelyn Wynn, Mairead Hyland, The Lighthouse Lab in Alderley Park and Alex Alderton, Roberto Amato, Jeffrey Barrett, Sonia Goncalves, Ewan Harrison, David K. Jackson, Ian Johnston, Dominic Kwiatkowski, Cordelia Langford, John Sillitoe on behalf of the Wellcome Sanger Institute COVID-19 Surveillance Team |
| EPI_ISL_2352696, EPI_ISL_2352697 | Lighthouse Lab in Milton Keynes | Wellcome Sanger Institute for the COVID-19 Genomics UK (COG-UK) Consortium | The Lighthouse Lab in Milton Keynes and Alex Alderton, Roberto Amato, Jeffrey Barrett, Sonia Goncalves, Ewan Harrison, David K. Jackson, Ian Johnston, Dominic Kwiatkowski, Cordelia Langford, John Sillitoe on behalf of the Wellcome Sanger Institute COVID-19 Surveillance Team |
| EPI_ISL_2352698 | Lighthouse Lab in Alderley Park | Wellcome Sanger Institute for the COVID-19 Genomics UK (COG-UK) Consortium | Jacquelyn Wynn, Mairead Hyland, The Lighthouse Lab in Alderley Park and Alex Alderton, Roberto Amato, Jeffrey Barrett, Sonia Goncalves, Ewan Harrison, David K. Jackson, Ian Johnston, Dominic Kwiatkowski, Cordelia Langford, John Sillitoe on behalf of the Wellcome Sanger Institute COVID-19 Surveillance Team |
| EPI_ISL_2352699, EPI_ISL_2352700 | Lighthouse Lab in Milton Keynes | Wellcome Sanger Institute for the COVID-19 Genomics UK (COG-UK) Consortium | The Lighthouse Lab in Milton Keynes and Alex Alderton, Roberto Amato, Jeffrey Barrett, Sonia Goncalves, Ewan Harrison, David K. Jackson, Ian Johnston, Dominic Kwiatkowski, Cordelia Langford, John Sillitoe on behalf of the Wellcome Sanger Institute COVID-19 Surveillance Team |
| EPI_ISL_2352701, EPI_ISL_2352702 | Lighthouse Lab in Alderley Park | Wellcome Sanger Institute for the COVID-19 Genomics UK (COG-UK) Consortium | Jacquelyn Wynn, Mairead Hyland, The Lighthouse Lab in Alderley Park and Alex Alderton, Roberto Amato, Jeffrey Barrett, Sonia Goncalves, Ewan Harrison, David K. Jackson, Ian Johnston, Dominic Kwiatkowski, Cordelia Langford, John Sillitoe on behalf of the Wellcome Sanger Institute COVID-19 Surveillance Team |
| EPI_ISL_2352703, EPI_ISL_2352704 | Lighthouse Lab in Milton Keynes | Wellcome Sanger Institute for the COVID-19 Genomics UK (COG-UK) Consortium | The Lighthouse Lab in Milton Keynes and Alex Alderton, Roberto Amato, Jeffrey Barrett, Sonia Goncalves, Ewan Harrison, David K. Jackson, Ian Johnston, Dominic Kwiatkowski, Cordelia Langford, John Sillitoe on behalf of the Wellcome Sanger Institute COVID-19 Surveillance Team |
| EPI_ISL_2352705, EPI_ISL_2352706, EPI_ISL_2352707 | Lighthouse Lab in Alderley Park | Wellcome Sanger Institute for the COVID-19 Genomics UK (COG-UK) Consortium | Jacquelyn Wynn, Mairead Hyland, The Lighthouse Lab in Alderley Park and Alex Alderton, Roberto Amato, Jeffrey Barrett, Sonia Goncalves, Ewan Harrison, David K. Jackson, Ian Johnston, Dominic Kwiatkowski, Cordelia Langford, John Sillitoe on behalf of the Wellcome Sanger Institute COVID-19 Surveillance Team |
| EPI_ISL_2352708 | Lighthouse Lab in Milton Keynes | Wellcome Sanger Institute for the COVID-19 Genomics UK (COG-UK) Consortium | The Lighthouse Lab in Milton Keynes and Alex Alderton, Roberto Amato, Jeffrey Barrett, Sonia Goncalves, Ewan Harrison, David K. Jackson, Ian Johnston, Dominic Kwiatkowski, Cordelia Langford, John Sillitoe on behalf of the Wellcome Sanger Institute COVID-19 Surveillance Team |
| EPI_ISL_2352709 | Lighthouse Lab in Alderley Park | Wellcome Sanger Institute for the COVID-19 Genomics UK (COG-UK) Consortium | Jacquelyn Wynn, Mairead Hyland, The Lighthouse Lab in Alderley Park and Alex Alderton, Roberto Amato, Jeffrey Barrett, Sonia Goncalves, Ewan Harrison, David K. Jackson, Ian Johnston, Dominic Kwiatkowski, Cordelia Langford, John Sillitoe on behalf of the Wellcome Sanger Institute COVID-19 Surveillance Team |
| EPI_ISL_2352710 | Lighthouse Lab in Milton Keynes | Wellcome Sanger Institute for the COVID-19 Genomics UK (COG-UK) Consortium | The Lighthouse Lab in Milton Keynes and Alex Alderton, Roberto Amato, Jeffrey Barrett, Sonia Goncalves, Ewan Harrison, David K. Jackson, Ian Johnston, Dominic Kwiatkowski, Cordelia Langford, John Sillitoe on behalf of the Wellcome Sanger Institute COVID-19 Surveillance Team |
| EPI_ISL_2352711, EPI_ISL_2352712, EPI_ISL_2352713 | Lighthouse Lab in Alderley Park | Wellcome Sanger Institute for the COVID-19 Genomics UK (COG-UK) Consortium | Jacquelyn Wynn, Mairead Hyland, The Lighthouse Lab in Alderley Park and Alex Alderton, Roberto Amato, Jeffrey Barrett, Sonia Goncalves, Ewan Harrison, David K. Jackson, Ian Johnston, Dominic Kwiatkowski, Cordelia Langford, John Sillitoe on behalf of the Wellcome Sanger Institute COVID-19 Surveillance Team |
| EPI_ISL_2352714, EPI_ISL_2352715, EPI_ISL_2352716 | Lighthouse Lab in Milton Keynes | Wellcome Sanger Institute for the COVID-19 Genomics UK (COG-UK) Consortium | The Lighthouse Lab in Milton Keynes and Alex Alderton, Roberto Amato, Jeffrey Barrett, Sonia Goncalves, Ewan Harrison, David K. Jackson, Ian Johnston, Dominic Kwiatkowski, Cordelia Langford, John Sillitoe on behalf of the Wellcome Sanger Institute COVID-19 Surveillance Team |
| EPI_ISL_2352717 | Lighthouse Lab in Alderley Park | Wellcome Sanger Institute for the COVID-19 Genomics UK (COG-UK) Consortium | Jacquelyn Wynn, Mairead Hyland, The Lighthouse Lab in Alderley Park and Alex Alderton, Roberto Amato, Jeffrey Barrett, Sonia Goncalves, Ewan Harrison, David K. Jackson, Ian Johnston, Dominic Kwiatkowski, Cordelia Langford, John Sillitoe on behalf of the Wellcome Sanger Institute COVID-19 Surveillance Team |
| EPI_ISL_2352719 | Lighthouse Lab in Milton Keynes | Wellcome Sanger Institute for the COVID-19 Genomics UK (COG-UK) Consortium | The Lighthouse Lab in Milton Keynes and Alex Alderton, Roberto Amato, Jeffrey Barrett, Sonia Goncalves, Ewan Harrison, David K. Jackson, Ian Johnston, Dominic Kwiatkowski, Cordelia Langford, John Sillitoe on behalf of the Wellcome Sanger Institute COVID-19 Surveillance Team |
| EPI_ISL_2352720 | Lighthouse Lab in Glasgow | Wellcome Sanger Institute for the COVID-19 Genomics UK (COG-UK) Consortium | Harper VanSteenhouse, Yumi Kasai, David Gray, Carol Clugston, Anna Dominiczak and Alex Alderton, Roberto Amato, Jeffrey Barrett, Sonia Goncalves, Ewan Harrison, David K. Jackson, Ian Johnston, Dominic Kwiatkowski, Cordelia Langford, John Sillitoe on behalf of the Wellcome Sanger Institute COVID-19 Surveillance Team |
| EPI_ISL_2352723, EPI_ISL_2352726 | Lighthouse Lab in Milton Keynes | Wellcome Sanger Institute for the COVID-19 Genomics UK (COG-UK) Consortium | The Lighthouse Lab in Milton Keynes and Alex Alderton, Roberto Amato, Jeffrey Barrett, Sonia Goncalves, Ewan Harrison, David K. Jackson, Ian Johnston, Dominic Kwiatkowski, Cordelia Langford, John Sillitoe on behalf of the Wellcome Sanger Institute COVID-19 Surveillance Team |
| EPI_ISL_2352729, EPI_ISL_2352730, EPI_ISL_2352731, EPI_ISL_2352732 | Lighthouse Lab in Alderley Park | Wellcome Sanger Institute for the COVID-19 Genomics UK (COG-UK) Consortium | Jacquelyn Wynn, Mairead Hyland, The Lighthouse Lab in Alderley Park and Alex Alderton, Roberto Amato, Jeffrey Barrett, Sonia Goncalves, Ewan Harrison, David K. Jackson, Ian Johnston, Dominic Kwiatkowski, Cordelia Langford, John Sillitoe on behalf of the Wellcome Sanger Institute COVID-19 Surveillance Team |
| EPI_ISL_2352733 | Lighthouse Lab in Milton Keynes | Wellcome Sanger Institute for the COVID-19 Genomics UK (COG-UK) Consortium | The Lighthouse Lab in Milton Keynes and Alex Alderton, Roberto Amato, Jeffrey Barrett, Sonia Goncalves, Ewan Harrison, David K. Jackson, Ian Johnston, Dominic Kwiatkowski, Cordelia Langford, John Sillitoe on behalf of the Wellcome Sanger Institute COVID-19 Surveillance Team |
| EPI_ISL_2352734, EPI_ISL_2352735 | Lighthouse Lab in Alderley Park | Wellcome Sanger Institute for the COVID-19 Genomics UK (COG-UK) Consortium | Jacquelyn Wynn, Mairead Hyland, The Lighthouse Lab in Alderley Park and Alex Alderton, Roberto Amato, Jeffrey Barrett, Sonia Goncalves, Ewan Harrison, David K. Jackson, Ian Johnston, Dominic Kwiatkowski, Cordelia Langford, John Sillitoe on behalf of the Wellcome Sanger Institute COVID-19 Surveillance Team |
| EPI_ISL_2352736 | Lighthouse Lab in Milton Keynes | Wellcome Sanger Institute for the COVID-19 Genomics UK (COG-UK) Consortium | The Lighthouse Lab in Milton Keynes and Alex Alderton, Roberto Amato, Jeffrey Barrett, Sonia Goncalves, Ewan Harrison, David K. Jackson, Ian Johnston, Dominic Kwiatkowski, Cordelia Langford, John Sillitoe on behalf of the Wellcome Sanger Institute COVID-19 Surveillance Team |
| EPI_ISL_2352737 | Lighthouse Lab in Glasgow | Wellcome Sanger Institute for the COVID-19 Genomics UK (COG-UK) Consortium | Harper VanSteenhouse, Yumi Kasai, David Gray, Carol Clugston, Anna Dominiczak and Alex Alderton, Roberto Amato, Jeffrey Barrett, Sonia Goncalves, Ewan Harrison, David K. Jackson, Ian Johnston, Dominic Kwiatkowski, Cordelia Langford, John Sillitoe on behalf of the Wellcome Sanger Institute COVID-19 Surveillance Team |
| EPI_ISL_2352738 | Lighthouse Lab in Milton Keynes | Wellcome Sanger Institute for the COVID-19 Genomics UK (COG-UK) Consortium | The Lighthouse Lab in Milton Keynes and Alex Alderton, Roberto Amato, Jeffrey Barrett, Sonia Goncalves, Ewan Harrison, David K. Jackson, Ian Johnston, Dominic Kwiatkowski, Cordelia Langford, John Sillitoe on behalf of the Wellcome Sanger Institute COVID-19 Surveillance Team |
| EPI_ISL_2352739 | Lighthouse Lab in Glasgow | Wellcome Sanger Institute for the COVID-19 Genomics UK (COG-UK) Consortium | Harper VanSteenhouse, Yumi Kasai, David Gray, Carol Clugston, Anna Dominiczak and Alex Alderton, Roberto Amato, Jeffrey Barrett, Sonia Goncalves, Ewan Harrison, David K. Jackson, Ian Johnston, Dominic Kwiatkowski, Cordelia Langford, John Sillitoe on behalf of the Wellcome Sanger Institute COVID-19 Surveillance Team |
| EPI_ISL_2352740, EPI_ISL_2352741 | Lighthouse Lab in Alderley Park | Wellcome Sanger Institute for the COVID-19 Genomics UK (COG-UK) Consortium | Jacquelyn Wynn, Mairead Hyland, The Lighthouse Lab in Alderley Park and Alex Alderton, Roberto Amato, Jeffrey Barrett, Sonia Goncalves, Ewan Harrison, David K. Jackson, Ian Johnston, Dominic Kwiatkowski, Cordelia Langford, John Sillitoe on behalf of the Wellcome Sanger Institute COVID-19 Surveillance Team |
| EPI_ISL_2352742 | Lighthouse Lab in Glasgow | Wellcome Sanger Institute for the COVID-19 Genomics UK (COG-UK) Consortium | Harper VanSteenhouse, Yumi Kasai, David Gray, Carol Clugston, Anna Dominiczak and Alex Alderton, Roberto Amato, Jeffrey Barrett, Sonia Goncalves, Ewan Harrison, David K. Jackson, Ian Johnston, Dominic Kwiatkowski, Cordelia Langford, John Sillitoe on behalf of the Wellcome Sanger Institute COVID-19 Surveillance Team |
| EPI_ISL_2352743 | Lighthouse Lab in Alderley Park | Wellcome Sanger Institute for the COVID-19 Genomics UK (COG-UK) Consortium | Jacquelyn Wynn, Mairead Hyland, The Lighthouse Lab in Alderley Park and Alex Alderton, Roberto Amato, Jeffrey Barrett, Sonia Goncalves, Ewan Harrison, David K. Jackson, Ian Johnston, Dominic Kwiatkowski, Cordelia Langford, John Sillitoe on behalf of the Wellcome Sanger Institute COVID-19 Surveillance Team |
| EPI_ISL_2352745, EPI_ISL_2352746, EPI_ISL_2352748, EPI_ISL_2352749 | Lighthouse Lab in Milton Keynes | Wellcome Sanger Institute for the COVID-19 Genomics UK (COG-UK) Consortium | The Lighthouse Lab in Milton Keynes and Alex Alderton, Roberto Amato, Jeffrey Barrett, Sonia Goncalves, Ewan Harrison, David K. Jackson, Ian Johnston, Dominic Kwiatkowski, Cordelia Langford, John Sillitoe on behalf of the Wellcome Sanger Institute COVID-19 Surveillance Team |
| EPI_ISL_2352751, EPI_ISL_2352752, EPI_ISL_2352753, EPI_ISL_2352756, EPI_ISL_2352757 | Lighthouse Lab in Alderley Park | Wellcome Sanger Institute for the COVID-19 Genomics UK (COG-UK) Consortium | Jacquelyn Wynn, Mairead Hyland, The Lighthouse Lab in Alderley Park and Alex Alderton, Roberto Amato, Jeffrey Barrett, Sonia Goncalves, Ewan Harrison, David K. Jackson, Ian Johnston, Dominic Kwiatkowski, Cordelia Langford, John Sillitoe on behalf of the Wellcome Sanger Institute COVID-19 Surveillance Team |
| EPI_ISL_2352758, EPI_ISL_2352759, EPI_ISL_2352760 | Lighthouse Lab in Milton Keynes | Wellcome Sanger Institute for the COVID-19 Genomics UK (COG-UK) Consortium | The Lighthouse Lab in Milton Keynes and Alex Alderton, Roberto Amato, Jeffrey Barrett, Sonia Goncalves, Ewan Harrison, David K. Jackson, Ian Johnston, Dominic Kwiatkowski, Cordelia Langford, John Sillitoe on behalf of the Wellcome Sanger Institute COVID-19 Surveillance Team |
| EPI_ISL_2352761 | Lighthouse Lab in Alderley Park | Wellcome Sanger Institute for the COVID-19 Genomics UK (COG-UK) Consortium | Jacquelyn Wynn, Mairead Hyland, The Lighthouse Lab in Alderley Park and Alex Alderton, Roberto Amato, Jeffrey Barrett, Sonia Goncalves, Ewan Harrison, David K. Jackson, Ian Johnston, Dominic Kwiatkowski, Cordelia Langford, John Sillitoe on behalf of the Wellcome Sanger Institute COVID-19 Surveillance Team |
| EPI_ISL_2352763 | Lighthouse Lab in Milton Keynes | Wellcome Sanger Institute for the COVID-19 Genomics UK (COG-UK) Consortium | The Lighthouse Lab in Milton Keynes and Alex Alderton, Roberto Amato, Jeffrey Barrett, Sonia Goncalves, Ewan Harrison, David K. Jackson, Ian Johnston, Dominic Kwiatkowski, Cordelia Langford, John Sillitoe on behalf of the Wellcome Sanger Institute COVID-19 Surveillance Team |
| EPI_ISL_2352764, EPI_ISL_2352765 | Lighthouse Lab in Alderley Park | Wellcome Sanger Institute for the COVID-19 Genomics UK (COG-UK) Consortium | Jacquelyn Wynn, Mairead Hyland, The Lighthouse Lab in Alderley Park and Alex Alderton, Roberto Amato, Jeffrey Barrett, Sonia Goncalves, Ewan Harrison, David K. Jackson, Ian Johnston, Dominic Kwiatkowski, Cordelia Langford, John Sillitoe on behalf of the Wellcome Sanger Institute COVID-19 Surveillance Team |
| EPI_ISL_2352767, EPI_ISL_2352768 | Lighthouse Lab in Milton Keynes | Wellcome Sanger Institute for the COVID-19 Genomics UK (COG-UK) Consortium | The Lighthouse Lab in Milton Keynes and Alex Alderton, Roberto Amato, Jeffrey Barrett, Sonia Goncalves, Ewan Harrison, David K. Jackson, Ian Johnston, Dominic Kwiatkowski, Cordelia Langford, John Sillitoe on behalf of the Wellcome Sanger Institute COVID-19 Surveillance Team |
| EPI_ISL_2352769 | Lighthouse Lab in Alderley Park | Wellcome Sanger Institute for the COVID-19 Genomics UK (COG-UK) | Jacquelyn Wynn, Mairead Hyland, The Lighthouse Lab in Alderley Park and Alex Alderton, Roberto Amato, Jeffrey Barrett, Sonia Goncalves, Ewan Harrison, David K. Jackson, Ian Johnston, Dominic Kwiatkowski, Cordelia Langford, John Sillitoe on |

| | | | |
|---|---|---|---|
| | | Consortium | behalf of the Wellcome Sanger Institute COVID-19 Surveillance Team |
| EPI_ISL_2352770, EPI_ISL_2352771 | Lighthouse Lab in Milton Keynes | Wellcome Sanger Institute for the COVID-19 Genomics UK (COG-UK) Consortium | The Lighthouse Lab in Milton Keynes and Alex Alderton, Roberto Amato, Jeffrey Barrett, Sonia Goncalves, Ewan Harrison, David K. Jackson, Ian Johnston, Dominic Kwiatkowski, Cordelia Langford, John Sillitoe on behalf of the Wellcome Sanger Institute COVID-19 Surveillance Team |
| EPI_ISL_2352772, EPI_ISL_2352774, EPI_ISL_2352775 | Lighthouse Lab in Alderley Park | Wellcome Sanger Institute for the COVID-19 Genomics UK (COG-UK) Consortium | Jacquelyn Wynn, Mairead Hyland, The Lighthouse Lab in Alderley Park and Alex Alderton, Roberto Amato, Jeffrey Barrett, Sonia Goncalves, Ewan Harrison, David K. Jackson, Ian Johnston, Dominic Kwiatkowski, Cordelia Langford, John Sillitoe on behalf of the Wellcome Sanger Institute COVID-19 Surveillance Team |
| EPI_ISL_2352777, EPI_ISL_2352782, EPI_ISL_2352783 | Lighthouse Lab in Milton Keynes | Wellcome Sanger Institute for the COVID-19 Genomics UK (COG-UK) Consortium | The Lighthouse Lab in Milton Keynes and Alex Alderton, Roberto Amato, Jeffrey Barrett, Sonia Goncalves, Ewan Harrison, David K. Jackson, Ian Johnston, Dominic Kwiatkowski, Cordelia Langford, John Sillitoe on behalf of the Wellcome Sanger Institute COVID-19 Surveillance Team |
| EPI_ISL_2352784, EPI_ISL_2352785, EPI_ISL_2352786 | Lighthouse Lab in Alderley Park | Wellcome Sanger Institute for the COVID-19 Genomics UK (COG-UK) Consortium | Jacquelyn Wynn, Mairead Hyland, The Lighthouse Lab in Alderley Park and Alex Alderton, Roberto Amato, Jeffrey Barrett, Sonia Goncalves, Ewan Harrison, David K. Jackson, Ian Johnston, Dominic Kwiatkowski, Cordelia Langford, John Sillitoe on behalf of the Wellcome Sanger Institute COVID-19 Surveillance Team |
| EPI_ISL_2352787, EPI_ISL_2352788, EPI_ISL_2352789, EPI_ISL_2352790 | Lighthouse Lab in Milton Keynes | Wellcome Sanger Institute for the COVID-19 Genomics UK (COG-UK) Consortium | The Lighthouse Lab in Milton Keynes and Alex Alderton, Roberto Amato, Jeffrey Barrett, Sonia Goncalves, Ewan Harrison, David K. Jackson, Ian Johnston, Dominic Kwiatkowski, Cordelia Langford, John Sillitoe on behalf of the Wellcome Sanger Institute COVID-19 Surveillance Team |
| EPI_ISL_2352791 | Lighthouse Lab in Alderley Park | Wellcome Sanger Institute for the COVID-19 Genomics UK (COG-UK) Consortium | Jacquelyn Wynn, Mairead Hyland, The Lighthouse Lab in Alderley Park and Alex Alderton, Roberto Amato, Jeffrey Barrett, Sonia Goncalves, Ewan Harrison, David K. Jackson, Ian Johnston, Dominic Kwiatkowski, Cordelia Langford, John Sillitoe on behalf of the Wellcome Sanger Institute COVID-19 Surveillance Team |
| EPI_ISL_2352792, EPI_ISL_2352793, EPI_ISL_2352795, EPI_ISL_2352796, EPI_ISL_2352798 | Lighthouse Lab in Milton Keynes | Wellcome Sanger Institute for the COVID-19 Genomics UK (COG-UK) Consortium | The Lighthouse Lab in Milton Keynes and Alex Alderton, Roberto Amato, Jeffrey Barrett, Sonia Goncalves, Ewan Harrison, David K. Jackson, Ian Johnston, Dominic Kwiatkowski, Cordelia Langford, John Sillitoe on behalf of the Wellcome Sanger Institute COVID-19 Surveillance Team |
| EPI_ISL_2352799 | Lighthouse Lab in Alderley Park | Wellcome Sanger Institute for the COVID-19 Genomics UK (COG-UK) Consortium | Jacquelyn Wynn, Mairead Hyland, The Lighthouse Lab in Alderley Park and Alex Alderton, Roberto Amato, Jeffrey Barrett, Sonia Goncalves, Ewan Harrison, David K. Jackson, Ian Johnston, Dominic Kwiatkowski, Cordelia Langford, John Sillitoe on behalf of the Wellcome Sanger Institute COVID-19 Surveillance Team |
| EPI_ISL_2352800 | Lighthouse Lab in Glasgow | Wellcome Sanger Institute for the COVID-19 Genomics UK (COG-UK) Consortium | Harper VanSteenhouse, Yumi Kasai, David Gray, Carol Clugston, Anna Dominiczak and Alex Alderton, Roberto Amato, Jeffrey Barrett, Sonia Goncalves, Ewan Harrison, David K. Jackson, Ian Johnston, Dominic Kwiatkowski, Cordelia Langford, John Sillitoe on behalf of the Wellcome Sanger Institute COVID-19 Surveillance Team |
| EPI_ISL_2352801 | Lighthouse Lab in Alderley Park | Wellcome Sanger Institute for the COVID-19 Genomics UK (COG-UK) Consortium | Jacquelyn Wynn, Mairead Hyland, The Lighthouse Lab in Alderley Park and Alex Alderton, Roberto Amato, Jeffrey Barrett, Sonia Goncalves, Ewan Harrison, David K. Jackson, Ian Johnston, Dominic Kwiatkowski, Cordelia Langford, John Sillitoe on behalf of the Wellcome Sanger Institute COVID-19 Surveillance Team |
| EPI_ISL_2352802, EPI_ISL_2352803 | Lighthouse Lab in Milton Keynes | Wellcome Sanger Institute for the COVID-19 Genomics UK (COG-UK) Consortium | The Lighthouse Lab in Milton Keynes and Alex Alderton, Roberto Amato, Jeffrey Barrett, Sonia Goncalves, Ewan Harrison, David K. Jackson, Ian Johnston, Dominic Kwiatkowski, Cordelia Langford, John Sillitoe on behalf of the Wellcome Sanger Institute COVID-19 Surveillance Team |
| EPI_ISL_2352804 | Lighthouse Lab in Glasgow | Wellcome Sanger Institute for the COVID-19 Genomics UK (COG-UK) Consortium | Harper VanSteenhouse, Yumi Kasai, David Gray, Carol Clugston, Anna Dominiczak and Alex Alderton, Roberto Amato, Jeffrey Barrett, Sonia Goncalves, Ewan Harrison, David K. Jackson, Ian Johnston, Dominic Kwiatkowski, Cordelia Langford, John Sillitoe on behalf of the Wellcome Sanger Institute COVID-19 Surveillance Team |
| EPI_ISL_2352805 | Lighthouse Lab in Alderley Park | Wellcome Sanger Institute for the COVID-19 Genomics UK (COG-UK) Consortium | Jacquelyn Wynn, Mairead Hyland, The Lighthouse Lab in Alderley Park and Alex Alderton, Roberto Amato, Jeffrey Barrett, Sonia Goncalves, Ewan Harrison, David K. Jackson, Ian Johnston, Dominic Kwiatkowski, Cordelia Langford, John Sillitoe on behalf of the Wellcome Sanger Institute COVID-19 Surveillance Team |
| EPI_ISL_2352807 | Lighthouse Lab in Glasgow | Wellcome Sanger Institute for the COVID-19 Genomics UK (COG-UK) Consortium | Harper VanSteenhouse, Yumi Kasai, David Gray, Carol Clugston, Anna Dominiczak and Alex Alderton, Roberto Amato, Jeffrey Barrett, Sonia Goncalves, Ewan Harrison, David K. Jackson, Ian Johnston, Dominic Kwiatkowski, Cordelia Langford, John Sillitoe on behalf of the Wellcome Sanger Institute COVID-19 Surveillance Team |
| EPI_ISL_2352808 | Lighthouse Lab in Alderley Park | Wellcome Sanger Institute for the COVID-19 Genomics UK (COG-UK) Consortium | Jacquelyn Wynn, Mairead Hyland, The Lighthouse Lab in Alderley Park and Alex Alderton, Roberto Amato, Jeffrey Barrett, Sonia Goncalves, Ewan Harrison, David K. Jackson, Ian Johnston, Dominic Kwiatkowski, Cordelia Langford, John Sillitoe on behalf of the Wellcome Sanger Institute COVID-19 Surveillance Team |
| EPI_ISL_2352810, EPI_ISL_2352811, EPI_ISL_2352812 | Lighthouse Lab in Milton Keynes | Wellcome Sanger Institute for the COVID-19 Genomics UK (COG-UK) Consortium | The Lighthouse Lab in Milton Keynes and Alex Alderton, Roberto Amato, Jeffrey Barrett, Sonia Goncalves, Ewan Harrison, David K. Jackson, Ian Johnston, Dominic Kwiatkowski, Cordelia Langford, John Sillitoe on behalf of the Wellcome Sanger Institute COVID-19 Surveillance Team |
| EPI_ISL_2352813 | Lighthouse Lab in Alderley Park | Wellcome Sanger Institute for the COVID-19 Genomics UK (COG-UK) Consortium | Jacquelyn Wynn, Mairead Hyland, The Lighthouse Lab in Alderley Park and Alex Alderton, Roberto Amato, Jeffrey Barrett, Sonia Goncalves, Ewan Harrison, David K. Jackson, Ian Johnston, Dominic Kwiatkowski, Cordelia Langford, John Sillitoe on behalf of the Wellcome Sanger Institute COVID-19 Surveillance Team |
| EPI_ISL_2352814 | Lighthouse Lab in Glasgow | Wellcome Sanger Institute for the COVID-19 Genomics UK (COG-UK) Consortium | Harper VanSteenhouse, Yumi Kasai, David Gray, Carol Clugston, Anna Dominiczak and Alex Alderton, Roberto Amato, Jeffrey Barrett, Sonia Goncalves, Ewan Harrison, David K. Jackson, Ian Johnston, Dominic Kwiatkowski, Cordelia Langford, John Sillitoe on behalf of the Wellcome Sanger Institute COVID-19 Surveillance Team |
| EPI_ISL_2352815 | Lighthouse Lab in Alderley Park | Wellcome Sanger Institute for the COVID-19 Genomics UK (COG-UK) Consortium | Jacquelyn Wynn, Mairead Hyland, The Lighthouse Lab in Alderley Park and Alex Alderton, Roberto Amato, Jeffrey Barrett, Sonia Goncalves, Ewan Harrison, David K. Jackson, Ian Johnston, Dominic Kwiatkowski, Cordelia Langford, John Sillitoe on behalf of the Wellcome Sanger Institute COVID-19 Surveillance Team |
| EPI_ISL_2352816, EPI_ISL_2352817 | Lighthouse Lab in Milton Keynes | Wellcome Sanger Institute for the COVID-19 Genomics UK (COG-UK) Consortium | The Lighthouse Lab in Milton Keynes and Alex Alderton, Roberto Amato, Jeffrey Barrett, Sonia Goncalves, Ewan Harrison, David K. Jackson, Ian Johnston, Dominic Kwiatkowski, Cordelia Langford, John Sillitoe on behalf of the Wellcome Sanger Institute COVID-19 Surveillance Team |
| EPI_ISL_2352818 | Lighthouse Lab in Alderley Park | Wellcome Sanger Institute for the COVID-19 Genomics UK (COG-UK) Consortium | Jacquelyn Wynn, Mairead Hyland, The Lighthouse Lab in Alderley Park and Alex Alderton, Roberto Amato, Jeffrey Barrett, Sonia Goncalves, Ewan Harrison, David K. Jackson, Ian Johnston, Dominic Kwiatkowski, Cordelia Langford, John Sillitoe on behalf of the Wellcome Sanger Institute COVID-19 Surveillance Team |
| EPI_ISL_2352819 | Lighthouse Lab in Milton Keynes | Wellcome Sanger Institute for the COVID-19 Genomics UK (COG-UK) Consortium | The Lighthouse Lab in Milton Keynes and Alex Alderton, Roberto Amato, Jeffrey Barrett, Sonia Goncalves, Ewan Harrison, David K. Jackson, Ian Johnston, Dominic Kwiatkowski, Cordelia Langford, John Sillitoe on behalf of the Wellcome Sanger Institute COVID-19 Surveillance Team |
| EPI_ISL_2352821 | Lighthouse Lab in Glasgow | Wellcome Sanger Institute for the COVID-19 Genomics UK (COG-UK) Consortium | Harper VanSteenhouse, Yumi Kasai, David Gray, Carol Clugston, Anna Dominiczak and Alex Alderton, Roberto Amato, Jeffrey Barrett, Sonia Goncalves, Ewan Harrison, David K. Jackson, Ian Johnston, Dominic Kwiatkowski, Cordelia Langford, John Sillitoe on behalf of the Wellcome Sanger Institute COVID-19 Surveillance Team |
| EPI_ISL_2352824, EPI_ISL_2352825 | Lighthouse Lab in Milton Keynes | Wellcome Sanger Institute for the COVID-19 Genomics UK (COG-UK) Consortium | The Lighthouse Lab in Milton Keynes and Alex Alderton, Roberto Amato, Jeffrey Barrett, Sonia Goncalves, Ewan Harrison, David K. Jackson, Ian Johnston, Dominic Kwiatkowski, Cordelia Langford, John Sillitoe on behalf of the Wellcome Sanger Institute COVID-19 Surveillance Team |
| EPI_ISL_2352827 | Lighthouse Lab in Alderley Park | Wellcome Sanger Institute for the COVID-19 Genomics UK (COG-UK) Consortium | Jacquelyn Wynn, Mairead Hyland, The Lighthouse Lab in Alderley Park and Alex Alderton, Roberto Amato, Jeffrey Barrett, Sonia Goncalves, Ewan Harrison, David K. Jackson, Ian Johnston, Dominic Kwiatkowski, Cordelia Langford, John Sillitoe on behalf of the Wellcome Sanger Institute COVID-19 Surveillance Team |
| EPI_ISL_2352828, EPI_ISL_2352831, EPI_ISL_2352832 | Lighthouse Lab in Milton Keynes | Wellcome Sanger Institute for the COVID-19 Genomics UK (COG-UK) Consortium | The Lighthouse Lab in Milton Keynes and Alex Alderton, Roberto Amato, Jeffrey Barrett, Sonia Goncalves, Ewan Harrison, David K. Jackson, Ian Johnston, Dominic Kwiatkowski, Cordelia Langford, John Sillitoe on behalf of the Wellcome Sanger Institute COVID-19 Surveillance Team |
| EPI_ISL_2352833 | Lighthouse Lab in Alderley Park | Wellcome Sanger Institute for the COVID-19 Genomics UK (COG-UK) Consortium | Jacquelyn Wynn, Mairead Hyland, The Lighthouse Lab in Alderley Park and Alex Alderton, Roberto Amato, Jeffrey Barrett, Sonia Goncalves, Ewan Harrison, David K. Jackson, Ian Johnston, Dominic Kwiatkowski, Cordelia Langford, John Sillitoe on behalf of the Wellcome Sanger Institute COVID-19 Surveillance Team |
| EPI_ISL_2352834 | Lighthouse Lab in Glasgow | Wellcome Sanger Institute for the COVID-19 Genomics UK (COG-UK) Consortium | Harper VanSteenhouse, Yumi Kasai, David Gray, Carol Clugston, Anna Dominiczak and Alex Alderton, Roberto Amato, Jeffrey Barrett, Sonia Goncalves, Ewan Harrison, David K. Jackson, Ian Johnston, Dominic Kwiatkowski, Cordelia Langford, John Sillitoe on behalf of the Wellcome Sanger Institute COVID-19 Surveillance Team |
| EPI_ISL_2352835 | Lighthouse Lab in Milton Keynes | Wellcome Sanger Institute for the COVID-19 Genomics UK (COG-UK) Consortium | The Lighthouse Lab in Milton Keynes and Alex Alderton, Roberto Amato, Jeffrey Barrett, Sonia Goncalves, Ewan Harrison, David K. Jackson, Ian Johnston, Dominic Kwiatkowski, Cordelia Langford, John Sillitoe on behalf of the Wellcome Sanger Institute COVID-19 Surveillance Team |
| EPI_ISL_2352836 | Lighthouse Lab in Alderley Park | Wellcome Sanger Institute for the COVID-19 Genomics UK (COG-UK) Consortium | Jacquelyn Wynn, Mairead Hyland, The Lighthouse Lab in Alderley Park and Alex Alderton, Roberto Amato, Jeffrey Barrett, Sonia Goncalves, Ewan Harrison, David K. Jackson, Ian Johnston, Dominic Kwiatkowski, Cordelia Langford, John Sillitoe on behalf of the Wellcome Sanger Institute COVID-19 Surveillance Team |
| EPI_ISL_2352837, EPI_ISL_2352838, EPI_ISL_2352841, EPI_ISL_2352842 | Lighthouse Lab in Milton Keynes | Wellcome Sanger Institute for the COVID-19 Genomics UK (COG-UK) Consortium | The Lighthouse Lab in Milton Keynes and Alex Alderton, Roberto Amato, Jeffrey Barrett, Sonia Goncalves, Ewan Harrison, David K. Jackson, Ian Johnston, Dominic Kwiatkowski, Cordelia Langford, John Sillitoe on behalf of the Wellcome Sanger Institute COVID-19 Surveillance Team |
| EPI_ISL_2352843, EPI_ISL_2352844, EPI_ISL_2352845 | Lighthouse Lab in Alderley Park | Wellcome Sanger Institute for the COVID-19 Genomics UK (COG-UK) Consortium | Jacquelyn Wynn, Mairead Hyland, The Lighthouse Lab in Alderley Park and Alex Alderton, Roberto Amato, Jeffrey Barrett, Sonia Goncalves, Ewan Harrison, David K. Jackson, Ian Johnston, Dominic Kwiatkowski, Cordelia Langford, John Sillitoe on behalf of the Wellcome Sanger Institute COVID-19 Surveillance Team |
| EPI_ISL_2352846, EPI_ISL_2352847 | Lighthouse Lab in Milton Keynes | Wellcome Sanger Institute for the COVID-19 Genomics UK (COG-UK) Consortium | The Lighthouse Lab in Milton Keynes and Alex Alderton, Roberto Amato, Jeffrey Barrett, Sonia Goncalves, Ewan Harrison, David K. Jackson, Ian Johnston, Dominic Kwiatkowski, Cordelia Langford, John Sillitoe on behalf of the Wellcome Sanger Institute COVID-19 Surveillance Team |
| EPI_ISL_2352848 | Lighthouse Lab in Alderley Park | Wellcome Sanger Institute for the COVID-19 Genomics UK (COG-UK) Consortium | Jacquelyn Wynn, Mairead Hyland, The Lighthouse Lab in Alderley Park and Alex Alderton, Roberto Amato, Jeffrey Barrett, Sonia Goncalves, Ewan Harrison, David K. Jackson, Ian Johnston, Dominic Kwiatkowski, Cordelia Langford, John Sillitoe on behalf of the Wellcome Sanger Institute COVID-19 Surveillance Team |
| EPI_ISL_2352852 | Lighthouse Lab in Milton Keynes | Wellcome Sanger Institute for the COVID-19 Genomics UK (COG-UK) Consortium | The Lighthouse Lab in Milton Keynes and Alex Alderton, Roberto Amato, Jeffrey Barrett, Sonia Goncalves, Ewan Harrison, David K. Jackson, Ian Johnston, Dominic Kwiatkowski, Cordelia Langford, John Sillitoe on behalf of the Wellcome Sanger Institute COVID-19 Surveillance Team |
| EPI_ISL_2352853 | Lighthouse Lab in Alderley Park | Wellcome Sanger Institute for the COVID-19 Genomics UK (COG-UK) Consortium | Jacquelyn Wynn, Mairead Hyland, The Lighthouse Lab in Alderley Park and Alex Alderton, Roberto Amato, Jeffrey Barrett, Sonia Goncalves, Ewan Harrison, David K. Jackson, Ian Johnston, Dominic Kwiatkowski, Cordelia Langford, John Sillitoe on behalf of the Wellcome Sanger Institute COVID-19 Surveillance Team |
| EPI_ISL_2352856, EPI_ISL_2352857, EPI_ISL_2352859, EPI_ISL_2352860 | Lighthouse Lab in Milton Keynes | Wellcome Sanger Institute for the COVID-19 Genomics UK (COG-UK) Consortium | The Lighthouse Lab in Milton Keynes and Alex Alderton, Roberto Amato, Jeffrey Barrett, Sonia Goncalves, Ewan Harrison, David K. Jackson, Ian Johnston, Dominic Kwiatkowski, Cordelia Langford, John Sillitoe on behalf of the Wellcome Sanger Institute COVID-19 Surveillance Team |
| EPI_ISL_2352861 | Lighthouse Lab in Alderley Park | Wellcome Sanger Institute for the COVID-19 Genomics UK (COG-UK) Consortium | Jacquelyn Wynn, Mairead Hyland, The Lighthouse Lab in Alderley Park and Alex Alderton, Roberto Amato, Jeffrey Barrett, Sonia Goncalves, Ewan Harrison, David K. Jackson, Ian Johnston, Dominic Kwiatkowski, Cordelia Langford, John Sillitoe on behalf of the Wellcome Sanger Institute COVID-19 Surveillance Team |

| | | | |
|---|---|---|---|
| EPI_ISL_2352863 | Lighthouse Lab in Glasgow | Wellcome Sanger Institute for the COVID-19 Genomics UK (COG-UK) Consortium | Harper VanSteenhouse, Yumi Kasai, David Gray, Carol Clugston, Anna Dominiczak and Alex Alderton, Roberto Amato, Jeffrey Barrett, Sonia Goncalves, Ewan Harrison, David K. Jackson, Ian Johnston, Dominic Kwiatkowski, Cordelia Langford, John Sillitoe on behalf of the Wellcome Sanger Institute COVID-19 Surveillance Team |
| EPI_ISL_2352864 | Lighthouse Lab in Milton Keynes | Wellcome Sanger Institute for the COVID-19 Genomics UK (COG-UK) Consortium | The Lighthouse Lab in Milton Keynes and Alex Alderton, Roberto Amato, Jeffrey Barrett, Sonia Goncalves, Ewan Harrison, David K. Jackson, Ian Johnston, Dominic Kwiatkowski, Cordelia Langford, John Sillitoe on behalf of the Wellcome Sanger Institute COVID-19 Surveillance Team |
| EPI_ISL_2352865 | Lighthouse Lab in Glasgow | Wellcome Sanger Institute for the COVID-19 Genomics UK (COG-UK) Consortium | Harper VanSteenhouse, Yumi Kasai, David Gray, Carol Clugston, Anna Dominiczak and Alex Alderton, Roberto Amato, Jeffrey Barrett, Sonia Goncalves, Ewan Harrison, David K. Jackson, Ian Johnston, Dominic Kwiatkowski, Cordelia Langford, John Sillitoe on behalf of the Wellcome Sanger Institute COVID-19 Surveillance Team |
| EPI_ISL_2352866 | Lighthouse Lab in Alderley Park | Wellcome Sanger Institute for the COVID-19 Genomics UK (COG-UK) Consortium | Jacquelyn Wynn, Mairead Hyland, The Lighthouse Lab in Alderley Park and Alex Alderton, Roberto Amato, Jeffrey Barrett, Sonia Goncalves, Ewan Harrison, David K. Jackson, Ian Johnston, Dominic Kwiatkowski, Cordelia Langford, John Sillitoe on behalf of the Wellcome Sanger Institute COVID-19 Surveillance Team |
| EPI_ISL_2352867, EPI_ISL_2352868, EPI_ISL_2352869 | Lighthouse Lab in Milton Keynes | Wellcome Sanger Institute for the COVID-19 Genomics UK (COG-UK) Consortium | The Lighthouse Lab in Milton Keynes and Alex Alderton, Roberto Amato, Jeffrey Barrett, Sonia Goncalves, Ewan Harrison, David K. Jackson, Ian Johnston, Dominic Kwiatkowski, Cordelia Langford, John Sillitoe on behalf of the Wellcome Sanger Institute COVID-19 Surveillance Team |
| EPI_ISL_2352870 | Lighthouse Lab in Alderley Park | Wellcome Sanger Institute for the COVID-19 Genomics UK (COG-UK) Consortium | Jacquelyn Wynn, Mairead Hyland, The Lighthouse Lab in Alderley Park and Alex Alderton, Roberto Amato, Jeffrey Barrett, Sonia Goncalves, Ewan Harrison, David K. Jackson, Ian Johnston, Dominic Kwiatkowski, Cordelia Langford, John Sillitoe on behalf of the Wellcome Sanger Institute COVID-19 Surveillance Team |
| EPI_ISL_2352871 | Lighthouse Lab in Glasgow | Wellcome Sanger Institute for the COVID-19 Genomics UK (COG-UK) Consortium | Harper VanSteenhouse, Yumi Kasai, David Gray, Carol Clugston, Anna Dominiczak and Alex Alderton, Roberto Amato, Jeffrey Barrett, Sonia Goncalves, Ewan Harrison, David K. Jackson, Ian Johnston, Dominic Kwiatkowski, Cordelia Langford, John Sillitoe on behalf of the Wellcome Sanger Institute COVID-19 Surveillance Team |
| EPI_ISL_2352872, EPI_ISL_2352873, EPI_ISL_2352875, EPI_ISL_2352876 | Lighthouse Lab in Alderley Park | Wellcome Sanger Institute for the COVID-19 Genomics UK (COG-UK) Consortium | Jacquelyn Wynn, Mairead Hyland, The Lighthouse Lab in Alderley Park and Alex Alderton, Roberto Amato, Jeffrey Barrett, Sonia Goncalves, Ewan Harrison, David K. Jackson, Ian Johnston, Dominic Kwiatkowski, Cordelia Langford, John Sillitoe on behalf of the Wellcome Sanger Institute COVID-19 Surveillance Team |
| EPI_ISL_2352878, EPI_ISL_2352879, EPI_ISL_2352880, EPI_ISL_2352881 | Lighthouse Lab in Milton Keynes | Wellcome Sanger Institute for the COVID-19 Genomics UK (COG-UK) Consortium | The Lighthouse Lab in Milton Keynes and Alex Alderton, Roberto Amato, Jeffrey Barrett, Sonia Goncalves, Ewan Harrison, David K. Jackson, Ian Johnston, Dominic Kwiatkowski, Cordelia Langford, John Sillitoe on behalf of the Wellcome Sanger Institute COVID-19 Surveillance Team |
| EPI_ISL_2352882, EPI_ISL_2352884, EPI_ISL_2352885 | Lighthouse Lab in Alderley Park | Wellcome Sanger Institute for the COVID-19 Genomics UK (COG-UK) Consortium | Jacquelyn Wynn, Mairead Hyland, The Lighthouse Lab in Alderley Park and Alex Alderton, Roberto Amato, Jeffrey Barrett, Sonia Goncalves, Ewan Harrison, David K. Jackson, Ian Johnston, Dominic Kwiatkowski, Cordelia Langford, John Sillitoe on behalf of the Wellcome Sanger Institute COVID-19 Surveillance Team |
| EPI_ISL_2352886, EPI_ISL_2352887, EPI_ISL_2352888 | Lighthouse Lab in Milton Keynes | Wellcome Sanger Institute for the COVID-19 Genomics UK (COG-UK) Consortium | The Lighthouse Lab in Milton Keynes and Alex Alderton, Roberto Amato, Jeffrey Barrett, Sonia Goncalves, Ewan Harrison, David K. Jackson, Ian Johnston, Dominic Kwiatkowski, Cordelia Langford, John Sillitoe on behalf of the Wellcome Sanger Institute COVID-19 Surveillance Team |
| EPI_ISL_2352889 | Lighthouse Lab in Glasgow | Wellcome Sanger Institute for the COVID-19 Genomics UK (COG-UK) Consortium | Harper VanSteenhouse, Yumi Kasai, David Gray, Carol Clugston, Anna Dominiczak and Alex Alderton, Roberto Amato, Jeffrey Barrett, Sonia Goncalves, Ewan Harrison, David K. Jackson, Ian Johnston, Dominic Kwiatkowski, Cordelia Langford, John Sillitoe on behalf of the Wellcome Sanger Institute COVID-19 Surveillance Team |
| EPI_ISL_2352890, EPI_ISL_2352891 | Lighthouse Lab in Milton Keynes | Wellcome Sanger Institute for the COVID-19 Genomics UK (COG-UK) Consortium | The Lighthouse Lab in Milton Keynes and Alex Alderton, Roberto Amato, Jeffrey Barrett, Sonia Goncalves, Ewan Harrison, David K. Jackson, Ian Johnston, Dominic Kwiatkowski, Cordelia Langford, John Sillitoe on behalf of the Wellcome Sanger Institute COVID-19 Surveillance Team |
| EPI_ISL_2352892 | Lighthouse Lab in Glasgow | Wellcome Sanger Institute for the COVID-19 Genomics UK (COG-UK) Consortium | Harper VanSteenhouse, Yumi Kasai, David Gray, Carol Clugston, Anna Dominiczak and Alex Alderton, Roberto Amato, Jeffrey Barrett, Sonia Goncalves, Ewan Harrison, David K. Jackson, Ian Johnston, Dominic Kwiatkowski, Cordelia Langford, John Sillitoe on behalf of the Wellcome Sanger Institute COVID-19 Surveillance Team |
| EPI_ISL_2352894 | Lighthouse Lab in Alderley Park | Wellcome Sanger Institute for the COVID-19 Genomics UK (COG-UK) Consortium | Jacquelyn Wynn, Mairead Hyland, The Lighthouse Lab in Alderley Park and Alex Alderton, Roberto Amato, Jeffrey Barrett, Sonia Goncalves, Ewan Harrison, David K. Jackson, Ian Johnston, Dominic Kwiatkowski, Cordelia Langford, John Sillitoe on behalf of the Wellcome Sanger Institute COVID-19 Surveillance Team |
| EPI_ISL_2352895, EPI_ISL_2352896, EPI_ISL_2352897 | Lighthouse Lab in Milton Keynes | Wellcome Sanger Institute for the COVID-19 Genomics UK (COG-UK) Consortium | The Lighthouse Lab in Milton Keynes and Alex Alderton, Roberto Amato, Jeffrey Barrett, Sonia Goncalves, Ewan Harrison, David K. Jackson, Ian Johnston, Dominic Kwiatkowski, Cordelia Langford, John Sillitoe on behalf of the Wellcome Sanger Institute COVID-19 Surveillance Team |
| EPI_ISL_2352898 | Lighthouse Lab in Alderley Park | Wellcome Sanger Institute for the COVID-19 Genomics UK (COG-UK) Consortium | Jacquelyn Wynn, Mairead Hyland, The Lighthouse Lab in Alderley Park and Alex Alderton, Roberto Amato, Jeffrey Barrett, Sonia Goncalves, Ewan Harrison, David K. Jackson, Ian Johnston, Dominic Kwiatkowski, Cordelia Langford, John Sillitoe on behalf of the Wellcome Sanger Institute COVID-19 Surveillance Team |
| EPI_ISL_2352900, EPI_ISL_2352902 | Lighthouse Lab in Milton Keynes | Wellcome Sanger Institute for the COVID-19 Genomics UK (COG-UK) Consortium | The Lighthouse Lab in Milton Keynes and Alex Alderton, Roberto Amato, Jeffrey Barrett, Sonia Goncalves, Ewan Harrison, David K. Jackson, Ian Johnston, Dominic Kwiatkowski, Cordelia Langford, John Sillitoe on behalf of the Wellcome Sanger Institute COVID-19 Surveillance Team |
| EPI_ISL_2352903 | Lighthouse Lab in Alderley Park | Wellcome Sanger Institute for the COVID-19 Genomics UK (COG-UK) Consortium | Jacquelyn Wynn, Mairead Hyland, The Lighthouse Lab in Alderley Park and Alex Alderton, Roberto Amato, Jeffrey Barrett, Sonia Goncalves, Ewan Harrison, David K. Jackson, Ian Johnston, Dominic Kwiatkowski, Cordelia Langford, John Sillitoe on behalf of the Wellcome Sanger Institute COVID-19 Surveillance Team |
| EPI_ISL_2352905, EPI_ISL_2352906 | Lighthouse Lab in Milton Keynes | Wellcome Sanger Institute for the COVID-19 Genomics UK (COG-UK) Consortium | The Lighthouse Lab in Milton Keynes and Alex Alderton, Roberto Amato, Jeffrey Barrett, Sonia Goncalves, Ewan Harrison, David K. Jackson, Ian Johnston, Dominic Kwiatkowski, Cordelia Langford, John Sillitoe on behalf of the Wellcome Sanger Institute COVID-19 Surveillance Team |
| EPI_ISL_2352907 | Lighthouse Lab in Alderley Park | Wellcome Sanger Institute for the COVID-19 Genomics UK (COG-UK) Consortium | Jacquelyn Wynn, Mairead Hyland, The Lighthouse Lab in Alderley Park and Alex Alderton, Roberto Amato, Jeffrey Barrett, Sonia Goncalves, Ewan Harrison, David K. Jackson, Ian Johnston, Dominic Kwiatkowski, Cordelia Langford, John Sillitoe on behalf of the Wellcome Sanger Institute COVID-19 Surveillance Team |
| EPI_ISL_2352908, EPI_ISL_2352909, EPI_ISL_2352910, EPI_ISL_2352912 | Lighthouse Lab in Milton Keynes | Wellcome Sanger Institute for the COVID-19 Genomics UK (COG-UK) Consortium | The Lighthouse Lab in Milton Keynes and Alex Alderton, Roberto Amato, Jeffrey Barrett, Sonia Goncalves, Ewan Harrison, David K. Jackson, Ian Johnston, Dominic Kwiatkowski, Cordelia Langford, John Sillitoe on behalf of the Wellcome Sanger Institute COVID-19 Surveillance Team |
| EPI_ISL_2352913 | Lighthouse Lab in Alderley Park | Wellcome Sanger Institute for the COVID-19 Genomics UK (COG-UK) Consortium | Jacquelyn Wynn, Mairead Hyland, The Lighthouse Lab in Alderley Park and Alex Alderton, Roberto Amato, Jeffrey Barrett, Sonia Goncalves, Ewan Harrison, David K. Jackson, Ian Johnston, Dominic Kwiatkowski, Cordelia Langford, John Sillitoe on behalf of the Wellcome Sanger Institute COVID-19 Surveillance Team |
| EPI_ISL_2352914 | Lighthouse Lab in Milton Keynes | Wellcome Sanger Institute for the COVID-19 Genomics UK (COG-UK) Consortium | The Lighthouse Lab in Milton Keynes and Alex Alderton, Roberto Amato, Jeffrey Barrett, Sonia Goncalves, Ewan Harrison, David K. Jackson, Ian Johnston, Dominic Kwiatkowski, Cordelia Langford, John Sillitoe on behalf of the Wellcome Sanger Institute COVID-19 Surveillance Team |
| EPI_ISL_2352915, EPI_ISL_2352916, EPI_ISL_2352918 | Lighthouse Lab in Alderley Park | Wellcome Sanger Institute for the COVID-19 Genomics UK (COG-UK) Consortium | Jacquelyn Wynn, Mairead Hyland, The Lighthouse Lab in Alderley Park and Alex Alderton, Roberto Amato, Jeffrey Barrett, Sonia Goncalves, Ewan Harrison, David K. Jackson, Ian Johnston, Dominic Kwiatkowski, Cordelia Langford, John Sillitoe on behalf of the Wellcome Sanger Institute COVID-19 Surveillance Team |
| EPI_ISL_2352919 | Lighthouse Lab in Milton Keynes | Wellcome Sanger Institute for the COVID-19 Genomics UK (COG-UK) Consortium | The Lighthouse Lab in Milton Keynes and Alex Alderton, Roberto Amato, Jeffrey Barrett, Sonia Goncalves, Ewan Harrison, David K. Jackson, Ian Johnston, Dominic Kwiatkowski, Cordelia Langford, John Sillitoe on behalf of the Wellcome Sanger Institute COVID-19 Surveillance Team |
| EPI_ISL_2352920 | Lighthouse Lab in Glasgow | Wellcome Sanger Institute for the COVID-19 Genomics UK (COG-UK) Consortium | Harper VanSteenhouse, Yumi Kasai, David Gray, Carol Clugston, Anna Dominiczak and Alex Alderton, Roberto Amato, Jeffrey Barrett, Sonia Goncalves, Ewan Harrison, David K. Jackson, Ian Johnston, Dominic Kwiatkowski, Cordelia Langford, John Sillitoe on behalf of the Wellcome Sanger Institute COVID-19 Surveillance Team |
| EPI_ISL_2352921, EPI_ISL_2352923, EPI_ISL_2352925 | Lighthouse Lab in Milton Keynes | Wellcome Sanger Institute for the COVID-19 Genomics UK (COG-UK) Consortium | The Lighthouse Lab in Milton Keynes and Alex Alderton, Roberto Amato, Jeffrey Barrett, Sonia Goncalves, Ewan Harrison, David K. Jackson, Ian Johnston, Dominic Kwiatkowski, Cordelia Langford, John Sillitoe on behalf of the Wellcome Sanger Institute COVID-19 Surveillance Team |
| EPI_ISL_2352926 | Lighthouse Lab in Glasgow | Wellcome Sanger Institute for the COVID-19 Genomics UK (COG-UK) Consortium | Harper VanSteenhouse, Yumi Kasai, David Gray, Carol Clugston, Anna Dominiczak and Alex Alderton, Roberto Amato, Jeffrey Barrett, Sonia Goncalves, Ewan Harrison, David K. Jackson, Ian Johnston, Dominic Kwiatkowski, Cordelia Langford, John Sillitoe on behalf of the Wellcome Sanger Institute COVID-19 Surveillance Team |
| EPI_ISL_2352927, EPI_ISL_2352928, EPI_ISL_2352929 | Lighthouse Lab in Milton Keynes | Wellcome Sanger Institute for the COVID-19 Genomics UK (COG-UK) Consortium | The Lighthouse Lab in Milton Keynes and Alex Alderton, Roberto Amato, Jeffrey Barrett, Sonia Goncalves, Ewan Harrison, David K. Jackson, Ian Johnston, Dominic Kwiatkowski, Cordelia Langford, John Sillitoe on behalf of the Wellcome Sanger Institute COVID-19 Surveillance Team |
| EPI_ISL_2352930 | Lighthouse Lab in Alderley Park | Wellcome Sanger Institute for the COVID-19 Genomics UK (COG-UK) Consortium | Jacquelyn Wynn, Mairead Hyland, The Lighthouse Lab in Alderley Park and Alex Alderton, Roberto Amato, Jeffrey Barrett, Sonia Goncalves, Ewan Harrison, David K. Jackson, Ian Johnston, Dominic Kwiatkowski, Cordelia Langford, John Sillitoe on behalf of the Wellcome Sanger Institute COVID-19 Surveillance Team |
| EPI_ISL_2352931 | Lighthouse Lab in Glasgow | Wellcome Sanger Institute for the COVID-19 Genomics UK (COG-UK) Consortium | Harper VanSteenhouse, Yumi Kasai, David Gray, Carol Clugston, Anna Dominiczak and Alex Alderton, Roberto Amato, Jeffrey Barrett, Sonia Goncalves, Ewan Harrison, David K. Jackson, Ian Johnston, Dominic Kwiatkowski, Cordelia Langford, John Sillitoe on behalf of the Wellcome Sanger Institute COVID-19 Surveillance Team |
| EPI_ISL_2352932 | Lighthouse Lab in Alderley Park | Wellcome Sanger Institute for the COVID-19 Genomics UK (COG-UK) Consortium | Jacquelyn Wynn, Mairead Hyland, The Lighthouse Lab in Alderley Park and Alex Alderton, Roberto Amato, Jeffrey Barrett, Sonia Goncalves, Ewan Harrison, David K. Jackson, Ian Johnston, Dominic Kwiatkowski, Cordelia Langford, John Sillitoe on behalf of the Wellcome Sanger Institute COVID-19 Surveillance Team |
| EPI_ISL_2352933 | Lighthouse Lab in Milton Keynes | Wellcome Sanger Institute for the COVID-19 Genomics UK (COG-UK) Consortium | The Lighthouse Lab in Milton Keynes and Alex Alderton, Roberto Amato, Jeffrey Barrett, Sonia Goncalves, Ewan Harrison, David K. Jackson, Ian Johnston, Dominic Kwiatkowski, Cordelia Langford, John Sillitoe on behalf of the Wellcome Sanger Institute COVID-19 Surveillance Team |
| EPI_ISL_2352934, EPI_ISL_2352935, EPI_ISL_2352936, EPI_ISL_2352937 | Lighthouse Lab in Alderley Park | Wellcome Sanger Institute for the COVID-19 Genomics UK (COG-UK) Consortium | Jacquelyn Wynn, Mairead Hyland, The Lighthouse Lab in Alderley Park and Alex Alderton, Roberto Amato, Jeffrey Barrett, Sonia Goncalves, Ewan Harrison, David K. Jackson, Ian Johnston, Dominic Kwiatkowski, Cordelia Langford, John Sillitoe on behalf of the Wellcome Sanger Institute COVID-19 Surveillance Team |
| EPI_ISL_2352938 | Lighthouse Lab in Glasgow | Wellcome Sanger Institute for the COVID-19 Genomics UK (COG-UK) Consortium | Harper VanSteenhouse, Yumi Kasai, David Gray, Carol Clugston, Anna Dominiczak and Alex Alderton, Roberto Amato, Jeffrey Barrett, Sonia Goncalves, Ewan Harrison, David K. Jackson, Ian Johnston, Dominic Kwiatkowski, Cordelia Langford, John Sillitoe on behalf of the Wellcome Sanger Institute COVID-19 Surveillance Team |
| EPI_ISL_2352939, EPI_ISL_2352940 | Lighthouse Lab in Milton Keynes | Wellcome Sanger Institute for the COVID-19 Genomics UK (COG-UK) Consortium | The Lighthouse Lab in Milton Keynes and Alex Alderton, Roberto Amato, Jeffrey Barrett, Sonia Goncalves, Ewan Harrison, David K. Jackson, Ian Johnston, Dominic Kwiatkowski, Cordelia Langford, John Sillitoe on behalf of the Wellcome Sanger Institute COVID-19 Surveillance Team |
| EPI_ISL_2352941, EPI_ISL_2352942 | Lighthouse Lab in Alderley Park | Wellcome Sanger Institute for the COVID-19 Genomics UK (COG-UK) Consortium | Jacquelyn Wynn, Mairead Hyland, The Lighthouse Lab in Alderley Park and Alex Alderton, Roberto Amato, Jeffrey Barrett, Sonia Goncalves, Ewan Harrison, David K. Jackson, Ian Johnston, Dominic Kwiatkowski, Cordelia Langford, John Sillitoe on behalf of the Wellcome Sanger Institute COVID-19 Surveillance Team |
| EPI_ISL_2352943 | Lighthouse Lab in Milton Keynes | Wellcome Sanger Institute for the COVID-19 Genomics UK (COG-UK) Consortium | The Lighthouse Lab in Milton Keynes and Alex Alderton, Roberto Amato, Jeffrey Barrett, Sonia Goncalves, Ewan Harrison, David K. Jackson, Ian Johnston, Dominic Kwiatkowski, Cordelia Langford, John Sillitoe on behalf of the Wellcome Sanger Institute COVID-19 Surveillance Team |

| | | | |
|---|---|---|---|
| EPI_ISL_2352944 | Lighthouse Lab in Glasgow | Wellcome Sanger Institute for the COVID-19 Genomics UK (COG-UK) Consortium | Harper VanSteenhouse, Yumi Kasai, David Gray, Carol Clugston, Anna Dominiczak and Alex Alderton, Roberto Amato, Jeffrey Barrett, Sonia Goncalves, Ewan Harrison, David K. Jackson, Ian Johnston, Dominic Kwiatkowski, Cordelia Langford, John Sillitoe on behalf of the Wellcome Sanger Institute COVID-19 Surveillance Team |
| EPI_ISL_2352945, EPI_ISL_2352946 | Lighthouse Lab in Milton Keynes | Wellcome Sanger Institute for the COVID-19 Genomics UK (COG-UK) Consortium | The Lighthouse Lab in Milton Keynes and Alex Alderton, Roberto Amato, Jeffrey Barrett, Sonia Goncalves, Ewan Harrison, David K. Jackson, Ian Johnston, Dominic Kwiatkowski, Cordelia Langford, John Sillitoe on behalf of the Wellcome Sanger Institute COVID-19 Surveillance Team |
| EPI_ISL_2352947 | Lighthouse Lab in Alderley Park | Wellcome Sanger Institute for the COVID-19 Genomics UK (COG-UK) Consortium | Jacquelyn Wynn, Mairead Hyland, The Lighthouse Lab in Alderley Park and Alex Alderton, Roberto Amato, Jeffrey Barrett, Sonia Goncalves, Ewan Harrison, David K. Jackson, Ian Johnston, Dominic Kwiatkowski, Cordelia Langford, John Sillitoe on behalf of the Wellcome Sanger Institute COVID-19 Surveillance Team |
| EPI_ISL_2352948 | Lighthouse Lab in Glasgow | Wellcome Sanger Institute for the COVID-19 Genomics UK (COG-UK) Consortium | Harper VanSteenhouse, Yumi Kasai, David Gray, Carol Clugston, Anna Dominiczak and Alex Alderton, Roberto Amato, Jeffrey Barrett, Sonia Goncalves, Ewan Harrison, David K. Jackson, Ian Johnston, Dominic Kwiatkowski, Cordelia Langford, John Sillitoe on behalf of the Wellcome Sanger Institute COVID-19 Surveillance Team |
| EPI_ISL_2352950, EPI_ISL_2352951 | Lighthouse Lab in Alderley Park | Wellcome Sanger Institute for the COVID-19 Genomics UK (COG-UK) Consortium | Jacquelyn Wynn, Mairead Hyland, The Lighthouse Lab in Alderley Park and Alex Alderton, Roberto Amato, Jeffrey Barrett, Sonia Goncalves, Ewan Harrison, David K. Jackson, Ian Johnston, Dominic Kwiatkowski, Cordelia Langford, John Sillitoe on behalf of the Wellcome Sanger Institute COVID-19 Surveillance Team |
| EPI_ISL_2352952 | Lighthouse Lab in Glasgow | Wellcome Sanger Institute for the COVID-19 Genomics UK (COG-UK) Consortium | Harper VanSteenhouse, Yumi Kasai, David Gray, Carol Clugston, Anna Dominiczak and Alex Alderton, Roberto Amato, Jeffrey Barrett, Sonia Goncalves, Ewan Harrison, David K. Jackson, Ian Johnston, Dominic Kwiatkowski, Cordelia Langford, John Sillitoe on behalf of the Wellcome Sanger Institute COVID-19 Surveillance Team |
| EPI_ISL_2352953 | Lighthouse Lab in Alderley Park | Wellcome Sanger Institute for the COVID-19 Genomics UK (COG-UK) Consortium | Jacquelyn Wynn, Mairead Hyland, The Lighthouse Lab in Alderley Park and Alex Alderton, Roberto Amato, Jeffrey Barrett, Sonia Goncalves, Ewan Harrison, David K. Jackson, Ian Johnston, Dominic Kwiatkowski, Cordelia Langford, John Sillitoe on behalf of the Wellcome Sanger Institute COVID-19 Surveillance Team |
| EPI_ISL_2352954 | Lighthouse Lab in Glasgow | Wellcome Sanger Institute for the COVID-19 Genomics UK (COG-UK) Consortium | Harper VanSteenhouse, Yumi Kasai, David Gray, Carol Clugston, Anna Dominiczak and Alex Alderton, Roberto Amato, Jeffrey Barrett, Sonia Goncalves, Ewan Harrison, David K. Jackson, Ian Johnston, Dominic Kwiatkowski, Cordelia Langford, John Sillitoe on behalf of the Wellcome Sanger Institute COVID-19 Surveillance Team |
| EPI_ISL_2352956, EPI_ISL_2352957 | Lighthouse Lab in Milton Keynes | Wellcome Sanger Institute for the COVID-19 Genomics UK (COG-UK) Consortium | The Lighthouse Lab in Milton Keynes and Alex Alderton, Roberto Amato, Jeffrey Barrett, Sonia Goncalves, Ewan Harrison, David K. Jackson, Ian Johnston, Dominic Kwiatkowski, Cordelia Langford, John Sillitoe on behalf of the Wellcome Sanger Institute COVID-19 Surveillance Team |
| EPI_ISL_2352958 | Lighthouse Lab in Alderley Park | Wellcome Sanger Institute for the COVID-19 Genomics UK (COG-UK) Consortium | Jacquelyn Wynn, Mairead Hyland, The Lighthouse Lab in Alderley Park and Alex Alderton, Roberto Amato, Jeffrey Barrett, Sonia Goncalves, Ewan Harrison, David K. Jackson, Ian Johnston, Dominic Kwiatkowski, Cordelia Langford, John Sillitoe on behalf of the Wellcome Sanger Institute COVID-19 Surveillance Team |
| EPI_ISL_2352959 | Lighthouse Lab in Milton Keynes | Wellcome Sanger Institute for the COVID-19 Genomics UK (COG-UK) Consortium | The Lighthouse Lab in Milton Keynes and Alex Alderton, Roberto Amato, Jeffrey Barrett, Sonia Goncalves, Ewan Harrison, David K. Jackson, Ian Johnston, Dominic Kwiatkowski, Cordelia Langford, John Sillitoe on behalf of the Wellcome Sanger Institute COVID-19 Surveillance Team |
| EPI_ISL_2352960 | Lighthouse Lab in Alderley Park | Wellcome Sanger Institute for the COVID-19 Genomics UK (COG-UK) Consortium | Jacquelyn Wynn, Mairead Hyland, The Lighthouse Lab in Alderley Park and Alex Alderton, Roberto Amato, Jeffrey Barrett, Sonia Goncalves, Ewan Harrison, David K. Jackson, Ian Johnston, Dominic Kwiatkowski, Cordelia Langford, John Sillitoe on behalf of the Wellcome Sanger Institute COVID-19 Surveillance Team |
| EPI_ISL_2352961, EPI_ISL_2352962 | Lighthouse Lab in Milton Keynes | Wellcome Sanger Institute for the COVID-19 Genomics UK (COG-UK) Consortium | The Lighthouse Lab in Milton Keynes and Alex Alderton, Roberto Amato, Jeffrey Barrett, Sonia Goncalves, Ewan Harrison, David K. Jackson, Ian Johnston, Dominic Kwiatkowski, Cordelia Langford, John Sillitoe on behalf of the Wellcome Sanger Institute COVID-19 Surveillance Team |
| EPI_ISL_2352963, EPI_ISL_2352965 | Lighthouse Lab in Alderley Park | Wellcome Sanger Institute for the COVID-19 Genomics UK (COG-UK) Consortium | Jacquelyn Wynn, Mairead Hyland, The Lighthouse Lab in Alderley Park and Alex Alderton, Roberto Amato, Jeffrey Barrett, Sonia Goncalves, Ewan Harrison, David K. Jackson, Ian Johnston, Dominic Kwiatkowski, Cordelia Langford, John Sillitoe on behalf of the Wellcome Sanger Institute COVID-19 Surveillance Team |
| EPI_ISL_2352966 | Lighthouse Lab in Milton Keynes | Wellcome Sanger Institute for the COVID-19 Genomics UK (COG-UK) Consortium | The Lighthouse Lab in Milton Keynes and Alex Alderton, Roberto Amato, Jeffrey Barrett, Sonia Goncalves, Ewan Harrison, David K. Jackson, Ian Johnston, Dominic Kwiatkowski, Cordelia Langford, John Sillitoe on behalf of the Wellcome Sanger Institute COVID-19 Surveillance Team |
| EPI_ISL_2352968 | Lighthouse Lab in Alderley Park | Wellcome Sanger Institute for the COVID-19 Genomics UK (COG-UK) Consortium | Jacquelyn Wynn, Mairead Hyland, The Lighthouse Lab in Alderley Park and Alex Alderton, Roberto Amato, Jeffrey Barrett, Sonia Goncalves, Ewan Harrison, David K. Jackson, Ian Johnston, Dominic Kwiatkowski, Cordelia Langford, John Sillitoe on behalf of the Wellcome Sanger Institute COVID-19 Surveillance Team |
| EPI_ISL_2352969 | Lighthouse Lab in Milton Keynes | Wellcome Sanger Institute for the COVID-19 Genomics UK (COG-UK) Consortium | The Lighthouse Lab in Milton Keynes and Alex Alderton, Roberto Amato, Jeffrey Barrett, Sonia Goncalves, Ewan Harrison, David K. Jackson, Ian Johnston, Dominic Kwiatkowski, Cordelia Langford, John Sillitoe on behalf of the Wellcome Sanger Institute COVID-19 Surveillance Team |
| EPI_ISL_2352970 | Lighthouse Lab in Alderley Park | Wellcome Sanger Institute for the COVID-19 Genomics UK (COG-UK) Consortium | Jacquelyn Wynn, Mairead Hyland, The Lighthouse Lab in Alderley Park and Alex Alderton, Roberto Amato, Jeffrey Barrett, Sonia Goncalves, Ewan Harrison, David K. Jackson, Ian Johnston, Dominic Kwiatkowski, Cordelia Langford, John Sillitoe on behalf of the Wellcome Sanger Institute COVID-19 Surveillance Team |
| EPI_ISL_2352971, EPI_ISL_2352972 | Lighthouse Lab in Milton Keynes | Wellcome Sanger Institute for the COVID-19 Genomics UK (COG-UK) Consortium | The Lighthouse Lab in Milton Keynes and Alex Alderton, Roberto Amato, Jeffrey Barrett, Sonia Goncalves, Ewan Harrison, David K. Jackson, Ian Johnston, Dominic Kwiatkowski, Cordelia Langford, John Sillitoe on behalf of the Wellcome Sanger Institute COVID-19 Surveillance Team |
| EPI_ISL_2352974 | Lighthouse Lab in Alderley Park | Wellcome Sanger Institute for the COVID-19 Genomics UK (COG-UK) Consortium | Jacquelyn Wynn, Mairead Hyland, The Lighthouse Lab in Alderley Park and Alex Alderton, Roberto Amato, Jeffrey Barrett, Sonia Goncalves, Ewan Harrison, David K. Jackson, Ian Johnston, Dominic Kwiatkowski, Cordelia Langford, John Sillitoe on behalf of the Wellcome Sanger Institute COVID-19 Surveillance Team |
| EPI_ISL_2352975 | Lighthouse Lab in Milton Keynes | Wellcome Sanger Institute for the COVID-19 Genomics UK (COG-UK) Consortium | The Lighthouse Lab in Milton Keynes and Alex Alderton, Roberto Amato, Jeffrey Barrett, Sonia Goncalves, Ewan Harrison, David K. Jackson, Ian Johnston, Dominic Kwiatkowski, Cordelia Langford, John Sillitoe on behalf of the Wellcome Sanger Institute COVID-19 Surveillance Team |
| EPI_ISL_2352976 | Lighthouse Lab in Alderley Park | Wellcome Sanger Institute for the COVID-19 Genomics UK (COG-UK) Consortium | Jacquelyn Wynn, Mairead Hyland, The Lighthouse Lab in Alderley Park and Alex Alderton, Roberto Amato, Jeffrey Barrett, Sonia Goncalves, Ewan Harrison, David K. Jackson, Ian Johnston, Dominic Kwiatkowski, Cordelia Langford, John Sillitoe on behalf of the Wellcome Sanger Institute COVID-19 Surveillance Team |
| EPI_ISL_2352977, EPI_ISL_2352979 | Lighthouse Lab in Milton Keynes | Wellcome Sanger Institute for the COVID-19 Genomics UK (COG-UK) Consortium | The Lighthouse Lab in Milton Keynes and Alex Alderton, Roberto Amato, Jeffrey Barrett, Sonia Goncalves, Ewan Harrison, David K. Jackson, Ian Johnston, Dominic Kwiatkowski, Cordelia Langford, John Sillitoe on behalf of the Wellcome Sanger Institute COVID-19 Surveillance Team |
| EPI_ISL_2352980, EPI_ISL_2352981 | Lighthouse Lab in Alderley Park | Wellcome Sanger Institute for the COVID-19 Genomics UK (COG-UK) Consortium | Jacquelyn Wynn, Mairead Hyland, The Lighthouse Lab in Alderley Park and Alex Alderton, Roberto Amato, Jeffrey Barrett, Sonia Goncalves, Ewan Harrison, David K. Jackson, Ian Johnston, Dominic Kwiatkowski, Cordelia Langford, John Sillitoe on behalf of the Wellcome Sanger Institute COVID-19 Surveillance Team |
| EPI_ISL_2352982 | Lighthouse Lab in Milton Keynes | Wellcome Sanger Institute for the COVID-19 Genomics UK (COG-UK) Consortium | The Lighthouse Lab in Milton Keynes and Alex Alderton, Roberto Amato, Jeffrey Barrett, Sonia Goncalves, Ewan Harrison, David K. Jackson, Ian Johnston, Dominic Kwiatkowski, Cordelia Langford, John Sillitoe on behalf of the Wellcome Sanger Institute COVID-19 Surveillance Team |
| EPI_ISL_2352983, EPI_ISL_2352984 | Lighthouse Lab in Alderley Park | Wellcome Sanger Institute for the COVID-19 Genomics UK (COG-UK) Consortium | Jacquelyn Wynn, Mairead Hyland, The Lighthouse Lab in Alderley Park and Alex Alderton, Roberto Amato, Jeffrey Barrett, Sonia Goncalves, Ewan Harrison, David K. Jackson, Ian Johnston, Dominic Kwiatkowski, Cordelia Langford, John Sillitoe on behalf of the Wellcome Sanger Institute COVID-19 Surveillance Team |
| EPI_ISL_2352985, EPI_ISL_2352986, EPI_ISL_2352987, EPI_ISL_2352988 | Lighthouse Lab in Milton Keynes | Wellcome Sanger Institute for the COVID-19 Genomics UK (COG-UK) Consortium | The Lighthouse Lab in Milton Keynes and Alex Alderton, Roberto Amato, Jeffrey Barrett, Sonia Goncalves, Ewan Harrison, David K. Jackson, Ian Johnston, Dominic Kwiatkowski, Cordelia Langford, John Sillitoe on behalf of the Wellcome Sanger Institute COVID-19 Surveillance Team |
| EPI_ISL_2352989 | Lighthouse Lab in Glasgow | Wellcome Sanger Institute for the COVID-19 Genomics UK (COG-UK) Consortium | Harper VanSteenhouse, Yumi Kasai, David Gray, Carol Clugston, Anna Dominiczak and Alex Alderton, Roberto Amato, Jeffrey Barrett, Sonia Goncalves, Ewan Harrison, David K. Jackson, Ian Johnston, Dominic Kwiatkowski, Cordelia Langford, John Sillitoe on behalf of the Wellcome Sanger Institute COVID-19 Surveillance Team |
| EPI_ISL_2352990, EPI_ISL_2352991 | Lighthouse Lab in Alderley Park | Wellcome Sanger Institute for the COVID-19 Genomics UK (COG-UK) Consortium | Jacquelyn Wynn, Mairead Hyland, The Lighthouse Lab in Alderley Park and Alex Alderton, Roberto Amato, Jeffrey Barrett, Sonia Goncalves, Ewan Harrison, David K. Jackson, Ian Johnston, Dominic Kwiatkowski, Cordelia Langford, John Sillitoe on behalf of the Wellcome Sanger Institute COVID-19 Surveillance Team |
| EPI_ISL_2352992 | Lighthouse Lab in Milton Keynes | Wellcome Sanger Institute for the COVID-19 Genomics UK (COG-UK) Consortium | The Lighthouse Lab in Milton Keynes and Alex Alderton, Roberto Amato, Jeffrey Barrett, Sonia Goncalves, Ewan Harrison, David K. Jackson, Ian Johnston, Dominic Kwiatkowski, Cordelia Langford, John Sillitoe on behalf of the Wellcome Sanger Institute COVID-19 Surveillance Team |
| EPI_ISL_2352993 | Lighthouse Lab in Glasgow | Wellcome Sanger Institute for the COVID-19 Genomics UK (COG-UK) Consortium | Harper VanSteenhouse, Yumi Kasai, David Gray, Carol Clugston, Anna Dominiczak and Alex Alderton, Roberto Amato, Jeffrey Barrett, Sonia Goncalves, Ewan Harrison, David K. Jackson, Ian Johnston, Dominic Kwiatkowski, Cordelia Langford, John Sillitoe on behalf of the Wellcome Sanger Institute COVID-19 Surveillance Team |
| EPI_ISL_2352994 | Lighthouse Lab in Alderley Park | Wellcome Sanger Institute for the COVID-19 Genomics UK (COG-UK) Consortium | Jacquelyn Wynn, Mairead Hyland, The Lighthouse Lab in Alderley Park and Alex Alderton, Roberto Amato, Jeffrey Barrett, Sonia Goncalves, Ewan Harrison, David K. Jackson, Ian Johnston, Dominic Kwiatkowski, Cordelia Langford, John Sillitoe on behalf of the Wellcome Sanger Institute COVID-19 Surveillance Team |
| EPI_ISL_2352996 | Lighthouse Lab in Milton Keynes | Wellcome Sanger Institute for the COVID-19 Genomics UK (COG-UK) Consortium | The Lighthouse Lab in Milton Keynes and Alex Alderton, Roberto Amato, Jeffrey Barrett, Sonia Goncalves, Ewan Harrison, David K. Jackson, Ian Johnston, Dominic Kwiatkowski, Cordelia Langford, John Sillitoe on behalf of the Wellcome Sanger Institute COVID-19 Surveillance Team |
| EPI_ISL_2352997, EPI_ISL_2352998, EPI_ISL_2352999, EPI_ISL_2353000, EPI_ISL_2353001, EPI_ISL_2353005, EPI_ISL_2353006, EPI_ISL_2353007, EPI_ISL_2353008 | Lighthouse Lab in Alderley Park | Wellcome Sanger Institute for the COVID-19 Genomics UK (COG-UK) Consortium | Jacquelyn Wynn, Mairead Hyland, The Lighthouse Lab in Alderley Park and Alex Alderton, Roberto Amato, Jeffrey Barrett, Sonia Goncalves, Ewan Harrison, David K. Jackson, Ian Johnston, Dominic Kwiatkowski, Cordelia Langford, John Sillitoe on behalf of the Wellcome Sanger Institute COVID-19 Surveillance Team |
| EPI_ISL_2353009 | Lighthouse Lab in Milton Keynes | Wellcome Sanger Institute for the COVID-19 Genomics UK (COG-UK) Consortium | The Lighthouse Lab in Milton Keynes and Alex Alderton, Roberto Amato, Jeffrey Barrett, Sonia Goncalves, Ewan Harrison, David K. Jackson, Ian Johnston, Dominic Kwiatkowski, Cordelia Langford, John Sillitoe on behalf of the Wellcome Sanger Institute COVID-19 Surveillance Team |
| EPI_ISL_2353011, EPI_ISL_2353012 | Lighthouse Lab in Alderley Park | Wellcome Sanger Institute for the COVID-19 Genomics UK (COG-UK) Consortium | Jacquelyn Wynn, Mairead Hyland, The Lighthouse Lab in Alderley Park and Alex Alderton, Roberto Amato, Jeffrey Barrett, Sonia Goncalves, Ewan Harrison, David K. Jackson, Ian Johnston, Dominic Kwiatkowski, Cordelia Langford, John Sillitoe on behalf of the Wellcome Sanger Institute COVID-19 Surveillance Team |
| EPI_ISL_2353013, EPI_ISL_2353014 | Lighthouse Lab in Milton Keynes | Wellcome Sanger Institute for the COVID-19 Genomics UK (COG-UK) Consortium | The Lighthouse Lab in Milton Keynes and Alex Alderton, Roberto Amato, Jeffrey Barrett, Sonia Goncalves, Ewan Harrison, David K. Jackson, Ian Johnston, Dominic Kwiatkowski, Cordelia Langford, John Sillitoe on behalf of the Wellcome Sanger Institute COVID-19 Surveillance Team |
| EPI_ISL_2353015, EPI_ISL_2353016, EPI_ISL_2353017, EPI_ISL_2353018, EPI_ISL_2353019, EPI_ISL_2353020, EPI_ISL_2353022, EPI_ISL_2353023 | Lighthouse Lab in Alderley Park | Wellcome Sanger Institute for the COVID-19 Genomics UK (COG-UK) Consortium | Jacquelyn Wynn, Mairead Hyland, The Lighthouse Lab in Alderley Park and Alex Alderton, Roberto Amato, Jeffrey Barrett, Sonia Goncalves, Ewan Harrison, David K. Jackson, Ian Johnston, Dominic Kwiatkowski, Cordelia Langford, John Sillitoe on behalf of the Wellcome Sanger Institute COVID-19 Surveillance Team |
| EPI_ISL_2353024 | Lighthouse Lab in Milton Keynes | Wellcome Sanger Institute for the COVID-19 Genomics UK (COG-UK) Consortium | The Lighthouse Lab in Milton Keynes and Alex Alderton, Roberto Amato, Jeffrey Barrett, Sonia Goncalves, Ewan Harrison, David K. Jackson, Ian Johnston, Dominic Kwiatkowski, Cordelia Langford, John Sillitoe on behalf of the Wellcome Sanger Institute COVID-19 Surveillance Team |

| | | | |
|---|---|---|---|
| EPI_ISL_2353026, EPI_ISL_2353028, EPI_ISL_2353029 | Lighthouse Lab in Alderley Park | Wellcome Sanger Institute for the COVID-19 Genomics UK (COG-UK) Consortium | Jacquelyn Wynn, Mairead Hyland, The Lighthouse Lab in Alderley Park and Alex Alderton, Roberto Amato, Jeffrey Barrett, Sonia Goncalves, Ewan Harrison, David K. Jackson, Ian Johnston, Dominic Kwiatkowski, Cordelia Langford, John Sillitoe on behalf of the Wellcome Sanger Institute COVID-19 Surveillance Team |
| EPI_ISL_2353030 | Lighthouse Lab in Milton Keynes | Wellcome Sanger Institute for the COVID-19 Genomics UK (COG-UK) Consortium | The Lighthouse Lab in Milton Keynes and Alex Alderton, Roberto Amato, Jeffrey Barrett, Sonia Goncalves, Ewan Harrison, David K. Jackson, Ian Johnston, Dominic Kwiatkowski, Cordelia Langford, John Sillitoe on behalf of the Wellcome Sanger Institute COVID-19 Surveillance Team |
| EPI_ISL_2353032, EPI_ISL_2353034, EPI_ISL_2353035, EPI_ISL_2353036 | Lighthouse Lab in Alderley Park | Wellcome Sanger Institute for the COVID-19 Genomics UK (COG-UK) Consortium | Jacquelyn Wynn, Mairead Hyland, The Lighthouse Lab in Alderley Park and Alex Alderton, Roberto Amato, Jeffrey Barrett, Sonia Goncalves, Ewan Harrison, David K. Jackson, Ian Johnston, Dominic Kwiatkowski, Cordelia Langford, John Sillitoe on behalf of the Wellcome Sanger Institute COVID-19 Surveillance Team |
| EPI_ISL_2353037, EPI_ISL_2353039, EPI_ISL_2353040 | Lighthouse Lab in Milton Keynes | Wellcome Sanger Institute for the COVID-19 Genomics UK (COG-UK) Consortium | The Lighthouse Lab in Milton Keynes and Alex Alderton, Roberto Amato, Jeffrey Barrett, Sonia Goncalves, Ewan Harrison, David K. Jackson, Ian Johnston, Dominic Kwiatkowski, Cordelia Langford, John Sillitoe on behalf of the Wellcome Sanger Institute COVID-19 Surveillance Team |
| EPI_ISL_2353041, EPI_ISL_2353042, EPI_ISL_2353043, EPI_ISL_2353044, EPI_ISL_2353045, EPI_ISL_2353046, EPI_ISL_2353047, EPI_ISL_2353049, EPI_ISL_2353050, EPI_ISL_2353052 | Lighthouse Lab in Alderley Park | Wellcome Sanger Institute for the COVID-19 Genomics UK (COG-UK) Consortium | Jacquelyn Wynn, Mairead Hyland, The Lighthouse Lab in Alderley Park and Alex Alderton, Roberto Amato, Jeffrey Barrett, Sonia Goncalves, Ewan Harrison, David K. Jackson, Ian Johnston, Dominic Kwiatkowski, Cordelia Langford, John Sillitoe on behalf of the Wellcome Sanger Institute COVID-19 Surveillance Team |
| EPI_ISL_2353053 | Lighthouse Lab in Milton Keynes | Wellcome Sanger Institute for the COVID-19 Genomics UK (COG-UK) Consortium | The Lighthouse Lab in Milton Keynes and Alex Alderton, Roberto Amato, Jeffrey Barrett, Sonia Goncalves, Ewan Harrison, David K. Jackson, Ian Johnston, Dominic Kwiatkowski, Cordelia Langford, John Sillitoe on behalf of the Wellcome Sanger Institute COVID-19 Surveillance Team |
| EPI_ISL_2353055, EPI_ISL_2353056, EPI_ISL_2353059, EPI_ISL_2353060, EPI_ISL_2353061 | Lighthouse Lab in Alderley Park | Wellcome Sanger Institute for the COVID-19 Genomics UK (COG-UK) Consortium | Jacquelyn Wynn, Mairead Hyland, The Lighthouse Lab in Alderley Park and Alex Alderton, Roberto Amato, Jeffrey Barrett, Sonia Goncalves, Ewan Harrison, David K. Jackson, Ian Johnston, Dominic Kwiatkowski, Cordelia Langford, John Sillitoe on behalf of the Wellcome Sanger Institute COVID-19 Surveillance Team |
| EPI_ISL_2353064, EPI_ISL_2353066, EPI_ISL_2353067, EPI_ISL_2353068 | Lighthouse Lab in Milton Keynes | Wellcome Sanger Institute for the COVID-19 Genomics UK (COG-UK) Consortium | The Lighthouse Lab in Milton Keynes and Alex Alderton, Roberto Amato, Jeffrey Barrett, Sonia Goncalves, Ewan Harrison, David K. Jackson, Ian Johnston, Dominic Kwiatkowski, Cordelia Langford, John Sillitoe on behalf of the Wellcome Sanger Institute COVID-19 Surveillance Team |
| EPI_ISL_2353069, EPI_ISL_2353070 | Lighthouse Lab in Alderley Park | Wellcome Sanger Institute for the COVID-19 Genomics UK (COG-UK) Consortium | Jacquelyn Wynn, Mairead Hyland, The Lighthouse Lab in Alderley Park and Alex Alderton, Roberto Amato, Jeffrey Barrett, Sonia Goncalves, Ewan Harrison, David K. Jackson, Ian Johnston, Dominic Kwiatkowski, Cordelia Langford, John Sillitoe on behalf of the Wellcome Sanger Institute COVID-19 Surveillance Team |
| EPI_ISL_2353071, EPI_ISL_2353073 | Lighthouse Lab in Milton Keynes | Wellcome Sanger Institute for the COVID-19 Genomics UK (COG-UK) Consortium | The Lighthouse Lab in Milton Keynes and Alex Alderton, Roberto Amato, Jeffrey Barrett, Sonia Goncalves, Ewan Harrison, David K. Jackson, Ian Johnston, Dominic Kwiatkowski, Cordelia Langford, John Sillitoe on behalf of the Wellcome Sanger Institute COVID-19 Surveillance Team |
| EPI_ISL_2353075, EPI_ISL_2353076, EPI_ISL_2353077, EPI_ISL_2353078, EPI_ISL_2353080, EPI_ISL_2353081, EPI_ISL_2353082 | Lighthouse Lab in Alderley Park | Wellcome Sanger Institute for the COVID-19 Genomics UK (COG-UK) Consortium | Jacquelyn Wynn, Mairead Hyland, The Lighthouse Lab in Alderley Park and Alex Alderton, Roberto Amato, Jeffrey Barrett, Sonia Goncalves, Ewan Harrison, David K. Jackson, Ian Johnston, Dominic Kwiatkowski, Cordelia Langford, John Sillitoe on behalf of the Wellcome Sanger Institute COVID-19 Surveillance Team |
| EPI_ISL_2353084, EPI_ISL_2353085, EPI_ISL_2353086, EPI_ISL_2353087, EPI_ISL_2353089, EPI_ISL_2353090, EPI_ISL_2353091 | Lighthouse Lab in Milton Keynes | Wellcome Sanger Institute for the COVID-19 Genomics UK (COG-UK) Consortium | The Lighthouse Lab in Milton Keynes and Alex Alderton, Roberto Amato, Jeffrey Barrett, Sonia Goncalves, Ewan Harrison, David K. Jackson, Ian Johnston, Dominic Kwiatkowski, Cordelia Langford, John Sillitoe on behalf of the Wellcome Sanger Institute COVID-19 Surveillance Team |
| EPI_ISL_2353092, EPI_ISL_2353094, EPI_ISL_2353095, EPI_ISL_2353096 | Lighthouse Lab in Alderley Park | Wellcome Sanger Institute for the COVID-19 Genomics UK (COG-UK) Consortium | Jacquelyn Wynn, Mairead Hyland, The Lighthouse Lab in Alderley Park and Alex Alderton, Roberto Amato, Jeffrey Barrett, Sonia Goncalves, Ewan Harrison, David K. Jackson, Ian Johnston, Dominic Kwiatkowski, Cordelia Langford, John Sillitoe on behalf of the Wellcome Sanger Institute COVID-19 Surveillance Team |
| EPI_ISL_2353097 | Lighthouse Lab in Milton Keynes | Wellcome Sanger Institute for the COVID-19 Genomics UK (COG-UK) Consortium | The Lighthouse Lab in Milton Keynes and Alex Alderton, Roberto Amato, Jeffrey Barrett, Sonia Goncalves, Ewan Harrison, David K. Jackson, Ian Johnston, Dominic Kwiatkowski, Cordelia Langford, John Sillitoe on behalf of the Wellcome Sanger Institute COVID-19 Surveillance Team |
| EPI_ISL_2353098, EPI_ISL_2353099, EPI_ISL_2353100, EPI_ISL_2353101, EPI_ISL_2353102 | Lighthouse Lab in Alderley Park | Wellcome Sanger Institute for the COVID-19 Genomics UK (COG-UK) Consortium | Jacquelyn Wynn, Mairead Hyland, The Lighthouse Lab in Alderley Park and Alex Alderton, Roberto Amato, Jeffrey Barrett, Sonia Goncalves, Ewan Harrison, David K. Jackson, Ian Johnston, Dominic Kwiatkowski, Cordelia Langford, John Sillitoe on behalf of the Wellcome Sanger Institute COVID-19 Surveillance Team |
| EPI_ISL_2353103 | Lighthouse Lab in Milton Keynes | Wellcome Sanger Institute for the COVID-19 Genomics UK (COG-UK) Consortium | The Lighthouse Lab in Milton Keynes and Alex Alderton, Roberto Amato, Jeffrey Barrett, Sonia Goncalves, Ewan Harrison, David K. Jackson, Ian Johnston, Dominic Kwiatkowski, Cordelia Langford, John Sillitoe on behalf of the Wellcome Sanger Institute COVID-19 Surveillance Team |
| EPI_ISL_2353105 | Lighthouse Lab in Alderley Park | Wellcome Sanger Institute for the COVID-19 Genomics UK (COG-UK) Consortium | Jacquelyn Wynn, Mairead Hyland, The Lighthouse Lab in Alderley Park and Alex Alderton, Roberto Amato, Jeffrey Barrett, Sonia Goncalves, Ewan Harrison, David K. Jackson, Ian Johnston, Dominic Kwiatkowski, Cordelia Langford, John Sillitoe on behalf of the Wellcome Sanger Institute COVID-19 Surveillance Team |
| EPI_ISL_2353106 | Lighthouse Lab in Milton Keynes | Wellcome Sanger Institute for the COVID-19 Genomics UK (COG-UK) Consortium | The Lighthouse Lab in Milton Keynes and Alex Alderton, Roberto Amato, Jeffrey Barrett, Sonia Goncalves, Ewan Harrison, David K. Jackson, Ian Johnston, Dominic Kwiatkowski, Cordelia Langford, John Sillitoe on behalf of the Wellcome Sanger Institute COVID-19 Surveillance Team |
| EPI_ISL_2353107, EPI_ISL_2353108, EPI_ISL_2353109, EPI_ISL_2353110, EPI_ISL_2353111 | Lighthouse Lab in Alderley Park | Wellcome Sanger Institute for the COVID-19 Genomics UK (COG-UK) Consortium | Jacquelyn Wynn, Mairead Hyland, The Lighthouse Lab in Alderley Park and Alex Alderton, Roberto Amato, Jeffrey Barrett, Sonia Goncalves, Ewan Harrison, David K. Jackson, Ian Johnston, Dominic Kwiatkowski, Cordelia Langford, John Sillitoe on behalf of the Wellcome Sanger Institute COVID-19 Surveillance Team |
| EPI_ISL_2353114 | Lighthouse Lab in Milton Keynes | Wellcome Sanger Institute for the COVID-19 Genomics UK (COG-UK) Consortium | The Lighthouse Lab in Milton Keynes and Alex Alderton, Roberto Amato, Jeffrey Barrett, Sonia Goncalves, Ewan Harrison, David K. Jackson, Ian Johnston, Dominic Kwiatkowski, Cordelia Langford, John Sillitoe on behalf of the Wellcome Sanger Institute COVID-19 Surveillance Team |
| EPI_ISL_2353116, EPI_ISL_2353117, EPI_ISL_2353118, EPI_ISL_2353119, EPI_ISL_2353121, EPI_ISL_2353122 | Lighthouse Lab in Alderley Park | Wellcome Sanger Institute for the COVID-19 Genomics UK (COG-UK) Consortium | Jacquelyn Wynn, Mairead Hyland, The Lighthouse Lab in Alderley Park and Alex Alderton, Roberto Amato, Jeffrey Barrett, Sonia Goncalves, Ewan Harrison, David K. Jackson, Ian Johnston, Dominic Kwiatkowski, Cordelia Langford, John Sillitoe on behalf of the Wellcome Sanger Institute COVID-19 Surveillance Team |
| EPI_ISL_2353123 | Lighthouse Lab in Milton Keynes | Wellcome Sanger Institute for the COVID-19 Genomics UK (COG-UK) Consortium | The Lighthouse Lab in Milton Keynes and Alex Alderton, Roberto Amato, Jeffrey Barrett, Sonia Goncalves, Ewan Harrison, David K. Jackson, Ian Johnston, Dominic Kwiatkowski, Cordelia Langford, John Sillitoe on behalf of the Wellcome Sanger Institute COVID-19 Surveillance Team |
| EPI_ISL_2353124, EPI_ISL_2353125 | Lighthouse Lab in Alderley Park | Wellcome Sanger Institute for the COVID-19 Genomics UK (COG-UK) Consortium | Jacquelyn Wynn, Mairead Hyland, The Lighthouse Lab in Alderley Park and Alex Alderton, Roberto Amato, Jeffrey Barrett, Sonia Goncalves, Ewan Harrison, David K. Jackson, Ian Johnston, Dominic Kwiatkowski, Cordelia Langford, John Sillitoe on behalf of the Wellcome Sanger Institute COVID-19 Surveillance Team |
| EPI_ISL_2353126, EPI_ISL_2353127 | Lighthouse Lab in Milton Keynes | Wellcome Sanger Institute for the COVID-19 Genomics UK (COG-UK) Consortium | The Lighthouse Lab in Milton Keynes and Alex Alderton, Roberto Amato, Jeffrey Barrett, Sonia Goncalves, Ewan Harrison, David K. Jackson, Ian Johnston, Dominic Kwiatkowski, Cordelia Langford, John Sillitoe on behalf of the Wellcome Sanger Institute COVID-19 Surveillance Team |
| EPI_ISL_2353128, EPI_ISL_2353129, EPI_ISL_2353132 | Lighthouse Lab in Alderley Park | Wellcome Sanger Institute for the COVID-19 Genomics UK (COG-UK) Consortium | Jacquelyn Wynn, Mairead Hyland, The Lighthouse Lab in Alderley Park and Alex Alderton, Roberto Amato, Jeffrey Barrett, Sonia Goncalves, Ewan Harrison, David K. Jackson, Ian Johnston, Dominic Kwiatkowski, Cordelia Langford, John Sillitoe on behalf of the Wellcome Sanger Institute COVID-19 Surveillance Team |
| EPI_ISL_2353134, EPI_ISL_2353136 | Lighthouse Lab in Milton Keynes | Wellcome Sanger Institute for the COVID-19 Genomics UK (COG-UK) Consortium | The Lighthouse Lab in Milton Keynes and Alex Alderton, Roberto Amato, Jeffrey Barrett, Sonia Goncalves, Ewan Harrison, David K. Jackson, Ian Johnston, Dominic Kwiatkowski, Cordelia Langford, John Sillitoe on behalf of the Wellcome Sanger Institute COVID-19 Surveillance Team |
| EPI_ISL_2353137 | Lighthouse Lab in Alderley Park | Wellcome Sanger Institute for the COVID-19 Genomics UK (COG-UK) Consortium | Jacquelyn Wynn, Mairead Hyland, The Lighthouse Lab in Alderley Park and Alex Alderton, Roberto Amato, Jeffrey Barrett, Sonia Goncalves, Ewan Harrison, David K. Jackson, Ian Johnston, Dominic Kwiatkowski, Cordelia Langford, John Sillitoe on behalf of the Wellcome Sanger Institute COVID-19 Surveillance Team |
| EPI_ISL_2353138 | Lighthouse Lab in Milton Keynes | Wellcome Sanger Institute for the COVID-19 Genomics UK (COG-UK) Consortium | The Lighthouse Lab in Milton Keynes and Alex Alderton, Roberto Amato, Jeffrey Barrett, Sonia Goncalves, Ewan Harrison, David K. Jackson, Ian Johnston, Dominic Kwiatkowski, Cordelia Langford, John Sillitoe on behalf of the Wellcome Sanger Institute COVID-19 Surveillance Team |
| EPI_ISL_2353139 | Lighthouse Lab in Alderley Park | Wellcome Sanger Institute for the COVID-19 Genomics UK (COG-UK) Consortium | Jacquelyn Wynn, Mairead Hyland, The Lighthouse Lab in Alderley Park and Alex Alderton, Roberto Amato, Jeffrey Barrett, Sonia Goncalves, Ewan Harrison, David K. Jackson, Ian Johnston, Dominic Kwiatkowski, Cordelia Langford, John Sillitoe on behalf of the Wellcome Sanger Institute COVID-19 Surveillance Team |

| | | | |
|---|---|---|---|
| EPI_ISL_2353140 | Lighthouse Lab in Milton Keynes | Wellcome Sanger Institute for the COVID-19 Genomics UK (COG-UK) Consortium | The Lighthouse Lab in Milton Keynes and Alex Alderton, Roberto Amato, Jeffrey Barrett, Sonia Goncalves, Ewan Harrison, David K. Jackson, Ian Johnston, Dominic Kwiatkowski, Cordelia Langford, John Sillitoe on behalf of the Wellcome Sanger Institute COVID-19 Surveillance Team |
| EPI_ISL_2353141 | Lighthouse Lab in Alderley Park | Wellcome Sanger Institute for the COVID-19 Genomics UK (COG-UK) Consortium | Jacquelyn Wynn, Mairead Hyland, The Lighthouse Lab in Alderley Park and Alex Alderton, Roberto Amato, Jeffrey Barrett, Sonia Goncalves, Ewan Harrison, David K. Jackson, Ian Johnston, Dominic Kwiatkowski, Cordelia Langford, John Sillitoe on behalf of the Wellcome Sanger Institute COVID-19 Surveillance Team |
| EPI_ISL_2353143 | Lighthouse Lab in Milton Keynes | Wellcome Sanger Institute for the COVID-19 Genomics UK (COG-UK) Consortium | The Lighthouse Lab in Milton Keynes and Alex Alderton, Roberto Amato, Jeffrey Barrett, Sonia Goncalves, Ewan Harrison, David K. Jackson, Ian Johnston, Dominic Kwiatkowski, Cordelia Langford, John Sillitoe on behalf of the Wellcome Sanger Institute COVID-19 Surveillance Team |
| EPI_ISL_2353144 | Lighthouse Lab in Alderley Park | Wellcome Sanger Institute for the COVID-19 Genomics UK (COG-UK) Consortium | Jacquelyn Wynn, Mairead Hyland, The Lighthouse Lab in Alderley Park and Alex Alderton, Roberto Amato, Jeffrey Barrett, Sonia Goncalves, Ewan Harrison, David K. Jackson, Ian Johnston, Dominic Kwiatkowski, Cordelia Langford, John Sillitoe on behalf of the Wellcome Sanger Institute COVID-19 Surveillance Team |
| EPI_ISL_2353145 | Lighthouse Lab in Milton Keynes | Wellcome Sanger Institute for the COVID-19 Genomics UK (COG-UK) Consortium | The Lighthouse Lab in Milton Keynes and Alex Alderton, Roberto Amato, Jeffrey Barrett, Sonia Goncalves, Ewan Harrison, David K. Jackson, Ian Johnston, Dominic Kwiatkowski, Cordelia Langford, John Sillitoe on behalf of the Wellcome Sanger Institute COVID-19 Surveillance Team |
| EPI_ISL_2353146, EPI_ISL_2353147, EPI_ISL_2353149, EPI_ISL_2353150, EPI_ISL_2353151, EPI_ISL_2353152, EPI_ISL_2353154 | Lighthouse Lab in Alderley Park | Wellcome Sanger Institute for the COVID-19 Genomics UK (COG-UK) Consortium | Jacquelyn Wynn, Mairead Hyland, The Lighthouse Lab in Alderley Park and Alex Alderton, Roberto Amato, Jeffrey Barrett, Sonia Goncalves, Ewan Harrison, David K. Jackson, Ian Johnston, Dominic Kwiatkowski, Cordelia Langford, John Sillitoe on behalf of the Wellcome Sanger Institute COVID-19 Surveillance Team |
| EPI_ISL_2353157 | Lighthouse Lab in Milton Keynes | Wellcome Sanger Institute for the COVID-19 Genomics UK (COG-UK) Consortium | The Lighthouse Lab in Milton Keynes and Alex Alderton, Roberto Amato, Jeffrey Barrett, Sonia Goncalves, Ewan Harrison, David K. Jackson, Ian Johnston, Dominic Kwiatkowski, Cordelia Langford, John Sillitoe on behalf of the Wellcome Sanger Institute COVID-19 Surveillance Team |
| EPI_ISL_2353158, EPI_ISL_2353159 | Lighthouse Lab in Alderley Park | Wellcome Sanger Institute for the COVID-19 Genomics UK (COG-UK) Consortium | Jacquelyn Wynn, Mairead Hyland, The Lighthouse Lab in Alderley Park and Alex Alderton, Roberto Amato, Jeffrey Barrett, Sonia Goncalves, Ewan Harrison, David K. Jackson, Ian Johnston, Dominic Kwiatkowski, Cordelia Langford, John Sillitoe on behalf of the Wellcome Sanger Institute COVID-19 Surveillance Team |
| EPI_ISL_2353160 | Lighthouse Lab in Milton Keynes | Wellcome Sanger Institute for the COVID-19 Genomics UK (COG-UK) Consortium | The Lighthouse Lab in Milton Keynes and Alex Alderton, Roberto Amato, Jeffrey Barrett, Sonia Goncalves, Ewan Harrison, David K. Jackson, Ian Johnston, Dominic Kwiatkowski, Cordelia Langford, John Sillitoe on behalf of the Wellcome Sanger Institute COVID-19 Surveillance Team |
| EPI_ISL_2353161, EPI_ISL_2353164, EPI_ISL_2353165, EPI_ISL_2353166, EPI_ISL_2353170, EPI_ISL_2353171, EPI_ISL_2353172, EPI_ISL_2353173 | Lighthouse Lab in Alderley Park | Wellcome Sanger Institute for the COVID-19 Genomics UK (COG-UK) Consortium | Jacquelyn Wynn, Mairead Hyland, The Lighthouse Lab in Alderley Park and Alex Alderton, Roberto Amato, Jeffrey Barrett, Sonia Goncalves, Ewan Harrison, David K. Jackson, Ian Johnston, Dominic Kwiatkowski, Cordelia Langford, John Sillitoe on behalf of the Wellcome Sanger Institute COVID-19 Surveillance Team |
| EPI_ISL_2353175 | Lighthouse Lab in Milton Keynes | Wellcome Sanger Institute for the COVID-19 Genomics UK (COG-UK) Consortium | The Lighthouse Lab in Milton Keynes and Alex Alderton, Roberto Amato, Jeffrey Barrett, Sonia Goncalves, Ewan Harrison, David K. Jackson, Ian Johnston, Dominic Kwiatkowski, Cordelia Langford, John Sillitoe on behalf of the Wellcome Sanger Institute COVID-19 Surveillance Team |
| EPI_ISL_2353176 | Lighthouse Lab in Alderley Park | Wellcome Sanger Institute for the COVID-19 Genomics UK (COG-UK) Consortium | Jacquelyn Wynn, Mairead Hyland, The Lighthouse Lab in Alderley Park and Alex Alderton, Roberto Amato, Jeffrey Barrett, Sonia Goncalves, Ewan Harrison, David K. Jackson, Ian Johnston, Dominic Kwiatkowski, Cordelia Langford, John Sillitoe on behalf of the Wellcome Sanger Institute COVID-19 Surveillance Team |
| EPI_ISL_2353177, EPI_ISL_2353178, EPI_ISL_2353180 | Lighthouse Lab in Milton Keynes | Wellcome Sanger Institute for the COVID-19 Genomics UK (COG-UK) Consortium | The Lighthouse Lab in Milton Keynes and Alex Alderton, Roberto Amato, Jeffrey Barrett, Sonia Goncalves, Ewan Harrison, David K. Jackson, Ian Johnston, Dominic Kwiatkowski, Cordelia Langford, John Sillitoe on behalf of the Wellcome Sanger Institute COVID-19 Surveillance Team |
| EPI_ISL_2353182 | Lighthouse Lab in Alderley Park | Wellcome Sanger Institute for the COVID-19 Genomics UK (COG-UK) Consortium | Jacquelyn Wynn, Mairead Hyland, The Lighthouse Lab in Alderley Park and Alex Alderton, Roberto Amato, Jeffrey Barrett, Sonia Goncalves, Ewan Harrison, David K. Jackson, Ian Johnston, Dominic Kwiatkowski, Cordelia Langford, John Sillitoe on behalf of the Wellcome Sanger Institute COVID-19 Surveillance Team |
| EPI_ISL_2353184, EPI_ISL_2353185 | Lighthouse Lab in Milton Keynes | Wellcome Sanger Institute for the COVID-19 Genomics UK (COG-UK) Consortium | The Lighthouse Lab in Milton Keynes and Alex Alderton, Roberto Amato, Jeffrey Barrett, Sonia Goncalves, Ewan Harrison, David K. Jackson, Ian Johnston, Dominic Kwiatkowski, Cordelia Langford, John Sillitoe on behalf of the Wellcome Sanger Institute COVID-19 Surveillance Team |
| EPI_ISL_2353186, EPI_ISL_2353187 | Lighthouse Lab in Alderley Park | Wellcome Sanger Institute for the COVID-19 Genomics UK (COG-UK) Consortium | Jacquelyn Wynn, Mairead Hyland, The Lighthouse Lab in Alderley Park and Alex Alderton, Roberto Amato, Jeffrey Barrett, Sonia Goncalves, Ewan Harrison, David K. Jackson, Ian Johnston, Dominic Kwiatkowski, Cordelia Langford, John Sillitoe on behalf of the Wellcome Sanger Institute COVID-19 Surveillance Team |
| EPI_ISL_2353188 | Lighthouse Lab in Milton Keynes | Wellcome Sanger Institute for the COVID-19 Genomics UK (COG-UK) Consortium | The Lighthouse Lab in Milton Keynes and Alex Alderton, Roberto Amato, Jeffrey Barrett, Sonia Goncalves, Ewan Harrison, David K. Jackson, Ian Johnston, Dominic Kwiatkowski, Cordelia Langford, John Sillitoe on behalf of the Wellcome Sanger Institute COVID-19 Surveillance Team |
| EPI_ISL_2353190, EPI_ISL_2353192 | Lighthouse Lab in Alderley Park | Wellcome Sanger Institute for the COVID-19 Genomics UK (COG-UK) Consortium | Jacquelyn Wynn, Mairead Hyland, The Lighthouse Lab in Alderley Park and Alex Alderton, Roberto Amato, Jeffrey Barrett, Sonia Goncalves, Ewan Harrison, David K. Jackson, Ian Johnston, Dominic Kwiatkowski, Cordelia Langford, John Sillitoe on behalf of the Wellcome Sanger Institute COVID-19 Surveillance Team |
| EPI_ISL_2353193, EPI_ISL_2353194, EPI_ISL_2353196, EPI_ISL_2353198, EPI_ISL_2353199, EPI_ISL_2353200 | Lighthouse Lab in Milton Keynes | Wellcome Sanger Institute for the COVID-19 Genomics UK (COG-UK) Consortium | The Lighthouse Lab in Milton Keynes and Alex Alderton, Roberto Amato, Jeffrey Barrett, Sonia Goncalves, Ewan Harrison, David K. Jackson, Ian Johnston, Dominic Kwiatkowski, Cordelia Langford, John Sillitoe on behalf of the Wellcome Sanger Institute COVID-19 Surveillance Team |
| EPI_ISL_2353202, EPI_ISL_2353203 | Lighthouse Lab in Alderley Park | Wellcome Sanger Institute for the COVID-19 Genomics UK (COG-UK) Consortium | Jacquelyn Wynn, Mairead Hyland, The Lighthouse Lab in Alderley Park and Alex Alderton, Roberto Amato, Jeffrey Barrett, Sonia Goncalves, Ewan Harrison, David K. Jackson, Ian Johnston, Dominic Kwiatkowski, Cordelia Langford, John Sillitoe on behalf of the Wellcome Sanger Institute COVID-19 Surveillance Team |
| EPI_ISL_2353204, EPI_ISL_2353205 | Lighthouse Lab in Milton Keynes | Wellcome Sanger Institute for the COVID-19 Genomics UK (COG-UK) Consortium | The Lighthouse Lab in Milton Keynes and Alex Alderton, Roberto Amato, Jeffrey Barrett, Sonia Goncalves, Ewan Harrison, David K. Jackson, Ian Johnston, Dominic Kwiatkowski, Cordelia Langford, John Sillitoe on behalf of the Wellcome Sanger Institute COVID-19 Surveillance Team |
| EPI_ISL_2353206 | Lighthouse Lab in Alderley Park | Wellcome Sanger Institute for the COVID-19 Genomics UK (COG-UK) Consortium | Jacquelyn Wynn, Mairead Hyland, The Lighthouse Lab in Alderley Park and Alex Alderton, Roberto Amato, Jeffrey Barrett, Sonia Goncalves, Ewan Harrison, David K. Jackson, Ian Johnston, Dominic Kwiatkowski, Cordelia Langford, John Sillitoe on behalf of the Wellcome Sanger Institute COVID-19 Surveillance Team |
| EPI_ISL_2353209, EPI_ISL_2353211, EPI_ISL_2353213 | Lighthouse Lab in Milton Keynes | Wellcome Sanger Institute for the COVID-19 Genomics UK (COG-UK) Consortium | The Lighthouse Lab in Milton Keynes and Alex Alderton, Roberto Amato, Jeffrey Barrett, Sonia Goncalves, Ewan Harrison, David K. Jackson, Ian Johnston, Dominic Kwiatkowski, Cordelia Langford, John Sillitoe on behalf of the Wellcome Sanger Institute COVID-19 Surveillance Team |
| EPI_ISL_2353215, EPI_ISL_2353216, EPI_ISL_2353218 | Lighthouse Lab in Alderley Park | Wellcome Sanger Institute for the COVID-19 Genomics UK (COG-UK) Consortium | Jacquelyn Wynn, Mairead Hyland, The Lighthouse Lab in Alderley Park and Alex Alderton, Roberto Amato, Jeffrey Barrett, Sonia Goncalves, Ewan Harrison, David K. Jackson, Ian Johnston, Dominic Kwiatkowski, Cordelia Langford, John Sillitoe on behalf of the Wellcome Sanger Institute COVID-19 Surveillance Team |
| EPI_ISL_2353219 | Lighthouse Lab in Milton Keynes | Wellcome Sanger Institute for the COVID-19 Genomics UK (COG-UK) Consortium | The Lighthouse Lab in Milton Keynes and Alex Alderton, Roberto Amato, Jeffrey Barrett, Sonia Goncalves, Ewan Harrison, David K. Jackson, Ian Johnston, Dominic Kwiatkowski, Cordelia Langford, John Sillitoe on behalf of the Wellcome Sanger Institute COVID-19 Surveillance Team |
| EPI_ISL_2353220, EPI_ISL_2353221, EPI_ISL_2353222 | Lighthouse Lab in Alderley Park | Wellcome Sanger Institute for the COVID-19 Genomics UK (COG-UK) Consortium | Jacquelyn Wynn, Mairead Hyland, The Lighthouse Lab in Alderley Park and Alex Alderton, Roberto Amato, Jeffrey Barrett, Sonia Goncalves, Ewan Harrison, David K. Jackson, Ian Johnston, Dominic Kwiatkowski, Cordelia Langford, John Sillitoe on behalf of the Wellcome Sanger Institute COVID-19 Surveillance Team |
| EPI_ISL_2353223 | Lighthouse Lab in Milton Keynes | Wellcome Sanger Institute for the COVID-19 Genomics UK (COG-UK) Consortium | The Lighthouse Lab in Milton Keynes and Alex Alderton, Roberto Amato, Jeffrey Barrett, Sonia Goncalves, Ewan Harrison, David K. Jackson, Ian Johnston, Dominic Kwiatkowski, Cordelia Langford, John Sillitoe on behalf of the Wellcome Sanger Institute COVID-19 Surveillance Team |
| EPI_ISL_2353224, EPI_ISL_2353225, EPI_ISL_2353226, EPI_ISL_2353227, EPI_ISL_2353228 | Lighthouse Lab in Alderley Park | Wellcome Sanger Institute for the COVID-19 Genomics UK (COG-UK) Consortium | Jacquelyn Wynn, Mairead Hyland, The Lighthouse Lab in Alderley Park and Alex Alderton, Roberto Amato, Jeffrey Barrett, Sonia Goncalves, Ewan Harrison, David K. Jackson, Ian Johnston, Dominic Kwiatkowski, Cordelia Langford, John Sillitoe on behalf of the Wellcome Sanger Institute COVID-19 Surveillance Team |
| EPI_ISL_2353229 | Lighthouse Lab in Milton Keynes | Wellcome Sanger Institute for the COVID-19 Genomics UK (COG-UK) Consortium | The Lighthouse Lab in Milton Keynes and Alex Alderton, Roberto Amato, Jeffrey Barrett, Sonia Goncalves, Ewan Harrison, David K. Jackson, Ian Johnston, Dominic Kwiatkowski, Cordelia Langford, John Sillitoe on behalf of the Wellcome Sanger Institute COVID-19 Surveillance Team |
| EPI_ISL_2353230, EPI_ISL_2353231 | Lighthouse Lab in Alderley Park | Wellcome Sanger Institute for the COVID-19 Genomics UK (COG-UK) Consortium | Jacquelyn Wynn, Mairead Hyland, The Lighthouse Lab in Alderley Park and Alex Alderton, Roberto Amato, Jeffrey Barrett, Sonia Goncalves, Ewan Harrison, David K. Jackson, Ian Johnston, Dominic Kwiatkowski, Cordelia Langford, John Sillitoe on behalf of the Wellcome Sanger Institute COVID-19 Surveillance Team |
| EPI_ISL_2353234 | Lighthouse Lab in Milton Keynes | Wellcome Sanger Institute for the COVID-19 Genomics UK (COG-UK) Consortium | The Lighthouse Lab in Milton Keynes and Alex Alderton, Roberto Amato, Jeffrey Barrett, Sonia Goncalves, Ewan Harrison, David K. Jackson, Ian Johnston, Dominic Kwiatkowski, Cordelia Langford, John Sillitoe on behalf of the Wellcome Sanger Institute COVID-19 Surveillance Team |
| EPI_ISL_2353235, EPI_ISL_2353237, EPI_ISL_2353239 | Lighthouse Lab in Alderley Park | Wellcome Sanger Institute for the COVID-19 Genomics UK (COG-UK) Consortium | Jacquelyn Wynn, Mairead Hyland, The Lighthouse Lab in Alderley Park and Alex Alderton, Roberto Amato, Jeffrey Barrett, Sonia Goncalves, Ewan Harrison, David K. Jackson, Ian Johnston, Dominic Kwiatkowski, Cordelia Langford, John Sillitoe on behalf of the Wellcome Sanger Institute COVID-19 Surveillance Team |
| EPI_ISL_2353240 | Lighthouse Lab in Milton Keynes | Wellcome Sanger Institute for the COVID-19 Genomics UK (COG-UK) Consortium | The Lighthouse Lab in Milton Keynes and Alex Alderton, Roberto Amato, Jeffrey Barrett, Sonia Goncalves, Ewan Harrison, David K. Jackson, Ian Johnston, Dominic Kwiatkowski, Cordelia Langford, John Sillitoe on behalf of the Wellcome Sanger Institute COVID-19 Surveillance Team |
| EPI_ISL_2353241, EPI_ISL_2353242, EPI_ISL_2353243 | Lighthouse Lab in Alderley Park | Wellcome Sanger Institute for the COVID-19 Genomics UK (COG-UK) Consortium | Jacquelyn Wynn, Mairead Hyland, The Lighthouse Lab in Alderley Park and Alex Alderton, Roberto Amato, Jeffrey Barrett, Sonia Goncalves, Ewan Harrison, David K. Jackson, Ian Johnston, Dominic Kwiatkowski, Cordelia Langford, John Sillitoe on behalf of the Wellcome Sanger Institute COVID-19 Surveillance Team |
| EPI_ISL_2353244 | Lighthouse Lab in Milton Keynes | Wellcome Sanger Institute for the COVID-19 Genomics UK (COG-UK) Consortium | The Lighthouse Lab in Milton Keynes and Alex Alderton, Roberto Amato, Jeffrey Barrett, Sonia Goncalves, Ewan Harrison, David K. Jackson, Ian Johnston, Dominic Kwiatkowski, Cordelia Langford, John Sillitoe on behalf of the Wellcome Sanger Institute COVID-19 Surveillance Team |

| | | | |
|---|---|---|---|
| EPI_ISL_2353245 | Lighthouse Lab in Alderley Park | Wellcome Sanger Institute for the COVID-19 Genomics UK (COG-UK) Consortium | Jacquelyn Wynn, Mairead Hyland, The Lighthouse Lab in Alderley Park and Alex Alderton, Roberto Amato, Jeffrey Barrett, Sonia Goncalves, Ewan Harrison, David K. Jackson, Ian Johnston, Dominic Kwiatkowski, Cordelia Langford, John Sillitoe on behalf of the Wellcome Sanger Institute COVID-19 Surveillance Team |
| EPI_ISL_2353246 | Lighthouse Lab in Milton Keynes | Wellcome Sanger Institute for the COVID-19 Genomics UK (COG-UK) Consortium | The Lighthouse Lab in Milton Keynes and Alex Alderton, Roberto Amato, Jeffrey Barrett, Sonia Goncalves, Ewan Harrison, David K. Jackson, Ian Johnston, Dominic Kwiatkowski, Cordelia Langford, John Sillitoe on behalf of the Wellcome Sanger Institute COVID-19 Surveillance Team |
| EPI_ISL_2353248, EPI_ISL_2353249, EPI_ISL_2353250, EPI_ISL_2353251 | Lighthouse Lab in Alderley Park | Wellcome Sanger Institute for the COVID-19 Genomics UK (COG-UK) Consortium | Jacquelyn Wynn, Mairead Hyland, The Lighthouse Lab in Alderley Park and Alex Alderton, Roberto Amato, Jeffrey Barrett, Sonia Goncalves, Ewan Harrison, David K. Jackson, Ian Johnston, Dominic Kwiatkowski, Cordelia Langford, John Sillitoe on behalf of the Wellcome Sanger Institute COVID-19 Surveillance Team |
| EPI_ISL_2353252, EPI_ISL_2353253, EPI_ISL_2353255, EPI_ISL_2353256 | Lighthouse Lab in Milton Keynes | Wellcome Sanger Institute for the COVID-19 Genomics UK (COG-UK) Consortium | The Lighthouse Lab in Milton Keynes and Alex Alderton, Roberto Amato, Jeffrey Barrett, Sonia Goncalves, Ewan Harrison, David K. Jackson, Ian Johnston, Dominic Kwiatkowski, Cordelia Langford, John Sillitoe on behalf of the Wellcome Sanger Institute COVID-19 Surveillance Team |
| EPI_ISL_2353257 | Lighthouse Lab in Alderley Park | Wellcome Sanger Institute for the COVID-19 Genomics UK (COG-UK) Consortium | Jacquelyn Wynn, Mairead Hyland, The Lighthouse Lab in Alderley Park and Alex Alderton, Roberto Amato, Jeffrey Barrett, Sonia Goncalves, Ewan Harrison, David K. Jackson, Ian Johnston, Dominic Kwiatkowski, Cordelia Langford, John Sillitoe on behalf of the Wellcome Sanger Institute COVID-19 Surveillance Team |
| EPI_ISL_2353258 | Lighthouse Laboratory Plymouth | Wellcome Sanger Institute for the COVID-19 Genomics UK (COG-UK) Consortium | Lighthouse Laboratory Plymouth and Alex Alderton, Roberto Amato, Jeffrey Barrett, Sonia Goncalves, Ewan Harrison, David K. Jackson, Ian Johnston, Dominic Kwiatkowski, Cordelia Langford, John Sillitoe on behalf of the Wellcome Sanger Institute COVID-19 Surveillance Team |
| EPI_ISL_2353259, EPI_ISL_2353260, EPI_ISL_2353262 | Lighthouse Lab in Milton Keynes | Wellcome Sanger Institute for the COVID-19 Genomics UK (COG-UK) Consortium | The Lighthouse Lab in Milton Keynes and Alex Alderton, Roberto Amato, Jeffrey Barrett, Sonia Goncalves, Ewan Harrison, David K. Jackson, Ian Johnston, Dominic Kwiatkowski, Cordelia Langford, John Sillitoe on behalf of the Wellcome Sanger Institute COVID-19 Surveillance Team |
| EPI_ISL_2353264 | Lighthouse Laboratory Plymouth | Wellcome Sanger Institute for the COVID-19 Genomics UK (COG-UK) Consortium | Lighthouse Laboratory Plymouth and Alex Alderton, Roberto Amato, Jeffrey Barrett, Sonia Goncalves, Ewan Harrison, David K. Jackson, Ian Johnston, Dominic Kwiatkowski, Cordelia Langford, John Sillitoe on behalf of the Wellcome Sanger Institute COVID-19 Surveillance Team |
| EPI_ISL_2353265, EPI_ISL_2353266 | Lighthouse Lab in Alderley Park | Wellcome Sanger Institute for the COVID-19 Genomics UK (COG-UK) Consortium | Jacquelyn Wynn, Mairead Hyland, The Lighthouse Lab in Alderley Park and Alex Alderton, Roberto Amato, Jeffrey Barrett, Sonia Goncalves, Ewan Harrison, David K. Jackson, Ian Johnston, Dominic Kwiatkowski, Cordelia Langford, John Sillitoe on behalf of the Wellcome Sanger Institute COVID-19 Surveillance Team |
| EPI_ISL_2353268, EPI_ISL_2353269, EPI_ISL_2353274 | Lighthouse Lab in Milton Keynes | Wellcome Sanger Institute for the COVID-19 Genomics UK (COG-UK) Consortium | The Lighthouse Lab in Milton Keynes and Alex Alderton, Roberto Amato, Jeffrey Barrett, Sonia Goncalves, Ewan Harrison, David K. Jackson, Ian Johnston, Dominic Kwiatkowski, Cordelia Langford, John Sillitoe on behalf of the Wellcome Sanger Institute COVID-19 Surveillance Team |
| EPI_ISL_2353275, EPI_ISL_2353276 | Lighthouse Lab in Alderley Park | Wellcome Sanger Institute for the COVID-19 Genomics UK (COG-UK) Consortium | Jacquelyn Wynn, Mairead Hyland, The Lighthouse Lab in Alderley Park and Alex Alderton, Roberto Amato, Jeffrey Barrett, Sonia Goncalves, Ewan Harrison, David K. Jackson, Ian Johnston, Dominic Kwiatkowski, Cordelia Langford, John Sillitoe on behalf of the Wellcome Sanger Institute COVID-19 Surveillance Team |
| EPI_ISL_2353278, EPI_ISL_2353279 | Lighthouse Lab in Milton Keynes | Wellcome Sanger Institute for the COVID-19 Genomics UK (COG-UK) Consortium | The Lighthouse Lab in Milton Keynes and Alex Alderton, Roberto Amato, Jeffrey Barrett, Sonia Goncalves, Ewan Harrison, David K. Jackson, Ian Johnston, Dominic Kwiatkowski, Cordelia Langford, John Sillitoe on behalf of the Wellcome Sanger Institute COVID-19 Surveillance Team |
| EPI_ISL_2353281 | Lighthouse Lab in Alderley Park | Wellcome Sanger Institute for the COVID-19 Genomics UK (COG-UK) Consortium | Jacquelyn Wynn, Mairead Hyland, The Lighthouse Lab in Alderley Park and Alex Alderton, Roberto Amato, Jeffrey Barrett, Sonia Goncalves, Ewan Harrison, David K. Jackson, Ian Johnston, Dominic Kwiatkowski, Cordelia Langford, John Sillitoe on behalf of the Wellcome Sanger Institute COVID-19 Surveillance Team |
| EPI_ISL_2353282, EPI_ISL_2353287, EPI_ISL_2353288, EPI_ISL_2353289 | Lighthouse Lab in Milton Keynes | Wellcome Sanger Institute for the COVID-19 Genomics UK (COG-UK) Consortium | The Lighthouse Lab in Milton Keynes and Alex Alderton, Roberto Amato, Jeffrey Barrett, Sonia Goncalves, Ewan Harrison, David K. Jackson, Ian Johnston, Dominic Kwiatkowski, Cordelia Langford, John Sillitoe on behalf of the Wellcome Sanger Institute COVID-19 Surveillance Team |
| EPI_ISL_2353291 | Lighthouse Lab in Alderley Park | Wellcome Sanger Institute for the COVID-19 Genomics UK (COG-UK) Consortium | Jacquelyn Wynn, Mairead Hyland, The Lighthouse Lab in Alderley Park and Alex Alderton, Roberto Amato, Jeffrey Barrett, Sonia Goncalves, Ewan Harrison, David K. Jackson, Ian Johnston, Dominic Kwiatkowski, Cordelia Langford, John Sillitoe on behalf of the Wellcome Sanger Institute COVID-19 Surveillance Team |
| EPI_ISL_2353292, EPI_ISL_2353293, EPI_ISL_2353295, EPI_ISL_2353296 | Lighthouse Lab in Milton Keynes | Wellcome Sanger Institute for the COVID-19 Genomics UK (COG-UK) Consortium | The Lighthouse Lab in Milton Keynes and Alex Alderton, Roberto Amato, Jeffrey Barrett, Sonia Goncalves, Ewan Harrison, David K. Jackson, Ian Johnston, Dominic Kwiatkowski, Cordelia Langford, John Sillitoe on behalf of the Wellcome Sanger Institute COVID-19 Surveillance Team |
| EPI_ISL_2353297 | Lighthouse Lab in Alderley Park | Wellcome Sanger Institute for the COVID-19 Genomics UK (COG-UK) Consortium | Jacquelyn Wynn, Mairead Hyland, The Lighthouse Lab in Alderley Park and Alex Alderton, Roberto Amato, Jeffrey Barrett, Sonia Goncalves, Ewan Harrison, David K. Jackson, Ian Johnston, Dominic Kwiatkowski, Cordelia Langford, John Sillitoe on behalf of the Wellcome Sanger Institute COVID-19 Surveillance Team |
| EPI_ISL_2353298, EPI_ISL_2353300 | Lighthouse Lab in Milton Keynes | Wellcome Sanger Institute for the COVID-19 Genomics UK (COG-UK) Consortium | The Lighthouse Lab in Milton Keynes and Alex Alderton, Roberto Amato, Jeffrey Barrett, Sonia Goncalves, Ewan Harrison, David K. Jackson, Ian Johnston, Dominic Kwiatkowski, Cordelia Langford, John Sillitoe on behalf of the Wellcome Sanger Institute COVID-19 Surveillance Team |
| EPI_ISL_2353301, EPI_ISL_2353303 | Lighthouse Lab in Alderley Park | Wellcome Sanger Institute for the COVID-19 Genomics UK (COG-UK) Consortium | Jacquelyn Wynn, Mairead Hyland, The Lighthouse Lab in Alderley Park and Alex Alderton, Roberto Amato, Jeffrey Barrett, Sonia Goncalves, Ewan Harrison, David K. Jackson, Ian Johnston, Dominic Kwiatkowski, Cordelia Langford, John Sillitoe on behalf of the Wellcome Sanger Institute COVID-19 Surveillance Team |
| EPI_ISL_2353304 | Lighthouse Lab in Milton Keynes | Wellcome Sanger Institute for the COVID-19 Genomics UK (COG-UK) Consortium | The Lighthouse Lab in Milton Keynes and Alex Alderton, Roberto Amato, Jeffrey Barrett, Sonia Goncalves, Ewan Harrison, David K. Jackson, Ian Johnston, Dominic Kwiatkowski, Cordelia Langford, John Sillitoe on behalf of the Wellcome Sanger Institute COVID-19 Surveillance Team |
| EPI_ISL_2353306, EPI_ISL_2353307, EPI_ISL_2353309 | Lighthouse Lab in Alderley Park | Wellcome Sanger Institute for the COVID-19 Genomics UK (COG-UK) Consortium | Jacquelyn Wynn, Mairead Hyland, The Lighthouse Lab in Alderley Park and Alex Alderton, Roberto Amato, Jeffrey Barrett, Sonia Goncalves, Ewan Harrison, David K. Jackson, Ian Johnston, Dominic Kwiatkowski, Cordelia Langford, John Sillitoe on behalf of the Wellcome Sanger Institute COVID-19 Surveillance Team |
| EPI_ISL_2353314 | Lighthouse Lab in Milton Keynes | Wellcome Sanger Institute for the COVID-19 Genomics UK (COG-UK) Consortium | The Lighthouse Lab in Milton Keynes and Alex Alderton, Roberto Amato, Jeffrey Barrett, Sonia Goncalves, Ewan Harrison, David K. Jackson, Ian Johnston, Dominic Kwiatkowski, Cordelia Langford, John Sillitoe on behalf of the Wellcome Sanger Institute COVID-19 Surveillance Team |
| EPI_ISL_2353316 | Lighthouse Lab in Alderley Park | Wellcome Sanger Institute for the COVID-19 Genomics UK (COG-UK) Consortium | Jacquelyn Wynn, Mairead Hyland, The Lighthouse Lab in Alderley Park and Alex Alderton, Roberto Amato, Jeffrey Barrett, Sonia Goncalves, Ewan Harrison, David K. Jackson, Ian Johnston, Dominic Kwiatkowski, Cordelia Langford, John Sillitoe on behalf of the Wellcome Sanger Institute COVID-19 Surveillance Team |
| EPI_ISL_2353317 | Lighthouse Lab in Milton Keynes | Wellcome Sanger Institute for the COVID-19 Genomics UK (COG-UK) Consortium | The Lighthouse Lab in Milton Keynes and Alex Alderton, Roberto Amato, Jeffrey Barrett, Sonia Goncalves, Ewan Harrison, David K. Jackson, Ian Johnston, Dominic Kwiatkowski, Cordelia Langford, John Sillitoe on behalf of the Wellcome Sanger Institute COVID-19 Surveillance Team |
| EPI_ISL_2353318 | Lighthouse Laboratory Plymouth | Wellcome Sanger Institute for the COVID-19 Genomics UK (COG-UK) Consortium | Lighthouse Laboratory Plymouth and Alex Alderton, Roberto Amato, Jeffrey Barrett, Sonia Goncalves, Ewan Harrison, David K. Jackson, Ian Johnston, Dominic Kwiatkowski, Cordelia Langford, John Sillitoe on behalf of the Wellcome Sanger Institute COVID-19 Surveillance Team |
| EPI_ISL_2353319, EPI_ISL_2353320 | Lighthouse Lab in Milton Keynes | Wellcome Sanger Institute for the COVID-19 Genomics UK (COG-UK) Consortium | The Lighthouse Lab in Milton Keynes and Alex Alderton, Roberto Amato, Jeffrey Barrett, Sonia Goncalves, Ewan Harrison, David K. Jackson, Ian Johnston, Dominic Kwiatkowski, Cordelia Langford, John Sillitoe on behalf of the Wellcome Sanger Institute COVID-19 Surveillance Team |
| EPI_ISL_2353321 | Lighthouse Lab in Alderley Park | Wellcome Sanger Institute for the COVID-19 Genomics UK (COG-UK) Consortium | Jacquelyn Wynn, Mairead Hyland, The Lighthouse Lab in Alderley Park and Alex Alderton, Roberto Amato, Jeffrey Barrett, Sonia Goncalves, Ewan Harrison, David K. Jackson, Ian Johnston, Dominic Kwiatkowski, Cordelia Langford, John Sillitoe on behalf of the Wellcome Sanger Institute COVID-19 Surveillance Team |
| EPI_ISL_2353322, EPI_ISL_2353325, EPI_ISL_2353326 | Lighthouse Lab in Milton Keynes | Wellcome Sanger Institute for the COVID-19 Genomics UK (COG-UK) Consortium | The Lighthouse Lab in Milton Keynes and Alex Alderton, Roberto Amato, Jeffrey Barrett, Sonia Goncalves, Ewan Harrison, David K. Jackson, Ian Johnston, Dominic Kwiatkowski, Cordelia Langford, John Sillitoe on behalf of the Wellcome Sanger Institute COVID-19 Surveillance Team |
| EPI_ISL_2353327 | Lighthouse Lab in Alderley Park | Wellcome Sanger Institute for the COVID-19 Genomics UK (COG-UK) Consortium | Jacquelyn Wynn, Mairead Hyland, The Lighthouse Lab in Alderley Park and Alex Alderton, Roberto Amato, Jeffrey Barrett, Sonia Goncalves, Ewan Harrison, David K. Jackson, Ian Johnston, Dominic Kwiatkowski, Cordelia Langford, John Sillitoe on behalf of the Wellcome Sanger Institute COVID-19 Surveillance Team |
| EPI_ISL_2353328, EPI_ISL_2353329, EPI_ISL_2353330 | Lighthouse Lab in Milton Keynes | Wellcome Sanger Institute for the COVID-19 Genomics UK (COG-UK) Consortium | The Lighthouse Lab in Milton Keynes and Alex Alderton, Roberto Amato, Jeffrey Barrett, Sonia Goncalves, Ewan Harrison, David K. Jackson, Ian Johnston, Dominic Kwiatkowski, Cordelia Langford, John Sillitoe on behalf of the Wellcome Sanger Institute COVID-19 Surveillance Team |
| EPI_ISL_2353332 | Lighthouse Lab in Alderley Park | Wellcome Sanger Institute for the COVID-19 Genomics UK (COG-UK) Consortium | Jacquelyn Wynn, Mairead Hyland, The Lighthouse Lab in Alderley Park and Alex Alderton, Roberto Amato, Jeffrey Barrett, Sonia Goncalves, Ewan Harrison, David K. Jackson, Ian Johnston, Dominic Kwiatkowski, Cordelia Langford, John Sillitoe on behalf of the Wellcome Sanger Institute COVID-19 Surveillance Team |
| EPI_ISL_2353333 | Lighthouse Laboratory Plymouth | Wellcome Sanger Institute for the COVID-19 Genomics UK (COG-UK) Consortium | Lighthouse Laboratory Plymouth and Alex Alderton, Roberto Amato, Jeffrey Barrett, Sonia Goncalves, Ewan Harrison, David K. Jackson, Ian Johnston, Dominic Kwiatkowski, Cordelia Langford, John Sillitoe on behalf of the Wellcome Sanger Institute COVID-19 Surveillance Team |
| EPI_ISL_2353334 | Lighthouse Lab in Milton Keynes | Wellcome Sanger Institute for the COVID-19 Genomics UK (COG-UK) Consortium | The Lighthouse Lab in Milton Keynes and Alex Alderton, Roberto Amato, Jeffrey Barrett, Sonia Goncalves, Ewan Harrison, David K. Jackson, Ian Johnston, Dominic Kwiatkowski, Cordelia Langford, John Sillitoe on behalf of the Wellcome Sanger Institute COVID-19 Surveillance Team |
| EPI_ISL_2353335 | Lighthouse Laboratory Plymouth | Wellcome Sanger Institute for the COVID-19 Genomics UK (COG-UK) Consortium | Lighthouse Laboratory Plymouth and Alex Alderton, Roberto Amato, Jeffrey Barrett, Sonia Goncalves, Ewan Harrison, David K. Jackson, Ian Johnston, Dominic Kwiatkowski, Cordelia Langford, John Sillitoe on behalf of the Wellcome Sanger Institute COVID-19 Surveillance Team |
| EPI_ISL_2353336, EPI_ISL_2353337, EPI_ISL_2353339 | Lighthouse Lab in Alderley Park | Wellcome Sanger Institute for the COVID-19 Genomics UK (COG-UK) Consortium | Jacquelyn Wynn, Mairead Hyland, The Lighthouse Lab in Alderley Park and Alex Alderton, Roberto Amato, Jeffrey Barrett, Sonia Goncalves, Ewan Harrison, David K. Jackson, Ian Johnston, Dominic Kwiatkowski, Cordelia Langford, John Sillitoe on behalf of the Wellcome Sanger Institute COVID-19 Surveillance Team |
| EPI_ISL_2353340, EPI_ISL_2353341, EPI_ISL_2353342, EPI_ISL_2353343, EPI_ISL_2353344, EPI_ISL_2353345 | Lighthouse Lab in Milton Keynes | Wellcome Sanger Institute for the COVID-19 Genomics UK (COG-UK) Consortium | The Lighthouse Lab in Milton Keynes and Alex Alderton, Roberto Amato, Jeffrey Barrett, Sonia Goncalves, Ewan Harrison, David K. Jackson, Ian Johnston, Dominic Kwiatkowski, Cordelia Langford, John Sillitoe on behalf of the Wellcome Sanger Institute COVID-19 Surveillance Team |
| EPI_ISL_2353346, EPI_ISL_2353348, EPI_ISL_2353349 | Lighthouse Laboratory Plymouth | Wellcome Sanger Institute for the COVID-19 Genomics UK (COG-UK) Consortium | Lighthouse Laboratory Plymouth and Alex Alderton, Roberto Amato, Jeffrey Barrett, Sonia Goncalves, Ewan Harrison, David K. Jackson, Ian Johnston, Dominic Kwiatkowski, Cordelia Langford, John Sillitoe on behalf of the Wellcome Sanger Institute COVID-19 Surveillance Team |
| EPI_ISL_2353350 | Lighthouse Lab in Alderley Park | Wellcome Sanger Institute for the COVID-19 Genomics UK (COG-UK) Consortium | Jacquelyn Wynn, Mairead Hyland, The Lighthouse Lab in Alderley Park and Alex Alderton, Roberto Amato, Jeffrey Barrett, Sonia Goncalves, Ewan Harrison, David K. Jackson, Ian Johnston, Dominic Kwiatkowski, Cordelia Langford, John Sillitoe on |

| | | | |
|---|---|---|---|
| | | Consortium | behalf of the Wellcome Sanger Institute COVID-19 Surveillance Team |
| EPI_ISL_2353351, EPI_ISL_2353352, EPI_ISL_2353353 | Lighthouse Lab in Milton Keynes | Wellcome Sanger Institute for the COVID-19 Genomics UK (COG-UK) Consortium | The Lighthouse Lab in Milton Keynes and Alex Alderton, Roberto Amato, Jeffrey Barrett, Sonia Goncalves, Ewan Harrison, David K. Jackson, Ian Johnston, Dominic Kwiatkowski, Cordelia Langford, John Sillitoe on behalf of the Wellcome Sanger Institute COVID-19 Surveillance Team |
| EPI_ISL_2353354 | Lighthouse Lab in Alderley Park | Wellcome Sanger Institute for the COVID-19 Genomics UK (COG-UK) Consortium | Jacquelyn Wynn, Mairead Hyland, The Lighthouse Lab in Alderley Park and Alex Alderton, Roberto Amato, Jeffrey Barrett, Sonia Goncalves, Ewan Harrison, David K. Jackson, Ian Johnston, Dominic Kwiatkowski, Cordelia Langford, John Sillitoe on behalf of the Wellcome Sanger Institute COVID-19 Surveillance Team |
| EPI_ISL_2353356, EPI_ISL_2353357, EPI_ISL_2353359 | Lighthouse Lab in Milton Keynes | Wellcome Sanger Institute for the COVID-19 Genomics UK (COG-UK) Consortium | The Lighthouse Lab in Milton Keynes and Alex Alderton, Roberto Amato, Jeffrey Barrett, Sonia Goncalves, Ewan Harrison, David K. Jackson, Ian Johnston, Dominic Kwiatkowski, Cordelia Langford, John Sillitoe on behalf of the Wellcome Sanger Institute COVID-19 Surveillance Team |
| EPI_ISL_2353360 | Lighthouse Lab in Alderley Park | Wellcome Sanger Institute for the COVID-19 Genomics UK (COG-UK) Consortium | Jacquelyn Wynn, Mairead Hyland, The Lighthouse Lab in Alderley Park and Alex Alderton, Roberto Amato, Jeffrey Barrett, Sonia Goncalves, Ewan Harrison, David K. Jackson, Ian Johnston, Dominic Kwiatkowski, Cordelia Langford, John Sillitoe on behalf of the Wellcome Sanger Institute COVID-19 Surveillance Team |
| EPI_ISL_2353361 | Lighthouse Laboratory Plymouth | Wellcome Sanger Institute for the COVID-19 Genomics UK (COG-UK) Consortium | Lighthouse Laboratory Plymouth and Alex Alderton, Roberto Amato, Jeffrey Barrett, Sonia Goncalves, Ewan Harrison, David K. Jackson, Ian Johnston, Dominic Kwiatkowski, Cordelia Langford, John Sillitoe on behalf of the Wellcome Sanger Institute COVID-19 Surveillance Team |
| EPI_ISL_2353362, EPI_ISL_2353363 | Lighthouse Lab in Milton Keynes | Wellcome Sanger Institute for the COVID-19 Genomics UK (COG-UK) Consortium | The Lighthouse Lab in Milton Keynes and Alex Alderton, Roberto Amato, Jeffrey Barrett, Sonia Goncalves, Ewan Harrison, David K. Jackson, Ian Johnston, Dominic Kwiatkowski, Cordelia Langford, John Sillitoe on behalf of the Wellcome Sanger Institute COVID-19 Surveillance Team |
| EPI_ISL_2353364 | Lighthouse Lab in Alderley Park | Wellcome Sanger Institute for the COVID-19 Genomics UK (COG-UK) Consortium | Jacquelyn Wynn, Mairead Hyland, The Lighthouse Lab in Alderley Park and Alex Alderton, Roberto Amato, Jeffrey Barrett, Sonia Goncalves, Ewan Harrison, David K. Jackson, Ian Johnston, Dominic Kwiatkowski, Cordelia Langford, John Sillitoe on behalf of the Wellcome Sanger Institute COVID-19 Surveillance Team |
| EPI_ISL_2353365, EPI_ISL_2353366, EPI_ISL_2353367 | Lighthouse Lab in Milton Keynes | Wellcome Sanger Institute for the COVID-19 Genomics UK (COG-UK) Consortium | The Lighthouse Lab in Milton Keynes and Alex Alderton, Roberto Amato, Jeffrey Barrett, Sonia Goncalves, Ewan Harrison, David K. Jackson, Ian Johnston, Dominic Kwiatkowski, Cordelia Langford, John Sillitoe on behalf of the Wellcome Sanger Institute COVID-19 Surveillance Team |
| EPI_ISL_2353368 | Lighthouse Lab in Alderley Park | Wellcome Sanger Institute for the COVID-19 Genomics UK (COG-UK) Consortium | Jacquelyn Wynn, Mairead Hyland, The Lighthouse Lab in Alderley Park and Alex Alderton, Roberto Amato, Jeffrey Barrett, Sonia Goncalves, Ewan Harrison, David K. Jackson, Ian Johnston, Dominic Kwiatkowski, Cordelia Langford, John Sillitoe on behalf of the Wellcome Sanger Institute COVID-19 Surveillance Team |
| EPI_ISL_2353369 | Lighthouse Lab in Milton Keynes | Wellcome Sanger Institute for the COVID-19 Genomics UK (COG-UK) Consortium | The Lighthouse Lab in Milton Keynes and Alex Alderton, Roberto Amato, Jeffrey Barrett, Sonia Goncalves, Ewan Harrison, David K. Jackson, Ian Johnston, Dominic Kwiatkowski, Cordelia Langford, John Sillitoe on behalf of the Wellcome Sanger Institute COVID-19 Surveillance Team |
| EPI_ISL_2353370 | Lighthouse Laboratory Plymouth | Wellcome Sanger Institute for the COVID-19 Genomics UK (COG-UK) Consortium | Lighthouse Laboratory Plymouth and Alex Alderton, Roberto Amato, Jeffrey Barrett, Sonia Goncalves, Ewan Harrison, David K. Jackson, Ian Johnston, Dominic Kwiatkowski, Cordelia Langford, John Sillitoe on behalf of the Wellcome Sanger Institute COVID-19 Surveillance Team |
| EPI_ISL_2353372 | Lighthouse Lab in Alderley Park | Wellcome Sanger Institute for the COVID-19 Genomics UK (COG-UK) Consortium | Jacquelyn Wynn, Mairead Hyland, The Lighthouse Lab in Alderley Park and Alex Alderton, Roberto Amato, Jeffrey Barrett, Sonia Goncalves, Ewan Harrison, David K. Jackson, Ian Johnston, Dominic Kwiatkowski, Cordelia Langford, John Sillitoe on behalf of the Wellcome Sanger Institute COVID-19 Surveillance Team |
| EPI_ISL_2353373, EPI_ISL_2353375 | Lighthouse Lab in Milton Keynes | Wellcome Sanger Institute for the COVID-19 Genomics UK (COG-UK) Consortium | The Lighthouse Lab in Milton Keynes and Alex Alderton, Roberto Amato, Jeffrey Barrett, Sonia Goncalves, Ewan Harrison, David K. Jackson, Ian Johnston, Dominic Kwiatkowski, Cordelia Langford, John Sillitoe on behalf of the Wellcome Sanger Institute COVID-19 Surveillance Team |
| EPI_ISL_2353376 | Lighthouse Laboratory Plymouth | Wellcome Sanger Institute for the COVID-19 Genomics UK (COG-UK) Consortium | Lighthouse Laboratory Plymouth and Alex Alderton, Roberto Amato, Jeffrey Barrett, Sonia Goncalves, Ewan Harrison, David K. Jackson, Ian Johnston, Dominic Kwiatkowski, Cordelia Langford, John Sillitoe on behalf of the Wellcome Sanger Institute COVID-19 Surveillance Team |
| EPI_ISL_2353377 | Lighthouse Lab in Alderley Park | Wellcome Sanger Institute for the COVID-19 Genomics UK (COG-UK) Consortium | Jacquelyn Wynn, Mairead Hyland, The Lighthouse Lab in Alderley Park and Alex Alderton, Roberto Amato, Jeffrey Barrett, Sonia Goncalves, Ewan Harrison, David K. Jackson, Ian Johnston, Dominic Kwiatkowski, Cordelia Langford, John Sillitoe on behalf of the Wellcome Sanger Institute COVID-19 Surveillance Team |
| EPI_ISL_2353378 | Lighthouse Lab in Milton Keynes | Wellcome Sanger Institute for the COVID-19 Genomics UK (COG-UK) Consortium | The Lighthouse Lab in Milton Keynes and Alex Alderton, Roberto Amato, Jeffrey Barrett, Sonia Goncalves, Ewan Harrison, David K. Jackson, Ian Johnston, Dominic Kwiatkowski, Cordelia Langford, John Sillitoe on behalf of the Wellcome Sanger Institute COVID-19 Surveillance Team |
| EPI_ISL_2353379 | Lighthouse Lab in Alderley Park | Wellcome Sanger Institute for the COVID-19 Genomics UK (COG-UK) Consortium | Jacquelyn Wynn, Mairead Hyland, The Lighthouse Lab in Alderley Park and Alex Alderton, Roberto Amato, Jeffrey Barrett, Sonia Goncalves, Ewan Harrison, David K. Jackson, Ian Johnston, Dominic Kwiatkowski, Cordelia Langford, John Sillitoe on behalf of the Wellcome Sanger Institute COVID-19 Surveillance Team |
| EPI_ISL_2353380 | Lighthouse Laboratory Plymouth | Wellcome Sanger Institute for the COVID-19 Genomics UK (COG-UK) Consortium | Lighthouse Laboratory Plymouth and Alex Alderton, Roberto Amato, Jeffrey Barrett, Sonia Goncalves, Ewan Harrison, David K. Jackson, Ian Johnston, Dominic Kwiatkowski, Cordelia Langford, John Sillitoe on behalf of the Wellcome Sanger Institute COVID-19 Surveillance Team |
| EPI_ISL_2353381 | Lighthouse Lab in Alderley Park | Wellcome Sanger Institute for the COVID-19 Genomics UK (COG-UK) Consortium | Jacquelyn Wynn, Mairead Hyland, The Lighthouse Lab in Alderley Park and Alex Alderton, Roberto Amato, Jeffrey Barrett, Sonia Goncalves, Ewan Harrison, David K. Jackson, Ian Johnston, Dominic Kwiatkowski, Cordelia Langford, John Sillitoe on behalf of the Wellcome Sanger Institute COVID-19 Surveillance Team |
| EPI_ISL_2353382 | Lighthouse Laboratory Plymouth | Wellcome Sanger Institute for the COVID-19 Genomics UK (COG-UK) Consortium | Lighthouse Laboratory Plymouth and Alex Alderton, Roberto Amato, Jeffrey Barrett, Sonia Goncalves, Ewan Harrison, David K. Jackson, Ian Johnston, Dominic Kwiatkowski, Cordelia Langford, John Sillitoe on behalf of the Wellcome Sanger Institute COVID-19 Surveillance Team |
| EPI_ISL_2353383, EPI_ISL_2353384 | Lighthouse Lab in Milton Keynes | Wellcome Sanger Institute for the COVID-19 Genomics UK (COG-UK) Consortium | The Lighthouse Lab in Milton Keynes and Alex Alderton, Roberto Amato, Jeffrey Barrett, Sonia Goncalves, Ewan Harrison, David K. Jackson, Ian Johnston, Dominic Kwiatkowski, Cordelia Langford, John Sillitoe on behalf of the Wellcome Sanger Institute COVID-19 Surveillance Team |
| EPI_ISL_2353386, EPI_ISL_2353387 | Lighthouse Lab in Alderley Park | Wellcome Sanger Institute for the COVID-19 Genomics UK (COG-UK) Consortium | Jacquelyn Wynn, Mairead Hyland, The Lighthouse Lab in Alderley Park and Alex Alderton, Roberto Amato, Jeffrey Barrett, Sonia Goncalves, Ewan Harrison, David K. Jackson, Ian Johnston, Dominic Kwiatkowski, Cordelia Langford, John Sillitoe on behalf of the Wellcome Sanger Institute COVID-19 Surveillance Team |
| EPI_ISL_2353388, EPI_ISL_2353389, EPI_ISL_2353391 | Lighthouse Lab in Milton Keynes | Wellcome Sanger Institute for the COVID-19 Genomics UK (COG-UK) Consortium | The Lighthouse Lab in Milton Keynes and Alex Alderton, Roberto Amato, Jeffrey Barrett, Sonia Goncalves, Ewan Harrison, David K. Jackson, Ian Johnston, Dominic Kwiatkowski, Cordelia Langford, John Sillitoe on behalf of the Wellcome Sanger Institute COVID-19 Surveillance Team |
| EPI_ISL_2353392 | Lighthouse Lab in Alderley Park | Wellcome Sanger Institute for the COVID-19 Genomics UK (COG-UK) Consortium | Jacquelyn Wynn, Mairead Hyland, The Lighthouse Lab in Alderley Park and Alex Alderton, Roberto Amato, Jeffrey Barrett, Sonia Goncalves, Ewan Harrison, David K. Jackson, Ian Johnston, Dominic Kwiatkowski, Cordelia Langford, John Sillitoe on behalf of the Wellcome Sanger Institute COVID-19 Surveillance Team |
| EPI_ISL_2353393 | Lighthouse Lab in Milton Keynes | Wellcome Sanger Institute for the COVID-19 Genomics UK (COG-UK) Consortium | The Lighthouse Lab in Milton Keynes and Alex Alderton, Roberto Amato, Jeffrey Barrett, Sonia Goncalves, Ewan Harrison, David K. Jackson, Ian Johnston, Dominic Kwiatkowski, Cordelia Langford, John Sillitoe on behalf of the Wellcome Sanger Institute COVID-19 Surveillance Team |
| EPI_ISL_2353394 | Lighthouse Lab in Alderley Park | Wellcome Sanger Institute for the COVID-19 Genomics UK (COG-UK) Consortium | Jacquelyn Wynn, Mairead Hyland, The Lighthouse Lab in Alderley Park and Alex Alderton, Roberto Amato, Jeffrey Barrett, Sonia Goncalves, Ewan Harrison, David K. Jackson, Ian Johnston, Dominic Kwiatkowski, Cordelia Langford, John Sillitoe on behalf of the Wellcome Sanger Institute COVID-19 Surveillance Team |
| EPI_ISL_2353395, EPI_ISL_2353397 | Lighthouse Lab in Milton Keynes | Wellcome Sanger Institute for the COVID-19 Genomics UK (COG-UK) Consortium | The Lighthouse Lab in Milton Keynes and Alex Alderton, Roberto Amato, Jeffrey Barrett, Sonia Goncalves, Ewan Harrison, David K. Jackson, Ian Johnston, Dominic Kwiatkowski, Cordelia Langford, John Sillitoe on behalf of the Wellcome Sanger Institute COVID-19 Surveillance Team |
| EPI_ISL_2353399 | Lighthouse Lab in Alderley Park | Wellcome Sanger Institute for the COVID-19 Genomics UK (COG-UK) Consortium | Jacquelyn Wynn, Mairead Hyland, The Lighthouse Lab in Alderley Park and Alex Alderton, Roberto Amato, Jeffrey Barrett, Sonia Goncalves, Ewan Harrison, David K. Jackson, Ian Johnston, Dominic Kwiatkowski, Cordelia Langford, John Sillitoe on behalf of the Wellcome Sanger Institute COVID-19 Surveillance Team |
| EPI_ISL_2353400, EPI_ISL_2353401 | Lighthouse Lab in Milton Keynes | Wellcome Sanger Institute for the COVID-19 Genomics UK (COG-UK) Consortium | The Lighthouse Lab in Milton Keynes and Alex Alderton, Roberto Amato, Jeffrey Barrett, Sonia Goncalves, Ewan Harrison, David K. Jackson, Ian Johnston, Dominic Kwiatkowski, Cordelia Langford, John Sillitoe on behalf of the Wellcome Sanger Institute COVID-19 Surveillance Team |
| EPI_ISL_2353402 | Lighthouse Laboratory Plymouth | Wellcome Sanger Institute for the COVID-19 Genomics UK (COG-UK) Consortium | Lighthouse Laboratory Plymouth and Alex Alderton, Roberto Amato, Jeffrey Barrett, Sonia Goncalves, Ewan Harrison, David K. Jackson, Ian Johnston, Dominic Kwiatkowski, Cordelia Langford, John Sillitoe on behalf of the Wellcome Sanger Institute COVID-19 Surveillance Team |
| EPI_ISL_2353403 | Lighthouse Lab in Milton Keynes | Wellcome Sanger Institute for the COVID-19 Genomics UK (COG-UK) Consortium | The Lighthouse Lab in Milton Keynes and Alex Alderton, Roberto Amato, Jeffrey Barrett, Sonia Goncalves, Ewan Harrison, David K. Jackson, Ian Johnston, Dominic Kwiatkowski, Cordelia Langford, John Sillitoe on behalf of the Wellcome Sanger Institute COVID-19 Surveillance Team |
| EPI_ISL_2353406, EPI_ISL_2353410, EPI_ISL_2353412 | Lighthouse Lab in Alderley Park | Wellcome Sanger Institute for the COVID-19 Genomics UK (COG-UK) Consortium | Jacquelyn Wynn, Mairead Hyland, The Lighthouse Lab in Alderley Park and Alex Alderton, Roberto Amato, Jeffrey Barrett, Sonia Goncalves, Ewan Harrison, David K. Jackson, Ian Johnston, Dominic Kwiatkowski, Cordelia Langford, John Sillitoe on behalf of the Wellcome Sanger Institute COVID-19 Surveillance Team |
| EPI_ISL_2353414, EPI_ISL_2353415 | Lighthouse Lab in Milton Keynes | Wellcome Sanger Institute for the COVID-19 Genomics UK (COG-UK) Consortium | The Lighthouse Lab in Milton Keynes and Alex Alderton, Roberto Amato, Jeffrey Barrett, Sonia Goncalves, Ewan Harrison, David K. Jackson, Ian Johnston, Dominic Kwiatkowski, Cordelia Langford, John Sillitoe on behalf of the Wellcome Sanger Institute COVID-19 Surveillance Team |
| EPI_ISL_2353417 | Lighthouse Laboratory Plymouth | Wellcome Sanger Institute for the COVID-19 Genomics UK (COG-UK) Consortium | Lighthouse Laboratory Plymouth and Alex Alderton, Roberto Amato, Jeffrey Barrett, Sonia Goncalves, Ewan Harrison, David K. Jackson, Ian Johnston, Dominic Kwiatkowski, Cordelia Langford, John Sillitoe on behalf of the Wellcome Sanger Institute COVID-19 Surveillance Team |
| EPI_ISL_2353418 | Lighthouse Lab in Alderley Park | Wellcome Sanger Institute for the COVID-19 Genomics UK (COG-UK) Consortium | Jacquelyn Wynn, Mairead Hyland, The Lighthouse Lab in Alderley Park and Alex Alderton, Roberto Amato, Jeffrey Barrett, Sonia Goncalves, Ewan Harrison, David K. Jackson, Ian Johnston, Dominic Kwiatkowski, Cordelia Langford, John Sillitoe on behalf of the Wellcome Sanger Institute COVID-19 Surveillance Team |
| EPI_ISL_2353419 | Lighthouse Lab in Milton Keynes | Wellcome Sanger Institute for the COVID-19 Genomics UK (COG-UK) Consortium | The Lighthouse Lab in Milton Keynes and Alex Alderton, Roberto Amato, Jeffrey Barrett, Sonia Goncalves, Ewan Harrison, David K. Jackson, Ian Johnston, Dominic Kwiatkowski, Cordelia Langford, John Sillitoe on behalf of the Wellcome Sanger Institute COVID-19 Surveillance Team |
| EPI_ISL_2353420 | Lighthouse Laboratory Plymouth | Wellcome Sanger Institute for the COVID-19 Genomics UK (COG-UK) Consortium | Lighthouse Laboratory Plymouth and Alex Alderton, Roberto Amato, Jeffrey Barrett, Sonia Goncalves, Ewan Harrison, David K. Jackson, Ian Johnston, Dominic Kwiatkowski, Cordelia Langford, John Sillitoe on behalf of the Wellcome Sanger Institute COVID-19 Surveillance Team |
| EPI_ISL_2353422, EPI_ISL_2353423, EPI_ISL_2353425, EPI_ISL_2353426, EPI_ISL_2353427 | Lighthouse Lab in Milton Keynes | Wellcome Sanger Institute for the COVID-19 Genomics UK (COG-UK) Consortium | The Lighthouse Lab in Milton Keynes and Alex Alderton, Roberto Amato, Jeffrey Barrett, Sonia Goncalves, Ewan Harrison, David K. Jackson, Ian Johnston, Dominic Kwiatkowski, Cordelia Langford, John Sillitoe on behalf of the Wellcome Sanger Institute COVID-19 Surveillance Team |
| EPI_ISL_2353428 | Lighthouse Lab in Alderley Park | Wellcome Sanger Institute for the COVID-19 Genomics UK (COG-UK) Consortium | Jacquelyn Wynn, Mairead Hyland, The Lighthouse Lab in Alderley Park and Alex Alderton, Roberto Amato, Jeffrey Barrett, Sonia Goncalves, Ewan Harrison, David K. Jackson, Ian Johnston, Dominic Kwiatkowski, Cordelia Langford, John Sillitoe on behalf of the Wellcome Sanger Institute COVID-19 Surveillance Team |
| EPI_ISL_2353429 | Lighthouse Laboratory Plymouth | Wellcome Sanger Institute for the COVID-19 Genomics UK (COG-UK) Consortium | Lighthouse Laboratory Plymouth and Alex Alderton, Roberto Amato, Jeffrey Barrett, Sonia Goncalves, Ewan Harrison, David K. Jackson, Ian Johnston, Dominic Kwiatkowski, Cordelia Langford, John Sillitoe on behalf of the Wellcome Sanger Institute COVID-19 Surveillance Team |
| EPI_ISL_2353431, EPI_ISL_2353433 | Lighthouse Lab in Alderley Park | Wellcome Sanger Institute for the COVID-19 Genomics UK (COG-UK) Consortium | Jacquelyn Wynn, Mairead Hyland, The Lighthouse Lab in Alderley Park and Alex Alderton, Roberto Amato, Jeffrey Barrett, Sonia Goncalves, Ewan Harrison, David K. Jackson, Ian Johnston, Dominic Kwiatkowski, Cordelia Langford, John Sillitoe on behalf of the Wellcome Sanger Institute COVID-19 Surveillance Team |

| | | | |
|---|---|---|---|
| EPI_ISL_2353434, EPI_ISL_2353435 | Lighthouse Lab in Milton Keynes | Wellcome Sanger Institute for the COVID-19 Genomics UK (COG-UK) Consortium | The Lighthouse Lab in Milton Keynes and Alex Alderton, Roberto Amato, Jeffrey Barrett, Sonia Goncalves, Ewan Harrison, David K. Jackson, Ian Johnston, Dominic Kwiatkowski, Cordelia Langford, John Sillitoe on behalf of the Wellcome Sanger Institute COVID-19 Surveillance Team |
| EPI_ISL_2353436, EPI_ISL_2353437 | Lighthouse Lab in Alderley Park | Wellcome Sanger Institute for the COVID-19 Genomics UK (COG-UK) Consortium | Jacquelyn Wynn, Mairead Hyland, The Lighthouse Lab in Alderley Park and Alex Alderton, Roberto Amato, Jeffrey Barrett, Sonia Goncalves, Ewan Harrison, David K. Jackson, Ian Johnston, Dominic Kwiatkowski, Cordelia Langford, John Sillitoe on behalf of the Wellcome Sanger Institute COVID-19 Surveillance Team |
| EPI_ISL_2353439 | Lighthouse Lab in Milton Keynes | Wellcome Sanger Institute for the COVID-19 Genomics UK (COG-UK) Consortium | The Lighthouse Lab in Milton Keynes and Alex Alderton, Roberto Amato, Jeffrey Barrett, Sonia Goncalves, Ewan Harrison, David K. Jackson, Ian Johnston, Dominic Kwiatkowski, Cordelia Langford, John Sillitoe on behalf of the Wellcome Sanger Institute COVID-19 Surveillance Team |
| EPI_ISL_2353440 | Lighthouse Lab in Alderley Park | Wellcome Sanger Institute for the COVID-19 Genomics UK (COG-UK) Consortium | Jacquelyn Wynn, Mairead Hyland, The Lighthouse Lab in Alderley Park and Alex Alderton, Roberto Amato, Jeffrey Barrett, Sonia Goncalves, Ewan Harrison, David K. Jackson, Ian Johnston, Dominic Kwiatkowski, Cordelia Langford, John Sillitoe on behalf of the Wellcome Sanger Institute COVID-19 Surveillance Team |
| EPI_ISL_2353441, EPI_ISL_2353443 | Lighthouse Lab in Milton Keynes | Wellcome Sanger Institute for the COVID-19 Genomics UK (COG-UK) Consortium | The Lighthouse Lab in Milton Keynes and Alex Alderton, Roberto Amato, Jeffrey Barrett, Sonia Goncalves, Ewan Harrison, David K. Jackson, Ian Johnston, Dominic Kwiatkowski, Cordelia Langford, John Sillitoe on behalf of the Wellcome Sanger Institute COVID-19 Surveillance Team |
| EPI_ISL_2353444 | Lighthouse Lab in Alderley Park | Wellcome Sanger Institute for the COVID-19 Genomics UK (COG-UK) Consortium | Jacquelyn Wynn, Mairead Hyland, The Lighthouse Lab in Alderley Park and Alex Alderton, Roberto Amato, Jeffrey Barrett, Sonia Goncalves, Ewan Harrison, David K. Jackson, Ian Johnston, Dominic Kwiatkowski, Cordelia Langford, John Sillitoe on behalf of the Wellcome Sanger Institute COVID-19 Surveillance Team |
| EPI_ISL_2353445, EPI_ISL_2353446 | Lighthouse Lab in Milton Keynes | Wellcome Sanger Institute for the COVID-19 Genomics UK (COG-UK) Consortium | The Lighthouse Lab in Milton Keynes and Alex Alderton, Roberto Amato, Jeffrey Barrett, Sonia Goncalves, Ewan Harrison, David K. Jackson, Ian Johnston, Dominic Kwiatkowski, Cordelia Langford, John Sillitoe on behalf of the Wellcome Sanger Institute COVID-19 Surveillance Team |
| EPI_ISL_2353447, EPI_ISL_2353448 | Lighthouse Laboratory Plymouth | Wellcome Sanger Institute for the COVID-19 Genomics UK (COG-UK) Consortium | Lighthouse Laboratory Plymouth and Alex Alderton, Roberto Amato, Jeffrey Barrett, Sonia Goncalves, Ewan Harrison, David K. Jackson, Ian Johnston, Dominic Kwiatkowski, Cordelia Langford, John Sillitoe on behalf of the Wellcome Sanger Institute COVID-19 Surveillance Team |
| EPI_ISL_2353449, EPI_ISL_2353451, EPI_ISL_2353453, EPI_ISL_2353456 | Lighthouse Lab in Milton Keynes | Wellcome Sanger Institute for the COVID-19 Genomics UK (COG-UK) Consortium | The Lighthouse Lab in Milton Keynes and Alex Alderton, Roberto Amato, Jeffrey Barrett, Sonia Goncalves, Ewan Harrison, David K. Jackson, Ian Johnston, Dominic Kwiatkowski, Cordelia Langford, John Sillitoe on behalf of the Wellcome Sanger Institute COVID-19 Surveillance Team |
| EPI_ISL_2353458 | Lighthouse Lab in Alderley Park | Wellcome Sanger Institute for the COVID-19 Genomics UK (COG-UK) Consortium | Jacquelyn Wynn, Mairead Hyland, The Lighthouse Lab in Alderley Park and Alex Alderton, Roberto Amato, Jeffrey Barrett, Sonia Goncalves, Ewan Harrison, David K. Jackson, Ian Johnston, Dominic Kwiatkowski, Cordelia Langford, John Sillitoe on behalf of the Wellcome Sanger Institute COVID-19 Surveillance Team |
| EPI_ISL_2353459 | Lighthouse Laboratory Plymouth | Wellcome Sanger Institute for the COVID-19 Genomics UK (COG-UK) Consortium | Lighthouse Laboratory Plymouth and Alex Alderton, Roberto Amato, Jeffrey Barrett, Sonia Goncalves, Ewan Harrison, David K. Jackson, Ian Johnston, Dominic Kwiatkowski, Cordelia Langford, John Sillitoe on behalf of the Wellcome Sanger Institute COVID-19 Surveillance Team |
| EPI_ISL_2353460 | Lighthouse Lab in Milton Keynes | Wellcome Sanger Institute for the COVID-19 Genomics UK (COG-UK) Consortium | The Lighthouse Lab in Milton Keynes and Alex Alderton, Roberto Amato, Jeffrey Barrett, Sonia Goncalves, Ewan Harrison, David K. Jackson, Ian Johnston, Dominic Kwiatkowski, Cordelia Langford, John Sillitoe on behalf of the Wellcome Sanger Institute COVID-19 Surveillance Team |
| EPI_ISL_2353461 | Lighthouse Lab in Alderley Park | Wellcome Sanger Institute for the COVID-19 Genomics UK (COG-UK) Consortium | Jacquelyn Wynn, Mairead Hyland, The Lighthouse Lab in Alderley Park and Alex Alderton, Roberto Amato, Jeffrey Barrett, Sonia Goncalves, Ewan Harrison, David K. Jackson, Ian Johnston, Dominic Kwiatkowski, Cordelia Langford, John Sillitoe on behalf of the Wellcome Sanger Institute COVID-19 Surveillance Team |
| EPI_ISL_2353462, EPI_ISL_2353464, EPI_ISL_2353465, EPI_ISL_2353466, EPI_ISL_2353469, EPI_ISL_2353470, EPI_ISL_2353473 | Lighthouse Lab in Milton Keynes | Wellcome Sanger Institute for the COVID-19 Genomics UK (COG-UK) Consortium | The Lighthouse Lab in Milton Keynes and Alex Alderton, Roberto Amato, Jeffrey Barrett, Sonia Goncalves, Ewan Harrison, David K. Jackson, Ian Johnston, Dominic Kwiatkowski, Cordelia Langford, John Sillitoe on behalf of the Wellcome Sanger Institute COVID-19 Surveillance Team |
| EPI_ISL_2353474 | Lighthouse Lab in Alderley Park | Wellcome Sanger Institute for the COVID-19 Genomics UK (COG-UK) Consortium | Jacquelyn Wynn, Mairead Hyland, The Lighthouse Lab in Alderley Park and Alex Alderton, Roberto Amato, Jeffrey Barrett, Sonia Goncalves, Ewan Harrison, David K. Jackson, Ian Johnston, Dominic Kwiatkowski, Cordelia Langford, John Sillitoe on behalf of the Wellcome Sanger Institute COVID-19 Surveillance Team |
| EPI_ISL_2353476, EPI_ISL_2353477 | Lighthouse Lab in Milton Keynes | Wellcome Sanger Institute for the COVID-19 Genomics UK (COG-UK) Consortium | The Lighthouse Lab in Milton Keynes and Alex Alderton, Roberto Amato, Jeffrey Barrett, Sonia Goncalves, Ewan Harrison, David K. Jackson, Ian Johnston, Dominic Kwiatkowski, Cordelia Langford, John Sillitoe on behalf of the Wellcome Sanger Institute COVID-19 Surveillance Team |
| EPI_ISL_2353478 | Lighthouse Lab in Alderley Park | Wellcome Sanger Institute for the COVID-19 Genomics UK (COG-UK) Consortium | Jacquelyn Wynn, Mairead Hyland, The Lighthouse Lab in Alderley Park and Alex Alderton, Roberto Amato, Jeffrey Barrett, Sonia Goncalves, Ewan Harrison, David K. Jackson, Ian Johnston, Dominic Kwiatkowski, Cordelia Langford, John Sillitoe on behalf of the Wellcome Sanger Institute COVID-19 Surveillance Team |
| EPI_ISL_2353479 | Lighthouse Laboratory Plymouth | Wellcome Sanger Institute for the COVID-19 Genomics UK (COG-UK) Consortium | Lighthouse Laboratory Plymouth and Alex Alderton, Roberto Amato, Jeffrey Barrett, Sonia Goncalves, Ewan Harrison, David K. Jackson, Ian Johnston, Dominic Kwiatkowski, Cordelia Langford, John Sillitoe on behalf of the Wellcome Sanger Institute COVID-19 Surveillance Team |
| EPI_ISL_2353481 | Lighthouse Lab in Milton Keynes | Wellcome Sanger Institute for the COVID-19 Genomics UK (COG-UK) Consortium | The Lighthouse Lab in Milton Keynes and Alex Alderton, Roberto Amato, Jeffrey Barrett, Sonia Goncalves, Ewan Harrison, David K. Jackson, Ian Johnston, Dominic Kwiatkowski, Cordelia Langford, John Sillitoe on behalf of the Wellcome Sanger Institute COVID-19 Surveillance Team |
| EPI_ISL_2353482 | Lighthouse Laboratory Plymouth | Wellcome Sanger Institute for the COVID-19 Genomics UK (COG-UK) Consortium | Lighthouse Laboratory Plymouth and Alex Alderton, Roberto Amato, Jeffrey Barrett, Sonia Goncalves, Ewan Harrison, David K. Jackson, Ian Johnston, Dominic Kwiatkowski, Cordelia Langford, John Sillitoe on behalf of the Wellcome Sanger Institute COVID-19 Surveillance Team |
| EPI_ISL_2353483 | Lighthouse Lab in Alderley Park | Wellcome Sanger Institute for the COVID-19 Genomics UK (COG-UK) Consortium | Jacquelyn Wynn, Mairead Hyland, The Lighthouse Lab in Alderley Park and Alex Alderton, Roberto Amato, Jeffrey Barrett, Sonia Goncalves, Ewan Harrison, David K. Jackson, Ian Johnston, Dominic Kwiatkowski, Cordelia Langford, John Sillitoe on behalf of the Wellcome Sanger Institute COVID-19 Surveillance Team |
| EPI_ISL_2353484, EPI_ISL_2353485 | Lighthouse Lab in Milton Keynes | Wellcome Sanger Institute for the COVID-19 Genomics UK (COG-UK) Consortium | The Lighthouse Lab in Milton Keynes and Alex Alderton, Roberto Amato, Jeffrey Barrett, Sonia Goncalves, Ewan Harrison, David K. Jackson, Ian Johnston, Dominic Kwiatkowski, Cordelia Langford, John Sillitoe on behalf of the Wellcome Sanger Institute COVID-19 Surveillance Team |
| EPI_ISL_2353486 | Lighthouse Lab in Alderley Park | Wellcome Sanger Institute for the COVID-19 Genomics UK (COG-UK) Consortium | Jacquelyn Wynn, Mairead Hyland, The Lighthouse Lab in Alderley Park and Alex Alderton, Roberto Amato, Jeffrey Barrett, Sonia Goncalves, Ewan Harrison, David K. Jackson, Ian Johnston, Dominic Kwiatkowski, Cordelia Langford, John Sillitoe on behalf of the Wellcome Sanger Institute COVID-19 Surveillance Team |
| EPI_ISL_2353487 | Lighthouse Lab in Milton Keynes | Wellcome Sanger Institute for the COVID-19 Genomics UK (COG-UK) Consortium | The Lighthouse Lab in Milton Keynes and Alex Alderton, Roberto Amato, Jeffrey Barrett, Sonia Goncalves, Ewan Harrison, David K. Jackson, Ian Johnston, Dominic Kwiatkowski, Cordelia Langford, John Sillitoe on behalf of the Wellcome Sanger Institute COVID-19 Surveillance Team |
| EPI_ISL_2353488 | Lighthouse Lab in Alderley Park | Wellcome Sanger Institute for the COVID-19 Genomics UK (COG-UK) Consortium | Jacquelyn Wynn, Mairead Hyland, The Lighthouse Lab in Alderley Park and Alex Alderton, Roberto Amato, Jeffrey Barrett, Sonia Goncalves, Ewan Harrison, David K. Jackson, Ian Johnston, Dominic Kwiatkowski, Cordelia Langford, John Sillitoe on behalf of the Wellcome Sanger Institute COVID-19 Surveillance Team |
| EPI_ISL_2353489 | Lighthouse Laboratory Plymouth | Wellcome Sanger Institute for the COVID-19 Genomics UK (COG-UK) Consortium | Lighthouse Laboratory Plymouth and Alex Alderton, Roberto Amato, Jeffrey Barrett, Sonia Goncalves, Ewan Harrison, David K. Jackson, Ian Johnston, Dominic Kwiatkowski, Cordelia Langford, John Sillitoe on behalf of the Wellcome Sanger Institute COVID-19 Surveillance Team |
| EPI_ISL_2353490 | Lighthouse Lab in Milton Keynes | Wellcome Sanger Institute for the COVID-19 Genomics UK (COG-UK) Consortium | The Lighthouse Lab in Milton Keynes and Alex Alderton, Roberto Amato, Jeffrey Barrett, Sonia Goncalves, Ewan Harrison, David K. Jackson, Ian Johnston, Dominic Kwiatkowski, Cordelia Langford, John Sillitoe on behalf of the Wellcome Sanger Institute COVID-19 Surveillance Team |
| EPI_ISL_2353491 | Lighthouse Laboratory Plymouth | Wellcome Sanger Institute for the COVID-19 Genomics UK (COG-UK) Consortium | Lighthouse Laboratory Plymouth and Alex Alderton, Roberto Amato, Jeffrey Barrett, Sonia Goncalves, Ewan Harrison, David K. Jackson, Ian Johnston, Dominic Kwiatkowski, Cordelia Langford, John Sillitoe on behalf of the Wellcome Sanger Institute COVID-19 Surveillance Team |
| EPI_ISL_2353492, EPI_ISL_2353493 | Lighthouse Lab in Milton Keynes | Wellcome Sanger Institute for the COVID-19 Genomics UK (COG-UK) Consortium | The Lighthouse Lab in Milton Keynes and Alex Alderton, Roberto Amato, Jeffrey Barrett, Sonia Goncalves, Ewan Harrison, David K. Jackson, Ian Johnston, Dominic Kwiatkowski, Cordelia Langford, John Sillitoe on behalf of the Wellcome Sanger Institute COVID-19 Surveillance Team |
| EPI_ISL_2353494 | Lighthouse Lab in Alderley Park | Wellcome Sanger Institute for the COVID-19 Genomics UK (COG-UK) Consortium | Jacquelyn Wynn, Mairead Hyland, The Lighthouse Lab in Alderley Park and Alex Alderton, Roberto Amato, Jeffrey Barrett, Sonia Goncalves, Ewan Harrison, David K. Jackson, Ian Johnston, Dominic Kwiatkowski, Cordelia Langford, John Sillitoe on behalf of the Wellcome Sanger Institute COVID-19 Surveillance Team |
| EPI_ISL_2353495 | Lighthouse Lab in Milton Keynes | Wellcome Sanger Institute for the COVID-19 Genomics UK (COG-UK) Consortium | The Lighthouse Lab in Milton Keynes and Alex Alderton, Roberto Amato, Jeffrey Barrett, Sonia Goncalves, Ewan Harrison, David K. Jackson, Ian Johnston, Dominic Kwiatkowski, Cordelia Langford, John Sillitoe on behalf of the Wellcome Sanger Institute COVID-19 Surveillance Team |
| EPI_ISL_2353496, EPI_ISL_2353497 | Lighthouse Lab in Alderley Park | Wellcome Sanger Institute for the COVID-19 Genomics UK (COG-UK) Consortium | Jacquelyn Wynn, Mairead Hyland, The Lighthouse Lab in Alderley Park and Alex Alderton, Roberto Amato, Jeffrey Barrett, Sonia Goncalves, Ewan Harrison, David K. Jackson, Ian Johnston, Dominic Kwiatkowski, Cordelia Langford, John Sillitoe on behalf of the Wellcome Sanger Institute COVID-19 Surveillance Team |
| EPI_ISL_2353498 | Lighthouse Lab in Milton Keynes | Wellcome Sanger Institute for the COVID-19 Genomics UK (COG-UK) Consortium | The Lighthouse Lab in Milton Keynes and Alex Alderton, Roberto Amato, Jeffrey Barrett, Sonia Goncalves, Ewan Harrison, David K. Jackson, Ian Johnston, Dominic Kwiatkowski, Cordelia Langford, John Sillitoe on behalf of the Wellcome Sanger Institute COVID-19 Surveillance Team |
| EPI_ISL_2353499 | Lighthouse Lab in Alderley Park | Wellcome Sanger Institute for the COVID-19 Genomics UK (COG-UK) Consortium | Jacquelyn Wynn, Mairead Hyland, The Lighthouse Lab in Alderley Park and Alex Alderton, Roberto Amato, Jeffrey Barrett, Sonia Goncalves, Ewan Harrison, David K. Jackson, Ian Johnston, Dominic Kwiatkowski, Cordelia Langford, John Sillitoe on behalf of the Wellcome Sanger Institute COVID-19 Surveillance Team |
| EPI_ISL_2353501, EPI_ISL_2353502 | Lighthouse Lab in Milton Keynes | Wellcome Sanger Institute for the COVID-19 Genomics UK (COG-UK) Consortium | The Lighthouse Lab in Milton Keynes and Alex Alderton, Roberto Amato, Jeffrey Barrett, Sonia Goncalves, Ewan Harrison, David K. Jackson, Ian Johnston, Dominic Kwiatkowski, Cordelia Langford, John Sillitoe on behalf of the Wellcome Sanger Institute COVID-19 Surveillance Team |
| EPI_ISL_2353504 | Lighthouse Laboratory Plymouth | Wellcome Sanger Institute for the COVID-19 Genomics UK (COG-UK) Consortium | Lighthouse Laboratory Plymouth and Alex Alderton, Roberto Amato, Jeffrey Barrett, Sonia Goncalves, Ewan Harrison, David K. Jackson, Ian Johnston, Dominic Kwiatkowski, Cordelia Langford, John Sillitoe on behalf of the Wellcome Sanger Institute COVID-19 Surveillance Team |
| EPI_ISL_2353505, EPI_ISL_2353506, EPI_ISL_2353507 | Lighthouse Lab in Milton Keynes | Wellcome Sanger Institute for the COVID-19 Genomics UK (COG-UK) Consortium | The Lighthouse Lab in Milton Keynes and Alex Alderton, Roberto Amato, Jeffrey Barrett, Sonia Goncalves, Ewan Harrison, David K. Jackson, Ian Johnston, Dominic Kwiatkowski, Cordelia Langford, John Sillitoe on behalf of the Wellcome Sanger Institute COVID-19 Surveillance Team |
| EPI_ISL_2353508 | Lighthouse Lab in Alderley Park | Wellcome Sanger Institute for the COVID-19 Genomics UK (COG-UK) Consortium | Jacquelyn Wynn, Mairead Hyland, The Lighthouse Lab in Alderley Park and Alex Alderton, Roberto Amato, Jeffrey Barrett, Sonia Goncalves, Ewan Harrison, David K. Jackson, Ian Johnston, Dominic Kwiatkowski, Cordelia Langford, John Sillitoe on behalf of the Wellcome Sanger Institute COVID-19 Surveillance Team |
| EPI_ISL_2353509 | Lighthouse Laboratory Plymouth | Wellcome Sanger Institute for the COVID-19 Genomics UK (COG-UK) Consortium | Lighthouse Laboratory Plymouth and Alex Alderton, Roberto Amato, Jeffrey Barrett, Sonia Goncalves, Ewan Harrison, David K. Jackson, Ian Johnston, Dominic Kwiatkowski, Cordelia Langford, John Sillitoe on behalf of the Wellcome Sanger Institute COVID-19 Surveillance Team |
| EPI_ISL_2353510, EPI_ISL_2353511, EPI_ISL_2353512, EPI_ISL_2353513 | Lighthouse Lab in Milton Keynes | Wellcome Sanger Institute for the COVID-19 Genomics UK (COG-UK) Consortium | The Lighthouse Lab in Milton Keynes and Alex Alderton, Roberto Amato, Jeffrey Barrett, Sonia Goncalves, Ewan Harrison, David K. Jackson, Ian Johnston, Dominic Kwiatkowski, Cordelia Langford, John Sillitoe on behalf of the Wellcome Sanger Institute COVID-19 Surveillance Team |
| EPI_ISL_2353514 | Lighthouse Lab in Alderley Park | Wellcome Sanger Institute for the COVID-19 Genomics UK (COG-UK) Consortium | Jacquelyn Wynn, Mairead Hyland, The Lighthouse Lab in Alderley Park and Alex Alderton, Roberto Amato, Jeffrey Barrett, Sonia Goncalves, Ewan Harrison, David K. Jackson, Ian Johnston, Dominic Kwiatkowski, Cordelia Langford, John Sillitoe on behalf of the Wellcome Sanger Institute COVID-19 Surveillance Team |

| | | | |
|---|---|---|---|
| EPI_ISL_2353515, EPI_ISL_2353516, EPI_ISL_2353519 | Lighthouse Lab in Milton Keynes | Wellcome Sanger Institute for the COVID-19 Genomics UK (COG-UK) Consortium | The Lighthouse Lab in Milton Keynes and Alex Alderton, Roberto Amato, Jeffrey Barrett, Sonia Goncalves, Ewan Harrison, David K. Jackson, Ian Johnston, Dominic Kwiatkowski, Cordelia Langford, John Sillitoe on behalf of the Wellcome Sanger Institute COVID-19 Surveillance Team |
| EPI_ISL_2353520 | Lighthouse Lab in Alderley Park | Wellcome Sanger Institute for the COVID-19 Genomics UK (COG-UK) Consortium | Jacquelyn Wynn, Mairead Hyland, The Lighthouse Lab in Alderley Park and Alex Alderton, Roberto Amato, Jeffrey Barrett, Sonia Goncalves, Ewan Harrison, David K. Jackson, Ian Johnston, Dominic Kwiatkowski, Cordelia Langford, John Sillitoe on behalf of the Wellcome Sanger Institute COVID-19 Surveillance Team |
| EPI_ISL_2353521, EPI_ISL_2353522 | Lighthouse Lab in Milton Keynes | Wellcome Sanger Institute for the COVID-19 Genomics UK (COG-UK) Consortium | The Lighthouse Lab in Milton Keynes and Alex Alderton, Roberto Amato, Jeffrey Barrett, Sonia Goncalves, Ewan Harrison, David K. Jackson, Ian Johnston, Dominic Kwiatkowski, Cordelia Langford, John Sillitoe on behalf of the Wellcome Sanger Institute COVID-19 Surveillance Team |
| EPI_ISL_2353524 | Lighthouse Lab in Alderley Park | Wellcome Sanger Institute for the COVID-19 Genomics UK (COG-UK) Consortium | Jacquelyn Wynn, Mairead Hyland, The Lighthouse Lab in Alderley Park and Alex Alderton, Roberto Amato, Jeffrey Barrett, Sonia Goncalves, Ewan Harrison, David K. Jackson, Ian Johnston, Dominic Kwiatkowski, Cordelia Langford, John Sillitoe on behalf of the Wellcome Sanger Institute COVID-19 Surveillance Team |
| EPI_ISL_2353525, EPI_ISL_2353527, EPI_ISL_2353528, EPI_ISL_2353529 | Lighthouse Lab in Milton Keynes | Wellcome Sanger Institute for the COVID-19 Genomics UK (COG-UK) Consortium | The Lighthouse Lab in Milton Keynes and Alex Alderton, Roberto Amato, Jeffrey Barrett, Sonia Goncalves, Ewan Harrison, David K. Jackson, Ian Johnston, Dominic Kwiatkowski, Cordelia Langford, John Sillitoe on behalf of the Wellcome Sanger Institute COVID-19 Surveillance Team |
| EPI_ISL_2353530, EPI_ISL_2353532, EPI_ISL_2353533 | Lighthouse Lab in Alderley Park | Wellcome Sanger Institute for the COVID-19 Genomics UK (COG-UK) Consortium | Jacquelyn Wynn, Mairead Hyland, The Lighthouse Lab in Alderley Park and Alex Alderton, Roberto Amato, Jeffrey Barrett, Sonia Goncalves, Ewan Harrison, David K. Jackson, Ian Johnston, Dominic Kwiatkowski, Cordelia Langford, John Sillitoe on behalf of the Wellcome Sanger Institute COVID-19 Surveillance Team |
| EPI_ISL_2353534 | Lighthouse Lab in Milton Keynes | Wellcome Sanger Institute for the COVID-19 Genomics UK (COG-UK) Consortium | The Lighthouse Lab in Milton Keynes and Alex Alderton, Roberto Amato, Jeffrey Barrett, Sonia Goncalves, Ewan Harrison, David K. Jackson, Ian Johnston, Dominic Kwiatkowski, Cordelia Langford, John Sillitoe on behalf of the Wellcome Sanger Institute COVID-19 Surveillance Team |
| EPI_ISL_2353535 | Lighthouse Lab in Alderley Park | Wellcome Sanger Institute for the COVID-19 Genomics UK (COG-UK) Consortium | Jacquelyn Wynn, Mairead Hyland, The Lighthouse Lab in Alderley Park and Alex Alderton, Roberto Amato, Jeffrey Barrett, Sonia Goncalves, Ewan Harrison, David K. Jackson, Ian Johnston, Dominic Kwiatkowski, Cordelia Langford, John Sillitoe on behalf of the Wellcome Sanger Institute COVID-19 Surveillance Team |
| EPI_ISL_2353536, EPI_ISL_2353537 | Lighthouse Lab in Milton Keynes | Wellcome Sanger Institute for the COVID-19 Genomics UK (COG-UK) Consortium | The Lighthouse Lab in Milton Keynes and Alex Alderton, Roberto Amato, Jeffrey Barrett, Sonia Goncalves, Ewan Harrison, David K. Jackson, Ian Johnston, Dominic Kwiatkowski, Cordelia Langford, John Sillitoe on behalf of the Wellcome Sanger Institute COVID-19 Surveillance Team |
| EPI_ISL_2353538 | Lighthouse Lab in Alderley Park | Wellcome Sanger Institute for the COVID-19 Genomics UK (COG-UK) Consortium | Jacquelyn Wynn, Mairead Hyland, The Lighthouse Lab in Alderley Park and Alex Alderton, Roberto Amato, Jeffrey Barrett, Sonia Goncalves, Ewan Harrison, David K. Jackson, Ian Johnston, Dominic Kwiatkowski, Cordelia Langford, John Sillitoe on behalf of the Wellcome Sanger Institute COVID-19 Surveillance Team |
| EPI_ISL_2353539 | Lighthouse Lab in Milton Keynes | Wellcome Sanger Institute for the COVID-19 Genomics UK (COG-UK) Consortium | The Lighthouse Lab in Milton Keynes and Alex Alderton, Roberto Amato, Jeffrey Barrett, Sonia Goncalves, Ewan Harrison, David K. Jackson, Ian Johnston, Dominic Kwiatkowski, Cordelia Langford, John Sillitoe on behalf of the Wellcome Sanger Institute COVID-19 Surveillance Team |
| EPI_ISL_2353540 | Lighthouse Lab in Alderley Park | Wellcome Sanger Institute for the COVID-19 Genomics UK (COG-UK) Consortium | Jacquelyn Wynn, Mairead Hyland, The Lighthouse Lab in Alderley Park and Alex Alderton, Roberto Amato, Jeffrey Barrett, Sonia Goncalves, Ewan Harrison, David K. Jackson, Ian Johnston, Dominic Kwiatkowski, Cordelia Langford, John Sillitoe on behalf of the Wellcome Sanger Institute COVID-19 Surveillance Team |
| EPI_ISL_2353541, EPI_ISL_2353543, EPI_ISL_2353545 | Lighthouse Laboratory Plymouth | Wellcome Sanger Institute for the COVID-19 Genomics UK (COG-UK) Consortium | Lighthouse Laboratory Plymouth and Alex Alderton, Roberto Amato, Jeffrey Barrett, Sonia Goncalves, Ewan Harrison, David K. Jackson, Ian Johnston, Dominic Kwiatkowski, Cordelia Langford, John Sillitoe on behalf of the Wellcome Sanger Institute COVID-19 Surveillance Team |
| EPI_ISL_2353546 | Lighthouse Lab in Milton Keynes | Wellcome Sanger Institute for the COVID-19 Genomics UK (COG-UK) Consortium | The Lighthouse Lab in Milton Keynes and Alex Alderton, Roberto Amato, Jeffrey Barrett, Sonia Goncalves, Ewan Harrison, David K. Jackson, Ian Johnston, Dominic Kwiatkowski, Cordelia Langford, John Sillitoe on behalf of the Wellcome Sanger Institute COVID-19 Surveillance Team |
| EPI_ISL_2353547, EPI_ISL_2353548 | Lighthouse Laboratory Plymouth | Wellcome Sanger Institute for the COVID-19 Genomics UK (COG-UK) Consortium | Lighthouse Laboratory Plymouth and Alex Alderton, Roberto Amato, Jeffrey Barrett, Sonia Goncalves, Ewan Harrison, David K. Jackson, Ian Johnston, Dominic Kwiatkowski, Cordelia Langford, John Sillitoe on behalf of the Wellcome Sanger Institute COVID-19 Surveillance Team |
| EPI_ISL_2353550 | Lighthouse Lab in Milton Keynes | Wellcome Sanger Institute for the COVID-19 Genomics UK (COG-UK) Consortium | The Lighthouse Lab in Milton Keynes and Alex Alderton, Roberto Amato, Jeffrey Barrett, Sonia Goncalves, Ewan Harrison, David K. Jackson, Ian Johnston, Dominic Kwiatkowski, Cordelia Langford, John Sillitoe on behalf of the Wellcome Sanger Institute COVID-19 Surveillance Team |
| EPI_ISL_2353551 | Lighthouse Lab in Alderley Park | Wellcome Sanger Institute for the COVID-19 Genomics UK (COG-UK) Consortium | Jacquelyn Wynn, Mairead Hyland, The Lighthouse Lab in Alderley Park and Alex Alderton, Roberto Amato, Jeffrey Barrett, Sonia Goncalves, Ewan Harrison, David K. Jackson, Ian Johnston, Dominic Kwiatkowski, Cordelia Langford, John Sillitoe on behalf of the Wellcome Sanger Institute COVID-19 Surveillance Team |
| EPI_ISL_2353552, EPI_ISL_2353553 | Lighthouse Lab in Milton Keynes | Wellcome Sanger Institute for the COVID-19 Genomics UK (COG-UK) Consortium | The Lighthouse Lab in Milton Keynes and Alex Alderton, Roberto Amato, Jeffrey Barrett, Sonia Goncalves, Ewan Harrison, David K. Jackson, Ian Johnston, Dominic Kwiatkowski, Cordelia Langford, John Sillitoe on behalf of the Wellcome Sanger Institute COVID-19 Surveillance Team |
| EPI_ISL_2353554 | Lighthouse Lab in Alderley Park | Wellcome Sanger Institute for the COVID-19 Genomics UK (COG-UK) Consortium | Jacquelyn Wynn, Mairead Hyland, The Lighthouse Lab in Alderley Park and Alex Alderton, Roberto Amato, Jeffrey Barrett, Sonia Goncalves, Ewan Harrison, David K. Jackson, Ian Johnston, Dominic Kwiatkowski, Cordelia Langford, John Sillitoe on behalf of the Wellcome Sanger Institute COVID-19 Surveillance Team |
| EPI_ISL_2353555 | Lighthouse Lab in Milton Keynes | Wellcome Sanger Institute for the COVID-19 Genomics UK (COG-UK) Consortium | The Lighthouse Lab in Milton Keynes and Alex Alderton, Roberto Amato, Jeffrey Barrett, Sonia Goncalves, Ewan Harrison, David K. Jackson, Ian Johnston, Dominic Kwiatkowski, Cordelia Langford, John Sillitoe on behalf of the Wellcome Sanger Institute COVID-19 Surveillance Team |
| EPI_ISL_2353556 | Lighthouse Lab in Alderley Park | Wellcome Sanger Institute for the COVID-19 Genomics UK (COG-UK) Consortium | Jacquelyn Wynn, Mairead Hyland, The Lighthouse Lab in Alderley Park and Alex Alderton, Roberto Amato, Jeffrey Barrett, Sonia Goncalves, Ewan Harrison, David K. Jackson, Ian Johnston, Dominic Kwiatkowski, Cordelia Langford, John Sillitoe on behalf of the Wellcome Sanger Institute COVID-19 Surveillance Team |
| EPI_ISL_2353557 | Lighthouse Laboratory Plymouth | Wellcome Sanger Institute for the COVID-19 Genomics UK (COG-UK) Consortium | Lighthouse Laboratory Plymouth and Alex Alderton, Roberto Amato, Jeffrey Barrett, Sonia Goncalves, Ewan Harrison, David K. Jackson, Ian Johnston, Dominic Kwiatkowski, Cordelia Langford, John Sillitoe on behalf of the Wellcome Sanger Institute COVID-19 Surveillance Team |
| EPI_ISL_2353559, EPI_ISL_2353560 | Lighthouse Lab in Milton Keynes | Wellcome Sanger Institute for the COVID-19 Genomics UK (COG-UK) Consortium | The Lighthouse Lab in Milton Keynes and Alex Alderton, Roberto Amato, Jeffrey Barrett, Sonia Goncalves, Ewan Harrison, David K. Jackson, Ian Johnston, Dominic Kwiatkowski, Cordelia Langford, John Sillitoe on behalf of the Wellcome Sanger Institute COVID-19 Surveillance Team |
| EPI_ISL_2353561 | Lighthouse Laboratory Plymouth | Wellcome Sanger Institute for the COVID-19 Genomics UK (COG-UK) Consortium | Lighthouse Laboratory Plymouth and Alex Alderton, Roberto Amato, Jeffrey Barrett, Sonia Goncalves, Ewan Harrison, David K. Jackson, Ian Johnston, Dominic Kwiatkowski, Cordelia Langford, John Sillitoe on behalf of the Wellcome Sanger Institute COVID-19 Surveillance Team |
| EPI_ISL_2353564 | Lighthouse Lab in Milton Keynes | Wellcome Sanger Institute for the COVID-19 Genomics UK (COG-UK) Consortium | The Lighthouse Lab in Milton Keynes and Alex Alderton, Roberto Amato, Jeffrey Barrett, Sonia Goncalves, Ewan Harrison, David K. Jackson, Ian Johnston, Dominic Kwiatkowski, Cordelia Langford, John Sillitoe on behalf of the Wellcome Sanger Institute COVID-19 Surveillance Team |
| EPI_ISL_2353566 | Lighthouse Lab in Alderley Park | Wellcome Sanger Institute for the COVID-19 Genomics UK (COG-UK) Consortium | Jacquelyn Wynn, Mairead Hyland, The Lighthouse Lab in Alderley Park and Alex Alderton, Roberto Amato, Jeffrey Barrett, Sonia Goncalves, Ewan Harrison, David K. Jackson, Ian Johnston, Dominic Kwiatkowski, Cordelia Langford, John Sillitoe on behalf of the Wellcome Sanger Institute COVID-19 Surveillance Team |
| EPI_ISL_2353567 | Lighthouse Laboratory Plymouth | Wellcome Sanger Institute for the COVID-19 Genomics UK (COG-UK) Consortium | Lighthouse Laboratory Plymouth and Alex Alderton, Roberto Amato, Jeffrey Barrett, Sonia Goncalves, Ewan Harrison, David K. Jackson, Ian Johnston, Dominic Kwiatkowski, Cordelia Langford, John Sillitoe on behalf of the Wellcome Sanger Institute COVID-19 Surveillance Team |
| EPI_ISL_2353569 | Lighthouse Lab in Alderley Park | Wellcome Sanger Institute for the COVID-19 Genomics UK (COG-UK) Consortium | Jacquelyn Wynn, Mairead Hyland, The Lighthouse Lab in Alderley Park and Alex Alderton, Roberto Amato, Jeffrey Barrett, Sonia Goncalves, Ewan Harrison, David K. Jackson, Ian Johnston, Dominic Kwiatkowski, Cordelia Langford, John Sillitoe on behalf of the Wellcome Sanger Institute COVID-19 Surveillance Team |
| EPI_ISL_2353570, EPI_ISL_2353571, EPI_ISL_2353573, EPI_ISL_2353575 | Lighthouse Lab in Milton Keynes | Wellcome Sanger Institute for the COVID-19 Genomics UK (COG-UK) Consortium | The Lighthouse Lab in Milton Keynes and Alex Alderton, Roberto Amato, Jeffrey Barrett, Sonia Goncalves, Ewan Harrison, David K. Jackson, Ian Johnston, Dominic Kwiatkowski, Cordelia Langford, John Sillitoe on behalf of the Wellcome Sanger Institute COVID-19 Surveillance Team |
| EPI_ISL_2353576 | Lighthouse Laboratory Plymouth | Wellcome Sanger Institute for the COVID-19 Genomics UK (COG-UK) Consortium | Lighthouse Laboratory Plymouth and Alex Alderton, Roberto Amato, Jeffrey Barrett, Sonia Goncalves, Ewan Harrison, David K. Jackson, Ian Johnston, Dominic Kwiatkowski, Cordelia Langford, John Sillitoe on behalf of the Wellcome Sanger Institute COVID-19 Surveillance Team |
| EPI_ISL_2353577 | Lighthouse Lab in Milton Keynes | Wellcome Sanger Institute for the COVID-19 Genomics UK (COG-UK) Consortium | The Lighthouse Lab in Milton Keynes and Alex Alderton, Roberto Amato, Jeffrey Barrett, Sonia Goncalves, Ewan Harrison, David K. Jackson, Ian Johnston, Dominic Kwiatkowski, Cordelia Langford, John Sillitoe on behalf of the Wellcome Sanger Institute COVID-19 Surveillance Team |
| EPI_ISL_2353578 | Lighthouse Laboratory Plymouth | Wellcome Sanger Institute for the COVID-19 Genomics UK (COG-UK) Consortium | Lighthouse Laboratory Plymouth and Alex Alderton, Roberto Amato, Jeffrey Barrett, Sonia Goncalves, Ewan Harrison, David K. Jackson, Ian Johnston, Dominic Kwiatkowski, Cordelia Langford, John Sillitoe on behalf of the Wellcome Sanger Institute COVID-19 Surveillance Team |
| EPI_ISL_2353579 | Lighthouse Lab in Alderley Park | Wellcome Sanger Institute for the COVID-19 Genomics UK (COG-UK) Consortium | Jacquelyn Wynn, Mairead Hyland, The Lighthouse Lab in Alderley Park and Alex Alderton, Roberto Amato, Jeffrey Barrett, Sonia Goncalves, Ewan Harrison, David K. Jackson, Ian Johnston, Dominic Kwiatkowski, Cordelia Langford, John Sillitoe on behalf of the Wellcome Sanger Institute COVID-19 Surveillance Team |
| EPI_ISL_2353581 | Lighthouse Lab in Milton Keynes | Wellcome Sanger Institute for the COVID-19 Genomics UK (COG-UK) Consortium | The Lighthouse Lab in Milton Keynes and Alex Alderton, Roberto Amato, Jeffrey Barrett, Sonia Goncalves, Ewan Harrison, David K. Jackson, Ian Johnston, Dominic Kwiatkowski, Cordelia Langford, John Sillitoe on behalf of the Wellcome Sanger Institute COVID-19 Surveillance Team |
| EPI_ISL_2353582 | Lighthouse Lab in Alderley Park | Wellcome Sanger Institute for the COVID-19 Genomics UK (COG-UK) Consortium | Jacquelyn Wynn, Mairead Hyland, The Lighthouse Lab in Alderley Park and Alex Alderton, Roberto Amato, Jeffrey Barrett, Sonia Goncalves, Ewan Harrison, David K. Jackson, Ian Johnston, Dominic Kwiatkowski, Cordelia Langford, John Sillitoe on behalf of the Wellcome Sanger Institute COVID-19 Surveillance Team |
| EPI_ISL_2353583 | Lighthouse Laboratory Plymouth | Wellcome Sanger Institute for the COVID-19 Genomics UK (COG-UK) Consortium | Lighthouse Laboratory Plymouth and Alex Alderton, Roberto Amato, Jeffrey Barrett, Sonia Goncalves, Ewan Harrison, David K. Jackson, Ian Johnston, Dominic Kwiatkowski, Cordelia Langford, John Sillitoe on behalf of the Wellcome Sanger Institute COVID-19 Surveillance Team |
| EPI_ISL_2353584 | Lighthouse Lab in Alderley Park | Wellcome Sanger Institute for the COVID-19 Genomics UK (COG-UK) Consortium | Jacquelyn Wynn, Mairead Hyland, The Lighthouse Lab in Alderley Park and Alex Alderton, Roberto Amato, Jeffrey Barrett, Sonia Goncalves, Ewan Harrison, David K. Jackson, Ian Johnston, Dominic Kwiatkowski, Cordelia Langford, John Sillitoe on behalf of the Wellcome Sanger Institute COVID-19 Surveillance Team |
| EPI_ISL_2353585 | Lighthouse Lab in Milton Keynes | Wellcome Sanger Institute for the COVID-19 Genomics UK (COG-UK) Consortium | The Lighthouse Lab in Milton Keynes and Alex Alderton, Roberto Amato, Jeffrey Barrett, Sonia Goncalves, Ewan Harrison, David K. Jackson, Ian Johnston, Dominic Kwiatkowski, Cordelia Langford, John Sillitoe on behalf of the Wellcome Sanger Institute COVID-19 Surveillance Team |
| EPI_ISL_2353586, EPI_ISL_2353587, EPI_ISL_2353588 | Lighthouse Lab in Alderley Park | Wellcome Sanger Institute for the COVID-19 Genomics UK (COG-UK) Consortium | Jacquelyn Wynn, Mairead Hyland, The Lighthouse Lab in Alderley Park and Alex Alderton, Roberto Amato, Jeffrey Barrett, Sonia Goncalves, Ewan Harrison, David K. Jackson, Ian Johnston, Dominic Kwiatkowski, Cordelia Langford, John Sillitoe on behalf of the Wellcome Sanger Institute COVID-19 Surveillance Team |
| EPI_ISL_2353589 | Lighthouse Lab in Milton Keynes | Wellcome Sanger Institute for the COVID-19 Genomics UK (COG-UK) Consortium | The Lighthouse Lab in Milton Keynes and Alex Alderton, Roberto Amato, Jeffrey Barrett, Sonia Goncalves, Ewan Harrison, David K. Jackson, Ian Johnston, Dominic Kwiatkowski, Cordelia Langford, John Sillitoe on behalf of the Wellcome Sanger Institute COVID-19 Surveillance Team |
| EPI_ISL_2353590 | Lighthouse Lab in Alderley Park | Wellcome Sanger Institute for the COVID-19 Genomics UK (COG-UK) Consortium | Jacquelyn Wynn, Mairead Hyland, The Lighthouse Lab in Alderley Park and Alex Alderton, Roberto Amato, Jeffrey Barrett, Sonia Goncalves, Ewan Harrison, David K. Jackson, Ian Johnston, Dominic Kwiatkowski, Cordelia Langford, John Sillitoe on behalf of the Wellcome Sanger Institute COVID-19 Surveillance Team |
| EPI_ISL_2353591 | Lighthouse Lab in Milton Keynes | Wellcome Sanger Institute for the COVID-19 Genomics UK (COG-UK) Consortium | The Lighthouse Lab in Milton Keynes and Alex Alderton, Roberto Amato, Jeffrey Barrett, Sonia Goncalves, Ewan Harrison, David K. Jackson, Ian Johnston, Dominic Kwiatkowski, Cordelia Langford, John Sillitoe on behalf of the Wellcome Sanger Institute COVID-19 Surveillance Team |

| | | | |
|---|---|---|---|
| EPI_ISL_2353592 | Lighthouse Lab in Glasgow | Wellcome Sanger Institute for the COVID-19 Genomics UK (COG-UK) Consortium | Harper VanSteenhouse, Yumi Kasai, David Gray, Carol Clugston, Anna Dominiczak and Alex Alderton, Roberto Amato, Jeffrey Barrett, Sonia Goncalves, Ewan Harrison, David K. Jackson, Ian Johnston, Dominic Kwiatkowski, Cordelia Langford, John Sillitoe on behalf of the Wellcome Sanger Institute COVID-19 Surveillance Team |
| EPI_ISL_2353593, EPI_ISL_2353595, EPI_ISL_2353596, EPI_ISL_2353597, EPI_ISL_2353598 | Lighthouse Lab in Alderley Park | Wellcome Sanger Institute for the COVID-19 Genomics UK (COG-UK) Consortium | Jacquelyn Wynn, Mairead Hyland, The Lighthouse Lab in Alderley Park and Alex Alderton, Roberto Amato, Jeffrey Barrett, Sonia Goncalves, Ewan Harrison, David K. Jackson, Ian Johnston, Dominic Kwiatkowski, Cordelia Langford, John Sillitoe on behalf of the Wellcome Sanger Institute COVID-19 Surveillance Team |
| EPI_ISL_2353600, EPI_ISL_2353601 | Lighthouse Lab in Milton Keynes | Wellcome Sanger Institute for the COVID-19 Genomics UK (COG-UK) Consortium | The Lighthouse Lab in Milton Keynes and Alex Alderton, Roberto Amato, Jeffrey Barrett, Sonia Goncalves, Ewan Harrison, David K. Jackson, Ian Johnston, Dominic Kwiatkowski, Cordelia Langford, John Sillitoe on behalf of the Wellcome Sanger Institute COVID-19 Surveillance Team |
| EPI_ISL_2353602 | Lighthouse Lab in Alderley Park | Wellcome Sanger Institute for the COVID-19 Genomics UK (COG-UK) Consortium | Jacquelyn Wynn, Mairead Hyland, The Lighthouse Lab in Alderley Park and Alex Alderton, Roberto Amato, Jeffrey Barrett, Sonia Goncalves, Ewan Harrison, David K. Jackson, Ian Johnston, Dominic Kwiatkowski, Cordelia Langford, John Sillitoe on behalf of the Wellcome Sanger Institute COVID-19 Surveillance Team |
| EPI_ISL_2353604 | Lighthouse Lab in Milton Keynes | Wellcome Sanger Institute for the COVID-19 Genomics UK (COG-UK) Consortium | The Lighthouse Lab in Milton Keynes and Alex Alderton, Roberto Amato, Jeffrey Barrett, Sonia Goncalves, Ewan Harrison, David K. Jackson, Ian Johnston, Dominic Kwiatkowski, Cordelia Langford, John Sillitoe on behalf of the Wellcome Sanger Institute COVID-19 Surveillance Team |
| EPI_ISL_2353605, EPI_ISL_2353606, EPI_ISL_2353607, EPI_ISL_2353608, EPI_ISL_2353609, EPI_ISL_2353610, EPI_ISL_2353611, EPI_ISL_2353612 | Lighthouse Lab in Alderley Park | Wellcome Sanger Institute for the COVID-19 Genomics UK (COG-UK) Consortium | Jacquelyn Wynn, Mairead Hyland, The Lighthouse Lab in Alderley Park and Alex Alderton, Roberto Amato, Jeffrey Barrett, Sonia Goncalves, Ewan Harrison, David K. Jackson, Ian Johnston, Dominic Kwiatkowski, Cordelia Langford, John Sillitoe on behalf of the Wellcome Sanger Institute COVID-19 Surveillance Team |
| EPI_ISL_2353613 | Lighthouse Lab in Milton Keynes | Wellcome Sanger Institute for the COVID-19 Genomics UK (COG-UK) Consortium | The Lighthouse Lab in Milton Keynes and Alex Alderton, Roberto Amato, Jeffrey Barrett, Sonia Goncalves, Ewan Harrison, David K. Jackson, Ian Johnston, Dominic Kwiatkowski, Cordelia Langford, John Sillitoe on behalf of the Wellcome Sanger Institute COVID-19 Surveillance Team |
| EPI_ISL_2353614 | Lighthouse Lab in Glasgow | Wellcome Sanger Institute for the COVID-19 Genomics UK (COG-UK) Consortium | Harper VanSteenhouse, Yumi Kasai, David Gray, Carol Clugston, Anna Dominiczak and Alex Alderton, Roberto Amato, Jeffrey Barrett, Sonia Goncalves, Ewan Harrison, David K. Jackson, Ian Johnston, Dominic Kwiatkowski, Cordelia Langford, John Sillitoe on behalf of the Wellcome Sanger Institute COVID-19 Surveillance Team |
| EPI_ISL_2353617 | Lighthouse Lab in Milton Keynes | Wellcome Sanger Institute for the COVID-19 Genomics UK (COG-UK) Consortium | The Lighthouse Lab in Milton Keynes and Alex Alderton, Roberto Amato, Jeffrey Barrett, Sonia Goncalves, Ewan Harrison, David K. Jackson, Ian Johnston, Dominic Kwiatkowski, Cordelia Langford, John Sillitoe on behalf of the Wellcome Sanger Institute COVID-19 Surveillance Team |
| EPI_ISL_2353619 | Lighthouse Lab in Alderley Park | Wellcome Sanger Institute for the COVID-19 Genomics UK (COG-UK) Consortium | Jacquelyn Wynn, Mairead Hyland, The Lighthouse Lab in Alderley Park and Alex Alderton, Roberto Amato, Jeffrey Barrett, Sonia Goncalves, Ewan Harrison, David K. Jackson, Ian Johnston, Dominic Kwiatkowski, Cordelia Langford, John Sillitoe on behalf of the Wellcome Sanger Institute COVID-19 Surveillance Team |
| EPI_ISL_2353620 | Lighthouse Lab in Milton Keynes | Wellcome Sanger Institute for the COVID-19 Genomics UK (COG-UK) Consortium | The Lighthouse Lab in Milton Keynes and Alex Alderton, Roberto Amato, Jeffrey Barrett, Sonia Goncalves, Ewan Harrison, David K. Jackson, Ian Johnston, Dominic Kwiatkowski, Cordelia Langford, John Sillitoe on behalf of the Wellcome Sanger Institute COVID-19 Surveillance Team |
| EPI_ISL_2353622 | Lighthouse Lab in Glasgow | Wellcome Sanger Institute for the COVID-19 Genomics UK (COG-UK) Consortium | Harper VanSteenhouse, Yumi Kasai, David Gray, Carol Clugston, Anna Dominiczak and Alex Alderton, Roberto Amato, Jeffrey Barrett, Sonia Goncalves, Ewan Harrison, David K. Jackson, Ian Johnston, Dominic Kwiatkowski, Cordelia Langford, John Sillitoe on behalf of the Wellcome Sanger Institute COVID-19 Surveillance Team |
| EPI_ISL_2353624, EPI_ISL_2353625, EPI_ISL_2353626, EPI_ISL_2353627 | Lighthouse Lab in Alderley Park | Wellcome Sanger Institute for the COVID-19 Genomics UK (COG-UK) Consortium | Jacquelyn Wynn, Mairead Hyland, The Lighthouse Lab in Alderley Park and Alex Alderton, Roberto Amato, Jeffrey Barrett, Sonia Goncalves, Ewan Harrison, David K. Jackson, Ian Johnston, Dominic Kwiatkowski, Cordelia Langford, John Sillitoe on behalf of the Wellcome Sanger Institute COVID-19 Surveillance Team |
| EPI_ISL_2353629 | Lighthouse Lab in Milton Keynes | Wellcome Sanger Institute for the COVID-19 Genomics UK (COG-UK) Consortium | The Lighthouse Lab in Milton Keynes and Alex Alderton, Roberto Amato, Jeffrey Barrett, Sonia Goncalves, Ewan Harrison, David K. Jackson, Ian Johnston, Dominic Kwiatkowski, Cordelia Langford, John Sillitoe on behalf of the Wellcome Sanger Institute COVID-19 Surveillance Team |
| EPI_ISL_2353631, EPI_ISL_2353632 | Lighthouse Lab in Alderley Park | Wellcome Sanger Institute for the COVID-19 Genomics UK (COG-UK) Consortium | Jacquelyn Wynn, Mairead Hyland, The Lighthouse Lab in Alderley Park and Alex Alderton, Roberto Amato, Jeffrey Barrett, Sonia Goncalves, Ewan Harrison, David K. Jackson, Ian Johnston, Dominic Kwiatkowski, Cordelia Langford, John Sillitoe on behalf of the Wellcome Sanger Institute COVID-19 Surveillance Team |
| EPI_ISL_2353634 | Lighthouse Lab in Milton Keynes | Wellcome Sanger Institute for the COVID-19 Genomics UK (COG-UK) Consortium | The Lighthouse Lab in Milton Keynes and Alex Alderton, Roberto Amato, Jeffrey Barrett, Sonia Goncalves, Ewan Harrison, David K. Jackson, Ian Johnston, Dominic Kwiatkowski, Cordelia Langford, John Sillitoe on behalf of the Wellcome Sanger Institute COVID-19 Surveillance Team |
| EPI_ISL_2353635, EPI_ISL_2353636 | Lighthouse Lab in Alderley Park | Wellcome Sanger Institute for the COVID-19 Genomics UK (COG-UK) Consortium | Jacquelyn Wynn, Mairead Hyland, The Lighthouse Lab in Alderley Park and Alex Alderton, Roberto Amato, Jeffrey Barrett, Sonia Goncalves, Ewan Harrison, David K. Jackson, Ian Johnston, Dominic Kwiatkowski, Cordelia Langford, John Sillitoe on behalf of the Wellcome Sanger Institute COVID-19 Surveillance Team |
| EPI_ISL_2353638, EPI_ISL_2353640 | Lighthouse Lab in Milton Keynes | Wellcome Sanger Institute for the COVID-19 Genomics UK (COG-UK) Consortium | The Lighthouse Lab in Milton Keynes and Alex Alderton, Roberto Amato, Jeffrey Barrett, Sonia Goncalves, Ewan Harrison, David K. Jackson, Ian Johnston, Dominic Kwiatkowski, Cordelia Langford, John Sillitoe on behalf of the Wellcome Sanger Institute COVID-19 Surveillance Team |
| EPI_ISL_2353641, EPI_ISL_2353642, EPI_ISL_2353646, EPI_ISL_2353647 | Lighthouse Lab in Alderley Park | Wellcome Sanger Institute for the COVID-19 Genomics UK (COG-UK) Consortium | Jacquelyn Wynn, Mairead Hyland, The Lighthouse Lab in Alderley Park and Alex Alderton, Roberto Amato, Jeffrey Barrett, Sonia Goncalves, Ewan Harrison, David K. Jackson, Ian Johnston, Dominic Kwiatkowski, Cordelia Langford, John Sillitoe on behalf of the Wellcome Sanger Institute COVID-19 Surveillance Team |
| EPI_ISL_2353648 | Lighthouse Lab in Milton Keynes | Wellcome Sanger Institute for the COVID-19 Genomics UK (COG-UK) Consortium | The Lighthouse Lab in Milton Keynes and Alex Alderton, Roberto Amato, Jeffrey Barrett, Sonia Goncalves, Ewan Harrison, David K. Jackson, Ian Johnston, Dominic Kwiatkowski, Cordelia Langford, John Sillitoe on behalf of the Wellcome Sanger Institute COVID-19 Surveillance Team |
| EPI_ISL_2353649, EPI_ISL_2353650, EPI_ISL_2353651, EPI_ISL_2353653, EPI_ISL_2353654 | Lighthouse Lab in Alderley Park | Wellcome Sanger Institute for the COVID-19 Genomics UK (COG-UK) Consortium | Jacquelyn Wynn, Mairead Hyland, The Lighthouse Lab in Alderley Park and Alex Alderton, Roberto Amato, Jeffrey Barrett, Sonia Goncalves, Ewan Harrison, David K. Jackson, Ian Johnston, Dominic Kwiatkowski, Cordelia Langford, John Sillitoe on behalf of the Wellcome Sanger Institute COVID-19 Surveillance Team |
| EPI_ISL_2353655 | Lighthouse Lab in Milton Keynes | Wellcome Sanger Institute for the COVID-19 Genomics UK (COG-UK) Consortium | The Lighthouse Lab in Milton Keynes and Alex Alderton, Roberto Amato, Jeffrey Barrett, Sonia Goncalves, Ewan Harrison, David K. Jackson, Ian Johnston, Dominic Kwiatkowski, Cordelia Langford, John Sillitoe on behalf of the Wellcome Sanger Institute COVID-19 Surveillance Team |
| EPI_ISL_2353656, EPI_ISL_2353657, EPI_ISL_2353660 | Lighthouse Lab in Alderley Park | Wellcome Sanger Institute for the COVID-19 Genomics UK (COG-UK) Consortium | Jacquelyn Wynn, Mairead Hyland, The Lighthouse Lab in Alderley Park and Alex Alderton, Roberto Amato, Jeffrey Barrett, Sonia Goncalves, Ewan Harrison, David K. Jackson, Ian Johnston, Dominic Kwiatkowski, Cordelia Langford, John Sillitoe on behalf of the Wellcome Sanger Institute COVID-19 Surveillance Team |
| EPI_ISL_2353661 | Lighthouse Lab in Milton Keynes | Wellcome Sanger Institute for the COVID-19 Genomics UK (COG-UK) Consortium | The Lighthouse Lab in Milton Keynes and Alex Alderton, Roberto Amato, Jeffrey Barrett, Sonia Goncalves, Ewan Harrison, David K. Jackson, Ian Johnston, Dominic Kwiatkowski, Cordelia Langford, John Sillitoe on behalf of the Wellcome Sanger Institute COVID-19 Surveillance Team |
| EPI_ISL_2353663, EPI_ISL_2353664, EPI_ISL_2353665, EPI_ISL_2353667 | Lighthouse Lab in Alderley Park | Wellcome Sanger Institute for the COVID-19 Genomics UK (COG-UK) Consortium | Jacquelyn Wynn, Mairead Hyland, The Lighthouse Lab in Alderley Park and Alex Alderton, Roberto Amato, Jeffrey Barrett, Sonia Goncalves, Ewan Harrison, David K. Jackson, Ian Johnston, Dominic Kwiatkowski, Cordelia Langford, John Sillitoe on behalf of the Wellcome Sanger Institute COVID-19 Surveillance Team |
| EPI_ISL_2353668 | Lighthouse Lab in Glasgow | Wellcome Sanger Institute for the COVID-19 Genomics UK (COG-UK) Consortium | Harper VanSteenhouse, Yumi Kasai, David Gray, Carol Clugston, Anna Dominiczak and Alex Alderton, Roberto Amato, Jeffrey Barrett, Sonia Goncalves, Ewan Harrison, David K. Jackson, Ian Johnston, Dominic Kwiatkowski, Cordelia Langford, John Sillitoe on behalf of the Wellcome Sanger Institute COVID-19 Surveillance Team |
| EPI_ISL_2353671 | Lighthouse Lab in Milton Keynes | Wellcome Sanger Institute for the COVID-19 Genomics UK (COG-UK) Consortium | The Lighthouse Lab in Milton Keynes and Alex Alderton, Roberto Amato, Jeffrey Barrett, Sonia Goncalves, Ewan Harrison, David K. Jackson, Ian Johnston, Dominic Kwiatkowski, Cordelia Langford, John Sillitoe on behalf of the Wellcome Sanger Institute COVID-19 Surveillance Team |
| EPI_ISL_2353672, EPI_ISL_2353673 | Lighthouse Lab in Alderley Park | Wellcome Sanger Institute for the COVID-19 Genomics UK (COG-UK) Consortium | Jacquelyn Wynn, Mairead Hyland, The Lighthouse Lab in Alderley Park and Alex Alderton, Roberto Amato, Jeffrey Barrett, Sonia Goncalves, Ewan Harrison, David K. Jackson, Ian Johnston, Dominic Kwiatkowski, Cordelia Langford, John Sillitoe on behalf of the Wellcome Sanger Institute COVID-19 Surveillance Team |
| EPI_ISL_2353674 | Lighthouse Lab in Milton Keynes | Wellcome Sanger Institute for the COVID-19 Genomics UK (COG-UK) Consortium | The Lighthouse Lab in Milton Keynes and Alex Alderton, Roberto Amato, Jeffrey Barrett, Sonia Goncalves, Ewan Harrison, David K. Jackson, Ian Johnston, Dominic Kwiatkowski, Cordelia Langford, John Sillitoe on behalf of the Wellcome Sanger Institute COVID-19 Surveillance Team |
| EPI_ISL_2353676 | Lighthouse Lab in Alderley Park | Wellcome Sanger Institute for the COVID-19 Genomics UK (COG-UK) Consortium | Jacquelyn Wynn, Mairead Hyland, The Lighthouse Lab in Alderley Park and Alex Alderton, Roberto Amato, Jeffrey Barrett, Sonia Goncalves, Ewan Harrison, David K. Jackson, Ian Johnston, Dominic Kwiatkowski, Cordelia Langford, John Sillitoe on behalf of the Wellcome Sanger Institute COVID-19 Surveillance Team |
| EPI_ISL_2353678 | Lighthouse Lab in Glasgow | Wellcome Sanger Institute for the COVID-19 Genomics UK (COG-UK) Consortium | Harper VanSteenhouse, Yumi Kasai, David Gray, Carol Clugston, Anna Dominiczak and Alex Alderton, Roberto Amato, Jeffrey Barrett, Sonia Goncalves, Ewan Harrison, David K. Jackson, Ian Johnston, Dominic Kwiatkowski, Cordelia Langford, John Sillitoe on behalf of the Wellcome Sanger Institute COVID-19 Surveillance Team |
| EPI_ISL_2353679, EPI_ISL_2353680 | Lighthouse Lab in Alderley Park | Wellcome Sanger Institute for the COVID-19 Genomics UK (COG-UK) Consortium | Jacquelyn Wynn, Mairead Hyland, The Lighthouse Lab in Alderley Park and Alex Alderton, Roberto Amato, Jeffrey Barrett, Sonia Goncalves, Ewan Harrison, David K. Jackson, Ian Johnston, Dominic Kwiatkowski, Cordelia Langford, John Sillitoe on behalf of the Wellcome Sanger Institute COVID-19 Surveillance Team |
| EPI_ISL_2353681 | Lighthouse Lab in Milton Keynes | Wellcome Sanger Institute for the COVID-19 Genomics UK (COG-UK) Consortium | The Lighthouse Lab in Milton Keynes and Alex Alderton, Roberto Amato, Jeffrey Barrett, Sonia Goncalves, Ewan Harrison, David K. Jackson, Ian Johnston, Dominic Kwiatkowski, Cordelia Langford, John Sillitoe on behalf of the Wellcome Sanger Institute COVID-19 Surveillance Team |
| EPI_ISL_2353682, EPI_ISL_2353683 | Lighthouse Lab in Alderley Park | Wellcome Sanger Institute for the COVID-19 Genomics UK (COG-UK) Consortium | Jacquelyn Wynn, Mairead Hyland, The Lighthouse Lab in Alderley Park and Alex Alderton, Roberto Amato, Jeffrey Barrett, Sonia Goncalves, Ewan Harrison, David K. Jackson, Ian Johnston, Dominic Kwiatkowski, Cordelia Langford, John Sillitoe on behalf of the Wellcome Sanger Institute COVID-19 Surveillance Team |
| EPI_ISL_2353684, EPI_ISL_2353685, EPI_ISL_2353686 | Lighthouse Lab in Milton Keynes | Wellcome Sanger Institute for the COVID-19 Genomics UK (COG-UK) Consortium | The Lighthouse Lab in Milton Keynes and Alex Alderton, Roberto Amato, Jeffrey Barrett, Sonia Goncalves, Ewan Harrison, David K. Jackson, Ian Johnston, Dominic Kwiatkowski, Cordelia Langford, John Sillitoe on behalf of the Wellcome Sanger Institute COVID-19 Surveillance Team |
| EPI_ISL_2353687, EPI_ISL_2353689, EPI_ISL_2353690, EPI_ISL_2353691, EPI_ISL_2353693, | Lighthouse Lab in Alderley Park | Wellcome Sanger Institute for the COVID-19 Genomics UK (COG-UK) Consortium | Jacquelyn Wynn, Mairead Hyland, The Lighthouse Lab in Alderley Park and Alex Alderton, Roberto Amato, Jeffrey Barrett, Sonia Goncalves, Ewan Harrison, David K. Jackson, Ian Johnston, Dominic Kwiatkowski, Cordelia Langford, John Sillitoe on behalf of the Wellcome Sanger Institute COVID-19 Surveillance Team |

| | | | |
|---|---|---|---|
| EPI_ISL_2353694, EPI_ISL_2353696, EPI_ISL_2353697, EPI_ISL_2353698, EPI_ISL_2353699 | | | |
| EPI_ISL_2353700 | Lighthouse Lab in Glasgow | Wellcome Sanger Institute for the COVID-19 Genomics UK (COG-UK) Consortium | Harper VanSteenhouse, Yumi Kasai, David Gray, Carol Clugston, Anna Dominiczak and Alex Alderton, Roberto Amato, Jeffrey Barrett, Sonia Goncalves, Ewan Harrison, David K. Jackson, Ian Johnston, Dominic Kwiatkowski, Cordelia Langford, John Sillitoe on behalf of the Wellcome Sanger Institute COVID-19 Surveillance Team |
| EPI_ISL_2353702, EPI_ISL_2353703, EPI_ISL_2353704 | Lighthouse Lab in Milton Keynes | Wellcome Sanger Institute for the COVID-19 Genomics UK (COG-UK) Consortium | The Lighthouse Lab in Milton Keynes and Alex Alderton, Roberto Amato, Jeffrey Barrett, Sonia Goncalves, Ewan Harrison, David K. Jackson, Ian Johnston, Dominic Kwiatkowski, Cordelia Langford, John Sillitoe on behalf of the Wellcome Sanger Institute COVID-19 Surveillance Team |
| EPI_ISL_2353708 | Lighthouse Lab in Alderley Park | Wellcome Sanger Institute for the COVID-19 Genomics UK (COG-UK) Consortium | Jacquelyn Wynn, Mairead Hyland, The Lighthouse Lab in Alderley Park and Alex Alderton, Roberto Amato, Jeffrey Barrett, Sonia Goncalves, Ewan Harrison, David K. Jackson, Ian Johnston, Dominic Kwiatkowski, Cordelia Langford, John Sillitoe on behalf of the Wellcome Sanger Institute COVID-19 Surveillance Team |
| EPI_ISL_2353710, EPI_ISL_2353712 | Lighthouse Lab in Milton Keynes | Wellcome Sanger Institute for the COVID-19 Genomics UK (COG-UK) Consortium | The Lighthouse Lab in Milton Keynes and Alex Alderton, Roberto Amato, Jeffrey Barrett, Sonia Goncalves, Ewan Harrison, David K. Jackson, Ian Johnston, Dominic Kwiatkowski, Cordelia Langford, John Sillitoe on behalf of the Wellcome Sanger Institute COVID-19 Surveillance Team |
| EPI_ISL_2353714 | Lighthouse Lab in Alderley Park | Wellcome Sanger Institute for the COVID-19 Genomics UK (COG-UK) Consortium | Jacquelyn Wynn, Mairead Hyland, The Lighthouse Lab in Alderley Park and Alex Alderton, Roberto Amato, Jeffrey Barrett, Sonia Goncalves, Ewan Harrison, David K. Jackson, Ian Johnston, Dominic Kwiatkowski, Cordelia Langford, John Sillitoe on behalf of the Wellcome Sanger Institute COVID-19 Surveillance Team |
| EPI_ISL_2353715 | Lighthouse Lab in Glasgow | Wellcome Sanger Institute for the COVID-19 Genomics UK (COG-UK) Consortium | Harper VanSteenhouse, Yumi Kasai, David Gray, Carol Clugston, Anna Dominiczak and Alex Alderton, Roberto Amato, Jeffrey Barrett, Sonia Goncalves, Ewan Harrison, David K. Jackson, Ian Johnston, Dominic Kwiatkowski, Cordelia Langford, John Sillitoe on behalf of the Wellcome Sanger Institute COVID-19 Surveillance Team |
| EPI_ISL_2353717 | Lighthouse Lab in Alderley Park | Wellcome Sanger Institute for the COVID-19 Genomics UK (COG-UK) Consortium | Jacquelyn Wynn, Mairead Hyland, The Lighthouse Lab in Alderley Park and Alex Alderton, Roberto Amato, Jeffrey Barrett, Sonia Goncalves, Ewan Harrison, David K. Jackson, Ian Johnston, Dominic Kwiatkowski, Cordelia Langford, John Sillitoe on behalf of the Wellcome Sanger Institute COVID-19 Surveillance Team |
| EPI_ISL_2353718, EPI_ISL_2353719 | Lighthouse Lab in Milton Keynes | Wellcome Sanger Institute for the COVID-19 Genomics UK (COG-UK) Consortium | The Lighthouse Lab in Milton Keynes and Alex Alderton, Roberto Amato, Jeffrey Barrett, Sonia Goncalves, Ewan Harrison, David K. Jackson, Ian Johnston, Dominic Kwiatkowski, Cordelia Langford, John Sillitoe on behalf of the Wellcome Sanger Institute COVID-19 Surveillance Team |
| EPI_ISL_2353721 | Lighthouse Lab in Alderley Park | Wellcome Sanger Institute for the COVID-19 Genomics UK (COG-UK) Consortium | Jacquelyn Wynn, Mairead Hyland, The Lighthouse Lab in Alderley Park and Alex Alderton, Roberto Amato, Jeffrey Barrett, Sonia Goncalves, Ewan Harrison, David K. Jackson, Ian Johnston, Dominic Kwiatkowski, Cordelia Langford, John Sillitoe on behalf of the Wellcome Sanger Institute COVID-19 Surveillance Team |
| EPI_ISL_2353722 | Lighthouse Lab in Milton Keynes | Wellcome Sanger Institute for the COVID-19 Genomics UK (COG-UK) Consortium | The Lighthouse Lab in Milton Keynes and Alex Alderton, Roberto Amato, Jeffrey Barrett, Sonia Goncalves, Ewan Harrison, David K. Jackson, Ian Johnston, Dominic Kwiatkowski, Cordelia Langford, John Sillitoe on behalf of the Wellcome Sanger Institute COVID-19 Surveillance Team |
| EPI_ISL_2353723, EPI_ISL_2353724 | Lighthouse Lab in Alderley Park | Wellcome Sanger Institute for the COVID-19 Genomics UK (COG-UK) Consortium | Jacquelyn Wynn, Mairead Hyland, The Lighthouse Lab in Alderley Park and Alex Alderton, Roberto Amato, Jeffrey Barrett, Sonia Goncalves, Ewan Harrison, David K. Jackson, Ian Johnston, Dominic Kwiatkowski, Cordelia Langford, John Sillitoe on behalf of the Wellcome Sanger Institute COVID-19 Surveillance Team |
| EPI_ISL_2353727 | Lighthouse Lab in Glasgow | Wellcome Sanger Institute for the COVID-19 Genomics UK (COG-UK) Consortium | Harper VanSteenhouse, Yumi Kasai, David Gray, Carol Clugston, Anna Dominiczak and Alex Alderton, Roberto Amato, Jeffrey Barrett, Sonia Goncalves, Ewan Harrison, David K. Jackson, Ian Johnston, Dominic Kwiatkowski, Cordelia Langford, John Sillitoe on behalf of the Wellcome Sanger Institute COVID-19 Surveillance Team |
| EPI_ISL_2353729, EPI_ISL_2353730, EPI_ISL_2353731, EPI_ISL_2353732, EPI_ISL_2353733, EPI_ISL_2353734 | Lighthouse Lab in Alderley Park | Wellcome Sanger Institute for the COVID-19 Genomics UK (COG-UK) Consortium | Jacquelyn Wynn, Mairead Hyland, The Lighthouse Lab in Alderley Park and Alex Alderton, Roberto Amato, Jeffrey Barrett, Sonia Goncalves, Ewan Harrison, David K. Jackson, Ian Johnston, Dominic Kwiatkowski, Cordelia Langford, John Sillitoe on behalf of the Wellcome Sanger Institute COVID-19 Surveillance Team |
| EPI_ISL_2353737 | Lighthouse Lab in Milton Keynes | Wellcome Sanger Institute for the COVID-19 Genomics UK (COG-UK) Consortium | The Lighthouse Lab in Milton Keynes and Alex Alderton, Roberto Amato, Jeffrey Barrett, Sonia Goncalves, Ewan Harrison, David K. Jackson, Ian Johnston, Dominic Kwiatkowski, Cordelia Langford, John Sillitoe on behalf of the Wellcome Sanger Institute COVID-19 Surveillance Team |
| EPI_ISL_2353738 | Lighthouse Lab in Alderley Park | Wellcome Sanger Institute for the COVID-19 Genomics UK (COG-UK) Consortium | Jacquelyn Wynn, Mairead Hyland, The Lighthouse Lab in Alderley Park and Alex Alderton, Roberto Amato, Jeffrey Barrett, Sonia Goncalves, Ewan Harrison, David K. Jackson, Ian Johnston, Dominic Kwiatkowski, Cordelia Langford, John Sillitoe on behalf of the Wellcome Sanger Institute COVID-19 Surveillance Team |
| EPI_ISL_2353739 | Lighthouse Lab in Milton Keynes | Wellcome Sanger Institute for the COVID-19 Genomics UK (COG-UK) Consortium | The Lighthouse Lab in Milton Keynes and Alex Alderton, Roberto Amato, Jeffrey Barrett, Sonia Goncalves, Ewan Harrison, David K. Jackson, Ian Johnston, Dominic Kwiatkowski, Cordelia Langford, John Sillitoe on behalf of the Wellcome Sanger Institute COVID-19 Surveillance Team |
| EPI_ISL_2353741 | Lighthouse Lab in Alderley Park | Wellcome Sanger Institute for the COVID-19 Genomics UK (COG-UK) Consortium | Jacquelyn Wynn, Mairead Hyland, The Lighthouse Lab in Alderley Park and Alex Alderton, Roberto Amato, Jeffrey Barrett, Sonia Goncalves, Ewan Harrison, David K. Jackson, Ian Johnston, Dominic Kwiatkowski, Cordelia Langford, John Sillitoe on behalf of the Wellcome Sanger Institute COVID-19 Surveillance Team |
| EPI_ISL_2353743 | Lighthouse Lab in Milton Keynes | Wellcome Sanger Institute for the COVID-19 Genomics UK (COG-UK) Consortium | The Lighthouse Lab in Milton Keynes and Alex Alderton, Roberto Amato, Jeffrey Barrett, Sonia Goncalves, Ewan Harrison, David K. Jackson, Ian Johnston, Dominic Kwiatkowski, Cordelia Langford, John Sillitoe on behalf of the Wellcome Sanger Institute COVID-19 Surveillance Team |
| EPI_ISL_2353745, EPI_ISL_2353749 | Lighthouse Lab in Alderley Park | Wellcome Sanger Institute for the COVID-19 Genomics UK (COG-UK) Consortium | Jacquelyn Wynn, Mairead Hyland, The Lighthouse Lab in Alderley Park and Alex Alderton, Roberto Amato, Jeffrey Barrett, Sonia Goncalves, Ewan Harrison, David K. Jackson, Ian Johnston, Dominic Kwiatkowski, Cordelia Langford, John Sillitoe on behalf of the Wellcome Sanger Institute COVID-19 Surveillance Team |
| EPI_ISL_2353750 | Lighthouse Lab in Milton Keynes | Wellcome Sanger Institute for the COVID-19 Genomics UK (COG-UK) Consortium | The Lighthouse Lab in Milton Keynes and Alex Alderton, Roberto Amato, Jeffrey Barrett, Sonia Goncalves, Ewan Harrison, David K. Jackson, Ian Johnston, Dominic Kwiatkowski, Cordelia Langford, John Sillitoe on behalf of the Wellcome Sanger Institute COVID-19 Surveillance Team |
| EPI_ISL_2353752, EPI_ISL_2353753, EPI_ISL_2353755 | Lighthouse Lab in Alderley Park | Wellcome Sanger Institute for the COVID-19 Genomics UK (COG-UK) Consortium | Jacquelyn Wynn, Mairead Hyland, The Lighthouse Lab in Alderley Park and Alex Alderton, Roberto Amato, Jeffrey Barrett, Sonia Goncalves, Ewan Harrison, David K. Jackson, Ian Johnston, Dominic Kwiatkowski, Cordelia Langford, John Sillitoe on behalf of the Wellcome Sanger Institute COVID-19 Surveillance Team |
| EPI_ISL_2353756 | Lighthouse Lab in Milton Keynes | Wellcome Sanger Institute for the COVID-19 Genomics UK (COG-UK) Consortium | The Lighthouse Lab in Milton Keynes and Alex Alderton, Roberto Amato, Jeffrey Barrett, Sonia Goncalves, Ewan Harrison, David K. Jackson, Ian Johnston, Dominic Kwiatkowski, Cordelia Langford, John Sillitoe on behalf of the Wellcome Sanger Institute COVID-19 Surveillance Team |
| EPI_ISL_2353757, EPI_ISL_2353759, EPI_ISL_2353760, EPI_ISL_2353762 | Lighthouse Lab in Alderley Park | Wellcome Sanger Institute for the COVID-19 Genomics UK (COG-UK) Consortium | Jacquelyn Wynn, Mairead Hyland, The Lighthouse Lab in Alderley Park and Alex Alderton, Roberto Amato, Jeffrey Barrett, Sonia Goncalves, Ewan Harrison, David K. Jackson, Ian Johnston, Dominic Kwiatkowski, Cordelia Langford, John Sillitoe on behalf of the Wellcome Sanger Institute COVID-19 Surveillance Team |
| EPI_ISL_2353763 | Lighthouse Lab in Milton Keynes | Wellcome Sanger Institute for the COVID-19 Genomics UK (COG-UK) Consortium | The Lighthouse Lab in Milton Keynes and Alex Alderton, Roberto Amato, Jeffrey Barrett, Sonia Goncalves, Ewan Harrison, David K. Jackson, Ian Johnston, Dominic Kwiatkowski, Cordelia Langford, John Sillitoe on behalf of the Wellcome Sanger Institute COVID-19 Surveillance Team |
| EPI_ISL_2353764, EPI_ISL_2353765, EPI_ISL_2353766, EPI_ISL_2353768, EPI_ISL_2353770 | Lighthouse Lab in Alderley Park | Wellcome Sanger Institute for the COVID-19 Genomics UK (COG-UK) Consortium | Jacquelyn Wynn, Mairead Hyland, The Lighthouse Lab in Alderley Park and Alex Alderton, Roberto Amato, Jeffrey Barrett, Sonia Goncalves, Ewan Harrison, David K. Jackson, Ian Johnston, Dominic Kwiatkowski, Cordelia Langford, John Sillitoe on behalf of the Wellcome Sanger Institute COVID-19 Surveillance Team |
| EPI_ISL_2353771 | Lighthouse Lab in Milton Keynes | Wellcome Sanger Institute for the COVID-19 Genomics UK (COG-UK) Consortium | The Lighthouse Lab in Milton Keynes and Alex Alderton, Roberto Amato, Jeffrey Barrett, Sonia Goncalves, Ewan Harrison, David K. Jackson, Ian Johnston, Dominic Kwiatkowski, Cordelia Langford, John Sillitoe on behalf of the Wellcome Sanger Institute COVID-19 Surveillance Team |
| EPI_ISL_2353772, EPI_ISL_2353773, EPI_ISL_2353777, EPI_ISL_2353778 | Lighthouse Lab in Alderley Park | Wellcome Sanger Institute for the COVID-19 Genomics UK (COG-UK) Consortium | Jacquelyn Wynn, Mairead Hyland, The Lighthouse Lab in Alderley Park and Alex Alderton, Roberto Amato, Jeffrey Barrett, Sonia Goncalves, Ewan Harrison, David K. Jackson, Ian Johnston, Dominic Kwiatkowski, Cordelia Langford, John Sillitoe on behalf of the Wellcome Sanger Institute COVID-19 Surveillance Team |
| EPI_ISL_2353779 | Lighthouse Lab in Milton Keynes | Wellcome Sanger Institute for the COVID-19 Genomics UK (COG-UK) Consortium | The Lighthouse Lab in Milton Keynes and Alex Alderton, Roberto Amato, Jeffrey Barrett, Sonia Goncalves, Ewan Harrison, David K. Jackson, Ian Johnston, Dominic Kwiatkowski, Cordelia Langford, John Sillitoe on behalf of the Wellcome Sanger Institute COVID-19 Surveillance Team |
| EPI_ISL_2353780, EPI_ISL_2353781, EPI_ISL_2353782, EPI_ISL_2353783, EPI_ISL_2353784, EPI_ISL_2353785, EPI_ISL_2353786, EPI_ISL_2353787, EPI_ISL_2353788, EPI_ISL_2353789, EPI_ISL_2353790, EPI_ISL_2353791, EPI_ISL_2353792, EPI_ISL_2353793, EPI_ISL_2353794, EPI_ISL_2353795, EPI_ISL_2353797, EPI_ISL_2353798, EPI_ISL_2353800, EPI_ISL_2353802 | | | |
| see above | Lighthouse Lab in Alderley Park | Wellcome Sanger Institute for the COVID-19 Genomics UK (COG-UK) Consortium | Jacquelyn Wynn, Mairead Hyland, The Lighthouse Lab in Alderley Park and Alex Alderton, Roberto Amato, Jeffrey Barrett, Sonia Goncalves, Ewan Harrison, David K. Jackson, Ian Johnston, Dominic Kwiatkowski, Cordelia Langford, John Sillitoe on behalf of the Wellcome Sanger Institute COVID-19 Surveillance Team |
| EPI_ISL_2353803 | Lighthouse Lab in Milton Keynes | Wellcome Sanger Institute for the COVID-19 Genomics UK (COG-UK) Consortium | The Lighthouse Lab in Milton Keynes and Alex Alderton, Roberto Amato, Jeffrey Barrett, Sonia Goncalves, Ewan Harrison, David K. Jackson, Ian Johnston, Dominic Kwiatkowski, Cordelia Langford, John Sillitoe on behalf of the Wellcome Sanger Institute COVID-19 Surveillance Team |
| EPI_ISL_2353806 | Lighthouse Lab in Alderley Park | Wellcome Sanger Institute for the COVID-19 Genomics UK (COG-UK) Consortium | Jacquelyn Wynn, Mairead Hyland, The Lighthouse Lab in Alderley Park and Alex Alderton, Roberto Amato, Jeffrey Barrett, Sonia Goncalves, Ewan Harrison, David K. Jackson, Ian Johnston, Dominic Kwiatkowski, Cordelia Langford, John Sillitoe on behalf of the Wellcome Sanger Institute COVID-19 Surveillance Team |
| EPI_ISL_2353807, EPI_ISL_2353808 | Lighthouse Lab in Milton Keynes | Wellcome Sanger Institute for the COVID-19 Genomics UK (COG-UK) Consortium | The Lighthouse Lab in Milton Keynes and Alex Alderton, Roberto Amato, Jeffrey Barrett, Sonia Goncalves, Ewan Harrison, David K. Jackson, Ian Johnston, Dominic Kwiatkowski, Cordelia Langford, John Sillitoe on behalf of the Wellcome Sanger Institute COVID-19 Surveillance Team |
| EPI_ISL_2353809 | Lighthouse Lab in Alderley Park | Wellcome Sanger Institute for the COVID-19 Genomics UK (COG-UK) Consortium | Jacquelyn Wynn, Mairead Hyland, The Lighthouse Lab in Alderley Park and Alex Alderton, Roberto Amato, Jeffrey Barrett, Sonia Goncalves, Ewan Harrison, David K. Jackson, Ian Johnston, Dominic Kwiatkowski, Cordelia Langford, John Sillitoe on behalf of the Wellcome Sanger Institute COVID-19 Surveillance Team |
| EPI_ISL_2353810 | Lighthouse Lab in Milton Keynes | Wellcome Sanger Institute for the COVID-19 Genomics UK (COG-UK) Consortium | The Lighthouse Lab in Milton Keynes and Alex Alderton, Roberto Amato, Jeffrey Barrett, Sonia Goncalves, Ewan Harrison, David K. Jackson, Ian Johnston, Dominic Kwiatkowski, Cordelia Langford, John Sillitoe on behalf of the Wellcome Sanger Institute COVID-19 Surveillance Team |
| EPI_ISL_2353812 | Lighthouse Lab in Alderley Park | Wellcome Sanger Institute for the COVID-19 Genomics UK (COG-UK) Consortium | Jacquelyn Wynn, Mairead Hyland, The Lighthouse Lab in Alderley Park and Alex Alderton, Roberto Amato, Jeffrey Barrett, Sonia Goncalves, Ewan Harrison, David K. Jackson, Ian Johnston, Dominic Kwiatkowski, Cordelia Langford, John Sillitoe on behalf of the Wellcome Sanger Institute COVID-19 Surveillance Team |
| EPI_ISL_2353813, EPI_ISL_2353814 | Lighthouse Lab in Milton Keynes | Wellcome Sanger Institute for the COVID-19 Genomics UK (COG-UK) Consortium | The Lighthouse Lab in Milton Keynes and Alex Alderton, Roberto Amato, Jeffrey Barrett, Sonia Goncalves, Ewan Harrison, David K. Jackson, Ian Johnston, Dominic Kwiatkowski, Cordelia Langford, John Sillitoe on behalf of the Wellcome Sanger Institute COVID-19 Surveillance Team |
| EPI_ISL_2353815, EPI_ISL_2353816, | Lighthouse Lab in Alderley Park | Wellcome Sanger Institute for the COVID-19 Genomics UK (COG-UK) Consortium | Jacquelyn Wynn, Mairead Hyland, The Lighthouse Lab in Alderley Park and Alex Alderton, Roberto Amato, Jeffrey Barrett, Sonia Goncalves, Ewan Harrison, David K. Jackson, Ian Johnston, Dominic Kwiatkowski, Cordelia Langford, John Sillitoe on behalf of the Wellcome Sanger Institute COVID-19 Surveillance Team |

| | | | |
|---|---|---|---|
| EPI_ISL_2353817, EPI_ISL_2353818 | | | |
| EPI_ISL_2353819, EPI_ISL_2353821 | Lighthouse Lab in Milton Keynes | Wellcome Sanger Institute for the COVID-19 Genomics UK (COG-UK) Consortium | The Lighthouse Lab in Milton Keynes and Alex Alderton, Roberto Amato, Jeffrey Barrett, Sonia Goncalves, Ewan Harrison, David K. Jackson, Ian Johnston, Dominic Kwiatkowski, Cordelia Langford, John Sillitoe on behalf of the Wellcome Sanger Institute COVID-19 Surveillance Team |
| EPI_ISL_2353822, EPI_ISL_2353823, EPI_ISL_2353824 | Lighthouse Lab in Alderley Park | Wellcome Sanger Institute for the COVID-19 Genomics UK (COG-UK) Consortium | Jacquelyn Wynn, Mairead Hyland, The Lighthouse Lab in Alderley Park and Alex Alderton, Roberto Amato, Jeffrey Barrett, Sonia Goncalves, Ewan Harrison, David K. Jackson, Ian Johnston, Dominic Kwiatkowski, Cordelia Langford, John Sillitoe on behalf of the Wellcome Sanger Institute COVID-19 Surveillance Team |
| EPI_ISL_2353825 | Lighthouse Lab in Glasgow | Wellcome Sanger Institute for the COVID-19 Genomics UK (COG-UK) Consortium | Harper VanSteenhouse, Yumi Kasai, David Gray, Carol Clugston, Anna Dominiczak and Alex Alderton, Roberto Amato, Jeffrey Barrett, Sonia Goncalves, Ewan Harrison, David K. Jackson, Ian Johnston, Dominic Kwiatkowski, Cordelia Langford, John Sillitoe on behalf of the Wellcome Sanger Institute COVID-19 Surveillance Team |
| EPI_ISL_2353826 | Lighthouse Lab in Alderley Park | Wellcome Sanger Institute for the COVID-19 Genomics UK (COG-UK) Consortium | Jacquelyn Wynn, Mairead Hyland, The Lighthouse Lab in Alderley Park and Alex Alderton, Roberto Amato, Jeffrey Barrett, Sonia Goncalves, Ewan Harrison, David K. Jackson, Ian Johnston, Dominic Kwiatkowski, Cordelia Langford, John Sillitoe on behalf of the Wellcome Sanger Institute COVID-19 Surveillance Team |
| EPI_ISL_2353827 | Lighthouse Lab in Glasgow | Wellcome Sanger Institute for the COVID-19 Genomics UK (COG-UK) Consortium | Harper VanSteenhouse, Yumi Kasai, David Gray, Carol Clugston, Anna Dominiczak and Alex Alderton, Roberto Amato, Jeffrey Barrett, Sonia Goncalves, Ewan Harrison, David K. Jackson, Ian Johnston, Dominic Kwiatkowski, Cordelia Langford, John Sillitoe on behalf of the Wellcome Sanger Institute COVID-19 Surveillance Team |
| EPI_ISL_2353830, EPI_ISL_2353833 | Lighthouse Lab in Alderley Park | Wellcome Sanger Institute for the COVID-19 Genomics UK (COG-UK) Consortium | Jacquelyn Wynn, Mairead Hyland, The Lighthouse Lab in Alderley Park and Alex Alderton, Roberto Amato, Jeffrey Barrett, Sonia Goncalves, Ewan Harrison, David K. Jackson, Ian Johnston, Dominic Kwiatkowski, Cordelia Langford, John Sillitoe on behalf of the Wellcome Sanger Institute COVID-19 Surveillance Team |
| EPI_ISL_2353835 | Lighthouse Lab in Milton Keynes | Wellcome Sanger Institute for the COVID-19 Genomics UK (COG-UK) Consortium | The Lighthouse Lab in Milton Keynes and Alex Alderton, Roberto Amato, Jeffrey Barrett, Sonia Goncalves, Ewan Harrison, David K. Jackson, Ian Johnston, Dominic Kwiatkowski, Cordelia Langford, John Sillitoe on behalf of the Wellcome Sanger Institute COVID-19 Surveillance Team |
| EPI_ISL_2353838, EPI_ISL_2353839, EPI_ISL_2353840, EPI_ISL_2353841, EPI_ISL_2353842, EPI_ISL_2353843 | Lighthouse Lab in Alderley Park | Wellcome Sanger Institute for the COVID-19 Genomics UK (COG-UK) Consortium | Jacquelyn Wynn, Mairead Hyland, The Lighthouse Lab in Alderley Park and Alex Alderton, Roberto Amato, Jeffrey Barrett, Sonia Goncalves, Ewan Harrison, David K. Jackson, Ian Johnston, Dominic Kwiatkowski, Cordelia Langford, John Sillitoe on behalf of the Wellcome Sanger Institute COVID-19 Surveillance Team |
| EPI_ISL_2353844 | Lighthouse Lab in Milton Keynes | Wellcome Sanger Institute for the COVID-19 Genomics UK (COG-UK) Consortium | The Lighthouse Lab in Milton Keynes and Alex Alderton, Roberto Amato, Jeffrey Barrett, Sonia Goncalves, Ewan Harrison, David K. Jackson, Ian Johnston, Dominic Kwiatkowski, Cordelia Langford, John Sillitoe on behalf of the Wellcome Sanger Institute COVID-19 Surveillance Team |
| EPI_ISL_2353845 | Lighthouse Lab in Glasgow | Wellcome Sanger Institute for the COVID-19 Genomics UK (COG-UK) Consortium | Harper VanSteenhouse, Yumi Kasai, David Gray, Carol Clugston, Anna Dominiczak and Alex Alderton, Roberto Amato, Jeffrey Barrett, Sonia Goncalves, Ewan Harrison, David K. Jackson, Ian Johnston, Dominic Kwiatkowski, Cordelia Langford, John Sillitoe on behalf of the Wellcome Sanger Institute COVID-19 Surveillance Team |
| EPI_ISL_2353847, EPI_ISL_2353848, EPI_ISL_2353850 | Lighthouse Lab in Alderley Park | Wellcome Sanger Institute for the COVID-19 Genomics UK (COG-UK) Consortium | Jacquelyn Wynn, Mairead Hyland, The Lighthouse Lab in Alderley Park and Alex Alderton, Roberto Amato, Jeffrey Barrett, Sonia Goncalves, Ewan Harrison, David K. Jackson, Ian Johnston, Dominic Kwiatkowski, Cordelia Langford, John Sillitoe on behalf of the Wellcome Sanger Institute COVID-19 Surveillance Team |
| EPI_ISL_2353852, EPI_ISL_2353853 | Lighthouse Lab in Milton Keynes | Wellcome Sanger Institute for the COVID-19 Genomics UK (COG-UK) Consortium | The Lighthouse Lab in Milton Keynes and Alex Alderton, Roberto Amato, Jeffrey Barrett, Sonia Goncalves, Ewan Harrison, David K. Jackson, Ian Johnston, Dominic Kwiatkowski, Cordelia Langford, John Sillitoe on behalf of the Wellcome Sanger Institute COVID-19 Surveillance Team |
| EPI_ISL_2353854 | Lighthouse Lab in Alderley Park | Wellcome Sanger Institute for the COVID-19 Genomics UK (COG-UK) Consortium | Jacquelyn Wynn, Mairead Hyland, The Lighthouse Lab in Alderley Park and Alex Alderton, Roberto Amato, Jeffrey Barrett, Sonia Goncalves, Ewan Harrison, David K. Jackson, Ian Johnston, Dominic Kwiatkowski, Cordelia Langford, John Sillitoe on behalf of the Wellcome Sanger Institute COVID-19 Surveillance Team |
| EPI_ISL_2353856, EPI_ISL_2353858 | Lighthouse Lab in Milton Keynes | Wellcome Sanger Institute for the COVID-19 Genomics UK (COG-UK) Consortium | The Lighthouse Lab in Milton Keynes and Alex Alderton, Roberto Amato, Jeffrey Barrett, Sonia Goncalves, Ewan Harrison, David K. Jackson, Ian Johnston, Dominic Kwiatkowski, Cordelia Langford, John Sillitoe on behalf of the Wellcome Sanger Institute COVID-19 Surveillance Team |
| EPI_ISL_2353859 | Lighthouse Lab in Glasgow | Wellcome Sanger Institute for the COVID-19 Genomics UK (COG-UK) Consortium | Harper VanSteenhouse, Yumi Kasai, David Gray, Carol Clugston, Anna Dominiczak and Alex Alderton, Roberto Amato, Jeffrey Barrett, Sonia Goncalves, Ewan Harrison, David K. Jackson, Ian Johnston, Dominic Kwiatkowski, Cordelia Langford, John Sillitoe on behalf of the Wellcome Sanger Institute COVID-19 Surveillance Team |
| EPI_ISL_2353860, EPI_ISL_2353861, EPI_ISL_2353862 | Lighthouse Lab in Alderley Park | Wellcome Sanger Institute for the COVID-19 Genomics UK (COG-UK) Consortium | Jacquelyn Wynn, Mairead Hyland, The Lighthouse Lab in Alderley Park and Alex Alderton, Roberto Amato, Jeffrey Barrett, Sonia Goncalves, Ewan Harrison, David K. Jackson, Ian Johnston, Dominic Kwiatkowski, Cordelia Langford, John Sillitoe on behalf of the Wellcome Sanger Institute COVID-19 Surveillance Team |
| EPI_ISL_2353864, EPI_ISL_2353867, EPI_ISL_2353870, EPI_ISL_2353871, EPI_ISL_2353872, EPI_ISL_2353873, EPI_ISL_2353874, EPI_ISL_2353875, EPI_ISL_2353876, EPI_ISL_2353879, EPI_ISL_2353882, EPI_ISL_2353885, EPI_ISL_2353886, EPI_ISL_2353888, EPI_ISL_2353889, EPI_ISL_2353890, EPI_ISL_2353892, EPI_ISL_2353893, EPI_ISL_2353895, EPI_ISL_2353897, EPI_ISL_2353898, EPI_ISL_2353899, EPI_ISL_2353900, EPI_ISL_2353903, EPI_ISL_2353905, EPI_ISL_2353906, EPI_ISL_2353907, EPI_ISL_2353910, EPI_ISL_2353912, EPI_ISL_2353913, EPI_ISL_2353915, EPI_ISL_2353916, EPI_ISL_2353917, EPI_ISL_2353920, EPI_ISL_2353921, EPI_ISL_2353924, EPI_ISL_2353925, EPI_ISL_2353926, EPI_ISL_2353927, EPI_ISL_2353929, EPI_ISL_2353932, EPI_ISL_2353934, EPI_ISL_2353935, EPI_ISL_2353937, EPI_ISL_2353939 | | | |
| see above | Lighthouse Lab in Glasgow | Wellcome Sanger Institute for the COVID-19 Genomics UK (COG-UK) Consortium | Harper VanSteenhouse, Yumi Kasai, David Gray, Carol Clugston, Anna Dominiczak and Alex Alderton, Roberto Amato, Jeffrey Barrett, Sonia Goncalves, Ewan Harrison, David K. Jackson, Ian Johnston, Dominic Kwiatkowski, Cordelia Langford, John Sillitoe on behalf of the Wellcome Sanger Institute COVID-19 Surveillance Team |
| EPI_ISL_2353940 | Lighthouse Lab in Milton Keynes | Wellcome Sanger Institute for the COVID-19 Genomics UK (COG-UK) Consortium | The Lighthouse Lab in Milton Keynes and Alex Alderton, Roberto Amato, Jeffrey Barrett, Sonia Goncalves, Ewan Harrison, David K. Jackson, Ian Johnston, Dominic Kwiatkowski, Cordelia Langford, John Sillitoe on behalf of the Wellcome Sanger Institute COVID-19 Surveillance Team |
| EPI_ISL_2353943, EPI_ISL_2353944, EPI_ISL_2353947, EPI_ISL_2353948, EPI_ISL_2353951, EPI_ISL_2353952, EPI_ISL_2353954, EPI_ISL_2353956, EPI_ISL_2353957, EPI_ISL_2353958, EPI_ISL_2353959, EPI_ISL_2353961, EPI_ISL_2353965, EPI_ISL_2353967, EPI_ISL_2353968, EPI_ISL_2353975, EPI_ISL_2353976, EPI_ISL_2353977, EPI_ISL_2353978, EPI_ISL_2353980, EPI_ISL_2353981, EPI_ISL_2353982, EPI_ISL_2353987, EPI_ISL_2353988, EPI_ISL_2353989, EPI_ISL_2353991, EPI_ISL_2353992, EPI_ISL_2353998, EPI_ISL_2353999, EPI_ISL_2354000, EPI_ISL_2354003, EPI_ISL_2354004, EPI_ISL_2354005, EPI_ISL_2354006, EPI_ISL_2354007, EPI_ISL_2354010, EPI_ISL_2354011, EPI_ISL_2354016, EPI_ISL_2354017, EPI_ISL_2354020, EPI_ISL_2354021, EPI_ISL_2354023, EPI_ISL_2354024, EPI_ISL_2354025, EPI_ISL_2354029, EPI_ISL_2354030, EPI_ISL_2354034, EPI_ISL_2354035, EPI_ISL_2354036, EPI_ISL_2354038, EPI_ISL_2354040, EPI_ISL_2354041, EPI_ISL_2354043, EPI_ISL_2354044, EPI_ISL_2354045, EPI_ISL_2354046, EPI_ISL_2354048, EPI_ISL_2354051, EPI_ISL_2354054, EPI_ISL_2354056, EPI_ISL_2354057, EPI_ISL_2354058, EPI_ISL_2354059, EPI_ISL_2354061, EPI_ISL_2354062, EPI_ISL_2354064, EPI_ISL_2354066, EPI_ISL_2354068, EPI_ISL_2354069, EPI_ISL_2354070, EPI_ISL_2354071, EPI_ISL_2354072, EPI_ISL_2354073, EPI_ISL_2354076, EPI_ISL_2354077, EPI_ISL_2354078, EPI_ISL_2354079, EPI_ISL_2354081, EPI_ISL_2354082, EPI_ISL_2354083, EPI_ISL_2354084, EPI_ISL_2354085, EPI_ISL_2354086, EPI_ISL_2354087, EPI_ISL_2354089, EPI_ISL_2354091, EPI_ISL_2354092, EPI_ISL_2354096, EPI_ISL_2354097, EPI_ISL_2354099, EPI_ISL_2354100, EPI_ISL_2354101, EPI_ISL_2354103, EPI_ISL_2354109, EPI_ISL_2354111, EPI_ISL_2354113, EPI_ISL_2354114, EPI_ISL_2354115, EPI_ISL_2354116, EPI_ISL_2354118, EPI_ISL_2354119, EPI_ISL_2354122, EPI_ISL_2354125, EPI_ISL_2354127, EPI_ISL_2354128, EPI_ISL_2354129, EPI_ISL_2354130, EPI_ISL_2354131, EPI_ISL_2354132, EPI_ISL_2354133, EPI_ISL_2354134, EPI_ISL_2354137, EPI_ISL_2354138, EPI_ISL_2354139, EPI_ISL_2354141, EPI_ISL_2354142, EPI_ISL_2354143, EPI_ISL_2354144, EPI_ISL_2354145, EPI_ISL_2354146, EPI_ISL_2354148, EPI_ISL_2354150, EPI_ISL_2354154, EPI_ISL_2354155, EPI_ISL_2354158, EPI_ISL_2354160, EPI_ISL_2354161, EPI_ISL_2354163, EPI_ISL_2354164, EPI_ISL_2354165, EPI_ISL_2354167, EPI_ISL_2354169, EPI_ISL_2354172, EPI_ISL_2354173, EPI_ISL_2354175, EPI_ISL_2354177, EPI_ISL_2354178, EPI_ISL_2354181, EPI_ISL_2354183, EPI_ISL_2354187, EPI_ISL_2354191, EPI_ISL_2354193, EPI_ISL_2354199, EPI_ISL_2354202, EPI_ISL_2354204, EPI_ISL_2354207, EPI_ISL_2354208, EPI_ISL_2354209, EPI_ISL_2354212, EPI_ISL_2354214, EPI_ISL_2354216, EPI_ISL_2354219, EPI_ISL_2354220, EPI_ISL_2354222, EPI_ISL_2354224, EPI_ISL_2354226, EPI_ISL_2354227, EPI_ISL_2354229, EPI_ISL_2354233, EPI_ISL_2354234, EPI_ISL_2354236, EPI_ISL_2354239, EPI_ISL_2354240, EPI_ISL_2354241, EPI_ISL_2354244, EPI_ISL_2354245, EPI_ISL_2354247, EPI_ISL_2354251, EPI_ISL_2354252, EPI_ISL_2354254, EPI_ISL_2354256, EPI_ISL_2354257, EPI_ISL_2354259, EPI_ISL_2354260, EPI_ISL_2354262, EPI_ISL_2354263, EPI_ISL_2354264, EPI_ISL_2354265, EPI_ISL_2354267, EPI_ISL_2354268, EPI_ISL_2354271, EPI_ISL_2354274, EPI_ISL_2354275, EPI_ISL_2354277, EPI_ISL_2354279, EPI_ISL_2354284, EPI_ISL_2354285, EPI_ISL_2354293, EPI_ISL_2354298, EPI_ISL_2354299, EPI_ISL_2354300, EPI_ISL_2354301, EPI_ISL_2354302, EPI_ISL_2354303, EPI_ISL_2354304, EPI_ISL_2354305, EPI_ISL_2354306, EPI_ISL_2354307, EPI_ISL_2354308, EPI_ISL_2354310, EPI_ISL_2354311, EPI_ISL_2354312, EPI_ISL_2354313, EPI_ISL_2354315, EPI_ISL_2354318, EPI_ISL_2354320, EPI_ISL_2354321, EPI_ISL_2354323, EPI_ISL_2354330, EPI_ISL_2354332, EPI_ISL_2354333, EPI_ISL_2354335, EPI_ISL_2354336, EPI_ISL_2354337, EPI_ISL_2354339, EPI_ISL_2354343, EPI_ISL_2354344, EPI_ISL_2354345, EPI_ISL_2354349, EPI_ISL_2354350, EPI_ISL_2354352, EPI_ISL_2354353, EPI_ISL_2354354, EPI_ISL_2354356, EPI_ISL_2354359, EPI_ISL_2354362, EPI_ISL_2354364, EPI_ISL_2354367, EPI_ISL_2354370, EPI_ISL_2354371, EPI_ISL_2354372, EPI_ISL_2354374, EPI_ISL_2354375, EPI_ISL_2354376, EPI_ISL_2354377, EPI_ISL_2354379, EPI_ISL_2354380, EPI_ISL_2354382, EPI_ISL_2354384, EPI_ISL_2354388, EPI_ISL_2354390, EPI_ISL_2354391, EPI_ISL_2354392, EPI_ISL_2354397, EPI_ISL_2354398, EPI_ISL_2354399, EPI_ISL_2354403, EPI_ISL_2354405, EPI_ISL_2354406, EPI_ISL_2354407, EPI_ISL_2354408, EPI_ISL_2354409, EPI_ISL_2354410, EPI_ISL_2354411, EPI_ISL_2354412, EPI_ISL_2354413, EPI_ISL_2354414, EPI_ISL_2354415, EPI_ISL_2354417, EPI_ISL_2354418, EPI_ISL_2354420, EPI_ISL_2354421, EPI_ISL_2354423, EPI_ISL_2354424, EPI_ISL_2354427, EPI_ISL_2354429, EPI_ISL_2354430, EPI_ISL_2354431, EPI_ISL_2354432, EPI_ISL_2354434, EPI_ISL_2354436, EPI_ISL_2354437, EPI_ISL_2354438, EPI_ISL_2354439, EPI_ISL_2354443, EPI_ISL_2354445, EPI_ISL_2354446, EPI_ISL_2354448, EPI_ISL_2354449, EPI_ISL_2354451, EPI_ISL_2354454, EPI_ISL_2354455, EPI_ISL_2354456, EPI_ISL_2354457, EPI_ISL_2354459, EPI_ISL_2354462, EPI_ISL_2354463, EPI_ISL_2354465, EPI_ISL_2354467, EPI_ISL_2354471, EPI_ISL_2354472, EPI_ISL_2354473, EPI_ISL_2354475, EPI_ISL_2354476, EPI_ISL_2354477 | | | |
| see above | Lighthouse Lab in Glasgow | Wellcome Sanger Institute for the COVID-19 Genomics UK (COG-UK) Consortium | Harper VanSteenhouse, Yumi Kasai, David Gray, Carol Clugston, Anna Dominiczak and Alex Alderton, Roberto Amato, Jeffrey Barrett, Sonia Goncalves, Ewan Harrison, David K. Jackson, Ian Johnston, Dominic Kwiatkowski, Cordelia Langford, John Sillitoe on behalf of the Wellcome Sanger Institute COVID-19 Surveillance Team |
| EPI_ISL_2354489 | Microbiology Division, SC DHEC | Microbiology Division, SC DHEC | Flores,H., Freeman,J. |
| EPI_ISL_2354503, EPI_ISL_2354507, EPI_ISL_2354508, EPI_ISL_2354510, EPI_ISL_2354511, EPI_ISL_2354513, EPI_ISL_2354514, EPI_ISL_2354515, EPI_ISL_2354516, EPI_ISL_2354517, EPI_ISL_2354518, EPI_ISL_2354519, EPI_ISL_2354520, EPI_ISL_2354528, EPI_ISL_2354530, EPI_ISL_2354531, EPI_ISL_2354532 | | | |
| see above | University of Birmingham | COVID-19 Genomics UK (COG-UK) Consortium | Institute of Microbiology, University of Birmingham: Claire McMurray, Joanne Stockton, Samuel Nicholls, Radoslaw Poplawski, Will Rowe, Josh Quick, Nicholas Loman. University of Birmingham Testing Laboratory: Celina M Whalley, Andrew Bosworth, Charlotte Poxon, Kasun Wanigasooriya, Oliver Pickles, Mike Kidd, Alex Richter, Andrew D Beggs PHE Heartlands Lab: Husam Osman, Andrew Bosworth. Queen Elizabeth Hospital: Anna Casey |
| EPI_ISL_2354543, EPI_ISL_2354544, EPI_ISL_2354545 | Department of Pathology, University of Cambridge | COVID-19 Genomics UK (COG-UK) Consortium | Aminu S. Jahun, Yasmin Chaudhry, Iliana Georgana, Myra Hosmillo, Rhys Izuagbe, William L. Hamilton, Martin D. Curran, Surendra Parmar, Ian Goodfellow |
| EPI_ISL_2354550, EPI_ISL_2354560, EPI_ISL_2354562, EPI_ISL_2354563, EPI_ISL_2354564, EPI_ISL_2354569, EPI_ISL_2354576, EPI_ISL_2354577, EPI_ISL_2354581, EPI_ISL_2354587, EPI_ISL_2354589, EPI_ISL_2354590, EPI_ISL_2354591, EPI_ISL_2354592, EPI_ISL_2354594, EPI_ISL_2354598, EPI_ISL_2354599, EPI_ISL_2354600, EPI_ISL_2354601, EPI_ISL_2354603, EPI_ISL_2354604, EPI_ISL_2354605, EPI_ISL_2354606, EPI_ISL_2354607, EPI_ISL_2354608, EPI_ISL_2354611, EPI_ISL_2354614, EPI_ISL_2354615, EPI_ISL_2354616, EPI_ISL_2354617, EPI_ISL_2354618, EPI_ISL_2354619, EPI_ISL_2354620, EPI_ISL_2354621, EPI_ISL_2354622, EPI_ISL_2354623, EPI_ISL_2354626, EPI_ISL_2354628, EPI_ISL_2354629, EPI_ISL_2354630, EPI_ISL_2354632, EPI_ISL_2354633 | | | |
| see above | Virology Department, Royal Infirmary of Edinburgh, NHS Lothian / School of Biological Sciences, University of Edinburgh | COVID-19 Genomics UK (COG-UK) Consortium | McHugh M, Dewar R, Cotton S, Gallagher A, Maloney D, Rooke S, O'Toole Á, Scher E, Hill V, McCrone JT, Colquhoun R, Yu X, Jackson B, Rambaut A, Templeton K |
| EPI_ISL_2355346 | University Hospitals Of Leicester NHS Trust and DeepSeq Nottingham | COVID-19 Genomics UK (COG-UK) Consortium | Christopher Holmes, Paul Bird, Thomas Helmer, Karlie Fallon, Julian Tang, Jonathan Ball, Patrick McClure, Joeseph Chappell, Nadine Holmes, Matthew Carlisle, Christopher Moore, Fei Sang, Johnny Debebe, Victoria Wright, Matthew Loose |
| EPI_ISL_2355350, EPI_ISL_2355351, EPI_ISL_2355352, EPI_ISL_2355356, EPI_ISL_2355357, EPI_ISL_2355364, EPI_ISL_2355366, EPI_ISL_2355367, EPI_ISL_2355368, EPI_ISL_2355369, EPI_ISL_2355370, EPI_ISL_2355371 | | | |
| see above | Liverpool Clinical Laboratories | COVID-19 Genomics UK (COG-UK) Consortium | Sam Haldenby, Alistair Darby, Steve Paterson, Anita Lucaci, Julian Hiscox, M Almsaud, A Alrezaihi, Muhannad Alruwaili, Stuart D Armstrong, Jones Benjamin, Eleanor G Bentley, Anu Chawla, Jordan J Clark, Angela Cowell, Richard Eccles, Isabel García-Dorival, Matthew Gemmell, Alessandro Gerada, PKF Gilmore, Richard Gregory, Ximeng Han, Catherine Hartley, Margaret Hughes, Miren Iturriza-Gomara, James Johnson, L Luu, Jenifer Manson, Charlotte Nelson, Elaine O'Toole, Cassie Olateju, Rebekah Penrice-Randal , Lucille Rainbow, N.P Randle, Trevor Ian Robinson, Parul Sharma, Ghada T Shawli, James P Stewart, Neil Swainston, Ecaterina Vamos, Joanne Watts, Mark Whitehead, Hermione Webster |
| EPI_ISL_2355374, EPI_ISL_2355375 | Barts Health NHS Trust | COVID-19 Genomics UK (COG-UK) Consortium | CUTINO-MOGUEL, Maria-Teresa; HARRINGTON, David; OWOYEMI, Dola; KULASEGARAN-SHYLINI, Raghavendran; BROAD, Claire; KELE, Beatrix |
| EPI_ISL_2355376, EPI_ISL_2355378, EPI_ISL_2355379, EPI_ISL_2355382, EPI_ISL_2355383, EPI_ISL_2355384, EPI_ISL_2355386, EPI_ISL_2355388, EPI_ISL_2355389, EPI_ISL_2355390, EPI_ISL_2355391, EPI_ISL_2355392, EPI_ISL_2355394, EPI_ISL_2355395, EPI_ISL_2355396, EPI_ISL_2355397, EPI_ISL_2355398, EPI_ISL_2355402, EPI_ISL_2355403, EPI_ISL_2355404, EPI_ISL_2355405, EPI_ISL_2355406, EPI_ISL_2355407, EPI_ISL_2355408, EPI_ISL_2355409, EPI_ISL_2355410, EPI_ISL_2355411, EPI_ISL_2355412, EPI_ISL_2355413, EPI_ISL_2355414, EPI_ISL_2355416, EPI_ISL_2355417, EPI_ISL_2355418, EPI_ISL_2355419, EPI_ISL_2355420, EPI_ISL_2355421, EPI_ISL_2355422, EPI_ISL_2355424, EPI_ISL_2355427, EPI_ISL_2355429, EPI_ISL_2355433, EPI_ISL_2355435, EPI_ISL_2355436, EPI_ISL_2355437, EPI_ISL_2355438, EPI_ISL_2355439 | | | |
| see above | University College London, Great Ormond Street Hospital for Children NHS Foundation Trust, Imperial College Healthcare NHS Trust | COVID-19 Genomics UK (COG-UK) Consortium | Sergi Castellano, Rachel Williams, Mark Kristiansen, Paola Resende Silva, Sunando Roy, Tony Brooks, Helena Tutill, Paola Niola, Patricia Dyal, Charlotte Williams, Leysa Forrest, Yasmin Panchbhaya, Jacqueline Findlay, Samuel Weeks, Julianne Brown, Kathryn Harris, Paul Randell, James Price, Alison Holmes, Judith Breuer |
| EPI_ISL_2355441, EPI_ISL_2355442, EPI_ISL_2355445, EPI_ISL_2355455, EPI_ISL_2355466, EPI_ISL_2355477, EPI_ISL_2355479, EPI_ISL_2355481, EPI_ISL_2355485, EPI_ISL_2355486, EPI_ISL_2355487, EPI_ISL_2355491, EPI_ISL_2355497, EPI_ISL_2355498, EPI_ISL_2355499, EPI_ISL_2355502, EPI_ISL_2355505, EPI_ISL_2355513, EPI_ISL_2355519, EPI_ISL_2355526, EPI_ISL_2355532, EPI_ISL_2355540, EPI_ISL_2355541, EPI_ISL_2355544, EPI_ISL_2355551, EPI_ISL_2355554, EPI_ISL_2355562, EPI_ISL_2355565, EPI_ISL_2355577, EPI_ISL_2355578, EPI_ISL_2355596, EPI_ISL_2355598, EPI_ISL_2355600, EPI_ISL_2355601, EPI_ISL_2355606, EPI_ISL_2355618, EPI_ISL_2355629, EPI_ISL_2355632, EPI_ISL_2355634, EPI_ISL_2355642, EPI_ISL_2355643, EPI_ISL_2355645, EPI_ISL_2355650, EPI_ISL_2355652, EPI_ISL_2355655, EPI_ISL_2355658, EPI_ISL_2355664, EPI_ISL_2355665, EPI_ISL_2355671, EPI_ISL_2355674, EPI_ISL_2355675, EPI_ISL_2355680, EPI_ISL_2355688, EPI_ISL_2355694, EPI_ISL_2355696, EPI_ISL_2355697, EPI_ISL_2355703, EPI_ISL_2355706, EPI_ISL_2355713, EPI_ISL_2355716, EPI_ISL_2355719, EPI_ISL_2355721, EPI_ISL_2355725, EPI_ISL_2355727, EPI_ISL_2355732, EPI_ISL_2355735, EPI_ISL_2355738, EPI_ISL_2355739, EPI_ISL_2355740, EPI_ISL_2355741, EPI_ISL_2355746, EPI_ISL_2355754, EPI_ISL_2355759, EPI_ISL_2355760, EPI_ISL_2355765, EPI_ISL_2355767, EPI_ISL_2355769, EPI_ISL_2355773, EPI_ISL_2355774, EPI_ISL_2355777, EPI_ISL_2355778, EPI_ISL_2355785, EPI_ISL_2355787, EPI_ISL_2355791, EPI_ISL_2355792, EPI_ISL_2355793, EPI_ISL_2355794, EPI_ISL_2355804, EPI_ISL_2355819, EPI_ISL_2355827, EPI_ISL_2355828, EPI_ISL_2355837, EPI_ISL_2355839, EPI_ISL_2355861, EPI_ISL_2355896, EPI_ISL_2355899, EPI_ISL_2355908, EPI_ISL_2355931, EPI_ISL_2355932, EPI_ISL_2355946, EPI_ISL_2355947, EPI_ISL_2355955, EPI_ISL_2355956, EPI_ISL_2355958, EPI_ISL_2355962, EPI_ISL_2355964, EPI_ISL_2355972, EPI_ISL_2356000, EPI_ISL_2356011, EPI_ISL_2356019, EPI_ISL_2356020, EPI_ISL_2356025, EPI_ISL_2356031, EPI_ISL_2356035, EPI_ISL_2356046, EPI_ISL_2356057, EPI_ISL_2356060, EPI_ISL_2356069, EPI_ISL_2356072, EPI_ISL_2356088, EPI_ISL_2356091, EPI_ISL_2356096, EPI_ISL_2356102, EPI_ISL_2356108, EPI_ISL_2356109, EPI_ISL_2356128, EPI_ISL_2356134, EPI_ISL_2356136, EPI_ISL_2356141, EPI_ISL_2356143, EPI_ISL_2356149, EPI_ISL_2356150, | | | |

| | | | |
|---|---|---|---|
| EPI_ISL_2356156, EPI_ISL_2356162, EPI_ISL_2356169, EPI_ISL_2356170, EPI_ISL_2356185, EPI_ISL_2356189, EPI_ISL_2356196, EPI_ISL_2356202, EPI_ISL_2356203, EPI_ISL_2356206, EPI_ISL_2356230, EPI_ISL_2356248, EPI_ISL_2356254, EPI_ISL_2356255, EPI_ISL_2356256, EPI_ISL_2356258, EPI_ISL_2356268, EPI_ISL_2356272, EPI_ISL_2356281, EPI_ISL_2356282, EPI_ISL_2356287, EPI_ISL_2356290, EPI_ISL_2356291, EPI_ISL_2356299, EPI_ISL_2356301, EPI_ISL_2356304, EPI_ISL_2356308, EPI_ISL_2356317, EPI_ISL_2356320, EPI_ISL_2356321, EPI_ISL_2356323, EPI_ISL_2356336, EPI_ISL_2356342, EPI_ISL_2356343, EPI_ISL_2356349, EPI_ISL_2356351, EPI_ISL_2356361, EPI_ISL_2356362, EPI_ISL_2356368, EPI_ISL_2356375, EPI_ISL_2356378, EPI_ISL_2356379, EPI_ISL_2356381, EPI_ISL_2356382, EPI_ISL_2356383, EPI_ISL_2356384, EPI_ISL_2356398, EPI_ISL_2356399, EPI_ISL_2356401, EPI_ISL_2356406, EPI_ISL_2356407, EPI_ISL_2356414, EPI_ISL_2356419, EPI_ISL_2356448, EPI_ISL_2356456, EPI_ISL_2356462, EPI_ISL_2356467, EPI_ISL_2356476, EPI_ISL_2356486, EPI_ISL_2356494, EPI_ISL_2356497, EPI_ISL_2356498, EPI_ISL_2356501, EPI_ISL_2356505, EPI_ISL_2356507, EPI_ISL_2356508, EPI_ISL_2356509, EPI_ISL_2356512, EPI_ISL_2356514, EPI_ISL_2356515, EPI_ISL_2356517, EPI_ISL_2356518, EPI_ISL_2356522, EPI_ISL_2356527, EPI_ISL_2356530, EPI_ISL_2356539, EPI_ISL_2356544, EPI_ISL_2356546, EPI_ISL_2356550, EPI_ISL_2356559, EPI_ISL_2356560, EPI_ISL_2356571, EPI_ISL_2356575, EPI_ISL_2356581, EPI_ISL_2356597, EPI_ISL_2356613, EPI_ISL_2356614, EPI_ISL_2356616, EPI_ISL_2356619, EPI_ISL_2356626, EPI_ISL_2356627, EPI_ISL_2356632, EPI_ISL_2356638, EPI_ISL_2356641, EPI_ISL_2356643, EPI_ISL_2356647, EPI_ISL_2356652, EPI_ISL_2356669, EPI_ISL_2356670, EPI_ISL_2356672, EPI_ISL_2356673, EPI_ISL_2356676, EPI_ISL_2356679, EPI_ISL_2356681 | | | |
| see above | Northumbria University / South Tees Hospitals NHS Foundation Trust / North Cumbria Integrated Care NHS Foundation Trust / North Tees and Hartlepool NHS Foundation Trust / Newcastle Hospitals NHS Foundation Trust | COVID-19 Genomics UK (COG-UK) Consortium | Darren L Smith,Andrew Nelson,Matthew Bashton,Greg R Young,Joshua Loh,John Allan,Mohammad A Tariq,Giles S Holt,Gary Black,Wen C Yew,Lynn Dover,Paul Baker,Steve Liggett,Sarah Essex,Jane Greenaway,Debra Padgett,Clive Graham,Garren Scott,Edward Barton,Emma Swindells,Brendan Payne,Jennifer Collins,Yusri Taha,Gary Eltringham |
| EPI_ISL_2356695, EPI_ISL_2356696 | Quadram Institute Bioscience | COVID-19 Genomics UK (COG-UK) Consortium | Dave J. Baker, Gemma L. Kay, Alp Aydin, Thanh Le-Viet, Steven Rudder, Ana P. Tedim, Anastasia Kolyva, Maria Diaz, Leonardo de Oliveira Martins, Nabil-Fareed Alikhan, Lizzie Meadows, Rachael Stanley, Ngozi Elumogo, Muhammed Yasir, Nicholas M. Thomson, Alexander J Trotter, Rachel Gilroy, Samuel Bloomfield, Claire Stuart, Andrew Bell, Reenesh Prakash, Samir Dervisevic, Alison E. Mather, John Wain, Mark Webber, Andrew J. Page, Justin O'Grady |
| EPI_ISL_2356699, EPI_ISL_2356700, EPI_ISL_2356701, EPI_ISL_2356702, EPI_ISL_2356703, EPI_ISL_2356704, EPI_ISL_2356706 | Queens Medical Centre, Clinical Microbiology Department / DeepSeq Nottingham | COVID-19 Genomics UK (COG-UK) Consortium | Gemma Clark, Wendy Smith, Manjinder Khakh, Vicki M Fleming, Michelle M Lister, Hannah Howson-Wells, Jonathan Ball, Timothy Byaruhanga, Jayasree Dey, Emily Park, Jack Hill, Patrick McClure, Joseph Chappell, Theocharis Tsoleridis, Nadine Holmes, Matthew Carlisle, Christopher Moore, Fei Sang, Johnny Debebe, Victoria Wright, Matthew Loose |
| EPI_ISL_2356720, EPI_ISL_2356724, EPI_ISL_2356726, EPI_ISL_2356731, EPI_ISL_2356734, EPI_ISL_2356739, EPI_ISL_2356742, EPI_ISL_2356748, EPI_ISL_2356751, EPI_ISL_2356760, EPI_ISL_2356761, EPI_ISL_2356772, EPI_ISL_2356779, EPI_ISL_2356783, EPI_ISL_2356791, EPI_ISL_2356792, EPI_ISL_2356793, EPI_ISL_2356795, EPI_ISL_2356800, EPI_ISL_2356813, EPI_ISL_2356824, EPI_ISL_2356826, EPI_ISL_2356842, EPI_ISL_2356869, EPI_ISL_2356870, EPI_ISL_2356871 | | | |
| see above | Wales Specialist Virology Centre Sequencing lab: Pathogen Genomics Unit | Public Health Wales Microbiology Cardiff Wales Specialist Virology Centre | Catherine Moore, Johnathan Evans, Laura Gifford, Malorie Perry, Simon Cottrell, Angela Marchbank, Alec Birchley, Alexander Adams, Amy Gaskin, Bree Gatica-Wilcox, Jason Coombes, Joel Southgate, Lauren Gilbert, Lee Graham, Nicole Pacchiarini, Sara Kumziene-Summerhayes, Sarah Taylor, Sophie Jones, Sara Rey, Matthew Bull, Joanne Watkins, Sally Corden, Tom Connor |
| EPI_ISL_2356891 | Virology Department, Sheffield Teaching Hospitals NHS Foundation Trust/Department of Infection, Immunity and Cardiovascular Disease, The Medical School, University of Sheffield | COVID-19 Genomics UK (COG-UK) Consortium | Thushan de Silva, Matthew Parker, Nikki Smith, Adri Angyal, Rebecca Brown, Luke Green, Rachel Tucker, Paul Parsons, Danielle Groves, Katie Johnson, Laura Carrilero, Alex Keeley, Dave Partridge, Matthew Wyles, Benjamin Lindsey, Mehmet Yavuz, Mohammad Raza, Cariad Evans |
| EPI_ISL_2357738, EPI_ISL_2357742, EPI_ISL_2357743, EPI_ISL_2357744, EPI_ISL_2357745 | National Institute of Public Health | State Veterinary Institute Prague | Nagy,A;Jirincova,H;Suri,T;Trnka,D;Vecerova,J |
| EPI_ISL_2357759 | Furst Medical Laboratory | Norwegian Institute of Public Health, Department of Virology | Kathrine Stene-Johansen, Kamilla Heddeland Instefjord, Hilde Elshaug, Garcia Llorente Ignacio, Jon Bråte, Line Victoria Moen, Engebretsen Serina Beate,Pedersen Benedikte Nevjen, Debech Nadia, Atiya R Ali,Marie Paulsen Madsen, Rasmus Riis Kopperud, Hilde Vollan, Karoline Bragstad, Olav Hungnes |
| EPI_ISL_2357777 | Medical Microbiology Unit, Department for Laboratory Medicine, Drammen Hospital, Vestre Viken Health Trust, | Norwegian Institute of Public Health, Department of Virology | Kathrine Stene-Johansen, Kamilla Heddeland Instefjord, Hilde Elshaug, Garcia Llorente Ignacio, Jon Bråte, Line Victoria Moen, Engebretsen Serina Beate,Pedersen Benedikte Nevjen, Debech Nadia, Atiya R Ali,Marie Paulsen Madsen, Rasmus Riis Kopperud, Hilde Vollan, Karoline Bragstad, Olav Hungnes |
| EPI_ISL_2357849 | Akershus University Hospital, Department for Microbiology and Infectious Disease Control | Norwegian Institute of Public Health, Department of Virology | Kathrine Stene-Johansen, Kamilla Heddeland Instefjord, Hilde Elshaug, Garcia Llorente Ignacio, Jon Bråte, Engebretsen Serina Beate,Pedersen Benedikte Nevjen, Debech Nadia, Atiya R Ali,Marie Paulsen Madsen, Rasmus Riis Kopperud, Hilde Vollan, Karoline Bragstad, Olav Hungnes |
| EPI_ISL_2357867 | Furst Medical Laboratory | Norwegian Institute of Public Health, Department of Virology | Kathrine Stene-Johansen, Kamilla Heddeland Instefjord, Hilde Elshaug, Garcia Llorente Ignacio, Jon Bråte, Engebretsen Serina Beate,Pedersen Benedikte Nevjen, Debech Nadia, Atiya R Ali,Marie Paulsen Madsen, Rasmus Riis Kopperud, Hilde Vollan, Karoline Bragstad, Olav Hungnes |
| EPI_ISL_2357962, EPI_ISL_2357963, EPI_ISL_2357964, EPI_ISL_2357998, EPI_ISL_2357999, EPI_ISL_2358012 | Dept. of Microbiology and Infection Control, Akershus University Hospital HF | Dept. of Microbiology and Infection Control, Akershus University Hospital HF | Hege Vangstein Aamot, Alexander Hesselberg Løvestad |
| EPI_ISL_2358230 | Algemeen Medisch Labo | Labo Klinische Biologie, UZA | Marie Le Mercier, Jasmine Coppens, Basil Britto Xavier, Christine Lammens, Veerle Matheeussen, Herman Goossens |
| EPI_ISL_2360239 | UniversitätsSpital Zürich 009 | Institute of Medical Virology | Verena Kufner, Gabriela Ziltener, Maryam Zaheri, Stefan Schmutz, Annette Audigé, Maria Grünberg, Kevin Steiner, Jon Huder, Cyril Shah, Riccarda Capaul, Guido Bloemberg, Jürg Böni, Michael Huber, Alexandra Trkola |
| EPI_ISL_2360241 | Stadtspital Triemli | Institute of Medical Virology | Verena Kufner, Gabriela Ziltener, Maryam Zaheri, Stefan Schmutz, Annette Audigé, Maria Grünberg, Kevin Steiner, Jon Huder, Cyril Shah, Riccarda Capaul, Guido Bloemberg, Jürg Böni, Michael Huber, Alexandra Trkola |
| EPI_ISL_2360242 | Medica | Institute of Medical Virology | Verena Kufner, Gabriela Ziltener, Maryam Zaheri, Stefan Schmutz, Annette Audigé, Maria Grünberg, Kevin Steiner, Jon Huder, Cyril Shah, Riccarda Capaul, Guido Bloemberg, Jürg Böni, Michael Huber, Alexandra Trkola |
| EPI_ISL_2360252 | Shahid Ghodossi Laboratory | National Influenza Center | J Yavarian,K Sadeghi, NZ Shafiei Jandaghi, V Salimi, A Nejati, N Ghavvami,F Ajaminejad and T Mokhtari Azad |
| EPI_ISL_2360253 | Maku Molecular Laboratory | National Influenza Center | A Nejati, J Yavarian,K Sadeghi, NZ Shafiei Jandaghi, V Salimi, N Ghavvami,F Ajaminejad and T Mokhtari Azad |
| EPI_ISL_2360254 | Shahid Ghodossi Laboratory | National Influenza Center | K Sadeghi, A Nejati, J Yavarian, NZ Shafiei Jandaghi, V Salimi,F Ajaminejad,N Ghavvami and T Mokhtari Azad |
| EPI_ISL_2360256 | Boshehr University of Medical Sciences | National Influenza Center | NZ Shafiei Jandaghi, V Salimi, A Nejati, K Sadeghi, J Yavarian, Mostafa Salehi-Vaziri, F Ajaminejad,N Ghavvami and T Mokhtari Azad |
| EPI_ISL_2360257 | Boshehr University of Medical Sciences | National Influenza Center | V Salimi,NZ Shafiei Jandaghi, J Yavarian, A Nejati, K Sadeghi,Mostafa Salehi-Vaziri , N Ghavvami,F Ajaminejad and T Mokhtari Azad |
| EPI_ISL_2361439 | IN State Department of Health Laboratory Services | IN State Department of Health Laboratory Services | Cassandra Campion, Jamie Yeadon, Brian Pope, Lixia Liu, Kyle Brownlee, Melissa Hindenlang, Mark Glazier |
| EPI_ISL_2361493, EPI_ISL_2361494 | Laboratorio HUB -Azienda Ospedaliero Universitaria - AOU – Cagliari | Laboratorio SPOKE Biologia Molecolare -Azienda Ospedaliero Universitaria - AOU – Cagliari | Germano Orrù, Sara Fais, Valentina Medda, Alessandra Scano, Miriam Loddo, Riccardo Cappai, Ferdinando Coghe |
| EPI_ISL_2361830, EPI_ISL_2361831, EPI_ISL_2361832, EPI_ISL_2361833, EPI_ISL_2361834, EPI_ISL_2361835, EPI_ISL_2361836, EPI_ISL_2361837, EPI_ISL_2361838, EPI_ISL_2361840, EPI_ISL_2361841, EPI_ISL_2361843, EPI_ISL_2361844, EPI_ISL_2361845, EPI_ISL_2361846, EPI_ISL_2361847, EPI_ISL_2361848, EPI_ISL_2361849, EPI_ISL_2361850, EPI_ISL_2361851, EPI_ISL_2361852, EPI_ISL_2361853, EPI_ISL_2361854, EPI_ISL_2361855, EPI_ISL_2361856, EPI_ISL_2361857, EPI_ISL_2361858, EPI_ISL_2361859, EPI_ISL_2361860, EPI_ISL_2361862, EPI_ISL_2361864, EPI_ISL_2361865, EPI_ISL_2361866, EPI_ISL_2361867, EPI_ISL_2361868, EPI_ISL_2361869, EPI_ISL_2361871, EPI_ISL_2361872, EPI_ISL_2361873, EPI_ISL_2361874, EPI_ISL_2361875, EPI_ISL_2361876, EPI_ISL_2361877, EPI_ISL_2361878 | | | |
| see above | ICMR-National Institute for Research In Tuberculosis | NIV Influenza | Dr Padmapriyadarsini.C, Dr Luke Elizabeth Hanna, Dr S Siva Kumar, Dr Nandhini Palani, Mr Anbazagan Selvaraj |
| EPI_ISL_2361925, EPI_ISL_2361946 | HOSPITAL UNIVERSITARIO SON ESPASES | HOSPITAL UNIVERSITARIO SON ESPASES | Carla López-Causapé, Pablo Fraile-Ribot, Antonio Oliver, SeqCovid |
| EPI_ISL_2362018, EPI_ISL_2362019, EPI_ISL_2362024, EPI_ISL_2362025, EPI_ISL_2362033 | Virginia Division of Consolidated Laboratory Services | Virginia Division of Consolidated Laboratory Services | Virginia DCLS |
| EPI_ISL_2362054, EPI_ISL_2362062 | FUNDACIO ALTHAIA-MANRESA | Banc de Sang i Teixits | Francisco Vidal, Irene Corrales, Nina Borràs, Carlos Hobeich, Natália Comes, Lorena Ramírez, Noemí Gonzalez, Silvia Sauleda, Maria Glòria Soria |
| EPI_ISL_2362074 | CAP TORELLÓ | Banc de Sang i Teixits | Francisco Vidal, Irene Corrales, Nina Borràs, Carlos Hobeich, Natália Comes, Lorena Ramírez, Noemí Gonzalez, Silvia Sauleda, Maria Glòria Soria |
| EPI_ISL_2362089, EPI_ISL_2362090, EPI_ISL_2362091, EPI_ISL_2362092, EPI_ISL_2362093, EPI_ISL_2362094, EPI_ISL_2362095, EPI_ISL_2362096, EPI_ISL_2362097, EPI_ISL_2362098, EPI_ISL_2362099, EPI_ISL_2362100, EPI_ISL_2362101, EPI_ISL_2362102, EPI_ISL_2362103, EPI_ISL_2362104, EPI_ISL_2362106, EPI_ISL_2362107, EPI_ISL_2362108, EPI_ISL_2362109, EPI_ISL_2362112, EPI_ISL_2362113, EPI_ISL_2362115, EPI_ISL_2362116, EPI_ISL_2362117, EPI_ISL_2362118, EPI_ISL_2362119 | | | |
| see above | CHLC | Instituto Nacional de Saude (INSA) | Borges et al |
| EPI_ISL_2362360 | Massachusetts General Hospital | Infectious Disease Program, Broad Institute of Harvard and MIT | Lemieux,J.E., Siddle,K.J., Shaw,B., Adams,G., Pierce,V., Turbett,S., Anahtar,M., Branda,J., Slater,D., Harris,J., Lin,A.E., Gladden-Young,A., Lagerborg,K., Rudy,M., DeRuff,K., Carter,A., Normandin,E., Bauer,M., Reilly,S., Tomkins-Tinch,C., Loreth,C., Chaluvadi,S., Neumann,A., Cusick,C., Chapman,S.B., Gnirke,A., Flowers,K., Cerrato,F., Birren,B.W., Gallagher,G., Smole,S., Park,D.J., MacInnis,B.L., Ryan,E., LaRocque,R., Rosenberg,E. and Sabeti,P.C. |
| EPI_ISL_2362528, EPI_ISL_2362530 | St James's Hospital, Dublin 8, Ireland. | ELDA biotech | Deirdre M. Broderick, Peter R. Flanagan, Brendan Crowley |
| EPI_ISL_2362614, EPI_ISL_2362616, EPI_ISL_2362624, EPI_ISL_2362625 | Platform BIS UZA/UAntwerpen | Labo Klinische Biologie, UZA | Marie Le Mercier, Jasmine Coppens, Basil Britto Xavier, Christine Lammens, Veerle Matheeussen, Herman Goossens |
| EPI_ISL_2362674, EPI_ISL_2362675, EPI_ISL_2362678, EPI_ISL_2362679, EPI_ISL_2362680, EPI_ISL_2362683, EPI_ISL_2362684, EPI_ISL_2362685, EPI_ISL_2362686 | Microbiology and Virology Unit, Florence Careggi University Hospital | Microbiology and Virology Unit, Florence Careggi University Hospital | Vincenzo Di Pilato, Marco Coppi, Fabio Morecchiato, Noemi Aiezza, Ilaria Baccani, Nicla Giovacchini, Alberto Antonelli, Emanuele Gori, Gian Maria Rossolini |
| EPI_ISL_2362860, EPI_ISL_2362935, EPI_ISL_2362951, EPI_ISL_2362955 | Lab voor klinische biologie | Lab voor klinische biologie | Marija Janevska, Hannelore Hamerlinck, Bruno Verhasselt |

| | | | |
|---|---|---|---|
| EPI_ISL_2363042, EPI_ISL_2363072, EPI_ISL_2363077, EPI_ISL_2363178, EPI_ISL_2363194, EPI_ISL_2363196, EPI_ISL_2363199, EPI_ISL_2363211, EPI_ISL_2363283, EPI_ISL_2363288, EPI_ISL_2363291, EPI_ISL_2363297, EPI_ISL_2363307, EPI_ISL_2363322, EPI_ISL_2363324 | | | |
| see above | National Virus Reference Laboratory | National Virus Reference Laboratory | Zoe Yandle, Charlene Bennett, Gabriel Gonzalez, Michael Carr, Jonathan Dean, Cillian F De Gascun |
| EPI_ISL_2363328, EPI_ISL_2363379, EPI_ISL_2363380 | National Virus Reference Laboratory | National Virus Reference Laboratory | Fiona Crispie, Calum Walsh, Matthew McCabe, Zoe Yandle, Charlene Bennet, Gabriel Gonzalez, Michael Carr, Jonathan Dean, Paul Cotter, Cillian F De Gascun |
| EPI_ISL_2363818, EPI_ISL_2363821 | Vita Laboratoriot Oy | Expert Microbiology, National Institute for Health and Welfare | Soile Blomqvist, Jani Halkilahti, Kirsi Liitsola, Haider al-Hello, Päivi Laurila, Erika Lindh, Teemu Smura, Sari Hannula, Pekka Ellonen, Niina Ikonen, Carita Savolainen-Kopra |
| EPI_ISL_2363870, EPI_ISL_2363874 | Fimlab Laboratoriot Oy Tampere | Expert Microbiology, National Institute for Health and Welfare | Soile Blomqvist, Jani Halkilahti, Kirsi Liitsola, Haider al-Hello, Päivi Laurila, Erika Lindh, Teemu Smura, Sari Hannula, Pekka Ellonen, Niina Ikonen, Carita Savolainen-Kopra |
| EPI_ISL_2363996, EPI_ISL_2363998, EPI_ISL_2364003, EPI_ISL_2364004, EPI_ISL_2364006 | TYKS, Kliininen mikrobiologia | Expert Microbiology, National Institute for Health and Welfare | Soile Blomqvist, Jani Halkilahti, Kirsi Liitsola, Haider al-Hello, Päivi Laurila, Erika Lindh, Teemu Smura, Sari Hannula, Pekka Ellonen, Niina Ikonen, Carita Savolainen-Kopra |
| EPI_ISL_2364992 | Greek Genome Center, Biomedical Research Foundation of the Academy of Athens (BRFAA) | Greek Genome Center, Biomedical Research Foundation of the Academy of Athens (BRFAA) | Emmanouil Athanasiadis, Giannis Vatsellas, Theodoros Loupis, Katerina Zoi, Dimitrios Thanos |
| EPI_ISL_2365681 | WSSE w Gdańsku | 1. National Institute of Public Health - National Institute of Hygiene, Warsaw, Poland 2. Biobank Lab, University of Lodz 3. Laboratory of Respiratory Viruses, Teaching and Clinical Center of the Medical University of Lodz | Dominik Strapagiel, Marta Sobalska-Kwapis, Klaudyna Królikowska, Jakub Lach, Marcin Słomka, Tomasz Płoszaj, Magdalena Traczyk-Borszyńska, Izabela Dróżdż, Maciej Borowiec, Katarzyna Zacharczuk, Magdalena Nowakowska, Małgorzata Sadkowska-Todys, Tomasz Wołkowicz |
| EPI_ISL_2365687 | Invicta | 1. National Institute of Public Health - National Institute of Hygiene, Warsaw, Poland 2. Biobank Lab, University of Lodz 3. Laboratory of Respiratory Viruses, Teaching and Clinical Center of the Medical University of Lodz | Dominik Strapagiel, Marta Sobalska-Kwapis, Klaudyna Królikowska, Jakub Lach, Marcin Słomka, Tomasz Płoszaj, Magdalena Traczyk-Borszyńska, Izabela Dróżdż, Maciej Borowiec, Katarzyna Zacharczuk, Magdalena Nowakowska, Małgorzata Sadkowska-Todys, Tomasz Wołkowicz |
| EPI_ISL_2365706 | Cell-T | 1. National Institute of Public Health - National Institute of Hygiene, Warsaw, Poland 2. Biobank Lab, University of Lodz 3. Laboratory of Respiratory Viruses, Teaching and Clinical Center of the Medical University of Lodz | Dominik Strapagiel, Marta Sobalska-Kwapis, Klaudyna Królikowska, Jakub Lach, Marcin Słomka, Tomasz Płoszaj, Magdalena Traczyk-Borszyńska, Izabela Dróżdż, Maciej Borowiec, Katarzyna Zacharczuk, Magdalena Nowakowska, Małgorzata Sadkowska-Todys, Tomasz Wołkowicz |
| EPI_ISL_2365789 | Diagnostyka Sp. z o.o. | 1. National Institute of Public Health - National Institute of Hygiene, Warsaw, Poland 2. Biobank Lab, University of Lodz 3. Laboratory of Respiratory Viruses, Teaching and Clinical Center of the Medical University of Lodz | Dominik Strapagiel, Marta Sobalska-Kwapis, Klaudyna Królikowska, Jakub Lach, Marcin Słomka, Tomasz Płoszaj, Magdalena Traczyk-Borszyńska, Izabela Dróżdż, Maciej Borowiec, Katarzyna Zacharczuk, Magdalena Nowakowska, Małgorzata Sadkowska-Todys, Tomasz Wołkowicz |
| EPI_ISL_2365935, EPI_ISL_2365936, EPI_ISL_2365945 | University of Bari Biomedical Sciences and Human Oncology-Policlinico | University of Bari Biomedical Sciences and Human Oncology | Chironna M., Sallustio A., Loconsole D., Accogli M. |
| EPI_ISL_2365971 | Lighthouse Lab in Milton Keynes | Wellcome Sanger Institute for the COVID-19 Genomics UK (COG-UK) Consortium | The Lighthouse Lab in Milton Keynes and Alex Alderton, Roberto Amato, Jeffrey Barrett, Sonia Goncalves, Ewan Harrison, David K. Jackson, Ian Johnston, Dominic Kwiatkowski, Cordelia Langford, John Sillitoe on behalf of the Wellcome Sanger Institute COVID-19 Surveillance Team |
| EPI_ISL_2365972, EPI_ISL_2365973, EPI_ISL_2365974, EPI_ISL_2365975, EPI_ISL_2365976 | Lighthouse Lab in Alderley Park | Wellcome Sanger Institute for the COVID-19 Genomics UK (COG-UK) Consortium | Jacquelyn Wynn, Mairead Hyland, The Lighthouse Lab in Alderley Park and Alex Alderton, Roberto Amato, Jeffrey Barrett, Sonia Goncalves, Ewan Harrison, David K. Jackson, Ian Johnston, Dominic Kwiatkowski, Cordelia Langford, John Sillitoe on behalf of the Wellcome Sanger Institute COVID-19 Surveillance Team |
| EPI_ISL_2365977 | Lighthouse Lab in Milton Keynes | Wellcome Sanger Institute for the COVID-19 Genomics UK (COG-UK) Consortium | The Lighthouse Lab in Milton Keynes and Alex Alderton, Roberto Amato, Jeffrey Barrett, Sonia Goncalves, Ewan Harrison, David K. Jackson, Ian Johnston, Dominic Kwiatkowski, Cordelia Langford, John Sillitoe on behalf of the Wellcome Sanger Institute COVID-19 Surveillance Team |
| EPI_ISL_2365979, EPI_ISL_2365981, EPI_ISL_2365985, EPI_ISL_2365986 | Lighthouse Lab in Alderley Park | Wellcome Sanger Institute for the COVID-19 Genomics UK (COG-UK) Consortium | Jacquelyn Wynn, Mairead Hyland, The Lighthouse Lab in Alderley Park and Alex Alderton, Roberto Amato, Jeffrey Barrett, Sonia Goncalves, Ewan Harrison, David K. Jackson, Ian Johnston, Dominic Kwiatkowski, Cordelia Langford, John Sillitoe on behalf of the Wellcome Sanger Institute COVID-19 Surveillance Team |
| EPI_ISL_2365987 | Lighthouse Lab in Milton Keynes | Wellcome Sanger Institute for the COVID-19 Genomics UK (COG-UK) Consortium | The Lighthouse Lab in Milton Keynes and Alex Alderton, Roberto Amato, Jeffrey Barrett, Sonia Goncalves, Ewan Harrison, David K. Jackson, Ian Johnston, Dominic Kwiatkowski, Cordelia Langford, John Sillitoe on behalf of the Wellcome Sanger Institute COVID-19 Surveillance Team |
| EPI_ISL_2365989, EPI_ISL_2365990, EPI_ISL_2365991, EPI_ISL_2365992, EPI_ISL_2365993, EPI_ISL_2365994 | Lighthouse Lab in Alderley Park | Wellcome Sanger Institute for the COVID-19 Genomics UK (COG-UK) Consortium | Jacquelyn Wynn, Mairead Hyland, The Lighthouse Lab in Alderley Park and Alex Alderton, Roberto Amato, Jeffrey Barrett, Sonia Goncalves, Ewan Harrison, David K. Jackson, Ian Johnston, Dominic Kwiatkowski, Cordelia Langford, John Sillitoe on behalf of the Wellcome Sanger Institute COVID-19 Surveillance Team |
| EPI_ISL_2365995 | Lighthouse Lab in Milton Keynes | Wellcome Sanger Institute for the COVID-19 Genomics UK (COG-UK) Consortium | The Lighthouse Lab in Milton Keynes and Alex Alderton, Roberto Amato, Jeffrey Barrett, Sonia Goncalves, Ewan Harrison, David K. Jackson, Ian Johnston, Dominic Kwiatkowski, Cordelia Langford, John Sillitoe on behalf of the Wellcome Sanger Institute COVID-19 Surveillance Team |
| EPI_ISL_2365996, EPI_ISL_2365997, EPI_ISL_2365998, EPI_ISL_2366000, EPI_ISL_2366001, EPI_ISL_2366003 | Lighthouse Lab in Alderley Park | Wellcome Sanger Institute for the COVID-19 Genomics UK (COG-UK) Consortium | Jacquelyn Wynn, Mairead Hyland, The Lighthouse Lab in Alderley Park and Alex Alderton, Roberto Amato, Jeffrey Barrett, Sonia Goncalves, Ewan Harrison, David K. Jackson, Ian Johnston, Dominic Kwiatkowski, Cordelia Langford, John Sillitoe on behalf of the Wellcome Sanger Institute COVID-19 Surveillance Team |
| EPI_ISL_2366004 | Lighthouse Lab in Milton Keynes | Wellcome Sanger Institute for the COVID-19 Genomics UK (COG-UK) Consortium | The Lighthouse Lab in Milton Keynes and Alex Alderton, Roberto Amato, Jeffrey Barrett, Sonia Goncalves, Ewan Harrison, David K. Jackson, Ian Johnston, Dominic Kwiatkowski, Cordelia Langford, John Sillitoe on behalf of the Wellcome Sanger Institute COVID-19 Surveillance Team |
| EPI_ISL_2366005 | Lighthouse Lab in Alderley Park | Wellcome Sanger Institute for the COVID-19 Genomics UK (COG-UK) Consortium | Jacquelyn Wynn, Mairead Hyland, The Lighthouse Lab in Alderley Park and Alex Alderton, Roberto Amato, Jeffrey Barrett, Sonia Goncalves, Ewan Harrison, David K. Jackson, Ian Johnston, Dominic Kwiatkowski, Cordelia Langford, John Sillitoe on behalf of the Wellcome Sanger Institute COVID-19 Surveillance Team |
| EPI_ISL_2366006, EPI_ISL_2366007 | Lighthouse Lab in Milton Keynes | Wellcome Sanger Institute for the COVID-19 Genomics UK (COG-UK) Consortium | The Lighthouse Lab in Milton Keynes and Alex Alderton, Roberto Amato, Jeffrey Barrett, Sonia Goncalves, Ewan Harrison, David K. Jackson, Ian Johnston, Dominic Kwiatkowski, Cordelia Langford, John Sillitoe on behalf of the Wellcome Sanger Institute COVID-19 Surveillance Team |
| EPI_ISL_2366009 | Lighthouse Lab in Alderley Park | Wellcome Sanger Institute for the COVID-19 Genomics UK (COG-UK) Consortium | Jacquelyn Wynn, Mairead Hyland, The Lighthouse Lab in Alderley Park and Alex Alderton, Roberto Amato, Jeffrey Barrett, Sonia Goncalves, Ewan Harrison, David K. Jackson, Ian Johnston, Dominic Kwiatkowski, Cordelia Langford, John Sillitoe on behalf of the Wellcome Sanger Institute COVID-19 Surveillance Team |
| EPI_ISL_2366011, EPI_ISL_2366012 | Lighthouse Lab in Milton Keynes | Wellcome Sanger Institute for the COVID-19 Genomics UK (COG-UK) Consortium | The Lighthouse Lab in Milton Keynes and Alex Alderton, Roberto Amato, Jeffrey Barrett, Sonia Goncalves, Ewan Harrison, David K. Jackson, Ian Johnston, Dominic Kwiatkowski, Cordelia Langford, John Sillitoe on behalf of the Wellcome Sanger Institute COVID-19 Surveillance Team |
| EPI_ISL_2366013, EPI_ISL_2366014, EPI_ISL_2366016, EPI_ISL_2366018, EPI_ISL_2366019, EPI_ISL_2366020 | Lighthouse Lab in Alderley Park | Wellcome Sanger Institute for the COVID-19 Genomics UK (COG-UK) Consortium | Jacquelyn Wynn, Mairead Hyland, The Lighthouse Lab in Alderley Park and Alex Alderton, Roberto Amato, Jeffrey Barrett, Sonia Goncalves, Ewan Harrison, David K. Jackson, Ian Johnston, Dominic Kwiatkowski, Cordelia Langford, John Sillitoe on behalf of the Wellcome Sanger Institute COVID-19 Surveillance Team |
| EPI_ISL_2366023 | Lighthouse Lab in Milton Keynes | Wellcome Sanger Institute for the COVID-19 Genomics UK (COG-UK) Consortium | The Lighthouse Lab in Milton Keynes and Alex Alderton, Roberto Amato, Jeffrey Barrett, Sonia Goncalves, Ewan Harrison, David K. Jackson, Ian Johnston, Dominic Kwiatkowski, Cordelia Langford, John Sillitoe on behalf of the Wellcome Sanger Institute COVID-19 Surveillance Team |
| EPI_ISL_2366024, EPI_ISL_2366025, EPI_ISL_2366027, EPI_ISL_2366029, EPI_ISL_2366031, EPI_ISL_2366033, EPI_ISL_2366034, EPI_ISL_2366035, EPI_ISL_2366036 | Lighthouse Lab in Alderley Park | Wellcome Sanger Institute for the COVID-19 Genomics UK (COG-UK) Consortium | Jacquelyn Wynn, Mairead Hyland, The Lighthouse Lab in Alderley Park and Alex Alderton, Roberto Amato, Jeffrey Barrett, Sonia Goncalves, Ewan Harrison, David K. Jackson, Ian Johnston, Dominic Kwiatkowski, Cordelia Langford, John Sillitoe on behalf of the Wellcome Sanger Institute COVID-19 Surveillance Team |
| EPI_ISL_2366038, EPI_ISL_2366040, EPI_ISL_2366041, EPI_ISL_2366042 | Lighthouse Lab in Milton Keynes | Wellcome Sanger Institute for the COVID-19 Genomics UK (COG-UK) Consortium | The Lighthouse Lab in Milton Keynes and Alex Alderton, Roberto Amato, Jeffrey Barrett, Sonia Goncalves, Ewan Harrison, David K. Jackson, Ian Johnston, Dominic Kwiatkowski, Cordelia Langford, John Sillitoe on behalf of the Wellcome Sanger Institute COVID-19 Surveillance Team |
| EPI_ISL_2366043, | Lighthouse Lab in Alderley Park | Wellcome Sanger Institute for the COVID-19 Genomics UK (COG-UK) Consortium | Jacquelyn Wynn, Mairead Hyland, The Lighthouse Lab in Alderley Park and Alex Alderton, Roberto Amato, Jeffrey Barrett, Sonia Goncalves, Ewan Harrison, David K. Jackson, Ian Johnston, Dominic Kwiatkowski, Cordelia Langford, John Sillitoe on |

| | | | |
|---|---|---|---|
| EPI_ISL_2366044, EPI_ISL_2366045, EPI_ISL_2366046 | | Consortium | behalf of the Wellcome Sanger Institute COVID-19 Surveillance Team |
| EPI_ISL_2366047 | Lighthouse Lab in Milton Keynes | Wellcome Sanger Institute for the COVID-19 Genomics UK (COG-UK) Consortium | The Lighthouse Lab in Milton Keynes and Alex Alderton, Roberto Amato, Jeffrey Barrett, Sonia Goncalves, Ewan Harrison, David K. Jackson, Ian Johnston, Dominic Kwiatkowski, Cordelia Langford, John Sillitoe on behalf of the Wellcome Sanger Institute COVID-19 Surveillance Team |
| EPI_ISL_2366048 | Lighthouse Lab in Alderley Park | Wellcome Sanger Institute for the COVID-19 Genomics UK (COG-UK) Consortium | Jacquelyn Wynn, Mairead Hyland, The Lighthouse Lab in Alderley Park and Alex Alderton, Roberto Amato, Jeffrey Barrett, Sonia Goncalves, Ewan Harrison, David K. Jackson, Ian Johnston, Dominic Kwiatkowski, Cordelia Langford, John Sillitoe on behalf of the Wellcome Sanger Institute COVID-19 Surveillance Team |
| EPI_ISL_2366049, EPI_ISL_2366050, EPI_ISL_2366051 | Lighthouse Lab in Milton Keynes | Wellcome Sanger Institute for the COVID-19 Genomics UK (COG-UK) Consortium | The Lighthouse Lab in Milton Keynes and Alex Alderton, Roberto Amato, Jeffrey Barrett, Sonia Goncalves, Ewan Harrison, David K. Jackson, Ian Johnston, Dominic Kwiatkowski, Cordelia Langford, John Sillitoe on behalf of the Wellcome Sanger Institute COVID-19 Surveillance Team |
| EPI_ISL_2366053, EPI_ISL_2366054, EPI_ISL_2366056, EPI_ISL_2366057, EPI_ISL_2366061, EPI_ISL_2366062, EPI_ISL_2366064, EPI_ISL_2366069, EPI_ISL_2366070, EPI_ISL_2366071, EPI_ISL_2366073, EPI_ISL_2366074, EPI_ISL_2366075, EPI_ISL_2366076, EPI_ISL_2366077, EPI_ISL_2366078, EPI_ISL_2366079, EPI_ISL_2366080, EPI_ISL_2366082, EPI_ISL_2366083, EPI_ISL_2366084, EPI_ISL_2366085, EPI_ISL_2366089, EPI_ISL_2366092, EPI_ISL_2366093 | | | |
| see above | Lighthouse Lab in Alderley Park | Wellcome Sanger Institute for the COVID-19 Genomics UK (COG-UK) Consortium | Jacquelyn Wynn, Mairead Hyland, The Lighthouse Lab in Alderley Park and Alex Alderton, Roberto Amato, Jeffrey Barrett, Sonia Goncalves, Ewan Harrison, David K. Jackson, Ian Johnston, Dominic Kwiatkowski, Cordelia Langford, John Sillitoe on behalf of the Wellcome Sanger Institute COVID-19 Surveillance Team |
| EPI_ISL_2366094 | Lighthouse Lab in Milton Keynes | Wellcome Sanger Institute for the COVID-19 Genomics UK (COG-UK) Consortium | The Lighthouse Lab in Milton Keynes and Alex Alderton, Roberto Amato, Jeffrey Barrett, Sonia Goncalves, Ewan Harrison, David K. Jackson, Ian Johnston, Dominic Kwiatkowski, Cordelia Langford, John Sillitoe on behalf of the Wellcome Sanger Institute COVID-19 Surveillance Team |
| EPI_ISL_2366095, EPI_ISL_2366096, EPI_ISL_2366097, EPI_ISL_2366098 | Lighthouse Lab in Alderley Park | Wellcome Sanger Institute for the COVID-19 Genomics UK (COG-UK) Consortium | Jacquelyn Wynn, Mairead Hyland, The Lighthouse Lab in Alderley Park and Alex Alderton, Roberto Amato, Jeffrey Barrett, Sonia Goncalves, Ewan Harrison, David K. Jackson, Ian Johnston, Dominic Kwiatkowski, Cordelia Langford, John Sillitoe on behalf of the Wellcome Sanger Institute COVID-19 Surveillance Team |
| EPI_ISL_2366100 | Lighthouse Lab in Milton Keynes | Wellcome Sanger Institute for the COVID-19 Genomics UK (COG-UK) Consortium | The Lighthouse Lab in Milton Keynes and Alex Alderton, Roberto Amato, Jeffrey Barrett, Sonia Goncalves, Ewan Harrison, David K. Jackson, Ian Johnston, Dominic Kwiatkowski, Cordelia Langford, John Sillitoe on behalf of the Wellcome Sanger Institute COVID-19 Surveillance Team |
| EPI_ISL_2366102, EPI_ISL_2366103 | Lighthouse Lab in Alderley Park | Wellcome Sanger Institute for the COVID-19 Genomics UK (COG-UK) Consortium | Jacquelyn Wynn, Mairead Hyland, The Lighthouse Lab in Alderley Park and Alex Alderton, Roberto Amato, Jeffrey Barrett, Sonia Goncalves, Ewan Harrison, David K. Jackson, Ian Johnston, Dominic Kwiatkowski, Cordelia Langford, John Sillitoe on behalf of the Wellcome Sanger Institute COVID-19 Surveillance Team |
| EPI_ISL_2366105 | Lighthouse Lab in Milton Keynes | Wellcome Sanger Institute for the COVID-19 Genomics UK (COG-UK) Consortium | The Lighthouse Lab in Milton Keynes and Alex Alderton, Roberto Amato, Jeffrey Barrett, Sonia Goncalves, Ewan Harrison, David K. Jackson, Ian Johnston, Dominic Kwiatkowski, Cordelia Langford, John Sillitoe on behalf of the Wellcome Sanger Institute COVID-19 Surveillance Team |
| EPI_ISL_2366107, EPI_ISL_2366108, EPI_ISL_2366109 | Lighthouse Lab in Alderley Park | Wellcome Sanger Institute for the COVID-19 Genomics UK (COG-UK) Consortium | Jacquelyn Wynn, Mairead Hyland, The Lighthouse Lab in Alderley Park and Alex Alderton, Roberto Amato, Jeffrey Barrett, Sonia Goncalves, Ewan Harrison, David K. Jackson, Ian Johnston, Dominic Kwiatkowski, Cordelia Langford, John Sillitoe on behalf of the Wellcome Sanger Institute COVID-19 Surveillance Team |
| EPI_ISL_2366110, EPI_ISL_2366112, EPI_ISL_2366113 | Lighthouse Lab in Milton Keynes | Wellcome Sanger Institute for the COVID-19 Genomics UK (COG-UK) Consortium | The Lighthouse Lab in Milton Keynes and Alex Alderton, Roberto Amato, Jeffrey Barrett, Sonia Goncalves, Ewan Harrison, David K. Jackson, Ian Johnston, Dominic Kwiatkowski, Cordelia Langford, John Sillitoe on behalf of the Wellcome Sanger Institute COVID-19 Surveillance Team |
| EPI_ISL_2366114, EPI_ISL_2366115, EPI_ISL_2366116 | Lighthouse Lab in Alderley Park | Wellcome Sanger Institute for the COVID-19 Genomics UK (COG-UK) Consortium | Jacquelyn Wynn, Mairead Hyland, The Lighthouse Lab in Alderley Park and Alex Alderton, Roberto Amato, Jeffrey Barrett, Sonia Goncalves, Ewan Harrison, David K. Jackson, Ian Johnston, Dominic Kwiatkowski, Cordelia Langford, John Sillitoe on behalf of the Wellcome Sanger Institute COVID-19 Surveillance Team |
| EPI_ISL_2366117 | Lighthouse Lab in Milton Keynes | Wellcome Sanger Institute for the COVID-19 Genomics UK (COG-UK) Consortium | The Lighthouse Lab in Milton Keynes and Alex Alderton, Roberto Amato, Jeffrey Barrett, Sonia Goncalves, Ewan Harrison, David K. Jackson, Ian Johnston, Dominic Kwiatkowski, Cordelia Langford, John Sillitoe on behalf of the Wellcome Sanger Institute COVID-19 Surveillance Team |
| EPI_ISL_2366119 | Lighthouse Lab in Alderley Park | Wellcome Sanger Institute for the COVID-19 Genomics UK (COG-UK) Consortium | Jacquelyn Wynn, Mairead Hyland, The Lighthouse Lab in Alderley Park and Alex Alderton, Roberto Amato, Jeffrey Barrett, Sonia Goncalves, Ewan Harrison, David K. Jackson, Ian Johnston, Dominic Kwiatkowski, Cordelia Langford, John Sillitoe on behalf of the Wellcome Sanger Institute COVID-19 Surveillance Team |
| EPI_ISL_2366120 | Lighthouse Lab in Milton Keynes | Wellcome Sanger Institute for the COVID-19 Genomics UK (COG-UK) Consortium | The Lighthouse Lab in Milton Keynes and Alex Alderton, Roberto Amato, Jeffrey Barrett, Sonia Goncalves, Ewan Harrison, David K. Jackson, Ian Johnston, Dominic Kwiatkowski, Cordelia Langford, John Sillitoe on behalf of the Wellcome Sanger Institute COVID-19 Surveillance Team |
| EPI_ISL_2366121 | Lighthouse Lab in Alderley Park | Wellcome Sanger Institute for the COVID-19 Genomics UK (COG-UK) Consortium | Jacquelyn Wynn, Mairead Hyland, The Lighthouse Lab in Alderley Park and Alex Alderton, Roberto Amato, Jeffrey Barrett, Sonia Goncalves, Ewan Harrison, David K. Jackson, Ian Johnston, Dominic Kwiatkowski, Cordelia Langford, John Sillitoe on behalf of the Wellcome Sanger Institute COVID-19 Surveillance Team |
| EPI_ISL_2366122 | Lighthouse Lab in Milton Keynes | Wellcome Sanger Institute for the COVID-19 Genomics UK (COG-UK) Consortium | The Lighthouse Lab in Milton Keynes and Alex Alderton, Roberto Amato, Jeffrey Barrett, Sonia Goncalves, Ewan Harrison, David K. Jackson, Ian Johnston, Dominic Kwiatkowski, Cordelia Langford, John Sillitoe on behalf of the Wellcome Sanger Institute COVID-19 Surveillance Team |
| EPI_ISL_2366124 | Lighthouse Lab in Alderley Park | Wellcome Sanger Institute for the COVID-19 Genomics UK (COG-UK) Consortium | Jacquelyn Wynn, Mairead Hyland, The Lighthouse Lab in Alderley Park and Alex Alderton, Roberto Amato, Jeffrey Barrett, Sonia Goncalves, Ewan Harrison, David K. Jackson, Ian Johnston, Dominic Kwiatkowski, Cordelia Langford, John Sillitoe on behalf of the Wellcome Sanger Institute COVID-19 Surveillance Team |
| EPI_ISL_2366125 | Lighthouse Lab in Milton Keynes | Wellcome Sanger Institute for the COVID-19 Genomics UK (COG-UK) Consortium | The Lighthouse Lab in Milton Keynes and Alex Alderton, Roberto Amato, Jeffrey Barrett, Sonia Goncalves, Ewan Harrison, David K. Jackson, Ian Johnston, Dominic Kwiatkowski, Cordelia Langford, John Sillitoe on behalf of the Wellcome Sanger Institute COVID-19 Surveillance Team |
| EPI_ISL_2366126, EPI_ISL_2366127, EPI_ISL_2366129, EPI_ISL_2366130, EPI_ISL_2366131, EPI_ISL_2366132 | Lighthouse Lab in Alderley Park | Wellcome Sanger Institute for the COVID-19 Genomics UK (COG-UK) Consortium | Jacquelyn Wynn, Mairead Hyland, The Lighthouse Lab in Alderley Park and Alex Alderton, Roberto Amato, Jeffrey Barrett, Sonia Goncalves, Ewan Harrison, David K. Jackson, Ian Johnston, Dominic Kwiatkowski, Cordelia Langford, John Sillitoe on behalf of the Wellcome Sanger Institute COVID-19 Surveillance Team |
| EPI_ISL_2366133, EPI_ISL_2366134 | Lighthouse Lab in Milton Keynes | Wellcome Sanger Institute for the COVID-19 Genomics UK (COG-UK) Consortium | The Lighthouse Lab in Milton Keynes and Alex Alderton, Roberto Amato, Jeffrey Barrett, Sonia Goncalves, Ewan Harrison, David K. Jackson, Ian Johnston, Dominic Kwiatkowski, Cordelia Langford, John Sillitoe on behalf of the Wellcome Sanger Institute COVID-19 Surveillance Team |
| EPI_ISL_2366135, EPI_ISL_2366136, EPI_ISL_2366137, EPI_ISL_2366138 | Lighthouse Lab in Alderley Park | Wellcome Sanger Institute for the COVID-19 Genomics UK (COG-UK) Consortium | Jacquelyn Wynn, Mairead Hyland, The Lighthouse Lab in Alderley Park and Alex Alderton, Roberto Amato, Jeffrey Barrett, Sonia Goncalves, Ewan Harrison, David K. Jackson, Ian Johnston, Dominic Kwiatkowski, Cordelia Langford, John Sillitoe on behalf of the Wellcome Sanger Institute COVID-19 Surveillance Team |
| EPI_ISL_2366140 | Lighthouse Lab in Milton Keynes | Wellcome Sanger Institute for the COVID-19 Genomics UK (COG-UK) Consortium | The Lighthouse Lab in Milton Keynes and Alex Alderton, Roberto Amato, Jeffrey Barrett, Sonia Goncalves, Ewan Harrison, David K. Jackson, Ian Johnston, Dominic Kwiatkowski, Cordelia Langford, John Sillitoe on behalf of the Wellcome Sanger Institute COVID-19 Surveillance Team |
| EPI_ISL_2366141, EPI_ISL_2366142 | Lighthouse Lab in Alderley Park | Wellcome Sanger Institute for the COVID-19 Genomics UK (COG-UK) Consortium | Jacquelyn Wynn, Mairead Hyland, The Lighthouse Lab in Alderley Park and Alex Alderton, Roberto Amato, Jeffrey Barrett, Sonia Goncalves, Ewan Harrison, David K. Jackson, Ian Johnston, Dominic Kwiatkowski, Cordelia Langford, John Sillitoe on behalf of the Wellcome Sanger Institute COVID-19 Surveillance Team |
| EPI_ISL_2366143 | Lighthouse Lab in Milton Keynes | Wellcome Sanger Institute for the COVID-19 Genomics UK (COG-UK) Consortium | The Lighthouse Lab in Milton Keynes and Alex Alderton, Roberto Amato, Jeffrey Barrett, Sonia Goncalves, Ewan Harrison, David K. Jackson, Ian Johnston, Dominic Kwiatkowski, Cordelia Langford, John Sillitoe on behalf of the Wellcome Sanger Institute COVID-19 Surveillance Team |
| EPI_ISL_2366144, EPI_ISL_2366145, EPI_ISL_2366146 | Lighthouse Lab in Alderley Park | Wellcome Sanger Institute for the COVID-19 Genomics UK (COG-UK) Consortium | Jacquelyn Wynn, Mairead Hyland, The Lighthouse Lab in Alderley Park and Alex Alderton, Roberto Amato, Jeffrey Barrett, Sonia Goncalves, Ewan Harrison, David K. Jackson, Ian Johnston, Dominic Kwiatkowski, Cordelia Langford, John Sillitoe on behalf of the Wellcome Sanger Institute COVID-19 Surveillance Team |
| EPI_ISL_2366148 | Lighthouse Lab in Milton Keynes | Wellcome Sanger Institute for the COVID-19 Genomics UK (COG-UK) Consortium | The Lighthouse Lab in Milton Keynes and Alex Alderton, Roberto Amato, Jeffrey Barrett, Sonia Goncalves, Ewan Harrison, David K. Jackson, Ian Johnston, Dominic Kwiatkowski, Cordelia Langford, John Sillitoe on behalf of the Wellcome Sanger Institute COVID-19 Surveillance Team |
| EPI_ISL_2366149, EPI_ISL_2366150, EPI_ISL_2366151 | Lighthouse Lab in Alderley Park | Wellcome Sanger Institute for the COVID-19 Genomics UK (COG-UK) Consortium | Jacquelyn Wynn, Mairead Hyland, The Lighthouse Lab in Alderley Park and Alex Alderton, Roberto Amato, Jeffrey Barrett, Sonia Goncalves, Ewan Harrison, David K. Jackson, Ian Johnston, Dominic Kwiatkowski, Cordelia Langford, John Sillitoe on behalf of the Wellcome Sanger Institute COVID-19 Surveillance Team |
| EPI_ISL_2366152 | Lighthouse Lab in Milton Keynes | Wellcome Sanger Institute for the COVID-19 Genomics UK (COG-UK) Consortium | The Lighthouse Lab in Milton Keynes and Alex Alderton, Roberto Amato, Jeffrey Barrett, Sonia Goncalves, Ewan Harrison, David K. Jackson, Ian Johnston, Dominic Kwiatkowski, Cordelia Langford, John Sillitoe on behalf of the Wellcome Sanger Institute COVID-19 Surveillance Team |
| EPI_ISL_2366154, EPI_ISL_2366155 | Lighthouse Lab in Alderley Park | Wellcome Sanger Institute for the COVID-19 Genomics UK (COG-UK) Consortium | Jacquelyn Wynn, Mairead Hyland, The Lighthouse Lab in Alderley Park and Alex Alderton, Roberto Amato, Jeffrey Barrett, Sonia Goncalves, Ewan Harrison, David K. Jackson, Ian Johnston, Dominic Kwiatkowski, Cordelia Langford, John Sillitoe on behalf of the Wellcome Sanger Institute COVID-19 Surveillance Team |
| EPI_ISL_2366157 | Lighthouse Lab in Milton Keynes | Wellcome Sanger Institute for the COVID-19 Genomics UK (COG-UK) Consortium | The Lighthouse Lab in Milton Keynes and Alex Alderton, Roberto Amato, Jeffrey Barrett, Sonia Goncalves, Ewan Harrison, David K. Jackson, Ian Johnston, Dominic Kwiatkowski, Cordelia Langford, John Sillitoe on behalf of the Wellcome Sanger Institute COVID-19 Surveillance Team |
| EPI_ISL_2366159, EPI_ISL_2366160, EPI_ISL_2366161 | Lighthouse Lab in Alderley Park | Wellcome Sanger Institute for the COVID-19 Genomics UK (COG-UK) Consortium | Jacquelyn Wynn, Mairead Hyland, The Lighthouse Lab in Alderley Park and Alex Alderton, Roberto Amato, Jeffrey Barrett, Sonia Goncalves, Ewan Harrison, David K. Jackson, Ian Johnston, Dominic Kwiatkowski, Cordelia Langford, John Sillitoe on behalf of the Wellcome Sanger Institute COVID-19 Surveillance Team |
| EPI_ISL_2366162, EPI_ISL_2366163, EPI_ISL_2366166 | Lighthouse Lab in Milton Keynes | Wellcome Sanger Institute for the COVID-19 Genomics UK (COG-UK) Consortium | The Lighthouse Lab in Milton Keynes and Alex Alderton, Roberto Amato, Jeffrey Barrett, Sonia Goncalves, Ewan Harrison, David K. Jackson, Ian Johnston, Dominic Kwiatkowski, Cordelia Langford, John Sillitoe on behalf of the Wellcome Sanger Institute COVID-19 Surveillance Team |
| EPI_ISL_2366170, EPI_ISL_2366171, EPI_ISL_2366172, EPI_ISL_2366173 | Lighthouse Lab in Alderley Park | Wellcome Sanger Institute for the COVID-19 Genomics UK (COG-UK) Consortium | Jacquelyn Wynn, Mairead Hyland, The Lighthouse Lab in Alderley Park and Alex Alderton, Roberto Amato, Jeffrey Barrett, Sonia Goncalves, Ewan Harrison, David K. Jackson, Ian Johnston, Dominic Kwiatkowski, Cordelia Langford, John Sillitoe on behalf of the Wellcome Sanger Institute COVID-19 Surveillance Team |
| EPI_ISL_2366176 | Lighthouse Lab in Milton Keynes | Wellcome Sanger Institute for the COVID-19 Genomics UK (COG-UK) Consortium | The Lighthouse Lab in Milton Keynes and Alex Alderton, Roberto Amato, Jeffrey Barrett, Sonia Goncalves, Ewan Harrison, David K. Jackson, Ian Johnston, Dominic Kwiatkowski, Cordelia Langford, John Sillitoe on behalf of the Wellcome Sanger Institute COVID-19 Surveillance Team |
| EPI_ISL_2366177, | Lighthouse Lab in Alderley Park | Wellcome Sanger Institute for the COVID-19 Genomics UK (COG-UK) | Jacquelyn Wynn, Mairead Hyland, The Lighthouse Lab in Alderley Park and Alex Alderton, Roberto Amato, Jeffrey Barrett, Sonia Goncalves, Ewan Harrison, David K. Jackson, Ian Johnston, Dominic Kwiatkowski, Cordelia Langford, John Sillitoe on |

| | | | |
|---|---|---|---|
| EPI_ISL_2366178, EPI_ISL_2366181, EPI_ISL_2366184, EPI_ISL_2366185, EPI_ISL_2366186, EPI_ISL_2366187, EPI_ISL_2366188, EPI_ISL_2366191, EPI_ISL_2366192 | | Consortium | behalf of the Wellcome Sanger Institute COVID-19 Surveillance Team |
| EPI_ISL_2366193, EPI_ISL_2366194, EPI_ISL_2366196 | Lighthouse Lab in Milton Keynes | Wellcome Sanger Institute for the COVID-19 Genomics UK (COG-UK) Consortium | The Lighthouse Lab in Milton Keynes and Alex Alderton, Roberto Amato, Jeffrey Barrett, Sonia Goncalves, Ewan Harrison, David K. Jackson, Ian Johnston, Dominic Kwiatkowski, Cordelia Langford, John Sillitoe on behalf of the Wellcome Sanger Institute COVID-19 Surveillance Team |
| EPI_ISL_2366197, EPI_ISL_2366198, EPI_ISL_2366199, EPI_ISL_2366200, EPI_ISL_2366202 | Lighthouse Lab in Alderley Park | Wellcome Sanger Institute for the COVID-19 Genomics UK (COG-UK) Consortium | Jacquelyn Wynn, Mairead Hyland, The Lighthouse Lab in Alderley Park and Alex Alderton, Roberto Amato, Jeffrey Barrett, Sonia Goncalves, Ewan Harrison, David K. Jackson, Ian Johnston, Dominic Kwiatkowski, Cordelia Langford, John Sillitoe on behalf of the Wellcome Sanger Institute COVID-19 Surveillance Team |
| EPI_ISL_2366203 | Lighthouse Lab in Milton Keynes | Wellcome Sanger Institute for the COVID-19 Genomics UK (COG-UK) Consortium | The Lighthouse Lab in Milton Keynes and Alex Alderton, Roberto Amato, Jeffrey Barrett, Sonia Goncalves, Ewan Harrison, David K. Jackson, Ian Johnston, Dominic Kwiatkowski, Cordelia Langford, John Sillitoe on behalf of the Wellcome Sanger Institute COVID-19 Surveillance Team |
| EPI_ISL_2366206, EPI_ISL_2366207, EPI_ISL_2366208, EPI_ISL_2366209, EPI_ISL_2366210, EPI_ISL_2366211, EPI_ISL_2366213, EPI_ISL_2366214 | Lighthouse Lab in Alderley Park | Wellcome Sanger Institute for the COVID-19 Genomics UK (COG-UK) Consortium | Jacquelyn Wynn, Mairead Hyland, The Lighthouse Lab in Alderley Park and Alex Alderton, Roberto Amato, Jeffrey Barrett, Sonia Goncalves, Ewan Harrison, David K. Jackson, Ian Johnston, Dominic Kwiatkowski, Cordelia Langford, John Sillitoe on behalf of the Wellcome Sanger Institute COVID-19 Surveillance Team |
| EPI_ISL_2366215, EPI_ISL_2366216 | Lighthouse Lab in Milton Keynes | Wellcome Sanger Institute for the COVID-19 Genomics UK (COG-UK) Consortium | The Lighthouse Lab in Milton Keynes and Alex Alderton, Roberto Amato, Jeffrey Barrett, Sonia Goncalves, Ewan Harrison, David K. Jackson, Ian Johnston, Dominic Kwiatkowski, Cordelia Langford, John Sillitoe on behalf of the Wellcome Sanger Institute COVID-19 Surveillance Team |
| EPI_ISL_2366219, EPI_ISL_2366220 | Lighthouse Lab in Alderley Park | Wellcome Sanger Institute for the COVID-19 Genomics UK (COG-UK) Consortium | Jacquelyn Wynn, Mairead Hyland, The Lighthouse Lab in Alderley Park and Alex Alderton, Roberto Amato, Jeffrey Barrett, Sonia Goncalves, Ewan Harrison, David K. Jackson, Ian Johnston, Dominic Kwiatkowski, Cordelia Langford, John Sillitoe on behalf of the Wellcome Sanger Institute COVID-19 Surveillance Team |
| EPI_ISL_2366221 | Lighthouse Lab in Milton Keynes | Wellcome Sanger Institute for the COVID-19 Genomics UK (COG-UK) Consortium | The Lighthouse Lab in Milton Keynes and Alex Alderton, Roberto Amato, Jeffrey Barrett, Sonia Goncalves, Ewan Harrison, David K. Jackson, Ian Johnston, Dominic Kwiatkowski, Cordelia Langford, John Sillitoe on behalf of the Wellcome Sanger Institute COVID-19 Surveillance Team |
| EPI_ISL_2366222, EPI_ISL_2366224, EPI_ISL_2366225 | Lighthouse Lab in Alderley Park | Wellcome Sanger Institute for the COVID-19 Genomics UK (COG-UK) Consortium | Jacquelyn Wynn, Mairead Hyland, The Lighthouse Lab in Alderley Park and Alex Alderton, Roberto Amato, Jeffrey Barrett, Sonia Goncalves, Ewan Harrison, David K. Jackson, Ian Johnston, Dominic Kwiatkowski, Cordelia Langford, John Sillitoe on behalf of the Wellcome Sanger Institute COVID-19 Surveillance Team |
| EPI_ISL_2366226, EPI_ISL_2366228, EPI_ISL_2366229 | Lighthouse Lab in Milton Keynes | Wellcome Sanger Institute for the COVID-19 Genomics UK (COG-UK) Consortium | The Lighthouse Lab in Milton Keynes and Alex Alderton, Roberto Amato, Jeffrey Barrett, Sonia Goncalves, Ewan Harrison, David K. Jackson, Ian Johnston, Dominic Kwiatkowski, Cordelia Langford, John Sillitoe on behalf of the Wellcome Sanger Institute COVID-19 Surveillance Team |
| EPI_ISL_2366231, EPI_ISL_2366232 | Lighthouse Lab in Alderley Park | Wellcome Sanger Institute for the COVID-19 Genomics UK (COG-UK) Consortium | Jacquelyn Wynn, Mairead Hyland, The Lighthouse Lab in Alderley Park and Alex Alderton, Roberto Amato, Jeffrey Barrett, Sonia Goncalves, Ewan Harrison, David K. Jackson, Ian Johnston, Dominic Kwiatkowski, Cordelia Langford, John Sillitoe on behalf of the Wellcome Sanger Institute COVID-19 Surveillance Team |
| EPI_ISL_2366233, EPI_ISL_2366234, EPI_ISL_2366235 | Lighthouse Lab in Milton Keynes | Wellcome Sanger Institute for the COVID-19 Genomics UK (COG-UK) Consortium | The Lighthouse Lab in Milton Keynes and Alex Alderton, Roberto Amato, Jeffrey Barrett, Sonia Goncalves, Ewan Harrison, David K. Jackson, Ian Johnston, Dominic Kwiatkowski, Cordelia Langford, John Sillitoe on behalf of the Wellcome Sanger Institute COVID-19 Surveillance Team |
| EPI_ISL_2366236, EPI_ISL_2366237 | Lighthouse Lab in Alderley Park | Wellcome Sanger Institute for the COVID-19 Genomics UK (COG-UK) Consortium | Jacquelyn Wynn, Mairead Hyland, The Lighthouse Lab in Alderley Park and Alex Alderton, Roberto Amato, Jeffrey Barrett, Sonia Goncalves, Ewan Harrison, David K. Jackson, Ian Johnston, Dominic Kwiatkowski, Cordelia Langford, John Sillitoe on behalf of the Wellcome Sanger Institute COVID-19 Surveillance Team |
| EPI_ISL_2366239 | Lighthouse Lab in Milton Keynes | Wellcome Sanger Institute for the COVID-19 Genomics UK (COG-UK) Consortium | The Lighthouse Lab in Milton Keynes and Alex Alderton, Roberto Amato, Jeffrey Barrett, Sonia Goncalves, Ewan Harrison, David K. Jackson, Ian Johnston, Dominic Kwiatkowski, Cordelia Langford, John Sillitoe on behalf of the Wellcome Sanger Institute COVID-19 Surveillance Team |
| EPI_ISL_2366241 | Lighthouse Lab in Alderley Park | Wellcome Sanger Institute for the COVID-19 Genomics UK (COG-UK) Consortium | Jacquelyn Wynn, Mairead Hyland, The Lighthouse Lab in Alderley Park and Alex Alderton, Roberto Amato, Jeffrey Barrett, Sonia Goncalves, Ewan Harrison, David K. Jackson, Ian Johnston, Dominic Kwiatkowski, Cordelia Langford, John Sillitoe on behalf of the Wellcome Sanger Institute COVID-19 Surveillance Team |
| EPI_ISL_2366243 | Lighthouse Lab in Milton Keynes | Wellcome Sanger Institute for the COVID-19 Genomics UK (COG-UK) Consortium | The Lighthouse Lab in Milton Keynes and Alex Alderton, Roberto Amato, Jeffrey Barrett, Sonia Goncalves, Ewan Harrison, David K. Jackson, Ian Johnston, Dominic Kwiatkowski, Cordelia Langford, John Sillitoe on behalf of the Wellcome Sanger Institute COVID-19 Surveillance Team |
| EPI_ISL_2366245, EPI_ISL_2366246, EPI_ISL_2366247 | Lighthouse Lab in Alderley Park | Wellcome Sanger Institute for the COVID-19 Genomics UK (COG-UK) Consortium | Jacquelyn Wynn, Mairead Hyland, The Lighthouse Lab in Alderley Park and Alex Alderton, Roberto Amato, Jeffrey Barrett, Sonia Goncalves, Ewan Harrison, David K. Jackson, Ian Johnston, Dominic Kwiatkowski, Cordelia Langford, John Sillitoe on behalf of the Wellcome Sanger Institute COVID-19 Surveillance Team |
| EPI_ISL_2366249 | Lighthouse Lab in Milton Keynes | Wellcome Sanger Institute for the COVID-19 Genomics UK (COG-UK) Consortium | The Lighthouse Lab in Milton Keynes and Alex Alderton, Roberto Amato, Jeffrey Barrett, Sonia Goncalves, Ewan Harrison, David K. Jackson, Ian Johnston, Dominic Kwiatkowski, Cordelia Langford, John Sillitoe on behalf of the Wellcome Sanger Institute COVID-19 Surveillance Team |
| EPI_ISL_2366250, EPI_ISL_2366252, EPI_ISL_2366253 | Lighthouse Lab in Alderley Park | Wellcome Sanger Institute for the COVID-19 Genomics UK (COG-UK) Consortium | Jacquelyn Wynn, Mairead Hyland, The Lighthouse Lab in Alderley Park and Alex Alderton, Roberto Amato, Jeffrey Barrett, Sonia Goncalves, Ewan Harrison, David K. Jackson, Ian Johnston, Dominic Kwiatkowski, Cordelia Langford, John Sillitoe on behalf of the Wellcome Sanger Institute COVID-19 Surveillance Team |
| EPI_ISL_2366254 | Lighthouse Lab in Milton Keynes | Wellcome Sanger Institute for the COVID-19 Genomics UK (COG-UK) Consortium | The Lighthouse Lab in Milton Keynes and Alex Alderton, Roberto Amato, Jeffrey Barrett, Sonia Goncalves, Ewan Harrison, David K. Jackson, Ian Johnston, Dominic Kwiatkowski, Cordelia Langford, John Sillitoe on behalf of the Wellcome Sanger Institute COVID-19 Surveillance Team |
| EPI_ISL_2366255, EPI_ISL_2366256, EPI_ISL_2366257, EPI_ISL_2366258 | Lighthouse Lab in Alderley Park | Wellcome Sanger Institute for the COVID-19 Genomics UK (COG-UK) Consortium | Jacquelyn Wynn, Mairead Hyland, The Lighthouse Lab in Alderley Park and Alex Alderton, Roberto Amato, Jeffrey Barrett, Sonia Goncalves, Ewan Harrison, David K. Jackson, Ian Johnston, Dominic Kwiatkowski, Cordelia Langford, John Sillitoe on behalf of the Wellcome Sanger Institute COVID-19 Surveillance Team |
| EPI_ISL_2366261 | Lighthouse Lab in Milton Keynes | Wellcome Sanger Institute for the COVID-19 Genomics UK (COG-UK) Consortium | The Lighthouse Lab in Milton Keynes and Alex Alderton, Roberto Amato, Jeffrey Barrett, Sonia Goncalves, Ewan Harrison, David K. Jackson, Ian Johnston, Dominic Kwiatkowski, Cordelia Langford, John Sillitoe on behalf of the Wellcome Sanger Institute COVID-19 Surveillance Team |
| EPI_ISL_2366262 | Lighthouse Lab in Alderley Park | Wellcome Sanger Institute for the COVID-19 Genomics UK (COG-UK) Consortium | Jacquelyn Wynn, Mairead Hyland, The Lighthouse Lab in Alderley Park and Alex Alderton, Roberto Amato, Jeffrey Barrett, Sonia Goncalves, Ewan Harrison, David K. Jackson, Ian Johnston, Dominic Kwiatkowski, Cordelia Langford, John Sillitoe on behalf of the Wellcome Sanger Institute COVID-19 Surveillance Team |
| EPI_ISL_2366264 | Lighthouse Lab in Milton Keynes | Wellcome Sanger Institute for the COVID-19 Genomics UK (COG-UK) Consortium | The Lighthouse Lab in Milton Keynes and Alex Alderton, Roberto Amato, Jeffrey Barrett, Sonia Goncalves, Ewan Harrison, David K. Jackson, Ian Johnston, Dominic Kwiatkowski, Cordelia Langford, John Sillitoe on behalf of the Wellcome Sanger Institute COVID-19 Surveillance Team |
| EPI_ISL_2366265, EPI_ISL_2366266 | Lighthouse Lab in Alderley Park | Wellcome Sanger Institute for the COVID-19 Genomics UK (COG-UK) Consortium | Jacquelyn Wynn, Mairead Hyland, The Lighthouse Lab in Alderley Park and Alex Alderton, Roberto Amato, Jeffrey Barrett, Sonia Goncalves, Ewan Harrison, David K. Jackson, Ian Johnston, Dominic Kwiatkowski, Cordelia Langford, John Sillitoe on behalf of the Wellcome Sanger Institute COVID-19 Surveillance Team |
| EPI_ISL_2366269 | Lighthouse Lab in Milton Keynes | Wellcome Sanger Institute for the COVID-19 Genomics UK (COG-UK) Consortium | The Lighthouse Lab in Milton Keynes and Alex Alderton, Roberto Amato, Jeffrey Barrett, Sonia Goncalves, Ewan Harrison, David K. Jackson, Ian Johnston, Dominic Kwiatkowski, Cordelia Langford, John Sillitoe on behalf of the Wellcome Sanger Institute COVID-19 Surveillance Team |
| EPI_ISL_2366270 | Lighthouse Lab in Glasgow | Wellcome Sanger Institute for the COVID-19 Genomics UK (COG-UK) Consortium | Harper VanSteenhouse, Yumi Kasai, David Gray, Carol Clugston, Anna Dominiczak and Alex Alderton, Roberto Amato, Jeffrey Barrett, Sonia Goncalves, Ewan Harrison, David K. Jackson, Ian Johnston, Dominic Kwiatkowski, Cordelia Langford, John Sillitoe on behalf of the Wellcome Sanger Institute COVID-19 Surveillance Team |
| EPI_ISL_2366275, EPI_ISL_2366276, EPI_ISL_2366278, EPI_ISL_2366279, EPI_ISL_2366280, EPI_ISL_2366281, EPI_ISL_2366282, EPI_ISL_2366283, EPI_ISL_2366284, EPI_ISL_2366285, EPI_ISL_2366286, EPI_ISL_2366288, EPI_ISL_2366289, EPI_ISL_2366290, EPI_ISL_2366292, EPI_ISL_2366294, EPI_ISL_2366295 | | | |
| see above | Lighthouse Lab in Milton Keynes | Wellcome Sanger Institute for the COVID-19 Genomics UK (COG-UK) Consortium | The Lighthouse Lab in Milton Keynes and Alex Alderton, Roberto Amato, Jeffrey Barrett, Sonia Goncalves, Ewan Harrison, David K. Jackson, Ian Johnston, Dominic Kwiatkowski, Cordelia Langford, John Sillitoe on behalf of the Wellcome Sanger Institute COVID-19 Surveillance Team |
| EPI_ISL_2366296, EPI_ISL_2366298, EPI_ISL_2366299 | Lighthouse Lab in Glasgow | Wellcome Sanger Institute for the COVID-19 Genomics UK (COG-UK) Consortium | Harper VanSteenhouse, Yumi Kasai, David Gray, Carol Clugston, Anna Dominiczak and Alex Alderton, Roberto Amato, Jeffrey Barrett, Sonia Goncalves, Ewan Harrison, David K. Jackson, Ian Johnston, Dominic Kwiatkowski, Cordelia Langford, John Sillitoe on behalf of the Wellcome Sanger Institute COVID-19 Surveillance Team |
| EPI_ISL_2366301 | Lighthouse Lab in Milton Keynes | Wellcome Sanger Institute for the COVID-19 Genomics UK (COG-UK) Consortium | The Lighthouse Lab in Milton Keynes and Alex Alderton, Roberto Amato, Jeffrey Barrett, Sonia Goncalves, Ewan Harrison, David K. Jackson, Ian Johnston, Dominic Kwiatkowski, Cordelia Langford, John Sillitoe on behalf of the Wellcome Sanger Institute COVID-19 Surveillance Team |
| EPI_ISL_2366302 | Lighthouse Lab in Alderley Park | Wellcome Sanger Institute for the COVID-19 Genomics UK (COG-UK) Consortium | Jacquelyn Wynn, Mairead Hyland, The Lighthouse Lab in Alderley Park and Alex Alderton, Roberto Amato, Jeffrey Barrett, Sonia Goncalves, Ewan Harrison, David K. Jackson, Ian Johnston, Dominic Kwiatkowski, Cordelia Langford, John Sillitoe on behalf of the Wellcome Sanger Institute COVID-19 Surveillance Team |
| EPI_ISL_2366303 | Lighthouse Lab in Glasgow | Wellcome Sanger Institute for the COVID-19 Genomics UK (COG-UK) Consortium | Harper VanSteenhouse, Yumi Kasai, David Gray, Carol Clugston, Anna Dominiczak and Alex Alderton, Roberto Amato, Jeffrey Barrett, Sonia Goncalves, Ewan Harrison, David K. Jackson, Ian Johnston, Dominic Kwiatkowski, Cordelia Langford, John Sillitoe on behalf of the Wellcome Sanger Institute COVID-19 Surveillance Team |
| EPI_ISL_2366305 | Lighthouse Lab in Alderley Park | Wellcome Sanger Institute for the COVID-19 Genomics UK (COG-UK) Consortium | Jacquelyn Wynn, Mairead Hyland, The Lighthouse Lab in Alderley Park and Alex Alderton, Roberto Amato, Jeffrey Barrett, Sonia Goncalves, Ewan Harrison, David K. Jackson, Ian Johnston, Dominic Kwiatkowski, Cordelia Langford, John Sillitoe on behalf of the Wellcome Sanger Institute COVID-19 Surveillance Team |
| EPI_ISL_2366306, EPI_ISL_2366307 | Lighthouse Lab in Milton Keynes | Wellcome Sanger Institute for the COVID-19 Genomics UK (COG-UK) Consortium | The Lighthouse Lab in Milton Keynes and Alex Alderton, Roberto Amato, Jeffrey Barrett, Sonia Goncalves, Ewan Harrison, David K. Jackson, Ian Johnston, Dominic Kwiatkowski, Cordelia Langford, John Sillitoe on behalf of the Wellcome Sanger Institute COVID-19 Surveillance Team |
| EPI_ISL_2366308 | Lighthouse Lab in Alderley Park | Wellcome Sanger Institute for the COVID-19 Genomics UK (COG-UK) | Jacquelyn Wynn, Mairead Hyland, The Lighthouse Lab in Alderley Park and Alex Alderton, Roberto Amato, Jeffrey Barrett, Sonia Goncalves, Ewan Harrison, David K. Jackson, Ian Johnston, Dominic Kwiatkowski, Cordelia Langford, John Sillitoe on |

| | | | |
|---|---|---|---|
| | | Consortium | behalf of the Wellcome Sanger Institute COVID-19 Surveillance Team |
| EPI_ISL_2366310 | Lighthouse Lab in Glasgow | Wellcome Sanger Institute for the COVID-19 Genomics UK (COG-UK) Consortium | Harper VanSteenhouse, Yumi Kasai, David Gray, Carol Clugston, Anna Dominiczak and Alex Alderton, Roberto Amato, Jeffrey Barrett, Sonia Goncalves, Ewan Harrison, David K. Jackson, Ian Johnston, Dominic Kwiatkowski, Cordelia Langford, John Sillitoe on behalf of the Wellcome Sanger Institute COVID-19 Surveillance Team |
| EPI_ISL_2366313 | Lighthouse Lab in Alderley Park | Wellcome Sanger Institute for the COVID-19 Genomics UK (COG-UK) Consortium | Jacquelyn Wynn, Mairead Hyland, The Lighthouse Lab in Alderley Park and Alex Alderton, Roberto Amato, Jeffrey Barrett, Sonia Goncalves, Ewan Harrison, David K. Jackson, Ian Johnston, Dominic Kwiatkowski, Cordelia Langford, John Sillitoe on behalf of the Wellcome Sanger Institute COVID-19 Surveillance Team |
| EPI_ISL_2366314 | Lighthouse Lab in Glasgow | Wellcome Sanger Institute for the COVID-19 Genomics UK (COG-UK) Consortium | Harper VanSteenhouse, Yumi Kasai, David Gray, Carol Clugston, Anna Dominiczak and Alex Alderton, Roberto Amato, Jeffrey Barrett, Sonia Goncalves, Ewan Harrison, David K. Jackson, Ian Johnston, Dominic Kwiatkowski, Cordelia Langford, John Sillitoe on behalf of the Wellcome Sanger Institute COVID-19 Surveillance Team |
| EPI_ISL_2366315, EPI_ISL_2366317 | Lighthouse Lab in Milton Keynes | Wellcome Sanger Institute for the COVID-19 Genomics UK (COG-UK) Consortium | The Lighthouse Lab in Milton Keynes and Alex Alderton, Roberto Amato, Jeffrey Barrett, Sonia Goncalves, Ewan Harrison, David K. Jackson, Ian Johnston, Dominic Kwiatkowski, Cordelia Langford, John Sillitoe on behalf of the Wellcome Sanger Institute COVID-19 Surveillance Team |
| EPI_ISL_2366318 | Lighthouse Lab in Alderley Park | Wellcome Sanger Institute for the COVID-19 Genomics UK (COG-UK) Consortium | Jacquelyn Wynn, Mairead Hyland, The Lighthouse Lab in Alderley Park and Alex Alderton, Roberto Amato, Jeffrey Barrett, Sonia Goncalves, Ewan Harrison, David K. Jackson, Ian Johnston, Dominic Kwiatkowski, Cordelia Langford, John Sillitoe on behalf of the Wellcome Sanger Institute COVID-19 Surveillance Team |
| EPI_ISL_2366319 | Lighthouse Lab in Milton Keynes | Wellcome Sanger Institute for the COVID-19 Genomics UK (COG-UK) Consortium | The Lighthouse Lab in Milton Keynes and Alex Alderton, Roberto Amato, Jeffrey Barrett, Sonia Goncalves, Ewan Harrison, David K. Jackson, Ian Johnston, Dominic Kwiatkowski, Cordelia Langford, John Sillitoe on behalf of the Wellcome Sanger Institute COVID-19 Surveillance Team |
| EPI_ISL_2366321 | Lighthouse Lab in Glasgow | Wellcome Sanger Institute for the COVID-19 Genomics UK (COG-UK) Consortium | Harper VanSteenhouse, Yumi Kasai, David Gray, Carol Clugston, Anna Dominiczak and Alex Alderton, Roberto Amato, Jeffrey Barrett, Sonia Goncalves, Ewan Harrison, David K. Jackson, Ian Johnston, Dominic Kwiatkowski, Cordelia Langford, John Sillitoe on behalf of the Wellcome Sanger Institute COVID-19 Surveillance Team |
| EPI_ISL_2366323, EPI_ISL_2366324 | Lighthouse Lab in Milton Keynes | Wellcome Sanger Institute for the COVID-19 Genomics UK (COG-UK) Consortium | The Lighthouse Lab in Milton Keynes and Alex Alderton, Roberto Amato, Jeffrey Barrett, Sonia Goncalves, Ewan Harrison, David K. Jackson, Ian Johnston, Dominic Kwiatkowski, Cordelia Langford, John Sillitoe on behalf of the Wellcome Sanger Institute COVID-19 Surveillance Team |
| EPI_ISL_2366326 | Lighthouse Lab in Alderley Park | Wellcome Sanger Institute for the COVID-19 Genomics UK (COG-UK) Consortium | Jacquelyn Wynn, Mairead Hyland, The Lighthouse Lab in Alderley Park and Alex Alderton, Roberto Amato, Jeffrey Barrett, Sonia Goncalves, Ewan Harrison, David K. Jackson, Ian Johnston, Dominic Kwiatkowski, Cordelia Langford, John Sillitoe on behalf of the Wellcome Sanger Institute COVID-19 Surveillance Team |
| EPI_ISL_2366327 | Lighthouse Lab in Milton Keynes | Wellcome Sanger Institute for the COVID-19 Genomics UK (COG-UK) Consortium | The Lighthouse Lab in Milton Keynes and Alex Alderton, Roberto Amato, Jeffrey Barrett, Sonia Goncalves, Ewan Harrison, David K. Jackson, Ian Johnston, Dominic Kwiatkowski, Cordelia Langford, John Sillitoe on behalf of the Wellcome Sanger Institute COVID-19 Surveillance Team |
| EPI_ISL_2366330, EPI_ISL_2366331 | Lighthouse Lab in Alderley Park | Wellcome Sanger Institute for the COVID-19 Genomics UK (COG-UK) Consortium | Jacquelyn Wynn, Mairead Hyland, The Lighthouse Lab in Alderley Park and Alex Alderton, Roberto Amato, Jeffrey Barrett, Sonia Goncalves, Ewan Harrison, David K. Jackson, Ian Johnston, Dominic Kwiatkowski, Cordelia Langford, John Sillitoe on behalf of the Wellcome Sanger Institute COVID-19 Surveillance Team |
| EPI_ISL_2366332, EPI_ISL_2366333, EPI_ISL_2366334, EPI_ISL_2366336, EPI_ISL_2366337, EPI_ISL_2366338 | Lighthouse Lab in Milton Keynes | Wellcome Sanger Institute for the COVID-19 Genomics UK (COG-UK) Consortium | The Lighthouse Lab in Milton Keynes and Alex Alderton, Roberto Amato, Jeffrey Barrett, Sonia Goncalves, Ewan Harrison, David K. Jackson, Ian Johnston, Dominic Kwiatkowski, Cordelia Langford, John Sillitoe on behalf of the Wellcome Sanger Institute COVID-19 Surveillance Team |
| EPI_ISL_2366339 | Lighthouse Lab in Alderley Park | Wellcome Sanger Institute for the COVID-19 Genomics UK (COG-UK) Consortium | Jacquelyn Wynn, Mairead Hyland, The Lighthouse Lab in Alderley Park and Alex Alderton, Roberto Amato, Jeffrey Barrett, Sonia Goncalves, Ewan Harrison, David K. Jackson, Ian Johnston, Dominic Kwiatkowski, Cordelia Langford, John Sillitoe on behalf of the Wellcome Sanger Institute COVID-19 Surveillance Team |
| EPI_ISL_2366340 | Lighthouse Lab in Glasgow | Wellcome Sanger Institute for the COVID-19 Genomics UK (COG-UK) Consortium | Harper VanSteenhouse, Yumi Kasai, David Gray, Carol Clugston, Anna Dominiczak and Alex Alderton, Roberto Amato, Jeffrey Barrett, Sonia Goncalves, Ewan Harrison, David K. Jackson, Ian Johnston, Dominic Kwiatkowski, Cordelia Langford, John Sillitoe on behalf of the Wellcome Sanger Institute COVID-19 Surveillance Team |
| EPI_ISL_2366341, EPI_ISL_2366342 | Lighthouse Lab in Milton Keynes | Wellcome Sanger Institute for the COVID-19 Genomics UK (COG-UK) Consortium | The Lighthouse Lab in Milton Keynes and Alex Alderton, Roberto Amato, Jeffrey Barrett, Sonia Goncalves, Ewan Harrison, David K. Jackson, Ian Johnston, Dominic Kwiatkowski, Cordelia Langford, John Sillitoe on behalf of the Wellcome Sanger Institute COVID-19 Surveillance Team |
| EPI_ISL_2366343 | Lighthouse Lab in Alderley Park | Wellcome Sanger Institute for the COVID-19 Genomics UK (COG-UK) Consortium | Jacquelyn Wynn, Mairead Hyland, The Lighthouse Lab in Alderley Park and Alex Alderton, Roberto Amato, Jeffrey Barrett, Sonia Goncalves, Ewan Harrison, David K. Jackson, Ian Johnston, Dominic Kwiatkowski, Cordelia Langford, John Sillitoe on behalf of the Wellcome Sanger Institute COVID-19 Surveillance Team |
| EPI_ISL_2366344 | Lighthouse Lab in Glasgow | Wellcome Sanger Institute for the COVID-19 Genomics UK (COG-UK) Consortium | Harper VanSteenhouse, Yumi Kasai, David Gray, Carol Clugston, Anna Dominiczak and Alex Alderton, Roberto Amato, Jeffrey Barrett, Sonia Goncalves, Ewan Harrison, David K. Jackson, Ian Johnston, Dominic Kwiatkowski, Cordelia Langford, John Sillitoe on behalf of the Wellcome Sanger Institute COVID-19 Surveillance Team |
| EPI_ISL_2366345, EPI_ISL_2366346, EPI_ISL_2366347, EPI_ISL_2366349, EPI_ISL_2366350 | Lighthouse Lab in Milton Keynes | Wellcome Sanger Institute for the COVID-19 Genomics UK (COG-UK) Consortium | The Lighthouse Lab in Milton Keynes and Alex Alderton, Roberto Amato, Jeffrey Barrett, Sonia Goncalves, Ewan Harrison, David K. Jackson, Ian Johnston, Dominic Kwiatkowski, Cordelia Langford, John Sillitoe on behalf of the Wellcome Sanger Institute COVID-19 Surveillance Team |
| EPI_ISL_2366352 | Lighthouse Lab in Alderley Park | Wellcome Sanger Institute for the COVID-19 Genomics UK (COG-UK) Consortium | Jacquelyn Wynn, Mairead Hyland, The Lighthouse Lab in Alderley Park and Alex Alderton, Roberto Amato, Jeffrey Barrett, Sonia Goncalves, Ewan Harrison, David K. Jackson, Ian Johnston, Dominic Kwiatkowski, Cordelia Langford, John Sillitoe on behalf of the Wellcome Sanger Institute COVID-19 Surveillance Team |
| EPI_ISL_2366353 | Lighthouse Lab in Glasgow | Wellcome Sanger Institute for the COVID-19 Genomics UK (COG-UK) Consortium | Harper VanSteenhouse, Yumi Kasai, David Gray, Carol Clugston, Anna Dominiczak and Alex Alderton, Roberto Amato, Jeffrey Barrett, Sonia Goncalves, Ewan Harrison, David K. Jackson, Ian Johnston, Dominic Kwiatkowski, Cordelia Langford, John Sillitoe on behalf of the Wellcome Sanger Institute COVID-19 Surveillance Team |
| EPI_ISL_2366354, EPI_ISL_2366355 | Lighthouse Lab in Milton Keynes | Wellcome Sanger Institute for the COVID-19 Genomics UK (COG-UK) Consortium | The Lighthouse Lab in Milton Keynes and Alex Alderton, Roberto Amato, Jeffrey Barrett, Sonia Goncalves, Ewan Harrison, David K. Jackson, Ian Johnston, Dominic Kwiatkowski, Cordelia Langford, John Sillitoe on behalf of the Wellcome Sanger Institute COVID-19 Surveillance Team |
| EPI_ISL_2366356, EPI_ISL_2366357 | Lighthouse Lab in Alderley Park | Wellcome Sanger Institute for the COVID-19 Genomics UK (COG-UK) Consortium | Jacquelyn Wynn, Mairead Hyland, The Lighthouse Lab in Alderley Park and Alex Alderton, Roberto Amato, Jeffrey Barrett, Sonia Goncalves, Ewan Harrison, David K. Jackson, Ian Johnston, Dominic Kwiatkowski, Cordelia Langford, John Sillitoe on behalf of the Wellcome Sanger Institute COVID-19 Surveillance Team |
| EPI_ISL_2366358 | Lighthouse Lab in Milton Keynes | Wellcome Sanger Institute for the COVID-19 Genomics UK (COG-UK) Consortium | The Lighthouse Lab in Milton Keynes and Alex Alderton, Roberto Amato, Jeffrey Barrett, Sonia Goncalves, Ewan Harrison, David K. Jackson, Ian Johnston, Dominic Kwiatkowski, Cordelia Langford, John Sillitoe on behalf of the Wellcome Sanger Institute COVID-19 Surveillance Team |
| EPI_ISL_2366359 | Lighthouse Lab in Alderley Park | Wellcome Sanger Institute for the COVID-19 Genomics UK (COG-UK) Consortium | Jacquelyn Wynn, Mairead Hyland, The Lighthouse Lab in Alderley Park and Alex Alderton, Roberto Amato, Jeffrey Barrett, Sonia Goncalves, Ewan Harrison, David K. Jackson, Ian Johnston, Dominic Kwiatkowski, Cordelia Langford, John Sillitoe on behalf of the Wellcome Sanger Institute COVID-19 Surveillance Team |
| EPI_ISL_2366360 | Lighthouse Lab in Milton Keynes | Wellcome Sanger Institute for the COVID-19 Genomics UK (COG-UK) Consortium | The Lighthouse Lab in Milton Keynes and Alex Alderton, Roberto Amato, Jeffrey Barrett, Sonia Goncalves, Ewan Harrison, David K. Jackson, Ian Johnston, Dominic Kwiatkowski, Cordelia Langford, John Sillitoe on behalf of the Wellcome Sanger Institute COVID-19 Surveillance Team |
| EPI_ISL_2366361, EPI_ISL_2366362 | Lighthouse Lab in Alderley Park | Wellcome Sanger Institute for the COVID-19 Genomics UK (COG-UK) Consortium | Jacquelyn Wynn, Mairead Hyland, The Lighthouse Lab in Alderley Park and Alex Alderton, Roberto Amato, Jeffrey Barrett, Sonia Goncalves, Ewan Harrison, David K. Jackson, Ian Johnston, Dominic Kwiatkowski, Cordelia Langford, John Sillitoe on behalf of the Wellcome Sanger Institute COVID-19 Surveillance Team |
| EPI_ISL_2366363, EPI_ISL_2366365 | Lighthouse Lab in Milton Keynes | Wellcome Sanger Institute for the COVID-19 Genomics UK (COG-UK) Consortium | The Lighthouse Lab in Milton Keynes and Alex Alderton, Roberto Amato, Jeffrey Barrett, Sonia Goncalves, Ewan Harrison, David K. Jackson, Ian Johnston, Dominic Kwiatkowski, Cordelia Langford, John Sillitoe on behalf of the Wellcome Sanger Institute COVID-19 Surveillance Team |
| EPI_ISL_2366366 | Lighthouse Lab in Alderley Park | Wellcome Sanger Institute for the COVID-19 Genomics UK (COG-UK) Consortium | Jacquelyn Wynn, Mairead Hyland, The Lighthouse Lab in Alderley Park and Alex Alderton, Roberto Amato, Jeffrey Barrett, Sonia Goncalves, Ewan Harrison, David K. Jackson, Ian Johnston, Dominic Kwiatkowski, Cordelia Langford, John Sillitoe on behalf of the Wellcome Sanger Institute COVID-19 Surveillance Team |
| EPI_ISL_2366368, EPI_ISL_2366369, EPI_ISL_2366370, EPI_ISL_2366371, EPI_ISL_2366372, EPI_ISL_2366373, EPI_ISL_2366374 | Lighthouse Lab in Milton Keynes | Wellcome Sanger Institute for the COVID-19 Genomics UK (COG-UK) Consortium | The Lighthouse Lab in Milton Keynes and Alex Alderton, Roberto Amato, Jeffrey Barrett, Sonia Goncalves, Ewan Harrison, David K. Jackson, Ian Johnston, Dominic Kwiatkowski, Cordelia Langford, John Sillitoe on behalf of the Wellcome Sanger Institute COVID-19 Surveillance Team |
| EPI_ISL_2366375, EPI_ISL_2366377 | Lighthouse Lab in Alderley Park | Wellcome Sanger Institute for the COVID-19 Genomics UK (COG-UK) Consortium | Jacquelyn Wynn, Mairead Hyland, The Lighthouse Lab in Alderley Park and Alex Alderton, Roberto Amato, Jeffrey Barrett, Sonia Goncalves, Ewan Harrison, David K. Jackson, Ian Johnston, Dominic Kwiatkowski, Cordelia Langford, John Sillitoe on behalf of the Wellcome Sanger Institute COVID-19 Surveillance Team |
| EPI_ISL_2366378, EPI_ISL_2366379, EPI_ISL_2366380 | Lighthouse Lab in Milton Keynes | Wellcome Sanger Institute for the COVID-19 Genomics UK (COG-UK) Consortium | The Lighthouse Lab in Milton Keynes and Alex Alderton, Roberto Amato, Jeffrey Barrett, Sonia Goncalves, Ewan Harrison, David K. Jackson, Ian Johnston, Dominic Kwiatkowski, Cordelia Langford, John Sillitoe on behalf of the Wellcome Sanger Institute COVID-19 Surveillance Team |
| EPI_ISL_2366381 | Lighthouse Lab in Glasgow | Wellcome Sanger Institute for the COVID-19 Genomics UK (COG-UK) Consortium | Harper VanSteenhouse, Yumi Kasai, David Gray, Carol Clugston, Anna Dominiczak and Alex Alderton, Roberto Amato, Jeffrey Barrett, Sonia Goncalves, Ewan Harrison, David K. Jackson, Ian Johnston, Dominic Kwiatkowski, Cordelia Langford, John Sillitoe on behalf of the Wellcome Sanger Institute COVID-19 Surveillance Team |
| EPI_ISL_2366383 | Lighthouse Lab in Milton Keynes | Wellcome Sanger Institute for the COVID-19 Genomics UK (COG-UK) Consortium | The Lighthouse Lab in Milton Keynes and Alex Alderton, Roberto Amato, Jeffrey Barrett, Sonia Goncalves, Ewan Harrison, David K. Jackson, Ian Johnston, Dominic Kwiatkowski, Cordelia Langford, John Sillitoe on behalf of the Wellcome Sanger Institute COVID-19 Surveillance Team |
| EPI_ISL_2366384 | Lighthouse Lab in Alderley Park | Wellcome Sanger Institute for the COVID-19 Genomics UK (COG-UK) Consortium | Jacquelyn Wynn, Mairead Hyland, The Lighthouse Lab in Alderley Park and Alex Alderton, Roberto Amato, Jeffrey Barrett, Sonia Goncalves, Ewan Harrison, David K. Jackson, Ian Johnston, Dominic Kwiatkowski, Cordelia Langford, John Sillitoe on behalf of the Wellcome Sanger Institute COVID-19 Surveillance Team |
| EPI_ISL_2366386, EPI_ISL_2366387 | Lighthouse Lab in Milton Keynes | Wellcome Sanger Institute for the COVID-19 Genomics UK (COG-UK) Consortium | The Lighthouse Lab in Milton Keynes and Alex Alderton, Roberto Amato, Jeffrey Barrett, Sonia Goncalves, Ewan Harrison, David K. Jackson, Ian Johnston, Dominic Kwiatkowski, Cordelia Langford, John Sillitoe on behalf of the Wellcome Sanger Institute COVID-19 Surveillance Team |
| EPI_ISL_2366389, EPI_ISL_2366392 | Lighthouse Lab in Alderley Park | Wellcome Sanger Institute for the COVID-19 Genomics UK (COG-UK) Consortium | Jacquelyn Wynn, Mairead Hyland, The Lighthouse Lab in Alderley Park and Alex Alderton, Roberto Amato, Jeffrey Barrett, Sonia Goncalves, Ewan Harrison, David K. Jackson, Ian Johnston, Dominic Kwiatkowski, Cordelia Langford, John Sillitoe on behalf of the Wellcome Sanger Institute COVID-19 Surveillance Team |
| EPI_ISL_2366393, EPI_ISL_2366394 | Lighthouse Lab in Milton Keynes | Wellcome Sanger Institute for the COVID-19 Genomics UK (COG-UK) Consortium | The Lighthouse Lab in Milton Keynes and Alex Alderton, Roberto Amato, Jeffrey Barrett, Sonia Goncalves, Ewan Harrison, David K. Jackson, Ian Johnston, Dominic Kwiatkowski, Cordelia Langford, John Sillitoe on behalf of the Wellcome Sanger Institute COVID-19 Surveillance Team |
| EPI_ISL_2366395 | Lighthouse Lab in Alderley Park | Wellcome Sanger Institute for the COVID-19 Genomics UK (COG-UK) Consortium | Jacquelyn Wynn, Mairead Hyland, The Lighthouse Lab in Alderley Park and Alex Alderton, Roberto Amato, Jeffrey Barrett, Sonia Goncalves, Ewan Harrison, David K. Jackson, Ian Johnston, Dominic Kwiatkowski, Cordelia Langford, John Sillitoe on behalf of the Wellcome Sanger Institute COVID-19 Surveillance Team |
| EPI_ISL_2366396 | Lighthouse Lab in Milton Keynes | Wellcome Sanger Institute for the COVID-19 Genomics UK (COG-UK) Consortium | The Lighthouse Lab in Milton Keynes and Alex Alderton, Roberto Amato, Jeffrey Barrett, Sonia Goncalves, Ewan Harrison, David K. Jackson, Ian Johnston, Dominic Kwiatkowski, Cordelia Langford, John Sillitoe on behalf of the Wellcome Sanger Institute COVID-19 Surveillance Team |
| EPI_ISL_2366397, | Lighthouse Lab in Alderley Park | Wellcome Sanger Institute for the COVID-19 Genomics UK (COG-UK) | Jacquelyn Wynn, Mairead Hyland, The Lighthouse Lab in Alderley Park and Alex Alderton, Roberto Amato, Jeffrey Barrett, Sonia Goncalves, Ewan Harrison, David K. Jackson, Ian Johnston, Dominic Kwiatkowski, Cordelia Langford, John Sillitoe on |

| | | | |
|---|---|---|---|
| EPI_ISL_2366398 | | Consortium | behalf of the Wellcome Sanger Institute COVID-19 Surveillance Team |
| EPI_ISL_2366399, EPI_ISL_2366400, EPI_ISL_2366402 | Lighthouse Lab in Milton Keynes | Wellcome Sanger Institute for the COVID-19 Genomics UK (COG-UK) Consortium | The Lighthouse Lab in Milton Keynes and Alex Alderton, Roberto Amato, Jeffrey Barrett, Sonia Goncalves, Ewan Harrison, David K. Jackson, Ian Johnston, Dominic Kwiatkowski, Cordelia Langford, John Sillitoe on behalf of the Wellcome Sanger Institute COVID-19 Surveillance Team |
| EPI_ISL_2366404 | Lighthouse Lab in Alderley Park | Wellcome Sanger Institute for the COVID-19 Genomics UK (COG-UK) Consortium | Jacquelyn Wynn, Mairead Hyland, The Lighthouse Lab in Alderley Park and Alex Alderton, Roberto Amato, Jeffrey Barrett, Sonia Goncalves, Ewan Harrison, David K. Jackson, Ian Johnston, Dominic Kwiatkowski, Cordelia Langford, John Sillitoe on behalf of the Wellcome Sanger Institute COVID-19 Surveillance Team |
| EPI_ISL_2366405 | Lighthouse Lab in Glasgow | Wellcome Sanger Institute for the COVID-19 Genomics UK (COG-UK) Consortium | Harper VanSteenhouse, Yumi Kasai, David Gray, Carol Clugston, Anna Dominiczak and Alex Alderton, Roberto Amato, Jeffrey Barrett, Sonia Goncalves, Ewan Harrison, David K. Jackson, Ian Johnston, Dominic Kwiatkowski, Cordelia Langford, John Sillitoe on behalf of the Wellcome Sanger Institute COVID-19 Surveillance Team |
| EPI_ISL_2366408, EPI_ISL_2366409 | Lighthouse Lab in Milton Keynes | Wellcome Sanger Institute for the COVID-19 Genomics UK (COG-UK) Consortium | The Lighthouse Lab in Milton Keynes and Alex Alderton, Roberto Amato, Jeffrey Barrett, Sonia Goncalves, Ewan Harrison, David K. Jackson, Ian Johnston, Dominic Kwiatkowski, Cordelia Langford, John Sillitoe on behalf of the Wellcome Sanger Institute COVID-19 Surveillance Team |
| EPI_ISL_2366411 | Lighthouse Lab in Alderley Park | Wellcome Sanger Institute for the COVID-19 Genomics UK (COG-UK) Consortium | Jacquelyn Wynn, Mairead Hyland, The Lighthouse Lab in Alderley Park and Alex Alderton, Roberto Amato, Jeffrey Barrett, Sonia Goncalves, Ewan Harrison, David K. Jackson, Ian Johnston, Dominic Kwiatkowski, Cordelia Langford, John Sillitoe on behalf of the Wellcome Sanger Institute COVID-19 Surveillance Team |
| EPI_ISL_2366415 | Lighthouse Lab in Milton Keynes | Wellcome Sanger Institute for the COVID-19 Genomics UK (COG-UK) Consortium | The Lighthouse Lab in Milton Keynes and Alex Alderton, Roberto Amato, Jeffrey Barrett, Sonia Goncalves, Ewan Harrison, David K. Jackson, Ian Johnston, Dominic Kwiatkowski, Cordelia Langford, John Sillitoe on behalf of the Wellcome Sanger Institute COVID-19 Surveillance Team |
| EPI_ISL_2366416, EPI_ISL_2366417 | Lighthouse Lab in Alderley Park | Wellcome Sanger Institute for the COVID-19 Genomics UK (COG-UK) Consortium | Jacquelyn Wynn, Mairead Hyland, The Lighthouse Lab in Alderley Park and Alex Alderton, Roberto Amato, Jeffrey Barrett, Sonia Goncalves, Ewan Harrison, David K. Jackson, Ian Johnston, Dominic Kwiatkowski, Cordelia Langford, John Sillitoe on behalf of the Wellcome Sanger Institute COVID-19 Surveillance Team |
| EPI_ISL_2366421, EPI_ISL_2366422 | Lighthouse Lab in Milton Keynes | Wellcome Sanger Institute for the COVID-19 Genomics UK (COG-UK) Consortium | The Lighthouse Lab in Milton Keynes and Alex Alderton, Roberto Amato, Jeffrey Barrett, Sonia Goncalves, Ewan Harrison, David K. Jackson, Ian Johnston, Dominic Kwiatkowski, Cordelia Langford, John Sillitoe on behalf of the Wellcome Sanger Institute COVID-19 Surveillance Team |
| EPI_ISL_2366426 | Lighthouse Lab in Alderley Park | Wellcome Sanger Institute for the COVID-19 Genomics UK (COG-UK) Consortium | Jacquelyn Wynn, Mairead Hyland, The Lighthouse Lab in Alderley Park and Alex Alderton, Roberto Amato, Jeffrey Barrett, Sonia Goncalves, Ewan Harrison, David K. Jackson, Ian Johnston, Dominic Kwiatkowski, Cordelia Langford, John Sillitoe on behalf of the Wellcome Sanger Institute COVID-19 Surveillance Team |
| EPI_ISL_2366429, EPI_ISL_2366431, EPI_ISL_2366432 | Lighthouse Lab in Milton Keynes | Wellcome Sanger Institute for the COVID-19 Genomics UK (COG-UK) Consortium | The Lighthouse Lab in Milton Keynes and Alex Alderton, Roberto Amato, Jeffrey Barrett, Sonia Goncalves, Ewan Harrison, David K. Jackson, Ian Johnston, Dominic Kwiatkowski, Cordelia Langford, John Sillitoe on behalf of the Wellcome Sanger Institute COVID-19 Surveillance Team |
| EPI_ISL_2366433 | Lighthouse Lab in Alderley Park | Wellcome Sanger Institute for the COVID-19 Genomics UK (COG-UK) Consortium | Jacquelyn Wynn, Mairead Hyland, The Lighthouse Lab in Alderley Park and Alex Alderton, Roberto Amato, Jeffrey Barrett, Sonia Goncalves, Ewan Harrison, David K. Jackson, Ian Johnston, Dominic Kwiatkowski, Cordelia Langford, John Sillitoe on behalf of the Wellcome Sanger Institute COVID-19 Surveillance Team |
| EPI_ISL_2366435, EPI_ISL_2366436, EPI_ISL_2366437 | Lighthouse Lab in Milton Keynes | Wellcome Sanger Institute for the COVID-19 Genomics UK (COG-UK) Consortium | The Lighthouse Lab in Milton Keynes and Alex Alderton, Roberto Amato, Jeffrey Barrett, Sonia Goncalves, Ewan Harrison, David K. Jackson, Ian Johnston, Dominic Kwiatkowski, Cordelia Langford, John Sillitoe on behalf of the Wellcome Sanger Institute COVID-19 Surveillance Team |
| EPI_ISL_2366439 | Lighthouse Lab in Alderley Park | Wellcome Sanger Institute for the COVID-19 Genomics UK (COG-UK) Consortium | Jacquelyn Wynn, Mairead Hyland, The Lighthouse Lab in Alderley Park and Alex Alderton, Roberto Amato, Jeffrey Barrett, Sonia Goncalves, Ewan Harrison, David K. Jackson, Ian Johnston, Dominic Kwiatkowski, Cordelia Langford, John Sillitoe on behalf of the Wellcome Sanger Institute COVID-19 Surveillance Team |
| EPI_ISL_2366440 | Lighthouse Lab in Milton Keynes | Wellcome Sanger Institute for the COVID-19 Genomics UK (COG-UK) Consortium | The Lighthouse Lab in Milton Keynes and Alex Alderton, Roberto Amato, Jeffrey Barrett, Sonia Goncalves, Ewan Harrison, David K. Jackson, Ian Johnston, Dominic Kwiatkowski, Cordelia Langford, John Sillitoe on behalf of the Wellcome Sanger Institute COVID-19 Surveillance Team |
| EPI_ISL_2366441 | Lighthouse Lab in Alderley Park | Wellcome Sanger Institute for the COVID-19 Genomics UK (COG-UK) Consortium | Jacquelyn Wynn, Mairead Hyland, The Lighthouse Lab in Alderley Park and Alex Alderton, Roberto Amato, Jeffrey Barrett, Sonia Goncalves, Ewan Harrison, David K. Jackson, Ian Johnston, Dominic Kwiatkowski, Cordelia Langford, John Sillitoe on behalf of the Wellcome Sanger Institute COVID-19 Surveillance Team |
| EPI_ISL_2366442 | Lighthouse Lab in Milton Keynes | Wellcome Sanger Institute for the COVID-19 Genomics UK (COG-UK) Consortium | The Lighthouse Lab in Milton Keynes and Alex Alderton, Roberto Amato, Jeffrey Barrett, Sonia Goncalves, Ewan Harrison, David K. Jackson, Ian Johnston, Dominic Kwiatkowski, Cordelia Langford, John Sillitoe on behalf of the Wellcome Sanger Institute COVID-19 Surveillance Team |
| EPI_ISL_2366443 | Lighthouse Lab in Alderley Park | Wellcome Sanger Institute for the COVID-19 Genomics UK (COG-UK) Consortium | Jacquelyn Wynn, Mairead Hyland, The Lighthouse Lab in Alderley Park and Alex Alderton, Roberto Amato, Jeffrey Barrett, Sonia Goncalves, Ewan Harrison, David K. Jackson, Ian Johnston, Dominic Kwiatkowski, Cordelia Langford, John Sillitoe on behalf of the Wellcome Sanger Institute COVID-19 Surveillance Team |
| EPI_ISL_2366444, EPI_ISL_2366445 | Lighthouse Lab in Milton Keynes | Wellcome Sanger Institute for the COVID-19 Genomics UK (COG-UK) Consortium | The Lighthouse Lab in Milton Keynes and Alex Alderton, Roberto Amato, Jeffrey Barrett, Sonia Goncalves, Ewan Harrison, David K. Jackson, Ian Johnston, Dominic Kwiatkowski, Cordelia Langford, John Sillitoe on behalf of the Wellcome Sanger Institute COVID-19 Surveillance Team |
| EPI_ISL_2366446, EPI_ISL_2366447 | Lighthouse Lab in Alderley Park | Wellcome Sanger Institute for the COVID-19 Genomics UK (COG-UK) Consortium | Jacquelyn Wynn, Mairead Hyland, The Lighthouse Lab in Alderley Park and Alex Alderton, Roberto Amato, Jeffrey Barrett, Sonia Goncalves, Ewan Harrison, David K. Jackson, Ian Johnston, Dominic Kwiatkowski, Cordelia Langford, John Sillitoe on behalf of the Wellcome Sanger Institute COVID-19 Surveillance Team |
| EPI_ISL_2366450 | Lighthouse Lab in Milton Keynes | Wellcome Sanger Institute for the COVID-19 Genomics UK (COG-UK) Consortium | The Lighthouse Lab in Milton Keynes and Alex Alderton, Roberto Amato, Jeffrey Barrett, Sonia Goncalves, Ewan Harrison, David K. Jackson, Ian Johnston, Dominic Kwiatkowski, Cordelia Langford, John Sillitoe on behalf of the Wellcome Sanger Institute COVID-19 Surveillance Team |
| EPI_ISL_2366451 | Lighthouse Lab in Glasgow | Wellcome Sanger Institute for the COVID-19 Genomics UK (COG-UK) Consortium | Harper VanSteenhouse, Yumi Kasai, David Gray, Carol Clugston, Anna Dominiczak and Alex Alderton, Roberto Amato, Jeffrey Barrett, Sonia Goncalves, Ewan Harrison, David K. Jackson, Ian Johnston, Dominic Kwiatkowski, Cordelia Langford, John Sillitoe on behalf of the Wellcome Sanger Institute COVID-19 Surveillance Team |
| EPI_ISL_2366454, EPI_ISL_2366455, EPI_ISL_2366456 | Lighthouse Lab in Milton Keynes | Wellcome Sanger Institute for the COVID-19 Genomics UK (COG-UK) Consortium | The Lighthouse Lab in Milton Keynes and Alex Alderton, Roberto Amato, Jeffrey Barrett, Sonia Goncalves, Ewan Harrison, David K. Jackson, Ian Johnston, Dominic Kwiatkowski, Cordelia Langford, John Sillitoe on behalf of the Wellcome Sanger Institute COVID-19 Surveillance Team |
| EPI_ISL_2366457 | Lighthouse Lab in Alderley Park | Wellcome Sanger Institute for the COVID-19 Genomics UK (COG-UK) Consortium | Jacquelyn Wynn, Mairead Hyland, The Lighthouse Lab in Alderley Park and Alex Alderton, Roberto Amato, Jeffrey Barrett, Sonia Goncalves, Ewan Harrison, David K. Jackson, Ian Johnston, Dominic Kwiatkowski, Cordelia Langford, John Sillitoe on behalf of the Wellcome Sanger Institute COVID-19 Surveillance Team |
| EPI_ISL_2366458, EPI_ISL_2366462, EPI_ISL_2366463 | Lighthouse Lab in Milton Keynes | Wellcome Sanger Institute for the COVID-19 Genomics UK (COG-UK) Consortium | The Lighthouse Lab in Milton Keynes and Alex Alderton, Roberto Amato, Jeffrey Barrett, Sonia Goncalves, Ewan Harrison, David K. Jackson, Ian Johnston, Dominic Kwiatkowski, Cordelia Langford, John Sillitoe on behalf of the Wellcome Sanger Institute COVID-19 Surveillance Team |
| EPI_ISL_2366465 | Lighthouse Lab in Alderley Park | Wellcome Sanger Institute for the COVID-19 Genomics UK (COG-UK) Consortium | Jacquelyn Wynn, Mairead Hyland, The Lighthouse Lab in Alderley Park and Alex Alderton, Roberto Amato, Jeffrey Barrett, Sonia Goncalves, Ewan Harrison, David K. Jackson, Ian Johnston, Dominic Kwiatkowski, Cordelia Langford, John Sillitoe on behalf of the Wellcome Sanger Institute COVID-19 Surveillance Team |
| EPI_ISL_2366466 | Lighthouse Lab in Milton Keynes | Wellcome Sanger Institute for the COVID-19 Genomics UK (COG-UK) Consortium | The Lighthouse Lab in Milton Keynes and Alex Alderton, Roberto Amato, Jeffrey Barrett, Sonia Goncalves, Ewan Harrison, David K. Jackson, Ian Johnston, Dominic Kwiatkowski, Cordelia Langford, John Sillitoe on behalf of the Wellcome Sanger Institute COVID-19 Surveillance Team |
| EPI_ISL_2366468, EPI_ISL_2366469 | Lighthouse Lab in Alderley Park | Wellcome Sanger Institute for the COVID-19 Genomics UK (COG-UK) Consortium | Jacquelyn Wynn, Mairead Hyland, The Lighthouse Lab in Alderley Park and Alex Alderton, Roberto Amato, Jeffrey Barrett, Sonia Goncalves, Ewan Harrison, David K. Jackson, Ian Johnston, Dominic Kwiatkowski, Cordelia Langford, John Sillitoe on behalf of the Wellcome Sanger Institute COVID-19 Surveillance Team |
| EPI_ISL_2366471 | Lighthouse Lab in Milton Keynes | Wellcome Sanger Institute for the COVID-19 Genomics UK (COG-UK) Consortium | The Lighthouse Lab in Milton Keynes and Alex Alderton, Roberto Amato, Jeffrey Barrett, Sonia Goncalves, Ewan Harrison, David K. Jackson, Ian Johnston, Dominic Kwiatkowski, Cordelia Langford, John Sillitoe on behalf of the Wellcome Sanger Institute COVID-19 Surveillance Team |
| EPI_ISL_2366472 | Lighthouse Lab in Alderley Park | Wellcome Sanger Institute for the COVID-19 Genomics UK (COG-UK) Consortium | Jacquelyn Wynn, Mairead Hyland, The Lighthouse Lab in Alderley Park and Alex Alderton, Roberto Amato, Jeffrey Barrett, Sonia Goncalves, Ewan Harrison, David K. Jackson, Ian Johnston, Dominic Kwiatkowski, Cordelia Langford, John Sillitoe on behalf of the Wellcome Sanger Institute COVID-19 Surveillance Team |
| EPI_ISL_2366474, EPI_ISL_2366475 | Lighthouse Lab in Milton Keynes | Wellcome Sanger Institute for the COVID-19 Genomics UK (COG-UK) Consortium | The Lighthouse Lab in Milton Keynes and Alex Alderton, Roberto Amato, Jeffrey Barrett, Sonia Goncalves, Ewan Harrison, David K. Jackson, Ian Johnston, Dominic Kwiatkowski, Cordelia Langford, John Sillitoe on behalf of the Wellcome Sanger Institute COVID-19 Surveillance Team |
| EPI_ISL_2366477 | Lighthouse Lab in Alderley Park | Wellcome Sanger Institute for the COVID-19 Genomics UK (COG-UK) Consortium | Jacquelyn Wynn, Mairead Hyland, The Lighthouse Lab in Alderley Park and Alex Alderton, Roberto Amato, Jeffrey Barrett, Sonia Goncalves, Ewan Harrison, David K. Jackson, Ian Johnston, Dominic Kwiatkowski, Cordelia Langford, John Sillitoe on behalf of the Wellcome Sanger Institute COVID-19 Surveillance Team |
| EPI_ISL_2366479 | Lighthouse Lab in Glasgow | Wellcome Sanger Institute for the COVID-19 Genomics UK (COG-UK) Consortium | Harper VanSteenhouse, Yumi Kasai, David Gray, Carol Clugston, Anna Dominiczak and Alex Alderton, Roberto Amato, Jeffrey Barrett, Sonia Goncalves, Ewan Harrison, David K. Jackson, Ian Johnston, Dominic Kwiatkowski, Cordelia Langford, John Sillitoe on behalf of the Wellcome Sanger Institute COVID-19 Surveillance Team |
| EPI_ISL_2366482, EPI_ISL_2366483 | Lighthouse Lab in Milton Keynes | Wellcome Sanger Institute for the COVID-19 Genomics UK (COG-UK) Consortium | The Lighthouse Lab in Milton Keynes and Alex Alderton, Roberto Amato, Jeffrey Barrett, Sonia Goncalves, Ewan Harrison, David K. Jackson, Ian Johnston, Dominic Kwiatkowski, Cordelia Langford, John Sillitoe on behalf of the Wellcome Sanger Institute COVID-19 Surveillance Team |
| EPI_ISL_2366484, EPI_ISL_2366485 | Lighthouse Lab in Alderley Park | Wellcome Sanger Institute for the COVID-19 Genomics UK (COG-UK) Consortium | Jacquelyn Wynn, Mairead Hyland, The Lighthouse Lab in Alderley Park and Alex Alderton, Roberto Amato, Jeffrey Barrett, Sonia Goncalves, Ewan Harrison, David K. Jackson, Ian Johnston, Dominic Kwiatkowski, Cordelia Langford, John Sillitoe on behalf of the Wellcome Sanger Institute COVID-19 Surveillance Team |
| EPI_ISL_2366487, EPI_ISL_2366488, EPI_ISL_2366489, EPI_ISL_2366490 | Lighthouse Lab in Milton Keynes | Wellcome Sanger Institute for the COVID-19 Genomics UK (COG-UK) Consortium | The Lighthouse Lab in Milton Keynes and Alex Alderton, Roberto Amato, Jeffrey Barrett, Sonia Goncalves, Ewan Harrison, David K. Jackson, Ian Johnston, Dominic Kwiatkowski, Cordelia Langford, John Sillitoe on behalf of the Wellcome Sanger Institute COVID-19 Surveillance Team |
| EPI_ISL_2366491 | Lighthouse Lab in Glasgow | Wellcome Sanger Institute for the COVID-19 Genomics UK (COG-UK) Consortium | Harper VanSteenhouse, Yumi Kasai, David Gray, Carol Clugston, Anna Dominiczak and Alex Alderton, Roberto Amato, Jeffrey Barrett, Sonia Goncalves, Ewan Harrison, David K. Jackson, Ian Johnston, Dominic Kwiatkowski, Cordelia Langford, John Sillitoe on behalf of the Wellcome Sanger Institute COVID-19 Surveillance Team |
| EPI_ISL_2366493 | Lighthouse Lab in Alderley Park | Wellcome Sanger Institute for the COVID-19 Genomics UK (COG-UK) Consortium | Jacquelyn Wynn, Mairead Hyland, The Lighthouse Lab in Alderley Park and Alex Alderton, Roberto Amato, Jeffrey Barrett, Sonia Goncalves, Ewan Harrison, David K. Jackson, Ian Johnston, Dominic Kwiatkowski, Cordelia Langford, John Sillitoe on behalf of the Wellcome Sanger Institute COVID-19 Surveillance Team |
| EPI_ISL_2366494, EPI_ISL_2366495, EPI_ISL_2366496, EPI_ISL_2366497, EPI_ISL_2366498 | Lighthouse Lab in Milton Keynes | Wellcome Sanger Institute for the COVID-19 Genomics UK (COG-UK) Consortium | The Lighthouse Lab in Milton Keynes and Alex Alderton, Roberto Amato, Jeffrey Barrett, Sonia Goncalves, Ewan Harrison, David K. Jackson, Ian Johnston, Dominic Kwiatkowski, Cordelia Langford, John Sillitoe on behalf of the Wellcome Sanger Institute COVID-19 Surveillance Team |
| EPI_ISL_2366500 | Lighthouse Lab in Alderley Park | Wellcome Sanger Institute for the COVID-19 Genomics UK (COG-UK) Consortium | Jacquelyn Wynn, Mairead Hyland, The Lighthouse Lab in Alderley Park and Alex Alderton, Roberto Amato, Jeffrey Barrett, Sonia Goncalves, Ewan Harrison, David K. Jackson, Ian Johnston, Dominic Kwiatkowski, Cordelia Langford, John Sillitoe on behalf of the Wellcome Sanger Institute COVID-19 Surveillance Team |
| EPI_ISL_2366502 | Lighthouse Lab in Milton Keynes | Wellcome Sanger Institute for the COVID-19 Genomics UK (COG-UK) Consortium | The Lighthouse Lab in Milton Keynes and Alex Alderton, Roberto Amato, Jeffrey Barrett, Sonia Goncalves, Ewan Harrison, David K. Jackson, Ian Johnston, Dominic Kwiatkowski, Cordelia Langford, John Sillitoe on behalf of the Wellcome Sanger Institute COVID-19 Surveillance Team |
| EPI_ISL_2366503 | Lighthouse Lab in Glasgow | Wellcome Sanger Institute for the COVID-19 Genomics UK (COG-UK) | Harper VanSteenhouse, Yumi Kasai, David Gray, Carol Clugston, Anna Dominiczak and Alex Alderton, Roberto Amato, Jeffrey Barrett, Sonia Goncalves, Ewan Harrison, David K. Jackson, Ian Johnston, Dominic Kwiatkowski, Cordelia Langford, |

| | | | |
|---|---|---|---|
| | | Consortium | John Sillitoe on behalf of the Wellcome Sanger Institute COVID-19 Surveillance Team |
| EPI_ISL_2366504 | Lighthouse Lab in Alderley Park | Wellcome Sanger Institute for the COVID-19 Genomics UK (COG-UK) Consortium | Jacquelyn Wynn, Mairead Hyland, The Lighthouse Lab in Alderley Park and Alex Alderton, Roberto Amato, Jeffrey Barrett, Sonia Goncalves, Ewan Harrison, David K. Jackson, Ian Johnston, Dominic Kwiatkowski, Cordelia Langford, John Sillitoe on behalf of the Wellcome Sanger Institute COVID-19 Surveillance Team |
| EPI_ISL_2366506, EPI_ISL_2366507, EPI_ISL_2366508, EPI_ISL_2366509, EPI_ISL_2366511 | Lighthouse Lab in Milton Keynes | Wellcome Sanger Institute for the COVID-19 Genomics UK (COG-UK) Consortium | The Lighthouse Lab in Milton Keynes and Alex Alderton, Roberto Amato, Jeffrey Barrett, Sonia Goncalves, Ewan Harrison, David K. Jackson, Ian Johnston, Dominic Kwiatkowski, Cordelia Langford, John Sillitoe on behalf of the Wellcome Sanger Institute COVID-19 Surveillance Team |
| EPI_ISL_2366513 | Lighthouse Lab in Alderley Park | Wellcome Sanger Institute for the COVID-19 Genomics UK (COG-UK) Consortium | Jacquelyn Wynn, Mairead Hyland, The Lighthouse Lab in Alderley Park and Alex Alderton, Roberto Amato, Jeffrey Barrett, Sonia Goncalves, Ewan Harrison, David K. Jackson, Ian Johnston, Dominic Kwiatkowski, Cordelia Langford, John Sillitoe on behalf of the Wellcome Sanger Institute COVID-19 Surveillance Team |
| EPI_ISL_2366514 | Lighthouse Lab in Milton Keynes | Wellcome Sanger Institute for the COVID-19 Genomics UK (COG-UK) Consortium | The Lighthouse Lab in Milton Keynes and Alex Alderton, Roberto Amato, Jeffrey Barrett, Sonia Goncalves, Ewan Harrison, David K. Jackson, Ian Johnston, Dominic Kwiatkowski, Cordelia Langford, John Sillitoe on behalf of the Wellcome Sanger Institute COVID-19 Surveillance Team |
| EPI_ISL_2366516 | Lighthouse Lab in Alderley Park | Wellcome Sanger Institute for the COVID-19 Genomics UK (COG-UK) Consortium | Jacquelyn Wynn, Mairead Hyland, The Lighthouse Lab in Alderley Park and Alex Alderton, Roberto Amato, Jeffrey Barrett, Sonia Goncalves, Ewan Harrison, David K. Jackson, Ian Johnston, Dominic Kwiatkowski, Cordelia Langford, John Sillitoe on behalf of the Wellcome Sanger Institute COVID-19 Surveillance Team |
| EPI_ISL_2366517, EPI_ISL_2366519, EPI_ISL_2366521 | Lighthouse Lab in Milton Keynes | Wellcome Sanger Institute for the COVID-19 Genomics UK (COG-UK) Consortium | The Lighthouse Lab in Milton Keynes and Alex Alderton, Roberto Amato, Jeffrey Barrett, Sonia Goncalves, Ewan Harrison, David K. Jackson, Ian Johnston, Dominic Kwiatkowski, Cordelia Langford, John Sillitoe on behalf of the Wellcome Sanger Institute COVID-19 Surveillance Team |
| EPI_ISL_2366523 | Lighthouse Lab in Alderley Park | Wellcome Sanger Institute for the COVID-19 Genomics UK (COG-UK) Consortium | Jacquelyn Wynn, Mairead Hyland, The Lighthouse Lab in Alderley Park and Alex Alderton, Roberto Amato, Jeffrey Barrett, Sonia Goncalves, Ewan Harrison, David K. Jackson, Ian Johnston, Dominic Kwiatkowski, Cordelia Langford, John Sillitoe on behalf of the Wellcome Sanger Institute COVID-19 Surveillance Team |
| EPI_ISL_2366524 | Lighthouse Lab in Milton Keynes | Wellcome Sanger Institute for the COVID-19 Genomics UK (COG-UK) Consortium | The Lighthouse Lab in Milton Keynes and Alex Alderton, Roberto Amato, Jeffrey Barrett, Sonia Goncalves, Ewan Harrison, David K. Jackson, Ian Johnston, Dominic Kwiatkowski, Cordelia Langford, John Sillitoe on behalf of the Wellcome Sanger Institute COVID-19 Surveillance Team |
| EPI_ISL_2366525 | Lighthouse Lab in Alderley Park | Wellcome Sanger Institute for the COVID-19 Genomics UK (COG-UK) Consortium | Jacquelyn Wynn, Mairead Hyland, The Lighthouse Lab in Alderley Park and Alex Alderton, Roberto Amato, Jeffrey Barrett, Sonia Goncalves, Ewan Harrison, David K. Jackson, Ian Johnston, Dominic Kwiatkowski, Cordelia Langford, John Sillitoe on behalf of the Wellcome Sanger Institute COVID-19 Surveillance Team |
| EPI_ISL_2366526 | Lighthouse Lab in Milton Keynes | Wellcome Sanger Institute for the COVID-19 Genomics UK (COG-UK) Consortium | The Lighthouse Lab in Milton Keynes and Alex Alderton, Roberto Amato, Jeffrey Barrett, Sonia Goncalves, Ewan Harrison, David K. Jackson, Ian Johnston, Dominic Kwiatkowski, Cordelia Langford, John Sillitoe on behalf of the Wellcome Sanger Institute COVID-19 Surveillance Team |
| EPI_ISL_2366528 | Lighthouse Lab in Alderley Park | Wellcome Sanger Institute for the COVID-19 Genomics UK (COG-UK) Consortium | Jacquelyn Wynn, Mairead Hyland, The Lighthouse Lab in Alderley Park and Alex Alderton, Roberto Amato, Jeffrey Barrett, Sonia Goncalves, Ewan Harrison, David K. Jackson, Ian Johnston, Dominic Kwiatkowski, Cordelia Langford, John Sillitoe on behalf of the Wellcome Sanger Institute COVID-19 Surveillance Team |
| EPI_ISL_2366529, EPI_ISL_2366530 | Lighthouse Lab in Glasgow | Wellcome Sanger Institute for the COVID-19 Genomics UK (COG-UK) Consortium | Harper VanSteenhouse, Yumi Kasai, David Gray, Carol Clugston, Anna Dominiczak and Alex Alderton, Roberto Amato, Jeffrey Barrett, Sonia Goncalves, Ewan Harrison, David K. Jackson, Ian Johnston, Dominic Kwiatkowski, Cordelia Langford, John Sillitoe on behalf of the Wellcome Sanger Institute COVID-19 Surveillance Team |
| EPI_ISL_2366534 | Lighthouse Lab in Milton Keynes | Wellcome Sanger Institute for the COVID-19 Genomics UK (COG-UK) Consortium | The Lighthouse Lab in Milton Keynes and Alex Alderton, Roberto Amato, Jeffrey Barrett, Sonia Goncalves, Ewan Harrison, David K. Jackson, Ian Johnston, Dominic Kwiatkowski, Cordelia Langford, John Sillitoe on behalf of the Wellcome Sanger Institute COVID-19 Surveillance Team |
| EPI_ISL_2366535 | Lighthouse Lab in Alderley Park | Wellcome Sanger Institute for the COVID-19 Genomics UK (COG-UK) Consortium | Jacquelyn Wynn, Mairead Hyland, The Lighthouse Lab in Alderley Park and Alex Alderton, Roberto Amato, Jeffrey Barrett, Sonia Goncalves, Ewan Harrison, David K. Jackson, Ian Johnston, Dominic Kwiatkowski, Cordelia Langford, John Sillitoe on behalf of the Wellcome Sanger Institute COVID-19 Surveillance Team |
| EPI_ISL_2366536, EPI_ISL_2366537, EPI_ISL_2366539, EPI_ISL_2366540 | Lighthouse Lab in Milton Keynes | Wellcome Sanger Institute for the COVID-19 Genomics UK (COG-UK) Consortium | The Lighthouse Lab in Milton Keynes and Alex Alderton, Roberto Amato, Jeffrey Barrett, Sonia Goncalves, Ewan Harrison, David K. Jackson, Ian Johnston, Dominic Kwiatkowski, Cordelia Langford, John Sillitoe on behalf of the Wellcome Sanger Institute COVID-19 Surveillance Team |
| EPI_ISL_2366541 | Lighthouse Lab in Alderley Park | Wellcome Sanger Institute for the COVID-19 Genomics UK (COG-UK) Consortium | Jacquelyn Wynn, Mairead Hyland, The Lighthouse Lab in Alderley Park and Alex Alderton, Roberto Amato, Jeffrey Barrett, Sonia Goncalves, Ewan Harrison, David K. Jackson, Ian Johnston, Dominic Kwiatkowski, Cordelia Langford, John Sillitoe on behalf of the Wellcome Sanger Institute COVID-19 Surveillance Team |
| EPI_ISL_2366542, EPI_ISL_2366544, EPI_ISL_2366545 | Lighthouse Lab in Milton Keynes | Wellcome Sanger Institute for the COVID-19 Genomics UK (COG-UK) Consortium | The Lighthouse Lab in Milton Keynes and Alex Alderton, Roberto Amato, Jeffrey Barrett, Sonia Goncalves, Ewan Harrison, David K. Jackson, Ian Johnston, Dominic Kwiatkowski, Cordelia Langford, John Sillitoe on behalf of the Wellcome Sanger Institute COVID-19 Surveillance Team |
| EPI_ISL_2366548 | Lighthouse Lab in Alderley Park | Wellcome Sanger Institute for the COVID-19 Genomics UK (COG-UK) Consortium | Jacquelyn Wynn, Mairead Hyland, The Lighthouse Lab in Alderley Park and Alex Alderton, Roberto Amato, Jeffrey Barrett, Sonia Goncalves, Ewan Harrison, David K. Jackson, Ian Johnston, Dominic Kwiatkowski, Cordelia Langford, John Sillitoe on behalf of the Wellcome Sanger Institute COVID-19 Surveillance Team |
| EPI_ISL_2366549, EPI_ISL_2366550, EPI_ISL_2366551 | Lighthouse Lab in Milton Keynes | Wellcome Sanger Institute for the COVID-19 Genomics UK (COG-UK) Consortium | The Lighthouse Lab in Milton Keynes and Alex Alderton, Roberto Amato, Jeffrey Barrett, Sonia Goncalves, Ewan Harrison, David K. Jackson, Ian Johnston, Dominic Kwiatkowski, Cordelia Langford, John Sillitoe on behalf of the Wellcome Sanger Institute COVID-19 Surveillance Team |
| EPI_ISL_2366552 | Lighthouse Lab in Glasgow | Wellcome Sanger Institute for the COVID-19 Genomics UK (COG-UK) Consortium | Harper VanSteenhouse, Yumi Kasai, David Gray, Carol Clugston, Anna Dominiczak and Alex Alderton, Roberto Amato, Jeffrey Barrett, Sonia Goncalves, Ewan Harrison, David K. Jackson, Ian Johnston, Dominic Kwiatkowski, Cordelia Langford, John Sillitoe on behalf of the Wellcome Sanger Institute COVID-19 Surveillance Team |
| EPI_ISL_2366555, EPI_ISL_2366556, EPI_ISL_2366558, EPI_ISL_2366559 | Lighthouse Lab in Milton Keynes | Wellcome Sanger Institute for the COVID-19 Genomics UK (COG-UK) Consortium | The Lighthouse Lab in Milton Keynes and Alex Alderton, Roberto Amato, Jeffrey Barrett, Sonia Goncalves, Ewan Harrison, David K. Jackson, Ian Johnston, Dominic Kwiatkowski, Cordelia Langford, John Sillitoe on behalf of the Wellcome Sanger Institute COVID-19 Surveillance Team |
| EPI_ISL_2366561, EPI_ISL_2366562 | Lighthouse Lab in Alderley Park | Wellcome Sanger Institute for the COVID-19 Genomics UK (COG-UK) Consortium | Jacquelyn Wynn, Mairead Hyland, The Lighthouse Lab in Alderley Park and Alex Alderton, Roberto Amato, Jeffrey Barrett, Sonia Goncalves, Ewan Harrison, David K. Jackson, Ian Johnston, Dominic Kwiatkowski, Cordelia Langford, John Sillitoe on behalf of the Wellcome Sanger Institute COVID-19 Surveillance Team |
| EPI_ISL_2366563, EPI_ISL_2366566, EPI_ISL_2366568 | Lighthouse Lab in Milton Keynes | Wellcome Sanger Institute for the COVID-19 Genomics UK (COG-UK) Consortium | The Lighthouse Lab in Milton Keynes and Alex Alderton, Roberto Amato, Jeffrey Barrett, Sonia Goncalves, Ewan Harrison, David K. Jackson, Ian Johnston, Dominic Kwiatkowski, Cordelia Langford, John Sillitoe on behalf of the Wellcome Sanger Institute COVID-19 Surveillance Team |
| EPI_ISL_2366569 | Lighthouse Lab in Alderley Park | Wellcome Sanger Institute for the COVID-19 Genomics UK (COG-UK) Consortium | Jacquelyn Wynn, Mairead Hyland, The Lighthouse Lab in Alderley Park and Alex Alderton, Roberto Amato, Jeffrey Barrett, Sonia Goncalves, Ewan Harrison, David K. Jackson, Ian Johnston, Dominic Kwiatkowski, Cordelia Langford, John Sillitoe on behalf of the Wellcome Sanger Institute COVID-19 Surveillance Team |
| EPI_ISL_2366570, EPI_ISL_2366572, EPI_ISL_2366573, EPI_ISL_2366574, EPI_ISL_2366575, EPI_ISL_2366576, EPI_ISL_2366577 | Lighthouse Lab in Milton Keynes | Wellcome Sanger Institute for the COVID-19 Genomics UK (COG-UK) Consortium | The Lighthouse Lab in Milton Keynes and Alex Alderton, Roberto Amato, Jeffrey Barrett, Sonia Goncalves, Ewan Harrison, David K. Jackson, Ian Johnston, Dominic Kwiatkowski, Cordelia Langford, John Sillitoe on behalf of the Wellcome Sanger Institute COVID-19 Surveillance Team |
| EPI_ISL_2366578 | Lighthouse Lab in Glasgow | Wellcome Sanger Institute for the COVID-19 Genomics UK (COG-UK) Consortium | Harper VanSteenhouse, Yumi Kasai, David Gray, Carol Clugston, Anna Dominiczak and Alex Alderton, Roberto Amato, Jeffrey Barrett, Sonia Goncalves, Ewan Harrison, David K. Jackson, Ian Johnston, Dominic Kwiatkowski, Cordelia Langford, John Sillitoe on behalf of the Wellcome Sanger Institute COVID-19 Surveillance Team |
| EPI_ISL_2366579, EPI_ISL_2366580 | Lighthouse Lab in Milton Keynes | Wellcome Sanger Institute for the COVID-19 Genomics UK (COG-UK) Consortium | The Lighthouse Lab in Milton Keynes and Alex Alderton, Roberto Amato, Jeffrey Barrett, Sonia Goncalves, Ewan Harrison, David K. Jackson, Ian Johnston, Dominic Kwiatkowski, Cordelia Langford, John Sillitoe on behalf of the Wellcome Sanger Institute COVID-19 Surveillance Team |
| EPI_ISL_2366581 | Lighthouse Lab in Alderley Park | Wellcome Sanger Institute for the COVID-19 Genomics UK (COG-UK) Consortium | Jacquelyn Wynn, Mairead Hyland, The Lighthouse Lab in Alderley Park and Alex Alderton, Roberto Amato, Jeffrey Barrett, Sonia Goncalves, Ewan Harrison, David K. Jackson, Ian Johnston, Dominic Kwiatkowski, Cordelia Langford, John Sillitoe on behalf of the Wellcome Sanger Institute COVID-19 Surveillance Team |
| EPI_ISL_2366582, EPI_ISL_2366583 | Lighthouse Lab in Milton Keynes | Wellcome Sanger Institute for the COVID-19 Genomics UK (COG-UK) Consortium | The Lighthouse Lab in Milton Keynes and Alex Alderton, Roberto Amato, Jeffrey Barrett, Sonia Goncalves, Ewan Harrison, David K. Jackson, Ian Johnston, Dominic Kwiatkowski, Cordelia Langford, John Sillitoe on behalf of the Wellcome Sanger Institute COVID-19 Surveillance Team |
| EPI_ISL_2366584 | Lighthouse Lab in Alderley Park | Wellcome Sanger Institute for the COVID-19 Genomics UK (COG-UK) Consortium | Jacquelyn Wynn, Mairead Hyland, The Lighthouse Lab in Alderley Park and Alex Alderton, Roberto Amato, Jeffrey Barrett, Sonia Goncalves, Ewan Harrison, David K. Jackson, Ian Johnston, Dominic Kwiatkowski, Cordelia Langford, John Sillitoe on behalf of the Wellcome Sanger Institute COVID-19 Surveillance Team |
| EPI_ISL_2366585, EPI_ISL_2366586, EPI_ISL_2366587 | Lighthouse Lab in Milton Keynes | Wellcome Sanger Institute for the COVID-19 Genomics UK (COG-UK) Consortium | The Lighthouse Lab in Milton Keynes and Alex Alderton, Roberto Amato, Jeffrey Barrett, Sonia Goncalves, Ewan Harrison, David K. Jackson, Ian Johnston, Dominic Kwiatkowski, Cordelia Langford, John Sillitoe on behalf of the Wellcome Sanger Institute COVID-19 Surveillance Team |
| EPI_ISL_2366588 | Lighthouse Lab in Glasgow | Wellcome Sanger Institute for the COVID-19 Genomics UK (COG-UK) Consortium | Harper VanSteenhouse, Yumi Kasai, David Gray, Carol Clugston, Anna Dominiczak and Alex Alderton, Roberto Amato, Jeffrey Barrett, Sonia Goncalves, Ewan Harrison, David K. Jackson, Ian Johnston, Dominic Kwiatkowski, Cordelia Langford, John Sillitoe on behalf of the Wellcome Sanger Institute COVID-19 Surveillance Team |
| EPI_ISL_2366589, EPI_ISL_2366590, EPI_ISL_2366592 | Lighthouse Lab in Milton Keynes | Wellcome Sanger Institute for the COVID-19 Genomics UK (COG-UK) Consortium | The Lighthouse Lab in Milton Keynes and Alex Alderton, Roberto Amato, Jeffrey Barrett, Sonia Goncalves, Ewan Harrison, David K. Jackson, Ian Johnston, Dominic Kwiatkowski, Cordelia Langford, John Sillitoe on behalf of the Wellcome Sanger Institute COVID-19 Surveillance Team |
| EPI_ISL_2366593, EPI_ISL_2366595, EPI_ISL_2366596, EPI_ISL_2366597 | Lighthouse Lab in Alderley Park | Wellcome Sanger Institute for the COVID-19 Genomics UK (COG-UK) Consortium | Jacquelyn Wynn, Mairead Hyland, The Lighthouse Lab in Alderley Park and Alex Alderton, Roberto Amato, Jeffrey Barrett, Sonia Goncalves, Ewan Harrison, David K. Jackson, Ian Johnston, Dominic Kwiatkowski, Cordelia Langford, John Sillitoe on behalf of the Wellcome Sanger Institute COVID-19 Surveillance Team |
| EPI_ISL_2366598, EPI_ISL_2366599, EPI_ISL_2366600, EPI_ISL_2366602 | Lighthouse Lab in Milton Keynes | Wellcome Sanger Institute for the COVID-19 Genomics UK (COG-UK) Consortium | The Lighthouse Lab in Milton Keynes and Alex Alderton, Roberto Amato, Jeffrey Barrett, Sonia Goncalves, Ewan Harrison, David K. Jackson, Ian Johnston, Dominic Kwiatkowski, Cordelia Langford, John Sillitoe on behalf of the Wellcome Sanger Institute COVID-19 Surveillance Team |
| EPI_ISL_2366603 | Lighthouse Lab in Alderley Park | Wellcome Sanger Institute for the COVID-19 Genomics UK (COG-UK) | Jacquelyn Wynn, Mairead Hyland, The Lighthouse Lab in Alderley Park and Alex Alderton, Roberto Amato, Jeffrey Barrett, Sonia Goncalves, Ewan Harrison, David K. Jackson, Ian Johnston, Dominic Kwiatkowski, Cordelia Langford, John Sillitoe on |

| | | | |
|---|---|---|---|
| | | Consortium | behalf of the Wellcome Sanger Institute COVID-19 Surveillance Team |
| EPI_ISL_2366605, EPI_ISL_2366606 | Lighthouse Lab in Milton Keynes | Wellcome Sanger Institute for the COVID-19 Genomics UK (COG-UK) Consortium | The Lighthouse Lab in Milton Keynes and Alex Alderton, Roberto Amato, Jeffrey Barrett, Sonia Goncalves, Ewan Harrison, David K. Jackson, Ian Johnston, Dominic Kwiatkowski, Cordelia Langford, John Sillitoe on behalf of the Wellcome Sanger Institute COVID-19 Surveillance Team |
| EPI_ISL_2366607, EPI_ISL_2366608, EPI_ISL_2366609, EPI_ISL_2366611, EPI_ISL_2366612, EPI_ISL_2366613, EPI_ISL_2366614, EPI_ISL_2366615, EPI_ISL_2366617, EPI_ISL_2366618, EPI_ISL_2366619, EPI_ISL_2366620 | | | |
| see above | Lighthouse Lab in Alderley Park | Wellcome Sanger Institute for the COVID-19 Genomics UK (COG-UK) Consortium | Jacquelyn Wynn, Mairead Hyland, The Lighthouse Lab in Alderley Park and Alex Alderton, Roberto Amato, Jeffrey Barrett, Sonia Goncalves, Ewan Harrison, David K. Jackson, Ian Johnston, Dominic Kwiatkowski, Cordelia Langford, John Sillitoe on behalf of the Wellcome Sanger Institute COVID-19 Surveillance Team |
| EPI_ISL_2367243 | Clinical Virology | Clinical Bacteriology | Tim Roloff, Fanny Wegner, Helena MB Seth-Smith, Alfredo Mari, Karoline Leuzinger, Julia Bielicki, Manuel Battegay, Hans Hirsch, Adrian Egli |
| EPI_ISL_2367643 | Helix/Illumina | Centers for Disease Control and Prevention Division of Viral Diseases, Pathogen Discovery | Dakota Howard, Dhwani Batra, Peter W. Cook, Kara Moser, Adrian Paskey, Jason Caravas, Benjamin Rambo-Martin, Shatavia Morrison, Christopher Gulvick, Scott Sammons, Yvette Unoarumhi, Darlene Wagner, Matthew Schmerer, Eileen de Feo, Jan Antico, Christine Tran, Matthew Tolentino, Shannon Wickline, Kim Gietzen, Brad Sickler, Jingtao Liu, Eric Allen, Phil Febbo, Nicole L. Washington, Simon White, Geraint Levan, Kelly Schiabor Barrett, Elizabeth Cirulli, Alexandre Bolze, Ary Ascencio, Charlotte Rivera-Garcia, Ryan Cho, Jason Nguyen, Sherry Wang, Jimmy Ramirez, Tyler Cassens, Efren Sandoval, Magnus Isaksson, William Lee, David Becker, Marc Laurent, James Lu, Clinton R. Paden, Duncan MacCannell |
| EPI_ISL_2367692, EPI_ISL_2367713, EPI_ISL_2367802, EPI_ISL_2367836, EPI_ISL_2367897, EPI_ISL_2367929, EPI_ISL_2367941, EPI_ISL_2367983 | Quest Diagnostics Incorporated | Centers for Disease Control and Prevention Division of Viral Diseases, Pathogen Discovery | Dakota Howard, Dhwani Batra, Peter W. Cook, Kara Moser, Adrian Paskey, Jason Caravas, Benjamin Rambo-Martin, Shatavia Morrison, Christopher Gulvick, Scott Sammons, Yvette Unoarumhi, Darlene Wagner, Matthew Schmerer, S. H. Rosenthal, A. Gerasimova, R. M. Kagan, B. Anderson, M. Hua, Y. Liu, L.E. Bernstein, K.E. Livingston, A. Perez, I. A. Shlyakhter, R. V. Rolando, R. Owen, P. Tanpaiboon, F. Lacbawan, Clinton R. Paden, Duncan MacCannell |
| EPI_ISL_2368081, EPI_ISL_2368082, EPI_ISL_2368083, EPI_ISL_2368103, EPI_ISL_2368104, EPI_ISL_2368105, EPI_ISL_2368106, EPI_ISL_2368107, EPI_ISL_2368108, EPI_ISL_2368109, EPI_ISL_2368111, EPI_ISL_2368121, EPI_ISL_2368123, EPI_ISL_2368124, EPI_ISL_2368128, EPI_ISL_2368130, EPI_ISL_2368172 | | | |
| see above | Fulgent Genetics | Centers for Disease Control and Prevention Division of Viral Diseases, Pathogen Discovery | Dakota Howard, Dhwani Batra, Peter W. Cook, Kara Moser, Adrian Paskey, Jason Caravas, Benjamin Rambo-Martin, Shatavia Morrison, Christopher Gulvick, Scott Sammons, Yvette Unoarumhi, Darlene Wagner, Matthew Schmerer, Harry Gao, Mickey Li, John Gao, Joseph Fierro, Benafsh Sapra, Becky Tsai, Yan Meng, Doreen Ng, James Xie, Clinton R. Paden, Duncan MacCannell |
| EPI_ISL_2368243, EPI_ISL_2368393, EPI_ISL_2368418, EPI_ISL_2368430, EPI_ISL_2368447, EPI_ISL_2368448, EPI_ISL_2368463, EPI_ISL_2368498, EPI_ISL_2368666, EPI_ISL_2368756, EPI_ISL_2368826, EPI_ISL_2368827, EPI_ISL_2368887, EPI_ISL_2368903 | | | |
| see above | Aegis Sciences Corporation | Centers for Disease Control and Prevention Division of Viral Diseases, Pathogen Discovery | Dakota Howard, Dhwani Batra, Peter W. Cook, Kara Moser, Adrian Paskey, Jason Caravas, Benjamin Rambo-Martin, Shatavia Morrison, Christopher Gulvick, Scott Sammons, Yvette Unoarumhi, Darlene Wagner, Matthew Schmerer, Cyndi Clark, Patrick Campbell, Rob Case, Vikramsinha Ghorpade, Holly Houdeshell, Ola Kvalvaag, Dillon Nall, Ethan Sanders, Alec Vest, Shaun Westlund, Matthew Hardison, Clinton R. Paden, Duncan MacCannell |
| EPI_ISL_2369020, EPI_ISL_2369239, EPI_ISL_2369242 | Quest Diagnostics Incorporated | Centers for Disease Control and Prevention Division of Viral Diseases, Pathogen Discovery | Dakota Howard, Dhwani Batra, Peter W. Cook, Kara Moser, Adrian Paskey, Jason Caravas, Benjamin Rambo-Martin, Shatavia Morrison, Christopher Gulvick, Scott Sammons, Yvette Unoarumhi, Darlene Wagner, Matthew Schmerer, S. H. Rosenthal, A. Gerasimova, R. M. Kagan, B. Anderson, M. Hua, Y. Liu, L.E. Bernstein, K.E. Livingston, A. Perez, I. A. Shlyakhter, R. V. Rolando, R. Owen, P. Tanpaiboon, F. Lacbawan, Clinton R. Paden, Duncan MacCannell |
| EPI_ISL_2369968, EPI_ISL_2370046, EPI_ISL_2370135 | Aegis Sciences Corporation | Centers for Disease Control and Prevention Division of Viral Diseases, Pathogen Discovery | Dakota Howard, Dhwani Batra, Peter W. Cook, Kara Moser, Adrian Paskey, Jason Caravas, Benjamin Rambo-Martin, Shatavia Morrison, Christopher Gulvick, Scott Sammons, Yvette Unoarumhi, Darlene Wagner, Matthew Schmerer, Cyndi Clark, Patrick Campbell, Rob Case, Vikramsinha Ghorpade, Holly Houdeshell, Ola Kvalvaag, Dillon Nall, Ethan Sanders, Alec Vest, Shaun Westlund, Matthew Hardison, Clinton R. Paden, Duncan MacCannell |
| EPI_ISL_2370147 | Greek Genome Center, Biomedical Research Foundation of the Academy of Athens (BRFAA) | Greek Genome Center, Biomedical Research Foundation of the Academy of Athens (BRFAA) | Emmanouil Athanasiadis, Giannis Vatsellas, Theodoros Loupis, Katerina Zoi, Dimitrios Thanos |
| EPI_ISL_2370180, EPI_ISL_2370194, EPI_ISL_2370200, EPI_ISL_2370217, EPI_ISL_2370237, EPI_ISL_2370334, EPI_ISL_2370372, EPI_ISL_2370411, EPI_ISL_2370420, EPI_ISL_2370480, EPI_ISL_2370507, EPI_ISL_2370533 | | | |
| see above | Aegis Sciences Corporation | Centers for Disease Control and Prevention Division of Viral Diseases, Pathogen Discovery | Dakota Howard, Dhwani Batra, Peter W. Cook, Kara Moser, Adrian Paskey, Jason Caravas, Benjamin Rambo-Martin, Shatavia Morrison, Christopher Gulvick, Scott Sammons, Yvette Unoarumhi, Darlene Wagner, Matthew Schmerer, Cyndi Clark, Patrick Campbell, Rob Case, Vikramsinha Ghorpade, Holly Houdeshell, Ola Kvalvaag, Dillon Nall, Ethan Sanders, Alec Vest, Shaun Westlund, Matthew Hardison, Clinton R. Paden, Duncan MacCannell |
| EPI_ISL_2371543, EPI_ISL_2371546, EPI_ISL_2371552, EPI_ISL_2371553, EPI_ISL_2371557, EPI_ISL_2371562, EPI_ISL_2371569, EPI_ISL_2371604, EPI_ISL_2371607, EPI_ISL_2371621, EPI_ISL_2371622, EPI_ISL_2371627, EPI_ISL_2371642, EPI_ISL_2371666 | | | |
| see above | Fulgent Genetics | Centers for Disease Control and Prevention Division of Viral Diseases, Pathogen Discovery | Dakota Howard, Dhwani Batra, Peter W. Cook, Kara Moser, Adrian Paskey, Jason Caravas, Benjamin Rambo-Martin, Shatavia Morrison, Christopher Gulvick, Scott Sammons, Yvette Unoarumhi, Darlene Wagner, Matthew Schmerer, Harry Gao, Mickey Li, John Gao, Joseph Fierro, Benafsh Sapra, Becky Tsai, Yan Meng, Doreen Ng, James Xie, Clinton R. Paden, Duncan MacCannell |
| EPI_ISL_2371742, EPI_ISL_2371804, EPI_ISL_2371814, EPI_ISL_2371831, EPI_ISL_2371884, EPI_ISL_2371903, EPI_ISL_2371914, EPI_ISL_2371921, EPI_ISL_2371942, EPI_ISL_2372010, EPI_ISL_2372036, EPI_ISL_2372111, EPI_ISL_2372175, EPI_ISL_2372177, EPI_ISL_2372198, EPI_ISL_2372203, EPI_ISL_2372248 | | | |
| see above | Quest Diagnostics Incorporated | Centers for Disease Control and Prevention Division of Viral Diseases, Pathogen Discovery | Dakota Howard, Dhwani Batra, Peter W. Cook, Kara Moser, Adrian Paskey, Jason Caravas, Benjamin Rambo-Martin, Shatavia Morrison, Christopher Gulvick, Scott Sammons, Yvette Unoarumhi, Darlene Wagner, Matthew Schmerer, S. H. Rosenthal, A. Gerasimova, R. M. Kagan, B. Anderson, M. Hua, Y. Liu, L.E. Bernstein, K.E. Livingston, A. Perez, I. A. Shlyakhter, R. V. Rolando, R. Owen, P. Tanpaiboon, F. Lacbawan, Clinton R. Paden, Duncan MacCannell |
| EPI_ISL_2372266, EPI_ISL_2372267 | OUCRU | OUCRU | Nguyen Van Vinh Chau, Nguyen To Anh, Nguyen Thi thu Hong, Nghiem My Ngoc, Huynh Trung Trieu, Le Nguyen Truc Nhu, Nguyen Thi Han Ny, Lam Minh Yen, Ngo Ngoc Quang Minh, Nguyen Thanh Phong, Nguyen Thanh Truong, Le Thi Thu Huong, Tran Nguyen Hoang Tu, Le Manh Hung, Tran Tan Thanh, Nguyen Thanh Dung, Nguyen Tri Dung, Guy Thwaites, Le Van Tan |
| EPI_ISL_2372341, EPI_ISL_2372359, EPI_ISL_2372460 | Aegis Sciences Corporation | Centers for Disease Control and Prevention Division of Viral Diseases, Pathogen Discovery | Dakota Howard, Dhwani Batra, Peter W. Cook, Kara Moser, Adrian Paskey, Jason Caravas, Benjamin Rambo-Martin, Shatavia Morrison, Christopher Gulvick, Scott Sammons, Yvette Unoarumhi, Darlene Wagner, Matthew Schmerer, Cyndi Clark, Patrick Campbell, Rob Case, Vikramsinha Ghorpade, Holly Houdeshell, Ola Kvalvaag, Dillon Nall, Ethan Sanders, Alec Vest, Shaun Westlund, Matthew Hardison, Clinton R. Paden, Duncan MacCannell |
| EPI_ISL_2372720 | Microbiology Department, Laboratori Clínic Metropolitana Nord. Hospital Universitari Germans Trias i Pujol | Can Ruti SARS-CoV-2 Sequencing Hub (HUGTiP/IrsiCaixa/IGTP) | Marc Noguera-Julian, Pilar Armengol, Ignacio Blanco, Antoni E Bordoy, Francesc Catala-Moll, Pere-Joan Cardona, Maria Casadellà, Cristina Casañ, Gemma Clara, Bonaventura Clotet, Cristina Esteban, Montserrat Giménez, Mercedes Guerrero, Anna Not, Roger Pared |
| EPI_ISL_2372725 | Microbiologia CATLAB | Can Ruti SARS-CoV-2 Sequencing Hub (HUGTiP/IrsiCaixa/IGTP) | Marc Noguera-Julian, Pilar Armengol, Ignacio Blanco, Antoni E Bordoy, Francesc Catala-Moll, Pere-Joan Cardona, Maria Casadellà, Cristina Casañ, Gemma Clara, Bonaventura Clotet, Cristina Esteban, Montserrat Giménez, Mercedes Guerrero, Anna Not, Roger Pared |
| EPI_ISL_2372733 | Microbiology Department, Laboratori Clínic Metropolitana Nord. Hospital Universitari Germans Trias i Pujol | Can Ruti SARS-CoV-2 Sequencing Hub (HUGTiP/IrsiCaixa/IGTP) | Marc Noguera-Julian, Pilar Armengol, Ignacio Blanco, Antoni E Bordoy, Francesc Catala-Moll, Pere-Joan Cardona, Maria Casadellà, Cristina Casañ, Gemma Clara, Bonaventura Clotet, Cristina Esteban, Montserrat Giménez, Mercedes Guerrero, Anna Not, Roger Pared |
| EPI_ISL_2372737 | Microbiologia CATLAB | Can Ruti SARS-CoV-2 Sequencing Hub (HUGTiP/IrsiCaixa/IGTP) | Marc Noguera-Julian, Pilar Armengol, Ignacio Blanco, Antoni E Bordoy, Francesc Catala-Moll, Pere-Joan Cardona, Maria Casadellà, Cristina Casañ, Gemma Clara, Bonaventura Clotet, Cristina Esteban, Montserrat Giménez, Mercedes Guerrero, Anna Not, Roger Pared |
| EPI_ISL_2372744 | Microbiology Department, Laboratori Clínic Metropolitana Nord. Hospital Universitari Germans Trias i Pujol | Can Ruti SARS-CoV-2 Sequencing Hub (HUGTiP/IrsiCaixa/IGTP) | Marc Noguera-Julian, Pilar Armengol, Ignacio Blanco, Antoni E Bordoy, Francesc Catala-Moll, Pere-Joan Cardona, Maria Casadellà, Cristina Casañ, Gemma Clara, Bonaventura Clotet, Cristina Esteban, Montserrat Giménez, Mercedes Guerrero, Anna Not, Roger Pared |
| EPI_ISL_2372749 | Microbiologia CATLAB | Can Ruti SARS-CoV-2 Sequencing Hub (HUGTiP/IrsiCaixa/IGTP) | Marc Noguera-Julian, Pilar Armengol, Ignacio Blanco, Antoni E Bordoy, Francesc Catala-Moll, Pere-Joan Cardona, Maria Casadellà, Cristina Casañ, Gemma Clara, Bonaventura Clotet, Cristina Esteban, Montserrat Giménez, Mercedes Guerrero, Anna Not, Roger Pared |
| EPI_ISL_2372758 | Microbiology Department, Laboratori Clínic Metropolitana Nord. Hospital Universitari Germans Trias i Pujol | Can Ruti SARS-CoV-2 Sequencing Hub (HUGTiP/IrsiCaixa/IGTP) | Marc Noguera-Julian, Pilar Armengol, Ignacio Blanco, Antoni E Bordoy, Francesc Catala-Moll, Pere-Joan Cardona, Maria Casadellà, Cristina Casañ, Gemma Clara, Bonaventura Clotet, Cristina Esteban, Montserrat Giménez, Mercedes Guerrero, Anna Not, Roger Pared |
| EPI_ISL_2372761, EPI_ISL_2372772 | Microbiologia CATLAB | Can Ruti SARS-CoV-2 Sequencing Hub (HUGTiP/IrsiCaixa/IGTP) | Marc Noguera-Julian, Pilar Armengol, Ignacio Blanco, Antoni E Bordoy, Francesc Catala-Moll, Pere-Joan Cardona, Maria Casadellà, Cristina Casañ, Gemma Clara, Bonaventura Clotet, Cristina Esteban, Montserrat Giménez, Mercedes Guerrero, Anna Not, Roger Pared |
| EPI_ISL_2372781 | Microbiology Department, Laboratori Clínic Metropolitana Nord. Hospital Universitari Germans Trias i Pujol | Can Ruti SARS-CoV-2 Sequencing Hub (HUGTiP/IrsiCaixa/IGTP) | Marc Noguera-Julian, Pilar Armengol, Ignacio Blanco, Antoni E Bordoy, Francesc Catala-Moll, Pere-Joan Cardona, Maria Casadellà, Cristina Casañ, Gemma Clara, Bonaventura Clotet, Cristina Esteban, Montserrat Giménez, Mercedes Guerrero, Anna Not, Roger Pared |
| EPI_ISL_2372806, EPI_ISL_2372814, EPI_ISL_2372815, EPI_ISL_2372817, EPI_ISL_2372818, EPI_ISL_2372819, EPI_ISL_2372821, EPI_ISL_2372823, EPI_ISL_2372824, EPI_ISL_2372829, EPI_ISL_2372844, EPI_ISL_2372847, EPI_ISL_2372849, EPI_ISL_2372853, EPI_ISL_2372856, EPI_ISL_2372857, EPI_ISL_2372859, EPI_ISL_2372884, EPI_ISL_2372888, EPI_ISL_2372894 | | | |
| see above | Azienda Sanitaria dell'Alto Adige Laboratorio Aziendale di Microbiologia e Virologia | Istituto di Genomica Applicata | Elisabetta Pagani, Irene Bianconi, Elisabetta Giacobazzi, Elisa Masi, Stefanie Wieser, Irena Jurman, Vera Vendramin, Gabriele Magris, Eleonora Paparelli, Davide Scaglione, Michele Morgante |
| EPI_ISL_2372961, EPI_ISL_2373021, EPI_ISL_2373038 | Northwestern Memorial Hospital | Northwestern University - Ozer Lab | Ramon Lorenzo-Redondo, Lacy M. Simons, Taylor J. Dean, Chad J. Achenbach, Lawrence J. Jennings, Chao Qi, Michael G. Ison, Judd F. Hultquist, Egon A. Ozer |
| EPI_ISL_2373108 | National Influenza Center, National Institute of Hygiene and Epidemiology (NIHE) | National Influenza Center, National Institute of Hygiene and Epidemiology (NIHE) | Le Quynh Mai, Taichiro Takemura, Nguyen Le Khanh Hang, Hoang Vu Mai Phuong, Ung Thi Hong Trang, Le Thi Thanh, Nguyen Vu Son, Vuong Duc Cuong, Pham Thi Hien, Tran Thu Huong, Nguyen Phuong Anh, Nguyen Co Thach, Tran Thi Hien, Vũ Hiếu, Nguyen Thi Kieu Anh, Dang Duc Anh |
| EPI_ISL_2373109 | National Influenza Center, National Institute of Hygiene and Epidemiology (NIHE) | National Influenza Center, National Institute of Hygiene and Epidemiology (NIHE) | Le Quynh Mai, Taichiro Takemura, Nguyen Le Khanh Hang, Hoang Vu Mai Phuong, Ung Thi Hong Trang, Le Thi Thanh, Nguyen Vu Son, Vuong Duc Cuong, Pham Thi Hien, Tran Thu Huong, Nguyen Phuong Anh, Nguyen Co Thach, Tran Thi Hien, Vũ Hiếu, Nguyen Thi Quyen, Dang Duc Anh |
| EPI_ISL_2373110, EPI_ISL_2373111 | National Influenza Center, National Institute of Hygiene and Epidemiology (NIHE) | National Influenza Center, National Institute of Hygiene and Epidemiology (NIHE) | Le Quynh Mai, Taichiro Takemura, Nguyen Le Khanh Hang, Hoang Vu Mai Phuong, Ung Thi Hong Trang, Le Thi Thanh, Nguyen Vu Son, Vuong Duc Cuong, Pham Thi Hien, Tran Thu Huong, Nguyen Phuong Anh, Nguyen Co Thach, Tran Thi Hien, Vũ Hiếu, Nguyen Thi Kieu Anh, Dang Duc Anh |
| EPI_ISL_2373137, EPI_ISL_2373138, EPI_ISL_2373141, EPI_ISL_2373142, EPI_ISL_2373143, EPI_ISL_2373144, EPI_ISL_2373145, EPI_ISL_2373146, EPI_ISL_2373147, EPI_ISL_2373148, EPI_ISL_2373149, EPI_ISL_2373150, EPI_ISL_2373151, EPI_ISL_2373152, EPI_ISL_2373153 | | | |
| see above | AP SSO | CSIR-Centre for Cellular and Molecular Biology-INSACOG | Lamuk Zaveri,Tulasi Nagabandi, Ara Sreenivas,Shreekant Verma, Amareshwar Vodapalli ,Blessy B John,Viswagithe S L,B Himasri,Valli Nagalakshmi Undamatla,Payel Mukherjee,Sofia Banu,Archana Bharadwaj Siva,Sharath Chandra Thota,Karthik Bharadwaj Tallapaka,Rakesh K Mishra,Divya Tej Sowpati |
| EPI_ISL_2373154, EPI_ISL_2373156, EPI_ISL_2373157, EPI_ISL_2373158, EPI_ISL_2373160, EPI_ISL_2373161, EPI_ISL_2373162, EPI_ISL_2373163, EPI_ISL_2373164, EPI_ISL_2373165, EPI_ISL_2373167, EPI_ISL_2373168, EPI_ISL_2373169, EPI_ISL_2373170, EPI_ISL_2373172, EPI_ISL_2373173, EPI_ISL_2373174, EPI_ISL_2373175, EPI_ISL_2373176, EPI_ISL_2373177, EPI_ISL_2373178, EPI_ISL_2373179, EPI_ISL_2373180, EPI_ISL_2373182 | | | |
| see above | AP SSO | CSIR-Centre for Cellular and Molecular Biology-INSACOG | Tulasi Nagabandi, Lamuk Zaveri,Ara Sreenivas,Shreekant Verma, Amareshwar Vodapalli ,Blessy B John,Viswagithe S L,B Himasri,Valli Nagalakshmi Undamatla,Payel Mukherjee,Sofia Banu,Archana Bharadwaj Siva,Sharath Chandra Thota,Karthik Bharadwaj Tallapaka,Rakesh K Mishra,Divya Tej Sowpati |
| EPI_ISL_2373183, EPI_ISL_2373185, EPI_ISL_2373186, EPI_ISL_2373187, EPI_ISL_2373188, EPI_ISL_2373189, EPI_ISL_2373190, EPI_ISL_2373191, EPI_ISL_2373193, EPI_ISL_2373195, EPI_ISL_2373196, EPI_ISL_2373197, EPI_ISL_2373198, EPI_ISL_2373199, EPI_ISL_2373200, EPI_ISL_2373201, EPI_ISL_2373202, EPI_ISL_2373203, EPI_ISL_2373204, EPI_ISL_2373205, EPI_ISL_2373206, EPI_ISL_2373207, EPI_ISL_2373208 | | | |
| see above | AP SSO | CSIR-Centre for Cellular and Molecular Biology-INSACOG | Blessy B John,Viswagithe S L,Tulasi Nagabandi, Lamuk Zaveri,Ara Sreenivas,Shreekant Verma, Amareshwar Vodapalli ,B Himasri,Valli Nagalakshmi Undamatla,Payel Mukherjee,Sofia Banu,Archana Bharadwaj Siva,Sharath Chandra Thota,Karthik Bharadwaj Tallapaka,Rakesh K Mishra,Divya Tej Sowpati |
| EPI_ISL_2373210, EPI_ISL_2373211, EPI_ISL_2373212, EPI_ISL_2373213, | AP SSO | CSIR-Centre for Cellular and Molecular Biology-INSACOG | Tulasi Nagabandi, Lamuk Zaveri,Ara Sreenivas,Shreekant Verma, Amareshwar Vodapalli ,Blessy B John,Viswagithe S L,B Himasri,Valli Nagalakshmi Undamatla,Payel Mukherjee,Sofia Banu,Archana Bharadwaj Siva,Sharath Chandra Thota,Karthik Bharadwaj Tallapaka,Rakesh K Mishra,Divya Tej Sowpati |

| | | | |
|---|---|---|---|
| EPI_ISL_2373215 | | | |
| EPI_ISL_2373216, EPI_ISL_2373217, EPI_ISL_2373218, EPI_ISL_2373219, EPI_ISL_2373220, EPI_ISL_2373221 | AP SSO | CSIR-Centre for Cellular and Molecular Biology-INSACOG | Blessy B John,Viswagithe S L,Tulasi Nagabandi, Lamuk Zaveri,Ara Sreenivas,Shreekant Verma, Amareshwar Vodapalli ,B Himasri,Valli Nagalakshmi Undamatla,Payel Mukherjee,Sofia Banu,Archana Bharadwaj Siva,Sharath Chandra Thota,Karthik Bharadwaj Tallapaka,Rakesh K Mishra,Divya Tej Sowpati |
| EPI_ISL_2373222, EPI_ISL_2373223, EPI_ISL_2373224, EPI_ISL_2373225, EPI_ISL_2373226, EPI_ISL_2373227, EPI_ISL_2373228, EPI_ISL_2373229, EPI_ISL_2373230, EPI_ISL_2373231, EPI_ISL_2373232, EPI_ISL_2373233, EPI_ISL_2373234, EPI_ISL_2373235, EPI_ISL_2373236, EPI_ISL_2373237, EPI_ISL_2373238, EPI_ISL_2373239, EPI_ISL_2373241 | | | |
| see above | AP SSO | CSIR-Centre for Cellular and Molecular Biology-INSACOG | Ara Sreenivas,Shreekant Verma,Tulasi Nagabandi, Lamuk Zaveri, Amareshwar Vodapalli ,Blessy B John,Viswagithe S L,B Himasri,Valli Nagalakshmi Undamatla,Payel Mukherjee,Sofia Banu,Archana Bharadwaj Siva,Sharath Chandra Thota,Karthik Bharadwaj Tallapaka,Rakesh K Mishra,Divya Tej Sowpati |
| EPI_ISL_2373243, EPI_ISL_2373246, EPI_ISL_2373247, EPI_ISL_2373248, EPI_ISL_2373249, EPI_ISL_2373251, EPI_ISL_2373252, EPI_ISL_2373253, EPI_ISL_2373254 | CSIR-Centre for Cellular and Molecular Biology | CSIR-Centre for Cellular and Molecular Biology-INSACOG | Ara Sreenivas,Shreekant Verma,Tulasi Nagabandi, Lamuk Zaveri, Amareshwar Vodapalli ,Blessy B John,Viswagithe S L,B Himasri,Valli Nagalakshmi Undamatla,Payel Mukherjee,Sofia Banu,Archana Bharadwaj Siva,Sharath Chandra Thota,Karthik Bharadwaj Tallapaka,Rakesh K Mishra,Divya Tej Sowpati |
| EPI_ISL_2373256, EPI_ISL_2373257, EPI_ISL_2373262, EPI_ISL_2373263, EPI_ISL_2373264, EPI_ISL_2373265, EPI_ISL_2373266, EPI_ISL_2373269, EPI_ISL_2373271, EPI_ISL_2373272, EPI_ISL_2373273, EPI_ISL_2373274, EPI_ISL_2373275 | | | |
| see above | CSIR-Centre for Cellular and Molecular Biology | CSIR-Centre for Cellular and Molecular Biology-INSACOG | Lamuk Zaveri,Tulasi Nagabandi, Ara Sreenivas,Shreekant Verma, Amareshwar Vodapalli ,Blessy B John,Viswagithe S L,B Himasri,Valli Nagalakshmi Undamatla,Payel Mukherjee,Sofia Banu,Archana Bharadwaj Siva,Sharath Chandra Thota,Karthik Bharadwaj Tallapaka,Rakesh K Mishra,Divya Tej Sowpati |
| EPI_ISL_2373276, EPI_ISL_2373277, EPI_ISL_2373278, EPI_ISL_2373279, EPI_ISL_2373280, EPI_ISL_2373281, EPI_ISL_2373282, EPI_ISL_2373286, EPI_ISL_2373287, EPI_ISL_2373288, EPI_ISL_2373289, EPI_ISL_2373290 | | | |
| see above | CSIR-Centre for Cellular and Molecular Biology | CSIR-Centre for Cellular and Molecular Biology-INSACOG | Blessy B John,Viswagithe S L,Tulasi Nagabandi, Lamuk Zaveri,Ara Sreenivas,Shreekant Verma, Amareshwar Vodapalli ,B Himasri,Valli Nagalakshmi Undamatla,Payel Mukherjee,Sofia Banu,Archana Bharadwaj Siva,Sharath Chandra Thota,Karthik Bharadwaj Tallapaka,Rakesh K Mishra,Divya Tej Sowpati |
| EPI_ISL_2373291, EPI_ISL_2373292, EPI_ISL_2373293, EPI_ISL_2373294, EPI_ISL_2373295, EPI_ISL_2373296, EPI_ISL_2373297, EPI_ISL_2373298, EPI_ISL_2373299, EPI_ISL_2373300, EPI_ISL_2373301, EPI_ISL_2373302, EPI_ISL_2373303, EPI_ISL_2373304, EPI_ISL_2373305, EPI_ISL_2373306 | | | |
| see above | CSIR-Centre for Cellular and Molecular Biology | CSIR-Centre for Cellular and Molecular Biology-INSACOG | Ara Sreenivas,Shreekant Verma,Tulasi Nagabandi, Lamuk Zaveri, Amareshwar Vodapalli ,Blessy B John,Viswagithe S L,B Himasri,Valli Nagalakshmi Undamatla,Payel Mukherjee,Sofia Banu,Archana Bharadwaj Siva,Sharath Chandra Thota,Karthik Bharadwaj Tallapaka,Rakesh K Mishra,Divya Tej Sowpati |
| EPI_ISL_2373320, EPI_ISL_2373323 | Mapmygenome | CSIR-Centre for Cellular and Molecular Biology-INSACOG | B Himasri,Valli Nagalakshmi Undamatla,Lamuk Zaveri,Tulasi Nagabandi, Ara Sreenivas,Shreekant Verma, Amareshwar Vodapalli ,Blessy B John,Viswagithe S L,Payel Mukherjee,Sofia Banu,Archana Bharadwaj Siva,Sharath Chandra Thota,Karthik Bharadwaj Tallapaka,Rakesh K Mishra,Divya Tej Sowpati |
| EPI_ISL_2373331, EPI_ISL_2373340 | Mapmygenome | CSIR-Centre for Cellular and Molecular Biology-INSACOG | Tulasi Nagabandi, Lamuk Zaveri,Ara Sreenivas,Shreekant Verma, Amareshwar Vodapalli ,Blessy B John,Viswagithe S L,B Himasri,Valli Nagalakshmi Undamatla,Payel Mukherjee,Sofia Banu,Archana Bharadwaj Siva,Sharath Chandra Thota,Karthik Bharadwaj Tallapaka,Rakesh K Mishra,Divya Tej Sowpati |
| EPI_ISL_2373341, EPI_ISL_2373342 | CSIR-Centre for Cellular and Molecular Biology | CSIR-Centre for Cellular and Molecular Biology-INSACOG | Tulasi Nagabandi, Lamuk Zaveri,Ara Sreenivas,Shreekant Verma, Amareshwar Vodapalli ,Blessy B John,Viswagithe S L,B Himasri,Valli Nagalakshmi Undamatla,Payel Mukherjee,Sofia Banu,Archana Bharadwaj Siva,Sharath Chandra Thota,Karthik Bharadwaj Tallapaka,Rakesh K Mishra,Divya Tej Sowpati |
| EPI_ISL_2373349 | CSIR-Centre for Cellular and Molecular Biology | CSIR-Centre for Cellular and Molecular Biology-INSACOG | Payel Mukherjee,Lamuk Zaveri,Tulasi Nagabandi, Ara Sreenivas,Shreekant Verma, Amareshwar Vodapalli ,Blessy B John,Viswagithe S L,B Himasri,Valli Nagalakshmi Undamatla,Sofia Banu,Archana Bharadwaj Siva,Sharath Chandra Thota,Karthik Bharadwaj Tallapaka,Rakesh K Mishra,Divya Tej Sowpati |
| EPI_ISL_2373360, EPI_ISL_2373367, EPI_ISL_2373369 | Mapmygenome | CSIR-Centre for Cellular and Molecular Biology-INSACOG | Payel Mukherjee,Lamuk Zaveri,Tulasi Nagabandi, Ara Sreenivas,Shreekant Verma, Amareshwar Vodapalli ,Blessy B John,Viswagithe S L,B Himasri,Valli Nagalakshmi Undamatla,Sofia Banu,Archana Bharadwaj Siva,Sharath Chandra Thota,Karthik Bharadwaj Tallapaka,Rakesh K Mishra,Divya Tej Sowpati |
| EPI_ISL_2373377, EPI_ISL_2373378, EPI_ISL_2373379 | CSIR-Centre for Cellular and Molecular Biology | CSIR-Centre for Cellular and Molecular Biology-INSACOG | B Himasri,Valli Nagalakshmi Undamatla,Lamuk Zaveri,Tulasi Nagabandi, Ara Sreenivas,Shreekant Verma, Amareshwar Vodapalli ,Blessy B John,Viswagithe S L,Payel Mukherjee,Sofia Banu,Archana Bharadwaj Siva,Sharath Chandra Thota,Karthik Bharadwaj Tallapaka,Rakesh K Mishra,Divya Tej Sowpati |
| EPI_ISL_2373380, EPI_ISL_2373381, EPI_ISL_2373384, EPI_ISL_2373385, EPI_ISL_2373386, EPI_ISL_2373389, EPI_ISL_2373390, EPI_ISL_2373391, EPI_ISL_2373392, EPI_ISL_2373393, EPI_ISL_2373394, EPI_ISL_2373395, EPI_ISL_2373396 | | | |
| see above | CSIR-Centre for Cellular and Molecular Biology | CSIR-Centre for Cellular and Molecular Biology-INSACOG | Lamuk Zaveri,Tulasi Nagabandi, Ara Sreenivas,Shreekant Verma, Amareshwar Vodapalli ,Blessy B John,Viswagithe S L,B Himasri,Valli Nagalakshmi Undamatla,Payel Mukherjee,Sofia Banu,Archana Bharadwaj Siva,Sharath Chandra Thota,Karthik Bharadwaj Tallapaka,Rakesh K Mishra,Divya Tej Sowpati |
| EPI_ISL_2373398, EPI_ISL_2373399, EPI_ISL_2373400, EPI_ISL_2373401, EPI_ISL_2373402, EPI_ISL_2373403, EPI_ISL_2373404, EPI_ISL_2373405, EPI_ISL_2373406, EPI_ISL_2373407, EPI_ISL_2373408, EPI_ISL_2373409, EPI_ISL_2373410, EPI_ISL_2373412 | | | |
| see above | CSIR-Centre for Cellular and Molecular Biology | CSIR-Centre for Cellular and Molecular Biology-INSACOG | Blessy B John,Viswagithe S L,Tulasi Nagabandi, Lamuk Zaveri,Ara Sreenivas,Shreekant Verma, Amareshwar Vodapalli ,B Himasri,Valli Nagalakshmi Undamatla,Payel Mukherjee,Sofia Banu,Archana Bharadwaj Siva,Sharath Chandra Thota,Karthik Bharadwaj Tallapaka,Rakesh K Mishra,Divya Tej Sowpati |
| EPI_ISL_2373413, EPI_ISL_2373415, EPI_ISL_2373416, EPI_ISL_2373418, EPI_ISL_2373419, EPI_ISL_2373421, EPI_ISL_2373422, EPI_ISL_2373423, EPI_ISL_2373424, EPI_ISL_2373425, EPI_ISL_2373426, EPI_ISL_2373427, EPI_ISL_2373428, EPI_ISL_2373429 | | | |
| see above | CSIR-Centre for Cellular and Molecular Biology | CSIR-Centre for Cellular and Molecular Biology-INSACOG | Ara Sreenivas,Shreekant Verma,Tulasi Nagabandi, Lamuk Zaveri, Amareshwar Vodapalli ,Blessy B John,Viswagithe S L,B Himasri,Valli Nagalakshmi Undamatla,Payel Mukherjee,Sofia Banu,Archana Bharadwaj Siva,Sharath Chandra Thota,Karthik Bharadwaj Tallapaka,Rakesh K Mishra,Divya Tej Sowpati |
| EPI_ISL_2373430, EPI_ISL_2373431, EPI_ISL_2373432, EPI_ISL_2373433, EPI_ISL_2373434, EPI_ISL_2373435, EPI_ISL_2373436, EPI_ISL_2373437, EPI_ISL_2373438, EPI_ISL_2373439 | CSIR-Centre for Cellular and Molecular Biology | CSIR-Centre for Cellular and Molecular Biology-INSACOG | Lamuk Zaveri,Tulasi Nagabandi, Ara Sreenivas,Shreekant Verma, Amareshwar Vodapalli ,Blessy B John,Viswagithe S L,B Himasri,Valli Nagalakshmi Undamatla,Payel Mukherjee,Sofia Banu,Archana Bharadwaj Siva,Sharath Chandra Thota,Karthik Bharadwaj Tallapaka,Rakesh K Mishra,Divya Tej Sowpati |
| EPI_ISL_2373440, EPI_ISL_2373441, EPI_ISL_2373442, EPI_ISL_2373443, EPI_ISL_2373444, EPI_ISL_2373445, EPI_ISL_2373446, EPI_ISL_2373448, EPI_ISL_2373449, EPI_ISL_2373450, EPI_ISL_2373451, EPI_ISL_2373452, EPI_ISL_2373453 | | | |
| see above | CSIR-Centre for Cellular and Molecular Biology | CSIR-Centre for Cellular and Molecular Biology-INSACOG | Tulasi Nagabandi, Lamuk Zaveri,Ara Sreenivas,Shreekant Verma, Amareshwar Vodapalli ,Blessy B John,Viswagithe S L,B Himasri,Valli Nagalakshmi Undamatla,Payel Mukherjee,Sofia Banu,Archana Bharadwaj Siva,Sharath Chandra Thota,Karthik Bharadwaj Tallapaka,Rakesh K Mishra,Divya Tej Sowpati |
| EPI_ISL_2373454, EPI_ISL_2373455, EPI_ISL_2373456, EPI_ISL_2373457, EPI_ISL_2373458, EPI_ISL_2373459, EPI_ISL_2373460, EPI_ISL_2373461, EPI_ISL_2373462, EPI_ISL_2373463 | CSIR-Centre for Cellular and Molecular Biology | CSIR-Centre for Cellular and Molecular Biology-INSACOG | Payel Mukherjee,Lamuk Zaveri,Tulasi Nagabandi, Ara Sreenivas,Shreekant Verma, Amareshwar Vodapalli ,Blessy B John,Viswagithe S L,B Himasri,Valli Nagalakshmi Undamatla,Sofia Banu,Archana Bharadwaj Siva,Sharath Chandra Thota,Karthik Bharadwaj Tallapaka,Rakesh K Mishra,Divya Tej Sowpati |
| EPI_ISL_2373464, EPI_ISL_2373465, EPI_ISL_2373466, EPI_ISL_2373467, EPI_ISL_2373468, EPI_ISL_2373469, EPI_ISL_2373470, EPI_ISL_2373471, EPI_ISL_2373472, EPI_ISL_2373473, EPI_ISL_2373474, EPI_ISL_2373475, EPI_ISL_2373476, EPI_ISL_2373477, EPI_ISL_2373478 | | | |
| see above | CSIR-Centre for Cellular and Molecular Biology | CSIR-Centre for Cellular and Molecular Biology-INSACOG | Tulasi Nagabandi, Lamuk Zaveri,Ara Sreenivas,Shreekant Verma, Amareshwar Vodapalli ,Blessy B John,Viswagithe S L,B Himasri,Valli Nagalakshmi Undamatla,Payel Mukherjee,Sofia Banu,Archana Bharadwaj Siva,Sharath Chandra Thota,Karthik Bharadwaj Tallapaka,Rakesh K Mishra,Divya Tej Sowpati |
| EPI_ISL_2373479, EPI_ISL_2373480, EPI_ISL_2373481, EPI_ISL_2373482, EPI_ISL_2373483, EPI_ISL_2373484, EPI_ISL_2373485, EPI_ISL_2373486 | CSIR-Centre for Cellular and Molecular Biology | CSIR-Centre for Cellular and Molecular Biology-INSACOG | Payel Mukherjee,Lamuk Zaveri,Tulasi Nagabandi, Ara Sreenivas,Shreekant Verma, Amareshwar Vodapalli ,Blessy B John,Viswagithe S L,B Himasri,Valli Nagalakshmi Undamatla,Sofia Banu,Archana Bharadwaj Siva,Sharath Chandra Thota,Karthik Bharadwaj Tallapaka,Rakesh K Mishra,Divya Tej Sowpati |
| EPI_ISL_2373487 | CSIR-Centre for Cellular and Molecular Biology | CSIR-Centre for Cellular and Molecular Biology-INSACOG | B Himasri,Valli Nagalakshmi Undamatla,Lamuk Zaveri,Tulasi Nagabandi, Ara Sreenivas,Shreekant Verma, Amareshwar Vodapalli ,Blessy B John,Viswagithe S L,Payel Mukherjee,Sofia Banu,Archana Bharadwaj Siva,Sharath Chandra Thota,Karthik Bharadwaj Tallapaka,Rakesh K Mishra,Divya Tej Sowpati |
| EPI_ISL_2373489, EPI_ISL_2373490, EPI_ISL_2373491, EPI_ISL_2373492, EPI_ISL_2373493, EPI_ISL_2373494, EPI_ISL_2373495, EPI_ISL_2373496, EPI_ISL_2373497, EPI_ISL_2373498, EPI_ISL_2373499, EPI_ISL_2373500 | | | |
| see above | CSIR-Centre for Cellular and Molecular Biology | CSIR-Centre for Cellular and Molecular Biology-INSACOG | Payel Mukherjee,Lamuk Zaveri,Tulasi Nagabandi, Ara Sreenivas,Shreekant Verma, Amareshwar Vodapalli ,Blessy B John,Viswagithe S L,B Himasri,Valli Nagalakshmi Undamatla,Sofia Banu,Archana Bharadwaj Siva,Sharath Chandra Thota,Karthik Bharadwaj Tallapaka,Rakesh K Mishra,Divya Tej Sowpati |
| EPI_ISL_2373501 | Banaras Hindu University | CSIR-Centre for Cellular and Molecular Biology-INSACOG | Royana Singh, Saurabh Singh, Jaya Chakraborty, Priyoneel Basu, Chetan Sahni, Deepa Devadas, Saumya Singh, Manpreet Kaur, Nitish Kumar Singh, Abhay Kumar Yadav, Ajay Kumar Yadav, Ashish, Shani Vishwakarma,Payel Mukherjee,Lamuk Zaveri,Tulasi Nagabandi, ,Blessy B John,Viswagithe S L,B Himasri,Sofia Banu,Archana Bharadwaj Siva,Sharath Chandra Thota,Gyaneshwer Chaubey,Karthik Bharadwaj Tallapaka,Rakesh K Mishra,Divya Tej Sowpati |
| EPI_ISL_2373502 | Banaras Hindu University | CSIR-Centre for Cellular and Molecular Biology-INSACOG | Royana Singh,Saumya Singh, Manpreet Kaur, Nitish Kumar Singh, Abhay Kumar Yadav, Ajay Kumar Yadav, Ashish, Shani Vishwakarma,B Himasri,Amareshwar Vodapalli ,Blessy B John,Viswagithe S L,Payel Mukherjee,Sofia Banu,Archana Bharadwaj Siva,Sharath Chandra Thota,Gyaneshwer Chaubey,Karthik Bharadwaj Tallapaka,Rakesh K Mishra,Divya Tej Sowpati |
| EPI_ISL_2373503, EPI_ISL_2373504, EPI_ISL_2373505, EPI_ISL_2373506, EPI_ISL_2373509, EPI_ISL_2373510, EPI_ISL_2373512, EPI_ISL_2373514, EPI_ISL_2373515, EPI_ISL_2373517, EPI_ISL_2373519 | | | |

| | | | |
|---|---|---|---|
| see above | Banaras Hindu University | CSIR-Centre for Cellular and Molecular Biology-INSACOG | Royana Singh, Saurabh Singh, Jaya Chakraborty, Priyoneel Basu, Chetan Sahni, Deepa Devadas, Saumya Singh, Manpreet Kaur, Nitish Kumar Singh, Abhay Kumar Yadav, Ajay Kumar Yadav, Ashish, Shani Vishwakarma,Payel Mukherjee,Lamuk Zaveri,Tulasi Nagabandi, ,Blessy B John,Viswagithe S L,B Himasri,Sofia Banu,Archana Bharadwaj Siva,Sharath Chandra Thota,Gyaneshwer Chaubey,Karthik Bharadwaj Tallapaka,Rakesh K Mishra,Divya Tej Sowpati |
| EPI_ISL_2373521, EPI_ISL_2373522, EPI_ISL_2373523, EPI_ISL_2373524, EPI_ISL_2373525, EPI_ISL_2373526, EPI_ISL_2373527, EPI_ISL_2373529, EPI_ISL_2373530, EPI_ISL_2373531, EPI_ISL_2373532, EPI_ISL_2373533, EPI_ISL_2373534, EPI_ISL_2373535, EPI_ISL_2373536, EPI_ISL_2373545 | | | |
| see above | Banaras Hindu University | CSIR-Centre for Cellular and Molecular Biology-INSACOG | Surendra Pratap Mishra, Royana Singh,Shivani Mishra, Shivam Tiwari, Gunjan Rai, Umesh Choudhary, Maneesha Upadhyay,Lamuk Zaveri,Tulasi Nagabandi, Ara Sreenivas,Shreekant Verma, Amareshwar Vodapalli ,Valli Nagalakshmi Undamatla,Payel Mukherjee,Archana Bharadwaj Siva,Sharath Chandra Thota,Gyaneshwer Chaubey,Karthik Bharadwaj Tallapaka,Rakesh K Mishra,Divya Tej Sowpati |
| EPI_ISL_2373546, EPI_ISL_2373547, EPI_ISL_2373554, EPI_ISL_2373559, EPI_ISL_2373561, EPI_ISL_2373562, EPI_ISL_2373567 | Banaras Hindu University | CSIR-Centre for Cellular and Molecular Biology-INSACOG | Prajjval Pratap Singh ,Yashvant Patel, Royana Singh,Debashruti Das , Abhishek Pathak , Tribhuwan Mohan Mohapatra, Vijay Kumar Shukla,Tulasi Nagabandi, Lamuk Zaveri, Amareshwar Vodapalli ,Blessy B John,Viswagithe S L,Payel Mukherjee,Sofia Banu,Archana Bharadwaj Siva,Sharath Chandra Thota,Gyaneshwer Chaubey,Karthik Bharadwaj Tallapaka,Rakesh K Mishra,Divya Tej Sowpati |
| EPI_ISL_2373576, EPI_ISL_2373590, EPI_ISL_2373593, EPI_ISL_2373601 | Banaras Hindu University | CSIR-Centre for Cellular and Molecular Biology-INSACOG | Royana Singh,Saumya Singh, Manpreet Kaur, Nitish Kumar Singh, Abhay Kumar Yadav, Ajay Kumar Yadav, Ashish, Shani Vishwakarma,B Himasri,Amareshwar Vodapalli ,Blessy B John,Viswagithe S L,Payel Mukherjee,Sofia Banu,Archana Bharadwaj Siva,Sharath Chandra Thota,Gyaneshwer Chaubey,Karthik Bharadwaj Tallapaka,Rakesh K Mishra,Divya Tej Sowpati |
| EPI_ISL_2373603 | Banaras Hindu University | CSIR-Centre for Cellular and Molecular Biology-INSACOG | Royana Singh, Saurabh Singh, Jaya Chakraborty, Priyoneel Basu, Chetan Sahni, Deepa Devadas, Saumya Singh, Manpreet Kaur, Nitish Kumar Singh, Abhay Kumar Yadav, Ajay Kumar Yadav, Ashish, Shani Vishwakarma,Payel Mukherjee,Lamuk Zaveri,Tulasi Nagabandi, ,Blessy B John,Viswagithe S L,B Himasri,Sofia Banu,Archana Bharadwaj Siva,Sharath Chandra Thota,Gyaneshwer Chaubey,Karthik Bharadwaj Tallapaka,Rakesh K Mishra,Divya Tej Sowpati |
| EPI_ISL_2373615, EPI_ISL_2373620 | Mapmygenome | CSIR-Centre for Cellular and Molecular Biology - INSACOG | Lamuk Zaveri, Ara Sreenivas, Onkar Kulkarni, Sofia Banu, Shreekant Verma, Amareshwar Vodapalli , Viswagithe S L, B Himasri, Sharath Chandra Thota, Karthik Bharadwaj Tallapaka, Rakesh K Mishra,Divya Tej Sowpati |
| EPI_ISL_2373628, EPI_ISL_2373635 | Mapmygenome | CSIR-Centre for Cellular and Molecular Biology - INSACOG | Onkar Kulkarni, Sofia Banu, Lamuk Zaveri, Ara Sreenivas, Shreekant Verma, Amareshwar Vodapalli , Viswagithe S L, B Himasri, Sharath Chandra Thota, Karthik Bharadwaj Tallapaka, Rakesh K Mishra,Divya Tej Sowpati |
| EPI_ISL_2373640, EPI_ISL_2373643 | Mapmygenome | CSIR-Centre for Cellular and Molecular Biology - INSACOG | Ara Sreenivas, Onkar Kulkarni, Lamuk Zaveri, Sofia Banu, Shreekant Verma, Amareshwar Vodapalli , Viswagithe S L, B Himasri, Sharath Chandra Thota, Karthik Bharadwaj Tallapaka, Rakesh K Mishra,Divya Tej Sowpati |
| EPI_ISL_2373743, EPI_ISL_2373746 | AX BIO OCEAN | CNR Virus des Infections Respiratoires - France SUD | Antonin Bal, Gregory Destras, Gwendolyne Burfin, Hadrien Regue, Quentin Semanas, Martine Valette, Bruno Lina, Laurence Josset |
| EPI_ISL_2373878, EPI_ISL_2373880, EPI_ISL_2373881 | MIRIALIS CLUSES BECHET | CNR Virus des Infections Respiratoires - France SUD | Antonin Bal, Gregory Destras, Gwendolyne Burfin, Hadrien Regue, Quentin Semanas, Martine Valette, Bruno Lina, Laurence Josset |
| EPI_ISL_2374497, EPI_ISL_2374508, EPI_ISL_2374509, EPI_ISL_2374510, EPI_ISL_2374511, EPI_ISL_2374513 | Furst Medical Laboratory | Norwegian Institute of Public Health, Department of Virology | Kathrine Stene-Johansen, Kamilla Heddeland Instefjord, Hilde Elshaug, Garcia Llorente Ignacio, Jon Bråte, Line Victoria Moen, Engebretsen Serina Beate,Pedersen Benedikte Nevjen, Debech Nadia, Atiya R Ali,Marie Paulsen Madsen, Rasmus Riis Kopperud, Hilde Vollan, Karoline Bragstad, Olav Hungnes |
| EPI_ISL_2374542 | Akershus University Hospital, Department for Microbiology and Infectious Disease Control | Norwegian Institute of Public Health, Department of Virology | Kathrine Stene-Johansen, Kamilla Heddeland Instefjord, Hilde Elshaug, Garcia Llorente Ignacio, Jon Bråte, Line Victoria Moen, Engebretsen Serina Beate,Pedersen Benedikte Nevjen, Debech Nadia, Atiya R Ali,Marie Paulsen Madsen, Rasmus Riis Kopperud, Hilde Vollan, Karoline Bragstad, Olav Hungnes |
| EPI_ISL_2375015, EPI_ISL_2375060 | Viollier AG | Department of Biosystems Science and Engineering, ETH Zürich | Chaoran Chen, Sarah Nadeau, Catharine Aquino, Ivan Topolsky, Philipp Jablonski, Lara Fuhrmann, David Dreifuss, Katharina Jahn, Daniel Ehrsam, Isabel Stürmer, Andreia Cabral de Gouvea, Maria Domenica Moccia, Simon Grüter, Timothy Sykes, Lennart Opitz, Griffin White, Laura Neff, Doris Popovic, Andrea Patrignani, Jay Tracy, Ralph Schlapbach, Christiane Beckmann, Maurice Redondo, Olivier Kobel, Christoph Noppen, Sophie Seidel, Noemie Santamaria de Souza, Niko Beerenwinkel, Tanja Stadler |
| EPI_ISL_2375080, EPI_ISL_2375122, EPI_ISL_2375177, EPI_ISL_2375211, EPI_ISL_2375282 | Viollier AG | Department of Biosystems Science and Engineering, ETH Zürich | Christian Beisel, Sarah Nadeau, Chaoran Chen, Ivan Topolsky, Philipp Jablonski, Lara Fuhrmann, David Dreifuss, Katharina Jahn, Rebecca Denes, Mirjam Feldkamp, Ina Nissen, Natascha Santacroce, Elodie Burcklen, Christiane Beckmann, Maurice Redondo, Olivier Kobel, Christoph Noppen, Sophie Seidel, Noemie Santamaria de Souza, Niko Beerenwinkel, Tanja Stadler |
| EPI_ISL_2376038 | University of Michigan Clinical Microbiology Laboratory | Lauring Lab, University of Michigan, Department of Microbiology and Immunology | Gilbert |
| EPI_ISL_2376127 | NJDOH, Public Health and Environmental Laboratories | NJ_PHEL | Lindsey Bodnar, Shiv K. Verma, Jacquelyn Deverell, Dana Woell, Allison Roder, Byeong Jeong |
| EPI_ISL_2376218 | Heilig hart Lier | Imeda Hospital | Johan Frans, Dagmar Obbels, Hanne Valgaeren |
| EPI_ISL_2376285, EPI_ISL_2376288 | ULSS 4 Veneto Orientale | Istituto Zooprofilattico Sperimentale delle Venezie | Adelaide Milani, Alessia Schivo, Annalisa Salviato, Elisa Palumbo, Erika Giorgia Quaranta, Luca Tassoni, Ambra Pastori, Edoardo Giussani, Alice Fusaro, Isabella Monne, Calogero Terregino, Antonia Ricci |
| EPI_ISL_2376289, EPI_ISL_2376290, EPI_ISL_2376291 | ULSS 3 Venezia | Istituto Zooprofilattico Sperimentale delle Venezie | Adelaide Milani, Alessia Schivo, Annalisa Salviato, Elisa Palumbo, Erika Giorgia Quaranta, Luca Tassoni, Ambra Pastori, Edoardo Giussani, Alice Fusaro, Isabella Monne, Calogero Terregino, Antonia Ricci |
| EPI_ISL_2376292, EPI_ISL_2376294 | ULSS 2 Marca Trevigiana | Istituto Zooprofilattico Sperimentale delle Venezie | Adelaide Milani, Alessia Schivo, Annalisa Salviato, Elisa Palumbo, Erika Giorgia Quaranta, Luca Tassoni, Ambra Pastori, Edoardo Giussani, Alice Fusaro, Isabella Monne, Calogero Terregino, Antonia Ricci |
| EPI_ISL_2376296, EPI_ISL_2376297 | ULSS 8 Berica | Istituto Zooprofilattico Sperimentale delle Venezie | Adelaide Milani, Alessia Schivo, Annalisa Salviato, Elisa Palumbo, Erika Giorgia Quaranta, Luca Tassoni, Ambra Pastori, Edoardo Giussani, Alice Fusaro, Isabella Monne, Calogero Terregino, Antonia Ricci |
| EPI_ISL_2376379, EPI_ISL_2376380 | ULSS 3 Venezia | Istituto Zooprofilattico Sperimentale delle Venezie | Adelaide Milani, Alessia Schivo, Annalisa Salviato, Elisa Palumbo, Erika Giorgia Quaranta, Luca Tassoni, Ambra Pastori, Edoardo Giussani, Alice Fusaro, Isabella Monne, Calogero Terregino, Antonia Ricci |
| EPI_ISL_2376402, EPI_ISL_2376404, EPI_ISL_2376405, EPI_ISL_2376406, EPI_ISL_2376407, EPI_ISL_2376408, EPI_ISL_2376409, EPI_ISL_2376410 | ULSS 2 Marca Trevigiana | Istituto Zooprofilattico Sperimentale delle Venezie | Adelaide Milani, Alessia Schivo, Annalisa Salviato, Elisa Palumbo, Erika Giorgia Quaranta, Luca Tassoni, Ambra Pastori, Edoardo Giussani, Alice Fusaro, Isabella Monne, Calogero Terregino, Antonia Ricci |
| EPI_ISL_2376420 | Ospedale Magalini | Istituto Zooprofilattico Sperimentale delle Venezie | Adelaide Milani, Alessia Schivo, Annalisa Salviato, Elisa Palumbo, Erika Giorgia Quaranta, Luca Tassoni, Ambra Pastori, Edoardo Giussani, Alice Fusaro, Isabella Monne, Calogero Terregino, Antonia Ricci |
| EPI_ISL_2376421 | CHLC | Instituto Nacional de Saude (INSA) | Borges et al |
| EPI_ISL_2376424 | GA Department of Public Health Laboratory | Genomics and Discovery, Respiratory Viruses Branch, Division of Viral Diseases, Centers for Disease Control and Prevention | Anna Kelleher, Ying Tao, Yan Li, Jing Zhang, Brian Lynch, Krista Queen, Anna Uehara, Peter Cook, Han Jia Justin Ng, Rachel Marine, Clinton R. Paden, Dhwani Batra, Haibin Wang, Tara Coalter, Jasmine Padilla, Morgan Davis, Mili Sheth, Sarah Nobles, Mark Burroughs, Justin Lee, Adam Retchless, Suxiang Tong |
| EPI_ISL_2377203, EPI_ISL_2377204, EPI_ISL_2377205, EPI_ISL_2377206, EPI_ISL_2377207, EPI_ISL_2377208 | WSSE Katowice | 1. National Institute of Public Health - National Institute of Hygiene; 2. Eurofins Genomics Europe Sequencing GmbH | Wołkowicz Tomasz, Zacharczuk Katarzyna, Sadkowska-Todys Małgorzata, Gierczyńki Rafał, Eurofins Genomics Europe Sequencing Team, ECDC COVID-19 WGS support team |
| EPI_ISL_2377636 | Maryland Genomics, Institute for Genome Sciences, University of Maryland School of Medicine | Maryland Genomics, Institute for Genome Sciences, University of Maryland School of Medicine | Tallon, Luke J; Sadzewicz, Lisa D; Humphrys, Mike; Ott, Sandra; Roussey, Holly; Mehta, Aditya; Vavikolanu, Kranthi; Fraser, Claire M; Ravel, Jacques |
| EPI_ISL_2377736 | Texas Department of State Health Services (TXDSHS) | Texas Department of State Health Services (TXDSHS) | Rashmi Tuladhar, Bonnie Oh, Jenny Zhang, Maliha Rahman, Mayela Pedrueza, Anita Pokharel, Karen Bobier, Lorraine Rodriguez, Myong Koag, Chun Wang, Rachel Lee, Grace Kubin |
| EPI_ISL_2377818, EPI_ISL_2377852 | UCK | 1. Tricity SARS-CoV-2 sequencing consortium: University of Gdansk, Medical University of Gdansk, Vaxican Ltd., Invicta Ltd. 2. National Institute of Public Health - National Institute of Hygiene, Warsaw, Poland | Maciej Kosinski, Celina Cybulska, Krystyna Bienkowska Szewczyk, Maciej Grzybek, Karolina Gackowska, Marcin Lubocki, Katarzyna Groth, Lukasz Rabalski, Katarzyna Zacharczuk, Magdalena Nowakowska, Małgorzata Sadkowska-Todys, Tomasz Wołkowicz |
| EPI_ISL_2377869, EPI_ISL_2377875 | WSSE w Gdańsku | 1. Tricity SARS-CoV-2 sequencing consortium: University of Gdansk, Medical University of Gdansk, Vaxican Ltd., Invicta Ltd. 2. National Institute of Public Health - National Institute of Hygiene, Warsaw, Poland | Maciej Kosinski, Celina Cybulska, Krystyna Bienkowska Szewczyk, Maciej Grzybek, Karolina Gackowska, Marcin Lubocki, Katarzyna Groth, Lukasz Rabalski, Katarzyna Zacharczuk, Magdalena Nowakowska, Małgorzata Sadkowska-Todys, Tomasz Wołkowicz |
| EPI_ISL_2377955, EPI_ISL_2377967, EPI_ISL_2378174, EPI_ISL_2378500 | State Testing Facility | Altius Institute for Biomedical Research | Daniel Bates, Rebecca Bruders, Michael Buckley, Mark Frerker, Amanda Gale, Clem Green, Muhammad Halimun, Kneshay Harper, Matt Hartman, Alex Isner, Audra Johnson, Jessica Kunder, Lauren Mitchell, Jemma Nelson, Alex Nguyen, Sofia Olsson, Sadie Patraw, Tobias Ragoczy, Joshua Richards, Jean Robinston, Jacob Rodriguez, John Stamatoyannopoulos, Eric Thorland, Julia Wald |
| EPI_ISL_2378732 | Hiroyuki Asakura Tokyo Metropolitan Institute of Public Health Department of Microbiology | Hiroyuki Asakura Tokyo Metropolitan Institute of Public Health Department of Microbiology | Asakura,H., Kumagai,R., Yoshida,I., Fujiwara,T., Miyake,H.,Nagashima,M., Suzuki,J. and Sadamasu,K. |

| | | | |
|---|---|---|---|
| EPI_ISL_2378856, EPI_ISL_2378863 | Washington State Department of Health Public Health Laboratories | Washington State Department of Health Public Health Laboratories | Drew MacKellar, Philip Dykema, Denny Russell, Joenice Gonzalez, Hannah Gray, Geoff Melly, Vanessa De Los Santos, Darren Lucas, JohnAric Peterson, Avi Singh, Rebecca Cao |
| EPI_ISL_2379203 | National Center of Infectious and Parasitic Diseases | National Center of Infectious and Parasitic Diseases | Alexiev et al |
| EPI_ISL_2379315, EPI_ISL_2379316, EPI_ISL_2379318, EPI_ISL_2379320, EPI_ISL_2379321 | RAILWAY | INSACOG-KA, NIMHANS | Chitra Pattabiraman, Anson Kunjumon George,Darshan Sreenivas,Pramada Prasad, Ananthapadmanabha Kotambail, Chetan G K, Gautham Arunachal Udupi, Anita S Desai, V Ravi |
| EPI_ISL_2379322, EPI_ISL_2379323, EPI_ISL_2379324, EPI_ISL_2379325, EPI_ISL_2379326, EPI_ISL_2379327, EPI_ISL_2379328, EPI_ISL_2379329, EPI_ISL_2379330 | AGRAHARA PHC | INSACOG-KA, NIMHANS | Chitra Pattabiraman, Anson Kunjumon George,Darshan Sreenivas,Pramada Prasad, Ananthapadmanabha Kotambail, Chetan G K, Gautham Arunachal Udupi, Anita S Desai, V Ravi |
| EPI_ISL_2379347 | RAILWAY H | INSACOG-KA, NIMHANS | Chitra Pattabiraman, Anson Kunjumon George,Darshan Sreenivas,Pramada Prasad, Ananthapadmanabha Kotambail, Chetan G K, Gautham Arunachal Udupi, Anita S Desai, V Ravi |
| EPI_ISL_2379349, EPI_ISL_2379350 | HOSA HALLI | INSACOG-KA, NIMHANS | Chitra Pattabiraman, Anson Kunjumon George,Darshan Sreenivas,Pramada Prasad, Ananthapadmanabha Kotambail, Chetan G K, Gautham Arunachal Udupi, Anita S Desai, V Ravi |
| EPI_ISL_2379353 | GOVIDRAJ NAGAR | INSACOG-KA, NIMHANS | Chitra Pattabiraman, Anson Kunjumon George,Darshan Sreenivas,Pramada Prasad, Ananthapadmanabha Kotambail, Chetan G K, Gautham Arunachal Udupi, Anita S Desai, V Ravi |
| EPI_ISL_2379356 | AD HALLI | INSACOG-KA, NIMHANS | Chitra Pattabiraman, Anson Kunjumon George,Darshan Sreenivas,Pramada Prasad, Ananthapadmanabha Kotambail, Chetan G K, Gautham Arunachal Udupi, Anita S Desai, V Ravi |
| EPI_ISL_2379357 | NIMHANS | INSACOG-KA, NIMHANS | Chitra Pattabiraman, Anson Kunjumon George,Darshan Sreenivas,Pramada Prasad, Ananthapadmanabha Kotambail, Chetan G K, Gautham Arunachal Udupi, Anita S Desai, V Ravi |
| EPI_ISL_2379358, EPI_ISL_2379359 | AD HALLI | INSACOG-KA, NIMHANS | Chitra Pattabiraman, Anson Kunjumon George,Darshan Sreenivas,Pramada Prasad, Ananthapadmanabha Kotambail, Chetan G K, Gautham Arunachal Udupi, Anita S Desai, V Ravi |
| EPI_ISL_2379360 | NIMHANS | INSACOG-KA, NIMHANS | Chitra Pattabiraman, Anson Kunjumon George,Darshan Sreenivas,Pramada Prasad, Ananthapadmanabha Kotambail, Chetan G K, Gautham Arunachal Udupi, Anita S Desai, V Ravi |
| EPI_ISL_2379361, EPI_ISL_2379362 | AD HALLI | INSACOG-KA, NIMHANS | Chitra Pattabiraman, Anson Kunjumon George,Darshan Sreenivas,Pramada Prasad, Ananthapadmanabha Kotambail, Chetan G K, Gautham Arunachal Udupi, Anita S Desai, V Ravi |
| EPI_ISL_2379363, EPI_ISL_2379364, EPI_ISL_2379365, EPI_ISL_2379366 | HOSAHALLI | INSACOG-KA, NIMHANS | Chitra Pattabiraman, Anson Kunjumon George,Darshan Sreenivas,Pramada Prasad, Ananthapadmanabha Kotambail, Chetan G K, Gautham Arunachal Udupi, Anita S Desai, V Ravi |
| EPI_ISL_2379367, EPI_ISL_2379368, EPI_ISL_2379369, EPI_ISL_2379370, EPI_ISL_2379371 | MOODALAPALYA | INSACOG-KA, NIMHANS | Chitra Pattabiraman, Anson Kunjumon George,Darshan Sreenivas,Pramada Prasad, Ananthapadmanabha Kotambail, Chetan G K, Gautham Arunachal Udupi, Anita S Desai, V Ravi |
| EPI_ISL_2379372, EPI_ISL_2379373, EPI_ISL_2379374 | GOVIDRAJNAGAR | INSACOG-KA, NIMHANS | Chitra Pattabiraman, Anson Kunjumon George,Darshan Sreenivas,Pramada Prasad, Ananthapadmanabha Kotambail, Chetan G K, Gautham Arunachal Udupi, Anita S Desai, V Ravi |
| EPI_ISL_2379375 | RAILWAY H | INSACOG-KA, NIMHANS | Chitra Pattabiraman, Anson Kunjumon George,Darshan Sreenivas,Pramada Prasad, Ananthapadmanabha Kotambail, Chetan G K, Gautham Arunachal Udupi, Anita S Desai, V Ravi |
| EPI_ISL_2379376 | JAYDEVA | INSACOG-KA, NIMHANS | Chitra Pattabiraman, Anson Kunjumon George,Darshan Sreenivas,Pramada Prasad, Ananthapadmanabha Kotambail, Chetan G K, Gautham Arunachal Udupi, Anita S Desai, V Ravi |
| EPI_ISL_2379377 | RAILWAY H | INSACOG-KA, NIMHANS | Chitra Pattabiraman, Anson Kunjumon George,Darshan Sreenivas,Pramada Prasad, Ananthapadmanabha Kotambail, Chetan G K, Gautham Arunachal Udupi, Anita S Desai, V Ravi |
| EPI_ISL_2379378 | AD HALLI | INSACOG-KA, NIMHANS | Chitra Pattabiraman, Anson Kunjumon George,Darshan Sreenivas,Pramada Prasad, Ananthapadmanabha Kotambail, Chetan G K, Gautham Arunachal Udupi, Anita S Desai, V Ravi |
| EPI_ISL_2379379 | PANTHRPALYA | INSACOG-KA, NIMHANS | Chitra Pattabiraman, Anson Kunjumon George,Darshan Sreenivas,Pramada Prasad, Ananthapadmanabha Kotambail, Chetan G K, Gautham Arunachal Udupi, Anita S Desai, V Ravi |
| EPI_ISL_2379380 | HOSAHALLI | INSACOG-KA, NIMHANS | Chitra Pattabiraman, Anson Kunjumon George,Darshan Sreenivas,Pramada Prasad, Ananthapadmanabha Kotambail, Chetan G K, Gautham Arunachal Udupi, Anita S Desai, V Ravi |
| EPI_ISL_2379381, EPI_ISL_2379382 | MOODALAPALYA | INSACOG-KA, NIMHANS | Chitra Pattabiraman, Anson Kunjumon George,Darshan Sreenivas,Pramada Prasad, Ananthapadmanabha Kotambail, Chetan G K, Gautham Arunachal Udupi, Anita S Desai, V Ravi |
| EPI_ISL_2379383 | NETHAJI CIRCLE | INSACOG-KA, NIMHANS | Chitra Pattabiraman, Anson Kunjumon George,Darshan Sreenivas,Pramada Prasad, Ananthapadmanabha Kotambail, Chetan G K, Gautham Arunachal Udupi, Anita S Desai, V Ravi |
| EPI_ISL_2379384, EPI_ISL_2379385, EPI_ISL_2379386 | GANDHIGRAMA | INSACOG-KA, NIMHANS | Chitra Pattabiraman, Anson Kunjumon George,Darshan Sreenivas,Pramada Prasad, Ananthapadmanabha Kotambail, Chetan G K, Gautham Arunachal Udupi, Anita S Desai, V Ravi |
| EPI_ISL_2379387 | NAGAPPA BLOCK | INSACOG-KA, NIMHANS | Chitra Pattabiraman, Anson Kunjumon George,Darshan Sreenivas,Pramada Prasad, Ananthapadmanabha Kotambail, Chetan G K, Gautham Arunachal Udupi, Anita S Desai, V Ravi |
| EPI_ISL_2379388 | NIMHANS | INSACOG-KA, NIMHANS | Chitra Pattabiraman, Anson Kunjumon George,Darshan Sreenivas,Pramada Prasad, Ananthapadmanabha Kotambail, Chetan G K, Gautham Arunachal Udupi, Anita S Desai, V Ravi |
| EPI_ISL_2379389 | RAILWAY H | INSACOG-KA, NIMHANS | Chitra Pattabiraman, Anson Kunjumon George,Darshan Sreenivas,Pramada Prasad, Ananthapadmanabha Kotambail, Chetan G K, Gautham Arunachal Udupi, Anita S Desai, V Ravi |
| EPI_ISL_2379390 | AD HALLI | INSACOG-KA, NIMHANS | Chitra Pattabiraman, Anson Kunjumon George,Darshan Sreenivas,Pramada Prasad, Ananthapadmanabha Kotambail, Chetan G K, Gautham Arunachal Udupi, Anita S Desai, V Ravi |
| EPI_ISL_2379391 | SHANKARNAGAR | INSACOG-KA, NIMHANS | Chitra Pattabiraman, Anson Kunjumon George,Darshan Sreenivas,Pramada Prasad, Ananthapadmanabha Kotambail, Chetan G K, Gautham Arunachal Udupi, Anita S Desai, V Ravi |
| EPI_ISL_2379393 | PEENYA KAMALANAGAR | INSACOG-KA, NIMHANS | Chitra Pattabiraman, Anson Kunjumon George,Darshan Sreenivas,Pramada Prasad, Ananthapadmanabha Kotambail, Chetan G K, Gautham Arunachal Udupi, Anita S Desai, V Ravi |
| EPI_ISL_2379394, EPI_ISL_2379395 | ASHOKPURAM | INSACOG-KA, NIMHANS | Chitra Pattabiraman, Anson Kunjumon George,Darshan Sreenivas,Pramada Prasad, Ananthapadmanabha Kotambail, Chetan G K, Gautham Arunachal Udupi, Anita S Desai, V Ravi |
| EPI_ISL_2379397 | RAILWAY | INSACOG-KA, NIMHANS | Chitra Pattabiraman, Anson Kunjumon George,Darshan Sreenivas,Pramada Prasad, Ananthapadmanabha Kotambail, Chetan G K, Gautham Arunachal Udupi, Anita S Desai, V Ravi |
| EPI_ISL_2379398 | NIMHANS | INSACOG-KA, NIMHANS | Chitra Pattabiraman, Anson Kunjumon George,Darshan Sreenivas,Pramada Prasad, Ananthapadmanabha Kotambail, Chetan G K, Gautham Arunachal Udupi, Anita S Desai, V Ravi |
| EPI_ISL_2379400 | PEENYA KAMALA NAGAR | INSACOG-KA, NIMHANS | Chitra Pattabiraman, Anson Kunjumon George,Darshan Sreenivas,Pramada Prasad, Ananthapadmanabha Kotambail, Chetan G K, Gautham Arunachal Udupi, Anita S Desai, V Ravi |
| EPI_ISL_2379401 | ASHOKPURAM | INSACOG-KA, NIMHANS | Chitra Pattabiraman, Anson Kunjumon George,Darshan Sreenivas,Pramada Prasad, Ananthapadmanabha Kotambail, Chetan G K, Gautham Arunachal Udupi, Anita S Desai, V Ravi |
| EPI_ISL_2379402, EPI_ISL_2379403 | MAHALAKSHMI LAYOUT | INSACOG-KA, NIMHANS | Chitra Pattabiraman, Anson Kunjumon George,Darshan Sreenivas,Pramada Prasad, Ananthapadmanabha Kotambail, Chetan G K, Gautham Arunachal Udupi, Anita S Desai, V Ravi |
| EPI_ISL_2379404, EPI_ISL_2379405, EPI_ISL_2379406, EPI_ISL_2379407, EPI_ISL_2379408, EPI_ISL_2379409, EPI_ISL_2379410 | DSU MYSURU | INSACOG-KA, NIMHANS | Chitra Pattabiraman, Anson Kunjumon George,Darshan Sreenivas,Pramada Prasad, Ananthapadmanabha Kotambail, Chetan G K, Gautham Arunachal Udupi, Anita S Desai, V Ravi |
| EPI_ISL_2379412, EPI_ISL_2379413 | DSU MANDYA | INSACOG-KA, NIMHANS | Chitra Pattabiraman, Anson Kunjumon George,Darshan Sreenivas,Pramada Prasad, Ananthapadmanabha Kotambail, Chetan G K, Gautham Arunachal Udupi, Anita S Desai, V Ravi |
| EPI_ISL_2379414 | SHANKARNAGR | INSACOG-KA, NIMHANS | Chitra Pattabiraman, Anson Kunjumon George,Darshan Sreenivas,Pramada Prasad, Ananthapadmanabha Kotambail, Chetan G K, Gautham Arunachal Udupi, Anita S Desai, V Ravi |
| EPI_ISL_2379415 | PEENYA KAMALANAGAR | INSACOG-KA, NIMHANS | Chitra Pattabiraman, Anson Kunjumon George,Darshan Sreenivas,Pramada Prasad, Ananthapadmanabha Kotambail, Chetan G K, Gautham Arunachal Udupi, Anita S Desai, V Ravi |
| EPI_ISL_2379416, EPI_ISL_2379417 | DSU MANDYA | INSACOG-KA, NIMHANS | Chitra Pattabiraman, Anson Kunjumon George,Darshan Sreenivas,Pramada Prasad, Ananthapadmanabha Kotambail, Chetan G K, Gautham Arunachal Udupi, Anita S Desai, V Ravi |
| EPI_ISL_2379418, EPI_ISL_2379419, EPI_ISL_2379420, EPI_ISL_2379421, EPI_ISL_2379422 | DSU CHAMRAJANAGARA | INSACOG-KA, NIMHANS | Chitra Pattabiraman, Anson Kunjumon George,Darshan Sreenivas,Pramada Prasad, Ananthapadmanabha Kotambail, Chetan G K, Gautham Arunachal Udupi, Anita S Desai, V Ravi |
| EPI_ISL_2379423, EPI_ISL_2379424, EPI_ISL_2379425 | DSU KODAGU | INSACOG-KA, NIMHANS | Chitra Pattabiraman, Anson Kunjumon George,Darshan Sreenivas,Pramada Prasad, Ananthapadmanabha Kotambail, Chetan G K, Gautham Arunachal Udupi, Anita S Desai, V Ravi |
| EPI_ISL_2379427, EPI_ISL_2379428 | MAHALAKSHMI LAYOUT | INSACOG-KA, NIMHANS | Chitra Pattabiraman, Anson Kunjumon George,Darshan Sreenivas,Pramada Prasad, Ananthapadmanabha Kotambail, Chetan G K, Gautham Arunachal Udupi, Anita S Desai, V Ravi |
| EPI_ISL_2379429 | DSU KODAGU | INSACOG-KA, NIMHANS | Chitra Pattabiraman, Anson Kunjumon George,Darshan Sreenivas,Pramada Prasad, Ananthapadmanabha Kotambail, Chetan G K, Gautham Arunachal Udupi, Anita S Desai, V Ravi |

| | | | |
|---|---|---|---|
| EPI_ISL_2379430, EPI_ISL_2379431 | RAILWAY H | INSACOG-KA, NIMHANS | Chitra Pattabiraman, Anson Kunjumon George,Darshan Sreenivas,Pramada Prasad, Ananthapadmanabha Kotambail, Chetan G K, Gautham Arunachal Udupi, Anita S Desai, V Ravi |
| EPI_ISL_2379433, EPI_ISL_2379443 | Institute for Medical Research, Infectious Disease Research Centre, National Institutes of Health, Ministry of Health Malaysia | Institute for Medical Research, Infectious Disease Research Centre, National Institutes of Health, Ministry of Health Malaysia | Suppiah J, Kamel K, Mohd Zawawi Z, Azizan MA, Ramly N, Robert F, Thayan R |
| EPI_ISL_2379462, EPI_ISL_2379464, EPI_ISL_2379465, EPI_ISL_2379466, EPI_ISL_2379467, EPI_ISL_2379468, EPI_ISL_2379469, EPI_ISL_2379470, EPI_ISL_2379471, EPI_ISL_2379472, EPI_ISL_2379473, EPI_ISL_2379474, EPI_ISL_2379475, EPI_ISL_2379476, EPI_ISL_2379477, EPI_ISL_2379478, EPI_ISL_2379479, EPI_ISL_2379480, EPI_ISL_2379481, EPI_ISL_2379482, EPI_ISL_2379483, EPI_ISL_2379484, EPI_ISL_2379485, EPI_ISL_2379486, EPI_ISL_2379488, EPI_ISL_2379489, EPI_ISL_2379490, EPI_ISL_2379491, EPI_ISL_2379493, EPI_ISL_2379494, EPI_ISL_2379496, EPI_ISL_2379497, EPI_ISL_2379498, EPI_ISL_2379499, EPI_ISL_2379500, EPI_ISL_2379501, EPI_ISL_2379502, EPI_ISL_2379504, EPI_ISL_2379505, EPI_ISL_2379506, EPI_ISL_2379507, EPI_ISL_2379508, EPI_ISL_2379509, EPI_ISL_2379510, EPI_ISL_2379511, EPI_ISL_2379513, EPI_ISL_2379514, EPI_ISL_2379515, EPI_ISL_2379518, EPI_ISL_2379519, EPI_ISL_2379520, EPI_ISL_2379521, EPI_ISL_2379522, EPI_ISL_2379523, EPI_ISL_2379526, EPI_ISL_2379527, EPI_ISL_2379528, EPI_ISL_2379529, EPI_ISL_2379530, EPI_ISL_2379531, EPI_ISL_2379532, EPI_ISL_2379533, EPI_ISL_2379535, EPI_ISL_2379536, EPI_ISL_2379537, EPI_ISL_2379538, EPI_ISL_2379539, EPI_ISL_2379540, EPI_ISL_2379541, EPI_ISL_2379542, EPI_ISL_2379543, EPI_ISL_2379544, EPI_ISL_2379545, EPI_ISL_2379546, EPI_ISL_2379547, EPI_ISL_2379548, EPI_ISL_2379549, EPI_ISL_2379550, EPI_ISL_2379551, EPI_ISL_2379552, EPI_ISL_2379553, EPI_ISL_2379554, EPI_ISL_2379555, EPI_ISL_2379556, EPI_ISL_2379557, EPI_ISL_2379558, EPI_ISL_2379559, EPI_ISL_2379560, EPI_ISL_2379561, EPI_ISL_2379562, EPI_ISL_2379563, EPI_ISL_2379564, EPI_ISL_2379565, EPI_ISL_2379566, EPI_ISL_2379567, EPI_ISL_2379568, EPI_ISL_2379569, EPI_ISL_2379570, EPI_ISL_2379571, EPI_ISL_2379572, EPI_ISL_2379573, EPI_ISL_2379575, EPI_ISL_2379577, EPI_ISL_2379578, EPI_ISL_2379579, EPI_ISL_2379580, EPI_ISL_2379581, EPI_ISL_2379582, EPI_ISL_2379583, EPI_ISL_2379584, EPI_ISL_2379585, EPI_ISL_2379586, EPI_ISL_2379587, EPI_ISL_2379588, EPI_ISL_2379589, EPI_ISL_2379590, EPI_ISL_2379591, EPI_ISL_2379592, EPI_ISL_2379593, EPI_ISL_2379594, EPI_ISL_2379595, EPI_ISL_2379596, EPI_ISL_2379598, EPI_ISL_2379599, EPI_ISL_2379600, EPI_ISL_2379601, EPI_ISL_2379602, EPI_ISL_2379603, EPI_ISL_2379604, EPI_ISL_2379606, EPI_ISL_2379607, EPI_ISL_2379608, EPI_ISL_2379610, EPI_ISL_2379611, EPI_ISL_2379612, EPI_ISL_2379613, EPI_ISL_2379614, EPI_ISL_2379615, EPI_ISL_2379616, EPI_ISL_2379617, EPI_ISL_2379618, EPI_ISL_2379619, EPI_ISL_2379620, EPI_ISL_2379621, EPI_ISL_2379622, EPI_ISL_2379623, EPI_ISL_2379624, EPI_ISL_2379625, EPI_ISL_2379627, EPI_ISL_2379628, EPI_ISL_2379629, EPI_ISL_2379630, EPI_ISL_2379631, EPI_ISL_2379632, EPI_ISL_2379633, EPI_ISL_2379634, EPI_ISL_2379636, EPI_ISL_2379637, EPI_ISL_2379638, EPI_ISL_2379639, EPI_ISL_2379640, EPI_ISL_2379641, EPI_ISL_2379642, EPI_ISL_2379643, EPI_ISL_2379644, EPI_ISL_2379645, EPI_ISL_2379646, EPI_ISL_2379647, EPI_ISL_2379648, EPI_ISL_2379649 | | | |
| see above | Dr S Raju, Director of Public Heath and Preventive Medicine | inStem NCBS - INSACOG | Uma Ramakrishnan Dasaradhi Palakodeti Aswin SaiNarain |
| EPI_ISL_2379650 | Institute for Medical Research, Infectious Disease Research Centre, National Institutes of Health, Ministry of Health Malaysia | Institute for Medical Research, Infectious Disease Research Centre, National Institutes of Health, Ministry of Health Malaysia | Suppiah J, Kamel K, Mohd Zawawi Z, Azizan MA, Ramly N, Robert F, Thayan R |
| EPI_ISL_2379667 | Valais Hospital, Central Institute | Valais Hospital, Central Institute | Alexis Dumoulin, Lorenzo Cerutti, Henri Pegeot, Melyssa Elies, Deborah Penet, Keith Harshman, Ioannis Xenarios, Emmanouil Dermitzakis |
| EPI_ISL_2379748 | National Heart Institute | Institute for Medical Research, Infectious Disease Research Centre, National Institutes of Health, Ministry of Health Malaysia | Suppiah J, Kamel K, Mohd Zawawi Z, Azizan MA, Ramly N, Robert F, Thayan R |
| EPI_ISL_2380047 | Microbiology Department, Laboratori Clínic Metropolitana Nord. Hospital Universitari Germans Trias i Pujol. | Can Ruti SARS-CoV-2 Sequencing Hub (HUGTiP/IrsiCaixa/IGTP) | Marc Noguera-Julian, Pilar Armengol, Ignacio Blanco, Antoni E Bordoy, Francesc Catala-Moll, Pere-Joan Cardona, Maria Casadellà, Cristina Casañ, Gemma Clara, Bonaventura Clotet, Cristina Esteban, Montserrat Giménez, Mercedes Guerrero, Anna Not, Roger Pared |
| EPI_ISL_2380495 | Medical Laboratories Duesseldorf | Center of Medical Microbiology, Virology, and Hospital Hygiene, University of Duesseldorf | Maximilian Damagnez;Alexander Dilthey;Angelika Helmer;Torsten Houwaart;Lisanna Hülse;Christian Lange;Malte Kohns Vasconcelos;Nadine Lübke;Jessica Nicolai;Klaus Pfeffer;Daniel Strelow;Jörg Timm;Andreas Walker;Tobias Wienemann |
| EPI_ISL_2380551 | Institute for Infectious Diseases | Institute for Infectious Diseases | Stefan Neuenschwander, Christian Baumann, Miguel A Terrazos Miani, Cora Sägesser, Pascal Bittel, Peter Keller, Franziska Suter-Riniker, Stephen L Leib, Alban Ramette |
| EPI_ISL_2380647, EPI_ISL_2380766, EPI_ISL_2380767, EPI_ISL_2381523, EPI_ISL_2381664, EPI_ISL_2381735, EPI_ISL_2381736 | ZOTZ KLIMAS MVZ Düsseldorf-Centrum GbR ÜBAG für Labormedizin, Genetik, Zytologie, Pathologie | Center of Medical Microbiology, Virology, and Hospital Hygiene, University of Duesseldorf | Maximilian Damagnez;Alexander Dilthey;Patrick Finzer;Katrin Hoffmann;Torsten Houwaart;Lisanna Hülse;Malte Kohns Vasconcelos;Nadine Lübke;Jessica Nicolai;Klaus Pfeffer;Daniel Strelow;Jörg Timm;Andreas Walker;Tobias Wienemann;Rainer Zotz |
| EPI_ISL_2382091, EPI_ISL_2382124, EPI_ISL_2382145, EPI_ISL_2382151, EPI_ISL_2382176 | Mako Medical | Centers for Disease Control and Prevention Division of Viral Diseases, Pathogen Discovery | Dakota Howard, Dhwani Batra, Peter W. Cook, Kara Moser, Adrian Paskey, Jason Caravas, Benjamin Rambo-Martin, Shatavia Morrison, Christopher Gulvick, Scott Sammons, Yvette Unoarumhi, Darlene Wagner, Matthew Schmerer,Matthew Tugwell, Lauren Moon, Clinton R. Paden, Duncan MacCannell |
| EPI_ISL_2382576, EPI_ISL_2382590, EPI_ISL_2382618, EPI_ISL_2382619 | Labor Berlin Charité Vivantes GmbH / Institut für Virologie | Charité Universitätsmedizin Berlin, Institut für Virologie/Labor Berlin | Peter Menzel, Christine Stephan, Rolf Schwarzer, Victor M Corman, Barbara Mühlemann, Terry Jones, Christian Drosten |
| EPI_ISL_2382812 | SYNLAB | GIGA Medical Genomics | Keith Durkin, Maria Artesi, Sébastien Bontems, Raphaël Boreux, Bouchra Boujemla, Nathalie Renotte, Cécile Meex, Pierrette Melin, Marie-Pierre Hayette, Vincent Bours |
| EPI_ISL_2382824 | Labo Luc Olivier | GIGA Medical Genomics | Keith Durkin, Maria Artesi, Sébastien Bontems, Raphaël Boreux, Bouchra Boujemla, Nathalie Renotte, Cécile Meex, Pierrette Melin, Marie-Pierre Hayette, Vincent Bours |
| EPI_ISL_2382848, EPI_ISL_2382863, EPI_ISL_2382870 | University of Liège COVID-19 testing center | GIGA Medical Genomics | Keith Durkin, Maria Artesi, Bouchra Boujemla, Nathalie Renotte, Cécile Meex, Sébastien Bontems, Fabrice Bureau, Laurent Gillet, Wouter Coppieters, Marie-Pierre Hayette, Vincent Bours |
| EPI_ISL_2382915, EPI_ISL_2382917, EPI_ISL_2382918, EPI_ISL_2382919, EPI_ISL_2382920, EPI_ISL_2382922, EPI_ISL_2382923, EPI_ISL_2382924, EPI_ISL_2382925, EPI_ISL_2382926, EPI_ISL_2382927, EPI_ISL_2382928, EPI_ISL_2382929, EPI_ISL_2382930, EPI_ISL_2382931, EPI_ISL_2382932, EPI_ISL_2382933, EPI_ISL_2382936, EPI_ISL_2382937, EPI_ISL_2382938, EPI_ISL_2382939, EPI_ISL_2382940, EPI_ISL_2382941, EPI_ISL_2382942, EPI_ISL_2382943, EPI_ISL_2382946, EPI_ISL_2382947, EPI_ISL_2382948, EPI_ISL_2382954, EPI_ISL_2382955, EPI_ISL_2382956, EPI_ISL_2382957, EPI_ISL_2382958, EPI_ISL_2382959, EPI_ISL_2382960, EPI_ISL_2382961, EPI_ISL_2382962, EPI_ISL_2382963, EPI_ISL_2382964, EPI_ISL_2382965, EPI_ISL_2382966, EPI_ISL_2382969, EPI_ISL_2382970, EPI_ISL_2382971, EPI_ISL_2382972, EPI_ISL_2382973, EPI_ISL_2382974, EPI_ISL_2382975, EPI_ISL_2382977, EPI_ISL_2382979, EPI_ISL_2382980, EPI_ISL_2382981, EPI_ISL_2382982, EPI_ISL_2382983, EPI_ISL_2382988, EPI_ISL_2382996, EPI_ISL_2382998, EPI_ISL_2383004, EPI_ISL_2383005 | | | |
| see above | National Centre For Cell Science | National Centre For Cell Science-INSACOG | Dhiraj Paul, Shivang P. Bhanushali, Mohak P Gujare, Mitali Inamdar, Manoj Kumar Bhat, Ajay Pillai, INSACOG Consortium team, Yogesh Shouche |
| EPI_ISL_2383022, EPI_ISL_2383027, EPI_ISL_2383028, EPI_ISL_2383029, EPI_ISL_2383030, EPI_ISL_2383032 | Wichita State University- Molecular Diagonstic lab | Kansas Health and Environmental Lab | Katherine Wiggins, Mike Grose, Jonathan Barnell, Ben Olsen, and Phil Adam |
| EPI_ISL_2383132, EPI_ISL_2383133, EPI_ISL_2383134, EPI_ISL_2383135, EPI_ISL_2383136, EPI_ISL_2383148, EPI_ISL_2383151, EPI_ISL_2383152 | Wyoming Public Health Laboratory | Wyoming Public Health Laboratory | Jim Mildenberger, Wanda Manley, Noah Hull, Taylor Fearing, Lynette Gumbleton, Channing Weber, Ashley Norberg, Chayse Rowley, Marley Goetz, Brian Dominguez, Elliot Thomasson, Sam Britz, Cari Sloma, and Rob Christensen |
| EPI_ISL_2383189 | LSUHS Emerging Viral Threat Laboratory | LSUHS Emerging Viral Threat Laboratory | Gregory L. Ware, Alexander Mijalis, Jeremy P. Kamil, Jennifer L. Carroll, Maarten Van Diest, Rona S. Scott, Andrew D. Yurochko, Christopher G. Kevil, John A. Vanchiere |
| EPI_ISL_2383315 | Servicio de Microbiología. Hospital Clínico Universitario de Valencia | SeqCOVID-SPAIN consortium/IBV(CSIC) | David Navarro Ortega, Eliseo Albert Vicent, Ignacio Torres and SeqCOVID-SPAIN consortium |
| EPI_ISL_2383603, EPI_ISL_2383639, EPI_ISL_2383844, EPI_ISL_2383872, EPI_ISL_2383876 | DPHL | Delaware Public Health Lab | Rebecca Savage |
| EPI_ISL_2383898 | WVDHHR - Office of Laboratory Services | Centers for Disease Control and Prevention Division of Viral Diseases, Pathogen Discovery | Mili Sheth, Sarah Nobles, Jasmine Padilla, Mark Burroughs, Shoshona Le, Katie Dillon, Peter Cook, Clinton R. Paden, Dhwani Batra, Krista Queen, Kristen Knipe, Dakota Howard, Yvette Unoarumhi, Darlene Wagner, Matthew Schmerer, Ben L. Rambo-Martin, Kristine Lacek, Sam Shepard, Alison Laufer Halpin, Dave Wentworth, Vivien Dugan, Suxiang Tong, Justin Lee |
| EPI_ISL_2383982, EPI_ISL_2383984 | AL Dept. of Public Health Bureau of Clinical Laboratories | Centers for Disease Control and Prevention Division of Viral Diseases, Pathogen Discovery | Mili Sheth, Sarah Nobles, Jasmine Padilla, Mark Burroughs, Shoshona Le, Katie Dillon, Peter Cook, Clinton R. Paden, Dhwani Batra, Krista Queen, Kristen Knipe, Dakota Howard, Yvette Unoarumhi, Darlene Wagner, Matthew Schmerer, Ben L. Rambo-Martin, Kristine Lacek, Sam Shepard, Alison Laufer Halpin, Dave Wentworth, Vivien Dugan, Suxiang Tong, Justin Lee |
| EPI_ISL_2384050 | IA State Hygienic Laboratory | Centers for Disease Control and Prevention Division of Viral Diseases, Pathogen Discovery | Mili Sheth, Sarah Nobles, Jasmine Padilla, Mark Burroughs, Shoshona Le, Katie Dillon, Peter Cook, Clinton R. Paden, Dhwani Batra, Krista Queen, Kristen Knipe, Dakota Howard, Yvette Unoarumhi, Darlene Wagner, Matthew Schmerer, Ben L. Rambo-Martin, Kristine Lacek, Sam Shepard, Alison Laufer Halpin, Dave Wentworth, Vivien Dugan, Suxiang Tong, Justin Lee |
| EPI_ISL_2384069 | CT-Dr. Katherine A. Kelley State Public Health Lab | Centers for Disease Control and Prevention Division of Viral Diseases, Pathogen Discovery | Mili Sheth, Sarah Nobles, Jasmine Padilla, Mark Burroughs, Shoshona Le, Katie Dillon, Peter Cook, Clinton R. Paden, Dhwani Batra, Krista Queen, Kristen Knipe, Dakota Howard, Yvette Unoarumhi, Darlene Wagner, Matthew Schmerer, Ben L. Rambo-Martin, Kristine Lacek, Sam Shepard, Alison Laufer Halpin, Dave Wentworth, Vivien Dugan, Suxiang Tong, Justin Lee |
| EPI_ISL_2384139 | Alaska State Virology Lab, Alaska State Public Health Labs | Alaska State Virology Lab, Alaska State Public Health Labs | DeRonde,S., House,E., Zidek,J., Redlinger,M., Smith,L., Chen,J. |
| EPI_ISL_2384664 | Labor Dr. Fenner und Kollegen | Heinrich Pette Institute, Leibniz Institute for Experimental Virology | Alexis Robitaille, Thomas Günther, Johannes Knobloch, Martin Aepfelbacher, Nicole Fischer, Adam Grundhoff |
| EPI_ISL_2385077, EPI_ISL_2385078, EPI_ISL_2385081, EPI_ISL_2385085, EPI_ISL_2385090, EPI_ISL_2385091, EPI_ISL_2385095, EPI_ISL_2385096, EPI_ISL_2385100, EPI_ISL_2385101, EPI_ISL_2385113, EPI_ISL_2385114, EPI_ISL_2385116, EPI_ISL_2385119, EPI_ISL_2385121, EPI_ISL_2385123, EPI_ISL_2385124 | | | |
| see above | WHO National Influenza Centre Russian Federation | WHO National Influenza Centre Russian Federation | Andrey Komissarov, Artem Fadeev, Kseniya Komissarova, Oula Masour, Kirill Varchenko, Mikhail Bakaev, Tamila Musaeva, Maria Timofeeva, Veronika Eder, Maria Pisareva, Nikita Yolshin, Daria Danilenko, Ksenia Safina, Elena Nabieva, Georgii Bazykin, Dmitry Lioznov |
| EPI_ISL_2385125, EPI_ISL_2385126, EPI_ISL_2385128, EPI_ISL_2385129, EPI_ISL_2385135, EPI_ISL_2385138, EPI_ISL_2385139, EPI_ISL_2385142, EPI_ISL_2385143, EPI_ISL_2385144, EPI_ISL_2385145, EPI_ISL_2385146, EPI_ISL_2385147, EPI_ISL_2385148, EPI_ISL_2385149, EPI_ISL_2385150, EPI_ISL_2385152, EPI_ISL_2385153, EPI_ISL_2385158, EPI_ISL_2385159, EPI_ISL_2385161, EPI_ISL_2385162, EPI_ISL_2385163, EPI_ISL_2385165, EPI_ISL_2385166, EPI_ISL_2385167, EPI_ISL_2385169, EPI_ISL_2385170, EPI_ISL_2385172, EPI_ISL_2385173, EPI_ISL_2385174, EPI_ISL_2385175, EPI_ISL_2385176, EPI_ISL_2385177, EPI_ISL_2385178, EPI_ISL_2385179, EPI_ISL_2385181, EPI_ISL_2385184, EPI_ISL_2385185, EPI_ISL_2385187, EPI_ISL_2385188, EPI_ISL_2385189, EPI_ISL_2385190, EPI_ISL_2385193, EPI_ISL_2385194, EPI_ISL_2385195, EPI_ISL_2385204, EPI_ISL_2385206, EPI_ISL_2385207, EPI_ISL_2385208, EPI_ISL_2385211 | | | |
| see above | WHO National Influenza Centre Russian Federation | WHO National Influenza Centre Russian Federation | Andrey Komissarov, Artem Fadeev, Kseniya Komissarova, Oula Masour, Kirill Varchenko, Mikhail Bakaev, Tamila Musaeva, Maria Timofeeva, Veronika Eder, Maria Pisareva, Nikita Yolshin, Daria Danilenko, Maria Baturova, Alexey Masharsky, Ksenia Safina, Elena Nabieva, Georgii Bazykin, Dmitry Lioznov |
| EPI_ISL_2385214, EPI_ISL_2385226, EPI_ISL_2385230, EPI_ISL_2385233, EPI_ISL_2385242, EPI_ISL_2385245, EPI_ISL_2385266, EPI_ISL_2385267, EPI_ISL_2385279, EPI_ISL_2385280, EPI_ISL_2385284, EPI_ISL_2385288, EPI_ISL_2385292, EPI_ISL_2385308 | | | |
| see above | HELIX LLC | WHO National Influenza Centre Russian Federation | Andrey Komissarov, Artem Fadeev, Kseniya Komissarova, Oula Masour, Kirill Varchenko, Mikhail Bakaev, Tamila Musaeva, Maria Timofeeva, Veronika Eder, Maria Pisareva, Nikita Yolshin, Daria Danilenko, Ksenia Safina, Elena Nabieva, Georgii Bazykin, Dmitry Lioznov |
| EPI_ISL_2385311, EPI_ISL_2385319, EPI_ISL_2385323, EPI_ISL_2385324, | WHO National Influenza Centre Russian Federation | WHO National Influenza Centre Russian Federation | Andrey Komissarov, Artem Fadeev, Kseniya Komissarova, Oula Masour, Kirill Varchenko, Mikhail Bakaev, Tamila Musaeva, Maria Timofeeva, Veronika Eder, Maria Pisareva, Nikita Yolshin, Daria Danilenko, Ksenia Safina, Elena Nabieva, Georgii Bazykin, Dmitry Lioznov |

| | | | |
|---|---|---|---|
| EPI_ISL_2385333 | | | |
| EPI_ISL_2385428 | WHO National Influenza Centre Russian Federation | WHO National Influenza Centre Russian Federation | Andrey Komissarov, Artem Fadeev, Kseniya Komissarova, Oula Masour, Kirill Varchenko, Mikhail Bakaev, Tamila Musaeva, Maria Timofeeva, Veronika Eder, Maria Pisareva, Nikita Yolshin, Daria Danilenko, Maria Baturova, Alexey Masharsky, Ksenia Safina, Elena Nabieva, Georgii Bazykin, Dmitry Lioznov |
| EPI_ISL_2385968, EPI_ISL_2385969, EPI_ISL_2385972, EPI_ISL_2385973 | Queensland Health Forensic and Scientific Services | Queensland Health Forensic and Scientific Services | Son Nguyen |
| EPI_ISL_2386191 | Medizinisches Labor Ostsachsen MVZ GbR Nebenbetriebsstätte Dresden | Robert Koch Institute | unknown |
| EPI_ISL_2386212 | Limbach - MVZ Labor Eveld & Kollegen Essen | Robert Koch Institute | unknown |
| EPI_ISL_2386317 | SYNLAB MVZ Weiden | Robert Koch Institute | unknown |
| EPI_ISL_2386337 | SYNLAB MVZ Trier | Robert Koch Institute | unknown |
| EPI_ISL_2386345, EPI_ISL_2386346, EPI_ISL_2386361 | SYNLAB MVZ Weiden | Robert Koch Institute | unknown |
| EPI_ISL_2386368 | SYNLAB MVZ Trier | Robert Koch Institute | unknown |
| EPI_ISL_2386403, EPI_ISL_2386424 | SYNLAB MVZ Weiden | Robert Koch Institute | unknown |
| EPI_ISL_2386460, EPI_ISL_2386472, EPI_ISL_2386509 | SYNLAB MVZ Leverkusen | Robert Koch Institute | unknown |
| EPI_ISL_2386672, EPI_ISL_2386685, EPI_ISL_2386700, EPI_ISL_2386730, EPI_ISL_2386764 | Labor ZOTZ\|KLIMAS; MVZ Düsseldorf-Centrum | Robert Koch Institute | unknown |
| EPI_ISL_2386808, EPI_ISL_2386838, EPI_ISL_2386840 | Labor Becker & Kollegen (Standort München) | Robert Koch Institute | unknown |
| EPI_ISL_2386874 | Dianovis GmbH Greiz | Robert Koch Institute | unknown |
| EPI_ISL_2386977, EPI_ISL_2386995, EPI_ISL_2387037, EPI_ISL_2387041, EPI_ISL_2387091, EPI_ISL_2387108, EPI_ISL_2387117, EPI_ISL_2387165 | CENTOGENE Frankfurt Laboratory: Niederlassung Industriepark Höchst | Robert Koch Institute | unknown |
| EPI_ISL_2387354, EPI_ISL_2387384, EPI_ISL_2387404, EPI_ISL_2387435, EPI_ISL_2387445 | MVZ Dr. Eberhard & Partner Dortmund | Robert Koch Institute | unknown |
| EPI_ISL_2387466, EPI_ISL_2387472 | SYNLAB MVZ Weiden | Robert Koch Institute | unknown |
| EPI_ISL_2387517 | Synlab MVZ Augsburg | Robert Koch Institute | unknown |
| EPI_ISL_2387530 | SYNLAB MVZ Leverkusen | Robert Koch Institute | unknown |
| EPI_ISL_2387546 | SYNLAB MVZ Leinfelden-Echterdingen | Robert Koch Institute | unknown |
| EPI_ISL_2387806 | SYNLAB MVZ Trier | Robert Koch Institute | unknown |
| EPI_ISL_2387857, EPI_ISL_2387863 | SYNLAB MVZ Weiden | Robert Koch Institute | unknown |
| EPI_ISL_2387983 | SYNLAB MVZ Leinfelden-Echterdingen | Robert Koch Institute | unknown |
| EPI_ISL_2387999 | SYNLAB MVZ Leverkusen | Robert Koch Institute | unknown |
| EPI_ISL_2388120, EPI_ISL_2388139, EPI_ISL_2388142, EPI_ISL_2388159 | MVZ Labor Dr. Limbach & Kollegen GbR | Robert Koch Institute | unknown |
| EPI_ISL_2388189 | MVZ Dr. Eberhard & Partner Dortmund | Robert Koch Institute | unknown |
| EPI_ISL_2388326, EPI_ISL_2388343, EPI_ISL_2388386, EPI_ISL_2388428, EPI_ISL_2388504, EPI_ISL_2388581, EPI_ISL_2388583, EPI_ISL_2388592 | Sonic - MVZ Medizinisches Labor Bremen GmbH | Robert Koch Institute | unknown |
| EPI_ISL_2388611 | Labor 28 MVZ GmbH | Robert Koch Institute | unknown |
| EPI_ISL_2388659, EPI_ISL_2388709, EPI_ISL_2388738, EPI_ISL_2388740, EPI_ISL_2388803 | Sonic - MVZ Medizinisches Labor Bremen GmbH | Robert Koch Institute | unknown |
| EPI_ISL_2388844 | Bioscientia Labor Wermsdorf | Robert Koch Institute | unknown |
| EPI_ISL_2388867 | Sonic - MVZ Medizinisches Labor Bremen GmbH | Robert Koch Institute | unknown |
| EPI_ISL_2388942, EPI_ISL_2388951, EPI_ISL_2388983, EPI_ISL_2388984, EPI_ISL_2389026 | Bioscientia Labor Wermsdorf | Robert Koch Institute | unknown |
| EPI_ISL_2389082, EPI_ISL_2389083, EPI_ISL_2389084, EPI_ISL_2389085, EPI_ISL_2389086, EPI_ISL_2389088, EPI_ISL_2389093, EPI_ISL_2389094, EPI_ISL_2389096, EPI_ISL_2389104, EPI_ISL_2389107, EPI_ISL_2389111, EPI_ISL_2389112, EPI_ISL_2389116, EPI_ISL_2389117, EPI_ISL_2389119, EPI_ISL_2389124, EPI_ISL_2389125 | | | |
| see above | Medizinische Laboratorien Düsseldorf | Robert Koch Institute | unknown |
| EPI_ISL_2389337, EPI_ISL_2389352 | Labor Dr. Heidrich & Kollegen MVZ GmbH Hamburg | Robert Koch Institute | unknown |
| EPI_ISL_2389360 | Med. Labor Prof. Schenk Dr. Ansorge & Kollegen | Robert Koch Institute | unknown |
| EPI_ISL_2389371, EPI_ISL_2389377 | Robert Koch-Institut ZBS1 | Robert Koch Institute | unknown |
| EPI_ISL_2389440 | Universitätsklinikum des Saarlandes - Institut für Virologie | Robert Koch Institute | unknown |
| EPI_ISL_2389514 | SYNLAB MVZ Heidelberg | Robert Koch Institute | unknown |
| EPI_ISL_2389603 | SYNLAB MVZ Trier | Robert Koch Institute | unknown |
| EPI_ISL_2389632 | B.J. Medical College and Civil Hospital, Ahmedabad | Gujarat Biotechnology Research Centre | Zuber Saiyed,Ramesh Pandit,Janvi Raval,Zarna Patel,Nitin Savaliya,Dinesh Kumar,Twinkle Soni,Sonal Sharma ,Pranay Shah,Umang Mishra,Nitesh Shah,Chaitanya Joshi,Madhvi Joshi |
| EPI_ISL_2389634 | B.J. Medical College and Civil Hospital, Ahmedabad | Gujarat Biotechnology Research Centre | Ramesh Pandit,Janvi Raval,Zarna Patel,Nitin Savaliya,Dinesh Kumar,Twinkle Soni,Sonal Sharma ,Zuber Saiyed,Pranay Shah,Umang Mishra,Nitesh Shah,Chaitanya Joshi,Madhvi Joshi |

| | | | |
|---|---|---|---|
| EPI_ISL_2389636 | B.J. Medical College and Civil Hospital, Ahmedabad | Gujarat Biotechnology Research Centre | Janvi Raval,Zarna Patel,Nitin Savaliya,Dinesh Kumar,Twinkle Soni,Sonal Sharma ,Zuber Saiyed,Ramesh Pandit,Pranay Shah,Umang Mishra,Nitesh Shah,Chaitanya Joshi,Madhvi Joshi |
| EPI_ISL_2389638 | B.J. Medical College and Civil Hospital, Ahmedabad | Gujarat Biotechnology Research Centre | Zarna Patel,Nitin Savaliya,Dinesh Kumar,Twinkle Soni,Sonal Sharma ,Zuber Saiyed,Ramesh Pandit,Janvi Raval,Pranay Shah,Umang Mishra,Nitesh Shah,Chaitanya Joshi,Madhvi Joshi |
| EPI_ISL_2389639 | Praxisgemeinschaft für Laboratoriumsmedizin Labor Blumenstraße; Praxis Dr. E. Wietschel & Dr. F. Wietschel | Robert Koch Institute | unknown |
| EPI_ISL_2389640 | B.J. Medical College and Civil Hospital, Ahmedabad | Gujarat Biotechnology Research Centre | Nitin Savaliya,Dinesh Kumar,Twinkle Soni,Sonal Sharma ,Zuber Saiyed,Ramesh Pandit,Janvi Raval,Zarna Patel,Pranay Shah,Umang Mishra,Nitesh Shah,Chaitanya Joshi,Madhvi Joshi |
| EPI_ISL_2389641 | B.J. Medical College and Civil Hospital, Ahmedabad | Gujarat Biotechnology Research Centre | Dinesh Kumar,Twinkle Soni,Sonal Sharma ,Zuber Saiyed,Ramesh Pandit,Janvi Raval,Zarna Patel,Nitin Savaliya,Pranay Shah,Umang Mishra,Nitesh Shah,Chaitanya Joshi,Madhvi Joshi |
| EPI_ISL_2389645 | B.J. Medical College and Civil Hospital, Ahmedabad | Gujarat Biotechnology Research Centre | Sonal Sharma ,Zuber Saiyed,Ramesh Pandit,Janvi Raval,Zarna Patel,Nitin Savaliya,Dinesh Kumar,Twinkle Soni,Pranay Shah,Umang Mishra,Nitesh Shah,Chaitanya Joshi,Madhvi Joshi |
| EPI_ISL_2389647 | B.J. Medical College and Civil Hospital, Ahmedabad | Gujarat Biotechnology Research Centre | Zuber Saiyed,Ramesh Pandit,Janvi Raval,Zarna Patel,Nitin Savaliya,Dinesh Kumar,Twinkle Soni,Sonal Sharma ,Pranay Shah,Umang Mishra,Nitesh Shah,Chaitanya Joshi,Madhvi Joshi |
| EPI_ISL_2389649 | B.J. Medical College and Civil Hospital, Ahmedabad | Gujarat Biotechnology Research Centre | Ramesh Pandit,Janvi Raval,Zarna Patel,Nitin Savaliya,Dinesh Kumar,Twinkle Soni,Sonal Sharma ,Zuber Saiyed,Pranay Shah,Umang Mishra,Nitesh Shah,Chaitanya Joshi,Madhvi Joshi |
| EPI_ISL_2389651 | B.J. Medical College and Civil Hospital, Ahmedabad | Gujarat Biotechnology Research Centre | Janvi Raval,Zarna Patel,Nitin Savaliya,Dinesh Kumar,Twinkle Soni,Sonal Sharma ,Zuber Saiyed,Ramesh Pandit,Pranay Shah,Umang Mishra,Nitesh Shah,Chaitanya Joshi,Madhvi Joshi |
| EPI_ISL_2389653 | B.J. Medical College and Civil Hospital, Ahmedabad | Gujarat Biotechnology Research Centre | Zarna Patel,Nitin Savaliya,Dinesh Kumar,Twinkle Soni,Sonal Sharma ,Zuber Saiyed,Ramesh Pandit,Janvi Raval,Pranay Shah,Umang Mishra,Nitesh Shah,Chaitanya Joshi,Madhvi Joshi |
| EPI_ISL_2389655 | B.J. Medical College and Civil Hospital, Ahmedabad | Gujarat Biotechnology Research Centre | Nitin Savaliya,Dinesh Kumar,Twinkle Soni,Sonal Sharma ,Zuber Saiyed,Ramesh Pandit,Janvi Raval,Zarna Patel,Pranay Shah,Umang Mishra,Nitesh Shah,Chaitanya Joshi,Madhvi Joshi |
| EPI_ISL_2389656 | Praxisgemeinschaft für Laboratoriumsmedizin Labor Blumenstraße; Praxis Dr. E. Wietschel & Dr. F. Wietschel | Robert Koch Institute | unknown |
| EPI_ISL_2389657 | B.J. Medical College and Civil Hospital, Ahmedabad | Gujarat Biotechnology Research Centre | Dinesh Kumar,Twinkle Soni,Sonal Sharma ,Zuber Saiyed,Ramesh Pandit,Janvi Raval,Zarna Patel,Nitin Savaliya,Pranay Shah,Umang Mishra,Nitesh Shah,Chaitanya Joshi,Madhvi Joshi |
| EPI_ISL_2389708, EPI_ISL_2389712, EPI_ISL_2389719, EPI_ISL_2389726, EPI_ISL_2389768 | SYNLAB MVZ Weiden | Robert Koch Institute | unknown |
| EPI_ISL_2389792 | SYNLAB MVZ Leverkusen | Robert Koch Institute | unknown |
| EPI_ISL_2389829 | SYNLAB MVZ Weiden | Robert Koch Institute | unknown |
| EPI_ISL_2389852, EPI_ISL_2389853 | Eurofins MVZ Labor Gelsenkirchen | Robert Koch Institute | unknown |
| EPI_ISL_2389915, EPI_ISL_2389924, EPI_ISL_2389927, EPI_ISL_2389938, EPI_ISL_2389939, EPI_ISL_2389949 | Labor Becker & Kollegen (Standort München) | Robert Koch Institute | unknown |
| EPI_ISL_2389988 | IMD - Labor Oderland | Robert Koch Institute | unknown |
| EPI_ISL_2390048 | MVZ Labor Dr. Quade & Kollegen GmbH | Robert Koch Institute | unknown |
| EPI_ISL_2390099 | MVZ Labor Dr. Limbach & Kollegen GbR | Robert Koch Institute | unknown |
| EPI_ISL_2390131, EPI_ISL_2390150 | MVZ Labor Dr. Fenner und Kollegen (Standort Hamburg) | Robert Koch Institute | unknown |
| EPI_ISL_2390182, EPI_ISL_2390183, EPI_ISL_2390189, EPI_ISL_2390192, EPI_ISL_2390193, EPI_ISL_2390194, EPI_ISL_2390195 | Universitätsklinikum Bonn - Institut für Virologie | Robert Koch Institute | unknown |
| EPI_ISL_2390229 | MVZ für Laboratoriumsmedizin und Mikrobiologie Koblenz-Mittelrhein (Labor Koblenz) | Robert Koch Institute | unknown |
| EPI_ISL_2390316 | MVZ Labor Dr. Limbach & Kollegen GbR | Robert Koch Institute | unknown |
| EPI_ISL_2390335 | Uniklinikum Carl Gustav Carus an der TU Dresden; Institut für Virologie | Robert Koch Institute | unknown |
| EPI_ISL_2390358 | Städtisches Klinikum Dresden Institut für Klinische Chemie und Laboratoriumsmedizin | Robert Koch Institute | unknown |
| EPI_ISL_2390360 | Uniklinikum Carl Gustav Carus an der TU Dresden; Institut für Virologie | Robert Koch Institute | unknown |
| EPI_ISL_2390450 | Fraunhofer-Institut für Zelltherapie und Immunologie IZI AG Next-Generation Diagnostics | Robert Koch Institute | unknown |
| EPI_ISL_2390617, EPI_ISL_2390621, EPI_ISL_2390624, EPI_ISL_2390625, EPI_ISL_2390636, EPI_ISL_2390639, EPI_ISL_2390645, EPI_ISL_2390646, EPI_ISL_2390656, EPI_ISL_2390662, EPI_ISL_2390692, EPI_ISL_2390716, EPI_ISL_2390725 | | | |
| see above | Labor ZOTZ\|KLIMAS; MVZ Düsseldorf-Centrum | Robert Koch Institute | unknown |
| EPI_ISL_2390753, EPI_ISL_2390762 | MVZ Labor Krone GbR | Robert Koch Institute | unknown |
| EPI_ISL_2391477 | N.H.L Municipal Medical College, Ahmedabad | Gujarat Biotechnology Research Centre | Twinkle Soni,Sonal Sharma ,Zuber Saiyed,Ramesh Pandit,Janvi Raval,Zarna Patel,Nitin Savaliya,Dinesh Kumar,Jayshri Pethani,Umang Mishra,Nitesh Shah,Chaitanya Joshi,Madhvi Joshi |
| EPI_ISL_2391478 | N.H.L Municipal Medical College, Ahmedabad | Gujarat Biotechnology Research Centre | Sonal Sharma ,Zuber Saiyed,Ramesh Pandit,Janvi Raval,Zarna Patel,Nitin Savaliya,Dinesh Kumar,Twinkle Soni,Jayshri Pethani,Umang Mishra,Nitesh Shah,Chaitanya Joshi,Madhvi Joshi |
| EPI_ISL_2391481 | N.H.L Municipal Medical College, Ahmedabad | Gujarat Biotechnology Research Centre | Janvi Raval,Zarna Patel,Nitin Savaliya,Dinesh Kumar,Twinkle Soni,Sonal Sharma ,Zuber Saiyed,Ramesh Pandit,Jayshri Pethani,Umang Mishra,Nitesh Shah,Chaitanya Joshi,Madhvi Joshi |
| EPI_ISL_2391482 | N.H.L Municipal Medical College, Ahmedabad | Gujarat Biotechnology Research Centre | Zarna Patel,Nitin Savaliya,Dinesh Kumar,Twinkle Soni,Sonal Sharma ,Zuber Saiyed,Ramesh Pandit,Janvi Raval,Jayshri Pethani,Umang Mishra,Nitesh Shah,Chaitanya Joshi,Madhvi Joshi |
| EPI_ISL_2391483 | N.H.L Municipal Medical College, Ahmedabad | Gujarat Biotechnology Research Centre | Nitin Savaliya,Dinesh Kumar,Twinkle Soni,Sonal Sharma ,Zuber Saiyed,Ramesh Pandit,Janvi Raval,Zarna Patel,Jayshri Pethani,Umang Mishra,Nitesh Shah,Chaitanya Joshi,Madhvi Joshi |
| EPI_ISL_2391484 | N.H.L Municipal Medical College, Ahmedabad | Gujarat Biotechnology Research Centre | Dinesh Kumar,Twinkle Soni,Sonal Sharma ,Zuber Saiyed,Ramesh Pandit,Janvi Raval,Zarna Patel,Nitin Savaliya,Jayshri Pethani,Umang Mishra,Nitesh Shah,Chaitanya Joshi,Madhvi Joshi |
| EPI_ISL_2391486 | N.H.L Municipal Medical College, Ahmedabad | Gujarat Biotechnology Research Centre | Sonal Sharma ,Zuber Saiyed,Ramesh Pandit,Janvi Raval,Zarna Patel,Nitin Savaliya,Dinesh Kumar,Twinkle Soni,Jayshri Pethani,Umang Mishra,Nitesh Shah,Chaitanya Joshi,Madhvi Joshi |
| EPI_ISL_2391487 | N.H.L Municipal Medical College, Ahmedabad | Gujarat Biotechnology Research Centre | Zuber Saiyed,Ramesh Pandit,Janvi Raval,Zarna Patel,Nitin Savaliya,Dinesh Kumar,Twinkle Soni,Sonal Sharma ,Jayshri Pethani,Umang Mishra,Nitesh Shah,Chaitanya Joshi,Madhvi Joshi |
| EPI_ISL_2391488 | BJ Medical College and Civil Hospital, Ahmedabad | Gujarat Biotechnology Research Centre | Janvi Raval,Zarna Patel,Nitin Savaliya,Dinesh Kumar,Twinkle Soni,Sonal Sharma ,Zuber Saiyed,Ramesh Pandit,Pranay Shah,Umang Mishra,Nitesh Shah,Chaitanya Joshi,Madhvi Joshi |
| EPI_ISL_2391489 | GMERS, Government Medical College (GMC), Gotri | Gujarat Biotechnology Research Centre | Zarna Patel,Nitin Savaliya,Dinesh Kumar,Twinkle Soni,Sonal Sharma ,Zuber Saiyed,Ramesh Pandit,Janvi Raval,Bithika Duttaroy,Umang Mishra,Nitesh Shah,Chaitanya Joshi,Madhvi Joshi |
| EPI_ISL_2391490 | GMERS, Government Medical College (GMC), Gotri | Gujarat Biotechnology Research Centre | Nitin Savaliya,Dinesh Kumar,Twinkle Soni,Sonal Sharma ,Zuber Saiyed,Ramesh Pandit,Janvi Raval,Zarna Patel,Bithika Duttaroy,Umang Mishra,Nitesh Shah,Chaitanya Joshi,Madhvi Joshi |
| EPI_ISL_2391491 | GMERS, Government Medical College (GMC), Gotri | Gujarat Biotechnology Research Centre | Dinesh Kumar,Twinkle Soni,Sonal Sharma ,Zuber Saiyed,Ramesh Pandit,Janvi Raval,Zarna Patel,Nitin Savaliya,Bithika Duttaroy,Umang Mishra,Nitesh Shah,Chaitanya Joshi,Madhvi Joshi |
| EPI_ISL_2391492 | GMERS, Government Medical College (GMC), Gotri | Gujarat Biotechnology Research Centre | Twinkle Soni,Sonal Sharma ,Zuber Saiyed,Ramesh Pandit,Janvi Raval,Zarna Patel,Nitin Savaliya,Dinesh Kumar,Bithika Duttaroy,Umang Mishra,Nitesh Shah,Chaitanya Joshi,Madhvi Joshi |
| EPI_ISL_2391493 | GMERS, Government Medical College (GMC), Gotri | Gujarat Biotechnology Research Centre | Sonal Sharma ,Zuber Saiyed,Ramesh Pandit,Janvi Raval,Zarna Patel,Nitin Savaliya,Dinesh Kumar,Twinkle Soni,Bithika Duttaroy,Umang Mishra,Nitesh Shah,Chaitanya Joshi,Madhvi Joshi |
| EPI_ISL_2391494 | GMERS, Government Medical College (GMC), Gotri | Gujarat Biotechnology Research Centre | Zuber Saiyed,Ramesh Pandit,Janvi Raval,Zarna Patel,Nitin Savaliya,Dinesh Kumar,Twinkle Soni,Sonal Sharma ,Bithika Duttaroy,Umang Mishra,Nitesh Shah,Chaitanya Joshi,Madhvi Joshi |
| EPI_ISL_2391495 | GMERS, Government Medical College (GMC), Gotri | Gujarat Biotechnology Research Centre | Ramesh Pandit,Janvi Raval,Zarna Patel,Nitin Savaliya,Dinesh Kumar,Twinkle Soni,Sonal Sharma ,Zuber Saiyed,Bithika Duttaroy,Umang Mishra,Nitesh Shah,Chaitanya Joshi,Madhvi Joshi |
| EPI_ISL_2391496 | GMERS, Government Medical College (GMC), Gotri | Gujarat Biotechnology Research Centre | Janvi Raval,Zarna Patel,Nitin Savaliya,Dinesh Kumar,Twinkle Soni,Sonal Sharma ,Zuber Saiyed,Ramesh Pandit,Bithika Duttaroy,Umang Mishra,Nitesh Shah,Chaitanya Joshi,Madhvi Joshi |
| EPI_ISL_2391497 | GMERS, Government Medical College (GMC), Gotri | Gujarat Biotechnology Research Centre | Zarna Patel,Nitin Savaliya,Dinesh Kumar,Twinkle Soni,Sonal Sharma ,Zuber Saiyed,Ramesh Pandit,Janvi Raval,Bithika Duttaroy,Umang Mishra,Nitesh Shah,Chaitanya Joshi,Madhvi Joshi |
| EPI_ISL_2391498 | GMERS, Government Medical College (GMC), Gotri | Gujarat Biotechnology Research Centre | Nitin Savaliya,Dinesh Kumar,Twinkle Soni,Sonal Sharma ,Zuber Saiyed,Ramesh Pandit,Janvi Raval,Zarna Patel,Bithika Duttaroy,Umang Mishra,Nitesh Shah,Chaitanya Joshi,Madhvi Joshi |
| EPI_ISL_2391501 | GMERS, Government Medical College (GMC), Gotri | Gujarat Biotechnology Research Centre | Sonal Sharma ,Zuber Saiyed,Ramesh Pandit,Janvi Raval,Zarna Patel,Nitin Savaliya,Dinesh Kumar,Twinkle Soni,Bithika Duttaroy,Umang Mishra,Nitesh Shah,Chaitanya Joshi,Madhvi Joshi |
| EPI_ISL_2391502 | GMERS, Government Medical College (GMC), Gotri | Gujarat Biotechnology Research Centre | Zuber Saiyed,Ramesh Pandit,Janvi Raval,Zarna Patel,Nitin Savaliya,Dinesh Kumar,Twinkle Soni,Sonal Sharma ,Bithika Duttaroy,Umang Mishra,Nitesh Shah,Chaitanya Joshi,Madhvi Joshi |
| EPI_ISL_2391503 | GMERS, Government Medical College (GMC), Gotri | Gujarat Biotechnology Research Centre | Ramesh Pandit,Janvi Raval,Zarna Patel,Nitin Savaliya,Dinesh Kumar,Twinkle Soni,Sonal Sharma ,Zuber Saiyed,Bithika Duttaroy,Umang Mishra,Nitesh Shah,Chaitanya Joshi,Madhvi Joshi |
| EPI_ISL_2391788 | Huntington Memorial Hospital | Los Angeles County Public Health Laboratories | P. Hemarajata et al. |
| EPI_ISL_2391792 | Providence Saint Joseph Medical Center | Los Angeles County Public Health Laboratories | P. Hemarajata et al. |
| EPI_ISL_2391807 | Huntington Memorial Hospital | Los Angeles County Public Health Laboratories | P. Hemarajata et al. |
| EPI_ISL_2391814, EPI_ISL_2391843, | Hospital Universitari Vall d'Hebron - Vall d'Hebron Institut de Recerca | Hospital Universitari Vall d'Hebron - Vall d'Hebron Institut de Recerca | Cristina Andrés, Maria Piñana, Alejandra González-Sánchez, Damir Garcia-Cehic, Ariadna Rando, Juliana Esperalba, Maria Gema Codina, Carla Castillo, Maria Carmen Martin, Tomàs Pumarola, Josep Quer, Andrés Antón |

| | | | |
|---|---|---|---|
| EPI_ISL_2391845, EPI_ISL_2391899, EPI_ISL_2391912, EPI_ISL_2391913, EPI_ISL_2391926 | | | |
| EPI_ISL_2392108 | ULSS 3 Venezia | Istituto Zooprofilattico Sperimentale delle Venezie | Adelaide Milani, Alessia Schivo, Annalisa Salviato, Elisa Palumbo, Erika Giorgia Quaranta, Luca Tassoni, Ambra Pastori, Edoardo Giussani, Alice Fusaro, Isabella Monne, Calogero Terregino, Antonia Ricci |
| EPI_ISL_2392196 | Lighthouse Lab in Milton Keynes | Wellcome Sanger Institute for the COVID-19 Genomics UK (COG-UK) Consortium | The Lighthouse Lab in Milton Keynes and Alex Alderton, Roberto Amato, Jeffrey Barrett, Sonia Goncalves, Ewan Harrison, David K. Jackson, Ian Johnston, Dominic Kwiatkowski, Cordelia Langford, John Sillitoe on behalf of the Wellcome Sanger Institute COVID-19 Surveillance Team |
| EPI_ISL_2392197 | Lighthouse Lab in Alderley Park | Wellcome Sanger Institute for the COVID-19 Genomics UK (COG-UK) Consortium | Jacquelyn Wynn, Mairead Hyland, The Lighthouse Lab in Alderley Park and Alex Alderton, Roberto Amato, Jeffrey Barrett, Sonia Goncalves, Ewan Harrison, David K. Jackson, Ian Johnston, Dominic Kwiatkowski, Cordelia Langford, John Sillitoe on behalf of the Wellcome Sanger Institute COVID-19 Surveillance Team |
| EPI_ISL_2392198, EPI_ISL_2392199 | Lighthouse Lab in Milton Keynes | Wellcome Sanger Institute for the COVID-19 Genomics UK (COG-UK) Consortium | The Lighthouse Lab in Milton Keynes and Alex Alderton, Roberto Amato, Jeffrey Barrett, Sonia Goncalves, Ewan Harrison, David K. Jackson, Ian Johnston, Dominic Kwiatkowski, Cordelia Langford, John Sillitoe on behalf of the Wellcome Sanger Institute COVID-19 Surveillance Team |
| EPI_ISL_2392200, EPI_ISL_2392202, EPI_ISL_2392203, EPI_ISL_2392208, EPI_ISL_2392209 | Lighthouse Lab in Alderley Park | Wellcome Sanger Institute for the COVID-19 Genomics UK (COG-UK) Consortium | Jacquelyn Wynn, Mairead Hyland, The Lighthouse Lab in Alderley Park and Alex Alderton, Roberto Amato, Jeffrey Barrett, Sonia Goncalves, Ewan Harrison, David K. Jackson, Ian Johnston, Dominic Kwiatkowski, Cordelia Langford, John Sillitoe on behalf of the Wellcome Sanger Institute COVID-19 Surveillance Team |
| EPI_ISL_2392210, EPI_ISL_2392211 | Lighthouse Lab in Milton Keynes | Wellcome Sanger Institute for the COVID-19 Genomics UK (COG-UK) Consortium | The Lighthouse Lab in Milton Keynes and Alex Alderton, Roberto Amato, Jeffrey Barrett, Sonia Goncalves, Ewan Harrison, David K. Jackson, Ian Johnston, Dominic Kwiatkowski, Cordelia Langford, John Sillitoe on behalf of the Wellcome Sanger Institute COVID-19 Surveillance Team |
| EPI_ISL_2392213, EPI_ISL_2392214, EPI_ISL_2392215, EPI_ISL_2392216, EPI_ISL_2392218, EPI_ISL_2392219 | Lighthouse Lab in Alderley Park | Wellcome Sanger Institute for the COVID-19 Genomics UK (COG-UK) Consortium | Jacquelyn Wynn, Mairead Hyland, The Lighthouse Lab in Alderley Park and Alex Alderton, Roberto Amato, Jeffrey Barrett, Sonia Goncalves, Ewan Harrison, David K. Jackson, Ian Johnston, Dominic Kwiatkowski, Cordelia Langford, John Sillitoe on behalf of the Wellcome Sanger Institute COVID-19 Surveillance Team |
| EPI_ISL_2392220 | Lighthouse Lab in Milton Keynes | Wellcome Sanger Institute for the COVID-19 Genomics UK (COG-UK) Consortium | The Lighthouse Lab in Milton Keynes and Alex Alderton, Roberto Amato, Jeffrey Barrett, Sonia Goncalves, Ewan Harrison, David K. Jackson, Ian Johnston, Dominic Kwiatkowski, Cordelia Langford, John Sillitoe on behalf of the Wellcome Sanger Institute COVID-19 Surveillance Team |
| EPI_ISL_2392221, EPI_ISL_2392222 | Lighthouse Lab in Alderley Park | Wellcome Sanger Institute for the COVID-19 Genomics UK (COG-UK) Consortium | Jacquelyn Wynn, Mairead Hyland, The Lighthouse Lab in Alderley Park and Alex Alderton, Roberto Amato, Jeffrey Barrett, Sonia Goncalves, Ewan Harrison, David K. Jackson, Ian Johnston, Dominic Kwiatkowski, Cordelia Langford, John Sillitoe on behalf of the Wellcome Sanger Institute COVID-19 Surveillance Team |
| EPI_ISL_2392223, EPI_ISL_2392224, EPI_ISL_2392225, EPI_ISL_2392227 | Lighthouse Lab in Milton Keynes | Wellcome Sanger Institute for the COVID-19 Genomics UK (COG-UK) Consortium | The Lighthouse Lab in Milton Keynes and Alex Alderton, Roberto Amato, Jeffrey Barrett, Sonia Goncalves, Ewan Harrison, David K. Jackson, Ian Johnston, Dominic Kwiatkowski, Cordelia Langford, John Sillitoe on behalf of the Wellcome Sanger Institute COVID-19 Surveillance Team |
| EPI_ISL_2392228, EPI_ISL_2392229, EPI_ISL_2392232, EPI_ISL_2392233, EPI_ISL_2392234 | Lighthouse Lab in Alderley Park | Wellcome Sanger Institute for the COVID-19 Genomics UK (COG-UK) Consortium | Jacquelyn Wynn, Mairead Hyland, The Lighthouse Lab in Alderley Park and Alex Alderton, Roberto Amato, Jeffrey Barrett, Sonia Goncalves, Ewan Harrison, David K. Jackson, Ian Johnston, Dominic Kwiatkowski, Cordelia Langford, John Sillitoe on behalf of the Wellcome Sanger Institute COVID-19 Surveillance Team |
| EPI_ISL_2392235, EPI_ISL_2392236 | Lighthouse Lab in Milton Keynes | Wellcome Sanger Institute for the COVID-19 Genomics UK (COG-UK) Consortium | The Lighthouse Lab in Milton Keynes and Alex Alderton, Roberto Amato, Jeffrey Barrett, Sonia Goncalves, Ewan Harrison, David K. Jackson, Ian Johnston, Dominic Kwiatkowski, Cordelia Langford, John Sillitoe on behalf of the Wellcome Sanger Institute COVID-19 Surveillance Team |
| EPI_ISL_2392237, EPI_ISL_2392238, EPI_ISL_2392239 | Lighthouse Lab in Alderley Park | Wellcome Sanger Institute for the COVID-19 Genomics UK (COG-UK) Consortium | Jacquelyn Wynn, Mairead Hyland, The Lighthouse Lab in Alderley Park and Alex Alderton, Roberto Amato, Jeffrey Barrett, Sonia Goncalves, Ewan Harrison, David K. Jackson, Ian Johnston, Dominic Kwiatkowski, Cordelia Langford, John Sillitoe on behalf of the Wellcome Sanger Institute COVID-19 Surveillance Team |
| EPI_ISL_2392242, EPI_ISL_2392243 | Lighthouse Lab in Milton Keynes | Wellcome Sanger Institute for the COVID-19 Genomics UK (COG-UK) Consortium | The Lighthouse Lab in Milton Keynes and Alex Alderton, Roberto Amato, Jeffrey Barrett, Sonia Goncalves, Ewan Harrison, David K. Jackson, Ian Johnston, Dominic Kwiatkowski, Cordelia Langford, John Sillitoe on behalf of the Wellcome Sanger Institute COVID-19 Surveillance Team |
| EPI_ISL_2392244, EPI_ISL_2392245, EPI_ISL_2392247, EPI_ISL_2392250, EPI_ISL_2392252, EPI_ISL_2392253, EPI_ISL_2392254, EPI_ISL_2392256, EPI_ISL_2392257, EPI_ISL_2392258, EPI_ISL_2392260, EPI_ISL_2392261, EPI_ISL_2392262 | | | |
| see above | Lighthouse Lab in Alderley Park | Wellcome Sanger Institute for the COVID-19 Genomics UK (COG-UK) Consortium | Jacquelyn Wynn, Mairead Hyland, The Lighthouse Lab in Alderley Park and Alex Alderton, Roberto Amato, Jeffrey Barrett, Sonia Goncalves, Ewan Harrison, David K. Jackson, Ian Johnston, Dominic Kwiatkowski, Cordelia Langford, John Sillitoe on behalf of the Wellcome Sanger Institute COVID-19 Surveillance Team |
| EPI_ISL_2392263 | Lighthouse Lab in Milton Keynes | Wellcome Sanger Institute for the COVID-19 Genomics UK (COG-UK) Consortium | The Lighthouse Lab in Milton Keynes and Alex Alderton, Roberto Amato, Jeffrey Barrett, Sonia Goncalves, Ewan Harrison, David K. Jackson, Ian Johnston, Dominic Kwiatkowski, Cordelia Langford, John Sillitoe on behalf of the Wellcome Sanger Institute COVID-19 Surveillance Team |
| EPI_ISL_2392264 | Lighthouse Lab in Alderley Park | Wellcome Sanger Institute for the COVID-19 Genomics UK (COG-UK) Consortium | Jacquelyn Wynn, Mairead Hyland, The Lighthouse Lab in Alderley Park and Alex Alderton, Roberto Amato, Jeffrey Barrett, Sonia Goncalves, Ewan Harrison, David K. Jackson, Ian Johnston, Dominic Kwiatkowski, Cordelia Langford, John Sillitoe on behalf of the Wellcome Sanger Institute COVID-19 Surveillance Team |
| EPI_ISL_2392265, EPI_ISL_2392266 | Lighthouse Lab in Milton Keynes | Wellcome Sanger Institute for the COVID-19 Genomics UK (COG-UK) Consortium | The Lighthouse Lab in Milton Keynes and Alex Alderton, Roberto Amato, Jeffrey Barrett, Sonia Goncalves, Ewan Harrison, David K. Jackson, Ian Johnston, Dominic Kwiatkowski, Cordelia Langford, John Sillitoe on behalf of the Wellcome Sanger Institute COVID-19 Surveillance Team |
| EPI_ISL_2392267, EPI_ISL_2392268, EPI_ISL_2392270, EPI_ISL_2392271, EPI_ISL_2392272, EPI_ISL_2392273, EPI_ISL_2392274, EPI_ISL_2392275, EPI_ISL_2392276 | Lighthouse Lab in Alderley Park | Wellcome Sanger Institute for the COVID-19 Genomics UK (COG-UK) Consortium | Jacquelyn Wynn, Mairead Hyland, The Lighthouse Lab in Alderley Park and Alex Alderton, Roberto Amato, Jeffrey Barrett, Sonia Goncalves, Ewan Harrison, David K. Jackson, Ian Johnston, Dominic Kwiatkowski, Cordelia Langford, John Sillitoe on behalf of the Wellcome Sanger Institute COVID-19 Surveillance Team |
| EPI_ISL_2392277 | Lighthouse Lab in Milton Keynes | Wellcome Sanger Institute for the COVID-19 Genomics UK (COG-UK) Consortium | The Lighthouse Lab in Milton Keynes and Alex Alderton, Roberto Amato, Jeffrey Barrett, Sonia Goncalves, Ewan Harrison, David K. Jackson, Ian Johnston, Dominic Kwiatkowski, Cordelia Langford, John Sillitoe on behalf of the Wellcome Sanger Institute COVID-19 Surveillance Team |
| EPI_ISL_2392278, EPI_ISL_2392280, EPI_ISL_2392281, EPI_ISL_2392282, EPI_ISL_2392285, EPI_ISL_2392286, EPI_ISL_2392287, EPI_ISL_2392288, EPI_ISL_2392291 | Lighthouse Lab in Alderley Park | Wellcome Sanger Institute for the COVID-19 Genomics UK (COG-UK) Consortium | Jacquelyn Wynn, Mairead Hyland, The Lighthouse Lab in Alderley Park and Alex Alderton, Roberto Amato, Jeffrey Barrett, Sonia Goncalves, Ewan Harrison, David K. Jackson, Ian Johnston, Dominic Kwiatkowski, Cordelia Langford, John Sillitoe on behalf of the Wellcome Sanger Institute COVID-19 Surveillance Team |
| EPI_ISL_2392292 | Lighthouse Lab in Milton Keynes | Wellcome Sanger Institute for the COVID-19 Genomics UK (COG-UK) Consortium | The Lighthouse Lab in Milton Keynes and Alex Alderton, Roberto Amato, Jeffrey Barrett, Sonia Goncalves, Ewan Harrison, David K. Jackson, Ian Johnston, Dominic Kwiatkowski, Cordelia Langford, John Sillitoe on behalf of the Wellcome Sanger Institute COVID-19 Surveillance Team |
| EPI_ISL_2392293 | Lighthouse Lab in Alderley Park | Wellcome Sanger Institute for the COVID-19 Genomics UK (COG-UK) Consortium | Jacquelyn Wynn, Mairead Hyland, The Lighthouse Lab in Alderley Park and Alex Alderton, Roberto Amato, Jeffrey Barrett, Sonia Goncalves, Ewan Harrison, David K. Jackson, Ian Johnston, Dominic Kwiatkowski, Cordelia Langford, John Sillitoe on behalf of the Wellcome Sanger Institute COVID-19 Surveillance Team |
| EPI_ISL_2392294 | Lighthouse Lab in Milton Keynes | Wellcome Sanger Institute for the COVID-19 Genomics UK (COG-UK) Consortium | The Lighthouse Lab in Milton Keynes and Alex Alderton, Roberto Amato, Jeffrey Barrett, Sonia Goncalves, Ewan Harrison, David K. Jackson, Ian Johnston, Dominic Kwiatkowski, Cordelia Langford, John Sillitoe on behalf of the Wellcome Sanger Institute COVID-19 Surveillance Team |
| EPI_ISL_2392295, EPI_ISL_2392296, EPI_ISL_2392297, EPI_ISL_2392298, EPI_ISL_2392299, EPI_ISL_2392300, EPI_ISL_2392302, EPI_ISL_2392303 | Lighthouse Lab in Alderley Park | Wellcome Sanger Institute for the COVID-19 Genomics UK (COG-UK) Consortium | Jacquelyn Wynn, Mairead Hyland, The Lighthouse Lab in Alderley Park and Alex Alderton, Roberto Amato, Jeffrey Barrett, Sonia Goncalves, Ewan Harrison, David K. Jackson, Ian Johnston, Dominic Kwiatkowski, Cordelia Langford, John Sillitoe on behalf of the Wellcome Sanger Institute COVID-19 Surveillance Team |
| EPI_ISL_2392304 | Lighthouse Lab in Milton Keynes | Wellcome Sanger Institute for the COVID-19 Genomics UK (COG-UK) Consortium | The Lighthouse Lab in Milton Keynes and Alex Alderton, Roberto Amato, Jeffrey Barrett, Sonia Goncalves, Ewan Harrison, David K. Jackson, Ian Johnston, Dominic Kwiatkowski, Cordelia Langford, John Sillitoe on behalf of the Wellcome Sanger Institute COVID-19 Surveillance Team |
| EPI_ISL_2392305, EPI_ISL_2392307, EPI_ISL_2392308, EPI_ISL_2392309 | Lighthouse Lab in Alderley Park | Wellcome Sanger Institute for the COVID-19 Genomics UK (COG-UK) Consortium | Jacquelyn Wynn, Mairead Hyland, The Lighthouse Lab in Alderley Park and Alex Alderton, Roberto Amato, Jeffrey Barrett, Sonia Goncalves, Ewan Harrison, David K. Jackson, Ian Johnston, Dominic Kwiatkowski, Cordelia Langford, John Sillitoe on behalf of the Wellcome Sanger Institute COVID-19 Surveillance Team |
| EPI_ISL_2392310 | Lighthouse Lab in Milton Keynes | Wellcome Sanger Institute for the COVID-19 Genomics UK (COG-UK) Consortium | The Lighthouse Lab in Milton Keynes and Alex Alderton, Roberto Amato, Jeffrey Barrett, Sonia Goncalves, Ewan Harrison, David K. Jackson, Ian Johnston, Dominic Kwiatkowski, Cordelia Langford, John Sillitoe on behalf of the Wellcome Sanger Institute COVID-19 Surveillance Team |

| | | | |
|---|---|---|---|
| EPI_ISL_2392311, EPI_ISL_2392312, EPI_ISL_2392313, EPI_ISL_2392314, EPI_ISL_2392315 | Lighthouse Lab in Alderley Park | Wellcome Sanger Institute for the COVID-19 Genomics UK (COG-UK) Consortium | Jacquelyn Wynn, Mairead Hyland, The Lighthouse Lab in Alderley Park and Alex Alderton, Roberto Amato, Jeffrey Barrett, Sonia Goncalves, Ewan Harrison, David K. Jackson, Ian Johnston, Dominic Kwiatkowski, Cordelia Langford, John Sillitoe on behalf of the Wellcome Sanger Institute COVID-19 Surveillance Team |
| EPI_ISL_2392316 | Lighthouse Lab in Milton Keynes | Wellcome Sanger Institute for the COVID-19 Genomics UK (COG-UK) Consortium | The Lighthouse Lab in Milton Keynes and Alex Alderton, Roberto Amato, Jeffrey Barrett, Sonia Goncalves, Ewan Harrison, David K. Jackson, Ian Johnston, Dominic Kwiatkowski, Cordelia Langford, John Sillitoe on behalf of the Wellcome Sanger Institute COVID-19 Surveillance Team |
| EPI_ISL_2392318, EPI_ISL_2392319, EPI_ISL_2392320, EPI_ISL_2392321, EPI_ISL_2392322, EPI_ISL_2392324, EPI_ISL_2392325, EPI_ISL_2392326, EPI_ISL_2392327, EPI_ISL_2392328, EPI_ISL_2392329, EPI_ISL_2392330, EPI_ISL_2392331, EPI_ISL_2392332, EPI_ISL_2392334, EPI_ISL_2392335 | | | |
| see above | Lighthouse Lab in Alderley Park | Wellcome Sanger Institute for the COVID-19 Genomics UK (COG-UK) Consortium | Jacquelyn Wynn, Mairead Hyland, The Lighthouse Lab in Alderley Park and Alex Alderton, Roberto Amato, Jeffrey Barrett, Sonia Goncalves, Ewan Harrison, David K. Jackson, Ian Johnston, Dominic Kwiatkowski, Cordelia Langford, John Sillitoe on behalf of the Wellcome Sanger Institute COVID-19 Surveillance Team |
| EPI_ISL_2392336 | Lighthouse Lab in Milton Keynes | Wellcome Sanger Institute for the COVID-19 Genomics UK (COG-UK) Consortium | The Lighthouse Lab in Milton Keynes and Alex Alderton, Roberto Amato, Jeffrey Barrett, Sonia Goncalves, Ewan Harrison, David K. Jackson, Ian Johnston, Dominic Kwiatkowski, Cordelia Langford, John Sillitoe on behalf of the Wellcome Sanger Institute COVID-19 Surveillance Team |
| EPI_ISL_2392337 | Lighthouse Lab in Alderley Park | Wellcome Sanger Institute for the COVID-19 Genomics UK (COG-UK) Consortium | Jacquelyn Wynn, Mairead Hyland, The Lighthouse Lab in Alderley Park and Alex Alderton, Roberto Amato, Jeffrey Barrett, Sonia Goncalves, Ewan Harrison, David K. Jackson, Ian Johnston, Dominic Kwiatkowski, Cordelia Langford, John Sillitoe on behalf of the Wellcome Sanger Institute COVID-19 Surveillance Team |
| EPI_ISL_2392338 | Lighthouse Lab in Milton Keynes | Wellcome Sanger Institute for the COVID-19 Genomics UK (COG-UK) Consortium | The Lighthouse Lab in Milton Keynes and Alex Alderton, Roberto Amato, Jeffrey Barrett, Sonia Goncalves, Ewan Harrison, David K. Jackson, Ian Johnston, Dominic Kwiatkowski, Cordelia Langford, John Sillitoe on behalf of the Wellcome Sanger Institute COVID-19 Surveillance Team |
| EPI_ISL_2392339, EPI_ISL_2392340, EPI_ISL_2392341, EPI_ISL_2392342, EPI_ISL_2392343, EPI_ISL_2392344 | Lighthouse Lab in Alderley Park | Wellcome Sanger Institute for the COVID-19 Genomics UK (COG-UK) Consortium | Jacquelyn Wynn, Mairead Hyland, The Lighthouse Lab in Alderley Park and Alex Alderton, Roberto Amato, Jeffrey Barrett, Sonia Goncalves, Ewan Harrison, David K. Jackson, Ian Johnston, Dominic Kwiatkowski, Cordelia Langford, John Sillitoe on behalf of the Wellcome Sanger Institute COVID-19 Surveillance Team |
| EPI_ISL_2392345 | Lighthouse Lab in Milton Keynes | Wellcome Sanger Institute for the COVID-19 Genomics UK (COG-UK) Consortium | The Lighthouse Lab in Milton Keynes and Alex Alderton, Roberto Amato, Jeffrey Barrett, Sonia Goncalves, Ewan Harrison, David K. Jackson, Ian Johnston, Dominic Kwiatkowski, Cordelia Langford, John Sillitoe on behalf of the Wellcome Sanger Institute COVID-19 Surveillance Team |
| EPI_ISL_2392346, EPI_ISL_2392347, EPI_ISL_2392348 | Lighthouse Lab in Alderley Park | Wellcome Sanger Institute for the COVID-19 Genomics UK (COG-UK) Consortium | Jacquelyn Wynn, Mairead Hyland, The Lighthouse Lab in Alderley Park and Alex Alderton, Roberto Amato, Jeffrey Barrett, Sonia Goncalves, Ewan Harrison, David K. Jackson, Ian Johnston, Dominic Kwiatkowski, Cordelia Langford, John Sillitoe on behalf of the Wellcome Sanger Institute COVID-19 Surveillance Team |
| EPI_ISL_2392349 | Lighthouse Lab in Milton Keynes | Wellcome Sanger Institute for the COVID-19 Genomics UK (COG-UK) Consortium | The Lighthouse Lab in Milton Keynes and Alex Alderton, Roberto Amato, Jeffrey Barrett, Sonia Goncalves, Ewan Harrison, David K. Jackson, Ian Johnston, Dominic Kwiatkowski, Cordelia Langford, John Sillitoe on behalf of the Wellcome Sanger Institute COVID-19 Surveillance Team |
| EPI_ISL_2392350, EPI_ISL_2392351, EPI_ISL_2392352 | Lighthouse Lab in Alderley Park | Wellcome Sanger Institute for the COVID-19 Genomics UK (COG-UK) Consortium | Jacquelyn Wynn, Mairead Hyland, The Lighthouse Lab in Alderley Park and Alex Alderton, Roberto Amato, Jeffrey Barrett, Sonia Goncalves, Ewan Harrison, David K. Jackson, Ian Johnston, Dominic Kwiatkowski, Cordelia Langford, John Sillitoe on behalf of the Wellcome Sanger Institute COVID-19 Surveillance Team |
| EPI_ISL_2392353, EPI_ISL_2392355, EPI_ISL_2392356 | Lighthouse Lab in Milton Keynes | Wellcome Sanger Institute for the COVID-19 Genomics UK (COG-UK) Consortium | The Lighthouse Lab in Milton Keynes and Alex Alderton, Roberto Amato, Jeffrey Barrett, Sonia Goncalves, Ewan Harrison, David K. Jackson, Ian Johnston, Dominic Kwiatkowski, Cordelia Langford, John Sillitoe on behalf of the Wellcome Sanger Institute COVID-19 Surveillance Team |
| EPI_ISL_2392357, EPI_ISL_2392359, EPI_ISL_2392360 | Lighthouse Lab in Alderley Park | Wellcome Sanger Institute for the COVID-19 Genomics UK (COG-UK) Consortium | Jacquelyn Wynn, Mairead Hyland, The Lighthouse Lab in Alderley Park and Alex Alderton, Roberto Amato, Jeffrey Barrett, Sonia Goncalves, Ewan Harrison, David K. Jackson, Ian Johnston, Dominic Kwiatkowski, Cordelia Langford, John Sillitoe on behalf of the Wellcome Sanger Institute COVID-19 Surveillance Team |
| EPI_ISL_2392362 | Lighthouse Lab in Milton Keynes | Wellcome Sanger Institute for the COVID-19 Genomics UK (COG-UK) Consortium | The Lighthouse Lab in Milton Keynes and Alex Alderton, Roberto Amato, Jeffrey Barrett, Sonia Goncalves, Ewan Harrison, David K. Jackson, Ian Johnston, Dominic Kwiatkowski, Cordelia Langford, John Sillitoe on behalf of the Wellcome Sanger Institute COVID-19 Surveillance Team |
| EPI_ISL_2392364, EPI_ISL_2392365, EPI_ISL_2392366, EPI_ISL_2392367 | Lighthouse Lab in Alderley Park | Wellcome Sanger Institute for the COVID-19 Genomics UK (COG-UK) Consortium | Jacquelyn Wynn, Mairead Hyland, The Lighthouse Lab in Alderley Park and Alex Alderton, Roberto Amato, Jeffrey Barrett, Sonia Goncalves, Ewan Harrison, David K. Jackson, Ian Johnston, Dominic Kwiatkowski, Cordelia Langford, John Sillitoe on behalf of the Wellcome Sanger Institute COVID-19 Surveillance Team |
| EPI_ISL_2392368 | Lighthouse Lab in Milton Keynes | Wellcome Sanger Institute for the COVID-19 Genomics UK (COG-UK) Consortium | The Lighthouse Lab in Milton Keynes and Alex Alderton, Roberto Amato, Jeffrey Barrett, Sonia Goncalves, Ewan Harrison, David K. Jackson, Ian Johnston, Dominic Kwiatkowski, Cordelia Langford, John Sillitoe on behalf of the Wellcome Sanger Institute COVID-19 Surveillance Team |
| EPI_ISL_2392369, EPI_ISL_2392371, EPI_ISL_2392372, EPI_ISL_2392373, EPI_ISL_2392374, EPI_ISL_2392375, EPI_ISL_2392376, EPI_ISL_2392378, EPI_ISL_2392380 | Lighthouse Lab in Alderley Park | Wellcome Sanger Institute for the COVID-19 Genomics UK (COG-UK) Consortium | Jacquelyn Wynn, Mairead Hyland, The Lighthouse Lab in Alderley Park and Alex Alderton, Roberto Amato, Jeffrey Barrett, Sonia Goncalves, Ewan Harrison, David K. Jackson, Ian Johnston, Dominic Kwiatkowski, Cordelia Langford, John Sillitoe on behalf of the Wellcome Sanger Institute COVID-19 Surveillance Team |
| EPI_ISL_2392381 | Lighthouse Lab in Milton Keynes | Wellcome Sanger Institute for the COVID-19 Genomics UK (COG-UK) Consortium | The Lighthouse Lab in Milton Keynes and Alex Alderton, Roberto Amato, Jeffrey Barrett, Sonia Goncalves, Ewan Harrison, David K. Jackson, Ian Johnston, Dominic Kwiatkowski, Cordelia Langford, John Sillitoe on behalf of the Wellcome Sanger Institute COVID-19 Surveillance Team |
| EPI_ISL_2392382, EPI_ISL_2392383, EPI_ISL_2392384, EPI_ISL_2392385 | Lighthouse Lab in Alderley Park | Wellcome Sanger Institute for the COVID-19 Genomics UK (COG-UK) Consortium | Jacquelyn Wynn, Mairead Hyland, The Lighthouse Lab in Alderley Park and Alex Alderton, Roberto Amato, Jeffrey Barrett, Sonia Goncalves, Ewan Harrison, David K. Jackson, Ian Johnston, Dominic Kwiatkowski, Cordelia Langford, John Sillitoe on behalf of the Wellcome Sanger Institute COVID-19 Surveillance Team |
| EPI_ISL_2392387 | Lighthouse Lab in Milton Keynes | Wellcome Sanger Institute for the COVID-19 Genomics UK (COG-UK) Consortium | The Lighthouse Lab in Milton Keynes and Alex Alderton, Roberto Amato, Jeffrey Barrett, Sonia Goncalves, Ewan Harrison, David K. Jackson, Ian Johnston, Dominic Kwiatkowski, Cordelia Langford, John Sillitoe on behalf of the Wellcome Sanger Institute COVID-19 Surveillance Team |
| EPI_ISL_2392388 | Lighthouse Lab in Alderley Park | Wellcome Sanger Institute for the COVID-19 Genomics UK (COG-UK) Consortium | Jacquelyn Wynn, Mairead Hyland, The Lighthouse Lab in Alderley Park and Alex Alderton, Roberto Amato, Jeffrey Barrett, Sonia Goncalves, Ewan Harrison, David K. Jackson, Ian Johnston, Dominic Kwiatkowski, Cordelia Langford, John Sillitoe on behalf of the Wellcome Sanger Institute COVID-19 Surveillance Team |
| EPI_ISL_2392389 | Lighthouse Lab in Glasgow | Wellcome Sanger Institute for the COVID-19 Genomics UK (COG-UK) Consortium | Harper VanSteenhouse, Yumi Kasai, David Gray, Carol Clugston, Anna Dominiczak and Alex Alderton, Roberto Amato, Jeffrey Barrett, Sonia Goncalves, Ewan Harrison, David K. Jackson, Ian Johnston, Dominic Kwiatkowski, Cordelia Langford, John Sillitoe on behalf of the Wellcome Sanger Institute COVID-19 Surveillance Team |
| EPI_ISL_2392390, EPI_ISL_2392391 | Lighthouse Lab in Milton Keynes | Wellcome Sanger Institute for the COVID-19 Genomics UK (COG-UK) Consortium | The Lighthouse Lab in Milton Keynes and Alex Alderton, Roberto Amato, Jeffrey Barrett, Sonia Goncalves, Ewan Harrison, David K. Jackson, Ian Johnston, Dominic Kwiatkowski, Cordelia Langford, John Sillitoe on behalf of the Wellcome Sanger Institute COVID-19 Surveillance Team |
| EPI_ISL_2392392 | Lighthouse Lab in Alderley Park | Wellcome Sanger Institute for the COVID-19 Genomics UK (COG-UK) Consortium | Jacquelyn Wynn, Mairead Hyland, The Lighthouse Lab in Alderley Park and Alex Alderton, Roberto Amato, Jeffrey Barrett, Sonia Goncalves, Ewan Harrison, David K. Jackson, Ian Johnston, Dominic Kwiatkowski, Cordelia Langford, John Sillitoe on behalf of the Wellcome Sanger Institute COVID-19 Surveillance Team |
| EPI_ISL_2392393 | Lighthouse Lab in Milton Keynes | Wellcome Sanger Institute for the COVID-19 Genomics UK (COG-UK) Consortium | The Lighthouse Lab in Milton Keynes and Alex Alderton, Roberto Amato, Jeffrey Barrett, Sonia Goncalves, Ewan Harrison, David K. Jackson, Ian Johnston, Dominic Kwiatkowski, Cordelia Langford, John Sillitoe on behalf of the Wellcome Sanger Institute COVID-19 Surveillance Team |
| EPI_ISL_2392394, EPI_ISL_2392395 | Lighthouse Lab in Alderley Park | Wellcome Sanger Institute for the COVID-19 Genomics UK (COG-UK) Consortium | Jacquelyn Wynn, Mairead Hyland, The Lighthouse Lab in Alderley Park and Alex Alderton, Roberto Amato, Jeffrey Barrett, Sonia Goncalves, Ewan Harrison, David K. Jackson, Ian Johnston, Dominic Kwiatkowski, Cordelia Langford, John Sillitoe on behalf of the Wellcome Sanger Institute COVID-19 Surveillance Team |
| EPI_ISL_2392397, EPI_ISL_2392398 | Lighthouse Lab in Milton Keynes | Wellcome Sanger Institute for the COVID-19 Genomics UK (COG-UK) Consortium | The Lighthouse Lab in Milton Keynes and Alex Alderton, Roberto Amato, Jeffrey Barrett, Sonia Goncalves, Ewan Harrison, David K. Jackson, Ian Johnston, Dominic Kwiatkowski, Cordelia Langford, John Sillitoe on behalf of the Wellcome Sanger Institute COVID-19 Surveillance Team |
| EPI_ISL_2392399, EPI_ISL_2392400, EPI_ISL_2392401, EPI_ISL_2392402, EPI_ISL_2392403 | Lighthouse Lab in Alderley Park | Wellcome Sanger Institute for the COVID-19 Genomics UK (COG-UK) Consortium | Jacquelyn Wynn, Mairead Hyland, The Lighthouse Lab in Alderley Park and Alex Alderton, Roberto Amato, Jeffrey Barrett, Sonia Goncalves, Ewan Harrison, David K. Jackson, Ian Johnston, Dominic Kwiatkowski, Cordelia Langford, John Sillitoe on behalf of the Wellcome Sanger Institute COVID-19 Surveillance Team |
| EPI_ISL_2392404, EPI_ISL_2392405 | Lighthouse Lab in Milton Keynes | Wellcome Sanger Institute for the COVID-19 Genomics UK (COG-UK) Consortium | The Lighthouse Lab in Milton Keynes and Alex Alderton, Roberto Amato, Jeffrey Barrett, Sonia Goncalves, Ewan Harrison, David K. Jackson, Ian Johnston, Dominic Kwiatkowski, Cordelia Langford, John Sillitoe on behalf of the Wellcome Sanger Institute COVID-19 Surveillance Team |
| EPI_ISL_2392407, EPI_ISL_2392408, EPI_ISL_2392409, EPI_ISL_2392410, EPI_ISL_2392411, EPI_ISL_2392412, EPI_ISL_2392413, EPI_ISL_2392414, EPI_ISL_2392415, EPI_ISL_2392418 | Lighthouse Lab in Alderley Park | Wellcome Sanger Institute for the COVID-19 Genomics UK (COG-UK) Consortium | Jacquelyn Wynn, Mairead Hyland, The Lighthouse Lab in Alderley Park and Alex Alderton, Roberto Amato, Jeffrey Barrett, Sonia Goncalves, Ewan Harrison, David K. Jackson, Ian Johnston, Dominic Kwiatkowski, Cordelia Langford, John Sillitoe on behalf of the Wellcome Sanger Institute COVID-19 Surveillance Team |
| EPI_ISL_2392421 | Lighthouse Lab in Milton Keynes | Wellcome Sanger Institute for the COVID-19 Genomics UK (COG-UK) Consortium | The Lighthouse Lab in Milton Keynes and Alex Alderton, Roberto Amato, Jeffrey Barrett, Sonia Goncalves, Ewan Harrison, David K. Jackson, Ian Johnston, Dominic Kwiatkowski, Cordelia Langford, John Sillitoe on behalf of the Wellcome Sanger Institute COVID-19 Surveillance Team |

| | | | |
|---|---|---|---|
| EPI_ISL_2392422, EPI_ISL_2392423, EPI_ISL_2392425, EPI_ISL_2392426, EPI_ISL_2392427, EPI_ISL_2392428, EPI_ISL_2392429 | Lighthouse Lab in Alderley Park | Wellcome Sanger Institute for the COVID-19 Genomics UK (COG-UK) Consortium | Jacquelyn Wynn, Mairead Hyland, The Lighthouse Lab in Alderley Park and Alex Alderton, Roberto Amato, Jeffrey Barrett, Sonia Goncalves, Ewan Harrison, David K. Jackson, Ian Johnston, Dominic Kwiatkowski, Cordelia Langford, John Sillitoe on behalf of the Wellcome Sanger Institute COVID-19 Surveillance Team |
| EPI_ISL_2392430 | Lighthouse Lab in Milton Keynes | Wellcome Sanger Institute for the COVID-19 Genomics UK (COG-UK) Consortium | The Lighthouse Lab in Milton Keynes and Alex Alderton, Roberto Amato, Jeffrey Barrett, Sonia Goncalves, Ewan Harrison, David K. Jackson, Ian Johnston, Dominic Kwiatkowski, Cordelia Langford, John Sillitoe on behalf of the Wellcome Sanger Institute COVID-19 Surveillance Team |
| EPI_ISL_2392431 | Lighthouse Lab in Alderley Park | Wellcome Sanger Institute for the COVID-19 Genomics UK (COG-UK) Consortium | Jacquelyn Wynn, Mairead Hyland, The Lighthouse Lab in Alderley Park and Alex Alderton, Roberto Amato, Jeffrey Barrett, Sonia Goncalves, Ewan Harrison, David K. Jackson, Ian Johnston, Dominic Kwiatkowski, Cordelia Langford, John Sillitoe on behalf of the Wellcome Sanger Institute COVID-19 Surveillance Team |
| EPI_ISL_2392433 | Lighthouse Lab in Milton Keynes | Wellcome Sanger Institute for the COVID-19 Genomics UK (COG-UK) Consortium | The Lighthouse Lab in Milton Keynes and Alex Alderton, Roberto Amato, Jeffrey Barrett, Sonia Goncalves, Ewan Harrison, David K. Jackson, Ian Johnston, Dominic Kwiatkowski, Cordelia Langford, John Sillitoe on behalf of the Wellcome Sanger Institute COVID-19 Surveillance Team |
| EPI_ISL_2392434, EPI_ISL_2392435 | Lighthouse Lab in Alderley Park | Wellcome Sanger Institute for the COVID-19 Genomics UK (COG-UK) Consortium | Jacquelyn Wynn, Mairead Hyland, The Lighthouse Lab in Alderley Park and Alex Alderton, Roberto Amato, Jeffrey Barrett, Sonia Goncalves, Ewan Harrison, David K. Jackson, Ian Johnston, Dominic Kwiatkowski, Cordelia Langford, John Sillitoe on behalf of the Wellcome Sanger Institute COVID-19 Surveillance Team |
| EPI_ISL_2392436, EPI_ISL_2392438 | Lighthouse Lab in Milton Keynes | Wellcome Sanger Institute for the COVID-19 Genomics UK (COG-UK) Consortium | The Lighthouse Lab in Milton Keynes and Alex Alderton, Roberto Amato, Jeffrey Barrett, Sonia Goncalves, Ewan Harrison, David K. Jackson, Ian Johnston, Dominic Kwiatkowski, Cordelia Langford, John Sillitoe on behalf of the Wellcome Sanger Institute COVID-19 Surveillance Team |
| EPI_ISL_2392439, EPI_ISL_2392440 | Lighthouse Lab in Alderley Park | Wellcome Sanger Institute for the COVID-19 Genomics UK (COG-UK) Consortium | Jacquelyn Wynn, Mairead Hyland, The Lighthouse Lab in Alderley Park and Alex Alderton, Roberto Amato, Jeffrey Barrett, Sonia Goncalves, Ewan Harrison, David K. Jackson, Ian Johnston, Dominic Kwiatkowski, Cordelia Langford, John Sillitoe on behalf of the Wellcome Sanger Institute COVID-19 Surveillance Team |
| EPI_ISL_2392441 | Lighthouse Lab in Milton Keynes | Wellcome Sanger Institute for the COVID-19 Genomics UK (COG-UK) Consortium | The Lighthouse Lab in Milton Keynes and Alex Alderton, Roberto Amato, Jeffrey Barrett, Sonia Goncalves, Ewan Harrison, David K. Jackson, Ian Johnston, Dominic Kwiatkowski, Cordelia Langford, John Sillitoe on behalf of the Wellcome Sanger Institute COVID-19 Surveillance Team |
| EPI_ISL_2392443, EPI_ISL_2392444 | Lighthouse Lab in Alderley Park | Wellcome Sanger Institute for the COVID-19 Genomics UK (COG-UK) Consortium | Jacquelyn Wynn, Mairead Hyland, The Lighthouse Lab in Alderley Park and Alex Alderton, Roberto Amato, Jeffrey Barrett, Sonia Goncalves, Ewan Harrison, David K. Jackson, Ian Johnston, Dominic Kwiatkowski, Cordelia Langford, John Sillitoe on behalf of the Wellcome Sanger Institute COVID-19 Surveillance Team |
| EPI_ISL_2392445 | Lighthouse Lab in Milton Keynes | Wellcome Sanger Institute for the COVID-19 Genomics UK (COG-UK) Consortium | The Lighthouse Lab in Milton Keynes and Alex Alderton, Roberto Amato, Jeffrey Barrett, Sonia Goncalves, Ewan Harrison, David K. Jackson, Ian Johnston, Dominic Kwiatkowski, Cordelia Langford, John Sillitoe on behalf of the Wellcome Sanger Institute COVID-19 Surveillance Team |
| EPI_ISL_2392446, EPI_ISL_2392447 | Lighthouse Lab in Alderley Park | Wellcome Sanger Institute for the COVID-19 Genomics UK (COG-UK) Consortium | Jacquelyn Wynn, Mairead Hyland, The Lighthouse Lab in Alderley Park and Alex Alderton, Roberto Amato, Jeffrey Barrett, Sonia Goncalves, Ewan Harrison, David K. Jackson, Ian Johnston, Dominic Kwiatkowski, Cordelia Langford, John Sillitoe on behalf of the Wellcome Sanger Institute COVID-19 Surveillance Team |
| EPI_ISL_2392448 | Lighthouse Lab in Milton Keynes | Wellcome Sanger Institute for the COVID-19 Genomics UK (COG-UK) Consortium | The Lighthouse Lab in Milton Keynes and Alex Alderton, Roberto Amato, Jeffrey Barrett, Sonia Goncalves, Ewan Harrison, David K. Jackson, Ian Johnston, Dominic Kwiatkowski, Cordelia Langford, John Sillitoe on behalf of the Wellcome Sanger Institute COVID-19 Surveillance Team |
| EPI_ISL_2392449, EPI_ISL_2392450, EPI_ISL_2392451, EPI_ISL_2392452 | Lighthouse Lab in Alderley Park | Wellcome Sanger Institute for the COVID-19 Genomics UK (COG-UK) Consortium | Jacquelyn Wynn, Mairead Hyland, The Lighthouse Lab in Alderley Park and Alex Alderton, Roberto Amato, Jeffrey Barrett, Sonia Goncalves, Ewan Harrison, David K. Jackson, Ian Johnston, Dominic Kwiatkowski, Cordelia Langford, John Sillitoe on behalf of the Wellcome Sanger Institute COVID-19 Surveillance Team |
| EPI_ISL_2392454 | Lighthouse Lab in Milton Keynes | Wellcome Sanger Institute for the COVID-19 Genomics UK (COG-UK) Consortium | The Lighthouse Lab in Milton Keynes and Alex Alderton, Roberto Amato, Jeffrey Barrett, Sonia Goncalves, Ewan Harrison, David K. Jackson, Ian Johnston, Dominic Kwiatkowski, Cordelia Langford, John Sillitoe on behalf of the Wellcome Sanger Institute COVID-19 Surveillance Team |
| EPI_ISL_2392455 | Lighthouse Lab in Alderley Park | Wellcome Sanger Institute for the COVID-19 Genomics UK (COG-UK) Consortium | Jacquelyn Wynn, Mairead Hyland, The Lighthouse Lab in Alderley Park and Alex Alderton, Roberto Amato, Jeffrey Barrett, Sonia Goncalves, Ewan Harrison, David K. Jackson, Ian Johnston, Dominic Kwiatkowski, Cordelia Langford, John Sillitoe on behalf of the Wellcome Sanger Institute COVID-19 Surveillance Team |
| EPI_ISL_2392456, EPI_ISL_2392457 | Lighthouse Lab in Milton Keynes | Wellcome Sanger Institute for the COVID-19 Genomics UK (COG-UK) Consortium | The Lighthouse Lab in Milton Keynes and Alex Alderton, Roberto Amato, Jeffrey Barrett, Sonia Goncalves, Ewan Harrison, David K. Jackson, Ian Johnston, Dominic Kwiatkowski, Cordelia Langford, John Sillitoe on behalf of the Wellcome Sanger Institute COVID-19 Surveillance Team |
| EPI_ISL_2392458, EPI_ISL_2392459, EPI_ISL_2392460, EPI_ISL_2392461 | Lighthouse Lab in Alderley Park | Wellcome Sanger Institute for the COVID-19 Genomics UK (COG-UK) Consortium | Jacquelyn Wynn, Mairead Hyland, The Lighthouse Lab in Alderley Park and Alex Alderton, Roberto Amato, Jeffrey Barrett, Sonia Goncalves, Ewan Harrison, David K. Jackson, Ian Johnston, Dominic Kwiatkowski, Cordelia Langford, John Sillitoe on behalf of the Wellcome Sanger Institute COVID-19 Surveillance Team |
| EPI_ISL_2392462 | Lighthouse Lab in Milton Keynes | Wellcome Sanger Institute for the COVID-19 Genomics UK (COG-UK) Consortium | The Lighthouse Lab in Milton Keynes and Alex Alderton, Roberto Amato, Jeffrey Barrett, Sonia Goncalves, Ewan Harrison, David K. Jackson, Ian Johnston, Dominic Kwiatkowski, Cordelia Langford, John Sillitoe on behalf of the Wellcome Sanger Institute COVID-19 Surveillance Team |
| EPI_ISL_2392463, EPI_ISL_2392464, EPI_ISL_2392465, EPI_ISL_2392467 | Lighthouse Lab in Alderley Park | Wellcome Sanger Institute for the COVID-19 Genomics UK (COG-UK) Consortium | Jacquelyn Wynn, Mairead Hyland, The Lighthouse Lab in Alderley Park and Alex Alderton, Roberto Amato, Jeffrey Barrett, Sonia Goncalves, Ewan Harrison, David K. Jackson, Ian Johnston, Dominic Kwiatkowski, Cordelia Langford, John Sillitoe on behalf of the Wellcome Sanger Institute COVID-19 Surveillance Team |
| EPI_ISL_2392468 | Lighthouse Lab in Milton Keynes | Wellcome Sanger Institute for the COVID-19 Genomics UK (COG-UK) Consortium | The Lighthouse Lab in Milton Keynes and Alex Alderton, Roberto Amato, Jeffrey Barrett, Sonia Goncalves, Ewan Harrison, David K. Jackson, Ian Johnston, Dominic Kwiatkowski, Cordelia Langford, John Sillitoe on behalf of the Wellcome Sanger Institute COVID-19 Surveillance Team |
| EPI_ISL_2392469, EPI_ISL_2392470, EPI_ISL_2392471, EPI_ISL_2392472, EPI_ISL_2392473, EPI_ISL_2392474, EPI_ISL_2392475, EPI_ISL_2392476, EPI_ISL_2392477, EPI_ISL_2392478, EPI_ISL_2392479, EPI_ISL_2392480 | | | |
| see above | Lighthouse Lab in Alderley Park | Wellcome Sanger Institute for the COVID-19 Genomics UK (COG-UK) Consortium | Jacquelyn Wynn, Mairead Hyland, The Lighthouse Lab in Alderley Park and Alex Alderton, Roberto Amato, Jeffrey Barrett, Sonia Goncalves, Ewan Harrison, David K. Jackson, Ian Johnston, Dominic Kwiatkowski, Cordelia Langford, John Sillitoe on behalf of the Wellcome Sanger Institute COVID-19 Surveillance Team |
| EPI_ISL_2392481 | Lighthouse Lab in Milton Keynes | Wellcome Sanger Institute for the COVID-19 Genomics UK (COG-UK) Consortium | The Lighthouse Lab in Milton Keynes and Alex Alderton, Roberto Amato, Jeffrey Barrett, Sonia Goncalves, Ewan Harrison, David K. Jackson, Ian Johnston, Dominic Kwiatkowski, Cordelia Langford, John Sillitoe on behalf of the Wellcome Sanger Institute COVID-19 Surveillance Team |
| EPI_ISL_2392482, EPI_ISL_2392483, EPI_ISL_2392484, EPI_ISL_2392485, EPI_ISL_2392486, EPI_ISL_2392487, EPI_ISL_2392489, EPI_ISL_2392490, EPI_ISL_2392491, EPI_ISL_2392492, EPI_ISL_2392493, EPI_ISL_2392494, EPI_ISL_2392495, EPI_ISL_2392496, EPI_ISL_2392497, EPI_ISL_2392498, EPI_ISL_2392499, EPI_ISL_2392500 | | | |
| see above | Lighthouse Lab in Alderley Park | Wellcome Sanger Institute for the COVID-19 Genomics UK (COG-UK) Consortium | Jacquelyn Wynn, Mairead Hyland, The Lighthouse Lab in Alderley Park and Alex Alderton, Roberto Amato, Jeffrey Barrett, Sonia Goncalves, Ewan Harrison, David K. Jackson, Ian Johnston, Dominic Kwiatkowski, Cordelia Langford, John Sillitoe on behalf of the Wellcome Sanger Institute COVID-19 Surveillance Team |
| EPI_ISL_2392501 | Lighthouse Lab in Milton Keynes | Wellcome Sanger Institute for the COVID-19 Genomics UK (COG-UK) Consortium | The Lighthouse Lab in Milton Keynes and Alex Alderton, Roberto Amato, Jeffrey Barrett, Sonia Goncalves, Ewan Harrison, David K. Jackson, Ian Johnston, Dominic Kwiatkowski, Cordelia Langford, John Sillitoe on behalf of the Wellcome Sanger Institute COVID-19 Surveillance Team |
| EPI_ISL_2392502, EPI_ISL_2392503 | Lighthouse Lab in Alderley Park | Wellcome Sanger Institute for the COVID-19 Genomics UK (COG-UK) Consortium | Jacquelyn Wynn, Mairead Hyland, The Lighthouse Lab in Alderley Park and Alex Alderton, Roberto Amato, Jeffrey Barrett, Sonia Goncalves, Ewan Harrison, David K. Jackson, Ian Johnston, Dominic Kwiatkowski, Cordelia Langford, John Sillitoe on behalf of the Wellcome Sanger Institute COVID-19 Surveillance Team |
| EPI_ISL_2392504 | Lighthouse Lab in Milton Keynes | Wellcome Sanger Institute for the COVID-19 Genomics UK (COG-UK) Consortium | The Lighthouse Lab in Milton Keynes and Alex Alderton, Roberto Amato, Jeffrey Barrett, Sonia Goncalves, Ewan Harrison, David K. Jackson, Ian Johnston, Dominic Kwiatkowski, Cordelia Langford, John Sillitoe on behalf of the Wellcome Sanger Institute COVID-19 Surveillance Team |
| EPI_ISL_2392505, EPI_ISL_2392506, EPI_ISL_2392507, EPI_ISL_2392508, EPI_ISL_2392509 | Lighthouse Lab in Alderley Park | Wellcome Sanger Institute for the COVID-19 Genomics UK (COG-UK) Consortium | Jacquelyn Wynn, Mairead Hyland, The Lighthouse Lab in Alderley Park and Alex Alderton, Roberto Amato, Jeffrey Barrett, Sonia Goncalves, Ewan Harrison, David K. Jackson, Ian Johnston, Dominic Kwiatkowski, Cordelia Langford, John Sillitoe on behalf of the Wellcome Sanger Institute COVID-19 Surveillance Team |
| EPI_ISL_2392510 | Lighthouse Lab in Milton Keynes | Wellcome Sanger Institute for the COVID-19 Genomics UK (COG-UK) Consortium | The Lighthouse Lab in Milton Keynes and Alex Alderton, Roberto Amato, Jeffrey Barrett, Sonia Goncalves, Ewan Harrison, David K. Jackson, Ian Johnston, Dominic Kwiatkowski, Cordelia Langford, John Sillitoe on behalf of the Wellcome Sanger Institute COVID-19 Surveillance Team |
| EPI_ISL_2392511, EPI_ISL_2392512 | Lighthouse Lab in Alderley Park | Wellcome Sanger Institute for the COVID-19 Genomics UK (COG-UK) Consortium | Jacquelyn Wynn, Mairead Hyland, The Lighthouse Lab in Alderley Park and Alex Alderton, Roberto Amato, Jeffrey Barrett, Sonia Goncalves, Ewan Harrison, David K. Jackson, Ian Johnston, Dominic Kwiatkowski, Cordelia Langford, John Sillitoe on behalf of the Wellcome Sanger Institute COVID-19 Surveillance Team |
| EPI_ISL_2392513 | Lighthouse Lab in Milton Keynes | Wellcome Sanger Institute for the COVID-19 Genomics UK (COG-UK) Consortium | The Lighthouse Lab in Milton Keynes and Alex Alderton, Roberto Amato, Jeffrey Barrett, Sonia Goncalves, Ewan Harrison, David K. Jackson, Ian Johnston, Dominic Kwiatkowski, Cordelia Langford, John Sillitoe on behalf of the Wellcome Sanger Institute COVID-19 Surveillance Team |
| EPI_ISL_2392514, EPI_ISL_2392515, EPI_ISL_2392516, EPI_ISL_2392517, EPI_ISL_2392518, EPI_ISL_2392519, EPI_ISL_2392520, EPI_ISL_2392521 | Lighthouse Lab in Alderley Park | Wellcome Sanger Institute for the COVID-19 Genomics UK (COG-UK) Consortium | Jacquelyn Wynn, Mairead Hyland, The Lighthouse Lab in Alderley Park and Alex Alderton, Roberto Amato, Jeffrey Barrett, Sonia Goncalves, Ewan Harrison, David K. Jackson, Ian Johnston, Dominic Kwiatkowski, Cordelia Langford, John Sillitoe on behalf of the Wellcome Sanger Institute COVID-19 Surveillance Team |
| EPI_ISL_2392523, EPI_ISL_2392524, EPI_ISL_2392525 | Lighthouse Lab in Milton Keynes | Wellcome Sanger Institute for the COVID-19 Genomics UK (COG-UK) Consortium | The Lighthouse Lab in Milton Keynes and Alex Alderton, Roberto Amato, Jeffrey Barrett, Sonia Goncalves, Ewan Harrison, David K. Jackson, Ian Johnston, Dominic Kwiatkowski, Cordelia Langford, John Sillitoe on behalf of the Wellcome Sanger Institute COVID-19 Surveillance Team |
| EPI_ISL_2392526, EPI_ISL_2392527, EPI_ISL_2392529 | Lighthouse Lab in Alderley Park | Wellcome Sanger Institute for the COVID-19 Genomics UK (COG-UK) Consortium | Jacquelyn Wynn, Mairead Hyland, The Lighthouse Lab in Alderley Park and Alex Alderton, Roberto Amato, Jeffrey Barrett, Sonia Goncalves, Ewan Harrison, David K. Jackson, Ian Johnston, Dominic Kwiatkowski, Cordelia Langford, John Sillitoe on behalf of the Wellcome Sanger Institute COVID-19 Surveillance Team |
| EPI_ISL_2392531, EPI_ISL_2392532, EPI_ISL_2392533 | Lighthouse Lab in Milton Keynes | Wellcome Sanger Institute for the COVID-19 Genomics UK (COG-UK) Consortium | The Lighthouse Lab in Milton Keynes and Alex Alderton, Roberto Amato, Jeffrey Barrett, Sonia Goncalves, Ewan Harrison, David K. Jackson, Ian Johnston, Dominic Kwiatkowski, Cordelia Langford, John Sillitoe on behalf of the Wellcome Sanger Institute COVID-19 Surveillance Team |
| EPI_ISL_2392534 | Lighthouse Lab in Alderley Park | Wellcome Sanger Institute for the COVID-19 Genomics UK (COG-UK) | Jacquelyn Wynn, Mairead Hyland, The Lighthouse Lab in Alderley Park and Alex Alderton, Roberto Amato, Jeffrey Barrett, Sonia Goncalves, Ewan Harrison, David K. Jackson, Ian Johnston, Dominic Kwiatkowski, Cordelia Langford, John Sillitoe on |

| | | | |
|---|---|---|---|
| | | Consortium | behalf of the Wellcome Sanger Institute COVID-19 Surveillance Team |
| EPI_ISL_2392535 | Lighthouse Lab in Milton Keynes | Wellcome Sanger Institute for the COVID-19 Genomics UK (COG-UK) Consortium | The Lighthouse Lab in Milton Keynes and Alex Alderton, Roberto Amato, Jeffrey Barrett, Sonia Goncalves, Ewan Harrison, David K. Jackson, Ian Johnston, Dominic Kwiatkowski, Cordelia Langford, John Sillitoe on behalf of the Wellcome Sanger Institute COVID-19 Surveillance Team |
| EPI_ISL_2392536, EPI_ISL_2392537, EPI_ISL_2392538 | Lighthouse Lab in Alderley Park | Wellcome Sanger Institute for the COVID-19 Genomics UK (COG-UK) Consortium | Jacquelyn Wynn, Mairead Hyland, The Lighthouse Lab in Alderley Park and Alex Alderton, Roberto Amato, Jeffrey Barrett, Sonia Goncalves, Ewan Harrison, David K. Jackson, Ian Johnston, Dominic Kwiatkowski, Cordelia Langford, John Sillitoe on behalf of the Wellcome Sanger Institute COVID-19 Surveillance Team |
| EPI_ISL_2392540 | Lighthouse Lab in Milton Keynes | Wellcome Sanger Institute for the COVID-19 Genomics UK (COG-UK) Consortium | The Lighthouse Lab in Milton Keynes and Alex Alderton, Roberto Amato, Jeffrey Barrett, Sonia Goncalves, Ewan Harrison, David K. Jackson, Ian Johnston, Dominic Kwiatkowski, Cordelia Langford, John Sillitoe on behalf of the Wellcome Sanger Institute COVID-19 Surveillance Team |
| EPI_ISL_2392541, EPI_ISL_2392542 | Lighthouse Lab in Alderley Park | Wellcome Sanger Institute for the COVID-19 Genomics UK (COG-UK) Consortium | Jacquelyn Wynn, Mairead Hyland, The Lighthouse Lab in Alderley Park and Alex Alderton, Roberto Amato, Jeffrey Barrett, Sonia Goncalves, Ewan Harrison, David K. Jackson, Ian Johnston, Dominic Kwiatkowski, Cordelia Langford, John Sillitoe on behalf of the Wellcome Sanger Institute COVID-19 Surveillance Team |
| EPI_ISL_2392543 | Lighthouse Lab in Milton Keynes | Wellcome Sanger Institute for the COVID-19 Genomics UK (COG-UK) Consortium | The Lighthouse Lab in Milton Keynes and Alex Alderton, Roberto Amato, Jeffrey Barrett, Sonia Goncalves, Ewan Harrison, David K. Jackson, Ian Johnston, Dominic Kwiatkowski, Cordelia Langford, John Sillitoe on behalf of the Wellcome Sanger Institute COVID-19 Surveillance Team |
| EPI_ISL_2392545, EPI_ISL_2392547, EPI_ISL_2392548, EPI_ISL_2392549 | Lighthouse Lab in Alderley Park | Wellcome Sanger Institute for the COVID-19 Genomics UK (COG-UK) Consortium | Jacquelyn Wynn, Mairead Hyland, The Lighthouse Lab in Alderley Park and Alex Alderton, Roberto Amato, Jeffrey Barrett, Sonia Goncalves, Ewan Harrison, David K. Jackson, Ian Johnston, Dominic Kwiatkowski, Cordelia Langford, John Sillitoe on behalf of the Wellcome Sanger Institute COVID-19 Surveillance Team |
| EPI_ISL_2392550, EPI_ISL_2392551, EPI_ISL_2392552, EPI_ISL_2392553, EPI_ISL_2392554, EPI_ISL_2392555 | Lighthouse Lab in Milton Keynes | Wellcome Sanger Institute for the COVID-19 Genomics UK (COG-UK) Consortium | The Lighthouse Lab in Milton Keynes and Alex Alderton, Roberto Amato, Jeffrey Barrett, Sonia Goncalves, Ewan Harrison, David K. Jackson, Ian Johnston, Dominic Kwiatkowski, Cordelia Langford, John Sillitoe on behalf of the Wellcome Sanger Institute COVID-19 Surveillance Team |
| EPI_ISL_2392556, EPI_ISL_2392557 | Lighthouse Lab in Alderley Park | Wellcome Sanger Institute for the COVID-19 Genomics UK (COG-UK) Consortium | Jacquelyn Wynn, Mairead Hyland, The Lighthouse Lab in Alderley Park and Alex Alderton, Roberto Amato, Jeffrey Barrett, Sonia Goncalves, Ewan Harrison, David K. Jackson, Ian Johnston, Dominic Kwiatkowski, Cordelia Langford, John Sillitoe on behalf of the Wellcome Sanger Institute COVID-19 Surveillance Team |
| EPI_ISL_2392558, EPI_ISL_2392559, EPI_ISL_2392560, EPI_ISL_2392561 | Lighthouse Lab in Milton Keynes | Wellcome Sanger Institute for the COVID-19 Genomics UK (COG-UK) Consortium | The Lighthouse Lab in Milton Keynes and Alex Alderton, Roberto Amato, Jeffrey Barrett, Sonia Goncalves, Ewan Harrison, David K. Jackson, Ian Johnston, Dominic Kwiatkowski, Cordelia Langford, John Sillitoe on behalf of the Wellcome Sanger Institute COVID-19 Surveillance Team |
| EPI_ISL_2392562 | Lighthouse Lab in Alderley Park | Wellcome Sanger Institute for the COVID-19 Genomics UK (COG-UK) Consortium | Jacquelyn Wynn, Mairead Hyland, The Lighthouse Lab in Alderley Park and Alex Alderton, Roberto Amato, Jeffrey Barrett, Sonia Goncalves, Ewan Harrison, David K. Jackson, Ian Johnston, Dominic Kwiatkowski, Cordelia Langford, John Sillitoe on behalf of the Wellcome Sanger Institute COVID-19 Surveillance Team |
| EPI_ISL_2392563 | Lighthouse Lab in Milton Keynes | Wellcome Sanger Institute for the COVID-19 Genomics UK (COG-UK) Consortium | The Lighthouse Lab in Milton Keynes and Alex Alderton, Roberto Amato, Jeffrey Barrett, Sonia Goncalves, Ewan Harrison, David K. Jackson, Ian Johnston, Dominic Kwiatkowski, Cordelia Langford, John Sillitoe on behalf of the Wellcome Sanger Institute COVID-19 Surveillance Team |
| EPI_ISL_2392564, EPI_ISL_2392565 | Lighthouse Lab in Alderley Park | Wellcome Sanger Institute for the COVID-19 Genomics UK (COG-UK) Consortium | Jacquelyn Wynn, Mairead Hyland, The Lighthouse Lab in Alderley Park and Alex Alderton, Roberto Amato, Jeffrey Barrett, Sonia Goncalves, Ewan Harrison, David K. Jackson, Ian Johnston, Dominic Kwiatkowski, Cordelia Langford, John Sillitoe on behalf of the Wellcome Sanger Institute COVID-19 Surveillance Team |
| EPI_ISL_2392567, EPI_ISL_2392568 | Lighthouse Lab in Milton Keynes | Wellcome Sanger Institute for the COVID-19 Genomics UK (COG-UK) Consortium | The Lighthouse Lab in Milton Keynes and Alex Alderton, Roberto Amato, Jeffrey Barrett, Sonia Goncalves, Ewan Harrison, David K. Jackson, Ian Johnston, Dominic Kwiatkowski, Cordelia Langford, John Sillitoe on behalf of the Wellcome Sanger Institute COVID-19 Surveillance Team |
| EPI_ISL_2392569 | Lighthouse Lab in Alderley Park | Wellcome Sanger Institute for the COVID-19 Genomics UK (COG-UK) Consortium | Jacquelyn Wynn, Mairead Hyland, The Lighthouse Lab in Alderley Park and Alex Alderton, Roberto Amato, Jeffrey Barrett, Sonia Goncalves, Ewan Harrison, David K. Jackson, Ian Johnston, Dominic Kwiatkowski, Cordelia Langford, John Sillitoe on behalf of the Wellcome Sanger Institute COVID-19 Surveillance Team |
| EPI_ISL_2392570, EPI_ISL_2392571 | Lighthouse Lab in Milton Keynes | Wellcome Sanger Institute for the COVID-19 Genomics UK (COG-UK) Consortium | The Lighthouse Lab in Milton Keynes and Alex Alderton, Roberto Amato, Jeffrey Barrett, Sonia Goncalves, Ewan Harrison, David K. Jackson, Ian Johnston, Dominic Kwiatkowski, Cordelia Langford, John Sillitoe on behalf of the Wellcome Sanger Institute COVID-19 Surveillance Team |
| EPI_ISL_2392573, EPI_ISL_2392574, EPI_ISL_2392575 | Lighthouse Lab in Alderley Park | Wellcome Sanger Institute for the COVID-19 Genomics UK (COG-UK) Consortium | Jacquelyn Wynn, Mairead Hyland, The Lighthouse Lab in Alderley Park and Alex Alderton, Roberto Amato, Jeffrey Barrett, Sonia Goncalves, Ewan Harrison, David K. Jackson, Ian Johnston, Dominic Kwiatkowski, Cordelia Langford, John Sillitoe on behalf of the Wellcome Sanger Institute COVID-19 Surveillance Team |
| EPI_ISL_2392576, EPI_ISL_2392577, EPI_ISL_2392578 | Lighthouse Lab in Milton Keynes | Wellcome Sanger Institute for the COVID-19 Genomics UK (COG-UK) Consortium | The Lighthouse Lab in Milton Keynes and Alex Alderton, Roberto Amato, Jeffrey Barrett, Sonia Goncalves, Ewan Harrison, David K. Jackson, Ian Johnston, Dominic Kwiatkowski, Cordelia Langford, John Sillitoe on behalf of the Wellcome Sanger Institute COVID-19 Surveillance Team |
| EPI_ISL_2392579 | Lighthouse Lab in Glasgow | Wellcome Sanger Institute for the COVID-19 Genomics UK (COG-UK) Consortium | Harper VanSteenhouse, Yumi Kasai, David Gray, Carol Clugston, Anna Dominiczak and Alex Alderton, Roberto Amato, Jeffrey Barrett, Sonia Goncalves, Ewan Harrison, David K. Jackson, Ian Johnston, Dominic Kwiatkowski, Cordelia Langford, John Sillitoe on behalf of the Wellcome Sanger Institute COVID-19 Surveillance Team |
| EPI_ISL_2392580 | Lighthouse Lab in Alderley Park | Wellcome Sanger Institute for the COVID-19 Genomics UK (COG-UK) Consortium | Jacquelyn Wynn, Mairead Hyland, The Lighthouse Lab in Alderley Park and Alex Alderton, Roberto Amato, Jeffrey Barrett, Sonia Goncalves, Ewan Harrison, David K. Jackson, Ian Johnston, Dominic Kwiatkowski, Cordelia Langford, John Sillitoe on behalf of the Wellcome Sanger Institute COVID-19 Surveillance Team |
| EPI_ISL_2392581 | Lighthouse Lab in Milton Keynes | Wellcome Sanger Institute for the COVID-19 Genomics UK (COG-UK) Consortium | The Lighthouse Lab in Milton Keynes and Alex Alderton, Roberto Amato, Jeffrey Barrett, Sonia Goncalves, Ewan Harrison, David K. Jackson, Ian Johnston, Dominic Kwiatkowski, Cordelia Langford, John Sillitoe on behalf of the Wellcome Sanger Institute COVID-19 Surveillance Team |
| EPI_ISL_2392582 | Lighthouse Lab in Glasgow | Wellcome Sanger Institute for the COVID-19 Genomics UK (COG-UK) Consortium | Harper VanSteenhouse, Yumi Kasai, David Gray, Carol Clugston, Anna Dominiczak and Alex Alderton, Roberto Amato, Jeffrey Barrett, Sonia Goncalves, Ewan Harrison, David K. Jackson, Ian Johnston, Dominic Kwiatkowski, Cordelia Langford, John Sillitoe on behalf of the Wellcome Sanger Institute COVID-19 Surveillance Team |
| EPI_ISL_2392583 | Lighthouse Lab in Milton Keynes | Wellcome Sanger Institute for the COVID-19 Genomics UK (COG-UK) Consortium | The Lighthouse Lab in Milton Keynes and Alex Alderton, Roberto Amato, Jeffrey Barrett, Sonia Goncalves, Ewan Harrison, David K. Jackson, Ian Johnston, Dominic Kwiatkowski, Cordelia Langford, John Sillitoe on behalf of the Wellcome Sanger Institute COVID-19 Surveillance Team |
| EPI_ISL_2392584, EPI_ISL_2392585 | Lighthouse Lab in Alderley Park | Wellcome Sanger Institute for the COVID-19 Genomics UK (COG-UK) Consortium | Jacquelyn Wynn, Mairead Hyland, The Lighthouse Lab in Alderley Park and Alex Alderton, Roberto Amato, Jeffrey Barrett, Sonia Goncalves, Ewan Harrison, David K. Jackson, Ian Johnston, Dominic Kwiatkowski, Cordelia Langford, John Sillitoe on behalf of the Wellcome Sanger Institute COVID-19 Surveillance Team |
| EPI_ISL_2392586 | Lighthouse Lab in Milton Keynes | Wellcome Sanger Institute for the COVID-19 Genomics UK (COG-UK) Consortium | The Lighthouse Lab in Milton Keynes and Alex Alderton, Roberto Amato, Jeffrey Barrett, Sonia Goncalves, Ewan Harrison, David K. Jackson, Ian Johnston, Dominic Kwiatkowski, Cordelia Langford, John Sillitoe on behalf of the Wellcome Sanger Institute COVID-19 Surveillance Team |
| EPI_ISL_2392587 | Lighthouse Lab in Alderley Park | Wellcome Sanger Institute for the COVID-19 Genomics UK (COG-UK) Consortium | Jacquelyn Wynn, Mairead Hyland, The Lighthouse Lab in Alderley Park and Alex Alderton, Roberto Amato, Jeffrey Barrett, Sonia Goncalves, Ewan Harrison, David K. Jackson, Ian Johnston, Dominic Kwiatkowski, Cordelia Langford, John Sillitoe on behalf of the Wellcome Sanger Institute COVID-19 Surveillance Team |
| EPI_ISL_2392588, EPI_ISL_2392589, EPI_ISL_2392590, EPI_ISL_2392591 | Lighthouse Lab in Milton Keynes | Wellcome Sanger Institute for the COVID-19 Genomics UK (COG-UK) Consortium | The Lighthouse Lab in Milton Keynes and Alex Alderton, Roberto Amato, Jeffrey Barrett, Sonia Goncalves, Ewan Harrison, David K. Jackson, Ian Johnston, Dominic Kwiatkowski, Cordelia Langford, John Sillitoe on behalf of the Wellcome Sanger Institute COVID-19 Surveillance Team |
| EPI_ISL_2392594, EPI_ISL_2392596, EPI_ISL_2392597, EPI_ISL_2392598 | Lighthouse Lab in Alderley Park | Wellcome Sanger Institute for the COVID-19 Genomics UK (COG-UK) Consortium | Jacquelyn Wynn, Mairead Hyland, The Lighthouse Lab in Alderley Park and Alex Alderton, Roberto Amato, Jeffrey Barrett, Sonia Goncalves, Ewan Harrison, David K. Jackson, Ian Johnston, Dominic Kwiatkowski, Cordelia Langford, John Sillitoe on behalf of the Wellcome Sanger Institute COVID-19 Surveillance Team |
| EPI_ISL_2392599 | Lighthouse Lab in Milton Keynes | Wellcome Sanger Institute for the COVID-19 Genomics UK (COG-UK) Consortium | The Lighthouse Lab in Milton Keynes and Alex Alderton, Roberto Amato, Jeffrey Barrett, Sonia Goncalves, Ewan Harrison, David K. Jackson, Ian Johnston, Dominic Kwiatkowski, Cordelia Langford, John Sillitoe on behalf of the Wellcome Sanger Institute COVID-19 Surveillance Team |
| EPI_ISL_2392600, EPI_ISL_2392601 | Lighthouse Lab in Alderley Park | Wellcome Sanger Institute for the COVID-19 Genomics UK (COG-UK) Consortium | Jacquelyn Wynn, Mairead Hyland, The Lighthouse Lab in Alderley Park and Alex Alderton, Roberto Amato, Jeffrey Barrett, Sonia Goncalves, Ewan Harrison, David K. Jackson, Ian Johnston, Dominic Kwiatkowski, Cordelia Langford, John Sillitoe on behalf of the Wellcome Sanger Institute COVID-19 Surveillance Team |
| EPI_ISL_2392602 | Lighthouse Lab in Milton Keynes | Wellcome Sanger Institute for the COVID-19 Genomics UK (COG-UK) Consortium | The Lighthouse Lab in Milton Keynes and Alex Alderton, Roberto Amato, Jeffrey Barrett, Sonia Goncalves, Ewan Harrison, David K. Jackson, Ian Johnston, Dominic Kwiatkowski, Cordelia Langford, John Sillitoe on behalf of the Wellcome Sanger Institute COVID-19 Surveillance Team |
| EPI_ISL_2392603 | Lighthouse Lab in Alderley Park | Wellcome Sanger Institute for the COVID-19 Genomics UK (COG-UK) Consortium | Jacquelyn Wynn, Mairead Hyland, The Lighthouse Lab in Alderley Park and Alex Alderton, Roberto Amato, Jeffrey Barrett, Sonia Goncalves, Ewan Harrison, David K. Jackson, Ian Johnston, Dominic Kwiatkowski, Cordelia Langford, John Sillitoe on behalf of the Wellcome Sanger Institute COVID-19 Surveillance Team |
| EPI_ISL_2392604 | Lighthouse Lab in Milton Keynes | Wellcome Sanger Institute for the COVID-19 Genomics UK (COG-UK) Consortium | The Lighthouse Lab in Milton Keynes and Alex Alderton, Roberto Amato, Jeffrey Barrett, Sonia Goncalves, Ewan Harrison, David K. Jackson, Ian Johnston, Dominic Kwiatkowski, Cordelia Langford, John Sillitoe on behalf of the Wellcome Sanger Institute COVID-19 Surveillance Team |
| EPI_ISL_2392605 | Lighthouse Lab in Alderley Park | Wellcome Sanger Institute for the COVID-19 Genomics UK (COG-UK) Consortium | Jacquelyn Wynn, Mairead Hyland, The Lighthouse Lab in Alderley Park and Alex Alderton, Roberto Amato, Jeffrey Barrett, Sonia Goncalves, Ewan Harrison, David K. Jackson, Ian Johnston, Dominic Kwiatkowski, Cordelia Langford, John Sillitoe on behalf of the Wellcome Sanger Institute COVID-19 Surveillance Team |
| EPI_ISL_2392606 | Lighthouse Lab in Milton Keynes | Wellcome Sanger Institute for the COVID-19 Genomics UK (COG-UK) Consortium | The Lighthouse Lab in Milton Keynes and Alex Alderton, Roberto Amato, Jeffrey Barrett, Sonia Goncalves, Ewan Harrison, David K. Jackson, Ian Johnston, Dominic Kwiatkowski, Cordelia Langford, John Sillitoe on behalf of the Wellcome Sanger Institute COVID-19 Surveillance Team |
| EPI_ISL_2392607, EPI_ISL_2392608 | Lighthouse Lab in Alderley Park | Wellcome Sanger Institute for the COVID-19 Genomics UK (COG-UK) Consortium | Jacquelyn Wynn, Mairead Hyland, The Lighthouse Lab in Alderley Park and Alex Alderton, Roberto Amato, Jeffrey Barrett, Sonia Goncalves, Ewan Harrison, David K. Jackson, Ian Johnston, Dominic Kwiatkowski, Cordelia Langford, John Sillitoe on behalf of the Wellcome Sanger Institute COVID-19 Surveillance Team |
| EPI_ISL_2392609 | Lighthouse Lab in Milton Keynes | Wellcome Sanger Institute for the COVID-19 Genomics UK (COG-UK) Consortium | The Lighthouse Lab in Milton Keynes and Alex Alderton, Roberto Amato, Jeffrey Barrett, Sonia Goncalves, Ewan Harrison, David K. Jackson, Ian Johnston, Dominic Kwiatkowski, Cordelia Langford, John Sillitoe on behalf of the Wellcome Sanger Institute COVID-19 Surveillance Team |
| EPI_ISL_2392610 | Lighthouse Lab in Glasgow | Wellcome Sanger Institute for the COVID-19 Genomics UK (COG-UK) Consortium | Harper VanSteenhouse, Yumi Kasai, David Gray, Carol Clugston, Anna Dominiczak and Alex Alderton, Roberto Amato, Jeffrey Barrett, Sonia Goncalves, Ewan Harrison, David K. Jackson, Ian Johnston, Dominic Kwiatkowski, Cordelia Langford, John Sillitoe on behalf of the Wellcome Sanger Institute COVID-19 Surveillance Team |
| EPI_ISL_2392611 | Lighthouse Lab in Alderley Park | Wellcome Sanger Institute for the COVID-19 Genomics UK (COG-UK) Consortium | Jacquelyn Wynn, Mairead Hyland, The Lighthouse Lab in Alderley Park and Alex Alderton, Roberto Amato, Jeffrey Barrett, Sonia Goncalves, Ewan Harrison, David K. Jackson, Ian Johnston, Dominic Kwiatkowski, Cordelia Langford, John Sillitoe on behalf of the Wellcome Sanger Institute COVID-19 Surveillance Team |
| EPI_ISL_2392612 | Lighthouse Lab in Milton Keynes | Wellcome Sanger Institute for the COVID-19 Genomics UK (COG-UK) | The Lighthouse Lab in Milton Keynes and Alex Alderton, Roberto Amato, Jeffrey Barrett, Sonia Goncalves, Ewan Harrison, David K. Jackson, Ian Johnston, Dominic Kwiatkowski, Cordelia Langford, John Sillitoe on behalf of the Wellcome Sanger |

| | | | |
|---|---|---|---|
| | | Consortium | Institute COVID-19 Surveillance Team |
| EPI_ISL_2392613, EPI_ISL_2392614 | Lighthouse Lab in Alderley Park | Wellcome Sanger Institute for the COVID-19 Genomics UK (COG-UK) Consortium | Jacquelyn Wynn, Mairead Hyland, The Lighthouse Lab in Alderley Park and Alex Alderton, Roberto Amato, Jeffrey Barrett, Sonia Goncalves, Ewan Harrison, David K. Jackson, Ian Johnston, Dominic Kwiatkowski, Cordelia Langford, John Sillitoe on behalf of the Wellcome Sanger Institute COVID-19 Surveillance Team |
| EPI_ISL_2392615, EPI_ISL_2392616 | Lighthouse Lab in Milton Keynes | Wellcome Sanger Institute for the COVID-19 Genomics UK (COG-UK) Consortium | The Lighthouse Lab in Milton Keynes and Alex Alderton, Roberto Amato, Jeffrey Barrett, Sonia Goncalves, Ewan Harrison, David K. Jackson, Ian Johnston, Dominic Kwiatkowski, Cordelia Langford, John Sillitoe on behalf of the Wellcome Sanger Institute COVID-19 Surveillance Team |
| EPI_ISL_2392617, EPI_ISL_2392620, EPI_ISL_2392621, EPI_ISL_2392622 | Lighthouse Lab in Alderley Park | Wellcome Sanger Institute for the COVID-19 Genomics UK (COG-UK) Consortium | Jacquelyn Wynn, Mairead Hyland, The Lighthouse Lab in Alderley Park and Alex Alderton, Roberto Amato, Jeffrey Barrett, Sonia Goncalves, Ewan Harrison, David K. Jackson, Ian Johnston, Dominic Kwiatkowski, Cordelia Langford, John Sillitoe on behalf of the Wellcome Sanger Institute COVID-19 Surveillance Team |
| EPI_ISL_2392623, EPI_ISL_2392624 | Lighthouse Lab in Milton Keynes | Wellcome Sanger Institute for the COVID-19 Genomics UK (COG-UK) Consortium | The Lighthouse Lab in Milton Keynes and Alex Alderton, Roberto Amato, Jeffrey Barrett, Sonia Goncalves, Ewan Harrison, David K. Jackson, Ian Johnston, Dominic Kwiatkowski, Cordelia Langford, John Sillitoe on behalf of the Wellcome Sanger Institute COVID-19 Surveillance Team |
| EPI_ISL_2392625, EPI_ISL_2392628 | Lighthouse Lab in Alderley Park | Wellcome Sanger Institute for the COVID-19 Genomics UK (COG-UK) Consortium | Jacquelyn Wynn, Mairead Hyland, The Lighthouse Lab in Alderley Park and Alex Alderton, Roberto Amato, Jeffrey Barrett, Sonia Goncalves, Ewan Harrison, David K. Jackson, Ian Johnston, Dominic Kwiatkowski, Cordelia Langford, John Sillitoe on behalf of the Wellcome Sanger Institute COVID-19 Surveillance Team |
| EPI_ISL_2392629, EPI_ISL_2392631 | Lighthouse Lab in Milton Keynes | Wellcome Sanger Institute for the COVID-19 Genomics UK (COG-UK) Consortium | The Lighthouse Lab in Milton Keynes and Alex Alderton, Roberto Amato, Jeffrey Barrett, Sonia Goncalves, Ewan Harrison, David K. Jackson, Ian Johnston, Dominic Kwiatkowski, Cordelia Langford, John Sillitoe on behalf of the Wellcome Sanger Institute COVID-19 Surveillance Team |
| EPI_ISL_2392632, EPI_ISL_2392633, EPI_ISL_2392634, EPI_ISL_2392635, EPI_ISL_2392636 | Lighthouse Lab in Alderley Park | Wellcome Sanger Institute for the COVID-19 Genomics UK (COG-UK) Consortium | Jacquelyn Wynn, Mairead Hyland, The Lighthouse Lab in Alderley Park and Alex Alderton, Roberto Amato, Jeffrey Barrett, Sonia Goncalves, Ewan Harrison, David K. Jackson, Ian Johnston, Dominic Kwiatkowski, Cordelia Langford, John Sillitoe on behalf of the Wellcome Sanger Institute COVID-19 Surveillance Team |
| EPI_ISL_2392637 | Lighthouse Lab in Milton Keynes | Wellcome Sanger Institute for the COVID-19 Genomics UK (COG-UK) Consortium | The Lighthouse Lab in Milton Keynes and Alex Alderton, Roberto Amato, Jeffrey Barrett, Sonia Goncalves, Ewan Harrison, David K. Jackson, Ian Johnston, Dominic Kwiatkowski, Cordelia Langford, John Sillitoe on behalf of the Wellcome Sanger Institute COVID-19 Surveillance Team |
| EPI_ISL_2392638 | Lighthouse Lab in Alderley Park | Wellcome Sanger Institute for the COVID-19 Genomics UK (COG-UK) Consortium | Jacquelyn Wynn, Mairead Hyland, The Lighthouse Lab in Alderley Park and Alex Alderton, Roberto Amato, Jeffrey Barrett, Sonia Goncalves, Ewan Harrison, David K. Jackson, Ian Johnston, Dominic Kwiatkowski, Cordelia Langford, John Sillitoe on behalf of the Wellcome Sanger Institute COVID-19 Surveillance Team |
| EPI_ISL_2392639, EPI_ISL_2392640 | Lighthouse Lab in Glasgow | Wellcome Sanger Institute for the COVID-19 Genomics UK (COG-UK) Consortium | Harper VanSteenhouse, Yumi Kasai, David Gray, Carol Clugston, Anna Dominiczak and Alex Alderton, Roberto Amato, Jeffrey Barrett, Sonia Goncalves, Ewan Harrison, David K. Jackson, Ian Johnston, Dominic Kwiatkowski, Cordelia Langford, John Sillitoe on behalf of the Wellcome Sanger Institute COVID-19 Surveillance Team |
| EPI_ISL_2392642 | Lighthouse Lab in Alderley Park | Wellcome Sanger Institute for the COVID-19 Genomics UK (COG-UK) Consortium | Jacquelyn Wynn, Mairead Hyland, The Lighthouse Lab in Alderley Park and Alex Alderton, Roberto Amato, Jeffrey Barrett, Sonia Goncalves, Ewan Harrison, David K. Jackson, Ian Johnston, Dominic Kwiatkowski, Cordelia Langford, John Sillitoe on behalf of the Wellcome Sanger Institute COVID-19 Surveillance Team |
| EPI_ISL_2392643 | Lighthouse Lab in Milton Keynes | Wellcome Sanger Institute for the COVID-19 Genomics UK (COG-UK) Consortium | The Lighthouse Lab in Milton Keynes and Alex Alderton, Roberto Amato, Jeffrey Barrett, Sonia Goncalves, Ewan Harrison, David K. Jackson, Ian Johnston, Dominic Kwiatkowski, Cordelia Langford, John Sillitoe on behalf of the Wellcome Sanger Institute COVID-19 Surveillance Team |
| EPI_ISL_2392644, EPI_ISL_2392645, EPI_ISL_2392646, EPI_ISL_2392647, EPI_ISL_2392648 | Lighthouse Lab in Alderley Park | Wellcome Sanger Institute for the COVID-19 Genomics UK (COG-UK) Consortium | Jacquelyn Wynn, Mairead Hyland, The Lighthouse Lab in Alderley Park and Alex Alderton, Roberto Amato, Jeffrey Barrett, Sonia Goncalves, Ewan Harrison, David K. Jackson, Ian Johnston, Dominic Kwiatkowski, Cordelia Langford, John Sillitoe on behalf of the Wellcome Sanger Institute COVID-19 Surveillance Team |
| EPI_ISL_2392649 | Lighthouse Lab in Milton Keynes | Wellcome Sanger Institute for the COVID-19 Genomics UK (COG-UK) Consortium | The Lighthouse Lab in Milton Keynes and Alex Alderton, Roberto Amato, Jeffrey Barrett, Sonia Goncalves, Ewan Harrison, David K. Jackson, Ian Johnston, Dominic Kwiatkowski, Cordelia Langford, John Sillitoe on behalf of the Wellcome Sanger Institute COVID-19 Surveillance Team |
| EPI_ISL_2392651, EPI_ISL_2392652, EPI_ISL_2392653, EPI_ISL_2392654, EPI_ISL_2392656, EPI_ISL_2392657, EPI_ISL_2392658, EPI_ISL_2392659 | Lighthouse Lab in Alderley Park | Wellcome Sanger Institute for the COVID-19 Genomics UK (COG-UK) Consortium | Jacquelyn Wynn, Mairead Hyland, The Lighthouse Lab in Alderley Park and Alex Alderton, Roberto Amato, Jeffrey Barrett, Sonia Goncalves, Ewan Harrison, David K. Jackson, Ian Johnston, Dominic Kwiatkowski, Cordelia Langford, John Sillitoe on behalf of the Wellcome Sanger Institute COVID-19 Surveillance Team |
| EPI_ISL_2392660 | Lighthouse Lab in Milton Keynes | Wellcome Sanger Institute for the COVID-19 Genomics UK (COG-UK) Consortium | The Lighthouse Lab in Milton Keynes and Alex Alderton, Roberto Amato, Jeffrey Barrett, Sonia Goncalves, Ewan Harrison, David K. Jackson, Ian Johnston, Dominic Kwiatkowski, Cordelia Langford, John Sillitoe on behalf of the Wellcome Sanger Institute COVID-19 Surveillance Team |
| EPI_ISL_2392661, EPI_ISL_2392662, EPI_ISL_2392663 | Lighthouse Lab in Alderley Park | Wellcome Sanger Institute for the COVID-19 Genomics UK (COG-UK) Consortium | Jacquelyn Wynn, Mairead Hyland, The Lighthouse Lab in Alderley Park and Alex Alderton, Roberto Amato, Jeffrey Barrett, Sonia Goncalves, Ewan Harrison, David K. Jackson, Ian Johnston, Dominic Kwiatkowski, Cordelia Langford, John Sillitoe on behalf of the Wellcome Sanger Institute COVID-19 Surveillance Team |
| EPI_ISL_2392664 | Lighthouse Lab in Glasgow | Wellcome Sanger Institute for the COVID-19 Genomics UK (COG-UK) Consortium | Harper VanSteenhouse, Yumi Kasai, David Gray, Carol Clugston, Anna Dominiczak and Alex Alderton, Roberto Amato, Jeffrey Barrett, Sonia Goncalves, Ewan Harrison, David K. Jackson, Ian Johnston, Dominic Kwiatkowski, Cordelia Langford, John Sillitoe on behalf of the Wellcome Sanger Institute COVID-19 Surveillance Team |
| EPI_ISL_2392665 | Lighthouse Lab in Alderley Park | Wellcome Sanger Institute for the COVID-19 Genomics UK (COG-UK) Consortium | Jacquelyn Wynn, Mairead Hyland, The Lighthouse Lab in Alderley Park and Alex Alderton, Roberto Amato, Jeffrey Barrett, Sonia Goncalves, Ewan Harrison, David K. Jackson, Ian Johnston, Dominic Kwiatkowski, Cordelia Langford, John Sillitoe on behalf of the Wellcome Sanger Institute COVID-19 Surveillance Team |
| EPI_ISL_2392666 | Lighthouse Lab in Glasgow | Wellcome Sanger Institute for the COVID-19 Genomics UK (COG-UK) Consortium | Harper VanSteenhouse, Yumi Kasai, David Gray, Carol Clugston, Anna Dominiczak and Alex Alderton, Roberto Amato, Jeffrey Barrett, Sonia Goncalves, Ewan Harrison, David K. Jackson, Ian Johnston, Dominic Kwiatkowski, Cordelia Langford, John Sillitoe on behalf of the Wellcome Sanger Institute COVID-19 Surveillance Team |
| EPI_ISL_2392668, EPI_ISL_2392669, EPI_ISL_2392670, EPI_ISL_2392671 | Lighthouse Lab in Alderley Park | Wellcome Sanger Institute for the COVID-19 Genomics UK (COG-UK) Consortium | Jacquelyn Wynn, Mairead Hyland, The Lighthouse Lab in Alderley Park and Alex Alderton, Roberto Amato, Jeffrey Barrett, Sonia Goncalves, Ewan Harrison, David K. Jackson, Ian Johnston, Dominic Kwiatkowski, Cordelia Langford, John Sillitoe on behalf of the Wellcome Sanger Institute COVID-19 Surveillance Team |
| EPI_ISL_2392672, EPI_ISL_2392673 | Lighthouse Lab in Glasgow | Wellcome Sanger Institute for the COVID-19 Genomics UK (COG-UK) Consortium | Harper VanSteenhouse, Yumi Kasai, David Gray, Carol Clugston, Anna Dominiczak and Alex Alderton, Roberto Amato, Jeffrey Barrett, Sonia Goncalves, Ewan Harrison, David K. Jackson, Ian Johnston, Dominic Kwiatkowski, Cordelia Langford, John Sillitoe on behalf of the Wellcome Sanger Institute COVID-19 Surveillance Team |
| EPI_ISL_2392674, EPI_ISL_2392675, EPI_ISL_2392676 | Lighthouse Lab in Alderley Park | Wellcome Sanger Institute for the COVID-19 Genomics UK (COG-UK) Consortium | Jacquelyn Wynn, Mairead Hyland, The Lighthouse Lab in Alderley Park and Alex Alderton, Roberto Amato, Jeffrey Barrett, Sonia Goncalves, Ewan Harrison, David K. Jackson, Ian Johnston, Dominic Kwiatkowski, Cordelia Langford, John Sillitoe on behalf of the Wellcome Sanger Institute COVID-19 Surveillance Team |
| EPI_ISL_2392677 | Lighthouse Lab in Milton Keynes | Wellcome Sanger Institute for the COVID-19 Genomics UK (COG-UK) Consortium | The Lighthouse Lab in Milton Keynes and Alex Alderton, Roberto Amato, Jeffrey Barrett, Sonia Goncalves, Ewan Harrison, David K. Jackson, Ian Johnston, Dominic Kwiatkowski, Cordelia Langford, John Sillitoe on behalf of the Wellcome Sanger Institute COVID-19 Surveillance Team |
| EPI_ISL_2392678 | Lighthouse Lab in Glasgow | Wellcome Sanger Institute for the COVID-19 Genomics UK (COG-UK) Consortium | Harper VanSteenhouse, Yumi Kasai, David Gray, Carol Clugston, Anna Dominiczak and Alex Alderton, Roberto Amato, Jeffrey Barrett, Sonia Goncalves, Ewan Harrison, David K. Jackson, Ian Johnston, Dominic Kwiatkowski, Cordelia Langford, John Sillitoe on behalf of the Wellcome Sanger Institute COVID-19 Surveillance Team |
| EPI_ISL_2392679 | Lighthouse Lab in Milton Keynes | Wellcome Sanger Institute for the COVID-19 Genomics UK (COG-UK) Consortium | The Lighthouse Lab in Milton Keynes and Alex Alderton, Roberto Amato, Jeffrey Barrett, Sonia Goncalves, Ewan Harrison, David K. Jackson, Ian Johnston, Dominic Kwiatkowski, Cordelia Langford, John Sillitoe on behalf of the Wellcome Sanger Institute COVID-19 Surveillance Team |
| EPI_ISL_2392680 | Lighthouse Lab in Alderley Park | Wellcome Sanger Institute for the COVID-19 Genomics UK (COG-UK) Consortium | Jacquelyn Wynn, Mairead Hyland, The Lighthouse Lab in Alderley Park and Alex Alderton, Roberto Amato, Jeffrey Barrett, Sonia Goncalves, Ewan Harrison, David K. Jackson, Ian Johnston, Dominic Kwiatkowski, Cordelia Langford, John Sillitoe on behalf of the Wellcome Sanger Institute COVID-19 Surveillance Team |
| EPI_ISL_2392681 | Lighthouse Lab in Milton Keynes | Wellcome Sanger Institute for the COVID-19 Genomics UK (COG-UK) Consortium | The Lighthouse Lab in Milton Keynes and Alex Alderton, Roberto Amato, Jeffrey Barrett, Sonia Goncalves, Ewan Harrison, David K. Jackson, Ian Johnston, Dominic Kwiatkowski, Cordelia Langford, John Sillitoe on behalf of the Wellcome Sanger Institute COVID-19 Surveillance Team |
| EPI_ISL_2392682 | Lighthouse Lab in Alderley Park | Wellcome Sanger Institute for the COVID-19 Genomics UK (COG-UK) Consortium | Jacquelyn Wynn, Mairead Hyland, The Lighthouse Lab in Alderley Park and Alex Alderton, Roberto Amato, Jeffrey Barrett, Sonia Goncalves, Ewan Harrison, David K. Jackson, Ian Johnston, Dominic Kwiatkowski, Cordelia Langford, John Sillitoe on behalf of the Wellcome Sanger Institute COVID-19 Surveillance Team |
| EPI_ISL_2392683 | Lighthouse Lab in Milton Keynes | Wellcome Sanger Institute for the COVID-19 Genomics UK (COG-UK) Consortium | The Lighthouse Lab in Milton Keynes and Alex Alderton, Roberto Amato, Jeffrey Barrett, Sonia Goncalves, Ewan Harrison, David K. Jackson, Ian Johnston, Dominic Kwiatkowski, Cordelia Langford, John Sillitoe on behalf of the Wellcome Sanger Institute COVID-19 Surveillance Team |
| EPI_ISL_2392684 | Lighthouse Lab in Alderley Park | Wellcome Sanger Institute for the COVID-19 Genomics UK (COG-UK) Consortium | Jacquelyn Wynn, Mairead Hyland, The Lighthouse Lab in Alderley Park and Alex Alderton, Roberto Amato, Jeffrey Barrett, Sonia Goncalves, Ewan Harrison, David K. Jackson, Ian Johnston, Dominic Kwiatkowski, Cordelia Langford, John Sillitoe on behalf of the Wellcome Sanger Institute COVID-19 Surveillance Team |
| EPI_ISL_2392685 | Lighthouse Lab in Glasgow | Wellcome Sanger Institute for the COVID-19 Genomics UK (COG-UK) Consortium | Harper VanSteenhouse, Yumi Kasai, David Gray, Carol Clugston, Anna Dominiczak and Alex Alderton, Roberto Amato, Jeffrey Barrett, Sonia Goncalves, Ewan Harrison, David K. Jackson, Ian Johnston, Dominic Kwiatkowski, Cordelia Langford, John Sillitoe on behalf of the Wellcome Sanger Institute COVID-19 Surveillance Team |
| EPI_ISL_2392686, EPI_ISL_2392687 | Lighthouse Lab in Milton Keynes | Wellcome Sanger Institute for the COVID-19 Genomics UK (COG-UK) Consortium | The Lighthouse Lab in Milton Keynes and Alex Alderton, Roberto Amato, Jeffrey Barrett, Sonia Goncalves, Ewan Harrison, David K. Jackson, Ian Johnston, Dominic Kwiatkowski, Cordelia Langford, John Sillitoe on behalf of the Wellcome Sanger Institute COVID-19 Surveillance Team |
| EPI_ISL_2392689, EPI_ISL_2392690, EPI_ISL_2392691, EPI_ISL_2392692, EPI_ISL_2392693, EPI_ISL_2392694 | Lighthouse Lab in Alderley Park | Wellcome Sanger Institute for the COVID-19 Genomics UK (COG-UK) Consortium | Jacquelyn Wynn, Mairead Hyland, The Lighthouse Lab in Alderley Park and Alex Alderton, Roberto Amato, Jeffrey Barrett, Sonia Goncalves, Ewan Harrison, David K. Jackson, Ian Johnston, Dominic Kwiatkowski, Cordelia Langford, John Sillitoe on behalf of the Wellcome Sanger Institute COVID-19 Surveillance Team |
| EPI_ISL_2392695 | Lighthouse Lab in Glasgow | Wellcome Sanger Institute for the COVID-19 Genomics UK (COG-UK) Consortium | Harper VanSteenhouse, Yumi Kasai, David Gray, Carol Clugston, Anna Dominiczak and Alex Alderton, Roberto Amato, Jeffrey Barrett, Sonia Goncalves, Ewan Harrison, David K. Jackson, Ian Johnston, Dominic Kwiatkowski, Cordelia Langford, John Sillitoe on behalf of the Wellcome Sanger Institute COVID-19 Surveillance Team |
| EPI_ISL_2392696 | Lighthouse Lab in Alderley Park | Wellcome Sanger Institute for the COVID-19 Genomics UK (COG-UK) | Jacquelyn Wynn, Mairead Hyland, The Lighthouse Lab in Alderley Park and Alex Alderton, Roberto Amato, Jeffrey Barrett, Sonia Goncalves, Ewan Harrison, David K. Jackson, Ian Johnston, Dominic Kwiatkowski, Cordelia Langford, John Sillitoe on |

| | | | |
|---|---|---|---|
| | | Consortium | behalf of the Wellcome Sanger Institute COVID-19 Surveillance Team |
| EPI_ISL_2392697 | Lighthouse Lab in Milton Keynes | Wellcome Sanger Institute for the COVID-19 Genomics UK (COG-UK) Consortium | The Lighthouse Lab in Milton Keynes and Alex Alderton, Roberto Amato, Jeffrey Barrett, Sonia Goncalves, Ewan Harrison, David K. Jackson, Ian Johnston, Dominic Kwiatkowski, Cordelia Langford, John Sillitoe on behalf of the Wellcome Sanger Institute COVID-19 Surveillance Team |
| EPI_ISL_2392699 | Lighthouse Lab in Alderley Park | Wellcome Sanger Institute for the COVID-19 Genomics UK (COG-UK) Consortium | Jacquelyn Wynn, Mairead Hyland, The Lighthouse Lab in Alderley Park and Alex Alderton, Roberto Amato, Jeffrey Barrett, Sonia Goncalves, Ewan Harrison, David K. Jackson, Ian Johnston, Dominic Kwiatkowski, Cordelia Langford, John Sillitoe on behalf of the Wellcome Sanger Institute COVID-19 Surveillance Team |
| EPI_ISL_2392700 | Lighthouse Lab in Milton Keynes | Wellcome Sanger Institute for the COVID-19 Genomics UK (COG-UK) Consortium | The Lighthouse Lab in Milton Keynes and Alex Alderton, Roberto Amato, Jeffrey Barrett, Sonia Goncalves, Ewan Harrison, David K. Jackson, Ian Johnston, Dominic Kwiatkowski, Cordelia Langford, John Sillitoe on behalf of the Wellcome Sanger Institute COVID-19 Surveillance Team |
| EPI_ISL_2392701, EPI_ISL_2392702 | Lighthouse Lab in Alderley Park | Wellcome Sanger Institute for the COVID-19 Genomics UK (COG-UK) Consortium | Jacquelyn Wynn, Mairead Hyland, The Lighthouse Lab in Alderley Park and Alex Alderton, Roberto Amato, Jeffrey Barrett, Sonia Goncalves, Ewan Harrison, David K. Jackson, Ian Johnston, Dominic Kwiatkowski, Cordelia Langford, John Sillitoe on behalf of the Wellcome Sanger Institute COVID-19 Surveillance Team |
| EPI_ISL_2392703, EPI_ISL_2392704 | Lighthouse Lab in Milton Keynes | Wellcome Sanger Institute for the COVID-19 Genomics UK (COG-UK) Consortium | The Lighthouse Lab in Milton Keynes and Alex Alderton, Roberto Amato, Jeffrey Barrett, Sonia Goncalves, Ewan Harrison, David K. Jackson, Ian Johnston, Dominic Kwiatkowski, Cordelia Langford, John Sillitoe on behalf of the Wellcome Sanger Institute COVID-19 Surveillance Team |
| EPI_ISL_2392706, EPI_ISL_2392707, EPI_ISL_2392708 | Lighthouse Lab in Alderley Park | Wellcome Sanger Institute for the COVID-19 Genomics UK (COG-UK) Consortium | Jacquelyn Wynn, Mairead Hyland, The Lighthouse Lab in Alderley Park and Alex Alderton, Roberto Amato, Jeffrey Barrett, Sonia Goncalves, Ewan Harrison, David K. Jackson, Ian Johnston, Dominic Kwiatkowski, Cordelia Langford, John Sillitoe on behalf of the Wellcome Sanger Institute COVID-19 Surveillance Team |
| EPI_ISL_2392709, EPI_ISL_2392711 | Lighthouse Lab in Milton Keynes | Wellcome Sanger Institute for the COVID-19 Genomics UK (COG-UK) Consortium | The Lighthouse Lab in Milton Keynes and Alex Alderton, Roberto Amato, Jeffrey Barrett, Sonia Goncalves, Ewan Harrison, David K. Jackson, Ian Johnston, Dominic Kwiatkowski, Cordelia Langford, John Sillitoe on behalf of the Wellcome Sanger Institute COVID-19 Surveillance Team |
| EPI_ISL_2392712, EPI_ISL_2392713, EPI_ISL_2392715 | Lighthouse Lab in Alderley Park | Wellcome Sanger Institute for the COVID-19 Genomics UK (COG-UK) Consortium | Jacquelyn Wynn, Mairead Hyland, The Lighthouse Lab in Alderley Park and Alex Alderton, Roberto Amato, Jeffrey Barrett, Sonia Goncalves, Ewan Harrison, David K. Jackson, Ian Johnston, Dominic Kwiatkowski, Cordelia Langford, John Sillitoe on behalf of the Wellcome Sanger Institute COVID-19 Surveillance Team |
| EPI_ISL_2392717 | Lighthouse Lab in Milton Keynes | Wellcome Sanger Institute for the COVID-19 Genomics UK (COG-UK) Consortium | The Lighthouse Lab in Milton Keynes and Alex Alderton, Roberto Amato, Jeffrey Barrett, Sonia Goncalves, Ewan Harrison, David K. Jackson, Ian Johnston, Dominic Kwiatkowski, Cordelia Langford, John Sillitoe on behalf of the Wellcome Sanger Institute COVID-19 Surveillance Team |
| EPI_ISL_2392718 | Lighthouse Lab in Alderley Park | Wellcome Sanger Institute for the COVID-19 Genomics UK (COG-UK) Consortium | Jacquelyn Wynn, Mairead Hyland, The Lighthouse Lab in Alderley Park and Alex Alderton, Roberto Amato, Jeffrey Barrett, Sonia Goncalves, Ewan Harrison, David K. Jackson, Ian Johnston, Dominic Kwiatkowski, Cordelia Langford, John Sillitoe on behalf of the Wellcome Sanger Institute COVID-19 Surveillance Team |
| EPI_ISL_2392719 | Lighthouse Lab in Milton Keynes | Wellcome Sanger Institute for the COVID-19 Genomics UK (COG-UK) Consortium | The Lighthouse Lab in Milton Keynes and Alex Alderton, Roberto Amato, Jeffrey Barrett, Sonia Goncalves, Ewan Harrison, David K. Jackson, Ian Johnston, Dominic Kwiatkowski, Cordelia Langford, John Sillitoe on behalf of the Wellcome Sanger Institute COVID-19 Surveillance Team |
| EPI_ISL_2392720, EPI_ISL_2392721 | Lighthouse Lab in Alderley Park | Wellcome Sanger Institute for the COVID-19 Genomics UK (COG-UK) Consortium | Jacquelyn Wynn, Mairead Hyland, The Lighthouse Lab in Alderley Park and Alex Alderton, Roberto Amato, Jeffrey Barrett, Sonia Goncalves, Ewan Harrison, David K. Jackson, Ian Johnston, Dominic Kwiatkowski, Cordelia Langford, John Sillitoe on behalf of the Wellcome Sanger Institute COVID-19 Surveillance Team |
| EPI_ISL_2392722 | Lighthouse Lab in Milton Keynes | Wellcome Sanger Institute for the COVID-19 Genomics UK (COG-UK) Consortium | The Lighthouse Lab in Milton Keynes and Alex Alderton, Roberto Amato, Jeffrey Barrett, Sonia Goncalves, Ewan Harrison, David K. Jackson, Ian Johnston, Dominic Kwiatkowski, Cordelia Langford, John Sillitoe on behalf of the Wellcome Sanger Institute COVID-19 Surveillance Team |
| EPI_ISL_2392723 | Lighthouse Lab in Alderley Park | Wellcome Sanger Institute for the COVID-19 Genomics UK (COG-UK) Consortium | Jacquelyn Wynn, Mairead Hyland, The Lighthouse Lab in Alderley Park and Alex Alderton, Roberto Amato, Jeffrey Barrett, Sonia Goncalves, Ewan Harrison, David K. Jackson, Ian Johnston, Dominic Kwiatkowski, Cordelia Langford, John Sillitoe on behalf of the Wellcome Sanger Institute COVID-19 Surveillance Team |
| EPI_ISL_2392724 | Lighthouse Lab in Glasgow | Wellcome Sanger Institute for the COVID-19 Genomics UK (COG-UK) Consortium | Harper VanSteenhouse, Yumi Kasai, David Gray, Carol Clugston, Anna Dominiczak and Alex Alderton, Roberto Amato, Jeffrey Barrett, Sonia Goncalves, Ewan Harrison, David K. Jackson, Ian Johnston, Dominic Kwiatkowski, Cordelia Langford, John Sillitoe on behalf of the Wellcome Sanger Institute COVID-19 Surveillance Team |
| EPI_ISL_2392725, EPI_ISL_2392726 | Lighthouse Lab in Alderley Park | Wellcome Sanger Institute for the COVID-19 Genomics UK (COG-UK) Consortium | Jacquelyn Wynn, Mairead Hyland, The Lighthouse Lab in Alderley Park and Alex Alderton, Roberto Amato, Jeffrey Barrett, Sonia Goncalves, Ewan Harrison, David K. Jackson, Ian Johnston, Dominic Kwiatkowski, Cordelia Langford, John Sillitoe on behalf of the Wellcome Sanger Institute COVID-19 Surveillance Team |
| EPI_ISL_2392727 | Lighthouse Lab in Milton Keynes | Wellcome Sanger Institute for the COVID-19 Genomics UK (COG-UK) Consortium | The Lighthouse Lab in Milton Keynes and Alex Alderton, Roberto Amato, Jeffrey Barrett, Sonia Goncalves, Ewan Harrison, David K. Jackson, Ian Johnston, Dominic Kwiatkowski, Cordelia Langford, John Sillitoe on behalf of the Wellcome Sanger Institute COVID-19 Surveillance Team |
| EPI_ISL_2392728, EPI_ISL_2392729, EPI_ISL_2392731 | Lighthouse Lab in Alderley Park | Wellcome Sanger Institute for the COVID-19 Genomics UK (COG-UK) Consortium | Jacquelyn Wynn, Mairead Hyland, The Lighthouse Lab in Alderley Park and Alex Alderton, Roberto Amato, Jeffrey Barrett, Sonia Goncalves, Ewan Harrison, David K. Jackson, Ian Johnston, Dominic Kwiatkowski, Cordelia Langford, John Sillitoe on behalf of the Wellcome Sanger Institute COVID-19 Surveillance Team |
| EPI_ISL_2392732 | Lighthouse Lab in Glasgow | Wellcome Sanger Institute for the COVID-19 Genomics UK (COG-UK) Consortium | Harper VanSteenhouse, Yumi Kasai, David Gray, Carol Clugston, Anna Dominiczak and Alex Alderton, Roberto Amato, Jeffrey Barrett, Sonia Goncalves, Ewan Harrison, David K. Jackson, Ian Johnston, Dominic Kwiatkowski, Cordelia Langford, John Sillitoe on behalf of the Wellcome Sanger Institute COVID-19 Surveillance Team |
| EPI_ISL_2392733, EPI_ISL_2392734 | Lighthouse Lab in Milton Keynes | Wellcome Sanger Institute for the COVID-19 Genomics UK (COG-UK) Consortium | The Lighthouse Lab in Milton Keynes and Alex Alderton, Roberto Amato, Jeffrey Barrett, Sonia Goncalves, Ewan Harrison, David K. Jackson, Ian Johnston, Dominic Kwiatkowski, Cordelia Langford, John Sillitoe on behalf of the Wellcome Sanger Institute COVID-19 Surveillance Team |
| EPI_ISL_2392735, EPI_ISL_2392738 | Lighthouse Lab in Alderley Park | Wellcome Sanger Institute for the COVID-19 Genomics UK (COG-UK) Consortium | Jacquelyn Wynn, Mairead Hyland, The Lighthouse Lab in Alderley Park and Alex Alderton, Roberto Amato, Jeffrey Barrett, Sonia Goncalves, Ewan Harrison, David K. Jackson, Ian Johnston, Dominic Kwiatkowski, Cordelia Langford, John Sillitoe on behalf of the Wellcome Sanger Institute COVID-19 Surveillance Team |
| EPI_ISL_2392739, EPI_ISL_2392740, EPI_ISL_2392741 | Lighthouse Lab in Milton Keynes | Wellcome Sanger Institute for the COVID-19 Genomics UK (COG-UK) Consortium | The Lighthouse Lab in Milton Keynes and Alex Alderton, Roberto Amato, Jeffrey Barrett, Sonia Goncalves, Ewan Harrison, David K. Jackson, Ian Johnston, Dominic Kwiatkowski, Cordelia Langford, John Sillitoe on behalf of the Wellcome Sanger Institute COVID-19 Surveillance Team |
| EPI_ISL_2392742 | Lighthouse Lab in Alderley Park | Wellcome Sanger Institute for the COVID-19 Genomics UK (COG-UK) Consortium | Jacquelyn Wynn, Mairead Hyland, The Lighthouse Lab in Alderley Park and Alex Alderton, Roberto Amato, Jeffrey Barrett, Sonia Goncalves, Ewan Harrison, David K. Jackson, Ian Johnston, Dominic Kwiatkowski, Cordelia Langford, John Sillitoe on behalf of the Wellcome Sanger Institute COVID-19 Surveillance Team |
| EPI_ISL_2392743, EPI_ISL_2392744 | Lighthouse Lab in Milton Keynes | Wellcome Sanger Institute for the COVID-19 Genomics UK (COG-UK) Consortium | The Lighthouse Lab in Milton Keynes and Alex Alderton, Roberto Amato, Jeffrey Barrett, Sonia Goncalves, Ewan Harrison, David K. Jackson, Ian Johnston, Dominic Kwiatkowski, Cordelia Langford, John Sillitoe on behalf of the Wellcome Sanger Institute COVID-19 Surveillance Team |
| EPI_ISL_2392745 | Lighthouse Lab in Alderley Park | Wellcome Sanger Institute for the COVID-19 Genomics UK (COG-UK) Consortium | Jacquelyn Wynn, Mairead Hyland, The Lighthouse Lab in Alderley Park and Alex Alderton, Roberto Amato, Jeffrey Barrett, Sonia Goncalves, Ewan Harrison, David K. Jackson, Ian Johnston, Dominic Kwiatkowski, Cordelia Langford, John Sillitoe on behalf of the Wellcome Sanger Institute COVID-19 Surveillance Team |
| EPI_ISL_2392746, EPI_ISL_2392747, EPI_ISL_2392748, EPI_ISL_2392749 | Lighthouse Lab in Milton Keynes | Wellcome Sanger Institute for the COVID-19 Genomics UK (COG-UK) Consortium | The Lighthouse Lab in Milton Keynes and Alex Alderton, Roberto Amato, Jeffrey Barrett, Sonia Goncalves, Ewan Harrison, David K. Jackson, Ian Johnston, Dominic Kwiatkowski, Cordelia Langford, John Sillitoe on behalf of the Wellcome Sanger Institute COVID-19 Surveillance Team |
| EPI_ISL_2392751 | Lighthouse Lab in Alderley Park | Wellcome Sanger Institute for the COVID-19 Genomics UK (COG-UK) Consortium | Jacquelyn Wynn, Mairead Hyland, The Lighthouse Lab in Alderley Park and Alex Alderton, Roberto Amato, Jeffrey Barrett, Sonia Goncalves, Ewan Harrison, David K. Jackson, Ian Johnston, Dominic Kwiatkowski, Cordelia Langford, John Sillitoe on behalf of the Wellcome Sanger Institute COVID-19 Surveillance Team |
| EPI_ISL_2392752 | Lighthouse Lab in Milton Keynes | Wellcome Sanger Institute for the COVID-19 Genomics UK (COG-UK) Consortium | The Lighthouse Lab in Milton Keynes and Alex Alderton, Roberto Amato, Jeffrey Barrett, Sonia Goncalves, Ewan Harrison, David K. Jackson, Ian Johnston, Dominic Kwiatkowski, Cordelia Langford, John Sillitoe on behalf of the Wellcome Sanger Institute COVID-19 Surveillance Team |
| EPI_ISL_2392753, EPI_ISL_2392755, EPI_ISL_2392756 | Lighthouse Lab in Alderley Park | Wellcome Sanger Institute for the COVID-19 Genomics UK (COG-UK) Consortium | Jacquelyn Wynn, Mairead Hyland, The Lighthouse Lab in Alderley Park and Alex Alderton, Roberto Amato, Jeffrey Barrett, Sonia Goncalves, Ewan Harrison, David K. Jackson, Ian Johnston, Dominic Kwiatkowski, Cordelia Langford, John Sillitoe on behalf of the Wellcome Sanger Institute COVID-19 Surveillance Team |
| EPI_ISL_2392757, EPI_ISL_2392758 | Lighthouse Lab in Milton Keynes | Wellcome Sanger Institute for the COVID-19 Genomics UK (COG-UK) Consortium | The Lighthouse Lab in Milton Keynes and Alex Alderton, Roberto Amato, Jeffrey Barrett, Sonia Goncalves, Ewan Harrison, David K. Jackson, Ian Johnston, Dominic Kwiatkowski, Cordelia Langford, John Sillitoe on behalf of the Wellcome Sanger Institute COVID-19 Surveillance Team |
| EPI_ISL_2392760, EPI_ISL_2392761 | Lighthouse Lab in Alderley Park | Wellcome Sanger Institute for the COVID-19 Genomics UK (COG-UK) Consortium | Jacquelyn Wynn, Mairead Hyland, The Lighthouse Lab in Alderley Park and Alex Alderton, Roberto Amato, Jeffrey Barrett, Sonia Goncalves, Ewan Harrison, David K. Jackson, Ian Johnston, Dominic Kwiatkowski, Cordelia Langford, John Sillitoe on behalf of the Wellcome Sanger Institute COVID-19 Surveillance Team |
| EPI_ISL_2392762 | Lighthouse Lab in Milton Keynes | Wellcome Sanger Institute for the COVID-19 Genomics UK (COG-UK) Consortium | The Lighthouse Lab in Milton Keynes and Alex Alderton, Roberto Amato, Jeffrey Barrett, Sonia Goncalves, Ewan Harrison, David K. Jackson, Ian Johnston, Dominic Kwiatkowski, Cordelia Langford, John Sillitoe on behalf of the Wellcome Sanger Institute COVID-19 Surveillance Team |
| EPI_ISL_2392763 | Lighthouse Lab in Alderley Park | Wellcome Sanger Institute for the COVID-19 Genomics UK (COG-UK) Consortium | Jacquelyn Wynn, Mairead Hyland, The Lighthouse Lab in Alderley Park and Alex Alderton, Roberto Amato, Jeffrey Barrett, Sonia Goncalves, Ewan Harrison, David K. Jackson, Ian Johnston, Dominic Kwiatkowski, Cordelia Langford, John Sillitoe on behalf of the Wellcome Sanger Institute COVID-19 Surveillance Team |
| EPI_ISL_2392764, EPI_ISL_2392765, EPI_ISL_2392766, EPI_ISL_2392767, EPI_ISL_2392768, EPI_ISL_2392769, EPI_ISL_2392770 | Lighthouse Lab in Milton Keynes | Wellcome Sanger Institute for the COVID-19 Genomics UK (COG-UK) Consortium | The Lighthouse Lab in Milton Keynes and Alex Alderton, Roberto Amato, Jeffrey Barrett, Sonia Goncalves, Ewan Harrison, David K. Jackson, Ian Johnston, Dominic Kwiatkowski, Cordelia Langford, John Sillitoe on behalf of the Wellcome Sanger Institute COVID-19 Surveillance Team |
| EPI_ISL_2392772 | Lighthouse Lab in Glasgow | Wellcome Sanger Institute for the COVID-19 Genomics UK (COG-UK) Consortium | Harper VanSteenhouse, Yumi Kasai, David Gray, Carol Clugston, Anna Dominiczak and Alex Alderton, Roberto Amato, Jeffrey Barrett, Sonia Goncalves, Ewan Harrison, David K. Jackson, Ian Johnston, Dominic Kwiatkowski, Cordelia Langford, John Sillitoe on behalf of the Wellcome Sanger Institute COVID-19 Surveillance Team |
| EPI_ISL_2392773, EPI_ISL_2392774, EPI_ISL_2392775, EPI_ISL_2392776, EPI_ISL_2392777, EPI_ISL_2392778, EPI_ISL_2392779, EPI_ISL_2392783, EPI_ISL_2392784, EPI_ISL_2392786, EPI_ISL_2392787, EPI_ISL_2392788, EPI_ISL_2392789, EPI_ISL_2392790, EPI_ISL_2392791, EPI_ISL_2392792, EPI_ISL_2392793, EPI_ISL_2392795, EPI_ISL_2392796, EPI_ISL_2392797 | | | |
| see above | Lighthouse Lab in Milton Keynes | Wellcome Sanger Institute for the COVID-19 Genomics UK (COG-UK) Consortium | The Lighthouse Lab in Milton Keynes and Alex Alderton, Roberto Amato, Jeffrey Barrett, Sonia Goncalves, Ewan Harrison, David K. Jackson, Ian Johnston, Dominic Kwiatkowski, Cordelia Langford, John Sillitoe on behalf of the Wellcome Sanger Institute COVID-19 Surveillance Team |
| EPI_ISL_2392798 | Lighthouse Lab in Glasgow | Wellcome Sanger Institute for the COVID-19 Genomics UK (COG-UK) Consortium | Harper VanSteenhouse, Yumi Kasai, David Gray, Carol Clugston, Anna Dominiczak and Alex Alderton, Roberto Amato, Jeffrey Barrett, Sonia Goncalves, Ewan Harrison, David K. Jackson, Ian Johnston, Dominic Kwiatkowski, Cordelia Langford, John Sillitoe on behalf of the Wellcome Sanger Institute COVID-19 Surveillance Team |
| EPI_ISL_2392799, EPI_ISL_2392800, EPI_ISL_2392801, EPI_ISL_2392802, EPI_ISL_2392803, EPI_ISL_2392804, EPI_ISL_2392805, EPI_ISL_2392806, EPI_ISL_2392807, EPI_ISL_2392808, EPI_ISL_2392809, EPI_ISL_2392811, EPI_ISL_2392812, EPI_ISL_2392813, EPI_ISL_2392814, EPI_ISL_2392815, EPI_ISL_2392816, EPI_ISL_2392820, EPI_ISL_2392821, EPI_ISL_2392823, EPI_ISL_2392824, EPI_ISL_2392825, EPI_ISL_2392827, EPI_ISL_2392828, EPI_ISL_2392829, EPI_ISL_2392830, EPI_ISL_2392831, EPI_ISL_2392833, EPI_ISL_2392836, EPI_ISL_2392837, EPI_ISL_2392838 | | | |
| see above | Lighthouse Lab in Milton Keynes | Wellcome Sanger Institute for the COVID-19 Genomics UK (COG-UK) Consortium | The Lighthouse Lab in Milton Keynes and Alex Alderton, Roberto Amato, Jeffrey Barrett, Sonia Goncalves, Ewan Harrison, David K. Jackson, Ian Johnston, Dominic Kwiatkowski, Cordelia Langford, John Sillitoe on behalf of the Wellcome Sanger Institute COVID-19 Surveillance Team |

| | | | |
|---|---|---|---|
| EPI_ISL_2392840 | Lighthouse Lab in Glasgow | Wellcome Sanger Institute for the COVID-19 Genomics UK (COG-UK) Consortium | Harper VanSteenhouse, Yumi Kasai, David Gray, Carol Clugston, Anna Dominiczak and Alex Alderton, Roberto Amato, Jeffrey Barrett, Sonia Goncalves, Ewan Harrison, David K. Jackson, Ian Johnston, Dominic Kwiatkowski, Cordelia Langford, John Sillitoe on behalf of the Wellcome Sanger Institute COVID-19 Surveillance Team |
| EPI_ISL_2392841, EPI_ISL_2392842, EPI_ISL_2392843 | Lighthouse Lab in Milton Keynes | Wellcome Sanger Institute for the COVID-19 Genomics UK (COG-UK) Consortium | The Lighthouse Lab in Milton Keynes and Alex Alderton, Roberto Amato, Jeffrey Barrett, Sonia Goncalves, Ewan Harrison, David K. Jackson, Ian Johnston, Dominic Kwiatkowski, Cordelia Langford, John Sillitoe on behalf of the Wellcome Sanger Institute COVID-19 Surveillance Team |
| EPI_ISL_2392844 | Lighthouse Lab in Glasgow | Wellcome Sanger Institute for the COVID-19 Genomics UK (COG-UK) Consortium | Harper VanSteenhouse, Yumi Kasai, David Gray, Carol Clugston, Anna Dominiczak and Alex Alderton, Roberto Amato, Jeffrey Barrett, Sonia Goncalves, Ewan Harrison, David K. Jackson, Ian Johnston, Dominic Kwiatkowski, Cordelia Langford, John Sillitoe on behalf of the Wellcome Sanger Institute COVID-19 Surveillance Team |
| EPI_ISL_2392845, EPI_ISL_2392846, EPI_ISL_2392847, EPI_ISL_2392848, EPI_ISL_2392849, EPI_ISL_2392850, EPI_ISL_2392853, EPI_ISL_2392854, EPI_ISL_2392856, EPI_ISL_2392857, EPI_ISL_2392858, EPI_ISL_2392860, EPI_ISL_2392862, EPI_ISL_2392863, EPI_ISL_2392864, EPI_ISL_2392865, EPI_ISL_2392866, EPI_ISL_2392867, EPI_ISL_2392868, EPI_ISL_2392869, EPI_ISL_2392870, EPI_ISL_2392871, EPI_ISL_2392872, EPI_ISL_2392873, EPI_ISL_2392874, EPI_ISL_2392875, EPI_ISL_2392876, EPI_ISL_2392877, EPI_ISL_2392878, EPI_ISL_2392880, EPI_ISL_2392881, EPI_ISL_2392882, EPI_ISL_2392883, EPI_ISL_2392884, EPI_ISL_2392885, EPI_ISL_2392886, EPI_ISL_2392887, EPI_ISL_2392888 | | | |
| see above | Lighthouse Lab in Milton Keynes | Wellcome Sanger Institute for the COVID-19 Genomics UK (COG-UK) Consortium | The Lighthouse Lab in Milton Keynes and Alex Alderton, Roberto Amato, Jeffrey Barrett, Sonia Goncalves, Ewan Harrison, David K. Jackson, Ian Johnston, Dominic Kwiatkowski, Cordelia Langford, John Sillitoe on behalf of the Wellcome Sanger Institute COVID-19 Surveillance Team |
| EPI_ISL_2392889 | Lighthouse Lab in Glasgow | Wellcome Sanger Institute for the COVID-19 Genomics UK (COG-UK) Consortium | Harper VanSteenhouse, Yumi Kasai, David Gray, Carol Clugston, Anna Dominiczak and Alex Alderton, Roberto Amato, Jeffrey Barrett, Sonia Goncalves, Ewan Harrison, David K. Jackson, Ian Johnston, Dominic Kwiatkowski, Cordelia Langford, John Sillitoe on behalf of the Wellcome Sanger Institute COVID-19 Surveillance Team |
| EPI_ISL_2392890, EPI_ISL_2392891, EPI_ISL_2392892, EPI_ISL_2392893, EPI_ISL_2392894, EPI_ISL_2392895, EPI_ISL_2392896, EPI_ISL_2392897, EPI_ISL_2392899, EPI_ISL_2392900, EPI_ISL_2392901, EPI_ISL_2392902, EPI_ISL_2392903 | | | |
| see above | Lighthouse Lab in Milton Keynes | Wellcome Sanger Institute for the COVID-19 Genomics UK (COG-UK) Consortium | The Lighthouse Lab in Milton Keynes and Alex Alderton, Roberto Amato, Jeffrey Barrett, Sonia Goncalves, Ewan Harrison, David K. Jackson, Ian Johnston, Dominic Kwiatkowski, Cordelia Langford, John Sillitoe on behalf of the Wellcome Sanger Institute COVID-19 Surveillance Team |
| EPI_ISL_2392904 | Lighthouse Lab in Glasgow | Wellcome Sanger Institute for the COVID-19 Genomics UK (COG-UK) Consortium | Harper VanSteenhouse, Yumi Kasai, David Gray, Carol Clugston, Anna Dominiczak and Alex Alderton, Roberto Amato, Jeffrey Barrett, Sonia Goncalves, Ewan Harrison, David K. Jackson, Ian Johnston, Dominic Kwiatkowski, Cordelia Langford, John Sillitoe on behalf of the Wellcome Sanger Institute COVID-19 Surveillance Team |
| EPI_ISL_2392905, EPI_ISL_2392907, EPI_ISL_2392908, EPI_ISL_2392910, EPI_ISL_2392911, EPI_ISL_2392912, EPI_ISL_2392913, EPI_ISL_2392914, EPI_ISL_2392915, EPI_ISL_2392916, EPI_ISL_2392918 | | | |
| see above | Lighthouse Lab in Milton Keynes | Wellcome Sanger Institute for the COVID-19 Genomics UK (COG-UK) Consortium | The Lighthouse Lab in Milton Keynes and Alex Alderton, Roberto Amato, Jeffrey Barrett, Sonia Goncalves, Ewan Harrison, David K. Jackson, Ian Johnston, Dominic Kwiatkowski, Cordelia Langford, John Sillitoe on behalf of the Wellcome Sanger Institute COVID-19 Surveillance Team |
| EPI_ISL_2392919 | Lighthouse Lab in Glasgow | Wellcome Sanger Institute for the COVID-19 Genomics UK (COG-UK) Consortium | Harper VanSteenhouse, Yumi Kasai, David Gray, Carol Clugston, Anna Dominiczak and Alex Alderton, Roberto Amato, Jeffrey Barrett, Sonia Goncalves, Ewan Harrison, David K. Jackson, Ian Johnston, Dominic Kwiatkowski, Cordelia Langford, John Sillitoe on behalf of the Wellcome Sanger Institute COVID-19 Surveillance Team |
| EPI_ISL_2392920, EPI_ISL_2392921, EPI_ISL_2392923, EPI_ISL_2392924, EPI_ISL_2392925, EPI_ISL_2392926, EPI_ISL_2392927 | Lighthouse Lab in Milton Keynes | Wellcome Sanger Institute for the COVID-19 Genomics UK (COG-UK) Consortium | The Lighthouse Lab in Milton Keynes and Alex Alderton, Roberto Amato, Jeffrey Barrett, Sonia Goncalves, Ewan Harrison, David K. Jackson, Ian Johnston, Dominic Kwiatkowski, Cordelia Langford, John Sillitoe on behalf of the Wellcome Sanger Institute COVID-19 Surveillance Team |
| EPI_ISL_2392928 | Lighthouse Lab in Glasgow | Wellcome Sanger Institute for the COVID-19 Genomics UK (COG-UK) Consortium | Harper VanSteenhouse, Yumi Kasai, David Gray, Carol Clugston, Anna Dominiczak and Alex Alderton, Roberto Amato, Jeffrey Barrett, Sonia Goncalves, Ewan Harrison, David K. Jackson, Ian Johnston, Dominic Kwiatkowski, Cordelia Langford, John Sillitoe on behalf of the Wellcome Sanger Institute COVID-19 Surveillance Team |
| EPI_ISL_2392930, EPI_ISL_2392931, EPI_ISL_2392932, EPI_ISL_2392933, EPI_ISL_2392934, EPI_ISL_2392938, EPI_ISL_2392939, EPI_ISL_2392940, EPI_ISL_2392941, EPI_ISL_2392943 | Lighthouse Lab in Milton Keynes | Wellcome Sanger Institute for the COVID-19 Genomics UK (COG-UK) Consortium | The Lighthouse Lab in Milton Keynes and Alex Alderton, Roberto Amato, Jeffrey Barrett, Sonia Goncalves, Ewan Harrison, David K. Jackson, Ian Johnston, Dominic Kwiatkowski, Cordelia Langford, John Sillitoe on behalf of the Wellcome Sanger Institute COVID-19 Surveillance Team |
| EPI_ISL_2392944, EPI_ISL_2392945, EPI_ISL_2392946, EPI_ISL_2392947 | Lighthouse Lab in Alderley Park | Wellcome Sanger Institute for the COVID-19 Genomics UK (COG-UK) Consortium | Jacquelyn Wynn, Mairead Hyland, The Lighthouse Lab in Alderley Park and Alex Alderton, Roberto Amato, Jeffrey Barrett, Sonia Goncalves, Ewan Harrison, David K. Jackson, Ian Johnston, Dominic Kwiatkowski, Cordelia Langford, John Sillitoe on behalf of the Wellcome Sanger Institute COVID-19 Surveillance Team |
| EPI_ISL_2392948 | Lighthouse Lab in Glasgow | Wellcome Sanger Institute for the COVID-19 Genomics UK (COG-UK) Consortium | Harper VanSteenhouse, Yumi Kasai, David Gray, Carol Clugston, Anna Dominiczak and Alex Alderton, Roberto Amato, Jeffrey Barrett, Sonia Goncalves, Ewan Harrison, David K. Jackson, Ian Johnston, Dominic Kwiatkowski, Cordelia Langford, John Sillitoe on behalf of the Wellcome Sanger Institute COVID-19 Surveillance Team |
| EPI_ISL_2392949, EPI_ISL_2392950, EPI_ISL_2392951, EPI_ISL_2392953, EPI_ISL_2392954, EPI_ISL_2392956, EPI_ISL_2392957 | Lighthouse Lab in Alderley Park | Wellcome Sanger Institute for the COVID-19 Genomics UK (COG-UK) Consortium | Jacquelyn Wynn, Mairead Hyland, The Lighthouse Lab in Alderley Park and Alex Alderton, Roberto Amato, Jeffrey Barrett, Sonia Goncalves, Ewan Harrison, David K. Jackson, Ian Johnston, Dominic Kwiatkowski, Cordelia Langford, John Sillitoe on behalf of the Wellcome Sanger Institute COVID-19 Surveillance Team |
| EPI_ISL_2392958 | Lighthouse Lab in Glasgow | Wellcome Sanger Institute for the COVID-19 Genomics UK (COG-UK) Consortium | Harper VanSteenhouse, Yumi Kasai, David Gray, Carol Clugston, Anna Dominiczak and Alex Alderton, Roberto Amato, Jeffrey Barrett, Sonia Goncalves, Ewan Harrison, David K. Jackson, Ian Johnston, Dominic Kwiatkowski, Cordelia Langford, John Sillitoe on behalf of the Wellcome Sanger Institute COVID-19 Surveillance Team |
| EPI_ISL_2392959, EPI_ISL_2392960, EPI_ISL_2392961, EPI_ISL_2392962, EPI_ISL_2392963, EPI_ISL_2392964, EPI_ISL_2392965, EPI_ISL_2392969, EPI_ISL_2392970, EPI_ISL_2392971, EPI_ISL_2392973, EPI_ISL_2392974, EPI_ISL_2392975, EPI_ISL_2392977, EPI_ISL_2392978, EPI_ISL_2392979, EPI_ISL_2392980, EPI_ISL_2392981, EPI_ISL_2392982, EPI_ISL_2392984, EPI_ISL_2392985, EPI_ISL_2392987, EPI_ISL_2392988, EPI_ISL_2392990, EPI_ISL_2392991, EPI_ISL_2392992, EPI_ISL_2392993, EPI_ISL_2392994, EPI_ISL_2392996, EPI_ISL_2392997, EPI_ISL_2392999, EPI_ISL_2393001, EPI_ISL_2393002, EPI_ISL_2393003 | | | |
| see above | Lighthouse Lab in Alderley Park | Wellcome Sanger Institute for the COVID-19 Genomics UK (COG-UK) Consortium | Jacquelyn Wynn, Mairead Hyland, The Lighthouse Lab in Alderley Park and Alex Alderton, Roberto Amato, Jeffrey Barrett, Sonia Goncalves, Ewan Harrison, David K. Jackson, Ian Johnston, Dominic Kwiatkowski, Cordelia Langford, John Sillitoe on behalf of the Wellcome Sanger Institute COVID-19 Surveillance Team |
| EPI_ISL_2393004 | Lighthouse Lab in Glasgow | Wellcome Sanger Institute for the COVID-19 Genomics UK (COG-UK) Consortium | Harper VanSteenhouse, Yumi Kasai, David Gray, Carol Clugston, Anna Dominiczak and Alex Alderton, Roberto Amato, Jeffrey Barrett, Sonia Goncalves, Ewan Harrison, David K. Jackson, Ian Johnston, Dominic Kwiatkowski, Cordelia Langford, John Sillitoe on behalf of the Wellcome Sanger Institute COVID-19 Surveillance Team |
| EPI_ISL_2393005, EPI_ISL_2393006, EPI_ISL_2393007, EPI_ISL_2393009, EPI_ISL_2393010, EPI_ISL_2393012, EPI_ISL_2393013, EPI_ISL_2393014, EPI_ISL_2393015, EPI_ISL_2393016, EPI_ISL_2393017, EPI_ISL_2393018, EPI_ISL_2393019, EPI_ISL_2393020 | | | |
| see above | Lighthouse Lab in Alderley Park | Wellcome Sanger Institute for the COVID-19 Genomics UK (COG-UK) Consortium | Jacquelyn Wynn, Mairead Hyland, The Lighthouse Lab in Alderley Park and Alex Alderton, Roberto Amato, Jeffrey Barrett, Sonia Goncalves, Ewan Harrison, David K. Jackson, Ian Johnston, Dominic Kwiatkowski, Cordelia Langford, John Sillitoe on behalf of the Wellcome Sanger Institute COVID-19 Surveillance Team |
| EPI_ISL_2393021 | Lighthouse Lab in Glasgow | Wellcome Sanger Institute for the COVID-19 Genomics UK (COG-UK) Consortium | Harper VanSteenhouse, Yumi Kasai, David Gray, Carol Clugston, Anna Dominiczak and Alex Alderton, Roberto Amato, Jeffrey Barrett, Sonia Goncalves, Ewan Harrison, David K. Jackson, Ian Johnston, Dominic Kwiatkowski, Cordelia Langford, John Sillitoe on behalf of the Wellcome Sanger Institute COVID-19 Surveillance Team |
| EPI_ISL_2393022, EPI_ISL_2393023, EPI_ISL_2393024, EPI_ISL_2393025, EPI_ISL_2393027, EPI_ISL_2393028, EPI_ISL_2393029, EPI_ISL_2393031, EPI_ISL_2393032, EPI_ISL_2393033, EPI_ISL_2393034, EPI_ISL_2393036, EPI_ISL_2393037, EPI_ISL_2393039, EPI_ISL_2393040, EPI_ISL_2393041, EPI_ISL_2393042, EPI_ISL_2393043, EPI_ISL_2393044, EPI_ISL_2393046, EPI_ISL_2393047, EPI_ISL_2393048, EPI_ISL_2393049, EPI_ISL_2393050, EPI_ISL_2393051, EPI_ISL_2393053, EPI_ISL_2393054, EPI_ISL_2393055, EPI_ISL_2393056, EPI_ISL_2393057, EPI_ISL_2393058, EPI_ISL_2393059, EPI_ISL_2393060, EPI_ISL_2393061, EPI_ISL_2393062, EPI_ISL_2393063, EPI_ISL_2393064, EPI_ISL_2393065, EPI_ISL_2393066, EPI_ISL_2393067, EPI_ISL_2393068, EPI_ISL_2393069, EPI_ISL_2393070, EPI_ISL_2393071, EPI_ISL_2393073, EPI_ISL_2393074, EPI_ISL_2393075, EPI_ISL_2393077, EPI_ISL_2393079, EPI_ISL_2393080, EPI_ISL_2393081, EPI_ISL_2393082, EPI_ISL_2393084 | | | |
| see above | Lighthouse Lab in Alderley Park | Wellcome Sanger Institute for the COVID-19 Genomics UK (COG-UK) Consortium | Jacquelyn Wynn, Mairead Hyland, The Lighthouse Lab in Alderley Park and Alex Alderton, Roberto Amato, Jeffrey Barrett, Sonia Goncalves, Ewan Harrison, David K. Jackson, Ian Johnston, Dominic Kwiatkowski, Cordelia Langford, John Sillitoe on behalf of the Wellcome Sanger Institute COVID-19 Surveillance Team |
| EPI_ISL_2393085 | Lighthouse Lab in Glasgow | Wellcome Sanger Institute for the COVID-19 Genomics UK (COG-UK) Consortium | Harper VanSteenhouse, Yumi Kasai, David Gray, Carol Clugston, Anna Dominiczak and Alex Alderton, Roberto Amato, Jeffrey Barrett, Sonia Goncalves, Ewan Harrison, David K. Jackson, Ian Johnston, Dominic Kwiatkowski, Cordelia Langford, John Sillitoe on behalf of the Wellcome Sanger Institute COVID-19 Surveillance Team |
| EPI_ISL_2393086, EPI_ISL_2393087, EPI_ISL_2393088, EPI_ISL_2393089, EPI_ISL_2393090, EPI_ISL_2393091, EPI_ISL_2393093, EPI_ISL_2393094, EPI_ISL_2393096, EPI_ISL_2393097, EPI_ISL_2393098 | | | |
| see above | Lighthouse Lab in Alderley Park | Wellcome Sanger Institute for the COVID-19 Genomics UK (COG-UK) Consortium | Jacquelyn Wynn, Mairead Hyland, The Lighthouse Lab in Alderley Park and Alex Alderton, Roberto Amato, Jeffrey Barrett, Sonia Goncalves, Ewan Harrison, David K. Jackson, Ian Johnston, Dominic Kwiatkowski, Cordelia Langford, John Sillitoe on behalf of the Wellcome Sanger Institute COVID-19 Surveillance Team |
| EPI_ISL_2393100 | Lighthouse Lab in Glasgow | Wellcome Sanger Institute for the COVID-19 Genomics UK (COG-UK) Consortium | Harper VanSteenhouse, Yumi Kasai, David Gray, Carol Clugston, Anna Dominiczak and Alex Alderton, Roberto Amato, Jeffrey Barrett, Sonia Goncalves, Ewan Harrison, David K. Jackson, Ian Johnston, Dominic Kwiatkowski, Cordelia Langford, John Sillitoe on behalf of the Wellcome Sanger Institute COVID-19 Surveillance Team |
| EPI_ISL_2393101, EPI_ISL_2393102, EPI_ISL_2393103, EPI_ISL_2393104, EPI_ISL_2393105, EPI_ISL_2393107, EPI_ISL_2393108, EPI_ISL_2393109, EPI_ISL_2393110, EPI_ISL_2393111, EPI_ISL_2393112, EPI_ISL_2393113, EPI_ISL_2393115, EPI_ISL_2393116, EPI_ISL_2393117, EPI_ISL_2393119, EPI_ISL_2393121, EPI_ISL_2393122, EPI_ISL_2393123, EPI_ISL_2393124, EPI_ISL_2393125, EPI_ISL_2393126, EPI_ISL_2393127, EPI_ISL_2393129, EPI_ISL_2393130 | | | |
| see above | Lighthouse Lab in Alderley Park | Wellcome Sanger Institute for the COVID-19 Genomics UK (COG-UK) Consortium | Jacquelyn Wynn, Mairead Hyland, The Lighthouse Lab in Alderley Park and Alex Alderton, Roberto Amato, Jeffrey Barrett, Sonia Goncalves, Ewan Harrison, David K. Jackson, Ian Johnston, Dominic Kwiatkowski, Cordelia Langford, John Sillitoe on behalf of the Wellcome Sanger Institute COVID-19 Surveillance Team |
| EPI_ISL_2393131 | Lighthouse Lab in Glasgow | Wellcome Sanger Institute for the COVID-19 Genomics UK (COG-UK) Consortium | Harper VanSteenhouse, Yumi Kasai, David Gray, Carol Clugston, Anna Dominiczak and Alex Alderton, Roberto Amato, Jeffrey Barrett, Sonia Goncalves, Ewan Harrison, David K. Jackson, Ian Johnston, Dominic Kwiatkowski, Cordelia Langford, John Sillitoe on behalf of the Wellcome Sanger Institute COVID-19 Surveillance Team |
| EPI_ISL_2393132, EPI_ISL_2393133, EPI_ISL_2393134 | Lighthouse Lab in Alderley Park | Wellcome Sanger Institute for the COVID-19 Genomics UK (COG-UK) Consortium | Jacquelyn Wynn, Mairead Hyland, The Lighthouse Lab in Alderley Park and Alex Alderton, Roberto Amato, Jeffrey Barrett, Sonia Goncalves, Ewan Harrison, David K. Jackson, Ian Johnston, Dominic Kwiatkowski, Cordelia Langford, John Sillitoe on behalf of the Wellcome Sanger Institute COVID-19 Surveillance Team |
| EPI_ISL_2393135 | Lighthouse Lab in Milton Keynes | Wellcome Sanger Institute for the COVID-19 Genomics UK (COG-UK) Consortium | The Lighthouse Lab in Milton Keynes and Alex Alderton, Roberto Amato, Jeffrey Barrett, Sonia Goncalves, Ewan Harrison, David K. Jackson, Ian Johnston, Dominic Kwiatkowski, Cordelia Langford, John Sillitoe on behalf of the Wellcome Sanger Institute COVID-19 Surveillance Team |
| EPI_ISL_2393137 | Lighthouse Laboratory Plymouth | Wellcome Sanger Institute for the COVID-19 Genomics UK (COG-UK) Consortium | Lighthouse Laboratory Plymouth and Alex Alderton, Roberto Amato, Jeffrey Barrett, Sonia Goncalves, Ewan Harrison, David K. Jackson, Ian Johnston, Dominic Kwiatkowski, Cordelia Langford, John Sillitoe on behalf of the Wellcome Sanger Institute COVID-19 Surveillance Team |

| | | | |
|---|---|---|---|
| EPI_ISL_2393138, EPI_ISL_2393139 | Lighthouse Lab in Milton Keynes | Wellcome Sanger Institute for the COVID-19 Genomics UK (COG-UK) Consortium | The Lighthouse Lab in Milton Keynes and Alex Alderton, Roberto Amato, Jeffrey Barrett, Sonia Goncalves, Ewan Harrison, David K. Jackson, Ian Johnston, Dominic Kwiatkowski, Cordelia Langford, John Sillitoe on behalf of the Wellcome Sanger Institute COVID-19 Surveillance Team |
| EPI_ISL_2393140 | Lighthouse Laboratory Plymouth | Wellcome Sanger Institute for the COVID-19 Genomics UK (COG-UK) Consortium | Lighthouse Laboratory Plymouth and Alex Alderton, Roberto Amato, Jeffrey Barrett, Sonia Goncalves, Ewan Harrison, David K. Jackson, Ian Johnston, Dominic Kwiatkowski, Cordelia Langford, John Sillitoe on behalf of the Wellcome Sanger Institute COVID-19 Surveillance Team |
| EPI_ISL_2393143 | Lighthouse Lab in Alderley Park | Wellcome Sanger Institute for the COVID-19 Genomics UK (COG-UK) Consortium | Jacquelyn Wynn, Mairead Hyland, The Lighthouse Lab in Alderley Park and Alex Alderton, Roberto Amato, Jeffrey Barrett, Sonia Goncalves, Ewan Harrison, David K. Jackson, Ian Johnston, Dominic Kwiatkowski, Cordelia Langford, John Sillitoe on behalf of the Wellcome Sanger Institute COVID-19 Surveillance Team |
| EPI_ISL_2393144, EPI_ISL_2393145, EPI_ISL_2393146 | Lighthouse Lab in Milton Keynes | Wellcome Sanger Institute for the COVID-19 Genomics UK (COG-UK) Consortium | The Lighthouse Lab in Milton Keynes and Alex Alderton, Roberto Amato, Jeffrey Barrett, Sonia Goncalves, Ewan Harrison, David K. Jackson, Ian Johnston, Dominic Kwiatkowski, Cordelia Langford, John Sillitoe on behalf of the Wellcome Sanger Institute COVID-19 Surveillance Team |
| EPI_ISL_2393147 | Lighthouse Lab in Alderley Park | Wellcome Sanger Institute for the COVID-19 Genomics UK (COG-UK) Consortium | Jacquelyn Wynn, Mairead Hyland, The Lighthouse Lab in Alderley Park and Alex Alderton, Roberto Amato, Jeffrey Barrett, Sonia Goncalves, Ewan Harrison, David K. Jackson, Ian Johnston, Dominic Kwiatkowski, Cordelia Langford, John Sillitoe on behalf of the Wellcome Sanger Institute COVID-19 Surveillance Team |
| EPI_ISL_2393148 | Lighthouse Lab in Milton Keynes | Wellcome Sanger Institute for the COVID-19 Genomics UK (COG-UK) Consortium | The Lighthouse Lab in Milton Keynes and Alex Alderton, Roberto Amato, Jeffrey Barrett, Sonia Goncalves, Ewan Harrison, David K. Jackson, Ian Johnston, Dominic Kwiatkowski, Cordelia Langford, John Sillitoe on behalf of the Wellcome Sanger Institute COVID-19 Surveillance Team |
| EPI_ISL_2393149 | Lighthouse Lab in Alderley Park | Wellcome Sanger Institute for the COVID-19 Genomics UK (COG-UK) Consortium | Jacquelyn Wynn, Mairead Hyland, The Lighthouse Lab in Alderley Park and Alex Alderton, Roberto Amato, Jeffrey Barrett, Sonia Goncalves, Ewan Harrison, David K. Jackson, Ian Johnston, Dominic Kwiatkowski, Cordelia Langford, John Sillitoe on behalf of the Wellcome Sanger Institute COVID-19 Surveillance Team |
| EPI_ISL_2393150 | Lighthouse Laboratory Plymouth | Wellcome Sanger Institute for the COVID-19 Genomics UK (COG-UK) Consortium | Lighthouse Laboratory Plymouth and Alex Alderton, Roberto Amato, Jeffrey Barrett, Sonia Goncalves, Ewan Harrison, David K. Jackson, Ian Johnston, Dominic Kwiatkowski, Cordelia Langford, John Sillitoe on behalf of the Wellcome Sanger Institute COVID-19 Surveillance Team |
| EPI_ISL_2393151, EPI_ISL_2393153 | Lighthouse Lab in Alderley Park | Wellcome Sanger Institute for the COVID-19 Genomics UK (COG-UK) Consortium | Jacquelyn Wynn, Mairead Hyland, The Lighthouse Lab in Alderley Park and Alex Alderton, Roberto Amato, Jeffrey Barrett, Sonia Goncalves, Ewan Harrison, David K. Jackson, Ian Johnston, Dominic Kwiatkowski, Cordelia Langford, John Sillitoe on behalf of the Wellcome Sanger Institute COVID-19 Surveillance Team |
| EPI_ISL_2393154, EPI_ISL_2393155, EPI_ISL_2393156 | Lighthouse Lab in Milton Keynes | Wellcome Sanger Institute for the COVID-19 Genomics UK (COG-UK) Consortium | The Lighthouse Lab in Milton Keynes and Alex Alderton, Roberto Amato, Jeffrey Barrett, Sonia Goncalves, Ewan Harrison, David K. Jackson, Ian Johnston, Dominic Kwiatkowski, Cordelia Langford, John Sillitoe on behalf of the Wellcome Sanger Institute COVID-19 Surveillance Team |
| EPI_ISL_2393157 | Lighthouse Laboratory Plymouth | Wellcome Sanger Institute for the COVID-19 Genomics UK (COG-UK) Consortium | Lighthouse Laboratory Plymouth and Alex Alderton, Roberto Amato, Jeffrey Barrett, Sonia Goncalves, Ewan Harrison, David K. Jackson, Ian Johnston, Dominic Kwiatkowski, Cordelia Langford, John Sillitoe on behalf of the Wellcome Sanger Institute COVID-19 Surveillance Team |
| EPI_ISL_2393161, EPI_ISL_2393163, EPI_ISL_2393164, EPI_ISL_2393165 | Lighthouse Lab in Alderley Park | Wellcome Sanger Institute for the COVID-19 Genomics UK (COG-UK) Consortium | Jacquelyn Wynn, Mairead Hyland, The Lighthouse Lab in Alderley Park and Alex Alderton, Roberto Amato, Jeffrey Barrett, Sonia Goncalves, Ewan Harrison, David K. Jackson, Ian Johnston, Dominic Kwiatkowski, Cordelia Langford, John Sillitoe on behalf of the Wellcome Sanger Institute COVID-19 Surveillance Team |
| EPI_ISL_2393168 | Lighthouse Lab in Milton Keynes | Wellcome Sanger Institute for the COVID-19 Genomics UK (COG-UK) Consortium | The Lighthouse Lab in Milton Keynes and Alex Alderton, Roberto Amato, Jeffrey Barrett, Sonia Goncalves, Ewan Harrison, David K. Jackson, Ian Johnston, Dominic Kwiatkowski, Cordelia Langford, John Sillitoe on behalf of the Wellcome Sanger Institute COVID-19 Surveillance Team |
| EPI_ISL_2393169, EPI_ISL_2393170 | Lighthouse Lab in Alderley Park | Wellcome Sanger Institute for the COVID-19 Genomics UK (COG-UK) Consortium | Jacquelyn Wynn, Mairead Hyland, The Lighthouse Lab in Alderley Park and Alex Alderton, Roberto Amato, Jeffrey Barrett, Sonia Goncalves, Ewan Harrison, David K. Jackson, Ian Johnston, Dominic Kwiatkowski, Cordelia Langford, John Sillitoe on behalf of the Wellcome Sanger Institute COVID-19 Surveillance Team |
| EPI_ISL_2393171, EPI_ISL_2393172 | Lighthouse Lab in Milton Keynes | Wellcome Sanger Institute for the COVID-19 Genomics UK (COG-UK) Consortium | The Lighthouse Lab in Milton Keynes and Alex Alderton, Roberto Amato, Jeffrey Barrett, Sonia Goncalves, Ewan Harrison, David K. Jackson, Ian Johnston, Dominic Kwiatkowski, Cordelia Langford, John Sillitoe on behalf of the Wellcome Sanger Institute COVID-19 Surveillance Team |
| EPI_ISL_2393174, EPI_ISL_2393175, EPI_ISL_2393176 | Lighthouse Lab in Alderley Park | Wellcome Sanger Institute for the COVID-19 Genomics UK (COG-UK) Consortium | Jacquelyn Wynn, Mairead Hyland, The Lighthouse Lab in Alderley Park and Alex Alderton, Roberto Amato, Jeffrey Barrett, Sonia Goncalves, Ewan Harrison, David K. Jackson, Ian Johnston, Dominic Kwiatkowski, Cordelia Langford, John Sillitoe on behalf of the Wellcome Sanger Institute COVID-19 Surveillance Team |
| EPI_ISL_2393177 | Lighthouse Laboratory Plymouth | Wellcome Sanger Institute for the COVID-19 Genomics UK (COG-UK) Consortium | Lighthouse Laboratory Plymouth and Alex Alderton, Roberto Amato, Jeffrey Barrett, Sonia Goncalves, Ewan Harrison, David K. Jackson, Ian Johnston, Dominic Kwiatkowski, Cordelia Langford, John Sillitoe on behalf of the Wellcome Sanger Institute COVID-19 Surveillance Team |
| EPI_ISL_2393178, EPI_ISL_2393179 | Lighthouse Lab in Alderley Park | Wellcome Sanger Institute for the COVID-19 Genomics UK (COG-UK) Consortium | Jacquelyn Wynn, Mairead Hyland, The Lighthouse Lab in Alderley Park and Alex Alderton, Roberto Amato, Jeffrey Barrett, Sonia Goncalves, Ewan Harrison, David K. Jackson, Ian Johnston, Dominic Kwiatkowski, Cordelia Langford, John Sillitoe on behalf of the Wellcome Sanger Institute COVID-19 Surveillance Team |
| EPI_ISL_2393180 | Lighthouse Laboratory Plymouth | Wellcome Sanger Institute for the COVID-19 Genomics UK (COG-UK) Consortium | Lighthouse Laboratory Plymouth and Alex Alderton, Roberto Amato, Jeffrey Barrett, Sonia Goncalves, Ewan Harrison, David K. Jackson, Ian Johnston, Dominic Kwiatkowski, Cordelia Langford, John Sillitoe on behalf of the Wellcome Sanger Institute COVID-19 Surveillance Team |
| EPI_ISL_2393181, EPI_ISL_2393182, EPI_ISL_2393183, EPI_ISL_2393184, EPI_ISL_2393186 | Lighthouse Lab in Alderley Park | Wellcome Sanger Institute for the COVID-19 Genomics UK (COG-UK) Consortium | Jacquelyn Wynn, Mairead Hyland, The Lighthouse Lab in Alderley Park and Alex Alderton, Roberto Amato, Jeffrey Barrett, Sonia Goncalves, Ewan Harrison, David K. Jackson, Ian Johnston, Dominic Kwiatkowski, Cordelia Langford, John Sillitoe on behalf of the Wellcome Sanger Institute COVID-19 Surveillance Team |
| EPI_ISL_2393187, EPI_ISL_2393188 | Lighthouse Lab in Milton Keynes | Wellcome Sanger Institute for the COVID-19 Genomics UK (COG-UK) Consortium | The Lighthouse Lab in Milton Keynes and Alex Alderton, Roberto Amato, Jeffrey Barrett, Sonia Goncalves, Ewan Harrison, David K. Jackson, Ian Johnston, Dominic Kwiatkowski, Cordelia Langford, John Sillitoe on behalf of the Wellcome Sanger Institute COVID-19 Surveillance Team |
| EPI_ISL_2393190, EPI_ISL_2393192, EPI_ISL_2393194 | Lighthouse Lab in Alderley Park | Wellcome Sanger Institute for the COVID-19 Genomics UK (COG-UK) Consortium | Jacquelyn Wynn, Mairead Hyland, The Lighthouse Lab in Alderley Park and Alex Alderton, Roberto Amato, Jeffrey Barrett, Sonia Goncalves, Ewan Harrison, David K. Jackson, Ian Johnston, Dominic Kwiatkowski, Cordelia Langford, John Sillitoe on behalf of the Wellcome Sanger Institute COVID-19 Surveillance Team |
| EPI_ISL_2393195 | Lighthouse Laboratory Plymouth | Wellcome Sanger Institute for the COVID-19 Genomics UK (COG-UK) Consortium | Lighthouse Laboratory Plymouth and Alex Alderton, Roberto Amato, Jeffrey Barrett, Sonia Goncalves, Ewan Harrison, David K. Jackson, Ian Johnston, Dominic Kwiatkowski, Cordelia Langford, John Sillitoe on behalf of the Wellcome Sanger Institute COVID-19 Surveillance Team |
| EPI_ISL_2393197, EPI_ISL_2393198 | Lighthouse Lab in Alderley Park | Wellcome Sanger Institute for the COVID-19 Genomics UK (COG-UK) Consortium | Jacquelyn Wynn, Mairead Hyland, The Lighthouse Lab in Alderley Park and Alex Alderton, Roberto Amato, Jeffrey Barrett, Sonia Goncalves, Ewan Harrison, David K. Jackson, Ian Johnston, Dominic Kwiatkowski, Cordelia Langford, John Sillitoe on behalf of the Wellcome Sanger Institute COVID-19 Surveillance Team |
| EPI_ISL_2393199 | Lighthouse Laboratory Plymouth | Wellcome Sanger Institute for the COVID-19 Genomics UK (COG-UK) Consortium | Lighthouse Laboratory Plymouth and Alex Alderton, Roberto Amato, Jeffrey Barrett, Sonia Goncalves, Ewan Harrison, David K. Jackson, Ian Johnston, Dominic Kwiatkowski, Cordelia Langford, John Sillitoe on behalf of the Wellcome Sanger Institute COVID-19 Surveillance Team |
| EPI_ISL_2393200 | Lighthouse Lab in Milton Keynes | Wellcome Sanger Institute for the COVID-19 Genomics UK (COG-UK) Consortium | The Lighthouse Lab in Milton Keynes and Alex Alderton, Roberto Amato, Jeffrey Barrett, Sonia Goncalves, Ewan Harrison, David K. Jackson, Ian Johnston, Dominic Kwiatkowski, Cordelia Langford, John Sillitoe on behalf of the Wellcome Sanger Institute COVID-19 Surveillance Team |
| EPI_ISL_2393201, EPI_ISL_2393202, EPI_ISL_2393203, EPI_ISL_2393204 | Lighthouse Lab in Alderley Park | Wellcome Sanger Institute for the COVID-19 Genomics UK (COG-UK) Consortium | Jacquelyn Wynn, Mairead Hyland, The Lighthouse Lab in Alderley Park and Alex Alderton, Roberto Amato, Jeffrey Barrett, Sonia Goncalves, Ewan Harrison, David K. Jackson, Ian Johnston, Dominic Kwiatkowski, Cordelia Langford, John Sillitoe on behalf of the Wellcome Sanger Institute COVID-19 Surveillance Team |
| EPI_ISL_2393206 | Lighthouse Lab in Milton Keynes | Wellcome Sanger Institute for the COVID-19 Genomics UK (COG-UK) Consortium | The Lighthouse Lab in Milton Keynes and Alex Alderton, Roberto Amato, Jeffrey Barrett, Sonia Goncalves, Ewan Harrison, David K. Jackson, Ian Johnston, Dominic Kwiatkowski, Cordelia Langford, John Sillitoe on behalf of the Wellcome Sanger Institute COVID-19 Surveillance Team |
| EPI_ISL_2393207, EPI_ISL_2393213, EPI_ISL_2393216, EPI_ISL_2393217 | Lighthouse Lab in Alderley Park | Wellcome Sanger Institute for the COVID-19 Genomics UK (COG-UK) Consortium | Jacquelyn Wynn, Mairead Hyland, The Lighthouse Lab in Alderley Park and Alex Alderton, Roberto Amato, Jeffrey Barrett, Sonia Goncalves, Ewan Harrison, David K. Jackson, Ian Johnston, Dominic Kwiatkowski, Cordelia Langford, John Sillitoe on behalf of the Wellcome Sanger Institute COVID-19 Surveillance Team |
| EPI_ISL_2393218 | Lighthouse Lab in Milton Keynes | Wellcome Sanger Institute for the COVID-19 Genomics UK (COG-UK) Consortium | The Lighthouse Lab in Milton Keynes and Alex Alderton, Roberto Amato, Jeffrey Barrett, Sonia Goncalves, Ewan Harrison, David K. Jackson, Ian Johnston, Dominic Kwiatkowski, Cordelia Langford, John Sillitoe on behalf of the Wellcome Sanger Institute COVID-19 Surveillance Team |
| EPI_ISL_2393220 | Lighthouse Lab in Alderley Park | Wellcome Sanger Institute for the COVID-19 Genomics UK (COG-UK) Consortium | Jacquelyn Wynn, Mairead Hyland, The Lighthouse Lab in Alderley Park and Alex Alderton, Roberto Amato, Jeffrey Barrett, Sonia Goncalves, Ewan Harrison, David K. Jackson, Ian Johnston, Dominic Kwiatkowski, Cordelia Langford, John Sillitoe on behalf of the Wellcome Sanger Institute COVID-19 Surveillance Team |
| EPI_ISL_2393221 | Lighthouse Lab in Milton Keynes | Wellcome Sanger Institute for the COVID-19 Genomics UK (COG-UK) Consortium | The Lighthouse Lab in Milton Keynes and Alex Alderton, Roberto Amato, Jeffrey Barrett, Sonia Goncalves, Ewan Harrison, David K. Jackson, Ian Johnston, Dominic Kwiatkowski, Cordelia Langford, John Sillitoe on behalf of the Wellcome Sanger Institute COVID-19 Surveillance Team |
| EPI_ISL_2393222, EPI_ISL_2393223, EPI_ISL_2393224, EPI_ISL_2393225 | Lighthouse Lab in Alderley Park | Wellcome Sanger Institute for the COVID-19 Genomics UK (COG-UK) Consortium | Jacquelyn Wynn, Mairead Hyland, The Lighthouse Lab in Alderley Park and Alex Alderton, Roberto Amato, Jeffrey Barrett, Sonia Goncalves, Ewan Harrison, David K. Jackson, Ian Johnston, Dominic Kwiatkowski, Cordelia Langford, John Sillitoe on behalf of the Wellcome Sanger Institute COVID-19 Surveillance Team |
| EPI_ISL_2393226 | Lighthouse Laboratory Plymouth | Wellcome Sanger Institute for the COVID-19 Genomics UK (COG-UK) Consortium | Lighthouse Laboratory Plymouth and Alex Alderton, Roberto Amato, Jeffrey Barrett, Sonia Goncalves, Ewan Harrison, David K. Jackson, Ian Johnston, Dominic Kwiatkowski, Cordelia Langford, John Sillitoe on behalf of the Wellcome Sanger Institute COVID-19 Surveillance Team |
| EPI_ISL_2393227, EPI_ISL_2393230, EPI_ISL_2393231, EPI_ISL_2393232 | Lighthouse Lab in Alderley Park | Wellcome Sanger Institute for the COVID-19 Genomics UK (COG-UK) Consortium | Jacquelyn Wynn, Mairead Hyland, The Lighthouse Lab in Alderley Park and Alex Alderton, Roberto Amato, Jeffrey Barrett, Sonia Goncalves, Ewan Harrison, David K. Jackson, Ian Johnston, Dominic Kwiatkowski, Cordelia Langford, John Sillitoe on behalf of the Wellcome Sanger Institute COVID-19 Surveillance Team |
| EPI_ISL_2393233 | Lighthouse Laboratory Plymouth | Wellcome Sanger Institute for the COVID-19 Genomics UK (COG-UK) Consortium | Lighthouse Laboratory Plymouth and Alex Alderton, Roberto Amato, Jeffrey Barrett, Sonia Goncalves, Ewan Harrison, David K. Jackson, Ian Johnston, Dominic Kwiatkowski, Cordelia Langford, John Sillitoe on behalf of the Wellcome Sanger Institute COVID-19 Surveillance Team |
| EPI_ISL_2393234, EPI_ISL_2393235, EPI_ISL_2393237, EPI_ISL_2393238, | Lighthouse Lab in Alderley Park | Wellcome Sanger Institute for the COVID-19 Genomics UK (COG-UK) Consortium | Jacquelyn Wynn, Mairead Hyland, The Lighthouse Lab in Alderley Park and Alex Alderton, Roberto Amato, Jeffrey Barrett, Sonia Goncalves, Ewan Harrison, David K. Jackson, Ian Johnston, Dominic Kwiatkowski, Cordelia Langford, John Sillitoe on behalf of the Wellcome Sanger Institute COVID-19 Surveillance Team |

| | | | |
|---|---|---|---|
| EPI_ISL_2393239, EPI_ISL_2393241, EPI_ISL_2393242, EPI_ISL_2393246 | | | |
| EPI_ISL_2393250, EPI_ISL_2393251 | Lighthouse Laboratory Plymouth | Wellcome Sanger Institute for the COVID-19 Genomics UK (COG-UK) Consortium | Lighthouse Laboratory Plymouth and Alex Alderton, Roberto Amato, Jeffrey Barrett, Sonia Goncalves, Ewan Harrison, David K. Jackson, Ian Johnston, Dominic Kwiatkowski, Cordelia Langford, John Sillitoe on behalf of the Wellcome Sanger Institute COVID-19 Surveillance Team |
| EPI_ISL_2393252, EPI_ISL_2393253, EPI_ISL_2393254, EPI_ISL_2393255, EPI_ISL_2393258 | Lighthouse Lab in Alderley Park | Wellcome Sanger Institute for the COVID-19 Genomics UK (COG-UK) Consortium | Jacquelyn Wynn, Mairead Hyland, The Lighthouse Lab in Alderley Park and Alex Alderton, Roberto Amato, Jeffrey Barrett, Sonia Goncalves, Ewan Harrison, David K. Jackson, Ian Johnston, Dominic Kwiatkowski, Cordelia Langford, John Sillitoe on behalf of the Wellcome Sanger Institute COVID-19 Surveillance Team |
| EPI_ISL_2393259 | Lighthouse Lab in Milton Keynes | Wellcome Sanger Institute for the COVID-19 Genomics UK (COG-UK) Consortium | The Lighthouse Lab in Milton Keynes and Alex Alderton, Roberto Amato, Jeffrey Barrett, Sonia Goncalves, Ewan Harrison, David K. Jackson, Ian Johnston, Dominic Kwiatkowski, Cordelia Langford, John Sillitoe on behalf of the Wellcome Sanger Institute COVID-19 Surveillance Team |
| EPI_ISL_2393260, EPI_ISL_2393261, EPI_ISL_2393262, EPI_ISL_2393263 | Lighthouse Lab in Alderley Park | Wellcome Sanger Institute for the COVID-19 Genomics UK (COG-UK) Consortium | Jacquelyn Wynn, Mairead Hyland, The Lighthouse Lab in Alderley Park and Alex Alderton, Roberto Amato, Jeffrey Barrett, Sonia Goncalves, Ewan Harrison, David K. Jackson, Ian Johnston, Dominic Kwiatkowski, Cordelia Langford, John Sillitoe on behalf of the Wellcome Sanger Institute COVID-19 Surveillance Team |
| EPI_ISL_2393264 | Lighthouse Laboratory Plymouth | Wellcome Sanger Institute for the COVID-19 Genomics UK (COG-UK) Consortium | Lighthouse Laboratory Plymouth and Alex Alderton, Roberto Amato, Jeffrey Barrett, Sonia Goncalves, Ewan Harrison, David K. Jackson, Ian Johnston, Dominic Kwiatkowski, Cordelia Langford, John Sillitoe on behalf of the Wellcome Sanger Institute COVID-19 Surveillance Team |
| EPI_ISL_2393265, EPI_ISL_2393266, EPI_ISL_2393268, EPI_ISL_2393269, EPI_ISL_2393271, EPI_ISL_2393272, EPI_ISL_2393274, EPI_ISL_2393275 | Lighthouse Lab in Alderley Park | Wellcome Sanger Institute for the COVID-19 Genomics UK (COG-UK) Consortium | Jacquelyn Wynn, Mairead Hyland, The Lighthouse Lab in Alderley Park and Alex Alderton, Roberto Amato, Jeffrey Barrett, Sonia Goncalves, Ewan Harrison, David K. Jackson, Ian Johnston, Dominic Kwiatkowski, Cordelia Langford, John Sillitoe on behalf of the Wellcome Sanger Institute COVID-19 Surveillance Team |
| EPI_ISL_2393276 | Lighthouse Lab in Milton Keynes | Wellcome Sanger Institute for the COVID-19 Genomics UK (COG-UK) Consortium | The Lighthouse Lab in Milton Keynes and Alex Alderton, Roberto Amato, Jeffrey Barrett, Sonia Goncalves, Ewan Harrison, David K. Jackson, Ian Johnston, Dominic Kwiatkowski, Cordelia Langford, John Sillitoe on behalf of the Wellcome Sanger Institute COVID-19 Surveillance Team |
| EPI_ISL_2393277 | Lighthouse Lab in Alderley Park | Wellcome Sanger Institute for the COVID-19 Genomics UK (COG-UK) Consortium | Jacquelyn Wynn, Mairead Hyland, The Lighthouse Lab in Alderley Park and Alex Alderton, Roberto Amato, Jeffrey Barrett, Sonia Goncalves, Ewan Harrison, David K. Jackson, Ian Johnston, Dominic Kwiatkowski, Cordelia Langford, John Sillitoe on behalf of the Wellcome Sanger Institute COVID-19 Surveillance Team |
| EPI_ISL_2393278 | Lighthouse Laboratory Plymouth | Wellcome Sanger Institute for the COVID-19 Genomics UK (COG-UK) Consortium | Lighthouse Laboratory Plymouth and Alex Alderton, Roberto Amato, Jeffrey Barrett, Sonia Goncalves, Ewan Harrison, David K. Jackson, Ian Johnston, Dominic Kwiatkowski, Cordelia Langford, John Sillitoe on behalf of the Wellcome Sanger Institute COVID-19 Surveillance Team |
| EPI_ISL_2393280 | Lighthouse Lab in Alderley Park | Wellcome Sanger Institute for the COVID-19 Genomics UK (COG-UK) Consortium | Jacquelyn Wynn, Mairead Hyland, The Lighthouse Lab in Alderley Park and Alex Alderton, Roberto Amato, Jeffrey Barrett, Sonia Goncalves, Ewan Harrison, David K. Jackson, Ian Johnston, Dominic Kwiatkowski, Cordelia Langford, John Sillitoe on behalf of the Wellcome Sanger Institute COVID-19 Surveillance Team |
| EPI_ISL_2393281 | Lighthouse Laboratory Plymouth | Wellcome Sanger Institute for the COVID-19 Genomics UK (COG-UK) Consortium | Lighthouse Laboratory Plymouth and Alex Alderton, Roberto Amato, Jeffrey Barrett, Sonia Goncalves, Ewan Harrison, David K. Jackson, Ian Johnston, Dominic Kwiatkowski, Cordelia Langford, John Sillitoe on behalf of the Wellcome Sanger Institute COVID-19 Surveillance Team |
| EPI_ISL_2393282 | Lighthouse Lab in Alderley Park | Wellcome Sanger Institute for the COVID-19 Genomics UK (COG-UK) Consortium | Jacquelyn Wynn, Mairead Hyland, The Lighthouse Lab in Alderley Park and Alex Alderton, Roberto Amato, Jeffrey Barrett, Sonia Goncalves, Ewan Harrison, David K. Jackson, Ian Johnston, Dominic Kwiatkowski, Cordelia Langford, John Sillitoe on behalf of the Wellcome Sanger Institute COVID-19 Surveillance Team |
| EPI_ISL_2393284 | Lighthouse Laboratory Plymouth | Wellcome Sanger Institute for the COVID-19 Genomics UK (COG-UK) Consortium | Lighthouse Laboratory Plymouth and Alex Alderton, Roberto Amato, Jeffrey Barrett, Sonia Goncalves, Ewan Harrison, David K. Jackson, Ian Johnston, Dominic Kwiatkowski, Cordelia Langford, John Sillitoe on behalf of the Wellcome Sanger Institute COVID-19 Surveillance Team |
| EPI_ISL_2393285, EPI_ISL_2393286 | Lighthouse Lab in Alderley Park | Wellcome Sanger Institute for the COVID-19 Genomics UK (COG-UK) Consortium | Jacquelyn Wynn, Mairead Hyland, The Lighthouse Lab in Alderley Park and Alex Alderton, Roberto Amato, Jeffrey Barrett, Sonia Goncalves, Ewan Harrison, David K. Jackson, Ian Johnston, Dominic Kwiatkowski, Cordelia Langford, John Sillitoe on behalf of the Wellcome Sanger Institute COVID-19 Surveillance Team |
| EPI_ISL_2393288 | Lighthouse Lab in Milton Keynes | Wellcome Sanger Institute for the COVID-19 Genomics UK (COG-UK) Consortium | The Lighthouse Lab in Milton Keynes and Alex Alderton, Roberto Amato, Jeffrey Barrett, Sonia Goncalves, Ewan Harrison, David K. Jackson, Ian Johnston, Dominic Kwiatkowski, Cordelia Langford, John Sillitoe on behalf of the Wellcome Sanger Institute COVID-19 Surveillance Team |
| EPI_ISL_2393289, EPI_ISL_2393290, EPI_ISL_2393291, EPI_ISL_2393292 | Lighthouse Lab in Alderley Park | Wellcome Sanger Institute for the COVID-19 Genomics UK (COG-UK) Consortium | Jacquelyn Wynn, Mairead Hyland, The Lighthouse Lab in Alderley Park and Alex Alderton, Roberto Amato, Jeffrey Barrett, Sonia Goncalves, Ewan Harrison, David K. Jackson, Ian Johnston, Dominic Kwiatkowski, Cordelia Langford, John Sillitoe on behalf of the Wellcome Sanger Institute COVID-19 Surveillance Team |
| EPI_ISL_2393294 | Lighthouse Lab in Milton Keynes | Wellcome Sanger Institute for the COVID-19 Genomics UK (COG-UK) Consortium | The Lighthouse Lab in Milton Keynes and Alex Alderton, Roberto Amato, Jeffrey Barrett, Sonia Goncalves, Ewan Harrison, David K. Jackson, Ian Johnston, Dominic Kwiatkowski, Cordelia Langford, John Sillitoe on behalf of the Wellcome Sanger Institute COVID-19 Surveillance Team |
| EPI_ISL_2393295 | Lighthouse Laboratory Plymouth | Wellcome Sanger Institute for the COVID-19 Genomics UK (COG-UK) Consortium | Lighthouse Laboratory Plymouth and Alex Alderton, Roberto Amato, Jeffrey Barrett, Sonia Goncalves, Ewan Harrison, David K. Jackson, Ian Johnston, Dominic Kwiatkowski, Cordelia Langford, John Sillitoe on behalf of the Wellcome Sanger Institute COVID-19 Surveillance Team |
| EPI_ISL_2393297, EPI_ISL_2393299, EPI_ISL_2393300, EPI_ISL_2393301, EPI_ISL_2393302 | Lighthouse Lab in Alderley Park | Wellcome Sanger Institute for the COVID-19 Genomics UK (COG-UK) Consortium | Jacquelyn Wynn, Mairead Hyland, The Lighthouse Lab in Alderley Park and Alex Alderton, Roberto Amato, Jeffrey Barrett, Sonia Goncalves, Ewan Harrison, David K. Jackson, Ian Johnston, Dominic Kwiatkowski, Cordelia Langford, John Sillitoe on behalf of the Wellcome Sanger Institute COVID-19 Surveillance Team |
| EPI_ISL_2393305, EPI_ISL_2393308, EPI_ISL_2393309 | Lighthouse Lab in Milton Keynes | Wellcome Sanger Institute for the COVID-19 Genomics UK (COG-UK) Consortium | The Lighthouse Lab in Milton Keynes and Alex Alderton, Roberto Amato, Jeffrey Barrett, Sonia Goncalves, Ewan Harrison, David K. Jackson, Ian Johnston, Dominic Kwiatkowski, Cordelia Langford, John Sillitoe on behalf of the Wellcome Sanger Institute COVID-19 Surveillance Team |
| EPI_ISL_2393310 | Lighthouse Lab in Alderley Park | Wellcome Sanger Institute for the COVID-19 Genomics UK (COG-UK) Consortium | Jacquelyn Wynn, Mairead Hyland, The Lighthouse Lab in Alderley Park and Alex Alderton, Roberto Amato, Jeffrey Barrett, Sonia Goncalves, Ewan Harrison, David K. Jackson, Ian Johnston, Dominic Kwiatkowski, Cordelia Langford, John Sillitoe on behalf of the Wellcome Sanger Institute COVID-19 Surveillance Team |
| EPI_ISL_2393311 | Lighthouse Laboratory Plymouth | Wellcome Sanger Institute for the COVID-19 Genomics UK (COG-UK) Consortium | Lighthouse Laboratory Plymouth and Alex Alderton, Roberto Amato, Jeffrey Barrett, Sonia Goncalves, Ewan Harrison, David K. Jackson, Ian Johnston, Dominic Kwiatkowski, Cordelia Langford, John Sillitoe on behalf of the Wellcome Sanger Institute COVID-19 Surveillance Team |
| EPI_ISL_2393312, EPI_ISL_2393313, EPI_ISL_2393314 | Lighthouse Lab in Alderley Park | Wellcome Sanger Institute for the COVID-19 Genomics UK (COG-UK) Consortium | Jacquelyn Wynn, Mairead Hyland, The Lighthouse Lab in Alderley Park and Alex Alderton, Roberto Amato, Jeffrey Barrett, Sonia Goncalves, Ewan Harrison, David K. Jackson, Ian Johnston, Dominic Kwiatkowski, Cordelia Langford, John Sillitoe on behalf of the Wellcome Sanger Institute COVID-19 Surveillance Team |
| EPI_ISL_2393318 | Lighthouse Laboratory Plymouth | Wellcome Sanger Institute for the COVID-19 Genomics UK (COG-UK) Consortium | Lighthouse Laboratory Plymouth and Alex Alderton, Roberto Amato, Jeffrey Barrett, Sonia Goncalves, Ewan Harrison, David K. Jackson, Ian Johnston, Dominic Kwiatkowski, Cordelia Langford, John Sillitoe on behalf of the Wellcome Sanger Institute COVID-19 Surveillance Team |
| EPI_ISL_2393319, EPI_ISL_2393320, EPI_ISL_2393321 | Lighthouse Lab in Milton Keynes | Wellcome Sanger Institute for the COVID-19 Genomics UK (COG-UK) Consortium | The Lighthouse Lab in Milton Keynes and Alex Alderton, Roberto Amato, Jeffrey Barrett, Sonia Goncalves, Ewan Harrison, David K. Jackson, Ian Johnston, Dominic Kwiatkowski, Cordelia Langford, John Sillitoe on behalf of the Wellcome Sanger Institute COVID-19 Surveillance Team |
| EPI_ISL_2393323 | Lighthouse Laboratory Plymouth | Wellcome Sanger Institute for the COVID-19 Genomics UK (COG-UK) Consortium | Lighthouse Laboratory Plymouth and Alex Alderton, Roberto Amato, Jeffrey Barrett, Sonia Goncalves, Ewan Harrison, David K. Jackson, Ian Johnston, Dominic Kwiatkowski, Cordelia Langford, John Sillitoe on behalf of the Wellcome Sanger Institute COVID-19 Surveillance Team |
| EPI_ISL_2393326 | Lighthouse Lab in Alderley Park | Wellcome Sanger Institute for the COVID-19 Genomics UK (COG-UK) Consortium | Jacquelyn Wynn, Mairead Hyland, The Lighthouse Lab in Alderley Park and Alex Alderton, Roberto Amato, Jeffrey Barrett, Sonia Goncalves, Ewan Harrison, David K. Jackson, Ian Johnston, Dominic Kwiatkowski, Cordelia Langford, John Sillitoe on behalf of the Wellcome Sanger Institute COVID-19 Surveillance Team |
| EPI_ISL_2393327 | Lighthouse Lab in Milton Keynes | Wellcome Sanger Institute for the COVID-19 Genomics UK (COG-UK) Consortium | The Lighthouse Lab in Milton Keynes and Alex Alderton, Roberto Amato, Jeffrey Barrett, Sonia Goncalves, Ewan Harrison, David K. Jackson, Ian Johnston, Dominic Kwiatkowski, Cordelia Langford, John Sillitoe on behalf of the Wellcome Sanger Institute COVID-19 Surveillance Team |
| EPI_ISL_2393328 | Lighthouse Laboratory Plymouth | Wellcome Sanger Institute for the COVID-19 Genomics UK (COG-UK) Consortium | Lighthouse Laboratory Plymouth and Alex Alderton, Roberto Amato, Jeffrey Barrett, Sonia Goncalves, Ewan Harrison, David K. Jackson, Ian Johnston, Dominic Kwiatkowski, Cordelia Langford, John Sillitoe on behalf of the Wellcome Sanger Institute COVID-19 Surveillance Team |
| EPI_ISL_2393329, EPI_ISL_2393330, EPI_ISL_2393331 | Lighthouse Lab in Alderley Park | Wellcome Sanger Institute for the COVID-19 Genomics UK (COG-UK) Consortium | Jacquelyn Wynn, Mairead Hyland, The Lighthouse Lab in Alderley Park and Alex Alderton, Roberto Amato, Jeffrey Barrett, Sonia Goncalves, Ewan Harrison, David K. Jackson, Ian Johnston, Dominic Kwiatkowski, Cordelia Langford, John Sillitoe on behalf of the Wellcome Sanger Institute COVID-19 Surveillance Team |
| EPI_ISL_2393332 | Lighthouse Laboratory Plymouth | Wellcome Sanger Institute for the COVID-19 Genomics UK (COG-UK) Consortium | Lighthouse Laboratory Plymouth and Alex Alderton, Roberto Amato, Jeffrey Barrett, Sonia Goncalves, Ewan Harrison, David K. Jackson, Ian Johnston, Dominic Kwiatkowski, Cordelia Langford, John Sillitoe on behalf of the Wellcome Sanger Institute COVID-19 Surveillance Team |
| EPI_ISL_2393333, EPI_ISL_2393334, EPI_ISL_2393335 | Lighthouse Lab in Alderley Park | Wellcome Sanger Institute for the COVID-19 Genomics UK (COG-UK) Consortium | Jacquelyn Wynn, Mairead Hyland, The Lighthouse Lab in Alderley Park and Alex Alderton, Roberto Amato, Jeffrey Barrett, Sonia Goncalves, Ewan Harrison, David K. Jackson, Ian Johnston, Dominic Kwiatkowski, Cordelia Langford, John Sillitoe on behalf of the Wellcome Sanger Institute COVID-19 Surveillance Team |
| EPI_ISL_2393338 | Lighthouse Lab in Milton Keynes | Wellcome Sanger Institute for the COVID-19 Genomics UK (COG-UK) Consortium | The Lighthouse Lab in Milton Keynes and Alex Alderton, Roberto Amato, Jeffrey Barrett, Sonia Goncalves, Ewan Harrison, David K. Jackson, Ian Johnston, Dominic Kwiatkowski, Cordelia Langford, John Sillitoe on behalf of the Wellcome Sanger Institute COVID-19 Surveillance Team |
| EPI_ISL_2393339, EPI_ISL_2393340, EPI_ISL_2393341 | Lighthouse Lab in Alderley Park | Wellcome Sanger Institute for the COVID-19 Genomics UK (COG-UK) Consortium | Jacquelyn Wynn, Mairead Hyland, The Lighthouse Lab in Alderley Park and Alex Alderton, Roberto Amato, Jeffrey Barrett, Sonia Goncalves, Ewan Harrison, David K. Jackson, Ian Johnston, Dominic Kwiatkowski, Cordelia Langford, John Sillitoe on behalf of the Wellcome Sanger Institute COVID-19 Surveillance Team |
| EPI_ISL_2393343 | Lighthouse Laboratory Plymouth | Wellcome Sanger Institute for the COVID-19 Genomics UK (COG-UK) | Lighthouse Laboratory Plymouth and Alex Alderton, Roberto Amato, Jeffrey Barrett, Sonia Goncalves, Ewan Harrison, David K. Jackson, Ian Johnston, Dominic Kwiatkowski, Cordelia Langford, John Sillitoe on behalf of the Wellcome Sanger |

| | | | |
|---|---|---|---|
| | | Consortium | Institute COVID-19 Surveillance Team |
| EPI_ISL_2393346, EPI_ISL_2393348 | Lighthouse Lab in Alderley Park | Wellcome Sanger Institute for the COVID-19 Genomics UK (COG-UK) Consortium | Jacquelyn Wynn, Mairead Hyland, The Lighthouse Lab in Alderley Park and Alex Alderton, Roberto Amato, Jeffrey Barrett, Sonia Goncalves, Ewan Harrison, David K. Jackson, Ian Johnston, Dominic Kwiatkowski, Cordelia Langford, John Sillitoe on behalf of the Wellcome Sanger Institute COVID-19 Surveillance Team |
| EPI_ISL_2393349, EPI_ISL_2393350 | Lighthouse Lab in Milton Keynes | Wellcome Sanger Institute for the COVID-19 Genomics UK (COG-UK) Consortium | The Lighthouse Lab in Milton Keynes and Alex Alderton, Roberto Amato, Jeffrey Barrett, Sonia Goncalves, Ewan Harrison, David K. Jackson, Ian Johnston, Dominic Kwiatkowski, Cordelia Langford, John Sillitoe on behalf of the Wellcome Sanger Institute COVID-19 Surveillance Team |
| EPI_ISL_2393351 | Lighthouse Laboratory Plymouth | Wellcome Sanger Institute for the COVID-19 Genomics UK (COG-UK) Consortium | Lighthouse Laboratory Plymouth and Alex Alderton, Roberto Amato, Jeffrey Barrett, Sonia Goncalves, Ewan Harrison, David K. Jackson, Ian Johnston, Dominic Kwiatkowski, Cordelia Langford, John Sillitoe on behalf of the Wellcome Sanger Institute COVID-19 Surveillance Team |
| EPI_ISL_2393352 | Lighthouse Lab in Milton Keynes | Wellcome Sanger Institute for the COVID-19 Genomics UK (COG-UK) Consortium | The Lighthouse Lab in Milton Keynes and Alex Alderton, Roberto Amato, Jeffrey Barrett, Sonia Goncalves, Ewan Harrison, David K. Jackson, Ian Johnston, Dominic Kwiatkowski, Cordelia Langford, John Sillitoe on behalf of the Wellcome Sanger Institute COVID-19 Surveillance Team |
| EPI_ISL_2393354, EPI_ISL_2393355, EPI_ISL_2393356 | Lighthouse Lab in Alderley Park | Wellcome Sanger Institute for the COVID-19 Genomics UK (COG-UK) Consortium | Jacquelyn Wynn, Mairead Hyland, The Lighthouse Lab in Alderley Park and Alex Alderton, Roberto Amato, Jeffrey Barrett, Sonia Goncalves, Ewan Harrison, David K. Jackson, Ian Johnston, Dominic Kwiatkowski, Cordelia Langford, John Sillitoe on behalf of the Wellcome Sanger Institute COVID-19 Surveillance Team |
| EPI_ISL_2393357 | Lighthouse Lab in Milton Keynes | Wellcome Sanger Institute for the COVID-19 Genomics UK (COG-UK) Consortium | The Lighthouse Lab in Milton Keynes and Alex Alderton, Roberto Amato, Jeffrey Barrett, Sonia Goncalves, Ewan Harrison, David K. Jackson, Ian Johnston, Dominic Kwiatkowski, Cordelia Langford, John Sillitoe on behalf of the Wellcome Sanger Institute COVID-19 Surveillance Team |
| EPI_ISL_2393358 | Lighthouse Laboratory Plymouth | Wellcome Sanger Institute for the COVID-19 Genomics UK (COG-UK) Consortium | Lighthouse Laboratory Plymouth and Alex Alderton, Roberto Amato, Jeffrey Barrett, Sonia Goncalves, Ewan Harrison, David K. Jackson, Ian Johnston, Dominic Kwiatkowski, Cordelia Langford, John Sillitoe on behalf of the Wellcome Sanger Institute COVID-19 Surveillance Team |
| EPI_ISL_2393359 | Lighthouse Lab in Milton Keynes | Wellcome Sanger Institute for the COVID-19 Genomics UK (COG-UK) Consortium | The Lighthouse Lab in Milton Keynes and Alex Alderton, Roberto Amato, Jeffrey Barrett, Sonia Goncalves, Ewan Harrison, David K. Jackson, Ian Johnston, Dominic Kwiatkowski, Cordelia Langford, John Sillitoe on behalf of the Wellcome Sanger Institute COVID-19 Surveillance Team |
| EPI_ISL_2393360 | Lighthouse Laboratory Plymouth | Wellcome Sanger Institute for the COVID-19 Genomics UK (COG-UK) Consortium | Lighthouse Laboratory Plymouth and Alex Alderton, Roberto Amato, Jeffrey Barrett, Sonia Goncalves, Ewan Harrison, David K. Jackson, Ian Johnston, Dominic Kwiatkowski, Cordelia Langford, John Sillitoe on behalf of the Wellcome Sanger Institute COVID-19 Surveillance Team |
| EPI_ISL_2393362, EPI_ISL_2393363, EPI_ISL_2393364, EPI_ISL_2393365, EPI_ISL_2393366 | Lighthouse Lab in Alderley Park | Wellcome Sanger Institute for the COVID-19 Genomics UK (COG-UK) Consortium | Jacquelyn Wynn, Mairead Hyland, The Lighthouse Lab in Alderley Park and Alex Alderton, Roberto Amato, Jeffrey Barrett, Sonia Goncalves, Ewan Harrison, David K. Jackson, Ian Johnston, Dominic Kwiatkowski, Cordelia Langford, John Sillitoe on behalf of the Wellcome Sanger Institute COVID-19 Surveillance Team |
| EPI_ISL_2393367 | Lighthouse Lab in Milton Keynes | Wellcome Sanger Institute for the COVID-19 Genomics UK (COG-UK) Consortium | The Lighthouse Lab in Milton Keynes and Alex Alderton, Roberto Amato, Jeffrey Barrett, Sonia Goncalves, Ewan Harrison, David K. Jackson, Ian Johnston, Dominic Kwiatkowski, Cordelia Langford, John Sillitoe on behalf of the Wellcome Sanger Institute COVID-19 Surveillance Team |
| EPI_ISL_2393369 | Lighthouse Lab in Alderley Park | Wellcome Sanger Institute for the COVID-19 Genomics UK (COG-UK) Consortium | Jacquelyn Wynn, Mairead Hyland, The Lighthouse Lab in Alderley Park and Alex Alderton, Roberto Amato, Jeffrey Barrett, Sonia Goncalves, Ewan Harrison, David K. Jackson, Ian Johnston, Dominic Kwiatkowski, Cordelia Langford, John Sillitoe on behalf of the Wellcome Sanger Institute COVID-19 Surveillance Team |
| EPI_ISL_2393370 | Lighthouse Lab in Milton Keynes | Wellcome Sanger Institute for the COVID-19 Genomics UK (COG-UK) Consortium | The Lighthouse Lab in Milton Keynes and Alex Alderton, Roberto Amato, Jeffrey Barrett, Sonia Goncalves, Ewan Harrison, David K. Jackson, Ian Johnston, Dominic Kwiatkowski, Cordelia Langford, John Sillitoe on behalf of the Wellcome Sanger Institute COVID-19 Surveillance Team |
| EPI_ISL_2393371 | Lighthouse Lab in Alderley Park | Wellcome Sanger Institute for the COVID-19 Genomics UK (COG-UK) Consortium | Jacquelyn Wynn, Mairead Hyland, The Lighthouse Lab in Alderley Park and Alex Alderton, Roberto Amato, Jeffrey Barrett, Sonia Goncalves, Ewan Harrison, David K. Jackson, Ian Johnston, Dominic Kwiatkowski, Cordelia Langford, John Sillitoe on behalf of the Wellcome Sanger Institute COVID-19 Surveillance Team |
| EPI_ISL_2393372 | Lighthouse Laboratory Plymouth | Wellcome Sanger Institute for the COVID-19 Genomics UK (COG-UK) Consortium | Lighthouse Laboratory Plymouth and Alex Alderton, Roberto Amato, Jeffrey Barrett, Sonia Goncalves, Ewan Harrison, David K. Jackson, Ian Johnston, Dominic Kwiatkowski, Cordelia Langford, John Sillitoe on behalf of the Wellcome Sanger Institute COVID-19 Surveillance Team |
| EPI_ISL_2393373 | Lighthouse Lab in Alderley Park | Wellcome Sanger Institute for the COVID-19 Genomics UK (COG-UK) Consortium | Jacquelyn Wynn, Mairead Hyland, The Lighthouse Lab in Alderley Park and Alex Alderton, Roberto Amato, Jeffrey Barrett, Sonia Goncalves, Ewan Harrison, David K. Jackson, Ian Johnston, Dominic Kwiatkowski, Cordelia Langford, John Sillitoe on behalf of the Wellcome Sanger Institute COVID-19 Surveillance Team |
| EPI_ISL_2393374 | Lighthouse Laboratory Plymouth | Wellcome Sanger Institute for the COVID-19 Genomics UK (COG-UK) Consortium | Lighthouse Laboratory Plymouth and Alex Alderton, Roberto Amato, Jeffrey Barrett, Sonia Goncalves, Ewan Harrison, David K. Jackson, Ian Johnston, Dominic Kwiatkowski, Cordelia Langford, John Sillitoe on behalf of the Wellcome Sanger Institute COVID-19 Surveillance Team |
| EPI_ISL_2393375 | Lighthouse Lab in Milton Keynes | Wellcome Sanger Institute for the COVID-19 Genomics UK (COG-UK) Consortium | The Lighthouse Lab in Milton Keynes and Alex Alderton, Roberto Amato, Jeffrey Barrett, Sonia Goncalves, Ewan Harrison, David K. Jackson, Ian Johnston, Dominic Kwiatkowski, Cordelia Langford, John Sillitoe on behalf of the Wellcome Sanger Institute COVID-19 Surveillance Team |
| EPI_ISL_2393376 | Lighthouse Laboratory Plymouth | Wellcome Sanger Institute for the COVID-19 Genomics UK (COG-UK) Consortium | Lighthouse Laboratory Plymouth and Alex Alderton, Roberto Amato, Jeffrey Barrett, Sonia Goncalves, Ewan Harrison, David K. Jackson, Ian Johnston, Dominic Kwiatkowski, Cordelia Langford, John Sillitoe on behalf of the Wellcome Sanger Institute COVID-19 Surveillance Team |
| EPI_ISL_2393377, EPI_ISL_2393378, EPI_ISL_2393379, EPI_ISL_2393381, EPI_ISL_2393382 | Lighthouse Lab in Alderley Park | Wellcome Sanger Institute for the COVID-19 Genomics UK (COG-UK) Consortium | Jacquelyn Wynn, Mairead Hyland, The Lighthouse Lab in Alderley Park and Alex Alderton, Roberto Amato, Jeffrey Barrett, Sonia Goncalves, Ewan Harrison, David K. Jackson, Ian Johnston, Dominic Kwiatkowski, Cordelia Langford, John Sillitoe on behalf of the Wellcome Sanger Institute COVID-19 Surveillance Team |
| EPI_ISL_2393387 | Lighthouse Laboratory Plymouth | Wellcome Sanger Institute for the COVID-19 Genomics UK (COG-UK) Consortium | Lighthouse Laboratory Plymouth and Alex Alderton, Roberto Amato, Jeffrey Barrett, Sonia Goncalves, Ewan Harrison, David K. Jackson, Ian Johnston, Dominic Kwiatkowski, Cordelia Langford, John Sillitoe on behalf of the Wellcome Sanger Institute COVID-19 Surveillance Team |
| EPI_ISL_2393389 | Lighthouse Lab in Milton Keynes | Wellcome Sanger Institute for the COVID-19 Genomics UK (COG-UK) Consortium | The Lighthouse Lab in Milton Keynes and Alex Alderton, Roberto Amato, Jeffrey Barrett, Sonia Goncalves, Ewan Harrison, David K. Jackson, Ian Johnston, Dominic Kwiatkowski, Cordelia Langford, John Sillitoe on behalf of the Wellcome Sanger Institute COVID-19 Surveillance Team |
| EPI_ISL_2393390 | Lighthouse Lab in Alderley Park | Wellcome Sanger Institute for the COVID-19 Genomics UK (COG-UK) Consortium | Jacquelyn Wynn, Mairead Hyland, The Lighthouse Lab in Alderley Park and Alex Alderton, Roberto Amato, Jeffrey Barrett, Sonia Goncalves, Ewan Harrison, David K. Jackson, Ian Johnston, Dominic Kwiatkowski, Cordelia Langford, John Sillitoe on behalf of the Wellcome Sanger Institute COVID-19 Surveillance Team |
| EPI_ISL_2393392, EPI_ISL_2393395, EPI_ISL_2393396, EPI_ISL_2393397 | Lighthouse Lab in Milton Keynes | Wellcome Sanger Institute for the COVID-19 Genomics UK (COG-UK) Consortium | The Lighthouse Lab in Milton Keynes and Alex Alderton, Roberto Amato, Jeffrey Barrett, Sonia Goncalves, Ewan Harrison, David K. Jackson, Ian Johnston, Dominic Kwiatkowski, Cordelia Langford, John Sillitoe on behalf of the Wellcome Sanger Institute COVID-19 Surveillance Team |
| EPI_ISL_2393398, EPI_ISL_2393400, EPI_ISL_2393401, EPI_ISL_2393402, EPI_ISL_2393403 | Lighthouse Lab in Alderley Park | Wellcome Sanger Institute for the COVID-19 Genomics UK (COG-UK) Consortium | Jacquelyn Wynn, Mairead Hyland, The Lighthouse Lab in Alderley Park and Alex Alderton, Roberto Amato, Jeffrey Barrett, Sonia Goncalves, Ewan Harrison, David K. Jackson, Ian Johnston, Dominic Kwiatkowski, Cordelia Langford, John Sillitoe on behalf of the Wellcome Sanger Institute COVID-19 Surveillance Team |
| EPI_ISL_2393405 | Lighthouse Laboratory Plymouth | Wellcome Sanger Institute for the COVID-19 Genomics UK (COG-UK) Consortium | Lighthouse Laboratory Plymouth and Alex Alderton, Roberto Amato, Jeffrey Barrett, Sonia Goncalves, Ewan Harrison, David K. Jackson, Ian Johnston, Dominic Kwiatkowski, Cordelia Langford, John Sillitoe on behalf of the Wellcome Sanger Institute COVID-19 Surveillance Team |
| EPI_ISL_2393407 | Lighthouse Lab in Milton Keynes | Wellcome Sanger Institute for the COVID-19 Genomics UK (COG-UK) Consortium | The Lighthouse Lab in Milton Keynes and Alex Alderton, Roberto Amato, Jeffrey Barrett, Sonia Goncalves, Ewan Harrison, David K. Jackson, Ian Johnston, Dominic Kwiatkowski, Cordelia Langford, John Sillitoe on behalf of the Wellcome Sanger Institute COVID-19 Surveillance Team |
| EPI_ISL_2393408 | Lighthouse Lab in Alderley Park | Wellcome Sanger Institute for the COVID-19 Genomics UK (COG-UK) Consortium | Jacquelyn Wynn, Mairead Hyland, The Lighthouse Lab in Alderley Park and Alex Alderton, Roberto Amato, Jeffrey Barrett, Sonia Goncalves, Ewan Harrison, David K. Jackson, Ian Johnston, Dominic Kwiatkowski, Cordelia Langford, John Sillitoe on behalf of the Wellcome Sanger Institute COVID-19 Surveillance Team |
| EPI_ISL_2393410 | Lighthouse Lab in Milton Keynes | Wellcome Sanger Institute for the COVID-19 Genomics UK (COG-UK) Consortium | The Lighthouse Lab in Milton Keynes and Alex Alderton, Roberto Amato, Jeffrey Barrett, Sonia Goncalves, Ewan Harrison, David K. Jackson, Ian Johnston, Dominic Kwiatkowski, Cordelia Langford, John Sillitoe on behalf of the Wellcome Sanger Institute COVID-19 Surveillance Team |
| EPI_ISL_2393411, EPI_ISL_2393412 | Lighthouse Lab in Alderley Park | Wellcome Sanger Institute for the COVID-19 Genomics UK (COG-UK) Consortium | Jacquelyn Wynn, Mairead Hyland, The Lighthouse Lab in Alderley Park and Alex Alderton, Roberto Amato, Jeffrey Barrett, Sonia Goncalves, Ewan Harrison, David K. Jackson, Ian Johnston, Dominic Kwiatkowski, Cordelia Langford, John Sillitoe on behalf of the Wellcome Sanger Institute COVID-19 Surveillance Team |
| EPI_ISL_2393414 | Lighthouse Lab in Milton Keynes | Wellcome Sanger Institute for the COVID-19 Genomics UK (COG-UK) Consortium | The Lighthouse Lab in Milton Keynes and Alex Alderton, Roberto Amato, Jeffrey Barrett, Sonia Goncalves, Ewan Harrison, David K. Jackson, Ian Johnston, Dominic Kwiatkowski, Cordelia Langford, John Sillitoe on behalf of the Wellcome Sanger Institute COVID-19 Surveillance Team |
| EPI_ISL_2393416, EPI_ISL_2393417, EPI_ISL_2393418 | Lighthouse Lab in Alderley Park | Wellcome Sanger Institute for the COVID-19 Genomics UK (COG-UK) Consortium | Jacquelyn Wynn, Mairead Hyland, The Lighthouse Lab in Alderley Park and Alex Alderton, Roberto Amato, Jeffrey Barrett, Sonia Goncalves, Ewan Harrison, David K. Jackson, Ian Johnston, Dominic Kwiatkowski, Cordelia Langford, John Sillitoe on behalf of the Wellcome Sanger Institute COVID-19 Surveillance Team |
| EPI_ISL_2393419 | Lighthouse Lab in Milton Keynes | Wellcome Sanger Institute for the COVID-19 Genomics UK (COG-UK) Consortium | The Lighthouse Lab in Milton Keynes and Alex Alderton, Roberto Amato, Jeffrey Barrett, Sonia Goncalves, Ewan Harrison, David K. Jackson, Ian Johnston, Dominic Kwiatkowski, Cordelia Langford, John Sillitoe on behalf of the Wellcome Sanger Institute COVID-19 Surveillance Team |
| EPI_ISL_2393420 | Lighthouse Laboratory Plymouth | Wellcome Sanger Institute for the COVID-19 Genomics UK (COG-UK) Consortium | Lighthouse Laboratory Plymouth and Alex Alderton, Roberto Amato, Jeffrey Barrett, Sonia Goncalves, Ewan Harrison, David K. Jackson, Ian Johnston, Dominic Kwiatkowski, Cordelia Langford, John Sillitoe on behalf of the Wellcome Sanger Institute COVID-19 Surveillance Team |
| EPI_ISL_2393421, EPI_ISL_2393422, EPI_ISL_2393423 | Lighthouse Lab in Alderley Park | Wellcome Sanger Institute for the COVID-19 Genomics UK (COG-UK) Consortium | Jacquelyn Wynn, Mairead Hyland, The Lighthouse Lab in Alderley Park and Alex Alderton, Roberto Amato, Jeffrey Barrett, Sonia Goncalves, Ewan Harrison, David K. Jackson, Ian Johnston, Dominic Kwiatkowski, Cordelia Langford, John Sillitoe on behalf of the Wellcome Sanger Institute COVID-19 Surveillance Team |
| EPI_ISL_2393424 | Lighthouse Laboratory Plymouth | Wellcome Sanger Institute for the COVID-19 Genomics UK (COG-UK) Consortium | Lighthouse Laboratory Plymouth and Alex Alderton, Roberto Amato, Jeffrey Barrett, Sonia Goncalves, Ewan Harrison, David K. Jackson, Ian Johnston, Dominic Kwiatkowski, Cordelia Langford, John Sillitoe on behalf of the Wellcome Sanger Institute COVID-19 Surveillance Team |
| EPI_ISL_2393426 | Lighthouse Lab in Alderley Park | Wellcome Sanger Institute for the COVID-19 Genomics UK (COG-UK) Consortium | Jacquelyn Wynn, Mairead Hyland, The Lighthouse Lab in Alderley Park and Alex Alderton, Roberto Amato, Jeffrey Barrett, Sonia Goncalves, Ewan Harrison, David K. Jackson, Ian Johnston, Dominic Kwiatkowski, Cordelia Langford, John Sillitoe on behalf of the Wellcome Sanger Institute COVID-19 Surveillance Team |
| EPI_ISL_2393428 | Lighthouse Laboratory Plymouth | Wellcome Sanger Institute for the COVID-19 Genomics UK (COG-UK) Consortium | Lighthouse Laboratory Plymouth and Alex Alderton, Roberto Amato, Jeffrey Barrett, Sonia Goncalves, Ewan Harrison, David K. Jackson, Ian Johnston, Dominic Kwiatkowski, Cordelia Langford, John Sillitoe on behalf of the Wellcome Sanger Institute COVID-19 Surveillance Team |
| EPI_ISL_2393429 | Lighthouse Lab in Alderley Park | Wellcome Sanger Institute for the COVID-19 Genomics UK (COG-UK) Consortium | Jacquelyn Wynn, Mairead Hyland, The Lighthouse Lab in Alderley Park and Alex Alderton, Roberto Amato, Jeffrey Barrett, Sonia Goncalves, Ewan Harrison, David K. Jackson, Ian Johnston, Dominic Kwiatkowski, Cordelia Langford, John Sillitoe on behalf of the Wellcome Sanger Institute COVID-19 Surveillance Team |

| | | | |
|---|---|---|---|
| EPI_ISL_2393431 | Lighthouse Laboratory Plymouth | Wellcome Sanger Institute for the COVID-19 Genomics UK (COG-UK) Consortium | Lighthouse Laboratory Plymouth and Alex Alderton, Roberto Amato, Jeffrey Barrett, Sonia Goncalves, Ewan Harrison, David K. Jackson, Ian Johnston, Dominic Kwiatkowski, Cordelia Langford, John Sillitoe on behalf of the Wellcome Sanger Institute COVID-19 Surveillance Team |
| EPI_ISL_2393432, EPI_ISL_2393433 | Lighthouse Lab in Alderley Park | Wellcome Sanger Institute for the COVID-19 Genomics UK (COG-UK) Consortium | Jacquelyn Wynn, Mairead Hyland, The Lighthouse Lab in Alderley Park and Alex Alderton, Roberto Amato, Jeffrey Barrett, Sonia Goncalves, Ewan Harrison, David K. Jackson, Ian Johnston, Dominic Kwiatkowski, Cordelia Langford, John Sillitoe on behalf of the Wellcome Sanger Institute COVID-19 Surveillance Team |
| EPI_ISL_2393434 | Lighthouse Lab in Milton Keynes | Wellcome Sanger Institute for the COVID-19 Genomics UK (COG-UK) Consortium | The Lighthouse Lab in Milton Keynes and Alex Alderton, Roberto Amato, Jeffrey Barrett, Sonia Goncalves, Ewan Harrison, David K. Jackson, Ian Johnston, Dominic Kwiatkowski, Cordelia Langford, John Sillitoe on behalf of the Wellcome Sanger Institute COVID-19 Surveillance Team |
| EPI_ISL_2393435 | Lighthouse Laboratory Plymouth | Wellcome Sanger Institute for the COVID-19 Genomics UK (COG-UK) Consortium | Lighthouse Laboratory Plymouth and Alex Alderton, Roberto Amato, Jeffrey Barrett, Sonia Goncalves, Ewan Harrison, David K. Jackson, Ian Johnston, Dominic Kwiatkowski, Cordelia Langford, John Sillitoe on behalf of the Wellcome Sanger Institute COVID-19 Surveillance Team |
| EPI_ISL_2393436, EPI_ISL_2393438 | Lighthouse Lab in Alderley Park | Wellcome Sanger Institute for the COVID-19 Genomics UK (COG-UK) Consortium | Jacquelyn Wynn, Mairead Hyland, The Lighthouse Lab in Alderley Park and Alex Alderton, Roberto Amato, Jeffrey Barrett, Sonia Goncalves, Ewan Harrison, David K. Jackson, Ian Johnston, Dominic Kwiatkowski, Cordelia Langford, John Sillitoe on behalf of the Wellcome Sanger Institute COVID-19 Surveillance Team |
| EPI_ISL_2393440 | Lighthouse Lab in Milton Keynes | Wellcome Sanger Institute for the COVID-19 Genomics UK (COG-UK) Consortium | The Lighthouse Lab in Milton Keynes and Alex Alderton, Roberto Amato, Jeffrey Barrett, Sonia Goncalves, Ewan Harrison, David K. Jackson, Ian Johnston, Dominic Kwiatkowski, Cordelia Langford, John Sillitoe on behalf of the Wellcome Sanger Institute COVID-19 Surveillance Team |
| EPI_ISL_2393441 | Lighthouse Laboratory Plymouth | Wellcome Sanger Institute for the COVID-19 Genomics UK (COG-UK) Consortium | Lighthouse Laboratory Plymouth and Alex Alderton, Roberto Amato, Jeffrey Barrett, Sonia Goncalves, Ewan Harrison, David K. Jackson, Ian Johnston, Dominic Kwiatkowski, Cordelia Langford, John Sillitoe on behalf of the Wellcome Sanger Institute COVID-19 Surveillance Team |
| EPI_ISL_2393442 | Lighthouse Lab in Milton Keynes | Wellcome Sanger Institute for the COVID-19 Genomics UK (COG-UK) Consortium | The Lighthouse Lab in Milton Keynes and Alex Alderton, Roberto Amato, Jeffrey Barrett, Sonia Goncalves, Ewan Harrison, David K. Jackson, Ian Johnston, Dominic Kwiatkowski, Cordelia Langford, John Sillitoe on behalf of the Wellcome Sanger Institute COVID-19 Surveillance Team |
| EPI_ISL_2393444 | Lighthouse Lab in Alderley Park | Wellcome Sanger Institute for the COVID-19 Genomics UK (COG-UK) Consortium | Jacquelyn Wynn, Mairead Hyland, The Lighthouse Lab in Alderley Park and Alex Alderton, Roberto Amato, Jeffrey Barrett, Sonia Goncalves, Ewan Harrison, David K. Jackson, Ian Johnston, Dominic Kwiatkowski, Cordelia Langford, John Sillitoe on behalf of the Wellcome Sanger Institute COVID-19 Surveillance Team |
| EPI_ISL_2393445, EPI_ISL_2393446 | Lighthouse Laboratory Plymouth | Wellcome Sanger Institute for the COVID-19 Genomics UK (COG-UK) Consortium | Lighthouse Laboratory Plymouth and Alex Alderton, Roberto Amato, Jeffrey Barrett, Sonia Goncalves, Ewan Harrison, David K. Jackson, Ian Johnston, Dominic Kwiatkowski, Cordelia Langford, John Sillitoe on behalf of the Wellcome Sanger Institute COVID-19 Surveillance Team |
| EPI_ISL_2393447 | Lighthouse Lab in Alderley Park | Wellcome Sanger Institute for the COVID-19 Genomics UK (COG-UK) Consortium | Jacquelyn Wynn, Mairead Hyland, The Lighthouse Lab in Alderley Park and Alex Alderton, Roberto Amato, Jeffrey Barrett, Sonia Goncalves, Ewan Harrison, David K. Jackson, Ian Johnston, Dominic Kwiatkowski, Cordelia Langford, John Sillitoe on behalf of the Wellcome Sanger Institute COVID-19 Surveillance Team |
| EPI_ISL_2393448, EPI_ISL_2393450, EPI_ISL_2393451, EPI_ISL_2393452 | Lighthouse Lab in Milton Keynes | Wellcome Sanger Institute for the COVID-19 Genomics UK (COG-UK) Consortium | The Lighthouse Lab in Milton Keynes and Alex Alderton, Roberto Amato, Jeffrey Barrett, Sonia Goncalves, Ewan Harrison, David K. Jackson, Ian Johnston, Dominic Kwiatkowski, Cordelia Langford, John Sillitoe on behalf of the Wellcome Sanger Institute COVID-19 Surveillance Team |
| EPI_ISL_2393453 | Lighthouse Laboratory Plymouth | Wellcome Sanger Institute for the COVID-19 Genomics UK (COG-UK) Consortium | Lighthouse Laboratory Plymouth and Alex Alderton, Roberto Amato, Jeffrey Barrett, Sonia Goncalves, Ewan Harrison, David K. Jackson, Ian Johnston, Dominic Kwiatkowski, Cordelia Langford, John Sillitoe on behalf of the Wellcome Sanger Institute COVID-19 Surveillance Team |
| EPI_ISL_2393454 | Lighthouse Lab in Milton Keynes | Wellcome Sanger Institute for the COVID-19 Genomics UK (COG-UK) Consortium | The Lighthouse Lab in Milton Keynes and Alex Alderton, Roberto Amato, Jeffrey Barrett, Sonia Goncalves, Ewan Harrison, David K. Jackson, Ian Johnston, Dominic Kwiatkowski, Cordelia Langford, John Sillitoe on behalf of the Wellcome Sanger Institute COVID-19 Surveillance Team |
| EPI_ISL_2393455, EPI_ISL_2393456, EPI_ISL_2393457 | Lighthouse Lab in Alderley Park | Wellcome Sanger Institute for the COVID-19 Genomics UK (COG-UK) Consortium | Jacquelyn Wynn, Mairead Hyland, The Lighthouse Lab in Alderley Park and Alex Alderton, Roberto Amato, Jeffrey Barrett, Sonia Goncalves, Ewan Harrison, David K. Jackson, Ian Johnston, Dominic Kwiatkowski, Cordelia Langford, John Sillitoe on behalf of the Wellcome Sanger Institute COVID-19 Surveillance Team |
| EPI_ISL_2393458 | Lighthouse Lab in Milton Keynes | Wellcome Sanger Institute for the COVID-19 Genomics UK (COG-UK) Consortium | The Lighthouse Lab in Milton Keynes and Alex Alderton, Roberto Amato, Jeffrey Barrett, Sonia Goncalves, Ewan Harrison, David K. Jackson, Ian Johnston, Dominic Kwiatkowski, Cordelia Langford, John Sillitoe on behalf of the Wellcome Sanger Institute COVID-19 Surveillance Team |
| EPI_ISL_2393459 | Lighthouse Lab in Alderley Park | Wellcome Sanger Institute for the COVID-19 Genomics UK (COG-UK) Consortium | Jacquelyn Wynn, Mairead Hyland, The Lighthouse Lab in Alderley Park and Alex Alderton, Roberto Amato, Jeffrey Barrett, Sonia Goncalves, Ewan Harrison, David K. Jackson, Ian Johnston, Dominic Kwiatkowski, Cordelia Langford, John Sillitoe on behalf of the Wellcome Sanger Institute COVID-19 Surveillance Team |
| EPI_ISL_2393460 | Lighthouse Lab in Milton Keynes | Wellcome Sanger Institute for the COVID-19 Genomics UK (COG-UK) Consortium | The Lighthouse Lab in Milton Keynes and Alex Alderton, Roberto Amato, Jeffrey Barrett, Sonia Goncalves, Ewan Harrison, David K. Jackson, Ian Johnston, Dominic Kwiatkowski, Cordelia Langford, John Sillitoe on behalf of the Wellcome Sanger Institute COVID-19 Surveillance Team |
| EPI_ISL_2393461 | Lighthouse Lab in Alderley Park | Wellcome Sanger Institute for the COVID-19 Genomics UK (COG-UK) Consortium | Jacquelyn Wynn, Mairead Hyland, The Lighthouse Lab in Alderley Park and Alex Alderton, Roberto Amato, Jeffrey Barrett, Sonia Goncalves, Ewan Harrison, David K. Jackson, Ian Johnston, Dominic Kwiatkowski, Cordelia Langford, John Sillitoe on behalf of the Wellcome Sanger Institute COVID-19 Surveillance Team |
| EPI_ISL_2393462 | Lighthouse Lab in Milton Keynes | Wellcome Sanger Institute for the COVID-19 Genomics UK (COG-UK) Consortium | The Lighthouse Lab in Milton Keynes and Alex Alderton, Roberto Amato, Jeffrey Barrett, Sonia Goncalves, Ewan Harrison, David K. Jackson, Ian Johnston, Dominic Kwiatkowski, Cordelia Langford, John Sillitoe on behalf of the Wellcome Sanger Institute COVID-19 Surveillance Team |
| EPI_ISL_2393463, EPI_ISL_2393464, EPI_ISL_2393465, EPI_ISL_2393466 | Lighthouse Lab in Alderley Park | Wellcome Sanger Institute for the COVID-19 Genomics UK (COG-UK) Consortium | Jacquelyn Wynn, Mairead Hyland, The Lighthouse Lab in Alderley Park and Alex Alderton, Roberto Amato, Jeffrey Barrett, Sonia Goncalves, Ewan Harrison, David K. Jackson, Ian Johnston, Dominic Kwiatkowski, Cordelia Langford, John Sillitoe on behalf of the Wellcome Sanger Institute COVID-19 Surveillance Team |
| EPI_ISL_2393467 | Lighthouse Lab in Milton Keynes | Wellcome Sanger Institute for the COVID-19 Genomics UK (COG-UK) Consortium | The Lighthouse Lab in Milton Keynes and Alex Alderton, Roberto Amato, Jeffrey Barrett, Sonia Goncalves, Ewan Harrison, David K. Jackson, Ian Johnston, Dominic Kwiatkowski, Cordelia Langford, John Sillitoe on behalf of the Wellcome Sanger Institute COVID-19 Surveillance Team |
| EPI_ISL_2393468 | Lighthouse Lab in Alderley Park | Wellcome Sanger Institute for the COVID-19 Genomics UK (COG-UK) Consortium | Jacquelyn Wynn, Mairead Hyland, The Lighthouse Lab in Alderley Park and Alex Alderton, Roberto Amato, Jeffrey Barrett, Sonia Goncalves, Ewan Harrison, David K. Jackson, Ian Johnston, Dominic Kwiatkowski, Cordelia Langford, John Sillitoe on behalf of the Wellcome Sanger Institute COVID-19 Surveillance Team |
| EPI_ISL_2393469 | Lighthouse Lab in Milton Keynes | Wellcome Sanger Institute for the COVID-19 Genomics UK (COG-UK) Consortium | The Lighthouse Lab in Milton Keynes and Alex Alderton, Roberto Amato, Jeffrey Barrett, Sonia Goncalves, Ewan Harrison, David K. Jackson, Ian Johnston, Dominic Kwiatkowski, Cordelia Langford, John Sillitoe on behalf of the Wellcome Sanger Institute COVID-19 Surveillance Team |
| EPI_ISL_2393470 | Lighthouse Laboratory Plymouth | Wellcome Sanger Institute for the COVID-19 Genomics UK (COG-UK) Consortium | Lighthouse Laboratory Plymouth and Alex Alderton, Roberto Amato, Jeffrey Barrett, Sonia Goncalves, Ewan Harrison, David K. Jackson, Ian Johnston, Dominic Kwiatkowski, Cordelia Langford, John Sillitoe on behalf of the Wellcome Sanger Institute COVID-19 Surveillance Team |
| EPI_ISL_2393472 | Lighthouse Lab in Alderley Park | Wellcome Sanger Institute for the COVID-19 Genomics UK (COG-UK) Consortium | Jacquelyn Wynn, Mairead Hyland, The Lighthouse Lab in Alderley Park and Alex Alderton, Roberto Amato, Jeffrey Barrett, Sonia Goncalves, Ewan Harrison, David K. Jackson, Ian Johnston, Dominic Kwiatkowski, Cordelia Langford, John Sillitoe on behalf of the Wellcome Sanger Institute COVID-19 Surveillance Team |
| EPI_ISL_2393473 | Lighthouse Lab in Milton Keynes | Wellcome Sanger Institute for the COVID-19 Genomics UK (COG-UK) Consortium | The Lighthouse Lab in Milton Keynes and Alex Alderton, Roberto Amato, Jeffrey Barrett, Sonia Goncalves, Ewan Harrison, David K. Jackson, Ian Johnston, Dominic Kwiatkowski, Cordelia Langford, John Sillitoe on behalf of the Wellcome Sanger Institute COVID-19 Surveillance Team |
| EPI_ISL_2393474, EPI_ISL_2393476 | Lighthouse Lab in Alderley Park | Wellcome Sanger Institute for the COVID-19 Genomics UK (COG-UK) Consortium | Jacquelyn Wynn, Mairead Hyland, The Lighthouse Lab in Alderley Park and Alex Alderton, Roberto Amato, Jeffrey Barrett, Sonia Goncalves, Ewan Harrison, David K. Jackson, Ian Johnston, Dominic Kwiatkowski, Cordelia Langford, John Sillitoe on behalf of the Wellcome Sanger Institute COVID-19 Surveillance Team |
| EPI_ISL_2393477 | Lighthouse Lab in Milton Keynes | Wellcome Sanger Institute for the COVID-19 Genomics UK (COG-UK) Consortium | The Lighthouse Lab in Milton Keynes and Alex Alderton, Roberto Amato, Jeffrey Barrett, Sonia Goncalves, Ewan Harrison, David K. Jackson, Ian Johnston, Dominic Kwiatkowski, Cordelia Langford, John Sillitoe on behalf of the Wellcome Sanger Institute COVID-19 Surveillance Team |
| EPI_ISL_2393478, EPI_ISL_2393479, EPI_ISL_2393480 | Lighthouse Lab in Alderley Park | Wellcome Sanger Institute for the COVID-19 Genomics UK (COG-UK) Consortium | Jacquelyn Wynn, Mairead Hyland, The Lighthouse Lab in Alderley Park and Alex Alderton, Roberto Amato, Jeffrey Barrett, Sonia Goncalves, Ewan Harrison, David K. Jackson, Ian Johnston, Dominic Kwiatkowski, Cordelia Langford, John Sillitoe on behalf of the Wellcome Sanger Institute COVID-19 Surveillance Team |
| EPI_ISL_2393481 | Lighthouse Lab in Milton Keynes | Wellcome Sanger Institute for the COVID-19 Genomics UK (COG-UK) Consortium | The Lighthouse Lab in Milton Keynes and Alex Alderton, Roberto Amato, Jeffrey Barrett, Sonia Goncalves, Ewan Harrison, David K. Jackson, Ian Johnston, Dominic Kwiatkowski, Cordelia Langford, John Sillitoe on behalf of the Wellcome Sanger Institute COVID-19 Surveillance Team |
| EPI_ISL_2393482 | Lighthouse Lab in Alderley Park | Wellcome Sanger Institute for the COVID-19 Genomics UK (COG-UK) Consortium | Jacquelyn Wynn, Mairead Hyland, The Lighthouse Lab in Alderley Park and Alex Alderton, Roberto Amato, Jeffrey Barrett, Sonia Goncalves, Ewan Harrison, David K. Jackson, Ian Johnston, Dominic Kwiatkowski, Cordelia Langford, John Sillitoe on behalf of the Wellcome Sanger Institute COVID-19 Surveillance Team |
| EPI_ISL_2393483, EPI_ISL_2393484 | Lighthouse Laboratory Plymouth | Wellcome Sanger Institute for the COVID-19 Genomics UK (COG-UK) Consortium | Lighthouse Laboratory Plymouth and Alex Alderton, Roberto Amato, Jeffrey Barrett, Sonia Goncalves, Ewan Harrison, David K. Jackson, Ian Johnston, Dominic Kwiatkowski, Cordelia Langford, John Sillitoe on behalf of the Wellcome Sanger Institute COVID-19 Surveillance Team |
| EPI_ISL_2393485, EPI_ISL_2393487, EPI_ISL_2393488, EPI_ISL_2393489 | Lighthouse Lab in Alderley Park | Wellcome Sanger Institute for the COVID-19 Genomics UK (COG-UK) Consortium | Jacquelyn Wynn, Mairead Hyland, The Lighthouse Lab in Alderley Park and Alex Alderton, Roberto Amato, Jeffrey Barrett, Sonia Goncalves, Ewan Harrison, David K. Jackson, Ian Johnston, Dominic Kwiatkowski, Cordelia Langford, John Sillitoe on behalf of the Wellcome Sanger Institute COVID-19 Surveillance Team |
| EPI_ISL_2393490 | Lighthouse Lab in Milton Keynes | Wellcome Sanger Institute for the COVID-19 Genomics UK (COG-UK) Consortium | The Lighthouse Lab in Milton Keynes and Alex Alderton, Roberto Amato, Jeffrey Barrett, Sonia Goncalves, Ewan Harrison, David K. Jackson, Ian Johnston, Dominic Kwiatkowski, Cordelia Langford, John Sillitoe on behalf of the Wellcome Sanger Institute COVID-19 Surveillance Team |
| EPI_ISL_2393491, EPI_ISL_2393492 | Lighthouse Lab in Glasgow | Wellcome Sanger Institute for the COVID-19 Genomics UK (COG-UK) Consortium | Harper VanSteenhouse, Yumi Kasai, David Gray, Carol Clugston, Anna Dominiczak and Alex Alderton, Roberto Amato, Jeffrey Barrett, Sonia Goncalves, Ewan Harrison, David K. Jackson, Ian Johnston, Dominic Kwiatkowski, Cordelia Langford, John Sillitoe on behalf of the Wellcome Sanger Institute COVID-19 Surveillance Team |
| EPI_ISL_2393493, EPI_ISL_2393494, EPI_ISL_2393495, EPI_ISL_2393496, EPI_ISL_2393497 | Lighthouse Lab in Alderley Park | Wellcome Sanger Institute for the COVID-19 Genomics UK (COG-UK) Consortium | Jacquelyn Wynn, Mairead Hyland, The Lighthouse Lab in Alderley Park and Alex Alderton, Roberto Amato, Jeffrey Barrett, Sonia Goncalves, Ewan Harrison, David K. Jackson, Ian Johnston, Dominic Kwiatkowski, Cordelia Langford, John Sillitoe on behalf of the Wellcome Sanger Institute COVID-19 Surveillance Team |
| EPI_ISL_2393498 | Lighthouse Lab in Milton Keynes | Wellcome Sanger Institute for the COVID-19 Genomics UK (COG-UK) Consortium | The Lighthouse Lab in Milton Keynes and Alex Alderton, Roberto Amato, Jeffrey Barrett, Sonia Goncalves, Ewan Harrison, David K. Jackson, Ian Johnston, Dominic Kwiatkowski, Cordelia Langford, John Sillitoe on behalf of the Wellcome Sanger Institute COVID-19 Surveillance Team |
| EPI_ISL_2393499 | Lighthouse Lab in Alderley Park | Wellcome Sanger Institute for the COVID-19 Genomics UK (COG-UK) Consortium | Jacquelyn Wynn, Mairead Hyland, The Lighthouse Lab in Alderley Park and Alex Alderton, Roberto Amato, Jeffrey Barrett, Sonia Goncalves, Ewan Harrison, David K. Jackson, Ian Johnston, Dominic Kwiatkowski, Cordelia Langford, John Sillitoe on behalf of the Wellcome Sanger Institute COVID-19 Surveillance Team |
| EPI_ISL_2393500 | Lighthouse Lab in Milton Keynes | Wellcome Sanger Institute for the COVID-19 Genomics UK (COG-UK) Consortium | The Lighthouse Lab in Milton Keynes and Alex Alderton, Roberto Amato, Jeffrey Barrett, Sonia Goncalves, Ewan Harrison, David K. Jackson, Ian Johnston, Dominic Kwiatkowski, Cordelia Langford, John Sillitoe on behalf of the Wellcome Sanger Institute COVID-19 Surveillance Team |
| EPI_ISL_2393501, | Lighthouse Lab in Alderley Park | Wellcome Sanger Institute for the COVID-19 Genomics UK (COG-UK) | Jacquelyn Wynn, Mairead Hyland, The Lighthouse Lab in Alderley Park and Alex Alderton, Roberto Amato, Jeffrey Barrett, Sonia Goncalves, Ewan Harrison, David K. Jackson, Ian Johnston, Dominic Kwiatkowski, Cordelia Langford, John Sillitoe on |

| | | | |
|---|---|---|---|
| EPI_ISL_2393502 | | Consortium | behalf of the Wellcome Sanger Institute COVID-19 Surveillance Team |
| EPI_ISL_2393503 | Lighthouse Lab in Milton Keynes | Wellcome Sanger Institute for the COVID-19 Genomics UK (COG-UK) Consortium | The Lighthouse Lab in Milton Keynes and Alex Alderton, Roberto Amato, Jeffrey Barrett, Sonia Goncalves, Ewan Harrison, David K. Jackson, Ian Johnston, Dominic Kwiatkowski, Cordelia Langford, John Sillitoe on behalf of the Wellcome Sanger Institute COVID-19 Surveillance Team |
| EPI_ISL_2393504, EPI_ISL_2393505, EPI_ISL_2393506, EPI_ISL_2393507, EPI_ISL_2393508, EPI_ISL_2393509, EPI_ISL_2393510, EPI_ISL_2393511, EPI_ISL_2393512, EPI_ISL_2393513 | Lighthouse Lab in Alderley Park | Wellcome Sanger Institute for the COVID-19 Genomics UK (COG-UK) Consortium | Jacquelyn Wynn, Mairead Hyland, The Lighthouse Lab in Alderley Park and Alex Alderton, Roberto Amato, Jeffrey Barrett, Sonia Goncalves, Ewan Harrison, David K. Jackson, Ian Johnston, Dominic Kwiatkowski, Cordelia Langford, John Sillitoe on behalf of the Wellcome Sanger Institute COVID-19 Surveillance Team |
| EPI_ISL_2393514 | Lighthouse Lab in Milton Keynes | Wellcome Sanger Institute for the COVID-19 Genomics UK (COG-UK) Consortium | The Lighthouse Lab in Milton Keynes and Alex Alderton, Roberto Amato, Jeffrey Barrett, Sonia Goncalves, Ewan Harrison, David K. Jackson, Ian Johnston, Dominic Kwiatkowski, Cordelia Langford, John Sillitoe on behalf of the Wellcome Sanger Institute COVID-19 Surveillance Team |
| EPI_ISL_2393515, EPI_ISL_2393516, EPI_ISL_2393517, EPI_ISL_2393518 | Lighthouse Lab in Alderley Park | Wellcome Sanger Institute for the COVID-19 Genomics UK (COG-UK) Consortium | Jacquelyn Wynn, Mairead Hyland, The Lighthouse Lab in Alderley Park and Alex Alderton, Roberto Amato, Jeffrey Barrett, Sonia Goncalves, Ewan Harrison, David K. Jackson, Ian Johnston, Dominic Kwiatkowski, Cordelia Langford, John Sillitoe on behalf of the Wellcome Sanger Institute COVID-19 Surveillance Team |
| EPI_ISL_2393519, EPI_ISL_2393520 | Lighthouse Lab in Milton Keynes | Wellcome Sanger Institute for the COVID-19 Genomics UK (COG-UK) Consortium | The Lighthouse Lab in Milton Keynes and Alex Alderton, Roberto Amato, Jeffrey Barrett, Sonia Goncalves, Ewan Harrison, David K. Jackson, Ian Johnston, Dominic Kwiatkowski, Cordelia Langford, John Sillitoe on behalf of the Wellcome Sanger Institute COVID-19 Surveillance Team |
| EPI_ISL_2393521, EPI_ISL_2393522 | Lighthouse Lab in Alderley Park | Wellcome Sanger Institute for the COVID-19 Genomics UK (COG-UK) Consortium | Jacquelyn Wynn, Mairead Hyland, The Lighthouse Lab in Alderley Park and Alex Alderton, Roberto Amato, Jeffrey Barrett, Sonia Goncalves, Ewan Harrison, David K. Jackson, Ian Johnston, Dominic Kwiatkowski, Cordelia Langford, John Sillitoe on behalf of the Wellcome Sanger Institute COVID-19 Surveillance Team |
| EPI_ISL_2393523, EPI_ISL_2393524 | Lighthouse Lab in Milton Keynes | Wellcome Sanger Institute for the COVID-19 Genomics UK (COG-UK) Consortium | The Lighthouse Lab in Milton Keynes and Alex Alderton, Roberto Amato, Jeffrey Barrett, Sonia Goncalves, Ewan Harrison, David K. Jackson, Ian Johnston, Dominic Kwiatkowski, Cordelia Langford, John Sillitoe on behalf of the Wellcome Sanger Institute COVID-19 Surveillance Team |
| EPI_ISL_2393525, EPI_ISL_2393526, EPI_ISL_2393527, EPI_ISL_2393528, EPI_ISL_2393529, EPI_ISL_2393530, EPI_ISL_2393531, EPI_ISL_2393532 | Lighthouse Lab in Alderley Park | Wellcome Sanger Institute for the COVID-19 Genomics UK (COG-UK) Consortium | Jacquelyn Wynn, Mairead Hyland, The Lighthouse Lab in Alderley Park and Alex Alderton, Roberto Amato, Jeffrey Barrett, Sonia Goncalves, Ewan Harrison, David K. Jackson, Ian Johnston, Dominic Kwiatkowski, Cordelia Langford, John Sillitoe on behalf of the Wellcome Sanger Institute COVID-19 Surveillance Team |
| EPI_ISL_2393533 | Lighthouse Lab in Milton Keynes | Wellcome Sanger Institute for the COVID-19 Genomics UK (COG-UK) Consortium | The Lighthouse Lab in Milton Keynes and Alex Alderton, Roberto Amato, Jeffrey Barrett, Sonia Goncalves, Ewan Harrison, David K. Jackson, Ian Johnston, Dominic Kwiatkowski, Cordelia Langford, John Sillitoe on behalf of the Wellcome Sanger Institute COVID-19 Surveillance Team |
| EPI_ISL_2393534 | Lighthouse Lab in Alderley Park | Wellcome Sanger Institute for the COVID-19 Genomics UK (COG-UK) Consortium | Jacquelyn Wynn, Mairead Hyland, The Lighthouse Lab in Alderley Park and Alex Alderton, Roberto Amato, Jeffrey Barrett, Sonia Goncalves, Ewan Harrison, David K. Jackson, Ian Johnston, Dominic Kwiatkowski, Cordelia Langford, John Sillitoe on behalf of the Wellcome Sanger Institute COVID-19 Surveillance Team |
| EPI_ISL_2393536 | Lighthouse Lab in Milton Keynes | Wellcome Sanger Institute for the COVID-19 Genomics UK (COG-UK) Consortium | The Lighthouse Lab in Milton Keynes and Alex Alderton, Roberto Amato, Jeffrey Barrett, Sonia Goncalves, Ewan Harrison, David K. Jackson, Ian Johnston, Dominic Kwiatkowski, Cordelia Langford, John Sillitoe on behalf of the Wellcome Sanger Institute COVID-19 Surveillance Team |
| EPI_ISL_2393538, EPI_ISL_2393539, EPI_ISL_2393540, EPI_ISL_2393541, EPI_ISL_2393542 | Lighthouse Lab in Alderley Park | Wellcome Sanger Institute for the COVID-19 Genomics UK (COG-UK) Consortium | Jacquelyn Wynn, Mairead Hyland, The Lighthouse Lab in Alderley Park and Alex Alderton, Roberto Amato, Jeffrey Barrett, Sonia Goncalves, Ewan Harrison, David K. Jackson, Ian Johnston, Dominic Kwiatkowski, Cordelia Langford, John Sillitoe on behalf of the Wellcome Sanger Institute COVID-19 Surveillance Team |
| EPI_ISL_2393543 | Lighthouse Lab in Milton Keynes | Wellcome Sanger Institute for the COVID-19 Genomics UK (COG-UK) Consortium | The Lighthouse Lab in Milton Keynes and Alex Alderton, Roberto Amato, Jeffrey Barrett, Sonia Goncalves, Ewan Harrison, David K. Jackson, Ian Johnston, Dominic Kwiatkowski, Cordelia Langford, John Sillitoe on behalf of the Wellcome Sanger Institute COVID-19 Surveillance Team |
| EPI_ISL_2393544, EPI_ISL_2393545, EPI_ISL_2393546, EPI_ISL_2393548, EPI_ISL_2393549, EPI_ISL_2393550, EPI_ISL_2393551 | Lighthouse Lab in Alderley Park | Wellcome Sanger Institute for the COVID-19 Genomics UK (COG-UK) Consortium | Jacquelyn Wynn, Mairead Hyland, The Lighthouse Lab in Alderley Park and Alex Alderton, Roberto Amato, Jeffrey Barrett, Sonia Goncalves, Ewan Harrison, David K. Jackson, Ian Johnston, Dominic Kwiatkowski, Cordelia Langford, John Sillitoe on behalf of the Wellcome Sanger Institute COVID-19 Surveillance Team |
| EPI_ISL_2393552 | Lighthouse Lab in Milton Keynes | Wellcome Sanger Institute for the COVID-19 Genomics UK (COG-UK) Consortium | The Lighthouse Lab in Milton Keynes and Alex Alderton, Roberto Amato, Jeffrey Barrett, Sonia Goncalves, Ewan Harrison, David K. Jackson, Ian Johnston, Dominic Kwiatkowski, Cordelia Langford, John Sillitoe on behalf of the Wellcome Sanger Institute COVID-19 Surveillance Team |
| EPI_ISL_2393553 | Lighthouse Lab in Glasgow | Wellcome Sanger Institute for the COVID-19 Genomics UK (COG-UK) Consortium | Harper VanSteenhouse, Yumi Kasai, David Gray, Carol Clugston, Anna Dominiczak and Alex Alderton, Roberto Amato, Jeffrey Barrett, Sonia Goncalves, Ewan Harrison, David K. Jackson, Ian Johnston, Dominic Kwiatkowski, Cordelia Langford, John Sillitoe on behalf of the Wellcome Sanger Institute COVID-19 Surveillance Team |
| EPI_ISL_2393554, EPI_ISL_2393556, EPI_ISL_2393557, EPI_ISL_2393558 | Lighthouse Lab in Alderley Park | Wellcome Sanger Institute for the COVID-19 Genomics UK (COG-UK) Consortium | Jacquelyn Wynn, Mairead Hyland, The Lighthouse Lab in Alderley Park and Alex Alderton, Roberto Amato, Jeffrey Barrett, Sonia Goncalves, Ewan Harrison, David K. Jackson, Ian Johnston, Dominic Kwiatkowski, Cordelia Langford, John Sillitoe on behalf of the Wellcome Sanger Institute COVID-19 Surveillance Team |
| EPI_ISL_2393560 | Lighthouse Lab in Milton Keynes | Wellcome Sanger Institute for the COVID-19 Genomics UK (COG-UK) Consortium | The Lighthouse Lab in Milton Keynes and Alex Alderton, Roberto Amato, Jeffrey Barrett, Sonia Goncalves, Ewan Harrison, David K. Jackson, Ian Johnston, Dominic Kwiatkowski, Cordelia Langford, John Sillitoe on behalf of the Wellcome Sanger Institute COVID-19 Surveillance Team |
| EPI_ISL_2393561 | Lighthouse Lab in Alderley Park | Wellcome Sanger Institute for the COVID-19 Genomics UK (COG-UK) Consortium | Jacquelyn Wynn, Mairead Hyland, The Lighthouse Lab in Alderley Park and Alex Alderton, Roberto Amato, Jeffrey Barrett, Sonia Goncalves, Ewan Harrison, David K. Jackson, Ian Johnston, Dominic Kwiatkowski, Cordelia Langford, John Sillitoe on behalf of the Wellcome Sanger Institute COVID-19 Surveillance Team |
| EPI_ISL_2393562 | Lighthouse Lab in Milton Keynes | Wellcome Sanger Institute for the COVID-19 Genomics UK (COG-UK) Consortium | The Lighthouse Lab in Milton Keynes and Alex Alderton, Roberto Amato, Jeffrey Barrett, Sonia Goncalves, Ewan Harrison, David K. Jackson, Ian Johnston, Dominic Kwiatkowski, Cordelia Langford, John Sillitoe on behalf of the Wellcome Sanger Institute COVID-19 Surveillance Team |
| EPI_ISL_2393563, EPI_ISL_2393564, EPI_ISL_2393567, EPI_ISL_2393568, EPI_ISL_2393570, EPI_ISL_2393571 | Lighthouse Lab in Alderley Park | Wellcome Sanger Institute for the COVID-19 Genomics UK (COG-UK) Consortium | Jacquelyn Wynn, Mairead Hyland, The Lighthouse Lab in Alderley Park and Alex Alderton, Roberto Amato, Jeffrey Barrett, Sonia Goncalves, Ewan Harrison, David K. Jackson, Ian Johnston, Dominic Kwiatkowski, Cordelia Langford, John Sillitoe on behalf of the Wellcome Sanger Institute COVID-19 Surveillance Team |
| EPI_ISL_2393572, EPI_ISL_2393573 | Lighthouse Lab in Milton Keynes | Wellcome Sanger Institute for the COVID-19 Genomics UK (COG-UK) Consortium | The Lighthouse Lab in Milton Keynes and Alex Alderton, Roberto Amato, Jeffrey Barrett, Sonia Goncalves, Ewan Harrison, David K. Jackson, Ian Johnston, Dominic Kwiatkowski, Cordelia Langford, John Sillitoe on behalf of the Wellcome Sanger Institute COVID-19 Surveillance Team |
| EPI_ISL_2393574, EPI_ISL_2393575 | Lighthouse Lab in Alderley Park | Wellcome Sanger Institute for the COVID-19 Genomics UK (COG-UK) Consortium | Jacquelyn Wynn, Mairead Hyland, The Lighthouse Lab in Alderley Park and Alex Alderton, Roberto Amato, Jeffrey Barrett, Sonia Goncalves, Ewan Harrison, David K. Jackson, Ian Johnston, Dominic Kwiatkowski, Cordelia Langford, John Sillitoe on behalf of the Wellcome Sanger Institute COVID-19 Surveillance Team |
| EPI_ISL_2393576 | Lighthouse Lab in Milton Keynes | Wellcome Sanger Institute for the COVID-19 Genomics UK (COG-UK) Consortium | The Lighthouse Lab in Milton Keynes and Alex Alderton, Roberto Amato, Jeffrey Barrett, Sonia Goncalves, Ewan Harrison, David K. Jackson, Ian Johnston, Dominic Kwiatkowski, Cordelia Langford, John Sillitoe on behalf of the Wellcome Sanger Institute COVID-19 Surveillance Team |
| EPI_ISL_2393577, EPI_ISL_2393578, EPI_ISL_2393579, EPI_ISL_2393581, EPI_ISL_2393582, EPI_ISL_2393583 | Lighthouse Lab in Alderley Park | Wellcome Sanger Institute for the COVID-19 Genomics UK (COG-UK) Consortium | Jacquelyn Wynn, Mairead Hyland, The Lighthouse Lab in Alderley Park and Alex Alderton, Roberto Amato, Jeffrey Barrett, Sonia Goncalves, Ewan Harrison, David K. Jackson, Ian Johnston, Dominic Kwiatkowski, Cordelia Langford, John Sillitoe on behalf of the Wellcome Sanger Institute COVID-19 Surveillance Team |
| EPI_ISL_2393584 | Lighthouse Lab in Milton Keynes | Wellcome Sanger Institute for the COVID-19 Genomics UK (COG-UK) Consortium | The Lighthouse Lab in Milton Keynes and Alex Alderton, Roberto Amato, Jeffrey Barrett, Sonia Goncalves, Ewan Harrison, David K. Jackson, Ian Johnston, Dominic Kwiatkowski, Cordelia Langford, John Sillitoe on behalf of the Wellcome Sanger Institute COVID-19 Surveillance Team |
| EPI_ISL_2393585 | Lighthouse Lab in Alderley Park | Wellcome Sanger Institute for the COVID-19 Genomics UK (COG-UK) Consortium | Jacquelyn Wynn, Mairead Hyland, The Lighthouse Lab in Alderley Park and Alex Alderton, Roberto Amato, Jeffrey Barrett, Sonia Goncalves, Ewan Harrison, David K. Jackson, Ian Johnston, Dominic Kwiatkowski, Cordelia Langford, John Sillitoe on behalf of the Wellcome Sanger Institute COVID-19 Surveillance Team |
| EPI_ISL_2393586 | Lighthouse Lab in Glasgow | Wellcome Sanger Institute for the COVID-19 Genomics UK (COG-UK) Consortium | Harper VanSteenhouse, Yumi Kasai, David Gray, Carol Clugston, Anna Dominiczak and Alex Alderton, Roberto Amato, Jeffrey Barrett, Sonia Goncalves, Ewan Harrison, David K. Jackson, Ian Johnston, Dominic Kwiatkowski, Cordelia Langford, John Sillitoe on behalf of the Wellcome Sanger Institute COVID-19 Surveillance Team |
| EPI_ISL_2393587, EPI_ISL_2393588 | Lighthouse Lab in Alderley Park | Wellcome Sanger Institute for the COVID-19 Genomics UK (COG-UK) Consortium | Jacquelyn Wynn, Mairead Hyland, The Lighthouse Lab in Alderley Park and Alex Alderton, Roberto Amato, Jeffrey Barrett, Sonia Goncalves, Ewan Harrison, David K. Jackson, Ian Johnston, Dominic Kwiatkowski, Cordelia Langford, John Sillitoe on behalf of the Wellcome Sanger Institute COVID-19 Surveillance Team |
| EPI_ISL_2393589 | Lighthouse Lab in Glasgow | Wellcome Sanger Institute for the COVID-19 Genomics UK (COG-UK) Consortium | Harper VanSteenhouse, Yumi Kasai, David Gray, Carol Clugston, Anna Dominiczak and Alex Alderton, Roberto Amato, Jeffrey Barrett, Sonia Goncalves, Ewan Harrison, David K. Jackson, Ian Johnston, Dominic Kwiatkowski, Cordelia Langford, John Sillitoe on behalf of the Wellcome Sanger Institute COVID-19 Surveillance Team |
| EPI_ISL_2393590, | Lighthouse Lab in Alderley Park | Wellcome Sanger Institute for the COVID-19 Genomics UK (COG-UK) | Jacquelyn Wynn, Mairead Hyland, The Lighthouse Lab in Alderley Park and Alex Alderton, Roberto Amato, Jeffrey Barrett, Sonia Goncalves, Ewan Harrison, David K. Jackson, Ian Johnston, Dominic Kwiatkowski, Cordelia Langford, John Sillitoe on |

| | | | |
|---|---|---|---|
| EPI_ISL_2393591, EPI_ISL_2393593 | | Consortium | behalf of the Wellcome Sanger Institute COVID-19 Surveillance Team |
| EPI_ISL_2393594 | Lighthouse Lab in Milton Keynes | Wellcome Sanger Institute for the COVID-19 Genomics UK (COG-UK) Consortium | The Lighthouse Lab in Milton Keynes and Alex Alderton, Roberto Amato, Jeffrey Barrett, Sonia Goncalves, Ewan Harrison, David K. Jackson, Ian Johnston, Dominic Kwiatkowski, Cordelia Langford, John Sillitoe on behalf of the Wellcome Sanger Institute COVID-19 Surveillance Team |
| EPI_ISL_2393595 | Lighthouse Lab in Alderley Park | Wellcome Sanger Institute for the COVID-19 Genomics UK (COG-UK) Consortium | Jacquelyn Wynn, Mairead Hyland, The Lighthouse Lab in Alderley Park and Alex Alderton, Roberto Amato, Jeffrey Barrett, Sonia Goncalves, Ewan Harrison, David K. Jackson, Ian Johnston, Dominic Kwiatkowski, Cordelia Langford, John Sillitoe on behalf of the Wellcome Sanger Institute COVID-19 Surveillance Team |
| EPI_ISL_2393597 | Lighthouse Lab in Glasgow | Wellcome Sanger Institute for the COVID-19 Genomics UK (COG-UK) Consortium | Harper VanSteenhouse, Yumi Kasai, David Gray, Carol Clugston, Anna Dominiczak and Alex Alderton, Roberto Amato, Jeffrey Barrett, Sonia Goncalves, Ewan Harrison, David K. Jackson, Ian Johnston, Dominic Kwiatkowski, Cordelia Langford, John Sillitoe on behalf of the Wellcome Sanger Institute COVID-19 Surveillance Team |
| EPI_ISL_2393598, EPI_ISL_2393600 | Lighthouse Lab in Milton Keynes | Wellcome Sanger Institute for the COVID-19 Genomics UK (COG-UK) Consortium | The Lighthouse Lab in Milton Keynes and Alex Alderton, Roberto Amato, Jeffrey Barrett, Sonia Goncalves, Ewan Harrison, David K. Jackson, Ian Johnston, Dominic Kwiatkowski, Cordelia Langford, John Sillitoe on behalf of the Wellcome Sanger Institute COVID-19 Surveillance Team |
| EPI_ISL_2393601, EPI_ISL_2393602, EPI_ISL_2393603, EPI_ISL_2393604, EPI_ISL_2393605, EPI_ISL_2393606, EPI_ISL_2393607, EPI_ISL_2393608, EPI_ISL_2393610 | Lighthouse Lab in Alderley Park | Wellcome Sanger Institute for the COVID-19 Genomics UK (COG-UK) Consortium | Jacquelyn Wynn, Mairead Hyland, The Lighthouse Lab in Alderley Park and Alex Alderton, Roberto Amato, Jeffrey Barrett, Sonia Goncalves, Ewan Harrison, David K. Jackson, Ian Johnston, Dominic Kwiatkowski, Cordelia Langford, John Sillitoe on behalf of the Wellcome Sanger Institute COVID-19 Surveillance Team |
| EPI_ISL_2393611 | Lighthouse Lab in Glasgow | Wellcome Sanger Institute for the COVID-19 Genomics UK (COG-UK) Consortium | Harper VanSteenhouse, Yumi Kasai, David Gray, Carol Clugston, Anna Dominiczak and Alex Alderton, Roberto Amato, Jeffrey Barrett, Sonia Goncalves, Ewan Harrison, David K. Jackson, Ian Johnston, Dominic Kwiatkowski, Cordelia Langford, John Sillitoe on behalf of the Wellcome Sanger Institute COVID-19 Surveillance Team |
| EPI_ISL_2393612 | Lighthouse Lab in Alderley Park | Wellcome Sanger Institute for the COVID-19 Genomics UK (COG-UK) Consortium | Jacquelyn Wynn, Mairead Hyland, The Lighthouse Lab in Alderley Park and Alex Alderton, Roberto Amato, Jeffrey Barrett, Sonia Goncalves, Ewan Harrison, David K. Jackson, Ian Johnston, Dominic Kwiatkowski, Cordelia Langford, John Sillitoe on behalf of the Wellcome Sanger Institute COVID-19 Surveillance Team |
| EPI_ISL_2393613 | Lighthouse Lab in Milton Keynes | Wellcome Sanger Institute for the COVID-19 Genomics UK (COG-UK) Consortium | The Lighthouse Lab in Milton Keynes and Alex Alderton, Roberto Amato, Jeffrey Barrett, Sonia Goncalves, Ewan Harrison, David K. Jackson, Ian Johnston, Dominic Kwiatkowski, Cordelia Langford, John Sillitoe on behalf of the Wellcome Sanger Institute COVID-19 Surveillance Team |
| EPI_ISL_2393614, EPI_ISL_2393615 | Lighthouse Lab in Alderley Park | Wellcome Sanger Institute for the COVID-19 Genomics UK (COG-UK) Consortium | Jacquelyn Wynn, Mairead Hyland, The Lighthouse Lab in Alderley Park and Alex Alderton, Roberto Amato, Jeffrey Barrett, Sonia Goncalves, Ewan Harrison, David K. Jackson, Ian Johnston, Dominic Kwiatkowski, Cordelia Langford, John Sillitoe on behalf of the Wellcome Sanger Institute COVID-19 Surveillance Team |
| EPI_ISL_2393616, EPI_ISL_2393618, EPI_ISL_2393619, EPI_ISL_2393622 | Lighthouse Lab in Milton Keynes | Wellcome Sanger Institute for the COVID-19 Genomics UK (COG-UK) Consortium | The Lighthouse Lab in Milton Keynes and Alex Alderton, Roberto Amato, Jeffrey Barrett, Sonia Goncalves, Ewan Harrison, David K. Jackson, Ian Johnston, Dominic Kwiatkowski, Cordelia Langford, John Sillitoe on behalf of the Wellcome Sanger Institute COVID-19 Surveillance Team |
| EPI_ISL_2393623, EPI_ISL_2393624 | Lighthouse Lab in Alderley Park | Wellcome Sanger Institute for the COVID-19 Genomics UK (COG-UK) Consortium | Jacquelyn Wynn, Mairead Hyland, The Lighthouse Lab in Alderley Park and Alex Alderton, Roberto Amato, Jeffrey Barrett, Sonia Goncalves, Ewan Harrison, David K. Jackson, Ian Johnston, Dominic Kwiatkowski, Cordelia Langford, John Sillitoe on behalf of the Wellcome Sanger Institute COVID-19 Surveillance Team |
| EPI_ISL_2393625, EPI_ISL_2393626 | Lighthouse Lab in Milton Keynes | Wellcome Sanger Institute for the COVID-19 Genomics UK (COG-UK) Consortium | The Lighthouse Lab in Milton Keynes and Alex Alderton, Roberto Amato, Jeffrey Barrett, Sonia Goncalves, Ewan Harrison, David K. Jackson, Ian Johnston, Dominic Kwiatkowski, Cordelia Langford, John Sillitoe on behalf of the Wellcome Sanger Institute COVID-19 Surveillance Team |
| EPI_ISL_2393627, EPI_ISL_2393628, EPI_ISL_2393629, EPI_ISL_2393630, EPI_ISL_2393631, EPI_ISL_2393633, EPI_ISL_2393634, EPI_ISL_2393635, EPI_ISL_2393636, EPI_ISL_2393638, EPI_ISL_2393639, EPI_ISL_2393640, EPI_ISL_2393641, EPI_ISL_2393642 | | | |
| see above | Lighthouse Lab in Alderley Park | Wellcome Sanger Institute for the COVID-19 Genomics UK (COG-UK) Consortium | Jacquelyn Wynn, Mairead Hyland, The Lighthouse Lab in Alderley Park and Alex Alderton, Roberto Amato, Jeffrey Barrett, Sonia Goncalves, Ewan Harrison, David K. Jackson, Ian Johnston, Dominic Kwiatkowski, Cordelia Langford, John Sillitoe on behalf of the Wellcome Sanger Institute COVID-19 Surveillance Team |
| EPI_ISL_2393644 | Lighthouse Lab in Milton Keynes | Wellcome Sanger Institute for the COVID-19 Genomics UK (COG-UK) Consortium | The Lighthouse Lab in Milton Keynes and Alex Alderton, Roberto Amato, Jeffrey Barrett, Sonia Goncalves, Ewan Harrison, David K. Jackson, Ian Johnston, Dominic Kwiatkowski, Cordelia Langford, John Sillitoe on behalf of the Wellcome Sanger Institute COVID-19 Surveillance Team |
| EPI_ISL_2393645, EPI_ISL_2393646 | Lighthouse Lab in Alderley Park | Wellcome Sanger Institute for the COVID-19 Genomics UK (COG-UK) Consortium | Jacquelyn Wynn, Mairead Hyland, The Lighthouse Lab in Alderley Park and Alex Alderton, Roberto Amato, Jeffrey Barrett, Sonia Goncalves, Ewan Harrison, David K. Jackson, Ian Johnston, Dominic Kwiatkowski, Cordelia Langford, John Sillitoe on behalf of the Wellcome Sanger Institute COVID-19 Surveillance Team |
| EPI_ISL_2393647 | Lighthouse Lab in Milton Keynes | Wellcome Sanger Institute for the COVID-19 Genomics UK (COG-UK) Consortium | The Lighthouse Lab in Milton Keynes and Alex Alderton, Roberto Amato, Jeffrey Barrett, Sonia Goncalves, Ewan Harrison, David K. Jackson, Ian Johnston, Dominic Kwiatkowski, Cordelia Langford, John Sillitoe on behalf of the Wellcome Sanger Institute COVID-19 Surveillance Team |
| EPI_ISL_2393649 | Lighthouse Lab in Alderley Park | Wellcome Sanger Institute for the COVID-19 Genomics UK (COG-UK) Consortium | Jacquelyn Wynn, Mairead Hyland, The Lighthouse Lab in Alderley Park and Alex Alderton, Roberto Amato, Jeffrey Barrett, Sonia Goncalves, Ewan Harrison, David K. Jackson, Ian Johnston, Dominic Kwiatkowski, Cordelia Langford, John Sillitoe on behalf of the Wellcome Sanger Institute COVID-19 Surveillance Team |
| EPI_ISL_2393650 | Lighthouse Lab in Milton Keynes | Wellcome Sanger Institute for the COVID-19 Genomics UK (COG-UK) Consortium | The Lighthouse Lab in Milton Keynes and Alex Alderton, Roberto Amato, Jeffrey Barrett, Sonia Goncalves, Ewan Harrison, David K. Jackson, Ian Johnston, Dominic Kwiatkowski, Cordelia Langford, John Sillitoe on behalf of the Wellcome Sanger Institute COVID-19 Surveillance Team |
| EPI_ISL_2393651 | Lighthouse Lab in Alderley Park | Wellcome Sanger Institute for the COVID-19 Genomics UK (COG-UK) Consortium | Jacquelyn Wynn, Mairead Hyland, The Lighthouse Lab in Alderley Park and Alex Alderton, Roberto Amato, Jeffrey Barrett, Sonia Goncalves, Ewan Harrison, David K. Jackson, Ian Johnston, Dominic Kwiatkowski, Cordelia Langford, John Sillitoe on behalf of the Wellcome Sanger Institute COVID-19 Surveillance Team |
| EPI_ISL_2393652, EPI_ISL_2393653, EPI_ISL_2393654 | Lighthouse Lab in Milton Keynes | Wellcome Sanger Institute for the COVID-19 Genomics UK (COG-UK) Consortium | The Lighthouse Lab in Milton Keynes and Alex Alderton, Roberto Amato, Jeffrey Barrett, Sonia Goncalves, Ewan Harrison, David K. Jackson, Ian Johnston, Dominic Kwiatkowski, Cordelia Langford, John Sillitoe on behalf of the Wellcome Sanger Institute COVID-19 Surveillance Team |
| EPI_ISL_2393655 | Lighthouse Lab in Glasgow | Wellcome Sanger Institute for the COVID-19 Genomics UK (COG-UK) Consortium | Harper VanSteenhouse, Yumi Kasai, David Gray, Carol Clugston, Anna Dominiczak and Alex Alderton, Roberto Amato, Jeffrey Barrett, Sonia Goncalves, Ewan Harrison, David K. Jackson, Ian Johnston, Dominic Kwiatkowski, Cordelia Langford, John Sillitoe on behalf of the Wellcome Sanger Institute COVID-19 Surveillance Team |
| EPI_ISL_2393656 | Lighthouse Lab in Milton Keynes | Wellcome Sanger Institute for the COVID-19 Genomics UK (COG-UK) Consortium | The Lighthouse Lab in Milton Keynes and Alex Alderton, Roberto Amato, Jeffrey Barrett, Sonia Goncalves, Ewan Harrison, David K. Jackson, Ian Johnston, Dominic Kwiatkowski, Cordelia Langford, John Sillitoe on behalf of the Wellcome Sanger Institute COVID-19 Surveillance Team |
| EPI_ISL_2393657, EPI_ISL_2393658, EPI_ISL_2393659 | Lighthouse Lab in Alderley Park | Wellcome Sanger Institute for the COVID-19 Genomics UK (COG-UK) Consortium | Jacquelyn Wynn, Mairead Hyland, The Lighthouse Lab in Alderley Park and Alex Alderton, Roberto Amato, Jeffrey Barrett, Sonia Goncalves, Ewan Harrison, David K. Jackson, Ian Johnston, Dominic Kwiatkowski, Cordelia Langford, John Sillitoe on behalf of the Wellcome Sanger Institute COVID-19 Surveillance Team |
| EPI_ISL_2393660 | Lighthouse Lab in Milton Keynes | Wellcome Sanger Institute for the COVID-19 Genomics UK (COG-UK) Consortium | The Lighthouse Lab in Milton Keynes and Alex Alderton, Roberto Amato, Jeffrey Barrett, Sonia Goncalves, Ewan Harrison, David K. Jackson, Ian Johnston, Dominic Kwiatkowski, Cordelia Langford, John Sillitoe on behalf of the Wellcome Sanger Institute COVID-19 Surveillance Team |
| EPI_ISL_2393661, EPI_ISL_2393662, EPI_ISL_2393664 | Lighthouse Lab in Alderley Park | Wellcome Sanger Institute for the COVID-19 Genomics UK (COG-UK) Consortium | Jacquelyn Wynn, Mairead Hyland, The Lighthouse Lab in Alderley Park and Alex Alderton, Roberto Amato, Jeffrey Barrett, Sonia Goncalves, Ewan Harrison, David K. Jackson, Ian Johnston, Dominic Kwiatkowski, Cordelia Langford, John Sillitoe on behalf of the Wellcome Sanger Institute COVID-19 Surveillance Team |
| EPI_ISL_2393665 | Lighthouse Lab in Milton Keynes | Wellcome Sanger Institute for the COVID-19 Genomics UK (COG-UK) Consortium | The Lighthouse Lab in Milton Keynes and Alex Alderton, Roberto Amato, Jeffrey Barrett, Sonia Goncalves, Ewan Harrison, David K. Jackson, Ian Johnston, Dominic Kwiatkowski, Cordelia Langford, John Sillitoe on behalf of the Wellcome Sanger Institute COVID-19 Surveillance Team |
| EPI_ISL_2393667 | Lighthouse Lab in Alderley Park | Wellcome Sanger Institute for the COVID-19 Genomics UK (COG-UK) Consortium | Jacquelyn Wynn, Mairead Hyland, The Lighthouse Lab in Alderley Park and Alex Alderton, Roberto Amato, Jeffrey Barrett, Sonia Goncalves, Ewan Harrison, David K. Jackson, Ian Johnston, Dominic Kwiatkowski, Cordelia Langford, John Sillitoe on behalf of the Wellcome Sanger Institute COVID-19 Surveillance Team |
| EPI_ISL_2393668 | Lighthouse Lab in Milton Keynes | Wellcome Sanger Institute for the COVID-19 Genomics UK (COG-UK) Consortium | The Lighthouse Lab in Milton Keynes and Alex Alderton, Roberto Amato, Jeffrey Barrett, Sonia Goncalves, Ewan Harrison, David K. Jackson, Ian Johnston, Dominic Kwiatkowski, Cordelia Langford, John Sillitoe on behalf of the Wellcome Sanger Institute COVID-19 Surveillance Team |
| EPI_ISL_2393669, EPI_ISL_2393670, EPI_ISL_2393671, EPI_ISL_2393672, EPI_ISL_2393673 | Lighthouse Lab in Alderley Park | Wellcome Sanger Institute for the COVID-19 Genomics UK (COG-UK) Consortium | Jacquelyn Wynn, Mairead Hyland, The Lighthouse Lab in Alderley Park and Alex Alderton, Roberto Amato, Jeffrey Barrett, Sonia Goncalves, Ewan Harrison, David K. Jackson, Ian Johnston, Dominic Kwiatkowski, Cordelia Langford, John Sillitoe on behalf of the Wellcome Sanger Institute COVID-19 Surveillance Team |
| EPI_ISL_2393674 | Lighthouse Lab in Milton Keynes | Wellcome Sanger Institute for the COVID-19 Genomics UK (COG-UK) Consortium | The Lighthouse Lab in Milton Keynes and Alex Alderton, Roberto Amato, Jeffrey Barrett, Sonia Goncalves, Ewan Harrison, David K. Jackson, Ian Johnston, Dominic Kwiatkowski, Cordelia Langford, John Sillitoe on behalf of the Wellcome Sanger Institute COVID-19 Surveillance Team |
| EPI_ISL_2393676 | Lighthouse Lab in Alderley Park | Wellcome Sanger Institute for the COVID-19 Genomics UK (COG-UK) Consortium | Jacquelyn Wynn, Mairead Hyland, The Lighthouse Lab in Alderley Park and Alex Alderton, Roberto Amato, Jeffrey Barrett, Sonia Goncalves, Ewan Harrison, David K. Jackson, Ian Johnston, Dominic Kwiatkowski, Cordelia Langford, John Sillitoe on behalf of the Wellcome Sanger Institute COVID-19 Surveillance Team |
| EPI_ISL_2393677 | Lighthouse Lab in Milton Keynes | Wellcome Sanger Institute for the COVID-19 Genomics UK (COG-UK) Consortium | The Lighthouse Lab in Milton Keynes and Alex Alderton, Roberto Amato, Jeffrey Barrett, Sonia Goncalves, Ewan Harrison, David K. Jackson, Ian Johnston, Dominic Kwiatkowski, Cordelia Langford, John Sillitoe on behalf of the Wellcome Sanger Institute COVID-19 Surveillance Team |
| EPI_ISL_2393678 | Lighthouse Lab in Glasgow | Wellcome Sanger Institute for the COVID-19 Genomics UK (COG-UK) Consortium | Harper VanSteenhouse, Yumi Kasai, David Gray, Carol Clugston, Anna Dominiczak and Alex Alderton, Roberto Amato, Jeffrey Barrett, Sonia Goncalves, Ewan Harrison, David K. Jackson, Ian Johnston, Dominic Kwiatkowski, Cordelia Langford, John Sillitoe on behalf of the Wellcome Sanger Institute COVID-19 Surveillance Team |
| EPI_ISL_2393679, EPI_ISL_2393680 | Lighthouse Lab in Milton Keynes | Wellcome Sanger Institute for the COVID-19 Genomics UK (COG-UK) Consortium | The Lighthouse Lab in Milton Keynes and Alex Alderton, Roberto Amato, Jeffrey Barrett, Sonia Goncalves, Ewan Harrison, David K. Jackson, Ian Johnston, Dominic Kwiatkowski, Cordelia Langford, John Sillitoe on behalf of the Wellcome Sanger Institute COVID-19 Surveillance Team |
| EPI_ISL_2393681, EPI_ISL_2393682, EPI_ISL_2393684, EPI_ISL_2393685 | Lighthouse Lab in Alderley Park | Wellcome Sanger Institute for the COVID-19 Genomics UK (COG-UK) Consortium | Jacquelyn Wynn, Mairead Hyland, The Lighthouse Lab in Alderley Park and Alex Alderton, Roberto Amato, Jeffrey Barrett, Sonia Goncalves, Ewan Harrison, David K. Jackson, Ian Johnston, Dominic Kwiatkowski, Cordelia Langford, John Sillitoe on behalf of the Wellcome Sanger Institute COVID-19 Surveillance Team |
| EPI_ISL_2393686 | Lighthouse Lab in Milton Keynes | Wellcome Sanger Institute for the COVID-19 Genomics UK (COG-UK) Consortium | The Lighthouse Lab in Milton Keynes and Alex Alderton, Roberto Amato, Jeffrey Barrett, Sonia Goncalves, Ewan Harrison, David K. Jackson, Ian Johnston, Dominic Kwiatkowski, Cordelia Langford, John Sillitoe on behalf of the Wellcome Sanger Institute COVID-19 Surveillance Team |
| EPI_ISL_2393687, | Lighthouse Lab in Alderley Park | Wellcome Sanger Institute for the COVID-19 Genomics UK (COG-UK) | Jacquelyn Wynn, Mairead Hyland, The Lighthouse Lab in Alderley Park and Alex Alderton, Roberto Amato, Jeffrey Barrett, Sonia Goncalves, Ewan Harrison, David K. Jackson, Ian Johnston, Dominic Kwiatkowski, Cordelia Langford, John Sillitoe on |

| | | | |
|---|---|---|---|
| EPI_ISL_2393688, EPI_ISL_2393689, EPI_ISL_2393690, EPI_ISL_2393691, EPI_ISL_2393692 | | Consortium | behalf of the Wellcome Sanger Institute COVID-19 Surveillance Team |
| EPI_ISL_2393693 | Lighthouse Lab in Glasgow | Wellcome Sanger Institute for the COVID-19 Genomics UK (COG-UK) Consortium | Harper VanSteenhouse, Yumi Kasai, David Gray, Carol Clugston, Anna Dominiczak and Alex Alderton, Roberto Amato, Jeffrey Barrett, Sonia Goncalves, Ewan Harrison, David K. Jackson, Ian Johnston, Dominic Kwiatkowski, Cordelia Langford, John Sillitoe on behalf of the Wellcome Sanger Institute COVID-19 Surveillance Team |
| EPI_ISL_2393694 | Lighthouse Lab in Alderley Park | Wellcome Sanger Institute for the COVID-19 Genomics UK (COG-UK) Consortium | Jacquelyn Wynn, Mairead Hyland, The Lighthouse Lab in Alderley Park and Alex Alderton, Roberto Amato, Jeffrey Barrett, Sonia Goncalves, Ewan Harrison, David K. Jackson, Ian Johnston, Dominic Kwiatkowski, Cordelia Langford, John Sillitoe on behalf of the Wellcome Sanger Institute COVID-19 Surveillance Team |
| EPI_ISL_2393695 | Lighthouse Lab in Milton Keynes | Wellcome Sanger Institute for the COVID-19 Genomics UK (COG-UK) Consortium | The Lighthouse Lab in Milton Keynes and Alex Alderton, Roberto Amato, Jeffrey Barrett, Sonia Goncalves, Ewan Harrison, David K. Jackson, Ian Johnston, Dominic Kwiatkowski, Cordelia Langford, John Sillitoe on behalf of the Wellcome Sanger Institute COVID-19 Surveillance Team |
| EPI_ISL_2393696, EPI_ISL_2393697, EPI_ISL_2393699 | Lighthouse Lab in Alderley Park | Wellcome Sanger Institute for the COVID-19 Genomics UK (COG-UK) Consortium | Jacquelyn Wynn, Mairead Hyland, The Lighthouse Lab in Alderley Park and Alex Alderton, Roberto Amato, Jeffrey Barrett, Sonia Goncalves, Ewan Harrison, David K. Jackson, Ian Johnston, Dominic Kwiatkowski, Cordelia Langford, John Sillitoe on behalf of the Wellcome Sanger Institute COVID-19 Surveillance Team |
| EPI_ISL_2393700 | Lighthouse Lab in Milton Keynes | Wellcome Sanger Institute for the COVID-19 Genomics UK (COG-UK) Consortium | The Lighthouse Lab in Milton Keynes and Alex Alderton, Roberto Amato, Jeffrey Barrett, Sonia Goncalves, Ewan Harrison, David K. Jackson, Ian Johnston, Dominic Kwiatkowski, Cordelia Langford, John Sillitoe on behalf of the Wellcome Sanger Institute COVID-19 Surveillance Team |
| EPI_ISL_2393702, EPI_ISL_2393704, EPI_ISL_2393705, EPI_ISL_2393707 | Lighthouse Lab in Alderley Park | Wellcome Sanger Institute for the COVID-19 Genomics UK (COG-UK) Consortium | Jacquelyn Wynn, Mairead Hyland, The Lighthouse Lab in Alderley Park and Alex Alderton, Roberto Amato, Jeffrey Barrett, Sonia Goncalves, Ewan Harrison, David K. Jackson, Ian Johnston, Dominic Kwiatkowski, Cordelia Langford, John Sillitoe on behalf of the Wellcome Sanger Institute COVID-19 Surveillance Team |
| EPI_ISL_2393708 | Lighthouse Lab in Milton Keynes | Wellcome Sanger Institute for the COVID-19 Genomics UK (COG-UK) Consortium | The Lighthouse Lab in Milton Keynes and Alex Alderton, Roberto Amato, Jeffrey Barrett, Sonia Goncalves, Ewan Harrison, David K. Jackson, Ian Johnston, Dominic Kwiatkowski, Cordelia Langford, John Sillitoe on behalf of the Wellcome Sanger Institute COVID-19 Surveillance Team |
| EPI_ISL_2393709 | Lighthouse Lab in Alderley Park | Wellcome Sanger Institute for the COVID-19 Genomics UK (COG-UK) Consortium | Jacquelyn Wynn, Mairead Hyland, The Lighthouse Lab in Alderley Park and Alex Alderton, Roberto Amato, Jeffrey Barrett, Sonia Goncalves, Ewan Harrison, David K. Jackson, Ian Johnston, Dominic Kwiatkowski, Cordelia Langford, John Sillitoe on behalf of the Wellcome Sanger Institute COVID-19 Surveillance Team |
| EPI_ISL_2393711, EPI_ISL_2393712 | Lighthouse Lab in Milton Keynes | Wellcome Sanger Institute for the COVID-19 Genomics UK (COG-UK) Consortium | The Lighthouse Lab in Milton Keynes and Alex Alderton, Roberto Amato, Jeffrey Barrett, Sonia Goncalves, Ewan Harrison, David K. Jackson, Ian Johnston, Dominic Kwiatkowski, Cordelia Langford, John Sillitoe on behalf of the Wellcome Sanger Institute COVID-19 Surveillance Team |
| EPI_ISL_2393713 | Lighthouse Lab in Alderley Park | Wellcome Sanger Institute for the COVID-19 Genomics UK (COG-UK) Consortium | Jacquelyn Wynn, Mairead Hyland, The Lighthouse Lab in Alderley Park and Alex Alderton, Roberto Amato, Jeffrey Barrett, Sonia Goncalves, Ewan Harrison, David K. Jackson, Ian Johnston, Dominic Kwiatkowski, Cordelia Langford, John Sillitoe on behalf of the Wellcome Sanger Institute COVID-19 Surveillance Team |
| EPI_ISL_2393714 | Lighthouse Lab in Milton Keynes | Wellcome Sanger Institute for the COVID-19 Genomics UK (COG-UK) Consortium | The Lighthouse Lab in Milton Keynes and Alex Alderton, Roberto Amato, Jeffrey Barrett, Sonia Goncalves, Ewan Harrison, David K. Jackson, Ian Johnston, Dominic Kwiatkowski, Cordelia Langford, John Sillitoe on behalf of the Wellcome Sanger Institute COVID-19 Surveillance Team |
| EPI_ISL_2393715, EPI_ISL_2393717 | Lighthouse Lab in Alderley Park | Wellcome Sanger Institute for the COVID-19 Genomics UK (COG-UK) Consortium | Jacquelyn Wynn, Mairead Hyland, The Lighthouse Lab in Alderley Park and Alex Alderton, Roberto Amato, Jeffrey Barrett, Sonia Goncalves, Ewan Harrison, David K. Jackson, Ian Johnston, Dominic Kwiatkowski, Cordelia Langford, John Sillitoe on behalf of the Wellcome Sanger Institute COVID-19 Surveillance Team |
| EPI_ISL_2393718 | Lighthouse Lab in Milton Keynes | Wellcome Sanger Institute for the COVID-19 Genomics UK (COG-UK) Consortium | The Lighthouse Lab in Milton Keynes and Alex Alderton, Roberto Amato, Jeffrey Barrett, Sonia Goncalves, Ewan Harrison, David K. Jackson, Ian Johnston, Dominic Kwiatkowski, Cordelia Langford, John Sillitoe on behalf of the Wellcome Sanger Institute COVID-19 Surveillance Team |
| EPI_ISL_2393721, EPI_ISL_2393722 | Lighthouse Lab in Alderley Park | Wellcome Sanger Institute for the COVID-19 Genomics UK (COG-UK) Consortium | Jacquelyn Wynn, Mairead Hyland, The Lighthouse Lab in Alderley Park and Alex Alderton, Roberto Amato, Jeffrey Barrett, Sonia Goncalves, Ewan Harrison, David K. Jackson, Ian Johnston, Dominic Kwiatkowski, Cordelia Langford, John Sillitoe on behalf of the Wellcome Sanger Institute COVID-19 Surveillance Team |
| EPI_ISL_2393723, EPI_ISL_2393724 | Lighthouse Lab in Milton Keynes | Wellcome Sanger Institute for the COVID-19 Genomics UK (COG-UK) Consortium | The Lighthouse Lab in Milton Keynes and Alex Alderton, Roberto Amato, Jeffrey Barrett, Sonia Goncalves, Ewan Harrison, David K. Jackson, Ian Johnston, Dominic Kwiatkowski, Cordelia Langford, John Sillitoe on behalf of the Wellcome Sanger Institute COVID-19 Surveillance Team |
| EPI_ISL_2393725 | Lighthouse Lab in Alderley Park | Wellcome Sanger Institute for the COVID-19 Genomics UK (COG-UK) Consortium | Jacquelyn Wynn, Mairead Hyland, The Lighthouse Lab in Alderley Park and Alex Alderton, Roberto Amato, Jeffrey Barrett, Sonia Goncalves, Ewan Harrison, David K. Jackson, Ian Johnston, Dominic Kwiatkowski, Cordelia Langford, John Sillitoe on behalf of the Wellcome Sanger Institute COVID-19 Surveillance Team |
| EPI_ISL_2393729 | Lighthouse Lab in Glasgow | Wellcome Sanger Institute for the COVID-19 Genomics UK (COG-UK) Consortium | Harper VanSteenhouse, Yumi Kasai, David Gray, Carol Clugston, Anna Dominiczak and Alex Alderton, Roberto Amato, Jeffrey Barrett, Sonia Goncalves, Ewan Harrison, David K. Jackson, Ian Johnston, Dominic Kwiatkowski, Cordelia Langford, John Sillitoe on behalf of the Wellcome Sanger Institute COVID-19 Surveillance Team |
| EPI_ISL_2393730, EPI_ISL_2393731 | Lighthouse Lab in Alderley Park | Wellcome Sanger Institute for the COVID-19 Genomics UK (COG-UK) Consortium | Jacquelyn Wynn, Mairead Hyland, The Lighthouse Lab in Alderley Park and Alex Alderton, Roberto Amato, Jeffrey Barrett, Sonia Goncalves, Ewan Harrison, David K. Jackson, Ian Johnston, Dominic Kwiatkowski, Cordelia Langford, John Sillitoe on behalf of the Wellcome Sanger Institute COVID-19 Surveillance Team |
| EPI_ISL_2393733 | Lighthouse Lab in Milton Keynes | Wellcome Sanger Institute for the COVID-19 Genomics UK (COG-UK) Consortium | The Lighthouse Lab in Milton Keynes and Alex Alderton, Roberto Amato, Jeffrey Barrett, Sonia Goncalves, Ewan Harrison, David K. Jackson, Ian Johnston, Dominic Kwiatkowski, Cordelia Langford, John Sillitoe on behalf of the Wellcome Sanger Institute COVID-19 Surveillance Team |
| EPI_ISL_2393734, EPI_ISL_2393735, EPI_ISL_2393736, EPI_ISL_2393737, EPI_ISL_2393738, EPI_ISL_2393739 | Lighthouse Lab in Alderley Park | Wellcome Sanger Institute for the COVID-19 Genomics UK (COG-UK) Consortium | Jacquelyn Wynn, Mairead Hyland, The Lighthouse Lab in Alderley Park and Alex Alderton, Roberto Amato, Jeffrey Barrett, Sonia Goncalves, Ewan Harrison, David K. Jackson, Ian Johnston, Dominic Kwiatkowski, Cordelia Langford, John Sillitoe on behalf of the Wellcome Sanger Institute COVID-19 Surveillance Team |
| EPI_ISL_2393740 | Lighthouse Lab in Milton Keynes | Wellcome Sanger Institute for the COVID-19 Genomics UK (COG-UK) Consortium | The Lighthouse Lab in Milton Keynes and Alex Alderton, Roberto Amato, Jeffrey Barrett, Sonia Goncalves, Ewan Harrison, David K. Jackson, Ian Johnston, Dominic Kwiatkowski, Cordelia Langford, John Sillitoe on behalf of the Wellcome Sanger Institute COVID-19 Surveillance Team |
| EPI_ISL_2393741 | Lighthouse Lab in Alderley Park | Wellcome Sanger Institute for the COVID-19 Genomics UK (COG-UK) Consortium | Jacquelyn Wynn, Mairead Hyland, The Lighthouse Lab in Alderley Park and Alex Alderton, Roberto Amato, Jeffrey Barrett, Sonia Goncalves, Ewan Harrison, David K. Jackson, Ian Johnston, Dominic Kwiatkowski, Cordelia Langford, John Sillitoe on behalf of the Wellcome Sanger Institute COVID-19 Surveillance Team |
| EPI_ISL_2393742 | Lighthouse Lab in Milton Keynes | Wellcome Sanger Institute for the COVID-19 Genomics UK (COG-UK) Consortium | The Lighthouse Lab in Milton Keynes and Alex Alderton, Roberto Amato, Jeffrey Barrett, Sonia Goncalves, Ewan Harrison, David K. Jackson, Ian Johnston, Dominic Kwiatkowski, Cordelia Langford, John Sillitoe on behalf of the Wellcome Sanger Institute COVID-19 Surveillance Team |
| EPI_ISL_2393743, EPI_ISL_2393745 | Lighthouse Lab in Alderley Park | Wellcome Sanger Institute for the COVID-19 Genomics UK (COG-UK) Consortium | Jacquelyn Wynn, Mairead Hyland, The Lighthouse Lab in Alderley Park and Alex Alderton, Roberto Amato, Jeffrey Barrett, Sonia Goncalves, Ewan Harrison, David K. Jackson, Ian Johnston, Dominic Kwiatkowski, Cordelia Langford, John Sillitoe on behalf of the Wellcome Sanger Institute COVID-19 Surveillance Team |
| EPI_ISL_2393747, EPI_ISL_2393748, EPI_ISL_2393749 | Lighthouse Lab in Milton Keynes | Wellcome Sanger Institute for the COVID-19 Genomics UK (COG-UK) Consortium | The Lighthouse Lab in Milton Keynes and Alex Alderton, Roberto Amato, Jeffrey Barrett, Sonia Goncalves, Ewan Harrison, David K. Jackson, Ian Johnston, Dominic Kwiatkowski, Cordelia Langford, John Sillitoe on behalf of the Wellcome Sanger Institute COVID-19 Surveillance Team |
| EPI_ISL_2393750, EPI_ISL_2393751, EPI_ISL_2393752, EPI_ISL_2393753 | Lighthouse Lab in Alderley Park | Wellcome Sanger Institute for the COVID-19 Genomics UK (COG-UK) Consortium | Jacquelyn Wynn, Mairead Hyland, The Lighthouse Lab in Alderley Park and Alex Alderton, Roberto Amato, Jeffrey Barrett, Sonia Goncalves, Ewan Harrison, David K. Jackson, Ian Johnston, Dominic Kwiatkowski, Cordelia Langford, John Sillitoe on behalf of the Wellcome Sanger Institute COVID-19 Surveillance Team |
| EPI_ISL_2393754 | Lighthouse Lab in Milton Keynes | Wellcome Sanger Institute for the COVID-19 Genomics UK (COG-UK) Consortium | The Lighthouse Lab in Milton Keynes and Alex Alderton, Roberto Amato, Jeffrey Barrett, Sonia Goncalves, Ewan Harrison, David K. Jackson, Ian Johnston, Dominic Kwiatkowski, Cordelia Langford, John Sillitoe on behalf of the Wellcome Sanger Institute COVID-19 Surveillance Team |
| EPI_ISL_2393755, EPI_ISL_2393756, EPI_ISL_2393757, EPI_ISL_2393758, EPI_ISL_2393759, EPI_ISL_2393761, EPI_ISL_2393762, EPI_ISL_2393764 | Lighthouse Lab in Alderley Park | Wellcome Sanger Institute for the COVID-19 Genomics UK (COG-UK) Consortium | Jacquelyn Wynn, Mairead Hyland, The Lighthouse Lab in Alderley Park and Alex Alderton, Roberto Amato, Jeffrey Barrett, Sonia Goncalves, Ewan Harrison, David K. Jackson, Ian Johnston, Dominic Kwiatkowski, Cordelia Langford, John Sillitoe on behalf of the Wellcome Sanger Institute COVID-19 Surveillance Team |
| EPI_ISL_2393766, EPI_ISL_2393767 | Lighthouse Lab in Milton Keynes | Wellcome Sanger Institute for the COVID-19 Genomics UK (COG-UK) Consortium | The Lighthouse Lab in Milton Keynes and Alex Alderton, Roberto Amato, Jeffrey Barrett, Sonia Goncalves, Ewan Harrison, David K. Jackson, Ian Johnston, Dominic Kwiatkowski, Cordelia Langford, John Sillitoe on behalf of the Wellcome Sanger Institute COVID-19 Surveillance Team |
| EPI_ISL_2393768, EPI_ISL_2393769, EPI_ISL_2393770 | Lighthouse Lab in Alderley Park | Wellcome Sanger Institute for the COVID-19 Genomics UK (COG-UK) Consortium | Jacquelyn Wynn, Mairead Hyland, The Lighthouse Lab in Alderley Park and Alex Alderton, Roberto Amato, Jeffrey Barrett, Sonia Goncalves, Ewan Harrison, David K. Jackson, Ian Johnston, Dominic Kwiatkowski, Cordelia Langford, John Sillitoe on behalf of the Wellcome Sanger Institute COVID-19 Surveillance Team |
| EPI_ISL_2393771, EPI_ISL_2393772 | Lighthouse Lab in Milton Keynes | Wellcome Sanger Institute for the COVID-19 Genomics UK (COG-UK) Consortium | The Lighthouse Lab in Milton Keynes and Alex Alderton, Roberto Amato, Jeffrey Barrett, Sonia Goncalves, Ewan Harrison, David K. Jackson, Ian Johnston, Dominic Kwiatkowski, Cordelia Langford, John Sillitoe on behalf of the Wellcome Sanger Institute COVID-19 Surveillance Team |
| EPI_ISL_2393775 | Lighthouse Lab in Alderley Park | Wellcome Sanger Institute for the COVID-19 Genomics UK (COG-UK) Consortium | Jacquelyn Wynn, Mairead Hyland, The Lighthouse Lab in Alderley Park and Alex Alderton, Roberto Amato, Jeffrey Barrett, Sonia Goncalves, Ewan Harrison, David K. Jackson, Ian Johnston, Dominic Kwiatkowski, Cordelia Langford, John Sillitoe on behalf of the Wellcome Sanger Institute COVID-19 Surveillance Team |
| EPI_ISL_2393776, EPI_ISL_2393778, EPI_ISL_2393779 | Lighthouse Lab in Milton Keynes | Wellcome Sanger Institute for the COVID-19 Genomics UK (COG-UK) Consortium | The Lighthouse Lab in Milton Keynes and Alex Alderton, Roberto Amato, Jeffrey Barrett, Sonia Goncalves, Ewan Harrison, David K. Jackson, Ian Johnston, Dominic Kwiatkowski, Cordelia Langford, John Sillitoe on behalf of the Wellcome Sanger Institute COVID-19 Surveillance Team |
| EPI_ISL_2393780, EPI_ISL_2393781, EPI_ISL_2393783 | Lighthouse Lab in Alderley Park | Wellcome Sanger Institute for the COVID-19 Genomics UK (COG-UK) Consortium | Jacquelyn Wynn, Mairead Hyland, The Lighthouse Lab in Alderley Park and Alex Alderton, Roberto Amato, Jeffrey Barrett, Sonia Goncalves, Ewan Harrison, David K. Jackson, Ian Johnston, Dominic Kwiatkowski, Cordelia Langford, John Sillitoe on behalf of the Wellcome Sanger Institute COVID-19 Surveillance Team |
| EPI_ISL_2393784, EPI_ISL_2393785 | Lighthouse Lab in Milton Keynes | Wellcome Sanger Institute for the COVID-19 Genomics UK (COG-UK) Consortium | The Lighthouse Lab in Milton Keynes and Alex Alderton, Roberto Amato, Jeffrey Barrett, Sonia Goncalves, Ewan Harrison, David K. Jackson, Ian Johnston, Dominic Kwiatkowski, Cordelia Langford, John Sillitoe on behalf of the Wellcome Sanger Institute COVID-19 Surveillance Team |

| | | | |
|---|---|---|---|
| EPI_ISL_2393789 | Lighthouse Lab in Alderley Park | Wellcome Sanger Institute for the COVID-19 Genomics UK (COG-UK) Consortium | Jacquelyn Wynn, Mairead Hyland, The Lighthouse Lab in Alderley Park and Alex Alderton, Roberto Amato, Jeffrey Barrett, Sonia Goncalves, Ewan Harrison, David K. Jackson, Ian Johnston, Dominic Kwiatkowski, Cordelia Langford, John Sillitoe on behalf of the Wellcome Sanger Institute COVID-19 Surveillance Team |
| EPI_ISL_2393790 | Lighthouse Lab in Milton Keynes | Wellcome Sanger Institute for the COVID-19 Genomics UK (COG-UK) Consortium | The Lighthouse Lab in Milton Keynes and Alex Alderton, Roberto Amato, Jeffrey Barrett, Sonia Goncalves, Ewan Harrison, David K. Jackson, Ian Johnston, Dominic Kwiatkowski, Cordelia Langford, John Sillitoe on behalf of the Wellcome Sanger Institute COVID-19 Surveillance Team |
| EPI_ISL_2393791, EPI_ISL_2393792, EPI_ISL_2393794, EPI_ISL_2393795 | Lighthouse Lab in Alderley Park | Wellcome Sanger Institute for the COVID-19 Genomics UK (COG-UK) Consortium | Jacquelyn Wynn, Mairead Hyland, The Lighthouse Lab in Alderley Park and Alex Alderton, Roberto Amato, Jeffrey Barrett, Sonia Goncalves, Ewan Harrison, David K. Jackson, Ian Johnston, Dominic Kwiatkowski, Cordelia Langford, John Sillitoe on behalf of the Wellcome Sanger Institute COVID-19 Surveillance Team |
| EPI_ISL_2393796, EPI_ISL_2393799 | Lighthouse Lab in Milton Keynes | Wellcome Sanger Institute for the COVID-19 Genomics UK (COG-UK) Consortium | The Lighthouse Lab in Milton Keynes and Alex Alderton, Roberto Amato, Jeffrey Barrett, Sonia Goncalves, Ewan Harrison, David K. Jackson, Ian Johnston, Dominic Kwiatkowski, Cordelia Langford, John Sillitoe on behalf of the Wellcome Sanger Institute COVID-19 Surveillance Team |
| EPI_ISL_2393800, EPI_ISL_2393801, EPI_ISL_2393802, EPI_ISL_2393803 | Lighthouse Lab in Alderley Park | Wellcome Sanger Institute for the COVID-19 Genomics UK (COG-UK) Consortium | Jacquelyn Wynn, Mairead Hyland, The Lighthouse Lab in Alderley Park and Alex Alderton, Roberto Amato, Jeffrey Barrett, Sonia Goncalves, Ewan Harrison, David K. Jackson, Ian Johnston, Dominic Kwiatkowski, Cordelia Langford, John Sillitoe on behalf of the Wellcome Sanger Institute COVID-19 Surveillance Team |
| EPI_ISL_2393804 | Lighthouse Lab in Milton Keynes | Wellcome Sanger Institute for the COVID-19 Genomics UK (COG-UK) Consortium | The Lighthouse Lab in Milton Keynes and Alex Alderton, Roberto Amato, Jeffrey Barrett, Sonia Goncalves, Ewan Harrison, David K. Jackson, Ian Johnston, Dominic Kwiatkowski, Cordelia Langford, John Sillitoe on behalf of the Wellcome Sanger Institute COVID-19 Surveillance Team |
| EPI_ISL_2393805, EPI_ISL_2393806, EPI_ISL_2393807, EPI_ISL_2393808, EPI_ISL_2393809 | Lighthouse Lab in Alderley Park | Wellcome Sanger Institute for the COVID-19 Genomics UK (COG-UK) Consortium | Jacquelyn Wynn, Mairead Hyland, The Lighthouse Lab in Alderley Park and Alex Alderton, Roberto Amato, Jeffrey Barrett, Sonia Goncalves, Ewan Harrison, David K. Jackson, Ian Johnston, Dominic Kwiatkowski, Cordelia Langford, John Sillitoe on behalf of the Wellcome Sanger Institute COVID-19 Surveillance Team |
| EPI_ISL_2393810 | Lighthouse Lab in Milton Keynes | Wellcome Sanger Institute for the COVID-19 Genomics UK (COG-UK) Consortium | The Lighthouse Lab in Milton Keynes and Alex Alderton, Roberto Amato, Jeffrey Barrett, Sonia Goncalves, Ewan Harrison, David K. Jackson, Ian Johnston, Dominic Kwiatkowski, Cordelia Langford, John Sillitoe on behalf of the Wellcome Sanger Institute COVID-19 Surveillance Team |
| EPI_ISL_2393812, EPI_ISL_2393813 | Lighthouse Lab in Alderley Park | Wellcome Sanger Institute for the COVID-19 Genomics UK (COG-UK) Consortium | Jacquelyn Wynn, Mairead Hyland, The Lighthouse Lab in Alderley Park and Alex Alderton, Roberto Amato, Jeffrey Barrett, Sonia Goncalves, Ewan Harrison, David K. Jackson, Ian Johnston, Dominic Kwiatkowski, Cordelia Langford, John Sillitoe on behalf of the Wellcome Sanger Institute COVID-19 Surveillance Team |
| EPI_ISL_2393814 | Lighthouse Lab in Milton Keynes | Wellcome Sanger Institute for the COVID-19 Genomics UK (COG-UK) Consortium | The Lighthouse Lab in Milton Keynes and Alex Alderton, Roberto Amato, Jeffrey Barrett, Sonia Goncalves, Ewan Harrison, David K. Jackson, Ian Johnston, Dominic Kwiatkowski, Cordelia Langford, John Sillitoe on behalf of the Wellcome Sanger Institute COVID-19 Surveillance Team |
| EPI_ISL_2393815 | Lighthouse Lab in Alderley Park | Wellcome Sanger Institute for the COVID-19 Genomics UK (COG-UK) Consortium | Jacquelyn Wynn, Mairead Hyland, The Lighthouse Lab in Alderley Park and Alex Alderton, Roberto Amato, Jeffrey Barrett, Sonia Goncalves, Ewan Harrison, David K. Jackson, Ian Johnston, Dominic Kwiatkowski, Cordelia Langford, John Sillitoe on behalf of the Wellcome Sanger Institute COVID-19 Surveillance Team |
| EPI_ISL_2393816, EPI_ISL_2393817, EPI_ISL_2393818, EPI_ISL_2393819, EPI_ISL_2393820, EPI_ISL_2393821 | Lighthouse Lab in Milton Keynes | Wellcome Sanger Institute for the COVID-19 Genomics UK (COG-UK) Consortium | The Lighthouse Lab in Milton Keynes and Alex Alderton, Roberto Amato, Jeffrey Barrett, Sonia Goncalves, Ewan Harrison, David K. Jackson, Ian Johnston, Dominic Kwiatkowski, Cordelia Langford, John Sillitoe on behalf of the Wellcome Sanger Institute COVID-19 Surveillance Team |
| EPI_ISL_2393822, EPI_ISL_2393823 | Lighthouse Lab in Alderley Park | Wellcome Sanger Institute for the COVID-19 Genomics UK (COG-UK) Consortium | Jacquelyn Wynn, Mairead Hyland, The Lighthouse Lab in Alderley Park and Alex Alderton, Roberto Amato, Jeffrey Barrett, Sonia Goncalves, Ewan Harrison, David K. Jackson, Ian Johnston, Dominic Kwiatkowski, Cordelia Langford, John Sillitoe on behalf of the Wellcome Sanger Institute COVID-19 Surveillance Team |
| EPI_ISL_2393827, EPI_ISL_2393828, EPI_ISL_2393829, EPI_ISL_2393830 | Lighthouse Lab in Milton Keynes | Wellcome Sanger Institute for the COVID-19 Genomics UK (COG-UK) Consortium | The Lighthouse Lab in Milton Keynes and Alex Alderton, Roberto Amato, Jeffrey Barrett, Sonia Goncalves, Ewan Harrison, David K. Jackson, Ian Johnston, Dominic Kwiatkowski, Cordelia Langford, John Sillitoe on behalf of the Wellcome Sanger Institute COVID-19 Surveillance Team |
| EPI_ISL_2393831 | Lighthouse Lab in Glasgow | Wellcome Sanger Institute for the COVID-19 Genomics UK (COG-UK) Consortium | Harper VanSteenhouse, Yumi Kasai, David Gray, Carol Clugston, Anna Dominiczak and Alex Alderton, Roberto Amato, Jeffrey Barrett, Sonia Goncalves, Ewan Harrison, David K. Jackson, Ian Johnston, Dominic Kwiatkowski, Cordelia Langford, John Sillitoe on behalf of the Wellcome Sanger Institute COVID-19 Surveillance Team |
| EPI_ISL_2393832, EPI_ISL_2393833 | Lighthouse Lab in Alderley Park | Wellcome Sanger Institute for the COVID-19 Genomics UK (COG-UK) Consortium | Jacquelyn Wynn, Mairead Hyland, The Lighthouse Lab in Alderley Park and Alex Alderton, Roberto Amato, Jeffrey Barrett, Sonia Goncalves, Ewan Harrison, David K. Jackson, Ian Johnston, Dominic Kwiatkowski, Cordelia Langford, John Sillitoe on behalf of the Wellcome Sanger Institute COVID-19 Surveillance Team |
| EPI_ISL_2393834 | Lighthouse Lab in Milton Keynes | Wellcome Sanger Institute for the COVID-19 Genomics UK (COG-UK) Consortium | The Lighthouse Lab in Milton Keynes and Alex Alderton, Roberto Amato, Jeffrey Barrett, Sonia Goncalves, Ewan Harrison, David K. Jackson, Ian Johnston, Dominic Kwiatkowski, Cordelia Langford, John Sillitoe on behalf of the Wellcome Sanger Institute COVID-19 Surveillance Team |
| EPI_ISL_2393835 | Lighthouse Lab in Alderley Park | Wellcome Sanger Institute for the COVID-19 Genomics UK (COG-UK) Consortium | Jacquelyn Wynn, Mairead Hyland, The Lighthouse Lab in Alderley Park and Alex Alderton, Roberto Amato, Jeffrey Barrett, Sonia Goncalves, Ewan Harrison, David K. Jackson, Ian Johnston, Dominic Kwiatkowski, Cordelia Langford, John Sillitoe on behalf of the Wellcome Sanger Institute COVID-19 Surveillance Team |
| EPI_ISL_2393836, EPI_ISL_2393837 | Lighthouse Lab in Milton Keynes | Wellcome Sanger Institute for the COVID-19 Genomics UK (COG-UK) Consortium | The Lighthouse Lab in Milton Keynes and Alex Alderton, Roberto Amato, Jeffrey Barrett, Sonia Goncalves, Ewan Harrison, David K. Jackson, Ian Johnston, Dominic Kwiatkowski, Cordelia Langford, John Sillitoe on behalf of the Wellcome Sanger Institute COVID-19 Surveillance Team |
| EPI_ISL_2393838 | Lighthouse Lab in Glasgow | Wellcome Sanger Institute for the COVID-19 Genomics UK (COG-UK) Consortium | Harper VanSteenhouse, Yumi Kasai, David Gray, Carol Clugston, Anna Dominiczak and Alex Alderton, Roberto Amato, Jeffrey Barrett, Sonia Goncalves, Ewan Harrison, David K. Jackson, Ian Johnston, Dominic Kwiatkowski, Cordelia Langford, John Sillitoe on behalf of the Wellcome Sanger Institute COVID-19 Surveillance Team |
| EPI_ISL_2393839 | Lighthouse Lab in Milton Keynes | Wellcome Sanger Institute for the COVID-19 Genomics UK (COG-UK) Consortium | The Lighthouse Lab in Milton Keynes and Alex Alderton, Roberto Amato, Jeffrey Barrett, Sonia Goncalves, Ewan Harrison, David K. Jackson, Ian Johnston, Dominic Kwiatkowski, Cordelia Langford, John Sillitoe on behalf of the Wellcome Sanger Institute COVID-19 Surveillance Team |
| EPI_ISL_2393840 | Lighthouse Lab in Alderley Park | Wellcome Sanger Institute for the COVID-19 Genomics UK (COG-UK) Consortium | Jacquelyn Wynn, Mairead Hyland, The Lighthouse Lab in Alderley Park and Alex Alderton, Roberto Amato, Jeffrey Barrett, Sonia Goncalves, Ewan Harrison, David K. Jackson, Ian Johnston, Dominic Kwiatkowski, Cordelia Langford, John Sillitoe on behalf of the Wellcome Sanger Institute COVID-19 Surveillance Team |
| EPI_ISL_2393841, EPI_ISL_2393842 | Lighthouse Lab in Milton Keynes | Wellcome Sanger Institute for the COVID-19 Genomics UK (COG-UK) Consortium | The Lighthouse Lab in Milton Keynes and Alex Alderton, Roberto Amato, Jeffrey Barrett, Sonia Goncalves, Ewan Harrison, David K. Jackson, Ian Johnston, Dominic Kwiatkowski, Cordelia Langford, John Sillitoe on behalf of the Wellcome Sanger Institute COVID-19 Surveillance Team |
| EPI_ISL_2393844, EPI_ISL_2393845 | Lighthouse Lab in Alderley Park | Wellcome Sanger Institute for the COVID-19 Genomics UK (COG-UK) Consortium | Jacquelyn Wynn, Mairead Hyland, The Lighthouse Lab in Alderley Park and Alex Alderton, Roberto Amato, Jeffrey Barrett, Sonia Goncalves, Ewan Harrison, David K. Jackson, Ian Johnston, Dominic Kwiatkowski, Cordelia Langford, John Sillitoe on behalf of the Wellcome Sanger Institute COVID-19 Surveillance Team |
| EPI_ISL_2393846 | Lighthouse Lab in Milton Keynes | Wellcome Sanger Institute for the COVID-19 Genomics UK (COG-UK) Consortium | The Lighthouse Lab in Milton Keynes and Alex Alderton, Roberto Amato, Jeffrey Barrett, Sonia Goncalves, Ewan Harrison, David K. Jackson, Ian Johnston, Dominic Kwiatkowski, Cordelia Langford, John Sillitoe on behalf of the Wellcome Sanger Institute COVID-19 Surveillance Team |
| EPI_ISL_2393848, EPI_ISL_2393850, EPI_ISL_2393851, EPI_ISL_2393853 | Lighthouse Lab in Alderley Park | Wellcome Sanger Institute for the COVID-19 Genomics UK (COG-UK) Consortium | Jacquelyn Wynn, Mairead Hyland, The Lighthouse Lab in Alderley Park and Alex Alderton, Roberto Amato, Jeffrey Barrett, Sonia Goncalves, Ewan Harrison, David K. Jackson, Ian Johnston, Dominic Kwiatkowski, Cordelia Langford, John Sillitoe on behalf of the Wellcome Sanger Institute COVID-19 Surveillance Team |
| EPI_ISL_2393854 | Lighthouse Lab in Milton Keynes | Wellcome Sanger Institute for the COVID-19 Genomics UK (COG-UK) Consortium | The Lighthouse Lab in Milton Keynes and Alex Alderton, Roberto Amato, Jeffrey Barrett, Sonia Goncalves, Ewan Harrison, David K. Jackson, Ian Johnston, Dominic Kwiatkowski, Cordelia Langford, John Sillitoe on behalf of the Wellcome Sanger Institute COVID-19 Surveillance Team |
| EPI_ISL_2393855, EPI_ISL_2393856, EPI_ISL_2393857 | Lighthouse Lab in Alderley Park | Wellcome Sanger Institute for the COVID-19 Genomics UK (COG-UK) Consortium | Jacquelyn Wynn, Mairead Hyland, The Lighthouse Lab in Alderley Park and Alex Alderton, Roberto Amato, Jeffrey Barrett, Sonia Goncalves, Ewan Harrison, David K. Jackson, Ian Johnston, Dominic Kwiatkowski, Cordelia Langford, John Sillitoe on behalf of the Wellcome Sanger Institute COVID-19 Surveillance Team |
| EPI_ISL_2393858, EPI_ISL_2393859, EPI_ISL_2393860 | Lighthouse Lab in Milton Keynes | Wellcome Sanger Institute for the COVID-19 Genomics UK (COG-UK) Consortium | The Lighthouse Lab in Milton Keynes and Alex Alderton, Roberto Amato, Jeffrey Barrett, Sonia Goncalves, Ewan Harrison, David K. Jackson, Ian Johnston, Dominic Kwiatkowski, Cordelia Langford, John Sillitoe on behalf of the Wellcome Sanger Institute COVID-19 Surveillance Team |
| EPI_ISL_2393863 | Lighthouse Lab in Alderley Park | Wellcome Sanger Institute for the COVID-19 Genomics UK (COG-UK) Consortium | Jacquelyn Wynn, Mairead Hyland, The Lighthouse Lab in Alderley Park and Alex Alderton, Roberto Amato, Jeffrey Barrett, Sonia Goncalves, Ewan Harrison, David K. Jackson, Ian Johnston, Dominic Kwiatkowski, Cordelia Langford, John Sillitoe on behalf of the Wellcome Sanger Institute COVID-19 Surveillance Team |
| EPI_ISL_2393864 | Lighthouse Lab in Milton Keynes | Wellcome Sanger Institute for the COVID-19 Genomics UK (COG-UK) Consortium | The Lighthouse Lab in Milton Keynes and Alex Alderton, Roberto Amato, Jeffrey Barrett, Sonia Goncalves, Ewan Harrison, David K. Jackson, Ian Johnston, Dominic Kwiatkowski, Cordelia Langford, John Sillitoe on behalf of the Wellcome Sanger Institute COVID-19 Surveillance Team |
| EPI_ISL_2393867, EPI_ISL_2393868 | Lighthouse Lab in Alderley Park | Wellcome Sanger Institute for the COVID-19 Genomics UK (COG-UK) Consortium | Jacquelyn Wynn, Mairead Hyland, The Lighthouse Lab in Alderley Park and Alex Alderton, Roberto Amato, Jeffrey Barrett, Sonia Goncalves, Ewan Harrison, David K. Jackson, Ian Johnston, Dominic Kwiatkowski, Cordelia Langford, John Sillitoe on behalf of the Wellcome Sanger Institute COVID-19 Surveillance Team |
| EPI_ISL_2393869 | Lighthouse Lab in Milton Keynes | Wellcome Sanger Institute for the COVID-19 Genomics UK (COG-UK) Consortium | The Lighthouse Lab in Milton Keynes and Alex Alderton, Roberto Amato, Jeffrey Barrett, Sonia Goncalves, Ewan Harrison, David K. Jackson, Ian Johnston, Dominic Kwiatkowski, Cordelia Langford, John Sillitoe on behalf of the Wellcome Sanger Institute COVID-19 Surveillance Team |
| EPI_ISL_2393870 | Lighthouse Lab in Alderley Park | Wellcome Sanger Institute for the COVID-19 Genomics UK (COG-UK) Consortium | Jacquelyn Wynn, Mairead Hyland, The Lighthouse Lab in Alderley Park and Alex Alderton, Roberto Amato, Jeffrey Barrett, Sonia Goncalves, Ewan Harrison, David K. Jackson, Ian Johnston, Dominic Kwiatkowski, Cordelia Langford, John Sillitoe on behalf of the Wellcome Sanger Institute COVID-19 Surveillance Team |
| EPI_ISL_2393871 | Lighthouse Lab in Milton Keynes | Wellcome Sanger Institute for the COVID-19 Genomics UK (COG-UK) Consortium | The Lighthouse Lab in Milton Keynes and Alex Alderton, Roberto Amato, Jeffrey Barrett, Sonia Goncalves, Ewan Harrison, David K. Jackson, Ian Johnston, Dominic Kwiatkowski, Cordelia Langford, John Sillitoe on behalf of the Wellcome Sanger Institute COVID-19 Surveillance Team |
| EPI_ISL_2393872 | Lighthouse Lab in Alderley Park | Wellcome Sanger Institute for the COVID-19 Genomics UK (COG-UK) Consortium | Jacquelyn Wynn, Mairead Hyland, The Lighthouse Lab in Alderley Park and Alex Alderton, Roberto Amato, Jeffrey Barrett, Sonia Goncalves, Ewan Harrison, David K. Jackson, Ian Johnston, Dominic Kwiatkowski, Cordelia Langford, John Sillitoe on behalf of the Wellcome Sanger Institute COVID-19 Surveillance Team |
| EPI_ISL_2393873, EPI_ISL_2393874 | Lighthouse Lab in Milton Keynes | Wellcome Sanger Institute for the COVID-19 Genomics UK (COG-UK) Consortium | The Lighthouse Lab in Milton Keynes and Alex Alderton, Roberto Amato, Jeffrey Barrett, Sonia Goncalves, Ewan Harrison, David K. Jackson, Ian Johnston, Dominic Kwiatkowski, Cordelia Langford, John Sillitoe on behalf of the Wellcome Sanger Institute COVID-19 Surveillance Team |
| EPI_ISL_2393875, EPI_ISL_2393876 | Lighthouse Lab in Alderley Park | Wellcome Sanger Institute for the COVID-19 Genomics UK (COG-UK) Consortium | Jacquelyn Wynn, Mairead Hyland, The Lighthouse Lab in Alderley Park and Alex Alderton, Roberto Amato, Jeffrey Barrett, Sonia Goncalves, Ewan Harrison, David K. Jackson, Ian Johnston, Dominic Kwiatkowski, Cordelia Langford, John Sillitoe on behalf of the Wellcome Sanger Institute COVID-19 Surveillance Team |

| | | | |
|---|---|---|---|
| EPI_ISL_2393877, EPI_ISL_2393879 | Lighthouse Lab in Milton Keynes | Wellcome Sanger Institute for the COVID-19 Genomics UK (COG-UK) Consortium | The Lighthouse Lab in Milton Keynes and Alex Alderton, Roberto Amato, Jeffrey Barrett, Sonia Goncalves, Ewan Harrison, David K. Jackson, Ian Johnston, Dominic Kwiatkowski, Cordelia Langford, John Sillitoe on behalf of the Wellcome Sanger Institute COVID-19 Surveillance Team |
| EPI_ISL_2393880 | Lighthouse Lab in Alderley Park | Wellcome Sanger Institute for the COVID-19 Genomics UK (COG-UK) Consortium | Jacquelyn Wynn, Mairead Hyland, The Lighthouse Lab in Alderley Park and Alex Alderton, Roberto Amato, Jeffrey Barrett, Sonia Goncalves, Ewan Harrison, David K. Jackson, Ian Johnston, Dominic Kwiatkowski, Cordelia Langford, John Sillitoe on behalf of the Wellcome Sanger Institute COVID-19 Surveillance Team |
| EPI_ISL_2393881, EPI_ISL_2393882, EPI_ISL_2393883 | Lighthouse Lab in Milton Keynes | Wellcome Sanger Institute for the COVID-19 Genomics UK (COG-UK) Consortium | The Lighthouse Lab in Milton Keynes and Alex Alderton, Roberto Amato, Jeffrey Barrett, Sonia Goncalves, Ewan Harrison, David K. Jackson, Ian Johnston, Dominic Kwiatkowski, Cordelia Langford, John Sillitoe on behalf of the Wellcome Sanger Institute COVID-19 Surveillance Team |
| EPI_ISL_2393884, EPI_ISL_2393885, EPI_ISL_2393887 | Lighthouse Lab in Alderley Park | Wellcome Sanger Institute for the COVID-19 Genomics UK (COG-UK) Consortium | Jacquelyn Wynn, Mairead Hyland, The Lighthouse Lab in Alderley Park and Alex Alderton, Roberto Amato, Jeffrey Barrett, Sonia Goncalves, Ewan Harrison, David K. Jackson, Ian Johnston, Dominic Kwiatkowski, Cordelia Langford, John Sillitoe on behalf of the Wellcome Sanger Institute COVID-19 Surveillance Team |
| EPI_ISL_2393888 | Lighthouse Lab in Milton Keynes | Wellcome Sanger Institute for the COVID-19 Genomics UK (COG-UK) Consortium | The Lighthouse Lab in Milton Keynes and Alex Alderton, Roberto Amato, Jeffrey Barrett, Sonia Goncalves, Ewan Harrison, David K. Jackson, Ian Johnston, Dominic Kwiatkowski, Cordelia Langford, John Sillitoe on behalf of the Wellcome Sanger Institute COVID-19 Surveillance Team |
| EPI_ISL_2393890 | Lighthouse Lab in Alderley Park | Wellcome Sanger Institute for the COVID-19 Genomics UK (COG-UK) Consortium | Jacquelyn Wynn, Mairead Hyland, The Lighthouse Lab in Alderley Park and Alex Alderton, Roberto Amato, Jeffrey Barrett, Sonia Goncalves, Ewan Harrison, David K. Jackson, Ian Johnston, Dominic Kwiatkowski, Cordelia Langford, John Sillitoe on behalf of the Wellcome Sanger Institute COVID-19 Surveillance Team |
| EPI_ISL_2393891 | Lighthouse Lab in Milton Keynes | Wellcome Sanger Institute for the COVID-19 Genomics UK (COG-UK) Consortium | The Lighthouse Lab in Milton Keynes and Alex Alderton, Roberto Amato, Jeffrey Barrett, Sonia Goncalves, Ewan Harrison, David K. Jackson, Ian Johnston, Dominic Kwiatkowski, Cordelia Langford, John Sillitoe on behalf of the Wellcome Sanger Institute COVID-19 Surveillance Team |
| EPI_ISL_2393892 | Lighthouse Lab in Alderley Park | Wellcome Sanger Institute for the COVID-19 Genomics UK (COG-UK) Consortium | Jacquelyn Wynn, Mairead Hyland, The Lighthouse Lab in Alderley Park and Alex Alderton, Roberto Amato, Jeffrey Barrett, Sonia Goncalves, Ewan Harrison, David K. Jackson, Ian Johnston, Dominic Kwiatkowski, Cordelia Langford, John Sillitoe on behalf of the Wellcome Sanger Institute COVID-19 Surveillance Team |
| EPI_ISL_2393893, EPI_ISL_2393894 | Lighthouse Lab in Milton Keynes | Wellcome Sanger Institute for the COVID-19 Genomics UK (COG-UK) Consortium | The Lighthouse Lab in Milton Keynes and Alex Alderton, Roberto Amato, Jeffrey Barrett, Sonia Goncalves, Ewan Harrison, David K. Jackson, Ian Johnston, Dominic Kwiatkowski, Cordelia Langford, John Sillitoe on behalf of the Wellcome Sanger Institute COVID-19 Surveillance Team |
| EPI_ISL_2393895 | Lighthouse Lab in Alderley Park | Wellcome Sanger Institute for the COVID-19 Genomics UK (COG-UK) Consortium | Jacquelyn Wynn, Mairead Hyland, The Lighthouse Lab in Alderley Park and Alex Alderton, Roberto Amato, Jeffrey Barrett, Sonia Goncalves, Ewan Harrison, David K. Jackson, Ian Johnston, Dominic Kwiatkowski, Cordelia Langford, John Sillitoe on behalf of the Wellcome Sanger Institute COVID-19 Surveillance Team |
| EPI_ISL_2393897 | Lighthouse Lab in Milton Keynes | Wellcome Sanger Institute for the COVID-19 Genomics UK (COG-UK) Consortium | The Lighthouse Lab in Milton Keynes and Alex Alderton, Roberto Amato, Jeffrey Barrett, Sonia Goncalves, Ewan Harrison, David K. Jackson, Ian Johnston, Dominic Kwiatkowski, Cordelia Langford, John Sillitoe on behalf of the Wellcome Sanger Institute COVID-19 Surveillance Team |
| EPI_ISL_2393899 | Lighthouse Lab in Glasgow | Wellcome Sanger Institute for the COVID-19 Genomics UK (COG-UK) Consortium | Harper VanSteenhouse, Yumi Kasai, David Gray, Carol Clugston, Anna Dominiczak and Alex Alderton, Roberto Amato, Jeffrey Barrett, Sonia Goncalves, Ewan Harrison, David K. Jackson, Ian Johnston, Dominic Kwiatkowski, Cordelia Langford, John Sillitoe on behalf of the Wellcome Sanger Institute COVID-19 Surveillance Team |
| EPI_ISL_2393900, EPI_ISL_2393901 | Lighthouse Lab in Alderley Park | Wellcome Sanger Institute for the COVID-19 Genomics UK (COG-UK) Consortium | Jacquelyn Wynn, Mairead Hyland, The Lighthouse Lab in Alderley Park and Alex Alderton, Roberto Amato, Jeffrey Barrett, Sonia Goncalves, Ewan Harrison, David K. Jackson, Ian Johnston, Dominic Kwiatkowski, Cordelia Langford, John Sillitoe on behalf of the Wellcome Sanger Institute COVID-19 Surveillance Team |
| EPI_ISL_2393902 | Lighthouse Lab in Glasgow | Wellcome Sanger Institute for the COVID-19 Genomics UK (COG-UK) Consortium | Harper VanSteenhouse, Yumi Kasai, David Gray, Carol Clugston, Anna Dominiczak and Alex Alderton, Roberto Amato, Jeffrey Barrett, Sonia Goncalves, Ewan Harrison, David K. Jackson, Ian Johnston, Dominic Kwiatkowski, Cordelia Langford, John Sillitoe on behalf of the Wellcome Sanger Institute COVID-19 Surveillance Team |
| EPI_ISL_2393904 | Lighthouse Lab in Milton Keynes | Wellcome Sanger Institute for the COVID-19 Genomics UK (COG-UK) Consortium | The Lighthouse Lab in Milton Keynes and Alex Alderton, Roberto Amato, Jeffrey Barrett, Sonia Goncalves, Ewan Harrison, David K. Jackson, Ian Johnston, Dominic Kwiatkowski, Cordelia Langford, John Sillitoe on behalf of the Wellcome Sanger Institute COVID-19 Surveillance Team |
| EPI_ISL_2393905, EPI_ISL_2393906 | Lighthouse Lab in Alderley Park | Wellcome Sanger Institute for the COVID-19 Genomics UK (COG-UK) Consortium | Jacquelyn Wynn, Mairead Hyland, The Lighthouse Lab in Alderley Park and Alex Alderton, Roberto Amato, Jeffrey Barrett, Sonia Goncalves, Ewan Harrison, David K. Jackson, Ian Johnston, Dominic Kwiatkowski, Cordelia Langford, John Sillitoe on behalf of the Wellcome Sanger Institute COVID-19 Surveillance Team |
| EPI_ISL_2393907 | Lighthouse Lab in Milton Keynes | Wellcome Sanger Institute for the COVID-19 Genomics UK (COG-UK) Consortium | The Lighthouse Lab in Milton Keynes and Alex Alderton, Roberto Amato, Jeffrey Barrett, Sonia Goncalves, Ewan Harrison, David K. Jackson, Ian Johnston, Dominic Kwiatkowski, Cordelia Langford, John Sillitoe on behalf of the Wellcome Sanger Institute COVID-19 Surveillance Team |
| EPI_ISL_2393908, EPI_ISL_2393909, EPI_ISL_2393910 | Lighthouse Lab in Alderley Park | Wellcome Sanger Institute for the COVID-19 Genomics UK (COG-UK) Consortium | Jacquelyn Wynn, Mairead Hyland, The Lighthouse Lab in Alderley Park and Alex Alderton, Roberto Amato, Jeffrey Barrett, Sonia Goncalves, Ewan Harrison, David K. Jackson, Ian Johnston, Dominic Kwiatkowski, Cordelia Langford, John Sillitoe on behalf of the Wellcome Sanger Institute COVID-19 Surveillance Team |
| EPI_ISL_2393911, EPI_ISL_2393912 | Lighthouse Lab in Milton Keynes | Wellcome Sanger Institute for the COVID-19 Genomics UK (COG-UK) Consortium | The Lighthouse Lab in Milton Keynes and Alex Alderton, Roberto Amato, Jeffrey Barrett, Sonia Goncalves, Ewan Harrison, David K. Jackson, Ian Johnston, Dominic Kwiatkowski, Cordelia Langford, John Sillitoe on behalf of the Wellcome Sanger Institute COVID-19 Surveillance Team |
| EPI_ISL_2393914, EPI_ISL_2393915, EPI_ISL_2393918 | Lighthouse Lab in Alderley Park | Wellcome Sanger Institute for the COVID-19 Genomics UK (COG-UK) Consortium | Jacquelyn Wynn, Mairead Hyland, The Lighthouse Lab in Alderley Park and Alex Alderton, Roberto Amato, Jeffrey Barrett, Sonia Goncalves, Ewan Harrison, David K. Jackson, Ian Johnston, Dominic Kwiatkowski, Cordelia Langford, John Sillitoe on behalf of the Wellcome Sanger Institute COVID-19 Surveillance Team |
| EPI_ISL_2393919, EPI_ISL_2393921 | Lighthouse Lab in Milton Keynes | Wellcome Sanger Institute for the COVID-19 Genomics UK (COG-UK) Consortium | The Lighthouse Lab in Milton Keynes and Alex Alderton, Roberto Amato, Jeffrey Barrett, Sonia Goncalves, Ewan Harrison, David K. Jackson, Ian Johnston, Dominic Kwiatkowski, Cordelia Langford, John Sillitoe on behalf of the Wellcome Sanger Institute COVID-19 Surveillance Team |
| EPI_ISL_2393922, EPI_ISL_2393923, EPI_ISL_2393924 | Lighthouse Lab in Alderley Park | Wellcome Sanger Institute for the COVID-19 Genomics UK (COG-UK) Consortium | Jacquelyn Wynn, Mairead Hyland, The Lighthouse Lab in Alderley Park and Alex Alderton, Roberto Amato, Jeffrey Barrett, Sonia Goncalves, Ewan Harrison, David K. Jackson, Ian Johnston, Dominic Kwiatkowski, Cordelia Langford, John Sillitoe on behalf of the Wellcome Sanger Institute COVID-19 Surveillance Team |
| EPI_ISL_2393925 | Lighthouse Lab in Milton Keynes | Wellcome Sanger Institute for the COVID-19 Genomics UK (COG-UK) Consortium | The Lighthouse Lab in Milton Keynes and Alex Alderton, Roberto Amato, Jeffrey Barrett, Sonia Goncalves, Ewan Harrison, David K. Jackson, Ian Johnston, Dominic Kwiatkowski, Cordelia Langford, John Sillitoe on behalf of the Wellcome Sanger Institute COVID-19 Surveillance Team |
| EPI_ISL_2393926 | Lighthouse Lab in Glasgow | Wellcome Sanger Institute for the COVID-19 Genomics UK (COG-UK) Consortium | Harper VanSteenhouse, Yumi Kasai, David Gray, Carol Clugston, Anna Dominiczak and Alex Alderton, Roberto Amato, Jeffrey Barrett, Sonia Goncalves, Ewan Harrison, David K. Jackson, Ian Johnston, Dominic Kwiatkowski, Cordelia Langford, John Sillitoe on behalf of the Wellcome Sanger Institute COVID-19 Surveillance Team |
| EPI_ISL_2393928 | Lighthouse Lab in Milton Keynes | Wellcome Sanger Institute for the COVID-19 Genomics UK (COG-UK) Consortium | The Lighthouse Lab in Milton Keynes and Alex Alderton, Roberto Amato, Jeffrey Barrett, Sonia Goncalves, Ewan Harrison, David K. Jackson, Ian Johnston, Dominic Kwiatkowski, Cordelia Langford, John Sillitoe on behalf of the Wellcome Sanger Institute COVID-19 Surveillance Team |
| EPI_ISL_2393930 | Lighthouse Lab in Alderley Park | Wellcome Sanger Institute for the COVID-19 Genomics UK (COG-UK) Consortium | Jacquelyn Wynn, Mairead Hyland, The Lighthouse Lab in Alderley Park and Alex Alderton, Roberto Amato, Jeffrey Barrett, Sonia Goncalves, Ewan Harrison, David K. Jackson, Ian Johnston, Dominic Kwiatkowski, Cordelia Langford, John Sillitoe on behalf of the Wellcome Sanger Institute COVID-19 Surveillance Team |
| EPI_ISL_2393931 | Lighthouse Lab in Milton Keynes | Wellcome Sanger Institute for the COVID-19 Genomics UK (COG-UK) Consortium | The Lighthouse Lab in Milton Keynes and Alex Alderton, Roberto Amato, Jeffrey Barrett, Sonia Goncalves, Ewan Harrison, David K. Jackson, Ian Johnston, Dominic Kwiatkowski, Cordelia Langford, John Sillitoe on behalf of the Wellcome Sanger Institute COVID-19 Surveillance Team |
| EPI_ISL_2393932 | Lighthouse Lab in Alderley Park | Wellcome Sanger Institute for the COVID-19 Genomics UK (COG-UK) Consortium | Jacquelyn Wynn, Mairead Hyland, The Lighthouse Lab in Alderley Park and Alex Alderton, Roberto Amato, Jeffrey Barrett, Sonia Goncalves, Ewan Harrison, David K. Jackson, Ian Johnston, Dominic Kwiatkowski, Cordelia Langford, John Sillitoe on behalf of the Wellcome Sanger Institute COVID-19 Surveillance Team |
| EPI_ISL_2393933 | Lighthouse Lab in Milton Keynes | Wellcome Sanger Institute for the COVID-19 Genomics UK (COG-UK) Consortium | The Lighthouse Lab in Milton Keynes and Alex Alderton, Roberto Amato, Jeffrey Barrett, Sonia Goncalves, Ewan Harrison, David K. Jackson, Ian Johnston, Dominic Kwiatkowski, Cordelia Langford, John Sillitoe on behalf of the Wellcome Sanger Institute COVID-19 Surveillance Team |
| EPI_ISL_2393934 | Lighthouse Lab in Alderley Park | Wellcome Sanger Institute for the COVID-19 Genomics UK (COG-UK) Consortium | Jacquelyn Wynn, Mairead Hyland, The Lighthouse Lab in Alderley Park and Alex Alderton, Roberto Amato, Jeffrey Barrett, Sonia Goncalves, Ewan Harrison, David K. Jackson, Ian Johnston, Dominic Kwiatkowski, Cordelia Langford, John Sillitoe on behalf of the Wellcome Sanger Institute COVID-19 Surveillance Team |
| EPI_ISL_2393937 | Lighthouse Lab in Milton Keynes | Wellcome Sanger Institute for the COVID-19 Genomics UK (COG-UK) Consortium | The Lighthouse Lab in Milton Keynes and Alex Alderton, Roberto Amato, Jeffrey Barrett, Sonia Goncalves, Ewan Harrison, David K. Jackson, Ian Johnston, Dominic Kwiatkowski, Cordelia Langford, John Sillitoe on behalf of the Wellcome Sanger Institute COVID-19 Surveillance Team |
| EPI_ISL_2393938 | Lighthouse Lab in Alderley Park | Wellcome Sanger Institute for the COVID-19 Genomics UK (COG-UK) Consortium | Jacquelyn Wynn, Mairead Hyland, The Lighthouse Lab in Alderley Park and Alex Alderton, Roberto Amato, Jeffrey Barrett, Sonia Goncalves, Ewan Harrison, David K. Jackson, Ian Johnston, Dominic Kwiatkowski, Cordelia Langford, John Sillitoe on behalf of the Wellcome Sanger Institute COVID-19 Surveillance Team |
| EPI_ISL_2393941 | Lighthouse Lab in Milton Keynes | Wellcome Sanger Institute for the COVID-19 Genomics UK (COG-UK) Consortium | The Lighthouse Lab in Milton Keynes and Alex Alderton, Roberto Amato, Jeffrey Barrett, Sonia Goncalves, Ewan Harrison, David K. Jackson, Ian Johnston, Dominic Kwiatkowski, Cordelia Langford, John Sillitoe on behalf of the Wellcome Sanger Institute COVID-19 Surveillance Team |
| EPI_ISL_2393942 | Lighthouse Lab in Glasgow | Wellcome Sanger Institute for the COVID-19 Genomics UK (COG-UK) Consortium | Harper VanSteenhouse, Yumi Kasai, David Gray, Carol Clugston, Anna Dominiczak and Alex Alderton, Roberto Amato, Jeffrey Barrett, Sonia Goncalves, Ewan Harrison, David K. Jackson, Ian Johnston, Dominic Kwiatkowski, Cordelia Langford, John Sillitoe on behalf of the Wellcome Sanger Institute COVID-19 Surveillance Team |
| EPI_ISL_2393943 | Lighthouse Lab in Alderley Park | Wellcome Sanger Institute for the COVID-19 Genomics UK (COG-UK) Consortium | Jacquelyn Wynn, Mairead Hyland, The Lighthouse Lab in Alderley Park and Alex Alderton, Roberto Amato, Jeffrey Barrett, Sonia Goncalves, Ewan Harrison, David K. Jackson, Ian Johnston, Dominic Kwiatkowski, Cordelia Langford, John Sillitoe on behalf of the Wellcome Sanger Institute COVID-19 Surveillance Team |
| EPI_ISL_2393945 | Lighthouse Lab in Glasgow | Wellcome Sanger Institute for the COVID-19 Genomics UK (COG-UK) Consortium | Harper VanSteenhouse, Yumi Kasai, David Gray, Carol Clugston, Anna Dominiczak and Alex Alderton, Roberto Amato, Jeffrey Barrett, Sonia Goncalves, Ewan Harrison, David K. Jackson, Ian Johnston, Dominic Kwiatkowski, Cordelia Langford, John Sillitoe on behalf of the Wellcome Sanger Institute COVID-19 Surveillance Team |
| EPI_ISL_2393948 | Lighthouse Lab in Alderley Park | Wellcome Sanger Institute for the COVID-19 Genomics UK (COG-UK) Consortium | Jacquelyn Wynn, Mairead Hyland, The Lighthouse Lab in Alderley Park and Alex Alderton, Roberto Amato, Jeffrey Barrett, Sonia Goncalves, Ewan Harrison, David K. Jackson, Ian Johnston, Dominic Kwiatkowski, Cordelia Langford, John Sillitoe on behalf of the Wellcome Sanger Institute COVID-19 Surveillance Team |
| EPI_ISL_2393949 | Lighthouse Lab in Milton Keynes | Wellcome Sanger Institute for the COVID-19 Genomics UK (COG-UK) Consortium | The Lighthouse Lab in Milton Keynes and Alex Alderton, Roberto Amato, Jeffrey Barrett, Sonia Goncalves, Ewan Harrison, David K. Jackson, Ian Johnston, Dominic Kwiatkowski, Cordelia Langford, John Sillitoe on behalf of the Wellcome Sanger Institute COVID-19 Surveillance Team |
| EPI_ISL_2393950 | Lighthouse Lab in Alderley Park | Wellcome Sanger Institute for the COVID-19 Genomics UK (COG-UK) Consortium | Jacquelyn Wynn, Mairead Hyland, The Lighthouse Lab in Alderley Park and Alex Alderton, Roberto Amato, Jeffrey Barrett, Sonia Goncalves, Ewan Harrison, David K. Jackson, Ian Johnston, Dominic Kwiatkowski, Cordelia Langford, John Sillitoe on behalf of the Wellcome Sanger Institute COVID-19 Surveillance Team |
| EPI_ISL_2393951, EPI_ISL_2393953, EPI_ISL_2393955, EPI_ISL_2393956 | Lighthouse Lab in Milton Keynes | Wellcome Sanger Institute for the COVID-19 Genomics UK (COG-UK) Consortium | The Lighthouse Lab in Milton Keynes and Alex Alderton, Roberto Amato, Jeffrey Barrett, Sonia Goncalves, Ewan Harrison, David K. Jackson, Ian Johnston, Dominic Kwiatkowski, Cordelia Langford, John Sillitoe on behalf of the Wellcome Sanger Institute COVID-19 Surveillance Team |
| EPI_ISL_2393957 | Lighthouse Lab in Alderley Park | Wellcome Sanger Institute for the COVID-19 Genomics UK (COG-UK) Consortium | Jacquelyn Wynn, Mairead Hyland, The Lighthouse Lab in Alderley Park and Alex Alderton, Roberto Amato, Jeffrey Barrett, Sonia Goncalves, Ewan Harrison, David K. Jackson, Ian Johnston, Dominic Kwiatkowski, Cordelia Langford, John Sillitoe on behalf of the Wellcome Sanger Institute COVID-19 Surveillance Team |
| EPI_ISL_2393958 | Lighthouse Lab in Glasgow | Wellcome Sanger Institute for the COVID-19 Genomics UK (COG-UK) Consortium | Harper VanSteenhouse, Yumi Kasai, David Gray, Carol Clugston, Anna Dominiczak and Alex Alderton, Roberto Amato, Jeffrey Barrett, Sonia Goncalves, Ewan Harrison, David K. Jackson, Ian Johnston, Dominic Kwiatkowski, Cordelia Langford, John Sillitoe on behalf of the Wellcome Sanger Institute COVID-19 Surveillance Team |

| | | | |
|---|---|---|---|
| EPI_ISL_2393959, EPI_ISL_2393960, EPI_ISL_2393961 | Lighthouse Lab in Alderley Park | Wellcome Sanger Institute for the COVID-19 Genomics UK (COG-UK) Consortium | Jacquelyn Wynn, Mairead Hyland, The Lighthouse Lab in Alderley Park and Alex Alderton, Roberto Amato, Jeffrey Barrett, Sonia Goncalves, Ewan Harrison, David K. Jackson, Ian Johnston, Dominic Kwiatkowski, Cordelia Langford, John Sillitoe on behalf of the Wellcome Sanger Institute COVID-19 Surveillance Team |
| EPI_ISL_2393962 | Lighthouse Lab in Milton Keynes | Wellcome Sanger Institute for the COVID-19 Genomics UK (COG-UK) Consortium | The Lighthouse Lab in Milton Keynes and Alex Alderton, Roberto Amato, Jeffrey Barrett, Sonia Goncalves, Ewan Harrison, David K. Jackson, Ian Johnston, Dominic Kwiatkowski, Cordelia Langford, John Sillitoe on behalf of the Wellcome Sanger Institute COVID-19 Surveillance Team |
| EPI_ISL_2393963 | Lighthouse Lab in Alderley Park | Wellcome Sanger Institute for the COVID-19 Genomics UK (COG-UK) Consortium | Jacquelyn Wynn, Mairead Hyland, The Lighthouse Lab in Alderley Park and Alex Alderton, Roberto Amato, Jeffrey Barrett, Sonia Goncalves, Ewan Harrison, David K. Jackson, Ian Johnston, Dominic Kwiatkowski, Cordelia Langford, John Sillitoe on behalf of the Wellcome Sanger Institute COVID-19 Surveillance Team |
| EPI_ISL_2393965 | Lighthouse Lab in Glasgow | Wellcome Sanger Institute for the COVID-19 Genomics UK (COG-UK) Consortium | Harper VanSteenhouse, Yumi Kasai, David Gray, Carol Clugston, Anna Dominiczak and Alex Alderton, Roberto Amato, Jeffrey Barrett, Sonia Goncalves, Ewan Harrison, David K. Jackson, Ian Johnston, Dominic Kwiatkowski, Cordelia Langford, John Sillitoe on behalf of the Wellcome Sanger Institute COVID-19 Surveillance Team |
| EPI_ISL_2393966 | Lighthouse Lab in Alderley Park | Wellcome Sanger Institute for the COVID-19 Genomics UK (COG-UK) Consortium | Jacquelyn Wynn, Mairead Hyland, The Lighthouse Lab in Alderley Park and Alex Alderton, Roberto Amato, Jeffrey Barrett, Sonia Goncalves, Ewan Harrison, David K. Jackson, Ian Johnston, Dominic Kwiatkowski, Cordelia Langford, John Sillitoe on behalf of the Wellcome Sanger Institute COVID-19 Surveillance Team |
| EPI_ISL_2393967 | Lighthouse Lab in Milton Keynes | Wellcome Sanger Institute for the COVID-19 Genomics UK (COG-UK) Consortium | The Lighthouse Lab in Milton Keynes and Alex Alderton, Roberto Amato, Jeffrey Barrett, Sonia Goncalves, Ewan Harrison, David K. Jackson, Ian Johnston, Dominic Kwiatkowski, Cordelia Langford, John Sillitoe on behalf of the Wellcome Sanger Institute COVID-19 Surveillance Team |
| EPI_ISL_2393968 | Lighthouse Lab in Alderley Park | Wellcome Sanger Institute for the COVID-19 Genomics UK (COG-UK) Consortium | Jacquelyn Wynn, Mairead Hyland, The Lighthouse Lab in Alderley Park and Alex Alderton, Roberto Amato, Jeffrey Barrett, Sonia Goncalves, Ewan Harrison, David K. Jackson, Ian Johnston, Dominic Kwiatkowski, Cordelia Langford, John Sillitoe on behalf of the Wellcome Sanger Institute COVID-19 Surveillance Team |
| EPI_ISL_2393970 | Lighthouse Lab in Milton Keynes | Wellcome Sanger Institute for the COVID-19 Genomics UK (COG-UK) Consortium | The Lighthouse Lab in Milton Keynes and Alex Alderton, Roberto Amato, Jeffrey Barrett, Sonia Goncalves, Ewan Harrison, David K. Jackson, Ian Johnston, Dominic Kwiatkowski, Cordelia Langford, John Sillitoe on behalf of the Wellcome Sanger Institute COVID-19 Surveillance Team |
| EPI_ISL_2393971 | Lighthouse Lab in Alderley Park | Wellcome Sanger Institute for the COVID-19 Genomics UK (COG-UK) Consortium | Jacquelyn Wynn, Mairead Hyland, The Lighthouse Lab in Alderley Park and Alex Alderton, Roberto Amato, Jeffrey Barrett, Sonia Goncalves, Ewan Harrison, David K. Jackson, Ian Johnston, Dominic Kwiatkowski, Cordelia Langford, John Sillitoe on behalf of the Wellcome Sanger Institute COVID-19 Surveillance Team |
| EPI_ISL_2393972 | Lighthouse Lab in Glasgow | Wellcome Sanger Institute for the COVID-19 Genomics UK (COG-UK) Consortium | Harper VanSteenhouse, Yumi Kasai, David Gray, Carol Clugston, Anna Dominiczak and Alex Alderton, Roberto Amato, Jeffrey Barrett, Sonia Goncalves, Ewan Harrison, David K. Jackson, Ian Johnston, Dominic Kwiatkowski, Cordelia Langford, John Sillitoe on behalf of the Wellcome Sanger Institute COVID-19 Surveillance Team |
| EPI_ISL_2393973, EPI_ISL_2393974 | Lighthouse Lab in Alderley Park | Wellcome Sanger Institute for the COVID-19 Genomics UK (COG-UK) Consortium | Jacquelyn Wynn, Mairead Hyland, The Lighthouse Lab in Alderley Park and Alex Alderton, Roberto Amato, Jeffrey Barrett, Sonia Goncalves, Ewan Harrison, David K. Jackson, Ian Johnston, Dominic Kwiatkowski, Cordelia Langford, John Sillitoe on behalf of the Wellcome Sanger Institute COVID-19 Surveillance Team |
| EPI_ISL_2393975, EPI_ISL_2393976, EPI_ISL_2393978 | Lighthouse Lab in Milton Keynes | Wellcome Sanger Institute for the COVID-19 Genomics UK (COG-UK) Consortium | The Lighthouse Lab in Milton Keynes and Alex Alderton, Roberto Amato, Jeffrey Barrett, Sonia Goncalves, Ewan Harrison, David K. Jackson, Ian Johnston, Dominic Kwiatkowski, Cordelia Langford, John Sillitoe on behalf of the Wellcome Sanger Institute COVID-19 Surveillance Team |
| EPI_ISL_2393979, EPI_ISL_2393980, EPI_ISL_2393981 | Lighthouse Lab in Alderley Park | Wellcome Sanger Institute for the COVID-19 Genomics UK (COG-UK) Consortium | Jacquelyn Wynn, Mairead Hyland, The Lighthouse Lab in Alderley Park and Alex Alderton, Roberto Amato, Jeffrey Barrett, Sonia Goncalves, Ewan Harrison, David K. Jackson, Ian Johnston, Dominic Kwiatkowski, Cordelia Langford, John Sillitoe on behalf of the Wellcome Sanger Institute COVID-19 Surveillance Team |
| EPI_ISL_2393982, EPI_ISL_2393983 | Lighthouse Lab in Milton Keynes | Wellcome Sanger Institute for the COVID-19 Genomics UK (COG-UK) Consortium | The Lighthouse Lab in Milton Keynes and Alex Alderton, Roberto Amato, Jeffrey Barrett, Sonia Goncalves, Ewan Harrison, David K. Jackson, Ian Johnston, Dominic Kwiatkowski, Cordelia Langford, John Sillitoe on behalf of the Wellcome Sanger Institute COVID-19 Surveillance Team |
| EPI_ISL_2393984, EPI_ISL_2393985, EPI_ISL_2393986 | Lighthouse Lab in Alderley Park | Wellcome Sanger Institute for the COVID-19 Genomics UK (COG-UK) Consortium | Jacquelyn Wynn, Mairead Hyland, The Lighthouse Lab in Alderley Park and Alex Alderton, Roberto Amato, Jeffrey Barrett, Sonia Goncalves, Ewan Harrison, David K. Jackson, Ian Johnston, Dominic Kwiatkowski, Cordelia Langford, John Sillitoe on behalf of the Wellcome Sanger Institute COVID-19 Surveillance Team |
| EPI_ISL_2393988 | Lighthouse Lab in Glasgow | Wellcome Sanger Institute for the COVID-19 Genomics UK (COG-UK) Consortium | Harper VanSteenhouse, Yumi Kasai, David Gray, Carol Clugston, Anna Dominiczak and Alex Alderton, Roberto Amato, Jeffrey Barrett, Sonia Goncalves, Ewan Harrison, David K. Jackson, Ian Johnston, Dominic Kwiatkowski, Cordelia Langford, John Sillitoe on behalf of the Wellcome Sanger Institute COVID-19 Surveillance Team |
| EPI_ISL_2393991, EPI_ISL_2393992 | Lighthouse Lab in Milton Keynes | Wellcome Sanger Institute for the COVID-19 Genomics UK (COG-UK) Consortium | The Lighthouse Lab in Milton Keynes and Alex Alderton, Roberto Amato, Jeffrey Barrett, Sonia Goncalves, Ewan Harrison, David K. Jackson, Ian Johnston, Dominic Kwiatkowski, Cordelia Langford, John Sillitoe on behalf of the Wellcome Sanger Institute COVID-19 Surveillance Team |
| EPI_ISL_2393993, EPI_ISL_2393994, EPI_ISL_2393995 | Lighthouse Lab in Alderley Park | Wellcome Sanger Institute for the COVID-19 Genomics UK (COG-UK) Consortium | Jacquelyn Wynn, Mairead Hyland, The Lighthouse Lab in Alderley Park and Alex Alderton, Roberto Amato, Jeffrey Barrett, Sonia Goncalves, Ewan Harrison, David K. Jackson, Ian Johnston, Dominic Kwiatkowski, Cordelia Langford, John Sillitoe on behalf of the Wellcome Sanger Institute COVID-19 Surveillance Team |
| EPI_ISL_2393996, EPI_ISL_2393997 | Lighthouse Lab in Glasgow | Wellcome Sanger Institute for the COVID-19 Genomics UK (COG-UK) Consortium | Harper VanSteenhouse, Yumi Kasai, David Gray, Carol Clugston, Anna Dominiczak and Alex Alderton, Roberto Amato, Jeffrey Barrett, Sonia Goncalves, Ewan Harrison, David K. Jackson, Ian Johnston, Dominic Kwiatkowski, Cordelia Langford, John Sillitoe on behalf of the Wellcome Sanger Institute COVID-19 Surveillance Team |
| EPI_ISL_2393998, EPI_ISL_2393999 | Lighthouse Lab in Milton Keynes | Wellcome Sanger Institute for the COVID-19 Genomics UK (COG-UK) Consortium | The Lighthouse Lab in Milton Keynes and Alex Alderton, Roberto Amato, Jeffrey Barrett, Sonia Goncalves, Ewan Harrison, David K. Jackson, Ian Johnston, Dominic Kwiatkowski, Cordelia Langford, John Sillitoe on behalf of the Wellcome Sanger Institute COVID-19 Surveillance Team |
| EPI_ISL_2394000 | Lighthouse Lab in Alderley Park | Wellcome Sanger Institute for the COVID-19 Genomics UK (COG-UK) Consortium | Jacquelyn Wynn, Mairead Hyland, The Lighthouse Lab in Alderley Park and Alex Alderton, Roberto Amato, Jeffrey Barrett, Sonia Goncalves, Ewan Harrison, David K. Jackson, Ian Johnston, Dominic Kwiatkowski, Cordelia Langford, John Sillitoe on behalf of the Wellcome Sanger Institute COVID-19 Surveillance Team |
| EPI_ISL_2394001 | Lighthouse Lab in Glasgow | Wellcome Sanger Institute for the COVID-19 Genomics UK (COG-UK) Consortium | Harper VanSteenhouse, Yumi Kasai, David Gray, Carol Clugston, Anna Dominiczak and Alex Alderton, Roberto Amato, Jeffrey Barrett, Sonia Goncalves, Ewan Harrison, David K. Jackson, Ian Johnston, Dominic Kwiatkowski, Cordelia Langford, John Sillitoe on behalf of the Wellcome Sanger Institute COVID-19 Surveillance Team |
| EPI_ISL_2394002 | Lighthouse Lab in Alderley Park | Wellcome Sanger Institute for the COVID-19 Genomics UK (COG-UK) Consortium | Jacquelyn Wynn, Mairead Hyland, The Lighthouse Lab in Alderley Park and Alex Alderton, Roberto Amato, Jeffrey Barrett, Sonia Goncalves, Ewan Harrison, David K. Jackson, Ian Johnston, Dominic Kwiatkowski, Cordelia Langford, John Sillitoe on behalf of the Wellcome Sanger Institute COVID-19 Surveillance Team |
| EPI_ISL_2394003, EPI_ISL_2394004, EPI_ISL_2394006 | Lighthouse Lab in Milton Keynes | Wellcome Sanger Institute for the COVID-19 Genomics UK (COG-UK) Consortium | The Lighthouse Lab in Milton Keynes and Alex Alderton, Roberto Amato, Jeffrey Barrett, Sonia Goncalves, Ewan Harrison, David K. Jackson, Ian Johnston, Dominic Kwiatkowski, Cordelia Langford, John Sillitoe on behalf of the Wellcome Sanger Institute COVID-19 Surveillance Team |
| EPI_ISL_2394007, EPI_ISL_2394009, EPI_ISL_2394010, EPI_ISL_2394011, EPI_ISL_2394012, EPI_ISL_2394013, EPI_ISL_2394014 | Lighthouse Lab in Alderley Park | Wellcome Sanger Institute for the COVID-19 Genomics UK (COG-UK) Consortium | Jacquelyn Wynn, Mairead Hyland, The Lighthouse Lab in Alderley Park and Alex Alderton, Roberto Amato, Jeffrey Barrett, Sonia Goncalves, Ewan Harrison, David K. Jackson, Ian Johnston, Dominic Kwiatkowski, Cordelia Langford, John Sillitoe on behalf of the Wellcome Sanger Institute COVID-19 Surveillance Team |
| EPI_ISL_2394015, EPI_ISL_2394017 | Lighthouse Lab in Milton Keynes | Wellcome Sanger Institute for the COVID-19 Genomics UK (COG-UK) Consortium | The Lighthouse Lab in Milton Keynes and Alex Alderton, Roberto Amato, Jeffrey Barrett, Sonia Goncalves, Ewan Harrison, David K. Jackson, Ian Johnston, Dominic Kwiatkowski, Cordelia Langford, John Sillitoe on behalf of the Wellcome Sanger Institute COVID-19 Surveillance Team |
| EPI_ISL_2394018, EPI_ISL_2394019 | Lighthouse Lab in Alderley Park | Wellcome Sanger Institute for the COVID-19 Genomics UK (COG-UK) Consortium | Jacquelyn Wynn, Mairead Hyland, The Lighthouse Lab in Alderley Park and Alex Alderton, Roberto Amato, Jeffrey Barrett, Sonia Goncalves, Ewan Harrison, David K. Jackson, Ian Johnston, Dominic Kwiatkowski, Cordelia Langford, John Sillitoe on behalf of the Wellcome Sanger Institute COVID-19 Surveillance Team |
| EPI_ISL_2394020 | Lighthouse Lab in Milton Keynes | Wellcome Sanger Institute for the COVID-19 Genomics UK (COG-UK) Consortium | The Lighthouse Lab in Milton Keynes and Alex Alderton, Roberto Amato, Jeffrey Barrett, Sonia Goncalves, Ewan Harrison, David K. Jackson, Ian Johnston, Dominic Kwiatkowski, Cordelia Langford, John Sillitoe on behalf of the Wellcome Sanger Institute COVID-19 Surveillance Team |
| EPI_ISL_2394021, EPI_ISL_2394022, EPI_ISL_2394023 | Lighthouse Lab in Alderley Park | Wellcome Sanger Institute for the COVID-19 Genomics UK (COG-UK) Consortium | Jacquelyn Wynn, Mairead Hyland, The Lighthouse Lab in Alderley Park and Alex Alderton, Roberto Amato, Jeffrey Barrett, Sonia Goncalves, Ewan Harrison, David K. Jackson, Ian Johnston, Dominic Kwiatkowski, Cordelia Langford, John Sillitoe on behalf of the Wellcome Sanger Institute COVID-19 Surveillance Team |
| EPI_ISL_2394024, EPI_ISL_2394025, EPI_ISL_2394027 | Lighthouse Lab in Milton Keynes | Wellcome Sanger Institute for the COVID-19 Genomics UK (COG-UK) Consortium | The Lighthouse Lab in Milton Keynes and Alex Alderton, Roberto Amato, Jeffrey Barrett, Sonia Goncalves, Ewan Harrison, David K. Jackson, Ian Johnston, Dominic Kwiatkowski, Cordelia Langford, John Sillitoe on behalf of the Wellcome Sanger Institute COVID-19 Surveillance Team |
| EPI_ISL_2394029, EPI_ISL_2394030 | Lighthouse Lab in Alderley Park | Wellcome Sanger Institute for the COVID-19 Genomics UK (COG-UK) Consortium | Jacquelyn Wynn, Mairead Hyland, The Lighthouse Lab in Alderley Park and Alex Alderton, Roberto Amato, Jeffrey Barrett, Sonia Goncalves, Ewan Harrison, David K. Jackson, Ian Johnston, Dominic Kwiatkowski, Cordelia Langford, John Sillitoe on behalf of the Wellcome Sanger Institute COVID-19 Surveillance Team |
| EPI_ISL_2394031 | Lighthouse Lab in Glasgow | Wellcome Sanger Institute for the COVID-19 Genomics UK (COG-UK) Consortium | Harper VanSteenhouse, Yumi Kasai, David Gray, Carol Clugston, Anna Dominiczak and Alex Alderton, Roberto Amato, Jeffrey Barrett, Sonia Goncalves, Ewan Harrison, David K. Jackson, Ian Johnston, Dominic Kwiatkowski, Cordelia Langford, John Sillitoe on behalf of the Wellcome Sanger Institute COVID-19 Surveillance Team |
| EPI_ISL_2394032 | Lighthouse Lab in Milton Keynes | Wellcome Sanger Institute for the COVID-19 Genomics UK (COG-UK) Consortium | The Lighthouse Lab in Milton Keynes and Alex Alderton, Roberto Amato, Jeffrey Barrett, Sonia Goncalves, Ewan Harrison, David K. Jackson, Ian Johnston, Dominic Kwiatkowski, Cordelia Langford, John Sillitoe on behalf of the Wellcome Sanger Institute COVID-19 Surveillance Team |
| EPI_ISL_2394033 | Lighthouse Lab in Glasgow | Wellcome Sanger Institute for the COVID-19 Genomics UK (COG-UK) Consortium | Harper VanSteenhouse, Yumi Kasai, David Gray, Carol Clugston, Anna Dominiczak and Alex Alderton, Roberto Amato, Jeffrey Barrett, Sonia Goncalves, Ewan Harrison, David K. Jackson, Ian Johnston, Dominic Kwiatkowski, Cordelia Langford, John Sillitoe on behalf of the Wellcome Sanger Institute COVID-19 Surveillance Team |
| EPI_ISL_2394034 | Lighthouse Lab in Milton Keynes | Wellcome Sanger Institute for the COVID-19 Genomics UK (COG-UK) Consortium | The Lighthouse Lab in Milton Keynes and Alex Alderton, Roberto Amato, Jeffrey Barrett, Sonia Goncalves, Ewan Harrison, David K. Jackson, Ian Johnston, Dominic Kwiatkowski, Cordelia Langford, John Sillitoe on behalf of the Wellcome Sanger Institute COVID-19 Surveillance Team |
| EPI_ISL_2394035, EPI_ISL_2394036, EPI_ISL_2394039, EPI_ISL_2394040, EPI_ISL_2394041, EPI_ISL_2394042, EPI_ISL_2394043, EPI_ISL_2394046, EPI_ISL_2394047, EPI_ISL_2394048 | Lighthouse Lab in Alderley Park | Wellcome Sanger Institute for the COVID-19 Genomics UK (COG-UK) Consortium | Jacquelyn Wynn, Mairead Hyland, The Lighthouse Lab in Alderley Park and Alex Alderton, Roberto Amato, Jeffrey Barrett, Sonia Goncalves, Ewan Harrison, David K. Jackson, Ian Johnston, Dominic Kwiatkowski, Cordelia Langford, John Sillitoe on behalf of the Wellcome Sanger Institute COVID-19 Surveillance Team |
| EPI_ISL_2394050 | Lighthouse Lab in Glasgow | Wellcome Sanger Institute for the COVID-19 Genomics UK (COG-UK) | Harper VanSteenhouse, Yumi Kasai, David Gray, Carol Clugston, Anna Dominiczak and Alex Alderton, Roberto Amato, Jeffrey Barrett, Sonia Goncalves, Ewan Harrison, David K. Jackson, Ian Johnston, Dominic Kwiatkowski, Cordelia Langford, |

| | | | |
|---|---|---|---|
| | | Consortium | John Sillitoe on behalf of the Wellcome Sanger Institute COVID-19 Surveillance Team |
| EPI_ISL_2394051 | Lighthouse Lab in Alderley Park | Wellcome Sanger Institute for the COVID-19 Genomics UK (COG-UK) Consortium | Jacquelyn Wynn, Mairead Hyland, The Lighthouse Lab in Alderley Park and Alex Alderton, Roberto Amato, Jeffrey Barrett, Sonia Goncalves, Ewan Harrison, David K. Jackson, Ian Johnston, Dominic Kwiatkowski, Cordelia Langford, John Sillitoe on behalf of the Wellcome Sanger Institute COVID-19 Surveillance Team |
| EPI_ISL_2394052 | Lighthouse Lab in Glasgow | Wellcome Sanger Institute for the COVID-19 Genomics UK (COG-UK) Consortium | Harper VanSteenhouse, Yumi Kasai, David Gray, Carol Clugston, Anna Dominiczak and Alex Alderton, Roberto Amato, Jeffrey Barrett, Sonia Goncalves, Ewan Harrison, David K. Jackson, Ian Johnston, Dominic Kwiatkowski, Cordelia Langford, John Sillitoe on behalf of the Wellcome Sanger Institute COVID-19 Surveillance Team |
| EPI_ISL_2394053, EPI_ISL_2394054, EPI_ISL_2394055, EPI_ISL_2394058, EPI_ISL_2394059, EPI_ISL_2394060, EPI_ISL_2394061, EPI_ISL_2394062, EPI_ISL_2394063, EPI_ISL_2394064, EPI_ISL_2394065, EPI_ISL_2394066, EPI_ISL_2394067, EPI_ISL_2394068, EPI_ISL_2394070, EPI_ISL_2394071, EPI_ISL_2394072, EPI_ISL_2394073, EPI_ISL_2394074, EPI_ISL_2394075, EPI_ISL_2394076, EPI_ISL_2394077, EPI_ISL_2394079, EPI_ISL_2394080, EPI_ISL_2394081, EPI_ISL_2394082, EPI_ISL_2394083, EPI_ISL_2394085, EPI_ISL_2394086, EPI_ISL_2394087, EPI_ISL_2394088, EPI_ISL_2394089 | | | |
| see above | Lighthouse Lab in Alderley Park | Wellcome Sanger Institute for the COVID-19 Genomics UK (COG-UK) Consortium | Jacquelyn Wynn, Mairead Hyland, The Lighthouse Lab in Alderley Park and Alex Alderton, Roberto Amato, Jeffrey Barrett, Sonia Goncalves, Ewan Harrison, David K. Jackson, Ian Johnston, Dominic Kwiatkowski, Cordelia Langford, John Sillitoe on behalf of the Wellcome Sanger Institute COVID-19 Surveillance Team |
| EPI_ISL_2394090 | Lighthouse Lab in Glasgow | Wellcome Sanger Institute for the COVID-19 Genomics UK (COG-UK) Consortium | Harper VanSteenhouse, Yumi Kasai, David Gray, Carol Clugston, Anna Dominiczak and Alex Alderton, Roberto Amato, Jeffrey Barrett, Sonia Goncalves, Ewan Harrison, David K. Jackson, Ian Johnston, Dominic Kwiatkowski, Cordelia Langford, John Sillitoe on behalf of the Wellcome Sanger Institute COVID-19 Surveillance Team |
| EPI_ISL_2394091, EPI_ISL_2394092, EPI_ISL_2394093, EPI_ISL_2394095 | Lighthouse Lab in Alderley Park | Wellcome Sanger Institute for the COVID-19 Genomics UK (COG-UK) Consortium | Jacquelyn Wynn, Mairead Hyland, The Lighthouse Lab in Alderley Park and Alex Alderton, Roberto Amato, Jeffrey Barrett, Sonia Goncalves, Ewan Harrison, David K. Jackson, Ian Johnston, Dominic Kwiatkowski, Cordelia Langford, John Sillitoe on behalf of the Wellcome Sanger Institute COVID-19 Surveillance Team |
| EPI_ISL_2394097 | Lighthouse Lab in Glasgow | Wellcome Sanger Institute for the COVID-19 Genomics UK (COG-UK) Consortium | Harper VanSteenhouse, Yumi Kasai, David Gray, Carol Clugston, Anna Dominiczak and Alex Alderton, Roberto Amato, Jeffrey Barrett, Sonia Goncalves, Ewan Harrison, David K. Jackson, Ian Johnston, Dominic Kwiatkowski, Cordelia Langford, John Sillitoe on behalf of the Wellcome Sanger Institute COVID-19 Surveillance Team |
| EPI_ISL_2394098, EPI_ISL_2394099, EPI_ISL_2394102, EPI_ISL_2394104, EPI_ISL_2394106, EPI_ISL_2394107, EPI_ISL_2394109, EPI_ISL_2394110, EPI_ISL_2394111, EPI_ISL_2394112, EPI_ISL_2394113, EPI_ISL_2394114 | | | |
| see above | Lighthouse Lab in Alderley Park | Wellcome Sanger Institute for the COVID-19 Genomics UK (COG-UK) Consortium | Jacquelyn Wynn, Mairead Hyland, The Lighthouse Lab in Alderley Park and Alex Alderton, Roberto Amato, Jeffrey Barrett, Sonia Goncalves, Ewan Harrison, David K. Jackson, Ian Johnston, Dominic Kwiatkowski, Cordelia Langford, John Sillitoe on behalf of the Wellcome Sanger Institute COVID-19 Surveillance Team |
| EPI_ISL_2394115 | Lighthouse Lab in Glasgow | Wellcome Sanger Institute for the COVID-19 Genomics UK (COG-UK) Consortium | Harper VanSteenhouse, Yumi Kasai, David Gray, Carol Clugston, Anna Dominiczak and Alex Alderton, Roberto Amato, Jeffrey Barrett, Sonia Goncalves, Ewan Harrison, David K. Jackson, Ian Johnston, Dominic Kwiatkowski, Cordelia Langford, John Sillitoe on behalf of the Wellcome Sanger Institute COVID-19 Surveillance Team |
| EPI_ISL_2394116, EPI_ISL_2394117, EPI_ISL_2394119, EPI_ISL_2394120, EPI_ISL_2394121, EPI_ISL_2394122, EPI_ISL_2394123, EPI_ISL_2394124, EPI_ISL_2394125, EPI_ISL_2394126, EPI_ISL_2394127, EPI_ISL_2394128, EPI_ISL_2394129, EPI_ISL_2394131, EPI_ISL_2394132, EPI_ISL_2394133, EPI_ISL_2394134, EPI_ISL_2394135, EPI_ISL_2394136 | | | |
| see above | Lighthouse Lab in Alderley Park | Wellcome Sanger Institute for the COVID-19 Genomics UK (COG-UK) Consortium | Jacquelyn Wynn, Mairead Hyland, The Lighthouse Lab in Alderley Park and Alex Alderton, Roberto Amato, Jeffrey Barrett, Sonia Goncalves, Ewan Harrison, David K. Jackson, Ian Johnston, Dominic Kwiatkowski, Cordelia Langford, John Sillitoe on behalf of the Wellcome Sanger Institute COVID-19 Surveillance Team |
| EPI_ISL_2394137 | Lighthouse Lab in Glasgow | Wellcome Sanger Institute for the COVID-19 Genomics UK (COG-UK) Consortium | Harper VanSteenhouse, Yumi Kasai, David Gray, Carol Clugston, Anna Dominiczak and Alex Alderton, Roberto Amato, Jeffrey Barrett, Sonia Goncalves, Ewan Harrison, David K. Jackson, Ian Johnston, Dominic Kwiatkowski, Cordelia Langford, John Sillitoe on behalf of the Wellcome Sanger Institute COVID-19 Surveillance Team |
| EPI_ISL_2394140, EPI_ISL_2394141, EPI_ISL_2394142, EPI_ISL_2394143, EPI_ISL_2394144, EPI_ISL_2394145 | Lighthouse Lab in Alderley Park | Wellcome Sanger Institute for the COVID-19 Genomics UK (COG-UK) Consortium | Jacquelyn Wynn, Mairead Hyland, The Lighthouse Lab in Alderley Park and Alex Alderton, Roberto Amato, Jeffrey Barrett, Sonia Goncalves, Ewan Harrison, David K. Jackson, Ian Johnston, Dominic Kwiatkowski, Cordelia Langford, John Sillitoe on behalf of the Wellcome Sanger Institute COVID-19 Surveillance Team |
| EPI_ISL_2394146 | Lighthouse Lab in Glasgow | Wellcome Sanger Institute for the COVID-19 Genomics UK (COG-UK) Consortium | Harper VanSteenhouse, Yumi Kasai, David Gray, Carol Clugston, Anna Dominiczak and Alex Alderton, Roberto Amato, Jeffrey Barrett, Sonia Goncalves, Ewan Harrison, David K. Jackson, Ian Johnston, Dominic Kwiatkowski, Cordelia Langford, John Sillitoe on behalf of the Wellcome Sanger Institute COVID-19 Surveillance Team |
| EPI_ISL_2394147, EPI_ISL_2394148, EPI_ISL_2394149, EPI_ISL_2394150, EPI_ISL_2394151, EPI_ISL_2394152, EPI_ISL_2394154, EPI_ISL_2394155, EPI_ISL_2394156, EPI_ISL_2394157, EPI_ISL_2394158, EPI_ISL_2394159, EPI_ISL_2394160, EPI_ISL_2394161, EPI_ISL_2394162, EPI_ISL_2394165, EPI_ISL_2394166, EPI_ISL_2394168, EPI_ISL_2394169 | | | |
| see above | Lighthouse Lab in Alderley Park | Wellcome Sanger Institute for the COVID-19 Genomics UK (COG-UK) Consortium | Jacquelyn Wynn, Mairead Hyland, The Lighthouse Lab in Alderley Park and Alex Alderton, Roberto Amato, Jeffrey Barrett, Sonia Goncalves, Ewan Harrison, David K. Jackson, Ian Johnston, Dominic Kwiatkowski, Cordelia Langford, John Sillitoe on behalf of the Wellcome Sanger Institute COVID-19 Surveillance Team |
| EPI_ISL_2394170 | Lighthouse Lab in Glasgow | Wellcome Sanger Institute for the COVID-19 Genomics UK (COG-UK) Consortium | Harper VanSteenhouse, Yumi Kasai, David Gray, Carol Clugston, Anna Dominiczak and Alex Alderton, Roberto Amato, Jeffrey Barrett, Sonia Goncalves, Ewan Harrison, David K. Jackson, Ian Johnston, Dominic Kwiatkowski, Cordelia Langford, John Sillitoe on behalf of the Wellcome Sanger Institute COVID-19 Surveillance Team |
| EPI_ISL_2394172, EPI_ISL_2394173, EPI_ISL_2394174, EPI_ISL_2394175 | Lighthouse Lab in Alderley Park | Wellcome Sanger Institute for the COVID-19 Genomics UK (COG-UK) Consortium | Jacquelyn Wynn, Mairead Hyland, The Lighthouse Lab in Alderley Park and Alex Alderton, Roberto Amato, Jeffrey Barrett, Sonia Goncalves, Ewan Harrison, David K. Jackson, Ian Johnston, Dominic Kwiatkowski, Cordelia Langford, John Sillitoe on behalf of the Wellcome Sanger Institute COVID-19 Surveillance Team |
| EPI_ISL_2394176 | Lighthouse Lab in Glasgow | Wellcome Sanger Institute for the COVID-19 Genomics UK (COG-UK) Consortium | Harper VanSteenhouse, Yumi Kasai, David Gray, Carol Clugston, Anna Dominiczak and Alex Alderton, Roberto Amato, Jeffrey Barrett, Sonia Goncalves, Ewan Harrison, David K. Jackson, Ian Johnston, Dominic Kwiatkowski, Cordelia Langford, John Sillitoe on behalf of the Wellcome Sanger Institute COVID-19 Surveillance Team |
| EPI_ISL_2394177, EPI_ISL_2394178, EPI_ISL_2394179 | Lighthouse Lab in Alderley Park | Wellcome Sanger Institute for the COVID-19 Genomics UK (COG-UK) Consortium | Jacquelyn Wynn, Mairead Hyland, The Lighthouse Lab in Alderley Park and Alex Alderton, Roberto Amato, Jeffrey Barrett, Sonia Goncalves, Ewan Harrison, David K. Jackson, Ian Johnston, Dominic Kwiatkowski, Cordelia Langford, John Sillitoe on behalf of the Wellcome Sanger Institute COVID-19 Surveillance Team |
| EPI_ISL_2394180 | Lighthouse Lab in Glasgow | Wellcome Sanger Institute for the COVID-19 Genomics UK (COG-UK) Consortium | Harper VanSteenhouse, Yumi Kasai, David Gray, Carol Clugston, Anna Dominiczak and Alex Alderton, Roberto Amato, Jeffrey Barrett, Sonia Goncalves, Ewan Harrison, David K. Jackson, Ian Johnston, Dominic Kwiatkowski, Cordelia Langford, John Sillitoe on behalf of the Wellcome Sanger Institute COVID-19 Surveillance Team |
| EPI_ISL_2394181, EPI_ISL_2394182, EPI_ISL_2394183, EPI_ISL_2394184, EPI_ISL_2394185, EPI_ISL_2394186, EPI_ISL_2394187, EPI_ISL_2394188, EPI_ISL_2394189, EPI_ISL_2394190, EPI_ISL_2394191, EPI_ISL_2394192, EPI_ISL_2394193, EPI_ISL_2394194, EPI_ISL_2394195, EPI_ISL_2394196, EPI_ISL_2394197, EPI_ISL_2394198, EPI_ISL_2394200, EPI_ISL_2394201, EPI_ISL_2394202, EPI_ISL_2394204, EPI_ISL_2394205, EPI_ISL_2394206, EPI_ISL_2394207, EPI_ISL_2394208, EPI_ISL_2394209, EPI_ISL_2394210, EPI_ISL_2394211, EPI_ISL_2394212, EPI_ISL_2394213, EPI_ISL_2394214, EPI_ISL_2394215 | | | |
| see above | Lighthouse Lab in Alderley Park | Wellcome Sanger Institute for the COVID-19 Genomics UK (COG-UK) Consortium | Jacquelyn Wynn, Mairead Hyland, The Lighthouse Lab in Alderley Park and Alex Alderton, Roberto Amato, Jeffrey Barrett, Sonia Goncalves, Ewan Harrison, David K. Jackson, Ian Johnston, Dominic Kwiatkowski, Cordelia Langford, John Sillitoe on behalf of the Wellcome Sanger Institute COVID-19 Surveillance Team |
| EPI_ISL_2394216 | Lighthouse Lab in Glasgow | Wellcome Sanger Institute for the COVID-19 Genomics UK (COG-UK) Consortium | Harper VanSteenhouse, Yumi Kasai, David Gray, Carol Clugston, Anna Dominiczak and Alex Alderton, Roberto Amato, Jeffrey Barrett, Sonia Goncalves, Ewan Harrison, David K. Jackson, Ian Johnston, Dominic Kwiatkowski, Cordelia Langford, John Sillitoe on behalf of the Wellcome Sanger Institute COVID-19 Surveillance Team |
| EPI_ISL_2394217, EPI_ISL_2394218, EPI_ISL_2394219, EPI_ISL_2394223, EPI_ISL_2394224, EPI_ISL_2394225, EPI_ISL_2394226, EPI_ISL_2394227, EPI_ISL_2394228, EPI_ISL_2394230, EPI_ISL_2394231, EPI_ISL_2394232, EPI_ISL_2394233, EPI_ISL_2394234, EPI_ISL_2394235, EPI_ISL_2394236, EPI_ISL_2394237, EPI_ISL_2394238, EPI_ISL_2394239, EPI_ISL_2394240, EPI_ISL_2394241 | | | |
| see above | Lighthouse Lab in Alderley Park | Wellcome Sanger Institute for the COVID-19 Genomics UK (COG-UK) Consortium | Jacquelyn Wynn, Mairead Hyland, The Lighthouse Lab in Alderley Park and Alex Alderton, Roberto Amato, Jeffrey Barrett, Sonia Goncalves, Ewan Harrison, David K. Jackson, Ian Johnston, Dominic Kwiatkowski, Cordelia Langford, John Sillitoe on behalf of the Wellcome Sanger Institute COVID-19 Surveillance Team |
| EPI_ISL_2394242 | Lighthouse Lab in Glasgow | Wellcome Sanger Institute for the COVID-19 Genomics UK (COG-UK) Consortium | Harper VanSteenhouse, Yumi Kasai, David Gray, Carol Clugston, Anna Dominiczak and Alex Alderton, Roberto Amato, Jeffrey Barrett, Sonia Goncalves, Ewan Harrison, David K. Jackson, Ian Johnston, Dominic Kwiatkowski, Cordelia Langford, John Sillitoe on behalf of the Wellcome Sanger Institute COVID-19 Surveillance Team |
| EPI_ISL_2394243, EPI_ISL_2394244, EPI_ISL_2394245 | Lighthouse Lab in Alderley Park | Wellcome Sanger Institute for the COVID-19 Genomics UK (COG-UK) Consortium | Jacquelyn Wynn, Mairead Hyland, The Lighthouse Lab in Alderley Park and Alex Alderton, Roberto Amato, Jeffrey Barrett, Sonia Goncalves, Ewan Harrison, David K. Jackson, Ian Johnston, Dominic Kwiatkowski, Cordelia Langford, John Sillitoe on behalf of the Wellcome Sanger Institute COVID-19 Surveillance Team |
| EPI_ISL_2394246, EPI_ISL_2394247 | Lighthouse Lab in Glasgow | Wellcome Sanger Institute for the COVID-19 Genomics UK (COG-UK) Consortium | Harper VanSteenhouse, Yumi Kasai, David Gray, Carol Clugston, Anna Dominiczak and Alex Alderton, Roberto Amato, Jeffrey Barrett, Sonia Goncalves, Ewan Harrison, David K. Jackson, Ian Johnston, Dominic Kwiatkowski, Cordelia Langford, John Sillitoe on behalf of the Wellcome Sanger Institute COVID-19 Surveillance Team |
| EPI_ISL_2394248, EPI_ISL_2394249, EPI_ISL_2394250, EPI_ISL_2394251, EPI_ISL_2394253, EPI_ISL_2394254, EPI_ISL_2394255 | Lighthouse Lab in Alderley Park | Wellcome Sanger Institute for the COVID-19 Genomics UK (COG-UK) Consortium | Jacquelyn Wynn, Mairead Hyland, The Lighthouse Lab in Alderley Park and Alex Alderton, Roberto Amato, Jeffrey Barrett, Sonia Goncalves, Ewan Harrison, David K. Jackson, Ian Johnston, Dominic Kwiatkowski, Cordelia Langford, John Sillitoe on behalf of the Wellcome Sanger Institute COVID-19 Surveillance Team |
| EPI_ISL_2394256 | Lighthouse Lab in Glasgow | Wellcome Sanger Institute for the COVID-19 Genomics UK (COG-UK) Consortium | Harper VanSteenhouse, Yumi Kasai, David Gray, Carol Clugston, Anna Dominiczak and Alex Alderton, Roberto Amato, Jeffrey Barrett, Sonia Goncalves, Ewan Harrison, David K. Jackson, Ian Johnston, Dominic Kwiatkowski, Cordelia Langford, John Sillitoe on behalf of the Wellcome Sanger Institute COVID-19 Surveillance Team |
| EPI_ISL_2394257, EPI_ISL_2394258, EPI_ISL_2394259 | Lighthouse Lab in Alderley Park | Wellcome Sanger Institute for the COVID-19 Genomics UK (COG-UK) Consortium | Jacquelyn Wynn, Mairead Hyland, The Lighthouse Lab in Alderley Park and Alex Alderton, Roberto Amato, Jeffrey Barrett, Sonia Goncalves, Ewan Harrison, David K. Jackson, Ian Johnston, Dominic Kwiatkowski, Cordelia Langford, John Sillitoe on behalf of the Wellcome Sanger Institute COVID-19 Surveillance Team |
| EPI_ISL_2394260 | Lighthouse Lab in Glasgow | Wellcome Sanger Institute for the COVID-19 Genomics UK (COG-UK) Consortium | Harper VanSteenhouse, Yumi Kasai, David Gray, Carol Clugston, Anna Dominiczak and Alex Alderton, Roberto Amato, Jeffrey Barrett, Sonia Goncalves, Ewan Harrison, David K. Jackson, Ian Johnston, Dominic Kwiatkowski, Cordelia Langford, John Sillitoe on behalf of the Wellcome Sanger Institute COVID-19 Surveillance Team |
| EPI_ISL_2394261, EPI_ISL_2394262, EPI_ISL_2394264, EPI_ISL_2394265, EPI_ISL_2394266, EPI_ISL_2394267, EPI_ISL_2394268, EPI_ISL_2394269, EPI_ISL_2394270, EPI_ISL_2394271 | Lighthouse Lab in Alderley Park | Wellcome Sanger Institute for the COVID-19 Genomics UK (COG-UK) Consortium | Jacquelyn Wynn, Mairead Hyland, The Lighthouse Lab in Alderley Park and Alex Alderton, Roberto Amato, Jeffrey Barrett, Sonia Goncalves, Ewan Harrison, David K. Jackson, Ian Johnston, Dominic Kwiatkowski, Cordelia Langford, John Sillitoe on behalf of the Wellcome Sanger Institute COVID-19 Surveillance Team |
| EPI_ISL_2394272 | Lighthouse Lab in Glasgow | Wellcome Sanger Institute for the COVID-19 Genomics UK (COG-UK) Consortium | Harper VanSteenhouse, Yumi Kasai, David Gray, Carol Clugston, Anna Dominiczak and Alex Alderton, Roberto Amato, Jeffrey Barrett, Sonia Goncalves, Ewan Harrison, David K. Jackson, Ian Johnston, Dominic Kwiatkowski, Cordelia Langford, John Sillitoe on behalf of the Wellcome Sanger Institute COVID-19 Surveillance Team |
| EPI_ISL_2394273, | Lighthouse Lab in Alderley Park | Wellcome Sanger Institute for the COVID-19 Genomics UK (COG-UK) | Jacquelyn Wynn, Mairead Hyland, The Lighthouse Lab in Alderley Park and Alex Alderton, Roberto Amato, Jeffrey Barrett, Sonia Goncalves, Ewan Harrison, David K. Jackson, Ian Johnston, Dominic Kwiatkowski, Cordelia Langford, John Sillitoe on |

| | | | |
|---|---|---|---|
| EPI_ISL_2394274, EPI_ISL_2394275, EPI_ISL_2394276, EPI_ISL_2394277, EPI_ISL_2394278, EPI_ISL_2394279, EPI_ISL_2394280, EPI_ISL_2394281 | | Consortium | behalf of the Wellcome Sanger Institute COVID-19 Surveillance Team |
| EPI_ISL_2394282 | Lighthouse Lab in Glasgow | Wellcome Sanger Institute for the COVID-19 Genomics UK (COG-UK) Consortium | Harper VanSteenhouse, Yumi Kasai, David Gray, Carol Clugston, Anna Dominiczak and Alex Alderton, Roberto Amato, Jeffrey Barrett, Sonia Goncalves, Ewan Harrison, David K. Jackson, Ian Johnston, Dominic Kwiatkowski, Cordelia Langford, John Sillitoe on behalf of the Wellcome Sanger Institute COVID-19 Surveillance Team |
| EPI_ISL_2394283, EPI_ISL_2394284, EPI_ISL_2394285, EPI_ISL_2394286, EPI_ISL_2394287, EPI_ISL_2394288, EPI_ISL_2394289, EPI_ISL_2394290, EPI_ISL_2394291, EPI_ISL_2394294, EPI_ISL_2394295, EPI_ISL_2394296, EPI_ISL_2394297, EPI_ISL_2394299, EPI_ISL_2394300, EPI_ISL_2394301, EPI_ISL_2394302 | | | |
| see above | Lighthouse Lab in Alderley Park | Wellcome Sanger Institute for the COVID-19 Genomics UK (COG-UK) Consortium | Jacquelyn Wynn, Mairead Hyland, The Lighthouse Lab in Alderley Park and Alex Alderton, Roberto Amato, Jeffrey Barrett, Sonia Goncalves, Ewan Harrison, David K. Jackson, Ian Johnston, Dominic Kwiatkowski, Cordelia Langford, John Sillitoe on behalf of the Wellcome Sanger Institute COVID-19 Surveillance Team |
| EPI_ISL_2394303 | Lighthouse Lab in Glasgow | Wellcome Sanger Institute for the COVID-19 Genomics UK (COG-UK) Consortium | Harper VanSteenhouse, Yumi Kasai, David Gray, Carol Clugston, Anna Dominiczak and Alex Alderton, Roberto Amato, Jeffrey Barrett, Sonia Goncalves, Ewan Harrison, David K. Jackson, Ian Johnston, Dominic Kwiatkowski, Cordelia Langford, John Sillitoe on behalf of the Wellcome Sanger Institute COVID-19 Surveillance Team |
| EPI_ISL_2394304, EPI_ISL_2394305, EPI_ISL_2394306, EPI_ISL_2394307, EPI_ISL_2394308, EPI_ISL_2394309, EPI_ISL_2394310, EPI_ISL_2394311 | Lighthouse Lab in Alderley Park | Wellcome Sanger Institute for the COVID-19 Genomics UK (COG-UK) Consortium | Jacquelyn Wynn, Mairead Hyland, The Lighthouse Lab in Alderley Park and Alex Alderton, Roberto Amato, Jeffrey Barrett, Sonia Goncalves, Ewan Harrison, David K. Jackson, Ian Johnston, Dominic Kwiatkowski, Cordelia Langford, John Sillitoe on behalf of the Wellcome Sanger Institute COVID-19 Surveillance Team |
| EPI_ISL_2394312 | Lighthouse Lab in Glasgow | Wellcome Sanger Institute for the COVID-19 Genomics UK (COG-UK) Consortium | Harper VanSteenhouse, Yumi Kasai, David Gray, Carol Clugston, Anna Dominiczak and Alex Alderton, Roberto Amato, Jeffrey Barrett, Sonia Goncalves, Ewan Harrison, David K. Jackson, Ian Johnston, Dominic Kwiatkowski, Cordelia Langford, John Sillitoe on behalf of the Wellcome Sanger Institute COVID-19 Surveillance Team |
| EPI_ISL_2394313, EPI_ISL_2394314, EPI_ISL_2394315, EPI_ISL_2394317, EPI_ISL_2394318, EPI_ISL_2394319, EPI_ISL_2394320 | Lighthouse Lab in Alderley Park | Wellcome Sanger Institute for the COVID-19 Genomics UK (COG-UK) Consortium | Jacquelyn Wynn, Mairead Hyland, The Lighthouse Lab in Alderley Park and Alex Alderton, Roberto Amato, Jeffrey Barrett, Sonia Goncalves, Ewan Harrison, David K. Jackson, Ian Johnston, Dominic Kwiatkowski, Cordelia Langford, John Sillitoe on behalf of the Wellcome Sanger Institute COVID-19 Surveillance Team |
| EPI_ISL_2394321, EPI_ISL_2394322, EPI_ISL_2394323 | Lighthouse Lab in Glasgow | Wellcome Sanger Institute for the COVID-19 Genomics UK (COG-UK) Consortium | Harper VanSteenhouse, Yumi Kasai, David Gray, Carol Clugston, Anna Dominiczak and Alex Alderton, Roberto Amato, Jeffrey Barrett, Sonia Goncalves, Ewan Harrison, David K. Jackson, Ian Johnston, Dominic Kwiatkowski, Cordelia Langford, John Sillitoe on behalf of the Wellcome Sanger Institute COVID-19 Surveillance Team |
| EPI_ISL_2394324, EPI_ISL_2394325, EPI_ISL_2394326 | Lighthouse Lab in Alderley Park | Wellcome Sanger Institute for the COVID-19 Genomics UK (COG-UK) Consortium | Jacquelyn Wynn, Mairead Hyland, The Lighthouse Lab in Alderley Park and Alex Alderton, Roberto Amato, Jeffrey Barrett, Sonia Goncalves, Ewan Harrison, David K. Jackson, Ian Johnston, Dominic Kwiatkowski, Cordelia Langford, John Sillitoe on behalf of the Wellcome Sanger Institute COVID-19 Surveillance Team |
| EPI_ISL_2394327, EPI_ISL_2394328 | Lighthouse Lab in Glasgow | Wellcome Sanger Institute for the COVID-19 Genomics UK (COG-UK) Consortium | Harper VanSteenhouse, Yumi Kasai, David Gray, Carol Clugston, Anna Dominiczak and Alex Alderton, Roberto Amato, Jeffrey Barrett, Sonia Goncalves, Ewan Harrison, David K. Jackson, Ian Johnston, Dominic Kwiatkowski, Cordelia Langford, John Sillitoe on behalf of the Wellcome Sanger Institute COVID-19 Surveillance Team |
| EPI_ISL_2394329, EPI_ISL_2394330, EPI_ISL_2394331, EPI_ISL_2394332, EPI_ISL_2394333 | Lighthouse Lab in Alderley Park | Wellcome Sanger Institute for the COVID-19 Genomics UK (COG-UK) Consortium | Jacquelyn Wynn, Mairead Hyland, The Lighthouse Lab in Alderley Park and Alex Alderton, Roberto Amato, Jeffrey Barrett, Sonia Goncalves, Ewan Harrison, David K. Jackson, Ian Johnston, Dominic Kwiatkowski, Cordelia Langford, John Sillitoe on behalf of the Wellcome Sanger Institute COVID-19 Surveillance Team |
| EPI_ISL_2394334 | Lighthouse Lab in Glasgow | Wellcome Sanger Institute for the COVID-19 Genomics UK (COG-UK) Consortium | Harper VanSteenhouse, Yumi Kasai, David Gray, Carol Clugston, Anna Dominiczak and Alex Alderton, Roberto Amato, Jeffrey Barrett, Sonia Goncalves, Ewan Harrison, David K. Jackson, Ian Johnston, Dominic Kwiatkowski, Cordelia Langford, John Sillitoe on behalf of the Wellcome Sanger Institute COVID-19 Surveillance Team |
| EPI_ISL_2394335 | Lighthouse Lab in Alderley Park | Wellcome Sanger Institute for the COVID-19 Genomics UK (COG-UK) Consortium | Jacquelyn Wynn, Mairead Hyland, The Lighthouse Lab in Alderley Park and Alex Alderton, Roberto Amato, Jeffrey Barrett, Sonia Goncalves, Ewan Harrison, David K. Jackson, Ian Johnston, Dominic Kwiatkowski, Cordelia Langford, John Sillitoe on behalf of the Wellcome Sanger Institute COVID-19 Surveillance Team |
| EPI_ISL_2394336 | Lighthouse Lab in Glasgow | Wellcome Sanger Institute for the COVID-19 Genomics UK (COG-UK) Consortium | Harper VanSteenhouse, Yumi Kasai, David Gray, Carol Clugston, Anna Dominiczak and Alex Alderton, Roberto Amato, Jeffrey Barrett, Sonia Goncalves, Ewan Harrison, David K. Jackson, Ian Johnston, Dominic Kwiatkowski, Cordelia Langford, John Sillitoe on behalf of the Wellcome Sanger Institute COVID-19 Surveillance Team |
| EPI_ISL_2394337, EPI_ISL_2394339, EPI_ISL_2394340 | Lighthouse Lab in Alderley Park | Wellcome Sanger Institute for the COVID-19 Genomics UK (COG-UK) Consortium | Jacquelyn Wynn, Mairead Hyland, The Lighthouse Lab in Alderley Park and Alex Alderton, Roberto Amato, Jeffrey Barrett, Sonia Goncalves, Ewan Harrison, David K. Jackson, Ian Johnston, Dominic Kwiatkowski, Cordelia Langford, John Sillitoe on behalf of the Wellcome Sanger Institute COVID-19 Surveillance Team |
| EPI_ISL_2394341, EPI_ISL_2394342 | Lighthouse Lab in Glasgow | Wellcome Sanger Institute for the COVID-19 Genomics UK (COG-UK) Consortium | Harper VanSteenhouse, Yumi Kasai, David Gray, Carol Clugston, Anna Dominiczak and Alex Alderton, Roberto Amato, Jeffrey Barrett, Sonia Goncalves, Ewan Harrison, David K. Jackson, Ian Johnston, Dominic Kwiatkowski, Cordelia Langford, John Sillitoe on behalf of the Wellcome Sanger Institute COVID-19 Surveillance Team |
| EPI_ISL_2394343 | Lighthouse Lab in Alderley Park | Wellcome Sanger Institute for the COVID-19 Genomics UK (COG-UK) Consortium | Jacquelyn Wynn, Mairead Hyland, The Lighthouse Lab in Alderley Park and Alex Alderton, Roberto Amato, Jeffrey Barrett, Sonia Goncalves, Ewan Harrison, David K. Jackson, Ian Johnston, Dominic Kwiatkowski, Cordelia Langford, John Sillitoe on behalf of the Wellcome Sanger Institute COVID-19 Surveillance Team |
| EPI_ISL_2394346 | Lighthouse Lab in Glasgow | Wellcome Sanger Institute for the COVID-19 Genomics UK (COG-UK) Consortium | Harper VanSteenhouse, Yumi Kasai, David Gray, Carol Clugston, Anna Dominiczak and Alex Alderton, Roberto Amato, Jeffrey Barrett, Sonia Goncalves, Ewan Harrison, David K. Jackson, Ian Johnston, Dominic Kwiatkowski, Cordelia Langford, John Sillitoe on behalf of the Wellcome Sanger Institute COVID-19 Surveillance Team |
| EPI_ISL_2394348, EPI_ISL_2394349 | Lighthouse Lab in Alderley Park | Wellcome Sanger Institute for the COVID-19 Genomics UK (COG-UK) Consortium | Jacquelyn Wynn, Mairead Hyland, The Lighthouse Lab in Alderley Park and Alex Alderton, Roberto Amato, Jeffrey Barrett, Sonia Goncalves, Ewan Harrison, David K. Jackson, Ian Johnston, Dominic Kwiatkowski, Cordelia Langford, John Sillitoe on behalf of the Wellcome Sanger Institute COVID-19 Surveillance Team |
| EPI_ISL_2394350 | Lighthouse Lab in Glasgow | Wellcome Sanger Institute for the COVID-19 Genomics UK (COG-UK) Consortium | Harper VanSteenhouse, Yumi Kasai, David Gray, Carol Clugston, Anna Dominiczak and Alex Alderton, Roberto Amato, Jeffrey Barrett, Sonia Goncalves, Ewan Harrison, David K. Jackson, Ian Johnston, Dominic Kwiatkowski, Cordelia Langford, John Sillitoe on behalf of the Wellcome Sanger Institute COVID-19 Surveillance Team |
| EPI_ISL_2394351 | Lighthouse Lab in Alderley Park | Wellcome Sanger Institute for the COVID-19 Genomics UK (COG-UK) Consortium | Jacquelyn Wynn, Mairead Hyland, The Lighthouse Lab in Alderley Park and Alex Alderton, Roberto Amato, Jeffrey Barrett, Sonia Goncalves, Ewan Harrison, David K. Jackson, Ian Johnston, Dominic Kwiatkowski, Cordelia Langford, John Sillitoe on behalf of the Wellcome Sanger Institute COVID-19 Surveillance Team |
| EPI_ISL_2394353, EPI_ISL_2394354 | Lighthouse Lab in Glasgow | Wellcome Sanger Institute for the COVID-19 Genomics UK (COG-UK) Consortium | Harper VanSteenhouse, Yumi Kasai, David Gray, Carol Clugston, Anna Dominiczak and Alex Alderton, Roberto Amato, Jeffrey Barrett, Sonia Goncalves, Ewan Harrison, David K. Jackson, Ian Johnston, Dominic Kwiatkowski, Cordelia Langford, John Sillitoe on behalf of the Wellcome Sanger Institute COVID-19 Surveillance Team |
| EPI_ISL_2394355 | Lighthouse Lab in Alderley Park | Wellcome Sanger Institute for the COVID-19 Genomics UK (COG-UK) Consortium | Jacquelyn Wynn, Mairead Hyland, The Lighthouse Lab in Alderley Park and Alex Alderton, Roberto Amato, Jeffrey Barrett, Sonia Goncalves, Ewan Harrison, David K. Jackson, Ian Johnston, Dominic Kwiatkowski, Cordelia Langford, John Sillitoe on behalf of the Wellcome Sanger Institute COVID-19 Surveillance Team |
| EPI_ISL_2394356 | Lighthouse Lab in Glasgow | Wellcome Sanger Institute for the COVID-19 Genomics UK (COG-UK) Consortium | Harper VanSteenhouse, Yumi Kasai, David Gray, Carol Clugston, Anna Dominiczak and Alex Alderton, Roberto Amato, Jeffrey Barrett, Sonia Goncalves, Ewan Harrison, David K. Jackson, Ian Johnston, Dominic Kwiatkowski, Cordelia Langford, John Sillitoe on behalf of the Wellcome Sanger Institute COVID-19 Surveillance Team |
| EPI_ISL_2394357, EPI_ISL_2394358, EPI_ISL_2394359 | Lighthouse Lab in Alderley Park | Wellcome Sanger Institute for the COVID-19 Genomics UK (COG-UK) Consortium | Jacquelyn Wynn, Mairead Hyland, The Lighthouse Lab in Alderley Park and Alex Alderton, Roberto Amato, Jeffrey Barrett, Sonia Goncalves, Ewan Harrison, David K. Jackson, Ian Johnston, Dominic Kwiatkowski, Cordelia Langford, John Sillitoe on behalf of the Wellcome Sanger Institute COVID-19 Surveillance Team |
| EPI_ISL_2394360 | Lighthouse Lab in Glasgow | Wellcome Sanger Institute for the COVID-19 Genomics UK (COG-UK) Consortium | Harper VanSteenhouse, Yumi Kasai, David Gray, Carol Clugston, Anna Dominiczak and Alex Alderton, Roberto Amato, Jeffrey Barrett, Sonia Goncalves, Ewan Harrison, David K. Jackson, Ian Johnston, Dominic Kwiatkowski, Cordelia Langford, John Sillitoe on behalf of the Wellcome Sanger Institute COVID-19 Surveillance Team |
| EPI_ISL_2394361, EPI_ISL_2394362 | Lighthouse Lab in Alderley Park | Wellcome Sanger Institute for the COVID-19 Genomics UK (COG-UK) Consortium | Jacquelyn Wynn, Mairead Hyland, The Lighthouse Lab in Alderley Park and Alex Alderton, Roberto Amato, Jeffrey Barrett, Sonia Goncalves, Ewan Harrison, David K. Jackson, Ian Johnston, Dominic Kwiatkowski, Cordelia Langford, John Sillitoe on behalf of the Wellcome Sanger Institute COVID-19 Surveillance Team |
| EPI_ISL_2394363 | Lighthouse Lab in Glasgow | Wellcome Sanger Institute for the COVID-19 Genomics UK (COG-UK) Consortium | Harper VanSteenhouse, Yumi Kasai, David Gray, Carol Clugston, Anna Dominiczak and Alex Alderton, Roberto Amato, Jeffrey Barrett, Sonia Goncalves, Ewan Harrison, David K. Jackson, Ian Johnston, Dominic Kwiatkowski, Cordelia Langford, John Sillitoe on behalf of the Wellcome Sanger Institute COVID-19 Surveillance Team |
| EPI_ISL_2394364, EPI_ISL_2394365, EPI_ISL_2394366, EPI_ISL_2394368 | Lighthouse Lab in Alderley Park | Wellcome Sanger Institute for the COVID-19 Genomics UK (COG-UK) Consortium | Jacquelyn Wynn, Mairead Hyland, The Lighthouse Lab in Alderley Park and Alex Alderton, Roberto Amato, Jeffrey Barrett, Sonia Goncalves, Ewan Harrison, David K. Jackson, Ian Johnston, Dominic Kwiatkowski, Cordelia Langford, John Sillitoe on behalf of the Wellcome Sanger Institute COVID-19 Surveillance Team |
| EPI_ISL_2394369 | Lighthouse Lab in Glasgow | Wellcome Sanger Institute for the COVID-19 Genomics UK (COG-UK) Consortium | Harper VanSteenhouse, Yumi Kasai, David Gray, Carol Clugston, Anna Dominiczak and Alex Alderton, Roberto Amato, Jeffrey Barrett, Sonia Goncalves, Ewan Harrison, David K. Jackson, Ian Johnston, Dominic Kwiatkowski, Cordelia Langford, John Sillitoe on behalf of the Wellcome Sanger Institute COVID-19 Surveillance Team |
| EPI_ISL_2394370, EPI_ISL_2394371 | Lighthouse Lab in Alderley Park | Wellcome Sanger Institute for the COVID-19 Genomics UK (COG-UK) Consortium | Jacquelyn Wynn, Mairead Hyland, The Lighthouse Lab in Alderley Park and Alex Alderton, Roberto Amato, Jeffrey Barrett, Sonia Goncalves, Ewan Harrison, David K. Jackson, Ian Johnston, Dominic Kwiatkowski, Cordelia Langford, John Sillitoe on behalf of the Wellcome Sanger Institute COVID-19 Surveillance Team |
| EPI_ISL_2394372 | Lighthouse Lab in Glasgow | Wellcome Sanger Institute for the COVID-19 Genomics UK (COG-UK) Consortium | Harper VanSteenhouse, Yumi Kasai, David Gray, Carol Clugston, Anna Dominiczak and Alex Alderton, Roberto Amato, Jeffrey Barrett, Sonia Goncalves, Ewan Harrison, David K. Jackson, Ian Johnston, Dominic Kwiatkowski, Cordelia Langford, John Sillitoe on behalf of the Wellcome Sanger Institute COVID-19 Surveillance Team |
| EPI_ISL_2394373 | Lighthouse Lab in Alderley Park | Wellcome Sanger Institute for the COVID-19 Genomics UK (COG-UK) Consortium | Jacquelyn Wynn, Mairead Hyland, The Lighthouse Lab in Alderley Park and Alex Alderton, Roberto Amato, Jeffrey Barrett, Sonia Goncalves, Ewan Harrison, David K. Jackson, Ian Johnston, Dominic Kwiatkowski, Cordelia Langford, John Sillitoe on behalf of the Wellcome Sanger Institute COVID-19 Surveillance Team |
| EPI_ISL_2394374 | Lighthouse Lab in Glasgow | Wellcome Sanger Institute for the COVID-19 Genomics UK (COG-UK) Consortium | Harper VanSteenhouse, Yumi Kasai, David Gray, Carol Clugston, Anna Dominiczak and Alex Alderton, Roberto Amato, Jeffrey Barrett, Sonia Goncalves, Ewan Harrison, David K. Jackson, Ian Johnston, Dominic Kwiatkowski, Cordelia Langford, John Sillitoe on behalf of the Wellcome Sanger Institute COVID-19 Surveillance Team |

| | | | |
|---|---|---|---|
| EPI_ISL_2394375, EPI_ISL_2394376, EPI_ISL_2394377, EPI_ISL_2394378, EPI_ISL_2394379, EPI_ISL_2394381, EPI_ISL_2394382 | Lighthouse Lab in Alderley Park | Wellcome Sanger Institute for the COVID-19 Genomics UK (COG-UK) Consortium | Jacquelyn Wynn, Mairead Hyland, The Lighthouse Lab in Alderley Park and Alex Alderton, Roberto Amato, Jeffrey Barrett, Sonia Goncalves, Ewan Harrison, David K. Jackson, Ian Johnston, Dominic Kwiatkowski, Cordelia Langford, John Sillitoe on behalf of the Wellcome Sanger Institute COVID-19 Surveillance Team |
| EPI_ISL_2394383, EPI_ISL_2394384 | Lighthouse Lab in Glasgow | Wellcome Sanger Institute for the COVID-19 Genomics UK (COG-UK) Consortium | Harper VanSteenhouse, Yumi Kasai, David Gray, Carol Clugston, Anna Dominiczak and Alex Alderton, Roberto Amato, Jeffrey Barrett, Sonia Goncalves, Ewan Harrison, David K. Jackson, Ian Johnston, Dominic Kwiatkowski, Cordelia Langford, John Sillitoe on behalf of the Wellcome Sanger Institute COVID-19 Surveillance Team |
| EPI_ISL_2394385, EPI_ISL_2394387, EPI_ISL_2394388, EPI_ISL_2394389, EPI_ISL_2394390, EPI_ISL_2394392, EPI_ISL_2394393, EPI_ISL_2394394, EPI_ISL_2394395, EPI_ISL_2394396, EPI_ISL_2394397, EPI_ISL_2394398, EPI_ISL_2394399, EPI_ISL_2394400, EPI_ISL_2394402, EPI_ISL_2394403, EPI_ISL_2394404, EPI_ISL_2394405, EPI_ISL_2394406, EPI_ISL_2394407 | | | |
| see above | Lighthouse Lab in Alderley Park | Wellcome Sanger Institute for the COVID-19 Genomics UK (COG-UK) Consortium | Jacquelyn Wynn, Mairead Hyland, The Lighthouse Lab in Alderley Park and Alex Alderton, Roberto Amato, Jeffrey Barrett, Sonia Goncalves, Ewan Harrison, David K. Jackson, Ian Johnston, Dominic Kwiatkowski, Cordelia Langford, John Sillitoe on behalf of the Wellcome Sanger Institute COVID-19 Surveillance Team |
| EPI_ISL_2394408 | Lighthouse Lab in Glasgow | Wellcome Sanger Institute for the COVID-19 Genomics UK (COG-UK) Consortium | Harper VanSteenhouse, Yumi Kasai, David Gray, Carol Clugston, Anna Dominiczak and Alex Alderton, Roberto Amato, Jeffrey Barrett, Sonia Goncalves, Ewan Harrison, David K. Jackson, Ian Johnston, Dominic Kwiatkowski, Cordelia Langford, John Sillitoe on behalf of the Wellcome Sanger Institute COVID-19 Surveillance Team |
| EPI_ISL_2394409, EPI_ISL_2394410, EPI_ISL_2394411, EPI_ISL_2394413 | Lighthouse Lab in Alderley Park | Wellcome Sanger Institute for the COVID-19 Genomics UK (COG-UK) Consortium | Jacquelyn Wynn, Mairead Hyland, The Lighthouse Lab in Alderley Park and Alex Alderton, Roberto Amato, Jeffrey Barrett, Sonia Goncalves, Ewan Harrison, David K. Jackson, Ian Johnston, Dominic Kwiatkowski, Cordelia Langford, John Sillitoe on behalf of the Wellcome Sanger Institute COVID-19 Surveillance Team |
| EPI_ISL_2394414, EPI_ISL_2394415 | Lighthouse Lab in Glasgow | Wellcome Sanger Institute for the COVID-19 Genomics UK (COG-UK) Consortium | Harper VanSteenhouse, Yumi Kasai, David Gray, Carol Clugston, Anna Dominiczak and Alex Alderton, Roberto Amato, Jeffrey Barrett, Sonia Goncalves, Ewan Harrison, David K. Jackson, Ian Johnston, Dominic Kwiatkowski, Cordelia Langford, John Sillitoe on behalf of the Wellcome Sanger Institute COVID-19 Surveillance Team |
| EPI_ISL_2394418 | Lighthouse Lab in Alderley Park | Wellcome Sanger Institute for the COVID-19 Genomics UK (COG-UK) Consortium | Jacquelyn Wynn, Mairead Hyland, The Lighthouse Lab in Alderley Park and Alex Alderton, Roberto Amato, Jeffrey Barrett, Sonia Goncalves, Ewan Harrison, David K. Jackson, Ian Johnston, Dominic Kwiatkowski, Cordelia Langford, John Sillitoe on behalf of the Wellcome Sanger Institute COVID-19 Surveillance Team |
| EPI_ISL_2394419 | Lighthouse Lab in Glasgow | Wellcome Sanger Institute for the COVID-19 Genomics UK (COG-UK) Consortium | Harper VanSteenhouse, Yumi Kasai, David Gray, Carol Clugston, Anna Dominiczak and Alex Alderton, Roberto Amato, Jeffrey Barrett, Sonia Goncalves, Ewan Harrison, David K. Jackson, Ian Johnston, Dominic Kwiatkowski, Cordelia Langford, John Sillitoe on behalf of the Wellcome Sanger Institute COVID-19 Surveillance Team |
| EPI_ISL_2394420 | Lighthouse Lab in Alderley Park | Wellcome Sanger Institute for the COVID-19 Genomics UK (COG-UK) Consortium | Jacquelyn Wynn, Mairead Hyland, The Lighthouse Lab in Alderley Park and Alex Alderton, Roberto Amato, Jeffrey Barrett, Sonia Goncalves, Ewan Harrison, David K. Jackson, Ian Johnston, Dominic Kwiatkowski, Cordelia Langford, John Sillitoe on behalf of the Wellcome Sanger Institute COVID-19 Surveillance Team |
| EPI_ISL_2394421 | Lighthouse Lab in Glasgow | Wellcome Sanger Institute for the COVID-19 Genomics UK (COG-UK) Consortium | Harper VanSteenhouse, Yumi Kasai, David Gray, Carol Clugston, Anna Dominiczak and Alex Alderton, Roberto Amato, Jeffrey Barrett, Sonia Goncalves, Ewan Harrison, David K. Jackson, Ian Johnston, Dominic Kwiatkowski, Cordelia Langford, John Sillitoe on behalf of the Wellcome Sanger Institute COVID-19 Surveillance Team |
| EPI_ISL_2394422, EPI_ISL_2394423, EPI_ISL_2394424 | Lighthouse Lab in Alderley Park | Wellcome Sanger Institute for the COVID-19 Genomics UK (COG-UK) Consortium | Jacquelyn Wynn, Mairead Hyland, The Lighthouse Lab in Alderley Park and Alex Alderton, Roberto Amato, Jeffrey Barrett, Sonia Goncalves, Ewan Harrison, David K. Jackson, Ian Johnston, Dominic Kwiatkowski, Cordelia Langford, John Sillitoe on behalf of the Wellcome Sanger Institute COVID-19 Surveillance Team |
| EPI_ISL_2394425 | Lighthouse Lab in Glasgow | Wellcome Sanger Institute for the COVID-19 Genomics UK (COG-UK) Consortium | Harper VanSteenhouse, Yumi Kasai, David Gray, Carol Clugston, Anna Dominiczak and Alex Alderton, Roberto Amato, Jeffrey Barrett, Sonia Goncalves, Ewan Harrison, David K. Jackson, Ian Johnston, Dominic Kwiatkowski, Cordelia Langford, John Sillitoe on behalf of the Wellcome Sanger Institute COVID-19 Surveillance Team |
| EPI_ISL_2394426 | Lighthouse Lab in Alderley Park | Wellcome Sanger Institute for the COVID-19 Genomics UK (COG-UK) Consortium | Jacquelyn Wynn, Mairead Hyland, The Lighthouse Lab in Alderley Park and Alex Alderton, Roberto Amato, Jeffrey Barrett, Sonia Goncalves, Ewan Harrison, David K. Jackson, Ian Johnston, Dominic Kwiatkowski, Cordelia Langford, John Sillitoe on behalf of the Wellcome Sanger Institute COVID-19 Surveillance Team |
| EPI_ISL_2394427, EPI_ISL_2394428 | Lighthouse Lab in Glasgow | Wellcome Sanger Institute for the COVID-19 Genomics UK (COG-UK) Consortium | Harper VanSteenhouse, Yumi Kasai, David Gray, Carol Clugston, Anna Dominiczak and Alex Alderton, Roberto Amato, Jeffrey Barrett, Sonia Goncalves, Ewan Harrison, David K. Jackson, Ian Johnston, Dominic Kwiatkowski, Cordelia Langford, John Sillitoe on behalf of the Wellcome Sanger Institute COVID-19 Surveillance Team |
| EPI_ISL_2394430 | Lighthouse Lab in Alderley Park | Wellcome Sanger Institute for the COVID-19 Genomics UK (COG-UK) Consortium | Jacquelyn Wynn, Mairead Hyland, The Lighthouse Lab in Alderley Park and Alex Alderton, Roberto Amato, Jeffrey Barrett, Sonia Goncalves, Ewan Harrison, David K. Jackson, Ian Johnston, Dominic Kwiatkowski, Cordelia Langford, John Sillitoe on behalf of the Wellcome Sanger Institute COVID-19 Surveillance Team |
| EPI_ISL_2394431 | Lighthouse Lab in Glasgow | Wellcome Sanger Institute for the COVID-19 Genomics UK (COG-UK) Consortium | Harper VanSteenhouse, Yumi Kasai, David Gray, Carol Clugston, Anna Dominiczak and Alex Alderton, Roberto Amato, Jeffrey Barrett, Sonia Goncalves, Ewan Harrison, David K. Jackson, Ian Johnston, Dominic Kwiatkowski, Cordelia Langford, John Sillitoe on behalf of the Wellcome Sanger Institute COVID-19 Surveillance Team |
| EPI_ISL_2394432, EPI_ISL_2394433, EPI_ISL_2394435, EPI_ISL_2394436, EPI_ISL_2394437, EPI_ISL_2394438, EPI_ISL_2394439 | Lighthouse Lab in Alderley Park | Wellcome Sanger Institute for the COVID-19 Genomics UK (COG-UK) Consortium | Jacquelyn Wynn, Mairead Hyland, The Lighthouse Lab in Alderley Park and Alex Alderton, Roberto Amato, Jeffrey Barrett, Sonia Goncalves, Ewan Harrison, David K. Jackson, Ian Johnston, Dominic Kwiatkowski, Cordelia Langford, John Sillitoe on behalf of the Wellcome Sanger Institute COVID-19 Surveillance Team |
| EPI_ISL_2394442 | Lighthouse Lab in Glasgow | Wellcome Sanger Institute for the COVID-19 Genomics UK (COG-UK) Consortium | Harper VanSteenhouse, Yumi Kasai, David Gray, Carol Clugston, Anna Dominiczak and Alex Alderton, Roberto Amato, Jeffrey Barrett, Sonia Goncalves, Ewan Harrison, David K. Jackson, Ian Johnston, Dominic Kwiatkowski, Cordelia Langford, John Sillitoe on behalf of the Wellcome Sanger Institute COVID-19 Surveillance Team |
| EPI_ISL_2394443, EPI_ISL_2394444, EPI_ISL_2394445, EPI_ISL_2394446 | Lighthouse Lab in Alderley Park | Wellcome Sanger Institute for the COVID-19 Genomics UK (COG-UK) Consortium | Jacquelyn Wynn, Mairead Hyland, The Lighthouse Lab in Alderley Park and Alex Alderton, Roberto Amato, Jeffrey Barrett, Sonia Goncalves, Ewan Harrison, David K. Jackson, Ian Johnston, Dominic Kwiatkowski, Cordelia Langford, John Sillitoe on behalf of the Wellcome Sanger Institute COVID-19 Surveillance Team |
| EPI_ISL_2394447 | Lighthouse Lab in Glasgow | Wellcome Sanger Institute for the COVID-19 Genomics UK (COG-UK) Consortium | Harper VanSteenhouse, Yumi Kasai, David Gray, Carol Clugston, Anna Dominiczak and Alex Alderton, Roberto Amato, Jeffrey Barrett, Sonia Goncalves, Ewan Harrison, David K. Jackson, Ian Johnston, Dominic Kwiatkowski, Cordelia Langford, John Sillitoe on behalf of the Wellcome Sanger Institute COVID-19 Surveillance Team |
| EPI_ISL_2394448 | Lighthouse Lab in Alderley Park | Wellcome Sanger Institute for the COVID-19 Genomics UK (COG-UK) Consortium | Jacquelyn Wynn, Mairead Hyland, The Lighthouse Lab in Alderley Park and Alex Alderton, Roberto Amato, Jeffrey Barrett, Sonia Goncalves, Ewan Harrison, David K. Jackson, Ian Johnston, Dominic Kwiatkowski, Cordelia Langford, John Sillitoe on behalf of the Wellcome Sanger Institute COVID-19 Surveillance Team |
| EPI_ISL_2394449, EPI_ISL_2394450 | Lighthouse Lab in Glasgow | Wellcome Sanger Institute for the COVID-19 Genomics UK (COG-UK) Consortium | Harper VanSteenhouse, Yumi Kasai, David Gray, Carol Clugston, Anna Dominiczak and Alex Alderton, Roberto Amato, Jeffrey Barrett, Sonia Goncalves, Ewan Harrison, David K. Jackson, Ian Johnston, Dominic Kwiatkowski, Cordelia Langford, John Sillitoe on behalf of the Wellcome Sanger Institute COVID-19 Surveillance Team |
| EPI_ISL_2394451 | Lighthouse Lab in Alderley Park | Wellcome Sanger Institute for the COVID-19 Genomics UK (COG-UK) Consortium | Jacquelyn Wynn, Mairead Hyland, The Lighthouse Lab in Alderley Park and Alex Alderton, Roberto Amato, Jeffrey Barrett, Sonia Goncalves, Ewan Harrison, David K. Jackson, Ian Johnston, Dominic Kwiatkowski, Cordelia Langford, John Sillitoe on behalf of the Wellcome Sanger Institute COVID-19 Surveillance Team |
| EPI_ISL_2394452 | Lighthouse Lab in Glasgow | Wellcome Sanger Institute for the COVID-19 Genomics UK (COG-UK) Consortium | Harper VanSteenhouse, Yumi Kasai, David Gray, Carol Clugston, Anna Dominiczak and Alex Alderton, Roberto Amato, Jeffrey Barrett, Sonia Goncalves, Ewan Harrison, David K. Jackson, Ian Johnston, Dominic Kwiatkowski, Cordelia Langford, John Sillitoe on behalf of the Wellcome Sanger Institute COVID-19 Surveillance Team |
| EPI_ISL_2394453 | Lighthouse Lab in Alderley Park | Wellcome Sanger Institute for the COVID-19 Genomics UK (COG-UK) Consortium | Jacquelyn Wynn, Mairead Hyland, The Lighthouse Lab in Alderley Park and Alex Alderton, Roberto Amato, Jeffrey Barrett, Sonia Goncalves, Ewan Harrison, David K. Jackson, Ian Johnston, Dominic Kwiatkowski, Cordelia Langford, John Sillitoe on behalf of the Wellcome Sanger Institute COVID-19 Surveillance Team |
| EPI_ISL_2394454 | Lighthouse Lab in Glasgow | Wellcome Sanger Institute for the COVID-19 Genomics UK (COG-UK) Consortium | Harper VanSteenhouse, Yumi Kasai, David Gray, Carol Clugston, Anna Dominiczak and Alex Alderton, Roberto Amato, Jeffrey Barrett, Sonia Goncalves, Ewan Harrison, David K. Jackson, Ian Johnston, Dominic Kwiatkowski, Cordelia Langford, John Sillitoe on behalf of the Wellcome Sanger Institute COVID-19 Surveillance Team |
| EPI_ISL_2394455, EPI_ISL_2394456, EPI_ISL_2394457, EPI_ISL_2394458 | Lighthouse Lab in Alderley Park | Wellcome Sanger Institute for the COVID-19 Genomics UK (COG-UK) Consortium | Jacquelyn Wynn, Mairead Hyland, The Lighthouse Lab in Alderley Park and Alex Alderton, Roberto Amato, Jeffrey Barrett, Sonia Goncalves, Ewan Harrison, David K. Jackson, Ian Johnston, Dominic Kwiatkowski, Cordelia Langford, John Sillitoe on behalf of the Wellcome Sanger Institute COVID-19 Surveillance Team |
| EPI_ISL_2394459 | Lighthouse Lab in Glasgow | Wellcome Sanger Institute for the COVID-19 Genomics UK (COG-UK) Consortium | Harper VanSteenhouse, Yumi Kasai, David Gray, Carol Clugston, Anna Dominiczak and Alex Alderton, Roberto Amato, Jeffrey Barrett, Sonia Goncalves, Ewan Harrison, David K. Jackson, Ian Johnston, Dominic Kwiatkowski, Cordelia Langford, John Sillitoe on behalf of the Wellcome Sanger Institute COVID-19 Surveillance Team |
| EPI_ISL_2394460, EPI_ISL_2394461 | Lighthouse Lab in Alderley Park | Wellcome Sanger Institute for the COVID-19 Genomics UK (COG-UK) Consortium | Jacquelyn Wynn, Mairead Hyland, The Lighthouse Lab in Alderley Park and Alex Alderton, Roberto Amato, Jeffrey Barrett, Sonia Goncalves, Ewan Harrison, David K. Jackson, Ian Johnston, Dominic Kwiatkowski, Cordelia Langford, John Sillitoe on behalf of the Wellcome Sanger Institute COVID-19 Surveillance Team |
| EPI_ISL_2394463 | Lighthouse Lab in Glasgow | Wellcome Sanger Institute for the COVID-19 Genomics UK (COG-UK) Consortium | Harper VanSteenhouse, Yumi Kasai, David Gray, Carol Clugston, Anna Dominiczak and Alex Alderton, Roberto Amato, Jeffrey Barrett, Sonia Goncalves, Ewan Harrison, David K. Jackson, Ian Johnston, Dominic Kwiatkowski, Cordelia Langford, John Sillitoe on behalf of the Wellcome Sanger Institute COVID-19 Surveillance Team |
| EPI_ISL_2394464 | Lighthouse Lab in Alderley Park | Wellcome Sanger Institute for the COVID-19 Genomics UK (COG-UK) Consortium | Jacquelyn Wynn, Mairead Hyland, The Lighthouse Lab in Alderley Park and Alex Alderton, Roberto Amato, Jeffrey Barrett, Sonia Goncalves, Ewan Harrison, David K. Jackson, Ian Johnston, Dominic Kwiatkowski, Cordelia Langford, John Sillitoe on behalf of the Wellcome Sanger Institute COVID-19 Surveillance Team |
| EPI_ISL_2394465 | Lighthouse Lab in Glasgow | Wellcome Sanger Institute for the COVID-19 Genomics UK (COG-UK) Consortium | Harper VanSteenhouse, Yumi Kasai, David Gray, Carol Clugston, Anna Dominiczak and Alex Alderton, Roberto Amato, Jeffrey Barrett, Sonia Goncalves, Ewan Harrison, David K. Jackson, Ian Johnston, Dominic Kwiatkowski, Cordelia Langford, John Sillitoe on behalf of the Wellcome Sanger Institute COVID-19 Surveillance Team |
| EPI_ISL_2394466, EPI_ISL_2394468 | Lighthouse Lab in Alderley Park | Wellcome Sanger Institute for the COVID-19 Genomics UK (COG-UK) Consortium | Jacquelyn Wynn, Mairead Hyland, The Lighthouse Lab in Alderley Park and Alex Alderton, Roberto Amato, Jeffrey Barrett, Sonia Goncalves, Ewan Harrison, David K. Jackson, Ian Johnston, Dominic Kwiatkowski, Cordelia Langford, John Sillitoe on behalf of the Wellcome Sanger Institute COVID-19 Surveillance Team |
| EPI_ISL_2394469, EPI_ISL_2394470 | Lighthouse Lab in Glasgow | Wellcome Sanger Institute for the COVID-19 Genomics UK (COG-UK) Consortium | Harper VanSteenhouse, Yumi Kasai, David Gray, Carol Clugston, Anna Dominiczak and Alex Alderton, Roberto Amato, Jeffrey Barrett, Sonia Goncalves, Ewan Harrison, David K. Jackson, Ian Johnston, Dominic Kwiatkowski, Cordelia Langford, John Sillitoe on behalf of the Wellcome Sanger Institute COVID-19 Surveillance Team |
| EPI_ISL_2394471 | Lighthouse Lab in Alderley Park | Wellcome Sanger Institute for the COVID-19 Genomics UK (COG-UK) Consortium | Jacquelyn Wynn, Mairead Hyland, The Lighthouse Lab in Alderley Park and Alex Alderton, Roberto Amato, Jeffrey Barrett, Sonia Goncalves, Ewan Harrison, David K. Jackson, Ian Johnston, Dominic Kwiatkowski, Cordelia Langford, John Sillitoe on behalf of the Wellcome Sanger Institute COVID-19 Surveillance Team |
| EPI_ISL_2394473, EPI_ISL_2394474 | Lighthouse Lab in Glasgow | Wellcome Sanger Institute for the COVID-19 Genomics UK (COG-UK) Consortium | Harper VanSteenhouse, Yumi Kasai, David Gray, Carol Clugston, Anna Dominiczak and Alex Alderton, Roberto Amato, Jeffrey Barrett, Sonia Goncalves, Ewan Harrison, David K. Jackson, Ian Johnston, Dominic Kwiatkowski, Cordelia Langford, John Sillitoe on behalf of the Wellcome Sanger Institute COVID-19 Surveillance Team |
| EPI_ISL_2394475, EPI_ISL_2394476, EPI_ISL_2394477, | Lighthouse Lab in Alderley Park | Wellcome Sanger Institute for the COVID-19 Genomics UK (COG-UK) Consortium | Jacquelyn Wynn, Mairead Hyland, The Lighthouse Lab in Alderley Park and Alex Alderton, Roberto Amato, Jeffrey Barrett, Sonia Goncalves, Ewan Harrison, David K. Jackson, Ian Johnston, Dominic Kwiatkowski, Cordelia Langford, John Sillitoe on behalf of the Wellcome Sanger Institute COVID-19 Surveillance Team |

| | | | |
|---|---|---|---|
| EPI_ISL_2394478, EPI_ISL_2394479, EPI_ISL_2394480 | | | |
| EPI_ISL_2394481 | Lighthouse Lab in Glasgow | Wellcome Sanger Institute for the COVID-19 Genomics UK (COG-UK) Consortium | Harper VanSteenhouse, Yumi Kasai, David Gray, Carol Clugston, Anna Dominiczak and Alex Alderton, Roberto Amato, Jeffrey Barrett, Sonia Goncalves, Ewan Harrison, David K. Jackson, Ian Johnston, Dominic Kwiatkowski, Cordelia Langford, John Sillitoe on behalf of the Wellcome Sanger Institute COVID-19 Surveillance Team |
| EPI_ISL_2394482, EPI_ISL_2394483 | Lighthouse Lab in Alderley Park | Wellcome Sanger Institute for the COVID-19 Genomics UK (COG-UK) Consortium | Jacquelyn Wynn, Mairead Hyland, The Lighthouse Lab in Alderley Park and Alex Alderton, Roberto Amato, Jeffrey Barrett, Sonia Goncalves, Ewan Harrison, David K. Jackson, Ian Johnston, Dominic Kwiatkowski, Cordelia Langford, John Sillitoe on behalf of the Wellcome Sanger Institute COVID-19 Surveillance Team |
| EPI_ISL_2394484 | Lighthouse Lab in Glasgow | Wellcome Sanger Institute for the COVID-19 Genomics UK (COG-UK) Consortium | Harper VanSteenhouse, Yumi Kasai, David Gray, Carol Clugston, Anna Dominiczak and Alex Alderton, Roberto Amato, Jeffrey Barrett, Sonia Goncalves, Ewan Harrison, David K. Jackson, Ian Johnston, Dominic Kwiatkowski, Cordelia Langford, John Sillitoe on behalf of the Wellcome Sanger Institute COVID-19 Surveillance Team |
| EPI_ISL_2394485, EPI_ISL_2394486, EPI_ISL_2394487 | Lighthouse Lab in Alderley Park | Wellcome Sanger Institute for the COVID-19 Genomics UK (COG-UK) Consortium | Jacquelyn Wynn, Mairead Hyland, The Lighthouse Lab in Alderley Park and Alex Alderton, Roberto Amato, Jeffrey Barrett, Sonia Goncalves, Ewan Harrison, David K. Jackson, Ian Johnston, Dominic Kwiatkowski, Cordelia Langford, John Sillitoe on behalf of the Wellcome Sanger Institute COVID-19 Surveillance Team |
| EPI_ISL_2394488, EPI_ISL_2394489, EPI_ISL_2394490 | Lighthouse Lab in Glasgow | Wellcome Sanger Institute for the COVID-19 Genomics UK (COG-UK) Consortium | Harper VanSteenhouse, Yumi Kasai, David Gray, Carol Clugston, Anna Dominiczak and Alex Alderton, Roberto Amato, Jeffrey Barrett, Sonia Goncalves, Ewan Harrison, David K. Jackson, Ian Johnston, Dominic Kwiatkowski, Cordelia Langford, John Sillitoe on behalf of the Wellcome Sanger Institute COVID-19 Surveillance Team |
| EPI_ISL_2394491 | Lighthouse Lab in Alderley Park | Wellcome Sanger Institute for the COVID-19 Genomics UK (COG-UK) Consortium | Jacquelyn Wynn, Mairead Hyland, The Lighthouse Lab in Alderley Park and Alex Alderton, Roberto Amato, Jeffrey Barrett, Sonia Goncalves, Ewan Harrison, David K. Jackson, Ian Johnston, Dominic Kwiatkowski, Cordelia Langford, John Sillitoe on behalf of the Wellcome Sanger Institute COVID-19 Surveillance Team |
| EPI_ISL_2394492, EPI_ISL_2394493 | Lighthouse Lab in Glasgow | Wellcome Sanger Institute for the COVID-19 Genomics UK (COG-UK) Consortium | Harper VanSteenhouse, Yumi Kasai, David Gray, Carol Clugston, Anna Dominiczak and Alex Alderton, Roberto Amato, Jeffrey Barrett, Sonia Goncalves, Ewan Harrison, David K. Jackson, Ian Johnston, Dominic Kwiatkowski, Cordelia Langford, John Sillitoe on behalf of the Wellcome Sanger Institute COVID-19 Surveillance Team |
| EPI_ISL_2394494, EPI_ISL_2394495, EPI_ISL_2394496, EPI_ISL_2394497, EPI_ISL_2394498 | Lighthouse Lab in Alderley Park | Wellcome Sanger Institute for the COVID-19 Genomics UK (COG-UK) Consortium | Jacquelyn Wynn, Mairead Hyland, The Lighthouse Lab in Alderley Park and Alex Alderton, Roberto Amato, Jeffrey Barrett, Sonia Goncalves, Ewan Harrison, David K. Jackson, Ian Johnston, Dominic Kwiatkowski, Cordelia Langford, John Sillitoe on behalf of the Wellcome Sanger Institute COVID-19 Surveillance Team |
| EPI_ISL_2394499 | Lighthouse Lab in Glasgow | Wellcome Sanger Institute for the COVID-19 Genomics UK (COG-UK) Consortium | Harper VanSteenhouse, Yumi Kasai, David Gray, Carol Clugston, Anna Dominiczak and Alex Alderton, Roberto Amato, Jeffrey Barrett, Sonia Goncalves, Ewan Harrison, David K. Jackson, Ian Johnston, Dominic Kwiatkowski, Cordelia Langford, John Sillitoe on behalf of the Wellcome Sanger Institute COVID-19 Surveillance Team |
| EPI_ISL_2394500, EPI_ISL_2394501, EPI_ISL_2394502, EPI_ISL_2394503, EPI_ISL_2394504, EPI_ISL_2394505, EPI_ISL_2394506, EPI_ISL_2394507, EPI_ISL_2394508 | Lighthouse Lab in Alderley Park | Wellcome Sanger Institute for the COVID-19 Genomics UK (COG-UK) Consortium | Jacquelyn Wynn, Mairead Hyland, The Lighthouse Lab in Alderley Park and Alex Alderton, Roberto Amato, Jeffrey Barrett, Sonia Goncalves, Ewan Harrison, David K. Jackson, Ian Johnston, Dominic Kwiatkowski, Cordelia Langford, John Sillitoe on behalf of the Wellcome Sanger Institute COVID-19 Surveillance Team |
| EPI_ISL_2394509, EPI_ISL_2394511 | Lighthouse Lab in Glasgow | Wellcome Sanger Institute for the COVID-19 Genomics UK (COG-UK) Consortium | Harper VanSteenhouse, Yumi Kasai, David Gray, Carol Clugston, Anna Dominiczak and Alex Alderton, Roberto Amato, Jeffrey Barrett, Sonia Goncalves, Ewan Harrison, David K. Jackson, Ian Johnston, Dominic Kwiatkowski, Cordelia Langford, John Sillitoe on behalf of the Wellcome Sanger Institute COVID-19 Surveillance Team |
| EPI_ISL_2394512, EPI_ISL_2394513, EPI_ISL_2394514, EPI_ISL_2394515, EPI_ISL_2394516, EPI_ISL_2394517, EPI_ISL_2394518 | Lighthouse Lab in Alderley Park | Wellcome Sanger Institute for the COVID-19 Genomics UK (COG-UK) Consortium | Jacquelyn Wynn, Mairead Hyland, The Lighthouse Lab in Alderley Park and Alex Alderton, Roberto Amato, Jeffrey Barrett, Sonia Goncalves, Ewan Harrison, David K. Jackson, Ian Johnston, Dominic Kwiatkowski, Cordelia Langford, John Sillitoe on behalf of the Wellcome Sanger Institute COVID-19 Surveillance Team |
| EPI_ISL_2394519, EPI_ISL_2394520 | Lighthouse Lab in Glasgow | Wellcome Sanger Institute for the COVID-19 Genomics UK (COG-UK) Consortium | Harper VanSteenhouse, Yumi Kasai, David Gray, Carol Clugston, Anna Dominiczak and Alex Alderton, Roberto Amato, Jeffrey Barrett, Sonia Goncalves, Ewan Harrison, David K. Jackson, Ian Johnston, Dominic Kwiatkowski, Cordelia Langford, John Sillitoe on behalf of the Wellcome Sanger Institute COVID-19 Surveillance Team |
| EPI_ISL_2394521, EPI_ISL_2394522, EPI_ISL_2394523, EPI_ISL_2394524, EPI_ISL_2394526, EPI_ISL_2394527 | Lighthouse Lab in Alderley Park | Wellcome Sanger Institute for the COVID-19 Genomics UK (COG-UK) Consortium | Jacquelyn Wynn, Mairead Hyland, The Lighthouse Lab in Alderley Park and Alex Alderton, Roberto Amato, Jeffrey Barrett, Sonia Goncalves, Ewan Harrison, David K. Jackson, Ian Johnston, Dominic Kwiatkowski, Cordelia Langford, John Sillitoe on behalf of the Wellcome Sanger Institute COVID-19 Surveillance Team |
| EPI_ISL_2394528 | Lighthouse Lab in Glasgow | Wellcome Sanger Institute for the COVID-19 Genomics UK (COG-UK) Consortium | Harper VanSteenhouse, Yumi Kasai, David Gray, Carol Clugston, Anna Dominiczak and Alex Alderton, Roberto Amato, Jeffrey Barrett, Sonia Goncalves, Ewan Harrison, David K. Jackson, Ian Johnston, Dominic Kwiatkowski, Cordelia Langford, John Sillitoe on behalf of the Wellcome Sanger Institute COVID-19 Surveillance Team |
| EPI_ISL_2394529, EPI_ISL_2394530, EPI_ISL_2394531 | Lighthouse Lab in Alderley Park | Wellcome Sanger Institute for the COVID-19 Genomics UK (COG-UK) Consortium | Jacquelyn Wynn, Mairead Hyland, The Lighthouse Lab in Alderley Park and Alex Alderton, Roberto Amato, Jeffrey Barrett, Sonia Goncalves, Ewan Harrison, David K. Jackson, Ian Johnston, Dominic Kwiatkowski, Cordelia Langford, John Sillitoe on behalf of the Wellcome Sanger Institute COVID-19 Surveillance Team |
| EPI_ISL_2394533, EPI_ISL_2394534 | Lighthouse Lab in Glasgow | Wellcome Sanger Institute for the COVID-19 Genomics UK (COG-UK) Consortium | Harper VanSteenhouse, Yumi Kasai, David Gray, Carol Clugston, Anna Dominiczak and Alex Alderton, Roberto Amato, Jeffrey Barrett, Sonia Goncalves, Ewan Harrison, David K. Jackson, Ian Johnston, Dominic Kwiatkowski, Cordelia Langford, John Sillitoe on behalf of the Wellcome Sanger Institute COVID-19 Surveillance Team |
| EPI_ISL_2394535, EPI_ISL_2394536 | Lighthouse Lab in Alderley Park | Wellcome Sanger Institute for the COVID-19 Genomics UK (COG-UK) Consortium | Jacquelyn Wynn, Mairead Hyland, The Lighthouse Lab in Alderley Park and Alex Alderton, Roberto Amato, Jeffrey Barrett, Sonia Goncalves, Ewan Harrison, David K. Jackson, Ian Johnston, Dominic Kwiatkowski, Cordelia Langford, John Sillitoe on behalf of the Wellcome Sanger Institute COVID-19 Surveillance Team |
| EPI_ISL_2394537 | Lighthouse Lab in Glasgow | Wellcome Sanger Institute for the COVID-19 Genomics UK (COG-UK) Consortium | Harper VanSteenhouse, Yumi Kasai, David Gray, Carol Clugston, Anna Dominiczak and Alex Alderton, Roberto Amato, Jeffrey Barrett, Sonia Goncalves, Ewan Harrison, David K. Jackson, Ian Johnston, Dominic Kwiatkowski, Cordelia Langford, John Sillitoe on behalf of the Wellcome Sanger Institute COVID-19 Surveillance Team |
| EPI_ISL_2394538, EPI_ISL_2394539 | Lighthouse Lab in Alderley Park | Wellcome Sanger Institute for the COVID-19 Genomics UK (COG-UK) Consortium | Jacquelyn Wynn, Mairead Hyland, The Lighthouse Lab in Alderley Park and Alex Alderton, Roberto Amato, Jeffrey Barrett, Sonia Goncalves, Ewan Harrison, David K. Jackson, Ian Johnston, Dominic Kwiatkowski, Cordelia Langford, John Sillitoe on behalf of the Wellcome Sanger Institute COVID-19 Surveillance Team |
| EPI_ISL_2394540, EPI_ISL_2394542 | Lighthouse Lab in Glasgow | Wellcome Sanger Institute for the COVID-19 Genomics UK (COG-UK) Consortium | Harper VanSteenhouse, Yumi Kasai, David Gray, Carol Clugston, Anna Dominiczak and Alex Alderton, Roberto Amato, Jeffrey Barrett, Sonia Goncalves, Ewan Harrison, David K. Jackson, Ian Johnston, Dominic Kwiatkowski, Cordelia Langford, John Sillitoe on behalf of the Wellcome Sanger Institute COVID-19 Surveillance Team |
| EPI_ISL_2394543, EPI_ISL_2394544, EPI_ISL_2394546 | Lighthouse Lab in Alderley Park | Wellcome Sanger Institute for the COVID-19 Genomics UK (COG-UK) Consortium | Jacquelyn Wynn, Mairead Hyland, The Lighthouse Lab in Alderley Park and Alex Alderton, Roberto Amato, Jeffrey Barrett, Sonia Goncalves, Ewan Harrison, David K. Jackson, Ian Johnston, Dominic Kwiatkowski, Cordelia Langford, John Sillitoe on behalf of the Wellcome Sanger Institute COVID-19 Surveillance Team |
| EPI_ISL_2394547, EPI_ISL_2394548, EPI_ISL_2394549, EPI_ISL_2394550, EPI_ISL_2394551 | Lighthouse Lab in Glasgow | Wellcome Sanger Institute for the COVID-19 Genomics UK (COG-UK) Consortium | Harper VanSteenhouse, Yumi Kasai, David Gray, Carol Clugston, Anna Dominiczak and Alex Alderton, Roberto Amato, Jeffrey Barrett, Sonia Goncalves, Ewan Harrison, David K. Jackson, Ian Johnston, Dominic Kwiatkowski, Cordelia Langford, John Sillitoe on behalf of the Wellcome Sanger Institute COVID-19 Surveillance Team |
| EPI_ISL_2394552, EPI_ISL_2394553, EPI_ISL_2394554, EPI_ISL_2394555, EPI_ISL_2394556 | Lighthouse Lab in Alderley Park | Wellcome Sanger Institute for the COVID-19 Genomics UK (COG-UK) Consortium | Jacquelyn Wynn, Mairead Hyland, The Lighthouse Lab in Alderley Park and Alex Alderton, Roberto Amato, Jeffrey Barrett, Sonia Goncalves, Ewan Harrison, David K. Jackson, Ian Johnston, Dominic Kwiatkowski, Cordelia Langford, John Sillitoe on behalf of the Wellcome Sanger Institute COVID-19 Surveillance Team |
| EPI_ISL_2394557 | Lighthouse Lab in Glasgow | Wellcome Sanger Institute for the COVID-19 Genomics UK (COG-UK) Consortium | Harper VanSteenhouse, Yumi Kasai, David Gray, Carol Clugston, Anna Dominiczak and Alex Alderton, Roberto Amato, Jeffrey Barrett, Sonia Goncalves, Ewan Harrison, David K. Jackson, Ian Johnston, Dominic Kwiatkowski, Cordelia Langford, John Sillitoe on behalf of the Wellcome Sanger Institute COVID-19 Surveillance Team |
| EPI_ISL_2394558 | Lighthouse Lab in Alderley Park | Wellcome Sanger Institute for the COVID-19 Genomics UK (COG-UK) Consortium | Jacquelyn Wynn, Mairead Hyland, The Lighthouse Lab in Alderley Park and Alex Alderton, Roberto Amato, Jeffrey Barrett, Sonia Goncalves, Ewan Harrison, David K. Jackson, Ian Johnston, Dominic Kwiatkowski, Cordelia Langford, John Sillitoe on behalf of the Wellcome Sanger Institute COVID-19 Surveillance Team |
| EPI_ISL_2394559 | Lighthouse Lab in Glasgow | Wellcome Sanger Institute for the COVID-19 Genomics UK (COG-UK) Consortium | Harper VanSteenhouse, Yumi Kasai, David Gray, Carol Clugston, Anna Dominiczak and Alex Alderton, Roberto Amato, Jeffrey Barrett, Sonia Goncalves, Ewan Harrison, David K. Jackson, Ian Johnston, Dominic Kwiatkowski, Cordelia Langford, John Sillitoe on behalf of the Wellcome Sanger Institute COVID-19 Surveillance Team |
| EPI_ISL_2394560 | Lighthouse Lab in Alderley Park | Wellcome Sanger Institute for the COVID-19 Genomics UK (COG-UK) Consortium | Jacquelyn Wynn, Mairead Hyland, The Lighthouse Lab in Alderley Park and Alex Alderton, Roberto Amato, Jeffrey Barrett, Sonia Goncalves, Ewan Harrison, David K. Jackson, Ian Johnston, Dominic Kwiatkowski, Cordelia Langford, John Sillitoe on behalf of the Wellcome Sanger Institute COVID-19 Surveillance Team |
| EPI_ISL_2394561 | Lighthouse Lab in Glasgow | Wellcome Sanger Institute for the COVID-19 Genomics UK (COG-UK) Consortium | Harper VanSteenhouse, Yumi Kasai, David Gray, Carol Clugston, Anna Dominiczak and Alex Alderton, Roberto Amato, Jeffrey Barrett, Sonia Goncalves, Ewan Harrison, David K. Jackson, Ian Johnston, Dominic Kwiatkowski, Cordelia Langford, John Sillitoe on behalf of the Wellcome Sanger Institute COVID-19 Surveillance Team |
| EPI_ISL_2394563, EPI_ISL_2394565 | Lighthouse Lab in Alderley Park | Wellcome Sanger Institute for the COVID-19 Genomics UK (COG-UK) Consortium | Jacquelyn Wynn, Mairead Hyland, The Lighthouse Lab in Alderley Park and Alex Alderton, Roberto Amato, Jeffrey Barrett, Sonia Goncalves, Ewan Harrison, David K. Jackson, Ian Johnston, Dominic Kwiatkowski, Cordelia Langford, John Sillitoe on behalf of the Wellcome Sanger Institute COVID-19 Surveillance Team |
| EPI_ISL_2394566 | Lighthouse Lab in Glasgow | Wellcome Sanger Institute for the COVID-19 Genomics UK (COG-UK) Consortium | Harper VanSteenhouse, Yumi Kasai, David Gray, Carol Clugston, Anna Dominiczak and Alex Alderton, Roberto Amato, Jeffrey Barrett, Sonia Goncalves, Ewan Harrison, David K. Jackson, Ian Johnston, Dominic Kwiatkowski, Cordelia Langford, John Sillitoe on behalf of the Wellcome Sanger Institute COVID-19 Surveillance Team |
| EPI_ISL_2394567, EPI_ISL_2394568 | Lighthouse Lab in Alderley Park | Wellcome Sanger Institute for the COVID-19 Genomics UK (COG-UK) Consortium | Jacquelyn Wynn, Mairead Hyland, The Lighthouse Lab in Alderley Park and Alex Alderton, Roberto Amato, Jeffrey Barrett, Sonia Goncalves, Ewan Harrison, David K. Jackson, Ian Johnston, Dominic Kwiatkowski, Cordelia Langford, John Sillitoe on behalf of the Wellcome Sanger Institute COVID-19 Surveillance Team |

| | | | |
|---|---|---|---|
| EPI_ISL_2394569 | Lighthouse Lab in Glasgow | Wellcome Sanger Institute for the COVID-19 Genomics UK (COG-UK) Consortium | Harper VanSteenhouse, Yumi Kasai, David Gray, Carol Clugston, Anna Dominiczak and Alex Alderton, Roberto Amato, Jeffrey Barrett, Sonia Goncalves, Ewan Harrison, David K. Jackson, Ian Johnston, Dominic Kwiatkowski, Cordelia Langford, John Sillitoe on behalf of the Wellcome Sanger Institute COVID-19 Surveillance Team |
| EPI_ISL_2394570 | Lighthouse Lab in Alderley Park | Wellcome Sanger Institute for the COVID-19 Genomics UK (COG-UK) Consortium | Jacquelyn Wynn, Mairead Hyland, The Lighthouse Lab in Alderley Park and Alex Alderton, Roberto Amato, Jeffrey Barrett, Sonia Goncalves, Ewan Harrison, David K. Jackson, Ian Johnston, Dominic Kwiatkowski, Cordelia Langford, John Sillitoe on behalf of the Wellcome Sanger Institute COVID-19 Surveillance Team |
| EPI_ISL_2394571, EPI_ISL_2394572 | Lighthouse Lab in Glasgow | Wellcome Sanger Institute for the COVID-19 Genomics UK (COG-UK) Consortium | Harper VanSteenhouse, Yumi Kasai, David Gray, Carol Clugston, Anna Dominiczak and Alex Alderton, Roberto Amato, Jeffrey Barrett, Sonia Goncalves, Ewan Harrison, David K. Jackson, Ian Johnston, Dominic Kwiatkowski, Cordelia Langford, John Sillitoe on behalf of the Wellcome Sanger Institute COVID-19 Surveillance Team |
| EPI_ISL_2394573, EPI_ISL_2394574 | Lighthouse Lab in Alderley Park | Wellcome Sanger Institute for the COVID-19 Genomics UK (COG-UK) Consortium | Jacquelyn Wynn, Mairead Hyland, The Lighthouse Lab in Alderley Park and Alex Alderton, Roberto Amato, Jeffrey Barrett, Sonia Goncalves, Ewan Harrison, David K. Jackson, Ian Johnston, Dominic Kwiatkowski, Cordelia Langford, John Sillitoe on behalf of the Wellcome Sanger Institute COVID-19 Surveillance Team |
| EPI_ISL_2394576 | Lighthouse Lab in Glasgow | Wellcome Sanger Institute for the COVID-19 Genomics UK (COG-UK) Consortium | Harper VanSteenhouse, Yumi Kasai, David Gray, Carol Clugston, Anna Dominiczak and Alex Alderton, Roberto Amato, Jeffrey Barrett, Sonia Goncalves, Ewan Harrison, David K. Jackson, Ian Johnston, Dominic Kwiatkowski, Cordelia Langford, John Sillitoe on behalf of the Wellcome Sanger Institute COVID-19 Surveillance Team |
| EPI_ISL_2394577, EPI_ISL_2394578, EPI_ISL_2394580, EPI_ISL_2394581, EPI_ISL_2394582, EPI_ISL_2394583, EPI_ISL_2394584 | Lighthouse Lab in Alderley Park | Wellcome Sanger Institute for the COVID-19 Genomics UK (COG-UK) Consortium | Jacquelyn Wynn, Mairead Hyland, The Lighthouse Lab in Alderley Park and Alex Alderton, Roberto Amato, Jeffrey Barrett, Sonia Goncalves, Ewan Harrison, David K. Jackson, Ian Johnston, Dominic Kwiatkowski, Cordelia Langford, John Sillitoe on behalf of the Wellcome Sanger Institute COVID-19 Surveillance Team |
| EPI_ISL_2394585 | Lighthouse Lab in Glasgow | Wellcome Sanger Institute for the COVID-19 Genomics UK (COG-UK) Consortium | Harper VanSteenhouse, Yumi Kasai, David Gray, Carol Clugston, Anna Dominiczak and Alex Alderton, Roberto Amato, Jeffrey Barrett, Sonia Goncalves, Ewan Harrison, David K. Jackson, Ian Johnston, Dominic Kwiatkowski, Cordelia Langford, John Sillitoe on behalf of the Wellcome Sanger Institute COVID-19 Surveillance Team |
| EPI_ISL_2394586 | Health Services Laboratories | Wellcome Sanger Institute for the COVID-19 Genomics UK (COG-UK) Consortium | Health Services Laboratories and Alex Alderton, Roberto Amato, Jeffrey Barrett, Sonia Goncalves, Ewan Harrison, David K. Jackson, Ian Johnston, Dominic Kwiatkowski, Cordelia Langford, John Sillitoe on behalf of the Wellcome Sanger Institute COVID-19 Surveillance Team |
| EPI_ISL_2394588, EPI_ISL_2394589, EPI_ISL_2394590, EPI_ISL_2394591, EPI_ISL_2394592, EPI_ISL_2394593, EPI_ISL_2394595, EPI_ISL_2394596, EPI_ISL_2394597, EPI_ISL_2394598, EPI_ISL_2394600, EPI_ISL_2394602, EPI_ISL_2394603, EPI_ISL_2394604, EPI_ISL_2394605, EPI_ISL_2394606 | | | |
| see above | Lighthouse Lab in Alderley Park | Wellcome Sanger Institute for the COVID-19 Genomics UK (COG-UK) Consortium | Jacquelyn Wynn, Mairead Hyland, The Lighthouse Lab in Alderley Park and Alex Alderton, Roberto Amato, Jeffrey Barrett, Sonia Goncalves, Ewan Harrison, David K. Jackson, Ian Johnston, Dominic Kwiatkowski, Cordelia Langford, John Sillitoe on behalf of the Wellcome Sanger Institute COVID-19 Surveillance Team |
| EPI_ISL_2394607 | Health Services Laboratories | Wellcome Sanger Institute for the COVID-19 Genomics UK (COG-UK) Consortium | Health Services Laboratories and Alex Alderton, Roberto Amato, Jeffrey Barrett, Sonia Goncalves, Ewan Harrison, David K. Jackson, Ian Johnston, Dominic Kwiatkowski, Cordelia Langford, John Sillitoe on behalf of the Wellcome Sanger Institute COVID-19 Surveillance Team |
| EPI_ISL_2394608, EPI_ISL_2394609, EPI_ISL_2394610, EPI_ISL_2394611 | Lighthouse Lab in Alderley Park | Wellcome Sanger Institute for the COVID-19 Genomics UK (COG-UK) Consortium | Jacquelyn Wynn, Mairead Hyland, The Lighthouse Lab in Alderley Park and Alex Alderton, Roberto Amato, Jeffrey Barrett, Sonia Goncalves, Ewan Harrison, David K. Jackson, Ian Johnston, Dominic Kwiatkowski, Cordelia Langford, John Sillitoe on behalf of the Wellcome Sanger Institute COVID-19 Surveillance Team |
| EPI_ISL_2394612 | Health Services Laboratories | Wellcome Sanger Institute for the COVID-19 Genomics UK (COG-UK) Consortium | Health Services Laboratories and Alex Alderton, Roberto Amato, Jeffrey Barrett, Sonia Goncalves, Ewan Harrison, David K. Jackson, Ian Johnston, Dominic Kwiatkowski, Cordelia Langford, John Sillitoe on behalf of the Wellcome Sanger Institute COVID-19 Surveillance Team |
| EPI_ISL_2394613 | Lighthouse Lab in Alderley Park | Wellcome Sanger Institute for the COVID-19 Genomics UK (COG-UK) Consortium | Jacquelyn Wynn, Mairead Hyland, The Lighthouse Lab in Alderley Park and Alex Alderton, Roberto Amato, Jeffrey Barrett, Sonia Goncalves, Ewan Harrison, David K. Jackson, Ian Johnston, Dominic Kwiatkowski, Cordelia Langford, John Sillitoe on behalf of the Wellcome Sanger Institute COVID-19 Surveillance Team |
| EPI_ISL_2394614 | Health Services Laboratories | Wellcome Sanger Institute for the COVID-19 Genomics UK (COG-UK) Consortium | Health Services Laboratories and Alex Alderton, Roberto Amato, Jeffrey Barrett, Sonia Goncalves, Ewan Harrison, David K. Jackson, Ian Johnston, Dominic Kwiatkowski, Cordelia Langford, John Sillitoe on behalf of the Wellcome Sanger Institute COVID-19 Surveillance Team |
| EPI_ISL_2394615, EPI_ISL_2394616, EPI_ISL_2394617 | Lighthouse Lab in Alderley Park | Wellcome Sanger Institute for the COVID-19 Genomics UK (COG-UK) Consortium | Jacquelyn Wynn, Mairead Hyland, The Lighthouse Lab in Alderley Park and Alex Alderton, Roberto Amato, Jeffrey Barrett, Sonia Goncalves, Ewan Harrison, David K. Jackson, Ian Johnston, Dominic Kwiatkowski, Cordelia Langford, John Sillitoe on behalf of the Wellcome Sanger Institute COVID-19 Surveillance Team |
| EPI_ISL_2394618 | Lighthouse Lab in Glasgow | Wellcome Sanger Institute for the COVID-19 Genomics UK (COG-UK) Consortium | Harper VanSteenhouse, Yumi Kasai, David Gray, Carol Clugston, Anna Dominiczak and Alex Alderton, Roberto Amato, Jeffrey Barrett, Sonia Goncalves, Ewan Harrison, David K. Jackson, Ian Johnston, Dominic Kwiatkowski, Cordelia Langford, John Sillitoe on behalf of the Wellcome Sanger Institute COVID-19 Surveillance Team |
| EPI_ISL_2394619, EPI_ISL_2394620, EPI_ISL_2394623 | Lighthouse Lab in Alderley Park | Wellcome Sanger Institute for the COVID-19 Genomics UK (COG-UK) Consortium | Jacquelyn Wynn, Mairead Hyland, The Lighthouse Lab in Alderley Park and Alex Alderton, Roberto Amato, Jeffrey Barrett, Sonia Goncalves, Ewan Harrison, David K. Jackson, Ian Johnston, Dominic Kwiatkowski, Cordelia Langford, John Sillitoe on behalf of the Wellcome Sanger Institute COVID-19 Surveillance Team |
| EPI_ISL_2394624 | Health Services Laboratories | Wellcome Sanger Institute for the COVID-19 Genomics UK (COG-UK) Consortium | Health Services Laboratories and Alex Alderton, Roberto Amato, Jeffrey Barrett, Sonia Goncalves, Ewan Harrison, David K. Jackson, Ian Johnston, Dominic Kwiatkowski, Cordelia Langford, John Sillitoe on behalf of the Wellcome Sanger Institute COVID-19 Surveillance Team |
| EPI_ISL_2394625, EPI_ISL_2394626, EPI_ISL_2394628, EPI_ISL_2394629, EPI_ISL_2394631, EPI_ISL_2394632, EPI_ISL_2394633, EPI_ISL_2394634, EPI_ISL_2394635, EPI_ISL_2394636, EPI_ISL_2394637 | | | |
| see above | Lighthouse Lab in Alderley Park | Wellcome Sanger Institute for the COVID-19 Genomics UK (COG-UK) Consortium | Jacquelyn Wynn, Mairead Hyland, The Lighthouse Lab in Alderley Park and Alex Alderton, Roberto Amato, Jeffrey Barrett, Sonia Goncalves, Ewan Harrison, David K. Jackson, Ian Johnston, Dominic Kwiatkowski, Cordelia Langford, John Sillitoe on behalf of the Wellcome Sanger Institute COVID-19 Surveillance Team |
| EPI_ISL_2394638 | Health Services Laboratories | Wellcome Sanger Institute for the COVID-19 Genomics UK (COG-UK) Consortium | Health Services Laboratories and Alex Alderton, Roberto Amato, Jeffrey Barrett, Sonia Goncalves, Ewan Harrison, David K. Jackson, Ian Johnston, Dominic Kwiatkowski, Cordelia Langford, John Sillitoe on behalf of the Wellcome Sanger Institute COVID-19 Surveillance Team |
| EPI_ISL_2394639 | Lighthouse Lab in Alderley Park | Wellcome Sanger Institute for the COVID-19 Genomics UK (COG-UK) Consortium | Jacquelyn Wynn, Mairead Hyland, The Lighthouse Lab in Alderley Park and Alex Alderton, Roberto Amato, Jeffrey Barrett, Sonia Goncalves, Ewan Harrison, David K. Jackson, Ian Johnston, Dominic Kwiatkowski, Cordelia Langford, John Sillitoe on behalf of the Wellcome Sanger Institute COVID-19 Surveillance Team |
| EPI_ISL_2394640 | Health Services Laboratories | Wellcome Sanger Institute for the COVID-19 Genomics UK (COG-UK) Consortium | Health Services Laboratories and Alex Alderton, Roberto Amato, Jeffrey Barrett, Sonia Goncalves, Ewan Harrison, David K. Jackson, Ian Johnston, Dominic Kwiatkowski, Cordelia Langford, John Sillitoe on behalf of the Wellcome Sanger Institute COVID-19 Surveillance Team |
| EPI_ISL_2394641 | Lighthouse Lab in Alderley Park | Wellcome Sanger Institute for the COVID-19 Genomics UK (COG-UK) Consortium | Jacquelyn Wynn, Mairead Hyland, The Lighthouse Lab in Alderley Park and Alex Alderton, Roberto Amato, Jeffrey Barrett, Sonia Goncalves, Ewan Harrison, David K. Jackson, Ian Johnston, Dominic Kwiatkowski, Cordelia Langford, John Sillitoe on behalf of the Wellcome Sanger Institute COVID-19 Surveillance Team |
| EPI_ISL_2394642 | Health Services Laboratories | Wellcome Sanger Institute for the COVID-19 Genomics UK (COG-UK) Consortium | Health Services Laboratories and Alex Alderton, Roberto Amato, Jeffrey Barrett, Sonia Goncalves, Ewan Harrison, David K. Jackson, Ian Johnston, Dominic Kwiatkowski, Cordelia Langford, John Sillitoe on behalf of the Wellcome Sanger Institute COVID-19 Surveillance Team |
| EPI_ISL_2394643, EPI_ISL_2394644, EPI_ISL_2394645, EPI_ISL_2394647, EPI_ISL_2394648, EPI_ISL_2394649, EPI_ISL_2394650, EPI_ISL_2394651, EPI_ISL_2394652, EPI_ISL_2394653, EPI_ISL_2394654, EPI_ISL_2394655, EPI_ISL_2394658, EPI_ISL_2394659, EPI_ISL_2394660, EPI_ISL_2394661, EPI_ISL_2394662, EPI_ISL_2394663, EPI_ISL_2394664, EPI_ISL_2394665, EPI_ISL_2394667, EPI_ISL_2394669, EPI_ISL_2394670, EPI_ISL_2394671, EPI_ISL_2394672, EPI_ISL_2394673, EPI_ISL_2394674, EPI_ISL_2394676, EPI_ISL_2394677, EPI_ISL_2394678, EPI_ISL_2394679, EPI_ISL_2394680, EPI_ISL_2394681, EPI_ISL_2394682, EPI_ISL_2394683, EPI_ISL_2394684, EPI_ISL_2394685, EPI_ISL_2394686 | | | |
| see above | Lighthouse Lab in Alderley Park | Wellcome Sanger Institute for the COVID-19 Genomics UK (COG-UK) Consortium | Jacquelyn Wynn, Mairead Hyland, The Lighthouse Lab in Alderley Park and Alex Alderton, Roberto Amato, Jeffrey Barrett, Sonia Goncalves, Ewan Harrison, David K. Jackson, Ian Johnston, Dominic Kwiatkowski, Cordelia Langford, John Sillitoe on behalf of the Wellcome Sanger Institute COVID-19 Surveillance Team |
| EPI_ISL_2394687 | Health Services Laboratories | Wellcome Sanger Institute for the COVID-19 Genomics UK (COG-UK) Consortium | Health Services Laboratories and Alex Alderton, Roberto Amato, Jeffrey Barrett, Sonia Goncalves, Ewan Harrison, David K. Jackson, Ian Johnston, Dominic Kwiatkowski, Cordelia Langford, John Sillitoe on behalf of the Wellcome Sanger Institute COVID-19 Surveillance Team |
| EPI_ISL_2394688 | Lighthouse Lab in Alderley Park | Wellcome Sanger Institute for the COVID-19 Genomics UK (COG-UK) Consortium | Jacquelyn Wynn, Mairead Hyland, The Lighthouse Lab in Alderley Park and Alex Alderton, Roberto Amato, Jeffrey Barrett, Sonia Goncalves, Ewan Harrison, David K. Jackson, Ian Johnston, Dominic Kwiatkowski, Cordelia Langford, John Sillitoe on behalf of the Wellcome Sanger Institute COVID-19 Surveillance Team |
| EPI_ISL_2394689 | Lighthouse Lab in Glasgow | Wellcome Sanger Institute for the COVID-19 Genomics UK (COG-UK) Consortium | Harper VanSteenhouse, Yumi Kasai, David Gray, Carol Clugston, Anna Dominiczak and Alex Alderton, Roberto Amato, Jeffrey Barrett, Sonia Goncalves, Ewan Harrison, David K. Jackson, Ian Johnston, Dominic Kwiatkowski, Cordelia Langford, John Sillitoe on behalf of the Wellcome Sanger Institute COVID-19 Surveillance Team |
| EPI_ISL_2394690, EPI_ISL_2394691, EPI_ISL_2394692, EPI_ISL_2394693 | Lighthouse Lab in Alderley Park | Wellcome Sanger Institute for the COVID-19 Genomics UK (COG-UK) Consortium | Jacquelyn Wynn, Mairead Hyland, The Lighthouse Lab in Alderley Park and Alex Alderton, Roberto Amato, Jeffrey Barrett, Sonia Goncalves, Ewan Harrison, David K. Jackson, Ian Johnston, Dominic Kwiatkowski, Cordelia Langford, John Sillitoe on behalf of the Wellcome Sanger Institute COVID-19 Surveillance Team |
| EPI_ISL_2394694 | Health Services Laboratories | Wellcome Sanger Institute for the COVID-19 Genomics UK (COG-UK) Consortium | Health Services Laboratories and Alex Alderton, Roberto Amato, Jeffrey Barrett, Sonia Goncalves, Ewan Harrison, David K. Jackson, Ian Johnston, Dominic Kwiatkowski, Cordelia Langford, John Sillitoe on behalf of the Wellcome Sanger Institute COVID-19 Surveillance Team |
| EPI_ISL_2394695, EPI_ISL_2394696, EPI_ISL_2394697, EPI_ISL_2394698, EPI_ISL_2394699 | Lighthouse Lab in Alderley Park | Wellcome Sanger Institute for the COVID-19 Genomics UK (COG-UK) Consortium | Jacquelyn Wynn, Mairead Hyland, The Lighthouse Lab in Alderley Park and Alex Alderton, Roberto Amato, Jeffrey Barrett, Sonia Goncalves, Ewan Harrison, David K. Jackson, Ian Johnston, Dominic Kwiatkowski, Cordelia Langford, John Sillitoe on behalf of the Wellcome Sanger Institute COVID-19 Surveillance Team |
| EPI_ISL_2394700 | Lighthouse Lab in Glasgow | Wellcome Sanger Institute for the COVID-19 Genomics UK (COG-UK) Consortium | Harper VanSteenhouse, Yumi Kasai, David Gray, Carol Clugston, Anna Dominiczak and Alex Alderton, Roberto Amato, Jeffrey Barrett, Sonia Goncalves, Ewan Harrison, David K. Jackson, Ian Johnston, Dominic Kwiatkowski, Cordelia Langford, John Sillitoe on behalf of the Wellcome Sanger Institute COVID-19 Surveillance Team |
| EPI_ISL_2394701, EPI_ISL_2394702, EPI_ISL_2394704, EPI_ISL_2394705, EPI_ISL_2394706 | Lighthouse Lab in Alderley Park | Wellcome Sanger Institute for the COVID-19 Genomics UK (COG-UK) Consortium | Jacquelyn Wynn, Mairead Hyland, The Lighthouse Lab in Alderley Park and Alex Alderton, Roberto Amato, Jeffrey Barrett, Sonia Goncalves, Ewan Harrison, David K. Jackson, Ian Johnston, Dominic Kwiatkowski, Cordelia Langford, John Sillitoe on behalf of the Wellcome Sanger Institute COVID-19 Surveillance Team |
| EPI_ISL_2394707 | Lighthouse Lab in Glasgow | Wellcome Sanger Institute for the COVID-19 Genomics UK (COG-UK) Consortium | Harper VanSteenhouse, Yumi Kasai, David Gray, Carol Clugston, Anna Dominiczak and Alex Alderton, Roberto Amato, Jeffrey Barrett, Sonia Goncalves, Ewan Harrison, David K. Jackson, Ian Johnston, Dominic Kwiatkowski, Cordelia Langford, John Sillitoe on behalf of the Wellcome Sanger Institute COVID-19 Surveillance Team |
| EPI_ISL_2394708 | Lighthouse Lab in Alderley Park | Wellcome Sanger Institute for the COVID-19 Genomics UK (COG-UK) Consortium | Jacquelyn Wynn, Mairead Hyland, The Lighthouse Lab in Alderley Park and Alex Alderton, Roberto Amato, Jeffrey Barrett, Sonia Goncalves, Ewan Harrison, David K. Jackson, Ian Johnston, Dominic Kwiatkowski, Cordelia Langford, John Sillitoe on behalf of the Wellcome Sanger Institute COVID-19 Surveillance Team |
| EPI_ISL_2394710 | Lighthouse Lab in Glasgow | Wellcome Sanger Institute for the COVID-19 Genomics UK (COG-UK) Consortium | Harper VanSteenhouse, Yumi Kasai, David Gray, Carol Clugston, Anna Dominiczak and Alex Alderton, Roberto Amato, Jeffrey Barrett, Sonia Goncalves, Ewan Harrison, David K. Jackson, Ian Johnston, Dominic Kwiatkowski, Cordelia Langford, John Sillitoe on behalf of the Wellcome Sanger Institute COVID-19 Surveillance Team |

| | | | |
|---|---|---|---|
| EPI_ISL_2394711, EPI_ISL_2394713, EPI_ISL_2394714, EPI_ISL_2394715, EPI_ISL_2394716 | Lighthouse Lab in Alderley Park | Wellcome Sanger Institute for the COVID-19 Genomics UK (COG-UK) Consortium | Jacquelyn Wynn, Mairead Hyland, The Lighthouse Lab in Alderley Park and Alex Alderton, Roberto Amato, Jeffrey Barrett, Sonia Goncalves, Ewan Harrison, David K. Jackson, Ian Johnston, Dominic Kwiatkowski, Cordelia Langford, John Sillitoe on behalf of the Wellcome Sanger Institute COVID-19 Surveillance Team |
| EPI_ISL_2394717 | Health Services Laboratories | Wellcome Sanger Institute for the COVID-19 Genomics UK (COG-UK) Consortium | Health Services Laboratories and Alex Alderton, Roberto Amato, Jeffrey Barrett, Sonia Goncalves, Ewan Harrison, David K. Jackson, Ian Johnston, Dominic Kwiatkowski, Cordelia Langford, John Sillitoe on behalf of the Wellcome Sanger Institute COVID-19 Surveillance Team |
| EPI_ISL_2394718 | Lighthouse Lab in Alderley Park | Wellcome Sanger Institute for the COVID-19 Genomics UK (COG-UK) Consortium | Jacquelyn Wynn, Mairead Hyland, The Lighthouse Lab in Alderley Park and Alex Alderton, Roberto Amato, Jeffrey Barrett, Sonia Goncalves, Ewan Harrison, David K. Jackson, Ian Johnston, Dominic Kwiatkowski, Cordelia Langford, John Sillitoe on behalf of the Wellcome Sanger Institute COVID-19 Surveillance Team |
| EPI_ISL_2394719 | Health Services Laboratories | Wellcome Sanger Institute for the COVID-19 Genomics UK (COG-UK) Consortium | Health Services Laboratories and Alex Alderton, Roberto Amato, Jeffrey Barrett, Sonia Goncalves, Ewan Harrison, David K. Jackson, Ian Johnston, Dominic Kwiatkowski, Cordelia Langford, John Sillitoe on behalf of the Wellcome Sanger Institute COVID-19 Surveillance Team |
| EPI_ISL_2394720, EPI_ISL_2394721, EPI_ISL_2394722, EPI_ISL_2394723, EPI_ISL_2394724, EPI_ISL_2394725, EPI_ISL_2394726, EPI_ISL_2394727, EPI_ISL_2394728, EPI_ISL_2394729, EPI_ISL_2394730, EPI_ISL_2394731, EPI_ISL_2394732, EPI_ISL_2394733, EPI_ISL_2394734, EPI_ISL_2394735, EPI_ISL_2394736, EPI_ISL_2394737 | | | |
| see above | Lighthouse Lab in Alderley Park | Wellcome Sanger Institute for the COVID-19 Genomics UK (COG-UK) Consortium | Jacquelyn Wynn, Mairead Hyland, The Lighthouse Lab in Alderley Park and Alex Alderton, Roberto Amato, Jeffrey Barrett, Sonia Goncalves, Ewan Harrison, David K. Jackson, Ian Johnston, Dominic Kwiatkowski, Cordelia Langford, John Sillitoe on behalf of the Wellcome Sanger Institute COVID-19 Surveillance Team |
| EPI_ISL_2394738 | Health Services Laboratories | Wellcome Sanger Institute for the COVID-19 Genomics UK (COG-UK) Consortium | Health Services Laboratories and Alex Alderton, Roberto Amato, Jeffrey Barrett, Sonia Goncalves, Ewan Harrison, David K. Jackson, Ian Johnston, Dominic Kwiatkowski, Cordelia Langford, John Sillitoe on behalf of the Wellcome Sanger Institute COVID-19 Surveillance Team |
| EPI_ISL_2394739, EPI_ISL_2394740, EPI_ISL_2394741 | Lighthouse Lab in Alderley Park | Wellcome Sanger Institute for the COVID-19 Genomics UK (COG-UK) Consortium | Jacquelyn Wynn, Mairead Hyland, The Lighthouse Lab in Alderley Park and Alex Alderton, Roberto Amato, Jeffrey Barrett, Sonia Goncalves, Ewan Harrison, David K. Jackson, Ian Johnston, Dominic Kwiatkowski, Cordelia Langford, John Sillitoe on behalf of the Wellcome Sanger Institute COVID-19 Surveillance Team |
| EPI_ISL_2394742 | Lighthouse Lab in Glasgow | Wellcome Sanger Institute for the COVID-19 Genomics UK (COG-UK) Consortium | Harper VanSteenhouse, Yumi Kasai, David Gray, Carol Clugston, Anna Dominiczak and Alex Alderton, Roberto Amato, Jeffrey Barrett, Sonia Goncalves, Ewan Harrison, David K. Jackson, Ian Johnston, Dominic Kwiatkowski, Cordelia Langford, John Sillitoe on behalf of the Wellcome Sanger Institute COVID-19 Surveillance Team |
| EPI_ISL_2394744, EPI_ISL_2394746 | Lighthouse Lab in Alderley Park | Wellcome Sanger Institute for the COVID-19 Genomics UK (COG-UK) Consortium | Jacquelyn Wynn, Mairead Hyland, The Lighthouse Lab in Alderley Park and Alex Alderton, Roberto Amato, Jeffrey Barrett, Sonia Goncalves, Ewan Harrison, David K. Jackson, Ian Johnston, Dominic Kwiatkowski, Cordelia Langford, John Sillitoe on behalf of the Wellcome Sanger Institute COVID-19 Surveillance Team |
| EPI_ISL_2394747 | Health Services Laboratories | Wellcome Sanger Institute for the COVID-19 Genomics UK (COG-UK) Consortium | Health Services Laboratories and Alex Alderton, Roberto Amato, Jeffrey Barrett, Sonia Goncalves, Ewan Harrison, David K. Jackson, Ian Johnston, Dominic Kwiatkowski, Cordelia Langford, John Sillitoe on behalf of the Wellcome Sanger Institute COVID-19 Surveillance Team |
| EPI_ISL_2394748, EPI_ISL_2394749, EPI_ISL_2394750, EPI_ISL_2394751, EPI_ISL_2394753, EPI_ISL_2394754 | Lighthouse Lab in Alderley Park | Wellcome Sanger Institute for the COVID-19 Genomics UK (COG-UK) Consortium | Jacquelyn Wynn, Mairead Hyland, The Lighthouse Lab in Alderley Park and Alex Alderton, Roberto Amato, Jeffrey Barrett, Sonia Goncalves, Ewan Harrison, David K. Jackson, Ian Johnston, Dominic Kwiatkowski, Cordelia Langford, John Sillitoe on behalf of the Wellcome Sanger Institute COVID-19 Surveillance Team |
| EPI_ISL_2394755 | Health Services Laboratories | Wellcome Sanger Institute for the COVID-19 Genomics UK (COG-UK) Consortium | Health Services Laboratories and Alex Alderton, Roberto Amato, Jeffrey Barrett, Sonia Goncalves, Ewan Harrison, David K. Jackson, Ian Johnston, Dominic Kwiatkowski, Cordelia Langford, John Sillitoe on behalf of the Wellcome Sanger Institute COVID-19 Surveillance Team |
| EPI_ISL_2394756, EPI_ISL_2394757, EPI_ISL_2394758, EPI_ISL_2394759 | Lighthouse Lab in Alderley Park | Wellcome Sanger Institute for the COVID-19 Genomics UK (COG-UK) Consortium | Jacquelyn Wynn, Mairead Hyland, The Lighthouse Lab in Alderley Park and Alex Alderton, Roberto Amato, Jeffrey Barrett, Sonia Goncalves, Ewan Harrison, David K. Jackson, Ian Johnston, Dominic Kwiatkowski, Cordelia Langford, John Sillitoe on behalf of the Wellcome Sanger Institute COVID-19 Surveillance Team |
| EPI_ISL_2394760 | Health Services Laboratories | Wellcome Sanger Institute for the COVID-19 Genomics UK (COG-UK) Consortium | Health Services Laboratories and Alex Alderton, Roberto Amato, Jeffrey Barrett, Sonia Goncalves, Ewan Harrison, David K. Jackson, Ian Johnston, Dominic Kwiatkowski, Cordelia Langford, John Sillitoe on behalf of the Wellcome Sanger Institute COVID-19 Surveillance Team |
| EPI_ISL_2394761 | Lighthouse Lab in Alderley Park | Wellcome Sanger Institute for the COVID-19 Genomics UK (COG-UK) Consortium | Jacquelyn Wynn, Mairead Hyland, The Lighthouse Lab in Alderley Park and Alex Alderton, Roberto Amato, Jeffrey Barrett, Sonia Goncalves, Ewan Harrison, David K. Jackson, Ian Johnston, Dominic Kwiatkowski, Cordelia Langford, John Sillitoe on behalf of the Wellcome Sanger Institute COVID-19 Surveillance Team |
| EPI_ISL_2394762 | Lighthouse Lab in Glasgow | Wellcome Sanger Institute for the COVID-19 Genomics UK (COG-UK) Consortium | Harper VanSteenhouse, Yumi Kasai, David Gray, Carol Clugston, Anna Dominiczak and Alex Alderton, Roberto Amato, Jeffrey Barrett, Sonia Goncalves, Ewan Harrison, David K. Jackson, Ian Johnston, Dominic Kwiatkowski, Cordelia Langford, John Sillitoe on behalf of the Wellcome Sanger Institute COVID-19 Surveillance Team |
| EPI_ISL_2394763, EPI_ISL_2394764, EPI_ISL_2394765, EPI_ISL_2394766 | Lighthouse Lab in Alderley Park | Wellcome Sanger Institute for the COVID-19 Genomics UK (COG-UK) Consortium | Jacquelyn Wynn, Mairead Hyland, The Lighthouse Lab in Alderley Park and Alex Alderton, Roberto Amato, Jeffrey Barrett, Sonia Goncalves, Ewan Harrison, David K. Jackson, Ian Johnston, Dominic Kwiatkowski, Cordelia Langford, John Sillitoe on behalf of the Wellcome Sanger Institute COVID-19 Surveillance Team |
| EPI_ISL_2394767 | Lighthouse Lab in Glasgow | Wellcome Sanger Institute for the COVID-19 Genomics UK (COG-UK) Consortium | Harper VanSteenhouse, Yumi Kasai, David Gray, Carol Clugston, Anna Dominiczak and Alex Alderton, Roberto Amato, Jeffrey Barrett, Sonia Goncalves, Ewan Harrison, David K. Jackson, Ian Johnston, Dominic Kwiatkowski, Cordelia Langford, John Sillitoe on behalf of the Wellcome Sanger Institute COVID-19 Surveillance Team |
| EPI_ISL_2394768, EPI_ISL_2394769, EPI_ISL_2394772, EPI_ISL_2394773, EPI_ISL_2394774, EPI_ISL_2394776, EPI_ISL_2394778, EPI_ISL_2394779 | Lighthouse Lab in Alderley Park | Wellcome Sanger Institute for the COVID-19 Genomics UK (COG-UK) Consortium | Jacquelyn Wynn, Mairead Hyland, The Lighthouse Lab in Alderley Park and Alex Alderton, Roberto Amato, Jeffrey Barrett, Sonia Goncalves, Ewan Harrison, David K. Jackson, Ian Johnston, Dominic Kwiatkowski, Cordelia Langford, John Sillitoe on behalf of the Wellcome Sanger Institute COVID-19 Surveillance Team |
| EPI_ISL_2394780 | Lighthouse Lab in Glasgow | Wellcome Sanger Institute for the COVID-19 Genomics UK (COG-UK) Consortium | Harper VanSteenhouse, Yumi Kasai, David Gray, Carol Clugston, Anna Dominiczak and Alex Alderton, Roberto Amato, Jeffrey Barrett, Sonia Goncalves, Ewan Harrison, David K. Jackson, Ian Johnston, Dominic Kwiatkowski, Cordelia Langford, John Sillitoe on behalf of the Wellcome Sanger Institute COVID-19 Surveillance Team |
| EPI_ISL_2394781, EPI_ISL_2394782, EPI_ISL_2394783 | Lighthouse Lab in Alderley Park | Wellcome Sanger Institute for the COVID-19 Genomics UK (COG-UK) Consortium | Jacquelyn Wynn, Mairead Hyland, The Lighthouse Lab in Alderley Park and Alex Alderton, Roberto Amato, Jeffrey Barrett, Sonia Goncalves, Ewan Harrison, David K. Jackson, Ian Johnston, Dominic Kwiatkowski, Cordelia Langford, John Sillitoe on behalf of the Wellcome Sanger Institute COVID-19 Surveillance Team |
| EPI_ISL_2394784 | Health Services Laboratories | Wellcome Sanger Institute for the COVID-19 Genomics UK (COG-UK) Consortium | Health Services Laboratories and Alex Alderton, Roberto Amato, Jeffrey Barrett, Sonia Goncalves, Ewan Harrison, David K. Jackson, Ian Johnston, Dominic Kwiatkowski, Cordelia Langford, John Sillitoe on behalf of the Wellcome Sanger Institute COVID-19 Surveillance Team |
| EPI_ISL_2394785, EPI_ISL_2394786, EPI_ISL_2394787, EPI_ISL_2394788, EPI_ISL_2394789, EPI_ISL_2394791, EPI_ISL_2394792, EPI_ISL_2394793, EPI_ISL_2394794, EPI_ISL_2394795 | Lighthouse Lab in Alderley Park | Wellcome Sanger Institute for the COVID-19 Genomics UK (COG-UK) Consortium | Jacquelyn Wynn, Mairead Hyland, The Lighthouse Lab in Alderley Park and Alex Alderton, Roberto Amato, Jeffrey Barrett, Sonia Goncalves, Ewan Harrison, David K. Jackson, Ian Johnston, Dominic Kwiatkowski, Cordelia Langford, John Sillitoe on behalf of the Wellcome Sanger Institute COVID-19 Surveillance Team |
| EPI_ISL_2394797 | Health Services Laboratories | Wellcome Sanger Institute for the COVID-19 Genomics UK (COG-UK) Consortium | Health Services Laboratories and Alex Alderton, Roberto Amato, Jeffrey Barrett, Sonia Goncalves, Ewan Harrison, David K. Jackson, Ian Johnston, Dominic Kwiatkowski, Cordelia Langford, John Sillitoe on behalf of the Wellcome Sanger Institute COVID-19 Surveillance Team |
| EPI_ISL_2394799, EPI_ISL_2394800, EPI_ISL_2394802 | Lighthouse Lab in Alderley Park | Wellcome Sanger Institute for the COVID-19 Genomics UK (COG-UK) Consortium | Jacquelyn Wynn, Mairead Hyland, The Lighthouse Lab in Alderley Park and Alex Alderton, Roberto Amato, Jeffrey Barrett, Sonia Goncalves, Ewan Harrison, David K. Jackson, Ian Johnston, Dominic Kwiatkowski, Cordelia Langford, John Sillitoe on behalf of the Wellcome Sanger Institute COVID-19 Surveillance Team |
| EPI_ISL_2394803 | Lighthouse Lab in Glasgow | Wellcome Sanger Institute for the COVID-19 Genomics UK (COG-UK) Consortium | Harper VanSteenhouse, Yumi Kasai, David Gray, Carol Clugston, Anna Dominiczak and Alex Alderton, Roberto Amato, Jeffrey Barrett, Sonia Goncalves, Ewan Harrison, David K. Jackson, Ian Johnston, Dominic Kwiatkowski, Cordelia Langford, John Sillitoe on behalf of the Wellcome Sanger Institute COVID-19 Surveillance Team |
| EPI_ISL_2394804 | Health Services Laboratories | Wellcome Sanger Institute for the COVID-19 Genomics UK (COG-UK) Consortium | Health Services Laboratories and Alex Alderton, Roberto Amato, Jeffrey Barrett, Sonia Goncalves, Ewan Harrison, David K. Jackson, Ian Johnston, Dominic Kwiatkowski, Cordelia Langford, John Sillitoe on behalf of the Wellcome Sanger Institute COVID-19 Surveillance Team |
| EPI_ISL_2394805, EPI_ISL_2394806 | Lighthouse Lab in Alderley Park | Wellcome Sanger Institute for the COVID-19 Genomics UK (COG-UK) Consortium | Jacquelyn Wynn, Mairead Hyland, The Lighthouse Lab in Alderley Park and Alex Alderton, Roberto Amato, Jeffrey Barrett, Sonia Goncalves, Ewan Harrison, David K. Jackson, Ian Johnston, Dominic Kwiatkowski, Cordelia Langford, John Sillitoe on behalf of the Wellcome Sanger Institute COVID-19 Surveillance Team |
| EPI_ISL_2394807 | Health Services Laboratories | Wellcome Sanger Institute for the COVID-19 Genomics UK (COG-UK) Consortium | Health Services Laboratories and Alex Alderton, Roberto Amato, Jeffrey Barrett, Sonia Goncalves, Ewan Harrison, David K. Jackson, Ian Johnston, Dominic Kwiatkowski, Cordelia Langford, John Sillitoe on behalf of the Wellcome Sanger Institute COVID-19 Surveillance Team |
| EPI_ISL_2394808, EPI_ISL_2394809, EPI_ISL_2394810, EPI_ISL_2394811, EPI_ISL_2394813, EPI_ISL_2394814, EPI_ISL_2394815, EPI_ISL_2394816, EPI_ISL_2394817, EPI_ISL_2394818, EPI_ISL_2394819, EPI_ISL_2394820 | | | |
| see above | Lighthouse Lab in Alderley Park | Wellcome Sanger Institute for the COVID-19 Genomics UK (COG-UK) Consortium | Jacquelyn Wynn, Mairead Hyland, The Lighthouse Lab in Alderley Park and Alex Alderton, Roberto Amato, Jeffrey Barrett, Sonia Goncalves, Ewan Harrison, David K. Jackson, Ian Johnston, Dominic Kwiatkowski, Cordelia Langford, John Sillitoe on behalf of the Wellcome Sanger Institute COVID-19 Surveillance Team |
| EPI_ISL_2394821 | Health Services Laboratories | Wellcome Sanger Institute for the COVID-19 Genomics UK (COG-UK) Consortium | Health Services Laboratories and Alex Alderton, Roberto Amato, Jeffrey Barrett, Sonia Goncalves, Ewan Harrison, David K. Jackson, Ian Johnston, Dominic Kwiatkowski, Cordelia Langford, John Sillitoe on behalf of the Wellcome Sanger Institute COVID-19 Surveillance Team |
| EPI_ISL_2394823, EPI_ISL_2394824, EPI_ISL_2394826, EPI_ISL_2394828, EPI_ISL_2394829, EPI_ISL_2394830, EPI_ISL_2394831, EPI_ISL_2394832, EPI_ISL_2394833, EPI_ISL_2394834, EPI_ISL_2394835, EPI_ISL_2394836, EPI_ISL_2394838, EPI_ISL_2394839 | | | |
| see above | Lighthouse Lab in Alderley Park | Wellcome Sanger Institute for the COVID-19 Genomics UK (COG-UK) Consortium | Jacquelyn Wynn, Mairead Hyland, The Lighthouse Lab in Alderley Park and Alex Alderton, Roberto Amato, Jeffrey Barrett, Sonia Goncalves, Ewan Harrison, David K. Jackson, Ian Johnston, Dominic Kwiatkowski, Cordelia Langford, John Sillitoe on behalf of the Wellcome Sanger Institute COVID-19 Surveillance Team |

| | | | |
|---|---|---|---|
| EPI_ISL_2394840 | Health Services Laboratories | Wellcome Sanger Institute for the COVID-19 Genomics UK (COG-UK) Consortium | Health Services Laboratories and Alex Alderton, Roberto Amato, Jeffrey Barrett, Sonia Goncalves, Ewan Harrison, David K. Jackson, Ian Johnston, Dominic Kwiatkowski, Cordelia Langford, John Sillitoe on behalf of the Wellcome Sanger Institute COVID-19 Surveillance Team |
| EPI_ISL_2394841, EPI_ISL_2394843 | Lighthouse Lab in Alderley Park | Wellcome Sanger Institute for the COVID-19 Genomics UK (COG-UK) Consortium | Jacquelyn Wynn, Mairead Hyland, The Lighthouse Lab in Alderley Park and Alex Alderton, Roberto Amato, Jeffrey Barrett, Sonia Goncalves, Ewan Harrison, David K. Jackson, Ian Johnston, Dominic Kwiatkowski, Cordelia Langford, John Sillitoe on behalf of the Wellcome Sanger Institute COVID-19 Surveillance Team |
| EPI_ISL_2394844 | Health Services Laboratories | Wellcome Sanger Institute for the COVID-19 Genomics UK (COG-UK) Consortium | Health Services Laboratories and Alex Alderton, Roberto Amato, Jeffrey Barrett, Sonia Goncalves, Ewan Harrison, David K. Jackson, Ian Johnston, Dominic Kwiatkowski, Cordelia Langford, John Sillitoe on behalf of the Wellcome Sanger Institute COVID-19 Surveillance Team |
| EPI_ISL_2394845 | Lighthouse Lab in Alderley Park | Wellcome Sanger Institute for the COVID-19 Genomics UK (COG-UK) Consortium | Jacquelyn Wynn, Mairead Hyland, The Lighthouse Lab in Alderley Park and Alex Alderton, Roberto Amato, Jeffrey Barrett, Sonia Goncalves, Ewan Harrison, David K. Jackson, Ian Johnston, Dominic Kwiatkowski, Cordelia Langford, John Sillitoe on behalf of the Wellcome Sanger Institute COVID-19 Surveillance Team |
| EPI_ISL_2394846 | Lighthouse Lab in Glasgow | Wellcome Sanger Institute for the COVID-19 Genomics UK (COG-UK) Consortium | Harper VanSteenhouse, Yumi Kasai, David Gray, Carol Clugston, Anna Dominiczak and Alex Alderton, Roberto Amato, Jeffrey Barrett, Sonia Goncalves, Ewan Harrison, David K. Jackson, Ian Johnston, Dominic Kwiatkowski, Cordelia Langford, John Sillitoe on behalf of the Wellcome Sanger Institute COVID-19 Surveillance Team |
| EPI_ISL_2394847, EPI_ISL_2394848, EPI_ISL_2394849 | Lighthouse Lab in Alderley Park | Wellcome Sanger Institute for the COVID-19 Genomics UK (COG-UK) Consortium | Jacquelyn Wynn, Mairead Hyland, The Lighthouse Lab in Alderley Park and Alex Alderton, Roberto Amato, Jeffrey Barrett, Sonia Goncalves, Ewan Harrison, David K. Jackson, Ian Johnston, Dominic Kwiatkowski, Cordelia Langford, John Sillitoe on behalf of the Wellcome Sanger Institute COVID-19 Surveillance Team |
| EPI_ISL_2394850 | Health Services Laboratories | Wellcome Sanger Institute for the COVID-19 Genomics UK (COG-UK) Consortium | Health Services Laboratories and Alex Alderton, Roberto Amato, Jeffrey Barrett, Sonia Goncalves, Ewan Harrison, David K. Jackson, Ian Johnston, Dominic Kwiatkowski, Cordelia Langford, John Sillitoe on behalf of the Wellcome Sanger Institute COVID-19 Surveillance Team |
| EPI_ISL_2394851, EPI_ISL_2394852, EPI_ISL_2394853 | Lighthouse Lab in Alderley Park | Wellcome Sanger Institute for the COVID-19 Genomics UK (COG-UK) Consortium | Jacquelyn Wynn, Mairead Hyland, The Lighthouse Lab in Alderley Park and Alex Alderton, Roberto Amato, Jeffrey Barrett, Sonia Goncalves, Ewan Harrison, David K. Jackson, Ian Johnston, Dominic Kwiatkowski, Cordelia Langford, John Sillitoe on behalf of the Wellcome Sanger Institute COVID-19 Surveillance Team |
| EPI_ISL_2394856 | Lighthouse Lab in Glasgow | Wellcome Sanger Institute for the COVID-19 Genomics UK (COG-UK) Consortium | Harper VanSteenhouse, Yumi Kasai, David Gray, Carol Clugston, Anna Dominiczak and Alex Alderton, Roberto Amato, Jeffrey Barrett, Sonia Goncalves, Ewan Harrison, David K. Jackson, Ian Johnston, Dominic Kwiatkowski, Cordelia Langford, John Sillitoe on behalf of the Wellcome Sanger Institute COVID-19 Surveillance Team |
| EPI_ISL_2394857 | Lighthouse Lab in Alderley Park | Wellcome Sanger Institute for the COVID-19 Genomics UK (COG-UK) Consortium | Jacquelyn Wynn, Mairead Hyland, The Lighthouse Lab in Alderley Park and Alex Alderton, Roberto Amato, Jeffrey Barrett, Sonia Goncalves, Ewan Harrison, David K. Jackson, Ian Johnston, Dominic Kwiatkowski, Cordelia Langford, John Sillitoe on behalf of the Wellcome Sanger Institute COVID-19 Surveillance Team |
| EPI_ISL_2394858 | Health Services Laboratories | Wellcome Sanger Institute for the COVID-19 Genomics UK (COG-UK) Consortium | Health Services Laboratories and Alex Alderton, Roberto Amato, Jeffrey Barrett, Sonia Goncalves, Ewan Harrison, David K. Jackson, Ian Johnston, Dominic Kwiatkowski, Cordelia Langford, John Sillitoe on behalf of the Wellcome Sanger Institute COVID-19 Surveillance Team |
| EPI_ISL_2394859, EPI_ISL_2394860, EPI_ISL_2394861, EPI_ISL_2394863 | Lighthouse Lab in Alderley Park | Wellcome Sanger Institute for the COVID-19 Genomics UK (COG-UK) Consortium | Jacquelyn Wynn, Mairead Hyland, The Lighthouse Lab in Alderley Park and Alex Alderton, Roberto Amato, Jeffrey Barrett, Sonia Goncalves, Ewan Harrison, David K. Jackson, Ian Johnston, Dominic Kwiatkowski, Cordelia Langford, John Sillitoe on behalf of the Wellcome Sanger Institute COVID-19 Surveillance Team |
| EPI_ISL_2394865 | Lighthouse Lab in Milton Keynes | Wellcome Sanger Institute for the COVID-19 Genomics UK (COG-UK) Consortium | The Lighthouse Lab in Milton Keynes and Alex Alderton, Roberto Amato, Jeffrey Barrett, Sonia Goncalves, Ewan Harrison, David K. Jackson, Ian Johnston, Dominic Kwiatkowski, Cordelia Langford, John Sillitoe on behalf of the Wellcome Sanger Institute COVID-19 Surveillance Team |
| EPI_ISL_2394866 | Lighthouse Lab in Alderley Park | Wellcome Sanger Institute for the COVID-19 Genomics UK (COG-UK) Consortium | Jacquelyn Wynn, Mairead Hyland, The Lighthouse Lab in Alderley Park and Alex Alderton, Roberto Amato, Jeffrey Barrett, Sonia Goncalves, Ewan Harrison, David K. Jackson, Ian Johnston, Dominic Kwiatkowski, Cordelia Langford, John Sillitoe on behalf of the Wellcome Sanger Institute COVID-19 Surveillance Team |
| EPI_ISL_2394867, EPI_ISL_2394868, EPI_ISL_2394869, EPI_ISL_2394870 | Lighthouse Lab in Milton Keynes | Wellcome Sanger Institute for the COVID-19 Genomics UK (COG-UK) Consortium | The Lighthouse Lab in Milton Keynes and Alex Alderton, Roberto Amato, Jeffrey Barrett, Sonia Goncalves, Ewan Harrison, David K. Jackson, Ian Johnston, Dominic Kwiatkowski, Cordelia Langford, John Sillitoe on behalf of the Wellcome Sanger Institute COVID-19 Surveillance Team |
| EPI_ISL_2394871 | Lighthouse Lab in Alderley Park | Wellcome Sanger Institute for the COVID-19 Genomics UK (COG-UK) Consortium | Jacquelyn Wynn, Mairead Hyland, The Lighthouse Lab in Alderley Park and Alex Alderton, Roberto Amato, Jeffrey Barrett, Sonia Goncalves, Ewan Harrison, David K. Jackson, Ian Johnston, Dominic Kwiatkowski, Cordelia Langford, John Sillitoe on behalf of the Wellcome Sanger Institute COVID-19 Surveillance Team |
| EPI_ISL_2394872 | Lighthouse Lab in Milton Keynes | Wellcome Sanger Institute for the COVID-19 Genomics UK (COG-UK) Consortium | The Lighthouse Lab in Milton Keynes and Alex Alderton, Roberto Amato, Jeffrey Barrett, Sonia Goncalves, Ewan Harrison, David K. Jackson, Ian Johnston, Dominic Kwiatkowski, Cordelia Langford, John Sillitoe on behalf of the Wellcome Sanger Institute COVID-19 Surveillance Team |
| EPI_ISL_2394873 | Lighthouse Lab in Alderley Park | Wellcome Sanger Institute for the COVID-19 Genomics UK (COG-UK) Consortium | Jacquelyn Wynn, Mairead Hyland, The Lighthouse Lab in Alderley Park and Alex Alderton, Roberto Amato, Jeffrey Barrett, Sonia Goncalves, Ewan Harrison, David K. Jackson, Ian Johnston, Dominic Kwiatkowski, Cordelia Langford, John Sillitoe on behalf of the Wellcome Sanger Institute COVID-19 Surveillance Team |
| EPI_ISL_2394874, EPI_ISL_2394875, EPI_ISL_2394877 | Lighthouse Lab in Milton Keynes | Wellcome Sanger Institute for the COVID-19 Genomics UK (COG-UK) Consortium | The Lighthouse Lab in Milton Keynes and Alex Alderton, Roberto Amato, Jeffrey Barrett, Sonia Goncalves, Ewan Harrison, David K. Jackson, Ian Johnston, Dominic Kwiatkowski, Cordelia Langford, John Sillitoe on behalf of the Wellcome Sanger Institute COVID-19 Surveillance Team |
| EPI_ISL_2394878 | Lighthouse Lab in Glasgow | Wellcome Sanger Institute for the COVID-19 Genomics UK (COG-UK) Consortium | Harper VanSteenhouse, Yumi Kasai, David Gray, Carol Clugston, Anna Dominiczak and Alex Alderton, Roberto Amato, Jeffrey Barrett, Sonia Goncalves, Ewan Harrison, David K. Jackson, Ian Johnston, Dominic Kwiatkowski, Cordelia Langford, John Sillitoe on behalf of the Wellcome Sanger Institute COVID-19 Surveillance Team |
| EPI_ISL_2394879 | Lighthouse Lab in Alderley Park | Wellcome Sanger Institute for the COVID-19 Genomics UK (COG-UK) Consortium | Jacquelyn Wynn, Mairead Hyland, The Lighthouse Lab in Alderley Park and Alex Alderton, Roberto Amato, Jeffrey Barrett, Sonia Goncalves, Ewan Harrison, David K. Jackson, Ian Johnston, Dominic Kwiatkowski, Cordelia Langford, John Sillitoe on behalf of the Wellcome Sanger Institute COVID-19 Surveillance Team |
| EPI_ISL_2394880 | Lighthouse Lab in Milton Keynes | Wellcome Sanger Institute for the COVID-19 Genomics UK (COG-UK) Consortium | The Lighthouse Lab in Milton Keynes and Alex Alderton, Roberto Amato, Jeffrey Barrett, Sonia Goncalves, Ewan Harrison, David K. Jackson, Ian Johnston, Dominic Kwiatkowski, Cordelia Langford, John Sillitoe on behalf of the Wellcome Sanger Institute COVID-19 Surveillance Team |
| EPI_ISL_2394881 | Lighthouse Lab in Alderley Park | Wellcome Sanger Institute for the COVID-19 Genomics UK (COG-UK) Consortium | Jacquelyn Wynn, Mairead Hyland, The Lighthouse Lab in Alderley Park and Alex Alderton, Roberto Amato, Jeffrey Barrett, Sonia Goncalves, Ewan Harrison, David K. Jackson, Ian Johnston, Dominic Kwiatkowski, Cordelia Langford, John Sillitoe on behalf of the Wellcome Sanger Institute COVID-19 Surveillance Team |
| EPI_ISL_2394882, EPI_ISL_2394883 | Lighthouse Lab in Milton Keynes | Wellcome Sanger Institute for the COVID-19 Genomics UK (COG-UK) Consortium | The Lighthouse Lab in Milton Keynes and Alex Alderton, Roberto Amato, Jeffrey Barrett, Sonia Goncalves, Ewan Harrison, David K. Jackson, Ian Johnston, Dominic Kwiatkowski, Cordelia Langford, John Sillitoe on behalf of the Wellcome Sanger Institute COVID-19 Surveillance Team |
| EPI_ISL_2394884, EPI_ISL_2394885 | Lighthouse Lab in Alderley Park | Wellcome Sanger Institute for the COVID-19 Genomics UK (COG-UK) Consortium | Jacquelyn Wynn, Mairead Hyland, The Lighthouse Lab in Alderley Park and Alex Alderton, Roberto Amato, Jeffrey Barrett, Sonia Goncalves, Ewan Harrison, David K. Jackson, Ian Johnston, Dominic Kwiatkowski, Cordelia Langford, John Sillitoe on behalf of the Wellcome Sanger Institute COVID-19 Surveillance Team |
| EPI_ISL_2394886 | Lighthouse Lab in Milton Keynes | Wellcome Sanger Institute for the COVID-19 Genomics UK (COG-UK) Consortium | The Lighthouse Lab in Milton Keynes and Alex Alderton, Roberto Amato, Jeffrey Barrett, Sonia Goncalves, Ewan Harrison, David K. Jackson, Ian Johnston, Dominic Kwiatkowski, Cordelia Langford, John Sillitoe on behalf of the Wellcome Sanger Institute COVID-19 Surveillance Team |
| EPI_ISL_2394887, EPI_ISL_2394888, EPI_ISL_2394889, EPI_ISL_2394890, EPI_ISL_2394891 | Lighthouse Lab in Alderley Park | Wellcome Sanger Institute for the COVID-19 Genomics UK (COG-UK) Consortium | Jacquelyn Wynn, Mairead Hyland, The Lighthouse Lab in Alderley Park and Alex Alderton, Roberto Amato, Jeffrey Barrett, Sonia Goncalves, Ewan Harrison, David K. Jackson, Ian Johnston, Dominic Kwiatkowski, Cordelia Langford, John Sillitoe on behalf of the Wellcome Sanger Institute COVID-19 Surveillance Team |
| EPI_ISL_2394892 | Lighthouse Lab in Glasgow | Wellcome Sanger Institute for the COVID-19 Genomics UK (COG-UK) Consortium | Harper VanSteenhouse, Yumi Kasai, David Gray, Carol Clugston, Anna Dominiczak and Alex Alderton, Roberto Amato, Jeffrey Barrett, Sonia Goncalves, Ewan Harrison, David K. Jackson, Ian Johnston, Dominic Kwiatkowski, Cordelia Langford, John Sillitoe on behalf of the Wellcome Sanger Institute COVID-19 Surveillance Team |
| EPI_ISL_2394894, EPI_ISL_2394896 | Lighthouse Lab in Milton Keynes | Wellcome Sanger Institute for the COVID-19 Genomics UK (COG-UK) Consortium | The Lighthouse Lab in Milton Keynes and Alex Alderton, Roberto Amato, Jeffrey Barrett, Sonia Goncalves, Ewan Harrison, David K. Jackson, Ian Johnston, Dominic Kwiatkowski, Cordelia Langford, John Sillitoe on behalf of the Wellcome Sanger Institute COVID-19 Surveillance Team |
| EPI_ISL_2394898, EPI_ISL_2394899, EPI_ISL_2394900 | Lighthouse Lab in Alderley Park | Wellcome Sanger Institute for the COVID-19 Genomics UK (COG-UK) Consortium | Jacquelyn Wynn, Mairead Hyland, The Lighthouse Lab in Alderley Park and Alex Alderton, Roberto Amato, Jeffrey Barrett, Sonia Goncalves, Ewan Harrison, David K. Jackson, Ian Johnston, Dominic Kwiatkowski, Cordelia Langford, John Sillitoe on behalf of the Wellcome Sanger Institute COVID-19 Surveillance Team |
| EPI_ISL_2394902, EPI_ISL_2394904, EPI_ISL_2394905 | Lighthouse Lab in Milton Keynes | Wellcome Sanger Institute for the COVID-19 Genomics UK (COG-UK) Consortium | The Lighthouse Lab in Milton Keynes and Alex Alderton, Roberto Amato, Jeffrey Barrett, Sonia Goncalves, Ewan Harrison, David K. Jackson, Ian Johnston, Dominic Kwiatkowski, Cordelia Langford, John Sillitoe on behalf of the Wellcome Sanger Institute COVID-19 Surveillance Team |
| EPI_ISL_2394906 | Lighthouse Lab in Alderley Park | Wellcome Sanger Institute for the COVID-19 Genomics UK (COG-UK) Consortium | Jacquelyn Wynn, Mairead Hyland, The Lighthouse Lab in Alderley Park and Alex Alderton, Roberto Amato, Jeffrey Barrett, Sonia Goncalves, Ewan Harrison, David K. Jackson, Ian Johnston, Dominic Kwiatkowski, Cordelia Langford, John Sillitoe on behalf of the Wellcome Sanger Institute COVID-19 Surveillance Team |
| EPI_ISL_2394907, EPI_ISL_2394908 | Lighthouse Lab in Milton Keynes | Wellcome Sanger Institute for the COVID-19 Genomics UK (COG-UK) Consortium | The Lighthouse Lab in Milton Keynes and Alex Alderton, Roberto Amato, Jeffrey Barrett, Sonia Goncalves, Ewan Harrison, David K. Jackson, Ian Johnston, Dominic Kwiatkowski, Cordelia Langford, John Sillitoe on behalf of the Wellcome Sanger Institute COVID-19 Surveillance Team |
| EPI_ISL_2394909 | Lighthouse Lab in Alderley Park | Wellcome Sanger Institute for the COVID-19 Genomics UK (COG-UK) Consortium | Jacquelyn Wynn, Mairead Hyland, The Lighthouse Lab in Alderley Park and Alex Alderton, Roberto Amato, Jeffrey Barrett, Sonia Goncalves, Ewan Harrison, David K. Jackson, Ian Johnston, Dominic Kwiatkowski, Cordelia Langford, John Sillitoe on behalf of the Wellcome Sanger Institute COVID-19 Surveillance Team |
| EPI_ISL_2394910, EPI_ISL_2394911 | Lighthouse Lab in Milton Keynes | Wellcome Sanger Institute for the COVID-19 Genomics UK (COG-UK) Consortium | The Lighthouse Lab in Milton Keynes and Alex Alderton, Roberto Amato, Jeffrey Barrett, Sonia Goncalves, Ewan Harrison, David K. Jackson, Ian Johnston, Dominic Kwiatkowski, Cordelia Langford, John Sillitoe on behalf of the Wellcome Sanger Institute COVID-19 Surveillance Team |
| EPI_ISL_2394912 | Lighthouse Lab in Alderley Park | Wellcome Sanger Institute for the COVID-19 Genomics UK (COG-UK) Consortium | Jacquelyn Wynn, Mairead Hyland, The Lighthouse Lab in Alderley Park and Alex Alderton, Roberto Amato, Jeffrey Barrett, Sonia Goncalves, Ewan Harrison, David K. Jackson, Ian Johnston, Dominic Kwiatkowski, Cordelia Langford, John Sillitoe on behalf of the Wellcome Sanger Institute COVID-19 Surveillance Team |
| EPI_ISL_2394913 | Lighthouse Lab in Milton Keynes | Wellcome Sanger Institute for the COVID-19 Genomics UK (COG-UK) Consortium | The Lighthouse Lab in Milton Keynes and Alex Alderton, Roberto Amato, Jeffrey Barrett, Sonia Goncalves, Ewan Harrison, David K. Jackson, Ian Johnston, Dominic Kwiatkowski, Cordelia Langford, John Sillitoe on behalf of the Wellcome Sanger Institute COVID-19 Surveillance Team |
| EPI_ISL_2394914, EPI_ISL_2394916 | Lighthouse Lab in Alderley Park | Wellcome Sanger Institute for the COVID-19 Genomics UK (COG-UK) Consortium | Jacquelyn Wynn, Mairead Hyland, The Lighthouse Lab in Alderley Park and Alex Alderton, Roberto Amato, Jeffrey Barrett, Sonia Goncalves, Ewan Harrison, David K. Jackson, Ian Johnston, Dominic Kwiatkowski, Cordelia Langford, John Sillitoe on behalf of the Wellcome Sanger Institute COVID-19 Surveillance Team |
| EPI_ISL_2394917, EPI_ISL_2394918 | Lighthouse Lab in Milton Keynes | Wellcome Sanger Institute for the COVID-19 Genomics UK (COG-UK) Consortium | The Lighthouse Lab in Milton Keynes and Alex Alderton, Roberto Amato, Jeffrey Barrett, Sonia Goncalves, Ewan Harrison, David K. Jackson, Ian Johnston, Dominic Kwiatkowski, Cordelia Langford, John Sillitoe on behalf of the Wellcome Sanger Institute COVID-19 Surveillance Team |
| EPI_ISL_2394919 | Lighthouse Lab in Glasgow | Wellcome Sanger Institute for the COVID-19 Genomics UK (COG-UK) Consortium | Harper VanSteenhouse, Yumi Kasai, David Gray, Carol Clugston, Anna Dominiczak and Alex Alderton, Roberto Amato, Jeffrey Barrett, Sonia Goncalves, Ewan Harrison, David K. Jackson, Ian Johnston, Dominic Kwiatkowski, Cordelia Langford, John Sillitoe on behalf of the Wellcome Sanger Institute COVID-19 Surveillance Team |
| EPI_ISL_2394920 | Lighthouse Lab in Milton Keynes | Wellcome Sanger Institute for the COVID-19 Genomics UK (COG-UK) | The Lighthouse Lab in Milton Keynes and Alex Alderton, Roberto Amato, Jeffrey Barrett, Sonia Goncalves, Ewan Harrison, David K. Jackson, Ian Johnston, Dominic Kwiatkowski, Cordelia Langford, John Sillitoe on behalf of the Wellcome Sanger |

| | | | |
|---|---|---|---|
| | | Consortium | Institute COVID-19 Surveillance Team |
| EPI_ISL_2394922 | Lighthouse Lab in Alderley Park | Wellcome Sanger Institute for the COVID-19 Genomics UK (COG-UK) Consortium | Jacquelyn Wynn, Mairead Hyland, The Lighthouse Lab in Alderley Park and Alex Alderton, Roberto Amato, Jeffrey Barrett, Sonia Goncalves, Ewan Harrison, David K. Jackson, Ian Johnston, Dominic Kwiatkowski, Cordelia Langford, John Sillitoe on behalf of the Wellcome Sanger Institute COVID-19 Surveillance Team |
| EPI_ISL_2394923 | Lighthouse Lab in Milton Keynes | Wellcome Sanger Institute for the COVID-19 Genomics UK (COG-UK) Consortium | The Lighthouse Lab in Milton Keynes and Alex Alderton, Roberto Amato, Jeffrey Barrett, Sonia Goncalves, Ewan Harrison, David K. Jackson, Ian Johnston, Dominic Kwiatkowski, Cordelia Langford, John Sillitoe on behalf of the Wellcome Sanger Institute COVID-19 Surveillance Team |
| EPI_ISL_2394924 | Lighthouse Lab in Alderley Park | Wellcome Sanger Institute for the COVID-19 Genomics UK (COG-UK) Consortium | Jacquelyn Wynn, Mairead Hyland, The Lighthouse Lab in Alderley Park and Alex Alderton, Roberto Amato, Jeffrey Barrett, Sonia Goncalves, Ewan Harrison, David K. Jackson, Ian Johnston, Dominic Kwiatkowski, Cordelia Langford, John Sillitoe on behalf of the Wellcome Sanger Institute COVID-19 Surveillance Team |
| EPI_ISL_2394925, EPI_ISL_2394926 | Lighthouse Lab in Milton Keynes | Wellcome Sanger Institute for the COVID-19 Genomics UK (COG-UK) Consortium | The Lighthouse Lab in Milton Keynes and Alex Alderton, Roberto Amato, Jeffrey Barrett, Sonia Goncalves, Ewan Harrison, David K. Jackson, Ian Johnston, Dominic Kwiatkowski, Cordelia Langford, John Sillitoe on behalf of the Wellcome Sanger Institute COVID-19 Surveillance Team |
| EPI_ISL_2394928 | Lighthouse Lab in Alderley Park | Wellcome Sanger Institute for the COVID-19 Genomics UK (COG-UK) Consortium | Jacquelyn Wynn, Mairead Hyland, The Lighthouse Lab in Alderley Park and Alex Alderton, Roberto Amato, Jeffrey Barrett, Sonia Goncalves, Ewan Harrison, David K. Jackson, Ian Johnston, Dominic Kwiatkowski, Cordelia Langford, John Sillitoe on behalf of the Wellcome Sanger Institute COVID-19 Surveillance Team |
| EPI_ISL_2394929, EPI_ISL_2394931 | Lighthouse Lab in Milton Keynes | Wellcome Sanger Institute for the COVID-19 Genomics UK (COG-UK) Consortium | The Lighthouse Lab in Milton Keynes and Alex Alderton, Roberto Amato, Jeffrey Barrett, Sonia Goncalves, Ewan Harrison, David K. Jackson, Ian Johnston, Dominic Kwiatkowski, Cordelia Langford, John Sillitoe on behalf of the Wellcome Sanger Institute COVID-19 Surveillance Team |
| EPI_ISL_2394932 | Lighthouse Lab in Alderley Park | Wellcome Sanger Institute for the COVID-19 Genomics UK (COG-UK) Consortium | Jacquelyn Wynn, Mairead Hyland, The Lighthouse Lab in Alderley Park and Alex Alderton, Roberto Amato, Jeffrey Barrett, Sonia Goncalves, Ewan Harrison, David K. Jackson, Ian Johnston, Dominic Kwiatkowski, Cordelia Langford, John Sillitoe on behalf of the Wellcome Sanger Institute COVID-19 Surveillance Team |
| EPI_ISL_2394933, EPI_ISL_2394934 | Lighthouse Lab in Milton Keynes | Wellcome Sanger Institute for the COVID-19 Genomics UK (COG-UK) Consortium | The Lighthouse Lab in Milton Keynes and Alex Alderton, Roberto Amato, Jeffrey Barrett, Sonia Goncalves, Ewan Harrison, David K. Jackson, Ian Johnston, Dominic Kwiatkowski, Cordelia Langford, John Sillitoe on behalf of the Wellcome Sanger Institute COVID-19 Surveillance Team |
| EPI_ISL_2394938, EPI_ISL_2394939, EPI_ISL_2394940 | Lighthouse Lab in Alderley Park | Wellcome Sanger Institute for the COVID-19 Genomics UK (COG-UK) Consortium | Jacquelyn Wynn, Mairead Hyland, The Lighthouse Lab in Alderley Park and Alex Alderton, Roberto Amato, Jeffrey Barrett, Sonia Goncalves, Ewan Harrison, David K. Jackson, Ian Johnston, Dominic Kwiatkowski, Cordelia Langford, John Sillitoe on behalf of the Wellcome Sanger Institute COVID-19 Surveillance Team |
| EPI_ISL_2394943 | Lighthouse Lab in Milton Keynes | Wellcome Sanger Institute for the COVID-19 Genomics UK (COG-UK) Consortium | The Lighthouse Lab in Milton Keynes and Alex Alderton, Roberto Amato, Jeffrey Barrett, Sonia Goncalves, Ewan Harrison, David K. Jackson, Ian Johnston, Dominic Kwiatkowski, Cordelia Langford, John Sillitoe on behalf of the Wellcome Sanger Institute COVID-19 Surveillance Team |
| EPI_ISL_2394945, EPI_ISL_2394946, EPI_ISL_2394947, EPI_ISL_2394948, EPI_ISL_2394949 | Lighthouse Lab in Alderley Park | Wellcome Sanger Institute for the COVID-19 Genomics UK (COG-UK) Consortium | Jacquelyn Wynn, Mairead Hyland, The Lighthouse Lab in Alderley Park and Alex Alderton, Roberto Amato, Jeffrey Barrett, Sonia Goncalves, Ewan Harrison, David K. Jackson, Ian Johnston, Dominic Kwiatkowski, Cordelia Langford, John Sillitoe on behalf of the Wellcome Sanger Institute COVID-19 Surveillance Team |
| EPI_ISL_2394950 | Lighthouse Lab in Milton Keynes | Wellcome Sanger Institute for the COVID-19 Genomics UK (COG-UK) Consortium | The Lighthouse Lab in Milton Keynes and Alex Alderton, Roberto Amato, Jeffrey Barrett, Sonia Goncalves, Ewan Harrison, David K. Jackson, Ian Johnston, Dominic Kwiatkowski, Cordelia Langford, John Sillitoe on behalf of the Wellcome Sanger Institute COVID-19 Surveillance Team |
| EPI_ISL_2394951, EPI_ISL_2394952, EPI_ISL_2394954, EPI_ISL_2394955 | Lighthouse Lab in Alderley Park | Wellcome Sanger Institute for the COVID-19 Genomics UK (COG-UK) Consortium | Jacquelyn Wynn, Mairead Hyland, The Lighthouse Lab in Alderley Park and Alex Alderton, Roberto Amato, Jeffrey Barrett, Sonia Goncalves, Ewan Harrison, David K. Jackson, Ian Johnston, Dominic Kwiatkowski, Cordelia Langford, John Sillitoe on behalf of the Wellcome Sanger Institute COVID-19 Surveillance Team |
| EPI_ISL_2394956, EPI_ISL_2394957 | Lighthouse Lab in Milton Keynes | Wellcome Sanger Institute for the COVID-19 Genomics UK (COG-UK) Consortium | The Lighthouse Lab in Milton Keynes and Alex Alderton, Roberto Amato, Jeffrey Barrett, Sonia Goncalves, Ewan Harrison, David K. Jackson, Ian Johnston, Dominic Kwiatkowski, Cordelia Langford, John Sillitoe on behalf of the Wellcome Sanger Institute COVID-19 Surveillance Team |
| EPI_ISL_2394958, EPI_ISL_2394959, EPI_ISL_2394960 | Lighthouse Lab in Alderley Park | Wellcome Sanger Institute for the COVID-19 Genomics UK (COG-UK) Consortium | Jacquelyn Wynn, Mairead Hyland, The Lighthouse Lab in Alderley Park and Alex Alderton, Roberto Amato, Jeffrey Barrett, Sonia Goncalves, Ewan Harrison, David K. Jackson, Ian Johnston, Dominic Kwiatkowski, Cordelia Langford, John Sillitoe on behalf of the Wellcome Sanger Institute COVID-19 Surveillance Team |
| EPI_ISL_2394961 | Lighthouse Lab in Milton Keynes | Wellcome Sanger Institute for the COVID-19 Genomics UK (COG-UK) Consortium | The Lighthouse Lab in Milton Keynes and Alex Alderton, Roberto Amato, Jeffrey Barrett, Sonia Goncalves, Ewan Harrison, David K. Jackson, Ian Johnston, Dominic Kwiatkowski, Cordelia Langford, John Sillitoe on behalf of the Wellcome Sanger Institute COVID-19 Surveillance Team |
| EPI_ISL_2394962 | Lighthouse Lab in Alderley Park | Wellcome Sanger Institute for the COVID-19 Genomics UK (COG-UK) Consortium | Jacquelyn Wynn, Mairead Hyland, The Lighthouse Lab in Alderley Park and Alex Alderton, Roberto Amato, Jeffrey Barrett, Sonia Goncalves, Ewan Harrison, David K. Jackson, Ian Johnston, Dominic Kwiatkowski, Cordelia Langford, John Sillitoe on behalf of the Wellcome Sanger Institute COVID-19 Surveillance Team |
| EPI_ISL_2394963 | Lighthouse Lab in Milton Keynes | Wellcome Sanger Institute for the COVID-19 Genomics UK (COG-UK) Consortium | The Lighthouse Lab in Milton Keynes and Alex Alderton, Roberto Amato, Jeffrey Barrett, Sonia Goncalves, Ewan Harrison, David K. Jackson, Ian Johnston, Dominic Kwiatkowski, Cordelia Langford, John Sillitoe on behalf of the Wellcome Sanger Institute COVID-19 Surveillance Team |
| EPI_ISL_2394964 | Lighthouse Lab in Alderley Park | Wellcome Sanger Institute for the COVID-19 Genomics UK (COG-UK) Consortium | Jacquelyn Wynn, Mairead Hyland, The Lighthouse Lab in Alderley Park and Alex Alderton, Roberto Amato, Jeffrey Barrett, Sonia Goncalves, Ewan Harrison, David K. Jackson, Ian Johnston, Dominic Kwiatkowski, Cordelia Langford, John Sillitoe on behalf of the Wellcome Sanger Institute COVID-19 Surveillance Team |
| EPI_ISL_2394965, EPI_ISL_2394966 | Lighthouse Lab in Milton Keynes | Wellcome Sanger Institute for the COVID-19 Genomics UK (COG-UK) Consortium | The Lighthouse Lab in Milton Keynes and Alex Alderton, Roberto Amato, Jeffrey Barrett, Sonia Goncalves, Ewan Harrison, David K. Jackson, Ian Johnston, Dominic Kwiatkowski, Cordelia Langford, John Sillitoe on behalf of the Wellcome Sanger Institute COVID-19 Surveillance Team |
| EPI_ISL_2394967, EPI_ISL_2394969, EPI_ISL_2394970, EPI_ISL_2394971 | Lighthouse Lab in Alderley Park | Wellcome Sanger Institute for the COVID-19 Genomics UK (COG-UK) Consortium | Jacquelyn Wynn, Mairead Hyland, The Lighthouse Lab in Alderley Park and Alex Alderton, Roberto Amato, Jeffrey Barrett, Sonia Goncalves, Ewan Harrison, David K. Jackson, Ian Johnston, Dominic Kwiatkowski, Cordelia Langford, John Sillitoe on behalf of the Wellcome Sanger Institute COVID-19 Surveillance Team |
| EPI_ISL_2394972 | Lighthouse Lab in Milton Keynes | Wellcome Sanger Institute for the COVID-19 Genomics UK (COG-UK) Consortium | The Lighthouse Lab in Milton Keynes and Alex Alderton, Roberto Amato, Jeffrey Barrett, Sonia Goncalves, Ewan Harrison, David K. Jackson, Ian Johnston, Dominic Kwiatkowski, Cordelia Langford, John Sillitoe on behalf of the Wellcome Sanger Institute COVID-19 Surveillance Team |
| EPI_ISL_2394973 | Lighthouse Lab in Alderley Park | Wellcome Sanger Institute for the COVID-19 Genomics UK (COG-UK) Consortium | Jacquelyn Wynn, Mairead Hyland, The Lighthouse Lab in Alderley Park and Alex Alderton, Roberto Amato, Jeffrey Barrett, Sonia Goncalves, Ewan Harrison, David K. Jackson, Ian Johnston, Dominic Kwiatkowski, Cordelia Langford, John Sillitoe on behalf of the Wellcome Sanger Institute COVID-19 Surveillance Team |
| EPI_ISL_2394974 | Lighthouse Lab in Milton Keynes | Wellcome Sanger Institute for the COVID-19 Genomics UK (COG-UK) Consortium | The Lighthouse Lab in Milton Keynes and Alex Alderton, Roberto Amato, Jeffrey Barrett, Sonia Goncalves, Ewan Harrison, David K. Jackson, Ian Johnston, Dominic Kwiatkowski, Cordelia Langford, John Sillitoe on behalf of the Wellcome Sanger Institute COVID-19 Surveillance Team |
| EPI_ISL_2394975 | Lighthouse Lab in Glasgow | Wellcome Sanger Institute for the COVID-19 Genomics UK (COG-UK) Consortium | Harper VanSteenhouse, Yumi Kasai, David Gray, Carol Clugston, Anna Dominiczak and Alex Alderton, Roberto Amato, Jeffrey Barrett, Sonia Goncalves, Ewan Harrison, David K. Jackson, Ian Johnston, Dominic Kwiatkowski, Cordelia Langford, John Sillitoe on behalf of the Wellcome Sanger Institute COVID-19 Surveillance Team |
| EPI_ISL_2394976, EPI_ISL_2394977, EPI_ISL_2394979, EPI_ISL_2394980 | Lighthouse Lab in Alderley Park | Wellcome Sanger Institute for the COVID-19 Genomics UK (COG-UK) Consortium | Jacquelyn Wynn, Mairead Hyland, The Lighthouse Lab in Alderley Park and Alex Alderton, Roberto Amato, Jeffrey Barrett, Sonia Goncalves, Ewan Harrison, David K. Jackson, Ian Johnston, Dominic Kwiatkowski, Cordelia Langford, John Sillitoe on behalf of the Wellcome Sanger Institute COVID-19 Surveillance Team |
| EPI_ISL_2394981, EPI_ISL_2394982, EPI_ISL_2394983, EPI_ISL_2394984 | Lighthouse Lab in Milton Keynes | Wellcome Sanger Institute for the COVID-19 Genomics UK (COG-UK) Consortium | The Lighthouse Lab in Milton Keynes and Alex Alderton, Roberto Amato, Jeffrey Barrett, Sonia Goncalves, Ewan Harrison, David K. Jackson, Ian Johnston, Dominic Kwiatkowski, Cordelia Langford, John Sillitoe on behalf of the Wellcome Sanger Institute COVID-19 Surveillance Team |
| EPI_ISL_2394985, EPI_ISL_2394986 | Lighthouse Lab in Alderley Park | Wellcome Sanger Institute for the COVID-19 Genomics UK (COG-UK) Consortium | Jacquelyn Wynn, Mairead Hyland, The Lighthouse Lab in Alderley Park and Alex Alderton, Roberto Amato, Jeffrey Barrett, Sonia Goncalves, Ewan Harrison, David K. Jackson, Ian Johnston, Dominic Kwiatkowski, Cordelia Langford, John Sillitoe on behalf of the Wellcome Sanger Institute COVID-19 Surveillance Team |
| EPI_ISL_2394987 | Lighthouse Lab in Milton Keynes | Wellcome Sanger Institute for the COVID-19 Genomics UK (COG-UK) Consortium | The Lighthouse Lab in Milton Keynes and Alex Alderton, Roberto Amato, Jeffrey Barrett, Sonia Goncalves, Ewan Harrison, David K. Jackson, Ian Johnston, Dominic Kwiatkowski, Cordelia Langford, John Sillitoe on behalf of the Wellcome Sanger Institute COVID-19 Surveillance Team |
| EPI_ISL_2394988 | Lighthouse Lab in Alderley Park | Wellcome Sanger Institute for the COVID-19 Genomics UK (COG-UK) Consortium | Jacquelyn Wynn, Mairead Hyland, The Lighthouse Lab in Alderley Park and Alex Alderton, Roberto Amato, Jeffrey Barrett, Sonia Goncalves, Ewan Harrison, David K. Jackson, Ian Johnston, Dominic Kwiatkowski, Cordelia Langford, John Sillitoe on behalf of the Wellcome Sanger Institute COVID-19 Surveillance Team |
| EPI_ISL_2394990, EPI_ISL_2394992, EPI_ISL_2394993 | Lighthouse Lab in Milton Keynes | Wellcome Sanger Institute for the COVID-19 Genomics UK (COG-UK) Consortium | The Lighthouse Lab in Milton Keynes and Alex Alderton, Roberto Amato, Jeffrey Barrett, Sonia Goncalves, Ewan Harrison, David K. Jackson, Ian Johnston, Dominic Kwiatkowski, Cordelia Langford, John Sillitoe on behalf of the Wellcome Sanger Institute COVID-19 Surveillance Team |
| EPI_ISL_2394995 | Lighthouse Lab in Alderley Park | Wellcome Sanger Institute for the COVID-19 Genomics UK (COG-UK) Consortium | Jacquelyn Wynn, Mairead Hyland, The Lighthouse Lab in Alderley Park and Alex Alderton, Roberto Amato, Jeffrey Barrett, Sonia Goncalves, Ewan Harrison, David K. Jackson, Ian Johnston, Dominic Kwiatkowski, Cordelia Langford, John Sillitoe on behalf of the Wellcome Sanger Institute COVID-19 Surveillance Team |
| EPI_ISL_2394996 | Lighthouse Lab in Milton Keynes | Wellcome Sanger Institute for the COVID-19 Genomics UK (COG-UK) Consortium | The Lighthouse Lab in Milton Keynes and Alex Alderton, Roberto Amato, Jeffrey Barrett, Sonia Goncalves, Ewan Harrison, David K. Jackson, Ian Johnston, Dominic Kwiatkowski, Cordelia Langford, John Sillitoe on behalf of the Wellcome Sanger Institute COVID-19 Surveillance Team |
| EPI_ISL_2394997, EPI_ISL_2394998 | Lighthouse Lab in Alderley Park | Wellcome Sanger Institute for the COVID-19 Genomics UK (COG-UK) Consortium | Jacquelyn Wynn, Mairead Hyland, The Lighthouse Lab in Alderley Park and Alex Alderton, Roberto Amato, Jeffrey Barrett, Sonia Goncalves, Ewan Harrison, David K. Jackson, Ian Johnston, Dominic Kwiatkowski, Cordelia Langford, John Sillitoe on behalf of the Wellcome Sanger Institute COVID-19 Surveillance Team |
| EPI_ISL_2394999 | Lighthouse Lab in Milton Keynes | Wellcome Sanger Institute for the COVID-19 Genomics UK (COG-UK) Consortium | The Lighthouse Lab in Milton Keynes and Alex Alderton, Roberto Amato, Jeffrey Barrett, Sonia Goncalves, Ewan Harrison, David K. Jackson, Ian Johnston, Dominic Kwiatkowski, Cordelia Langford, John Sillitoe on behalf of the Wellcome Sanger Institute COVID-19 Surveillance Team |
| EPI_ISL_2395000, EPI_ISL_2395002, EPI_ISL_2395003, EPI_ISL_2395004 | Lighthouse Lab in Alderley Park | Wellcome Sanger Institute for the COVID-19 Genomics UK (COG-UK) Consortium | Jacquelyn Wynn, Mairead Hyland, The Lighthouse Lab in Alderley Park and Alex Alderton, Roberto Amato, Jeffrey Barrett, Sonia Goncalves, Ewan Harrison, David K. Jackson, Ian Johnston, Dominic Kwiatkowski, Cordelia Langford, John Sillitoe on behalf of the Wellcome Sanger Institute COVID-19 Surveillance Team |
| EPI_ISL_2395005, EPI_ISL_2395007 | Lighthouse Lab in Milton Keynes | Wellcome Sanger Institute for the COVID-19 Genomics UK (COG-UK) Consortium | The Lighthouse Lab in Milton Keynes and Alex Alderton, Roberto Amato, Jeffrey Barrett, Sonia Goncalves, Ewan Harrison, David K. Jackson, Ian Johnston, Dominic Kwiatkowski, Cordelia Langford, John Sillitoe on behalf of the Wellcome Sanger Institute COVID-19 Surveillance Team |
| EPI_ISL_2395009, EPI_ISL_2395011 | Lighthouse Lab in Alderley Park | Wellcome Sanger Institute for the COVID-19 Genomics UK (COG-UK) Consortium | Jacquelyn Wynn, Mairead Hyland, The Lighthouse Lab in Alderley Park and Alex Alderton, Roberto Amato, Jeffrey Barrett, Sonia Goncalves, Ewan Harrison, David K. Jackson, Ian Johnston, Dominic Kwiatkowski, Cordelia Langford, John Sillitoe on behalf of the Wellcome Sanger Institute COVID-19 Surveillance Team |

| | | | |
|---|---|---|---|
| EPI_ISL_2395012 | Lighthouse Lab in Milton Keynes | Wellcome Sanger Institute for the COVID-19 Genomics UK (COG-UK) Consortium | The Lighthouse Lab in Milton Keynes and Alex Alderton, Roberto Amato, Jeffrey Barrett, Sonia Goncalves, Ewan Harrison, David K. Jackson, Ian Johnston, Dominic Kwiatkowski, Cordelia Langford, John Sillitoe on behalf of the Wellcome Sanger Institute COVID-19 Surveillance Team |
| EPI_ISL_2395014 | Lighthouse Lab in Alderley Park | Wellcome Sanger Institute for the COVID-19 Genomics UK (COG-UK) Consortium | Jacquelyn Wynn, Mairead Hyland, The Lighthouse Lab in Alderley Park and Alex Alderton, Roberto Amato, Jeffrey Barrett, Sonia Goncalves, Ewan Harrison, David K. Jackson, Ian Johnston, Dominic Kwiatkowski, Cordelia Langford, John Sillitoe on behalf of the Wellcome Sanger Institute COVID-19 Surveillance Team |
| EPI_ISL_2395015 | Lighthouse Lab in Glasgow | Wellcome Sanger Institute for the COVID-19 Genomics UK (COG-UK) Consortium | Harper VanSteenhouse, Yumi Kasai, David Gray, Carol Clugston, Anna Dominiczak and Alex Alderton, Roberto Amato, Jeffrey Barrett, Sonia Goncalves, Ewan Harrison, David K. Jackson, Ian Johnston, Dominic Kwiatkowski, Cordelia Langford, John Sillitoe on behalf of the Wellcome Sanger Institute COVID-19 Surveillance Team |
| EPI_ISL_2395016, EPI_ISL_2395018, EPI_ISL_2395019 | Lighthouse Lab in Alderley Park | Wellcome Sanger Institute for the COVID-19 Genomics UK (COG-UK) Consortium | Jacquelyn Wynn, Mairead Hyland, The Lighthouse Lab in Alderley Park and Alex Alderton, Roberto Amato, Jeffrey Barrett, Sonia Goncalves, Ewan Harrison, David K. Jackson, Ian Johnston, Dominic Kwiatkowski, Cordelia Langford, John Sillitoe on behalf of the Wellcome Sanger Institute COVID-19 Surveillance Team |
| EPI_ISL_2395022 | Lighthouse Lab in Milton Keynes | Wellcome Sanger Institute for the COVID-19 Genomics UK (COG-UK) Consortium | The Lighthouse Lab in Milton Keynes and Alex Alderton, Roberto Amato, Jeffrey Barrett, Sonia Goncalves, Ewan Harrison, David K. Jackson, Ian Johnston, Dominic Kwiatkowski, Cordelia Langford, John Sillitoe on behalf of the Wellcome Sanger Institute COVID-19 Surveillance Team |
| EPI_ISL_2395023, EPI_ISL_2395025 | Lighthouse Lab in Alderley Park | Wellcome Sanger Institute for the COVID-19 Genomics UK (COG-UK) Consortium | Jacquelyn Wynn, Mairead Hyland, The Lighthouse Lab in Alderley Park and Alex Alderton, Roberto Amato, Jeffrey Barrett, Sonia Goncalves, Ewan Harrison, David K. Jackson, Ian Johnston, Dominic Kwiatkowski, Cordelia Langford, John Sillitoe on behalf of the Wellcome Sanger Institute COVID-19 Surveillance Team |
| EPI_ISL_2395028 | Lighthouse Lab in Milton Keynes | Wellcome Sanger Institute for the COVID-19 Genomics UK (COG-UK) Consortium | The Lighthouse Lab in Milton Keynes and Alex Alderton, Roberto Amato, Jeffrey Barrett, Sonia Goncalves, Ewan Harrison, David K. Jackson, Ian Johnston, Dominic Kwiatkowski, Cordelia Langford, John Sillitoe on behalf of the Wellcome Sanger Institute COVID-19 Surveillance Team |
| EPI_ISL_2395030 | Lighthouse Lab in Alderley Park | Wellcome Sanger Institute for the COVID-19 Genomics UK (COG-UK) Consortium | Jacquelyn Wynn, Mairead Hyland, The Lighthouse Lab in Alderley Park and Alex Alderton, Roberto Amato, Jeffrey Barrett, Sonia Goncalves, Ewan Harrison, David K. Jackson, Ian Johnston, Dominic Kwiatkowski, Cordelia Langford, John Sillitoe on behalf of the Wellcome Sanger Institute COVID-19 Surveillance Team |
| EPI_ISL_2395031 | Lighthouse Lab in Milton Keynes | Wellcome Sanger Institute for the COVID-19 Genomics UK (COG-UK) Consortium | The Lighthouse Lab in Milton Keynes and Alex Alderton, Roberto Amato, Jeffrey Barrett, Sonia Goncalves, Ewan Harrison, David K. Jackson, Ian Johnston, Dominic Kwiatkowski, Cordelia Langford, John Sillitoe on behalf of the Wellcome Sanger Institute COVID-19 Surveillance Team |
| EPI_ISL_2395032 | Lighthouse Lab in Alderley Park | Wellcome Sanger Institute for the COVID-19 Genomics UK (COG-UK) Consortium | Jacquelyn Wynn, Mairead Hyland, The Lighthouse Lab in Alderley Park and Alex Alderton, Roberto Amato, Jeffrey Barrett, Sonia Goncalves, Ewan Harrison, David K. Jackson, Ian Johnston, Dominic Kwiatkowski, Cordelia Langford, John Sillitoe on behalf of the Wellcome Sanger Institute COVID-19 Surveillance Team |
| EPI_ISL_2395033, EPI_ISL_2395035 | Lighthouse Lab in Milton Keynes | Wellcome Sanger Institute for the COVID-19 Genomics UK (COG-UK) Consortium | The Lighthouse Lab in Milton Keynes and Alex Alderton, Roberto Amato, Jeffrey Barrett, Sonia Goncalves, Ewan Harrison, David K. Jackson, Ian Johnston, Dominic Kwiatkowski, Cordelia Langford, John Sillitoe on behalf of the Wellcome Sanger Institute COVID-19 Surveillance Team |
| EPI_ISL_2395036 | Lighthouse Lab in Alderley Park | Wellcome Sanger Institute for the COVID-19 Genomics UK (COG-UK) Consortium | Jacquelyn Wynn, Mairead Hyland, The Lighthouse Lab in Alderley Park and Alex Alderton, Roberto Amato, Jeffrey Barrett, Sonia Goncalves, Ewan Harrison, David K. Jackson, Ian Johnston, Dominic Kwiatkowski, Cordelia Langford, John Sillitoe on behalf of the Wellcome Sanger Institute COVID-19 Surveillance Team |
| EPI_ISL_2395037 | Lighthouse Lab in Milton Keynes | Wellcome Sanger Institute for the COVID-19 Genomics UK (COG-UK) Consortium | The Lighthouse Lab in Milton Keynes and Alex Alderton, Roberto Amato, Jeffrey Barrett, Sonia Goncalves, Ewan Harrison, David K. Jackson, Ian Johnston, Dominic Kwiatkowski, Cordelia Langford, John Sillitoe on behalf of the Wellcome Sanger Institute COVID-19 Surveillance Team |
| EPI_ISL_2395038, EPI_ISL_2395039 | Lighthouse Lab in Alderley Park | Wellcome Sanger Institute for the COVID-19 Genomics UK (COG-UK) Consortium | Jacquelyn Wynn, Mairead Hyland, The Lighthouse Lab in Alderley Park and Alex Alderton, Roberto Amato, Jeffrey Barrett, Sonia Goncalves, Ewan Harrison, David K. Jackson, Ian Johnston, Dominic Kwiatkowski, Cordelia Langford, John Sillitoe on behalf of the Wellcome Sanger Institute COVID-19 Surveillance Team |
| EPI_ISL_2395040 | Lighthouse Lab in Milton Keynes | Wellcome Sanger Institute for the COVID-19 Genomics UK (COG-UK) Consortium | The Lighthouse Lab in Milton Keynes and Alex Alderton, Roberto Amato, Jeffrey Barrett, Sonia Goncalves, Ewan Harrison, David K. Jackson, Ian Johnston, Dominic Kwiatkowski, Cordelia Langford, John Sillitoe on behalf of the Wellcome Sanger Institute COVID-19 Surveillance Team |
| EPI_ISL_2395041, EPI_ISL_2395042 | Lighthouse Lab in Alderley Park | Wellcome Sanger Institute for the COVID-19 Genomics UK (COG-UK) Consortium | Jacquelyn Wynn, Mairead Hyland, The Lighthouse Lab in Alderley Park and Alex Alderton, Roberto Amato, Jeffrey Barrett, Sonia Goncalves, Ewan Harrison, David K. Jackson, Ian Johnston, Dominic Kwiatkowski, Cordelia Langford, John Sillitoe on behalf of the Wellcome Sanger Institute COVID-19 Surveillance Team |
| EPI_ISL_2395043, EPI_ISL_2395044 | Lighthouse Lab in Glasgow | Wellcome Sanger Institute for the COVID-19 Genomics UK (COG-UK) Consortium | Harper VanSteenhouse, Yumi Kasai, David Gray, Carol Clugston, Anna Dominiczak and Alex Alderton, Roberto Amato, Jeffrey Barrett, Sonia Goncalves, Ewan Harrison, David K. Jackson, Ian Johnston, Dominic Kwiatkowski, Cordelia Langford, John Sillitoe on behalf of the Wellcome Sanger Institute COVID-19 Surveillance Team |
| EPI_ISL_2395045, EPI_ISL_2395046, EPI_ISL_2395047, EPI_ISL_2395048, EPI_ISL_2395049, EPI_ISL_2395050, EPI_ISL_2395051, EPI_ISL_2395052, EPI_ISL_2395053, EPI_ISL_2395054, EPI_ISL_2395055, EPI_ISL_2395056, EPI_ISL_2395057, EPI_ISL_2395058, EPI_ISL_2395059, EPI_ISL_2395060, EPI_ISL_2395061, EPI_ISL_2395062, EPI_ISL_2395063, EPI_ISL_2395064, EPI_ISL_2395065, EPI_ISL_2395066, EPI_ISL_2395067, EPI_ISL_2395069, EPI_ISL_2395070, EPI_ISL_2395071, EPI_ISL_2395072 | | | |
| see above | Lighthouse Lab in Alderley Park | Wellcome Sanger Institute for the COVID-19 Genomics UK (COG-UK) Consortium | Jacquelyn Wynn, Mairead Hyland, The Lighthouse Lab in Alderley Park and Alex Alderton, Roberto Amato, Jeffrey Barrett, Sonia Goncalves, Ewan Harrison, David K. Jackson, Ian Johnston, Dominic Kwiatkowski, Cordelia Langford, John Sillitoe on behalf of the Wellcome Sanger Institute COVID-19 Surveillance Team |
| EPI_ISL_2395073 | Lighthouse Lab in Glasgow | Wellcome Sanger Institute for the COVID-19 Genomics UK (COG-UK) Consortium | Harper VanSteenhouse, Yumi Kasai, David Gray, Carol Clugston, Anna Dominiczak and Alex Alderton, Roberto Amato, Jeffrey Barrett, Sonia Goncalves, Ewan Harrison, David K. Jackson, Ian Johnston, Dominic Kwiatkowski, Cordelia Langford, John Sillitoe on behalf of the Wellcome Sanger Institute COVID-19 Surveillance Team |
| EPI_ISL_2395074, EPI_ISL_2395075, EPI_ISL_2395076, EPI_ISL_2395077, EPI_ISL_2395078, EPI_ISL_2395079, EPI_ISL_2395080, EPI_ISL_2395081 | Lighthouse Lab in Alderley Park | Wellcome Sanger Institute for the COVID-19 Genomics UK (COG-UK) Consortium | Jacquelyn Wynn, Mairead Hyland, The Lighthouse Lab in Alderley Park and Alex Alderton, Roberto Amato, Jeffrey Barrett, Sonia Goncalves, Ewan Harrison, David K. Jackson, Ian Johnston, Dominic Kwiatkowski, Cordelia Langford, John Sillitoe on behalf of the Wellcome Sanger Institute COVID-19 Surveillance Team |
| EPI_ISL_2395082 | Lighthouse Lab in Glasgow | Wellcome Sanger Institute for the COVID-19 Genomics UK (COG-UK) Consortium | Harper VanSteenhouse, Yumi Kasai, David Gray, Carol Clugston, Anna Dominiczak and Alex Alderton, Roberto Amato, Jeffrey Barrett, Sonia Goncalves, Ewan Harrison, David K. Jackson, Ian Johnston, Dominic Kwiatkowski, Cordelia Langford, John Sillitoe on behalf of the Wellcome Sanger Institute COVID-19 Surveillance Team |
| EPI_ISL_2395083, EPI_ISL_2395084, EPI_ISL_2395085, EPI_ISL_2395086, EPI_ISL_2395088, EPI_ISL_2395089, EPI_ISL_2395090, EPI_ISL_2395091, EPI_ISL_2395093, EPI_ISL_2395095, EPI_ISL_2395096, EPI_ISL_2395097, EPI_ISL_2395098, EPI_ISL_2395099, EPI_ISL_2395100, EPI_ISL_2395101, EPI_ISL_2395103, EPI_ISL_2395104, EPI_ISL_2395105, EPI_ISL_2395108, EPI_ISL_2395110, EPI_ISL_2395111 | | | |
| see above | Lighthouse Lab in Alderley Park | Wellcome Sanger Institute for the COVID-19 Genomics UK (COG-UK) Consortium | Jacquelyn Wynn, Mairead Hyland, The Lighthouse Lab in Alderley Park and Alex Alderton, Roberto Amato, Jeffrey Barrett, Sonia Goncalves, Ewan Harrison, David K. Jackson, Ian Johnston, Dominic Kwiatkowski, Cordelia Langford, John Sillitoe on behalf of the Wellcome Sanger Institute COVID-19 Surveillance Team |
| EPI_ISL_2395112 | Lighthouse Lab in Glasgow | Wellcome Sanger Institute for the COVID-19 Genomics UK (COG-UK) Consortium | Harper VanSteenhouse, Yumi Kasai, David Gray, Carol Clugston, Anna Dominiczak and Alex Alderton, Roberto Amato, Jeffrey Barrett, Sonia Goncalves, Ewan Harrison, David K. Jackson, Ian Johnston, Dominic Kwiatkowski, Cordelia Langford, John Sillitoe on behalf of the Wellcome Sanger Institute COVID-19 Surveillance Team |
| EPI_ISL_2395113, EPI_ISL_2395114, EPI_ISL_2395115, EPI_ISL_2395116, EPI_ISL_2395117, EPI_ISL_2395118, EPI_ISL_2395119, EPI_ISL_2395120, EPI_ISL_2395121, EPI_ISL_2395122 | Lighthouse Lab in Alderley Park | Wellcome Sanger Institute for the COVID-19 Genomics UK (COG-UK) Consortium | Jacquelyn Wynn, Mairead Hyland, The Lighthouse Lab in Alderley Park and Alex Alderton, Roberto Amato, Jeffrey Barrett, Sonia Goncalves, Ewan Harrison, David K. Jackson, Ian Johnston, Dominic Kwiatkowski, Cordelia Langford, John Sillitoe on behalf of the Wellcome Sanger Institute COVID-19 Surveillance Team |
| EPI_ISL_2395123 | Lighthouse Lab in Glasgow | Wellcome Sanger Institute for the COVID-19 Genomics UK (COG-UK) Consortium | Harper VanSteenhouse, Yumi Kasai, David Gray, Carol Clugston, Anna Dominiczak and Alex Alderton, Roberto Amato, Jeffrey Barrett, Sonia Goncalves, Ewan Harrison, David K. Jackson, Ian Johnston, Dominic Kwiatkowski, Cordelia Langford, John Sillitoe on behalf of the Wellcome Sanger Institute COVID-19 Surveillance Team |
| EPI_ISL_2395124, EPI_ISL_2395126, EPI_ISL_2395128, EPI_ISL_2395129, EPI_ISL_2395130, EPI_ISL_2395131 | Lighthouse Lab in Alderley Park | Wellcome Sanger Institute for the COVID-19 Genomics UK (COG-UK) Consortium | Jacquelyn Wynn, Mairead Hyland, The Lighthouse Lab in Alderley Park and Alex Alderton, Roberto Amato, Jeffrey Barrett, Sonia Goncalves, Ewan Harrison, David K. Jackson, Ian Johnston, Dominic Kwiatkowski, Cordelia Langford, John Sillitoe on behalf of the Wellcome Sanger Institute COVID-19 Surveillance Team |
| EPI_ISL_2395132 | Lighthouse Lab in Glasgow | Wellcome Sanger Institute for the COVID-19 Genomics UK (COG-UK) Consortium | Harper VanSteenhouse, Yumi Kasai, David Gray, Carol Clugston, Anna Dominiczak and Alex Alderton, Roberto Amato, Jeffrey Barrett, Sonia Goncalves, Ewan Harrison, David K. Jackson, Ian Johnston, Dominic Kwiatkowski, Cordelia Langford, John Sillitoe on behalf of the Wellcome Sanger Institute COVID-19 Surveillance Team |
| EPI_ISL_2395133, EPI_ISL_2395134, EPI_ISL_2395137, EPI_ISL_2395138, EPI_ISL_2395139, EPI_ISL_2395140, EPI_ISL_2395141, EPI_ISL_2395142, EPI_ISL_2395143, EPI_ISL_2395146, EPI_ISL_2395147, EPI_ISL_2395148, EPI_ISL_2395151, EPI_ISL_2395152 | | | |
| see above | Lighthouse Lab in Alderley Park | Wellcome Sanger Institute for the COVID-19 Genomics UK (COG-UK) Consortium | Jacquelyn Wynn, Mairead Hyland, The Lighthouse Lab in Alderley Park and Alex Alderton, Roberto Amato, Jeffrey Barrett, Sonia Goncalves, Ewan Harrison, David K. Jackson, Ian Johnston, Dominic Kwiatkowski, Cordelia Langford, John Sillitoe on behalf of the Wellcome Sanger Institute COVID-19 Surveillance Team |
| EPI_ISL_2395153 | Lighthouse Lab in Glasgow | Wellcome Sanger Institute for the COVID-19 Genomics UK (COG-UK) Consortium | Harper VanSteenhouse, Yumi Kasai, David Gray, Carol Clugston, Anna Dominiczak and Alex Alderton, Roberto Amato, Jeffrey Barrett, Sonia Goncalves, Ewan Harrison, David K. Jackson, Ian Johnston, Dominic Kwiatkowski, Cordelia Langford, John Sillitoe on behalf of the Wellcome Sanger Institute COVID-19 Surveillance Team |
| EPI_ISL_2395154, EPI_ISL_2395155, EPI_ISL_2395156, EPI_ISL_2395157, EPI_ISL_2395158, EPI_ISL_2395159, EPI_ISL_2395161, EPI_ISL_2395162, EPI_ISL_2395163, EPI_ISL_2395164, EPI_ISL_2395165, EPI_ISL_2395166, EPI_ISL_2395167, EPI_ISL_2395168, EPI_ISL_2395169, EPI_ISL_2395170, EPI_ISL_2395171 | | | |
| see above | Lighthouse Lab in Alderley Park | Wellcome Sanger Institute for the COVID-19 Genomics UK (COG-UK) Consortium | Jacquelyn Wynn, Mairead Hyland, The Lighthouse Lab in Alderley Park and Alex Alderton, Roberto Amato, Jeffrey Barrett, Sonia Goncalves, Ewan Harrison, David K. Jackson, Ian Johnston, Dominic Kwiatkowski, Cordelia Langford, John Sillitoe on behalf of the Wellcome Sanger Institute COVID-19 Surveillance Team |
| EPI_ISL_2395172 | Lighthouse Lab in Glasgow | Wellcome Sanger Institute for the COVID-19 Genomics UK (COG-UK) Consortium | Harper VanSteenhouse, Yumi Kasai, David Gray, Carol Clugston, Anna Dominiczak and Alex Alderton, Roberto Amato, Jeffrey Barrett, Sonia Goncalves, Ewan Harrison, David K. Jackson, Ian Johnston, Dominic Kwiatkowski, Cordelia Langford, John Sillitoe on behalf of the Wellcome Sanger Institute COVID-19 Surveillance Team |
| EPI_ISL_2395173, EPI_ISL_2395174, EPI_ISL_2395175 | Lighthouse Lab in Alderley Park | Wellcome Sanger Institute for the COVID-19 Genomics UK (COG-UK) Consortium | Jacquelyn Wynn, Mairead Hyland, The Lighthouse Lab in Alderley Park and Alex Alderton, Roberto Amato, Jeffrey Barrett, Sonia Goncalves, Ewan Harrison, David K. Jackson, Ian Johnston, Dominic Kwiatkowski, Cordelia Langford, John Sillitoe on behalf of the Wellcome Sanger Institute COVID-19 Surveillance Team |
| EPI_ISL_2395176 | Lighthouse Lab in Glasgow | Wellcome Sanger Institute for the COVID-19 Genomics UK (COG-UK) Consortium | Harper VanSteenhouse, Yumi Kasai, David Gray, Carol Clugston, Anna Dominiczak and Alex Alderton, Roberto Amato, Jeffrey Barrett, Sonia Goncalves, Ewan Harrison, David K. Jackson, Ian Johnston, Dominic Kwiatkowski, Cordelia Langford, John Sillitoe on behalf of the Wellcome Sanger Institute COVID-19 Surveillance Team |
| EPI_ISL_2395177, EPI_ISL_2395178, | Lighthouse Lab in Alderley Park | Wellcome Sanger Institute for the COVID-19 Genomics UK (COG-UK) Consortium | Jacquelyn Wynn, Mairead Hyland, The Lighthouse Lab in Alderley Park and Alex Alderton, Roberto Amato, Jeffrey Barrett, Sonia Goncalves, Ewan Harrison, David K. Jackson, Ian Johnston, Dominic Kwiatkowski, Cordelia Langford, John Sillitoe on behalf of the Wellcome Sanger Institute COVID-19 Surveillance Team |

| | | | |
|---|---|---|---|
| EPI_ISL_2395179, EPI_ISL_2395180, EPI_ISL_2395181, EPI_ISL_2395182, EPI_ISL_2395183 | | | |
| EPI_ISL_2395184 | Lighthouse Lab in Glasgow | Wellcome Sanger Institute for the COVID-19 Genomics UK (COG-UK) Consortium | Harper VanSteenhouse, Yumi Kasai, David Gray, Carol Clugston, Anna Dominiczak and Alex Alderton, Roberto Amato, Jeffrey Barrett, Sonia Goncalves, Ewan Harrison, David K. Jackson, Ian Johnston, Dominic Kwiatkowski, Cordelia Langford, John Sillitoe on behalf of the Wellcome Sanger Institute COVID-19 Surveillance Team |
| EPI_ISL_2395185, EPI_ISL_2395186, EPI_ISL_2395187, EPI_ISL_2395188, EPI_ISL_2395189, EPI_ISL_2395190, EPI_ISL_2395191, EPI_ISL_2395192, EPI_ISL_2395193, EPI_ISL_2395194 | Lighthouse Lab in Alderley Park | Wellcome Sanger Institute for the COVID-19 Genomics UK (COG-UK) Consortium | Jacquelyn Wynn, Mairead Hyland, The Lighthouse Lab in Alderley Park and Alex Alderton, Roberto Amato, Jeffrey Barrett, Sonia Goncalves, Ewan Harrison, David K. Jackson, Ian Johnston, Dominic Kwiatkowski, Cordelia Langford, John Sillitoe on behalf of the Wellcome Sanger Institute COVID-19 Surveillance Team |
| EPI_ISL_2395195 | Lighthouse Lab in Glasgow | Wellcome Sanger Institute for the COVID-19 Genomics UK (COG-UK) Consortium | Harper VanSteenhouse, Yumi Kasai, David Gray, Carol Clugston, Anna Dominiczak and Alex Alderton, Roberto Amato, Jeffrey Barrett, Sonia Goncalves, Ewan Harrison, David K. Jackson, Ian Johnston, Dominic Kwiatkowski, Cordelia Langford, John Sillitoe on behalf of the Wellcome Sanger Institute COVID-19 Surveillance Team |
| EPI_ISL_2395196, EPI_ISL_2395197, EPI_ISL_2395198, EPI_ISL_2395199, EPI_ISL_2395200, EPI_ISL_2395201, EPI_ISL_2395202, EPI_ISL_2395203, EPI_ISL_2395204, EPI_ISL_2395205, EPI_ISL_2395206, EPI_ISL_2395207, EPI_ISL_2395208 | | | |
| see above | Lighthouse Lab in Alderley Park | Wellcome Sanger Institute for the COVID-19 Genomics UK (COG-UK) Consortium | Jacquelyn Wynn, Mairead Hyland, The Lighthouse Lab in Alderley Park and Alex Alderton, Roberto Amato, Jeffrey Barrett, Sonia Goncalves, Ewan Harrison, David K. Jackson, Ian Johnston, Dominic Kwiatkowski, Cordelia Langford, John Sillitoe on behalf of the Wellcome Sanger Institute COVID-19 Surveillance Team |
| EPI_ISL_2395209 | Lighthouse Lab in Glasgow | Wellcome Sanger Institute for the COVID-19 Genomics UK (COG-UK) Consortium | Harper VanSteenhouse, Yumi Kasai, David Gray, Carol Clugston, Anna Dominiczak and Alex Alderton, Roberto Amato, Jeffrey Barrett, Sonia Goncalves, Ewan Harrison, David K. Jackson, Ian Johnston, Dominic Kwiatkowski, Cordelia Langford, John Sillitoe on behalf of the Wellcome Sanger Institute COVID-19 Surveillance Team |
| EPI_ISL_2395210, EPI_ISL_2395211 | Lighthouse Lab in Alderley Park | Wellcome Sanger Institute for the COVID-19 Genomics UK (COG-UK) Consortium | Jacquelyn Wynn, Mairead Hyland, The Lighthouse Lab in Alderley Park and Alex Alderton, Roberto Amato, Jeffrey Barrett, Sonia Goncalves, Ewan Harrison, David K. Jackson, Ian Johnston, Dominic Kwiatkowski, Cordelia Langford, John Sillitoe on behalf of the Wellcome Sanger Institute COVID-19 Surveillance Team |
| EPI_ISL_2395212, EPI_ISL_2395213 | Lighthouse Lab in Glasgow | Wellcome Sanger Institute for the COVID-19 Genomics UK (COG-UK) Consortium | Harper VanSteenhouse, Yumi Kasai, David Gray, Carol Clugston, Anna Dominiczak and Alex Alderton, Roberto Amato, Jeffrey Barrett, Sonia Goncalves, Ewan Harrison, David K. Jackson, Ian Johnston, Dominic Kwiatkowski, Cordelia Langford, John Sillitoe on behalf of the Wellcome Sanger Institute COVID-19 Surveillance Team |
| EPI_ISL_2395215, EPI_ISL_2395216, EPI_ISL_2395217, EPI_ISL_2395218, EPI_ISL_2395219, EPI_ISL_2395221, EPI_ISL_2395222 | Lighthouse Lab in Alderley Park | Wellcome Sanger Institute for the COVID-19 Genomics UK (COG-UK) Consortium | Jacquelyn Wynn, Mairead Hyland, The Lighthouse Lab in Alderley Park and Alex Alderton, Roberto Amato, Jeffrey Barrett, Sonia Goncalves, Ewan Harrison, David K. Jackson, Ian Johnston, Dominic Kwiatkowski, Cordelia Langford, John Sillitoe on behalf of the Wellcome Sanger Institute COVID-19 Surveillance Team |
| EPI_ISL_2395223 | Lighthouse Lab in Glasgow | Wellcome Sanger Institute for the COVID-19 Genomics UK (COG-UK) Consortium | Harper VanSteenhouse, Yumi Kasai, David Gray, Carol Clugston, Anna Dominiczak and Alex Alderton, Roberto Amato, Jeffrey Barrett, Sonia Goncalves, Ewan Harrison, David K. Jackson, Ian Johnston, Dominic Kwiatkowski, Cordelia Langford, John Sillitoe on behalf of the Wellcome Sanger Institute COVID-19 Surveillance Team |
| EPI_ISL_2395224, EPI_ISL_2395225, EPI_ISL_2395226, EPI_ISL_2395227 | Lighthouse Lab in Alderley Park | Wellcome Sanger Institute for the COVID-19 Genomics UK (COG-UK) Consortium | Jacquelyn Wynn, Mairead Hyland, The Lighthouse Lab in Alderley Park and Alex Alderton, Roberto Amato, Jeffrey Barrett, Sonia Goncalves, Ewan Harrison, David K. Jackson, Ian Johnston, Dominic Kwiatkowski, Cordelia Langford, John Sillitoe on behalf of the Wellcome Sanger Institute COVID-19 Surveillance Team |
| EPI_ISL_2395228 | Lighthouse Lab in Glasgow | Wellcome Sanger Institute for the COVID-19 Genomics UK (COG-UK) Consortium | Harper VanSteenhouse, Yumi Kasai, David Gray, Carol Clugston, Anna Dominiczak and Alex Alderton, Roberto Amato, Jeffrey Barrett, Sonia Goncalves, Ewan Harrison, David K. Jackson, Ian Johnston, Dominic Kwiatkowski, Cordelia Langford, John Sillitoe on behalf of the Wellcome Sanger Institute COVID-19 Surveillance Team |
| EPI_ISL_2395229, EPI_ISL_2395230, EPI_ISL_2395231, EPI_ISL_2395232, EPI_ISL_2395234, EPI_ISL_2395235, EPI_ISL_2395236 | Lighthouse Lab in Alderley Park | Wellcome Sanger Institute for the COVID-19 Genomics UK (COG-UK) Consortium | Jacquelyn Wynn, Mairead Hyland, The Lighthouse Lab in Alderley Park and Alex Alderton, Roberto Amato, Jeffrey Barrett, Sonia Goncalves, Ewan Harrison, David K. Jackson, Ian Johnston, Dominic Kwiatkowski, Cordelia Langford, John Sillitoe on behalf of the Wellcome Sanger Institute COVID-19 Surveillance Team |
| EPI_ISL_2395238 | Lighthouse Lab in Glasgow | Wellcome Sanger Institute for the COVID-19 Genomics UK (COG-UK) Consortium | Harper VanSteenhouse, Yumi Kasai, David Gray, Carol Clugston, Anna Dominiczak and Alex Alderton, Roberto Amato, Jeffrey Barrett, Sonia Goncalves, Ewan Harrison, David K. Jackson, Ian Johnston, Dominic Kwiatkowski, Cordelia Langford, John Sillitoe on behalf of the Wellcome Sanger Institute COVID-19 Surveillance Team |
| EPI_ISL_2395239, EPI_ISL_2395240, EPI_ISL_2395242, EPI_ISL_2395243, EPI_ISL_2395245, EPI_ISL_2395246, EPI_ISL_2395247 | Lighthouse Lab in Alderley Park | Wellcome Sanger Institute for the COVID-19 Genomics UK (COG-UK) Consortium | Jacquelyn Wynn, Mairead Hyland, The Lighthouse Lab in Alderley Park and Alex Alderton, Roberto Amato, Jeffrey Barrett, Sonia Goncalves, Ewan Harrison, David K. Jackson, Ian Johnston, Dominic Kwiatkowski, Cordelia Langford, John Sillitoe on behalf of the Wellcome Sanger Institute COVID-19 Surveillance Team |
| EPI_ISL_2395248, EPI_ISL_2395249 | Lighthouse Lab in Glasgow | Wellcome Sanger Institute for the COVID-19 Genomics UK (COG-UK) Consortium | Harper VanSteenhouse, Yumi Kasai, David Gray, Carol Clugston, Anna Dominiczak and Alex Alderton, Roberto Amato, Jeffrey Barrett, Sonia Goncalves, Ewan Harrison, David K. Jackson, Ian Johnston, Dominic Kwiatkowski, Cordelia Langford, John Sillitoe on behalf of the Wellcome Sanger Institute COVID-19 Surveillance Team |
| EPI_ISL_2395250 | Lighthouse Lab in Alderley Park | Wellcome Sanger Institute for the COVID-19 Genomics UK (COG-UK) Consortium | Jacquelyn Wynn, Mairead Hyland, The Lighthouse Lab in Alderley Park and Alex Alderton, Roberto Amato, Jeffrey Barrett, Sonia Goncalves, Ewan Harrison, David K. Jackson, Ian Johnston, Dominic Kwiatkowski, Cordelia Langford, John Sillitoe on behalf of the Wellcome Sanger Institute COVID-19 Surveillance Team |
| EPI_ISL_2395251 | Lighthouse Lab in Glasgow | Wellcome Sanger Institute for the COVID-19 Genomics UK (COG-UK) Consortium | Harper VanSteenhouse, Yumi Kasai, David Gray, Carol Clugston, Anna Dominiczak and Alex Alderton, Roberto Amato, Jeffrey Barrett, Sonia Goncalves, Ewan Harrison, David K. Jackson, Ian Johnston, Dominic Kwiatkowski, Cordelia Langford, John Sillitoe on behalf of the Wellcome Sanger Institute COVID-19 Surveillance Team |
| EPI_ISL_2395252, EPI_ISL_2395253, EPI_ISL_2395254, EPI_ISL_2395255, EPI_ISL_2395257, EPI_ISL_2395258, EPI_ISL_2395259, EPI_ISL_2395260, EPI_ISL_2395261, EPI_ISL_2395262, EPI_ISL_2395263, EPI_ISL_2395264, EPI_ISL_2395266, EPI_ISL_2395268, EPI_ISL_2395269, EPI_ISL_2395270, EPI_ISL_2395271, EPI_ISL_2395272, EPI_ISL_2395274, EPI_ISL_2395276, EPI_ISL_2395277, EPI_ISL_2395278, EPI_ISL_2395279 | | | |
| see above | Lighthouse Lab in Alderley Park | Wellcome Sanger Institute for the COVID-19 Genomics UK (COG-UK) Consortium | Jacquelyn Wynn, Mairead Hyland, The Lighthouse Lab in Alderley Park and Alex Alderton, Roberto Amato, Jeffrey Barrett, Sonia Goncalves, Ewan Harrison, David K. Jackson, Ian Johnston, Dominic Kwiatkowski, Cordelia Langford, John Sillitoe on behalf of the Wellcome Sanger Institute COVID-19 Surveillance Team |
| EPI_ISL_2395280 | Lighthouse Lab in Glasgow | Wellcome Sanger Institute for the COVID-19 Genomics UK (COG-UK) Consortium | Harper VanSteenhouse, Yumi Kasai, David Gray, Carol Clugston, Anna Dominiczak and Alex Alderton, Roberto Amato, Jeffrey Barrett, Sonia Goncalves, Ewan Harrison, David K. Jackson, Ian Johnston, Dominic Kwiatkowski, Cordelia Langford, John Sillitoe on behalf of the Wellcome Sanger Institute COVID-19 Surveillance Team |
| EPI_ISL_2395281, EPI_ISL_2395282, EPI_ISL_2395283, EPI_ISL_2395284, EPI_ISL_2395285, EPI_ISL_2395286, EPI_ISL_2395287, EPI_ISL_2395288, EPI_ISL_2395289, EPI_ISL_2395290, EPI_ISL_2395291, EPI_ISL_2395292, EPI_ISL_2395293, EPI_ISL_2395295, EPI_ISL_2395296, EPI_ISL_2395297, EPI_ISL_2395298, EPI_ISL_2395299, EPI_ISL_2395300, EPI_ISL_2395301, EPI_ISL_2395302, EPI_ISL_2395303, EPI_ISL_2395304, EPI_ISL_2395305 | | | |
| see above | Lighthouse Lab in Alderley Park | Wellcome Sanger Institute for the COVID-19 Genomics UK (COG-UK) Consortium | Jacquelyn Wynn, Mairead Hyland, The Lighthouse Lab in Alderley Park and Alex Alderton, Roberto Amato, Jeffrey Barrett, Sonia Goncalves, Ewan Harrison, David K. Jackson, Ian Johnston, Dominic Kwiatkowski, Cordelia Langford, John Sillitoe on behalf of the Wellcome Sanger Institute COVID-19 Surveillance Team |
| EPI_ISL_2395307, EPI_ISL_2395308 | Lighthouse Lab in Glasgow | Wellcome Sanger Institute for the COVID-19 Genomics UK (COG-UK) Consortium | Harper VanSteenhouse, Yumi Kasai, David Gray, Carol Clugston, Anna Dominiczak and Alex Alderton, Roberto Amato, Jeffrey Barrett, Sonia Goncalves, Ewan Harrison, David K. Jackson, Ian Johnston, Dominic Kwiatkowski, Cordelia Langford, John Sillitoe on behalf of the Wellcome Sanger Institute COVID-19 Surveillance Team |
| EPI_ISL_2395310, EPI_ISL_2395311, EPI_ISL_2395313, EPI_ISL_2395314, EPI_ISL_2395315, EPI_ISL_2395316 | Lighthouse Lab in Alderley Park | Wellcome Sanger Institute for the COVID-19 Genomics UK (COG-UK) Consortium | Jacquelyn Wynn, Mairead Hyland, The Lighthouse Lab in Alderley Park and Alex Alderton, Roberto Amato, Jeffrey Barrett, Sonia Goncalves, Ewan Harrison, David K. Jackson, Ian Johnston, Dominic Kwiatkowski, Cordelia Langford, John Sillitoe on behalf of the Wellcome Sanger Institute COVID-19 Surveillance Team |
| EPI_ISL_2395318, EPI_ISL_2395319 | Lighthouse Lab in Glasgow | Wellcome Sanger Institute for the COVID-19 Genomics UK (COG-UK) Consortium | Harper VanSteenhouse, Yumi Kasai, David Gray, Carol Clugston, Anna Dominiczak and Alex Alderton, Roberto Amato, Jeffrey Barrett, Sonia Goncalves, Ewan Harrison, David K. Jackson, Ian Johnston, Dominic Kwiatkowski, Cordelia Langford, John Sillitoe on behalf of the Wellcome Sanger Institute COVID-19 Surveillance Team |
| EPI_ISL_2395320, EPI_ISL_2395321, EPI_ISL_2395323, EPI_ISL_2395324, EPI_ISL_2395325, EPI_ISL_2395326, EPI_ISL_2395327, EPI_ISL_2395328, EPI_ISL_2395329, EPI_ISL_2395330, EPI_ISL_2395331, EPI_ISL_2395332, EPI_ISL_2395333, EPI_ISL_2395334, EPI_ISL_2395335, EPI_ISL_2395336 | | | |
| see above | Lighthouse Lab in Alderley Park | Wellcome Sanger Institute for the COVID-19 Genomics UK (COG-UK) Consortium | Jacquelyn Wynn, Mairead Hyland, The Lighthouse Lab in Alderley Park and Alex Alderton, Roberto Amato, Jeffrey Barrett, Sonia Goncalves, Ewan Harrison, David K. Jackson, Ian Johnston, Dominic Kwiatkowski, Cordelia Langford, John Sillitoe on behalf of the Wellcome Sanger Institute COVID-19 Surveillance Team |
| EPI_ISL_2395338, EPI_ISL_2395339 | Lighthouse Lab in Milton Keynes | Wellcome Sanger Institute for the COVID-19 Genomics UK (COG-UK) Consortium | The Lighthouse Lab in Milton Keynes and Alex Alderton, Roberto Amato, Jeffrey Barrett, Sonia Goncalves, Ewan Harrison, David K. Jackson, Ian Johnston, Dominic Kwiatkowski, Cordelia Langford, John Sillitoe on behalf of the Wellcome Sanger Institute COVID-19 Surveillance Team |
| EPI_ISL_2395340 | Lighthouse Lab in Glasgow | Wellcome Sanger Institute for the COVID-19 Genomics UK (COG-UK) Consortium | Harper VanSteenhouse, Yumi Kasai, David Gray, Carol Clugston, Anna Dominiczak and Alex Alderton, Roberto Amato, Jeffrey Barrett, Sonia Goncalves, Ewan Harrison, David K. Jackson, Ian Johnston, Dominic Kwiatkowski, Cordelia Langford, John Sillitoe on behalf of the Wellcome Sanger Institute COVID-19 Surveillance Team |
| EPI_ISL_2395341 | Lighthouse Lab in Milton Keynes | Wellcome Sanger Institute for the COVID-19 Genomics UK (COG-UK) Consortium | The Lighthouse Lab in Milton Keynes and Alex Alderton, Roberto Amato, Jeffrey Barrett, Sonia Goncalves, Ewan Harrison, David K. Jackson, Ian Johnston, Dominic Kwiatkowski, Cordelia Langford, John Sillitoe on behalf of the Wellcome Sanger Institute COVID-19 Surveillance Team |
| EPI_ISL_2395342, | Lighthouse Lab in Alderley Park | Wellcome Sanger Institute for the COVID-19 Genomics UK (COG-UK) | Jacquelyn Wynn, Mairead Hyland, The Lighthouse Lab in Alderley Park and Alex Alderton, Roberto Amato, Jeffrey Barrett, Sonia Goncalves, Ewan Harrison, David K. Jackson, Ian Johnston, Dominic Kwiatkowski, Cordelia Langford, John Sillitoe on |

| | | | |
|---|---|---|---|
| EPI_ISL_2395343 | | Consortium | behalf of the Wellcome Sanger Institute COVID-19 Surveillance Team |
| EPI_ISL_2395344 | Lighthouse Lab in Glasgow | Wellcome Sanger Institute for the COVID-19 Genomics UK (COG-UK) Consortium | Harper VanSteenhouse, Yumi Kasai, David Gray, Carol Clugston, Anna Dominiczak and Alex Alderton, Roberto Amato, Jeffrey Barrett, Sonia Goncalves, Ewan Harrison, David K. Jackson, Ian Johnston, Dominic Kwiatkowski, Cordelia Langford, John Sillitoe on behalf of the Wellcome Sanger Institute COVID-19 Surveillance Team |
| EPI_ISL_2395345, EPI_ISL_2395346 | Lighthouse Lab in Milton Keynes | Wellcome Sanger Institute for the COVID-19 Genomics UK (COG-UK) Consortium | The Lighthouse Lab in Milton Keynes and Alex Alderton, Roberto Amato, Jeffrey Barrett, Sonia Goncalves, Ewan Harrison, David K. Jackson, Ian Johnston, Dominic Kwiatkowski, Cordelia Langford, John Sillitoe on behalf of the Wellcome Sanger Institute COVID-19 Surveillance Team |
| EPI_ISL_2395348, EPI_ISL_2395350 | Lighthouse Lab in Glasgow | Wellcome Sanger Institute for the COVID-19 Genomics UK (COG-UK) Consortium | Harper VanSteenhouse, Yumi Kasai, David Gray, Carol Clugston, Anna Dominiczak and Alex Alderton, Roberto Amato, Jeffrey Barrett, Sonia Goncalves, Ewan Harrison, David K. Jackson, Ian Johnston, Dominic Kwiatkowski, Cordelia Langford, John Sillitoe on behalf of the Wellcome Sanger Institute COVID-19 Surveillance Team |
| EPI_ISL_2395352, EPI_ISL_2395353, EPI_ISL_2395354, EPI_ISL_2395356, EPI_ISL_2395357, EPI_ISL_2395358, EPI_ISL_2395359 | Lighthouse Lab in Milton Keynes | Wellcome Sanger Institute for the COVID-19 Genomics UK (COG-UK) Consortium | The Lighthouse Lab in Milton Keynes and Alex Alderton, Roberto Amato, Jeffrey Barrett, Sonia Goncalves, Ewan Harrison, David K. Jackson, Ian Johnston, Dominic Kwiatkowski, Cordelia Langford, John Sillitoe on behalf of the Wellcome Sanger Institute COVID-19 Surveillance Team |
| EPI_ISL_2395360, EPI_ISL_2395361 | Lighthouse Lab in Alderley Park | Wellcome Sanger Institute for the COVID-19 Genomics UK (COG-UK) Consortium | Jacquelyn Wynn, Mairead Hyland, The Lighthouse Lab in Alderley Park and Alex Alderton, Roberto Amato, Jeffrey Barrett, Sonia Goncalves, Ewan Harrison, David K. Jackson, Ian Johnston, Dominic Kwiatkowski, Cordelia Langford, John Sillitoe on behalf of the Wellcome Sanger Institute COVID-19 Surveillance Team |
| EPI_ISL_2395362 | Lighthouse Lab in Glasgow | Wellcome Sanger Institute for the COVID-19 Genomics UK (COG-UK) Consortium | Harper VanSteenhouse, Yumi Kasai, David Gray, Carol Clugston, Anna Dominiczak and Alex Alderton, Roberto Amato, Jeffrey Barrett, Sonia Goncalves, Ewan Harrison, David K. Jackson, Ian Johnston, Dominic Kwiatkowski, Cordelia Langford, John Sillitoe on behalf of the Wellcome Sanger Institute COVID-19 Surveillance Team |
| EPI_ISL_2395363, EPI_ISL_2395364 | Lighthouse Lab in Alderley Park | Wellcome Sanger Institute for the COVID-19 Genomics UK (COG-UK) Consortium | Jacquelyn Wynn, Mairead Hyland, The Lighthouse Lab in Alderley Park and Alex Alderton, Roberto Amato, Jeffrey Barrett, Sonia Goncalves, Ewan Harrison, David K. Jackson, Ian Johnston, Dominic Kwiatkowski, Cordelia Langford, John Sillitoe on behalf of the Wellcome Sanger Institute COVID-19 Surveillance Team |
| EPI_ISL_2395365 | Lighthouse Lab in Glasgow | Wellcome Sanger Institute for the COVID-19 Genomics UK (COG-UK) Consortium | Harper VanSteenhouse, Yumi Kasai, David Gray, Carol Clugston, Anna Dominiczak and Alex Alderton, Roberto Amato, Jeffrey Barrett, Sonia Goncalves, Ewan Harrison, David K. Jackson, Ian Johnston, Dominic Kwiatkowski, Cordelia Langford, John Sillitoe on behalf of the Wellcome Sanger Institute COVID-19 Surveillance Team |
| EPI_ISL_2395366 | Lighthouse Lab in Milton Keynes | Wellcome Sanger Institute for the COVID-19 Genomics UK (COG-UK) Consortium | The Lighthouse Lab in Milton Keynes and Alex Alderton, Roberto Amato, Jeffrey Barrett, Sonia Goncalves, Ewan Harrison, David K. Jackson, Ian Johnston, Dominic Kwiatkowski, Cordelia Langford, John Sillitoe on behalf of the Wellcome Sanger Institute COVID-19 Surveillance Team |
| EPI_ISL_2395367 | Lighthouse Lab in Alderley Park | Wellcome Sanger Institute for the COVID-19 Genomics UK (COG-UK) Consortium | Jacquelyn Wynn, Mairead Hyland, The Lighthouse Lab in Alderley Park and Alex Alderton, Roberto Amato, Jeffrey Barrett, Sonia Goncalves, Ewan Harrison, David K. Jackson, Ian Johnston, Dominic Kwiatkowski, Cordelia Langford, John Sillitoe on behalf of the Wellcome Sanger Institute COVID-19 Surveillance Team |
| EPI_ISL_2395372 | Lighthouse Lab in Milton Keynes | Wellcome Sanger Institute for the COVID-19 Genomics UK (COG-UK) Consortium | The Lighthouse Lab in Milton Keynes and Alex Alderton, Roberto Amato, Jeffrey Barrett, Sonia Goncalves, Ewan Harrison, David K. Jackson, Ian Johnston, Dominic Kwiatkowski, Cordelia Langford, John Sillitoe on behalf of the Wellcome Sanger Institute COVID-19 Surveillance Team |
| EPI_ISL_2395374 | Lighthouse Lab in Glasgow | Wellcome Sanger Institute for the COVID-19 Genomics UK (COG-UK) Consortium | Harper VanSteenhouse, Yumi Kasai, David Gray, Carol Clugston, Anna Dominiczak and Alex Alderton, Roberto Amato, Jeffrey Barrett, Sonia Goncalves, Ewan Harrison, David K. Jackson, Ian Johnston, Dominic Kwiatkowski, Cordelia Langford, John Sillitoe on behalf of the Wellcome Sanger Institute COVID-19 Surveillance Team |
| EPI_ISL_2395375, EPI_ISL_2395376 | Lighthouse Lab in Milton Keynes | Wellcome Sanger Institute for the COVID-19 Genomics UK (COG-UK) Consortium | The Lighthouse Lab in Milton Keynes and Alex Alderton, Roberto Amato, Jeffrey Barrett, Sonia Goncalves, Ewan Harrison, David K. Jackson, Ian Johnston, Dominic Kwiatkowski, Cordelia Langford, John Sillitoe on behalf of the Wellcome Sanger Institute COVID-19 Surveillance Team |
| EPI_ISL_2395377, EPI_ISL_2395378 | Lighthouse Lab in Glasgow | Wellcome Sanger Institute for the COVID-19 Genomics UK (COG-UK) Consortium | Harper VanSteenhouse, Yumi Kasai, David Gray, Carol Clugston, Anna Dominiczak and Alex Alderton, Roberto Amato, Jeffrey Barrett, Sonia Goncalves, Ewan Harrison, David K. Jackson, Ian Johnston, Dominic Kwiatkowski, Cordelia Langford, John Sillitoe on behalf of the Wellcome Sanger Institute COVID-19 Surveillance Team |
| EPI_ISL_2395379, EPI_ISL_2395380 | Lighthouse Lab in Alderley Park | Wellcome Sanger Institute for the COVID-19 Genomics UK (COG-UK) Consortium | Jacquelyn Wynn, Mairead Hyland, The Lighthouse Lab in Alderley Park and Alex Alderton, Roberto Amato, Jeffrey Barrett, Sonia Goncalves, Ewan Harrison, David K. Jackson, Ian Johnston, Dominic Kwiatkowski, Cordelia Langford, John Sillitoe on behalf of the Wellcome Sanger Institute COVID-19 Surveillance Team |
| EPI_ISL_2395382, EPI_ISL_2395383 | Lighthouse Lab in Milton Keynes | Wellcome Sanger Institute for the COVID-19 Genomics UK (COG-UK) Consortium | The Lighthouse Lab in Milton Keynes and Alex Alderton, Roberto Amato, Jeffrey Barrett, Sonia Goncalves, Ewan Harrison, David K. Jackson, Ian Johnston, Dominic Kwiatkowski, Cordelia Langford, John Sillitoe on behalf of the Wellcome Sanger Institute COVID-19 Surveillance Team |
| EPI_ISL_2395386 | Lighthouse Lab in Glasgow | Wellcome Sanger Institute for the COVID-19 Genomics UK (COG-UK) Consortium | Harper VanSteenhouse, Yumi Kasai, David Gray, Carol Clugston, Anna Dominiczak and Alex Alderton, Roberto Amato, Jeffrey Barrett, Sonia Goncalves, Ewan Harrison, David K. Jackson, Ian Johnston, Dominic Kwiatkowski, Cordelia Langford, John Sillitoe on behalf of the Wellcome Sanger Institute COVID-19 Surveillance Team |
| EPI_ISL_2395387 | Lighthouse Lab in Alderley Park | Wellcome Sanger Institute for the COVID-19 Genomics UK (COG-UK) Consortium | Jacquelyn Wynn, Mairead Hyland, The Lighthouse Lab in Alderley Park and Alex Alderton, Roberto Amato, Jeffrey Barrett, Sonia Goncalves, Ewan Harrison, David K. Jackson, Ian Johnston, Dominic Kwiatkowski, Cordelia Langford, John Sillitoe on behalf of the Wellcome Sanger Institute COVID-19 Surveillance Team |
| EPI_ISL_2395388 | Lighthouse Lab in Glasgow | Wellcome Sanger Institute for the COVID-19 Genomics UK (COG-UK) Consortium | Harper VanSteenhouse, Yumi Kasai, David Gray, Carol Clugston, Anna Dominiczak and Alex Alderton, Roberto Amato, Jeffrey Barrett, Sonia Goncalves, Ewan Harrison, David K. Jackson, Ian Johnston, Dominic Kwiatkowski, Cordelia Langford, John Sillitoe on behalf of the Wellcome Sanger Institute COVID-19 Surveillance Team |
| EPI_ISL_2395389 | Lighthouse Lab in Milton Keynes | Wellcome Sanger Institute for the COVID-19 Genomics UK (COG-UK) Consortium | The Lighthouse Lab in Milton Keynes and Alex Alderton, Roberto Amato, Jeffrey Barrett, Sonia Goncalves, Ewan Harrison, David K. Jackson, Ian Johnston, Dominic Kwiatkowski, Cordelia Langford, John Sillitoe on behalf of the Wellcome Sanger Institute COVID-19 Surveillance Team |
| EPI_ISL_2395390, EPI_ISL_2395391 | Lighthouse Lab in Alderley Park | Wellcome Sanger Institute for the COVID-19 Genomics UK (COG-UK) Consortium | Jacquelyn Wynn, Mairead Hyland, The Lighthouse Lab in Alderley Park and Alex Alderton, Roberto Amato, Jeffrey Barrett, Sonia Goncalves, Ewan Harrison, David K. Jackson, Ian Johnston, Dominic Kwiatkowski, Cordelia Langford, John Sillitoe on behalf of the Wellcome Sanger Institute COVID-19 Surveillance Team |
| EPI_ISL_2395392 | Lighthouse Lab in Milton Keynes | Wellcome Sanger Institute for the COVID-19 Genomics UK (COG-UK) Consortium | The Lighthouse Lab in Milton Keynes and Alex Alderton, Roberto Amato, Jeffrey Barrett, Sonia Goncalves, Ewan Harrison, David K. Jackson, Ian Johnston, Dominic Kwiatkowski, Cordelia Langford, John Sillitoe on behalf of the Wellcome Sanger Institute COVID-19 Surveillance Team |
| EPI_ISL_2395393, EPI_ISL_2395394 | Lighthouse Lab in Glasgow | Wellcome Sanger Institute for the COVID-19 Genomics UK (COG-UK) Consortium | Harper VanSteenhouse, Yumi Kasai, David Gray, Carol Clugston, Anna Dominiczak and Alex Alderton, Roberto Amato, Jeffrey Barrett, Sonia Goncalves, Ewan Harrison, David K. Jackson, Ian Johnston, Dominic Kwiatkowski, Cordelia Langford, John Sillitoe on behalf of the Wellcome Sanger Institute COVID-19 Surveillance Team |
| EPI_ISL_2395396 | Lighthouse Lab in Milton Keynes | Wellcome Sanger Institute for the COVID-19 Genomics UK (COG-UK) Consortium | The Lighthouse Lab in Milton Keynes and Alex Alderton, Roberto Amato, Jeffrey Barrett, Sonia Goncalves, Ewan Harrison, David K. Jackson, Ian Johnston, Dominic Kwiatkowski, Cordelia Langford, John Sillitoe on behalf of the Wellcome Sanger Institute COVID-19 Surveillance Team |
| EPI_ISL_2395398, EPI_ISL_2395399, EPI_ISL_2395401 | Lighthouse Lab in Glasgow | Wellcome Sanger Institute for the COVID-19 Genomics UK (COG-UK) Consortium | Harper VanSteenhouse, Yumi Kasai, David Gray, Carol Clugston, Anna Dominiczak and Alex Alderton, Roberto Amato, Jeffrey Barrett, Sonia Goncalves, Ewan Harrison, David K. Jackson, Ian Johnston, Dominic Kwiatkowski, Cordelia Langford, John Sillitoe on behalf of the Wellcome Sanger Institute COVID-19 Surveillance Team |
| EPI_ISL_2395402 | Lighthouse Lab in Milton Keynes | Wellcome Sanger Institute for the COVID-19 Genomics UK (COG-UK) Consortium | The Lighthouse Lab in Milton Keynes and Alex Alderton, Roberto Amato, Jeffrey Barrett, Sonia Goncalves, Ewan Harrison, David K. Jackson, Ian Johnston, Dominic Kwiatkowski, Cordelia Langford, John Sillitoe on behalf of the Wellcome Sanger Institute COVID-19 Surveillance Team |
| EPI_ISL_2395403 | Lighthouse Lab in Glasgow | Wellcome Sanger Institute for the COVID-19 Genomics UK (COG-UK) Consortium | Harper VanSteenhouse, Yumi Kasai, David Gray, Carol Clugston, Anna Dominiczak and Alex Alderton, Roberto Amato, Jeffrey Barrett, Sonia Goncalves, Ewan Harrison, David K. Jackson, Ian Johnston, Dominic Kwiatkowski, Cordelia Langford, John Sillitoe on behalf of the Wellcome Sanger Institute COVID-19 Surveillance Team |
| EPI_ISL_2395404, EPI_ISL_2395405, EPI_ISL_2395407, EPI_ISL_2395408, EPI_ISL_2395409, EPI_ISL_2395410 | Lighthouse Lab in Milton Keynes | Wellcome Sanger Institute for the COVID-19 Genomics UK (COG-UK) Consortium | The Lighthouse Lab in Milton Keynes and Alex Alderton, Roberto Amato, Jeffrey Barrett, Sonia Goncalves, Ewan Harrison, David K. Jackson, Ian Johnston, Dominic Kwiatkowski, Cordelia Langford, John Sillitoe on behalf of the Wellcome Sanger Institute COVID-19 Surveillance Team |
| EPI_ISL_2395411 | Lighthouse Lab in Alderley Park | Wellcome Sanger Institute for the COVID-19 Genomics UK (COG-UK) Consortium | Jacquelyn Wynn, Mairead Hyland, The Lighthouse Lab in Alderley Park and Alex Alderton, Roberto Amato, Jeffrey Barrett, Sonia Goncalves, Ewan Harrison, David K. Jackson, Ian Johnston, Dominic Kwiatkowski, Cordelia Langford, John Sillitoe on behalf of the Wellcome Sanger Institute COVID-19 Surveillance Team |
| EPI_ISL_2395413 | Lighthouse Lab in Glasgow | Wellcome Sanger Institute for the COVID-19 Genomics UK (COG-UK) Consortium | Harper VanSteenhouse, Yumi Kasai, David Gray, Carol Clugston, Anna Dominiczak and Alex Alderton, Roberto Amato, Jeffrey Barrett, Sonia Goncalves, Ewan Harrison, David K. Jackson, Ian Johnston, Dominic Kwiatkowski, Cordelia Langford, John Sillitoe on behalf of the Wellcome Sanger Institute COVID-19 Surveillance Team |
| EPI_ISL_2395414, EPI_ISL_2395417, EPI_ISL_2395418, EPI_ISL_2395419, EPI_ISL_2395420 | Lighthouse Lab in Milton Keynes | Wellcome Sanger Institute for the COVID-19 Genomics UK (COG-UK) Consortium | The Lighthouse Lab in Milton Keynes and Alex Alderton, Roberto Amato, Jeffrey Barrett, Sonia Goncalves, Ewan Harrison, David K. Jackson, Ian Johnston, Dominic Kwiatkowski, Cordelia Langford, John Sillitoe on behalf of the Wellcome Sanger Institute COVID-19 Surveillance Team |
| EPI_ISL_2395422 | Lighthouse Lab in Alderley Park | Wellcome Sanger Institute for the COVID-19 Genomics UK (COG-UK) Consortium | Jacquelyn Wynn, Mairead Hyland, The Lighthouse Lab in Alderley Park and Alex Alderton, Roberto Amato, Jeffrey Barrett, Sonia Goncalves, Ewan Harrison, David K. Jackson, Ian Johnston, Dominic Kwiatkowski, Cordelia Langford, John Sillitoe on behalf of the Wellcome Sanger Institute COVID-19 Surveillance Team |
| EPI_ISL_2395423 | Lighthouse Lab in Glasgow | Wellcome Sanger Institute for the COVID-19 Genomics UK (COG-UK) Consortium | Harper VanSteenhouse, Yumi Kasai, David Gray, Carol Clugston, Anna Dominiczak and Alex Alderton, Roberto Amato, Jeffrey Barrett, Sonia Goncalves, Ewan Harrison, David K. Jackson, Ian Johnston, Dominic Kwiatkowski, Cordelia Langford, John Sillitoe on behalf of the Wellcome Sanger Institute COVID-19 Surveillance Team |
| EPI_ISL_2395425 | Lighthouse Lab in Milton Keynes | Wellcome Sanger Institute for the COVID-19 Genomics UK (COG-UK) Consortium | The Lighthouse Lab in Milton Keynes and Alex Alderton, Roberto Amato, Jeffrey Barrett, Sonia Goncalves, Ewan Harrison, David K. Jackson, Ian Johnston, Dominic Kwiatkowski, Cordelia Langford, John Sillitoe on behalf of the Wellcome Sanger Institute COVID-19 Surveillance Team |
| EPI_ISL_2395426 | Lighthouse Lab in Alderley Park | Wellcome Sanger Institute for the COVID-19 Genomics UK (COG-UK) Consortium | Jacquelyn Wynn, Mairead Hyland, The Lighthouse Lab in Alderley Park and Alex Alderton, Roberto Amato, Jeffrey Barrett, Sonia Goncalves, Ewan Harrison, David K. Jackson, Ian Johnston, Dominic Kwiatkowski, Cordelia Langford, John Sillitoe on behalf of the Wellcome Sanger Institute COVID-19 Surveillance Team |
| EPI_ISL_2395427, EPI_ISL_2395428 | Lighthouse Lab in Milton Keynes | Wellcome Sanger Institute for the COVID-19 Genomics UK (COG-UK) Consortium | The Lighthouse Lab in Milton Keynes and Alex Alderton, Roberto Amato, Jeffrey Barrett, Sonia Goncalves, Ewan Harrison, David K. Jackson, Ian Johnston, Dominic Kwiatkowski, Cordelia Langford, John Sillitoe on behalf of the Wellcome Sanger Institute COVID-19 Surveillance Team |
| EPI_ISL_2395429, EPI_ISL_2395430, EPI_ISL_2395432 | Lighthouse Lab in Glasgow | Wellcome Sanger Institute for the COVID-19 Genomics UK (COG-UK) Consortium | Harper VanSteenhouse, Yumi Kasai, David Gray, Carol Clugston, Anna Dominiczak and Alex Alderton, Roberto Amato, Jeffrey Barrett, Sonia Goncalves, Ewan Harrison, David K. Jackson, Ian Johnston, Dominic Kwiatkowski, Cordelia Langford, John Sillitoe on behalf of the Wellcome Sanger Institute COVID-19 Surveillance Team |
| EPI_ISL_2395433 | Lighthouse Lab in Milton Keynes | Wellcome Sanger Institute for the COVID-19 Genomics UK (COG-UK) Consortium | The Lighthouse Lab in Milton Keynes and Alex Alderton, Roberto Amato, Jeffrey Barrett, Sonia Goncalves, Ewan Harrison, David K. Jackson, Ian Johnston, Dominic Kwiatkowski, Cordelia Langford, John Sillitoe on behalf of the Wellcome Sanger Institute COVID-19 Surveillance Team |
| EPI_ISL_2395435 | Lighthouse Lab in Glasgow | Wellcome Sanger Institute for the COVID-19 Genomics UK (COG-UK) Consortium | Harper VanSteenhouse, Yumi Kasai, David Gray, Carol Clugston, Anna Dominiczak and Alex Alderton, Roberto Amato, Jeffrey Barrett, Sonia Goncalves, Ewan Harrison, David K. Jackson, Ian Johnston, Dominic Kwiatkowski, Cordelia Langford, John Sillitoe on behalf of the Wellcome Sanger Institute COVID-19 Surveillance Team |

| | | | |
|---|---|---|---|
| EPI_ISL_2395436 | Lighthouse Lab in Milton Keynes | Wellcome Sanger Institute for the COVID-19 Genomics UK (COG-UK) Consortium | The Lighthouse Lab in Milton Keynes and Alex Alderton, Roberto Amato, Jeffrey Barrett, Sonia Goncalves, Ewan Harrison, David K. Jackson, Ian Johnston, Dominic Kwiatkowski, Cordelia Langford, John Sillitoe on behalf of the Wellcome Sanger Institute COVID-19 Surveillance Team |
| EPI_ISL_2395437 | Lighthouse Lab in Alderley Park | Wellcome Sanger Institute for the COVID-19 Genomics UK (COG-UK) Consortium | Jacquelyn Wynn, Mairead Hyland, The Lighthouse Lab in Alderley Park and Alex Alderton, Roberto Amato, Jeffrey Barrett, Sonia Goncalves, Ewan Harrison, David K. Jackson, Ian Johnston, Dominic Kwiatkowski, Cordelia Langford, John Sillitoe on behalf of the Wellcome Sanger Institute COVID-19 Surveillance Team |
| EPI_ISL_2395438, EPI_ISL_2395439 | Lighthouse Lab in Milton Keynes | Wellcome Sanger Institute for the COVID-19 Genomics UK (COG-UK) Consortium | The Lighthouse Lab in Milton Keynes and Alex Alderton, Roberto Amato, Jeffrey Barrett, Sonia Goncalves, Ewan Harrison, David K. Jackson, Ian Johnston, Dominic Kwiatkowski, Cordelia Langford, John Sillitoe on behalf of the Wellcome Sanger Institute COVID-19 Surveillance Team |
| EPI_ISL_2395440 | Lighthouse Lab in Glasgow | Wellcome Sanger Institute for the COVID-19 Genomics UK (COG-UK) Consortium | Harper VanSteenhouse, Yumi Kasai, David Gray, Carol Clugston, Anna Dominiczak and Alex Alderton, Roberto Amato, Jeffrey Barrett, Sonia Goncalves, Ewan Harrison, David K. Jackson, Ian Johnston, Dominic Kwiatkowski, Cordelia Langford, John Sillitoe on behalf of the Wellcome Sanger Institute COVID-19 Surveillance Team |
| EPI_ISL_2395441 | Lighthouse Lab in Milton Keynes | Wellcome Sanger Institute for the COVID-19 Genomics UK (COG-UK) Consortium | The Lighthouse Lab in Milton Keynes and Alex Alderton, Roberto Amato, Jeffrey Barrett, Sonia Goncalves, Ewan Harrison, David K. Jackson, Ian Johnston, Dominic Kwiatkowski, Cordelia Langford, John Sillitoe on behalf of the Wellcome Sanger Institute COVID-19 Surveillance Team |
| EPI_ISL_2395442 | Lighthouse Lab in Glasgow | Wellcome Sanger Institute for the COVID-19 Genomics UK (COG-UK) Consortium | Harper VanSteenhouse, Yumi Kasai, David Gray, Carol Clugston, Anna Dominiczak and Alex Alderton, Roberto Amato, Jeffrey Barrett, Sonia Goncalves, Ewan Harrison, David K. Jackson, Ian Johnston, Dominic Kwiatkowski, Cordelia Langford, John Sillitoe on behalf of the Wellcome Sanger Institute COVID-19 Surveillance Team |
| EPI_ISL_2395444, EPI_ISL_2395446 | Lighthouse Lab in Milton Keynes | Wellcome Sanger Institute for the COVID-19 Genomics UK (COG-UK) Consortium | The Lighthouse Lab in Milton Keynes and Alex Alderton, Roberto Amato, Jeffrey Barrett, Sonia Goncalves, Ewan Harrison, David K. Jackson, Ian Johnston, Dominic Kwiatkowski, Cordelia Langford, John Sillitoe on behalf of the Wellcome Sanger Institute COVID-19 Surveillance Team |
| EPI_ISL_2395449, EPI_ISL_2395450 | Lighthouse Lab in Glasgow | Wellcome Sanger Institute for the COVID-19 Genomics UK (COG-UK) Consortium | Harper VanSteenhouse, Yumi Kasai, David Gray, Carol Clugston, Anna Dominiczak and Alex Alderton, Roberto Amato, Jeffrey Barrett, Sonia Goncalves, Ewan Harrison, David K. Jackson, Ian Johnston, Dominic Kwiatkowski, Cordelia Langford, John Sillitoe on behalf of the Wellcome Sanger Institute COVID-19 Surveillance Team |
| EPI_ISL_2395451, EPI_ISL_2395453 | Lighthouse Lab in Milton Keynes | Wellcome Sanger Institute for the COVID-19 Genomics UK (COG-UK) Consortium | The Lighthouse Lab in Milton Keynes and Alex Alderton, Roberto Amato, Jeffrey Barrett, Sonia Goncalves, Ewan Harrison, David K. Jackson, Ian Johnston, Dominic Kwiatkowski, Cordelia Langford, John Sillitoe on behalf of the Wellcome Sanger Institute COVID-19 Surveillance Team |
| EPI_ISL_2395455 | Lighthouse Lab in Glasgow | Wellcome Sanger Institute for the COVID-19 Genomics UK (COG-UK) Consortium | Harper VanSteenhouse, Yumi Kasai, David Gray, Carol Clugston, Anna Dominiczak and Alex Alderton, Roberto Amato, Jeffrey Barrett, Sonia Goncalves, Ewan Harrison, David K. Jackson, Ian Johnston, Dominic Kwiatkowski, Cordelia Langford, John Sillitoe on behalf of the Wellcome Sanger Institute COVID-19 Surveillance Team |
| EPI_ISL_2395457 | Lighthouse Lab in Milton Keynes | Wellcome Sanger Institute for the COVID-19 Genomics UK (COG-UK) Consortium | The Lighthouse Lab in Milton Keynes and Alex Alderton, Roberto Amato, Jeffrey Barrett, Sonia Goncalves, Ewan Harrison, David K. Jackson, Ian Johnston, Dominic Kwiatkowski, Cordelia Langford, John Sillitoe on behalf of the Wellcome Sanger Institute COVID-19 Surveillance Team |
| EPI_ISL_2395458, EPI_ISL_2395459 | Lighthouse Lab in Glasgow | Wellcome Sanger Institute for the COVID-19 Genomics UK (COG-UK) Consortium | Harper VanSteenhouse, Yumi Kasai, David Gray, Carol Clugston, Anna Dominiczak and Alex Alderton, Roberto Amato, Jeffrey Barrett, Sonia Goncalves, Ewan Harrison, David K. Jackson, Ian Johnston, Dominic Kwiatkowski, Cordelia Langford, John Sillitoe on behalf of the Wellcome Sanger Institute COVID-19 Surveillance Team |
| EPI_ISL_2395460, EPI_ISL_2395461, EPI_ISL_2395462 | Lighthouse Lab in Alderley Park | Wellcome Sanger Institute for the COVID-19 Genomics UK (COG-UK) Consortium | Jacquelyn Wynn, Mairead Hyland, The Lighthouse Lab in Alderley Park and Alex Alderton, Roberto Amato, Jeffrey Barrett, Sonia Goncalves, Ewan Harrison, David K. Jackson, Ian Johnston, Dominic Kwiatkowski, Cordelia Langford, John Sillitoe on behalf of the Wellcome Sanger Institute COVID-19 Surveillance Team |
| EPI_ISL_2395463, EPI_ISL_2395464 | Lighthouse Lab in Milton Keynes | Wellcome Sanger Institute for the COVID-19 Genomics UK (COG-UK) Consortium | The Lighthouse Lab in Milton Keynes and Alex Alderton, Roberto Amato, Jeffrey Barrett, Sonia Goncalves, Ewan Harrison, David K. Jackson, Ian Johnston, Dominic Kwiatkowski, Cordelia Langford, John Sillitoe on behalf of the Wellcome Sanger Institute COVID-19 Surveillance Team |
| EPI_ISL_2395465 | Lighthouse Lab in Glasgow | Wellcome Sanger Institute for the COVID-19 Genomics UK (COG-UK) Consortium | Harper VanSteenhouse, Yumi Kasai, David Gray, Carol Clugston, Anna Dominiczak and Alex Alderton, Roberto Amato, Jeffrey Barrett, Sonia Goncalves, Ewan Harrison, David K. Jackson, Ian Johnston, Dominic Kwiatkowski, Cordelia Langford, John Sillitoe on behalf of the Wellcome Sanger Institute COVID-19 Surveillance Team |
| EPI_ISL_2395466 | Lighthouse Lab in Milton Keynes | Wellcome Sanger Institute for the COVID-19 Genomics UK (COG-UK) Consortium | The Lighthouse Lab in Milton Keynes and Alex Alderton, Roberto Amato, Jeffrey Barrett, Sonia Goncalves, Ewan Harrison, David K. Jackson, Ian Johnston, Dominic Kwiatkowski, Cordelia Langford, John Sillitoe on behalf of the Wellcome Sanger Institute COVID-19 Surveillance Team |
| EPI_ISL_2395467 | Lighthouse Lab in Alderley Park | Wellcome Sanger Institute for the COVID-19 Genomics UK (COG-UK) Consortium | Jacquelyn Wynn, Mairead Hyland, The Lighthouse Lab in Alderley Park and Alex Alderton, Roberto Amato, Jeffrey Barrett, Sonia Goncalves, Ewan Harrison, David K. Jackson, Ian Johnston, Dominic Kwiatkowski, Cordelia Langford, John Sillitoe on behalf of the Wellcome Sanger Institute COVID-19 Surveillance Team |
| EPI_ISL_2395468 | Lighthouse Lab in Milton Keynes | Wellcome Sanger Institute for the COVID-19 Genomics UK (COG-UK) Consortium | The Lighthouse Lab in Milton Keynes and Alex Alderton, Roberto Amato, Jeffrey Barrett, Sonia Goncalves, Ewan Harrison, David K. Jackson, Ian Johnston, Dominic Kwiatkowski, Cordelia Langford, John Sillitoe on behalf of the Wellcome Sanger Institute COVID-19 Surveillance Team |
| EPI_ISL_2395469 | Lighthouse Lab in Alderley Park | Wellcome Sanger Institute for the COVID-19 Genomics UK (COG-UK) Consortium | Jacquelyn Wynn, Mairead Hyland, The Lighthouse Lab in Alderley Park and Alex Alderton, Roberto Amato, Jeffrey Barrett, Sonia Goncalves, Ewan Harrison, David K. Jackson, Ian Johnston, Dominic Kwiatkowski, Cordelia Langford, John Sillitoe on behalf of the Wellcome Sanger Institute COVID-19 Surveillance Team |
| EPI_ISL_2395470 | Lighthouse Lab in Milton Keynes | Wellcome Sanger Institute for the COVID-19 Genomics UK (COG-UK) Consortium | The Lighthouse Lab in Milton Keynes and Alex Alderton, Roberto Amato, Jeffrey Barrett, Sonia Goncalves, Ewan Harrison, David K. Jackson, Ian Johnston, Dominic Kwiatkowski, Cordelia Langford, John Sillitoe on behalf of the Wellcome Sanger Institute COVID-19 Surveillance Team |
| EPI_ISL_2395471, EPI_ISL_2395472, EPI_ISL_2395473, EPI_ISL_2395474 | Lighthouse Lab in Glasgow | Wellcome Sanger Institute for the COVID-19 Genomics UK (COG-UK) Consortium | Harper VanSteenhouse, Yumi Kasai, David Gray, Carol Clugston, Anna Dominiczak and Alex Alderton, Roberto Amato, Jeffrey Barrett, Sonia Goncalves, Ewan Harrison, David K. Jackson, Ian Johnston, Dominic Kwiatkowski, Cordelia Langford, John Sillitoe on behalf of the Wellcome Sanger Institute COVID-19 Surveillance Team |
| EPI_ISL_2395475, EPI_ISL_2395476, EPI_ISL_2395477, EPI_ISL_2395480 | Lighthouse Lab in Milton Keynes | Wellcome Sanger Institute for the COVID-19 Genomics UK (COG-UK) Consortium | The Lighthouse Lab in Milton Keynes and Alex Alderton, Roberto Amato, Jeffrey Barrett, Sonia Goncalves, Ewan Harrison, David K. Jackson, Ian Johnston, Dominic Kwiatkowski, Cordelia Langford, John Sillitoe on behalf of the Wellcome Sanger Institute COVID-19 Surveillance Team |
| EPI_ISL_2395481 | Lighthouse Lab in Glasgow | Wellcome Sanger Institute for the COVID-19 Genomics UK (COG-UK) Consortium | Harper VanSteenhouse, Yumi Kasai, David Gray, Carol Clugston, Anna Dominiczak and Alex Alderton, Roberto Amato, Jeffrey Barrett, Sonia Goncalves, Ewan Harrison, David K. Jackson, Ian Johnston, Dominic Kwiatkowski, Cordelia Langford, John Sillitoe on behalf of the Wellcome Sanger Institute COVID-19 Surveillance Team |
| EPI_ISL_2395482 | Lighthouse Lab in Milton Keynes | Wellcome Sanger Institute for the COVID-19 Genomics UK (COG-UK) Consortium | The Lighthouse Lab in Milton Keynes and Alex Alderton, Roberto Amato, Jeffrey Barrett, Sonia Goncalves, Ewan Harrison, David K. Jackson, Ian Johnston, Dominic Kwiatkowski, Cordelia Langford, John Sillitoe on behalf of the Wellcome Sanger Institute COVID-19 Surveillance Team |
| EPI_ISL_2395484, EPI_ISL_2395485, EPI_ISL_2395486 | Lighthouse Lab in Alderley Park | Wellcome Sanger Institute for the COVID-19 Genomics UK (COG-UK) Consortium | Jacquelyn Wynn, Mairead Hyland, The Lighthouse Lab in Alderley Park and Alex Alderton, Roberto Amato, Jeffrey Barrett, Sonia Goncalves, Ewan Harrison, David K. Jackson, Ian Johnston, Dominic Kwiatkowski, Cordelia Langford, John Sillitoe on behalf of the Wellcome Sanger Institute COVID-19 Surveillance Team |
| EPI_ISL_2395488, EPI_ISL_2395490 | Lighthouse Lab in Milton Keynes | Wellcome Sanger Institute for the COVID-19 Genomics UK (COG-UK) Consortium | The Lighthouse Lab in Milton Keynes and Alex Alderton, Roberto Amato, Jeffrey Barrett, Sonia Goncalves, Ewan Harrison, David K. Jackson, Ian Johnston, Dominic Kwiatkowski, Cordelia Langford, John Sillitoe on behalf of the Wellcome Sanger Institute COVID-19 Surveillance Team |
| EPI_ISL_2395492 | Lighthouse Lab in Glasgow | Wellcome Sanger Institute for the COVID-19 Genomics UK (COG-UK) Consortium | Harper VanSteenhouse, Yumi Kasai, David Gray, Carol Clugston, Anna Dominiczak and Alex Alderton, Roberto Amato, Jeffrey Barrett, Sonia Goncalves, Ewan Harrison, David K. Jackson, Ian Johnston, Dominic Kwiatkowski, Cordelia Langford, John Sillitoe on behalf of the Wellcome Sanger Institute COVID-19 Surveillance Team |
| EPI_ISL_2395493 | Lighthouse Lab in Milton Keynes | Wellcome Sanger Institute for the COVID-19 Genomics UK (COG-UK) Consortium | The Lighthouse Lab in Milton Keynes and Alex Alderton, Roberto Amato, Jeffrey Barrett, Sonia Goncalves, Ewan Harrison, David K. Jackson, Ian Johnston, Dominic Kwiatkowski, Cordelia Langford, John Sillitoe on behalf of the Wellcome Sanger Institute COVID-19 Surveillance Team |
| EPI_ISL_2395494 | Lighthouse Lab in Alderley Park | Wellcome Sanger Institute for the COVID-19 Genomics UK (COG-UK) Consortium | Jacquelyn Wynn, Mairead Hyland, The Lighthouse Lab in Alderley Park and Alex Alderton, Roberto Amato, Jeffrey Barrett, Sonia Goncalves, Ewan Harrison, David K. Jackson, Ian Johnston, Dominic Kwiatkowski, Cordelia Langford, John Sillitoe on behalf of the Wellcome Sanger Institute COVID-19 Surveillance Team |
| EPI_ISL_2395495, EPI_ISL_2395496 | Lighthouse Lab in Milton Keynes | Wellcome Sanger Institute for the COVID-19 Genomics UK (COG-UK) Consortium | The Lighthouse Lab in Milton Keynes and Alex Alderton, Roberto Amato, Jeffrey Barrett, Sonia Goncalves, Ewan Harrison, David K. Jackson, Ian Johnston, Dominic Kwiatkowski, Cordelia Langford, John Sillitoe on behalf of the Wellcome Sanger Institute COVID-19 Surveillance Team |
| EPI_ISL_2395497 | Lighthouse Lab in Alderley Park | Wellcome Sanger Institute for the COVID-19 Genomics UK (COG-UK) Consortium | Jacquelyn Wynn, Mairead Hyland, The Lighthouse Lab in Alderley Park and Alex Alderton, Roberto Amato, Jeffrey Barrett, Sonia Goncalves, Ewan Harrison, David K. Jackson, Ian Johnston, Dominic Kwiatkowski, Cordelia Langford, John Sillitoe on behalf of the Wellcome Sanger Institute COVID-19 Surveillance Team |
| EPI_ISL_2395498, EPI_ISL_2395499, EPI_ISL_2395500 | Lighthouse Lab in Milton Keynes | Wellcome Sanger Institute for the COVID-19 Genomics UK (COG-UK) Consortium | The Lighthouse Lab in Milton Keynes and Alex Alderton, Roberto Amato, Jeffrey Barrett, Sonia Goncalves, Ewan Harrison, David K. Jackson, Ian Johnston, Dominic Kwiatkowski, Cordelia Langford, John Sillitoe on behalf of the Wellcome Sanger Institute COVID-19 Surveillance Team |
| EPI_ISL_2395502 | Lighthouse Lab in Alderley Park | Wellcome Sanger Institute for the COVID-19 Genomics UK (COG-UK) Consortium | Jacquelyn Wynn, Mairead Hyland, The Lighthouse Lab in Alderley Park and Alex Alderton, Roberto Amato, Jeffrey Barrett, Sonia Goncalves, Ewan Harrison, David K. Jackson, Ian Johnston, Dominic Kwiatkowski, Cordelia Langford, John Sillitoe on behalf of the Wellcome Sanger Institute COVID-19 Surveillance Team |
| EPI_ISL_2395503 | Lighthouse Lab in Milton Keynes | Wellcome Sanger Institute for the COVID-19 Genomics UK (COG-UK) Consortium | The Lighthouse Lab in Milton Keynes and Alex Alderton, Roberto Amato, Jeffrey Barrett, Sonia Goncalves, Ewan Harrison, David K. Jackson, Ian Johnston, Dominic Kwiatkowski, Cordelia Langford, John Sillitoe on behalf of the Wellcome Sanger Institute COVID-19 Surveillance Team |
| EPI_ISL_2395504 | Lighthouse Lab in Glasgow | Wellcome Sanger Institute for the COVID-19 Genomics UK (COG-UK) Consortium | Harper VanSteenhouse, Yumi Kasai, David Gray, Carol Clugston, Anna Dominiczak and Alex Alderton, Roberto Amato, Jeffrey Barrett, Sonia Goncalves, Ewan Harrison, David K. Jackson, Ian Johnston, Dominic Kwiatkowski, Cordelia Langford, John Sillitoe on behalf of the Wellcome Sanger Institute COVID-19 Surveillance Team |
| EPI_ISL_2395505 | Lighthouse Lab in Alderley Park | Wellcome Sanger Institute for the COVID-19 Genomics UK (COG-UK) Consortium | Jacquelyn Wynn, Mairead Hyland, The Lighthouse Lab in Alderley Park and Alex Alderton, Roberto Amato, Jeffrey Barrett, Sonia Goncalves, Ewan Harrison, David K. Jackson, Ian Johnston, Dominic Kwiatkowski, Cordelia Langford, John Sillitoe on behalf of the Wellcome Sanger Institute COVID-19 Surveillance Team |
| EPI_ISL_2395506, EPI_ISL_2395507 | Lighthouse Lab in Milton Keynes | Wellcome Sanger Institute for the COVID-19 Genomics UK (COG-UK) Consortium | The Lighthouse Lab in Milton Keynes and Alex Alderton, Roberto Amato, Jeffrey Barrett, Sonia Goncalves, Ewan Harrison, David K. Jackson, Ian Johnston, Dominic Kwiatkowski, Cordelia Langford, John Sillitoe on behalf of the Wellcome Sanger Institute COVID-19 Surveillance Team |
| EPI_ISL_2395509, EPI_ISL_2395510 | Lighthouse Lab in Glasgow | Wellcome Sanger Institute for the COVID-19 Genomics UK (COG-UK) Consortium | Harper VanSteenhouse, Yumi Kasai, David Gray, Carol Clugston, Anna Dominiczak and Alex Alderton, Roberto Amato, Jeffrey Barrett, Sonia Goncalves, Ewan Harrison, David K. Jackson, Ian Johnston, Dominic Kwiatkowski, Cordelia Langford, John Sillitoe on behalf of the Wellcome Sanger Institute COVID-19 Surveillance Team |
| EPI_ISL_2395511, EPI_ISL_2395513, EPI_ISL_2395514 | Lighthouse Lab in Milton Keynes | Wellcome Sanger Institute for the COVID-19 Genomics UK (COG-UK) Consortium | The Lighthouse Lab in Milton Keynes and Alex Alderton, Roberto Amato, Jeffrey Barrett, Sonia Goncalves, Ewan Harrison, David K. Jackson, Ian Johnston, Dominic Kwiatkowski, Cordelia Langford, John Sillitoe on behalf of the Wellcome Sanger Institute COVID-19 Surveillance Team |
| EPI_ISL_2395515 | Lighthouse Lab in Glasgow | Wellcome Sanger Institute for the COVID-19 Genomics UK (COG-UK) Consortium | Harper VanSteenhouse, Yumi Kasai, David Gray, Carol Clugston, Anna Dominiczak and Alex Alderton, Roberto Amato, Jeffrey Barrett, Sonia Goncalves, Ewan Harrison, David K. Jackson, Ian Johnston, Dominic Kwiatkowski, Cordelia Langford, John Sillitoe on behalf of the Wellcome Sanger Institute COVID-19 Surveillance Team |
| EPI_ISL_2395516 | Lighthouse Lab in Milton Keynes | Wellcome Sanger Institute for the COVID-19 Genomics UK (COG-UK) Consortium | The Lighthouse Lab in Milton Keynes and Alex Alderton, Roberto Amato, Jeffrey Barrett, Sonia Goncalves, Ewan Harrison, David K. Jackson, Ian Johnston, Dominic Kwiatkowski, Cordelia Langford, John Sillitoe on behalf of the Wellcome Sanger Institute COVID-19 Surveillance Team |
| EPI_ISL_2395517 | Lighthouse Lab in Glasgow | Wellcome Sanger Institute for the COVID-19 Genomics UK (COG-UK) Consortium | Harper VanSteenhouse, Yumi Kasai, David Gray, Carol Clugston, Anna Dominiczak and Alex Alderton, Roberto Amato, Jeffrey Barrett, Sonia Goncalves, Ewan Harrison, David K. Jackson, Ian Johnston, Dominic Kwiatkowski, Cordelia Langford, John Sillitoe on behalf of the Wellcome Sanger Institute COVID-19 Surveillance Team |

| | | | |
|---|---|---|---|
| EPI_ISL_2395518, EPI_ISL_2395519, EPI_ISL_2395520 | Lighthouse Lab in Milton Keynes | Wellcome Sanger Institute for the COVID-19 Genomics UK (COG-UK) Consortium | The Lighthouse Lab in Milton Keynes and Alex Alderton, Roberto Amato, Jeffrey Barrett, Sonia Goncalves, Ewan Harrison, David K. Jackson, Ian Johnston, Dominic Kwiatkowski, Cordelia Langford, John Sillitoe on behalf of the Wellcome Sanger Institute COVID-19 Surveillance Team |
| EPI_ISL_2395521 | Lighthouse Lab in Alderley Park | Wellcome Sanger Institute for the COVID-19 Genomics UK (COG-UK) Consortium | Jacquelyn Wynn, Mairead Hyland, The Lighthouse Lab in Alderley Park and Alex Alderton, Roberto Amato, Jeffrey Barrett, Sonia Goncalves, Ewan Harrison, David K. Jackson, Ian Johnston, Dominic Kwiatkowski, Cordelia Langford, John Sillitoe on behalf of the Wellcome Sanger Institute COVID-19 Surveillance Team |
| EPI_ISL_2395522 | Lighthouse Lab in Milton Keynes | Wellcome Sanger Institute for the COVID-19 Genomics UK (COG-UK) Consortium | The Lighthouse Lab in Milton Keynes and Alex Alderton, Roberto Amato, Jeffrey Barrett, Sonia Goncalves, Ewan Harrison, David K. Jackson, Ian Johnston, Dominic Kwiatkowski, Cordelia Langford, John Sillitoe on behalf of the Wellcome Sanger Institute COVID-19 Surveillance Team |
| EPI_ISL_2395524 | Lighthouse Lab in Alderley Park | Wellcome Sanger Institute for the COVID-19 Genomics UK (COG-UK) Consortium | Jacquelyn Wynn, Mairead Hyland, The Lighthouse Lab in Alderley Park and Alex Alderton, Roberto Amato, Jeffrey Barrett, Sonia Goncalves, Ewan Harrison, David K. Jackson, Ian Johnston, Dominic Kwiatkowski, Cordelia Langford, John Sillitoe on behalf of the Wellcome Sanger Institute COVID-19 Surveillance Team |
| EPI_ISL_2395525 | Lighthouse Lab in Milton Keynes | Wellcome Sanger Institute for the COVID-19 Genomics UK (COG-UK) Consortium | The Lighthouse Lab in Milton Keynes and Alex Alderton, Roberto Amato, Jeffrey Barrett, Sonia Goncalves, Ewan Harrison, David K. Jackson, Ian Johnston, Dominic Kwiatkowski, Cordelia Langford, John Sillitoe on behalf of the Wellcome Sanger Institute COVID-19 Surveillance Team |
| EPI_ISL_2395526, EPI_ISL_2395527, EPI_ISL_2395528 | Lighthouse Lab in Glasgow | Wellcome Sanger Institute for the COVID-19 Genomics UK (COG-UK) Consortium | Harper VanSteenhouse, Yumi Kasai, David Gray, Carol Clugston, Anna Dominiczak and Alex Alderton, Roberto Amato, Jeffrey Barrett, Sonia Goncalves, Ewan Harrison, David K. Jackson, Ian Johnston, Dominic Kwiatkowski, Cordelia Langford, John Sillitoe on behalf of the Wellcome Sanger Institute COVID-19 Surveillance Team |
| EPI_ISL_2395529 | Lighthouse Lab in Milton Keynes | Wellcome Sanger Institute for the COVID-19 Genomics UK (COG-UK) Consortium | The Lighthouse Lab in Milton Keynes and Alex Alderton, Roberto Amato, Jeffrey Barrett, Sonia Goncalves, Ewan Harrison, David K. Jackson, Ian Johnston, Dominic Kwiatkowski, Cordelia Langford, John Sillitoe on behalf of the Wellcome Sanger Institute COVID-19 Surveillance Team |
| EPI_ISL_2395530 | Lighthouse Lab in Alderley Park | Wellcome Sanger Institute for the COVID-19 Genomics UK (COG-UK) Consortium | Jacquelyn Wynn, Mairead Hyland, The Lighthouse Lab in Alderley Park and Alex Alderton, Roberto Amato, Jeffrey Barrett, Sonia Goncalves, Ewan Harrison, David K. Jackson, Ian Johnston, Dominic Kwiatkowski, Cordelia Langford, John Sillitoe on behalf of the Wellcome Sanger Institute COVID-19 Surveillance Team |
| EPI_ISL_2395532, EPI_ISL_2395533 | Lighthouse Lab in Milton Keynes | Wellcome Sanger Institute for the COVID-19 Genomics UK (COG-UK) Consortium | The Lighthouse Lab in Milton Keynes and Alex Alderton, Roberto Amato, Jeffrey Barrett, Sonia Goncalves, Ewan Harrison, David K. Jackson, Ian Johnston, Dominic Kwiatkowski, Cordelia Langford, John Sillitoe on behalf of the Wellcome Sanger Institute COVID-19 Surveillance Team |
| EPI_ISL_2395535 | Lighthouse Lab in Alderley Park | Wellcome Sanger Institute for the COVID-19 Genomics UK (COG-UK) Consortium | Jacquelyn Wynn, Mairead Hyland, The Lighthouse Lab in Alderley Park and Alex Alderton, Roberto Amato, Jeffrey Barrett, Sonia Goncalves, Ewan Harrison, David K. Jackson, Ian Johnston, Dominic Kwiatkowski, Cordelia Langford, John Sillitoe on behalf of the Wellcome Sanger Institute COVID-19 Surveillance Team |
| EPI_ISL_2395536 | Lighthouse Lab in Milton Keynes | Wellcome Sanger Institute for the COVID-19 Genomics UK (COG-UK) Consortium | The Lighthouse Lab in Milton Keynes and Alex Alderton, Roberto Amato, Jeffrey Barrett, Sonia Goncalves, Ewan Harrison, David K. Jackson, Ian Johnston, Dominic Kwiatkowski, Cordelia Langford, John Sillitoe on behalf of the Wellcome Sanger Institute COVID-19 Surveillance Team |
| EPI_ISL_2395538 | Lighthouse Lab in Glasgow | Wellcome Sanger Institute for the COVID-19 Genomics UK (COG-UK) Consortium | Harper VanSteenhouse, Yumi Kasai, David Gray, Carol Clugston, Anna Dominiczak and Alex Alderton, Roberto Amato, Jeffrey Barrett, Sonia Goncalves, Ewan Harrison, David K. Jackson, Ian Johnston, Dominic Kwiatkowski, Cordelia Langford, John Sillitoe on behalf of the Wellcome Sanger Institute COVID-19 Surveillance Team |
| EPI_ISL_2395541, EPI_ISL_2395543 | Lighthouse Lab in Milton Keynes | Wellcome Sanger Institute for the COVID-19 Genomics UK (COG-UK) Consortium | The Lighthouse Lab in Milton Keynes and Alex Alderton, Roberto Amato, Jeffrey Barrett, Sonia Goncalves, Ewan Harrison, David K. Jackson, Ian Johnston, Dominic Kwiatkowski, Cordelia Langford, John Sillitoe on behalf of the Wellcome Sanger Institute COVID-19 Surveillance Team |
| EPI_ISL_2395544 | Lighthouse Lab in Glasgow | Wellcome Sanger Institute for the COVID-19 Genomics UK (COG-UK) Consortium | Harper VanSteenhouse, Yumi Kasai, David Gray, Carol Clugston, Anna Dominiczak and Alex Alderton, Roberto Amato, Jeffrey Barrett, Sonia Goncalves, Ewan Harrison, David K. Jackson, Ian Johnston, Dominic Kwiatkowski, Cordelia Langford, John Sillitoe on behalf of the Wellcome Sanger Institute COVID-19 Surveillance Team |
| EPI_ISL_2395545, EPI_ISL_2395546 | Lighthouse Lab in Milton Keynes | Wellcome Sanger Institute for the COVID-19 Genomics UK (COG-UK) Consortium | The Lighthouse Lab in Milton Keynes and Alex Alderton, Roberto Amato, Jeffrey Barrett, Sonia Goncalves, Ewan Harrison, David K. Jackson, Ian Johnston, Dominic Kwiatkowski, Cordelia Langford, John Sillitoe on behalf of the Wellcome Sanger Institute COVID-19 Surveillance Team |
| EPI_ISL_2395547 | Lighthouse Lab in Glasgow | Wellcome Sanger Institute for the COVID-19 Genomics UK (COG-UK) Consortium | Harper VanSteenhouse, Yumi Kasai, David Gray, Carol Clugston, Anna Dominiczak and Alex Alderton, Roberto Amato, Jeffrey Barrett, Sonia Goncalves, Ewan Harrison, David K. Jackson, Ian Johnston, Dominic Kwiatkowski, Cordelia Langford, John Sillitoe on behalf of the Wellcome Sanger Institute COVID-19 Surveillance Team |
| EPI_ISL_2395548, EPI_ISL_2395549 | Lighthouse Lab in Milton Keynes | Wellcome Sanger Institute for the COVID-19 Genomics UK (COG-UK) Consortium | The Lighthouse Lab in Milton Keynes and Alex Alderton, Roberto Amato, Jeffrey Barrett, Sonia Goncalves, Ewan Harrison, David K. Jackson, Ian Johnston, Dominic Kwiatkowski, Cordelia Langford, John Sillitoe on behalf of the Wellcome Sanger Institute COVID-19 Surveillance Team |
| EPI_ISL_2395551 | Lighthouse Lab in Alderley Park | Wellcome Sanger Institute for the COVID-19 Genomics UK (COG-UK) Consortium | Jacquelyn Wynn, Mairead Hyland, The Lighthouse Lab in Alderley Park and Alex Alderton, Roberto Amato, Jeffrey Barrett, Sonia Goncalves, Ewan Harrison, David K. Jackson, Ian Johnston, Dominic Kwiatkowski, Cordelia Langford, John Sillitoe on behalf of the Wellcome Sanger Institute COVID-19 Surveillance Team |
| EPI_ISL_2395552, EPI_ISL_2395553, EPI_ISL_2395554 | Lighthouse Lab in Milton Keynes | Wellcome Sanger Institute for the COVID-19 Genomics UK (COG-UK) Consortium | The Lighthouse Lab in Milton Keynes and Alex Alderton, Roberto Amato, Jeffrey Barrett, Sonia Goncalves, Ewan Harrison, David K. Jackson, Ian Johnston, Dominic Kwiatkowski, Cordelia Langford, John Sillitoe on behalf of the Wellcome Sanger Institute COVID-19 Surveillance Team |
| EPI_ISL_2395555 | Lighthouse Lab in Glasgow | Wellcome Sanger Institute for the COVID-19 Genomics UK (COG-UK) Consortium | Harper VanSteenhouse, Yumi Kasai, David Gray, Carol Clugston, Anna Dominiczak and Alex Alderton, Roberto Amato, Jeffrey Barrett, Sonia Goncalves, Ewan Harrison, David K. Jackson, Ian Johnston, Dominic Kwiatkowski, Cordelia Langford, John Sillitoe on behalf of the Wellcome Sanger Institute COVID-19 Surveillance Team |
| EPI_ISL_2395556 | Lighthouse Lab in Milton Keynes | Wellcome Sanger Institute for the COVID-19 Genomics UK (COG-UK) Consortium | The Lighthouse Lab in Milton Keynes and Alex Alderton, Roberto Amato, Jeffrey Barrett, Sonia Goncalves, Ewan Harrison, David K. Jackson, Ian Johnston, Dominic Kwiatkowski, Cordelia Langford, John Sillitoe on behalf of the Wellcome Sanger Institute COVID-19 Surveillance Team |
| EPI_ISL_2395557 | Lighthouse Lab in Glasgow | Wellcome Sanger Institute for the COVID-19 Genomics UK (COG-UK) Consortium | Harper VanSteenhouse, Yumi Kasai, David Gray, Carol Clugston, Anna Dominiczak and Alex Alderton, Roberto Amato, Jeffrey Barrett, Sonia Goncalves, Ewan Harrison, David K. Jackson, Ian Johnston, Dominic Kwiatkowski, Cordelia Langford, John Sillitoe on behalf of the Wellcome Sanger Institute COVID-19 Surveillance Team |
| EPI_ISL_2395559, EPI_ISL_2395560, EPI_ISL_2395561 | Lighthouse Lab in Milton Keynes | Wellcome Sanger Institute for the COVID-19 Genomics UK (COG-UK) Consortium | The Lighthouse Lab in Milton Keynes and Alex Alderton, Roberto Amato, Jeffrey Barrett, Sonia Goncalves, Ewan Harrison, David K. Jackson, Ian Johnston, Dominic Kwiatkowski, Cordelia Langford, John Sillitoe on behalf of the Wellcome Sanger Institute COVID-19 Surveillance Team |
| EPI_ISL_2395562 | Lighthouse Lab in Glasgow | Wellcome Sanger Institute for the COVID-19 Genomics UK (COG-UK) Consortium | Harper VanSteenhouse, Yumi Kasai, David Gray, Carol Clugston, Anna Dominiczak and Alex Alderton, Roberto Amato, Jeffrey Barrett, Sonia Goncalves, Ewan Harrison, David K. Jackson, Ian Johnston, Dominic Kwiatkowski, Cordelia Langford, John Sillitoe on behalf of the Wellcome Sanger Institute COVID-19 Surveillance Team |
| EPI_ISL_2395563, EPI_ISL_2395564 | Lighthouse Lab in Milton Keynes | Wellcome Sanger Institute for the COVID-19 Genomics UK (COG-UK) Consortium | The Lighthouse Lab in Milton Keynes and Alex Alderton, Roberto Amato, Jeffrey Barrett, Sonia Goncalves, Ewan Harrison, David K. Jackson, Ian Johnston, Dominic Kwiatkowski, Cordelia Langford, John Sillitoe on behalf of the Wellcome Sanger Institute COVID-19 Surveillance Team |
| EPI_ISL_2395566 | Lighthouse Lab in Glasgow | Wellcome Sanger Institute for the COVID-19 Genomics UK (COG-UK) Consortium | Harper VanSteenhouse, Yumi Kasai, David Gray, Carol Clugston, Anna Dominiczak and Alex Alderton, Roberto Amato, Jeffrey Barrett, Sonia Goncalves, Ewan Harrison, David K. Jackson, Ian Johnston, Dominic Kwiatkowski, Cordelia Langford, John Sillitoe on behalf of the Wellcome Sanger Institute COVID-19 Surveillance Team |
| EPI_ISL_2395567, EPI_ISL_2395568, EPI_ISL_2395569 | Lighthouse Lab in Alderley Park | Wellcome Sanger Institute for the COVID-19 Genomics UK (COG-UK) Consortium | Jacquelyn Wynn, Mairead Hyland, The Lighthouse Lab in Alderley Park and Alex Alderton, Roberto Amato, Jeffrey Barrett, Sonia Goncalves, Ewan Harrison, David K. Jackson, Ian Johnston, Dominic Kwiatkowski, Cordelia Langford, John Sillitoe on behalf of the Wellcome Sanger Institute COVID-19 Surveillance Team |
| EPI_ISL_2395571, EPI_ISL_2395572, EPI_ISL_2395573 | Lighthouse Lab in Milton Keynes | Wellcome Sanger Institute for the COVID-19 Genomics UK (COG-UK) Consortium | The Lighthouse Lab in Milton Keynes and Alex Alderton, Roberto Amato, Jeffrey Barrett, Sonia Goncalves, Ewan Harrison, David K. Jackson, Ian Johnston, Dominic Kwiatkowski, Cordelia Langford, John Sillitoe on behalf of the Wellcome Sanger Institute COVID-19 Surveillance Team |
| EPI_ISL_2395575, EPI_ISL_2395576 | Lighthouse Lab in Alderley Park | Wellcome Sanger Institute for the COVID-19 Genomics UK (COG-UK) Consortium | Jacquelyn Wynn, Mairead Hyland, The Lighthouse Lab in Alderley Park and Alex Alderton, Roberto Amato, Jeffrey Barrett, Sonia Goncalves, Ewan Harrison, David K. Jackson, Ian Johnston, Dominic Kwiatkowski, Cordelia Langford, John Sillitoe on behalf of the Wellcome Sanger Institute COVID-19 Surveillance Team |
| EPI_ISL_2395578, EPI_ISL_2395579 | Lighthouse Lab in Milton Keynes | Wellcome Sanger Institute for the COVID-19 Genomics UK (COG-UK) Consortium | The Lighthouse Lab in Milton Keynes and Alex Alderton, Roberto Amato, Jeffrey Barrett, Sonia Goncalves, Ewan Harrison, David K. Jackson, Ian Johnston, Dominic Kwiatkowski, Cordelia Langford, John Sillitoe on behalf of the Wellcome Sanger Institute COVID-19 Surveillance Team |
| EPI_ISL_2395580 | Lighthouse Lab in Glasgow | Wellcome Sanger Institute for the COVID-19 Genomics UK (COG-UK) Consortium | Harper VanSteenhouse, Yumi Kasai, David Gray, Carol Clugston, Anna Dominiczak and Alex Alderton, Roberto Amato, Jeffrey Barrett, Sonia Goncalves, Ewan Harrison, David K. Jackson, Ian Johnston, Dominic Kwiatkowski, Cordelia Langford, John Sillitoe on behalf of the Wellcome Sanger Institute COVID-19 Surveillance Team |
| EPI_ISL_2395581 | Lighthouse Lab in Milton Keynes | Wellcome Sanger Institute for the COVID-19 Genomics UK (COG-UK) Consortium | The Lighthouse Lab in Milton Keynes and Alex Alderton, Roberto Amato, Jeffrey Barrett, Sonia Goncalves, Ewan Harrison, David K. Jackson, Ian Johnston, Dominic Kwiatkowski, Cordelia Langford, John Sillitoe on behalf of the Wellcome Sanger Institute COVID-19 Surveillance Team |
| EPI_ISL_2395584, EPI_ISL_2395588, EPI_ISL_2395589 | Lighthouse Lab in Alderley Park | Wellcome Sanger Institute for the COVID-19 Genomics UK (COG-UK) Consortium | Jacquelyn Wynn, Mairead Hyland, The Lighthouse Lab in Alderley Park and Alex Alderton, Roberto Amato, Jeffrey Barrett, Sonia Goncalves, Ewan Harrison, David K. Jackson, Ian Johnston, Dominic Kwiatkowski, Cordelia Langford, John Sillitoe on behalf of the Wellcome Sanger Institute COVID-19 Surveillance Team |
| EPI_ISL_2395592, EPI_ISL_2395594 | Lighthouse Lab in Milton Keynes | Wellcome Sanger Institute for the COVID-19 Genomics UK (COG-UK) Consortium | The Lighthouse Lab in Milton Keynes and Alex Alderton, Roberto Amato, Jeffrey Barrett, Sonia Goncalves, Ewan Harrison, David K. Jackson, Ian Johnston, Dominic Kwiatkowski, Cordelia Langford, John Sillitoe on behalf of the Wellcome Sanger Institute COVID-19 Surveillance Team |
| EPI_ISL_2395595, EPI_ISL_2395596 | Lighthouse Lab in Alderley Park | Wellcome Sanger Institute for the COVID-19 Genomics UK (COG-UK) Consortium | Jacquelyn Wynn, Mairead Hyland, The Lighthouse Lab in Alderley Park and Alex Alderton, Roberto Amato, Jeffrey Barrett, Sonia Goncalves, Ewan Harrison, David K. Jackson, Ian Johnston, Dominic Kwiatkowski, Cordelia Langford, John Sillitoe on behalf of the Wellcome Sanger Institute COVID-19 Surveillance Team |
| EPI_ISL_2395597 | Lighthouse Lab in Glasgow | Wellcome Sanger Institute for the COVID-19 Genomics UK (COG-UK) Consortium | Harper VanSteenhouse, Yumi Kasai, David Gray, Carol Clugston, Anna Dominiczak and Alex Alderton, Roberto Amato, Jeffrey Barrett, Sonia Goncalves, Ewan Harrison, David K. Jackson, Ian Johnston, Dominic Kwiatkowski, Cordelia Langford, John Sillitoe on behalf of the Wellcome Sanger Institute COVID-19 Surveillance Team |
| EPI_ISL_2395598 | Lighthouse Lab in Milton Keynes | Wellcome Sanger Institute for the COVID-19 Genomics UK (COG-UK) Consortium | The Lighthouse Lab in Milton Keynes and Alex Alderton, Roberto Amato, Jeffrey Barrett, Sonia Goncalves, Ewan Harrison, David K. Jackson, Ian Johnston, Dominic Kwiatkowski, Cordelia Langford, John Sillitoe on behalf of the Wellcome Sanger Institute COVID-19 Surveillance Team |
| EPI_ISL_2395599, EPI_ISL_2395600 | Lighthouse Lab in Alderley Park | Wellcome Sanger Institute for the COVID-19 Genomics UK (COG-UK) Consortium | Jacquelyn Wynn, Mairead Hyland, The Lighthouse Lab in Alderley Park and Alex Alderton, Roberto Amato, Jeffrey Barrett, Sonia Goncalves, Ewan Harrison, David K. Jackson, Ian Johnston, Dominic Kwiatkowski, Cordelia Langford, John Sillitoe on behalf of the Wellcome Sanger Institute COVID-19 Surveillance Team |
| EPI_ISL_2395601 | Lighthouse Lab in Glasgow | Wellcome Sanger Institute for the COVID-19 Genomics UK (COG-UK) Consortium | Harper VanSteenhouse, Yumi Kasai, David Gray, Carol Clugston, Anna Dominiczak and Alex Alderton, Roberto Amato, Jeffrey Barrett, Sonia Goncalves, Ewan Harrison, David K. Jackson, Ian Johnston, Dominic Kwiatkowski, Cordelia Langford, John Sillitoe on behalf of the Wellcome Sanger Institute COVID-19 Surveillance Team |
| EPI_ISL_2395603, EPI_ISL_2395604 | Lighthouse Lab in Alderley Park | Wellcome Sanger Institute for the COVID-19 Genomics UK (COG-UK) Consortium | Jacquelyn Wynn, Mairead Hyland, The Lighthouse Lab in Alderley Park and Alex Alderton, Roberto Amato, Jeffrey Barrett, Sonia Goncalves, Ewan Harrison, David K. Jackson, Ian Johnston, Dominic Kwiatkowski, Cordelia Langford, John Sillitoe on behalf of the Wellcome Sanger Institute COVID-19 Surveillance Team |
| EPI_ISL_2395605, EPI_ISL_2395607, EPI_ISL_2395608, EPI_ISL_2395609 | Lighthouse Lab in Milton Keynes | Wellcome Sanger Institute for the COVID-19 Genomics UK (COG-UK) Consortium | The Lighthouse Lab in Milton Keynes and Alex Alderton, Roberto Amato, Jeffrey Barrett, Sonia Goncalves, Ewan Harrison, David K. Jackson, Ian Johnston, Dominic Kwiatkowski, Cordelia Langford, John Sillitoe on behalf of the Wellcome Sanger Institute COVID-19 Surveillance Team |
| EPI_ISL_2395610 | Lighthouse Lab in Glasgow | Wellcome Sanger Institute for the COVID-19 Genomics UK (COG-UK) | Harper VanSteenhouse, Yumi Kasai, David Gray, Carol Clugston, Anna Dominiczak and Alex Alderton, Roberto Amato, Jeffrey Barrett, Sonia Goncalves, Ewan Harrison, David K. Jackson, Ian Johnston, Dominic Kwiatkowski, Cordelia Langford, |

| | | | |
|---|---|---|---|
| | | Consortium | John Sillitoe on behalf of the Wellcome Sanger Institute COVID-19 Surveillance Team |
| EPI_ISL_2395611, EPI_ISL_2395612, EPI_ISL_2395613, EPI_ISL_2395614, EPI_ISL_2395615, EPI_ISL_2395616, EPI_ISL_2395617 | Lighthouse Lab in Milton Keynes | Wellcome Sanger Institute for the COVID-19 Genomics UK (COG-UK) Consortium | The Lighthouse Lab in Milton Keynes and Alex Alderton, Roberto Amato, Jeffrey Barrett, Sonia Goncalves, Ewan Harrison, David K. Jackson, Ian Johnston, Dominic Kwiatkowski, Cordelia Langford, John Sillitoe on behalf of the Wellcome Sanger Institute COVID-19 Surveillance Team |
| EPI_ISL_2395618 | Lighthouse Lab in Alderley Park | Wellcome Sanger Institute for the COVID-19 Genomics UK (COG-UK) Consortium | Jacquelyn Wynn, Mairead Hyland, The Lighthouse Lab in Alderley Park and Alex Alderton, Roberto Amato, Jeffrey Barrett, Sonia Goncalves, Ewan Harrison, David K. Jackson, Ian Johnston, Dominic Kwiatkowski, Cordelia Langford, John Sillitoe on behalf of the Wellcome Sanger Institute COVID-19 Surveillance Team |
| EPI_ISL_2395620 | Lighthouse Lab in Milton Keynes | Wellcome Sanger Institute for the COVID-19 Genomics UK (COG-UK) Consortium | The Lighthouse Lab in Milton Keynes and Alex Alderton, Roberto Amato, Jeffrey Barrett, Sonia Goncalves, Ewan Harrison, David K. Jackson, Ian Johnston, Dominic Kwiatkowski, Cordelia Langford, John Sillitoe on behalf of the Wellcome Sanger Institute COVID-19 Surveillance Team |
| EPI_ISL_2395621 | Lighthouse Lab in Glasgow | Wellcome Sanger Institute for the COVID-19 Genomics UK (COG-UK) Consortium | Harper VanSteenhouse, Yumi Kasai, David Gray, Carol Clugston, Anna Dominiczak and Alex Alderton, Roberto Amato, Jeffrey Barrett, Sonia Goncalves, Ewan Harrison, David K. Jackson, Ian Johnston, Dominic Kwiatkowski, Cordelia Langford, John Sillitoe on behalf of the Wellcome Sanger Institute COVID-19 Surveillance Team |
| EPI_ISL_2395622 | Lighthouse Lab in Alderley Park | Wellcome Sanger Institute for the COVID-19 Genomics UK (COG-UK) Consortium | Jacquelyn Wynn, Mairead Hyland, The Lighthouse Lab in Alderley Park and Alex Alderton, Roberto Amato, Jeffrey Barrett, Sonia Goncalves, Ewan Harrison, David K. Jackson, Ian Johnston, Dominic Kwiatkowski, Cordelia Langford, John Sillitoe on behalf of the Wellcome Sanger Institute COVID-19 Surveillance Team |
| EPI_ISL_2395623 | Lighthouse Lab in Milton Keynes | Wellcome Sanger Institute for the COVID-19 Genomics UK (COG-UK) Consortium | The Lighthouse Lab in Milton Keynes and Alex Alderton, Roberto Amato, Jeffrey Barrett, Sonia Goncalves, Ewan Harrison, David K. Jackson, Ian Johnston, Dominic Kwiatkowski, Cordelia Langford, John Sillitoe on behalf of the Wellcome Sanger Institute COVID-19 Surveillance Team |
| EPI_ISL_2395624 | Lighthouse Lab in Alderley Park | Wellcome Sanger Institute for the COVID-19 Genomics UK (COG-UK) Consortium | Jacquelyn Wynn, Mairead Hyland, The Lighthouse Lab in Alderley Park and Alex Alderton, Roberto Amato, Jeffrey Barrett, Sonia Goncalves, Ewan Harrison, David K. Jackson, Ian Johnston, Dominic Kwiatkowski, Cordelia Langford, John Sillitoe on behalf of the Wellcome Sanger Institute COVID-19 Surveillance Team |
| EPI_ISL_2395625 | Lighthouse Lab in Milton Keynes | Wellcome Sanger Institute for the COVID-19 Genomics UK (COG-UK) Consortium | The Lighthouse Lab in Milton Keynes and Alex Alderton, Roberto Amato, Jeffrey Barrett, Sonia Goncalves, Ewan Harrison, David K. Jackson, Ian Johnston, Dominic Kwiatkowski, Cordelia Langford, John Sillitoe on behalf of the Wellcome Sanger Institute COVID-19 Surveillance Team |
| EPI_ISL_2395626 | Lighthouse Lab in Glasgow | Wellcome Sanger Institute for the COVID-19 Genomics UK (COG-UK) Consortium | Harper VanSteenhouse, Yumi Kasai, David Gray, Carol Clugston, Anna Dominiczak and Alex Alderton, Roberto Amato, Jeffrey Barrett, Sonia Goncalves, Ewan Harrison, David K. Jackson, Ian Johnston, Dominic Kwiatkowski, Cordelia Langford, John Sillitoe on behalf of the Wellcome Sanger Institute COVID-19 Surveillance Team |
| EPI_ISL_2395627 | Lighthouse Lab in Milton Keynes | Wellcome Sanger Institute for the COVID-19 Genomics UK (COG-UK) Consortium | The Lighthouse Lab in Milton Keynes and Alex Alderton, Roberto Amato, Jeffrey Barrett, Sonia Goncalves, Ewan Harrison, David K. Jackson, Ian Johnston, Dominic Kwiatkowski, Cordelia Langford, John Sillitoe on behalf of the Wellcome Sanger Institute COVID-19 Surveillance Team |
| EPI_ISL_2395628 | Lighthouse Lab in Glasgow | Wellcome Sanger Institute for the COVID-19 Genomics UK (COG-UK) Consortium | Harper VanSteenhouse, Yumi Kasai, David Gray, Carol Clugston, Anna Dominiczak and Alex Alderton, Roberto Amato, Jeffrey Barrett, Sonia Goncalves, Ewan Harrison, David K. Jackson, Ian Johnston, Dominic Kwiatkowski, Cordelia Langford, John Sillitoe on behalf of the Wellcome Sanger Institute COVID-19 Surveillance Team |
| EPI_ISL_2395629 | Lighthouse Lab in Alderley Park | Wellcome Sanger Institute for the COVID-19 Genomics UK (COG-UK) Consortium | Jacquelyn Wynn, Mairead Hyland, The Lighthouse Lab in Alderley Park and Alex Alderton, Roberto Amato, Jeffrey Barrett, Sonia Goncalves, Ewan Harrison, David K. Jackson, Ian Johnston, Dominic Kwiatkowski, Cordelia Langford, John Sillitoe on behalf of the Wellcome Sanger Institute COVID-19 Surveillance Team |
| EPI_ISL_2395631 | Lighthouse Lab in Milton Keynes | Wellcome Sanger Institute for the COVID-19 Genomics UK (COG-UK) Consortium | The Lighthouse Lab in Milton Keynes and Alex Alderton, Roberto Amato, Jeffrey Barrett, Sonia Goncalves, Ewan Harrison, David K. Jackson, Ian Johnston, Dominic Kwiatkowski, Cordelia Langford, John Sillitoe on behalf of the Wellcome Sanger Institute COVID-19 Surveillance Team |
| EPI_ISL_2395632 | Lighthouse Lab in Glasgow | Wellcome Sanger Institute for the COVID-19 Genomics UK (COG-UK) Consortium | Harper VanSteenhouse, Yumi Kasai, David Gray, Carol Clugston, Anna Dominiczak and Alex Alderton, Roberto Amato, Jeffrey Barrett, Sonia Goncalves, Ewan Harrison, David K. Jackson, Ian Johnston, Dominic Kwiatkowski, Cordelia Langford, John Sillitoe on behalf of the Wellcome Sanger Institute COVID-19 Surveillance Team |
| EPI_ISL_2395633 | Lighthouse Lab in Milton Keynes | Wellcome Sanger Institute for the COVID-19 Genomics UK (COG-UK) Consortium | The Lighthouse Lab in Milton Keynes and Alex Alderton, Roberto Amato, Jeffrey Barrett, Sonia Goncalves, Ewan Harrison, David K. Jackson, Ian Johnston, Dominic Kwiatkowski, Cordelia Langford, John Sillitoe on behalf of the Wellcome Sanger Institute COVID-19 Surveillance Team |
| EPI_ISL_2395634 | Lighthouse Lab in Alderley Park | Wellcome Sanger Institute for the COVID-19 Genomics UK (COG-UK) Consortium | Jacquelyn Wynn, Mairead Hyland, The Lighthouse Lab in Alderley Park and Alex Alderton, Roberto Amato, Jeffrey Barrett, Sonia Goncalves, Ewan Harrison, David K. Jackson, Ian Johnston, Dominic Kwiatkowski, Cordelia Langford, John Sillitoe on behalf of the Wellcome Sanger Institute COVID-19 Surveillance Team |
| EPI_ISL_2395635, EPI_ISL_2395636, EPI_ISL_2395637 | Lighthouse Lab in Glasgow | Wellcome Sanger Institute for the COVID-19 Genomics UK (COG-UK) Consortium | Harper VanSteenhouse, Yumi Kasai, David Gray, Carol Clugston, Anna Dominiczak and Alex Alderton, Roberto Amato, Jeffrey Barrett, Sonia Goncalves, Ewan Harrison, David K. Jackson, Ian Johnston, Dominic Kwiatkowski, Cordelia Langford, John Sillitoe on behalf of the Wellcome Sanger Institute COVID-19 Surveillance Team |
| EPI_ISL_2395638, EPI_ISL_2395639 | Lighthouse Lab in Milton Keynes | Wellcome Sanger Institute for the COVID-19 Genomics UK (COG-UK) Consortium | The Lighthouse Lab in Milton Keynes and Alex Alderton, Roberto Amato, Jeffrey Barrett, Sonia Goncalves, Ewan Harrison, David K. Jackson, Ian Johnston, Dominic Kwiatkowski, Cordelia Langford, John Sillitoe on behalf of the Wellcome Sanger Institute COVID-19 Surveillance Team |
| EPI_ISL_2395640 | Lighthouse Lab in Alderley Park | Wellcome Sanger Institute for the COVID-19 Genomics UK (COG-UK) Consortium | Jacquelyn Wynn, Mairead Hyland, The Lighthouse Lab in Alderley Park and Alex Alderton, Roberto Amato, Jeffrey Barrett, Sonia Goncalves, Ewan Harrison, David K. Jackson, Ian Johnston, Dominic Kwiatkowski, Cordelia Langford, John Sillitoe on behalf of the Wellcome Sanger Institute COVID-19 Surveillance Team |
| EPI_ISL_2395641, EPI_ISL_2395642 | Lighthouse Lab in Milton Keynes | Wellcome Sanger Institute for the COVID-19 Genomics UK (COG-UK) Consortium | The Lighthouse Lab in Milton Keynes and Alex Alderton, Roberto Amato, Jeffrey Barrett, Sonia Goncalves, Ewan Harrison, David K. Jackson, Ian Johnston, Dominic Kwiatkowski, Cordelia Langford, John Sillitoe on behalf of the Wellcome Sanger Institute COVID-19 Surveillance Team |
| EPI_ISL_2395643 | Lighthouse Lab in Alderley Park | Wellcome Sanger Institute for the COVID-19 Genomics UK (COG-UK) Consortium | Jacquelyn Wynn, Mairead Hyland, The Lighthouse Lab in Alderley Park and Alex Alderton, Roberto Amato, Jeffrey Barrett, Sonia Goncalves, Ewan Harrison, David K. Jackson, Ian Johnston, Dominic Kwiatkowski, Cordelia Langford, John Sillitoe on behalf of the Wellcome Sanger Institute COVID-19 Surveillance Team |
| EPI_ISL_2395645 | Lighthouse Lab in Milton Keynes | Wellcome Sanger Institute for the COVID-19 Genomics UK (COG-UK) Consortium | The Lighthouse Lab in Milton Keynes and Alex Alderton, Roberto Amato, Jeffrey Barrett, Sonia Goncalves, Ewan Harrison, David K. Jackson, Ian Johnston, Dominic Kwiatkowski, Cordelia Langford, John Sillitoe on behalf of the Wellcome Sanger Institute COVID-19 Surveillance Team |
| EPI_ISL_2395646 | Lighthouse Lab in Alderley Park | Wellcome Sanger Institute for the COVID-19 Genomics UK (COG-UK) Consortium | Jacquelyn Wynn, Mairead Hyland, The Lighthouse Lab in Alderley Park and Alex Alderton, Roberto Amato, Jeffrey Barrett, Sonia Goncalves, Ewan Harrison, David K. Jackson, Ian Johnston, Dominic Kwiatkowski, Cordelia Langford, John Sillitoe on behalf of the Wellcome Sanger Institute COVID-19 Surveillance Team |
| EPI_ISL_2395647 | Lighthouse Lab in Milton Keynes | Wellcome Sanger Institute for the COVID-19 Genomics UK (COG-UK) Consortium | The Lighthouse Lab in Milton Keynes and Alex Alderton, Roberto Amato, Jeffrey Barrett, Sonia Goncalves, Ewan Harrison, David K. Jackson, Ian Johnston, Dominic Kwiatkowski, Cordelia Langford, John Sillitoe on behalf of the Wellcome Sanger Institute COVID-19 Surveillance Team |
| EPI_ISL_2395648 | Lighthouse Lab in Alderley Park | Wellcome Sanger Institute for the COVID-19 Genomics UK (COG-UK) Consortium | Jacquelyn Wynn, Mairead Hyland, The Lighthouse Lab in Alderley Park and Alex Alderton, Roberto Amato, Jeffrey Barrett, Sonia Goncalves, Ewan Harrison, David K. Jackson, Ian Johnston, Dominic Kwiatkowski, Cordelia Langford, John Sillitoe on behalf of the Wellcome Sanger Institute COVID-19 Surveillance Team |
| EPI_ISL_2395649, EPI_ISL_2395651, EPI_ISL_2395652, EPI_ISL_2395653 | Lighthouse Lab in Milton Keynes | Wellcome Sanger Institute for the COVID-19 Genomics UK (COG-UK) Consortium | The Lighthouse Lab in Milton Keynes and Alex Alderton, Roberto Amato, Jeffrey Barrett, Sonia Goncalves, Ewan Harrison, David K. Jackson, Ian Johnston, Dominic Kwiatkowski, Cordelia Langford, John Sillitoe on behalf of the Wellcome Sanger Institute COVID-19 Surveillance Team |
| EPI_ISL_2395654, EPI_ISL_2395656 | Lighthouse Lab in Glasgow | Wellcome Sanger Institute for the COVID-19 Genomics UK (COG-UK) Consortium | Harper VanSteenhouse, Yumi Kasai, David Gray, Carol Clugston, Anna Dominiczak and Alex Alderton, Roberto Amato, Jeffrey Barrett, Sonia Goncalves, Ewan Harrison, David K. Jackson, Ian Johnston, Dominic Kwiatkowski, Cordelia Langford, John Sillitoe on behalf of the Wellcome Sanger Institute COVID-19 Surveillance Team |
| EPI_ISL_2395657 | Lighthouse Lab in Milton Keynes | Wellcome Sanger Institute for the COVID-19 Genomics UK (COG-UK) Consortium | The Lighthouse Lab in Milton Keynes and Alex Alderton, Roberto Amato, Jeffrey Barrett, Sonia Goncalves, Ewan Harrison, David K. Jackson, Ian Johnston, Dominic Kwiatkowski, Cordelia Langford, John Sillitoe on behalf of the Wellcome Sanger Institute COVID-19 Surveillance Team |
| EPI_ISL_2395658, EPI_ISL_2395659 | Lighthouse Lab in Alderley Park | Wellcome Sanger Institute for the COVID-19 Genomics UK (COG-UK) Consortium | Jacquelyn Wynn, Mairead Hyland, The Lighthouse Lab in Alderley Park and Alex Alderton, Roberto Amato, Jeffrey Barrett, Sonia Goncalves, Ewan Harrison, David K. Jackson, Ian Johnston, Dominic Kwiatkowski, Cordelia Langford, John Sillitoe on behalf of the Wellcome Sanger Institute COVID-19 Surveillance Team |
| EPI_ISL_2395662 | Lighthouse Lab in Milton Keynes | Wellcome Sanger Institute for the COVID-19 Genomics UK (COG-UK) Consortium | The Lighthouse Lab in Milton Keynes and Alex Alderton, Roberto Amato, Jeffrey Barrett, Sonia Goncalves, Ewan Harrison, David K. Jackson, Ian Johnston, Dominic Kwiatkowski, Cordelia Langford, John Sillitoe on behalf of the Wellcome Sanger Institute COVID-19 Surveillance Team |
| EPI_ISL_2395664 | Lighthouse Lab in Alderley Park | Wellcome Sanger Institute for the COVID-19 Genomics UK (COG-UK) Consortium | Jacquelyn Wynn, Mairead Hyland, The Lighthouse Lab in Alderley Park and Alex Alderton, Roberto Amato, Jeffrey Barrett, Sonia Goncalves, Ewan Harrison, David K. Jackson, Ian Johnston, Dominic Kwiatkowski, Cordelia Langford, John Sillitoe on behalf of the Wellcome Sanger Institute COVID-19 Surveillance Team |
| EPI_ISL_2395665 | Lighthouse Lab in Milton Keynes | Wellcome Sanger Institute for the COVID-19 Genomics UK (COG-UK) Consortium | The Lighthouse Lab in Milton Keynes and Alex Alderton, Roberto Amato, Jeffrey Barrett, Sonia Goncalves, Ewan Harrison, David K. Jackson, Ian Johnston, Dominic Kwiatkowski, Cordelia Langford, John Sillitoe on behalf of the Wellcome Sanger Institute COVID-19 Surveillance Team |
| EPI_ISL_2395666 | Lighthouse Lab in Glasgow | Wellcome Sanger Institute for the COVID-19 Genomics UK (COG-UK) Consortium | Harper VanSteenhouse, Yumi Kasai, David Gray, Carol Clugston, Anna Dominiczak and Alex Alderton, Roberto Amato, Jeffrey Barrett, Sonia Goncalves, Ewan Harrison, David K. Jackson, Ian Johnston, Dominic Kwiatkowski, Cordelia Langford, John Sillitoe on behalf of the Wellcome Sanger Institute COVID-19 Surveillance Team |
| EPI_ISL_2395668, EPI_ISL_2395669 | Lighthouse Lab in Milton Keynes | Wellcome Sanger Institute for the COVID-19 Genomics UK (COG-UK) Consortium | The Lighthouse Lab in Milton Keynes and Alex Alderton, Roberto Amato, Jeffrey Barrett, Sonia Goncalves, Ewan Harrison, David K. Jackson, Ian Johnston, Dominic Kwiatkowski, Cordelia Langford, John Sillitoe on behalf of the Wellcome Sanger Institute COVID-19 Surveillance Team |
| EPI_ISL_2395670, EPI_ISL_2395671 | Lighthouse Lab in Glasgow | Wellcome Sanger Institute for the COVID-19 Genomics UK (COG-UK) Consortium | Harper VanSteenhouse, Yumi Kasai, David Gray, Carol Clugston, Anna Dominiczak and Alex Alderton, Roberto Amato, Jeffrey Barrett, Sonia Goncalves, Ewan Harrison, David K. Jackson, Ian Johnston, Dominic Kwiatkowski, Cordelia Langford, John Sillitoe on behalf of the Wellcome Sanger Institute COVID-19 Surveillance Team |
| EPI_ISL_2395672 | Lighthouse Lab in Milton Keynes | Wellcome Sanger Institute for the COVID-19 Genomics UK (COG-UK) Consortium | The Lighthouse Lab in Milton Keynes and Alex Alderton, Roberto Amato, Jeffrey Barrett, Sonia Goncalves, Ewan Harrison, David K. Jackson, Ian Johnston, Dominic Kwiatkowski, Cordelia Langford, John Sillitoe on behalf of the Wellcome Sanger Institute COVID-19 Surveillance Team |
| EPI_ISL_2395674 | Lighthouse Lab in Alderley Park | Wellcome Sanger Institute for the COVID-19 Genomics UK (COG-UK) Consortium | Jacquelyn Wynn, Mairead Hyland, The Lighthouse Lab in Alderley Park and Alex Alderton, Roberto Amato, Jeffrey Barrett, Sonia Goncalves, Ewan Harrison, David K. Jackson, Ian Johnston, Dominic Kwiatkowski, Cordelia Langford, John Sillitoe on behalf of the Wellcome Sanger Institute COVID-19 Surveillance Team |
| EPI_ISL_2395676 | Lighthouse Lab in Glasgow | Wellcome Sanger Institute for the COVID-19 Genomics UK (COG-UK) Consortium | Harper VanSteenhouse, Yumi Kasai, David Gray, Carol Clugston, Anna Dominiczak and Alex Alderton, Roberto Amato, Jeffrey Barrett, Sonia Goncalves, Ewan Harrison, David K. Jackson, Ian Johnston, Dominic Kwiatkowski, Cordelia Langford, John Sillitoe on behalf of the Wellcome Sanger Institute COVID-19 Surveillance Team |
| EPI_ISL_2395677 | Lighthouse Lab in Milton Keynes | Wellcome Sanger Institute for the COVID-19 Genomics UK (COG-UK) Consortium | The Lighthouse Lab in Milton Keynes and Alex Alderton, Roberto Amato, Jeffrey Barrett, Sonia Goncalves, Ewan Harrison, David K. Jackson, Ian Johnston, Dominic Kwiatkowski, Cordelia Langford, John Sillitoe on behalf of the Wellcome Sanger Institute COVID-19 Surveillance Team |
| EPI_ISL_2395678 | Lighthouse Lab in Glasgow | Wellcome Sanger Institute for the COVID-19 Genomics UK (COG-UK) Consortium | Harper VanSteenhouse, Yumi Kasai, David Gray, Carol Clugston, Anna Dominiczak and Alex Alderton, Roberto Amato, Jeffrey Barrett, Sonia Goncalves, Ewan Harrison, David K. Jackson, Ian Johnston, Dominic Kwiatkowski, Cordelia Langford, John Sillitoe on behalf of the Wellcome Sanger Institute COVID-19 Surveillance Team |
| EPI_ISL_2395679 | Lighthouse Lab in Alderley Park | Wellcome Sanger Institute for the COVID-19 Genomics UK (COG-UK) Consortium | Jacquelyn Wynn, Mairead Hyland, The Lighthouse Lab in Alderley Park and Alex Alderton, Roberto Amato, Jeffrey Barrett, Sonia Goncalves, Ewan Harrison, David K. Jackson, Ian Johnston, Dominic Kwiatkowski, Cordelia Langford, John Sillitoe on behalf of the Wellcome Sanger Institute COVID-19 Surveillance Team |

| | | | |
|---|---|---|---|
| EPI_ISL_2395680, EPI_ISL_2395681 | Lighthouse Lab in Glasgow | Wellcome Sanger Institute for the COVID-19 Genomics UK (COG-UK) Consortium | Harper VanSteenhouse, Yumi Kasai, David Gray, Carol Clugston, Anna Dominiczak and Alex Alderton, Roberto Amato, Jeffrey Barrett, Sonia Goncalves, Ewan Harrison, David K. Jackson, Ian Johnston, Dominic Kwiatkowski, Cordelia Langford, John Sillitoe on behalf of the Wellcome Sanger Institute COVID-19 Surveillance Team |
| EPI_ISL_2395682, EPI_ISL_2395683 | Lighthouse Lab in Alderley Park | Wellcome Sanger Institute for the COVID-19 Genomics UK (COG-UK) Consortium | Jacquelyn Wynn, Mairead Hyland, The Lighthouse Lab in Alderley Park and Alex Alderton, Roberto Amato, Jeffrey Barrett, Sonia Goncalves, Ewan Harrison, David K. Jackson, Ian Johnston, Dominic Kwiatkowski, Cordelia Langford, John Sillitoe on behalf of the Wellcome Sanger Institute COVID-19 Surveillance Team |
| EPI_ISL_2395687, EPI_ISL_2395688 | Lighthouse Lab in Glasgow | Wellcome Sanger Institute for the COVID-19 Genomics UK (COG-UK) Consortium | Harper VanSteenhouse, Yumi Kasai, David Gray, Carol Clugston, Anna Dominiczak and Alex Alderton, Roberto Amato, Jeffrey Barrett, Sonia Goncalves, Ewan Harrison, David K. Jackson, Ian Johnston, Dominic Kwiatkowski, Cordelia Langford, John Sillitoe on behalf of the Wellcome Sanger Institute COVID-19 Surveillance Team |
| EPI_ISL_2395691 | Lighthouse Lab in Milton Keynes | Wellcome Sanger Institute for the COVID-19 Genomics UK (COG-UK) Consortium | The Lighthouse Lab in Milton Keynes and Alex Alderton, Roberto Amato, Jeffrey Barrett, Sonia Goncalves, Ewan Harrison, David K. Jackson, Ian Johnston, Dominic Kwiatkowski, Cordelia Langford, John Sillitoe on behalf of the Wellcome Sanger Institute COVID-19 Surveillance Team |
| EPI_ISL_2395693 | Lighthouse Lab in Glasgow | Wellcome Sanger Institute for the COVID-19 Genomics UK (COG-UK) Consortium | Harper VanSteenhouse, Yumi Kasai, David Gray, Carol Clugston, Anna Dominiczak and Alex Alderton, Roberto Amato, Jeffrey Barrett, Sonia Goncalves, Ewan Harrison, David K. Jackson, Ian Johnston, Dominic Kwiatkowski, Cordelia Langford, John Sillitoe on behalf of the Wellcome Sanger Institute COVID-19 Surveillance Team |
| EPI_ISL_2395696, EPI_ISL_2395697 | Lighthouse Lab in Milton Keynes | Wellcome Sanger Institute for the COVID-19 Genomics UK (COG-UK) Consortium | The Lighthouse Lab in Milton Keynes and Alex Alderton, Roberto Amato, Jeffrey Barrett, Sonia Goncalves, Ewan Harrison, David K. Jackson, Ian Johnston, Dominic Kwiatkowski, Cordelia Langford, John Sillitoe on behalf of the Wellcome Sanger Institute COVID-19 Surveillance Team |
| EPI_ISL_2395700 | Lighthouse Lab in Glasgow | Wellcome Sanger Institute for the COVID-19 Genomics UK (COG-UK) Consortium | Harper VanSteenhouse, Yumi Kasai, David Gray, Carol Clugston, Anna Dominiczak and Alex Alderton, Roberto Amato, Jeffrey Barrett, Sonia Goncalves, Ewan Harrison, David K. Jackson, Ian Johnston, Dominic Kwiatkowski, Cordelia Langford, John Sillitoe on behalf of the Wellcome Sanger Institute COVID-19 Surveillance Team |
| EPI_ISL_2395702, EPI_ISL_2395703 | Lighthouse Lab in Milton Keynes | Wellcome Sanger Institute for the COVID-19 Genomics UK (COG-UK) Consortium | The Lighthouse Lab in Milton Keynes and Alex Alderton, Roberto Amato, Jeffrey Barrett, Sonia Goncalves, Ewan Harrison, David K. Jackson, Ian Johnston, Dominic Kwiatkowski, Cordelia Langford, John Sillitoe on behalf of the Wellcome Sanger Institute COVID-19 Surveillance Team |
| EPI_ISL_2395704 | Lighthouse Lab in Glasgow | Wellcome Sanger Institute for the COVID-19 Genomics UK (COG-UK) Consortium | Harper VanSteenhouse, Yumi Kasai, David Gray, Carol Clugston, Anna Dominiczak and Alex Alderton, Roberto Amato, Jeffrey Barrett, Sonia Goncalves, Ewan Harrison, David K. Jackson, Ian Johnston, Dominic Kwiatkowski, Cordelia Langford, John Sillitoe on behalf of the Wellcome Sanger Institute COVID-19 Surveillance Team |
| EPI_ISL_2395705 | Lighthouse Lab in Milton Keynes | Wellcome Sanger Institute for the COVID-19 Genomics UK (COG-UK) Consortium | The Lighthouse Lab in Milton Keynes and Alex Alderton, Roberto Amato, Jeffrey Barrett, Sonia Goncalves, Ewan Harrison, David K. Jackson, Ian Johnston, Dominic Kwiatkowski, Cordelia Langford, John Sillitoe on behalf of the Wellcome Sanger Institute COVID-19 Surveillance Team |
| EPI_ISL_2395707, EPI_ISL_2395712 | Lighthouse Lab in Glasgow | Wellcome Sanger Institute for the COVID-19 Genomics UK (COG-UK) Consortium | Harper VanSteenhouse, Yumi Kasai, David Gray, Carol Clugston, Anna Dominiczak and Alex Alderton, Roberto Amato, Jeffrey Barrett, Sonia Goncalves, Ewan Harrison, David K. Jackson, Ian Johnston, Dominic Kwiatkowski, Cordelia Langford, John Sillitoe on behalf of the Wellcome Sanger Institute COVID-19 Surveillance Team |
| EPI_ISL_2395714 | Lighthouse Lab in Milton Keynes | Wellcome Sanger Institute for the COVID-19 Genomics UK (COG-UK) Consortium | The Lighthouse Lab in Milton Keynes and Alex Alderton, Roberto Amato, Jeffrey Barrett, Sonia Goncalves, Ewan Harrison, David K. Jackson, Ian Johnston, Dominic Kwiatkowski, Cordelia Langford, John Sillitoe on behalf of the Wellcome Sanger Institute COVID-19 Surveillance Team |
| EPI_ISL_2395716 | Lighthouse Lab in Glasgow | Wellcome Sanger Institute for the COVID-19 Genomics UK (COG-UK) Consortium | Harper VanSteenhouse, Yumi Kasai, David Gray, Carol Clugston, Anna Dominiczak and Alex Alderton, Roberto Amato, Jeffrey Barrett, Sonia Goncalves, Ewan Harrison, David K. Jackson, Ian Johnston, Dominic Kwiatkowski, Cordelia Langford, John Sillitoe on behalf of the Wellcome Sanger Institute COVID-19 Surveillance Team |
| EPI_ISL_2395717 | Lighthouse Lab in Milton Keynes | Wellcome Sanger Institute for the COVID-19 Genomics UK (COG-UK) Consortium | The Lighthouse Lab in Milton Keynes and Alex Alderton, Roberto Amato, Jeffrey Barrett, Sonia Goncalves, Ewan Harrison, David K. Jackson, Ian Johnston, Dominic Kwiatkowski, Cordelia Langford, John Sillitoe on behalf of the Wellcome Sanger Institute COVID-19 Surveillance Team |
| EPI_ISL_2395718 | Lighthouse Lab in Glasgow | Wellcome Sanger Institute for the COVID-19 Genomics UK (COG-UK) Consortium | Harper VanSteenhouse, Yumi Kasai, David Gray, Carol Clugston, Anna Dominiczak and Alex Alderton, Roberto Amato, Jeffrey Barrett, Sonia Goncalves, Ewan Harrison, David K. Jackson, Ian Johnston, Dominic Kwiatkowski, Cordelia Langford, John Sillitoe on behalf of the Wellcome Sanger Institute COVID-19 Surveillance Team |
| EPI_ISL_2395719, EPI_ISL_2395720, EPI_ISL_2395721 | Lighthouse Lab in Milton Keynes | Wellcome Sanger Institute for the COVID-19 Genomics UK (COG-UK) Consortium | The Lighthouse Lab in Milton Keynes and Alex Alderton, Roberto Amato, Jeffrey Barrett, Sonia Goncalves, Ewan Harrison, David K. Jackson, Ian Johnston, Dominic Kwiatkowski, Cordelia Langford, John Sillitoe on behalf of the Wellcome Sanger Institute COVID-19 Surveillance Team |
| EPI_ISL_2395723, EPI_ISL_2395724, EPI_ISL_2395726 | Lighthouse Lab in Glasgow | Wellcome Sanger Institute for the COVID-19 Genomics UK (COG-UK) Consortium | Harper VanSteenhouse, Yumi Kasai, David Gray, Carol Clugston, Anna Dominiczak and Alex Alderton, Roberto Amato, Jeffrey Barrett, Sonia Goncalves, Ewan Harrison, David K. Jackson, Ian Johnston, Dominic Kwiatkowski, Cordelia Langford, John Sillitoe on behalf of the Wellcome Sanger Institute COVID-19 Surveillance Team |
| EPI_ISL_2395727, EPI_ISL_2395731 | Lighthouse Lab in Milton Keynes | Wellcome Sanger Institute for the COVID-19 Genomics UK (COG-UK) Consortium | The Lighthouse Lab in Milton Keynes and Alex Alderton, Roberto Amato, Jeffrey Barrett, Sonia Goncalves, Ewan Harrison, David K. Jackson, Ian Johnston, Dominic Kwiatkowski, Cordelia Langford, John Sillitoe on behalf of the Wellcome Sanger Institute COVID-19 Surveillance Team |
| EPI_ISL_2395732, EPI_ISL_2395733, EPI_ISL_2395734 | Lighthouse Lab in Glasgow | Wellcome Sanger Institute for the COVID-19 Genomics UK (COG-UK) Consortium | Harper VanSteenhouse, Yumi Kasai, David Gray, Carol Clugston, Anna Dominiczak and Alex Alderton, Roberto Amato, Jeffrey Barrett, Sonia Goncalves, Ewan Harrison, David K. Jackson, Ian Johnston, Dominic Kwiatkowski, Cordelia Langford, John Sillitoe on behalf of the Wellcome Sanger Institute COVID-19 Surveillance Team |
| EPI_ISL_2395736 | Lighthouse Lab in Milton Keynes | Wellcome Sanger Institute for the COVID-19 Genomics UK (COG-UK) Consortium | The Lighthouse Lab in Milton Keynes and Alex Alderton, Roberto Amato, Jeffrey Barrett, Sonia Goncalves, Ewan Harrison, David K. Jackson, Ian Johnston, Dominic Kwiatkowski, Cordelia Langford, John Sillitoe on behalf of the Wellcome Sanger Institute COVID-19 Surveillance Team |
| EPI_ISL_2395737, EPI_ISL_2395738, EPI_ISL_2395739, EPI_ISL_2395740 | Lighthouse Lab in Glasgow | Wellcome Sanger Institute for the COVID-19 Genomics UK (COG-UK) Consortium | Harper VanSteenhouse, Yumi Kasai, David Gray, Carol Clugston, Anna Dominiczak and Alex Alderton, Roberto Amato, Jeffrey Barrett, Sonia Goncalves, Ewan Harrison, David K. Jackson, Ian Johnston, Dominic Kwiatkowski, Cordelia Langford, John Sillitoe on behalf of the Wellcome Sanger Institute COVID-19 Surveillance Team |
| EPI_ISL_2395741 | Lighthouse Lab in Milton Keynes | Wellcome Sanger Institute for the COVID-19 Genomics UK (COG-UK) Consortium | The Lighthouse Lab in Milton Keynes and Alex Alderton, Roberto Amato, Jeffrey Barrett, Sonia Goncalves, Ewan Harrison, David K. Jackson, Ian Johnston, Dominic Kwiatkowski, Cordelia Langford, John Sillitoe on behalf of the Wellcome Sanger Institute COVID-19 Surveillance Team |
| EPI_ISL_2395742, EPI_ISL_2395743 | Lighthouse Lab in Glasgow | Wellcome Sanger Institute for the COVID-19 Genomics UK (COG-UK) Consortium | Harper VanSteenhouse, Yumi Kasai, David Gray, Carol Clugston, Anna Dominiczak and Alex Alderton, Roberto Amato, Jeffrey Barrett, Sonia Goncalves, Ewan Harrison, David K. Jackson, Ian Johnston, Dominic Kwiatkowski, Cordelia Langford, John Sillitoe on behalf of the Wellcome Sanger Institute COVID-19 Surveillance Team |
| EPI_ISL_2395744, EPI_ISL_2395748, EPI_ISL_2395749, EPI_ISL_2395751 | Lighthouse Lab in Milton Keynes | Wellcome Sanger Institute for the COVID-19 Genomics UK (COG-UK) Consortium | The Lighthouse Lab in Milton Keynes and Alex Alderton, Roberto Amato, Jeffrey Barrett, Sonia Goncalves, Ewan Harrison, David K. Jackson, Ian Johnston, Dominic Kwiatkowski, Cordelia Langford, John Sillitoe on behalf of the Wellcome Sanger Institute COVID-19 Surveillance Team |
| EPI_ISL_2395753 | Lighthouse Lab in Glasgow | Wellcome Sanger Institute for the COVID-19 Genomics UK (COG-UK) Consortium | Harper VanSteenhouse, Yumi Kasai, David Gray, Carol Clugston, Anna Dominiczak and Alex Alderton, Roberto Amato, Jeffrey Barrett, Sonia Goncalves, Ewan Harrison, David K. Jackson, Ian Johnston, Dominic Kwiatkowski, Cordelia Langford, John Sillitoe on behalf of the Wellcome Sanger Institute COVID-19 Surveillance Team |
| EPI_ISL_2395754 | Lighthouse Lab in Milton Keynes | Wellcome Sanger Institute for the COVID-19 Genomics UK (COG-UK) Consortium | The Lighthouse Lab in Milton Keynes and Alex Alderton, Roberto Amato, Jeffrey Barrett, Sonia Goncalves, Ewan Harrison, David K. Jackson, Ian Johnston, Dominic Kwiatkowski, Cordelia Langford, John Sillitoe on behalf of the Wellcome Sanger Institute COVID-19 Surveillance Team |
| EPI_ISL_2395755, EPI_ISL_2395756 | Lighthouse Lab in Glasgow | Wellcome Sanger Institute for the COVID-19 Genomics UK (COG-UK) Consortium | Harper VanSteenhouse, Yumi Kasai, David Gray, Carol Clugston, Anna Dominiczak and Alex Alderton, Roberto Amato, Jeffrey Barrett, Sonia Goncalves, Ewan Harrison, David K. Jackson, Ian Johnston, Dominic Kwiatkowski, Cordelia Langford, John Sillitoe on behalf of the Wellcome Sanger Institute COVID-19 Surveillance Team |
| EPI_ISL_2395757, EPI_ISL_2395759 | Lighthouse Lab in Milton Keynes | Wellcome Sanger Institute for the COVID-19 Genomics UK (COG-UK) Consortium | The Lighthouse Lab in Milton Keynes and Alex Alderton, Roberto Amato, Jeffrey Barrett, Sonia Goncalves, Ewan Harrison, David K. Jackson, Ian Johnston, Dominic Kwiatkowski, Cordelia Langford, John Sillitoe on behalf of the Wellcome Sanger Institute COVID-19 Surveillance Team |
| EPI_ISL_2395760 | Lighthouse Lab in Glasgow | Wellcome Sanger Institute for the COVID-19 Genomics UK (COG-UK) Consortium | Harper VanSteenhouse, Yumi Kasai, David Gray, Carol Clugston, Anna Dominiczak and Alex Alderton, Roberto Amato, Jeffrey Barrett, Sonia Goncalves, Ewan Harrison, David K. Jackson, Ian Johnston, Dominic Kwiatkowski, Cordelia Langford, John Sillitoe on behalf of the Wellcome Sanger Institute COVID-19 Surveillance Team |
| EPI_ISL_2395762 | Lighthouse Lab in Milton Keynes | Wellcome Sanger Institute for the COVID-19 Genomics UK (COG-UK) Consortium | The Lighthouse Lab in Milton Keynes and Alex Alderton, Roberto Amato, Jeffrey Barrett, Sonia Goncalves, Ewan Harrison, David K. Jackson, Ian Johnston, Dominic Kwiatkowski, Cordelia Langford, John Sillitoe on behalf of the Wellcome Sanger Institute COVID-19 Surveillance Team |
| EPI_ISL_2395763 | Lighthouse Lab in Glasgow | Wellcome Sanger Institute for the COVID-19 Genomics UK (COG-UK) Consortium | Harper VanSteenhouse, Yumi Kasai, David Gray, Carol Clugston, Anna Dominiczak and Alex Alderton, Roberto Amato, Jeffrey Barrett, Sonia Goncalves, Ewan Harrison, David K. Jackson, Ian Johnston, Dominic Kwiatkowski, Cordelia Langford, John Sillitoe on behalf of the Wellcome Sanger Institute COVID-19 Surveillance Team |
| EPI_ISL_2395765 | Lighthouse Lab in Milton Keynes | Wellcome Sanger Institute for the COVID-19 Genomics UK (COG-UK) Consortium | The Lighthouse Lab in Milton Keynes and Alex Alderton, Roberto Amato, Jeffrey Barrett, Sonia Goncalves, Ewan Harrison, David K. Jackson, Ian Johnston, Dominic Kwiatkowski, Cordelia Langford, John Sillitoe on behalf of the Wellcome Sanger Institute COVID-19 Surveillance Team |
| EPI_ISL_2395767, EPI_ISL_2395769, EPI_ISL_2395770 | Lighthouse Lab in Glasgow | Wellcome Sanger Institute for the COVID-19 Genomics UK (COG-UK) Consortium | Harper VanSteenhouse, Yumi Kasai, David Gray, Carol Clugston, Anna Dominiczak and Alex Alderton, Roberto Amato, Jeffrey Barrett, Sonia Goncalves, Ewan Harrison, David K. Jackson, Ian Johnston, Dominic Kwiatkowski, Cordelia Langford, John Sillitoe on behalf of the Wellcome Sanger Institute COVID-19 Surveillance Team |
| EPI_ISL_2395772 | Lighthouse Lab in Milton Keynes | Wellcome Sanger Institute for the COVID-19 Genomics UK (COG-UK) Consortium | The Lighthouse Lab in Milton Keynes and Alex Alderton, Roberto Amato, Jeffrey Barrett, Sonia Goncalves, Ewan Harrison, David K. Jackson, Ian Johnston, Dominic Kwiatkowski, Cordelia Langford, John Sillitoe on behalf of the Wellcome Sanger Institute COVID-19 Surveillance Team |
| EPI_ISL_2395774 | Lighthouse Lab in Glasgow | Wellcome Sanger Institute for the COVID-19 Genomics UK (COG-UK) Consortium | Harper VanSteenhouse, Yumi Kasai, David Gray, Carol Clugston, Anna Dominiczak and Alex Alderton, Roberto Amato, Jeffrey Barrett, Sonia Goncalves, Ewan Harrison, David K. Jackson, Ian Johnston, Dominic Kwiatkowski, Cordelia Langford, John Sillitoe on behalf of the Wellcome Sanger Institute COVID-19 Surveillance Team |
| EPI_ISL_2395776, EPI_ISL_2395777 | Lighthouse Lab in Milton Keynes | Wellcome Sanger Institute for the COVID-19 Genomics UK (COG-UK) Consortium | The Lighthouse Lab in Milton Keynes and Alex Alderton, Roberto Amato, Jeffrey Barrett, Sonia Goncalves, Ewan Harrison, David K. Jackson, Ian Johnston, Dominic Kwiatkowski, Cordelia Langford, John Sillitoe on behalf of the Wellcome Sanger Institute COVID-19 Surveillance Team |
| EPI_ISL_2395779 | Lighthouse Lab in Glasgow | Wellcome Sanger Institute for the COVID-19 Genomics UK (COG-UK) Consortium | Harper VanSteenhouse, Yumi Kasai, David Gray, Carol Clugston, Anna Dominiczak and Alex Alderton, Roberto Amato, Jeffrey Barrett, Sonia Goncalves, Ewan Harrison, David K. Jackson, Ian Johnston, Dominic Kwiatkowski, Cordelia Langford, John Sillitoe on behalf of the Wellcome Sanger Institute COVID-19 Surveillance Team |
| EPI_ISL_2395780, EPI_ISL_2395781, EPI_ISL_2395782 | Lighthouse Lab in Milton Keynes | Wellcome Sanger Institute for the COVID-19 Genomics UK (COG-UK) Consortium | The Lighthouse Lab in Milton Keynes and Alex Alderton, Roberto Amato, Jeffrey Barrett, Sonia Goncalves, Ewan Harrison, David K. Jackson, Ian Johnston, Dominic Kwiatkowski, Cordelia Langford, John Sillitoe on behalf of the Wellcome Sanger Institute COVID-19 Surveillance Team |
| EPI_ISL_2395784, EPI_ISL_2395786 | Lighthouse Lab in Glasgow | Wellcome Sanger Institute for the COVID-19 Genomics UK (COG-UK) Consortium | Harper VanSteenhouse, Yumi Kasai, David Gray, Carol Clugston, Anna Dominiczak and Alex Alderton, Roberto Amato, Jeffrey Barrett, Sonia Goncalves, Ewan Harrison, David K. Jackson, Ian Johnston, Dominic Kwiatkowski, Cordelia Langford, John Sillitoe on behalf of the Wellcome Sanger Institute COVID-19 Surveillance Team |
| EPI_ISL_2395789 | Lighthouse Lab in Milton Keynes | Wellcome Sanger Institute for the COVID-19 Genomics UK (COG-UK) Consortium | The Lighthouse Lab in Milton Keynes and Alex Alderton, Roberto Amato, Jeffrey Barrett, Sonia Goncalves, Ewan Harrison, David K. Jackson, Ian Johnston, Dominic Kwiatkowski, Cordelia Langford, John Sillitoe on behalf of the Wellcome Sanger Institute COVID-19 Surveillance Team |
| EPI_ISL_2395791, EPI_ISL_2395795, EPI_ISL_2395796 | Lighthouse Lab in Glasgow | Wellcome Sanger Institute for the COVID-19 Genomics UK (COG-UK) Consortium | Harper VanSteenhouse, Yumi Kasai, David Gray, Carol Clugston, Anna Dominiczak and Alex Alderton, Roberto Amato, Jeffrey Barrett, Sonia Goncalves, Ewan Harrison, David K. Jackson, Ian Johnston, Dominic Kwiatkowski, Cordelia Langford, John Sillitoe on behalf of the Wellcome Sanger Institute COVID-19 Surveillance Team |

| | | | |
|---|---|---|---|
| EPI_ISL_2395797, EPI_ISL_2395798 | Lighthouse Lab in Milton Keynes | Wellcome Sanger Institute for the COVID-19 Genomics UK (COG-UK) Consortium | The Lighthouse Lab in Milton Keynes and Alex Alderton, Roberto Amato, Jeffrey Barrett, Sonia Goncalves, Ewan Harrison, David K. Jackson, Ian Johnston, Dominic Kwiatkowski, Cordelia Langford, John Sillitoe on behalf of the Wellcome Sanger Institute COVID-19 Surveillance Team |
| EPI_ISL_2395799, EPI_ISL_2395801 | Lighthouse Lab in Glasgow | Wellcome Sanger Institute for the COVID-19 Genomics UK (COG-UK) Consortium | Harper VanSteenhouse, Yumi Kasai, David Gray, Carol Clugston, Anna Dominiczak and Alex Alderton, Roberto Amato, Jeffrey Barrett, Sonia Goncalves, Ewan Harrison, David K. Jackson, Ian Johnston, Dominic Kwiatkowski, Cordelia Langford, John Sillitoe on behalf of the Wellcome Sanger Institute COVID-19 Surveillance Team |
| EPI_ISL_2395803 | Lighthouse Lab in Milton Keynes | Wellcome Sanger Institute for the COVID-19 Genomics UK (COG-UK) Consortium | The Lighthouse Lab in Milton Keynes and Alex Alderton, Roberto Amato, Jeffrey Barrett, Sonia Goncalves, Ewan Harrison, David K. Jackson, Ian Johnston, Dominic Kwiatkowski, Cordelia Langford, John Sillitoe on behalf of the Wellcome Sanger Institute COVID-19 Surveillance Team |
| EPI_ISL_2395804 | Lighthouse Lab in Glasgow | Wellcome Sanger Institute for the COVID-19 Genomics UK (COG-UK) Consortium | Harper VanSteenhouse, Yumi Kasai, David Gray, Carol Clugston, Anna Dominiczak and Alex Alderton, Roberto Amato, Jeffrey Barrett, Sonia Goncalves, Ewan Harrison, David K. Jackson, Ian Johnston, Dominic Kwiatkowski, Cordelia Langford, John Sillitoe on behalf of the Wellcome Sanger Institute COVID-19 Surveillance Team |
| EPI_ISL_2395807, EPI_ISL_2395811 | Lighthouse Lab in Milton Keynes | Wellcome Sanger Institute for the COVID-19 Genomics UK (COG-UK) Consortium | The Lighthouse Lab in Milton Keynes and Alex Alderton, Roberto Amato, Jeffrey Barrett, Sonia Goncalves, Ewan Harrison, David K. Jackson, Ian Johnston, Dominic Kwiatkowski, Cordelia Langford, John Sillitoe on behalf of the Wellcome Sanger Institute COVID-19 Surveillance Team |
| EPI_ISL_2395812, EPI_ISL_2395813 | Lighthouse Lab in Glasgow | Wellcome Sanger Institute for the COVID-19 Genomics UK (COG-UK) Consortium | Harper VanSteenhouse, Yumi Kasai, David Gray, Carol Clugston, Anna Dominiczak and Alex Alderton, Roberto Amato, Jeffrey Barrett, Sonia Goncalves, Ewan Harrison, David K. Jackson, Ian Johnston, Dominic Kwiatkowski, Cordelia Langford, John Sillitoe on behalf of the Wellcome Sanger Institute COVID-19 Surveillance Team |
| EPI_ISL_2395814 | Lighthouse Lab in Milton Keynes | Wellcome Sanger Institute for the COVID-19 Genomics UK (COG-UK) Consortium | The Lighthouse Lab in Milton Keynes and Alex Alderton, Roberto Amato, Jeffrey Barrett, Sonia Goncalves, Ewan Harrison, David K. Jackson, Ian Johnston, Dominic Kwiatkowski, Cordelia Langford, John Sillitoe on behalf of the Wellcome Sanger Institute COVID-19 Surveillance Team |
| EPI_ISL_2395815, EPI_ISL_2395817, EPI_ISL_2395818, EPI_ISL_2395821 | Lighthouse Lab in Glasgow | Wellcome Sanger Institute for the COVID-19 Genomics UK (COG-UK) Consortium | Harper VanSteenhouse, Yumi Kasai, David Gray, Carol Clugston, Anna Dominiczak and Alex Alderton, Roberto Amato, Jeffrey Barrett, Sonia Goncalves, Ewan Harrison, David K. Jackson, Ian Johnston, Dominic Kwiatkowski, Cordelia Langford, John Sillitoe on behalf of the Wellcome Sanger Institute COVID-19 Surveillance Team |
| EPI_ISL_2395822 | Lighthouse Lab in Milton Keynes | Wellcome Sanger Institute for the COVID-19 Genomics UK (COG-UK) Consortium | The Lighthouse Lab in Milton Keynes and Alex Alderton, Roberto Amato, Jeffrey Barrett, Sonia Goncalves, Ewan Harrison, David K. Jackson, Ian Johnston, Dominic Kwiatkowski, Cordelia Langford, John Sillitoe on behalf of the Wellcome Sanger Institute COVID-19 Surveillance Team |
| EPI_ISL_2395824 | Lighthouse Lab in Glasgow | Wellcome Sanger Institute for the COVID-19 Genomics UK (COG-UK) Consortium | Harper VanSteenhouse, Yumi Kasai, David Gray, Carol Clugston, Anna Dominiczak and Alex Alderton, Roberto Amato, Jeffrey Barrett, Sonia Goncalves, Ewan Harrison, David K. Jackson, Ian Johnston, Dominic Kwiatkowski, Cordelia Langford, John Sillitoe on behalf of the Wellcome Sanger Institute COVID-19 Surveillance Team |
| EPI_ISL_2395825 | Lighthouse Lab in Milton Keynes | Wellcome Sanger Institute for the COVID-19 Genomics UK (COG-UK) Consortium | The Lighthouse Lab in Milton Keynes and Alex Alderton, Roberto Amato, Jeffrey Barrett, Sonia Goncalves, Ewan Harrison, David K. Jackson, Ian Johnston, Dominic Kwiatkowski, Cordelia Langford, John Sillitoe on behalf of the Wellcome Sanger Institute COVID-19 Surveillance Team |
| EPI_ISL_2395827 | Lighthouse Lab in Glasgow | Wellcome Sanger Institute for the COVID-19 Genomics UK (COG-UK) Consortium | Harper VanSteenhouse, Yumi Kasai, David Gray, Carol Clugston, Anna Dominiczak and Alex Alderton, Roberto Amato, Jeffrey Barrett, Sonia Goncalves, Ewan Harrison, David K. Jackson, Ian Johnston, Dominic Kwiatkowski, Cordelia Langford, John Sillitoe on behalf of the Wellcome Sanger Institute COVID-19 Surveillance Team |
| EPI_ISL_2395828 | Lighthouse Lab in Milton Keynes | Wellcome Sanger Institute for the COVID-19 Genomics UK (COG-UK) Consortium | The Lighthouse Lab in Milton Keynes and Alex Alderton, Roberto Amato, Jeffrey Barrett, Sonia Goncalves, Ewan Harrison, David K. Jackson, Ian Johnston, Dominic Kwiatkowski, Cordelia Langford, John Sillitoe on behalf of the Wellcome Sanger Institute COVID-19 Surveillance Team |
| EPI_ISL_2395830 | Lighthouse Lab in Glasgow | Wellcome Sanger Institute for the COVID-19 Genomics UK (COG-UK) Consortium | Harper VanSteenhouse, Yumi Kasai, David Gray, Carol Clugston, Anna Dominiczak and Alex Alderton, Roberto Amato, Jeffrey Barrett, Sonia Goncalves, Ewan Harrison, David K. Jackson, Ian Johnston, Dominic Kwiatkowski, Cordelia Langford, John Sillitoe on behalf of the Wellcome Sanger Institute COVID-19 Surveillance Team |
| EPI_ISL_2395831, EPI_ISL_2395832 | Lighthouse Lab in Milton Keynes | Wellcome Sanger Institute for the COVID-19 Genomics UK (COG-UK) Consortium | The Lighthouse Lab in Milton Keynes and Alex Alderton, Roberto Amato, Jeffrey Barrett, Sonia Goncalves, Ewan Harrison, David K. Jackson, Ian Johnston, Dominic Kwiatkowski, Cordelia Langford, John Sillitoe on behalf of the Wellcome Sanger Institute COVID-19 Surveillance Team |
| EPI_ISL_2395833 | Lighthouse Lab in Glasgow | Wellcome Sanger Institute for the COVID-19 Genomics UK (COG-UK) Consortium | Harper VanSteenhouse, Yumi Kasai, David Gray, Carol Clugston, Anna Dominiczak and Alex Alderton, Roberto Amato, Jeffrey Barrett, Sonia Goncalves, Ewan Harrison, David K. Jackson, Ian Johnston, Dominic Kwiatkowski, Cordelia Langford, John Sillitoe on behalf of the Wellcome Sanger Institute COVID-19 Surveillance Team |
| EPI_ISL_2395834 | Lighthouse Lab in Milton Keynes | Wellcome Sanger Institute for the COVID-19 Genomics UK (COG-UK) Consortium | The Lighthouse Lab in Milton Keynes and Alex Alderton, Roberto Amato, Jeffrey Barrett, Sonia Goncalves, Ewan Harrison, David K. Jackson, Ian Johnston, Dominic Kwiatkowski, Cordelia Langford, John Sillitoe on behalf of the Wellcome Sanger Institute COVID-19 Surveillance Team |
| EPI_ISL_2395835 | Lighthouse Lab in Glasgow | Wellcome Sanger Institute for the COVID-19 Genomics UK (COG-UK) Consortium | Harper VanSteenhouse, Yumi Kasai, David Gray, Carol Clugston, Anna Dominiczak and Alex Alderton, Roberto Amato, Jeffrey Barrett, Sonia Goncalves, Ewan Harrison, David K. Jackson, Ian Johnston, Dominic Kwiatkowski, Cordelia Langford, John Sillitoe on behalf of the Wellcome Sanger Institute COVID-19 Surveillance Team |
| EPI_ISL_2395836, EPI_ISL_2395837, EPI_ISL_2395839, EPI_ISL_2395840 | Lighthouse Lab in Milton Keynes | Wellcome Sanger Institute for the COVID-19 Genomics UK (COG-UK) Consortium | The Lighthouse Lab in Milton Keynes and Alex Alderton, Roberto Amato, Jeffrey Barrett, Sonia Goncalves, Ewan Harrison, David K. Jackson, Ian Johnston, Dominic Kwiatkowski, Cordelia Langford, John Sillitoe on behalf of the Wellcome Sanger Institute COVID-19 Surveillance Team |
| EPI_ISL_2395843 | Lighthouse Lab in Glasgow | Wellcome Sanger Institute for the COVID-19 Genomics UK (COG-UK) Consortium | Harper VanSteenhouse, Yumi Kasai, David Gray, Carol Clugston, Anna Dominiczak and Alex Alderton, Roberto Amato, Jeffrey Barrett, Sonia Goncalves, Ewan Harrison, David K. Jackson, Ian Johnston, Dominic Kwiatkowski, Cordelia Langford, John Sillitoe on behalf of the Wellcome Sanger Institute COVID-19 Surveillance Team |
| EPI_ISL_2395844, EPI_ISL_2395845, EPI_ISL_2395847 | Lighthouse Lab in Milton Keynes | Wellcome Sanger Institute for the COVID-19 Genomics UK (COG-UK) Consortium | The Lighthouse Lab in Milton Keynes and Alex Alderton, Roberto Amato, Jeffrey Barrett, Sonia Goncalves, Ewan Harrison, David K. Jackson, Ian Johnston, Dominic Kwiatkowski, Cordelia Langford, John Sillitoe on behalf of the Wellcome Sanger Institute COVID-19 Surveillance Team |
| EPI_ISL_2395848 | Lighthouse Lab in Glasgow | Wellcome Sanger Institute for the COVID-19 Genomics UK (COG-UK) Consortium | Harper VanSteenhouse, Yumi Kasai, David Gray, Carol Clugston, Anna Dominiczak and Alex Alderton, Roberto Amato, Jeffrey Barrett, Sonia Goncalves, Ewan Harrison, David K. Jackson, Ian Johnston, Dominic Kwiatkowski, Cordelia Langford, John Sillitoe on behalf of the Wellcome Sanger Institute COVID-19 Surveillance Team |
| EPI_ISL_2395849 | Lighthouse Lab in Milton Keynes | Wellcome Sanger Institute for the COVID-19 Genomics UK (COG-UK) Consortium | The Lighthouse Lab in Milton Keynes and Alex Alderton, Roberto Amato, Jeffrey Barrett, Sonia Goncalves, Ewan Harrison, David K. Jackson, Ian Johnston, Dominic Kwiatkowski, Cordelia Langford, John Sillitoe on behalf of the Wellcome Sanger Institute COVID-19 Surveillance Team |
| EPI_ISL_2395850 | Lighthouse Lab in Glasgow | Wellcome Sanger Institute for the COVID-19 Genomics UK (COG-UK) Consortium | Harper VanSteenhouse, Yumi Kasai, David Gray, Carol Clugston, Anna Dominiczak and Alex Alderton, Roberto Amato, Jeffrey Barrett, Sonia Goncalves, Ewan Harrison, David K. Jackson, Ian Johnston, Dominic Kwiatkowski, Cordelia Langford, John Sillitoe on behalf of the Wellcome Sanger Institute COVID-19 Surveillance Team |
| EPI_ISL_2395851 | Lighthouse Lab in Milton Keynes | Wellcome Sanger Institute for the COVID-19 Genomics UK (COG-UK) Consortium | The Lighthouse Lab in Milton Keynes and Alex Alderton, Roberto Amato, Jeffrey Barrett, Sonia Goncalves, Ewan Harrison, David K. Jackson, Ian Johnston, Dominic Kwiatkowski, Cordelia Langford, John Sillitoe on behalf of the Wellcome Sanger Institute COVID-19 Surveillance Team |
| EPI_ISL_2395853, EPI_ISL_2395854 | Lighthouse Lab in Glasgow | Wellcome Sanger Institute for the COVID-19 Genomics UK (COG-UK) Consortium | Harper VanSteenhouse, Yumi Kasai, David Gray, Carol Clugston, Anna Dominiczak and Alex Alderton, Roberto Amato, Jeffrey Barrett, Sonia Goncalves, Ewan Harrison, David K. Jackson, Ian Johnston, Dominic Kwiatkowski, Cordelia Langford, John Sillitoe on behalf of the Wellcome Sanger Institute COVID-19 Surveillance Team |
| EPI_ISL_2395856, EPI_ISL_2395857, EPI_ISL_2395858 | Lighthouse Lab in Milton Keynes | Wellcome Sanger Institute for the COVID-19 Genomics UK (COG-UK) Consortium | The Lighthouse Lab in Milton Keynes and Alex Alderton, Roberto Amato, Jeffrey Barrett, Sonia Goncalves, Ewan Harrison, David K. Jackson, Ian Johnston, Dominic Kwiatkowski, Cordelia Langford, John Sillitoe on behalf of the Wellcome Sanger Institute COVID-19 Surveillance Team |
| EPI_ISL_2395860, EPI_ISL_2395862, EPI_ISL_2395863 | Lighthouse Lab in Glasgow | Wellcome Sanger Institute for the COVID-19 Genomics UK (COG-UK) Consortium | Harper VanSteenhouse, Yumi Kasai, David Gray, Carol Clugston, Anna Dominiczak and Alex Alderton, Roberto Amato, Jeffrey Barrett, Sonia Goncalves, Ewan Harrison, David K. Jackson, Ian Johnston, Dominic Kwiatkowski, Cordelia Langford, John Sillitoe on behalf of the Wellcome Sanger Institute COVID-19 Surveillance Team |
| EPI_ISL_2395864 | Lighthouse Lab in Milton Keynes | Wellcome Sanger Institute for the COVID-19 Genomics UK (COG-UK) Consortium | The Lighthouse Lab in Milton Keynes and Alex Alderton, Roberto Amato, Jeffrey Barrett, Sonia Goncalves, Ewan Harrison, David K. Jackson, Ian Johnston, Dominic Kwiatkowski, Cordelia Langford, John Sillitoe on behalf of the Wellcome Sanger Institute COVID-19 Surveillance Team |
| EPI_ISL_2395868 | Lighthouse Lab in Glasgow | Wellcome Sanger Institute for the COVID-19 Genomics UK (COG-UK) Consortium | Harper VanSteenhouse, Yumi Kasai, David Gray, Carol Clugston, Anna Dominiczak and Alex Alderton, Roberto Amato, Jeffrey Barrett, Sonia Goncalves, Ewan Harrison, David K. Jackson, Ian Johnston, Dominic Kwiatkowski, Cordelia Langford, John Sillitoe on behalf of the Wellcome Sanger Institute COVID-19 Surveillance Team |
| EPI_ISL_2395869 | Lighthouse Lab in Milton Keynes | Wellcome Sanger Institute for the COVID-19 Genomics UK (COG-UK) Consortium | The Lighthouse Lab in Milton Keynes and Alex Alderton, Roberto Amato, Jeffrey Barrett, Sonia Goncalves, Ewan Harrison, David K. Jackson, Ian Johnston, Dominic Kwiatkowski, Cordelia Langford, John Sillitoe on behalf of the Wellcome Sanger Institute COVID-19 Surveillance Team |
| EPI_ISL_2395870, EPI_ISL_2395872 | Lighthouse Lab in Glasgow | Wellcome Sanger Institute for the COVID-19 Genomics UK (COG-UK) Consortium | Harper VanSteenhouse, Yumi Kasai, David Gray, Carol Clugston, Anna Dominiczak and Alex Alderton, Roberto Amato, Jeffrey Barrett, Sonia Goncalves, Ewan Harrison, David K. Jackson, Ian Johnston, Dominic Kwiatkowski, Cordelia Langford, John Sillitoe on behalf of the Wellcome Sanger Institute COVID-19 Surveillance Team |
| EPI_ISL_2395874, EPI_ISL_2395875, EPI_ISL_2395876 | Lighthouse Lab in Milton Keynes | Wellcome Sanger Institute for the COVID-19 Genomics UK (COG-UK) Consortium | The Lighthouse Lab in Milton Keynes and Alex Alderton, Roberto Amato, Jeffrey Barrett, Sonia Goncalves, Ewan Harrison, David K. Jackson, Ian Johnston, Dominic Kwiatkowski, Cordelia Langford, John Sillitoe on behalf of the Wellcome Sanger Institute COVID-19 Surveillance Team |
| EPI_ISL_2395877 | Lighthouse Lab in Glasgow | Wellcome Sanger Institute for the COVID-19 Genomics UK (COG-UK) Consortium | Harper VanSteenhouse, Yumi Kasai, David Gray, Carol Clugston, Anna Dominiczak and Alex Alderton, Roberto Amato, Jeffrey Barrett, Sonia Goncalves, Ewan Harrison, David K. Jackson, Ian Johnston, Dominic Kwiatkowski, Cordelia Langford, John Sillitoe on behalf of the Wellcome Sanger Institute COVID-19 Surveillance Team |
| EPI_ISL_2395878, EPI_ISL_2395880, EPI_ISL_2395881, EPI_ISL_2395882, EPI_ISL_2395883, EPI_ISL_2395884 | Lighthouse Lab in Milton Keynes | Wellcome Sanger Institute for the COVID-19 Genomics UK (COG-UK) Consortium | The Lighthouse Lab in Milton Keynes and Alex Alderton, Roberto Amato, Jeffrey Barrett, Sonia Goncalves, Ewan Harrison, David K. Jackson, Ian Johnston, Dominic Kwiatkowski, Cordelia Langford, John Sillitoe on behalf of the Wellcome Sanger Institute COVID-19 Surveillance Team |
| EPI_ISL_2395886 | Lighthouse Lab in Glasgow | Wellcome Sanger Institute for the COVID-19 Genomics UK (COG-UK) Consortium | Harper VanSteenhouse, Yumi Kasai, David Gray, Carol Clugston, Anna Dominiczak and Alex Alderton, Roberto Amato, Jeffrey Barrett, Sonia Goncalves, Ewan Harrison, David K. Jackson, Ian Johnston, Dominic Kwiatkowski, Cordelia Langford, John Sillitoe on behalf of the Wellcome Sanger Institute COVID-19 Surveillance Team |
| EPI_ISL_2395889, EPI_ISL_2395893 | Lighthouse Lab in Milton Keynes | Wellcome Sanger Institute for the COVID-19 Genomics UK (COG-UK) Consortium | The Lighthouse Lab in Milton Keynes and Alex Alderton, Roberto Amato, Jeffrey Barrett, Sonia Goncalves, Ewan Harrison, David K. Jackson, Ian Johnston, Dominic Kwiatkowski, Cordelia Langford, John Sillitoe on behalf of the Wellcome Sanger Institute COVID-19 Surveillance Team |
| EPI_ISL_2395895 | Lighthouse Lab in Glasgow | Wellcome Sanger Institute for the COVID-19 Genomics UK (COG-UK) Consortium | Harper VanSteenhouse, Yumi Kasai, David Gray, Carol Clugston, Anna Dominiczak and Alex Alderton, Roberto Amato, Jeffrey Barrett, Sonia Goncalves, Ewan Harrison, David K. Jackson, Ian Johnston, Dominic Kwiatkowski, Cordelia Langford, John Sillitoe on behalf of the Wellcome Sanger Institute COVID-19 Surveillance Team |
| EPI_ISL_2395897 | Lighthouse Lab in Milton Keynes | Wellcome Sanger Institute for the COVID-19 Genomics UK (COG-UK) Consortium | The Lighthouse Lab in Milton Keynes and Alex Alderton, Roberto Amato, Jeffrey Barrett, Sonia Goncalves, Ewan Harrison, David K. Jackson, Ian Johnston, Dominic Kwiatkowski, Cordelia Langford, John Sillitoe on behalf of the Wellcome Sanger Institute COVID-19 Surveillance Team |
| EPI_ISL_2395898, EPI_ISL_2395899 | Lighthouse Lab in Glasgow | Wellcome Sanger Institute for the COVID-19 Genomics UK (COG-UK) Consortium | Harper VanSteenhouse, Yumi Kasai, David Gray, Carol Clugston, Anna Dominiczak and Alex Alderton, Roberto Amato, Jeffrey Barrett, Sonia Goncalves, Ewan Harrison, David K. Jackson, Ian Johnston, Dominic Kwiatkowski, Cordelia Langford, John Sillitoe on behalf of the Wellcome Sanger Institute COVID-19 Surveillance Team |

| | | | |
|---|---|---|---|
| EPI_ISL_2395902 | Lighthouse Lab in Milton Keynes | Wellcome Sanger Institute for the COVID-19 Genomics UK (COG-UK) Consortium | The Lighthouse Lab in Milton Keynes and Alex Alderton, Roberto Amato, Jeffrey Barrett, Sonia Goncalves, Ewan Harrison, David K. Jackson, Ian Johnston, Dominic Kwiatkowski, Cordelia Langford, John Sillitoe on behalf of the Wellcome Sanger Institute COVID-19 Surveillance Team |
| EPI_ISL_2395905 | Lighthouse Lab in Glasgow | Wellcome Sanger Institute for the COVID-19 Genomics UK (COG-UK) Consortium | Harper VanSteenhouse, Yumi Kasai, David Gray, Carol Clugston, Anna Dominiczak and Alex Alderton, Roberto Amato, Jeffrey Barrett, Sonia Goncalves, Ewan Harrison, David K. Jackson, Ian Johnston, Dominic Kwiatkowski, Cordelia Langford, John Sillitoe on behalf of the Wellcome Sanger Institute COVID-19 Surveillance Team |
| EPI_ISL_2395907 | Lighthouse Lab in Milton Keynes | Wellcome Sanger Institute for the COVID-19 Genomics UK (COG-UK) Consortium | The Lighthouse Lab in Milton Keynes and Alex Alderton, Roberto Amato, Jeffrey Barrett, Sonia Goncalves, Ewan Harrison, David K. Jackson, Ian Johnston, Dominic Kwiatkowski, Cordelia Langford, John Sillitoe on behalf of the Wellcome Sanger Institute COVID-19 Surveillance Team |
| EPI_ISL_2395908 | Lighthouse Lab in Glasgow | Wellcome Sanger Institute for the COVID-19 Genomics UK (COG-UK) Consortium | Harper VanSteenhouse, Yumi Kasai, David Gray, Carol Clugston, Anna Dominiczak and Alex Alderton, Roberto Amato, Jeffrey Barrett, Sonia Goncalves, Ewan Harrison, David K. Jackson, Ian Johnston, Dominic Kwiatkowski, Cordelia Langford, John Sillitoe on behalf of the Wellcome Sanger Institute COVID-19 Surveillance Team |
| EPI_ISL_2395909 | Lighthouse Lab in Milton Keynes | Wellcome Sanger Institute for the COVID-19 Genomics UK (COG-UK) Consortium | The Lighthouse Lab in Milton Keynes and Alex Alderton, Roberto Amato, Jeffrey Barrett, Sonia Goncalves, Ewan Harrison, David K. Jackson, Ian Johnston, Dominic Kwiatkowski, Cordelia Langford, John Sillitoe on behalf of the Wellcome Sanger Institute COVID-19 Surveillance Team |
| EPI_ISL_2395912, EPI_ISL_2395915 | Lighthouse Lab in Glasgow | Wellcome Sanger Institute for the COVID-19 Genomics UK (COG-UK) Consortium | Harper VanSteenhouse, Yumi Kasai, David Gray, Carol Clugston, Anna Dominiczak and Alex Alderton, Roberto Amato, Jeffrey Barrett, Sonia Goncalves, Ewan Harrison, David K. Jackson, Ian Johnston, Dominic Kwiatkowski, Cordelia Langford, John Sillitoe on behalf of the Wellcome Sanger Institute COVID-19 Surveillance Team |
| EPI_ISL_2395917 | Lighthouse Lab in Milton Keynes | Wellcome Sanger Institute for the COVID-19 Genomics UK (COG-UK) Consortium | The Lighthouse Lab in Milton Keynes and Alex Alderton, Roberto Amato, Jeffrey Barrett, Sonia Goncalves, Ewan Harrison, David K. Jackson, Ian Johnston, Dominic Kwiatkowski, Cordelia Langford, John Sillitoe on behalf of the Wellcome Sanger Institute COVID-19 Surveillance Team |
| EPI_ISL_2395918, EPI_ISL_2395922 | Lighthouse Lab in Glasgow | Wellcome Sanger Institute for the COVID-19 Genomics UK (COG-UK) Consortium | Harper VanSteenhouse, Yumi Kasai, David Gray, Carol Clugston, Anna Dominiczak and Alex Alderton, Roberto Amato, Jeffrey Barrett, Sonia Goncalves, Ewan Harrison, David K. Jackson, Ian Johnston, Dominic Kwiatkowski, Cordelia Langford, John Sillitoe on behalf of the Wellcome Sanger Institute COVID-19 Surveillance Team |
| EPI_ISL_2395923, EPI_ISL_2395924 | Lighthouse Lab in Milton Keynes | Wellcome Sanger Institute for the COVID-19 Genomics UK (COG-UK) Consortium | The Lighthouse Lab in Milton Keynes and Alex Alderton, Roberto Amato, Jeffrey Barrett, Sonia Goncalves, Ewan Harrison, David K. Jackson, Ian Johnston, Dominic Kwiatkowski, Cordelia Langford, John Sillitoe on behalf of the Wellcome Sanger Institute COVID-19 Surveillance Team |
| EPI_ISL_2395926 | Lighthouse Lab in Glasgow | Wellcome Sanger Institute for the COVID-19 Genomics UK (COG-UK) Consortium | Harper VanSteenhouse, Yumi Kasai, David Gray, Carol Clugston, Anna Dominiczak and Alex Alderton, Roberto Amato, Jeffrey Barrett, Sonia Goncalves, Ewan Harrison, David K. Jackson, Ian Johnston, Dominic Kwiatkowski, Cordelia Langford, John Sillitoe on behalf of the Wellcome Sanger Institute COVID-19 Surveillance Team |
| EPI_ISL_2395928, EPI_ISL_2395929 | Lighthouse Lab in Milton Keynes | Wellcome Sanger Institute for the COVID-19 Genomics UK (COG-UK) Consortium | The Lighthouse Lab in Milton Keynes and Alex Alderton, Roberto Amato, Jeffrey Barrett, Sonia Goncalves, Ewan Harrison, David K. Jackson, Ian Johnston, Dominic Kwiatkowski, Cordelia Langford, John Sillitoe on behalf of the Wellcome Sanger Institute COVID-19 Surveillance Team |
| EPI_ISL_2395930, EPI_ISL_2395933 | Lighthouse Lab in Glasgow | Wellcome Sanger Institute for the COVID-19 Genomics UK (COG-UK) Consortium | Harper VanSteenhouse, Yumi Kasai, David Gray, Carol Clugston, Anna Dominiczak and Alex Alderton, Roberto Amato, Jeffrey Barrett, Sonia Goncalves, Ewan Harrison, David K. Jackson, Ian Johnston, Dominic Kwiatkowski, Cordelia Langford, John Sillitoe on behalf of the Wellcome Sanger Institute COVID-19 Surveillance Team |
| EPI_ISL_2395934 | Lighthouse Lab in Milton Keynes | Wellcome Sanger Institute for the COVID-19 Genomics UK (COG-UK) Consortium | The Lighthouse Lab in Milton Keynes and Alex Alderton, Roberto Amato, Jeffrey Barrett, Sonia Goncalves, Ewan Harrison, David K. Jackson, Ian Johnston, Dominic Kwiatkowski, Cordelia Langford, John Sillitoe on behalf of the Wellcome Sanger Institute COVID-19 Surveillance Team |
| EPI_ISL_2395935 | Lighthouse Lab in Glasgow | Wellcome Sanger Institute for the COVID-19 Genomics UK (COG-UK) Consortium | Harper VanSteenhouse, Yumi Kasai, David Gray, Carol Clugston, Anna Dominiczak and Alex Alderton, Roberto Amato, Jeffrey Barrett, Sonia Goncalves, Ewan Harrison, David K. Jackson, Ian Johnston, Dominic Kwiatkowski, Cordelia Langford, John Sillitoe on behalf of the Wellcome Sanger Institute COVID-19 Surveillance Team |
| EPI_ISL_2395936 | Lighthouse Lab in Milton Keynes | Wellcome Sanger Institute for the COVID-19 Genomics UK (COG-UK) Consortium | The Lighthouse Lab in Milton Keynes and Alex Alderton, Roberto Amato, Jeffrey Barrett, Sonia Goncalves, Ewan Harrison, David K. Jackson, Ian Johnston, Dominic Kwiatkowski, Cordelia Langford, John Sillitoe on behalf of the Wellcome Sanger Institute COVID-19 Surveillance Team |
| EPI_ISL_2395937, EPI_ISL_2395938, EPI_ISL_2395939, EPI_ISL_2395940 | Lighthouse Lab in Glasgow | Wellcome Sanger Institute for the COVID-19 Genomics UK (COG-UK) Consortium | Harper VanSteenhouse, Yumi Kasai, David Gray, Carol Clugston, Anna Dominiczak and Alex Alderton, Roberto Amato, Jeffrey Barrett, Sonia Goncalves, Ewan Harrison, David K. Jackson, Ian Johnston, Dominic Kwiatkowski, Cordelia Langford, John Sillitoe on behalf of the Wellcome Sanger Institute COVID-19 Surveillance Team |
| EPI_ISL_2395941, EPI_ISL_2395943, EPI_ISL_2395944, EPI_ISL_2395945 | Lighthouse Lab in Milton Keynes | Wellcome Sanger Institute for the COVID-19 Genomics UK (COG-UK) Consortium | The Lighthouse Lab in Milton Keynes and Alex Alderton, Roberto Amato, Jeffrey Barrett, Sonia Goncalves, Ewan Harrison, David K. Jackson, Ian Johnston, Dominic Kwiatkowski, Cordelia Langford, John Sillitoe on behalf of the Wellcome Sanger Institute COVID-19 Surveillance Team |
| EPI_ISL_2395946 | Lighthouse Lab in Glasgow | Wellcome Sanger Institute for the COVID-19 Genomics UK (COG-UK) Consortium | Harper VanSteenhouse, Yumi Kasai, David Gray, Carol Clugston, Anna Dominiczak and Alex Alderton, Roberto Amato, Jeffrey Barrett, Sonia Goncalves, Ewan Harrison, David K. Jackson, Ian Johnston, Dominic Kwiatkowski, Cordelia Langford, John Sillitoe on behalf of the Wellcome Sanger Institute COVID-19 Surveillance Team |
| EPI_ISL_2395947, EPI_ISL_2395948, EPI_ISL_2395949 | Lighthouse Lab in Milton Keynes | Wellcome Sanger Institute for the COVID-19 Genomics UK (COG-UK) Consortium | The Lighthouse Lab in Milton Keynes and Alex Alderton, Roberto Amato, Jeffrey Barrett, Sonia Goncalves, Ewan Harrison, David K. Jackson, Ian Johnston, Dominic Kwiatkowski, Cordelia Langford, John Sillitoe on behalf of the Wellcome Sanger Institute COVID-19 Surveillance Team |
| EPI_ISL_2395950 | Lighthouse Lab in Glasgow | Wellcome Sanger Institute for the COVID-19 Genomics UK (COG-UK) Consortium | Harper VanSteenhouse, Yumi Kasai, David Gray, Carol Clugston, Anna Dominiczak and Alex Alderton, Roberto Amato, Jeffrey Barrett, Sonia Goncalves, Ewan Harrison, David K. Jackson, Ian Johnston, Dominic Kwiatkowski, Cordelia Langford, John Sillitoe on behalf of the Wellcome Sanger Institute COVID-19 Surveillance Team |
| EPI_ISL_2395951 | Lighthouse Lab in Milton Keynes | Wellcome Sanger Institute for the COVID-19 Genomics UK (COG-UK) Consortium | The Lighthouse Lab in Milton Keynes and Alex Alderton, Roberto Amato, Jeffrey Barrett, Sonia Goncalves, Ewan Harrison, David K. Jackson, Ian Johnston, Dominic Kwiatkowski, Cordelia Langford, John Sillitoe on behalf of the Wellcome Sanger Institute COVID-19 Surveillance Team |
| EPI_ISL_2395952 | Lighthouse Lab in Glasgow | Wellcome Sanger Institute for the COVID-19 Genomics UK (COG-UK) Consortium | Harper VanSteenhouse, Yumi Kasai, David Gray, Carol Clugston, Anna Dominiczak and Alex Alderton, Roberto Amato, Jeffrey Barrett, Sonia Goncalves, Ewan Harrison, David K. Jackson, Ian Johnston, Dominic Kwiatkowski, Cordelia Langford, John Sillitoe on behalf of the Wellcome Sanger Institute COVID-19 Surveillance Team |
| EPI_ISL_2395954, EPI_ISL_2395955, EPI_ISL_2395956, EPI_ISL_2395957, EPI_ISL_2395958, EPI_ISL_2395959, EPI_ISL_2395961, EPI_ISL_2395964, EPI_ISL_2395965, EPI_ISL_2395966, EPI_ISL_2395967 | | | |
| see above | Lighthouse Lab in Milton Keynes | Wellcome Sanger Institute for the COVID-19 Genomics UK (COG-UK) Consortium | The Lighthouse Lab in Milton Keynes and Alex Alderton, Roberto Amato, Jeffrey Barrett, Sonia Goncalves, Ewan Harrison, David K. Jackson, Ian Johnston, Dominic Kwiatkowski, Cordelia Langford, John Sillitoe on behalf of the Wellcome Sanger Institute COVID-19 Surveillance Team |
| EPI_ISL_2395968, EPI_ISL_2395969 | Lighthouse Lab in Glasgow | Wellcome Sanger Institute for the COVID-19 Genomics UK (COG-UK) Consortium | Harper VanSteenhouse, Yumi Kasai, David Gray, Carol Clugston, Anna Dominiczak and Alex Alderton, Roberto Amato, Jeffrey Barrett, Sonia Goncalves, Ewan Harrison, David K. Jackson, Ian Johnston, Dominic Kwiatkowski, Cordelia Langford, John Sillitoe on behalf of the Wellcome Sanger Institute COVID-19 Surveillance Team |
| EPI_ISL_2395971, EPI_ISL_2395972, EPI_ISL_2395973, EPI_ISL_2395974 | Lighthouse Lab in Milton Keynes | Wellcome Sanger Institute for the COVID-19 Genomics UK (COG-UK) Consortium | The Lighthouse Lab in Milton Keynes and Alex Alderton, Roberto Amato, Jeffrey Barrett, Sonia Goncalves, Ewan Harrison, David K. Jackson, Ian Johnston, Dominic Kwiatkowski, Cordelia Langford, John Sillitoe on behalf of the Wellcome Sanger Institute COVID-19 Surveillance Team |
| EPI_ISL_2395975, EPI_ISL_2395976 | Lighthouse Lab in Glasgow | Wellcome Sanger Institute for the COVID-19 Genomics UK (COG-UK) Consortium | Harper VanSteenhouse, Yumi Kasai, David Gray, Carol Clugston, Anna Dominiczak and Alex Alderton, Roberto Amato, Jeffrey Barrett, Sonia Goncalves, Ewan Harrison, David K. Jackson, Ian Johnston, Dominic Kwiatkowski, Cordelia Langford, John Sillitoe on behalf of the Wellcome Sanger Institute COVID-19 Surveillance Team |
| EPI_ISL_2395977 | Lighthouse Lab in Milton Keynes | Wellcome Sanger Institute for the COVID-19 Genomics UK (COG-UK) Consortium | The Lighthouse Lab in Milton Keynes and Alex Alderton, Roberto Amato, Jeffrey Barrett, Sonia Goncalves, Ewan Harrison, David K. Jackson, Ian Johnston, Dominic Kwiatkowski, Cordelia Langford, John Sillitoe on behalf of the Wellcome Sanger Institute COVID-19 Surveillance Team |
| EPI_ISL_2395981, EPI_ISL_2395982, EPI_ISL_2395983, EPI_ISL_2395984, EPI_ISL_2395985 | Lighthouse Lab in Glasgow | Wellcome Sanger Institute for the COVID-19 Genomics UK (COG-UK) Consortium | Harper VanSteenhouse, Yumi Kasai, David Gray, Carol Clugston, Anna Dominiczak and Alex Alderton, Roberto Amato, Jeffrey Barrett, Sonia Goncalves, Ewan Harrison, David K. Jackson, Ian Johnston, Dominic Kwiatkowski, Cordelia Langford, John Sillitoe on behalf of the Wellcome Sanger Institute COVID-19 Surveillance Team |
| EPI_ISL_2395986, EPI_ISL_2395987, EPI_ISL_2395988 | Lighthouse Lab in Milton Keynes | Wellcome Sanger Institute for the COVID-19 Genomics UK (COG-UK) Consortium | The Lighthouse Lab in Milton Keynes and Alex Alderton, Roberto Amato, Jeffrey Barrett, Sonia Goncalves, Ewan Harrison, David K. Jackson, Ian Johnston, Dominic Kwiatkowski, Cordelia Langford, John Sillitoe on behalf of the Wellcome Sanger Institute COVID-19 Surveillance Team |
| EPI_ISL_2395989 | Lighthouse Lab in Glasgow | Wellcome Sanger Institute for the COVID-19 Genomics UK (COG-UK) Consortium | Harper VanSteenhouse, Yumi Kasai, David Gray, Carol Clugston, Anna Dominiczak and Alex Alderton, Roberto Amato, Jeffrey Barrett, Sonia Goncalves, Ewan Harrison, David K. Jackson, Ian Johnston, Dominic Kwiatkowski, Cordelia Langford, John Sillitoe on behalf of the Wellcome Sanger Institute COVID-19 Surveillance Team |
| EPI_ISL_2395990, EPI_ISL_2395991, EPI_ISL_2395992, EPI_ISL_2395994, EPI_ISL_2395995, EPI_ISL_2395996 | Lighthouse Lab in Milton Keynes | Wellcome Sanger Institute for the COVID-19 Genomics UK (COG-UK) Consortium | The Lighthouse Lab in Milton Keynes and Alex Alderton, Roberto Amato, Jeffrey Barrett, Sonia Goncalves, Ewan Harrison, David K. Jackson, Ian Johnston, Dominic Kwiatkowski, Cordelia Langford, John Sillitoe on behalf of the Wellcome Sanger Institute COVID-19 Surveillance Team |
| EPI_ISL_2395999, EPI_ISL_2396002, EPI_ISL_2396003 | Lighthouse Lab in Glasgow | Wellcome Sanger Institute for the COVID-19 Genomics UK (COG-UK) Consortium | Harper VanSteenhouse, Yumi Kasai, David Gray, Carol Clugston, Anna Dominiczak and Alex Alderton, Roberto Amato, Jeffrey Barrett, Sonia Goncalves, Ewan Harrison, David K. Jackson, Ian Johnston, Dominic Kwiatkowski, Cordelia Langford, John Sillitoe on behalf of the Wellcome Sanger Institute COVID-19 Surveillance Team |
| EPI_ISL_2396005, EPI_ISL_2396006, EPI_ISL_2396009 | Lighthouse Lab in Milton Keynes | Wellcome Sanger Institute for the COVID-19 Genomics UK (COG-UK) Consortium | The Lighthouse Lab in Milton Keynes and Alex Alderton, Roberto Amato, Jeffrey Barrett, Sonia Goncalves, Ewan Harrison, David K. Jackson, Ian Johnston, Dominic Kwiatkowski, Cordelia Langford, John Sillitoe on behalf of the Wellcome Sanger Institute COVID-19 Surveillance Team |
| EPI_ISL_2396010 | Lighthouse Lab in Glasgow | Wellcome Sanger Institute for the COVID-19 Genomics UK (COG-UK) Consortium | Harper VanSteenhouse, Yumi Kasai, David Gray, Carol Clugston, Anna Dominiczak and Alex Alderton, Roberto Amato, Jeffrey Barrett, Sonia Goncalves, Ewan Harrison, David K. Jackson, Ian Johnston, Dominic Kwiatkowski, Cordelia Langford, John Sillitoe on behalf of the Wellcome Sanger Institute COVID-19 Surveillance Team |
| EPI_ISL_2396012, EPI_ISL_2396013 | Lighthouse Lab in Milton Keynes | Wellcome Sanger Institute for the COVID-19 Genomics UK (COG-UK) Consortium | The Lighthouse Lab in Milton Keynes and Alex Alderton, Roberto Amato, Jeffrey Barrett, Sonia Goncalves, Ewan Harrison, David K. Jackson, Ian Johnston, Dominic Kwiatkowski, Cordelia Langford, John Sillitoe on behalf of the Wellcome Sanger Institute COVID-19 Surveillance Team |
| EPI_ISL_2396015 | Lighthouse Lab in Glasgow | Wellcome Sanger Institute for the COVID-19 Genomics UK (COG-UK) Consortium | Harper VanSteenhouse, Yumi Kasai, David Gray, Carol Clugston, Anna Dominiczak and Alex Alderton, Roberto Amato, Jeffrey Barrett, Sonia Goncalves, Ewan Harrison, David K. Jackson, Ian Johnston, Dominic Kwiatkowski, Cordelia Langford, John Sillitoe on behalf of the Wellcome Sanger Institute COVID-19 Surveillance Team |
| EPI_ISL_2396016, EPI_ISL_2396017, EPI_ISL_2396018, EPI_ISL_2396019, EPI_ISL_2396020, EPI_ISL_2396021, EPI_ISL_2396022, EPI_ISL_2396023, EPI_ISL_2396024, EPI_ISL_2396025, EPI_ISL_2396026, EPI_ISL_2396027, EPI_ISL_2396028, EPI_ISL_2396031, EPI_ISL_2396033, EPI_ISL_2396034, EPI_ISL_2396036, EPI_ISL_2396037, EPI_ISL_2396038, EPI_ISL_2396039, EPI_ISL_2396040, EPI_ISL_2396041, EPI_ISL_2396042, EPI_ISL_2396043, EPI_ISL_2396044, EPI_ISL_2396045, EPI_ISL_2396047, EPI_ISL_2396048, EPI_ISL_2396049, EPI_ISL_2396050, EPI_ISL_2396051, EPI_ISL_2396052, EPI_ISL_2396053, EPI_ISL_2396054, EPI_ISL_2396055, EPI_ISL_2396056, EPI_ISL_2396057, EPI_ISL_2396058, EPI_ISL_2396059, EPI_ISL_2396060, EPI_ISL_2396061, EPI_ISL_2396063, EPI_ISL_2396065, EPI_ISL_2396066, EPI_ISL_2396067, EPI_ISL_2396068, EPI_ISL_2396069, EPI_ISL_2396070, EPI_ISL_2396071, EPI_ISL_2396072, EPI_ISL_2396073, EPI_ISL_2396074, EPI_ISL_2396077, EPI_ISL_2396078, EPI_ISL_2396080, EPI_ISL_2396081, EPI_ISL_2396084, EPI_ISL_2396085, EPI_ISL_2396086, EPI_ISL_2396087, EPI_ISL_2396088, EPI_ISL_2396089, EPI_ISL_2396090, EPI_ISL_2396091, EPI_ISL_2396092, EPI_ISL_2396093, | | | |

| | | | |
|---|---|---|---|
| EPI_ISL_2396094, EPI_ISL_2396095, EPI_ISL_2396096, EPI_ISL_2396097, EPI_ISL_2396098, EPI_ISL_2396099, EPI_ISL_2396100, EPI_ISL_2396101, EPI_ISL_2396102, EPI_ISL_2396103, EPI_ISL_2396104, EPI_ISL_2396105, EPI_ISL_2396106, EPI_ISL_2396108, EPI_ISL_2396109, EPI_ISL_2396110, EPI_ISL_2396111, EPI_ISL_2396112, EPI_ISL_2396113, EPI_ISL_2396114, EPI_ISL_2396116, EPI_ISL_2396117, EPI_ISL_2396118, EPI_ISL_2396119, EPI_ISL_2396120, EPI_ISL_2396121, EPI_ISL_2396122, EPI_ISL_2396123, EPI_ISL_2396124, EPI_ISL_2396126, EPI_ISL_2396129, EPI_ISL_2396130, EPI_ISL_2396131, EPI_ISL_2396132, EPI_ISL_2396134, EPI_ISL_2396135, EPI_ISL_2396136, EPI_ISL_2396138, EPI_ISL_2396139, EPI_ISL_2396140, EPI_ISL_2396144, EPI_ISL_2396145, EPI_ISL_2396148, EPI_ISL_2396149, EPI_ISL_2396150, EPI_ISL_2396151, EPI_ISL_2396152, EPI_ISL_2396154, EPI_ISL_2396155, EPI_ISL_2396156, EPI_ISL_2396158, EPI_ISL_2396159, EPI_ISL_2396160, EPI_ISL_2396162, EPI_ISL_2396163, EPI_ISL_2396166, EPI_ISL_2396167, EPI_ISL_2396168, EPI_ISL_2396170, EPI_ISL_2396171, EPI_ISL_2396172, EPI_ISL_2396173, EPI_ISL_2396174, EPI_ISL_2396178, EPI_ISL_2396180, EPI_ISL_2396181, EPI_ISL_2396182, EPI_ISL_2396183, EPI_ISL_2396186, EPI_ISL_2396187, EPI_ISL_2396188, EPI_ISL_2396189, EPI_ISL_2396190, EPI_ISL_2396191, EPI_ISL_2396192, EPI_ISL_2396193, EPI_ISL_2396194, EPI_ISL_2396197, EPI_ISL_2396198, EPI_ISL_2396199, EPI_ISL_2396201, EPI_ISL_2396202, EPI_ISL_2396204, EPI_ISL_2396206, EPI_ISL_2396207, EPI_ISL_2396208, EPI_ISL_2396209, EPI_ISL_2396211, EPI_ISL_2396213, EPI_ISL_2396214, EPI_ISL_2396215, EPI_ISL_2396216, EPI_ISL_2396218, EPI_ISL_2396219, EPI_ISL_2396220, EPI_ISL_2396221, EPI_ISL_2396222, EPI_ISL_2396224, EPI_ISL_2396225, EPI_ISL_2396226, EPI_ISL_2396228, EPI_ISL_2396229, EPI_ISL_2396230, EPI_ISL_2396231, EPI_ISL_2396234, EPI_ISL_2396235, EPI_ISL_2396236, EPI_ISL_2396239, EPI_ISL_2396240, EPI_ISL_2396241, EPI_ISL_2396242, EPI_ISL_2396243, EPI_ISL_2396244, EPI_ISL_2396245, EPI_ISL_2396246, EPI_ISL_2396247, EPI_ISL_2396249, EPI_ISL_2396250, EPI_ISL_2396251, EPI_ISL_2396252, EPI_ISL_2396254, EPI_ISL_2396255, EPI_ISL_2396257, EPI_ISL_2396258, EPI_ISL_2396260, EPI_ISL_2396261, EPI_ISL_2396262, EPI_ISL_2396264, EPI_ISL_2396265, EPI_ISL_2396266, EPI_ISL_2396267, EPI_ISL_2396268, EPI_ISL_2396269, EPI_ISL_2396271, EPI_ISL_2396272, EPI_ISL_2396275, EPI_ISL_2396277, EPI_ISL_2396279, EPI_ISL_2396281, EPI_ISL_2396283, EPI_ISL_2396284, EPI_ISL_2396285, EPI_ISL_2396286, EPI_ISL_2396287, EPI_ISL_2396288, EPI_ISL_2396289, EPI_ISL_2396291, EPI_ISL_2396292, EPI_ISL_2396293, EPI_ISL_2396294, EPI_ISL_2396295, EPI_ISL_2396298, EPI_ISL_2396299, EPI_ISL_2396300, EPI_ISL_2396301, EPI_ISL_2396303, EPI_ISL_2396304, EPI_ISL_2396306, EPI_ISL_2396307, EPI_ISL_2396308, EPI_ISL_2396310, EPI_ISL_2396311, EPI_ISL_2396312, EPI_ISL_2396313, EPI_ISL_2396315, EPI_ISL_2396316, EPI_ISL_2396317, EPI_ISL_2396318, EPI_ISL_2396320, EPI_ISL_2396323, EPI_ISL_2396325, EPI_ISL_2396326, EPI_ISL_2396327, EPI_ISL_2396328, EPI_ISL_2396329, EPI_ISL_2396330, EPI_ISL_2396331, EPI_ISL_2396332, EPI_ISL_2396333, EPI_ISL_2396335, EPI_ISL_2396337, EPI_ISL_2396338, EPI_ISL_2396339, EPI_ISL_2396340, EPI_ISL_2396341, EPI_ISL_2396343 | | | |
| see above | Lighthouse Lab in Alderley Park | Wellcome Sanger Institute for the COVID-19 Genomics UK (COG-UK) Consortium | Jacquelyn Wynn, Mairead Hyland, The Lighthouse Lab in Alderley Park and Alex Alderton, Roberto Amato, Jeffrey Barrett, Sonia Goncalves, Ewan Harrison, David K. Jackson, Ian Johnston, Dominic Kwiatkowski, Cordelia Langford, John Sillitoe on behalf of the Wellcome Sanger Institute COVID-19 Surveillance Team |
| EPI_ISL_2396355, EPI_ISL_2396357, EPI_ISL_2396358, EPI_ISL_2396360, EPI_ISL_2396365, EPI_ISL_2396366 | Lighthouse Lab in Glasgow | Wellcome Sanger Institute for the COVID-19 Genomics UK (COG-UK) Consortium | Harper VanSteenhouse, Yumi Kasai, David Gray, Carol Clugston, Anna Dominiczak and Alex Alderton, Roberto Amato, Jeffrey Barrett, Sonia Goncalves, Ewan Harrison, David K. Jackson, Ian Johnston, Dominic Kwiatkowski, Cordelia Langford, John Sillitoe on behalf of the Wellcome Sanger Institute COVID-19 Surveillance Team |
| EPI_ISL_2396370 | Lighthouse Lab in Milton Keynes | Wellcome Sanger Institute for the COVID-19 Genomics UK (COG-UK) Consortium | The Lighthouse Lab in Milton Keynes and Alex Alderton, Roberto Amato, Jeffrey Barrett, Sonia Goncalves, Ewan Harrison, David K. Jackson, Ian Johnston, Dominic Kwiatkowski, Cordelia Langford, John Sillitoe on behalf of the Wellcome Sanger Institute COVID-19 Surveillance Team |
| EPI_ISL_2396375, EPI_ISL_2396377, EPI_ISL_2396380, EPI_ISL_2396388 | Lighthouse Lab in Glasgow | Wellcome Sanger Institute for the COVID-19 Genomics UK (COG-UK) Consortium | Harper VanSteenhouse, Yumi Kasai, David Gray, Carol Clugston, Anna Dominiczak and Alex Alderton, Roberto Amato, Jeffrey Barrett, Sonia Goncalves, Ewan Harrison, David K. Jackson, Ian Johnston, Dominic Kwiatkowski, Cordelia Langford, John Sillitoe on behalf of the Wellcome Sanger Institute COVID-19 Surveillance Team |
| EPI_ISL_2396389 | Lighthouse Lab in Alderley Park | Wellcome Sanger Institute for the COVID-19 Genomics UK (COG-UK) Consortium | Jacquelyn Wynn, Mairead Hyland, The Lighthouse Lab in Alderley Park and Alex Alderton, Roberto Amato, Jeffrey Barrett, Sonia Goncalves, Ewan Harrison, David K. Jackson, Ian Johnston, Dominic Kwiatkowski, Cordelia Langford, John Sillitoe on behalf of the Wellcome Sanger Institute COVID-19 Surveillance Team |
| EPI_ISL_2396390, EPI_ISL_2396392, EPI_ISL_2396397, EPI_ISL_2396399, EPI_ISL_2396401, EPI_ISL_2396404, EPI_ISL_2396406, EPI_ISL_2396407, EPI_ISL_2396408, EPI_ISL_2396412, EPI_ISL_2396414, EPI_ISL_2396418, EPI_ISL_2396422, EPI_ISL_2396424, EPI_ISL_2396425, EPI_ISL_2396427, EPI_ISL_2396428, EPI_ISL_2396431, EPI_ISL_2396452, EPI_ISL_2396454, EPI_ISL_2396456, EPI_ISL_2396457, EPI_ISL_2396462, EPI_ISL_2396464, EPI_ISL_2396466, EPI_ISL_2396470, EPI_ISL_2396471, EPI_ISL_2396472, EPI_ISL_2396476, EPI_ISL_2396477, EPI_ISL_2396484, EPI_ISL_2396489, EPI_ISL_2396490, EPI_ISL_2396491, EPI_ISL_2396493, EPI_ISL_2396495, EPI_ISL_2396496, EPI_ISL_2396503, EPI_ISL_2396508, EPI_ISL_2396509, EPI_ISL_2396514, EPI_ISL_2396515, EPI_ISL_2396516, EPI_ISL_2396517, EPI_ISL_2396518, EPI_ISL_2396519, EPI_ISL_2396520, EPI_ISL_2396521, EPI_ISL_2396522, EPI_ISL_2396524, EPI_ISL_2396525, EPI_ISL_2396527, EPI_ISL_2396528, EPI_ISL_2396529, EPI_ISL_2396531, EPI_ISL_2396532, EPI_ISL_2396534, EPI_ISL_2396535, EPI_ISL_2396536, EPI_ISL_2396540, EPI_ISL_2396543, EPI_ISL_2396544, EPI_ISL_2396545, EPI_ISL_2396546, EPI_ISL_2396549, EPI_ISL_2396550, EPI_ISL_2396553, EPI_ISL_2396554, EPI_ISL_2396555, EPI_ISL_2396557, EPI_ISL_2396558, EPI_ISL_2396559, EPI_ISL_2396560, EPI_ISL_2396561, EPI_ISL_2396562, EPI_ISL_2396563, EPI_ISL_2396567, EPI_ISL_2396568, EPI_ISL_2396569, EPI_ISL_2396570, EPI_ISL_2396573, EPI_ISL_2396574, EPI_ISL_2396575, EPI_ISL_2396576, EPI_ISL_2396577, EPI_ISL_2396578, EPI_ISL_2396580, EPI_ISL_2396581, EPI_ISL_2396582, EPI_ISL_2396583, EPI_ISL_2396585, EPI_ISL_2396588, EPI_ISL_2396589, EPI_ISL_2396590, EPI_ISL_2396592, EPI_ISL_2396593, EPI_ISL_2396594, EPI_ISL_2396597, EPI_ISL_2396598, EPI_ISL_2396600, EPI_ISL_2396602, EPI_ISL_2396603, EPI_ISL_2396604, EPI_ISL_2396605, EPI_ISL_2396606, EPI_ISL_2396607, EPI_ISL_2396608, EPI_ISL_2396609, EPI_ISL_2396611, EPI_ISL_2396612, EPI_ISL_2396613, EPI_ISL_2396616, EPI_ISL_2396617, EPI_ISL_2396618, EPI_ISL_2396619, EPI_ISL_2396620, EPI_ISL_2396622, EPI_ISL_2396623, EPI_ISL_2396624, EPI_ISL_2396626, EPI_ISL_2396628, EPI_ISL_2396629, EPI_ISL_2396631, EPI_ISL_2396633, EPI_ISL_2396634, EPI_ISL_2396635, EPI_ISL_2396636, EPI_ISL_2396637, EPI_ISL_2396638, EPI_ISL_2396639, EPI_ISL_2396642, EPI_ISL_2396643, EPI_ISL_2396644, EPI_ISL_2396646, EPI_ISL_2396647, EPI_ISL_2396648, EPI_ISL_2396649, EPI_ISL_2396650, EPI_ISL_2396651, EPI_ISL_2396652, EPI_ISL_2396653, EPI_ISL_2396654, EPI_ISL_2396655, EPI_ISL_2396656, EPI_ISL_2396657, EPI_ISL_2396658, EPI_ISL_2396659, EPI_ISL_2396660, EPI_ISL_2396662, EPI_ISL_2396663, EPI_ISL_2396664, EPI_ISL_2396665, EPI_ISL_2396666, EPI_ISL_2396667, EPI_ISL_2396668, EPI_ISL_2396669, EPI_ISL_2396670, EPI_ISL_2396671, EPI_ISL_2396675, EPI_ISL_2396677, EPI_ISL_2396679, EPI_ISL_2396680, EPI_ISL_2396681, EPI_ISL_2396682, EPI_ISL_2396683, EPI_ISL_2396684, EPI_ISL_2396686, EPI_ISL_2396687, EPI_ISL_2396688, EPI_ISL_2396689, EPI_ISL_2396690, EPI_ISL_2396691, EPI_ISL_2396695, EPI_ISL_2396696, EPI_ISL_2396697, EPI_ISL_2396698, EPI_ISL_2396700, EPI_ISL_2396701, EPI_ISL_2396702, EPI_ISL_2396703, EPI_ISL_2396704, EPI_ISL_2396705, EPI_ISL_2396706, EPI_ISL_2396708, EPI_ISL_2396709, EPI_ISL_2396710 | | | |
| see above | Lighthouse Lab in Glasgow | Wellcome Sanger Institute for the COVID-19 Genomics UK (COG-UK) Consortium | Harper VanSteenhouse, Yumi Kasai, David Gray, Carol Clugston, Anna Dominiczak and Alex Alderton, Roberto Amato, Jeffrey Barrett, Sonia Goncalves, Ewan Harrison, David K. Jackson, Ian Johnston, Dominic Kwiatkowski, Cordelia Langford, John Sillitoe on behalf of the Wellcome Sanger Institute COVID-19 Surveillance Team |
| EPI_ISL_2396712 | Lighthouse Laboratory Plymouth | Wellcome Sanger Institute for the COVID-19 Genomics UK (COG-UK) Consortium | Lighthouse Laboratory Plymouth and Alex Alderton, Roberto Amato, Jeffrey Barrett, Sonia Goncalves, Ewan Harrison, David K. Jackson, Ian Johnston, Dominic Kwiatkowski, Cordelia Langford, John Sillitoe on behalf of the Wellcome Sanger Institute COVID-19 Surveillance Team |
| EPI_ISL_2396713, EPI_ISL_2396714, EPI_ISL_2396715, EPI_ISL_2396719, EPI_ISL_2396720, EPI_ISL_2396722, EPI_ISL_2396723, EPI_ISL_2396725, EPI_ISL_2396726, EPI_ISL_2396727, EPI_ISL_2396728, EPI_ISL_2396729, EPI_ISL_2396730, EPI_ISL_2396734, EPI_ISL_2396735, EPI_ISL_2396737, EPI_ISL_2396738, EPI_ISL_2396740, EPI_ISL_2396741, EPI_ISL_2396742, EPI_ISL_2396743, EPI_ISL_2396744, EPI_ISL_2396745, EPI_ISL_2396746, EPI_ISL_2396747, EPI_ISL_2396749, EPI_ISL_2396751, EPI_ISL_2396752 | | | |
| see above | Lighthouse Lab in Glasgow | Wellcome Sanger Institute for the COVID-19 Genomics UK (COG-UK) Consortium | Harper VanSteenhouse, Yumi Kasai, David Gray, Carol Clugston, Anna Dominiczak and Alex Alderton, Roberto Amato, Jeffrey Barrett, Sonia Goncalves, Ewan Harrison, David K. Jackson, Ian Johnston, Dominic Kwiatkowski, Cordelia Langford, John Sillitoe on behalf of the Wellcome Sanger Institute COVID-19 Surveillance Team |
| EPI_ISL_2396753 | Lighthouse Lab in Milton Keynes | Wellcome Sanger Institute for the COVID-19 Genomics UK (COG-UK) Consortium | The Lighthouse Lab in Milton Keynes and Alex Alderton, Roberto Amato, Jeffrey Barrett, Sonia Goncalves, Ewan Harrison, David K. Jackson, Ian Johnston, Dominic Kwiatkowski, Cordelia Langford, John Sillitoe on behalf of the Wellcome Sanger Institute COVID-19 Surveillance Team |
| EPI_ISL_2396754, EPI_ISL_2396756, EPI_ISL_2396761, EPI_ISL_2396762, EPI_ISL_2396763, EPI_ISL_2396765, EPI_ISL_2396766, EPI_ISL_2396768, EPI_ISL_2396769, EPI_ISL_2396771, EPI_ISL_2396772, EPI_ISL_2396778, EPI_ISL_2396779, EPI_ISL_2396786, EPI_ISL_2396787, EPI_ISL_2396790, EPI_ISL_2396791, EPI_ISL_2396792, EPI_ISL_2396794, EPI_ISL_2396797 | | | |
| see above | Lighthouse Lab in Glasgow | Wellcome Sanger Institute for the COVID-19 Genomics UK (COG-UK) Consortium | Harper VanSteenhouse, Yumi Kasai, David Gray, Carol Clugston, Anna Dominiczak and Alex Alderton, Roberto Amato, Jeffrey Barrett, Sonia Goncalves, Ewan Harrison, David K. Jackson, Ian Johnston, Dominic Kwiatkowski, Cordelia Langford, John Sillitoe on behalf of the Wellcome Sanger Institute COVID-19 Surveillance Team |
| EPI_ISL_2396799 | Lighthouse Lab in Alderley Park | Wellcome Sanger Institute for the COVID-19 Genomics UK (COG-UK) Consortium | Jacquelyn Wynn, Mairead Hyland, The Lighthouse Lab in Alderley Park and Alex Alderton, Roberto Amato, Jeffrey Barrett, Sonia Goncalves, Ewan Harrison, David K. Jackson, Ian Johnston, Dominic Kwiatkowski, Cordelia Langford, John Sillitoe on behalf of the Wellcome Sanger Institute COVID-19 Surveillance Team |
| EPI_ISL_2396803, EPI_ISL_2396804 | Randox Laboratories | Wellcome Sanger Institute for the COVID-19 Genomics UK (COG-UK) Consortium | Randox Laboratories and Alex Alderton, Roberto Amato, Jeffrey Barrett, Sonia Goncalves, Ewan Harrison, David K. Jackson, Ian Johnston, Dominic Kwiatkowski, Cordelia Langford, John Sillitoe on behalf of the Wellcome Sanger Institute COVID-19 Surveillance Team |
| EPI_ISL_2396812, EPI_ISL_2396813, EPI_ISL_2396814, EPI_ISL_2396815, EPI_ISL_2396816 | Lighthouse Lab in Milton Keynes | Wellcome Sanger Institute for the COVID-19 Genomics UK (COG-UK) Consortium | The Lighthouse Lab in Milton Keynes and Alex Alderton, Roberto Amato, Jeffrey Barrett, Sonia Goncalves, Ewan Harrison, David K. Jackson, Ian Johnston, Dominic Kwiatkowski, Cordelia Langford, John Sillitoe on behalf of the Wellcome Sanger Institute COVID-19 Surveillance Team |
| EPI_ISL_2396828, EPI_ISL_2396834, EPI_ISL_2396838 | Randox Laboratories | Wellcome Sanger Institute for the COVID-19 Genomics UK (COG-UK) Consortium | Randox Laboratories and Alex Alderton, Roberto Amato, Jeffrey Barrett, Sonia Goncalves, Ewan Harrison, David K. Jackson, Ian Johnston, Dominic Kwiatkowski, Cordelia Langford, John Sillitoe on behalf of the Wellcome Sanger Institute COVID-19 Surveillance Team |
| EPI_ISL_2397179, EPI_ISL_2397185, EPI_ISL_2397200 | Klinisch Laboratorium ZNA | Klinisch Laboratorium ZNA | Verstrepen et al. |
| EPI_ISL_2397205, EPI_ISL_2397207 | Plateforme de testing Namuroise | Plateforme de testing Namuroise | Lesly Nyinkeu Kemamen; Nicolas Gilliard; Otto Gaetan ; Denis Olivier ; Degosserie Jonathan ; Mullier François |
| EPI_ISL_2397364, EPI_ISL_2397375, EPI_ISL_2397387, EPI_ISL_2397390, EPI_ISL_2397398, EPI_ISL_2397424, EPI_ISL_2397470, EPI_ISL_2397521, EPI_ISL_2397578, EPI_ISL_2397580, EPI_ISL_2397590, EPI_ISL_2397597, EPI_ISL_2397614, EPI_ISL_2397641, EPI_ISL_2397654, EPI_ISL_2397661, EPI_ISL_2397672, EPI_ISL_2397677, EPI_ISL_2397699, EPI_ISL_2397700, EPI_ISL_2397723, EPI_ISL_2397725, EPI_ISL_2397733, EPI_ISL_2397740, EPI_ISL_2397758, EPI_ISL_2397779 | | | |
| see above | Quest Diagnostics Incorporated | Centers for Disease Control and Prevention Division of Viral Diseases, Pathogen Discovery | Dakota Howard, Dhwani Batra, Peter W. Cook, Kara Moser, Adrian Paskey, Jason Caravas, Benjamin Rambo-Martin, Shatavia Morrison, Christopher Gulvick, Scott Sammons, Yvette Unoarumhi, Darlene Wagner, Matthew Schmerer, S. H. Rosenthal, A. Gerasimova, R. M. Kagan, B. Anderson, M. Hua, Y. Liu, L.E. Bernstein, K.E. Livingston, A. Perez, I. A. Shlyakhter, R. V. Rolando, R. Owen, P. Tanpaiboon, F. Lacbawan, Clinton R. Paden, Duncan MacCannell |
| EPI_ISL_2397814, EPI_ISL_2397929, EPI_ISL_2397930, EPI_ISL_2398004, EPI_ISL_2398018, EPI_ISL_2398033, EPI_ISL_2398049, EPI_ISL_2398087, EPI_ISL_2398122, EPI_ISL_2398135, EPI_ISL_2398154, EPI_ISL_2398179, EPI_ISL_2398206, EPI_ISL_2398210, EPI_ISL_2398265, EPI_ISL_2398267, EPI_ISL_2398272, EPI_ISL_2398284, EPI_ISL_2398353, EPI_ISL_2398356, EPI_ISL_2398360, EPI_ISL_2398432, EPI_ISL_2398435, EPI_ISL_2398471, EPI_ISL_2398475, EPI_ISL_2398510, EPI_ISL_2398529, EPI_ISL_2398535, EPI_ISL_2398556, EPI_ISL_2398560, EPI_ISL_2398561, EPI_ISL_2398566, EPI_ISL_2398569, EPI_ISL_2398571, EPI_ISL_2398661, EPI_ISL_2398662, EPI_ISL_2398663, EPI_ISL_2398664, EPI_ISL_2398694, EPI_ISL_2398738, EPI_ISL_2398834, EPI_ISL_2398836, EPI_ISL_2398837, EPI_ISL_2398883, EPI_ISL_2398900, EPI_ISL_2398923, EPI_ISL_2398952, EPI_ISL_2398987, EPI_ISL_2398990, EPI_ISL_2399078, EPI_ISL_2399106, EPI_ISL_2399109, EPI_ISL_2399120, EPI_ISL_2399131, EPI_ISL_2399158, EPI_ISL_2399160, EPI_ISL_2399161, EPI_ISL_2399205, EPI_ISL_2399232, EPI_ISL_2399291, EPI_ISL_2399347, EPI_ISL_2399406, EPI_ISL_2399416 | | | |
| see above | Laboratory Corporation of America | Centers for Disease Control and Prevention Division of Viral Diseases, Pathogen Discovery | Dakota Howard, Dhwani Batra, Peter W. Cook, Kara Moser, Adrian Paskey, Jason Caravas, Benjamin Rambo-Martin, Shatavia Morrison, Christopher Gulvick, Scott Sammons, Yvette Unoarumhi, Darlene Wagner, Matthew Schmerer, Minoo Agarwal, Eyad Almasri, Debbie Boles, Ayla Burns, Nuthawin Charoensri, Oren Cohen, Susan Countryman, Mary Ann Cristobal, Bobbi Croy, Suzanne Dale, Hrushikesh Deshmukh, Amanda Douglas, Vincent Drouillon, Marcia Eisenberg, Howard Engler, Rama Ghatti, Prashant Gupta, Susan Hicks, Jake Humphrey, Lax Iyer, Lisa Pfefferle, Manoj Jain, Matthew Robinson, Mohan Kolli, Brian Krueger, Tim Kuphal, Stanley Letovsky, Michael Levandoski, Craig Lukasik, Jonathan Meltzer, Brian Norvell, Mindy Nye, Scott Parker, Christos Petropoulos, John Pruitt, Steven Ragan, Scott Ryan, Mike Sapeta, Jana Schroth, Suresh Babu Selvaraju, Goran Stevovic, Amanda Suchanek, Andrea Throop, Lyndon Tilson, Thomas Urban, Joe Voshell, Kimberly Wagner, Jonathan Williams, Mary Williamson, Qian Zeng, Tricia Zwiefelhofer, Clinton R. Paden, Duncan MacCannell |
| EPI_ISL_2399427 | UT-Unified State Labs: Public Health Utah DOH | Centers for Disease Control and Prevention Division of Viral Diseases, Pathogen Discovery | Mili Sheth, Sarah Nobles, Jasmine Padilla, Mark Burroughs, Shoshona Le, Katie Dillon, Peter Cook, Clinton R. Paden, Dhwani Batra, Krista Queen, Kristen Knipe, Dakota Howard, Yvette Unoarumhi, Darlene Wagner, Matthew Schmerer, Ben L. Rambo-Martin, Kristine Lacek, Sam Shepard, Alison Laufer Halpin, Dave Wentworth, Vivien Dugan, Suxiang Tong, Justin Lee |
| EPI_ISL_2399438 | Molecular Diagnostics Pathology Department Mater Dei Hospital Malta | Molecular Diagnostics Pathology Department Mater Dei Hospital Malta | G Zahra, R Borg, C Cilia, L Grech, M Briffa |
| EPI_ISL_2399440, EPI_ISL_2399441, EPI_ISL_2399442, EPI_ISL_2399443, EPI_ISL_2399444, EPI_ISL_2399445, EPI_ISL_2399446, EPI_ISL_2399447, EPI_ISL_2399448, EPI_ISL_2399449, EPI_ISL_2399450, EPI_ISL_2399451, EPI_ISL_2399452, EPI_ISL_2399453, EPI_ISL_2399454, EPI_ISL_2399455, EPI_ISL_2399456 | | | |
| see above | Institute of Epidemiology, Disease Control and Research (IEDCR) | IEDCR-ideSHi-icddr,b | Hassan Afrad, Sadia Rahman, Manjur Hossain Khan, Firdausi Qadri, Tahmina Shirin |
| EPI_ISL_2399793, EPI_ISL_2399958, EPI_ISL_2399961, EPI_ISL_2399962, EPI_ISL_2399963, EPI_ISL_2399964, EPI_ISL_2399974, EPI_ISL_2399977, EPI_ISL_2399978, EPI_ISL_2399979, EPI_ISL_2399991 | | | |
| see above | National Virus Reference Laboratory | National Virus Reference Laboratory | Zoe Yandle, Charlene Bennett, Gabriel Gonzalez, Michael Carr, Jonathan Dean, Cillian F De Gascun |
| EPI_ISL_2400078, EPI_ISL_2400079 | Medical Microbiology Unit, Department for Laboratory Medicine, Drammen Hospital, Vestre Viken Health Trust, | Norwegian Institute of Public Health, Department of Virology | Kathrine Stene-Johansen, Kamilla Heddeland Instefjord, Hilde Elshaug, Garcia Llorente Ignacio, Jon Bråte, Line Victoria Moen, Engebretsen Serina Beate,Pedersen Benedikte Nevjen, Debech Nadia, Atiya R Ali,Marie Paulsen Madsen, Rasmus Riis Kopperud, Hilde Vollan, Karoline Bragstad, Olav Hungnes |
| EPI_ISL_2400135, EPI_ISL_2400138, EPI_ISL_2400139, EPI_ISL_2400143, EPI_ISL_2400144, EPI_ISL_2400148, EPI_ISL_2400149, EPI_ISL_2400151, EPI_ISL_2400153, EPI_ISL_2400154, EPI_ISL_2400156, EPI_ISL_2400157, EPI_ISL_2400158, EPI_ISL_2400159, EPI_ISL_2400160, EPI_ISL_2400161, EPI_ISL_2400166, EPI_ISL_2400167, EPI_ISL_2400170, EPI_ISL_2400171, EPI_ISL_2400177, EPI_ISL_2400178, EPI_ISL_2400181, EPI_ISL_2400182, EPI_ISL_2400183, EPI_ISL_2400186, EPI_ISL_2400188, EPI_ISL_2400198, EPI_ISL_2400199, EPI_ISL_2400201, EPI_ISL_2400204, EPI_ISL_2400205, EPI_ISL_2400206, EPI_ISL_2400208, EPI_ISL_2400212, EPI_ISL_2400213, EPI_ISL_2400215, EPI_ISL_2400220, EPI_ISL_2400221, EPI_ISL_2400222, EPI_ISL_2400223, EPI_ISL_2400225 | | | |
| see above | INSACOG-WB | National Institute of Biomedical Genomics – INSACOG | Arindam Maitra, Bhaswati Bandyopadhyay, Nidhan Kumar Biswas, Tamal Ghosh, Sreedhar Chinnaswamy, Ajay Chakraborti, Saumitra Das |
| EPI_ISL_2400233, EPI_ISL_2400235, EPI_ISL_2400239, EPI_ISL_2400240 | Wichita State University - Molecular Diagnostics Lab | Kansas Health and Environmental Lab | Mike Grose, Katherine Wiggins, Jonathan Barnell, Ben Olsen, and Phil Adam |
| EPI_ISL_2400293, | Kansas Health and Environmental Lab | Kansas Health and Environmental Lab | Mike Grose, Katherine Wiggins, Jonathan Barnell, Ben Olsen, and Phil Adam |

| | | | |
|---|---|---|---|
| EPI_ISL_2400297, EPI_ISL_2400298, EPI_ISL_2400299 | | | |
| EPI_ISL_2400304, EPI_ISL_2400305 | Mapmygenome | CSIR-Centre for Cellular and Molecular Biology - INSACOG | Sofia Banu, Lamuk Zaveri, Ara Sreenivas, Onkar Kulkarni, Shreekant Verma, Amareshwar Vodapalli , Viswagithe S L, B Himasri, Sharath Chandra Thota, Karthik Bharadwaj Tallapaka, Rakesh K Mishra,Divya Tej Sowpati |
| EPI_ISL_2400307 | Mapmygenome | CSIR-Centre for Cellular and Molecular Biology - INSACOG | Onkar Kulkarni, Sofia Banu, Lamuk Zaveri, Ara Sreenivas, Shreekant Verma, Amareshwar Vodapalli , Viswagithe S L, B Himasri, Sharath Chandra Thota, Karthik Bharadwaj Tallapaka, Rakesh K Mishra,Divya Tej Sowpati |
| EPI_ISL_2400311 | Mapmygenome | CSIR-Centre for Cellular and Molecular Biology - INSACOG | Ara Sreenivas, Onkar Kulkarni, Lamuk Zaveri, Sofia Banu, Shreekant Verma, Amareshwar Vodapalli , Viswagithe S L, B Himasri, Sharath Chandra Thota, Karthik Bharadwaj Tallapaka, Rakesh K Mishra,Divya Tej Sowpati |
| EPI_ISL_2400319 | CSIR-Centre for Cellular and Molecular Biology | CSIR-Centre for Cellular and Molecular Biology-INSACOG | Ara Sreenivas,Shreekant Verma,Tulasi Nagabandi, Lamuk Zaveri, Amareshwar Vodapalli ,Blessy B John,Viswagithe S L,B Himasri,Valli Nagalakshmi Undamatla,Payel Mukherjee,Sofia Banu,Archana Bharadwaj Siva,Sharath Chandra Thota,Karthik Bharadwaj Tallapaka,Rakesh K Mishra,Divya Tej Sowpati |
| EPI_ISL_2400320 | Banaras Hindu University | CSIR-Centre for Cellular and Molecular Biology-INSACOG | Surendra Pratap Mishra, Royana Singh,Shivani Mishra, Shivam Tiwari, Gunjan Rai, Umesh Choudhary, Maneesha Upadhyay,Lamuk Zaveri,Tulasi Nagabandi, Ara Sreenivas,Shreekant Verma, Amareshwar Vodapalli ,Valli Nagalakshmi Undamatla,Payel Mukherjee,Archana Bharadwaj Siva,Sharath Chandra Thota,Gyaneshwer Chaubey,Karthik Bharadwaj Tallapaka,Rakesh K Mishra,Divya Tej Sowpati |
| EPI_ISL_2400322 | Banaras Hindu University | CSIR-Centre for Cellular and Molecular Biology-INSACOG | Royana Singh, Saurabh Singh, Jaya Chakraborty, Priyoneel Basu, Chetan Sahni, Deepa Devadas, Saumya Singh, Manpreet Kaur, Nitish Kumar Singh, Abhay Kumar Yadav, Ajay Kumar Yadav, Ashish, Shani Vishwakarma,Payel Mukherjee,Lamuk Zaveri,Tulasi Nagabandi, ,Blessy B John,Viswagithe S L,B Himasri,Sofia Banu,Archana Bharadwaj Siva,Sharath Chandra Thota,Gyaneshwer Chaubey,Karthik Bharadwaj Tallapaka,Rakesh K Mishra,Divya Tej Sowpati |
| EPI_ISL_2400474, EPI_ISL_2400478, EPI_ISL_2400492 | Maryland Genomics, Institute for Genome Sciences, University of Maryland School of Medicine | Maryland Genomics, Institute for Genome Sciences, University of Maryland School of Medicine | Tallon, Luke J; Sadzewicz, Lisa D; Humphrys, Mike; Ott, Sandra; Roussey, Holly; Mehta, Aditya; Vavikolanu, Kranthi; Fraser, Claire M; Ravel, Jacques |
| EPI_ISL_2400520, EPI_ISL_2400521 | Indian Council of Medical Research-National Institute of Virology, Microbial Containment Complex | Indian Council of Medical Research-National Institute of Virology, Microbial Containment Complex | Pragya D Yadav |
| EPI_ISL_2400536 | Hôpitaux Robert Schuman | Laboratoire national de sante, Microbiology, Microbial Genomics Platform | Anke Wienecke-Baldacchino, Catherine Ragimbeau,Jessica Tapp, Fatu Djabi, Lise Pignon, Raoul Salmon, Alain Hakim, Nawfal Faik, Tamir Abdelrahman |
| EPI_ISL_2400730, EPI_ISL_2400820, EPI_ISL_2400821, EPI_ISL_2400824, EPI_ISL_2400827, EPI_ISL_2400828, EPI_ISL_2400829, EPI_ISL_2400830 | Laboratoires d'analyses medicales - Ketterthill | Laboratoire national de sante, Microbiology, Microbial Genomics Platform | Anke Wienecke-Baldacchino, Catherine Ragimbeau,Jessica Tapp, Fatu Djabi, Lise Pignon, Raoul Salmon, Serge Vedy, Caroline Scheiber, Tamir Abdelrahman |
| EPI_ISL_2400874, EPI_ISL_2400877, EPI_ISL_2400918, EPI_ISL_2400942, EPI_ISL_2400960 | Laboratoires Reunis | Laboratoire national de sante, Microbiology, Microbial Genomics Platform | Anke Wienecke-Baldacchino, Catherine Ragimbeau,Jessica Tapp, Fatu Djabi, Lise Pignon, Raoul Salmon, Bernard Weber, Tamir Abdelrahman |
| EPI_ISL_2401121, EPI_ISL_2401137, EPI_ISL_2401138, EPI_ISL_2401139, EPI_ISL_2401152, EPI_ISL_2401177 | Laboratoire national de sante, Microbiology, Virology | Laboratoire national de sante, Microbiology, Microbial Genomics Platform | Anke Wienecke-Baldacchino, Catherine Ragimbeau,Jessica Tapp, Fatu Djabi, Lise Pignon, Raoul Salmon, Trung Nguyen Nguyen, Tamir Abdelrahman |
| EPI_ISL_2401412, EPI_ISL_2401414, EPI_ISL_2401509, EPI_ISL_2401510, EPI_ISL_2401513, EPI_ISL_2401514, EPI_ISL_2401515, EPI_ISL_2401516, EPI_ISL_2401517, EPI_ISL_2401526, EPI_ISL_2401576, EPI_ISL_2401615, EPI_ISL_2401625, EPI_ISL_2401633, EPI_ISL_2401651, EPI_ISL_2401653, EPI_ISL_2401661, EPI_ISL_2401662, EPI_ISL_2401678, EPI_ISL_2401687, EPI_ISL_2401698, EPI_ISL_2401704, EPI_ISL_2401708, EPI_ISL_2401709, EPI_ISL_2401720, EPI_ISL_2401729, EPI_ISL_2401748, EPI_ISL_2401749 | | | |
| see above | BioneXt Lab | Laboratoire national de sante, Microbiology, Microbial Genomics Platform | Anke Wienecke-Baldacchino, Catherine Ragimbeau,Jessica Tapp, Fatu Djabi, Lise Pignon, Raoul Salmon, Thibault Ferrandon, Tamir Abdelrahman |
| EPI_ISL_2401863 | Hospital Center Emile Mayrisch | Laboratoire national de sante, Microbiology, Microbial Genomics Platform | Anke Wienecke-Baldacchino, Catherine Ragimbeau,Jessica Tapp, Fatu Djabi, Lise Pignon, Raoul Salmon, Cynthia Oxacelay, Tamir Abdelrahman |
| EPI_ISL_2401928 | Hospital Center Luxembourg | Laboratoire national de sante, Microbiology, Microbial Genomics Platform | Anke Wienecke-Baldacchino, Catherine Ragimbeau,Jessica Tapp, Fatu Djabi, Lise Pignon, Raoul Salmon, Michel Kohnen, Jean-Hugues Francois, Tamir Abdelrahman |
| EPI_ISL_2402378, EPI_ISL_2402390, EPI_ISL_2402399, EPI_ISL_2402443 | SYNLAB | GIGA Medical Genomics | Keith Durkin, Maria Artesi, Sébastien Bontems, Raphaël Boreux, Bouchra Boujemla, Nathalie Renotte, Cécile Meex, Pierrette Melin, Marie-Pierre Hayette, Vincent Bours |
| EPI_ISL_2402461 | New South Wales Health Pathology Royal Prince Alfred Hospital | Microbiology RPAH | Foster, C.; Au, J.; Ruiz Silva, M.; Deveson, I.; Bull, R.; Van Hal, S.; Rawlinson, W. |
| EPI_ISL_2402477, EPI_ISL_2402478 | Hospital General Universitario Gregorio Marañón | Hospital General Universitario Gregorio Marañón | Sergio Buenestado Serrano, Pedro Sola Campoy, Laura Pérez-Lago, Cristina Rodriguez-Grande, Marta Herranz Martin, Victor Manuel de la Cueva, Julia Suárez, Pilar Catalán, Patricia Muñoz, Darío García de Viedma |
| EPI_ISL_2402535, EPI_ISL_2402541 | OLVZ Aalst | OLVZ Aalst | Astrid Holderbeke |
| EPI_ISL_2402558 | The Caribbean Public Health Agency | Carrington Lab, Department of PreClinical Sciences, Faculty of Medical Sciences, The University of the West Indies | Nikita S. D. Sahadeo, Arianne Brown-Jordan, Sarah Hill, Vernie Ramkissoon, Aisha Andrewin, Naresh Nandram, Avery Hinds, Karla Georges, Risha Singh, SueMin Nathaniel, Nuno Faria, Oliver Pybus, Christopher Oura, Gabriel Escobar, Christine V. F. Carrington |
| EPI_ISL_2402575, EPI_ISL_2402577 | Minnesota Department of Health, Public Health Laboratory | Minnesota Department of Health, Public Health Laboratory | Alexandra Lorentz, Jacob Garfin, Matt Plumb, and Xiong Wang |
| EPI_ISL_2402819 | Weill Cornell Medicine | New York Genome Center | Michael Zody, Andre Corvelo, Dayna M. Oschwald, Samantha Fennessey, Tom Maniatis, Melissa Cushing, Olivier Elemento, Margaret Elizabeth Ross, Chris Mason, Priya Velu, Hanna Rennert, Arryn Craney, Lars F Westblade |
| EPI_ISL_2403108 | Illinois Department of Public Health - Springfield Lab | Illinois Department of Public Health - Springfield Lab | Bryan Sim, Gordon McCall |
| EPI_ISL_2403370 | Ministry of Health Turkey | Ministry of Health Turkey | Fatma Bayrakdar, Yasemin Cosgun, Suleyman Yalcin, Gulay Korukluoglu |
| EPI_ISL_2404730 | Wyoming Public Health Laboratory | Wyoming Public Health Laboratory | Jim Mildenberger, Taylor Fearing, Channing Weber, Ashley Norberg, Chayse Rowley, Marley Goetz, Brian Dominguez, Elliot Thomasson, Brittany Oher, Sam Britz, Cari Sloma, and Rob Christensen |
| EPI_ISL_2404806, EPI_ISL_2404815, EPI_ISL_2404816, EPI_ISL_2404820, EPI_ISL_2404824 | VMC | Santa Clara County public Health Laboratory | Santa Clara County Public Health Department |
| EPI_ISL_2404826 | GOOD SAMARITAN HOSPTIAL | Santa Clara County public Health Laboratory | Santa Clara County Public Health Department |
| EPI_ISL_2404827 | PAMF PALO ALTO | Santa Clara County public Health Laboratory | Santa Clara County Public Health Department |
| EPI_ISL_2404828, EPI_ISL_2404829, EPI_ISL_2404838 | VMC | Santa Clara County public Health Laboratory | Santa Clara County Public Health Department |
| EPI_ISL_2404984, EPI_ISL_2404986, EPI_ISL_2404990, EPI_ISL_2404991 | Area of Virology, Serology and Virology Division (SAViD), New South Wales Health Pathology Randwick | Virology Research Laboratory; Area of Virology, Serology and Virology Division (SAViD), New South Wales Health Pathology Randwick | Foster, C.; Au, J.; Ruiz Silva, M.; Deveson, I.; Bull, R.; Van Hal, S.; Rawlinson, W. |
| EPI_ISL_2405107 | Molecular Biology Laboratory, Westchester Medical Center | Genomics Core Laboratory, New York Medical College | Weihua Huang, Utsav Pandey, Jian Zhuge, Changhong Yin |
| EPI_ISL_2405109, EPI_ISL_2405110 | South Eastern Area Laboratory Services (SEALS) | NSW Health Pathology - Institute of Clinical Pathology and Medical Research; Westmead Hospital; University of Sydney | CIDM-PH et al. |
| EPI_ISL_2405111, EPI_ISL_2405112 | Sydney South West Pathology Service (SSWPS) - Royal Prince Alfred Hospital - NSW Health Pathology | NSW Health Pathology - Institute of Clinical Pathology and Medical Research; Westmead Hospital; University of Sydney | CIDM-PH et al. |
| EPI_ISL_2405113, EPI_ISL_2405114, EPI_ISL_2405115, EPI_ISL_2405116, EPI_ISL_2405117 | South Eastern Area Laboratory Services (SEALS) | NSW Health Pathology - Institute of Clinical Pathology and Medical Research; Westmead Hospital; University of Sydney | CIDM-PH et al. |
| EPI_ISL_2405118 | N.H.L Municipal Medical College, Ahmedabad | Gujarat Biotechnology Research Centre | Janvi Raval,Zarna Patel,Nitin Savaliya,Dinesh Kumar,Twinkle Soni,Sonal Sharma ,Zuber Saiyed,Ramesh Pandit,Jayshri Pethani,Umang Mishra,Nitesh Shah,Chaitanya Joshi,Madhvi Joshi |
| EPI_ISL_2405119 | N.H.L Municipal Medical College, Ahmedabad | Gujarat Biotechnology Research Centre | Zarna Patel,Nitin Savaliya,Dinesh Kumar,Twinkle Soni,Sonal Sharma ,Zuber Saiyed,Ramesh Pandit,Janvi Raval,Jayshri Pethani,Umang Mishra,Nitesh Shah,Chaitanya Joshi,Madhvi Joshi |
| EPI_ISL_2405120 | N.H.L Municipal Medical College, Ahmedabad | Gujarat Biotechnology Research Centre | Nitin Savaliya,Dinesh Kumar,Twinkle Soni,Sonal Sharma ,Zuber Saiyed,Ramesh Pandit,Janvi Raval,Zarna Patel,Jayshri Pethani,Umang Mishra,Nitesh Shah,Chaitanya Joshi,Madhvi Joshi |
| EPI_ISL_2405121 | N.H.L Municipal Medical College, Ahmedabad | Gujarat Biotechnology Research Centre | Dinesh Kumar,Twinkle Soni,Sonal Sharma ,Zuber Saiyed,Ramesh Pandit,Janvi Raval,Zarna Patel,Nitin Savaliya,Jayshri Pethani,Umang Mishra,Nitesh Shah,Chaitanya Joshi,Madhvi Joshi |

| | | | |
|---|---|---|---|
| EPI_ISL_2405122 | N.H.L Municipal Medical College, Ahmedabad | Gujarat Biotechnology Research Centre | Twinkle Soni,Sonal Sharma ,Zuber Saiyed,Ramesh Pandit,Janvi Raval,Zarna Patel,Nitin Savaliya,Dinesh Kumar,Jayshri Pethani,Umang Mishra,Nitesh Shah,Chaitanya Joshi,Madhvi Joshi |
| EPI_ISL_2405123 | N.H.L Municipal Medical College, Ahmedabad | Gujarat Biotechnology Research Centre | Sonal Sharma ,Zuber Saiyed,Ramesh Pandit,Janvi Raval,Zarna Patel,Nitin Savaliya,Dinesh Kumar,Twinkle Soni,Jayshri Pethani,Umang Mishra,Nitesh Shah,Chaitanya Joshi,Madhvi Joshi |
| EPI_ISL_2405125 | GMERS Government Medical College (GMC), Gotri | Gujarat Biotechnology Research Centre | Ramesh Pandit,Janvi Raval,Zarna Patel,Nitin Savaliya,Dinesh Kumar,Twinkle Soni,Sonal Sharma ,Zuber Saiyed,Bithika Duttaroy,Umang Mishra,Nitesh Shah,Chaitanya Joshi,Madhvi Joshi |
| EPI_ISL_2405126 | GMERS Government Medical College (GMC), Gotri | Gujarat Biotechnology Research Centre | Janvi Raval,Zarna Patel,Nitin Savaliya,Dinesh Kumar,Twinkle Soni,Sonal Sharma ,Zuber Saiyed,Ramesh Pandit,Bithika Duttaroy,Umang Mishra,Nitesh Shah,Chaitanya Joshi,Madhvi Joshi |
| EPI_ISL_2405127 | BJ Medical College and Civil Hospital, Ahmedabad | Gujarat Biotechnology Research Centre | Zarna Patel,Nitin Savaliya,Dinesh Kumar,Twinkle Soni,Sonal Sharma ,Zuber Saiyed,Ramesh Pandit,Janvi Raval,Pranay Shah,Umang Mishra,Nitesh Shah,Chaitanya Joshi,Madhvi Joshi |
| EPI_ISL_2405128 | BJ Medical College and Civil Hospital, Ahmedabad | Gujarat Biotechnology Research Centre | Nitin Savaliya,Dinesh Kumar,Twinkle Soni,Sonal Sharma ,Zuber Saiyed,Ramesh Pandit,Janvi Raval,Zarna Patel,Pranay Shah,Umang Mishra,Nitesh Shah,Chaitanya Joshi,Madhvi Joshi |
| EPI_ISL_2405129 | BJ Medical College and Civil Hospital, Ahmedabad | Gujarat Biotechnology Research Centre | Dinesh Kumar,Twinkle Soni,Sonal Sharma ,Zuber Saiyed,Ramesh Pandit,Janvi Raval,Zarna Patel,Nitin Savaliya,Pranay Shah,Umang Mishra,Nitesh Shah,Chaitanya Joshi,Madhvi Joshi |
| EPI_ISL_2405130 | BJ Medical College and Civil Hospital, Ahmedabad | Gujarat Biotechnology Research Centre | Twinkle Soni,Sonal Sharma ,Zuber Saiyed,Ramesh Pandit,Janvi Raval,Zarna Patel,Nitin Savaliya,Dinesh Kumar,Pranay Shah,Umang Mishra,Nitesh Shah,Chaitanya Joshi,Madhvi Joshi |
| EPI_ISL_2405131 | BJ Medical College and Civil Hospital, Ahmedabad | Gujarat Biotechnology Research Centre | Sonal Sharma ,Zuber Saiyed,Ramesh Pandit,Janvi Raval,Zarna Patel,Nitin Savaliya,Dinesh Kumar,Twinkle Soni,Pranay Shah,Umang Mishra,Nitesh Shah,Chaitanya Joshi,Madhvi Joshi |
| EPI_ISL_2405132 | BJ Medical College and Civil Hospital, Ahmedabad | Gujarat Biotechnology Research Centre | Zuber Saiyed,Ramesh Pandit,Janvi Raval,Zarna Patel,Nitin Savaliya,Dinesh Kumar,Twinkle Soni,Sonal Sharma ,Pranay Shah,Umang Mishra,Nitesh Shah,Chaitanya Joshi,Madhvi Joshi |
| EPI_ISL_2405133 | BJ Medical College and Civil Hospital, Ahmedabad | Gujarat Biotechnology Research Centre | Ramesh Pandit,Janvi Raval,Zarna Patel,Nitin Savaliya,Dinesh Kumar,Twinkle Soni,Sonal Sharma ,Zuber Saiyed,Pranay Shah,Umang Mishra,Nitesh Shah,Chaitanya Joshi,Madhvi Joshi |
| EPI_ISL_2405134 | BJ Medical College and Civil Hospital, Ahmedabad | Gujarat Biotechnology Research Centre | Janvi Raval,Zarna Patel,Nitin Savaliya,Dinesh Kumar,Twinkle Soni,Sonal Sharma ,Zuber Saiyed,Ramesh Pandit,Pranay Shah,Umang Mishra,Nitesh Shah,Chaitanya Joshi,Madhvi Joshi |
| EPI_ISL_2405135 | BJ Medical College and Civil Hospital, Ahmedabad | Gujarat Biotechnology Research Centre | Zarna Patel,Nitin Savaliya,Dinesh Kumar,Twinkle Soni,Sonal Sharma ,Zuber Saiyed,Ramesh Pandit,Janvi Raval,Pranay Shah,Umang Mishra,Nitesh Shah,Chaitanya Joshi,Madhvi Joshi |
| EPI_ISL_2405136 | BJ Medical College and Civil Hospital, Ahmedabad | Gujarat Biotechnology Research Centre | Nitin Savaliya,Dinesh Kumar,Twinkle Soni,Sonal Sharma ,Zuber Saiyed,Ramesh Pandit,Janvi Raval,Zarna Patel,Pranay Shah,Umang Mishra,Nitesh Shah,Chaitanya Joshi,Madhvi Joshi |
| EPI_ISL_2405137 | BJ Medical College and Civil Hospital, Ahmedabad | Gujarat Biotechnology Research Centre | Dinesh Kumar,Twinkle Soni,Sonal Sharma ,Zuber Saiyed,Ramesh Pandit,Janvi Raval,Zarna Patel,Nitin Savaliya,Pranay Shah,Umang Mishra,Nitesh Shah,Chaitanya Joshi,Madhvi Joshi |
| EPI_ISL_2405138 | BJ Medical College and Civil Hospital, Ahmedabad | Gujarat Biotechnology Research Centre | Twinkle Soni,Sonal Sharma ,Zuber Saiyed,Ramesh Pandit,Janvi Raval,Zarna Patel,Nitin Savaliya,Dinesh Kumar,Pranay Shah,Umang Mishra,Nitesh Shah,Chaitanya Joshi,Madhvi Joshi |
| EPI_ISL_2405139 | BJ Medical College and Civil Hospital, Ahmedabad | Gujarat Biotechnology Research Centre | Sonal Sharma ,Zuber Saiyed,Ramesh Pandit,Janvi Raval,Zarna Patel,Nitin Savaliya,Dinesh Kumar,Twinkle Soni,Pranay Shah,Umang Mishra,Nitesh Shah,Chaitanya Joshi,Madhvi Joshi |
| EPI_ISL_2405140 | BJ Medical College and Civil Hospital, Ahmedabad | Gujarat Biotechnology Research Centre | Zuber Saiyed,Ramesh Pandit,Janvi Raval,Zarna Patel,Nitin Savaliya,Dinesh Kumar,Twinkle Soni,Sonal Sharma ,Pranay Shah,Umang Mishra,Nitesh Shah,Chaitanya Joshi,Madhvi Joshi |
| EPI_ISL_2405141 | BJ Medical College and Civil Hospital, Ahmedabad | Gujarat Biotechnology Research Centre | Ramesh Pandit,Janvi Raval,Zarna Patel,Nitin Savaliya,Dinesh Kumar,Twinkle Soni,Sonal Sharma ,Zuber Saiyed,Pranay Shah,Umang Mishra,Nitesh Shah,Chaitanya Joshi,Madhvi Joshi |
| EPI_ISL_2405142 | BJ Medical College and Civil Hospital, Ahmedabad | Gujarat Biotechnology Research Centre | Janvi Raval,Zarna Patel,Nitin Savaliya,Dinesh Kumar,Twinkle Soni,Sonal Sharma ,Zuber Saiyed,Ramesh Pandit,Pranay Shah,Umang Mishra,Nitesh Shah,Chaitanya Joshi,Madhvi Joshi |
| EPI_ISL_2405143 | BJ Medical College and Civil Hospital, Ahmedabad | Gujarat Biotechnology Research Centre | Zarna Patel,Nitin Savaliya,Dinesh Kumar,Twinkle Soni,Sonal Sharma ,Zuber Saiyed,Ramesh Pandit,Janvi Raval,Pranay Shah,Umang Mishra,Nitesh Shah,Chaitanya Joshi,Madhvi Joshi |
| EPI_ISL_2405144 | BJ Medical College and Civil Hospital, Ahmedabad | Gujarat Biotechnology Research Centre | Nitin Savaliya,Dinesh Kumar,Twinkle Soni,Sonal Sharma ,Zuber Saiyed,Ramesh Pandit,Janvi Raval,Zarna Patel,Pranay Shah,Umang Mishra,Nitesh Shah,Chaitanya Joshi,Madhvi Joshi |
| EPI_ISL_2405209, EPI_ISL_2405210 | Flow Diagnostics Lab | Los Angeles County Public Health Laboratories | P. Hemarajata et al. |
| EPI_ISL_2405251, EPI_ISL_2405325 | University Hospitals of Geneva, Laboratory of Virology | HUG, Laboratory of Virology and the Health2030 Genome Center | Samuel Cordey, Ana Rita Goncalves, Laurent Kaiser, Lorenzo Cerutti, Henri Pegeot, Melyssa Elies, Deborah Penet, Keith Harshman, Ioannis Xenarios, Emmanouil Dermitzakis |
| EPI_ISL_2405344, EPI_ISL_2405356 | Microbiological Diagnostic Unit - Public Health Laboratory (MDU-PHL) | MDU-PHL | Seemann T., Sait, M.L., Sherry, N.L. |
| EPI_ISL_2405621, EPI_ISL_2405646, EPI_ISL_2405815, EPI_ISL_2405912, EPI_ISL_2405937, EPI_ISL_2406370 | Dutch COVID-19 response team | National Institute for Public Health and the Environment (RIVM) | Adam Meijer, Harry Vennema, Dirk Eggink, Jeroen Cremer, Sharon van den Brink, Bas van der Veer, AnneMarie van den Brandt, Lisa Wijsman, Kim Freriks, Ryanne Jaarsma, Euníce Then, Lynn Aarts, Sanne Bos, Melissa van Tuil, Linda van de Nes, Sjoerd Kuiling, James Groot, Florian Zwagemaker, Dennis Schmitz, Annelies Kroneman, Karim Hajji, Chantal Reusken, on behalf of the national COVID-19 response team |
| EPI_ISL_2406459, EPI_ISL_2406460 | Battambang Provincial Laboratory | Virology Unit, Institut Pasteur du Cambodge | Cecile Troupin, Leakhena Pum, Jurre Y Siegers, Ly Sovann, Kraing Sidonn, Yi Sengdoeurn, Chin Savuth, Chau Darapheak, Veasna Duong, Erik A Karlsson |
| EPI_ISL_2406461 | Oddar Meanchey Rapid Response Team | Virology Unit, Institut Pasteur du Cambodge | Cecile Troupin, Leakhena Pum, Jurre Y Siegers, Ly Sovann, Kraing Sidonn, Yi Sengdoeurn, Chin Savuth, Chau Darapheak, Veasna Duong, Erik A Karlsson |
| EPI_ISL_2406490, EPI_ISL_2406495 | LabPLUS | Institute of Environmental Science and Research (ESR) | Rachel Boyle, SallyAnn Harbison, Olivia Stroeven, Xiaoyun Ren, Matt Storey, Nikki Freed, Muhammad Faisal, Jing Wang, Hermes Perez, Anja Werno, Antje van der Linden, Arlo Upton, Chris Mansell, David Hammer, Dragana Drinkovic, Gary McAuliffe, Hana Sofia Andersson, James Ussher, Jill Sherwood, Josh Freeman, Julia Howard, Juliet Elvy, Mary DeAlmeida, Matt Blakiston, Matthew Rogers, Max Bloomfield, Michael Addidle, Michelle Balm, Sally Roberts, Sarah Jefferies, Sharmini Muttaiyah, Susan Morpeth, Susan Taylor, Timothy Blackmore, Vani Sathyendran, Veronica Playle, Virginia Hope, Erasmus Smit, Lauren Jelly, Olin Silander, Joep de Ligt |
| EPI_ISL_2406506 | Laboratoire Virologie Saint Louis APHP | Laboratoire Virologie Saint Louis APHP | Maud Salmona, Marie Laure Chaix, Severine Mercier Delarue, Marie Laure Néré, Linda Feghoul, Jérome Le Goff, Constance Delauguerre, Sophia Achaibou |
| EPI_ISL_2406590, EPI_ISL_2406591, EPI_ISL_2406594, EPI_ISL_2406595, EPI_ISL_2406596, EPI_ISL_2406597, EPI_ISL_2406598, EPI_ISL_2406599, EPI_ISL_2406600, EPI_ISL_2406602, EPI_ISL_2406606, EPI_ISL_2406607, EPI_ISL_2406608, EPI_ISL_2406609, EPI_ISL_2406611, EPI_ISL_2406613, EPI_ISL_2406614, EPI_ISL_2406615, EPI_ISL_2406618, EPI_ISL_2406619 | | | |
| see above | Lighthouse Lab in Glasgow | Wellcome Sanger Institute for the COVID-19 Genomics UK (COG-UK) Consortium | Harper VanSteenhouse, Yumi Kasai, David Gray, Carol Clugston, Anna Dominiczak and Alex Alderton, Roberto Amato, Jeffrey Barrett, Sonia Goncalves, Ewan Harrison, David K. Jackson, Ian Johnston, Dominic Kwiatkowski, Cordelia Langford, John Sillitoe on behalf of the Wellcome Sanger Institute COVID-19 Surveillance Team |
| EPI_ISL_2406622 | Lighthouse Lab in Milton Keynes | Wellcome Sanger Institute for the COVID-19 Genomics UK (COG-UK) Consortium | The Lighthouse Lab in Milton Keynes and Alex Alderton, Roberto Amato, Jeffrey Barrett, Sonia Goncalves, Ewan Harrison, David K. Jackson, Ian Johnston, Dominic Kwiatkowski, Cordelia Langford, John Sillitoe on behalf of the Wellcome Sanger Institute COVID-19 Surveillance Team |
| EPI_ISL_2406623, EPI_ISL_2406624, EPI_ISL_2406625, EPI_ISL_2406626, EPI_ISL_2406627, EPI_ISL_2406631, EPI_ISL_2406635, EPI_ISL_2406637, EPI_ISL_2406638, EPI_ISL_2406639, EPI_ISL_2406641, EPI_ISL_2406642, EPI_ISL_2406645, EPI_ISL_2406646, EPI_ISL_2406648, EPI_ISL_2406651, EPI_ISL_2406653, EPI_ISL_2406654, EPI_ISL_2406656, EPI_ISL_2406658, EPI_ISL_2406659, EPI_ISL_2406661, EPI_ISL_2406662, EPI_ISL_2406663, EPI_ISL_2406664, EPI_ISL_2406665, EPI_ISL_2406666, EPI_ISL_2406667, EPI_ISL_2406669, EPI_ISL_2406670, EPI_ISL_2406672, EPI_ISL_2406675, EPI_ISL_2406679, EPI_ISL_2406681, EPI_ISL_2406686, EPI_ISL_2406688, EPI_ISL_2406689, EPI_ISL_2406690, EPI_ISL_2406691, EPI_ISL_2406693, EPI_ISL_2406694, EPI_ISL_2406695, EPI_ISL_2406697, EPI_ISL_2406699, EPI_ISL_2406700, EPI_ISL_2406702, EPI_ISL_2406706, EPI_ISL_2406707, EPI_ISL_2406709, EPI_ISL_2406711, EPI_ISL_2406716, EPI_ISL_2406717, EPI_ISL_2406719, EPI_ISL_2406720, EPI_ISL_2406721, EPI_ISL_2406722, EPI_ISL_2406723, EPI_ISL_2406724, EPI_ISL_2406725, EPI_ISL_2406727, EPI_ISL_2406729, EPI_ISL_2406730, EPI_ISL_2406731, EPI_ISL_2406732, EPI_ISL_2406736, EPI_ISL_2406738, EPI_ISL_2406739, EPI_ISL_2406740, EPI_ISL_2406742, EPI_ISL_2406743, EPI_ISL_2406744, EPI_ISL_2406745 | | | |
| see above | Lighthouse Lab in Glasgow | Wellcome Sanger Institute for the COVID-19 Genomics UK (COG-UK) Consortium | Harper VanSteenhouse, Yumi Kasai, David Gray, Carol Clugston, Anna Dominiczak and Alex Alderton, Roberto Amato, Jeffrey Barrett, Sonia Goncalves, Ewan Harrison, David K. Jackson, Ian Johnston, Dominic Kwiatkowski, Cordelia Langford, John Sillitoe on behalf of the Wellcome Sanger Institute COVID-19 Surveillance Team |
| EPI_ISL_2406746 | Lighthouse Lab in Milton Keynes | Wellcome Sanger Institute for the COVID-19 Genomics UK (COG-UK) Consortium | The Lighthouse Lab in Milton Keynes and Alex Alderton, Roberto Amato, Jeffrey Barrett, Sonia Goncalves, Ewan Harrison, David K. Jackson, Ian Johnston, Dominic Kwiatkowski, Cordelia Langford, John Sillitoe on behalf of the Wellcome Sanger Institute COVID-19 Surveillance Team |
| EPI_ISL_2406748, EPI_ISL_2406749, EPI_ISL_2406750, EPI_ISL_2406751, EPI_ISL_2406752, EPI_ISL_2406753, EPI_ISL_2406754, EPI_ISL_2406757, EPI_ISL_2406758, EPI_ISL_2406760, EPI_ISL_2406762, EPI_ISL_2406767, EPI_ISL_2406769, EPI_ISL_2406771, EPI_ISL_2406772, EPI_ISL_2406773, EPI_ISL_2406774, EPI_ISL_2406775, EPI_ISL_2406777, EPI_ISL_2406778, EPI_ISL_2406779, EPI_ISL_2406781, EPI_ISL_2406782, EPI_ISL_2406783, EPI_ISL_2406784, EPI_ISL_2406786, EPI_ISL_2406788, EPI_ISL_2406790, EPI_ISL_2406793, EPI_ISL_2406794, EPI_ISL_2406795, EPI_ISL_2406796, EPI_ISL_2406797, EPI_ISL_2406798, EPI_ISL_2406799, EPI_ISL_2406801, EPI_ISL_2406802, EPI_ISL_2406803, EPI_ISL_2406805, EPI_ISL_2406806, EPI_ISL_2406807, EPI_ISL_2406808, EPI_ISL_2406810, EPI_ISL_2406813, EPI_ISL_2406814, EPI_ISL_2406815, EPI_ISL_2406816, EPI_ISL_2406817, EPI_ISL_2406818, EPI_ISL_2406819, EPI_ISL_2406820, EPI_ISL_2406821, EPI_ISL_2406823, EPI_ISL_2406824, EPI_ISL_2406826, EPI_ISL_2406827, EPI_ISL_2406828, EPI_ISL_2406829, EPI_ISL_2406830, EPI_ISL_2406834, EPI_ISL_2406836, EPI_ISL_2406837, EPI_ISL_2406838, EPI_ISL_2406839, EPI_ISL_2406840, EPI_ISL_2406843, EPI_ISL_2406845, EPI_ISL_2406846, EPI_ISL_2406847, EPI_ISL_2406849, EPI_ISL_2406850, EPI_ISL_2406851, EPI_ISL_2406852 | | | |
| see above | Lighthouse Lab in Glasgow | Wellcome Sanger Institute for the COVID-19 Genomics UK (COG-UK) Consortium | Harper VanSteenhouse, Yumi Kasai, David Gray, Carol Clugston, Anna Dominiczak and Alex Alderton, Roberto Amato, Jeffrey Barrett, Sonia Goncalves, Ewan Harrison, David K. Jackson, Ian Johnston, Dominic Kwiatkowski, Cordelia Langford, John Sillitoe on behalf of the Wellcome Sanger Institute COVID-19 Surveillance Team |
| EPI_ISL_2406857 | Lighthouse Lab in Milton Keynes | Wellcome Sanger Institute for the COVID-19 Genomics UK (COG-UK) Consortium | The Lighthouse Lab in Milton Keynes and Alex Alderton, Roberto Amato, Jeffrey Barrett, Sonia Goncalves, Ewan Harrison, David K. Jackson, Ian Johnston, Dominic Kwiatkowski, Cordelia Langford, John Sillitoe on behalf of the Wellcome Sanger Institute COVID-19 Surveillance Team |
| EPI_ISL_2406860, EPI_ISL_2406862, EPI_ISL_2406864, EPI_ISL_2406865, EPI_ISL_2406866, EPI_ISL_2406867, EPI_ISL_2406870, EPI_ISL_2406872, EPI_ISL_2406874, EPI_ISL_2406875, EPI_ISL_2406876, EPI_ISL_2406877, EPI_ISL_2406880, EPI_ISL_2406881, EPI_ISL_2406883, EPI_ISL_2406886, EPI_ISL_2406887, EPI_ISL_2406888, EPI_ISL_2406889, EPI_ISL_2406894 | | | |
| see above | Lighthouse Lab in Glasgow | Wellcome Sanger Institute for the COVID-19 Genomics UK (COG-UK) Consortium | Harper VanSteenhouse, Yumi Kasai, David Gray, Carol Clugston, Anna Dominiczak and Alex Alderton, Roberto Amato, Jeffrey Barrett, Sonia Goncalves, Ewan Harrison, David K. Jackson, Ian Johnston, Dominic Kwiatkowski, Cordelia Langford, John Sillitoe on behalf of the Wellcome Sanger Institute COVID-19 Surveillance Team |
| EPI_ISL_2406895 | Lighthouse Lab in Alderley Park | Wellcome Sanger Institute for the COVID-19 Genomics UK (COG-UK) Consortium | Jacquelyn Wynn, Mairead Hyland, The Lighthouse Lab in Alderley Park and Alex Alderton, Roberto Amato, Jeffrey Barrett, Sonia Goncalves, Ewan Harrison, David K. Jackson, Ian Johnston, Dominic Kwiatkowski, Cordelia Langford, John Sillitoe on behalf of the Wellcome Sanger Institute COVID-19 Surveillance Team |
| EPI_ISL_2406896 | Lighthouse Lab in Milton Keynes | Wellcome Sanger Institute for the COVID-19 Genomics UK (COG-UK) Consortium | The Lighthouse Lab in Milton Keynes and Alex Alderton, Roberto Amato, Jeffrey Barrett, Sonia Goncalves, Ewan Harrison, David K. Jackson, Ian Johnston, Dominic Kwiatkowski, Cordelia Langford, John Sillitoe on behalf of the Wellcome Sanger Institute COVID-19 Surveillance Team |
| EPI_ISL_2406897 | Lighthouse Lab in Alderley Park | Wellcome Sanger Institute for the COVID-19 Genomics UK (COG-UK) Consortium | Jacquelyn Wynn, Mairead Hyland, The Lighthouse Lab in Alderley Park and Alex Alderton, Roberto Amato, Jeffrey Barrett, Sonia Goncalves, Ewan Harrison, David K. Jackson, Ian Johnston, Dominic Kwiatkowski, Cordelia Langford, John Sillitoe on behalf of the Wellcome Sanger Institute COVID-19 Surveillance Team |
| EPI_ISL_2406898 | Lighthouse Lab in Glasgow | Wellcome Sanger Institute for the COVID-19 Genomics UK (COG-UK) Consortium | Harper VanSteenhouse, Yumi Kasai, David Gray, Carol Clugston, Anna Dominiczak and Alex Alderton, Roberto Amato, Jeffrey Barrett, Sonia Goncalves, Ewan Harrison, David K. Jackson, Ian Johnston, Dominic Kwiatkowski, Cordelia Langford, John Sillitoe on behalf of the Wellcome Sanger Institute COVID-19 Surveillance Team |
| EPI_ISL_2406899, EPI_ISL_2406900 | Lighthouse Lab in Milton Keynes | Wellcome Sanger Institute for the COVID-19 Genomics UK (COG-UK) Consortium | The Lighthouse Lab in Milton Keynes and Alex Alderton, Roberto Amato, Jeffrey Barrett, Sonia Goncalves, Ewan Harrison, David K. Jackson, Ian Johnston, Dominic Kwiatkowski, Cordelia Langford, John Sillitoe on behalf of the Wellcome Sanger Institute COVID-19 Surveillance Team |
| EPI_ISL_2406901, EPI_ISL_2406902, EPI_ISL_2406903 | Lighthouse Lab in Alderley Park | Wellcome Sanger Institute for the COVID-19 Genomics UK (COG-UK) Consortium | Jacquelyn Wynn, Mairead Hyland, The Lighthouse Lab in Alderley Park and Alex Alderton, Roberto Amato, Jeffrey Barrett, Sonia Goncalves, Ewan Harrison, David K. Jackson, Ian Johnston, Dominic Kwiatkowski, Cordelia Langford, John Sillitoe on behalf of the Wellcome Sanger Institute COVID-19 Surveillance Team |
| EPI_ISL_2406904, EPI_ISL_2406905 | Lighthouse Lab in Milton Keynes | Wellcome Sanger Institute for the COVID-19 Genomics UK (COG-UK) Consortium | The Lighthouse Lab in Milton Keynes and Alex Alderton, Roberto Amato, Jeffrey Barrett, Sonia Goncalves, Ewan Harrison, David K. Jackson, Ian Johnston, Dominic Kwiatkowski, Cordelia Langford, John Sillitoe on behalf of the Wellcome Sanger Institute COVID-19 Surveillance Team |
| EPI_ISL_2406906 | Lighthouse Lab in Alderley Park | Wellcome Sanger Institute for the COVID-19 Genomics UK (COG-UK) Consortium | Jacquelyn Wynn, Mairead Hyland, The Lighthouse Lab in Alderley Park and Alex Alderton, Roberto Amato, Jeffrey Barrett, Sonia Goncalves, Ewan Harrison, David K. Jackson, Ian Johnston, Dominic Kwiatkowski, Cordelia Langford, John Sillitoe on behalf of the Wellcome Sanger Institute COVID-19 Surveillance Team |
| EPI_ISL_2406907 | Lighthouse Lab in Milton Keynes | Wellcome Sanger Institute for the COVID-19 Genomics UK (COG-UK) Consortium | The Lighthouse Lab in Milton Keynes and Alex Alderton, Roberto Amato, Jeffrey Barrett, Sonia Goncalves, Ewan Harrison, David K. Jackson, Ian Johnston, Dominic Kwiatkowski, Cordelia Langford, John Sillitoe on behalf of the Wellcome Sanger Institute COVID-19 Surveillance Team |
| EPI_ISL_2406908 | Lighthouse Lab in Alderley Park | Wellcome Sanger Institute for the COVID-19 Genomics UK (COG-UK) Consortium | Jacquelyn Wynn, Mairead Hyland, The Lighthouse Lab in Alderley Park and Alex Alderton, Roberto Amato, Jeffrey Barrett, Sonia Goncalves, Ewan Harrison, David K. Jackson, Ian Johnston, Dominic Kwiatkowski, Cordelia Langford, John Sillitoe on behalf of the Wellcome Sanger Institute COVID-19 Surveillance Team |
| EPI_ISL_2406909 | Lighthouse Lab in Glasgow | Wellcome Sanger Institute for the COVID-19 Genomics UK (COG-UK) Consortium | Harper VanSteenhouse, Yumi Kasai, David Gray, Carol Clugston, Anna Dominiczak and Alex Alderton, Roberto Amato, Jeffrey Barrett, Sonia Goncalves, Ewan Harrison, David K. Jackson, Ian Johnston, Dominic Kwiatkowski, Cordelia Langford, John Sillitoe on behalf of the Wellcome Sanger Institute COVID-19 Surveillance Team |

| | | | |
|---|---|---|---|
| EPI_ISL_2406911 | Lighthouse Lab in Alderley Park | Wellcome Sanger Institute for the COVID-19 Genomics UK (COG-UK) Consortium | Jacquelyn Wynn, Mairead Hyland, The Lighthouse Lab in Alderley Park and Alex Alderton, Roberto Amato, Jeffrey Barrett, Sonia Goncalves, Ewan Harrison, David K. Jackson, Ian Johnston, Dominic Kwiatkowski, Cordelia Langford, John Sillitoe on behalf of the Wellcome Sanger Institute COVID-19 Surveillance Team |
| EPI_ISL_2406912 | Lighthouse Lab in Glasgow | Wellcome Sanger Institute for the COVID-19 Genomics UK (COG-UK) Consortium | Harper VanSteenhouse, Yumi Kasai, David Gray, Carol Clugston, Anna Dominiczak and Alex Alderton, Roberto Amato, Jeffrey Barrett, Sonia Goncalves, Ewan Harrison, David K. Jackson, Ian Johnston, Dominic Kwiatkowski, Cordelia Langford, John Sillitoe on behalf of the Wellcome Sanger Institute COVID-19 Surveillance Team |
| EPI_ISL_2406913 | Lighthouse Lab in Milton Keynes | Wellcome Sanger Institute for the COVID-19 Genomics UK (COG-UK) Consortium | The Lighthouse Lab in Milton Keynes and Alex Alderton, Roberto Amato, Jeffrey Barrett, Sonia Goncalves, Ewan Harrison, David K. Jackson, Ian Johnston, Dominic Kwiatkowski, Cordelia Langford, John Sillitoe on behalf of the Wellcome Sanger Institute COVID-19 Surveillance Team |
| EPI_ISL_2406914, EPI_ISL_2406915, EPI_ISL_2406916 | Lighthouse Lab in Alderley Park | Wellcome Sanger Institute for the COVID-19 Genomics UK (COG-UK) Consortium | Jacquelyn Wynn, Mairead Hyland, The Lighthouse Lab in Alderley Park and Alex Alderton, Roberto Amato, Jeffrey Barrett, Sonia Goncalves, Ewan Harrison, David K. Jackson, Ian Johnston, Dominic Kwiatkowski, Cordelia Langford, John Sillitoe on behalf of the Wellcome Sanger Institute COVID-19 Surveillance Team |
| EPI_ISL_2406917 | Lighthouse Lab in Milton Keynes | Wellcome Sanger Institute for the COVID-19 Genomics UK (COG-UK) Consortium | The Lighthouse Lab in Milton Keynes and Alex Alderton, Roberto Amato, Jeffrey Barrett, Sonia Goncalves, Ewan Harrison, David K. Jackson, Ian Johnston, Dominic Kwiatkowski, Cordelia Langford, John Sillitoe on behalf of the Wellcome Sanger Institute COVID-19 Surveillance Team |
| EPI_ISL_2406919, EPI_ISL_2406920 | Lighthouse Lab in Alderley Park | Wellcome Sanger Institute for the COVID-19 Genomics UK (COG-UK) Consortium | Jacquelyn Wynn, Mairead Hyland, The Lighthouse Lab in Alderley Park and Alex Alderton, Roberto Amato, Jeffrey Barrett, Sonia Goncalves, Ewan Harrison, David K. Jackson, Ian Johnston, Dominic Kwiatkowski, Cordelia Langford, John Sillitoe on behalf of the Wellcome Sanger Institute COVID-19 Surveillance Team |
| EPI_ISL_2406921 | Lighthouse Lab in Milton Keynes | Wellcome Sanger Institute for the COVID-19 Genomics UK (COG-UK) Consortium | The Lighthouse Lab in Milton Keynes and Alex Alderton, Roberto Amato, Jeffrey Barrett, Sonia Goncalves, Ewan Harrison, David K. Jackson, Ian Johnston, Dominic Kwiatkowski, Cordelia Langford, John Sillitoe on behalf of the Wellcome Sanger Institute COVID-19 Surveillance Team |
| EPI_ISL_2406922 | Lighthouse Lab in Glasgow | Wellcome Sanger Institute for the COVID-19 Genomics UK (COG-UK) Consortium | Harper VanSteenhouse, Yumi Kasai, David Gray, Carol Clugston, Anna Dominiczak and Alex Alderton, Roberto Amato, Jeffrey Barrett, Sonia Goncalves, Ewan Harrison, David K. Jackson, Ian Johnston, Dominic Kwiatkowski, Cordelia Langford, John Sillitoe on behalf of the Wellcome Sanger Institute COVID-19 Surveillance Team |
| EPI_ISL_2406923 | Lighthouse Lab in Alderley Park | Wellcome Sanger Institute for the COVID-19 Genomics UK (COG-UK) Consortium | Jacquelyn Wynn, Mairead Hyland, The Lighthouse Lab in Alderley Park and Alex Alderton, Roberto Amato, Jeffrey Barrett, Sonia Goncalves, Ewan Harrison, David K. Jackson, Ian Johnston, Dominic Kwiatkowski, Cordelia Langford, John Sillitoe on behalf of the Wellcome Sanger Institute COVID-19 Surveillance Team |
| EPI_ISL_2406924 | Lighthouse Lab in Milton Keynes | Wellcome Sanger Institute for the COVID-19 Genomics UK (COG-UK) Consortium | The Lighthouse Lab in Milton Keynes and Alex Alderton, Roberto Amato, Jeffrey Barrett, Sonia Goncalves, Ewan Harrison, David K. Jackson, Ian Johnston, Dominic Kwiatkowski, Cordelia Langford, John Sillitoe on behalf of the Wellcome Sanger Institute COVID-19 Surveillance Team |
| EPI_ISL_2406925 | Lighthouse Lab in Alderley Park | Wellcome Sanger Institute for the COVID-19 Genomics UK (COG-UK) Consortium | Jacquelyn Wynn, Mairead Hyland, The Lighthouse Lab in Alderley Park and Alex Alderton, Roberto Amato, Jeffrey Barrett, Sonia Goncalves, Ewan Harrison, David K. Jackson, Ian Johnston, Dominic Kwiatkowski, Cordelia Langford, John Sillitoe on behalf of the Wellcome Sanger Institute COVID-19 Surveillance Team |
| EPI_ISL_2406926 | Lighthouse Lab in Milton Keynes | Wellcome Sanger Institute for the COVID-19 Genomics UK (COG-UK) Consortium | The Lighthouse Lab in Milton Keynes and Alex Alderton, Roberto Amato, Jeffrey Barrett, Sonia Goncalves, Ewan Harrison, David K. Jackson, Ian Johnston, Dominic Kwiatkowski, Cordelia Langford, John Sillitoe on behalf of the Wellcome Sanger Institute COVID-19 Surveillance Team |
| EPI_ISL_2406927 | Lighthouse Lab in Alderley Park | Wellcome Sanger Institute for the COVID-19 Genomics UK (COG-UK) Consortium | Jacquelyn Wynn, Mairead Hyland, The Lighthouse Lab in Alderley Park and Alex Alderton, Roberto Amato, Jeffrey Barrett, Sonia Goncalves, Ewan Harrison, David K. Jackson, Ian Johnston, Dominic Kwiatkowski, Cordelia Langford, John Sillitoe on behalf of the Wellcome Sanger Institute COVID-19 Surveillance Team |
| EPI_ISL_2406929, EPI_ISL_2406930, EPI_ISL_2406931 | Lighthouse Lab in Milton Keynes | Wellcome Sanger Institute for the COVID-19 Genomics UK (COG-UK) Consortium | The Lighthouse Lab in Milton Keynes and Alex Alderton, Roberto Amato, Jeffrey Barrett, Sonia Goncalves, Ewan Harrison, David K. Jackson, Ian Johnston, Dominic Kwiatkowski, Cordelia Langford, John Sillitoe on behalf of the Wellcome Sanger Institute COVID-19 Surveillance Team |
| EPI_ISL_2406932, EPI_ISL_2406933, EPI_ISL_2406934 | Lighthouse Lab in Alderley Park | Wellcome Sanger Institute for the COVID-19 Genomics UK (COG-UK) Consortium | Jacquelyn Wynn, Mairead Hyland, The Lighthouse Lab in Alderley Park and Alex Alderton, Roberto Amato, Jeffrey Barrett, Sonia Goncalves, Ewan Harrison, David K. Jackson, Ian Johnston, Dominic Kwiatkowski, Cordelia Langford, John Sillitoe on behalf of the Wellcome Sanger Institute COVID-19 Surveillance Team |
| EPI_ISL_2406935, EPI_ISL_2406937 | Lighthouse Lab in Milton Keynes | Wellcome Sanger Institute for the COVID-19 Genomics UK (COG-UK) Consortium | The Lighthouse Lab in Milton Keynes and Alex Alderton, Roberto Amato, Jeffrey Barrett, Sonia Goncalves, Ewan Harrison, David K. Jackson, Ian Johnston, Dominic Kwiatkowski, Cordelia Langford, John Sillitoe on behalf of the Wellcome Sanger Institute COVID-19 Surveillance Team |
| EPI_ISL_2406938 | Lighthouse Lab in Glasgow | Wellcome Sanger Institute for the COVID-19 Genomics UK (COG-UK) Consortium | Harper VanSteenhouse, Yumi Kasai, David Gray, Carol Clugston, Anna Dominiczak and Alex Alderton, Roberto Amato, Jeffrey Barrett, Sonia Goncalves, Ewan Harrison, David K. Jackson, Ian Johnston, Dominic Kwiatkowski, Cordelia Langford, John Sillitoe on behalf of the Wellcome Sanger Institute COVID-19 Surveillance Team |
| EPI_ISL_2406939 | Lighthouse Lab in Alderley Park | Wellcome Sanger Institute for the COVID-19 Genomics UK (COG-UK) Consortium | Jacquelyn Wynn, Mairead Hyland, The Lighthouse Lab in Alderley Park and Alex Alderton, Roberto Amato, Jeffrey Barrett, Sonia Goncalves, Ewan Harrison, David K. Jackson, Ian Johnston, Dominic Kwiatkowski, Cordelia Langford, John Sillitoe on behalf of the Wellcome Sanger Institute COVID-19 Surveillance Team |
| EPI_ISL_2406940 | Lighthouse Lab in Glasgow | Wellcome Sanger Institute for the COVID-19 Genomics UK (COG-UK) Consortium | Harper VanSteenhouse, Yumi Kasai, David Gray, Carol Clugston, Anna Dominiczak and Alex Alderton, Roberto Amato, Jeffrey Barrett, Sonia Goncalves, Ewan Harrison, David K. Jackson, Ian Johnston, Dominic Kwiatkowski, Cordelia Langford, John Sillitoe on behalf of the Wellcome Sanger Institute COVID-19 Surveillance Team |
| EPI_ISL_2406941 | Lighthouse Lab in Milton Keynes | Wellcome Sanger Institute for the COVID-19 Genomics UK (COG-UK) Consortium | The Lighthouse Lab in Milton Keynes and Alex Alderton, Roberto Amato, Jeffrey Barrett, Sonia Goncalves, Ewan Harrison, David K. Jackson, Ian Johnston, Dominic Kwiatkowski, Cordelia Langford, John Sillitoe on behalf of the Wellcome Sanger Institute COVID-19 Surveillance Team |
| EPI_ISL_2406942 | Lighthouse Lab in Glasgow | Wellcome Sanger Institute for the COVID-19 Genomics UK (COG-UK) Consortium | Harper VanSteenhouse, Yumi Kasai, David Gray, Carol Clugston, Anna Dominiczak and Alex Alderton, Roberto Amato, Jeffrey Barrett, Sonia Goncalves, Ewan Harrison, David K. Jackson, Ian Johnston, Dominic Kwiatkowski, Cordelia Langford, John Sillitoe on behalf of the Wellcome Sanger Institute COVID-19 Surveillance Team |
| EPI_ISL_2406943 | Lighthouse Lab in Alderley Park | Wellcome Sanger Institute for the COVID-19 Genomics UK (COG-UK) Consortium | Jacquelyn Wynn, Mairead Hyland, The Lighthouse Lab in Alderley Park and Alex Alderton, Roberto Amato, Jeffrey Barrett, Sonia Goncalves, Ewan Harrison, David K. Jackson, Ian Johnston, Dominic Kwiatkowski, Cordelia Langford, John Sillitoe on behalf of the Wellcome Sanger Institute COVID-19 Surveillance Team |
| EPI_ISL_2406944 | Lighthouse Lab in Milton Keynes | Wellcome Sanger Institute for the COVID-19 Genomics UK (COG-UK) Consortium | The Lighthouse Lab in Milton Keynes and Alex Alderton, Roberto Amato, Jeffrey Barrett, Sonia Goncalves, Ewan Harrison, David K. Jackson, Ian Johnston, Dominic Kwiatkowski, Cordelia Langford, John Sillitoe on behalf of the Wellcome Sanger Institute COVID-19 Surveillance Team |
| EPI_ISL_2406945 | Lighthouse Lab in Alderley Park | Wellcome Sanger Institute for the COVID-19 Genomics UK (COG-UK) Consortium | Jacquelyn Wynn, Mairead Hyland, The Lighthouse Lab in Alderley Park and Alex Alderton, Roberto Amato, Jeffrey Barrett, Sonia Goncalves, Ewan Harrison, David K. Jackson, Ian Johnston, Dominic Kwiatkowski, Cordelia Langford, John Sillitoe on behalf of the Wellcome Sanger Institute COVID-19 Surveillance Team |
| EPI_ISL_2406947 | Lighthouse Lab in Milton Keynes | Wellcome Sanger Institute for the COVID-19 Genomics UK (COG-UK) Consortium | The Lighthouse Lab in Milton Keynes and Alex Alderton, Roberto Amato, Jeffrey Barrett, Sonia Goncalves, Ewan Harrison, David K. Jackson, Ian Johnston, Dominic Kwiatkowski, Cordelia Langford, John Sillitoe on behalf of the Wellcome Sanger Institute COVID-19 Surveillance Team |
| EPI_ISL_2406948 | Lighthouse Lab in Glasgow | Wellcome Sanger Institute for the COVID-19 Genomics UK (COG-UK) Consortium | Harper VanSteenhouse, Yumi Kasai, David Gray, Carol Clugston, Anna Dominiczak and Alex Alderton, Roberto Amato, Jeffrey Barrett, Sonia Goncalves, Ewan Harrison, David K. Jackson, Ian Johnston, Dominic Kwiatkowski, Cordelia Langford, John Sillitoe on behalf of the Wellcome Sanger Institute COVID-19 Surveillance Team |
| EPI_ISL_2406949, EPI_ISL_2406950, EPI_ISL_2406951, EPI_ISL_2406953, EPI_ISL_2406954, EPI_ISL_2406956 | Lighthouse Lab in Alderley Park | Wellcome Sanger Institute for the COVID-19 Genomics UK (COG-UK) Consortium | Jacquelyn Wynn, Mairead Hyland, The Lighthouse Lab in Alderley Park and Alex Alderton, Roberto Amato, Jeffrey Barrett, Sonia Goncalves, Ewan Harrison, David K. Jackson, Ian Johnston, Dominic Kwiatkowski, Cordelia Langford, John Sillitoe on behalf of the Wellcome Sanger Institute COVID-19 Surveillance Team |
| EPI_ISL_2406957 | Lighthouse Lab in Glasgow | Wellcome Sanger Institute for the COVID-19 Genomics UK (COG-UK) Consortium | Harper VanSteenhouse, Yumi Kasai, David Gray, Carol Clugston, Anna Dominiczak and Alex Alderton, Roberto Amato, Jeffrey Barrett, Sonia Goncalves, Ewan Harrison, David K. Jackson, Ian Johnston, Dominic Kwiatkowski, Cordelia Langford, John Sillitoe on behalf of the Wellcome Sanger Institute COVID-19 Surveillance Team |
| EPI_ISL_2406958, EPI_ISL_2406960 | Lighthouse Lab in Milton Keynes | Wellcome Sanger Institute for the COVID-19 Genomics UK (COG-UK) Consortium | The Lighthouse Lab in Milton Keynes and Alex Alderton, Roberto Amato, Jeffrey Barrett, Sonia Goncalves, Ewan Harrison, David K. Jackson, Ian Johnston, Dominic Kwiatkowski, Cordelia Langford, John Sillitoe on behalf of the Wellcome Sanger Institute COVID-19 Surveillance Team |
| EPI_ISL_2406962, EPI_ISL_2406963 | Lighthouse Lab in Alderley Park | Wellcome Sanger Institute for the COVID-19 Genomics UK (COG-UK) Consortium | Jacquelyn Wynn, Mairead Hyland, The Lighthouse Lab in Alderley Park and Alex Alderton, Roberto Amato, Jeffrey Barrett, Sonia Goncalves, Ewan Harrison, David K. Jackson, Ian Johnston, Dominic Kwiatkowski, Cordelia Langford, John Sillitoe on behalf of the Wellcome Sanger Institute COVID-19 Surveillance Team |
| EPI_ISL_2406964 | Lighthouse Lab in Milton Keynes | Wellcome Sanger Institute for the COVID-19 Genomics UK (COG-UK) Consortium | The Lighthouse Lab in Milton Keynes and Alex Alderton, Roberto Amato, Jeffrey Barrett, Sonia Goncalves, Ewan Harrison, David K. Jackson, Ian Johnston, Dominic Kwiatkowski, Cordelia Langford, John Sillitoe on behalf of the Wellcome Sanger Institute COVID-19 Surveillance Team |
| EPI_ISL_2406965 | Lighthouse Lab in Alderley Park | Wellcome Sanger Institute for the COVID-19 Genomics UK (COG-UK) Consortium | Jacquelyn Wynn, Mairead Hyland, The Lighthouse Lab in Alderley Park and Alex Alderton, Roberto Amato, Jeffrey Barrett, Sonia Goncalves, Ewan Harrison, David K. Jackson, Ian Johnston, Dominic Kwiatkowski, Cordelia Langford, John Sillitoe on behalf of the Wellcome Sanger Institute COVID-19 Surveillance Team |
| EPI_ISL_2406968, EPI_ISL_2406969, EPI_ISL_2406970, EPI_ISL_2406971, EPI_ISL_2406972 | Lighthouse Lab in Milton Keynes | Wellcome Sanger Institute for the COVID-19 Genomics UK (COG-UK) Consortium | The Lighthouse Lab in Milton Keynes and Alex Alderton, Roberto Amato, Jeffrey Barrett, Sonia Goncalves, Ewan Harrison, David K. Jackson, Ian Johnston, Dominic Kwiatkowski, Cordelia Langford, John Sillitoe on behalf of the Wellcome Sanger Institute COVID-19 Surveillance Team |
| EPI_ISL_2406973 | Lighthouse Lab in Alderley Park | Wellcome Sanger Institute for the COVID-19 Genomics UK (COG-UK) Consortium | Jacquelyn Wynn, Mairead Hyland, The Lighthouse Lab in Alderley Park and Alex Alderton, Roberto Amato, Jeffrey Barrett, Sonia Goncalves, Ewan Harrison, David K. Jackson, Ian Johnston, Dominic Kwiatkowski, Cordelia Langford, John Sillitoe on behalf of the Wellcome Sanger Institute COVID-19 Surveillance Team |
| EPI_ISL_2406974 | Lighthouse Lab in Glasgow | Wellcome Sanger Institute for the COVID-19 Genomics UK (COG-UK) Consortium | Harper VanSteenhouse, Yumi Kasai, David Gray, Carol Clugston, Anna Dominiczak and Alex Alderton, Roberto Amato, Jeffrey Barrett, Sonia Goncalves, Ewan Harrison, David K. Jackson, Ian Johnston, Dominic Kwiatkowski, Cordelia Langford, John Sillitoe on behalf of the Wellcome Sanger Institute COVID-19 Surveillance Team |
| EPI_ISL_2406976, EPI_ISL_2406977 | Lighthouse Lab in Alderley Park | Wellcome Sanger Institute for the COVID-19 Genomics UK (COG-UK) Consortium | Jacquelyn Wynn, Mairead Hyland, The Lighthouse Lab in Alderley Park and Alex Alderton, Roberto Amato, Jeffrey Barrett, Sonia Goncalves, Ewan Harrison, David K. Jackson, Ian Johnston, Dominic Kwiatkowski, Cordelia Langford, John Sillitoe on behalf of the Wellcome Sanger Institute COVID-19 Surveillance Team |
| EPI_ISL_2406978 | Lighthouse Lab in Glasgow | Wellcome Sanger Institute for the COVID-19 Genomics UK (COG-UK) Consortium | Harper VanSteenhouse, Yumi Kasai, David Gray, Carol Clugston, Anna Dominiczak and Alex Alderton, Roberto Amato, Jeffrey Barrett, Sonia Goncalves, Ewan Harrison, David K. Jackson, Ian Johnston, Dominic Kwiatkowski, Cordelia Langford, John Sillitoe on behalf of the Wellcome Sanger Institute COVID-19 Surveillance Team |
| EPI_ISL_2406979, EPI_ISL_2406980, EPI_ISL_2406981, EPI_ISL_2406983 | Lighthouse Lab in Alderley Park | Wellcome Sanger Institute for the COVID-19 Genomics UK (COG-UK) Consortium | Jacquelyn Wynn, Mairead Hyland, The Lighthouse Lab in Alderley Park and Alex Alderton, Roberto Amato, Jeffrey Barrett, Sonia Goncalves, Ewan Harrison, David K. Jackson, Ian Johnston, Dominic Kwiatkowski, Cordelia Langford, John Sillitoe on behalf of the Wellcome Sanger Institute COVID-19 Surveillance Team |
| EPI_ISL_2406984 | Lighthouse Lab in Milton Keynes | Wellcome Sanger Institute for the COVID-19 Genomics UK (COG-UK) Consortium | The Lighthouse Lab in Milton Keynes and Alex Alderton, Roberto Amato, Jeffrey Barrett, Sonia Goncalves, Ewan Harrison, David K. Jackson, Ian Johnston, Dominic Kwiatkowski, Cordelia Langford, John Sillitoe on behalf of the Wellcome Sanger Institute COVID-19 Surveillance Team |
| EPI_ISL_2406985, EPI_ISL_2406986 | Lighthouse Lab in Alderley Park | Wellcome Sanger Institute for the COVID-19 Genomics UK (COG-UK) Consortium | Jacquelyn Wynn, Mairead Hyland, The Lighthouse Lab in Alderley Park and Alex Alderton, Roberto Amato, Jeffrey Barrett, Sonia Goncalves, Ewan Harrison, David K. Jackson, Ian Johnston, Dominic Kwiatkowski, Cordelia Langford, John Sillitoe on behalf of the Wellcome Sanger Institute COVID-19 Surveillance Team |

| | | | |
|---|---|---|---|
| EPI_ISL_2406987 | Lighthouse Lab in Milton Keynes | Wellcome Sanger Institute for the COVID-19 Genomics UK (COG-UK) Consortium | The Lighthouse Lab in Milton Keynes and Alex Alderton, Roberto Amato, Jeffrey Barrett, Sonia Goncalves, Ewan Harrison, David K. Jackson, Ian Johnston, Dominic Kwiatkowski, Cordelia Langford, John Sillitoe on behalf of the Wellcome Sanger Institute COVID-19 Surveillance Team |
| EPI_ISL_2406988, EPI_ISL_2406989 | Lighthouse Lab in Alderley Park | Wellcome Sanger Institute for the COVID-19 Genomics UK (COG-UK) Consortium | Jacquelyn Wynn, Mairead Hyland, The Lighthouse Lab in Alderley Park and Alex Alderton, Roberto Amato, Jeffrey Barrett, Sonia Goncalves, Ewan Harrison, David K. Jackson, Ian Johnston, Dominic Kwiatkowski, Cordelia Langford, John Sillitoe on behalf of the Wellcome Sanger Institute COVID-19 Surveillance Team |
| EPI_ISL_2406990, EPI_ISL_2406991 | Lighthouse Lab in Milton Keynes | Wellcome Sanger Institute for the COVID-19 Genomics UK (COG-UK) Consortium | The Lighthouse Lab in Milton Keynes and Alex Alderton, Roberto Amato, Jeffrey Barrett, Sonia Goncalves, Ewan Harrison, David K. Jackson, Ian Johnston, Dominic Kwiatkowski, Cordelia Langford, John Sillitoe on behalf of the Wellcome Sanger Institute COVID-19 Surveillance Team |
| EPI_ISL_2406992 | Lighthouse Lab in Alderley Park | Wellcome Sanger Institute for the COVID-19 Genomics UK (COG-UK) Consortium | Jacquelyn Wynn, Mairead Hyland, The Lighthouse Lab in Alderley Park and Alex Alderton, Roberto Amato, Jeffrey Barrett, Sonia Goncalves, Ewan Harrison, David K. Jackson, Ian Johnston, Dominic Kwiatkowski, Cordelia Langford, John Sillitoe on behalf of the Wellcome Sanger Institute COVID-19 Surveillance Team |
| EPI_ISL_2406993 | Lighthouse Lab in Glasgow | Wellcome Sanger Institute for the COVID-19 Genomics UK (COG-UK) Consortium | Harper VanSteenhouse, Yumi Kasai, David Gray, Carol Clugston, Anna Dominiczak and Alex Alderton, Roberto Amato, Jeffrey Barrett, Sonia Goncalves, Ewan Harrison, David K. Jackson, Ian Johnston, Dominic Kwiatkowski, Cordelia Langford, John Sillitoe on behalf of the Wellcome Sanger Institute COVID-19 Surveillance Team |
| EPI_ISL_2406994 | Lighthouse Lab in Alderley Park | Wellcome Sanger Institute for the COVID-19 Genomics UK (COG-UK) Consortium | Jacquelyn Wynn, Mairead Hyland, The Lighthouse Lab in Alderley Park and Alex Alderton, Roberto Amato, Jeffrey Barrett, Sonia Goncalves, Ewan Harrison, David K. Jackson, Ian Johnston, Dominic Kwiatkowski, Cordelia Langford, John Sillitoe on behalf of the Wellcome Sanger Institute COVID-19 Surveillance Team |
| EPI_ISL_2406995 | Lighthouse Lab in Glasgow | Wellcome Sanger Institute for the COVID-19 Genomics UK (COG-UK) Consortium | Harper VanSteenhouse, Yumi Kasai, David Gray, Carol Clugston, Anna Dominiczak and Alex Alderton, Roberto Amato, Jeffrey Barrett, Sonia Goncalves, Ewan Harrison, David K. Jackson, Ian Johnston, Dominic Kwiatkowski, Cordelia Langford, John Sillitoe on behalf of the Wellcome Sanger Institute COVID-19 Surveillance Team |
| EPI_ISL_2406996 | Lighthouse Lab in Alderley Park | Wellcome Sanger Institute for the COVID-19 Genomics UK (COG-UK) Consortium | Jacquelyn Wynn, Mairead Hyland, The Lighthouse Lab in Alderley Park and Alex Alderton, Roberto Amato, Jeffrey Barrett, Sonia Goncalves, Ewan Harrison, David K. Jackson, Ian Johnston, Dominic Kwiatkowski, Cordelia Langford, John Sillitoe on behalf of the Wellcome Sanger Institute COVID-19 Surveillance Team |
| EPI_ISL_2406997 | Lighthouse Lab in Glasgow | Wellcome Sanger Institute for the COVID-19 Genomics UK (COG-UK) Consortium | Harper VanSteenhouse, Yumi Kasai, David Gray, Carol Clugston, Anna Dominiczak and Alex Alderton, Roberto Amato, Jeffrey Barrett, Sonia Goncalves, Ewan Harrison, David K. Jackson, Ian Johnston, Dominic Kwiatkowski, Cordelia Langford, John Sillitoe on behalf of the Wellcome Sanger Institute COVID-19 Surveillance Team |
| EPI_ISL_2406998 | Lighthouse Lab in Alderley Park | Wellcome Sanger Institute for the COVID-19 Genomics UK (COG-UK) Consortium | Jacquelyn Wynn, Mairead Hyland, The Lighthouse Lab in Alderley Park and Alex Alderton, Roberto Amato, Jeffrey Barrett, Sonia Goncalves, Ewan Harrison, David K. Jackson, Ian Johnston, Dominic Kwiatkowski, Cordelia Langford, John Sillitoe on behalf of the Wellcome Sanger Institute COVID-19 Surveillance Team |
| EPI_ISL_2406999 | Lighthouse Lab in Glasgow | Wellcome Sanger Institute for the COVID-19 Genomics UK (COG-UK) Consortium | Harper VanSteenhouse, Yumi Kasai, David Gray, Carol Clugston, Anna Dominiczak and Alex Alderton, Roberto Amato, Jeffrey Barrett, Sonia Goncalves, Ewan Harrison, David K. Jackson, Ian Johnston, Dominic Kwiatkowski, Cordelia Langford, John Sillitoe on behalf of the Wellcome Sanger Institute COVID-19 Surveillance Team |
| EPI_ISL_2407000 | Lighthouse Lab in Alderley Park | Wellcome Sanger Institute for the COVID-19 Genomics UK (COG-UK) Consortium | Jacquelyn Wynn, Mairead Hyland, The Lighthouse Lab in Alderley Park and Alex Alderton, Roberto Amato, Jeffrey Barrett, Sonia Goncalves, Ewan Harrison, David K. Jackson, Ian Johnston, Dominic Kwiatkowski, Cordelia Langford, John Sillitoe on behalf of the Wellcome Sanger Institute COVID-19 Surveillance Team |
| EPI_ISL_2407001 | Lighthouse Lab in Glasgow | Wellcome Sanger Institute for the COVID-19 Genomics UK (COG-UK) Consortium | Harper VanSteenhouse, Yumi Kasai, David Gray, Carol Clugston, Anna Dominiczak and Alex Alderton, Roberto Amato, Jeffrey Barrett, Sonia Goncalves, Ewan Harrison, David K. Jackson, Ian Johnston, Dominic Kwiatkowski, Cordelia Langford, John Sillitoe on behalf of the Wellcome Sanger Institute COVID-19 Surveillance Team |
| EPI_ISL_2407002, EPI_ISL_2407003, EPI_ISL_2407004 | Lighthouse Lab in Milton Keynes | Wellcome Sanger Institute for the COVID-19 Genomics UK (COG-UK) Consortium | The Lighthouse Lab in Milton Keynes and Alex Alderton, Roberto Amato, Jeffrey Barrett, Sonia Goncalves, Ewan Harrison, David K. Jackson, Ian Johnston, Dominic Kwiatkowski, Cordelia Langford, John Sillitoe on behalf of the Wellcome Sanger Institute COVID-19 Surveillance Team |
| EPI_ISL_2407005 | Lighthouse Lab in Alderley Park | Wellcome Sanger Institute for the COVID-19 Genomics UK (COG-UK) Consortium | Jacquelyn Wynn, Mairead Hyland, The Lighthouse Lab in Alderley Park and Alex Alderton, Roberto Amato, Jeffrey Barrett, Sonia Goncalves, Ewan Harrison, David K. Jackson, Ian Johnston, Dominic Kwiatkowski, Cordelia Langford, John Sillitoe on behalf of the Wellcome Sanger Institute COVID-19 Surveillance Team |
| EPI_ISL_2407006 | Lighthouse Lab in Milton Keynes | Wellcome Sanger Institute for the COVID-19 Genomics UK (COG-UK) Consortium | The Lighthouse Lab in Milton Keynes and Alex Alderton, Roberto Amato, Jeffrey Barrett, Sonia Goncalves, Ewan Harrison, David K. Jackson, Ian Johnston, Dominic Kwiatkowski, Cordelia Langford, John Sillitoe on behalf of the Wellcome Sanger Institute COVID-19 Surveillance Team |
| EPI_ISL_2407007 | Lighthouse Lab in Alderley Park | Wellcome Sanger Institute for the COVID-19 Genomics UK (COG-UK) Consortium | Jacquelyn Wynn, Mairead Hyland, The Lighthouse Lab in Alderley Park and Alex Alderton, Roberto Amato, Jeffrey Barrett, Sonia Goncalves, Ewan Harrison, David K. Jackson, Ian Johnston, Dominic Kwiatkowski, Cordelia Langford, John Sillitoe on behalf of the Wellcome Sanger Institute COVID-19 Surveillance Team |
| EPI_ISL_2407008, EPI_ISL_2407009, EPI_ISL_2407011, EPI_ISL_2407012 | Lighthouse Lab in Milton Keynes | Wellcome Sanger Institute for the COVID-19 Genomics UK (COG-UK) Consortium | The Lighthouse Lab in Milton Keynes and Alex Alderton, Roberto Amato, Jeffrey Barrett, Sonia Goncalves, Ewan Harrison, David K. Jackson, Ian Johnston, Dominic Kwiatkowski, Cordelia Langford, John Sillitoe on behalf of the Wellcome Sanger Institute COVID-19 Surveillance Team |
| EPI_ISL_2407014, EPI_ISL_2407015 | Lighthouse Lab in Alderley Park | Wellcome Sanger Institute for the COVID-19 Genomics UK (COG-UK) Consortium | Jacquelyn Wynn, Mairead Hyland, The Lighthouse Lab in Alderley Park and Alex Alderton, Roberto Amato, Jeffrey Barrett, Sonia Goncalves, Ewan Harrison, David K. Jackson, Ian Johnston, Dominic Kwiatkowski, Cordelia Langford, John Sillitoe on behalf of the Wellcome Sanger Institute COVID-19 Surveillance Team |
| EPI_ISL_2407016 | Lighthouse Lab in Milton Keynes | Wellcome Sanger Institute for the COVID-19 Genomics UK (COG-UK) Consortium | The Lighthouse Lab in Milton Keynes and Alex Alderton, Roberto Amato, Jeffrey Barrett, Sonia Goncalves, Ewan Harrison, David K. Jackson, Ian Johnston, Dominic Kwiatkowski, Cordelia Langford, John Sillitoe on behalf of the Wellcome Sanger Institute COVID-19 Surveillance Team |
| EPI_ISL_2407017 | Lighthouse Lab in Glasgow | Wellcome Sanger Institute for the COVID-19 Genomics UK (COG-UK) Consortium | Harper VanSteenhouse, Yumi Kasai, David Gray, Carol Clugston, Anna Dominiczak and Alex Alderton, Roberto Amato, Jeffrey Barrett, Sonia Goncalves, Ewan Harrison, David K. Jackson, Ian Johnston, Dominic Kwiatkowski, Cordelia Langford, John Sillitoe on behalf of the Wellcome Sanger Institute COVID-19 Surveillance Team |
| EPI_ISL_2407020 | Lighthouse Lab in Alderley Park | Wellcome Sanger Institute for the COVID-19 Genomics UK (COG-UK) Consortium | Jacquelyn Wynn, Mairead Hyland, The Lighthouse Lab in Alderley Park and Alex Alderton, Roberto Amato, Jeffrey Barrett, Sonia Goncalves, Ewan Harrison, David K. Jackson, Ian Johnston, Dominic Kwiatkowski, Cordelia Langford, John Sillitoe on behalf of the Wellcome Sanger Institute COVID-19 Surveillance Team |
| EPI_ISL_2407021 | Lighthouse Lab in Milton Keynes | Wellcome Sanger Institute for the COVID-19 Genomics UK (COG-UK) Consortium | The Lighthouse Lab in Milton Keynes and Alex Alderton, Roberto Amato, Jeffrey Barrett, Sonia Goncalves, Ewan Harrison, David K. Jackson, Ian Johnston, Dominic Kwiatkowski, Cordelia Langford, John Sillitoe on behalf of the Wellcome Sanger Institute COVID-19 Surveillance Team |
| EPI_ISL_2407022 | Lighthouse Lab in Glasgow | Wellcome Sanger Institute for the COVID-19 Genomics UK (COG-UK) Consortium | Harper VanSteenhouse, Yumi Kasai, David Gray, Carol Clugston, Anna Dominiczak and Alex Alderton, Roberto Amato, Jeffrey Barrett, Sonia Goncalves, Ewan Harrison, David K. Jackson, Ian Johnston, Dominic Kwiatkowski, Cordelia Langford, John Sillitoe on behalf of the Wellcome Sanger Institute COVID-19 Surveillance Team |
| EPI_ISL_2407024 | Lighthouse Lab in Milton Keynes | Wellcome Sanger Institute for the COVID-19 Genomics UK (COG-UK) Consortium | The Lighthouse Lab in Milton Keynes and Alex Alderton, Roberto Amato, Jeffrey Barrett, Sonia Goncalves, Ewan Harrison, David K. Jackson, Ian Johnston, Dominic Kwiatkowski, Cordelia Langford, John Sillitoe on behalf of the Wellcome Sanger Institute COVID-19 Surveillance Team |
| EPI_ISL_2407025 | Lighthouse Lab in Alderley Park | Wellcome Sanger Institute for the COVID-19 Genomics UK (COG-UK) Consortium | Jacquelyn Wynn, Mairead Hyland, The Lighthouse Lab in Alderley Park and Alex Alderton, Roberto Amato, Jeffrey Barrett, Sonia Goncalves, Ewan Harrison, David K. Jackson, Ian Johnston, Dominic Kwiatkowski, Cordelia Langford, John Sillitoe on behalf of the Wellcome Sanger Institute COVID-19 Surveillance Team |
| EPI_ISL_2407026, EPI_ISL_2407027, EPI_ISL_2407028 | Lighthouse Lab in Glasgow | Wellcome Sanger Institute for the COVID-19 Genomics UK (COG-UK) Consortium | Harper VanSteenhouse, Yumi Kasai, David Gray, Carol Clugston, Anna Dominiczak and Alex Alderton, Roberto Amato, Jeffrey Barrett, Sonia Goncalves, Ewan Harrison, David K. Jackson, Ian Johnston, Dominic Kwiatkowski, Cordelia Langford, John Sillitoe on behalf of the Wellcome Sanger Institute COVID-19 Surveillance Team |
| EPI_ISL_2407030 | Lighthouse Lab in Alderley Park | Wellcome Sanger Institute for the COVID-19 Genomics UK (COG-UK) Consortium | Jacquelyn Wynn, Mairead Hyland, The Lighthouse Lab in Alderley Park and Alex Alderton, Roberto Amato, Jeffrey Barrett, Sonia Goncalves, Ewan Harrison, David K. Jackson, Ian Johnston, Dominic Kwiatkowski, Cordelia Langford, John Sillitoe on behalf of the Wellcome Sanger Institute COVID-19 Surveillance Team |
| EPI_ISL_2407031 | Lighthouse Lab in Milton Keynes | Wellcome Sanger Institute for the COVID-19 Genomics UK (COG-UK) Consortium | The Lighthouse Lab in Milton Keynes and Alex Alderton, Roberto Amato, Jeffrey Barrett, Sonia Goncalves, Ewan Harrison, David K. Jackson, Ian Johnston, Dominic Kwiatkowski, Cordelia Langford, John Sillitoe on behalf of the Wellcome Sanger Institute COVID-19 Surveillance Team |
| EPI_ISL_2407032, EPI_ISL_2407033 | Lighthouse Lab in Alderley Park | Wellcome Sanger Institute for the COVID-19 Genomics UK (COG-UK) Consortium | Jacquelyn Wynn, Mairead Hyland, The Lighthouse Lab in Alderley Park and Alex Alderton, Roberto Amato, Jeffrey Barrett, Sonia Goncalves, Ewan Harrison, David K. Jackson, Ian Johnston, Dominic Kwiatkowski, Cordelia Langford, John Sillitoe on behalf of the Wellcome Sanger Institute COVID-19 Surveillance Team |
| EPI_ISL_2407034 | Lighthouse Lab in Glasgow | Wellcome Sanger Institute for the COVID-19 Genomics UK (COG-UK) Consortium | Harper VanSteenhouse, Yumi Kasai, David Gray, Carol Clugston, Anna Dominiczak and Alex Alderton, Roberto Amato, Jeffrey Barrett, Sonia Goncalves, Ewan Harrison, David K. Jackson, Ian Johnston, Dominic Kwiatkowski, Cordelia Langford, John Sillitoe on behalf of the Wellcome Sanger Institute COVID-19 Surveillance Team |
| EPI_ISL_2407035 | Lighthouse Lab in Milton Keynes | Wellcome Sanger Institute for the COVID-19 Genomics UK (COG-UK) Consortium | The Lighthouse Lab in Milton Keynes and Alex Alderton, Roberto Amato, Jeffrey Barrett, Sonia Goncalves, Ewan Harrison, David K. Jackson, Ian Johnston, Dominic Kwiatkowski, Cordelia Langford, John Sillitoe on behalf of the Wellcome Sanger Institute COVID-19 Surveillance Team |
| EPI_ISL_2407037, EPI_ISL_2407038 | Lighthouse Lab in Alderley Park | Wellcome Sanger Institute for the COVID-19 Genomics UK (COG-UK) Consortium | Jacquelyn Wynn, Mairead Hyland, The Lighthouse Lab in Alderley Park and Alex Alderton, Roberto Amato, Jeffrey Barrett, Sonia Goncalves, Ewan Harrison, David K. Jackson, Ian Johnston, Dominic Kwiatkowski, Cordelia Langford, John Sillitoe on behalf of the Wellcome Sanger Institute COVID-19 Surveillance Team |
| EPI_ISL_2407039 | Lighthouse Lab in Glasgow | Wellcome Sanger Institute for the COVID-19 Genomics UK (COG-UK) Consortium | Harper VanSteenhouse, Yumi Kasai, David Gray, Carol Clugston, Anna Dominiczak and Alex Alderton, Roberto Amato, Jeffrey Barrett, Sonia Goncalves, Ewan Harrison, David K. Jackson, Ian Johnston, Dominic Kwiatkowski, Cordelia Langford, John Sillitoe on behalf of the Wellcome Sanger Institute COVID-19 Surveillance Team |
| EPI_ISL_2407040 | Lighthouse Lab in Alderley Park | Wellcome Sanger Institute for the COVID-19 Genomics UK (COG-UK) Consortium | Jacquelyn Wynn, Mairead Hyland, The Lighthouse Lab in Alderley Park and Alex Alderton, Roberto Amato, Jeffrey Barrett, Sonia Goncalves, Ewan Harrison, David K. Jackson, Ian Johnston, Dominic Kwiatkowski, Cordelia Langford, John Sillitoe on behalf of the Wellcome Sanger Institute COVID-19 Surveillance Team |
| EPI_ISL_2407041 | Lighthouse Lab in Milton Keynes | Wellcome Sanger Institute for the COVID-19 Genomics UK (COG-UK) Consortium | The Lighthouse Lab in Milton Keynes and Alex Alderton, Roberto Amato, Jeffrey Barrett, Sonia Goncalves, Ewan Harrison, David K. Jackson, Ian Johnston, Dominic Kwiatkowski, Cordelia Langford, John Sillitoe on behalf of the Wellcome Sanger Institute COVID-19 Surveillance Team |
| EPI_ISL_2407042 | Lighthouse Lab in Alderley Park | Wellcome Sanger Institute for the COVID-19 Genomics UK (COG-UK) Consortium | Jacquelyn Wynn, Mairead Hyland, The Lighthouse Lab in Alderley Park and Alex Alderton, Roberto Amato, Jeffrey Barrett, Sonia Goncalves, Ewan Harrison, David K. Jackson, Ian Johnston, Dominic Kwiatkowski, Cordelia Langford, John Sillitoe on behalf of the Wellcome Sanger Institute COVID-19 Surveillance Team |
| EPI_ISL_2407043 | Lighthouse Lab in Milton Keynes | Wellcome Sanger Institute for the COVID-19 Genomics UK (COG-UK) Consortium | The Lighthouse Lab in Milton Keynes and Alex Alderton, Roberto Amato, Jeffrey Barrett, Sonia Goncalves, Ewan Harrison, David K. Jackson, Ian Johnston, Dominic Kwiatkowski, Cordelia Langford, John Sillitoe on behalf of the Wellcome Sanger Institute COVID-19 Surveillance Team |
| EPI_ISL_2407044 | Lighthouse Lab in Glasgow | Wellcome Sanger Institute for the COVID-19 Genomics UK (COG-UK) Consortium | Harper VanSteenhouse, Yumi Kasai, David Gray, Carol Clugston, Anna Dominiczak and Alex Alderton, Roberto Amato, Jeffrey Barrett, Sonia Goncalves, Ewan Harrison, David K. Jackson, Ian Johnston, Dominic Kwiatkowski, Cordelia Langford, John Sillitoe on behalf of the Wellcome Sanger Institute COVID-19 Surveillance Team |
| EPI_ISL_2407045 | Lighthouse Lab in Milton Keynes | Wellcome Sanger Institute for the COVID-19 Genomics UK (COG-UK) Consortium | The Lighthouse Lab in Milton Keynes and Alex Alderton, Roberto Amato, Jeffrey Barrett, Sonia Goncalves, Ewan Harrison, David K. Jackson, Ian Johnston, Dominic Kwiatkowski, Cordelia Langford, John Sillitoe on behalf of the Wellcome Sanger Institute COVID-19 Surveillance Team |
| EPI_ISL_2407046 | Lighthouse Lab in Alderley Park | Wellcome Sanger Institute for the COVID-19 Genomics UK (COG-UK) Consortium | Jacquelyn Wynn, Mairead Hyland, The Lighthouse Lab in Alderley Park and Alex Alderton, Roberto Amato, Jeffrey Barrett, Sonia Goncalves, Ewan Harrison, David K. Jackson, Ian Johnston, Dominic Kwiatkowski, Cordelia Langford, John Sillitoe on behalf of the Wellcome Sanger Institute COVID-19 Surveillance Team |
| EPI_ISL_2407047, EPI_ISL_2407048, EPI_ISL_2407049 | Lighthouse Lab in Milton Keynes | Wellcome Sanger Institute for the COVID-19 Genomics UK (COG-UK) Consortium | The Lighthouse Lab in Milton Keynes and Alex Alderton, Roberto Amato, Jeffrey Barrett, Sonia Goncalves, Ewan Harrison, David K. Jackson, Ian Johnston, Dominic Kwiatkowski, Cordelia Langford, John Sillitoe on behalf of the Wellcome Sanger Institute COVID-19 Surveillance Team |
| EPI_ISL_2407051 | Lighthouse Lab in Alderley Park | Wellcome Sanger Institute for the COVID-19 Genomics UK (COG-UK) Consortium | Jacquelyn Wynn, Mairead Hyland, The Lighthouse Lab in Alderley Park and Alex Alderton, Roberto Amato, Jeffrey Barrett, Sonia Goncalves, Ewan Harrison, David K. Jackson, Ian Johnston, Dominic Kwiatkowski, Cordelia Langford, John Sillitoe on behalf of the Wellcome Sanger Institute COVID-19 Surveillance Team |

| | | | |
|---|---|---|---|
| EPI_ISL_2407052 | Lighthouse Lab in Milton Keynes | Wellcome Sanger Institute for the COVID-19 Genomics UK (COG-UK) Consortium | The Lighthouse Lab in Milton Keynes and Alex Alderton, Roberto Amato, Jeffrey Barrett, Sonia Goncalves, Ewan Harrison, David K. Jackson, Ian Johnston, Dominic Kwiatkowski, Cordelia Langford, John Sillitoe on behalf of the Wellcome Sanger Institute COVID-19 Surveillance Team |
| EPI_ISL_2407053 | Lighthouse Lab in Alderley Park | Wellcome Sanger Institute for the COVID-19 Genomics UK (COG-UK) Consortium | Jacquelyn Wynn, Mairead Hyland, The Lighthouse Lab in Alderley Park and Alex Alderton, Roberto Amato, Jeffrey Barrett, Sonia Goncalves, Ewan Harrison, David K. Jackson, Ian Johnston, Dominic Kwiatkowski, Cordelia Langford, John Sillitoe on behalf of the Wellcome Sanger Institute COVID-19 Surveillance Team |
| EPI_ISL_2407054 | Lighthouse Lab in Milton Keynes | Wellcome Sanger Institute for the COVID-19 Genomics UK (COG-UK) Consortium | The Lighthouse Lab in Milton Keynes and Alex Alderton, Roberto Amato, Jeffrey Barrett, Sonia Goncalves, Ewan Harrison, David K. Jackson, Ian Johnston, Dominic Kwiatkowski, Cordelia Langford, John Sillitoe on behalf of the Wellcome Sanger Institute COVID-19 Surveillance Team |
| EPI_ISL_2407056, EPI_ISL_2407057 | Lighthouse Lab in Glasgow | Wellcome Sanger Institute for the COVID-19 Genomics UK (COG-UK) Consortium | Harper VanSteenhouse, Yumi Kasai, David Gray, Carol Clugston, Anna Dominiczak and Alex Alderton, Roberto Amato, Jeffrey Barrett, Sonia Goncalves, Ewan Harrison, David K. Jackson, Ian Johnston, Dominic Kwiatkowski, Cordelia Langford, John Sillitoe on behalf of the Wellcome Sanger Institute COVID-19 Surveillance Team |
| EPI_ISL_2407058, EPI_ISL_2407060 | Lighthouse Lab in Milton Keynes | Wellcome Sanger Institute for the COVID-19 Genomics UK (COG-UK) Consortium | The Lighthouse Lab in Milton Keynes and Alex Alderton, Roberto Amato, Jeffrey Barrett, Sonia Goncalves, Ewan Harrison, David K. Jackson, Ian Johnston, Dominic Kwiatkowski, Cordelia Langford, John Sillitoe on behalf of the Wellcome Sanger Institute COVID-19 Surveillance Team |
| EPI_ISL_2407061 | Lighthouse Lab in Glasgow | Wellcome Sanger Institute for the COVID-19 Genomics UK (COG-UK) Consortium | Harper VanSteenhouse, Yumi Kasai, David Gray, Carol Clugston, Anna Dominiczak and Alex Alderton, Roberto Amato, Jeffrey Barrett, Sonia Goncalves, Ewan Harrison, David K. Jackson, Ian Johnston, Dominic Kwiatkowski, Cordelia Langford, John Sillitoe on behalf of the Wellcome Sanger Institute COVID-19 Surveillance Team |
| EPI_ISL_2407062 | Lighthouse Lab in Alderley Park | Wellcome Sanger Institute for the COVID-19 Genomics UK (COG-UK) Consortium | Jacquelyn Wynn, Mairead Hyland, The Lighthouse Lab in Alderley Park and Alex Alderton, Roberto Amato, Jeffrey Barrett, Sonia Goncalves, Ewan Harrison, David K. Jackson, Ian Johnston, Dominic Kwiatkowski, Cordelia Langford, John Sillitoe on behalf of the Wellcome Sanger Institute COVID-19 Surveillance Team |
| EPI_ISL_2407063 | Lighthouse Lab in Glasgow | Wellcome Sanger Institute for the COVID-19 Genomics UK (COG-UK) Consortium | Harper VanSteenhouse, Yumi Kasai, David Gray, Carol Clugston, Anna Dominiczak and Alex Alderton, Roberto Amato, Jeffrey Barrett, Sonia Goncalves, Ewan Harrison, David K. Jackson, Ian Johnston, Dominic Kwiatkowski, Cordelia Langford, John Sillitoe on behalf of the Wellcome Sanger Institute COVID-19 Surveillance Team |
| EPI_ISL_2407064, EPI_ISL_2407066, EPI_ISL_2407067 | Lighthouse Lab in Alderley Park | Wellcome Sanger Institute for the COVID-19 Genomics UK (COG-UK) Consortium | Jacquelyn Wynn, Mairead Hyland, The Lighthouse Lab in Alderley Park and Alex Alderton, Roberto Amato, Jeffrey Barrett, Sonia Goncalves, Ewan Harrison, David K. Jackson, Ian Johnston, Dominic Kwiatkowski, Cordelia Langford, John Sillitoe on behalf of the Wellcome Sanger Institute COVID-19 Surveillance Team |
| EPI_ISL_2407068, EPI_ISL_2407069, EPI_ISL_2407070, EPI_ISL_2407071 | Lighthouse Lab in Milton Keynes | Wellcome Sanger Institute for the COVID-19 Genomics UK (COG-UK) Consortium | The Lighthouse Lab in Milton Keynes and Alex Alderton, Roberto Amato, Jeffrey Barrett, Sonia Goncalves, Ewan Harrison, David K. Jackson, Ian Johnston, Dominic Kwiatkowski, Cordelia Langford, John Sillitoe on behalf of the Wellcome Sanger Institute COVID-19 Surveillance Team |
| EPI_ISL_2407072 | Lighthouse Lab in Glasgow | Wellcome Sanger Institute for the COVID-19 Genomics UK (COG-UK) Consortium | Harper VanSteenhouse, Yumi Kasai, David Gray, Carol Clugston, Anna Dominiczak and Alex Alderton, Roberto Amato, Jeffrey Barrett, Sonia Goncalves, Ewan Harrison, David K. Jackson, Ian Johnston, Dominic Kwiatkowski, Cordelia Langford, John Sillitoe on behalf of the Wellcome Sanger Institute COVID-19 Surveillance Team |
| EPI_ISL_2407073, EPI_ISL_2407074, EPI_ISL_2407075 | Lighthouse Lab in Milton Keynes | Wellcome Sanger Institute for the COVID-19 Genomics UK (COG-UK) Consortium | The Lighthouse Lab in Milton Keynes and Alex Alderton, Roberto Amato, Jeffrey Barrett, Sonia Goncalves, Ewan Harrison, David K. Jackson, Ian Johnston, Dominic Kwiatkowski, Cordelia Langford, John Sillitoe on behalf of the Wellcome Sanger Institute COVID-19 Surveillance Team |
| EPI_ISL_2407076 | Lighthouse Lab in Glasgow | Wellcome Sanger Institute for the COVID-19 Genomics UK (COG-UK) Consortium | Harper VanSteenhouse, Yumi Kasai, David Gray, Carol Clugston, Anna Dominiczak and Alex Alderton, Roberto Amato, Jeffrey Barrett, Sonia Goncalves, Ewan Harrison, David K. Jackson, Ian Johnston, Dominic Kwiatkowski, Cordelia Langford, John Sillitoe on behalf of the Wellcome Sanger Institute COVID-19 Surveillance Team |
| EPI_ISL_2407077, EPI_ISL_2407079, EPI_ISL_2407080, EPI_ISL_2407081 | Lighthouse Lab in Milton Keynes | Wellcome Sanger Institute for the COVID-19 Genomics UK (COG-UK) Consortium | The Lighthouse Lab in Milton Keynes and Alex Alderton, Roberto Amato, Jeffrey Barrett, Sonia Goncalves, Ewan Harrison, David K. Jackson, Ian Johnston, Dominic Kwiatkowski, Cordelia Langford, John Sillitoe on behalf of the Wellcome Sanger Institute COVID-19 Surveillance Team |
| EPI_ISL_2407082 | Lighthouse Lab in Glasgow | Wellcome Sanger Institute for the COVID-19 Genomics UK (COG-UK) Consortium | Harper VanSteenhouse, Yumi Kasai, David Gray, Carol Clugston, Anna Dominiczak and Alex Alderton, Roberto Amato, Jeffrey Barrett, Sonia Goncalves, Ewan Harrison, David K. Jackson, Ian Johnston, Dominic Kwiatkowski, Cordelia Langford, John Sillitoe on behalf of the Wellcome Sanger Institute COVID-19 Surveillance Team |
| EPI_ISL_2407083, EPI_ISL_2407084 | Lighthouse Lab in Alderley Park | Wellcome Sanger Institute for the COVID-19 Genomics UK (COG-UK) Consortium | Jacquelyn Wynn, Mairead Hyland, The Lighthouse Lab in Alderley Park and Alex Alderton, Roberto Amato, Jeffrey Barrett, Sonia Goncalves, Ewan Harrison, David K. Jackson, Ian Johnston, Dominic Kwiatkowski, Cordelia Langford, John Sillitoe on behalf of the Wellcome Sanger Institute COVID-19 Surveillance Team |
| EPI_ISL_2407085 | Lighthouse Lab in Glasgow | Wellcome Sanger Institute for the COVID-19 Genomics UK (COG-UK) Consortium | Harper VanSteenhouse, Yumi Kasai, David Gray, Carol Clugston, Anna Dominiczak and Alex Alderton, Roberto Amato, Jeffrey Barrett, Sonia Goncalves, Ewan Harrison, David K. Jackson, Ian Johnston, Dominic Kwiatkowski, Cordelia Langford, John Sillitoe on behalf of the Wellcome Sanger Institute COVID-19 Surveillance Team |
| EPI_ISL_2407086, EPI_ISL_2407087 | Lighthouse Lab in Alderley Park | Wellcome Sanger Institute for the COVID-19 Genomics UK (COG-UK) Consortium | Jacquelyn Wynn, Mairead Hyland, The Lighthouse Lab in Alderley Park and Alex Alderton, Roberto Amato, Jeffrey Barrett, Sonia Goncalves, Ewan Harrison, David K. Jackson, Ian Johnston, Dominic Kwiatkowski, Cordelia Langford, John Sillitoe on behalf of the Wellcome Sanger Institute COVID-19 Surveillance Team |
| EPI_ISL_2407088, EPI_ISL_2407089 | Lighthouse Lab in Glasgow | Wellcome Sanger Institute for the COVID-19 Genomics UK (COG-UK) Consortium | Harper VanSteenhouse, Yumi Kasai, David Gray, Carol Clugston, Anna Dominiczak and Alex Alderton, Roberto Amato, Jeffrey Barrett, Sonia Goncalves, Ewan Harrison, David K. Jackson, Ian Johnston, Dominic Kwiatkowski, Cordelia Langford, John Sillitoe on behalf of the Wellcome Sanger Institute COVID-19 Surveillance Team |
| EPI_ISL_2407090 | Lighthouse Lab in Milton Keynes | Wellcome Sanger Institute for the COVID-19 Genomics UK (COG-UK) Consortium | The Lighthouse Lab in Milton Keynes and Alex Alderton, Roberto Amato, Jeffrey Barrett, Sonia Goncalves, Ewan Harrison, David K. Jackson, Ian Johnston, Dominic Kwiatkowski, Cordelia Langford, John Sillitoe on behalf of the Wellcome Sanger Institute COVID-19 Surveillance Team |
| EPI_ISL_2407091 | Lighthouse Lab in Alderley Park | Wellcome Sanger Institute for the COVID-19 Genomics UK (COG-UK) Consortium | Jacquelyn Wynn, Mairead Hyland, The Lighthouse Lab in Alderley Park and Alex Alderton, Roberto Amato, Jeffrey Barrett, Sonia Goncalves, Ewan Harrison, David K. Jackson, Ian Johnston, Dominic Kwiatkowski, Cordelia Langford, John Sillitoe on behalf of the Wellcome Sanger Institute COVID-19 Surveillance Team |
| EPI_ISL_2407093 | Lighthouse Lab in Glasgow | Wellcome Sanger Institute for the COVID-19 Genomics UK (COG-UK) Consortium | Harper VanSteenhouse, Yumi Kasai, David Gray, Carol Clugston, Anna Dominiczak and Alex Alderton, Roberto Amato, Jeffrey Barrett, Sonia Goncalves, Ewan Harrison, David K. Jackson, Ian Johnston, Dominic Kwiatkowski, Cordelia Langford, John Sillitoe on behalf of the Wellcome Sanger Institute COVID-19 Surveillance Team |
| EPI_ISL_2407094, EPI_ISL_2407095, EPI_ISL_2407096 | Lighthouse Lab in Alderley Park | Wellcome Sanger Institute for the COVID-19 Genomics UK (COG-UK) Consortium | Jacquelyn Wynn, Mairead Hyland, The Lighthouse Lab in Alderley Park and Alex Alderton, Roberto Amato, Jeffrey Barrett, Sonia Goncalves, Ewan Harrison, David K. Jackson, Ian Johnston, Dominic Kwiatkowski, Cordelia Langford, John Sillitoe on behalf of the Wellcome Sanger Institute COVID-19 Surveillance Team |
| EPI_ISL_2407098, EPI_ISL_2407099, EPI_ISL_2407100 | Lighthouse Lab in Milton Keynes | Wellcome Sanger Institute for the COVID-19 Genomics UK (COG-UK) Consortium | The Lighthouse Lab in Milton Keynes and Alex Alderton, Roberto Amato, Jeffrey Barrett, Sonia Goncalves, Ewan Harrison, David K. Jackson, Ian Johnston, Dominic Kwiatkowski, Cordelia Langford, John Sillitoe on behalf of the Wellcome Sanger Institute COVID-19 Surveillance Team |
| EPI_ISL_2407101 | Lighthouse Lab in Alderley Park | Wellcome Sanger Institute for the COVID-19 Genomics UK (COG-UK) Consortium | Jacquelyn Wynn, Mairead Hyland, The Lighthouse Lab in Alderley Park and Alex Alderton, Roberto Amato, Jeffrey Barrett, Sonia Goncalves, Ewan Harrison, David K. Jackson, Ian Johnston, Dominic Kwiatkowski, Cordelia Langford, John Sillitoe on behalf of the Wellcome Sanger Institute COVID-19 Surveillance Team |
| EPI_ISL_2407102 | Lighthouse Lab in Milton Keynes | Wellcome Sanger Institute for the COVID-19 Genomics UK (COG-UK) Consortium | The Lighthouse Lab in Milton Keynes and Alex Alderton, Roberto Amato, Jeffrey Barrett, Sonia Goncalves, Ewan Harrison, David K. Jackson, Ian Johnston, Dominic Kwiatkowski, Cordelia Langford, John Sillitoe on behalf of the Wellcome Sanger Institute COVID-19 Surveillance Team |
| EPI_ISL_2407103, EPI_ISL_2407105, EPI_ISL_2407106 | Lighthouse Lab in Alderley Park | Wellcome Sanger Institute for the COVID-19 Genomics UK (COG-UK) Consortium | Jacquelyn Wynn, Mairead Hyland, The Lighthouse Lab in Alderley Park and Alex Alderton, Roberto Amato, Jeffrey Barrett, Sonia Goncalves, Ewan Harrison, David K. Jackson, Ian Johnston, Dominic Kwiatkowski, Cordelia Langford, John Sillitoe on behalf of the Wellcome Sanger Institute COVID-19 Surveillance Team |
| EPI_ISL_2407107 | Lighthouse Lab in Milton Keynes | Wellcome Sanger Institute for the COVID-19 Genomics UK (COG-UK) Consortium | The Lighthouse Lab in Milton Keynes and Alex Alderton, Roberto Amato, Jeffrey Barrett, Sonia Goncalves, Ewan Harrison, David K. Jackson, Ian Johnston, Dominic Kwiatkowski, Cordelia Langford, John Sillitoe on behalf of the Wellcome Sanger Institute COVID-19 Surveillance Team |
| EPI_ISL_2407108 | Lighthouse Lab in Alderley Park | Wellcome Sanger Institute for the COVID-19 Genomics UK (COG-UK) Consortium | Jacquelyn Wynn, Mairead Hyland, The Lighthouse Lab in Alderley Park and Alex Alderton, Roberto Amato, Jeffrey Barrett, Sonia Goncalves, Ewan Harrison, David K. Jackson, Ian Johnston, Dominic Kwiatkowski, Cordelia Langford, John Sillitoe on behalf of the Wellcome Sanger Institute COVID-19 Surveillance Team |
| EPI_ISL_2407109 | Lighthouse Lab in Milton Keynes | Wellcome Sanger Institute for the COVID-19 Genomics UK (COG-UK) Consortium | The Lighthouse Lab in Milton Keynes and Alex Alderton, Roberto Amato, Jeffrey Barrett, Sonia Goncalves, Ewan Harrison, David K. Jackson, Ian Johnston, Dominic Kwiatkowski, Cordelia Langford, John Sillitoe on behalf of the Wellcome Sanger Institute COVID-19 Surveillance Team |
| EPI_ISL_2407111 | Lighthouse Lab in Alderley Park | Wellcome Sanger Institute for the COVID-19 Genomics UK (COG-UK) Consortium | Jacquelyn Wynn, Mairead Hyland, The Lighthouse Lab in Alderley Park and Alex Alderton, Roberto Amato, Jeffrey Barrett, Sonia Goncalves, Ewan Harrison, David K. Jackson, Ian Johnston, Dominic Kwiatkowski, Cordelia Langford, John Sillitoe on behalf of the Wellcome Sanger Institute COVID-19 Surveillance Team |
| EPI_ISL_2407112, EPI_ISL_2407113, EPI_ISL_2407114 | Lighthouse Lab in Glasgow | Wellcome Sanger Institute for the COVID-19 Genomics UK (COG-UK) Consortium | Harper VanSteenhouse, Yumi Kasai, David Gray, Carol Clugston, Anna Dominiczak and Alex Alderton, Roberto Amato, Jeffrey Barrett, Sonia Goncalves, Ewan Harrison, David K. Jackson, Ian Johnston, Dominic Kwiatkowski, Cordelia Langford, John Sillitoe on behalf of the Wellcome Sanger Institute COVID-19 Surveillance Team |
| EPI_ISL_2407115, EPI_ISL_2407116, EPI_ISL_2407117, EPI_ISL_2407118, EPI_ISL_2407119, EPI_ISL_2407120, EPI_ISL_2407121, EPI_ISL_2407122, EPI_ISL_2407123, EPI_ISL_2407124, EPI_ISL_2407125, EPI_ISL_2407126, EPI_ISL_2407127, EPI_ISL_2407128 | | | |
| see above | Lighthouse Lab in Alderley Park | Wellcome Sanger Institute for the COVID-19 Genomics UK (COG-UK) Consortium | Jacquelyn Wynn, Mairead Hyland, The Lighthouse Lab in Alderley Park and Alex Alderton, Roberto Amato, Jeffrey Barrett, Sonia Goncalves, Ewan Harrison, David K. Jackson, Ian Johnston, Dominic Kwiatkowski, Cordelia Langford, John Sillitoe on behalf of the Wellcome Sanger Institute COVID-19 Surveillance Team |
| EPI_ISL_2407129 | Lighthouse Lab in Glasgow | Wellcome Sanger Institute for the COVID-19 Genomics UK (COG-UK) Consortium | Harper VanSteenhouse, Yumi Kasai, David Gray, Carol Clugston, Anna Dominiczak and Alex Alderton, Roberto Amato, Jeffrey Barrett, Sonia Goncalves, Ewan Harrison, David K. Jackson, Ian Johnston, Dominic Kwiatkowski, Cordelia Langford, John Sillitoe on behalf of the Wellcome Sanger Institute COVID-19 Surveillance Team |
| EPI_ISL_2407132 | Lighthouse Lab in Alderley Park | Wellcome Sanger Institute for the COVID-19 Genomics UK (COG-UK) Consortium | Jacquelyn Wynn, Mairead Hyland, The Lighthouse Lab in Alderley Park and Alex Alderton, Roberto Amato, Jeffrey Barrett, Sonia Goncalves, Ewan Harrison, David K. Jackson, Ian Johnston, Dominic Kwiatkowski, Cordelia Langford, John Sillitoe on behalf of the Wellcome Sanger Institute COVID-19 Surveillance Team |
| EPI_ISL_2407133, EPI_ISL_2407134 | Lighthouse Lab in Glasgow | Wellcome Sanger Institute for the COVID-19 Genomics UK (COG-UK) Consortium | Harper VanSteenhouse, Yumi Kasai, David Gray, Carol Clugston, Anna Dominiczak and Alex Alderton, Roberto Amato, Jeffrey Barrett, Sonia Goncalves, Ewan Harrison, David K. Jackson, Ian Johnston, Dominic Kwiatkowski, Cordelia Langford, John Sillitoe on behalf of the Wellcome Sanger Institute COVID-19 Surveillance Team |
| EPI_ISL_2407136, EPI_ISL_2407137, EPI_ISL_2407138, EPI_ISL_2407140, EPI_ISL_2407142, EPI_ISL_2407143, EPI_ISL_2407144, EPI_ISL_2407145, EPI_ISL_2407147, EPI_ISL_2407148 | Lighthouse Lab in Alderley Park | Wellcome Sanger Institute for the COVID-19 Genomics UK (COG-UK) Consortium | Jacquelyn Wynn, Mairead Hyland, The Lighthouse Lab in Alderley Park and Alex Alderton, Roberto Amato, Jeffrey Barrett, Sonia Goncalves, Ewan Harrison, David K. Jackson, Ian Johnston, Dominic Kwiatkowski, Cordelia Langford, John Sillitoe on behalf of the Wellcome Sanger Institute COVID-19 Surveillance Team |

| | | | |
|---|---|---|---|
| EPI_ISL_2407149, EPI_ISL_2407150 | Lighthouse Lab in Glasgow | Wellcome Sanger Institute for the COVID-19 Genomics UK (COG-UK) Consortium | Harper VanSteenhouse, Yumi Kasai, David Gray, Carol Clugston, Anna Dominiczak and Alex Alderton, Roberto Amato, Jeffrey Barrett, Sonia Goncalves, Ewan Harrison, David K. Jackson, Ian Johnston, Dominic Kwiatkowski, Cordelia Langford, John Sillitoe on behalf of the Wellcome Sanger Institute COVID-19 Surveillance Team |
| EPI_ISL_2407151, EPI_ISL_2407152, EPI_ISL_2407153, EPI_ISL_2407154, EPI_ISL_2407155, EPI_ISL_2407156, EPI_ISL_2407157, EPI_ISL_2407158, EPI_ISL_2407159, EPI_ISL_2407160 | Lighthouse Lab in Alderley Park | Wellcome Sanger Institute for the COVID-19 Genomics UK (COG-UK) Consortium | Jacquelyn Wynn, Mairead Hyland, The Lighthouse Lab in Alderley Park and Alex Alderton, Roberto Amato, Jeffrey Barrett, Sonia Goncalves, Ewan Harrison, David K. Jackson, Ian Johnston, Dominic Kwiatkowski, Cordelia Langford, John Sillitoe on behalf of the Wellcome Sanger Institute COVID-19 Surveillance Team |
| EPI_ISL_2407161 | Lighthouse Lab in Glasgow | Wellcome Sanger Institute for the COVID-19 Genomics UK (COG-UK) Consortium | Harper VanSteenhouse, Yumi Kasai, David Gray, Carol Clugston, Anna Dominiczak and Alex Alderton, Roberto Amato, Jeffrey Barrett, Sonia Goncalves, Ewan Harrison, David K. Jackson, Ian Johnston, Dominic Kwiatkowski, Cordelia Langford, John Sillitoe on behalf of the Wellcome Sanger Institute COVID-19 Surveillance Team |
| EPI_ISL_2407162, EPI_ISL_2407163, EPI_ISL_2407164, EPI_ISL_2407165, EPI_ISL_2407166, EPI_ISL_2407167, EPI_ISL_2407168, EPI_ISL_2407169, EPI_ISL_2407170, EPI_ISL_2407171, EPI_ISL_2407173 | | | |
| see above | Lighthouse Lab in Alderley Park | Wellcome Sanger Institute for the COVID-19 Genomics UK (COG-UK) Consortium | Jacquelyn Wynn, Mairead Hyland, The Lighthouse Lab in Alderley Park and Alex Alderton, Roberto Amato, Jeffrey Barrett, Sonia Goncalves, Ewan Harrison, David K. Jackson, Ian Johnston, Dominic Kwiatkowski, Cordelia Langford, John Sillitoe on behalf of the Wellcome Sanger Institute COVID-19 Surveillance Team |
| EPI_ISL_2407174 | Lighthouse Lab in Glasgow | Wellcome Sanger Institute for the COVID-19 Genomics UK (COG-UK) Consortium | Harper VanSteenhouse, Yumi Kasai, David Gray, Carol Clugston, Anna Dominiczak and Alex Alderton, Roberto Amato, Jeffrey Barrett, Sonia Goncalves, Ewan Harrison, David K. Jackson, Ian Johnston, Dominic Kwiatkowski, Cordelia Langford, John Sillitoe on behalf of the Wellcome Sanger Institute COVID-19 Surveillance Team |
| EPI_ISL_2407175 | Lighthouse Lab in Alderley Park | Wellcome Sanger Institute for the COVID-19 Genomics UK (COG-UK) Consortium | Jacquelyn Wynn, Mairead Hyland, The Lighthouse Lab in Alderley Park and Alex Alderton, Roberto Amato, Jeffrey Barrett, Sonia Goncalves, Ewan Harrison, David K. Jackson, Ian Johnston, Dominic Kwiatkowski, Cordelia Langford, John Sillitoe on behalf of the Wellcome Sanger Institute COVID-19 Surveillance Team |
| EPI_ISL_2407176 | Lighthouse Lab in Glasgow | Wellcome Sanger Institute for the COVID-19 Genomics UK (COG-UK) Consortium | Harper VanSteenhouse, Yumi Kasai, David Gray, Carol Clugston, Anna Dominiczak and Alex Alderton, Roberto Amato, Jeffrey Barrett, Sonia Goncalves, Ewan Harrison, David K. Jackson, Ian Johnston, Dominic Kwiatkowski, Cordelia Langford, John Sillitoe on behalf of the Wellcome Sanger Institute COVID-19 Surveillance Team |
| EPI_ISL_2407177, EPI_ISL_2407178, EPI_ISL_2407179, EPI_ISL_2407180, EPI_ISL_2407181, EPI_ISL_2407182, EPI_ISL_2407183, EPI_ISL_2407184, EPI_ISL_2407186, EPI_ISL_2407187, EPI_ISL_2407188, EPI_ISL_2407190, EPI_ISL_2407191, EPI_ISL_2407192, EPI_ISL_2407195, EPI_ISL_2407196, EPI_ISL_2407197, EPI_ISL_2407198, EPI_ISL_2407199, EPI_ISL_2407200, EPI_ISL_2407201, EPI_ISL_2407202, EPI_ISL_2407203, EPI_ISL_2407204, EPI_ISL_2407205, EPI_ISL_2407206, EPI_ISL_2407207 | | | |
| see above | Lighthouse Lab in Alderley Park | Wellcome Sanger Institute for the COVID-19 Genomics UK (COG-UK) Consortium | Jacquelyn Wynn, Mairead Hyland, The Lighthouse Lab in Alderley Park and Alex Alderton, Roberto Amato, Jeffrey Barrett, Sonia Goncalves, Ewan Harrison, David K. Jackson, Ian Johnston, Dominic Kwiatkowski, Cordelia Langford, John Sillitoe on behalf of the Wellcome Sanger Institute COVID-19 Surveillance Team |
| EPI_ISL_2407208 | Lighthouse Lab in Glasgow | Wellcome Sanger Institute for the COVID-19 Genomics UK (COG-UK) Consortium | Harper VanSteenhouse, Yumi Kasai, David Gray, Carol Clugston, Anna Dominiczak and Alex Alderton, Roberto Amato, Jeffrey Barrett, Sonia Goncalves, Ewan Harrison, David K. Jackson, Ian Johnston, Dominic Kwiatkowski, Cordelia Langford, John Sillitoe on behalf of the Wellcome Sanger Institute COVID-19 Surveillance Team |
| EPI_ISL_2407209, EPI_ISL_2407211, EPI_ISL_2407212, EPI_ISL_2407213, EPI_ISL_2407214, EPI_ISL_2407216, EPI_ISL_2407217 | Lighthouse Lab in Alderley Park | Wellcome Sanger Institute for the COVID-19 Genomics UK (COG-UK) Consortium | Jacquelyn Wynn, Mairead Hyland, The Lighthouse Lab in Alderley Park and Alex Alderton, Roberto Amato, Jeffrey Barrett, Sonia Goncalves, Ewan Harrison, David K. Jackson, Ian Johnston, Dominic Kwiatkowski, Cordelia Langford, John Sillitoe on behalf of the Wellcome Sanger Institute COVID-19 Surveillance Team |
| EPI_ISL_2407219 | Lighthouse Lab in Glasgow | Wellcome Sanger Institute for the COVID-19 Genomics UK (COG-UK) Consortium | Harper VanSteenhouse, Yumi Kasai, David Gray, Carol Clugston, Anna Dominiczak and Alex Alderton, Roberto Amato, Jeffrey Barrett, Sonia Goncalves, Ewan Harrison, David K. Jackson, Ian Johnston, Dominic Kwiatkowski, Cordelia Langford, John Sillitoe on behalf of the Wellcome Sanger Institute COVID-19 Surveillance Team |
| EPI_ISL_2407220, EPI_ISL_2407221, EPI_ISL_2407222, EPI_ISL_2407223, EPI_ISL_2407224, EPI_ISL_2407225, EPI_ISL_2407226, EPI_ISL_2407227, EPI_ISL_2407228, EPI_ISL_2407229, EPI_ISL_2407231, EPI_ISL_2407233, EPI_ISL_2407234, EPI_ISL_2407235, EPI_ISL_2407236, EPI_ISL_2407238, EPI_ISL_2407239 | | | |
| see above | Lighthouse Lab in Alderley Park | Wellcome Sanger Institute for the COVID-19 Genomics UK (COG-UK) Consortium | Jacquelyn Wynn, Mairead Hyland, The Lighthouse Lab in Alderley Park and Alex Alderton, Roberto Amato, Jeffrey Barrett, Sonia Goncalves, Ewan Harrison, David K. Jackson, Ian Johnston, Dominic Kwiatkowski, Cordelia Langford, John Sillitoe on behalf of the Wellcome Sanger Institute COVID-19 Surveillance Team |
| EPI_ISL_2407240, EPI_ISL_2407241 | Lighthouse Lab in Glasgow | Wellcome Sanger Institute for the COVID-19 Genomics UK (COG-UK) Consortium | Harper VanSteenhouse, Yumi Kasai, David Gray, Carol Clugston, Anna Dominiczak and Alex Alderton, Roberto Amato, Jeffrey Barrett, Sonia Goncalves, Ewan Harrison, David K. Jackson, Ian Johnston, Dominic Kwiatkowski, Cordelia Langford, John Sillitoe on behalf of the Wellcome Sanger Institute COVID-19 Surveillance Team |
| EPI_ISL_2407242, EPI_ISL_2407244, EPI_ISL_2407246, EPI_ISL_2407248 | Lighthouse Lab in Alderley Park | Wellcome Sanger Institute for the COVID-19 Genomics UK (COG-UK) Consortium | Jacquelyn Wynn, Mairead Hyland, The Lighthouse Lab in Alderley Park and Alex Alderton, Roberto Amato, Jeffrey Barrett, Sonia Goncalves, Ewan Harrison, David K. Jackson, Ian Johnston, Dominic Kwiatkowski, Cordelia Langford, John Sillitoe on behalf of the Wellcome Sanger Institute COVID-19 Surveillance Team |
| EPI_ISL_2407249 | Lighthouse Lab in Glasgow | Wellcome Sanger Institute for the COVID-19 Genomics UK (COG-UK) Consortium | Harper VanSteenhouse, Yumi Kasai, David Gray, Carol Clugston, Anna Dominiczak and Alex Alderton, Roberto Amato, Jeffrey Barrett, Sonia Goncalves, Ewan Harrison, David K. Jackson, Ian Johnston, Dominic Kwiatkowski, Cordelia Langford, John Sillitoe on behalf of the Wellcome Sanger Institute COVID-19 Surveillance Team |
| EPI_ISL_2407250, EPI_ISL_2407251, EPI_ISL_2407252, EPI_ISL_2407253, EPI_ISL_2407254 | Lighthouse Lab in Alderley Park | Wellcome Sanger Institute for the COVID-19 Genomics UK (COG-UK) Consortium | Jacquelyn Wynn, Mairead Hyland, The Lighthouse Lab in Alderley Park and Alex Alderton, Roberto Amato, Jeffrey Barrett, Sonia Goncalves, Ewan Harrison, David K. Jackson, Ian Johnston, Dominic Kwiatkowski, Cordelia Langford, John Sillitoe on behalf of the Wellcome Sanger Institute COVID-19 Surveillance Team |
| EPI_ISL_2407255 | Lighthouse Lab in Glasgow | Wellcome Sanger Institute for the COVID-19 Genomics UK (COG-UK) Consortium | Harper VanSteenhouse, Yumi Kasai, David Gray, Carol Clugston, Anna Dominiczak and Alex Alderton, Roberto Amato, Jeffrey Barrett, Sonia Goncalves, Ewan Harrison, David K. Jackson, Ian Johnston, Dominic Kwiatkowski, Cordelia Langford, John Sillitoe on behalf of the Wellcome Sanger Institute COVID-19 Surveillance Team |
| EPI_ISL_2407256 | Lighthouse Lab in Alderley Park | Wellcome Sanger Institute for the COVID-19 Genomics UK (COG-UK) Consortium | Jacquelyn Wynn, Mairead Hyland, The Lighthouse Lab in Alderley Park and Alex Alderton, Roberto Amato, Jeffrey Barrett, Sonia Goncalves, Ewan Harrison, David K. Jackson, Ian Johnston, Dominic Kwiatkowski, Cordelia Langford, John Sillitoe on behalf of the Wellcome Sanger Institute COVID-19 Surveillance Team |
| EPI_ISL_2407259 | Lighthouse Lab in Glasgow | Wellcome Sanger Institute for the COVID-19 Genomics UK (COG-UK) Consortium | Harper VanSteenhouse, Yumi Kasai, David Gray, Carol Clugston, Anna Dominiczak and Alex Alderton, Roberto Amato, Jeffrey Barrett, Sonia Goncalves, Ewan Harrison, David K. Jackson, Ian Johnston, Dominic Kwiatkowski, Cordelia Langford, John Sillitoe on behalf of the Wellcome Sanger Institute COVID-19 Surveillance Team |
| EPI_ISL_2407260 | Lighthouse Lab in Alderley Park | Wellcome Sanger Institute for the COVID-19 Genomics UK (COG-UK) Consortium | Jacquelyn Wynn, Mairead Hyland, The Lighthouse Lab in Alderley Park and Alex Alderton, Roberto Amato, Jeffrey Barrett, Sonia Goncalves, Ewan Harrison, David K. Jackson, Ian Johnston, Dominic Kwiatkowski, Cordelia Langford, John Sillitoe on behalf of the Wellcome Sanger Institute COVID-19 Surveillance Team |
| EPI_ISL_2407261 | Lighthouse Lab in Glasgow | Wellcome Sanger Institute for the COVID-19 Genomics UK (COG-UK) Consortium | Harper VanSteenhouse, Yumi Kasai, David Gray, Carol Clugston, Anna Dominiczak and Alex Alderton, Roberto Amato, Jeffrey Barrett, Sonia Goncalves, Ewan Harrison, David K. Jackson, Ian Johnston, Dominic Kwiatkowski, Cordelia Langford, John Sillitoe on behalf of the Wellcome Sanger Institute COVID-19 Surveillance Team |
| EPI_ISL_2407262, EPI_ISL_2407263, EPI_ISL_2407264, EPI_ISL_2407266, EPI_ISL_2407267, EPI_ISL_2407268, EPI_ISL_2407269, EPI_ISL_2407270, EPI_ISL_2407272, EPI_ISL_2407273, EPI_ISL_2407274, EPI_ISL_2407275, EPI_ISL_2407276, EPI_ISL_2407277, EPI_ISL_2407278, EPI_ISL_2407280, EPI_ISL_2407281, EPI_ISL_2407282, EPI_ISL_2407283, EPI_ISL_2407284 | | | |
| see above | Lighthouse Lab in Alderley Park | Wellcome Sanger Institute for the COVID-19 Genomics UK (COG-UK) Consortium | Jacquelyn Wynn, Mairead Hyland, The Lighthouse Lab in Alderley Park and Alex Alderton, Roberto Amato, Jeffrey Barrett, Sonia Goncalves, Ewan Harrison, David K. Jackson, Ian Johnston, Dominic Kwiatkowski, Cordelia Langford, John Sillitoe on behalf of the Wellcome Sanger Institute COVID-19 Surveillance Team |
| EPI_ISL_2407285, EPI_ISL_2407287, EPI_ISL_2407289, EPI_ISL_2407291, EPI_ISL_2407292, EPI_ISL_2407294, EPI_ISL_2407295, EPI_ISL_2407296, EPI_ISL_2407297, EPI_ISL_2407299, EPI_ISL_2407300, EPI_ISL_2407301, EPI_ISL_2407304, EPI_ISL_2407308, EPI_ISL_2407309, EPI_ISL_2407310, EPI_ISL_2407312, EPI_ISL_2407313, EPI_ISL_2407316, EPI_ISL_2407317, EPI_ISL_2407318, EPI_ISL_2407322, EPI_ISL_2407324, EPI_ISL_2407325, EPI_ISL_2407326, EPI_ISL_2407327, EPI_ISL_2407328, EPI_ISL_2407329, EPI_ISL_2407330, EPI_ISL_2407332, EPI_ISL_2407333, EPI_ISL_2407336, EPI_ISL_2407337, EPI_ISL_2407339, EPI_ISL_2407340, EPI_ISL_2407341, EPI_ISL_2407345, EPI_ISL_2407346, EPI_ISL_2407349, EPI_ISL_2407351, EPI_ISL_2407352, EPI_ISL_2407353, EPI_ISL_2407355, EPI_ISL_2407357, EPI_ISL_2407362, EPI_ISL_2407363, EPI_ISL_2407364, EPI_ISL_2407366, EPI_ISL_2407367, EPI_ISL_2407368, EPI_ISL_2407371, EPI_ISL_2407372, EPI_ISL_2407373, EPI_ISL_2407376, EPI_ISL_2407377, EPI_ISL_2407378, EPI_ISL_2407379, EPI_ISL_2407381, EPI_ISL_2407383, EPI_ISL_2407384, EPI_ISL_2407385, EPI_ISL_2407386, EPI_ISL_2407387, EPI_ISL_2407389, EPI_ISL_2407391, EPI_ISL_2407392, EPI_ISL_2407393, EPI_ISL_2407394, EPI_ISL_2407399, EPI_ISL_2407400, EPI_ISL_2407401, EPI_ISL_2407403, EPI_ISL_2407406, EPI_ISL_2407407, EPI_ISL_2407409, EPI_ISL_2407410, EPI_ISL_2407412, EPI_ISL_2407414, EPI_ISL_2407415, EPI_ISL_2407416, EPI_ISL_2407418, EPI_ISL_2407419, EPI_ISL_2407420, EPI_ISL_2407421, EPI_ISL_2407422, EPI_ISL_2407423, EPI_ISL_2407424, EPI_ISL_2407425, EPI_ISL_2407426, EPI_ISL_2407427, EPI_ISL_2407428, EPI_ISL_2407429, EPI_ISL_2407436, EPI_ISL_2407438, EPI_ISL_2407439, EPI_ISL_2407440, EPI_ISL_2407441, EPI_ISL_2407444, EPI_ISL_2407445, EPI_ISL_2407446, EPI_ISL_2407447, EPI_ISL_2407448, EPI_ISL_2407449, EPI_ISL_2407451, EPI_ISL_2407452, EPI_ISL_2407453, EPI_ISL_2407455, EPI_ISL_2407456, EPI_ISL_2407461, EPI_ISL_2407463, EPI_ISL_2407466 | | | |
| see above | Lighthouse Lab in Glasgow | Wellcome Sanger Institute for the COVID-19 Genomics UK (COG-UK) Consortium | Harper VanSteenhouse, Yumi Kasai, David Gray, Carol Clugston, Anna Dominiczak and Alex Alderton, Roberto Amato, Jeffrey Barrett, Sonia Goncalves, Ewan Harrison, David K. Jackson, Ian Johnston, Dominic Kwiatkowski, Cordelia Langford, John Sillitoe on behalf of the Wellcome Sanger Institute COVID-19 Surveillance Team |
| EPI_ISL_2407467 | Lighthouse Lab in Alderley Park | Wellcome Sanger Institute for the COVID-19 Genomics UK (COG-UK) Consortium | Jacquelyn Wynn, Mairead Hyland, The Lighthouse Lab in Alderley Park and Alex Alderton, Roberto Amato, Jeffrey Barrett, Sonia Goncalves, Ewan Harrison, David K. Jackson, Ian Johnston, Dominic Kwiatkowski, Cordelia Langford, John Sillitoe on behalf of the Wellcome Sanger Institute COVID-19 Surveillance Team |
| EPI_ISL_2407468, EPI_ISL_2407469, EPI_ISL_2407471, EPI_ISL_2407473, EPI_ISL_2407474, EPI_ISL_2407476, EPI_ISL_2407477, EPI_ISL_2407480, EPI_ISL_2407481, EPI_ISL_2407482, EPI_ISL_2407484, EPI_ISL_2407486, EPI_ISL_2407487, EPI_ISL_2407488, EPI_ISL_2407489, EPI_ISL_2407494, EPI_ISL_2407495, EPI_ISL_2407498, EPI_ISL_2407499, EPI_ISL_2407500, EPI_ISL_2407502, EPI_ISL_2407503, EPI_ISL_2407504, EPI_ISL_2407505, EPI_ISL_2407506, EPI_ISL_2407508, EPI_ISL_2407511, EPI_ISL_2407512, EPI_ISL_2407513, EPI_ISL_2407514, EPI_ISL_2407515, EPI_ISL_2407516, EPI_ISL_2407517, EPI_ISL_2407521, EPI_ISL_2407522, EPI_ISL_2407523, EPI_ISL_2407524, EPI_ISL_2407525, EPI_ISL_2407526, EPI_ISL_2407527, EPI_ISL_2407528, EPI_ISL_2407529, EPI_ISL_2407531, EPI_ISL_2407532, EPI_ISL_2407535, EPI_ISL_2407536, EPI_ISL_2407539, EPI_ISL_2407542, EPI_ISL_2407544, EPI_ISL_2407545, EPI_ISL_2407546, EPI_ISL_2407547, EPI_ISL_2407548, EPI_ISL_2407552, EPI_ISL_2407554, EPI_ISL_2407556, EPI_ISL_2407560, EPI_ISL_2407561, EPI_ISL_2407562, EPI_ISL_2407566, EPI_ISL_2407568, EPI_ISL_2407571, EPI_ISL_2407573, EPI_ISL_2407574, EPI_ISL_2407577, EPI_ISL_2407579, EPI_ISL_2407581, EPI_ISL_2407582, EPI_ISL_2407584, EPI_ISL_2407585, EPI_ISL_2407586, EPI_ISL_2407587, EPI_ISL_2407589, EPI_ISL_2407591, EPI_ISL_2407593, EPI_ISL_2407595, EPI_ISL_2407596, EPI_ISL_2407599, EPI_ISL_2407601, EPI_ISL_2407602, EPI_ISL_2407603, EPI_ISL_2407604, EPI_ISL_2407605, EPI_ISL_2407606, EPI_ISL_2407607 | | | |
| see above | Lighthouse Lab in Glasgow | Wellcome Sanger Institute for the COVID-19 Genomics UK (COG-UK) Consortium | Harper VanSteenhouse, Yumi Kasai, David Gray, Carol Clugston, Anna Dominiczak and Alex Alderton, Roberto Amato, Jeffrey Barrett, Sonia Goncalves, Ewan Harrison, David K. Jackson, Ian Johnston, Dominic Kwiatkowski, Cordelia Langford, John Sillitoe on behalf of the Wellcome Sanger Institute COVID-19 Surveillance Team |
| EPI_ISL_2407610 | Lighthouse Lab in Alderley Park | Wellcome Sanger Institute for the COVID-19 Genomics UK (COG-UK) Consortium | Jacquelyn Wynn, Mairead Hyland, The Lighthouse Lab in Alderley Park and Alex Alderton, Roberto Amato, Jeffrey Barrett, Sonia Goncalves, Ewan Harrison, David K. Jackson, Ian Johnston, Dominic Kwiatkowski, Cordelia Langford, John Sillitoe on behalf of the Wellcome Sanger Institute COVID-19 Surveillance Team |
| EPI_ISL_2407611, EPI_ISL_2407612, EPI_ISL_2407614, EPI_ISL_2407617, EPI_ISL_2407619, EPI_ISL_2407621, EPI_ISL_2407622, EPI_ISL_2407623, EPI_ISL_2407624, EPI_ISL_2407625, EPI_ISL_2407630, EPI_ISL_2407632, EPI_ISL_2407633, EPI_ISL_2407634, EPI_ISL_2407635, EPI_ISL_2407636, EPI_ISL_2407637, EPI_ISL_2407638, EPI_ISL_2407639 | | | |
| see above | Lighthouse Lab in Glasgow | Wellcome Sanger Institute for the COVID-19 Genomics UK (COG-UK) Consortium | Harper VanSteenhouse, Yumi Kasai, David Gray, Carol Clugston, Anna Dominiczak and Alex Alderton, Roberto Amato, Jeffrey Barrett, Sonia Goncalves, Ewan Harrison, David K. Jackson, Ian Johnston, Dominic Kwiatkowski, Cordelia Langford, John Sillitoe on behalf of the Wellcome Sanger Institute COVID-19 Surveillance Team |
| EPI_ISL_2407644 | Lighthouse Lab in Alderley Park | Wellcome Sanger Institute for the COVID-19 Genomics UK (COG-UK) Consortium | Jacquelyn Wynn, Mairead Hyland, The Lighthouse Lab in Alderley Park and Alex Alderton, Roberto Amato, Jeffrey Barrett, Sonia Goncalves, Ewan Harrison, David K. Jackson, Ian Johnston, Dominic Kwiatkowski, Cordelia Langford, John Sillitoe on behalf of the Wellcome Sanger Institute COVID-19 Surveillance Team |
| EPI_ISL_2407645, EPI_ISL_2407649, EPI_ISL_2407655, EPI_ISL_2407657, EPI_ISL_2407659, EPI_ISL_2407660, EPI_ISL_2407664, EPI_ISL_2407665, EPI_ISL_2407666, EPI_ISL_2407669, EPI_ISL_2407673, EPI_ISL_2407674, EPI_ISL_2407675, EPI_ISL_2407677, EPI_ISL_2407678, EPI_ISL_2407679, EPI_ISL_2407680, EPI_ISL_2407681, EPI_ISL_2407682, EPI_ISL_2407684, EPI_ISL_2407687, EPI_ISL_2407688, EPI_ISL_2407691, EPI_ISL_2407692, EPI_ISL_2407693, EPI_ISL_2407695, EPI_ISL_2407696, EPI_ISL_2407697, EPI_ISL_2407699, EPI_ISL_2407700, EPI_ISL_2407701, EPI_ISL_2407702, EPI_ISL_2407703, EPI_ISL_2407704, EPI_ISL_2407707, EPI_ISL_2407709, EPI_ISL_2407711, EPI_ISL_2407712, EPI_ISL_2407713, EPI_ISL_2407716, EPI_ISL_2407717, EPI_ISL_2407720, EPI_ISL_2407725, EPI_ISL_2407730, EPI_ISL_2407731, EPI_ISL_2407732, EPI_ISL_2407736, EPI_ISL_2407737, EPI_ISL_2407738, EPI_ISL_2407739, EPI_ISL_2407741, EPI_ISL_2407742, EPI_ISL_2407743, EPI_ISL_2407745, EPI_ISL_2407747, EPI_ISL_2407748, EPI_ISL_2407749, EPI_ISL_2407750, EPI_ISL_2407751, EPI_ISL_2407753, EPI_ISL_2407755, EPI_ISL_2407757, EPI_ISL_2407758, EPI_ISL_2407759, EPI_ISL_2407761, EPI_ISL_2407762, EPI_ISL_2407766, EPI_ISL_2407767, EPI_ISL_2407768, EPI_ISL_2407769, EPI_ISL_2407771, EPI_ISL_2407774, EPI_ISL_2407776, EPI_ISL_2407777, EPI_ISL_2407779, EPI_ISL_2407781, EPI_ISL_2407782, EPI_ISL_2407786, EPI_ISL_2407788, EPI_ISL_2407790, EPI_ISL_2407793, EPI_ISL_2407796, EPI_ISL_2407797, EPI_ISL_2407798, EPI_ISL_2407799, EPI_ISL_2407800, EPI_ISL_2407801, EPI_ISL_2407802, EPI_ISL_2407805, EPI_ISL_2407809, EPI_ISL_2407810, EPI_ISL_2407812, EPI_ISL_2407813, EPI_ISL_2407814, EPI_ISL_2407816, EPI_ISL_2407817, EPI_ISL_2407818, EPI_ISL_2407820, EPI_ISL_2407821, EPI_ISL_2407822, EPI_ISL_2407823, EPI_ISL_2407825, EPI_ISL_2407827, EPI_ISL_2407829, EPI_ISL_2407831, EPI_ISL_2407833, EPI_ISL_2407834, EPI_ISL_2407835, EPI_ISL_2407836, EPI_ISL_2407838, | | | |

| | | | |
|---|---|---|---|
| EPI_ISL_2407839, EPI_ISL_2407840, EPI_ISL_2407841, EPI_ISL_2407843, EPI_ISL_2407844, EPI_ISL_2407846, EPI_ISL_2407847, EPI_ISL_2407848, EPI_ISL_2407849, EPI_ISL_2407850, EPI_ISL_2407851, EPI_ISL_2407853, EPI_ISL_2407855, EPI_ISL_2407858, EPI_ISL_2407860, EPI_ISL_2407861, EPI_ISL_2407864, EPI_ISL_2407865, EPI_ISL_2407867, EPI_ISL_2407870, EPI_ISL_2407871, EPI_ISL_2407872, EPI_ISL_2407874, EPI_ISL_2407875, EPI_ISL_2407877, EPI_ISL_2407878, EPI_ISL_2407879, EPI_ISL_2407881, EPI_ISL_2407882, EPI_ISL_2407884, EPI_ISL_2407885, EPI_ISL_2407886, EPI_ISL_2407889, EPI_ISL_2407890, EPI_ISL_2407891, EPI_ISL_2407892, EPI_ISL_2407893, EPI_ISL_2407894, EPI_ISL_2407895, EPI_ISL_2407896, EPI_ISL_2407897, EPI_ISL_2407898, EPI_ISL_2407901, EPI_ISL_2407903, EPI_ISL_2407904, EPI_ISL_2407906, EPI_ISL_2407907, EPI_ISL_2407908, EPI_ISL_2407909, EPI_ISL_2407912, EPI_ISL_2407913, EPI_ISL_2407915, EPI_ISL_2407916, EPI_ISL_2407917, EPI_ISL_2407918, EPI_ISL_2407919, EPI_ISL_2407921, EPI_ISL_2407922, EPI_ISL_2407923, EPI_ISL_2407924, EPI_ISL_2407925, EPI_ISL_2407929, EPI_ISL_2407931, EPI_ISL_2407933, EPI_ISL_2407935, EPI_ISL_2407937, EPI_ISL_2407939, EPI_ISL_2407940, EPI_ISL_2407944, EPI_ISL_2407945, EPI_ISL_2407946, EPI_ISL_2407947, EPI_ISL_2407948, EPI_ISL_2407949, EPI_ISL_2407950, EPI_ISL_2407951, EPI_ISL_2407952, EPI_ISL_2407955, EPI_ISL_2407956, EPI_ISL_2407957, EPI_ISL_2407959, EPI_ISL_2407961, EPI_ISL_2407963, EPI_ISL_2407965, EPI_ISL_2407966, EPI_ISL_2407967, EPI_ISL_2407969, EPI_ISL_2407970, EPI_ISL_2407971, EPI_ISL_2407972, EPI_ISL_2407973, EPI_ISL_2407975, EPI_ISL_2407977, EPI_ISL_2407981, EPI_ISL_2407982, EPI_ISL_2407983, EPI_ISL_2407986, EPI_ISL_2407987, EPI_ISL_2407988, EPI_ISL_2407989, EPI_ISL_2407990, EPI_ISL_2407991, EPI_ISL_2407992, EPI_ISL_2407993, EPI_ISL_2407996, EPI_ISL_2407998, EPI_ISL_2408000, EPI_ISL_2408001, EPI_ISL_2408002, EPI_ISL_2408005, EPI_ISL_2408006, EPI_ISL_2408007, EPI_ISL_2408009, EPI_ISL_2408010, EPI_ISL_2408013, EPI_ISL_2408014, EPI_ISL_2408015, EPI_ISL_2408016, EPI_ISL_2408017, EPI_ISL_2408020, EPI_ISL_2408021, EPI_ISL_2408022, EPI_ISL_2408023, EPI_ISL_2408025, EPI_ISL_2408026, EPI_ISL_2408027, EPI_ISL_2408028, EPI_ISL_2408029, EPI_ISL_2408030, EPI_ISL_2408031, EPI_ISL_2408032, EPI_ISL_2408033, EPI_ISL_2408034, EPI_ISL_2408036, EPI_ISL_2408037, EPI_ISL_2408038, EPI_ISL_2408039, EPI_ISL_2408040, EPI_ISL_2408041, EPI_ISL_2408042, EPI_ISL_2408044, EPI_ISL_2408045, EPI_ISL_2408046, EPI_ISL_2408047, EPI_ISL_2408048 | | | |
| see above | Lighthouse Lab in Glasgow | Wellcome Sanger Institute for the COVID-19 Genomics UK (COG-UK) Consortium | Harper VanSteenhouse, Yumi Kasai, David Gray, Carol Clugston, Anna Dominiczak and Alex Alderton, Roberto Amato, Jeffrey Barrett, Sonia Goncalves, Ewan Harrison, David K. Jackson, Ian Johnston, Dominic Kwiatkowski, Cordelia Langford, John Sillitoe on behalf of the Wellcome Sanger Institute COVID-19 Surveillance Team |
| EPI_ISL_2408049 | Lighthouse Lab in Alderley Park | Wellcome Sanger Institute for the COVID-19 Genomics UK (COG-UK) Consortium | Jacquelyn Wynn, Mairead Hyland, The Lighthouse Lab in Alderley Park and Alex Alderton, Roberto Amato, Jeffrey Barrett, Sonia Goncalves, Ewan Harrison, David K. Jackson, Ian Johnston, Dominic Kwiatkowski, Cordelia Langford, John Sillitoe on behalf of the Wellcome Sanger Institute COVID-19 Surveillance Team |
| EPI_ISL_2408050, EPI_ISL_2408052, EPI_ISL_2408053, EPI_ISL_2408054, EPI_ISL_2408055, EPI_ISL_2408056 | Lighthouse Lab in Glasgow | Wellcome Sanger Institute for the COVID-19 Genomics UK (COG-UK) Consortium | Harper VanSteenhouse, Yumi Kasai, David Gray, Carol Clugston, Anna Dominiczak and Alex Alderton, Roberto Amato, Jeffrey Barrett, Sonia Goncalves, Ewan Harrison, David K. Jackson, Ian Johnston, Dominic Kwiatkowski, Cordelia Langford, John Sillitoe on behalf of the Wellcome Sanger Institute COVID-19 Surveillance Team |
| EPI_ISL_2408057, EPI_ISL_2408058 | Lighthouse Lab in Alderley Park | Wellcome Sanger Institute for the COVID-19 Genomics UK (COG-UK) Consortium | Jacquelyn Wynn, Mairead Hyland, The Lighthouse Lab in Alderley Park and Alex Alderton, Roberto Amato, Jeffrey Barrett, Sonia Goncalves, Ewan Harrison, David K. Jackson, Ian Johnston, Dominic Kwiatkowski, Cordelia Langford, John Sillitoe on behalf of the Wellcome Sanger Institute COVID-19 Surveillance Team |
| EPI_ISL_2408059, EPI_ISL_2408060 | Lighthouse Lab in Glasgow | Wellcome Sanger Institute for the COVID-19 Genomics UK (COG-UK) Consortium | Harper VanSteenhouse, Yumi Kasai, David Gray, Carol Clugston, Anna Dominiczak and Alex Alderton, Roberto Amato, Jeffrey Barrett, Sonia Goncalves, Ewan Harrison, David K. Jackson, Ian Johnston, Dominic Kwiatkowski, Cordelia Langford, John Sillitoe on behalf of the Wellcome Sanger Institute COVID-19 Surveillance Team |
| EPI_ISL_2408061 | Lighthouse Lab in Milton Keynes | Wellcome Sanger Institute for the COVID-19 Genomics UK (COG-UK) Consortium | The Lighthouse Lab in Milton Keynes and Alex Alderton, Roberto Amato, Jeffrey Barrett, Sonia Goncalves, Ewan Harrison, David K. Jackson, Ian Johnston, Dominic Kwiatkowski, Cordelia Langford, John Sillitoe on behalf of the Wellcome Sanger Institute COVID-19 Surveillance Team |
| EPI_ISL_2408255, EPI_ISL_2408260, EPI_ISL_2408314, EPI_ISL_2408315, EPI_ISL_2408316, EPI_ISL_2408321, EPI_ISL_2408323, EPI_ISL_2408326, EPI_ISL_2408327, EPI_ISL_2408328, EPI_ISL_2408330, EPI_ISL_2408338, EPI_ISL_2408340, EPI_ISL_2408352, EPI_ISL_2408358, EPI_ISL_2408359, EPI_ISL_2408360, EPI_ISL_2408364, EPI_ISL_2408368, EPI_ISL_2408384 | | | |
| see above | Ministry of Public Health / Hamad Medical Corporation | Biomedical Research Center (BRC), Qatar University / Qatar Genome Project (QGP) | BRC: Fatiha M. Benslimane, Heba A. Al-Khatib, Oal Al-Jamal, Dana Al-Batesh, Hadi M. Yassine, Asmaa A. Al-Thani. MOPH and HMC: Abdullatif Al-Khal, Muna A. S. Al-Maslamani, Mashael A. Al-Bader, Hamda Alromaihi, Roberto Bertollini, Peter V. Coyle, Einas A. E. Al-Kuwari, Hamad E. Al-Romaihi, Salih Al-Marri, Mohammed Al-Thani, Reham A. El-Kahlout. QBB: Tasneem Al-Hamad, Dina Elgakhlab QGP: Fatima H. Al-Kuwari, Chadi Saad |
| EPI_ISL_2408445 | Clinical Virology | Clinical Bacteriology | Tim Roloff, Fanny Wegner, Helena MB Seth-Smith, Alfredo Mari, Karoline Leuzinger, Julia Bielicki, Manuel Battegay, Hans Hirsch, Adrian Egli |
| EPI_ISL_2411693, EPI_ISL_2411761, EPI_ISL_2411877, EPI_ISL_2412260, EPI_ISL_2412644, EPI_ISL_2412739, EPI_ISL_2412850 | Department of Bacteria, Parasites and Fungi, Statens Serum Institut, Copenhagen, Denmark | Statens Serum Institut Bioinformatics and Microbial Genomics | Danish Covid-19 Genome Consortium |
| EPI_ISL_2414357, EPI_ISL_2414459, EPI_ISL_2414464 | Swedish national genomic surveillance program of SARS-CoV-2 | The Public Health Agency of Sweden | Maximilian Riess, Maria Lind Karlberg, Alma Brolund, Swedish national genomic surveillance program of SARS-CoV-2 |
| EPI_ISL_2414490 | Department of Bacteria, Parasites and Fungi, Statens Serum Institut, Copenhagen, Denmark | Statens Serum Institut Bioinformatics and Microbial Genomics | Danish Covid-19 Genome Consortium |
| EPI_ISL_2414793, EPI_ISL_2415198, EPI_ISL_2415482, EPI_ISL_2415484, EPI_ISL_2415611, EPI_ISL_2415935, EPI_ISL_2416022, EPI_ISL_2416025 | Swedish national genomic surveillance program of SARS-CoV-2 | The Public Health Agency of Sweden | Maximilian Riess, Maria Lind Karlberg, Alma Brolund, Swedish national genomic surveillance program of SARS-CoV-2 |
| EPI_ISL_2416932, EPI_ISL_2417283, EPI_ISL_2417828 | Department of Bacteria, Parasites and Fungi, Statens Serum Institut, Copenhagen, Denmark | Statens Serum Institut Bioinformatics and Microbial Genomics | Danish Covid-19 Genome Consortium |
| EPI_ISL_2418112, EPI_ISL_2418223, EPI_ISL_2418250, EPI_ISL_2418275, EPI_ISL_2418290, EPI_ISL_2418297, EPI_ISL_2418305, EPI_ISL_2418315, EPI_ISL_2418320, EPI_ISL_2418322, EPI_ISL_2418328, EPI_ISL_2418647, EPI_ISL_2419791 | | | |
| see above | Swedish national genomic surveillance program of SARS-CoV-2 | The Public Health Agency of Sweden | Maximilian Riess, Maria Lind Karlberg, Alma Brolund, Swedish national genomic surveillance program of SARS-CoV-2 |
| EPI_ISL_2420168 | IN State Department of Health Laboratory Services | IN State Department of Health Laboratory Services | Cassandra Campion, Jamie Yeadon, Brian Pope, Lixia Liu, Kyle Brownlee, Melissa Hindenlang, Mark Glazier |
| EPI_ISL_2420172, EPI_ISL_2420215, EPI_ISL_2420238 | Valais Hospital, Central Institute | Valais Hospital, Central Institute | Alexis Dumoulin, Lorenzo Cerutti, Henri Pegeot, Melyssa Elies, Deborah Penet, Keith Harshman, Ioannis Xenarios, Emmanouil Dermitzakis |
| EPI_ISL_2420245 | IN State Department of Health Laboratory Services | IN State Department of Health Laboratory Services | Cassandra Campion, Jamie Yeadon, Brian Pope, Lixia Liu, Kyle Brownlee, Melissa Hindenlang, Mark Glazier |
| EPI_ISL_2420342 | Laboratoire Virologie Saint Louis APHP | Laboratoire Virologie Saint Louis APHP | Maud Salmona, Marie Laure Chaix, Severine Mercier Delarue, Marie Laure Néré, Linda Feghoul, Jérome Le Goff, Constance Delauguerre, Sophia Achaibou |
| EPI_ISL_2420473 | San Gallicano Dermatological Institute I.F.O. | INMI Lazzaro Spallanzani IRCCS | G Bonfiglio, O Butera, CEM Gruber, F Santini, B Bartolini, E Giombini, F Messina, M Rueca, F Pimpinelli, F Ensoli, E Trento, MR Capobianchi, A Di Caro |
| EPI_ISL_2420474 | INMI Lazzaro Spallanzani IRCCS | INMI Lazzaro Spallanzani IRCCS | O Butera, CEM Gruber, F Santini, B Bartolini, E Giombini, F Messina, M Rueca, G Bonfiglio, A Di Caro, MR Capobianchi |
| EPI_ISL_2421240 | BOSTON HEART DIAGNOSTICS CORP | Wadsworth Center, New York State Department of Health | Kirsten St. George, Daryl M. Lamson, Alexis Russell, Matthew Shudt, Melissa A Leisner, Jonathan Plitnick, Catharine Prussing, Navjot Singh, John Kelly, Erasmus Schneider, Erica Lasek-Nesselquist |
| EPI_ISL_2421279, EPI_ISL_2421280, EPI_ISL_2421281, EPI_ISL_2421282 | Laboratori de Referencia de Catalunya | Laboratori de Referencia de Catalunya | Ramirez A., Bellosillo B., Padilla E., Vilas A., Hernandez JJ., Canal M. |
| EPI_ISL_2421819, EPI_ISL_2421902, EPI_ISL_2421958, EPI_ISL_2421966, EPI_ISL_2421970, EPI_ISL_2421971, EPI_ISL_2422090 | Aegis Sciences Corporation | Centers for Disease Control and Prevention Division of Viral Diseases, Pathogen Discovery | Dakota Howard, Dhwani Batra, Peter W. Cook, Kara Moser, Adrian Paskey, Jason Caravas, Benjamin Rambo-Martin, Shatavia Morrison, Christopher Gulvick, Scott Sammons, Yvette Unoarumhi, Darlene Wagner, Matthew Schmerer, Cyndi Clark, Patrick Campbell, Rob Case, Vikramsinha Ghorpade, Holly Houdeshell, Ola Kvalvaag, Dillon Nall, Ethan Sanders, Alec Vest, Shaun Westlund, Matthew Hardison, Clinton R. Paden, Duncan MacCannell |
| EPI_ISL_2422123 | Algemeen Medisch Labo | Labo Klinische Biologie, UZA | Marie Le Mercier, Jasmine Coppens, Basil Britto Xavier, Christine Lammens, Veerle Matheeussen, Herman Goossens |
| EPI_ISL_2422173, EPI_ISL_2422181 | Platform BIS UZA/UAntwerpen | Labo Klinische Biologie, UZA | Marie Le Mercier, Jasmine Coppens, Basil Britto Xavier, Christine Lammens, Veerle Matheeussen, Herman Goossens |
| EPI_ISL_2422226 | Labo Klinische Biologie, UZA | Labo Klinische Biologie, UZA | Marie Le Mercier, Jasmine Coppens, Basil Britto Xavier, Christine Lammens, Veerle Matheeussen, Herman Goossens |
| EPI_ISL_2422370, EPI_ISL_2422371, EPI_ISL_2422399 | Jessa | Jessa | Berden et al. on behalf of the Jessa_cmdLab |
| EPI_ISL_2422442 | AR Dept. of Health-PHL, Molecular Diagnostics | Genomics and Discovery, Respiratory Viruses Branch, Division of Viral Diseases, Centers for Disease Control and Prevention | Anna Kelleher, Ying Tao, Yan Li, Jing Zhang, Brian Lynch, Krista Queen, Anna Uehara, Peter Cook, Han Jia Justin Ng, Rachel Marine, Clinton R. Paden, Dhwani Batra, Haibin Wang, Tara Coalter, Jasmine Padilla, Morgan Davis, Mili Sheth, Sarah Nobles, Mark Burroughs, Justin Lee, Adam Retchless, Suxiang Tong |
| EPI_ISL_2422459 | GA Department of Public Health Laboratory | Genomics and Discovery, Respiratory Viruses Branch, Division of Viral Diseases, Centers for Disease Control and Prevention | Anna Kelleher, Ying Tao, Yan Li, Jing Zhang, Brian Lynch, Krista Queen, Anna Uehara, Peter Cook, Han Jia Justin Ng, Rachel Marine, Clinton R. Paden, Dhwani Batra, Haibin Wang, Tara Coalter, Jasmine Padilla, Morgan Davis, Mili Sheth, Sarah Nobles, Mark Burroughs, Justin Lee, Adam Retchless, Suxiang Tong |
| EPI_ISL_2423033 | Diagnostyka. Laboratoria Medyczne. | 1. ViroGenetics - BSL3 Laboratory of Virology, Małopolska Centre of Biotechnology, Jagiellonian University; 2. Diagtron Laboratoria Lukasz Rabalski | Rabalski L., Zielinska M, Gromowski,T., Rogalska-Kupiec M., Swadzba J., Wydmanski,W., Mazur-Panasiuk,N., Kowalski,M., Maciej Kosinski, Natalia Derewonko, Szulc,P., Sylwia Januszczak, Labaj,P.P., Foremny J, Pyrc,K. |
| EPI_ISL_2423090, EPI_ISL_2423128, EPI_ISL_2423135, EPI_ISL_2423139 | Diagnostyka. Laboratoria Medyczne. | 1. ViroGenetics - BSL3 Laboratory of Virology, Małopolska Centre of Biotechnology, Jagiellonian University; 2. genXone SA, Research & Development Laboratory | Mazur-Panasiuk,N., Nowicki G, Gromowski,T., Drweska-Matelska N, Grabowski J, Brylak A, Gidlewicz A, Szeszko K, Sykulski M, Krych L, Kowalski,M., Wydmanski W., Szulc,P., Januszczak S, Labaj,P.P., Kaszuba M, Pyrc,K. |
| EPI_ISL_2423473, EPI_ISL_2423486 | Diagnostyka. Laboratoria Medyczne. | 1. ViroGenetics - BSL3 Laboratory of Virology, Małopolska Centre of Biotechnology, Jagiellonian University; 2. Diagtron Laboratoria Lukasz Rabalski | Rabalski L., Gromowski,T., Rogalska-Kupiec M., Swadzba J., Wydmanski,W., Mazur-Panasiuk,N., Kowalski,M., Maciej Kosinski, Natalia Derewonko, Szulc,P., Sylwia Januszczak, Labaj,P.P., Pyrc,K. |
| EPI_ISL_2423549 | TXDSHS | TXDSHS | Rashmi Tuladhar, Bonnie Oh, Jenny Zhang, Maliha Rahman, Mayela Pedrueza, Anita Pokharel, Karen Bobier, Lorraine Rodriguez, Myong Koag, Chun Wang, Rachel Lee, Grace Kubin |

| | | | |
|---|---|---|---|
| EPI_ISL_2423826, EPI_ISL_2423911 | TGen North | TGen North | "Jolene Bowers, Heather Centner, Chris French, Hayley Yaglom, Ashlyn Pfeiffer, Darrin Lemmer, Dave Engelthaler, The Arizona COVID Genomics Union (ACGU)" |
| EPI_ISL_2424439, EPI_ISL_2424440, EPI_ISL_2424441, EPI_ISL_2424442, EPI_ISL_2424443, EPI_ISL_2424444, EPI_ISL_2425097, EPI_ISL_2425122, EPI_ISL_2425254, EPI_ISL_2425255, EPI_ISL_2425256, EPI_ISL_2425257, EPI_ISL_2425258, EPI_ISL_2425316, EPI_ISL_2425317, EPI_ISL_2425318, EPI_ISL_2425319, EPI_ISL_2425320, EPI_ISL_2425321, EPI_ISL_2425322 | | | |
| see above | KU Leuven, Rega Institute, Clinical and Epidemiological Virology | KU Leuven, Rega Institute, Clinical and Epidemiological Virology | Tony Wawina-Bokalanga, Bert Vanmechelen, Joan Marti-Carerras, Piet Maes |
| EPI_ISL_2425397 | M Health Fairview | Minnesota Department of Health, Public Health Laboratory | Alexandra Lorentz, Jacob Garfin, Matt Plumb, and Xiong Wang |
| EPI_ISL_2425410, EPI_ISL_2425414, EPI_ISL_2425415 | MD PHL | MD PHL | Maryland Department of Health Laboratories Administration |
| EPI_ISL_2425458, EPI_ISL_2425478 | Virginia Division of Consolidated Laboratory Services | Virginia Division of Consolidated Laboratory Services | Virginia DCLS |
| EPI_ISL_2425841 | Furst Medical Laboratory | Norwegian Institute of Public Health, Department of Virology | Kathrine Stene-Johansen, Kamilla Heddeland Instefjord, Hilde Elshaug, Garcia Llorente Ignacio, Jon Bråte, Engebretsen Serina Beate,Pedersen Benedikte Nevjen, Line Victoria Moen, Debech Nadia, Atiya R Ali,Marie Paulsen Madsen, Rasmus Riis Kopperud, Hilde Vollan, Karoline Bragstad, Olav Hungnes |
| EPI_ISL_2426102, EPI_ISL_2426114 | Yale Clinical Virology Lab | Grubaugh Lab - Yale School of Public Health | Joseph Fauver, Mallery Breban, Isabel Ott, Tara Alpert, Mary Petrone, Anderson Brito, Chantal Vogels, Annie Watkins, Chaney Kalinich, Jessica Rothman, Marie L. Landry, Nathan Grubaugh |
| EPI_ISL_2426148, EPI_ISL_2426149, EPI_ISL_2426151, EPI_ISL_2426153, EPI_ISL_2426154, EPI_ISL_2426155, EPI_ISL_2426156, EPI_ISL_2426159, EPI_ISL_2426160, EPI_ISL_2426161, EPI_ISL_2426162, EPI_ISL_2426163, EPI_ISL_2426164, EPI_ISL_2426165, EPI_ISL_2426166, EPI_ISL_2426167, EPI_ISL_2426168, EPI_ISL_2426170, EPI_ISL_2426171, EPI_ISL_2426175, EPI_ISL_2426177, EPI_ISL_2426178, EPI_ISL_2426179, EPI_ISL_2426180, EPI_ISL_2426182, EPI_ISL_2426183, EPI_ISL_2426186, EPI_ISL_2426187 | | | |
| see above | Clinical Virology Laboratory, Institute of Liver and Biliary Sciences | ILBS - IGIB | Ekta Gupta, Sheetalnath Rooge, Pratibha Kale, Amit Pandey, Chhagan Bihari Sharma, Partha Chattopadhyay, Priti Devi, Ranjeet Maurya, Neha Jha, Priyanka Mehta, Manish Kumar, Pooja Sharma, Sheeba Saifi, Aparna S, Sarfaraz Alam, Bharathram Uppili, Mohammed Faruq, Anurag Agrawal, Rajesh Pandey, Shiv Kumar Sarin |
| EPI_ISL_2426190, EPI_ISL_2426191, EPI_ISL_2426192, EPI_ISL_2426193, EPI_ISL_2426198, EPI_ISL_2426200, EPI_ISL_2426205, EPI_ISL_2426206, EPI_ISL_2426210, EPI_ISL_2426214, EPI_ISL_2426216, EPI_ISL_2426217, EPI_ISL_2426222, EPI_ISL_2426223, EPI_ISL_2426224, EPI_ISL_2426226, EPI_ISL_2426227, EPI_ISL_2426230, EPI_ISL_2426231, EPI_ISL_2426232, EPI_ISL_2426234, EPI_ISL_2426235, EPI_ISL_2426239, EPI_ISL_2426242, EPI_ISL_2426243, EPI_ISL_2426245, EPI_ISL_2426248, EPI_ISL_2426249, EPI_ISL_2426251, EPI_ISL_2426252 | | | |
| see above | All India Institute of Medical Sciences, Delhi | CSIR Institute of Genomics and Integrative Biology | Urvashi B Singh, Mercy Rophina, Rama Chaudhry, Vigneshwar Senthivel, Kiran Bala, Rahul C. Bhoyar, Bani Jolly, Nayer Jamshed, Mohamed Imran,Ritu Gupta, Praveen Aggarwal, Mohit Kumar Divakar, Subrata Sinha , Arvinden VR, Anjali Bajaj, Afra Shamnath, Abhinav Jain, COVID CBNAAT CORE GROUP , Vinod Scaria, Sridhar Sivasubbu, Randeep Guleria |
| EPI_ISL_2426255, EPI_ISL_2426256, EPI_ISL_2426257, EPI_ISL_2426258, EPI_ISL_2426259 | Thai Red Cross Emerging Infectious Diseases Clinical Center and Faculty of Medicine, Chulalongkorn University | Thai Red Cross Emerging Infectious Diseases Clinical Center and Faculty of Medicine, Chulalongkorn University | Opass Putcharoen, Stefan Fernandez, Leilani Paitoonpong, Anthony R. Jones, Gompol Suwanpimolkul, Chonticha Klungthong, Watsamon Jantarabenjakul, Piyawan Chinnawirotpisan, Torvorapanit Pattama, Khajohn Joonlasak, Wudtichai Manasatienkij, Supaporn Wacharapluesadee |
| EPI_ISL_2426266, EPI_ISL_2426300, EPI_ISL_2426303, EPI_ISL_2426340, EPI_ISL_2426352, EPI_ISL_2426372 | UW Virology Lab | UW Virology Lab | Pavitra Roychoudhury, Hong Xie, Lasata Shrestha, Tien V. Nguyen, Shah Mohamed Bakhash, Michelle Lin, Noah R. Baker, Ricardo Perez, Sean Ellis, Nathan Breit, Robert J. Livingston, Meei-Li Huang, Keith R Jerome, Patrick Mathias, Alexander Greninger |
| EPI_ISL_2426447, EPI_ISL_2426457, EPI_ISL_2426475, EPI_ISL_2426480, EPI_ISL_2426498, EPI_ISL_2426523, EPI_ISL_2426556 | UW Virology Lab | UW Virology Lab | Pavitra Roychoudhury, Hong Xie, Lasata Shrestha, Tien V. Nguyen, Shah Mohamed Bakhash, Michelle Lin, Noah R. Baker, Sean Ellis, Meei-Li Huang, Keith R Jerome, Alexander Greninger |
| EPI_ISL_2426583, EPI_ISL_2426585 | SYNLAB | GIGA Medical Genomics | Keith Durkin, Maria Artesi, Sébastien Bontems, Raphaël Boreux, Bouchra Boujemla, Nathalie Renotte, Cécile Meex, Pierrette Melin, Marie-Pierre Hayette, Vincent Bours |
| EPI_ISL_2426586 | Department of Clinical Microbiology | GIGA Medical Genomics | Keith Durkin, Maria Artesi, Sébastien Bontems, Raphaël Boreux, Bouchra Boujemla, Nathalie Renotte, Cécile Meex, Pierrette Melin, Marie-Pierre Hayette, Vincent Bours |
| EPI_ISL_2426596, EPI_ISL_2426602, EPI_ISL_2426608, EPI_ISL_2426614, EPI_ISL_2426618 | SYNLAB | GIGA Medical Genomics | Keith Durkin, Maria Artesi, Sébastien Bontems, Raphaël Boreux, Bouchra Boujemla, Nathalie Renotte, Cécile Meex, Pierrette Melin, Marie-Pierre Hayette, Vincent Bours |
| EPI_ISL_2426646, EPI_ISL_2426647 | Servicio de Microbiologia Hospital Ramon y Cajal | Servicio de Microbiologia Hospital Ramon y Cajal | Ponce M, Galan JC, Martinez L. Abreu M, y Gonzalez-Alba JM |
| EPI_ISL_2426721 | US Air Force School of Aerospace Medicine | US Air Force School of Aerospace Medicine | Anthony Fries, Jennifer Meyer, William Gruner, Amanda Javorina, Carol Garrett, Sarah Purves, Clarise Starr, Elizabeth Macias |
| EPI_ISL_2426989 | UC Davis Genome Center and Healthy Davis Together | UC Davis Genome Center | UC Davis |
| EPI_ISL_2427162 | Austrian Agency for Health and Food Safety (AGES) | Bergthaler laboratory, CeMM Research Center for Molecular Medicine of the Austrian Academy of Sciences | Lukas Endler, Anna Schedl, Fabian Amman, Petr Triska, Thomas Penz, Benedikt Agerer, Maelle Le Moing, Michael Schuster, Bekir Erguner, Jan Laine, Martin Senekowitsch, Christoph Bock, Andreas Bergthaler |
| EPI_ISL_2427343, EPI_ISL_2427353 | Cliniques universitaires Saint-Luc | UCLouvain/IREC/MBLG | Jean Ruelle, Ophélie Simon, Bertrand Bearzatto, Benoit Kabamba Mukadi |
| EPI_ISL_2427381, EPI_ISL_2427389, EPI_ISL_2427396, EPI_ISL_2427399 | Clinique Saint-Pierre Ottignies | UCLouvain/IREC/MBLG | Jean Ruelle, Ophélie Simon, Bertrand Bearzatto, Benoit Kabamba Mukadi |
| EPI_ISL_2427441, EPI_ISL_2427444, EPI_ISL_2427452, EPI_ISL_2427454, EPI_ISL_2427467, EPI_ISL_2427482 | NYU Langone Health | Departments of Pathology and Medicine, New York University School of Medicine | Adriana Heguy, Dacia Dimartino, Emily Guzman, Christian Marier, Peter Meyn, Sitharam Ramaswami, Gael Westby, Paul Zappile, Yutong Zhang, Paolo Cotzia, Guiqing Wang |
| EPI_ISL_2429136, EPI_ISL_2429137, EPI_ISL_2429138, EPI_ISL_2429141 | Institute for Medical Research, Infectious Disease Research Centre, National Institutes of Health, Ministry of Health Malaysia | Institute for Medical Research, Infectious Disease Research Centre, National Institutes of Health, Ministry of Health Malaysia | Suppiah J, Kamel K, Mohd Zawawi Z, Azizan MA, Ramly N, Robert F, Thayan R |
| EPI_ISL_2429651, EPI_ISL_2430149, EPI_ISL_2430527 | Department of Bacteria, Parasites and Fungi, Statens Serum Institut, Copenhagen, Denmark | Statens Serum Institut Bioinformatics and Microbial Genomics | Danish Covid-19 Genome Consortium |
| EPI_ISL_2432604, EPI_ISL_2432624, EPI_ISL_2432627, EPI_ISL_2432642, EPI_ISL_2432644, EPI_ISL_2432645, EPI_ISL_2432670, EPI_ISL_2432682, EPI_ISL_2432691, EPI_ISL_2432697, EPI_ISL_2432699, EPI_ISL_2432728, EPI_ISL_2432729, EPI_ISL_2432789, EPI_ISL_2432797, EPI_ISL_2432803, EPI_ISL_2432807, EPI_ISL_2432820, EPI_ISL_2432826 | | | |
| see above | UW Virology Lab | UW Virology Lab | Pavitra Roychoudhury, Hong Xie, Lasata Shrestha, Tien V. Nguyen, Shah Mohamed Bakhash, Michelle Lin, Noah R. Baker, Ricardo Perez, Sean Ellis, Nathan Breit, Robert J. Livingston, Meei-Li Huang, Keith R Jerome, Patrick Mathias, Alexander Greninger |
| EPI_ISL_2432843, EPI_ISL_2432850, EPI_ISL_2432873, EPI_ISL_2432879, EPI_ISL_2432910 | UW Virology Lab | UW Virology Lab | Pavitra Roychoudhury, Hong Xie, Lasata Shrestha, Tien V. Nguyen, Shah Mohamed Bakhash, Michelle Lin, Noah R. Baker, Sean Ellis, Meei-Li Huang, Keith R Jerome, Alexander Greninger |
| EPI_ISL_2433100, EPI_ISL_2433253, EPI_ISL_2433255 | Pathology and Laboratory Medicine Institute, Cleveland Clinic, Ohio, USA | Pathology and Laboratory Medicine Institute, Cleveland Clinic, Ohio, USA | Concetta Peck, Jennifer Starbuck, Joy Nakitandwe, Kristen McDonnell, David Plunkett, Zheng Jin Tu, Jay Brock, Yu-Wei Cheng, Gary Procop, Daniel Rhoads, Daniel H. Farkas, David Bosler |
| EPI_ISL_2433259, EPI_ISL_2433260, EPI_ISL_2433268, EPI_ISL_2433269, EPI_ISL_2433276 | INSACOG-Tripura | National Institute of Biomedical Genomics – INSACOG | Arindam Maitra, Tapan Majumdar, Nidhan Kumar Biswas, Saikat Majumder, Sreedhar Chinnaswamy, Apurba Sarkar, Saumitra Das |
| EPI_ISL_2433277, EPI_ISL_2433278, EPI_ISL_2433279 | INSACOG-WB | National Institute of Biomedical Genomics – INSACOG | Arindam Maitra, Bhaswati Bandyopadhyay, Nidhan Kumar Biswas, Tamal Ghosh, Sreedhar Chinnaswamy, Ajay Chakraborti, Saumitra Das |
| EPI_ISL_2433287, EPI_ISL_2433290 | Chulabhorn Hospital | Division of Genomic Medicine and Innovation support,Department of Medical Sciences, Ministry of Public Health, Thailand | Surakameth Mahasirimongkol,Nuanjun Wichukchinda,Archawin Rojanawiwat,Pilailuk Akkapaiboon Okada ,Waritta Sawaengdee,Penpitcha Thawong,Pundharika Piboonsiri,Jirapha Pakdee,Natthakul Bunneang |
| EPI_ISL_2433295 | Kasemrad international hospital | Division of Genomic Medicine and Innovation support,Department of | Surakameth Mahasirimongkol,Nuanjun Wichukchinda,Archawin Rojanawiwat,Pilailuk Akkapaiboon Okada ,Waritta Sawaengdee,Penpitcha Thawong,Pundharika Piboonsiri,Jirapha Pakdee,Natthakul Bunneang |

| | | | |
|---|---|---|---|
| | | Medical Sciences, Ministry of Public Health, Thailand | |
| EPI_ISL_2433305 | Chulabhorn Hospital | Division of Genomic Medicine and Innovation support,Department of Medical Sciences, Ministry of Public Health, Thailand | Surakameth Mahasirimongkol,Nuanjun Wichukchinda,Archawin Rojanawiwat,Pilailuk Akkapaiboon Okada ,Waritta Sawaengdee,Penpitcha Thawong,Pundharika Piboonsiri,Jirapha Pakdee,Natthakul Bunneang |
| EPI_ISL_2433313, EPI_ISL_2433572, EPI_ISL_2433573, EPI_ISL_2433574, EPI_ISL_2433575, EPI_ISL_2433576, EPI_ISL_2433577, EPI_ISL_2433578, EPI_ISL_2433579, EPI_ISL_2433580, EPI_ISL_2433581, EPI_ISL_2433582, EPI_ISL_2433583, EPI_ISL_2433584, EPI_ISL_2433585, EPI_ISL_2433586, EPI_ISL_2433587, EPI_ISL_2433588, EPI_ISL_2433589, EPI_ISL_2433590, EPI_ISL_2433591, EPI_ISL_2433592, EPI_ISL_2433593, EPI_ISL_2433594, EPI_ISL_2433595, EPI_ISL_2433596, EPI_ISL_2433597, EPI_ISL_2433598, EPI_ISL_2433599, EPI_ISL_2433600, EPI_ISL_2433601, EPI_ISL_2433602, EPI_ISL_2433603, EPI_ISL_2433604, EPI_ISL_2433605, EPI_ISL_2433606, EPI_ISL_2433607, EPI_ISL_2433608, EPI_ISL_2433609, EPI_ISL_2433610, EPI_ISL_2433611, EPI_ISL_2433612, EPI_ISL_2433613, EPI_ISL_2433614, EPI_ISL_2433615, EPI_ISL_2433616, EPI_ISL_2433617 | | | |
| see above | Kasemrad international hospital | Division of Genomic Medicine and Innovation support,Department of Medical Sciences, Ministry of Public Health, Thailand | Surakameth Mahasirimongkol,Nuanjun Wichukchinda,Archawin Rojanawiwat,Pilailuk Akkapaiboon Okada ,Waritta Sawaengdee,Penpitcha Thawong,Pundharika Piboonsiri,Jirapha Pakdee,Natthakul Bunneang |
| EPI_ISL_2433618 | Regional Medical Science Center 2 Phitsanulok | Division of Genomic Medicine and Innovation support,Department of Medical Sciences, Ministry of Public Health, Thailand | Surakameth Mahasirimongkol,Nuanjun Wichukchinda,Archawin Rojanawiwat,Pilailuk Akkapaiboon Okada ,Waritta Sawaengdee,Penpitcha Thawong,Pundharika Piboonsiri,Jirapha Pakdee,Natthakul Bunneang |
| EPI_ISL_2433619, EPI_ISL_2433620, EPI_ISL_2433621, EPI_ISL_2433622, EPI_ISL_2433623, EPI_ISL_2433624, EPI_ISL_2433625, EPI_ISL_2433626, EPI_ISL_2433627, EPI_ISL_2433628, EPI_ISL_2433629, EPI_ISL_2433630, EPI_ISL_2433631, EPI_ISL_2433632, EPI_ISL_2433633, EPI_ISL_2433634, EPI_ISL_2433635, EPI_ISL_2433636, EPI_ISL_2433637, EPI_ISL_2433638, EPI_ISL_2433639, EPI_ISL_2433640, EPI_ISL_2433641, EPI_ISL_2433642, EPI_ISL_2433643, EPI_ISL_2433644, EPI_ISL_2433645 | | | |
| see above | Chulabhorn Hospital | Division of Genomic Medicine and Innovation support,Department of Medical Sciences, Ministry of Public Health, Thailand | Surakameth Mahasirimongkol,Nuanjun Wichukchinda,Archawin Rojanawiwat,Pilailuk Akkapaiboon Okada ,Waritta Sawaengdee,Penpitcha Thawong,Pundharika Piboonsiri,Jirapha Pakdee,Natthakul Bunneang |
| EPI_ISL_2433662, EPI_ISL_2433688, EPI_ISL_2433691, EPI_ISL_2433694, EPI_ISL_2433696, EPI_ISL_2433700, EPI_ISL_2433702, EPI_ISL_2433704 | Dianalabs SA | Genesupport | Tanguy ARAUD |
| EPI_ISL_2433928 | Public Health Virology-Forensic and Scientific Services | Public Health Virology-Forensic and Scientific Services | Alyssa T. Pyke |
| EPI_ISL_2433938 | National Institute of Laboratory Medicine and Referral Center | Genomic Research Lab, BCSIR | Md. Murshed Hasan Sarkar, Abu Sayeed Mohammad Mahmud, Mohammad Samir Uzzaman, Eshrar Osman, Md. Ahasan Habib, Shahina Akter, Tanjina Akhter Banu, Barna Goswami, Iffat Jahan, Mohammad Mohi Uddin, Md. Kamrul Islam, Tasnim Nafisa, Md. Maruf Ahmed Molla, Mahmuda Yeasmin, Asish Kumar Ghosh, Arifa Akram, Md. Salim Khan |
| EPI_ISL_2433969 | Microbiologia CATLAB | Can Ruti SARS-CoV-2 Sequencing Hub (HUGTiP/IrsiCaixa/IGTP) | Marc Noguera-Julian, Pilar Armengol, Ignacio Blanco, Antoni E Bordoy, Francesc Catala-Moll, Pere-Joan Cardona, Maria Casadellà, Cristina Casañ, Gemma Clara, Bonaventura Clotet, Cristina Esteban, Montserrat Giménez, Mercedes Guerrero, Anna Not, Roger Paredes, Mariona Parera, Verónica Saludes, Alba Sánchez, and Elisa Martró on behalf of the Can Ruti SARS-CoV-2 Sequencing Hub. |
| EPI_ISL_2433972, EPI_ISL_2433975, EPI_ISL_2433976, EPI_ISL_2433984, EPI_ISL_2433987, EPI_ISL_2433990 | Microbiology Department, Laboratori Clínic Metropolitana Nord. Hospital Universitari Germans Trias i Pujol | Can Ruti SARS-CoV-2 Sequencing Hub (HUGTiP/IrsiCaixa/IGTP) | Marc Noguera-Julian, Pilar Armengol, Ignacio Blanco, Antoni E Bordoy, Francesc Catala-Moll, Pere-Joan Cardona, Maria Casadellà, Cristina Casañ, Gemma Clara, Bonaventura Clotet, Cristina Esteban, Montserrat Giménez, Mercedes Guerrero, Anna Not, Roger Paredes, Mariona Parera, Verónica Saludes, Alba Sánchez, and Elisa Martró on behalf of the Can Ruti SARS-CoV-2 Sequencing Hub. |
| EPI_ISL_2433994, EPI_ISL_2434005 | Microbiologia CATLAB | Can Ruti SARS-CoV-2 Sequencing Hub (HUGTiP/IrsiCaixa/IGTP) | Marc Noguera-Julian, Pilar Armengol, Ignacio Blanco, Antoni E Bordoy, Francesc Catala-Moll, Pere-Joan Cardona, Maria Casadellà, Cristina Casañ, Gemma Clara, Bonaventura Clotet, Cristina Esteban, Montserrat Giménez, Mercedes Guerrero, Anna Not, Roger Paredes, Mariona Parera, Verónica Saludes, Alba Sánchez, and Elisa Martró on behalf of the Can Ruti SARS-CoV-2 Sequencing Hub. |
| EPI_ISL_2434007, EPI_ISL_2434011 | Microbiology Department, Laboratori Clínic Metropolitana Nord. Hospital Universitari Germans Trias i Pujol | Can Ruti SARS-CoV-2 Sequencing Hub (HUGTiP/IrsiCaixa/IGTP) | Marc Noguera-Julian, Pilar Armengol, Ignacio Blanco, Antoni E Bordoy, Francesc Catala-Moll, Pere-Joan Cardona, Maria Casadellà, Cristina Casañ, Gemma Clara, Bonaventura Clotet, Cristina Esteban, Montserrat Giménez, Mercedes Guerrero, Anna Not, Roger Paredes, Mariona Parera, Verónica Saludes, Alba Sánchez, and Elisa Martró on behalf of the Can Ruti SARS-CoV-2 Sequencing Hub. |
| EPI_ISL_2434017 | Microbiologia CATLAB | Can Ruti SARS-CoV-2 Sequencing Hub (HUGTiP/IrsiCaixa/IGTP) | Marc Noguera-Julian, Pilar Armengol, Ignacio Blanco, Antoni E Bordoy, Francesc Catala-Moll, Pere-Joan Cardona, Maria Casadellà, Cristina Casañ, Gemma Clara, Bonaventura Clotet, Cristina Esteban, Montserrat Giménez, Mercedes Guerrero, Anna Not, Roger Paredes, Mariona Parera, Verónica Saludes, Alba Sánchez, and Elisa Martró on behalf of the Can Ruti SARS-CoV-2 Sequencing Hub. |
| EPI_ISL_2434018, EPI_ISL_2434019 | Microbiology Department, Laboratori Clínic Metropolitana Nord. Hospital Universitari Germans Trias i Pujol | Can Ruti SARS-CoV-2 Sequencing Hub (HUGTiP/IrsiCaixa/IGTP) | Marc Noguera-Julian, Pilar Armengol, Ignacio Blanco, Antoni E Bordoy, Francesc Catala-Moll, Pere-Joan Cardona, Maria Casadellà, Cristina Casañ, Gemma Clara, Bonaventura Clotet, Cristina Esteban, Montserrat Giménez, Mercedes Guerrero, Anna Not, Roger Paredes, Mariona Parera, Verónica Saludes, Alba Sánchez, and Elisa Martró on behalf of the Can Ruti SARS-CoV-2 Sequencing Hub. |
| EPI_ISL_2434028, EPI_ISL_2434029 | Microbiologia CATLAB | Can Ruti SARS-CoV-2 Sequencing Hub (HUGTiP/IrsiCaixa/IGTP) | Marc Noguera-Julian, Pilar Armengol, Ignacio Blanco, Antoni E Bordoy, Francesc Catala-Moll, Pere-Joan Cardona, Maria Casadellà, Cristina Casañ, Gemma Clara, Bonaventura Clotet, Cristina Esteban, Montserrat Giménez, Mercedes Guerrero, Anna Not, Roger Paredes, Mariona Parera, Verónica Saludes, Alba Sánchez, and Elisa Martró on behalf of the Can Ruti SARS-CoV-2 Sequencing Hub. |
| EPI_ISL_2434037 | Microbiology Department, Laboratori Clínic Metropolitana Nord. Hospital Universitari Germans Trias i Pujol | Can Ruti SARS-CoV-2 Sequencing Hub (HUGTiP/IrsiCaixa/IGTP) | Marc Noguera-Julian, Pilar Armengol, Ignacio Blanco, Antoni E Bordoy, Francesc Catala-Moll, Pere-Joan Cardona, Maria Casadellà, Cristina Casañ, Gemma Clara, Bonaventura Clotet, Cristina Esteban, Montserrat Giménez, Mercedes Guerrero, Anna Not, Roger Paredes, Mariona Parera, Verónica Saludes, Alba Sánchez, and Elisa Martró on behalf of the Can Ruti SARS-CoV-2 Sequencing Hub. |
| EPI_ISL_2434040 | Microbiologia CATLAB | Can Ruti SARS-CoV-2 Sequencing Hub (HUGTiP/IrsiCaixa/IGTP) | Marc Noguera-Julian, Pilar Armengol, Ignacio Blanco, Antoni E Bordoy, Francesc Catala-Moll, Pere-Joan Cardona, Maria Casadellà, Cristina Casañ, Gemma Clara, Bonaventura Clotet, Cristina Esteban, Montserrat Giménez, Mercedes Guerrero, Anna Not, Roger Paredes, Mariona Parera, Verónica Saludes, Alba Sánchez, and Elisa Martró on behalf of the Can Ruti SARS-CoV-2 Sequencing Hub. |
| EPI_ISL_2434047, EPI_ISL_2434048 | Microbiology Department, Laboratori Clínic Metropolitana Nord. Hospital Universitari Germans Trias i Pujol | Can Ruti SARS-CoV-2 Sequencing Hub (HUGTiP/IrsiCaixa/IGTP) | Marc Noguera-Julian, Pilar Armengol, Ignacio Blanco, Antoni E Bordoy, Francesc Catala-Moll, Pere-Joan Cardona, Maria Casadellà, Cristina Casañ, Gemma Clara, Bonaventura Clotet, Cristina Esteban, Montserrat Giménez, Mercedes Guerrero, Anna Not, Roger Paredes, Mariona Parera, Verónica Saludes, Alba Sánchez, and Elisa Martró on behalf of the Can Ruti SARS-CoV-2 Sequencing Hub. |
| EPI_ISL_2434050, EPI_ISL_2434051, EPI_ISL_2434052 | Microbiologia CATLAB | Can Ruti SARS-CoV-2 Sequencing Hub (HUGTiP/IrsiCaixa/IGTP) | Marc Noguera-Julian, Pilar Armengol, Ignacio Blanco, Antoni E Bordoy, Francesc Catala-Moll, Pere-Joan Cardona, Maria Casadellà, Cristina Casañ, Gemma Clara, Bonaventura Clotet, Cristina Esteban, Montserrat Giménez, Mercedes Guerrero, Anna Not, Roger Paredes, Mariona Parera, Verónica Saludes, Alba Sánchez, and Elisa Martró on behalf of the Can Ruti SARS-CoV-2 Sequencing Hub. |
| EPI_ISL_2434139, EPI_ISL_2434140, EPI_ISL_2434141, EPI_ISL_2434142, EPI_ISL_2434143 | Institute of Epidemiology, Disease Control and Research (IEDCR) | IEDCR-ideSHi-icddr,b | Hassan Afrad, Sadia Rahman, Manjur Hossain Khan, Firdausi Qadri, Tahmina Shirin |
| EPI_ISL_2434148, EPI_ISL_2434150 | AZ Klina | AZ Klina | Carl Vael - Lynsey Berckmans |
| EPI_ISL_2434158 | National Hospital for Tropical Diseases | Oxford University Clinical Research Unit, Hanoi | Pham Ngoc Thach, Nguyen Thi Tam, Van Dinh Trang, Le Van Duyet, Nguyen Thu Trang, Nguyen Thi Hong Thuong, Nguyen Thi Kim Chi, Phan Manh Cuong, Thomas Kesteman, H.Rogier van Doorn on behalf of the OUCRU COVID-19 research group |
| EPI_ISL_2434159 | AZ Klina | AZ Klina | Carl Vael - Lynsey Berckmans |
| EPI_ISL_2434161 | Genomik Solidaritas Indonesia Laboratorium | Genomik Solidaritas Indonesia Laboratorium | Alfin Mohammad Abdillah, Dhahlia Agustina Cahyono, Normastuti Adhini Tantyo, Annisa Muthiah Sukirman, Carissa Sintca Wijaya, Vania Gavrila Wikasa, Gracia Felias Enos Korompis, Meutia Ayuputeri Kumaheri, Anindya Pradipta Susanto, Ariel Pradipta |
| EPI_ISL_2434163 | AZ Klina | AZ Klina | Carl Vael - Lynsey Berckmans |
| EPI_ISL_2434174 | Department of Virology | Department of Virology | Massab Umair, Aamer Ikram, Muhammad Salman, Nazish Badar, Zaira Rehman, Muhammad Ammar, Syed Adnan Haider |
| EPI_ISL_2434176, EPI_ISL_2434177, EPI_ISL_2434178, EPI_ISL_2434181, EPI_ISL_2434182, EPI_ISL_2434184, EPI_ISL_2434189, EPI_ISL_2434190 | Quadram Institute Bioscience | COVID-19 Genomics UK (COG-UK) Consortium | Dave J. Baker, Gemma L. Kay, Alp Aydin, Thanh Le-Viet, Steven Rudder, Ana P. Tedim, Anastasia Kolyva, Maria Diaz, Leonardo de Oliveira Martins, Nabil-Fareed Alikhan, Lizzie Meadows, Rachael Stanley, Ngozi Elumogo, Muhammed Yasir, Nicholas M. Thomson, Alexander J Trotter, Rachel Gilroy, Samuel Bloomfield, Claire Stuart, Andrew Bell, Reenesh Prakash, Samir Dervisevic, Alison E. Mather, John Wain, Mark Webber, Andrew J. Page, Justin O'Grady |
| EPI_ISL_2434195, EPI_ISL_2434197, EPI_ISL_2434201 | West of Scotland Specialist Virology Centre, NHSGGC / MRC-University of Glasgow Centre for Virus Research | COVID-19 Genomics UK (COG-UK) Consortium | Ana da Silva Filipe, Natasha Johnson, Kathy Smollett, Daniel Mair, Stephen Carmichael, Alice Broos, Lily Tong, Jenna Nichols, Kyriaki Nomikou; Sarah McDonald; Ioulia Tsatsani, Richard Orton, Joseph Hughes, Sreenu Vattipally, David L Robertson; Alasdair MacLean, Rory Gunson; Sharif Shaaban, Matthew Holden; Rachel Blacow, Guy Mollett, Kathy Li, James Shepherd, Antonia Ho, Emma Thomson |
| EPI_ISL_2434218, EPI_ISL_2434220, EPI_ISL_2434222, EPI_ISL_2434223, EPI_ISL_2434224, EPI_ISL_2434226, EPI_ISL_2434236 | Virology Department, Royal Infirmary of Edinburgh, NHS Lothian / School of Biological Sciences, University of Edinburgh | COVID-19 Genomics UK (COG-UK) Consortium | McHugh M, Dewar R, Cotton S, Gallagher A, Maloney D, Rooke S, O'Toole Á, Scher E, Hill V, McCrone JT, Colquhoun R, Yu X, Jackson B, Rambaut A, Templeton K |
| EPI_ISL_2434248, EPI_ISL_2434249, EPI_ISL_2434252, EPI_ISL_2434253, EPI_ISL_2434254, EPI_ISL_2434255, EPI_ISL_2434257, EPI_ISL_2434258 | Liverpool Clinical Laboratories | COVID-19 Genomics UK (COG-UK) Consortium | Sam Haldenby, Alistair Darby, Steve Paterson, Anita Lucaci, Julian Hiscox, M Almsaud, A Alrezaihi, Muhannad Alruwaili, Stuart D Armstrong, Jones Benjamin, Eleanor G Bentley, Anu Chawla, Jordan J Clark, Angela Cowell, Richard Eccles, Isabel García-Dorival, Matthew Gemmell, Alessandro Gerada, PKF Gilmore, Richard Gregory, Ximeng Han, Catherine Hartley, Margaret Hughes, Miren Iturriza-Gomara, James Johnson, L Luu, Jenifer Manson, Charlotte Nelson, Elaine O'Toole, Cassie Olateju, Rebekah Penrice-Randal , Lucille Rainbow, N.P Randle, Trevor Ian Robinson, Parul Sharma, Ghada T Shawli, James P Stewart, Neil Swainston, Ecaterina Vamos, Joanne Watts, Mark Whitehead, Hermione Webster |
| EPI_ISL_2434259, EPI_ISL_2434260, EPI_ISL_2434261, EPI_ISL_2434262, EPI_ISL_2434264, EPI_ISL_2434266, EPI_ISL_2434267, EPI_ISL_2434268, EPI_ISL_2434269, EPI_ISL_2434270, EPI_ISL_2434271 | | | |
| see above | Barts Health NHS Trust | COVID-19 Genomics UK (COG-UK) Consortium | CUTINO-MOGUEL, Maria-Teresa; HARRINGTON, David; OWOYEMI, Dola; KULASEGARAN-SHYLINI, Raghavendran; BROAD, Claire; KELE, Beatrix |
| EPI_ISL_2434272, EPI_ISL_2434274, EPI_ISL_2434275, EPI_ISL_2434278, EPI_ISL_2434279, EPI_ISL_2434280, EPI_ISL_2434281, EPI_ISL_2434287, EPI_ISL_2434289, EPI_ISL_2434290, EPI_ISL_2434292, EPI_ISL_2434293, EPI_ISL_2434296, EPI_ISL_2434300, EPI_ISL_2434301, EPI_ISL_2434304, EPI_ISL_2434305, EPI_ISL_2434308, EPI_ISL_2434313, EPI_ISL_2434316, EPI_ISL_2434317, EPI_ISL_2434323, | | | |

| | | | |
|---|---|---|---|
| EPI_ISL_2434324, EPI_ISL_2434325, EPI_ISL_2434329, EPI_ISL_2434330, EPI_ISL_2434336, EPI_ISL_2434339, EPI_ISL_2434342, EPI_ISL_2434343, EPI_ISL_2434345, EPI_ISL_2434347, EPI_ISL_2434350, EPI_ISL_2434352, EPI_ISL_2434354, EPI_ISL_2434362, EPI_ISL_2434377, EPI_ISL_2434381, EPI_ISL_2434387, EPI_ISL_2434388, EPI_ISL_2434391, EPI_ISL_2434402, EPI_ISL_2434432, EPI_ISL_2434447, EPI_ISL_2434466, EPI_ISL_2434474, EPI_ISL_2434487, EPI_ISL_2434499, EPI_ISL_2434582, EPI_ISL_2434597, EPI_ISL_2434602, EPI_ISL_2434654, EPI_ISL_2434657, EPI_ISL_2434663, EPI_ISL_2434670, EPI_ISL_2434678, EPI_ISL_2434680, EPI_ISL_2434681, EPI_ISL_2434685, EPI_ISL_2434700, EPI_ISL_2434713, EPI_ISL_2434721, EPI_ISL_2434724, EPI_ISL_2434732, EPI_ISL_2434738, EPI_ISL_2434739, EPI_ISL_2434750, EPI_ISL_2434751, EPI_ISL_2434755 | | | |
| see above | Northumbria University / South Tees Hospitals NHS Foundation Trust / North Cumbria Integrated Care NHS Foundation Trust / North Tees and Hartlepool NHS Foundation Trust / Newcastle Hospitals NHS Foundation Trust | COVID-19 Genomics UK (COG-UK) Consortium | Darren L Smith,Andrew Nelson,Matthew Bashton,Greg R Young,Joshua Loh,John Allan,Mohammad A Tariq,Giles S Holt,Gary Black,Wen C Yew,Lynn Dover,Paul Baker,Steve Liggett,Sarah Essex,Jane Greenaway,Debra Padgett,Clive Graham,Garren Scott,Edward Barton,Emma Swindells,Brendan Payne,Jennifer Collins,Yusri Taha,Gary Eltringham |
| EPI_ISL_2434759, EPI_ISL_2434760 | Queens Medical Centre, Clinical Microbiology Department / DeepSeq Nottingham | COVID-19 Genomics UK (COG-UK) Consortium | Gemma Clark, Wendy Smith, Manjinder Khakh, Vicki M Fleming, Michelle M Lister, Hannah Howson-Wells, Jonathan Ball, Timothy Byaruhanga, Jayasree Dey, Emily Park, Jack Hill, Patrick McClure, Joseph Chappell, Theocharis Tsoleridis, Nadine Holmes, Matthew Carlisle, Christopher Moore, Fei Sang, Johnny Debebe, Victoria Wright, Matthew Loose |
| EPI_ISL_2434762, EPI_ISL_2434765, EPI_ISL_2434767, EPI_ISL_2434768, EPI_ISL_2434769, EPI_ISL_2434770, EPI_ISL_2434771, EPI_ISL_2434773, EPI_ISL_2434775, EPI_ISL_2434776, EPI_ISL_2434778, EPI_ISL_2434779 | | | |
| see above | Oxford Viromics, NDM, University of Oxford; Oxford University Hospitals; Basingstoke and North Hampshire Hospital | COVID-19 Genomics UK (COG-UK) Consortium | Tanya Golubchik, David Bonsall, George Macintyre, Amy Trebes, Mariateresa de Cesare, Catrin Moore, Alex Mobbs, Anita Justice, Robert Shaw, Monique Andersson, Timothy Peto, Emma Wise, Nathan Moore, Jessica Lynch, Nick Cortes, Matilde Mori, Stephen Kidd, David Buck, John Todd, Christophe Fraser |
| EPI_ISL_2434781 | Department of Virology | Department of Virology | Massab Umair, Aamer Ikram, Muhammad Salman, Nazish Badar, Zaira Rehman, Muhammad Ammar, Syed Adnan Haider |
| EPI_ISL_2434782, EPI_ISL_2434784, EPI_ISL_2434793, EPI_ISL_2434794, EPI_ISL_2434799, EPI_ISL_2434802, EPI_ISL_2434804, EPI_ISL_2434808, EPI_ISL_2434814, EPI_ISL_2434815, EPI_ISL_2434816, EPI_ISL_2434818, EPI_ISL_2434819, EPI_ISL_2434821, EPI_ISL_2434822, EPI_ISL_2434824, EPI_ISL_2434825, EPI_ISL_2434829, EPI_ISL_2434831, EPI_ISL_2434832, EPI_ISL_2434833, EPI_ISL_2434834, EPI_ISL_2434835, EPI_ISL_2434837, EPI_ISL_2434838, EPI_ISL_2434839, EPI_ISL_2434840, EPI_ISL_2434841, EPI_ISL_2434842, EPI_ISL_2434843, EPI_ISL_2434847, EPI_ISL_2434849, EPI_ISL_2434852, EPI_ISL_2434853, EPI_ISL_2434854, EPI_ISL_2434855, EPI_ISL_2434857, EPI_ISL_2434858, EPI_ISL_2434860, EPI_ISL_2434862, EPI_ISL_2434864, EPI_ISL_2434865, EPI_ISL_2434868, EPI_ISL_2434870, EPI_ISL_2434871, EPI_ISL_2434872, EPI_ISL_2434875, EPI_ISL_2434876, EPI_ISL_2434877, EPI_ISL_2434891, EPI_ISL_2434892, EPI_ISL_2434894, EPI_ISL_2434896, EPI_ISL_2434897, EPI_ISL_2434898, EPI_ISL_2434900, EPI_ISL_2434902, EPI_ISL_2434903, EPI_ISL_2434904, EPI_ISL_2434905, EPI_ISL_2434906, EPI_ISL_2434907, EPI_ISL_2434908, EPI_ISL_2434909, EPI_ISL_2434910, EPI_ISL_2434911, EPI_ISL_2434912, EPI_ISL_2434913, EPI_ISL_2434915, EPI_ISL_2434916, EPI_ISL_2434917, EPI_ISL_2434918, EPI_ISL_2434920, EPI_ISL_2434921, EPI_ISL_2434922, EPI_ISL_2434924, EPI_ISL_2434925, EPI_ISL_2434926, EPI_ISL_2434927, EPI_ISL_2434929, EPI_ISL_2434934, EPI_ISL_2434935, EPI_ISL_2434936, EPI_ISL_2434937, EPI_ISL_2434942, EPI_ISL_2434943, EPI_ISL_2434945, EPI_ISL_2434946, EPI_ISL_2434947, EPI_ISL_2434948, EPI_ISL_2434949, EPI_ISL_2434950, EPI_ISL_2434951, EPI_ISL_2434956, EPI_ISL_2434957, EPI_ISL_2434958, EPI_ISL_2434959, EPI_ISL_2434960, EPI_ISL_2434962, EPI_ISL_2434964 | | | |
| see above | Originating lab: Wales Specialist Virology Centre Sequencing lab: Pathogen Genomics Unit | Public Health Wales Microbiology Cardiff Wales Specialist Virology Centre | Catherine Moore, Johnathan Evans, Laura Gifford, Malorie Perry, Simon Cottrell, Angela Marchbank, Alec Birchley, Alexander Adams, Amy Gaskin, Bree Gatica-Wilcox, Jason Coombes, Joel Southgate, Lauren Gilbert, Lee Graham, Nicole Pacchiarini, Sara Kumziene-Summerhayes, Sarah Taylor, Sophie Jones, Sara Rey, Matthew Bull, Joanne Watkins, Sally Corden, Tom Connor |
| EPI_ISL_2434966, EPI_ISL_2434967, EPI_ISL_2434969 | Virology Department, Sheffield Teaching Hospitals NHS Foundation Trust/Department of Infection, Immunity and Cardiovascular Disease, The Medical School, University of Sheffield | COVID-19 Genomics UK (COG-UK) Consortium | Thushan de Silva, Matthew Parker, Nikki Smith, Adri Angyal, Rebecca Brown, Luke Green, Rachel Tucker, Paul Parsons, Danielle Groves, Katie Johnson, Laura Carrilero, Alex Keeley, Dave Partridge, Matthew Wyles, Benjamin Lindsey, Mehmet Yavuz, Mohammad Raza, Cariad Evans |
| EPI_ISL_2434972, EPI_ISL_2434976, EPI_ISL_2434982 | Department of Virology | Department of Virology | Massab Umair, Aamer Ikram, Muhammad Salman, Nazish Badar, Zaira Rehman, Muhammad Ammar, Syed Adnan Haider |
| EPI_ISL_2434988 | Genomik Solidaritas Indonesia Laboratorium | Genomik Solidaritas Indonesia Laboratorium | Alfin Mohammad Abdillah, Dhahlia Agustina Cahyono, Normastuti Adhini Tantyo, Annisa Muthiah Sukirman, Carissa Sintca Wijaya, Vania Gavrila Wikasa, Gracia Felias Enos Korompis, Meutia Ayuputeri Kumaheri, Anindya Pradipta Susanto, Ariel Pradipta |
| EPI_ISL_2434991 | Genomik Solidaritas Indonesia Laboratorium | Genomik Solidaritas Indonesia Laboratorium | Dhahlia Agustina Cahyono, Alfin Mohammad Abdillah, Normastuti Adhini Tantyo, Annisa Muthiah Sukirman, Carissa Sintca Wijaya, Vania Gavrila Wikasa, Gracia Felias Enos Korompis, Meutia Ayuputeri Kumaheri, Anindya Pradipta Susanto, Ariel Pradipta |
| EPI_ISL_2434992, EPI_ISL_2434993, EPI_ISL_2434994, EPI_ISL_2434995, EPI_ISL_2434997, EPI_ISL_2434998, EPI_ISL_2435000, EPI_ISL_2435002, EPI_ISL_2435003 | Lighthouse Lab in Alderley Park | Wellcome Sanger Institute for the COVID-19 Genomics UK (COG-UK) Consortium | Jacquelyn Wynn, Mairead Hyland, The Lighthouse Lab in Alderley Park and Alex Alderton, Roberto Amato, Jeffrey Barrett, Sonia Goncalves, Ewan Harrison, David K. Jackson, Ian Johnston, Dominic Kwiatkowski, Cordelia Langford, John Sillitoe on behalf of the Wellcome Sanger Institute COVID-19 Surveillance Team |
| EPI_ISL_2435004 | Lighthouse Lab in Glasgow | Wellcome Sanger Institute for the COVID-19 Genomics UK (COG-UK) Consortium | Harper VanSteenhouse, Yumi Kasai, David Gray, Carol Clugston, Anna Dominiczak and Alex Alderton, Roberto Amato, Jeffrey Barrett, Sonia Goncalves, Ewan Harrison, David K. Jackson, Ian Johnston, Dominic Kwiatkowski, Cordelia Langford, John Sillitoe on behalf of the Wellcome Sanger Institute COVID-19 Surveillance Team |
| EPI_ISL_2435007, EPI_ISL_2435009, EPI_ISL_2435010, EPI_ISL_2435011, EPI_ISL_2435012, EPI_ISL_2435013, EPI_ISL_2435014, EPI_ISL_2435016, EPI_ISL_2435017, EPI_ISL_2435018, EPI_ISL_2435019, EPI_ISL_2435020, EPI_ISL_2435021, EPI_ISL_2435022, EPI_ISL_2435023, EPI_ISL_2435025, EPI_ISL_2435026, EPI_ISL_2435027 | | | |
| see above | Lighthouse Lab in Alderley Park | Wellcome Sanger Institute for the COVID-19 Genomics UK (COG-UK) Consortium | Jacquelyn Wynn, Mairead Hyland, The Lighthouse Lab in Alderley Park and Alex Alderton, Roberto Amato, Jeffrey Barrett, Sonia Goncalves, Ewan Harrison, David K. Jackson, Ian Johnston, Dominic Kwiatkowski, Cordelia Langford, John Sillitoe on behalf of the Wellcome Sanger Institute COVID-19 Surveillance Team |
| EPI_ISL_2435029 | Lighthouse Lab in Glasgow | Wellcome Sanger Institute for the COVID-19 Genomics UK (COG-UK) Consortium | Harper VanSteenhouse, Yumi Kasai, David Gray, Carol Clugston, Anna Dominiczak and Alex Alderton, Roberto Amato, Jeffrey Barrett, Sonia Goncalves, Ewan Harrison, David K. Jackson, Ian Johnston, Dominic Kwiatkowski, Cordelia Langford, John Sillitoe on behalf of the Wellcome Sanger Institute COVID-19 Surveillance Team |
| EPI_ISL_2435031 | Lighthouse Lab in Alderley Park | Wellcome Sanger Institute for the COVID-19 Genomics UK (COG-UK) Consortium | Jacquelyn Wynn, Mairead Hyland, The Lighthouse Lab in Alderley Park and Alex Alderton, Roberto Amato, Jeffrey Barrett, Sonia Goncalves, Ewan Harrison, David K. Jackson, Ian Johnston, Dominic Kwiatkowski, Cordelia Langford, John Sillitoe on behalf of the Wellcome Sanger Institute COVID-19 Surveillance Team |
| EPI_ISL_2435032 | Lighthouse Lab in Glasgow | Wellcome Sanger Institute for the COVID-19 Genomics UK (COG-UK) Consortium | Harper VanSteenhouse, Yumi Kasai, David Gray, Carol Clugston, Anna Dominiczak and Alex Alderton, Roberto Amato, Jeffrey Barrett, Sonia Goncalves, Ewan Harrison, David K. Jackson, Ian Johnston, Dominic Kwiatkowski, Cordelia Langford, John Sillitoe on behalf of the Wellcome Sanger Institute COVID-19 Surveillance Team |
| EPI_ISL_2435033 | Lighthouse Lab in Alderley Park | Wellcome Sanger Institute for the COVID-19 Genomics UK (COG-UK) Consortium | Jacquelyn Wynn, Mairead Hyland, The Lighthouse Lab in Alderley Park and Alex Alderton, Roberto Amato, Jeffrey Barrett, Sonia Goncalves, Ewan Harrison, David K. Jackson, Ian Johnston, Dominic Kwiatkowski, Cordelia Langford, John Sillitoe on behalf of the Wellcome Sanger Institute COVID-19 Surveillance Team |
| EPI_ISL_2435034 | Lighthouse Lab in Glasgow | Wellcome Sanger Institute for the COVID-19 Genomics UK (COG-UK) Consortium | Harper VanSteenhouse, Yumi Kasai, David Gray, Carol Clugston, Anna Dominiczak and Alex Alderton, Roberto Amato, Jeffrey Barrett, Sonia Goncalves, Ewan Harrison, David K. Jackson, Ian Johnston, Dominic Kwiatkowski, Cordelia Langford, John Sillitoe on behalf of the Wellcome Sanger Institute COVID-19 Surveillance Team |
| EPI_ISL_2435035 | Lighthouse Lab in Alderley Park | Wellcome Sanger Institute for the COVID-19 Genomics UK (COG-UK) Consortium | Jacquelyn Wynn, Mairead Hyland, The Lighthouse Lab in Alderley Park and Alex Alderton, Roberto Amato, Jeffrey Barrett, Sonia Goncalves, Ewan Harrison, David K. Jackson, Ian Johnston, Dominic Kwiatkowski, Cordelia Langford, John Sillitoe on behalf of the Wellcome Sanger Institute COVID-19 Surveillance Team |
| EPI_ISL_2435036 | Lighthouse Lab in Glasgow | Wellcome Sanger Institute for the COVID-19 Genomics UK (COG-UK) Consortium | Harper VanSteenhouse, Yumi Kasai, David Gray, Carol Clugston, Anna Dominiczak and Alex Alderton, Roberto Amato, Jeffrey Barrett, Sonia Goncalves, Ewan Harrison, David K. Jackson, Ian Johnston, Dominic Kwiatkowski, Cordelia Langford, John Sillitoe on behalf of the Wellcome Sanger Institute COVID-19 Surveillance Team |
| EPI_ISL_2435037, EPI_ISL_2435040 | Lighthouse Lab in Alderley Park | Wellcome Sanger Institute for the COVID-19 Genomics UK (COG-UK) Consortium | Jacquelyn Wynn, Mairead Hyland, The Lighthouse Lab in Alderley Park and Alex Alderton, Roberto Amato, Jeffrey Barrett, Sonia Goncalves, Ewan Harrison, David K. Jackson, Ian Johnston, Dominic Kwiatkowski, Cordelia Langford, John Sillitoe on behalf of the Wellcome Sanger Institute COVID-19 Surveillance Team |
| EPI_ISL_2435041, EPI_ISL_2435042 | Lighthouse Lab in Glasgow | Wellcome Sanger Institute for the COVID-19 Genomics UK (COG-UK) Consortium | Harper VanSteenhouse, Yumi Kasai, David Gray, Carol Clugston, Anna Dominiczak and Alex Alderton, Roberto Amato, Jeffrey Barrett, Sonia Goncalves, Ewan Harrison, David K. Jackson, Ian Johnston, Dominic Kwiatkowski, Cordelia Langford, John Sillitoe on behalf of the Wellcome Sanger Institute COVID-19 Surveillance Team |
| EPI_ISL_2435043, EPI_ISL_2435046, EPI_ISL_2435047, EPI_ISL_2435048, EPI_ISL_2435049, EPI_ISL_2435050, EPI_ISL_2435051 | Lighthouse Lab in Alderley Park | Wellcome Sanger Institute for the COVID-19 Genomics UK (COG-UK) Consortium | Jacquelyn Wynn, Mairead Hyland, The Lighthouse Lab in Alderley Park and Alex Alderton, Roberto Amato, Jeffrey Barrett, Sonia Goncalves, Ewan Harrison, David K. Jackson, Ian Johnston, Dominic Kwiatkowski, Cordelia Langford, John Sillitoe on behalf of the Wellcome Sanger Institute COVID-19 Surveillance Team |
| EPI_ISL_2435052 | Lighthouse Lab in Glasgow | Wellcome Sanger Institute for the COVID-19 Genomics UK (COG-UK) Consortium | Harper VanSteenhouse, Yumi Kasai, David Gray, Carol Clugston, Anna Dominiczak and Alex Alderton, Roberto Amato, Jeffrey Barrett, Sonia Goncalves, Ewan Harrison, David K. Jackson, Ian Johnston, Dominic Kwiatkowski, Cordelia Langford, John Sillitoe on behalf of the Wellcome Sanger Institute COVID-19 Surveillance Team |
| EPI_ISL_2435053, EPI_ISL_2435054 | Lighthouse Lab in Alderley Park | Wellcome Sanger Institute for the COVID-19 Genomics UK (COG-UK) Consortium | Jacquelyn Wynn, Mairead Hyland, The Lighthouse Lab in Alderley Park and Alex Alderton, Roberto Amato, Jeffrey Barrett, Sonia Goncalves, Ewan Harrison, David K. Jackson, Ian Johnston, Dominic Kwiatkowski, Cordelia Langford, John Sillitoe on behalf of the Wellcome Sanger Institute COVID-19 Surveillance Team |
| EPI_ISL_2435055 | Lighthouse Lab in Glasgow | Wellcome Sanger Institute for the COVID-19 Genomics UK (COG-UK) Consortium | Harper VanSteenhouse, Yumi Kasai, David Gray, Carol Clugston, Anna Dominiczak and Alex Alderton, Roberto Amato, Jeffrey Barrett, Sonia Goncalves, Ewan Harrison, David K. Jackson, Ian Johnston, Dominic Kwiatkowski, Cordelia Langford, John Sillitoe on behalf of the Wellcome Sanger Institute COVID-19 Surveillance Team |
| EPI_ISL_2435056, EPI_ISL_2435057, EPI_ISL_2435058, EPI_ISL_2435059, EPI_ISL_2435060 | Lighthouse Lab in Alderley Park | Wellcome Sanger Institute for the COVID-19 Genomics UK (COG-UK) Consortium | Jacquelyn Wynn, Mairead Hyland, The Lighthouse Lab in Alderley Park and Alex Alderton, Roberto Amato, Jeffrey Barrett, Sonia Goncalves, Ewan Harrison, David K. Jackson, Ian Johnston, Dominic Kwiatkowski, Cordelia Langford, John Sillitoe on behalf of the Wellcome Sanger Institute COVID-19 Surveillance Team |
| EPI_ISL_2435061, EPI_ISL_2435062 | Lighthouse Lab in Glasgow | Wellcome Sanger Institute for the COVID-19 Genomics UK (COG-UK) Consortium | Harper VanSteenhouse, Yumi Kasai, David Gray, Carol Clugston, Anna Dominiczak and Alex Alderton, Roberto Amato, Jeffrey Barrett, Sonia Goncalves, Ewan Harrison, David K. Jackson, Ian Johnston, Dominic Kwiatkowski, Cordelia Langford, John Sillitoe on behalf of the Wellcome Sanger Institute COVID-19 Surveillance Team |
| EPI_ISL_2435063, EPI_ISL_2435064, EPI_ISL_2435065, EPI_ISL_2435066, EPI_ISL_2435067, EPI_ISL_2435068, EPI_ISL_2435069, EPI_ISL_2435070, EPI_ISL_2435071, EPI_ISL_2435072, EPI_ISL_2435073 | | | |
| see above | Lighthouse Lab in Alderley Park | Wellcome Sanger Institute for the COVID-19 Genomics UK (COG-UK) Consortium | Jacquelyn Wynn, Mairead Hyland, The Lighthouse Lab in Alderley Park and Alex Alderton, Roberto Amato, Jeffrey Barrett, Sonia Goncalves, Ewan Harrison, David K. Jackson, Ian Johnston, Dominic Kwiatkowski, Cordelia Langford, John Sillitoe on behalf of the Wellcome Sanger Institute COVID-19 Surveillance Team |
| EPI_ISL_2435074, EPI_ISL_2435075 | Lighthouse Lab in Glasgow | Wellcome Sanger Institute for the COVID-19 Genomics UK (COG-UK) Consortium | Harper VanSteenhouse, Yumi Kasai, David Gray, Carol Clugston, Anna Dominiczak and Alex Alderton, Roberto Amato, Jeffrey Barrett, Sonia Goncalves, Ewan Harrison, David K. Jackson, Ian Johnston, Dominic Kwiatkowski, Cordelia Langford, John Sillitoe on behalf of the Wellcome Sanger Institute COVID-19 Surveillance Team |
| EPI_ISL_2435076 | Lighthouse Lab in Alderley Park | Wellcome Sanger Institute for the COVID-19 Genomics UK (COG-UK) Consortium | Jacquelyn Wynn, Mairead Hyland, The Lighthouse Lab in Alderley Park and Alex Alderton, Roberto Amato, Jeffrey Barrett, Sonia Goncalves, Ewan Harrison, David K. Jackson, Ian Johnston, Dominic Kwiatkowski, Cordelia Langford, John Sillitoe on behalf of the Wellcome Sanger Institute COVID-19 Surveillance Team |
| EPI_ISL_2435077 | Lighthouse Lab in Glasgow | Wellcome Sanger Institute for the COVID-19 Genomics UK (COG-UK) Consortium | Harper VanSteenhouse, Yumi Kasai, David Gray, Carol Clugston, Anna Dominiczak and Alex Alderton, Roberto Amato, Jeffrey Barrett, Sonia Goncalves, Ewan Harrison, David K. Jackson, Ian Johnston, Dominic Kwiatkowski, Cordelia Langford, John Sillitoe on behalf of the Wellcome Sanger Institute COVID-19 Surveillance Team |
| EPI_ISL_2435078, EPI_ISL_2435079, | Lighthouse Lab in Alderley Park | Wellcome Sanger Institute for the COVID-19 Genomics UK (COG-UK) Consortium | Jacquelyn Wynn, Mairead Hyland, The Lighthouse Lab in Alderley Park and Alex Alderton, Roberto Amato, Jeffrey Barrett, Sonia Goncalves, Ewan Harrison, David K. Jackson, Ian Johnston, Dominic Kwiatkowski, Cordelia Langford, John Sillitoe on behalf of the Wellcome Sanger Institute COVID-19 Surveillance Team |

| | | | |
|---|---|---|---|
| EPI_ISL_2435080, EPI_ISL_2435081, EPI_ISL_2435082, EPI_ISL_2435083, EPI_ISL_2435084, EPI_ISL_2435085, EPI_ISL_2435087 | | | |
| EPI_ISL_2435088 | Lighthouse Lab in Glasgow | Wellcome Sanger Institute for the COVID-19 Genomics UK (COG-UK) Consortium | Harper VanSteenhouse, Yumi Kasai, David Gray, Carol Clugston, Anna Dominiczak and Alex Alderton, Roberto Amato, Jeffrey Barrett, Sonia Goncalves, Ewan Harrison, David K. Jackson, Ian Johnston, Dominic Kwiatkowski, Cordelia Langford, John Sillitoe on behalf of the Wellcome Sanger Institute COVID-19 Surveillance Team |
| EPI_ISL_2435089, EPI_ISL_2435090, EPI_ISL_2435092, EPI_ISL_2435093, EPI_ISL_2435094 | Lighthouse Lab in Alderley Park | Wellcome Sanger Institute for the COVID-19 Genomics UK (COG-UK) Consortium | Jacquelyn Wynn, Mairead Hyland, The Lighthouse Lab in Alderley Park and Alex Alderton, Roberto Amato, Jeffrey Barrett, Sonia Goncalves, Ewan Harrison, David K. Jackson, Ian Johnston, Dominic Kwiatkowski, Cordelia Langford, John Sillitoe on behalf of the Wellcome Sanger Institute COVID-19 Surveillance Team |
| EPI_ISL_2435095 | Lighthouse Lab in Glasgow | Wellcome Sanger Institute for the COVID-19 Genomics UK (COG-UK) Consortium | Harper VanSteenhouse, Yumi Kasai, David Gray, Carol Clugston, Anna Dominiczak and Alex Alderton, Roberto Amato, Jeffrey Barrett, Sonia Goncalves, Ewan Harrison, David K. Jackson, Ian Johnston, Dominic Kwiatkowski, Cordelia Langford, John Sillitoe on behalf of the Wellcome Sanger Institute COVID-19 Surveillance Team |
| EPI_ISL_2435097, EPI_ISL_2435098 | Lighthouse Lab in Alderley Park | Wellcome Sanger Institute for the COVID-19 Genomics UK (COG-UK) Consortium | Jacquelyn Wynn, Mairead Hyland, The Lighthouse Lab in Alderley Park and Alex Alderton, Roberto Amato, Jeffrey Barrett, Sonia Goncalves, Ewan Harrison, David K. Jackson, Ian Johnston, Dominic Kwiatkowski, Cordelia Langford, John Sillitoe on behalf of the Wellcome Sanger Institute COVID-19 Surveillance Team |
| EPI_ISL_2435099 | Lighthouse Lab in Glasgow | Wellcome Sanger Institute for the COVID-19 Genomics UK (COG-UK) Consortium | Harper VanSteenhouse, Yumi Kasai, David Gray, Carol Clugston, Anna Dominiczak and Alex Alderton, Roberto Amato, Jeffrey Barrett, Sonia Goncalves, Ewan Harrison, David K. Jackson, Ian Johnston, Dominic Kwiatkowski, Cordelia Langford, John Sillitoe on behalf of the Wellcome Sanger Institute COVID-19 Surveillance Team |
| EPI_ISL_2435100, EPI_ISL_2435101, EPI_ISL_2435102, EPI_ISL_2435103, EPI_ISL_2435104, EPI_ISL_2435105, EPI_ISL_2435106 | Lighthouse Lab in Alderley Park | Wellcome Sanger Institute for the COVID-19 Genomics UK (COG-UK) Consortium | Jacquelyn Wynn, Mairead Hyland, The Lighthouse Lab in Alderley Park and Alex Alderton, Roberto Amato, Jeffrey Barrett, Sonia Goncalves, Ewan Harrison, David K. Jackson, Ian Johnston, Dominic Kwiatkowski, Cordelia Langford, John Sillitoe on behalf of the Wellcome Sanger Institute COVID-19 Surveillance Team |
| EPI_ISL_2435107 | Lighthouse Lab in Glasgow | Wellcome Sanger Institute for the COVID-19 Genomics UK (COG-UK) Consortium | Harper VanSteenhouse, Yumi Kasai, David Gray, Carol Clugston, Anna Dominiczak and Alex Alderton, Roberto Amato, Jeffrey Barrett, Sonia Goncalves, Ewan Harrison, David K. Jackson, Ian Johnston, Dominic Kwiatkowski, Cordelia Langford, John Sillitoe on behalf of the Wellcome Sanger Institute COVID-19 Surveillance Team |
| EPI_ISL_2435109 | Lighthouse Lab in Alderley Park | Wellcome Sanger Institute for the COVID-19 Genomics UK (COG-UK) Consortium | Jacquelyn Wynn, Mairead Hyland, The Lighthouse Lab in Alderley Park and Alex Alderton, Roberto Amato, Jeffrey Barrett, Sonia Goncalves, Ewan Harrison, David K. Jackson, Ian Johnston, Dominic Kwiatkowski, Cordelia Langford, John Sillitoe on behalf of the Wellcome Sanger Institute COVID-19 Surveillance Team |
| EPI_ISL_2435110 | Lighthouse Lab in Glasgow | Wellcome Sanger Institute for the COVID-19 Genomics UK (COG-UK) Consortium | Harper VanSteenhouse, Yumi Kasai, David Gray, Carol Clugston, Anna Dominiczak and Alex Alderton, Roberto Amato, Jeffrey Barrett, Sonia Goncalves, Ewan Harrison, David K. Jackson, Ian Johnston, Dominic Kwiatkowski, Cordelia Langford, John Sillitoe on behalf of the Wellcome Sanger Institute COVID-19 Surveillance Team |
| EPI_ISL_2435111, EPI_ISL_2435112, EPI_ISL_2435113 | Lighthouse Lab in Alderley Park | Wellcome Sanger Institute for the COVID-19 Genomics UK (COG-UK) Consortium | Jacquelyn Wynn, Mairead Hyland, The Lighthouse Lab in Alderley Park and Alex Alderton, Roberto Amato, Jeffrey Barrett, Sonia Goncalves, Ewan Harrison, David K. Jackson, Ian Johnston, Dominic Kwiatkowski, Cordelia Langford, John Sillitoe on behalf of the Wellcome Sanger Institute COVID-19 Surveillance Team |
| EPI_ISL_2435114, EPI_ISL_2435115 | Lighthouse Lab in Glasgow | Wellcome Sanger Institute for the COVID-19 Genomics UK (COG-UK) Consortium | Harper VanSteenhouse, Yumi Kasai, David Gray, Carol Clugston, Anna Dominiczak and Alex Alderton, Roberto Amato, Jeffrey Barrett, Sonia Goncalves, Ewan Harrison, David K. Jackson, Ian Johnston, Dominic Kwiatkowski, Cordelia Langford, John Sillitoe on behalf of the Wellcome Sanger Institute COVID-19 Surveillance Team |
| EPI_ISL_2435116 | Lighthouse Lab in Alderley Park | Wellcome Sanger Institute for the COVID-19 Genomics UK (COG-UK) Consortium | Jacquelyn Wynn, Mairead Hyland, The Lighthouse Lab in Alderley Park and Alex Alderton, Roberto Amato, Jeffrey Barrett, Sonia Goncalves, Ewan Harrison, David K. Jackson, Ian Johnston, Dominic Kwiatkowski, Cordelia Langford, John Sillitoe on behalf of the Wellcome Sanger Institute COVID-19 Surveillance Team |
| EPI_ISL_2435117 | Lighthouse Lab in Glasgow | Wellcome Sanger Institute for the COVID-19 Genomics UK (COG-UK) Consortium | Harper VanSteenhouse, Yumi Kasai, David Gray, Carol Clugston, Anna Dominiczak and Alex Alderton, Roberto Amato, Jeffrey Barrett, Sonia Goncalves, Ewan Harrison, David K. Jackson, Ian Johnston, Dominic Kwiatkowski, Cordelia Langford, John Sillitoe on behalf of the Wellcome Sanger Institute COVID-19 Surveillance Team |
| EPI_ISL_2435118, EPI_ISL_2435119, EPI_ISL_2435120, EPI_ISL_2435121, EPI_ISL_2435122 | Lighthouse Lab in Alderley Park | Wellcome Sanger Institute for the COVID-19 Genomics UK (COG-UK) Consortium | Jacquelyn Wynn, Mairead Hyland, The Lighthouse Lab in Alderley Park and Alex Alderton, Roberto Amato, Jeffrey Barrett, Sonia Goncalves, Ewan Harrison, David K. Jackson, Ian Johnston, Dominic Kwiatkowski, Cordelia Langford, John Sillitoe on behalf of the Wellcome Sanger Institute COVID-19 Surveillance Team |
| EPI_ISL_2435124 | Lighthouse Lab in Glasgow | Wellcome Sanger Institute for the COVID-19 Genomics UK (COG-UK) Consortium | Harper VanSteenhouse, Yumi Kasai, David Gray, Carol Clugston, Anna Dominiczak and Alex Alderton, Roberto Amato, Jeffrey Barrett, Sonia Goncalves, Ewan Harrison, David K. Jackson, Ian Johnston, Dominic Kwiatkowski, Cordelia Langford, John Sillitoe on behalf of the Wellcome Sanger Institute COVID-19 Surveillance Team |
| EPI_ISL_2435125 | Lighthouse Lab in Alderley Park | Wellcome Sanger Institute for the COVID-19 Genomics UK (COG-UK) Consortium | Jacquelyn Wynn, Mairead Hyland, The Lighthouse Lab in Alderley Park and Alex Alderton, Roberto Amato, Jeffrey Barrett, Sonia Goncalves, Ewan Harrison, David K. Jackson, Ian Johnston, Dominic Kwiatkowski, Cordelia Langford, John Sillitoe on behalf of the Wellcome Sanger Institute COVID-19 Surveillance Team |
| EPI_ISL_2435126 | Lighthouse Lab in Glasgow | Wellcome Sanger Institute for the COVID-19 Genomics UK (COG-UK) Consortium | Harper VanSteenhouse, Yumi Kasai, David Gray, Carol Clugston, Anna Dominiczak and Alex Alderton, Roberto Amato, Jeffrey Barrett, Sonia Goncalves, Ewan Harrison, David K. Jackson, Ian Johnston, Dominic Kwiatkowski, Cordelia Langford, John Sillitoe on behalf of the Wellcome Sanger Institute COVID-19 Surveillance Team |
| EPI_ISL_2435127, EPI_ISL_2435128 | Lighthouse Lab in Alderley Park | Wellcome Sanger Institute for the COVID-19 Genomics UK (COG-UK) Consortium | Jacquelyn Wynn, Mairead Hyland, The Lighthouse Lab in Alderley Park and Alex Alderton, Roberto Amato, Jeffrey Barrett, Sonia Goncalves, Ewan Harrison, David K. Jackson, Ian Johnston, Dominic Kwiatkowski, Cordelia Langford, John Sillitoe on behalf of the Wellcome Sanger Institute COVID-19 Surveillance Team |
| EPI_ISL_2435129 | Lighthouse Lab in Glasgow | Wellcome Sanger Institute for the COVID-19 Genomics UK (COG-UK) Consortium | Harper VanSteenhouse, Yumi Kasai, David Gray, Carol Clugston, Anna Dominiczak and Alex Alderton, Roberto Amato, Jeffrey Barrett, Sonia Goncalves, Ewan Harrison, David K. Jackson, Ian Johnston, Dominic Kwiatkowski, Cordelia Langford, John Sillitoe on behalf of the Wellcome Sanger Institute COVID-19 Surveillance Team |
| EPI_ISL_2435130, EPI_ISL_2435131 | Lighthouse Lab in Alderley Park | Wellcome Sanger Institute for the COVID-19 Genomics UK (COG-UK) Consortium | Jacquelyn Wynn, Mairead Hyland, The Lighthouse Lab in Alderley Park and Alex Alderton, Roberto Amato, Jeffrey Barrett, Sonia Goncalves, Ewan Harrison, David K. Jackson, Ian Johnston, Dominic Kwiatkowski, Cordelia Langford, John Sillitoe on behalf of the Wellcome Sanger Institute COVID-19 Surveillance Team |
| EPI_ISL_2435132 | Lighthouse Lab in Glasgow | Wellcome Sanger Institute for the COVID-19 Genomics UK (COG-UK) Consortium | Harper VanSteenhouse, Yumi Kasai, David Gray, Carol Clugston, Anna Dominiczak and Alex Alderton, Roberto Amato, Jeffrey Barrett, Sonia Goncalves, Ewan Harrison, David K. Jackson, Ian Johnston, Dominic Kwiatkowski, Cordelia Langford, John Sillitoe on behalf of the Wellcome Sanger Institute COVID-19 Surveillance Team |
| EPI_ISL_2435133, EPI_ISL_2435134, EPI_ISL_2435135 | Lighthouse Lab in Alderley Park | Wellcome Sanger Institute for the COVID-19 Genomics UK (COG-UK) Consortium | Jacquelyn Wynn, Mairead Hyland, The Lighthouse Lab in Alderley Park and Alex Alderton, Roberto Amato, Jeffrey Barrett, Sonia Goncalves, Ewan Harrison, David K. Jackson, Ian Johnston, Dominic Kwiatkowski, Cordelia Langford, John Sillitoe on behalf of the Wellcome Sanger Institute COVID-19 Surveillance Team |
| EPI_ISL_2435136 | Lighthouse Lab in Glasgow | Wellcome Sanger Institute for the COVID-19 Genomics UK (COG-UK) Consortium | Harper VanSteenhouse, Yumi Kasai, David Gray, Carol Clugston, Anna Dominiczak and Alex Alderton, Roberto Amato, Jeffrey Barrett, Sonia Goncalves, Ewan Harrison, David K. Jackson, Ian Johnston, Dominic Kwiatkowski, Cordelia Langford, John Sillitoe on behalf of the Wellcome Sanger Institute COVID-19 Surveillance Team |
| EPI_ISL_2435137, EPI_ISL_2435140 | Lighthouse Lab in Alderley Park | Wellcome Sanger Institute for the COVID-19 Genomics UK (COG-UK) Consortium | Jacquelyn Wynn, Mairead Hyland, The Lighthouse Lab in Alderley Park and Alex Alderton, Roberto Amato, Jeffrey Barrett, Sonia Goncalves, Ewan Harrison, David K. Jackson, Ian Johnston, Dominic Kwiatkowski, Cordelia Langford, John Sillitoe on behalf of the Wellcome Sanger Institute COVID-19 Surveillance Team |
| EPI_ISL_2435141 | Lighthouse Lab in Glasgow | Wellcome Sanger Institute for the COVID-19 Genomics UK (COG-UK) Consortium | Harper VanSteenhouse, Yumi Kasai, David Gray, Carol Clugston, Anna Dominiczak and Alex Alderton, Roberto Amato, Jeffrey Barrett, Sonia Goncalves, Ewan Harrison, David K. Jackson, Ian Johnston, Dominic Kwiatkowski, Cordelia Langford, John Sillitoe on behalf of the Wellcome Sanger Institute COVID-19 Surveillance Team |
| EPI_ISL_2435142, EPI_ISL_2435143, EPI_ISL_2435144 | Lighthouse Lab in Alderley Park | Wellcome Sanger Institute for the COVID-19 Genomics UK (COG-UK) Consortium | Jacquelyn Wynn, Mairead Hyland, The Lighthouse Lab in Alderley Park and Alex Alderton, Roberto Amato, Jeffrey Barrett, Sonia Goncalves, Ewan Harrison, David K. Jackson, Ian Johnston, Dominic Kwiatkowski, Cordelia Langford, John Sillitoe on behalf of the Wellcome Sanger Institute COVID-19 Surveillance Team |
| EPI_ISL_2435145 | Lighthouse Lab in Glasgow | Wellcome Sanger Institute for the COVID-19 Genomics UK (COG-UK) Consortium | Harper VanSteenhouse, Yumi Kasai, David Gray, Carol Clugston, Anna Dominiczak and Alex Alderton, Roberto Amato, Jeffrey Barrett, Sonia Goncalves, Ewan Harrison, David K. Jackson, Ian Johnston, Dominic Kwiatkowski, Cordelia Langford, John Sillitoe on behalf of the Wellcome Sanger Institute COVID-19 Surveillance Team |
| EPI_ISL_2435146, EPI_ISL_2435147, EPI_ISL_2435148, EPI_ISL_2435149, EPI_ISL_2435150, EPI_ISL_2435151 | Lighthouse Lab in Alderley Park | Wellcome Sanger Institute for the COVID-19 Genomics UK (COG-UK) Consortium | Jacquelyn Wynn, Mairead Hyland, The Lighthouse Lab in Alderley Park and Alex Alderton, Roberto Amato, Jeffrey Barrett, Sonia Goncalves, Ewan Harrison, David K. Jackson, Ian Johnston, Dominic Kwiatkowski, Cordelia Langford, John Sillitoe on behalf of the Wellcome Sanger Institute COVID-19 Surveillance Team |
| EPI_ISL_2435152 | Lighthouse Lab in Glasgow | Wellcome Sanger Institute for the COVID-19 Genomics UK (COG-UK) Consortium | Harper VanSteenhouse, Yumi Kasai, David Gray, Carol Clugston, Anna Dominiczak and Alex Alderton, Roberto Amato, Jeffrey Barrett, Sonia Goncalves, Ewan Harrison, David K. Jackson, Ian Johnston, Dominic Kwiatkowski, Cordelia Langford, John Sillitoe on behalf of the Wellcome Sanger Institute COVID-19 Surveillance Team |
| EPI_ISL_2435153, EPI_ISL_2435154, EPI_ISL_2435155, EPI_ISL_2435156, EPI_ISL_2435157, EPI_ISL_2435158, EPI_ISL_2435159, EPI_ISL_2435160, EPI_ISL_2435161, EPI_ISL_2435162, EPI_ISL_2435165, EPI_ISL_2435166 | | | |
| see above | Lighthouse Lab in Alderley Park | Wellcome Sanger Institute for the COVID-19 Genomics UK (COG-UK) Consortium | Jacquelyn Wynn, Mairead Hyland, The Lighthouse Lab in Alderley Park and Alex Alderton, Roberto Amato, Jeffrey Barrett, Sonia Goncalves, Ewan Harrison, David K. Jackson, Ian Johnston, Dominic Kwiatkowski, Cordelia Langford, John Sillitoe on behalf of the Wellcome Sanger Institute COVID-19 Surveillance Team |
| EPI_ISL_2435168 | Lighthouse Lab in Glasgow | Wellcome Sanger Institute for the COVID-19 Genomics UK (COG-UK) Consortium | Harper VanSteenhouse, Yumi Kasai, David Gray, Carol Clugston, Anna Dominiczak and Alex Alderton, Roberto Amato, Jeffrey Barrett, Sonia Goncalves, Ewan Harrison, David K. Jackson, Ian Johnston, Dominic Kwiatkowski, Cordelia Langford, John Sillitoe on behalf of the Wellcome Sanger Institute COVID-19 Surveillance Team |
| EPI_ISL_2435169, EPI_ISL_2435171, EPI_ISL_2435174 | Lighthouse Lab in Alderley Park | Wellcome Sanger Institute for the COVID-19 Genomics UK (COG-UK) Consortium | Jacquelyn Wynn, Mairead Hyland, The Lighthouse Lab in Alderley Park and Alex Alderton, Roberto Amato, Jeffrey Barrett, Sonia Goncalves, Ewan Harrison, David K. Jackson, Ian Johnston, Dominic Kwiatkowski, Cordelia Langford, John Sillitoe on behalf of the Wellcome Sanger Institute COVID-19 Surveillance Team |
| EPI_ISL_2435175 | Lighthouse Lab in Glasgow | Wellcome Sanger Institute for the COVID-19 Genomics UK (COG-UK) Consortium | Harper VanSteenhouse, Yumi Kasai, David Gray, Carol Clugston, Anna Dominiczak and Alex Alderton, Roberto Amato, Jeffrey Barrett, Sonia Goncalves, Ewan Harrison, David K. Jackson, Ian Johnston, Dominic Kwiatkowski, Cordelia Langford, John Sillitoe on behalf of the Wellcome Sanger Institute COVID-19 Surveillance Team |
| EPI_ISL_2435176, EPI_ISL_2435177 | Lighthouse Lab in Alderley Park | Wellcome Sanger Institute for the COVID-19 Genomics UK (COG-UK) Consortium | Jacquelyn Wynn, Mairead Hyland, The Lighthouse Lab in Alderley Park and Alex Alderton, Roberto Amato, Jeffrey Barrett, Sonia Goncalves, Ewan Harrison, David K. Jackson, Ian Johnston, Dominic Kwiatkowski, Cordelia Langford, John Sillitoe on behalf of the Wellcome Sanger Institute COVID-19 Surveillance Team |

| | | | |
|---|---|---|---|
| EPI_ISL_2435178, EPI_ISL_2435179 | Lighthouse Lab in Glasgow | Wellcome Sanger Institute for the COVID-19 Genomics UK (COG-UK) Consortium | Harper VanSteenhouse, Yumi Kasai, David Gray, Carol Clugston, Anna Dominiczak and Alex Alderton, Roberto Amato, Jeffrey Barrett, Sonia Goncalves, Ewan Harrison, David K. Jackson, Ian Johnston, Dominic Kwiatkowski, Cordelia Langford, John Sillitoe on behalf of the Wellcome Sanger Institute COVID-19 Surveillance Team |
| EPI_ISL_2435180, EPI_ISL_2435181, EPI_ISL_2435182, EPI_ISL_2435183, EPI_ISL_2435184, EPI_ISL_2435185, EPI_ISL_2435186, EPI_ISL_2435187, EPI_ISL_2435188, EPI_ISL_2435189, EPI_ISL_2435190, EPI_ISL_2435191, EPI_ISL_2435192, EPI_ISL_2435193, EPI_ISL_2435194, EPI_ISL_2435195, EPI_ISL_2435196, EPI_ISL_2435197 | | | |
| see above | Lighthouse Lab in Alderley Park | Wellcome Sanger Institute for the COVID-19 Genomics UK (COG-UK) Consortium | Jacquelyn Wynn, Mairead Hyland, The Lighthouse Lab in Alderley Park and Alex Alderton, Roberto Amato, Jeffrey Barrett, Sonia Goncalves, Ewan Harrison, David K. Jackson, Ian Johnston, Dominic Kwiatkowski, Cordelia Langford, John Sillitoe on behalf of the Wellcome Sanger Institute COVID-19 Surveillance Team |
| EPI_ISL_2435198 | Lighthouse Lab in Glasgow | Wellcome Sanger Institute for the COVID-19 Genomics UK (COG-UK) Consortium | Harper VanSteenhouse, Yumi Kasai, David Gray, Carol Clugston, Anna Dominiczak and Alex Alderton, Roberto Amato, Jeffrey Barrett, Sonia Goncalves, Ewan Harrison, David K. Jackson, Ian Johnston, Dominic Kwiatkowski, Cordelia Langford, John Sillitoe on behalf of the Wellcome Sanger Institute COVID-19 Surveillance Team |
| EPI_ISL_2435199, EPI_ISL_2435200, EPI_ISL_2435201, EPI_ISL_2435202, EPI_ISL_2435203, EPI_ISL_2435204, EPI_ISL_2435205, EPI_ISL_2435206, EPI_ISL_2435207, EPI_ISL_2435208, EPI_ISL_2435211, EPI_ISL_2435212, EPI_ISL_2435213 | | | |
| see above | Lighthouse Lab in Alderley Park | Wellcome Sanger Institute for the COVID-19 Genomics UK (COG-UK) Consortium | Jacquelyn Wynn, Mairead Hyland, The Lighthouse Lab in Alderley Park and Alex Alderton, Roberto Amato, Jeffrey Barrett, Sonia Goncalves, Ewan Harrison, David K. Jackson, Ian Johnston, Dominic Kwiatkowski, Cordelia Langford, John Sillitoe on behalf of the Wellcome Sanger Institute COVID-19 Surveillance Team |
| EPI_ISL_2435214 | Lighthouse Lab in Glasgow | Wellcome Sanger Institute for the COVID-19 Genomics UK (COG-UK) Consortium | Harper VanSteenhouse, Yumi Kasai, David Gray, Carol Clugston, Anna Dominiczak and Alex Alderton, Roberto Amato, Jeffrey Barrett, Sonia Goncalves, Ewan Harrison, David K. Jackson, Ian Johnston, Dominic Kwiatkowski, Cordelia Langford, John Sillitoe on behalf of the Wellcome Sanger Institute COVID-19 Surveillance Team |
| EPI_ISL_2435215 | Lighthouse Lab in Alderley Park | Wellcome Sanger Institute for the COVID-19 Genomics UK (COG-UK) Consortium | Jacquelyn Wynn, Mairead Hyland, The Lighthouse Lab in Alderley Park and Alex Alderton, Roberto Amato, Jeffrey Barrett, Sonia Goncalves, Ewan Harrison, David K. Jackson, Ian Johnston, Dominic Kwiatkowski, Cordelia Langford, John Sillitoe on behalf of the Wellcome Sanger Institute COVID-19 Surveillance Team |
| EPI_ISL_2435216 | Lighthouse Lab in Glasgow | Wellcome Sanger Institute for the COVID-19 Genomics UK (COG-UK) Consortium | Harper VanSteenhouse, Yumi Kasai, David Gray, Carol Clugston, Anna Dominiczak and Alex Alderton, Roberto Amato, Jeffrey Barrett, Sonia Goncalves, Ewan Harrison, David K. Jackson, Ian Johnston, Dominic Kwiatkowski, Cordelia Langford, John Sillitoe on behalf of the Wellcome Sanger Institute COVID-19 Surveillance Team |
| EPI_ISL_2435217, EPI_ISL_2435218, EPI_ISL_2435219, EPI_ISL_2435221, EPI_ISL_2435223, EPI_ISL_2435224 | Lighthouse Lab in Alderley Park | Wellcome Sanger Institute for the COVID-19 Genomics UK (COG-UK) Consortium | Jacquelyn Wynn, Mairead Hyland, The Lighthouse Lab in Alderley Park and Alex Alderton, Roberto Amato, Jeffrey Barrett, Sonia Goncalves, Ewan Harrison, David K. Jackson, Ian Johnston, Dominic Kwiatkowski, Cordelia Langford, John Sillitoe on behalf of the Wellcome Sanger Institute COVID-19 Surveillance Team |
| EPI_ISL_2435225, EPI_ISL_2435226, EPI_ISL_2435227 | Lighthouse Lab in Glasgow | Wellcome Sanger Institute for the COVID-19 Genomics UK (COG-UK) Consortium | Harper VanSteenhouse, Yumi Kasai, David Gray, Carol Clugston, Anna Dominiczak and Alex Alderton, Roberto Amato, Jeffrey Barrett, Sonia Goncalves, Ewan Harrison, David K. Jackson, Ian Johnston, Dominic Kwiatkowski, Cordelia Langford, John Sillitoe on behalf of the Wellcome Sanger Institute COVID-19 Surveillance Team |
| EPI_ISL_2435228, EPI_ISL_2435229, EPI_ISL_2435230, EPI_ISL_2435231, EPI_ISL_2435232, EPI_ISL_2435233, EPI_ISL_2435234, EPI_ISL_2435235, EPI_ISL_2435236, EPI_ISL_2435237, EPI_ISL_2435238, EPI_ISL_2435239, EPI_ISL_2435240, EPI_ISL_2435241, EPI_ISL_2435242, EPI_ISL_2435243, EPI_ISL_2435244, EPI_ISL_2435245, EPI_ISL_2435246 | | | |
| see above | Lighthouse Lab in Alderley Park | Wellcome Sanger Institute for the COVID-19 Genomics UK (COG-UK) Consortium | Jacquelyn Wynn, Mairead Hyland, The Lighthouse Lab in Alderley Park and Alex Alderton, Roberto Amato, Jeffrey Barrett, Sonia Goncalves, Ewan Harrison, David K. Jackson, Ian Johnston, Dominic Kwiatkowski, Cordelia Langford, John Sillitoe on behalf of the Wellcome Sanger Institute COVID-19 Surveillance Team |
| EPI_ISL_2435247 | Lighthouse Lab in Glasgow | Wellcome Sanger Institute for the COVID-19 Genomics UK (COG-UK) Consortium | Harper VanSteenhouse, Yumi Kasai, David Gray, Carol Clugston, Anna Dominiczak and Alex Alderton, Roberto Amato, Jeffrey Barrett, Sonia Goncalves, Ewan Harrison, David K. Jackson, Ian Johnston, Dominic Kwiatkowski, Cordelia Langford, John Sillitoe on behalf of the Wellcome Sanger Institute COVID-19 Surveillance Team |
| EPI_ISL_2435248, EPI_ISL_2435249, EPI_ISL_2435250, EPI_ISL_2435251 | Lighthouse Lab in Alderley Park | Wellcome Sanger Institute for the COVID-19 Genomics UK (COG-UK) Consortium | Jacquelyn Wynn, Mairead Hyland, The Lighthouse Lab in Alderley Park and Alex Alderton, Roberto Amato, Jeffrey Barrett, Sonia Goncalves, Ewan Harrison, David K. Jackson, Ian Johnston, Dominic Kwiatkowski, Cordelia Langford, John Sillitoe on behalf of the Wellcome Sanger Institute COVID-19 Surveillance Team |
| EPI_ISL_2435252 | Lighthouse Lab in Glasgow | Wellcome Sanger Institute for the COVID-19 Genomics UK (COG-UK) Consortium | Harper VanSteenhouse, Yumi Kasai, David Gray, Carol Clugston, Anna Dominiczak and Alex Alderton, Roberto Amato, Jeffrey Barrett, Sonia Goncalves, Ewan Harrison, David K. Jackson, Ian Johnston, Dominic Kwiatkowski, Cordelia Langford, John Sillitoe on behalf of the Wellcome Sanger Institute COVID-19 Surveillance Team |
| EPI_ISL_2435253 | Lighthouse Lab in Alderley Park | Wellcome Sanger Institute for the COVID-19 Genomics UK (COG-UK) Consortium | Jacquelyn Wynn, Mairead Hyland, The Lighthouse Lab in Alderley Park and Alex Alderton, Roberto Amato, Jeffrey Barrett, Sonia Goncalves, Ewan Harrison, David K. Jackson, Ian Johnston, Dominic Kwiatkowski, Cordelia Langford, John Sillitoe on behalf of the Wellcome Sanger Institute COVID-19 Surveillance Team |
| EPI_ISL_2435254, EPI_ISL_2435255 | Lighthouse Lab in Glasgow | Wellcome Sanger Institute for the COVID-19 Genomics UK (COG-UK) Consortium | Harper VanSteenhouse, Yumi Kasai, David Gray, Carol Clugston, Anna Dominiczak and Alex Alderton, Roberto Amato, Jeffrey Barrett, Sonia Goncalves, Ewan Harrison, David K. Jackson, Ian Johnston, Dominic Kwiatkowski, Cordelia Langford, John Sillitoe on behalf of the Wellcome Sanger Institute COVID-19 Surveillance Team |
| EPI_ISL_2435256, EPI_ISL_2435257 | Lighthouse Lab in Alderley Park | Wellcome Sanger Institute for the COVID-19 Genomics UK (COG-UK) Consortium | Jacquelyn Wynn, Mairead Hyland, The Lighthouse Lab in Alderley Park and Alex Alderton, Roberto Amato, Jeffrey Barrett, Sonia Goncalves, Ewan Harrison, David K. Jackson, Ian Johnston, Dominic Kwiatkowski, Cordelia Langford, John Sillitoe on behalf of the Wellcome Sanger Institute COVID-19 Surveillance Team |
| EPI_ISL_2435258 | Lighthouse Lab in Glasgow | Wellcome Sanger Institute for the COVID-19 Genomics UK (COG-UK) Consortium | Harper VanSteenhouse, Yumi Kasai, David Gray, Carol Clugston, Anna Dominiczak and Alex Alderton, Roberto Amato, Jeffrey Barrett, Sonia Goncalves, Ewan Harrison, David K. Jackson, Ian Johnston, Dominic Kwiatkowski, Cordelia Langford, John Sillitoe on behalf of the Wellcome Sanger Institute COVID-19 Surveillance Team |
| EPI_ISL_2435259, EPI_ISL_2435260, EPI_ISL_2435261, EPI_ISL_2435262 | Lighthouse Lab in Alderley Park | Wellcome Sanger Institute for the COVID-19 Genomics UK (COG-UK) Consortium | Jacquelyn Wynn, Mairead Hyland, The Lighthouse Lab in Alderley Park and Alex Alderton, Roberto Amato, Jeffrey Barrett, Sonia Goncalves, Ewan Harrison, David K. Jackson, Ian Johnston, Dominic Kwiatkowski, Cordelia Langford, John Sillitoe on behalf of the Wellcome Sanger Institute COVID-19 Surveillance Team |
| EPI_ISL_2435263 | Lighthouse Lab in Glasgow | Wellcome Sanger Institute for the COVID-19 Genomics UK (COG-UK) Consortium | Harper VanSteenhouse, Yumi Kasai, David Gray, Carol Clugston, Anna Dominiczak and Alex Alderton, Roberto Amato, Jeffrey Barrett, Sonia Goncalves, Ewan Harrison, David K. Jackson, Ian Johnston, Dominic Kwiatkowski, Cordelia Langford, John Sillitoe on behalf of the Wellcome Sanger Institute COVID-19 Surveillance Team |
| EPI_ISL_2435264, EPI_ISL_2435265, EPI_ISL_2435266, EPI_ISL_2435267, EPI_ISL_2435268, EPI_ISL_2435269, EPI_ISL_2435270, EPI_ISL_2435271, EPI_ISL_2435272, EPI_ISL_2435273, EPI_ISL_2435274 | | | |
| see above | Lighthouse Lab in Alderley Park | Wellcome Sanger Institute for the COVID-19 Genomics UK (COG-UK) Consortium | Jacquelyn Wynn, Mairead Hyland, The Lighthouse Lab in Alderley Park and Alex Alderton, Roberto Amato, Jeffrey Barrett, Sonia Goncalves, Ewan Harrison, David K. Jackson, Ian Johnston, Dominic Kwiatkowski, Cordelia Langford, John Sillitoe on behalf of the Wellcome Sanger Institute COVID-19 Surveillance Team |
| EPI_ISL_2435275 | Lighthouse Lab in Glasgow | Wellcome Sanger Institute for the COVID-19 Genomics UK (COG-UK) Consortium | Harper VanSteenhouse, Yumi Kasai, David Gray, Carol Clugston, Anna Dominiczak and Alex Alderton, Roberto Amato, Jeffrey Barrett, Sonia Goncalves, Ewan Harrison, David K. Jackson, Ian Johnston, Dominic Kwiatkowski, Cordelia Langford, John Sillitoe on behalf of the Wellcome Sanger Institute COVID-19 Surveillance Team |
| EPI_ISL_2435276, EPI_ISL_2435277 | Lighthouse Lab in Alderley Park | Wellcome Sanger Institute for the COVID-19 Genomics UK (COG-UK) Consortium | Jacquelyn Wynn, Mairead Hyland, The Lighthouse Lab in Alderley Park and Alex Alderton, Roberto Amato, Jeffrey Barrett, Sonia Goncalves, Ewan Harrison, David K. Jackson, Ian Johnston, Dominic Kwiatkowski, Cordelia Langford, John Sillitoe on behalf of the Wellcome Sanger Institute COVID-19 Surveillance Team |
| EPI_ISL_2435278 | Lighthouse Lab in Glasgow | Wellcome Sanger Institute for the COVID-19 Genomics UK (COG-UK) Consortium | Harper VanSteenhouse, Yumi Kasai, David Gray, Carol Clugston, Anna Dominiczak and Alex Alderton, Roberto Amato, Jeffrey Barrett, Sonia Goncalves, Ewan Harrison, David K. Jackson, Ian Johnston, Dominic Kwiatkowski, Cordelia Langford, John Sillitoe on behalf of the Wellcome Sanger Institute COVID-19 Surveillance Team |
| EPI_ISL_2435279, EPI_ISL_2435280 | Lighthouse Lab in Alderley Park | Wellcome Sanger Institute for the COVID-19 Genomics UK (COG-UK) Consortium | Jacquelyn Wynn, Mairead Hyland, The Lighthouse Lab in Alderley Park and Alex Alderton, Roberto Amato, Jeffrey Barrett, Sonia Goncalves, Ewan Harrison, David K. Jackson, Ian Johnston, Dominic Kwiatkowski, Cordelia Langford, John Sillitoe on behalf of the Wellcome Sanger Institute COVID-19 Surveillance Team |
| EPI_ISL_2435281, EPI_ISL_2435282 | Lighthouse Lab in Glasgow | Wellcome Sanger Institute for the COVID-19 Genomics UK (COG-UK) Consortium | Harper VanSteenhouse, Yumi Kasai, David Gray, Carol Clugston, Anna Dominiczak and Alex Alderton, Roberto Amato, Jeffrey Barrett, Sonia Goncalves, Ewan Harrison, David K. Jackson, Ian Johnston, Dominic Kwiatkowski, Cordelia Langford, John Sillitoe on behalf of the Wellcome Sanger Institute COVID-19 Surveillance Team |
| EPI_ISL_2435283, EPI_ISL_2435284, EPI_ISL_2435285, EPI_ISL_2435287, EPI_ISL_2435288, EPI_ISL_2435289 | Lighthouse Lab in Alderley Park | Wellcome Sanger Institute for the COVID-19 Genomics UK (COG-UK) Consortium | Jacquelyn Wynn, Mairead Hyland, The Lighthouse Lab in Alderley Park and Alex Alderton, Roberto Amato, Jeffrey Barrett, Sonia Goncalves, Ewan Harrison, David K. Jackson, Ian Johnston, Dominic Kwiatkowski, Cordelia Langford, John Sillitoe on behalf of the Wellcome Sanger Institute COVID-19 Surveillance Team |
| EPI_ISL_2435292 | Lighthouse Lab in Glasgow | Wellcome Sanger Institute for the COVID-19 Genomics UK (COG-UK) Consortium | Harper VanSteenhouse, Yumi Kasai, David Gray, Carol Clugston, Anna Dominiczak and Alex Alderton, Roberto Amato, Jeffrey Barrett, Sonia Goncalves, Ewan Harrison, David K. Jackson, Ian Johnston, Dominic Kwiatkowski, Cordelia Langford, John Sillitoe on behalf of the Wellcome Sanger Institute COVID-19 Surveillance Team |
| EPI_ISL_2435293, EPI_ISL_2435294 | Lighthouse Lab in Alderley Park | Wellcome Sanger Institute for the COVID-19 Genomics UK (COG-UK) Consortium | Jacquelyn Wynn, Mairead Hyland, The Lighthouse Lab in Alderley Park and Alex Alderton, Roberto Amato, Jeffrey Barrett, Sonia Goncalves, Ewan Harrison, David K. Jackson, Ian Johnston, Dominic Kwiatkowski, Cordelia Langford, John Sillitoe on behalf of the Wellcome Sanger Institute COVID-19 Surveillance Team |
| EPI_ISL_2435295, EPI_ISL_2435296 | Lighthouse Lab in Glasgow | Wellcome Sanger Institute for the COVID-19 Genomics UK (COG-UK) Consortium | Harper VanSteenhouse, Yumi Kasai, David Gray, Carol Clugston, Anna Dominiczak and Alex Alderton, Roberto Amato, Jeffrey Barrett, Sonia Goncalves, Ewan Harrison, David K. Jackson, Ian Johnston, Dominic Kwiatkowski, Cordelia Langford, John Sillitoe on behalf of the Wellcome Sanger Institute COVID-19 Surveillance Team |
| EPI_ISL_2435297, EPI_ISL_2435298 | Lighthouse Lab in Alderley Park | Wellcome Sanger Institute for the COVID-19 Genomics UK (COG-UK) Consortium | Jacquelyn Wynn, Mairead Hyland, The Lighthouse Lab in Alderley Park and Alex Alderton, Roberto Amato, Jeffrey Barrett, Sonia Goncalves, Ewan Harrison, David K. Jackson, Ian Johnston, Dominic Kwiatkowski, Cordelia Langford, John Sillitoe on behalf of the Wellcome Sanger Institute COVID-19 Surveillance Team |
| EPI_ISL_2435299, EPI_ISL_2435300 | Lighthouse Lab in Glasgow | Wellcome Sanger Institute for the COVID-19 Genomics UK (COG-UK) Consortium | Harper VanSteenhouse, Yumi Kasai, David Gray, Carol Clugston, Anna Dominiczak and Alex Alderton, Roberto Amato, Jeffrey Barrett, Sonia Goncalves, Ewan Harrison, David K. Jackson, Ian Johnston, Dominic Kwiatkowski, Cordelia Langford, John Sillitoe on behalf of the Wellcome Sanger Institute COVID-19 Surveillance Team |
| EPI_ISL_2435301, EPI_ISL_2435302, EPI_ISL_2435303, EPI_ISL_2435304 | Lighthouse Lab in Alderley Park | Wellcome Sanger Institute for the COVID-19 Genomics UK (COG-UK) Consortium | Jacquelyn Wynn, Mairead Hyland, The Lighthouse Lab in Alderley Park and Alex Alderton, Roberto Amato, Jeffrey Barrett, Sonia Goncalves, Ewan Harrison, David K. Jackson, Ian Johnston, Dominic Kwiatkowski, Cordelia Langford, John Sillitoe on behalf of the Wellcome Sanger Institute COVID-19 Surveillance Team |
| EPI_ISL_2435305, EPI_ISL_2435306 | Lighthouse Lab in Glasgow | Wellcome Sanger Institute for the COVID-19 Genomics UK (COG-UK) Consortium | Harper VanSteenhouse, Yumi Kasai, David Gray, Carol Clugston, Anna Dominiczak and Alex Alderton, Roberto Amato, Jeffrey Barrett, Sonia Goncalves, Ewan Harrison, David K. Jackson, Ian Johnston, Dominic Kwiatkowski, Cordelia Langford, John Sillitoe on behalf of the Wellcome Sanger Institute COVID-19 Surveillance Team |
| EPI_ISL_2435307, EPI_ISL_2435308, EPI_ISL_2435309, EPI_ISL_2435310 | Lighthouse Lab in Alderley Park | Wellcome Sanger Institute for the COVID-19 Genomics UK (COG-UK) Consortium | Jacquelyn Wynn, Mairead Hyland, The Lighthouse Lab in Alderley Park and Alex Alderton, Roberto Amato, Jeffrey Barrett, Sonia Goncalves, Ewan Harrison, David K. Jackson, Ian Johnston, Dominic Kwiatkowski, Cordelia Langford, John Sillitoe on behalf of the Wellcome Sanger Institute COVID-19 Surveillance Team |
| EPI_ISL_2435311 | Lighthouse Lab in Glasgow | Wellcome Sanger Institute for the COVID-19 Genomics UK (COG-UK) Consortium | Harper VanSteenhouse, Yumi Kasai, David Gray, Carol Clugston, Anna Dominiczak and Alex Alderton, Roberto Amato, Jeffrey Barrett, Sonia Goncalves, Ewan Harrison, David K. Jackson, Ian Johnston, Dominic Kwiatkowski, Cordelia Langford, John Sillitoe on behalf of the Wellcome Sanger Institute COVID-19 Surveillance Team |
| EPI_ISL_2435312 | Lighthouse Lab in Alderley Park | Wellcome Sanger Institute for the COVID-19 Genomics UK (COG-UK) | Jacquelyn Wynn, Mairead Hyland, The Lighthouse Lab in Alderley Park and Alex Alderton, Roberto Amato, Jeffrey Barrett, Sonia Goncalves, Ewan Harrison, David K. Jackson, Ian Johnston, Dominic Kwiatkowski, Cordelia Langford, John Sillitoe on |

| | | | |
|---|---|---|---|
| | | Consortium | behalf of the Wellcome Sanger Institute COVID-19 Surveillance Team |
| EPI_ISL_2435313, EPI_ISL_2435315 | Lighthouse Lab in Glasgow | Wellcome Sanger Institute for the COVID-19 Genomics UK (COG-UK) Consortium | Harper VanSteenhouse, Yumi Kasai, David Gray, Carol Clugston, Anna Dominiczak and Alex Alderton, Roberto Amato, Jeffrey Barrett, Sonia Goncalves, Ewan Harrison, David K. Jackson, Ian Johnston, Dominic Kwiatkowski, Cordelia Langford, John Sillitoe on behalf of the Wellcome Sanger Institute COVID-19 Surveillance Team |
| EPI_ISL_2435316, EPI_ISL_2435317 | Lighthouse Lab in Alderley Park | Wellcome Sanger Institute for the COVID-19 Genomics UK (COG-UK) Consortium | Jacquelyn Wynn, Mairead Hyland, The Lighthouse Lab in Alderley Park and Alex Alderton, Roberto Amato, Jeffrey Barrett, Sonia Goncalves, Ewan Harrison, David K. Jackson, Ian Johnston, Dominic Kwiatkowski, Cordelia Langford, John Sillitoe on behalf of the Wellcome Sanger Institute COVID-19 Surveillance Team |
| EPI_ISL_2435318, EPI_ISL_2435319 | Lighthouse Lab in Glasgow | Wellcome Sanger Institute for the COVID-19 Genomics UK (COG-UK) Consortium | Harper VanSteenhouse, Yumi Kasai, David Gray, Carol Clugston, Anna Dominiczak and Alex Alderton, Roberto Amato, Jeffrey Barrett, Sonia Goncalves, Ewan Harrison, David K. Jackson, Ian Johnston, Dominic Kwiatkowski, Cordelia Langford, John Sillitoe on behalf of the Wellcome Sanger Institute COVID-19 Surveillance Team |
| EPI_ISL_2435320, EPI_ISL_2435321, EPI_ISL_2435323, EPI_ISL_2435324, EPI_ISL_2435327, EPI_ISL_2435328 | Lighthouse Lab in Alderley Park | Wellcome Sanger Institute for the COVID-19 Genomics UK (COG-UK) Consortium | Jacquelyn Wynn, Mairead Hyland, The Lighthouse Lab in Alderley Park and Alex Alderton, Roberto Amato, Jeffrey Barrett, Sonia Goncalves, Ewan Harrison, David K. Jackson, Ian Johnston, Dominic Kwiatkowski, Cordelia Langford, John Sillitoe on behalf of the Wellcome Sanger Institute COVID-19 Surveillance Team |
| EPI_ISL_2435330 | Lighthouse Lab in Glasgow | Wellcome Sanger Institute for the COVID-19 Genomics UK (COG-UK) Consortium | Harper VanSteenhouse, Yumi Kasai, David Gray, Carol Clugston, Anna Dominiczak and Alex Alderton, Roberto Amato, Jeffrey Barrett, Sonia Goncalves, Ewan Harrison, David K. Jackson, Ian Johnston, Dominic Kwiatkowski, Cordelia Langford, John Sillitoe on behalf of the Wellcome Sanger Institute COVID-19 Surveillance Team |
| EPI_ISL_2435331, EPI_ISL_2435332 | Lighthouse Lab in Alderley Park | Wellcome Sanger Institute for the COVID-19 Genomics UK (COG-UK) Consortium | Jacquelyn Wynn, Mairead Hyland, The Lighthouse Lab in Alderley Park and Alex Alderton, Roberto Amato, Jeffrey Barrett, Sonia Goncalves, Ewan Harrison, David K. Jackson, Ian Johnston, Dominic Kwiatkowski, Cordelia Langford, John Sillitoe on behalf of the Wellcome Sanger Institute COVID-19 Surveillance Team |
| EPI_ISL_2435333 | Lighthouse Lab in Glasgow | Wellcome Sanger Institute for the COVID-19 Genomics UK (COG-UK) Consortium | Harper VanSteenhouse, Yumi Kasai, David Gray, Carol Clugston, Anna Dominiczak and Alex Alderton, Roberto Amato, Jeffrey Barrett, Sonia Goncalves, Ewan Harrison, David K. Jackson, Ian Johnston, Dominic Kwiatkowski, Cordelia Langford, John Sillitoe on behalf of the Wellcome Sanger Institute COVID-19 Surveillance Team |
| EPI_ISL_2435334 | Lighthouse Lab in Alderley Park | Wellcome Sanger Institute for the COVID-19 Genomics UK (COG-UK) Consortium | Jacquelyn Wynn, Mairead Hyland, The Lighthouse Lab in Alderley Park and Alex Alderton, Roberto Amato, Jeffrey Barrett, Sonia Goncalves, Ewan Harrison, David K. Jackson, Ian Johnston, Dominic Kwiatkowski, Cordelia Langford, John Sillitoe on behalf of the Wellcome Sanger Institute COVID-19 Surveillance Team |
| EPI_ISL_2435335, EPI_ISL_2435337 | Lighthouse Lab in Glasgow | Wellcome Sanger Institute for the COVID-19 Genomics UK (COG-UK) Consortium | Harper VanSteenhouse, Yumi Kasai, David Gray, Carol Clugston, Anna Dominiczak and Alex Alderton, Roberto Amato, Jeffrey Barrett, Sonia Goncalves, Ewan Harrison, David K. Jackson, Ian Johnston, Dominic Kwiatkowski, Cordelia Langford, John Sillitoe on behalf of the Wellcome Sanger Institute COVID-19 Surveillance Team |
| EPI_ISL_2435338, EPI_ISL_2435339, EPI_ISL_2435340, EPI_ISL_2435341, EPI_ISL_2435342, EPI_ISL_2435344, EPI_ISL_2435345 | Lighthouse Lab in Alderley Park | Wellcome Sanger Institute for the COVID-19 Genomics UK (COG-UK) Consortium | Jacquelyn Wynn, Mairead Hyland, The Lighthouse Lab in Alderley Park and Alex Alderton, Roberto Amato, Jeffrey Barrett, Sonia Goncalves, Ewan Harrison, David K. Jackson, Ian Johnston, Dominic Kwiatkowski, Cordelia Langford, John Sillitoe on behalf of the Wellcome Sanger Institute COVID-19 Surveillance Team |
| EPI_ISL_2435346, EPI_ISL_2435347 | Lighthouse Lab in Milton Keynes | Wellcome Sanger Institute for the COVID-19 Genomics UK (COG-UK) Consortium | The Lighthouse Lab in Milton Keynes and Alex Alderton, Roberto Amato, Jeffrey Barrett, Sonia Goncalves, Ewan Harrison, David K. Jackson, Ian Johnston, Dominic Kwiatkowski, Cordelia Langford, John Sillitoe on behalf of the Wellcome Sanger Institute COVID-19 Surveillance Team |
| EPI_ISL_2435348 | Lighthouse Lab in Alderley Park | Wellcome Sanger Institute for the COVID-19 Genomics UK (COG-UK) Consortium | Jacquelyn Wynn, Mairead Hyland, The Lighthouse Lab in Alderley Park and Alex Alderton, Roberto Amato, Jeffrey Barrett, Sonia Goncalves, Ewan Harrison, David K. Jackson, Ian Johnston, Dominic Kwiatkowski, Cordelia Langford, John Sillitoe on behalf of the Wellcome Sanger Institute COVID-19 Surveillance Team |
| EPI_ISL_2435349 | Lighthouse Lab in Milton Keynes | Wellcome Sanger Institute for the COVID-19 Genomics UK (COG-UK) Consortium | The Lighthouse Lab in Milton Keynes and Alex Alderton, Roberto Amato, Jeffrey Barrett, Sonia Goncalves, Ewan Harrison, David K. Jackson, Ian Johnston, Dominic Kwiatkowski, Cordelia Langford, John Sillitoe on behalf of the Wellcome Sanger Institute COVID-19 Surveillance Team |
| EPI_ISL_2435350, EPI_ISL_2435351, EPI_ISL_2435352 | Lighthouse Lab in Alderley Park | Wellcome Sanger Institute for the COVID-19 Genomics UK (COG-UK) Consortium | Jacquelyn Wynn, Mairead Hyland, The Lighthouse Lab in Alderley Park and Alex Alderton, Roberto Amato, Jeffrey Barrett, Sonia Goncalves, Ewan Harrison, David K. Jackson, Ian Johnston, Dominic Kwiatkowski, Cordelia Langford, John Sillitoe on behalf of the Wellcome Sanger Institute COVID-19 Surveillance Team |
| EPI_ISL_2435354 | Lighthouse Lab in Glasgow | Wellcome Sanger Institute for the COVID-19 Genomics UK (COG-UK) Consortium | Harper VanSteenhouse, Yumi Kasai, David Gray, Carol Clugston, Anna Dominiczak and Alex Alderton, Roberto Amato, Jeffrey Barrett, Sonia Goncalves, Ewan Harrison, David K. Jackson, Ian Johnston, Dominic Kwiatkowski, Cordelia Langford, John Sillitoe on behalf of the Wellcome Sanger Institute COVID-19 Surveillance Team |
| EPI_ISL_2435355 | Lighthouse Lab in Alderley Park | Wellcome Sanger Institute for the COVID-19 Genomics UK (COG-UK) Consortium | Jacquelyn Wynn, Mairead Hyland, The Lighthouse Lab in Alderley Park and Alex Alderton, Roberto Amato, Jeffrey Barrett, Sonia Goncalves, Ewan Harrison, David K. Jackson, Ian Johnston, Dominic Kwiatkowski, Cordelia Langford, John Sillitoe on behalf of the Wellcome Sanger Institute COVID-19 Surveillance Team |
| EPI_ISL_2435356 | Lighthouse Lab in Milton Keynes | Wellcome Sanger Institute for the COVID-19 Genomics UK (COG-UK) Consortium | The Lighthouse Lab in Milton Keynes and Alex Alderton, Roberto Amato, Jeffrey Barrett, Sonia Goncalves, Ewan Harrison, David K. Jackson, Ian Johnston, Dominic Kwiatkowski, Cordelia Langford, John Sillitoe on behalf of the Wellcome Sanger Institute COVID-19 Surveillance Team |
| EPI_ISL_2435357 | Lighthouse Lab in Alderley Park | Wellcome Sanger Institute for the COVID-19 Genomics UK (COG-UK) Consortium | Jacquelyn Wynn, Mairead Hyland, The Lighthouse Lab in Alderley Park and Alex Alderton, Roberto Amato, Jeffrey Barrett, Sonia Goncalves, Ewan Harrison, David K. Jackson, Ian Johnston, Dominic Kwiatkowski, Cordelia Langford, John Sillitoe on behalf of the Wellcome Sanger Institute COVID-19 Surveillance Team |
| EPI_ISL_2435358 | Lighthouse Lab in Milton Keynes | Wellcome Sanger Institute for the COVID-19 Genomics UK (COG-UK) Consortium | The Lighthouse Lab in Milton Keynes and Alex Alderton, Roberto Amato, Jeffrey Barrett, Sonia Goncalves, Ewan Harrison, David K. Jackson, Ian Johnston, Dominic Kwiatkowski, Cordelia Langford, John Sillitoe on behalf of the Wellcome Sanger Institute COVID-19 Surveillance Team |
| EPI_ISL_2435359, EPI_ISL_2435360 | Lighthouse Lab in Alderley Park | Wellcome Sanger Institute for the COVID-19 Genomics UK (COG-UK) Consortium | Jacquelyn Wynn, Mairead Hyland, The Lighthouse Lab in Alderley Park and Alex Alderton, Roberto Amato, Jeffrey Barrett, Sonia Goncalves, Ewan Harrison, David K. Jackson, Ian Johnston, Dominic Kwiatkowski, Cordelia Langford, John Sillitoe on behalf of the Wellcome Sanger Institute COVID-19 Surveillance Team |
| EPI_ISL_2435361 | Lighthouse Lab in Milton Keynes | Wellcome Sanger Institute for the COVID-19 Genomics UK (COG-UK) Consortium | The Lighthouse Lab in Milton Keynes and Alex Alderton, Roberto Amato, Jeffrey Barrett, Sonia Goncalves, Ewan Harrison, David K. Jackson, Ian Johnston, Dominic Kwiatkowski, Cordelia Langford, John Sillitoe on behalf of the Wellcome Sanger Institute COVID-19 Surveillance Team |
| EPI_ISL_2435362, EPI_ISL_2435363, EPI_ISL_2435365 | Lighthouse Lab in Alderley Park | Wellcome Sanger Institute for the COVID-19 Genomics UK (COG-UK) Consortium | Jacquelyn Wynn, Mairead Hyland, The Lighthouse Lab in Alderley Park and Alex Alderton, Roberto Amato, Jeffrey Barrett, Sonia Goncalves, Ewan Harrison, David K. Jackson, Ian Johnston, Dominic Kwiatkowski, Cordelia Langford, John Sillitoe on behalf of the Wellcome Sanger Institute COVID-19 Surveillance Team |
| EPI_ISL_2435366, EPI_ISL_2435368 | Lighthouse Lab in Milton Keynes | Wellcome Sanger Institute for the COVID-19 Genomics UK (COG-UK) Consortium | The Lighthouse Lab in Milton Keynes and Alex Alderton, Roberto Amato, Jeffrey Barrett, Sonia Goncalves, Ewan Harrison, David K. Jackson, Ian Johnston, Dominic Kwiatkowski, Cordelia Langford, John Sillitoe on behalf of the Wellcome Sanger Institute COVID-19 Surveillance Team |
| EPI_ISL_2435369, EPI_ISL_2435370, EPI_ISL_2435371, EPI_ISL_2435372, EPI_ISL_2435373, EPI_ISL_2435374, EPI_ISL_2435376, EPI_ISL_2435377 | Lighthouse Lab in Alderley Park | Wellcome Sanger Institute for the COVID-19 Genomics UK (COG-UK) Consortium | Jacquelyn Wynn, Mairead Hyland, The Lighthouse Lab in Alderley Park and Alex Alderton, Roberto Amato, Jeffrey Barrett, Sonia Goncalves, Ewan Harrison, David K. Jackson, Ian Johnston, Dominic Kwiatkowski, Cordelia Langford, John Sillitoe on behalf of the Wellcome Sanger Institute COVID-19 Surveillance Team |
| EPI_ISL_2435378 | Lighthouse Lab in Milton Keynes | Wellcome Sanger Institute for the COVID-19 Genomics UK (COG-UK) Consortium | The Lighthouse Lab in Milton Keynes and Alex Alderton, Roberto Amato, Jeffrey Barrett, Sonia Goncalves, Ewan Harrison, David K. Jackson, Ian Johnston, Dominic Kwiatkowski, Cordelia Langford, John Sillitoe on behalf of the Wellcome Sanger Institute COVID-19 Surveillance Team |
| EPI_ISL_2435380, EPI_ISL_2435381 | Lighthouse Lab in Alderley Park | Wellcome Sanger Institute for the COVID-19 Genomics UK (COG-UK) Consortium | Jacquelyn Wynn, Mairead Hyland, The Lighthouse Lab in Alderley Park and Alex Alderton, Roberto Amato, Jeffrey Barrett, Sonia Goncalves, Ewan Harrison, David K. Jackson, Ian Johnston, Dominic Kwiatkowski, Cordelia Langford, John Sillitoe on behalf of the Wellcome Sanger Institute COVID-19 Surveillance Team |
| EPI_ISL_2435382, EPI_ISL_2435386 | Lighthouse Lab in Milton Keynes | Wellcome Sanger Institute for the COVID-19 Genomics UK (COG-UK) Consortium | The Lighthouse Lab in Milton Keynes and Alex Alderton, Roberto Amato, Jeffrey Barrett, Sonia Goncalves, Ewan Harrison, David K. Jackson, Ian Johnston, Dominic Kwiatkowski, Cordelia Langford, John Sillitoe on behalf of the Wellcome Sanger Institute COVID-19 Surveillance Team |
| EPI_ISL_2435387 | Lighthouse Lab in Alderley Park | Wellcome Sanger Institute for the COVID-19 Genomics UK (COG-UK) Consortium | Jacquelyn Wynn, Mairead Hyland, The Lighthouse Lab in Alderley Park and Alex Alderton, Roberto Amato, Jeffrey Barrett, Sonia Goncalves, Ewan Harrison, David K. Jackson, Ian Johnston, Dominic Kwiatkowski, Cordelia Langford, John Sillitoe on behalf of the Wellcome Sanger Institute COVID-19 Surveillance Team |
| EPI_ISL_2435388, EPI_ISL_2435389 | Lighthouse Lab in Milton Keynes | Wellcome Sanger Institute for the COVID-19 Genomics UK (COG-UK) Consortium | The Lighthouse Lab in Milton Keynes and Alex Alderton, Roberto Amato, Jeffrey Barrett, Sonia Goncalves, Ewan Harrison, David K. Jackson, Ian Johnston, Dominic Kwiatkowski, Cordelia Langford, John Sillitoe on behalf of the Wellcome Sanger Institute COVID-19 Surveillance Team |
| EPI_ISL_2435390 | Lighthouse Lab in Alderley Park | Wellcome Sanger Institute for the COVID-19 Genomics UK (COG-UK) Consortium | Jacquelyn Wynn, Mairead Hyland, The Lighthouse Lab in Alderley Park and Alex Alderton, Roberto Amato, Jeffrey Barrett, Sonia Goncalves, Ewan Harrison, David K. Jackson, Ian Johnston, Dominic Kwiatkowski, Cordelia Langford, John Sillitoe on behalf of the Wellcome Sanger Institute COVID-19 Surveillance Team |
| EPI_ISL_2435391 | Lighthouse Lab in Milton Keynes | Wellcome Sanger Institute for the COVID-19 Genomics UK (COG-UK) Consortium | The Lighthouse Lab in Milton Keynes and Alex Alderton, Roberto Amato, Jeffrey Barrett, Sonia Goncalves, Ewan Harrison, David K. Jackson, Ian Johnston, Dominic Kwiatkowski, Cordelia Langford, John Sillitoe on behalf of the Wellcome Sanger Institute COVID-19 Surveillance Team |
| EPI_ISL_2435392 | Lighthouse Lab in Glasgow | Wellcome Sanger Institute for the COVID-19 Genomics UK (COG-UK) Consortium | Harper VanSteenhouse, Yumi Kasai, David Gray, Carol Clugston, Anna Dominiczak and Alex Alderton, Roberto Amato, Jeffrey Barrett, Sonia Goncalves, Ewan Harrison, David K. Jackson, Ian Johnston, Dominic Kwiatkowski, Cordelia Langford, John Sillitoe on behalf of the Wellcome Sanger Institute COVID-19 Surveillance Team |
| EPI_ISL_2435393, EPI_ISL_2435394 | Lighthouse Lab in Alderley Park | Wellcome Sanger Institute for the COVID-19 Genomics UK (COG-UK) Consortium | Jacquelyn Wynn, Mairead Hyland, The Lighthouse Lab in Alderley Park and Alex Alderton, Roberto Amato, Jeffrey Barrett, Sonia Goncalves, Ewan Harrison, David K. Jackson, Ian Johnston, Dominic Kwiatkowski, Cordelia Langford, John Sillitoe on behalf of the Wellcome Sanger Institute COVID-19 Surveillance Team |
| EPI_ISL_2435395 | Lighthouse Lab in Milton Keynes | Wellcome Sanger Institute for the COVID-19 Genomics UK (COG-UK) Consortium | The Lighthouse Lab in Milton Keynes and Alex Alderton, Roberto Amato, Jeffrey Barrett, Sonia Goncalves, Ewan Harrison, David K. Jackson, Ian Johnston, Dominic Kwiatkowski, Cordelia Langford, John Sillitoe on behalf of the Wellcome Sanger Institute COVID-19 Surveillance Team |
| EPI_ISL_2435396 | Lighthouse Lab in Glasgow | Wellcome Sanger Institute for the COVID-19 Genomics UK (COG-UK) Consortium | Harper VanSteenhouse, Yumi Kasai, David Gray, Carol Clugston, Anna Dominiczak and Alex Alderton, Roberto Amato, Jeffrey Barrett, Sonia Goncalves, Ewan Harrison, David K. Jackson, Ian Johnston, Dominic Kwiatkowski, Cordelia Langford, John Sillitoe on behalf of the Wellcome Sanger Institute COVID-19 Surveillance Team |
| EPI_ISL_2435400 | Lighthouse Lab in Milton Keynes | Wellcome Sanger Institute for the COVID-19 Genomics UK (COG-UK) Consortium | The Lighthouse Lab in Milton Keynes and Alex Alderton, Roberto Amato, Jeffrey Barrett, Sonia Goncalves, Ewan Harrison, David K. Jackson, Ian Johnston, Dominic Kwiatkowski, Cordelia Langford, John Sillitoe on behalf of the Wellcome Sanger Institute COVID-19 Surveillance Team |
| EPI_ISL_2435401, EPI_ISL_2435402, EPI_ISL_2435403, | Lighthouse Lab in Alderley Park | Wellcome Sanger Institute for the COVID-19 Genomics UK (COG-UK) Consortium | Jacquelyn Wynn, Mairead Hyland, The Lighthouse Lab in Alderley Park and Alex Alderton, Roberto Amato, Jeffrey Barrett, Sonia Goncalves, Ewan Harrison, David K. Jackson, Ian Johnston, Dominic Kwiatkowski, Cordelia Langford, John Sillitoe on behalf of the Wellcome Sanger Institute COVID-19 Surveillance Team |

| | | | |
|---|---|---|---|
| EPI_ISL_2435404 | | | |
| EPI_ISL_2435405 | Lighthouse Lab in Milton Keynes | Wellcome Sanger Institute for the COVID-19 Genomics UK (COG-UK) Consortium | The Lighthouse Lab in Milton Keynes and Alex Alderton, Roberto Amato, Jeffrey Barrett, Sonia Goncalves, Ewan Harrison, David K. Jackson, Ian Johnston, Dominic Kwiatkowski, Cordelia Langford, John Sillitoe on behalf of the Wellcome Sanger Institute COVID-19 Surveillance Team |
| EPI_ISL_2435406, EPI_ISL_2435407, EPI_ISL_2435408 | Lighthouse Lab in Alderley Park | Wellcome Sanger Institute for the COVID-19 Genomics UK (COG-UK) Consortium | Jacquelyn Wynn, Mairead Hyland, The Lighthouse Lab in Alderley Park and Alex Alderton, Roberto Amato, Jeffrey Barrett, Sonia Goncalves, Ewan Harrison, David K. Jackson, Ian Johnston, Dominic Kwiatkowski, Cordelia Langford, John Sillitoe on behalf of the Wellcome Sanger Institute COVID-19 Surveillance Team |
| EPI_ISL_2435409 | Lighthouse Lab in Milton Keynes | Wellcome Sanger Institute for the COVID-19 Genomics UK (COG-UK) Consortium | The Lighthouse Lab in Milton Keynes and Alex Alderton, Roberto Amato, Jeffrey Barrett, Sonia Goncalves, Ewan Harrison, David K. Jackson, Ian Johnston, Dominic Kwiatkowski, Cordelia Langford, John Sillitoe on behalf of the Wellcome Sanger Institute COVID-19 Surveillance Team |
| EPI_ISL_2435410, EPI_ISL_2435411, EPI_ISL_2435412 | Lighthouse Lab in Alderley Park | Wellcome Sanger Institute for the COVID-19 Genomics UK (COG-UK) Consortium | Jacquelyn Wynn, Mairead Hyland, The Lighthouse Lab in Alderley Park and Alex Alderton, Roberto Amato, Jeffrey Barrett, Sonia Goncalves, Ewan Harrison, David K. Jackson, Ian Johnston, Dominic Kwiatkowski, Cordelia Langford, John Sillitoe on behalf of the Wellcome Sanger Institute COVID-19 Surveillance Team |
| EPI_ISL_2435413 | Lighthouse Lab in Milton Keynes | Wellcome Sanger Institute for the COVID-19 Genomics UK (COG-UK) Consortium | The Lighthouse Lab in Milton Keynes and Alex Alderton, Roberto Amato, Jeffrey Barrett, Sonia Goncalves, Ewan Harrison, David K. Jackson, Ian Johnston, Dominic Kwiatkowski, Cordelia Langford, John Sillitoe on behalf of the Wellcome Sanger Institute COVID-19 Surveillance Team |
| EPI_ISL_2435414 | Lighthouse Lab in Alderley Park | Wellcome Sanger Institute for the COVID-19 Genomics UK (COG-UK) Consortium | Jacquelyn Wynn, Mairead Hyland, The Lighthouse Lab in Alderley Park and Alex Alderton, Roberto Amato, Jeffrey Barrett, Sonia Goncalves, Ewan Harrison, David K. Jackson, Ian Johnston, Dominic Kwiatkowski, Cordelia Langford, John Sillitoe on behalf of the Wellcome Sanger Institute COVID-19 Surveillance Team |
| EPI_ISL_2435415 | Lighthouse Lab in Milton Keynes | Wellcome Sanger Institute for the COVID-19 Genomics UK (COG-UK) Consortium | The Lighthouse Lab in Milton Keynes and Alex Alderton, Roberto Amato, Jeffrey Barrett, Sonia Goncalves, Ewan Harrison, David K. Jackson, Ian Johnston, Dominic Kwiatkowski, Cordelia Langford, John Sillitoe on behalf of the Wellcome Sanger Institute COVID-19 Surveillance Team |
| EPI_ISL_2435416, EPI_ISL_2435417, EPI_ISL_2435418 | Lighthouse Lab in Alderley Park | Wellcome Sanger Institute for the COVID-19 Genomics UK (COG-UK) Consortium | Jacquelyn Wynn, Mairead Hyland, The Lighthouse Lab in Alderley Park and Alex Alderton, Roberto Amato, Jeffrey Barrett, Sonia Goncalves, Ewan Harrison, David K. Jackson, Ian Johnston, Dominic Kwiatkowski, Cordelia Langford, John Sillitoe on behalf of the Wellcome Sanger Institute COVID-19 Surveillance Team |
| EPI_ISL_2435419 | Lighthouse Lab in Milton Keynes | Wellcome Sanger Institute for the COVID-19 Genomics UK (COG-UK) Consortium | The Lighthouse Lab in Milton Keynes and Alex Alderton, Roberto Amato, Jeffrey Barrett, Sonia Goncalves, Ewan Harrison, David K. Jackson, Ian Johnston, Dominic Kwiatkowski, Cordelia Langford, John Sillitoe on behalf of the Wellcome Sanger Institute COVID-19 Surveillance Team |
| EPI_ISL_2435420, EPI_ISL_2435421, EPI_ISL_2435422, EPI_ISL_2435425, EPI_ISL_2435426, EPI_ISL_2435427 | Lighthouse Lab in Alderley Park | Wellcome Sanger Institute for the COVID-19 Genomics UK (COG-UK) Consortium | Jacquelyn Wynn, Mairead Hyland, The Lighthouse Lab in Alderley Park and Alex Alderton, Roberto Amato, Jeffrey Barrett, Sonia Goncalves, Ewan Harrison, David K. Jackson, Ian Johnston, Dominic Kwiatkowski, Cordelia Langford, John Sillitoe on behalf of the Wellcome Sanger Institute COVID-19 Surveillance Team |
| EPI_ISL_2435428, EPI_ISL_2435430 | Lighthouse Lab in Milton Keynes | Wellcome Sanger Institute for the COVID-19 Genomics UK (COG-UK) Consortium | The Lighthouse Lab in Milton Keynes and Alex Alderton, Roberto Amato, Jeffrey Barrett, Sonia Goncalves, Ewan Harrison, David K. Jackson, Ian Johnston, Dominic Kwiatkowski, Cordelia Langford, John Sillitoe on behalf of the Wellcome Sanger Institute COVID-19 Surveillance Team |
| EPI_ISL_2435431, EPI_ISL_2435434, EPI_ISL_2435435 | Lighthouse Lab in Alderley Park | Wellcome Sanger Institute for the COVID-19 Genomics UK (COG-UK) Consortium | Jacquelyn Wynn, Mairead Hyland, The Lighthouse Lab in Alderley Park and Alex Alderton, Roberto Amato, Jeffrey Barrett, Sonia Goncalves, Ewan Harrison, David K. Jackson, Ian Johnston, Dominic Kwiatkowski, Cordelia Langford, John Sillitoe on behalf of the Wellcome Sanger Institute COVID-19 Surveillance Team |
| EPI_ISL_2435436 | Lighthouse Lab in Milton Keynes | Wellcome Sanger Institute for the COVID-19 Genomics UK (COG-UK) Consortium | The Lighthouse Lab in Milton Keynes and Alex Alderton, Roberto Amato, Jeffrey Barrett, Sonia Goncalves, Ewan Harrison, David K. Jackson, Ian Johnston, Dominic Kwiatkowski, Cordelia Langford, John Sillitoe on behalf of the Wellcome Sanger Institute COVID-19 Surveillance Team |
| EPI_ISL_2435437 | Lighthouse Lab in Alderley Park | Wellcome Sanger Institute for the COVID-19 Genomics UK (COG-UK) Consortium | Jacquelyn Wynn, Mairead Hyland, The Lighthouse Lab in Alderley Park and Alex Alderton, Roberto Amato, Jeffrey Barrett, Sonia Goncalves, Ewan Harrison, David K. Jackson, Ian Johnston, Dominic Kwiatkowski, Cordelia Langford, John Sillitoe on behalf of the Wellcome Sanger Institute COVID-19 Surveillance Team |
| EPI_ISL_2435438 | Lighthouse Lab in Milton Keynes | Wellcome Sanger Institute for the COVID-19 Genomics UK (COG-UK) Consortium | The Lighthouse Lab in Milton Keynes and Alex Alderton, Roberto Amato, Jeffrey Barrett, Sonia Goncalves, Ewan Harrison, David K. Jackson, Ian Johnston, Dominic Kwiatkowski, Cordelia Langford, John Sillitoe on behalf of the Wellcome Sanger Institute COVID-19 Surveillance Team |
| EPI_ISL_2435439 | Lighthouse Lab in Glasgow | Wellcome Sanger Institute for the COVID-19 Genomics UK (COG-UK) Consortium | Harper VanSteenhouse, Yumi Kasai, David Gray, Carol Clugston, Anna Dominiczak and Alex Alderton, Roberto Amato, Jeffrey Barrett, Sonia Goncalves, Ewan Harrison, David K. Jackson, Ian Johnston, Dominic Kwiatkowski, Cordelia Langford, John Sillitoe on behalf of the Wellcome Sanger Institute COVID-19 Surveillance Team |
| EPI_ISL_2435440 | Lighthouse Lab in Alderley Park | Wellcome Sanger Institute for the COVID-19 Genomics UK (COG-UK) Consortium | Jacquelyn Wynn, Mairead Hyland, The Lighthouse Lab in Alderley Park and Alex Alderton, Roberto Amato, Jeffrey Barrett, Sonia Goncalves, Ewan Harrison, David K. Jackson, Ian Johnston, Dominic Kwiatkowski, Cordelia Langford, John Sillitoe on behalf of the Wellcome Sanger Institute COVID-19 Surveillance Team |
| EPI_ISL_2435441 | Lighthouse Lab in Milton Keynes | Wellcome Sanger Institute for the COVID-19 Genomics UK (COG-UK) Consortium | The Lighthouse Lab in Milton Keynes and Alex Alderton, Roberto Amato, Jeffrey Barrett, Sonia Goncalves, Ewan Harrison, David K. Jackson, Ian Johnston, Dominic Kwiatkowski, Cordelia Langford, John Sillitoe on behalf of the Wellcome Sanger Institute COVID-19 Surveillance Team |
| EPI_ISL_2435442 | Lighthouse Lab in Glasgow | Wellcome Sanger Institute for the COVID-19 Genomics UK (COG-UK) Consortium | Harper VanSteenhouse, Yumi Kasai, David Gray, Carol Clugston, Anna Dominiczak and Alex Alderton, Roberto Amato, Jeffrey Barrett, Sonia Goncalves, Ewan Harrison, David K. Jackson, Ian Johnston, Dominic Kwiatkowski, Cordelia Langford, John Sillitoe on behalf of the Wellcome Sanger Institute COVID-19 Surveillance Team |
| EPI_ISL_2435443, EPI_ISL_2435444, EPI_ISL_2435445, EPI_ISL_2435446 | Lighthouse Lab in Milton Keynes | Wellcome Sanger Institute for the COVID-19 Genomics UK (COG-UK) Consortium | The Lighthouse Lab in Milton Keynes and Alex Alderton, Roberto Amato, Jeffrey Barrett, Sonia Goncalves, Ewan Harrison, David K. Jackson, Ian Johnston, Dominic Kwiatkowski, Cordelia Langford, John Sillitoe on behalf of the Wellcome Sanger Institute COVID-19 Surveillance Team |
| EPI_ISL_2435447 | Lighthouse Lab in Glasgow | Wellcome Sanger Institute for the COVID-19 Genomics UK (COG-UK) Consortium | Harper VanSteenhouse, Yumi Kasai, David Gray, Carol Clugston, Anna Dominiczak and Alex Alderton, Roberto Amato, Jeffrey Barrett, Sonia Goncalves, Ewan Harrison, David K. Jackson, Ian Johnston, Dominic Kwiatkowski, Cordelia Langford, John Sillitoe on behalf of the Wellcome Sanger Institute COVID-19 Surveillance Team |
| EPI_ISL_2435448, EPI_ISL_2435449, EPI_ISL_2435450 | Lighthouse Lab in Milton Keynes | Wellcome Sanger Institute for the COVID-19 Genomics UK (COG-UK) Consortium | The Lighthouse Lab in Milton Keynes and Alex Alderton, Roberto Amato, Jeffrey Barrett, Sonia Goncalves, Ewan Harrison, David K. Jackson, Ian Johnston, Dominic Kwiatkowski, Cordelia Langford, John Sillitoe on behalf of the Wellcome Sanger Institute COVID-19 Surveillance Team |
| EPI_ISL_2435451, EPI_ISL_2435452 | Lighthouse Lab in Alderley Park | Wellcome Sanger Institute for the COVID-19 Genomics UK (COG-UK) Consortium | Jacquelyn Wynn, Mairead Hyland, The Lighthouse Lab in Alderley Park and Alex Alderton, Roberto Amato, Jeffrey Barrett, Sonia Goncalves, Ewan Harrison, David K. Jackson, Ian Johnston, Dominic Kwiatkowski, Cordelia Langford, John Sillitoe on behalf of the Wellcome Sanger Institute COVID-19 Surveillance Team |
| EPI_ISL_2435454, EPI_ISL_2435455 | Lighthouse Lab in Milton Keynes | Wellcome Sanger Institute for the COVID-19 Genomics UK (COG-UK) Consortium | The Lighthouse Lab in Milton Keynes and Alex Alderton, Roberto Amato, Jeffrey Barrett, Sonia Goncalves, Ewan Harrison, David K. Jackson, Ian Johnston, Dominic Kwiatkowski, Cordelia Langford, John Sillitoe on behalf of the Wellcome Sanger Institute COVID-19 Surveillance Team |
| EPI_ISL_2435456 | Lighthouse Lab in Alderley Park | Wellcome Sanger Institute for the COVID-19 Genomics UK (COG-UK) Consortium | Jacquelyn Wynn, Mairead Hyland, The Lighthouse Lab in Alderley Park and Alex Alderton, Roberto Amato, Jeffrey Barrett, Sonia Goncalves, Ewan Harrison, David K. Jackson, Ian Johnston, Dominic Kwiatkowski, Cordelia Langford, John Sillitoe on behalf of the Wellcome Sanger Institute COVID-19 Surveillance Team |
| EPI_ISL_2435457 | Lighthouse Lab in Milton Keynes | Wellcome Sanger Institute for the COVID-19 Genomics UK (COG-UK) Consortium | The Lighthouse Lab in Milton Keynes and Alex Alderton, Roberto Amato, Jeffrey Barrett, Sonia Goncalves, Ewan Harrison, David K. Jackson, Ian Johnston, Dominic Kwiatkowski, Cordelia Langford, John Sillitoe on behalf of the Wellcome Sanger Institute COVID-19 Surveillance Team |
| EPI_ISL_2435458 | Lighthouse Lab in Alderley Park | Wellcome Sanger Institute for the COVID-19 Genomics UK (COG-UK) Consortium | Jacquelyn Wynn, Mairead Hyland, The Lighthouse Lab in Alderley Park and Alex Alderton, Roberto Amato, Jeffrey Barrett, Sonia Goncalves, Ewan Harrison, David K. Jackson, Ian Johnston, Dominic Kwiatkowski, Cordelia Langford, John Sillitoe on behalf of the Wellcome Sanger Institute COVID-19 Surveillance Team |
| EPI_ISL_2435459 | Lighthouse Lab in Milton Keynes | Wellcome Sanger Institute for the COVID-19 Genomics UK (COG-UK) Consortium | The Lighthouse Lab in Milton Keynes and Alex Alderton, Roberto Amato, Jeffrey Barrett, Sonia Goncalves, Ewan Harrison, David K. Jackson, Ian Johnston, Dominic Kwiatkowski, Cordelia Langford, John Sillitoe on behalf of the Wellcome Sanger Institute COVID-19 Surveillance Team |
| EPI_ISL_2435460 | Lighthouse Lab in Glasgow | Wellcome Sanger Institute for the COVID-19 Genomics UK (COG-UK) Consortium | Harper VanSteenhouse, Yumi Kasai, David Gray, Carol Clugston, Anna Dominiczak and Alex Alderton, Roberto Amato, Jeffrey Barrett, Sonia Goncalves, Ewan Harrison, David K. Jackson, Ian Johnston, Dominic Kwiatkowski, Cordelia Langford, John Sillitoe on behalf of the Wellcome Sanger Institute COVID-19 Surveillance Team |
| EPI_ISL_2435461, EPI_ISL_2435463, EPI_ISL_2435464, EPI_ISL_2435465 | Lighthouse Lab in Milton Keynes | Wellcome Sanger Institute for the COVID-19 Genomics UK (COG-UK) Consortium | The Lighthouse Lab in Milton Keynes and Alex Alderton, Roberto Amato, Jeffrey Barrett, Sonia Goncalves, Ewan Harrison, David K. Jackson, Ian Johnston, Dominic Kwiatkowski, Cordelia Langford, John Sillitoe on behalf of the Wellcome Sanger Institute COVID-19 Surveillance Team |
| EPI_ISL_2435466, EPI_ISL_2435467, EPI_ISL_2435468, EPI_ISL_2435469 | Lighthouse Lab in Alderley Park | Wellcome Sanger Institute for the COVID-19 Genomics UK (COG-UK) Consortium | Jacquelyn Wynn, Mairead Hyland, The Lighthouse Lab in Alderley Park and Alex Alderton, Roberto Amato, Jeffrey Barrett, Sonia Goncalves, Ewan Harrison, David K. Jackson, Ian Johnston, Dominic Kwiatkowski, Cordelia Langford, John Sillitoe on behalf of the Wellcome Sanger Institute COVID-19 Surveillance Team |
| EPI_ISL_2435470 | Lighthouse Lab in Glasgow | Wellcome Sanger Institute for the COVID-19 Genomics UK (COG-UK) Consortium | Harper VanSteenhouse, Yumi Kasai, David Gray, Carol Clugston, Anna Dominiczak and Alex Alderton, Roberto Amato, Jeffrey Barrett, Sonia Goncalves, Ewan Harrison, David K. Jackson, Ian Johnston, Dominic Kwiatkowski, Cordelia Langford, John Sillitoe on behalf of the Wellcome Sanger Institute COVID-19 Surveillance Team |
| EPI_ISL_2435472 | Lighthouse Lab in Milton Keynes | Wellcome Sanger Institute for the COVID-19 Genomics UK (COG-UK) Consortium | The Lighthouse Lab in Milton Keynes and Alex Alderton, Roberto Amato, Jeffrey Barrett, Sonia Goncalves, Ewan Harrison, David K. Jackson, Ian Johnston, Dominic Kwiatkowski, Cordelia Langford, John Sillitoe on behalf of the Wellcome Sanger Institute COVID-19 Surveillance Team |
| EPI_ISL_2435474 | Lighthouse Lab in Alderley Park | Wellcome Sanger Institute for the COVID-19 Genomics UK (COG-UK) Consortium | Jacquelyn Wynn, Mairead Hyland, The Lighthouse Lab in Alderley Park and Alex Alderton, Roberto Amato, Jeffrey Barrett, Sonia Goncalves, Ewan Harrison, David K. Jackson, Ian Johnston, Dominic Kwiatkowski, Cordelia Langford, John Sillitoe on behalf of the Wellcome Sanger Institute COVID-19 Surveillance Team |
| EPI_ISL_2435475 | Lighthouse Lab in Milton Keynes | Wellcome Sanger Institute for the COVID-19 Genomics UK (COG-UK) Consortium | The Lighthouse Lab in Milton Keynes and Alex Alderton, Roberto Amato, Jeffrey Barrett, Sonia Goncalves, Ewan Harrison, David K. Jackson, Ian Johnston, Dominic Kwiatkowski, Cordelia Langford, John Sillitoe on behalf of the Wellcome Sanger Institute COVID-19 Surveillance Team |
| EPI_ISL_2435476, EPI_ISL_2435478, EPI_ISL_2435479, EPI_ISL_2435480, EPI_ISL_2435481 | Lighthouse Lab in Alderley Park | Wellcome Sanger Institute for the COVID-19 Genomics UK (COG-UK) Consortium | Jacquelyn Wynn, Mairead Hyland, The Lighthouse Lab in Alderley Park and Alex Alderton, Roberto Amato, Jeffrey Barrett, Sonia Goncalves, Ewan Harrison, David K. Jackson, Ian Johnston, Dominic Kwiatkowski, Cordelia Langford, John Sillitoe on behalf of the Wellcome Sanger Institute COVID-19 Surveillance Team |
| EPI_ISL_2435483 | Lighthouse Lab in Milton Keynes | Wellcome Sanger Institute for the COVID-19 Genomics UK (COG-UK) Consortium | The Lighthouse Lab in Milton Keynes and Alex Alderton, Roberto Amato, Jeffrey Barrett, Sonia Goncalves, Ewan Harrison, David K. Jackson, Ian Johnston, Dominic Kwiatkowski, Cordelia Langford, John Sillitoe on behalf of the Wellcome Sanger Institute COVID-19 Surveillance Team |

| | | | |
|---|---|---|---|
| EPI_ISL_2435484, EPI_ISL_2435486 | Lighthouse Lab in Alderley Park | Wellcome Sanger Institute for the COVID-19 Genomics UK (COG-UK) Consortium | Jacquelyn Wynn, Mairead Hyland, The Lighthouse Lab in Alderley Park and Alex Alderton, Roberto Amato, Jeffrey Barrett, Sonia Goncalves, Ewan Harrison, David K. Jackson, Ian Johnston, Dominic Kwiatkowski, Cordelia Langford, John Sillitoe on behalf of the Wellcome Sanger Institute COVID-19 Surveillance Team |
| EPI_ISL_2435487 | Lighthouse Lab in Milton Keynes | Wellcome Sanger Institute for the COVID-19 Genomics UK (COG-UK) Consortium | The Lighthouse Lab in Milton Keynes and Alex Alderton, Roberto Amato, Jeffrey Barrett, Sonia Goncalves, Ewan Harrison, David K. Jackson, Ian Johnston, Dominic Kwiatkowski, Cordelia Langford, John Sillitoe on behalf of the Wellcome Sanger Institute COVID-19 Surveillance Team |
| EPI_ISL_2435488, EPI_ISL_2435489, EPI_ISL_2435491 | Lighthouse Lab in Alderley Park | Wellcome Sanger Institute for the COVID-19 Genomics UK (COG-UK) Consortium | Jacquelyn Wynn, Mairead Hyland, The Lighthouse Lab in Alderley Park and Alex Alderton, Roberto Amato, Jeffrey Barrett, Sonia Goncalves, Ewan Harrison, David K. Jackson, Ian Johnston, Dominic Kwiatkowski, Cordelia Langford, John Sillitoe on behalf of the Wellcome Sanger Institute COVID-19 Surveillance Team |
| EPI_ISL_2435493 | Lighthouse Lab in Glasgow | Wellcome Sanger Institute for the COVID-19 Genomics UK (COG-UK) Consortium | Harper VanSteenhouse, Yumi Kasai, David Gray, Carol Clugston, Anna Dominiczak and Alex Alderton, Roberto Amato, Jeffrey Barrett, Sonia Goncalves, Ewan Harrison, David K. Jackson, Ian Johnston, Dominic Kwiatkowski, Cordelia Langford, John Sillitoe on behalf of the Wellcome Sanger Institute COVID-19 Surveillance Team |
| EPI_ISL_2435494, EPI_ISL_2435495 | Lighthouse Lab in Milton Keynes | Wellcome Sanger Institute for the COVID-19 Genomics UK (COG-UK) Consortium | The Lighthouse Lab in Milton Keynes and Alex Alderton, Roberto Amato, Jeffrey Barrett, Sonia Goncalves, Ewan Harrison, David K. Jackson, Ian Johnston, Dominic Kwiatkowski, Cordelia Langford, John Sillitoe on behalf of the Wellcome Sanger Institute COVID-19 Surveillance Team |
| EPI_ISL_2435498 | Lighthouse Lab in Alderley Park | Wellcome Sanger Institute for the COVID-19 Genomics UK (COG-UK) Consortium | Jacquelyn Wynn, Mairead Hyland, The Lighthouse Lab in Alderley Park and Alex Alderton, Roberto Amato, Jeffrey Barrett, Sonia Goncalves, Ewan Harrison, David K. Jackson, Ian Johnston, Dominic Kwiatkowski, Cordelia Langford, John Sillitoe on behalf of the Wellcome Sanger Institute COVID-19 Surveillance Team |
| EPI_ISL_2435499 | Lighthouse Lab in Milton Keynes | Wellcome Sanger Institute for the COVID-19 Genomics UK (COG-UK) Consortium | The Lighthouse Lab in Milton Keynes and Alex Alderton, Roberto Amato, Jeffrey Barrett, Sonia Goncalves, Ewan Harrison, David K. Jackson, Ian Johnston, Dominic Kwiatkowski, Cordelia Langford, John Sillitoe on behalf of the Wellcome Sanger Institute COVID-19 Surveillance Team |
| EPI_ISL_2435500 | Lighthouse Lab in Alderley Park | Wellcome Sanger Institute for the COVID-19 Genomics UK (COG-UK) Consortium | Jacquelyn Wynn, Mairead Hyland, The Lighthouse Lab in Alderley Park and Alex Alderton, Roberto Amato, Jeffrey Barrett, Sonia Goncalves, Ewan Harrison, David K. Jackson, Ian Johnston, Dominic Kwiatkowski, Cordelia Langford, John Sillitoe on behalf of the Wellcome Sanger Institute COVID-19 Surveillance Team |
| EPI_ISL_2435501, EPI_ISL_2435502 | Lighthouse Lab in Milton Keynes | Wellcome Sanger Institute for the COVID-19 Genomics UK (COG-UK) Consortium | The Lighthouse Lab in Milton Keynes and Alex Alderton, Roberto Amato, Jeffrey Barrett, Sonia Goncalves, Ewan Harrison, David K. Jackson, Ian Johnston, Dominic Kwiatkowski, Cordelia Langford, John Sillitoe on behalf of the Wellcome Sanger Institute COVID-19 Surveillance Team |
| EPI_ISL_2435504 | Lighthouse Lab in Alderley Park | Wellcome Sanger Institute for the COVID-19 Genomics UK (COG-UK) Consortium | Jacquelyn Wynn, Mairead Hyland, The Lighthouse Lab in Alderley Park and Alex Alderton, Roberto Amato, Jeffrey Barrett, Sonia Goncalves, Ewan Harrison, David K. Jackson, Ian Johnston, Dominic Kwiatkowski, Cordelia Langford, John Sillitoe on behalf of the Wellcome Sanger Institute COVID-19 Surveillance Team |
| EPI_ISL_2435505, EPI_ISL_2435506, EPI_ISL_2435507 | Lighthouse Lab in Milton Keynes | Wellcome Sanger Institute for the COVID-19 Genomics UK (COG-UK) Consortium | The Lighthouse Lab in Milton Keynes and Alex Alderton, Roberto Amato, Jeffrey Barrett, Sonia Goncalves, Ewan Harrison, David K. Jackson, Ian Johnston, Dominic Kwiatkowski, Cordelia Langford, John Sillitoe on behalf of the Wellcome Sanger Institute COVID-19 Surveillance Team |
| EPI_ISL_2435508, EPI_ISL_2435509, EPI_ISL_2435510 | Lighthouse Lab in Alderley Park | Wellcome Sanger Institute for the COVID-19 Genomics UK (COG-UK) Consortium | Jacquelyn Wynn, Mairead Hyland, The Lighthouse Lab in Alderley Park and Alex Alderton, Roberto Amato, Jeffrey Barrett, Sonia Goncalves, Ewan Harrison, David K. Jackson, Ian Johnston, Dominic Kwiatkowski, Cordelia Langford, John Sillitoe on behalf of the Wellcome Sanger Institute COVID-19 Surveillance Team |
| EPI_ISL_2435511 | Lighthouse Lab in Milton Keynes | Wellcome Sanger Institute for the COVID-19 Genomics UK (COG-UK) Consortium | The Lighthouse Lab in Milton Keynes and Alex Alderton, Roberto Amato, Jeffrey Barrett, Sonia Goncalves, Ewan Harrison, David K. Jackson, Ian Johnston, Dominic Kwiatkowski, Cordelia Langford, John Sillitoe on behalf of the Wellcome Sanger Institute COVID-19 Surveillance Team |
| EPI_ISL_2435512 | Lighthouse Lab in Alderley Park | Wellcome Sanger Institute for the COVID-19 Genomics UK (COG-UK) Consortium | Jacquelyn Wynn, Mairead Hyland, The Lighthouse Lab in Alderley Park and Alex Alderton, Roberto Amato, Jeffrey Barrett, Sonia Goncalves, Ewan Harrison, David K. Jackson, Ian Johnston, Dominic Kwiatkowski, Cordelia Langford, John Sillitoe on behalf of the Wellcome Sanger Institute COVID-19 Surveillance Team |
| EPI_ISL_2435513 | Lighthouse Lab in Milton Keynes | Wellcome Sanger Institute for the COVID-19 Genomics UK (COG-UK) Consortium | The Lighthouse Lab in Milton Keynes and Alex Alderton, Roberto Amato, Jeffrey Barrett, Sonia Goncalves, Ewan Harrison, David K. Jackson, Ian Johnston, Dominic Kwiatkowski, Cordelia Langford, John Sillitoe on behalf of the Wellcome Sanger Institute COVID-19 Surveillance Team |
| EPI_ISL_2435514, EPI_ISL_2435515, EPI_ISL_2435516, EPI_ISL_2435517, EPI_ISL_2435518, EPI_ISL_2435519 | Lighthouse Lab in Alderley Park | Wellcome Sanger Institute for the COVID-19 Genomics UK (COG-UK) Consortium | Jacquelyn Wynn, Mairead Hyland, The Lighthouse Lab in Alderley Park and Alex Alderton, Roberto Amato, Jeffrey Barrett, Sonia Goncalves, Ewan Harrison, David K. Jackson, Ian Johnston, Dominic Kwiatkowski, Cordelia Langford, John Sillitoe on behalf of the Wellcome Sanger Institute COVID-19 Surveillance Team |
| EPI_ISL_2435520, EPI_ISL_2435521 | Lighthouse Lab in Milton Keynes | Wellcome Sanger Institute for the COVID-19 Genomics UK (COG-UK) Consortium | The Lighthouse Lab in Milton Keynes and Alex Alderton, Roberto Amato, Jeffrey Barrett, Sonia Goncalves, Ewan Harrison, David K. Jackson, Ian Johnston, Dominic Kwiatkowski, Cordelia Langford, John Sillitoe on behalf of the Wellcome Sanger Institute COVID-19 Surveillance Team |
| EPI_ISL_2435522, EPI_ISL_2435524 | Lighthouse Lab in Alderley Park | Wellcome Sanger Institute for the COVID-19 Genomics UK (COG-UK) Consortium | Jacquelyn Wynn, Mairead Hyland, The Lighthouse Lab in Alderley Park and Alex Alderton, Roberto Amato, Jeffrey Barrett, Sonia Goncalves, Ewan Harrison, David K. Jackson, Ian Johnston, Dominic Kwiatkowski, Cordelia Langford, John Sillitoe on behalf of the Wellcome Sanger Institute COVID-19 Surveillance Team |
| EPI_ISL_2435525 | Lighthouse Lab in Milton Keynes | Wellcome Sanger Institute for the COVID-19 Genomics UK (COG-UK) Consortium | The Lighthouse Lab in Milton Keynes and Alex Alderton, Roberto Amato, Jeffrey Barrett, Sonia Goncalves, Ewan Harrison, David K. Jackson, Ian Johnston, Dominic Kwiatkowski, Cordelia Langford, John Sillitoe on behalf of the Wellcome Sanger Institute COVID-19 Surveillance Team |
| EPI_ISL_2435527 | Lighthouse Lab in Alderley Park | Wellcome Sanger Institute for the COVID-19 Genomics UK (COG-UK) Consortium | Jacquelyn Wynn, Mairead Hyland, The Lighthouse Lab in Alderley Park and Alex Alderton, Roberto Amato, Jeffrey Barrett, Sonia Goncalves, Ewan Harrison, David K. Jackson, Ian Johnston, Dominic Kwiatkowski, Cordelia Langford, John Sillitoe on behalf of the Wellcome Sanger Institute COVID-19 Surveillance Team |
| EPI_ISL_2435528, EPI_ISL_2435529 | Lighthouse Lab in Milton Keynes | Wellcome Sanger Institute for the COVID-19 Genomics UK (COG-UK) Consortium | The Lighthouse Lab in Milton Keynes and Alex Alderton, Roberto Amato, Jeffrey Barrett, Sonia Goncalves, Ewan Harrison, David K. Jackson, Ian Johnston, Dominic Kwiatkowski, Cordelia Langford, John Sillitoe on behalf of the Wellcome Sanger Institute COVID-19 Surveillance Team |
| EPI_ISL_2435530, EPI_ISL_2435531 | Lighthouse Lab in Alderley Park | Wellcome Sanger Institute for the COVID-19 Genomics UK (COG-UK) Consortium | Jacquelyn Wynn, Mairead Hyland, The Lighthouse Lab in Alderley Park and Alex Alderton, Roberto Amato, Jeffrey Barrett, Sonia Goncalves, Ewan Harrison, David K. Jackson, Ian Johnston, Dominic Kwiatkowski, Cordelia Langford, John Sillitoe on behalf of the Wellcome Sanger Institute COVID-19 Surveillance Team |
| EPI_ISL_2435532 | Lighthouse Lab in Glasgow | Wellcome Sanger Institute for the COVID-19 Genomics UK (COG-UK) Consortium | Harper VanSteenhouse, Yumi Kasai, David Gray, Carol Clugston, Anna Dominiczak and Alex Alderton, Roberto Amato, Jeffrey Barrett, Sonia Goncalves, Ewan Harrison, David K. Jackson, Ian Johnston, Dominic Kwiatkowski, Cordelia Langford, John Sillitoe on behalf of the Wellcome Sanger Institute COVID-19 Surveillance Team |
| EPI_ISL_2435533, EPI_ISL_2435534 | Lighthouse Lab in Alderley Park | Wellcome Sanger Institute for the COVID-19 Genomics UK (COG-UK) Consortium | Jacquelyn Wynn, Mairead Hyland, The Lighthouse Lab in Alderley Park and Alex Alderton, Roberto Amato, Jeffrey Barrett, Sonia Goncalves, Ewan Harrison, David K. Jackson, Ian Johnston, Dominic Kwiatkowski, Cordelia Langford, John Sillitoe on behalf of the Wellcome Sanger Institute COVID-19 Surveillance Team |
| EPI_ISL_2435535 | Lighthouse Lab in Milton Keynes | Wellcome Sanger Institute for the COVID-19 Genomics UK (COG-UK) Consortium | The Lighthouse Lab in Milton Keynes and Alex Alderton, Roberto Amato, Jeffrey Barrett, Sonia Goncalves, Ewan Harrison, David K. Jackson, Ian Johnston, Dominic Kwiatkowski, Cordelia Langford, John Sillitoe on behalf of the Wellcome Sanger Institute COVID-19 Surveillance Team |
| EPI_ISL_2435536, EPI_ISL_2435538, EPI_ISL_2435539, EPI_ISL_2435540, EPI_ISL_2435541 | Lighthouse Lab in Alderley Park | Wellcome Sanger Institute for the COVID-19 Genomics UK (COG-UK) Consortium | Jacquelyn Wynn, Mairead Hyland, The Lighthouse Lab in Alderley Park and Alex Alderton, Roberto Amato, Jeffrey Barrett, Sonia Goncalves, Ewan Harrison, David K. Jackson, Ian Johnston, Dominic Kwiatkowski, Cordelia Langford, John Sillitoe on behalf of the Wellcome Sanger Institute COVID-19 Surveillance Team |
| EPI_ISL_2435542 | Lighthouse Lab in Milton Keynes | Wellcome Sanger Institute for the COVID-19 Genomics UK (COG-UK) Consortium | The Lighthouse Lab in Milton Keynes and Alex Alderton, Roberto Amato, Jeffrey Barrett, Sonia Goncalves, Ewan Harrison, David K. Jackson, Ian Johnston, Dominic Kwiatkowski, Cordelia Langford, John Sillitoe on behalf of the Wellcome Sanger Institute COVID-19 Surveillance Team |
| EPI_ISL_2435543 | Lighthouse Lab in Alderley Park | Wellcome Sanger Institute for the COVID-19 Genomics UK (COG-UK) Consortium | Jacquelyn Wynn, Mairead Hyland, The Lighthouse Lab in Alderley Park and Alex Alderton, Roberto Amato, Jeffrey Barrett, Sonia Goncalves, Ewan Harrison, David K. Jackson, Ian Johnston, Dominic Kwiatkowski, Cordelia Langford, John Sillitoe on behalf of the Wellcome Sanger Institute COVID-19 Surveillance Team |
| EPI_ISL_2435544 | Lighthouse Lab in Milton Keynes | Wellcome Sanger Institute for the COVID-19 Genomics UK (COG-UK) Consortium | The Lighthouse Lab in Milton Keynes and Alex Alderton, Roberto Amato, Jeffrey Barrett, Sonia Goncalves, Ewan Harrison, David K. Jackson, Ian Johnston, Dominic Kwiatkowski, Cordelia Langford, John Sillitoe on behalf of the Wellcome Sanger Institute COVID-19 Surveillance Team |
| EPI_ISL_2435546 | Lighthouse Lab in Alderley Park | Wellcome Sanger Institute for the COVID-19 Genomics UK (COG-UK) Consortium | Jacquelyn Wynn, Mairead Hyland, The Lighthouse Lab in Alderley Park and Alex Alderton, Roberto Amato, Jeffrey Barrett, Sonia Goncalves, Ewan Harrison, David K. Jackson, Ian Johnston, Dominic Kwiatkowski, Cordelia Langford, John Sillitoe on behalf of the Wellcome Sanger Institute COVID-19 Surveillance Team |
| EPI_ISL_2435547 | Lighthouse Lab in Milton Keynes | Wellcome Sanger Institute for the COVID-19 Genomics UK (COG-UK) Consortium | The Lighthouse Lab in Milton Keynes and Alex Alderton, Roberto Amato, Jeffrey Barrett, Sonia Goncalves, Ewan Harrison, David K. Jackson, Ian Johnston, Dominic Kwiatkowski, Cordelia Langford, John Sillitoe on behalf of the Wellcome Sanger Institute COVID-19 Surveillance Team |
| EPI_ISL_2435548, EPI_ISL_2435550, EPI_ISL_2435551 | Lighthouse Lab in Alderley Park | Wellcome Sanger Institute for the COVID-19 Genomics UK (COG-UK) Consortium | Jacquelyn Wynn, Mairead Hyland, The Lighthouse Lab in Alderley Park and Alex Alderton, Roberto Amato, Jeffrey Barrett, Sonia Goncalves, Ewan Harrison, David K. Jackson, Ian Johnston, Dominic Kwiatkowski, Cordelia Langford, John Sillitoe on behalf of the Wellcome Sanger Institute COVID-19 Surveillance Team |
| EPI_ISL_2435552, EPI_ISL_2435553 | Lighthouse Lab in Milton Keynes | Wellcome Sanger Institute for the COVID-19 Genomics UK (COG-UK) Consortium | The Lighthouse Lab in Milton Keynes and Alex Alderton, Roberto Amato, Jeffrey Barrett, Sonia Goncalves, Ewan Harrison, David K. Jackson, Ian Johnston, Dominic Kwiatkowski, Cordelia Langford, John Sillitoe on behalf of the Wellcome Sanger Institute COVID-19 Surveillance Team |
| EPI_ISL_2435554, EPI_ISL_2435555 | Lighthouse Lab in Alderley Park | Wellcome Sanger Institute for the COVID-19 Genomics UK (COG-UK) Consortium | Jacquelyn Wynn, Mairead Hyland, The Lighthouse Lab in Alderley Park and Alex Alderton, Roberto Amato, Jeffrey Barrett, Sonia Goncalves, Ewan Harrison, David K. Jackson, Ian Johnston, Dominic Kwiatkowski, Cordelia Langford, John Sillitoe on behalf of the Wellcome Sanger Institute COVID-19 Surveillance Team |
| EPI_ISL_2435556 | Lighthouse Lab in Glasgow | Wellcome Sanger Institute for the COVID-19 Genomics UK (COG-UK) Consortium | Harper VanSteenhouse, Yumi Kasai, David Gray, Carol Clugston, Anna Dominiczak and Alex Alderton, Roberto Amato, Jeffrey Barrett, Sonia Goncalves, Ewan Harrison, David K. Jackson, Ian Johnston, Dominic Kwiatkowski, Cordelia Langford, John Sillitoe on behalf of the Wellcome Sanger Institute COVID-19 Surveillance Team |
| EPI_ISL_2435557 | Lighthouse Lab in Alderley Park | Wellcome Sanger Institute for the COVID-19 Genomics UK (COG-UK) Consortium | Jacquelyn Wynn, Mairead Hyland, The Lighthouse Lab in Alderley Park and Alex Alderton, Roberto Amato, Jeffrey Barrett, Sonia Goncalves, Ewan Harrison, David K. Jackson, Ian Johnston, Dominic Kwiatkowski, Cordelia Langford, John Sillitoe on behalf of the Wellcome Sanger Institute COVID-19 Surveillance Team |
| EPI_ISL_2435558 | Lighthouse Lab in Glasgow | Wellcome Sanger Institute for the COVID-19 Genomics UK (COG-UK) Consortium | Harper VanSteenhouse, Yumi Kasai, David Gray, Carol Clugston, Anna Dominiczak and Alex Alderton, Roberto Amato, Jeffrey Barrett, Sonia Goncalves, Ewan Harrison, David K. Jackson, Ian Johnston, Dominic Kwiatkowski, Cordelia Langford, John Sillitoe on behalf of the Wellcome Sanger Institute COVID-19 Surveillance Team |
| EPI_ISL_2435559, EPI_ISL_2435560 | Lighthouse Lab in Milton Keynes | Wellcome Sanger Institute for the COVID-19 Genomics UK (COG-UK) Consortium | The Lighthouse Lab in Milton Keynes and Alex Alderton, Roberto Amato, Jeffrey Barrett, Sonia Goncalves, Ewan Harrison, David K. Jackson, Ian Johnston, Dominic Kwiatkowski, Cordelia Langford, John Sillitoe on behalf of the Wellcome Sanger Institute COVID-19 Surveillance Team |
| EPI_ISL_2435562 | Lighthouse Lab in Alderley Park | Wellcome Sanger Institute for the COVID-19 Genomics UK (COG-UK) Consortium | Jacquelyn Wynn, Mairead Hyland, The Lighthouse Lab in Alderley Park and Alex Alderton, Roberto Amato, Jeffrey Barrett, Sonia Goncalves, Ewan Harrison, David K. Jackson, Ian Johnston, Dominic Kwiatkowski, Cordelia Langford, John Sillitoe on behalf of the Wellcome Sanger Institute COVID-19 Surveillance Team |
| EPI_ISL_2435563 | Lighthouse Lab in Milton Keynes | Wellcome Sanger Institute for the COVID-19 Genomics UK (COG-UK) Consortium | The Lighthouse Lab in Milton Keynes and Alex Alderton, Roberto Amato, Jeffrey Barrett, Sonia Goncalves, Ewan Harrison, David K. Jackson, Ian Johnston, Dominic Kwiatkowski, Cordelia Langford, John Sillitoe on behalf of the Wellcome Sanger Institute COVID-19 Surveillance Team |
| EPI_ISL_2435564 | Lighthouse Lab in Alderley Park | Wellcome Sanger Institute for the COVID-19 Genomics UK (COG-UK) Consortium | Jacquelyn Wynn, Mairead Hyland, The Lighthouse Lab in Alderley Park and Alex Alderton, Roberto Amato, Jeffrey Barrett, Sonia Goncalves, Ewan Harrison, David K. Jackson, Ian Johnston, Dominic Kwiatkowski, Cordelia Langford, John Sillitoe on behalf of the Wellcome Sanger Institute COVID-19 Surveillance Team |

| | | | |
|---|---|---|---|
| EPI_ISL_2435565, EPI_ISL_2435566 | Lighthouse Lab in Milton Keynes | Wellcome Sanger Institute for the COVID-19 Genomics UK (COG-UK) Consortium | The Lighthouse Lab in Milton Keynes and Alex Alderton, Roberto Amato, Jeffrey Barrett, Sonia Goncalves, Ewan Harrison, David K. Jackson, Ian Johnston, Dominic Kwiatkowski, Cordelia Langford, John Sillitoe on behalf of the Wellcome Sanger Institute COVID-19 Surveillance Team |

EPI_ISL_2435567, EPI_ISL_2435568, EPI_ISL_2435569, EPI_ISL_2435570, EPI_ISL_2435571, EPI_ISL_2435572, EPI_ISL_2435573, EPI_ISL_2435574, EPI_ISL_2435575, EPI_ISL_2435576, EPI_ISL_2435577, EPI_ISL_2435578, EPI_ISL_2435579

| | | | |
|---|---|---|---|
| see above | Lighthouse Lab in Alderley Park | Wellcome Sanger Institute for the COVID-19 Genomics UK (COG-UK) Consortium | Jacquelyn Wynn, Mairead Hyland, The Lighthouse Lab in Alderley Park and Alex Alderton, Roberto Amato, Jeffrey Barrett, Sonia Goncalves, Ewan Harrison, David K. Jackson, Ian Johnston, Dominic Kwiatkowski, Cordelia Langford, John Sillitoe on behalf of the Wellcome Sanger Institute COVID-19 Surveillance Team |
| EPI_ISL_2435580 | Lighthouse Lab in Milton Keynes | Wellcome Sanger Institute for the COVID-19 Genomics UK (COG-UK) Consortium | The Lighthouse Lab in Milton Keynes and Alex Alderton, Roberto Amato, Jeffrey Barrett, Sonia Goncalves, Ewan Harrison, David K. Jackson, Ian Johnston, Dominic Kwiatkowski, Cordelia Langford, John Sillitoe on behalf of the Wellcome Sanger Institute COVID-19 Surveillance Team |
| EPI_ISL_2435581, EPI_ISL_2435582, EPI_ISL_2435583, EPI_ISL_2435584, EPI_ISL_2435585 | Lighthouse Lab in Alderley Park | Wellcome Sanger Institute for the COVID-19 Genomics UK (COG-UK) Consortium | Jacquelyn Wynn, Mairead Hyland, The Lighthouse Lab in Alderley Park and Alex Alderton, Roberto Amato, Jeffrey Barrett, Sonia Goncalves, Ewan Harrison, David K. Jackson, Ian Johnston, Dominic Kwiatkowski, Cordelia Langford, John Sillitoe on behalf of the Wellcome Sanger Institute COVID-19 Surveillance Team |
| EPI_ISL_2435587 | Lighthouse Lab in Glasgow | Wellcome Sanger Institute for the COVID-19 Genomics UK (COG-UK) Consortium | Harper VanSteenhouse, Yumi Kasai, David Gray, Carol Clugston, Anna Dominiczak and Alex Alderton, Roberto Amato, Jeffrey Barrett, Sonia Goncalves, Ewan Harrison, David K. Jackson, Ian Johnston, Dominic Kwiatkowski, Cordelia Langford, John Sillitoe on behalf of the Wellcome Sanger Institute COVID-19 Surveillance Team |
| EPI_ISL_2435588 | Lighthouse Lab in Alderley Park | Wellcome Sanger Institute for the COVID-19 Genomics UK (COG-UK) Consortium | Jacquelyn Wynn, Mairead Hyland, The Lighthouse Lab in Alderley Park and Alex Alderton, Roberto Amato, Jeffrey Barrett, Sonia Goncalves, Ewan Harrison, David K. Jackson, Ian Johnston, Dominic Kwiatkowski, Cordelia Langford, John Sillitoe on behalf of the Wellcome Sanger Institute COVID-19 Surveillance Team |
| EPI_ISL_2435590 | Lighthouse Lab in Milton Keynes | Wellcome Sanger Institute for the COVID-19 Genomics UK (COG-UK) Consortium | The Lighthouse Lab in Milton Keynes and Alex Alderton, Roberto Amato, Jeffrey Barrett, Sonia Goncalves, Ewan Harrison, David K. Jackson, Ian Johnston, Dominic Kwiatkowski, Cordelia Langford, John Sillitoe on behalf of the Wellcome Sanger Institute COVID-19 Surveillance Team |
| EPI_ISL_2435591 | Lighthouse Lab in Alderley Park | Wellcome Sanger Institute for the COVID-19 Genomics UK (COG-UK) Consortium | Jacquelyn Wynn, Mairead Hyland, The Lighthouse Lab in Alderley Park and Alex Alderton, Roberto Amato, Jeffrey Barrett, Sonia Goncalves, Ewan Harrison, David K. Jackson, Ian Johnston, Dominic Kwiatkowski, Cordelia Langford, John Sillitoe on behalf of the Wellcome Sanger Institute COVID-19 Surveillance Team |
| EPI_ISL_2435592, EPI_ISL_2435593 | Lighthouse Lab in Milton Keynes | Wellcome Sanger Institute for the COVID-19 Genomics UK (COG-UK) Consortium | The Lighthouse Lab in Milton Keynes and Alex Alderton, Roberto Amato, Jeffrey Barrett, Sonia Goncalves, Ewan Harrison, David K. Jackson, Ian Johnston, Dominic Kwiatkowski, Cordelia Langford, John Sillitoe on behalf of the Wellcome Sanger Institute COVID-19 Surveillance Team |
| EPI_ISL_2435594, EPI_ISL_2435595, EPI_ISL_2435596 | Lighthouse Lab in Alderley Park | Wellcome Sanger Institute for the COVID-19 Genomics UK (COG-UK) Consortium | Jacquelyn Wynn, Mairead Hyland, The Lighthouse Lab in Alderley Park and Alex Alderton, Roberto Amato, Jeffrey Barrett, Sonia Goncalves, Ewan Harrison, David K. Jackson, Ian Johnston, Dominic Kwiatkowski, Cordelia Langford, John Sillitoe on behalf of the Wellcome Sanger Institute COVID-19 Surveillance Team |
| EPI_ISL_2435597 | Lighthouse Lab in Milton Keynes | Wellcome Sanger Institute for the COVID-19 Genomics UK (COG-UK) Consortium | The Lighthouse Lab in Milton Keynes and Alex Alderton, Roberto Amato, Jeffrey Barrett, Sonia Goncalves, Ewan Harrison, David K. Jackson, Ian Johnston, Dominic Kwiatkowski, Cordelia Langford, John Sillitoe on behalf of the Wellcome Sanger Institute COVID-19 Surveillance Team |
| EPI_ISL_2435599 | Lighthouse Lab in Alderley Park | Wellcome Sanger Institute for the COVID-19 Genomics UK (COG-UK) Consortium | Jacquelyn Wynn, Mairead Hyland, The Lighthouse Lab in Alderley Park and Alex Alderton, Roberto Amato, Jeffrey Barrett, Sonia Goncalves, Ewan Harrison, David K. Jackson, Ian Johnston, Dominic Kwiatkowski, Cordelia Langford, John Sillitoe on behalf of the Wellcome Sanger Institute COVID-19 Surveillance Team |
| EPI_ISL_2435600 | Lighthouse Lab in Milton Keynes | Wellcome Sanger Institute for the COVID-19 Genomics UK (COG-UK) Consortium | The Lighthouse Lab in Milton Keynes and Alex Alderton, Roberto Amato, Jeffrey Barrett, Sonia Goncalves, Ewan Harrison, David K. Jackson, Ian Johnston, Dominic Kwiatkowski, Cordelia Langford, John Sillitoe on behalf of the Wellcome Sanger Institute COVID-19 Surveillance Team |
| EPI_ISL_2435601 | Lighthouse Lab in Glasgow | Wellcome Sanger Institute for the COVID-19 Genomics UK (COG-UK) Consortium | Harper VanSteenhouse, Yumi Kasai, David Gray, Carol Clugston, Anna Dominiczak and Alex Alderton, Roberto Amato, Jeffrey Barrett, Sonia Goncalves, Ewan Harrison, David K. Jackson, Ian Johnston, Dominic Kwiatkowski, Cordelia Langford, John Sillitoe on behalf of the Wellcome Sanger Institute COVID-19 Surveillance Team |
| EPI_ISL_2435603 | Lighthouse Lab in Alderley Park | Wellcome Sanger Institute for the COVID-19 Genomics UK (COG-UK) Consortium | Jacquelyn Wynn, Mairead Hyland, The Lighthouse Lab in Alderley Park and Alex Alderton, Roberto Amato, Jeffrey Barrett, Sonia Goncalves, Ewan Harrison, David K. Jackson, Ian Johnston, Dominic Kwiatkowski, Cordelia Langford, John Sillitoe on behalf of the Wellcome Sanger Institute COVID-19 Surveillance Team |
| EPI_ISL_2435605 | Lighthouse Lab in Glasgow | Wellcome Sanger Institute for the COVID-19 Genomics UK (COG-UK) Consortium | Harper VanSteenhouse, Yumi Kasai, David Gray, Carol Clugston, Anna Dominiczak and Alex Alderton, Roberto Amato, Jeffrey Barrett, Sonia Goncalves, Ewan Harrison, David K. Jackson, Ian Johnston, Dominic Kwiatkowski, Cordelia Langford, John Sillitoe on behalf of the Wellcome Sanger Institute COVID-19 Surveillance Team |
| EPI_ISL_2435606, EPI_ISL_2435607, EPI_ISL_2435608, EPI_ISL_2435609, EPI_ISL_2435610, EPI_ISL_2435612, EPI_ISL_2435613, EPI_ISL_2435614, EPI_ISL_2435615, EPI_ISL_2435616 | Lighthouse Lab in Alderley Park | Wellcome Sanger Institute for the COVID-19 Genomics UK (COG-UK) Consortium | Jacquelyn Wynn, Mairead Hyland, The Lighthouse Lab in Alderley Park and Alex Alderton, Roberto Amato, Jeffrey Barrett, Sonia Goncalves, Ewan Harrison, David K. Jackson, Ian Johnston, Dominic Kwiatkowski, Cordelia Langford, John Sillitoe on behalf of the Wellcome Sanger Institute COVID-19 Surveillance Team |
| EPI_ISL_2435617 | Lighthouse Lab in Milton Keynes | Wellcome Sanger Institute for the COVID-19 Genomics UK (COG-UK) Consortium | The Lighthouse Lab in Milton Keynes and Alex Alderton, Roberto Amato, Jeffrey Barrett, Sonia Goncalves, Ewan Harrison, David K. Jackson, Ian Johnston, Dominic Kwiatkowski, Cordelia Langford, John Sillitoe on behalf of the Wellcome Sanger Institute COVID-19 Surveillance Team |
| EPI_ISL_2435618, EPI_ISL_2435620 | Lighthouse Lab in Alderley Park | Wellcome Sanger Institute for the COVID-19 Genomics UK (COG-UK) Consortium | Jacquelyn Wynn, Mairead Hyland, The Lighthouse Lab in Alderley Park and Alex Alderton, Roberto Amato, Jeffrey Barrett, Sonia Goncalves, Ewan Harrison, David K. Jackson, Ian Johnston, Dominic Kwiatkowski, Cordelia Langford, John Sillitoe on behalf of the Wellcome Sanger Institute COVID-19 Surveillance Team |
| EPI_ISL_2435621 | Lighthouse Lab in Milton Keynes | Wellcome Sanger Institute for the COVID-19 Genomics UK (COG-UK) Consortium | The Lighthouse Lab in Milton Keynes and Alex Alderton, Roberto Amato, Jeffrey Barrett, Sonia Goncalves, Ewan Harrison, David K. Jackson, Ian Johnston, Dominic Kwiatkowski, Cordelia Langford, John Sillitoe on behalf of the Wellcome Sanger Institute COVID-19 Surveillance Team |
| EPI_ISL_2435622, EPI_ISL_2435623, EPI_ISL_2435624, EPI_ISL_2435625, EPI_ISL_2435626, EPI_ISL_2435627 | Lighthouse Lab in Alderley Park | Wellcome Sanger Institute for the COVID-19 Genomics UK (COG-UK) Consortium | Jacquelyn Wynn, Mairead Hyland, The Lighthouse Lab in Alderley Park and Alex Alderton, Roberto Amato, Jeffrey Barrett, Sonia Goncalves, Ewan Harrison, David K. Jackson, Ian Johnston, Dominic Kwiatkowski, Cordelia Langford, John Sillitoe on behalf of the Wellcome Sanger Institute COVID-19 Surveillance Team |
| EPI_ISL_2435628 | Lighthouse Lab in Glasgow | Wellcome Sanger Institute for the COVID-19 Genomics UK (COG-UK) Consortium | Harper VanSteenhouse, Yumi Kasai, David Gray, Carol Clugston, Anna Dominiczak and Alex Alderton, Roberto Amato, Jeffrey Barrett, Sonia Goncalves, Ewan Harrison, David K. Jackson, Ian Johnston, Dominic Kwiatkowski, Cordelia Langford, John Sillitoe on behalf of the Wellcome Sanger Institute COVID-19 Surveillance Team |
| EPI_ISL_2435629, EPI_ISL_2435630, EPI_ISL_2435631, EPI_ISL_2435632, EPI_ISL_2435633, EPI_ISL_2435634 | Lighthouse Lab in Alderley Park | Wellcome Sanger Institute for the COVID-19 Genomics UK (COG-UK) Consortium | Jacquelyn Wynn, Mairead Hyland, The Lighthouse Lab in Alderley Park and Alex Alderton, Roberto Amato, Jeffrey Barrett, Sonia Goncalves, Ewan Harrison, David K. Jackson, Ian Johnston, Dominic Kwiatkowski, Cordelia Langford, John Sillitoe on behalf of the Wellcome Sanger Institute COVID-19 Surveillance Team |
| EPI_ISL_2435635 | Lighthouse Lab in Milton Keynes | Wellcome Sanger Institute for the COVID-19 Genomics UK (COG-UK) Consortium | The Lighthouse Lab in Milton Keynes and Alex Alderton, Roberto Amato, Jeffrey Barrett, Sonia Goncalves, Ewan Harrison, David K. Jackson, Ian Johnston, Dominic Kwiatkowski, Cordelia Langford, John Sillitoe on behalf of the Wellcome Sanger Institute COVID-19 Surveillance Team |
| EPI_ISL_2435636, EPI_ISL_2435637, EPI_ISL_2435638, EPI_ISL_2435639, EPI_ISL_2435640 | Lighthouse Lab in Alderley Park | Wellcome Sanger Institute for the COVID-19 Genomics UK (COG-UK) Consortium | Jacquelyn Wynn, Mairead Hyland, The Lighthouse Lab in Alderley Park and Alex Alderton, Roberto Amato, Jeffrey Barrett, Sonia Goncalves, Ewan Harrison, David K. Jackson, Ian Johnston, Dominic Kwiatkowski, Cordelia Langford, John Sillitoe on behalf of the Wellcome Sanger Institute COVID-19 Surveillance Team |
| EPI_ISL_2435641 | Lighthouse Lab in Milton Keynes | Wellcome Sanger Institute for the COVID-19 Genomics UK (COG-UK) Consortium | The Lighthouse Lab in Milton Keynes and Alex Alderton, Roberto Amato, Jeffrey Barrett, Sonia Goncalves, Ewan Harrison, David K. Jackson, Ian Johnston, Dominic Kwiatkowski, Cordelia Langford, John Sillitoe on behalf of the Wellcome Sanger Institute COVID-19 Surveillance Team |
| EPI_ISL_2435642 | Lighthouse Lab in Alderley Park | Wellcome Sanger Institute for the COVID-19 Genomics UK (COG-UK) Consortium | Jacquelyn Wynn, Mairead Hyland, The Lighthouse Lab in Alderley Park and Alex Alderton, Roberto Amato, Jeffrey Barrett, Sonia Goncalves, Ewan Harrison, David K. Jackson, Ian Johnston, Dominic Kwiatkowski, Cordelia Langford, John Sillitoe on behalf of the Wellcome Sanger Institute COVID-19 Surveillance Team |
| EPI_ISL_2435644, EPI_ISL_2435645 | Lighthouse Lab in Milton Keynes | Wellcome Sanger Institute for the COVID-19 Genomics UK (COG-UK) Consortium | The Lighthouse Lab in Milton Keynes and Alex Alderton, Roberto Amato, Jeffrey Barrett, Sonia Goncalves, Ewan Harrison, David K. Jackson, Ian Johnston, Dominic Kwiatkowski, Cordelia Langford, John Sillitoe on behalf of the Wellcome Sanger Institute COVID-19 Surveillance Team |
| EPI_ISL_2435646, EPI_ISL_2435649, EPI_ISL_2435650, EPI_ISL_2435654, EPI_ISL_2435655, EPI_ISL_2435656, EPI_ISL_2435659, EPI_ISL_2435660, EPI_ISL_2435661 | Lighthouse Lab in Alderley Park | Wellcome Sanger Institute for the COVID-19 Genomics UK (COG-UK) Consortium | Jacquelyn Wynn, Mairead Hyland, The Lighthouse Lab in Alderley Park and Alex Alderton, Roberto Amato, Jeffrey Barrett, Sonia Goncalves, Ewan Harrison, David K. Jackson, Ian Johnston, Dominic Kwiatkowski, Cordelia Langford, John Sillitoe on behalf of the Wellcome Sanger Institute COVID-19 Surveillance Team |
| EPI_ISL_2435662 | Lighthouse Lab in Milton Keynes | Wellcome Sanger Institute for the COVID-19 Genomics UK (COG-UK) Consortium | The Lighthouse Lab in Milton Keynes and Alex Alderton, Roberto Amato, Jeffrey Barrett, Sonia Goncalves, Ewan Harrison, David K. Jackson, Ian Johnston, Dominic Kwiatkowski, Cordelia Langford, John Sillitoe on behalf of the Wellcome Sanger Institute COVID-19 Surveillance Team |
| EPI_ISL_2435663, EPI_ISL_2435665 | Lighthouse Lab in Alderley Park | Wellcome Sanger Institute for the COVID-19 Genomics UK (COG-UK) Consortium | Jacquelyn Wynn, Mairead Hyland, The Lighthouse Lab in Alderley Park and Alex Alderton, Roberto Amato, Jeffrey Barrett, Sonia Goncalves, Ewan Harrison, David K. Jackson, Ian Johnston, Dominic Kwiatkowski, Cordelia Langford, John Sillitoe on behalf of the Wellcome Sanger Institute COVID-19 Surveillance Team |
| EPI_ISL_2435666 | Lighthouse Lab in Milton Keynes | Wellcome Sanger Institute for the COVID-19 Genomics UK (COG-UK) Consortium | The Lighthouse Lab in Milton Keynes and Alex Alderton, Roberto Amato, Jeffrey Barrett, Sonia Goncalves, Ewan Harrison, David K. Jackson, Ian Johnston, Dominic Kwiatkowski, Cordelia Langford, John Sillitoe on behalf of the Wellcome Sanger Institute COVID-19 Surveillance Team |

| | | | |
|---|---|---|---|
| EPI_ISL_2435667 | Lighthouse Lab in Glasgow | Wellcome Sanger Institute for the COVID-19 Genomics UK (COG-UK) Consortium | Harper VanSteenhouse, Yumi Kasai, David Gray, Carol Clugston, Anna Dominiczak and Alex Alderton, Roberto Amato, Jeffrey Barrett, Sonia Goncalves, Ewan Harrison, David K. Jackson, Ian Johnston, Dominic Kwiatkowski, Cordelia Langford, John Sillitoe on behalf of the Wellcome Sanger Institute COVID-19 Surveillance Team |
| EPI_ISL_2435669, EPI_ISL_2435672, EPI_ISL_2435673, EPI_ISL_2435674 | Lighthouse Lab in Alderley Park | Wellcome Sanger Institute for the COVID-19 Genomics UK (COG-UK) Consortium | Jacquelyn Wynn, Mairead Hyland, The Lighthouse Lab in Alderley Park and Alex Alderton, Roberto Amato, Jeffrey Barrett, Sonia Goncalves, Ewan Harrison, David K. Jackson, Ian Johnston, Dominic Kwiatkowski, Cordelia Langford, John Sillitoe on behalf of the Wellcome Sanger Institute COVID-19 Surveillance Team |
| EPI_ISL_2435675 | Lighthouse Lab in Milton Keynes | Wellcome Sanger Institute for the COVID-19 Genomics UK (COG-UK) Consortium | The Lighthouse Lab in Milton Keynes and Alex Alderton, Roberto Amato, Jeffrey Barrett, Sonia Goncalves, Ewan Harrison, David K. Jackson, Ian Johnston, Dominic Kwiatkowski, Cordelia Langford, John Sillitoe on behalf of the Wellcome Sanger Institute COVID-19 Surveillance Team |
| EPI_ISL_2435676, EPI_ISL_2435679, EPI_ISL_2435680, EPI_ISL_2435681, EPI_ISL_2435682, EPI_ISL_2435683, EPI_ISL_2435684, EPI_ISL_2435686, EPI_ISL_2435687, EPI_ISL_2435689 | Lighthouse Lab in Alderley Park | Wellcome Sanger Institute for the COVID-19 Genomics UK (COG-UK) Consortium | Jacquelyn Wynn, Mairead Hyland, The Lighthouse Lab in Alderley Park and Alex Alderton, Roberto Amato, Jeffrey Barrett, Sonia Goncalves, Ewan Harrison, David K. Jackson, Ian Johnston, Dominic Kwiatkowski, Cordelia Langford, John Sillitoe on behalf of the Wellcome Sanger Institute COVID-19 Surveillance Team |
| EPI_ISL_2435690 | Lighthouse Lab in Milton Keynes | Wellcome Sanger Institute for the COVID-19 Genomics UK (COG-UK) Consortium | The Lighthouse Lab in Milton Keynes and Alex Alderton, Roberto Amato, Jeffrey Barrett, Sonia Goncalves, Ewan Harrison, David K. Jackson, Ian Johnston, Dominic Kwiatkowski, Cordelia Langford, John Sillitoe on behalf of the Wellcome Sanger Institute COVID-19 Surveillance Team |
| EPI_ISL_2435691 | Lighthouse Lab in Alderley Park | Wellcome Sanger Institute for the COVID-19 Genomics UK (COG-UK) Consortium | Jacquelyn Wynn, Mairead Hyland, The Lighthouse Lab in Alderley Park and Alex Alderton, Roberto Amato, Jeffrey Barrett, Sonia Goncalves, Ewan Harrison, David K. Jackson, Ian Johnston, Dominic Kwiatkowski, Cordelia Langford, John Sillitoe on behalf of the Wellcome Sanger Institute COVID-19 Surveillance Team |
| EPI_ISL_2435692 | Lighthouse Lab in Milton Keynes | Wellcome Sanger Institute for the COVID-19 Genomics UK (COG-UK) Consortium | The Lighthouse Lab in Milton Keynes and Alex Alderton, Roberto Amato, Jeffrey Barrett, Sonia Goncalves, Ewan Harrison, David K. Jackson, Ian Johnston, Dominic Kwiatkowski, Cordelia Langford, John Sillitoe on behalf of the Wellcome Sanger Institute COVID-19 Surveillance Team |
| EPI_ISL_2435693 | Lighthouse Lab in Glasgow | Wellcome Sanger Institute for the COVID-19 Genomics UK (COG-UK) Consortium | Harper VanSteenhouse, Yumi Kasai, David Gray, Carol Clugston, Anna Dominiczak and Alex Alderton, Roberto Amato, Jeffrey Barrett, Sonia Goncalves, Ewan Harrison, David K. Jackson, Ian Johnston, Dominic Kwiatkowski, Cordelia Langford, John Sillitoe on behalf of the Wellcome Sanger Institute COVID-19 Surveillance Team |
| EPI_ISL_2435694, EPI_ISL_2435695 | Lighthouse Lab in Alderley Park | Wellcome Sanger Institute for the COVID-19 Genomics UK (COG-UK) Consortium | Jacquelyn Wynn, Mairead Hyland, The Lighthouse Lab in Alderley Park and Alex Alderton, Roberto Amato, Jeffrey Barrett, Sonia Goncalves, Ewan Harrison, David K. Jackson, Ian Johnston, Dominic Kwiatkowski, Cordelia Langford, John Sillitoe on behalf of the Wellcome Sanger Institute COVID-19 Surveillance Team |
| EPI_ISL_2435697 | Lighthouse Lab in Milton Keynes | Wellcome Sanger Institute for the COVID-19 Genomics UK (COG-UK) Consortium | The Lighthouse Lab in Milton Keynes and Alex Alderton, Roberto Amato, Jeffrey Barrett, Sonia Goncalves, Ewan Harrison, David K. Jackson, Ian Johnston, Dominic Kwiatkowski, Cordelia Langford, John Sillitoe on behalf of the Wellcome Sanger Institute COVID-19 Surveillance Team |
| EPI_ISL_2435698, EPI_ISL_2435699, EPI_ISL_2435700 | Lighthouse Lab in Alderley Park | Wellcome Sanger Institute for the COVID-19 Genomics UK (COG-UK) Consortium | Jacquelyn Wynn, Mairead Hyland, The Lighthouse Lab in Alderley Park and Alex Alderton, Roberto Amato, Jeffrey Barrett, Sonia Goncalves, Ewan Harrison, David K. Jackson, Ian Johnston, Dominic Kwiatkowski, Cordelia Langford, John Sillitoe on behalf of the Wellcome Sanger Institute COVID-19 Surveillance Team |
| EPI_ISL_2435701 | Lighthouse Lab in Glasgow | Wellcome Sanger Institute for the COVID-19 Genomics UK (COG-UK) Consortium | Harper VanSteenhouse, Yumi Kasai, David Gray, Carol Clugston, Anna Dominiczak and Alex Alderton, Roberto Amato, Jeffrey Barrett, Sonia Goncalves, Ewan Harrison, David K. Jackson, Ian Johnston, Dominic Kwiatkowski, Cordelia Langford, John Sillitoe on behalf of the Wellcome Sanger Institute COVID-19 Surveillance Team |
| EPI_ISL_2435702 | Lighthouse Lab in Alderley Park | Wellcome Sanger Institute for the COVID-19 Genomics UK (COG-UK) Consortium | Jacquelyn Wynn, Mairead Hyland, The Lighthouse Lab in Alderley Park and Alex Alderton, Roberto Amato, Jeffrey Barrett, Sonia Goncalves, Ewan Harrison, David K. Jackson, Ian Johnston, Dominic Kwiatkowski, Cordelia Langford, John Sillitoe on behalf of the Wellcome Sanger Institute COVID-19 Surveillance Team |
| EPI_ISL_2435703 | Lighthouse Lab in Glasgow | Wellcome Sanger Institute for the COVID-19 Genomics UK (COG-UK) Consortium | Harper VanSteenhouse, Yumi Kasai, David Gray, Carol Clugston, Anna Dominiczak and Alex Alderton, Roberto Amato, Jeffrey Barrett, Sonia Goncalves, Ewan Harrison, David K. Jackson, Ian Johnston, Dominic Kwiatkowski, Cordelia Langford, John Sillitoe on behalf of the Wellcome Sanger Institute COVID-19 Surveillance Team |
| EPI_ISL_2435704, EPI_ISL_2435705, EPI_ISL_2435706, EPI_ISL_2435707, EPI_ISL_2435709, EPI_ISL_2435711 | Lighthouse Lab in Alderley Park | Wellcome Sanger Institute for the COVID-19 Genomics UK (COG-UK) Consortium | Jacquelyn Wynn, Mairead Hyland, The Lighthouse Lab in Alderley Park and Alex Alderton, Roberto Amato, Jeffrey Barrett, Sonia Goncalves, Ewan Harrison, David K. Jackson, Ian Johnston, Dominic Kwiatkowski, Cordelia Langford, John Sillitoe on behalf of the Wellcome Sanger Institute COVID-19 Surveillance Team |
| EPI_ISL_2435712 | Lighthouse Lab in Milton Keynes | Wellcome Sanger Institute for the COVID-19 Genomics UK (COG-UK) Consortium | The Lighthouse Lab in Milton Keynes and Alex Alderton, Roberto Amato, Jeffrey Barrett, Sonia Goncalves, Ewan Harrison, David K. Jackson, Ian Johnston, Dominic Kwiatkowski, Cordelia Langford, John Sillitoe on behalf of the Wellcome Sanger Institute COVID-19 Surveillance Team |
| EPI_ISL_2435713, EPI_ISL_2435715 | Lighthouse Lab in Alderley Park | Wellcome Sanger Institute for the COVID-19 Genomics UK (COG-UK) Consortium | Jacquelyn Wynn, Mairead Hyland, The Lighthouse Lab in Alderley Park and Alex Alderton, Roberto Amato, Jeffrey Barrett, Sonia Goncalves, Ewan Harrison, David K. Jackson, Ian Johnston, Dominic Kwiatkowski, Cordelia Langford, John Sillitoe on behalf of the Wellcome Sanger Institute COVID-19 Surveillance Team |
| EPI_ISL_2435716 | Lighthouse Lab in Milton Keynes | Wellcome Sanger Institute for the COVID-19 Genomics UK (COG-UK) Consortium | The Lighthouse Lab in Milton Keynes and Alex Alderton, Roberto Amato, Jeffrey Barrett, Sonia Goncalves, Ewan Harrison, David K. Jackson, Ian Johnston, Dominic Kwiatkowski, Cordelia Langford, John Sillitoe on behalf of the Wellcome Sanger Institute COVID-19 Surveillance Team |
| EPI_ISL_2435717, EPI_ISL_2435718 | Lighthouse Lab in Alderley Park | Wellcome Sanger Institute for the COVID-19 Genomics UK (COG-UK) Consortium | Jacquelyn Wynn, Mairead Hyland, The Lighthouse Lab in Alderley Park and Alex Alderton, Roberto Amato, Jeffrey Barrett, Sonia Goncalves, Ewan Harrison, David K. Jackson, Ian Johnston, Dominic Kwiatkowski, Cordelia Langford, John Sillitoe on behalf of the Wellcome Sanger Institute COVID-19 Surveillance Team |
| EPI_ISL_2435719, EPI_ISL_2435720, EPI_ISL_2435721, EPI_ISL_2435722, EPI_ISL_2435723, EPI_ISL_2435724 | Lighthouse Lab in Milton Keynes | Wellcome Sanger Institute for the COVID-19 Genomics UK (COG-UK) Consortium | The Lighthouse Lab in Milton Keynes and Alex Alderton, Roberto Amato, Jeffrey Barrett, Sonia Goncalves, Ewan Harrison, David K. Jackson, Ian Johnston, Dominic Kwiatkowski, Cordelia Langford, John Sillitoe on behalf of the Wellcome Sanger Institute COVID-19 Surveillance Team |
| EPI_ISL_2435725, EPI_ISL_2435727, EPI_ISL_2435728, EPI_ISL_2435729, EPI_ISL_2435732 | Lighthouse Lab in Alderley Park | Wellcome Sanger Institute for the COVID-19 Genomics UK (COG-UK) Consortium | Jacquelyn Wynn, Mairead Hyland, The Lighthouse Lab in Alderley Park and Alex Alderton, Roberto Amato, Jeffrey Barrett, Sonia Goncalves, Ewan Harrison, David K. Jackson, Ian Johnston, Dominic Kwiatkowski, Cordelia Langford, John Sillitoe on behalf of the Wellcome Sanger Institute COVID-19 Surveillance Team |
| EPI_ISL_2435733 | Lighthouse Lab in Glasgow | Wellcome Sanger Institute for the COVID-19 Genomics UK (COG-UK) Consortium | Harper VanSteenhouse, Yumi Kasai, David Gray, Carol Clugston, Anna Dominiczak and Alex Alderton, Roberto Amato, Jeffrey Barrett, Sonia Goncalves, Ewan Harrison, David K. Jackson, Ian Johnston, Dominic Kwiatkowski, Cordelia Langford, John Sillitoe on behalf of the Wellcome Sanger Institute COVID-19 Surveillance Team |
| EPI_ISL_2435734, EPI_ISL_2435735 | Lighthouse Lab in Alderley Park | Wellcome Sanger Institute for the COVID-19 Genomics UK (COG-UK) Consortium | Jacquelyn Wynn, Mairead Hyland, The Lighthouse Lab in Alderley Park and Alex Alderton, Roberto Amato, Jeffrey Barrett, Sonia Goncalves, Ewan Harrison, David K. Jackson, Ian Johnston, Dominic Kwiatkowski, Cordelia Langford, John Sillitoe on behalf of the Wellcome Sanger Institute COVID-19 Surveillance Team |
| EPI_ISL_2435736 | Lighthouse Lab in Glasgow | Wellcome Sanger Institute for the COVID-19 Genomics UK (COG-UK) Consortium | Harper VanSteenhouse, Yumi Kasai, David Gray, Carol Clugston, Anna Dominiczak and Alex Alderton, Roberto Amato, Jeffrey Barrett, Sonia Goncalves, Ewan Harrison, David K. Jackson, Ian Johnston, Dominic Kwiatkowski, Cordelia Langford, John Sillitoe on behalf of the Wellcome Sanger Institute COVID-19 Surveillance Team |
| EPI_ISL_2435737, EPI_ISL_2435738, EPI_ISL_2435739 | Lighthouse Lab in Alderley Park | Wellcome Sanger Institute for the COVID-19 Genomics UK (COG-UK) Consortium | Jacquelyn Wynn, Mairead Hyland, The Lighthouse Lab in Alderley Park and Alex Alderton, Roberto Amato, Jeffrey Barrett, Sonia Goncalves, Ewan Harrison, David K. Jackson, Ian Johnston, Dominic Kwiatkowski, Cordelia Langford, John Sillitoe on behalf of the Wellcome Sanger Institute COVID-19 Surveillance Team |
| EPI_ISL_2435740 | Lighthouse Lab in Glasgow | Wellcome Sanger Institute for the COVID-19 Genomics UK (COG-UK) Consortium | Harper VanSteenhouse, Yumi Kasai, David Gray, Carol Clugston, Anna Dominiczak and Alex Alderton, Roberto Amato, Jeffrey Barrett, Sonia Goncalves, Ewan Harrison, David K. Jackson, Ian Johnston, Dominic Kwiatkowski, Cordelia Langford, John Sillitoe on behalf of the Wellcome Sanger Institute COVID-19 Surveillance Team |
| EPI_ISL_2435741, EPI_ISL_2435742, EPI_ISL_2435743, EPI_ISL_2435744, EPI_ISL_2435745, EPI_ISL_2435746 | Lighthouse Lab in Alderley Park | Wellcome Sanger Institute for the COVID-19 Genomics UK (COG-UK) Consortium | Jacquelyn Wynn, Mairead Hyland, The Lighthouse Lab in Alderley Park and Alex Alderton, Roberto Amato, Jeffrey Barrett, Sonia Goncalves, Ewan Harrison, David K. Jackson, Ian Johnston, Dominic Kwiatkowski, Cordelia Langford, John Sillitoe on behalf of the Wellcome Sanger Institute COVID-19 Surveillance Team |
| EPI_ISL_2435747 | Lighthouse Lab in Glasgow | Wellcome Sanger Institute for the COVID-19 Genomics UK (COG-UK) Consortium | Harper VanSteenhouse, Yumi Kasai, David Gray, Carol Clugston, Anna Dominiczak and Alex Alderton, Roberto Amato, Jeffrey Barrett, Sonia Goncalves, Ewan Harrison, David K. Jackson, Ian Johnston, Dominic Kwiatkowski, Cordelia Langford, John Sillitoe on behalf of the Wellcome Sanger Institute COVID-19 Surveillance Team |
| EPI_ISL_2435748, EPI_ISL_2435749 | Lighthouse Lab in Milton Keynes | Wellcome Sanger Institute for the COVID-19 Genomics UK (COG-UK) Consortium | The Lighthouse Lab in Milton Keynes and Alex Alderton, Roberto Amato, Jeffrey Barrett, Sonia Goncalves, Ewan Harrison, David K. Jackson, Ian Johnston, Dominic Kwiatkowski, Cordelia Langford, John Sillitoe on behalf of the Wellcome Sanger Institute COVID-19 Surveillance Team |
| EPI_ISL_2435750 | Lighthouse Lab in Glasgow | Wellcome Sanger Institute for the COVID-19 Genomics UK (COG-UK) Consortium | Harper VanSteenhouse, Yumi Kasai, David Gray, Carol Clugston, Anna Dominiczak and Alex Alderton, Roberto Amato, Jeffrey Barrett, Sonia Goncalves, Ewan Harrison, David K. Jackson, Ian Johnston, Dominic Kwiatkowski, Cordelia Langford, John Sillitoe on behalf of the Wellcome Sanger Institute COVID-19 Surveillance Team |
| EPI_ISL_2435752 | Lighthouse Lab in Alderley Park | Wellcome Sanger Institute for the COVID-19 Genomics UK (COG-UK) Consortium | Jacquelyn Wynn, Mairead Hyland, The Lighthouse Lab in Alderley Park and Alex Alderton, Roberto Amato, Jeffrey Barrett, Sonia Goncalves, Ewan Harrison, David K. Jackson, Ian Johnston, Dominic Kwiatkowski, Cordelia Langford, John Sillitoe on behalf of the Wellcome Sanger Institute COVID-19 Surveillance Team |
| EPI_ISL_2435753 | Lighthouse Lab in Glasgow | Wellcome Sanger Institute for the COVID-19 Genomics UK (COG-UK) Consortium | Harper VanSteenhouse, Yumi Kasai, David Gray, Carol Clugston, Anna Dominiczak and Alex Alderton, Roberto Amato, Jeffrey Barrett, Sonia Goncalves, Ewan Harrison, David K. Jackson, Ian Johnston, Dominic Kwiatkowski, Cordelia Langford, John Sillitoe on behalf of the Wellcome Sanger Institute COVID-19 Surveillance Team |
| EPI_ISL_2435755, EPI_ISL_2435756, EPI_ISL_2435757 | Lighthouse Lab in Alderley Park | Wellcome Sanger Institute for the COVID-19 Genomics UK (COG-UK) Consortium | Jacquelyn Wynn, Mairead Hyland, The Lighthouse Lab in Alderley Park and Alex Alderton, Roberto Amato, Jeffrey Barrett, Sonia Goncalves, Ewan Harrison, David K. Jackson, Ian Johnston, Dominic Kwiatkowski, Cordelia Langford, John Sillitoe on behalf of the Wellcome Sanger Institute COVID-19 Surveillance Team |
| EPI_ISL_2435758 | Lighthouse Lab in Milton Keynes | Wellcome Sanger Institute for the COVID-19 Genomics UK (COG-UK) | The Lighthouse Lab in Milton Keynes and Alex Alderton, Roberto Amato, Jeffrey Barrett, Sonia Goncalves, Ewan Harrison, David K. Jackson, Ian Johnston, Dominic Kwiatkowski, Cordelia Langford, John Sillitoe on behalf of the Wellcome Sanger |

| | | | |
|---|---|---|---|
| | | Consortium | Institute COVID-19 Surveillance Team |
| EPI_ISL_2435760, EPI_ISL_2435761 | Lighthouse Lab in Alderley Park | Wellcome Sanger Institute for the COVID-19 Genomics UK (COG-UK) Consortium | Jacquelyn Wynn, Mairead Hyland, The Lighthouse Lab in Alderley Park and Alex Alderton, Roberto Amato, Jeffrey Barrett, Sonia Goncalves, Ewan Harrison, David K. Jackson, Ian Johnston, Dominic Kwiatkowski, Cordelia Langford, John Sillitoe on behalf of the Wellcome Sanger Institute COVID-19 Surveillance Team |
| EPI_ISL_2435762 | Lighthouse Lab in Milton Keynes | Wellcome Sanger Institute for the COVID-19 Genomics UK (COG-UK) Consortium | The Lighthouse Lab in Milton Keynes and Alex Alderton, Roberto Amato, Jeffrey Barrett, Sonia Goncalves, Ewan Harrison, David K. Jackson, Ian Johnston, Dominic Kwiatkowski, Cordelia Langford, John Sillitoe on behalf of the Wellcome Sanger Institute COVID-19 Surveillance Team |
| EPI_ISL_2435763, EPI_ISL_2435764, EPI_ISL_2435765 | Lighthouse Lab in Alderley Park | Wellcome Sanger Institute for the COVID-19 Genomics UK (COG-UK) Consortium | Jacquelyn Wynn, Mairead Hyland, The Lighthouse Lab in Alderley Park and Alex Alderton, Roberto Amato, Jeffrey Barrett, Sonia Goncalves, Ewan Harrison, David K. Jackson, Ian Johnston, Dominic Kwiatkowski, Cordelia Langford, John Sillitoe on behalf of the Wellcome Sanger Institute COVID-19 Surveillance Team |
| EPI_ISL_2435766 | Lighthouse Lab in Milton Keynes | Wellcome Sanger Institute for the COVID-19 Genomics UK (COG-UK) Consortium | The Lighthouse Lab in Milton Keynes and Alex Alderton, Roberto Amato, Jeffrey Barrett, Sonia Goncalves, Ewan Harrison, David K. Jackson, Ian Johnston, Dominic Kwiatkowski, Cordelia Langford, John Sillitoe on behalf of the Wellcome Sanger Institute COVID-19 Surveillance Team |
| EPI_ISL_2435768, EPI_ISL_2435770, EPI_ISL_2435771, EPI_ISL_2435772, EPI_ISL_2435773 | Lighthouse Lab in Alderley Park | Wellcome Sanger Institute for the COVID-19 Genomics UK (COG-UK) Consortium | Jacquelyn Wynn, Mairead Hyland, The Lighthouse Lab in Alderley Park and Alex Alderton, Roberto Amato, Jeffrey Barrett, Sonia Goncalves, Ewan Harrison, David K. Jackson, Ian Johnston, Dominic Kwiatkowski, Cordelia Langford, John Sillitoe on behalf of the Wellcome Sanger Institute COVID-19 Surveillance Team |
| EPI_ISL_2435774 | Lighthouse Lab in Glasgow | Wellcome Sanger Institute for the COVID-19 Genomics UK (COG-UK) Consortium | Harper VanSteenhouse, Yumi Kasai, David Gray, Carol Clugston, Anna Dominiczak and Alex Alderton, Roberto Amato, Jeffrey Barrett, Sonia Goncalves, Ewan Harrison, David K. Jackson, Ian Johnston, Dominic Kwiatkowski, Cordelia Langford, John Sillitoe on behalf of the Wellcome Sanger Institute COVID-19 Surveillance Team |
| EPI_ISL_2435775, EPI_ISL_2435776, EPI_ISL_2435777, EPI_ISL_2435778 | Lighthouse Lab in Alderley Park | Wellcome Sanger Institute for the COVID-19 Genomics UK (COG-UK) Consortium | Jacquelyn Wynn, Mairead Hyland, The Lighthouse Lab in Alderley Park and Alex Alderton, Roberto Amato, Jeffrey Barrett, Sonia Goncalves, Ewan Harrison, David K. Jackson, Ian Johnston, Dominic Kwiatkowski, Cordelia Langford, John Sillitoe on behalf of the Wellcome Sanger Institute COVID-19 Surveillance Team |
| EPI_ISL_2435779 | Lighthouse Lab in Milton Keynes | Wellcome Sanger Institute for the COVID-19 Genomics UK (COG-UK) Consortium | The Lighthouse Lab in Milton Keynes and Alex Alderton, Roberto Amato, Jeffrey Barrett, Sonia Goncalves, Ewan Harrison, David K. Jackson, Ian Johnston, Dominic Kwiatkowski, Cordelia Langford, John Sillitoe on behalf of the Wellcome Sanger Institute COVID-19 Surveillance Team |
| EPI_ISL_2435780, EPI_ISL_2435781, EPI_ISL_2435782, EPI_ISL_2435783, EPI_ISL_2435784, EPI_ISL_2435785, EPI_ISL_2435786, EPI_ISL_2435787, EPI_ISL_2435788 | Lighthouse Lab in Alderley Park | Wellcome Sanger Institute for the COVID-19 Genomics UK (COG-UK) Consortium | Jacquelyn Wynn, Mairead Hyland, The Lighthouse Lab in Alderley Park and Alex Alderton, Roberto Amato, Jeffrey Barrett, Sonia Goncalves, Ewan Harrison, David K. Jackson, Ian Johnston, Dominic Kwiatkowski, Cordelia Langford, John Sillitoe on behalf of the Wellcome Sanger Institute COVID-19 Surveillance Team |
| EPI_ISL_2435789 | Lighthouse Lab in Milton Keynes | Wellcome Sanger Institute for the COVID-19 Genomics UK (COG-UK) Consortium | The Lighthouse Lab in Milton Keynes and Alex Alderton, Roberto Amato, Jeffrey Barrett, Sonia Goncalves, Ewan Harrison, David K. Jackson, Ian Johnston, Dominic Kwiatkowski, Cordelia Langford, John Sillitoe on behalf of the Wellcome Sanger Institute COVID-19 Surveillance Team |
| EPI_ISL_2435791 | Lighthouse Lab in Glasgow | Wellcome Sanger Institute for the COVID-19 Genomics UK (COG-UK) Consortium | Harper VanSteenhouse, Yumi Kasai, David Gray, Carol Clugston, Anna Dominiczak and Alex Alderton, Roberto Amato, Jeffrey Barrett, Sonia Goncalves, Ewan Harrison, David K. Jackson, Ian Johnston, Dominic Kwiatkowski, Cordelia Langford, John Sillitoe on behalf of the Wellcome Sanger Institute COVID-19 Surveillance Team |
| EPI_ISL_2435792 | Lighthouse Lab in Alderley Park | Wellcome Sanger Institute for the COVID-19 Genomics UK (COG-UK) Consortium | Jacquelyn Wynn, Mairead Hyland, The Lighthouse Lab in Alderley Park and Alex Alderton, Roberto Amato, Jeffrey Barrett, Sonia Goncalves, Ewan Harrison, David K. Jackson, Ian Johnston, Dominic Kwiatkowski, Cordelia Langford, John Sillitoe on behalf of the Wellcome Sanger Institute COVID-19 Surveillance Team |
| EPI_ISL_2435793 | Lighthouse Lab in Milton Keynes | Wellcome Sanger Institute for the COVID-19 Genomics UK (COG-UK) Consortium | The Lighthouse Lab in Milton Keynes and Alex Alderton, Roberto Amato, Jeffrey Barrett, Sonia Goncalves, Ewan Harrison, David K. Jackson, Ian Johnston, Dominic Kwiatkowski, Cordelia Langford, John Sillitoe on behalf of the Wellcome Sanger Institute COVID-19 Surveillance Team |
| EPI_ISL_2435794, EPI_ISL_2435795, EPI_ISL_2435796, EPI_ISL_2435797 | Lighthouse Lab in Alderley Park | Wellcome Sanger Institute for the COVID-19 Genomics UK (COG-UK) Consortium | Jacquelyn Wynn, Mairead Hyland, The Lighthouse Lab in Alderley Park and Alex Alderton, Roberto Amato, Jeffrey Barrett, Sonia Goncalves, Ewan Harrison, David K. Jackson, Ian Johnston, Dominic Kwiatkowski, Cordelia Langford, John Sillitoe on behalf of the Wellcome Sanger Institute COVID-19 Surveillance Team |
| EPI_ISL_2435798 | Lighthouse Lab in Milton Keynes | Wellcome Sanger Institute for the COVID-19 Genomics UK (COG-UK) Consortium | The Lighthouse Lab in Milton Keynes and Alex Alderton, Roberto Amato, Jeffrey Barrett, Sonia Goncalves, Ewan Harrison, David K. Jackson, Ian Johnston, Dominic Kwiatkowski, Cordelia Langford, John Sillitoe on behalf of the Wellcome Sanger Institute COVID-19 Surveillance Team |
| EPI_ISL_2435799 | Lighthouse Lab in Alderley Park | Wellcome Sanger Institute for the COVID-19 Genomics UK (COG-UK) Consortium | Jacquelyn Wynn, Mairead Hyland, The Lighthouse Lab in Alderley Park and Alex Alderton, Roberto Amato, Jeffrey Barrett, Sonia Goncalves, Ewan Harrison, David K. Jackson, Ian Johnston, Dominic Kwiatkowski, Cordelia Langford, John Sillitoe on behalf of the Wellcome Sanger Institute COVID-19 Surveillance Team |
| EPI_ISL_2435800 | Lighthouse Lab in Milton Keynes | Wellcome Sanger Institute for the COVID-19 Genomics UK (COG-UK) Consortium | The Lighthouse Lab in Milton Keynes and Alex Alderton, Roberto Amato, Jeffrey Barrett, Sonia Goncalves, Ewan Harrison, David K. Jackson, Ian Johnston, Dominic Kwiatkowski, Cordelia Langford, John Sillitoe on behalf of the Wellcome Sanger Institute COVID-19 Surveillance Team |
| EPI_ISL_2435801 | Lighthouse Lab in Glasgow | Wellcome Sanger Institute for the COVID-19 Genomics UK (COG-UK) Consortium | Harper VanSteenhouse, Yumi Kasai, David Gray, Carol Clugston, Anna Dominiczak and Alex Alderton, Roberto Amato, Jeffrey Barrett, Sonia Goncalves, Ewan Harrison, David K. Jackson, Ian Johnston, Dominic Kwiatkowski, Cordelia Langford, John Sillitoe on behalf of the Wellcome Sanger Institute COVID-19 Surveillance Team |
| EPI_ISL_2435802, EPI_ISL_2435803, EPI_ISL_2435804 | Lighthouse Lab in Alderley Park | Wellcome Sanger Institute for the COVID-19 Genomics UK (COG-UK) Consortium | Jacquelyn Wynn, Mairead Hyland, The Lighthouse Lab in Alderley Park and Alex Alderton, Roberto Amato, Jeffrey Barrett, Sonia Goncalves, Ewan Harrison, David K. Jackson, Ian Johnston, Dominic Kwiatkowski, Cordelia Langford, John Sillitoe on behalf of the Wellcome Sanger Institute COVID-19 Surveillance Team |
| EPI_ISL_2435805 | Lighthouse Lab in Glasgow | Wellcome Sanger Institute for the COVID-19 Genomics UK (COG-UK) Consortium | Harper VanSteenhouse, Yumi Kasai, David Gray, Carol Clugston, Anna Dominiczak and Alex Alderton, Roberto Amato, Jeffrey Barrett, Sonia Goncalves, Ewan Harrison, David K. Jackson, Ian Johnston, Dominic Kwiatkowski, Cordelia Langford, John Sillitoe on behalf of the Wellcome Sanger Institute COVID-19 Surveillance Team |
| EPI_ISL_2435806, EPI_ISL_2435808, EPI_ISL_2435809, EPI_ISL_2435810, EPI_ISL_2435811 | Lighthouse Lab in Alderley Park | Wellcome Sanger Institute for the COVID-19 Genomics UK (COG-UK) Consortium | Jacquelyn Wynn, Mairead Hyland, The Lighthouse Lab in Alderley Park and Alex Alderton, Roberto Amato, Jeffrey Barrett, Sonia Goncalves, Ewan Harrison, David K. Jackson, Ian Johnston, Dominic Kwiatkowski, Cordelia Langford, John Sillitoe on behalf of the Wellcome Sanger Institute COVID-19 Surveillance Team |
| EPI_ISL_2435812 | Lighthouse Lab in Glasgow | Wellcome Sanger Institute for the COVID-19 Genomics UK (COG-UK) Consortium | Harper VanSteenhouse, Yumi Kasai, David Gray, Carol Clugston, Anna Dominiczak and Alex Alderton, Roberto Amato, Jeffrey Barrett, Sonia Goncalves, Ewan Harrison, David K. Jackson, Ian Johnston, Dominic Kwiatkowski, Cordelia Langford, John Sillitoe on behalf of the Wellcome Sanger Institute COVID-19 Surveillance Team |
| EPI_ISL_2435813 | Lighthouse Lab in Alderley Park | Wellcome Sanger Institute for the COVID-19 Genomics UK (COG-UK) Consortium | Jacquelyn Wynn, Mairead Hyland, The Lighthouse Lab in Alderley Park and Alex Alderton, Roberto Amato, Jeffrey Barrett, Sonia Goncalves, Ewan Harrison, David K. Jackson, Ian Johnston, Dominic Kwiatkowski, Cordelia Langford, John Sillitoe on behalf of the Wellcome Sanger Institute COVID-19 Surveillance Team |
| EPI_ISL_2435814, EPI_ISL_2435815 | Lighthouse Lab in Milton Keynes | Wellcome Sanger Institute for the COVID-19 Genomics UK (COG-UK) Consortium | The Lighthouse Lab in Milton Keynes and Alex Alderton, Roberto Amato, Jeffrey Barrett, Sonia Goncalves, Ewan Harrison, David K. Jackson, Ian Johnston, Dominic Kwiatkowski, Cordelia Langford, John Sillitoe on behalf of the Wellcome Sanger Institute COVID-19 Surveillance Team |
| EPI_ISL_2435816 | Lighthouse Lab in Alderley Park | Wellcome Sanger Institute for the COVID-19 Genomics UK (COG-UK) Consortium | Jacquelyn Wynn, Mairead Hyland, The Lighthouse Lab in Alderley Park and Alex Alderton, Roberto Amato, Jeffrey Barrett, Sonia Goncalves, Ewan Harrison, David K. Jackson, Ian Johnston, Dominic Kwiatkowski, Cordelia Langford, John Sillitoe on behalf of the Wellcome Sanger Institute COVID-19 Surveillance Team |
| EPI_ISL_2435818 | Lighthouse Lab in Milton Keynes | Wellcome Sanger Institute for the COVID-19 Genomics UK (COG-UK) Consortium | The Lighthouse Lab in Milton Keynes and Alex Alderton, Roberto Amato, Jeffrey Barrett, Sonia Goncalves, Ewan Harrison, David K. Jackson, Ian Johnston, Dominic Kwiatkowski, Cordelia Langford, John Sillitoe on behalf of the Wellcome Sanger Institute COVID-19 Surveillance Team |
| EPI_ISL_2435819 | Lighthouse Lab in Glasgow | Wellcome Sanger Institute for the COVID-19 Genomics UK (COG-UK) Consortium | Harper VanSteenhouse, Yumi Kasai, David Gray, Carol Clugston, Anna Dominiczak and Alex Alderton, Roberto Amato, Jeffrey Barrett, Sonia Goncalves, Ewan Harrison, David K. Jackson, Ian Johnston, Dominic Kwiatkowski, Cordelia Langford, John Sillitoe on behalf of the Wellcome Sanger Institute COVID-19 Surveillance Team |
| EPI_ISL_2435820, EPI_ISL_2435822, EPI_ISL_2435823, EPI_ISL_2435824, EPI_ISL_2435825, EPI_ISL_2435826, EPI_ISL_2435827, EPI_ISL_2435828, EPI_ISL_2435829, EPI_ISL_2435832, EPI_ISL_2435834, EPI_ISL_2435835 | | | |
| see above | Lighthouse Lab in Alderley Park | Wellcome Sanger Institute for the COVID-19 Genomics UK (COG-UK) Consortium | Jacquelyn Wynn, Mairead Hyland, The Lighthouse Lab in Alderley Park and Alex Alderton, Roberto Amato, Jeffrey Barrett, Sonia Goncalves, Ewan Harrison, David K. Jackson, Ian Johnston, Dominic Kwiatkowski, Cordelia Langford, John Sillitoe on behalf of the Wellcome Sanger Institute COVID-19 Surveillance Team |
| EPI_ISL_2435837 | Lighthouse Lab in Glasgow | Wellcome Sanger Institute for the COVID-19 Genomics UK (COG-UK) Consortium | Harper VanSteenhouse, Yumi Kasai, David Gray, Carol Clugston, Anna Dominiczak and Alex Alderton, Roberto Amato, Jeffrey Barrett, Sonia Goncalves, Ewan Harrison, David K. Jackson, Ian Johnston, Dominic Kwiatkowski, Cordelia Langford, John Sillitoe on behalf of the Wellcome Sanger Institute COVID-19 Surveillance Team |
| EPI_ISL_2435838, EPI_ISL_2435839, EPI_ISL_2435840, EPI_ISL_2435841 | Lighthouse Lab in Alderley Park | Wellcome Sanger Institute for the COVID-19 Genomics UK (COG-UK) Consortium | Jacquelyn Wynn, Mairead Hyland, The Lighthouse Lab in Alderley Park and Alex Alderton, Roberto Amato, Jeffrey Barrett, Sonia Goncalves, Ewan Harrison, David K. Jackson, Ian Johnston, Dominic Kwiatkowski, Cordelia Langford, John Sillitoe on behalf of the Wellcome Sanger Institute COVID-19 Surveillance Team |
| EPI_ISL_2435842 | Lighthouse Lab in Glasgow | Wellcome Sanger Institute for the COVID-19 Genomics UK (COG-UK) Consortium | Harper VanSteenhouse, Yumi Kasai, David Gray, Carol Clugston, Anna Dominiczak and Alex Alderton, Roberto Amato, Jeffrey Barrett, Sonia Goncalves, Ewan Harrison, David K. Jackson, Ian Johnston, Dominic Kwiatkowski, Cordelia Langford, John Sillitoe on behalf of the Wellcome Sanger Institute COVID-19 Surveillance Team |
| EPI_ISL_2435843, EPI_ISL_2435844, EPI_ISL_2435845, EPI_ISL_2435846, EPI_ISL_2435847, EPI_ISL_2435849, EPI_ISL_2435850, EPI_ISL_2435851, EPI_ISL_2435852, EPI_ISL_2435853, EPI_ISL_2435855 | | | |
| see above | Lighthouse Lab in Alderley Park | Wellcome Sanger Institute for the COVID-19 Genomics UK (COG-UK) Consortium | Jacquelyn Wynn, Mairead Hyland, The Lighthouse Lab in Alderley Park and Alex Alderton, Roberto Amato, Jeffrey Barrett, Sonia Goncalves, Ewan Harrison, David K. Jackson, Ian Johnston, Dominic Kwiatkowski, Cordelia Langford, John Sillitoe on behalf of the Wellcome Sanger Institute COVID-19 Surveillance Team |
| EPI_ISL_2435856 | Lighthouse Lab in Milton Keynes | Wellcome Sanger Institute for the COVID-19 Genomics UK (COG-UK) Consortium | The Lighthouse Lab in Milton Keynes and Alex Alderton, Roberto Amato, Jeffrey Barrett, Sonia Goncalves, Ewan Harrison, David K. Jackson, Ian Johnston, Dominic Kwiatkowski, Cordelia Langford, John Sillitoe on behalf of the Wellcome Sanger Institute COVID-19 Surveillance Team |
| EPI_ISL_2435857, EPI_ISL_2435858 | Lighthouse Lab in Alderley Park | Wellcome Sanger Institute for the COVID-19 Genomics UK (COG-UK) Consortium | Jacquelyn Wynn, Mairead Hyland, The Lighthouse Lab in Alderley Park and Alex Alderton, Roberto Amato, Jeffrey Barrett, Sonia Goncalves, Ewan Harrison, David K. Jackson, Ian Johnston, Dominic Kwiatkowski, Cordelia Langford, John Sillitoe on behalf of the Wellcome Sanger Institute COVID-19 Surveillance Team |
| EPI_ISL_2435859 | Lighthouse Lab in Milton Keynes | Wellcome Sanger Institute for the COVID-19 Genomics UK (COG-UK) Consortium | The Lighthouse Lab in Milton Keynes and Alex Alderton, Roberto Amato, Jeffrey Barrett, Sonia Goncalves, Ewan Harrison, David K. Jackson, Ian Johnston, Dominic Kwiatkowski, Cordelia Langford, John Sillitoe on behalf of the Wellcome Sanger Institute COVID-19 Surveillance Team |

| | | | |
|---|---|---|---|
| EPI_ISL_2435862, EPI_ISL_2435863 | Lighthouse Lab in Glasgow | Wellcome Sanger Institute for the COVID-19 Genomics UK (COG-UK) Consortium | Harper VanSteenhouse, Yumi Kasai, David Gray, Carol Clugston, Anna Dominiczak and Alex Alderton, Roberto Amato, Jeffrey Barrett, Sonia Goncalves, Ewan Harrison, David K. Jackson, Ian Johnston, Dominic Kwiatkowski, Cordelia Langford, John Sillitoe on behalf of the Wellcome Sanger Institute COVID-19 Surveillance Team |
| EPI_ISL_2435865 | Lighthouse Lab in Alderley Park | Wellcome Sanger Institute for the COVID-19 Genomics UK (COG-UK) Consortium | Jacquelyn Wynn, Mairead Hyland, The Lighthouse Lab in Alderley Park and Alex Alderton, Roberto Amato, Jeffrey Barrett, Sonia Goncalves, Ewan Harrison, David K. Jackson, Ian Johnston, Dominic Kwiatkowski, Cordelia Langford, John Sillitoe on behalf of the Wellcome Sanger Institute COVID-19 Surveillance Team |
| EPI_ISL_2435866 | Lighthouse Lab in Milton Keynes | Wellcome Sanger Institute for the COVID-19 Genomics UK (COG-UK) Consortium | The Lighthouse Lab in Milton Keynes and Alex Alderton, Roberto Amato, Jeffrey Barrett, Sonia Goncalves, Ewan Harrison, David K. Jackson, Ian Johnston, Dominic Kwiatkowski, Cordelia Langford, John Sillitoe on behalf of the Wellcome Sanger Institute COVID-19 Surveillance Team |
| EPI_ISL_2435867, EPI_ISL_2435868, EPI_ISL_2435869, EPI_ISL_2435872, EPI_ISL_2435873, EPI_ISL_2435874, EPI_ISL_2435875, EPI_ISL_2435876, EPI_ISL_2435877, EPI_ISL_2435879, EPI_ISL_2435882, EPI_ISL_2435884, EPI_ISL_2435885, EPI_ISL_2435886, EPI_ISL_2435887, EPI_ISL_2435888, EPI_ISL_2435889, EPI_ISL_2435890, EPI_ISL_2435891, EPI_ISL_2435892, EPI_ISL_2435893, EPI_ISL_2435894, EPI_ISL_2435895, EPI_ISL_2435896, EPI_ISL_2435897, EPI_ISL_2435899, EPI_ISL_2435901, EPI_ISL_2435902, EPI_ISL_2435903, EPI_ISL_2435904, EPI_ISL_2435905, EPI_ISL_2435906, EPI_ISL_2435907, EPI_ISL_2435908, EPI_ISL_2435909, EPI_ISL_2435910, EPI_ISL_2435912, EPI_ISL_2435913, EPI_ISL_2435915, EPI_ISL_2435916, EPI_ISL_2435917, EPI_ISL_2435918, EPI_ISL_2435920, EPI_ISL_2435921, EPI_ISL_2435922, EPI_ISL_2435923, EPI_ISL_2435925, EPI_ISL_2435926, EPI_ISL_2435927, EPI_ISL_2435928, EPI_ISL_2435929, EPI_ISL_2435930, EPI_ISL_2435931, EPI_ISL_2435932, EPI_ISL_2435933, EPI_ISL_2435934, EPI_ISL_2435935, EPI_ISL_2435936, EPI_ISL_2435937, EPI_ISL_2435938, EPI_ISL_2435939, EPI_ISL_2435940, EPI_ISL_2435941, EPI_ISL_2435942, EPI_ISL_2435943, EPI_ISL_2435944, EPI_ISL_2435946, EPI_ISL_2435947, EPI_ISL_2435949, EPI_ISL_2435950, EPI_ISL_2435952, EPI_ISL_2435953, EPI_ISL_2435954, EPI_ISL_2435955, EPI_ISL_2435956, EPI_ISL_2435957, EPI_ISL_2435959, EPI_ISL_2435960, EPI_ISL_2435961, EPI_ISL_2435962, EPI_ISL_2435963, EPI_ISL_2435964, EPI_ISL_2435965, EPI_ISL_2435966, EPI_ISL_2435967, EPI_ISL_2435968, EPI_ISL_2435969, EPI_ISL_2435970, EPI_ISL_2435972, EPI_ISL_2435973, EPI_ISL_2435974, EPI_ISL_2435975, EPI_ISL_2435976, EPI_ISL_2435977, EPI_ISL_2435978, EPI_ISL_2435980, EPI_ISL_2435982, EPI_ISL_2435983, EPI_ISL_2435984, EPI_ISL_2435985, EPI_ISL_2435986, EPI_ISL_2435988, EPI_ISL_2435989, EPI_ISL_2435990, EPI_ISL_2435994, EPI_ISL_2435995, EPI_ISL_2435996, EPI_ISL_2435997, EPI_ISL_2435998, EPI_ISL_2435999, EPI_ISL_2436001, EPI_ISL_2436002, EPI_ISL_2436003, EPI_ISL_2436004, EPI_ISL_2436005, EPI_ISL_2436008, EPI_ISL_2436009, EPI_ISL_2436010, EPI_ISL_2436011, EPI_ISL_2436013, EPI_ISL_2436014, EPI_ISL_2436015, EPI_ISL_2436016, EPI_ISL_2436017, EPI_ISL_2436018, EPI_ISL_2436019, EPI_ISL_2436020, EPI_ISL_2436021, EPI_ISL_2436022, EPI_ISL_2436024, EPI_ISL_2436025, EPI_ISL_2436026, EPI_ISL_2436027, EPI_ISL_2436028, EPI_ISL_2436029, EPI_ISL_2436031, EPI_ISL_2436032, EPI_ISL_2436033, EPI_ISL_2436034, EPI_ISL_2436035, EPI_ISL_2436036, EPI_ISL_2436037, EPI_ISL_2436038, EPI_ISL_2436039, EPI_ISL_2436040, EPI_ISL_2436041, EPI_ISL_2436042, EPI_ISL_2436043, EPI_ISL_2436046, EPI_ISL_2436047, EPI_ISL_2436049, EPI_ISL_2436050, EPI_ISL_2436051, EPI_ISL_2436052, EPI_ISL_2436053, EPI_ISL_2436054, EPI_ISL_2436055, EPI_ISL_2436056, EPI_ISL_2436057, EPI_ISL_2436058, EPI_ISL_2436059, EPI_ISL_2436061, EPI_ISL_2436062, EPI_ISL_2436063, EPI_ISL_2436064, EPI_ISL_2436065, EPI_ISL_2436066, EPI_ISL_2436067, EPI_ISL_2436068, EPI_ISL_2436069, EPI_ISL_2436070, EPI_ISL_2436071, EPI_ISL_2436072, EPI_ISL_2436074, EPI_ISL_2436075, EPI_ISL_2436076, EPI_ISL_2436077, EPI_ISL_2436078, EPI_ISL_2436079, EPI_ISL_2436080, EPI_ISL_2436081, EPI_ISL_2436082, EPI_ISL_2436083, EPI_ISL_2436085, EPI_ISL_2436086, EPI_ISL_2436087, EPI_ISL_2436088, EPI_ISL_2436089, EPI_ISL_2436090, EPI_ISL_2436091, EPI_ISL_2436092, EPI_ISL_2436093, EPI_ISL_2436094, EPI_ISL_2436095, EPI_ISL_2436096, EPI_ISL_2436097, EPI_ISL_2436098, EPI_ISL_2436100, EPI_ISL_2436101, EPI_ISL_2436102, EPI_ISL_2436103, EPI_ISL_2436104, EPI_ISL_2436106, EPI_ISL_2436107, EPI_ISL_2436109, EPI_ISL_2436110, EPI_ISL_2436112, EPI_ISL_2436113, EPI_ISL_2436114, EPI_ISL_2436115, EPI_ISL_2436116, EPI_ISL_2436117, EPI_ISL_2436119, EPI_ISL_2436120, EPI_ISL_2436122, EPI_ISL_2436123, EPI_ISL_2436124, EPI_ISL_2436125, EPI_ISL_2436126, EPI_ISL_2436127, EPI_ISL_2436128, EPI_ISL_2436129, EPI_ISL_2436130, EPI_ISL_2436131, EPI_ISL_2436133, EPI_ISL_2436134, EPI_ISL_2436135, EPI_ISL_2436137, EPI_ISL_2436138, EPI_ISL_2436139, EPI_ISL_2436140, EPI_ISL_2436141, EPI_ISL_2436142, EPI_ISL_2436144, EPI_ISL_2436145, EPI_ISL_2436146, EPI_ISL_2436147, EPI_ISL_2436148, EPI_ISL_2436149, EPI_ISL_2436150, EPI_ISL_2436151, EPI_ISL_2436152, EPI_ISL_2436153, EPI_ISL_2436154, EPI_ISL_2436155, EPI_ISL_2436156, EPI_ISL_2436158, EPI_ISL_2436159, EPI_ISL_2436160, EPI_ISL_2436161, EPI_ISL_2436162, EPI_ISL_2436163, EPI_ISL_2436164, EPI_ISL_2436165, EPI_ISL_2436167, EPI_ISL_2436169, EPI_ISL_2436170, EPI_ISL_2436171, EPI_ISL_2436172, EPI_ISL_2436173, EPI_ISL_2436174, EPI_ISL_2436175, EPI_ISL_2436176, EPI_ISL_2436177, EPI_ISL_2436178, EPI_ISL_2436179, EPI_ISL_2436181, EPI_ISL_2436182, EPI_ISL_2436183, EPI_ISL_2436184, EPI_ISL_2436187, EPI_ISL_2436188, EPI_ISL_2436189, EPI_ISL_2436190, EPI_ISL_2436191, EPI_ISL_2436192, EPI_ISL_2436193, EPI_ISL_2436194, EPI_ISL_2436195, EPI_ISL_2436196, EPI_ISL_2436197, EPI_ISL_2436198, EPI_ISL_2436199, EPI_ISL_2436200, EPI_ISL_2436202, EPI_ISL_2436203, EPI_ISL_2436204, EPI_ISL_2436205, EPI_ISL_2436206, EPI_ISL_2436207, EPI_ISL_2436208, EPI_ISL_2436209, EPI_ISL_2436212, EPI_ISL_2436214, EPI_ISL_2436215, EPI_ISL_2436216, EPI_ISL_2436217, EPI_ISL_2436218, EPI_ISL_2436219, EPI_ISL_2436220 | | | |
| see above | Lighthouse Lab in Alderley Park | Wellcome Sanger Institute for the COVID-19 Genomics UK (COG-UK) Consortium | Jacquelyn Wynn, Mairead Hyland, The Lighthouse Lab in Alderley Park and Alex Alderton, Roberto Amato, Jeffrey Barrett, Sonia Goncalves, Ewan Harrison, David K. Jackson, Ian Johnston, Dominic Kwiatkowski, Cordelia Langford, John Sillitoe on behalf of the Wellcome Sanger Institute COVID-19 Surveillance Team |
| EPI_ISL_2436222 | Lighthouse Lab in Glasgow | Wellcome Sanger Institute for the COVID-19 Genomics UK (COG-UK) Consortium | Harper VanSteenhouse, Yumi Kasai, David Gray, Carol Clugston, Anna Dominiczak and Alex Alderton, Roberto Amato, Jeffrey Barrett, Sonia Goncalves, Ewan Harrison, David K. Jackson, Ian Johnston, Dominic Kwiatkowski, Cordelia Langford, John Sillitoe on behalf of the Wellcome Sanger Institute COVID-19 Surveillance Team |
| EPI_ISL_2436223 | Lighthouse Laboratory Plymouth | Wellcome Sanger Institute for the COVID-19 Genomics UK (COG-UK) Consortium | Lighthouse Laboratory Plymouth and Alex Alderton, Roberto Amato, Jeffrey Barrett, Sonia Goncalves, Ewan Harrison, David K. Jackson, Ian Johnston, Dominic Kwiatkowski, Cordelia Langford, John Sillitoe on behalf of the Wellcome Sanger Institute COVID-19 Surveillance Team |
| EPI_ISL_2436224, EPI_ISL_2436225 | Lighthouse Lab in Alderley Park | Wellcome Sanger Institute for the COVID-19 Genomics UK (COG-UK) Consortium | Jacquelyn Wynn, Mairead Hyland, The Lighthouse Lab in Alderley Park and Alex Alderton, Roberto Amato, Jeffrey Barrett, Sonia Goncalves, Ewan Harrison, David K. Jackson, Ian Johnston, Dominic Kwiatkowski, Cordelia Langford, John Sillitoe on behalf of the Wellcome Sanger Institute COVID-19 Surveillance Team |
| EPI_ISL_2436226 | Lighthouse Laboratory Plymouth | Wellcome Sanger Institute for the COVID-19 Genomics UK (COG-UK) Consortium | Lighthouse Laboratory Plymouth and Alex Alderton, Roberto Amato, Jeffrey Barrett, Sonia Goncalves, Ewan Harrison, David K. Jackson, Ian Johnston, Dominic Kwiatkowski, Cordelia Langford, John Sillitoe on behalf of the Wellcome Sanger Institute COVID-19 Surveillance Team |
| EPI_ISL_2436227 | Lighthouse Lab in Glasgow | Wellcome Sanger Institute for the COVID-19 Genomics UK (COG-UK) Consortium | Harper VanSteenhouse, Yumi Kasai, David Gray, Carol Clugston, Anna Dominiczak and Alex Alderton, Roberto Amato, Jeffrey Barrett, Sonia Goncalves, Ewan Harrison, David K. Jackson, Ian Johnston, Dominic Kwiatkowski, Cordelia Langford, John Sillitoe on behalf of the Wellcome Sanger Institute COVID-19 Surveillance Team |
| EPI_ISL_2436228 | Lighthouse Lab in Alderley Park | Wellcome Sanger Institute for the COVID-19 Genomics UK (COG-UK) Consortium | Jacquelyn Wynn, Mairead Hyland, The Lighthouse Lab in Alderley Park and Alex Alderton, Roberto Amato, Jeffrey Barrett, Sonia Goncalves, Ewan Harrison, David K. Jackson, Ian Johnston, Dominic Kwiatkowski, Cordelia Langford, John Sillitoe on behalf of the Wellcome Sanger Institute COVID-19 Surveillance Team |
| EPI_ISL_2436229 | Lighthouse Lab in Glasgow | Wellcome Sanger Institute for the COVID-19 Genomics UK (COG-UK) Consortium | Harper VanSteenhouse, Yumi Kasai, David Gray, Carol Clugston, Anna Dominiczak and Alex Alderton, Roberto Amato, Jeffrey Barrett, Sonia Goncalves, Ewan Harrison, David K. Jackson, Ian Johnston, Dominic Kwiatkowski, Cordelia Langford, John Sillitoe on behalf of the Wellcome Sanger Institute COVID-19 Surveillance Team |
| EPI_ISL_2436230 | Lighthouse Laboratory Plymouth | Wellcome Sanger Institute for the COVID-19 Genomics UK (COG-UK) Consortium | Lighthouse Laboratory Plymouth and Alex Alderton, Roberto Amato, Jeffrey Barrett, Sonia Goncalves, Ewan Harrison, David K. Jackson, Ian Johnston, Dominic Kwiatkowski, Cordelia Langford, John Sillitoe on behalf of the Wellcome Sanger Institute COVID-19 Surveillance Team |
| EPI_ISL_2436231 | Lighthouse Lab in Glasgow | Wellcome Sanger Institute for the COVID-19 Genomics UK (COG-UK) Consortium | Harper VanSteenhouse, Yumi Kasai, David Gray, Carol Clugston, Anna Dominiczak and Alex Alderton, Roberto Amato, Jeffrey Barrett, Sonia Goncalves, Ewan Harrison, David K. Jackson, Ian Johnston, Dominic Kwiatkowski, Cordelia Langford, John Sillitoe on behalf of the Wellcome Sanger Institute COVID-19 Surveillance Team |
| EPI_ISL_2436232 | Lighthouse Lab in Milton Keynes | Wellcome Sanger Institute for the COVID-19 Genomics UK (COG-UK) Consortium | The Lighthouse Lab in Milton Keynes and Alex Alderton, Roberto Amato, Jeffrey Barrett, Sonia Goncalves, Ewan Harrison, David K. Jackson, Ian Johnston, Dominic Kwiatkowski, Cordelia Langford, John Sillitoe on behalf of the Wellcome Sanger Institute COVID-19 Surveillance Team |
| EPI_ISL_2436233, EPI_ISL_2436234 | Lighthouse Lab in Glasgow | Wellcome Sanger Institute for the COVID-19 Genomics UK (COG-UK) Consortium | Harper VanSteenhouse, Yumi Kasai, David Gray, Carol Clugston, Anna Dominiczak and Alex Alderton, Roberto Amato, Jeffrey Barrett, Sonia Goncalves, Ewan Harrison, David K. Jackson, Ian Johnston, Dominic Kwiatkowski, Cordelia Langford, John Sillitoe on behalf of the Wellcome Sanger Institute COVID-19 Surveillance Team |
| EPI_ISL_2436235 | Lighthouse Lab in Milton Keynes | Wellcome Sanger Institute for the COVID-19 Genomics UK (COG-UK) Consortium | The Lighthouse Lab in Milton Keynes and Alex Alderton, Roberto Amato, Jeffrey Barrett, Sonia Goncalves, Ewan Harrison, David K. Jackson, Ian Johnston, Dominic Kwiatkowski, Cordelia Langford, John Sillitoe on behalf of the Wellcome Sanger Institute COVID-19 Surveillance Team |
| EPI_ISL_2436237, EPI_ISL_2436238, EPI_ISL_2436239, EPI_ISL_2436240 | Health Services Laboratories | Wellcome Sanger Institute for the COVID-19 Genomics UK (COG-UK) Consortium | Health Services Laboratories and Alex Alderton, Roberto Amato, Jeffrey Barrett, Sonia Goncalves, Ewan Harrison, David K. Jackson, Ian Johnston, Dominic Kwiatkowski, Cordelia Langford, John Sillitoe on behalf of the Wellcome Sanger Institute COVID-19 Surveillance Team |
| EPI_ISL_2436241, EPI_ISL_2436242 | Lighthouse Lab in Glasgow | Wellcome Sanger Institute for the COVID-19 Genomics UK (COG-UK) Consortium | Harper VanSteenhouse, Yumi Kasai, David Gray, Carol Clugston, Anna Dominiczak and Alex Alderton, Roberto Amato, Jeffrey Barrett, Sonia Goncalves, Ewan Harrison, David K. Jackson, Ian Johnston, Dominic Kwiatkowski, Cordelia Langford, John Sillitoe on behalf of the Wellcome Sanger Institute COVID-19 Surveillance Team |
| EPI_ISL_2436243 | Lighthouse Lab in Alderley Park | Wellcome Sanger Institute for the COVID-19 Genomics UK (COG-UK) Consortium | Jacquelyn Wynn, Mairead Hyland, The Lighthouse Lab in Alderley Park and Alex Alderton, Roberto Amato, Jeffrey Barrett, Sonia Goncalves, Ewan Harrison, David K. Jackson, Ian Johnston, Dominic Kwiatkowski, Cordelia Langford, John Sillitoe on behalf of the Wellcome Sanger Institute COVID-19 Surveillance Team |
| EPI_ISL_2436244 | Lighthouse Laboratory Plymouth | Wellcome Sanger Institute for the COVID-19 Genomics UK (COG-UK) Consortium | Lighthouse Laboratory Plymouth and Alex Alderton, Roberto Amato, Jeffrey Barrett, Sonia Goncalves, Ewan Harrison, David K. Jackson, Ian Johnston, Dominic Kwiatkowski, Cordelia Langford, John Sillitoe on behalf of the Wellcome Sanger Institute COVID-19 Surveillance Team |
| EPI_ISL_2436245, EPI_ISL_2436246 | Lighthouse Lab in Milton Keynes | Wellcome Sanger Institute for the COVID-19 Genomics UK (COG-UK) Consortium | The Lighthouse Lab in Milton Keynes and Alex Alderton, Roberto Amato, Jeffrey Barrett, Sonia Goncalves, Ewan Harrison, David K. Jackson, Ian Johnston, Dominic Kwiatkowski, Cordelia Langford, John Sillitoe on behalf of the Wellcome Sanger Institute COVID-19 Surveillance Team |
| EPI_ISL_2436247 | Lighthouse Laboratory Plymouth | Wellcome Sanger Institute for the COVID-19 Genomics UK (COG-UK) Consortium | Lighthouse Laboratory Plymouth and Alex Alderton, Roberto Amato, Jeffrey Barrett, Sonia Goncalves, Ewan Harrison, David K. Jackson, Ian Johnston, Dominic Kwiatkowski, Cordelia Langford, John Sillitoe on behalf of the Wellcome Sanger Institute COVID-19 Surveillance Team |
| EPI_ISL_2436248 | Lighthouse Lab in Alderley Park | Wellcome Sanger Institute for the COVID-19 Genomics UK (COG-UK) Consortium | Jacquelyn Wynn, Mairead Hyland, The Lighthouse Lab in Alderley Park and Alex Alderton, Roberto Amato, Jeffrey Barrett, Sonia Goncalves, Ewan Harrison, David K. Jackson, Ian Johnston, Dominic Kwiatkowski, Cordelia Langford, John Sillitoe on behalf of the Wellcome Sanger Institute COVID-19 Surveillance Team |
| EPI_ISL_2436249 | Lighthouse Lab in Milton Keynes | Wellcome Sanger Institute for the COVID-19 Genomics UK (COG-UK) Consortium | The Lighthouse Lab in Milton Keynes and Alex Alderton, Roberto Amato, Jeffrey Barrett, Sonia Goncalves, Ewan Harrison, David K. Jackson, Ian Johnston, Dominic Kwiatkowski, Cordelia Langford, John Sillitoe on behalf of the Wellcome Sanger Institute COVID-19 Surveillance Team |
| EPI_ISL_2436250 | Health Services Laboratories | Wellcome Sanger Institute for the COVID-19 Genomics UK (COG-UK) Consortium | Health Services Laboratories and Alex Alderton, Roberto Amato, Jeffrey Barrett, Sonia Goncalves, Ewan Harrison, David K. Jackson, Ian Johnston, Dominic Kwiatkowski, Cordelia Langford, John Sillitoe on behalf of the Wellcome Sanger Institute COVID-19 Surveillance Team |
| EPI_ISL_2436251 | Lighthouse Lab in Alderley Park | Wellcome Sanger Institute for the COVID-19 Genomics UK (COG-UK) Consortium | Jacquelyn Wynn, Mairead Hyland, The Lighthouse Lab in Alderley Park and Alex Alderton, Roberto Amato, Jeffrey Barrett, Sonia Goncalves, Ewan Harrison, David K. Jackson, Ian Johnston, Dominic Kwiatkowski, Cordelia Langford, John Sillitoe on behalf of the Wellcome Sanger Institute COVID-19 Surveillance Team |
| EPI_ISL_2436253 | Lighthouse Lab in Glasgow | Wellcome Sanger Institute for the COVID-19 Genomics UK (COG-UK) Consortium | Harper VanSteenhouse, Yumi Kasai, David Gray, Carol Clugston, Anna Dominiczak and Alex Alderton, Roberto Amato, Jeffrey Barrett, Sonia Goncalves, Ewan Harrison, David K. Jackson, Ian Johnston, Dominic Kwiatkowski, Cordelia Langford, John Sillitoe on behalf of the Wellcome Sanger Institute COVID-19 Surveillance Team |
| EPI_ISL_2436254 | Lighthouse Lab in Alderley Park | Wellcome Sanger Institute for the COVID-19 Genomics UK (COG-UK) Consortium | Jacquelyn Wynn, Mairead Hyland, The Lighthouse Lab in Alderley Park and Alex Alderton, Roberto Amato, Jeffrey Barrett, Sonia Goncalves, Ewan Harrison, David K. Jackson, Ian Johnston, Dominic Kwiatkowski, Cordelia Langford, John Sillitoe on behalf of the Wellcome Sanger Institute COVID-19 Surveillance Team |
| EPI_ISL_2436255, EPI_ISL_2436256 | Lighthouse Lab in Milton Keynes | Wellcome Sanger Institute for the COVID-19 Genomics UK (COG-UK) Consortium | The Lighthouse Lab in Milton Keynes and Alex Alderton, Roberto Amato, Jeffrey Barrett, Sonia Goncalves, Ewan Harrison, David K. Jackson, Ian Johnston, Dominic Kwiatkowski, Cordelia Langford, John Sillitoe on behalf of the Wellcome Sanger Institute COVID-19 Surveillance Team |
| EPI_ISL_2436257, EPI_ISL_2436258 | Lighthouse Lab in Glasgow | Wellcome Sanger Institute for the COVID-19 Genomics UK (COG-UK) Consortium | Harper VanSteenhouse, Yumi Kasai, David Gray, Carol Clugston, Anna Dominiczak and Alex Alderton, Roberto Amato, Jeffrey Barrett, Sonia Goncalves, Ewan Harrison, David K. Jackson, Ian Johnston, Dominic Kwiatkowski, Cordelia Langford, John Sillitoe on behalf of the Wellcome Sanger Institute COVID-19 Surveillance Team |
| EPI_ISL_2436260 | Lighthouse Lab in Alderley Park | Wellcome Sanger Institute for the COVID-19 Genomics UK (COG-UK) Consortium | Jacquelyn Wynn, Mairead Hyland, The Lighthouse Lab in Alderley Park and Alex Alderton, Roberto Amato, Jeffrey Barrett, Sonia Goncalves, Ewan Harrison, David K. Jackson, Ian Johnston, Dominic Kwiatkowski, Cordelia Langford, John Sillitoe on behalf of the Wellcome Sanger Institute COVID-19 Surveillance Team |
| EPI_ISL_2436261 | Health Services Laboratories | Wellcome Sanger Institute for the COVID-19 Genomics UK (COG-UK) Consortium | Health Services Laboratories and Alex Alderton, Roberto Amato, Jeffrey Barrett, Sonia Goncalves, Ewan Harrison, David K. Jackson, Ian Johnston, Dominic Kwiatkowski, Cordelia Langford, John Sillitoe on behalf of the Wellcome Sanger Institute COVID-19 Surveillance Team |
| EPI_ISL_2436263 | Lighthouse Lab in Alderley Park | Wellcome Sanger Institute for the COVID-19 Genomics UK (COG-UK) Consortium | Jacquelyn Wynn, Mairead Hyland, The Lighthouse Lab in Alderley Park and Alex Alderton, Roberto Amato, Jeffrey Barrett, Sonia Goncalves, Ewan Harrison, David K. Jackson, Ian Johnston, Dominic Kwiatkowski, Cordelia Langford, John Sillitoe on behalf of the Wellcome Sanger Institute COVID-19 Surveillance Team |
| EPI_ISL_2436264 | Health Services Laboratories | Wellcome Sanger Institute for the COVID-19 Genomics UK (COG-UK) Consortium | Health Services Laboratories and Alex Alderton, Roberto Amato, Jeffrey Barrett, Sonia Goncalves, Ewan Harrison, David K. Jackson, Ian Johnston, Dominic Kwiatkowski, Cordelia Langford, John Sillitoe on behalf of the Wellcome Sanger Institute COVID-19 Surveillance Team |
| EPI_ISL_2436267 | Lighthouse Lab in Glasgow | Wellcome Sanger Institute for the COVID-19 Genomics UK (COG-UK) Consortium | Harper VanSteenhouse, Yumi Kasai, David Gray, Carol Clugston, Anna Dominiczak and Alex Alderton, Roberto Amato, Jeffrey Barrett, Sonia Goncalves, Ewan Harrison, David K. Jackson, Ian Johnston, Dominic Kwiatkowski, Cordelia Langford, John Sillitoe on behalf of the Wellcome Sanger Institute COVID-19 Surveillance Team |
| EPI_ISL_2436268 | Lighthouse Laboratory Plymouth | Wellcome Sanger Institute for the COVID-19 Genomics UK (COG-UK) Consortium | Lighthouse Laboratory Plymouth and Alex Alderton, Roberto Amato, Jeffrey Barrett, Sonia Goncalves, Ewan Harrison, David K. Jackson, Ian Johnston, Dominic Kwiatkowski, Cordelia Langford, John Sillitoe on behalf of the Wellcome Sanger Institute COVID-19 Surveillance Team |
| EPI_ISL_2436269 | Lighthouse Lab in Glasgow | Wellcome Sanger Institute for the COVID-19 Genomics UK (COG-UK) Consortium | Harper VanSteenhouse, Yumi Kasai, David Gray, Carol Clugston, Anna Dominiczak and Alex Alderton, Roberto Amato, Jeffrey Barrett, Sonia Goncalves, Ewan Harrison, David K. Jackson, Ian Johnston, Dominic Kwiatkowski, Cordelia Langford, John Sillitoe on behalf of the Wellcome Sanger Institute COVID-19 Surveillance Team |
| EPI_ISL_2436270 | Health Services Laboratories | Wellcome Sanger Institute for the COVID-19 Genomics UK (COG-UK) Consortium | Health Services Laboratories and Alex Alderton, Roberto Amato, Jeffrey Barrett, Sonia Goncalves, Ewan Harrison, David K. Jackson, Ian Johnston, Dominic Kwiatkowski, Cordelia Langford, John Sillitoe on behalf of the Wellcome Sanger Institute COVID-19 Surveillance Team |

| | | | |
|---|---|---|---|
| EPI_ISL_2436273 | Lighthouse Lab in Glasgow | Wellcome Sanger Institute for the COVID-19 Genomics UK (COG-UK) Consortium | Harper VanSteenhouse, Yumi Kasai, David Gray, Carol Clugston, Anna Dominiczak and Alex Alderton, Roberto Amato, Jeffrey Barrett, Sonia Goncalves, Ewan Harrison, David K. Jackson, Ian Johnston, Dominic Kwiatkowski, Cordelia Langford, John Sillitoe on behalf of the Wellcome Sanger Institute COVID-19 Surveillance Team |
| EPI_ISL_2436274 | Health Services Laboratories | Wellcome Sanger Institute for the COVID-19 Genomics UK (COG-UK) Consortium | Health Services Laboratories and Alex Alderton, Roberto Amato, Jeffrey Barrett, Sonia Goncalves, Ewan Harrison, David K. Jackson, Ian Johnston, Dominic Kwiatkowski, Cordelia Langford, John Sillitoe on behalf of the Wellcome Sanger Institute COVID-19 Surveillance Team |
| EPI_ISL_2436275 | Lighthouse Lab in Glasgow | Wellcome Sanger Institute for the COVID-19 Genomics UK (COG-UK) Consortium | Harper VanSteenhouse, Yumi Kasai, David Gray, Carol Clugston, Anna Dominiczak and Alex Alderton, Roberto Amato, Jeffrey Barrett, Sonia Goncalves, Ewan Harrison, David K. Jackson, Ian Johnston, Dominic Kwiatkowski, Cordelia Langford, John Sillitoe on behalf of the Wellcome Sanger Institute COVID-19 Surveillance Team |
| EPI_ISL_2436276 | Lighthouse Laboratory Plymouth | Wellcome Sanger Institute for the COVID-19 Genomics UK (COG-UK) Consortium | Lighthouse Laboratory Plymouth and Alex Alderton, Roberto Amato, Jeffrey Barrett, Sonia Goncalves, Ewan Harrison, David K. Jackson, Ian Johnston, Dominic Kwiatkowski, Cordelia Langford, John Sillitoe on behalf of the Wellcome Sanger Institute COVID-19 Surveillance Team |
| EPI_ISL_2436277 | Lighthouse Lab in Glasgow | Wellcome Sanger Institute for the COVID-19 Genomics UK (COG-UK) Consortium | Harper VanSteenhouse, Yumi Kasai, David Gray, Carol Clugston, Anna Dominiczak and Alex Alderton, Roberto Amato, Jeffrey Barrett, Sonia Goncalves, Ewan Harrison, David K. Jackson, Ian Johnston, Dominic Kwiatkowski, Cordelia Langford, John Sillitoe on behalf of the Wellcome Sanger Institute COVID-19 Surveillance Team |
| EPI_ISL_2436278 | Health Services Laboratories | Wellcome Sanger Institute for the COVID-19 Genomics UK (COG-UK) Consortium | Health Services Laboratories and Alex Alderton, Roberto Amato, Jeffrey Barrett, Sonia Goncalves, Ewan Harrison, David K. Jackson, Ian Johnston, Dominic Kwiatkowski, Cordelia Langford, John Sillitoe on behalf of the Wellcome Sanger Institute COVID-19 Surveillance Team |
| EPI_ISL_2436279, EPI_ISL_2436281 | Lighthouse Lab in Milton Keynes | Wellcome Sanger Institute for the COVID-19 Genomics UK (COG-UK) Consortium | The Lighthouse Lab in Milton Keynes and Alex Alderton, Roberto Amato, Jeffrey Barrett, Sonia Goncalves, Ewan Harrison, David K. Jackson, Ian Johnston, Dominic Kwiatkowski, Cordelia Langford, John Sillitoe on behalf of the Wellcome Sanger Institute COVID-19 Surveillance Team |
| EPI_ISL_2436282 | Lighthouse Lab in Alderley Park | Wellcome Sanger Institute for the COVID-19 Genomics UK (COG-UK) Consortium | Jacquelyn Wynn, Mairead Hyland, The Lighthouse Lab in Alderley Park and Alex Alderton, Roberto Amato, Jeffrey Barrett, Sonia Goncalves, Ewan Harrison, David K. Jackson, Ian Johnston, Dominic Kwiatkowski, Cordelia Langford, John Sillitoe on behalf of the Wellcome Sanger Institute COVID-19 Surveillance Team |
| EPI_ISL_2436283 | Lighthouse Lab in Milton Keynes | Wellcome Sanger Institute for the COVID-19 Genomics UK (COG-UK) Consortium | The Lighthouse Lab in Milton Keynes and Alex Alderton, Roberto Amato, Jeffrey Barrett, Sonia Goncalves, Ewan Harrison, David K. Jackson, Ian Johnston, Dominic Kwiatkowski, Cordelia Langford, John Sillitoe on behalf of the Wellcome Sanger Institute COVID-19 Surveillance Team |
| EPI_ISL_2436284 | Lighthouse Lab in Alderley Park | Wellcome Sanger Institute for the COVID-19 Genomics UK (COG-UK) Consortium | Jacquelyn Wynn, Mairead Hyland, The Lighthouse Lab in Alderley Park and Alex Alderton, Roberto Amato, Jeffrey Barrett, Sonia Goncalves, Ewan Harrison, David K. Jackson, Ian Johnston, Dominic Kwiatkowski, Cordelia Langford, John Sillitoe on behalf of the Wellcome Sanger Institute COVID-19 Surveillance Team |
| EPI_ISL_2436286 | Health Services Laboratories | Wellcome Sanger Institute for the COVID-19 Genomics UK (COG-UK) Consortium | Health Services Laboratories and Alex Alderton, Roberto Amato, Jeffrey Barrett, Sonia Goncalves, Ewan Harrison, David K. Jackson, Ian Johnston, Dominic Kwiatkowski, Cordelia Langford, John Sillitoe on behalf of the Wellcome Sanger Institute COVID-19 Surveillance Team |
| EPI_ISL_2436287 | Lighthouse Lab in Glasgow | Wellcome Sanger Institute for the COVID-19 Genomics UK (COG-UK) Consortium | Harper VanSteenhouse, Yumi Kasai, David Gray, Carol Clugston, Anna Dominiczak and Alex Alderton, Roberto Amato, Jeffrey Barrett, Sonia Goncalves, Ewan Harrison, David K. Jackson, Ian Johnston, Dominic Kwiatkowski, Cordelia Langford, John Sillitoe on behalf of the Wellcome Sanger Institute COVID-19 Surveillance Team |
| EPI_ISL_2436288, EPI_ISL_2436290 | Lighthouse Laboratory Plymouth | Wellcome Sanger Institute for the COVID-19 Genomics UK (COG-UK) Consortium | Lighthouse Laboratory Plymouth and Alex Alderton, Roberto Amato, Jeffrey Barrett, Sonia Goncalves, Ewan Harrison, David K. Jackson, Ian Johnston, Dominic Kwiatkowski, Cordelia Langford, John Sillitoe on behalf of the Wellcome Sanger Institute COVID-19 Surveillance Team |
| EPI_ISL_2436291, EPI_ISL_2436292 | Lighthouse Lab in Alderley Park | Wellcome Sanger Institute for the COVID-19 Genomics UK (COG-UK) Consortium | Jacquelyn Wynn, Mairead Hyland, The Lighthouse Lab in Alderley Park and Alex Alderton, Roberto Amato, Jeffrey Barrett, Sonia Goncalves, Ewan Harrison, David K. Jackson, Ian Johnston, Dominic Kwiatkowski, Cordelia Langford, John Sillitoe on behalf of the Wellcome Sanger Institute COVID-19 Surveillance Team |
| EPI_ISL_2436293 | Lighthouse Lab in Milton Keynes | Wellcome Sanger Institute for the COVID-19 Genomics UK (COG-UK) Consortium | The Lighthouse Lab in Milton Keynes and Alex Alderton, Roberto Amato, Jeffrey Barrett, Sonia Goncalves, Ewan Harrison, David K. Jackson, Ian Johnston, Dominic Kwiatkowski, Cordelia Langford, John Sillitoe on behalf of the Wellcome Sanger Institute COVID-19 Surveillance Team |
| EPI_ISL_2436294 | Health Services Laboratories | Wellcome Sanger Institute for the COVID-19 Genomics UK (COG-UK) Consortium | Health Services Laboratories and Alex Alderton, Roberto Amato, Jeffrey Barrett, Sonia Goncalves, Ewan Harrison, David K. Jackson, Ian Johnston, Dominic Kwiatkowski, Cordelia Langford, John Sillitoe on behalf of the Wellcome Sanger Institute COVID-19 Surveillance Team |
| EPI_ISL_2436295 | Lighthouse Lab in Milton Keynes | Wellcome Sanger Institute for the COVID-19 Genomics UK (COG-UK) Consortium | The Lighthouse Lab in Milton Keynes and Alex Alderton, Roberto Amato, Jeffrey Barrett, Sonia Goncalves, Ewan Harrison, David K. Jackson, Ian Johnston, Dominic Kwiatkowski, Cordelia Langford, John Sillitoe on behalf of the Wellcome Sanger Institute COVID-19 Surveillance Team |
| EPI_ISL_2436296 | Lighthouse Lab in Glasgow | Wellcome Sanger Institute for the COVID-19 Genomics UK (COG-UK) Consortium | Harper VanSteenhouse, Yumi Kasai, David Gray, Carol Clugston, Anna Dominiczak and Alex Alderton, Roberto Amato, Jeffrey Barrett, Sonia Goncalves, Ewan Harrison, David K. Jackson, Ian Johnston, Dominic Kwiatkowski, Cordelia Langford, John Sillitoe on behalf of the Wellcome Sanger Institute COVID-19 Surveillance Team |
| EPI_ISL_2436298 | Health Services Laboratories | Wellcome Sanger Institute for the COVID-19 Genomics UK (COG-UK) Consortium | Health Services Laboratories and Alex Alderton, Roberto Amato, Jeffrey Barrett, Sonia Goncalves, Ewan Harrison, David K. Jackson, Ian Johnston, Dominic Kwiatkowski, Cordelia Langford, John Sillitoe on behalf of the Wellcome Sanger Institute COVID-19 Surveillance Team |
| EPI_ISL_2436299 | Lighthouse Lab in Milton Keynes | Wellcome Sanger Institute for the COVID-19 Genomics UK (COG-UK) Consortium | The Lighthouse Lab in Milton Keynes and Alex Alderton, Roberto Amato, Jeffrey Barrett, Sonia Goncalves, Ewan Harrison, David K. Jackson, Ian Johnston, Dominic Kwiatkowski, Cordelia Langford, John Sillitoe on behalf of the Wellcome Sanger Institute COVID-19 Surveillance Team |
| EPI_ISL_2436300 | Lighthouse Lab in Alderley Park | Wellcome Sanger Institute for the COVID-19 Genomics UK (COG-UK) Consortium | Jacquelyn Wynn, Mairead Hyland, The Lighthouse Lab in Alderley Park and Alex Alderton, Roberto Amato, Jeffrey Barrett, Sonia Goncalves, Ewan Harrison, David K. Jackson, Ian Johnston, Dominic Kwiatkowski, Cordelia Langford, John Sillitoe on behalf of the Wellcome Sanger Institute COVID-19 Surveillance Team |
| EPI_ISL_2436301 | Lighthouse Lab in Milton Keynes | Wellcome Sanger Institute for the COVID-19 Genomics UK (COG-UK) Consortium | The Lighthouse Lab in Milton Keynes and Alex Alderton, Roberto Amato, Jeffrey Barrett, Sonia Goncalves, Ewan Harrison, David K. Jackson, Ian Johnston, Dominic Kwiatkowski, Cordelia Langford, John Sillitoe on behalf of the Wellcome Sanger Institute COVID-19 Surveillance Team |
| EPI_ISL_2436303, EPI_ISL_2436304 | Lighthouse Lab in Glasgow | Wellcome Sanger Institute for the COVID-19 Genomics UK (COG-UK) Consortium | Harper VanSteenhouse, Yumi Kasai, David Gray, Carol Clugston, Anna Dominiczak and Alex Alderton, Roberto Amato, Jeffrey Barrett, Sonia Goncalves, Ewan Harrison, David K. Jackson, Ian Johnston, Dominic Kwiatkowski, Cordelia Langford, John Sillitoe on behalf of the Wellcome Sanger Institute COVID-19 Surveillance Team |
| EPI_ISL_2436305 | Lighthouse Lab in Milton Keynes | Wellcome Sanger Institute for the COVID-19 Genomics UK (COG-UK) Consortium | The Lighthouse Lab in Milton Keynes and Alex Alderton, Roberto Amato, Jeffrey Barrett, Sonia Goncalves, Ewan Harrison, David K. Jackson, Ian Johnston, Dominic Kwiatkowski, Cordelia Langford, John Sillitoe on behalf of the Wellcome Sanger Institute COVID-19 Surveillance Team |
| EPI_ISL_2436306 | Lighthouse Lab in Alderley Park | Wellcome Sanger Institute for the COVID-19 Genomics UK (COG-UK) Consortium | Jacquelyn Wynn, Mairead Hyland, The Lighthouse Lab in Alderley Park and Alex Alderton, Roberto Amato, Jeffrey Barrett, Sonia Goncalves, Ewan Harrison, David K. Jackson, Ian Johnston, Dominic Kwiatkowski, Cordelia Langford, John Sillitoe on behalf of the Wellcome Sanger Institute COVID-19 Surveillance Team |
| EPI_ISL_2436307, EPI_ISL_2436308, EPI_ISL_2436309 | Lighthouse Lab in Glasgow | Wellcome Sanger Institute for the COVID-19 Genomics UK (COG-UK) Consortium | Harper VanSteenhouse, Yumi Kasai, David Gray, Carol Clugston, Anna Dominiczak and Alex Alderton, Roberto Amato, Jeffrey Barrett, Sonia Goncalves, Ewan Harrison, David K. Jackson, Ian Johnston, Dominic Kwiatkowski, Cordelia Langford, John Sillitoe on behalf of the Wellcome Sanger Institute COVID-19 Surveillance Team |
| EPI_ISL_2436310 | Lighthouse Lab in Milton Keynes | Wellcome Sanger Institute for the COVID-19 Genomics UK (COG-UK) Consortium | The Lighthouse Lab in Milton Keynes and Alex Alderton, Roberto Amato, Jeffrey Barrett, Sonia Goncalves, Ewan Harrison, David K. Jackson, Ian Johnston, Dominic Kwiatkowski, Cordelia Langford, John Sillitoe on behalf of the Wellcome Sanger Institute COVID-19 Surveillance Team |
| EPI_ISL_2436311 | Lighthouse Lab in Glasgow | Wellcome Sanger Institute for the COVID-19 Genomics UK (COG-UK) Consortium | Harper VanSteenhouse, Yumi Kasai, David Gray, Carol Clugston, Anna Dominiczak and Alex Alderton, Roberto Amato, Jeffrey Barrett, Sonia Goncalves, Ewan Harrison, David K. Jackson, Ian Johnston, Dominic Kwiatkowski, Cordelia Langford, John Sillitoe on behalf of the Wellcome Sanger Institute COVID-19 Surveillance Team |
| EPI_ISL_2436312 | Lighthouse Laboratory Plymouth | Wellcome Sanger Institute for the COVID-19 Genomics UK (COG-UK) Consortium | Lighthouse Laboratory Plymouth and Alex Alderton, Roberto Amato, Jeffrey Barrett, Sonia Goncalves, Ewan Harrison, David K. Jackson, Ian Johnston, Dominic Kwiatkowski, Cordelia Langford, John Sillitoe on behalf of the Wellcome Sanger Institute COVID-19 Surveillance Team |
| EPI_ISL_2436313 | Lighthouse Lab in Milton Keynes | Wellcome Sanger Institute for the COVID-19 Genomics UK (COG-UK) Consortium | The Lighthouse Lab in Milton Keynes and Alex Alderton, Roberto Amato, Jeffrey Barrett, Sonia Goncalves, Ewan Harrison, David K. Jackson, Ian Johnston, Dominic Kwiatkowski, Cordelia Langford, John Sillitoe on behalf of the Wellcome Sanger Institute COVID-19 Surveillance Team |
| EPI_ISL_2436314 | Lighthouse Lab in Alderley Park | Wellcome Sanger Institute for the COVID-19 Genomics UK (COG-UK) Consortium | Jacquelyn Wynn, Mairead Hyland, The Lighthouse Lab in Alderley Park and Alex Alderton, Roberto Amato, Jeffrey Barrett, Sonia Goncalves, Ewan Harrison, David K. Jackson, Ian Johnston, Dominic Kwiatkowski, Cordelia Langford, John Sillitoe on behalf of the Wellcome Sanger Institute COVID-19 Surveillance Team |
| EPI_ISL_2436315, EPI_ISL_2436316 | Lighthouse Lab in Glasgow | Wellcome Sanger Institute for the COVID-19 Genomics UK (COG-UK) Consortium | Harper VanSteenhouse, Yumi Kasai, David Gray, Carol Clugston, Anna Dominiczak and Alex Alderton, Roberto Amato, Jeffrey Barrett, Sonia Goncalves, Ewan Harrison, David K. Jackson, Ian Johnston, Dominic Kwiatkowski, Cordelia Langford, John Sillitoe on behalf of the Wellcome Sanger Institute COVID-19 Surveillance Team |
| EPI_ISL_2436317 | Health Services Laboratories | Wellcome Sanger Institute for the COVID-19 Genomics UK (COG-UK) Consortium | Health Services Laboratories and Alex Alderton, Roberto Amato, Jeffrey Barrett, Sonia Goncalves, Ewan Harrison, David K. Jackson, Ian Johnston, Dominic Kwiatkowski, Cordelia Langford, John Sillitoe on behalf of the Wellcome Sanger Institute COVID-19 Surveillance Team |
| EPI_ISL_2436318 | Lighthouse Lab in Glasgow | Wellcome Sanger Institute for the COVID-19 Genomics UK (COG-UK) Consortium | Harper VanSteenhouse, Yumi Kasai, David Gray, Carol Clugston, Anna Dominiczak and Alex Alderton, Roberto Amato, Jeffrey Barrett, Sonia Goncalves, Ewan Harrison, David K. Jackson, Ian Johnston, Dominic Kwiatkowski, Cordelia Langford, John Sillitoe on behalf of the Wellcome Sanger Institute COVID-19 Surveillance Team |
| EPI_ISL_2436319, EPI_ISL_2436320 | Health Services Laboratories | Wellcome Sanger Institute for the COVID-19 Genomics UK (COG-UK) Consortium | Health Services Laboratories and Alex Alderton, Roberto Amato, Jeffrey Barrett, Sonia Goncalves, Ewan Harrison, David K. Jackson, Ian Johnston, Dominic Kwiatkowski, Cordelia Langford, John Sillitoe on behalf of the Wellcome Sanger Institute COVID-19 Surveillance Team |
| EPI_ISL_2436321, EPI_ISL_2436322 | Lighthouse Lab in Glasgow | Wellcome Sanger Institute for the COVID-19 Genomics UK (COG-UK) Consortium | Harper VanSteenhouse, Yumi Kasai, David Gray, Carol Clugston, Anna Dominiczak and Alex Alderton, Roberto Amato, Jeffrey Barrett, Sonia Goncalves, Ewan Harrison, David K. Jackson, Ian Johnston, Dominic Kwiatkowski, Cordelia Langford, John Sillitoe on behalf of the Wellcome Sanger Institute COVID-19 Surveillance Team |
| EPI_ISL_2436323 | Lighthouse Laboratory Plymouth | Wellcome Sanger Institute for the COVID-19 Genomics UK (COG-UK) Consortium | Lighthouse Laboratory Plymouth and Alex Alderton, Roberto Amato, Jeffrey Barrett, Sonia Goncalves, Ewan Harrison, David K. Jackson, Ian Johnston, Dominic Kwiatkowski, Cordelia Langford, John Sillitoe on behalf of the Wellcome Sanger Institute COVID-19 Surveillance Team |
| EPI_ISL_2436324, EPI_ISL_2436326 | Lighthouse Lab in Alderley Park | Wellcome Sanger Institute for the COVID-19 Genomics UK (COG-UK) Consortium | Jacquelyn Wynn, Mairead Hyland, The Lighthouse Lab in Alderley Park and Alex Alderton, Roberto Amato, Jeffrey Barrett, Sonia Goncalves, Ewan Harrison, David K. Jackson, Ian Johnston, Dominic Kwiatkowski, Cordelia Langford, John Sillitoe on behalf of the Wellcome Sanger Institute COVID-19 Surveillance Team |
| EPI_ISL_2436328 | Lighthouse Laboratory Plymouth | Wellcome Sanger Institute for the COVID-19 Genomics UK (COG-UK) Consortium | Lighthouse Laboratory Plymouth and Alex Alderton, Roberto Amato, Jeffrey Barrett, Sonia Goncalves, Ewan Harrison, David K. Jackson, Ian Johnston, Dominic Kwiatkowski, Cordelia Langford, John Sillitoe on behalf of the Wellcome Sanger Institute COVID-19 Surveillance Team |
| EPI_ISL_2436329, EPI_ISL_2436330, EPI_ISL_2436331, EPI_ISL_2436332, EPI_ISL_2436333 | Lighthouse Lab in Alderley Park | Wellcome Sanger Institute for the COVID-19 Genomics UK (COG-UK) Consortium | Jacquelyn Wynn, Mairead Hyland, The Lighthouse Lab in Alderley Park and Alex Alderton, Roberto Amato, Jeffrey Barrett, Sonia Goncalves, Ewan Harrison, David K. Jackson, Ian Johnston, Dominic Kwiatkowski, Cordelia Langford, John Sillitoe on behalf of the Wellcome Sanger Institute COVID-19 Surveillance Team |
| EPI_ISL_2436334, EPI_ISL_2436336 | Lighthouse Lab in Glasgow | Wellcome Sanger Institute for the COVID-19 Genomics UK (COG-UK) Consortium | Harper VanSteenhouse, Yumi Kasai, David Gray, Carol Clugston, Anna Dominiczak and Alex Alderton, Roberto Amato, Jeffrey Barrett, Sonia Goncalves, Ewan Harrison, David K. Jackson, Ian Johnston, Dominic Kwiatkowski, Cordelia Langford, John Sillitoe on behalf of the Wellcome Sanger Institute COVID-19 Surveillance Team |
| EPI_ISL_2436337 | Lighthouse Lab in Alderley Park | Wellcome Sanger Institute for the COVID-19 Genomics UK (COG-UK) Consortium | Jacquelyn Wynn, Mairead Hyland, The Lighthouse Lab in Alderley Park and Alex Alderton, Roberto Amato, Jeffrey Barrett, Sonia Goncalves, Ewan Harrison, David K. Jackson, Ian Johnston, Dominic Kwiatkowski, Cordelia Langford, John Sillitoe on behalf of the Wellcome Sanger Institute COVID-19 Surveillance Team |
| EPI_ISL_2436338, EPI_ISL_2436339 | Health Services Laboratories | Wellcome Sanger Institute for the COVID-19 Genomics UK (COG-UK) Consortium | Health Services Laboratories and Alex Alderton, Roberto Amato, Jeffrey Barrett, Sonia Goncalves, Ewan Harrison, David K. Jackson, Ian Johnston, Dominic Kwiatkowski, Cordelia Langford, John Sillitoe on behalf of the Wellcome Sanger Institute COVID-19 Surveillance Team |
| EPI_ISL_2436340 | Lighthouse Lab in Glasgow | Wellcome Sanger Institute for the COVID-19 Genomics UK (COG-UK) | Harper VanSteenhouse, Yumi Kasai, David Gray, Carol Clugston, Anna Dominiczak and Alex Alderton, Roberto Amato, Jeffrey Barrett, Sonia Goncalves, Ewan Harrison, David K. Jackson, Ian Johnston, Dominic Kwiatkowski, Cordelia Langford, |

| | | | |
|---|---|---|---|
| | | Consortium | John Sillitoe on behalf of the Wellcome Sanger Institute COVID-19 Surveillance Team |
| EPI_ISL_2436341 | Lighthouse Laboratory Plymouth | Wellcome Sanger Institute for the COVID-19 Genomics UK (COG-UK) Consortium | Lighthouse Laboratory Plymouth and Alex Alderton, Roberto Amato, Jeffrey Barrett, Sonia Goncalves, Ewan Harrison, David K. Jackson, Ian Johnston, Dominic Kwiatkowski, Cordelia Langford, John Sillitoe on behalf of the Wellcome Sanger Institute COVID-19 Surveillance Team |
| EPI_ISL_2436342 | Lighthouse Lab in Alderley Park | Wellcome Sanger Institute for the COVID-19 Genomics UK (COG-UK) Consortium | Jacquelyn Wynn, Mairead Hyland, The Lighthouse Lab in Alderley Park and Alex Alderton, Roberto Amato, Jeffrey Barrett, Sonia Goncalves, Ewan Harrison, David K. Jackson, Ian Johnston, Dominic Kwiatkowski, Cordelia Langford, John Sillitoe on behalf of the Wellcome Sanger Institute COVID-19 Surveillance Team |
| EPI_ISL_2436343, EPI_ISL_2436344 | Lighthouse Lab in Glasgow | Wellcome Sanger Institute for the COVID-19 Genomics UK (COG-UK) Consortium | Harper VanSteenhouse, Yumi Kasai, David Gray, Carol Clugston, Anna Dominiczak and Alex Alderton, Roberto Amato, Jeffrey Barrett, Sonia Goncalves, Ewan Harrison, David K. Jackson, Ian Johnston, Dominic Kwiatkowski, Cordelia Langford, John Sillitoe on behalf of the Wellcome Sanger Institute COVID-19 Surveillance Team |
| EPI_ISL_2436345 | Lighthouse Laboratory Plymouth | Wellcome Sanger Institute for the COVID-19 Genomics UK (COG-UK) Consortium | Lighthouse Laboratory Plymouth and Alex Alderton, Roberto Amato, Jeffrey Barrett, Sonia Goncalves, Ewan Harrison, David K. Jackson, Ian Johnston, Dominic Kwiatkowski, Cordelia Langford, John Sillitoe on behalf of the Wellcome Sanger Institute COVID-19 Surveillance Team |
| EPI_ISL_2436346 | Lighthouse Lab in Glasgow | Wellcome Sanger Institute for the COVID-19 Genomics UK (COG-UK) Consortium | Harper VanSteenhouse, Yumi Kasai, David Gray, Carol Clugston, Anna Dominiczak and Alex Alderton, Roberto Amato, Jeffrey Barrett, Sonia Goncalves, Ewan Harrison, David K. Jackson, Ian Johnston, Dominic Kwiatkowski, Cordelia Langford, John Sillitoe on behalf of the Wellcome Sanger Institute COVID-19 Surveillance Team |
| EPI_ISL_2436347 | Lighthouse Lab in Alderley Park | Wellcome Sanger Institute for the COVID-19 Genomics UK (COG-UK) Consortium | Jacquelyn Wynn, Mairead Hyland, The Lighthouse Lab in Alderley Park and Alex Alderton, Roberto Amato, Jeffrey Barrett, Sonia Goncalves, Ewan Harrison, David K. Jackson, Ian Johnston, Dominic Kwiatkowski, Cordelia Langford, John Sillitoe on behalf of the Wellcome Sanger Institute COVID-19 Surveillance Team |
| EPI_ISL_2436349, EPI_ISL_2436350 | Health Services Laboratories | Wellcome Sanger Institute for the COVID-19 Genomics UK (COG-UK) Consortium | Health Services Laboratories and Alex Alderton, Roberto Amato, Jeffrey Barrett, Sonia Goncalves, Ewan Harrison, David K. Jackson, Ian Johnston, Dominic Kwiatkowski, Cordelia Langford, John Sillitoe on behalf of the Wellcome Sanger Institute COVID-19 Surveillance Team |
| EPI_ISL_2436351, EPI_ISL_2436352 | Lighthouse Lab in Glasgow | Wellcome Sanger Institute for the COVID-19 Genomics UK (COG-UK) Consortium | Harper VanSteenhouse, Yumi Kasai, David Gray, Carol Clugston, Anna Dominiczak and Alex Alderton, Roberto Amato, Jeffrey Barrett, Sonia Goncalves, Ewan Harrison, David K. Jackson, Ian Johnston, Dominic Kwiatkowski, Cordelia Langford, John Sillitoe on behalf of the Wellcome Sanger Institute COVID-19 Surveillance Team |
| EPI_ISL_2436353 | Lighthouse Lab in Milton Keynes | Wellcome Sanger Institute for the COVID-19 Genomics UK (COG-UK) Consortium | The Lighthouse Lab in Milton Keynes and Alex Alderton, Roberto Amato, Jeffrey Barrett, Sonia Goncalves, Ewan Harrison, David K. Jackson, Ian Johnston, Dominic Kwiatkowski, Cordelia Langford, John Sillitoe on behalf of the Wellcome Sanger Institute COVID-19 Surveillance Team |
| EPI_ISL_2436354, EPI_ISL_2436355, EPI_ISL_2436356 | Lighthouse Lab in Glasgow | Wellcome Sanger Institute for the COVID-19 Genomics UK (COG-UK) Consortium | Harper VanSteenhouse, Yumi Kasai, David Gray, Carol Clugston, Anna Dominiczak and Alex Alderton, Roberto Amato, Jeffrey Barrett, Sonia Goncalves, Ewan Harrison, David K. Jackson, Ian Johnston, Dominic Kwiatkowski, Cordelia Langford, John Sillitoe on behalf of the Wellcome Sanger Institute COVID-19 Surveillance Team |
| EPI_ISL_2436357 | Lighthouse Lab in Alderley Park | Wellcome Sanger Institute for the COVID-19 Genomics UK (COG-UK) Consortium | Jacquelyn Wynn, Mairead Hyland, The Lighthouse Lab in Alderley Park and Alex Alderton, Roberto Amato, Jeffrey Barrett, Sonia Goncalves, Ewan Harrison, David K. Jackson, Ian Johnston, Dominic Kwiatkowski, Cordelia Langford, John Sillitoe on behalf of the Wellcome Sanger Institute COVID-19 Surveillance Team |
| EPI_ISL_2436358 | Health Services Laboratories | Wellcome Sanger Institute for the COVID-19 Genomics UK (COG-UK) Consortium | Health Services Laboratories and Alex Alderton, Roberto Amato, Jeffrey Barrett, Sonia Goncalves, Ewan Harrison, David K. Jackson, Ian Johnston, Dominic Kwiatkowski, Cordelia Langford, John Sillitoe on behalf of the Wellcome Sanger Institute COVID-19 Surveillance Team |
| EPI_ISL_2436359 | Lighthouse Lab in Milton Keynes | Wellcome Sanger Institute for the COVID-19 Genomics UK (COG-UK) Consortium | The Lighthouse Lab in Milton Keynes and Alex Alderton, Roberto Amato, Jeffrey Barrett, Sonia Goncalves, Ewan Harrison, David K. Jackson, Ian Johnston, Dominic Kwiatkowski, Cordelia Langford, John Sillitoe on behalf of the Wellcome Sanger Institute COVID-19 Surveillance Team |
| EPI_ISL_2436360, EPI_ISL_2436362 | Lighthouse Lab in Alderley Park | Wellcome Sanger Institute for the COVID-19 Genomics UK (COG-UK) Consortium | Jacquelyn Wynn, Mairead Hyland, The Lighthouse Lab in Alderley Park and Alex Alderton, Roberto Amato, Jeffrey Barrett, Sonia Goncalves, Ewan Harrison, David K. Jackson, Ian Johnston, Dominic Kwiatkowski, Cordelia Langford, John Sillitoe on behalf of the Wellcome Sanger Institute COVID-19 Surveillance Team |
| EPI_ISL_2436363 | Lighthouse Laboratory Plymouth | Wellcome Sanger Institute for the COVID-19 Genomics UK (COG-UK) Consortium | Lighthouse Laboratory Plymouth and Alex Alderton, Roberto Amato, Jeffrey Barrett, Sonia Goncalves, Ewan Harrison, David K. Jackson, Ian Johnston, Dominic Kwiatkowski, Cordelia Langford, John Sillitoe on behalf of the Wellcome Sanger Institute COVID-19 Surveillance Team |
| EPI_ISL_2436364 | Lighthouse Lab in Alderley Park | Wellcome Sanger Institute for the COVID-19 Genomics UK (COG-UK) Consortium | Jacquelyn Wynn, Mairead Hyland, The Lighthouse Lab in Alderley Park and Alex Alderton, Roberto Amato, Jeffrey Barrett, Sonia Goncalves, Ewan Harrison, David K. Jackson, Ian Johnston, Dominic Kwiatkowski, Cordelia Langford, John Sillitoe on behalf of the Wellcome Sanger Institute COVID-19 Surveillance Team |
| EPI_ISL_2436365 | Lighthouse Lab in Milton Keynes | Wellcome Sanger Institute for the COVID-19 Genomics UK (COG-UK) Consortium | The Lighthouse Lab in Milton Keynes and Alex Alderton, Roberto Amato, Jeffrey Barrett, Sonia Goncalves, Ewan Harrison, David K. Jackson, Ian Johnston, Dominic Kwiatkowski, Cordelia Langford, John Sillitoe on behalf of the Wellcome Sanger Institute COVID-19 Surveillance Team |
| EPI_ISL_2436366 | Lighthouse Laboratory Plymouth | Wellcome Sanger Institute for the COVID-19 Genomics UK (COG-UK) Consortium | Lighthouse Laboratory Plymouth and Alex Alderton, Roberto Amato, Jeffrey Barrett, Sonia Goncalves, Ewan Harrison, David K. Jackson, Ian Johnston, Dominic Kwiatkowski, Cordelia Langford, John Sillitoe on behalf of the Wellcome Sanger Institute COVID-19 Surveillance Team |
| EPI_ISL_2436368, EPI_ISL_2436370 | Lighthouse Lab in Glasgow | Wellcome Sanger Institute for the COVID-19 Genomics UK (COG-UK) Consortium | Harper VanSteenhouse, Yumi Kasai, David Gray, Carol Clugston, Anna Dominiczak and Alex Alderton, Roberto Amato, Jeffrey Barrett, Sonia Goncalves, Ewan Harrison, David K. Jackson, Ian Johnston, Dominic Kwiatkowski, Cordelia Langford, John Sillitoe on behalf of the Wellcome Sanger Institute COVID-19 Surveillance Team |
| EPI_ISL_2436371 | Lighthouse Lab in Alderley Park | Wellcome Sanger Institute for the COVID-19 Genomics UK (COG-UK) Consortium | Jacquelyn Wynn, Mairead Hyland, The Lighthouse Lab in Alderley Park and Alex Alderton, Roberto Amato, Jeffrey Barrett, Sonia Goncalves, Ewan Harrison, David K. Jackson, Ian Johnston, Dominic Kwiatkowski, Cordelia Langford, John Sillitoe on behalf of the Wellcome Sanger Institute COVID-19 Surveillance Team |
| EPI_ISL_2436372 | Health Services Laboratories | Wellcome Sanger Institute for the COVID-19 Genomics UK (COG-UK) Consortium | Health Services Laboratories and Alex Alderton, Roberto Amato, Jeffrey Barrett, Sonia Goncalves, Ewan Harrison, David K. Jackson, Ian Johnston, Dominic Kwiatkowski, Cordelia Langford, John Sillitoe on behalf of the Wellcome Sanger Institute COVID-19 Surveillance Team |
| EPI_ISL_2436373 | Lighthouse Lab in Milton Keynes | Wellcome Sanger Institute for the COVID-19 Genomics UK (COG-UK) Consortium | The Lighthouse Lab in Milton Keynes and Alex Alderton, Roberto Amato, Jeffrey Barrett, Sonia Goncalves, Ewan Harrison, David K. Jackson, Ian Johnston, Dominic Kwiatkowski, Cordelia Langford, John Sillitoe on behalf of the Wellcome Sanger Institute COVID-19 Surveillance Team |
| EPI_ISL_2436374, EPI_ISL_2436375 | Lighthouse Lab in Alderley Park | Wellcome Sanger Institute for the COVID-19 Genomics UK (COG-UK) Consortium | Jacquelyn Wynn, Mairead Hyland, The Lighthouse Lab in Alderley Park and Alex Alderton, Roberto Amato, Jeffrey Barrett, Sonia Goncalves, Ewan Harrison, David K. Jackson, Ian Johnston, Dominic Kwiatkowski, Cordelia Langford, John Sillitoe on behalf of the Wellcome Sanger Institute COVID-19 Surveillance Team |
| EPI_ISL_2436376 | Lighthouse Lab in Glasgow | Wellcome Sanger Institute for the COVID-19 Genomics UK (COG-UK) Consortium | Harper VanSteenhouse, Yumi Kasai, David Gray, Carol Clugston, Anna Dominiczak and Alex Alderton, Roberto Amato, Jeffrey Barrett, Sonia Goncalves, Ewan Harrison, David K. Jackson, Ian Johnston, Dominic Kwiatkowski, Cordelia Langford, John Sillitoe on behalf of the Wellcome Sanger Institute COVID-19 Surveillance Team |
| EPI_ISL_2436378 | Health Services Laboratories | Wellcome Sanger Institute for the COVID-19 Genomics UK (COG-UK) Consortium | Health Services Laboratories and Alex Alderton, Roberto Amato, Jeffrey Barrett, Sonia Goncalves, Ewan Harrison, David K. Jackson, Ian Johnston, Dominic Kwiatkowski, Cordelia Langford, John Sillitoe on behalf of the Wellcome Sanger Institute COVID-19 Surveillance Team |
| EPI_ISL_2436379, EPI_ISL_2436380 | Lighthouse Lab in Alderley Park | Wellcome Sanger Institute for the COVID-19 Genomics UK (COG-UK) Consortium | Jacquelyn Wynn, Mairead Hyland, The Lighthouse Lab in Alderley Park and Alex Alderton, Roberto Amato, Jeffrey Barrett, Sonia Goncalves, Ewan Harrison, David K. Jackson, Ian Johnston, Dominic Kwiatkowski, Cordelia Langford, John Sillitoe on behalf of the Wellcome Sanger Institute COVID-19 Surveillance Team |
| EPI_ISL_2436381 | Health Services Laboratories | Wellcome Sanger Institute for the COVID-19 Genomics UK (COG-UK) Consortium | Health Services Laboratories and Alex Alderton, Roberto Amato, Jeffrey Barrett, Sonia Goncalves, Ewan Harrison, David K. Jackson, Ian Johnston, Dominic Kwiatkowski, Cordelia Langford, John Sillitoe on behalf of the Wellcome Sanger Institute COVID-19 Surveillance Team |
| EPI_ISL_2436382 | Lighthouse Lab in Milton Keynes | Wellcome Sanger Institute for the COVID-19 Genomics UK (COG-UK) Consortium | The Lighthouse Lab in Milton Keynes and Alex Alderton, Roberto Amato, Jeffrey Barrett, Sonia Goncalves, Ewan Harrison, David K. Jackson, Ian Johnston, Dominic Kwiatkowski, Cordelia Langford, John Sillitoe on behalf of the Wellcome Sanger Institute COVID-19 Surveillance Team |
| EPI_ISL_2436383 | Lighthouse Lab in Alderley Park | Wellcome Sanger Institute for the COVID-19 Genomics UK (COG-UK) Consortium | Jacquelyn Wynn, Mairead Hyland, The Lighthouse Lab in Alderley Park and Alex Alderton, Roberto Amato, Jeffrey Barrett, Sonia Goncalves, Ewan Harrison, David K. Jackson, Ian Johnston, Dominic Kwiatkowski, Cordelia Langford, John Sillitoe on behalf of the Wellcome Sanger Institute COVID-19 Surveillance Team |
| EPI_ISL_2436384, EPI_ISL_2436385, EPI_ISL_2436386, EPI_ISL_2436387 | Lighthouse Lab in Milton Keynes | Wellcome Sanger Institute for the COVID-19 Genomics UK (COG-UK) Consortium | The Lighthouse Lab in Milton Keynes and Alex Alderton, Roberto Amato, Jeffrey Barrett, Sonia Goncalves, Ewan Harrison, David K. Jackson, Ian Johnston, Dominic Kwiatkowski, Cordelia Langford, John Sillitoe on behalf of the Wellcome Sanger Institute COVID-19 Surveillance Team |
| EPI_ISL_2436388, EPI_ISL_2436389 | Lighthouse Lab in Glasgow | Wellcome Sanger Institute for the COVID-19 Genomics UK (COG-UK) Consortium | Harper VanSteenhouse, Yumi Kasai, David Gray, Carol Clugston, Anna Dominiczak and Alex Alderton, Roberto Amato, Jeffrey Barrett, Sonia Goncalves, Ewan Harrison, David K. Jackson, Ian Johnston, Dominic Kwiatkowski, Cordelia Langford, John Sillitoe on behalf of the Wellcome Sanger Institute COVID-19 Surveillance Team |
| EPI_ISL_2436390 | Lighthouse Lab in Milton Keynes | Wellcome Sanger Institute for the COVID-19 Genomics UK (COG-UK) Consortium | The Lighthouse Lab in Milton Keynes and Alex Alderton, Roberto Amato, Jeffrey Barrett, Sonia Goncalves, Ewan Harrison, David K. Jackson, Ian Johnston, Dominic Kwiatkowski, Cordelia Langford, John Sillitoe on behalf of the Wellcome Sanger Institute COVID-19 Surveillance Team |
| EPI_ISL_2436392 | Lighthouse Lab in Alderley Park | Wellcome Sanger Institute for the COVID-19 Genomics UK (COG-UK) Consortium | Jacquelyn Wynn, Mairead Hyland, The Lighthouse Lab in Alderley Park and Alex Alderton, Roberto Amato, Jeffrey Barrett, Sonia Goncalves, Ewan Harrison, David K. Jackson, Ian Johnston, Dominic Kwiatkowski, Cordelia Langford, John Sillitoe on behalf of the Wellcome Sanger Institute COVID-19 Surveillance Team |
| EPI_ISL_2436393, EPI_ISL_2436394, EPI_ISL_2436395, EPI_ISL_2436396 | Lighthouse Lab in Milton Keynes | Wellcome Sanger Institute for the COVID-19 Genomics UK (COG-UK) Consortium | The Lighthouse Lab in Milton Keynes and Alex Alderton, Roberto Amato, Jeffrey Barrett, Sonia Goncalves, Ewan Harrison, David K. Jackson, Ian Johnston, Dominic Kwiatkowski, Cordelia Langford, John Sillitoe on behalf of the Wellcome Sanger Institute COVID-19 Surveillance Team |
| EPI_ISL_2436397, EPI_ISL_2436398 | Health Services Laboratories | Wellcome Sanger Institute for the COVID-19 Genomics UK (COG-UK) Consortium | Health Services Laboratories and Alex Alderton, Roberto Amato, Jeffrey Barrett, Sonia Goncalves, Ewan Harrison, David K. Jackson, Ian Johnston, Dominic Kwiatkowski, Cordelia Langford, John Sillitoe on behalf of the Wellcome Sanger Institute COVID-19 Surveillance Team |
| EPI_ISL_2436400 | Lighthouse Lab in Glasgow | Wellcome Sanger Institute for the COVID-19 Genomics UK (COG-UK) Consortium | Harper VanSteenhouse, Yumi Kasai, David Gray, Carol Clugston, Anna Dominiczak and Alex Alderton, Roberto Amato, Jeffrey Barrett, Sonia Goncalves, Ewan Harrison, David K. Jackson, Ian Johnston, Dominic Kwiatkowski, Cordelia Langford, John Sillitoe on behalf of the Wellcome Sanger Institute COVID-19 Surveillance Team |
| EPI_ISL_2436401 | Lighthouse Lab in Milton Keynes | Wellcome Sanger Institute for the COVID-19 Genomics UK (COG-UK) Consortium | The Lighthouse Lab in Milton Keynes and Alex Alderton, Roberto Amato, Jeffrey Barrett, Sonia Goncalves, Ewan Harrison, David K. Jackson, Ian Johnston, Dominic Kwiatkowski, Cordelia Langford, John Sillitoe on behalf of the Wellcome Sanger Institute COVID-19 Surveillance Team |
| EPI_ISL_2436402, EPI_ISL_2436403 | Lighthouse Lab in Alderley Park | Wellcome Sanger Institute for the COVID-19 Genomics UK (COG-UK) Consortium | Jacquelyn Wynn, Mairead Hyland, The Lighthouse Lab in Alderley Park and Alex Alderton, Roberto Amato, Jeffrey Barrett, Sonia Goncalves, Ewan Harrison, David K. Jackson, Ian Johnston, Dominic Kwiatkowski, Cordelia Langford, John Sillitoe on behalf of the Wellcome Sanger Institute COVID-19 Surveillance Team |
| EPI_ISL_2436404 | Health Services Laboratories | Wellcome Sanger Institute for the COVID-19 Genomics UK (COG-UK) Consortium | Health Services Laboratories and Alex Alderton, Roberto Amato, Jeffrey Barrett, Sonia Goncalves, Ewan Harrison, David K. Jackson, Ian Johnston, Dominic Kwiatkowski, Cordelia Langford, John Sillitoe on behalf of the Wellcome Sanger Institute COVID-19 Surveillance Team |
| EPI_ISL_2436405 | Lighthouse Lab in Milton Keynes | Wellcome Sanger Institute for the COVID-19 Genomics UK (COG-UK) Consortium | The Lighthouse Lab in Milton Keynes and Alex Alderton, Roberto Amato, Jeffrey Barrett, Sonia Goncalves, Ewan Harrison, David K. Jackson, Ian Johnston, Dominic Kwiatkowski, Cordelia Langford, John Sillitoe on behalf of the Wellcome Sanger Institute COVID-19 Surveillance Team |
| EPI_ISL_2436406, EPI_ISL_2436407, EPI_ISL_2436408 | Lighthouse Lab in Glasgow | Wellcome Sanger Institute for the COVID-19 Genomics UK (COG-UK) Consortium | Harper VanSteenhouse, Yumi Kasai, David Gray, Carol Clugston, Anna Dominiczak and Alex Alderton, Roberto Amato, Jeffrey Barrett, Sonia Goncalves, Ewan Harrison, David K. Jackson, Ian Johnston, Dominic Kwiatkowski, Cordelia Langford, John Sillitoe on behalf of the Wellcome Sanger Institute COVID-19 Surveillance Team |
| EPI_ISL_2436409 | Lighthouse Lab in Alderley Park | Wellcome Sanger Institute for the COVID-19 Genomics UK (COG-UK) Consortium | Jacquelyn Wynn, Mairead Hyland, The Lighthouse Lab in Alderley Park and Alex Alderton, Roberto Amato, Jeffrey Barrett, Sonia Goncalves, Ewan Harrison, David K. Jackson, Ian Johnston, Dominic Kwiatkowski, Cordelia Langford, John Sillitoe on behalf of the Wellcome Sanger Institute COVID-19 Surveillance Team |

| | | | |
|---|---|---|---|
| EPI_ISL_2436410 | Health Services Laboratories | Wellcome Sanger Institute for the COVID-19 Genomics UK (COG-UK) Consortium | Health Services Laboratories and Alex Alderton, Roberto Amato, Jeffrey Barrett, Sonia Goncalves, Ewan Harrison, David K. Jackson, Ian Johnston, Dominic Kwiatkowski, Cordelia Langford, John Sillitoe on behalf of the Wellcome Sanger Institute COVID-19 Surveillance Team |
| EPI_ISL_2436411 | Lighthouse Lab in Glasgow | Wellcome Sanger Institute for the COVID-19 Genomics UK (COG-UK) Consortium | Harper VanSteenhouse, Yumi Kasai, David Gray, Carol Clugston, Anna Dominiczak and Alex Alderton, Roberto Amato, Jeffrey Barrett, Sonia Goncalves, Ewan Harrison, David K. Jackson, Ian Johnston, Dominic Kwiatkowski, Cordelia Langford, John Sillitoe on behalf of the Wellcome Sanger Institute COVID-19 Surveillance Team |
| EPI_ISL_2436412 | Lighthouse Lab in Alderley Park | Wellcome Sanger Institute for the COVID-19 Genomics UK (COG-UK) Consortium | Jacquelyn Wynn, Mairead Hyland, The Lighthouse Lab in Alderley Park and Alex Alderton, Roberto Amato, Jeffrey Barrett, Sonia Goncalves, Ewan Harrison, David K. Jackson, Ian Johnston, Dominic Kwiatkowski, Cordelia Langford, John Sillitoe on behalf of the Wellcome Sanger Institute COVID-19 Surveillance Team |
| EPI_ISL_2436413 | Lighthouse Lab in Glasgow | Wellcome Sanger Institute for the COVID-19 Genomics UK (COG-UK) Consortium | Harper VanSteenhouse, Yumi Kasai, David Gray, Carol Clugston, Anna Dominiczak and Alex Alderton, Roberto Amato, Jeffrey Barrett, Sonia Goncalves, Ewan Harrison, David K. Jackson, Ian Johnston, Dominic Kwiatkowski, Cordelia Langford, John Sillitoe on behalf of the Wellcome Sanger Institute COVID-19 Surveillance Team |
| EPI_ISL_2436414 | Lighthouse Lab in Alderley Park | Wellcome Sanger Institute for the COVID-19 Genomics UK (COG-UK) Consortium | Jacquelyn Wynn, Mairead Hyland, The Lighthouse Lab in Alderley Park and Alex Alderton, Roberto Amato, Jeffrey Barrett, Sonia Goncalves, Ewan Harrison, David K. Jackson, Ian Johnston, Dominic Kwiatkowski, Cordelia Langford, John Sillitoe on behalf of the Wellcome Sanger Institute COVID-19 Surveillance Team |
| EPI_ISL_2436415 | Health Services Laboratories | Wellcome Sanger Institute for the COVID-19 Genomics UK (COG-UK) Consortium | Health Services Laboratories and Alex Alderton, Roberto Amato, Jeffrey Barrett, Sonia Goncalves, Ewan Harrison, David K. Jackson, Ian Johnston, Dominic Kwiatkowski, Cordelia Langford, John Sillitoe on behalf of the Wellcome Sanger Institute COVID-19 Surveillance Team |
| EPI_ISL_2436416 | Lighthouse Laboratory Plymouth | Wellcome Sanger Institute for the COVID-19 Genomics UK (COG-UK) Consortium | Lighthouse Laboratory Plymouth and Alex Alderton, Roberto Amato, Jeffrey Barrett, Sonia Goncalves, Ewan Harrison, David K. Jackson, Ian Johnston, Dominic Kwiatkowski, Cordelia Langford, John Sillitoe on behalf of the Wellcome Sanger Institute COVID-19 Surveillance Team |
| EPI_ISL_2436417 | Lighthouse Lab in Glasgow | Wellcome Sanger Institute for the COVID-19 Genomics UK (COG-UK) Consortium | Harper VanSteenhouse, Yumi Kasai, David Gray, Carol Clugston, Anna Dominiczak and Alex Alderton, Roberto Amato, Jeffrey Barrett, Sonia Goncalves, Ewan Harrison, David K. Jackson, Ian Johnston, Dominic Kwiatkowski, Cordelia Langford, John Sillitoe on behalf of the Wellcome Sanger Institute COVID-19 Surveillance Team |
| EPI_ISL_2436418 | Lighthouse Laboratory Plymouth | Wellcome Sanger Institute for the COVID-19 Genomics UK (COG-UK) Consortium | Lighthouse Laboratory Plymouth and Alex Alderton, Roberto Amato, Jeffrey Barrett, Sonia Goncalves, Ewan Harrison, David K. Jackson, Ian Johnston, Dominic Kwiatkowski, Cordelia Langford, John Sillitoe on behalf of the Wellcome Sanger Institute COVID-19 Surveillance Team |
| EPI_ISL_2436419 | Health Services Laboratories | Wellcome Sanger Institute for the COVID-19 Genomics UK (COG-UK) Consortium | Health Services Laboratories and Alex Alderton, Roberto Amato, Jeffrey Barrett, Sonia Goncalves, Ewan Harrison, David K. Jackson, Ian Johnston, Dominic Kwiatkowski, Cordelia Langford, John Sillitoe on behalf of the Wellcome Sanger Institute COVID-19 Surveillance Team |
| EPI_ISL_2436421 | Lighthouse Lab in Alderley Park | Wellcome Sanger Institute for the COVID-19 Genomics UK (COG-UK) Consortium | Jacquelyn Wynn, Mairead Hyland, The Lighthouse Lab in Alderley Park and Alex Alderton, Roberto Amato, Jeffrey Barrett, Sonia Goncalves, Ewan Harrison, David K. Jackson, Ian Johnston, Dominic Kwiatkowski, Cordelia Langford, John Sillitoe on behalf of the Wellcome Sanger Institute COVID-19 Surveillance Team |
| EPI_ISL_2436422 | Health Services Laboratories | Wellcome Sanger Institute for the COVID-19 Genomics UK (COG-UK) Consortium | Health Services Laboratories and Alex Alderton, Roberto Amato, Jeffrey Barrett, Sonia Goncalves, Ewan Harrison, David K. Jackson, Ian Johnston, Dominic Kwiatkowski, Cordelia Langford, John Sillitoe on behalf of the Wellcome Sanger Institute COVID-19 Surveillance Team |
| EPI_ISL_2436423 | Lighthouse Lab in Glasgow | Wellcome Sanger Institute for the COVID-19 Genomics UK (COG-UK) Consortium | Harper VanSteenhouse, Yumi Kasai, David Gray, Carol Clugston, Anna Dominiczak and Alex Alderton, Roberto Amato, Jeffrey Barrett, Sonia Goncalves, Ewan Harrison, David K. Jackson, Ian Johnston, Dominic Kwiatkowski, Cordelia Langford, John Sillitoe on behalf of the Wellcome Sanger Institute COVID-19 Surveillance Team |
| EPI_ISL_2436424 | Lighthouse Lab in Milton Keynes | Wellcome Sanger Institute for the COVID-19 Genomics UK (COG-UK) Consortium | The Lighthouse Lab in Milton Keynes and Alex Alderton, Roberto Amato, Jeffrey Barrett, Sonia Goncalves, Ewan Harrison, David K. Jackson, Ian Johnston, Dominic Kwiatkowski, Cordelia Langford, John Sillitoe on behalf of the Wellcome Sanger Institute COVID-19 Surveillance Team |
| EPI_ISL_2436425, EPI_ISL_2436426 | Lighthouse Lab in Alderley Park | Wellcome Sanger Institute for the COVID-19 Genomics UK (COG-UK) Consortium | Jacquelyn Wynn, Mairead Hyland, The Lighthouse Lab in Alderley Park and Alex Alderton, Roberto Amato, Jeffrey Barrett, Sonia Goncalves, Ewan Harrison, David K. Jackson, Ian Johnston, Dominic Kwiatkowski, Cordelia Langford, John Sillitoe on behalf of the Wellcome Sanger Institute COVID-19 Surveillance Team |
| EPI_ISL_2436427 | Lighthouse Lab in Milton Keynes | Wellcome Sanger Institute for the COVID-19 Genomics UK (COG-UK) Consortium | The Lighthouse Lab in Milton Keynes and Alex Alderton, Roberto Amato, Jeffrey Barrett, Sonia Goncalves, Ewan Harrison, David K. Jackson, Ian Johnston, Dominic Kwiatkowski, Cordelia Langford, John Sillitoe on behalf of the Wellcome Sanger Institute COVID-19 Surveillance Team |
| EPI_ISL_2436428 | Lighthouse Lab in Glasgow | Wellcome Sanger Institute for the COVID-19 Genomics UK (COG-UK) Consortium | Harper VanSteenhouse, Yumi Kasai, David Gray, Carol Clugston, Anna Dominiczak and Alex Alderton, Roberto Amato, Jeffrey Barrett, Sonia Goncalves, Ewan Harrison, David K. Jackson, Ian Johnston, Dominic Kwiatkowski, Cordelia Langford, John Sillitoe on behalf of the Wellcome Sanger Institute COVID-19 Surveillance Team |
| EPI_ISL_2436429 | Lighthouse Lab in Milton Keynes | Wellcome Sanger Institute for the COVID-19 Genomics UK (COG-UK) Consortium | The Lighthouse Lab in Milton Keynes and Alex Alderton, Roberto Amato, Jeffrey Barrett, Sonia Goncalves, Ewan Harrison, David K. Jackson, Ian Johnston, Dominic Kwiatkowski, Cordelia Langford, John Sillitoe on behalf of the Wellcome Sanger Institute COVID-19 Surveillance Team |
| EPI_ISL_2436430 | Health Services Laboratories | Wellcome Sanger Institute for the COVID-19 Genomics UK (COG-UK) Consortium | Health Services Laboratories and Alex Alderton, Roberto Amato, Jeffrey Barrett, Sonia Goncalves, Ewan Harrison, David K. Jackson, Ian Johnston, Dominic Kwiatkowski, Cordelia Langford, John Sillitoe on behalf of the Wellcome Sanger Institute COVID-19 Surveillance Team |
| EPI_ISL_2436431 | Lighthouse Lab in Alderley Park | Wellcome Sanger Institute for the COVID-19 Genomics UK (COG-UK) Consortium | Jacquelyn Wynn, Mairead Hyland, The Lighthouse Lab in Alderley Park and Alex Alderton, Roberto Amato, Jeffrey Barrett, Sonia Goncalves, Ewan Harrison, David K. Jackson, Ian Johnston, Dominic Kwiatkowski, Cordelia Langford, John Sillitoe on behalf of the Wellcome Sanger Institute COVID-19 Surveillance Team |
| EPI_ISL_2436432, EPI_ISL_2436433 | Lighthouse Lab in Milton Keynes | Wellcome Sanger Institute for the COVID-19 Genomics UK (COG-UK) Consortium | The Lighthouse Lab in Milton Keynes and Alex Alderton, Roberto Amato, Jeffrey Barrett, Sonia Goncalves, Ewan Harrison, David K. Jackson, Ian Johnston, Dominic Kwiatkowski, Cordelia Langford, John Sillitoe on behalf of the Wellcome Sanger Institute COVID-19 Surveillance Team |
| EPI_ISL_2436434 | Lighthouse Lab in Glasgow | Wellcome Sanger Institute for the COVID-19 Genomics UK (COG-UK) Consortium | Harper VanSteenhouse, Yumi Kasai, David Gray, Carol Clugston, Anna Dominiczak and Alex Alderton, Roberto Amato, Jeffrey Barrett, Sonia Goncalves, Ewan Harrison, David K. Jackson, Ian Johnston, Dominic Kwiatkowski, Cordelia Langford, John Sillitoe on behalf of the Wellcome Sanger Institute COVID-19 Surveillance Team |
| EPI_ISL_2436435 | Health Services Laboratories | Wellcome Sanger Institute for the COVID-19 Genomics UK (COG-UK) Consortium | Health Services Laboratories and Alex Alderton, Roberto Amato, Jeffrey Barrett, Sonia Goncalves, Ewan Harrison, David K. Jackson, Ian Johnston, Dominic Kwiatkowski, Cordelia Langford, John Sillitoe on behalf of the Wellcome Sanger Institute COVID-19 Surveillance Team |
| EPI_ISL_2436436 | Lighthouse Lab in Glasgow | Wellcome Sanger Institute for the COVID-19 Genomics UK (COG-UK) Consortium | Harper VanSteenhouse, Yumi Kasai, David Gray, Carol Clugston, Anna Dominiczak and Alex Alderton, Roberto Amato, Jeffrey Barrett, Sonia Goncalves, Ewan Harrison, David K. Jackson, Ian Johnston, Dominic Kwiatkowski, Cordelia Langford, John Sillitoe on behalf of the Wellcome Sanger Institute COVID-19 Surveillance Team |
| EPI_ISL_2436437, EPI_ISL_2436438 | Lighthouse Lab in Alderley Park | Wellcome Sanger Institute for the COVID-19 Genomics UK (COG-UK) Consortium | Jacquelyn Wynn, Mairead Hyland, The Lighthouse Lab in Alderley Park and Alex Alderton, Roberto Amato, Jeffrey Barrett, Sonia Goncalves, Ewan Harrison, David K. Jackson, Ian Johnston, Dominic Kwiatkowski, Cordelia Langford, John Sillitoe on behalf of the Wellcome Sanger Institute COVID-19 Surveillance Team |
| EPI_ISL_2436439, EPI_ISL_2436440 | Health Services Laboratories | Wellcome Sanger Institute for the COVID-19 Genomics UK (COG-UK) Consortium | Health Services Laboratories and Alex Alderton, Roberto Amato, Jeffrey Barrett, Sonia Goncalves, Ewan Harrison, David K. Jackson, Ian Johnston, Dominic Kwiatkowski, Cordelia Langford, John Sillitoe on behalf of the Wellcome Sanger Institute COVID-19 Surveillance Team |
| EPI_ISL_2436442 | Lighthouse Lab in Glasgow | Wellcome Sanger Institute for the COVID-19 Genomics UK (COG-UK) Consortium | Harper VanSteenhouse, Yumi Kasai, David Gray, Carol Clugston, Anna Dominiczak and Alex Alderton, Roberto Amato, Jeffrey Barrett, Sonia Goncalves, Ewan Harrison, David K. Jackson, Ian Johnston, Dominic Kwiatkowski, Cordelia Langford, John Sillitoe on behalf of the Wellcome Sanger Institute COVID-19 Surveillance Team |
| EPI_ISL_2436443 | Health Services Laboratories | Wellcome Sanger Institute for the COVID-19 Genomics UK (COG-UK) Consortium | Health Services Laboratories and Alex Alderton, Roberto Amato, Jeffrey Barrett, Sonia Goncalves, Ewan Harrison, David K. Jackson, Ian Johnston, Dominic Kwiatkowski, Cordelia Langford, John Sillitoe on behalf of the Wellcome Sanger Institute COVID-19 Surveillance Team |
| EPI_ISL_2436444 | Lighthouse Lab in Milton Keynes | Wellcome Sanger Institute for the COVID-19 Genomics UK (COG-UK) Consortium | The Lighthouse Lab in Milton Keynes and Alex Alderton, Roberto Amato, Jeffrey Barrett, Sonia Goncalves, Ewan Harrison, David K. Jackson, Ian Johnston, Dominic Kwiatkowski, Cordelia Langford, John Sillitoe on behalf of the Wellcome Sanger Institute COVID-19 Surveillance Team |
| EPI_ISL_2436445 | Lighthouse Lab in Glasgow | Wellcome Sanger Institute for the COVID-19 Genomics UK (COG-UK) Consortium | Harper VanSteenhouse, Yumi Kasai, David Gray, Carol Clugston, Anna Dominiczak and Alex Alderton, Roberto Amato, Jeffrey Barrett, Sonia Goncalves, Ewan Harrison, David K. Jackson, Ian Johnston, Dominic Kwiatkowski, Cordelia Langford, John Sillitoe on behalf of the Wellcome Sanger Institute COVID-19 Surveillance Team |
| EPI_ISL_2436446 | Lighthouse Lab in Alderley Park | Wellcome Sanger Institute for the COVID-19 Genomics UK (COG-UK) Consortium | Jacquelyn Wynn, Mairead Hyland, The Lighthouse Lab in Alderley Park and Alex Alderton, Roberto Amato, Jeffrey Barrett, Sonia Goncalves, Ewan Harrison, David K. Jackson, Ian Johnston, Dominic Kwiatkowski, Cordelia Langford, John Sillitoe on behalf of the Wellcome Sanger Institute COVID-19 Surveillance Team |
| EPI_ISL_2436447, EPI_ISL_2436449 | Lighthouse Lab in Glasgow | Wellcome Sanger Institute for the COVID-19 Genomics UK (COG-UK) Consortium | Harper VanSteenhouse, Yumi Kasai, David Gray, Carol Clugston, Anna Dominiczak and Alex Alderton, Roberto Amato, Jeffrey Barrett, Sonia Goncalves, Ewan Harrison, David K. Jackson, Ian Johnston, Dominic Kwiatkowski, Cordelia Langford, John Sillitoe on behalf of the Wellcome Sanger Institute COVID-19 Surveillance Team |
| EPI_ISL_2436450, EPI_ISL_2436451, EPI_ISL_2436452, EPI_ISL_2436453 | Lighthouse Lab in Alderley Park | Wellcome Sanger Institute for the COVID-19 Genomics UK (COG-UK) Consortium | Jacquelyn Wynn, Mairead Hyland, The Lighthouse Lab in Alderley Park and Alex Alderton, Roberto Amato, Jeffrey Barrett, Sonia Goncalves, Ewan Harrison, David K. Jackson, Ian Johnston, Dominic Kwiatkowski, Cordelia Langford, John Sillitoe on behalf of the Wellcome Sanger Institute COVID-19 Surveillance Team |
| EPI_ISL_2436455 | Lighthouse Lab in Milton Keynes | Wellcome Sanger Institute for the COVID-19 Genomics UK (COG-UK) Consortium | The Lighthouse Lab in Milton Keynes and Alex Alderton, Roberto Amato, Jeffrey Barrett, Sonia Goncalves, Ewan Harrison, David K. Jackson, Ian Johnston, Dominic Kwiatkowski, Cordelia Langford, John Sillitoe on behalf of the Wellcome Sanger Institute COVID-19 Surveillance Team |
| EPI_ISL_2436456, EPI_ISL_2436457 | Lighthouse Lab in Alderley Park | Wellcome Sanger Institute for the COVID-19 Genomics UK (COG-UK) Consortium | Jacquelyn Wynn, Mairead Hyland, The Lighthouse Lab in Alderley Park and Alex Alderton, Roberto Amato, Jeffrey Barrett, Sonia Goncalves, Ewan Harrison, David K. Jackson, Ian Johnston, Dominic Kwiatkowski, Cordelia Langford, John Sillitoe on behalf of the Wellcome Sanger Institute COVID-19 Surveillance Team |
| EPI_ISL_2436458 | Lighthouse Lab in Milton Keynes | Wellcome Sanger Institute for the COVID-19 Genomics UK (COG-UK) Consortium | The Lighthouse Lab in Milton Keynes and Alex Alderton, Roberto Amato, Jeffrey Barrett, Sonia Goncalves, Ewan Harrison, David K. Jackson, Ian Johnston, Dominic Kwiatkowski, Cordelia Langford, John Sillitoe on behalf of the Wellcome Sanger Institute COVID-19 Surveillance Team |
| EPI_ISL_2436459 | Lighthouse Laboratory Plymouth | Wellcome Sanger Institute for the COVID-19 Genomics UK (COG-UK) Consortium | Lighthouse Laboratory Plymouth and Alex Alderton, Roberto Amato, Jeffrey Barrett, Sonia Goncalves, Ewan Harrison, David K. Jackson, Ian Johnston, Dominic Kwiatkowski, Cordelia Langford, John Sillitoe on behalf of the Wellcome Sanger Institute COVID-19 Surveillance Team |
| EPI_ISL_2436460 | Lighthouse Lab in Milton Keynes | Wellcome Sanger Institute for the COVID-19 Genomics UK (COG-UK) Consortium | The Lighthouse Lab in Milton Keynes and Alex Alderton, Roberto Amato, Jeffrey Barrett, Sonia Goncalves, Ewan Harrison, David K. Jackson, Ian Johnston, Dominic Kwiatkowski, Cordelia Langford, John Sillitoe on behalf of the Wellcome Sanger Institute COVID-19 Surveillance Team |
| EPI_ISL_2436461 | Lighthouse Lab in Alderley Park | Wellcome Sanger Institute for the COVID-19 Genomics UK (COG-UK) Consortium | Jacquelyn Wynn, Mairead Hyland, The Lighthouse Lab in Alderley Park and Alex Alderton, Roberto Amato, Jeffrey Barrett, Sonia Goncalves, Ewan Harrison, David K. Jackson, Ian Johnston, Dominic Kwiatkowski, Cordelia Langford, John Sillitoe on behalf of the Wellcome Sanger Institute COVID-19 Surveillance Team |
| EPI_ISL_2436463 | Health Services Laboratories | Wellcome Sanger Institute for the COVID-19 Genomics UK (COG-UK) Consortium | Health Services Laboratories and Alex Alderton, Roberto Amato, Jeffrey Barrett, Sonia Goncalves, Ewan Harrison, David K. Jackson, Ian Johnston, Dominic Kwiatkowski, Cordelia Langford, John Sillitoe on behalf of the Wellcome Sanger Institute COVID-19 Surveillance Team |
| EPI_ISL_2436464, EPI_ISL_2436465 | Lighthouse Lab in Glasgow | Wellcome Sanger Institute for the COVID-19 Genomics UK (COG-UK) Consortium | Harper VanSteenhouse, Yumi Kasai, David Gray, Carol Clugston, Anna Dominiczak and Alex Alderton, Roberto Amato, Jeffrey Barrett, Sonia Goncalves, Ewan Harrison, David K. Jackson, Ian Johnston, Dominic Kwiatkowski, Cordelia Langford, John Sillitoe on behalf of the Wellcome Sanger Institute COVID-19 Surveillance Team |
| EPI_ISL_2436466 | Lighthouse Lab in Milton Keynes | Wellcome Sanger Institute for the COVID-19 Genomics UK (COG-UK) Consortium | The Lighthouse Lab in Milton Keynes and Alex Alderton, Roberto Amato, Jeffrey Barrett, Sonia Goncalves, Ewan Harrison, David K. Jackson, Ian Johnston, Dominic Kwiatkowski, Cordelia Langford, John Sillitoe on behalf of the Wellcome Sanger Institute COVID-19 Surveillance Team |
| EPI_ISL_2436467 | Lighthouse Laboratory Plymouth | Wellcome Sanger Institute for the COVID-19 Genomics UK (COG-UK) Consortium | Lighthouse Laboratory Plymouth and Alex Alderton, Roberto Amato, Jeffrey Barrett, Sonia Goncalves, Ewan Harrison, David K. Jackson, Ian Johnston, Dominic Kwiatkowski, Cordelia Langford, John Sillitoe on behalf of the Wellcome Sanger Institute COVID-19 Surveillance Team |
| EPI_ISL_2436468 | Lighthouse Lab in Alderley Park | Wellcome Sanger Institute for the COVID-19 Genomics UK (COG-UK) Consortium | Jacquelyn Wynn, Mairead Hyland, The Lighthouse Lab in Alderley Park and Alex Alderton, Roberto Amato, Jeffrey Barrett, Sonia Goncalves, Ewan Harrison, David K. Jackson, Ian Johnston, Dominic Kwiatkowski, Cordelia Langford, John Sillitoe on behalf of the Wellcome Sanger Institute COVID-19 Surveillance Team |

| | | | |
|---|---|---|---|
| EPI_ISL_2436470 | Lighthouse Lab in Glasgow | Wellcome Sanger Institute for the COVID-19 Genomics UK (COG-UK) Consortium | Harper VanSteenhouse, Yumi Kasai, David Gray, Carol Clugston, Anna Dominiczak and Alex Alderton, Roberto Amato, Jeffrey Barrett, Sonia Goncalves, Ewan Harrison, David K. Jackson, Ian Johnston, Dominic Kwiatkowski, Cordelia Langford, John Sillitoe on behalf of the Wellcome Sanger Institute COVID-19 Surveillance Team |
| EPI_ISL_2436472 | Health Services Laboratories | Wellcome Sanger Institute for the COVID-19 Genomics UK (COG-UK) Consortium | Health Services Laboratories and Alex Alderton, Roberto Amato, Jeffrey Barrett, Sonia Goncalves, Ewan Harrison, David K. Jackson, Ian Johnston, Dominic Kwiatkowski, Cordelia Langford, John Sillitoe on behalf of the Wellcome Sanger Institute COVID-19 Surveillance Team |
| EPI_ISL_2436473, EPI_ISL_2436474 | Lighthouse Lab in Alderley Park | Wellcome Sanger Institute for the COVID-19 Genomics UK (COG-UK) Consortium | Jacquelyn Wynn, Mairead Hyland, The Lighthouse Lab in Alderley Park and Alex Alderton, Roberto Amato, Jeffrey Barrett, Sonia Goncalves, Ewan Harrison, David K. Jackson, Ian Johnston, Dominic Kwiatkowski, Cordelia Langford, John Sillitoe on behalf of the Wellcome Sanger Institute COVID-19 Surveillance Team |
| EPI_ISL_2436475 | Health Services Laboratories | Wellcome Sanger Institute for the COVID-19 Genomics UK (COG-UK) Consortium | Health Services Laboratories and Alex Alderton, Roberto Amato, Jeffrey Barrett, Sonia Goncalves, Ewan Harrison, David K. Jackson, Ian Johnston, Dominic Kwiatkowski, Cordelia Langford, John Sillitoe on behalf of the Wellcome Sanger Institute COVID-19 Surveillance Team |
| EPI_ISL_2436476 | Lighthouse Lab in Alderley Park | Wellcome Sanger Institute for the COVID-19 Genomics UK (COG-UK) Consortium | Jacquelyn Wynn, Mairead Hyland, The Lighthouse Lab in Alderley Park and Alex Alderton, Roberto Amato, Jeffrey Barrett, Sonia Goncalves, Ewan Harrison, David K. Jackson, Ian Johnston, Dominic Kwiatkowski, Cordelia Langford, John Sillitoe on behalf of the Wellcome Sanger Institute COVID-19 Surveillance Team |
| EPI_ISL_2436477, EPI_ISL_2436478 | Health Services Laboratories | Wellcome Sanger Institute for the COVID-19 Genomics UK (COG-UK) Consortium | Health Services Laboratories and Alex Alderton, Roberto Amato, Jeffrey Barrett, Sonia Goncalves, Ewan Harrison, David K. Jackson, Ian Johnston, Dominic Kwiatkowski, Cordelia Langford, John Sillitoe on behalf of the Wellcome Sanger Institute COVID-19 Surveillance Team |
| EPI_ISL_2436479 | Lighthouse Lab in Alderley Park | Wellcome Sanger Institute for the COVID-19 Genomics UK (COG-UK) Consortium | Jacquelyn Wynn, Mairead Hyland, The Lighthouse Lab in Alderley Park and Alex Alderton, Roberto Amato, Jeffrey Barrett, Sonia Goncalves, Ewan Harrison, David K. Jackson, Ian Johnston, Dominic Kwiatkowski, Cordelia Langford, John Sillitoe on behalf of the Wellcome Sanger Institute COVID-19 Surveillance Team |
| EPI_ISL_2436480, EPI_ISL_2436481 | Lighthouse Lab in Milton Keynes | Wellcome Sanger Institute for the COVID-19 Genomics UK (COG-UK) Consortium | The Lighthouse Lab in Milton Keynes and Alex Alderton, Roberto Amato, Jeffrey Barrett, Sonia Goncalves, Ewan Harrison, David K. Jackson, Ian Johnston, Dominic Kwiatkowski, Cordelia Langford, John Sillitoe on behalf of the Wellcome Sanger Institute COVID-19 Surveillance Team |
| EPI_ISL_2436482 | Health Services Laboratories | Wellcome Sanger Institute for the COVID-19 Genomics UK (COG-UK) Consortium | Health Services Laboratories and Alex Alderton, Roberto Amato, Jeffrey Barrett, Sonia Goncalves, Ewan Harrison, David K. Jackson, Ian Johnston, Dominic Kwiatkowski, Cordelia Langford, John Sillitoe on behalf of the Wellcome Sanger Institute COVID-19 Surveillance Team |
| EPI_ISL_2436483 | Lighthouse Lab in Glasgow | Wellcome Sanger Institute for the COVID-19 Genomics UK (COG-UK) Consortium | Harper VanSteenhouse, Yumi Kasai, David Gray, Carol Clugston, Anna Dominiczak and Alex Alderton, Roberto Amato, Jeffrey Barrett, Sonia Goncalves, Ewan Harrison, David K. Jackson, Ian Johnston, Dominic Kwiatkowski, Cordelia Langford, John Sillitoe on behalf of the Wellcome Sanger Institute COVID-19 Surveillance Team |
| EPI_ISL_2436484, EPI_ISL_2436485, EPI_ISL_2436486, EPI_ISL_2436487 | Health Services Laboratories | Wellcome Sanger Institute for the COVID-19 Genomics UK (COG-UK) Consortium | Health Services Laboratories and Alex Alderton, Roberto Amato, Jeffrey Barrett, Sonia Goncalves, Ewan Harrison, David K. Jackson, Ian Johnston, Dominic Kwiatkowski, Cordelia Langford, John Sillitoe on behalf of the Wellcome Sanger Institute COVID-19 Surveillance Team |
| EPI_ISL_2436488 | Lighthouse Lab in Milton Keynes | Wellcome Sanger Institute for the COVID-19 Genomics UK (COG-UK) Consortium | The Lighthouse Lab in Milton Keynes and Alex Alderton, Roberto Amato, Jeffrey Barrett, Sonia Goncalves, Ewan Harrison, David K. Jackson, Ian Johnston, Dominic Kwiatkowski, Cordelia Langford, John Sillitoe on behalf of the Wellcome Sanger Institute COVID-19 Surveillance Team |
| EPI_ISL_2436490 | Lighthouse Laboratory Plymouth | Wellcome Sanger Institute for the COVID-19 Genomics UK (COG-UK) Consortium | Lighthouse Laboratory Plymouth and Alex Alderton, Roberto Amato, Jeffrey Barrett, Sonia Goncalves, Ewan Harrison, David K. Jackson, Ian Johnston, Dominic Kwiatkowski, Cordelia Langford, John Sillitoe on behalf of the Wellcome Sanger Institute COVID-19 Surveillance Team |
| EPI_ISL_2436491 | Lighthouse Lab in Alderley Park | Wellcome Sanger Institute for the COVID-19 Genomics UK (COG-UK) Consortium | Jacquelyn Wynn, Mairead Hyland, The Lighthouse Lab in Alderley Park and Alex Alderton, Roberto Amato, Jeffrey Barrett, Sonia Goncalves, Ewan Harrison, David K. Jackson, Ian Johnston, Dominic Kwiatkowski, Cordelia Langford, John Sillitoe on behalf of the Wellcome Sanger Institute COVID-19 Surveillance Team |
| EPI_ISL_2436492 | Lighthouse Lab in Milton Keynes | Wellcome Sanger Institute for the COVID-19 Genomics UK (COG-UK) Consortium | The Lighthouse Lab in Milton Keynes and Alex Alderton, Roberto Amato, Jeffrey Barrett, Sonia Goncalves, Ewan Harrison, David K. Jackson, Ian Johnston, Dominic Kwiatkowski, Cordelia Langford, John Sillitoe on behalf of the Wellcome Sanger Institute COVID-19 Surveillance Team |
| EPI_ISL_2436493 | Health Services Laboratories | Wellcome Sanger Institute for the COVID-19 Genomics UK (COG-UK) Consortium | Health Services Laboratories and Alex Alderton, Roberto Amato, Jeffrey Barrett, Sonia Goncalves, Ewan Harrison, David K. Jackson, Ian Johnston, Dominic Kwiatkowski, Cordelia Langford, John Sillitoe on behalf of the Wellcome Sanger Institute COVID-19 Surveillance Team |
| EPI_ISL_2436494, EPI_ISL_2436495, EPI_ISL_2436496, EPI_ISL_2436497, EPI_ISL_2436498 | Lighthouse Lab in Milton Keynes | Wellcome Sanger Institute for the COVID-19 Genomics UK (COG-UK) Consortium | The Lighthouse Lab in Milton Keynes and Alex Alderton, Roberto Amato, Jeffrey Barrett, Sonia Goncalves, Ewan Harrison, David K. Jackson, Ian Johnston, Dominic Kwiatkowski, Cordelia Langford, John Sillitoe on behalf of the Wellcome Sanger Institute COVID-19 Surveillance Team |
| EPI_ISL_2436499 | Lighthouse Laboratory Plymouth | Wellcome Sanger Institute for the COVID-19 Genomics UK (COG-UK) Consortium | Lighthouse Laboratory Plymouth and Alex Alderton, Roberto Amato, Jeffrey Barrett, Sonia Goncalves, Ewan Harrison, David K. Jackson, Ian Johnston, Dominic Kwiatkowski, Cordelia Langford, John Sillitoe on behalf of the Wellcome Sanger Institute COVID-19 Surveillance Team |
| EPI_ISL_2436501 | Health Services Laboratories | Wellcome Sanger Institute for the COVID-19 Genomics UK (COG-UK) Consortium | Health Services Laboratories and Alex Alderton, Roberto Amato, Jeffrey Barrett, Sonia Goncalves, Ewan Harrison, David K. Jackson, Ian Johnston, Dominic Kwiatkowski, Cordelia Langford, John Sillitoe on behalf of the Wellcome Sanger Institute COVID-19 Surveillance Team |
| EPI_ISL_2436503 | Lighthouse Lab in Milton Keynes | Wellcome Sanger Institute for the COVID-19 Genomics UK (COG-UK) Consortium | The Lighthouse Lab in Milton Keynes and Alex Alderton, Roberto Amato, Jeffrey Barrett, Sonia Goncalves, Ewan Harrison, David K. Jackson, Ian Johnston, Dominic Kwiatkowski, Cordelia Langford, John Sillitoe on behalf of the Wellcome Sanger Institute COVID-19 Surveillance Team |
| EPI_ISL_2436504, EPI_ISL_2436505 | Lighthouse Lab in Alderley Park | Wellcome Sanger Institute for the COVID-19 Genomics UK (COG-UK) Consortium | Jacquelyn Wynn, Mairead Hyland, The Lighthouse Lab in Alderley Park and Alex Alderton, Roberto Amato, Jeffrey Barrett, Sonia Goncalves, Ewan Harrison, David K. Jackson, Ian Johnston, Dominic Kwiatkowski, Cordelia Langford, John Sillitoe on behalf of the Wellcome Sanger Institute COVID-19 Surveillance Team |
| EPI_ISL_2436506, EPI_ISL_2436507, EPI_ISL_2436509 | Lighthouse Lab in Milton Keynes | Wellcome Sanger Institute for the COVID-19 Genomics UK (COG-UK) Consortium | The Lighthouse Lab in Milton Keynes and Alex Alderton, Roberto Amato, Jeffrey Barrett, Sonia Goncalves, Ewan Harrison, David K. Jackson, Ian Johnston, Dominic Kwiatkowski, Cordelia Langford, John Sillitoe on behalf of the Wellcome Sanger Institute COVID-19 Surveillance Team |
| EPI_ISL_2436511 | Lighthouse Lab in Glasgow | Wellcome Sanger Institute for the COVID-19 Genomics UK (COG-UK) Consortium | Harper VanSteenhouse, Yumi Kasai, David Gray, Carol Clugston, Anna Dominiczak and Alex Alderton, Roberto Amato, Jeffrey Barrett, Sonia Goncalves, Ewan Harrison, David K. Jackson, Ian Johnston, Dominic Kwiatkowski, Cordelia Langford, John Sillitoe on behalf of the Wellcome Sanger Institute COVID-19 Surveillance Team |
| EPI_ISL_2436512, EPI_ISL_2436513 | Lighthouse Lab in Milton Keynes | Wellcome Sanger Institute for the COVID-19 Genomics UK (COG-UK) Consortium | The Lighthouse Lab in Milton Keynes and Alex Alderton, Roberto Amato, Jeffrey Barrett, Sonia Goncalves, Ewan Harrison, David K. Jackson, Ian Johnston, Dominic Kwiatkowski, Cordelia Langford, John Sillitoe on behalf of the Wellcome Sanger Institute COVID-19 Surveillance Team |
| EPI_ISL_2436514 | Lighthouse Lab in Glasgow | Wellcome Sanger Institute for the COVID-19 Genomics UK (COG-UK) Consortium | Harper VanSteenhouse, Yumi Kasai, David Gray, Carol Clugston, Anna Dominiczak and Alex Alderton, Roberto Amato, Jeffrey Barrett, Sonia Goncalves, Ewan Harrison, David K. Jackson, Ian Johnston, Dominic Kwiatkowski, Cordelia Langford, John Sillitoe on behalf of the Wellcome Sanger Institute COVID-19 Surveillance Team |
| EPI_ISL_2436515 | Health Services Laboratories | Wellcome Sanger Institute for the COVID-19 Genomics UK (COG-UK) Consortium | Health Services Laboratories and Alex Alderton, Roberto Amato, Jeffrey Barrett, Sonia Goncalves, Ewan Harrison, David K. Jackson, Ian Johnston, Dominic Kwiatkowski, Cordelia Langford, John Sillitoe on behalf of the Wellcome Sanger Institute COVID-19 Surveillance Team |
| EPI_ISL_2436516, EPI_ISL_2436517 | Lighthouse Lab in Alderley Park | Wellcome Sanger Institute for the COVID-19 Genomics UK (COG-UK) Consortium | Jacquelyn Wynn, Mairead Hyland, The Lighthouse Lab in Alderley Park and Alex Alderton, Roberto Amato, Jeffrey Barrett, Sonia Goncalves, Ewan Harrison, David K. Jackson, Ian Johnston, Dominic Kwiatkowski, Cordelia Langford, John Sillitoe on behalf of the Wellcome Sanger Institute COVID-19 Surveillance Team |
| EPI_ISL_2436518, EPI_ISL_2436520 | Lighthouse Lab in Milton Keynes | Wellcome Sanger Institute for the COVID-19 Genomics UK (COG-UK) Consortium | The Lighthouse Lab in Milton Keynes and Alex Alderton, Roberto Amato, Jeffrey Barrett, Sonia Goncalves, Ewan Harrison, David K. Jackson, Ian Johnston, Dominic Kwiatkowski, Cordelia Langford, John Sillitoe on behalf of the Wellcome Sanger Institute COVID-19 Surveillance Team |
| EPI_ISL_2436521 | Lighthouse Lab in Glasgow | Wellcome Sanger Institute for the COVID-19 Genomics UK (COG-UK) Consortium | Harper VanSteenhouse, Yumi Kasai, David Gray, Carol Clugston, Anna Dominiczak and Alex Alderton, Roberto Amato, Jeffrey Barrett, Sonia Goncalves, Ewan Harrison, David K. Jackson, Ian Johnston, Dominic Kwiatkowski, Cordelia Langford, John Sillitoe on behalf of the Wellcome Sanger Institute COVID-19 Surveillance Team |
| EPI_ISL_2436522 | Lighthouse Lab in Alderley Park | Wellcome Sanger Institute for the COVID-19 Genomics UK (COG-UK) Consortium | Jacquelyn Wynn, Mairead Hyland, The Lighthouse Lab in Alderley Park and Alex Alderton, Roberto Amato, Jeffrey Barrett, Sonia Goncalves, Ewan Harrison, David K. Jackson, Ian Johnston, Dominic Kwiatkowski, Cordelia Langford, John Sillitoe on behalf of the Wellcome Sanger Institute COVID-19 Surveillance Team |
| EPI_ISL_2436523 | Lighthouse Lab in Milton Keynes | Wellcome Sanger Institute for the COVID-19 Genomics UK (COG-UK) Consortium | The Lighthouse Lab in Milton Keynes and Alex Alderton, Roberto Amato, Jeffrey Barrett, Sonia Goncalves, Ewan Harrison, David K. Jackson, Ian Johnston, Dominic Kwiatkowski, Cordelia Langford, John Sillitoe on behalf of the Wellcome Sanger Institute COVID-19 Surveillance Team |
| EPI_ISL_2436524 | Lighthouse Lab in Alderley Park | Wellcome Sanger Institute for the COVID-19 Genomics UK (COG-UK) Consortium | Jacquelyn Wynn, Mairead Hyland, The Lighthouse Lab in Alderley Park and Alex Alderton, Roberto Amato, Jeffrey Barrett, Sonia Goncalves, Ewan Harrison, David K. Jackson, Ian Johnston, Dominic Kwiatkowski, Cordelia Langford, John Sillitoe on behalf of the Wellcome Sanger Institute COVID-19 Surveillance Team |
| EPI_ISL_2436525 | Health Services Laboratories | Wellcome Sanger Institute for the COVID-19 Genomics UK (COG-UK) Consortium | Health Services Laboratories and Alex Alderton, Roberto Amato, Jeffrey Barrett, Sonia Goncalves, Ewan Harrison, David K. Jackson, Ian Johnston, Dominic Kwiatkowski, Cordelia Langford, John Sillitoe on behalf of the Wellcome Sanger Institute COVID-19 Surveillance Team |
| EPI_ISL_2436526 | Lighthouse Laboratory Plymouth | Wellcome Sanger Institute for the COVID-19 Genomics UK (COG-UK) Consortium | Lighthouse Laboratory Plymouth and Alex Alderton, Roberto Amato, Jeffrey Barrett, Sonia Goncalves, Ewan Harrison, David K. Jackson, Ian Johnston, Dominic Kwiatkowski, Cordelia Langford, John Sillitoe on behalf of the Wellcome Sanger Institute COVID-19 Surveillance Team |
| EPI_ISL_2436528 | Lighthouse Lab in Glasgow | Wellcome Sanger Institute for the COVID-19 Genomics UK (COG-UK) Consortium | Harper VanSteenhouse, Yumi Kasai, David Gray, Carol Clugston, Anna Dominiczak and Alex Alderton, Roberto Amato, Jeffrey Barrett, Sonia Goncalves, Ewan Harrison, David K. Jackson, Ian Johnston, Dominic Kwiatkowski, Cordelia Langford, John Sillitoe on behalf of the Wellcome Sanger Institute COVID-19 Surveillance Team |
| EPI_ISL_2436530, EPI_ISL_2436531 | Lighthouse Lab in Milton Keynes | Wellcome Sanger Institute for the COVID-19 Genomics UK (COG-UK) Consortium | The Lighthouse Lab in Milton Keynes and Alex Alderton, Roberto Amato, Jeffrey Barrett, Sonia Goncalves, Ewan Harrison, David K. Jackson, Ian Johnston, Dominic Kwiatkowski, Cordelia Langford, John Sillitoe on behalf of the Wellcome Sanger Institute COVID-19 Surveillance Team |
| EPI_ISL_2436532, EPI_ISL_2436533 | Lighthouse Lab in Alderley Park | Wellcome Sanger Institute for the COVID-19 Genomics UK (COG-UK) Consortium | Jacquelyn Wynn, Mairead Hyland, The Lighthouse Lab in Alderley Park and Alex Alderton, Roberto Amato, Jeffrey Barrett, Sonia Goncalves, Ewan Harrison, David K. Jackson, Ian Johnston, Dominic Kwiatkowski, Cordelia Langford, John Sillitoe on behalf of the Wellcome Sanger Institute COVID-19 Surveillance Team |
| EPI_ISL_2436535 | Health Services Laboratories | Wellcome Sanger Institute for the COVID-19 Genomics UK (COG-UK) Consortium | Health Services Laboratories and Alex Alderton, Roberto Amato, Jeffrey Barrett, Sonia Goncalves, Ewan Harrison, David K. Jackson, Ian Johnston, Dominic Kwiatkowski, Cordelia Langford, John Sillitoe on behalf of the Wellcome Sanger Institute COVID-19 Surveillance Team |
| EPI_ISL_2436536 | Lighthouse Lab in Alderley Park | Wellcome Sanger Institute for the COVID-19 Genomics UK (COG-UK) Consortium | Jacquelyn Wynn, Mairead Hyland, The Lighthouse Lab in Alderley Park and Alex Alderton, Roberto Amato, Jeffrey Barrett, Sonia Goncalves, Ewan Harrison, David K. Jackson, Ian Johnston, Dominic Kwiatkowski, Cordelia Langford, John Sillitoe on behalf of the Wellcome Sanger Institute COVID-19 Surveillance Team |
| EPI_ISL_2436537, EPI_ISL_2436539 | Lighthouse Laboratory Plymouth | Wellcome Sanger Institute for the COVID-19 Genomics UK (COG-UK) Consortium | Lighthouse Laboratory Plymouth and Alex Alderton, Roberto Amato, Jeffrey Barrett, Sonia Goncalves, Ewan Harrison, David K. Jackson, Ian Johnston, Dominic Kwiatkowski, Cordelia Langford, John Sillitoe on behalf of the Wellcome Sanger Institute COVID-19 Surveillance Team |
| EPI_ISL_2436541, EPI_ISL_2436542, EPI_ISL_2436543 | Lighthouse Lab in Alderley Park | Wellcome Sanger Institute for the COVID-19 Genomics UK (COG-UK) Consortium | Jacquelyn Wynn, Mairead Hyland, The Lighthouse Lab in Alderley Park and Alex Alderton, Roberto Amato, Jeffrey Barrett, Sonia Goncalves, Ewan Harrison, David K. Jackson, Ian Johnston, Dominic Kwiatkowski, Cordelia Langford, John Sillitoe on behalf of the Wellcome Sanger Institute COVID-19 Surveillance Team |
| EPI_ISL_2436544 | Lighthouse Laboratory Plymouth | Wellcome Sanger Institute for the COVID-19 Genomics UK (COG-UK) Consortium | Lighthouse Laboratory Plymouth and Alex Alderton, Roberto Amato, Jeffrey Barrett, Sonia Goncalves, Ewan Harrison, David K. Jackson, Ian Johnston, Dominic Kwiatkowski, Cordelia Langford, John Sillitoe on behalf of the Wellcome Sanger Institute COVID-19 Surveillance Team |
| EPI_ISL_2436545, | Lighthouse Lab in Milton Keynes | Wellcome Sanger Institute for the COVID-19 Genomics UK (COG-UK) | The Lighthouse Lab in Milton Keynes and Alex Alderton, Roberto Amato, Jeffrey Barrett, Sonia Goncalves, Ewan Harrison, David K. Jackson, Ian Johnston, Dominic Kwiatkowski, Cordelia Langford, John Sillitoe on behalf of the Wellcome Sanger |

| | | | |
|---|---|---|---|
| EPI_ISL_2436546 | | Consortium | Institute COVID-19 Surveillance Team |
| EPI_ISL_2436547 | Lighthouse Lab in Glasgow | Wellcome Sanger Institute for the COVID-19 Genomics UK (COG-UK) Consortium | Harper VanSteenhouse, Yumi Kasai, David Gray, Carol Clugston, Anna Dominiczak and Alex Alderton, Roberto Amato, Jeffrey Barrett, Sonia Goncalves, Ewan Harrison, David K. Jackson, Ian Johnston, Dominic Kwiatkowski, Cordelia Langford, John Sillitoe on behalf of the Wellcome Sanger Institute COVID-19 Surveillance Team |
| EPI_ISL_2436548, EPI_ISL_2436549 | Lighthouse Laboratory Plymouth | Wellcome Sanger Institute for the COVID-19 Genomics UK (COG-UK) Consortium | Lighthouse Laboratory Plymouth and Alex Alderton, Roberto Amato, Jeffrey Barrett, Sonia Goncalves, Ewan Harrison, David K. Jackson, Ian Johnston, Dominic Kwiatkowski, Cordelia Langford, John Sillitoe on behalf of the Wellcome Sanger Institute COVID-19 Surveillance Team |
| EPI_ISL_2436550 | Health Services Laboratories | Wellcome Sanger Institute for the COVID-19 Genomics UK (COG-UK) Consortium | Health Services Laboratories and Alex Alderton, Roberto Amato, Jeffrey Barrett, Sonia Goncalves, Ewan Harrison, David K. Jackson, Ian Johnston, Dominic Kwiatkowski, Cordelia Langford, John Sillitoe on behalf of the Wellcome Sanger Institute COVID-19 Surveillance Team |
| EPI_ISL_2436551 | Lighthouse Lab in Alderley Park | Wellcome Sanger Institute for the COVID-19 Genomics UK (COG-UK) Consortium | Jacquelyn Wynn, Mairead Hyland, The Lighthouse Lab in Alderley Park and Alex Alderton, Roberto Amato, Jeffrey Barrett, Sonia Goncalves, Ewan Harrison, David K. Jackson, Ian Johnston, Dominic Kwiatkowski, Cordelia Langford, John Sillitoe on behalf of the Wellcome Sanger Institute COVID-19 Surveillance Team |
| EPI_ISL_2436552 | Lighthouse Lab in Glasgow | Wellcome Sanger Institute for the COVID-19 Genomics UK (COG-UK) Consortium | Harper VanSteenhouse, Yumi Kasai, David Gray, Carol Clugston, Anna Dominiczak and Alex Alderton, Roberto Amato, Jeffrey Barrett, Sonia Goncalves, Ewan Harrison, David K. Jackson, Ian Johnston, Dominic Kwiatkowski, Cordelia Langford, John Sillitoe on behalf of the Wellcome Sanger Institute COVID-19 Surveillance Team |
| EPI_ISL_2436553 | Health Services Laboratories | Wellcome Sanger Institute for the COVID-19 Genomics UK (COG-UK) Consortium | Health Services Laboratories and Alex Alderton, Roberto Amato, Jeffrey Barrett, Sonia Goncalves, Ewan Harrison, David K. Jackson, Ian Johnston, Dominic Kwiatkowski, Cordelia Langford, John Sillitoe on behalf of the Wellcome Sanger Institute COVID-19 Surveillance Team |
| EPI_ISL_2436554 | Lighthouse Lab in Milton Keynes | Wellcome Sanger Institute for the COVID-19 Genomics UK (COG-UK) Consortium | The Lighthouse Lab in Milton Keynes and Alex Alderton, Roberto Amato, Jeffrey Barrett, Sonia Goncalves, Ewan Harrison, David K. Jackson, Ian Johnston, Dominic Kwiatkowski, Cordelia Langford, John Sillitoe on behalf of the Wellcome Sanger Institute COVID-19 Surveillance Team |
| EPI_ISL_2436556 | Health Services Laboratories | Wellcome Sanger Institute for the COVID-19 Genomics UK (COG-UK) Consortium | Health Services Laboratories and Alex Alderton, Roberto Amato, Jeffrey Barrett, Sonia Goncalves, Ewan Harrison, David K. Jackson, Ian Johnston, Dominic Kwiatkowski, Cordelia Langford, John Sillitoe on behalf of the Wellcome Sanger Institute COVID-19 Surveillance Team |
| EPI_ISL_2436557 | Lighthouse Lab in Alderley Park | Wellcome Sanger Institute for the COVID-19 Genomics UK (COG-UK) Consortium | Jacquelyn Wynn, Mairead Hyland, The Lighthouse Lab in Alderley Park and Alex Alderton, Roberto Amato, Jeffrey Barrett, Sonia Goncalves, Ewan Harrison, David K. Jackson, Ian Johnston, Dominic Kwiatkowski, Cordelia Langford, John Sillitoe on behalf of the Wellcome Sanger Institute COVID-19 Surveillance Team |
| EPI_ISL_2436558 | Lighthouse Lab in Milton Keynes | Wellcome Sanger Institute for the COVID-19 Genomics UK (COG-UK) Consortium | The Lighthouse Lab in Milton Keynes and Alex Alderton, Roberto Amato, Jeffrey Barrett, Sonia Goncalves, Ewan Harrison, David K. Jackson, Ian Johnston, Dominic Kwiatkowski, Cordelia Langford, John Sillitoe on behalf of the Wellcome Sanger Institute COVID-19 Surveillance Team |
| EPI_ISL_2436559 | Lighthouse Laboratory Plymouth | Wellcome Sanger Institute for the COVID-19 Genomics UK (COG-UK) Consortium | Lighthouse Laboratory Plymouth and Alex Alderton, Roberto Amato, Jeffrey Barrett, Sonia Goncalves, Ewan Harrison, David K. Jackson, Ian Johnston, Dominic Kwiatkowski, Cordelia Langford, John Sillitoe on behalf of the Wellcome Sanger Institute COVID-19 Surveillance Team |
| EPI_ISL_2436560 | Lighthouse Lab in Glasgow | Wellcome Sanger Institute for the COVID-19 Genomics UK (COG-UK) Consortium | Harper VanSteenhouse, Yumi Kasai, David Gray, Carol Clugston, Anna Dominiczak and Alex Alderton, Roberto Amato, Jeffrey Barrett, Sonia Goncalves, Ewan Harrison, David K. Jackson, Ian Johnston, Dominic Kwiatkowski, Cordelia Langford, John Sillitoe on behalf of the Wellcome Sanger Institute COVID-19 Surveillance Team |
| EPI_ISL_2436561 | Health Services Laboratories | Wellcome Sanger Institute for the COVID-19 Genomics UK (COG-UK) Consortium | Health Services Laboratories and Alex Alderton, Roberto Amato, Jeffrey Barrett, Sonia Goncalves, Ewan Harrison, David K. Jackson, Ian Johnston, Dominic Kwiatkowski, Cordelia Langford, John Sillitoe on behalf of the Wellcome Sanger Institute COVID-19 Surveillance Team |
| EPI_ISL_2436562 | Lighthouse Lab in Alderley Park | Wellcome Sanger Institute for the COVID-19 Genomics UK (COG-UK) Consortium | Jacquelyn Wynn, Mairead Hyland, The Lighthouse Lab in Alderley Park and Alex Alderton, Roberto Amato, Jeffrey Barrett, Sonia Goncalves, Ewan Harrison, David K. Jackson, Ian Johnston, Dominic Kwiatkowski, Cordelia Langford, John Sillitoe on behalf of the Wellcome Sanger Institute COVID-19 Surveillance Team |
| EPI_ISL_2436563 | Health Services Laboratories | Wellcome Sanger Institute for the COVID-19 Genomics UK (COG-UK) Consortium | Health Services Laboratories and Alex Alderton, Roberto Amato, Jeffrey Barrett, Sonia Goncalves, Ewan Harrison, David K. Jackson, Ian Johnston, Dominic Kwiatkowski, Cordelia Langford, John Sillitoe on behalf of the Wellcome Sanger Institute COVID-19 Surveillance Team |
| EPI_ISL_2436564 | Lighthouse Laboratory Plymouth | Wellcome Sanger Institute for the COVID-19 Genomics UK (COG-UK) Consortium | Lighthouse Laboratory Plymouth and Alex Alderton, Roberto Amato, Jeffrey Barrett, Sonia Goncalves, Ewan Harrison, David K. Jackson, Ian Johnston, Dominic Kwiatkowski, Cordelia Langford, John Sillitoe on behalf of the Wellcome Sanger Institute COVID-19 Surveillance Team |
| EPI_ISL_2436565 | Lighthouse Lab in Milton Keynes | Wellcome Sanger Institute for the COVID-19 Genomics UK (COG-UK) Consortium | The Lighthouse Lab in Milton Keynes and Alex Alderton, Roberto Amato, Jeffrey Barrett, Sonia Goncalves, Ewan Harrison, David K. Jackson, Ian Johnston, Dominic Kwiatkowski, Cordelia Langford, John Sillitoe on behalf of the Wellcome Sanger Institute COVID-19 Surveillance Team |
| EPI_ISL_2436566 | Lighthouse Lab in Alderley Park | Wellcome Sanger Institute for the COVID-19 Genomics UK (COG-UK) Consortium | Jacquelyn Wynn, Mairead Hyland, The Lighthouse Lab in Alderley Park and Alex Alderton, Roberto Amato, Jeffrey Barrett, Sonia Goncalves, Ewan Harrison, David K. Jackson, Ian Johnston, Dominic Kwiatkowski, Cordelia Langford, John Sillitoe on behalf of the Wellcome Sanger Institute COVID-19 Surveillance Team |
| EPI_ISL_2436567 | Lighthouse Laboratory Plymouth | Wellcome Sanger Institute for the COVID-19 Genomics UK (COG-UK) Consortium | Lighthouse Laboratory Plymouth and Alex Alderton, Roberto Amato, Jeffrey Barrett, Sonia Goncalves, Ewan Harrison, David K. Jackson, Ian Johnston, Dominic Kwiatkowski, Cordelia Langford, John Sillitoe on behalf of the Wellcome Sanger Institute COVID-19 Surveillance Team |
| EPI_ISL_2436568 | Lighthouse Lab in Glasgow | Wellcome Sanger Institute for the COVID-19 Genomics UK (COG-UK) Consortium | Harper VanSteenhouse, Yumi Kasai, David Gray, Carol Clugston, Anna Dominiczak and Alex Alderton, Roberto Amato, Jeffrey Barrett, Sonia Goncalves, Ewan Harrison, David K. Jackson, Ian Johnston, Dominic Kwiatkowski, Cordelia Langford, John Sillitoe on behalf of the Wellcome Sanger Institute COVID-19 Surveillance Team |
| EPI_ISL_2436569, EPI_ISL_2436570 | Lighthouse Lab in Alderley Park | Wellcome Sanger Institute for the COVID-19 Genomics UK (COG-UK) Consortium | Jacquelyn Wynn, Mairead Hyland, The Lighthouse Lab in Alderley Park and Alex Alderton, Roberto Amato, Jeffrey Barrett, Sonia Goncalves, Ewan Harrison, David K. Jackson, Ian Johnston, Dominic Kwiatkowski, Cordelia Langford, John Sillitoe on behalf of the Wellcome Sanger Institute COVID-19 Surveillance Team |
| EPI_ISL_2436571, EPI_ISL_2436572 | Lighthouse Lab in Milton Keynes | Wellcome Sanger Institute for the COVID-19 Genomics UK (COG-UK) Consortium | The Lighthouse Lab in Milton Keynes and Alex Alderton, Roberto Amato, Jeffrey Barrett, Sonia Goncalves, Ewan Harrison, David K. Jackson, Ian Johnston, Dominic Kwiatkowski, Cordelia Langford, John Sillitoe on behalf of the Wellcome Sanger Institute COVID-19 Surveillance Team |
| EPI_ISL_2436573 | Lighthouse Lab in Alderley Park | Wellcome Sanger Institute for the COVID-19 Genomics UK (COG-UK) Consortium | Jacquelyn Wynn, Mairead Hyland, The Lighthouse Lab in Alderley Park and Alex Alderton, Roberto Amato, Jeffrey Barrett, Sonia Goncalves, Ewan Harrison, David K. Jackson, Ian Johnston, Dominic Kwiatkowski, Cordelia Langford, John Sillitoe on behalf of the Wellcome Sanger Institute COVID-19 Surveillance Team |
| EPI_ISL_2436574 | Lighthouse Lab in Glasgow | Wellcome Sanger Institute for the COVID-19 Genomics UK (COG-UK) Consortium | Harper VanSteenhouse, Yumi Kasai, David Gray, Carol Clugston, Anna Dominiczak and Alex Alderton, Roberto Amato, Jeffrey Barrett, Sonia Goncalves, Ewan Harrison, David K. Jackson, Ian Johnston, Dominic Kwiatkowski, Cordelia Langford, John Sillitoe on behalf of the Wellcome Sanger Institute COVID-19 Surveillance Team |
| EPI_ISL_2436575, EPI_ISL_2436576 | Lighthouse Lab in Alderley Park | Wellcome Sanger Institute for the COVID-19 Genomics UK (COG-UK) Consortium | Jacquelyn Wynn, Mairead Hyland, The Lighthouse Lab in Alderley Park and Alex Alderton, Roberto Amato, Jeffrey Barrett, Sonia Goncalves, Ewan Harrison, David K. Jackson, Ian Johnston, Dominic Kwiatkowski, Cordelia Langford, John Sillitoe on behalf of the Wellcome Sanger Institute COVID-19 Surveillance Team |
| EPI_ISL_2436577 | Lighthouse Lab in Glasgow | Wellcome Sanger Institute for the COVID-19 Genomics UK (COG-UK) Consortium | Harper VanSteenhouse, Yumi Kasai, David Gray, Carol Clugston, Anna Dominiczak and Alex Alderton, Roberto Amato, Jeffrey Barrett, Sonia Goncalves, Ewan Harrison, David K. Jackson, Ian Johnston, Dominic Kwiatkowski, Cordelia Langford, John Sillitoe on behalf of the Wellcome Sanger Institute COVID-19 Surveillance Team |
| EPI_ISL_2436578, EPI_ISL_2436579, EPI_ISL_2436580, EPI_ISL_2436581 | Lighthouse Lab in Alderley Park | Wellcome Sanger Institute for the COVID-19 Genomics UK (COG-UK) Consortium | Jacquelyn Wynn, Mairead Hyland, The Lighthouse Lab in Alderley Park and Alex Alderton, Roberto Amato, Jeffrey Barrett, Sonia Goncalves, Ewan Harrison, David K. Jackson, Ian Johnston, Dominic Kwiatkowski, Cordelia Langford, John Sillitoe on behalf of the Wellcome Sanger Institute COVID-19 Surveillance Team |
| EPI_ISL_2436582 | Lighthouse Lab in Glasgow | Wellcome Sanger Institute for the COVID-19 Genomics UK (COG-UK) Consortium | Harper VanSteenhouse, Yumi Kasai, David Gray, Carol Clugston, Anna Dominiczak and Alex Alderton, Roberto Amato, Jeffrey Barrett, Sonia Goncalves, Ewan Harrison, David K. Jackson, Ian Johnston, Dominic Kwiatkowski, Cordelia Langford, John Sillitoe on behalf of the Wellcome Sanger Institute COVID-19 Surveillance Team |
| EPI_ISL_2436584 | Lighthouse Lab in Milton Keynes | Wellcome Sanger Institute for the COVID-19 Genomics UK (COG-UK) Consortium | The Lighthouse Lab in Milton Keynes and Alex Alderton, Roberto Amato, Jeffrey Barrett, Sonia Goncalves, Ewan Harrison, David K. Jackson, Ian Johnston, Dominic Kwiatkowski, Cordelia Langford, John Sillitoe on behalf of the Wellcome Sanger Institute COVID-19 Surveillance Team |
| EPI_ISL_2436585 | Lighthouse Lab in Alderley Park | Wellcome Sanger Institute for the COVID-19 Genomics UK (COG-UK) Consortium | Jacquelyn Wynn, Mairead Hyland, The Lighthouse Lab in Alderley Park and Alex Alderton, Roberto Amato, Jeffrey Barrett, Sonia Goncalves, Ewan Harrison, David K. Jackson, Ian Johnston, Dominic Kwiatkowski, Cordelia Langford, John Sillitoe on behalf of the Wellcome Sanger Institute COVID-19 Surveillance Team |
| EPI_ISL_2436586 | Lighthouse Lab in Glasgow | Wellcome Sanger Institute for the COVID-19 Genomics UK (COG-UK) Consortium | Harper VanSteenhouse, Yumi Kasai, David Gray, Carol Clugston, Anna Dominiczak and Alex Alderton, Roberto Amato, Jeffrey Barrett, Sonia Goncalves, Ewan Harrison, David K. Jackson, Ian Johnston, Dominic Kwiatkowski, Cordelia Langford, John Sillitoe on behalf of the Wellcome Sanger Institute COVID-19 Surveillance Team |
| EPI_ISL_2436587, EPI_ISL_2436589 | Lighthouse Lab in Milton Keynes | Wellcome Sanger Institute for the COVID-19 Genomics UK (COG-UK) Consortium | The Lighthouse Lab in Milton Keynes and Alex Alderton, Roberto Amato, Jeffrey Barrett, Sonia Goncalves, Ewan Harrison, David K. Jackson, Ian Johnston, Dominic Kwiatkowski, Cordelia Langford, John Sillitoe on behalf of the Wellcome Sanger Institute COVID-19 Surveillance Team |
| EPI_ISL_2436590, EPI_ISL_2436591 | Lighthouse Lab in Alderley Park | Wellcome Sanger Institute for the COVID-19 Genomics UK (COG-UK) Consortium | Jacquelyn Wynn, Mairead Hyland, The Lighthouse Lab in Alderley Park and Alex Alderton, Roberto Amato, Jeffrey Barrett, Sonia Goncalves, Ewan Harrison, David K. Jackson, Ian Johnston, Dominic Kwiatkowski, Cordelia Langford, John Sillitoe on behalf of the Wellcome Sanger Institute COVID-19 Surveillance Team |
| EPI_ISL_2436592, EPI_ISL_2436593 | Lighthouse Lab in Milton Keynes | Wellcome Sanger Institute for the COVID-19 Genomics UK (COG-UK) Consortium | The Lighthouse Lab in Milton Keynes and Alex Alderton, Roberto Amato, Jeffrey Barrett, Sonia Goncalves, Ewan Harrison, David K. Jackson, Ian Johnston, Dominic Kwiatkowski, Cordelia Langford, John Sillitoe on behalf of the Wellcome Sanger Institute COVID-19 Surveillance Team |
| EPI_ISL_2436594, EPI_ISL_2436596 | Lighthouse Lab in Alderley Park | Wellcome Sanger Institute for the COVID-19 Genomics UK (COG-UK) Consortium | Jacquelyn Wynn, Mairead Hyland, The Lighthouse Lab in Alderley Park and Alex Alderton, Roberto Amato, Jeffrey Barrett, Sonia Goncalves, Ewan Harrison, David K. Jackson, Ian Johnston, Dominic Kwiatkowski, Cordelia Langford, John Sillitoe on behalf of the Wellcome Sanger Institute COVID-19 Surveillance Team |
| EPI_ISL_2436597, EPI_ISL_2436598 | Lighthouse Lab in Glasgow | Wellcome Sanger Institute for the COVID-19 Genomics UK (COG-UK) Consortium | Harper VanSteenhouse, Yumi Kasai, David Gray, Carol Clugston, Anna Dominiczak and Alex Alderton, Roberto Amato, Jeffrey Barrett, Sonia Goncalves, Ewan Harrison, David K. Jackson, Ian Johnston, Dominic Kwiatkowski, Cordelia Langford, John Sillitoe on behalf of the Wellcome Sanger Institute COVID-19 Surveillance Team |
| EPI_ISL_2436599 | Lighthouse Lab in Alderley Park | Wellcome Sanger Institute for the COVID-19 Genomics UK (COG-UK) Consortium | Jacquelyn Wynn, Mairead Hyland, The Lighthouse Lab in Alderley Park and Alex Alderton, Roberto Amato, Jeffrey Barrett, Sonia Goncalves, Ewan Harrison, David K. Jackson, Ian Johnston, Dominic Kwiatkowski, Cordelia Langford, John Sillitoe on behalf of the Wellcome Sanger Institute COVID-19 Surveillance Team |
| EPI_ISL_2436600 | Lighthouse Lab in Milton Keynes | Wellcome Sanger Institute for the COVID-19 Genomics UK (COG-UK) Consortium | The Lighthouse Lab in Milton Keynes and Alex Alderton, Roberto Amato, Jeffrey Barrett, Sonia Goncalves, Ewan Harrison, David K. Jackson, Ian Johnston, Dominic Kwiatkowski, Cordelia Langford, John Sillitoe on behalf of the Wellcome Sanger Institute COVID-19 Surveillance Team |
| EPI_ISL_2436602, EPI_ISL_2436603, EPI_ISL_2436604, EPI_ISL_2436605, EPI_ISL_2436606 | Lighthouse Lab in Alderley Park | Wellcome Sanger Institute for the COVID-19 Genomics UK (COG-UK) Consortium | Jacquelyn Wynn, Mairead Hyland, The Lighthouse Lab in Alderley Park and Alex Alderton, Roberto Amato, Jeffrey Barrett, Sonia Goncalves, Ewan Harrison, David K. Jackson, Ian Johnston, Dominic Kwiatkowski, Cordelia Langford, John Sillitoe on behalf of the Wellcome Sanger Institute COVID-19 Surveillance Team |
| EPI_ISL_2436607 | Lighthouse Lab in Glasgow | Wellcome Sanger Institute for the COVID-19 Genomics UK (COG-UK) Consortium | Harper VanSteenhouse, Yumi Kasai, David Gray, Carol Clugston, Anna Dominiczak and Alex Alderton, Roberto Amato, Jeffrey Barrett, Sonia Goncalves, Ewan Harrison, David K. Jackson, Ian Johnston, Dominic Kwiatkowski, Cordelia Langford, John Sillitoe on behalf of the Wellcome Sanger Institute COVID-19 Surveillance Team |
| EPI_ISL_2436608 | Lighthouse Lab in Milton Keynes | Wellcome Sanger Institute for the COVID-19 Genomics UK (COG-UK) Consortium | The Lighthouse Lab in Milton Keynes and Alex Alderton, Roberto Amato, Jeffrey Barrett, Sonia Goncalves, Ewan Harrison, David K. Jackson, Ian Johnston, Dominic Kwiatkowski, Cordelia Langford, John Sillitoe on behalf of the Wellcome Sanger Institute COVID-19 Surveillance Team |
| EPI_ISL_2436609, EPI_ISL_2436610, EPI_ISL_2436611, EPI_ISL_2436612, | Lighthouse Lab in Alderley Park | Wellcome Sanger Institute for the COVID-19 Genomics UK (COG-UK) Consortium | Jacquelyn Wynn, Mairead Hyland, The Lighthouse Lab in Alderley Park and Alex Alderton, Roberto Amato, Jeffrey Barrett, Sonia Goncalves, Ewan Harrison, David K. Jackson, Ian Johnston, Dominic Kwiatkowski, Cordelia Langford, John Sillitoe on behalf of the Wellcome Sanger Institute COVID-19 Surveillance Team |

| | | | |
|---|---|---|---|
| EPI_ISL_2436614, EPI_ISL_2436615, EPI_ISL_2436616 | | | |
| EPI_ISL_2436617 | Lighthouse Lab in Milton Keynes | Wellcome Sanger Institute for the COVID-19 Genomics UK (COG-UK) Consortium | The Lighthouse Lab in Milton Keynes and Alex Alderton, Roberto Amato, Jeffrey Barrett, Sonia Goncalves, Ewan Harrison, David K. Jackson, Ian Johnston, Dominic Kwiatkowski, Cordelia Langford, John Sillitoe on behalf of the Wellcome Sanger Institute COVID-19 Surveillance Team |
| EPI_ISL_2436618, EPI_ISL_2436619 | Lighthouse Lab in Alderley Park | Wellcome Sanger Institute for the COVID-19 Genomics UK (COG-UK) Consortium | Jacquelyn Wynn, Mairead Hyland, The Lighthouse Lab in Alderley Park and Alex Alderton, Roberto Amato, Jeffrey Barrett, Sonia Goncalves, Ewan Harrison, David K. Jackson, Ian Johnston, Dominic Kwiatkowski, Cordelia Langford, John Sillitoe on behalf of the Wellcome Sanger Institute COVID-19 Surveillance Team |
| EPI_ISL_2436620 | Lighthouse Lab in Glasgow | Wellcome Sanger Institute for the COVID-19 Genomics UK (COG-UK) Consortium | Harper VanSteenhouse, Yumi Kasai, David Gray, Carol Clugston, Anna Dominiczak and Alex Alderton, Roberto Amato, Jeffrey Barrett, Sonia Goncalves, Ewan Harrison, David K. Jackson, Ian Johnston, Dominic Kwiatkowski, Cordelia Langford, John Sillitoe on behalf of the Wellcome Sanger Institute COVID-19 Surveillance Team |
| EPI_ISL_2436621 | Lighthouse Lab in Alderley Park | Wellcome Sanger Institute for the COVID-19 Genomics UK (COG-UK) Consortium | Jacquelyn Wynn, Mairead Hyland, The Lighthouse Lab in Alderley Park and Alex Alderton, Roberto Amato, Jeffrey Barrett, Sonia Goncalves, Ewan Harrison, David K. Jackson, Ian Johnston, Dominic Kwiatkowski, Cordelia Langford, John Sillitoe on behalf of the Wellcome Sanger Institute COVID-19 Surveillance Team |
| EPI_ISL_2436623 | Lighthouse Lab in Glasgow | Wellcome Sanger Institute for the COVID-19 Genomics UK (COG-UK) Consortium | Harper VanSteenhouse, Yumi Kasai, David Gray, Carol Clugston, Anna Dominiczak and Alex Alderton, Roberto Amato, Jeffrey Barrett, Sonia Goncalves, Ewan Harrison, David K. Jackson, Ian Johnston, Dominic Kwiatkowski, Cordelia Langford, John Sillitoe on behalf of the Wellcome Sanger Institute COVID-19 Surveillance Team |
| EPI_ISL_2436624 | Lighthouse Lab in Alderley Park | Wellcome Sanger Institute for the COVID-19 Genomics UK (COG-UK) Consortium | Jacquelyn Wynn, Mairead Hyland, The Lighthouse Lab in Alderley Park and Alex Alderton, Roberto Amato, Jeffrey Barrett, Sonia Goncalves, Ewan Harrison, David K. Jackson, Ian Johnston, Dominic Kwiatkowski, Cordelia Langford, John Sillitoe on behalf of the Wellcome Sanger Institute COVID-19 Surveillance Team |
| EPI_ISL_2436625 | Lighthouse Lab in Milton Keynes | Wellcome Sanger Institute for the COVID-19 Genomics UK (COG-UK) Consortium | The Lighthouse Lab in Milton Keynes and Alex Alderton, Roberto Amato, Jeffrey Barrett, Sonia Goncalves, Ewan Harrison, David K. Jackson, Ian Johnston, Dominic Kwiatkowski, Cordelia Langford, John Sillitoe on behalf of the Wellcome Sanger Institute COVID-19 Surveillance Team |
| EPI_ISL_2436627 | Lighthouse Lab in Alderley Park | Wellcome Sanger Institute for the COVID-19 Genomics UK (COG-UK) Consortium | Jacquelyn Wynn, Mairead Hyland, The Lighthouse Lab in Alderley Park and Alex Alderton, Roberto Amato, Jeffrey Barrett, Sonia Goncalves, Ewan Harrison, David K. Jackson, Ian Johnston, Dominic Kwiatkowski, Cordelia Langford, John Sillitoe on behalf of the Wellcome Sanger Institute COVID-19 Surveillance Team |
| EPI_ISL_2436628 | Lighthouse Lab in Milton Keynes | Wellcome Sanger Institute for the COVID-19 Genomics UK (COG-UK) Consortium | The Lighthouse Lab in Milton Keynes and Alex Alderton, Roberto Amato, Jeffrey Barrett, Sonia Goncalves, Ewan Harrison, David K. Jackson, Ian Johnston, Dominic Kwiatkowski, Cordelia Langford, John Sillitoe on behalf of the Wellcome Sanger Institute COVID-19 Surveillance Team |
| EPI_ISL_2436629, EPI_ISL_2436632, EPI_ISL_2436633 | Lighthouse Lab in Alderley Park | Wellcome Sanger Institute for the COVID-19 Genomics UK (COG-UK) Consortium | Jacquelyn Wynn, Mairead Hyland, The Lighthouse Lab in Alderley Park and Alex Alderton, Roberto Amato, Jeffrey Barrett, Sonia Goncalves, Ewan Harrison, David K. Jackson, Ian Johnston, Dominic Kwiatkowski, Cordelia Langford, John Sillitoe on behalf of the Wellcome Sanger Institute COVID-19 Surveillance Team |
| EPI_ISL_2436634, EPI_ISL_2436635, EPI_ISL_2436636 | Lighthouse Lab in Glasgow | Wellcome Sanger Institute for the COVID-19 Genomics UK (COG-UK) Consortium | Harper VanSteenhouse, Yumi Kasai, David Gray, Carol Clugston, Anna Dominiczak and Alex Alderton, Roberto Amato, Jeffrey Barrett, Sonia Goncalves, Ewan Harrison, David K. Jackson, Ian Johnston, Dominic Kwiatkowski, Cordelia Langford, John Sillitoe on behalf of the Wellcome Sanger Institute COVID-19 Surveillance Team |
| EPI_ISL_2436637, EPI_ISL_2436638 | Lighthouse Lab in Alderley Park | Wellcome Sanger Institute for the COVID-19 Genomics UK (COG-UK) Consortium | Jacquelyn Wynn, Mairead Hyland, The Lighthouse Lab in Alderley Park and Alex Alderton, Roberto Amato, Jeffrey Barrett, Sonia Goncalves, Ewan Harrison, David K. Jackson, Ian Johnston, Dominic Kwiatkowski, Cordelia Langford, John Sillitoe on behalf of the Wellcome Sanger Institute COVID-19 Surveillance Team |
| EPI_ISL_2436639 | Lighthouse Lab in Milton Keynes | Wellcome Sanger Institute for the COVID-19 Genomics UK (COG-UK) Consortium | The Lighthouse Lab in Milton Keynes and Alex Alderton, Roberto Amato, Jeffrey Barrett, Sonia Goncalves, Ewan Harrison, David K. Jackson, Ian Johnston, Dominic Kwiatkowski, Cordelia Langford, John Sillitoe on behalf of the Wellcome Sanger Institute COVID-19 Surveillance Team |
| EPI_ISL_2436640 | Lighthouse Lab in Alderley Park | Wellcome Sanger Institute for the COVID-19 Genomics UK (COG-UK) Consortium | Jacquelyn Wynn, Mairead Hyland, The Lighthouse Lab in Alderley Park and Alex Alderton, Roberto Amato, Jeffrey Barrett, Sonia Goncalves, Ewan Harrison, David K. Jackson, Ian Johnston, Dominic Kwiatkowski, Cordelia Langford, John Sillitoe on behalf of the Wellcome Sanger Institute COVID-19 Surveillance Team |
| EPI_ISL_2436641 | Lighthouse Lab in Glasgow | Wellcome Sanger Institute for the COVID-19 Genomics UK (COG-UK) Consortium | Harper VanSteenhouse, Yumi Kasai, David Gray, Carol Clugston, Anna Dominiczak and Alex Alderton, Roberto Amato, Jeffrey Barrett, Sonia Goncalves, Ewan Harrison, David K. Jackson, Ian Johnston, Dominic Kwiatkowski, Cordelia Langford, John Sillitoe on behalf of the Wellcome Sanger Institute COVID-19 Surveillance Team |
| EPI_ISL_2436643 | Lighthouse Lab in Alderley Park | Wellcome Sanger Institute for the COVID-19 Genomics UK (COG-UK) Consortium | Jacquelyn Wynn, Mairead Hyland, The Lighthouse Lab in Alderley Park and Alex Alderton, Roberto Amato, Jeffrey Barrett, Sonia Goncalves, Ewan Harrison, David K. Jackson, Ian Johnston, Dominic Kwiatkowski, Cordelia Langford, John Sillitoe on behalf of the Wellcome Sanger Institute COVID-19 Surveillance Team |
| EPI_ISL_2436644 | Lighthouse Lab in Milton Keynes | Wellcome Sanger Institute for the COVID-19 Genomics UK (COG-UK) Consortium | The Lighthouse Lab in Milton Keynes and Alex Alderton, Roberto Amato, Jeffrey Barrett, Sonia Goncalves, Ewan Harrison, David K. Jackson, Ian Johnston, Dominic Kwiatkowski, Cordelia Langford, John Sillitoe on behalf of the Wellcome Sanger Institute COVID-19 Surveillance Team |
| EPI_ISL_2436645, EPI_ISL_2436646, EPI_ISL_2436647 | Lighthouse Lab in Alderley Park | Wellcome Sanger Institute for the COVID-19 Genomics UK (COG-UK) Consortium | Jacquelyn Wynn, Mairead Hyland, The Lighthouse Lab in Alderley Park and Alex Alderton, Roberto Amato, Jeffrey Barrett, Sonia Goncalves, Ewan Harrison, David K. Jackson, Ian Johnston, Dominic Kwiatkowski, Cordelia Langford, John Sillitoe on behalf of the Wellcome Sanger Institute COVID-19 Surveillance Team |
| EPI_ISL_2436649 | Lighthouse Lab in Milton Keynes | Wellcome Sanger Institute for the COVID-19 Genomics UK (COG-UK) Consortium | The Lighthouse Lab in Milton Keynes and Alex Alderton, Roberto Amato, Jeffrey Barrett, Sonia Goncalves, Ewan Harrison, David K. Jackson, Ian Johnston, Dominic Kwiatkowski, Cordelia Langford, John Sillitoe on behalf of the Wellcome Sanger Institute COVID-19 Surveillance Team |
| EPI_ISL_2436650 | Lighthouse Lab in Glasgow | Wellcome Sanger Institute for the COVID-19 Genomics UK (COG-UK) Consortium | Harper VanSteenhouse, Yumi Kasai, David Gray, Carol Clugston, Anna Dominiczak and Alex Alderton, Roberto Amato, Jeffrey Barrett, Sonia Goncalves, Ewan Harrison, David K. Jackson, Ian Johnston, Dominic Kwiatkowski, Cordelia Langford, John Sillitoe on behalf of the Wellcome Sanger Institute COVID-19 Surveillance Team |
| EPI_ISL_2436651 | Lighthouse Lab in Milton Keynes | Wellcome Sanger Institute for the COVID-19 Genomics UK (COG-UK) Consortium | The Lighthouse Lab in Milton Keynes and Alex Alderton, Roberto Amato, Jeffrey Barrett, Sonia Goncalves, Ewan Harrison, David K. Jackson, Ian Johnston, Dominic Kwiatkowski, Cordelia Langford, John Sillitoe on behalf of the Wellcome Sanger Institute COVID-19 Surveillance Team |
| EPI_ISL_2436652, EPI_ISL_2436653, EPI_ISL_2436654, EPI_ISL_2436655 | Lighthouse Lab in Alderley Park | Wellcome Sanger Institute for the COVID-19 Genomics UK (COG-UK) Consortium | Jacquelyn Wynn, Mairead Hyland, The Lighthouse Lab in Alderley Park and Alex Alderton, Roberto Amato, Jeffrey Barrett, Sonia Goncalves, Ewan Harrison, David K. Jackson, Ian Johnston, Dominic Kwiatkowski, Cordelia Langford, John Sillitoe on behalf of the Wellcome Sanger Institute COVID-19 Surveillance Team |
| EPI_ISL_2436657 | Lighthouse Lab in Milton Keynes | Wellcome Sanger Institute for the COVID-19 Genomics UK (COG-UK) Consortium | The Lighthouse Lab in Milton Keynes and Alex Alderton, Roberto Amato, Jeffrey Barrett, Sonia Goncalves, Ewan Harrison, David K. Jackson, Ian Johnston, Dominic Kwiatkowski, Cordelia Langford, John Sillitoe on behalf of the Wellcome Sanger Institute COVID-19 Surveillance Team |
| EPI_ISL_2436658, EPI_ISL_2436659, EPI_ISL_2436660, EPI_ISL_2436661 | Lighthouse Lab in Alderley Park | Wellcome Sanger Institute for the COVID-19 Genomics UK (COG-UK) Consortium | Jacquelyn Wynn, Mairead Hyland, The Lighthouse Lab in Alderley Park and Alex Alderton, Roberto Amato, Jeffrey Barrett, Sonia Goncalves, Ewan Harrison, David K. Jackson, Ian Johnston, Dominic Kwiatkowski, Cordelia Langford, John Sillitoe on behalf of the Wellcome Sanger Institute COVID-19 Surveillance Team |
| EPI_ISL_2436662 | Lighthouse Lab in Glasgow | Wellcome Sanger Institute for the COVID-19 Genomics UK (COG-UK) Consortium | Harper VanSteenhouse, Yumi Kasai, David Gray, Carol Clugston, Anna Dominiczak and Alex Alderton, Roberto Amato, Jeffrey Barrett, Sonia Goncalves, Ewan Harrison, David K. Jackson, Ian Johnston, Dominic Kwiatkowski, Cordelia Langford, John Sillitoe on behalf of the Wellcome Sanger Institute COVID-19 Surveillance Team |
| EPI_ISL_2436663 | Lighthouse Lab in Alderley Park | Wellcome Sanger Institute for the COVID-19 Genomics UK (COG-UK) Consortium | Jacquelyn Wynn, Mairead Hyland, The Lighthouse Lab in Alderley Park and Alex Alderton, Roberto Amato, Jeffrey Barrett, Sonia Goncalves, Ewan Harrison, David K. Jackson, Ian Johnston, Dominic Kwiatkowski, Cordelia Langford, John Sillitoe on behalf of the Wellcome Sanger Institute COVID-19 Surveillance Team |
| EPI_ISL_2436664 | Lighthouse Lab in Glasgow | Wellcome Sanger Institute for the COVID-19 Genomics UK (COG-UK) Consortium | Harper VanSteenhouse, Yumi Kasai, David Gray, Carol Clugston, Anna Dominiczak and Alex Alderton, Roberto Amato, Jeffrey Barrett, Sonia Goncalves, Ewan Harrison, David K. Jackson, Ian Johnston, Dominic Kwiatkowski, Cordelia Langford, John Sillitoe on behalf of the Wellcome Sanger Institute COVID-19 Surveillance Team |
| EPI_ISL_2436665 | Lighthouse Lab in Milton Keynes | Wellcome Sanger Institute for the COVID-19 Genomics UK (COG-UK) Consortium | The Lighthouse Lab in Milton Keynes and Alex Alderton, Roberto Amato, Jeffrey Barrett, Sonia Goncalves, Ewan Harrison, David K. Jackson, Ian Johnston, Dominic Kwiatkowski, Cordelia Langford, John Sillitoe on behalf of the Wellcome Sanger Institute COVID-19 Surveillance Team |
| EPI_ISL_2436666, EPI_ISL_2436667 | Lighthouse Lab in Alderley Park | Wellcome Sanger Institute for the COVID-19 Genomics UK (COG-UK) Consortium | Jacquelyn Wynn, Mairead Hyland, The Lighthouse Lab in Alderley Park and Alex Alderton, Roberto Amato, Jeffrey Barrett, Sonia Goncalves, Ewan Harrison, David K. Jackson, Ian Johnston, Dominic Kwiatkowski, Cordelia Langford, John Sillitoe on behalf of the Wellcome Sanger Institute COVID-19 Surveillance Team |
| EPI_ISL_2436669, EPI_ISL_2436670, EPI_ISL_2436672 | Lighthouse Lab in Milton Keynes | Wellcome Sanger Institute for the COVID-19 Genomics UK (COG-UK) Consortium | The Lighthouse Lab in Milton Keynes and Alex Alderton, Roberto Amato, Jeffrey Barrett, Sonia Goncalves, Ewan Harrison, David K. Jackson, Ian Johnston, Dominic Kwiatkowski, Cordelia Langford, John Sillitoe on behalf of the Wellcome Sanger Institute COVID-19 Surveillance Team |
| EPI_ISL_2436673 | Lighthouse Lab in Alderley Park | Wellcome Sanger Institute for the COVID-19 Genomics UK (COG-UK) Consortium | Jacquelyn Wynn, Mairead Hyland, The Lighthouse Lab in Alderley Park and Alex Alderton, Roberto Amato, Jeffrey Barrett, Sonia Goncalves, Ewan Harrison, David K. Jackson, Ian Johnston, Dominic Kwiatkowski, Cordelia Langford, John Sillitoe on behalf of the Wellcome Sanger Institute COVID-19 Surveillance Team |
| EPI_ISL_2436674 | Lighthouse Lab in Milton Keynes | Wellcome Sanger Institute for the COVID-19 Genomics UK (COG-UK) Consortium | The Lighthouse Lab in Milton Keynes and Alex Alderton, Roberto Amato, Jeffrey Barrett, Sonia Goncalves, Ewan Harrison, David K. Jackson, Ian Johnston, Dominic Kwiatkowski, Cordelia Langford, John Sillitoe on behalf of the Wellcome Sanger Institute COVID-19 Surveillance Team |
| EPI_ISL_2436675 | Lighthouse Lab in Alderley Park | Wellcome Sanger Institute for the COVID-19 Genomics UK (COG-UK) Consortium | Jacquelyn Wynn, Mairead Hyland, The Lighthouse Lab in Alderley Park and Alex Alderton, Roberto Amato, Jeffrey Barrett, Sonia Goncalves, Ewan Harrison, David K. Jackson, Ian Johnston, Dominic Kwiatkowski, Cordelia Langford, John Sillitoe on behalf of the Wellcome Sanger Institute COVID-19 Surveillance Team |
| EPI_ISL_2436676 | Lighthouse Lab in Milton Keynes | Wellcome Sanger Institute for the COVID-19 Genomics UK (COG-UK) Consortium | The Lighthouse Lab in Milton Keynes and Alex Alderton, Roberto Amato, Jeffrey Barrett, Sonia Goncalves, Ewan Harrison, David K. Jackson, Ian Johnston, Dominic Kwiatkowski, Cordelia Langford, John Sillitoe on behalf of the Wellcome Sanger Institute COVID-19 Surveillance Team |
| EPI_ISL_2436677 | Lighthouse Lab in Alderley Park | Wellcome Sanger Institute for the COVID-19 Genomics UK (COG-UK) Consortium | Jacquelyn Wynn, Mairead Hyland, The Lighthouse Lab in Alderley Park and Alex Alderton, Roberto Amato, Jeffrey Barrett, Sonia Goncalves, Ewan Harrison, David K. Jackson, Ian Johnston, Dominic Kwiatkowski, Cordelia Langford, John Sillitoe on behalf of the Wellcome Sanger Institute COVID-19 Surveillance Team |
| EPI_ISL_2436678 | Lighthouse Lab in Milton Keynes | Wellcome Sanger Institute for the COVID-19 Genomics UK (COG-UK) Consortium | The Lighthouse Lab in Milton Keynes and Alex Alderton, Roberto Amato, Jeffrey Barrett, Sonia Goncalves, Ewan Harrison, David K. Jackson, Ian Johnston, Dominic Kwiatkowski, Cordelia Langford, John Sillitoe on behalf of the Wellcome Sanger Institute COVID-19 Surveillance Team |
| EPI_ISL_2436679 | Lighthouse Lab in Alderley Park | Wellcome Sanger Institute for the COVID-19 Genomics UK (COG-UK) Consortium | Jacquelyn Wynn, Mairead Hyland, The Lighthouse Lab in Alderley Park and Alex Alderton, Roberto Amato, Jeffrey Barrett, Sonia Goncalves, Ewan Harrison, David K. Jackson, Ian Johnston, Dominic Kwiatkowski, Cordelia Langford, John Sillitoe on behalf of the Wellcome Sanger Institute COVID-19 Surveillance Team |
| EPI_ISL_2436680 | Lighthouse Lab in Milton Keynes | Wellcome Sanger Institute for the COVID-19 Genomics UK (COG-UK) Consortium | The Lighthouse Lab in Milton Keynes and Alex Alderton, Roberto Amato, Jeffrey Barrett, Sonia Goncalves, Ewan Harrison, David K. Jackson, Ian Johnston, Dominic Kwiatkowski, Cordelia Langford, John Sillitoe on behalf of the Wellcome Sanger Institute COVID-19 Surveillance Team |
| EPI_ISL_2436681 | Lighthouse Lab in Alderley Park | Wellcome Sanger Institute for the COVID-19 Genomics UK (COG-UK) Consortium | Jacquelyn Wynn, Mairead Hyland, The Lighthouse Lab in Alderley Park and Alex Alderton, Roberto Amato, Jeffrey Barrett, Sonia Goncalves, Ewan Harrison, David K. Jackson, Ian Johnston, Dominic Kwiatkowski, Cordelia Langford, John Sillitoe on behalf of the Wellcome Sanger Institute COVID-19 Surveillance Team |
| EPI_ISL_2436683, EPI_ISL_2436684 | Lighthouse Lab in Milton Keynes | Wellcome Sanger Institute for the COVID-19 Genomics UK (COG-UK) Consortium | The Lighthouse Lab in Milton Keynes and Alex Alderton, Roberto Amato, Jeffrey Barrett, Sonia Goncalves, Ewan Harrison, David K. Jackson, Ian Johnston, Dominic Kwiatkowski, Cordelia Langford, John Sillitoe on behalf of the Wellcome Sanger Institute COVID-19 Surveillance Team |

| | | | |
|---|---|---|---|
| EPI_ISL_2436685 | Lighthouse Lab in Alderley Park | Wellcome Sanger Institute for the COVID-19 Genomics UK (COG-UK) Consortium | Jacquelyn Wynn, Mairead Hyland, The Lighthouse Lab in Alderley Park and Alex Alderton, Roberto Amato, Jeffrey Barrett, Sonia Goncalves, Ewan Harrison, David K. Jackson, Ian Johnston, Dominic Kwiatkowski, Cordelia Langford, John Sillitoe on behalf of the Wellcome Sanger Institute COVID-19 Surveillance Team |
| EPI_ISL_2436686 | Lighthouse Lab in Glasgow | Wellcome Sanger Institute for the COVID-19 Genomics UK (COG-UK) Consortium | Harper VanSteenhouse, Yumi Kasai, David Gray, Carol Clugston, Anna Dominiczak and Alex Alderton, Roberto Amato, Jeffrey Barrett, Sonia Goncalves, Ewan Harrison, David K. Jackson, Ian Johnston, Dominic Kwiatkowski, Cordelia Langford, John Sillitoe on behalf of the Wellcome Sanger Institute COVID-19 Surveillance Team |
| EPI_ISL_2436687, EPI_ISL_2436688, EPI_ISL_2436690, EPI_ISL_2436691 | Lighthouse Lab in Alderley Park | Wellcome Sanger Institute for the COVID-19 Genomics UK (COG-UK) Consortium | Jacquelyn Wynn, Mairead Hyland, The Lighthouse Lab in Alderley Park and Alex Alderton, Roberto Amato, Jeffrey Barrett, Sonia Goncalves, Ewan Harrison, David K. Jackson, Ian Johnston, Dominic Kwiatkowski, Cordelia Langford, John Sillitoe on behalf of the Wellcome Sanger Institute COVID-19 Surveillance Team |
| EPI_ISL_2436692 | Lighthouse Lab in Milton Keynes | Wellcome Sanger Institute for the COVID-19 Genomics UK (COG-UK) Consortium | The Lighthouse Lab in Milton Keynes and Alex Alderton, Roberto Amato, Jeffrey Barrett, Sonia Goncalves, Ewan Harrison, David K. Jackson, Ian Johnston, Dominic Kwiatkowski, Cordelia Langford, John Sillitoe on behalf of the Wellcome Sanger Institute COVID-19 Surveillance Team |
| EPI_ISL_2436693, EPI_ISL_2436694, EPI_ISL_2436695 | Lighthouse Lab in Alderley Park | Wellcome Sanger Institute for the COVID-19 Genomics UK (COG-UK) Consortium | Jacquelyn Wynn, Mairead Hyland, The Lighthouse Lab in Alderley Park and Alex Alderton, Roberto Amato, Jeffrey Barrett, Sonia Goncalves, Ewan Harrison, David K. Jackson, Ian Johnston, Dominic Kwiatkowski, Cordelia Langford, John Sillitoe on behalf of the Wellcome Sanger Institute COVID-19 Surveillance Team |
| EPI_ISL_2436696 | Lighthouse Lab in Milton Keynes | Wellcome Sanger Institute for the COVID-19 Genomics UK (COG-UK) Consortium | The Lighthouse Lab in Milton Keynes and Alex Alderton, Roberto Amato, Jeffrey Barrett, Sonia Goncalves, Ewan Harrison, David K. Jackson, Ian Johnston, Dominic Kwiatkowski, Cordelia Langford, John Sillitoe on behalf of the Wellcome Sanger Institute COVID-19 Surveillance Team |
| EPI_ISL_2436697 | Lighthouse Lab in Glasgow | Wellcome Sanger Institute for the COVID-19 Genomics UK (COG-UK) Consortium | Harper VanSteenhouse, Yumi Kasai, David Gray, Carol Clugston, Anna Dominiczak and Alex Alderton, Roberto Amato, Jeffrey Barrett, Sonia Goncalves, Ewan Harrison, David K. Jackson, Ian Johnston, Dominic Kwiatkowski, Cordelia Langford, John Sillitoe on behalf of the Wellcome Sanger Institute COVID-19 Surveillance Team |
| EPI_ISL_2436698 | Lighthouse Lab in Milton Keynes | Wellcome Sanger Institute for the COVID-19 Genomics UK (COG-UK) Consortium | The Lighthouse Lab in Milton Keynes and Alex Alderton, Roberto Amato, Jeffrey Barrett, Sonia Goncalves, Ewan Harrison, David K. Jackson, Ian Johnston, Dominic Kwiatkowski, Cordelia Langford, John Sillitoe on behalf of the Wellcome Sanger Institute COVID-19 Surveillance Team |
| EPI_ISL_2436699 | Lighthouse Lab in Glasgow | Wellcome Sanger Institute for the COVID-19 Genomics UK (COG-UK) Consortium | Harper VanSteenhouse, Yumi Kasai, David Gray, Carol Clugston, Anna Dominiczak and Alex Alderton, Roberto Amato, Jeffrey Barrett, Sonia Goncalves, Ewan Harrison, David K. Jackson, Ian Johnston, Dominic Kwiatkowski, Cordelia Langford, John Sillitoe on behalf of the Wellcome Sanger Institute COVID-19 Surveillance Team |
| EPI_ISL_2436700, EPI_ISL_2436701, EPI_ISL_2436702, EPI_ISL_2436703, EPI_ISL_2436704 | Lighthouse Lab in Alderley Park | Wellcome Sanger Institute for the COVID-19 Genomics UK (COG-UK) Consortium | Jacquelyn Wynn, Mairead Hyland, The Lighthouse Lab in Alderley Park and Alex Alderton, Roberto Amato, Jeffrey Barrett, Sonia Goncalves, Ewan Harrison, David K. Jackson, Ian Johnston, Dominic Kwiatkowski, Cordelia Langford, John Sillitoe on behalf of the Wellcome Sanger Institute COVID-19 Surveillance Team |
| EPI_ISL_2436710 | Lighthouse Lab in Milton Keynes | Wellcome Sanger Institute for the COVID-19 Genomics UK (COG-UK) Consortium | The Lighthouse Lab in Milton Keynes and Alex Alderton, Roberto Amato, Jeffrey Barrett, Sonia Goncalves, Ewan Harrison, David K. Jackson, Ian Johnston, Dominic Kwiatkowski, Cordelia Langford, John Sillitoe on behalf of the Wellcome Sanger Institute COVID-19 Surveillance Team |
| EPI_ISL_2436711 | Lighthouse Lab in Alderley Park | Wellcome Sanger Institute for the COVID-19 Genomics UK (COG-UK) Consortium | Jacquelyn Wynn, Mairead Hyland, The Lighthouse Lab in Alderley Park and Alex Alderton, Roberto Amato, Jeffrey Barrett, Sonia Goncalves, Ewan Harrison, David K. Jackson, Ian Johnston, Dominic Kwiatkowski, Cordelia Langford, John Sillitoe on behalf of the Wellcome Sanger Institute COVID-19 Surveillance Team |
| EPI_ISL_2436714 | Lighthouse Lab in Milton Keynes | Wellcome Sanger Institute for the COVID-19 Genomics UK (COG-UK) Consortium | The Lighthouse Lab in Milton Keynes and Alex Alderton, Roberto Amato, Jeffrey Barrett, Sonia Goncalves, Ewan Harrison, David K. Jackson, Ian Johnston, Dominic Kwiatkowski, Cordelia Langford, John Sillitoe on behalf of the Wellcome Sanger Institute COVID-19 Surveillance Team |
| EPI_ISL_2436716, EPI_ISL_2436717, EPI_ISL_2436718, EPI_ISL_2436719 | Lighthouse Lab in Alderley Park | Wellcome Sanger Institute for the COVID-19 Genomics UK (COG-UK) Consortium | Jacquelyn Wynn, Mairead Hyland, The Lighthouse Lab in Alderley Park and Alex Alderton, Roberto Amato, Jeffrey Barrett, Sonia Goncalves, Ewan Harrison, David K. Jackson, Ian Johnston, Dominic Kwiatkowski, Cordelia Langford, John Sillitoe on behalf of the Wellcome Sanger Institute COVID-19 Surveillance Team |
| EPI_ISL_2436720 | Lighthouse Lab in Milton Keynes | Wellcome Sanger Institute for the COVID-19 Genomics UK (COG-UK) Consortium | The Lighthouse Lab in Milton Keynes and Alex Alderton, Roberto Amato, Jeffrey Barrett, Sonia Goncalves, Ewan Harrison, David K. Jackson, Ian Johnston, Dominic Kwiatkowski, Cordelia Langford, John Sillitoe on behalf of the Wellcome Sanger Institute COVID-19 Surveillance Team |
| EPI_ISL_2436721 | Lighthouse Lab in Glasgow | Wellcome Sanger Institute for the COVID-19 Genomics UK (COG-UK) Consortium | Harper VanSteenhouse, Yumi Kasai, David Gray, Carol Clugston, Anna Dominiczak and Alex Alderton, Roberto Amato, Jeffrey Barrett, Sonia Goncalves, Ewan Harrison, David K. Jackson, Ian Johnston, Dominic Kwiatkowski, Cordelia Langford, John Sillitoe on behalf of the Wellcome Sanger Institute COVID-19 Surveillance Team |
| EPI_ISL_2436722 | Lighthouse Lab in Alderley Park | Wellcome Sanger Institute for the COVID-19 Genomics UK (COG-UK) Consortium | Jacquelyn Wynn, Mairead Hyland, The Lighthouse Lab in Alderley Park and Alex Alderton, Roberto Amato, Jeffrey Barrett, Sonia Goncalves, Ewan Harrison, David K. Jackson, Ian Johnston, Dominic Kwiatkowski, Cordelia Langford, John Sillitoe on behalf of the Wellcome Sanger Institute COVID-19 Surveillance Team |
| EPI_ISL_2436723 | Lighthouse Lab in Glasgow | Wellcome Sanger Institute for the COVID-19 Genomics UK (COG-UK) Consortium | Harper VanSteenhouse, Yumi Kasai, David Gray, Carol Clugston, Anna Dominiczak and Alex Alderton, Roberto Amato, Jeffrey Barrett, Sonia Goncalves, Ewan Harrison, David K. Jackson, Ian Johnston, Dominic Kwiatkowski, Cordelia Langford, John Sillitoe on behalf of the Wellcome Sanger Institute COVID-19 Surveillance Team |
| EPI_ISL_2436724, EPI_ISL_2436725 | Lighthouse Lab in Alderley Park | Wellcome Sanger Institute for the COVID-19 Genomics UK (COG-UK) Consortium | Jacquelyn Wynn, Mairead Hyland, The Lighthouse Lab in Alderley Park and Alex Alderton, Roberto Amato, Jeffrey Barrett, Sonia Goncalves, Ewan Harrison, David K. Jackson, Ian Johnston, Dominic Kwiatkowski, Cordelia Langford, John Sillitoe on behalf of the Wellcome Sanger Institute COVID-19 Surveillance Team |
| EPI_ISL_2436727 | Lighthouse Lab in Glasgow | Wellcome Sanger Institute for the COVID-19 Genomics UK (COG-UK) Consortium | Harper VanSteenhouse, Yumi Kasai, David Gray, Carol Clugston, Anna Dominiczak and Alex Alderton, Roberto Amato, Jeffrey Barrett, Sonia Goncalves, Ewan Harrison, David K. Jackson, Ian Johnston, Dominic Kwiatkowski, Cordelia Langford, John Sillitoe on behalf of the Wellcome Sanger Institute COVID-19 Surveillance Team |
| EPI_ISL_2436728 | Lighthouse Lab in Milton Keynes | Wellcome Sanger Institute for the COVID-19 Genomics UK (COG-UK) Consortium | The Lighthouse Lab in Milton Keynes and Alex Alderton, Roberto Amato, Jeffrey Barrett, Sonia Goncalves, Ewan Harrison, David K. Jackson, Ian Johnston, Dominic Kwiatkowski, Cordelia Langford, John Sillitoe on behalf of the Wellcome Sanger Institute COVID-19 Surveillance Team |
| EPI_ISL_2436730 | Lighthouse Lab in Alderley Park | Wellcome Sanger Institute for the COVID-19 Genomics UK (COG-UK) Consortium | Jacquelyn Wynn, Mairead Hyland, The Lighthouse Lab in Alderley Park and Alex Alderton, Roberto Amato, Jeffrey Barrett, Sonia Goncalves, Ewan Harrison, David K. Jackson, Ian Johnston, Dominic Kwiatkowski, Cordelia Langford, John Sillitoe on behalf of the Wellcome Sanger Institute COVID-19 Surveillance Team |
| EPI_ISL_2436732 | Lighthouse Lab in Milton Keynes | Wellcome Sanger Institute for the COVID-19 Genomics UK (COG-UK) Consortium | The Lighthouse Lab in Milton Keynes and Alex Alderton, Roberto Amato, Jeffrey Barrett, Sonia Goncalves, Ewan Harrison, David K. Jackson, Ian Johnston, Dominic Kwiatkowski, Cordelia Langford, John Sillitoe on behalf of the Wellcome Sanger Institute COVID-19 Surveillance Team |
| EPI_ISL_2436733 | Lighthouse Lab in Glasgow | Wellcome Sanger Institute for the COVID-19 Genomics UK (COG-UK) Consortium | Harper VanSteenhouse, Yumi Kasai, David Gray, Carol Clugston, Anna Dominiczak and Alex Alderton, Roberto Amato, Jeffrey Barrett, Sonia Goncalves, Ewan Harrison, David K. Jackson, Ian Johnston, Dominic Kwiatkowski, Cordelia Langford, John Sillitoe on behalf of the Wellcome Sanger Institute COVID-19 Surveillance Team |
| EPI_ISL_2436734 | Lighthouse Lab in Alderley Park | Wellcome Sanger Institute for the COVID-19 Genomics UK (COG-UK) Consortium | Jacquelyn Wynn, Mairead Hyland, The Lighthouse Lab in Alderley Park and Alex Alderton, Roberto Amato, Jeffrey Barrett, Sonia Goncalves, Ewan Harrison, David K. Jackson, Ian Johnston, Dominic Kwiatkowski, Cordelia Langford, John Sillitoe on behalf of the Wellcome Sanger Institute COVID-19 Surveillance Team |
| EPI_ISL_2436735, EPI_ISL_2436736 | Lighthouse Lab in Milton Keynes | Wellcome Sanger Institute for the COVID-19 Genomics UK (COG-UK) Consortium | The Lighthouse Lab in Milton Keynes and Alex Alderton, Roberto Amato, Jeffrey Barrett, Sonia Goncalves, Ewan Harrison, David K. Jackson, Ian Johnston, Dominic Kwiatkowski, Cordelia Langford, John Sillitoe on behalf of the Wellcome Sanger Institute COVID-19 Surveillance Team |
| EPI_ISL_2436737 | Lighthouse Lab in Glasgow | Wellcome Sanger Institute for the COVID-19 Genomics UK (COG-UK) Consortium | Harper VanSteenhouse, Yumi Kasai, David Gray, Carol Clugston, Anna Dominiczak and Alex Alderton, Roberto Amato, Jeffrey Barrett, Sonia Goncalves, Ewan Harrison, David K. Jackson, Ian Johnston, Dominic Kwiatkowski, Cordelia Langford, John Sillitoe on behalf of the Wellcome Sanger Institute COVID-19 Surveillance Team |
| EPI_ISL_2436738, EPI_ISL_2436739, EPI_ISL_2436740 | Lighthouse Lab in Milton Keynes | Wellcome Sanger Institute for the COVID-19 Genomics UK (COG-UK) Consortium | The Lighthouse Lab in Milton Keynes and Alex Alderton, Roberto Amato, Jeffrey Barrett, Sonia Goncalves, Ewan Harrison, David K. Jackson, Ian Johnston, Dominic Kwiatkowski, Cordelia Langford, John Sillitoe on behalf of the Wellcome Sanger Institute COVID-19 Surveillance Team |
| EPI_ISL_2436741, EPI_ISL_2436742 | Lighthouse Lab in Alderley Park | Wellcome Sanger Institute for the COVID-19 Genomics UK (COG-UK) Consortium | Jacquelyn Wynn, Mairead Hyland, The Lighthouse Lab in Alderley Park and Alex Alderton, Roberto Amato, Jeffrey Barrett, Sonia Goncalves, Ewan Harrison, David K. Jackson, Ian Johnston, Dominic Kwiatkowski, Cordelia Langford, John Sillitoe on behalf of the Wellcome Sanger Institute COVID-19 Surveillance Team |
| EPI_ISL_2436744 | Lighthouse Lab in Milton Keynes | Wellcome Sanger Institute for the COVID-19 Genomics UK (COG-UK) Consortium | The Lighthouse Lab in Milton Keynes and Alex Alderton, Roberto Amato, Jeffrey Barrett, Sonia Goncalves, Ewan Harrison, David K. Jackson, Ian Johnston, Dominic Kwiatkowski, Cordelia Langford, John Sillitoe on behalf of the Wellcome Sanger Institute COVID-19 Surveillance Team |
| EPI_ISL_2436745 | Lighthouse Lab in Alderley Park | Wellcome Sanger Institute for the COVID-19 Genomics UK (COG-UK) Consortium | Jacquelyn Wynn, Mairead Hyland, The Lighthouse Lab in Alderley Park and Alex Alderton, Roberto Amato, Jeffrey Barrett, Sonia Goncalves, Ewan Harrison, David K. Jackson, Ian Johnston, Dominic Kwiatkowski, Cordelia Langford, John Sillitoe on behalf of the Wellcome Sanger Institute COVID-19 Surveillance Team |
| EPI_ISL_2436746 | Lighthouse Lab in Milton Keynes | Wellcome Sanger Institute for the COVID-19 Genomics UK (COG-UK) Consortium | The Lighthouse Lab in Milton Keynes and Alex Alderton, Roberto Amato, Jeffrey Barrett, Sonia Goncalves, Ewan Harrison, David K. Jackson, Ian Johnston, Dominic Kwiatkowski, Cordelia Langford, John Sillitoe on behalf of the Wellcome Sanger Institute COVID-19 Surveillance Team |
| EPI_ISL_2436747, EPI_ISL_2436749, EPI_ISL_2436750, EPI_ISL_2436751 | Lighthouse Lab in Alderley Park | Wellcome Sanger Institute for the COVID-19 Genomics UK (COG-UK) Consortium | Jacquelyn Wynn, Mairead Hyland, The Lighthouse Lab in Alderley Park and Alex Alderton, Roberto Amato, Jeffrey Barrett, Sonia Goncalves, Ewan Harrison, David K. Jackson, Ian Johnston, Dominic Kwiatkowski, Cordelia Langford, John Sillitoe on behalf of the Wellcome Sanger Institute COVID-19 Surveillance Team |
| EPI_ISL_2436753 | Lighthouse Lab in Glasgow | Wellcome Sanger Institute for the COVID-19 Genomics UK (COG-UK) Consortium | Harper VanSteenhouse, Yumi Kasai, David Gray, Carol Clugston, Anna Dominiczak and Alex Alderton, Roberto Amato, Jeffrey Barrett, Sonia Goncalves, Ewan Harrison, David K. Jackson, Ian Johnston, Dominic Kwiatkowski, Cordelia Langford, John Sillitoe on behalf of the Wellcome Sanger Institute COVID-19 Surveillance Team |
| EPI_ISL_2436754, EPI_ISL_2436755, EPI_ISL_2436756 | Lighthouse Lab in Alderley Park | Wellcome Sanger Institute for the COVID-19 Genomics UK (COG-UK) Consortium | Jacquelyn Wynn, Mairead Hyland, The Lighthouse Lab in Alderley Park and Alex Alderton, Roberto Amato, Jeffrey Barrett, Sonia Goncalves, Ewan Harrison, David K. Jackson, Ian Johnston, Dominic Kwiatkowski, Cordelia Langford, John Sillitoe on behalf of the Wellcome Sanger Institute COVID-19 Surveillance Team |
| EPI_ISL_2436757 | Lighthouse Lab in Milton Keynes | Wellcome Sanger Institute for the COVID-19 Genomics UK (COG-UK) Consortium | The Lighthouse Lab in Milton Keynes and Alex Alderton, Roberto Amato, Jeffrey Barrett, Sonia Goncalves, Ewan Harrison, David K. Jackson, Ian Johnston, Dominic Kwiatkowski, Cordelia Langford, John Sillitoe on behalf of the Wellcome Sanger Institute COVID-19 Surveillance Team |
| EPI_ISL_2436758 | Lighthouse Lab in Alderley Park | Wellcome Sanger Institute for the COVID-19 Genomics UK (COG-UK) Consortium | Jacquelyn Wynn, Mairead Hyland, The Lighthouse Lab in Alderley Park and Alex Alderton, Roberto Amato, Jeffrey Barrett, Sonia Goncalves, Ewan Harrison, David K. Jackson, Ian Johnston, Dominic Kwiatkowski, Cordelia Langford, John Sillitoe on behalf of the Wellcome Sanger Institute COVID-19 Surveillance Team |
| EPI_ISL_2436759 | Lighthouse Lab in Glasgow | Wellcome Sanger Institute for the COVID-19 Genomics UK (COG-UK) Consortium | Harper VanSteenhouse, Yumi Kasai, David Gray, Carol Clugston, Anna Dominiczak and Alex Alderton, Roberto Amato, Jeffrey Barrett, Sonia Goncalves, Ewan Harrison, David K. Jackson, Ian Johnston, Dominic Kwiatkowski, Cordelia Langford, John Sillitoe on behalf of the Wellcome Sanger Institute COVID-19 Surveillance Team |
| EPI_ISL_2436761 | Lighthouse Lab in Milton Keynes | Wellcome Sanger Institute for the COVID-19 Genomics UK (COG-UK) Consortium | The Lighthouse Lab in Milton Keynes and Alex Alderton, Roberto Amato, Jeffrey Barrett, Sonia Goncalves, Ewan Harrison, David K. Jackson, Ian Johnston, Dominic Kwiatkowski, Cordelia Langford, John Sillitoe on behalf of the Wellcome Sanger Institute COVID-19 Surveillance Team |
| EPI_ISL_2436762 | Lighthouse Lab in Glasgow | Wellcome Sanger Institute for the COVID-19 Genomics UK (COG-UK) Consortium | Harper VanSteenhouse, Yumi Kasai, David Gray, Carol Clugston, Anna Dominiczak and Alex Alderton, Roberto Amato, Jeffrey Barrett, Sonia Goncalves, Ewan Harrison, David K. Jackson, Ian Johnston, Dominic Kwiatkowski, Cordelia Langford, John Sillitoe on behalf of the Wellcome Sanger Institute COVID-19 Surveillance Team |
| EPI_ISL_2436763 | Lighthouse Lab in Alderley Park | Wellcome Sanger Institute for the COVID-19 Genomics UK (COG-UK) | Jacquelyn Wynn, Mairead Hyland, The Lighthouse Lab in Alderley Park and Alex Alderton, Roberto Amato, Jeffrey Barrett, Sonia Goncalves, Ewan Harrison, David K. Jackson, Ian Johnston, Dominic Kwiatkowski, Cordelia Langford, John Sillitoe on |

| | | | |
|---|---|---|---|
| | | Consortium | behalf of the Wellcome Sanger Institute COVID-19 Surveillance Team |
| EPI_ISL_2436765, EPI_ISL_2436766 | Lighthouse Lab in Milton Keynes | Wellcome Sanger Institute for the COVID-19 Genomics UK (COG-UK) Consortium | The Lighthouse Lab in Milton Keynes and Alex Alderton, Roberto Amato, Jeffrey Barrett, Sonia Goncalves, Ewan Harrison, David K. Jackson, Ian Johnston, Dominic Kwiatkowski, Cordelia Langford, John Sillitoe on behalf of the Wellcome Sanger Institute COVID-19 Surveillance Team |
| EPI_ISL_2436767 | Lighthouse Lab in Glasgow | Wellcome Sanger Institute for the COVID-19 Genomics UK (COG-UK) Consortium | Harper VanSteenhouse, Yumi Kasai, David Gray, Carol Clugston, Anna Dominiczak and Alex Alderton, Roberto Amato, Jeffrey Barrett, Sonia Goncalves, Ewan Harrison, David K. Jackson, Ian Johnston, Dominic Kwiatkowski, Cordelia Langford, John Sillitoe on behalf of the Wellcome Sanger Institute COVID-19 Surveillance Team |
| EPI_ISL_2436768 | Lighthouse Lab in Milton Keynes | Wellcome Sanger Institute for the COVID-19 Genomics UK (COG-UK) Consortium | The Lighthouse Lab in Milton Keynes and Alex Alderton, Roberto Amato, Jeffrey Barrett, Sonia Goncalves, Ewan Harrison, David K. Jackson, Ian Johnston, Dominic Kwiatkowski, Cordelia Langford, John Sillitoe on behalf of the Wellcome Sanger Institute COVID-19 Surveillance Team |
| EPI_ISL_2436769 | Lighthouse Lab in Glasgow | Wellcome Sanger Institute for the COVID-19 Genomics UK (COG-UK) Consortium | Harper VanSteenhouse, Yumi Kasai, David Gray, Carol Clugston, Anna Dominiczak and Alex Alderton, Roberto Amato, Jeffrey Barrett, Sonia Goncalves, Ewan Harrison, David K. Jackson, Ian Johnston, Dominic Kwiatkowski, Cordelia Langford, John Sillitoe on behalf of the Wellcome Sanger Institute COVID-19 Surveillance Team |
| EPI_ISL_2436770 | Lighthouse Lab in Milton Keynes | Wellcome Sanger Institute for the COVID-19 Genomics UK (COG-UK) Consortium | The Lighthouse Lab in Milton Keynes and Alex Alderton, Roberto Amato, Jeffrey Barrett, Sonia Goncalves, Ewan Harrison, David K. Jackson, Ian Johnston, Dominic Kwiatkowski, Cordelia Langford, John Sillitoe on behalf of the Wellcome Sanger Institute COVID-19 Surveillance Team |
| EPI_ISL_2436773 | Lighthouse Lab in Glasgow | Wellcome Sanger Institute for the COVID-19 Genomics UK (COG-UK) Consortium | Harper VanSteenhouse, Yumi Kasai, David Gray, Carol Clugston, Anna Dominiczak and Alex Alderton, Roberto Amato, Jeffrey Barrett, Sonia Goncalves, Ewan Harrison, David K. Jackson, Ian Johnston, Dominic Kwiatkowski, Cordelia Langford, John Sillitoe on behalf of the Wellcome Sanger Institute COVID-19 Surveillance Team |
| EPI_ISL_2436774, EPI_ISL_2436775 | Lighthouse Lab in Alderley Park | Wellcome Sanger Institute for the COVID-19 Genomics UK (COG-UK) Consortium | Jacquelyn Wynn, Mairead Hyland, The Lighthouse Lab in Alderley Park and Alex Alderton, Roberto Amato, Jeffrey Barrett, Sonia Goncalves, Ewan Harrison, David K. Jackson, Ian Johnston, Dominic Kwiatkowski, Cordelia Langford, John Sillitoe on behalf of the Wellcome Sanger Institute COVID-19 Surveillance Team |
| EPI_ISL_2436776 | Lighthouse Lab in Milton Keynes | Wellcome Sanger Institute for the COVID-19 Genomics UK (COG-UK) Consortium | The Lighthouse Lab in Milton Keynes and Alex Alderton, Roberto Amato, Jeffrey Barrett, Sonia Goncalves, Ewan Harrison, David K. Jackson, Ian Johnston, Dominic Kwiatkowski, Cordelia Langford, John Sillitoe on behalf of the Wellcome Sanger Institute COVID-19 Surveillance Team |
| EPI_ISL_2436777 | Lighthouse Lab in Alderley Park | Wellcome Sanger Institute for the COVID-19 Genomics UK (COG-UK) Consortium | Jacquelyn Wynn, Mairead Hyland, The Lighthouse Lab in Alderley Park and Alex Alderton, Roberto Amato, Jeffrey Barrett, Sonia Goncalves, Ewan Harrison, David K. Jackson, Ian Johnston, Dominic Kwiatkowski, Cordelia Langford, John Sillitoe on behalf of the Wellcome Sanger Institute COVID-19 Surveillance Team |
| EPI_ISL_2436778, EPI_ISL_2436779 | Lighthouse Lab in Milton Keynes | Wellcome Sanger Institute for the COVID-19 Genomics UK (COG-UK) Consortium | The Lighthouse Lab in Milton Keynes and Alex Alderton, Roberto Amato, Jeffrey Barrett, Sonia Goncalves, Ewan Harrison, David K. Jackson, Ian Johnston, Dominic Kwiatkowski, Cordelia Langford, John Sillitoe on behalf of the Wellcome Sanger Institute COVID-19 Surveillance Team |
| EPI_ISL_2436780 | Lighthouse Lab in Alderley Park | Wellcome Sanger Institute for the COVID-19 Genomics UK (COG-UK) Consortium | Jacquelyn Wynn, Mairead Hyland, The Lighthouse Lab in Alderley Park and Alex Alderton, Roberto Amato, Jeffrey Barrett, Sonia Goncalves, Ewan Harrison, David K. Jackson, Ian Johnston, Dominic Kwiatkowski, Cordelia Langford, John Sillitoe on behalf of the Wellcome Sanger Institute COVID-19 Surveillance Team |
| EPI_ISL_2436781 | Lighthouse Lab in Glasgow | Wellcome Sanger Institute for the COVID-19 Genomics UK (COG-UK) Consortium | Harper VanSteenhouse, Yumi Kasai, David Gray, Carol Clugston, Anna Dominiczak and Alex Alderton, Roberto Amato, Jeffrey Barrett, Sonia Goncalves, Ewan Harrison, David K. Jackson, Ian Johnston, Dominic Kwiatkowski, Cordelia Langford, John Sillitoe on behalf of the Wellcome Sanger Institute COVID-19 Surveillance Team |
| EPI_ISL_2436782 | Lighthouse Lab in Alderley Park | Wellcome Sanger Institute for the COVID-19 Genomics UK (COG-UK) Consortium | Jacquelyn Wynn, Mairead Hyland, The Lighthouse Lab in Alderley Park and Alex Alderton, Roberto Amato, Jeffrey Barrett, Sonia Goncalves, Ewan Harrison, David K. Jackson, Ian Johnston, Dominic Kwiatkowski, Cordelia Langford, John Sillitoe on behalf of the Wellcome Sanger Institute COVID-19 Surveillance Team |
| EPI_ISL_2436783 | Lighthouse Lab in Milton Keynes | Wellcome Sanger Institute for the COVID-19 Genomics UK (COG-UK) Consortium | The Lighthouse Lab in Milton Keynes and Alex Alderton, Roberto Amato, Jeffrey Barrett, Sonia Goncalves, Ewan Harrison, David K. Jackson, Ian Johnston, Dominic Kwiatkowski, Cordelia Langford, John Sillitoe on behalf of the Wellcome Sanger Institute COVID-19 Surveillance Team |
| EPI_ISL_2436784, EPI_ISL_2436785, EPI_ISL_2436786 | Lighthouse Lab in Alderley Park | Wellcome Sanger Institute for the COVID-19 Genomics UK (COG-UK) Consortium | Jacquelyn Wynn, Mairead Hyland, The Lighthouse Lab in Alderley Park and Alex Alderton, Roberto Amato, Jeffrey Barrett, Sonia Goncalves, Ewan Harrison, David K. Jackson, Ian Johnston, Dominic Kwiatkowski, Cordelia Langford, John Sillitoe on behalf of the Wellcome Sanger Institute COVID-19 Surveillance Team |
| EPI_ISL_2436791, EPI_ISL_2436793 | Lighthouse Lab in Milton Keynes | Wellcome Sanger Institute for the COVID-19 Genomics UK (COG-UK) Consortium | The Lighthouse Lab in Milton Keynes and Alex Alderton, Roberto Amato, Jeffrey Barrett, Sonia Goncalves, Ewan Harrison, David K. Jackson, Ian Johnston, Dominic Kwiatkowski, Cordelia Langford, John Sillitoe on behalf of the Wellcome Sanger Institute COVID-19 Surveillance Team |
| EPI_ISL_2436794 | Lighthouse Lab in Alderley Park | Wellcome Sanger Institute for the COVID-19 Genomics UK (COG-UK) Consortium | Jacquelyn Wynn, Mairead Hyland, The Lighthouse Lab in Alderley Park and Alex Alderton, Roberto Amato, Jeffrey Barrett, Sonia Goncalves, Ewan Harrison, David K. Jackson, Ian Johnston, Dominic Kwiatkowski, Cordelia Langford, John Sillitoe on behalf of the Wellcome Sanger Institute COVID-19 Surveillance Team |
| EPI_ISL_2436795 | Lighthouse Lab in Milton Keynes | Wellcome Sanger Institute for the COVID-19 Genomics UK (COG-UK) Consortium | The Lighthouse Lab in Milton Keynes and Alex Alderton, Roberto Amato, Jeffrey Barrett, Sonia Goncalves, Ewan Harrison, David K. Jackson, Ian Johnston, Dominic Kwiatkowski, Cordelia Langford, John Sillitoe on behalf of the Wellcome Sanger Institute COVID-19 Surveillance Team |
| EPI_ISL_2436796, EPI_ISL_2436797, EPI_ISL_2436798 | Lighthouse Lab in Glasgow | Wellcome Sanger Institute for the COVID-19 Genomics UK (COG-UK) Consortium | Harper VanSteenhouse, Yumi Kasai, David Gray, Carol Clugston, Anna Dominiczak and Alex Alderton, Roberto Amato, Jeffrey Barrett, Sonia Goncalves, Ewan Harrison, David K. Jackson, Ian Johnston, Dominic Kwiatkowski, Cordelia Langford, John Sillitoe on behalf of the Wellcome Sanger Institute COVID-19 Surveillance Team |
| EPI_ISL_2436799, EPI_ISL_2436800, EPI_ISL_2436803 | Lighthouse Lab in Alderley Park | Wellcome Sanger Institute for the COVID-19 Genomics UK (COG-UK) Consortium | Jacquelyn Wynn, Mairead Hyland, The Lighthouse Lab in Alderley Park and Alex Alderton, Roberto Amato, Jeffrey Barrett, Sonia Goncalves, Ewan Harrison, David K. Jackson, Ian Johnston, Dominic Kwiatkowski, Cordelia Langford, John Sillitoe on behalf of the Wellcome Sanger Institute COVID-19 Surveillance Team |
| EPI_ISL_2436804, EPI_ISL_2436806 | Lighthouse Lab in Glasgow | Wellcome Sanger Institute for the COVID-19 Genomics UK (COG-UK) Consortium | Harper VanSteenhouse, Yumi Kasai, David Gray, Carol Clugston, Anna Dominiczak and Alex Alderton, Roberto Amato, Jeffrey Barrett, Sonia Goncalves, Ewan Harrison, David K. Jackson, Ian Johnston, Dominic Kwiatkowski, Cordelia Langford, John Sillitoe on behalf of the Wellcome Sanger Institute COVID-19 Surveillance Team |
| EPI_ISL_2436807 | Lighthouse Lab in Alderley Park | Wellcome Sanger Institute for the COVID-19 Genomics UK (COG-UK) Consortium | Jacquelyn Wynn, Mairead Hyland, The Lighthouse Lab in Alderley Park and Alex Alderton, Roberto Amato, Jeffrey Barrett, Sonia Goncalves, Ewan Harrison, David K. Jackson, Ian Johnston, Dominic Kwiatkowski, Cordelia Langford, John Sillitoe on behalf of the Wellcome Sanger Institute COVID-19 Surveillance Team |
| EPI_ISL_2436809 | Lighthouse Lab in Milton Keynes | Wellcome Sanger Institute for the COVID-19 Genomics UK (COG-UK) Consortium | The Lighthouse Lab in Milton Keynes and Alex Alderton, Roberto Amato, Jeffrey Barrett, Sonia Goncalves, Ewan Harrison, David K. Jackson, Ian Johnston, Dominic Kwiatkowski, Cordelia Langford, John Sillitoe on behalf of the Wellcome Sanger Institute COVID-19 Surveillance Team |
| EPI_ISL_2436810, EPI_ISL_2436811 | Lighthouse Lab in Alderley Park | Wellcome Sanger Institute for the COVID-19 Genomics UK (COG-UK) Consortium | Jacquelyn Wynn, Mairead Hyland, The Lighthouse Lab in Alderley Park and Alex Alderton, Roberto Amato, Jeffrey Barrett, Sonia Goncalves, Ewan Harrison, David K. Jackson, Ian Johnston, Dominic Kwiatkowski, Cordelia Langford, John Sillitoe on behalf of the Wellcome Sanger Institute COVID-19 Surveillance Team |
| EPI_ISL_2436813 | Lighthouse Lab in Glasgow | Wellcome Sanger Institute for the COVID-19 Genomics UK (COG-UK) Consortium | Harper VanSteenhouse, Yumi Kasai, David Gray, Carol Clugston, Anna Dominiczak and Alex Alderton, Roberto Amato, Jeffrey Barrett, Sonia Goncalves, Ewan Harrison, David K. Jackson, Ian Johnston, Dominic Kwiatkowski, Cordelia Langford, John Sillitoe on behalf of the Wellcome Sanger Institute COVID-19 Surveillance Team |
| EPI_ISL_2436814, EPI_ISL_2436817 | Lighthouse Lab in Alderley Park | Wellcome Sanger Institute for the COVID-19 Genomics UK (COG-UK) Consortium | Jacquelyn Wynn, Mairead Hyland, The Lighthouse Lab in Alderley Park and Alex Alderton, Roberto Amato, Jeffrey Barrett, Sonia Goncalves, Ewan Harrison, David K. Jackson, Ian Johnston, Dominic Kwiatkowski, Cordelia Langford, John Sillitoe on behalf of the Wellcome Sanger Institute COVID-19 Surveillance Team |
| EPI_ISL_2436818 | Lighthouse Lab in Milton Keynes | Wellcome Sanger Institute for the COVID-19 Genomics UK (COG-UK) Consortium | The Lighthouse Lab in Milton Keynes and Alex Alderton, Roberto Amato, Jeffrey Barrett, Sonia Goncalves, Ewan Harrison, David K. Jackson, Ian Johnston, Dominic Kwiatkowski, Cordelia Langford, John Sillitoe on behalf of the Wellcome Sanger Institute COVID-19 Surveillance Team |
| EPI_ISL_2436819 | Lighthouse Lab in Glasgow | Wellcome Sanger Institute for the COVID-19 Genomics UK (COG-UK) Consortium | Harper VanSteenhouse, Yumi Kasai, David Gray, Carol Clugston, Anna Dominiczak and Alex Alderton, Roberto Amato, Jeffrey Barrett, Sonia Goncalves, Ewan Harrison, David K. Jackson, Ian Johnston, Dominic Kwiatkowski, Cordelia Langford, John Sillitoe on behalf of the Wellcome Sanger Institute COVID-19 Surveillance Team |
| EPI_ISL_2436820, EPI_ISL_2436821, EPI_ISL_2436822 | Lighthouse Lab in Alderley Park | Wellcome Sanger Institute for the COVID-19 Genomics UK (COG-UK) Consortium | Jacquelyn Wynn, Mairead Hyland, The Lighthouse Lab in Alderley Park and Alex Alderton, Roberto Amato, Jeffrey Barrett, Sonia Goncalves, Ewan Harrison, David K. Jackson, Ian Johnston, Dominic Kwiatkowski, Cordelia Langford, John Sillitoe on behalf of the Wellcome Sanger Institute COVID-19 Surveillance Team |
| EPI_ISL_2436823 | Lighthouse Lab in Milton Keynes | Wellcome Sanger Institute for the COVID-19 Genomics UK (COG-UK) Consortium | The Lighthouse Lab in Milton Keynes and Alex Alderton, Roberto Amato, Jeffrey Barrett, Sonia Goncalves, Ewan Harrison, David K. Jackson, Ian Johnston, Dominic Kwiatkowski, Cordelia Langford, John Sillitoe on behalf of the Wellcome Sanger Institute COVID-19 Surveillance Team |
| EPI_ISL_2436825 | Lighthouse Lab in Alderley Park | Wellcome Sanger Institute for the COVID-19 Genomics UK (COG-UK) Consortium | Jacquelyn Wynn, Mairead Hyland, The Lighthouse Lab in Alderley Park and Alex Alderton, Roberto Amato, Jeffrey Barrett, Sonia Goncalves, Ewan Harrison, David K. Jackson, Ian Johnston, Dominic Kwiatkowski, Cordelia Langford, John Sillitoe on behalf of the Wellcome Sanger Institute COVID-19 Surveillance Team |
| EPI_ISL_2436827 | Lighthouse Lab in Glasgow | Wellcome Sanger Institute for the COVID-19 Genomics UK (COG-UK) Consortium | Harper VanSteenhouse, Yumi Kasai, David Gray, Carol Clugston, Anna Dominiczak and Alex Alderton, Roberto Amato, Jeffrey Barrett, Sonia Goncalves, Ewan Harrison, David K. Jackson, Ian Johnston, Dominic Kwiatkowski, Cordelia Langford, John Sillitoe on behalf of the Wellcome Sanger Institute COVID-19 Surveillance Team |
| EPI_ISL_2436829 | Lighthouse Lab in Alderley Park | Wellcome Sanger Institute for the COVID-19 Genomics UK (COG-UK) Consortium | Jacquelyn Wynn, Mairead Hyland, The Lighthouse Lab in Alderley Park and Alex Alderton, Roberto Amato, Jeffrey Barrett, Sonia Goncalves, Ewan Harrison, David K. Jackson, Ian Johnston, Dominic Kwiatkowski, Cordelia Langford, John Sillitoe on behalf of the Wellcome Sanger Institute COVID-19 Surveillance Team |
| EPI_ISL_2436830 | Lighthouse Lab in Glasgow | Wellcome Sanger Institute for the COVID-19 Genomics UK (COG-UK) Consortium | Harper VanSteenhouse, Yumi Kasai, David Gray, Carol Clugston, Anna Dominiczak and Alex Alderton, Roberto Amato, Jeffrey Barrett, Sonia Goncalves, Ewan Harrison, David K. Jackson, Ian Johnston, Dominic Kwiatkowski, Cordelia Langford, John Sillitoe on behalf of the Wellcome Sanger Institute COVID-19 Surveillance Team |
| EPI_ISL_2436831 | Lighthouse Lab in Alderley Park | Wellcome Sanger Institute for the COVID-19 Genomics UK (COG-UK) Consortium | Jacquelyn Wynn, Mairead Hyland, The Lighthouse Lab in Alderley Park and Alex Alderton, Roberto Amato, Jeffrey Barrett, Sonia Goncalves, Ewan Harrison, David K. Jackson, Ian Johnston, Dominic Kwiatkowski, Cordelia Langford, John Sillitoe on behalf of the Wellcome Sanger Institute COVID-19 Surveillance Team |
| EPI_ISL_2436835 | Lighthouse Lab in Glasgow | Wellcome Sanger Institute for the COVID-19 Genomics UK (COG-UK) Consortium | Harper VanSteenhouse, Yumi Kasai, David Gray, Carol Clugston, Anna Dominiczak and Alex Alderton, Roberto Amato, Jeffrey Barrett, Sonia Goncalves, Ewan Harrison, David K. Jackson, Ian Johnston, Dominic Kwiatkowski, Cordelia Langford, John Sillitoe on behalf of the Wellcome Sanger Institute COVID-19 Surveillance Team |
| EPI_ISL_2436836, EPI_ISL_2436837, EPI_ISL_2436839, EPI_ISL_2436840, EPI_ISL_2436841, EPI_ISL_2436842 | Lighthouse Lab in Alderley Park | Wellcome Sanger Institute for the COVID-19 Genomics UK (COG-UK) Consortium | Jacquelyn Wynn, Mairead Hyland, The Lighthouse Lab in Alderley Park and Alex Alderton, Roberto Amato, Jeffrey Barrett, Sonia Goncalves, Ewan Harrison, David K. Jackson, Ian Johnston, Dominic Kwiatkowski, Cordelia Langford, John Sillitoe on behalf of the Wellcome Sanger Institute COVID-19 Surveillance Team |
| EPI_ISL_2436843 | Lighthouse Lab in Glasgow | Wellcome Sanger Institute for the COVID-19 Genomics UK (COG-UK) Consortium | Harper VanSteenhouse, Yumi Kasai, David Gray, Carol Clugston, Anna Dominiczak and Alex Alderton, Roberto Amato, Jeffrey Barrett, Sonia Goncalves, Ewan Harrison, David K. Jackson, Ian Johnston, Dominic Kwiatkowski, Cordelia Langford, John Sillitoe on behalf of the Wellcome Sanger Institute COVID-19 Surveillance Team |
| EPI_ISL_2436844 | Lighthouse Lab in Alderley Park | Wellcome Sanger Institute for the COVID-19 Genomics UK (COG-UK) Consortium | Jacquelyn Wynn, Mairead Hyland, The Lighthouse Lab in Alderley Park and Alex Alderton, Roberto Amato, Jeffrey Barrett, Sonia Goncalves, Ewan Harrison, David K. Jackson, Ian Johnston, Dominic Kwiatkowski, Cordelia Langford, John Sillitoe on behalf of the Wellcome Sanger Institute COVID-19 Surveillance Team |
| EPI_ISL_2436845 | Lighthouse Lab in Milton Keynes | Wellcome Sanger Institute for the COVID-19 Genomics UK (COG-UK) Consortium | The Lighthouse Lab in Milton Keynes and Alex Alderton, Roberto Amato, Jeffrey Barrett, Sonia Goncalves, Ewan Harrison, David K. Jackson, Ian Johnston, Dominic Kwiatkowski, Cordelia Langford, John Sillitoe on behalf of the Wellcome Sanger Institute COVID-19 Surveillance Team |
| EPI_ISL_2436846, EPI_ISL_2436847 | Lighthouse Lab in Alderley Park | Wellcome Sanger Institute for the COVID-19 Genomics UK (COG-UK) Consortium | Jacquelyn Wynn, Mairead Hyland, The Lighthouse Lab in Alderley Park and Alex Alderton, Roberto Amato, Jeffrey Barrett, Sonia Goncalves, Ewan Harrison, David K. Jackson, Ian Johnston, Dominic Kwiatkowski, Cordelia Langford, John Sillitoe on behalf of the Wellcome Sanger Institute COVID-19 Surveillance Team |

| | | | |
|---|---|---|---|
| EPI_ISL_2436848 | Lighthouse Lab in Glasgow | Wellcome Sanger Institute for the COVID-19 Genomics UK (COG-UK) Consortium | Harper VanSteenhouse, Yumi Kasai, David Gray, Carol Clugston, Anna Dominiczak and Alex Alderton, Roberto Amato, Jeffrey Barrett, Sonia Goncalves, Ewan Harrison, David K. Jackson, Ian Johnston, Dominic Kwiatkowski, Cordelia Langford, John Sillitoe on behalf of the Wellcome Sanger Institute COVID-19 Surveillance Team |
| EPI_ISL_2436849 | Lighthouse Lab in Alderley Park | Wellcome Sanger Institute for the COVID-19 Genomics UK (COG-UK) Consortium | Jacquelyn Wynn, Mairead Hyland, The Lighthouse Lab in Alderley Park and Alex Alderton, Roberto Amato, Jeffrey Barrett, Sonia Goncalves, Ewan Harrison, David K. Jackson, Ian Johnston, Dominic Kwiatkowski, Cordelia Langford, John Sillitoe on behalf of the Wellcome Sanger Institute COVID-19 Surveillance Team |
| EPI_ISL_2436850 | Lighthouse Lab in Glasgow | Wellcome Sanger Institute for the COVID-19 Genomics UK (COG-UK) Consortium | Harper VanSteenhouse, Yumi Kasai, David Gray, Carol Clugston, Anna Dominiczak and Alex Alderton, Roberto Amato, Jeffrey Barrett, Sonia Goncalves, Ewan Harrison, David K. Jackson, Ian Johnston, Dominic Kwiatkowski, Cordelia Langford, John Sillitoe on behalf of the Wellcome Sanger Institute COVID-19 Surveillance Team |
| EPI_ISL_2436851, EPI_ISL_2436852 | Lighthouse Lab in Milton Keynes | Wellcome Sanger Institute for the COVID-19 Genomics UK (COG-UK) Consortium | The Lighthouse Lab in Milton Keynes and Alex Alderton, Roberto Amato, Jeffrey Barrett, Sonia Goncalves, Ewan Harrison, David K. Jackson, Ian Johnston, Dominic Kwiatkowski, Cordelia Langford, John Sillitoe on behalf of the Wellcome Sanger Institute COVID-19 Surveillance Team |
| EPI_ISL_2436853 | Lighthouse Lab in Alderley Park | Wellcome Sanger Institute for the COVID-19 Genomics UK (COG-UK) Consortium | Jacquelyn Wynn, Mairead Hyland, The Lighthouse Lab in Alderley Park and Alex Alderton, Roberto Amato, Jeffrey Barrett, Sonia Goncalves, Ewan Harrison, David K. Jackson, Ian Johnston, Dominic Kwiatkowski, Cordelia Langford, John Sillitoe on behalf of the Wellcome Sanger Institute COVID-19 Surveillance Team |
| EPI_ISL_2436854, EPI_ISL_2436855 | Lighthouse Lab in Milton Keynes | Wellcome Sanger Institute for the COVID-19 Genomics UK (COG-UK) Consortium | The Lighthouse Lab in Milton Keynes and Alex Alderton, Roberto Amato, Jeffrey Barrett, Sonia Goncalves, Ewan Harrison, David K. Jackson, Ian Johnston, Dominic Kwiatkowski, Cordelia Langford, John Sillitoe on behalf of the Wellcome Sanger Institute COVID-19 Surveillance Team |
| EPI_ISL_2436856 | Lighthouse Lab in Glasgow | Wellcome Sanger Institute for the COVID-19 Genomics UK (COG-UK) Consortium | Harper VanSteenhouse, Yumi Kasai, David Gray, Carol Clugston, Anna Dominiczak and Alex Alderton, Roberto Amato, Jeffrey Barrett, Sonia Goncalves, Ewan Harrison, David K. Jackson, Ian Johnston, Dominic Kwiatkowski, Cordelia Langford, John Sillitoe on behalf of the Wellcome Sanger Institute COVID-19 Surveillance Team |
| EPI_ISL_2436857, EPI_ISL_2436858 | Lighthouse Lab in Milton Keynes | Wellcome Sanger Institute for the COVID-19 Genomics UK (COG-UK) Consortium | The Lighthouse Lab in Milton Keynes and Alex Alderton, Roberto Amato, Jeffrey Barrett, Sonia Goncalves, Ewan Harrison, David K. Jackson, Ian Johnston, Dominic Kwiatkowski, Cordelia Langford, John Sillitoe on behalf of the Wellcome Sanger Institute COVID-19 Surveillance Team |
| EPI_ISL_2436859, EPI_ISL_2436860 | Lighthouse Lab in Glasgow | Wellcome Sanger Institute for the COVID-19 Genomics UK (COG-UK) Consortium | Harper VanSteenhouse, Yumi Kasai, David Gray, Carol Clugston, Anna Dominiczak and Alex Alderton, Roberto Amato, Jeffrey Barrett, Sonia Goncalves, Ewan Harrison, David K. Jackson, Ian Johnston, Dominic Kwiatkowski, Cordelia Langford, John Sillitoe on behalf of the Wellcome Sanger Institute COVID-19 Surveillance Team |
| EPI_ISL_2436862, EPI_ISL_2436863 | Lighthouse Lab in Alderley Park | Wellcome Sanger Institute for the COVID-19 Genomics UK (COG-UK) Consortium | Jacquelyn Wynn, Mairead Hyland, The Lighthouse Lab in Alderley Park and Alex Alderton, Roberto Amato, Jeffrey Barrett, Sonia Goncalves, Ewan Harrison, David K. Jackson, Ian Johnston, Dominic Kwiatkowski, Cordelia Langford, John Sillitoe on behalf of the Wellcome Sanger Institute COVID-19 Surveillance Team |
| EPI_ISL_2436864 | Lighthouse Lab in Glasgow | Wellcome Sanger Institute for the COVID-19 Genomics UK (COG-UK) Consortium | Harper VanSteenhouse, Yumi Kasai, David Gray, Carol Clugston, Anna Dominiczak and Alex Alderton, Roberto Amato, Jeffrey Barrett, Sonia Goncalves, Ewan Harrison, David K. Jackson, Ian Johnston, Dominic Kwiatkowski, Cordelia Langford, John Sillitoe on behalf of the Wellcome Sanger Institute COVID-19 Surveillance Team |
| EPI_ISL_2436865 | Lighthouse Lab in Milton Keynes | Wellcome Sanger Institute for the COVID-19 Genomics UK (COG-UK) Consortium | The Lighthouse Lab in Milton Keynes and Alex Alderton, Roberto Amato, Jeffrey Barrett, Sonia Goncalves, Ewan Harrison, David K. Jackson, Ian Johnston, Dominic Kwiatkowski, Cordelia Langford, John Sillitoe on behalf of the Wellcome Sanger Institute COVID-19 Surveillance Team |
| EPI_ISL_2436866 | Lighthouse Lab in Alderley Park | Wellcome Sanger Institute for the COVID-19 Genomics UK (COG-UK) Consortium | Jacquelyn Wynn, Mairead Hyland, The Lighthouse Lab in Alderley Park and Alex Alderton, Roberto Amato, Jeffrey Barrett, Sonia Goncalves, Ewan Harrison, David K. Jackson, Ian Johnston, Dominic Kwiatkowski, Cordelia Langford, John Sillitoe on behalf of the Wellcome Sanger Institute COVID-19 Surveillance Team |
| EPI_ISL_2436867 | Lighthouse Lab in Milton Keynes | Wellcome Sanger Institute for the COVID-19 Genomics UK (COG-UK) Consortium | The Lighthouse Lab in Milton Keynes and Alex Alderton, Roberto Amato, Jeffrey Barrett, Sonia Goncalves, Ewan Harrison, David K. Jackson, Ian Johnston, Dominic Kwiatkowski, Cordelia Langford, John Sillitoe on behalf of the Wellcome Sanger Institute COVID-19 Surveillance Team |
| EPI_ISL_2436868 | Lighthouse Lab in Glasgow | Wellcome Sanger Institute for the COVID-19 Genomics UK (COG-UK) Consortium | Harper VanSteenhouse, Yumi Kasai, David Gray, Carol Clugston, Anna Dominiczak and Alex Alderton, Roberto Amato, Jeffrey Barrett, Sonia Goncalves, Ewan Harrison, David K. Jackson, Ian Johnston, Dominic Kwiatkowski, Cordelia Langford, John Sillitoe on behalf of the Wellcome Sanger Institute COVID-19 Surveillance Team |
| EPI_ISL_2436869, EPI_ISL_2436870, EPI_ISL_2436872 | Lighthouse Lab in Alderley Park | Wellcome Sanger Institute for the COVID-19 Genomics UK (COG-UK) Consortium | Jacquelyn Wynn, Mairead Hyland, The Lighthouse Lab in Alderley Park and Alex Alderton, Roberto Amato, Jeffrey Barrett, Sonia Goncalves, Ewan Harrison, David K. Jackson, Ian Johnston, Dominic Kwiatkowski, Cordelia Langford, John Sillitoe on behalf of the Wellcome Sanger Institute COVID-19 Surveillance Team |
| EPI_ISL_2436873 | Lighthouse Lab in Glasgow | Wellcome Sanger Institute for the COVID-19 Genomics UK (COG-UK) Consortium | Harper VanSteenhouse, Yumi Kasai, David Gray, Carol Clugston, Anna Dominiczak and Alex Alderton, Roberto Amato, Jeffrey Barrett, Sonia Goncalves, Ewan Harrison, David K. Jackson, Ian Johnston, Dominic Kwiatkowski, Cordelia Langford, John Sillitoe on behalf of the Wellcome Sanger Institute COVID-19 Surveillance Team |
| EPI_ISL_2436874, EPI_ISL_2436875, EPI_ISL_2436876, EPI_ISL_2436878 | Lighthouse Lab in Alderley Park | Wellcome Sanger Institute for the COVID-19 Genomics UK (COG-UK) Consortium | Jacquelyn Wynn, Mairead Hyland, The Lighthouse Lab in Alderley Park and Alex Alderton, Roberto Amato, Jeffrey Barrett, Sonia Goncalves, Ewan Harrison, David K. Jackson, Ian Johnston, Dominic Kwiatkowski, Cordelia Langford, John Sillitoe on behalf of the Wellcome Sanger Institute COVID-19 Surveillance Team |
| EPI_ISL_2436879 | Lighthouse Lab in Milton Keynes | Wellcome Sanger Institute for the COVID-19 Genomics UK (COG-UK) Consortium | The Lighthouse Lab in Milton Keynes and Alex Alderton, Roberto Amato, Jeffrey Barrett, Sonia Goncalves, Ewan Harrison, David K. Jackson, Ian Johnston, Dominic Kwiatkowski, Cordelia Langford, John Sillitoe on behalf of the Wellcome Sanger Institute COVID-19 Surveillance Team |
| EPI_ISL_2436880 | Lighthouse Lab in Alderley Park | Wellcome Sanger Institute for the COVID-19 Genomics UK (COG-UK) Consortium | Jacquelyn Wynn, Mairead Hyland, The Lighthouse Lab in Alderley Park and Alex Alderton, Roberto Amato, Jeffrey Barrett, Sonia Goncalves, Ewan Harrison, David K. Jackson, Ian Johnston, Dominic Kwiatkowski, Cordelia Langford, John Sillitoe on behalf of the Wellcome Sanger Institute COVID-19 Surveillance Team |
| EPI_ISL_2436881, EPI_ISL_2436883 | Lighthouse Lab in Milton Keynes | Wellcome Sanger Institute for the COVID-19 Genomics UK (COG-UK) Consortium | The Lighthouse Lab in Milton Keynes and Alex Alderton, Roberto Amato, Jeffrey Barrett, Sonia Goncalves, Ewan Harrison, David K. Jackson, Ian Johnston, Dominic Kwiatkowski, Cordelia Langford, John Sillitoe on behalf of the Wellcome Sanger Institute COVID-19 Surveillance Team |
| EPI_ISL_2436884 | Lighthouse Lab in Alderley Park | Wellcome Sanger Institute for the COVID-19 Genomics UK (COG-UK) Consortium | Jacquelyn Wynn, Mairead Hyland, The Lighthouse Lab in Alderley Park and Alex Alderton, Roberto Amato, Jeffrey Barrett, Sonia Goncalves, Ewan Harrison, David K. Jackson, Ian Johnston, Dominic Kwiatkowski, Cordelia Langford, John Sillitoe on behalf of the Wellcome Sanger Institute COVID-19 Surveillance Team |
| EPI_ISL_2436887 | Lighthouse Lab in Glasgow | Wellcome Sanger Institute for the COVID-19 Genomics UK (COG-UK) Consortium | Harper VanSteenhouse, Yumi Kasai, David Gray, Carol Clugston, Anna Dominiczak and Alex Alderton, Roberto Amato, Jeffrey Barrett, Sonia Goncalves, Ewan Harrison, David K. Jackson, Ian Johnston, Dominic Kwiatkowski, Cordelia Langford, John Sillitoe on behalf of the Wellcome Sanger Institute COVID-19 Surveillance Team |
| EPI_ISL_2436888 | Lighthouse Lab in Alderley Park | Wellcome Sanger Institute for the COVID-19 Genomics UK (COG-UK) Consortium | Jacquelyn Wynn, Mairead Hyland, The Lighthouse Lab in Alderley Park and Alex Alderton, Roberto Amato, Jeffrey Barrett, Sonia Goncalves, Ewan Harrison, David K. Jackson, Ian Johnston, Dominic Kwiatkowski, Cordelia Langford, John Sillitoe on behalf of the Wellcome Sanger Institute COVID-19 Surveillance Team |
| EPI_ISL_2436889 | Lighthouse Lab in Glasgow | Wellcome Sanger Institute for the COVID-19 Genomics UK (COG-UK) Consortium | Harper VanSteenhouse, Yumi Kasai, David Gray, Carol Clugston, Anna Dominiczak and Alex Alderton, Roberto Amato, Jeffrey Barrett, Sonia Goncalves, Ewan Harrison, David K. Jackson, Ian Johnston, Dominic Kwiatkowski, Cordelia Langford, John Sillitoe on behalf of the Wellcome Sanger Institute COVID-19 Surveillance Team |
| EPI_ISL_2436890 | Lighthouse Lab in Milton Keynes | Wellcome Sanger Institute for the COVID-19 Genomics UK (COG-UK) Consortium | The Lighthouse Lab in Milton Keynes and Alex Alderton, Roberto Amato, Jeffrey Barrett, Sonia Goncalves, Ewan Harrison, David K. Jackson, Ian Johnston, Dominic Kwiatkowski, Cordelia Langford, John Sillitoe on behalf of the Wellcome Sanger Institute COVID-19 Surveillance Team |
| EPI_ISL_2436892, EPI_ISL_2436893 | Lighthouse Lab in Alderley Park | Wellcome Sanger Institute for the COVID-19 Genomics UK (COG-UK) Consortium | Jacquelyn Wynn, Mairead Hyland, The Lighthouse Lab in Alderley Park and Alex Alderton, Roberto Amato, Jeffrey Barrett, Sonia Goncalves, Ewan Harrison, David K. Jackson, Ian Johnston, Dominic Kwiatkowski, Cordelia Langford, John Sillitoe on behalf of the Wellcome Sanger Institute COVID-19 Surveillance Team |
| EPI_ISL_2436894 | Lighthouse Lab in Milton Keynes | Wellcome Sanger Institute for the COVID-19 Genomics UK (COG-UK) Consortium | The Lighthouse Lab in Milton Keynes and Alex Alderton, Roberto Amato, Jeffrey Barrett, Sonia Goncalves, Ewan Harrison, David K. Jackson, Ian Johnston, Dominic Kwiatkowski, Cordelia Langford, John Sillitoe on behalf of the Wellcome Sanger Institute COVID-19 Surveillance Team |
| EPI_ISL_2436895 | Lighthouse Lab in Alderley Park | Wellcome Sanger Institute for the COVID-19 Genomics UK (COG-UK) Consortium | Jacquelyn Wynn, Mairead Hyland, The Lighthouse Lab in Alderley Park and Alex Alderton, Roberto Amato, Jeffrey Barrett, Sonia Goncalves, Ewan Harrison, David K. Jackson, Ian Johnston, Dominic Kwiatkowski, Cordelia Langford, John Sillitoe on behalf of the Wellcome Sanger Institute COVID-19 Surveillance Team |
| EPI_ISL_2436896 | Lighthouse Lab in Milton Keynes | Wellcome Sanger Institute for the COVID-19 Genomics UK (COG-UK) Consortium | The Lighthouse Lab in Milton Keynes and Alex Alderton, Roberto Amato, Jeffrey Barrett, Sonia Goncalves, Ewan Harrison, David K. Jackson, Ian Johnston, Dominic Kwiatkowski, Cordelia Langford, John Sillitoe on behalf of the Wellcome Sanger Institute COVID-19 Surveillance Team |
| EPI_ISL_2436897, EPI_ISL_2436899 | Lighthouse Lab in Glasgow | Wellcome Sanger Institute for the COVID-19 Genomics UK (COG-UK) Consortium | Harper VanSteenhouse, Yumi Kasai, David Gray, Carol Clugston, Anna Dominiczak and Alex Alderton, Roberto Amato, Jeffrey Barrett, Sonia Goncalves, Ewan Harrison, David K. Jackson, Ian Johnston, Dominic Kwiatkowski, Cordelia Langford, John Sillitoe on behalf of the Wellcome Sanger Institute COVID-19 Surveillance Team |
| EPI_ISL_2436900 | Lighthouse Lab in Alderley Park | Wellcome Sanger Institute for the COVID-19 Genomics UK (COG-UK) Consortium | Jacquelyn Wynn, Mairead Hyland, The Lighthouse Lab in Alderley Park and Alex Alderton, Roberto Amato, Jeffrey Barrett, Sonia Goncalves, Ewan Harrison, David K. Jackson, Ian Johnston, Dominic Kwiatkowski, Cordelia Langford, John Sillitoe on behalf of the Wellcome Sanger Institute COVID-19 Surveillance Team |
| EPI_ISL_2436901, EPI_ISL_2436902 | Lighthouse Lab in Glasgow | Wellcome Sanger Institute for the COVID-19 Genomics UK (COG-UK) Consortium | Harper VanSteenhouse, Yumi Kasai, David Gray, Carol Clugston, Anna Dominiczak and Alex Alderton, Roberto Amato, Jeffrey Barrett, Sonia Goncalves, Ewan Harrison, David K. Jackson, Ian Johnston, Dominic Kwiatkowski, Cordelia Langford, John Sillitoe on behalf of the Wellcome Sanger Institute COVID-19 Surveillance Team |
| EPI_ISL_2436903 | Lighthouse Lab in Alderley Park | Wellcome Sanger Institute for the COVID-19 Genomics UK (COG-UK) Consortium | Jacquelyn Wynn, Mairead Hyland, The Lighthouse Lab in Alderley Park and Alex Alderton, Roberto Amato, Jeffrey Barrett, Sonia Goncalves, Ewan Harrison, David K. Jackson, Ian Johnston, Dominic Kwiatkowski, Cordelia Langford, John Sillitoe on behalf of the Wellcome Sanger Institute COVID-19 Surveillance Team |
| EPI_ISL_2436904 | Lighthouse Lab in Glasgow | Wellcome Sanger Institute for the COVID-19 Genomics UK (COG-UK) Consortium | Harper VanSteenhouse, Yumi Kasai, David Gray, Carol Clugston, Anna Dominiczak and Alex Alderton, Roberto Amato, Jeffrey Barrett, Sonia Goncalves, Ewan Harrison, David K. Jackson, Ian Johnston, Dominic Kwiatkowski, Cordelia Langford, John Sillitoe on behalf of the Wellcome Sanger Institute COVID-19 Surveillance Team |
| EPI_ISL_2436905 | Lighthouse Lab in Milton Keynes | Wellcome Sanger Institute for the COVID-19 Genomics UK (COG-UK) Consortium | The Lighthouse Lab in Milton Keynes and Alex Alderton, Roberto Amato, Jeffrey Barrett, Sonia Goncalves, Ewan Harrison, David K. Jackson, Ian Johnston, Dominic Kwiatkowski, Cordelia Langford, John Sillitoe on behalf of the Wellcome Sanger Institute COVID-19 Surveillance Team |
| EPI_ISL_2436906 | Lighthouse Lab in Alderley Park | Wellcome Sanger Institute for the COVID-19 Genomics UK (COG-UK) Consortium | Jacquelyn Wynn, Mairead Hyland, The Lighthouse Lab in Alderley Park and Alex Alderton, Roberto Amato, Jeffrey Barrett, Sonia Goncalves, Ewan Harrison, David K. Jackson, Ian Johnston, Dominic Kwiatkowski, Cordelia Langford, John Sillitoe on behalf of the Wellcome Sanger Institute COVID-19 Surveillance Team |
| EPI_ISL_2436907 | Lighthouse Lab in Milton Keynes | Wellcome Sanger Institute for the COVID-19 Genomics UK (COG-UK) Consortium | The Lighthouse Lab in Milton Keynes and Alex Alderton, Roberto Amato, Jeffrey Barrett, Sonia Goncalves, Ewan Harrison, David K. Jackson, Ian Johnston, Dominic Kwiatkowski, Cordelia Langford, John Sillitoe on behalf of the Wellcome Sanger Institute COVID-19 Surveillance Team |
| EPI_ISL_2436909, EPI_ISL_2436910, EPI_ISL_2436911, EPI_ISL_2436912, EPI_ISL_2436913, EPI_ISL_2436914, EPI_ISL_2436915, EPI_ISL_2436916, EPI_ISL_2436917, EPI_ISL_2436918, EPI_ISL_2436919, EPI_ISL_2436920, EPI_ISL_2436921 | | | |
| see above | Lighthouse Lab in Alderley Park | Wellcome Sanger Institute for the COVID-19 Genomics UK (COG-UK) Consortium | Jacquelyn Wynn, Mairead Hyland, The Lighthouse Lab in Alderley Park and Alex Alderton, Roberto Amato, Jeffrey Barrett, Sonia Goncalves, Ewan Harrison, David K. Jackson, Ian Johnston, Dominic Kwiatkowski, Cordelia Langford, John Sillitoe on behalf of the Wellcome Sanger Institute COVID-19 Surveillance Team |
| EPI_ISL_2436922 | Lighthouse Lab in Glasgow | Wellcome Sanger Institute for the COVID-19 Genomics UK (COG-UK) Consortium | Harper VanSteenhouse, Yumi Kasai, David Gray, Carol Clugston, Anna Dominiczak and Alex Alderton, Roberto Amato, Jeffrey Barrett, Sonia Goncalves, Ewan Harrison, David K. Jackson, Ian Johnston, Dominic Kwiatkowski, Cordelia Langford, John Sillitoe on behalf of the Wellcome Sanger Institute COVID-19 Surveillance Team |
| EPI_ISL_2436924, EPI_ISL_2436925 | Lighthouse Lab in Alderley Park | Wellcome Sanger Institute for the COVID-19 Genomics UK (COG-UK) Consortium | Jacquelyn Wynn, Mairead Hyland, The Lighthouse Lab in Alderley Park and Alex Alderton, Roberto Amato, Jeffrey Barrett, Sonia Goncalves, Ewan Harrison, David K. Jackson, Ian Johnston, Dominic Kwiatkowski, Cordelia Langford, John Sillitoe on behalf of the Wellcome Sanger Institute COVID-19 Surveillance Team |
| EPI_ISL_2436926 | Lighthouse Lab in Milton Keynes | Wellcome Sanger Institute for the COVID-19 Genomics UK (COG-UK) Consortium | The Lighthouse Lab in Milton Keynes and Alex Alderton, Roberto Amato, Jeffrey Barrett, Sonia Goncalves, Ewan Harrison, David K. Jackson, Ian Johnston, Dominic Kwiatkowski, Cordelia Langford, John Sillitoe on behalf of the Wellcome Sanger Institute COVID-19 Surveillance Team |
| EPI_ISL_2436928, EPI_ISL_2436929 | Lighthouse Lab in Alderley Park | Wellcome Sanger Institute for the COVID-19 Genomics UK (COG-UK) Consortium | Jacquelyn Wynn, Mairead Hyland, The Lighthouse Lab in Alderley Park and Alex Alderton, Roberto Amato, Jeffrey Barrett, Sonia Goncalves, Ewan Harrison, David K. Jackson, Ian Johnston, Dominic Kwiatkowski, Cordelia Langford, John Sillitoe on behalf of the Wellcome Sanger Institute COVID-19 Surveillance Team |

| | | | |
|---|---|---|---|
| EPI_ISL_2436930 | Lighthouse Lab in Milton Keynes | Wellcome Sanger Institute for the COVID-19 Genomics UK (COG-UK) Consortium | The Lighthouse Lab in Milton Keynes and Alex Alderton, Roberto Amato, Jeffrey Barrett, Sonia Goncalves, Ewan Harrison, David K. Jackson, Ian Johnston, Dominic Kwiatkowski, Cordelia Langford, John Sillitoe on behalf of the Wellcome Sanger Institute COVID-19 Surveillance Team |
| EPI_ISL_2436931 | Lighthouse Lab in Alderley Park | Wellcome Sanger Institute for the COVID-19 Genomics UK (COG-UK) Consortium | Jacquelyn Wynn, Mairead Hyland, The Lighthouse Lab in Alderley Park and Alex Alderton, Roberto Amato, Jeffrey Barrett, Sonia Goncalves, Ewan Harrison, David K. Jackson, Ian Johnston, Dominic Kwiatkowski, Cordelia Langford, John Sillitoe on behalf of the Wellcome Sanger Institute COVID-19 Surveillance Team |
| EPI_ISL_2436932, EPI_ISL_2436934 | Lighthouse Lab in Milton Keynes | Wellcome Sanger Institute for the COVID-19 Genomics UK (COG-UK) Consortium | The Lighthouse Lab in Milton Keynes and Alex Alderton, Roberto Amato, Jeffrey Barrett, Sonia Goncalves, Ewan Harrison, David K. Jackson, Ian Johnston, Dominic Kwiatkowski, Cordelia Langford, John Sillitoe on behalf of the Wellcome Sanger Institute COVID-19 Surveillance Team |
| EPI_ISL_2436935, EPI_ISL_2436936 | Lighthouse Lab in Alderley Park | Wellcome Sanger Institute for the COVID-19 Genomics UK (COG-UK) Consortium | Jacquelyn Wynn, Mairead Hyland, The Lighthouse Lab in Alderley Park and Alex Alderton, Roberto Amato, Jeffrey Barrett, Sonia Goncalves, Ewan Harrison, David K. Jackson, Ian Johnston, Dominic Kwiatkowski, Cordelia Langford, John Sillitoe on behalf of the Wellcome Sanger Institute COVID-19 Surveillance Team |
| EPI_ISL_2436937 | Health Services Laboratories | Wellcome Sanger Institute for the COVID-19 Genomics UK (COG-UK) Consortium | Health Services Laboratories and Alex Alderton, Roberto Amato, Jeffrey Barrett, Sonia Goncalves, Ewan Harrison, David K. Jackson, Ian Johnston, Dominic Kwiatkowski, Cordelia Langford, John Sillitoe on behalf of the Wellcome Sanger Institute COVID-19 Surveillance Team |
| EPI_ISL_2436938, EPI_ISL_2436939 | Lighthouse Lab in Milton Keynes | Wellcome Sanger Institute for the COVID-19 Genomics UK (COG-UK) Consortium | The Lighthouse Lab in Milton Keynes and Alex Alderton, Roberto Amato, Jeffrey Barrett, Sonia Goncalves, Ewan Harrison, David K. Jackson, Ian Johnston, Dominic Kwiatkowski, Cordelia Langford, John Sillitoe on behalf of the Wellcome Sanger Institute COVID-19 Surveillance Team |
| EPI_ISL_2436940 | Health Services Laboratories | Wellcome Sanger Institute for the COVID-19 Genomics UK (COG-UK) Consortium | Health Services Laboratories and Alex Alderton, Roberto Amato, Jeffrey Barrett, Sonia Goncalves, Ewan Harrison, David K. Jackson, Ian Johnston, Dominic Kwiatkowski, Cordelia Langford, John Sillitoe on behalf of the Wellcome Sanger Institute COVID-19 Surveillance Team |
| EPI_ISL_2436941, EPI_ISL_2436942, EPI_ISL_2436943, EPI_ISL_2436944, EPI_ISL_2436946 | Lighthouse Lab in Alderley Park | Wellcome Sanger Institute for the COVID-19 Genomics UK (COG-UK) Consortium | Jacquelyn Wynn, Mairead Hyland, The Lighthouse Lab in Alderley Park and Alex Alderton, Roberto Amato, Jeffrey Barrett, Sonia Goncalves, Ewan Harrison, David K. Jackson, Ian Johnston, Dominic Kwiatkowski, Cordelia Langford, John Sillitoe on behalf of the Wellcome Sanger Institute COVID-19 Surveillance Team |
| EPI_ISL_2436947 | Lighthouse Lab in Milton Keynes | Wellcome Sanger Institute for the COVID-19 Genomics UK (COG-UK) Consortium | The Lighthouse Lab in Milton Keynes and Alex Alderton, Roberto Amato, Jeffrey Barrett, Sonia Goncalves, Ewan Harrison, David K. Jackson, Ian Johnston, Dominic Kwiatkowski, Cordelia Langford, John Sillitoe on behalf of the Wellcome Sanger Institute COVID-19 Surveillance Team |
| EPI_ISL_2436949 | Health Services Laboratories | Wellcome Sanger Institute for the COVID-19 Genomics UK (COG-UK) Consortium | Health Services Laboratories and Alex Alderton, Roberto Amato, Jeffrey Barrett, Sonia Goncalves, Ewan Harrison, David K. Jackson, Ian Johnston, Dominic Kwiatkowski, Cordelia Langford, John Sillitoe on behalf of the Wellcome Sanger Institute COVID-19 Surveillance Team |
| EPI_ISL_2436952, EPI_ISL_2436954 | Lighthouse Lab in Alderley Park | Wellcome Sanger Institute for the COVID-19 Genomics UK (COG-UK) Consortium | Jacquelyn Wynn, Mairead Hyland, The Lighthouse Lab in Alderley Park and Alex Alderton, Roberto Amato, Jeffrey Barrett, Sonia Goncalves, Ewan Harrison, David K. Jackson, Ian Johnston, Dominic Kwiatkowski, Cordelia Langford, John Sillitoe on behalf of the Wellcome Sanger Institute COVID-19 Surveillance Team |
| EPI_ISL_2436955 | Lighthouse Lab in Milton Keynes | Wellcome Sanger Institute for the COVID-19 Genomics UK (COG-UK) Consortium | The Lighthouse Lab in Milton Keynes and Alex Alderton, Roberto Amato, Jeffrey Barrett, Sonia Goncalves, Ewan Harrison, David K. Jackson, Ian Johnston, Dominic Kwiatkowski, Cordelia Langford, John Sillitoe on behalf of the Wellcome Sanger Institute COVID-19 Surveillance Team |
| EPI_ISL_2436956, EPI_ISL_2436957, EPI_ISL_2436958 | Lighthouse Lab in Alderley Park | Wellcome Sanger Institute for the COVID-19 Genomics UK (COG-UK) Consortium | Jacquelyn Wynn, Mairead Hyland, The Lighthouse Lab in Alderley Park and Alex Alderton, Roberto Amato, Jeffrey Barrett, Sonia Goncalves, Ewan Harrison, David K. Jackson, Ian Johnston, Dominic Kwiatkowski, Cordelia Langford, John Sillitoe on behalf of the Wellcome Sanger Institute COVID-19 Surveillance Team |
| EPI_ISL_2436960 | Lighthouse Lab in Milton Keynes | Wellcome Sanger Institute for the COVID-19 Genomics UK (COG-UK) Consortium | The Lighthouse Lab in Milton Keynes and Alex Alderton, Roberto Amato, Jeffrey Barrett, Sonia Goncalves, Ewan Harrison, David K. Jackson, Ian Johnston, Dominic Kwiatkowski, Cordelia Langford, John Sillitoe on behalf of the Wellcome Sanger Institute COVID-19 Surveillance Team |
| EPI_ISL_2436961 | Health Services Laboratories | Wellcome Sanger Institute for the COVID-19 Genomics UK (COG-UK) Consortium | Health Services Laboratories and Alex Alderton, Roberto Amato, Jeffrey Barrett, Sonia Goncalves, Ewan Harrison, David K. Jackson, Ian Johnston, Dominic Kwiatkowski, Cordelia Langford, John Sillitoe on behalf of the Wellcome Sanger Institute COVID-19 Surveillance Team |
| EPI_ISL_2436962, EPI_ISL_2436963, EPI_ISL_2436964, EPI_ISL_2436965, EPI_ISL_2436966 | Lighthouse Lab in Alderley Park | Wellcome Sanger Institute for the COVID-19 Genomics UK (COG-UK) Consortium | Jacquelyn Wynn, Mairead Hyland, The Lighthouse Lab in Alderley Park and Alex Alderton, Roberto Amato, Jeffrey Barrett, Sonia Goncalves, Ewan Harrison, David K. Jackson, Ian Johnston, Dominic Kwiatkowski, Cordelia Langford, John Sillitoe on behalf of the Wellcome Sanger Institute COVID-19 Surveillance Team |
| EPI_ISL_2436968 | Lighthouse Lab in Milton Keynes | Wellcome Sanger Institute for the COVID-19 Genomics UK (COG-UK) Consortium | The Lighthouse Lab in Milton Keynes and Alex Alderton, Roberto Amato, Jeffrey Barrett, Sonia Goncalves, Ewan Harrison, David K. Jackson, Ian Johnston, Dominic Kwiatkowski, Cordelia Langford, John Sillitoe on behalf of the Wellcome Sanger Institute COVID-19 Surveillance Team |
| EPI_ISL_2436969, EPI_ISL_2436971, EPI_ISL_2436973 | Lighthouse Lab in Alderley Park | Wellcome Sanger Institute for the COVID-19 Genomics UK (COG-UK) Consortium | Jacquelyn Wynn, Mairead Hyland, The Lighthouse Lab in Alderley Park and Alex Alderton, Roberto Amato, Jeffrey Barrett, Sonia Goncalves, Ewan Harrison, David K. Jackson, Ian Johnston, Dominic Kwiatkowski, Cordelia Langford, John Sillitoe on behalf of the Wellcome Sanger Institute COVID-19 Surveillance Team |
| EPI_ISL_2436975 | Lighthouse Lab in Milton Keynes | Wellcome Sanger Institute for the COVID-19 Genomics UK (COG-UK) Consortium | The Lighthouse Lab in Milton Keynes and Alex Alderton, Roberto Amato, Jeffrey Barrett, Sonia Goncalves, Ewan Harrison, David K. Jackson, Ian Johnston, Dominic Kwiatkowski, Cordelia Langford, John Sillitoe on behalf of the Wellcome Sanger Institute COVID-19 Surveillance Team |
| EPI_ISL_2436977, EPI_ISL_2436978 | Lighthouse Lab in Alderley Park | Wellcome Sanger Institute for the COVID-19 Genomics UK (COG-UK) Consortium | Jacquelyn Wynn, Mairead Hyland, The Lighthouse Lab in Alderley Park and Alex Alderton, Roberto Amato, Jeffrey Barrett, Sonia Goncalves, Ewan Harrison, David K. Jackson, Ian Johnston, Dominic Kwiatkowski, Cordelia Langford, John Sillitoe on behalf of the Wellcome Sanger Institute COVID-19 Surveillance Team |
| EPI_ISL_2436979 | Health Services Laboratories | Wellcome Sanger Institute for the COVID-19 Genomics UK (COG-UK) Consortium | Health Services Laboratories and Alex Alderton, Roberto Amato, Jeffrey Barrett, Sonia Goncalves, Ewan Harrison, David K. Jackson, Ian Johnston, Dominic Kwiatkowski, Cordelia Langford, John Sillitoe on behalf of the Wellcome Sanger Institute COVID-19 Surveillance Team |
| EPI_ISL_2436980 | Lighthouse Lab in Alderley Park | Wellcome Sanger Institute for the COVID-19 Genomics UK (COG-UK) Consortium | Jacquelyn Wynn, Mairead Hyland, The Lighthouse Lab in Alderley Park and Alex Alderton, Roberto Amato, Jeffrey Barrett, Sonia Goncalves, Ewan Harrison, David K. Jackson, Ian Johnston, Dominic Kwiatkowski, Cordelia Langford, John Sillitoe on behalf of the Wellcome Sanger Institute COVID-19 Surveillance Team |
| EPI_ISL_2436981 | Health Services Laboratories | Wellcome Sanger Institute for the COVID-19 Genomics UK (COG-UK) Consortium | Health Services Laboratories and Alex Alderton, Roberto Amato, Jeffrey Barrett, Sonia Goncalves, Ewan Harrison, David K. Jackson, Ian Johnston, Dominic Kwiatkowski, Cordelia Langford, John Sillitoe on behalf of the Wellcome Sanger Institute COVID-19 Surveillance Team |
| EPI_ISL_2436982, EPI_ISL_2436986 | Lighthouse Lab in Alderley Park | Wellcome Sanger Institute for the COVID-19 Genomics UK (COG-UK) Consortium | Jacquelyn Wynn, Mairead Hyland, The Lighthouse Lab in Alderley Park and Alex Alderton, Roberto Amato, Jeffrey Barrett, Sonia Goncalves, Ewan Harrison, David K. Jackson, Ian Johnston, Dominic Kwiatkowski, Cordelia Langford, John Sillitoe on behalf of the Wellcome Sanger Institute COVID-19 Surveillance Team |
| EPI_ISL_2436988 | Lighthouse Lab in Milton Keynes | Wellcome Sanger Institute for the COVID-19 Genomics UK (COG-UK) Consortium | The Lighthouse Lab in Milton Keynes and Alex Alderton, Roberto Amato, Jeffrey Barrett, Sonia Goncalves, Ewan Harrison, David K. Jackson, Ian Johnston, Dominic Kwiatkowski, Cordelia Langford, John Sillitoe on behalf of the Wellcome Sanger Institute COVID-19 Surveillance Team |
| EPI_ISL_2436989 | Health Services Laboratories | Wellcome Sanger Institute for the COVID-19 Genomics UK (COG-UK) Consortium | Health Services Laboratories and Alex Alderton, Roberto Amato, Jeffrey Barrett, Sonia Goncalves, Ewan Harrison, David K. Jackson, Ian Johnston, Dominic Kwiatkowski, Cordelia Langford, John Sillitoe on behalf of the Wellcome Sanger Institute COVID-19 Surveillance Team |
| EPI_ISL_2436990, EPI_ISL_2436991 | Lighthouse Lab in Alderley Park | Wellcome Sanger Institute for the COVID-19 Genomics UK (COG-UK) Consortium | Jacquelyn Wynn, Mairead Hyland, The Lighthouse Lab in Alderley Park and Alex Alderton, Roberto Amato, Jeffrey Barrett, Sonia Goncalves, Ewan Harrison, David K. Jackson, Ian Johnston, Dominic Kwiatkowski, Cordelia Langford, John Sillitoe on behalf of the Wellcome Sanger Institute COVID-19 Surveillance Team |
| EPI_ISL_2436992 | Lighthouse Lab in Milton Keynes | Wellcome Sanger Institute for the COVID-19 Genomics UK (COG-UK) Consortium | The Lighthouse Lab in Milton Keynes and Alex Alderton, Roberto Amato, Jeffrey Barrett, Sonia Goncalves, Ewan Harrison, David K. Jackson, Ian Johnston, Dominic Kwiatkowski, Cordelia Langford, John Sillitoe on behalf of the Wellcome Sanger Institute COVID-19 Surveillance Team |
| EPI_ISL_2436993, EPI_ISL_2436994 | Lighthouse Lab in Alderley Park | Wellcome Sanger Institute for the COVID-19 Genomics UK (COG-UK) Consortium | Jacquelyn Wynn, Mairead Hyland, The Lighthouse Lab in Alderley Park and Alex Alderton, Roberto Amato, Jeffrey Barrett, Sonia Goncalves, Ewan Harrison, David K. Jackson, Ian Johnston, Dominic Kwiatkowski, Cordelia Langford, John Sillitoe on behalf of the Wellcome Sanger Institute COVID-19 Surveillance Team |
| EPI_ISL_2436995 | Health Services Laboratories | Wellcome Sanger Institute for the COVID-19 Genomics UK (COG-UK) Consortium | Health Services Laboratories and Alex Alderton, Roberto Amato, Jeffrey Barrett, Sonia Goncalves, Ewan Harrison, David K. Jackson, Ian Johnston, Dominic Kwiatkowski, Cordelia Langford, John Sillitoe on behalf of the Wellcome Sanger Institute COVID-19 Surveillance Team |
| EPI_ISL_2436996 | Lighthouse Lab in Alderley Park | Wellcome Sanger Institute for the COVID-19 Genomics UK (COG-UK) Consortium | Jacquelyn Wynn, Mairead Hyland, The Lighthouse Lab in Alderley Park and Alex Alderton, Roberto Amato, Jeffrey Barrett, Sonia Goncalves, Ewan Harrison, David K. Jackson, Ian Johnston, Dominic Kwiatkowski, Cordelia Langford, John Sillitoe on behalf of the Wellcome Sanger Institute COVID-19 Surveillance Team |
| EPI_ISL_2436997 | Lighthouse Lab in Milton Keynes | Wellcome Sanger Institute for the COVID-19 Genomics UK (COG-UK) Consortium | The Lighthouse Lab in Milton Keynes and Alex Alderton, Roberto Amato, Jeffrey Barrett, Sonia Goncalves, Ewan Harrison, David K. Jackson, Ian Johnston, Dominic Kwiatkowski, Cordelia Langford, John Sillitoe on behalf of the Wellcome Sanger Institute COVID-19 Surveillance Team |
| EPI_ISL_2436999 | Health Services Laboratories | Wellcome Sanger Institute for the COVID-19 Genomics UK (COG-UK) Consortium | Health Services Laboratories and Alex Alderton, Roberto Amato, Jeffrey Barrett, Sonia Goncalves, Ewan Harrison, David K. Jackson, Ian Johnston, Dominic Kwiatkowski, Cordelia Langford, John Sillitoe on behalf of the Wellcome Sanger Institute COVID-19 Surveillance Team |
| EPI_ISL_2437000 | Lighthouse Lab in Milton Keynes | Wellcome Sanger Institute for the COVID-19 Genomics UK (COG-UK) Consortium | The Lighthouse Lab in Milton Keynes and Alex Alderton, Roberto Amato, Jeffrey Barrett, Sonia Goncalves, Ewan Harrison, David K. Jackson, Ian Johnston, Dominic Kwiatkowski, Cordelia Langford, John Sillitoe on behalf of the Wellcome Sanger Institute COVID-19 Surveillance Team |
| EPI_ISL_2437001, EPI_ISL_2437002 | Lighthouse Lab in Alderley Park | Wellcome Sanger Institute for the COVID-19 Genomics UK (COG-UK) Consortium | Jacquelyn Wynn, Mairead Hyland, The Lighthouse Lab in Alderley Park and Alex Alderton, Roberto Amato, Jeffrey Barrett, Sonia Goncalves, Ewan Harrison, David K. Jackson, Ian Johnston, Dominic Kwiatkowski, Cordelia Langford, John Sillitoe on behalf of the Wellcome Sanger Institute COVID-19 Surveillance Team |
| EPI_ISL_2437003, EPI_ISL_2437004, EPI_ISL_2437006 | Health Services Laboratories | Wellcome Sanger Institute for the COVID-19 Genomics UK (COG-UK) Consortium | Health Services Laboratories and Alex Alderton, Roberto Amato, Jeffrey Barrett, Sonia Goncalves, Ewan Harrison, David K. Jackson, Ian Johnston, Dominic Kwiatkowski, Cordelia Langford, John Sillitoe on behalf of the Wellcome Sanger Institute COVID-19 Surveillance Team |
| EPI_ISL_2437007, EPI_ISL_2437008 | Lighthouse Lab in Milton Keynes | Wellcome Sanger Institute for the COVID-19 Genomics UK (COG-UK) Consortium | The Lighthouse Lab in Milton Keynes and Alex Alderton, Roberto Amato, Jeffrey Barrett, Sonia Goncalves, Ewan Harrison, David K. Jackson, Ian Johnston, Dominic Kwiatkowski, Cordelia Langford, John Sillitoe on behalf of the Wellcome Sanger Institute COVID-19 Surveillance Team |
| EPI_ISL_2437010 | Health Services Laboratories | Wellcome Sanger Institute for the COVID-19 Genomics UK (COG-UK) Consortium | Health Services Laboratories and Alex Alderton, Roberto Amato, Jeffrey Barrett, Sonia Goncalves, Ewan Harrison, David K. Jackson, Ian Johnston, Dominic Kwiatkowski, Cordelia Langford, John Sillitoe on behalf of the Wellcome Sanger Institute COVID-19 Surveillance Team |
| EPI_ISL_2437011, EPI_ISL_2437012 | Lighthouse Lab in Alderley Park | Wellcome Sanger Institute for the COVID-19 Genomics UK (COG-UK) Consortium | Jacquelyn Wynn, Mairead Hyland, The Lighthouse Lab in Alderley Park and Alex Alderton, Roberto Amato, Jeffrey Barrett, Sonia Goncalves, Ewan Harrison, David K. Jackson, Ian Johnston, Dominic Kwiatkowski, Cordelia Langford, John Sillitoe on behalf of the Wellcome Sanger Institute COVID-19 Surveillance Team |
| EPI_ISL_2437013, EPI_ISL_2437014, EPI_ISL_2437016 | Lighthouse Lab in Milton Keynes | Wellcome Sanger Institute for the COVID-19 Genomics UK (COG-UK) Consortium | The Lighthouse Lab in Milton Keynes and Alex Alderton, Roberto Amato, Jeffrey Barrett, Sonia Goncalves, Ewan Harrison, David K. Jackson, Ian Johnston, Dominic Kwiatkowski, Cordelia Langford, John Sillitoe on behalf of the Wellcome Sanger Institute COVID-19 Surveillance Team |
| EPI_ISL_2437017 | Health Services Laboratories | Wellcome Sanger Institute for the COVID-19 Genomics UK (COG-UK) Consortium | Health Services Laboratories and Alex Alderton, Roberto Amato, Jeffrey Barrett, Sonia Goncalves, Ewan Harrison, David K. Jackson, Ian Johnston, Dominic Kwiatkowski, Cordelia Langford, John Sillitoe on behalf of the Wellcome Sanger Institute COVID-19 Surveillance Team |

| | | | |
|---|---|---|---|
| EPI_ISL_2437019 | Lighthouse Lab in Alderley Park | Wellcome Sanger Institute for the COVID-19 Genomics UK (COG-UK) Consortium | Jacquelyn Wynn, Mairead Hyland, The Lighthouse Lab in Alderley Park and Alex Alderton, Roberto Amato, Jeffrey Barrett, Sonia Goncalves, Ewan Harrison, David K. Jackson, Ian Johnston, Dominic Kwiatkowski, Cordelia Langford, John Sillitoe on behalf of the Wellcome Sanger Institute COVID-19 Surveillance Team |
| EPI_ISL_2437022, EPI_ISL_2437023, EPI_ISL_2437025 | Health Services Laboratories | Wellcome Sanger Institute for the COVID-19 Genomics UK (COG-UK) Consortium | Health Services Laboratories and Alex Alderton, Roberto Amato, Jeffrey Barrett, Sonia Goncalves, Ewan Harrison, David K. Jackson, Ian Johnston, Dominic Kwiatkowski, Cordelia Langford, John Sillitoe on behalf of the Wellcome Sanger Institute COVID-19 Surveillance Team |
| EPI_ISL_2437026 | Lighthouse Lab in Milton Keynes | Wellcome Sanger Institute for the COVID-19 Genomics UK (COG-UK) Consortium | The Lighthouse Lab in Milton Keynes and Alex Alderton, Roberto Amato, Jeffrey Barrett, Sonia Goncalves, Ewan Harrison, David K. Jackson, Ian Johnston, Dominic Kwiatkowski, Cordelia Langford, John Sillitoe on behalf of the Wellcome Sanger Institute COVID-19 Surveillance Team |
| EPI_ISL_2437027, EPI_ISL_2437028 | Lighthouse Lab in Alderley Park | Wellcome Sanger Institute for the COVID-19 Genomics UK (COG-UK) Consortium | Jacquelyn Wynn, Mairead Hyland, The Lighthouse Lab in Alderley Park and Alex Alderton, Roberto Amato, Jeffrey Barrett, Sonia Goncalves, Ewan Harrison, David K. Jackson, Ian Johnston, Dominic Kwiatkowski, Cordelia Langford, John Sillitoe on behalf of the Wellcome Sanger Institute COVID-19 Surveillance Team |
| EPI_ISL_2437029, EPI_ISL_2437030 | Lighthouse Lab in Milton Keynes | Wellcome Sanger Institute for the COVID-19 Genomics UK (COG-UK) Consortium | The Lighthouse Lab in Milton Keynes and Alex Alderton, Roberto Amato, Jeffrey Barrett, Sonia Goncalves, Ewan Harrison, David K. Jackson, Ian Johnston, Dominic Kwiatkowski, Cordelia Langford, John Sillitoe on behalf of the Wellcome Sanger Institute COVID-19 Surveillance Team |
| EPI_ISL_2437032, EPI_ISL_2437033 | Lighthouse Lab in Alderley Park | Wellcome Sanger Institute for the COVID-19 Genomics UK (COG-UK) Consortium | Jacquelyn Wynn, Mairead Hyland, The Lighthouse Lab in Alderley Park and Alex Alderton, Roberto Amato, Jeffrey Barrett, Sonia Goncalves, Ewan Harrison, David K. Jackson, Ian Johnston, Dominic Kwiatkowski, Cordelia Langford, John Sillitoe on behalf of the Wellcome Sanger Institute COVID-19 Surveillance Team |
| EPI_ISL_2437034 | Lighthouse Lab in Milton Keynes | Wellcome Sanger Institute for the COVID-19 Genomics UK (COG-UK) Consortium | The Lighthouse Lab in Milton Keynes and Alex Alderton, Roberto Amato, Jeffrey Barrett, Sonia Goncalves, Ewan Harrison, David K. Jackson, Ian Johnston, Dominic Kwiatkowski, Cordelia Langford, John Sillitoe on behalf of the Wellcome Sanger Institute COVID-19 Surveillance Team |
| EPI_ISL_2437035 | Lighthouse Lab in Alderley Park | Wellcome Sanger Institute for the COVID-19 Genomics UK (COG-UK) Consortium | Jacquelyn Wynn, Mairead Hyland, The Lighthouse Lab in Alderley Park and Alex Alderton, Roberto Amato, Jeffrey Barrett, Sonia Goncalves, Ewan Harrison, David K. Jackson, Ian Johnston, Dominic Kwiatkowski, Cordelia Langford, John Sillitoe on behalf of the Wellcome Sanger Institute COVID-19 Surveillance Team |
| EPI_ISL_2437036 | Lighthouse Lab in Milton Keynes | Wellcome Sanger Institute for the COVID-19 Genomics UK (COG-UK) Consortium | The Lighthouse Lab in Milton Keynes and Alex Alderton, Roberto Amato, Jeffrey Barrett, Sonia Goncalves, Ewan Harrison, David K. Jackson, Ian Johnston, Dominic Kwiatkowski, Cordelia Langford, John Sillitoe on behalf of the Wellcome Sanger Institute COVID-19 Surveillance Team |
| EPI_ISL_2437037, EPI_ISL_2437038 | Lighthouse Lab in Alderley Park | Wellcome Sanger Institute for the COVID-19 Genomics UK (COG-UK) Consortium | Jacquelyn Wynn, Mairead Hyland, The Lighthouse Lab in Alderley Park and Alex Alderton, Roberto Amato, Jeffrey Barrett, Sonia Goncalves, Ewan Harrison, David K. Jackson, Ian Johnston, Dominic Kwiatkowski, Cordelia Langford, John Sillitoe on behalf of the Wellcome Sanger Institute COVID-19 Surveillance Team |
| EPI_ISL_2437039 | Health Services Laboratories | Wellcome Sanger Institute for the COVID-19 Genomics UK (COG-UK) Consortium | Health Services Laboratories and Alex Alderton, Roberto Amato, Jeffrey Barrett, Sonia Goncalves, Ewan Harrison, David K. Jackson, Ian Johnston, Dominic Kwiatkowski, Cordelia Langford, John Sillitoe on behalf of the Wellcome Sanger Institute COVID-19 Surveillance Team |
| EPI_ISL_2437040, EPI_ISL_2437041, EPI_ISL_2437042, EPI_ISL_2437043 | Lighthouse Lab in Alderley Park | Wellcome Sanger Institute for the COVID-19 Genomics UK (COG-UK) Consortium | Jacquelyn Wynn, Mairead Hyland, The Lighthouse Lab in Alderley Park and Alex Alderton, Roberto Amato, Jeffrey Barrett, Sonia Goncalves, Ewan Harrison, David K. Jackson, Ian Johnston, Dominic Kwiatkowski, Cordelia Langford, John Sillitoe on behalf of the Wellcome Sanger Institute COVID-19 Surveillance Team |
| EPI_ISL_2437044 | Health Services Laboratories | Wellcome Sanger Institute for the COVID-19 Genomics UK (COG-UK) Consortium | Health Services Laboratories and Alex Alderton, Roberto Amato, Jeffrey Barrett, Sonia Goncalves, Ewan Harrison, David K. Jackson, Ian Johnston, Dominic Kwiatkowski, Cordelia Langford, John Sillitoe on behalf of the Wellcome Sanger Institute COVID-19 Surveillance Team |
| EPI_ISL_2437046, EPI_ISL_2437047, EPI_ISL_2437048 | Lighthouse Lab in Alderley Park | Wellcome Sanger Institute for the COVID-19 Genomics UK (COG-UK) Consortium | Jacquelyn Wynn, Mairead Hyland, The Lighthouse Lab in Alderley Park and Alex Alderton, Roberto Amato, Jeffrey Barrett, Sonia Goncalves, Ewan Harrison, David K. Jackson, Ian Johnston, Dominic Kwiatkowski, Cordelia Langford, John Sillitoe on behalf of the Wellcome Sanger Institute COVID-19 Surveillance Team |
| EPI_ISL_2437049 | Health Services Laboratories | Wellcome Sanger Institute for the COVID-19 Genomics UK (COG-UK) Consortium | Health Services Laboratories and Alex Alderton, Roberto Amato, Jeffrey Barrett, Sonia Goncalves, Ewan Harrison, David K. Jackson, Ian Johnston, Dominic Kwiatkowski, Cordelia Langford, John Sillitoe on behalf of the Wellcome Sanger Institute COVID-19 Surveillance Team |
| EPI_ISL_2437050 | Lighthouse Lab in Milton Keynes | Wellcome Sanger Institute for the COVID-19 Genomics UK (COG-UK) Consortium | The Lighthouse Lab in Milton Keynes and Alex Alderton, Roberto Amato, Jeffrey Barrett, Sonia Goncalves, Ewan Harrison, David K. Jackson, Ian Johnston, Dominic Kwiatkowski, Cordelia Langford, John Sillitoe on behalf of the Wellcome Sanger Institute COVID-19 Surveillance Team |
| EPI_ISL_2437051, EPI_ISL_2437052, EPI_ISL_2437053, EPI_ISL_2437054, EPI_ISL_2437055, EPI_ISL_2437056, EPI_ISL_2437057 | Lighthouse Lab in Alderley Park | Wellcome Sanger Institute for the COVID-19 Genomics UK (COG-UK) Consortium | Jacquelyn Wynn, Mairead Hyland, The Lighthouse Lab in Alderley Park and Alex Alderton, Roberto Amato, Jeffrey Barrett, Sonia Goncalves, Ewan Harrison, David K. Jackson, Ian Johnston, Dominic Kwiatkowski, Cordelia Langford, John Sillitoe on behalf of the Wellcome Sanger Institute COVID-19 Surveillance Team |
| EPI_ISL_2437058 | Lighthouse Lab in Milton Keynes | Wellcome Sanger Institute for the COVID-19 Genomics UK (COG-UK) Consortium | The Lighthouse Lab in Milton Keynes and Alex Alderton, Roberto Amato, Jeffrey Barrett, Sonia Goncalves, Ewan Harrison, David K. Jackson, Ian Johnston, Dominic Kwiatkowski, Cordelia Langford, John Sillitoe on behalf of the Wellcome Sanger Institute COVID-19 Surveillance Team |
| EPI_ISL_2437059, EPI_ISL_2437060, EPI_ISL_2437062, EPI_ISL_2437063 | Lighthouse Lab in Alderley Park | Wellcome Sanger Institute for the COVID-19 Genomics UK (COG-UK) Consortium | Jacquelyn Wynn, Mairead Hyland, The Lighthouse Lab in Alderley Park and Alex Alderton, Roberto Amato, Jeffrey Barrett, Sonia Goncalves, Ewan Harrison, David K. Jackson, Ian Johnston, Dominic Kwiatkowski, Cordelia Langford, John Sillitoe on behalf of the Wellcome Sanger Institute COVID-19 Surveillance Team |
| EPI_ISL_2437064 | Health Services Laboratories | Wellcome Sanger Institute for the COVID-19 Genomics UK (COG-UK) Consortium | Health Services Laboratories and Alex Alderton, Roberto Amato, Jeffrey Barrett, Sonia Goncalves, Ewan Harrison, David K. Jackson, Ian Johnston, Dominic Kwiatkowski, Cordelia Langford, John Sillitoe on behalf of the Wellcome Sanger Institute COVID-19 Surveillance Team |
| EPI_ISL_2437065 | Lighthouse Lab in Alderley Park | Wellcome Sanger Institute for the COVID-19 Genomics UK (COG-UK) Consortium | Jacquelyn Wynn, Mairead Hyland, The Lighthouse Lab in Alderley Park and Alex Alderton, Roberto Amato, Jeffrey Barrett, Sonia Goncalves, Ewan Harrison, David K. Jackson, Ian Johnston, Dominic Kwiatkowski, Cordelia Langford, John Sillitoe on behalf of the Wellcome Sanger Institute COVID-19 Surveillance Team |
| EPI_ISL_2437067 | Health Services Laboratories | Wellcome Sanger Institute for the COVID-19 Genomics UK (COG-UK) Consortium | Health Services Laboratories and Alex Alderton, Roberto Amato, Jeffrey Barrett, Sonia Goncalves, Ewan Harrison, David K. Jackson, Ian Johnston, Dominic Kwiatkowski, Cordelia Langford, John Sillitoe on behalf of the Wellcome Sanger Institute COVID-19 Surveillance Team |
| EPI_ISL_2437068 | Lighthouse Lab in Milton Keynes | Wellcome Sanger Institute for the COVID-19 Genomics UK (COG-UK) Consortium | The Lighthouse Lab in Milton Keynes and Alex Alderton, Roberto Amato, Jeffrey Barrett, Sonia Goncalves, Ewan Harrison, David K. Jackson, Ian Johnston, Dominic Kwiatkowski, Cordelia Langford, John Sillitoe on behalf of the Wellcome Sanger Institute COVID-19 Surveillance Team |
| EPI_ISL_2437069 | Lighthouse Lab in Alderley Park | Wellcome Sanger Institute for the COVID-19 Genomics UK (COG-UK) Consortium | Jacquelyn Wynn, Mairead Hyland, The Lighthouse Lab in Alderley Park and Alex Alderton, Roberto Amato, Jeffrey Barrett, Sonia Goncalves, Ewan Harrison, David K. Jackson, Ian Johnston, Dominic Kwiatkowski, Cordelia Langford, John Sillitoe on behalf of the Wellcome Sanger Institute COVID-19 Surveillance Team |
| EPI_ISL_2437070 | Lighthouse Lab in Milton Keynes | Wellcome Sanger Institute for the COVID-19 Genomics UK (COG-UK) Consortium | The Lighthouse Lab in Milton Keynes and Alex Alderton, Roberto Amato, Jeffrey Barrett, Sonia Goncalves, Ewan Harrison, David K. Jackson, Ian Johnston, Dominic Kwiatkowski, Cordelia Langford, John Sillitoe on behalf of the Wellcome Sanger Institute COVID-19 Surveillance Team |
| EPI_ISL_2437071, EPI_ISL_2437072, EPI_ISL_2437073 | Lighthouse Lab in Alderley Park | Wellcome Sanger Institute for the COVID-19 Genomics UK (COG-UK) Consortium | Jacquelyn Wynn, Mairead Hyland, The Lighthouse Lab in Alderley Park and Alex Alderton, Roberto Amato, Jeffrey Barrett, Sonia Goncalves, Ewan Harrison, David K. Jackson, Ian Johnston, Dominic Kwiatkowski, Cordelia Langford, John Sillitoe on behalf of the Wellcome Sanger Institute COVID-19 Surveillance Team |
| EPI_ISL_2437074 | Lighthouse Lab in Milton Keynes | Wellcome Sanger Institute for the COVID-19 Genomics UK (COG-UK) Consortium | The Lighthouse Lab in Milton Keynes and Alex Alderton, Roberto Amato, Jeffrey Barrett, Sonia Goncalves, Ewan Harrison, David K. Jackson, Ian Johnston, Dominic Kwiatkowski, Cordelia Langford, John Sillitoe on behalf of the Wellcome Sanger Institute COVID-19 Surveillance Team |
| EPI_ISL_2437075 | Health Services Laboratories | Wellcome Sanger Institute for the COVID-19 Genomics UK (COG-UK) Consortium | Health Services Laboratories and Alex Alderton, Roberto Amato, Jeffrey Barrett, Sonia Goncalves, Ewan Harrison, David K. Jackson, Ian Johnston, Dominic Kwiatkowski, Cordelia Langford, John Sillitoe on behalf of the Wellcome Sanger Institute COVID-19 Surveillance Team |
| EPI_ISL_2437076, EPI_ISL_2437077 | Lighthouse Lab in Alderley Park | Wellcome Sanger Institute for the COVID-19 Genomics UK (COG-UK) Consortium | Jacquelyn Wynn, Mairead Hyland, The Lighthouse Lab in Alderley Park and Alex Alderton, Roberto Amato, Jeffrey Barrett, Sonia Goncalves, Ewan Harrison, David K. Jackson, Ian Johnston, Dominic Kwiatkowski, Cordelia Langford, John Sillitoe on behalf of the Wellcome Sanger Institute COVID-19 Surveillance Team |
| EPI_ISL_2437078 | Health Services Laboratories | Wellcome Sanger Institute for the COVID-19 Genomics UK (COG-UK) Consortium | Health Services Laboratories and Alex Alderton, Roberto Amato, Jeffrey Barrett, Sonia Goncalves, Ewan Harrison, David K. Jackson, Ian Johnston, Dominic Kwiatkowski, Cordelia Langford, John Sillitoe on behalf of the Wellcome Sanger Institute COVID-19 Surveillance Team |
| EPI_ISL_2437079, EPI_ISL_2437080, EPI_ISL_2437081, EPI_ISL_2437082, EPI_ISL_2437084, EPI_ISL_2437085, EPI_ISL_2437086 | Lighthouse Lab in Alderley Park | Wellcome Sanger Institute for the COVID-19 Genomics UK (COG-UK) Consortium | Jacquelyn Wynn, Mairead Hyland, The Lighthouse Lab in Alderley Park and Alex Alderton, Roberto Amato, Jeffrey Barrett, Sonia Goncalves, Ewan Harrison, David K. Jackson, Ian Johnston, Dominic Kwiatkowski, Cordelia Langford, John Sillitoe on behalf of the Wellcome Sanger Institute COVID-19 Surveillance Team |
| EPI_ISL_2437089 | Lighthouse Lab in Milton Keynes | Wellcome Sanger Institute for the COVID-19 Genomics UK (COG-UK) Consortium | The Lighthouse Lab in Milton Keynes and Alex Alderton, Roberto Amato, Jeffrey Barrett, Sonia Goncalves, Ewan Harrison, David K. Jackson, Ian Johnston, Dominic Kwiatkowski, Cordelia Langford, John Sillitoe on behalf of the Wellcome Sanger Institute COVID-19 Surveillance Team |
| EPI_ISL_2437090 | Health Services Laboratories | Wellcome Sanger Institute for the COVID-19 Genomics UK (COG-UK) Consortium | Health Services Laboratories and Alex Alderton, Roberto Amato, Jeffrey Barrett, Sonia Goncalves, Ewan Harrison, David K. Jackson, Ian Johnston, Dominic Kwiatkowski, Cordelia Langford, John Sillitoe on behalf of the Wellcome Sanger Institute COVID-19 Surveillance Team |
| EPI_ISL_2437091 | Lighthouse Lab in Alderley Park | Wellcome Sanger Institute for the COVID-19 Genomics UK (COG-UK) Consortium | Jacquelyn Wynn, Mairead Hyland, The Lighthouse Lab in Alderley Park and Alex Alderton, Roberto Amato, Jeffrey Barrett, Sonia Goncalves, Ewan Harrison, David K. Jackson, Ian Johnston, Dominic Kwiatkowski, Cordelia Langford, John Sillitoe on behalf of the Wellcome Sanger Institute COVID-19 Surveillance Team |
| EPI_ISL_2437092 | Health Services Laboratories | Wellcome Sanger Institute for the COVID-19 Genomics UK (COG-UK) Consortium | Health Services Laboratories and Alex Alderton, Roberto Amato, Jeffrey Barrett, Sonia Goncalves, Ewan Harrison, David K. Jackson, Ian Johnston, Dominic Kwiatkowski, Cordelia Langford, John Sillitoe on behalf of the Wellcome Sanger Institute COVID-19 Surveillance Team |
| EPI_ISL_2437094 | Lighthouse Lab in Milton Keynes | Wellcome Sanger Institute for the COVID-19 Genomics UK (COG-UK) Consortium | The Lighthouse Lab in Milton Keynes and Alex Alderton, Roberto Amato, Jeffrey Barrett, Sonia Goncalves, Ewan Harrison, David K. Jackson, Ian Johnston, Dominic Kwiatkowski, Cordelia Langford, John Sillitoe on behalf of the Wellcome Sanger Institute COVID-19 Surveillance Team |
| EPI_ISL_2437095 | Health Services Laboratories | Wellcome Sanger Institute for the COVID-19 Genomics UK (COG-UK) Consortium | Health Services Laboratories and Alex Alderton, Roberto Amato, Jeffrey Barrett, Sonia Goncalves, Ewan Harrison, David K. Jackson, Ian Johnston, Dominic Kwiatkowski, Cordelia Langford, John Sillitoe on behalf of the Wellcome Sanger Institute COVID-19 Surveillance Team |
| EPI_ISL_2437096, EPI_ISL_2437097 | Lighthouse Lab in Alderley Park | Wellcome Sanger Institute for the COVID-19 Genomics UK (COG-UK) Consortium | Jacquelyn Wynn, Mairead Hyland, The Lighthouse Lab in Alderley Park and Alex Alderton, Roberto Amato, Jeffrey Barrett, Sonia Goncalves, Ewan Harrison, David K. Jackson, Ian Johnston, Dominic Kwiatkowski, Cordelia Langford, John Sillitoe on behalf of the Wellcome Sanger Institute COVID-19 Surveillance Team |

| | | | |
|---|---|---|---|
| EPI_ISL_2437098 | Lighthouse Lab in Milton Keynes | Wellcome Sanger Institute for the COVID-19 Genomics UK (COG-UK) Consortium | The Lighthouse Lab in Milton Keynes and Alex Alderton, Roberto Amato, Jeffrey Barrett, Sonia Goncalves, Ewan Harrison, David K. Jackson, Ian Johnston, Dominic Kwiatkowski, Cordelia Langford, John Sillitoe on behalf of the Wellcome Sanger Institute COVID-19 Surveillance Team |
| EPI_ISL_2437099 | Lighthouse Lab in Alderley Park | Wellcome Sanger Institute for the COVID-19 Genomics UK (COG-UK) Consortium | Jacquelyn Wynn, Mairead Hyland, The Lighthouse Lab in Alderley Park and Alex Alderton, Roberto Amato, Jeffrey Barrett, Sonia Goncalves, Ewan Harrison, David K. Jackson, Ian Johnston, Dominic Kwiatkowski, Cordelia Langford, John Sillitoe on behalf of the Wellcome Sanger Institute COVID-19 Surveillance Team |
| EPI_ISL_2437100, EPI_ISL_2437101 | Health Services Laboratories | Wellcome Sanger Institute for the COVID-19 Genomics UK (COG-UK) Consortium | Health Services Laboratories and Alex Alderton, Roberto Amato, Jeffrey Barrett, Sonia Goncalves, Ewan Harrison, David K. Jackson, Ian Johnston, Dominic Kwiatkowski, Cordelia Langford, John Sillitoe on behalf of the Wellcome Sanger Institute COVID-19 Surveillance Team |
| EPI_ISL_2437103 | Lighthouse Lab in Alderley Park | Wellcome Sanger Institute for the COVID-19 Genomics UK (COG-UK) Consortium | Jacquelyn Wynn, Mairead Hyland, The Lighthouse Lab in Alderley Park and Alex Alderton, Roberto Amato, Jeffrey Barrett, Sonia Goncalves, Ewan Harrison, David K. Jackson, Ian Johnston, Dominic Kwiatkowski, Cordelia Langford, John Sillitoe on behalf of the Wellcome Sanger Institute COVID-19 Surveillance Team |
| EPI_ISL_2437104 | Health Services Laboratories | Wellcome Sanger Institute for the COVID-19 Genomics UK (COG-UK) Consortium | Health Services Laboratories and Alex Alderton, Roberto Amato, Jeffrey Barrett, Sonia Goncalves, Ewan Harrison, David K. Jackson, Ian Johnston, Dominic Kwiatkowski, Cordelia Langford, John Sillitoe on behalf of the Wellcome Sanger Institute COVID-19 Surveillance Team |
| EPI_ISL_2437105, EPI_ISL_2437106, EPI_ISL_2437107 | Lighthouse Lab in Alderley Park | Wellcome Sanger Institute for the COVID-19 Genomics UK (COG-UK) Consortium | Jacquelyn Wynn, Mairead Hyland, The Lighthouse Lab in Alderley Park and Alex Alderton, Roberto Amato, Jeffrey Barrett, Sonia Goncalves, Ewan Harrison, David K. Jackson, Ian Johnston, Dominic Kwiatkowski, Cordelia Langford, John Sillitoe on behalf of the Wellcome Sanger Institute COVID-19 Surveillance Team |
| EPI_ISL_2437109 | Health Services Laboratories | Wellcome Sanger Institute for the COVID-19 Genomics UK (COG-UK) Consortium | Health Services Laboratories and Alex Alderton, Roberto Amato, Jeffrey Barrett, Sonia Goncalves, Ewan Harrison, David K. Jackson, Ian Johnston, Dominic Kwiatkowski, Cordelia Langford, John Sillitoe on behalf of the Wellcome Sanger Institute COVID-19 Surveillance Team |
| EPI_ISL_2437110, EPI_ISL_2437111, EPI_ISL_2437112, EPI_ISL_2437113 | Lighthouse Lab in Alderley Park | Wellcome Sanger Institute for the COVID-19 Genomics UK (COG-UK) Consortium | Jacquelyn Wynn, Mairead Hyland, The Lighthouse Lab in Alderley Park and Alex Alderton, Roberto Amato, Jeffrey Barrett, Sonia Goncalves, Ewan Harrison, David K. Jackson, Ian Johnston, Dominic Kwiatkowski, Cordelia Langford, John Sillitoe on behalf of the Wellcome Sanger Institute COVID-19 Surveillance Team |
| EPI_ISL_2437114 | Health Services Laboratories | Wellcome Sanger Institute for the COVID-19 Genomics UK (COG-UK) Consortium | Health Services Laboratories and Alex Alderton, Roberto Amato, Jeffrey Barrett, Sonia Goncalves, Ewan Harrison, David K. Jackson, Ian Johnston, Dominic Kwiatkowski, Cordelia Langford, John Sillitoe on behalf of the Wellcome Sanger Institute COVID-19 Surveillance Team |
| EPI_ISL_2437115 | Lighthouse Lab in Alderley Park | Wellcome Sanger Institute for the COVID-19 Genomics UK (COG-UK) Consortium | Jacquelyn Wynn, Mairead Hyland, The Lighthouse Lab in Alderley Park and Alex Alderton, Roberto Amato, Jeffrey Barrett, Sonia Goncalves, Ewan Harrison, David K. Jackson, Ian Johnston, Dominic Kwiatkowski, Cordelia Langford, John Sillitoe on behalf of the Wellcome Sanger Institute COVID-19 Surveillance Team |
| EPI_ISL_2437116 | Lighthouse Lab in Milton Keynes | Wellcome Sanger Institute for the COVID-19 Genomics UK (COG-UK) Consortium | The Lighthouse Lab in Milton Keynes and Alex Alderton, Roberto Amato, Jeffrey Barrett, Sonia Goncalves, Ewan Harrison, David K. Jackson, Ian Johnston, Dominic Kwiatkowski, Cordelia Langford, John Sillitoe on behalf of the Wellcome Sanger Institute COVID-19 Surveillance Team |
| EPI_ISL_2437117 | Health Services Laboratories | Wellcome Sanger Institute for the COVID-19 Genomics UK (COG-UK) Consortium | Health Services Laboratories and Alex Alderton, Roberto Amato, Jeffrey Barrett, Sonia Goncalves, Ewan Harrison, David K. Jackson, Ian Johnston, Dominic Kwiatkowski, Cordelia Langford, John Sillitoe on behalf of the Wellcome Sanger Institute COVID-19 Surveillance Team |
| EPI_ISL_2437119, EPI_ISL_2437120, EPI_ISL_2437121, EPI_ISL_2437124, EPI_ISL_2437125 | Lighthouse Lab in Alderley Park | Wellcome Sanger Institute for the COVID-19 Genomics UK (COG-UK) Consortium | Jacquelyn Wynn, Mairead Hyland, The Lighthouse Lab in Alderley Park and Alex Alderton, Roberto Amato, Jeffrey Barrett, Sonia Goncalves, Ewan Harrison, David K. Jackson, Ian Johnston, Dominic Kwiatkowski, Cordelia Langford, John Sillitoe on behalf of the Wellcome Sanger Institute COVID-19 Surveillance Team |
| EPI_ISL_2437127 | Lighthouse Lab in Milton Keynes | Wellcome Sanger Institute for the COVID-19 Genomics UK (COG-UK) Consortium | The Lighthouse Lab in Milton Keynes and Alex Alderton, Roberto Amato, Jeffrey Barrett, Sonia Goncalves, Ewan Harrison, David K. Jackson, Ian Johnston, Dominic Kwiatkowski, Cordelia Langford, John Sillitoe on behalf of the Wellcome Sanger Institute COVID-19 Surveillance Team |
| EPI_ISL_2437129, EPI_ISL_2437130 | Lighthouse Lab in Alderley Park | Wellcome Sanger Institute for the COVID-19 Genomics UK (COG-UK) Consortium | Jacquelyn Wynn, Mairead Hyland, The Lighthouse Lab in Alderley Park and Alex Alderton, Roberto Amato, Jeffrey Barrett, Sonia Goncalves, Ewan Harrison, David K. Jackson, Ian Johnston, Dominic Kwiatkowski, Cordelia Langford, John Sillitoe on behalf of the Wellcome Sanger Institute COVID-19 Surveillance Team |
| EPI_ISL_2437131 | Health Services Laboratories | Wellcome Sanger Institute for the COVID-19 Genomics UK (COG-UK) Consortium | Health Services Laboratories and Alex Alderton, Roberto Amato, Jeffrey Barrett, Sonia Goncalves, Ewan Harrison, David K. Jackson, Ian Johnston, Dominic Kwiatkowski, Cordelia Langford, John Sillitoe on behalf of the Wellcome Sanger Institute COVID-19 Surveillance Team |
| EPI_ISL_2437132 | Lighthouse Lab in Milton Keynes | Wellcome Sanger Institute for the COVID-19 Genomics UK (COG-UK) Consortium | The Lighthouse Lab in Milton Keynes and Alex Alderton, Roberto Amato, Jeffrey Barrett, Sonia Goncalves, Ewan Harrison, David K. Jackson, Ian Johnston, Dominic Kwiatkowski, Cordelia Langford, John Sillitoe on behalf of the Wellcome Sanger Institute COVID-19 Surveillance Team |
| EPI_ISL_2437133, EPI_ISL_2437134, EPI_ISL_2437135, EPI_ISL_2437140 | Lighthouse Lab in Alderley Park | Wellcome Sanger Institute for the COVID-19 Genomics UK (COG-UK) Consortium | Jacquelyn Wynn, Mairead Hyland, The Lighthouse Lab in Alderley Park and Alex Alderton, Roberto Amato, Jeffrey Barrett, Sonia Goncalves, Ewan Harrison, David K. Jackson, Ian Johnston, Dominic Kwiatkowski, Cordelia Langford, John Sillitoe on behalf of the Wellcome Sanger Institute COVID-19 Surveillance Team |
| EPI_ISL_2437142 | Health Services Laboratories | Wellcome Sanger Institute for the COVID-19 Genomics UK (COG-UK) Consortium | Health Services Laboratories and Alex Alderton, Roberto Amato, Jeffrey Barrett, Sonia Goncalves, Ewan Harrison, David K. Jackson, Ian Johnston, Dominic Kwiatkowski, Cordelia Langford, John Sillitoe on behalf of the Wellcome Sanger Institute COVID-19 Surveillance Team |
| EPI_ISL_2437143, EPI_ISL_2437144 | Lighthouse Lab in Milton Keynes | Wellcome Sanger Institute for the COVID-19 Genomics UK (COG-UK) Consortium | The Lighthouse Lab in Milton Keynes and Alex Alderton, Roberto Amato, Jeffrey Barrett, Sonia Goncalves, Ewan Harrison, David K. Jackson, Ian Johnston, Dominic Kwiatkowski, Cordelia Langford, John Sillitoe on behalf of the Wellcome Sanger Institute COVID-19 Surveillance Team |
| EPI_ISL_2437145 | Lighthouse Lab in Alderley Park | Wellcome Sanger Institute for the COVID-19 Genomics UK (COG-UK) Consortium | Jacquelyn Wynn, Mairead Hyland, The Lighthouse Lab in Alderley Park and Alex Alderton, Roberto Amato, Jeffrey Barrett, Sonia Goncalves, Ewan Harrison, David K. Jackson, Ian Johnston, Dominic Kwiatkowski, Cordelia Langford, John Sillitoe on behalf of the Wellcome Sanger Institute COVID-19 Surveillance Team |
| EPI_ISL_2437146, EPI_ISL_2437147 | Health Services Laboratories | Wellcome Sanger Institute for the COVID-19 Genomics UK (COG-UK) Consortium | Health Services Laboratories and Alex Alderton, Roberto Amato, Jeffrey Barrett, Sonia Goncalves, Ewan Harrison, David K. Jackson, Ian Johnston, Dominic Kwiatkowski, Cordelia Langford, John Sillitoe on behalf of the Wellcome Sanger Institute COVID-19 Surveillance Team |
| EPI_ISL_2437148 | Lighthouse Lab in Alderley Park | Wellcome Sanger Institute for the COVID-19 Genomics UK (COG-UK) Consortium | Jacquelyn Wynn, Mairead Hyland, The Lighthouse Lab in Alderley Park and Alex Alderton, Roberto Amato, Jeffrey Barrett, Sonia Goncalves, Ewan Harrison, David K. Jackson, Ian Johnston, Dominic Kwiatkowski, Cordelia Langford, John Sillitoe on behalf of the Wellcome Sanger Institute COVID-19 Surveillance Team |
| EPI_ISL_2437150 | Lighthouse Lab in Milton Keynes | Wellcome Sanger Institute for the COVID-19 Genomics UK (COG-UK) Consortium | The Lighthouse Lab in Milton Keynes and Alex Alderton, Roberto Amato, Jeffrey Barrett, Sonia Goncalves, Ewan Harrison, David K. Jackson, Ian Johnston, Dominic Kwiatkowski, Cordelia Langford, John Sillitoe on behalf of the Wellcome Sanger Institute COVID-19 Surveillance Team |
| EPI_ISL_2437151 | Health Services Laboratories | Wellcome Sanger Institute for the COVID-19 Genomics UK (COG-UK) Consortium | Health Services Laboratories and Alex Alderton, Roberto Amato, Jeffrey Barrett, Sonia Goncalves, Ewan Harrison, David K. Jackson, Ian Johnston, Dominic Kwiatkowski, Cordelia Langford, John Sillitoe on behalf of the Wellcome Sanger Institute COVID-19 Surveillance Team |
| EPI_ISL_2437152, EPI_ISL_2437153, EPI_ISL_2437154, EPI_ISL_2437155 | Lighthouse Lab in Alderley Park | Wellcome Sanger Institute for the COVID-19 Genomics UK (COG-UK) Consortium | Jacquelyn Wynn, Mairead Hyland, The Lighthouse Lab in Alderley Park and Alex Alderton, Roberto Amato, Jeffrey Barrett, Sonia Goncalves, Ewan Harrison, David K. Jackson, Ian Johnston, Dominic Kwiatkowski, Cordelia Langford, John Sillitoe on behalf of the Wellcome Sanger Institute COVID-19 Surveillance Team |
| EPI_ISL_2437156, EPI_ISL_2437157 | Lighthouse Lab in Milton Keynes | Wellcome Sanger Institute for the COVID-19 Genomics UK (COG-UK) Consortium | The Lighthouse Lab in Milton Keynes and Alex Alderton, Roberto Amato, Jeffrey Barrett, Sonia Goncalves, Ewan Harrison, David K. Jackson, Ian Johnston, Dominic Kwiatkowski, Cordelia Langford, John Sillitoe on behalf of the Wellcome Sanger Institute COVID-19 Surveillance Team |
| EPI_ISL_2437158, EPI_ISL_2437159, EPI_ISL_2437161, EPI_ISL_2437162, EPI_ISL_2437163 | Lighthouse Lab in Alderley Park | Wellcome Sanger Institute for the COVID-19 Genomics UK (COG-UK) Consortium | Jacquelyn Wynn, Mairead Hyland, The Lighthouse Lab in Alderley Park and Alex Alderton, Roberto Amato, Jeffrey Barrett, Sonia Goncalves, Ewan Harrison, David K. Jackson, Ian Johnston, Dominic Kwiatkowski, Cordelia Langford, John Sillitoe on behalf of the Wellcome Sanger Institute COVID-19 Surveillance Team |
| EPI_ISL_2437164, EPI_ISL_2437165 | Lighthouse Lab in Milton Keynes | Wellcome Sanger Institute for the COVID-19 Genomics UK (COG-UK) Consortium | The Lighthouse Lab in Milton Keynes and Alex Alderton, Roberto Amato, Jeffrey Barrett, Sonia Goncalves, Ewan Harrison, David K. Jackson, Ian Johnston, Dominic Kwiatkowski, Cordelia Langford, John Sillitoe on behalf of the Wellcome Sanger Institute COVID-19 Surveillance Team |
| EPI_ISL_2437166 | Health Services Laboratories | Wellcome Sanger Institute for the COVID-19 Genomics UK (COG-UK) Consortium | Health Services Laboratories and Alex Alderton, Roberto Amato, Jeffrey Barrett, Sonia Goncalves, Ewan Harrison, David K. Jackson, Ian Johnston, Dominic Kwiatkowski, Cordelia Langford, John Sillitoe on behalf of the Wellcome Sanger Institute COVID-19 Surveillance Team |
| EPI_ISL_2437167 | Lighthouse Lab in Alderley Park | Wellcome Sanger Institute for the COVID-19 Genomics UK (COG-UK) Consortium | Jacquelyn Wynn, Mairead Hyland, The Lighthouse Lab in Alderley Park and Alex Alderton, Roberto Amato, Jeffrey Barrett, Sonia Goncalves, Ewan Harrison, David K. Jackson, Ian Johnston, Dominic Kwiatkowski, Cordelia Langford, John Sillitoe on behalf of the Wellcome Sanger Institute COVID-19 Surveillance Team |
| EPI_ISL_2437168 | Health Services Laboratories | Wellcome Sanger Institute for the COVID-19 Genomics UK (COG-UK) Consortium | Health Services Laboratories and Alex Alderton, Roberto Amato, Jeffrey Barrett, Sonia Goncalves, Ewan Harrison, David K. Jackson, Ian Johnston, Dominic Kwiatkowski, Cordelia Langford, John Sillitoe on behalf of the Wellcome Sanger Institute COVID-19 Surveillance Team |
| EPI_ISL_2437169 | Lighthouse Lab in Alderley Park | Wellcome Sanger Institute for the COVID-19 Genomics UK (COG-UK) Consortium | Jacquelyn Wynn, Mairead Hyland, The Lighthouse Lab in Alderley Park and Alex Alderton, Roberto Amato, Jeffrey Barrett, Sonia Goncalves, Ewan Harrison, David K. Jackson, Ian Johnston, Dominic Kwiatkowski, Cordelia Langford, John Sillitoe on behalf of the Wellcome Sanger Institute COVID-19 Surveillance Team |
| EPI_ISL_2437170 | Health Services Laboratories | Wellcome Sanger Institute for the COVID-19 Genomics UK (COG-UK) Consortium | Health Services Laboratories and Alex Alderton, Roberto Amato, Jeffrey Barrett, Sonia Goncalves, Ewan Harrison, David K. Jackson, Ian Johnston, Dominic Kwiatkowski, Cordelia Langford, John Sillitoe on behalf of the Wellcome Sanger Institute COVID-19 Surveillance Team |
| EPI_ISL_2437171, EPI_ISL_2437172, EPI_ISL_2437173, EPI_ISL_2437179, EPI_ISL_2437180 | Lighthouse Lab in Alderley Park | Wellcome Sanger Institute for the COVID-19 Genomics UK (COG-UK) Consortium | Jacquelyn Wynn, Mairead Hyland, The Lighthouse Lab in Alderley Park and Alex Alderton, Roberto Amato, Jeffrey Barrett, Sonia Goncalves, Ewan Harrison, David K. Jackson, Ian Johnston, Dominic Kwiatkowski, Cordelia Langford, John Sillitoe on behalf of the Wellcome Sanger Institute COVID-19 Surveillance Team |
| EPI_ISL_2437181 | Health Services Laboratories | Wellcome Sanger Institute for the COVID-19 Genomics UK (COG-UK) Consortium | Health Services Laboratories and Alex Alderton, Roberto Amato, Jeffrey Barrett, Sonia Goncalves, Ewan Harrison, David K. Jackson, Ian Johnston, Dominic Kwiatkowski, Cordelia Langford, John Sillitoe on behalf of the Wellcome Sanger Institute COVID-19 Surveillance Team |
| EPI_ISL_2437184 | Lighthouse Lab in Alderley Park | Wellcome Sanger Institute for the COVID-19 Genomics UK (COG-UK) Consortium | Jacquelyn Wynn, Mairead Hyland, The Lighthouse Lab in Alderley Park and Alex Alderton, Roberto Amato, Jeffrey Barrett, Sonia Goncalves, Ewan Harrison, David K. Jackson, Ian Johnston, Dominic Kwiatkowski, Cordelia Langford, John Sillitoe on behalf of the Wellcome Sanger Institute COVID-19 Surveillance Team |
| EPI_ISL_2437185 | Lighthouse Lab in Milton Keynes | Wellcome Sanger Institute for the COVID-19 Genomics UK (COG-UK) Consortium | The Lighthouse Lab in Milton Keynes and Alex Alderton, Roberto Amato, Jeffrey Barrett, Sonia Goncalves, Ewan Harrison, David K. Jackson, Ian Johnston, Dominic Kwiatkowski, Cordelia Langford, John Sillitoe on behalf of the Wellcome Sanger Institute COVID-19 Surveillance Team |
| EPI_ISL_2437186, | Lighthouse Lab in Alderley Park | Wellcome Sanger Institute for the COVID-19 Genomics UK (COG-UK) | Jacquelyn Wynn, Mairead Hyland, The Lighthouse Lab in Alderley Park and Alex Alderton, Roberto Amato, Jeffrey Barrett, Sonia Goncalves, Ewan Harrison, David K. Jackson, Ian Johnston, Dominic Kwiatkowski, Cordelia Langford, John Sillitoe on |

| | | | |
|---|---|---|---|
| EPI_ISL_2437187 | | Consortium | behalf of the Wellcome Sanger Institute COVID-19 Surveillance Team |
| EPI_ISL_2437188 | Health Services Laboratories | Wellcome Sanger Institute for the COVID-19 Genomics UK (COG-UK) Consortium | Health Services Laboratories and Alex Alderton, Roberto Amato, Jeffrey Barrett, Sonia Goncalves, Ewan Harrison, David K. Jackson, Ian Johnston, Dominic Kwiatkowski, Cordelia Langford, John Sillitoe on behalf of the Wellcome Sanger Institute COVID-19 Surveillance Team |
| EPI_ISL_2437189, EPI_ISL_2437190, EPI_ISL_2437191 | Lighthouse Lab in Alderley Park | Wellcome Sanger Institute for the COVID-19 Genomics UK (COG-UK) Consortium | Jacquelyn Wynn, Mairead Hyland, The Lighthouse Lab in Alderley Park and Alex Alderton, Roberto Amato, Jeffrey Barrett, Sonia Goncalves, Ewan Harrison, David K. Jackson, Ian Johnston, Dominic Kwiatkowski, Cordelia Langford, John Sillitoe on behalf of the Wellcome Sanger Institute COVID-19 Surveillance Team |
| EPI_ISL_2437192 | Health Services Laboratories | Wellcome Sanger Institute for the COVID-19 Genomics UK (COG-UK) Consortium | Health Services Laboratories and Alex Alderton, Roberto Amato, Jeffrey Barrett, Sonia Goncalves, Ewan Harrison, David K. Jackson, Ian Johnston, Dominic Kwiatkowski, Cordelia Langford, John Sillitoe on behalf of the Wellcome Sanger Institute COVID-19 Surveillance Team |
| EPI_ISL_2437193, EPI_ISL_2437194 | Lighthouse Lab in Alderley Park | Wellcome Sanger Institute for the COVID-19 Genomics UK (COG-UK) Consortium | Jacquelyn Wynn, Mairead Hyland, The Lighthouse Lab in Alderley Park and Alex Alderton, Roberto Amato, Jeffrey Barrett, Sonia Goncalves, Ewan Harrison, David K. Jackson, Ian Johnston, Dominic Kwiatkowski, Cordelia Langford, John Sillitoe on behalf of the Wellcome Sanger Institute COVID-19 Surveillance Team |
| EPI_ISL_2437195, EPI_ISL_2437197 | Lighthouse Lab in Milton Keynes | Wellcome Sanger Institute for the COVID-19 Genomics UK (COG-UK) Consortium | The Lighthouse Lab in Milton Keynes and Alex Alderton, Roberto Amato, Jeffrey Barrett, Sonia Goncalves, Ewan Harrison, David K. Jackson, Ian Johnston, Dominic Kwiatkowski, Cordelia Langford, John Sillitoe on behalf of the Wellcome Sanger Institute COVID-19 Surveillance Team |
| EPI_ISL_2437198 | Lighthouse Lab in Alderley Park | Wellcome Sanger Institute for the COVID-19 Genomics UK (COG-UK) Consortium | Jacquelyn Wynn, Mairead Hyland, The Lighthouse Lab in Alderley Park and Alex Alderton, Roberto Amato, Jeffrey Barrett, Sonia Goncalves, Ewan Harrison, David K. Jackson, Ian Johnston, Dominic Kwiatkowski, Cordelia Langford, John Sillitoe on behalf of the Wellcome Sanger Institute COVID-19 Surveillance Team |
| EPI_ISL_2437199, EPI_ISL_2437201, EPI_ISL_2437203 | Lighthouse Lab in Milton Keynes | Wellcome Sanger Institute for the COVID-19 Genomics UK (COG-UK) Consortium | The Lighthouse Lab in Milton Keynes and Alex Alderton, Roberto Amato, Jeffrey Barrett, Sonia Goncalves, Ewan Harrison, David K. Jackson, Ian Johnston, Dominic Kwiatkowski, Cordelia Langford, John Sillitoe on behalf of the Wellcome Sanger Institute COVID-19 Surveillance Team |
| EPI_ISL_2437205, EPI_ISL_2437207, EPI_ISL_2437208, EPI_ISL_2437211, EPI_ISL_2437212, EPI_ISL_2437213, EPI_ISL_2437215 | Lighthouse Lab in Alderley Park | Wellcome Sanger Institute for the COVID-19 Genomics UK (COG-UK) Consortium | Jacquelyn Wynn, Mairead Hyland, The Lighthouse Lab in Alderley Park and Alex Alderton, Roberto Amato, Jeffrey Barrett, Sonia Goncalves, Ewan Harrison, David K. Jackson, Ian Johnston, Dominic Kwiatkowski, Cordelia Langford, John Sillitoe on behalf of the Wellcome Sanger Institute COVID-19 Surveillance Team |
| EPI_ISL_2437216, EPI_ISL_2437217 | Lighthouse Lab in Milton Keynes | Wellcome Sanger Institute for the COVID-19 Genomics UK (COG-UK) Consortium | The Lighthouse Lab in Milton Keynes and Alex Alderton, Roberto Amato, Jeffrey Barrett, Sonia Goncalves, Ewan Harrison, David K. Jackson, Ian Johnston, Dominic Kwiatkowski, Cordelia Langford, John Sillitoe on behalf of the Wellcome Sanger Institute COVID-19 Surveillance Team |
| EPI_ISL_2437219, EPI_ISL_2437220, EPI_ISL_2437221 | Lighthouse Lab in Alderley Park | Wellcome Sanger Institute for the COVID-19 Genomics UK (COG-UK) Consortium | Jacquelyn Wynn, Mairead Hyland, The Lighthouse Lab in Alderley Park and Alex Alderton, Roberto Amato, Jeffrey Barrett, Sonia Goncalves, Ewan Harrison, David K. Jackson, Ian Johnston, Dominic Kwiatkowski, Cordelia Langford, John Sillitoe on behalf of the Wellcome Sanger Institute COVID-19 Surveillance Team |
| EPI_ISL_2437223 | Health Services Laboratories | Wellcome Sanger Institute for the COVID-19 Genomics UK (COG-UK) Consortium | Health Services Laboratories and Alex Alderton, Roberto Amato, Jeffrey Barrett, Sonia Goncalves, Ewan Harrison, David K. Jackson, Ian Johnston, Dominic Kwiatkowski, Cordelia Langford, John Sillitoe on behalf of the Wellcome Sanger Institute COVID-19 Surveillance Team |
| EPI_ISL_2437225, EPI_ISL_2437226, EPI_ISL_2437227, EPI_ISL_2437228, EPI_ISL_2437229, EPI_ISL_2437230 | Lighthouse Lab in Alderley Park | Wellcome Sanger Institute for the COVID-19 Genomics UK (COG-UK) Consortium | Jacquelyn Wynn, Mairead Hyland, The Lighthouse Lab in Alderley Park and Alex Alderton, Roberto Amato, Jeffrey Barrett, Sonia Goncalves, Ewan Harrison, David K. Jackson, Ian Johnston, Dominic Kwiatkowski, Cordelia Langford, John Sillitoe on behalf of the Wellcome Sanger Institute COVID-19 Surveillance Team |
| EPI_ISL_2437231 | Lighthouse Lab in Milton Keynes | Wellcome Sanger Institute for the COVID-19 Genomics UK (COG-UK) Consortium | The Lighthouse Lab in Milton Keynes and Alex Alderton, Roberto Amato, Jeffrey Barrett, Sonia Goncalves, Ewan Harrison, David K. Jackson, Ian Johnston, Dominic Kwiatkowski, Cordelia Langford, John Sillitoe on behalf of the Wellcome Sanger Institute COVID-19 Surveillance Team |
| EPI_ISL_2437232 | Lighthouse Lab in Alderley Park | Wellcome Sanger Institute for the COVID-19 Genomics UK (COG-UK) Consortium | Jacquelyn Wynn, Mairead Hyland, The Lighthouse Lab in Alderley Park and Alex Alderton, Roberto Amato, Jeffrey Barrett, Sonia Goncalves, Ewan Harrison, David K. Jackson, Ian Johnston, Dominic Kwiatkowski, Cordelia Langford, John Sillitoe on behalf of the Wellcome Sanger Institute COVID-19 Surveillance Team |
| EPI_ISL_2437234, EPI_ISL_2437235 | Lighthouse Lab in Milton Keynes | Wellcome Sanger Institute for the COVID-19 Genomics UK (COG-UK) Consortium | The Lighthouse Lab in Milton Keynes and Alex Alderton, Roberto Amato, Jeffrey Barrett, Sonia Goncalves, Ewan Harrison, David K. Jackson, Ian Johnston, Dominic Kwiatkowski, Cordelia Langford, John Sillitoe on behalf of the Wellcome Sanger Institute COVID-19 Surveillance Team |
| EPI_ISL_2437236 | Health Services Laboratories | Wellcome Sanger Institute for the COVID-19 Genomics UK (COG-UK) Consortium | Health Services Laboratories and Alex Alderton, Roberto Amato, Jeffrey Barrett, Sonia Goncalves, Ewan Harrison, David K. Jackson, Ian Johnston, Dominic Kwiatkowski, Cordelia Langford, John Sillitoe on behalf of the Wellcome Sanger Institute COVID-19 Surveillance Team |
| EPI_ISL_2437238, EPI_ISL_2437239 | Lighthouse Lab in Alderley Park | Wellcome Sanger Institute for the COVID-19 Genomics UK (COG-UK) Consortium | Jacquelyn Wynn, Mairead Hyland, The Lighthouse Lab in Alderley Park and Alex Alderton, Roberto Amato, Jeffrey Barrett, Sonia Goncalves, Ewan Harrison, David K. Jackson, Ian Johnston, Dominic Kwiatkowski, Cordelia Langford, John Sillitoe on behalf of the Wellcome Sanger Institute COVID-19 Surveillance Team |
| EPI_ISL_2437240 | Lighthouse Lab in Milton Keynes | Wellcome Sanger Institute for the COVID-19 Genomics UK (COG-UK) Consortium | The Lighthouse Lab in Milton Keynes and Alex Alderton, Roberto Amato, Jeffrey Barrett, Sonia Goncalves, Ewan Harrison, David K. Jackson, Ian Johnston, Dominic Kwiatkowski, Cordelia Langford, John Sillitoe on behalf of the Wellcome Sanger Institute COVID-19 Surveillance Team |
| EPI_ISL_2437244 | Lighthouse Lab in Alderley Park | Wellcome Sanger Institute for the COVID-19 Genomics UK (COG-UK) Consortium | Jacquelyn Wynn, Mairead Hyland, The Lighthouse Lab in Alderley Park and Alex Alderton, Roberto Amato, Jeffrey Barrett, Sonia Goncalves, Ewan Harrison, David K. Jackson, Ian Johnston, Dominic Kwiatkowski, Cordelia Langford, John Sillitoe on behalf of the Wellcome Sanger Institute COVID-19 Surveillance Team |
| EPI_ISL_2437245 | Lighthouse Lab in Milton Keynes | Wellcome Sanger Institute for the COVID-19 Genomics UK (COG-UK) Consortium | The Lighthouse Lab in Milton Keynes and Alex Alderton, Roberto Amato, Jeffrey Barrett, Sonia Goncalves, Ewan Harrison, David K. Jackson, Ian Johnston, Dominic Kwiatkowski, Cordelia Langford, John Sillitoe on behalf of the Wellcome Sanger Institute COVID-19 Surveillance Team |
| EPI_ISL_2437246 | Lighthouse Lab in Alderley Park | Wellcome Sanger Institute for the COVID-19 Genomics UK (COG-UK) Consortium | Jacquelyn Wynn, Mairead Hyland, The Lighthouse Lab in Alderley Park and Alex Alderton, Roberto Amato, Jeffrey Barrett, Sonia Goncalves, Ewan Harrison, David K. Jackson, Ian Johnston, Dominic Kwiatkowski, Cordelia Langford, John Sillitoe on behalf of the Wellcome Sanger Institute COVID-19 Surveillance Team |
| EPI_ISL_2437247 | Health Services Laboratories | Wellcome Sanger Institute for the COVID-19 Genomics UK (COG-UK) Consortium | Health Services Laboratories and Alex Alderton, Roberto Amato, Jeffrey Barrett, Sonia Goncalves, Ewan Harrison, David K. Jackson, Ian Johnston, Dominic Kwiatkowski, Cordelia Langford, John Sillitoe on behalf of the Wellcome Sanger Institute COVID-19 Surveillance Team |
| EPI_ISL_2437248, EPI_ISL_2437249, EPI_ISL_2437251, EPI_ISL_2437252 | Lighthouse Lab in Alderley Park | Wellcome Sanger Institute for the COVID-19 Genomics UK (COG-UK) Consortium | Jacquelyn Wynn, Mairead Hyland, The Lighthouse Lab in Alderley Park and Alex Alderton, Roberto Amato, Jeffrey Barrett, Sonia Goncalves, Ewan Harrison, David K. Jackson, Ian Johnston, Dominic Kwiatkowski, Cordelia Langford, John Sillitoe on behalf of the Wellcome Sanger Institute COVID-19 Surveillance Team |
| EPI_ISL_2437253 | Lighthouse Lab in Milton Keynes | Wellcome Sanger Institute for the COVID-19 Genomics UK (COG-UK) Consortium | The Lighthouse Lab in Milton Keynes and Alex Alderton, Roberto Amato, Jeffrey Barrett, Sonia Goncalves, Ewan Harrison, David K. Jackson, Ian Johnston, Dominic Kwiatkowski, Cordelia Langford, John Sillitoe on behalf of the Wellcome Sanger Institute COVID-19 Surveillance Team |
| EPI_ISL_2437254 | Lighthouse Lab in Alderley Park | Wellcome Sanger Institute for the COVID-19 Genomics UK (COG-UK) Consortium | Jacquelyn Wynn, Mairead Hyland, The Lighthouse Lab in Alderley Park and Alex Alderton, Roberto Amato, Jeffrey Barrett, Sonia Goncalves, Ewan Harrison, David K. Jackson, Ian Johnston, Dominic Kwiatkowski, Cordelia Langford, John Sillitoe on behalf of the Wellcome Sanger Institute COVID-19 Surveillance Team |
| EPI_ISL_2437255 | Health Services Laboratories | Wellcome Sanger Institute for the COVID-19 Genomics UK (COG-UK) Consortium | Health Services Laboratories and Alex Alderton, Roberto Amato, Jeffrey Barrett, Sonia Goncalves, Ewan Harrison, David K. Jackson, Ian Johnston, Dominic Kwiatkowski, Cordelia Langford, John Sillitoe on behalf of the Wellcome Sanger Institute COVID-19 Surveillance Team |
| EPI_ISL_2437256, EPI_ISL_2437257, EPI_ISL_2437258, EPI_ISL_2437259 | Lighthouse Lab in Alderley Park | Wellcome Sanger Institute for the COVID-19 Genomics UK (COG-UK) Consortium | Jacquelyn Wynn, Mairead Hyland, The Lighthouse Lab in Alderley Park and Alex Alderton, Roberto Amato, Jeffrey Barrett, Sonia Goncalves, Ewan Harrison, David K. Jackson, Ian Johnston, Dominic Kwiatkowski, Cordelia Langford, John Sillitoe on behalf of the Wellcome Sanger Institute COVID-19 Surveillance Team |
| EPI_ISL_2437260 | Lighthouse Lab in Milton Keynes | Wellcome Sanger Institute for the COVID-19 Genomics UK (COG-UK) Consortium | The Lighthouse Lab in Milton Keynes and Alex Alderton, Roberto Amato, Jeffrey Barrett, Sonia Goncalves, Ewan Harrison, David K. Jackson, Ian Johnston, Dominic Kwiatkowski, Cordelia Langford, John Sillitoe on behalf of the Wellcome Sanger Institute COVID-19 Surveillance Team |
| EPI_ISL_2437263, EPI_ISL_2437264 | Health Services Laboratories | Wellcome Sanger Institute for the COVID-19 Genomics UK (COG-UK) Consortium | Health Services Laboratories and Alex Alderton, Roberto Amato, Jeffrey Barrett, Sonia Goncalves, Ewan Harrison, David K. Jackson, Ian Johnston, Dominic Kwiatkowski, Cordelia Langford, John Sillitoe on behalf of the Wellcome Sanger Institute COVID-19 Surveillance Team |
| EPI_ISL_2437265, EPI_ISL_2437266, EPI_ISL_2437267, EPI_ISL_2437268, EPI_ISL_2437269, EPI_ISL_2437270, EPI_ISL_2437271 | Lighthouse Lab in Alderley Park | Wellcome Sanger Institute for the COVID-19 Genomics UK (COG-UK) Consortium | Jacquelyn Wynn, Mairead Hyland, The Lighthouse Lab in Alderley Park and Alex Alderton, Roberto Amato, Jeffrey Barrett, Sonia Goncalves, Ewan Harrison, David K. Jackson, Ian Johnston, Dominic Kwiatkowski, Cordelia Langford, John Sillitoe on behalf of the Wellcome Sanger Institute COVID-19 Surveillance Team |
| EPI_ISL_2437272, EPI_ISL_2437274 | Health Services Laboratories | Wellcome Sanger Institute for the COVID-19 Genomics UK (COG-UK) Consortium | Health Services Laboratories and Alex Alderton, Roberto Amato, Jeffrey Barrett, Sonia Goncalves, Ewan Harrison, David K. Jackson, Ian Johnston, Dominic Kwiatkowski, Cordelia Langford, John Sillitoe on behalf of the Wellcome Sanger Institute COVID-19 Surveillance Team |
| EPI_ISL_2437275, EPI_ISL_2437277 | Lighthouse Lab in Alderley Park | Wellcome Sanger Institute for the COVID-19 Genomics UK (COG-UK) Consortium | Jacquelyn Wynn, Mairead Hyland, The Lighthouse Lab in Alderley Park and Alex Alderton, Roberto Amato, Jeffrey Barrett, Sonia Goncalves, Ewan Harrison, David K. Jackson, Ian Johnston, Dominic Kwiatkowski, Cordelia Langford, John Sillitoe on behalf of the Wellcome Sanger Institute COVID-19 Surveillance Team |
| EPI_ISL_2437278 | Lighthouse Lab in Glasgow | Wellcome Sanger Institute for the COVID-19 Genomics UK (COG-UK) Consortium | Harper VanSteenhouse, Yumi Kasai, David Gray, Carol Clugston, Anna Dominiczak and Alex Alderton, Roberto Amato, Jeffrey Barrett, Sonia Goncalves, Ewan Harrison, David K. Jackson, Ian Johnston, Dominic Kwiatkowski, Cordelia Langford, John Sillitoe on behalf of the Wellcome Sanger Institute COVID-19 Surveillance Team |
| EPI_ISL_2437279, EPI_ISL_2437281, EPI_ISL_2437284, EPI_ISL_2437285, EPI_ISL_2437286, EPI_ISL_2437287, EPI_ISL_2437288, | Lighthouse Lab in Alderley Park | Wellcome Sanger Institute for the COVID-19 Genomics UK (COG-UK) Consortium | Jacquelyn Wynn, Mairead Hyland, The Lighthouse Lab in Alderley Park and Alex Alderton, Roberto Amato, Jeffrey Barrett, Sonia Goncalves, Ewan Harrison, David K. Jackson, Ian Johnston, Dominic Kwiatkowski, Cordelia Langford, John Sillitoe on behalf of the Wellcome Sanger Institute COVID-19 Surveillance Team |

| | | | |
|---|---|---|---|
| EPI_ISL_2437289 | | | |
| EPI_ISL_2437290 | Lighthouse Lab in Glasgow | Wellcome Sanger Institute for the COVID-19 Genomics UK (COG-UK) Consortium | Harper VanSteenhouse, Yumi Kasai, David Gray, Carol Clugston, Anna Dominiczak and Alex Alderton, Roberto Amato, Jeffrey Barrett, Sonia Goncalves, Ewan Harrison, David K. Jackson, Ian Johnston, Dominic Kwiatkowski, Cordelia Langford, John Sillitoe on behalf of the Wellcome Sanger Institute COVID-19 Surveillance Team |
| EPI_ISL_2437291 | Lighthouse Laboratory Plymouth | Wellcome Sanger Institute for the COVID-19 Genomics UK (COG-UK) Consortium | Lighthouse Laboratory Plymouth and Alex Alderton, Roberto Amato, Jeffrey Barrett, Sonia Goncalves, Ewan Harrison, David K. Jackson, Ian Johnston, Dominic Kwiatkowski, Cordelia Langford, John Sillitoe on behalf of the Wellcome Sanger Institute COVID-19 Surveillance Team |
| EPI_ISL_2437292, EPI_ISL_2437293 | Lighthouse Lab in Alderley Park | Wellcome Sanger Institute for the COVID-19 Genomics UK (COG-UK) Consortium | Jacquelyn Wynn, Mairead Hyland, The Lighthouse Lab in Alderley Park and Alex Alderton, Roberto Amato, Jeffrey Barrett, Sonia Goncalves, Ewan Harrison, David K. Jackson, Ian Johnston, Dominic Kwiatkowski, Cordelia Langford, John Sillitoe on behalf of the Wellcome Sanger Institute COVID-19 Surveillance Team |
| EPI_ISL_2437294 | Lighthouse Laboratory Plymouth | Wellcome Sanger Institute for the COVID-19 Genomics UK (COG-UK) Consortium | Lighthouse Laboratory Plymouth and Alex Alderton, Roberto Amato, Jeffrey Barrett, Sonia Goncalves, Ewan Harrison, David K. Jackson, Ian Johnston, Dominic Kwiatkowski, Cordelia Langford, John Sillitoe on behalf of the Wellcome Sanger Institute COVID-19 Surveillance Team |
| EPI_ISL_2437295 | Lighthouse Lab in Glasgow | Wellcome Sanger Institute for the COVID-19 Genomics UK (COG-UK) Consortium | Harper VanSteenhouse, Yumi Kasai, David Gray, Carol Clugston, Anna Dominiczak and Alex Alderton, Roberto Amato, Jeffrey Barrett, Sonia Goncalves, Ewan Harrison, David K. Jackson, Ian Johnston, Dominic Kwiatkowski, Cordelia Langford, John Sillitoe on behalf of the Wellcome Sanger Institute COVID-19 Surveillance Team |
| EPI_ISL_2437296 | Lighthouse Laboratory Plymouth | Wellcome Sanger Institute for the COVID-19 Genomics UK (COG-UK) Consortium | Lighthouse Laboratory Plymouth and Alex Alderton, Roberto Amato, Jeffrey Barrett, Sonia Goncalves, Ewan Harrison, David K. Jackson, Ian Johnston, Dominic Kwiatkowski, Cordelia Langford, John Sillitoe on behalf of the Wellcome Sanger Institute COVID-19 Surveillance Team |
| EPI_ISL_2437297, EPI_ISL_2437302 | Lighthouse Lab in Alderley Park | Wellcome Sanger Institute for the COVID-19 Genomics UK (COG-UK) Consortium | Jacquelyn Wynn, Mairead Hyland, The Lighthouse Lab in Alderley Park and Alex Alderton, Roberto Amato, Jeffrey Barrett, Sonia Goncalves, Ewan Harrison, David K. Jackson, Ian Johnston, Dominic Kwiatkowski, Cordelia Langford, John Sillitoe on behalf of the Wellcome Sanger Institute COVID-19 Surveillance Team |
| EPI_ISL_2437305 | Lighthouse Laboratory Plymouth | Wellcome Sanger Institute for the COVID-19 Genomics UK (COG-UK) Consortium | Lighthouse Laboratory Plymouth and Alex Alderton, Roberto Amato, Jeffrey Barrett, Sonia Goncalves, Ewan Harrison, David K. Jackson, Ian Johnston, Dominic Kwiatkowski, Cordelia Langford, John Sillitoe on behalf of the Wellcome Sanger Institute COVID-19 Surveillance Team |
| EPI_ISL_2437306, EPI_ISL_2437307, EPI_ISL_2437309, EPI_ISL_2437310, EPI_ISL_2437311 | Lighthouse Lab in Alderley Park | Wellcome Sanger Institute for the COVID-19 Genomics UK (COG-UK) Consortium | Jacquelyn Wynn, Mairead Hyland, The Lighthouse Lab in Alderley Park and Alex Alderton, Roberto Amato, Jeffrey Barrett, Sonia Goncalves, Ewan Harrison, David K. Jackson, Ian Johnston, Dominic Kwiatkowski, Cordelia Langford, John Sillitoe on behalf of the Wellcome Sanger Institute COVID-19 Surveillance Team |
| EPI_ISL_2437312 | Lighthouse Laboratory Plymouth | Wellcome Sanger Institute for the COVID-19 Genomics UK (COG-UK) Consortium | Lighthouse Laboratory Plymouth and Alex Alderton, Roberto Amato, Jeffrey Barrett, Sonia Goncalves, Ewan Harrison, David K. Jackson, Ian Johnston, Dominic Kwiatkowski, Cordelia Langford, John Sillitoe on behalf of the Wellcome Sanger Institute COVID-19 Surveillance Team |
| EPI_ISL_2437314 | Lighthouse Lab in Alderley Park | Wellcome Sanger Institute for the COVID-19 Genomics UK (COG-UK) Consortium | Jacquelyn Wynn, Mairead Hyland, The Lighthouse Lab in Alderley Park and Alex Alderton, Roberto Amato, Jeffrey Barrett, Sonia Goncalves, Ewan Harrison, David K. Jackson, Ian Johnston, Dominic Kwiatkowski, Cordelia Langford, John Sillitoe on behalf of the Wellcome Sanger Institute COVID-19 Surveillance Team |
| EPI_ISL_2437315, EPI_ISL_2437316 | Lighthouse Lab in Glasgow | Wellcome Sanger Institute for the COVID-19 Genomics UK (COG-UK) Consortium | Harper VanSteenhouse, Yumi Kasai, David Gray, Carol Clugston, Anna Dominiczak and Alex Alderton, Roberto Amato, Jeffrey Barrett, Sonia Goncalves, Ewan Harrison, David K. Jackson, Ian Johnston, Dominic Kwiatkowski, Cordelia Langford, John Sillitoe on behalf of the Wellcome Sanger Institute COVID-19 Surveillance Team |
| EPI_ISL_2437317 | Lighthouse Lab in Milton Keynes | Wellcome Sanger Institute for the COVID-19 Genomics UK (COG-UK) Consortium | The Lighthouse Lab in Milton Keynes and Alex Alderton, Roberto Amato, Jeffrey Barrett, Sonia Goncalves, Ewan Harrison, David K. Jackson, Ian Johnston, Dominic Kwiatkowski, Cordelia Langford, John Sillitoe on behalf of the Wellcome Sanger Institute COVID-19 Surveillance Team |
| EPI_ISL_2437318, EPI_ISL_2437319, EPI_ISL_2437321 | Lighthouse Lab in Alderley Park | Wellcome Sanger Institute for the COVID-19 Genomics UK (COG-UK) Consortium | Jacquelyn Wynn, Mairead Hyland, The Lighthouse Lab in Alderley Park and Alex Alderton, Roberto Amato, Jeffrey Barrett, Sonia Goncalves, Ewan Harrison, David K. Jackson, Ian Johnston, Dominic Kwiatkowski, Cordelia Langford, John Sillitoe on behalf of the Wellcome Sanger Institute COVID-19 Surveillance Team |
| EPI_ISL_2437322 | Lighthouse Lab in Milton Keynes | Wellcome Sanger Institute for the COVID-19 Genomics UK (COG-UK) Consortium | The Lighthouse Lab in Milton Keynes and Alex Alderton, Roberto Amato, Jeffrey Barrett, Sonia Goncalves, Ewan Harrison, David K. Jackson, Ian Johnston, Dominic Kwiatkowski, Cordelia Langford, John Sillitoe on behalf of the Wellcome Sanger Institute COVID-19 Surveillance Team |
| EPI_ISL_2437323 | Lighthouse Laboratory Plymouth | Wellcome Sanger Institute for the COVID-19 Genomics UK (COG-UK) Consortium | Lighthouse Laboratory Plymouth and Alex Alderton, Roberto Amato, Jeffrey Barrett, Sonia Goncalves, Ewan Harrison, David K. Jackson, Ian Johnston, Dominic Kwiatkowski, Cordelia Langford, John Sillitoe on behalf of the Wellcome Sanger Institute COVID-19 Surveillance Team |
| EPI_ISL_2437324 | Lighthouse Lab in Alderley Park | Wellcome Sanger Institute for the COVID-19 Genomics UK (COG-UK) Consortium | Jacquelyn Wynn, Mairead Hyland, The Lighthouse Lab in Alderley Park and Alex Alderton, Roberto Amato, Jeffrey Barrett, Sonia Goncalves, Ewan Harrison, David K. Jackson, Ian Johnston, Dominic Kwiatkowski, Cordelia Langford, John Sillitoe on behalf of the Wellcome Sanger Institute COVID-19 Surveillance Team |
| EPI_ISL_2437325 | Lighthouse Lab in Milton Keynes | Wellcome Sanger Institute for the COVID-19 Genomics UK (COG-UK) Consortium | The Lighthouse Lab in Milton Keynes and Alex Alderton, Roberto Amato, Jeffrey Barrett, Sonia Goncalves, Ewan Harrison, David K. Jackson, Ian Johnston, Dominic Kwiatkowski, Cordelia Langford, John Sillitoe on behalf of the Wellcome Sanger Institute COVID-19 Surveillance Team |
| EPI_ISL_2437326 | Health Services Laboratories | Wellcome Sanger Institute for the COVID-19 Genomics UK (COG-UK) Consortium | Health Services Laboratories and Alex Alderton, Roberto Amato, Jeffrey Barrett, Sonia Goncalves, Ewan Harrison, David K. Jackson, Ian Johnston, Dominic Kwiatkowski, Cordelia Langford, John Sillitoe on behalf of the Wellcome Sanger Institute COVID-19 Surveillance Team |
| EPI_ISL_2437327, EPI_ISL_2437328 | Lighthouse Lab in Alderley Park | Wellcome Sanger Institute for the COVID-19 Genomics UK (COG-UK) Consortium | Jacquelyn Wynn, Mairead Hyland, The Lighthouse Lab in Alderley Park and Alex Alderton, Roberto Amato, Jeffrey Barrett, Sonia Goncalves, Ewan Harrison, David K. Jackson, Ian Johnston, Dominic Kwiatkowski, Cordelia Langford, John Sillitoe on behalf of the Wellcome Sanger Institute COVID-19 Surveillance Team |
| EPI_ISL_2437329 | Health Services Laboratories | Wellcome Sanger Institute for the COVID-19 Genomics UK (COG-UK) Consortium | Health Services Laboratories and Alex Alderton, Roberto Amato, Jeffrey Barrett, Sonia Goncalves, Ewan Harrison, David K. Jackson, Ian Johnston, Dominic Kwiatkowski, Cordelia Langford, John Sillitoe on behalf of the Wellcome Sanger Institute COVID-19 Surveillance Team |
| EPI_ISL_2437330 | Lighthouse Lab in Glasgow | Wellcome Sanger Institute for the COVID-19 Genomics UK (COG-UK) Consortium | Harper VanSteenhouse, Yumi Kasai, David Gray, Carol Clugston, Anna Dominiczak and Alex Alderton, Roberto Amato, Jeffrey Barrett, Sonia Goncalves, Ewan Harrison, David K. Jackson, Ian Johnston, Dominic Kwiatkowski, Cordelia Langford, John Sillitoe on behalf of the Wellcome Sanger Institute COVID-19 Surveillance Team |
| EPI_ISL_2437333, EPI_ISL_2437334 | Lighthouse Lab in Alderley Park | Wellcome Sanger Institute for the COVID-19 Genomics UK (COG-UK) Consortium | Jacquelyn Wynn, Mairead Hyland, The Lighthouse Lab in Alderley Park and Alex Alderton, Roberto Amato, Jeffrey Barrett, Sonia Goncalves, Ewan Harrison, David K. Jackson, Ian Johnston, Dominic Kwiatkowski, Cordelia Langford, John Sillitoe on behalf of the Wellcome Sanger Institute COVID-19 Surveillance Team |
| EPI_ISL_2437335 | Lighthouse Lab in Milton Keynes | Wellcome Sanger Institute for the COVID-19 Genomics UK (COG-UK) Consortium | The Lighthouse Lab in Milton Keynes and Alex Alderton, Roberto Amato, Jeffrey Barrett, Sonia Goncalves, Ewan Harrison, David K. Jackson, Ian Johnston, Dominic Kwiatkowski, Cordelia Langford, John Sillitoe on behalf of the Wellcome Sanger Institute COVID-19 Surveillance Team |
| EPI_ISL_2437336 | Lighthouse Lab in Alderley Park | Wellcome Sanger Institute for the COVID-19 Genomics UK (COG-UK) Consortium | Jacquelyn Wynn, Mairead Hyland, The Lighthouse Lab in Alderley Park and Alex Alderton, Roberto Amato, Jeffrey Barrett, Sonia Goncalves, Ewan Harrison, David K. Jackson, Ian Johnston, Dominic Kwiatkowski, Cordelia Langford, John Sillitoe on behalf of the Wellcome Sanger Institute COVID-19 Surveillance Team |
| EPI_ISL_2437337 | Lighthouse Laboratory Plymouth | Wellcome Sanger Institute for the COVID-19 Genomics UK (COG-UK) Consortium | Lighthouse Laboratory Plymouth and Alex Alderton, Roberto Amato, Jeffrey Barrett, Sonia Goncalves, Ewan Harrison, David K. Jackson, Ian Johnston, Dominic Kwiatkowski, Cordelia Langford, John Sillitoe on behalf of the Wellcome Sanger Institute COVID-19 Surveillance Team |
| EPI_ISL_2437338, EPI_ISL_2437339, EPI_ISL_2437340 | Lighthouse Lab in Alderley Park | Wellcome Sanger Institute for the COVID-19 Genomics UK (COG-UK) Consortium | Jacquelyn Wynn, Mairead Hyland, The Lighthouse Lab in Alderley Park and Alex Alderton, Roberto Amato, Jeffrey Barrett, Sonia Goncalves, Ewan Harrison, David K. Jackson, Ian Johnston, Dominic Kwiatkowski, Cordelia Langford, John Sillitoe on behalf of the Wellcome Sanger Institute COVID-19 Surveillance Team |
| EPI_ISL_2437342 | Lighthouse Lab in Milton Keynes | Wellcome Sanger Institute for the COVID-19 Genomics UK (COG-UK) Consortium | The Lighthouse Lab in Milton Keynes and Alex Alderton, Roberto Amato, Jeffrey Barrett, Sonia Goncalves, Ewan Harrison, David K. Jackson, Ian Johnston, Dominic Kwiatkowski, Cordelia Langford, John Sillitoe on behalf of the Wellcome Sanger Institute COVID-19 Surveillance Team |
| EPI_ISL_2437343 | Lighthouse Lab in Alderley Park | Wellcome Sanger Institute for the COVID-19 Genomics UK (COG-UK) Consortium | Jacquelyn Wynn, Mairead Hyland, The Lighthouse Lab in Alderley Park and Alex Alderton, Roberto Amato, Jeffrey Barrett, Sonia Goncalves, Ewan Harrison, David K. Jackson, Ian Johnston, Dominic Kwiatkowski, Cordelia Langford, John Sillitoe on behalf of the Wellcome Sanger Institute COVID-19 Surveillance Team |
| EPI_ISL_2437344 | Lighthouse Laboratory Plymouth | Wellcome Sanger Institute for the COVID-19 Genomics UK (COG-UK) Consortium | Lighthouse Laboratory Plymouth and Alex Alderton, Roberto Amato, Jeffrey Barrett, Sonia Goncalves, Ewan Harrison, David K. Jackson, Ian Johnston, Dominic Kwiatkowski, Cordelia Langford, John Sillitoe on behalf of the Wellcome Sanger Institute COVID-19 Surveillance Team |
| EPI_ISL_2437345 | Lighthouse Lab in Glasgow | Wellcome Sanger Institute for the COVID-19 Genomics UK (COG-UK) Consortium | Harper VanSteenhouse, Yumi Kasai, David Gray, Carol Clugston, Anna Dominiczak and Alex Alderton, Roberto Amato, Jeffrey Barrett, Sonia Goncalves, Ewan Harrison, David K. Jackson, Ian Johnston, Dominic Kwiatkowski, Cordelia Langford, John Sillitoe on behalf of the Wellcome Sanger Institute COVID-19 Surveillance Team |
| EPI_ISL_2437346 | Lighthouse Lab in Alderley Park | Wellcome Sanger Institute for the COVID-19 Genomics UK (COG-UK) Consortium | Jacquelyn Wynn, Mairead Hyland, The Lighthouse Lab in Alderley Park and Alex Alderton, Roberto Amato, Jeffrey Barrett, Sonia Goncalves, Ewan Harrison, David K. Jackson, Ian Johnston, Dominic Kwiatkowski, Cordelia Langford, John Sillitoe on behalf of the Wellcome Sanger Institute COVID-19 Surveillance Team |
| EPI_ISL_2437347 | Health Services Laboratories | Wellcome Sanger Institute for the COVID-19 Genomics UK (COG-UK) Consortium | Health Services Laboratories and Alex Alderton, Roberto Amato, Jeffrey Barrett, Sonia Goncalves, Ewan Harrison, David K. Jackson, Ian Johnston, Dominic Kwiatkowski, Cordelia Langford, John Sillitoe on behalf of the Wellcome Sanger Institute COVID-19 Surveillance Team |
| EPI_ISL_2437348, EPI_ISL_2437350 | Lighthouse Lab in Alderley Park | Wellcome Sanger Institute for the COVID-19 Genomics UK (COG-UK) Consortium | Jacquelyn Wynn, Mairead Hyland, The Lighthouse Lab in Alderley Park and Alex Alderton, Roberto Amato, Jeffrey Barrett, Sonia Goncalves, Ewan Harrison, David K. Jackson, Ian Johnston, Dominic Kwiatkowski, Cordelia Langford, John Sillitoe on behalf of the Wellcome Sanger Institute COVID-19 Surveillance Team |
| EPI_ISL_2437351 | Lighthouse Laboratory Plymouth | Wellcome Sanger Institute for the COVID-19 Genomics UK (COG-UK) Consortium | Lighthouse Laboratory Plymouth and Alex Alderton, Roberto Amato, Jeffrey Barrett, Sonia Goncalves, Ewan Harrison, David K. Jackson, Ian Johnston, Dominic Kwiatkowski, Cordelia Langford, John Sillitoe on behalf of the Wellcome Sanger Institute COVID-19 Surveillance Team |
| EPI_ISL_2437352 | Lighthouse Lab in Milton Keynes | Wellcome Sanger Institute for the COVID-19 Genomics UK (COG-UK) Consortium | The Lighthouse Lab in Milton Keynes and Alex Alderton, Roberto Amato, Jeffrey Barrett, Sonia Goncalves, Ewan Harrison, David K. Jackson, Ian Johnston, Dominic Kwiatkowski, Cordelia Langford, John Sillitoe on behalf of the Wellcome Sanger Institute COVID-19 Surveillance Team |
| EPI_ISL_2437353 | Lighthouse Laboratory Plymouth | Wellcome Sanger Institute for the COVID-19 Genomics UK (COG-UK) Consortium | Lighthouse Laboratory Plymouth and Alex Alderton, Roberto Amato, Jeffrey Barrett, Sonia Goncalves, Ewan Harrison, David K. Jackson, Ian Johnston, Dominic Kwiatkowski, Cordelia Langford, John Sillitoe on behalf of the Wellcome Sanger Institute COVID-19 Surveillance Team |
| EPI_ISL_2437354 | Lighthouse Lab in Glasgow | Wellcome Sanger Institute for the COVID-19 Genomics UK (COG-UK) Consortium | Harper VanSteenhouse, Yumi Kasai, David Gray, Carol Clugston, Anna Dominiczak and Alex Alderton, Roberto Amato, Jeffrey Barrett, Sonia Goncalves, Ewan Harrison, David K. Jackson, Ian Johnston, Dominic Kwiatkowski, Cordelia Langford, John Sillitoe on behalf of the Wellcome Sanger Institute COVID-19 Surveillance Team |
| EPI_ISL_2437357, EPI_ISL_2437358, EPI_ISL_2437359 | Lighthouse Lab in Alderley Park | Wellcome Sanger Institute for the COVID-19 Genomics UK (COG-UK) Consortium | Jacquelyn Wynn, Mairead Hyland, The Lighthouse Lab in Alderley Park and Alex Alderton, Roberto Amato, Jeffrey Barrett, Sonia Goncalves, Ewan Harrison, David K. Jackson, Ian Johnston, Dominic Kwiatkowski, Cordelia Langford, John Sillitoe on behalf of the Wellcome Sanger Institute COVID-19 Surveillance Team |
| EPI_ISL_2437360 | Lighthouse Lab in Milton Keynes | Wellcome Sanger Institute for the COVID-19 Genomics UK (COG-UK) Consortium | The Lighthouse Lab in Milton Keynes and Alex Alderton, Roberto Amato, Jeffrey Barrett, Sonia Goncalves, Ewan Harrison, David K. Jackson, Ian Johnston, Dominic Kwiatkowski, Cordelia Langford, John Sillitoe on behalf of the Wellcome Sanger Institute COVID-19 Surveillance Team |
| EPI_ISL_2437361 | Lighthouse Lab in Alderley Park | Wellcome Sanger Institute for the COVID-19 Genomics UK (COG-UK) Consortium | Jacquelyn Wynn, Mairead Hyland, The Lighthouse Lab in Alderley Park and Alex Alderton, Roberto Amato, Jeffrey Barrett, Sonia Goncalves, Ewan Harrison, David K. Jackson, Ian Johnston, Dominic Kwiatkowski, Cordelia Langford, John Sillitoe on behalf of the Wellcome Sanger Institute COVID-19 Surveillance Team |
| EPI_ISL_2437362 | Health Services Laboratories | Wellcome Sanger Institute for the COVID-19 Genomics UK (COG-UK) Consortium | Health Services Laboratories and Alex Alderton, Roberto Amato, Jeffrey Barrett, Sonia Goncalves, Ewan Harrison, David K. Jackson, Ian Johnston, Dominic Kwiatkowski, Cordelia Langford, John Sillitoe on behalf of the Wellcome Sanger Institute COVID-19 Surveillance Team |

| | | | |
|---|---|---|---|
| EPI_ISL_2437363, EPI_ISL_2437364, EPI_ISL_2437365, EPI_ISL_2437366, EPI_ISL_2437367, EPI_ISL_2437368, EPI_ISL_2437369 | Lighthouse Lab in Alderley Park | Wellcome Sanger Institute for the COVID-19 Genomics UK (COG-UK) Consortium | Jacquelyn Wynn, Mairead Hyland, The Lighthouse Lab in Alderley Park and Alex Alderton, Roberto Amato, Jeffrey Barrett, Sonia Goncalves, Ewan Harrison, David K. Jackson, Ian Johnston, Dominic Kwiatkowski, Cordelia Langford, John Sillitoe on behalf of the Wellcome Sanger Institute COVID-19 Surveillance Team |
| EPI_ISL_2437370, EPI_ISL_2437371 | Lighthouse Lab in Milton Keynes | Wellcome Sanger Institute for the COVID-19 Genomics UK (COG-UK) Consortium | The Lighthouse Lab in Milton Keynes and Alex Alderton, Roberto Amato, Jeffrey Barrett, Sonia Goncalves, Ewan Harrison, David K. Jackson, Ian Johnston, Dominic Kwiatkowski, Cordelia Langford, John Sillitoe on behalf of the Wellcome Sanger Institute COVID-19 Surveillance Team |
| EPI_ISL_2437372 | Lighthouse Lab in Alderley Park | Wellcome Sanger Institute for the COVID-19 Genomics UK (COG-UK) Consortium | Jacquelyn Wynn, Mairead Hyland, The Lighthouse Lab in Alderley Park and Alex Alderton, Roberto Amato, Jeffrey Barrett, Sonia Goncalves, Ewan Harrison, David K. Jackson, Ian Johnston, Dominic Kwiatkowski, Cordelia Langford, John Sillitoe on behalf of the Wellcome Sanger Institute COVID-19 Surveillance Team |
| EPI_ISL_2437375 | Lighthouse Lab in Glasgow | Wellcome Sanger Institute for the COVID-19 Genomics UK (COG-UK) Consortium | Harper VanSteenhouse, Yumi Kasai, David Gray, Carol Clugston, Anna Dominiczak and Alex Alderton, Roberto Amato, Jeffrey Barrett, Sonia Goncalves, Ewan Harrison, David K. Jackson, Ian Johnston, Dominic Kwiatkowski, Cordelia Langford, John Sillitoe on behalf of the Wellcome Sanger Institute COVID-19 Surveillance Team |
| EPI_ISL_2437376, EPI_ISL_2437377 | Lighthouse Lab in Alderley Park | Wellcome Sanger Institute for the COVID-19 Genomics UK (COG-UK) Consortium | Jacquelyn Wynn, Mairead Hyland, The Lighthouse Lab in Alderley Park and Alex Alderton, Roberto Amato, Jeffrey Barrett, Sonia Goncalves, Ewan Harrison, David K. Jackson, Ian Johnston, Dominic Kwiatkowski, Cordelia Langford, John Sillitoe on behalf of the Wellcome Sanger Institute COVID-19 Surveillance Team |
| EPI_ISL_2437378, EPI_ISL_2437379 | Lighthouse Laboratory Plymouth | Wellcome Sanger Institute for the COVID-19 Genomics UK (COG-UK) Consortium | Lighthouse Laboratory Plymouth and Alex Alderton, Roberto Amato, Jeffrey Barrett, Sonia Goncalves, Ewan Harrison, David K. Jackson, Ian Johnston, Dominic Kwiatkowski, Cordelia Langford, John Sillitoe on behalf of the Wellcome Sanger Institute COVID-19 Surveillance Team |
| EPI_ISL_2437380, EPI_ISL_2437381, EPI_ISL_2437383, EPI_ISL_2437384, EPI_ISL_2437385, EPI_ISL_2437386, EPI_ISL_2437387 | Lighthouse Lab in Alderley Park | Wellcome Sanger Institute for the COVID-19 Genomics UK (COG-UK) Consortium | Jacquelyn Wynn, Mairead Hyland, The Lighthouse Lab in Alderley Park and Alex Alderton, Roberto Amato, Jeffrey Barrett, Sonia Goncalves, Ewan Harrison, David K. Jackson, Ian Johnston, Dominic Kwiatkowski, Cordelia Langford, John Sillitoe on behalf of the Wellcome Sanger Institute COVID-19 Surveillance Team |
| EPI_ISL_2437389 | Lighthouse Laboratory Plymouth | Wellcome Sanger Institute for the COVID-19 Genomics UK (COG-UK) Consortium | Lighthouse Laboratory Plymouth and Alex Alderton, Roberto Amato, Jeffrey Barrett, Sonia Goncalves, Ewan Harrison, David K. Jackson, Ian Johnston, Dominic Kwiatkowski, Cordelia Langford, John Sillitoe on behalf of the Wellcome Sanger Institute COVID-19 Surveillance Team |
| EPI_ISL_2437390, EPI_ISL_2437391, EPI_ISL_2437392, EPI_ISL_2437393, EPI_ISL_2437394, EPI_ISL_2437396, EPI_ISL_2437397 | Lighthouse Lab in Alderley Park | Wellcome Sanger Institute for the COVID-19 Genomics UK (COG-UK) Consortium | Jacquelyn Wynn, Mairead Hyland, The Lighthouse Lab in Alderley Park and Alex Alderton, Roberto Amato, Jeffrey Barrett, Sonia Goncalves, Ewan Harrison, David K. Jackson, Ian Johnston, Dominic Kwiatkowski, Cordelia Langford, John Sillitoe on behalf of the Wellcome Sanger Institute COVID-19 Surveillance Team |
| EPI_ISL_2437398 | Lighthouse Lab in Glasgow | Wellcome Sanger Institute for the COVID-19 Genomics UK (COG-UK) Consortium | Harper VanSteenhouse, Yumi Kasai, David Gray, Carol Clugston, Anna Dominiczak and Alex Alderton, Roberto Amato, Jeffrey Barrett, Sonia Goncalves, Ewan Harrison, David K. Jackson, Ian Johnston, Dominic Kwiatkowski, Cordelia Langford, John Sillitoe on behalf of the Wellcome Sanger Institute COVID-19 Surveillance Team |
| EPI_ISL_2437399 | Health Services Laboratories | Wellcome Sanger Institute for the COVID-19 Genomics UK (COG-UK) Consortium | Health Services Laboratories and Alex Alderton, Roberto Amato, Jeffrey Barrett, Sonia Goncalves, Ewan Harrison, David K. Jackson, Ian Johnston, Dominic Kwiatkowski, Cordelia Langford, John Sillitoe on behalf of the Wellcome Sanger Institute COVID-19 Surveillance Team |
| EPI_ISL_2437401 | Lighthouse Lab in Alderley Park | Wellcome Sanger Institute for the COVID-19 Genomics UK (COG-UK) Consortium | Jacquelyn Wynn, Mairead Hyland, The Lighthouse Lab in Alderley Park and Alex Alderton, Roberto Amato, Jeffrey Barrett, Sonia Goncalves, Ewan Harrison, David K. Jackson, Ian Johnston, Dominic Kwiatkowski, Cordelia Langford, John Sillitoe on behalf of the Wellcome Sanger Institute COVID-19 Surveillance Team |
| EPI_ISL_2437403 | Lighthouse Lab in Glasgow | Wellcome Sanger Institute for the COVID-19 Genomics UK (COG-UK) Consortium | Harper VanSteenhouse, Yumi Kasai, David Gray, Carol Clugston, Anna Dominiczak and Alex Alderton, Roberto Amato, Jeffrey Barrett, Sonia Goncalves, Ewan Harrison, David K. Jackson, Ian Johnston, Dominic Kwiatkowski, Cordelia Langford, John Sillitoe on behalf of the Wellcome Sanger Institute COVID-19 Surveillance Team |
| EPI_ISL_2437404 | Health Services Laboratories | Wellcome Sanger Institute for the COVID-19 Genomics UK (COG-UK) Consortium | Health Services Laboratories and Alex Alderton, Roberto Amato, Jeffrey Barrett, Sonia Goncalves, Ewan Harrison, David K. Jackson, Ian Johnston, Dominic Kwiatkowski, Cordelia Langford, John Sillitoe on behalf of the Wellcome Sanger Institute COVID-19 Surveillance Team |
| EPI_ISL_2437406 | Lighthouse Lab in Alderley Park | Wellcome Sanger Institute for the COVID-19 Genomics UK (COG-UK) Consortium | Jacquelyn Wynn, Mairead Hyland, The Lighthouse Lab in Alderley Park and Alex Alderton, Roberto Amato, Jeffrey Barrett, Sonia Goncalves, Ewan Harrison, David K. Jackson, Ian Johnston, Dominic Kwiatkowski, Cordelia Langford, John Sillitoe on behalf of the Wellcome Sanger Institute COVID-19 Surveillance Team |
| EPI_ISL_2437408 | Lighthouse Lab in Milton Keynes | Wellcome Sanger Institute for the COVID-19 Genomics UK (COG-UK) Consortium | The Lighthouse Lab in Milton Keynes and Alex Alderton, Roberto Amato, Jeffrey Barrett, Sonia Goncalves, Ewan Harrison, David K. Jackson, Ian Johnston, Dominic Kwiatkowski, Cordelia Langford, John Sillitoe on behalf of the Wellcome Sanger Institute COVID-19 Surveillance Team |
| EPI_ISL_2437409 | Lighthouse Laboratory Plymouth | Wellcome Sanger Institute for the COVID-19 Genomics UK (COG-UK) Consortium | Lighthouse Laboratory Plymouth and Alex Alderton, Roberto Amato, Jeffrey Barrett, Sonia Goncalves, Ewan Harrison, David K. Jackson, Ian Johnston, Dominic Kwiatkowski, Cordelia Langford, John Sillitoe on behalf of the Wellcome Sanger Institute COVID-19 Surveillance Team |
| EPI_ISL_2437410, EPI_ISL_2437411 | Lighthouse Lab in Alderley Park | Wellcome Sanger Institute for the COVID-19 Genomics UK (COG-UK) Consortium | Jacquelyn Wynn, Mairead Hyland, The Lighthouse Lab in Alderley Park and Alex Alderton, Roberto Amato, Jeffrey Barrett, Sonia Goncalves, Ewan Harrison, David K. Jackson, Ian Johnston, Dominic Kwiatkowski, Cordelia Langford, John Sillitoe on behalf of the Wellcome Sanger Institute COVID-19 Surveillance Team |
| EPI_ISL_2437412 | Lighthouse Lab in Glasgow | Wellcome Sanger Institute for the COVID-19 Genomics UK (COG-UK) Consortium | Harper VanSteenhouse, Yumi Kasai, David Gray, Carol Clugston, Anna Dominiczak and Alex Alderton, Roberto Amato, Jeffrey Barrett, Sonia Goncalves, Ewan Harrison, David K. Jackson, Ian Johnston, Dominic Kwiatkowski, Cordelia Langford, John Sillitoe on behalf of the Wellcome Sanger Institute COVID-19 Surveillance Team |
| EPI_ISL_2437413, EPI_ISL_2437414 | Lighthouse Lab in Alderley Park | Wellcome Sanger Institute for the COVID-19 Genomics UK (COG-UK) Consortium | Jacquelyn Wynn, Mairead Hyland, The Lighthouse Lab in Alderley Park and Alex Alderton, Roberto Amato, Jeffrey Barrett, Sonia Goncalves, Ewan Harrison, David K. Jackson, Ian Johnston, Dominic Kwiatkowski, Cordelia Langford, John Sillitoe on behalf of the Wellcome Sanger Institute COVID-19 Surveillance Team |
| EPI_ISL_2437415 | Lighthouse Lab in Glasgow | Wellcome Sanger Institute for the COVID-19 Genomics UK (COG-UK) Consortium | Harper VanSteenhouse, Yumi Kasai, David Gray, Carol Clugston, Anna Dominiczak and Alex Alderton, Roberto Amato, Jeffrey Barrett, Sonia Goncalves, Ewan Harrison, David K. Jackson, Ian Johnston, Dominic Kwiatkowski, Cordelia Langford, John Sillitoe on behalf of the Wellcome Sanger Institute COVID-19 Surveillance Team |
| EPI_ISL_2437416, EPI_ISL_2437417, EPI_ISL_2437418, EPI_ISL_2437419, EPI_ISL_2437420 | Lighthouse Lab in Alderley Park | Wellcome Sanger Institute for the COVID-19 Genomics UK (COG-UK) Consortium | Jacquelyn Wynn, Mairead Hyland, The Lighthouse Lab in Alderley Park and Alex Alderton, Roberto Amato, Jeffrey Barrett, Sonia Goncalves, Ewan Harrison, David K. Jackson, Ian Johnston, Dominic Kwiatkowski, Cordelia Langford, John Sillitoe on behalf of the Wellcome Sanger Institute COVID-19 Surveillance Team |
| EPI_ISL_2437422 | Lighthouse Laboratory Plymouth | Wellcome Sanger Institute for the COVID-19 Genomics UK (COG-UK) Consortium | Lighthouse Laboratory Plymouth and Alex Alderton, Roberto Amato, Jeffrey Barrett, Sonia Goncalves, Ewan Harrison, David K. Jackson, Ian Johnston, Dominic Kwiatkowski, Cordelia Langford, John Sillitoe on behalf of the Wellcome Sanger Institute COVID-19 Surveillance Team |
| EPI_ISL_2437423 | Lighthouse Lab in Glasgow | Wellcome Sanger Institute for the COVID-19 Genomics UK (COG-UK) Consortium | Harper VanSteenhouse, Yumi Kasai, David Gray, Carol Clugston, Anna Dominiczak and Alex Alderton, Roberto Amato, Jeffrey Barrett, Sonia Goncalves, Ewan Harrison, David K. Jackson, Ian Johnston, Dominic Kwiatkowski, Cordelia Langford, John Sillitoe on behalf of the Wellcome Sanger Institute COVID-19 Surveillance Team |
| EPI_ISL_2437424 | Lighthouse Laboratory Plymouth | Wellcome Sanger Institute for the COVID-19 Genomics UK (COG-UK) Consortium | Lighthouse Laboratory Plymouth and Alex Alderton, Roberto Amato, Jeffrey Barrett, Sonia Goncalves, Ewan Harrison, David K. Jackson, Ian Johnston, Dominic Kwiatkowski, Cordelia Langford, John Sillitoe on behalf of the Wellcome Sanger Institute COVID-19 Surveillance Team |
| EPI_ISL_2437425 | Lighthouse Lab in Alderley Park | Wellcome Sanger Institute for the COVID-19 Genomics UK (COG-UK) Consortium | Jacquelyn Wynn, Mairead Hyland, The Lighthouse Lab in Alderley Park and Alex Alderton, Roberto Amato, Jeffrey Barrett, Sonia Goncalves, Ewan Harrison, David K. Jackson, Ian Johnston, Dominic Kwiatkowski, Cordelia Langford, John Sillitoe on behalf of the Wellcome Sanger Institute COVID-19 Surveillance Team |
| EPI_ISL_2437426 | Health Services Laboratories | Wellcome Sanger Institute for the COVID-19 Genomics UK (COG-UK) Consortium | Health Services Laboratories and Alex Alderton, Roberto Amato, Jeffrey Barrett, Sonia Goncalves, Ewan Harrison, David K. Jackson, Ian Johnston, Dominic Kwiatkowski, Cordelia Langford, John Sillitoe on behalf of the Wellcome Sanger Institute COVID-19 Surveillance Team |
| EPI_ISL_2437427 | Lighthouse Lab in Glasgow | Wellcome Sanger Institute for the COVID-19 Genomics UK (COG-UK) Consortium | Harper VanSteenhouse, Yumi Kasai, David Gray, Carol Clugston, Anna Dominiczak and Alex Alderton, Roberto Amato, Jeffrey Barrett, Sonia Goncalves, Ewan Harrison, David K. Jackson, Ian Johnston, Dominic Kwiatkowski, Cordelia Langford, John Sillitoe on behalf of the Wellcome Sanger Institute COVID-19 Surveillance Team |
| EPI_ISL_2437428, EPI_ISL_2437429 | Lighthouse Lab in Alderley Park | Wellcome Sanger Institute for the COVID-19 Genomics UK (COG-UK) Consortium | Jacquelyn Wynn, Mairead Hyland, The Lighthouse Lab in Alderley Park and Alex Alderton, Roberto Amato, Jeffrey Barrett, Sonia Goncalves, Ewan Harrison, David K. Jackson, Ian Johnston, Dominic Kwiatkowski, Cordelia Langford, John Sillitoe on behalf of the Wellcome Sanger Institute COVID-19 Surveillance Team |
| EPI_ISL_2437430 | Lighthouse Lab in Glasgow | Wellcome Sanger Institute for the COVID-19 Genomics UK (COG-UK) Consortium | Harper VanSteenhouse, Yumi Kasai, David Gray, Carol Clugston, Anna Dominiczak and Alex Alderton, Roberto Amato, Jeffrey Barrett, Sonia Goncalves, Ewan Harrison, David K. Jackson, Ian Johnston, Dominic Kwiatkowski, Cordelia Langford, John Sillitoe on behalf of the Wellcome Sanger Institute COVID-19 Surveillance Team |
| EPI_ISL_2437431, EPI_ISL_2437433 | Lighthouse Lab in Milton Keynes | Wellcome Sanger Institute for the COVID-19 Genomics UK (COG-UK) Consortium | The Lighthouse Lab in Milton Keynes and Alex Alderton, Roberto Amato, Jeffrey Barrett, Sonia Goncalves, Ewan Harrison, David K. Jackson, Ian Johnston, Dominic Kwiatkowski, Cordelia Langford, John Sillitoe on behalf of the Wellcome Sanger Institute COVID-19 Surveillance Team |
| EPI_ISL_2437434 | Lighthouse Lab in Alderley Park | Wellcome Sanger Institute for the COVID-19 Genomics UK (COG-UK) Consortium | Jacquelyn Wynn, Mairead Hyland, The Lighthouse Lab in Alderley Park and Alex Alderton, Roberto Amato, Jeffrey Barrett, Sonia Goncalves, Ewan Harrison, David K. Jackson, Ian Johnston, Dominic Kwiatkowski, Cordelia Langford, John Sillitoe on behalf of the Wellcome Sanger Institute COVID-19 Surveillance Team |
| EPI_ISL_2437435, EPI_ISL_2437436 | Lighthouse Lab in Milton Keynes | Wellcome Sanger Institute for the COVID-19 Genomics UK (COG-UK) Consortium | The Lighthouse Lab in Milton Keynes and Alex Alderton, Roberto Amato, Jeffrey Barrett, Sonia Goncalves, Ewan Harrison, David K. Jackson, Ian Johnston, Dominic Kwiatkowski, Cordelia Langford, John Sillitoe on behalf of the Wellcome Sanger Institute COVID-19 Surveillance Team |
| EPI_ISL_2437437 | Lighthouse Lab in Glasgow | Wellcome Sanger Institute for the COVID-19 Genomics UK (COG-UK) Consortium | Harper VanSteenhouse, Yumi Kasai, David Gray, Carol Clugston, Anna Dominiczak and Alex Alderton, Roberto Amato, Jeffrey Barrett, Sonia Goncalves, Ewan Harrison, David K. Jackson, Ian Johnston, Dominic Kwiatkowski, Cordelia Langford, John Sillitoe on behalf of the Wellcome Sanger Institute COVID-19 Surveillance Team |
| EPI_ISL_2437438, EPI_ISL_2437441 | Lighthouse Lab in Alderley Park | Wellcome Sanger Institute for the COVID-19 Genomics UK (COG-UK) Consortium | Jacquelyn Wynn, Mairead Hyland, The Lighthouse Lab in Alderley Park and Alex Alderton, Roberto Amato, Jeffrey Barrett, Sonia Goncalves, Ewan Harrison, David K. Jackson, Ian Johnston, Dominic Kwiatkowski, Cordelia Langford, John Sillitoe on behalf of the Wellcome Sanger Institute COVID-19 Surveillance Team |
| EPI_ISL_2437442, EPI_ISL_2437443 | Lighthouse Laboratory Plymouth | Wellcome Sanger Institute for the COVID-19 Genomics UK (COG-UK) Consortium | Lighthouse Laboratory Plymouth and Alex Alderton, Roberto Amato, Jeffrey Barrett, Sonia Goncalves, Ewan Harrison, David K. Jackson, Ian Johnston, Dominic Kwiatkowski, Cordelia Langford, John Sillitoe on behalf of the Wellcome Sanger Institute COVID-19 Surveillance Team |
| EPI_ISL_2437444, EPI_ISL_2437445, EPI_ISL_2437446 | Lighthouse Lab in Alderley Park | Wellcome Sanger Institute for the COVID-19 Genomics UK (COG-UK) Consortium | Jacquelyn Wynn, Mairead Hyland, The Lighthouse Lab in Alderley Park and Alex Alderton, Roberto Amato, Jeffrey Barrett, Sonia Goncalves, Ewan Harrison, David K. Jackson, Ian Johnston, Dominic Kwiatkowski, Cordelia Langford, John Sillitoe on behalf of the Wellcome Sanger Institute COVID-19 Surveillance Team |

| | | | |
|---|---|---|---|
| EPI_ISL_2437447 | Lighthouse Lab in Milton Keynes | Wellcome Sanger Institute for the COVID-19 Genomics UK (COG-UK) Consortium | The Lighthouse Lab in Milton Keynes and Alex Alderton, Roberto Amato, Jeffrey Barrett, Sonia Goncalves, Ewan Harrison, David K. Jackson, Ian Johnston, Dominic Kwiatkowski, Cordelia Langford, John Sillitoe on behalf of the Wellcome Sanger Institute COVID-19 Surveillance Team |
| EPI_ISL_2437449, EPI_ISL_2437450, EPI_ISL_2437451 | Lighthouse Lab in Alderley Park | Wellcome Sanger Institute for the COVID-19 Genomics UK (COG-UK) Consortium | Jacquelyn Wynn, Mairead Hyland, The Lighthouse Lab in Alderley Park and Alex Alderton, Roberto Amato, Jeffrey Barrett, Sonia Goncalves, Ewan Harrison, David K. Jackson, Ian Johnston, Dominic Kwiatkowski, Cordelia Langford, John Sillitoe on behalf of the Wellcome Sanger Institute COVID-19 Surveillance Team |
| EPI_ISL_2437452 | Lighthouse Lab in Milton Keynes | Wellcome Sanger Institute for the COVID-19 Genomics UK (COG-UK) Consortium | The Lighthouse Lab in Milton Keynes and Alex Alderton, Roberto Amato, Jeffrey Barrett, Sonia Goncalves, Ewan Harrison, David K. Jackson, Ian Johnston, Dominic Kwiatkowski, Cordelia Langford, John Sillitoe on behalf of the Wellcome Sanger Institute COVID-19 Surveillance Team |
| EPI_ISL_2437454 | Lighthouse Lab in Alderley Park | Wellcome Sanger Institute for the COVID-19 Genomics UK (COG-UK) Consortium | Jacquelyn Wynn, Mairead Hyland, The Lighthouse Lab in Alderley Park and Alex Alderton, Roberto Amato, Jeffrey Barrett, Sonia Goncalves, Ewan Harrison, David K. Jackson, Ian Johnston, Dominic Kwiatkowski, Cordelia Langford, John Sillitoe on behalf of the Wellcome Sanger Institute COVID-19 Surveillance Team |
| EPI_ISL_2437456 | Lighthouse Lab in Milton Keynes | Wellcome Sanger Institute for the COVID-19 Genomics UK (COG-UK) Consortium | The Lighthouse Lab in Milton Keynes and Alex Alderton, Roberto Amato, Jeffrey Barrett, Sonia Goncalves, Ewan Harrison, David K. Jackson, Ian Johnston, Dominic Kwiatkowski, Cordelia Langford, John Sillitoe on behalf of the Wellcome Sanger Institute COVID-19 Surveillance Team |
| EPI_ISL_2437457 | Lighthouse Lab in Glasgow | Wellcome Sanger Institute for the COVID-19 Genomics UK (COG-UK) Consortium | Harper VanSteenhouse, Yumi Kasai, David Gray, Carol Clugston, Anna Dominiczak and Alex Alderton, Roberto Amato, Jeffrey Barrett, Sonia Goncalves, Ewan Harrison, David K. Jackson, Ian Johnston, Dominic Kwiatkowski, Cordelia Langford, John Sillitoe on behalf of the Wellcome Sanger Institute COVID-19 Surveillance Team |
| EPI_ISL_2437458, EPI_ISL_2437459 | Lighthouse Lab in Milton Keynes | Wellcome Sanger Institute for the COVID-19 Genomics UK (COG-UK) Consortium | The Lighthouse Lab in Milton Keynes and Alex Alderton, Roberto Amato, Jeffrey Barrett, Sonia Goncalves, Ewan Harrison, David K. Jackson, Ian Johnston, Dominic Kwiatkowski, Cordelia Langford, John Sillitoe on behalf of the Wellcome Sanger Institute COVID-19 Surveillance Team |
| EPI_ISL_2437460, EPI_ISL_2437461 | Lighthouse Lab in Alderley Park | Wellcome Sanger Institute for the COVID-19 Genomics UK (COG-UK) Consortium | Jacquelyn Wynn, Mairead Hyland, The Lighthouse Lab in Alderley Park and Alex Alderton, Roberto Amato, Jeffrey Barrett, Sonia Goncalves, Ewan Harrison, David K. Jackson, Ian Johnston, Dominic Kwiatkowski, Cordelia Langford, John Sillitoe on behalf of the Wellcome Sanger Institute COVID-19 Surveillance Team |
| EPI_ISL_2437462 | Lighthouse Laboratory Plymouth | Wellcome Sanger Institute for the COVID-19 Genomics UK (COG-UK) Consortium | Lighthouse Laboratory Plymouth and Alex Alderton, Roberto Amato, Jeffrey Barrett, Sonia Goncalves, Ewan Harrison, David K. Jackson, Ian Johnston, Dominic Kwiatkowski, Cordelia Langford, John Sillitoe on behalf of the Wellcome Sanger Institute COVID-19 Surveillance Team |
| EPI_ISL_2437464 | Lighthouse Lab in Alderley Park | Wellcome Sanger Institute for the COVID-19 Genomics UK (COG-UK) Consortium | Jacquelyn Wynn, Mairead Hyland, The Lighthouse Lab in Alderley Park and Alex Alderton, Roberto Amato, Jeffrey Barrett, Sonia Goncalves, Ewan Harrison, David K. Jackson, Ian Johnston, Dominic Kwiatkowski, Cordelia Langford, John Sillitoe on behalf of the Wellcome Sanger Institute COVID-19 Surveillance Team |
| EPI_ISL_2437466, EPI_ISL_2437468 | Lighthouse Lab in Glasgow | Wellcome Sanger Institute for the COVID-19 Genomics UK (COG-UK) Consortium | Harper VanSteenhouse, Yumi Kasai, David Gray, Carol Clugston, Anna Dominiczak and Alex Alderton, Roberto Amato, Jeffrey Barrett, Sonia Goncalves, Ewan Harrison, David K. Jackson, Ian Johnston, Dominic Kwiatkowski, Cordelia Langford, John Sillitoe on behalf of the Wellcome Sanger Institute COVID-19 Surveillance Team |
| EPI_ISL_2437469 | Lighthouse Lab in Alderley Park | Wellcome Sanger Institute for the COVID-19 Genomics UK (COG-UK) Consortium | Jacquelyn Wynn, Mairead Hyland, The Lighthouse Lab in Alderley Park and Alex Alderton, Roberto Amato, Jeffrey Barrett, Sonia Goncalves, Ewan Harrison, David K. Jackson, Ian Johnston, Dominic Kwiatkowski, Cordelia Langford, John Sillitoe on behalf of the Wellcome Sanger Institute COVID-19 Surveillance Team |
| EPI_ISL_2437471 | Lighthouse Lab in Milton Keynes | Wellcome Sanger Institute for the COVID-19 Genomics UK (COG-UK) Consortium | The Lighthouse Lab in Milton Keynes and Alex Alderton, Roberto Amato, Jeffrey Barrett, Sonia Goncalves, Ewan Harrison, David K. Jackson, Ian Johnston, Dominic Kwiatkowski, Cordelia Langford, John Sillitoe on behalf of the Wellcome Sanger Institute COVID-19 Surveillance Team |
| EPI_ISL_2437472 | Health Services Laboratories | Wellcome Sanger Institute for the COVID-19 Genomics UK (COG-UK) Consortium | Health Services Laboratories and Alex Alderton, Roberto Amato, Jeffrey Barrett, Sonia Goncalves, Ewan Harrison, David K. Jackson, Ian Johnston, Dominic Kwiatkowski, Cordelia Langford, John Sillitoe on behalf of the Wellcome Sanger Institute COVID-19 Surveillance Team |
| EPI_ISL_2437473 | Lighthouse Laboratory Plymouth | Wellcome Sanger Institute for the COVID-19 Genomics UK (COG-UK) Consortium | Lighthouse Laboratory Plymouth and Alex Alderton, Roberto Amato, Jeffrey Barrett, Sonia Goncalves, Ewan Harrison, David K. Jackson, Ian Johnston, Dominic Kwiatkowski, Cordelia Langford, John Sillitoe on behalf of the Wellcome Sanger Institute COVID-19 Surveillance Team |
| EPI_ISL_2437476 | Lighthouse Lab in Alderley Park | Wellcome Sanger Institute for the COVID-19 Genomics UK (COG-UK) Consortium | Jacquelyn Wynn, Mairead Hyland, The Lighthouse Lab in Alderley Park and Alex Alderton, Roberto Amato, Jeffrey Barrett, Sonia Goncalves, Ewan Harrison, David K. Jackson, Ian Johnston, Dominic Kwiatkowski, Cordelia Langford, John Sillitoe on behalf of the Wellcome Sanger Institute COVID-19 Surveillance Team |
| EPI_ISL_2437477 | Lighthouse Lab in Milton Keynes | Wellcome Sanger Institute for the COVID-19 Genomics UK (COG-UK) Consortium | The Lighthouse Lab in Milton Keynes and Alex Alderton, Roberto Amato, Jeffrey Barrett, Sonia Goncalves, Ewan Harrison, David K. Jackson, Ian Johnston, Dominic Kwiatkowski, Cordelia Langford, John Sillitoe on behalf of the Wellcome Sanger Institute COVID-19 Surveillance Team |
| EPI_ISL_2437478, EPI_ISL_2437480 | Lighthouse Lab in Alderley Park | Wellcome Sanger Institute for the COVID-19 Genomics UK (COG-UK) Consortium | Jacquelyn Wynn, Mairead Hyland, The Lighthouse Lab in Alderley Park and Alex Alderton, Roberto Amato, Jeffrey Barrett, Sonia Goncalves, Ewan Harrison, David K. Jackson, Ian Johnston, Dominic Kwiatkowski, Cordelia Langford, John Sillitoe on behalf of the Wellcome Sanger Institute COVID-19 Surveillance Team |
| EPI_ISL_2437481 | Lighthouse Lab in Glasgow | Wellcome Sanger Institute for the COVID-19 Genomics UK (COG-UK) Consortium | Harper VanSteenhouse, Yumi Kasai, David Gray, Carol Clugston, Anna Dominiczak and Alex Alderton, Roberto Amato, Jeffrey Barrett, Sonia Goncalves, Ewan Harrison, David K. Jackson, Ian Johnston, Dominic Kwiatkowski, Cordelia Langford, John Sillitoe on behalf of the Wellcome Sanger Institute COVID-19 Surveillance Team |
| EPI_ISL_2437483 | Lighthouse Lab in Alderley Park | Wellcome Sanger Institute for the COVID-19 Genomics UK (COG-UK) Consortium | Jacquelyn Wynn, Mairead Hyland, The Lighthouse Lab in Alderley Park and Alex Alderton, Roberto Amato, Jeffrey Barrett, Sonia Goncalves, Ewan Harrison, David K. Jackson, Ian Johnston, Dominic Kwiatkowski, Cordelia Langford, John Sillitoe on behalf of the Wellcome Sanger Institute COVID-19 Surveillance Team |
| EPI_ISL_2437484, EPI_ISL_2437485 | Lighthouse Lab in Glasgow | Wellcome Sanger Institute for the COVID-19 Genomics UK (COG-UK) Consortium | Harper VanSteenhouse, Yumi Kasai, David Gray, Carol Clugston, Anna Dominiczak and Alex Alderton, Roberto Amato, Jeffrey Barrett, Sonia Goncalves, Ewan Harrison, David K. Jackson, Ian Johnston, Dominic Kwiatkowski, Cordelia Langford, John Sillitoe on behalf of the Wellcome Sanger Institute COVID-19 Surveillance Team |
| EPI_ISL_2437486, EPI_ISL_2437487 | Lighthouse Lab in Alderley Park | Wellcome Sanger Institute for the COVID-19 Genomics UK (COG-UK) Consortium | Jacquelyn Wynn, Mairead Hyland, The Lighthouse Lab in Alderley Park and Alex Alderton, Roberto Amato, Jeffrey Barrett, Sonia Goncalves, Ewan Harrison, David K. Jackson, Ian Johnston, Dominic Kwiatkowski, Cordelia Langford, John Sillitoe on behalf of the Wellcome Sanger Institute COVID-19 Surveillance Team |
| EPI_ISL_2437488 | Lighthouse Laboratory Plymouth | Wellcome Sanger Institute for the COVID-19 Genomics UK (COG-UK) Consortium | Lighthouse Laboratory Plymouth and Alex Alderton, Roberto Amato, Jeffrey Barrett, Sonia Goncalves, Ewan Harrison, David K. Jackson, Ian Johnston, Dominic Kwiatkowski, Cordelia Langford, John Sillitoe on behalf of the Wellcome Sanger Institute COVID-19 Surveillance Team |
| EPI_ISL_2437489 | Lighthouse Lab in Milton Keynes | Wellcome Sanger Institute for the COVID-19 Genomics UK (COG-UK) Consortium | The Lighthouse Lab in Milton Keynes and Alex Alderton, Roberto Amato, Jeffrey Barrett, Sonia Goncalves, Ewan Harrison, David K. Jackson, Ian Johnston, Dominic Kwiatkowski, Cordelia Langford, John Sillitoe on behalf of the Wellcome Sanger Institute COVID-19 Surveillance Team |
| EPI_ISL_2437490, EPI_ISL_2437491 | Lighthouse Lab in Glasgow | Wellcome Sanger Institute for the COVID-19 Genomics UK (COG-UK) Consortium | Harper VanSteenhouse, Yumi Kasai, David Gray, Carol Clugston, Anna Dominiczak and Alex Alderton, Roberto Amato, Jeffrey Barrett, Sonia Goncalves, Ewan Harrison, David K. Jackson, Ian Johnston, Dominic Kwiatkowski, Cordelia Langford, John Sillitoe on behalf of the Wellcome Sanger Institute COVID-19 Surveillance Team |
| EPI_ISL_2437492, EPI_ISL_2437493 | Lighthouse Lab in Alderley Park | Wellcome Sanger Institute for the COVID-19 Genomics UK (COG-UK) Consortium | Jacquelyn Wynn, Mairead Hyland, The Lighthouse Lab in Alderley Park and Alex Alderton, Roberto Amato, Jeffrey Barrett, Sonia Goncalves, Ewan Harrison, David K. Jackson, Ian Johnston, Dominic Kwiatkowski, Cordelia Langford, John Sillitoe on behalf of the Wellcome Sanger Institute COVID-19 Surveillance Team |
| EPI_ISL_2437494 | Health Services Laboratories | Wellcome Sanger Institute for the COVID-19 Genomics UK (COG-UK) Consortium | Health Services Laboratories and Alex Alderton, Roberto Amato, Jeffrey Barrett, Sonia Goncalves, Ewan Harrison, David K. Jackson, Ian Johnston, Dominic Kwiatkowski, Cordelia Langford, John Sillitoe on behalf of the Wellcome Sanger Institute COVID-19 Surveillance Team |
| EPI_ISL_2437495, EPI_ISL_2437496 | Lighthouse Lab in Alderley Park | Wellcome Sanger Institute for the COVID-19 Genomics UK (COG-UK) Consortium | Jacquelyn Wynn, Mairead Hyland, The Lighthouse Lab in Alderley Park and Alex Alderton, Roberto Amato, Jeffrey Barrett, Sonia Goncalves, Ewan Harrison, David K. Jackson, Ian Johnston, Dominic Kwiatkowski, Cordelia Langford, John Sillitoe on behalf of the Wellcome Sanger Institute COVID-19 Surveillance Team |
| EPI_ISL_2437497 | Health Services Laboratories | Wellcome Sanger Institute for the COVID-19 Genomics UK (COG-UK) Consortium | Health Services Laboratories and Alex Alderton, Roberto Amato, Jeffrey Barrett, Sonia Goncalves, Ewan Harrison, David K. Jackson, Ian Johnston, Dominic Kwiatkowski, Cordelia Langford, John Sillitoe on behalf of the Wellcome Sanger Institute COVID-19 Surveillance Team |
| EPI_ISL_2437498 | Lighthouse Laboratory Plymouth | Wellcome Sanger Institute for the COVID-19 Genomics UK (COG-UK) Consortium | Lighthouse Laboratory Plymouth and Alex Alderton, Roberto Amato, Jeffrey Barrett, Sonia Goncalves, Ewan Harrison, David K. Jackson, Ian Johnston, Dominic Kwiatkowski, Cordelia Langford, John Sillitoe on behalf of the Wellcome Sanger Institute COVID-19 Surveillance Team |
| EPI_ISL_2437499, EPI_ISL_2437500, EPI_ISL_2437501, EPI_ISL_2437502 | Lighthouse Lab in Alderley Park | Wellcome Sanger Institute for the COVID-19 Genomics UK (COG-UK) Consortium | Jacquelyn Wynn, Mairead Hyland, The Lighthouse Lab in Alderley Park and Alex Alderton, Roberto Amato, Jeffrey Barrett, Sonia Goncalves, Ewan Harrison, David K. Jackson, Ian Johnston, Dominic Kwiatkowski, Cordelia Langford, John Sillitoe on behalf of the Wellcome Sanger Institute COVID-19 Surveillance Team |
| EPI_ISL_2437503 | Lighthouse Lab in Milton Keynes | Wellcome Sanger Institute for the COVID-19 Genomics UK (COG-UK) Consortium | The Lighthouse Lab in Milton Keynes and Alex Alderton, Roberto Amato, Jeffrey Barrett, Sonia Goncalves, Ewan Harrison, David K. Jackson, Ian Johnston, Dominic Kwiatkowski, Cordelia Langford, John Sillitoe on behalf of the Wellcome Sanger Institute COVID-19 Surveillance Team |
| EPI_ISL_2437504, EPI_ISL_2437505, EPI_ISL_2437506, EPI_ISL_2437507, EPI_ISL_2437508, EPI_ISL_2437510 | Lighthouse Lab in Alderley Park | Wellcome Sanger Institute for the COVID-19 Genomics UK (COG-UK) Consortium | Jacquelyn Wynn, Mairead Hyland, The Lighthouse Lab in Alderley Park and Alex Alderton, Roberto Amato, Jeffrey Barrett, Sonia Goncalves, Ewan Harrison, David K. Jackson, Ian Johnston, Dominic Kwiatkowski, Cordelia Langford, John Sillitoe on behalf of the Wellcome Sanger Institute COVID-19 Surveillance Team |
| EPI_ISL_2437513 | Health Services Laboratories | Wellcome Sanger Institute for the COVID-19 Genomics UK (COG-UK) Consortium | Health Services Laboratories and Alex Alderton, Roberto Amato, Jeffrey Barrett, Sonia Goncalves, Ewan Harrison, David K. Jackson, Ian Johnston, Dominic Kwiatkowski, Cordelia Langford, John Sillitoe on behalf of the Wellcome Sanger Institute COVID-19 Surveillance Team |
| EPI_ISL_2437514 | Lighthouse Lab in Alderley Park | Wellcome Sanger Institute for the COVID-19 Genomics UK (COG-UK) Consortium | Jacquelyn Wynn, Mairead Hyland, The Lighthouse Lab in Alderley Park and Alex Alderton, Roberto Amato, Jeffrey Barrett, Sonia Goncalves, Ewan Harrison, David K. Jackson, Ian Johnston, Dominic Kwiatkowski, Cordelia Langford, John Sillitoe on behalf of the Wellcome Sanger Institute COVID-19 Surveillance Team |
| EPI_ISL_2437516 | Lighthouse Lab in Milton Keynes | Wellcome Sanger Institute for the COVID-19 Genomics UK (COG-UK) Consortium | The Lighthouse Lab in Milton Keynes and Alex Alderton, Roberto Amato, Jeffrey Barrett, Sonia Goncalves, Ewan Harrison, David K. Jackson, Ian Johnston, Dominic Kwiatkowski, Cordelia Langford, John Sillitoe on behalf of the Wellcome Sanger Institute COVID-19 Surveillance Team |
| EPI_ISL_2437519 | Lighthouse Lab in Glasgow | Wellcome Sanger Institute for the COVID-19 Genomics UK (COG-UK) Consortium | Harper VanSteenhouse, Yumi Kasai, David Gray, Carol Clugston, Anna Dominiczak and Alex Alderton, Roberto Amato, Jeffrey Barrett, Sonia Goncalves, Ewan Harrison, David K. Jackson, Ian Johnston, Dominic Kwiatkowski, Cordelia Langford, John Sillitoe on behalf of the Wellcome Sanger Institute COVID-19 Surveillance Team |
| EPI_ISL_2437520 | Lighthouse Lab in Alderley Park | Wellcome Sanger Institute for the COVID-19 Genomics UK (COG-UK) Consortium | Jacquelyn Wynn, Mairead Hyland, The Lighthouse Lab in Alderley Park and Alex Alderton, Roberto Amato, Jeffrey Barrett, Sonia Goncalves, Ewan Harrison, David K. Jackson, Ian Johnston, Dominic Kwiatkowski, Cordelia Langford, John Sillitoe on behalf of the Wellcome Sanger Institute COVID-19 Surveillance Team |
| EPI_ISL_2437521 | Lighthouse Lab in Glasgow | Wellcome Sanger Institute for the COVID-19 Genomics UK (COG-UK) Consortium | Harper VanSteenhouse, Yumi Kasai, David Gray, Carol Clugston, Anna Dominiczak and Alex Alderton, Roberto Amato, Jeffrey Barrett, Sonia Goncalves, Ewan Harrison, David K. Jackson, Ian Johnston, Dominic Kwiatkowski, Cordelia Langford, John Sillitoe on behalf of the Wellcome Sanger Institute COVID-19 Surveillance Team |
| EPI_ISL_2437522, EPI_ISL_2437523 | Lighthouse Lab in Alderley Park | Wellcome Sanger Institute for the COVID-19 Genomics UK (COG-UK) Consortium | Jacquelyn Wynn, Mairead Hyland, The Lighthouse Lab in Alderley Park and Alex Alderton, Roberto Amato, Jeffrey Barrett, Sonia Goncalves, Ewan Harrison, David K. Jackson, Ian Johnston, Dominic Kwiatkowski, Cordelia Langford, John Sillitoe on behalf of the Wellcome Sanger Institute COVID-19 Surveillance Team |
| EPI_ISL_2437524, EPI_ISL_2437525 | Lighthouse Lab in Glasgow | Wellcome Sanger Institute for the COVID-19 Genomics UK (COG-UK) Consortium | Harper VanSteenhouse, Yumi Kasai, David Gray, Carol Clugston, Anna Dominiczak and Alex Alderton, Roberto Amato, Jeffrey Barrett, Sonia Goncalves, Ewan Harrison, David K. Jackson, Ian Johnston, Dominic Kwiatkowski, Cordelia Langford, John Sillitoe on behalf of the Wellcome Sanger Institute COVID-19 Surveillance Team |
| EPI_ISL_2437526, EPI_ISL_2437527, EPI_ISL_2437529, | Lighthouse Lab in Alderley Park | Wellcome Sanger Institute for the COVID-19 Genomics UK (COG-UK) Consortium | Jacquelyn Wynn, Mairead Hyland, The Lighthouse Lab in Alderley Park and Alex Alderton, Roberto Amato, Jeffrey Barrett, Sonia Goncalves, Ewan Harrison, David K. Jackson, Ian Johnston, Dominic Kwiatkowski, Cordelia Langford, John Sillitoe on behalf of the Wellcome Sanger Institute COVID-19 Surveillance Team |

| | | | |
|---|---|---|---|
| EPI_ISL_2437531 | | | |
| EPI_ISL_2437533 | Lighthouse Lab in Milton Keynes | Wellcome Sanger Institute for the COVID-19 Genomics UK (COG-UK) Consortium | The Lighthouse Lab in Milton Keynes and Alex Alderton, Roberto Amato, Jeffrey Barrett, Sonia Goncalves, Ewan Harrison, David K. Jackson, Ian Johnston, Dominic Kwiatkowski, Cordelia Langford, John Sillitoe on behalf of the Wellcome Sanger Institute COVID-19 Surveillance Team |
| EPI_ISL_2437534, EPI_ISL_2437535 | Lighthouse Lab in Alderley Park | Wellcome Sanger Institute for the COVID-19 Genomics UK (COG-UK) Consortium | Jacquelyn Wynn, Mairead Hyland, The Lighthouse Lab in Alderley Park and Alex Alderton, Roberto Amato, Jeffrey Barrett, Sonia Goncalves, Ewan Harrison, David K. Jackson, Ian Johnston, Dominic Kwiatkowski, Cordelia Langford, John Sillitoe on behalf of the Wellcome Sanger Institute COVID-19 Surveillance Team |
| EPI_ISL_2437536 | Health Services Laboratories | Wellcome Sanger Institute for the COVID-19 Genomics UK (COG-UK) Consortium | Health Services Laboratories and Alex Alderton, Roberto Amato, Jeffrey Barrett, Sonia Goncalves, Ewan Harrison, David K. Jackson, Ian Johnston, Dominic Kwiatkowski, Cordelia Langford, John Sillitoe on behalf of the Wellcome Sanger Institute COVID-19 Surveillance Team |
| EPI_ISL_2437537, EPI_ISL_2437538 | Lighthouse Lab in Alderley Park | Wellcome Sanger Institute for the COVID-19 Genomics UK (COG-UK) Consortium | Jacquelyn Wynn, Mairead Hyland, The Lighthouse Lab in Alderley Park and Alex Alderton, Roberto Amato, Jeffrey Barrett, Sonia Goncalves, Ewan Harrison, David K. Jackson, Ian Johnston, Dominic Kwiatkowski, Cordelia Langford, John Sillitoe on behalf of the Wellcome Sanger Institute COVID-19 Surveillance Team |
| EPI_ISL_2437539 | Lighthouse Lab in Milton Keynes | Wellcome Sanger Institute for the COVID-19 Genomics UK (COG-UK) Consortium | The Lighthouse Lab in Milton Keynes and Alex Alderton, Roberto Amato, Jeffrey Barrett, Sonia Goncalves, Ewan Harrison, David K. Jackson, Ian Johnston, Dominic Kwiatkowski, Cordelia Langford, John Sillitoe on behalf of the Wellcome Sanger Institute COVID-19 Surveillance Team |
| EPI_ISL_2437541, EPI_ISL_2437543, EPI_ISL_2437544 | Lighthouse Lab in Alderley Park | Wellcome Sanger Institute for the COVID-19 Genomics UK (COG-UK) Consortium | Jacquelyn Wynn, Mairead Hyland, The Lighthouse Lab in Alderley Park and Alex Alderton, Roberto Amato, Jeffrey Barrett, Sonia Goncalves, Ewan Harrison, David K. Jackson, Ian Johnston, Dominic Kwiatkowski, Cordelia Langford, John Sillitoe on behalf of the Wellcome Sanger Institute COVID-19 Surveillance Team |
| EPI_ISL_2437545 | Lighthouse Laboratory Plymouth | Wellcome Sanger Institute for the COVID-19 Genomics UK (COG-UK) Consortium | Lighthouse Laboratory Plymouth and Alex Alderton, Roberto Amato, Jeffrey Barrett, Sonia Goncalves, Ewan Harrison, David K. Jackson, Ian Johnston, Dominic Kwiatkowski, Cordelia Langford, John Sillitoe on behalf of the Wellcome Sanger Institute COVID-19 Surveillance Team |
| EPI_ISL_2437546 | Lighthouse Lab in Milton Keynes | Wellcome Sanger Institute for the COVID-19 Genomics UK (COG-UK) Consortium | The Lighthouse Lab in Milton Keynes and Alex Alderton, Roberto Amato, Jeffrey Barrett, Sonia Goncalves, Ewan Harrison, David K. Jackson, Ian Johnston, Dominic Kwiatkowski, Cordelia Langford, John Sillitoe on behalf of the Wellcome Sanger Institute COVID-19 Surveillance Team |
| EPI_ISL_2437547 | Health Services Laboratories | Wellcome Sanger Institute for the COVID-19 Genomics UK (COG-UK) Consortium | Health Services Laboratories and Alex Alderton, Roberto Amato, Jeffrey Barrett, Sonia Goncalves, Ewan Harrison, David K. Jackson, Ian Johnston, Dominic Kwiatkowski, Cordelia Langford, John Sillitoe on behalf of the Wellcome Sanger Institute COVID-19 Surveillance Team |
| EPI_ISL_2437548 | Lighthouse Lab in Glasgow | Wellcome Sanger Institute for the COVID-19 Genomics UK (COG-UK) Consortium | Harper VanSteenhouse, Yumi Kasai, David Gray, Carol Clugston, Anna Dominiczak and Alex Alderton, Roberto Amato, Jeffrey Barrett, Sonia Goncalves, Ewan Harrison, David K. Jackson, Ian Johnston, Dominic Kwiatkowski, Cordelia Langford, John Sillitoe on behalf of the Wellcome Sanger Institute COVID-19 Surveillance Team |
| EPI_ISL_2437550, EPI_ISL_2437551 | Lighthouse Lab in Alderley Park | Wellcome Sanger Institute for the COVID-19 Genomics UK (COG-UK) Consortium | Jacquelyn Wynn, Mairead Hyland, The Lighthouse Lab in Alderley Park and Alex Alderton, Roberto Amato, Jeffrey Barrett, Sonia Goncalves, Ewan Harrison, David K. Jackson, Ian Johnston, Dominic Kwiatkowski, Cordelia Langford, John Sillitoe on behalf of the Wellcome Sanger Institute COVID-19 Surveillance Team |
| EPI_ISL_2437552 | Lighthouse Lab in Glasgow | Wellcome Sanger Institute for the COVID-19 Genomics UK (COG-UK) Consortium | Harper VanSteenhouse, Yumi Kasai, David Gray, Carol Clugston, Anna Dominiczak and Alex Alderton, Roberto Amato, Jeffrey Barrett, Sonia Goncalves, Ewan Harrison, David K. Jackson, Ian Johnston, Dominic Kwiatkowski, Cordelia Langford, John Sillitoe on behalf of the Wellcome Sanger Institute COVID-19 Surveillance Team |
| EPI_ISL_2437553 | Lighthouse Lab in Milton Keynes | Wellcome Sanger Institute for the COVID-19 Genomics UK (COG-UK) Consortium | The Lighthouse Lab in Milton Keynes and Alex Alderton, Roberto Amato, Jeffrey Barrett, Sonia Goncalves, Ewan Harrison, David K. Jackson, Ian Johnston, Dominic Kwiatkowski, Cordelia Langford, John Sillitoe on behalf of the Wellcome Sanger Institute COVID-19 Surveillance Team |
| EPI_ISL_2437554, EPI_ISL_2437555, EPI_ISL_2437558, EPI_ISL_2437560 | Lighthouse Lab in Alderley Park | Wellcome Sanger Institute for the COVID-19 Genomics UK (COG-UK) Consortium | Jacquelyn Wynn, Mairead Hyland, The Lighthouse Lab in Alderley Park and Alex Alderton, Roberto Amato, Jeffrey Barrett, Sonia Goncalves, Ewan Harrison, David K. Jackson, Ian Johnston, Dominic Kwiatkowski, Cordelia Langford, John Sillitoe on behalf of the Wellcome Sanger Institute COVID-19 Surveillance Team |
| EPI_ISL_2437561 | Lighthouse Lab in Milton Keynes | Wellcome Sanger Institute for the COVID-19 Genomics UK (COG-UK) Consortium | The Lighthouse Lab in Milton Keynes and Alex Alderton, Roberto Amato, Jeffrey Barrett, Sonia Goncalves, Ewan Harrison, David K. Jackson, Ian Johnston, Dominic Kwiatkowski, Cordelia Langford, John Sillitoe on behalf of the Wellcome Sanger Institute COVID-19 Surveillance Team |
| EPI_ISL_2437562 | Lighthouse Lab in Alderley Park | Wellcome Sanger Institute for the COVID-19 Genomics UK (COG-UK) Consortium | Jacquelyn Wynn, Mairead Hyland, The Lighthouse Lab in Alderley Park and Alex Alderton, Roberto Amato, Jeffrey Barrett, Sonia Goncalves, Ewan Harrison, David K. Jackson, Ian Johnston, Dominic Kwiatkowski, Cordelia Langford, John Sillitoe on behalf of the Wellcome Sanger Institute COVID-19 Surveillance Team |
| EPI_ISL_2437563 | Lighthouse Lab in Glasgow | Wellcome Sanger Institute for the COVID-19 Genomics UK (COG-UK) Consortium | Harper VanSteenhouse, Yumi Kasai, David Gray, Carol Clugston, Anna Dominiczak and Alex Alderton, Roberto Amato, Jeffrey Barrett, Sonia Goncalves, Ewan Harrison, David K. Jackson, Ian Johnston, Dominic Kwiatkowski, Cordelia Langford, John Sillitoe on behalf of the Wellcome Sanger Institute COVID-19 Surveillance Team |
| EPI_ISL_2437564 | Lighthouse Lab in Milton Keynes | Wellcome Sanger Institute for the COVID-19 Genomics UK (COG-UK) Consortium | The Lighthouse Lab in Milton Keynes and Alex Alderton, Roberto Amato, Jeffrey Barrett, Sonia Goncalves, Ewan Harrison, David K. Jackson, Ian Johnston, Dominic Kwiatkowski, Cordelia Langford, John Sillitoe on behalf of the Wellcome Sanger Institute COVID-19 Surveillance Team |
| EPI_ISL_2437565 | Lighthouse Laboratory Plymouth | Wellcome Sanger Institute for the COVID-19 Genomics UK (COG-UK) Consortium | Lighthouse Laboratory Plymouth and Alex Alderton, Roberto Amato, Jeffrey Barrett, Sonia Goncalves, Ewan Harrison, David K. Jackson, Ian Johnston, Dominic Kwiatkowski, Cordelia Langford, John Sillitoe on behalf of the Wellcome Sanger Institute COVID-19 Surveillance Team |
| EPI_ISL_2437566 | Lighthouse Lab in Glasgow | Wellcome Sanger Institute for the COVID-19 Genomics UK (COG-UK) Consortium | Harper VanSteenhouse, Yumi Kasai, David Gray, Carol Clugston, Anna Dominiczak and Alex Alderton, Roberto Amato, Jeffrey Barrett, Sonia Goncalves, Ewan Harrison, David K. Jackson, Ian Johnston, Dominic Kwiatkowski, Cordelia Langford, John Sillitoe on behalf of the Wellcome Sanger Institute COVID-19 Surveillance Team |
| EPI_ISL_2437567, EPI_ISL_2437568, EPI_ISL_2437569 | Lighthouse Lab in Alderley Park | Wellcome Sanger Institute for the COVID-19 Genomics UK (COG-UK) Consortium | Jacquelyn Wynn, Mairead Hyland, The Lighthouse Lab in Alderley Park and Alex Alderton, Roberto Amato, Jeffrey Barrett, Sonia Goncalves, Ewan Harrison, David K. Jackson, Ian Johnston, Dominic Kwiatkowski, Cordelia Langford, John Sillitoe on behalf of the Wellcome Sanger Institute COVID-19 Surveillance Team |
| EPI_ISL_2437572, EPI_ISL_2437573 | Lighthouse Lab in Glasgow | Wellcome Sanger Institute for the COVID-19 Genomics UK (COG-UK) Consortium | Harper VanSteenhouse, Yumi Kasai, David Gray, Carol Clugston, Anna Dominiczak and Alex Alderton, Roberto Amato, Jeffrey Barrett, Sonia Goncalves, Ewan Harrison, David K. Jackson, Ian Johnston, Dominic Kwiatkowski, Cordelia Langford, John Sillitoe on behalf of the Wellcome Sanger Institute COVID-19 Surveillance Team |
| EPI_ISL_2437574 | Lighthouse Lab in Alderley Park | Wellcome Sanger Institute for the COVID-19 Genomics UK (COG-UK) Consortium | Jacquelyn Wynn, Mairead Hyland, The Lighthouse Lab in Alderley Park and Alex Alderton, Roberto Amato, Jeffrey Barrett, Sonia Goncalves, Ewan Harrison, David K. Jackson, Ian Johnston, Dominic Kwiatkowski, Cordelia Langford, John Sillitoe on behalf of the Wellcome Sanger Institute COVID-19 Surveillance Team |
| EPI_ISL_2437575 | Lighthouse Lab in Milton Keynes | Wellcome Sanger Institute for the COVID-19 Genomics UK (COG-UK) Consortium | The Lighthouse Lab in Milton Keynes and Alex Alderton, Roberto Amato, Jeffrey Barrett, Sonia Goncalves, Ewan Harrison, David K. Jackson, Ian Johnston, Dominic Kwiatkowski, Cordelia Langford, John Sillitoe on behalf of the Wellcome Sanger Institute COVID-19 Surveillance Team |
| EPI_ISL_2437576, EPI_ISL_2437577 | Lighthouse Lab in Alderley Park | Wellcome Sanger Institute for the COVID-19 Genomics UK (COG-UK) Consortium | Jacquelyn Wynn, Mairead Hyland, The Lighthouse Lab in Alderley Park and Alex Alderton, Roberto Amato, Jeffrey Barrett, Sonia Goncalves, Ewan Harrison, David K. Jackson, Ian Johnston, Dominic Kwiatkowski, Cordelia Langford, John Sillitoe on behalf of the Wellcome Sanger Institute COVID-19 Surveillance Team |
| EPI_ISL_2437578 | Lighthouse Laboratory Plymouth | Wellcome Sanger Institute for the COVID-19 Genomics UK (COG-UK) Consortium | Lighthouse Laboratory Plymouth and Alex Alderton, Roberto Amato, Jeffrey Barrett, Sonia Goncalves, Ewan Harrison, David K. Jackson, Ian Johnston, Dominic Kwiatkowski, Cordelia Langford, John Sillitoe on behalf of the Wellcome Sanger Institute COVID-19 Surveillance Team |
| EPI_ISL_2437580, EPI_ISL_2437581, EPI_ISL_2437582 | Lighthouse Lab in Alderley Park | Wellcome Sanger Institute for the COVID-19 Genomics UK (COG-UK) Consortium | Jacquelyn Wynn, Mairead Hyland, The Lighthouse Lab in Alderley Park and Alex Alderton, Roberto Amato, Jeffrey Barrett, Sonia Goncalves, Ewan Harrison, David K. Jackson, Ian Johnston, Dominic Kwiatkowski, Cordelia Langford, John Sillitoe on behalf of the Wellcome Sanger Institute COVID-19 Surveillance Team |
| EPI_ISL_2437583 | Lighthouse Lab in Glasgow | Wellcome Sanger Institute for the COVID-19 Genomics UK (COG-UK) Consortium | Harper VanSteenhouse, Yumi Kasai, David Gray, Carol Clugston, Anna Dominiczak and Alex Alderton, Roberto Amato, Jeffrey Barrett, Sonia Goncalves, Ewan Harrison, David K. Jackson, Ian Johnston, Dominic Kwiatkowski, Cordelia Langford, John Sillitoe on behalf of the Wellcome Sanger Institute COVID-19 Surveillance Team |
| EPI_ISL_2437584, EPI_ISL_2437585 | Lighthouse Lab in Milton Keynes | Wellcome Sanger Institute for the COVID-19 Genomics UK (COG-UK) Consortium | The Lighthouse Lab in Milton Keynes and Alex Alderton, Roberto Amato, Jeffrey Barrett, Sonia Goncalves, Ewan Harrison, David K. Jackson, Ian Johnston, Dominic Kwiatkowski, Cordelia Langford, John Sillitoe on behalf of the Wellcome Sanger Institute COVID-19 Surveillance Team |
| EPI_ISL_2437586 | Lighthouse Lab in Glasgow | Wellcome Sanger Institute for the COVID-19 Genomics UK (COG-UK) Consortium | Harper VanSteenhouse, Yumi Kasai, David Gray, Carol Clugston, Anna Dominiczak and Alex Alderton, Roberto Amato, Jeffrey Barrett, Sonia Goncalves, Ewan Harrison, David K. Jackson, Ian Johnston, Dominic Kwiatkowski, Cordelia Langford, John Sillitoe on behalf of the Wellcome Sanger Institute COVID-19 Surveillance Team |
| EPI_ISL_2437587 | Lighthouse Lab in Alderley Park | Wellcome Sanger Institute for the COVID-19 Genomics UK (COG-UK) Consortium | Jacquelyn Wynn, Mairead Hyland, The Lighthouse Lab in Alderley Park and Alex Alderton, Roberto Amato, Jeffrey Barrett, Sonia Goncalves, Ewan Harrison, David K. Jackson, Ian Johnston, Dominic Kwiatkowski, Cordelia Langford, John Sillitoe on behalf of the Wellcome Sanger Institute COVID-19 Surveillance Team |
| EPI_ISL_2437588, EPI_ISL_2437589, EPI_ISL_2437590 | Lighthouse Laboratory Plymouth | Wellcome Sanger Institute for the COVID-19 Genomics UK (COG-UK) Consortium | Lighthouse Laboratory Plymouth and Alex Alderton, Roberto Amato, Jeffrey Barrett, Sonia Goncalves, Ewan Harrison, David K. Jackson, Ian Johnston, Dominic Kwiatkowski, Cordelia Langford, John Sillitoe on behalf of the Wellcome Sanger Institute COVID-19 Surveillance Team |
| EPI_ISL_2437592, EPI_ISL_2437593 | Lighthouse Lab in Alderley Park | Wellcome Sanger Institute for the COVID-19 Genomics UK (COG-UK) Consortium | Jacquelyn Wynn, Mairead Hyland, The Lighthouse Lab in Alderley Park and Alex Alderton, Roberto Amato, Jeffrey Barrett, Sonia Goncalves, Ewan Harrison, David K. Jackson, Ian Johnston, Dominic Kwiatkowski, Cordelia Langford, John Sillitoe on behalf of the Wellcome Sanger Institute COVID-19 Surveillance Team |
| EPI_ISL_2437594 | Lighthouse Laboratory Plymouth | Wellcome Sanger Institute for the COVID-19 Genomics UK (COG-UK) Consortium | Lighthouse Laboratory Plymouth and Alex Alderton, Roberto Amato, Jeffrey Barrett, Sonia Goncalves, Ewan Harrison, David K. Jackson, Ian Johnston, Dominic Kwiatkowski, Cordelia Langford, John Sillitoe on behalf of the Wellcome Sanger Institute COVID-19 Surveillance Team |
| EPI_ISL_2437595 | Lighthouse Lab in Alderley Park | Wellcome Sanger Institute for the COVID-19 Genomics UK (COG-UK) Consortium | Jacquelyn Wynn, Mairead Hyland, The Lighthouse Lab in Alderley Park and Alex Alderton, Roberto Amato, Jeffrey Barrett, Sonia Goncalves, Ewan Harrison, David K. Jackson, Ian Johnston, Dominic Kwiatkowski, Cordelia Langford, John Sillitoe on behalf of the Wellcome Sanger Institute COVID-19 Surveillance Team |
| EPI_ISL_2437596 | Health Services Laboratories | Wellcome Sanger Institute for the COVID-19 Genomics UK (COG-UK) Consortium | Health Services Laboratories and Alex Alderton, Roberto Amato, Jeffrey Barrett, Sonia Goncalves, Ewan Harrison, David K. Jackson, Ian Johnston, Dominic Kwiatkowski, Cordelia Langford, John Sillitoe on behalf of the Wellcome Sanger Institute COVID-19 Surveillance Team |
| EPI_ISL_2437597, EPI_ISL_2437599 | Lighthouse Lab in Alderley Park | Wellcome Sanger Institute for the COVID-19 Genomics UK (COG-UK) Consortium | Jacquelyn Wynn, Mairead Hyland, The Lighthouse Lab in Alderley Park and Alex Alderton, Roberto Amato, Jeffrey Barrett, Sonia Goncalves, Ewan Harrison, David K. Jackson, Ian Johnston, Dominic Kwiatkowski, Cordelia Langford, John Sillitoe on behalf of the Wellcome Sanger Institute COVID-19 Surveillance Team |
| EPI_ISL_2437600 | Lighthouse Lab in Glasgow | Wellcome Sanger Institute for the COVID-19 Genomics UK (COG-UK) Consortium | Harper VanSteenhouse, Yumi Kasai, David Gray, Carol Clugston, Anna Dominiczak and Alex Alderton, Roberto Amato, Jeffrey Barrett, Sonia Goncalves, Ewan Harrison, David K. Jackson, Ian Johnston, Dominic Kwiatkowski, Cordelia Langford, John Sillitoe on behalf of the Wellcome Sanger Institute COVID-19 Surveillance Team |
| EPI_ISL_2437601 | Lighthouse Lab in Alderley Park | Wellcome Sanger Institute for the COVID-19 Genomics UK (COG-UK) Consortium | Jacquelyn Wynn, Mairead Hyland, The Lighthouse Lab in Alderley Park and Alex Alderton, Roberto Amato, Jeffrey Barrett, Sonia Goncalves, Ewan Harrison, David K. Jackson, Ian Johnston, Dominic Kwiatkowski, Cordelia Langford, John Sillitoe on behalf of the Wellcome Sanger Institute COVID-19 Surveillance Team |
| EPI_ISL_2437604 | Lighthouse Lab in Milton Keynes | Wellcome Sanger Institute for the COVID-19 Genomics UK (COG-UK) Consortium | The Lighthouse Lab in Milton Keynes and Alex Alderton, Roberto Amato, Jeffrey Barrett, Sonia Goncalves, Ewan Harrison, David K. Jackson, Ian Johnston, Dominic Kwiatkowski, Cordelia Langford, John Sillitoe on behalf of the Wellcome Sanger Institute COVID-19 Surveillance Team |
| EPI_ISL_2437605, EPI_ISL_2437606 | Lighthouse Lab in Alderley Park | Wellcome Sanger Institute for the COVID-19 Genomics UK (COG-UK) Consortium | Jacquelyn Wynn, Mairead Hyland, The Lighthouse Lab in Alderley Park and Alex Alderton, Roberto Amato, Jeffrey Barrett, Sonia Goncalves, Ewan Harrison, David K. Jackson, Ian Johnston, Dominic Kwiatkowski, Cordelia Langford, John Sillitoe on behalf of the Wellcome Sanger Institute COVID-19 Surveillance Team |
| EPI_ISL_2437607 | Lighthouse Lab in Milton Keynes | Wellcome Sanger Institute for the COVID-19 Genomics UK (COG-UK) Consortium | The Lighthouse Lab in Milton Keynes and Alex Alderton, Roberto Amato, Jeffrey Barrett, Sonia Goncalves, Ewan Harrison, David K. Jackson, Ian Johnston, Dominic Kwiatkowski, Cordelia Langford, John Sillitoe on behalf of the Wellcome Sanger Institute COVID-19 Surveillance Team |

| | | | |
|---|---|---|---|
| EPI_ISL_2437609 | Lighthouse Lab in Alderley Park | Wellcome Sanger Institute for the COVID-19 Genomics UK (COG-UK) Consortium | Jacquelyn Wynn, Mairead Hyland, The Lighthouse Lab in Alderley Park and Alex Alderton, Roberto Amato, Jeffrey Barrett, Sonia Goncalves, Ewan Harrison, David K. Jackson, Ian Johnston, Dominic Kwiatkowski, Cordelia Langford, John Sillitoe on behalf of the Wellcome Sanger Institute COVID-19 Surveillance Team |
| EPI_ISL_2437610 | Lighthouse Lab in Glasgow | Wellcome Sanger Institute for the COVID-19 Genomics UK (COG-UK) Consortium | Harper VanSteenhouse, Yumi Kasai, David Gray, Carol Clugston, Anna Dominiczak and Alex Alderton, Roberto Amato, Jeffrey Barrett, Sonia Goncalves, Ewan Harrison, David K. Jackson, Ian Johnston, Dominic Kwiatkowski, Cordelia Langford, John Sillitoe on behalf of the Wellcome Sanger Institute COVID-19 Surveillance Team |
| EPI_ISL_2437611 | Lighthouse Laboratory Plymouth | Wellcome Sanger Institute for the COVID-19 Genomics UK (COG-UK) Consortium | Lighthouse Laboratory Plymouth and Alex Alderton, Roberto Amato, Jeffrey Barrett, Sonia Goncalves, Ewan Harrison, David K. Jackson, Ian Johnston, Dominic Kwiatkowski, Cordelia Langford, John Sillitoe on behalf of the Wellcome Sanger Institute COVID-19 Surveillance Team |
| EPI_ISL_2437612, EPI_ISL_2437613, EPI_ISL_2437614 | Lighthouse Lab in Alderley Park | Wellcome Sanger Institute for the COVID-19 Genomics UK (COG-UK) Consortium | Jacquelyn Wynn, Mairead Hyland, The Lighthouse Lab in Alderley Park and Alex Alderton, Roberto Amato, Jeffrey Barrett, Sonia Goncalves, Ewan Harrison, David K. Jackson, Ian Johnston, Dominic Kwiatkowski, Cordelia Langford, John Sillitoe on behalf of the Wellcome Sanger Institute COVID-19 Surveillance Team |
| EPI_ISL_2437615, EPI_ISL_2437616 | Lighthouse Lab in Milton Keynes | Wellcome Sanger Institute for the COVID-19 Genomics UK (COG-UK) Consortium | The Lighthouse Lab in Milton Keynes and Alex Alderton, Roberto Amato, Jeffrey Barrett, Sonia Goncalves, Ewan Harrison, David K. Jackson, Ian Johnston, Dominic Kwiatkowski, Cordelia Langford, John Sillitoe on behalf of the Wellcome Sanger Institute COVID-19 Surveillance Team |
| EPI_ISL_2437617 | Lighthouse Lab in Alderley Park | Wellcome Sanger Institute for the COVID-19 Genomics UK (COG-UK) Consortium | Jacquelyn Wynn, Mairead Hyland, The Lighthouse Lab in Alderley Park and Alex Alderton, Roberto Amato, Jeffrey Barrett, Sonia Goncalves, Ewan Harrison, David K. Jackson, Ian Johnston, Dominic Kwiatkowski, Cordelia Langford, John Sillitoe on behalf of the Wellcome Sanger Institute COVID-19 Surveillance Team |
| EPI_ISL_2437618, EPI_ISL_2437619 | Lighthouse Lab in Milton Keynes | Wellcome Sanger Institute for the COVID-19 Genomics UK (COG-UK) Consortium | The Lighthouse Lab in Milton Keynes and Alex Alderton, Roberto Amato, Jeffrey Barrett, Sonia Goncalves, Ewan Harrison, David K. Jackson, Ian Johnston, Dominic Kwiatkowski, Cordelia Langford, John Sillitoe on behalf of the Wellcome Sanger Institute COVID-19 Surveillance Team |
| EPI_ISL_2437620 | Lighthouse Lab in Glasgow | Wellcome Sanger Institute for the COVID-19 Genomics UK (COG-UK) Consortium | Harper VanSteenhouse, Yumi Kasai, David Gray, Carol Clugston, Anna Dominiczak and Alex Alderton, Roberto Amato, Jeffrey Barrett, Sonia Goncalves, Ewan Harrison, David K. Jackson, Ian Johnston, Dominic Kwiatkowski, Cordelia Langford, John Sillitoe on behalf of the Wellcome Sanger Institute COVID-19 Surveillance Team |
| EPI_ISL_2437621, EPI_ISL_2437622 | Lighthouse Lab in Alderley Park | Wellcome Sanger Institute for the COVID-19 Genomics UK (COG-UK) Consortium | Jacquelyn Wynn, Mairead Hyland, The Lighthouse Lab in Alderley Park and Alex Alderton, Roberto Amato, Jeffrey Barrett, Sonia Goncalves, Ewan Harrison, David K. Jackson, Ian Johnston, Dominic Kwiatkowski, Cordelia Langford, John Sillitoe on behalf of the Wellcome Sanger Institute COVID-19 Surveillance Team |
| EPI_ISL_2437623, EPI_ISL_2437624, EPI_ISL_2437625, EPI_ISL_2437626, EPI_ISL_2437627, EPI_ISL_2437628, EPI_ISL_2437629, EPI_ISL_2437631 | Lighthouse Lab in Milton Keynes | Wellcome Sanger Institute for the COVID-19 Genomics UK (COG-UK) Consortium | The Lighthouse Lab in Milton Keynes and Alex Alderton, Roberto Amato, Jeffrey Barrett, Sonia Goncalves, Ewan Harrison, David K. Jackson, Ian Johnston, Dominic Kwiatkowski, Cordelia Langford, John Sillitoe on behalf of the Wellcome Sanger Institute COVID-19 Surveillance Team |
| EPI_ISL_2437632 | Lighthouse Lab in Alderley Park | Wellcome Sanger Institute for the COVID-19 Genomics UK (COG-UK) Consortium | Jacquelyn Wynn, Mairead Hyland, The Lighthouse Lab in Alderley Park and Alex Alderton, Roberto Amato, Jeffrey Barrett, Sonia Goncalves, Ewan Harrison, David K. Jackson, Ian Johnston, Dominic Kwiatkowski, Cordelia Langford, John Sillitoe on behalf of the Wellcome Sanger Institute COVID-19 Surveillance Team |
| EPI_ISL_2437633 | Lighthouse Lab in Milton Keynes | Wellcome Sanger Institute for the COVID-19 Genomics UK (COG-UK) Consortium | The Lighthouse Lab in Milton Keynes and Alex Alderton, Roberto Amato, Jeffrey Barrett, Sonia Goncalves, Ewan Harrison, David K. Jackson, Ian Johnston, Dominic Kwiatkowski, Cordelia Langford, John Sillitoe on behalf of the Wellcome Sanger Institute COVID-19 Surveillance Team |
| EPI_ISL_2437634, EPI_ISL_2437635 | Lighthouse Lab in Glasgow | Wellcome Sanger Institute for the COVID-19 Genomics UK (COG-UK) Consortium | Harper VanSteenhouse, Yumi Kasai, David Gray, Carol Clugston, Anna Dominiczak and Alex Alderton, Roberto Amato, Jeffrey Barrett, Sonia Goncalves, Ewan Harrison, David K. Jackson, Ian Johnston, Dominic Kwiatkowski, Cordelia Langford, John Sillitoe on behalf of the Wellcome Sanger Institute COVID-19 Surveillance Team |
| EPI_ISL_2437636 | Lighthouse Lab in Milton Keynes | Wellcome Sanger Institute for the COVID-19 Genomics UK (COG-UK) Consortium | The Lighthouse Lab in Milton Keynes and Alex Alderton, Roberto Amato, Jeffrey Barrett, Sonia Goncalves, Ewan Harrison, David K. Jackson, Ian Johnston, Dominic Kwiatkowski, Cordelia Langford, John Sillitoe on behalf of the Wellcome Sanger Institute COVID-19 Surveillance Team |
| EPI_ISL_2437637 | Lighthouse Lab in Alderley Park | Wellcome Sanger Institute for the COVID-19 Genomics UK (COG-UK) Consortium | Jacquelyn Wynn, Mairead Hyland, The Lighthouse Lab in Alderley Park and Alex Alderton, Roberto Amato, Jeffrey Barrett, Sonia Goncalves, Ewan Harrison, David K. Jackson, Ian Johnston, Dominic Kwiatkowski, Cordelia Langford, John Sillitoe on behalf of the Wellcome Sanger Institute COVID-19 Surveillance Team |
| EPI_ISL_2437638, EPI_ISL_2437640 | Lighthouse Lab in Glasgow | Wellcome Sanger Institute for the COVID-19 Genomics UK (COG-UK) Consortium | Harper VanSteenhouse, Yumi Kasai, David Gray, Carol Clugston, Anna Dominiczak and Alex Alderton, Roberto Amato, Jeffrey Barrett, Sonia Goncalves, Ewan Harrison, David K. Jackson, Ian Johnston, Dominic Kwiatkowski, Cordelia Langford, John Sillitoe on behalf of the Wellcome Sanger Institute COVID-19 Surveillance Team |
| EPI_ISL_2437641, EPI_ISL_2437642 | Lighthouse Lab in Alderley Park | Wellcome Sanger Institute for the COVID-19 Genomics UK (COG-UK) Consortium | Jacquelyn Wynn, Mairead Hyland, The Lighthouse Lab in Alderley Park and Alex Alderton, Roberto Amato, Jeffrey Barrett, Sonia Goncalves, Ewan Harrison, David K. Jackson, Ian Johnston, Dominic Kwiatkowski, Cordelia Langford, John Sillitoe on behalf of the Wellcome Sanger Institute COVID-19 Surveillance Team |
| EPI_ISL_2437643 | Lighthouse Lab in Milton Keynes | Wellcome Sanger Institute for the COVID-19 Genomics UK (COG-UK) Consortium | The Lighthouse Lab in Milton Keynes and Alex Alderton, Roberto Amato, Jeffrey Barrett, Sonia Goncalves, Ewan Harrison, David K. Jackson, Ian Johnston, Dominic Kwiatkowski, Cordelia Langford, John Sillitoe on behalf of the Wellcome Sanger Institute COVID-19 Surveillance Team |
| EPI_ISL_2437645 | Lighthouse Lab in Glasgow | Wellcome Sanger Institute for the COVID-19 Genomics UK (COG-UK) Consortium | Harper VanSteenhouse, Yumi Kasai, David Gray, Carol Clugston, Anna Dominiczak and Alex Alderton, Roberto Amato, Jeffrey Barrett, Sonia Goncalves, Ewan Harrison, David K. Jackson, Ian Johnston, Dominic Kwiatkowski, Cordelia Langford, John Sillitoe on behalf of the Wellcome Sanger Institute COVID-19 Surveillance Team |
| EPI_ISL_2437646, EPI_ISL_2437647 | Lighthouse Lab in Milton Keynes | Wellcome Sanger Institute for the COVID-19 Genomics UK (COG-UK) Consortium | The Lighthouse Lab in Milton Keynes and Alex Alderton, Roberto Amato, Jeffrey Barrett, Sonia Goncalves, Ewan Harrison, David K. Jackson, Ian Johnston, Dominic Kwiatkowski, Cordelia Langford, John Sillitoe on behalf of the Wellcome Sanger Institute COVID-19 Surveillance Team |
| EPI_ISL_2437648, EPI_ISL_2437650 | Lighthouse Lab in Glasgow | Wellcome Sanger Institute for the COVID-19 Genomics UK (COG-UK) Consortium | Harper VanSteenhouse, Yumi Kasai, David Gray, Carol Clugston, Anna Dominiczak and Alex Alderton, Roberto Amato, Jeffrey Barrett, Sonia Goncalves, Ewan Harrison, David K. Jackson, Ian Johnston, Dominic Kwiatkowski, Cordelia Langford, John Sillitoe on behalf of the Wellcome Sanger Institute COVID-19 Surveillance Team |
| EPI_ISL_2437651 | Lighthouse Lab in Alderley Park | Wellcome Sanger Institute for the COVID-19 Genomics UK (COG-UK) Consortium | Jacquelyn Wynn, Mairead Hyland, The Lighthouse Lab in Alderley Park and Alex Alderton, Roberto Amato, Jeffrey Barrett, Sonia Goncalves, Ewan Harrison, David K. Jackson, Ian Johnston, Dominic Kwiatkowski, Cordelia Langford, John Sillitoe on behalf of the Wellcome Sanger Institute COVID-19 Surveillance Team |
| EPI_ISL_2437652 | Lighthouse Lab in Glasgow | Wellcome Sanger Institute for the COVID-19 Genomics UK (COG-UK) Consortium | Harper VanSteenhouse, Yumi Kasai, David Gray, Carol Clugston, Anna Dominiczak and Alex Alderton, Roberto Amato, Jeffrey Barrett, Sonia Goncalves, Ewan Harrison, David K. Jackson, Ian Johnston, Dominic Kwiatkowski, Cordelia Langford, John Sillitoe on behalf of the Wellcome Sanger Institute COVID-19 Surveillance Team |
| EPI_ISL_2437653 | Lighthouse Lab in Milton Keynes | Wellcome Sanger Institute for the COVID-19 Genomics UK (COG-UK) Consortium | The Lighthouse Lab in Milton Keynes and Alex Alderton, Roberto Amato, Jeffrey Barrett, Sonia Goncalves, Ewan Harrison, David K. Jackson, Ian Johnston, Dominic Kwiatkowski, Cordelia Langford, John Sillitoe on behalf of the Wellcome Sanger Institute COVID-19 Surveillance Team |
| EPI_ISL_2437654 | Lighthouse Lab in Alderley Park | Wellcome Sanger Institute for the COVID-19 Genomics UK (COG-UK) Consortium | Jacquelyn Wynn, Mairead Hyland, The Lighthouse Lab in Alderley Park and Alex Alderton, Roberto Amato, Jeffrey Barrett, Sonia Goncalves, Ewan Harrison, David K. Jackson, Ian Johnston, Dominic Kwiatkowski, Cordelia Langford, John Sillitoe on behalf of the Wellcome Sanger Institute COVID-19 Surveillance Team |
| EPI_ISL_2437655 | Lighthouse Lab in Milton Keynes | Wellcome Sanger Institute for the COVID-19 Genomics UK (COG-UK) Consortium | The Lighthouse Lab in Milton Keynes and Alex Alderton, Roberto Amato, Jeffrey Barrett, Sonia Goncalves, Ewan Harrison, David K. Jackson, Ian Johnston, Dominic Kwiatkowski, Cordelia Langford, John Sillitoe on behalf of the Wellcome Sanger Institute COVID-19 Surveillance Team |
| EPI_ISL_2437656, EPI_ISL_2437657 | Lighthouse Lab in Glasgow | Wellcome Sanger Institute for the COVID-19 Genomics UK (COG-UK) Consortium | Harper VanSteenhouse, Yumi Kasai, David Gray, Carol Clugston, Anna Dominiczak and Alex Alderton, Roberto Amato, Jeffrey Barrett, Sonia Goncalves, Ewan Harrison, David K. Jackson, Ian Johnston, Dominic Kwiatkowski, Cordelia Langford, John Sillitoe on behalf of the Wellcome Sanger Institute COVID-19 Surveillance Team |
| EPI_ISL_2437658 | Lighthouse Lab in Milton Keynes | Wellcome Sanger Institute for the COVID-19 Genomics UK (COG-UK) Consortium | The Lighthouse Lab in Milton Keynes and Alex Alderton, Roberto Amato, Jeffrey Barrett, Sonia Goncalves, Ewan Harrison, David K. Jackson, Ian Johnston, Dominic Kwiatkowski, Cordelia Langford, John Sillitoe on behalf of the Wellcome Sanger Institute COVID-19 Surveillance Team |
| EPI_ISL_2437659, EPI_ISL_2437660 | Lighthouse Lab in Glasgow | Wellcome Sanger Institute for the COVID-19 Genomics UK (COG-UK) Consortium | Harper VanSteenhouse, Yumi Kasai, David Gray, Carol Clugston, Anna Dominiczak and Alex Alderton, Roberto Amato, Jeffrey Barrett, Sonia Goncalves, Ewan Harrison, David K. Jackson, Ian Johnston, Dominic Kwiatkowski, Cordelia Langford, John Sillitoe on behalf of the Wellcome Sanger Institute COVID-19 Surveillance Team |
| EPI_ISL_2437662, EPI_ISL_2437663 | Lighthouse Lab in Milton Keynes | Wellcome Sanger Institute for the COVID-19 Genomics UK (COG-UK) Consortium | The Lighthouse Lab in Milton Keynes and Alex Alderton, Roberto Amato, Jeffrey Barrett, Sonia Goncalves, Ewan Harrison, David K. Jackson, Ian Johnston, Dominic Kwiatkowski, Cordelia Langford, John Sillitoe on behalf of the Wellcome Sanger Institute COVID-19 Surveillance Team |
| EPI_ISL_2437664 | Lighthouse Lab in Glasgow | Wellcome Sanger Institute for the COVID-19 Genomics UK (COG-UK) Consortium | Harper VanSteenhouse, Yumi Kasai, David Gray, Carol Clugston, Anna Dominiczak and Alex Alderton, Roberto Amato, Jeffrey Barrett, Sonia Goncalves, Ewan Harrison, David K. Jackson, Ian Johnston, Dominic Kwiatkowski, Cordelia Langford, John Sillitoe on behalf of the Wellcome Sanger Institute COVID-19 Surveillance Team |
| EPI_ISL_2437665, EPI_ISL_2437666 | Lighthouse Lab in Milton Keynes | Wellcome Sanger Institute for the COVID-19 Genomics UK (COG-UK) Consortium | The Lighthouse Lab in Milton Keynes and Alex Alderton, Roberto Amato, Jeffrey Barrett, Sonia Goncalves, Ewan Harrison, David K. Jackson, Ian Johnston, Dominic Kwiatkowski, Cordelia Langford, John Sillitoe on behalf of the Wellcome Sanger Institute COVID-19 Surveillance Team |
| EPI_ISL_2437667 | Lighthouse Lab in Glasgow | Wellcome Sanger Institute for the COVID-19 Genomics UK (COG-UK) Consortium | Harper VanSteenhouse, Yumi Kasai, David Gray, Carol Clugston, Anna Dominiczak and Alex Alderton, Roberto Amato, Jeffrey Barrett, Sonia Goncalves, Ewan Harrison, David K. Jackson, Ian Johnston, Dominic Kwiatkowski, Cordelia Langford, John Sillitoe on behalf of the Wellcome Sanger Institute COVID-19 Surveillance Team |
| EPI_ISL_2437668 | Lighthouse Lab in Alderley Park | Wellcome Sanger Institute for the COVID-19 Genomics UK (COG-UK) Consortium | Jacquelyn Wynn, Mairead Hyland, The Lighthouse Lab in Alderley Park and Alex Alderton, Roberto Amato, Jeffrey Barrett, Sonia Goncalves, Ewan Harrison, David K. Jackson, Ian Johnston, Dominic Kwiatkowski, Cordelia Langford, John Sillitoe on behalf of the Wellcome Sanger Institute COVID-19 Surveillance Team |
| EPI_ISL_2437669, EPI_ISL_2437670 | Lighthouse Lab in Glasgow | Wellcome Sanger Institute for the COVID-19 Genomics UK (COG-UK) Consortium | Harper VanSteenhouse, Yumi Kasai, David Gray, Carol Clugston, Anna Dominiczak and Alex Alderton, Roberto Amato, Jeffrey Barrett, Sonia Goncalves, Ewan Harrison, David K. Jackson, Ian Johnston, Dominic Kwiatkowski, Cordelia Langford, John Sillitoe on behalf of the Wellcome Sanger Institute COVID-19 Surveillance Team |
| EPI_ISL_2437671 | Lighthouse Lab in Alderley Park | Wellcome Sanger Institute for the COVID-19 Genomics UK (COG-UK) Consortium | Jacquelyn Wynn, Mairead Hyland, The Lighthouse Lab in Alderley Park and Alex Alderton, Roberto Amato, Jeffrey Barrett, Sonia Goncalves, Ewan Harrison, David K. Jackson, Ian Johnston, Dominic Kwiatkowski, Cordelia Langford, John Sillitoe on behalf of the Wellcome Sanger Institute COVID-19 Surveillance Team |
| EPI_ISL_2437672 | Lighthouse Lab in Glasgow | Wellcome Sanger Institute for the COVID-19 Genomics UK (COG-UK) Consortium | Harper VanSteenhouse, Yumi Kasai, David Gray, Carol Clugston, Anna Dominiczak and Alex Alderton, Roberto Amato, Jeffrey Barrett, Sonia Goncalves, Ewan Harrison, David K. Jackson, Ian Johnston, Dominic Kwiatkowski, Cordelia Langford, John Sillitoe on behalf of the Wellcome Sanger Institute COVID-19 Surveillance Team |
| EPI_ISL_2437673, EPI_ISL_2437674 | Lighthouse Lab in Milton Keynes | Wellcome Sanger Institute for the COVID-19 Genomics UK (COG-UK) Consortium | The Lighthouse Lab in Milton Keynes and Alex Alderton, Roberto Amato, Jeffrey Barrett, Sonia Goncalves, Ewan Harrison, David K. Jackson, Ian Johnston, Dominic Kwiatkowski, Cordelia Langford, John Sillitoe on behalf of the Wellcome Sanger Institute COVID-19 Surveillance Team |
| EPI_ISL_2437676 | Lighthouse Lab in Alderley Park | Wellcome Sanger Institute for the COVID-19 Genomics UK (COG-UK) Consortium | Jacquelyn Wynn, Mairead Hyland, The Lighthouse Lab in Alderley Park and Alex Alderton, Roberto Amato, Jeffrey Barrett, Sonia Goncalves, Ewan Harrison, David K. Jackson, Ian Johnston, Dominic Kwiatkowski, Cordelia Langford, John Sillitoe on behalf of the Wellcome Sanger Institute COVID-19 Surveillance Team |
| EPI_ISL_2437678, EPI_ISL_2437679, EPI_ISL_2437680, EPI_ISL_2437681 | Lighthouse Lab in Milton Keynes | Wellcome Sanger Institute for the COVID-19 Genomics UK (COG-UK) Consortium | The Lighthouse Lab in Milton Keynes and Alex Alderton, Roberto Amato, Jeffrey Barrett, Sonia Goncalves, Ewan Harrison, David K. Jackson, Ian Johnston, Dominic Kwiatkowski, Cordelia Langford, John Sillitoe on behalf of the Wellcome Sanger Institute COVID-19 Surveillance Team |
| EPI_ISL_2437682 | Lighthouse Lab in Alderley Park | Wellcome Sanger Institute for the COVID-19 Genomics UK (COG-UK) Consortium | Jacquelyn Wynn, Mairead Hyland, The Lighthouse Lab in Alderley Park and Alex Alderton, Roberto Amato, Jeffrey Barrett, Sonia Goncalves, Ewan Harrison, David K. Jackson, Ian Johnston, Dominic Kwiatkowski, Cordelia Langford, John Sillitoe on behalf of the Wellcome Sanger Institute COVID-19 Surveillance Team |

| | | | |
|---|---|---|---|
| EPI_ISL_2437683, EPI_ISL_2437684, EPI_ISL_2437685 | Lighthouse Lab in Milton Keynes | Wellcome Sanger Institute for the COVID-19 Genomics UK (COG-UK) Consortium | The Lighthouse Lab in Milton Keynes and Alex Alderton, Roberto Amato, Jeffrey Barrett, Sonia Goncalves, Ewan Harrison, David K. Jackson, Ian Johnston, Dominic Kwiatkowski, Cordelia Langford, John Sillitoe on behalf of the Wellcome Sanger Institute COVID-19 Surveillance Team |
| EPI_ISL_2437686 | Lighthouse Lab in Alderley Park | Wellcome Sanger Institute for the COVID-19 Genomics UK (COG-UK) Consortium | Jacquelyn Wynn, Mairead Hyland, The Lighthouse Lab in Alderley Park and Alex Alderton, Roberto Amato, Jeffrey Barrett, Sonia Goncalves, Ewan Harrison, David K. Jackson, Ian Johnston, Dominic Kwiatkowski, Cordelia Langford, John Sillitoe on behalf of the Wellcome Sanger Institute COVID-19 Surveillance Team |
| EPI_ISL_2437687 | Lighthouse Lab in Glasgow | Wellcome Sanger Institute for the COVID-19 Genomics UK (COG-UK) Consortium | Harper VanSteenhouse, Yumi Kasai, David Gray, Carol Clugston, Anna Dominiczak and Alex Alderton, Roberto Amato, Jeffrey Barrett, Sonia Goncalves, Ewan Harrison, David K. Jackson, Ian Johnston, Dominic Kwiatkowski, Cordelia Langford, John Sillitoe on behalf of the Wellcome Sanger Institute COVID-19 Surveillance Team |
| EPI_ISL_2437688 | Lighthouse Lab in Milton Keynes | Wellcome Sanger Institute for the COVID-19 Genomics UK (COG-UK) Consortium | The Lighthouse Lab in Milton Keynes and Alex Alderton, Roberto Amato, Jeffrey Barrett, Sonia Goncalves, Ewan Harrison, David K. Jackson, Ian Johnston, Dominic Kwiatkowski, Cordelia Langford, John Sillitoe on behalf of the Wellcome Sanger Institute COVID-19 Surveillance Team |
| EPI_ISL_2437689 | Lighthouse Lab in Glasgow | Wellcome Sanger Institute for the COVID-19 Genomics UK (COG-UK) Consortium | Harper VanSteenhouse, Yumi Kasai, David Gray, Carol Clugston, Anna Dominiczak and Alex Alderton, Roberto Amato, Jeffrey Barrett, Sonia Goncalves, Ewan Harrison, David K. Jackson, Ian Johnston, Dominic Kwiatkowski, Cordelia Langford, John Sillitoe on behalf of the Wellcome Sanger Institute COVID-19 Surveillance Team |
| EPI_ISL_2437690 | Lighthouse Lab in Milton Keynes | Wellcome Sanger Institute for the COVID-19 Genomics UK (COG-UK) Consortium | The Lighthouse Lab in Milton Keynes and Alex Alderton, Roberto Amato, Jeffrey Barrett, Sonia Goncalves, Ewan Harrison, David K. Jackson, Ian Johnston, Dominic Kwiatkowski, Cordelia Langford, John Sillitoe on behalf of the Wellcome Sanger Institute COVID-19 Surveillance Team |
| EPI_ISL_2437692 | Lighthouse Lab in Glasgow | Wellcome Sanger Institute for the COVID-19 Genomics UK (COG-UK) Consortium | Harper VanSteenhouse, Yumi Kasai, David Gray, Carol Clugston, Anna Dominiczak and Alex Alderton, Roberto Amato, Jeffrey Barrett, Sonia Goncalves, Ewan Harrison, David K. Jackson, Ian Johnston, Dominic Kwiatkowski, Cordelia Langford, John Sillitoe on behalf of the Wellcome Sanger Institute COVID-19 Surveillance Team |
| EPI_ISL_2437693 | Lighthouse Lab in Milton Keynes | Wellcome Sanger Institute for the COVID-19 Genomics UK (COG-UK) Consortium | The Lighthouse Lab in Milton Keynes and Alex Alderton, Roberto Amato, Jeffrey Barrett, Sonia Goncalves, Ewan Harrison, David K. Jackson, Ian Johnston, Dominic Kwiatkowski, Cordelia Langford, John Sillitoe on behalf of the Wellcome Sanger Institute COVID-19 Surveillance Team |
| EPI_ISL_2437694 | Lighthouse Lab in Glasgow | Wellcome Sanger Institute for the COVID-19 Genomics UK (COG-UK) Consortium | Harper VanSteenhouse, Yumi Kasai, David Gray, Carol Clugston, Anna Dominiczak and Alex Alderton, Roberto Amato, Jeffrey Barrett, Sonia Goncalves, Ewan Harrison, David K. Jackson, Ian Johnston, Dominic Kwiatkowski, Cordelia Langford, John Sillitoe on behalf of the Wellcome Sanger Institute COVID-19 Surveillance Team |
| EPI_ISL_2437695, EPI_ISL_2437697, EPI_ISL_2437699 | Lighthouse Lab in Milton Keynes | Wellcome Sanger Institute for the COVID-19 Genomics UK (COG-UK) Consortium | The Lighthouse Lab in Milton Keynes and Alex Alderton, Roberto Amato, Jeffrey Barrett, Sonia Goncalves, Ewan Harrison, David K. Jackson, Ian Johnston, Dominic Kwiatkowski, Cordelia Langford, John Sillitoe on behalf of the Wellcome Sanger Institute COVID-19 Surveillance Team |
| EPI_ISL_2437700 | Lighthouse Lab in Glasgow | Wellcome Sanger Institute for the COVID-19 Genomics UK (COG-UK) Consortium | Harper VanSteenhouse, Yumi Kasai, David Gray, Carol Clugston, Anna Dominiczak and Alex Alderton, Roberto Amato, Jeffrey Barrett, Sonia Goncalves, Ewan Harrison, David K. Jackson, Ian Johnston, Dominic Kwiatkowski, Cordelia Langford, John Sillitoe on behalf of the Wellcome Sanger Institute COVID-19 Surveillance Team |
| EPI_ISL_2437701 | Lighthouse Lab in Milton Keynes | Wellcome Sanger Institute for the COVID-19 Genomics UK (COG-UK) Consortium | The Lighthouse Lab in Milton Keynes and Alex Alderton, Roberto Amato, Jeffrey Barrett, Sonia Goncalves, Ewan Harrison, David K. Jackson, Ian Johnston, Dominic Kwiatkowski, Cordelia Langford, John Sillitoe on behalf of the Wellcome Sanger Institute COVID-19 Surveillance Team |
| EPI_ISL_2437702 | Lighthouse Lab in Glasgow | Wellcome Sanger Institute for the COVID-19 Genomics UK (COG-UK) Consortium | Harper VanSteenhouse, Yumi Kasai, David Gray, Carol Clugston, Anna Dominiczak and Alex Alderton, Roberto Amato, Jeffrey Barrett, Sonia Goncalves, Ewan Harrison, David K. Jackson, Ian Johnston, Dominic Kwiatkowski, Cordelia Langford, John Sillitoe on behalf of the Wellcome Sanger Institute COVID-19 Surveillance Team |
| EPI_ISL_2437703 | Lighthouse Lab in Alderley Park | Wellcome Sanger Institute for the COVID-19 Genomics UK (COG-UK) Consortium | Jacquelyn Wynn, Mairead Hyland, The Lighthouse Lab in Alderley Park and Alex Alderton, Roberto Amato, Jeffrey Barrett, Sonia Goncalves, Ewan Harrison, David K. Jackson, Ian Johnston, Dominic Kwiatkowski, Cordelia Langford, John Sillitoe on behalf of the Wellcome Sanger Institute COVID-19 Surveillance Team |
| EPI_ISL_2437704 | Lighthouse Lab in Milton Keynes | Wellcome Sanger Institute for the COVID-19 Genomics UK (COG-UK) Consortium | The Lighthouse Lab in Milton Keynes and Alex Alderton, Roberto Amato, Jeffrey Barrett, Sonia Goncalves, Ewan Harrison, David K. Jackson, Ian Johnston, Dominic Kwiatkowski, Cordelia Langford, John Sillitoe on behalf of the Wellcome Sanger Institute COVID-19 Surveillance Team |
| EPI_ISL_2437705 | Lighthouse Lab in Glasgow | Wellcome Sanger Institute for the COVID-19 Genomics UK (COG-UK) Consortium | Harper VanSteenhouse, Yumi Kasai, David Gray, Carol Clugston, Anna Dominiczak and Alex Alderton, Roberto Amato, Jeffrey Barrett, Sonia Goncalves, Ewan Harrison, David K. Jackson, Ian Johnston, Dominic Kwiatkowski, Cordelia Langford, John Sillitoe on behalf of the Wellcome Sanger Institute COVID-19 Surveillance Team |
| EPI_ISL_2437707, EPI_ISL_2437708 | Lighthouse Lab in Milton Keynes | Wellcome Sanger Institute for the COVID-19 Genomics UK (COG-UK) Consortium | The Lighthouse Lab in Milton Keynes and Alex Alderton, Roberto Amato, Jeffrey Barrett, Sonia Goncalves, Ewan Harrison, David K. Jackson, Ian Johnston, Dominic Kwiatkowski, Cordelia Langford, John Sillitoe on behalf of the Wellcome Sanger Institute COVID-19 Surveillance Team |
| EPI_ISL_2437709 | Lighthouse Lab in Glasgow | Wellcome Sanger Institute for the COVID-19 Genomics UK (COG-UK) Consortium | Harper VanSteenhouse, Yumi Kasai, David Gray, Carol Clugston, Anna Dominiczak and Alex Alderton, Roberto Amato, Jeffrey Barrett, Sonia Goncalves, Ewan Harrison, David K. Jackson, Ian Johnston, Dominic Kwiatkowski, Cordelia Langford, John Sillitoe on behalf of the Wellcome Sanger Institute COVID-19 Surveillance Team |
| EPI_ISL_2437711, EPI_ISL_2437712, EPI_ISL_2437713 | Lighthouse Lab in Milton Keynes | Wellcome Sanger Institute for the COVID-19 Genomics UK (COG-UK) Consortium | The Lighthouse Lab in Milton Keynes and Alex Alderton, Roberto Amato, Jeffrey Barrett, Sonia Goncalves, Ewan Harrison, David K. Jackson, Ian Johnston, Dominic Kwiatkowski, Cordelia Langford, John Sillitoe on behalf of the Wellcome Sanger Institute COVID-19 Surveillance Team |
| EPI_ISL_2437714 | Lighthouse Lab in Alderley Park | Wellcome Sanger Institute for the COVID-19 Genomics UK (COG-UK) Consortium | Jacquelyn Wynn, Mairead Hyland, The Lighthouse Lab in Alderley Park and Alex Alderton, Roberto Amato, Jeffrey Barrett, Sonia Goncalves, Ewan Harrison, David K. Jackson, Ian Johnston, Dominic Kwiatkowski, Cordelia Langford, John Sillitoe on behalf of the Wellcome Sanger Institute COVID-19 Surveillance Team |
| EPI_ISL_2437715 | Lighthouse Lab in Glasgow | Wellcome Sanger Institute for the COVID-19 Genomics UK (COG-UK) Consortium | Harper VanSteenhouse, Yumi Kasai, David Gray, Carol Clugston, Anna Dominiczak and Alex Alderton, Roberto Amato, Jeffrey Barrett, Sonia Goncalves, Ewan Harrison, David K. Jackson, Ian Johnston, Dominic Kwiatkowski, Cordelia Langford, John Sillitoe on behalf of the Wellcome Sanger Institute COVID-19 Surveillance Team |
| EPI_ISL_2437716 | Lighthouse Lab in Milton Keynes | Wellcome Sanger Institute for the COVID-19 Genomics UK (COG-UK) Consortium | The Lighthouse Lab in Milton Keynes and Alex Alderton, Roberto Amato, Jeffrey Barrett, Sonia Goncalves, Ewan Harrison, David K. Jackson, Ian Johnston, Dominic Kwiatkowski, Cordelia Langford, John Sillitoe on behalf of the Wellcome Sanger Institute COVID-19 Surveillance Team |
| EPI_ISL_2437718 | Lighthouse Lab in Alderley Park | Wellcome Sanger Institute for the COVID-19 Genomics UK (COG-UK) Consortium | Jacquelyn Wynn, Mairead Hyland, The Lighthouse Lab in Alderley Park and Alex Alderton, Roberto Amato, Jeffrey Barrett, Sonia Goncalves, Ewan Harrison, David K. Jackson, Ian Johnston, Dominic Kwiatkowski, Cordelia Langford, John Sillitoe on behalf of the Wellcome Sanger Institute COVID-19 Surveillance Team |
| EPI_ISL_2437719 | Lighthouse Lab in Glasgow | Wellcome Sanger Institute for the COVID-19 Genomics UK (COG-UK) Consortium | Harper VanSteenhouse, Yumi Kasai, David Gray, Carol Clugston, Anna Dominiczak and Alex Alderton, Roberto Amato, Jeffrey Barrett, Sonia Goncalves, Ewan Harrison, David K. Jackson, Ian Johnston, Dominic Kwiatkowski, Cordelia Langford, John Sillitoe on behalf of the Wellcome Sanger Institute COVID-19 Surveillance Team |
| EPI_ISL_2437720, EPI_ISL_2437722 | Lighthouse Lab in Alderley Park | Wellcome Sanger Institute for the COVID-19 Genomics UK (COG-UK) Consortium | Jacquelyn Wynn, Mairead Hyland, The Lighthouse Lab in Alderley Park and Alex Alderton, Roberto Amato, Jeffrey Barrett, Sonia Goncalves, Ewan Harrison, David K. Jackson, Ian Johnston, Dominic Kwiatkowski, Cordelia Langford, John Sillitoe on behalf of the Wellcome Sanger Institute COVID-19 Surveillance Team |
| EPI_ISL_2437723 | Lighthouse Lab in Glasgow | Wellcome Sanger Institute for the COVID-19 Genomics UK (COG-UK) Consortium | Harper VanSteenhouse, Yumi Kasai, David Gray, Carol Clugston, Anna Dominiczak and Alex Alderton, Roberto Amato, Jeffrey Barrett, Sonia Goncalves, Ewan Harrison, David K. Jackson, Ian Johnston, Dominic Kwiatkowski, Cordelia Langford, John Sillitoe on behalf of the Wellcome Sanger Institute COVID-19 Surveillance Team |
| EPI_ISL_2437724, EPI_ISL_2437725, EPI_ISL_2437726 | Lighthouse Lab in Milton Keynes | Wellcome Sanger Institute for the COVID-19 Genomics UK (COG-UK) Consortium | The Lighthouse Lab in Milton Keynes and Alex Alderton, Roberto Amato, Jeffrey Barrett, Sonia Goncalves, Ewan Harrison, David K. Jackson, Ian Johnston, Dominic Kwiatkowski, Cordelia Langford, John Sillitoe on behalf of the Wellcome Sanger Institute COVID-19 Surveillance Team |
| EPI_ISL_2437727 | Lighthouse Lab in Alderley Park | Wellcome Sanger Institute for the COVID-19 Genomics UK (COG-UK) Consortium | Jacquelyn Wynn, Mairead Hyland, The Lighthouse Lab in Alderley Park and Alex Alderton, Roberto Amato, Jeffrey Barrett, Sonia Goncalves, Ewan Harrison, David K. Jackson, Ian Johnston, Dominic Kwiatkowski, Cordelia Langford, John Sillitoe on behalf of the Wellcome Sanger Institute COVID-19 Surveillance Team |
| EPI_ISL_2437728, EPI_ISL_2437729 | Lighthouse Lab in Milton Keynes | Wellcome Sanger Institute for the COVID-19 Genomics UK (COG-UK) Consortium | The Lighthouse Lab in Milton Keynes and Alex Alderton, Roberto Amato, Jeffrey Barrett, Sonia Goncalves, Ewan Harrison, David K. Jackson, Ian Johnston, Dominic Kwiatkowski, Cordelia Langford, John Sillitoe on behalf of the Wellcome Sanger Institute COVID-19 Surveillance Team |
| EPI_ISL_2437731 | Lighthouse Lab in Alderley Park | Wellcome Sanger Institute for the COVID-19 Genomics UK (COG-UK) Consortium | Jacquelyn Wynn, Mairead Hyland, The Lighthouse Lab in Alderley Park and Alex Alderton, Roberto Amato, Jeffrey Barrett, Sonia Goncalves, Ewan Harrison, David K. Jackson, Ian Johnston, Dominic Kwiatkowski, Cordelia Langford, John Sillitoe on behalf of the Wellcome Sanger Institute COVID-19 Surveillance Team |
| EPI_ISL_2437732, EPI_ISL_2437733, EPI_ISL_2437734 | Lighthouse Lab in Milton Keynes | Wellcome Sanger Institute for the COVID-19 Genomics UK (COG-UK) Consortium | The Lighthouse Lab in Milton Keynes and Alex Alderton, Roberto Amato, Jeffrey Barrett, Sonia Goncalves, Ewan Harrison, David K. Jackson, Ian Johnston, Dominic Kwiatkowski, Cordelia Langford, John Sillitoe on behalf of the Wellcome Sanger Institute COVID-19 Surveillance Team |
| EPI_ISL_2437735 | Lighthouse Lab in Alderley Park | Wellcome Sanger Institute for the COVID-19 Genomics UK (COG-UK) Consortium | Jacquelyn Wynn, Mairead Hyland, The Lighthouse Lab in Alderley Park and Alex Alderton, Roberto Amato, Jeffrey Barrett, Sonia Goncalves, Ewan Harrison, David K. Jackson, Ian Johnston, Dominic Kwiatkowski, Cordelia Langford, John Sillitoe on behalf of the Wellcome Sanger Institute COVID-19 Surveillance Team |
| EPI_ISL_2437736 | Lighthouse Lab in Milton Keynes | Wellcome Sanger Institute for the COVID-19 Genomics UK (COG-UK) Consortium | The Lighthouse Lab in Milton Keynes and Alex Alderton, Roberto Amato, Jeffrey Barrett, Sonia Goncalves, Ewan Harrison, David K. Jackson, Ian Johnston, Dominic Kwiatkowski, Cordelia Langford, John Sillitoe on behalf of the Wellcome Sanger Institute COVID-19 Surveillance Team |
| EPI_ISL_2437737 | Lighthouse Lab in Glasgow | Wellcome Sanger Institute for the COVID-19 Genomics UK (COG-UK) Consortium | Harper VanSteenhouse, Yumi Kasai, David Gray, Carol Clugston, Anna Dominiczak and Alex Alderton, Roberto Amato, Jeffrey Barrett, Sonia Goncalves, Ewan Harrison, David K. Jackson, Ian Johnston, Dominic Kwiatkowski, Cordelia Langford, John Sillitoe on behalf of the Wellcome Sanger Institute COVID-19 Surveillance Team |
| EPI_ISL_2437738 | Lighthouse Lab in Alderley Park | Wellcome Sanger Institute for the COVID-19 Genomics UK (COG-UK) Consortium | Jacquelyn Wynn, Mairead Hyland, The Lighthouse Lab in Alderley Park and Alex Alderton, Roberto Amato, Jeffrey Barrett, Sonia Goncalves, Ewan Harrison, David K. Jackson, Ian Johnston, Dominic Kwiatkowski, Cordelia Langford, John Sillitoe on behalf of the Wellcome Sanger Institute COVID-19 Surveillance Team |
| EPI_ISL_2437739, EPI_ISL_2437740 | Lighthouse Lab in Milton Keynes | Wellcome Sanger Institute for the COVID-19 Genomics UK (COG-UK) Consortium | The Lighthouse Lab in Milton Keynes and Alex Alderton, Roberto Amato, Jeffrey Barrett, Sonia Goncalves, Ewan Harrison, David K. Jackson, Ian Johnston, Dominic Kwiatkowski, Cordelia Langford, John Sillitoe on behalf of the Wellcome Sanger Institute COVID-19 Surveillance Team |
| EPI_ISL_2437741 | Lighthouse Lab in Glasgow | Wellcome Sanger Institute for the COVID-19 Genomics UK (COG-UK) Consortium | Harper VanSteenhouse, Yumi Kasai, David Gray, Carol Clugston, Anna Dominiczak and Alex Alderton, Roberto Amato, Jeffrey Barrett, Sonia Goncalves, Ewan Harrison, David K. Jackson, Ian Johnston, Dominic Kwiatkowski, Cordelia Langford, John Sillitoe on behalf of the Wellcome Sanger Institute COVID-19 Surveillance Team |
| EPI_ISL_2437742 | Lighthouse Lab in Milton Keynes | Wellcome Sanger Institute for the COVID-19 Genomics UK (COG-UK) Consortium | The Lighthouse Lab in Milton Keynes and Alex Alderton, Roberto Amato, Jeffrey Barrett, Sonia Goncalves, Ewan Harrison, David K. Jackson, Ian Johnston, Dominic Kwiatkowski, Cordelia Langford, John Sillitoe on behalf of the Wellcome Sanger Institute COVID-19 Surveillance Team |
| EPI_ISL_2437743 | Lighthouse Lab in Alderley Park | Wellcome Sanger Institute for the COVID-19 Genomics UK (COG-UK) Consortium | Jacquelyn Wynn, Mairead Hyland, The Lighthouse Lab in Alderley Park and Alex Alderton, Roberto Amato, Jeffrey Barrett, Sonia Goncalves, Ewan Harrison, David K. Jackson, Ian Johnston, Dominic Kwiatkowski, Cordelia Langford, John Sillitoe on behalf of the Wellcome Sanger Institute COVID-19 Surveillance Team |
| EPI_ISL_2437745 | Lighthouse Lab in Milton Keynes | Wellcome Sanger Institute for the COVID-19 Genomics UK (COG-UK) Consortium | The Lighthouse Lab in Milton Keynes and Alex Alderton, Roberto Amato, Jeffrey Barrett, Sonia Goncalves, Ewan Harrison, David K. Jackson, Ian Johnston, Dominic Kwiatkowski, Cordelia Langford, John Sillitoe on behalf of the Wellcome Sanger Institute COVID-19 Surveillance Team |
| EPI_ISL_2437746 | Lighthouse Lab in Glasgow | Wellcome Sanger Institute for the COVID-19 Genomics UK (COG-UK) Consortium | Harper VanSteenhouse, Yumi Kasai, David Gray, Carol Clugston, Anna Dominiczak and Alex Alderton, Roberto Amato, Jeffrey Barrett, Sonia Goncalves, Ewan Harrison, David K. Jackson, Ian Johnston, Dominic Kwiatkowski, Cordelia Langford, John Sillitoe on behalf of the Wellcome Sanger Institute COVID-19 Surveillance Team |
| EPI_ISL_2437747, EPI_ISL_2437748 | Lighthouse Lab in Milton Keynes | Wellcome Sanger Institute for the COVID-19 Genomics UK (COG-UK) Consortium | The Lighthouse Lab in Milton Keynes and Alex Alderton, Roberto Amato, Jeffrey Barrett, Sonia Goncalves, Ewan Harrison, David K. Jackson, Ian Johnston, Dominic Kwiatkowski, Cordelia Langford, John Sillitoe on behalf of the Wellcome Sanger Institute COVID-19 Surveillance Team |
| EPI_ISL_2437749, EPI_ISL_2437750 | Lighthouse Lab in Glasgow | Wellcome Sanger Institute for the COVID-19 Genomics UK (COG-UK) Consortium | Harper VanSteenhouse, Yumi Kasai, David Gray, Carol Clugston, Anna Dominiczak and Alex Alderton, Roberto Amato, Jeffrey Barrett, Sonia Goncalves, Ewan Harrison, David K. Jackson, Ian Johnston, Dominic Kwiatkowski, Cordelia Langford, John Sillitoe on behalf of the Wellcome Sanger Institute COVID-19 Surveillance Team |

| | | | |
|---|---|---|---|
| EPI_ISL_2437751, EPI_ISL_2437752, EPI_ISL_2437753, EPI_ISL_2437754, EPI_ISL_2437755, EPI_ISL_2437757 | Lighthouse Lab in Milton Keynes | Wellcome Sanger Institute for the COVID-19 Genomics UK (COG-UK) Consortium | The Lighthouse Lab in Milton Keynes and Alex Alderton, Roberto Amato, Jeffrey Barrett, Sonia Goncalves, Ewan Harrison, David K. Jackson, Ian Johnston, Dominic Kwiatkowski, Cordelia Langford, John Sillitoe on behalf of the Wellcome Sanger Institute COVID-19 Surveillance Team |
| EPI_ISL_2437758, EPI_ISL_2437759 | Lighthouse Lab in Alderley Park | Wellcome Sanger Institute for the COVID-19 Genomics UK (COG-UK) Consortium | Jacquelyn Wynn, Mairead Hyland, The Lighthouse Lab in Alderley Park and Alex Alderton, Roberto Amato, Jeffrey Barrett, Sonia Goncalves, Ewan Harrison, David K. Jackson, Ian Johnston, Dominic Kwiatkowski, Cordelia Langford, John Sillitoe on behalf of the Wellcome Sanger Institute COVID-19 Surveillance Team |
| EPI_ISL_2437760 | Lighthouse Lab in Milton Keynes | Wellcome Sanger Institute for the COVID-19 Genomics UK (COG-UK) Consortium | The Lighthouse Lab in Milton Keynes and Alex Alderton, Roberto Amato, Jeffrey Barrett, Sonia Goncalves, Ewan Harrison, David K. Jackson, Ian Johnston, Dominic Kwiatkowski, Cordelia Langford, John Sillitoe on behalf of the Wellcome Sanger Institute COVID-19 Surveillance Team |
| EPI_ISL_2437761 | Lighthouse Lab in Glasgow | Wellcome Sanger Institute for the COVID-19 Genomics UK (COG-UK) Consortium | Harper VanSteenhouse, Yumi Kasai, David Gray, Carol Clugston, Anna Dominiczak and Alex Alderton, Roberto Amato, Jeffrey Barrett, Sonia Goncalves, Ewan Harrison, David K. Jackson, Ian Johnston, Dominic Kwiatkowski, Cordelia Langford, John Sillitoe on behalf of the Wellcome Sanger Institute COVID-19 Surveillance Team |
| EPI_ISL_2437762 | Lighthouse Lab in Milton Keynes | Wellcome Sanger Institute for the COVID-19 Genomics UK (COG-UK) Consortium | The Lighthouse Lab in Milton Keynes and Alex Alderton, Roberto Amato, Jeffrey Barrett, Sonia Goncalves, Ewan Harrison, David K. Jackson, Ian Johnston, Dominic Kwiatkowski, Cordelia Langford, John Sillitoe on behalf of the Wellcome Sanger Institute COVID-19 Surveillance Team |
| EPI_ISL_2437763 | Lighthouse Lab in Glasgow | Wellcome Sanger Institute for the COVID-19 Genomics UK (COG-UK) Consortium | Harper VanSteenhouse, Yumi Kasai, David Gray, Carol Clugston, Anna Dominiczak and Alex Alderton, Roberto Amato, Jeffrey Barrett, Sonia Goncalves, Ewan Harrison, David K. Jackson, Ian Johnston, Dominic Kwiatkowski, Cordelia Langford, John Sillitoe on behalf of the Wellcome Sanger Institute COVID-19 Surveillance Team |
| EPI_ISL_2437764, EPI_ISL_2437766, EPI_ISL_2437767 | Lighthouse Lab in Milton Keynes | Wellcome Sanger Institute for the COVID-19 Genomics UK (COG-UK) Consortium | The Lighthouse Lab in Milton Keynes and Alex Alderton, Roberto Amato, Jeffrey Barrett, Sonia Goncalves, Ewan Harrison, David K. Jackson, Ian Johnston, Dominic Kwiatkowski, Cordelia Langford, John Sillitoe on behalf of the Wellcome Sanger Institute COVID-19 Surveillance Team |
| EPI_ISL_2437769 | Lighthouse Lab in Alderley Park | Wellcome Sanger Institute for the COVID-19 Genomics UK (COG-UK) Consortium | Jacquelyn Wynn, Mairead Hyland, The Lighthouse Lab in Alderley Park and Alex Alderton, Roberto Amato, Jeffrey Barrett, Sonia Goncalves, Ewan Harrison, David K. Jackson, Ian Johnston, Dominic Kwiatkowski, Cordelia Langford, John Sillitoe on behalf of the Wellcome Sanger Institute COVID-19 Surveillance Team |
| EPI_ISL_2437771, EPI_ISL_2437772 | Lighthouse Lab in Milton Keynes | Wellcome Sanger Institute for the COVID-19 Genomics UK (COG-UK) Consortium | The Lighthouse Lab in Milton Keynes and Alex Alderton, Roberto Amato, Jeffrey Barrett, Sonia Goncalves, Ewan Harrison, David K. Jackson, Ian Johnston, Dominic Kwiatkowski, Cordelia Langford, John Sillitoe on behalf of the Wellcome Sanger Institute COVID-19 Surveillance Team |
| EPI_ISL_2437774 | Lighthouse Lab in Alderley Park | Wellcome Sanger Institute for the COVID-19 Genomics UK (COG-UK) Consortium | Jacquelyn Wynn, Mairead Hyland, The Lighthouse Lab in Alderley Park and Alex Alderton, Roberto Amato, Jeffrey Barrett, Sonia Goncalves, Ewan Harrison, David K. Jackson, Ian Johnston, Dominic Kwiatkowski, Cordelia Langford, John Sillitoe on behalf of the Wellcome Sanger Institute COVID-19 Surveillance Team |
| EPI_ISL_2437775 | Lighthouse Lab in Glasgow | Wellcome Sanger Institute for the COVID-19 Genomics UK (COG-UK) Consortium | Harper VanSteenhouse, Yumi Kasai, David Gray, Carol Clugston, Anna Dominiczak and Alex Alderton, Roberto Amato, Jeffrey Barrett, Sonia Goncalves, Ewan Harrison, David K. Jackson, Ian Johnston, Dominic Kwiatkowski, Cordelia Langford, John Sillitoe on behalf of the Wellcome Sanger Institute COVID-19 Surveillance Team |
| EPI_ISL_2437776 | Lighthouse Lab in Alderley Park | Wellcome Sanger Institute for the COVID-19 Genomics UK (COG-UK) Consortium | Jacquelyn Wynn, Mairead Hyland, The Lighthouse Lab in Alderley Park and Alex Alderton, Roberto Amato, Jeffrey Barrett, Sonia Goncalves, Ewan Harrison, David K. Jackson, Ian Johnston, Dominic Kwiatkowski, Cordelia Langford, John Sillitoe on behalf of the Wellcome Sanger Institute COVID-19 Surveillance Team |
| EPI_ISL_2437777 | Lighthouse Lab in Milton Keynes | Wellcome Sanger Institute for the COVID-19 Genomics UK (COG-UK) Consortium | The Lighthouse Lab in Milton Keynes and Alex Alderton, Roberto Amato, Jeffrey Barrett, Sonia Goncalves, Ewan Harrison, David K. Jackson, Ian Johnston, Dominic Kwiatkowski, Cordelia Langford, John Sillitoe on behalf of the Wellcome Sanger Institute COVID-19 Surveillance Team |
| EPI_ISL_2437778 | Lighthouse Lab in Glasgow | Wellcome Sanger Institute for the COVID-19 Genomics UK (COG-UK) Consortium | Harper VanSteenhouse, Yumi Kasai, David Gray, Carol Clugston, Anna Dominiczak and Alex Alderton, Roberto Amato, Jeffrey Barrett, Sonia Goncalves, Ewan Harrison, David K. Jackson, Ian Johnston, Dominic Kwiatkowski, Cordelia Langford, John Sillitoe on behalf of the Wellcome Sanger Institute COVID-19 Surveillance Team |
| EPI_ISL_2437779 | Lighthouse Lab in Alderley Park | Wellcome Sanger Institute for the COVID-19 Genomics UK (COG-UK) Consortium | Jacquelyn Wynn, Mairead Hyland, The Lighthouse Lab in Alderley Park and Alex Alderton, Roberto Amato, Jeffrey Barrett, Sonia Goncalves, Ewan Harrison, David K. Jackson, Ian Johnston, Dominic Kwiatkowski, Cordelia Langford, John Sillitoe on behalf of the Wellcome Sanger Institute COVID-19 Surveillance Team |
| EPI_ISL_2437780 | Lighthouse Lab in Glasgow | Wellcome Sanger Institute for the COVID-19 Genomics UK (COG-UK) Consortium | Harper VanSteenhouse, Yumi Kasai, David Gray, Carol Clugston, Anna Dominiczak and Alex Alderton, Roberto Amato, Jeffrey Barrett, Sonia Goncalves, Ewan Harrison, David K. Jackson, Ian Johnston, Dominic Kwiatkowski, Cordelia Langford, John Sillitoe on behalf of the Wellcome Sanger Institute COVID-19 Surveillance Team |
| EPI_ISL_2437781, EPI_ISL_2437782, EPI_ISL_2437783, EPI_ISL_2437784, EPI_ISL_2437785 | Lighthouse Lab in Milton Keynes | Wellcome Sanger Institute for the COVID-19 Genomics UK (COG-UK) Consortium | The Lighthouse Lab in Milton Keynes and Alex Alderton, Roberto Amato, Jeffrey Barrett, Sonia Goncalves, Ewan Harrison, David K. Jackson, Ian Johnston, Dominic Kwiatkowski, Cordelia Langford, John Sillitoe on behalf of the Wellcome Sanger Institute COVID-19 Surveillance Team |
| EPI_ISL_2437786, EPI_ISL_2437787 | Lighthouse Lab in Alderley Park | Wellcome Sanger Institute for the COVID-19 Genomics UK (COG-UK) Consortium | Jacquelyn Wynn, Mairead Hyland, The Lighthouse Lab in Alderley Park and Alex Alderton, Roberto Amato, Jeffrey Barrett, Sonia Goncalves, Ewan Harrison, David K. Jackson, Ian Johnston, Dominic Kwiatkowski, Cordelia Langford, John Sillitoe on behalf of the Wellcome Sanger Institute COVID-19 Surveillance Team |
| EPI_ISL_2437788 | Lighthouse Lab in Glasgow | Wellcome Sanger Institute for the COVID-19 Genomics UK (COG-UK) Consortium | Harper VanSteenhouse, Yumi Kasai, David Gray, Carol Clugston, Anna Dominiczak and Alex Alderton, Roberto Amato, Jeffrey Barrett, Sonia Goncalves, Ewan Harrison, David K. Jackson, Ian Johnston, Dominic Kwiatkowski, Cordelia Langford, John Sillitoe on behalf of the Wellcome Sanger Institute COVID-19 Surveillance Team |
| EPI_ISL_2437790 | Lighthouse Lab in Milton Keynes | Wellcome Sanger Institute for the COVID-19 Genomics UK (COG-UK) Consortium | The Lighthouse Lab in Milton Keynes and Alex Alderton, Roberto Amato, Jeffrey Barrett, Sonia Goncalves, Ewan Harrison, David K. Jackson, Ian Johnston, Dominic Kwiatkowski, Cordelia Langford, John Sillitoe on behalf of the Wellcome Sanger Institute COVID-19 Surveillance Team |
| EPI_ISL_2437792 | Lighthouse Lab in Glasgow | Wellcome Sanger Institute for the COVID-19 Genomics UK (COG-UK) Consortium | Harper VanSteenhouse, Yumi Kasai, David Gray, Carol Clugston, Anna Dominiczak and Alex Alderton, Roberto Amato, Jeffrey Barrett, Sonia Goncalves, Ewan Harrison, David K. Jackson, Ian Johnston, Dominic Kwiatkowski, Cordelia Langford, John Sillitoe on behalf of the Wellcome Sanger Institute COVID-19 Surveillance Team |
| EPI_ISL_2437793 | Lighthouse Lab in Alderley Park | Wellcome Sanger Institute for the COVID-19 Genomics UK (COG-UK) Consortium | Jacquelyn Wynn, Mairead Hyland, The Lighthouse Lab in Alderley Park and Alex Alderton, Roberto Amato, Jeffrey Barrett, Sonia Goncalves, Ewan Harrison, David K. Jackson, Ian Johnston, Dominic Kwiatkowski, Cordelia Langford, John Sillitoe on behalf of the Wellcome Sanger Institute COVID-19 Surveillance Team |
| EPI_ISL_2437794, EPI_ISL_2437795, EPI_ISL_2437796, EPI_ISL_2437797 | Lighthouse Lab in Milton Keynes | Wellcome Sanger Institute for the COVID-19 Genomics UK (COG-UK) Consortium | The Lighthouse Lab in Milton Keynes and Alex Alderton, Roberto Amato, Jeffrey Barrett, Sonia Goncalves, Ewan Harrison, David K. Jackson, Ian Johnston, Dominic Kwiatkowski, Cordelia Langford, John Sillitoe on behalf of the Wellcome Sanger Institute COVID-19 Surveillance Team |
| EPI_ISL_2437798 | Lighthouse Lab in Glasgow | Wellcome Sanger Institute for the COVID-19 Genomics UK (COG-UK) Consortium | Harper VanSteenhouse, Yumi Kasai, David Gray, Carol Clugston, Anna Dominiczak and Alex Alderton, Roberto Amato, Jeffrey Barrett, Sonia Goncalves, Ewan Harrison, David K. Jackson, Ian Johnston, Dominic Kwiatkowski, Cordelia Langford, John Sillitoe on behalf of the Wellcome Sanger Institute COVID-19 Surveillance Team |
| EPI_ISL_2437799 | Lighthouse Lab in Milton Keynes | Wellcome Sanger Institute for the COVID-19 Genomics UK (COG-UK) Consortium | The Lighthouse Lab in Milton Keynes and Alex Alderton, Roberto Amato, Jeffrey Barrett, Sonia Goncalves, Ewan Harrison, David K. Jackson, Ian Johnston, Dominic Kwiatkowski, Cordelia Langford, John Sillitoe on behalf of the Wellcome Sanger Institute COVID-19 Surveillance Team |
| EPI_ISL_2437800 | Lighthouse Lab in Alderley Park | Wellcome Sanger Institute for the COVID-19 Genomics UK (COG-UK) Consortium | Jacquelyn Wynn, Mairead Hyland, The Lighthouse Lab in Alderley Park and Alex Alderton, Roberto Amato, Jeffrey Barrett, Sonia Goncalves, Ewan Harrison, David K. Jackson, Ian Johnston, Dominic Kwiatkowski, Cordelia Langford, John Sillitoe on behalf of the Wellcome Sanger Institute COVID-19 Surveillance Team |
| EPI_ISL_2437801 | Lighthouse Lab in Milton Keynes | Wellcome Sanger Institute for the COVID-19 Genomics UK (COG-UK) Consortium | The Lighthouse Lab in Milton Keynes and Alex Alderton, Roberto Amato, Jeffrey Barrett, Sonia Goncalves, Ewan Harrison, David K. Jackson, Ian Johnston, Dominic Kwiatkowski, Cordelia Langford, John Sillitoe on behalf of the Wellcome Sanger Institute COVID-19 Surveillance Team |
| EPI_ISL_2437802, EPI_ISL_2437803 | Lighthouse Lab in Glasgow | Wellcome Sanger Institute for the COVID-19 Genomics UK (COG-UK) Consortium | Harper VanSteenhouse, Yumi Kasai, David Gray, Carol Clugston, Anna Dominiczak and Alex Alderton, Roberto Amato, Jeffrey Barrett, Sonia Goncalves, Ewan Harrison, David K. Jackson, Ian Johnston, Dominic Kwiatkowski, Cordelia Langford, John Sillitoe on behalf of the Wellcome Sanger Institute COVID-19 Surveillance Team |
| EPI_ISL_2437804, EPI_ISL_2437805 | Lighthouse Lab in Milton Keynes | Wellcome Sanger Institute for the COVID-19 Genomics UK (COG-UK) Consortium | The Lighthouse Lab in Milton Keynes and Alex Alderton, Roberto Amato, Jeffrey Barrett, Sonia Goncalves, Ewan Harrison, David K. Jackson, Ian Johnston, Dominic Kwiatkowski, Cordelia Langford, John Sillitoe on behalf of the Wellcome Sanger Institute COVID-19 Surveillance Team |
| EPI_ISL_2437806 | Lighthouse Lab in Glasgow | Wellcome Sanger Institute for the COVID-19 Genomics UK (COG-UK) Consortium | Harper VanSteenhouse, Yumi Kasai, David Gray, Carol Clugston, Anna Dominiczak and Alex Alderton, Roberto Amato, Jeffrey Barrett, Sonia Goncalves, Ewan Harrison, David K. Jackson, Ian Johnston, Dominic Kwiatkowski, Cordelia Langford, John Sillitoe on behalf of the Wellcome Sanger Institute COVID-19 Surveillance Team |
| EPI_ISL_2437807 | Lighthouse Lab in Alderley Park | Wellcome Sanger Institute for the COVID-19 Genomics UK (COG-UK) Consortium | Jacquelyn Wynn, Mairead Hyland, The Lighthouse Lab in Alderley Park and Alex Alderton, Roberto Amato, Jeffrey Barrett, Sonia Goncalves, Ewan Harrison, David K. Jackson, Ian Johnston, Dominic Kwiatkowski, Cordelia Langford, John Sillitoe on behalf of the Wellcome Sanger Institute COVID-19 Surveillance Team |
| EPI_ISL_2437808, EPI_ISL_2437809 | Lighthouse Lab in Milton Keynes | Wellcome Sanger Institute for the COVID-19 Genomics UK (COG-UK) Consortium | The Lighthouse Lab in Milton Keynes and Alex Alderton, Roberto Amato, Jeffrey Barrett, Sonia Goncalves, Ewan Harrison, David K. Jackson, Ian Johnston, Dominic Kwiatkowski, Cordelia Langford, John Sillitoe on behalf of the Wellcome Sanger Institute COVID-19 Surveillance Team |
| EPI_ISL_2437810 | Lighthouse Lab in Alderley Park | Wellcome Sanger Institute for the COVID-19 Genomics UK (COG-UK) Consortium | Jacquelyn Wynn, Mairead Hyland, The Lighthouse Lab in Alderley Park and Alex Alderton, Roberto Amato, Jeffrey Barrett, Sonia Goncalves, Ewan Harrison, David K. Jackson, Ian Johnston, Dominic Kwiatkowski, Cordelia Langford, John Sillitoe on behalf of the Wellcome Sanger Institute COVID-19 Surveillance Team |
| EPI_ISL_2437811 | Lighthouse Lab in Milton Keynes | Wellcome Sanger Institute for the COVID-19 Genomics UK (COG-UK) Consortium | The Lighthouse Lab in Milton Keynes and Alex Alderton, Roberto Amato, Jeffrey Barrett, Sonia Goncalves, Ewan Harrison, David K. Jackson, Ian Johnston, Dominic Kwiatkowski, Cordelia Langford, John Sillitoe on behalf of the Wellcome Sanger Institute COVID-19 Surveillance Team |
| EPI_ISL_2437812 | Lighthouse Lab in Glasgow | Wellcome Sanger Institute for the COVID-19 Genomics UK (COG-UK) Consortium | Harper VanSteenhouse, Yumi Kasai, David Gray, Carol Clugston, Anna Dominiczak and Alex Alderton, Roberto Amato, Jeffrey Barrett, Sonia Goncalves, Ewan Harrison, David K. Jackson, Ian Johnston, Dominic Kwiatkowski, Cordelia Langford, John Sillitoe on behalf of the Wellcome Sanger Institute COVID-19 Surveillance Team |
| EPI_ISL_2437813, EPI_ISL_2437814 | Lighthouse Lab in Milton Keynes | Wellcome Sanger Institute for the COVID-19 Genomics UK (COG-UK) Consortium | The Lighthouse Lab in Milton Keynes and Alex Alderton, Roberto Amato, Jeffrey Barrett, Sonia Goncalves, Ewan Harrison, David K. Jackson, Ian Johnston, Dominic Kwiatkowski, Cordelia Langford, John Sillitoe on behalf of the Wellcome Sanger Institute COVID-19 Surveillance Team |
| EPI_ISL_2437815 | Lighthouse Lab in Glasgow | Wellcome Sanger Institute for the COVID-19 Genomics UK (COG-UK) Consortium | Harper VanSteenhouse, Yumi Kasai, David Gray, Carol Clugston, Anna Dominiczak and Alex Alderton, Roberto Amato, Jeffrey Barrett, Sonia Goncalves, Ewan Harrison, David K. Jackson, Ian Johnston, Dominic Kwiatkowski, Cordelia Langford, John Sillitoe on behalf of the Wellcome Sanger Institute COVID-19 Surveillance Team |
| EPI_ISL_2437817 | Lighthouse Lab in Alderley Park | Wellcome Sanger Institute for the COVID-19 Genomics UK (COG-UK) Consortium | Jacquelyn Wynn, Mairead Hyland, The Lighthouse Lab in Alderley Park and Alex Alderton, Roberto Amato, Jeffrey Barrett, Sonia Goncalves, Ewan Harrison, David K. Jackson, Ian Johnston, Dominic Kwiatkowski, Cordelia Langford, John Sillitoe on behalf of the Wellcome Sanger Institute COVID-19 Surveillance Team |
| EPI_ISL_2437818 | Lighthouse Lab in Glasgow | Wellcome Sanger Institute for the COVID-19 Genomics UK (COG-UK) Consortium | Harper VanSteenhouse, Yumi Kasai, David Gray, Carol Clugston, Anna Dominiczak and Alex Alderton, Roberto Amato, Jeffrey Barrett, Sonia Goncalves, Ewan Harrison, David K. Jackson, Ian Johnston, Dominic Kwiatkowski, Cordelia Langford, John Sillitoe on behalf of the Wellcome Sanger Institute COVID-19 Surveillance Team |
| EPI_ISL_2437820 | Lighthouse Lab in Alderley Park | Wellcome Sanger Institute for the COVID-19 Genomics UK (COG-UK) Consortium | Jacquelyn Wynn, Mairead Hyland, The Lighthouse Lab in Alderley Park and Alex Alderton, Roberto Amato, Jeffrey Barrett, Sonia Goncalves, Ewan Harrison, David K. Jackson, Ian Johnston, Dominic Kwiatkowski, Cordelia Langford, John Sillitoe on behalf of the Wellcome Sanger Institute COVID-19 Surveillance Team |
| EPI_ISL_2437821, EPI_ISL_2437822, | Lighthouse Lab in Milton Keynes | Wellcome Sanger Institute for the COVID-19 Genomics UK (COG-UK) Consortium | The Lighthouse Lab in Milton Keynes and Alex Alderton, Roberto Amato, Jeffrey Barrett, Sonia Goncalves, Ewan Harrison, David K. Jackson, Ian Johnston, Dominic Kwiatkowski, Cordelia Langford, John Sillitoe on behalf of the Wellcome Sanger Institute COVID-19 Surveillance Team |

| | | | |
|---|---|---|---|
| EPI_ISL_2437823 | | | |
| EPI_ISL_2437824 | Lighthouse Lab in Alderley Park | Wellcome Sanger Institute for the COVID-19 Genomics UK (COG-UK) Consortium | Jacquelyn Wynn, Mairead Hyland, The Lighthouse Lab in Alderley Park and Alex Alderton, Roberto Amato, Jeffrey Barrett, Sonia Goncalves, Ewan Harrison, David K. Jackson, Ian Johnston, Dominic Kwiatkowski, Cordelia Langford, John Sillitoe on behalf of the Wellcome Sanger Institute COVID-19 Surveillance Team |
| EPI_ISL_2437825 | Lighthouse Lab in Milton Keynes | Wellcome Sanger Institute for the COVID-19 Genomics UK (COG-UK) Consortium | The Lighthouse Lab in Milton Keynes and Alex Alderton, Roberto Amato, Jeffrey Barrett, Sonia Goncalves, Ewan Harrison, David K. Jackson, Ian Johnston, Dominic Kwiatkowski, Cordelia Langford, John Sillitoe on behalf of the Wellcome Sanger Institute COVID-19 Surveillance Team |
| EPI_ISL_2437826, EPI_ISL_2437827, EPI_ISL_2437828, EPI_ISL_2437829, EPI_ISL_2437831, EPI_ISL_2437833, EPI_ISL_2437834 | Lighthouse Lab in Alderley Park | Wellcome Sanger Institute for the COVID-19 Genomics UK (COG-UK) Consortium | Jacquelyn Wynn, Mairead Hyland, The Lighthouse Lab in Alderley Park and Alex Alderton, Roberto Amato, Jeffrey Barrett, Sonia Goncalves, Ewan Harrison, David K. Jackson, Ian Johnston, Dominic Kwiatkowski, Cordelia Langford, John Sillitoe on behalf of the Wellcome Sanger Institute COVID-19 Surveillance Team |
| EPI_ISL_2437835, EPI_ISL_2437836 | Lighthouse Lab in Milton Keynes | Wellcome Sanger Institute for the COVID-19 Genomics UK (COG-UK) Consortium | The Lighthouse Lab in Milton Keynes and Alex Alderton, Roberto Amato, Jeffrey Barrett, Sonia Goncalves, Ewan Harrison, David K. Jackson, Ian Johnston, Dominic Kwiatkowski, Cordelia Langford, John Sillitoe on behalf of the Wellcome Sanger Institute COVID-19 Surveillance Team |
| EPI_ISL_2437837 | Lighthouse Lab in Alderley Park | Wellcome Sanger Institute for the COVID-19 Genomics UK (COG-UK) Consortium | Jacquelyn Wynn, Mairead Hyland, The Lighthouse Lab in Alderley Park and Alex Alderton, Roberto Amato, Jeffrey Barrett, Sonia Goncalves, Ewan Harrison, David K. Jackson, Ian Johnston, Dominic Kwiatkowski, Cordelia Langford, John Sillitoe on behalf of the Wellcome Sanger Institute COVID-19 Surveillance Team |
| EPI_ISL_2437838 | Lighthouse Lab in Glasgow | Wellcome Sanger Institute for the COVID-19 Genomics UK (COG-UK) Consortium | Harper VanSteenhouse, Yumi Kasai, David Gray, Carol Clugston, Anna Dominiczak and Alex Alderton, Roberto Amato, Jeffrey Barrett, Sonia Goncalves, Ewan Harrison, David K. Jackson, Ian Johnston, Dominic Kwiatkowski, Cordelia Langford, John Sillitoe on behalf of the Wellcome Sanger Institute COVID-19 Surveillance Team |
| EPI_ISL_2437839 | Lighthouse Lab in Alderley Park | Wellcome Sanger Institute for the COVID-19 Genomics UK (COG-UK) Consortium | Jacquelyn Wynn, Mairead Hyland, The Lighthouse Lab in Alderley Park and Alex Alderton, Roberto Amato, Jeffrey Barrett, Sonia Goncalves, Ewan Harrison, David K. Jackson, Ian Johnston, Dominic Kwiatkowski, Cordelia Langford, John Sillitoe on behalf of the Wellcome Sanger Institute COVID-19 Surveillance Team |
| EPI_ISL_2437840 | Lighthouse Lab in Milton Keynes | Wellcome Sanger Institute for the COVID-19 Genomics UK (COG-UK) Consortium | The Lighthouse Lab in Milton Keynes and Alex Alderton, Roberto Amato, Jeffrey Barrett, Sonia Goncalves, Ewan Harrison, David K. Jackson, Ian Johnston, Dominic Kwiatkowski, Cordelia Langford, John Sillitoe on behalf of the Wellcome Sanger Institute COVID-19 Surveillance Team |
| EPI_ISL_2437841 | Lighthouse Lab in Alderley Park | Wellcome Sanger Institute for the COVID-19 Genomics UK (COG-UK) Consortium | Jacquelyn Wynn, Mairead Hyland, The Lighthouse Lab in Alderley Park and Alex Alderton, Roberto Amato, Jeffrey Barrett, Sonia Goncalves, Ewan Harrison, David K. Jackson, Ian Johnston, Dominic Kwiatkowski, Cordelia Langford, John Sillitoe on behalf of the Wellcome Sanger Institute COVID-19 Surveillance Team |
| EPI_ISL_2437843, EPI_ISL_2437844, EPI_ISL_2437845, EPI_ISL_2437846 | Lighthouse Lab in Milton Keynes | Wellcome Sanger Institute for the COVID-19 Genomics UK (COG-UK) Consortium | The Lighthouse Lab in Milton Keynes and Alex Alderton, Roberto Amato, Jeffrey Barrett, Sonia Goncalves, Ewan Harrison, David K. Jackson, Ian Johnston, Dominic Kwiatkowski, Cordelia Langford, John Sillitoe on behalf of the Wellcome Sanger Institute COVID-19 Surveillance Team |
| EPI_ISL_2437847 | Lighthouse Lab in Alderley Park | Wellcome Sanger Institute for the COVID-19 Genomics UK (COG-UK) Consortium | Jacquelyn Wynn, Mairead Hyland, The Lighthouse Lab in Alderley Park and Alex Alderton, Roberto Amato, Jeffrey Barrett, Sonia Goncalves, Ewan Harrison, David K. Jackson, Ian Johnston, Dominic Kwiatkowski, Cordelia Langford, John Sillitoe on behalf of the Wellcome Sanger Institute COVID-19 Surveillance Team |
| EPI_ISL_2437848 | Lighthouse Lab in Glasgow | Wellcome Sanger Institute for the COVID-19 Genomics UK (COG-UK) Consortium | Harper VanSteenhouse, Yumi Kasai, David Gray, Carol Clugston, Anna Dominiczak and Alex Alderton, Roberto Amato, Jeffrey Barrett, Sonia Goncalves, Ewan Harrison, David K. Jackson, Ian Johnston, Dominic Kwiatkowski, Cordelia Langford, John Sillitoe on behalf of the Wellcome Sanger Institute COVID-19 Surveillance Team |
| EPI_ISL_2437849, EPI_ISL_2437850 | Lighthouse Lab in Milton Keynes | Wellcome Sanger Institute for the COVID-19 Genomics UK (COG-UK) Consortium | The Lighthouse Lab in Milton Keynes and Alex Alderton, Roberto Amato, Jeffrey Barrett, Sonia Goncalves, Ewan Harrison, David K. Jackson, Ian Johnston, Dominic Kwiatkowski, Cordelia Langford, John Sillitoe on behalf of the Wellcome Sanger Institute COVID-19 Surveillance Team |
| EPI_ISL_2437851 | Lighthouse Lab in Alderley Park | Wellcome Sanger Institute for the COVID-19 Genomics UK (COG-UK) Consortium | Jacquelyn Wynn, Mairead Hyland, The Lighthouse Lab in Alderley Park and Alex Alderton, Roberto Amato, Jeffrey Barrett, Sonia Goncalves, Ewan Harrison, David K. Jackson, Ian Johnston, Dominic Kwiatkowski, Cordelia Langford, John Sillitoe on behalf of the Wellcome Sanger Institute COVID-19 Surveillance Team |
| EPI_ISL_2437852, EPI_ISL_2437853 | Lighthouse Lab in Glasgow | Wellcome Sanger Institute for the COVID-19 Genomics UK (COG-UK) Consortium | Harper VanSteenhouse, Yumi Kasai, David Gray, Carol Clugston, Anna Dominiczak and Alex Alderton, Roberto Amato, Jeffrey Barrett, Sonia Goncalves, Ewan Harrison, David K. Jackson, Ian Johnston, Dominic Kwiatkowski, Cordelia Langford, John Sillitoe on behalf of the Wellcome Sanger Institute COVID-19 Surveillance Team |
| EPI_ISL_2437854, EPI_ISL_2437855, EPI_ISL_2437857 | Lighthouse Lab in Milton Keynes | Wellcome Sanger Institute for the COVID-19 Genomics UK (COG-UK) Consortium | The Lighthouse Lab in Milton Keynes and Alex Alderton, Roberto Amato, Jeffrey Barrett, Sonia Goncalves, Ewan Harrison, David K. Jackson, Ian Johnston, Dominic Kwiatkowski, Cordelia Langford, John Sillitoe on behalf of the Wellcome Sanger Institute COVID-19 Surveillance Team |
| EPI_ISL_2437858, EPI_ISL_2437859 | Lighthouse Lab in Alderley Park | Wellcome Sanger Institute for the COVID-19 Genomics UK (COG-UK) Consortium | Jacquelyn Wynn, Mairead Hyland, The Lighthouse Lab in Alderley Park and Alex Alderton, Roberto Amato, Jeffrey Barrett, Sonia Goncalves, Ewan Harrison, David K. Jackson, Ian Johnston, Dominic Kwiatkowski, Cordelia Langford, John Sillitoe on behalf of the Wellcome Sanger Institute COVID-19 Surveillance Team |
| EPI_ISL_2437860, EPI_ISL_2437861, EPI_ISL_2437862 | Lighthouse Lab in Glasgow | Wellcome Sanger Institute for the COVID-19 Genomics UK (COG-UK) Consortium | Harper VanSteenhouse, Yumi Kasai, David Gray, Carol Clugston, Anna Dominiczak and Alex Alderton, Roberto Amato, Jeffrey Barrett, Sonia Goncalves, Ewan Harrison, David K. Jackson, Ian Johnston, Dominic Kwiatkowski, Cordelia Langford, John Sillitoe on behalf of the Wellcome Sanger Institute COVID-19 Surveillance Team |
| EPI_ISL_2437863, EPI_ISL_2437864 | Lighthouse Lab in Milton Keynes | Wellcome Sanger Institute for the COVID-19 Genomics UK (COG-UK) Consortium | The Lighthouse Lab in Milton Keynes and Alex Alderton, Roberto Amato, Jeffrey Barrett, Sonia Goncalves, Ewan Harrison, David K. Jackson, Ian Johnston, Dominic Kwiatkowski, Cordelia Langford, John Sillitoe on behalf of the Wellcome Sanger Institute COVID-19 Surveillance Team |
| EPI_ISL_2437865 | Lighthouse Lab in Glasgow | Wellcome Sanger Institute for the COVID-19 Genomics UK (COG-UK) Consortium | Harper VanSteenhouse, Yumi Kasai, David Gray, Carol Clugston, Anna Dominiczak and Alex Alderton, Roberto Amato, Jeffrey Barrett, Sonia Goncalves, Ewan Harrison, David K. Jackson, Ian Johnston, Dominic Kwiatkowski, Cordelia Langford, John Sillitoe on behalf of the Wellcome Sanger Institute COVID-19 Surveillance Team |
| EPI_ISL_2437866, EPI_ISL_2437867 | Lighthouse Lab in Alderley Park | Wellcome Sanger Institute for the COVID-19 Genomics UK (COG-UK) Consortium | Jacquelyn Wynn, Mairead Hyland, The Lighthouse Lab in Alderley Park and Alex Alderton, Roberto Amato, Jeffrey Barrett, Sonia Goncalves, Ewan Harrison, David K. Jackson, Ian Johnston, Dominic Kwiatkowski, Cordelia Langford, John Sillitoe on behalf of the Wellcome Sanger Institute COVID-19 Surveillance Team |
| EPI_ISL_2437868, EPI_ISL_2437869, EPI_ISL_2437870 | Lighthouse Lab in Milton Keynes | Wellcome Sanger Institute for the COVID-19 Genomics UK (COG-UK) Consortium | The Lighthouse Lab in Milton Keynes and Alex Alderton, Roberto Amato, Jeffrey Barrett, Sonia Goncalves, Ewan Harrison, David K. Jackson, Ian Johnston, Dominic Kwiatkowski, Cordelia Langford, John Sillitoe on behalf of the Wellcome Sanger Institute COVID-19 Surveillance Team |
| EPI_ISL_2437871 | Lighthouse Lab in Alderley Park | Wellcome Sanger Institute for the COVID-19 Genomics UK (COG-UK) Consortium | Jacquelyn Wynn, Mairead Hyland, The Lighthouse Lab in Alderley Park and Alex Alderton, Roberto Amato, Jeffrey Barrett, Sonia Goncalves, Ewan Harrison, David K. Jackson, Ian Johnston, Dominic Kwiatkowski, Cordelia Langford, John Sillitoe on behalf of the Wellcome Sanger Institute COVID-19 Surveillance Team |
| EPI_ISL_2437872 | Lighthouse Lab in Glasgow | Wellcome Sanger Institute for the COVID-19 Genomics UK (COG-UK) Consortium | Harper VanSteenhouse, Yumi Kasai, David Gray, Carol Clugston, Anna Dominiczak and Alex Alderton, Roberto Amato, Jeffrey Barrett, Sonia Goncalves, Ewan Harrison, David K. Jackson, Ian Johnston, Dominic Kwiatkowski, Cordelia Langford, John Sillitoe on behalf of the Wellcome Sanger Institute COVID-19 Surveillance Team |
| EPI_ISL_2437874, EPI_ISL_2437876 | Lighthouse Lab in Milton Keynes | Wellcome Sanger Institute for the COVID-19 Genomics UK (COG-UK) Consortium | The Lighthouse Lab in Milton Keynes and Alex Alderton, Roberto Amato, Jeffrey Barrett, Sonia Goncalves, Ewan Harrison, David K. Jackson, Ian Johnston, Dominic Kwiatkowski, Cordelia Langford, John Sillitoe on behalf of the Wellcome Sanger Institute COVID-19 Surveillance Team |
| EPI_ISL_2437877 | Lighthouse Lab in Alderley Park | Wellcome Sanger Institute for the COVID-19 Genomics UK (COG-UK) Consortium | Jacquelyn Wynn, Mairead Hyland, The Lighthouse Lab in Alderley Park and Alex Alderton, Roberto Amato, Jeffrey Barrett, Sonia Goncalves, Ewan Harrison, David K. Jackson, Ian Johnston, Dominic Kwiatkowski, Cordelia Langford, John Sillitoe on behalf of the Wellcome Sanger Institute COVID-19 Surveillance Team |
| EPI_ISL_2437880 | Lighthouse Lab in Glasgow | Wellcome Sanger Institute for the COVID-19 Genomics UK (COG-UK) Consortium | Harper VanSteenhouse, Yumi Kasai, David Gray, Carol Clugston, Anna Dominiczak and Alex Alderton, Roberto Amato, Jeffrey Barrett, Sonia Goncalves, Ewan Harrison, David K. Jackson, Ian Johnston, Dominic Kwiatkowski, Cordelia Langford, John Sillitoe on behalf of the Wellcome Sanger Institute COVID-19 Surveillance Team |
| EPI_ISL_2437881 | Lighthouse Lab in Milton Keynes | Wellcome Sanger Institute for the COVID-19 Genomics UK (COG-UK) Consortium | The Lighthouse Lab in Milton Keynes and Alex Alderton, Roberto Amato, Jeffrey Barrett, Sonia Goncalves, Ewan Harrison, David K. Jackson, Ian Johnston, Dominic Kwiatkowski, Cordelia Langford, John Sillitoe on behalf of the Wellcome Sanger Institute COVID-19 Surveillance Team |
| EPI_ISL_2437882 | Lighthouse Lab in Alderley Park | Wellcome Sanger Institute for the COVID-19 Genomics UK (COG-UK) Consortium | Jacquelyn Wynn, Mairead Hyland, The Lighthouse Lab in Alderley Park and Alex Alderton, Roberto Amato, Jeffrey Barrett, Sonia Goncalves, Ewan Harrison, David K. Jackson, Ian Johnston, Dominic Kwiatkowski, Cordelia Langford, John Sillitoe on behalf of the Wellcome Sanger Institute COVID-19 Surveillance Team |
| EPI_ISL_2437883, EPI_ISL_2437886 | Lighthouse Lab in Glasgow | Wellcome Sanger Institute for the COVID-19 Genomics UK (COG-UK) Consortium | Harper VanSteenhouse, Yumi Kasai, David Gray, Carol Clugston, Anna Dominiczak and Alex Alderton, Roberto Amato, Jeffrey Barrett, Sonia Goncalves, Ewan Harrison, David K. Jackson, Ian Johnston, Dominic Kwiatkowski, Cordelia Langford, John Sillitoe on behalf of the Wellcome Sanger Institute COVID-19 Surveillance Team |
| EPI_ISL_2437887, EPI_ISL_2437888 | Lighthouse Lab in Milton Keynes | Wellcome Sanger Institute for the COVID-19 Genomics UK (COG-UK) Consortium | The Lighthouse Lab in Milton Keynes and Alex Alderton, Roberto Amato, Jeffrey Barrett, Sonia Goncalves, Ewan Harrison, David K. Jackson, Ian Johnston, Dominic Kwiatkowski, Cordelia Langford, John Sillitoe on behalf of the Wellcome Sanger Institute COVID-19 Surveillance Team |
| EPI_ISL_2437889 | Lighthouse Lab in Alderley Park | Wellcome Sanger Institute for the COVID-19 Genomics UK (COG-UK) Consortium | Jacquelyn Wynn, Mairead Hyland, The Lighthouse Lab in Alderley Park and Alex Alderton, Roberto Amato, Jeffrey Barrett, Sonia Goncalves, Ewan Harrison, David K. Jackson, Ian Johnston, Dominic Kwiatkowski, Cordelia Langford, John Sillitoe on behalf of the Wellcome Sanger Institute COVID-19 Surveillance Team |
| EPI_ISL_2437891 | Lighthouse Lab in Milton Keynes | Wellcome Sanger Institute for the COVID-19 Genomics UK (COG-UK) Consortium | The Lighthouse Lab in Milton Keynes and Alex Alderton, Roberto Amato, Jeffrey Barrett, Sonia Goncalves, Ewan Harrison, David K. Jackson, Ian Johnston, Dominic Kwiatkowski, Cordelia Langford, John Sillitoe on behalf of the Wellcome Sanger Institute COVID-19 Surveillance Team |
| EPI_ISL_2437892, EPI_ISL_2437893 | Lighthouse Lab in Glasgow | Wellcome Sanger Institute for the COVID-19 Genomics UK (COG-UK) Consortium | Harper VanSteenhouse, Yumi Kasai, David Gray, Carol Clugston, Anna Dominiczak and Alex Alderton, Roberto Amato, Jeffrey Barrett, Sonia Goncalves, Ewan Harrison, David K. Jackson, Ian Johnston, Dominic Kwiatkowski, Cordelia Langford, John Sillitoe on behalf of the Wellcome Sanger Institute COVID-19 Surveillance Team |
| EPI_ISL_2437894 | Lighthouse Lab in Milton Keynes | Wellcome Sanger Institute for the COVID-19 Genomics UK (COG-UK) Consortium | The Lighthouse Lab in Milton Keynes and Alex Alderton, Roberto Amato, Jeffrey Barrett, Sonia Goncalves, Ewan Harrison, David K. Jackson, Ian Johnston, Dominic Kwiatkowski, Cordelia Langford, John Sillitoe on behalf of the Wellcome Sanger Institute COVID-19 Surveillance Team |
| EPI_ISL_2437895, EPI_ISL_2437896 | Lighthouse Lab in Alderley Park | Wellcome Sanger Institute for the COVID-19 Genomics UK (COG-UK) Consortium | Jacquelyn Wynn, Mairead Hyland, The Lighthouse Lab in Alderley Park and Alex Alderton, Roberto Amato, Jeffrey Barrett, Sonia Goncalves, Ewan Harrison, David K. Jackson, Ian Johnston, Dominic Kwiatkowski, Cordelia Langford, John Sillitoe on behalf of the Wellcome Sanger Institute COVID-19 Surveillance Team |
| EPI_ISL_2437897 | Lighthouse Lab in Glasgow | Wellcome Sanger Institute for the COVID-19 Genomics UK (COG-UK) Consortium | Harper VanSteenhouse, Yumi Kasai, David Gray, Carol Clugston, Anna Dominiczak and Alex Alderton, Roberto Amato, Jeffrey Barrett, Sonia Goncalves, Ewan Harrison, David K. Jackson, Ian Johnston, Dominic Kwiatkowski, Cordelia Langford, John Sillitoe on behalf of the Wellcome Sanger Institute COVID-19 Surveillance Team |
| EPI_ISL_2437898, EPI_ISL_2437899, EPI_ISL_2437900 | Lighthouse Lab in Alderley Park | Wellcome Sanger Institute for the COVID-19 Genomics UK (COG-UK) Consortium | Jacquelyn Wynn, Mairead Hyland, The Lighthouse Lab in Alderley Park and Alex Alderton, Roberto Amato, Jeffrey Barrett, Sonia Goncalves, Ewan Harrison, David K. Jackson, Ian Johnston, Dominic Kwiatkowski, Cordelia Langford, John Sillitoe on behalf of the Wellcome Sanger Institute COVID-19 Surveillance Team |
| EPI_ISL_2437901 | Lighthouse Lab in Milton Keynes | Wellcome Sanger Institute for the COVID-19 Genomics UK (COG-UK) Consortium | The Lighthouse Lab in Milton Keynes and Alex Alderton, Roberto Amato, Jeffrey Barrett, Sonia Goncalves, Ewan Harrison, David K. Jackson, Ian Johnston, Dominic Kwiatkowski, Cordelia Langford, John Sillitoe on behalf of the Wellcome Sanger Institute COVID-19 Surveillance Team |
| EPI_ISL_2437902 | Lighthouse Lab in Alderley Park | Wellcome Sanger Institute for the COVID-19 Genomics UK (COG-UK) Consortium | Jacquelyn Wynn, Mairead Hyland, The Lighthouse Lab in Alderley Park and Alex Alderton, Roberto Amato, Jeffrey Barrett, Sonia Goncalves, Ewan Harrison, David K. Jackson, Ian Johnston, Dominic Kwiatkowski, Cordelia Langford, John Sillitoe on behalf of the Wellcome Sanger Institute COVID-19 Surveillance Team |

| | | | |
|---|---|---|---|
| EPI_ISL_2437904, EPI_ISL_2437905 | Lighthouse Lab in Milton Keynes | Wellcome Sanger Institute for the COVID-19 Genomics UK (COG-UK) Consortium | The Lighthouse Lab in Milton Keynes and Alex Alderton, Roberto Amato, Jeffrey Barrett, Sonia Goncalves, Ewan Harrison, David K. Jackson, Ian Johnston, Dominic Kwiatkowski, Cordelia Langford, John Sillitoe on behalf of the Wellcome Sanger Institute COVID-19 Surveillance Team |
| EPI_ISL_2437906 | Lighthouse Lab in Alderley Park | Wellcome Sanger Institute for the COVID-19 Genomics UK (COG-UK) Consortium | Jacquelyn Wynn, Mairead Hyland, The Lighthouse Lab in Alderley Park and Alex Alderton, Roberto Amato, Jeffrey Barrett, Sonia Goncalves, Ewan Harrison, David K. Jackson, Ian Johnston, Dominic Kwiatkowski, Cordelia Langford, John Sillitoe on behalf of the Wellcome Sanger Institute COVID-19 Surveillance Team |
| EPI_ISL_2437907 | Lighthouse Lab in Glasgow | Wellcome Sanger Institute for the COVID-19 Genomics UK (COG-UK) Consortium | Harper VanSteenhouse, Yumi Kasai, David Gray, Carol Clugston, Anna Dominiczak and Alex Alderton, Roberto Amato, Jeffrey Barrett, Sonia Goncalves, Ewan Harrison, David K. Jackson, Ian Johnston, Dominic Kwiatkowski, Cordelia Langford, John Sillitoe on behalf of the Wellcome Sanger Institute COVID-19 Surveillance Team |
| EPI_ISL_2437908, EPI_ISL_2437909 | Lighthouse Lab in Milton Keynes | Wellcome Sanger Institute for the COVID-19 Genomics UK (COG-UK) Consortium | The Lighthouse Lab in Milton Keynes and Alex Alderton, Roberto Amato, Jeffrey Barrett, Sonia Goncalves, Ewan Harrison, David K. Jackson, Ian Johnston, Dominic Kwiatkowski, Cordelia Langford, John Sillitoe on behalf of the Wellcome Sanger Institute COVID-19 Surveillance Team |
| EPI_ISL_2437910 | Lighthouse Lab in Glasgow | Wellcome Sanger Institute for the COVID-19 Genomics UK (COG-UK) Consortium | Harper VanSteenhouse, Yumi Kasai, David Gray, Carol Clugston, Anna Dominiczak and Alex Alderton, Roberto Amato, Jeffrey Barrett, Sonia Goncalves, Ewan Harrison, David K. Jackson, Ian Johnston, Dominic Kwiatkowski, Cordelia Langford, John Sillitoe on behalf of the Wellcome Sanger Institute COVID-19 Surveillance Team |
| EPI_ISL_2437911 | Lighthouse Lab in Milton Keynes | Wellcome Sanger Institute for the COVID-19 Genomics UK (COG-UK) Consortium | The Lighthouse Lab in Milton Keynes and Alex Alderton, Roberto Amato, Jeffrey Barrett, Sonia Goncalves, Ewan Harrison, David K. Jackson, Ian Johnston, Dominic Kwiatkowski, Cordelia Langford, John Sillitoe on behalf of the Wellcome Sanger Institute COVID-19 Surveillance Team |
| EPI_ISL_2437912 | Lighthouse Lab in Alderley Park | Wellcome Sanger Institute for the COVID-19 Genomics UK (COG-UK) Consortium | Jacquelyn Wynn, Mairead Hyland, The Lighthouse Lab in Alderley Park and Alex Alderton, Roberto Amato, Jeffrey Barrett, Sonia Goncalves, Ewan Harrison, David K. Jackson, Ian Johnston, Dominic Kwiatkowski, Cordelia Langford, John Sillitoe on behalf of the Wellcome Sanger Institute COVID-19 Surveillance Team |
| EPI_ISL_2437913 | Lighthouse Lab in Milton Keynes | Wellcome Sanger Institute for the COVID-19 Genomics UK (COG-UK) Consortium | The Lighthouse Lab in Milton Keynes and Alex Alderton, Roberto Amato, Jeffrey Barrett, Sonia Goncalves, Ewan Harrison, David K. Jackson, Ian Johnston, Dominic Kwiatkowski, Cordelia Langford, John Sillitoe on behalf of the Wellcome Sanger Institute COVID-19 Surveillance Team |
| EPI_ISL_2437914, EPI_ISL_2437915, EPI_ISL_2437917 | Lighthouse Lab in Glasgow | Wellcome Sanger Institute for the COVID-19 Genomics UK (COG-UK) Consortium | Harper VanSteenhouse, Yumi Kasai, David Gray, Carol Clugston, Anna Dominiczak and Alex Alderton, Roberto Amato, Jeffrey Barrett, Sonia Goncalves, Ewan Harrison, David K. Jackson, Ian Johnston, Dominic Kwiatkowski, Cordelia Langford, John Sillitoe on behalf of the Wellcome Sanger Institute COVID-19 Surveillance Team |
| EPI_ISL_2437919, EPI_ISL_2437921 | Lighthouse Lab in Alderley Park | Wellcome Sanger Institute for the COVID-19 Genomics UK (COG-UK) Consortium | Jacquelyn Wynn, Mairead Hyland, The Lighthouse Lab in Alderley Park and Alex Alderton, Roberto Amato, Jeffrey Barrett, Sonia Goncalves, Ewan Harrison, David K. Jackson, Ian Johnston, Dominic Kwiatkowski, Cordelia Langford, John Sillitoe on behalf of the Wellcome Sanger Institute COVID-19 Surveillance Team |
| EPI_ISL_2437922, EPI_ISL_2437923 | Lighthouse Lab in Milton Keynes | Wellcome Sanger Institute for the COVID-19 Genomics UK (COG-UK) Consortium | The Lighthouse Lab in Milton Keynes and Alex Alderton, Roberto Amato, Jeffrey Barrett, Sonia Goncalves, Ewan Harrison, David K. Jackson, Ian Johnston, Dominic Kwiatkowski, Cordelia Langford, John Sillitoe on behalf of the Wellcome Sanger Institute COVID-19 Surveillance Team |
| EPI_ISL_2437924 | Lighthouse Lab in Glasgow | Wellcome Sanger Institute for the COVID-19 Genomics UK (COG-UK) Consortium | Harper VanSteenhouse, Yumi Kasai, David Gray, Carol Clugston, Anna Dominiczak and Alex Alderton, Roberto Amato, Jeffrey Barrett, Sonia Goncalves, Ewan Harrison, David K. Jackson, Ian Johnston, Dominic Kwiatkowski, Cordelia Langford, John Sillitoe on behalf of the Wellcome Sanger Institute COVID-19 Surveillance Team |
| EPI_ISL_2437925 | Lighthouse Lab in Alderley Park | Wellcome Sanger Institute for the COVID-19 Genomics UK (COG-UK) Consortium | Jacquelyn Wynn, Mairead Hyland, The Lighthouse Lab in Alderley Park and Alex Alderton, Roberto Amato, Jeffrey Barrett, Sonia Goncalves, Ewan Harrison, David K. Jackson, Ian Johnston, Dominic Kwiatkowski, Cordelia Langford, John Sillitoe on behalf of the Wellcome Sanger Institute COVID-19 Surveillance Team |
| EPI_ISL_2437926 | Lighthouse Lab in Glasgow | Wellcome Sanger Institute for the COVID-19 Genomics UK (COG-UK) Consortium | Harper VanSteenhouse, Yumi Kasai, David Gray, Carol Clugston, Anna Dominiczak and Alex Alderton, Roberto Amato, Jeffrey Barrett, Sonia Goncalves, Ewan Harrison, David K. Jackson, Ian Johnston, Dominic Kwiatkowski, Cordelia Langford, John Sillitoe on behalf of the Wellcome Sanger Institute COVID-19 Surveillance Team |
| EPI_ISL_2437927, EPI_ISL_2437929 | Lighthouse Lab in Alderley Park | Wellcome Sanger Institute for the COVID-19 Genomics UK (COG-UK) Consortium | Jacquelyn Wynn, Mairead Hyland, The Lighthouse Lab in Alderley Park and Alex Alderton, Roberto Amato, Jeffrey Barrett, Sonia Goncalves, Ewan Harrison, David K. Jackson, Ian Johnston, Dominic Kwiatkowski, Cordelia Langford, John Sillitoe on behalf of the Wellcome Sanger Institute COVID-19 Surveillance Team |
| EPI_ISL_2437930, EPI_ISL_2437931 | Lighthouse Lab in Milton Keynes | Wellcome Sanger Institute for the COVID-19 Genomics UK (COG-UK) Consortium | The Lighthouse Lab in Milton Keynes and Alex Alderton, Roberto Amato, Jeffrey Barrett, Sonia Goncalves, Ewan Harrison, David K. Jackson, Ian Johnston, Dominic Kwiatkowski, Cordelia Langford, John Sillitoe on behalf of the Wellcome Sanger Institute COVID-19 Surveillance Team |
| EPI_ISL_2437932 | Lighthouse Lab in Alderley Park | Wellcome Sanger Institute for the COVID-19 Genomics UK (COG-UK) Consortium | Jacquelyn Wynn, Mairead Hyland, The Lighthouse Lab in Alderley Park and Alex Alderton, Roberto Amato, Jeffrey Barrett, Sonia Goncalves, Ewan Harrison, David K. Jackson, Ian Johnston, Dominic Kwiatkowski, Cordelia Langford, John Sillitoe on behalf of the Wellcome Sanger Institute COVID-19 Surveillance Team |
| EPI_ISL_2437934 | Lighthouse Lab in Glasgow | Wellcome Sanger Institute for the COVID-19 Genomics UK (COG-UK) Consortium | Harper VanSteenhouse, Yumi Kasai, David Gray, Carol Clugston, Anna Dominiczak and Alex Alderton, Roberto Amato, Jeffrey Barrett, Sonia Goncalves, Ewan Harrison, David K. Jackson, Ian Johnston, Dominic Kwiatkowski, Cordelia Langford, John Sillitoe on behalf of the Wellcome Sanger Institute COVID-19 Surveillance Team |
| EPI_ISL_2437935 | Lighthouse Lab in Alderley Park | Wellcome Sanger Institute for the COVID-19 Genomics UK (COG-UK) Consortium | Jacquelyn Wynn, Mairead Hyland, The Lighthouse Lab in Alderley Park and Alex Alderton, Roberto Amato, Jeffrey Barrett, Sonia Goncalves, Ewan Harrison, David K. Jackson, Ian Johnston, Dominic Kwiatkowski, Cordelia Langford, John Sillitoe on behalf of the Wellcome Sanger Institute COVID-19 Surveillance Team |
| EPI_ISL_2437937 | Lighthouse Lab in Glasgow | Wellcome Sanger Institute for the COVID-19 Genomics UK (COG-UK) Consortium | Harper VanSteenhouse, Yumi Kasai, David Gray, Carol Clugston, Anna Dominiczak and Alex Alderton, Roberto Amato, Jeffrey Barrett, Sonia Goncalves, Ewan Harrison, David K. Jackson, Ian Johnston, Dominic Kwiatkowski, Cordelia Langford, John Sillitoe on behalf of the Wellcome Sanger Institute COVID-19 Surveillance Team |
| EPI_ISL_2437938 | Lighthouse Lab in Alderley Park | Wellcome Sanger Institute for the COVID-19 Genomics UK (COG-UK) Consortium | Jacquelyn Wynn, Mairead Hyland, The Lighthouse Lab in Alderley Park and Alex Alderton, Roberto Amato, Jeffrey Barrett, Sonia Goncalves, Ewan Harrison, David K. Jackson, Ian Johnston, Dominic Kwiatkowski, Cordelia Langford, John Sillitoe on behalf of the Wellcome Sanger Institute COVID-19 Surveillance Team |
| EPI_ISL_2437939, EPI_ISL_2437940 | Lighthouse Lab in Milton Keynes | Wellcome Sanger Institute for the COVID-19 Genomics UK (COG-UK) Consortium | The Lighthouse Lab in Milton Keynes and Alex Alderton, Roberto Amato, Jeffrey Barrett, Sonia Goncalves, Ewan Harrison, David K. Jackson, Ian Johnston, Dominic Kwiatkowski, Cordelia Langford, John Sillitoe on behalf of the Wellcome Sanger Institute COVID-19 Surveillance Team |
| EPI_ISL_2437941, EPI_ISL_2437942 | Lighthouse Lab in Alderley Park | Wellcome Sanger Institute for the COVID-19 Genomics UK (COG-UK) Consortium | Jacquelyn Wynn, Mairead Hyland, The Lighthouse Lab in Alderley Park and Alex Alderton, Roberto Amato, Jeffrey Barrett, Sonia Goncalves, Ewan Harrison, David K. Jackson, Ian Johnston, Dominic Kwiatkowski, Cordelia Langford, John Sillitoe on behalf of the Wellcome Sanger Institute COVID-19 Surveillance Team |
| EPI_ISL_2437943 | Lighthouse Lab in Milton Keynes | Wellcome Sanger Institute for the COVID-19 Genomics UK (COG-UK) Consortium | The Lighthouse Lab in Milton Keynes and Alex Alderton, Roberto Amato, Jeffrey Barrett, Sonia Goncalves, Ewan Harrison, David K. Jackson, Ian Johnston, Dominic Kwiatkowski, Cordelia Langford, John Sillitoe on behalf of the Wellcome Sanger Institute COVID-19 Surveillance Team |
| EPI_ISL_2437944 | Lighthouse Lab in Alderley Park | Wellcome Sanger Institute for the COVID-19 Genomics UK (COG-UK) Consortium | Jacquelyn Wynn, Mairead Hyland, The Lighthouse Lab in Alderley Park and Alex Alderton, Roberto Amato, Jeffrey Barrett, Sonia Goncalves, Ewan Harrison, David K. Jackson, Ian Johnston, Dominic Kwiatkowski, Cordelia Langford, John Sillitoe on behalf of the Wellcome Sanger Institute COVID-19 Surveillance Team |
| EPI_ISL_2437945, EPI_ISL_2437946 | Lighthouse Lab in Glasgow | Wellcome Sanger Institute for the COVID-19 Genomics UK (COG-UK) Consortium | Harper VanSteenhouse, Yumi Kasai, David Gray, Carol Clugston, Anna Dominiczak and Alex Alderton, Roberto Amato, Jeffrey Barrett, Sonia Goncalves, Ewan Harrison, David K. Jackson, Ian Johnston, Dominic Kwiatkowski, Cordelia Langford, John Sillitoe on behalf of the Wellcome Sanger Institute COVID-19 Surveillance Team |
| EPI_ISL_2437947 | Lighthouse Lab in Milton Keynes | Wellcome Sanger Institute for the COVID-19 Genomics UK (COG-UK) Consortium | The Lighthouse Lab in Milton Keynes and Alex Alderton, Roberto Amato, Jeffrey Barrett, Sonia Goncalves, Ewan Harrison, David K. Jackson, Ian Johnston, Dominic Kwiatkowski, Cordelia Langford, John Sillitoe on behalf of the Wellcome Sanger Institute COVID-19 Surveillance Team |
| EPI_ISL_2437948 | Lighthouse Lab in Glasgow | Wellcome Sanger Institute for the COVID-19 Genomics UK (COG-UK) Consortium | Harper VanSteenhouse, Yumi Kasai, David Gray, Carol Clugston, Anna Dominiczak and Alex Alderton, Roberto Amato, Jeffrey Barrett, Sonia Goncalves, Ewan Harrison, David K. Jackson, Ian Johnston, Dominic Kwiatkowski, Cordelia Langford, John Sillitoe on behalf of the Wellcome Sanger Institute COVID-19 Surveillance Team |
| EPI_ISL_2437949, EPI_ISL_2437950, EPI_ISL_2437951, EPI_ISL_2437952, EPI_ISL_2437953, EPI_ISL_2437954 | Lighthouse Lab in Milton Keynes | Wellcome Sanger Institute for the COVID-19 Genomics UK (COG-UK) Consortium | The Lighthouse Lab in Milton Keynes and Alex Alderton, Roberto Amato, Jeffrey Barrett, Sonia Goncalves, Ewan Harrison, David K. Jackson, Ian Johnston, Dominic Kwiatkowski, Cordelia Langford, John Sillitoe on behalf of the Wellcome Sanger Institute COVID-19 Surveillance Team |
| EPI_ISL_2437958 | Lighthouse Lab in Alderley Park | Wellcome Sanger Institute for the COVID-19 Genomics UK (COG-UK) Consortium | Jacquelyn Wynn, Mairead Hyland, The Lighthouse Lab in Alderley Park and Alex Alderton, Roberto Amato, Jeffrey Barrett, Sonia Goncalves, Ewan Harrison, David K. Jackson, Ian Johnston, Dominic Kwiatkowski, Cordelia Langford, John Sillitoe on behalf of the Wellcome Sanger Institute COVID-19 Surveillance Team |
| EPI_ISL_2437959, EPI_ISL_2437960 | Lighthouse Lab in Milton Keynes | Wellcome Sanger Institute for the COVID-19 Genomics UK (COG-UK) Consortium | The Lighthouse Lab in Milton Keynes and Alex Alderton, Roberto Amato, Jeffrey Barrett, Sonia Goncalves, Ewan Harrison, David K. Jackson, Ian Johnston, Dominic Kwiatkowski, Cordelia Langford, John Sillitoe on behalf of the Wellcome Sanger Institute COVID-19 Surveillance Team |
| EPI_ISL_2437961 | Lighthouse Lab in Alderley Park | Wellcome Sanger Institute for the COVID-19 Genomics UK (COG-UK) Consortium | Jacquelyn Wynn, Mairead Hyland, The Lighthouse Lab in Alderley Park and Alex Alderton, Roberto Amato, Jeffrey Barrett, Sonia Goncalves, Ewan Harrison, David K. Jackson, Ian Johnston, Dominic Kwiatkowski, Cordelia Langford, John Sillitoe on behalf of the Wellcome Sanger Institute COVID-19 Surveillance Team |
| EPI_ISL_2437962 | Lighthouse Lab in Milton Keynes | Wellcome Sanger Institute for the COVID-19 Genomics UK (COG-UK) Consortium | The Lighthouse Lab in Milton Keynes and Alex Alderton, Roberto Amato, Jeffrey Barrett, Sonia Goncalves, Ewan Harrison, David K. Jackson, Ian Johnston, Dominic Kwiatkowski, Cordelia Langford, John Sillitoe on behalf of the Wellcome Sanger Institute COVID-19 Surveillance Team |
| EPI_ISL_2437963, EPI_ISL_2437964 | Lighthouse Lab in Glasgow | Wellcome Sanger Institute for the COVID-19 Genomics UK (COG-UK) Consortium | Harper VanSteenhouse, Yumi Kasai, David Gray, Carol Clugston, Anna Dominiczak and Alex Alderton, Roberto Amato, Jeffrey Barrett, Sonia Goncalves, Ewan Harrison, David K. Jackson, Ian Johnston, Dominic Kwiatkowski, Cordelia Langford, John Sillitoe on behalf of the Wellcome Sanger Institute COVID-19 Surveillance Team |
| EPI_ISL_2437967 | Lighthouse Lab in Milton Keynes | Wellcome Sanger Institute for the COVID-19 Genomics UK (COG-UK) Consortium | The Lighthouse Lab in Milton Keynes and Alex Alderton, Roberto Amato, Jeffrey Barrett, Sonia Goncalves, Ewan Harrison, David K. Jackson, Ian Johnston, Dominic Kwiatkowski, Cordelia Langford, John Sillitoe on behalf of the Wellcome Sanger Institute COVID-19 Surveillance Team |
| EPI_ISL_2437969 | Lighthouse Lab in Alderley Park | Wellcome Sanger Institute for the COVID-19 Genomics UK (COG-UK) Consortium | Jacquelyn Wynn, Mairead Hyland, The Lighthouse Lab in Alderley Park and Alex Alderton, Roberto Amato, Jeffrey Barrett, Sonia Goncalves, Ewan Harrison, David K. Jackson, Ian Johnston, Dominic Kwiatkowski, Cordelia Langford, John Sillitoe on behalf of the Wellcome Sanger Institute COVID-19 Surveillance Team |
| EPI_ISL_2437970 | Lighthouse Lab in Glasgow | Wellcome Sanger Institute for the COVID-19 Genomics UK (COG-UK) Consortium | Harper VanSteenhouse, Yumi Kasai, David Gray, Carol Clugston, Anna Dominiczak and Alex Alderton, Roberto Amato, Jeffrey Barrett, Sonia Goncalves, Ewan Harrison, David K. Jackson, Ian Johnston, Dominic Kwiatkowski, Cordelia Langford, John Sillitoe on behalf of the Wellcome Sanger Institute COVID-19 Surveillance Team |
| EPI_ISL_2437971 | Lighthouse Lab in Milton Keynes | Wellcome Sanger Institute for the COVID-19 Genomics UK (COG-UK) Consortium | The Lighthouse Lab in Milton Keynes and Alex Alderton, Roberto Amato, Jeffrey Barrett, Sonia Goncalves, Ewan Harrison, David K. Jackson, Ian Johnston, Dominic Kwiatkowski, Cordelia Langford, John Sillitoe on behalf of the Wellcome Sanger Institute COVID-19 Surveillance Team |
| EPI_ISL_2437972, EPI_ISL_2437973, EPI_ISL_2437974 | Lighthouse Lab in Glasgow | Wellcome Sanger Institute for the COVID-19 Genomics UK (COG-UK) Consortium | Harper VanSteenhouse, Yumi Kasai, David Gray, Carol Clugston, Anna Dominiczak and Alex Alderton, Roberto Amato, Jeffrey Barrett, Sonia Goncalves, Ewan Harrison, David K. Jackson, Ian Johnston, Dominic Kwiatkowski, Cordelia Langford, John Sillitoe on behalf of the Wellcome Sanger Institute COVID-19 Surveillance Team |
| EPI_ISL_2437975, EPI_ISL_2437976 | Lighthouse Lab in Alderley Park | Wellcome Sanger Institute for the COVID-19 Genomics UK (COG-UK) Consortium | Jacquelyn Wynn, Mairead Hyland, The Lighthouse Lab in Alderley Park and Alex Alderton, Roberto Amato, Jeffrey Barrett, Sonia Goncalves, Ewan Harrison, David K. Jackson, Ian Johnston, Dominic Kwiatkowski, Cordelia Langford, John Sillitoe on behalf of the Wellcome Sanger Institute COVID-19 Surveillance Team |
| EPI_ISL_2437977 | Lighthouse Lab in Milton Keynes | Wellcome Sanger Institute for the COVID-19 Genomics UK (COG-UK) Consortium | The Lighthouse Lab in Milton Keynes and Alex Alderton, Roberto Amato, Jeffrey Barrett, Sonia Goncalves, Ewan Harrison, David K. Jackson, Ian Johnston, Dominic Kwiatkowski, Cordelia Langford, John Sillitoe on behalf of the Wellcome Sanger Institute COVID-19 Surveillance Team |
| EPI_ISL_2437978, EPI_ISL_2437979 | Lighthouse Lab in Alderley Park | Wellcome Sanger Institute for the COVID-19 Genomics UK (COG-UK) Consortium | Jacquelyn Wynn, Mairead Hyland, The Lighthouse Lab in Alderley Park and Alex Alderton, Roberto Amato, Jeffrey Barrett, Sonia Goncalves, Ewan Harrison, David K. Jackson, Ian Johnston, Dominic Kwiatkowski, Cordelia Langford, John Sillitoe on behalf of the Wellcome Sanger Institute COVID-19 Surveillance Team |
| EPI_ISL_2437980 | Lighthouse Lab in Milton Keynes | Wellcome Sanger Institute for the COVID-19 Genomics UK (COG-UK) Consortium | The Lighthouse Lab in Milton Keynes and Alex Alderton, Roberto Amato, Jeffrey Barrett, Sonia Goncalves, Ewan Harrison, David K. Jackson, Ian Johnston, Dominic Kwiatkowski, Cordelia Langford, John Sillitoe on behalf of the Wellcome Sanger Institute COVID-19 Surveillance Team |

| | | | |
|---|---|---|---|
| EPI_ISL_2437981, EPI_ISL_2437982 | Lighthouse Lab in Glasgow | Wellcome Sanger Institute for the COVID-19 Genomics UK (COG-UK) Consortium | Harper VanSteenhouse, Yumi Kasai, David Gray, Carol Clugston, Anna Dominiczak and Alex Alderton, Roberto Amato, Jeffrey Barrett, Sonia Goncalves, Ewan Harrison, David K. Jackson, Ian Johnston, Dominic Kwiatkowski, Cordelia Langford, John Sillitoe on behalf of the Wellcome Sanger Institute COVID-19 Surveillance Team |
| EPI_ISL_2437983, EPI_ISL_2437984 | Lighthouse Lab in Alderley Park | Wellcome Sanger Institute for the COVID-19 Genomics UK (COG-UK) Consortium | Jacquelyn Wynn, Mairead Hyland, The Lighthouse Lab in Alderley Park and Alex Alderton, Roberto Amato, Jeffrey Barrett, Sonia Goncalves, Ewan Harrison, David K. Jackson, Ian Johnston, Dominic Kwiatkowski, Cordelia Langford, John Sillitoe on behalf of the Wellcome Sanger Institute COVID-19 Surveillance Team |
| EPI_ISL_2437985 | Lighthouse Lab in Glasgow | Wellcome Sanger Institute for the COVID-19 Genomics UK (COG-UK) Consortium | Harper VanSteenhouse, Yumi Kasai, David Gray, Carol Clugston, Anna Dominiczak and Alex Alderton, Roberto Amato, Jeffrey Barrett, Sonia Goncalves, Ewan Harrison, David K. Jackson, Ian Johnston, Dominic Kwiatkowski, Cordelia Langford, John Sillitoe on behalf of the Wellcome Sanger Institute COVID-19 Surveillance Team |
| EPI_ISL_2437987 | Lighthouse Lab in Alderley Park | Wellcome Sanger Institute for the COVID-19 Genomics UK (COG-UK) Consortium | Jacquelyn Wynn, Mairead Hyland, The Lighthouse Lab in Alderley Park and Alex Alderton, Roberto Amato, Jeffrey Barrett, Sonia Goncalves, Ewan Harrison, David K. Jackson, Ian Johnston, Dominic Kwiatkowski, Cordelia Langford, John Sillitoe on behalf of the Wellcome Sanger Institute COVID-19 Surveillance Team |
| EPI_ISL_2437988 | Lighthouse Lab in Milton Keynes | Wellcome Sanger Institute for the COVID-19 Genomics UK (COG-UK) Consortium | The Lighthouse Lab in Milton Keynes and Alex Alderton, Roberto Amato, Jeffrey Barrett, Sonia Goncalves, Ewan Harrison, David K. Jackson, Ian Johnston, Dominic Kwiatkowski, Cordelia Langford, John Sillitoe on behalf of the Wellcome Sanger Institute COVID-19 Surveillance Team |
| EPI_ISL_2437990 | Lighthouse Lab in Glasgow | Wellcome Sanger Institute for the COVID-19 Genomics UK (COG-UK) Consortium | Harper VanSteenhouse, Yumi Kasai, David Gray, Carol Clugston, Anna Dominiczak and Alex Alderton, Roberto Amato, Jeffrey Barrett, Sonia Goncalves, Ewan Harrison, David K. Jackson, Ian Johnston, Dominic Kwiatkowski, Cordelia Langford, John Sillitoe on behalf of the Wellcome Sanger Institute COVID-19 Surveillance Team |
| EPI_ISL_2437991 | Lighthouse Lab in Alderley Park | Wellcome Sanger Institute for the COVID-19 Genomics UK (COG-UK) Consortium | Jacquelyn Wynn, Mairead Hyland, The Lighthouse Lab in Alderley Park and Alex Alderton, Roberto Amato, Jeffrey Barrett, Sonia Goncalves, Ewan Harrison, David K. Jackson, Ian Johnston, Dominic Kwiatkowski, Cordelia Langford, John Sillitoe on behalf of the Wellcome Sanger Institute COVID-19 Surveillance Team |
| EPI_ISL_2437993, EPI_ISL_2438037, EPI_ISL_2438038, EPI_ISL_2438040, EPI_ISL_2438042, EPI_ISL_2438043, EPI_ISL_2438044 | Lighthouse Lab in Glasgow | Wellcome Sanger Institute for the COVID-19 Genomics UK (COG-UK) Consortium | Harper VanSteenhouse, Yumi Kasai, David Gray, Carol Clugston, Anna Dominiczak and Alex Alderton, Roberto Amato, Jeffrey Barrett, Sonia Goncalves, Ewan Harrison, David K. Jackson, Ian Johnston, Dominic Kwiatkowski, Cordelia Langford, John Sillitoe on behalf of the Wellcome Sanger Institute COVID-19 Surveillance Team |
| EPI_ISL_2438046 | Lighthouse Lab in Alderley Park | Wellcome Sanger Institute for the COVID-19 Genomics UK (COG-UK) Consortium | Jacquelyn Wynn, Mairead Hyland, The Lighthouse Lab in Alderley Park and Alex Alderton, Roberto Amato, Jeffrey Barrett, Sonia Goncalves, Ewan Harrison, David K. Jackson, Ian Johnston, Dominic Kwiatkowski, Cordelia Langford, John Sillitoe on behalf of the Wellcome Sanger Institute COVID-19 Surveillance Team |
| EPI_ISL_2438049 | Lighthouse Lab in Glasgow | Wellcome Sanger Institute for the COVID-19 Genomics UK (COG-UK) Consortium | Harper VanSteenhouse, Yumi Kasai, David Gray, Carol Clugston, Anna Dominiczak and Alex Alderton, Roberto Amato, Jeffrey Barrett, Sonia Goncalves, Ewan Harrison, David K. Jackson, Ian Johnston, Dominic Kwiatkowski, Cordelia Langford, John Sillitoe on behalf of the Wellcome Sanger Institute COVID-19 Surveillance Team |
| EPI_ISL_2438051, EPI_ISL_2438053 | Lighthouse Lab in Alderley Park | Wellcome Sanger Institute for the COVID-19 Genomics UK (COG-UK) Consortium | Jacquelyn Wynn, Mairead Hyland, The Lighthouse Lab in Alderley Park and Alex Alderton, Roberto Amato, Jeffrey Barrett, Sonia Goncalves, Ewan Harrison, David K. Jackson, Ian Johnston, Dominic Kwiatkowski, Cordelia Langford, John Sillitoe on behalf of the Wellcome Sanger Institute COVID-19 Surveillance Team |
| EPI_ISL_2438054 | Lighthouse Lab in Milton Keynes | Wellcome Sanger Institute for the COVID-19 Genomics UK (COG-UK) Consortium | The Lighthouse Lab in Milton Keynes and Alex Alderton, Roberto Amato, Jeffrey Barrett, Sonia Goncalves, Ewan Harrison, David K. Jackson, Ian Johnston, Dominic Kwiatkowski, Cordelia Langford, John Sillitoe on behalf of the Wellcome Sanger Institute COVID-19 Surveillance Team |
| EPI_ISL_2438056, EPI_ISL_2438058 | Lighthouse Lab in Alderley Park | Wellcome Sanger Institute for the COVID-19 Genomics UK (COG-UK) Consortium | Jacquelyn Wynn, Mairead Hyland, The Lighthouse Lab in Alderley Park and Alex Alderton, Roberto Amato, Jeffrey Barrett, Sonia Goncalves, Ewan Harrison, David K. Jackson, Ian Johnston, Dominic Kwiatkowski, Cordelia Langford, John Sillitoe on behalf of the Wellcome Sanger Institute COVID-19 Surveillance Team |
| EPI_ISL_2438060, EPI_ISL_2438061 | Lighthouse Lab in Milton Keynes | Wellcome Sanger Institute for the COVID-19 Genomics UK (COG-UK) Consortium | The Lighthouse Lab in Milton Keynes and Alex Alderton, Roberto Amato, Jeffrey Barrett, Sonia Goncalves, Ewan Harrison, David K. Jackson, Ian Johnston, Dominic Kwiatkowski, Cordelia Langford, John Sillitoe on behalf of the Wellcome Sanger Institute COVID-19 Surveillance Team |
| EPI_ISL_2438063 | Lighthouse Lab in Glasgow | Wellcome Sanger Institute for the COVID-19 Genomics UK (COG-UK) Consortium | Harper VanSteenhouse, Yumi Kasai, David Gray, Carol Clugston, Anna Dominiczak and Alex Alderton, Roberto Amato, Jeffrey Barrett, Sonia Goncalves, Ewan Harrison, David K. Jackson, Ian Johnston, Dominic Kwiatkowski, Cordelia Langford, John Sillitoe on behalf of the Wellcome Sanger Institute COVID-19 Surveillance Team |
| EPI_ISL_2438066 | Lighthouse Lab in Milton Keynes | Wellcome Sanger Institute for the COVID-19 Genomics UK (COG-UK) Consortium | The Lighthouse Lab in Milton Keynes and Alex Alderton, Roberto Amato, Jeffrey Barrett, Sonia Goncalves, Ewan Harrison, David K. Jackson, Ian Johnston, Dominic Kwiatkowski, Cordelia Langford, John Sillitoe on behalf of the Wellcome Sanger Institute COVID-19 Surveillance Team |
| EPI_ISL_2438067 | Lighthouse Lab in Glasgow | Wellcome Sanger Institute for the COVID-19 Genomics UK (COG-UK) Consortium | Harper VanSteenhouse, Yumi Kasai, David Gray, Carol Clugston, Anna Dominiczak and Alex Alderton, Roberto Amato, Jeffrey Barrett, Sonia Goncalves, Ewan Harrison, David K. Jackson, Ian Johnston, Dominic Kwiatkowski, Cordelia Langford, John Sillitoe on behalf of the Wellcome Sanger Institute COVID-19 Surveillance Team |
| EPI_ISL_2438070, EPI_ISL_2438072 | Lighthouse Lab in Milton Keynes | Wellcome Sanger Institute for the COVID-19 Genomics UK (COG-UK) Consortium | The Lighthouse Lab in Milton Keynes and Alex Alderton, Roberto Amato, Jeffrey Barrett, Sonia Goncalves, Ewan Harrison, David K. Jackson, Ian Johnston, Dominic Kwiatkowski, Cordelia Langford, John Sillitoe on behalf of the Wellcome Sanger Institute COVID-19 Surveillance Team |
| EPI_ISL_2438074, EPI_ISL_2438075 | Lighthouse Lab in Glasgow | Wellcome Sanger Institute for the COVID-19 Genomics UK (COG-UK) Consortium | Harper VanSteenhouse, Yumi Kasai, David Gray, Carol Clugston, Anna Dominiczak and Alex Alderton, Roberto Amato, Jeffrey Barrett, Sonia Goncalves, Ewan Harrison, David K. Jackson, Ian Johnston, Dominic Kwiatkowski, Cordelia Langford, John Sillitoe on behalf of the Wellcome Sanger Institute COVID-19 Surveillance Team |
| EPI_ISL_2438076, EPI_ISL_2438081 | Lighthouse Lab in Milton Keynes | Wellcome Sanger Institute for the COVID-19 Genomics UK (COG-UK) Consortium | The Lighthouse Lab in Milton Keynes and Alex Alderton, Roberto Amato, Jeffrey Barrett, Sonia Goncalves, Ewan Harrison, David K. Jackson, Ian Johnston, Dominic Kwiatkowski, Cordelia Langford, John Sillitoe on behalf of the Wellcome Sanger Institute COVID-19 Surveillance Team |
| EPI_ISL_2438084 | Lighthouse Lab in Glasgow | Wellcome Sanger Institute for the COVID-19 Genomics UK (COG-UK) Consortium | Harper VanSteenhouse, Yumi Kasai, David Gray, Carol Clugston, Anna Dominiczak and Alex Alderton, Roberto Amato, Jeffrey Barrett, Sonia Goncalves, Ewan Harrison, David K. Jackson, Ian Johnston, Dominic Kwiatkowski, Cordelia Langford, John Sillitoe on behalf of the Wellcome Sanger Institute COVID-19 Surveillance Team |
| EPI_ISL_2438085 | Lighthouse Lab in Alderley Park | Wellcome Sanger Institute for the COVID-19 Genomics UK (COG-UK) Consortium | Jacquelyn Wynn, Mairead Hyland, The Lighthouse Lab in Alderley Park and Alex Alderton, Roberto Amato, Jeffrey Barrett, Sonia Goncalves, Ewan Harrison, David K. Jackson, Ian Johnston, Dominic Kwiatkowski, Cordelia Langford, John Sillitoe on behalf of the Wellcome Sanger Institute COVID-19 Surveillance Team |
| EPI_ISL_2438087 | Lighthouse Lab in Glasgow | Wellcome Sanger Institute for the COVID-19 Genomics UK (COG-UK) Consortium | Harper VanSteenhouse, Yumi Kasai, David Gray, Carol Clugston, Anna Dominiczak and Alex Alderton, Roberto Amato, Jeffrey Barrett, Sonia Goncalves, Ewan Harrison, David K. Jackson, Ian Johnston, Dominic Kwiatkowski, Cordelia Langford, John Sillitoe on behalf of the Wellcome Sanger Institute COVID-19 Surveillance Team |
| EPI_ISL_2438090 | Lighthouse Lab in Milton Keynes | Wellcome Sanger Institute for the COVID-19 Genomics UK (COG-UK) Consortium | The Lighthouse Lab in Milton Keynes and Alex Alderton, Roberto Amato, Jeffrey Barrett, Sonia Goncalves, Ewan Harrison, David K. Jackson, Ian Johnston, Dominic Kwiatkowski, Cordelia Langford, John Sillitoe on behalf of the Wellcome Sanger Institute COVID-19 Surveillance Team |
| EPI_ISL_2438095, EPI_ISL_2438096 | Lighthouse Lab in Alderley Park | Wellcome Sanger Institute for the COVID-19 Genomics UK (COG-UK) Consortium | Jacquelyn Wynn, Mairead Hyland, The Lighthouse Lab in Alderley Park and Alex Alderton, Roberto Amato, Jeffrey Barrett, Sonia Goncalves, Ewan Harrison, David K. Jackson, Ian Johnston, Dominic Kwiatkowski, Cordelia Langford, John Sillitoe on behalf of the Wellcome Sanger Institute COVID-19 Surveillance Team |
| EPI_ISL_2438097, EPI_ISL_2438103, EPI_ISL_2438105, EPI_ISL_2438106 | Lighthouse Lab in Glasgow | Wellcome Sanger Institute for the COVID-19 Genomics UK (COG-UK) Consortium | Harper VanSteenhouse, Yumi Kasai, David Gray, Carol Clugston, Anna Dominiczak and Alex Alderton, Roberto Amato, Jeffrey Barrett, Sonia Goncalves, Ewan Harrison, David K. Jackson, Ian Johnston, Dominic Kwiatkowski, Cordelia Langford, John Sillitoe on behalf of the Wellcome Sanger Institute COVID-19 Surveillance Team |
| EPI_ISL_2438107, EPI_ISL_2438108, EPI_ISL_2438109 | Lighthouse Lab in Alderley Park | Wellcome Sanger Institute for the COVID-19 Genomics UK (COG-UK) Consortium | Jacquelyn Wynn, Mairead Hyland, The Lighthouse Lab in Alderley Park and Alex Alderton, Roberto Amato, Jeffrey Barrett, Sonia Goncalves, Ewan Harrison, David K. Jackson, Ian Johnston, Dominic Kwiatkowski, Cordelia Langford, John Sillitoe on behalf of the Wellcome Sanger Institute COVID-19 Surveillance Team |
| EPI_ISL_2438110 | Lighthouse Lab in Glasgow | Wellcome Sanger Institute for the COVID-19 Genomics UK (COG-UK) Consortium | Harper VanSteenhouse, Yumi Kasai, David Gray, Carol Clugston, Anna Dominiczak and Alex Alderton, Roberto Amato, Jeffrey Barrett, Sonia Goncalves, Ewan Harrison, David K. Jackson, Ian Johnston, Dominic Kwiatkowski, Cordelia Langford, John Sillitoe on behalf of the Wellcome Sanger Institute COVID-19 Surveillance Team |
| EPI_ISL_2438111 | Lighthouse Lab in Alderley Park | Wellcome Sanger Institute for the COVID-19 Genomics UK (COG-UK) Consortium | Jacquelyn Wynn, Mairead Hyland, The Lighthouse Lab in Alderley Park and Alex Alderton, Roberto Amato, Jeffrey Barrett, Sonia Goncalves, Ewan Harrison, David K. Jackson, Ian Johnston, Dominic Kwiatkowski, Cordelia Langford, John Sillitoe on behalf of the Wellcome Sanger Institute COVID-19 Surveillance Team |
| EPI_ISL_2438112 | Lighthouse Lab in Glasgow | Wellcome Sanger Institute for the COVID-19 Genomics UK (COG-UK) Consortium | Harper VanSteenhouse, Yumi Kasai, David Gray, Carol Clugston, Anna Dominiczak and Alex Alderton, Roberto Amato, Jeffrey Barrett, Sonia Goncalves, Ewan Harrison, David K. Jackson, Ian Johnston, Dominic Kwiatkowski, Cordelia Langford, John Sillitoe on behalf of the Wellcome Sanger Institute COVID-19 Surveillance Team |
| EPI_ISL_2438113 | Lighthouse Lab in Alderley Park | Wellcome Sanger Institute for the COVID-19 Genomics UK (COG-UK) Consortium | Jacquelyn Wynn, Mairead Hyland, The Lighthouse Lab in Alderley Park and Alex Alderton, Roberto Amato, Jeffrey Barrett, Sonia Goncalves, Ewan Harrison, David K. Jackson, Ian Johnston, Dominic Kwiatkowski, Cordelia Langford, John Sillitoe on behalf of the Wellcome Sanger Institute COVID-19 Surveillance Team |
| EPI_ISL_2438115 | Lighthouse Lab in Glasgow | Wellcome Sanger Institute for the COVID-19 Genomics UK (COG-UK) Consortium | Harper VanSteenhouse, Yumi Kasai, David Gray, Carol Clugston, Anna Dominiczak and Alex Alderton, Roberto Amato, Jeffrey Barrett, Sonia Goncalves, Ewan Harrison, David K. Jackson, Ian Johnston, Dominic Kwiatkowski, Cordelia Langford, John Sillitoe on behalf of the Wellcome Sanger Institute COVID-19 Surveillance Team |
| EPI_ISL_2438116 | Lighthouse Lab in Alderley Park | Wellcome Sanger Institute for the COVID-19 Genomics UK (COG-UK) Consortium | Jacquelyn Wynn, Mairead Hyland, The Lighthouse Lab in Alderley Park and Alex Alderton, Roberto Amato, Jeffrey Barrett, Sonia Goncalves, Ewan Harrison, David K. Jackson, Ian Johnston, Dominic Kwiatkowski, Cordelia Langford, John Sillitoe on behalf of the Wellcome Sanger Institute COVID-19 Surveillance Team |
| EPI_ISL_2438117 | Lighthouse Lab in Glasgow | Wellcome Sanger Institute for the COVID-19 Genomics UK (COG-UK) Consortium | Harper VanSteenhouse, Yumi Kasai, David Gray, Carol Clugston, Anna Dominiczak and Alex Alderton, Roberto Amato, Jeffrey Barrett, Sonia Goncalves, Ewan Harrison, David K. Jackson, Ian Johnston, Dominic Kwiatkowski, Cordelia Langford, John Sillitoe on behalf of the Wellcome Sanger Institute COVID-19 Surveillance Team |
| EPI_ISL_2438118, EPI_ISL_2438120 | Lighthouse Lab in Alderley Park | Wellcome Sanger Institute for the COVID-19 Genomics UK (COG-UK) Consortium | Jacquelyn Wynn, Mairead Hyland, The Lighthouse Lab in Alderley Park and Alex Alderton, Roberto Amato, Jeffrey Barrett, Sonia Goncalves, Ewan Harrison, David K. Jackson, Ian Johnston, Dominic Kwiatkowski, Cordelia Langford, John Sillitoe on behalf of the Wellcome Sanger Institute COVID-19 Surveillance Team |
| EPI_ISL_2438121, EPI_ISL_2438122 | Lighthouse Lab in Glasgow | Wellcome Sanger Institute for the COVID-19 Genomics UK (COG-UK) Consortium | Harper VanSteenhouse, Yumi Kasai, David Gray, Carol Clugston, Anna Dominiczak and Alex Alderton, Roberto Amato, Jeffrey Barrett, Sonia Goncalves, Ewan Harrison, David K. Jackson, Ian Johnston, Dominic Kwiatkowski, Cordelia Langford, John Sillitoe on behalf of the Wellcome Sanger Institute COVID-19 Surveillance Team |
| EPI_ISL_2438123 | Lighthouse Lab in Milton Keynes | Wellcome Sanger Institute for the COVID-19 Genomics UK (COG-UK) Consortium | The Lighthouse Lab in Milton Keynes and Alex Alderton, Roberto Amato, Jeffrey Barrett, Sonia Goncalves, Ewan Harrison, David K. Jackson, Ian Johnston, Dominic Kwiatkowski, Cordelia Langford, John Sillitoe on behalf of the Wellcome Sanger Institute COVID-19 Surveillance Team |
| EPI_ISL_2438124 | Lighthouse Lab in Alderley Park | Wellcome Sanger Institute for the COVID-19 Genomics UK (COG-UK) Consortium | Jacquelyn Wynn, Mairead Hyland, The Lighthouse Lab in Alderley Park and Alex Alderton, Roberto Amato, Jeffrey Barrett, Sonia Goncalves, Ewan Harrison, David K. Jackson, Ian Johnston, Dominic Kwiatkowski, Cordelia Langford, John Sillitoe on behalf of the Wellcome Sanger Institute COVID-19 Surveillance Team |
| EPI_ISL_2438125 | Lighthouse Lab in Glasgow | Wellcome Sanger Institute for the COVID-19 Genomics UK (COG-UK) Consortium | Harper VanSteenhouse, Yumi Kasai, David Gray, Carol Clugston, Anna Dominiczak and Alex Alderton, Roberto Amato, Jeffrey Barrett, Sonia Goncalves, Ewan Harrison, David K. Jackson, Ian Johnston, Dominic Kwiatkowski, Cordelia Langford, John Sillitoe on behalf of the Wellcome Sanger Institute COVID-19 Surveillance Team |
| EPI_ISL_2438126, EPI_ISL_2438128 | Lighthouse Lab in Milton Keynes | Wellcome Sanger Institute for the COVID-19 Genomics UK (COG-UK) Consortium | The Lighthouse Lab in Milton Keynes and Alex Alderton, Roberto Amato, Jeffrey Barrett, Sonia Goncalves, Ewan Harrison, David K. Jackson, Ian Johnston, Dominic Kwiatkowski, Cordelia Langford, John Sillitoe on behalf of the Wellcome Sanger Institute COVID-19 Surveillance Team |
| EPI_ISL_2438129 | Lighthouse Lab in Glasgow | Wellcome Sanger Institute for the COVID-19 Genomics UK (COG-UK) Consortium | Harper VanSteenhouse, Yumi Kasai, David Gray, Carol Clugston, Anna Dominiczak and Alex Alderton, Roberto Amato, Jeffrey Barrett, Sonia Goncalves, Ewan Harrison, David K. Jackson, Ian Johnston, Dominic Kwiatkowski, Cordelia Langford, John Sillitoe on behalf of the Wellcome Sanger Institute COVID-19 Surveillance Team |
| EPI_ISL_2438131, EPI_ISL_2438132 | Lighthouse Lab in Alderley Park | Wellcome Sanger Institute for the COVID-19 Genomics UK (COG-UK) Consortium | Jacquelyn Wynn, Mairead Hyland, The Lighthouse Lab in Alderley Park and Alex Alderton, Roberto Amato, Jeffrey Barrett, Sonia Goncalves, Ewan Harrison, David K. Jackson, Ian Johnston, Dominic Kwiatkowski, Cordelia Langford, John Sillitoe on behalf of the Wellcome Sanger Institute COVID-19 Surveillance Team |

| | | | |
|---|---|---|---|
| EPI_ISL_2438133, EPI_ISL_2438134, EPI_ISL_2438135 | Lighthouse Lab in Glasgow | Wellcome Sanger Institute for the COVID-19 Genomics UK (COG-UK) Consortium | Harper VanSteenhouse, Yumi Kasai, David Gray, Carol Clugston, Anna Dominiczak and Alex Alderton, Roberto Amato, Jeffrey Barrett, Sonia Goncalves, Ewan Harrison, David K. Jackson, Ian Johnston, Dominic Kwiatkowski, Cordelia Langford, John Sillitoe on behalf of the Wellcome Sanger Institute COVID-19 Surveillance Team |
| EPI_ISL_2438136 | Lighthouse Lab in Alderley Park | Wellcome Sanger Institute for the COVID-19 Genomics UK (COG-UK) Consortium | Jacquelyn Wynn, Mairead Hyland, The Lighthouse Lab in Alderley Park and Alex Alderton, Roberto Amato, Jeffrey Barrett, Sonia Goncalves, Ewan Harrison, David K. Jackson, Ian Johnston, Dominic Kwiatkowski, Cordelia Langford, John Sillitoe on behalf of the Wellcome Sanger Institute COVID-19 Surveillance Team |
| EPI_ISL_2438137, EPI_ISL_2438138 | Lighthouse Lab in Milton Keynes | Wellcome Sanger Institute for the COVID-19 Genomics UK (COG-UK) Consortium | The Lighthouse Lab in Milton Keynes and Alex Alderton, Roberto Amato, Jeffrey Barrett, Sonia Goncalves, Ewan Harrison, David K. Jackson, Ian Johnston, Dominic Kwiatkowski, Cordelia Langford, John Sillitoe on behalf of the Wellcome Sanger Institute COVID-19 Surveillance Team |
| EPI_ISL_2438139 | Lighthouse Lab in Glasgow | Wellcome Sanger Institute for the COVID-19 Genomics UK (COG-UK) Consortium | Harper VanSteenhouse, Yumi Kasai, David Gray, Carol Clugston, Anna Dominiczak and Alex Alderton, Roberto Amato, Jeffrey Barrett, Sonia Goncalves, Ewan Harrison, David K. Jackson, Ian Johnston, Dominic Kwiatkowski, Cordelia Langford, John Sillitoe on behalf of the Wellcome Sanger Institute COVID-19 Surveillance Team |
| EPI_ISL_2438140, EPI_ISL_2438141, EPI_ISL_2438142 | Lighthouse Lab in Milton Keynes | Wellcome Sanger Institute for the COVID-19 Genomics UK (COG-UK) Consortium | The Lighthouse Lab in Milton Keynes and Alex Alderton, Roberto Amato, Jeffrey Barrett, Sonia Goncalves, Ewan Harrison, David K. Jackson, Ian Johnston, Dominic Kwiatkowski, Cordelia Langford, John Sillitoe on behalf of the Wellcome Sanger Institute COVID-19 Surveillance Team |
| EPI_ISL_2438143 | Lighthouse Lab in Glasgow | Wellcome Sanger Institute for the COVID-19 Genomics UK (COG-UK) Consortium | Harper VanSteenhouse, Yumi Kasai, David Gray, Carol Clugston, Anna Dominiczak and Alex Alderton, Roberto Amato, Jeffrey Barrett, Sonia Goncalves, Ewan Harrison, David K. Jackson, Ian Johnston, Dominic Kwiatkowski, Cordelia Langford, John Sillitoe on behalf of the Wellcome Sanger Institute COVID-19 Surveillance Team |
| EPI_ISL_2438145, EPI_ISL_2438146 | Lighthouse Lab in Milton Keynes | Wellcome Sanger Institute for the COVID-19 Genomics UK (COG-UK) Consortium | The Lighthouse Lab in Milton Keynes and Alex Alderton, Roberto Amato, Jeffrey Barrett, Sonia Goncalves, Ewan Harrison, David K. Jackson, Ian Johnston, Dominic Kwiatkowski, Cordelia Langford, John Sillitoe on behalf of the Wellcome Sanger Institute COVID-19 Surveillance Team |
| EPI_ISL_2438147 | Lighthouse Lab in Glasgow | Wellcome Sanger Institute for the COVID-19 Genomics UK (COG-UK) Consortium | Harper VanSteenhouse, Yumi Kasai, David Gray, Carol Clugston, Anna Dominiczak and Alex Alderton, Roberto Amato, Jeffrey Barrett, Sonia Goncalves, Ewan Harrison, David K. Jackson, Ian Johnston, Dominic Kwiatkowski, Cordelia Langford, John Sillitoe on behalf of the Wellcome Sanger Institute COVID-19 Surveillance Team |
| EPI_ISL_2438148 | Lighthouse Lab in Alderley Park | Wellcome Sanger Institute for the COVID-19 Genomics UK (COG-UK) Consortium | Jacquelyn Wynn, Mairead Hyland, The Lighthouse Lab in Alderley Park and Alex Alderton, Roberto Amato, Jeffrey Barrett, Sonia Goncalves, Ewan Harrison, David K. Jackson, Ian Johnston, Dominic Kwiatkowski, Cordelia Langford, John Sillitoe on behalf of the Wellcome Sanger Institute COVID-19 Surveillance Team |
| EPI_ISL_2438149, EPI_ISL_2438150, EPI_ISL_2438151, EPI_ISL_2438152, EPI_ISL_2438154 | Lighthouse Lab in Glasgow | Wellcome Sanger Institute for the COVID-19 Genomics UK (COG-UK) Consortium | Harper VanSteenhouse, Yumi Kasai, David Gray, Carol Clugston, Anna Dominiczak and Alex Alderton, Roberto Amato, Jeffrey Barrett, Sonia Goncalves, Ewan Harrison, David K. Jackson, Ian Johnston, Dominic Kwiatkowski, Cordelia Langford, John Sillitoe on behalf of the Wellcome Sanger Institute COVID-19 Surveillance Team |
| EPI_ISL_2438155 | Lighthouse Lab in Milton Keynes | Wellcome Sanger Institute for the COVID-19 Genomics UK (COG-UK) Consortium | The Lighthouse Lab in Milton Keynes and Alex Alderton, Roberto Amato, Jeffrey Barrett, Sonia Goncalves, Ewan Harrison, David K. Jackson, Ian Johnston, Dominic Kwiatkowski, Cordelia Langford, John Sillitoe on behalf of the Wellcome Sanger Institute COVID-19 Surveillance Team |
| EPI_ISL_2438156 | Lighthouse Lab in Alderley Park | Wellcome Sanger Institute for the COVID-19 Genomics UK (COG-UK) Consortium | Jacquelyn Wynn, Mairead Hyland, The Lighthouse Lab in Alderley Park and Alex Alderton, Roberto Amato, Jeffrey Barrett, Sonia Goncalves, Ewan Harrison, David K. Jackson, Ian Johnston, Dominic Kwiatkowski, Cordelia Langford, John Sillitoe on behalf of the Wellcome Sanger Institute COVID-19 Surveillance Team |
| EPI_ISL_2438157 | Lighthouse Lab in Glasgow | Wellcome Sanger Institute for the COVID-19 Genomics UK (COG-UK) Consortium | Harper VanSteenhouse, Yumi Kasai, David Gray, Carol Clugston, Anna Dominiczak and Alex Alderton, Roberto Amato, Jeffrey Barrett, Sonia Goncalves, Ewan Harrison, David K. Jackson, Ian Johnston, Dominic Kwiatkowski, Cordelia Langford, John Sillitoe on behalf of the Wellcome Sanger Institute COVID-19 Surveillance Team |
| EPI_ISL_2438159 | Lighthouse Lab in Milton Keynes | Wellcome Sanger Institute for the COVID-19 Genomics UK (COG-UK) Consortium | The Lighthouse Lab in Milton Keynes and Alex Alderton, Roberto Amato, Jeffrey Barrett, Sonia Goncalves, Ewan Harrison, David K. Jackson, Ian Johnston, Dominic Kwiatkowski, Cordelia Langford, John Sillitoe on behalf of the Wellcome Sanger Institute COVID-19 Surveillance Team |
| EPI_ISL_2438160, EPI_ISL_2438161 | Lighthouse Lab in Glasgow | Wellcome Sanger Institute for the COVID-19 Genomics UK (COG-UK) Consortium | Harper VanSteenhouse, Yumi Kasai, David Gray, Carol Clugston, Anna Dominiczak and Alex Alderton, Roberto Amato, Jeffrey Barrett, Sonia Goncalves, Ewan Harrison, David K. Jackson, Ian Johnston, Dominic Kwiatkowski, Cordelia Langford, John Sillitoe on behalf of the Wellcome Sanger Institute COVID-19 Surveillance Team |
| EPI_ISL_2438162, EPI_ISL_2438165 | Lighthouse Lab in Milton Keynes | Wellcome Sanger Institute for the COVID-19 Genomics UK (COG-UK) Consortium | The Lighthouse Lab in Milton Keynes and Alex Alderton, Roberto Amato, Jeffrey Barrett, Sonia Goncalves, Ewan Harrison, David K. Jackson, Ian Johnston, Dominic Kwiatkowski, Cordelia Langford, John Sillitoe on behalf of the Wellcome Sanger Institute COVID-19 Surveillance Team |
| EPI_ISL_2438166, EPI_ISL_2438167 | Lighthouse Lab in Alderley Park | Wellcome Sanger Institute for the COVID-19 Genomics UK (COG-UK) Consortium | Jacquelyn Wynn, Mairead Hyland, The Lighthouse Lab in Alderley Park and Alex Alderton, Roberto Amato, Jeffrey Barrett, Sonia Goncalves, Ewan Harrison, David K. Jackson, Ian Johnston, Dominic Kwiatkowski, Cordelia Langford, John Sillitoe on behalf of the Wellcome Sanger Institute COVID-19 Surveillance Team |
| EPI_ISL_2438168, EPI_ISL_2438169, EPI_ISL_2438170 | Lighthouse Lab in Glasgow | Wellcome Sanger Institute for the COVID-19 Genomics UK (COG-UK) Consortium | Harper VanSteenhouse, Yumi Kasai, David Gray, Carol Clugston, Anna Dominiczak and Alex Alderton, Roberto Amato, Jeffrey Barrett, Sonia Goncalves, Ewan Harrison, David K. Jackson, Ian Johnston, Dominic Kwiatkowski, Cordelia Langford, John Sillitoe on behalf of the Wellcome Sanger Institute COVID-19 Surveillance Team |
| EPI_ISL_2438171 | Lighthouse Lab in Alderley Park | Wellcome Sanger Institute for the COVID-19 Genomics UK (COG-UK) Consortium | Jacquelyn Wynn, Mairead Hyland, The Lighthouse Lab in Alderley Park and Alex Alderton, Roberto Amato, Jeffrey Barrett, Sonia Goncalves, Ewan Harrison, David K. Jackson, Ian Johnston, Dominic Kwiatkowski, Cordelia Langford, John Sillitoe on behalf of the Wellcome Sanger Institute COVID-19 Surveillance Team |
| EPI_ISL_2438172, EPI_ISL_2438174 | Lighthouse Lab in Glasgow | Wellcome Sanger Institute for the COVID-19 Genomics UK (COG-UK) Consortium | Harper VanSteenhouse, Yumi Kasai, David Gray, Carol Clugston, Anna Dominiczak and Alex Alderton, Roberto Amato, Jeffrey Barrett, Sonia Goncalves, Ewan Harrison, David K. Jackson, Ian Johnston, Dominic Kwiatkowski, Cordelia Langford, John Sillitoe on behalf of the Wellcome Sanger Institute COVID-19 Surveillance Team |
| EPI_ISL_2438175, EPI_ISL_2438176 | Lighthouse Lab in Alderley Park | Wellcome Sanger Institute for the COVID-19 Genomics UK (COG-UK) Consortium | Jacquelyn Wynn, Mairead Hyland, The Lighthouse Lab in Alderley Park and Alex Alderton, Roberto Amato, Jeffrey Barrett, Sonia Goncalves, Ewan Harrison, David K. Jackson, Ian Johnston, Dominic Kwiatkowski, Cordelia Langford, John Sillitoe on behalf of the Wellcome Sanger Institute COVID-19 Surveillance Team |
| EPI_ISL_2438177 | Lighthouse Lab in Milton Keynes | Wellcome Sanger Institute for the COVID-19 Genomics UK (COG-UK) Consortium | The Lighthouse Lab in Milton Keynes and Alex Alderton, Roberto Amato, Jeffrey Barrett, Sonia Goncalves, Ewan Harrison, David K. Jackson, Ian Johnston, Dominic Kwiatkowski, Cordelia Langford, John Sillitoe on behalf of the Wellcome Sanger Institute COVID-19 Surveillance Team |
| EPI_ISL_2438178 | Lighthouse Lab in Alderley Park | Wellcome Sanger Institute for the COVID-19 Genomics UK (COG-UK) Consortium | Jacquelyn Wynn, Mairead Hyland, The Lighthouse Lab in Alderley Park and Alex Alderton, Roberto Amato, Jeffrey Barrett, Sonia Goncalves, Ewan Harrison, David K. Jackson, Ian Johnston, Dominic Kwiatkowski, Cordelia Langford, John Sillitoe on behalf of the Wellcome Sanger Institute COVID-19 Surveillance Team |
| EPI_ISL_2438179 | Lighthouse Lab in Glasgow | Wellcome Sanger Institute for the COVID-19 Genomics UK (COG-UK) Consortium | Harper VanSteenhouse, Yumi Kasai, David Gray, Carol Clugston, Anna Dominiczak and Alex Alderton, Roberto Amato, Jeffrey Barrett, Sonia Goncalves, Ewan Harrison, David K. Jackson, Ian Johnston, Dominic Kwiatkowski, Cordelia Langford, John Sillitoe on behalf of the Wellcome Sanger Institute COVID-19 Surveillance Team |
| EPI_ISL_2438180 | Lighthouse Lab in Milton Keynes | Wellcome Sanger Institute for the COVID-19 Genomics UK (COG-UK) Consortium | The Lighthouse Lab in Milton Keynes and Alex Alderton, Roberto Amato, Jeffrey Barrett, Sonia Goncalves, Ewan Harrison, David K. Jackson, Ian Johnston, Dominic Kwiatkowski, Cordelia Langford, John Sillitoe on behalf of the Wellcome Sanger Institute COVID-19 Surveillance Team |
| EPI_ISL_2438181 | Lighthouse Lab in Glasgow | Wellcome Sanger Institute for the COVID-19 Genomics UK (COG-UK) Consortium | Harper VanSteenhouse, Yumi Kasai, David Gray, Carol Clugston, Anna Dominiczak and Alex Alderton, Roberto Amato, Jeffrey Barrett, Sonia Goncalves, Ewan Harrison, David K. Jackson, Ian Johnston, Dominic Kwiatkowski, Cordelia Langford, John Sillitoe on behalf of the Wellcome Sanger Institute COVID-19 Surveillance Team |
| EPI_ISL_2438182 | Lighthouse Lab in Alderley Park | Wellcome Sanger Institute for the COVID-19 Genomics UK (COG-UK) Consortium | Jacquelyn Wynn, Mairead Hyland, The Lighthouse Lab in Alderley Park and Alex Alderton, Roberto Amato, Jeffrey Barrett, Sonia Goncalves, Ewan Harrison, David K. Jackson, Ian Johnston, Dominic Kwiatkowski, Cordelia Langford, John Sillitoe on behalf of the Wellcome Sanger Institute COVID-19 Surveillance Team |
| EPI_ISL_2438184, EPI_ISL_2438185, EPI_ISL_2438186, EPI_ISL_2438187 | Lighthouse Lab in Glasgow | Wellcome Sanger Institute for the COVID-19 Genomics UK (COG-UK) Consortium | Harper VanSteenhouse, Yumi Kasai, David Gray, Carol Clugston, Anna Dominiczak and Alex Alderton, Roberto Amato, Jeffrey Barrett, Sonia Goncalves, Ewan Harrison, David K. Jackson, Ian Johnston, Dominic Kwiatkowski, Cordelia Langford, John Sillitoe on behalf of the Wellcome Sanger Institute COVID-19 Surveillance Team |
| EPI_ISL_2438188 | Lighthouse Lab in Milton Keynes | Wellcome Sanger Institute for the COVID-19 Genomics UK (COG-UK) Consortium | The Lighthouse Lab in Milton Keynes and Alex Alderton, Roberto Amato, Jeffrey Barrett, Sonia Goncalves, Ewan Harrison, David K. Jackson, Ian Johnston, Dominic Kwiatkowski, Cordelia Langford, John Sillitoe on behalf of the Wellcome Sanger Institute COVID-19 Surveillance Team |
| EPI_ISL_2438189 | Lighthouse Lab in Alderley Park | Wellcome Sanger Institute for the COVID-19 Genomics UK (COG-UK) Consortium | Jacquelyn Wynn, Mairead Hyland, The Lighthouse Lab in Alderley Park and Alex Alderton, Roberto Amato, Jeffrey Barrett, Sonia Goncalves, Ewan Harrison, David K. Jackson, Ian Johnston, Dominic Kwiatkowski, Cordelia Langford, John Sillitoe on behalf of the Wellcome Sanger Institute COVID-19 Surveillance Team |
| EPI_ISL_2438190 | Lighthouse Lab in Glasgow | Wellcome Sanger Institute for the COVID-19 Genomics UK (COG-UK) Consortium | Harper VanSteenhouse, Yumi Kasai, David Gray, Carol Clugston, Anna Dominiczak and Alex Alderton, Roberto Amato, Jeffrey Barrett, Sonia Goncalves, Ewan Harrison, David K. Jackson, Ian Johnston, Dominic Kwiatkowski, Cordelia Langford, John Sillitoe on behalf of the Wellcome Sanger Institute COVID-19 Surveillance Team |
| EPI_ISL_2438191 | Lighthouse Lab in Milton Keynes | Wellcome Sanger Institute for the COVID-19 Genomics UK (COG-UK) Consortium | The Lighthouse Lab in Milton Keynes and Alex Alderton, Roberto Amato, Jeffrey Barrett, Sonia Goncalves, Ewan Harrison, David K. Jackson, Ian Johnston, Dominic Kwiatkowski, Cordelia Langford, John Sillitoe on behalf of the Wellcome Sanger Institute COVID-19 Surveillance Team |
| EPI_ISL_2438192 | Lighthouse Lab in Glasgow | Wellcome Sanger Institute for the COVID-19 Genomics UK (COG-UK) Consortium | Harper VanSteenhouse, Yumi Kasai, David Gray, Carol Clugston, Anna Dominiczak and Alex Alderton, Roberto Amato, Jeffrey Barrett, Sonia Goncalves, Ewan Harrison, David K. Jackson, Ian Johnston, Dominic Kwiatkowski, Cordelia Langford, John Sillitoe on behalf of the Wellcome Sanger Institute COVID-19 Surveillance Team |
| EPI_ISL_2438193, EPI_ISL_2438194, EPI_ISL_2438196 | Lighthouse Lab in Milton Keynes | Wellcome Sanger Institute for the COVID-19 Genomics UK (COG-UK) Consortium | The Lighthouse Lab in Milton Keynes and Alex Alderton, Roberto Amato, Jeffrey Barrett, Sonia Goncalves, Ewan Harrison, David K. Jackson, Ian Johnston, Dominic Kwiatkowski, Cordelia Langford, John Sillitoe on behalf of the Wellcome Sanger Institute COVID-19 Surveillance Team |
| EPI_ISL_2438198 | Lighthouse Lab in Alderley Park | Wellcome Sanger Institute for the COVID-19 Genomics UK (COG-UK) Consortium | Jacquelyn Wynn, Mairead Hyland, The Lighthouse Lab in Alderley Park and Alex Alderton, Roberto Amato, Jeffrey Barrett, Sonia Goncalves, Ewan Harrison, David K. Jackson, Ian Johnston, Dominic Kwiatkowski, Cordelia Langford, John Sillitoe on behalf of the Wellcome Sanger Institute COVID-19 Surveillance Team |
| EPI_ISL_2438199 | Lighthouse Lab in Milton Keynes | Wellcome Sanger Institute for the COVID-19 Genomics UK (COG-UK) Consortium | The Lighthouse Lab in Milton Keynes and Alex Alderton, Roberto Amato, Jeffrey Barrett, Sonia Goncalves, Ewan Harrison, David K. Jackson, Ian Johnston, Dominic Kwiatkowski, Cordelia Langford, John Sillitoe on behalf of the Wellcome Sanger Institute COVID-19 Surveillance Team |
| EPI_ISL_2438202 | Lighthouse Lab in Glasgow | Wellcome Sanger Institute for the COVID-19 Genomics UK (COG-UK) Consortium | Harper VanSteenhouse, Yumi Kasai, David Gray, Carol Clugston, Anna Dominiczak and Alex Alderton, Roberto Amato, Jeffrey Barrett, Sonia Goncalves, Ewan Harrison, David K. Jackson, Ian Johnston, Dominic Kwiatkowski, Cordelia Langford, John Sillitoe on behalf of the Wellcome Sanger Institute COVID-19 Surveillance Team |
| EPI_ISL_2438203 | Lighthouse Lab in Alderley Park | Wellcome Sanger Institute for the COVID-19 Genomics UK (COG-UK) Consortium | Jacquelyn Wynn, Mairead Hyland, The Lighthouse Lab in Alderley Park and Alex Alderton, Roberto Amato, Jeffrey Barrett, Sonia Goncalves, Ewan Harrison, David K. Jackson, Ian Johnston, Dominic Kwiatkowski, Cordelia Langford, John Sillitoe on behalf of the Wellcome Sanger Institute COVID-19 Surveillance Team |
| EPI_ISL_2438204 | Lighthouse Lab in Glasgow | Wellcome Sanger Institute for the COVID-19 Genomics UK (COG-UK) Consortium | Harper VanSteenhouse, Yumi Kasai, David Gray, Carol Clugston, Anna Dominiczak and Alex Alderton, Roberto Amato, Jeffrey Barrett, Sonia Goncalves, Ewan Harrison, David K. Jackson, Ian Johnston, Dominic Kwiatkowski, Cordelia Langford, John Sillitoe on behalf of the Wellcome Sanger Institute COVID-19 Surveillance Team |
| EPI_ISL_2438205 | Lighthouse Lab in Alderley Park | Wellcome Sanger Institute for the COVID-19 Genomics UK (COG-UK) Consortium | Jacquelyn Wynn, Mairead Hyland, The Lighthouse Lab in Alderley Park and Alex Alderton, Roberto Amato, Jeffrey Barrett, Sonia Goncalves, Ewan Harrison, David K. Jackson, Ian Johnston, Dominic Kwiatkowski, Cordelia Langford, John Sillitoe on behalf of the Wellcome Sanger Institute COVID-19 Surveillance Team |
| EPI_ISL_2438206 | Lighthouse Lab in Milton Keynes | Wellcome Sanger Institute for the COVID-19 Genomics UK (COG-UK) Consortium | The Lighthouse Lab in Milton Keynes and Alex Alderton, Roberto Amato, Jeffrey Barrett, Sonia Goncalves, Ewan Harrison, David K. Jackson, Ian Johnston, Dominic Kwiatkowski, Cordelia Langford, John Sillitoe on behalf of the Wellcome Sanger Institute COVID-19 Surveillance Team |

| | | | |
|---|---|---|---|
| EPI_ISL_2438207 | Lighthouse Lab in Alderley Park | Wellcome Sanger Institute for the COVID-19 Genomics UK (COG-UK) Consortium | Jacquelyn Wynn, Mairead Hyland, The Lighthouse Lab in Alderley Park and Alex Alderton, Roberto Amato, Jeffrey Barrett, Sonia Goncalves, Ewan Harrison, David K. Jackson, Ian Johnston, Dominic Kwiatkowski, Cordelia Langford, John Sillitoe on behalf of the Wellcome Sanger Institute COVID-19 Surveillance Team |
| EPI_ISL_2438208 | Lighthouse Lab in Milton Keynes | Wellcome Sanger Institute for the COVID-19 Genomics UK (COG-UK) Consortium | The Lighthouse Lab in Milton Keynes and Alex Alderton, Roberto Amato, Jeffrey Barrett, Sonia Goncalves, Ewan Harrison, David K. Jackson, Ian Johnston, Dominic Kwiatkowski, Cordelia Langford, John Sillitoe on behalf of the Wellcome Sanger Institute COVID-19 Surveillance Team |
| EPI_ISL_2438209, EPI_ISL_2438210, EPI_ISL_2438212 | Lighthouse Lab in Glasgow | Wellcome Sanger Institute for the COVID-19 Genomics UK (COG-UK) Consortium | Harper VanSteenhouse, Yumi Kasai, David Gray, Carol Clugston, Anna Dominiczak and Alex Alderton, Roberto Amato, Jeffrey Barrett, Sonia Goncalves, Ewan Harrison, David K. Jackson, Ian Johnston, Dominic Kwiatkowski, Cordelia Langford, John Sillitoe on behalf of the Wellcome Sanger Institute COVID-19 Surveillance Team |
| EPI_ISL_2438213 | Lighthouse Lab in Alderley Park | Wellcome Sanger Institute for the COVID-19 Genomics UK (COG-UK) Consortium | Jacquelyn Wynn, Mairead Hyland, The Lighthouse Lab in Alderley Park and Alex Alderton, Roberto Amato, Jeffrey Barrett, Sonia Goncalves, Ewan Harrison, David K. Jackson, Ian Johnston, Dominic Kwiatkowski, Cordelia Langford, John Sillitoe on behalf of the Wellcome Sanger Institute COVID-19 Surveillance Team |
| EPI_ISL_2438214 | Lighthouse Lab in Glasgow | Wellcome Sanger Institute for the COVID-19 Genomics UK (COG-UK) Consortium | Harper VanSteenhouse, Yumi Kasai, David Gray, Carol Clugston, Anna Dominiczak and Alex Alderton, Roberto Amato, Jeffrey Barrett, Sonia Goncalves, Ewan Harrison, David K. Jackson, Ian Johnston, Dominic Kwiatkowski, Cordelia Langford, John Sillitoe on behalf of the Wellcome Sanger Institute COVID-19 Surveillance Team |
| EPI_ISL_2438215 | Lighthouse Lab in Milton Keynes | Wellcome Sanger Institute for the COVID-19 Genomics UK (COG-UK) Consortium | The Lighthouse Lab in Milton Keynes and Alex Alderton, Roberto Amato, Jeffrey Barrett, Sonia Goncalves, Ewan Harrison, David K. Jackson, Ian Johnston, Dominic Kwiatkowski, Cordelia Langford, John Sillitoe on behalf of the Wellcome Sanger Institute COVID-19 Surveillance Team |
| EPI_ISL_2438217, EPI_ISL_2438218 | Lighthouse Lab in Alderley Park | Wellcome Sanger Institute for the COVID-19 Genomics UK (COG-UK) Consortium | Jacquelyn Wynn, Mairead Hyland, The Lighthouse Lab in Alderley Park and Alex Alderton, Roberto Amato, Jeffrey Barrett, Sonia Goncalves, Ewan Harrison, David K. Jackson, Ian Johnston, Dominic Kwiatkowski, Cordelia Langford, John Sillitoe on behalf of the Wellcome Sanger Institute COVID-19 Surveillance Team |
| EPI_ISL_2438219 | Lighthouse Lab in Glasgow | Wellcome Sanger Institute for the COVID-19 Genomics UK (COG-UK) Consortium | Harper VanSteenhouse, Yumi Kasai, David Gray, Carol Clugston, Anna Dominiczak and Alex Alderton, Roberto Amato, Jeffrey Barrett, Sonia Goncalves, Ewan Harrison, David K. Jackson, Ian Johnston, Dominic Kwiatkowski, Cordelia Langford, John Sillitoe on behalf of the Wellcome Sanger Institute COVID-19 Surveillance Team |
| EPI_ISL_2438220 | Lighthouse Lab in Milton Keynes | Wellcome Sanger Institute for the COVID-19 Genomics UK (COG-UK) Consortium | The Lighthouse Lab in Milton Keynes and Alex Alderton, Roberto Amato, Jeffrey Barrett, Sonia Goncalves, Ewan Harrison, David K. Jackson, Ian Johnston, Dominic Kwiatkowski, Cordelia Langford, John Sillitoe on behalf of the Wellcome Sanger Institute COVID-19 Surveillance Team |
| EPI_ISL_2438221 | Lighthouse Lab in Alderley Park | Wellcome Sanger Institute for the COVID-19 Genomics UK (COG-UK) Consortium | Jacquelyn Wynn, Mairead Hyland, The Lighthouse Lab in Alderley Park and Alex Alderton, Roberto Amato, Jeffrey Barrett, Sonia Goncalves, Ewan Harrison, David K. Jackson, Ian Johnston, Dominic Kwiatkowski, Cordelia Langford, John Sillitoe on behalf of the Wellcome Sanger Institute COVID-19 Surveillance Team |
| EPI_ISL_2438222 | Lighthouse Lab in Glasgow | Wellcome Sanger Institute for the COVID-19 Genomics UK (COG-UK) Consortium | Harper VanSteenhouse, Yumi Kasai, David Gray, Carol Clugston, Anna Dominiczak and Alex Alderton, Roberto Amato, Jeffrey Barrett, Sonia Goncalves, Ewan Harrison, David K. Jackson, Ian Johnston, Dominic Kwiatkowski, Cordelia Langford, John Sillitoe on behalf of the Wellcome Sanger Institute COVID-19 Surveillance Team |
| EPI_ISL_2438223, EPI_ISL_2438224, EPI_ISL_2438225 | Lighthouse Lab in Milton Keynes | Wellcome Sanger Institute for the COVID-19 Genomics UK (COG-UK) Consortium | The Lighthouse Lab in Milton Keynes and Alex Alderton, Roberto Amato, Jeffrey Barrett, Sonia Goncalves, Ewan Harrison, David K. Jackson, Ian Johnston, Dominic Kwiatkowski, Cordelia Langford, John Sillitoe on behalf of the Wellcome Sanger Institute COVID-19 Surveillance Team |
| EPI_ISL_2438226 | Lighthouse Lab in Alderley Park | Wellcome Sanger Institute for the COVID-19 Genomics UK (COG-UK) Consortium | Jacquelyn Wynn, Mairead Hyland, The Lighthouse Lab in Alderley Park and Alex Alderton, Roberto Amato, Jeffrey Barrett, Sonia Goncalves, Ewan Harrison, David K. Jackson, Ian Johnston, Dominic Kwiatkowski, Cordelia Langford, John Sillitoe on behalf of the Wellcome Sanger Institute COVID-19 Surveillance Team |
| EPI_ISL_2438227, EPI_ISL_2438228 | Lighthouse Lab in Glasgow | Wellcome Sanger Institute for the COVID-19 Genomics UK (COG-UK) Consortium | Harper VanSteenhouse, Yumi Kasai, David Gray, Carol Clugston, Anna Dominiczak and Alex Alderton, Roberto Amato, Jeffrey Barrett, Sonia Goncalves, Ewan Harrison, David K. Jackson, Ian Johnston, Dominic Kwiatkowski, Cordelia Langford, John Sillitoe on behalf of the Wellcome Sanger Institute COVID-19 Surveillance Team |
| EPI_ISL_2438229, EPI_ISL_2438230 | Lighthouse Lab in Alderley Park | Wellcome Sanger Institute for the COVID-19 Genomics UK (COG-UK) Consortium | Jacquelyn Wynn, Mairead Hyland, The Lighthouse Lab in Alderley Park and Alex Alderton, Roberto Amato, Jeffrey Barrett, Sonia Goncalves, Ewan Harrison, David K. Jackson, Ian Johnston, Dominic Kwiatkowski, Cordelia Langford, John Sillitoe on behalf of the Wellcome Sanger Institute COVID-19 Surveillance Team |
| EPI_ISL_2438231 | Lighthouse Lab in Milton Keynes | Wellcome Sanger Institute for the COVID-19 Genomics UK (COG-UK) Consortium | The Lighthouse Lab in Milton Keynes and Alex Alderton, Roberto Amato, Jeffrey Barrett, Sonia Goncalves, Ewan Harrison, David K. Jackson, Ian Johnston, Dominic Kwiatkowski, Cordelia Langford, John Sillitoe on behalf of the Wellcome Sanger Institute COVID-19 Surveillance Team |
| EPI_ISL_2438232 | Lighthouse Lab in Glasgow | Wellcome Sanger Institute for the COVID-19 Genomics UK (COG-UK) Consortium | Harper VanSteenhouse, Yumi Kasai, David Gray, Carol Clugston, Anna Dominiczak and Alex Alderton, Roberto Amato, Jeffrey Barrett, Sonia Goncalves, Ewan Harrison, David K. Jackson, Ian Johnston, Dominic Kwiatkowski, Cordelia Langford, John Sillitoe on behalf of the Wellcome Sanger Institute COVID-19 Surveillance Team |
| EPI_ISL_2438233 | Lighthouse Lab in Milton Keynes | Wellcome Sanger Institute for the COVID-19 Genomics UK (COG-UK) Consortium | The Lighthouse Lab in Milton Keynes and Alex Alderton, Roberto Amato, Jeffrey Barrett, Sonia Goncalves, Ewan Harrison, David K. Jackson, Ian Johnston, Dominic Kwiatkowski, Cordelia Langford, John Sillitoe on behalf of the Wellcome Sanger Institute COVID-19 Surveillance Team |
| EPI_ISL_2438234 | Lighthouse Lab in Glasgow | Wellcome Sanger Institute for the COVID-19 Genomics UK (COG-UK) Consortium | Harper VanSteenhouse, Yumi Kasai, David Gray, Carol Clugston, Anna Dominiczak and Alex Alderton, Roberto Amato, Jeffrey Barrett, Sonia Goncalves, Ewan Harrison, David K. Jackson, Ian Johnston, Dominic Kwiatkowski, Cordelia Langford, John Sillitoe on behalf of the Wellcome Sanger Institute COVID-19 Surveillance Team |
| EPI_ISL_2438235, EPI_ISL_2438238 | Lighthouse Lab in Alderley Park | Wellcome Sanger Institute for the COVID-19 Genomics UK (COG-UK) Consortium | Jacquelyn Wynn, Mairead Hyland, The Lighthouse Lab in Alderley Park and Alex Alderton, Roberto Amato, Jeffrey Barrett, Sonia Goncalves, Ewan Harrison, David K. Jackson, Ian Johnston, Dominic Kwiatkowski, Cordelia Langford, John Sillitoe on behalf of the Wellcome Sanger Institute COVID-19 Surveillance Team |
| EPI_ISL_2438240 | Lighthouse Lab in Glasgow | Wellcome Sanger Institute for the COVID-19 Genomics UK (COG-UK) Consortium | Harper VanSteenhouse, Yumi Kasai, David Gray, Carol Clugston, Anna Dominiczak and Alex Alderton, Roberto Amato, Jeffrey Barrett, Sonia Goncalves, Ewan Harrison, David K. Jackson, Ian Johnston, Dominic Kwiatkowski, Cordelia Langford, John Sillitoe on behalf of the Wellcome Sanger Institute COVID-19 Surveillance Team |
| EPI_ISL_2438241, EPI_ISL_2438244 | Lighthouse Lab in Milton Keynes | Wellcome Sanger Institute for the COVID-19 Genomics UK (COG-UK) Consortium | The Lighthouse Lab in Milton Keynes and Alex Alderton, Roberto Amato, Jeffrey Barrett, Sonia Goncalves, Ewan Harrison, David K. Jackson, Ian Johnston, Dominic Kwiatkowski, Cordelia Langford, John Sillitoe on behalf of the Wellcome Sanger Institute COVID-19 Surveillance Team |
| EPI_ISL_2438245, EPI_ISL_2438246, EPI_ISL_2438247 | Lighthouse Lab in Alderley Park | Wellcome Sanger Institute for the COVID-19 Genomics UK (COG-UK) Consortium | Jacquelyn Wynn, Mairead Hyland, The Lighthouse Lab in Alderley Park and Alex Alderton, Roberto Amato, Jeffrey Barrett, Sonia Goncalves, Ewan Harrison, David K. Jackson, Ian Johnston, Dominic Kwiatkowski, Cordelia Langford, John Sillitoe on behalf of the Wellcome Sanger Institute COVID-19 Surveillance Team |
| EPI_ISL_2438249 | Lighthouse Lab in Glasgow | Wellcome Sanger Institute for the COVID-19 Genomics UK (COG-UK) Consortium | Harper VanSteenhouse, Yumi Kasai, David Gray, Carol Clugston, Anna Dominiczak and Alex Alderton, Roberto Amato, Jeffrey Barrett, Sonia Goncalves, Ewan Harrison, David K. Jackson, Ian Johnston, Dominic Kwiatkowski, Cordelia Langford, John Sillitoe on behalf of the Wellcome Sanger Institute COVID-19 Surveillance Team |
| EPI_ISL_2438251 | Lighthouse Lab in Alderley Park | Wellcome Sanger Institute for the COVID-19 Genomics UK (COG-UK) Consortium | Jacquelyn Wynn, Mairead Hyland, The Lighthouse Lab in Alderley Park and Alex Alderton, Roberto Amato, Jeffrey Barrett, Sonia Goncalves, Ewan Harrison, David K. Jackson, Ian Johnston, Dominic Kwiatkowski, Cordelia Langford, John Sillitoe on behalf of the Wellcome Sanger Institute COVID-19 Surveillance Team |
| EPI_ISL_2438253 | Lighthouse Lab in Glasgow | Wellcome Sanger Institute for the COVID-19 Genomics UK (COG-UK) Consortium | Harper VanSteenhouse, Yumi Kasai, David Gray, Carol Clugston, Anna Dominiczak and Alex Alderton, Roberto Amato, Jeffrey Barrett, Sonia Goncalves, Ewan Harrison, David K. Jackson, Ian Johnston, Dominic Kwiatkowski, Cordelia Langford, John Sillitoe on behalf of the Wellcome Sanger Institute COVID-19 Surveillance Team |
| EPI_ISL_2438255 | Lighthouse Lab in Milton Keynes | Wellcome Sanger Institute for the COVID-19 Genomics UK (COG-UK) Consortium | The Lighthouse Lab in Milton Keynes and Alex Alderton, Roberto Amato, Jeffrey Barrett, Sonia Goncalves, Ewan Harrison, David K. Jackson, Ian Johnston, Dominic Kwiatkowski, Cordelia Langford, John Sillitoe on behalf of the Wellcome Sanger Institute COVID-19 Surveillance Team |
| EPI_ISL_2438257, EPI_ISL_2438258, EPI_ISL_2438259 | Lighthouse Lab in Glasgow | Wellcome Sanger Institute for the COVID-19 Genomics UK (COG-UK) Consortium | Harper VanSteenhouse, Yumi Kasai, David Gray, Carol Clugston, Anna Dominiczak and Alex Alderton, Roberto Amato, Jeffrey Barrett, Sonia Goncalves, Ewan Harrison, David K. Jackson, Ian Johnston, Dominic Kwiatkowski, Cordelia Langford, John Sillitoe on behalf of the Wellcome Sanger Institute COVID-19 Surveillance Team |
| EPI_ISL_2438264 | Lighthouse Lab in Milton Keynes | Wellcome Sanger Institute for the COVID-19 Genomics UK (COG-UK) Consortium | The Lighthouse Lab in Milton Keynes and Alex Alderton, Roberto Amato, Jeffrey Barrett, Sonia Goncalves, Ewan Harrison, David K. Jackson, Ian Johnston, Dominic Kwiatkowski, Cordelia Langford, John Sillitoe on behalf of the Wellcome Sanger Institute COVID-19 Surveillance Team |
| EPI_ISL_2438265 | Lighthouse Lab in Glasgow | Wellcome Sanger Institute for the COVID-19 Genomics UK (COG-UK) Consortium | Harper VanSteenhouse, Yumi Kasai, David Gray, Carol Clugston, Anna Dominiczak and Alex Alderton, Roberto Amato, Jeffrey Barrett, Sonia Goncalves, Ewan Harrison, David K. Jackson, Ian Johnston, Dominic Kwiatkowski, Cordelia Langford, John Sillitoe on behalf of the Wellcome Sanger Institute COVID-19 Surveillance Team |
| EPI_ISL_2438266 | Lighthouse Lab in Milton Keynes | Wellcome Sanger Institute for the COVID-19 Genomics UK (COG-UK) Consortium | The Lighthouse Lab in Milton Keynes and Alex Alderton, Roberto Amato, Jeffrey Barrett, Sonia Goncalves, Ewan Harrison, David K. Jackson, Ian Johnston, Dominic Kwiatkowski, Cordelia Langford, John Sillitoe on behalf of the Wellcome Sanger Institute COVID-19 Surveillance Team |
| EPI_ISL_2438268 | Lighthouse Lab in Glasgow | Wellcome Sanger Institute for the COVID-19 Genomics UK (COG-UK) Consortium | Harper VanSteenhouse, Yumi Kasai, David Gray, Carol Clugston, Anna Dominiczak and Alex Alderton, Roberto Amato, Jeffrey Barrett, Sonia Goncalves, Ewan Harrison, David K. Jackson, Ian Johnston, Dominic Kwiatkowski, Cordelia Langford, John Sillitoe on behalf of the Wellcome Sanger Institute COVID-19 Surveillance Team |
| EPI_ISL_2438269 | Lighthouse Lab in Alderley Park | Wellcome Sanger Institute for the COVID-19 Genomics UK (COG-UK) Consortium | Jacquelyn Wynn, Mairead Hyland, The Lighthouse Lab in Alderley Park and Alex Alderton, Roberto Amato, Jeffrey Barrett, Sonia Goncalves, Ewan Harrison, David K. Jackson, Ian Johnston, Dominic Kwiatkowski, Cordelia Langford, John Sillitoe on behalf of the Wellcome Sanger Institute COVID-19 Surveillance Team |
| EPI_ISL_2438270, EPI_ISL_2438271 | Lighthouse Lab in Milton Keynes | Wellcome Sanger Institute for the COVID-19 Genomics UK (COG-UK) Consortium | The Lighthouse Lab in Milton Keynes and Alex Alderton, Roberto Amato, Jeffrey Barrett, Sonia Goncalves, Ewan Harrison, David K. Jackson, Ian Johnston, Dominic Kwiatkowski, Cordelia Langford, John Sillitoe on behalf of the Wellcome Sanger Institute COVID-19 Surveillance Team |
| EPI_ISL_2438272 | Lighthouse Lab in Glasgow | Wellcome Sanger Institute for the COVID-19 Genomics UK (COG-UK) Consortium | Harper VanSteenhouse, Yumi Kasai, David Gray, Carol Clugston, Anna Dominiczak and Alex Alderton, Roberto Amato, Jeffrey Barrett, Sonia Goncalves, Ewan Harrison, David K. Jackson, Ian Johnston, Dominic Kwiatkowski, Cordelia Langford, John Sillitoe on behalf of the Wellcome Sanger Institute COVID-19 Surveillance Team |
| EPI_ISL_2438274 | Lighthouse Lab in Milton Keynes | Wellcome Sanger Institute for the COVID-19 Genomics UK (COG-UK) Consortium | The Lighthouse Lab in Milton Keynes and Alex Alderton, Roberto Amato, Jeffrey Barrett, Sonia Goncalves, Ewan Harrison, David K. Jackson, Ian Johnston, Dominic Kwiatkowski, Cordelia Langford, John Sillitoe on behalf of the Wellcome Sanger Institute COVID-19 Surveillance Team |
| EPI_ISL_2438275 | Lighthouse Lab in Glasgow | Wellcome Sanger Institute for the COVID-19 Genomics UK (COG-UK) Consortium | Harper VanSteenhouse, Yumi Kasai, David Gray, Carol Clugston, Anna Dominiczak and Alex Alderton, Roberto Amato, Jeffrey Barrett, Sonia Goncalves, Ewan Harrison, David K. Jackson, Ian Johnston, Dominic Kwiatkowski, Cordelia Langford, John Sillitoe on behalf of the Wellcome Sanger Institute COVID-19 Surveillance Team |
| EPI_ISL_2438276 | Lighthouse Lab in Milton Keynes | Wellcome Sanger Institute for the COVID-19 Genomics UK (COG-UK) Consortium | The Lighthouse Lab in Milton Keynes and Alex Alderton, Roberto Amato, Jeffrey Barrett, Sonia Goncalves, Ewan Harrison, David K. Jackson, Ian Johnston, Dominic Kwiatkowski, Cordelia Langford, John Sillitoe on behalf of the Wellcome Sanger Institute COVID-19 Surveillance Team |
| EPI_ISL_2438278 | Lighthouse Lab in Glasgow | Wellcome Sanger Institute for the COVID-19 Genomics UK (COG-UK) Consortium | Harper VanSteenhouse, Yumi Kasai, David Gray, Carol Clugston, Anna Dominiczak and Alex Alderton, Roberto Amato, Jeffrey Barrett, Sonia Goncalves, Ewan Harrison, David K. Jackson, Ian Johnston, Dominic Kwiatkowski, Cordelia Langford, John Sillitoe on behalf of the Wellcome Sanger Institute COVID-19 Surveillance Team |
| EPI_ISL_2438279 | Lighthouse Lab in Milton Keynes | Wellcome Sanger Institute for the COVID-19 Genomics UK (COG-UK) Consortium | The Lighthouse Lab in Milton Keynes and Alex Alderton, Roberto Amato, Jeffrey Barrett, Sonia Goncalves, Ewan Harrison, David K. Jackson, Ian Johnston, Dominic Kwiatkowski, Cordelia Langford, John Sillitoe on behalf of the Wellcome Sanger Institute COVID-19 Surveillance Team |
| EPI_ISL_2438280 | Lighthouse Lab in Glasgow | Wellcome Sanger Institute for the COVID-19 Genomics UK (COG-UK) Consortium | Harper VanSteenhouse, Yumi Kasai, David Gray, Carol Clugston, Anna Dominiczak and Alex Alderton, Roberto Amato, Jeffrey Barrett, Sonia Goncalves, Ewan Harrison, David K. Jackson, Ian Johnston, Dominic Kwiatkowski, Cordelia Langford, John Sillitoe on behalf of the Wellcome Sanger Institute COVID-19 Surveillance Team |
| EPI_ISL_2438281 | Lighthouse Lab in Milton Keynes | Wellcome Sanger Institute for the COVID-19 Genomics UK (COG-UK) Consortium | The Lighthouse Lab in Milton Keynes and Alex Alderton, Roberto Amato, Jeffrey Barrett, Sonia Goncalves, Ewan Harrison, David K. Jackson, Ian Johnston, Dominic Kwiatkowski, Cordelia Langford, John Sillitoe on behalf of the Wellcome Sanger Institute COVID-19 Surveillance Team |
| EPI_ISL_2438282 | Lighthouse Lab in Alderley Park | Wellcome Sanger Institute for the COVID-19 Genomics UK (COG-UK) Consortium | Jacquelyn Wynn, Mairead Hyland, The Lighthouse Lab in Alderley Park and Alex Alderton, Roberto Amato, Jeffrey Barrett, Sonia Goncalves, Ewan Harrison, David K. Jackson, Ian Johnston, Dominic Kwiatkowski, Cordelia Langford, John Sillitoe on behalf of the Wellcome Sanger Institute COVID-19 Surveillance Team |
| EPI_ISL_2438283 | Lighthouse Lab in Milton Keynes | Wellcome Sanger Institute for the COVID-19 Genomics UK (COG-UK) | The Lighthouse Lab in Milton Keynes and Alex Alderton, Roberto Amato, Jeffrey Barrett, Sonia Goncalves, Ewan Harrison, David K. Jackson, Ian Johnston, Dominic Kwiatkowski, Cordelia Langford, John Sillitoe on behalf of the Wellcome Sanger |

| | | | |
|---|---|---|---|
| | | Consortium | Institute COVID-19 Surveillance Team |
| EPI_ISL_2438284 | Lighthouse Lab in Glasgow | Wellcome Sanger Institute for the COVID-19 Genomics UK (COG-UK) Consortium | Harper VanSteenhouse, Yumi Kasai, David Gray, Carol Clugston, Anna Dominiczak and Alex Alderton, Roberto Amato, Jeffrey Barrett, Sonia Goncalves, Ewan Harrison, David K. Jackson, Ian Johnston, Dominic Kwiatkowski, Cordelia Langford, John Sillitoe on behalf of the Wellcome Sanger Institute COVID-19 Surveillance Team |
| EPI_ISL_2438287, EPI_ISL_2438289 | Lighthouse Lab in Alderley Park | Wellcome Sanger Institute for the COVID-19 Genomics UK (COG-UK) Consortium | Jacquelyn Wynn, Mairead Hyland, The Lighthouse Lab in Alderley Park and Alex Alderton, Roberto Amato, Jeffrey Barrett, Sonia Goncalves, Ewan Harrison, David K. Jackson, Ian Johnston, Dominic Kwiatkowski, Cordelia Langford, John Sillitoe on behalf of the Wellcome Sanger Institute COVID-19 Surveillance Team |
| EPI_ISL_2438290 | Lighthouse Lab in Milton Keynes | Wellcome Sanger Institute for the COVID-19 Genomics UK (COG-UK) Consortium | The Lighthouse Lab in Milton Keynes and Alex Alderton, Roberto Amato, Jeffrey Barrett, Sonia Goncalves, Ewan Harrison, David K. Jackson, Ian Johnston, Dominic Kwiatkowski, Cordelia Langford, John Sillitoe on behalf of the Wellcome Sanger Institute COVID-19 Surveillance Team |
| EPI_ISL_2438291 | Lighthouse Lab in Glasgow | Wellcome Sanger Institute for the COVID-19 Genomics UK (COG-UK) Consortium | Harper VanSteenhouse, Yumi Kasai, David Gray, Carol Clugston, Anna Dominiczak and Alex Alderton, Roberto Amato, Jeffrey Barrett, Sonia Goncalves, Ewan Harrison, David K. Jackson, Ian Johnston, Dominic Kwiatkowski, Cordelia Langford, John Sillitoe on behalf of the Wellcome Sanger Institute COVID-19 Surveillance Team |
| EPI_ISL_2438292 | Lighthouse Lab in Milton Keynes | Wellcome Sanger Institute for the COVID-19 Genomics UK (COG-UK) Consortium | The Lighthouse Lab in Milton Keynes and Alex Alderton, Roberto Amato, Jeffrey Barrett, Sonia Goncalves, Ewan Harrison, David K. Jackson, Ian Johnston, Dominic Kwiatkowski, Cordelia Langford, John Sillitoe on behalf of the Wellcome Sanger Institute COVID-19 Surveillance Team |
| EPI_ISL_2438293, EPI_ISL_2438294 | Lighthouse Lab in Alderley Park | Wellcome Sanger Institute for the COVID-19 Genomics UK (COG-UK) Consortium | Jacquelyn Wynn, Mairead Hyland, The Lighthouse Lab in Alderley Park and Alex Alderton, Roberto Amato, Jeffrey Barrett, Sonia Goncalves, Ewan Harrison, David K. Jackson, Ian Johnston, Dominic Kwiatkowski, Cordelia Langford, John Sillitoe on behalf of the Wellcome Sanger Institute COVID-19 Surveillance Team |
| EPI_ISL_2438296 | Lighthouse Lab in Glasgow | Wellcome Sanger Institute for the COVID-19 Genomics UK (COG-UK) Consortium | Harper VanSteenhouse, Yumi Kasai, David Gray, Carol Clugston, Anna Dominiczak and Alex Alderton, Roberto Amato, Jeffrey Barrett, Sonia Goncalves, Ewan Harrison, David K. Jackson, Ian Johnston, Dominic Kwiatkowski, Cordelia Langford, John Sillitoe on behalf of the Wellcome Sanger Institute COVID-19 Surveillance Team |
| EPI_ISL_2438298 | Lighthouse Lab in Alderley Park | Wellcome Sanger Institute for the COVID-19 Genomics UK (COG-UK) Consortium | Jacquelyn Wynn, Mairead Hyland, The Lighthouse Lab in Alderley Park and Alex Alderton, Roberto Amato, Jeffrey Barrett, Sonia Goncalves, Ewan Harrison, David K. Jackson, Ian Johnston, Dominic Kwiatkowski, Cordelia Langford, John Sillitoe on behalf of the Wellcome Sanger Institute COVID-19 Surveillance Team |
| EPI_ISL_2438299, EPI_ISL_2438300 | Lighthouse Lab in Glasgow | Wellcome Sanger Institute for the COVID-19 Genomics UK (COG-UK) Consortium | Harper VanSteenhouse, Yumi Kasai, David Gray, Carol Clugston, Anna Dominiczak and Alex Alderton, Roberto Amato, Jeffrey Barrett, Sonia Goncalves, Ewan Harrison, David K. Jackson, Ian Johnston, Dominic Kwiatkowski, Cordelia Langford, John Sillitoe on behalf of the Wellcome Sanger Institute COVID-19 Surveillance Team |
| EPI_ISL_2438301 | Lighthouse Lab in Alderley Park | Wellcome Sanger Institute for the COVID-19 Genomics UK (COG-UK) Consortium | Jacquelyn Wynn, Mairead Hyland, The Lighthouse Lab in Alderley Park and Alex Alderton, Roberto Amato, Jeffrey Barrett, Sonia Goncalves, Ewan Harrison, David K. Jackson, Ian Johnston, Dominic Kwiatkowski, Cordelia Langford, John Sillitoe on behalf of the Wellcome Sanger Institute COVID-19 Surveillance Team |
| EPI_ISL_2438302 | Lighthouse Lab in Milton Keynes | Wellcome Sanger Institute for the COVID-19 Genomics UK (COG-UK) Consortium | The Lighthouse Lab in Milton Keynes and Alex Alderton, Roberto Amato, Jeffrey Barrett, Sonia Goncalves, Ewan Harrison, David K. Jackson, Ian Johnston, Dominic Kwiatkowski, Cordelia Langford, John Sillitoe on behalf of the Wellcome Sanger Institute COVID-19 Surveillance Team |
| EPI_ISL_2438303, EPI_ISL_2438304, EPI_ISL_2438308 | Lighthouse Lab in Glasgow | Wellcome Sanger Institute for the COVID-19 Genomics UK (COG-UK) Consortium | Harper VanSteenhouse, Yumi Kasai, David Gray, Carol Clugston, Anna Dominiczak and Alex Alderton, Roberto Amato, Jeffrey Barrett, Sonia Goncalves, Ewan Harrison, David K. Jackson, Ian Johnston, Dominic Kwiatkowski, Cordelia Langford, John Sillitoe on behalf of the Wellcome Sanger Institute COVID-19 Surveillance Team |
| EPI_ISL_2438309, EPI_ISL_2438310 | Lighthouse Lab in Alderley Park | Wellcome Sanger Institute for the COVID-19 Genomics UK (COG-UK) Consortium | Jacquelyn Wynn, Mairead Hyland, The Lighthouse Lab in Alderley Park and Alex Alderton, Roberto Amato, Jeffrey Barrett, Sonia Goncalves, Ewan Harrison, David K. Jackson, Ian Johnston, Dominic Kwiatkowski, Cordelia Langford, John Sillitoe on behalf of the Wellcome Sanger Institute COVID-19 Surveillance Team |
| EPI_ISL_2438311 | Lighthouse Lab in Glasgow | Wellcome Sanger Institute for the COVID-19 Genomics UK (COG-UK) Consortium | Harper VanSteenhouse, Yumi Kasai, David Gray, Carol Clugston, Anna Dominiczak and Alex Alderton, Roberto Amato, Jeffrey Barrett, Sonia Goncalves, Ewan Harrison, David K. Jackson, Ian Johnston, Dominic Kwiatkowski, Cordelia Langford, John Sillitoe on behalf of the Wellcome Sanger Institute COVID-19 Surveillance Team |
| EPI_ISL_2438312, EPI_ISL_2438313, EPI_ISL_2438314 | Lighthouse Lab in Milton Keynes | Wellcome Sanger Institute for the COVID-19 Genomics UK (COG-UK) Consortium | The Lighthouse Lab in Milton Keynes and Alex Alderton, Roberto Amato, Jeffrey Barrett, Sonia Goncalves, Ewan Harrison, David K. Jackson, Ian Johnston, Dominic Kwiatkowski, Cordelia Langford, John Sillitoe on behalf of the Wellcome Sanger Institute COVID-19 Surveillance Team |
| EPI_ISL_2438316 | Lighthouse Lab in Alderley Park | Wellcome Sanger Institute for the COVID-19 Genomics UK (COG-UK) Consortium | Jacquelyn Wynn, Mairead Hyland, The Lighthouse Lab in Alderley Park and Alex Alderton, Roberto Amato, Jeffrey Barrett, Sonia Goncalves, Ewan Harrison, David K. Jackson, Ian Johnston, Dominic Kwiatkowski, Cordelia Langford, John Sillitoe on behalf of the Wellcome Sanger Institute COVID-19 Surveillance Team |
| EPI_ISL_2438317, EPI_ISL_2438318, EPI_ISL_2438319, EPI_ISL_2438320, EPI_ISL_2438321, EPI_ISL_2438322 | Lighthouse Lab in Glasgow | Wellcome Sanger Institute for the COVID-19 Genomics UK (COG-UK) Consortium | Harper VanSteenhouse, Yumi Kasai, David Gray, Carol Clugston, Anna Dominiczak and Alex Alderton, Roberto Amato, Jeffrey Barrett, Sonia Goncalves, Ewan Harrison, David K. Jackson, Ian Johnston, Dominic Kwiatkowski, Cordelia Langford, John Sillitoe on behalf of the Wellcome Sanger Institute COVID-19 Surveillance Team |
| EPI_ISL_2438323, EPI_ISL_2438324 | Lighthouse Lab in Alderley Park | Wellcome Sanger Institute for the COVID-19 Genomics UK (COG-UK) Consortium | Jacquelyn Wynn, Mairead Hyland, The Lighthouse Lab in Alderley Park and Alex Alderton, Roberto Amato, Jeffrey Barrett, Sonia Goncalves, Ewan Harrison, David K. Jackson, Ian Johnston, Dominic Kwiatkowski, Cordelia Langford, John Sillitoe on behalf of the Wellcome Sanger Institute COVID-19 Surveillance Team |
| EPI_ISL_2438326 | Lighthouse Lab in Milton Keynes | Wellcome Sanger Institute for the COVID-19 Genomics UK (COG-UK) Consortium | The Lighthouse Lab in Milton Keynes and Alex Alderton, Roberto Amato, Jeffrey Barrett, Sonia Goncalves, Ewan Harrison, David K. Jackson, Ian Johnston, Dominic Kwiatkowski, Cordelia Langford, John Sillitoe on behalf of the Wellcome Sanger Institute COVID-19 Surveillance Team |
| EPI_ISL_2438327, EPI_ISL_2438328 | Lighthouse Lab in Glasgow | Wellcome Sanger Institute for the COVID-19 Genomics UK (COG-UK) Consortium | Harper VanSteenhouse, Yumi Kasai, David Gray, Carol Clugston, Anna Dominiczak and Alex Alderton, Roberto Amato, Jeffrey Barrett, Sonia Goncalves, Ewan Harrison, David K. Jackson, Ian Johnston, Dominic Kwiatkowski, Cordelia Langford, John Sillitoe on behalf of the Wellcome Sanger Institute COVID-19 Surveillance Team |
| EPI_ISL_2438329 | Lighthouse Lab in Alderley Park | Wellcome Sanger Institute for the COVID-19 Genomics UK (COG-UK) Consortium | Jacquelyn Wynn, Mairead Hyland, The Lighthouse Lab in Alderley Park and Alex Alderton, Roberto Amato, Jeffrey Barrett, Sonia Goncalves, Ewan Harrison, David K. Jackson, Ian Johnston, Dominic Kwiatkowski, Cordelia Langford, John Sillitoe on behalf of the Wellcome Sanger Institute COVID-19 Surveillance Team |
| EPI_ISL_2438330, EPI_ISL_2438331, EPI_ISL_2438332, EPI_ISL_2438334 | Lighthouse Lab in Glasgow | Wellcome Sanger Institute for the COVID-19 Genomics UK (COG-UK) Consortium | Harper VanSteenhouse, Yumi Kasai, David Gray, Carol Clugston, Anna Dominiczak and Alex Alderton, Roberto Amato, Jeffrey Barrett, Sonia Goncalves, Ewan Harrison, David K. Jackson, Ian Johnston, Dominic Kwiatkowski, Cordelia Langford, John Sillitoe on behalf of the Wellcome Sanger Institute COVID-19 Surveillance Team |
| EPI_ISL_2438335 | Lighthouse Lab in Alderley Park | Wellcome Sanger Institute for the COVID-19 Genomics UK (COG-UK) Consortium | Jacquelyn Wynn, Mairead Hyland, The Lighthouse Lab in Alderley Park and Alex Alderton, Roberto Amato, Jeffrey Barrett, Sonia Goncalves, Ewan Harrison, David K. Jackson, Ian Johnston, Dominic Kwiatkowski, Cordelia Langford, John Sillitoe on behalf of the Wellcome Sanger Institute COVID-19 Surveillance Team |
| EPI_ISL_2438337 | Lighthouse Lab in Glasgow | Wellcome Sanger Institute for the COVID-19 Genomics UK (COG-UK) Consortium | Harper VanSteenhouse, Yumi Kasai, David Gray, Carol Clugston, Anna Dominiczak and Alex Alderton, Roberto Amato, Jeffrey Barrett, Sonia Goncalves, Ewan Harrison, David K. Jackson, Ian Johnston, Dominic Kwiatkowski, Cordelia Langford, John Sillitoe on behalf of the Wellcome Sanger Institute COVID-19 Surveillance Team |
| EPI_ISL_2438338 | Lighthouse Lab in Alderley Park | Wellcome Sanger Institute for the COVID-19 Genomics UK (COG-UK) Consortium | Jacquelyn Wynn, Mairead Hyland, The Lighthouse Lab in Alderley Park and Alex Alderton, Roberto Amato, Jeffrey Barrett, Sonia Goncalves, Ewan Harrison, David K. Jackson, Ian Johnston, Dominic Kwiatkowski, Cordelia Langford, John Sillitoe on behalf of the Wellcome Sanger Institute COVID-19 Surveillance Team |
| EPI_ISL_2438340, EPI_ISL_2438341, EPI_ISL_2438342, EPI_ISL_2438343 | Lighthouse Lab in Milton Keynes | Wellcome Sanger Institute for the COVID-19 Genomics UK (COG-UK) Consortium | The Lighthouse Lab in Milton Keynes and Alex Alderton, Roberto Amato, Jeffrey Barrett, Sonia Goncalves, Ewan Harrison, David K. Jackson, Ian Johnston, Dominic Kwiatkowski, Cordelia Langford, John Sillitoe on behalf of the Wellcome Sanger Institute COVID-19 Surveillance Team |
| EPI_ISL_2438344 | Lighthouse Lab in Glasgow | Wellcome Sanger Institute for the COVID-19 Genomics UK (COG-UK) Consortium | Harper VanSteenhouse, Yumi Kasai, David Gray, Carol Clugston, Anna Dominiczak and Alex Alderton, Roberto Amato, Jeffrey Barrett, Sonia Goncalves, Ewan Harrison, David K. Jackson, Ian Johnston, Dominic Kwiatkowski, Cordelia Langford, John Sillitoe on behalf of the Wellcome Sanger Institute COVID-19 Surveillance Team |
| EPI_ISL_2438346 | Lighthouse Lab in Alderley Park | Wellcome Sanger Institute for the COVID-19 Genomics UK (COG-UK) Consortium | Jacquelyn Wynn, Mairead Hyland, The Lighthouse Lab in Alderley Park and Alex Alderton, Roberto Amato, Jeffrey Barrett, Sonia Goncalves, Ewan Harrison, David K. Jackson, Ian Johnston, Dominic Kwiatkowski, Cordelia Langford, John Sillitoe on behalf of the Wellcome Sanger Institute COVID-19 Surveillance Team |
| EPI_ISL_2438348, EPI_ISL_2438349, EPI_ISL_2438350, EPI_ISL_2438351, EPI_ISL_2438353, EPI_ISL_2438354, EPI_ISL_2438355, EPI_ISL_2438356 | Lighthouse Lab in Glasgow | Wellcome Sanger Institute for the COVID-19 Genomics UK (COG-UK) Consortium | Harper VanSteenhouse, Yumi Kasai, David Gray, Carol Clugston, Anna Dominiczak and Alex Alderton, Roberto Amato, Jeffrey Barrett, Sonia Goncalves, Ewan Harrison, David K. Jackson, Ian Johnston, Dominic Kwiatkowski, Cordelia Langford, John Sillitoe on behalf of the Wellcome Sanger Institute COVID-19 Surveillance Team |
| EPI_ISL_2438360, EPI_ISL_2438361 | Lighthouse Lab in Milton Keynes | Wellcome Sanger Institute for the COVID-19 Genomics UK (COG-UK) Consortium | The Lighthouse Lab in Milton Keynes and Alex Alderton, Roberto Amato, Jeffrey Barrett, Sonia Goncalves, Ewan Harrison, David K. Jackson, Ian Johnston, Dominic Kwiatkowski, Cordelia Langford, John Sillitoe on behalf of the Wellcome Sanger Institute COVID-19 Surveillance Team |
| EPI_ISL_2438363 | Lighthouse Lab in Alderley Park | Wellcome Sanger Institute for the COVID-19 Genomics UK (COG-UK) Consortium | Jacquelyn Wynn, Mairead Hyland, The Lighthouse Lab in Alderley Park and Alex Alderton, Roberto Amato, Jeffrey Barrett, Sonia Goncalves, Ewan Harrison, David K. Jackson, Ian Johnston, Dominic Kwiatkowski, Cordelia Langford, John Sillitoe on behalf of the Wellcome Sanger Institute COVID-19 Surveillance Team |
| EPI_ISL_2438364 | Lighthouse Lab in Glasgow | Wellcome Sanger Institute for the COVID-19 Genomics UK (COG-UK) Consortium | Harper VanSteenhouse, Yumi Kasai, David Gray, Carol Clugston, Anna Dominiczak and Alex Alderton, Roberto Amato, Jeffrey Barrett, Sonia Goncalves, Ewan Harrison, David K. Jackson, Ian Johnston, Dominic Kwiatkowski, Cordelia Langford, John Sillitoe on behalf of the Wellcome Sanger Institute COVID-19 Surveillance Team |
| EPI_ISL_2438365, EPI_ISL_2438367 | Lighthouse Lab in Milton Keynes | Wellcome Sanger Institute for the COVID-19 Genomics UK (COG-UK) Consortium | The Lighthouse Lab in Milton Keynes and Alex Alderton, Roberto Amato, Jeffrey Barrett, Sonia Goncalves, Ewan Harrison, David K. Jackson, Ian Johnston, Dominic Kwiatkowski, Cordelia Langford, John Sillitoe on behalf of the Wellcome Sanger Institute COVID-19 Surveillance Team |
| EPI_ISL_2438368 | Lighthouse Lab in Alderley Park | Wellcome Sanger Institute for the COVID-19 Genomics UK (COG-UK) Consortium | Jacquelyn Wynn, Mairead Hyland, The Lighthouse Lab in Alderley Park and Alex Alderton, Roberto Amato, Jeffrey Barrett, Sonia Goncalves, Ewan Harrison, David K. Jackson, Ian Johnston, Dominic Kwiatkowski, Cordelia Langford, John Sillitoe on behalf of the Wellcome Sanger Institute COVID-19 Surveillance Team |
| EPI_ISL_2438369 | Lighthouse Lab in Glasgow | Wellcome Sanger Institute for the COVID-19 Genomics UK (COG-UK) Consortium | Harper VanSteenhouse, Yumi Kasai, David Gray, Carol Clugston, Anna Dominiczak and Alex Alderton, Roberto Amato, Jeffrey Barrett, Sonia Goncalves, Ewan Harrison, David K. Jackson, Ian Johnston, Dominic Kwiatkowski, Cordelia Langford, John Sillitoe on behalf of the Wellcome Sanger Institute COVID-19 Surveillance Team |
| EPI_ISL_2438370, EPI_ISL_2438371 | Lighthouse Lab in Milton Keynes | Wellcome Sanger Institute for the COVID-19 Genomics UK (COG-UK) Consortium | The Lighthouse Lab in Milton Keynes and Alex Alderton, Roberto Amato, Jeffrey Barrett, Sonia Goncalves, Ewan Harrison, David K. Jackson, Ian Johnston, Dominic Kwiatkowski, Cordelia Langford, John Sillitoe on behalf of the Wellcome Sanger Institute COVID-19 Surveillance Team |
| EPI_ISL_2438372 | Lighthouse Lab in Alderley Park | Wellcome Sanger Institute for the COVID-19 Genomics UK (COG-UK) Consortium | Jacquelyn Wynn, Mairead Hyland, The Lighthouse Lab in Alderley Park and Alex Alderton, Roberto Amato, Jeffrey Barrett, Sonia Goncalves, Ewan Harrison, David K. Jackson, Ian Johnston, Dominic Kwiatkowski, Cordelia Langford, John Sillitoe on behalf of the Wellcome Sanger Institute COVID-19 Surveillance Team |
| EPI_ISL_2438373 | Lighthouse Lab in Glasgow | Wellcome Sanger Institute for the COVID-19 Genomics UK (COG-UK) Consortium | Harper VanSteenhouse, Yumi Kasai, David Gray, Carol Clugston, Anna Dominiczak and Alex Alderton, Roberto Amato, Jeffrey Barrett, Sonia Goncalves, Ewan Harrison, David K. Jackson, Ian Johnston, Dominic Kwiatkowski, Cordelia Langford, John Sillitoe on behalf of the Wellcome Sanger Institute COVID-19 Surveillance Team |

| | | | |
|---|---|---|---|
| EPI_ISL_2438374, EPI_ISL_2438376 | Lighthouse Lab in Alderley Park | Wellcome Sanger Institute for the COVID-19 Genomics UK (COG-UK) Consortium | Jacquelyn Wynn, Mairead Hyland, The Lighthouse Lab in Alderley Park and Alex Alderton, Roberto Amato, Jeffrey Barrett, Sonia Goncalves, Ewan Harrison, David K. Jackson, Ian Johnston, Dominic Kwiatkowski, Cordelia Langford, John Sillitoe on behalf of the Wellcome Sanger Institute COVID-19 Surveillance Team |
| EPI_ISL_2438377 | Lighthouse Lab in Milton Keynes | Wellcome Sanger Institute for the COVID-19 Genomics UK (COG-UK) Consortium | The Lighthouse Lab in Milton Keynes and Alex Alderton, Roberto Amato, Jeffrey Barrett, Sonia Goncalves, Ewan Harrison, David K. Jackson, Ian Johnston, Dominic Kwiatkowski, Cordelia Langford, John Sillitoe on behalf of the Wellcome Sanger Institute COVID-19 Surveillance Team |
| EPI_ISL_2438378, EPI_ISL_2438379, EPI_ISL_2438380, EPI_ISL_2438383, EPI_ISL_2438386, EPI_ISL_2438388, EPI_ISL_2438389, EPI_ISL_2438390, EPI_ISL_2438395, EPI_ISL_2438396, EPI_ISL_2438398, EPI_ISL_2438399 | | | |
| see above | Lighthouse Lab in Glasgow | Wellcome Sanger Institute for the COVID-19 Genomics UK (COG-UK) Consortium | Harper VanSteenhouse, Yumi Kasai, David Gray, Carol Clugston, Anna Dominiczak and Alex Alderton, Roberto Amato, Jeffrey Barrett, Sonia Goncalves, Ewan Harrison, David K. Jackson, Ian Johnston, Dominic Kwiatkowski, Cordelia Langford, John Sillitoe on behalf of the Wellcome Sanger Institute COVID-19 Surveillance Team |
| EPI_ISL_2438402 | Lighthouse Lab in Alderley Park | Wellcome Sanger Institute for the COVID-19 Genomics UK (COG-UK) Consortium | Jacquelyn Wynn, Mairead Hyland, The Lighthouse Lab in Alderley Park and Alex Alderton, Roberto Amato, Jeffrey Barrett, Sonia Goncalves, Ewan Harrison, David K. Jackson, Ian Johnston, Dominic Kwiatkowski, Cordelia Langford, John Sillitoe on behalf of the Wellcome Sanger Institute COVID-19 Surveillance Team |
| EPI_ISL_2438403, EPI_ISL_2438405, EPI_ISL_2438407, EPI_ISL_2438408, EPI_ISL_2438409, EPI_ISL_2438410, EPI_ISL_2438411, EPI_ISL_2438412, EPI_ISL_2438414, EPI_ISL_2438415, EPI_ISL_2438417, EPI_ISL_2438418, EPI_ISL_2438419, EPI_ISL_2438420, EPI_ISL_2438421, EPI_ISL_2438423, EPI_ISL_2438425, EPI_ISL_2438426 | | | |
| see above | Lighthouse Lab in Glasgow | Wellcome Sanger Institute for the COVID-19 Genomics UK (COG-UK) Consortium | Harper VanSteenhouse, Yumi Kasai, David Gray, Carol Clugston, Anna Dominiczak and Alex Alderton, Roberto Amato, Jeffrey Barrett, Sonia Goncalves, Ewan Harrison, David K. Jackson, Ian Johnston, Dominic Kwiatkowski, Cordelia Langford, John Sillitoe on behalf of the Wellcome Sanger Institute COVID-19 Surveillance Team |
| EPI_ISL_2438427 | Lighthouse Lab in Milton Keynes | Wellcome Sanger Institute for the COVID-19 Genomics UK (COG-UK) Consortium | The Lighthouse Lab in Milton Keynes and Alex Alderton, Roberto Amato, Jeffrey Barrett, Sonia Goncalves, Ewan Harrison, David K. Jackson, Ian Johnston, Dominic Kwiatkowski, Cordelia Langford, John Sillitoe on behalf of the Wellcome Sanger Institute COVID-19 Surveillance Team |
| EPI_ISL_2438429, EPI_ISL_2438430, EPI_ISL_2438431, EPI_ISL_2438432, EPI_ISL_2438434, EPI_ISL_2438435, EPI_ISL_2438439, EPI_ISL_2438440, EPI_ISL_2438442, EPI_ISL_2438444, EPI_ISL_2438446, EPI_ISL_2438447, EPI_ISL_2438448, EPI_ISL_2438449, EPI_ISL_2438450, EPI_ISL_2438451, EPI_ISL_2438452, EPI_ISL_2438456, EPI_ISL_2438457, EPI_ISL_2438458, EPI_ISL_2438459, EPI_ISL_2438462, EPI_ISL_2438463, EPI_ISL_2438464, EPI_ISL_2438465, EPI_ISL_2438466, EPI_ISL_2438468, EPI_ISL_2438470, EPI_ISL_2438472, EPI_ISL_2438475, EPI_ISL_2438476, EPI_ISL_2438477, EPI_ISL_2438478, EPI_ISL_2438479, EPI_ISL_2438481, EPI_ISL_2438483, EPI_ISL_2438484, EPI_ISL_2438485, EPI_ISL_2438486, EPI_ISL_2438489, EPI_ISL_2438493, EPI_ISL_2438494, EPI_ISL_2438495, EPI_ISL_2438496, EPI_ISL_2438497, EPI_ISL_2438499, EPI_ISL_2438502, EPI_ISL_2438503, EPI_ISL_2438505, EPI_ISL_2438506, EPI_ISL_2438511, EPI_ISL_2438512, EPI_ISL_2438514, EPI_ISL_2438515, EPI_ISL_2438517, EPI_ISL_2438518 | | | |
| see above | Lighthouse Lab in Glasgow | Wellcome Sanger Institute for the COVID-19 Genomics UK (COG-UK) Consortium | Harper VanSteenhouse, Yumi Kasai, David Gray, Carol Clugston, Anna Dominiczak and Alex Alderton, Roberto Amato, Jeffrey Barrett, Sonia Goncalves, Ewan Harrison, David K. Jackson, Ian Johnston, Dominic Kwiatkowski, Cordelia Langford, John Sillitoe on behalf of the Wellcome Sanger Institute COVID-19 Surveillance Team |
| EPI_ISL_2438519 | Lighthouse Lab in Alderley Park | Wellcome Sanger Institute for the COVID-19 Genomics UK (COG-UK) Consortium | Jacquelyn Wynn, Mairead Hyland, The Lighthouse Lab in Alderley Park and Alex Alderton, Roberto Amato, Jeffrey Barrett, Sonia Goncalves, Ewan Harrison, David K. Jackson, Ian Johnston, Dominic Kwiatkowski, Cordelia Langford, John Sillitoe on behalf of the Wellcome Sanger Institute COVID-19 Surveillance Team |
| EPI_ISL_2438520, EPI_ISL_2438521, EPI_ISL_2438522, EPI_ISL_2438523, EPI_ISL_2438524, EPI_ISL_2438527, EPI_ISL_2438528, EPI_ISL_2438529, EPI_ISL_2438530, EPI_ISL_2438532, EPI_ISL_2438533, EPI_ISL_2438534, EPI_ISL_2438536, EPI_ISL_2438539, EPI_ISL_2438541 | | | |
| see above | Lighthouse Lab in Glasgow | Wellcome Sanger Institute for the COVID-19 Genomics UK (COG-UK) Consortium | Harper VanSteenhouse, Yumi Kasai, David Gray, Carol Clugston, Anna Dominiczak and Alex Alderton, Roberto Amato, Jeffrey Barrett, Sonia Goncalves, Ewan Harrison, David K. Jackson, Ian Johnston, Dominic Kwiatkowski, Cordelia Langford, John Sillitoe on behalf of the Wellcome Sanger Institute COVID-19 Surveillance Team |
| EPI_ISL_2438547, EPI_ISL_2438666 | Department of Virology | Department of Virology | Massab Umair, Aamer Ikram, Muhammad Salman, Nazish Badar, Zaira Rehman, Muhammad Ammar, Syed Adnan Haider |
| EPI_ISL_2438689 | HOSPITAL CAN MISSES | HOSPITAL UNIVERSITARIO SON ESPASES | Servicio de Microbiología, Hospital Universitario Son Espases |
| EPI_ISL_2438777, EPI_ISL_2438780, EPI_ISL_2438781, EPI_ISL_2438787, EPI_ISL_2438788, EPI_ISL_2438789, EPI_ISL_2438790, EPI_ISL_2438791, EPI_ISL_2438792, EPI_ISL_2438795, EPI_ISL_2438796, EPI_ISL_2438797, EPI_ISL_2438798, EPI_ISL_2438800 | | | |
| see above | Dept. of Microbiology and Infection Control, Akershus University Hospital HF | Dept. of Microbiology and Infection Control, Akershus University Hospital HF | Hege Vangstein Aamot, Alexander Hesselberg Løvestad |
| EPI_ISL_2438849, EPI_ISL_2438850, EPI_ISL_2438851, EPI_ISL_2438852, EPI_ISL_2438853 | Tokyo Metropolitan Institute of Public Health | Tokyo Metropolitan Institute of Public Health | Masaki Hayashi, Takako Yamazaki, Akane Negishi,Hiroshi Koike, Hirofumi Miyake, Takushi Fujiwara, Mami Nagashima, Jun Suzuki, Kenji Sadamasu |
| EPI_ISL_2439199 | Servicio Microbiología Hospital La Paz | Servicio Microbiología Hospital La Paz | Fernando Lázaro, Rubén Cáceres, Jesús Mingorance Cruz, Elie Dahdouh |
| EPI_ISL_2439298, EPI_ISL_2439305, EPI_ISL_2439307, EPI_ISL_2439313, EPI_ISL_2439321, EPI_ISL_2439339, EPI_ISL_2439349, EPI_ISL_2439351, EPI_ISL_2439373, EPI_ISL_2439375, EPI_ISL_2439390, EPI_ISL_2439407, EPI_ISL_2439424, EPI_ISL_2439432, EPI_ISL_2439438, EPI_ISL_2439444, EPI_ISL_2439476, EPI_ISL_2439483, EPI_ISL_2439491, EPI_ISL_2439499, EPI_ISL_2439505, EPI_ISL_2439511, EPI_ISL_2439527, EPI_ISL_2439530, EPI_ISL_2439539, EPI_ISL_2439540, EPI_ISL_2439547, EPI_ISL_2439549, EPI_ISL_2439552, EPI_ISL_2439558, EPI_ISL_2439579, EPI_ISL_2439581, EPI_ISL_2439636, EPI_ISL_2439685, EPI_ISL_2439725, EPI_ISL_2439768, EPI_ISL_2439778, EPI_ISL_2439788, EPI_ISL_2439805, EPI_ISL_2439808, EPI_ISL_2439836, EPI_ISL_2439876 | | | |
| see above | Quest Diagnostics Incorporated | Centers for Disease Control and Prevention Division of Viral Diseases, Pathogen Discovery | Dakota Howard, Dhwani Batra, Peter W. Cook, Kara Moser, Adrian Paskey, Jason Caravas, Benjamin Rambo-Martin, Shatavia Morrison, Christopher Gulvick, Scott Sammons, Yvette Unoarumhi, Darlene Wagner, Matthew Schmerer, S. H. Rosenthal, A. Gerasimova, R. M. Kagan, B. Anderson, M. Hua, Y. Liu, L.E. Bernstein, K.E. Livingston, A. Perez, I. A. Shlyakhter, R. V. Rolando, R. Owen, P. Tanpaiboon, F. Lacbawan, Clinton R. Paden, Duncan MacCannell |
| EPI_ISL_2440079, EPI_ISL_2440107, EPI_ISL_2440204, EPI_ISL_2440260, EPI_ISL_2440304, EPI_ISL_2440312, EPI_ISL_2440322, EPI_ISL_2440474, EPI_ISL_2440479, EPI_ISL_2440528 | Helix/Illumina | Centers for Disease Control and Prevention Division of Viral Diseases, Pathogen Discovery | Dakota Howard, Dhwani Batra, Peter W. Cook, Kara Moser, Adrian Paskey, Jason Caravas, Benjamin Rambo-Martin, Shatavia Morrison, Christopher Gulvick, Scott Sammons, Yvette Unoarumhi, Darlene Wagner, Matthew Schmerer, Eileen de Feo, Jan Antico, Christine Tran, Matthew Tolentino, Shannon Wickline, Kim Gietzen, Brad Sickler, Jingtao Liu, Eric Allen, Phil Febbo, Nicole L. Washington, Simon White, Geraint Levan, Kelly Schiabor Barrett, Elizabeth Cirulli, Alexandre Bolze, Ary Ascencio, Charlotte Rivera-Garcia, Ryan Cho, Jason Nguyen, Sherry Wang, Jimmy Ramirez, Tyler Cassens, Efren Sandoval, Magnus Isaksson, William Lee, David Becker, Marc Laurent, James Lu, Clinton R. Paden, Duncan MacCannell |
| EPI_ISL_2440539, EPI_ISL_2440541, EPI_ISL_2440576, EPI_ISL_2440580, EPI_ISL_2440585, EPI_ISL_2440590, EPI_ISL_2440626, EPI_ISL_2440656, EPI_ISL_2440658, EPI_ISL_2440670, EPI_ISL_2440688, EPI_ISL_2440689, EPI_ISL_2440711, EPI_ISL_2440739, EPI_ISL_2440794, EPI_ISL_2440795, EPI_ISL_2440832, EPI_ISL_2440841, EPI_ISL_2440845, EPI_ISL_2440848, EPI_ISL_2440856, EPI_ISL_2440858, EPI_ISL_2440888, EPI_ISL_2440893 | | | |
| see above | Quest Diagnostics Incorporated | Centers for Disease Control and Prevention Division of Viral Diseases, Pathogen Discovery | Dakota Howard, Dhwani Batra, Peter W. Cook, Kara Moser, Adrian Paskey, Jason Caravas, Benjamin Rambo-Martin, Shatavia Morrison, Christopher Gulvick, Scott Sammons, Yvette Unoarumhi, Darlene Wagner, Matthew Schmerer, S. H. Rosenthal, A. Gerasimova, R. M. Kagan, B. Anderson, M. Hua, Y. Liu, L.E. Bernstein, K.E. Livingston, A. Perez, I. A. Shlyakhter, R. V. Rolando, R. Owen, P. Tanpaiboon, F. Lacbawan, Clinton R. Paden, Duncan MacCannell |
| EPI_ISL_2440909, EPI_ISL_2440918, EPI_ISL_2440945, EPI_ISL_2440947, EPI_ISL_2440965, EPI_ISL_2440967, EPI_ISL_2440981, EPI_ISL_2441017, EPI_ISL_2441019, EPI_ISL_2441042, EPI_ISL_2441069 | | | |
| see above | Mako Medical | Centers for Disease Control and Prevention Division of Viral Diseases, Pathogen Discovery | Dakota Howard, Dhwani Batra, Peter W. Cook, Kara Moser, Adrian Paskey, Jason Caravas, Benjamin Rambo-Martin, Shatavia Morrison, Christopher Gulvick, Scott Sammons, Yvette Unoarumhi, Darlene Wagner, Matthew Schmerer,Matthew Tugwell, Lauren Moon, Clinton R. Paden, Duncan MacCannell |
| EPI_ISL_2441106, EPI_ISL_2441108, EPI_ISL_2441109, EPI_ISL_2441110, EPI_ISL_2441111, EPI_ISL_2441112, EPI_ISL_2441113, EPI_ISL_2441114 | AP SSO | CSIR-Centre for Cellular and Molecular Biology-INSACOG | Lamuk Zaveri,Tulasi Nagabandi,Payel Mukherjee,Onkar Kulkarni, Sofia Banu, Ara Sreenivas,Shreekant Verma, Amareshwar Vodapalli ,B Himasri,Valli Nagalakshmi Undamatla, Sumedha Avadhanula, Priya Nurkurthy, Vidhyadhari Methuku, Archana Bharadwaj Siva, Karthik Bharadwaj Tallapaka,Rakesh K Mishra,Divya Tej Sowpati |
| EPI_ISL_2441115, EPI_ISL_2441116, EPI_ISL_2441117, EPI_ISL_2441118, EPI_ISL_2441119, EPI_ISL_2441120, EPI_ISL_2441121, EPI_ISL_2441122, EPI_ISL_2441123, EPI_ISL_2441124, EPI_ISL_2441125, EPI_ISL_2441126, EPI_ISL_2441127, EPI_ISL_2441128 | | | |
| see above | AP SSO | CSIR-Centre for Cellular and Molecular Biology-INSACOG | Tulasi Nagabandi,Lamuk Zaveri,Payel Mukherjee,Onkar Kulkarni, Sofia Banu, Ara Sreenivas,Shreekant Verma, Amareshwar Vodapalli ,B Himasri,Valli Nagalakshmi Undamatla, Priya Nurkurthy,Sumedha Avadhanula, Vidhyadhari Methuku, Archana Bharadwaj Siva, Karthik Bharadwaj Tallapaka,Rakesh K Mishra,Divya Tej Sowpati |
| EPI_ISL_2441129, EPI_ISL_2441132, EPI_ISL_2441133, EPI_ISL_2441134, EPI_ISL_2441135, EPI_ISL_2441136, EPI_ISL_2441137, EPI_ISL_2441138, EPI_ISL_2441139 | AP SSO | CSIR-Centre for Cellular and Molecular Biology-INSACOG | Priya Nurkurthy,Ara Sreenivas,Shreekant Verma, Amareshwar Vodapalli ,Lamuk Zaveri,Tulasi Nagabandi,B Himasri,Valli Nagalakshmi Undamatla, Sumedha Avadhanula, Vidhyadhari Methuku,Payel Mukherjee,Onkar Kulkarni, Sofia Banu, Archana Bharadwaj Siva, Karthik Bharadwaj Tallapaka,Rakesh K Mishra,Divya Tej Sowpati |
| EPI_ISL_2441140, EPI_ISL_2441141, EPI_ISL_2441142, EPI_ISL_2441143, EPI_ISL_2441144, EPI_ISL_2441145, EPI_ISL_2441146, EPI_ISL_2441147, EPI_ISL_2441148, EPI_ISL_2441149, EPI_ISL_2441150 | | | |
| see above | AP SSO | CSIR-Centre for Cellular and Molecular Biology-INSACOG | Ara Sreenivas,Shreekant Verma, Amareshwar Vodapalli ,B Himasri,Valli Nagalakshmi Undamatla,Payel Mukherjee, Lamuk Zaveri, Onkar Kulkarni,Tulasi Nagabandi, Sofia Banu, ,Priya Nurkurthy, Vidhyadhari Methuku, Sumedha Avadhanula, Archana Bharadwaj Siva, Karthik Bharadwaj Tallapaka,Rakesh K Mishra,Divya Tej Sowpati |
| EPI_ISL_2441151, EPI_ISL_2441152, EPI_ISL_2441153, EPI_ISL_2441154, EPI_ISL_2441155, EPI_ISL_2441156, EPI_ISL_2441157, EPI_ISL_2441158, EPI_ISL_2441159, EPI_ISL_2441160, EPI_ISL_2441162, EPI_ISL_2441163, EPI_ISL_2441164, EPI_ISL_2441165, EPI_ISL_2441166, EPI_ISL_2441167, EPI_ISL_2441168 | | | |
| see above | AP SSO | CSIR-Centre for Cellular and Molecular Biology-INSACOG | Shreekant Verma, Amareshwar Vodapalli ,Ara Sreenivas,B Himasri,Valli Nagalakshmi Undamatla,Payel Mukherjee, Lamuk Zaveri, Onkar Kulkarni,Tulasi Nagabandi, Sofia Banu, Sumedha Avadhanula, Priya Nurkurthy, Vidhyadhari Methuku, Archana Bharadwaj Siva, Karthik Bharadwaj Tallapaka,Rakesh K Mishra,Divya Tej Sowpati |
| EPI_ISL_2441169, EPI_ISL_2441170, EPI_ISL_2441171, EPI_ISL_2441172, EPI_ISL_2441173, EPI_ISL_2441174, EPI_ISL_2441175, EPI_ISL_2441176, EPI_ISL_2441177, EPI_ISL_2441178, EPI_ISL_2441179, EPI_ISL_2441180, EPI_ISL_2441181, EPI_ISL_2441182, EPI_ISL_2441183, EPI_ISL_2441184, EPI_ISL_2441185, EPI_ISL_2441186, EPI_ISL_2441187, EPI_ISL_2441188, EPI_ISL_2441189, EPI_ISL_2441190, EPI_ISL_2441191 | | | |
| see above | AP SSO | CSIR-Centre for Cellular and Molecular Biology-INSACOG | Amareshwar Vodapalli,Ara Sreenivas,Shreekant Verma, B Himasri,Valli Nagalakshmi Undamatla,Payel Mukherjee, Lamuk Zaveri, Onkar Kulkarni,Tulasi Nagabandi, Sofia Banu, ,Priya Nurkurthy, Vidhyadhari Methuku, Sumedha Avadhanula, Archana Bharadwaj Siva, Karthik Bharadwaj Tallapaka,Rakesh K Mishra,Divya Tej Sowpati |
| EPI_ISL_2441192, EPI_ISL_2441193, EPI_ISL_2441194, EPI_ISL_2441195, EPI_ISL_2441196, EPI_ISL_2441197, EPI_ISL_2441198, EPI_ISL_2441199, EPI_ISL_2441200, EPI_ISL_2441201, EPI_ISL_2441202, EPI_ISL_2441203, EPI_ISL_2441204, EPI_ISL_2441205, EPI_ISL_2441206, EPI_ISL_2441207, EPI_ISL_2441208, EPI_ISL_2441209, EPI_ISL_2441210, EPI_ISL_2441211, EPI_ISL_2441212 | | | |
| see above | AP SSO | CSIR-Centre for Cellular and Molecular Biology-INSACOG | B Himasri,Amareshwar Vodapalli,Ara Sreenivas,Shreekant Verma,Valli Nagalakshmi Undamatla,Payel Mukherjee, Lamuk Zaveri, Onkar Kulkarni,Tulasi Nagabandi, Sofia Banu, Vidhyadhari Methuku,Priya Nurkurthy, Sumedha Avadhanula, Archana Bharadwaj Siva, Karthik Bharadwaj Tallapaka,Rakesh K Mishra,Divya Tej Sowpati |
| EPI_ISL_2441213, EPI_ISL_2441214, EPI_ISL_2441215, | AP SSO | CSIR-Centre for Cellular and Molecular Biology-INSACOG | Payel Mukherjee, Lamuk Zaveri, Onkar Kulkarni,Tulasi Nagabandi, Sofia Banu, Ara Sreenivas,Shreekant Verma, Amareshwar Vodapalli ,B Himasri,Valli Nagalakshmi Undamatla, Sumedha Avadhanula, Vidhyadhari Methuku, Priya Nurkurthy, Archana Bharadwaj Siva, Karthik Bharadwaj Tallapaka,Rakesh K Mishra,Divya Tej Sowpati |

| | | | |
|---|---|---|---|
| EPI_ISL_2441216, EPI_ISL_2441217, EPI_ISL_2441218, EPI_ISL_2441220, EPI_ISL_2441221 | | | |
| EPI_ISL_2441222, EPI_ISL_2441223, EPI_ISL_2441224, EPI_ISL_2441225, EPI_ISL_2441226, EPI_ISL_2441227, EPI_ISL_2441228, EPI_ISL_2441229 | AP SSO | CSIR-Centre for Cellular and Molecular Biology-INSACOG | Priya Nurkurthy,Ara Sreenivas,Shreekant Verma, Amareshwar Vodapalli ,Lamuk Zaveri,Tulasi Nagabandi,B Himasri,Valli Nagalakshmi Undamatla, Sumedha Avadhanula, Vidhyadhari Methuku,Payel Mukherjee,Onkar Kulkarni, Sofia Banu, Archana Bharadwaj Siva, Karthik Bharadwaj Tallapaka,Rakesh K Mishra,Divya Tej Sowpati |
| EPI_ISL_2441230, EPI_ISL_2441231, EPI_ISL_2441232, EPI_ISL_2441233, EPI_ISL_2441234, EPI_ISL_2441235, EPI_ISL_2441236, EPI_ISL_2441237, EPI_ISL_2441238, EPI_ISL_2441239, EPI_ISL_2441240, EPI_ISL_2441241 | | | |
| see above | AP SSO | CSIR-Centre for Cellular and Molecular Biology-INSACOG | Sofia Banu,Payel Mukherjee, Lamuk Zaveri, Onkar Kulkarni,Tulasi Nagabandi, Ara Sreenivas,Shreekant Verma, Amareshwar Vodapalli ,B Himasri,Valli Nagalakshmi Undamatla, Sumedha Avadhanula, Vidhyadhari Methuku, Priya Nurkurthy, Archana Bharadwaj Siva, Karthik Bharadwaj Tallapaka,Rakesh K Mishra,Divya Tej Sowpati |
| EPI_ISL_2441242, EPI_ISL_2441243, EPI_ISL_2441245, EPI_ISL_2441246, EPI_ISL_2441254, EPI_ISL_2441255, EPI_ISL_2441256, EPI_ISL_2441257 | AP SSO | CSIR-Centre for Cellular and Molecular Biology-INSACOG | Lamuk Zaveri,Tulasi Nagabandi,Payel Mukherjee,Onkar Kulkarni, Sofia Banu, Ara Sreenivas,Shreekant Verma, Amareshwar Vodapalli ,B Himasri,Valli Nagalakshmi Undamatla, Sumedha Avadhanula, Priya Nurkurthy, Vidhyadhari Methuku, Archana Bharadwaj Siva, Karthik Bharadwaj Tallapaka,Rakesh K Mishra,Divya Tej Sowpati |
| EPI_ISL_2441355, EPI_ISL_2441358, EPI_ISL_2441365, EPI_ISL_2441366 | Mapmygenome | CSIR-Centre for Cellular and Molecular Biology-INSACOG | Amareshwar Vodapalli,Ara Sreenivas,Shreekant Verma, B Himasri,Valli Nagalakshmi Undamatla,Payel Mukherjee, Lamuk Zaveri, Onkar Kulkarni,Tulasi Nagabandi, Sofia Banu, ,Priya Nurkurthy, Vidhyadhari Methuku, Sumedha Avadhanula, Archana Bharadwaj Siva, Karthik Bharadwaj Tallapaka,Rakesh K Mishra,Divya Tej Sowpati |
| EPI_ISL_2441367, EPI_ISL_2441368, EPI_ISL_2441369, EPI_ISL_2441370, EPI_ISL_2441371 | CSIR-Centre for Cellular and Molecular Biology | CSIR-Centre for Cellular and Molecular Biology-INSACOG | Amareshwar Vodapalli,Ara Sreenivas,Shreekant Verma, B Himasri,Valli Nagalakshmi Undamatla,Payel Mukherjee, Lamuk Zaveri, Onkar Kulkarni,Tulasi Nagabandi, Sofia Banu, ,Priya Nurkurthy, Vidhyadhari Methuku, Sumedha Avadhanula, Archana Bharadwaj Siva, Karthik Bharadwaj Tallapaka,Rakesh K Mishra,Divya Tej Sowpati |
| EPI_ISL_2441372, EPI_ISL_2441385, EPI_ISL_2441386, EPI_ISL_2441391, EPI_ISL_2441396, EPI_ISL_2441400, EPI_ISL_2441401, EPI_ISL_2441402, EPI_ISL_2441403, EPI_ISL_2441404, EPI_ISL_2441405, EPI_ISL_2441406, EPI_ISL_2441407, EPI_ISL_2441408, EPI_ISL_2441409, EPI_ISL_2441410, EPI_ISL_2441411, EPI_ISL_2441412, EPI_ISL_2441413, EPI_ISL_2441414, EPI_ISL_2441416, EPI_ISL_2441417, EPI_ISL_2441418, EPI_ISL_2441419, EPI_ISL_2441420, EPI_ISL_2441421, EPI_ISL_2441423, EPI_ISL_2441424, EPI_ISL_2441425, EPI_ISL_2441427, EPI_ISL_2441428, EPI_ISL_2441429, EPI_ISL_2441430, EPI_ISL_2441431, EPI_ISL_2441432, EPI_ISL_2441433, EPI_ISL_2441434, EPI_ISL_2441435, EPI_ISL_2441436, EPI_ISL_2441437, EPI_ISL_2441438, EPI_ISL_2441439, EPI_ISL_2441441, EPI_ISL_2441442, EPI_ISL_2441443, EPI_ISL_2441445, EPI_ISL_2441446, EPI_ISL_2441447, EPI_ISL_2441448, EPI_ISL_2441451, EPI_ISL_2441452, EPI_ISL_2441453, EPI_ISL_2441454, EPI_ISL_2441455, EPI_ISL_2441456, EPI_ISL_2441458, EPI_ISL_2441459, EPI_ISL_2441460, EPI_ISL_2441461, EPI_ISL_2441462, EPI_ISL_2441463, EPI_ISL_2441464, EPI_ISL_2441465, EPI_ISL_2441466, EPI_ISL_2441467, EPI_ISL_2441468, EPI_ISL_2441469, EPI_ISL_2441470, EPI_ISL_2441471 | | | |
| see above | CSIR-National Environmental Engineering Research Institute | CSIR-Centre for Cellular and Molecular Biology-INSACOG | Payel Mukherjee, Lamuk Zaveri, Onkar Kulkarni,Tulasi Nagabandi, Sofia Banu, Ara Sreenivas,Shreekant Verma, Amareshwar Vodapalli ,B Himasri,Valli Nagalakshmi Undamatla, Sumedha Avadhanula, Vidhyadhari Methuku, Priya Nurkurthy, Archana Bharadwaj Siva, Karthik Bharadwaj Tallapaka,Krishna Khairnar,Rakesh K Mishra,Divya Tej Sowpati |
| EPI_ISL_2441481, EPI_ISL_2441482, EPI_ISL_2441483, EPI_ISL_2441484, EPI_ISL_2441485, EPI_ISL_2441486, EPI_ISL_2441487, EPI_ISL_2441488, EPI_ISL_2441489, EPI_ISL_2441490 | CSIR-Centre for Cellular and Molecular Biology | CSIR-Centre for Cellular and Molecular Biology-INSACOG | B Himasri,Amareshwar Vodapalli,Ara Sreenivas,Shreekant Verma,Valli Nagalakshmi Undamatla,Payel Mukherjee, Lamuk Zaveri, Onkar Kulkarni,Tulasi Nagabandi, Sofia Banu, Vidhyadhari Methuku,Priya Nurkurthy, Sumedha Avadhanula, Archana Bharadwaj Siva, Karthik Bharadwaj Tallapaka,Rakesh K Mishra,Divya Tej Sowpati |
| EPI_ISL_2441491, EPI_ISL_2441492 | CSIR-Centre for Cellular and Molecular Biology | CSIR-Centre for Cellular and Molecular Biology-INSACOG | Vidhyadhari Methuku,Tulasi Nagabandi,Lamuk Zaveri, Ara Sreenivas,Shreekant Verma, Amareshwar Vodapalli ,B Himasri,Valli Nagalakshmi Undamatla, Priya Nurkurthy,Sumedha Avadhanula, Archana Bharadwaj Siva,Payel Mukherjee,Onkar Kulkarni, Sofia Banu, Karthik Bharadwaj Tallapaka,Rakesh K Mishra,Divya Tej Sowpati |
| EPI_ISL_2441493, EPI_ISL_2441494, EPI_ISL_2441495, EPI_ISL_2441496, EPI_ISL_2441497, EPI_ISL_2441498, EPI_ISL_2441499, EPI_ISL_2441500 | CSIR-Centre for Cellular and Molecular Biology | CSIR-Centre for Cellular and Molecular Biology-INSACOG | Payel Mukherjee, Lamuk Zaveri, Onkar Kulkarni,Tulasi Nagabandi, Sofia Banu, Ara Sreenivas,Shreekant Verma, Amareshwar Vodapalli ,B Himasri,Valli Nagalakshmi Undamatla, Sumedha Avadhanula, Vidhyadhari Methuku, Priya Nurkurthy, Archana Bharadwaj Siva, Karthik Bharadwaj Tallapaka,Rakesh K Mishra,Divya Tej Sowpati |
| EPI_ISL_2441501, EPI_ISL_2441502, EPI_ISL_2441503, EPI_ISL_2441504, EPI_ISL_2441505, EPI_ISL_2441506, EPI_ISL_2441507, EPI_ISL_2441508, EPI_ISL_2441509, EPI_ISL_2441510, EPI_ISL_2441511, EPI_ISL_2441512, EPI_ISL_2441513 | | | |
| see above | CSIR-Centre for Cellular and Molecular Biology | CSIR-Centre for Cellular and Molecular Biology-INSACOG | Lamuk Zaveri,Tulasi Nagabandi,Payel Mukherjee,Onkar Kulkarni, Sofia Banu, Ara Sreenivas,Shreekant Verma, Amareshwar Vodapalli ,B Himasri,Valli Nagalakshmi Undamatla, Sumedha Avadhanula, Priya Nurkurthy, Vidhyadhari Methuku, Archana Bharadwaj Siva, Karthik Bharadwaj Tallapaka,Rakesh K Mishra,Divya Tej Sowpati |
| EPI_ISL_2441514, EPI_ISL_2441515, EPI_ISL_2441516, EPI_ISL_2441517, EPI_ISL_2441518, EPI_ISL_2441520, EPI_ISL_2441521, EPI_ISL_2441522, EPI_ISL_2441523, EPI_ISL_2441524, EPI_ISL_2441525, EPI_ISL_2441526, EPI_ISL_2441527 | | | |
| see above | CSIR-Centre for Cellular and Molecular Biology | CSIR-Centre for Cellular and Molecular Biology-INSACOG | Tulasi Nagabandi,Lamuk Zaveri,Payel Mukherjee,Onkar Kulkarni, Sofia Banu, Ara Sreenivas,Shreekant Verma, Amareshwar Vodapalli ,B Himasri,Valli Nagalakshmi Undamatla, Priya Nurkurthy,Sumedha Avadhanula, Vidhyadhari Methuku, Archana Bharadwaj Siva, Karthik Bharadwaj Tallapaka,Rakesh K Mishra,Divya Tej Sowpati |
| EPI_ISL_2441528, EPI_ISL_2441529, EPI_ISL_2441530, EPI_ISL_2441531, EPI_ISL_2441532, EPI_ISL_2441533, EPI_ISL_2441534, EPI_ISL_2441535, EPI_ISL_2441536, EPI_ISL_2441537, EPI_ISL_2441538, EPI_ISL_2441539, EPI_ISL_2441540, EPI_ISL_2441541, EPI_ISL_2441542, EPI_ISL_2441543, EPI_ISL_2441544 | | | |
| see above | CSIR-Centre for Cellular and Molecular Biology | CSIR-Centre for Cellular and Molecular Biology-INSACOG | Ara Sreenivas,Shreekant Verma, Amareshwar Vodapalli ,B Himasri,Valli Nagalakshmi Undamatla,Payel Mukherjee, Lamuk Zaveri, Onkar Kulkarni,Tulasi Nagabandi, Sofia Banu, ,Priya Nurkurthy, Vidhyadhari Methuku, Sumedha Avadhanula, Archana Bharadwaj Siva, Karthik Bharadwaj Tallapaka,Rakesh K Mishra,Divya Tej Sowpati |
| EPI_ISL_2441545, EPI_ISL_2441546, EPI_ISL_2441547, EPI_ISL_2441548, EPI_ISL_2441549, EPI_ISL_2441550, EPI_ISL_2441551, EPI_ISL_2441552, EPI_ISL_2441553, EPI_ISL_2441556, EPI_ISL_2441557, EPI_ISL_2441558, EPI_ISL_2441559, EPI_ISL_2441560, EPI_ISL_2441561, EPI_ISL_2441562, EPI_ISL_2441563, EPI_ISL_2441564 | | | |
| see above | CSIR-Centre for Cellular and Molecular Biology | CSIR-Centre for Cellular and Molecular Biology-INSACOG | Shreekant Verma, Amareshwar Vodapalli ,Ara Sreenivas,B Himasri,Valli Nagalakshmi Undamatla,Payel Mukherjee, Lamuk Zaveri, Onkar Kulkarni,Tulasi Nagabandi, Sofia Banu, Sumedha Avadhanula, Priya Nurkurthy, Vidhyadhari Methuku, Archana Bharadwaj Siva, Karthik Bharadwaj Tallapaka,Rakesh K Mishra,Divya Tej Sowpati |
| EPI_ISL_2441565, EPI_ISL_2441566, EPI_ISL_2441567, EPI_ISL_2441568, EPI_ISL_2441569, EPI_ISL_2441570, EPI_ISL_2441571, EPI_ISL_2441572, EPI_ISL_2441573, EPI_ISL_2441574, EPI_ISL_2441575, EPI_ISL_2441576, EPI_ISL_2441577, EPI_ISL_2441578, EPI_ISL_2441580, EPI_ISL_2441581, EPI_ISL_2441582 | | | |
| see above | CSIR-Centre for Cellular and Molecular Biology | CSIR-Centre for Cellular and Molecular Biology-INSACOG | Amareshwar Vodapalli,Ara Sreenivas,Shreekant Verma, B Himasri,Valli Nagalakshmi Undamatla,Payel Mukherjee, Lamuk Zaveri, Onkar Kulkarni,Tulasi Nagabandi, Sofia Banu, ,Priya Nurkurthy, Vidhyadhari Methuku, Sumedha Avadhanula, Archana Bharadwaj Siva, Karthik Bharadwaj Tallapaka,Rakesh K Mishra,Divya Tej Sowpati |
| EPI_ISL_2441583, EPI_ISL_2441584, EPI_ISL_2441586, EPI_ISL_2441587, EPI_ISL_2441588, EPI_ISL_2441589, EPI_ISL_2441590, EPI_ISL_2441591, EPI_ISL_2441592, EPI_ISL_2441595 | CSIR-Centre for Cellular and Molecular Biology | CSIR-Centre for Cellular and Molecular Biology-INSACOG | B Himasri,Amareshwar Vodapalli,Ara Sreenivas,Shreekant Verma,Valli Nagalakshmi Undamatla,Payel Mukherjee, Lamuk Zaveri, Onkar Kulkarni,Tulasi Nagabandi, Sofia Banu, Vidhyadhari Methuku,Priya Nurkurthy, Sumedha Avadhanula, Archana Bharadwaj Siva, Karthik Bharadwaj Tallapaka,Rakesh K Mishra,Divya Tej Sowpati |
| EPI_ISL_2441598, EPI_ISL_2441599, EPI_ISL_2441600, EPI_ISL_2441601, EPI_ISL_2441602, EPI_ISL_2441603, EPI_ISL_2441604, EPI_ISL_2441605, EPI_ISL_2441606, EPI_ISL_2441607, EPI_ISL_2441608, EPI_ISL_2441609, EPI_ISL_2441610, EPI_ISL_2441611 | | | |
| see above | CSIR-Centre for Cellular and Molecular Biology | CSIR-Centre for Cellular and Molecular Biology-INSACOG | Sofia Banu,Payel Mukherjee, Lamuk Zaveri, Onkar Kulkarni,Tulasi Nagabandi, Ara Sreenivas,Shreekant Verma, Amareshwar Vodapalli ,B Himasri,Valli Nagalakshmi Undamatla, Sumedha Avadhanula, Vidhyadhari Methuku, Priya Nurkurthy, Archana Bharadwaj Siva, Karthik Bharadwaj Tallapaka,Rakesh K Mishra,Divya Tej Sowpati |
| EPI_ISL_2441612, EPI_ISL_2441613, EPI_ISL_2441614, EPI_ISL_2441615, EPI_ISL_2441616, EPI_ISL_2441617, EPI_ISL_2441618 | CSIR-Centre for Cellular and Molecular Biology | CSIR-Centre for Cellular and Molecular Biology-INSACOG | Payel Mukherjee, Lamuk Zaveri, Onkar Kulkarni,Tulasi Nagabandi, Sofia Banu, Ara Sreenivas,Shreekant Verma, Amareshwar Vodapalli ,B Himasri,Valli Nagalakshmi Undamatla, Sumedha Avadhanula, Vidhyadhari Methuku, Priya Nurkurthy, Archana Bharadwaj Siva, Karthik Bharadwaj Tallapaka,Rakesh K Mishra,Divya Tej Sowpati |
| EPI_ISL_2441619, EPI_ISL_2441620, EPI_ISL_2441621, EPI_ISL_2441622, EPI_ISL_2441623, EPI_ISL_2441624, EPI_ISL_2441625, EPI_ISL_2441626, EPI_ISL_2441627, EPI_ISL_2441628, EPI_ISL_2441629 | | | |
| see above | CSIR-Centre for Cellular and Molecular Biology | CSIR-Centre for Cellular and Molecular Biology-INSACOG | Sofia Banu,Payel Mukherjee, Lamuk Zaveri, Onkar Kulkarni,Tulasi Nagabandi, Ara Sreenivas,Shreekant Verma, Amareshwar Vodapalli ,B Himasri,Valli Nagalakshmi Undamatla, Sumedha Avadhanula, Vidhyadhari Methuku, Priya Nurkurthy, Archana Bharadwaj Siva, Karthik Bharadwaj Tallapaka,Rakesh K Mishra,Divya Tej Sowpati |

| | | | |
|---|---|---|---|
| EPI_ISL_2441634, EPI_ISL_2441635, EPI_ISL_2441636, EPI_ISL_2441637, EPI_ISL_2441638, EPI_ISL_2441639, EPI_ISL_2441640, EPI_ISL_2441641, EPI_ISL_2441642, EPI_ISL_2441643, EPI_ISL_2441644, EPI_ISL_2441645, EPI_ISL_2441646, EPI_ISL_2441647, EPI_ISL_2441648 | | | |
| see above | CSIR-Centre for Cellular and Molecular Biology | CSIR-Centre for Cellular and Molecular Biology-INSACOG | Lamuk Zaveri,Tulasi Nagabandi,Payel Mukherjee,Onkar Kulkarni, Sofia Banu, Ara Sreenivas,Shreekant Verma, Amareshwar Vodapalli ,B Himasri,Valli Nagalakshmi Undamatla, Sumedha Avadhanula, Priya Nurkurthy, Vidhyadhari Methuku, Archana Bharadwaj Siva, Karthik Bharadwaj Tallapaka,Rakesh K Mishra,Divya Tej Sowpati |
| EPI_ISL_2441649, EPI_ISL_2441650, EPI_ISL_2441651, EPI_ISL_2441652, EPI_ISL_2441653, EPI_ISL_2441654, EPI_ISL_2441655, EPI_ISL_2441656, EPI_ISL_2441657, EPI_ISL_2441658, EPI_ISL_2441659, EPI_ISL_2441660, EPI_ISL_2441661, EPI_ISL_2441662, EPI_ISL_2441663, EPI_ISL_2441664, EPI_ISL_2441665, EPI_ISL_2441666 | | | |
| see above | CSIR-Centre for Cellular and Molecular Biology | CSIR-Centre for Cellular and Molecular Biology-INSACOG | Tulasi Nagabandi,Lamuk Zaveri,Payel Mukherjee,Onkar Kulkarni, Sofia Banu, Ara Sreenivas,Shreekant Verma, Amareshwar Vodapalli ,B Himasri,Valli Nagalakshmi Undamatla, Priya Nurkurthy,Sumedha Avadhanula, Vidhyadhari Methuku, Archana Bharadwaj Siva, Karthik Bharadwaj Tallapaka,Rakesh K Mishra,Divya Tej Sowpati |
| EPI_ISL_2441667, EPI_ISL_2441668, EPI_ISL_2441669, EPI_ISL_2441673, EPI_ISL_2441674, EPI_ISL_2441675, EPI_ISL_2441676, EPI_ISL_2441677, EPI_ISL_2441678, EPI_ISL_2441679, EPI_ISL_2441680, EPI_ISL_2441681, EPI_ISL_2441682, EPI_ISL_2441683, EPI_ISL_2441684 | | | |
| see above | CSIR-Centre for Cellular and Molecular Biology | CSIR-Centre for Cellular and Molecular Biology-INSACOG | Ara Sreenivas,Shreekant Verma, Amareshwar Vodapalli ,B Himasri,Valli Nagalakshmi Undamatla,Payel Mukherjee, Lamuk Zaveri, Onkar Kulkarni,Tulasi Nagabandi, Sofia Banu, ,Priya Nurkurthy, Vidhyadhari Methuku, Sumedha Avadhanula, Archana Bharadwaj Siva, Karthik Bharadwaj Tallapaka,Rakesh K Mishra,Divya Tej Sowpati |
| EPI_ISL_2441685, EPI_ISL_2441686, EPI_ISL_2441687, EPI_ISL_2441688, EPI_ISL_2441689, EPI_ISL_2441690, EPI_ISL_2441691, EPI_ISL_2441692, EPI_ISL_2441693, EPI_ISL_2441694, EPI_ISL_2441695, EPI_ISL_2441696, EPI_ISL_2441697, EPI_ISL_2441698, EPI_ISL_2441699, EPI_ISL_2441700 | | | |
| see above | CSIR-Centre for Cellular and Molecular Biology | CSIR-Centre for Cellular and Molecular Biology-INSACOG | Shreekant Verma, Amareshwar Vodapalli ,Ara Sreenivas,B Himasri,Valli Nagalakshmi Undamatla,Payel Mukherjee, Lamuk Zaveri, Onkar Kulkarni,Tulasi Nagabandi, Sofia Banu, Sumedha Avadhanula, Priya Nurkurthy, Vidhyadhari Methuku, Archana Bharadwaj Siva, Karthik Bharadwaj Tallapaka,Rakesh K Mishra,Divya Tej Sowpati |
| EPI_ISL_2441701, EPI_ISL_2441702, EPI_ISL_2441703, EPI_ISL_2441704 | CSIR-Centre for Cellular and Molecular Biology | CSIR-Centre for Cellular and Molecular Biology-INSACOG | Sumedha Avadhanula, Lamuk Zaveri, Tulasi Nagabandi, Ara Sreenivas,Shreekant Verma, Amareshwar Vodapalli ,B Himasri,Valli Nagalakshmi Undamatla, Vidhyadhari Methuku, Priya Nurkurthy, Payel Mukherjee,Onkar Kulkarni, Sofia Banu, Archana Bharadwaj Siva, Karthik Bharadwaj Tallapaka,Rakesh K Mishra,Divya Tej Sowpati |
| EPI_ISL_2441705, EPI_ISL_2441706, EPI_ISL_2441707, EPI_ISL_2441708, EPI_ISL_2441709, EPI_ISL_2441710, EPI_ISL_2441711, EPI_ISL_2441712, EPI_ISL_2441715, EPI_ISL_2441716, EPI_ISL_2441717, EPI_ISL_2441718, EPI_ISL_2441720, EPI_ISL_2441721, EPI_ISL_2441722, EPI_ISL_2441723, EPI_ISL_2441724, EPI_ISL_2441725 | | | |
| see above | CSIR-Centre for Cellular and Molecular Biology | CSIR-Centre for Cellular and Molecular Biology-INSACOG | Amareshwar Vodapalli,Ara Sreenivas,Shreekant Verma, B Himasri,Valli Nagalakshmi Undamatla,Payel Mukherjee, Lamuk Zaveri, Onkar Kulkarni,Tulasi Nagabandi, Sofia Banu, ,Priya Nurkurthy, Vidhyadhari Methuku, Sumedha Avadhanula, Archana Bharadwaj Siva, Karthik Bharadwaj Tallapaka,Rakesh K Mishra,Divya Tej Sowpati |
| EPI_ISL_2441726, EPI_ISL_2441727, EPI_ISL_2441728, EPI_ISL_2441729, EPI_ISL_2441730, EPI_ISL_2441731, EPI_ISL_2441732, EPI_ISL_2441733, EPI_ISL_2441734, EPI_ISL_2441735, EPI_ISL_2441736, EPI_ISL_2441737 | | | |
| see above | CSIR-Centre for Cellular and Molecular Biology | CSIR-Centre for Cellular and Molecular Biology-INSACOG | B Himasri,Amareshwar Vodapalli,Ara Sreenivas,Shreekant Verma,Valli Nagalakshmi Undamatla,Payel Mukherjee, Lamuk Zaveri, Onkar Kulkarni,Tulasi Nagabandi, Sofia Banu, Vidhyadhari Methuku,Priya Nurkurthy, Sumedha Avadhanula, Archana Bharadwaj Siva, Karthik Bharadwaj Tallapaka,Rakesh K Mishra,Divya Tej Sowpati |
| EPI_ISL_2441738, EPI_ISL_2441740, EPI_ISL_2441741, EPI_ISL_2441742, EPI_ISL_2441743, EPI_ISL_2441744 | AP SSO | CSIR-Centre for Cellular and Molecular Biology-INSACOG | Lamuk Zaveri,Tulasi Nagabandi,Payel Mukherjee,Onkar Kulkarni, Sofia Banu, Ara Sreenivas,Shreekant Verma, Amareshwar Vodapalli ,B Himasri,Valli Nagalakshmi Undamatla, Sumedha Avadhanula, Priya Nurkurthy, Vidhyadhari Methuku, Archana Bharadwaj Siva, Karthik Bharadwaj Tallapaka,Rakesh K Mishra,Divya Tej Sowpati |
| EPI_ISL_2441745, EPI_ISL_2441746, EPI_ISL_2441747, EPI_ISL_2441748, EPI_ISL_2441749, EPI_ISL_2441750, EPI_ISL_2441751, EPI_ISL_2441752, EPI_ISL_2441753, EPI_ISL_2441754, EPI_ISL_2441755, EPI_ISL_2441756, EPI_ISL_2441757, EPI_ISL_2441758, EPI_ISL_2441759, EPI_ISL_2441760 | | | |
| see above | AP SSO | CSIR-Centre for Cellular and Molecular Biology-INSACOG | Tulasi Nagabandi,Lamuk Zaveri,Payel Mukherjee,Onkar Kulkarni, Sofia Banu, Ara Sreenivas,Shreekant Verma, Amareshwar Vodapalli ,B Himasri,Valli Nagalakshmi Undamatla, Priya Nurkurthy,Sumedha Avadhanula, Vidhyadhari Methuku, Archana Bharadwaj Siva, Karthik Bharadwaj Tallapaka,Rakesh K Mishra,Divya Tej Sowpati |
| EPI_ISL_2441761, EPI_ISL_2441762, EPI_ISL_2441763 | AP SSO | CSIR-Centre for Cellular and Molecular Biology-INSACOG | Ara Sreenivas,Shreekant Verma, Amareshwar Vodapalli ,B Himasri,Valli Nagalakshmi Undamatla,Payel Mukherjee, Lamuk Zaveri, Onkar Kulkarni,Tulasi Nagabandi, Sofia Banu, ,Priya Nurkurthy, Vidhyadhari Methuku, Sumedha Avadhanula, Archana Bharadwaj Siva, Karthik Bharadwaj Tallapaka,Rakesh K Mishra,Divya Tej Sowpati |
| EPI_ISL_2441764, EPI_ISL_2441765, EPI_ISL_2441766 | AP SSO | CSIR-Centre for Cellular and Molecular Biology-INSACOG | Shreekant Verma, Amareshwar Vodapalli ,Ara Sreenivas,B Himasri,Valli Nagalakshmi Undamatla,Payel Mukherjee, Lamuk Zaveri, Onkar Kulkarni,Tulasi Nagabandi, Sofia Banu, Sumedha Avadhanula, Priya Nurkurthy, Vidhyadhari Methuku, Archana Bharadwaj Siva, Karthik Bharadwaj Tallapaka,Rakesh K Mishra,Divya Tej Sowpati |
| EPI_ISL_2441767, EPI_ISL_2441768, EPI_ISL_2441769 | AP SSO | CSIR-Centre for Cellular and Molecular Biology-INSACOG | Amareshwar Vodapalli,Ara Sreenivas,Shreekant Verma, B Himasri,Valli Nagalakshmi Undamatla,Payel Mukherjee, Lamuk Zaveri, Onkar Kulkarni,Tulasi Nagabandi, Sofia Banu, ,Priya Nurkurthy, Vidhyadhari Methuku, Sumedha Avadhanula, Archana Bharadwaj Siva, Karthik Bharadwaj Tallapaka,Rakesh K Mishra,Divya Tej Sowpati |
| EPI_ISL_2441770, EPI_ISL_2441771 | AP SSO | CSIR-Centre for Cellular and Molecular Biology-INSACOG | B Himasri,Amareshwar Vodapalli,Ara Sreenivas,Shreekant Verma,Valli Nagalakshmi Undamatla,Payel Mukherjee, Lamuk Zaveri, Onkar Kulkarni,Tulasi Nagabandi, Sofia Banu, Vidhyadhari Methuku,Priya Nurkurthy, Sumedha Avadhanula, Archana Bharadwaj Siva, Karthik Bharadwaj Tallapaka,Rakesh K Mishra,Divya Tej Sowpati |
| EPI_ISL_2441808 | CHU de la Réunion - Nord | Laboratoire de virologie, CNR arbovirus Associé, Chu de la Réunion | Etienne Frumence, Marie-Christine Jaffar Bandjee, Nicolas Traversier, Sabrina Petit Genet |
| EPI_ISL_2441860, EPI_ISL_2441861, EPI_ISL_2441862, EPI_ISL_2441863, EPI_ISL_2441872, EPI_ISL_2441873, EPI_ISL_2441874, EPI_ISL_2441889, EPI_ISL_2441890, EPI_ISL_2441891, EPI_ISL_2441893, EPI_ISL_2441894, EPI_ISL_2441896, EPI_ISL_2441897, EPI_ISL_2441898, EPI_ISL_2441924, EPI_ISL_2441932, EPI_ISL_2441936, EPI_ISL_2441937, EPI_ISL_2441938, EPI_ISL_2441939, EPI_ISL_2441941, EPI_ISL_2441952, EPI_ISL_2441957 | | | |
| see above | Fulgent Genetics | Centers for Disease Control and Prevention Division of Viral Diseases, Pathogen Discovery | Dakota Howard, Dhwani Batra, Peter W. Cook, Kara Moser, Adrian Paskey, Jason Caravas, Benjamin Rambo-Martin, Shatavia Morrison, Christopher Gulvick, Scott Sammons, Yvette Unoarumhi, Darlene Wagner, Matthew Schmerer, Harry Gao, Mickey Li, John Gao, Joseph Fierro, Benafsh Sapra, Becky Tsai, Yan Meng, Doreen Ng, James Xie, Clinton R. Paden, Duncan MacCannell |
| EPI_ISL_2442341, EPI_ISL_2442343, EPI_ISL_2442347 | Wyoming Public Health Laboratory | Wyoming Public Health Laboratory | Jim Mildenberger, Taylor Fearing, Channing Weber, Ashley Norberg, Chayse Rowley, Marley Goetz, Brian Dominguez, Elliot Thomasson, Brittany Oher, Sam Britz, Cari Sloma, and Rob Christensen |
| EPI_ISL_2443048 | LESP Ciudad de Mexico | Instituto de Diagnostico y Referencia Epidemiologicos (INDRE) | Claudia Wong-Arambula, Abril Rodriguez-Maldonado, Vanessa Rivero-Arredondo, Ariadna Medina-Benitez, Joaquin Quiroz-Mercado, Sergio Rangel-Guerrero, Natividad Cruz-Ortiz, Tatiana Nunez-Garcia, Gisela Barrera-Badillo, Lucia Hernandez-Rivas, Irma Lopez-Martinez, Ernesto Ramirez-Gonzalez. |
| EPI_ISL_2443049 | LESP Quintana Roo | Instituto de Diagnostico y Referencia Epidemiologicos (INDRE) | Claudia Wong-Arambula, Abril Rodriguez-Maldonado, Vanessa Rivero-Arredondo, Ariadna Medina-Benitez, Joaquin Quiroz-Mercado, Sergio Rangel-Guerrero, Natividad Cruz-Ortiz, Tatiana Nunez-Garcia, Gisela Barrera-Badillo, Lucia Hernandez-Rivas, Irma Lopez-Martinez, Ernesto Ramirez-Gonzalez. |
| EPI_ISL_2443050 | LESP Michoacan | Instituto de Diagnostico y Referencia Epidemiologicos (INDRE) | Claudia Wong-Arambula, Abril Rodriguez-Maldonado, Vanessa Rivero-Arredondo, Ariadna Medina-Benitez, Joaquin Quiroz-Mercado, Sergio Rangel-Guerrero, Natividad Cruz-Ortiz, Tatiana Nunez-Garcia, Gisela Barrera-Badillo, Lucia Hernandez-Rivas, Irma Lopez-Martinez, Ernesto Ramirez-Gonzalez. |
| EPI_ISL_2443087, EPI_ISL_2443090, EPI_ISL_2443101, EPI_ISL_2443107, EPI_ISL_2443108, EPI_ISL_2443109 | NJDOH, Public Health and Environmental Laboratories | NJ_PHEL | Lindsey Bodnar, Shiv K. Verma, Ryan Pachucki, Chelsea San Filippo, Dana Woell, Allison Roder, Byeong Jeong |
| EPI_ISL_2443132, EPI_ISL_2443134, EPI_ISL_2443136, EPI_ISL_2443196, EPI_ISL_2443197, EPI_ISL_2443198, EPI_ISL_2443201 | Kansas Health and Environmental Lab | Kansas Health and Environmental Lab | Katherine Wiggins, Mike Grose, Jonathan Barnell, Ben Olsen, and Phil Adam |
| EPI_ISL_2443229, EPI_ISL_2443230, EPI_ISL_2443232, EPI_ISL_2443234, EPI_ISL_2443235, EPI_ISL_2443236, EPI_ISL_2443237, EPI_ISL_2443238, EPI_ISL_2443239, EPI_ISL_2443240, EPI_ISL_2443242, EPI_ISL_2443243, EPI_ISL_2443244, EPI_ISL_2443245, EPI_ISL_2443246, EPI_ISL_2443247, EPI_ISL_2443248, EPI_ISL_2443249, EPI_ISL_2443250, EPI_ISL_2443251, EPI_ISL_2443252, EPI_ISL_2443253, EPI_ISL_2443254, EPI_ISL_2443255, EPI_ISL_2443256, EPI_ISL_2443258, EPI_ISL_2443259, EPI_ISL_2443260, EPI_ISL_2443261, EPI_ISL_2443262, EPI_ISL_2443263, EPI_ISL_2443264, EPI_ISL_2443265, EPI_ISL_2443266, EPI_ISL_2443267, EPI_ISL_2443268, EPI_ISL_2443269, EPI_ISL_2443270, EPI_ISL_2443271, EPI_ISL_2443273, EPI_ISL_2443274, EPI_ISL_2443275, EPI_ISL_2443276, EPI_ISL_2443277, EPI_ISL_2443278, EPI_ISL_2443279, EPI_ISL_2443280, EPI_ISL_2443281, EPI_ISL_2443282, EPI_ISL_2443283, EPI_ISL_2443284, EPI_ISL_2443285, EPI_ISL_2443286, EPI_ISL_2443287 | | | |
| see above | Indian Council of Medical Research-National Institute of Virology, Microbial Containment Complex | Indian Council of Medical Research-National Institute of Virology, Microbial Containment Complex | Pragya D Yadav |
| EPI_ISL_2443295, EPI_ISL_2443296, EPI_ISL_2443300, EPI_ISL_2443304, EPI_ISL_2443319, EPI_ISL_2443323 | TXDSHS | TXDSHS | Rashmi Tuladhar, Bonnie Oh, Jenny Zhang, Maliha Rahman, Mayela Pedrueza, Anita Pokharel, Karen Bobier, Lorraine Rodriguez, Myong Koag, Chun Wang, Rachel Lee, Grace Kubin |
| EPI_ISL_2443326, EPI_ISL_2443327 | National Institute of Public Health | State Veterinary Institute Prague | Nagy,A;Jirincova,H;Suri,T;Trnka,D;Vecerova,J |
| EPI_ISL_2443381, EPI_ISL_2443382 | Indian Council of Medical Research-National Institute of Virology, Microbial Containment Complex | Indian Council of Medical Research-National Institute of Virology, Microbial Containment Complex | Pragya D. Yadav |
| EPI_ISL_2443384 | Platform BIS UZA/UAntwerpen | Labo Klinische Biologie, UZA | Marie Le Mercier, Jasmine Coppens, Basil Britto Xavier, Christine Lammens, Veerle Matheeussen, Herman Goossens |
| EPI_ISL_2443500, EPI_ISL_2443501, EPI_ISL_2443516, EPI_ISL_2443525 | Baylor Scott & White-Temple | Baylor Scott & White-Temple | Ari Rao, Linden Morales, Kimberly Walker, Marcus Volz, Shelby Johnson |
| EPI_ISL_2443545 | Lboratorio Central de Saude Publica do Estado do Parana (LACEN/PR) | Laboratory of Respiratory Viruses and Measles, Oswaldo Cruz Institute, FIOCRUZ | Paola Resende, Luciana Appolinario, Fernando Motta, Anna Carolina Paixao, Ana Carolina Mendonca, Alice Sampaio Rocha, Taina Venas, Elisa Cavalcante Pereira, Renata Serrano Lopes, Irina Riediger, Marilda Siqueira on behalf of the Fiocruz COVID-19 Genomic Surveillance Network |
| EPI_ISL_2443798 | M Health Fairview | Minnesota Department of Health, Public Health Laboratory | Alexandra Lorentz, Jacob Garfin, Matt Plumb, and Xiong Wang |
| EPI_ISL_2443802, | Minnesota Department of Health, Public Health Laboratory | Minnesota Department of Health, Public Health Laboratory | Alexandra Lorentz, Jacob Garfin, Matt Plumb, and Xiong Wang |

| | | | |
|---|---|---|---|
| EPI_ISL_2443838 | | | |
| EPI_ISL_2443850 | M Health Fairview | Minnesota Department of Health, Public Health Laboratory | Alexandra Lorentz, Jacob Garfin, Matt Plumb, and Xiong Wang |
| EPI_ISL_2444363 | CHU LILLE | CHU Lille - Laboratoire de Virologie | AIT YAHYA Emilie, ALIDJINOU Enagnon Kazali, BOCKET Laurence, CREPIN Michel, DEMAY Christophe, ENGELMANN Ilka, GEFFROY Sandrine, GUIGON Aurélie, LAMBERT Valérie, LAZREK Mouna, NOBILLIAUX Florian, PREVOST Brigitte, THUILLIER Caroline, TINEZ Claire |
| EPI_ISL_2444444, EPI_ISL_2444455, EPI_ISL_2444483, EPI_ISL_2444549 | Salud Digna | Instituto Nacional de Medicina Genomica | Hidalgo-Miranda A, Cedro-Tanda A, Mendoza-Vargas A, Reyes-Grajeda JP, Abraham Campos-Romero, Moreno-Camacho José Luis, Rodriguez-Gallegos Jorge, Luna-Ruiz Marco, Gonzalez-Barrera D, Rangel-DeLeon D, Munguia-Garza P, Ramirez-Vega O, Escobar-Arrazola, M, Herrera-Montalvo LA. |
| EPI_ISL_2444923, EPI_ISL_2444966 | Vitomed Sp. z o.o. | 1. Academic Center for Pathomorphological and Genetic-Molecular Diagnostics ltd, Bialystok, Poland 2. National Institute of Public Health - National Institute of Hygiene, Warsaw, Poland | Radosław Charkiewicz, Jacek Nikliński, Przemysław Biecek. Joanna Reszeć, Piotr Majewski, Anetta Sulewska, Piotr Karabowicz, Joanna Kiśluk, Janusz Dzięcioł, Konrad Raczkowski, Katarzyna Zacharczuk, Magdalena Nowakowska, Małgorzata Sadkowska-Todys, Tomasz Wołkowicz |
| EPI_ISL_2445707, EPI_ISL_2445709 | CEDOC | Instituto Nacional de Saude (INSA) | Borges et al |
| EPI_ISL_2445735 | CH Porto - H Sto Antonio | Instituto Nacional de Saude (INSA) | Borges et al |
| EPI_ISL_2445928 | H Santarem | Instituto Nacional de Saude (INSA) | Borges et al |
| EPI_ISL_2445929, EPI_ISL_2445930 | Cintramedica | Instituto Nacional de Saude (INSA) | Borges et al |
| EPI_ISL_2445931 | Germano de Sousa | Instituto Nacional de Saude (INSA) | Borges et al |
| EPI_ISL_2445935, EPI_ISL_2445936 | ULS Litoral Alentejano | Instituto Nacional de Saude (INSA) | Borges et al |
| EPI_ISL_2445954, EPI_ISL_2445955, EPI_ISL_2445956, EPI_ISL_2445957, EPI_ISL_2445958 | Affidea | Instituto Nacional de Saude (INSA) | Borges et al |
| EPI_ISL_2445962, EPI_ISL_2445963, EPI_ISL_2445964, EPI_ISL_2445966 | SYNLAB | Instituto Nacional de Saude (INSA) | Borges et al |
| EPI_ISL_2446053, EPI_ISL_2446059, EPI_ISL_2446066, EPI_ISL_2446068, EPI_ISL_2446086, EPI_ISL_2446087 | Pandemic Response Lab - NYC | Pandemic Response Lab, R&D | Henry Lee, Michael Hammerling, Melissa Hopkins, Cybill del Castillo, Shinyoung Clair Kang, William Ward, Pradeep Bugga, Sol Rey, Dylan Law, Katharine Nelson, Haiping Hao, Jon Laurent |
| EPI_ISL_2446095 | Lighthouse Lab in Alderley Park | Wellcome Sanger Institute for the COVID-19 Genomics UK (COG-UK) Consortium | Jacquelyn Wynn, Mairead Hyland, The Lighthouse Lab in Alderley Park and Alex Alderton, Roberto Amato, Jeffrey Barrett, Sonia Goncalves, Ewan Harrison, David K. Jackson, Ian Johnston, Dominic Kwiatkowski, Cordelia Langford, John Sillitoe on behalf of the Wellcome Sanger Institute COVID-19 Surveillance Team |
| EPI_ISL_2446097, EPI_ISL_2446098 | Lighthouse Lab in Milton Keynes | Wellcome Sanger Institute for the COVID-19 Genomics UK (COG-UK) Consortium | The Lighthouse Lab in Milton Keynes and Alex Alderton, Roberto Amato, Jeffrey Barrett, Sonia Goncalves, Ewan Harrison, David K. Jackson, Ian Johnston, Dominic Kwiatkowski, Cordelia Langford, John Sillitoe on behalf of the Wellcome Sanger Institute COVID-19 Surveillance Team |
| EPI_ISL_2446099 | Pandemic Response Lab - NYC | Pandemic Response Lab, R&D | Henry Lee, Michael Hammerling, Melissa Hopkins, Cybill del Castillo, Shinyoung Clair Kang, William Ward, Pradeep Bugga, Sol Rey, Dylan Law, Katharine Nelson, Haiping Hao, Jon Laurent |
| EPI_ISL_2446100 | Lighthouse Lab in Milton Keynes | Wellcome Sanger Institute for the COVID-19 Genomics UK (COG-UK) Consortium | The Lighthouse Lab in Milton Keynes and Alex Alderton, Roberto Amato, Jeffrey Barrett, Sonia Goncalves, Ewan Harrison, David K. Jackson, Ian Johnston, Dominic Kwiatkowski, Cordelia Langford, John Sillitoe on behalf of the Wellcome Sanger Institute COVID-19 Surveillance Team |
| EPI_ISL_2446101, EPI_ISL_2446103, EPI_ISL_2446104 | Lighthouse Lab in Alderley Park | Wellcome Sanger Institute for the COVID-19 Genomics UK (COG-UK) Consortium | Jacquelyn Wynn, Mairead Hyland, The Lighthouse Lab in Alderley Park and Alex Alderton, Roberto Amato, Jeffrey Barrett, Sonia Goncalves, Ewan Harrison, David K. Jackson, Ian Johnston, Dominic Kwiatkowski, Cordelia Langford, John Sillitoe on behalf of the Wellcome Sanger Institute COVID-19 Surveillance Team |
| EPI_ISL_2446105 | Pandemic Response Lab - NYC | Pandemic Response Lab, R&D | Henry Lee, Michael Hammerling, Melissa Hopkins, Cybill del Castillo, Shinyoung Clair Kang, William Ward, Pradeep Bugga, Sol Rey, Dylan Law, Katharine Nelson, Haiping Hao, Jon Laurent |
| EPI_ISL_2446106 | Lighthouse Lab in Glasgow | Wellcome Sanger Institute for the COVID-19 Genomics UK (COG-UK) Consortium | Harper VanSteenhouse, Yumi Kasai, David Gray, Carol Clugston, Anna Dominiczak and Alex Alderton, Roberto Amato, Jeffrey Barrett, Sonia Goncalves, Ewan Harrison, David K. Jackson, Ian Johnston, Dominic Kwiatkowski, Cordelia Langford, John Sillitoe on behalf of the Wellcome Sanger Institute COVID-19 Surveillance Team |
| EPI_ISL_2446107 | Pandemic Response Lab - NYC | Pandemic Response Lab, R&D | Henry Lee, Michael Hammerling, Melissa Hopkins, Cybill del Castillo, Shinyoung Clair Kang, William Ward, Pradeep Bugga, Sol Rey, Dylan Law, Katharine Nelson, Haiping Hao, Jon Laurent |
| EPI_ISL_2446110, EPI_ISL_2446112 | Lighthouse Lab in Glasgow | Wellcome Sanger Institute for the COVID-19 Genomics UK (COG-UK) Consortium | Harper VanSteenhouse, Yumi Kasai, David Gray, Carol Clugston, Anna Dominiczak and Alex Alderton, Roberto Amato, Jeffrey Barrett, Sonia Goncalves, Ewan Harrison, David K. Jackson, Ian Johnston, Dominic Kwiatkowski, Cordelia Langford, John Sillitoe on behalf of the Wellcome Sanger Institute COVID-19 Surveillance Team |
| EPI_ISL_2446113 | Lighthouse Lab in Milton Keynes | Wellcome Sanger Institute for the COVID-19 Genomics UK (COG-UK) Consortium | The Lighthouse Lab in Milton Keynes and Alex Alderton, Roberto Amato, Jeffrey Barrett, Sonia Goncalves, Ewan Harrison, David K. Jackson, Ian Johnston, Dominic Kwiatkowski, Cordelia Langford, John Sillitoe on behalf of the Wellcome Sanger Institute COVID-19 Surveillance Team |
| EPI_ISL_2446115 | Lighthouse Lab in Glasgow | Wellcome Sanger Institute for the COVID-19 Genomics UK (COG-UK) Consortium | Harper VanSteenhouse, Yumi Kasai, David Gray, Carol Clugston, Anna Dominiczak and Alex Alderton, Roberto Amato, Jeffrey Barrett, Sonia Goncalves, Ewan Harrison, David K. Jackson, Ian Johnston, Dominic Kwiatkowski, Cordelia Langford, John Sillitoe on behalf of the Wellcome Sanger Institute COVID-19 Surveillance Team |
| EPI_ISL_2446118 | Lighthouse Lab in Alderley Park | Wellcome Sanger Institute for the COVID-19 Genomics UK (COG-UK) Consortium | Jacquelyn Wynn, Mairead Hyland, The Lighthouse Lab in Alderley Park and Alex Alderton, Roberto Amato, Jeffrey Barrett, Sonia Goncalves, Ewan Harrison, David K. Jackson, Ian Johnston, Dominic Kwiatkowski, Cordelia Langford, John Sillitoe on behalf of the Wellcome Sanger Institute COVID-19 Surveillance Team |
| EPI_ISL_2446119 | Lighthouse Lab in Milton Keynes | Wellcome Sanger Institute for the COVID-19 Genomics UK (COG-UK) Consortium | The Lighthouse Lab in Milton Keynes and Alex Alderton, Roberto Amato, Jeffrey Barrett, Sonia Goncalves, Ewan Harrison, David K. Jackson, Ian Johnston, Dominic Kwiatkowski, Cordelia Langford, John Sillitoe on behalf of the Wellcome Sanger Institute COVID-19 Surveillance Team |
| EPI_ISL_2446121, EPI_ISL_2446122 | Lighthouse Lab in Alderley Park | Wellcome Sanger Institute for the COVID-19 Genomics UK (COG-UK) Consortium | Jacquelyn Wynn, Mairead Hyland, The Lighthouse Lab in Alderley Park and Alex Alderton, Roberto Amato, Jeffrey Barrett, Sonia Goncalves, Ewan Harrison, David K. Jackson, Ian Johnston, Dominic Kwiatkowski, Cordelia Langford, John Sillitoe on behalf of the Wellcome Sanger Institute COVID-19 Surveillance Team |
| EPI_ISL_2446123, EPI_ISL_2446125 | Lighthouse Lab in Glasgow | Wellcome Sanger Institute for the COVID-19 Genomics UK (COG-UK) Consortium | Harper VanSteenhouse, Yumi Kasai, David Gray, Carol Clugston, Anna Dominiczak and Alex Alderton, Roberto Amato, Jeffrey Barrett, Sonia Goncalves, Ewan Harrison, David K. Jackson, Ian Johnston, Dominic Kwiatkowski, Cordelia Langford, John Sillitoe on behalf of the Wellcome Sanger Institute COVID-19 Surveillance Team |
| EPI_ISL_2446126 | Lighthouse Lab in Milton Keynes | Wellcome Sanger Institute for the COVID-19 Genomics UK (COG-UK) Consortium | The Lighthouse Lab in Milton Keynes and Alex Alderton, Roberto Amato, Jeffrey Barrett, Sonia Goncalves, Ewan Harrison, David K. Jackson, Ian Johnston, Dominic Kwiatkowski, Cordelia Langford, John Sillitoe on behalf of the Wellcome Sanger Institute COVID-19 Surveillance Team |
| EPI_ISL_2446127, EPI_ISL_2446128 | Lighthouse Lab in Alderley Park | Wellcome Sanger Institute for the COVID-19 Genomics UK (COG-UK) Consortium | Jacquelyn Wynn, Mairead Hyland, The Lighthouse Lab in Alderley Park and Alex Alderton, Roberto Amato, Jeffrey Barrett, Sonia Goncalves, Ewan Harrison, David K. Jackson, Ian Johnston, Dominic Kwiatkowski, Cordelia Langford, John Sillitoe on behalf of the Wellcome Sanger Institute COVID-19 Surveillance Team |
| EPI_ISL_2446129 | Lighthouse Lab in Milton Keynes | Wellcome Sanger Institute for the COVID-19 Genomics UK (COG-UK) Consortium | The Lighthouse Lab in Milton Keynes and Alex Alderton, Roberto Amato, Jeffrey Barrett, Sonia Goncalves, Ewan Harrison, David K. Jackson, Ian Johnston, Dominic Kwiatkowski, Cordelia Langford, John Sillitoe on behalf of the Wellcome Sanger Institute COVID-19 Surveillance Team |
| EPI_ISL_2446130 | Lighthouse Lab in Alderley Park | Wellcome Sanger Institute for the COVID-19 Genomics UK (COG-UK) Consortium | Jacquelyn Wynn, Mairead Hyland, The Lighthouse Lab in Alderley Park and Alex Alderton, Roberto Amato, Jeffrey Barrett, Sonia Goncalves, Ewan Harrison, David K. Jackson, Ian Johnston, Dominic Kwiatkowski, Cordelia Langford, John Sillitoe on behalf of the Wellcome Sanger Institute COVID-19 Surveillance Team |
| EPI_ISL_2446131 | Lighthouse Lab in Glasgow | Wellcome Sanger Institute for the COVID-19 Genomics UK (COG-UK) Consortium | Harper VanSteenhouse, Yumi Kasai, David Gray, Carol Clugston, Anna Dominiczak and Alex Alderton, Roberto Amato, Jeffrey Barrett, Sonia Goncalves, Ewan Harrison, David K. Jackson, Ian Johnston, Dominic Kwiatkowski, Cordelia Langford, John Sillitoe on behalf of the Wellcome Sanger Institute COVID-19 Surveillance Team |
| EPI_ISL_2446132, EPI_ISL_2446133 | Lighthouse Lab in Milton Keynes | Wellcome Sanger Institute for the COVID-19 Genomics UK (COG-UK) Consortium | The Lighthouse Lab in Milton Keynes and Alex Alderton, Roberto Amato, Jeffrey Barrett, Sonia Goncalves, Ewan Harrison, David K. Jackson, Ian Johnston, Dominic Kwiatkowski, Cordelia Langford, John Sillitoe on behalf of the Wellcome Sanger Institute COVID-19 Surveillance Team |
| EPI_ISL_2446134 | Lighthouse Lab in Glasgow | Wellcome Sanger Institute for the COVID-19 Genomics UK (COG-UK) Consortium | Harper VanSteenhouse, Yumi Kasai, David Gray, Carol Clugston, Anna Dominiczak and Alex Alderton, Roberto Amato, Jeffrey Barrett, Sonia Goncalves, Ewan Harrison, David K. Jackson, Ian Johnston, Dominic Kwiatkowski, Cordelia Langford, John Sillitoe on behalf of the Wellcome Sanger Institute COVID-19 Surveillance Team |
| EPI_ISL_2446135 | Lighthouse Lab in Alderley Park | Wellcome Sanger Institute for the COVID-19 Genomics UK (COG-UK) Consortium | Jacquelyn Wynn, Mairead Hyland, The Lighthouse Lab in Alderley Park and Alex Alderton, Roberto Amato, Jeffrey Barrett, Sonia Goncalves, Ewan Harrison, David K. Jackson, Ian Johnston, Dominic Kwiatkowski, Cordelia Langford, John Sillitoe on behalf of the Wellcome Sanger Institute COVID-19 Surveillance Team |
| EPI_ISL_2446136 | Lighthouse Lab in Milton Keynes | Wellcome Sanger Institute for the COVID-19 Genomics UK (COG-UK) Consortium | The Lighthouse Lab in Milton Keynes and Alex Alderton, Roberto Amato, Jeffrey Barrett, Sonia Goncalves, Ewan Harrison, David K. Jackson, Ian Johnston, Dominic Kwiatkowski, Cordelia Langford, John Sillitoe on behalf of the Wellcome Sanger Institute COVID-19 Surveillance Team |
| EPI_ISL_2446137 | Lighthouse Lab in Alderley Park | Wellcome Sanger Institute for the COVID-19 Genomics UK (COG-UK) Consortium | Jacquelyn Wynn, Mairead Hyland, The Lighthouse Lab in Alderley Park and Alex Alderton, Roberto Amato, Jeffrey Barrett, Sonia Goncalves, Ewan Harrison, David K. Jackson, Ian Johnston, Dominic Kwiatkowski, Cordelia Langford, John Sillitoe on behalf of the Wellcome Sanger Institute COVID-19 Surveillance Team |
| EPI_ISL_2446138, EPI_ISL_2446139, EPI_ISL_2446140 | Lighthouse Lab in Milton Keynes | Wellcome Sanger Institute for the COVID-19 Genomics UK (COG-UK) Consortium | The Lighthouse Lab in Milton Keynes and Alex Alderton, Roberto Amato, Jeffrey Barrett, Sonia Goncalves, Ewan Harrison, David K. Jackson, Ian Johnston, Dominic Kwiatkowski, Cordelia Langford, John Sillitoe on behalf of the Wellcome Sanger Institute COVID-19 Surveillance Team |
| EPI_ISL_2446141 | Lighthouse Lab in Glasgow | Wellcome Sanger Institute for the COVID-19 Genomics UK (COG-UK) Consortium | Harper VanSteenhouse, Yumi Kasai, David Gray, Carol Clugston, Anna Dominiczak and Alex Alderton, Roberto Amato, Jeffrey Barrett, Sonia Goncalves, Ewan Harrison, David K. Jackson, Ian Johnston, Dominic Kwiatkowski, Cordelia Langford, John Sillitoe on behalf of the Wellcome Sanger Institute COVID-19 Surveillance Team |
| EPI_ISL_2446142, EPI_ISL_2446143, EPI_ISL_2446144 | Lighthouse Lab in Alderley Park | Wellcome Sanger Institute for the COVID-19 Genomics UK (COG-UK) Consortium | Jacquelyn Wynn, Mairead Hyland, The Lighthouse Lab in Alderley Park and Alex Alderton, Roberto Amato, Jeffrey Barrett, Sonia Goncalves, Ewan Harrison, David K. Jackson, Ian Johnston, Dominic Kwiatkowski, Cordelia Langford, John Sillitoe on behalf of the Wellcome Sanger Institute COVID-19 Surveillance Team |

| | | | |
|---|---|---|---|
| EPI_ISL_2446145, EPI_ISL_2446146 | Lighthouse Lab in Milton Keynes | Wellcome Sanger Institute for the COVID-19 Genomics UK (COG-UK) Consortium | The Lighthouse Lab in Milton Keynes and Alex Alderton, Roberto Amato, Jeffrey Barrett, Sonia Goncalves, Ewan Harrison, David K. Jackson, Ian Johnston, Dominic Kwiatkowski, Cordelia Langford, John Sillitoe on behalf of the Wellcome Sanger Institute COVID-19 Surveillance Team |
| EPI_ISL_2446147 | Lighthouse Lab in Glasgow | Wellcome Sanger Institute for the COVID-19 Genomics UK (COG-UK) Consortium | Harper VanSteenhouse, Yumi Kasai, David Gray, Carol Clugston, Anna Dominiczak and Alex Alderton, Roberto Amato, Jeffrey Barrett, Sonia Goncalves, Ewan Harrison, David K. Jackson, Ian Johnston, Dominic Kwiatkowski, Cordelia Langford, John Sillitoe on behalf of the Wellcome Sanger Institute COVID-19 Surveillance Team |
| EPI_ISL_2446148 | Lighthouse Lab in Milton Keynes | Wellcome Sanger Institute for the COVID-19 Genomics UK (COG-UK) Consortium | The Lighthouse Lab in Milton Keynes and Alex Alderton, Roberto Amato, Jeffrey Barrett, Sonia Goncalves, Ewan Harrison, David K. Jackson, Ian Johnston, Dominic Kwiatkowski, Cordelia Langford, John Sillitoe on behalf of the Wellcome Sanger Institute COVID-19 Surveillance Team |
| EPI_ISL_2446149 | Lighthouse Lab in Alderley Park | Wellcome Sanger Institute for the COVID-19 Genomics UK (COG-UK) Consortium | Jacquelyn Wynn, Mairead Hyland, The Lighthouse Lab in Alderley Park and Alex Alderton, Roberto Amato, Jeffrey Barrett, Sonia Goncalves, Ewan Harrison, David K. Jackson, Ian Johnston, Dominic Kwiatkowski, Cordelia Langford, John Sillitoe on behalf of the Wellcome Sanger Institute COVID-19 Surveillance Team |
| EPI_ISL_2446150, EPI_ISL_2446151 | Lighthouse Lab in Milton Keynes | Wellcome Sanger Institute for the COVID-19 Genomics UK (COG-UK) Consortium | The Lighthouse Lab in Milton Keynes and Alex Alderton, Roberto Amato, Jeffrey Barrett, Sonia Goncalves, Ewan Harrison, David K. Jackson, Ian Johnston, Dominic Kwiatkowski, Cordelia Langford, John Sillitoe on behalf of the Wellcome Sanger Institute COVID-19 Surveillance Team |
| EPI_ISL_2446152, EPI_ISL_2446153, EPI_ISL_2446154 | Lighthouse Lab in Alderley Park | Wellcome Sanger Institute for the COVID-19 Genomics UK (COG-UK) Consortium | Jacquelyn Wynn, Mairead Hyland, The Lighthouse Lab in Alderley Park and Alex Alderton, Roberto Amato, Jeffrey Barrett, Sonia Goncalves, Ewan Harrison, David K. Jackson, Ian Johnston, Dominic Kwiatkowski, Cordelia Langford, John Sillitoe on behalf of the Wellcome Sanger Institute COVID-19 Surveillance Team |
| EPI_ISL_2446155, EPI_ISL_2446156 | Lighthouse Lab in Milton Keynes | Wellcome Sanger Institute for the COVID-19 Genomics UK (COG-UK) Consortium | The Lighthouse Lab in Milton Keynes and Alex Alderton, Roberto Amato, Jeffrey Barrett, Sonia Goncalves, Ewan Harrison, David K. Jackson, Ian Johnston, Dominic Kwiatkowski, Cordelia Langford, John Sillitoe on behalf of the Wellcome Sanger Institute COVID-19 Surveillance Team |
| EPI_ISL_2446157 | Lighthouse Lab in Glasgow | Wellcome Sanger Institute for the COVID-19 Genomics UK (COG-UK) Consortium | Harper VanSteenhouse, Yumi Kasai, David Gray, Carol Clugston, Anna Dominiczak and Alex Alderton, Roberto Amato, Jeffrey Barrett, Sonia Goncalves, Ewan Harrison, David K. Jackson, Ian Johnston, Dominic Kwiatkowski, Cordelia Langford, John Sillitoe on behalf of the Wellcome Sanger Institute COVID-19 Surveillance Team |
| EPI_ISL_2446158 | Health Services Laboratories | Wellcome Sanger Institute for the COVID-19 Genomics UK (COG-UK) Consortium | Health Services Laboratories and Alex Alderton, Roberto Amato, Jeffrey Barrett, Sonia Goncalves, Ewan Harrison, David K. Jackson, Ian Johnston, Dominic Kwiatkowski, Cordelia Langford, John Sillitoe on behalf of the Wellcome Sanger Institute COVID-19 Surveillance Team |
| EPI_ISL_2446159 | Lighthouse Lab in Alderley Park | Wellcome Sanger Institute for the COVID-19 Genomics UK (COG-UK) Consortium | Jacquelyn Wynn, Mairead Hyland, The Lighthouse Lab in Alderley Park and Alex Alderton, Roberto Amato, Jeffrey Barrett, Sonia Goncalves, Ewan Harrison, David K. Jackson, Ian Johnston, Dominic Kwiatkowski, Cordelia Langford, John Sillitoe on behalf of the Wellcome Sanger Institute COVID-19 Surveillance Team |
| EPI_ISL_2446160, EPI_ISL_2446161 | Lighthouse Lab in Milton Keynes | Wellcome Sanger Institute for the COVID-19 Genomics UK (COG-UK) Consortium | The Lighthouse Lab in Milton Keynes and Alex Alderton, Roberto Amato, Jeffrey Barrett, Sonia Goncalves, Ewan Harrison, David K. Jackson, Ian Johnston, Dominic Kwiatkowski, Cordelia Langford, John Sillitoe on behalf of the Wellcome Sanger Institute COVID-19 Surveillance Team |
| EPI_ISL_2446162 | Lighthouse Lab in Alderley Park | Wellcome Sanger Institute for the COVID-19 Genomics UK (COG-UK) Consortium | Jacquelyn Wynn, Mairead Hyland, The Lighthouse Lab in Alderley Park and Alex Alderton, Roberto Amato, Jeffrey Barrett, Sonia Goncalves, Ewan Harrison, David K. Jackson, Ian Johnston, Dominic Kwiatkowski, Cordelia Langford, John Sillitoe on behalf of the Wellcome Sanger Institute COVID-19 Surveillance Team |
| EPI_ISL_2446163, EPI_ISL_2446164, EPI_ISL_2446165, EPI_ISL_2446166 | Lighthouse Lab in Milton Keynes | Wellcome Sanger Institute for the COVID-19 Genomics UK (COG-UK) Consortium | The Lighthouse Lab in Milton Keynes and Alex Alderton, Roberto Amato, Jeffrey Barrett, Sonia Goncalves, Ewan Harrison, David K. Jackson, Ian Johnston, Dominic Kwiatkowski, Cordelia Langford, John Sillitoe on behalf of the Wellcome Sanger Institute COVID-19 Surveillance Team |
| EPI_ISL_2446167 | Lighthouse Lab in Alderley Park | Wellcome Sanger Institute for the COVID-19 Genomics UK (COG-UK) Consortium | Jacquelyn Wynn, Mairead Hyland, The Lighthouse Lab in Alderley Park and Alex Alderton, Roberto Amato, Jeffrey Barrett, Sonia Goncalves, Ewan Harrison, David K. Jackson, Ian Johnston, Dominic Kwiatkowski, Cordelia Langford, John Sillitoe on behalf of the Wellcome Sanger Institute COVID-19 Surveillance Team |
| EPI_ISL_2446168 | Lighthouse Lab in Glasgow | Wellcome Sanger Institute for the COVID-19 Genomics UK (COG-UK) Consortium | Harper VanSteenhouse, Yumi Kasai, David Gray, Carol Clugston, Anna Dominiczak and Alex Alderton, Roberto Amato, Jeffrey Barrett, Sonia Goncalves, Ewan Harrison, David K. Jackson, Ian Johnston, Dominic Kwiatkowski, Cordelia Langford, John Sillitoe on behalf of the Wellcome Sanger Institute COVID-19 Surveillance Team |
| EPI_ISL_2446169, EPI_ISL_2446170, EPI_ISL_2446171 | Lighthouse Lab in Alderley Park | Wellcome Sanger Institute for the COVID-19 Genomics UK (COG-UK) Consortium | Jacquelyn Wynn, Mairead Hyland, The Lighthouse Lab in Alderley Park and Alex Alderton, Roberto Amato, Jeffrey Barrett, Sonia Goncalves, Ewan Harrison, David K. Jackson, Ian Johnston, Dominic Kwiatkowski, Cordelia Langford, John Sillitoe on behalf of the Wellcome Sanger Institute COVID-19 Surveillance Team |
| EPI_ISL_2446172, EPI_ISL_2446173, EPI_ISL_2446174 | Lighthouse Lab in Milton Keynes | Wellcome Sanger Institute for the COVID-19 Genomics UK (COG-UK) Consortium | The Lighthouse Lab in Milton Keynes and Alex Alderton, Roberto Amato, Jeffrey Barrett, Sonia Goncalves, Ewan Harrison, David K. Jackson, Ian Johnston, Dominic Kwiatkowski, Cordelia Langford, John Sillitoe on behalf of the Wellcome Sanger Institute COVID-19 Surveillance Team |
| EPI_ISL_2446175 | Lighthouse Lab in Alderley Park | Wellcome Sanger Institute for the COVID-19 Genomics UK (COG-UK) Consortium | Jacquelyn Wynn, Mairead Hyland, The Lighthouse Lab in Alderley Park and Alex Alderton, Roberto Amato, Jeffrey Barrett, Sonia Goncalves, Ewan Harrison, David K. Jackson, Ian Johnston, Dominic Kwiatkowski, Cordelia Langford, John Sillitoe on behalf of the Wellcome Sanger Institute COVID-19 Surveillance Team |
| EPI_ISL_2446176, EPI_ISL_2446177, EPI_ISL_2446178 | Lighthouse Lab in Milton Keynes | Wellcome Sanger Institute for the COVID-19 Genomics UK (COG-UK) Consortium | The Lighthouse Lab in Milton Keynes and Alex Alderton, Roberto Amato, Jeffrey Barrett, Sonia Goncalves, Ewan Harrison, David K. Jackson, Ian Johnston, Dominic Kwiatkowski, Cordelia Langford, John Sillitoe on behalf of the Wellcome Sanger Institute COVID-19 Surveillance Team |
| EPI_ISL_2446179 | Lighthouse Lab in Glasgow | Wellcome Sanger Institute for the COVID-19 Genomics UK (COG-UK) Consortium | Harper VanSteenhouse, Yumi Kasai, David Gray, Carol Clugston, Anna Dominiczak and Alex Alderton, Roberto Amato, Jeffrey Barrett, Sonia Goncalves, Ewan Harrison, David K. Jackson, Ian Johnston, Dominic Kwiatkowski, Cordelia Langford, John Sillitoe on behalf of the Wellcome Sanger Institute COVID-19 Surveillance Team |
| EPI_ISL_2446180 | Lighthouse Lab in Milton Keynes | Wellcome Sanger Institute for the COVID-19 Genomics UK (COG-UK) Consortium | The Lighthouse Lab in Milton Keynes and Alex Alderton, Roberto Amato, Jeffrey Barrett, Sonia Goncalves, Ewan Harrison, David K. Jackson, Ian Johnston, Dominic Kwiatkowski, Cordelia Langford, John Sillitoe on behalf of the Wellcome Sanger Institute COVID-19 Surveillance Team |
| EPI_ISL_2446181 | Lighthouse Lab in Glasgow | Wellcome Sanger Institute for the COVID-19 Genomics UK (COG-UK) Consortium | Harper VanSteenhouse, Yumi Kasai, David Gray, Carol Clugston, Anna Dominiczak and Alex Alderton, Roberto Amato, Jeffrey Barrett, Sonia Goncalves, Ewan Harrison, David K. Jackson, Ian Johnston, Dominic Kwiatkowski, Cordelia Langford, John Sillitoe on behalf of the Wellcome Sanger Institute COVID-19 Surveillance Team |
| EPI_ISL_2446182, EPI_ISL_2446183 | Lighthouse Lab in Milton Keynes | Wellcome Sanger Institute for the COVID-19 Genomics UK (COG-UK) Consortium | The Lighthouse Lab in Milton Keynes and Alex Alderton, Roberto Amato, Jeffrey Barrett, Sonia Goncalves, Ewan Harrison, David K. Jackson, Ian Johnston, Dominic Kwiatkowski, Cordelia Langford, John Sillitoe on behalf of the Wellcome Sanger Institute COVID-19 Surveillance Team |
| EPI_ISL_2446184 | Health Services Laboratories | Wellcome Sanger Institute for the COVID-19 Genomics UK (COG-UK) Consortium | Health Services Laboratories and Alex Alderton, Roberto Amato, Jeffrey Barrett, Sonia Goncalves, Ewan Harrison, David K. Jackson, Ian Johnston, Dominic Kwiatkowski, Cordelia Langford, John Sillitoe on behalf of the Wellcome Sanger Institute COVID-19 Surveillance Team |
| EPI_ISL_2446185, EPI_ISL_2446186 | Lighthouse Lab in Alderley Park | Wellcome Sanger Institute for the COVID-19 Genomics UK (COG-UK) Consortium | Jacquelyn Wynn, Mairead Hyland, The Lighthouse Lab in Alderley Park and Alex Alderton, Roberto Amato, Jeffrey Barrett, Sonia Goncalves, Ewan Harrison, David K. Jackson, Ian Johnston, Dominic Kwiatkowski, Cordelia Langford, John Sillitoe on behalf of the Wellcome Sanger Institute COVID-19 Surveillance Team |
| EPI_ISL_2446187 | Health Services Laboratories | Wellcome Sanger Institute for the COVID-19 Genomics UK (COG-UK) Consortium | Health Services Laboratories and Alex Alderton, Roberto Amato, Jeffrey Barrett, Sonia Goncalves, Ewan Harrison, David K. Jackson, Ian Johnston, Dominic Kwiatkowski, Cordelia Langford, John Sillitoe on behalf of the Wellcome Sanger Institute COVID-19 Surveillance Team |
| EPI_ISL_2446188, EPI_ISL_2446189 | Lighthouse Lab in Milton Keynes | Wellcome Sanger Institute for the COVID-19 Genomics UK (COG-UK) Consortium | The Lighthouse Lab in Milton Keynes and Alex Alderton, Roberto Amato, Jeffrey Barrett, Sonia Goncalves, Ewan Harrison, David K. Jackson, Ian Johnston, Dominic Kwiatkowski, Cordelia Langford, John Sillitoe on behalf of the Wellcome Sanger Institute COVID-19 Surveillance Team |
| EPI_ISL_2446190 | Lighthouse Lab in Alderley Park | Wellcome Sanger Institute for the COVID-19 Genomics UK (COG-UK) Consortium | Jacquelyn Wynn, Mairead Hyland, The Lighthouse Lab in Alderley Park and Alex Alderton, Roberto Amato, Jeffrey Barrett, Sonia Goncalves, Ewan Harrison, David K. Jackson, Ian Johnston, Dominic Kwiatkowski, Cordelia Langford, John Sillitoe on behalf of the Wellcome Sanger Institute COVID-19 Surveillance Team |
| EPI_ISL_2446191, EPI_ISL_2446192 | Lighthouse Lab in Milton Keynes | Wellcome Sanger Institute for the COVID-19 Genomics UK (COG-UK) Consortium | The Lighthouse Lab in Milton Keynes and Alex Alderton, Roberto Amato, Jeffrey Barrett, Sonia Goncalves, Ewan Harrison, David K. Jackson, Ian Johnston, Dominic Kwiatkowski, Cordelia Langford, John Sillitoe on behalf of the Wellcome Sanger Institute COVID-19 Surveillance Team |
| EPI_ISL_2446193 | Lighthouse Lab in Alderley Park | Wellcome Sanger Institute for the COVID-19 Genomics UK (COG-UK) Consortium | Jacquelyn Wynn, Mairead Hyland, The Lighthouse Lab in Alderley Park and Alex Alderton, Roberto Amato, Jeffrey Barrett, Sonia Goncalves, Ewan Harrison, David K. Jackson, Ian Johnston, Dominic Kwiatkowski, Cordelia Langford, John Sillitoe on behalf of the Wellcome Sanger Institute COVID-19 Surveillance Team |
| EPI_ISL_2446194 | Lighthouse Lab in Glasgow | Wellcome Sanger Institute for the COVID-19 Genomics UK (COG-UK) Consortium | Harper VanSteenhouse, Yumi Kasai, David Gray, Carol Clugston, Anna Dominiczak and Alex Alderton, Roberto Amato, Jeffrey Barrett, Sonia Goncalves, Ewan Harrison, David K. Jackson, Ian Johnston, Dominic Kwiatkowski, Cordelia Langford, John Sillitoe on behalf of the Wellcome Sanger Institute COVID-19 Surveillance Team |
| EPI_ISL_2446195 | Lighthouse Lab in Alderley Park | Wellcome Sanger Institute for the COVID-19 Genomics UK (COG-UK) Consortium | Jacquelyn Wynn, Mairead Hyland, The Lighthouse Lab in Alderley Park and Alex Alderton, Roberto Amato, Jeffrey Barrett, Sonia Goncalves, Ewan Harrison, David K. Jackson, Ian Johnston, Dominic Kwiatkowski, Cordelia Langford, John Sillitoe on behalf of the Wellcome Sanger Institute COVID-19 Surveillance Team |
| EPI_ISL_2446197, EPI_ISL_2446198 | Lighthouse Lab in Milton Keynes | Wellcome Sanger Institute for the COVID-19 Genomics UK (COG-UK) Consortium | The Lighthouse Lab in Milton Keynes and Alex Alderton, Roberto Amato, Jeffrey Barrett, Sonia Goncalves, Ewan Harrison, David K. Jackson, Ian Johnston, Dominic Kwiatkowski, Cordelia Langford, John Sillitoe on behalf of the Wellcome Sanger Institute COVID-19 Surveillance Team |
| EPI_ISL_2446199 | Lighthouse Lab in Alderley Park | Wellcome Sanger Institute for the COVID-19 Genomics UK (COG-UK) Consortium | Jacquelyn Wynn, Mairead Hyland, The Lighthouse Lab in Alderley Park and Alex Alderton, Roberto Amato, Jeffrey Barrett, Sonia Goncalves, Ewan Harrison, David K. Jackson, Ian Johnston, Dominic Kwiatkowski, Cordelia Langford, John Sillitoe on behalf of the Wellcome Sanger Institute COVID-19 Surveillance Team |
| EPI_ISL_2446200 | Lighthouse Lab in Glasgow | Wellcome Sanger Institute for the COVID-19 Genomics UK (COG-UK) Consortium | Harper VanSteenhouse, Yumi Kasai, David Gray, Carol Clugston, Anna Dominiczak and Alex Alderton, Roberto Amato, Jeffrey Barrett, Sonia Goncalves, Ewan Harrison, David K. Jackson, Ian Johnston, Dominic Kwiatkowski, Cordelia Langford, John Sillitoe on behalf of the Wellcome Sanger Institute COVID-19 Surveillance Team |
| EPI_ISL_2446201 | Lighthouse Lab in Milton Keynes | Wellcome Sanger Institute for the COVID-19 Genomics UK (COG-UK) Consortium | The Lighthouse Lab in Milton Keynes and Alex Alderton, Roberto Amato, Jeffrey Barrett, Sonia Goncalves, Ewan Harrison, David K. Jackson, Ian Johnston, Dominic Kwiatkowski, Cordelia Langford, John Sillitoe on behalf of the Wellcome Sanger Institute COVID-19 Surveillance Team |
| EPI_ISL_2446202 | Lighthouse Lab in Alderley Park | Wellcome Sanger Institute for the COVID-19 Genomics UK (COG-UK) Consortium | Jacquelyn Wynn, Mairead Hyland, The Lighthouse Lab in Alderley Park and Alex Alderton, Roberto Amato, Jeffrey Barrett, Sonia Goncalves, Ewan Harrison, David K. Jackson, Ian Johnston, Dominic Kwiatkowski, Cordelia Langford, John Sillitoe on behalf of the Wellcome Sanger Institute COVID-19 Surveillance Team |
| EPI_ISL_2446203 | Lighthouse Lab in Milton Keynes | Wellcome Sanger Institute for the COVID-19 Genomics UK (COG-UK) Consortium | The Lighthouse Lab in Milton Keynes and Alex Alderton, Roberto Amato, Jeffrey Barrett, Sonia Goncalves, Ewan Harrison, David K. Jackson, Ian Johnston, Dominic Kwiatkowski, Cordelia Langford, John Sillitoe on behalf of the Wellcome Sanger Institute COVID-19 Surveillance Team |
| EPI_ISL_2446204, EPI_ISL_2446205, EPI_ISL_2446206 | Lighthouse Lab in Alderley Park | Wellcome Sanger Institute for the COVID-19 Genomics UK (COG-UK) Consortium | Jacquelyn Wynn, Mairead Hyland, The Lighthouse Lab in Alderley Park and Alex Alderton, Roberto Amato, Jeffrey Barrett, Sonia Goncalves, Ewan Harrison, David K. Jackson, Ian Johnston, Dominic Kwiatkowski, Cordelia Langford, John Sillitoe on behalf of the Wellcome Sanger Institute COVID-19 Surveillance Team |
| EPI_ISL_2446207 | Lighthouse Lab in Milton Keynes | Wellcome Sanger Institute for the COVID-19 Genomics UK (COG-UK) Consortium | The Lighthouse Lab in Milton Keynes and Alex Alderton, Roberto Amato, Jeffrey Barrett, Sonia Goncalves, Ewan Harrison, David K. Jackson, Ian Johnston, Dominic Kwiatkowski, Cordelia Langford, John Sillitoe on behalf of the Wellcome Sanger Institute COVID-19 Surveillance Team |
| EPI_ISL_2446208, EPI_ISL_2446209 | Lighthouse Lab in Alderley Park | Wellcome Sanger Institute for the COVID-19 Genomics UK (COG-UK) Consortium | Jacquelyn Wynn, Mairead Hyland, The Lighthouse Lab in Alderley Park and Alex Alderton, Roberto Amato, Jeffrey Barrett, Sonia Goncalves, Ewan Harrison, David K. Jackson, Ian Johnston, Dominic Kwiatkowski, Cordelia Langford, John Sillitoe on behalf of the Wellcome Sanger Institute COVID-19 Surveillance Team |
| EPI_ISL_2446211 | Lighthouse Lab in Glasgow | Wellcome Sanger Institute for the COVID-19 Genomics UK (COG-UK) Consortium | Harper VanSteenhouse, Yumi Kasai, David Gray, Carol Clugston, Anna Dominiczak and Alex Alderton, Roberto Amato, Jeffrey Barrett, Sonia Goncalves, Ewan Harrison, David K. Jackson, Ian Johnston, Dominic Kwiatkowski, Cordelia Langford, John Sillitoe on behalf of the Wellcome Sanger Institute COVID-19 Surveillance Team |

| | | | |
|---|---|---|---|
| EPI_ISL_2446212 | Lighthouse Lab in Alderley Park | Wellcome Sanger Institute for the COVID-19 Genomics UK (COG-UK) Consortium | Jacquelyn Wynn, Mairead Hyland, The Lighthouse Lab in Alderley Park and Alex Alderton, Roberto Amato, Jeffrey Barrett, Sonia Goncalves, Ewan Harrison, David K. Jackson, Ian Johnston, Dominic Kwiatkowski, Cordelia Langford, John Sillitoe on behalf of the Wellcome Sanger Institute COVID-19 Surveillance Team |
| EPI_ISL_2446213 | Lighthouse Lab in Milton Keynes | Wellcome Sanger Institute for the COVID-19 Genomics UK (COG-UK) Consortium | The Lighthouse Lab in Milton Keynes and Alex Alderton, Roberto Amato, Jeffrey Barrett, Sonia Goncalves, Ewan Harrison, David K. Jackson, Ian Johnston, Dominic Kwiatkowski, Cordelia Langford, John Sillitoe on behalf of the Wellcome Sanger Institute COVID-19 Surveillance Team |
| EPI_ISL_2446214 | Health Services Laboratories | Wellcome Sanger Institute for the COVID-19 Genomics UK (COG-UK) Consortium | Health Services Laboratories and Alex Alderton, Roberto Amato, Jeffrey Barrett, Sonia Goncalves, Ewan Harrison, David K. Jackson, Ian Johnston, Dominic Kwiatkowski, Cordelia Langford, John Sillitoe on behalf of the Wellcome Sanger Institute COVID-19 Surveillance Team |
| EPI_ISL_2446215, EPI_ISL_2446216 | Lighthouse Lab in Alderley Park | Wellcome Sanger Institute for the COVID-19 Genomics UK (COG-UK) Consortium | Jacquelyn Wynn, Mairead Hyland, The Lighthouse Lab in Alderley Park and Alex Alderton, Roberto Amato, Jeffrey Barrett, Sonia Goncalves, Ewan Harrison, David K. Jackson, Ian Johnston, Dominic Kwiatkowski, Cordelia Langford, John Sillitoe on behalf of the Wellcome Sanger Institute COVID-19 Surveillance Team |
| EPI_ISL_2446217 | Lighthouse Lab in Milton Keynes | Wellcome Sanger Institute for the COVID-19 Genomics UK (COG-UK) Consortium | The Lighthouse Lab in Milton Keynes and Alex Alderton, Roberto Amato, Jeffrey Barrett, Sonia Goncalves, Ewan Harrison, David K. Jackson, Ian Johnston, Dominic Kwiatkowski, Cordelia Langford, John Sillitoe on behalf of the Wellcome Sanger Institute COVID-19 Surveillance Team |
| EPI_ISL_2446218 | Lighthouse Lab in Alderley Park | Wellcome Sanger Institute for the COVID-19 Genomics UK (COG-UK) Consortium | Jacquelyn Wynn, Mairead Hyland, The Lighthouse Lab in Alderley Park and Alex Alderton, Roberto Amato, Jeffrey Barrett, Sonia Goncalves, Ewan Harrison, David K. Jackson, Ian Johnston, Dominic Kwiatkowski, Cordelia Langford, John Sillitoe on behalf of the Wellcome Sanger Institute COVID-19 Surveillance Team |
| EPI_ISL_2446219 | Health Services Laboratories | Wellcome Sanger Institute for the COVID-19 Genomics UK (COG-UK) Consortium | Health Services Laboratories and Alex Alderton, Roberto Amato, Jeffrey Barrett, Sonia Goncalves, Ewan Harrison, David K. Jackson, Ian Johnston, Dominic Kwiatkowski, Cordelia Langford, John Sillitoe on behalf of the Wellcome Sanger Institute COVID-19 Surveillance Team |
| EPI_ISL_2446220 | Lighthouse Lab in Milton Keynes | Wellcome Sanger Institute for the COVID-19 Genomics UK (COG-UK) Consortium | The Lighthouse Lab in Milton Keynes and Alex Alderton, Roberto Amato, Jeffrey Barrett, Sonia Goncalves, Ewan Harrison, David K. Jackson, Ian Johnston, Dominic Kwiatkowski, Cordelia Langford, John Sillitoe on behalf of the Wellcome Sanger Institute COVID-19 Surveillance Team |
| EPI_ISL_2446221, EPI_ISL_2446222 | Lighthouse Lab in Alderley Park | Wellcome Sanger Institute for the COVID-19 Genomics UK (COG-UK) Consortium | Jacquelyn Wynn, Mairead Hyland, The Lighthouse Lab in Alderley Park and Alex Alderton, Roberto Amato, Jeffrey Barrett, Sonia Goncalves, Ewan Harrison, David K. Jackson, Ian Johnston, Dominic Kwiatkowski, Cordelia Langford, John Sillitoe on behalf of the Wellcome Sanger Institute COVID-19 Surveillance Team |
| EPI_ISL_2446224, EPI_ISL_2446225, EPI_ISL_2446226 | Lighthouse Lab in Milton Keynes | Wellcome Sanger Institute for the COVID-19 Genomics UK (COG-UK) Consortium | The Lighthouse Lab in Milton Keynes and Alex Alderton, Roberto Amato, Jeffrey Barrett, Sonia Goncalves, Ewan Harrison, David K. Jackson, Ian Johnston, Dominic Kwiatkowski, Cordelia Langford, John Sillitoe on behalf of the Wellcome Sanger Institute COVID-19 Surveillance Team |
| EPI_ISL_2446228, EPI_ISL_2446229 | Lighthouse Lab in Glasgow | Wellcome Sanger Institute for the COVID-19 Genomics UK (COG-UK) Consortium | Harper VanSteenhouse, Yumi Kasai, David Gray, Carol Clugston, Anna Dominiczak and Alex Alderton, Roberto Amato, Jeffrey Barrett, Sonia Goncalves, Ewan Harrison, David K. Jackson, Ian Johnston, Dominic Kwiatkowski, Cordelia Langford, John Sillitoe on behalf of the Wellcome Sanger Institute COVID-19 Surveillance Team |
| EPI_ISL_2446230 | Health Services Laboratories | Wellcome Sanger Institute for the COVID-19 Genomics UK (COG-UK) Consortium | Health Services Laboratories and Alex Alderton, Roberto Amato, Jeffrey Barrett, Sonia Goncalves, Ewan Harrison, David K. Jackson, Ian Johnston, Dominic Kwiatkowski, Cordelia Langford, John Sillitoe on behalf of the Wellcome Sanger Institute COVID-19 Surveillance Team |
| EPI_ISL_2446231 | Lighthouse Lab in Alderley Park | Wellcome Sanger Institute for the COVID-19 Genomics UK (COG-UK) Consortium | Jacquelyn Wynn, Mairead Hyland, The Lighthouse Lab in Alderley Park and Alex Alderton, Roberto Amato, Jeffrey Barrett, Sonia Goncalves, Ewan Harrison, David K. Jackson, Ian Johnston, Dominic Kwiatkowski, Cordelia Langford, John Sillitoe on behalf of the Wellcome Sanger Institute COVID-19 Surveillance Team |
| EPI_ISL_2446233, EPI_ISL_2446234, EPI_ISL_2446235 | Lighthouse Lab in Milton Keynes | Wellcome Sanger Institute for the COVID-19 Genomics UK (COG-UK) Consortium | The Lighthouse Lab in Milton Keynes and Alex Alderton, Roberto Amato, Jeffrey Barrett, Sonia Goncalves, Ewan Harrison, David K. Jackson, Ian Johnston, Dominic Kwiatkowski, Cordelia Langford, John Sillitoe on behalf of the Wellcome Sanger Institute COVID-19 Surveillance Team |
| EPI_ISL_2446236 | Lighthouse Lab in Glasgow | Wellcome Sanger Institute for the COVID-19 Genomics UK (COG-UK) Consortium | Harper VanSteenhouse, Yumi Kasai, David Gray, Carol Clugston, Anna Dominiczak and Alex Alderton, Roberto Amato, Jeffrey Barrett, Sonia Goncalves, Ewan Harrison, David K. Jackson, Ian Johnston, Dominic Kwiatkowski, Cordelia Langford, John Sillitoe on behalf of the Wellcome Sanger Institute COVID-19 Surveillance Team |
| EPI_ISL_2446237, EPI_ISL_2446239 | Lighthouse Lab in Milton Keynes | Wellcome Sanger Institute for the COVID-19 Genomics UK (COG-UK) Consortium | The Lighthouse Lab in Milton Keynes and Alex Alderton, Roberto Amato, Jeffrey Barrett, Sonia Goncalves, Ewan Harrison, David K. Jackson, Ian Johnston, Dominic Kwiatkowski, Cordelia Langford, John Sillitoe on behalf of the Wellcome Sanger Institute COVID-19 Surveillance Team |
| EPI_ISL_2446240 | Lighthouse Lab in Glasgow | Wellcome Sanger Institute for the COVID-19 Genomics UK (COG-UK) Consortium | Harper VanSteenhouse, Yumi Kasai, David Gray, Carol Clugston, Anna Dominiczak and Alex Alderton, Roberto Amato, Jeffrey Barrett, Sonia Goncalves, Ewan Harrison, David K. Jackson, Ian Johnston, Dominic Kwiatkowski, Cordelia Langford, John Sillitoe on behalf of the Wellcome Sanger Institute COVID-19 Surveillance Team |
| EPI_ISL_2446241, EPI_ISL_2446242 | Lighthouse Lab in Alderley Park | Wellcome Sanger Institute for the COVID-19 Genomics UK (COG-UK) Consortium | Jacquelyn Wynn, Mairead Hyland, The Lighthouse Lab in Alderley Park and Alex Alderton, Roberto Amato, Jeffrey Barrett, Sonia Goncalves, Ewan Harrison, David K. Jackson, Ian Johnston, Dominic Kwiatkowski, Cordelia Langford, John Sillitoe on behalf of the Wellcome Sanger Institute COVID-19 Surveillance Team |
| EPI_ISL_2446243 | Lighthouse Lab in Milton Keynes | Wellcome Sanger Institute for the COVID-19 Genomics UK (COG-UK) Consortium | The Lighthouse Lab in Milton Keynes and Alex Alderton, Roberto Amato, Jeffrey Barrett, Sonia Goncalves, Ewan Harrison, David K. Jackson, Ian Johnston, Dominic Kwiatkowski, Cordelia Langford, John Sillitoe on behalf of the Wellcome Sanger Institute COVID-19 Surveillance Team |
| EPI_ISL_2446244, EPI_ISL_2446247 | Lighthouse Lab in Alderley Park | Wellcome Sanger Institute for the COVID-19 Genomics UK (COG-UK) Consortium | Jacquelyn Wynn, Mairead Hyland, The Lighthouse Lab in Alderley Park and Alex Alderton, Roberto Amato, Jeffrey Barrett, Sonia Goncalves, Ewan Harrison, David K. Jackson, Ian Johnston, Dominic Kwiatkowski, Cordelia Langford, John Sillitoe on behalf of the Wellcome Sanger Institute COVID-19 Surveillance Team |
| EPI_ISL_2446248 | Lighthouse Lab in Glasgow | Wellcome Sanger Institute for the COVID-19 Genomics UK (COG-UK) Consortium | Harper VanSteenhouse, Yumi Kasai, David Gray, Carol Clugston, Anna Dominiczak and Alex Alderton, Roberto Amato, Jeffrey Barrett, Sonia Goncalves, Ewan Harrison, David K. Jackson, Ian Johnston, Dominic Kwiatkowski, Cordelia Langford, John Sillitoe on behalf of the Wellcome Sanger Institute COVID-19 Surveillance Team |
| EPI_ISL_2446249 | Lighthouse Lab in Milton Keynes | Wellcome Sanger Institute for the COVID-19 Genomics UK (COG-UK) Consortium | The Lighthouse Lab in Milton Keynes and Alex Alderton, Roberto Amato, Jeffrey Barrett, Sonia Goncalves, Ewan Harrison, David K. Jackson, Ian Johnston, Dominic Kwiatkowski, Cordelia Langford, John Sillitoe on behalf of the Wellcome Sanger Institute COVID-19 Surveillance Team |
| EPI_ISL_2446250 | Health Services Laboratories | Wellcome Sanger Institute for the COVID-19 Genomics UK (COG-UK) Consortium | Health Services Laboratories and Alex Alderton, Roberto Amato, Jeffrey Barrett, Sonia Goncalves, Ewan Harrison, David K. Jackson, Ian Johnston, Dominic Kwiatkowski, Cordelia Langford, John Sillitoe on behalf of the Wellcome Sanger Institute COVID-19 Surveillance Team |
| EPI_ISL_2446251 | Lighthouse Lab in Alderley Park | Wellcome Sanger Institute for the COVID-19 Genomics UK (COG-UK) Consortium | Jacquelyn Wynn, Mairead Hyland, The Lighthouse Lab in Alderley Park and Alex Alderton, Roberto Amato, Jeffrey Barrett, Sonia Goncalves, Ewan Harrison, David K. Jackson, Ian Johnston, Dominic Kwiatkowski, Cordelia Langford, John Sillitoe on behalf of the Wellcome Sanger Institute COVID-19 Surveillance Team |
| EPI_ISL_2446252 | Lighthouse Lab in Milton Keynes | Wellcome Sanger Institute for the COVID-19 Genomics UK (COG-UK) Consortium | The Lighthouse Lab in Milton Keynes and Alex Alderton, Roberto Amato, Jeffrey Barrett, Sonia Goncalves, Ewan Harrison, David K. Jackson, Ian Johnston, Dominic Kwiatkowski, Cordelia Langford, John Sillitoe on behalf of the Wellcome Sanger Institute COVID-19 Surveillance Team |
| EPI_ISL_2446253 | Lighthouse Lab in Alderley Park | Wellcome Sanger Institute for the COVID-19 Genomics UK (COG-UK) Consortium | Jacquelyn Wynn, Mairead Hyland, The Lighthouse Lab in Alderley Park and Alex Alderton, Roberto Amato, Jeffrey Barrett, Sonia Goncalves, Ewan Harrison, David K. Jackson, Ian Johnston, Dominic Kwiatkowski, Cordelia Langford, John Sillitoe on behalf of the Wellcome Sanger Institute COVID-19 Surveillance Team |
| EPI_ISL_2446254 | Lighthouse Lab in Glasgow | Wellcome Sanger Institute for the COVID-19 Genomics UK (COG-UK) Consortium | Harper VanSteenhouse, Yumi Kasai, David Gray, Carol Clugston, Anna Dominiczak and Alex Alderton, Roberto Amato, Jeffrey Barrett, Sonia Goncalves, Ewan Harrison, David K. Jackson, Ian Johnston, Dominic Kwiatkowski, Cordelia Langford, John Sillitoe on behalf of the Wellcome Sanger Institute COVID-19 Surveillance Team |
| EPI_ISL_2446255 | Lighthouse Lab in Milton Keynes | Wellcome Sanger Institute for the COVID-19 Genomics UK (COG-UK) Consortium | The Lighthouse Lab in Milton Keynes and Alex Alderton, Roberto Amato, Jeffrey Barrett, Sonia Goncalves, Ewan Harrison, David K. Jackson, Ian Johnston, Dominic Kwiatkowski, Cordelia Langford, John Sillitoe on behalf of the Wellcome Sanger Institute COVID-19 Surveillance Team |
| EPI_ISL_2446256 | Lighthouse Lab in Alderley Park | Wellcome Sanger Institute for the COVID-19 Genomics UK (COG-UK) Consortium | Jacquelyn Wynn, Mairead Hyland, The Lighthouse Lab in Alderley Park and Alex Alderton, Roberto Amato, Jeffrey Barrett, Sonia Goncalves, Ewan Harrison, David K. Jackson, Ian Johnston, Dominic Kwiatkowski, Cordelia Langford, John Sillitoe on behalf of the Wellcome Sanger Institute COVID-19 Surveillance Team |
| EPI_ISL_2446257 | Lighthouse Lab in Glasgow | Wellcome Sanger Institute for the COVID-19 Genomics UK (COG-UK) Consortium | Harper VanSteenhouse, Yumi Kasai, David Gray, Carol Clugston, Anna Dominiczak and Alex Alderton, Roberto Amato, Jeffrey Barrett, Sonia Goncalves, Ewan Harrison, David K. Jackson, Ian Johnston, Dominic Kwiatkowski, Cordelia Langford, John Sillitoe on behalf of the Wellcome Sanger Institute COVID-19 Surveillance Team |
| EPI_ISL_2446259, EPI_ISL_2446260 | Lighthouse Lab in Alderley Park | Wellcome Sanger Institute for the COVID-19 Genomics UK (COG-UK) Consortium | Jacquelyn Wynn, Mairead Hyland, The Lighthouse Lab in Alderley Park and Alex Alderton, Roberto Amato, Jeffrey Barrett, Sonia Goncalves, Ewan Harrison, David K. Jackson, Ian Johnston, Dominic Kwiatkowski, Cordelia Langford, John Sillitoe on behalf of the Wellcome Sanger Institute COVID-19 Surveillance Team |
| EPI_ISL_2446261 | Lighthouse Lab in Milton Keynes | Wellcome Sanger Institute for the COVID-19 Genomics UK (COG-UK) Consortium | The Lighthouse Lab in Milton Keynes and Alex Alderton, Roberto Amato, Jeffrey Barrett, Sonia Goncalves, Ewan Harrison, David K. Jackson, Ian Johnston, Dominic Kwiatkowski, Cordelia Langford, John Sillitoe on behalf of the Wellcome Sanger Institute COVID-19 Surveillance Team |
| EPI_ISL_2446262, EPI_ISL_2446263 | Lighthouse Lab in Alderley Park | Wellcome Sanger Institute for the COVID-19 Genomics UK (COG-UK) Consortium | Jacquelyn Wynn, Mairead Hyland, The Lighthouse Lab in Alderley Park and Alex Alderton, Roberto Amato, Jeffrey Barrett, Sonia Goncalves, Ewan Harrison, David K. Jackson, Ian Johnston, Dominic Kwiatkowski, Cordelia Langford, John Sillitoe on behalf of the Wellcome Sanger Institute COVID-19 Surveillance Team |
| EPI_ISL_2446264 | Lighthouse Lab in Milton Keynes | Wellcome Sanger Institute for the COVID-19 Genomics UK (COG-UK) Consortium | The Lighthouse Lab in Milton Keynes and Alex Alderton, Roberto Amato, Jeffrey Barrett, Sonia Goncalves, Ewan Harrison, David K. Jackson, Ian Johnston, Dominic Kwiatkowski, Cordelia Langford, John Sillitoe on behalf of the Wellcome Sanger Institute COVID-19 Surveillance Team |
| EPI_ISL_2446265, EPI_ISL_2446266, EPI_ISL_2446268 | Lighthouse Lab in Alderley Park | Wellcome Sanger Institute for the COVID-19 Genomics UK (COG-UK) Consortium | Jacquelyn Wynn, Mairead Hyland, The Lighthouse Lab in Alderley Park and Alex Alderton, Roberto Amato, Jeffrey Barrett, Sonia Goncalves, Ewan Harrison, David K. Jackson, Ian Johnston, Dominic Kwiatkowski, Cordelia Langford, John Sillitoe on behalf of the Wellcome Sanger Institute COVID-19 Surveillance Team |
| EPI_ISL_2446269, EPI_ISL_2446270, EPI_ISL_2446272, EPI_ISL_2446273 | Lighthouse Lab in Milton Keynes | Wellcome Sanger Institute for the COVID-19 Genomics UK (COG-UK) Consortium | The Lighthouse Lab in Milton Keynes and Alex Alderton, Roberto Amato, Jeffrey Barrett, Sonia Goncalves, Ewan Harrison, David K. Jackson, Ian Johnston, Dominic Kwiatkowski, Cordelia Langford, John Sillitoe on behalf of the Wellcome Sanger Institute COVID-19 Surveillance Team |
| EPI_ISL_2446274, EPI_ISL_2446275 | Lighthouse Lab in Alderley Park | Wellcome Sanger Institute for the COVID-19 Genomics UK (COG-UK) Consortium | Jacquelyn Wynn, Mairead Hyland, The Lighthouse Lab in Alderley Park and Alex Alderton, Roberto Amato, Jeffrey Barrett, Sonia Goncalves, Ewan Harrison, David K. Jackson, Ian Johnston, Dominic Kwiatkowski, Cordelia Langford, John Sillitoe on behalf of the Wellcome Sanger Institute COVID-19 Surveillance Team |
| EPI_ISL_2446276 | Lighthouse Lab in Milton Keynes | Wellcome Sanger Institute for the COVID-19 Genomics UK (COG-UK) Consortium | The Lighthouse Lab in Milton Keynes and Alex Alderton, Roberto Amato, Jeffrey Barrett, Sonia Goncalves, Ewan Harrison, David K. Jackson, Ian Johnston, Dominic Kwiatkowski, Cordelia Langford, John Sillitoe on behalf of the Wellcome Sanger Institute COVID-19 Surveillance Team |
| EPI_ISL_2446278 | Lighthouse Lab in Glasgow | Wellcome Sanger Institute for the COVID-19 Genomics UK (COG-UK) Consortium | Harper VanSteenhouse, Yumi Kasai, David Gray, Carol Clugston, Anna Dominiczak and Alex Alderton, Roberto Amato, Jeffrey Barrett, Sonia Goncalves, Ewan Harrison, David K. Jackson, Ian Johnston, Dominic Kwiatkowski, Cordelia Langford, John Sillitoe on behalf of the Wellcome Sanger Institute COVID-19 Surveillance Team |
| EPI_ISL_2446279, EPI_ISL_2446280 | Lighthouse Lab in Alderley Park | Wellcome Sanger Institute for the COVID-19 Genomics UK (COG-UK) Consortium | Jacquelyn Wynn, Mairead Hyland, The Lighthouse Lab in Alderley Park and Alex Alderton, Roberto Amato, Jeffrey Barrett, Sonia Goncalves, Ewan Harrison, David K. Jackson, Ian Johnston, Dominic Kwiatkowski, Cordelia Langford, John Sillitoe on behalf of the Wellcome Sanger Institute COVID-19 Surveillance Team |
| EPI_ISL_2446281 | Lighthouse Lab in Milton Keynes | Wellcome Sanger Institute for the COVID-19 Genomics UK (COG-UK) Consortium | The Lighthouse Lab in Milton Keynes and Alex Alderton, Roberto Amato, Jeffrey Barrett, Sonia Goncalves, Ewan Harrison, David K. Jackson, Ian Johnston, Dominic Kwiatkowski, Cordelia Langford, John Sillitoe on behalf of the Wellcome Sanger Institute COVID-19 Surveillance Team |
| EPI_ISL_2446282 | Lighthouse Lab in Alderley Park | Wellcome Sanger Institute for the COVID-19 Genomics UK (COG-UK) Consortium | Jacquelyn Wynn, Mairead Hyland, The Lighthouse Lab in Alderley Park and Alex Alderton, Roberto Amato, Jeffrey Barrett, Sonia Goncalves, Ewan Harrison, David K. Jackson, Ian Johnston, Dominic Kwiatkowski, Cordelia Langford, John Sillitoe on behalf of the Wellcome Sanger Institute COVID-19 Surveillance Team |
| EPI_ISL_2446283 | Lighthouse Lab in Milton Keynes | Wellcome Sanger Institute for the COVID-19 Genomics UK (COG-UK) Consortium | The Lighthouse Lab in Milton Keynes and Alex Alderton, Roberto Amato, Jeffrey Barrett, Sonia Goncalves, Ewan Harrison, David K. Jackson, Ian Johnston, Dominic Kwiatkowski, Cordelia Langford, John Sillitoe on behalf of the Wellcome Sanger Institute COVID-19 Surveillance Team |

| | | | |
|---|---|---|---|
| EPI_ISL_2446284 | Lighthouse Lab in Alderley Park | Wellcome Sanger Institute for the COVID-19 Genomics UK (COG-UK) Consortium | Jacquelyn Wynn, Mairead Hyland, The Lighthouse Lab in Alderley Park and Alex Alderton, Roberto Amato, Jeffrey Barrett, Sonia Goncalves, Ewan Harrison, David K. Jackson, Ian Johnston, Dominic Kwiatkowski, Cordelia Langford, John Sillitoe on behalf of the Wellcome Sanger Institute COVID-19 Surveillance Team |
| EPI_ISL_2446286 | Lighthouse Lab in Milton Keynes | Wellcome Sanger Institute for the COVID-19 Genomics UK (COG-UK) Consortium | The Lighthouse Lab in Milton Keynes and Alex Alderton, Roberto Amato, Jeffrey Barrett, Sonia Goncalves, Ewan Harrison, David K. Jackson, Ian Johnston, Dominic Kwiatkowski, Cordelia Langford, John Sillitoe on behalf of the Wellcome Sanger Institute COVID-19 Surveillance Team |
| EPI_ISL_2446287 | Health Services Laboratories | Wellcome Sanger Institute for the COVID-19 Genomics UK (COG-UK) Consortium | Health Services Laboratories and Alex Alderton, Roberto Amato, Jeffrey Barrett, Sonia Goncalves, Ewan Harrison, David K. Jackson, Ian Johnston, Dominic Kwiatkowski, Cordelia Langford, John Sillitoe on behalf of the Wellcome Sanger Institute COVID-19 Surveillance Team |
| EPI_ISL_2446288 | Lighthouse Lab in Alderley Park | Wellcome Sanger Institute for the COVID-19 Genomics UK (COG-UK) Consortium | Jacquelyn Wynn, Mairead Hyland, The Lighthouse Lab in Alderley Park and Alex Alderton, Roberto Amato, Jeffrey Barrett, Sonia Goncalves, Ewan Harrison, David K. Jackson, Ian Johnston, Dominic Kwiatkowski, Cordelia Langford, John Sillitoe on behalf of the Wellcome Sanger Institute COVID-19 Surveillance Team |
| EPI_ISL_2446289 | Lighthouse Lab in Glasgow | Wellcome Sanger Institute for the COVID-19 Genomics UK (COG-UK) Consortium | Harper VanSteenhouse, Yumi Kasai, David Gray, Carol Clugston, Anna Dominiczak and Alex Alderton, Roberto Amato, Jeffrey Barrett, Sonia Goncalves, Ewan Harrison, David K. Jackson, Ian Johnston, Dominic Kwiatkowski, Cordelia Langford, John Sillitoe on behalf of the Wellcome Sanger Institute COVID-19 Surveillance Team |
| EPI_ISL_2446290, EPI_ISL_2446291 | Lighthouse Lab in Alderley Park | Wellcome Sanger Institute for the COVID-19 Genomics UK (COG-UK) Consortium | Jacquelyn Wynn, Mairead Hyland, The Lighthouse Lab in Alderley Park and Alex Alderton, Roberto Amato, Jeffrey Barrett, Sonia Goncalves, Ewan Harrison, David K. Jackson, Ian Johnston, Dominic Kwiatkowski, Cordelia Langford, John Sillitoe on behalf of the Wellcome Sanger Institute COVID-19 Surveillance Team |
| EPI_ISL_2446293 | Lighthouse Lab in Milton Keynes | Wellcome Sanger Institute for the COVID-19 Genomics UK (COG-UK) Consortium | The Lighthouse Lab in Milton Keynes and Alex Alderton, Roberto Amato, Jeffrey Barrett, Sonia Goncalves, Ewan Harrison, David K. Jackson, Ian Johnston, Dominic Kwiatkowski, Cordelia Langford, John Sillitoe on behalf of the Wellcome Sanger Institute COVID-19 Surveillance Team |
| EPI_ISL_2446294 | Lighthouse Lab in Alderley Park | Wellcome Sanger Institute for the COVID-19 Genomics UK (COG-UK) Consortium | Jacquelyn Wynn, Mairead Hyland, The Lighthouse Lab in Alderley Park and Alex Alderton, Roberto Amato, Jeffrey Barrett, Sonia Goncalves, Ewan Harrison, David K. Jackson, Ian Johnston, Dominic Kwiatkowski, Cordelia Langford, John Sillitoe on behalf of the Wellcome Sanger Institute COVID-19 Surveillance Team |
| EPI_ISL_2446295, EPI_ISL_2446296, EPI_ISL_2446297 | Lighthouse Lab in Milton Keynes | Wellcome Sanger Institute for the COVID-19 Genomics UK (COG-UK) Consortium | The Lighthouse Lab in Milton Keynes and Alex Alderton, Roberto Amato, Jeffrey Barrett, Sonia Goncalves, Ewan Harrison, David K. Jackson, Ian Johnston, Dominic Kwiatkowski, Cordelia Langford, John Sillitoe on behalf of the Wellcome Sanger Institute COVID-19 Surveillance Team |
| EPI_ISL_2446298 | Lighthouse Lab in Glasgow | Wellcome Sanger Institute for the COVID-19 Genomics UK (COG-UK) Consortium | Harper VanSteenhouse, Yumi Kasai, David Gray, Carol Clugston, Anna Dominiczak and Alex Alderton, Roberto Amato, Jeffrey Barrett, Sonia Goncalves, Ewan Harrison, David K. Jackson, Ian Johnston, Dominic Kwiatkowski, Cordelia Langford, John Sillitoe on behalf of the Wellcome Sanger Institute COVID-19 Surveillance Team |
| EPI_ISL_2446300 | Lighthouse Lab in Milton Keynes | Wellcome Sanger Institute for the COVID-19 Genomics UK (COG-UK) Consortium | The Lighthouse Lab in Milton Keynes and Alex Alderton, Roberto Amato, Jeffrey Barrett, Sonia Goncalves, Ewan Harrison, David K. Jackson, Ian Johnston, Dominic Kwiatkowski, Cordelia Langford, John Sillitoe on behalf of the Wellcome Sanger Institute COVID-19 Surveillance Team |
| EPI_ISL_2446301 | Lighthouse Lab in Alderley Park | Wellcome Sanger Institute for the COVID-19 Genomics UK (COG-UK) Consortium | Jacquelyn Wynn, Mairead Hyland, The Lighthouse Lab in Alderley Park and Alex Alderton, Roberto Amato, Jeffrey Barrett, Sonia Goncalves, Ewan Harrison, David K. Jackson, Ian Johnston, Dominic Kwiatkowski, Cordelia Langford, John Sillitoe on behalf of the Wellcome Sanger Institute COVID-19 Surveillance Team |
| EPI_ISL_2446302 | Lighthouse Lab in Milton Keynes | Wellcome Sanger Institute for the COVID-19 Genomics UK (COG-UK) Consortium | The Lighthouse Lab in Milton Keynes and Alex Alderton, Roberto Amato, Jeffrey Barrett, Sonia Goncalves, Ewan Harrison, David K. Jackson, Ian Johnston, Dominic Kwiatkowski, Cordelia Langford, John Sillitoe on behalf of the Wellcome Sanger Institute COVID-19 Surveillance Team |
| EPI_ISL_2446303, EPI_ISL_2446304, EPI_ISL_2446305, EPI_ISL_2446306 | Lighthouse Lab in Alderley Park | Wellcome Sanger Institute for the COVID-19 Genomics UK (COG-UK) Consortium | Jacquelyn Wynn, Mairead Hyland, The Lighthouse Lab in Alderley Park and Alex Alderton, Roberto Amato, Jeffrey Barrett, Sonia Goncalves, Ewan Harrison, David K. Jackson, Ian Johnston, Dominic Kwiatkowski, Cordelia Langford, John Sillitoe on behalf of the Wellcome Sanger Institute COVID-19 Surveillance Team |
| EPI_ISL_2446307 | Lighthouse Lab in Glasgow | Wellcome Sanger Institute for the COVID-19 Genomics UK (COG-UK) Consortium | Harper VanSteenhouse, Yumi Kasai, David Gray, Carol Clugston, Anna Dominiczak and Alex Alderton, Roberto Amato, Jeffrey Barrett, Sonia Goncalves, Ewan Harrison, David K. Jackson, Ian Johnston, Dominic Kwiatkowski, Cordelia Langford, John Sillitoe on behalf of the Wellcome Sanger Institute COVID-19 Surveillance Team |
| EPI_ISL_2446308, EPI_ISL_2446309, EPI_ISL_2446310 | Lighthouse Lab in Alderley Park | Wellcome Sanger Institute for the COVID-19 Genomics UK (COG-UK) Consortium | Jacquelyn Wynn, Mairead Hyland, The Lighthouse Lab in Alderley Park and Alex Alderton, Roberto Amato, Jeffrey Barrett, Sonia Goncalves, Ewan Harrison, David K. Jackson, Ian Johnston, Dominic Kwiatkowski, Cordelia Langford, John Sillitoe on behalf of the Wellcome Sanger Institute COVID-19 Surveillance Team |
| EPI_ISL_2446311, EPI_ISL_2446312 | Lighthouse Lab in Milton Keynes | Wellcome Sanger Institute for the COVID-19 Genomics UK (COG-UK) Consortium | The Lighthouse Lab in Milton Keynes and Alex Alderton, Roberto Amato, Jeffrey Barrett, Sonia Goncalves, Ewan Harrison, David K. Jackson, Ian Johnston, Dominic Kwiatkowski, Cordelia Langford, John Sillitoe on behalf of the Wellcome Sanger Institute COVID-19 Surveillance Team |
| EPI_ISL_2446313 | Lighthouse Lab in Alderley Park | Wellcome Sanger Institute for the COVID-19 Genomics UK (COG-UK) Consortium | Jacquelyn Wynn, Mairead Hyland, The Lighthouse Lab in Alderley Park and Alex Alderton, Roberto Amato, Jeffrey Barrett, Sonia Goncalves, Ewan Harrison, David K. Jackson, Ian Johnston, Dominic Kwiatkowski, Cordelia Langford, John Sillitoe on behalf of the Wellcome Sanger Institute COVID-19 Surveillance Team |
| EPI_ISL_2446314 | Lighthouse Lab in Glasgow | Wellcome Sanger Institute for the COVID-19 Genomics UK (COG-UK) Consortium | Harper VanSteenhouse, Yumi Kasai, David Gray, Carol Clugston, Anna Dominiczak and Alex Alderton, Roberto Amato, Jeffrey Barrett, Sonia Goncalves, Ewan Harrison, David K. Jackson, Ian Johnston, Dominic Kwiatkowski, Cordelia Langford, John Sillitoe on behalf of the Wellcome Sanger Institute COVID-19 Surveillance Team |
| EPI_ISL_2446316 | Lighthouse Lab in Milton Keynes | Wellcome Sanger Institute for the COVID-19 Genomics UK (COG-UK) Consortium | The Lighthouse Lab in Milton Keynes and Alex Alderton, Roberto Amato, Jeffrey Barrett, Sonia Goncalves, Ewan Harrison, David K. Jackson, Ian Johnston, Dominic Kwiatkowski, Cordelia Langford, John Sillitoe on behalf of the Wellcome Sanger Institute COVID-19 Surveillance Team |
| EPI_ISL_2446317, EPI_ISL_2446318 | Lighthouse Lab in Alderley Park | Wellcome Sanger Institute for the COVID-19 Genomics UK (COG-UK) Consortium | Jacquelyn Wynn, Mairead Hyland, The Lighthouse Lab in Alderley Park and Alex Alderton, Roberto Amato, Jeffrey Barrett, Sonia Goncalves, Ewan Harrison, David K. Jackson, Ian Johnston, Dominic Kwiatkowski, Cordelia Langford, John Sillitoe on behalf of the Wellcome Sanger Institute COVID-19 Surveillance Team |
| EPI_ISL_2446319, EPI_ISL_2446320 | Lighthouse Lab in Milton Keynes | Wellcome Sanger Institute for the COVID-19 Genomics UK (COG-UK) Consortium | The Lighthouse Lab in Milton Keynes and Alex Alderton, Roberto Amato, Jeffrey Barrett, Sonia Goncalves, Ewan Harrison, David K. Jackson, Ian Johnston, Dominic Kwiatkowski, Cordelia Langford, John Sillitoe on behalf of the Wellcome Sanger Institute COVID-19 Surveillance Team |
| EPI_ISL_2446321 | Lighthouse Lab in Alderley Park | Wellcome Sanger Institute for the COVID-19 Genomics UK (COG-UK) Consortium | Jacquelyn Wynn, Mairead Hyland, The Lighthouse Lab in Alderley Park and Alex Alderton, Roberto Amato, Jeffrey Barrett, Sonia Goncalves, Ewan Harrison, David K. Jackson, Ian Johnston, Dominic Kwiatkowski, Cordelia Langford, John Sillitoe on behalf of the Wellcome Sanger Institute COVID-19 Surveillance Team |
| EPI_ISL_2446322, EPI_ISL_2446323, EPI_ISL_2446324, EPI_ISL_2446325 | Lighthouse Lab in Milton Keynes | Wellcome Sanger Institute for the COVID-19 Genomics UK (COG-UK) Consortium | The Lighthouse Lab in Milton Keynes and Alex Alderton, Roberto Amato, Jeffrey Barrett, Sonia Goncalves, Ewan Harrison, David K. Jackson, Ian Johnston, Dominic Kwiatkowski, Cordelia Langford, John Sillitoe on behalf of the Wellcome Sanger Institute COVID-19 Surveillance Team |
| EPI_ISL_2446326, EPI_ISL_2446327 | Lighthouse Lab in Alderley Park | Wellcome Sanger Institute for the COVID-19 Genomics UK (COG-UK) Consortium | Jacquelyn Wynn, Mairead Hyland, The Lighthouse Lab in Alderley Park and Alex Alderton, Roberto Amato, Jeffrey Barrett, Sonia Goncalves, Ewan Harrison, David K. Jackson, Ian Johnston, Dominic Kwiatkowski, Cordelia Langford, John Sillitoe on behalf of the Wellcome Sanger Institute COVID-19 Surveillance Team |
| EPI_ISL_2446328 | Lighthouse Lab in Milton Keynes | Wellcome Sanger Institute for the COVID-19 Genomics UK (COG-UK) Consortium | The Lighthouse Lab in Milton Keynes and Alex Alderton, Roberto Amato, Jeffrey Barrett, Sonia Goncalves, Ewan Harrison, David K. Jackson, Ian Johnston, Dominic Kwiatkowski, Cordelia Langford, John Sillitoe on behalf of the Wellcome Sanger Institute COVID-19 Surveillance Team |
| EPI_ISL_2446330 | Lighthouse Lab in Alderley Park | Wellcome Sanger Institute for the COVID-19 Genomics UK (COG-UK) Consortium | Jacquelyn Wynn, Mairead Hyland, The Lighthouse Lab in Alderley Park and Alex Alderton, Roberto Amato, Jeffrey Barrett, Sonia Goncalves, Ewan Harrison, David K. Jackson, Ian Johnston, Dominic Kwiatkowski, Cordelia Langford, John Sillitoe on behalf of the Wellcome Sanger Institute COVID-19 Surveillance Team |
| EPI_ISL_2446331, EPI_ISL_2446332 | Lighthouse Lab in Milton Keynes | Wellcome Sanger Institute for the COVID-19 Genomics UK (COG-UK) Consortium | The Lighthouse Lab in Milton Keynes and Alex Alderton, Roberto Amato, Jeffrey Barrett, Sonia Goncalves, Ewan Harrison, David K. Jackson, Ian Johnston, Dominic Kwiatkowski, Cordelia Langford, John Sillitoe on behalf of the Wellcome Sanger Institute COVID-19 Surveillance Team |
| EPI_ISL_2446333 | Lighthouse Lab in Glasgow | Wellcome Sanger Institute for the COVID-19 Genomics UK (COG-UK) Consortium | Harper VanSteenhouse, Yumi Kasai, David Gray, Carol Clugston, Anna Dominiczak and Alex Alderton, Roberto Amato, Jeffrey Barrett, Sonia Goncalves, Ewan Harrison, David K. Jackson, Ian Johnston, Dominic Kwiatkowski, Cordelia Langford, John Sillitoe on behalf of the Wellcome Sanger Institute COVID-19 Surveillance Team |
| EPI_ISL_2446334, EPI_ISL_2446335 | Lighthouse Lab in Alderley Park | Wellcome Sanger Institute for the COVID-19 Genomics UK (COG-UK) Consortium | Jacquelyn Wynn, Mairead Hyland, The Lighthouse Lab in Alderley Park and Alex Alderton, Roberto Amato, Jeffrey Barrett, Sonia Goncalves, Ewan Harrison, David K. Jackson, Ian Johnston, Dominic Kwiatkowski, Cordelia Langford, John Sillitoe on behalf of the Wellcome Sanger Institute COVID-19 Surveillance Team |
| EPI_ISL_2446336, EPI_ISL_2446337, EPI_ISL_2446338, EPI_ISL_2446339 | Lighthouse Lab in Milton Keynes | Wellcome Sanger Institute for the COVID-19 Genomics UK (COG-UK) Consortium | The Lighthouse Lab in Milton Keynes and Alex Alderton, Roberto Amato, Jeffrey Barrett, Sonia Goncalves, Ewan Harrison, David K. Jackson, Ian Johnston, Dominic Kwiatkowski, Cordelia Langford, John Sillitoe on behalf of the Wellcome Sanger Institute COVID-19 Surveillance Team |
| EPI_ISL_2446340 | Lighthouse Lab in Alderley Park | Wellcome Sanger Institute for the COVID-19 Genomics UK (COG-UK) Consortium | Jacquelyn Wynn, Mairead Hyland, The Lighthouse Lab in Alderley Park and Alex Alderton, Roberto Amato, Jeffrey Barrett, Sonia Goncalves, Ewan Harrison, David K. Jackson, Ian Johnston, Dominic Kwiatkowski, Cordelia Langford, John Sillitoe on behalf of the Wellcome Sanger Institute COVID-19 Surveillance Team |
| EPI_ISL_2446341 | Lighthouse Lab in Glasgow | Wellcome Sanger Institute for the COVID-19 Genomics UK (COG-UK) Consortium | Harper VanSteenhouse, Yumi Kasai, David Gray, Carol Clugston, Anna Dominiczak and Alex Alderton, Roberto Amato, Jeffrey Barrett, Sonia Goncalves, Ewan Harrison, David K. Jackson, Ian Johnston, Dominic Kwiatkowski, Cordelia Langford, John Sillitoe on behalf of the Wellcome Sanger Institute COVID-19 Surveillance Team |
| EPI_ISL_2446342, EPI_ISL_2446343 | Lighthouse Lab in Milton Keynes | Wellcome Sanger Institute for the COVID-19 Genomics UK (COG-UK) Consortium | The Lighthouse Lab in Milton Keynes and Alex Alderton, Roberto Amato, Jeffrey Barrett, Sonia Goncalves, Ewan Harrison, David K. Jackson, Ian Johnston, Dominic Kwiatkowski, Cordelia Langford, John Sillitoe on behalf of the Wellcome Sanger Institute COVID-19 Surveillance Team |
| EPI_ISL_2446344 | Lighthouse Lab in Alderley Park | Wellcome Sanger Institute for the COVID-19 Genomics UK (COG-UK) Consortium | Jacquelyn Wynn, Mairead Hyland, The Lighthouse Lab in Alderley Park and Alex Alderton, Roberto Amato, Jeffrey Barrett, Sonia Goncalves, Ewan Harrison, David K. Jackson, Ian Johnston, Dominic Kwiatkowski, Cordelia Langford, John Sillitoe on behalf of the Wellcome Sanger Institute COVID-19 Surveillance Team |
| EPI_ISL_2446345 | Lighthouse Lab in Milton Keynes | Wellcome Sanger Institute for the COVID-19 Genomics UK (COG-UK) Consortium | The Lighthouse Lab in Milton Keynes and Alex Alderton, Roberto Amato, Jeffrey Barrett, Sonia Goncalves, Ewan Harrison, David K. Jackson, Ian Johnston, Dominic Kwiatkowski, Cordelia Langford, John Sillitoe on behalf of the Wellcome Sanger Institute COVID-19 Surveillance Team |
| EPI_ISL_2446346 | Lighthouse Lab in Alderley Park | Wellcome Sanger Institute for the COVID-19 Genomics UK (COG-UK) Consortium | Jacquelyn Wynn, Mairead Hyland, The Lighthouse Lab in Alderley Park and Alex Alderton, Roberto Amato, Jeffrey Barrett, Sonia Goncalves, Ewan Harrison, David K. Jackson, Ian Johnston, Dominic Kwiatkowski, Cordelia Langford, John Sillitoe on behalf of the Wellcome Sanger Institute COVID-19 Surveillance Team |
| EPI_ISL_2446347 | Lighthouse Lab in Milton Keynes | Wellcome Sanger Institute for the COVID-19 Genomics UK (COG-UK) Consortium | The Lighthouse Lab in Milton Keynes and Alex Alderton, Roberto Amato, Jeffrey Barrett, Sonia Goncalves, Ewan Harrison, David K. Jackson, Ian Johnston, Dominic Kwiatkowski, Cordelia Langford, John Sillitoe on behalf of the Wellcome Sanger Institute COVID-19 Surveillance Team |
| EPI_ISL_2446349 | Lighthouse Lab in Alderley Park | Wellcome Sanger Institute for the COVID-19 Genomics UK (COG-UK) Consortium | Jacquelyn Wynn, Mairead Hyland, The Lighthouse Lab in Alderley Park and Alex Alderton, Roberto Amato, Jeffrey Barrett, Sonia Goncalves, Ewan Harrison, David K. Jackson, Ian Johnston, Dominic Kwiatkowski, Cordelia Langford, John Sillitoe on behalf of the Wellcome Sanger Institute COVID-19 Surveillance Team |
| EPI_ISL_2446350 | Lighthouse Lab in Milton Keynes | Wellcome Sanger Institute for the COVID-19 Genomics UK (COG-UK) Consortium | The Lighthouse Lab in Milton Keynes and Alex Alderton, Roberto Amato, Jeffrey Barrett, Sonia Goncalves, Ewan Harrison, David K. Jackson, Ian Johnston, Dominic Kwiatkowski, Cordelia Langford, John Sillitoe on behalf of the Wellcome Sanger Institute COVID-19 Surveillance Team |
| EPI_ISL_2446352 | Lighthouse Lab in Alderley Park | Wellcome Sanger Institute for the COVID-19 Genomics UK (COG-UK) Consortium | Jacquelyn Wynn, Mairead Hyland, The Lighthouse Lab in Alderley Park and Alex Alderton, Roberto Amato, Jeffrey Barrett, Sonia Goncalves, Ewan Harrison, David K. Jackson, Ian Johnston, Dominic Kwiatkowski, Cordelia Langford, John Sillitoe on behalf of the Wellcome Sanger Institute COVID-19 Surveillance Team |
| EPI_ISL_2446353, | Lighthouse Lab in Milton Keynes | Wellcome Sanger Institute for the COVID-19 Genomics UK (COG-UK) | The Lighthouse Lab in Milton Keynes and Alex Alderton, Roberto Amato, Jeffrey Barrett, Sonia Goncalves, Ewan Harrison, David K. Jackson, Ian Johnston, Dominic Kwiatkowski, Cordelia Langford, John Sillitoe on behalf of the Wellcome Sanger |

| | | | |
|---|---|---|---|
| EPI_ISL_2446354 | | Consortium | Institute COVID-19 Surveillance Team |
| EPI_ISL_2446355 | Lighthouse Lab in Alderley Park | Wellcome Sanger Institute for the COVID-19 Genomics UK (COG-UK) Consortium | Jacquelyn Wynn, Mairead Hyland, The Lighthouse Lab in Alderley Park and Alex Alderton, Roberto Amato, Jeffrey Barrett, Sonia Goncalves, Ewan Harrison, David K. Jackson, Ian Johnston, Dominic Kwiatkowski, Cordelia Langford, John Sillitoe on behalf of the Wellcome Sanger Institute COVID-19 Surveillance Team |
| EPI_ISL_2446356 | Lighthouse Lab in Milton Keynes | Wellcome Sanger Institute for the COVID-19 Genomics UK (COG-UK) Consortium | The Lighthouse Lab in Milton Keynes and Alex Alderton, Roberto Amato, Jeffrey Barrett, Sonia Goncalves, Ewan Harrison, David K. Jackson, Ian Johnston, Dominic Kwiatkowski, Cordelia Langford, John Sillitoe on behalf of the Wellcome Sanger Institute COVID-19 Surveillance Team |
| EPI_ISL_2446357 | Lighthouse Lab in Alderley Park | Wellcome Sanger Institute for the COVID-19 Genomics UK (COG-UK) Consortium | Jacquelyn Wynn, Mairead Hyland, The Lighthouse Lab in Alderley Park and Alex Alderton, Roberto Amato, Jeffrey Barrett, Sonia Goncalves, Ewan Harrison, David K. Jackson, Ian Johnston, Dominic Kwiatkowski, Cordelia Langford, John Sillitoe on behalf of the Wellcome Sanger Institute COVID-19 Surveillance Team |
| EPI_ISL_2446358 | Lighthouse Lab in Milton Keynes | Wellcome Sanger Institute for the COVID-19 Genomics UK (COG-UK) Consortium | The Lighthouse Lab in Milton Keynes and Alex Alderton, Roberto Amato, Jeffrey Barrett, Sonia Goncalves, Ewan Harrison, David K. Jackson, Ian Johnston, Dominic Kwiatkowski, Cordelia Langford, John Sillitoe on behalf of the Wellcome Sanger Institute COVID-19 Surveillance Team |
| EPI_ISL_2446359, EPI_ISL_2446360 | Lighthouse Lab in Alderley Park | Wellcome Sanger Institute for the COVID-19 Genomics UK (COG-UK) Consortium | Jacquelyn Wynn, Mairead Hyland, The Lighthouse Lab in Alderley Park and Alex Alderton, Roberto Amato, Jeffrey Barrett, Sonia Goncalves, Ewan Harrison, David K. Jackson, Ian Johnston, Dominic Kwiatkowski, Cordelia Langford, John Sillitoe on behalf of the Wellcome Sanger Institute COVID-19 Surveillance Team |
| EPI_ISL_2446361 | Lighthouse Lab in Glasgow | Wellcome Sanger Institute for the COVID-19 Genomics UK (COG-UK) Consortium | Harper VanSteenhouse, Yumi Kasai, David Gray, Carol Clugston, Anna Dominiczak and Alex Alderton, Roberto Amato, Jeffrey Barrett, Sonia Goncalves, Ewan Harrison, David K. Jackson, Ian Johnston, Dominic Kwiatkowski, Cordelia Langford, John Sillitoe on behalf of the Wellcome Sanger Institute COVID-19 Surveillance Team |
| EPI_ISL_2446362 | Lighthouse Lab in Alderley Park | Wellcome Sanger Institute for the COVID-19 Genomics UK (COG-UK) Consortium | Jacquelyn Wynn, Mairead Hyland, The Lighthouse Lab in Alderley Park and Alex Alderton, Roberto Amato, Jeffrey Barrett, Sonia Goncalves, Ewan Harrison, David K. Jackson, Ian Johnston, Dominic Kwiatkowski, Cordelia Langford, John Sillitoe on behalf of the Wellcome Sanger Institute COVID-19 Surveillance Team |
| EPI_ISL_2446363 | Lighthouse Lab in Glasgow | Wellcome Sanger Institute for the COVID-19 Genomics UK (COG-UK) Consortium | Harper VanSteenhouse, Yumi Kasai, David Gray, Carol Clugston, Anna Dominiczak and Alex Alderton, Roberto Amato, Jeffrey Barrett, Sonia Goncalves, Ewan Harrison, David K. Jackson, Ian Johnston, Dominic Kwiatkowski, Cordelia Langford, John Sillitoe on behalf of the Wellcome Sanger Institute COVID-19 Surveillance Team |
| EPI_ISL_2446364 | Lighthouse Lab in Milton Keynes | Wellcome Sanger Institute for the COVID-19 Genomics UK (COG-UK) Consortium | The Lighthouse Lab in Milton Keynes and Alex Alderton, Roberto Amato, Jeffrey Barrett, Sonia Goncalves, Ewan Harrison, David K. Jackson, Ian Johnston, Dominic Kwiatkowski, Cordelia Langford, John Sillitoe on behalf of the Wellcome Sanger Institute COVID-19 Surveillance Team |
| EPI_ISL_2446365 | Lighthouse Lab in Glasgow | Wellcome Sanger Institute for the COVID-19 Genomics UK (COG-UK) Consortium | Harper VanSteenhouse, Yumi Kasai, David Gray, Carol Clugston, Anna Dominiczak and Alex Alderton, Roberto Amato, Jeffrey Barrett, Sonia Goncalves, Ewan Harrison, David K. Jackson, Ian Johnston, Dominic Kwiatkowski, Cordelia Langford, John Sillitoe on behalf of the Wellcome Sanger Institute COVID-19 Surveillance Team |
| EPI_ISL_2446366 | Lighthouse Lab in Milton Keynes | Wellcome Sanger Institute for the COVID-19 Genomics UK (COG-UK) Consortium | The Lighthouse Lab in Milton Keynes and Alex Alderton, Roberto Amato, Jeffrey Barrett, Sonia Goncalves, Ewan Harrison, David K. Jackson, Ian Johnston, Dominic Kwiatkowski, Cordelia Langford, John Sillitoe on behalf of the Wellcome Sanger Institute COVID-19 Surveillance Team |
| EPI_ISL_2446367, EPI_ISL_2446368 | Lighthouse Lab in Glasgow | Wellcome Sanger Institute for the COVID-19 Genomics UK (COG-UK) Consortium | Harper VanSteenhouse, Yumi Kasai, David Gray, Carol Clugston, Anna Dominiczak and Alex Alderton, Roberto Amato, Jeffrey Barrett, Sonia Goncalves, Ewan Harrison, David K. Jackson, Ian Johnston, Dominic Kwiatkowski, Cordelia Langford, John Sillitoe on behalf of the Wellcome Sanger Institute COVID-19 Surveillance Team |
| EPI_ISL_2446369 | Lighthouse Lab in Milton Keynes | Wellcome Sanger Institute for the COVID-19 Genomics UK (COG-UK) Consortium | The Lighthouse Lab in Milton Keynes and Alex Alderton, Roberto Amato, Jeffrey Barrett, Sonia Goncalves, Ewan Harrison, David K. Jackson, Ian Johnston, Dominic Kwiatkowski, Cordelia Langford, John Sillitoe on behalf of the Wellcome Sanger Institute COVID-19 Surveillance Team |
| EPI_ISL_2446370 | Lighthouse Lab in Alderley Park | Wellcome Sanger Institute for the COVID-19 Genomics UK (COG-UK) Consortium | Jacquelyn Wynn, Mairead Hyland, The Lighthouse Lab in Alderley Park and Alex Alderton, Roberto Amato, Jeffrey Barrett, Sonia Goncalves, Ewan Harrison, David K. Jackson, Ian Johnston, Dominic Kwiatkowski, Cordelia Langford, John Sillitoe on behalf of the Wellcome Sanger Institute COVID-19 Surveillance Team |
| EPI_ISL_2446371 | Lighthouse Lab in Milton Keynes | Wellcome Sanger Institute for the COVID-19 Genomics UK (COG-UK) Consortium | The Lighthouse Lab in Milton Keynes and Alex Alderton, Roberto Amato, Jeffrey Barrett, Sonia Goncalves, Ewan Harrison, David K. Jackson, Ian Johnston, Dominic Kwiatkowski, Cordelia Langford, John Sillitoe on behalf of the Wellcome Sanger Institute COVID-19 Surveillance Team |
| EPI_ISL_2446372, EPI_ISL_2446373, EPI_ISL_2446374 | Lighthouse Lab in Alderley Park | Wellcome Sanger Institute for the COVID-19 Genomics UK (COG-UK) Consortium | Jacquelyn Wynn, Mairead Hyland, The Lighthouse Lab in Alderley Park and Alex Alderton, Roberto Amato, Jeffrey Barrett, Sonia Goncalves, Ewan Harrison, David K. Jackson, Ian Johnston, Dominic Kwiatkowski, Cordelia Langford, John Sillitoe on behalf of the Wellcome Sanger Institute COVID-19 Surveillance Team |
| EPI_ISL_2446375 | Lighthouse Lab in Milton Keynes | Wellcome Sanger Institute for the COVID-19 Genomics UK (COG-UK) Consortium | The Lighthouse Lab in Milton Keynes and Alex Alderton, Roberto Amato, Jeffrey Barrett, Sonia Goncalves, Ewan Harrison, David K. Jackson, Ian Johnston, Dominic Kwiatkowski, Cordelia Langford, John Sillitoe on behalf of the Wellcome Sanger Institute COVID-19 Surveillance Team |
| EPI_ISL_2446376 | Lighthouse Lab in Glasgow | Wellcome Sanger Institute for the COVID-19 Genomics UK (COG-UK) Consortium | Harper VanSteenhouse, Yumi Kasai, David Gray, Carol Clugston, Anna Dominiczak and Alex Alderton, Roberto Amato, Jeffrey Barrett, Sonia Goncalves, Ewan Harrison, David K. Jackson, Ian Johnston, Dominic Kwiatkowski, Cordelia Langford, John Sillitoe on behalf of the Wellcome Sanger Institute COVID-19 Surveillance Team |
| EPI_ISL_2446378, EPI_ISL_2446379 | Lighthouse Lab in Alderley Park | Wellcome Sanger Institute for the COVID-19 Genomics UK (COG-UK) Consortium | Jacquelyn Wynn, Mairead Hyland, The Lighthouse Lab in Alderley Park and Alex Alderton, Roberto Amato, Jeffrey Barrett, Sonia Goncalves, Ewan Harrison, David K. Jackson, Ian Johnston, Dominic Kwiatkowski, Cordelia Langford, John Sillitoe on behalf of the Wellcome Sanger Institute COVID-19 Surveillance Team |
| EPI_ISL_2446380 | Lighthouse Lab in Glasgow | Wellcome Sanger Institute for the COVID-19 Genomics UK (COG-UK) Consortium | Harper VanSteenhouse, Yumi Kasai, David Gray, Carol Clugston, Anna Dominiczak and Alex Alderton, Roberto Amato, Jeffrey Barrett, Sonia Goncalves, Ewan Harrison, David K. Jackson, Ian Johnston, Dominic Kwiatkowski, Cordelia Langford, John Sillitoe on behalf of the Wellcome Sanger Institute COVID-19 Surveillance Team |
| EPI_ISL_2446381, EPI_ISL_2446382, EPI_ISL_2446383 | Lighthouse Lab in Alderley Park | Wellcome Sanger Institute for the COVID-19 Genomics UK (COG-UK) Consortium | Jacquelyn Wynn, Mairead Hyland, The Lighthouse Lab in Alderley Park and Alex Alderton, Roberto Amato, Jeffrey Barrett, Sonia Goncalves, Ewan Harrison, David K. Jackson, Ian Johnston, Dominic Kwiatkowski, Cordelia Langford, John Sillitoe on behalf of the Wellcome Sanger Institute COVID-19 Surveillance Team |
| EPI_ISL_2446384 | Lighthouse Lab in Milton Keynes | Wellcome Sanger Institute for the COVID-19 Genomics UK (COG-UK) Consortium | The Lighthouse Lab in Milton Keynes and Alex Alderton, Roberto Amato, Jeffrey Barrett, Sonia Goncalves, Ewan Harrison, David K. Jackson, Ian Johnston, Dominic Kwiatkowski, Cordelia Langford, John Sillitoe on behalf of the Wellcome Sanger Institute COVID-19 Surveillance Team |
| EPI_ISL_2446385 | Lighthouse Lab in Glasgow | Wellcome Sanger Institute for the COVID-19 Genomics UK (COG-UK) Consortium | Harper VanSteenhouse, Yumi Kasai, David Gray, Carol Clugston, Anna Dominiczak and Alex Alderton, Roberto Amato, Jeffrey Barrett, Sonia Goncalves, Ewan Harrison, David K. Jackson, Ian Johnston, Dominic Kwiatkowski, Cordelia Langford, John Sillitoe on behalf of the Wellcome Sanger Institute COVID-19 Surveillance Team |
| EPI_ISL_2446386, EPI_ISL_2446387, EPI_ISL_2446388 | Lighthouse Lab in Alderley Park | Wellcome Sanger Institute for the COVID-19 Genomics UK (COG-UK) Consortium | Jacquelyn Wynn, Mairead Hyland, The Lighthouse Lab in Alderley Park and Alex Alderton, Roberto Amato, Jeffrey Barrett, Sonia Goncalves, Ewan Harrison, David K. Jackson, Ian Johnston, Dominic Kwiatkowski, Cordelia Langford, John Sillitoe on behalf of the Wellcome Sanger Institute COVID-19 Surveillance Team |
| EPI_ISL_2446389, EPI_ISL_2446390 | Lighthouse Lab in Glasgow | Wellcome Sanger Institute for the COVID-19 Genomics UK (COG-UK) Consortium | Harper VanSteenhouse, Yumi Kasai, David Gray, Carol Clugston, Anna Dominiczak and Alex Alderton, Roberto Amato, Jeffrey Barrett, Sonia Goncalves, Ewan Harrison, David K. Jackson, Ian Johnston, Dominic Kwiatkowski, Cordelia Langford, John Sillitoe on behalf of the Wellcome Sanger Institute COVID-19 Surveillance Team |
| EPI_ISL_2446392, EPI_ISL_2446393 | Lighthouse Lab in Alderley Park | Wellcome Sanger Institute for the COVID-19 Genomics UK (COG-UK) Consortium | Jacquelyn Wynn, Mairead Hyland, The Lighthouse Lab in Alderley Park and Alex Alderton, Roberto Amato, Jeffrey Barrett, Sonia Goncalves, Ewan Harrison, David K. Jackson, Ian Johnston, Dominic Kwiatkowski, Cordelia Langford, John Sillitoe on behalf of the Wellcome Sanger Institute COVID-19 Surveillance Team |
| EPI_ISL_2446394, EPI_ISL_2446395 | Lighthouse Lab in Glasgow | Wellcome Sanger Institute for the COVID-19 Genomics UK (COG-UK) Consortium | Harper VanSteenhouse, Yumi Kasai, David Gray, Carol Clugston, Anna Dominiczak and Alex Alderton, Roberto Amato, Jeffrey Barrett, Sonia Goncalves, Ewan Harrison, David K. Jackson, Ian Johnston, Dominic Kwiatkowski, Cordelia Langford, John Sillitoe on behalf of the Wellcome Sanger Institute COVID-19 Surveillance Team |
| EPI_ISL_2446396, EPI_ISL_2446397, EPI_ISL_2446398, EPI_ISL_2446399 | Lighthouse Lab in Alderley Park | Wellcome Sanger Institute for the COVID-19 Genomics UK (COG-UK) Consortium | Jacquelyn Wynn, Mairead Hyland, The Lighthouse Lab in Alderley Park and Alex Alderton, Roberto Amato, Jeffrey Barrett, Sonia Goncalves, Ewan Harrison, David K. Jackson, Ian Johnston, Dominic Kwiatkowski, Cordelia Langford, John Sillitoe on behalf of the Wellcome Sanger Institute COVID-19 Surveillance Team |
| EPI_ISL_2446401 | Lighthouse Lab in Milton Keynes | Wellcome Sanger Institute for the COVID-19 Genomics UK (COG-UK) Consortium | The Lighthouse Lab in Milton Keynes and Alex Alderton, Roberto Amato, Jeffrey Barrett, Sonia Goncalves, Ewan Harrison, David K. Jackson, Ian Johnston, Dominic Kwiatkowski, Cordelia Langford, John Sillitoe on behalf of the Wellcome Sanger Institute COVID-19 Surveillance Team |
| EPI_ISL_2446402 | Health Services Laboratories | Wellcome Sanger Institute for the COVID-19 Genomics UK (COG-UK) Consortium | Health Services Laboratories and Alex Alderton, Roberto Amato, Jeffrey Barrett, Sonia Goncalves, Ewan Harrison, David K. Jackson, Ian Johnston, Dominic Kwiatkowski, Cordelia Langford, John Sillitoe on behalf of the Wellcome Sanger Institute COVID-19 Surveillance Team |
| EPI_ISL_2446404 | Lighthouse Lab in Glasgow | Wellcome Sanger Institute for the COVID-19 Genomics UK (COG-UK) Consortium | Harper VanSteenhouse, Yumi Kasai, David Gray, Carol Clugston, Anna Dominiczak and Alex Alderton, Roberto Amato, Jeffrey Barrett, Sonia Goncalves, Ewan Harrison, David K. Jackson, Ian Johnston, Dominic Kwiatkowski, Cordelia Langford, John Sillitoe on behalf of the Wellcome Sanger Institute COVID-19 Surveillance Team |
| EPI_ISL_2446405 | Lighthouse Lab in Milton Keynes | Wellcome Sanger Institute for the COVID-19 Genomics UK (COG-UK) Consortium | The Lighthouse Lab in Milton Keynes and Alex Alderton, Roberto Amato, Jeffrey Barrett, Sonia Goncalves, Ewan Harrison, David K. Jackson, Ian Johnston, Dominic Kwiatkowski, Cordelia Langford, John Sillitoe on behalf of the Wellcome Sanger Institute COVID-19 Surveillance Team |
| EPI_ISL_2446406 | Lighthouse Lab in Alderley Park | Wellcome Sanger Institute for the COVID-19 Genomics UK (COG-UK) Consortium | Jacquelyn Wynn, Mairead Hyland, The Lighthouse Lab in Alderley Park and Alex Alderton, Roberto Amato, Jeffrey Barrett, Sonia Goncalves, Ewan Harrison, David K. Jackson, Ian Johnston, Dominic Kwiatkowski, Cordelia Langford, John Sillitoe on behalf of the Wellcome Sanger Institute COVID-19 Surveillance Team |
| EPI_ISL_2446407 | Lighthouse Lab in Glasgow | Wellcome Sanger Institute for the COVID-19 Genomics UK (COG-UK) Consortium | Harper VanSteenhouse, Yumi Kasai, David Gray, Carol Clugston, Anna Dominiczak and Alex Alderton, Roberto Amato, Jeffrey Barrett, Sonia Goncalves, Ewan Harrison, David K. Jackson, Ian Johnston, Dominic Kwiatkowski, Cordelia Langford, John Sillitoe on behalf of the Wellcome Sanger Institute COVID-19 Surveillance Team |
| EPI_ISL_2446408 | Lighthouse Lab in Milton Keynes | Wellcome Sanger Institute for the COVID-19 Genomics UK (COG-UK) Consortium | The Lighthouse Lab in Milton Keynes and Alex Alderton, Roberto Amato, Jeffrey Barrett, Sonia Goncalves, Ewan Harrison, David K. Jackson, Ian Johnston, Dominic Kwiatkowski, Cordelia Langford, John Sillitoe on behalf of the Wellcome Sanger Institute COVID-19 Surveillance Team |
| EPI_ISL_2446409, EPI_ISL_2446410, EPI_ISL_2446411, EPI_ISL_2446412, EPI_ISL_2446413 | Lighthouse Lab in Alderley Park | Wellcome Sanger Institute for the COVID-19 Genomics UK (COG-UK) Consortium | Jacquelyn Wynn, Mairead Hyland, The Lighthouse Lab in Alderley Park and Alex Alderton, Roberto Amato, Jeffrey Barrett, Sonia Goncalves, Ewan Harrison, David K. Jackson, Ian Johnston, Dominic Kwiatkowski, Cordelia Langford, John Sillitoe on behalf of the Wellcome Sanger Institute COVID-19 Surveillance Team |
| EPI_ISL_2446417, EPI_ISL_2446418 | Lighthouse Lab in Milton Keynes | Wellcome Sanger Institute for the COVID-19 Genomics UK (COG-UK) Consortium | The Lighthouse Lab in Milton Keynes and Alex Alderton, Roberto Amato, Jeffrey Barrett, Sonia Goncalves, Ewan Harrison, David K. Jackson, Ian Johnston, Dominic Kwiatkowski, Cordelia Langford, John Sillitoe on behalf of the Wellcome Sanger Institute COVID-19 Surveillance Team |
| EPI_ISL_2446420, EPI_ISL_2446421, EPI_ISL_2446422, EPI_ISL_2446423, EPI_ISL_2446424 | Lighthouse Lab in Alderley Park | Wellcome Sanger Institute for the COVID-19 Genomics UK (COG-UK) Consortium | Jacquelyn Wynn, Mairead Hyland, The Lighthouse Lab in Alderley Park and Alex Alderton, Roberto Amato, Jeffrey Barrett, Sonia Goncalves, Ewan Harrison, David K. Jackson, Ian Johnston, Dominic Kwiatkowski, Cordelia Langford, John Sillitoe on behalf of the Wellcome Sanger Institute COVID-19 Surveillance Team |
| EPI_ISL_2446425 | Lighthouse Lab in Milton Keynes | Wellcome Sanger Institute for the COVID-19 Genomics UK (COG-UK) Consortium | The Lighthouse Lab in Milton Keynes and Alex Alderton, Roberto Amato, Jeffrey Barrett, Sonia Goncalves, Ewan Harrison, David K. Jackson, Ian Johnston, Dominic Kwiatkowski, Cordelia Langford, John Sillitoe on behalf of the Wellcome Sanger Institute COVID-19 Surveillance Team |
| EPI_ISL_2446426, EPI_ISL_2446427 | Lighthouse Lab in Alderley Park | Wellcome Sanger Institute for the COVID-19 Genomics UK (COG-UK) Consortium | Jacquelyn Wynn, Mairead Hyland, The Lighthouse Lab in Alderley Park and Alex Alderton, Roberto Amato, Jeffrey Barrett, Sonia Goncalves, Ewan Harrison, David K. Jackson, Ian Johnston, Dominic Kwiatkowski, Cordelia Langford, John Sillitoe on behalf of the Wellcome Sanger Institute COVID-19 Surveillance Team |

| | | | |
|---|---|---|---|
| EPI_ISL_2446428 | Lighthouse Lab in Milton Keynes | Wellcome Sanger Institute for the COVID-19 Genomics UK (COG-UK) Consortium | The Lighthouse Lab in Milton Keynes and Alex Alderton, Roberto Amato, Jeffrey Barrett, Sonia Goncalves, Ewan Harrison, David K. Jackson, Ian Johnston, Dominic Kwiatkowski, Cordelia Langford, John Sillitoe on behalf of the Wellcome Sanger Institute COVID-19 Surveillance Team |
| EPI_ISL_2446429, EPI_ISL_2446431 | Lighthouse Lab in Alderley Park | Wellcome Sanger Institute for the COVID-19 Genomics UK (COG-UK) Consortium | Jacquelyn Wynn, Mairead Hyland, The Lighthouse Lab in Alderley Park and Alex Alderton, Roberto Amato, Jeffrey Barrett, Sonia Goncalves, Ewan Harrison, David K. Jackson, Ian Johnston, Dominic Kwiatkowski, Cordelia Langford, John Sillitoe on behalf of the Wellcome Sanger Institute COVID-19 Surveillance Team |
| EPI_ISL_2446433 | Lighthouse Lab in Milton Keynes | Wellcome Sanger Institute for the COVID-19 Genomics UK (COG-UK) Consortium | The Lighthouse Lab in Milton Keynes and Alex Alderton, Roberto Amato, Jeffrey Barrett, Sonia Goncalves, Ewan Harrison, David K. Jackson, Ian Johnston, Dominic Kwiatkowski, Cordelia Langford, John Sillitoe on behalf of the Wellcome Sanger Institute COVID-19 Surveillance Team |
| EPI_ISL_2446434, EPI_ISL_2446435, EPI_ISL_2446436 | Lighthouse Lab in Alderley Park | Wellcome Sanger Institute for the COVID-19 Genomics UK (COG-UK) Consortium | Jacquelyn Wynn, Mairead Hyland, The Lighthouse Lab in Alderley Park and Alex Alderton, Roberto Amato, Jeffrey Barrett, Sonia Goncalves, Ewan Harrison, David K. Jackson, Ian Johnston, Dominic Kwiatkowski, Cordelia Langford, John Sillitoe on behalf of the Wellcome Sanger Institute COVID-19 Surveillance Team |
| EPI_ISL_2446437, EPI_ISL_2446438 | Lighthouse Lab in Milton Keynes | Wellcome Sanger Institute for the COVID-19 Genomics UK (COG-UK) Consortium | The Lighthouse Lab in Milton Keynes and Alex Alderton, Roberto Amato, Jeffrey Barrett, Sonia Goncalves, Ewan Harrison, David K. Jackson, Ian Johnston, Dominic Kwiatkowski, Cordelia Langford, John Sillitoe on behalf of the Wellcome Sanger Institute COVID-19 Surveillance Team |
| EPI_ISL_2446439, EPI_ISL_2446440, EPI_ISL_2446443, EPI_ISL_2446444, EPI_ISL_2446445, EPI_ISL_2446448, EPI_ISL_2446449, EPI_ISL_2446450, EPI_ISL_2446451, EPI_ISL_2446452 | Lighthouse Lab in Alderley Park | Wellcome Sanger Institute for the COVID-19 Genomics UK (COG-UK) Consortium | Jacquelyn Wynn, Mairead Hyland, The Lighthouse Lab in Alderley Park and Alex Alderton, Roberto Amato, Jeffrey Barrett, Sonia Goncalves, Ewan Harrison, David K. Jackson, Ian Johnston, Dominic Kwiatkowski, Cordelia Langford, John Sillitoe on behalf of the Wellcome Sanger Institute COVID-19 Surveillance Team |
| EPI_ISL_2446453 | Lighthouse Lab in Milton Keynes | Wellcome Sanger Institute for the COVID-19 Genomics UK (COG-UK) Consortium | The Lighthouse Lab in Milton Keynes and Alex Alderton, Roberto Amato, Jeffrey Barrett, Sonia Goncalves, Ewan Harrison, David K. Jackson, Ian Johnston, Dominic Kwiatkowski, Cordelia Langford, John Sillitoe on behalf of the Wellcome Sanger Institute COVID-19 Surveillance Team |
| EPI_ISL_2446454, EPI_ISL_2446455 | Lighthouse Lab in Alderley Park | Wellcome Sanger Institute for the COVID-19 Genomics UK (COG-UK) Consortium | Jacquelyn Wynn, Mairead Hyland, The Lighthouse Lab in Alderley Park and Alex Alderton, Roberto Amato, Jeffrey Barrett, Sonia Goncalves, Ewan Harrison, David K. Jackson, Ian Johnston, Dominic Kwiatkowski, Cordelia Langford, John Sillitoe on behalf of the Wellcome Sanger Institute COVID-19 Surveillance Team |
| EPI_ISL_2446456, EPI_ISL_2446458 | Lighthouse Lab in Milton Keynes | Wellcome Sanger Institute for the COVID-19 Genomics UK (COG-UK) Consortium | The Lighthouse Lab in Milton Keynes and Alex Alderton, Roberto Amato, Jeffrey Barrett, Sonia Goncalves, Ewan Harrison, David K. Jackson, Ian Johnston, Dominic Kwiatkowski, Cordelia Langford, John Sillitoe on behalf of the Wellcome Sanger Institute COVID-19 Surveillance Team |
| EPI_ISL_2446459 | Lighthouse Lab in Alderley Park | Wellcome Sanger Institute for the COVID-19 Genomics UK (COG-UK) Consortium | Jacquelyn Wynn, Mairead Hyland, The Lighthouse Lab in Alderley Park and Alex Alderton, Roberto Amato, Jeffrey Barrett, Sonia Goncalves, Ewan Harrison, David K. Jackson, Ian Johnston, Dominic Kwiatkowski, Cordelia Langford, John Sillitoe on behalf of the Wellcome Sanger Institute COVID-19 Surveillance Team |
| EPI_ISL_2446460, EPI_ISL_2446461 | Lighthouse Lab in Milton Keynes | Wellcome Sanger Institute for the COVID-19 Genomics UK (COG-UK) Consortium | The Lighthouse Lab in Milton Keynes and Alex Alderton, Roberto Amato, Jeffrey Barrett, Sonia Goncalves, Ewan Harrison, David K. Jackson, Ian Johnston, Dominic Kwiatkowski, Cordelia Langford, John Sillitoe on behalf of the Wellcome Sanger Institute COVID-19 Surveillance Team |
| EPI_ISL_2446462, EPI_ISL_2446463, EPI_ISL_2446464, EPI_ISL_2446465 | Lighthouse Lab in Alderley Park | Wellcome Sanger Institute for the COVID-19 Genomics UK (COG-UK) Consortium | Jacquelyn Wynn, Mairead Hyland, The Lighthouse Lab in Alderley Park and Alex Alderton, Roberto Amato, Jeffrey Barrett, Sonia Goncalves, Ewan Harrison, David K. Jackson, Ian Johnston, Dominic Kwiatkowski, Cordelia Langford, John Sillitoe on behalf of the Wellcome Sanger Institute COVID-19 Surveillance Team |
| EPI_ISL_2446466 | Lighthouse Lab in Milton Keynes | Wellcome Sanger Institute for the COVID-19 Genomics UK (COG-UK) Consortium | The Lighthouse Lab in Milton Keynes and Alex Alderton, Roberto Amato, Jeffrey Barrett, Sonia Goncalves, Ewan Harrison, David K. Jackson, Ian Johnston, Dominic Kwiatkowski, Cordelia Langford, John Sillitoe on behalf of the Wellcome Sanger Institute COVID-19 Surveillance Team |
| EPI_ISL_2446467, EPI_ISL_2446468, EPI_ISL_2446469, EPI_ISL_2446470, EPI_ISL_2446472 | Lighthouse Lab in Alderley Park | Wellcome Sanger Institute for the COVID-19 Genomics UK (COG-UK) Consortium | Jacquelyn Wynn, Mairead Hyland, The Lighthouse Lab in Alderley Park and Alex Alderton, Roberto Amato, Jeffrey Barrett, Sonia Goncalves, Ewan Harrison, David K. Jackson, Ian Johnston, Dominic Kwiatkowski, Cordelia Langford, John Sillitoe on behalf of the Wellcome Sanger Institute COVID-19 Surveillance Team |
| EPI_ISL_2446473 | Lighthouse Lab in Milton Keynes | Wellcome Sanger Institute for the COVID-19 Genomics UK (COG-UK) Consortium | The Lighthouse Lab in Milton Keynes and Alex Alderton, Roberto Amato, Jeffrey Barrett, Sonia Goncalves, Ewan Harrison, David K. Jackson, Ian Johnston, Dominic Kwiatkowski, Cordelia Langford, John Sillitoe on behalf of the Wellcome Sanger Institute COVID-19 Surveillance Team |
| EPI_ISL_2446474 | Lighthouse Lab in Alderley Park | Wellcome Sanger Institute for the COVID-19 Genomics UK (COG-UK) Consortium | Jacquelyn Wynn, Mairead Hyland, The Lighthouse Lab in Alderley Park and Alex Alderton, Roberto Amato, Jeffrey Barrett, Sonia Goncalves, Ewan Harrison, David K. Jackson, Ian Johnston, Dominic Kwiatkowski, Cordelia Langford, John Sillitoe on behalf of the Wellcome Sanger Institute COVID-19 Surveillance Team |
| EPI_ISL_2446475 | Lighthouse Lab in Milton Keynes | Wellcome Sanger Institute for the COVID-19 Genomics UK (COG-UK) Consortium | The Lighthouse Lab in Milton Keynes and Alex Alderton, Roberto Amato, Jeffrey Barrett, Sonia Goncalves, Ewan Harrison, David K. Jackson, Ian Johnston, Dominic Kwiatkowski, Cordelia Langford, John Sillitoe on behalf of the Wellcome Sanger Institute COVID-19 Surveillance Team |
| EPI_ISL_2446476 | Lighthouse Lab in Alderley Park | Wellcome Sanger Institute for the COVID-19 Genomics UK (COG-UK) Consortium | Jacquelyn Wynn, Mairead Hyland, The Lighthouse Lab in Alderley Park and Alex Alderton, Roberto Amato, Jeffrey Barrett, Sonia Goncalves, Ewan Harrison, David K. Jackson, Ian Johnston, Dominic Kwiatkowski, Cordelia Langford, John Sillitoe on behalf of the Wellcome Sanger Institute COVID-19 Surveillance Team |
| EPI_ISL_2446477 | Lighthouse Lab in Milton Keynes | Wellcome Sanger Institute for the COVID-19 Genomics UK (COG-UK) Consortium | The Lighthouse Lab in Milton Keynes and Alex Alderton, Roberto Amato, Jeffrey Barrett, Sonia Goncalves, Ewan Harrison, David K. Jackson, Ian Johnston, Dominic Kwiatkowski, Cordelia Langford, John Sillitoe on behalf of the Wellcome Sanger Institute COVID-19 Surveillance Team |
| EPI_ISL_2446478 | Lighthouse Lab in Alderley Park | Wellcome Sanger Institute for the COVID-19 Genomics UK (COG-UK) Consortium | Jacquelyn Wynn, Mairead Hyland, The Lighthouse Lab in Alderley Park and Alex Alderton, Roberto Amato, Jeffrey Barrett, Sonia Goncalves, Ewan Harrison, David K. Jackson, Ian Johnston, Dominic Kwiatkowski, Cordelia Langford, John Sillitoe on behalf of the Wellcome Sanger Institute COVID-19 Surveillance Team |
| EPI_ISL_2446480, EPI_ISL_2446481 | Lighthouse Lab in Milton Keynes | Wellcome Sanger Institute for the COVID-19 Genomics UK (COG-UK) Consortium | The Lighthouse Lab in Milton Keynes and Alex Alderton, Roberto Amato, Jeffrey Barrett, Sonia Goncalves, Ewan Harrison, David K. Jackson, Ian Johnston, Dominic Kwiatkowski, Cordelia Langford, John Sillitoe on behalf of the Wellcome Sanger Institute COVID-19 Surveillance Team |
| EPI_ISL_2446482, EPI_ISL_2446483, EPI_ISL_2446485, EPI_ISL_2446486, EPI_ISL_2446487, EPI_ISL_2446488, EPI_ISL_2446490, EPI_ISL_2446491, EPI_ISL_2446492, EPI_ISL_2446493, EPI_ISL_2446494 | | | |
| see above | Lighthouse Lab in Alderley Park | Wellcome Sanger Institute for the COVID-19 Genomics UK (COG-UK) Consortium | Jacquelyn Wynn, Mairead Hyland, The Lighthouse Lab in Alderley Park and Alex Alderton, Roberto Amato, Jeffrey Barrett, Sonia Goncalves, Ewan Harrison, David K. Jackson, Ian Johnston, Dominic Kwiatkowski, Cordelia Langford, John Sillitoe on behalf of the Wellcome Sanger Institute COVID-19 Surveillance Team |
| EPI_ISL_2446495 | Lighthouse Lab in Milton Keynes | Wellcome Sanger Institute for the COVID-19 Genomics UK (COG-UK) Consortium | The Lighthouse Lab in Milton Keynes and Alex Alderton, Roberto Amato, Jeffrey Barrett, Sonia Goncalves, Ewan Harrison, David K. Jackson, Ian Johnston, Dominic Kwiatkowski, Cordelia Langford, John Sillitoe on behalf of the Wellcome Sanger Institute COVID-19 Surveillance Team |
| EPI_ISL_2446496, EPI_ISL_2446497 | Lighthouse Lab in Alderley Park | Wellcome Sanger Institute for the COVID-19 Genomics UK (COG-UK) Consortium | Jacquelyn Wynn, Mairead Hyland, The Lighthouse Lab in Alderley Park and Alex Alderton, Roberto Amato, Jeffrey Barrett, Sonia Goncalves, Ewan Harrison, David K. Jackson, Ian Johnston, Dominic Kwiatkowski, Cordelia Langford, John Sillitoe on behalf of the Wellcome Sanger Institute COVID-19 Surveillance Team |
| EPI_ISL_2446498 | Lighthouse Lab in Milton Keynes | Wellcome Sanger Institute for the COVID-19 Genomics UK (COG-UK) Consortium | The Lighthouse Lab in Milton Keynes and Alex Alderton, Roberto Amato, Jeffrey Barrett, Sonia Goncalves, Ewan Harrison, David K. Jackson, Ian Johnston, Dominic Kwiatkowski, Cordelia Langford, John Sillitoe on behalf of the Wellcome Sanger Institute COVID-19 Surveillance Team |
| EPI_ISL_2446499, EPI_ISL_2446501, EPI_ISL_2446502, EPI_ISL_2446503, EPI_ISL_2446504, EPI_ISL_2446505 | Lighthouse Lab in Alderley Park | Wellcome Sanger Institute for the COVID-19 Genomics UK (COG-UK) Consortium | Jacquelyn Wynn, Mairead Hyland, The Lighthouse Lab in Alderley Park and Alex Alderton, Roberto Amato, Jeffrey Barrett, Sonia Goncalves, Ewan Harrison, David K. Jackson, Ian Johnston, Dominic Kwiatkowski, Cordelia Langford, John Sillitoe on behalf of the Wellcome Sanger Institute COVID-19 Surveillance Team |
| EPI_ISL_2446507 | Lighthouse Lab in Milton Keynes | Wellcome Sanger Institute for the COVID-19 Genomics UK (COG-UK) Consortium | The Lighthouse Lab in Milton Keynes and Alex Alderton, Roberto Amato, Jeffrey Barrett, Sonia Goncalves, Ewan Harrison, David K. Jackson, Ian Johnston, Dominic Kwiatkowski, Cordelia Langford, John Sillitoe on behalf of the Wellcome Sanger Institute COVID-19 Surveillance Team |
| EPI_ISL_2446508, EPI_ISL_2446509, EPI_ISL_2446510, EPI_ISL_2446511, EPI_ISL_2446512, EPI_ISL_2446513 | Lighthouse Lab in Alderley Park | Wellcome Sanger Institute for the COVID-19 Genomics UK (COG-UK) Consortium | Jacquelyn Wynn, Mairead Hyland, The Lighthouse Lab in Alderley Park and Alex Alderton, Roberto Amato, Jeffrey Barrett, Sonia Goncalves, Ewan Harrison, David K. Jackson, Ian Johnston, Dominic Kwiatkowski, Cordelia Langford, John Sillitoe on behalf of the Wellcome Sanger Institute COVID-19 Surveillance Team |
| EPI_ISL_2446514 | Lighthouse Lab in Milton Keynes | Wellcome Sanger Institute for the COVID-19 Genomics UK (COG-UK) Consortium | The Lighthouse Lab in Milton Keynes and Alex Alderton, Roberto Amato, Jeffrey Barrett, Sonia Goncalves, Ewan Harrison, David K. Jackson, Ian Johnston, Dominic Kwiatkowski, Cordelia Langford, John Sillitoe on behalf of the Wellcome Sanger Institute COVID-19 Surveillance Team |
| EPI_ISL_2446515, EPI_ISL_2446516, EPI_ISL_2446517, EPI_ISL_2446518 | Lighthouse Lab in Alderley Park | Wellcome Sanger Institute for the COVID-19 Genomics UK (COG-UK) Consortium | Jacquelyn Wynn, Mairead Hyland, The Lighthouse Lab in Alderley Park and Alex Alderton, Roberto Amato, Jeffrey Barrett, Sonia Goncalves, Ewan Harrison, David K. Jackson, Ian Johnston, Dominic Kwiatkowski, Cordelia Langford, John Sillitoe on behalf of the Wellcome Sanger Institute COVID-19 Surveillance Team |
| EPI_ISL_2446519, EPI_ISL_2446520, EPI_ISL_2446521 | Lighthouse Lab in Milton Keynes | Wellcome Sanger Institute for the COVID-19 Genomics UK (COG-UK) Consortium | The Lighthouse Lab in Milton Keynes and Alex Alderton, Roberto Amato, Jeffrey Barrett, Sonia Goncalves, Ewan Harrison, David K. Jackson, Ian Johnston, Dominic Kwiatkowski, Cordelia Langford, John Sillitoe on behalf of the Wellcome Sanger Institute COVID-19 Surveillance Team |
| EPI_ISL_2446522, EPI_ISL_2446524, EPI_ISL_2446525 | Lighthouse Lab in Alderley Park | Wellcome Sanger Institute for the COVID-19 Genomics UK (COG-UK) Consortium | Jacquelyn Wynn, Mairead Hyland, The Lighthouse Lab in Alderley Park and Alex Alderton, Roberto Amato, Jeffrey Barrett, Sonia Goncalves, Ewan Harrison, David K. Jackson, Ian Johnston, Dominic Kwiatkowski, Cordelia Langford, John Sillitoe on behalf of the Wellcome Sanger Institute COVID-19 Surveillance Team |
| EPI_ISL_2446526 | Lighthouse Lab in Milton Keynes | Wellcome Sanger Institute for the COVID-19 Genomics UK (COG-UK) Consortium | The Lighthouse Lab in Milton Keynes and Alex Alderton, Roberto Amato, Jeffrey Barrett, Sonia Goncalves, Ewan Harrison, David K. Jackson, Ian Johnston, Dominic Kwiatkowski, Cordelia Langford, John Sillitoe on behalf of the Wellcome Sanger Institute COVID-19 Surveillance Team |
| EPI_ISL_2446527 | Lighthouse Lab in Alderley Park | Wellcome Sanger Institute for the COVID-19 Genomics UK (COG-UK) Consortium | Jacquelyn Wynn, Mairead Hyland, The Lighthouse Lab in Alderley Park and Alex Alderton, Roberto Amato, Jeffrey Barrett, Sonia Goncalves, Ewan Harrison, David K. Jackson, Ian Johnston, Dominic Kwiatkowski, Cordelia Langford, John Sillitoe on behalf of the Wellcome Sanger Institute COVID-19 Surveillance Team |

| | | | |
|---|---|---|---|
| EPI_ISL_2446529 | Lighthouse Lab in Milton Keynes | Wellcome Sanger Institute for the COVID-19 Genomics UK (COG-UK) Consortium | The Lighthouse Lab in Milton Keynes and Alex Alderton, Roberto Amato, Jeffrey Barrett, Sonia Goncalves, Ewan Harrison, David K. Jackson, Ian Johnston, Dominic Kwiatkowski, Cordelia Langford, John Sillitoe on behalf of the Wellcome Sanger Institute COVID-19 Surveillance Team |
| EPI_ISL_2446532, EPI_ISL_2446534, EPI_ISL_2446535 | Lighthouse Lab in Alderley Park | Wellcome Sanger Institute for the COVID-19 Genomics UK (COG-UK) Consortium | Jacquelyn Wynn, Mairead Hyland, The Lighthouse Lab in Alderley Park and Alex Alderton, Roberto Amato, Jeffrey Barrett, Sonia Goncalves, Ewan Harrison, David K. Jackson, Ian Johnston, Dominic Kwiatkowski, Cordelia Langford, John Sillitoe on behalf of the Wellcome Sanger Institute COVID-19 Surveillance Team |
| EPI_ISL_2446537 | Lighthouse Lab in Milton Keynes | Wellcome Sanger Institute for the COVID-19 Genomics UK (COG-UK) Consortium | The Lighthouse Lab in Milton Keynes and Alex Alderton, Roberto Amato, Jeffrey Barrett, Sonia Goncalves, Ewan Harrison, David K. Jackson, Ian Johnston, Dominic Kwiatkowski, Cordelia Langford, John Sillitoe on behalf of the Wellcome Sanger Institute COVID-19 Surveillance Team |
| EPI_ISL_2446538 | Lighthouse Lab in Alderley Park | Wellcome Sanger Institute for the COVID-19 Genomics UK (COG-UK) Consortium | Jacquelyn Wynn, Mairead Hyland, The Lighthouse Lab in Alderley Park and Alex Alderton, Roberto Amato, Jeffrey Barrett, Sonia Goncalves, Ewan Harrison, David K. Jackson, Ian Johnston, Dominic Kwiatkowski, Cordelia Langford, John Sillitoe on behalf of the Wellcome Sanger Institute COVID-19 Surveillance Team |
| EPI_ISL_2446539 | Lighthouse Lab in Milton Keynes | Wellcome Sanger Institute for the COVID-19 Genomics UK (COG-UK) Consortium | The Lighthouse Lab in Milton Keynes and Alex Alderton, Roberto Amato, Jeffrey Barrett, Sonia Goncalves, Ewan Harrison, David K. Jackson, Ian Johnston, Dominic Kwiatkowski, Cordelia Langford, John Sillitoe on behalf of the Wellcome Sanger Institute COVID-19 Surveillance Team |
| EPI_ISL_2446540, EPI_ISL_2446541, EPI_ISL_2446542, EPI_ISL_2446543 | Lighthouse Lab in Alderley Park | Wellcome Sanger Institute for the COVID-19 Genomics UK (COG-UK) Consortium | Jacquelyn Wynn, Mairead Hyland, The Lighthouse Lab in Alderley Park and Alex Alderton, Roberto Amato, Jeffrey Barrett, Sonia Goncalves, Ewan Harrison, David K. Jackson, Ian Johnston, Dominic Kwiatkowski, Cordelia Langford, John Sillitoe on behalf of the Wellcome Sanger Institute COVID-19 Surveillance Team |
| EPI_ISL_2446546 | Lighthouse Lab in Milton Keynes | Wellcome Sanger Institute for the COVID-19 Genomics UK (COG-UK) Consortium | The Lighthouse Lab in Milton Keynes and Alex Alderton, Roberto Amato, Jeffrey Barrett, Sonia Goncalves, Ewan Harrison, David K. Jackson, Ian Johnston, Dominic Kwiatkowski, Cordelia Langford, John Sillitoe on behalf of the Wellcome Sanger Institute COVID-19 Surveillance Team |
| EPI_ISL_2446547, EPI_ISL_2446548, EPI_ISL_2446551, EPI_ISL_2446552, EPI_ISL_2446553, EPI_ISL_2446554, EPI_ISL_2446556, EPI_ISL_2446557 | Lighthouse Lab in Alderley Park | Wellcome Sanger Institute for the COVID-19 Genomics UK (COG-UK) Consortium | Jacquelyn Wynn, Mairead Hyland, The Lighthouse Lab in Alderley Park and Alex Alderton, Roberto Amato, Jeffrey Barrett, Sonia Goncalves, Ewan Harrison, David K. Jackson, Ian Johnston, Dominic Kwiatkowski, Cordelia Langford, John Sillitoe on behalf of the Wellcome Sanger Institute COVID-19 Surveillance Team |
| EPI_ISL_2446559 | Lighthouse Lab in Milton Keynes | Wellcome Sanger Institute for the COVID-19 Genomics UK (COG-UK) Consortium | The Lighthouse Lab in Milton Keynes and Alex Alderton, Roberto Amato, Jeffrey Barrett, Sonia Goncalves, Ewan Harrison, David K. Jackson, Ian Johnston, Dominic Kwiatkowski, Cordelia Langford, John Sillitoe on behalf of the Wellcome Sanger Institute COVID-19 Surveillance Team |
| EPI_ISL_2446560, EPI_ISL_2446561, EPI_ISL_2446562, EPI_ISL_2446563, EPI_ISL_2446565, EPI_ISL_2446566, EPI_ISL_2446567 | Lighthouse Lab in Alderley Park | Wellcome Sanger Institute for the COVID-19 Genomics UK (COG-UK) Consortium | Jacquelyn Wynn, Mairead Hyland, The Lighthouse Lab in Alderley Park and Alex Alderton, Roberto Amato, Jeffrey Barrett, Sonia Goncalves, Ewan Harrison, David K. Jackson, Ian Johnston, Dominic Kwiatkowski, Cordelia Langford, John Sillitoe on behalf of the Wellcome Sanger Institute COVID-19 Surveillance Team |
| EPI_ISL_2446569, EPI_ISL_2446570 | Lighthouse Lab in Milton Keynes | Wellcome Sanger Institute for the COVID-19 Genomics UK (COG-UK) Consortium | The Lighthouse Lab in Milton Keynes and Alex Alderton, Roberto Amato, Jeffrey Barrett, Sonia Goncalves, Ewan Harrison, David K. Jackson, Ian Johnston, Dominic Kwiatkowski, Cordelia Langford, John Sillitoe on behalf of the Wellcome Sanger Institute COVID-19 Surveillance Team |
| EPI_ISL_2446571, EPI_ISL_2446572 | Lighthouse Lab in Alderley Park | Wellcome Sanger Institute for the COVID-19 Genomics UK (COG-UK) Consortium | Jacquelyn Wynn, Mairead Hyland, The Lighthouse Lab in Alderley Park and Alex Alderton, Roberto Amato, Jeffrey Barrett, Sonia Goncalves, Ewan Harrison, David K. Jackson, Ian Johnston, Dominic Kwiatkowski, Cordelia Langford, John Sillitoe on behalf of the Wellcome Sanger Institute COVID-19 Surveillance Team |
| EPI_ISL_2446573, EPI_ISL_2446574 | Lighthouse Lab in Milton Keynes | Wellcome Sanger Institute for the COVID-19 Genomics UK (COG-UK) Consortium | The Lighthouse Lab in Milton Keynes and Alex Alderton, Roberto Amato, Jeffrey Barrett, Sonia Goncalves, Ewan Harrison, David K. Jackson, Ian Johnston, Dominic Kwiatkowski, Cordelia Langford, John Sillitoe on behalf of the Wellcome Sanger Institute COVID-19 Surveillance Team |
| EPI_ISL_2446575, EPI_ISL_2446576, EPI_ISL_2446577 | Lighthouse Lab in Alderley Park | Wellcome Sanger Institute for the COVID-19 Genomics UK (COG-UK) Consortium | Jacquelyn Wynn, Mairead Hyland, The Lighthouse Lab in Alderley Park and Alex Alderton, Roberto Amato, Jeffrey Barrett, Sonia Goncalves, Ewan Harrison, David K. Jackson, Ian Johnston, Dominic Kwiatkowski, Cordelia Langford, John Sillitoe on behalf of the Wellcome Sanger Institute COVID-19 Surveillance Team |
| EPI_ISL_2446578 | Lighthouse Lab in Milton Keynes | Wellcome Sanger Institute for the COVID-19 Genomics UK (COG-UK) Consortium | The Lighthouse Lab in Milton Keynes and Alex Alderton, Roberto Amato, Jeffrey Barrett, Sonia Goncalves, Ewan Harrison, David K. Jackson, Ian Johnston, Dominic Kwiatkowski, Cordelia Langford, John Sillitoe on behalf of the Wellcome Sanger Institute COVID-19 Surveillance Team |
| EPI_ISL_2446579, EPI_ISL_2446580, EPI_ISL_2446581, EPI_ISL_2446582, EPI_ISL_2446583, EPI_ISL_2446584, EPI_ISL_2446585, EPI_ISL_2446586, EPI_ISL_2446587, EPI_ISL_2446588 | Lighthouse Lab in Alderley Park | Wellcome Sanger Institute for the COVID-19 Genomics UK (COG-UK) Consortium | Jacquelyn Wynn, Mairead Hyland, The Lighthouse Lab in Alderley Park and Alex Alderton, Roberto Amato, Jeffrey Barrett, Sonia Goncalves, Ewan Harrison, David K. Jackson, Ian Johnston, Dominic Kwiatkowski, Cordelia Langford, John Sillitoe on behalf of the Wellcome Sanger Institute COVID-19 Surveillance Team |
| EPI_ISL_2446589 | Lighthouse Lab in Milton Keynes | Wellcome Sanger Institute for the COVID-19 Genomics UK (COG-UK) Consortium | The Lighthouse Lab in Milton Keynes and Alex Alderton, Roberto Amato, Jeffrey Barrett, Sonia Goncalves, Ewan Harrison, David K. Jackson, Ian Johnston, Dominic Kwiatkowski, Cordelia Langford, John Sillitoe on behalf of the Wellcome Sanger Institute COVID-19 Surveillance Team |
| EPI_ISL_2446590, EPI_ISL_2446591, EPI_ISL_2446592, EPI_ISL_2446593, EPI_ISL_2446594, EPI_ISL_2446595, EPI_ISL_2446596, EPI_ISL_2446597 | Lighthouse Lab in Alderley Park | Wellcome Sanger Institute for the COVID-19 Genomics UK (COG-UK) Consortium | Jacquelyn Wynn, Mairead Hyland, The Lighthouse Lab in Alderley Park and Alex Alderton, Roberto Amato, Jeffrey Barrett, Sonia Goncalves, Ewan Harrison, David K. Jackson, Ian Johnston, Dominic Kwiatkowski, Cordelia Langford, John Sillitoe on behalf of the Wellcome Sanger Institute COVID-19 Surveillance Team |
| EPI_ISL_2446598 | Lighthouse Lab in Milton Keynes | Wellcome Sanger Institute for the COVID-19 Genomics UK (COG-UK) Consortium | The Lighthouse Lab in Milton Keynes and Alex Alderton, Roberto Amato, Jeffrey Barrett, Sonia Goncalves, Ewan Harrison, David K. Jackson, Ian Johnston, Dominic Kwiatkowski, Cordelia Langford, John Sillitoe on behalf of the Wellcome Sanger Institute COVID-19 Surveillance Team |
| EPI_ISL_2446599, EPI_ISL_2446600 | Lighthouse Lab in Alderley Park | Wellcome Sanger Institute for the COVID-19 Genomics UK (COG-UK) Consortium | Jacquelyn Wynn, Mairead Hyland, The Lighthouse Lab in Alderley Park and Alex Alderton, Roberto Amato, Jeffrey Barrett, Sonia Goncalves, Ewan Harrison, David K. Jackson, Ian Johnston, Dominic Kwiatkowski, Cordelia Langford, John Sillitoe on behalf of the Wellcome Sanger Institute COVID-19 Surveillance Team |
| EPI_ISL_2446601 | Lighthouse Lab in Milton Keynes | Wellcome Sanger Institute for the COVID-19 Genomics UK (COG-UK) Consortium | The Lighthouse Lab in Milton Keynes and Alex Alderton, Roberto Amato, Jeffrey Barrett, Sonia Goncalves, Ewan Harrison, David K. Jackson, Ian Johnston, Dominic Kwiatkowski, Cordelia Langford, John Sillitoe on behalf of the Wellcome Sanger Institute COVID-19 Surveillance Team |
| EPI_ISL_2446602, EPI_ISL_2446603, EPI_ISL_2446604 | Lighthouse Lab in Alderley Park | Wellcome Sanger Institute for the COVID-19 Genomics UK (COG-UK) Consortium | Jacquelyn Wynn, Mairead Hyland, The Lighthouse Lab in Alderley Park and Alex Alderton, Roberto Amato, Jeffrey Barrett, Sonia Goncalves, Ewan Harrison, David K. Jackson, Ian Johnston, Dominic Kwiatkowski, Cordelia Langford, John Sillitoe on behalf of the Wellcome Sanger Institute COVID-19 Surveillance Team |
| EPI_ISL_2446605 | Lighthouse Lab in Milton Keynes | Wellcome Sanger Institute for the COVID-19 Genomics UK (COG-UK) Consortium | The Lighthouse Lab in Milton Keynes and Alex Alderton, Roberto Amato, Jeffrey Barrett, Sonia Goncalves, Ewan Harrison, David K. Jackson, Ian Johnston, Dominic Kwiatkowski, Cordelia Langford, John Sillitoe on behalf of the Wellcome Sanger Institute COVID-19 Surveillance Team |
| EPI_ISL_2446606, EPI_ISL_2446607, EPI_ISL_2446608, EPI_ISL_2446609, EPI_ISL_2446610, EPI_ISL_2446612, EPI_ISL_2446613, EPI_ISL_2446614, EPI_ISL_2446615, EPI_ISL_2446618 | Lighthouse Lab in Alderley Park | Wellcome Sanger Institute for the COVID-19 Genomics UK (COG-UK) Consortium | Jacquelyn Wynn, Mairead Hyland, The Lighthouse Lab in Alderley Park and Alex Alderton, Roberto Amato, Jeffrey Barrett, Sonia Goncalves, Ewan Harrison, David K. Jackson, Ian Johnston, Dominic Kwiatkowski, Cordelia Langford, John Sillitoe on behalf of the Wellcome Sanger Institute COVID-19 Surveillance Team |
| EPI_ISL_2446619 | Lighthouse Lab in Milton Keynes | Wellcome Sanger Institute for the COVID-19 Genomics UK (COG-UK) Consortium | The Lighthouse Lab in Milton Keynes and Alex Alderton, Roberto Amato, Jeffrey Barrett, Sonia Goncalves, Ewan Harrison, David K. Jackson, Ian Johnston, Dominic Kwiatkowski, Cordelia Langford, John Sillitoe on behalf of the Wellcome Sanger Institute COVID-19 Surveillance Team |
| EPI_ISL_2446621, EPI_ISL_2446622, EPI_ISL_2446623, EPI_ISL_2446624 | Lighthouse Lab in Alderley Park | Wellcome Sanger Institute for the COVID-19 Genomics UK (COG-UK) Consortium | Jacquelyn Wynn, Mairead Hyland, The Lighthouse Lab in Alderley Park and Alex Alderton, Roberto Amato, Jeffrey Barrett, Sonia Goncalves, Ewan Harrison, David K. Jackson, Ian Johnston, Dominic Kwiatkowski, Cordelia Langford, John Sillitoe on behalf of the Wellcome Sanger Institute COVID-19 Surveillance Team |
| EPI_ISL_2446625 | Lighthouse Lab in Milton Keynes | Wellcome Sanger Institute for the COVID-19 Genomics UK (COG-UK) Consortium | The Lighthouse Lab in Milton Keynes and Alex Alderton, Roberto Amato, Jeffrey Barrett, Sonia Goncalves, Ewan Harrison, David K. Jackson, Ian Johnston, Dominic Kwiatkowski, Cordelia Langford, John Sillitoe on behalf of the Wellcome Sanger Institute COVID-19 Surveillance Team |
| EPI_ISL_2446626, EPI_ISL_2446627 | Lighthouse Lab in Alderley Park | Wellcome Sanger Institute for the COVID-19 Genomics UK (COG-UK) Consortium | Jacquelyn Wynn, Mairead Hyland, The Lighthouse Lab in Alderley Park and Alex Alderton, Roberto Amato, Jeffrey Barrett, Sonia Goncalves, Ewan Harrison, David K. Jackson, Ian Johnston, Dominic Kwiatkowski, Cordelia Langford, John Sillitoe on behalf of the Wellcome Sanger Institute COVID-19 Surveillance Team |
| EPI_ISL_2446628, | Lighthouse Lab in Milton Keynes | Wellcome Sanger Institute for the COVID-19 Genomics UK (COG-UK) | The Lighthouse Lab in Milton Keynes and Alex Alderton, Roberto Amato, Jeffrey Barrett, Sonia Goncalves, Ewan Harrison, David K. Jackson, Ian Johnston, Dominic Kwiatkowski, Cordelia Langford, John Sillitoe on behalf of the Wellcome Sanger |

| | | | |
|---|---|---|---|
| EPI_ISL_2446629, EPI_ISL_2446630 | | Consortium | Institute COVID-19 Surveillance Team |
| EPI_ISL_2446631, EPI_ISL_2446632, EPI_ISL_2446634, EPI_ISL_2446636, EPI_ISL_2446637 | Lighthouse Lab in Alderley Park | Wellcome Sanger Institute for the COVID-19 Genomics UK (COG-UK) Consortium | Jacquelyn Wynn, Mairead Hyland, The Lighthouse Lab in Alderley Park and Alex Alderton, Roberto Amato, Jeffrey Barrett, Sonia Goncalves, Ewan Harrison, David K. Jackson, Ian Johnston, Dominic Kwiatkowski, Cordelia Langford, John Sillitoe on behalf of the Wellcome Sanger Institute COVID-19 Surveillance Team |
| EPI_ISL_2446638 | Lighthouse Lab in Milton Keynes | Wellcome Sanger Institute for the COVID-19 Genomics UK (COG-UK) Consortium | The Lighthouse Lab in Milton Keynes and Alex Alderton, Roberto Amato, Jeffrey Barrett, Sonia Goncalves, Ewan Harrison, David K. Jackson, Ian Johnston, Dominic Kwiatkowski, Cordelia Langford, John Sillitoe on behalf of the Wellcome Sanger Institute COVID-19 Surveillance Team |
| EPI_ISL_2446639 | Lighthouse Lab in Alderley Park | Wellcome Sanger Institute for the COVID-19 Genomics UK (COG-UK) Consortium | Jacquelyn Wynn, Mairead Hyland, The Lighthouse Lab in Alderley Park and Alex Alderton, Roberto Amato, Jeffrey Barrett, Sonia Goncalves, Ewan Harrison, David K. Jackson, Ian Johnston, Dominic Kwiatkowski, Cordelia Langford, John Sillitoe on behalf of the Wellcome Sanger Institute COVID-19 Surveillance Team |
| EPI_ISL_2446640 | Lighthouse Lab in Milton Keynes | Wellcome Sanger Institute for the COVID-19 Genomics UK (COG-UK) Consortium | The Lighthouse Lab in Milton Keynes and Alex Alderton, Roberto Amato, Jeffrey Barrett, Sonia Goncalves, Ewan Harrison, David K. Jackson, Ian Johnston, Dominic Kwiatkowski, Cordelia Langford, John Sillitoe on behalf of the Wellcome Sanger Institute COVID-19 Surveillance Team |
| EPI_ISL_2446641, EPI_ISL_2446642, EPI_ISL_2446644 | Lighthouse Lab in Alderley Park | Wellcome Sanger Institute for the COVID-19 Genomics UK (COG-UK) Consortium | Jacquelyn Wynn, Mairead Hyland, The Lighthouse Lab in Alderley Park and Alex Alderton, Roberto Amato, Jeffrey Barrett, Sonia Goncalves, Ewan Harrison, David K. Jackson, Ian Johnston, Dominic Kwiatkowski, Cordelia Langford, John Sillitoe on behalf of the Wellcome Sanger Institute COVID-19 Surveillance Team |
| EPI_ISL_2446645, EPI_ISL_2446646 | Lighthouse Lab in Milton Keynes | Wellcome Sanger Institute for the COVID-19 Genomics UK (COG-UK) Consortium | The Lighthouse Lab in Milton Keynes and Alex Alderton, Roberto Amato, Jeffrey Barrett, Sonia Goncalves, Ewan Harrison, David K. Jackson, Ian Johnston, Dominic Kwiatkowski, Cordelia Langford, John Sillitoe on behalf of the Wellcome Sanger Institute COVID-19 Surveillance Team |
| EPI_ISL_2446647, EPI_ISL_2446648, EPI_ISL_2446649, EPI_ISL_2446650, EPI_ISL_2446651, EPI_ISL_2446652, EPI_ISL_2446653, EPI_ISL_2446654, EPI_ISL_2446655, EPI_ISL_2446656, EPI_ISL_2446658 | | | |
| see above | Lighthouse Lab in Alderley Park | Wellcome Sanger Institute for the COVID-19 Genomics UK (COG-UK) Consortium | Jacquelyn Wynn, Mairead Hyland, The Lighthouse Lab in Alderley Park and Alex Alderton, Roberto Amato, Jeffrey Barrett, Sonia Goncalves, Ewan Harrison, David K. Jackson, Ian Johnston, Dominic Kwiatkowski, Cordelia Langford, John Sillitoe on behalf of the Wellcome Sanger Institute COVID-19 Surveillance Team |
| EPI_ISL_2446660 | Lighthouse Lab in Milton Keynes | Wellcome Sanger Institute for the COVID-19 Genomics UK (COG-UK) Consortium | The Lighthouse Lab in Milton Keynes and Alex Alderton, Roberto Amato, Jeffrey Barrett, Sonia Goncalves, Ewan Harrison, David K. Jackson, Ian Johnston, Dominic Kwiatkowski, Cordelia Langford, John Sillitoe on behalf of the Wellcome Sanger Institute COVID-19 Surveillance Team |
| EPI_ISL_2446661 | Lighthouse Lab in Alderley Park | Wellcome Sanger Institute for the COVID-19 Genomics UK (COG-UK) Consortium | Jacquelyn Wynn, Mairead Hyland, The Lighthouse Lab in Alderley Park and Alex Alderton, Roberto Amato, Jeffrey Barrett, Sonia Goncalves, Ewan Harrison, David K. Jackson, Ian Johnston, Dominic Kwiatkowski, Cordelia Langford, John Sillitoe on behalf of the Wellcome Sanger Institute COVID-19 Surveillance Team |
| EPI_ISL_2446662 | Lighthouse Lab in Milton Keynes | Wellcome Sanger Institute for the COVID-19 Genomics UK (COG-UK) Consortium | The Lighthouse Lab in Milton Keynes and Alex Alderton, Roberto Amato, Jeffrey Barrett, Sonia Goncalves, Ewan Harrison, David K. Jackson, Ian Johnston, Dominic Kwiatkowski, Cordelia Langford, John Sillitoe on behalf of the Wellcome Sanger Institute COVID-19 Surveillance Team |
| EPI_ISL_2446663, EPI_ISL_2446664, EPI_ISL_2446666, EPI_ISL_2446667, EPI_ISL_2446668, EPI_ISL_2446669, EPI_ISL_2446670, EPI_ISL_2446671, EPI_ISL_2446672 | Lighthouse Lab in Alderley Park | Wellcome Sanger Institute for the COVID-19 Genomics UK (COG-UK) Consortium | Jacquelyn Wynn, Mairead Hyland, The Lighthouse Lab in Alderley Park and Alex Alderton, Roberto Amato, Jeffrey Barrett, Sonia Goncalves, Ewan Harrison, David K. Jackson, Ian Johnston, Dominic Kwiatkowski, Cordelia Langford, John Sillitoe on behalf of the Wellcome Sanger Institute COVID-19 Surveillance Team |
| EPI_ISL_2446673, EPI_ISL_2446676 | Lighthouse Lab in Milton Keynes | Wellcome Sanger Institute for the COVID-19 Genomics UK (COG-UK) Consortium | The Lighthouse Lab in Milton Keynes and Alex Alderton, Roberto Amato, Jeffrey Barrett, Sonia Goncalves, Ewan Harrison, David K. Jackson, Ian Johnston, Dominic Kwiatkowski, Cordelia Langford, John Sillitoe on behalf of the Wellcome Sanger Institute COVID-19 Surveillance Team |
| EPI_ISL_2446677 | Lighthouse Lab in Alderley Park | Wellcome Sanger Institute for the COVID-19 Genomics UK (COG-UK) Consortium | Jacquelyn Wynn, Mairead Hyland, The Lighthouse Lab in Alderley Park and Alex Alderton, Roberto Amato, Jeffrey Barrett, Sonia Goncalves, Ewan Harrison, David K. Jackson, Ian Johnston, Dominic Kwiatkowski, Cordelia Langford, John Sillitoe on behalf of the Wellcome Sanger Institute COVID-19 Surveillance Team |
| EPI_ISL_2446678 | Lighthouse Lab in Milton Keynes | Wellcome Sanger Institute for the COVID-19 Genomics UK (COG-UK) Consortium | The Lighthouse Lab in Milton Keynes and Alex Alderton, Roberto Amato, Jeffrey Barrett, Sonia Goncalves, Ewan Harrison, David K. Jackson, Ian Johnston, Dominic Kwiatkowski, Cordelia Langford, John Sillitoe on behalf of the Wellcome Sanger Institute COVID-19 Surveillance Team |
| EPI_ISL_2446679 | Lighthouse Lab in Alderley Park | Wellcome Sanger Institute for the COVID-19 Genomics UK (COG-UK) Consortium | Jacquelyn Wynn, Mairead Hyland, The Lighthouse Lab in Alderley Park and Alex Alderton, Roberto Amato, Jeffrey Barrett, Sonia Goncalves, Ewan Harrison, David K. Jackson, Ian Johnston, Dominic Kwiatkowski, Cordelia Langford, John Sillitoe on behalf of the Wellcome Sanger Institute COVID-19 Surveillance Team |
| EPI_ISL_2446680, EPI_ISL_2446681 | Lighthouse Lab in Milton Keynes | Wellcome Sanger Institute for the COVID-19 Genomics UK (COG-UK) Consortium | The Lighthouse Lab in Milton Keynes and Alex Alderton, Roberto Amato, Jeffrey Barrett, Sonia Goncalves, Ewan Harrison, David K. Jackson, Ian Johnston, Dominic Kwiatkowski, Cordelia Langford, John Sillitoe on behalf of the Wellcome Sanger Institute COVID-19 Surveillance Team |
| EPI_ISL_2446682, EPI_ISL_2446683, EPI_ISL_2446684, EPI_ISL_2446685 | Lighthouse Lab in Alderley Park | Wellcome Sanger Institute for the COVID-19 Genomics UK (COG-UK) Consortium | Jacquelyn Wynn, Mairead Hyland, The Lighthouse Lab in Alderley Park and Alex Alderton, Roberto Amato, Jeffrey Barrett, Sonia Goncalves, Ewan Harrison, David K. Jackson, Ian Johnston, Dominic Kwiatkowski, Cordelia Langford, John Sillitoe on behalf of the Wellcome Sanger Institute COVID-19 Surveillance Team |
| EPI_ISL_2446686, EPI_ISL_2446687 | Lighthouse Lab in Milton Keynes | Wellcome Sanger Institute for the COVID-19 Genomics UK (COG-UK) Consortium | The Lighthouse Lab in Milton Keynes and Alex Alderton, Roberto Amato, Jeffrey Barrett, Sonia Goncalves, Ewan Harrison, David K. Jackson, Ian Johnston, Dominic Kwiatkowski, Cordelia Langford, John Sillitoe on behalf of the Wellcome Sanger Institute COVID-19 Surveillance Team |
| EPI_ISL_2446688 | Lighthouse Lab in Alderley Park | Wellcome Sanger Institute for the COVID-19 Genomics UK (COG-UK) Consortium | Jacquelyn Wynn, Mairead Hyland, The Lighthouse Lab in Alderley Park and Alex Alderton, Roberto Amato, Jeffrey Barrett, Sonia Goncalves, Ewan Harrison, David K. Jackson, Ian Johnston, Dominic Kwiatkowski, Cordelia Langford, John Sillitoe on behalf of the Wellcome Sanger Institute COVID-19 Surveillance Team |
| EPI_ISL_2446690, EPI_ISL_2446692 | Lighthouse Lab in Milton Keynes | Wellcome Sanger Institute for the COVID-19 Genomics UK (COG-UK) Consortium | The Lighthouse Lab in Milton Keynes and Alex Alderton, Roberto Amato, Jeffrey Barrett, Sonia Goncalves, Ewan Harrison, David K. Jackson, Ian Johnston, Dominic Kwiatkowski, Cordelia Langford, John Sillitoe on behalf of the Wellcome Sanger Institute COVID-19 Surveillance Team |
| EPI_ISL_2446693, EPI_ISL_2446695, EPI_ISL_2446696, EPI_ISL_2446697, EPI_ISL_2446698, EPI_ISL_2446699 | Lighthouse Lab in Alderley Park | Wellcome Sanger Institute for the COVID-19 Genomics UK (COG-UK) Consortium | Jacquelyn Wynn, Mairead Hyland, The Lighthouse Lab in Alderley Park and Alex Alderton, Roberto Amato, Jeffrey Barrett, Sonia Goncalves, Ewan Harrison, David K. Jackson, Ian Johnston, Dominic Kwiatkowski, Cordelia Langford, John Sillitoe on behalf of the Wellcome Sanger Institute COVID-19 Surveillance Team |
| EPI_ISL_2446700 | Lighthouse Lab in Milton Keynes | Wellcome Sanger Institute for the COVID-19 Genomics UK (COG-UK) Consortium | The Lighthouse Lab in Milton Keynes and Alex Alderton, Roberto Amato, Jeffrey Barrett, Sonia Goncalves, Ewan Harrison, David K. Jackson, Ian Johnston, Dominic Kwiatkowski, Cordelia Langford, John Sillitoe on behalf of the Wellcome Sanger Institute COVID-19 Surveillance Team |
| EPI_ISL_2446702, EPI_ISL_2446703, EPI_ISL_2446704, EPI_ISL_2446706, EPI_ISL_2446707, EPI_ISL_2446708, EPI_ISL_2446709, EPI_ISL_2446711, EPI_ISL_2446712 | Lighthouse Lab in Alderley Park | Wellcome Sanger Institute for the COVID-19 Genomics UK (COG-UK) Consortium | Jacquelyn Wynn, Mairead Hyland, The Lighthouse Lab in Alderley Park and Alex Alderton, Roberto Amato, Jeffrey Barrett, Sonia Goncalves, Ewan Harrison, David K. Jackson, Ian Johnston, Dominic Kwiatkowski, Cordelia Langford, John Sillitoe on behalf of the Wellcome Sanger Institute COVID-19 Surveillance Team |
| EPI_ISL_2446713 | Lighthouse Lab in Milton Keynes | Wellcome Sanger Institute for the COVID-19 Genomics UK (COG-UK) Consortium | The Lighthouse Lab in Milton Keynes and Alex Alderton, Roberto Amato, Jeffrey Barrett, Sonia Goncalves, Ewan Harrison, David K. Jackson, Ian Johnston, Dominic Kwiatkowski, Cordelia Langford, John Sillitoe on behalf of the Wellcome Sanger Institute COVID-19 Surveillance Team |
| EPI_ISL_2446714, EPI_ISL_2446715, EPI_ISL_2446716, EPI_ISL_2446717, EPI_ISL_2446718, EPI_ISL_2446719, EPI_ISL_2446720, EPI_ISL_2446722 | Lighthouse Lab in Alderley Park | Wellcome Sanger Institute for the COVID-19 Genomics UK (COG-UK) Consortium | Jacquelyn Wynn, Mairead Hyland, The Lighthouse Lab in Alderley Park and Alex Alderton, Roberto Amato, Jeffrey Barrett, Sonia Goncalves, Ewan Harrison, David K. Jackson, Ian Johnston, Dominic Kwiatkowski, Cordelia Langford, John Sillitoe on behalf of the Wellcome Sanger Institute COVID-19 Surveillance Team |
| EPI_ISL_2446723 | Lighthouse Lab in Milton Keynes | Wellcome Sanger Institute for the COVID-19 Genomics UK (COG-UK) Consortium | The Lighthouse Lab in Milton Keynes and Alex Alderton, Roberto Amato, Jeffrey Barrett, Sonia Goncalves, Ewan Harrison, David K. Jackson, Ian Johnston, Dominic Kwiatkowski, Cordelia Langford, John Sillitoe on behalf of the Wellcome Sanger Institute COVID-19 Surveillance Team |
| EPI_ISL_2446725, EPI_ISL_2446726, EPI_ISL_2446727, EPI_ISL_2446728, EPI_ISL_2446729, EPI_ISL_2446730, EPI_ISL_2446731, EPI_ISL_2446732, EPI_ISL_2446733, EPI_ISL_2446734, EPI_ISL_2446735 | | | |
| see above | Lighthouse Lab in Alderley Park | Wellcome Sanger Institute for the COVID-19 Genomics UK (COG-UK) Consortium | Jacquelyn Wynn, Mairead Hyland, The Lighthouse Lab in Alderley Park and Alex Alderton, Roberto Amato, Jeffrey Barrett, Sonia Goncalves, Ewan Harrison, David K. Jackson, Ian Johnston, Dominic Kwiatkowski, Cordelia Langford, John Sillitoe on behalf of the Wellcome Sanger Institute COVID-19 Surveillance Team |
| EPI_ISL_2446736 | Lighthouse Lab in Milton Keynes | Wellcome Sanger Institute for the COVID-19 Genomics UK (COG-UK) Consortium | The Lighthouse Lab in Milton Keynes and Alex Alderton, Roberto Amato, Jeffrey Barrett, Sonia Goncalves, Ewan Harrison, David K. Jackson, Ian Johnston, Dominic Kwiatkowski, Cordelia Langford, John Sillitoe on behalf of the Wellcome Sanger Institute COVID-19 Surveillance Team |
| EPI_ISL_2446737, EPI_ISL_2446738, EPI_ISL_2446739, EPI_ISL_2446742, EPI_ISL_2446743 | Lighthouse Lab in Alderley Park | Wellcome Sanger Institute for the COVID-19 Genomics UK (COG-UK) Consortium | Jacquelyn Wynn, Mairead Hyland, The Lighthouse Lab in Alderley Park and Alex Alderton, Roberto Amato, Jeffrey Barrett, Sonia Goncalves, Ewan Harrison, David K. Jackson, Ian Johnston, Dominic Kwiatkowski, Cordelia Langford, John Sillitoe on behalf of the Wellcome Sanger Institute COVID-19 Surveillance Team |
| EPI_ISL_2446744, | Lighthouse Lab in Milton Keynes | Wellcome Sanger Institute for the COVID-19 Genomics UK (COG-UK) | The Lighthouse Lab in Milton Keynes and Alex Alderton, Roberto Amato, Jeffrey Barrett, Sonia Goncalves, Ewan Harrison, David K. Jackson, Ian Johnston, Dominic Kwiatkowski, Cordelia Langford, John Sillitoe on behalf of the Wellcome Sanger |

| | | | |
|---|---|---|---|
| EPI_ISL_2446745 | | Consortium | Institute COVID-19 Surveillance Team |
| EPI_ISL_2446746, EPI_ISL_2446747, EPI_ISL_2446748 | Lighthouse Lab in Alderley Park | Wellcome Sanger Institute for the COVID-19 Genomics UK (COG-UK) Consortium | Jacquelyn Wynn, Mairead Hyland, The Lighthouse Lab in Alderley Park and Alex Alderton, Roberto Amato, Jeffrey Barrett, Sonia Goncalves, Ewan Harrison, David K. Jackson, Ian Johnston, Dominic Kwiatkowski, Cordelia Langford, John Sillitoe on behalf of the Wellcome Sanger Institute COVID-19 Surveillance Team |
| EPI_ISL_2446749, EPI_ISL_2446750 | Lighthouse Lab in Milton Keynes | Wellcome Sanger Institute for the COVID-19 Genomics UK (COG-UK) Consortium | The Lighthouse Lab in Milton Keynes and Alex Alderton, Roberto Amato, Jeffrey Barrett, Sonia Goncalves, Ewan Harrison, David K. Jackson, Ian Johnston, Dominic Kwiatkowski, Cordelia Langford, John Sillitoe on behalf of the Wellcome Sanger Institute COVID-19 Surveillance Team |
| EPI_ISL_2446751 | Lighthouse Lab in Alderley Park | Wellcome Sanger Institute for the COVID-19 Genomics UK (COG-UK) Consortium | Jacquelyn Wynn, Mairead Hyland, The Lighthouse Lab in Alderley Park and Alex Alderton, Roberto Amato, Jeffrey Barrett, Sonia Goncalves, Ewan Harrison, David K. Jackson, Ian Johnston, Dominic Kwiatkowski, Cordelia Langford, John Sillitoe on behalf of the Wellcome Sanger Institute COVID-19 Surveillance Team |
| EPI_ISL_2446754, EPI_ISL_2446755, EPI_ISL_2446756 | Lighthouse Lab in Milton Keynes | Wellcome Sanger Institute for the COVID-19 Genomics UK (COG-UK) Consortium | The Lighthouse Lab in Milton Keynes and Alex Alderton, Roberto Amato, Jeffrey Barrett, Sonia Goncalves, Ewan Harrison, David K. Jackson, Ian Johnston, Dominic Kwiatkowski, Cordelia Langford, John Sillitoe on behalf of the Wellcome Sanger Institute COVID-19 Surveillance Team |
| EPI_ISL_2446757, EPI_ISL_2446758 | Lighthouse Lab in Alderley Park | Wellcome Sanger Institute for the COVID-19 Genomics UK (COG-UK) Consortium | Jacquelyn Wynn, Mairead Hyland, The Lighthouse Lab in Alderley Park and Alex Alderton, Roberto Amato, Jeffrey Barrett, Sonia Goncalves, Ewan Harrison, David K. Jackson, Ian Johnston, Dominic Kwiatkowski, Cordelia Langford, John Sillitoe on behalf of the Wellcome Sanger Institute COVID-19 Surveillance Team |
| EPI_ISL_2446760, EPI_ISL_2446761, EPI_ISL_2446762 | Lighthouse Lab in Milton Keynes | Wellcome Sanger Institute for the COVID-19 Genomics UK (COG-UK) Consortium | The Lighthouse Lab in Milton Keynes and Alex Alderton, Roberto Amato, Jeffrey Barrett, Sonia Goncalves, Ewan Harrison, David K. Jackson, Ian Johnston, Dominic Kwiatkowski, Cordelia Langford, John Sillitoe on behalf of the Wellcome Sanger Institute COVID-19 Surveillance Team |
| EPI_ISL_2446763 | Lighthouse Lab in Alderley Park | Wellcome Sanger Institute for the COVID-19 Genomics UK (COG-UK) Consortium | Jacquelyn Wynn, Mairead Hyland, The Lighthouse Lab in Alderley Park and Alex Alderton, Roberto Amato, Jeffrey Barrett, Sonia Goncalves, Ewan Harrison, David K. Jackson, Ian Johnston, Dominic Kwiatkowski, Cordelia Langford, John Sillitoe on behalf of the Wellcome Sanger Institute COVID-19 Surveillance Team |
| EPI_ISL_2446764, EPI_ISL_2446765, EPI_ISL_2446766, EPI_ISL_2446767, EPI_ISL_2446768, EPI_ISL_2446770 | Lighthouse Lab in Milton Keynes | Wellcome Sanger Institute for the COVID-19 Genomics UK (COG-UK) Consortium | The Lighthouse Lab in Milton Keynes and Alex Alderton, Roberto Amato, Jeffrey Barrett, Sonia Goncalves, Ewan Harrison, David K. Jackson, Ian Johnston, Dominic Kwiatkowski, Cordelia Langford, John Sillitoe on behalf of the Wellcome Sanger Institute COVID-19 Surveillance Team |
| EPI_ISL_2446771 | Lighthouse Lab in Alderley Park | Wellcome Sanger Institute for the COVID-19 Genomics UK (COG-UK) Consortium | Jacquelyn Wynn, Mairead Hyland, The Lighthouse Lab in Alderley Park and Alex Alderton, Roberto Amato, Jeffrey Barrett, Sonia Goncalves, Ewan Harrison, David K. Jackson, Ian Johnston, Dominic Kwiatkowski, Cordelia Langford, John Sillitoe on behalf of the Wellcome Sanger Institute COVID-19 Surveillance Team |
| EPI_ISL_2446772, EPI_ISL_2446773, EPI_ISL_2446774, EPI_ISL_2446775, EPI_ISL_2446776 | Lighthouse Lab in Milton Keynes | Wellcome Sanger Institute for the COVID-19 Genomics UK (COG-UK) Consortium | The Lighthouse Lab in Milton Keynes and Alex Alderton, Roberto Amato, Jeffrey Barrett, Sonia Goncalves, Ewan Harrison, David K. Jackson, Ian Johnston, Dominic Kwiatkowski, Cordelia Langford, John Sillitoe on behalf of the Wellcome Sanger Institute COVID-19 Surveillance Team |
| EPI_ISL_2446777 | Lighthouse Lab in Alderley Park | Wellcome Sanger Institute for the COVID-19 Genomics UK (COG-UK) Consortium | Jacquelyn Wynn, Mairead Hyland, The Lighthouse Lab in Alderley Park and Alex Alderton, Roberto Amato, Jeffrey Barrett, Sonia Goncalves, Ewan Harrison, David K. Jackson, Ian Johnston, Dominic Kwiatkowski, Cordelia Langford, John Sillitoe on behalf of the Wellcome Sanger Institute COVID-19 Surveillance Team |
| EPI_ISL_2446778, EPI_ISL_2446780 | Lighthouse Lab in Milton Keynes | Wellcome Sanger Institute for the COVID-19 Genomics UK (COG-UK) Consortium | The Lighthouse Lab in Milton Keynes and Alex Alderton, Roberto Amato, Jeffrey Barrett, Sonia Goncalves, Ewan Harrison, David K. Jackson, Ian Johnston, Dominic Kwiatkowski, Cordelia Langford, John Sillitoe on behalf of the Wellcome Sanger Institute COVID-19 Surveillance Team |
| EPI_ISL_2446781, EPI_ISL_2446782 | Lighthouse Lab in Alderley Park | Wellcome Sanger Institute for the COVID-19 Genomics UK (COG-UK) Consortium | Jacquelyn Wynn, Mairead Hyland, The Lighthouse Lab in Alderley Park and Alex Alderton, Roberto Amato, Jeffrey Barrett, Sonia Goncalves, Ewan Harrison, David K. Jackson, Ian Johnston, Dominic Kwiatkowski, Cordelia Langford, John Sillitoe on behalf of the Wellcome Sanger Institute COVID-19 Surveillance Team |
| EPI_ISL_2446783 | Lighthouse Lab in Milton Keynes | Wellcome Sanger Institute for the COVID-19 Genomics UK (COG-UK) Consortium | The Lighthouse Lab in Milton Keynes and Alex Alderton, Roberto Amato, Jeffrey Barrett, Sonia Goncalves, Ewan Harrison, David K. Jackson, Ian Johnston, Dominic Kwiatkowski, Cordelia Langford, John Sillitoe on behalf of the Wellcome Sanger Institute COVID-19 Surveillance Team |
| EPI_ISL_2446784, EPI_ISL_2446785 | Lighthouse Lab in Alderley Park | Wellcome Sanger Institute for the COVID-19 Genomics UK (COG-UK) Consortium | Jacquelyn Wynn, Mairead Hyland, The Lighthouse Lab in Alderley Park and Alex Alderton, Roberto Amato, Jeffrey Barrett, Sonia Goncalves, Ewan Harrison, David K. Jackson, Ian Johnston, Dominic Kwiatkowski, Cordelia Langford, John Sillitoe on behalf of the Wellcome Sanger Institute COVID-19 Surveillance Team |
| EPI_ISL_2446786 | Lighthouse Lab in Milton Keynes | Wellcome Sanger Institute for the COVID-19 Genomics UK (COG-UK) Consortium | The Lighthouse Lab in Milton Keynes and Alex Alderton, Roberto Amato, Jeffrey Barrett, Sonia Goncalves, Ewan Harrison, David K. Jackson, Ian Johnston, Dominic Kwiatkowski, Cordelia Langford, John Sillitoe on behalf of the Wellcome Sanger Institute COVID-19 Surveillance Team |
| EPI_ISL_2446787, EPI_ISL_2446789 | Lighthouse Lab in Alderley Park | Wellcome Sanger Institute for the COVID-19 Genomics UK (COG-UK) Consortium | Jacquelyn Wynn, Mairead Hyland, The Lighthouse Lab in Alderley Park and Alex Alderton, Roberto Amato, Jeffrey Barrett, Sonia Goncalves, Ewan Harrison, David K. Jackson, Ian Johnston, Dominic Kwiatkowski, Cordelia Langford, John Sillitoe on behalf of the Wellcome Sanger Institute COVID-19 Surveillance Team |
| EPI_ISL_2446790, EPI_ISL_2446792, EPI_ISL_2446793 | Lighthouse Lab in Milton Keynes | Wellcome Sanger Institute for the COVID-19 Genomics UK (COG-UK) Consortium | The Lighthouse Lab in Milton Keynes and Alex Alderton, Roberto Amato, Jeffrey Barrett, Sonia Goncalves, Ewan Harrison, David K. Jackson, Ian Johnston, Dominic Kwiatkowski, Cordelia Langford, John Sillitoe on behalf of the Wellcome Sanger Institute COVID-19 Surveillance Team |
| EPI_ISL_2446794 | Lighthouse Lab in Alderley Park | Wellcome Sanger Institute for the COVID-19 Genomics UK (COG-UK) Consortium | Jacquelyn Wynn, Mairead Hyland, The Lighthouse Lab in Alderley Park and Alex Alderton, Roberto Amato, Jeffrey Barrett, Sonia Goncalves, Ewan Harrison, David K. Jackson, Ian Johnston, Dominic Kwiatkowski, Cordelia Langford, John Sillitoe on behalf of the Wellcome Sanger Institute COVID-19 Surveillance Team |
| EPI_ISL_2446795, EPI_ISL_2446796 | Lighthouse Lab in Milton Keynes | Wellcome Sanger Institute for the COVID-19 Genomics UK (COG-UK) Consortium | The Lighthouse Lab in Milton Keynes and Alex Alderton, Roberto Amato, Jeffrey Barrett, Sonia Goncalves, Ewan Harrison, David K. Jackson, Ian Johnston, Dominic Kwiatkowski, Cordelia Langford, John Sillitoe on behalf of the Wellcome Sanger Institute COVID-19 Surveillance Team |
| EPI_ISL_2446797, EPI_ISL_2446798 | Lighthouse Lab in Alderley Park | Wellcome Sanger Institute for the COVID-19 Genomics UK (COG-UK) Consortium | Jacquelyn Wynn, Mairead Hyland, The Lighthouse Lab in Alderley Park and Alex Alderton, Roberto Amato, Jeffrey Barrett, Sonia Goncalves, Ewan Harrison, David K. Jackson, Ian Johnston, Dominic Kwiatkowski, Cordelia Langford, John Sillitoe on behalf of the Wellcome Sanger Institute COVID-19 Surveillance Team |
| EPI_ISL_2446800, EPI_ISL_2446801, EPI_ISL_2446802, EPI_ISL_2446803, EPI_ISL_2446804, EPI_ISL_2446805, EPI_ISL_2446806, EPI_ISL_2446807, EPI_ISL_2446808 | Lighthouse Lab in Milton Keynes | Wellcome Sanger Institute for the COVID-19 Genomics UK (COG-UK) Consortium | The Lighthouse Lab in Milton Keynes and Alex Alderton, Roberto Amato, Jeffrey Barrett, Sonia Goncalves, Ewan Harrison, David K. Jackson, Ian Johnston, Dominic Kwiatkowski, Cordelia Langford, John Sillitoe on behalf of the Wellcome Sanger Institute COVID-19 Surveillance Team |
| EPI_ISL_2446810 | Lighthouse Lab in Alderley Park | Wellcome Sanger Institute for the COVID-19 Genomics UK (COG-UK) Consortium | Jacquelyn Wynn, Mairead Hyland, The Lighthouse Lab in Alderley Park and Alex Alderton, Roberto Amato, Jeffrey Barrett, Sonia Goncalves, Ewan Harrison, David K. Jackson, Ian Johnston, Dominic Kwiatkowski, Cordelia Langford, John Sillitoe on behalf of the Wellcome Sanger Institute COVID-19 Surveillance Team |
| EPI_ISL_2446811 | Lighthouse Lab in Milton Keynes | Wellcome Sanger Institute for the COVID-19 Genomics UK (COG-UK) Consortium | The Lighthouse Lab in Milton Keynes and Alex Alderton, Roberto Amato, Jeffrey Barrett, Sonia Goncalves, Ewan Harrison, David K. Jackson, Ian Johnston, Dominic Kwiatkowski, Cordelia Langford, John Sillitoe on behalf of the Wellcome Sanger Institute COVID-19 Surveillance Team |
| EPI_ISL_2446812 | Lighthouse Lab in Alderley Park | Wellcome Sanger Institute for the COVID-19 Genomics UK (COG-UK) Consortium | Jacquelyn Wynn, Mairead Hyland, The Lighthouse Lab in Alderley Park and Alex Alderton, Roberto Amato, Jeffrey Barrett, Sonia Goncalves, Ewan Harrison, David K. Jackson, Ian Johnston, Dominic Kwiatkowski, Cordelia Langford, John Sillitoe on behalf of the Wellcome Sanger Institute COVID-19 Surveillance Team |
| EPI_ISL_2446813, EPI_ISL_2446814, EPI_ISL_2446815, EPI_ISL_2446817 | Lighthouse Lab in Milton Keynes | Wellcome Sanger Institute for the COVID-19 Genomics UK (COG-UK) Consortium | The Lighthouse Lab in Milton Keynes and Alex Alderton, Roberto Amato, Jeffrey Barrett, Sonia Goncalves, Ewan Harrison, David K. Jackson, Ian Johnston, Dominic Kwiatkowski, Cordelia Langford, John Sillitoe on behalf of the Wellcome Sanger Institute COVID-19 Surveillance Team |
| EPI_ISL_2446818, EPI_ISL_2446819 | Lighthouse Lab in Alderley Park | Wellcome Sanger Institute for the COVID-19 Genomics UK (COG-UK) Consortium | Jacquelyn Wynn, Mairead Hyland, The Lighthouse Lab in Alderley Park and Alex Alderton, Roberto Amato, Jeffrey Barrett, Sonia Goncalves, Ewan Harrison, David K. Jackson, Ian Johnston, Dominic Kwiatkowski, Cordelia Langford, John Sillitoe on behalf of the Wellcome Sanger Institute COVID-19 Surveillance Team |
| EPI_ISL_2446820, EPI_ISL_2446821, EPI_ISL_2446823 | Lighthouse Lab in Milton Keynes | Wellcome Sanger Institute for the COVID-19 Genomics UK (COG-UK) Consortium | The Lighthouse Lab in Milton Keynes and Alex Alderton, Roberto Amato, Jeffrey Barrett, Sonia Goncalves, Ewan Harrison, David K. Jackson, Ian Johnston, Dominic Kwiatkowski, Cordelia Langford, John Sillitoe on behalf of the Wellcome Sanger Institute COVID-19 Surveillance Team |
| EPI_ISL_2446824 | Lighthouse Lab in Alderley Park | Wellcome Sanger Institute for the COVID-19 Genomics UK (COG-UK) Consortium | Jacquelyn Wynn, Mairead Hyland, The Lighthouse Lab in Alderley Park and Alex Alderton, Roberto Amato, Jeffrey Barrett, Sonia Goncalves, Ewan Harrison, David K. Jackson, Ian Johnston, Dominic Kwiatkowski, Cordelia Langford, John Sillitoe on behalf of the Wellcome Sanger Institute COVID-19 Surveillance Team |
| EPI_ISL_2446825 | Lighthouse Lab in Milton Keynes | Wellcome Sanger Institute for the COVID-19 Genomics UK (COG-UK) Consortium | The Lighthouse Lab in Milton Keynes and Alex Alderton, Roberto Amato, Jeffrey Barrett, Sonia Goncalves, Ewan Harrison, David K. Jackson, Ian Johnston, Dominic Kwiatkowski, Cordelia Langford, John Sillitoe on behalf of the Wellcome Sanger Institute COVID-19 Surveillance Team |
| EPI_ISL_2446826 | Lighthouse Lab in Alderley Park | Wellcome Sanger Institute for the COVID-19 Genomics UK (COG-UK) Consortium | Jacquelyn Wynn, Mairead Hyland, The Lighthouse Lab in Alderley Park and Alex Alderton, Roberto Amato, Jeffrey Barrett, Sonia Goncalves, Ewan Harrison, David K. Jackson, Ian Johnston, Dominic Kwiatkowski, Cordelia Langford, John Sillitoe on behalf of the Wellcome Sanger Institute COVID-19 Surveillance Team |
| EPI_ISL_2446827, EPI_ISL_2446828, EPI_ISL_2446829 | Lighthouse Lab in Milton Keynes | Wellcome Sanger Institute for the COVID-19 Genomics UK (COG-UK) Consortium | The Lighthouse Lab in Milton Keynes and Alex Alderton, Roberto Amato, Jeffrey Barrett, Sonia Goncalves, Ewan Harrison, David K. Jackson, Ian Johnston, Dominic Kwiatkowski, Cordelia Langford, John Sillitoe on behalf of the Wellcome Sanger Institute COVID-19 Surveillance Team |
| EPI_ISL_2446830 | Lighthouse Lab in Alderley Park | Wellcome Sanger Institute for the COVID-19 Genomics UK (COG-UK) Consortium | Jacquelyn Wynn, Mairead Hyland, The Lighthouse Lab in Alderley Park and Alex Alderton, Roberto Amato, Jeffrey Barrett, Sonia Goncalves, Ewan Harrison, David K. Jackson, Ian Johnston, Dominic Kwiatkowski, Cordelia Langford, John Sillitoe on behalf of the Wellcome Sanger Institute COVID-19 Surveillance Team |
| EPI_ISL_2446831 | Lighthouse Lab in Milton Keynes | Wellcome Sanger Institute for the COVID-19 Genomics UK (COG-UK) Consortium | The Lighthouse Lab in Milton Keynes and Alex Alderton, Roberto Amato, Jeffrey Barrett, Sonia Goncalves, Ewan Harrison, David K. Jackson, Ian Johnston, Dominic Kwiatkowski, Cordelia Langford, John Sillitoe on behalf of the Wellcome Sanger Institute COVID-19 Surveillance Team |
| EPI_ISL_2446832 | Lighthouse Lab in Alderley Park | Wellcome Sanger Institute for the COVID-19 Genomics UK (COG-UK) Consortium | Jacquelyn Wynn, Mairead Hyland, The Lighthouse Lab in Alderley Park and Alex Alderton, Roberto Amato, Jeffrey Barrett, Sonia Goncalves, Ewan Harrison, David K. Jackson, Ian Johnston, Dominic Kwiatkowski, Cordelia Langford, John Sillitoe on behalf of the Wellcome Sanger Institute COVID-19 Surveillance Team |
| EPI_ISL_2446833 | Lighthouse Lab in Milton Keynes | Wellcome Sanger Institute for the COVID-19 Genomics UK (COG-UK) | The Lighthouse Lab in Milton Keynes and Alex Alderton, Roberto Amato, Jeffrey Barrett, Sonia Goncalves, Ewan Harrison, David K. Jackson, Ian Johnston, Dominic Kwiatkowski, Cordelia Langford, John Sillitoe on behalf of the Wellcome Sanger |

| | | | |
|---|---|---|---|
| | | Consortium | Institute COVID-19 Surveillance Team |
| EPI_ISL_2446834 | Lighthouse Lab in Alderley Park | Wellcome Sanger Institute for the COVID-19 Genomics UK (COG-UK) Consortium | Jacquelyn Wynn, Mairead Hyland, The Lighthouse Lab in Alderley Park and Alex Alderton, Roberto Amato, Jeffrey Barrett, Sonia Goncalves, Ewan Harrison, David K. Jackson, Ian Johnston, Dominic Kwiatkowski, Cordelia Langford, John Sillitoe on behalf of the Wellcome Sanger Institute COVID-19 Surveillance Team |
| EPI_ISL_2446835 | Lighthouse Lab in Milton Keynes | Wellcome Sanger Institute for the COVID-19 Genomics UK (COG-UK) Consortium | The Lighthouse Lab in Milton Keynes and Alex Alderton, Roberto Amato, Jeffrey Barrett, Sonia Goncalves, Ewan Harrison, David K. Jackson, Ian Johnston, Dominic Kwiatkowski, Cordelia Langford, John Sillitoe on behalf of the Wellcome Sanger Institute COVID-19 Surveillance Team |
| EPI_ISL_2446837 | Lighthouse Lab in Alderley Park | Wellcome Sanger Institute for the COVID-19 Genomics UK (COG-UK) Consortium | Jacquelyn Wynn, Mairead Hyland, The Lighthouse Lab in Alderley Park and Alex Alderton, Roberto Amato, Jeffrey Barrett, Sonia Goncalves, Ewan Harrison, David K. Jackson, Ian Johnston, Dominic Kwiatkowski, Cordelia Langford, John Sillitoe on behalf of the Wellcome Sanger Institute COVID-19 Surveillance Team |
| EPI_ISL_2446838 | Lighthouse Lab in Milton Keynes | Wellcome Sanger Institute for the COVID-19 Genomics UK (COG-UK) Consortium | The Lighthouse Lab in Milton Keynes and Alex Alderton, Roberto Amato, Jeffrey Barrett, Sonia Goncalves, Ewan Harrison, David K. Jackson, Ian Johnston, Dominic Kwiatkowski, Cordelia Langford, John Sillitoe on behalf of the Wellcome Sanger Institute COVID-19 Surveillance Team |
| EPI_ISL_2446839 | Lighthouse Lab in Alderley Park | Wellcome Sanger Institute for the COVID-19 Genomics UK (COG-UK) Consortium | Jacquelyn Wynn, Mairead Hyland, The Lighthouse Lab in Alderley Park and Alex Alderton, Roberto Amato, Jeffrey Barrett, Sonia Goncalves, Ewan Harrison, David K. Jackson, Ian Johnston, Dominic Kwiatkowski, Cordelia Langford, John Sillitoe on behalf of the Wellcome Sanger Institute COVID-19 Surveillance Team |
| EPI_ISL_2446840 | Lighthouse Lab in Milton Keynes | Wellcome Sanger Institute for the COVID-19 Genomics UK (COG-UK) Consortium | The Lighthouse Lab in Milton Keynes and Alex Alderton, Roberto Amato, Jeffrey Barrett, Sonia Goncalves, Ewan Harrison, David K. Jackson, Ian Johnston, Dominic Kwiatkowski, Cordelia Langford, John Sillitoe on behalf of the Wellcome Sanger Institute COVID-19 Surveillance Team |
| EPI_ISL_2446841, EPI_ISL_2446842 | Lighthouse Lab in Alderley Park | Wellcome Sanger Institute for the COVID-19 Genomics UK (COG-UK) Consortium | Jacquelyn Wynn, Mairead Hyland, The Lighthouse Lab in Alderley Park and Alex Alderton, Roberto Amato, Jeffrey Barrett, Sonia Goncalves, Ewan Harrison, David K. Jackson, Ian Johnston, Dominic Kwiatkowski, Cordelia Langford, John Sillitoe on behalf of the Wellcome Sanger Institute COVID-19 Surveillance Team |
| EPI_ISL_2446843, EPI_ISL_2446844 | Lighthouse Lab in Glasgow | Wellcome Sanger Institute for the COVID-19 Genomics UK (COG-UK) Consortium | Harper VanSteenhouse, Yumi Kasai, David Gray, Carol Clugston, Anna Dominiczak and Alex Alderton, Roberto Amato, Jeffrey Barrett, Sonia Goncalves, Ewan Harrison, David K. Jackson, Ian Johnston, Dominic Kwiatkowski, Cordelia Langford, John Sillitoe on behalf of the Wellcome Sanger Institute COVID-19 Surveillance Team |
| EPI_ISL_2446845, EPI_ISL_2446846, EPI_ISL_2446847 | Lighthouse Lab in Milton Keynes | Wellcome Sanger Institute for the COVID-19 Genomics UK (COG-UK) Consortium | The Lighthouse Lab in Milton Keynes and Alex Alderton, Roberto Amato, Jeffrey Barrett, Sonia Goncalves, Ewan Harrison, David K. Jackson, Ian Johnston, Dominic Kwiatkowski, Cordelia Langford, John Sillitoe on behalf of the Wellcome Sanger Institute COVID-19 Surveillance Team |
| EPI_ISL_2446848 | Lighthouse Lab in Alderley Park | Wellcome Sanger Institute for the COVID-19 Genomics UK (COG-UK) Consortium | Jacquelyn Wynn, Mairead Hyland, The Lighthouse Lab in Alderley Park and Alex Alderton, Roberto Amato, Jeffrey Barrett, Sonia Goncalves, Ewan Harrison, David K. Jackson, Ian Johnston, Dominic Kwiatkowski, Cordelia Langford, John Sillitoe on behalf of the Wellcome Sanger Institute COVID-19 Surveillance Team |
| EPI_ISL_2446849 | Lighthouse Lab in Milton Keynes | Wellcome Sanger Institute for the COVID-19 Genomics UK (COG-UK) Consortium | The Lighthouse Lab in Milton Keynes and Alex Alderton, Roberto Amato, Jeffrey Barrett, Sonia Goncalves, Ewan Harrison, David K. Jackson, Ian Johnston, Dominic Kwiatkowski, Cordelia Langford, John Sillitoe on behalf of the Wellcome Sanger Institute COVID-19 Surveillance Team |
| EPI_ISL_2446850 | Lighthouse Lab in Alderley Park | Wellcome Sanger Institute for the COVID-19 Genomics UK (COG-UK) Consortium | Jacquelyn Wynn, Mairead Hyland, The Lighthouse Lab in Alderley Park and Alex Alderton, Roberto Amato, Jeffrey Barrett, Sonia Goncalves, Ewan Harrison, David K. Jackson, Ian Johnston, Dominic Kwiatkowski, Cordelia Langford, John Sillitoe on behalf of the Wellcome Sanger Institute COVID-19 Surveillance Team |
| EPI_ISL_2446854 | Lighthouse Lab in Glasgow | Wellcome Sanger Institute for the COVID-19 Genomics UK (COG-UK) Consortium | Harper VanSteenhouse, Yumi Kasai, David Gray, Carol Clugston, Anna Dominiczak and Alex Alderton, Roberto Amato, Jeffrey Barrett, Sonia Goncalves, Ewan Harrison, David K. Jackson, Ian Johnston, Dominic Kwiatkowski, Cordelia Langford, John Sillitoe on behalf of the Wellcome Sanger Institute COVID-19 Surveillance Team |
| EPI_ISL_2446855 | Lighthouse Lab in Milton Keynes | Wellcome Sanger Institute for the COVID-19 Genomics UK (COG-UK) Consortium | The Lighthouse Lab in Milton Keynes and Alex Alderton, Roberto Amato, Jeffrey Barrett, Sonia Goncalves, Ewan Harrison, David K. Jackson, Ian Johnston, Dominic Kwiatkowski, Cordelia Langford, John Sillitoe on behalf of the Wellcome Sanger Institute COVID-19 Surveillance Team |
| EPI_ISL_2446856, EPI_ISL_2446857 | Lighthouse Lab in Alderley Park | Wellcome Sanger Institute for the COVID-19 Genomics UK (COG-UK) Consortium | Jacquelyn Wynn, Mairead Hyland, The Lighthouse Lab in Alderley Park and Alex Alderton, Roberto Amato, Jeffrey Barrett, Sonia Goncalves, Ewan Harrison, David K. Jackson, Ian Johnston, Dominic Kwiatkowski, Cordelia Langford, John Sillitoe on behalf of the Wellcome Sanger Institute COVID-19 Surveillance Team |
| EPI_ISL_2446858 | Lighthouse Lab in Glasgow | Wellcome Sanger Institute for the COVID-19 Genomics UK (COG-UK) Consortium | Harper VanSteenhouse, Yumi Kasai, David Gray, Carol Clugston, Anna Dominiczak and Alex Alderton, Roberto Amato, Jeffrey Barrett, Sonia Goncalves, Ewan Harrison, David K. Jackson, Ian Johnston, Dominic Kwiatkowski, Cordelia Langford, John Sillitoe on behalf of the Wellcome Sanger Institute COVID-19 Surveillance Team |
| EPI_ISL_2446859 | Lighthouse Lab in Alderley Park | Wellcome Sanger Institute for the COVID-19 Genomics UK (COG-UK) Consortium | Jacquelyn Wynn, Mairead Hyland, The Lighthouse Lab in Alderley Park and Alex Alderton, Roberto Amato, Jeffrey Barrett, Sonia Goncalves, Ewan Harrison, David K. Jackson, Ian Johnston, Dominic Kwiatkowski, Cordelia Langford, John Sillitoe on behalf of the Wellcome Sanger Institute COVID-19 Surveillance Team |
| EPI_ISL_2446861 | Lighthouse Lab in Milton Keynes | Wellcome Sanger Institute for the COVID-19 Genomics UK (COG-UK) Consortium | The Lighthouse Lab in Milton Keynes and Alex Alderton, Roberto Amato, Jeffrey Barrett, Sonia Goncalves, Ewan Harrison, David K. Jackson, Ian Johnston, Dominic Kwiatkowski, Cordelia Langford, John Sillitoe on behalf of the Wellcome Sanger Institute COVID-19 Surveillance Team |
| EPI_ISL_2446862 | Lighthouse Lab in Alderley Park | Wellcome Sanger Institute for the COVID-19 Genomics UK (COG-UK) Consortium | Jacquelyn Wynn, Mairead Hyland, The Lighthouse Lab in Alderley Park and Alex Alderton, Roberto Amato, Jeffrey Barrett, Sonia Goncalves, Ewan Harrison, David K. Jackson, Ian Johnston, Dominic Kwiatkowski, Cordelia Langford, John Sillitoe on behalf of the Wellcome Sanger Institute COVID-19 Surveillance Team |
| EPI_ISL_2446863 | Lighthouse Lab in Milton Keynes | Wellcome Sanger Institute for the COVID-19 Genomics UK (COG-UK) Consortium | The Lighthouse Lab in Milton Keynes and Alex Alderton, Roberto Amato, Jeffrey Barrett, Sonia Goncalves, Ewan Harrison, David K. Jackson, Ian Johnston, Dominic Kwiatkowski, Cordelia Langford, John Sillitoe on behalf of the Wellcome Sanger Institute COVID-19 Surveillance Team |
| EPI_ISL_2446864 | Lighthouse Lab in Alderley Park | Wellcome Sanger Institute for the COVID-19 Genomics UK (COG-UK) Consortium | Jacquelyn Wynn, Mairead Hyland, The Lighthouse Lab in Alderley Park and Alex Alderton, Roberto Amato, Jeffrey Barrett, Sonia Goncalves, Ewan Harrison, David K. Jackson, Ian Johnston, Dominic Kwiatkowski, Cordelia Langford, John Sillitoe on behalf of the Wellcome Sanger Institute COVID-19 Surveillance Team |
| EPI_ISL_2446865, EPI_ISL_2446866 | Lighthouse Lab in Milton Keynes | Wellcome Sanger Institute for the COVID-19 Genomics UK (COG-UK) Consortium | The Lighthouse Lab in Milton Keynes and Alex Alderton, Roberto Amato, Jeffrey Barrett, Sonia Goncalves, Ewan Harrison, David K. Jackson, Ian Johnston, Dominic Kwiatkowski, Cordelia Langford, John Sillitoe on behalf of the Wellcome Sanger Institute COVID-19 Surveillance Team |
| EPI_ISL_2446869 | Lighthouse Lab in Alderley Park | Wellcome Sanger Institute for the COVID-19 Genomics UK (COG-UK) Consortium | Jacquelyn Wynn, Mairead Hyland, The Lighthouse Lab in Alderley Park and Alex Alderton, Roberto Amato, Jeffrey Barrett, Sonia Goncalves, Ewan Harrison, David K. Jackson, Ian Johnston, Dominic Kwiatkowski, Cordelia Langford, John Sillitoe on behalf of the Wellcome Sanger Institute COVID-19 Surveillance Team |
| EPI_ISL_2446870 | Lighthouse Lab in Milton Keynes | Wellcome Sanger Institute for the COVID-19 Genomics UK (COG-UK) Consortium | The Lighthouse Lab in Milton Keynes and Alex Alderton, Roberto Amato, Jeffrey Barrett, Sonia Goncalves, Ewan Harrison, David K. Jackson, Ian Johnston, Dominic Kwiatkowski, Cordelia Langford, John Sillitoe on behalf of the Wellcome Sanger Institute COVID-19 Surveillance Team |
| EPI_ISL_2446871, EPI_ISL_2446872, EPI_ISL_2446875 | Lighthouse Lab in Alderley Park | Wellcome Sanger Institute for the COVID-19 Genomics UK (COG-UK) Consortium | Jacquelyn Wynn, Mairead Hyland, The Lighthouse Lab in Alderley Park and Alex Alderton, Roberto Amato, Jeffrey Barrett, Sonia Goncalves, Ewan Harrison, David K. Jackson, Ian Johnston, Dominic Kwiatkowski, Cordelia Langford, John Sillitoe on behalf of the Wellcome Sanger Institute COVID-19 Surveillance Team |
| EPI_ISL_2446876 | Lighthouse Lab in Milton Keynes | Wellcome Sanger Institute for the COVID-19 Genomics UK (COG-UK) Consortium | The Lighthouse Lab in Milton Keynes and Alex Alderton, Roberto Amato, Jeffrey Barrett, Sonia Goncalves, Ewan Harrison, David K. Jackson, Ian Johnston, Dominic Kwiatkowski, Cordelia Langford, John Sillitoe on behalf of the Wellcome Sanger Institute COVID-19 Surveillance Team |
| EPI_ISL_2446877 | Lighthouse Lab in Alderley Park | Wellcome Sanger Institute for the COVID-19 Genomics UK (COG-UK) Consortium | Jacquelyn Wynn, Mairead Hyland, The Lighthouse Lab in Alderley Park and Alex Alderton, Roberto Amato, Jeffrey Barrett, Sonia Goncalves, Ewan Harrison, David K. Jackson, Ian Johnston, Dominic Kwiatkowski, Cordelia Langford, John Sillitoe on behalf of the Wellcome Sanger Institute COVID-19 Surveillance Team |
| EPI_ISL_2446878, EPI_ISL_2446879, EPI_ISL_2446880 | Lighthouse Lab in Milton Keynes | Wellcome Sanger Institute for the COVID-19 Genomics UK (COG-UK) Consortium | The Lighthouse Lab in Milton Keynes and Alex Alderton, Roberto Amato, Jeffrey Barrett, Sonia Goncalves, Ewan Harrison, David K. Jackson, Ian Johnston, Dominic Kwiatkowski, Cordelia Langford, John Sillitoe on behalf of the Wellcome Sanger Institute COVID-19 Surveillance Team |
| EPI_ISL_2446881, EPI_ISL_2446882 | Lighthouse Lab in Alderley Park | Wellcome Sanger Institute for the COVID-19 Genomics UK (COG-UK) Consortium | Jacquelyn Wynn, Mairead Hyland, The Lighthouse Lab in Alderley Park and Alex Alderton, Roberto Amato, Jeffrey Barrett, Sonia Goncalves, Ewan Harrison, David K. Jackson, Ian Johnston, Dominic Kwiatkowski, Cordelia Langford, John Sillitoe on behalf of the Wellcome Sanger Institute COVID-19 Surveillance Team |
| EPI_ISL_2446883, EPI_ISL_2446884, EPI_ISL_2446885 | Lighthouse Lab in Milton Keynes | Wellcome Sanger Institute for the COVID-19 Genomics UK (COG-UK) Consortium | The Lighthouse Lab in Milton Keynes and Alex Alderton, Roberto Amato, Jeffrey Barrett, Sonia Goncalves, Ewan Harrison, David K. Jackson, Ian Johnston, Dominic Kwiatkowski, Cordelia Langford, John Sillitoe on behalf of the Wellcome Sanger Institute COVID-19 Surveillance Team |
| EPI_ISL_2446886 | Lighthouse Lab in Alderley Park | Wellcome Sanger Institute for the COVID-19 Genomics UK (COG-UK) Consortium | Jacquelyn Wynn, Mairead Hyland, The Lighthouse Lab in Alderley Park and Alex Alderton, Roberto Amato, Jeffrey Barrett, Sonia Goncalves, Ewan Harrison, David K. Jackson, Ian Johnston, Dominic Kwiatkowski, Cordelia Langford, John Sillitoe on behalf of the Wellcome Sanger Institute COVID-19 Surveillance Team |
| EPI_ISL_2446887, EPI_ISL_2446888, EPI_ISL_2446889, EPI_ISL_2446890, EPI_ISL_2446891, EPI_ISL_2446892 | Lighthouse Lab in Milton Keynes | Wellcome Sanger Institute for the COVID-19 Genomics UK (COG-UK) Consortium | The Lighthouse Lab in Milton Keynes and Alex Alderton, Roberto Amato, Jeffrey Barrett, Sonia Goncalves, Ewan Harrison, David K. Jackson, Ian Johnston, Dominic Kwiatkowski, Cordelia Langford, John Sillitoe on behalf of the Wellcome Sanger Institute COVID-19 Surveillance Team |
| EPI_ISL_2446893 | Lighthouse Lab in Alderley Park | Wellcome Sanger Institute for the COVID-19 Genomics UK (COG-UK) Consortium | Jacquelyn Wynn, Mairead Hyland, The Lighthouse Lab in Alderley Park and Alex Alderton, Roberto Amato, Jeffrey Barrett, Sonia Goncalves, Ewan Harrison, David K. Jackson, Ian Johnston, Dominic Kwiatkowski, Cordelia Langford, John Sillitoe on behalf of the Wellcome Sanger Institute COVID-19 Surveillance Team |
| EPI_ISL_2446894, EPI_ISL_2446895, EPI_ISL_2446896, EPI_ISL_2446897, EPI_ISL_2446898 | Lighthouse Lab in Milton Keynes | Wellcome Sanger Institute for the COVID-19 Genomics UK (COG-UK) Consortium | The Lighthouse Lab in Milton Keynes and Alex Alderton, Roberto Amato, Jeffrey Barrett, Sonia Goncalves, Ewan Harrison, David K. Jackson, Ian Johnston, Dominic Kwiatkowski, Cordelia Langford, John Sillitoe on behalf of the Wellcome Sanger Institute COVID-19 Surveillance Team |
| EPI_ISL_2446900 | Lighthouse Lab in Glasgow | Wellcome Sanger Institute for the COVID-19 Genomics UK (COG-UK) Consortium | Harper VanSteenhouse, Yumi Kasai, David Gray, Carol Clugston, Anna Dominiczak and Alex Alderton, Roberto Amato, Jeffrey Barrett, Sonia Goncalves, Ewan Harrison, David K. Jackson, Ian Johnston, Dominic Kwiatkowski, Cordelia Langford, John Sillitoe on behalf of the Wellcome Sanger Institute COVID-19 Surveillance Team |
| EPI_ISL_2446901, EPI_ISL_2446902, EPI_ISL_2446903 | Lighthouse Lab in Milton Keynes | Wellcome Sanger Institute for the COVID-19 Genomics UK (COG-UK) Consortium | The Lighthouse Lab in Milton Keynes and Alex Alderton, Roberto Amato, Jeffrey Barrett, Sonia Goncalves, Ewan Harrison, David K. Jackson, Ian Johnston, Dominic Kwiatkowski, Cordelia Langford, John Sillitoe on behalf of the Wellcome Sanger Institute COVID-19 Surveillance Team |
| EPI_ISL_2446904 | Lighthouse Lab in Alderley Park | Wellcome Sanger Institute for the COVID-19 Genomics UK (COG-UK) Consortium | Jacquelyn Wynn, Mairead Hyland, The Lighthouse Lab in Alderley Park and Alex Alderton, Roberto Amato, Jeffrey Barrett, Sonia Goncalves, Ewan Harrison, David K. Jackson, Ian Johnston, Dominic Kwiatkowski, Cordelia Langford, John Sillitoe on behalf of the Wellcome Sanger Institute COVID-19 Surveillance Team |
| EPI_ISL_2446905, EPI_ISL_2446906, EPI_ISL_2446907, EPI_ISL_2446908, EPI_ISL_2446909, EPI_ISL_2446910, | Lighthouse Lab in Milton Keynes | Wellcome Sanger Institute for the COVID-19 Genomics UK (COG-UK) Consortium | The Lighthouse Lab in Milton Keynes and Alex Alderton, Roberto Amato, Jeffrey Barrett, Sonia Goncalves, Ewan Harrison, David K. Jackson, Ian Johnston, Dominic Kwiatkowski, Cordelia Langford, John Sillitoe on behalf of the Wellcome Sanger Institute COVID-19 Surveillance Team |

| | | | |
|---|---|---|---|
| EPI_ISL_2446911 | | | |
| EPI_ISL_2446912, EPI_ISL_2446913 | Lighthouse Lab in Alderley Park | Wellcome Sanger Institute for the COVID-19 Genomics UK (COG-UK) Consortium | Jacquelyn Wynn, Mairead Hyland, The Lighthouse Lab in Alderley Park and Alex Alderton, Roberto Amato, Jeffrey Barrett, Sonia Goncalves, Ewan Harrison, David K. Jackson, Ian Johnston, Dominic Kwiatkowski, Cordelia Langford, John Sillitoe on behalf of the Wellcome Sanger Institute COVID-19 Surveillance Team |
| EPI_ISL_2446914 | Lighthouse Lab in Milton Keynes | Wellcome Sanger Institute for the COVID-19 Genomics UK (COG-UK) Consortium | The Lighthouse Lab in Milton Keynes and Alex Alderton, Roberto Amato, Jeffrey Barrett, Sonia Goncalves, Ewan Harrison, David K. Jackson, Ian Johnston, Dominic Kwiatkowski, Cordelia Langford, John Sillitoe on behalf of the Wellcome Sanger Institute COVID-19 Surveillance Team |
| EPI_ISL_2446915 | Lighthouse Lab in Alderley Park | Wellcome Sanger Institute for the COVID-19 Genomics UK (COG-UK) Consortium | Jacquelyn Wynn, Mairead Hyland, The Lighthouse Lab in Alderley Park and Alex Alderton, Roberto Amato, Jeffrey Barrett, Sonia Goncalves, Ewan Harrison, David K. Jackson, Ian Johnston, Dominic Kwiatkowski, Cordelia Langford, John Sillitoe on behalf of the Wellcome Sanger Institute COVID-19 Surveillance Team |
| EPI_ISL_2446916 | Lighthouse Lab in Milton Keynes | Wellcome Sanger Institute for the COVID-19 Genomics UK (COG-UK) Consortium | The Lighthouse Lab in Milton Keynes and Alex Alderton, Roberto Amato, Jeffrey Barrett, Sonia Goncalves, Ewan Harrison, David K. Jackson, Ian Johnston, Dominic Kwiatkowski, Cordelia Langford, John Sillitoe on behalf of the Wellcome Sanger Institute COVID-19 Surveillance Team |
| EPI_ISL_2446917 | Lighthouse Lab in Glasgow | Wellcome Sanger Institute for the COVID-19 Genomics UK (COG-UK) Consortium | Harper VanSteenhouse, Yumi Kasai, David Gray, Carol Clugston, Anna Dominiczak and Alex Alderton, Roberto Amato, Jeffrey Barrett, Sonia Goncalves, Ewan Harrison, David K. Jackson, Ian Johnston, Dominic Kwiatkowski, Cordelia Langford, John Sillitoe on behalf of the Wellcome Sanger Institute COVID-19 Surveillance Team |
| EPI_ISL_2446918, EPI_ISL_2446919, EPI_ISL_2446921, EPI_ISL_2446922, EPI_ISL_2446923 | Lighthouse Lab in Milton Keynes | Wellcome Sanger Institute for the COVID-19 Genomics UK (COG-UK) Consortium | The Lighthouse Lab in Milton Keynes and Alex Alderton, Roberto Amato, Jeffrey Barrett, Sonia Goncalves, Ewan Harrison, David K. Jackson, Ian Johnston, Dominic Kwiatkowski, Cordelia Langford, John Sillitoe on behalf of the Wellcome Sanger Institute COVID-19 Surveillance Team |
| EPI_ISL_2446924 | Lighthouse Lab in Alderley Park | Wellcome Sanger Institute for the COVID-19 Genomics UK (COG-UK) Consortium | Jacquelyn Wynn, Mairead Hyland, The Lighthouse Lab in Alderley Park and Alex Alderton, Roberto Amato, Jeffrey Barrett, Sonia Goncalves, Ewan Harrison, David K. Jackson, Ian Johnston, Dominic Kwiatkowski, Cordelia Langford, John Sillitoe on behalf of the Wellcome Sanger Institute COVID-19 Surveillance Team |
| EPI_ISL_2446925, EPI_ISL_2446926, EPI_ISL_2446927, EPI_ISL_2446928 | Lighthouse Lab in Milton Keynes | Wellcome Sanger Institute for the COVID-19 Genomics UK (COG-UK) Consortium | The Lighthouse Lab in Milton Keynes and Alex Alderton, Roberto Amato, Jeffrey Barrett, Sonia Goncalves, Ewan Harrison, David K. Jackson, Ian Johnston, Dominic Kwiatkowski, Cordelia Langford, John Sillitoe on behalf of the Wellcome Sanger Institute COVID-19 Surveillance Team |
| EPI_ISL_2446929 | Lighthouse Lab in Alderley Park | Wellcome Sanger Institute for the COVID-19 Genomics UK (COG-UK) Consortium | Jacquelyn Wynn, Mairead Hyland, The Lighthouse Lab in Alderley Park and Alex Alderton, Roberto Amato, Jeffrey Barrett, Sonia Goncalves, Ewan Harrison, David K. Jackson, Ian Johnston, Dominic Kwiatkowski, Cordelia Langford, John Sillitoe on behalf of the Wellcome Sanger Institute COVID-19 Surveillance Team |
| EPI_ISL_2446930, EPI_ISL_2446931 | Lighthouse Lab in Milton Keynes | Wellcome Sanger Institute for the COVID-19 Genomics UK (COG-UK) Consortium | The Lighthouse Lab in Milton Keynes and Alex Alderton, Roberto Amato, Jeffrey Barrett, Sonia Goncalves, Ewan Harrison, David K. Jackson, Ian Johnston, Dominic Kwiatkowski, Cordelia Langford, John Sillitoe on behalf of the Wellcome Sanger Institute COVID-19 Surveillance Team |
| EPI_ISL_2446933, EPI_ISL_2446934, EPI_ISL_2446935, EPI_ISL_2446936 | Lighthouse Lab in Alderley Park | Wellcome Sanger Institute for the COVID-19 Genomics UK (COG-UK) Consortium | Jacquelyn Wynn, Mairead Hyland, The Lighthouse Lab in Alderley Park and Alex Alderton, Roberto Amato, Jeffrey Barrett, Sonia Goncalves, Ewan Harrison, David K. Jackson, Ian Johnston, Dominic Kwiatkowski, Cordelia Langford, John Sillitoe on behalf of the Wellcome Sanger Institute COVID-19 Surveillance Team |
| EPI_ISL_2446937, EPI_ISL_2446938, EPI_ISL_2446939, EPI_ISL_2446940 | Lighthouse Lab in Milton Keynes | Wellcome Sanger Institute for the COVID-19 Genomics UK (COG-UK) Consortium | The Lighthouse Lab in Milton Keynes and Alex Alderton, Roberto Amato, Jeffrey Barrett, Sonia Goncalves, Ewan Harrison, David K. Jackson, Ian Johnston, Dominic Kwiatkowski, Cordelia Langford, John Sillitoe on behalf of the Wellcome Sanger Institute COVID-19 Surveillance Team |
| EPI_ISL_2446941 | Lighthouse Lab in Alderley Park | Wellcome Sanger Institute for the COVID-19 Genomics UK (COG-UK) Consortium | Jacquelyn Wynn, Mairead Hyland, The Lighthouse Lab in Alderley Park and Alex Alderton, Roberto Amato, Jeffrey Barrett, Sonia Goncalves, Ewan Harrison, David K. Jackson, Ian Johnston, Dominic Kwiatkowski, Cordelia Langford, John Sillitoe on behalf of the Wellcome Sanger Institute COVID-19 Surveillance Team |
| EPI_ISL_2446942, EPI_ISL_2446943, EPI_ISL_2446944, EPI_ISL_2446945, EPI_ISL_2446946 | Lighthouse Lab in Milton Keynes | Wellcome Sanger Institute for the COVID-19 Genomics UK (COG-UK) Consortium | The Lighthouse Lab in Milton Keynes and Alex Alderton, Roberto Amato, Jeffrey Barrett, Sonia Goncalves, Ewan Harrison, David K. Jackson, Ian Johnston, Dominic Kwiatkowski, Cordelia Langford, John Sillitoe on behalf of the Wellcome Sanger Institute COVID-19 Surveillance Team |
| EPI_ISL_2446947 | Lighthouse Lab in Alderley Park | Wellcome Sanger Institute for the COVID-19 Genomics UK (COG-UK) Consortium | Jacquelyn Wynn, Mairead Hyland, The Lighthouse Lab in Alderley Park and Alex Alderton, Roberto Amato, Jeffrey Barrett, Sonia Goncalves, Ewan Harrison, David K. Jackson, Ian Johnston, Dominic Kwiatkowski, Cordelia Langford, John Sillitoe on behalf of the Wellcome Sanger Institute COVID-19 Surveillance Team |
| EPI_ISL_2446948, EPI_ISL_2446949 | Lighthouse Lab in Milton Keynes | Wellcome Sanger Institute for the COVID-19 Genomics UK (COG-UK) Consortium | The Lighthouse Lab in Milton Keynes and Alex Alderton, Roberto Amato, Jeffrey Barrett, Sonia Goncalves, Ewan Harrison, David K. Jackson, Ian Johnston, Dominic Kwiatkowski, Cordelia Langford, John Sillitoe on behalf of the Wellcome Sanger Institute COVID-19 Surveillance Team |
| EPI_ISL_2446950 | Lighthouse Lab in Alderley Park | Wellcome Sanger Institute for the COVID-19 Genomics UK (COG-UK) Consortium | Jacquelyn Wynn, Mairead Hyland, The Lighthouse Lab in Alderley Park and Alex Alderton, Roberto Amato, Jeffrey Barrett, Sonia Goncalves, Ewan Harrison, David K. Jackson, Ian Johnston, Dominic Kwiatkowski, Cordelia Langford, John Sillitoe on behalf of the Wellcome Sanger Institute COVID-19 Surveillance Team |
| EPI_ISL_2446951 | Lighthouse Lab in Milton Keynes | Wellcome Sanger Institute for the COVID-19 Genomics UK (COG-UK) Consortium | The Lighthouse Lab in Milton Keynes and Alex Alderton, Roberto Amato, Jeffrey Barrett, Sonia Goncalves, Ewan Harrison, David K. Jackson, Ian Johnston, Dominic Kwiatkowski, Cordelia Langford, John Sillitoe on behalf of the Wellcome Sanger Institute COVID-19 Surveillance Team |
| EPI_ISL_2446952 | Lighthouse Lab in Glasgow | Wellcome Sanger Institute for the COVID-19 Genomics UK (COG-UK) Consortium | Harper VanSteenhouse, Yumi Kasai, David Gray, Carol Clugston, Anna Dominiczak and Alex Alderton, Roberto Amato, Jeffrey Barrett, Sonia Goncalves, Ewan Harrison, David K. Jackson, Ian Johnston, Dominic Kwiatkowski, Cordelia Langford, John Sillitoe on behalf of the Wellcome Sanger Institute COVID-19 Surveillance Team |
| EPI_ISL_2446953 | Lighthouse Lab in Alderley Park | Wellcome Sanger Institute for the COVID-19 Genomics UK (COG-UK) Consortium | Jacquelyn Wynn, Mairead Hyland, The Lighthouse Lab in Alderley Park and Alex Alderton, Roberto Amato, Jeffrey Barrett, Sonia Goncalves, Ewan Harrison, David K. Jackson, Ian Johnston, Dominic Kwiatkowski, Cordelia Langford, John Sillitoe on behalf of the Wellcome Sanger Institute COVID-19 Surveillance Team |
| EPI_ISL_2446954, EPI_ISL_2446955, EPI_ISL_2446956 | Lighthouse Lab in Milton Keynes | Wellcome Sanger Institute for the COVID-19 Genomics UK (COG-UK) Consortium | The Lighthouse Lab in Milton Keynes and Alex Alderton, Roberto Amato, Jeffrey Barrett, Sonia Goncalves, Ewan Harrison, David K. Jackson, Ian Johnston, Dominic Kwiatkowski, Cordelia Langford, John Sillitoe on behalf of the Wellcome Sanger Institute COVID-19 Surveillance Team |
| EPI_ISL_2446957, EPI_ISL_2446958 | Lighthouse Lab in Alderley Park | Wellcome Sanger Institute for the COVID-19 Genomics UK (COG-UK) Consortium | Jacquelyn Wynn, Mairead Hyland, The Lighthouse Lab in Alderley Park and Alex Alderton, Roberto Amato, Jeffrey Barrett, Sonia Goncalves, Ewan Harrison, David K. Jackson, Ian Johnston, Dominic Kwiatkowski, Cordelia Langford, John Sillitoe on behalf of the Wellcome Sanger Institute COVID-19 Surveillance Team |
| EPI_ISL_2446960 | Lighthouse Lab in Milton Keynes | Wellcome Sanger Institute for the COVID-19 Genomics UK (COG-UK) Consortium | The Lighthouse Lab in Milton Keynes and Alex Alderton, Roberto Amato, Jeffrey Barrett, Sonia Goncalves, Ewan Harrison, David K. Jackson, Ian Johnston, Dominic Kwiatkowski, Cordelia Langford, John Sillitoe on behalf of the Wellcome Sanger Institute COVID-19 Surveillance Team |
| EPI_ISL_2446961, EPI_ISL_2446962 | Lighthouse Lab in Alderley Park | Wellcome Sanger Institute for the COVID-19 Genomics UK (COG-UK) Consortium | Jacquelyn Wynn, Mairead Hyland, The Lighthouse Lab in Alderley Park and Alex Alderton, Roberto Amato, Jeffrey Barrett, Sonia Goncalves, Ewan Harrison, David K. Jackson, Ian Johnston, Dominic Kwiatkowski, Cordelia Langford, John Sillitoe on behalf of the Wellcome Sanger Institute COVID-19 Surveillance Team |
| EPI_ISL_2446964, EPI_ISL_2446965, EPI_ISL_2446966, EPI_ISL_2446967 | Lighthouse Lab in Milton Keynes | Wellcome Sanger Institute for the COVID-19 Genomics UK (COG-UK) Consortium | The Lighthouse Lab in Milton Keynes and Alex Alderton, Roberto Amato, Jeffrey Barrett, Sonia Goncalves, Ewan Harrison, David K. Jackson, Ian Johnston, Dominic Kwiatkowski, Cordelia Langford, John Sillitoe on behalf of the Wellcome Sanger Institute COVID-19 Surveillance Team |
| EPI_ISL_2446968 | Lighthouse Lab in Alderley Park | Wellcome Sanger Institute for the COVID-19 Genomics UK (COG-UK) Consortium | Jacquelyn Wynn, Mairead Hyland, The Lighthouse Lab in Alderley Park and Alex Alderton, Roberto Amato, Jeffrey Barrett, Sonia Goncalves, Ewan Harrison, David K. Jackson, Ian Johnston, Dominic Kwiatkowski, Cordelia Langford, John Sillitoe on behalf of the Wellcome Sanger Institute COVID-19 Surveillance Team |
| EPI_ISL_2446969 | Lighthouse Lab in Milton Keynes | Wellcome Sanger Institute for the COVID-19 Genomics UK (COG-UK) Consortium | The Lighthouse Lab in Milton Keynes and Alex Alderton, Roberto Amato, Jeffrey Barrett, Sonia Goncalves, Ewan Harrison, David K. Jackson, Ian Johnston, Dominic Kwiatkowski, Cordelia Langford, John Sillitoe on behalf of the Wellcome Sanger Institute COVID-19 Surveillance Team |
| EPI_ISL_2446970 | Lighthouse Lab in Alderley Park | Wellcome Sanger Institute for the COVID-19 Genomics UK (COG-UK) Consortium | Jacquelyn Wynn, Mairead Hyland, The Lighthouse Lab in Alderley Park and Alex Alderton, Roberto Amato, Jeffrey Barrett, Sonia Goncalves, Ewan Harrison, David K. Jackson, Ian Johnston, Dominic Kwiatkowski, Cordelia Langford, John Sillitoe on behalf of the Wellcome Sanger Institute COVID-19 Surveillance Team |
| EPI_ISL_2446972, EPI_ISL_2446973 | Lighthouse Lab in Milton Keynes | Wellcome Sanger Institute for the COVID-19 Genomics UK (COG-UK) Consortium | The Lighthouse Lab in Milton Keynes and Alex Alderton, Roberto Amato, Jeffrey Barrett, Sonia Goncalves, Ewan Harrison, David K. Jackson, Ian Johnston, Dominic Kwiatkowski, Cordelia Langford, John Sillitoe on behalf of the Wellcome Sanger Institute COVID-19 Surveillance Team |
| EPI_ISL_2446974 | Lighthouse Lab in Alderley Park | Wellcome Sanger Institute for the COVID-19 Genomics UK (COG-UK) Consortium | Jacquelyn Wynn, Mairead Hyland, The Lighthouse Lab in Alderley Park and Alex Alderton, Roberto Amato, Jeffrey Barrett, Sonia Goncalves, Ewan Harrison, David K. Jackson, Ian Johnston, Dominic Kwiatkowski, Cordelia Langford, John Sillitoe on behalf of the Wellcome Sanger Institute COVID-19 Surveillance Team |
| EPI_ISL_2446975, EPI_ISL_2446976 | Lighthouse Lab in Milton Keynes | Wellcome Sanger Institute for the COVID-19 Genomics UK (COG-UK) Consortium | The Lighthouse Lab in Milton Keynes and Alex Alderton, Roberto Amato, Jeffrey Barrett, Sonia Goncalves, Ewan Harrison, David K. Jackson, Ian Johnston, Dominic Kwiatkowski, Cordelia Langford, John Sillitoe on behalf of the Wellcome Sanger Institute COVID-19 Surveillance Team |
| EPI_ISL_2446977 | Lighthouse Lab in Alderley Park | Wellcome Sanger Institute for the COVID-19 Genomics UK (COG-UK) Consortium | Jacquelyn Wynn, Mairead Hyland, The Lighthouse Lab in Alderley Park and Alex Alderton, Roberto Amato, Jeffrey Barrett, Sonia Goncalves, Ewan Harrison, David K. Jackson, Ian Johnston, Dominic Kwiatkowski, Cordelia Langford, John Sillitoe on behalf of the Wellcome Sanger Institute COVID-19 Surveillance Team |
| EPI_ISL_2446978 | Lighthouse Lab in Glasgow | Wellcome Sanger Institute for the COVID-19 Genomics UK (COG-UK) Consortium | Harper VanSteenhouse, Yumi Kasai, David Gray, Carol Clugston, Anna Dominiczak and Alex Alderton, Roberto Amato, Jeffrey Barrett, Sonia Goncalves, Ewan Harrison, David K. Jackson, Ian Johnston, Dominic Kwiatkowski, Cordelia Langford, John Sillitoe on behalf of the Wellcome Sanger Institute COVID-19 Surveillance Team |
| EPI_ISL_2446979, EPI_ISL_2446980 | Lighthouse Lab in Milton Keynes | Wellcome Sanger Institute for the COVID-19 Genomics UK (COG-UK) Consortium | The Lighthouse Lab in Milton Keynes and Alex Alderton, Roberto Amato, Jeffrey Barrett, Sonia Goncalves, Ewan Harrison, David K. Jackson, Ian Johnston, Dominic Kwiatkowski, Cordelia Langford, John Sillitoe on behalf of the Wellcome Sanger Institute COVID-19 Surveillance Team |
| EPI_ISL_2446981, EPI_ISL_2446982, EPI_ISL_2446983, EPI_ISL_2446984, EPI_ISL_2446985 | Lighthouse Lab in Alderley Park | Wellcome Sanger Institute for the COVID-19 Genomics UK (COG-UK) Consortium | Jacquelyn Wynn, Mairead Hyland, The Lighthouse Lab in Alderley Park and Alex Alderton, Roberto Amato, Jeffrey Barrett, Sonia Goncalves, Ewan Harrison, David K. Jackson, Ian Johnston, Dominic Kwiatkowski, Cordelia Langford, John Sillitoe on behalf of the Wellcome Sanger Institute COVID-19 Surveillance Team |
| EPI_ISL_2446986, EPI_ISL_2446987 | Lighthouse Lab in Milton Keynes | Wellcome Sanger Institute for the COVID-19 Genomics UK (COG-UK) Consortium | The Lighthouse Lab in Milton Keynes and Alex Alderton, Roberto Amato, Jeffrey Barrett, Sonia Goncalves, Ewan Harrison, David K. Jackson, Ian Johnston, Dominic Kwiatkowski, Cordelia Langford, John Sillitoe on behalf of the Wellcome Sanger Institute COVID-19 Surveillance Team |
| EPI_ISL_2446988 | Lighthouse Lab in Alderley Park | Wellcome Sanger Institute for the COVID-19 Genomics UK (COG-UK) Consortium | Jacquelyn Wynn, Mairead Hyland, The Lighthouse Lab in Alderley Park and Alex Alderton, Roberto Amato, Jeffrey Barrett, Sonia Goncalves, Ewan Harrison, David K. Jackson, Ian Johnston, Dominic Kwiatkowski, Cordelia Langford, John Sillitoe on behalf of the Wellcome Sanger Institute COVID-19 Surveillance Team |

| | | | |
|---|---|---|---|
| EPI_ISL_2446989 | Lighthouse Lab in Milton Keynes | Wellcome Sanger Institute for the COVID-19 Genomics UK (COG-UK) Consortium | The Lighthouse Lab in Milton Keynes and Alex Alderton, Roberto Amato, Jeffrey Barrett, Sonia Goncalves, Ewan Harrison, David K. Jackson, Ian Johnston, Dominic Kwiatkowski, Cordelia Langford, John Sillitoe on behalf of the Wellcome Sanger Institute COVID-19 Surveillance Team |
| EPI_ISL_2446990 | Lighthouse Lab in Alderley Park | Wellcome Sanger Institute for the COVID-19 Genomics UK (COG-UK) Consortium | Jacquelyn Wynn, Mairead Hyland, The Lighthouse Lab in Alderley Park and Alex Alderton, Roberto Amato, Jeffrey Barrett, Sonia Goncalves, Ewan Harrison, David K. Jackson, Ian Johnston, Dominic Kwiatkowski, Cordelia Langford, John Sillitoe on behalf of the Wellcome Sanger Institute COVID-19 Surveillance Team |
| EPI_ISL_2446991, EPI_ISL_2446992 | Lighthouse Lab in Milton Keynes | Wellcome Sanger Institute for the COVID-19 Genomics UK (COG-UK) Consortium | The Lighthouse Lab in Milton Keynes and Alex Alderton, Roberto Amato, Jeffrey Barrett, Sonia Goncalves, Ewan Harrison, David K. Jackson, Ian Johnston, Dominic Kwiatkowski, Cordelia Langford, John Sillitoe on behalf of the Wellcome Sanger Institute COVID-19 Surveillance Team |
| EPI_ISL_2446993, EPI_ISL_2446994 | Lighthouse Lab in Alderley Park | Wellcome Sanger Institute for the COVID-19 Genomics UK (COG-UK) Consortium | Jacquelyn Wynn, Mairead Hyland, The Lighthouse Lab in Alderley Park and Alex Alderton, Roberto Amato, Jeffrey Barrett, Sonia Goncalves, Ewan Harrison, David K. Jackson, Ian Johnston, Dominic Kwiatkowski, Cordelia Langford, John Sillitoe on behalf of the Wellcome Sanger Institute COVID-19 Surveillance Team |
| EPI_ISL_2446995 | Lighthouse Lab in Glasgow | Wellcome Sanger Institute for the COVID-19 Genomics UK (COG-UK) Consortium | Harper VanSteenhouse, Yumi Kasai, David Gray, Carol Clugston, Anna Dominiczak and Alex Alderton, Roberto Amato, Jeffrey Barrett, Sonia Goncalves, Ewan Harrison, David K. Jackson, Ian Johnston, Dominic Kwiatkowski, Cordelia Langford, John Sillitoe on behalf of the Wellcome Sanger Institute COVID-19 Surveillance Team |
| EPI_ISL_2446996 | Lighthouse Lab in Milton Keynes | Wellcome Sanger Institute for the COVID-19 Genomics UK (COG-UK) Consortium | The Lighthouse Lab in Milton Keynes and Alex Alderton, Roberto Amato, Jeffrey Barrett, Sonia Goncalves, Ewan Harrison, David K. Jackson, Ian Johnston, Dominic Kwiatkowski, Cordelia Langford, John Sillitoe on behalf of the Wellcome Sanger Institute COVID-19 Surveillance Team |
| EPI_ISL_2446997 | Lighthouse Lab in Alderley Park | Wellcome Sanger Institute for the COVID-19 Genomics UK (COG-UK) Consortium | Jacquelyn Wynn, Mairead Hyland, The Lighthouse Lab in Alderley Park and Alex Alderton, Roberto Amato, Jeffrey Barrett, Sonia Goncalves, Ewan Harrison, David K. Jackson, Ian Johnston, Dominic Kwiatkowski, Cordelia Langford, John Sillitoe on behalf of the Wellcome Sanger Institute COVID-19 Surveillance Team |
| EPI_ISL_2446998, EPI_ISL_2447000, EPI_ISL_2447001, EPI_ISL_2447002, EPI_ISL_2447003 | Lighthouse Lab in Milton Keynes | Wellcome Sanger Institute for the COVID-19 Genomics UK (COG-UK) Consortium | The Lighthouse Lab in Milton Keynes and Alex Alderton, Roberto Amato, Jeffrey Barrett, Sonia Goncalves, Ewan Harrison, David K. Jackson, Ian Johnston, Dominic Kwiatkowski, Cordelia Langford, John Sillitoe on behalf of the Wellcome Sanger Institute COVID-19 Surveillance Team |
| EPI_ISL_2447004 | Lighthouse Lab in Alderley Park | Wellcome Sanger Institute for the COVID-19 Genomics UK (COG-UK) Consortium | Jacquelyn Wynn, Mairead Hyland, The Lighthouse Lab in Alderley Park and Alex Alderton, Roberto Amato, Jeffrey Barrett, Sonia Goncalves, Ewan Harrison, David K. Jackson, Ian Johnston, Dominic Kwiatkowski, Cordelia Langford, John Sillitoe on behalf of the Wellcome Sanger Institute COVID-19 Surveillance Team |
| EPI_ISL_2447006 | Lighthouse Lab in Milton Keynes | Wellcome Sanger Institute for the COVID-19 Genomics UK (COG-UK) Consortium | The Lighthouse Lab in Milton Keynes and Alex Alderton, Roberto Amato, Jeffrey Barrett, Sonia Goncalves, Ewan Harrison, David K. Jackson, Ian Johnston, Dominic Kwiatkowski, Cordelia Langford, John Sillitoe on behalf of the Wellcome Sanger Institute COVID-19 Surveillance Team |
| EPI_ISL_2447007, EPI_ISL_2447008 | Lighthouse Lab in Alderley Park | Wellcome Sanger Institute for the COVID-19 Genomics UK (COG-UK) Consortium | Jacquelyn Wynn, Mairead Hyland, The Lighthouse Lab in Alderley Park and Alex Alderton, Roberto Amato, Jeffrey Barrett, Sonia Goncalves, Ewan Harrison, David K. Jackson, Ian Johnston, Dominic Kwiatkowski, Cordelia Langford, John Sillitoe on behalf of the Wellcome Sanger Institute COVID-19 Surveillance Team |
| EPI_ISL_2447009, EPI_ISL_2447010 | Lighthouse Lab in Milton Keynes | Wellcome Sanger Institute for the COVID-19 Genomics UK (COG-UK) Consortium | The Lighthouse Lab in Milton Keynes and Alex Alderton, Roberto Amato, Jeffrey Barrett, Sonia Goncalves, Ewan Harrison, David K. Jackson, Ian Johnston, Dominic Kwiatkowski, Cordelia Langford, John Sillitoe on behalf of the Wellcome Sanger Institute COVID-19 Surveillance Team |
| EPI_ISL_2447011 | Lighthouse Lab in Alderley Park | Wellcome Sanger Institute for the COVID-19 Genomics UK (COG-UK) Consortium | Jacquelyn Wynn, Mairead Hyland, The Lighthouse Lab in Alderley Park and Alex Alderton, Roberto Amato, Jeffrey Barrett, Sonia Goncalves, Ewan Harrison, David K. Jackson, Ian Johnston, Dominic Kwiatkowski, Cordelia Langford, John Sillitoe on behalf of the Wellcome Sanger Institute COVID-19 Surveillance Team |
| EPI_ISL_2447012, EPI_ISL_2447013, EPI_ISL_2447015, EPI_ISL_2447016 | Lighthouse Lab in Milton Keynes | Wellcome Sanger Institute for the COVID-19 Genomics UK (COG-UK) Consortium | The Lighthouse Lab in Milton Keynes and Alex Alderton, Roberto Amato, Jeffrey Barrett, Sonia Goncalves, Ewan Harrison, David K. Jackson, Ian Johnston, Dominic Kwiatkowski, Cordelia Langford, John Sillitoe on behalf of the Wellcome Sanger Institute COVID-19 Surveillance Team |
| EPI_ISL_2447017, EPI_ISL_2447019, EPI_ISL_2447020 | Lighthouse Lab in Alderley Park | Wellcome Sanger Institute for the COVID-19 Genomics UK (COG-UK) Consortium | Jacquelyn Wynn, Mairead Hyland, The Lighthouse Lab in Alderley Park and Alex Alderton, Roberto Amato, Jeffrey Barrett, Sonia Goncalves, Ewan Harrison, David K. Jackson, Ian Johnston, Dominic Kwiatkowski, Cordelia Langford, John Sillitoe on behalf of the Wellcome Sanger Institute COVID-19 Surveillance Team |
| EPI_ISL_2447021 | Lighthouse Lab in Milton Keynes | Wellcome Sanger Institute for the COVID-19 Genomics UK (COG-UK) Consortium | The Lighthouse Lab in Milton Keynes and Alex Alderton, Roberto Amato, Jeffrey Barrett, Sonia Goncalves, Ewan Harrison, David K. Jackson, Ian Johnston, Dominic Kwiatkowski, Cordelia Langford, John Sillitoe on behalf of the Wellcome Sanger Institute COVID-19 Surveillance Team |
| EPI_ISL_2447022, EPI_ISL_2447023 | Lighthouse Lab in Alderley Park | Wellcome Sanger Institute for the COVID-19 Genomics UK (COG-UK) Consortium | Jacquelyn Wynn, Mairead Hyland, The Lighthouse Lab in Alderley Park and Alex Alderton, Roberto Amato, Jeffrey Barrett, Sonia Goncalves, Ewan Harrison, David K. Jackson, Ian Johnston, Dominic Kwiatkowski, Cordelia Langford, John Sillitoe on behalf of the Wellcome Sanger Institute COVID-19 Surveillance Team |
| EPI_ISL_2447024 | Lighthouse Lab in Milton Keynes | Wellcome Sanger Institute for the COVID-19 Genomics UK (COG-UK) Consortium | The Lighthouse Lab in Milton Keynes and Alex Alderton, Roberto Amato, Jeffrey Barrett, Sonia Goncalves, Ewan Harrison, David K. Jackson, Ian Johnston, Dominic Kwiatkowski, Cordelia Langford, John Sillitoe on behalf of the Wellcome Sanger Institute COVID-19 Surveillance Team |
| EPI_ISL_2447025 | Lighthouse Lab in Alderley Park | Wellcome Sanger Institute for the COVID-19 Genomics UK (COG-UK) Consortium | Jacquelyn Wynn, Mairead Hyland, The Lighthouse Lab in Alderley Park and Alex Alderton, Roberto Amato, Jeffrey Barrett, Sonia Goncalves, Ewan Harrison, David K. Jackson, Ian Johnston, Dominic Kwiatkowski, Cordelia Langford, John Sillitoe on behalf of the Wellcome Sanger Institute COVID-19 Surveillance Team |
| EPI_ISL_2447026 | Lighthouse Lab in Glasgow | Wellcome Sanger Institute for the COVID-19 Genomics UK (COG-UK) Consortium | Harper VanSteenhouse, Yumi Kasai, David Gray, Carol Clugston, Anna Dominiczak and Alex Alderton, Roberto Amato, Jeffrey Barrett, Sonia Goncalves, Ewan Harrison, David K. Jackson, Ian Johnston, Dominic Kwiatkowski, Cordelia Langford, John Sillitoe on behalf of the Wellcome Sanger Institute COVID-19 Surveillance Team |
| EPI_ISL_2447027 | Lighthouse Lab in Milton Keynes | Wellcome Sanger Institute for the COVID-19 Genomics UK (COG-UK) Consortium | The Lighthouse Lab in Milton Keynes and Alex Alderton, Roberto Amato, Jeffrey Barrett, Sonia Goncalves, Ewan Harrison, David K. Jackson, Ian Johnston, Dominic Kwiatkowski, Cordelia Langford, John Sillitoe on behalf of the Wellcome Sanger Institute COVID-19 Surveillance Team |
| EPI_ISL_2447028 | Lighthouse Lab in Glasgow | Wellcome Sanger Institute for the COVID-19 Genomics UK (COG-UK) Consortium | Harper VanSteenhouse, Yumi Kasai, David Gray, Carol Clugston, Anna Dominiczak and Alex Alderton, Roberto Amato, Jeffrey Barrett, Sonia Goncalves, Ewan Harrison, David K. Jackson, Ian Johnston, Dominic Kwiatkowski, Cordelia Langford, John Sillitoe on behalf of the Wellcome Sanger Institute COVID-19 Surveillance Team |
| EPI_ISL_2447029, EPI_ISL_2447030, EPI_ISL_2447031 | Lighthouse Lab in Alderley Park | Wellcome Sanger Institute for the COVID-19 Genomics UK (COG-UK) Consortium | Jacquelyn Wynn, Mairead Hyland, The Lighthouse Lab in Alderley Park and Alex Alderton, Roberto Amato, Jeffrey Barrett, Sonia Goncalves, Ewan Harrison, David K. Jackson, Ian Johnston, Dominic Kwiatkowski, Cordelia Langford, John Sillitoe on behalf of the Wellcome Sanger Institute COVID-19 Surveillance Team |
| EPI_ISL_2447033, EPI_ISL_2447034 | Lighthouse Lab in Milton Keynes | Wellcome Sanger Institute for the COVID-19 Genomics UK (COG-UK) Consortium | The Lighthouse Lab in Milton Keynes and Alex Alderton, Roberto Amato, Jeffrey Barrett, Sonia Goncalves, Ewan Harrison, David K. Jackson, Ian Johnston, Dominic Kwiatkowski, Cordelia Langford, John Sillitoe on behalf of the Wellcome Sanger Institute COVID-19 Surveillance Team |
| EPI_ISL_2447035, EPI_ISL_2447036 | Lighthouse Lab in Alderley Park | Wellcome Sanger Institute for the COVID-19 Genomics UK (COG-UK) Consortium | Jacquelyn Wynn, Mairead Hyland, The Lighthouse Lab in Alderley Park and Alex Alderton, Roberto Amato, Jeffrey Barrett, Sonia Goncalves, Ewan Harrison, David K. Jackson, Ian Johnston, Dominic Kwiatkowski, Cordelia Langford, John Sillitoe on behalf of the Wellcome Sanger Institute COVID-19 Surveillance Team |
| EPI_ISL_2447037 | Lighthouse Lab in Milton Keynes | Wellcome Sanger Institute for the COVID-19 Genomics UK (COG-UK) Consortium | The Lighthouse Lab in Milton Keynes and Alex Alderton, Roberto Amato, Jeffrey Barrett, Sonia Goncalves, Ewan Harrison, David K. Jackson, Ian Johnston, Dominic Kwiatkowski, Cordelia Langford, John Sillitoe on behalf of the Wellcome Sanger Institute COVID-19 Surveillance Team |
| EPI_ISL_2447038 | Lighthouse Lab in Glasgow | Wellcome Sanger Institute for the COVID-19 Genomics UK (COG-UK) Consortium | Harper VanSteenhouse, Yumi Kasai, David Gray, Carol Clugston, Anna Dominiczak and Alex Alderton, Roberto Amato, Jeffrey Barrett, Sonia Goncalves, Ewan Harrison, David K. Jackson, Ian Johnston, Dominic Kwiatkowski, Cordelia Langford, John Sillitoe on behalf of the Wellcome Sanger Institute COVID-19 Surveillance Team |
| EPI_ISL_2447039 | Lighthouse Lab in Alderley Park | Wellcome Sanger Institute for the COVID-19 Genomics UK (COG-UK) Consortium | Jacquelyn Wynn, Mairead Hyland, The Lighthouse Lab in Alderley Park and Alex Alderton, Roberto Amato, Jeffrey Barrett, Sonia Goncalves, Ewan Harrison, David K. Jackson, Ian Johnston, Dominic Kwiatkowski, Cordelia Langford, John Sillitoe on behalf of the Wellcome Sanger Institute COVID-19 Surveillance Team |
| EPI_ISL_2447040 | Lighthouse Lab in Glasgow | Wellcome Sanger Institute for the COVID-19 Genomics UK (COG-UK) Consortium | Harper VanSteenhouse, Yumi Kasai, David Gray, Carol Clugston, Anna Dominiczak and Alex Alderton, Roberto Amato, Jeffrey Barrett, Sonia Goncalves, Ewan Harrison, David K. Jackson, Ian Johnston, Dominic Kwiatkowski, Cordelia Langford, John Sillitoe on behalf of the Wellcome Sanger Institute COVID-19 Surveillance Team |
| EPI_ISL_2447041 | Lighthouse Lab in Milton Keynes | Wellcome Sanger Institute for the COVID-19 Genomics UK (COG-UK) Consortium | The Lighthouse Lab in Milton Keynes and Alex Alderton, Roberto Amato, Jeffrey Barrett, Sonia Goncalves, Ewan Harrison, David K. Jackson, Ian Johnston, Dominic Kwiatkowski, Cordelia Langford, John Sillitoe on behalf of the Wellcome Sanger Institute COVID-19 Surveillance Team |
| EPI_ISL_2447042 | Lighthouse Lab in Glasgow | Wellcome Sanger Institute for the COVID-19 Genomics UK (COG-UK) Consortium | Harper VanSteenhouse, Yumi Kasai, David Gray, Carol Clugston, Anna Dominiczak and Alex Alderton, Roberto Amato, Jeffrey Barrett, Sonia Goncalves, Ewan Harrison, David K. Jackson, Ian Johnston, Dominic Kwiatkowski, Cordelia Langford, John Sillitoe on behalf of the Wellcome Sanger Institute COVID-19 Surveillance Team |
| EPI_ISL_2447043, EPI_ISL_2447044 | Lighthouse Lab in Milton Keynes | Wellcome Sanger Institute for the COVID-19 Genomics UK (COG-UK) Consortium | The Lighthouse Lab in Milton Keynes and Alex Alderton, Roberto Amato, Jeffrey Barrett, Sonia Goncalves, Ewan Harrison, David K. Jackson, Ian Johnston, Dominic Kwiatkowski, Cordelia Langford, John Sillitoe on behalf of the Wellcome Sanger Institute COVID-19 Surveillance Team |
| EPI_ISL_2447045 | Lighthouse Lab in Alderley Park | Wellcome Sanger Institute for the COVID-19 Genomics UK (COG-UK) Consortium | Jacquelyn Wynn, Mairead Hyland, The Lighthouse Lab in Alderley Park and Alex Alderton, Roberto Amato, Jeffrey Barrett, Sonia Goncalves, Ewan Harrison, David K. Jackson, Ian Johnston, Dominic Kwiatkowski, Cordelia Langford, John Sillitoe on behalf of the Wellcome Sanger Institute COVID-19 Surveillance Team |
| EPI_ISL_2447046, EPI_ISL_2447047 | Lighthouse Lab in Milton Keynes | Wellcome Sanger Institute for the COVID-19 Genomics UK (COG-UK) Consortium | The Lighthouse Lab in Milton Keynes and Alex Alderton, Roberto Amato, Jeffrey Barrett, Sonia Goncalves, Ewan Harrison, David K. Jackson, Ian Johnston, Dominic Kwiatkowski, Cordelia Langford, John Sillitoe on behalf of the Wellcome Sanger Institute COVID-19 Surveillance Team |
| EPI_ISL_2447048 | Lighthouse Lab in Glasgow | Wellcome Sanger Institute for the COVID-19 Genomics UK (COG-UK) Consortium | Harper VanSteenhouse, Yumi Kasai, David Gray, Carol Clugston, Anna Dominiczak and Alex Alderton, Roberto Amato, Jeffrey Barrett, Sonia Goncalves, Ewan Harrison, David K. Jackson, Ian Johnston, Dominic Kwiatkowski, Cordelia Langford, John Sillitoe on behalf of the Wellcome Sanger Institute COVID-19 Surveillance Team |
| EPI_ISL_2447049 | Lighthouse Lab in Alderley Park | Wellcome Sanger Institute for the COVID-19 Genomics UK (COG-UK) Consortium | Jacquelyn Wynn, Mairead Hyland, The Lighthouse Lab in Alderley Park and Alex Alderton, Roberto Amato, Jeffrey Barrett, Sonia Goncalves, Ewan Harrison, David K. Jackson, Ian Johnston, Dominic Kwiatkowski, Cordelia Langford, John Sillitoe on behalf of the Wellcome Sanger Institute COVID-19 Surveillance Team |
| EPI_ISL_2447050 | Lighthouse Lab in Milton Keynes | Wellcome Sanger Institute for the COVID-19 Genomics UK (COG-UK) Consortium | The Lighthouse Lab in Milton Keynes and Alex Alderton, Roberto Amato, Jeffrey Barrett, Sonia Goncalves, Ewan Harrison, David K. Jackson, Ian Johnston, Dominic Kwiatkowski, Cordelia Langford, John Sillitoe on behalf of the Wellcome Sanger Institute COVID-19 Surveillance Team |
| EPI_ISL_2447051, EPI_ISL_2447052 | Lighthouse Lab in Alderley Park | Wellcome Sanger Institute for the COVID-19 Genomics UK (COG-UK) Consortium | Jacquelyn Wynn, Mairead Hyland, The Lighthouse Lab in Alderley Park and Alex Alderton, Roberto Amato, Jeffrey Barrett, Sonia Goncalves, Ewan Harrison, David K. Jackson, Ian Johnston, Dominic Kwiatkowski, Cordelia Langford, John Sillitoe on behalf of the Wellcome Sanger Institute COVID-19 Surveillance Team |
| EPI_ISL_2447053 | Lighthouse Lab in Glasgow | Wellcome Sanger Institute for the COVID-19 Genomics UK (COG-UK) Consortium | Harper VanSteenhouse, Yumi Kasai, David Gray, Carol Clugston, Anna Dominiczak and Alex Alderton, Roberto Amato, Jeffrey Barrett, Sonia Goncalves, Ewan Harrison, David K. Jackson, Ian Johnston, Dominic Kwiatkowski, Cordelia Langford, John Sillitoe on behalf of the Wellcome Sanger Institute COVID-19 Surveillance Team |
| EPI_ISL_2447054, EPI_ISL_2447055, EPI_ISL_2447056 | Lighthouse Lab in Milton Keynes | Wellcome Sanger Institute for the COVID-19 Genomics UK (COG-UK) Consortium | The Lighthouse Lab in Milton Keynes and Alex Alderton, Roberto Amato, Jeffrey Barrett, Sonia Goncalves, Ewan Harrison, David K. Jackson, Ian Johnston, Dominic Kwiatkowski, Cordelia Langford, John Sillitoe on behalf of the Wellcome Sanger Institute COVID-19 Surveillance Team |
| EPI_ISL_2447057 | Lighthouse Lab in Alderley Park | Wellcome Sanger Institute for the COVID-19 Genomics UK (COG-UK) Consortium | Jacquelyn Wynn, Mairead Hyland, The Lighthouse Lab in Alderley Park and Alex Alderton, Roberto Amato, Jeffrey Barrett, Sonia Goncalves, Ewan Harrison, David K. Jackson, Ian Johnston, Dominic Kwiatkowski, Cordelia Langford, John Sillitoe on behalf of the Wellcome Sanger Institute COVID-19 Surveillance Team |
| EPI_ISL_2447058 | Lighthouse Lab in Glasgow | Wellcome Sanger Institute for the COVID-19 Genomics UK (COG-UK) Consortium | Harper VanSteenhouse, Yumi Kasai, David Gray, Carol Clugston, Anna Dominiczak and Alex Alderton, Roberto Amato, Jeffrey Barrett, Sonia Goncalves, Ewan Harrison, David K. Jackson, Ian Johnston, Dominic Kwiatkowski, Cordelia Langford, John Sillitoe on behalf of the Wellcome Sanger Institute COVID-19 Surveillance Team |

| | | | |
|---|---|---|---|
| EPI_ISL_2447059, EPI_ISL_2447060, EPI_ISL_2447061, EPI_ISL_2447062 | Lighthouse Lab in Milton Keynes | Wellcome Sanger Institute for the COVID-19 Genomics UK (COG-UK) Consortium | The Lighthouse Lab in Milton Keynes and Alex Alderton, Roberto Amato, Jeffrey Barrett, Sonia Goncalves, Ewan Harrison, David K. Jackson, Ian Johnston, Dominic Kwiatkowski, Cordelia Langford, John Sillitoe on behalf of the Wellcome Sanger Institute COVID-19 Surveillance Team |
| EPI_ISL_2447063 | Lighthouse Lab in Alderley Park | Wellcome Sanger Institute for the COVID-19 Genomics UK (COG-UK) Consortium | Jacquelyn Wynn, Mairead Hyland, The Lighthouse Lab in Alderley Park and Alex Alderton, Roberto Amato, Jeffrey Barrett, Sonia Goncalves, Ewan Harrison, David K. Jackson, Ian Johnston, Dominic Kwiatkowski, Cordelia Langford, John Sillitoe on behalf of the Wellcome Sanger Institute COVID-19 Surveillance Team |
| EPI_ISL_2447064 | Lighthouse Lab in Glasgow | Wellcome Sanger Institute for the COVID-19 Genomics UK (COG-UK) Consortium | Harper VanSteenhouse, Yumi Kasai, David Gray, Carol Clugston, Anna Dominiczak and Alex Alderton, Roberto Amato, Jeffrey Barrett, Sonia Goncalves, Ewan Harrison, David K. Jackson, Ian Johnston, Dominic Kwiatkowski, Cordelia Langford, John Sillitoe on behalf of the Wellcome Sanger Institute COVID-19 Surveillance Team |
| EPI_ISL_2447066, EPI_ISL_2447067 | Lighthouse Lab in Alderley Park | Wellcome Sanger Institute for the COVID-19 Genomics UK (COG-UK) Consortium | Jacquelyn Wynn, Mairead Hyland, The Lighthouse Lab in Alderley Park and Alex Alderton, Roberto Amato, Jeffrey Barrett, Sonia Goncalves, Ewan Harrison, David K. Jackson, Ian Johnston, Dominic Kwiatkowski, Cordelia Langford, John Sillitoe on behalf of the Wellcome Sanger Institute COVID-19 Surveillance Team |
| EPI_ISL_2447068 | Lighthouse Lab in Milton Keynes | Wellcome Sanger Institute for the COVID-19 Genomics UK (COG-UK) Consortium | The Lighthouse Lab in Milton Keynes and Alex Alderton, Roberto Amato, Jeffrey Barrett, Sonia Goncalves, Ewan Harrison, David K. Jackson, Ian Johnston, Dominic Kwiatkowski, Cordelia Langford, John Sillitoe on behalf of the Wellcome Sanger Institute COVID-19 Surveillance Team |
| EPI_ISL_2447069 | Lighthouse Lab in Alderley Park | Wellcome Sanger Institute for the COVID-19 Genomics UK (COG-UK) Consortium | Jacquelyn Wynn, Mairead Hyland, The Lighthouse Lab in Alderley Park and Alex Alderton, Roberto Amato, Jeffrey Barrett, Sonia Goncalves, Ewan Harrison, David K. Jackson, Ian Johnston, Dominic Kwiatkowski, Cordelia Langford, John Sillitoe on behalf of the Wellcome Sanger Institute COVID-19 Surveillance Team |
| EPI_ISL_2447070 | Lighthouse Lab in Glasgow | Wellcome Sanger Institute for the COVID-19 Genomics UK (COG-UK) Consortium | Harper VanSteenhouse, Yumi Kasai, David Gray, Carol Clugston, Anna Dominiczak and Alex Alderton, Roberto Amato, Jeffrey Barrett, Sonia Goncalves, Ewan Harrison, David K. Jackson, Ian Johnston, Dominic Kwiatkowski, Cordelia Langford, John Sillitoe on behalf of the Wellcome Sanger Institute COVID-19 Surveillance Team |
| EPI_ISL_2447071 | Lighthouse Lab in Alderley Park | Wellcome Sanger Institute for the COVID-19 Genomics UK (COG-UK) Consortium | Jacquelyn Wynn, Mairead Hyland, The Lighthouse Lab in Alderley Park and Alex Alderton, Roberto Amato, Jeffrey Barrett, Sonia Goncalves, Ewan Harrison, David K. Jackson, Ian Johnston, Dominic Kwiatkowski, Cordelia Langford, John Sillitoe on behalf of the Wellcome Sanger Institute COVID-19 Surveillance Team |
| EPI_ISL_2447072, EPI_ISL_2447073 | Lighthouse Lab in Milton Keynes | Wellcome Sanger Institute for the COVID-19 Genomics UK (COG-UK) Consortium | The Lighthouse Lab in Milton Keynes and Alex Alderton, Roberto Amato, Jeffrey Barrett, Sonia Goncalves, Ewan Harrison, David K. Jackson, Ian Johnston, Dominic Kwiatkowski, Cordelia Langford, John Sillitoe on behalf of the Wellcome Sanger Institute COVID-19 Surveillance Team |
| EPI_ISL_2447075, EPI_ISL_2447076 | Lighthouse Lab in Alderley Park | Wellcome Sanger Institute for the COVID-19 Genomics UK (COG-UK) Consortium | Jacquelyn Wynn, Mairead Hyland, The Lighthouse Lab in Alderley Park and Alex Alderton, Roberto Amato, Jeffrey Barrett, Sonia Goncalves, Ewan Harrison, David K. Jackson, Ian Johnston, Dominic Kwiatkowski, Cordelia Langford, John Sillitoe on behalf of the Wellcome Sanger Institute COVID-19 Surveillance Team |
| EPI_ISL_2447077, EPI_ISL_2447079 | Lighthouse Lab in Glasgow | Wellcome Sanger Institute for the COVID-19 Genomics UK (COG-UK) Consortium | Harper VanSteenhouse, Yumi Kasai, David Gray, Carol Clugston, Anna Dominiczak and Alex Alderton, Roberto Amato, Jeffrey Barrett, Sonia Goncalves, Ewan Harrison, David K. Jackson, Ian Johnston, Dominic Kwiatkowski, Cordelia Langford, John Sillitoe on behalf of the Wellcome Sanger Institute COVID-19 Surveillance Team |
| EPI_ISL_2447080 | Lighthouse Lab in Alderley Park | Wellcome Sanger Institute for the COVID-19 Genomics UK (COG-UK) Consortium | Jacquelyn Wynn, Mairead Hyland, The Lighthouse Lab in Alderley Park and Alex Alderton, Roberto Amato, Jeffrey Barrett, Sonia Goncalves, Ewan Harrison, David K. Jackson, Ian Johnston, Dominic Kwiatkowski, Cordelia Langford, John Sillitoe on behalf of the Wellcome Sanger Institute COVID-19 Surveillance Team |
| EPI_ISL_2447081 | Lighthouse Lab in Milton Keynes | Wellcome Sanger Institute for the COVID-19 Genomics UK (COG-UK) Consortium | The Lighthouse Lab in Milton Keynes and Alex Alderton, Roberto Amato, Jeffrey Barrett, Sonia Goncalves, Ewan Harrison, David K. Jackson, Ian Johnston, Dominic Kwiatkowski, Cordelia Langford, John Sillitoe on behalf of the Wellcome Sanger Institute COVID-19 Surveillance Team |
| EPI_ISL_2447082 | Lighthouse Lab in Glasgow | Wellcome Sanger Institute for the COVID-19 Genomics UK (COG-UK) Consortium | Harper VanSteenhouse, Yumi Kasai, David Gray, Carol Clugston, Anna Dominiczak and Alex Alderton, Roberto Amato, Jeffrey Barrett, Sonia Goncalves, Ewan Harrison, David K. Jackson, Ian Johnston, Dominic Kwiatkowski, Cordelia Langford, John Sillitoe on behalf of the Wellcome Sanger Institute COVID-19 Surveillance Team |
| EPI_ISL_2447083 | Lighthouse Lab in Milton Keynes | Wellcome Sanger Institute for the COVID-19 Genomics UK (COG-UK) Consortium | The Lighthouse Lab in Milton Keynes and Alex Alderton, Roberto Amato, Jeffrey Barrett, Sonia Goncalves, Ewan Harrison, David K. Jackson, Ian Johnston, Dominic Kwiatkowski, Cordelia Langford, John Sillitoe on behalf of the Wellcome Sanger Institute COVID-19 Surveillance Team |
| EPI_ISL_2447084 | Lighthouse Lab in Alderley Park | Wellcome Sanger Institute for the COVID-19 Genomics UK (COG-UK) Consortium | Jacquelyn Wynn, Mairead Hyland, The Lighthouse Lab in Alderley Park and Alex Alderton, Roberto Amato, Jeffrey Barrett, Sonia Goncalves, Ewan Harrison, David K. Jackson, Ian Johnston, Dominic Kwiatkowski, Cordelia Langford, John Sillitoe on behalf of the Wellcome Sanger Institute COVID-19 Surveillance Team |
| EPI_ISL_2447085 | Lighthouse Lab in Glasgow | Wellcome Sanger Institute for the COVID-19 Genomics UK (COG-UK) Consortium | Harper VanSteenhouse, Yumi Kasai, David Gray, Carol Clugston, Anna Dominiczak and Alex Alderton, Roberto Amato, Jeffrey Barrett, Sonia Goncalves, Ewan Harrison, David K. Jackson, Ian Johnston, Dominic Kwiatkowski, Cordelia Langford, John Sillitoe on behalf of the Wellcome Sanger Institute COVID-19 Surveillance Team |
| EPI_ISL_2447086 | Lighthouse Lab in Milton Keynes | Wellcome Sanger Institute for the COVID-19 Genomics UK (COG-UK) Consortium | The Lighthouse Lab in Milton Keynes and Alex Alderton, Roberto Amato, Jeffrey Barrett, Sonia Goncalves, Ewan Harrison, David K. Jackson, Ian Johnston, Dominic Kwiatkowski, Cordelia Langford, John Sillitoe on behalf of the Wellcome Sanger Institute COVID-19 Surveillance Team |
| EPI_ISL_2447088 | Lighthouse Lab in Alderley Park | Wellcome Sanger Institute for the COVID-19 Genomics UK (COG-UK) Consortium | Jacquelyn Wynn, Mairead Hyland, The Lighthouse Lab in Alderley Park and Alex Alderton, Roberto Amato, Jeffrey Barrett, Sonia Goncalves, Ewan Harrison, David K. Jackson, Ian Johnston, Dominic Kwiatkowski, Cordelia Langford, John Sillitoe on behalf of the Wellcome Sanger Institute COVID-19 Surveillance Team |
| EPI_ISL_2447089 | Lighthouse Lab in Glasgow | Wellcome Sanger Institute for the COVID-19 Genomics UK (COG-UK) Consortium | Harper VanSteenhouse, Yumi Kasai, David Gray, Carol Clugston, Anna Dominiczak and Alex Alderton, Roberto Amato, Jeffrey Barrett, Sonia Goncalves, Ewan Harrison, David K. Jackson, Ian Johnston, Dominic Kwiatkowski, Cordelia Langford, John Sillitoe on behalf of the Wellcome Sanger Institute COVID-19 Surveillance Team |
| EPI_ISL_2447090 | Lighthouse Lab in Alderley Park | Wellcome Sanger Institute for the COVID-19 Genomics UK (COG-UK) Consortium | Jacquelyn Wynn, Mairead Hyland, The Lighthouse Lab in Alderley Park and Alex Alderton, Roberto Amato, Jeffrey Barrett, Sonia Goncalves, Ewan Harrison, David K. Jackson, Ian Johnston, Dominic Kwiatkowski, Cordelia Langford, John Sillitoe on behalf of the Wellcome Sanger Institute COVID-19 Surveillance Team |
| EPI_ISL_2447091, EPI_ISL_2447093 | Lighthouse Lab in Milton Keynes | Wellcome Sanger Institute for the COVID-19 Genomics UK (COG-UK) Consortium | The Lighthouse Lab in Milton Keynes and Alex Alderton, Roberto Amato, Jeffrey Barrett, Sonia Goncalves, Ewan Harrison, David K. Jackson, Ian Johnston, Dominic Kwiatkowski, Cordelia Langford, John Sillitoe on behalf of the Wellcome Sanger Institute COVID-19 Surveillance Team |
| EPI_ISL_2447094 | Lighthouse Lab in Alderley Park | Wellcome Sanger Institute for the COVID-19 Genomics UK (COG-UK) Consortium | Jacquelyn Wynn, Mairead Hyland, The Lighthouse Lab in Alderley Park and Alex Alderton, Roberto Amato, Jeffrey Barrett, Sonia Goncalves, Ewan Harrison, David K. Jackson, Ian Johnston, Dominic Kwiatkowski, Cordelia Langford, John Sillitoe on behalf of the Wellcome Sanger Institute COVID-19 Surveillance Team |
| EPI_ISL_2447095, EPI_ISL_2447096 | Lighthouse Lab in Milton Keynes | Wellcome Sanger Institute for the COVID-19 Genomics UK (COG-UK) Consortium | The Lighthouse Lab in Milton Keynes and Alex Alderton, Roberto Amato, Jeffrey Barrett, Sonia Goncalves, Ewan Harrison, David K. Jackson, Ian Johnston, Dominic Kwiatkowski, Cordelia Langford, John Sillitoe on behalf of the Wellcome Sanger Institute COVID-19 Surveillance Team |
| EPI_ISL_2447097, EPI_ISL_2447099 | Lighthouse Lab in Glasgow | Wellcome Sanger Institute for the COVID-19 Genomics UK (COG-UK) Consortium | Harper VanSteenhouse, Yumi Kasai, David Gray, Carol Clugston, Anna Dominiczak and Alex Alderton, Roberto Amato, Jeffrey Barrett, Sonia Goncalves, Ewan Harrison, David K. Jackson, Ian Johnston, Dominic Kwiatkowski, Cordelia Langford, John Sillitoe on behalf of the Wellcome Sanger Institute COVID-19 Surveillance Team |
| EPI_ISL_2447100 | Lighthouse Lab in Alderley Park | Wellcome Sanger Institute for the COVID-19 Genomics UK (COG-UK) Consortium | Jacquelyn Wynn, Mairead Hyland, The Lighthouse Lab in Alderley Park and Alex Alderton, Roberto Amato, Jeffrey Barrett, Sonia Goncalves, Ewan Harrison, David K. Jackson, Ian Johnston, Dominic Kwiatkowski, Cordelia Langford, John Sillitoe on behalf of the Wellcome Sanger Institute COVID-19 Surveillance Team |
| EPI_ISL_2447101 | Lighthouse Lab in Milton Keynes | Wellcome Sanger Institute for the COVID-19 Genomics UK (COG-UK) Consortium | The Lighthouse Lab in Milton Keynes and Alex Alderton, Roberto Amato, Jeffrey Barrett, Sonia Goncalves, Ewan Harrison, David K. Jackson, Ian Johnston, Dominic Kwiatkowski, Cordelia Langford, John Sillitoe on behalf of the Wellcome Sanger Institute COVID-19 Surveillance Team |
| EPI_ISL_2447102, EPI_ISL_2447103, EPI_ISL_2447104 | Lighthouse Lab in Alderley Park | Wellcome Sanger Institute for the COVID-19 Genomics UK (COG-UK) Consortium | Jacquelyn Wynn, Mairead Hyland, The Lighthouse Lab in Alderley Park and Alex Alderton, Roberto Amato, Jeffrey Barrett, Sonia Goncalves, Ewan Harrison, David K. Jackson, Ian Johnston, Dominic Kwiatkowski, Cordelia Langford, John Sillitoe on behalf of the Wellcome Sanger Institute COVID-19 Surveillance Team |
| EPI_ISL_2447105 | Lighthouse Lab in Milton Keynes | Wellcome Sanger Institute for the COVID-19 Genomics UK (COG-UK) Consortium | The Lighthouse Lab in Milton Keynes and Alex Alderton, Roberto Amato, Jeffrey Barrett, Sonia Goncalves, Ewan Harrison, David K. Jackson, Ian Johnston, Dominic Kwiatkowski, Cordelia Langford, John Sillitoe on behalf of the Wellcome Sanger Institute COVID-19 Surveillance Team |
| EPI_ISL_2447106, EPI_ISL_2447107, EPI_ISL_2447108 | Lighthouse Lab in Alderley Park | Wellcome Sanger Institute for the COVID-19 Genomics UK (COG-UK) Consortium | Jacquelyn Wynn, Mairead Hyland, The Lighthouse Lab in Alderley Park and Alex Alderton, Roberto Amato, Jeffrey Barrett, Sonia Goncalves, Ewan Harrison, David K. Jackson, Ian Johnston, Dominic Kwiatkowski, Cordelia Langford, John Sillitoe on behalf of the Wellcome Sanger Institute COVID-19 Surveillance Team |
| EPI_ISL_2447109 | Lighthouse Lab in Milton Keynes | Wellcome Sanger Institute for the COVID-19 Genomics UK (COG-UK) Consortium | The Lighthouse Lab in Milton Keynes and Alex Alderton, Roberto Amato, Jeffrey Barrett, Sonia Goncalves, Ewan Harrison, David K. Jackson, Ian Johnston, Dominic Kwiatkowski, Cordelia Langford, John Sillitoe on behalf of the Wellcome Sanger Institute COVID-19 Surveillance Team |
| EPI_ISL_2447110 | Lighthouse Lab in Alderley Park | Wellcome Sanger Institute for the COVID-19 Genomics UK (COG-UK) Consortium | Jacquelyn Wynn, Mairead Hyland, The Lighthouse Lab in Alderley Park and Alex Alderton, Roberto Amato, Jeffrey Barrett, Sonia Goncalves, Ewan Harrison, David K. Jackson, Ian Johnston, Dominic Kwiatkowski, Cordelia Langford, John Sillitoe on behalf of the Wellcome Sanger Institute COVID-19 Surveillance Team |
| EPI_ISL_2447111 | Lighthouse Lab in Milton Keynes | Wellcome Sanger Institute for the COVID-19 Genomics UK (COG-UK) Consortium | The Lighthouse Lab in Milton Keynes and Alex Alderton, Roberto Amato, Jeffrey Barrett, Sonia Goncalves, Ewan Harrison, David K. Jackson, Ian Johnston, Dominic Kwiatkowski, Cordelia Langford, John Sillitoe on behalf of the Wellcome Sanger Institute COVID-19 Surveillance Team |
| EPI_ISL_2447112 | Lighthouse Lab in Alderley Park | Wellcome Sanger Institute for the COVID-19 Genomics UK (COG-UK) Consortium | Jacquelyn Wynn, Mairead Hyland, The Lighthouse Lab in Alderley Park and Alex Alderton, Roberto Amato, Jeffrey Barrett, Sonia Goncalves, Ewan Harrison, David K. Jackson, Ian Johnston, Dominic Kwiatkowski, Cordelia Langford, John Sillitoe on behalf of the Wellcome Sanger Institute COVID-19 Surveillance Team |
| EPI_ISL_2447113 | Lighthouse Lab in Milton Keynes | Wellcome Sanger Institute for the COVID-19 Genomics UK (COG-UK) Consortium | The Lighthouse Lab in Milton Keynes and Alex Alderton, Roberto Amato, Jeffrey Barrett, Sonia Goncalves, Ewan Harrison, David K. Jackson, Ian Johnston, Dominic Kwiatkowski, Cordelia Langford, John Sillitoe on behalf of the Wellcome Sanger Institute COVID-19 Surveillance Team |
| EPI_ISL_2447114 | Lighthouse Lab in Glasgow | Wellcome Sanger Institute for the COVID-19 Genomics UK (COG-UK) Consortium | Harper VanSteenhouse, Yumi Kasai, David Gray, Carol Clugston, Anna Dominiczak and Alex Alderton, Roberto Amato, Jeffrey Barrett, Sonia Goncalves, Ewan Harrison, David K. Jackson, Ian Johnston, Dominic Kwiatkowski, Cordelia Langford, John Sillitoe on behalf of the Wellcome Sanger Institute COVID-19 Surveillance Team |
| EPI_ISL_2447115 | Lighthouse Lab in Milton Keynes | Wellcome Sanger Institute for the COVID-19 Genomics UK (COG-UK) Consortium | The Lighthouse Lab in Milton Keynes and Alex Alderton, Roberto Amato, Jeffrey Barrett, Sonia Goncalves, Ewan Harrison, David K. Jackson, Ian Johnston, Dominic Kwiatkowski, Cordelia Langford, John Sillitoe on behalf of the Wellcome Sanger Institute COVID-19 Surveillance Team |
| EPI_ISL_2447116 | Lighthouse Lab in Glasgow | Wellcome Sanger Institute for the COVID-19 Genomics UK (COG-UK) Consortium | Harper VanSteenhouse, Yumi Kasai, David Gray, Carol Clugston, Anna Dominiczak and Alex Alderton, Roberto Amato, Jeffrey Barrett, Sonia Goncalves, Ewan Harrison, David K. Jackson, Ian Johnston, Dominic Kwiatkowski, Cordelia Langford, John Sillitoe on behalf of the Wellcome Sanger Institute COVID-19 Surveillance Team |
| EPI_ISL_2447118 | Lighthouse Lab in Alderley Park | Wellcome Sanger Institute for the COVID-19 Genomics UK (COG-UK) Consortium | Jacquelyn Wynn, Mairead Hyland, The Lighthouse Lab in Alderley Park and Alex Alderton, Roberto Amato, Jeffrey Barrett, Sonia Goncalves, Ewan Harrison, David K. Jackson, Ian Johnston, Dominic Kwiatkowski, Cordelia Langford, John Sillitoe on behalf of the Wellcome Sanger Institute COVID-19 Surveillance Team |
| EPI_ISL_2447119 | Lighthouse Lab in Milton Keynes | Wellcome Sanger Institute for the COVID-19 Genomics UK (COG-UK) Consortium | The Lighthouse Lab in Milton Keynes and Alex Alderton, Roberto Amato, Jeffrey Barrett, Sonia Goncalves, Ewan Harrison, David K. Jackson, Ian Johnston, Dominic Kwiatkowski, Cordelia Langford, John Sillitoe on behalf of the Wellcome Sanger Institute COVID-19 Surveillance Team |
| EPI_ISL_2447121 | Lighthouse Lab in Alderley Park | Wellcome Sanger Institute for the COVID-19 Genomics UK (COG-UK) Consortium | Jacquelyn Wynn, Mairead Hyland, The Lighthouse Lab in Alderley Park and Alex Alderton, Roberto Amato, Jeffrey Barrett, Sonia Goncalves, Ewan Harrison, David K. Jackson, Ian Johnston, Dominic Kwiatkowski, Cordelia Langford, John Sillitoe on behalf of the Wellcome Sanger Institute COVID-19 Surveillance Team |
| EPI_ISL_2447122, EPI_ISL_2447124 | Lighthouse Lab in Milton Keynes | Wellcome Sanger Institute for the COVID-19 Genomics UK (COG-UK) Consortium | The Lighthouse Lab in Milton Keynes and Alex Alderton, Roberto Amato, Jeffrey Barrett, Sonia Goncalves, Ewan Harrison, David K. Jackson, Ian Johnston, Dominic Kwiatkowski, Cordelia Langford, John Sillitoe on behalf of the Wellcome Sanger Institute COVID-19 Surveillance Team |
| EPI_ISL_2447125 | Lighthouse Lab in Glasgow | Wellcome Sanger Institute for the COVID-19 Genomics UK (COG-UK) Consortium | Harper VanSteenhouse, Yumi Kasai, David Gray, Carol Clugston, Anna Dominiczak and Alex Alderton, Roberto Amato, Jeffrey Barrett, Sonia Goncalves, Ewan Harrison, David K. Jackson, Ian Johnston, Dominic Kwiatkowski, Cordelia Langford, John Sillitoe on behalf of the Wellcome Sanger Institute COVID-19 Surveillance Team |

| | | | |
|---|---|---|---|
| EPI_ISL_2447126, EPI_ISL_2447128, EPI_ISL_2447129 | Lighthouse Lab in Alderley Park | Wellcome Sanger Institute for the COVID-19 Genomics UK (COG-UK) Consortium | Jacquelyn Wynn, Mairead Hyland, The Lighthouse Lab in Alderley Park and Alex Alderton, Roberto Amato, Jeffrey Barrett, Sonia Goncalves, Ewan Harrison, David K. Jackson, Ian Johnston, Dominic Kwiatkowski, Cordelia Langford, John Sillitoe on behalf of the Wellcome Sanger Institute COVID-19 Surveillance Team |
| EPI_ISL_2447130 | Lighthouse Lab in Glasgow | Wellcome Sanger Institute for the COVID-19 Genomics UK (COG-UK) Consortium | Harper VanSteenhouse, Yumi Kasai, David Gray, Carol Clugston, Anna Dominiczak and Alex Alderton, Roberto Amato, Jeffrey Barrett, Sonia Goncalves, Ewan Harrison, David K. Jackson, Ian Johnston, Dominic Kwiatkowski, Cordelia Langford, John Sillitoe on behalf of the Wellcome Sanger Institute COVID-19 Surveillance Team |
| EPI_ISL_2447131 | Lighthouse Lab in Milton Keynes | Wellcome Sanger Institute for the COVID-19 Genomics UK (COG-UK) Consortium | The Lighthouse Lab in Milton Keynes and Alex Alderton, Roberto Amato, Jeffrey Barrett, Sonia Goncalves, Ewan Harrison, David K. Jackson, Ian Johnston, Dominic Kwiatkowski, Cordelia Langford, John Sillitoe on behalf of the Wellcome Sanger Institute COVID-19 Surveillance Team |
| EPI_ISL_2447132 | Lighthouse Lab in Glasgow | Wellcome Sanger Institute for the COVID-19 Genomics UK (COG-UK) Consortium | Harper VanSteenhouse, Yumi Kasai, David Gray, Carol Clugston, Anna Dominiczak and Alex Alderton, Roberto Amato, Jeffrey Barrett, Sonia Goncalves, Ewan Harrison, David K. Jackson, Ian Johnston, Dominic Kwiatkowski, Cordelia Langford, John Sillitoe on behalf of the Wellcome Sanger Institute COVID-19 Surveillance Team |
| EPI_ISL_2447133, EPI_ISL_2447134, EPI_ISL_2447135 | Lighthouse Lab in Milton Keynes | Wellcome Sanger Institute for the COVID-19 Genomics UK (COG-UK) Consortium | The Lighthouse Lab in Milton Keynes and Alex Alderton, Roberto Amato, Jeffrey Barrett, Sonia Goncalves, Ewan Harrison, David K. Jackson, Ian Johnston, Dominic Kwiatkowski, Cordelia Langford, John Sillitoe on behalf of the Wellcome Sanger Institute COVID-19 Surveillance Team |
| EPI_ISL_2447136 | Lighthouse Lab in Alderley Park | Wellcome Sanger Institute for the COVID-19 Genomics UK (COG-UK) Consortium | Jacquelyn Wynn, Mairead Hyland, The Lighthouse Lab in Alderley Park and Alex Alderton, Roberto Amato, Jeffrey Barrett, Sonia Goncalves, Ewan Harrison, David K. Jackson, Ian Johnston, Dominic Kwiatkowski, Cordelia Langford, John Sillitoe on behalf of the Wellcome Sanger Institute COVID-19 Surveillance Team |
| EPI_ISL_2447137, EPI_ISL_2447138, EPI_ISL_2447139, EPI_ISL_2447140 | Lighthouse Lab in Milton Keynes | Wellcome Sanger Institute for the COVID-19 Genomics UK (COG-UK) Consortium | The Lighthouse Lab in Milton Keynes and Alex Alderton, Roberto Amato, Jeffrey Barrett, Sonia Goncalves, Ewan Harrison, David K. Jackson, Ian Johnston, Dominic Kwiatkowski, Cordelia Langford, John Sillitoe on behalf of the Wellcome Sanger Institute COVID-19 Surveillance Team |
| EPI_ISL_2447141, EPI_ISL_2447142, EPI_ISL_2447143 | Lighthouse Lab in Alderley Park | Wellcome Sanger Institute for the COVID-19 Genomics UK (COG-UK) Consortium | Jacquelyn Wynn, Mairead Hyland, The Lighthouse Lab in Alderley Park and Alex Alderton, Roberto Amato, Jeffrey Barrett, Sonia Goncalves, Ewan Harrison, David K. Jackson, Ian Johnston, Dominic Kwiatkowski, Cordelia Langford, John Sillitoe on behalf of the Wellcome Sanger Institute COVID-19 Surveillance Team |
| EPI_ISL_2447144, EPI_ISL_2447145 | Lighthouse Lab in Milton Keynes | Wellcome Sanger Institute for the COVID-19 Genomics UK (COG-UK) Consortium | The Lighthouse Lab in Milton Keynes and Alex Alderton, Roberto Amato, Jeffrey Barrett, Sonia Goncalves, Ewan Harrison, David K. Jackson, Ian Johnston, Dominic Kwiatkowski, Cordelia Langford, John Sillitoe on behalf of the Wellcome Sanger Institute COVID-19 Surveillance Team |
| EPI_ISL_2447148 | Lighthouse Lab in Alderley Park | Wellcome Sanger Institute for the COVID-19 Genomics UK (COG-UK) Consortium | Jacquelyn Wynn, Mairead Hyland, The Lighthouse Lab in Alderley Park and Alex Alderton, Roberto Amato, Jeffrey Barrett, Sonia Goncalves, Ewan Harrison, David K. Jackson, Ian Johnston, Dominic Kwiatkowski, Cordelia Langford, John Sillitoe on behalf of the Wellcome Sanger Institute COVID-19 Surveillance Team |
| EPI_ISL_2447149, EPI_ISL_2447150 | Lighthouse Lab in Milton Keynes | Wellcome Sanger Institute for the COVID-19 Genomics UK (COG-UK) Consortium | The Lighthouse Lab in Milton Keynes and Alex Alderton, Roberto Amato, Jeffrey Barrett, Sonia Goncalves, Ewan Harrison, David K. Jackson, Ian Johnston, Dominic Kwiatkowski, Cordelia Langford, John Sillitoe on behalf of the Wellcome Sanger Institute COVID-19 Surveillance Team |
| EPI_ISL_2447151 | Lighthouse Lab in Glasgow | Wellcome Sanger Institute for the COVID-19 Genomics UK (COG-UK) Consortium | Harper VanSteenhouse, Yumi Kasai, David Gray, Carol Clugston, Anna Dominiczak and Alex Alderton, Roberto Amato, Jeffrey Barrett, Sonia Goncalves, Ewan Harrison, David K. Jackson, Ian Johnston, Dominic Kwiatkowski, Cordelia Langford, John Sillitoe on behalf of the Wellcome Sanger Institute COVID-19 Surveillance Team |
| EPI_ISL_2447152, EPI_ISL_2447153, EPI_ISL_2447154, EPI_ISL_2447155 | Lighthouse Lab in Alderley Park | Wellcome Sanger Institute for the COVID-19 Genomics UK (COG-UK) Consortium | Jacquelyn Wynn, Mairead Hyland, The Lighthouse Lab in Alderley Park and Alex Alderton, Roberto Amato, Jeffrey Barrett, Sonia Goncalves, Ewan Harrison, David K. Jackson, Ian Johnston, Dominic Kwiatkowski, Cordelia Langford, John Sillitoe on behalf of the Wellcome Sanger Institute COVID-19 Surveillance Team |
| EPI_ISL_2447156 | Lighthouse Lab in Milton Keynes | Wellcome Sanger Institute for the COVID-19 Genomics UK (COG-UK) Consortium | The Lighthouse Lab in Milton Keynes and Alex Alderton, Roberto Amato, Jeffrey Barrett, Sonia Goncalves, Ewan Harrison, David K. Jackson, Ian Johnston, Dominic Kwiatkowski, Cordelia Langford, John Sillitoe on behalf of the Wellcome Sanger Institute COVID-19 Surveillance Team |
| EPI_ISL_2447157 | Lighthouse Lab in Alderley Park | Wellcome Sanger Institute for the COVID-19 Genomics UK (COG-UK) Consortium | Jacquelyn Wynn, Mairead Hyland, The Lighthouse Lab in Alderley Park and Alex Alderton, Roberto Amato, Jeffrey Barrett, Sonia Goncalves, Ewan Harrison, David K. Jackson, Ian Johnston, Dominic Kwiatkowski, Cordelia Langford, John Sillitoe on behalf of the Wellcome Sanger Institute COVID-19 Surveillance Team |
| EPI_ISL_2447158, EPI_ISL_2447159 | Lighthouse Lab in Milton Keynes | Wellcome Sanger Institute for the COVID-19 Genomics UK (COG-UK) Consortium | The Lighthouse Lab in Milton Keynes and Alex Alderton, Roberto Amato, Jeffrey Barrett, Sonia Goncalves, Ewan Harrison, David K. Jackson, Ian Johnston, Dominic Kwiatkowski, Cordelia Langford, John Sillitoe on behalf of the Wellcome Sanger Institute COVID-19 Surveillance Team |
| EPI_ISL_2447160 | Lighthouse Lab in Alderley Park | Wellcome Sanger Institute for the COVID-19 Genomics UK (COG-UK) Consortium | Jacquelyn Wynn, Mairead Hyland, The Lighthouse Lab in Alderley Park and Alex Alderton, Roberto Amato, Jeffrey Barrett, Sonia Goncalves, Ewan Harrison, David K. Jackson, Ian Johnston, Dominic Kwiatkowski, Cordelia Langford, John Sillitoe on behalf of the Wellcome Sanger Institute COVID-19 Surveillance Team |
| EPI_ISL_2447161 | Lighthouse Lab in Milton Keynes | Wellcome Sanger Institute for the COVID-19 Genomics UK (COG-UK) Consortium | The Lighthouse Lab in Milton Keynes and Alex Alderton, Roberto Amato, Jeffrey Barrett, Sonia Goncalves, Ewan Harrison, David K. Jackson, Ian Johnston, Dominic Kwiatkowski, Cordelia Langford, John Sillitoe on behalf of the Wellcome Sanger Institute COVID-19 Surveillance Team |
| EPI_ISL_2447162 | Lighthouse Lab in Glasgow | Wellcome Sanger Institute for the COVID-19 Genomics UK (COG-UK) Consortium | Harper VanSteenhouse, Yumi Kasai, David Gray, Carol Clugston, Anna Dominiczak and Alex Alderton, Roberto Amato, Jeffrey Barrett, Sonia Goncalves, Ewan Harrison, David K. Jackson, Ian Johnston, Dominic Kwiatkowski, Cordelia Langford, John Sillitoe on behalf of the Wellcome Sanger Institute COVID-19 Surveillance Team |
| EPI_ISL_2447163 | Lighthouse Lab in Milton Keynes | Wellcome Sanger Institute for the COVID-19 Genomics UK (COG-UK) Consortium | The Lighthouse Lab in Milton Keynes and Alex Alderton, Roberto Amato, Jeffrey Barrett, Sonia Goncalves, Ewan Harrison, David K. Jackson, Ian Johnston, Dominic Kwiatkowski, Cordelia Langford, John Sillitoe on behalf of the Wellcome Sanger Institute COVID-19 Surveillance Team |
| EPI_ISL_2447164 | Lighthouse Lab in Alderley Park | Wellcome Sanger Institute for the COVID-19 Genomics UK (COG-UK) Consortium | Jacquelyn Wynn, Mairead Hyland, The Lighthouse Lab in Alderley Park and Alex Alderton, Roberto Amato, Jeffrey Barrett, Sonia Goncalves, Ewan Harrison, David K. Jackson, Ian Johnston, Dominic Kwiatkowski, Cordelia Langford, John Sillitoe on behalf of the Wellcome Sanger Institute COVID-19 Surveillance Team |
| EPI_ISL_2447165 | Lighthouse Lab in Glasgow | Wellcome Sanger Institute for the COVID-19 Genomics UK (COG-UK) Consortium | Harper VanSteenhouse, Yumi Kasai, David Gray, Carol Clugston, Anna Dominiczak and Alex Alderton, Roberto Amato, Jeffrey Barrett, Sonia Goncalves, Ewan Harrison, David K. Jackson, Ian Johnston, Dominic Kwiatkowski, Cordelia Langford, John Sillitoe on behalf of the Wellcome Sanger Institute COVID-19 Surveillance Team |
| EPI_ISL_2447166 | Lighthouse Lab in Milton Keynes | Wellcome Sanger Institute for the COVID-19 Genomics UK (COG-UK) Consortium | The Lighthouse Lab in Milton Keynes and Alex Alderton, Roberto Amato, Jeffrey Barrett, Sonia Goncalves, Ewan Harrison, David K. Jackson, Ian Johnston, Dominic Kwiatkowski, Cordelia Langford, John Sillitoe on behalf of the Wellcome Sanger Institute COVID-19 Surveillance Team |
| EPI_ISL_2447168, EPI_ISL_2447169 | Lighthouse Lab in Alderley Park | Wellcome Sanger Institute for the COVID-19 Genomics UK (COG-UK) Consortium | Jacquelyn Wynn, Mairead Hyland, The Lighthouse Lab in Alderley Park and Alex Alderton, Roberto Amato, Jeffrey Barrett, Sonia Goncalves, Ewan Harrison, David K. Jackson, Ian Johnston, Dominic Kwiatkowski, Cordelia Langford, John Sillitoe on behalf of the Wellcome Sanger Institute COVID-19 Surveillance Team |
| EPI_ISL_2447170, EPI_ISL_2447171 | Lighthouse Lab in Milton Keynes | Wellcome Sanger Institute for the COVID-19 Genomics UK (COG-UK) Consortium | The Lighthouse Lab in Milton Keynes and Alex Alderton, Roberto Amato, Jeffrey Barrett, Sonia Goncalves, Ewan Harrison, David K. Jackson, Ian Johnston, Dominic Kwiatkowski, Cordelia Langford, John Sillitoe on behalf of the Wellcome Sanger Institute COVID-19 Surveillance Team |
| EPI_ISL_2447172 | Lighthouse Lab in Alderley Park | Wellcome Sanger Institute for the COVID-19 Genomics UK (COG-UK) Consortium | Jacquelyn Wynn, Mairead Hyland, The Lighthouse Lab in Alderley Park and Alex Alderton, Roberto Amato, Jeffrey Barrett, Sonia Goncalves, Ewan Harrison, David K. Jackson, Ian Johnston, Dominic Kwiatkowski, Cordelia Langford, John Sillitoe on behalf of the Wellcome Sanger Institute COVID-19 Surveillance Team |
| EPI_ISL_2447173 | Lighthouse Lab in Glasgow | Wellcome Sanger Institute for the COVID-19 Genomics UK (COG-UK) Consortium | Harper VanSteenhouse, Yumi Kasai, David Gray, Carol Clugston, Anna Dominiczak and Alex Alderton, Roberto Amato, Jeffrey Barrett, Sonia Goncalves, Ewan Harrison, David K. Jackson, Ian Johnston, Dominic Kwiatkowski, Cordelia Langford, John Sillitoe on behalf of the Wellcome Sanger Institute COVID-19 Surveillance Team |
| EPI_ISL_2447174, EPI_ISL_2447175 | Lighthouse Lab in Milton Keynes | Wellcome Sanger Institute for the COVID-19 Genomics UK (COG-UK) Consortium | The Lighthouse Lab in Milton Keynes and Alex Alderton, Roberto Amato, Jeffrey Barrett, Sonia Goncalves, Ewan Harrison, David K. Jackson, Ian Johnston, Dominic Kwiatkowski, Cordelia Langford, John Sillitoe on behalf of the Wellcome Sanger Institute COVID-19 Surveillance Team |
| EPI_ISL_2447176, EPI_ISL_2447177 | Lighthouse Lab in Glasgow | Wellcome Sanger Institute for the COVID-19 Genomics UK (COG-UK) Consortium | Harper VanSteenhouse, Yumi Kasai, David Gray, Carol Clugston, Anna Dominiczak and Alex Alderton, Roberto Amato, Jeffrey Barrett, Sonia Goncalves, Ewan Harrison, David K. Jackson, Ian Johnston, Dominic Kwiatkowski, Cordelia Langford, John Sillitoe on behalf of the Wellcome Sanger Institute COVID-19 Surveillance Team |
| EPI_ISL_2447179 | Lighthouse Lab in Milton Keynes | Wellcome Sanger Institute for the COVID-19 Genomics UK (COG-UK) Consortium | The Lighthouse Lab in Milton Keynes and Alex Alderton, Roberto Amato, Jeffrey Barrett, Sonia Goncalves, Ewan Harrison, David K. Jackson, Ian Johnston, Dominic Kwiatkowski, Cordelia Langford, John Sillitoe on behalf of the Wellcome Sanger Institute COVID-19 Surveillance Team |
| EPI_ISL_2447180 | Lighthouse Lab in Alderley Park | Wellcome Sanger Institute for the COVID-19 Genomics UK (COG-UK) Consortium | Jacquelyn Wynn, Mairead Hyland, The Lighthouse Lab in Alderley Park and Alex Alderton, Roberto Amato, Jeffrey Barrett, Sonia Goncalves, Ewan Harrison, David K. Jackson, Ian Johnston, Dominic Kwiatkowski, Cordelia Langford, John Sillitoe on behalf of the Wellcome Sanger Institute COVID-19 Surveillance Team |
| EPI_ISL_2447182 | Lighthouse Lab in Milton Keynes | Wellcome Sanger Institute for the COVID-19 Genomics UK (COG-UK) Consortium | The Lighthouse Lab in Milton Keynes and Alex Alderton, Roberto Amato, Jeffrey Barrett, Sonia Goncalves, Ewan Harrison, David K. Jackson, Ian Johnston, Dominic Kwiatkowski, Cordelia Langford, John Sillitoe on behalf of the Wellcome Sanger Institute COVID-19 Surveillance Team |
| EPI_ISL_2447183 | Lighthouse Lab in Alderley Park | Wellcome Sanger Institute for the COVID-19 Genomics UK (COG-UK) Consortium | Jacquelyn Wynn, Mairead Hyland, The Lighthouse Lab in Alderley Park and Alex Alderton, Roberto Amato, Jeffrey Barrett, Sonia Goncalves, Ewan Harrison, David K. Jackson, Ian Johnston, Dominic Kwiatkowski, Cordelia Langford, John Sillitoe on behalf of the Wellcome Sanger Institute COVID-19 Surveillance Team |
| EPI_ISL_2447184 | Lighthouse Lab in Glasgow | Wellcome Sanger Institute for the COVID-19 Genomics UK (COG-UK) Consortium | Harper VanSteenhouse, Yumi Kasai, David Gray, Carol Clugston, Anna Dominiczak and Alex Alderton, Roberto Amato, Jeffrey Barrett, Sonia Goncalves, Ewan Harrison, David K. Jackson, Ian Johnston, Dominic Kwiatkowski, Cordelia Langford, John Sillitoe on behalf of the Wellcome Sanger Institute COVID-19 Surveillance Team |
| EPI_ISL_2447185, EPI_ISL_2447186, EPI_ISL_2447187 | Lighthouse Lab in Milton Keynes | Wellcome Sanger Institute for the COVID-19 Genomics UK (COG-UK) Consortium | The Lighthouse Lab in Milton Keynes and Alex Alderton, Roberto Amato, Jeffrey Barrett, Sonia Goncalves, Ewan Harrison, David K. Jackson, Ian Johnston, Dominic Kwiatkowski, Cordelia Langford, John Sillitoe on behalf of the Wellcome Sanger Institute COVID-19 Surveillance Team |
| EPI_ISL_2447188, EPI_ISL_2447189 | Lighthouse Lab in Alderley Park | Wellcome Sanger Institute for the COVID-19 Genomics UK (COG-UK) Consortium | Jacquelyn Wynn, Mairead Hyland, The Lighthouse Lab in Alderley Park and Alex Alderton, Roberto Amato, Jeffrey Barrett, Sonia Goncalves, Ewan Harrison, David K. Jackson, Ian Johnston, Dominic Kwiatkowski, Cordelia Langford, John Sillitoe on behalf of the Wellcome Sanger Institute COVID-19 Surveillance Team |
| EPI_ISL_2447190 | Lighthouse Lab in Milton Keynes | Wellcome Sanger Institute for the COVID-19 Genomics UK (COG-UK) Consortium | The Lighthouse Lab in Milton Keynes and Alex Alderton, Roberto Amato, Jeffrey Barrett, Sonia Goncalves, Ewan Harrison, David K. Jackson, Ian Johnston, Dominic Kwiatkowski, Cordelia Langford, John Sillitoe on behalf of the Wellcome Sanger Institute COVID-19 Surveillance Team |
| EPI_ISL_2447191 | Lighthouse Lab in Alderley Park | Wellcome Sanger Institute for the COVID-19 Genomics UK (COG-UK) Consortium | Jacquelyn Wynn, Mairead Hyland, The Lighthouse Lab in Alderley Park and Alex Alderton, Roberto Amato, Jeffrey Barrett, Sonia Goncalves, Ewan Harrison, David K. Jackson, Ian Johnston, Dominic Kwiatkowski, Cordelia Langford, John Sillitoe on behalf of the Wellcome Sanger Institute COVID-19 Surveillance Team |
| EPI_ISL_2447194 | Lighthouse Lab in Milton Keynes | Wellcome Sanger Institute for the COVID-19 Genomics UK (COG-UK) Consortium | The Lighthouse Lab in Milton Keynes and Alex Alderton, Roberto Amato, Jeffrey Barrett, Sonia Goncalves, Ewan Harrison, David K. Jackson, Ian Johnston, Dominic Kwiatkowski, Cordelia Langford, John Sillitoe on behalf of the Wellcome Sanger Institute COVID-19 Surveillance Team |
| EPI_ISL_2447195, EPI_ISL_2447198, EPI_ISL_2447199 | Lighthouse Lab in Alderley Park | Wellcome Sanger Institute for the COVID-19 Genomics UK (COG-UK) Consortium | Jacquelyn Wynn, Mairead Hyland, The Lighthouse Lab in Alderley Park and Alex Alderton, Roberto Amato, Jeffrey Barrett, Sonia Goncalves, Ewan Harrison, David K. Jackson, Ian Johnston, Dominic Kwiatkowski, Cordelia Langford, John Sillitoe on behalf of the Wellcome Sanger Institute COVID-19 Surveillance Team |
| EPI_ISL_2447200 | Lighthouse Lab in Glasgow | Wellcome Sanger Institute for the COVID-19 Genomics UK (COG-UK) Consortium | Harper VanSteenhouse, Yumi Kasai, David Gray, Carol Clugston, Anna Dominiczak and Alex Alderton, Roberto Amato, Jeffrey Barrett, Sonia Goncalves, Ewan Harrison, David K. Jackson, Ian Johnston, Dominic Kwiatkowski, Cordelia Langford, John Sillitoe on behalf of the Wellcome Sanger Institute COVID-19 Surveillance Team |
| EPI_ISL_2447201, | Lighthouse Lab in Alderley Park | Wellcome Sanger Institute for the COVID-19 Genomics UK (COG-UK) | Jacquelyn Wynn, Mairead Hyland, The Lighthouse Lab in Alderley Park and Alex Alderton, Roberto Amato, Jeffrey Barrett, Sonia Goncalves, Ewan Harrison, David K. Jackson, Ian Johnston, Dominic Kwiatkowski, Cordelia Langford, John Sillitoe on |

| | | | |
|---|---|---|---|
| EPI_ISL_2447202 | | Consortium | behalf of the Wellcome Sanger Institute COVID-19 Surveillance Team |
| EPI_ISL_2447203 | Lighthouse Lab in Milton Keynes | Wellcome Sanger Institute for the COVID-19 Genomics UK (COG-UK) Consortium | The Lighthouse Lab in Milton Keynes and Alex Alderton, Roberto Amato, Jeffrey Barrett, Sonia Goncalves, Ewan Harrison, David K. Jackson, Ian Johnston, Dominic Kwiatkowski, Cordelia Langford, John Sillitoe on behalf of the Wellcome Sanger Institute COVID-19 Surveillance Team |
| EPI_ISL_2447204, EPI_ISL_2447205 | Lighthouse Lab in Alderley Park | Wellcome Sanger Institute for the COVID-19 Genomics UK (COG-UK) Consortium | Jacquelyn Wynn, Mairead Hyland, The Lighthouse Lab in Alderley Park and Alex Alderton, Roberto Amato, Jeffrey Barrett, Sonia Goncalves, Ewan Harrison, David K. Jackson, Ian Johnston, Dominic Kwiatkowski, Cordelia Langford, John Sillitoe on behalf of the Wellcome Sanger Institute COVID-19 Surveillance Team |
| EPI_ISL_2447206, EPI_ISL_2447207 | Lighthouse Lab in Milton Keynes | Wellcome Sanger Institute for the COVID-19 Genomics UK (COG-UK) Consortium | The Lighthouse Lab in Milton Keynes and Alex Alderton, Roberto Amato, Jeffrey Barrett, Sonia Goncalves, Ewan Harrison, David K. Jackson, Ian Johnston, Dominic Kwiatkowski, Cordelia Langford, John Sillitoe on behalf of the Wellcome Sanger Institute COVID-19 Surveillance Team |
| EPI_ISL_2447208 | Lighthouse Lab in Glasgow | Wellcome Sanger Institute for the COVID-19 Genomics UK (COG-UK) Consortium | Harper VanSteenhouse, Yumi Kasai, David Gray, Carol Clugston, Anna Dominiczak and Alex Alderton, Roberto Amato, Jeffrey Barrett, Sonia Goncalves, Ewan Harrison, David K. Jackson, Ian Johnston, Dominic Kwiatkowski, Cordelia Langford, John Sillitoe on behalf of the Wellcome Sanger Institute COVID-19 Surveillance Team |
| EPI_ISL_2447209 | Lighthouse Lab in Alderley Park | Wellcome Sanger Institute for the COVID-19 Genomics UK (COG-UK) Consortium | Jacquelyn Wynn, Mairead Hyland, The Lighthouse Lab in Alderley Park and Alex Alderton, Roberto Amato, Jeffrey Barrett, Sonia Goncalves, Ewan Harrison, David K. Jackson, Ian Johnston, Dominic Kwiatkowski, Cordelia Langford, John Sillitoe on behalf of the Wellcome Sanger Institute COVID-19 Surveillance Team |
| EPI_ISL_2447210 | Lighthouse Lab in Milton Keynes | Wellcome Sanger Institute for the COVID-19 Genomics UK (COG-UK) Consortium | The Lighthouse Lab in Milton Keynes and Alex Alderton, Roberto Amato, Jeffrey Barrett, Sonia Goncalves, Ewan Harrison, David K. Jackson, Ian Johnston, Dominic Kwiatkowski, Cordelia Langford, John Sillitoe on behalf of the Wellcome Sanger Institute COVID-19 Surveillance Team |
| EPI_ISL_2447211 | Lighthouse Lab in Alderley Park | Wellcome Sanger Institute for the COVID-19 Genomics UK (COG-UK) Consortium | Jacquelyn Wynn, Mairead Hyland, The Lighthouse Lab in Alderley Park and Alex Alderton, Roberto Amato, Jeffrey Barrett, Sonia Goncalves, Ewan Harrison, David K. Jackson, Ian Johnston, Dominic Kwiatkowski, Cordelia Langford, John Sillitoe on behalf of the Wellcome Sanger Institute COVID-19 Surveillance Team |
| EPI_ISL_2447212 | Lighthouse Lab in Glasgow | Wellcome Sanger Institute for the COVID-19 Genomics UK (COG-UK) Consortium | Harper VanSteenhouse, Yumi Kasai, David Gray, Carol Clugston, Anna Dominiczak and Alex Alderton, Roberto Amato, Jeffrey Barrett, Sonia Goncalves, Ewan Harrison, David K. Jackson, Ian Johnston, Dominic Kwiatkowski, Cordelia Langford, John Sillitoe on behalf of the Wellcome Sanger Institute COVID-19 Surveillance Team |
| EPI_ISL_2447213, EPI_ISL_2447214 | Lighthouse Lab in Alderley Park | Wellcome Sanger Institute for the COVID-19 Genomics UK (COG-UK) Consortium | Jacquelyn Wynn, Mairead Hyland, The Lighthouse Lab in Alderley Park and Alex Alderton, Roberto Amato, Jeffrey Barrett, Sonia Goncalves, Ewan Harrison, David K. Jackson, Ian Johnston, Dominic Kwiatkowski, Cordelia Langford, John Sillitoe on behalf of the Wellcome Sanger Institute COVID-19 Surveillance Team |
| EPI_ISL_2447215, EPI_ISL_2447216, EPI_ISL_2447217 | Lighthouse Lab in Milton Keynes | Wellcome Sanger Institute for the COVID-19 Genomics UK (COG-UK) Consortium | The Lighthouse Lab in Milton Keynes and Alex Alderton, Roberto Amato, Jeffrey Barrett, Sonia Goncalves, Ewan Harrison, David K. Jackson, Ian Johnston, Dominic Kwiatkowski, Cordelia Langford, John Sillitoe on behalf of the Wellcome Sanger Institute COVID-19 Surveillance Team |
| EPI_ISL_2447218, EPI_ISL_2447219, EPI_ISL_2447220, EPI_ISL_2447221, EPI_ISL_2447222, EPI_ISL_2447223, EPI_ISL_2447225 | Lighthouse Lab in Alderley Park | Wellcome Sanger Institute for the COVID-19 Genomics UK (COG-UK) Consortium | Jacquelyn Wynn, Mairead Hyland, The Lighthouse Lab in Alderley Park and Alex Alderton, Roberto Amato, Jeffrey Barrett, Sonia Goncalves, Ewan Harrison, David K. Jackson, Ian Johnston, Dominic Kwiatkowski, Cordelia Langford, John Sillitoe on behalf of the Wellcome Sanger Institute COVID-19 Surveillance Team |
| EPI_ISL_2447226 | Lighthouse Lab in Milton Keynes | Wellcome Sanger Institute for the COVID-19 Genomics UK (COG-UK) Consortium | The Lighthouse Lab in Milton Keynes and Alex Alderton, Roberto Amato, Jeffrey Barrett, Sonia Goncalves, Ewan Harrison, David K. Jackson, Ian Johnston, Dominic Kwiatkowski, Cordelia Langford, John Sillitoe on behalf of the Wellcome Sanger Institute COVID-19 Surveillance Team |
| EPI_ISL_2447227, EPI_ISL_2447228 | Lighthouse Lab in Alderley Park | Wellcome Sanger Institute for the COVID-19 Genomics UK (COG-UK) Consortium | Jacquelyn Wynn, Mairead Hyland, The Lighthouse Lab in Alderley Park and Alex Alderton, Roberto Amato, Jeffrey Barrett, Sonia Goncalves, Ewan Harrison, David K. Jackson, Ian Johnston, Dominic Kwiatkowski, Cordelia Langford, John Sillitoe on behalf of the Wellcome Sanger Institute COVID-19 Surveillance Team |
| EPI_ISL_2447229, EPI_ISL_2447230, EPI_ISL_2447231 | Lighthouse Lab in Milton Keynes | Wellcome Sanger Institute for the COVID-19 Genomics UK (COG-UK) Consortium | The Lighthouse Lab in Milton Keynes and Alex Alderton, Roberto Amato, Jeffrey Barrett, Sonia Goncalves, Ewan Harrison, David K. Jackson, Ian Johnston, Dominic Kwiatkowski, Cordelia Langford, John Sillitoe on behalf of the Wellcome Sanger Institute COVID-19 Surveillance Team |
| EPI_ISL_2447232 | Lighthouse Lab in Alderley Park | Wellcome Sanger Institute for the COVID-19 Genomics UK (COG-UK) Consortium | Jacquelyn Wynn, Mairead Hyland, The Lighthouse Lab in Alderley Park and Alex Alderton, Roberto Amato, Jeffrey Barrett, Sonia Goncalves, Ewan Harrison, David K. Jackson, Ian Johnston, Dominic Kwiatkowski, Cordelia Langford, John Sillitoe on behalf of the Wellcome Sanger Institute COVID-19 Surveillance Team |
| EPI_ISL_2447234, EPI_ISL_2447235 | Lighthouse Laboratory Plymouth | Wellcome Sanger Institute for the COVID-19 Genomics UK (COG-UK) Consortium | Lighthouse Laboratory Plymouth and Alex Alderton, Roberto Amato, Jeffrey Barrett, Sonia Goncalves, Ewan Harrison, David K. Jackson, Ian Johnston, Dominic Kwiatkowski, Cordelia Langford, John Sillitoe on behalf of the Wellcome Sanger Institute COVID-19 Surveillance Team |
| EPI_ISL_2447237, EPI_ISL_2447239, EPI_ISL_2447244, EPI_ISL_2447249 | Health Services Laboratories | Wellcome Sanger Institute for the COVID-19 Genomics UK (COG-UK) Consortium | Health Services Laboratories and Alex Alderton, Roberto Amato, Jeffrey Barrett, Sonia Goncalves, Ewan Harrison, David K. Jackson, Ian Johnston, Dominic Kwiatkowski, Cordelia Langford, John Sillitoe on behalf of the Wellcome Sanger Institute COVID-19 Surveillance Team |
| EPI_ISL_2447252 | Lighthouse Laboratory Plymouth | Wellcome Sanger Institute for the COVID-19 Genomics UK (COG-UK) Consortium | Lighthouse Laboratory Plymouth and Alex Alderton, Roberto Amato, Jeffrey Barrett, Sonia Goncalves, Ewan Harrison, David K. Jackson, Ian Johnston, Dominic Kwiatkowski, Cordelia Langford, John Sillitoe on behalf of the Wellcome Sanger Institute COVID-19 Surveillance Team |
| EPI_ISL_2447253 | Health Services Laboratories | Wellcome Sanger Institute for the COVID-19 Genomics UK (COG-UK) Consortium | Health Services Laboratories and Alex Alderton, Roberto Amato, Jeffrey Barrett, Sonia Goncalves, Ewan Harrison, David K. Jackson, Ian Johnston, Dominic Kwiatkowski, Cordelia Langford, John Sillitoe on behalf of the Wellcome Sanger Institute COVID-19 Surveillance Team |
| EPI_ISL_2447254 | Lighthouse Laboratory Plymouth | Wellcome Sanger Institute for the COVID-19 Genomics UK (COG-UK) Consortium | Lighthouse Laboratory Plymouth and Alex Alderton, Roberto Amato, Jeffrey Barrett, Sonia Goncalves, Ewan Harrison, David K. Jackson, Ian Johnston, Dominic Kwiatkowski, Cordelia Langford, John Sillitoe on behalf of the Wellcome Sanger Institute COVID-19 Surveillance Team |
| EPI_ISL_2447258 | Health Services Laboratories | Wellcome Sanger Institute for the COVID-19 Genomics UK (COG-UK) Consortium | Health Services Laboratories and Alex Alderton, Roberto Amato, Jeffrey Barrett, Sonia Goncalves, Ewan Harrison, David K. Jackson, Ian Johnston, Dominic Kwiatkowski, Cordelia Langford, John Sillitoe on behalf of the Wellcome Sanger Institute COVID-19 Surveillance Team |
| EPI_ISL_2447262 | Lighthouse Laboratory Plymouth | Wellcome Sanger Institute for the COVID-19 Genomics UK (COG-UK) Consortium | Lighthouse Laboratory Plymouth and Alex Alderton, Roberto Amato, Jeffrey Barrett, Sonia Goncalves, Ewan Harrison, David K. Jackson, Ian Johnston, Dominic Kwiatkowski, Cordelia Langford, John Sillitoe on behalf of the Wellcome Sanger Institute COVID-19 Surveillance Team |
| EPI_ISL_2447264 | Health Services Laboratories | Wellcome Sanger Institute for the COVID-19 Genomics UK (COG-UK) Consortium | Health Services Laboratories and Alex Alderton, Roberto Amato, Jeffrey Barrett, Sonia Goncalves, Ewan Harrison, David K. Jackson, Ian Johnston, Dominic Kwiatkowski, Cordelia Langford, John Sillitoe on behalf of the Wellcome Sanger Institute COVID-19 Surveillance Team |
| EPI_ISL_2447265 | Lighthouse Laboratory Plymouth | Wellcome Sanger Institute for the COVID-19 Genomics UK (COG-UK) Consortium | Lighthouse Laboratory Plymouth and Alex Alderton, Roberto Amato, Jeffrey Barrett, Sonia Goncalves, Ewan Harrison, David K. Jackson, Ian Johnston, Dominic Kwiatkowski, Cordelia Langford, John Sillitoe on behalf of the Wellcome Sanger Institute COVID-19 Surveillance Team |
| EPI_ISL_2447266 | Health Services Laboratories | Wellcome Sanger Institute for the COVID-19 Genomics UK (COG-UK) Consortium | Health Services Laboratories and Alex Alderton, Roberto Amato, Jeffrey Barrett, Sonia Goncalves, Ewan Harrison, David K. Jackson, Ian Johnston, Dominic Kwiatkowski, Cordelia Langford, John Sillitoe on behalf of the Wellcome Sanger Institute COVID-19 Surveillance Team |
| EPI_ISL_2447267 | Lighthouse Laboratory Plymouth | Wellcome Sanger Institute for the COVID-19 Genomics UK (COG-UK) Consortium | Lighthouse Laboratory Plymouth and Alex Alderton, Roberto Amato, Jeffrey Barrett, Sonia Goncalves, Ewan Harrison, David K. Jackson, Ian Johnston, Dominic Kwiatkowski, Cordelia Langford, John Sillitoe on behalf of the Wellcome Sanger Institute COVID-19 Surveillance Team |
| EPI_ISL_2447268, EPI_ISL_2447271 | Health Services Laboratories | Wellcome Sanger Institute for the COVID-19 Genomics UK (COG-UK) Consortium | Health Services Laboratories and Alex Alderton, Roberto Amato, Jeffrey Barrett, Sonia Goncalves, Ewan Harrison, David K. Jackson, Ian Johnston, Dominic Kwiatkowski, Cordelia Langford, John Sillitoe on behalf of the Wellcome Sanger Institute COVID-19 Surveillance Team |
| EPI_ISL_2447276, EPI_ISL_2447279, EPI_ISL_2447283 | Lighthouse Laboratory Plymouth | Wellcome Sanger Institute for the COVID-19 Genomics UK (COG-UK) Consortium | Lighthouse Laboratory Plymouth and Alex Alderton, Roberto Amato, Jeffrey Barrett, Sonia Goncalves, Ewan Harrison, David K. Jackson, Ian Johnston, Dominic Kwiatkowski, Cordelia Langford, John Sillitoe on behalf of the Wellcome Sanger Institute COVID-19 Surveillance Team |
| EPI_ISL_2447285 | Health Services Laboratories | Wellcome Sanger Institute for the COVID-19 Genomics UK (COG-UK) Consortium | Health Services Laboratories and Alex Alderton, Roberto Amato, Jeffrey Barrett, Sonia Goncalves, Ewan Harrison, David K. Jackson, Ian Johnston, Dominic Kwiatkowski, Cordelia Langford, John Sillitoe on behalf of the Wellcome Sanger Institute COVID-19 Surveillance Team |
| EPI_ISL_2447286, EPI_ISL_2447288 | Lighthouse Lab in Glasgow | Wellcome Sanger Institute for the COVID-19 Genomics UK (COG-UK) Consortium | Harper VanSteenhouse, Yumi Kasai, David Gray, Carol Clugston, Anna Dominiczak and Alex Alderton, Roberto Amato, Jeffrey Barrett, Sonia Goncalves, Ewan Harrison, David K. Jackson, Ian Johnston, Dominic Kwiatkowski, Cordelia Langford, John Sillitoe on behalf of the Wellcome Sanger Institute COVID-19 Surveillance Team |
| EPI_ISL_2447297, EPI_ISL_2447304 | Health Services Laboratories | Wellcome Sanger Institute for the COVID-19 Genomics UK (COG-UK) Consortium | Health Services Laboratories and Alex Alderton, Roberto Amato, Jeffrey Barrett, Sonia Goncalves, Ewan Harrison, David K. Jackson, Ian Johnston, Dominic Kwiatkowski, Cordelia Langford, John Sillitoe on behalf of the Wellcome Sanger Institute COVID-19 Surveillance Team |
| EPI_ISL_2447307 | Lighthouse Laboratory Plymouth | Wellcome Sanger Institute for the COVID-19 Genomics UK (COG-UK) Consortium | Lighthouse Laboratory Plymouth and Alex Alderton, Roberto Amato, Jeffrey Barrett, Sonia Goncalves, Ewan Harrison, David K. Jackson, Ian Johnston, Dominic Kwiatkowski, Cordelia Langford, John Sillitoe on behalf of the Wellcome Sanger Institute COVID-19 Surveillance Team |
| EPI_ISL_2447308 | Health Services Laboratories | Wellcome Sanger Institute for the COVID-19 Genomics UK (COG-UK) Consortium | Health Services Laboratories and Alex Alderton, Roberto Amato, Jeffrey Barrett, Sonia Goncalves, Ewan Harrison, David K. Jackson, Ian Johnston, Dominic Kwiatkowski, Cordelia Langford, John Sillitoe on behalf of the Wellcome Sanger Institute COVID-19 Surveillance Team |
| EPI_ISL_2447318, EPI_ISL_2447319 | Lighthouse Laboratory Plymouth | Wellcome Sanger Institute for the COVID-19 Genomics UK (COG-UK) Consortium | Lighthouse Laboratory Plymouth and Alex Alderton, Roberto Amato, Jeffrey Barrett, Sonia Goncalves, Ewan Harrison, David K. Jackson, Ian Johnston, Dominic Kwiatkowski, Cordelia Langford, John Sillitoe on behalf of the Wellcome Sanger Institute COVID-19 Surveillance Team |
| EPI_ISL_2447323 | Lighthouse Lab in Glasgow | Wellcome Sanger Institute for the COVID-19 Genomics UK (COG-UK) Consortium | Harper VanSteenhouse, Yumi Kasai, David Gray, Carol Clugston, Anna Dominiczak and Alex Alderton, Roberto Amato, Jeffrey Barrett, Sonia Goncalves, Ewan Harrison, David K. Jackson, Ian Johnston, Dominic Kwiatkowski, Cordelia Langford, John Sillitoe on behalf of the Wellcome Sanger Institute COVID-19 Surveillance Team |
| EPI_ISL_2447325, EPI_ISL_2447330 | Lighthouse Laboratory Plymouth | Wellcome Sanger Institute for the COVID-19 Genomics UK (COG-UK) Consortium | Lighthouse Laboratory Plymouth and Alex Alderton, Roberto Amato, Jeffrey Barrett, Sonia Goncalves, Ewan Harrison, David K. Jackson, Ian Johnston, Dominic Kwiatkowski, Cordelia Langford, John Sillitoe on behalf of the Wellcome Sanger Institute COVID-19 Surveillance Team |
| EPI_ISL_2447332 | Lighthouse Lab in Glasgow | Wellcome Sanger Institute for the COVID-19 Genomics UK (COG-UK) Consortium | Harper VanSteenhouse, Yumi Kasai, David Gray, Carol Clugston, Anna Dominiczak and Alex Alderton, Roberto Amato, Jeffrey Barrett, Sonia Goncalves, Ewan Harrison, David K. Jackson, Ian Johnston, Dominic Kwiatkowski, Cordelia Langford, John Sillitoe on behalf of the Wellcome Sanger Institute COVID-19 Surveillance Team |
| EPI_ISL_2447334 | Lighthouse Laboratory Plymouth | Wellcome Sanger Institute for the COVID-19 Genomics UK (COG-UK) Consortium | Lighthouse Laboratory Plymouth and Alex Alderton, Roberto Amato, Jeffrey Barrett, Sonia Goncalves, Ewan Harrison, David K. Jackson, Ian Johnston, Dominic Kwiatkowski, Cordelia Langford, John Sillitoe on behalf of the Wellcome Sanger Institute COVID-19 Surveillance Team |
| EPI_ISL_2447336 | Lighthouse Lab in Glasgow | Wellcome Sanger Institute for the COVID-19 Genomics UK (COG-UK) Consortium | Harper VanSteenhouse, Yumi Kasai, David Gray, Carol Clugston, Anna Dominiczak and Alex Alderton, Roberto Amato, Jeffrey Barrett, Sonia Goncalves, Ewan Harrison, David K. Jackson, Ian Johnston, Dominic Kwiatkowski, Cordelia Langford, John Sillitoe on behalf of the Wellcome Sanger Institute COVID-19 Surveillance Team |
| EPI_ISL_2447338 | Lighthouse Laboratory Plymouth | Wellcome Sanger Institute for the COVID-19 Genomics UK (COG-UK) Consortium | Lighthouse Laboratory Plymouth and Alex Alderton, Roberto Amato, Jeffrey Barrett, Sonia Goncalves, Ewan Harrison, David K. Jackson, Ian Johnston, Dominic Kwiatkowski, Cordelia Langford, John Sillitoe on behalf of the Wellcome Sanger Institute COVID-19 Surveillance Team |

| | | | |
|---|---|---|---|
| EPI_ISL_2447340 | Health Services Laboratories | Wellcome Sanger Institute for the COVID-19 Genomics UK (COG-UK) Consortium | Health Services Laboratories and Alex Alderton, Roberto Amato, Jeffrey Barrett, Sonia Goncalves, Ewan Harrison, David K. Jackson, Ian Johnston, Dominic Kwiatkowski, Cordelia Langford, John Sillitoe on behalf of the Wellcome Sanger Institute COVID-19 Surveillance Team |
| EPI_ISL_2447341 | Lighthouse Laboratory Plymouth | Wellcome Sanger Institute for the COVID-19 Genomics UK (COG-UK) Consortium | Lighthouse Laboratory Plymouth and Alex Alderton, Roberto Amato, Jeffrey Barrett, Sonia Goncalves, Ewan Harrison, David K. Jackson, Ian Johnston, Dominic Kwiatkowski, Cordelia Langford, John Sillitoe on behalf of the Wellcome Sanger Institute COVID-19 Surveillance Team |
| EPI_ISL_2447343 | Health Services Laboratories | Wellcome Sanger Institute for the COVID-19 Genomics UK (COG-UK) Consortium | Health Services Laboratories and Alex Alderton, Roberto Amato, Jeffrey Barrett, Sonia Goncalves, Ewan Harrison, David K. Jackson, Ian Johnston, Dominic Kwiatkowski, Cordelia Langford, John Sillitoe on behalf of the Wellcome Sanger Institute COVID-19 Surveillance Team |
| EPI_ISL_2447346, EPI_ISL_2447348 | Lighthouse Laboratory Plymouth | Wellcome Sanger Institute for the COVID-19 Genomics UK (COG-UK) Consortium | Lighthouse Laboratory Plymouth and Alex Alderton, Roberto Amato, Jeffrey Barrett, Sonia Goncalves, Ewan Harrison, David K. Jackson, Ian Johnston, Dominic Kwiatkowski, Cordelia Langford, John Sillitoe on behalf of the Wellcome Sanger Institute COVID-19 Surveillance Team |
| EPI_ISL_2447349 | Health Services Laboratories | Wellcome Sanger Institute for the COVID-19 Genomics UK (COG-UK) Consortium | Health Services Laboratories and Alex Alderton, Roberto Amato, Jeffrey Barrett, Sonia Goncalves, Ewan Harrison, David K. Jackson, Ian Johnston, Dominic Kwiatkowski, Cordelia Langford, John Sillitoe on behalf of the Wellcome Sanger Institute COVID-19 Surveillance Team |
| EPI_ISL_2447350, EPI_ISL_2447355, EPI_ISL_2447356 | Lighthouse Laboratory Plymouth | Wellcome Sanger Institute for the COVID-19 Genomics UK (COG-UK) Consortium | Lighthouse Laboratory Plymouth and Alex Alderton, Roberto Amato, Jeffrey Barrett, Sonia Goncalves, Ewan Harrison, David K. Jackson, Ian Johnston, Dominic Kwiatkowski, Cordelia Langford, John Sillitoe on behalf of the Wellcome Sanger Institute COVID-19 Surveillance Team |
| EPI_ISL_2447358 | Health Services Laboratories | Wellcome Sanger Institute for the COVID-19 Genomics UK (COG-UK) Consortium | Health Services Laboratories and Alex Alderton, Roberto Amato, Jeffrey Barrett, Sonia Goncalves, Ewan Harrison, David K. Jackson, Ian Johnston, Dominic Kwiatkowski, Cordelia Langford, John Sillitoe on behalf of the Wellcome Sanger Institute COVID-19 Surveillance Team |
| EPI_ISL_2447359 | Lighthouse Laboratory Plymouth | Wellcome Sanger Institute for the COVID-19 Genomics UK (COG-UK) Consortium | Lighthouse Laboratory Plymouth and Alex Alderton, Roberto Amato, Jeffrey Barrett, Sonia Goncalves, Ewan Harrison, David K. Jackson, Ian Johnston, Dominic Kwiatkowski, Cordelia Langford, John Sillitoe on behalf of the Wellcome Sanger Institute COVID-19 Surveillance Team |
| EPI_ISL_2447361, EPI_ISL_2447362 | Health Services Laboratories | Wellcome Sanger Institute for the COVID-19 Genomics UK (COG-UK) Consortium | Health Services Laboratories and Alex Alderton, Roberto Amato, Jeffrey Barrett, Sonia Goncalves, Ewan Harrison, David K. Jackson, Ian Johnston, Dominic Kwiatkowski, Cordelia Langford, John Sillitoe on behalf of the Wellcome Sanger Institute COVID-19 Surveillance Team |
| EPI_ISL_2447363 | Lighthouse Lab in Milton Keynes | Wellcome Sanger Institute for the COVID-19 Genomics UK (COG-UK) Consortium | The Lighthouse Lab in Milton Keynes and Alex Alderton, Roberto Amato, Jeffrey Barrett, Sonia Goncalves, Ewan Harrison, David K. Jackson, Ian Johnston, Dominic Kwiatkowski, Cordelia Langford, John Sillitoe on behalf of the Wellcome Sanger Institute COVID-19 Surveillance Team |
| EPI_ISL_2447364 | Health Services Laboratories | Wellcome Sanger Institute for the COVID-19 Genomics UK (COG-UK) Consortium | Health Services Laboratories and Alex Alderton, Roberto Amato, Jeffrey Barrett, Sonia Goncalves, Ewan Harrison, David K. Jackson, Ian Johnston, Dominic Kwiatkowski, Cordelia Langford, John Sillitoe on behalf of the Wellcome Sanger Institute COVID-19 Surveillance Team |
| EPI_ISL_2447365, EPI_ISL_2447366 | Lighthouse Lab in Alderley Park | Wellcome Sanger Institute for the COVID-19 Genomics UK (COG-UK) Consortium | Jacquelyn Wynn, Mairead Hyland, The Lighthouse Lab in Alderley Park and Alex Alderton, Roberto Amato, Jeffrey Barrett, Sonia Goncalves, Ewan Harrison, David K. Jackson, Ian Johnston, Dominic Kwiatkowski, Cordelia Langford, John Sillitoe on behalf of the Wellcome Sanger Institute COVID-19 Surveillance Team |
| EPI_ISL_2447368 | Health Services Laboratories | Wellcome Sanger Institute for the COVID-19 Genomics UK (COG-UK) Consortium | Health Services Laboratories and Alex Alderton, Roberto Amato, Jeffrey Barrett, Sonia Goncalves, Ewan Harrison, David K. Jackson, Ian Johnston, Dominic Kwiatkowski, Cordelia Langford, John Sillitoe on behalf of the Wellcome Sanger Institute COVID-19 Surveillance Team |
| EPI_ISL_2447369 | Lighthouse Lab in Milton Keynes | Wellcome Sanger Institute for the COVID-19 Genomics UK (COG-UK) Consortium | The Lighthouse Lab in Milton Keynes and Alex Alderton, Roberto Amato, Jeffrey Barrett, Sonia Goncalves, Ewan Harrison, David K. Jackson, Ian Johnston, Dominic Kwiatkowski, Cordelia Langford, John Sillitoe on behalf of the Wellcome Sanger Institute COVID-19 Surveillance Team |
| EPI_ISL_2447370 | Lighthouse Lab in Alderley Park | Wellcome Sanger Institute for the COVID-19 Genomics UK (COG-UK) Consortium | Jacquelyn Wynn, Mairead Hyland, The Lighthouse Lab in Alderley Park and Alex Alderton, Roberto Amato, Jeffrey Barrett, Sonia Goncalves, Ewan Harrison, David K. Jackson, Ian Johnston, Dominic Kwiatkowski, Cordelia Langford, John Sillitoe on behalf of the Wellcome Sanger Institute COVID-19 Surveillance Team |
| EPI_ISL_2447371, EPI_ISL_2447372 | Lighthouse Lab in Milton Keynes | Wellcome Sanger Institute for the COVID-19 Genomics UK (COG-UK) Consortium | The Lighthouse Lab in Milton Keynes and Alex Alderton, Roberto Amato, Jeffrey Barrett, Sonia Goncalves, Ewan Harrison, David K. Jackson, Ian Johnston, Dominic Kwiatkowski, Cordelia Langford, John Sillitoe on behalf of the Wellcome Sanger Institute COVID-19 Surveillance Team |
| EPI_ISL_2447373 | Lighthouse Lab in Alderley Park | Wellcome Sanger Institute for the COVID-19 Genomics UK (COG-UK) Consortium | Jacquelyn Wynn, Mairead Hyland, The Lighthouse Lab in Alderley Park and Alex Alderton, Roberto Amato, Jeffrey Barrett, Sonia Goncalves, Ewan Harrison, David K. Jackson, Ian Johnston, Dominic Kwiatkowski, Cordelia Langford, John Sillitoe on behalf of the Wellcome Sanger Institute COVID-19 Surveillance Team |
| EPI_ISL_2447374 | Lighthouse Lab in Milton Keynes | Wellcome Sanger Institute for the COVID-19 Genomics UK (COG-UK) Consortium | The Lighthouse Lab in Milton Keynes and Alex Alderton, Roberto Amato, Jeffrey Barrett, Sonia Goncalves, Ewan Harrison, David K. Jackson, Ian Johnston, Dominic Kwiatkowski, Cordelia Langford, John Sillitoe on behalf of the Wellcome Sanger Institute COVID-19 Surveillance Team |
| EPI_ISL_2447375 | Lighthouse Lab in Alderley Park | Wellcome Sanger Institute for the COVID-19 Genomics UK (COG-UK) Consortium | Jacquelyn Wynn, Mairead Hyland, The Lighthouse Lab in Alderley Park and Alex Alderton, Roberto Amato, Jeffrey Barrett, Sonia Goncalves, Ewan Harrison, David K. Jackson, Ian Johnston, Dominic Kwiatkowski, Cordelia Langford, John Sillitoe on behalf of the Wellcome Sanger Institute COVID-19 Surveillance Team |
| EPI_ISL_2447376 | Lighthouse Lab in Milton Keynes | Wellcome Sanger Institute for the COVID-19 Genomics UK (COG-UK) Consortium | The Lighthouse Lab in Milton Keynes and Alex Alderton, Roberto Amato, Jeffrey Barrett, Sonia Goncalves, Ewan Harrison, David K. Jackson, Ian Johnston, Dominic Kwiatkowski, Cordelia Langford, John Sillitoe on behalf of the Wellcome Sanger Institute COVID-19 Surveillance Team |
| EPI_ISL_2447377, EPI_ISL_2447378, EPI_ISL_2447379, EPI_ISL_2447380, EPI_ISL_2447381 | Lighthouse Lab in Alderley Park | Wellcome Sanger Institute for the COVID-19 Genomics UK (COG-UK) Consortium | Jacquelyn Wynn, Mairead Hyland, The Lighthouse Lab in Alderley Park and Alex Alderton, Roberto Amato, Jeffrey Barrett, Sonia Goncalves, Ewan Harrison, David K. Jackson, Ian Johnston, Dominic Kwiatkowski, Cordelia Langford, John Sillitoe on behalf of the Wellcome Sanger Institute COVID-19 Surveillance Team |
| EPI_ISL_2447382 | Lighthouse Laboratory Plymouth | Wellcome Sanger Institute for the COVID-19 Genomics UK (COG-UK) Consortium | Lighthouse Laboratory Plymouth and Alex Alderton, Roberto Amato, Jeffrey Barrett, Sonia Goncalves, Ewan Harrison, David K. Jackson, Ian Johnston, Dominic Kwiatkowski, Cordelia Langford, John Sillitoe on behalf of the Wellcome Sanger Institute COVID-19 Surveillance Team |
| EPI_ISL_2447383 | Lighthouse Lab in Alderley Park | Wellcome Sanger Institute for the COVID-19 Genomics UK (COG-UK) Consortium | Jacquelyn Wynn, Mairead Hyland, The Lighthouse Lab in Alderley Park and Alex Alderton, Roberto Amato, Jeffrey Barrett, Sonia Goncalves, Ewan Harrison, David K. Jackson, Ian Johnston, Dominic Kwiatkowski, Cordelia Langford, John Sillitoe on behalf of the Wellcome Sanger Institute COVID-19 Surveillance Team |
| EPI_ISL_2447384 | Health Services Laboratories | Wellcome Sanger Institute for the COVID-19 Genomics UK (COG-UK) Consortium | Health Services Laboratories and Alex Alderton, Roberto Amato, Jeffrey Barrett, Sonia Goncalves, Ewan Harrison, David K. Jackson, Ian Johnston, Dominic Kwiatkowski, Cordelia Langford, John Sillitoe on behalf of the Wellcome Sanger Institute COVID-19 Surveillance Team |
| EPI_ISL_2447385, EPI_ISL_2447386, EPI_ISL_2447387, EPI_ISL_2447388 | Lighthouse Lab in Alderley Park | Wellcome Sanger Institute for the COVID-19 Genomics UK (COG-UK) Consortium | Jacquelyn Wynn, Mairead Hyland, The Lighthouse Lab in Alderley Park and Alex Alderton, Roberto Amato, Jeffrey Barrett, Sonia Goncalves, Ewan Harrison, David K. Jackson, Ian Johnston, Dominic Kwiatkowski, Cordelia Langford, John Sillitoe on behalf of the Wellcome Sanger Institute COVID-19 Surveillance Team |
| EPI_ISL_2447389, EPI_ISL_2447390 | Lighthouse Lab in Milton Keynes | Wellcome Sanger Institute for the COVID-19 Genomics UK (COG-UK) Consortium | The Lighthouse Lab in Milton Keynes and Alex Alderton, Roberto Amato, Jeffrey Barrett, Sonia Goncalves, Ewan Harrison, David K. Jackson, Ian Johnston, Dominic Kwiatkowski, Cordelia Langford, John Sillitoe on behalf of the Wellcome Sanger Institute COVID-19 Surveillance Team |
| EPI_ISL_2447391 | Lighthouse Lab in Alderley Park | Wellcome Sanger Institute for the COVID-19 Genomics UK (COG-UK) Consortium | Jacquelyn Wynn, Mairead Hyland, The Lighthouse Lab in Alderley Park and Alex Alderton, Roberto Amato, Jeffrey Barrett, Sonia Goncalves, Ewan Harrison, David K. Jackson, Ian Johnston, Dominic Kwiatkowski, Cordelia Langford, John Sillitoe on behalf of the Wellcome Sanger Institute COVID-19 Surveillance Team |
| EPI_ISL_2447392 | Lighthouse Lab in Milton Keynes | Wellcome Sanger Institute for the COVID-19 Genomics UK (COG-UK) Consortium | The Lighthouse Lab in Milton Keynes and Alex Alderton, Roberto Amato, Jeffrey Barrett, Sonia Goncalves, Ewan Harrison, David K. Jackson, Ian Johnston, Dominic Kwiatkowski, Cordelia Langford, John Sillitoe on behalf of the Wellcome Sanger Institute COVID-19 Surveillance Team |
| EPI_ISL_2447394, EPI_ISL_2447396, EPI_ISL_2447397 | Lighthouse Lab in Alderley Park | Wellcome Sanger Institute for the COVID-19 Genomics UK (COG-UK) Consortium | Jacquelyn Wynn, Mairead Hyland, The Lighthouse Lab in Alderley Park and Alex Alderton, Roberto Amato, Jeffrey Barrett, Sonia Goncalves, Ewan Harrison, David K. Jackson, Ian Johnston, Dominic Kwiatkowski, Cordelia Langford, John Sillitoe on behalf of the Wellcome Sanger Institute COVID-19 Surveillance Team |
| EPI_ISL_2447398, EPI_ISL_2447399, EPI_ISL_2447400 | Lighthouse Lab in Milton Keynes | Wellcome Sanger Institute for the COVID-19 Genomics UK (COG-UK) Consortium | The Lighthouse Lab in Milton Keynes and Alex Alderton, Roberto Amato, Jeffrey Barrett, Sonia Goncalves, Ewan Harrison, David K. Jackson, Ian Johnston, Dominic Kwiatkowski, Cordelia Langford, John Sillitoe on behalf of the Wellcome Sanger Institute COVID-19 Surveillance Team |
| EPI_ISL_2447401, EPI_ISL_2447402 | Lighthouse Lab in Alderley Park | Wellcome Sanger Institute for the COVID-19 Genomics UK (COG-UK) Consortium | Jacquelyn Wynn, Mairead Hyland, The Lighthouse Lab in Alderley Park and Alex Alderton, Roberto Amato, Jeffrey Barrett, Sonia Goncalves, Ewan Harrison, David K. Jackson, Ian Johnston, Dominic Kwiatkowski, Cordelia Langford, John Sillitoe on behalf of the Wellcome Sanger Institute COVID-19 Surveillance Team |
| EPI_ISL_2447404 | Lighthouse Laboratory Plymouth | Wellcome Sanger Institute for the COVID-19 Genomics UK (COG-UK) Consortium | Lighthouse Laboratory Plymouth and Alex Alderton, Roberto Amato, Jeffrey Barrett, Sonia Goncalves, Ewan Harrison, David K. Jackson, Ian Johnston, Dominic Kwiatkowski, Cordelia Langford, John Sillitoe on behalf of the Wellcome Sanger Institute COVID-19 Surveillance Team |
| EPI_ISL_2447405, EPI_ISL_2447407 | Lighthouse Lab in Alderley Park | Wellcome Sanger Institute for the COVID-19 Genomics UK (COG-UK) Consortium | Jacquelyn Wynn, Mairead Hyland, The Lighthouse Lab in Alderley Park and Alex Alderton, Roberto Amato, Jeffrey Barrett, Sonia Goncalves, Ewan Harrison, David K. Jackson, Ian Johnston, Dominic Kwiatkowski, Cordelia Langford, John Sillitoe on behalf of the Wellcome Sanger Institute COVID-19 Surveillance Team |
| EPI_ISL_2447408 | Health Services Laboratories | Wellcome Sanger Institute for the COVID-19 Genomics UK (COG-UK) Consortium | Health Services Laboratories and Alex Alderton, Roberto Amato, Jeffrey Barrett, Sonia Goncalves, Ewan Harrison, David K. Jackson, Ian Johnston, Dominic Kwiatkowski, Cordelia Langford, John Sillitoe on behalf of the Wellcome Sanger Institute COVID-19 Surveillance Team |
| EPI_ISL_2447409 | Lighthouse Lab in Alderley Park | Wellcome Sanger Institute for the COVID-19 Genomics UK (COG-UK) Consortium | Jacquelyn Wynn, Mairead Hyland, The Lighthouse Lab in Alderley Park and Alex Alderton, Roberto Amato, Jeffrey Barrett, Sonia Goncalves, Ewan Harrison, David K. Jackson, Ian Johnston, Dominic Kwiatkowski, Cordelia Langford, John Sillitoe on behalf of the Wellcome Sanger Institute COVID-19 Surveillance Team |
| EPI_ISL_2447410 | Lighthouse Laboratory Plymouth | Wellcome Sanger Institute for the COVID-19 Genomics UK (COG-UK) Consortium | Lighthouse Laboratory Plymouth and Alex Alderton, Roberto Amato, Jeffrey Barrett, Sonia Goncalves, Ewan Harrison, David K. Jackson, Ian Johnston, Dominic Kwiatkowski, Cordelia Langford, John Sillitoe on behalf of the Wellcome Sanger Institute COVID-19 Surveillance Team |
| EPI_ISL_2447411 | Lighthouse Lab in Milton Keynes | Wellcome Sanger Institute for the COVID-19 Genomics UK (COG-UK) Consortium | The Lighthouse Lab in Milton Keynes and Alex Alderton, Roberto Amato, Jeffrey Barrett, Sonia Goncalves, Ewan Harrison, David K. Jackson, Ian Johnston, Dominic Kwiatkowski, Cordelia Langford, John Sillitoe on behalf of the Wellcome Sanger Institute COVID-19 Surveillance Team |
| EPI_ISL_2447412 | Lighthouse Lab in Alderley Park | Wellcome Sanger Institute for the COVID-19 Genomics UK (COG-UK) Consortium | Jacquelyn Wynn, Mairead Hyland, The Lighthouse Lab in Alderley Park and Alex Alderton, Roberto Amato, Jeffrey Barrett, Sonia Goncalves, Ewan Harrison, David K. Jackson, Ian Johnston, Dominic Kwiatkowski, Cordelia Langford, John Sillitoe on behalf of the Wellcome Sanger Institute COVID-19 Surveillance Team |
| EPI_ISL_2447414, EPI_ISL_2447415 | Lighthouse Lab in Milton Keynes | Wellcome Sanger Institute for the COVID-19 Genomics UK (COG-UK) Consortium | The Lighthouse Lab in Milton Keynes and Alex Alderton, Roberto Amato, Jeffrey Barrett, Sonia Goncalves, Ewan Harrison, David K. Jackson, Ian Johnston, Dominic Kwiatkowski, Cordelia Langford, John Sillitoe on behalf of the Wellcome Sanger Institute COVID-19 Surveillance Team |
| EPI_ISL_2447416 | Lighthouse Lab in Alderley Park | Wellcome Sanger Institute for the COVID-19 Genomics UK (COG-UK) Consortium | Jacquelyn Wynn, Mairead Hyland, The Lighthouse Lab in Alderley Park and Alex Alderton, Roberto Amato, Jeffrey Barrett, Sonia Goncalves, Ewan Harrison, David K. Jackson, Ian Johnston, Dominic Kwiatkowski, Cordelia Langford, John Sillitoe on behalf of the Wellcome Sanger Institute COVID-19 Surveillance Team |
| EPI_ISL_2447417, EPI_ISL_2447418 | Lighthouse Lab in Milton Keynes | Wellcome Sanger Institute for the COVID-19 Genomics UK (COG-UK) Consortium | The Lighthouse Lab in Milton Keynes and Alex Alderton, Roberto Amato, Jeffrey Barrett, Sonia Goncalves, Ewan Harrison, David K. Jackson, Ian Johnston, Dominic Kwiatkowski, Cordelia Langford, John Sillitoe on behalf of the Wellcome Sanger Institute COVID-19 Surveillance Team |
| EPI_ISL_2447419 | Lighthouse Laboratory Plymouth | Wellcome Sanger Institute for the COVID-19 Genomics UK (COG-UK) | Lighthouse Laboratory Plymouth and Alex Alderton, Roberto Amato, Jeffrey Barrett, Sonia Goncalves, Ewan Harrison, David K. Jackson, Ian Johnston, Dominic Kwiatkowski, Cordelia Langford, John Sillitoe on behalf of the Wellcome Sanger |

| | | | |
|---|---|---|---|
| | | Consortium | Institute COVID-19 Surveillance Team |
| EPI_ISL_2447420, EPI_ISL_2447421, EPI_ISL_2447422 | Lighthouse Lab in Alderley Park | Wellcome Sanger Institute for the COVID-19 Genomics UK (COG-UK) Consortium | Jacquelyn Wynn, Mairead Hyland, The Lighthouse Lab in Alderley Park and Alex Alderton, Roberto Amato, Jeffrey Barrett, Sonia Goncalves, Ewan Harrison, David K. Jackson, Ian Johnston, Dominic Kwiatkowski, Cordelia Langford, John Sillitoe on behalf of the Wellcome Sanger Institute COVID-19 Surveillance Team |
| EPI_ISL_2447423, EPI_ISL_2447424 | Lighthouse Lab in Milton Keynes | Wellcome Sanger Institute for the COVID-19 Genomics UK (COG-UK) Consortium | The Lighthouse Lab in Milton Keynes and Alex Alderton, Roberto Amato, Jeffrey Barrett, Sonia Goncalves, Ewan Harrison, David K. Jackson, Ian Johnston, Dominic Kwiatkowski, Cordelia Langford, John Sillitoe on behalf of the Wellcome Sanger Institute COVID-19 Surveillance Team |
| EPI_ISL_2447425, EPI_ISL_2447426, EPI_ISL_2447427, EPI_ISL_2447428, EPI_ISL_2447429 | Lighthouse Lab in Alderley Park | Wellcome Sanger Institute for the COVID-19 Genomics UK (COG-UK) Consortium | Jacquelyn Wynn, Mairead Hyland, The Lighthouse Lab in Alderley Park and Alex Alderton, Roberto Amato, Jeffrey Barrett, Sonia Goncalves, Ewan Harrison, David K. Jackson, Ian Johnston, Dominic Kwiatkowski, Cordelia Langford, John Sillitoe on behalf of the Wellcome Sanger Institute COVID-19 Surveillance Team |
| EPI_ISL_2447430 | Lighthouse Lab in Milton Keynes | Wellcome Sanger Institute for the COVID-19 Genomics UK (COG-UK) Consortium | The Lighthouse Lab in Milton Keynes and Alex Alderton, Roberto Amato, Jeffrey Barrett, Sonia Goncalves, Ewan Harrison, David K. Jackson, Ian Johnston, Dominic Kwiatkowski, Cordelia Langford, John Sillitoe on behalf of the Wellcome Sanger Institute COVID-19 Surveillance Team |
| EPI_ISL_2447431 | Lighthouse Lab in Alderley Park | Wellcome Sanger Institute for the COVID-19 Genomics UK (COG-UK) Consortium | Jacquelyn Wynn, Mairead Hyland, The Lighthouse Lab in Alderley Park and Alex Alderton, Roberto Amato, Jeffrey Barrett, Sonia Goncalves, Ewan Harrison, David K. Jackson, Ian Johnston, Dominic Kwiatkowski, Cordelia Langford, John Sillitoe on behalf of the Wellcome Sanger Institute COVID-19 Surveillance Team |
| EPI_ISL_2447432 | Lighthouse Lab in Milton Keynes | Wellcome Sanger Institute for the COVID-19 Genomics UK (COG-UK) Consortium | The Lighthouse Lab in Milton Keynes and Alex Alderton, Roberto Amato, Jeffrey Barrett, Sonia Goncalves, Ewan Harrison, David K. Jackson, Ian Johnston, Dominic Kwiatkowski, Cordelia Langford, John Sillitoe on behalf of the Wellcome Sanger Institute COVID-19 Surveillance Team |
| EPI_ISL_2447433 | Lighthouse Laboratory Plymouth | Wellcome Sanger Institute for the COVID-19 Genomics UK (COG-UK) Consortium | Lighthouse Laboratory Plymouth and Alex Alderton, Roberto Amato, Jeffrey Barrett, Sonia Goncalves, Ewan Harrison, David K. Jackson, Ian Johnston, Dominic Kwiatkowski, Cordelia Langford, John Sillitoe on behalf of the Wellcome Sanger Institute COVID-19 Surveillance Team |
| EPI_ISL_2447434 | Lighthouse Lab in Milton Keynes | Wellcome Sanger Institute for the COVID-19 Genomics UK (COG-UK) Consortium | The Lighthouse Lab in Milton Keynes and Alex Alderton, Roberto Amato, Jeffrey Barrett, Sonia Goncalves, Ewan Harrison, David K. Jackson, Ian Johnston, Dominic Kwiatkowski, Cordelia Langford, John Sillitoe on behalf of the Wellcome Sanger Institute COVID-19 Surveillance Team |
| EPI_ISL_2447435 | Lighthouse Laboratory Plymouth | Wellcome Sanger Institute for the COVID-19 Genomics UK (COG-UK) Consortium | Lighthouse Laboratory Plymouth and Alex Alderton, Roberto Amato, Jeffrey Barrett, Sonia Goncalves, Ewan Harrison, David K. Jackson, Ian Johnston, Dominic Kwiatkowski, Cordelia Langford, John Sillitoe on behalf of the Wellcome Sanger Institute COVID-19 Surveillance Team |
| EPI_ISL_2447436, EPI_ISL_2447437 | Lighthouse Lab in Milton Keynes | Wellcome Sanger Institute for the COVID-19 Genomics UK (COG-UK) Consortium | The Lighthouse Lab in Milton Keynes and Alex Alderton, Roberto Amato, Jeffrey Barrett, Sonia Goncalves, Ewan Harrison, David K. Jackson, Ian Johnston, Dominic Kwiatkowski, Cordelia Langford, John Sillitoe on behalf of the Wellcome Sanger Institute COVID-19 Surveillance Team |
| EPI_ISL_2447438, EPI_ISL_2447439 | Lighthouse Lab in Alderley Park | Wellcome Sanger Institute for the COVID-19 Genomics UK (COG-UK) Consortium | Jacquelyn Wynn, Mairead Hyland, The Lighthouse Lab in Alderley Park and Alex Alderton, Roberto Amato, Jeffrey Barrett, Sonia Goncalves, Ewan Harrison, David K. Jackson, Ian Johnston, Dominic Kwiatkowski, Cordelia Langford, John Sillitoe on behalf of the Wellcome Sanger Institute COVID-19 Surveillance Team |
| EPI_ISL_2447440, EPI_ISL_2447441 | Lighthouse Laboratory Plymouth | Wellcome Sanger Institute for the COVID-19 Genomics UK (COG-UK) Consortium | Lighthouse Laboratory Plymouth and Alex Alderton, Roberto Amato, Jeffrey Barrett, Sonia Goncalves, Ewan Harrison, David K. Jackson, Ian Johnston, Dominic Kwiatkowski, Cordelia Langford, John Sillitoe on behalf of the Wellcome Sanger Institute COVID-19 Surveillance Team |
| EPI_ISL_2447442, EPI_ISL_2447443 | Lighthouse Lab in Alderley Park | Wellcome Sanger Institute for the COVID-19 Genomics UK (COG-UK) Consortium | Jacquelyn Wynn, Mairead Hyland, The Lighthouse Lab in Alderley Park and Alex Alderton, Roberto Amato, Jeffrey Barrett, Sonia Goncalves, Ewan Harrison, David K. Jackson, Ian Johnston, Dominic Kwiatkowski, Cordelia Langford, John Sillitoe on behalf of the Wellcome Sanger Institute COVID-19 Surveillance Team |
| EPI_ISL_2447444, EPI_ISL_2447445 | Health Services Laboratories | Wellcome Sanger Institute for the COVID-19 Genomics UK (COG-UK) Consortium | Health Services Laboratories and Alex Alderton, Roberto Amato, Jeffrey Barrett, Sonia Goncalves, Ewan Harrison, David K. Jackson, Ian Johnston, Dominic Kwiatkowski, Cordelia Langford, John Sillitoe on behalf of the Wellcome Sanger Institute COVID-19 Surveillance Team |
| EPI_ISL_2447446, EPI_ISL_2447447 | Lighthouse Lab in Alderley Park | Wellcome Sanger Institute for the COVID-19 Genomics UK (COG-UK) Consortium | Jacquelyn Wynn, Mairead Hyland, The Lighthouse Lab in Alderley Park and Alex Alderton, Roberto Amato, Jeffrey Barrett, Sonia Goncalves, Ewan Harrison, David K. Jackson, Ian Johnston, Dominic Kwiatkowski, Cordelia Langford, John Sillitoe on behalf of the Wellcome Sanger Institute COVID-19 Surveillance Team |
| EPI_ISL_2447448 | Lighthouse Lab in Milton Keynes | Wellcome Sanger Institute for the COVID-19 Genomics UK (COG-UK) Consortium | The Lighthouse Lab in Milton Keynes and Alex Alderton, Roberto Amato, Jeffrey Barrett, Sonia Goncalves, Ewan Harrison, David K. Jackson, Ian Johnston, Dominic Kwiatkowski, Cordelia Langford, John Sillitoe on behalf of the Wellcome Sanger Institute COVID-19 Surveillance Team |
| EPI_ISL_2447449 | Lighthouse Lab in Alderley Park | Wellcome Sanger Institute for the COVID-19 Genomics UK (COG-UK) Consortium | Jacquelyn Wynn, Mairead Hyland, The Lighthouse Lab in Alderley Park and Alex Alderton, Roberto Amato, Jeffrey Barrett, Sonia Goncalves, Ewan Harrison, David K. Jackson, Ian Johnston, Dominic Kwiatkowski, Cordelia Langford, John Sillitoe on behalf of the Wellcome Sanger Institute COVID-19 Surveillance Team |
| EPI_ISL_2447450 | Lighthouse Lab in Milton Keynes | Wellcome Sanger Institute for the COVID-19 Genomics UK (COG-UK) Consortium | The Lighthouse Lab in Milton Keynes and Alex Alderton, Roberto Amato, Jeffrey Barrett, Sonia Goncalves, Ewan Harrison, David K. Jackson, Ian Johnston, Dominic Kwiatkowski, Cordelia Langford, John Sillitoe on behalf of the Wellcome Sanger Institute COVID-19 Surveillance Team |
| EPI_ISL_2447451, EPI_ISL_2447452 | Lighthouse Lab in Alderley Park | Wellcome Sanger Institute for the COVID-19 Genomics UK (COG-UK) Consortium | Jacquelyn Wynn, Mairead Hyland, The Lighthouse Lab in Alderley Park and Alex Alderton, Roberto Amato, Jeffrey Barrett, Sonia Goncalves, Ewan Harrison, David K. Jackson, Ian Johnston, Dominic Kwiatkowski, Cordelia Langford, John Sillitoe on behalf of the Wellcome Sanger Institute COVID-19 Surveillance Team |
| EPI_ISL_2447453, EPI_ISL_2447455 | Lighthouse Laboratory Plymouth | Wellcome Sanger Institute for the COVID-19 Genomics UK (COG-UK) Consortium | Lighthouse Laboratory Plymouth and Alex Alderton, Roberto Amato, Jeffrey Barrett, Sonia Goncalves, Ewan Harrison, David K. Jackson, Ian Johnston, Dominic Kwiatkowski, Cordelia Langford, John Sillitoe on behalf of the Wellcome Sanger Institute COVID-19 Surveillance Team |
| EPI_ISL_2447456, EPI_ISL_2447457, EPI_ISL_2447458, EPI_ISL_2447459 | Lighthouse Lab in Milton Keynes | Wellcome Sanger Institute for the COVID-19 Genomics UK (COG-UK) Consortium | The Lighthouse Lab in Milton Keynes and Alex Alderton, Roberto Amato, Jeffrey Barrett, Sonia Goncalves, Ewan Harrison, David K. Jackson, Ian Johnston, Dominic Kwiatkowski, Cordelia Langford, John Sillitoe on behalf of the Wellcome Sanger Institute COVID-19 Surveillance Team |
| EPI_ISL_2447460 | Lighthouse Lab in Alderley Park | Wellcome Sanger Institute for the COVID-19 Genomics UK (COG-UK) Consortium | Jacquelyn Wynn, Mairead Hyland, The Lighthouse Lab in Alderley Park and Alex Alderton, Roberto Amato, Jeffrey Barrett, Sonia Goncalves, Ewan Harrison, David K. Jackson, Ian Johnston, Dominic Kwiatkowski, Cordelia Langford, John Sillitoe on behalf of the Wellcome Sanger Institute COVID-19 Surveillance Team |
| EPI_ISL_2447461, EPI_ISL_2447462, EPI_ISL_2447464 | Lighthouse Lab in Milton Keynes | Wellcome Sanger Institute for the COVID-19 Genomics UK (COG-UK) Consortium | The Lighthouse Lab in Milton Keynes and Alex Alderton, Roberto Amato, Jeffrey Barrett, Sonia Goncalves, Ewan Harrison, David K. Jackson, Ian Johnston, Dominic Kwiatkowski, Cordelia Langford, John Sillitoe on behalf of the Wellcome Sanger Institute COVID-19 Surveillance Team |
| EPI_ISL_2447466 | Health Services Laboratories | Wellcome Sanger Institute for the COVID-19 Genomics UK (COG-UK) Consortium | Health Services Laboratories and Alex Alderton, Roberto Amato, Jeffrey Barrett, Sonia Goncalves, Ewan Harrison, David K. Jackson, Ian Johnston, Dominic Kwiatkowski, Cordelia Langford, John Sillitoe on behalf of the Wellcome Sanger Institute COVID-19 Surveillance Team |
| EPI_ISL_2447467 | Lighthouse Lab in Alderley Park | Wellcome Sanger Institute for the COVID-19 Genomics UK (COG-UK) Consortium | Jacquelyn Wynn, Mairead Hyland, The Lighthouse Lab in Alderley Park and Alex Alderton, Roberto Amato, Jeffrey Barrett, Sonia Goncalves, Ewan Harrison, David K. Jackson, Ian Johnston, Dominic Kwiatkowski, Cordelia Langford, John Sillitoe on behalf of the Wellcome Sanger Institute COVID-19 Surveillance Team |
| EPI_ISL_2447468 | Lighthouse Lab in Milton Keynes | Wellcome Sanger Institute for the COVID-19 Genomics UK (COG-UK) Consortium | The Lighthouse Lab in Milton Keynes and Alex Alderton, Roberto Amato, Jeffrey Barrett, Sonia Goncalves, Ewan Harrison, David K. Jackson, Ian Johnston, Dominic Kwiatkowski, Cordelia Langford, John Sillitoe on behalf of the Wellcome Sanger Institute COVID-19 Surveillance Team |
| EPI_ISL_2447469 | Lighthouse Laboratory Plymouth | Wellcome Sanger Institute for the COVID-19 Genomics UK (COG-UK) Consortium | Lighthouse Laboratory Plymouth and Alex Alderton, Roberto Amato, Jeffrey Barrett, Sonia Goncalves, Ewan Harrison, David K. Jackson, Ian Johnston, Dominic Kwiatkowski, Cordelia Langford, John Sillitoe on behalf of the Wellcome Sanger Institute COVID-19 Surveillance Team |
| EPI_ISL_2447470 | Lighthouse Lab in Alderley Park | Wellcome Sanger Institute for the COVID-19 Genomics UK (COG-UK) Consortium | Jacquelyn Wynn, Mairead Hyland, The Lighthouse Lab in Alderley Park and Alex Alderton, Roberto Amato, Jeffrey Barrett, Sonia Goncalves, Ewan Harrison, David K. Jackson, Ian Johnston, Dominic Kwiatkowski, Cordelia Langford, John Sillitoe on behalf of the Wellcome Sanger Institute COVID-19 Surveillance Team |
| EPI_ISL_2447473, EPI_ISL_2447474 | Lighthouse Lab in Milton Keynes | Wellcome Sanger Institute for the COVID-19 Genomics UK (COG-UK) Consortium | The Lighthouse Lab in Milton Keynes and Alex Alderton, Roberto Amato, Jeffrey Barrett, Sonia Goncalves, Ewan Harrison, David K. Jackson, Ian Johnston, Dominic Kwiatkowski, Cordelia Langford, John Sillitoe on behalf of the Wellcome Sanger Institute COVID-19 Surveillance Team |
| EPI_ISL_2447475 | Lighthouse Lab in Alderley Park | Wellcome Sanger Institute for the COVID-19 Genomics UK (COG-UK) Consortium | Jacquelyn Wynn, Mairead Hyland, The Lighthouse Lab in Alderley Park and Alex Alderton, Roberto Amato, Jeffrey Barrett, Sonia Goncalves, Ewan Harrison, David K. Jackson, Ian Johnston, Dominic Kwiatkowski, Cordelia Langford, John Sillitoe on behalf of the Wellcome Sanger Institute COVID-19 Surveillance Team |
| EPI_ISL_2447476, EPI_ISL_2447477, EPI_ISL_2447478, EPI_ISL_2447479 | Lighthouse Lab in Milton Keynes | Wellcome Sanger Institute for the COVID-19 Genomics UK (COG-UK) Consortium | The Lighthouse Lab in Milton Keynes and Alex Alderton, Roberto Amato, Jeffrey Barrett, Sonia Goncalves, Ewan Harrison, David K. Jackson, Ian Johnston, Dominic Kwiatkowski, Cordelia Langford, John Sillitoe on behalf of the Wellcome Sanger Institute COVID-19 Surveillance Team |
| EPI_ISL_2447480, EPI_ISL_2447482, EPI_ISL_2447483 | Lighthouse Lab in Alderley Park | Wellcome Sanger Institute for the COVID-19 Genomics UK (COG-UK) Consortium | Jacquelyn Wynn, Mairead Hyland, The Lighthouse Lab in Alderley Park and Alex Alderton, Roberto Amato, Jeffrey Barrett, Sonia Goncalves, Ewan Harrison, David K. Jackson, Ian Johnston, Dominic Kwiatkowski, Cordelia Langford, John Sillitoe on behalf of the Wellcome Sanger Institute COVID-19 Surveillance Team |
| EPI_ISL_2447485 | Lighthouse Lab in Milton Keynes | Wellcome Sanger Institute for the COVID-19 Genomics UK (COG-UK) Consortium | The Lighthouse Lab in Milton Keynes and Alex Alderton, Roberto Amato, Jeffrey Barrett, Sonia Goncalves, Ewan Harrison, David K. Jackson, Ian Johnston, Dominic Kwiatkowski, Cordelia Langford, John Sillitoe on behalf of the Wellcome Sanger Institute COVID-19 Surveillance Team |
| EPI_ISL_2447487, EPI_ISL_2447488 | Lighthouse Lab in Alderley Park | Wellcome Sanger Institute for the COVID-19 Genomics UK (COG-UK) Consortium | Jacquelyn Wynn, Mairead Hyland, The Lighthouse Lab in Alderley Park and Alex Alderton, Roberto Amato, Jeffrey Barrett, Sonia Goncalves, Ewan Harrison, David K. Jackson, Ian Johnston, Dominic Kwiatkowski, Cordelia Langford, John Sillitoe on behalf of the Wellcome Sanger Institute COVID-19 Surveillance Team |
| EPI_ISL_2447489 | Lighthouse Laboratory Plymouth | Wellcome Sanger Institute for the COVID-19 Genomics UK (COG-UK) Consortium | Lighthouse Laboratory Plymouth and Alex Alderton, Roberto Amato, Jeffrey Barrett, Sonia Goncalves, Ewan Harrison, David K. Jackson, Ian Johnston, Dominic Kwiatkowski, Cordelia Langford, John Sillitoe on behalf of the Wellcome Sanger Institute COVID-19 Surveillance Team |
| EPI_ISL_2447490, EPI_ISL_2447491 | Lighthouse Lab in Alderley Park | Wellcome Sanger Institute for the COVID-19 Genomics UK (COG-UK) Consortium | Jacquelyn Wynn, Mairead Hyland, The Lighthouse Lab in Alderley Park and Alex Alderton, Roberto Amato, Jeffrey Barrett, Sonia Goncalves, Ewan Harrison, David K. Jackson, Ian Johnston, Dominic Kwiatkowski, Cordelia Langford, John Sillitoe on behalf of the Wellcome Sanger Institute COVID-19 Surveillance Team |
| EPI_ISL_2447492 | Lighthouse Lab in Milton Keynes | Wellcome Sanger Institute for the COVID-19 Genomics UK (COG-UK) Consortium | The Lighthouse Lab in Milton Keynes and Alex Alderton, Roberto Amato, Jeffrey Barrett, Sonia Goncalves, Ewan Harrison, David K. Jackson, Ian Johnston, Dominic Kwiatkowski, Cordelia Langford, John Sillitoe on behalf of the Wellcome Sanger Institute COVID-19 Surveillance Team |
| EPI_ISL_2447493, EPI_ISL_2447495, EPI_ISL_2447496 | Health Services Laboratories | Wellcome Sanger Institute for the COVID-19 Genomics UK (COG-UK) Consortium | Health Services Laboratories and Alex Alderton, Roberto Amato, Jeffrey Barrett, Sonia Goncalves, Ewan Harrison, David K. Jackson, Ian Johnston, Dominic Kwiatkowski, Cordelia Langford, John Sillitoe on behalf of the Wellcome Sanger Institute COVID-19 Surveillance Team |
| EPI_ISL_2447497, EPI_ISL_2447499 | Lighthouse Lab in Milton Keynes | Wellcome Sanger Institute for the COVID-19 Genomics UK (COG-UK) Consortium | The Lighthouse Lab in Milton Keynes and Alex Alderton, Roberto Amato, Jeffrey Barrett, Sonia Goncalves, Ewan Harrison, David K. Jackson, Ian Johnston, Dominic Kwiatkowski, Cordelia Langford, John Sillitoe on behalf of the Wellcome Sanger Institute COVID-19 Surveillance Team |
| EPI_ISL_2447500 | Lighthouse Lab in Alderley Park | Wellcome Sanger Institute for the COVID-19 Genomics UK (COG-UK) Consortium | Jacquelyn Wynn, Mairead Hyland, The Lighthouse Lab in Alderley Park and Alex Alderton, Roberto Amato, Jeffrey Barrett, Sonia Goncalves, Ewan Harrison, David K. Jackson, Ian Johnston, Dominic Kwiatkowski, Cordelia Langford, John Sillitoe on behalf of the Wellcome Sanger Institute COVID-19 Surveillance Team |

| | | | |
|---|---|---|---|
| EPI_ISL_2447501 | Lighthouse Lab in Milton Keynes | Wellcome Sanger Institute for the COVID-19 Genomics UK (COG-UK) Consortium | The Lighthouse Lab in Milton Keynes and Alex Alderton, Roberto Amato, Jeffrey Barrett, Sonia Goncalves, Ewan Harrison, David K. Jackson, Ian Johnston, Dominic Kwiatkowski, Cordelia Langford, John Sillitoe on behalf of the Wellcome Sanger Institute COVID-19 Surveillance Team |
| EPI_ISL_2447504, EPI_ISL_2447505, EPI_ISL_2447506 | Lighthouse Lab in Alderley Park | Wellcome Sanger Institute for the COVID-19 Genomics UK (COG-UK) Consortium | Jacquelyn Wynn, Mairead Hyland, The Lighthouse Lab in Alderley Park and Alex Alderton, Roberto Amato, Jeffrey Barrett, Sonia Goncalves, Ewan Harrison, David K. Jackson, Ian Johnston, Dominic Kwiatkowski, Cordelia Langford, John Sillitoe on behalf of the Wellcome Sanger Institute COVID-19 Surveillance Team |
| EPI_ISL_2447507 | Health Services Laboratories | Wellcome Sanger Institute for the COVID-19 Genomics UK (COG-UK) Consortium | Health Services Laboratories and Alex Alderton, Roberto Amato, Jeffrey Barrett, Sonia Goncalves, Ewan Harrison, David K. Jackson, Ian Johnston, Dominic Kwiatkowski, Cordelia Langford, John Sillitoe on behalf of the Wellcome Sanger Institute COVID-19 Surveillance Team |
| EPI_ISL_2447510 | Lighthouse Lab in Alderley Park | Wellcome Sanger Institute for the COVID-19 Genomics UK (COG-UK) Consortium | Jacquelyn Wynn, Mairead Hyland, The Lighthouse Lab in Alderley Park and Alex Alderton, Roberto Amato, Jeffrey Barrett, Sonia Goncalves, Ewan Harrison, David K. Jackson, Ian Johnston, Dominic Kwiatkowski, Cordelia Langford, John Sillitoe on behalf of the Wellcome Sanger Institute COVID-19 Surveillance Team |
| EPI_ISL_2447511 | Lighthouse Lab in Milton Keynes | Wellcome Sanger Institute for the COVID-19 Genomics UK (COG-UK) Consortium | The Lighthouse Lab in Milton Keynes and Alex Alderton, Roberto Amato, Jeffrey Barrett, Sonia Goncalves, Ewan Harrison, David K. Jackson, Ian Johnston, Dominic Kwiatkowski, Cordelia Langford, John Sillitoe on behalf of the Wellcome Sanger Institute COVID-19 Surveillance Team |
| EPI_ISL_2447513 | Health Services Laboratories | Wellcome Sanger Institute for the COVID-19 Genomics UK (COG-UK) Consortium | Health Services Laboratories and Alex Alderton, Roberto Amato, Jeffrey Barrett, Sonia Goncalves, Ewan Harrison, David K. Jackson, Ian Johnston, Dominic Kwiatkowski, Cordelia Langford, John Sillitoe on behalf of the Wellcome Sanger Institute COVID-19 Surveillance Team |
| EPI_ISL_2447515, EPI_ISL_2447516, EPI_ISL_2447517, EPI_ISL_2447518, EPI_ISL_2447520 | Lighthouse Lab in Alderley Park | Wellcome Sanger Institute for the COVID-19 Genomics UK (COG-UK) Consortium | Jacquelyn Wynn, Mairead Hyland, The Lighthouse Lab in Alderley Park and Alex Alderton, Roberto Amato, Jeffrey Barrett, Sonia Goncalves, Ewan Harrison, David K. Jackson, Ian Johnston, Dominic Kwiatkowski, Cordelia Langford, John Sillitoe on behalf of the Wellcome Sanger Institute COVID-19 Surveillance Team |
| EPI_ISL_2447521, EPI_ISL_2447522 | Lighthouse Laboratory Plymouth | Wellcome Sanger Institute for the COVID-19 Genomics UK (COG-UK) Consortium | Lighthouse Laboratory Plymouth and Alex Alderton, Roberto Amato, Jeffrey Barrett, Sonia Goncalves, Ewan Harrison, David K. Jackson, Ian Johnston, Dominic Kwiatkowski, Cordelia Langford, John Sillitoe on behalf of the Wellcome Sanger Institute COVID-19 Surveillance Team |
| EPI_ISL_2447523, EPI_ISL_2447524 | Lighthouse Lab in Milton Keynes | Wellcome Sanger Institute for the COVID-19 Genomics UK (COG-UK) Consortium | The Lighthouse Lab in Milton Keynes and Alex Alderton, Roberto Amato, Jeffrey Barrett, Sonia Goncalves, Ewan Harrison, David K. Jackson, Ian Johnston, Dominic Kwiatkowski, Cordelia Langford, John Sillitoe on behalf of the Wellcome Sanger Institute COVID-19 Surveillance Team |
| EPI_ISL_2447525, EPI_ISL_2447526 | Lighthouse Lab in Alderley Park | Wellcome Sanger Institute for the COVID-19 Genomics UK (COG-UK) Consortium | Jacquelyn Wynn, Mairead Hyland, The Lighthouse Lab in Alderley Park and Alex Alderton, Roberto Amato, Jeffrey Barrett, Sonia Goncalves, Ewan Harrison, David K. Jackson, Ian Johnston, Dominic Kwiatkowski, Cordelia Langford, John Sillitoe on behalf of the Wellcome Sanger Institute COVID-19 Surveillance Team |
| EPI_ISL_2447528 | Lighthouse Lab in Milton Keynes | Wellcome Sanger Institute for the COVID-19 Genomics UK (COG-UK) Consortium | The Lighthouse Lab in Milton Keynes and Alex Alderton, Roberto Amato, Jeffrey Barrett, Sonia Goncalves, Ewan Harrison, David K. Jackson, Ian Johnston, Dominic Kwiatkowski, Cordelia Langford, John Sillitoe on behalf of the Wellcome Sanger Institute COVID-19 Surveillance Team |
| EPI_ISL_2447529 | Health Services Laboratories | Wellcome Sanger Institute for the COVID-19 Genomics UK (COG-UK) Consortium | Health Services Laboratories and Alex Alderton, Roberto Amato, Jeffrey Barrett, Sonia Goncalves, Ewan Harrison, David K. Jackson, Ian Johnston, Dominic Kwiatkowski, Cordelia Langford, John Sillitoe on behalf of the Wellcome Sanger Institute COVID-19 Surveillance Team |
| EPI_ISL_2447530, EPI_ISL_2447531, EPI_ISL_2447532 | Lighthouse Lab in Alderley Park | Wellcome Sanger Institute for the COVID-19 Genomics UK (COG-UK) Consortium | Jacquelyn Wynn, Mairead Hyland, The Lighthouse Lab in Alderley Park and Alex Alderton, Roberto Amato, Jeffrey Barrett, Sonia Goncalves, Ewan Harrison, David K. Jackson, Ian Johnston, Dominic Kwiatkowski, Cordelia Langford, John Sillitoe on behalf of the Wellcome Sanger Institute COVID-19 Surveillance Team |
| EPI_ISL_2447533 | Lighthouse Lab in Milton Keynes | Wellcome Sanger Institute for the COVID-19 Genomics UK (COG-UK) Consortium | The Lighthouse Lab in Milton Keynes and Alex Alderton, Roberto Amato, Jeffrey Barrett, Sonia Goncalves, Ewan Harrison, David K. Jackson, Ian Johnston, Dominic Kwiatkowski, Cordelia Langford, John Sillitoe on behalf of the Wellcome Sanger Institute COVID-19 Surveillance Team |
| EPI_ISL_2447534, EPI_ISL_2447535 | Lighthouse Lab in Alderley Park | Wellcome Sanger Institute for the COVID-19 Genomics UK (COG-UK) Consortium | Jacquelyn Wynn, Mairead Hyland, The Lighthouse Lab in Alderley Park and Alex Alderton, Roberto Amato, Jeffrey Barrett, Sonia Goncalves, Ewan Harrison, David K. Jackson, Ian Johnston, Dominic Kwiatkowski, Cordelia Langford, John Sillitoe on behalf of the Wellcome Sanger Institute COVID-19 Surveillance Team |
| EPI_ISL_2447536 | Lighthouse Lab in Milton Keynes | Wellcome Sanger Institute for the COVID-19 Genomics UK (COG-UK) Consortium | The Lighthouse Lab in Milton Keynes and Alex Alderton, Roberto Amato, Jeffrey Barrett, Sonia Goncalves, Ewan Harrison, David K. Jackson, Ian Johnston, Dominic Kwiatkowski, Cordelia Langford, John Sillitoe on behalf of the Wellcome Sanger Institute COVID-19 Surveillance Team |
| EPI_ISL_2447537 | Lighthouse Laboratory Plymouth | Wellcome Sanger Institute for the COVID-19 Genomics UK (COG-UK) Consortium | Lighthouse Laboratory Plymouth and Alex Alderton, Roberto Amato, Jeffrey Barrett, Sonia Goncalves, Ewan Harrison, David K. Jackson, Ian Johnston, Dominic Kwiatkowski, Cordelia Langford, John Sillitoe on behalf of the Wellcome Sanger Institute COVID-19 Surveillance Team |
| EPI_ISL_2447538, EPI_ISL_2447539, EPI_ISL_2447540 | Lighthouse Lab in Alderley Park | Wellcome Sanger Institute for the COVID-19 Genomics UK (COG-UK) Consortium | Jacquelyn Wynn, Mairead Hyland, The Lighthouse Lab in Alderley Park and Alex Alderton, Roberto Amato, Jeffrey Barrett, Sonia Goncalves, Ewan Harrison, David K. Jackson, Ian Johnston, Dominic Kwiatkowski, Cordelia Langford, John Sillitoe on behalf of the Wellcome Sanger Institute COVID-19 Surveillance Team |
| EPI_ISL_2447541 | Health Services Laboratories | Wellcome Sanger Institute for the COVID-19 Genomics UK (COG-UK) Consortium | Health Services Laboratories and Alex Alderton, Roberto Amato, Jeffrey Barrett, Sonia Goncalves, Ewan Harrison, David K. Jackson, Ian Johnston, Dominic Kwiatkowski, Cordelia Langford, John Sillitoe on behalf of the Wellcome Sanger Institute COVID-19 Surveillance Team |
| EPI_ISL_2447542 | Lighthouse Lab in Milton Keynes | Wellcome Sanger Institute for the COVID-19 Genomics UK (COG-UK) Consortium | The Lighthouse Lab in Milton Keynes and Alex Alderton, Roberto Amato, Jeffrey Barrett, Sonia Goncalves, Ewan Harrison, David K. Jackson, Ian Johnston, Dominic Kwiatkowski, Cordelia Langford, John Sillitoe on behalf of the Wellcome Sanger Institute COVID-19 Surveillance Team |
| EPI_ISL_2447543 | Lighthouse Lab in Alderley Park | Wellcome Sanger Institute for the COVID-19 Genomics UK (COG-UK) Consortium | Jacquelyn Wynn, Mairead Hyland, The Lighthouse Lab in Alderley Park and Alex Alderton, Roberto Amato, Jeffrey Barrett, Sonia Goncalves, Ewan Harrison, David K. Jackson, Ian Johnston, Dominic Kwiatkowski, Cordelia Langford, John Sillitoe on behalf of the Wellcome Sanger Institute COVID-19 Surveillance Team |
| EPI_ISL_2447544 | Lighthouse Lab in Milton Keynes | Wellcome Sanger Institute for the COVID-19 Genomics UK (COG-UK) Consortium | The Lighthouse Lab in Milton Keynes and Alex Alderton, Roberto Amato, Jeffrey Barrett, Sonia Goncalves, Ewan Harrison, David K. Jackson, Ian Johnston, Dominic Kwiatkowski, Cordelia Langford, John Sillitoe on behalf of the Wellcome Sanger Institute COVID-19 Surveillance Team |
| EPI_ISL_2447545 | Health Services Laboratories | Wellcome Sanger Institute for the COVID-19 Genomics UK (COG-UK) Consortium | Health Services Laboratories and Alex Alderton, Roberto Amato, Jeffrey Barrett, Sonia Goncalves, Ewan Harrison, David K. Jackson, Ian Johnston, Dominic Kwiatkowski, Cordelia Langford, John Sillitoe on behalf of the Wellcome Sanger Institute COVID-19 Surveillance Team |
| EPI_ISL_2447547, EPI_ISL_2447548, EPI_ISL_2447549 | Lighthouse Lab in Alderley Park | Wellcome Sanger Institute for the COVID-19 Genomics UK (COG-UK) Consortium | Jacquelyn Wynn, Mairead Hyland, The Lighthouse Lab in Alderley Park and Alex Alderton, Roberto Amato, Jeffrey Barrett, Sonia Goncalves, Ewan Harrison, David K. Jackson, Ian Johnston, Dominic Kwiatkowski, Cordelia Langford, John Sillitoe on behalf of the Wellcome Sanger Institute COVID-19 Surveillance Team |
| EPI_ISL_2447550, EPI_ISL_2447551 | Health Services Laboratories | Wellcome Sanger Institute for the COVID-19 Genomics UK (COG-UK) Consortium | Health Services Laboratories and Alex Alderton, Roberto Amato, Jeffrey Barrett, Sonia Goncalves, Ewan Harrison, David K. Jackson, Ian Johnston, Dominic Kwiatkowski, Cordelia Langford, John Sillitoe on behalf of the Wellcome Sanger Institute COVID-19 Surveillance Team |
| EPI_ISL_2447552 | Lighthouse Lab in Milton Keynes | Wellcome Sanger Institute for the COVID-19 Genomics UK (COG-UK) Consortium | The Lighthouse Lab in Milton Keynes and Alex Alderton, Roberto Amato, Jeffrey Barrett, Sonia Goncalves, Ewan Harrison, David K. Jackson, Ian Johnston, Dominic Kwiatkowski, Cordelia Langford, John Sillitoe on behalf of the Wellcome Sanger Institute COVID-19 Surveillance Team |
| EPI_ISL_2447553 | Lighthouse Lab in Alderley Park | Wellcome Sanger Institute for the COVID-19 Genomics UK (COG-UK) Consortium | Jacquelyn Wynn, Mairead Hyland, The Lighthouse Lab in Alderley Park and Alex Alderton, Roberto Amato, Jeffrey Barrett, Sonia Goncalves, Ewan Harrison, David K. Jackson, Ian Johnston, Dominic Kwiatkowski, Cordelia Langford, John Sillitoe on behalf of the Wellcome Sanger Institute COVID-19 Surveillance Team |
| EPI_ISL_2447554 | Lighthouse Laboratory Plymouth | Wellcome Sanger Institute for the COVID-19 Genomics UK (COG-UK) Consortium | Lighthouse Laboratory Plymouth and Alex Alderton, Roberto Amato, Jeffrey Barrett, Sonia Goncalves, Ewan Harrison, David K. Jackson, Ian Johnston, Dominic Kwiatkowski, Cordelia Langford, John Sillitoe on behalf of the Wellcome Sanger Institute COVID-19 Surveillance Team |
| EPI_ISL_2447555, EPI_ISL_2447556 | Lighthouse Lab in Alderley Park | Wellcome Sanger Institute for the COVID-19 Genomics UK (COG-UK) Consortium | Jacquelyn Wynn, Mairead Hyland, The Lighthouse Lab in Alderley Park and Alex Alderton, Roberto Amato, Jeffrey Barrett, Sonia Goncalves, Ewan Harrison, David K. Jackson, Ian Johnston, Dominic Kwiatkowski, Cordelia Langford, John Sillitoe on behalf of the Wellcome Sanger Institute COVID-19 Surveillance Team |
| EPI_ISL_2447557 | Lighthouse Lab in Milton Keynes | Wellcome Sanger Institute for the COVID-19 Genomics UK (COG-UK) Consortium | The Lighthouse Lab in Milton Keynes and Alex Alderton, Roberto Amato, Jeffrey Barrett, Sonia Goncalves, Ewan Harrison, David K. Jackson, Ian Johnston, Dominic Kwiatkowski, Cordelia Langford, John Sillitoe on behalf of the Wellcome Sanger Institute COVID-19 Surveillance Team |
| EPI_ISL_2447558 | Lighthouse Laboratory Plymouth | Wellcome Sanger Institute for the COVID-19 Genomics UK (COG-UK) Consortium | Lighthouse Laboratory Plymouth and Alex Alderton, Roberto Amato, Jeffrey Barrett, Sonia Goncalves, Ewan Harrison, David K. Jackson, Ian Johnston, Dominic Kwiatkowski, Cordelia Langford, John Sillitoe on behalf of the Wellcome Sanger Institute COVID-19 Surveillance Team |
| EPI_ISL_2447559, EPI_ISL_2447560 | Lighthouse Lab in Alderley Park | Wellcome Sanger Institute for the COVID-19 Genomics UK (COG-UK) Consortium | Jacquelyn Wynn, Mairead Hyland, The Lighthouse Lab in Alderley Park and Alex Alderton, Roberto Amato, Jeffrey Barrett, Sonia Goncalves, Ewan Harrison, David K. Jackson, Ian Johnston, Dominic Kwiatkowski, Cordelia Langford, John Sillitoe on behalf of the Wellcome Sanger Institute COVID-19 Surveillance Team |
| EPI_ISL_2447561 | Health Services Laboratories | Wellcome Sanger Institute for the COVID-19 Genomics UK (COG-UK) Consortium | Health Services Laboratories and Alex Alderton, Roberto Amato, Jeffrey Barrett, Sonia Goncalves, Ewan Harrison, David K. Jackson, Ian Johnston, Dominic Kwiatkowski, Cordelia Langford, John Sillitoe on behalf of the Wellcome Sanger Institute COVID-19 Surveillance Team |
| EPI_ISL_2447562 | Lighthouse Lab in Milton Keynes | Wellcome Sanger Institute for the COVID-19 Genomics UK (COG-UK) Consortium | The Lighthouse Lab in Milton Keynes and Alex Alderton, Roberto Amato, Jeffrey Barrett, Sonia Goncalves, Ewan Harrison, David K. Jackson, Ian Johnston, Dominic Kwiatkowski, Cordelia Langford, John Sillitoe on behalf of the Wellcome Sanger Institute COVID-19 Surveillance Team |
| EPI_ISL_2447563, EPI_ISL_2447564 | Health Services Laboratories | Wellcome Sanger Institute for the COVID-19 Genomics UK (COG-UK) Consortium | Health Services Laboratories and Alex Alderton, Roberto Amato, Jeffrey Barrett, Sonia Goncalves, Ewan Harrison, David K. Jackson, Ian Johnston, Dominic Kwiatkowski, Cordelia Langford, John Sillitoe on behalf of the Wellcome Sanger Institute COVID-19 Surveillance Team |
| EPI_ISL_2447565 | Lighthouse Lab in Alderley Park | Wellcome Sanger Institute for the COVID-19 Genomics UK (COG-UK) Consortium | Jacquelyn Wynn, Mairead Hyland, The Lighthouse Lab in Alderley Park and Alex Alderton, Roberto Amato, Jeffrey Barrett, Sonia Goncalves, Ewan Harrison, David K. Jackson, Ian Johnston, Dominic Kwiatkowski, Cordelia Langford, John Sillitoe on behalf of the Wellcome Sanger Institute COVID-19 Surveillance Team |
| EPI_ISL_2447566 | Lighthouse Lab in Milton Keynes | Wellcome Sanger Institute for the COVID-19 Genomics UK (COG-UK) Consortium | The Lighthouse Lab in Milton Keynes and Alex Alderton, Roberto Amato, Jeffrey Barrett, Sonia Goncalves, Ewan Harrison, David K. Jackson, Ian Johnston, Dominic Kwiatkowski, Cordelia Langford, John Sillitoe on behalf of the Wellcome Sanger Institute COVID-19 Surveillance Team |
| EPI_ISL_2447567 | Lighthouse Lab in Alderley Park | Wellcome Sanger Institute for the COVID-19 Genomics UK (COG-UK) Consortium | Jacquelyn Wynn, Mairead Hyland, The Lighthouse Lab in Alderley Park and Alex Alderton, Roberto Amato, Jeffrey Barrett, Sonia Goncalves, Ewan Harrison, David K. Jackson, Ian Johnston, Dominic Kwiatkowski, Cordelia Langford, John Sillitoe on behalf of the Wellcome Sanger Institute COVID-19 Surveillance Team |
| EPI_ISL_2447568 | Health Services Laboratories | Wellcome Sanger Institute for the COVID-19 Genomics UK (COG-UK) Consortium | Health Services Laboratories and Alex Alderton, Roberto Amato, Jeffrey Barrett, Sonia Goncalves, Ewan Harrison, David K. Jackson, Ian Johnston, Dominic Kwiatkowski, Cordelia Langford, John Sillitoe on behalf of the Wellcome Sanger Institute COVID-19 Surveillance Team |
| EPI_ISL_2447569, EPI_ISL_2447570, EPI_ISL_2447571 | Lighthouse Lab in Alderley Park | Wellcome Sanger Institute for the COVID-19 Genomics UK (COG-UK) Consortium | Jacquelyn Wynn, Mairead Hyland, The Lighthouse Lab in Alderley Park and Alex Alderton, Roberto Amato, Jeffrey Barrett, Sonia Goncalves, Ewan Harrison, David K. Jackson, Ian Johnston, Dominic Kwiatkowski, Cordelia Langford, John Sillitoe on behalf of the Wellcome Sanger Institute COVID-19 Surveillance Team |
| EPI_ISL_2447572 | Lighthouse Lab in Milton Keynes | Wellcome Sanger Institute for the COVID-19 Genomics UK (COG-UK) Consortium | The Lighthouse Lab in Milton Keynes and Alex Alderton, Roberto Amato, Jeffrey Barrett, Sonia Goncalves, Ewan Harrison, David K. Jackson, Ian Johnston, Dominic Kwiatkowski, Cordelia Langford, John Sillitoe on behalf of the Wellcome Sanger Institute COVID-19 Surveillance Team |
| EPI_ISL_2447573 | Lighthouse Lab in Alderley Park | Wellcome Sanger Institute for the COVID-19 Genomics UK (COG-UK) Consortium | Jacquelyn Wynn, Mairead Hyland, The Lighthouse Lab in Alderley Park and Alex Alderton, Roberto Amato, Jeffrey Barrett, Sonia Goncalves, Ewan Harrison, David K. Jackson, Ian Johnston, Dominic Kwiatkowski, Cordelia Langford, John Sillitoe on behalf of the Wellcome Sanger Institute COVID-19 Surveillance Team |

| | | | |
|---|---|---|---|
| EPI_ISL_2447575 | Lighthouse Lab in Milton Keynes | Wellcome Sanger Institute for the COVID-19 Genomics UK (COG-UK) Consortium | The Lighthouse Lab in Milton Keynes and Alex Alderton, Roberto Amato, Jeffrey Barrett, Sonia Goncalves, Ewan Harrison, David K. Jackson, Ian Johnston, Dominic Kwiatkowski, Cordelia Langford, John Sillitoe on behalf of the Wellcome Sanger Institute COVID-19 Surveillance Team |
| EPI_ISL_2447576 | Health Services Laboratories | Wellcome Sanger Institute for the COVID-19 Genomics UK (COG-UK) Consortium | Health Services Laboratories and Alex Alderton, Roberto Amato, Jeffrey Barrett, Sonia Goncalves, Ewan Harrison, David K. Jackson, Ian Johnston, Dominic Kwiatkowski, Cordelia Langford, John Sillitoe on behalf of the Wellcome Sanger Institute COVID-19 Surveillance Team |
| EPI_ISL_2447577, EPI_ISL_2447578, EPI_ISL_2447579 | Lighthouse Lab in Alderley Park | Wellcome Sanger Institute for the COVID-19 Genomics UK (COG-UK) Consortium | Jacquelyn Wynn, Mairead Hyland, The Lighthouse Lab in Alderley Park and Alex Alderton, Roberto Amato, Jeffrey Barrett, Sonia Goncalves, Ewan Harrison, David K. Jackson, Ian Johnston, Dominic Kwiatkowski, Cordelia Langford, John Sillitoe on behalf of the Wellcome Sanger Institute COVID-19 Surveillance Team |
| EPI_ISL_2447580 | Health Services Laboratories | Wellcome Sanger Institute for the COVID-19 Genomics UK (COG-UK) Consortium | Health Services Laboratories and Alex Alderton, Roberto Amato, Jeffrey Barrett, Sonia Goncalves, Ewan Harrison, David K. Jackson, Ian Johnston, Dominic Kwiatkowski, Cordelia Langford, John Sillitoe on behalf of the Wellcome Sanger Institute COVID-19 Surveillance Team |
| EPI_ISL_2447581 | Lighthouse Lab in Alderley Park | Wellcome Sanger Institute for the COVID-19 Genomics UK (COG-UK) Consortium | Jacquelyn Wynn, Mairead Hyland, The Lighthouse Lab in Alderley Park and Alex Alderton, Roberto Amato, Jeffrey Barrett, Sonia Goncalves, Ewan Harrison, David K. Jackson, Ian Johnston, Dominic Kwiatkowski, Cordelia Langford, John Sillitoe on behalf of the Wellcome Sanger Institute COVID-19 Surveillance Team |
| EPI_ISL_2447582 | Lighthouse Lab in Milton Keynes | Wellcome Sanger Institute for the COVID-19 Genomics UK (COG-UK) Consortium | The Lighthouse Lab in Milton Keynes and Alex Alderton, Roberto Amato, Jeffrey Barrett, Sonia Goncalves, Ewan Harrison, David K. Jackson, Ian Johnston, Dominic Kwiatkowski, Cordelia Langford, John Sillitoe on behalf of the Wellcome Sanger Institute COVID-19 Surveillance Team |
| EPI_ISL_2447583 | Lighthouse Laboratory Plymouth | Wellcome Sanger Institute for the COVID-19 Genomics UK (COG-UK) Consortium | Lighthouse Laboratory Plymouth and Alex Alderton, Roberto Amato, Jeffrey Barrett, Sonia Goncalves, Ewan Harrison, David K. Jackson, Ian Johnston, Dominic Kwiatkowski, Cordelia Langford, John Sillitoe on behalf of the Wellcome Sanger Institute COVID-19 Surveillance Team |
| EPI_ISL_2447584, EPI_ISL_2447585, EPI_ISL_2447586, EPI_ISL_2447588, EPI_ISL_2447589, EPI_ISL_2447590, EPI_ISL_2447591 | Lighthouse Lab in Alderley Park | Wellcome Sanger Institute for the COVID-19 Genomics UK (COG-UK) Consortium | Jacquelyn Wynn, Mairead Hyland, The Lighthouse Lab in Alderley Park and Alex Alderton, Roberto Amato, Jeffrey Barrett, Sonia Goncalves, Ewan Harrison, David K. Jackson, Ian Johnston, Dominic Kwiatkowski, Cordelia Langford, John Sillitoe on behalf of the Wellcome Sanger Institute COVID-19 Surveillance Team |
| EPI_ISL_2447592 | Lighthouse Laboratory Plymouth | Wellcome Sanger Institute for the COVID-19 Genomics UK (COG-UK) Consortium | Lighthouse Laboratory Plymouth and Alex Alderton, Roberto Amato, Jeffrey Barrett, Sonia Goncalves, Ewan Harrison, David K. Jackson, Ian Johnston, Dominic Kwiatkowski, Cordelia Langford, John Sillitoe on behalf of the Wellcome Sanger Institute COVID-19 Surveillance Team |
| EPI_ISL_2447593, EPI_ISL_2447594, EPI_ISL_2447595, EPI_ISL_2447596, EPI_ISL_2447597 | Lighthouse Lab in Alderley Park | Wellcome Sanger Institute for the COVID-19 Genomics UK (COG-UK) Consortium | Jacquelyn Wynn, Mairead Hyland, The Lighthouse Lab in Alderley Park and Alex Alderton, Roberto Amato, Jeffrey Barrett, Sonia Goncalves, Ewan Harrison, David K. Jackson, Ian Johnston, Dominic Kwiatkowski, Cordelia Langford, John Sillitoe on behalf of the Wellcome Sanger Institute COVID-19 Surveillance Team |
| EPI_ISL_2447598, EPI_ISL_2447599 | Lighthouse Lab in Milton Keynes | Wellcome Sanger Institute for the COVID-19 Genomics UK (COG-UK) Consortium | The Lighthouse Lab in Milton Keynes and Alex Alderton, Roberto Amato, Jeffrey Barrett, Sonia Goncalves, Ewan Harrison, David K. Jackson, Ian Johnston, Dominic Kwiatkowski, Cordelia Langford, John Sillitoe on behalf of the Wellcome Sanger Institute COVID-19 Surveillance Team |
| EPI_ISL_2447600 | Health Services Laboratories | Wellcome Sanger Institute for the COVID-19 Genomics UK (COG-UK) Consortium | Health Services Laboratories and Alex Alderton, Roberto Amato, Jeffrey Barrett, Sonia Goncalves, Ewan Harrison, David K. Jackson, Ian Johnston, Dominic Kwiatkowski, Cordelia Langford, John Sillitoe on behalf of the Wellcome Sanger Institute COVID-19 Surveillance Team |
| EPI_ISL_2447601, EPI_ISL_2447602, EPI_ISL_2447603 | Lighthouse Lab in Alderley Park | Wellcome Sanger Institute for the COVID-19 Genomics UK (COG-UK) Consortium | Jacquelyn Wynn, Mairead Hyland, The Lighthouse Lab in Alderley Park and Alex Alderton, Roberto Amato, Jeffrey Barrett, Sonia Goncalves, Ewan Harrison, David K. Jackson, Ian Johnston, Dominic Kwiatkowski, Cordelia Langford, John Sillitoe on behalf of the Wellcome Sanger Institute COVID-19 Surveillance Team |
| EPI_ISL_2447607, EPI_ISL_2447613 | SARS-CoV-2 testing team, National Institute of Infectious Diseases | Pathogen Genomics Center, National Institute of Infectious Diseases | Tsuyoshi Sekizuka, Kentaro Itokawa, Rina Tanaka, Masanori Hashino, Nozomu Hanaoka, Masumichi Saito, Naomi Nojiri, Hazuka Y Furihata, Sana Uchikoba, Tsuguto Fujimoto, Makoto Kuroda |
| EPI_ISL_2447614 | Lighthouse Lab in Milton Keynes | Wellcome Sanger Institute for the COVID-19 Genomics UK (COG-UK) Consortium | The Lighthouse Lab in Milton Keynes and Alex Alderton, Roberto Amato, Jeffrey Barrett, Sonia Goncalves, Ewan Harrison, David K. Jackson, Ian Johnston, Dominic Kwiatkowski, Cordelia Langford, John Sillitoe on behalf of the Wellcome Sanger Institute COVID-19 Surveillance Team |
| EPI_ISL_2447615 | SARS-CoV-2 testing team, National Institute of Infectious Diseases | Pathogen Genomics Center, National Institute of Infectious Diseases | Tsuyoshi Sekizuka, Kentaro Itokawa, Rina Tanaka, Masanori Hashino, Nozomu Hanaoka, Masumichi Saito, Naomi Nojiri, Hazuka Y Furihata, Sana Uchikoba, Tsuguto Fujimoto, Makoto Kuroda |
| EPI_ISL_2447616 | Health Services Laboratories | Wellcome Sanger Institute for the COVID-19 Genomics UK (COG-UK) Consortium | Health Services Laboratories and Alex Alderton, Roberto Amato, Jeffrey Barrett, Sonia Goncalves, Ewan Harrison, David K. Jackson, Ian Johnston, Dominic Kwiatkowski, Cordelia Langford, John Sillitoe on behalf of the Wellcome Sanger Institute COVID-19 Surveillance Team |
| EPI_ISL_2447617 | SARS-CoV-2 testing team, National Institute of Infectious Diseases | Pathogen Genomics Center, National Institute of Infectious Diseases | Tsuyoshi Sekizuka, Kentaro Itokawa, Rina Tanaka, Masanori Hashino, Nozomu Hanaoka, Masumichi Saito, Naomi Nojiri, Hazuka Y Furihata, Sana Uchikoba, Tsuguto Fujimoto, Makoto Kuroda |
| EPI_ISL_2447618, EPI_ISL_2447619 | Lighthouse Lab in Alderley Park | Wellcome Sanger Institute for the COVID-19 Genomics UK (COG-UK) Consortium | Jacquelyn Wynn, Mairead Hyland, The Lighthouse Lab in Alderley Park and Alex Alderton, Roberto Amato, Jeffrey Barrett, Sonia Goncalves, Ewan Harrison, David K. Jackson, Ian Johnston, Dominic Kwiatkowski, Cordelia Langford, John Sillitoe on behalf of the Wellcome Sanger Institute COVID-19 Surveillance Team |
| EPI_ISL_2447621 | Lighthouse Lab in Milton Keynes | Wellcome Sanger Institute for the COVID-19 Genomics UK (COG-UK) Consortium | The Lighthouse Lab in Milton Keynes and Alex Alderton, Roberto Amato, Jeffrey Barrett, Sonia Goncalves, Ewan Harrison, David K. Jackson, Ian Johnston, Dominic Kwiatkowski, Cordelia Langford, John Sillitoe on behalf of the Wellcome Sanger Institute COVID-19 Surveillance Team |
| EPI_ISL_2447622 | Lighthouse Lab in Alderley Park | Wellcome Sanger Institute for the COVID-19 Genomics UK (COG-UK) Consortium | Jacquelyn Wynn, Mairead Hyland, The Lighthouse Lab in Alderley Park and Alex Alderton, Roberto Amato, Jeffrey Barrett, Sonia Goncalves, Ewan Harrison, David K. Jackson, Ian Johnston, Dominic Kwiatkowski, Cordelia Langford, John Sillitoe on behalf of the Wellcome Sanger Institute COVID-19 Surveillance Team |
| EPI_ISL_2447624 | Lighthouse Laboratory Plymouth | Wellcome Sanger Institute for the COVID-19 Genomics UK (COG-UK) Consortium | Lighthouse Laboratory Plymouth and Alex Alderton, Roberto Amato, Jeffrey Barrett, Sonia Goncalves, Ewan Harrison, David K. Jackson, Ian Johnston, Dominic Kwiatkowski, Cordelia Langford, John Sillitoe on behalf of the Wellcome Sanger Institute COVID-19 Surveillance Team |
| EPI_ISL_2447625 | SARS-CoV-2 testing team, National Institute of Infectious Diseases | Pathogen Genomics Center, National Institute of Infectious Diseases | Tsuyoshi Sekizuka, Kentaro Itokawa, Rina Tanaka, Masanori Hashino, Nozomu Hanaoka, Masumichi Saito, Naomi Nojiri, Hazuka Y Furihata, Sana Uchikoba, Tsuguto Fujimoto, Makoto Kuroda |
| EPI_ISL_2447626, EPI_ISL_2447627 | Lighthouse Lab in Alderley Park | Wellcome Sanger Institute for the COVID-19 Genomics UK (COG-UK) Consortium | Jacquelyn Wynn, Mairead Hyland, The Lighthouse Lab in Alderley Park and Alex Alderton, Roberto Amato, Jeffrey Barrett, Sonia Goncalves, Ewan Harrison, David K. Jackson, Ian Johnston, Dominic Kwiatkowski, Cordelia Langford, John Sillitoe on behalf of the Wellcome Sanger Institute COVID-19 Surveillance Team |
| EPI_ISL_2447629 | Lighthouse Laboratory Plymouth | Wellcome Sanger Institute for the COVID-19 Genomics UK (COG-UK) Consortium | Lighthouse Laboratory Plymouth and Alex Alderton, Roberto Amato, Jeffrey Barrett, Sonia Goncalves, Ewan Harrison, David K. Jackson, Ian Johnston, Dominic Kwiatkowski, Cordelia Langford, John Sillitoe on behalf of the Wellcome Sanger Institute COVID-19 Surveillance Team |
| EPI_ISL_2447630 | Lighthouse Lab in Alderley Park | Wellcome Sanger Institute for the COVID-19 Genomics UK (COG-UK) Consortium | Jacquelyn Wynn, Mairead Hyland, The Lighthouse Lab in Alderley Park and Alex Alderton, Roberto Amato, Jeffrey Barrett, Sonia Goncalves, Ewan Harrison, David K. Jackson, Ian Johnston, Dominic Kwiatkowski, Cordelia Langford, John Sillitoe on behalf of the Wellcome Sanger Institute COVID-19 Surveillance Team |
| EPI_ISL_2447631 | SARS-CoV-2 testing team, National Institute of Infectious Diseases | Pathogen Genomics Center, National Institute of Infectious Diseases | Tsuyoshi Sekizuka, Kentaro Itokawa, Rina Tanaka, Masanori Hashino, Nozomu Hanaoka, Masumichi Saito, Naomi Nojiri, Hazuka Y Furihata, Sana Uchikoba, Tsuguto Fujimoto, Makoto Kuroda |
| EPI_ISL_2447632 | Health Services Laboratories | Wellcome Sanger Institute for the COVID-19 Genomics UK (COG-UK) Consortium | Health Services Laboratories and Alex Alderton, Roberto Amato, Jeffrey Barrett, Sonia Goncalves, Ewan Harrison, David K. Jackson, Ian Johnston, Dominic Kwiatkowski, Cordelia Langford, John Sillitoe on behalf of the Wellcome Sanger Institute COVID-19 Surveillance Team |
| EPI_ISL_2447633, EPI_ISL_2447635, EPI_ISL_2447637, EPI_ISL_2447638, EPI_ISL_2447641 | Lighthouse Lab in Alderley Park | Wellcome Sanger Institute for the COVID-19 Genomics UK (COG-UK) Consortium | Jacquelyn Wynn, Mairead Hyland, The Lighthouse Lab in Alderley Park and Alex Alderton, Roberto Amato, Jeffrey Barrett, Sonia Goncalves, Ewan Harrison, David K. Jackson, Ian Johnston, Dominic Kwiatkowski, Cordelia Langford, John Sillitoe on behalf of the Wellcome Sanger Institute COVID-19 Surveillance Team |
| EPI_ISL_2447643 | Lighthouse Lab in Milton Keynes | Wellcome Sanger Institute for the COVID-19 Genomics UK (COG-UK) Consortium | The Lighthouse Lab in Milton Keynes and Alex Alderton, Roberto Amato, Jeffrey Barrett, Sonia Goncalves, Ewan Harrison, David K. Jackson, Ian Johnston, Dominic Kwiatkowski, Cordelia Langford, John Sillitoe on behalf of the Wellcome Sanger Institute COVID-19 Surveillance Team |
| EPI_ISL_2447645, EPI_ISL_2447646 | Lighthouse Lab in Alderley Park | Wellcome Sanger Institute for the COVID-19 Genomics UK (COG-UK) Consortium | Jacquelyn Wynn, Mairead Hyland, The Lighthouse Lab in Alderley Park and Alex Alderton, Roberto Amato, Jeffrey Barrett, Sonia Goncalves, Ewan Harrison, David K. Jackson, Ian Johnston, Dominic Kwiatkowski, Cordelia Langford, John Sillitoe on behalf of the Wellcome Sanger Institute COVID-19 Surveillance Team |
| EPI_ISL_2447647 | Lighthouse Lab in Milton Keynes | Wellcome Sanger Institute for the COVID-19 Genomics UK (COG-UK) Consortium | The Lighthouse Lab in Milton Keynes and Alex Alderton, Roberto Amato, Jeffrey Barrett, Sonia Goncalves, Ewan Harrison, David K. Jackson, Ian Johnston, Dominic Kwiatkowski, Cordelia Langford, John Sillitoe on behalf of the Wellcome Sanger Institute COVID-19 Surveillance Team |
| EPI_ISL_2447648 | SARS-CoV-2 testing team, National Institute of Infectious Diseases | Pathogen Genomics Center, National Institute of Infectious Diseases | Tsuyoshi Sekizuka, Kentaro Itokawa, Rina Tanaka, Masanori Hashino, Nozomu Hanaoka, Masumichi Saito, Naomi Nojiri, Hazuka Y Furihata, Sana Uchikoba, Tsuguto Fujimoto, Makoto Kuroda |
| EPI_ISL_2447651 | Health Services Laboratories | Wellcome Sanger Institute for the COVID-19 Genomics UK (COG-UK) Consortium | Health Services Laboratories and Alex Alderton, Roberto Amato, Jeffrey Barrett, Sonia Goncalves, Ewan Harrison, David K. Jackson, Ian Johnston, Dominic Kwiatkowski, Cordelia Langford, John Sillitoe on behalf of the Wellcome Sanger Institute COVID-19 Surveillance Team |
| EPI_ISL_2447652 | Lighthouse Lab in Alderley Park | Wellcome Sanger Institute for the COVID-19 Genomics UK (COG-UK) Consortium | Jacquelyn Wynn, Mairead Hyland, The Lighthouse Lab in Alderley Park and Alex Alderton, Roberto Amato, Jeffrey Barrett, Sonia Goncalves, Ewan Harrison, David K. Jackson, Ian Johnston, Dominic Kwiatkowski, Cordelia Langford, John Sillitoe on behalf of the Wellcome Sanger Institute COVID-19 Surveillance Team |
| EPI_ISL_2447654 | Lighthouse Laboratory Plymouth | Wellcome Sanger Institute for the COVID-19 Genomics UK (COG-UK) Consortium | Lighthouse Laboratory Plymouth and Alex Alderton, Roberto Amato, Jeffrey Barrett, Sonia Goncalves, Ewan Harrison, David K. Jackson, Ian Johnston, Dominic Kwiatkowski, Cordelia Langford, John Sillitoe on behalf of the Wellcome Sanger Institute COVID-19 Surveillance Team |
| EPI_ISL_2447655 | Lighthouse Lab in Alderley Park | Wellcome Sanger Institute for the COVID-19 Genomics UK (COG-UK) Consortium | Jacquelyn Wynn, Mairead Hyland, The Lighthouse Lab in Alderley Park and Alex Alderton, Roberto Amato, Jeffrey Barrett, Sonia Goncalves, Ewan Harrison, David K. Jackson, Ian Johnston, Dominic Kwiatkowski, Cordelia Langford, John Sillitoe on behalf of the Wellcome Sanger Institute COVID-19 Surveillance Team |
| EPI_ISL_2447656 | SARS-CoV-2 testing team, National Institute of Infectious Diseases | Pathogen Genomics Center, National Institute of Infectious Diseases | Tsuyoshi Sekizuka, Kentaro Itokawa, Rina Tanaka, Masanori Hashino, Nozomu Hanaoka, Masumichi Saito, Naomi Nojiri, Hazuka Y Furihata, Sana Uchikoba, Tsuguto Fujimoto, Makoto Kuroda |
| EPI_ISL_2447657 | Lighthouse Lab in Milton Keynes | Wellcome Sanger Institute for the COVID-19 Genomics UK (COG-UK) Consortium | The Lighthouse Lab in Milton Keynes and Alex Alderton, Roberto Amato, Jeffrey Barrett, Sonia Goncalves, Ewan Harrison, David K. Jackson, Ian Johnston, Dominic Kwiatkowski, Cordelia Langford, John Sillitoe on behalf of the Wellcome Sanger Institute COVID-19 Surveillance Team |
| EPI_ISL_2447660 | Lighthouse Lab in Alderley Park | Wellcome Sanger Institute for the COVID-19 Genomics UK (COG-UK) | Jacquelyn Wynn, Mairead Hyland, The Lighthouse Lab in Alderley Park and Alex Alderton, Roberto Amato, Jeffrey Barrett, Sonia Goncalves, Ewan Harrison, David K. Jackson, Ian Johnston, Dominic Kwiatkowski, Cordelia Langford, John Sillitoe on |

| | | | |
|---|---|---|---|
| | | Consortium | behalf of the Wellcome Sanger Institute COVID-19 Surveillance Team |
| EPI_ISL_2447663 | Lighthouse Laboratory Plymouth | Wellcome Sanger Institute for the COVID-19 Genomics UK (COG-UK) Consortium | Lighthouse Laboratory Plymouth and Alex Alderton, Roberto Amato, Jeffrey Barrett, Sonia Goncalves, Ewan Harrison, David K. Jackson, Ian Johnston, Dominic Kwiatkowski, Cordelia Langford, John Sillitoe on behalf of the Wellcome Sanger Institute COVID-19 Surveillance Team |
| EPI_ISL_2447664 | SARS-CoV-2 testing team, National Institute of Infectious Diseases | Pathogen Genomics Center, National Institute of Infectious Diseases | Tsuyoshi Sekizuka, Kentaro Itokawa, Rina Tanaka, Masanori Hashino, Nozomu Hanaoka, Masumichi Saito, Naomi Nojiri, Hazuka Y Furihata, Sana Uchikoba, Tsuguto Fujimoto, Makoto Kuroda |
| EPI_ISL_2447665 | Lighthouse Lab in Milton Keynes | Wellcome Sanger Institute for the COVID-19 Genomics UK (COG-UK) Consortium | The Lighthouse Lab in Milton Keynes and Alex Alderton, Roberto Amato, Jeffrey Barrett, Sonia Goncalves, Ewan Harrison, David K. Jackson, Ian Johnston, Dominic Kwiatkowski, Cordelia Langford, John Sillitoe on behalf of the Wellcome Sanger Institute COVID-19 Surveillance Team |
| EPI_ISL_2447666 | Lighthouse Lab in Alderley Park | Wellcome Sanger Institute for the COVID-19 Genomics UK (COG-UK) Consortium | Jacquelyn Wynn, Mairead Hyland, The Lighthouse Lab in Alderley Park and Alex Alderton, Roberto Amato, Jeffrey Barrett, Sonia Goncalves, Ewan Harrison, David K. Jackson, Ian Johnston, Dominic Kwiatkowski, Cordelia Langford, John Sillitoe on behalf of the Wellcome Sanger Institute COVID-19 Surveillance Team |
| EPI_ISL_2447667 | SARS-CoV-2 testing team, National Institute of Infectious Diseases | Pathogen Genomics Center, National Institute of Infectious Diseases | Tsuyoshi Sekizuka, Kentaro Itokawa, Rina Tanaka, Masanori Hashino, Nozomu Hanaoka, Masumichi Saito, Naomi Nojiri, Hazuka Y Furihata, Sana Uchikoba, Tsuguto Fujimoto, Makoto Kuroda |
| EPI_ISL_2447668, EPI_ISL_2447671 | Lighthouse Lab in Alderley Park | Wellcome Sanger Institute for the COVID-19 Genomics UK (COG-UK) Consortium | Jacquelyn Wynn, Mairead Hyland, The Lighthouse Lab in Alderley Park and Alex Alderton, Roberto Amato, Jeffrey Barrett, Sonia Goncalves, Ewan Harrison, David K. Jackson, Ian Johnston, Dominic Kwiatkowski, Cordelia Langford, John Sillitoe on behalf of the Wellcome Sanger Institute COVID-19 Surveillance Team |
| EPI_ISL_2447672 | Lighthouse Laboratory Plymouth | Wellcome Sanger Institute for the COVID-19 Genomics UK (COG-UK) Consortium | Lighthouse Laboratory Plymouth and Alex Alderton, Roberto Amato, Jeffrey Barrett, Sonia Goncalves, Ewan Harrison, David K. Jackson, Ian Johnston, Dominic Kwiatkowski, Cordelia Langford, John Sillitoe on behalf of the Wellcome Sanger Institute COVID-19 Surveillance Team |
| EPI_ISL_2447673 | SARS-CoV-2 testing team, National Institute of Infectious Diseases | Pathogen Genomics Center, National Institute of Infectious Diseases | Tsuyoshi Sekizuka, Kentaro Itokawa, Rina Tanaka, Masanori Hashino, Nozomu Hanaoka, Masumichi Saito, Naomi Nojiri, Hazuka Y Furihata, Sana Uchikoba, Tsuguto Fujimoto, Makoto Kuroda |
| EPI_ISL_2447674 | Lighthouse Lab in Alderley Park | Wellcome Sanger Institute for the COVID-19 Genomics UK (COG-UK) Consortium | Jacquelyn Wynn, Mairead Hyland, The Lighthouse Lab in Alderley Park and Alex Alderton, Roberto Amato, Jeffrey Barrett, Sonia Goncalves, Ewan Harrison, David K. Jackson, Ian Johnston, Dominic Kwiatkowski, Cordelia Langford, John Sillitoe on behalf of the Wellcome Sanger Institute COVID-19 Surveillance Team |
| EPI_ISL_2447675 | Health Services Laboratories | Wellcome Sanger Institute for the COVID-19 Genomics UK (COG-UK) Consortium | Health Services Laboratories and Alex Alderton, Roberto Amato, Jeffrey Barrett, Sonia Goncalves, Ewan Harrison, David K. Jackson, Ian Johnston, Dominic Kwiatkowski, Cordelia Langford, John Sillitoe on behalf of the Wellcome Sanger Institute COVID-19 Surveillance Team |
| EPI_ISL_2447676 | SARS-CoV-2 testing team, National Institute of Infectious Diseases | Pathogen Genomics Center, National Institute of Infectious Diseases | Tsuyoshi Sekizuka, Kentaro Itokawa, Rina Tanaka, Masanori Hashino, Nozomu Hanaoka, Masumichi Saito, Naomi Nojiri, Hazuka Y Furihata, Sana Uchikoba, Tsuguto Fujimoto, Makoto Kuroda |
| EPI_ISL_2447677 | Lighthouse Lab in Alderley Park | Wellcome Sanger Institute for the COVID-19 Genomics UK (COG-UK) Consortium | Jacquelyn Wynn, Mairead Hyland, The Lighthouse Lab in Alderley Park and Alex Alderton, Roberto Amato, Jeffrey Barrett, Sonia Goncalves, Ewan Harrison, David K. Jackson, Ian Johnston, Dominic Kwiatkowski, Cordelia Langford, John Sillitoe on behalf of the Wellcome Sanger Institute COVID-19 Surveillance Team |
| EPI_ISL_2447678, EPI_ISL_2447680 | SARS-CoV-2 testing team, National Institute of Infectious Diseases | Pathogen Genomics Center, National Institute of Infectious Diseases | Tsuyoshi Sekizuka, Kentaro Itokawa, Rina Tanaka, Masanori Hashino, Nozomu Hanaoka, Masumichi Saito, Naomi Nojiri, Hazuka Y Furihata, Sana Uchikoba, Tsuguto Fujimoto, Makoto Kuroda |
| EPI_ISL_2447681 | Lighthouse Laboratory Plymouth | Wellcome Sanger Institute for the COVID-19 Genomics UK (COG-UK) Consortium | Lighthouse Laboratory Plymouth and Alex Alderton, Roberto Amato, Jeffrey Barrett, Sonia Goncalves, Ewan Harrison, David K. Jackson, Ian Johnston, Dominic Kwiatkowski, Cordelia Langford, John Sillitoe on behalf of the Wellcome Sanger Institute COVID-19 Surveillance Team |
| EPI_ISL_2447682, EPI_ISL_2447684, EPI_ISL_2447685, EPI_ISL_2447688, EPI_ISL_2447690 | Lighthouse Lab in Alderley Park | Wellcome Sanger Institute for the COVID-19 Genomics UK (COG-UK) Consortium | Jacquelyn Wynn, Mairead Hyland, The Lighthouse Lab in Alderley Park and Alex Alderton, Roberto Amato, Jeffrey Barrett, Sonia Goncalves, Ewan Harrison, David K. Jackson, Ian Johnston, Dominic Kwiatkowski, Cordelia Langford, John Sillitoe on behalf of the Wellcome Sanger Institute COVID-19 Surveillance Team |
| EPI_ISL_2447691 | Lighthouse Lab in Milton Keynes | Wellcome Sanger Institute for the COVID-19 Genomics UK (COG-UK) Consortium | The Lighthouse Lab in Milton Keynes and Alex Alderton, Roberto Amato, Jeffrey Barrett, Sonia Goncalves, Ewan Harrison, David K. Jackson, Ian Johnston, Dominic Kwiatkowski, Cordelia Langford, John Sillitoe on behalf of the Wellcome Sanger Institute COVID-19 Surveillance Team |
| EPI_ISL_2447693 | Lighthouse Lab in Alderley Park | Wellcome Sanger Institute for the COVID-19 Genomics UK (COG-UK) Consortium | Jacquelyn Wynn, Mairead Hyland, The Lighthouse Lab in Alderley Park and Alex Alderton, Roberto Amato, Jeffrey Barrett, Sonia Goncalves, Ewan Harrison, David K. Jackson, Ian Johnston, Dominic Kwiatkowski, Cordelia Langford, John Sillitoe on behalf of the Wellcome Sanger Institute COVID-19 Surveillance Team |
| EPI_ISL_2447694 | SARS-CoV-2 testing team, National Institute of Infectious Diseases | Pathogen Genomics Center, National Institute of Infectious Diseases | Tsuyoshi Sekizuka, Kentaro Itokawa, Rina Tanaka, Masanori Hashino, Nozomu Hanaoka, Masumichi Saito, Naomi Nojiri, Hazuka Y Furihata, Sana Uchikoba, Tsuguto Fujimoto, Makoto Kuroda |
| EPI_ISL_2447695 | Lighthouse Laboratory Plymouth | Wellcome Sanger Institute for the COVID-19 Genomics UK (COG-UK) Consortium | Lighthouse Laboratory Plymouth and Alex Alderton, Roberto Amato, Jeffrey Barrett, Sonia Goncalves, Ewan Harrison, David K. Jackson, Ian Johnston, Dominic Kwiatkowski, Cordelia Langford, John Sillitoe on behalf of the Wellcome Sanger Institute COVID-19 Surveillance Team |
| EPI_ISL_2447696 | Lighthouse Lab in Alderley Park | Wellcome Sanger Institute for the COVID-19 Genomics UK (COG-UK) Consortium | Jacquelyn Wynn, Mairead Hyland, The Lighthouse Lab in Alderley Park and Alex Alderton, Roberto Amato, Jeffrey Barrett, Sonia Goncalves, Ewan Harrison, David K. Jackson, Ian Johnston, Dominic Kwiatkowski, Cordelia Langford, John Sillitoe on behalf of the Wellcome Sanger Institute COVID-19 Surveillance Team |
| EPI_ISL_2447698, EPI_ISL_2447699 | Lighthouse Lab in Milton Keynes | Wellcome Sanger Institute for the COVID-19 Genomics UK (COG-UK) Consortium | The Lighthouse Lab in Milton Keynes and Alex Alderton, Roberto Amato, Jeffrey Barrett, Sonia Goncalves, Ewan Harrison, David K. Jackson, Ian Johnston, Dominic Kwiatkowski, Cordelia Langford, John Sillitoe on behalf of the Wellcome Sanger Institute COVID-19 Surveillance Team |
| EPI_ISL_2447701 | Lighthouse Lab in Alderley Park | Wellcome Sanger Institute for the COVID-19 Genomics UK (COG-UK) Consortium | Jacquelyn Wynn, Mairead Hyland, The Lighthouse Lab in Alderley Park and Alex Alderton, Roberto Amato, Jeffrey Barrett, Sonia Goncalves, Ewan Harrison, David K. Jackson, Ian Johnston, Dominic Kwiatkowski, Cordelia Langford, John Sillitoe on behalf of the Wellcome Sanger Institute COVID-19 Surveillance Team |
| EPI_ISL_2447702 | Lighthouse Lab in Milton Keynes | Wellcome Sanger Institute for the COVID-19 Genomics UK (COG-UK) Consortium | The Lighthouse Lab in Milton Keynes and Alex Alderton, Roberto Amato, Jeffrey Barrett, Sonia Goncalves, Ewan Harrison, David K. Jackson, Ian Johnston, Dominic Kwiatkowski, Cordelia Langford, John Sillitoe on behalf of the Wellcome Sanger Institute COVID-19 Surveillance Team |
| EPI_ISL_2447703 | Lighthouse Lab in Alderley Park | Wellcome Sanger Institute for the COVID-19 Genomics UK (COG-UK) Consortium | Jacquelyn Wynn, Mairead Hyland, The Lighthouse Lab in Alderley Park and Alex Alderton, Roberto Amato, Jeffrey Barrett, Sonia Goncalves, Ewan Harrison, David K. Jackson, Ian Johnston, Dominic Kwiatkowski, Cordelia Langford, John Sillitoe on behalf of the Wellcome Sanger Institute COVID-19 Surveillance Team |
| EPI_ISL_2447704 | Lighthouse Laboratory Plymouth | Wellcome Sanger Institute for the COVID-19 Genomics UK (COG-UK) Consortium | Lighthouse Laboratory Plymouth and Alex Alderton, Roberto Amato, Jeffrey Barrett, Sonia Goncalves, Ewan Harrison, David K. Jackson, Ian Johnston, Dominic Kwiatkowski, Cordelia Langford, John Sillitoe on behalf of the Wellcome Sanger Institute COVID-19 Surveillance Team |
| EPI_ISL_2447705 | Health Services Laboratories | Wellcome Sanger Institute for the COVID-19 Genomics UK (COG-UK) Consortium | Health Services Laboratories and Alex Alderton, Roberto Amato, Jeffrey Barrett, Sonia Goncalves, Ewan Harrison, David K. Jackson, Ian Johnston, Dominic Kwiatkowski, Cordelia Langford, John Sillitoe on behalf of the Wellcome Sanger Institute COVID-19 Surveillance Team |
| EPI_ISL_2447706, EPI_ISL_2447707 | Lighthouse Lab in Alderley Park | Wellcome Sanger Institute for the COVID-19 Genomics UK (COG-UK) Consortium | Jacquelyn Wynn, Mairead Hyland, The Lighthouse Lab in Alderley Park and Alex Alderton, Roberto Amato, Jeffrey Barrett, Sonia Goncalves, Ewan Harrison, David K. Jackson, Ian Johnston, Dominic Kwiatkowski, Cordelia Langford, John Sillitoe on behalf of the Wellcome Sanger Institute COVID-19 Surveillance Team |
| EPI_ISL_2447709 | Health Services Laboratories | Wellcome Sanger Institute for the COVID-19 Genomics UK (COG-UK) Consortium | Health Services Laboratories and Alex Alderton, Roberto Amato, Jeffrey Barrett, Sonia Goncalves, Ewan Harrison, David K. Jackson, Ian Johnston, Dominic Kwiatkowski, Cordelia Langford, John Sillitoe on behalf of the Wellcome Sanger Institute COVID-19 Surveillance Team |
| EPI_ISL_2447710 | Lighthouse Lab in Alderley Park | Wellcome Sanger Institute for the COVID-19 Genomics UK (COG-UK) Consortium | Jacquelyn Wynn, Mairead Hyland, The Lighthouse Lab in Alderley Park and Alex Alderton, Roberto Amato, Jeffrey Barrett, Sonia Goncalves, Ewan Harrison, David K. Jackson, Ian Johnston, Dominic Kwiatkowski, Cordelia Langford, John Sillitoe on behalf of the Wellcome Sanger Institute COVID-19 Surveillance Team |
| EPI_ISL_2447711 | Lighthouse Lab in Milton Keynes | Wellcome Sanger Institute for the COVID-19 Genomics UK (COG-UK) Consortium | The Lighthouse Lab in Milton Keynes and Alex Alderton, Roberto Amato, Jeffrey Barrett, Sonia Goncalves, Ewan Harrison, David K. Jackson, Ian Johnston, Dominic Kwiatkowski, Cordelia Langford, John Sillitoe on behalf of the Wellcome Sanger Institute COVID-19 Surveillance Team |
| EPI_ISL_2447712 | Lighthouse Laboratory Plymouth | Wellcome Sanger Institute for the COVID-19 Genomics UK (COG-UK) Consortium | Lighthouse Laboratory Plymouth and Alex Alderton, Roberto Amato, Jeffrey Barrett, Sonia Goncalves, Ewan Harrison, David K. Jackson, Ian Johnston, Dominic Kwiatkowski, Cordelia Langford, John Sillitoe on behalf of the Wellcome Sanger Institute COVID-19 Surveillance Team |
| EPI_ISL_2447713 | Lighthouse Lab in Alderley Park | Wellcome Sanger Institute for the COVID-19 Genomics UK (COG-UK) Consortium | Jacquelyn Wynn, Mairead Hyland, The Lighthouse Lab in Alderley Park and Alex Alderton, Roberto Amato, Jeffrey Barrett, Sonia Goncalves, Ewan Harrison, David K. Jackson, Ian Johnston, Dominic Kwiatkowski, Cordelia Langford, John Sillitoe on behalf of the Wellcome Sanger Institute COVID-19 Surveillance Team |
| EPI_ISL_2447715, EPI_ISL_2447716 | Lighthouse Laboratory Plymouth | Wellcome Sanger Institute for the COVID-19 Genomics UK (COG-UK) Consortium | Lighthouse Laboratory Plymouth and Alex Alderton, Roberto Amato, Jeffrey Barrett, Sonia Goncalves, Ewan Harrison, David K. Jackson, Ian Johnston, Dominic Kwiatkowski, Cordelia Langford, John Sillitoe on behalf of the Wellcome Sanger Institute COVID-19 Surveillance Team |
| EPI_ISL_2447717 | Lighthouse Lab in Alderley Park | Wellcome Sanger Institute for the COVID-19 Genomics UK (COG-UK) Consortium | Jacquelyn Wynn, Mairead Hyland, The Lighthouse Lab in Alderley Park and Alex Alderton, Roberto Amato, Jeffrey Barrett, Sonia Goncalves, Ewan Harrison, David K. Jackson, Ian Johnston, Dominic Kwiatkowski, Cordelia Langford, John Sillitoe on behalf of the Wellcome Sanger Institute COVID-19 Surveillance Team |
| EPI_ISL_2447718 | Lighthouse Laboratory Plymouth | Wellcome Sanger Institute for the COVID-19 Genomics UK (COG-UK) Consortium | Lighthouse Laboratory Plymouth and Alex Alderton, Roberto Amato, Jeffrey Barrett, Sonia Goncalves, Ewan Harrison, David K. Jackson, Ian Johnston, Dominic Kwiatkowski, Cordelia Langford, John Sillitoe on behalf of the Wellcome Sanger Institute COVID-19 Surveillance Team |
| EPI_ISL_2447719 | Lighthouse Lab in Alderley Park | Wellcome Sanger Institute for the COVID-19 Genomics UK (COG-UK) Consortium | Jacquelyn Wynn, Mairead Hyland, The Lighthouse Lab in Alderley Park and Alex Alderton, Roberto Amato, Jeffrey Barrett, Sonia Goncalves, Ewan Harrison, David K. Jackson, Ian Johnston, Dominic Kwiatkowski, Cordelia Langford, John Sillitoe on behalf of the Wellcome Sanger Institute COVID-19 Surveillance Team |
| EPI_ISL_2447720, EPI_ISL_2447721 | Lighthouse Laboratory Plymouth | Wellcome Sanger Institute for the COVID-19 Genomics UK (COG-UK) Consortium | Lighthouse Laboratory Plymouth and Alex Alderton, Roberto Amato, Jeffrey Barrett, Sonia Goncalves, Ewan Harrison, David K. Jackson, Ian Johnston, Dominic Kwiatkowski, Cordelia Langford, John Sillitoe on behalf of the Wellcome Sanger Institute COVID-19 Surveillance Team |
| EPI_ISL_2447722 | Lighthouse Lab in Alderley Park | Wellcome Sanger Institute for the COVID-19 Genomics UK (COG-UK) Consortium | Jacquelyn Wynn, Mairead Hyland, The Lighthouse Lab in Alderley Park and Alex Alderton, Roberto Amato, Jeffrey Barrett, Sonia Goncalves, Ewan Harrison, David K. Jackson, Ian Johnston, Dominic Kwiatkowski, Cordelia Langford, John Sillitoe on behalf of the Wellcome Sanger Institute COVID-19 Surveillance Team |
| EPI_ISL_2447723 | Lighthouse Laboratory Plymouth | Wellcome Sanger Institute for the COVID-19 Genomics UK (COG-UK) Consortium | Lighthouse Laboratory Plymouth and Alex Alderton, Roberto Amato, Jeffrey Barrett, Sonia Goncalves, Ewan Harrison, David K. Jackson, Ian Johnston, Dominic Kwiatkowski, Cordelia Langford, John Sillitoe on behalf of the Wellcome Sanger Institute COVID-19 Surveillance Team |
| EPI_ISL_2447724, EPI_ISL_2447725 | Lighthouse Lab in Alderley Park | Wellcome Sanger Institute for the COVID-19 Genomics UK (COG-UK) Consortium | Jacquelyn Wynn, Mairead Hyland, The Lighthouse Lab in Alderley Park and Alex Alderton, Roberto Amato, Jeffrey Barrett, Sonia Goncalves, Ewan Harrison, David K. Jackson, Ian Johnston, Dominic Kwiatkowski, Cordelia Langford, John Sillitoe on behalf of the Wellcome Sanger Institute COVID-19 Surveillance Team |
| EPI_ISL_2447727 | Health Services Laboratories | Wellcome Sanger Institute for the COVID-19 Genomics UK (COG-UK) Consortium | Health Services Laboratories and Alex Alderton, Roberto Amato, Jeffrey Barrett, Sonia Goncalves, Ewan Harrison, David K. Jackson, Ian Johnston, Dominic Kwiatkowski, Cordelia Langford, John Sillitoe on behalf of the Wellcome Sanger Institute COVID-19 Surveillance Team |
| EPI_ISL_2447728 | Lighthouse Lab in Alderley Park | Wellcome Sanger Institute for the COVID-19 Genomics UK (COG-UK) Consortium | Jacquelyn Wynn, Mairead Hyland, The Lighthouse Lab in Alderley Park and Alex Alderton, Roberto Amato, Jeffrey Barrett, Sonia Goncalves, Ewan Harrison, David K. Jackson, Ian Johnston, Dominic Kwiatkowski, Cordelia Langford, John Sillitoe on behalf of the Wellcome Sanger Institute COVID-19 Surveillance Team |
| EPI_ISL_2447729 | Health Services Laboratories | Wellcome Sanger Institute for the COVID-19 Genomics UK (COG-UK) Consortium | Health Services Laboratories and Alex Alderton, Roberto Amato, Jeffrey Barrett, Sonia Goncalves, Ewan Harrison, David K. Jackson, Ian Johnston, Dominic Kwiatkowski, Cordelia Langford, John Sillitoe on behalf of the Wellcome Sanger Institute COVID-19 Surveillance Team |

| | | | |
|---|---|---|---|
| EPI_ISL_2447730, EPI_ISL_2447731 | Lighthouse Lab in Alderley Park | Wellcome Sanger Institute for the COVID-19 Genomics UK (COG-UK) Consortium | Jacquelyn Wynn, Mairead Hyland, The Lighthouse Lab in Alderley Park and Alex Alderton, Roberto Amato, Jeffrey Barrett, Sonia Goncalves, Ewan Harrison, David K. Jackson, Ian Johnston, Dominic Kwiatkowski, Cordelia Langford, John Sillitoe on behalf of the Wellcome Sanger Institute COVID-19 Surveillance Team |
| EPI_ISL_2447733 | Lighthouse Lab in Milton Keynes | Wellcome Sanger Institute for the COVID-19 Genomics UK (COG-UK) Consortium | The Lighthouse Lab in Milton Keynes and Alex Alderton, Roberto Amato, Jeffrey Barrett, Sonia Goncalves, Ewan Harrison, David K. Jackson, Ian Johnston, Dominic Kwiatkowski, Cordelia Langford, John Sillitoe on behalf of the Wellcome Sanger Institute COVID-19 Surveillance Team |
| EPI_ISL_2447734 | Lighthouse Laboratory Plymouth | Wellcome Sanger Institute for the COVID-19 Genomics UK (COG-UK) Consortium | Lighthouse Laboratory Plymouth and Alex Alderton, Roberto Amato, Jeffrey Barrett, Sonia Goncalves, Ewan Harrison, David K. Jackson, Ian Johnston, Dominic Kwiatkowski, Cordelia Langford, John Sillitoe on behalf of the Wellcome Sanger Institute COVID-19 Surveillance Team |
| EPI_ISL_2447735 | Lighthouse Lab in Milton Keynes | Wellcome Sanger Institute for the COVID-19 Genomics UK (COG-UK) Consortium | The Lighthouse Lab in Milton Keynes and Alex Alderton, Roberto Amato, Jeffrey Barrett, Sonia Goncalves, Ewan Harrison, David K. Jackson, Ian Johnston, Dominic Kwiatkowski, Cordelia Langford, John Sillitoe on behalf of the Wellcome Sanger Institute COVID-19 Surveillance Team |
| EPI_ISL_2447736 | Lighthouse Lab in Alderley Park | Wellcome Sanger Institute for the COVID-19 Genomics UK (COG-UK) Consortium | Jacquelyn Wynn, Mairead Hyland, The Lighthouse Lab in Alderley Park and Alex Alderton, Roberto Amato, Jeffrey Barrett, Sonia Goncalves, Ewan Harrison, David K. Jackson, Ian Johnston, Dominic Kwiatkowski, Cordelia Langford, John Sillitoe on behalf of the Wellcome Sanger Institute COVID-19 Surveillance Team |
| EPI_ISL_2447737 | Lighthouse Lab in Milton Keynes | Wellcome Sanger Institute for the COVID-19 Genomics UK (COG-UK) Consortium | The Lighthouse Lab in Milton Keynes and Alex Alderton, Roberto Amato, Jeffrey Barrett, Sonia Goncalves, Ewan Harrison, David K. Jackson, Ian Johnston, Dominic Kwiatkowski, Cordelia Langford, John Sillitoe on behalf of the Wellcome Sanger Institute COVID-19 Surveillance Team |
| EPI_ISL_2447738 | Lighthouse Lab in Alderley Park | Wellcome Sanger Institute for the COVID-19 Genomics UK (COG-UK) Consortium | Jacquelyn Wynn, Mairead Hyland, The Lighthouse Lab in Alderley Park and Alex Alderton, Roberto Amato, Jeffrey Barrett, Sonia Goncalves, Ewan Harrison, David K. Jackson, Ian Johnston, Dominic Kwiatkowski, Cordelia Langford, John Sillitoe on behalf of the Wellcome Sanger Institute COVID-19 Surveillance Team |
| EPI_ISL_2447739 | Lighthouse Laboratory Plymouth | Wellcome Sanger Institute for the COVID-19 Genomics UK (COG-UK) Consortium | Lighthouse Laboratory Plymouth and Alex Alderton, Roberto Amato, Jeffrey Barrett, Sonia Goncalves, Ewan Harrison, David K. Jackson, Ian Johnston, Dominic Kwiatkowski, Cordelia Langford, John Sillitoe on behalf of the Wellcome Sanger Institute COVID-19 Surveillance Team |
| EPI_ISL_2447740 | Lighthouse Lab in Alderley Park | Wellcome Sanger Institute for the COVID-19 Genomics UK (COG-UK) Consortium | Jacquelyn Wynn, Mairead Hyland, The Lighthouse Lab in Alderley Park and Alex Alderton, Roberto Amato, Jeffrey Barrett, Sonia Goncalves, Ewan Harrison, David K. Jackson, Ian Johnston, Dominic Kwiatkowski, Cordelia Langford, John Sillitoe on behalf of the Wellcome Sanger Institute COVID-19 Surveillance Team |
| EPI_ISL_2447742, EPI_ISL_2447743 | Lighthouse Lab in Milton Keynes | Wellcome Sanger Institute for the COVID-19 Genomics UK (COG-UK) Consortium | The Lighthouse Lab in Milton Keynes and Alex Alderton, Roberto Amato, Jeffrey Barrett, Sonia Goncalves, Ewan Harrison, David K. Jackson, Ian Johnston, Dominic Kwiatkowski, Cordelia Langford, John Sillitoe on behalf of the Wellcome Sanger Institute COVID-19 Surveillance Team |
| EPI_ISL_2447744 | Lighthouse Laboratory Plymouth | Wellcome Sanger Institute for the COVID-19 Genomics UK (COG-UK) Consortium | Lighthouse Laboratory Plymouth and Alex Alderton, Roberto Amato, Jeffrey Barrett, Sonia Goncalves, Ewan Harrison, David K. Jackson, Ian Johnston, Dominic Kwiatkowski, Cordelia Langford, John Sillitoe on behalf of the Wellcome Sanger Institute COVID-19 Surveillance Team |
| EPI_ISL_2447745, EPI_ISL_2447746, EPI_ISL_2447748 | Lighthouse Lab in Alderley Park | Wellcome Sanger Institute for the COVID-19 Genomics UK (COG-UK) Consortium | Jacquelyn Wynn, Mairead Hyland, The Lighthouse Lab in Alderley Park and Alex Alderton, Roberto Amato, Jeffrey Barrett, Sonia Goncalves, Ewan Harrison, David K. Jackson, Ian Johnston, Dominic Kwiatkowski, Cordelia Langford, John Sillitoe on behalf of the Wellcome Sanger Institute COVID-19 Surveillance Team |
| EPI_ISL_2447749 | Health Services Laboratories | Wellcome Sanger Institute for the COVID-19 Genomics UK (COG-UK) Consortium | Health Services Laboratories and Alex Alderton, Roberto Amato, Jeffrey Barrett, Sonia Goncalves, Ewan Harrison, David K. Jackson, Ian Johnston, Dominic Kwiatkowski, Cordelia Langford, John Sillitoe on behalf of the Wellcome Sanger Institute COVID-19 Surveillance Team |
| EPI_ISL_2447750 | Lighthouse Lab in Milton Keynes | Wellcome Sanger Institute for the COVID-19 Genomics UK (COG-UK) Consortium | The Lighthouse Lab in Milton Keynes and Alex Alderton, Roberto Amato, Jeffrey Barrett, Sonia Goncalves, Ewan Harrison, David K. Jackson, Ian Johnston, Dominic Kwiatkowski, Cordelia Langford, John Sillitoe on behalf of the Wellcome Sanger Institute COVID-19 Surveillance Team |
| EPI_ISL_2447751, EPI_ISL_2447752, EPI_ISL_2447753 | Lighthouse Lab in Alderley Park | Wellcome Sanger Institute for the COVID-19 Genomics UK (COG-UK) Consortium | Jacquelyn Wynn, Mairead Hyland, The Lighthouse Lab in Alderley Park and Alex Alderton, Roberto Amato, Jeffrey Barrett, Sonia Goncalves, Ewan Harrison, David K. Jackson, Ian Johnston, Dominic Kwiatkowski, Cordelia Langford, John Sillitoe on behalf of the Wellcome Sanger Institute COVID-19 Surveillance Team |
| EPI_ISL_2447754 | Lighthouse Lab in Milton Keynes | Wellcome Sanger Institute for the COVID-19 Genomics UK (COG-UK) Consortium | The Lighthouse Lab in Milton Keynes and Alex Alderton, Roberto Amato, Jeffrey Barrett, Sonia Goncalves, Ewan Harrison, David K. Jackson, Ian Johnston, Dominic Kwiatkowski, Cordelia Langford, John Sillitoe on behalf of the Wellcome Sanger Institute COVID-19 Surveillance Team |
| EPI_ISL_2447755, EPI_ISL_2447756, EPI_ISL_2447757 | Lighthouse Lab in Alderley Park | Wellcome Sanger Institute for the COVID-19 Genomics UK (COG-UK) Consortium | Jacquelyn Wynn, Mairead Hyland, The Lighthouse Lab in Alderley Park and Alex Alderton, Roberto Amato, Jeffrey Barrett, Sonia Goncalves, Ewan Harrison, David K. Jackson, Ian Johnston, Dominic Kwiatkowski, Cordelia Langford, John Sillitoe on behalf of the Wellcome Sanger Institute COVID-19 Surveillance Team |
| EPI_ISL_2447759 | Health Services Laboratories | Wellcome Sanger Institute for the COVID-19 Genomics UK (COG-UK) Consortium | Health Services Laboratories and Alex Alderton, Roberto Amato, Jeffrey Barrett, Sonia Goncalves, Ewan Harrison, David K. Jackson, Ian Johnston, Dominic Kwiatkowski, Cordelia Langford, John Sillitoe on behalf of the Wellcome Sanger Institute COVID-19 Surveillance Team |
| EPI_ISL_2447760 | Lighthouse Lab in Alderley Park | Wellcome Sanger Institute for the COVID-19 Genomics UK (COG-UK) Consortium | Jacquelyn Wynn, Mairead Hyland, The Lighthouse Lab in Alderley Park and Alex Alderton, Roberto Amato, Jeffrey Barrett, Sonia Goncalves, Ewan Harrison, David K. Jackson, Ian Johnston, Dominic Kwiatkowski, Cordelia Langford, John Sillitoe on behalf of the Wellcome Sanger Institute COVID-19 Surveillance Team |
| EPI_ISL_2447770 | Lighthouse Lab in Milton Keynes | Wellcome Sanger Institute for the COVID-19 Genomics UK (COG-UK) Consortium | The Lighthouse Lab in Milton Keynes and Alex Alderton, Roberto Amato, Jeffrey Barrett, Sonia Goncalves, Ewan Harrison, David K. Jackson, Ian Johnston, Dominic Kwiatkowski, Cordelia Langford, John Sillitoe on behalf of the Wellcome Sanger Institute COVID-19 Surveillance Team |
| EPI_ISL_2447771 | Randox Laboratories | Wellcome Sanger Institute for the COVID-19 Genomics UK (COG-UK) Consortium | Randox Laboratories and Alex Alderton, Roberto Amato, Jeffrey Barrett, Sonia Goncalves, Ewan Harrison, David K. Jackson, Ian Johnston, Dominic Kwiatkowski, Cordelia Langford, John Sillitoe on behalf of the Wellcome Sanger Institute COVID-19 Surveillance Team |
| EPI_ISL_2447907, EPI_ISL_2447918, EPI_ISL_2447924 | Virology, Universitätsklinikum des Saarlandes | Epigenetics, Saarland University | Kathrin Kattler, Nastasja Seiwert, Stefan Lohse, Sascha Tierling, Thorsten Pfuhl, Sigrun Smola, Jörn Walter |
| EPI_ISL_2448019 | Microbiology Department, Laboratori Clínic Metropolitana Nord. Hospital Universitari Germans Trias i Pujol | Can Ruti SARS-CoV-2 Sequencing Hub (HUGTiP/IrsiCaixa/IGTP) | Marc Noguera-Julian, Pilar Armengol, Ignacio Blanco, Antoni E Bordoy, Francesc Catala-Moll, Pere-Joan Cardona, Maria Casadell, Cristina Casa, Gemma Clara, Bonaventura Clotet, Cristina Esteban, Montserrat Gimnez, Mercedes Guerrero, Anna Not, Roger Paredes, Mariona Parera, Vernica Saludes, Alba Snchez, and Elisa Martr on behalf of the Can Ruti SARS-CoV-2 Sequencing Hub. |
| EPI_ISL_2448020 | Microbiologia CATLAB | Can Ruti SARS-CoV-2 Sequencing Hub (HUGTiP/IrsiCaixa/IGTP) | Marc Noguera-Julian, Pilar Armengol, Ignacio Blanco, Antoni E Bordoy, Francesc Catala-Moll, Pere-Joan Cardona, Maria Casadell, Cristina Casa, Gemma Clara, Bonaventura Clotet, Cristina Esteban, Montserrat Gimnez, Mercedes Guerrero, Anna Not, Roger Paredes, Mariona Parera, Vernica Saludes, Alba Snchez, and Elisa Martr on behalf of the Can Ruti SARS-CoV-2 Sequencing Hub. |
| EPI_ISL_2448021, EPI_ISL_2448027, EPI_ISL_2448029 | Microbiology Department, Laboratori Clínic Metropolitana Nord. Hospital Universitari Germans Trias i Pujol | Can Ruti SARS-CoV-2 Sequencing Hub (HUGTiP/IrsiCaixa/IGTP) | Marc Noguera-Julian, Pilar Armengol, Ignacio Blanco, Antoni E Bordoy, Francesc Catala-Moll, Pere-Joan Cardona, Maria Casadell, Cristina Casa, Gemma Clara, Bonaventura Clotet, Cristina Esteban, Montserrat Gimnez, Mercedes Guerrero, Anna Not, Roger Paredes, Mariona Parera, Vernica Saludes, Alba Snchez, and Elisa Martr on behalf of the Can Ruti SARS-CoV-2 Sequencing Hub. |
| EPI_ISL_2448031 | Microbiologia CATLAB | Can Ruti SARS-CoV-2 Sequencing Hub (HUGTiP/IrsiCaixa/IGTP) | Marc Noguera-Julian, Pilar Armengol, Ignacio Blanco, Antoni E Bordoy, Francesc Catala-Moll, Pere-Joan Cardona, Maria Casadell, Cristina Casa, Gemma Clara, Bonaventura Clotet, Cristina Esteban, Montserrat Gimnez, Mercedes Guerrero, Anna Not, Roger Paredes, Mariona Parera, Vernica Saludes, Alba Snchez, and Elisa Martr on behalf of the Can Ruti SARS-CoV-2 Sequencing Hub. |
| EPI_ISL_2448035 | Microbiology Department, Laboratori Clínic Metropolitana Nord. Hospital Universitari Germans Trias i Pujol | Can Ruti SARS-CoV-2 Sequencing Hub (HUGTiP/IrsiCaixa/IGTP) | Marc Noguera-Julian, Pilar Armengol, Ignacio Blanco, Antoni E Bordoy, Francesc Catala-Moll, Pere-Joan Cardona, Maria Casadell, Cristina Casa, Gemma Clara, Bonaventura Clotet, Cristina Esteban, Montserrat Gimnez, Mercedes Guerrero, Anna Not, Roger Paredes, Mariona Parera, Vernica Saludes, Alba Snchez, and Elisa Martr on behalf of the Can Ruti SARS-CoV-2 Sequencing Hub. |
| EPI_ISL_2448042 | Microbiologia CATLAB | Can Ruti SARS-CoV-2 Sequencing Hub (HUGTiP/IrsiCaixa/IGTP) | Marc Noguera-Julian, Pilar Armengol, Ignacio Blanco, Antoni E Bordoy, Francesc Catala-Moll, Pere-Joan Cardona, Maria Casadell, Cristina Casa, Gemma Clara, Bonaventura Clotet, Cristina Esteban, Montserrat Gimnez, Mercedes Guerrero, Anna Not, Roger Paredes, Mariona Parera, Vernica Saludes, Alba Snchez, and Elisa Martr on behalf of the Can Ruti SARS-CoV-2 Sequencing Hub. |
| EPI_ISL_2448043 | Microbiology Department, Laboratori Clínic Metropolitana Nord. Hospital Universitari Germans Trias i Pujol | Can Ruti SARS-CoV-2 Sequencing Hub (HUGTiP/IrsiCaixa/IGTP) | Marc Noguera-Julian, Pilar Armengol, Ignacio Blanco, Antoni E Bordoy, Francesc Catala-Moll, Pere-Joan Cardona, Maria Casadell, Cristina Casa, Gemma Clara, Bonaventura Clotet, Cristina Esteban, Montserrat Gimnez, Mercedes Guerrero, Anna Not, Roger Paredes, Mariona Parera, Vernica Saludes, Alba Snchez, and Elisa Martr on behalf of the Can Ruti SARS-CoV-2 Sequencing Hub. |
| EPI_ISL_2448053 | Microbiologia CATLAB | Can Ruti SARS-CoV-2 Sequencing Hub (HUGTiP/IrsiCaixa/IGTP) | Marc Noguera-Julian, Pilar Armengol, Ignacio Blanco, Antoni E Bordoy, Francesc Catala-Moll, Pere-Joan Cardona, Maria Casadell, Cristina Casa, Gemma Clara, Bonaventura Clotet, Cristina Esteban, Montserrat Gimnez, Mercedes Guerrero, Anna Not, Roger Paredes, Mariona Parera, Vernica Saludes, Alba Snchez, and Elisa Martr on behalf of the Can Ruti SARS-CoV-2 Sequencing Hub. |
| EPI_ISL_2448056, EPI_ISL_2448058 | Microbiology Department, Laboratori Clínic Metropolitana Nord. Hospital Universitari Germans Trias i Pujol | Can Ruti SARS-CoV-2 Sequencing Hub (HUGTiP/IrsiCaixa/IGTP) | Marc Noguera-Julian, Pilar Armengol, Ignacio Blanco, Antoni E Bordoy, Francesc Catala-Moll, Pere-Joan Cardona, Maria Casadell, Cristina Casa, Gemma Clara, Bonaventura Clotet, Cristina Esteban, Montserrat Gimnez, Mercedes Guerrero, Anna Not, Roger Paredes, Mariona Parera, Vernica Saludes, Alba Snchez, and Elisa Martr on behalf of the Can Ruti SARS-CoV-2 Sequencing Hub. |
| EPI_ISL_2448065 | Microbiologia CATLAB | Can Ruti SARS-CoV-2 Sequencing Hub (HUGTiP/IrsiCaixa/IGTP) | Marc Noguera-Julian, Pilar Armengol, Ignacio Blanco, Antoni E Bordoy, Francesc Catala-Moll, Pere-Joan Cardona, Maria Casadell, Cristina Casa, Gemma Clara, Bonaventura Clotet, Cristina Esteban, Montserrat Gimnez, Mercedes Guerrero, Anna Not, Roger Paredes, Mariona Parera, Vernica Saludes, Alba Snchez, and Elisa Martr on behalf of the Can Ruti SARS-CoV-2 Sequencing Hub. |
| EPI_ISL_2448066, EPI_ISL_2448067, EPI_ISL_2448072 | Microbiology Department, Laboratori Clínic Metropolitana Nord. Hospital Universitari Germans Trias i Pujol | Can Ruti SARS-CoV-2 Sequencing Hub (HUGTiP/IrsiCaixa/IGTP) | Marc Noguera-Julian, Pilar Armengol, Ignacio Blanco, Antoni E Bordoy, Francesc Catala-Moll, Pere-Joan Cardona, Maria Casadell, Cristina Casa, Gemma Clara, Bonaventura Clotet, Cristina Esteban, Montserrat Gimnez, Mercedes Guerrero, Anna Not, Roger Paredes, Mariona Parera, Vernica Saludes, Alba Snchez, and Elisa Martr on behalf of the Can Ruti SARS-CoV-2 Sequencing Hub. |
| EPI_ISL_2448075, EPI_ISL_2448076 | Microbiologia CATLAB | Can Ruti SARS-CoV-2 Sequencing Hub (HUGTiP/IrsiCaixa/IGTP) | Marc Noguera-Julian, Pilar Armengol, Ignacio Blanco, Antoni E Bordoy, Francesc Catala-Moll, Pere-Joan Cardona, Maria Casadell, Cristina Casa, Gemma Clara, Bonaventura Clotet, Cristina Esteban, Montserrat Gimnez, Mercedes Guerrero, Anna Not, Roger Paredes, Mariona Parera, Vernica Saludes, Alba Snchez, and Elisa Martr on behalf of the Can Ruti SARS-CoV-2 Sequencing Hub. |
| EPI_ISL_2448077 | Microbiology Department, Laboratori Clínic Metropolitana Nord. Hospital Universitari Germans Trias i Pujol | Can Ruti SARS-CoV-2 Sequencing Hub (HUGTiP/IrsiCaixa/IGTP) | Marc Noguera-Julian, Pilar Armengol, Ignacio Blanco, Antoni E Bordoy, Francesc Catala-Moll, Pere-Joan Cardona, Maria Casadell, Cristina Casa, Gemma Clara, Bonaventura Clotet, Cristina Esteban, Montserrat Gimnez, Mercedes Guerrero, Anna Not, Roger Paredes, Mariona Parera, Vernica Saludes, Alba Snchez, and Elisa Martr on behalf of the Can Ruti SARS-CoV-2 Sequencing Hub. |
| EPI_ISL_2448087, EPI_ISL_2448088 | Microbiologia CATLAB | Can Ruti SARS-CoV-2 Sequencing Hub (HUGTiP/IrsiCaixa/IGTP) | Marc Noguera-Julian, Pilar Armengol, Ignacio Blanco, Antoni E Bordoy, Francesc Catala-Moll, Pere-Joan Cardona, Maria Casadell, Cristina Casa, Gemma Clara, Bonaventura Clotet, Cristina Esteban, Montserrat Gimnez, Mercedes Guerrero, Anna Not, Roger Paredes, Mariona Parera, Vernica Saludes, Alba Snchez, and Elisa Martr on behalf of the Can Ruti SARS-CoV-2 Sequencing Hub. |
| EPI_ISL_2448095, EPI_ISL_2448097 | Microbiology Department, Laboratori Clínic Metropolitana Nord. Hospital Universitari Germans Trias i Pujol | Can Ruti SARS-CoV-2 Sequencing Hub (HUGTiP/IrsiCaixa/IGTP) | Marc Noguera-Julian, Pilar Armengol, Ignacio Blanco, Antoni E Bordoy, Francesc Catala-Moll, Pere-Joan Cardona, Maria Casadell, Cristina Casa, Gemma Clara, Bonaventura Clotet, Cristina Esteban, Montserrat Gimnez, Mercedes Guerrero, Anna Not, Roger Paredes, Mariona Parera, Vernica Saludes, Alba Snchez, and Elisa Martr on behalf of the Can Ruti SARS-CoV-2 Sequencing Hub. |
| EPI_ISL_2448098 | Microbiologia CATLAB | Can Ruti SARS-CoV-2 Sequencing Hub (HUGTiP/IrsiCaixa/IGTP) | Marc Noguera-Julian, Pilar Armengol, Ignacio Blanco, Antoni E Bordoy, Francesc Catala-Moll, Pere-Joan Cardona, Maria Casadell, Cristina Casa, Gemma Clara, Bonaventura Clotet, Cristina Esteban, Montserrat Gimnez, Mercedes Guerrero, Anna Not, Roger Paredes, Mariona Parera, Vernica Saludes, Alba Snchez, and Elisa Martr on behalf of the Can Ruti SARS-CoV-2 Sequencing Hub. |
| EPI_ISL_2448101, EPI_ISL_2448104 | Microbiology Department, Laboratori Clínic Metropolitana Nord. Hospital Universitari Germans Trias i Pujol | Can Ruti SARS-CoV-2 Sequencing Hub (HUGTiP/IrsiCaixa/IGTP) | Marc Noguera-Julian, Pilar Armengol, Ignacio Blanco, Antoni E Bordoy, Francesc Catala-Moll, Pere-Joan Cardona, Maria Casadell, Cristina Casa, Gemma Clara, Bonaventura Clotet, Cristina Esteban, Montserrat Gimnez, Mercedes Guerrero, Anna Not, Roger Paredes, Mariona Parera, Vernica Saludes, Alba Snchez, and Elisa Martr on behalf of the Can Ruti SARS-CoV-2 Sequencing Hub. |
| EPI_ISL_2448369, EPI_ISL_2448370, EPI_ISL_2448371, EPI_ISL_2448372, EPI_ISL_2448374, EPI_ISL_2448376 | Virology, Universitätsklinikum des Saarlandes | Epigenetics, Saarland University | Kathrin Kattler, Stefan Lohse, Sascha Tierling, Thorsten Pfuhl, Sigrun Smola, Jörn Walter |
| EPI_ISL_2448428, EPI_ISL_2448429, | ULSS 8 Berica | Istituto Zooprofilattico Sperimentale delle Venezie | Adelaide Milani, Alessia Schivo, Annalisa Salviato, Elisa Palumbo, Erika Giorgia Quaranta, Luca Tassoni, Ambra Pastori, Edoardo Giussani, Alice Fusaro, Isabella Monne, Calogero Terregino, Antonia Ricci |

| | | | |
|---|---|---|---|
| EPI_ISL_2448430, EPI_ISL_2448521 | | | |
| EPI_ISL_2448534 | ULSS 7 Pedemontana - Distretto 2 | Istituto Zooprofilattico Sperimentale delle Venezie | Adelaide Milani, Alessia Schivo, Annalisa Salviato, Elisa Palumbo, Erika Giorgia Quaranta, Luca Tassoni, Ambra Pastori, Edoardo Giussani, Alice Fusaro, Isabella Monne, Calogero Terregino, Antonia Ricci |
| EPI_ISL_2448617 | Microbiologia e Virologia | Istituto Zooprofilattico Sperimentale delle Venezie | Adelaide Milani, Alessia Schivo, Annalisa Salviato, Elisa Palumbo, Erika Giorgia Quaranta, Luca Tassoni, Ambra Pastori, Edoardo Giussani, Alice Fusaro, Isabella Monne, Calogero Terregino, Antonia Ricci |
| EPI_ISL_2448618, EPI_ISL_2448619, EPI_ISL_2448620 | ULSS 8 Berica | Istituto Zooprofilattico Sperimentale delle Venezie | Adelaide Milani, Alessia Schivo, Annalisa Salviato, Elisa Palumbo, Erika Giorgia Quaranta, Luca Tassoni, Ambra Pastori, Edoardo Giussani, Alice Fusaro, Isabella Monne, Calogero Terregino, Antonia Ricci |
| EPI_ISL_2448636 | Laboffice | CHU Poitiers | Caroline MICHAUD, Maxime PICHON, Manon PRAT, Valentin BON-BARET, Birama N'DIAYE, Magali GARCIA, Agnes BEBY-DEFAUX, Nicolas LEVEQUE |
| EPI_ISL_2448883 | UCK | 1. National Institute of Public Health - National Institute of Hygiene, Warsaw, Poland 2. Biobank Lab, University of Lodz 3. Laboratory of Respiratory Viruses, Teaching and Clinical Center of the Medical University of Lodz | Dominik Strapagiel, Marta Sobalska-Kwapis, Klaudyna Królikowska, Jakub Lach, Marcin Słomka, Tomasz Płoszaj, Magdalena Traczyk-Borszyńska, Izabela Dróżdż, Maciej Borowiec, Katarzyna Zacharczuk, Magdalena Nowakowska, Małgorzata Sadkowska-Todys, Tomasz Wołkowicz |
| EPI_ISL_2448918, EPI_ISL_2448927, EPI_ISL_2448928, EPI_ISL_2448929 | WSSE w Gdańsku | 1. National Institute of Public Health - National Institute of Hygiene, Warsaw, Poland 2. Biobank Lab, University of Lodz 3. Laboratory of Respiratory Viruses, Teaching and Clinical Center of the Medical University of Lodz | Dominik Strapagiel, Marta Sobalska-Kwapis, Klaudyna Królikowska, Jakub Lach, Marcin Słomka, Tomasz Płoszaj, Magdalena Traczyk-Borszyńska, Izabela Dróżdż, Maciej Borowiec, Katarzyna Zacharczuk, Magdalena Nowakowska, Małgorzata Sadkowska-Todys, Tomasz Wołkowicz |
| EPI_ISL_2448957 | NZOZ Medyczne Laboratorium Diagnostyczne | 1. National Institute of Public Health - National Institute of Hygiene, Warsaw, Poland 2. Biobank Lab, University of Lodz 3. Laboratory of Respiratory Viruses, Teaching and Clinical Center of the Medical University of Lodz | Dominik Strapagiel, Marta Sobalska-Kwapis, Klaudyna Królikowska, Jakub Lach, Marcin Słomka, Tomasz Płoszaj, Magdalena Traczyk-Borszyńska, Izabela Dróżdż, Maciej Borowiec, Katarzyna Zacharczuk, Magdalena Nowakowska, Małgorzata Sadkowska-Todys, Tomasz Wołkowicz |
| EPI_ISL_2449868 | Spital MŠnnedorf AG | Institute of Medical Virology | Daniel Ehrsam, Isabel StŸrmer, Catharine Aquino, Joel Wirz, Weihong Qi, Hubert Rehrauer, Verena Kufner, Gabriela Ziltener, Maryam Zaheri, Stefan Schmutz, Annette AudigŽ, Maria GrŸnberg, Kevin Steiner, Jon Huder, Cyril Shah, Riccarda Capaul, Guido Bloemberg, JŸrg BŠni, Michael Huber, Alexandra Trkola |
| EPI_ISL_2449950, EPI_ISL_2449965, EPI_ISL_2449966, EPI_ISL_2449967, EPI_ISL_2449987, EPI_ISL_2449988, EPI_ISL_2449989, EPI_ISL_2449990, EPI_ISL_2449994 | Dutch COVID-19 response team | Erasmus Medical Center | Bas Oude Munnink, Reina Sikkema, David Nieuwenhuijse, Irina Chestakova, Anne van der Linden, Marjan Boter, Emmanuelle Munger, Corine GeurtsvanKessel, Annemiek van der Eijk, Richard Molenkamp, Marion Koopmans, on behalf of the Dutch national COVID-19 respo |
| EPI_ISL_2450030 | Center for Laboratory Medicine | Center for Laboratory Medicine | Yannick Gerth |
| EPI_ISL_2450238, EPI_ISL_2450254, EPI_ISL_2450255, EPI_ISL_2450267, EPI_ISL_2450268, EPI_ISL_2450270, EPI_ISL_2450277, EPI_ISL_2450280, EPI_ISL_2450292, EPI_ISL_2450295, EPI_ISL_2450297, EPI_ISL_2450300, EPI_ISL_2450315, EPI_ISL_2450323, EPI_ISL_2450325, EPI_ISL_2450326 | | | |
| see above | Max von Pettenkofer Institute, Virology, National Reference Center for Retroviruses, LMU Munich | Laboratory for Functional Genome Analysis; Dept. Genomics; Gene Center of the LMU Munich | Max Muenchhoff; Stefan Krebs; Alexander Graf; Oliver Keppler; Helmut Blum |
| EPI_ISL_2450328, EPI_ISL_2450329 | NIMHANS | INSACOG-KA, NIMHANS | Chitra Pattabiraman, Anson Kunjumon George,Darshan Sreenivas, Ananthapadmanabha Kotambail, Chetan G K, Gautham Arunachal Udupi, Anita S Desai, V Ravi |
| EPI_ISL_2450330, EPI_ISL_2450331 | RH HOSAHALLI | INSACOG-KA, NIMHANS | Chitra Pattabiraman, Anson Kunjumon George,Darshan Sreenivas, Ananthapadmanabha Kotambail, Chetan G K, Gautham Arunachal Udupi, Anita S Desai, V Ravi |
| EPI_ISL_2450332 | UPHC GANGONDANAHALLI | INSACOG-KA, NIMHANS | Chitra Pattabiraman, Anson Kunjumon George,Darshan Sreenivas, Ananthapadmanabha Kotambail, Chetan G K, Gautham Arunachal Udupi, Anita S Desai, V Ravi |
| EPI_ISL_2450333 | JJR NAGAR UPHC | INSACOG-KA, NIMHANS | Chitra Pattabiraman, Anson Kunjumon George,Darshan Sreenivas, Ananthapadmanabha Kotambail, Chetan G K, Gautham Arunachal Udupi, Anita S Desai, V Ravi |
| EPI_ISL_2450334 | NIMHANS | INSACOG-KA, NIMHANS | Chitra Pattabiraman, Anson Kunjumon George,Darshan Sreenivas, Ananthapadmanabha Kotambail, Chetan G K, Gautham Arunachal Udupi, Anita S Desai, V Ravi |
| EPI_ISL_2450337 | SANJAY GANDHI INSTITUTE OF TRAUMA AND ORTHOPAEDICS | INSACOG-KA, NIMHANS | Chitra Pattabiraman, Anson Kunjumon George,Darshan Sreenivas, Ananthapadmanabha Kotambail, Chetan G K, Gautham Arunachal Udupi, Anita S Desai, V Ravi |
| EPI_ISL_2450338 | RAILWAY HOSPITAL | INSACOG-KA, NIMHANS | Chitra Pattabiraman, Anson Kunjumon George,Darshan Sreenivas, Ananthapadmanabha Kotambail, Chetan G K, Gautham Arunachal Udupi, Anita S Desai, V Ravi |
| EPI_ISL_2450339 | NIMHANS | INSACOG-KA, NIMHANS | Chitra Pattabiraman, Anson Kunjumon George,Darshan Sreenivas, Ananthapadmanabha Kotambail, Chetan G K, Gautham Arunachal Udupi, Anita S Desai, V Ravi |
| EPI_ISL_2450340, EPI_ISL_2450341, EPI_ISL_2450342 | RAILWAY | INSACOG-KA, NIMHANS | Chitra Pattabiraman, Anson Kunjumon George,Darshan Sreenivas, Ananthapadmanabha Kotambail, Chetan G K, Gautham Arunachal Udupi, Anita S Desai, V Ravi |
| EPI_ISL_2450343, EPI_ISL_2450344, EPI_ISL_2450345, EPI_ISL_2450346, EPI_ISL_2450347 | DSU KOLAR | INSACOG-KA, NIMHANS | Chitra Pattabiraman, Anson Kunjumon George,Darshan Sreenivas, Ananthapadmanabha Kotambail, Chetan G K, Gautham Arunachal Udupi, Anita S Desai, V Ravi |
| EPI_ISL_2450349, EPI_ISL_2450350, EPI_ISL_2450352 | THE KOLAR MOBILE TEAM | INSACOG-KA, NIMHANS | Chitra Pattabiraman, Anson Kunjumon George,Darshan Sreenivas, Ananthapadmanabha Kotambail, Chetan G K, Gautham Arunachal Udupi, Anita S Desai, V Ravi |
| EPI_ISL_2450354, EPI_ISL_2450356 | NIMHANS | INSACOG-KA, NIMHANS | Chitra Pattabiraman, Anson Kunjumon George,Darshan Sreenivas, Ananthapadmanabha Kotambail, Chetan G K, Gautham Arunachal Udupi, Anita S Desai, V Ravi |
| EPI_ISL_2450357, EPI_ISL_2450358 | UPHC A D HALLI | INSACOG-KA, NIMHANS | Chitra Pattabiraman, Anson Kunjumon George,Darshan Sreenivas, Ananthapadmanabha Kotambail, Chetan G K, Gautham Arunachal Udupi, Anita S Desai, V Ravi |
| EPI_ISL_2450359, EPI_ISL_2450360, EPI_ISL_2450361, EPI_ISL_2450362, EPI_ISL_2450363, EPI_ISL_2450364 | RAILWAY HOSPITAL | INSACOG-KA, NIMHANS | Chitra Pattabiraman, Anson Kunjumon George,Darshan Sreenivas, Ananthapadmanabha Kotambail, Chetan G K, Gautham Arunachal Udupi, Anita S Desai, V Ravi |
| EPI_ISL_2450365 | SRI JAYADEVA INSTITUTE OF CARDIOLOGY | INSACOG-KA, NIMHANS | Chitra Pattabiraman, Anson Kunjumon George,Darshan Sreenivas, Ananthapadmanabha Kotambail, Chetan G K, Gautham Arunachal Udupi, Anita S Desai, V Ravi |
| EPI_ISL_2450376 | Ospedale di Genzano - ASL RM 6 | INMI Lazzaro Spallanzani IRCCS | F Santini, B Bartolini, E Giombini, F Messina, M Rueca, G Bonfiglio, O Butera, CEM Gruber, G Tramini, E Conti, MR Capobianchi, A Di Caro |
| EPI_ISL_2450455, EPI_ISL_2450456, EPI_ISL_2450457, EPI_ISL_2450458, EPI_ISL_2450460, EPI_ISL_2450461, EPI_ISL_2450462, EPI_ISL_2450463, EPI_ISL_2450465, EPI_ISL_2450466, EPI_ISL_2450467, EPI_ISL_2450469, EPI_ISL_2450470, EPI_ISL_2450471, EPI_ISL_2450472, EPI_ISL_2450473, EPI_ISL_2450474, EPI_ISL_2450475, EPI_ISL_2450476, EPI_ISL_2450477, EPI_ISL_2450478, EPI_ISL_2450479, EPI_ISL_2450480, EPI_ISL_2450481, EPI_ISL_2450482, EPI_ISL_2450483, EPI_ISL_2450484, EPI_ISL_2450485, EPI_ISL_2450487, EPI_ISL_2450488, EPI_ISL_2450489, EPI_ISL_2450490, EPI_ISL_2450496, EPI_ISL_2450497, EPI_ISL_2450498, EPI_ISL_2450499, EPI_ISL_2450501, EPI_ISL_2450502, EPI_ISL_2450503, EPI_ISL_2450504, EPI_ISL_2450505, EPI_ISL_2450506, EPI_ISL_2450508, EPI_ISL_2450509, EPI_ISL_2450510, EPI_ISL_2450511, EPI_ISL_2450512, EPI_ISL_2450513, EPI_ISL_2450514, EPI_ISL_2450518, EPI_ISL_2450519, EPI_ISL_2450520, EPI_ISL_2450521, EPI_ISL_2450522, EPI_ISL_2450523, EPI_ISL_2450524, EPI_ISL_2450525, EPI_ISL_2450526, EPI_ISL_2450527, EPI_ISL_2450528, EPI_ISL_2450529, EPI_ISL_2450530, EPI_ISL_2450531, EPI_ISL_2450535, EPI_ISL_2450536, EPI_ISL_2450538, EPI_ISL_2450540, EPI_ISL_2450541, EPI_ISL_2450542 | | | |
| see above | WHO National Influenza Centre Russian Federation | WHO National Influenza Centre Russian Federation | Andrey Komissarov, Artem Fadeev, Kseniya Komissarova, Oula Mansour, Kirill Varchenko, Mikhail Bakaev, Tamila Musaeva, Maria Timofeeva, Veronika Eder, Maria Pisareva, Nikita Yolshin, Daria Danilenko, Ksenia Safina, Elena Nabieva, Georgii Bazykin, Dmitry Lioznov |
| EPI_ISL_2450544, EPI_ISL_2450550, EPI_ISL_2450551, EPI_ISL_2450553, EPI_ISL_2450554, EPI_ISL_2450559, EPI_ISL_2450560, EPI_ISL_2450561, EPI_ISL_2450563, EPI_ISL_2450565 | City Hospital No 40 | WHO National Influenza Centre Russian Federation | Andrey Komissarov, Artem Fadeev, Kseniya Komissarova, Oula Mansour, Kirill Varchenko, Mikhail Bakaev, Tamila Musaeva, Maria Timofeeva, Veronika Eder, Maria Pisareva, Nikita Yolshin, Daria Danilenko, Olga Shneider, Sergey Scherbak, Ksenia Safina, Elena Nabieva, Georgii Bazykin, Dmitry Lioznov |
| EPI_ISL_2450597, EPI_ISL_2450599, EPI_ISL_2450600, EPI_ISL_2450601, EPI_ISL_2450603, EPI_ISL_2450604 | WHO National Influenza Centre Russian Federation | WHO National Influenza Centre Russian Federation | Andrey Komissarov, Artem Fadeev, Kseniya Komissarova, Oula Mansour, Kirill Varchenko, Mikhail Bakaev, Tamila Musaeva, Maria Timofeeva, Veronika Eder, Maria Pisareva, Nikita Yolshin, Daria Danilenko, Ksenia Safina, Elena Nabieva, Georgii Bazykin, Dmitry Lioznov |
| EPI_ISL_2450606 | HELIX LLC | WHO National Influenza Centre Russian Federation | Andrey Komissarov, Artem Fadeev, Kseniya Komissarova, Oula Mansour, Kirill Varchenko, Mikhail Bakaev, Tamila Musaeva, Maria Timofeeva, Veronika Eder, Maria Pisareva, Nikita Yolshin, Daria Danilenko, Ksenia Safina, Elena Nabieva, Georgii Bazykin, Dmitry Lioznov |
| EPI_ISL_2450694 | CAP TORELLÓ | Banc de Sang i Teixits | Francisco Vidal, Irene Corrales, Nina Borràs, Carlos Hobeich, Natália Comes, Lorena Ramírez, Noemí Gonzalez, Silvia Sauleda, Maria Glòria Soria |
| EPI_ISL_2450716 | CAP SANT VICENÇ DE TORELLÓ | Banc de Sang i Teixits | Francisco Vidal, Irene Corrales, Nina Borràs, Carlos Hobeich, Natália Comes, Lorena Ramírez, Noemí Gonzalez, Silvia Sauleda, Maria Glòria Soria |

| | | | |
|---|---|---|---|
| EPI_ISL_2450972, EPI_ISL_2450977, EPI_ISL_2450983, EPI_ISL_2451007, EPI_ISL_2451045, EPI_ISL_2451072, EPI_ISL_2451135, EPI_ISL_2451139, EPI_ISL_2451145, EPI_ISL_2451147, EPI_ISL_2451168, EPI_ISL_2451260 | | | |
| see above | Mako Medical | Centers for Disease Control and Prevention Division of Viral Diseases, Pathogen Discovery | Dakota Howard, Dhwani Batra, Peter W. Cook, Kara Moser, Adrian Paskey, Jason Caravas, Benjamin Rambo-Martin, Shatavia Morrison, Christopher Gulvick, Scott Sammons, Yvette Unoarumhi, Darlene Wagner, Matthew Schmerer,Matthew Tugwell, Lauren Moon, Clinton R. Paden, Duncan MacCannell |
| EPI_ISL_2451293, EPI_ISL_2451302, EPI_ISL_2451329, EPI_ISL_2451336, EPI_ISL_2451351, EPI_ISL_2451356, EPI_ISL_2451361 | Quest Diagnostics Incorporated | Centers for Disease Control and Prevention Division of Viral Diseases, Pathogen Discovery | Dakota Howard, Dhwani Batra, Peter W. Cook, Kara Moser, Adrian Paskey, Jason Caravas, Benjamin Rambo-Martin, Shatavia Morrison, Christopher Gulvick, Scott Sammons, Yvette Unoarumhi, Darlene Wagner, Matthew Schmerer, S. H. Rosenthal, A. Gerasimova, R. M. Kagan, B. Anderson, M. Hua, Y. Liu, L.E. Bernstein, K.E. Livingston, A. Perez, I. A. Shlyakhter, R. V. Rolando, R. Owen, P. Tanpaiboon, F. Lacbawan, Clinton R. Paden, Duncan MacCannell |
| EPI_ISL_2451567 | MN PHL Division, Minnesota Department of Health | Centers for Disease Control and Prevention Division of Viral Diseases, Pathogen Discovery | Mili Sheth, Sarah Nobles, Jasmine Padilla, Mark Burroughs, Shoshona Le, Katie Dillon, Peter Cook, Clinton R. Paden, Dhwani Batra, Krista Queen, Kristen Knipe, Dakota Howard, Yvette Unoarumhi, Darlene Wagner, Matthew Schmerer, Ben L. Rambo-Martin, Kristine Lacek, Sam Shepard, Alison Laufer Halpin, Dave Wentworth, Vivien Dugan, Suxiang Tong, Justin Lee |
| EPI_ISL_2451575 | OK Public Health Laboratory, Oklahoma State DOH | Centers for Disease Control and Prevention Division of Viral Diseases, Pathogen Discovery | Mili Sheth, Sarah Nobles, Jasmine Padilla, Mark Burroughs, Shoshona Le, Katie Dillon, Peter Cook, Clinton R. Paden, Dhwani Batra, Krista Queen, Kristen Knipe, Dakota Howard, Yvette Unoarumhi, Darlene Wagner, Matthew Schmerer, Ben L. Rambo-Martin, Kristine Lacek, Sam Shepard, Alison Laufer Halpin, Dave Wentworth, Vivien Dugan, Suxiang Tong, Justin Lee |
| EPI_ISL_2451580 | KS Health and Environmental Laboratories | Centers for Disease Control and Prevention Division of Viral Diseases, Pathogen Discovery | Mili Sheth, Sarah Nobles, Jasmine Padilla, Mark Burroughs, Shoshona Le, Katie Dillon, Peter Cook, Clinton R. Paden, Dhwani Batra, Krista Queen, Kristen Knipe, Dakota Howard, Yvette Unoarumhi, Darlene Wagner, Matthew Schmerer, Ben L. Rambo-Martin, Kristine Lacek, Sam Shepard, Alison Laufer Halpin, Dave Wentworth, Vivien Dugan, Suxiang Tong, Justin Lee |
| EPI_ISL_2451626, EPI_ISL_2451628, EPI_ISL_2451661, EPI_ISL_2451663, EPI_ISL_2451758 | Public Health Ontario Laboratory | Public Health Ontario Laboratory | Vanessa G Allen, Philip Banh, Yao Chen, Richard de Borja, Alireza Eshaghi, Nahuel Fittipaldi, Christine Frantz, Jonathan B Gubbay, Jennifer L Guthrie, Lawrence Heisler, Esha Joshi, Michael Laszloffy, Aimin Li, Michael CY Li, Dean Maxwell, Sandeep Nagra, Samir N Patel, Jared Simpson, Karthikeyan Sivaraman, Ashleigh Sullivan, Yogi Sundaravadanam, Sarah Teatero, Andre Villegas, Matthew Watson, Sandra Zittermann |
| EPI_ISL_2451768 | DC Public Health Lab/ Dept. of Forensic Sciences | Centers for Disease Control and Prevention Division of Viral Diseases, Pathogen Discovery | Mili Sheth, Sarah Nobles, Jasmine Padilla, Mark Burroughs, Shoshona Le, Katie Dillon, Peter Cook, Clinton R. Paden, Dhwani Batra, Krista Queen, Kristen Knipe, Dakota Howard, Yvette Unoarumhi, Darlene Wagner, Matthew Schmerer, Ben L. Rambo-Martin, Kristine Lacek, Sam Shepard, Alison Laufer Halpin, Dave Wentworth, Vivien Dugan, Suxiang Tong, Justin Lee |
| EPI_ISL_2451780, EPI_ISL_2451784 | AR Dept. of Health-PHL, Molecular Diagnostics | Centers for Disease Control and Prevention Division of Viral Diseases, Pathogen Discovery | Mili Sheth, Sarah Nobles, Jasmine Padilla, Mark Burroughs, Shoshona Le, Katie Dillon, Peter Cook, Clinton R. Paden, Dhwani Batra, Krista Queen, Kristen Knipe, Dakota Howard, Yvette Unoarumhi, Darlene Wagner, Matthew Schmerer, Ben L. Rambo-Martin, Kristine Lacek, Sam Shepard, Alison Laufer Halpin, Dave Wentworth, Vivien Dugan, Suxiang Tong, Justin Lee |
| EPI_ISL_2451829 | MA State Public Health Laboratory | Centers for Disease Control and Prevention Division of Viral Diseases, Pathogen Discovery | Mili Sheth, Sarah Nobles, Jasmine Padilla, Mark Burroughs, Shoshona Le, Katie Dillon, Peter Cook, Clinton R. Paden, Dhwani Batra, Krista Queen, Kristen Knipe, Dakota Howard, Yvette Unoarumhi, Darlene Wagner, Matthew Schmerer, Ben L. Rambo-Martin, Kristine Lacek, Sam Shepard, Alison Laufer Halpin, Dave Wentworth, Vivien Dugan, Suxiang Tong, Justin Lee |
| EPI_ISL_2451890, EPI_ISL_2451893 | CA-Los Angeles County Public Health Laboratory | Centers for Disease Control and Prevention Division of Viral Diseases, Pathogen Discovery | Mili Sheth, Sarah Nobles, Jasmine Padilla, Mark Burroughs, Shoshona Le, Katie Dillon, Peter Cook, Clinton R. Paden, Dhwani Batra, Krista Queen, Kristen Knipe, Dakota Howard, Yvette Unoarumhi, Darlene Wagner, Matthew Schmerer, Ben L. Rambo-Martin, Kristine Lacek, Sam Shepard, Alison Laufer Halpin, Dave Wentworth, Vivien Dugan, Suxiang Tong, Justin Lee |
| EPI_ISL_2451907 | ND Dept. of Health Laboratory Services-Microbiology | Centers for Disease Control and Prevention Division of Viral Diseases, Pathogen Discovery | Mili Sheth, Sarah Nobles, Jasmine Padilla, Mark Burroughs, Shoshona Le, Katie Dillon, Peter Cook, Clinton R. Paden, Dhwani Batra, Krista Queen, Kristen Knipe, Dakota Howard, Yvette Unoarumhi, Darlene Wagner, Matthew Schmerer, Ben L. Rambo-Martin, Kristine Lacek, Sam Shepard, Alison Laufer Halpin, Dave Wentworth, Vivien Dugan, Suxiang Tong, Justin Lee |
| EPI_ISL_2451993, EPI_ISL_2452022 | Public Health Ontario Laboratory | Public Health Ontario Laboratory | Vanessa G Allen, Philip Banh, Yao Chen, Richard de Borja, Alireza Eshaghi, Nahuel Fittipaldi, Christine Frantz, Jonathan B Gubbay, Jennifer L Guthrie, Lawrence Heisler, Esha Joshi, Michael Laszloffy, Aimin Li, Michael CY Li, Dean Maxwell, Sandeep Nagra, Samir N Patel, Jared Simpson, Karthikeyan Sivaraman, Ashleigh Sullivan, Yogi Sundaravadanam, Sarah Teatero, Andre Villegas, Matthew Watson, Sandra Zittermann |
| EPI_ISL_2452031, EPI_ISL_2452037, EPI_ISL_2452040 | CDPH, Viral and Rickettsial Disease Laboratory | Centers for Disease Control and Prevention Division of Viral Diseases, Pathogen Discovery | Mili Sheth, Sarah Nobles, Jasmine Padilla, Mark Burroughs, Shoshona Le, Katie Dillon, Peter Cook, Clinton R. Paden, Dhwani Batra, Krista Queen, Kristen Knipe, Dakota Howard, Yvette Unoarumhi, Darlene Wagner, Matthew Schmerer, Ben L. Rambo-Martin, Kristine Lacek, Sam Shepard, Alison Laufer Halpin, Dave Wentworth, Vivien Dugan, Suxiang Tong, Justin Lee |
| EPI_ISL_2452090, EPI_ISL_2452092 | WY Public Health Laboratory | Centers for Disease Control and Prevention Division of Viral Diseases, Pathogen Discovery | Mili Sheth, Sarah Nobles, Jasmine Padilla, Mark Burroughs, Shoshona Le, Katie Dillon, Peter Cook, Clinton R. Paden, Dhwani Batra, Krista Queen, Kristen Knipe, Dakota Howard, Yvette Unoarumhi, Darlene Wagner, Matthew Schmerer, Ben L. Rambo-Martin, Kristine Lacek, Sam Shepard, Alison Laufer Halpin, Dave Wentworth, Vivien Dugan, Suxiang Tong, Justin Lee |
| EPI_ISL_2452150 | IN State Department of Health Laboratory Services | Centers for Disease Control and Prevention Division of Viral Diseases, Pathogen Discovery | Mili Sheth, Sarah Nobles, Jasmine Padilla, Mark Burroughs, Shoshona Le, Katie Dillon, Peter Cook, Clinton R. Paden, Dhwani Batra, Krista Queen, Kristen Knipe, Dakota Howard, Yvette Unoarumhi, Darlene Wagner, Matthew Schmerer, Ben L. Rambo-Martin, Kristine Lacek, Sam Shepard, Alison Laufer Halpin, Dave Wentworth, Vivien Dugan, Suxiang Tong, Justin Lee |
| EPI_ISL_2452165 | AL Dept. of Public Health Bureau of Clinical Laboratories | Centers for Disease Control and Prevention Division of Viral Diseases, Pathogen Discovery | Mili Sheth, Sarah Nobles, Jasmine Padilla, Mark Burroughs, Shoshona Le, Katie Dillon, Peter Cook, Clinton R. Paden, Dhwani Batra, Krista Queen, Kristen Knipe, Dakota Howard, Yvette Unoarumhi, Darlene Wagner, Matthew Schmerer, Ben L. Rambo-Martin, Kristine Lacek, Sam Shepard, Alison Laufer Halpin, Dave Wentworth, Vivien Dugan, Suxiang Tong, Justin Lee |
| EPI_ISL_2452177 | IA State Hygienic Laboratory | Centers for Disease Control and Prevention Division of Viral Diseases, Pathogen Discovery | Mili Sheth, Sarah Nobles, Jasmine Padilla, Mark Burroughs, Shoshona Le, Katie Dillon, Peter Cook, Clinton R. Paden, Dhwani Batra, Krista Queen, Kristen Knipe, Dakota Howard, Yvette Unoarumhi, Darlene Wagner, Matthew Schmerer, Ben L. Rambo-Martin, Kristine Lacek, Sam Shepard, Alison Laufer Halpin, Dave Wentworth, Vivien Dugan, Suxiang Tong, Justin Lee |
| EPI_ISL_2452282 | MD DOH Laboratories Administration | Centers for Disease Control and Prevention Division of Viral Diseases, Pathogen Discovery | Mili Sheth, Sarah Nobles, Jasmine Padilla, Mark Burroughs, Shoshona Le, Katie Dillon, Peter Cook, Clinton R. Paden, Dhwani Batra, Krista Queen, Kristen Knipe, Dakota Howard, Yvette Unoarumhi, Darlene Wagner, Matthew Schmerer, Ben L. Rambo-Martin, Kristine Lacek, Sam Shepard, Alison Laufer Halpin, Dave Wentworth, Vivien Dugan, Suxiang Tong, Justin Lee |
| EPI_ISL_2452319, EPI_ISL_2452333, EPI_ISL_2452516, EPI_ISL_2452523, EPI_ISL_2452525, EPI_ISL_2452526, EPI_ISL_2452547, EPI_ISL_2452584, EPI_ISL_2452593, EPI_ISL_2452601, EPI_ISL_2452625, EPI_ISL_2452654, EPI_ISL_2452659, EPI_ISL_2452673, EPI_ISL_2452686, EPI_ISL_2452695, EPI_ISL_2452704, EPI_ISL_2452710, EPI_ISL_2452717, EPI_ISL_2452720, EPI_ISL_2452725, EPI_ISL_2452728, EPI_ISL_2452730, EPI_ISL_2452732, EPI_ISL_2452734, EPI_ISL_2452735, EPI_ISL_2452737, EPI_ISL_2452738, EPI_ISL_2452739, EPI_ISL_2452741, EPI_ISL_2452742, EPI_ISL_2452743, EPI_ISL_2452744, EPI_ISL_2452746, EPI_ISL_2452747, EPI_ISL_2452748, EPI_ISL_2452749, EPI_ISL_2452755, EPI_ISL_2452793, EPI_ISL_2452799, EPI_ISL_2452802, EPI_ISL_2452804, EPI_ISL_2452806, EPI_ISL_2452807, EPI_ISL_2452811, EPI_ISL_2452815, EPI_ISL_2452816, EPI_ISL_2452818, EPI_ISL_2452820, EPI_ISL_2452821, EPI_ISL_2452823, EPI_ISL_2452824, EPI_ISL_2452827, EPI_ISL_2452828, EPI_ISL_2452829, EPI_ISL_2452832, EPI_ISL_2452844, EPI_ISL_2452867, EPI_ISL_2452868, EPI_ISL_2452884, EPI_ISL_2452885, EPI_ISL_2452886, EPI_ISL_2452891, EPI_ISL_2452899, EPI_ISL_2452901, EPI_ISL_2452917, EPI_ISL_2452921, EPI_ISL_2452944, EPI_ISL_2452945, EPI_ISL_2452951, EPI_ISL_2452953, EPI_ISL_2452955, EPI_ISL_2452958, EPI_ISL_2452966, EPI_ISL_2452967, EPI_ISL_2452978, EPI_ISL_2452987, EPI_ISL_2452995, EPI_ISL_2453000, EPI_ISL_2453008, EPI_ISL_2453009, EPI_ISL_2453010, EPI_ISL_2453013, EPI_ISL_2453024, EPI_ISL_2453029, EPI_ISL_2453036, EPI_ISL_2453045, EPI_ISL_2453061, EPI_ISL_2453062, EPI_ISL_2453066, EPI_ISL_2453079, EPI_ISL_2453100, EPI_ISL_2453101, EPI_ISL_2453105, EPI_ISL_2453108, EPI_ISL_2453116, EPI_ISL_2453136, EPI_ISL_2453148, EPI_ISL_2453151, EPI_ISL_2453152, EPI_ISL_2453155, EPI_ISL_2453158, EPI_ISL_2453162, EPI_ISL_2453174, EPI_ISL_2453182, EPI_ISL_2453184, EPI_ISL_2453197, EPI_ISL_2453205, EPI_ISL_2453208, EPI_ISL_2453209, EPI_ISL_2453213, EPI_ISL_2453214, EPI_ISL_2453215, EPI_ISL_2453216, EPI_ISL_2453217, EPI_ISL_2453219, EPI_ISL_2453220, EPI_ISL_2453221, EPI_ISL_2453223, EPI_ISL_2453248, EPI_ISL_2453253, EPI_ISL_2453269, EPI_ISL_2453280, EPI_ISL_2453282, EPI_ISL_2453283, EPI_ISL_2453284, EPI_ISL_2453291, EPI_ISL_2453299, EPI_ISL_2453304, EPI_ISL_2453314, EPI_ISL_2453316 | | | |
| see above | Public Health Ontario Laboratory | Public Health Ontario Laboratory | Vanessa G Allen, Philip Banh, Yao Chen, Richard de Borja, Alireza Eshaghi, Nahuel Fittipaldi, Christine Frantz, Jonathan B Gubbay, Jennifer L Guthrie, Lawrence Heisler, Esha Joshi, Michael Laszloffy, Aimin Li, Michael CY Li, Dean Maxwell, Sandeep Nagra, Samir N Patel, Jared Simpson, Karthikeyan Sivaraman, Ashleigh Sullivan, Yogi Sundaravadanam, Sarah Teatero, Andre Villegas, Matthew Watson, Sandra Zittermann |
| EPI_ISL_2453515, EPI_ISL_2453516, EPI_ISL_2453821, EPI_ISL_2453822, EPI_ISL_2453823, EPI_ISL_2453824, EPI_ISL_2453825, EPI_ISL_2453826, EPI_ISL_2453827, EPI_ISL_2453838, EPI_ISL_2453839, EPI_ISL_2453840, EPI_ISL_2453841, EPI_ISL_2453842, EPI_ISL_2453843, EPI_ISL_2453844, EPI_ISL_2453845, EPI_ISL_2453846, EPI_ISL_2453847, EPI_ISL_2453848, EPI_ISL_2453849, EPI_ISL_2453850, EPI_ISL_2453851, EPI_ISL_2453855, EPI_ISL_2453856, EPI_ISL_2453857, EPI_ISL_2453858, EPI_ISL_2453859, EPI_ISL_2453860, EPI_ISL_2453861, EPI_ISL_2453862, EPI_ISL_2453876, EPI_ISL_2453877, EPI_ISL_2453895, EPI_ISL_2454005 | | | |
| see above | MEPHI, Aix Marseille University | MEPHI, Aix Marseille University | Anthony LEVASSEUR |
| EPI_ISL_2454302 | Greek Genome Center, Biomedical Research Foundation of the Academy of Athens (BRFAA) | Greek Genome Center, Biomedical Research Foundation of the Academy of Athens (BRFAA) | Emmanouil Athanasiadis, Giannis Vatsellas, Theodoros Loupis, Katerina Zoi, Dimitrios Thanos |
| EPI_ISL_2454305, EPI_ISL_2454322 | TriCore Reference Laboratories | Center for Global Health, University of New Mexico Health Sciences Center | Daryl Domman, Kurt Schwalm, Valerie Morley, Cecilia Thompson, Kendra Pesko, Karissa Culbreath, Darrell Dinwiddie |
| EPI_ISL_2455082 | Houston Health Dept. | Houston Health Dept. | Ryker Penn, Pamela Brown, Adolpho Lara, Yanlai Lai |
| EPI_ISL_2455160 | Bruss | 1. Tricity SARS-CoV-2 sequencing consortium: University of Gdansk, Medical University of Gdansk, Vaxican Ltd., Invicta Ltd. 2. National Institute of Public Health - National Institute of Hygiene, Warsaw, Poland | Maciej Kosinski, Celina Cybulska, Krystyna Bienkowska Szewczyk, Maciej Grzybek, Karolina Gackowska, Marcin Lubocki, Katarzyna Groth, Lukasz Rabalski, Katarzyna Zacharczuk, Magdalena Nowakowska, Małgorzata Sadkowska-Todys, Tomasz Wołkowicz |
| EPI_ISL_2455193, EPI_ISL_2455195, EPI_ISL_2455203, EPI_ISL_2455205 | Diagnostyka | 1. Tricity SARS-CoV-2 sequencing consortium: University of Gdansk, Medical University of Gdansk, Vaxican Ltd., Invicta Ltd. 2. National Institute of Public Health - National Institute of Hygiene, Warsaw, Poland | Maciej Kosinski, Celina Cybulska, Krystyna Bienkowska Szewczyk, Maciej Grzybek, Karolina Gackowska, Marcin Lubocki, Katarzyna Groth, Lukasz Rabalski, Katarzyna Zacharczuk, Magdalena Nowakowska, Małgorzata Sadkowska-Todys, Tomasz Wołkowicz |
| EPI_ISL_2455220 | Area of Virology, Serology and Virology Division (SAViD), New South Wales Health Pathology Randwick | Virology Research Laboratory; Area of Virology, Serology and Virology Division (SAViD), New South Wales Health Pathology Randwick | Foster, C.; Au, J.; Ruiz Silva, M.; Deveson, I.; Bull, R.; Van Hal, S.; Rawlinson, W. |
| EPI_ISL_2455221, EPI_ISL_2455222, EPI_ISL_2455223, EPI_ISL_2455224 | National Hospital for Tropical Diseases | Oxford University Clinical Research Unit, Hanoi, Vietnam | Pham Ngoc Thach, Nguyen Thi Tam, Van Dinh Trang, Le Van Duyet, Nguyen Thu Trang, Nguyen Thi Hong Thuong, Nguyen Thi Kim Chi, Phan Manh Cuong, Thomas Kesteman, H.Rogier van Doorn on behalf of the OUCRU COVID-19 research group |
| EPI_ISL_2455227, EPI_ISL_2455231 | New South Wales Health Pathology Royal Prince Alfred Hospital | Microbiology RPAH | Foster, C.; Au, J.; Ruiz Silva, M.; Deveson, I.; Bull, R.; Van Hal, S.; Rawlinson, W. |
| EPI_ISL_2455233, EPI_ISL_2455234, EPI_ISL_2455235, EPI_ISL_2455238, EPI_ISL_2455239, EPI_ISL_2455240 | National Hospital for Tropical Diseases | Oxford University Clinical Research Unit, Hanoi, Vietnam | Pham Ngoc Thach, Nguyen Thi Tam, Van Dinh Trang, Le Van Duyet, Nguyen Thu Trang, Nguyen Thi Hong Thuong, Nguyen Thi Kim Chi, Phan Manh Cuong, Thomas Kesteman, H.Rogier van Doorn on behalf of the OUCRU COVID-19 research group |
| EPI_ISL_2455252, EPI_ISL_2455254, EPI_ISL_2455256 | Wyoming Public Health Laboratory | Wyoming Public Health Laboratory | Jim Mildenberger, Taylor Fearing, Channing Weber, Ashley Norberg, Chayse Rowley, Marley Goetz, Brian Dominguez, Elliot Thomasson, Brittany Oher, Sam Britz, Cari Sloma, and Rob Christensen |
| EPI_ISL_2455286 | Johns Hopkins Hospital Department of Pathology | Johns Hopkins Hospital Department of Pathology | C. Paul Morris, Chun Huai Luo, Adannaya Amadi, Matthew Schwartz, Nicholas Gallagher, Heba H. Mostafa |
| EPI_ISL_2455368 | Minnesota Department of Health, Public Health Laboratory | Minnesota Department of Health, Public Health Laboratory | Alexandra Lorentz, Jacob Garfin, Matt Plumb, and Xiong Wang |
| EPI_ISL_2455375, EPI_ISL_2455377, EPI_ISL_2455379 | HealthPartners Central Lab | Minnesota Department of Health, Public Health Laboratory | Alexandra Lorentz, Jacob Garfin, Matt Plumb, and Xiong Wang |
| EPI_ISL_2455398, | National Hospital for Tropical Diseases | Oxford University Clinical Research Unit, Hanoi, Vietnam | Pham Ngoc Thach, Nguyen Thi Tam, Van Dinh Trang, Le Van Duyet, Nguyen Thu Trang, Nguyen Thi Hong Thuong, Nguyen Thi Kim Chi, Phan Manh Cuong, Thomas Kesteman, H.Rogier van Doorn on behalf of the OUCRU COVID-19 research |

| | | | |
|---|---|---|---|
| EPI_ISL_2455399, EPI_ISL_2455415, EPI_ISL_2455416 | | | group |
| EPI_ISL_2455419, EPI_ISL_2455420 | Alaska State Virology Laboratory | Alaska State Virology Laboratory | Stephanie DeRonde, Elva House, Jacob Zidek, Lisa Smith, Ph.D., Jack Chen, Ph.D. |
| EPI_ISL_2455451 | National Hospital for Tropical Diseases | Oxford University Clinical Research Unit, Hanoi, Vietnam | Pham Ngoc Thach, Nguyen Thi Tam, Van Dinh Trang, Le Van Duyet, Nguyen Thu Trang, Nguyen Thi Hong Thuong, Nguyen Thi Kim Chi, Phan Manh Cuong, Thomas Kesteman, H.Rogier van Doorn on behalf of the OUCRU COVID-19 research group |
| EPI_ISL_2455453, EPI_ISL_2455456, EPI_ISL_2455464, EPI_ISL_2455466, EPI_ISL_2455467, EPI_ISL_2455468, EPI_ISL_2455469 | Kansas Health and Environmental Lab | Kansas Health and Environmental Lab | Katherine Wiggins, Mike Grose, Jonathan Barnell, Ben Olsen, and Phil Adam |
| EPI_ISL_2455479, EPI_ISL_2455480, EPI_ISL_2455481, EPI_ISL_2455482, EPI_ISL_2455483, EPI_ISL_2455484, EPI_ISL_2455485, EPI_ISL_2455486, EPI_ISL_2455487, EPI_ISL_2455488, EPI_ISL_2455489, EPI_ISL_2455490, EPI_ISL_2455491, EPI_ISL_2455492, EPI_ISL_2455494, EPI_ISL_2455495, EPI_ISL_2455496, EPI_ISL_2455497 | | | |
| see above | National Hospital for Tropical Diseases | Oxford University Clinical Research Unit, Hanoi, Vietnam | Pham Ngoc Thach, Nguyen Thi Tam, Van Dinh Trang, Le Van Duyet, Nguyen Thu Trang, Nguyen Thi Hong Thuong, Nguyen Thi Kim Chi, Phan Manh Cuong, Thomas Kesteman, H.Rogier van Doorn on behalf of the OUCRU COVID-19 research group |
| EPI_ISL_2455553 | LSUHS Emerging Viral Threat Laboratory | LSUHS Emerging Viral Threat Laboratory | Gregory L. Ware, Alexander Mijalis, Jeremy P. Kamil, Jennifer L. Carroll, Maarten Van Diest, Rona S. Scott, Andrew D. Yurochko, Christopher G. Kevil, John A. Vanchiere |
| EPI_ISL_2455554, EPI_ISL_2455559, EPI_ISL_2455560, EPI_ISL_2455561, EPI_ISL_2455583, EPI_ISL_2455585, EPI_ISL_2455587, EPI_ISL_2455677, EPI_ISL_2455895, EPI_ISL_2456004 | National Hospital for Tropical Diseases | Oxford University Clinical Research Unit, Hanoi, Vietnam | Pham Ngoc Thach, Nguyen Thi Tam, Van Dinh Trang, Le Van Duyet, Nguyen Thu Trang, Nguyen Thi Hong Thuong, Nguyen Thi Kim Chi, Phan Manh Cuong, Thomas Kesteman, H.Rogier van Doorn on behalf of the OUCRU COVID-19 research group |
| EPI_ISL_2456005, EPI_ISL_2456006 | Lighthouse Lab in Milton Keynes | Wellcome Sanger Institute for the COVID-19 Genomics UK (COG-UK) Consortium | The Lighthouse Lab in Milton Keynes and Alex Alderton, Roberto Amato, Jeffrey Barrett, Sonia Goncalves, Ewan Harrison, David K. Jackson, Ian Johnston, Dominic Kwiatkowski, Cordelia Langford, John Sillitoe on behalf of the Wellcome Sanger Institute COVID-19 Surveillance Team |
| EPI_ISL_2456008, EPI_ISL_2456009, EPI_ISL_2456010, EPI_ISL_2456011 | Lighthouse Lab in Alderley Park | Wellcome Sanger Institute for the COVID-19 Genomics UK (COG-UK) Consortium | Jacquelyn Wynn, Mairead Hyland, The Lighthouse Lab in Alderley Park and Alex Alderton, Roberto Amato, Jeffrey Barrett, Sonia Goncalves, Ewan Harrison, David K. Jackson, Ian Johnston, Dominic Kwiatkowski, Cordelia Langford, John Sillitoe on behalf of the Wellcome Sanger Institute COVID-19 Surveillance Team |
| EPI_ISL_2456012, EPI_ISL_2456013, EPI_ISL_2456014, EPI_ISL_2456015, EPI_ISL_2456016, EPI_ISL_2456017 | Lighthouse Lab in Milton Keynes | Wellcome Sanger Institute for the COVID-19 Genomics UK (COG-UK) Consortium | The Lighthouse Lab in Milton Keynes and Alex Alderton, Roberto Amato, Jeffrey Barrett, Sonia Goncalves, Ewan Harrison, David K. Jackson, Ian Johnston, Dominic Kwiatkowski, Cordelia Langford, John Sillitoe on behalf of the Wellcome Sanger Institute COVID-19 Surveillance Team |
| EPI_ISL_2456018 | Lighthouse Lab in Alderley Park | Wellcome Sanger Institute for the COVID-19 Genomics UK (COG-UK) Consortium | Jacquelyn Wynn, Mairead Hyland, The Lighthouse Lab in Alderley Park and Alex Alderton, Roberto Amato, Jeffrey Barrett, Sonia Goncalves, Ewan Harrison, David K. Jackson, Ian Johnston, Dominic Kwiatkowski, Cordelia Langford, John Sillitoe on behalf of the Wellcome Sanger Institute COVID-19 Surveillance Team |
| EPI_ISL_2456019, EPI_ISL_2456020, EPI_ISL_2456021, EPI_ISL_2456022, EPI_ISL_2456023, EPI_ISL_2456024, EPI_ISL_2456025, EPI_ISL_2456027, EPI_ISL_2456028, EPI_ISL_2456029, EPI_ISL_2456030, EPI_ISL_2456033, EPI_ISL_2456034 | | | |
| see above | Lighthouse Lab in Milton Keynes | Wellcome Sanger Institute for the COVID-19 Genomics UK (COG-UK) Consortium | The Lighthouse Lab in Milton Keynes and Alex Alderton, Roberto Amato, Jeffrey Barrett, Sonia Goncalves, Ewan Harrison, David K. Jackson, Ian Johnston, Dominic Kwiatkowski, Cordelia Langford, John Sillitoe on behalf of the Wellcome Sanger Institute COVID-19 Surveillance Team |
| EPI_ISL_2456035 | Lighthouse Lab in Alderley Park | Wellcome Sanger Institute for the COVID-19 Genomics UK (COG-UK) Consortium | Jacquelyn Wynn, Mairead Hyland, The Lighthouse Lab in Alderley Park and Alex Alderton, Roberto Amato, Jeffrey Barrett, Sonia Goncalves, Ewan Harrison, David K. Jackson, Ian Johnston, Dominic Kwiatkowski, Cordelia Langford, John Sillitoe on behalf of the Wellcome Sanger Institute COVID-19 Surveillance Team |
| EPI_ISL_2456036, EPI_ISL_2456037, EPI_ISL_2456038 | Lighthouse Lab in Milton Keynes | Wellcome Sanger Institute for the COVID-19 Genomics UK (COG-UK) Consortium | The Lighthouse Lab in Milton Keynes and Alex Alderton, Roberto Amato, Jeffrey Barrett, Sonia Goncalves, Ewan Harrison, David K. Jackson, Ian Johnston, Dominic Kwiatkowski, Cordelia Langford, John Sillitoe on behalf of the Wellcome Sanger Institute COVID-19 Surveillance Team |
| EPI_ISL_2456039 | Lighthouse Lab in Alderley Park | Wellcome Sanger Institute for the COVID-19 Genomics UK (COG-UK) Consortium | Jacquelyn Wynn, Mairead Hyland, The Lighthouse Lab in Alderley Park and Alex Alderton, Roberto Amato, Jeffrey Barrett, Sonia Goncalves, Ewan Harrison, David K. Jackson, Ian Johnston, Dominic Kwiatkowski, Cordelia Langford, John Sillitoe on behalf of the Wellcome Sanger Institute COVID-19 Surveillance Team |
| EPI_ISL_2456040, EPI_ISL_2456041, EPI_ISL_2456042, EPI_ISL_2456044 | Lighthouse Lab in Milton Keynes | Wellcome Sanger Institute for the COVID-19 Genomics UK (COG-UK) Consortium | The Lighthouse Lab in Milton Keynes and Alex Alderton, Roberto Amato, Jeffrey Barrett, Sonia Goncalves, Ewan Harrison, David K. Jackson, Ian Johnston, Dominic Kwiatkowski, Cordelia Langford, John Sillitoe on behalf of the Wellcome Sanger Institute COVID-19 Surveillance Team |
| EPI_ISL_2456045 | Lighthouse Lab in Alderley Park | Wellcome Sanger Institute for the COVID-19 Genomics UK (COG-UK) Consortium | Jacquelyn Wynn, Mairead Hyland, The Lighthouse Lab in Alderley Park and Alex Alderton, Roberto Amato, Jeffrey Barrett, Sonia Goncalves, Ewan Harrison, David K. Jackson, Ian Johnston, Dominic Kwiatkowski, Cordelia Langford, John Sillitoe on behalf of the Wellcome Sanger Institute COVID-19 Surveillance Team |
| EPI_ISL_2456046, EPI_ISL_2456047, EPI_ISL_2456048 | Lighthouse Lab in Milton Keynes | Wellcome Sanger Institute for the COVID-19 Genomics UK (COG-UK) Consortium | The Lighthouse Lab in Milton Keynes and Alex Alderton, Roberto Amato, Jeffrey Barrett, Sonia Goncalves, Ewan Harrison, David K. Jackson, Ian Johnston, Dominic Kwiatkowski, Cordelia Langford, John Sillitoe on behalf of the Wellcome Sanger Institute COVID-19 Surveillance Team |
| EPI_ISL_2456049 | Lighthouse Lab in Alderley Park | Wellcome Sanger Institute for the COVID-19 Genomics UK (COG-UK) Consortium | Jacquelyn Wynn, Mairead Hyland, The Lighthouse Lab in Alderley Park and Alex Alderton, Roberto Amato, Jeffrey Barrett, Sonia Goncalves, Ewan Harrison, David K. Jackson, Ian Johnston, Dominic Kwiatkowski, Cordelia Langford, John Sillitoe on behalf of the Wellcome Sanger Institute COVID-19 Surveillance Team |
| EPI_ISL_2456050, EPI_ISL_2456051, EPI_ISL_2456052 | Lighthouse Lab in Milton Keynes | Wellcome Sanger Institute for the COVID-19 Genomics UK (COG-UK) Consortium | The Lighthouse Lab in Milton Keynes and Alex Alderton, Roberto Amato, Jeffrey Barrett, Sonia Goncalves, Ewan Harrison, David K. Jackson, Ian Johnston, Dominic Kwiatkowski, Cordelia Langford, John Sillitoe on behalf of the Wellcome Sanger Institute COVID-19 Surveillance Team |
| EPI_ISL_2456054, EPI_ISL_2456055 | Lighthouse Lab in Alderley Park | Wellcome Sanger Institute for the COVID-19 Genomics UK (COG-UK) Consortium | Jacquelyn Wynn, Mairead Hyland, The Lighthouse Lab in Alderley Park and Alex Alderton, Roberto Amato, Jeffrey Barrett, Sonia Goncalves, Ewan Harrison, David K. Jackson, Ian Johnston, Dominic Kwiatkowski, Cordelia Langford, John Sillitoe on behalf of the Wellcome Sanger Institute COVID-19 Surveillance Team |
| EPI_ISL_2456056 | Lighthouse Lab in Milton Keynes | Wellcome Sanger Institute for the COVID-19 Genomics UK (COG-UK) Consortium | The Lighthouse Lab in Milton Keynes and Alex Alderton, Roberto Amato, Jeffrey Barrett, Sonia Goncalves, Ewan Harrison, David K. Jackson, Ian Johnston, Dominic Kwiatkowski, Cordelia Langford, John Sillitoe on behalf of the Wellcome Sanger Institute COVID-19 Surveillance Team |
| EPI_ISL_2456057 | Lighthouse Lab in Alderley Park | Wellcome Sanger Institute for the COVID-19 Genomics UK (COG-UK) Consortium | Jacquelyn Wynn, Mairead Hyland, The Lighthouse Lab in Alderley Park and Alex Alderton, Roberto Amato, Jeffrey Barrett, Sonia Goncalves, Ewan Harrison, David K. Jackson, Ian Johnston, Dominic Kwiatkowski, Cordelia Langford, John Sillitoe on behalf of the Wellcome Sanger Institute COVID-19 Surveillance Team |
| EPI_ISL_2456058, EPI_ISL_2456059 | Lighthouse Lab in Milton Keynes | Wellcome Sanger Institute for the COVID-19 Genomics UK (COG-UK) Consortium | The Lighthouse Lab in Milton Keynes and Alex Alderton, Roberto Amato, Jeffrey Barrett, Sonia Goncalves, Ewan Harrison, David K. Jackson, Ian Johnston, Dominic Kwiatkowski, Cordelia Langford, John Sillitoe on behalf of the Wellcome Sanger Institute COVID-19 Surveillance Team |
| EPI_ISL_2456060, EPI_ISL_2456061 | Lighthouse Lab in Alderley Park | Wellcome Sanger Institute for the COVID-19 Genomics UK (COG-UK) Consortium | Jacquelyn Wynn, Mairead Hyland, The Lighthouse Lab in Alderley Park and Alex Alderton, Roberto Amato, Jeffrey Barrett, Sonia Goncalves, Ewan Harrison, David K. Jackson, Ian Johnston, Dominic Kwiatkowski, Cordelia Langford, John Sillitoe on behalf of the Wellcome Sanger Institute COVID-19 Surveillance Team |
| EPI_ISL_2456062, EPI_ISL_2456063, EPI_ISL_2456065 | Lighthouse Lab in Milton Keynes | Wellcome Sanger Institute for the COVID-19 Genomics UK (COG-UK) Consortium | The Lighthouse Lab in Milton Keynes and Alex Alderton, Roberto Amato, Jeffrey Barrett, Sonia Goncalves, Ewan Harrison, David K. Jackson, Ian Johnston, Dominic Kwiatkowski, Cordelia Langford, John Sillitoe on behalf of the Wellcome Sanger Institute COVID-19 Surveillance Team |
| EPI_ISL_2456067 | Lighthouse Lab in Alderley Park | Wellcome Sanger Institute for the COVID-19 Genomics UK (COG-UK) Consortium | Jacquelyn Wynn, Mairead Hyland, The Lighthouse Lab in Alderley Park and Alex Alderton, Roberto Amato, Jeffrey Barrett, Sonia Goncalves, Ewan Harrison, David K. Jackson, Ian Johnston, Dominic Kwiatkowski, Cordelia Langford, John Sillitoe on behalf of the Wellcome Sanger Institute COVID-19 Surveillance Team |
| EPI_ISL_2456068, EPI_ISL_2456069 | Lighthouse Lab in Milton Keynes | Wellcome Sanger Institute for the COVID-19 Genomics UK (COG-UK) Consortium | The Lighthouse Lab in Milton Keynes and Alex Alderton, Roberto Amato, Jeffrey Barrett, Sonia Goncalves, Ewan Harrison, David K. Jackson, Ian Johnston, Dominic Kwiatkowski, Cordelia Langford, John Sillitoe on behalf of the Wellcome Sanger Institute COVID-19 Surveillance Team |
| EPI_ISL_2456070, EPI_ISL_2456071, EPI_ISL_2456072 | Lighthouse Lab in Alderley Park | Wellcome Sanger Institute for the COVID-19 Genomics UK (COG-UK) Consortium | Jacquelyn Wynn, Mairead Hyland, The Lighthouse Lab in Alderley Park and Alex Alderton, Roberto Amato, Jeffrey Barrett, Sonia Goncalves, Ewan Harrison, David K. Jackson, Ian Johnston, Dominic Kwiatkowski, Cordelia Langford, John Sillitoe on behalf of the Wellcome Sanger Institute COVID-19 Surveillance Team |
| EPI_ISL_2456073, EPI_ISL_2456074, EPI_ISL_2456075, EPI_ISL_2456076, EPI_ISL_2456077, EPI_ISL_2456078 | Lighthouse Lab in Milton Keynes | Wellcome Sanger Institute for the COVID-19 Genomics UK (COG-UK) Consortium | The Lighthouse Lab in Milton Keynes and Alex Alderton, Roberto Amato, Jeffrey Barrett, Sonia Goncalves, Ewan Harrison, David K. Jackson, Ian Johnston, Dominic Kwiatkowski, Cordelia Langford, John Sillitoe on behalf of the Wellcome Sanger Institute COVID-19 Surveillance Team |
| EPI_ISL_2456079 | Lighthouse Lab in Alderley Park | Wellcome Sanger Institute for the COVID-19 Genomics UK (COG-UK) Consortium | Jacquelyn Wynn, Mairead Hyland, The Lighthouse Lab in Alderley Park and Alex Alderton, Roberto Amato, Jeffrey Barrett, Sonia Goncalves, Ewan Harrison, David K. Jackson, Ian Johnston, Dominic Kwiatkowski, Cordelia Langford, John Sillitoe on behalf of the Wellcome Sanger Institute COVID-19 Surveillance Team |
| EPI_ISL_2456080, EPI_ISL_2456081, | Lighthouse Lab in Milton Keynes | Wellcome Sanger Institute for the COVID-19 Genomics UK (COG-UK) Consortium | The Lighthouse Lab in Milton Keynes and Alex Alderton, Roberto Amato, Jeffrey Barrett, Sonia Goncalves, Ewan Harrison, David K. Jackson, Ian Johnston, Dominic Kwiatkowski, Cordelia Langford, John Sillitoe on behalf of the Wellcome Sanger Institute COVID-19 Surveillance Team |

| | | | |
|---|---|---|---|
| EPI_ISL_2456082 | | | |
| EPI_ISL_2456083 | Lighthouse Lab in Alderley Park | Wellcome Sanger Institute for the COVID-19 Genomics UK (COG-UK) Consortium | Jacquelyn Wynn, Mairead Hyland, The Lighthouse Lab in Alderley Park and Alex Alderton, Roberto Amato, Jeffrey Barrett, Sonia Goncalves, Ewan Harrison, David K. Jackson, Ian Johnston, Dominic Kwiatkowski, Cordelia Langford, John Sillitoe on behalf of the Wellcome Sanger Institute COVID-19 Surveillance Team |
| EPI_ISL_2456084, EPI_ISL_2456085, EPI_ISL_2456086, EPI_ISL_2456087, EPI_ISL_2456088, EPI_ISL_2456089 | Lighthouse Lab in Milton Keynes | Wellcome Sanger Institute for the COVID-19 Genomics UK (COG-UK) Consortium | The Lighthouse Lab in Milton Keynes and Alex Alderton, Roberto Amato, Jeffrey Barrett, Sonia Goncalves, Ewan Harrison, David K. Jackson, Ian Johnston, Dominic Kwiatkowski, Cordelia Langford, John Sillitoe on behalf of the Wellcome Sanger Institute COVID-19 Surveillance Team |
| EPI_ISL_2456090 | Lighthouse Lab in Alderley Park | Wellcome Sanger Institute for the COVID-19 Genomics UK (COG-UK) Consortium | Jacquelyn Wynn, Mairead Hyland, The Lighthouse Lab in Alderley Park and Alex Alderton, Roberto Amato, Jeffrey Barrett, Sonia Goncalves, Ewan Harrison, David K. Jackson, Ian Johnston, Dominic Kwiatkowski, Cordelia Langford, John Sillitoe on behalf of the Wellcome Sanger Institute COVID-19 Surveillance Team |
| EPI_ISL_2456091 | Lighthouse Lab in Milton Keynes | Wellcome Sanger Institute for the COVID-19 Genomics UK (COG-UK) Consortium | The Lighthouse Lab in Milton Keynes and Alex Alderton, Roberto Amato, Jeffrey Barrett, Sonia Goncalves, Ewan Harrison, David K. Jackson, Ian Johnston, Dominic Kwiatkowski, Cordelia Langford, John Sillitoe on behalf of the Wellcome Sanger Institute COVID-19 Surveillance Team |
| EPI_ISL_2456092 | Lighthouse Lab in Alderley Park | Wellcome Sanger Institute for the COVID-19 Genomics UK (COG-UK) Consortium | Jacquelyn Wynn, Mairead Hyland, The Lighthouse Lab in Alderley Park and Alex Alderton, Roberto Amato, Jeffrey Barrett, Sonia Goncalves, Ewan Harrison, David K. Jackson, Ian Johnston, Dominic Kwiatkowski, Cordelia Langford, John Sillitoe on behalf of the Wellcome Sanger Institute COVID-19 Surveillance Team |
| EPI_ISL_2456093 | Lighthouse Lab in Milton Keynes | Wellcome Sanger Institute for the COVID-19 Genomics UK (COG-UK) Consortium | The Lighthouse Lab in Milton Keynes and Alex Alderton, Roberto Amato, Jeffrey Barrett, Sonia Goncalves, Ewan Harrison, David K. Jackson, Ian Johnston, Dominic Kwiatkowski, Cordelia Langford, John Sillitoe on behalf of the Wellcome Sanger Institute COVID-19 Surveillance Team |
| EPI_ISL_2456094, EPI_ISL_2456095, EPI_ISL_2456096, EPI_ISL_2456097 | Lighthouse Lab in Alderley Park | Wellcome Sanger Institute for the COVID-19 Genomics UK (COG-UK) Consortium | Jacquelyn Wynn, Mairead Hyland, The Lighthouse Lab in Alderley Park and Alex Alderton, Roberto Amato, Jeffrey Barrett, Sonia Goncalves, Ewan Harrison, David K. Jackson, Ian Johnston, Dominic Kwiatkowski, Cordelia Langford, John Sillitoe on behalf of the Wellcome Sanger Institute COVID-19 Surveillance Team |
| EPI_ISL_2456098, EPI_ISL_2456099, EPI_ISL_2456101, EPI_ISL_2456102, EPI_ISL_2456103, EPI_ISL_2456104, EPI_ISL_2456105, EPI_ISL_2456106, EPI_ISL_2456107, EPI_ISL_2456108 | Lighthouse Lab in Milton Keynes | Wellcome Sanger Institute for the COVID-19 Genomics UK (COG-UK) Consortium | The Lighthouse Lab in Milton Keynes and Alex Alderton, Roberto Amato, Jeffrey Barrett, Sonia Goncalves, Ewan Harrison, David K. Jackson, Ian Johnston, Dominic Kwiatkowski, Cordelia Langford, John Sillitoe on behalf of the Wellcome Sanger Institute COVID-19 Surveillance Team |
| EPI_ISL_2456109 | Lighthouse Lab in Alderley Park | Wellcome Sanger Institute for the COVID-19 Genomics UK (COG-UK) Consortium | Jacquelyn Wynn, Mairead Hyland, The Lighthouse Lab in Alderley Park and Alex Alderton, Roberto Amato, Jeffrey Barrett, Sonia Goncalves, Ewan Harrison, David K. Jackson, Ian Johnston, Dominic Kwiatkowski, Cordelia Langford, John Sillitoe on behalf of the Wellcome Sanger Institute COVID-19 Surveillance Team |
| EPI_ISL_2456110 | Lighthouse Lab in Milton Keynes | Wellcome Sanger Institute for the COVID-19 Genomics UK (COG-UK) Consortium | The Lighthouse Lab in Milton Keynes and Alex Alderton, Roberto Amato, Jeffrey Barrett, Sonia Goncalves, Ewan Harrison, David K. Jackson, Ian Johnston, Dominic Kwiatkowski, Cordelia Langford, John Sillitoe on behalf of the Wellcome Sanger Institute COVID-19 Surveillance Team |
| EPI_ISL_2456112 | Lighthouse Lab in Alderley Park | Wellcome Sanger Institute for the COVID-19 Genomics UK (COG-UK) Consortium | Jacquelyn Wynn, Mairead Hyland, The Lighthouse Lab in Alderley Park and Alex Alderton, Roberto Amato, Jeffrey Barrett, Sonia Goncalves, Ewan Harrison, David K. Jackson, Ian Johnston, Dominic Kwiatkowski, Cordelia Langford, John Sillitoe on behalf of the Wellcome Sanger Institute COVID-19 Surveillance Team |
| EPI_ISL_2456113, EPI_ISL_2456116 | Lighthouse Lab in Milton Keynes | Wellcome Sanger Institute for the COVID-19 Genomics UK (COG-UK) Consortium | The Lighthouse Lab in Milton Keynes and Alex Alderton, Roberto Amato, Jeffrey Barrett, Sonia Goncalves, Ewan Harrison, David K. Jackson, Ian Johnston, Dominic Kwiatkowski, Cordelia Langford, John Sillitoe on behalf of the Wellcome Sanger Institute COVID-19 Surveillance Team |
| EPI_ISL_2456117 | Utah Public Health Laboratory | Utah Public Health Laboratory | Erin L. Young, Kelly F. Oakeson, Tara Gallagher |
| EPI_ISL_2456118, EPI_ISL_2456121, EPI_ISL_2456122 | Lighthouse Lab in Milton Keynes | Wellcome Sanger Institute for the COVID-19 Genomics UK (COG-UK) Consortium | The Lighthouse Lab in Milton Keynes and Alex Alderton, Roberto Amato, Jeffrey Barrett, Sonia Goncalves, Ewan Harrison, David K. Jackson, Ian Johnston, Dominic Kwiatkowski, Cordelia Langford, John Sillitoe on behalf of the Wellcome Sanger Institute COVID-19 Surveillance Team |
| EPI_ISL_2456125 | Lighthouse Lab in Alderley Park | Wellcome Sanger Institute for the COVID-19 Genomics UK (COG-UK) Consortium | Jacquelyn Wynn, Mairead Hyland, The Lighthouse Lab in Alderley Park and Alex Alderton, Roberto Amato, Jeffrey Barrett, Sonia Goncalves, Ewan Harrison, David K. Jackson, Ian Johnston, Dominic Kwiatkowski, Cordelia Langford, John Sillitoe on behalf of the Wellcome Sanger Institute COVID-19 Surveillance Team |
| EPI_ISL_2456127 | Lighthouse Lab in Milton Keynes | Wellcome Sanger Institute for the COVID-19 Genomics UK (COG-UK) Consortium | The Lighthouse Lab in Milton Keynes and Alex Alderton, Roberto Amato, Jeffrey Barrett, Sonia Goncalves, Ewan Harrison, David K. Jackson, Ian Johnston, Dominic Kwiatkowski, Cordelia Langford, John Sillitoe on behalf of the Wellcome Sanger Institute COVID-19 Surveillance Team |
| EPI_ISL_2456128, EPI_ISL_2456130 | Lighthouse Lab in Alderley Park | Wellcome Sanger Institute for the COVID-19 Genomics UK (COG-UK) Consortium | Jacquelyn Wynn, Mairead Hyland, The Lighthouse Lab in Alderley Park and Alex Alderton, Roberto Amato, Jeffrey Barrett, Sonia Goncalves, Ewan Harrison, David K. Jackson, Ian Johnston, Dominic Kwiatkowski, Cordelia Langford, John Sillitoe on behalf of the Wellcome Sanger Institute COVID-19 Surveillance Team |
| EPI_ISL_2456131, EPI_ISL_2456133 | Lighthouse Lab in Milton Keynes | Wellcome Sanger Institute for the COVID-19 Genomics UK (COG-UK) Consortium | The Lighthouse Lab in Milton Keynes and Alex Alderton, Roberto Amato, Jeffrey Barrett, Sonia Goncalves, Ewan Harrison, David K. Jackson, Ian Johnston, Dominic Kwiatkowski, Cordelia Langford, John Sillitoe on behalf of the Wellcome Sanger Institute COVID-19 Surveillance Team |
| EPI_ISL_2456134 | Lighthouse Lab in Alderley Park | Wellcome Sanger Institute for the COVID-19 Genomics UK (COG-UK) Consortium | Jacquelyn Wynn, Mairead Hyland, The Lighthouse Lab in Alderley Park and Alex Alderton, Roberto Amato, Jeffrey Barrett, Sonia Goncalves, Ewan Harrison, David K. Jackson, Ian Johnston, Dominic Kwiatkowski, Cordelia Langford, John Sillitoe on behalf of the Wellcome Sanger Institute COVID-19 Surveillance Team |
| EPI_ISL_2456138 | Utah Public Health Laboratory | Utah Public Health Laboratory | Erin L. Young, Kelly F. Oakeson, Tara Gallagher |
| EPI_ISL_2456139 | Lighthouse Lab in Alderley Park | Wellcome Sanger Institute for the COVID-19 Genomics UK (COG-UK) Consortium | Jacquelyn Wynn, Mairead Hyland, The Lighthouse Lab in Alderley Park and Alex Alderton, Roberto Amato, Jeffrey Barrett, Sonia Goncalves, Ewan Harrison, David K. Jackson, Ian Johnston, Dominic Kwiatkowski, Cordelia Langford, John Sillitoe on behalf of the Wellcome Sanger Institute COVID-19 Surveillance Team |
| EPI_ISL_2456140 | Lighthouse Lab in Milton Keynes | Wellcome Sanger Institute for the COVID-19 Genomics UK (COG-UK) Consortium | The Lighthouse Lab in Milton Keynes and Alex Alderton, Roberto Amato, Jeffrey Barrett, Sonia Goncalves, Ewan Harrison, David K. Jackson, Ian Johnston, Dominic Kwiatkowski, Cordelia Langford, John Sillitoe on behalf of the Wellcome Sanger Institute COVID-19 Surveillance Team |
| EPI_ISL_2456141 | Utah Public Health Laboratory | Utah Public Health Laboratory | Erin L. Young, Kelly F. Oakeson, Tara Gallagher |
| EPI_ISL_2456142, EPI_ISL_2456144, EPI_ISL_2456145 | Lighthouse Lab in Alderley Park | Wellcome Sanger Institute for the COVID-19 Genomics UK (COG-UK) Consortium | Jacquelyn Wynn, Mairead Hyland, The Lighthouse Lab in Alderley Park and Alex Alderton, Roberto Amato, Jeffrey Barrett, Sonia Goncalves, Ewan Harrison, David K. Jackson, Ian Johnston, Dominic Kwiatkowski, Cordelia Langford, John Sillitoe on behalf of the Wellcome Sanger Institute COVID-19 Surveillance Team |
| EPI_ISL_2456148, EPI_ISL_2456150 | Lighthouse Lab in Milton Keynes | Wellcome Sanger Institute for the COVID-19 Genomics UK (COG-UK) Consortium | The Lighthouse Lab in Milton Keynes and Alex Alderton, Roberto Amato, Jeffrey Barrett, Sonia Goncalves, Ewan Harrison, David K. Jackson, Ian Johnston, Dominic Kwiatkowski, Cordelia Langford, John Sillitoe on behalf of the Wellcome Sanger Institute COVID-19 Surveillance Team |
| EPI_ISL_2456151 | Lighthouse Lab in Alderley Park | Wellcome Sanger Institute for the COVID-19 Genomics UK (COG-UK) Consortium | Jacquelyn Wynn, Mairead Hyland, The Lighthouse Lab in Alderley Park and Alex Alderton, Roberto Amato, Jeffrey Barrett, Sonia Goncalves, Ewan Harrison, David K. Jackson, Ian Johnston, Dominic Kwiatkowski, Cordelia Langford, John Sillitoe on behalf of the Wellcome Sanger Institute COVID-19 Surveillance Team |
| EPI_ISL_2456152, EPI_ISL_2456154, EPI_ISL_2456155 | Lighthouse Lab in Milton Keynes | Wellcome Sanger Institute for the COVID-19 Genomics UK (COG-UK) Consortium | The Lighthouse Lab in Milton Keynes and Alex Alderton, Roberto Amato, Jeffrey Barrett, Sonia Goncalves, Ewan Harrison, David K. Jackson, Ian Johnston, Dominic Kwiatkowski, Cordelia Langford, John Sillitoe on behalf of the Wellcome Sanger Institute COVID-19 Surveillance Team |
| EPI_ISL_2456156 | Utah Public Health Laboratory | Utah Public Health Laboratory | Erin L. Young, Kelly F. Oakeson, Tara Gallagher |
| EPI_ISL_2456157, EPI_ISL_2456158, EPI_ISL_2456159, EPI_ISL_2456161 | Lighthouse Lab in Milton Keynes | Wellcome Sanger Institute for the COVID-19 Genomics UK (COG-UK) Consortium | The Lighthouse Lab in Milton Keynes and Alex Alderton, Roberto Amato, Jeffrey Barrett, Sonia Goncalves, Ewan Harrison, David K. Jackson, Ian Johnston, Dominic Kwiatkowski, Cordelia Langford, John Sillitoe on behalf of the Wellcome Sanger Institute COVID-19 Surveillance Team |
| EPI_ISL_2456162, EPI_ISL_2456164, EPI_ISL_2456165 | Lighthouse Lab in Alderley Park | Wellcome Sanger Institute for the COVID-19 Genomics UK (COG-UK) Consortium | Jacquelyn Wynn, Mairead Hyland, The Lighthouse Lab in Alderley Park and Alex Alderton, Roberto Amato, Jeffrey Barrett, Sonia Goncalves, Ewan Harrison, David K. Jackson, Ian Johnston, Dominic Kwiatkowski, Cordelia Langford, John Sillitoe on behalf of the Wellcome Sanger Institute COVID-19 Surveillance Team |
| EPI_ISL_2456167, EPI_ISL_2456168 | Lighthouse Lab in Milton Keynes | Wellcome Sanger Institute for the COVID-19 Genomics UK (COG-UK) Consortium | The Lighthouse Lab in Milton Keynes and Alex Alderton, Roberto Amato, Jeffrey Barrett, Sonia Goncalves, Ewan Harrison, David K. Jackson, Ian Johnston, Dominic Kwiatkowski, Cordelia Langford, John Sillitoe on behalf of the Wellcome Sanger Institute COVID-19 Surveillance Team |
| EPI_ISL_2456170 | Lighthouse Lab in Alderley Park | Wellcome Sanger Institute for the COVID-19 Genomics UK (COG-UK) Consortium | Jacquelyn Wynn, Mairead Hyland, The Lighthouse Lab in Alderley Park and Alex Alderton, Roberto Amato, Jeffrey Barrett, Sonia Goncalves, Ewan Harrison, David K. Jackson, Ian Johnston, Dominic Kwiatkowski, Cordelia Langford, John Sillitoe on behalf of the Wellcome Sanger Institute COVID-19 Surveillance Team |
| EPI_ISL_2456171 | Lighthouse Lab in Milton Keynes | Wellcome Sanger Institute for the COVID-19 Genomics UK (COG-UK) Consortium | The Lighthouse Lab in Milton Keynes and Alex Alderton, Roberto Amato, Jeffrey Barrett, Sonia Goncalves, Ewan Harrison, David K. Jackson, Ian Johnston, Dominic Kwiatkowski, Cordelia Langford, John Sillitoe on behalf of the Wellcome Sanger Institute COVID-19 Surveillance Team |
| EPI_ISL_2456173 | Lighthouse Lab in Alderley Park | Wellcome Sanger Institute for the COVID-19 Genomics UK (COG-UK) Consortium | Jacquelyn Wynn, Mairead Hyland, The Lighthouse Lab in Alderley Park and Alex Alderton, Roberto Amato, Jeffrey Barrett, Sonia Goncalves, Ewan Harrison, David K. Jackson, Ian Johnston, Dominic Kwiatkowski, Cordelia Langford, John Sillitoe on behalf of the Wellcome Sanger Institute COVID-19 Surveillance Team |
| EPI_ISL_2456174, EPI_ISL_2456176, EPI_ISL_2456178, EPI_ISL_2456180, EPI_ISL_2456181, EPI_ISL_2456183, EPI_ISL_2456184, EPI_ISL_2456186, EPI_ISL_2456187, EPI_ISL_2456189, EPI_ISL_2456190 | | | |
| see above | Lighthouse Lab in Milton Keynes | Wellcome Sanger Institute for the COVID-19 Genomics UK (COG-UK) Consortium | The Lighthouse Lab in Milton Keynes and Alex Alderton, Roberto Amato, Jeffrey Barrett, Sonia Goncalves, Ewan Harrison, David K. Jackson, Ian Johnston, Dominic Kwiatkowski, Cordelia Langford, John Sillitoe on behalf of the Wellcome Sanger Institute COVID-19 Surveillance Team |
| EPI_ISL_2456192 | Lighthouse Lab in Alderley Park | Wellcome Sanger Institute for the COVID-19 Genomics UK (COG-UK) Consortium | Jacquelyn Wynn, Mairead Hyland, The Lighthouse Lab in Alderley Park and Alex Alderton, Roberto Amato, Jeffrey Barrett, Sonia Goncalves, Ewan Harrison, David K. Jackson, Ian Johnston, Dominic Kwiatkowski, Cordelia Langford, John Sillitoe on behalf of the Wellcome Sanger Institute COVID-19 Surveillance Team |
| EPI_ISL_2456195 | Lighthouse Lab in Milton Keynes | Wellcome Sanger Institute for the COVID-19 Genomics UK (COG-UK) Consortium | The Lighthouse Lab in Milton Keynes and Alex Alderton, Roberto Amato, Jeffrey Barrett, Sonia Goncalves, Ewan Harrison, David K. Jackson, Ian Johnston, Dominic Kwiatkowski, Cordelia Langford, John Sillitoe on behalf of the Wellcome Sanger Institute COVID-19 Surveillance Team |

| | | | |
|---|---|---|---|
| EPI_ISL_2456197 | Lighthouse Lab in Alderley Park | Wellcome Sanger Institute for the COVID-19 Genomics UK (COG-UK) Consortium | Jacquelyn Wynn, Mairead Hyland, The Lighthouse Lab in Alderley Park and Alex Alderton, Roberto Amato, Jeffrey Barrett, Sonia Goncalves, Ewan Harrison, David K. Jackson, Ian Johnston, Dominic Kwiatkowski, Cordelia Langford, John Sillitoe on behalf of the Wellcome Sanger Institute COVID-19 Surveillance Team |
| EPI_ISL_2456198, EPI_ISL_2456199 | Lighthouse Lab in Milton Keynes | Wellcome Sanger Institute for the COVID-19 Genomics UK (COG-UK) Consortium | The Lighthouse Lab in Milton Keynes and Alex Alderton, Roberto Amato, Jeffrey Barrett, Sonia Goncalves, Ewan Harrison, David K. Jackson, Ian Johnston, Dominic Kwiatkowski, Cordelia Langford, John Sillitoe on behalf of the Wellcome Sanger Institute COVID-19 Surveillance Team |
| EPI_ISL_2456201 | Lighthouse Lab in Alderley Park | Wellcome Sanger Institute for the COVID-19 Genomics UK (COG-UK) Consortium | Jacquelyn Wynn, Mairead Hyland, The Lighthouse Lab in Alderley Park and Alex Alderton, Roberto Amato, Jeffrey Barrett, Sonia Goncalves, Ewan Harrison, David K. Jackson, Ian Johnston, Dominic Kwiatkowski, Cordelia Langford, John Sillitoe on behalf of the Wellcome Sanger Institute COVID-19 Surveillance Team |
| EPI_ISL_2456202 | Lighthouse Lab in Milton Keynes | Wellcome Sanger Institute for the COVID-19 Genomics UK (COG-UK) Consortium | The Lighthouse Lab in Milton Keynes and Alex Alderton, Roberto Amato, Jeffrey Barrett, Sonia Goncalves, Ewan Harrison, David K. Jackson, Ian Johnston, Dominic Kwiatkowski, Cordelia Langford, John Sillitoe on behalf of the Wellcome Sanger Institute COVID-19 Surveillance Team |
| EPI_ISL_2456205 | Lighthouse Lab in Alderley Park | Wellcome Sanger Institute for the COVID-19 Genomics UK (COG-UK) Consortium | Jacquelyn Wynn, Mairead Hyland, The Lighthouse Lab in Alderley Park and Alex Alderton, Roberto Amato, Jeffrey Barrett, Sonia Goncalves, Ewan Harrison, David K. Jackson, Ian Johnston, Dominic Kwiatkowski, Cordelia Langford, John Sillitoe on behalf of the Wellcome Sanger Institute COVID-19 Surveillance Team |
| EPI_ISL_2456207 | Lighthouse Lab in Milton Keynes | Wellcome Sanger Institute for the COVID-19 Genomics UK (COG-UK) Consortium | The Lighthouse Lab in Milton Keynes and Alex Alderton, Roberto Amato, Jeffrey Barrett, Sonia Goncalves, Ewan Harrison, David K. Jackson, Ian Johnston, Dominic Kwiatkowski, Cordelia Langford, John Sillitoe on behalf of the Wellcome Sanger Institute COVID-19 Surveillance Team |
| EPI_ISL_2456208, EPI_ISL_2456210 | Lighthouse Lab in Alderley Park | Wellcome Sanger Institute for the COVID-19 Genomics UK (COG-UK) Consortium | Jacquelyn Wynn, Mairead Hyland, The Lighthouse Lab in Alderley Park and Alex Alderton, Roberto Amato, Jeffrey Barrett, Sonia Goncalves, Ewan Harrison, David K. Jackson, Ian Johnston, Dominic Kwiatkowski, Cordelia Langford, John Sillitoe on behalf of the Wellcome Sanger Institute COVID-19 Surveillance Team |
| EPI_ISL_2456211, EPI_ISL_2456213, EPI_ISL_2456214, EPI_ISL_2456216, EPI_ISL_2456217, EPI_ISL_2456219, EPI_ISL_2456220, EPI_ISL_2456221, EPI_ISL_2456223 | Lighthouse Lab in Milton Keynes | Wellcome Sanger Institute for the COVID-19 Genomics UK (COG-UK) Consortium | The Lighthouse Lab in Milton Keynes and Alex Alderton, Roberto Amato, Jeffrey Barrett, Sonia Goncalves, Ewan Harrison, David K. Jackson, Ian Johnston, Dominic Kwiatkowski, Cordelia Langford, John Sillitoe on behalf of the Wellcome Sanger Institute COVID-19 Surveillance Team |
| EPI_ISL_2456224 | Lighthouse Lab in Alderley Park | Wellcome Sanger Institute for the COVID-19 Genomics UK (COG-UK) Consortium | Jacquelyn Wynn, Mairead Hyland, The Lighthouse Lab in Alderley Park and Alex Alderton, Roberto Amato, Jeffrey Barrett, Sonia Goncalves, Ewan Harrison, David K. Jackson, Ian Johnston, Dominic Kwiatkowski, Cordelia Langford, John Sillitoe on behalf of the Wellcome Sanger Institute COVID-19 Surveillance Team |
| EPI_ISL_2456226 | Lighthouse Lab in Milton Keynes | Wellcome Sanger Institute for the COVID-19 Genomics UK (COG-UK) Consortium | The Lighthouse Lab in Milton Keynes and Alex Alderton, Roberto Amato, Jeffrey Barrett, Sonia Goncalves, Ewan Harrison, David K. Jackson, Ian Johnston, Dominic Kwiatkowski, Cordelia Langford, John Sillitoe on behalf of the Wellcome Sanger Institute COVID-19 Surveillance Team |
| EPI_ISL_2456227 | Lighthouse Lab in Alderley Park | Wellcome Sanger Institute for the COVID-19 Genomics UK (COG-UK) Consortium | Jacquelyn Wynn, Mairead Hyland, The Lighthouse Lab in Alderley Park and Alex Alderton, Roberto Amato, Jeffrey Barrett, Sonia Goncalves, Ewan Harrison, David K. Jackson, Ian Johnston, Dominic Kwiatkowski, Cordelia Langford, John Sillitoe on behalf of the Wellcome Sanger Institute COVID-19 Surveillance Team |
| EPI_ISL_2456229, EPI_ISL_2456232 | Lighthouse Lab in Milton Keynes | Wellcome Sanger Institute for the COVID-19 Genomics UK (COG-UK) Consortium | The Lighthouse Lab in Milton Keynes and Alex Alderton, Roberto Amato, Jeffrey Barrett, Sonia Goncalves, Ewan Harrison, David K. Jackson, Ian Johnston, Dominic Kwiatkowski, Cordelia Langford, John Sillitoe on behalf of the Wellcome Sanger Institute COVID-19 Surveillance Team |
| EPI_ISL_2456233, EPI_ISL_2456235, EPI_ISL_2456236 | Lighthouse Lab in Alderley Park | Wellcome Sanger Institute for the COVID-19 Genomics UK (COG-UK) Consortium | Jacquelyn Wynn, Mairead Hyland, The Lighthouse Lab in Alderley Park and Alex Alderton, Roberto Amato, Jeffrey Barrett, Sonia Goncalves, Ewan Harrison, David K. Jackson, Ian Johnston, Dominic Kwiatkowski, Cordelia Langford, John Sillitoe on behalf of the Wellcome Sanger Institute COVID-19 Surveillance Team |
| EPI_ISL_2456238 | Lighthouse Lab in Milton Keynes | Wellcome Sanger Institute for the COVID-19 Genomics UK (COG-UK) Consortium | The Lighthouse Lab in Milton Keynes and Alex Alderton, Roberto Amato, Jeffrey Barrett, Sonia Goncalves, Ewan Harrison, David K. Jackson, Ian Johnston, Dominic Kwiatkowski, Cordelia Langford, John Sillitoe on behalf of the Wellcome Sanger Institute COVID-19 Surveillance Team |
| EPI_ISL_2456239 | Lighthouse Lab in Alderley Park | Wellcome Sanger Institute for the COVID-19 Genomics UK (COG-UK) Consortium | Jacquelyn Wynn, Mairead Hyland, The Lighthouse Lab in Alderley Park and Alex Alderton, Roberto Amato, Jeffrey Barrett, Sonia Goncalves, Ewan Harrison, David K. Jackson, Ian Johnston, Dominic Kwiatkowski, Cordelia Langford, John Sillitoe on behalf of the Wellcome Sanger Institute COVID-19 Surveillance Team |
| EPI_ISL_2456241 | Lighthouse Lab in Milton Keynes | Wellcome Sanger Institute for the COVID-19 Genomics UK (COG-UK) Consortium | The Lighthouse Lab in Milton Keynes and Alex Alderton, Roberto Amato, Jeffrey Barrett, Sonia Goncalves, Ewan Harrison, David K. Jackson, Ian Johnston, Dominic Kwiatkowski, Cordelia Langford, John Sillitoe on behalf of the Wellcome Sanger Institute COVID-19 Surveillance Team |
| EPI_ISL_2456242 | Lighthouse Lab in Alderley Park | Wellcome Sanger Institute for the COVID-19 Genomics UK (COG-UK) Consortium | Jacquelyn Wynn, Mairead Hyland, The Lighthouse Lab in Alderley Park and Alex Alderton, Roberto Amato, Jeffrey Barrett, Sonia Goncalves, Ewan Harrison, David K. Jackson, Ian Johnston, Dominic Kwiatkowski, Cordelia Langford, John Sillitoe on behalf of the Wellcome Sanger Institute COVID-19 Surveillance Team |
| EPI_ISL_2456243, EPI_ISL_2456245, EPI_ISL_2456246, EPI_ISL_2456248 | Lighthouse Lab in Milton Keynes | Wellcome Sanger Institute for the COVID-19 Genomics UK (COG-UK) Consortium | The Lighthouse Lab in Milton Keynes and Alex Alderton, Roberto Amato, Jeffrey Barrett, Sonia Goncalves, Ewan Harrison, David K. Jackson, Ian Johnston, Dominic Kwiatkowski, Cordelia Langford, John Sillitoe on behalf of the Wellcome Sanger Institute COVID-19 Surveillance Team |
| EPI_ISL_2456249 | Lighthouse Lab in Alderley Park | Wellcome Sanger Institute for the COVID-19 Genomics UK (COG-UK) Consortium | Jacquelyn Wynn, Mairead Hyland, The Lighthouse Lab in Alderley Park and Alex Alderton, Roberto Amato, Jeffrey Barrett, Sonia Goncalves, Ewan Harrison, David K. Jackson, Ian Johnston, Dominic Kwiatkowski, Cordelia Langford, John Sillitoe on behalf of the Wellcome Sanger Institute COVID-19 Surveillance Team |
| EPI_ISL_2456251, EPI_ISL_2456252 | Lighthouse Lab in Milton Keynes | Wellcome Sanger Institute for the COVID-19 Genomics UK (COG-UK) Consortium | The Lighthouse Lab in Milton Keynes and Alex Alderton, Roberto Amato, Jeffrey Barrett, Sonia Goncalves, Ewan Harrison, David K. Jackson, Ian Johnston, Dominic Kwiatkowski, Cordelia Langford, John Sillitoe on behalf of the Wellcome Sanger Institute COVID-19 Surveillance Team |
| EPI_ISL_2456254 | Lighthouse Lab in Alderley Park | Wellcome Sanger Institute for the COVID-19 Genomics UK (COG-UK) Consortium | Jacquelyn Wynn, Mairead Hyland, The Lighthouse Lab in Alderley Park and Alex Alderton, Roberto Amato, Jeffrey Barrett, Sonia Goncalves, Ewan Harrison, David K. Jackson, Ian Johnston, Dominic Kwiatkowski, Cordelia Langford, John Sillitoe on behalf of the Wellcome Sanger Institute COVID-19 Surveillance Team |
| EPI_ISL_2456256, EPI_ISL_2456257, EPI_ISL_2456259, EPI_ISL_2456260, EPI_ISL_2456262 | Lighthouse Lab in Milton Keynes | Wellcome Sanger Institute for the COVID-19 Genomics UK (COG-UK) Consortium | The Lighthouse Lab in Milton Keynes and Alex Alderton, Roberto Amato, Jeffrey Barrett, Sonia Goncalves, Ewan Harrison, David K. Jackson, Ian Johnston, Dominic Kwiatkowski, Cordelia Langford, John Sillitoe on behalf of the Wellcome Sanger Institute COVID-19 Surveillance Team |
| EPI_ISL_2456263 | Lighthouse Lab in Alderley Park | Wellcome Sanger Institute for the COVID-19 Genomics UK (COG-UK) Consortium | Jacquelyn Wynn, Mairead Hyland, The Lighthouse Lab in Alderley Park and Alex Alderton, Roberto Amato, Jeffrey Barrett, Sonia Goncalves, Ewan Harrison, David K. Jackson, Ian Johnston, Dominic Kwiatkowski, Cordelia Langford, John Sillitoe on behalf of the Wellcome Sanger Institute COVID-19 Surveillance Team |
| EPI_ISL_2456265, EPI_ISL_2456266, EPI_ISL_2456268, EPI_ISL_2456269 | Lighthouse Lab in Milton Keynes | Wellcome Sanger Institute for the COVID-19 Genomics UK (COG-UK) Consortium | The Lighthouse Lab in Milton Keynes and Alex Alderton, Roberto Amato, Jeffrey Barrett, Sonia Goncalves, Ewan Harrison, David K. Jackson, Ian Johnston, Dominic Kwiatkowski, Cordelia Langford, John Sillitoe on behalf of the Wellcome Sanger Institute COVID-19 Surveillance Team |
| EPI_ISL_2456270 | Lighthouse Lab in Alderley Park | Wellcome Sanger Institute for the COVID-19 Genomics UK (COG-UK) Consortium | Jacquelyn Wynn, Mairead Hyland, The Lighthouse Lab in Alderley Park and Alex Alderton, Roberto Amato, Jeffrey Barrett, Sonia Goncalves, Ewan Harrison, David K. Jackson, Ian Johnston, Dominic Kwiatkowski, Cordelia Langford, John Sillitoe on behalf of the Wellcome Sanger Institute COVID-19 Surveillance Team |
| EPI_ISL_2456271 | Lighthouse Lab in Milton Keynes | Wellcome Sanger Institute for the COVID-19 Genomics UK (COG-UK) Consortium | The Lighthouse Lab in Milton Keynes and Alex Alderton, Roberto Amato, Jeffrey Barrett, Sonia Goncalves, Ewan Harrison, David K. Jackson, Ian Johnston, Dominic Kwiatkowski, Cordelia Langford, John Sillitoe on behalf of the Wellcome Sanger Institute COVID-19 Surveillance Team |
| EPI_ISL_2456272 | Lighthouse Lab in Alderley Park | Wellcome Sanger Institute for the COVID-19 Genomics UK (COG-UK) Consortium | Jacquelyn Wynn, Mairead Hyland, The Lighthouse Lab in Alderley Park and Alex Alderton, Roberto Amato, Jeffrey Barrett, Sonia Goncalves, Ewan Harrison, David K. Jackson, Ian Johnston, Dominic Kwiatkowski, Cordelia Langford, John Sillitoe on behalf of the Wellcome Sanger Institute COVID-19 Surveillance Team |
| EPI_ISL_2456274 | Lighthouse Lab in Milton Keynes | Wellcome Sanger Institute for the COVID-19 Genomics UK (COG-UK) Consortium | The Lighthouse Lab in Milton Keynes and Alex Alderton, Roberto Amato, Jeffrey Barrett, Sonia Goncalves, Ewan Harrison, David K. Jackson, Ian Johnston, Dominic Kwiatkowski, Cordelia Langford, John Sillitoe on behalf of the Wellcome Sanger Institute COVID-19 Surveillance Team |
| EPI_ISL_2456275, EPI_ISL_2456278 | Lighthouse Lab in Alderley Park | Wellcome Sanger Institute for the COVID-19 Genomics UK (COG-UK) Consortium | Jacquelyn Wynn, Mairead Hyland, The Lighthouse Lab in Alderley Park and Alex Alderton, Roberto Amato, Jeffrey Barrett, Sonia Goncalves, Ewan Harrison, David K. Jackson, Ian Johnston, Dominic Kwiatkowski, Cordelia Langford, John Sillitoe on behalf of the Wellcome Sanger Institute COVID-19 Surveillance Team |
| EPI_ISL_2456280, EPI_ISL_2456281, EPI_ISL_2456284, EPI_ISL_2456286 | Lighthouse Lab in Milton Keynes | Wellcome Sanger Institute for the COVID-19 Genomics UK (COG-UK) Consortium | The Lighthouse Lab in Milton Keynes and Alex Alderton, Roberto Amato, Jeffrey Barrett, Sonia Goncalves, Ewan Harrison, David K. Jackson, Ian Johnston, Dominic Kwiatkowski, Cordelia Langford, John Sillitoe on behalf of the Wellcome Sanger Institute COVID-19 Surveillance Team |
| EPI_ISL_2456287 | Lighthouse Lab in Alderley Park | Wellcome Sanger Institute for the COVID-19 Genomics UK (COG-UK) Consortium | Jacquelyn Wynn, Mairead Hyland, The Lighthouse Lab in Alderley Park and Alex Alderton, Roberto Amato, Jeffrey Barrett, Sonia Goncalves, Ewan Harrison, David K. Jackson, Ian Johnston, Dominic Kwiatkowski, Cordelia Langford, John Sillitoe on behalf of the Wellcome Sanger Institute COVID-19 Surveillance Team |
| EPI_ISL_2456289, EPI_ISL_2456290 | Lighthouse Lab in Milton Keynes | Wellcome Sanger Institute for the COVID-19 Genomics UK (COG-UK) Consortium | The Lighthouse Lab in Milton Keynes and Alex Alderton, Roberto Amato, Jeffrey Barrett, Sonia Goncalves, Ewan Harrison, David K. Jackson, Ian Johnston, Dominic Kwiatkowski, Cordelia Langford, John Sillitoe on behalf of the Wellcome Sanger Institute COVID-19 Surveillance Team |
| EPI_ISL_2456292 | Lighthouse Lab in Alderley Park | Wellcome Sanger Institute for the COVID-19 Genomics UK (COG-UK) Consortium | Jacquelyn Wynn, Mairead Hyland, The Lighthouse Lab in Alderley Park and Alex Alderton, Roberto Amato, Jeffrey Barrett, Sonia Goncalves, Ewan Harrison, David K. Jackson, Ian Johnston, Dominic Kwiatkowski, Cordelia Langford, John Sillitoe on behalf of the Wellcome Sanger Institute COVID-19 Surveillance Team |
| EPI_ISL_2456293, EPI_ISL_2456295, EPI_ISL_2456296, EPI_ISL_2456298 | Lighthouse Lab in Milton Keynes | Wellcome Sanger Institute for the COVID-19 Genomics UK (COG-UK) Consortium | The Lighthouse Lab in Milton Keynes and Alex Alderton, Roberto Amato, Jeffrey Barrett, Sonia Goncalves, Ewan Harrison, David K. Jackson, Ian Johnston, Dominic Kwiatkowski, Cordelia Langford, John Sillitoe on behalf of the Wellcome Sanger Institute COVID-19 Surveillance Team |
| EPI_ISL_2456299 | Lighthouse Lab in Alderley Park | Wellcome Sanger Institute for the COVID-19 Genomics UK (COG-UK) Consortium | Jacquelyn Wynn, Mairead Hyland, The Lighthouse Lab in Alderley Park and Alex Alderton, Roberto Amato, Jeffrey Barrett, Sonia Goncalves, Ewan Harrison, David K. Jackson, Ian Johnston, Dominic Kwiatkowski, Cordelia Langford, John Sillitoe on behalf of the Wellcome Sanger Institute COVID-19 Surveillance Team |
| EPI_ISL_2456301, EPI_ISL_2456302, EPI_ISL_2456304, EPI_ISL_2456305, EPI_ISL_2456306 | Lighthouse Lab in Milton Keynes | Wellcome Sanger Institute for the COVID-19 Genomics UK (COG-UK) Consortium | The Lighthouse Lab in Milton Keynes and Alex Alderton, Roberto Amato, Jeffrey Barrett, Sonia Goncalves, Ewan Harrison, David K. Jackson, Ian Johnston, Dominic Kwiatkowski, Cordelia Langford, John Sillitoe on behalf of the Wellcome Sanger Institute COVID-19 Surveillance Team |

| | | | |
|---|---|---|---|
| EPI_ISL_2456307 | Utah Public Health Laboratory | Utah Public Health Laboratory | Erin L. Young, Kelly F. Oakeson, Tara Gallagher |
| EPI_ISL_2456308, EPI_ISL_2456311, EPI_ISL_2456312, EPI_ISL_2456314, EPI_ISL_2456315, EPI_ISL_2456317, EPI_ISL_2456319, EPI_ISL_2456320 | Lighthouse Lab in Milton Keynes | Wellcome Sanger Institute for the COVID-19 Genomics UK (COG-UK) Consortium | The Lighthouse Lab in Milton Keynes and Alex Alderton, Roberto Amato, Jeffrey Barrett, Sonia Goncalves, Ewan Harrison, David K. Jackson, Ian Johnston, Dominic Kwiatkowski, Cordelia Langford, John Sillitoe on behalf of the Wellcome Sanger Institute COVID-19 Surveillance Team |
| EPI_ISL_2456322 | Lighthouse Lab in Alderley Park | Wellcome Sanger Institute for the COVID-19 Genomics UK (COG-UK) Consortium | Jacquelyn Wynn, Mairead Hyland, The Lighthouse Lab in Alderley Park and Alex Alderton, Roberto Amato, Jeffrey Barrett, Sonia Goncalves, Ewan Harrison, David K. Jackson, Ian Johnston, Dominic Kwiatkowski, Cordelia Langford, John Sillitoe on behalf of the Wellcome Sanger Institute COVID-19 Surveillance Team |
| EPI_ISL_2456323 | Lighthouse Lab in Milton Keynes | Wellcome Sanger Institute for the COVID-19 Genomics UK (COG-UK) Consortium | The Lighthouse Lab in Milton Keynes and Alex Alderton, Roberto Amato, Jeffrey Barrett, Sonia Goncalves, Ewan Harrison, David K. Jackson, Ian Johnston, Dominic Kwiatkowski, Cordelia Langford, John Sillitoe on behalf of the Wellcome Sanger Institute COVID-19 Surveillance Team |
| EPI_ISL_2456325, EPI_ISL_2456326, EPI_ISL_2456328 | Lighthouse Lab in Alderley Park | Wellcome Sanger Institute for the COVID-19 Genomics UK (COG-UK) Consortium | Jacquelyn Wynn, Mairead Hyland, The Lighthouse Lab in Alderley Park and Alex Alderton, Roberto Amato, Jeffrey Barrett, Sonia Goncalves, Ewan Harrison, David K. Jackson, Ian Johnston, Dominic Kwiatkowski, Cordelia Langford, John Sillitoe on behalf of the Wellcome Sanger Institute COVID-19 Surveillance Team |
| EPI_ISL_2456330 | National Hospital for Tropical Diseases | Oxford University Clinical Research Unit, Hanoi, Vietnam | Pham Ngoc Thach, Nguyen Thi Tam, Van Dinh Trang, Le Van Duyet, Nguyen Thu Trang, Nguyen Thi Hong Thuong, Nguyen Thi Kim Chi, Phan Manh Cuong, Thomas Kesteman, H.Rogier van Doorn on behalf of the OUCRU COVID-19 research group |
| EPI_ISL_2456332, EPI_ISL_2456333 | Lighthouse Lab in Milton Keynes | Wellcome Sanger Institute for the COVID-19 Genomics UK (COG-UK) Consortium | The Lighthouse Lab in Milton Keynes and Alex Alderton, Roberto Amato, Jeffrey Barrett, Sonia Goncalves, Ewan Harrison, David K. Jackson, Ian Johnston, Dominic Kwiatkowski, Cordelia Langford, John Sillitoe on behalf of the Wellcome Sanger Institute COVID-19 Surveillance Team |
| EPI_ISL_2456335 | Lighthouse Lab in Alderley Park | Wellcome Sanger Institute for the COVID-19 Genomics UK (COG-UK) Consortium | Jacquelyn Wynn, Mairead Hyland, The Lighthouse Lab in Alderley Park and Alex Alderton, Roberto Amato, Jeffrey Barrett, Sonia Goncalves, Ewan Harrison, David K. Jackson, Ian Johnston, Dominic Kwiatkowski, Cordelia Langford, John Sillitoe on behalf of the Wellcome Sanger Institute COVID-19 Surveillance Team |
| EPI_ISL_2456336, EPI_ISL_2456339, EPI_ISL_2456341, EPI_ISL_2456342, EPI_ISL_2456345, EPI_ISL_2456347, EPI_ISL_2456349, EPI_ISL_2456352, EPI_ISL_2456354, EPI_ISL_2456355, EPI_ISL_2456357, EPI_ISL_2456358, EPI_ISL_2456360 | | | |
| see above | Lighthouse Lab in Milton Keynes | Wellcome Sanger Institute for the COVID-19 Genomics UK (COG-UK) Consortium | The Lighthouse Lab in Milton Keynes and Alex Alderton, Roberto Amato, Jeffrey Barrett, Sonia Goncalves, Ewan Harrison, David K. Jackson, Ian Johnston, Dominic Kwiatkowski, Cordelia Langford, John Sillitoe on behalf of the Wellcome Sanger Institute COVID-19 Surveillance Team |
| EPI_ISL_2456361, EPI_ISL_2456363 | Lighthouse Lab in Alderley Park | Wellcome Sanger Institute for the COVID-19 Genomics UK (COG-UK) Consortium | Jacquelyn Wynn, Mairead Hyland, The Lighthouse Lab in Alderley Park and Alex Alderton, Roberto Amato, Jeffrey Barrett, Sonia Goncalves, Ewan Harrison, David K. Jackson, Ian Johnston, Dominic Kwiatkowski, Cordelia Langford, John Sillitoe on behalf of the Wellcome Sanger Institute COVID-19 Surveillance Team |
| EPI_ISL_2456364, EPI_ISL_2456366, EPI_ISL_2456368 | Lighthouse Lab in Milton Keynes | Wellcome Sanger Institute for the COVID-19 Genomics UK (COG-UK) Consortium | The Lighthouse Lab in Milton Keynes and Alex Alderton, Roberto Amato, Jeffrey Barrett, Sonia Goncalves, Ewan Harrison, David K. Jackson, Ian Johnston, Dominic Kwiatkowski, Cordelia Langford, John Sillitoe on behalf of the Wellcome Sanger Institute COVID-19 Surveillance Team |
| EPI_ISL_2456369 | Lighthouse Lab in Alderley Park | Wellcome Sanger Institute for the COVID-19 Genomics UK (COG-UK) Consortium | Jacquelyn Wynn, Mairead Hyland, The Lighthouse Lab in Alderley Park and Alex Alderton, Roberto Amato, Jeffrey Barrett, Sonia Goncalves, Ewan Harrison, David K. Jackson, Ian Johnston, Dominic Kwiatkowski, Cordelia Langford, John Sillitoe on behalf of the Wellcome Sanger Institute COVID-19 Surveillance Team |
| EPI_ISL_2456371, EPI_ISL_2456372, EPI_ISL_2456373, EPI_ISL_2456375, EPI_ISL_2456376 | Lighthouse Lab in Milton Keynes | Wellcome Sanger Institute for the COVID-19 Genomics UK (COG-UK) Consortium | The Lighthouse Lab in Milton Keynes and Alex Alderton, Roberto Amato, Jeffrey Barrett, Sonia Goncalves, Ewan Harrison, David K. Jackson, Ian Johnston, Dominic Kwiatkowski, Cordelia Langford, John Sillitoe on behalf of the Wellcome Sanger Institute COVID-19 Surveillance Team |
| EPI_ISL_2456378 | Lighthouse Lab in Alderley Park | Wellcome Sanger Institute for the COVID-19 Genomics UK (COG-UK) Consortium | Jacquelyn Wynn, Mairead Hyland, The Lighthouse Lab in Alderley Park and Alex Alderton, Roberto Amato, Jeffrey Barrett, Sonia Goncalves, Ewan Harrison, David K. Jackson, Ian Johnston, Dominic Kwiatkowski, Cordelia Langford, John Sillitoe on behalf of the Wellcome Sanger Institute COVID-19 Surveillance Team |
| EPI_ISL_2456379, EPI_ISL_2456381, EPI_ISL_2456382, EPI_ISL_2456384 | Lighthouse Lab in Milton Keynes | Wellcome Sanger Institute for the COVID-19 Genomics UK (COG-UK) Consortium | The Lighthouse Lab in Milton Keynes and Alex Alderton, Roberto Amato, Jeffrey Barrett, Sonia Goncalves, Ewan Harrison, David K. Jackson, Ian Johnston, Dominic Kwiatkowski, Cordelia Langford, John Sillitoe on behalf of the Wellcome Sanger Institute COVID-19 Surveillance Team |
| EPI_ISL_2456385, EPI_ISL_2456387 | Lighthouse Lab in Alderley Park | Wellcome Sanger Institute for the COVID-19 Genomics UK (COG-UK) Consortium | Jacquelyn Wynn, Mairead Hyland, The Lighthouse Lab in Alderley Park and Alex Alderton, Roberto Amato, Jeffrey Barrett, Sonia Goncalves, Ewan Harrison, David K. Jackson, Ian Johnston, Dominic Kwiatkowski, Cordelia Langford, John Sillitoe on behalf of the Wellcome Sanger Institute COVID-19 Surveillance Team |
| EPI_ISL_2456388, EPI_ISL_2456390, EPI_ISL_2456391, EPI_ISL_2456393 | Lighthouse Lab in Milton Keynes | Wellcome Sanger Institute for the COVID-19 Genomics UK (COG-UK) Consortium | The Lighthouse Lab in Milton Keynes and Alex Alderton, Roberto Amato, Jeffrey Barrett, Sonia Goncalves, Ewan Harrison, David K. Jackson, Ian Johnston, Dominic Kwiatkowski, Cordelia Langford, John Sillitoe on behalf of the Wellcome Sanger Institute COVID-19 Surveillance Team |
| EPI_ISL_2456394, EPI_ISL_2456396, EPI_ISL_2456397 | Lighthouse Lab in Alderley Park | Wellcome Sanger Institute for the COVID-19 Genomics UK (COG-UK) Consortium | Jacquelyn Wynn, Mairead Hyland, The Lighthouse Lab in Alderley Park and Alex Alderton, Roberto Amato, Jeffrey Barrett, Sonia Goncalves, Ewan Harrison, David K. Jackson, Ian Johnston, Dominic Kwiatkowski, Cordelia Langford, John Sillitoe on behalf of the Wellcome Sanger Institute COVID-19 Surveillance Team |
| EPI_ISL_2456400, EPI_ISL_2456402, EPI_ISL_2456403, EPI_ISL_2456405 | Lighthouse Lab in Milton Keynes | Wellcome Sanger Institute for the COVID-19 Genomics UK (COG-UK) Consortium | The Lighthouse Lab in Milton Keynes and Alex Alderton, Roberto Amato, Jeffrey Barrett, Sonia Goncalves, Ewan Harrison, David K. Jackson, Ian Johnston, Dominic Kwiatkowski, Cordelia Langford, John Sillitoe on behalf of the Wellcome Sanger Institute COVID-19 Surveillance Team |
| EPI_ISL_2456406 | Lighthouse Lab in Alderley Park | Wellcome Sanger Institute for the COVID-19 Genomics UK (COG-UK) Consortium | Jacquelyn Wynn, Mairead Hyland, The Lighthouse Lab in Alderley Park and Alex Alderton, Roberto Amato, Jeffrey Barrett, Sonia Goncalves, Ewan Harrison, David K. Jackson, Ian Johnston, Dominic Kwiatkowski, Cordelia Langford, John Sillitoe on behalf of the Wellcome Sanger Institute COVID-19 Surveillance Team |
| EPI_ISL_2456408, EPI_ISL_2456409, EPI_ISL_2456411, EPI_ISL_2456412, EPI_ISL_2456414, EPI_ISL_2456416, EPI_ISL_2456417 | Lighthouse Lab in Milton Keynes | Wellcome Sanger Institute for the COVID-19 Genomics UK (COG-UK) Consortium | The Lighthouse Lab in Milton Keynes and Alex Alderton, Roberto Amato, Jeffrey Barrett, Sonia Goncalves, Ewan Harrison, David K. Jackson, Ian Johnston, Dominic Kwiatkowski, Cordelia Langford, John Sillitoe on behalf of the Wellcome Sanger Institute COVID-19 Surveillance Team |
| EPI_ISL_2456419 | Lighthouse Lab in Alderley Park | Wellcome Sanger Institute for the COVID-19 Genomics UK (COG-UK) Consortium | Jacquelyn Wynn, Mairead Hyland, The Lighthouse Lab in Alderley Park and Alex Alderton, Roberto Amato, Jeffrey Barrett, Sonia Goncalves, Ewan Harrison, David K. Jackson, Ian Johnston, Dominic Kwiatkowski, Cordelia Langford, John Sillitoe on behalf of the Wellcome Sanger Institute COVID-19 Surveillance Team |
| EPI_ISL_2456421, EPI_ISL_2456423 | Lighthouse Lab in Milton Keynes | Wellcome Sanger Institute for the COVID-19 Genomics UK (COG-UK) Consortium | The Lighthouse Lab in Milton Keynes and Alex Alderton, Roberto Amato, Jeffrey Barrett, Sonia Goncalves, Ewan Harrison, David K. Jackson, Ian Johnston, Dominic Kwiatkowski, Cordelia Langford, John Sillitoe on behalf of the Wellcome Sanger Institute COVID-19 Surveillance Team |
| EPI_ISL_2456424, EPI_ISL_2456426 | Lighthouse Lab in Alderley Park | Wellcome Sanger Institute for the COVID-19 Genomics UK (COG-UK) Consortium | Jacquelyn Wynn, Mairead Hyland, The Lighthouse Lab in Alderley Park and Alex Alderton, Roberto Amato, Jeffrey Barrett, Sonia Goncalves, Ewan Harrison, David K. Jackson, Ian Johnston, Dominic Kwiatkowski, Cordelia Langford, John Sillitoe on behalf of the Wellcome Sanger Institute COVID-19 Surveillance Team |
| EPI_ISL_2456427 | Lighthouse Lab in Milton Keynes | Wellcome Sanger Institute for the COVID-19 Genomics UK (COG-UK) Consortium | The Lighthouse Lab in Milton Keynes and Alex Alderton, Roberto Amato, Jeffrey Barrett, Sonia Goncalves, Ewan Harrison, David K. Jackson, Ian Johnston, Dominic Kwiatkowski, Cordelia Langford, John Sillitoe on behalf of the Wellcome Sanger Institute COVID-19 Surveillance Team |
| EPI_ISL_2456429 | Lighthouse Lab in Alderley Park | Wellcome Sanger Institute for the COVID-19 Genomics UK (COG-UK) Consortium | Jacquelyn Wynn, Mairead Hyland, The Lighthouse Lab in Alderley Park and Alex Alderton, Roberto Amato, Jeffrey Barrett, Sonia Goncalves, Ewan Harrison, David K. Jackson, Ian Johnston, Dominic Kwiatkowski, Cordelia Langford, John Sillitoe on behalf of the Wellcome Sanger Institute COVID-19 Surveillance Team |
| EPI_ISL_2456431, EPI_ISL_2456432, EPI_ISL_2456433, EPI_ISL_2456435, EPI_ISL_2456436, EPI_ISL_2456437, EPI_ISL_2456440, EPI_ISL_2456442, EPI_ISL_2456443, EPI_ISL_2456445, EPI_ISL_2456446, EPI_ISL_2456448 | | | |
| see above | Lighthouse Lab in Milton Keynes | Wellcome Sanger Institute for the COVID-19 Genomics UK (COG-UK) Consortium | The Lighthouse Lab in Milton Keynes and Alex Alderton, Roberto Amato, Jeffrey Barrett, Sonia Goncalves, Ewan Harrison, David K. Jackson, Ian Johnston, Dominic Kwiatkowski, Cordelia Langford, John Sillitoe on behalf of the Wellcome Sanger Institute COVID-19 Surveillance Team |
| EPI_ISL_2456449 | Utah Public Health Laboratory | Utah Public Health Laboratory | Erin L. Young, Kelly F. Oakeson, Tara Gallagher |
| EPI_ISL_2456450 | Lighthouse Lab in Alderley Park | Wellcome Sanger Institute for the COVID-19 Genomics UK (COG-UK) Consortium | Jacquelyn Wynn, Mairead Hyland, The Lighthouse Lab in Alderley Park and Alex Alderton, Roberto Amato, Jeffrey Barrett, Sonia Goncalves, Ewan Harrison, David K. Jackson, Ian Johnston, Dominic Kwiatkowski, Cordelia Langford, John Sillitoe on behalf of the Wellcome Sanger Institute COVID-19 Surveillance Team |
| EPI_ISL_2456451, EPI_ISL_2456452, EPI_ISL_2456454, EPI_ISL_2456457, EPI_ISL_2456460, EPI_ISL_2456461, EPI_ISL_2456463, EPI_ISL_2456464, EPI_ISL_2456466, EPI_ISL_2456467, EPI_ISL_2456469, EPI_ISL_2456471, EPI_ISL_2456472, EPI_ISL_2456473, EPI_ISL_2456475, EPI_ISL_2456476, EPI_ISL_2456478, EPI_ISL_2456479, EPI_ISL_2456481, EPI_ISL_2456482, EPI_ISL_2456484, EPI_ISL_2456487, EPI_ISL_2456488, EPI_ISL_2456490, EPI_ISL_2456491, EPI_ISL_2456493, EPI_ISL_2456494 | | | |
| see above | Lighthouse Lab in Milton Keynes | Wellcome Sanger Institute for the COVID-19 Genomics UK (COG-UK) Consortium | The Lighthouse Lab in Milton Keynes and Alex Alderton, Roberto Amato, Jeffrey Barrett, Sonia Goncalves, Ewan Harrison, David K. Jackson, Ian Johnston, Dominic Kwiatkowski, Cordelia Langford, John Sillitoe on behalf of the Wellcome Sanger Institute COVID-19 Surveillance Team |
| EPI_ISL_2456497 | Lighthouse Lab in Alderley Park | Wellcome Sanger Institute for the COVID-19 Genomics UK (COG-UK) Consortium | Jacquelyn Wynn, Mairead Hyland, The Lighthouse Lab in Alderley Park and Alex Alderton, Roberto Amato, Jeffrey Barrett, Sonia Goncalves, Ewan Harrison, David K. Jackson, Ian Johnston, Dominic Kwiatkowski, Cordelia Langford, John Sillitoe on behalf of the Wellcome Sanger Institute COVID-19 Surveillance Team |
| EPI_ISL_2456498 | Lighthouse Lab in Milton Keynes | Wellcome Sanger Institute for the COVID-19 Genomics UK (COG-UK) Consortium | The Lighthouse Lab in Milton Keynes and Alex Alderton, Roberto Amato, Jeffrey Barrett, Sonia Goncalves, Ewan Harrison, David K. Jackson, Ian Johnston, Dominic Kwiatkowski, Cordelia Langford, John Sillitoe on behalf of the Wellcome Sanger Institute COVID-19 Surveillance Team |
| EPI_ISL_2456499 | Utah Public Health Laboratory | Utah Public Health Laboratory | Erin L. Young, Kelly F. Oakeson, Tara Gallagher |
| EPI_ISL_2456500, EPI_ISL_2456501, EPI_ISL_2456503, EPI_ISL_2456504, | Lighthouse Lab in Milton Keynes | Wellcome Sanger Institute for the COVID-19 Genomics UK (COG-UK) Consortium | The Lighthouse Lab in Milton Keynes and Alex Alderton, Roberto Amato, Jeffrey Barrett, Sonia Goncalves, Ewan Harrison, David K. Jackson, Ian Johnston, Dominic Kwiatkowski, Cordelia Langford, John Sillitoe on behalf of the Wellcome Sanger Institute COVID-19 Surveillance Team |

| | | | |
|---|---|---|---|
| EPI_ISL_2456506, EPI_ISL_2456507, EPI_ISL_2456511, EPI_ISL_2456512, EPI_ISL_2456514 | | | |
| EPI_ISL_2456515 | Lighthouse Lab in Alderley Park | Wellcome Sanger Institute for the COVID-19 Genomics UK (COG-UK) Consortium | Jacquelyn Wynn, Mairead Hyland, The Lighthouse Lab in Alderley Park and Alex Alderton, Roberto Amato, Jeffrey Barrett, Sonia Goncalves, Ewan Harrison, David K. Jackson, Ian Johnston, Dominic Kwiatkowski, Cordelia Langford, John Sillitoe on behalf of the Wellcome Sanger Institute COVID-19 Surveillance Team |
| EPI_ISL_2456517, EPI_ISL_2456518, EPI_ISL_2456520, EPI_ISL_2456521, EPI_ISL_2456523, EPI_ISL_2456524, EPI_ISL_2456526, EPI_ISL_2456529, EPI_ISL_2456530, EPI_ISL_2456532, EPI_ISL_2456533, EPI_ISL_2456535, EPI_ISL_2456536, EPI_ISL_2456539, EPI_ISL_2456540, EPI_ISL_2456542, EPI_ISL_2456543, EPI_ISL_2456545, EPI_ISL_2456548, EPI_ISL_2456549 | | | |
| see above | Lighthouse Lab in Milton Keynes | Wellcome Sanger Institute for the COVID-19 Genomics UK (COG-UK) Consortium | The Lighthouse Lab in Milton Keynes and Alex Alderton, Roberto Amato, Jeffrey Barrett, Sonia Goncalves, Ewan Harrison, David K. Jackson, Ian Johnston, Dominic Kwiatkowski, Cordelia Langford, John Sillitoe on behalf of the Wellcome Sanger Institute COVID-19 Surveillance Team |
| EPI_ISL_2456550 | Utah Public Health Laboratory | Utah Public Health Laboratory | Erin L. Young, Kelly F. Oakeson, Tara Gallagher |
| EPI_ISL_2456551, EPI_ISL_2456552, EPI_ISL_2456554, EPI_ISL_2456555, EPI_ISL_2456557, EPI_ISL_2456559, EPI_ISL_2456560, EPI_ISL_2456562, EPI_ISL_2456563, EPI_ISL_2456564, EPI_ISL_2456566, EPI_ISL_2456567, EPI_ISL_2456570, EPI_ISL_2456573 | | | |
| see above | Lighthouse Lab in Milton Keynes | Wellcome Sanger Institute for the COVID-19 Genomics UK (COG-UK) Consortium | The Lighthouse Lab in Milton Keynes and Alex Alderton, Roberto Amato, Jeffrey Barrett, Sonia Goncalves, Ewan Harrison, David K. Jackson, Ian Johnston, Dominic Kwiatkowski, Cordelia Langford, John Sillitoe on behalf of the Wellcome Sanger Institute COVID-19 Surveillance Team |
| EPI_ISL_2456574 | Utah Public Health Laboratory | Utah Public Health Laboratory | Erin L. Young, Kelly F. Oakeson, Tara Gallagher |
| EPI_ISL_2456575, EPI_ISL_2456576, EPI_ISL_2456578, EPI_ISL_2456579 | Lighthouse Lab in Milton Keynes | Wellcome Sanger Institute for the COVID-19 Genomics UK (COG-UK) Consortium | The Lighthouse Lab in Milton Keynes and Alex Alderton, Roberto Amato, Jeffrey Barrett, Sonia Goncalves, Ewan Harrison, David K. Jackson, Ian Johnston, Dominic Kwiatkowski, Cordelia Langford, John Sillitoe on behalf of the Wellcome Sanger Institute COVID-19 Surveillance Team |
| EPI_ISL_2456581 | Lighthouse Lab in Alderley Park | Wellcome Sanger Institute for the COVID-19 Genomics UK (COG-UK) Consortium | Jacquelyn Wynn, Mairead Hyland, The Lighthouse Lab in Alderley Park and Alex Alderton, Roberto Amato, Jeffrey Barrett, Sonia Goncalves, Ewan Harrison, David K. Jackson, Ian Johnston, Dominic Kwiatkowski, Cordelia Langford, John Sillitoe on behalf of the Wellcome Sanger Institute COVID-19 Surveillance Team |
| EPI_ISL_2456582 | Lighthouse Lab in Milton Keynes | Wellcome Sanger Institute for the COVID-19 Genomics UK (COG-UK) Consortium | The Lighthouse Lab in Milton Keynes and Alex Alderton, Roberto Amato, Jeffrey Barrett, Sonia Goncalves, Ewan Harrison, David K. Jackson, Ian Johnston, Dominic Kwiatkowski, Cordelia Langford, John Sillitoe on behalf of the Wellcome Sanger Institute COVID-19 Surveillance Team |
| EPI_ISL_2456583 | Utah Public Health Laboratory | Utah Public Health Laboratory | Erin L. Young, Kelly F. Oakeson, Tara Gallagher |
| EPI_ISL_2456584, EPI_ISL_2456585, EPI_ISL_2456587 | Lighthouse Lab in Milton Keynes | Wellcome Sanger Institute for the COVID-19 Genomics UK (COG-UK) Consortium | The Lighthouse Lab in Milton Keynes and Alex Alderton, Roberto Amato, Jeffrey Barrett, Sonia Goncalves, Ewan Harrison, David K. Jackson, Ian Johnston, Dominic Kwiatkowski, Cordelia Langford, John Sillitoe on behalf of the Wellcome Sanger Institute COVID-19 Surveillance Team |
| EPI_ISL_2456588 | Lighthouse Lab in Alderley Park | Wellcome Sanger Institute for the COVID-19 Genomics UK (COG-UK) Consortium | Jacquelyn Wynn, Mairead Hyland, The Lighthouse Lab in Alderley Park and Alex Alderton, Roberto Amato, Jeffrey Barrett, Sonia Goncalves, Ewan Harrison, David K. Jackson, Ian Johnston, Dominic Kwiatkowski, Cordelia Langford, John Sillitoe on behalf of the Wellcome Sanger Institute COVID-19 Surveillance Team |
| EPI_ISL_2456590, EPI_ISL_2456591, EPI_ISL_2456593, EPI_ISL_2456594, EPI_ISL_2456596, EPI_ISL_2456597, EPI_ISL_2456599, EPI_ISL_2456600 | Lighthouse Lab in Milton Keynes | Wellcome Sanger Institute for the COVID-19 Genomics UK (COG-UK) Consortium | The Lighthouse Lab in Milton Keynes and Alex Alderton, Roberto Amato, Jeffrey Barrett, Sonia Goncalves, Ewan Harrison, David K. Jackson, Ian Johnston, Dominic Kwiatkowski, Cordelia Langford, John Sillitoe on behalf of the Wellcome Sanger Institute COVID-19 Surveillance Team |
| EPI_ISL_2456601 | Utah Public Health Laboratory | Utah Public Health Laboratory | Erin L. Young, Kelly F. Oakeson, Tara Gallagher |
| EPI_ISL_2456602 | Lighthouse Lab in Milton Keynes | Wellcome Sanger Institute for the COVID-19 Genomics UK (COG-UK) Consortium | The Lighthouse Lab in Milton Keynes and Alex Alderton, Roberto Amato, Jeffrey Barrett, Sonia Goncalves, Ewan Harrison, David K. Jackson, Ian Johnston, Dominic Kwiatkowski, Cordelia Langford, John Sillitoe on behalf of the Wellcome Sanger Institute COVID-19 Surveillance Team |
| EPI_ISL_2456603 | Lighthouse Lab in Alderley Park | Wellcome Sanger Institute for the COVID-19 Genomics UK (COG-UK) Consortium | Jacquelyn Wynn, Mairead Hyland, The Lighthouse Lab in Alderley Park and Alex Alderton, Roberto Amato, Jeffrey Barrett, Sonia Goncalves, Ewan Harrison, David K. Jackson, Ian Johnston, Dominic Kwiatkowski, Cordelia Langford, John Sillitoe on behalf of the Wellcome Sanger Institute COVID-19 Surveillance Team |
| EPI_ISL_2456605, EPI_ISL_2456606 | Lighthouse Lab in Milton Keynes | Wellcome Sanger Institute for the COVID-19 Genomics UK (COG-UK) Consortium | The Lighthouse Lab in Milton Keynes and Alex Alderton, Roberto Amato, Jeffrey Barrett, Sonia Goncalves, Ewan Harrison, David K. Jackson, Ian Johnston, Dominic Kwiatkowski, Cordelia Langford, John Sillitoe on behalf of the Wellcome Sanger Institute COVID-19 Surveillance Team |
| EPI_ISL_2456607 | Utah Public Health Laboratory | Utah Public Health Laboratory | Erin L. Young, Kelly F. Oakeson, Tara Gallagher |
| EPI_ISL_2456608, EPI_ISL_2456610, EPI_ISL_2456611 | Lighthouse Lab in Milton Keynes | Wellcome Sanger Institute for the COVID-19 Genomics UK (COG-UK) Consortium | The Lighthouse Lab in Milton Keynes and Alex Alderton, Roberto Amato, Jeffrey Barrett, Sonia Goncalves, Ewan Harrison, David K. Jackson, Ian Johnston, Dominic Kwiatkowski, Cordelia Langford, John Sillitoe on behalf of the Wellcome Sanger Institute COVID-19 Surveillance Team |
| EPI_ISL_2456612 | Utah Public Health Laboratory | Utah Public Health Laboratory | Erin L. Young, Kelly F. Oakeson, Tara Gallagher |
| EPI_ISL_2456613, EPI_ISL_2456614, EPI_ISL_2456615, EPI_ISL_2456617, EPI_ISL_2456619, EPI_ISL_2456620, EPI_ISL_2456621, EPI_ISL_2456623, EPI_ISL_2456624 | Lighthouse Lab in Milton Keynes | Wellcome Sanger Institute for the COVID-19 Genomics UK (COG-UK) Consortium | The Lighthouse Lab in Milton Keynes and Alex Alderton, Roberto Amato, Jeffrey Barrett, Sonia Goncalves, Ewan Harrison, David K. Jackson, Ian Johnston, Dominic Kwiatkowski, Cordelia Langford, John Sillitoe on behalf of the Wellcome Sanger Institute COVID-19 Surveillance Team |
| EPI_ISL_2456625 | Utah Public Health Laboratory | Utah Public Health Laboratory | Erin L. Young, Kelly F. Oakeson, Tara Gallagher |
| EPI_ISL_2456626, EPI_ISL_2456628, EPI_ISL_2456630, EPI_ISL_2456633, EPI_ISL_2456634, EPI_ISL_2456636, EPI_ISL_2456637, EPI_ISL_2456640, EPI_ISL_2456642, EPI_ISL_2456643, EPI_ISL_2456644 | | | |
| see above | Lighthouse Lab in Milton Keynes | Wellcome Sanger Institute for the COVID-19 Genomics UK (COG-UK) Consortium | The Lighthouse Lab in Milton Keynes and Alex Alderton, Roberto Amato, Jeffrey Barrett, Sonia Goncalves, Ewan Harrison, David K. Jackson, Ian Johnston, Dominic Kwiatkowski, Cordelia Langford, John Sillitoe on behalf of the Wellcome Sanger Institute COVID-19 Surveillance Team |
| EPI_ISL_2456645 | Utah Public Health Laboratory | Utah Public Health Laboratory | Erin L. Young, Kelly F. Oakeson, Tara Gallagher |
| EPI_ISL_2456646, EPI_ISL_2456647, EPI_ISL_2456648, EPI_ISL_2456650, EPI_ISL_2456651, EPI_ISL_2456655, EPI_ISL_2456656, EPI_ISL_2456658, EPI_ISL_2456661, EPI_ISL_2456662, EPI_ISL_2456665 | | | |
| see above | Lighthouse Lab in Milton Keynes | Wellcome Sanger Institute for the COVID-19 Genomics UK (COG-UK) Consortium | The Lighthouse Lab in Milton Keynes and Alex Alderton, Roberto Amato, Jeffrey Barrett, Sonia Goncalves, Ewan Harrison, David K. Jackson, Ian Johnston, Dominic Kwiatkowski, Cordelia Langford, John Sillitoe on behalf of the Wellcome Sanger Institute COVID-19 Surveillance Team |
| EPI_ISL_2456666 | Utah Public Health Laboratory | Utah Public Health Laboratory | Erin L. Young, Kelly F. Oakeson, Tara Gallagher |
| EPI_ISL_2456667 | Lighthouse Lab in Alderley Park | Wellcome Sanger Institute for the COVID-19 Genomics UK (COG-UK) Consortium | Jacquelyn Wynn, Mairead Hyland, The Lighthouse Lab in Alderley Park and Alex Alderton, Roberto Amato, Jeffrey Barrett, Sonia Goncalves, Ewan Harrison, David K. Jackson, Ian Johnston, Dominic Kwiatkowski, Cordelia Langford, John Sillitoe on behalf of the Wellcome Sanger Institute COVID-19 Surveillance Team |
| EPI_ISL_2456668, EPI_ISL_2456671, EPI_ISL_2456672 | Lighthouse Lab in Milton Keynes | Wellcome Sanger Institute for the COVID-19 Genomics UK (COG-UK) Consortium | The Lighthouse Lab in Milton Keynes and Alex Alderton, Roberto Amato, Jeffrey Barrett, Sonia Goncalves, Ewan Harrison, David K. Jackson, Ian Johnston, Dominic Kwiatkowski, Cordelia Langford, John Sillitoe on behalf of the Wellcome Sanger Institute COVID-19 Surveillance Team |
| EPI_ISL_2456674 | Lighthouse Lab in Alderley Park | Wellcome Sanger Institute for the COVID-19 Genomics UK (COG-UK) Consortium | Jacquelyn Wynn, Mairead Hyland, The Lighthouse Lab in Alderley Park and Alex Alderton, Roberto Amato, Jeffrey Barrett, Sonia Goncalves, Ewan Harrison, David K. Jackson, Ian Johnston, Dominic Kwiatkowski, Cordelia Langford, John Sillitoe on behalf of the Wellcome Sanger Institute COVID-19 Surveillance Team |
| EPI_ISL_2456675, EPI_ISL_2456677 | Lighthouse Lab in Milton Keynes | Wellcome Sanger Institute for the COVID-19 Genomics UK (COG-UK) Consortium | The Lighthouse Lab in Milton Keynes and Alex Alderton, Roberto Amato, Jeffrey Barrett, Sonia Goncalves, Ewan Harrison, David K. Jackson, Ian Johnston, Dominic Kwiatkowski, Cordelia Langford, John Sillitoe on behalf of the Wellcome Sanger Institute COVID-19 Surveillance Team |
| EPI_ISL_2456678 | Utah Public Health Laboratory | Utah Public Health Laboratory | Erin L. Young, Kelly F. Oakeson, Tara Gallagher |
| EPI_ISL_2456679, EPI_ISL_2456680, EPI_ISL_2456682, EPI_ISL_2456683, EPI_ISL_2456684, EPI_ISL_2456686, EPI_ISL_2456687, EPI_ISL_2456689, EPI_ISL_2456690, EPI_ISL_2456692, EPI_ISL_2456693, EPI_ISL_2456695, EPI_ISL_2456696 | | | |
| see above | Lighthouse Lab in Milton Keynes | Wellcome Sanger Institute for the COVID-19 Genomics UK (COG-UK) Consortium | The Lighthouse Lab in Milton Keynes and Alex Alderton, Roberto Amato, Jeffrey Barrett, Sonia Goncalves, Ewan Harrison, David K. Jackson, Ian Johnston, Dominic Kwiatkowski, Cordelia Langford, John Sillitoe on behalf of the Wellcome Sanger Institute COVID-19 Surveillance Team |
| EPI_ISL_2456699, EPI_ISL_2456702, EPI_ISL_2456705, EPI_ISL_2456715, EPI_ISL_2456716, EPI_ISL_2456718, EPI_ISL_2456722, EPI_ISL_2456726, EPI_ISL_2456729, EPI_ISL_2456737 | Utah Public Health Laboratory | Utah Public Health Laboratory | Erin L. Young, Kelly F. Oakeson, Tara Gallagher |
| EPI_ISL_2456746 | Lighthouse Lab in Alderley Park | Wellcome Sanger Institute for the COVID-19 Genomics UK (COG-UK) Consortium | Jacquelyn Wynn, Mairead Hyland, The Lighthouse Lab in Alderley Park and Alex Alderton, Roberto Amato, Jeffrey Barrett, Sonia Goncalves, Ewan Harrison, David K. Jackson, Ian Johnston, Dominic Kwiatkowski, Cordelia Langford, John Sillitoe on behalf of the Wellcome Sanger Institute COVID-19 Surveillance Team |

| | | | |
|---|---|---|---|
| EPI_ISL_2456748, EPI_ISL_2456749, EPI_ISL_2456750 | Lighthouse Lab in Milton Keynes | Wellcome Sanger Institute for the COVID-19 Genomics UK (COG-UK) Consortium | The Lighthouse Lab in Milton Keynes and Alex Alderton, Roberto Amato, Jeffrey Barrett, Sonia Goncalves, Ewan Harrison, David K. Jackson, Ian Johnston, Dominic Kwiatkowski, Cordelia Langford, John Sillitoe on behalf of the Wellcome Sanger Institute COVID-19 Surveillance Team |
| EPI_ISL_2456752 | Lighthouse Lab in Glasgow | Wellcome Sanger Institute for the COVID-19 Genomics UK (COG-UK) Consortium | Harper VanSteenhouse, Yumi Kasai, David Gray, Carol Clugston, Anna Dominiczak and Alex Alderton, Roberto Amato, Jeffrey Barrett, Sonia Goncalves, Ewan Harrison, David K. Jackson, Ian Johnston, Dominic Kwiatkowski, Cordelia Langford, John Sillitoe on behalf of the Wellcome Sanger Institute COVID-19 Surveillance Team |
| EPI_ISL_2456753, EPI_ISL_2456754, EPI_ISL_2456755, EPI_ISL_2456757 | Lighthouse Lab in Milton Keynes | Wellcome Sanger Institute for the COVID-19 Genomics UK (COG-UK) Consortium | The Lighthouse Lab in Milton Keynes and Alex Alderton, Roberto Amato, Jeffrey Barrett, Sonia Goncalves, Ewan Harrison, David K. Jackson, Ian Johnston, Dominic Kwiatkowski, Cordelia Langford, John Sillitoe on behalf of the Wellcome Sanger Institute COVID-19 Surveillance Team |
| EPI_ISL_2456758 | Lighthouse Lab in Alderley Park | Wellcome Sanger Institute for the COVID-19 Genomics UK (COG-UK) Consortium | Jacquelyn Wynn, Mairead Hyland, The Lighthouse Lab in Alderley Park and Alex Alderton, Roberto Amato, Jeffrey Barrett, Sonia Goncalves, Ewan Harrison, David K. Jackson, Ian Johnston, Dominic Kwiatkowski, Cordelia Langford, John Sillitoe on behalf of the Wellcome Sanger Institute COVID-19 Surveillance Team |
| EPI_ISL_2456759, EPI_ISL_2456760, EPI_ISL_2456762, EPI_ISL_2456763, EPI_ISL_2456764, EPI_ISL_2456765, EPI_ISL_2456767 | Lighthouse Lab in Milton Keynes | Wellcome Sanger Institute for the COVID-19 Genomics UK (COG-UK) Consortium | The Lighthouse Lab in Milton Keynes and Alex Alderton, Roberto Amato, Jeffrey Barrett, Sonia Goncalves, Ewan Harrison, David K. Jackson, Ian Johnston, Dominic Kwiatkowski, Cordelia Langford, John Sillitoe on behalf of the Wellcome Sanger Institute COVID-19 Surveillance Team |
| EPI_ISL_2456768 | Lighthouse Lab in Alderley Park | Wellcome Sanger Institute for the COVID-19 Genomics UK (COG-UK) Consortium | Jacquelyn Wynn, Mairead Hyland, The Lighthouse Lab in Alderley Park and Alex Alderton, Roberto Amato, Jeffrey Barrett, Sonia Goncalves, Ewan Harrison, David K. Jackson, Ian Johnston, Dominic Kwiatkowski, Cordelia Langford, John Sillitoe on behalf of the Wellcome Sanger Institute COVID-19 Surveillance Team |
| EPI_ISL_2456769, EPI_ISL_2456771, EPI_ISL_2456772, EPI_ISL_2456774, EPI_ISL_2456776, EPI_ISL_2456777, EPI_ISL_2456778 | Lighthouse Lab in Milton Keynes | Wellcome Sanger Institute for the COVID-19 Genomics UK (COG-UK) Consortium | The Lighthouse Lab in Milton Keynes and Alex Alderton, Roberto Amato, Jeffrey Barrett, Sonia Goncalves, Ewan Harrison, David K. Jackson, Ian Johnston, Dominic Kwiatkowski, Cordelia Langford, John Sillitoe on behalf of the Wellcome Sanger Institute COVID-19 Surveillance Team |
| EPI_ISL_2456779 | Lighthouse Lab in Alderley Park | Wellcome Sanger Institute for the COVID-19 Genomics UK (COG-UK) Consortium | Jacquelyn Wynn, Mairead Hyland, The Lighthouse Lab in Alderley Park and Alex Alderton, Roberto Amato, Jeffrey Barrett, Sonia Goncalves, Ewan Harrison, David K. Jackson, Ian Johnston, Dominic Kwiatkowski, Cordelia Langford, John Sillitoe on behalf of the Wellcome Sanger Institute COVID-19 Surveillance Team |
| EPI_ISL_2456780, EPI_ISL_2456781, EPI_ISL_2456782, EPI_ISL_2456783, EPI_ISL_2456784, EPI_ISL_2456785, EPI_ISL_2456786, EPI_ISL_2456787, EPI_ISL_2456788, EPI_ISL_2456789, EPI_ISL_2456790, EPI_ISL_2456791, EPI_ISL_2456792, EPI_ISL_2456793, EPI_ISL_2456794, EPI_ISL_2456795, EPI_ISL_2456796, EPI_ISL_2456797, EPI_ISL_2456798, EPI_ISL_2456800, EPI_ISL_2456801, EPI_ISL_2456802, EPI_ISL_2456803, EPI_ISL_2456805, EPI_ISL_2456806, EPI_ISL_2456807, EPI_ISL_2456808, EPI_ISL_2456810, EPI_ISL_2456811, EPI_ISL_2456812, EPI_ISL_2456813, EPI_ISL_2456814, EPI_ISL_2456815, EPI_ISL_2456816 | | | |
| see above | Lighthouse Lab in Milton Keynes | Wellcome Sanger Institute for the COVID-19 Genomics UK (COG-UK) Consortium | The Lighthouse Lab in Milton Keynes and Alex Alderton, Roberto Amato, Jeffrey Barrett, Sonia Goncalves, Ewan Harrison, David K. Jackson, Ian Johnston, Dominic Kwiatkowski, Cordelia Langford, John Sillitoe on behalf of the Wellcome Sanger Institute COVID-19 Surveillance Team |
| EPI_ISL_2456817 | Lighthouse Lab in Alderley Park | Wellcome Sanger Institute for the COVID-19 Genomics UK (COG-UK) Consortium | Jacquelyn Wynn, Mairead Hyland, The Lighthouse Lab in Alderley Park and Alex Alderton, Roberto Amato, Jeffrey Barrett, Sonia Goncalves, Ewan Harrison, David K. Jackson, Ian Johnston, Dominic Kwiatkowski, Cordelia Langford, John Sillitoe on behalf of the Wellcome Sanger Institute COVID-19 Surveillance Team |
| EPI_ISL_2456818, EPI_ISL_2456819, EPI_ISL_2456820, EPI_ISL_2456821 | Lighthouse Lab in Milton Keynes | Wellcome Sanger Institute for the COVID-19 Genomics UK (COG-UK) Consortium | The Lighthouse Lab in Milton Keynes and Alex Alderton, Roberto Amato, Jeffrey Barrett, Sonia Goncalves, Ewan Harrison, David K. Jackson, Ian Johnston, Dominic Kwiatkowski, Cordelia Langford, John Sillitoe on behalf of the Wellcome Sanger Institute COVID-19 Surveillance Team |
| EPI_ISL_2456822 | Lighthouse Lab in Alderley Park | Wellcome Sanger Institute for the COVID-19 Genomics UK (COG-UK) Consortium | Jacquelyn Wynn, Mairead Hyland, The Lighthouse Lab in Alderley Park and Alex Alderton, Roberto Amato, Jeffrey Barrett, Sonia Goncalves, Ewan Harrison, David K. Jackson, Ian Johnston, Dominic Kwiatkowski, Cordelia Langford, John Sillitoe on behalf of the Wellcome Sanger Institute COVID-19 Surveillance Team |
| EPI_ISL_2456823, EPI_ISL_2456824, EPI_ISL_2456825, EPI_ISL_2456827, EPI_ISL_2456828, EPI_ISL_2456831, EPI_ISL_2456832 | Lighthouse Lab in Milton Keynes | Wellcome Sanger Institute for the COVID-19 Genomics UK (COG-UK) Consortium | The Lighthouse Lab in Milton Keynes and Alex Alderton, Roberto Amato, Jeffrey Barrett, Sonia Goncalves, Ewan Harrison, David K. Jackson, Ian Johnston, Dominic Kwiatkowski, Cordelia Langford, John Sillitoe on behalf of the Wellcome Sanger Institute COVID-19 Surveillance Team |
| EPI_ISL_2456833, EPI_ISL_2456834 | Lighthouse Lab in Alderley Park | Wellcome Sanger Institute for the COVID-19 Genomics UK (COG-UK) Consortium | Jacquelyn Wynn, Mairead Hyland, The Lighthouse Lab in Alderley Park and Alex Alderton, Roberto Amato, Jeffrey Barrett, Sonia Goncalves, Ewan Harrison, David K. Jackson, Ian Johnston, Dominic Kwiatkowski, Cordelia Langford, John Sillitoe on behalf of the Wellcome Sanger Institute COVID-19 Surveillance Team |
| EPI_ISL_2456835, EPI_ISL_2456836 | Lighthouse Lab in Milton Keynes | Wellcome Sanger Institute for the COVID-19 Genomics UK (COG-UK) Consortium | The Lighthouse Lab in Milton Keynes and Alex Alderton, Roberto Amato, Jeffrey Barrett, Sonia Goncalves, Ewan Harrison, David K. Jackson, Ian Johnston, Dominic Kwiatkowski, Cordelia Langford, John Sillitoe on behalf of the Wellcome Sanger Institute COVID-19 Surveillance Team |
| EPI_ISL_2456837, EPI_ISL_2456839 | Lighthouse Lab in Alderley Park | Wellcome Sanger Institute for the COVID-19 Genomics UK (COG-UK) Consortium | Jacquelyn Wynn, Mairead Hyland, The Lighthouse Lab in Alderley Park and Alex Alderton, Roberto Amato, Jeffrey Barrett, Sonia Goncalves, Ewan Harrison, David K. Jackson, Ian Johnston, Dominic Kwiatkowski, Cordelia Langford, John Sillitoe on behalf of the Wellcome Sanger Institute COVID-19 Surveillance Team |
| EPI_ISL_2456840, EPI_ISL_2456841 | Lighthouse Lab in Milton Keynes | Wellcome Sanger Institute for the COVID-19 Genomics UK (COG-UK) Consortium | The Lighthouse Lab in Milton Keynes and Alex Alderton, Roberto Amato, Jeffrey Barrett, Sonia Goncalves, Ewan Harrison, David K. Jackson, Ian Johnston, Dominic Kwiatkowski, Cordelia Langford, John Sillitoe on behalf of the Wellcome Sanger Institute COVID-19 Surveillance Team |
| EPI_ISL_2456842 | Health Services Laboratories | Wellcome Sanger Institute for the COVID-19 Genomics UK (COG-UK) Consortium | Health Services Laboratories and Alex Alderton, Roberto Amato, Jeffrey Barrett, Sonia Goncalves, Ewan Harrison, David K. Jackson, Ian Johnston, Dominic Kwiatkowski, Cordelia Langford, John Sillitoe on behalf of the Wellcome Sanger Institute COVID-19 Surveillance Team |
| EPI_ISL_2456843 | Lighthouse Lab in Milton Keynes | Wellcome Sanger Institute for the COVID-19 Genomics UK (COG-UK) Consortium | The Lighthouse Lab in Milton Keynes and Alex Alderton, Roberto Amato, Jeffrey Barrett, Sonia Goncalves, Ewan Harrison, David K. Jackson, Ian Johnston, Dominic Kwiatkowski, Cordelia Langford, John Sillitoe on behalf of the Wellcome Sanger Institute COVID-19 Surveillance Team |
| EPI_ISL_2456844 | Health Services Laboratories | Wellcome Sanger Institute for the COVID-19 Genomics UK (COG-UK) Consortium | Health Services Laboratories and Alex Alderton, Roberto Amato, Jeffrey Barrett, Sonia Goncalves, Ewan Harrison, David K. Jackson, Ian Johnston, Dominic Kwiatkowski, Cordelia Langford, John Sillitoe on behalf of the Wellcome Sanger Institute COVID-19 Surveillance Team |
| EPI_ISL_2456845, EPI_ISL_2456846 | Lighthouse Lab in Milton Keynes | Wellcome Sanger Institute for the COVID-19 Genomics UK (COG-UK) Consortium | The Lighthouse Lab in Milton Keynes and Alex Alderton, Roberto Amato, Jeffrey Barrett, Sonia Goncalves, Ewan Harrison, David K. Jackson, Ian Johnston, Dominic Kwiatkowski, Cordelia Langford, John Sillitoe on behalf of the Wellcome Sanger Institute COVID-19 Surveillance Team |
| EPI_ISL_2456847 | Lighthouse Lab in Alderley Park | Wellcome Sanger Institute for the COVID-19 Genomics UK (COG-UK) Consortium | Jacquelyn Wynn, Mairead Hyland, The Lighthouse Lab in Alderley Park and Alex Alderton, Roberto Amato, Jeffrey Barrett, Sonia Goncalves, Ewan Harrison, David K. Jackson, Ian Johnston, Dominic Kwiatkowski, Cordelia Langford, John Sillitoe on behalf of the Wellcome Sanger Institute COVID-19 Surveillance Team |
| EPI_ISL_2456850, EPI_ISL_2456851, EPI_ISL_2456852, EPI_ISL_2456853 | Lighthouse Lab in Milton Keynes | Wellcome Sanger Institute for the COVID-19 Genomics UK (COG-UK) Consortium | The Lighthouse Lab in Milton Keynes and Alex Alderton, Roberto Amato, Jeffrey Barrett, Sonia Goncalves, Ewan Harrison, David K. Jackson, Ian Johnston, Dominic Kwiatkowski, Cordelia Langford, John Sillitoe on behalf of the Wellcome Sanger Institute COVID-19 Surveillance Team |
| EPI_ISL_2456854, EPI_ISL_2456855, EPI_ISL_2456856 | Lighthouse Lab in Alderley Park | Wellcome Sanger Institute for the COVID-19 Genomics UK (COG-UK) Consortium | Jacquelyn Wynn, Mairead Hyland, The Lighthouse Lab in Alderley Park and Alex Alderton, Roberto Amato, Jeffrey Barrett, Sonia Goncalves, Ewan Harrison, David K. Jackson, Ian Johnston, Dominic Kwiatkowski, Cordelia Langford, John Sillitoe on behalf of the Wellcome Sanger Institute COVID-19 Surveillance Team |
| EPI_ISL_2456857 | Lighthouse Lab in Milton Keynes | Wellcome Sanger Institute for the COVID-19 Genomics UK (COG-UK) Consortium | The Lighthouse Lab in Milton Keynes and Alex Alderton, Roberto Amato, Jeffrey Barrett, Sonia Goncalves, Ewan Harrison, David K. Jackson, Ian Johnston, Dominic Kwiatkowski, Cordelia Langford, John Sillitoe on behalf of the Wellcome Sanger Institute COVID-19 Surveillance Team |
| EPI_ISL_2456858 | Lighthouse Lab in Alderley Park | Wellcome Sanger Institute for the COVID-19 Genomics UK (COG-UK) Consortium | Jacquelyn Wynn, Mairead Hyland, The Lighthouse Lab in Alderley Park and Alex Alderton, Roberto Amato, Jeffrey Barrett, Sonia Goncalves, Ewan Harrison, David K. Jackson, Ian Johnston, Dominic Kwiatkowski, Cordelia Langford, John Sillitoe on behalf of the Wellcome Sanger Institute COVID-19 Surveillance Team |
| EPI_ISL_2456859, EPI_ISL_2456861, EPI_ISL_2456862 | Lighthouse Lab in Milton Keynes | Wellcome Sanger Institute for the COVID-19 Genomics UK (COG-UK) Consortium | The Lighthouse Lab in Milton Keynes and Alex Alderton, Roberto Amato, Jeffrey Barrett, Sonia Goncalves, Ewan Harrison, David K. Jackson, Ian Johnston, Dominic Kwiatkowski, Cordelia Langford, John Sillitoe on behalf of the Wellcome Sanger Institute COVID-19 Surveillance Team |
| EPI_ISL_2456863 | Health Services Laboratories | Wellcome Sanger Institute for the COVID-19 Genomics UK (COG-UK) Consortium | Health Services Laboratories and Alex Alderton, Roberto Amato, Jeffrey Barrett, Sonia Goncalves, Ewan Harrison, David K. Jackson, Ian Johnston, Dominic Kwiatkowski, Cordelia Langford, John Sillitoe on behalf of the Wellcome Sanger Institute COVID-19 Surveillance Team |
| EPI_ISL_2456864, EPI_ISL_2456865 | Lighthouse Lab in Alderley Park | Wellcome Sanger Institute for the COVID-19 Genomics UK (COG-UK) Consortium | Jacquelyn Wynn, Mairead Hyland, The Lighthouse Lab in Alderley Park and Alex Alderton, Roberto Amato, Jeffrey Barrett, Sonia Goncalves, Ewan Harrison, David K. Jackson, Ian Johnston, Dominic Kwiatkowski, Cordelia Langford, John Sillitoe on behalf of the Wellcome Sanger Institute COVID-19 Surveillance Team |
| EPI_ISL_2456866 | Lighthouse Lab in Milton Keynes | Wellcome Sanger Institute for the COVID-19 Genomics UK (COG-UK) Consortium | The Lighthouse Lab in Milton Keynes and Alex Alderton, Roberto Amato, Jeffrey Barrett, Sonia Goncalves, Ewan Harrison, David K. Jackson, Ian Johnston, Dominic Kwiatkowski, Cordelia Langford, John Sillitoe on behalf of the Wellcome Sanger Institute COVID-19 Surveillance Team |
| EPI_ISL_2456867, EPI_ISL_2456868 | Lighthouse Lab in Alderley Park | Wellcome Sanger Institute for the COVID-19 Genomics UK (COG-UK) Consortium | Jacquelyn Wynn, Mairead Hyland, The Lighthouse Lab in Alderley Park and Alex Alderton, Roberto Amato, Jeffrey Barrett, Sonia Goncalves, Ewan Harrison, David K. Jackson, Ian Johnston, Dominic Kwiatkowski, Cordelia Langford, John Sillitoe on behalf of the Wellcome Sanger Institute COVID-19 Surveillance Team |
| EPI_ISL_2456869, EPI_ISL_2456870 | Lighthouse Lab in Milton Keynes | Wellcome Sanger Institute for the COVID-19 Genomics UK (COG-UK) Consortium | The Lighthouse Lab in Milton Keynes and Alex Alderton, Roberto Amato, Jeffrey Barrett, Sonia Goncalves, Ewan Harrison, David K. Jackson, Ian Johnston, Dominic Kwiatkowski, Cordelia Langford, John Sillitoe on behalf of the Wellcome Sanger Institute COVID-19 Surveillance Team |
| EPI_ISL_2456872, EPI_ISL_2456874 | Lighthouse Lab in Alderley Park | Wellcome Sanger Institute for the COVID-19 Genomics UK (COG-UK) Consortium | Jacquelyn Wynn, Mairead Hyland, The Lighthouse Lab in Alderley Park and Alex Alderton, Roberto Amato, Jeffrey Barrett, Sonia Goncalves, Ewan Harrison, David K. Jackson, Ian Johnston, Dominic Kwiatkowski, Cordelia Langford, John Sillitoe on behalf of the Wellcome Sanger Institute COVID-19 Surveillance Team |
| EPI_ISL_2456876, EPI_ISL_2456877 | Lighthouse Lab in Milton Keynes | Wellcome Sanger Institute for the COVID-19 Genomics UK (COG-UK) Consortium | The Lighthouse Lab in Milton Keynes and Alex Alderton, Roberto Amato, Jeffrey Barrett, Sonia Goncalves, Ewan Harrison, David K. Jackson, Ian Johnston, Dominic Kwiatkowski, Cordelia Langford, John Sillitoe on behalf of the Wellcome Sanger Institute COVID-19 Surveillance Team |
| EPI_ISL_2456878 | Health Services Laboratories | Wellcome Sanger Institute for the COVID-19 Genomics UK (COG-UK) | Health Services Laboratories and Alex Alderton, Roberto Amato, Jeffrey Barrett, Sonia Goncalves, Ewan Harrison, David K. Jackson, Ian Johnston, Dominic Kwiatkowski, Cordelia Langford, John Sillitoe on behalf of the Wellcome Sanger Institute |

| | | | |
|---|---|---|---|
| | | Consortium | COVID-19 Surveillance Team |
| EPI_ISL_2456879 | Lighthouse Lab in Milton Keynes | Wellcome Sanger Institute for the COVID-19 Genomics UK (COG-UK) Consortium | The Lighthouse Lab in Milton Keynes and Alex Alderton, Roberto Amato, Jeffrey Barrett, Sonia Goncalves, Ewan Harrison, David K. Jackson, Ian Johnston, Dominic Kwiatkowski, Cordelia Langford, John Sillitoe on behalf of the Wellcome Sanger Institute COVID-19 Surveillance Team |
| EPI_ISL_2456880, EPI_ISL_2456881 | Lighthouse Lab in Alderley Park | Wellcome Sanger Institute for the COVID-19 Genomics UK (COG-UK) Consortium | Jacquelyn Wynn, Mairead Hyland, The Lighthouse Lab in Alderley Park and Alex Alderton, Roberto Amato, Jeffrey Barrett, Sonia Goncalves, Ewan Harrison, David K. Jackson, Ian Johnston, Dominic Kwiatkowski, Cordelia Langford, John Sillitoe on behalf of the Wellcome Sanger Institute COVID-19 Surveillance Team |
| EPI_ISL_2456882 | Health Services Laboratories | Wellcome Sanger Institute for the COVID-19 Genomics UK (COG-UK) Consortium | Health Services Laboratories and Alex Alderton, Roberto Amato, Jeffrey Barrett, Sonia Goncalves, Ewan Harrison, David K. Jackson, Ian Johnston, Dominic Kwiatkowski, Cordelia Langford, John Sillitoe on behalf of the Wellcome Sanger Institute COVID-19 Surveillance Team |
| EPI_ISL_2456883 | Lighthouse Lab in Milton Keynes | Wellcome Sanger Institute for the COVID-19 Genomics UK (COG-UK) Consortium | The Lighthouse Lab in Milton Keynes and Alex Alderton, Roberto Amato, Jeffrey Barrett, Sonia Goncalves, Ewan Harrison, David K. Jackson, Ian Johnston, Dominic Kwiatkowski, Cordelia Langford, John Sillitoe on behalf of the Wellcome Sanger Institute COVID-19 Surveillance Team |
| EPI_ISL_2456884, EPI_ISL_2456885 | Lighthouse Lab in Alderley Park | Wellcome Sanger Institute for the COVID-19 Genomics UK (COG-UK) Consortium | Jacquelyn Wynn, Mairead Hyland, The Lighthouse Lab in Alderley Park and Alex Alderton, Roberto Amato, Jeffrey Barrett, Sonia Goncalves, Ewan Harrison, David K. Jackson, Ian Johnston, Dominic Kwiatkowski, Cordelia Langford, John Sillitoe on behalf of the Wellcome Sanger Institute COVID-19 Surveillance Team |
| EPI_ISL_2456886 | Health Services Laboratories | Wellcome Sanger Institute for the COVID-19 Genomics UK (COG-UK) Consortium | Health Services Laboratories and Alex Alderton, Roberto Amato, Jeffrey Barrett, Sonia Goncalves, Ewan Harrison, David K. Jackson, Ian Johnston, Dominic Kwiatkowski, Cordelia Langford, John Sillitoe on behalf of the Wellcome Sanger Institute COVID-19 Surveillance Team |
| EPI_ISL_2456887 | Lighthouse Lab in Alderley Park | Wellcome Sanger Institute for the COVID-19 Genomics UK (COG-UK) Consortium | Jacquelyn Wynn, Mairead Hyland, The Lighthouse Lab in Alderley Park and Alex Alderton, Roberto Amato, Jeffrey Barrett, Sonia Goncalves, Ewan Harrison, David K. Jackson, Ian Johnston, Dominic Kwiatkowski, Cordelia Langford, John Sillitoe on behalf of the Wellcome Sanger Institute COVID-19 Surveillance Team |
| EPI_ISL_2456889 | Lighthouse Lab in Milton Keynes | Wellcome Sanger Institute for the COVID-19 Genomics UK (COG-UK) Consortium | The Lighthouse Lab in Milton Keynes and Alex Alderton, Roberto Amato, Jeffrey Barrett, Sonia Goncalves, Ewan Harrison, David K. Jackson, Ian Johnston, Dominic Kwiatkowski, Cordelia Langford, John Sillitoe on behalf of the Wellcome Sanger Institute COVID-19 Surveillance Team |
| EPI_ISL_2456890 | Lighthouse Lab in Alderley Park | Wellcome Sanger Institute for the COVID-19 Genomics UK (COG-UK) Consortium | Jacquelyn Wynn, Mairead Hyland, The Lighthouse Lab in Alderley Park and Alex Alderton, Roberto Amato, Jeffrey Barrett, Sonia Goncalves, Ewan Harrison, David K. Jackson, Ian Johnston, Dominic Kwiatkowski, Cordelia Langford, John Sillitoe on behalf of the Wellcome Sanger Institute COVID-19 Surveillance Team |
| EPI_ISL_2456893, EPI_ISL_2456894, EPI_ISL_2456896 | Lighthouse Lab in Milton Keynes | Wellcome Sanger Institute for the COVID-19 Genomics UK (COG-UK) Consortium | The Lighthouse Lab in Milton Keynes and Alex Alderton, Roberto Amato, Jeffrey Barrett, Sonia Goncalves, Ewan Harrison, David K. Jackson, Ian Johnston, Dominic Kwiatkowski, Cordelia Langford, John Sillitoe on behalf of the Wellcome Sanger Institute COVID-19 Surveillance Team |
| EPI_ISL_2456897 | Lighthouse Lab in Alderley Park | Wellcome Sanger Institute for the COVID-19 Genomics UK (COG-UK) Consortium | Jacquelyn Wynn, Mairead Hyland, The Lighthouse Lab in Alderley Park and Alex Alderton, Roberto Amato, Jeffrey Barrett, Sonia Goncalves, Ewan Harrison, David K. Jackson, Ian Johnston, Dominic Kwiatkowski, Cordelia Langford, John Sillitoe on behalf of the Wellcome Sanger Institute COVID-19 Surveillance Team |
| EPI_ISL_2456898, EPI_ISL_2456899 | Lighthouse Lab in Milton Keynes | Wellcome Sanger Institute for the COVID-19 Genomics UK (COG-UK) Consortium | The Lighthouse Lab in Milton Keynes and Alex Alderton, Roberto Amato, Jeffrey Barrett, Sonia Goncalves, Ewan Harrison, David K. Jackson, Ian Johnston, Dominic Kwiatkowski, Cordelia Langford, John Sillitoe on behalf of the Wellcome Sanger Institute COVID-19 Surveillance Team |
| EPI_ISL_2456900, EPI_ISL_2456902, EPI_ISL_2456903, EPI_ISL_2456904, EPI_ISL_2456905 | Lighthouse Lab in Alderley Park | Wellcome Sanger Institute for the COVID-19 Genomics UK (COG-UK) Consortium | Jacquelyn Wynn, Mairead Hyland, The Lighthouse Lab in Alderley Park and Alex Alderton, Roberto Amato, Jeffrey Barrett, Sonia Goncalves, Ewan Harrison, David K. Jackson, Ian Johnston, Dominic Kwiatkowski, Cordelia Langford, John Sillitoe on behalf of the Wellcome Sanger Institute COVID-19 Surveillance Team |
| EPI_ISL_2456906 | Health Services Laboratories | Wellcome Sanger Institute for the COVID-19 Genomics UK (COG-UK) Consortium | Health Services Laboratories and Alex Alderton, Roberto Amato, Jeffrey Barrett, Sonia Goncalves, Ewan Harrison, David K. Jackson, Ian Johnston, Dominic Kwiatkowski, Cordelia Langford, John Sillitoe on behalf of the Wellcome Sanger Institute COVID-19 Surveillance Team |
| EPI_ISL_2456907 | Lighthouse Lab in Alderley Park | Wellcome Sanger Institute for the COVID-19 Genomics UK (COG-UK) Consortium | Jacquelyn Wynn, Mairead Hyland, The Lighthouse Lab in Alderley Park and Alex Alderton, Roberto Amato, Jeffrey Barrett, Sonia Goncalves, Ewan Harrison, David K. Jackson, Ian Johnston, Dominic Kwiatkowski, Cordelia Langford, John Sillitoe on behalf of the Wellcome Sanger Institute COVID-19 Surveillance Team |
| EPI_ISL_2456908 | Health Services Laboratories | Wellcome Sanger Institute for the COVID-19 Genomics UK (COG-UK) Consortium | Health Services Laboratories and Alex Alderton, Roberto Amato, Jeffrey Barrett, Sonia Goncalves, Ewan Harrison, David K. Jackson, Ian Johnston, Dominic Kwiatkowski, Cordelia Langford, John Sillitoe on behalf of the Wellcome Sanger Institute COVID-19 Surveillance Team |
| EPI_ISL_2456909, EPI_ISL_2456911 | Lighthouse Lab in Milton Keynes | Wellcome Sanger Institute for the COVID-19 Genomics UK (COG-UK) Consortium | The Lighthouse Lab in Milton Keynes and Alex Alderton, Roberto Amato, Jeffrey Barrett, Sonia Goncalves, Ewan Harrison, David K. Jackson, Ian Johnston, Dominic Kwiatkowski, Cordelia Langford, John Sillitoe on behalf of the Wellcome Sanger Institute COVID-19 Surveillance Team |
| EPI_ISL_2456912, EPI_ISL_2456913 | Lighthouse Lab in Alderley Park | Wellcome Sanger Institute for the COVID-19 Genomics UK (COG-UK) Consortium | Jacquelyn Wynn, Mairead Hyland, The Lighthouse Lab in Alderley Park and Alex Alderton, Roberto Amato, Jeffrey Barrett, Sonia Goncalves, Ewan Harrison, David K. Jackson, Ian Johnston, Dominic Kwiatkowski, Cordelia Langford, John Sillitoe on behalf of the Wellcome Sanger Institute COVID-19 Surveillance Team |
| EPI_ISL_2456915, EPI_ISL_2456917 | Lighthouse Lab in Milton Keynes | Wellcome Sanger Institute for the COVID-19 Genomics UK (COG-UK) Consortium | The Lighthouse Lab in Milton Keynes and Alex Alderton, Roberto Amato, Jeffrey Barrett, Sonia Goncalves, Ewan Harrison, David K. Jackson, Ian Johnston, Dominic Kwiatkowski, Cordelia Langford, John Sillitoe on behalf of the Wellcome Sanger Institute COVID-19 Surveillance Team |
| EPI_ISL_2456918, EPI_ISL_2456919, EPI_ISL_2456920, EPI_ISL_2456921 | Health Services Laboratories | Wellcome Sanger Institute for the COVID-19 Genomics UK (COG-UK) Consortium | Health Services Laboratories and Alex Alderton, Roberto Amato, Jeffrey Barrett, Sonia Goncalves, Ewan Harrison, David K. Jackson, Ian Johnston, Dominic Kwiatkowski, Cordelia Langford, John Sillitoe on behalf of the Wellcome Sanger Institute COVID-19 Surveillance Team |
| EPI_ISL_2456922 | Lighthouse Lab in Milton Keynes | Wellcome Sanger Institute for the COVID-19 Genomics UK (COG-UK) Consortium | The Lighthouse Lab in Milton Keynes and Alex Alderton, Roberto Amato, Jeffrey Barrett, Sonia Goncalves, Ewan Harrison, David K. Jackson, Ian Johnston, Dominic Kwiatkowski, Cordelia Langford, John Sillitoe on behalf of the Wellcome Sanger Institute COVID-19 Surveillance Team |
| EPI_ISL_2456923 | Lighthouse Lab in Alderley Park | Wellcome Sanger Institute for the COVID-19 Genomics UK (COG-UK) Consortium | Jacquelyn Wynn, Mairead Hyland, The Lighthouse Lab in Alderley Park and Alex Alderton, Roberto Amato, Jeffrey Barrett, Sonia Goncalves, Ewan Harrison, David K. Jackson, Ian Johnston, Dominic Kwiatkowski, Cordelia Langford, John Sillitoe on behalf of the Wellcome Sanger Institute COVID-19 Surveillance Team |
| EPI_ISL_2456924 | Health Services Laboratories | Wellcome Sanger Institute for the COVID-19 Genomics UK (COG-UK) Consortium | Health Services Laboratories and Alex Alderton, Roberto Amato, Jeffrey Barrett, Sonia Goncalves, Ewan Harrison, David K. Jackson, Ian Johnston, Dominic Kwiatkowski, Cordelia Langford, John Sillitoe on behalf of the Wellcome Sanger Institute COVID-19 Surveillance Team |
| EPI_ISL_2456925, EPI_ISL_2456926 | Lighthouse Lab in Milton Keynes | Wellcome Sanger Institute for the COVID-19 Genomics UK (COG-UK) Consortium | The Lighthouse Lab in Milton Keynes and Alex Alderton, Roberto Amato, Jeffrey Barrett, Sonia Goncalves, Ewan Harrison, David K. Jackson, Ian Johnston, Dominic Kwiatkowski, Cordelia Langford, John Sillitoe on behalf of the Wellcome Sanger Institute COVID-19 Surveillance Team |
| EPI_ISL_2456927, EPI_ISL_2456928 | Health Services Laboratories | Wellcome Sanger Institute for the COVID-19 Genomics UK (COG-UK) Consortium | Health Services Laboratories and Alex Alderton, Roberto Amato, Jeffrey Barrett, Sonia Goncalves, Ewan Harrison, David K. Jackson, Ian Johnston, Dominic Kwiatkowski, Cordelia Langford, John Sillitoe on behalf of the Wellcome Sanger Institute COVID-19 Surveillance Team |
| EPI_ISL_2456929 | Lighthouse Lab in Alderley Park | Wellcome Sanger Institute for the COVID-19 Genomics UK (COG-UK) Consortium | Jacquelyn Wynn, Mairead Hyland, The Lighthouse Lab in Alderley Park and Alex Alderton, Roberto Amato, Jeffrey Barrett, Sonia Goncalves, Ewan Harrison, David K. Jackson, Ian Johnston, Dominic Kwiatkowski, Cordelia Langford, John Sillitoe on behalf of the Wellcome Sanger Institute COVID-19 Surveillance Team |
| EPI_ISL_2456930 | Lighthouse Lab in Milton Keynes | Wellcome Sanger Institute for the COVID-19 Genomics UK (COG-UK) Consortium | The Lighthouse Lab in Milton Keynes and Alex Alderton, Roberto Amato, Jeffrey Barrett, Sonia Goncalves, Ewan Harrison, David K. Jackson, Ian Johnston, Dominic Kwiatkowski, Cordelia Langford, John Sillitoe on behalf of the Wellcome Sanger Institute COVID-19 Surveillance Team |
| EPI_ISL_2456933 | Lighthouse Lab in Alderley Park | Wellcome Sanger Institute for the COVID-19 Genomics UK (COG-UK) Consortium | Jacquelyn Wynn, Mairead Hyland, The Lighthouse Lab in Alderley Park and Alex Alderton, Roberto Amato, Jeffrey Barrett, Sonia Goncalves, Ewan Harrison, David K. Jackson, Ian Johnston, Dominic Kwiatkowski, Cordelia Langford, John Sillitoe on behalf of the Wellcome Sanger Institute COVID-19 Surveillance Team |
| EPI_ISL_2456934 | Lighthouse Lab in Milton Keynes | Wellcome Sanger Institute for the COVID-19 Genomics UK (COG-UK) Consortium | The Lighthouse Lab in Milton Keynes and Alex Alderton, Roberto Amato, Jeffrey Barrett, Sonia Goncalves, Ewan Harrison, David K. Jackson, Ian Johnston, Dominic Kwiatkowski, Cordelia Langford, John Sillitoe on behalf of the Wellcome Sanger Institute COVID-19 Surveillance Team |
| EPI_ISL_2456936, EPI_ISL_2456937 | Lighthouse Lab in Alderley Park | Wellcome Sanger Institute for the COVID-19 Genomics UK (COG-UK) Consortium | Jacquelyn Wynn, Mairead Hyland, The Lighthouse Lab in Alderley Park and Alex Alderton, Roberto Amato, Jeffrey Barrett, Sonia Goncalves, Ewan Harrison, David K. Jackson, Ian Johnston, Dominic Kwiatkowski, Cordelia Langford, John Sillitoe on behalf of the Wellcome Sanger Institute COVID-19 Surveillance Team |
| EPI_ISL_2456938, EPI_ISL_2456939 | Lighthouse Lab in Milton Keynes | Wellcome Sanger Institute for the COVID-19 Genomics UK (COG-UK) Consortium | The Lighthouse Lab in Milton Keynes and Alex Alderton, Roberto Amato, Jeffrey Barrett, Sonia Goncalves, Ewan Harrison, David K. Jackson, Ian Johnston, Dominic Kwiatkowski, Cordelia Langford, John Sillitoe on behalf of the Wellcome Sanger Institute COVID-19 Surveillance Team |
| EPI_ISL_2456940 | Lighthouse Lab in Alderley Park | Wellcome Sanger Institute for the COVID-19 Genomics UK (COG-UK) Consortium | Jacquelyn Wynn, Mairead Hyland, The Lighthouse Lab in Alderley Park and Alex Alderton, Roberto Amato, Jeffrey Barrett, Sonia Goncalves, Ewan Harrison, David K. Jackson, Ian Johnston, Dominic Kwiatkowski, Cordelia Langford, John Sillitoe on behalf of the Wellcome Sanger Institute COVID-19 Surveillance Team |
| EPI_ISL_2456941, EPI_ISL_2456942 | Lighthouse Lab in Milton Keynes | Wellcome Sanger Institute for the COVID-19 Genomics UK (COG-UK) Consortium | The Lighthouse Lab in Milton Keynes and Alex Alderton, Roberto Amato, Jeffrey Barrett, Sonia Goncalves, Ewan Harrison, David K. Jackson, Ian Johnston, Dominic Kwiatkowski, Cordelia Langford, John Sillitoe on behalf of the Wellcome Sanger Institute COVID-19 Surveillance Team |
| EPI_ISL_2456943, EPI_ISL_2456945, EPI_ISL_2456946 | Lighthouse Lab in Alderley Park | Wellcome Sanger Institute for the COVID-19 Genomics UK (COG-UK) Consortium | Jacquelyn Wynn, Mairead Hyland, The Lighthouse Lab in Alderley Park and Alex Alderton, Roberto Amato, Jeffrey Barrett, Sonia Goncalves, Ewan Harrison, David K. Jackson, Ian Johnston, Dominic Kwiatkowski, Cordelia Langford, John Sillitoe on behalf of the Wellcome Sanger Institute COVID-19 Surveillance Team |
| EPI_ISL_2456948 | Lighthouse Lab in Milton Keynes | Wellcome Sanger Institute for the COVID-19 Genomics UK (COG-UK) Consortium | The Lighthouse Lab in Milton Keynes and Alex Alderton, Roberto Amato, Jeffrey Barrett, Sonia Goncalves, Ewan Harrison, David K. Jackson, Ian Johnston, Dominic Kwiatkowski, Cordelia Langford, John Sillitoe on behalf of the Wellcome Sanger Institute COVID-19 Surveillance Team |
| EPI_ISL_2456949, EPI_ISL_2456950 | Lighthouse Lab in Alderley Park | Wellcome Sanger Institute for the COVID-19 Genomics UK (COG-UK) Consortium | Jacquelyn Wynn, Mairead Hyland, The Lighthouse Lab in Alderley Park and Alex Alderton, Roberto Amato, Jeffrey Barrett, Sonia Goncalves, Ewan Harrison, David K. Jackson, Ian Johnston, Dominic Kwiatkowski, Cordelia Langford, John Sillitoe on behalf of the Wellcome Sanger Institute COVID-19 Surveillance Team |
| EPI_ISL_2456951 | Health Services Laboratories | Wellcome Sanger Institute for the COVID-19 Genomics UK (COG-UK) Consortium | Health Services Laboratories and Alex Alderton, Roberto Amato, Jeffrey Barrett, Sonia Goncalves, Ewan Harrison, David K. Jackson, Ian Johnston, Dominic Kwiatkowski, Cordelia Langford, John Sillitoe on behalf of the Wellcome Sanger Institute COVID-19 Surveillance Team |
| EPI_ISL_2456953 | Lighthouse Lab in Alderley Park | Wellcome Sanger Institute for the COVID-19 Genomics UK (COG-UK) Consortium | Jacquelyn Wynn, Mairead Hyland, The Lighthouse Lab in Alderley Park and Alex Alderton, Roberto Amato, Jeffrey Barrett, Sonia Goncalves, Ewan Harrison, David K. Jackson, Ian Johnston, Dominic Kwiatkowski, Cordelia Langford, John Sillitoe on behalf of the Wellcome Sanger Institute COVID-19 Surveillance Team |
| EPI_ISL_2456955, EPI_ISL_2456956 | Lighthouse Lab in Milton Keynes | Wellcome Sanger Institute for the COVID-19 Genomics UK (COG-UK) Consortium | The Lighthouse Lab in Milton Keynes and Alex Alderton, Roberto Amato, Jeffrey Barrett, Sonia Goncalves, Ewan Harrison, David K. Jackson, Ian Johnston, Dominic Kwiatkowski, Cordelia Langford, John Sillitoe on behalf of the Wellcome Sanger Institute COVID-19 Surveillance Team |
| EPI_ISL_2456958, EPI_ISL_2456959 | Lighthouse Lab in Alderley Park | Wellcome Sanger Institute for the COVID-19 Genomics UK (COG-UK) Consortium | Jacquelyn Wynn, Mairead Hyland, The Lighthouse Lab in Alderley Park and Alex Alderton, Roberto Amato, Jeffrey Barrett, Sonia Goncalves, Ewan Harrison, David K. Jackson, Ian Johnston, Dominic Kwiatkowski, Cordelia Langford, John Sillitoe on behalf of the Wellcome Sanger Institute COVID-19 Surveillance Team |
| EPI_ISL_2456960 | Health Services Laboratories | Wellcome Sanger Institute for the COVID-19 Genomics UK (COG-UK) Consortium | Health Services Laboratories and Alex Alderton, Roberto Amato, Jeffrey Barrett, Sonia Goncalves, Ewan Harrison, David K. Jackson, Ian Johnston, Dominic Kwiatkowski, Cordelia Langford, John Sillitoe on behalf of the Wellcome Sanger Institute COVID-19 Surveillance Team |
| EPI_ISL_2456961 | Lighthouse Lab in Alderley Park | Wellcome Sanger Institute for the COVID-19 Genomics UK (COG-UK) Consortium | Jacquelyn Wynn, Mairead Hyland, The Lighthouse Lab in Alderley Park and Alex Alderton, Roberto Amato, Jeffrey Barrett, Sonia Goncalves, Ewan Harrison, David K. Jackson, Ian Johnston, Dominic Kwiatkowski, Cordelia Langford, John Sillitoe on behalf of the Wellcome Sanger Institute COVID-19 Surveillance Team |

| | | | |
|---|---|---|---|
| EPI_ISL_2456962 | Health Services Laboratories | Wellcome Sanger Institute for the COVID-19 Genomics UK (COG-UK) Consortium | Health Services Laboratories and Alex Alderton, Roberto Amato, Jeffrey Barrett, Sonia Goncalves, Ewan Harrison, David K. Jackson, Ian Johnston, Dominic Kwiatkowski, Cordelia Langford, John Sillitoe on behalf of the Wellcome Sanger Institute COVID-19 Surveillance Team |
| EPI_ISL_2456963 | Lighthouse Lab in Alderley Park | Wellcome Sanger Institute for the COVID-19 Genomics UK (COG-UK) Consortium | Jacquelyn Wynn, Mairead Hyland, The Lighthouse Lab in Alderley Park and Alex Alderton, Roberto Amato, Jeffrey Barrett, Sonia Goncalves, Ewan Harrison, David K. Jackson, Ian Johnston, Dominic Kwiatkowski, Cordelia Langford, John Sillitoe on behalf of the Wellcome Sanger Institute COVID-19 Surveillance Team |
| EPI_ISL_2456964 | Health Services Laboratories | Wellcome Sanger Institute for the COVID-19 Genomics UK (COG-UK) Consortium | Health Services Laboratories and Alex Alderton, Roberto Amato, Jeffrey Barrett, Sonia Goncalves, Ewan Harrison, David K. Jackson, Ian Johnston, Dominic Kwiatkowski, Cordelia Langford, John Sillitoe on behalf of the Wellcome Sanger Institute COVID-19 Surveillance Team |
| EPI_ISL_2456965 | Lighthouse Lab in Milton Keynes | Wellcome Sanger Institute for the COVID-19 Genomics UK (COG-UK) Consortium | The Lighthouse Lab in Milton Keynes and Alex Alderton, Roberto Amato, Jeffrey Barrett, Sonia Goncalves, Ewan Harrison, David K. Jackson, Ian Johnston, Dominic Kwiatkowski, Cordelia Langford, John Sillitoe on behalf of the Wellcome Sanger Institute COVID-19 Surveillance Team |
| EPI_ISL_2456966 | Lighthouse Lab in Alderley Park | Wellcome Sanger Institute for the COVID-19 Genomics UK (COG-UK) Consortium | Jacquelyn Wynn, Mairead Hyland, The Lighthouse Lab in Alderley Park and Alex Alderton, Roberto Amato, Jeffrey Barrett, Sonia Goncalves, Ewan Harrison, David K. Jackson, Ian Johnston, Dominic Kwiatkowski, Cordelia Langford, John Sillitoe on behalf of the Wellcome Sanger Institute COVID-19 Surveillance Team |
| EPI_ISL_2456967 | Lighthouse Lab in Milton Keynes | Wellcome Sanger Institute for the COVID-19 Genomics UK (COG-UK) Consortium | The Lighthouse Lab in Milton Keynes and Alex Alderton, Roberto Amato, Jeffrey Barrett, Sonia Goncalves, Ewan Harrison, David K. Jackson, Ian Johnston, Dominic Kwiatkowski, Cordelia Langford, John Sillitoe on behalf of the Wellcome Sanger Institute COVID-19 Surveillance Team |
| EPI_ISL_2456968, EPI_ISL_2456969 | Health Services Laboratories | Wellcome Sanger Institute for the COVID-19 Genomics UK (COG-UK) Consortium | Health Services Laboratories and Alex Alderton, Roberto Amato, Jeffrey Barrett, Sonia Goncalves, Ewan Harrison, David K. Jackson, Ian Johnston, Dominic Kwiatkowski, Cordelia Langford, John Sillitoe on behalf of the Wellcome Sanger Institute COVID-19 Surveillance Team |
| EPI_ISL_2456971 | Lighthouse Lab in Milton Keynes | Wellcome Sanger Institute for the COVID-19 Genomics UK (COG-UK) Consortium | The Lighthouse Lab in Milton Keynes and Alex Alderton, Roberto Amato, Jeffrey Barrett, Sonia Goncalves, Ewan Harrison, David K. Jackson, Ian Johnston, Dominic Kwiatkowski, Cordelia Langford, John Sillitoe on behalf of the Wellcome Sanger Institute COVID-19 Surveillance Team |
| EPI_ISL_2456972 | Lighthouse Lab in Alderley Park | Wellcome Sanger Institute for the COVID-19 Genomics UK (COG-UK) Consortium | Jacquelyn Wynn, Mairead Hyland, The Lighthouse Lab in Alderley Park and Alex Alderton, Roberto Amato, Jeffrey Barrett, Sonia Goncalves, Ewan Harrison, David K. Jackson, Ian Johnston, Dominic Kwiatkowski, Cordelia Langford, John Sillitoe on behalf of the Wellcome Sanger Institute COVID-19 Surveillance Team |
| EPI_ISL_2456973 | Lighthouse Lab in Milton Keynes | Wellcome Sanger Institute for the COVID-19 Genomics UK (COG-UK) Consortium | The Lighthouse Lab in Milton Keynes and Alex Alderton, Roberto Amato, Jeffrey Barrett, Sonia Goncalves, Ewan Harrison, David K. Jackson, Ian Johnston, Dominic Kwiatkowski, Cordelia Langford, John Sillitoe on behalf of the Wellcome Sanger Institute COVID-19 Surveillance Team |
| EPI_ISL_2456974 | Lighthouse Lab in Alderley Park | Wellcome Sanger Institute for the COVID-19 Genomics UK (COG-UK) Consortium | Jacquelyn Wynn, Mairead Hyland, The Lighthouse Lab in Alderley Park and Alex Alderton, Roberto Amato, Jeffrey Barrett, Sonia Goncalves, Ewan Harrison, David K. Jackson, Ian Johnston, Dominic Kwiatkowski, Cordelia Langford, John Sillitoe on behalf of the Wellcome Sanger Institute COVID-19 Surveillance Team |
| EPI_ISL_2456975 | Health Services Laboratories | Wellcome Sanger Institute for the COVID-19 Genomics UK (COG-UK) Consortium | Health Services Laboratories and Alex Alderton, Roberto Amato, Jeffrey Barrett, Sonia Goncalves, Ewan Harrison, David K. Jackson, Ian Johnston, Dominic Kwiatkowski, Cordelia Langford, John Sillitoe on behalf of the Wellcome Sanger Institute COVID-19 Surveillance Team |
| EPI_ISL_2456977, EPI_ISL_2456978 | Lighthouse Lab in Alderley Park | Wellcome Sanger Institute for the COVID-19 Genomics UK (COG-UK) Consortium | Jacquelyn Wynn, Mairead Hyland, The Lighthouse Lab in Alderley Park and Alex Alderton, Roberto Amato, Jeffrey Barrett, Sonia Goncalves, Ewan Harrison, David K. Jackson, Ian Johnston, Dominic Kwiatkowski, Cordelia Langford, John Sillitoe on behalf of the Wellcome Sanger Institute COVID-19 Surveillance Team |
| EPI_ISL_2456979 | Health Services Laboratories | Wellcome Sanger Institute for the COVID-19 Genomics UK (COG-UK) Consortium | Health Services Laboratories and Alex Alderton, Roberto Amato, Jeffrey Barrett, Sonia Goncalves, Ewan Harrison, David K. Jackson, Ian Johnston, Dominic Kwiatkowski, Cordelia Langford, John Sillitoe on behalf of the Wellcome Sanger Institute COVID-19 Surveillance Team |
| EPI_ISL_2456980, EPI_ISL_2456981 | Lighthouse Lab in Milton Keynes | Wellcome Sanger Institute for the COVID-19 Genomics UK (COG-UK) Consortium | The Lighthouse Lab in Milton Keynes and Alex Alderton, Roberto Amato, Jeffrey Barrett, Sonia Goncalves, Ewan Harrison, David K. Jackson, Ian Johnston, Dominic Kwiatkowski, Cordelia Langford, John Sillitoe on behalf of the Wellcome Sanger Institute COVID-19 Surveillance Team |
| EPI_ISL_2456982 | Health Services Laboratories | Wellcome Sanger Institute for the COVID-19 Genomics UK (COG-UK) Consortium | Health Services Laboratories and Alex Alderton, Roberto Amato, Jeffrey Barrett, Sonia Goncalves, Ewan Harrison, David K. Jackson, Ian Johnston, Dominic Kwiatkowski, Cordelia Langford, John Sillitoe on behalf of the Wellcome Sanger Institute COVID-19 Surveillance Team |
| EPI_ISL_2456983 | Lighthouse Lab in Milton Keynes | Wellcome Sanger Institute for the COVID-19 Genomics UK (COG-UK) Consortium | The Lighthouse Lab in Milton Keynes and Alex Alderton, Roberto Amato, Jeffrey Barrett, Sonia Goncalves, Ewan Harrison, David K. Jackson, Ian Johnston, Dominic Kwiatkowski, Cordelia Langford, John Sillitoe on behalf of the Wellcome Sanger Institute COVID-19 Surveillance Team |
| EPI_ISL_2456984 | Lighthouse Lab in Alderley Park | Wellcome Sanger Institute for the COVID-19 Genomics UK (COG-UK) Consortium | Jacquelyn Wynn, Mairead Hyland, The Lighthouse Lab in Alderley Park and Alex Alderton, Roberto Amato, Jeffrey Barrett, Sonia Goncalves, Ewan Harrison, David K. Jackson, Ian Johnston, Dominic Kwiatkowski, Cordelia Langford, John Sillitoe on behalf of the Wellcome Sanger Institute COVID-19 Surveillance Team |
| EPI_ISL_2456985, EPI_ISL_2456986 | Lighthouse Lab in Milton Keynes | Wellcome Sanger Institute for the COVID-19 Genomics UK (COG-UK) Consortium | The Lighthouse Lab in Milton Keynes and Alex Alderton, Roberto Amato, Jeffrey Barrett, Sonia Goncalves, Ewan Harrison, David K. Jackson, Ian Johnston, Dominic Kwiatkowski, Cordelia Langford, John Sillitoe on behalf of the Wellcome Sanger Institute COVID-19 Surveillance Team |
| EPI_ISL_2456987 | Lighthouse Lab in Alderley Park | Wellcome Sanger Institute for the COVID-19 Genomics UK (COG-UK) Consortium | Jacquelyn Wynn, Mairead Hyland, The Lighthouse Lab in Alderley Park and Alex Alderton, Roberto Amato, Jeffrey Barrett, Sonia Goncalves, Ewan Harrison, David K. Jackson, Ian Johnston, Dominic Kwiatkowski, Cordelia Langford, John Sillitoe on behalf of the Wellcome Sanger Institute COVID-19 Surveillance Team |
| EPI_ISL_2456988 | Health Services Laboratories | Wellcome Sanger Institute for the COVID-19 Genomics UK (COG-UK) Consortium | Health Services Laboratories and Alex Alderton, Roberto Amato, Jeffrey Barrett, Sonia Goncalves, Ewan Harrison, David K. Jackson, Ian Johnston, Dominic Kwiatkowski, Cordelia Langford, John Sillitoe on behalf of the Wellcome Sanger Institute COVID-19 Surveillance Team |
| EPI_ISL_2456990, EPI_ISL_2456991, EPI_ISL_2456992 | Lighthouse Lab in Milton Keynes | Wellcome Sanger Institute for the COVID-19 Genomics UK (COG-UK) Consortium | The Lighthouse Lab in Milton Keynes and Alex Alderton, Roberto Amato, Jeffrey Barrett, Sonia Goncalves, Ewan Harrison, David K. Jackson, Ian Johnston, Dominic Kwiatkowski, Cordelia Langford, John Sillitoe on behalf of the Wellcome Sanger Institute COVID-19 Surveillance Team |
| EPI_ISL_2456995 | Lighthouse Lab in Alderley Park | Wellcome Sanger Institute for the COVID-19 Genomics UK (COG-UK) Consortium | Jacquelyn Wynn, Mairead Hyland, The Lighthouse Lab in Alderley Park and Alex Alderton, Roberto Amato, Jeffrey Barrett, Sonia Goncalves, Ewan Harrison, David K. Jackson, Ian Johnston, Dominic Kwiatkowski, Cordelia Langford, John Sillitoe on behalf of the Wellcome Sanger Institute COVID-19 Surveillance Team |
| EPI_ISL_2456996 | Health Services Laboratories | Wellcome Sanger Institute for the COVID-19 Genomics UK (COG-UK) Consortium | Health Services Laboratories and Alex Alderton, Roberto Amato, Jeffrey Barrett, Sonia Goncalves, Ewan Harrison, David K. Jackson, Ian Johnston, Dominic Kwiatkowski, Cordelia Langford, John Sillitoe on behalf of the Wellcome Sanger Institute COVID-19 Surveillance Team |
| EPI_ISL_2456997 | Lighthouse Lab in Milton Keynes | Wellcome Sanger Institute for the COVID-19 Genomics UK (COG-UK) Consortium | The Lighthouse Lab in Milton Keynes and Alex Alderton, Roberto Amato, Jeffrey Barrett, Sonia Goncalves, Ewan Harrison, David K. Jackson, Ian Johnston, Dominic Kwiatkowski, Cordelia Langford, John Sillitoe on behalf of the Wellcome Sanger Institute COVID-19 Surveillance Team |
| EPI_ISL_2456998 | Health Services Laboratories | Wellcome Sanger Institute for the COVID-19 Genomics UK (COG-UK) Consortium | Health Services Laboratories and Alex Alderton, Roberto Amato, Jeffrey Barrett, Sonia Goncalves, Ewan Harrison, David K. Jackson, Ian Johnston, Dominic Kwiatkowski, Cordelia Langford, John Sillitoe on behalf of the Wellcome Sanger Institute COVID-19 Surveillance Team |
| EPI_ISL_2456999 | Lighthouse Lab in Alderley Park | Wellcome Sanger Institute for the COVID-19 Genomics UK (COG-UK) Consortium | Jacquelyn Wynn, Mairead Hyland, The Lighthouse Lab in Alderley Park and Alex Alderton, Roberto Amato, Jeffrey Barrett, Sonia Goncalves, Ewan Harrison, David K. Jackson, Ian Johnston, Dominic Kwiatkowski, Cordelia Langford, John Sillitoe on behalf of the Wellcome Sanger Institute COVID-19 Surveillance Team |
| EPI_ISL_2457001, EPI_ISL_2457002 | Lighthouse Lab in Milton Keynes | Wellcome Sanger Institute for the COVID-19 Genomics UK (COG-UK) Consortium | The Lighthouse Lab in Milton Keynes and Alex Alderton, Roberto Amato, Jeffrey Barrett, Sonia Goncalves, Ewan Harrison, David K. Jackson, Ian Johnston, Dominic Kwiatkowski, Cordelia Langford, John Sillitoe on behalf of the Wellcome Sanger Institute COVID-19 Surveillance Team |
| EPI_ISL_2457003 | Health Services Laboratories | Wellcome Sanger Institute for the COVID-19 Genomics UK (COG-UK) Consortium | Health Services Laboratories and Alex Alderton, Roberto Amato, Jeffrey Barrett, Sonia Goncalves, Ewan Harrison, David K. Jackson, Ian Johnston, Dominic Kwiatkowski, Cordelia Langford, John Sillitoe on behalf of the Wellcome Sanger Institute COVID-19 Surveillance Team |
| EPI_ISL_2457004, EPI_ISL_2457005, EPI_ISL_2457006 | Lighthouse Lab in Alderley Park | Wellcome Sanger Institute for the COVID-19 Genomics UK (COG-UK) Consortium | Jacquelyn Wynn, Mairead Hyland, The Lighthouse Lab in Alderley Park and Alex Alderton, Roberto Amato, Jeffrey Barrett, Sonia Goncalves, Ewan Harrison, David K. Jackson, Ian Johnston, Dominic Kwiatkowski, Cordelia Langford, John Sillitoe on behalf of the Wellcome Sanger Institute COVID-19 Surveillance Team |
| EPI_ISL_2457007 | Lighthouse Lab in Milton Keynes | Wellcome Sanger Institute for the COVID-19 Genomics UK (COG-UK) Consortium | The Lighthouse Lab in Milton Keynes and Alex Alderton, Roberto Amato, Jeffrey Barrett, Sonia Goncalves, Ewan Harrison, David K. Jackson, Ian Johnston, Dominic Kwiatkowski, Cordelia Langford, John Sillitoe on behalf of the Wellcome Sanger Institute COVID-19 Surveillance Team |
| EPI_ISL_2457008 | Lighthouse Lab in Alderley Park | Wellcome Sanger Institute for the COVID-19 Genomics UK (COG-UK) Consortium | Jacquelyn Wynn, Mairead Hyland, The Lighthouse Lab in Alderley Park and Alex Alderton, Roberto Amato, Jeffrey Barrett, Sonia Goncalves, Ewan Harrison, David K. Jackson, Ian Johnston, Dominic Kwiatkowski, Cordelia Langford, John Sillitoe on behalf of the Wellcome Sanger Institute COVID-19 Surveillance Team |
| EPI_ISL_2457009 | Lighthouse Lab in Milton Keynes | Wellcome Sanger Institute for the COVID-19 Genomics UK (COG-UK) Consortium | The Lighthouse Lab in Milton Keynes and Alex Alderton, Roberto Amato, Jeffrey Barrett, Sonia Goncalves, Ewan Harrison, David K. Jackson, Ian Johnston, Dominic Kwiatkowski, Cordelia Langford, John Sillitoe on behalf of the Wellcome Sanger Institute COVID-19 Surveillance Team |
| EPI_ISL_2457010, EPI_ISL_2457012, EPI_ISL_2457013 | Lighthouse Lab in Alderley Park | Wellcome Sanger Institute for the COVID-19 Genomics UK (COG-UK) Consortium | Jacquelyn Wynn, Mairead Hyland, The Lighthouse Lab in Alderley Park and Alex Alderton, Roberto Amato, Jeffrey Barrett, Sonia Goncalves, Ewan Harrison, David K. Jackson, Ian Johnston, Dominic Kwiatkowski, Cordelia Langford, John Sillitoe on behalf of the Wellcome Sanger Institute COVID-19 Surveillance Team |
| EPI_ISL_2457014 | Lighthouse Lab in Milton Keynes | Wellcome Sanger Institute for the COVID-19 Genomics UK (COG-UK) Consortium | The Lighthouse Lab in Milton Keynes and Alex Alderton, Roberto Amato, Jeffrey Barrett, Sonia Goncalves, Ewan Harrison, David K. Jackson, Ian Johnston, Dominic Kwiatkowski, Cordelia Langford, John Sillitoe on behalf of the Wellcome Sanger Institute COVID-19 Surveillance Team |
| EPI_ISL_2457015 | Lighthouse Lab in Alderley Park | Wellcome Sanger Institute for the COVID-19 Genomics UK (COG-UK) Consortium | Jacquelyn Wynn, Mairead Hyland, The Lighthouse Lab in Alderley Park and Alex Alderton, Roberto Amato, Jeffrey Barrett, Sonia Goncalves, Ewan Harrison, David K. Jackson, Ian Johnston, Dominic Kwiatkowski, Cordelia Langford, John Sillitoe on behalf of the Wellcome Sanger Institute COVID-19 Surveillance Team |
| EPI_ISL_2457016 | Lighthouse Lab in Milton Keynes | Wellcome Sanger Institute for the COVID-19 Genomics UK (COG-UK) Consortium | The Lighthouse Lab in Milton Keynes and Alex Alderton, Roberto Amato, Jeffrey Barrett, Sonia Goncalves, Ewan Harrison, David K. Jackson, Ian Johnston, Dominic Kwiatkowski, Cordelia Langford, John Sillitoe on behalf of the Wellcome Sanger Institute COVID-19 Surveillance Team |
| EPI_ISL_2457017 | Health Services Laboratories | Wellcome Sanger Institute for the COVID-19 Genomics UK (COG-UK) Consortium | Health Services Laboratories and Alex Alderton, Roberto Amato, Jeffrey Barrett, Sonia Goncalves, Ewan Harrison, David K. Jackson, Ian Johnston, Dominic Kwiatkowski, Cordelia Langford, John Sillitoe on behalf of the Wellcome Sanger Institute COVID-19 Surveillance Team |
| EPI_ISL_2457018 | Lighthouse Lab in Alderley Park | Wellcome Sanger Institute for the COVID-19 Genomics UK (COG-UK) Consortium | Jacquelyn Wynn, Mairead Hyland, The Lighthouse Lab in Alderley Park and Alex Alderton, Roberto Amato, Jeffrey Barrett, Sonia Goncalves, Ewan Harrison, David K. Jackson, Ian Johnston, Dominic Kwiatkowski, Cordelia Langford, John Sillitoe on behalf of the Wellcome Sanger Institute COVID-19 Surveillance Team |
| EPI_ISL_2457019 | Health Services Laboratories | Wellcome Sanger Institute for the COVID-19 Genomics UK (COG-UK) Consortium | Health Services Laboratories and Alex Alderton, Roberto Amato, Jeffrey Barrett, Sonia Goncalves, Ewan Harrison, David K. Jackson, Ian Johnston, Dominic Kwiatkowski, Cordelia Langford, John Sillitoe on behalf of the Wellcome Sanger Institute COVID-19 Surveillance Team |
| EPI_ISL_2457020, EPI_ISL_2457022 | Lighthouse Lab in Milton Keynes | Wellcome Sanger Institute for the COVID-19 Genomics UK (COG-UK) Consortium | The Lighthouse Lab in Milton Keynes and Alex Alderton, Roberto Amato, Jeffrey Barrett, Sonia Goncalves, Ewan Harrison, David K. Jackson, Ian Johnston, Dominic Kwiatkowski, Cordelia Langford, John Sillitoe on behalf of the Wellcome Sanger Institute COVID-19 Surveillance Team |
| EPI_ISL_2457023 | Health Services Laboratories | Wellcome Sanger Institute for the COVID-19 Genomics UK (COG-UK) Consortium | Health Services Laboratories and Alex Alderton, Roberto Amato, Jeffrey Barrett, Sonia Goncalves, Ewan Harrison, David K. Jackson, Ian Johnston, Dominic Kwiatkowski, Cordelia Langford, John Sillitoe on behalf of the Wellcome Sanger Institute COVID-19 Surveillance Team |
| EPI_ISL_2457024 | Lighthouse Lab in Milton Keynes | Wellcome Sanger Institute for the COVID-19 Genomics UK (COG-UK) Consortium | The Lighthouse Lab in Milton Keynes and Alex Alderton, Roberto Amato, Jeffrey Barrett, Sonia Goncalves, Ewan Harrison, David K. Jackson, Ian Johnston, Dominic Kwiatkowski, Cordelia Langford, John Sillitoe on behalf of the Wellcome Sanger Institute COVID-19 Surveillance Team |
| EPI_ISL_2457025, | Lighthouse Lab in Alderley Park | Wellcome Sanger Institute for the COVID-19 Genomics UK (COG-UK) | Jacquelyn Wynn, Mairead Hyland, The Lighthouse Lab in Alderley Park and Alex Alderton, Roberto Amato, Jeffrey Barrett, Sonia Goncalves, Ewan Harrison, David K. Jackson, Ian Johnston, Dominic Kwiatkowski, Cordelia Langford, John Sillitoe on |

| | | | |
|---|---|---|---|
| EPI_ISL_2457026, EPI_ISL_2457027, EPI_ISL_2457028 | | Consortium | behalf of the Wellcome Sanger Institute COVID-19 Surveillance Team |
| EPI_ISL_2457029 | Lighthouse Lab in Milton Keynes | Wellcome Sanger Institute for the COVID-19 Genomics UK (COG-UK) Consortium | The Lighthouse Lab in Milton Keynes and Alex Alderton, Roberto Amato, Jeffrey Barrett, Sonia Goncalves, Ewan Harrison, David K. Jackson, Ian Johnston, Dominic Kwiatkowski, Cordelia Langford, John Sillitoe on behalf of the Wellcome Sanger Institute COVID-19 Surveillance Team |
| EPI_ISL_2457030 | Health Services Laboratories | Wellcome Sanger Institute for the COVID-19 Genomics UK (COG-UK) Consortium | Health Services Laboratories and Alex Alderton, Roberto Amato, Jeffrey Barrett, Sonia Goncalves, Ewan Harrison, David K. Jackson, Ian Johnston, Dominic Kwiatkowski, Cordelia Langford, John Sillitoe on behalf of the Wellcome Sanger Institute COVID-19 Surveillance Team |
| EPI_ISL_2457031, EPI_ISL_2457032 | Lighthouse Lab in Alderley Park | Wellcome Sanger Institute for the COVID-19 Genomics UK (COG-UK) Consortium | Jacquelyn Wynn, Mairead Hyland, The Lighthouse Lab in Alderley Park and Alex Alderton, Roberto Amato, Jeffrey Barrett, Sonia Goncalves, Ewan Harrison, David K. Jackson, Ian Johnston, Dominic Kwiatkowski, Cordelia Langford, John Sillitoe on behalf of the Wellcome Sanger Institute COVID-19 Surveillance Team |
| EPI_ISL_2457033 | Lighthouse Lab in Milton Keynes | Wellcome Sanger Institute for the COVID-19 Genomics UK (COG-UK) Consortium | The Lighthouse Lab in Milton Keynes and Alex Alderton, Roberto Amato, Jeffrey Barrett, Sonia Goncalves, Ewan Harrison, David K. Jackson, Ian Johnston, Dominic Kwiatkowski, Cordelia Langford, John Sillitoe on behalf of the Wellcome Sanger Institute COVID-19 Surveillance Team |
| EPI_ISL_2457034 | Health Services Laboratories | Wellcome Sanger Institute for the COVID-19 Genomics UK (COG-UK) Consortium | Health Services Laboratories and Alex Alderton, Roberto Amato, Jeffrey Barrett, Sonia Goncalves, Ewan Harrison, David K. Jackson, Ian Johnston, Dominic Kwiatkowski, Cordelia Langford, John Sillitoe on behalf of the Wellcome Sanger Institute COVID-19 Surveillance Team |
| EPI_ISL_2457035, EPI_ISL_2457036 | Lighthouse Lab in Milton Keynes | Wellcome Sanger Institute for the COVID-19 Genomics UK (COG-UK) Consortium | The Lighthouse Lab in Milton Keynes and Alex Alderton, Roberto Amato, Jeffrey Barrett, Sonia Goncalves, Ewan Harrison, David K. Jackson, Ian Johnston, Dominic Kwiatkowski, Cordelia Langford, John Sillitoe on behalf of the Wellcome Sanger Institute COVID-19 Surveillance Team |
| EPI_ISL_2457038, EPI_ISL_2457039 | Lighthouse Lab in Alderley Park | Wellcome Sanger Institute for the COVID-19 Genomics UK (COG-UK) Consortium | Jacquelyn Wynn, Mairead Hyland, The Lighthouse Lab in Alderley Park and Alex Alderton, Roberto Amato, Jeffrey Barrett, Sonia Goncalves, Ewan Harrison, David K. Jackson, Ian Johnston, Dominic Kwiatkowski, Cordelia Langford, John Sillitoe on behalf of the Wellcome Sanger Institute COVID-19 Surveillance Team |
| EPI_ISL_2457040, EPI_ISL_2457041 | Health Services Laboratories | Wellcome Sanger Institute for the COVID-19 Genomics UK (COG-UK) Consortium | Health Services Laboratories and Alex Alderton, Roberto Amato, Jeffrey Barrett, Sonia Goncalves, Ewan Harrison, David K. Jackson, Ian Johnston, Dominic Kwiatkowski, Cordelia Langford, John Sillitoe on behalf of the Wellcome Sanger Institute COVID-19 Surveillance Team |
| EPI_ISL_2457042 | Lighthouse Lab in Alderley Park | Wellcome Sanger Institute for the COVID-19 Genomics UK (COG-UK) Consortium | Jacquelyn Wynn, Mairead Hyland, The Lighthouse Lab in Alderley Park and Alex Alderton, Roberto Amato, Jeffrey Barrett, Sonia Goncalves, Ewan Harrison, David K. Jackson, Ian Johnston, Dominic Kwiatkowski, Cordelia Langford, John Sillitoe on behalf of the Wellcome Sanger Institute COVID-19 Surveillance Team |
| EPI_ISL_2457043, EPI_ISL_2457044, EPI_ISL_2457045 | Lighthouse Lab in Milton Keynes | Wellcome Sanger Institute for the COVID-19 Genomics UK (COG-UK) Consortium | The Lighthouse Lab in Milton Keynes and Alex Alderton, Roberto Amato, Jeffrey Barrett, Sonia Goncalves, Ewan Harrison, David K. Jackson, Ian Johnston, Dominic Kwiatkowski, Cordelia Langford, John Sillitoe on behalf of the Wellcome Sanger Institute COVID-19 Surveillance Team |
| EPI_ISL_2457046 | Health Services Laboratories | Wellcome Sanger Institute for the COVID-19 Genomics UK (COG-UK) Consortium | Health Services Laboratories and Alex Alderton, Roberto Amato, Jeffrey Barrett, Sonia Goncalves, Ewan Harrison, David K. Jackson, Ian Johnston, Dominic Kwiatkowski, Cordelia Langford, John Sillitoe on behalf of the Wellcome Sanger Institute COVID-19 Surveillance Team |
| EPI_ISL_2457048 | Lighthouse Lab in Milton Keynes | Wellcome Sanger Institute for the COVID-19 Genomics UK (COG-UK) Consortium | The Lighthouse Lab in Milton Keynes and Alex Alderton, Roberto Amato, Jeffrey Barrett, Sonia Goncalves, Ewan Harrison, David K. Jackson, Ian Johnston, Dominic Kwiatkowski, Cordelia Langford, John Sillitoe on behalf of the Wellcome Sanger Institute COVID-19 Surveillance Team |
| EPI_ISL_2457049, EPI_ISL_2457051, EPI_ISL_2457052 | Lighthouse Lab in Alderley Park | Wellcome Sanger Institute for the COVID-19 Genomics UK (COG-UK) Consortium | Jacquelyn Wynn, Mairead Hyland, The Lighthouse Lab in Alderley Park and Alex Alderton, Roberto Amato, Jeffrey Barrett, Sonia Goncalves, Ewan Harrison, David K. Jackson, Ian Johnston, Dominic Kwiatkowski, Cordelia Langford, John Sillitoe on behalf of the Wellcome Sanger Institute COVID-19 Surveillance Team |
| EPI_ISL_2457054 | Lighthouse Lab in Milton Keynes | Wellcome Sanger Institute for the COVID-19 Genomics UK (COG-UK) Consortium | The Lighthouse Lab in Milton Keynes and Alex Alderton, Roberto Amato, Jeffrey Barrett, Sonia Goncalves, Ewan Harrison, David K. Jackson, Ian Johnston, Dominic Kwiatkowski, Cordelia Langford, John Sillitoe on behalf of the Wellcome Sanger Institute COVID-19 Surveillance Team |
| EPI_ISL_2457055 | Health Services Laboratories | Wellcome Sanger Institute for the COVID-19 Genomics UK (COG-UK) Consortium | Health Services Laboratories and Alex Alderton, Roberto Amato, Jeffrey Barrett, Sonia Goncalves, Ewan Harrison, David K. Jackson, Ian Johnston, Dominic Kwiatkowski, Cordelia Langford, John Sillitoe on behalf of the Wellcome Sanger Institute COVID-19 Surveillance Team |
| EPI_ISL_2457056 | Lighthouse Lab in Alderley Park | Wellcome Sanger Institute for the COVID-19 Genomics UK (COG-UK) Consortium | Jacquelyn Wynn, Mairead Hyland, The Lighthouse Lab in Alderley Park and Alex Alderton, Roberto Amato, Jeffrey Barrett, Sonia Goncalves, Ewan Harrison, David K. Jackson, Ian Johnston, Dominic Kwiatkowski, Cordelia Langford, John Sillitoe on behalf of the Wellcome Sanger Institute COVID-19 Surveillance Team |
| EPI_ISL_2457057, EPI_ISL_2457059, EPI_ISL_2457060 | Lighthouse Lab in Milton Keynes | Wellcome Sanger Institute for the COVID-19 Genomics UK (COG-UK) Consortium | The Lighthouse Lab in Milton Keynes and Alex Alderton, Roberto Amato, Jeffrey Barrett, Sonia Goncalves, Ewan Harrison, David K. Jackson, Ian Johnston, Dominic Kwiatkowski, Cordelia Langford, John Sillitoe on behalf of the Wellcome Sanger Institute COVID-19 Surveillance Team |
| EPI_ISL_2457061 | Stanford Health Care | Stanford University School of Medicine, Clinical Virology Laboratory | Daniel Solis, Mamdouh Sibai, Bernadette Troung, Becky Jiang, Michelle Verghese, ChunHong Huang, James Zehnder, Malaya K. Sahoo, and Benjamin A. Pinsky |
| EPI_ISL_2457063, EPI_ISL_2457064, EPI_ISL_2457065 | Lighthouse Lab in Milton Keynes | Wellcome Sanger Institute for the COVID-19 Genomics UK (COG-UK) Consortium | The Lighthouse Lab in Milton Keynes and Alex Alderton, Roberto Amato, Jeffrey Barrett, Sonia Goncalves, Ewan Harrison, David K. Jackson, Ian Johnston, Dominic Kwiatkowski, Cordelia Langford, John Sillitoe on behalf of the Wellcome Sanger Institute COVID-19 Surveillance Team |
| EPI_ISL_2457066, EPI_ISL_2457067, EPI_ISL_2457068 | Lighthouse Lab in Alderley Park | Wellcome Sanger Institute for the COVID-19 Genomics UK (COG-UK) Consortium | Jacquelyn Wynn, Mairead Hyland, The Lighthouse Lab in Alderley Park and Alex Alderton, Roberto Amato, Jeffrey Barrett, Sonia Goncalves, Ewan Harrison, David K. Jackson, Ian Johnston, Dominic Kwiatkowski, Cordelia Langford, John Sillitoe on behalf of the Wellcome Sanger Institute COVID-19 Surveillance Team |
| EPI_ISL_2457069, EPI_ISL_2457070, EPI_ISL_2457071, EPI_ISL_2457072 | Lighthouse Lab in Milton Keynes | Wellcome Sanger Institute for the COVID-19 Genomics UK (COG-UK) Consortium | The Lighthouse Lab in Milton Keynes and Alex Alderton, Roberto Amato, Jeffrey Barrett, Sonia Goncalves, Ewan Harrison, David K. Jackson, Ian Johnston, Dominic Kwiatkowski, Cordelia Langford, John Sillitoe on behalf of the Wellcome Sanger Institute COVID-19 Surveillance Team |
| EPI_ISL_2457073, EPI_ISL_2457074, EPI_ISL_2457076 | Lighthouse Lab in Alderley Park | Wellcome Sanger Institute for the COVID-19 Genomics UK (COG-UK) Consortium | Jacquelyn Wynn, Mairead Hyland, The Lighthouse Lab in Alderley Park and Alex Alderton, Roberto Amato, Jeffrey Barrett, Sonia Goncalves, Ewan Harrison, David K. Jackson, Ian Johnston, Dominic Kwiatkowski, Cordelia Langford, John Sillitoe on behalf of the Wellcome Sanger Institute COVID-19 Surveillance Team |
| EPI_ISL_2457077, EPI_ISL_2457078, EPI_ISL_2457080, EPI_ISL_2457082 | Lighthouse Lab in Milton Keynes | Wellcome Sanger Institute for the COVID-19 Genomics UK (COG-UK) Consortium | The Lighthouse Lab in Milton Keynes and Alex Alderton, Roberto Amato, Jeffrey Barrett, Sonia Goncalves, Ewan Harrison, David K. Jackson, Ian Johnston, Dominic Kwiatkowski, Cordelia Langford, John Sillitoe on behalf of the Wellcome Sanger Institute COVID-19 Surveillance Team |
| EPI_ISL_2457083 | Lighthouse Lab in Alderley Park | Wellcome Sanger Institute for the COVID-19 Genomics UK (COG-UK) Consortium | Jacquelyn Wynn, Mairead Hyland, The Lighthouse Lab in Alderley Park and Alex Alderton, Roberto Amato, Jeffrey Barrett, Sonia Goncalves, Ewan Harrison, David K. Jackson, Ian Johnston, Dominic Kwiatkowski, Cordelia Langford, John Sillitoe on behalf of the Wellcome Sanger Institute COVID-19 Surveillance Team |
| EPI_ISL_2457085 | Lighthouse Lab in Milton Keynes | Wellcome Sanger Institute for the COVID-19 Genomics UK (COG-UK) Consortium | The Lighthouse Lab in Milton Keynes and Alex Alderton, Roberto Amato, Jeffrey Barrett, Sonia Goncalves, Ewan Harrison, David K. Jackson, Ian Johnston, Dominic Kwiatkowski, Cordelia Langford, John Sillitoe on behalf of the Wellcome Sanger Institute COVID-19 Surveillance Team |
| EPI_ISL_2457086 | Lighthouse Lab in Alderley Park | Wellcome Sanger Institute for the COVID-19 Genomics UK (COG-UK) Consortium | Jacquelyn Wynn, Mairead Hyland, The Lighthouse Lab in Alderley Park and Alex Alderton, Roberto Amato, Jeffrey Barrett, Sonia Goncalves, Ewan Harrison, David K. Jackson, Ian Johnston, Dominic Kwiatkowski, Cordelia Langford, John Sillitoe on behalf of the Wellcome Sanger Institute COVID-19 Surveillance Team |
| EPI_ISL_2457087 | Health Services Laboratories | Wellcome Sanger Institute for the COVID-19 Genomics UK (COG-UK) Consortium | Health Services Laboratories and Alex Alderton, Roberto Amato, Jeffrey Barrett, Sonia Goncalves, Ewan Harrison, David K. Jackson, Ian Johnston, Dominic Kwiatkowski, Cordelia Langford, John Sillitoe on behalf of the Wellcome Sanger Institute COVID-19 Surveillance Team |
| EPI_ISL_2457088 | Lighthouse Lab in Alderley Park | Wellcome Sanger Institute for the COVID-19 Genomics UK (COG-UK) Consortium | Jacquelyn Wynn, Mairead Hyland, The Lighthouse Lab in Alderley Park and Alex Alderton, Roberto Amato, Jeffrey Barrett, Sonia Goncalves, Ewan Harrison, David K. Jackson, Ian Johnston, Dominic Kwiatkowski, Cordelia Langford, John Sillitoe on behalf of the Wellcome Sanger Institute COVID-19 Surveillance Team |
| EPI_ISL_2457089 | Lighthouse Lab in Milton Keynes | Wellcome Sanger Institute for the COVID-19 Genomics UK (COG-UK) Consortium | The Lighthouse Lab in Milton Keynes and Alex Alderton, Roberto Amato, Jeffrey Barrett, Sonia Goncalves, Ewan Harrison, David K. Jackson, Ian Johnston, Dominic Kwiatkowski, Cordelia Langford, John Sillitoe on behalf of the Wellcome Sanger Institute COVID-19 Surveillance Team |
| EPI_ISL_2457090 | Health Services Laboratories | Wellcome Sanger Institute for the COVID-19 Genomics UK (COG-UK) Consortium | Health Services Laboratories and Alex Alderton, Roberto Amato, Jeffrey Barrett, Sonia Goncalves, Ewan Harrison, David K. Jackson, Ian Johnston, Dominic Kwiatkowski, Cordelia Langford, John Sillitoe on behalf of the Wellcome Sanger Institute COVID-19 Surveillance Team |
| EPI_ISL_2457093 | Lighthouse Lab in Alderley Park | Wellcome Sanger Institute for the COVID-19 Genomics UK (COG-UK) Consortium | Jacquelyn Wynn, Mairead Hyland, The Lighthouse Lab in Alderley Park and Alex Alderton, Roberto Amato, Jeffrey Barrett, Sonia Goncalves, Ewan Harrison, David K. Jackson, Ian Johnston, Dominic Kwiatkowski, Cordelia Langford, John Sillitoe on behalf of the Wellcome Sanger Institute COVID-19 Surveillance Team |
| EPI_ISL_2457094 | Lighthouse Lab in Milton Keynes | Wellcome Sanger Institute for the COVID-19 Genomics UK (COG-UK) Consortium | The Lighthouse Lab in Milton Keynes and Alex Alderton, Roberto Amato, Jeffrey Barrett, Sonia Goncalves, Ewan Harrison, David K. Jackson, Ian Johnston, Dominic Kwiatkowski, Cordelia Langford, John Sillitoe on behalf of the Wellcome Sanger Institute COVID-19 Surveillance Team |
| EPI_ISL_2457095 | Lighthouse Lab in Alderley Park | Wellcome Sanger Institute for the COVID-19 Genomics UK (COG-UK) Consortium | Jacquelyn Wynn, Mairead Hyland, The Lighthouse Lab in Alderley Park and Alex Alderton, Roberto Amato, Jeffrey Barrett, Sonia Goncalves, Ewan Harrison, David K. Jackson, Ian Johnston, Dominic Kwiatkowski, Cordelia Langford, John Sillitoe on behalf of the Wellcome Sanger Institute COVID-19 Surveillance Team |
| EPI_ISL_2457096 | Lighthouse Lab in Milton Keynes | Wellcome Sanger Institute for the COVID-19 Genomics UK (COG-UK) Consortium | The Lighthouse Lab in Milton Keynes and Alex Alderton, Roberto Amato, Jeffrey Barrett, Sonia Goncalves, Ewan Harrison, David K. Jackson, Ian Johnston, Dominic Kwiatkowski, Cordelia Langford, John Sillitoe on behalf of the Wellcome Sanger Institute COVID-19 Surveillance Team |
| EPI_ISL_2457098 | Lighthouse Lab in Alderley Park | Wellcome Sanger Institute for the COVID-19 Genomics UK (COG-UK) Consortium | Jacquelyn Wynn, Mairead Hyland, The Lighthouse Lab in Alderley Park and Alex Alderton, Roberto Amato, Jeffrey Barrett, Sonia Goncalves, Ewan Harrison, David K. Jackson, Ian Johnston, Dominic Kwiatkowski, Cordelia Langford, John Sillitoe on behalf of the Wellcome Sanger Institute COVID-19 Surveillance Team |
| EPI_ISL_2457099 | Lighthouse Lab in Milton Keynes | Wellcome Sanger Institute for the COVID-19 Genomics UK (COG-UK) Consortium | The Lighthouse Lab in Milton Keynes and Alex Alderton, Roberto Amato, Jeffrey Barrett, Sonia Goncalves, Ewan Harrison, David K. Jackson, Ian Johnston, Dominic Kwiatkowski, Cordelia Langford, John Sillitoe on behalf of the Wellcome Sanger Institute COVID-19 Surveillance Team |
| EPI_ISL_2457100 | Health Services Laboratories | Wellcome Sanger Institute for the COVID-19 Genomics UK (COG-UK) Consortium | Health Services Laboratories and Alex Alderton, Roberto Amato, Jeffrey Barrett, Sonia Goncalves, Ewan Harrison, David K. Jackson, Ian Johnston, Dominic Kwiatkowski, Cordelia Langford, John Sillitoe on behalf of the Wellcome Sanger Institute COVID-19 Surveillance Team |
| EPI_ISL_2457101 | Lighthouse Lab in Alderley Park | Wellcome Sanger Institute for the COVID-19 Genomics UK (COG-UK) Consortium | Jacquelyn Wynn, Mairead Hyland, The Lighthouse Lab in Alderley Park and Alex Alderton, Roberto Amato, Jeffrey Barrett, Sonia Goncalves, Ewan Harrison, David K. Jackson, Ian Johnston, Dominic Kwiatkowski, Cordelia Langford, John Sillitoe on behalf of the Wellcome Sanger Institute COVID-19 Surveillance Team |
| EPI_ISL_2457102 | Lighthouse Lab in Milton Keynes | Wellcome Sanger Institute for the COVID-19 Genomics UK (COG-UK) Consortium | The Lighthouse Lab in Milton Keynes and Alex Alderton, Roberto Amato, Jeffrey Barrett, Sonia Goncalves, Ewan Harrison, David K. Jackson, Ian Johnston, Dominic Kwiatkowski, Cordelia Langford, John Sillitoe on behalf of the Wellcome Sanger Institute COVID-19 Surveillance Team |
| EPI_ISL_2457103 | Health Services Laboratories | Wellcome Sanger Institute for the COVID-19 Genomics UK (COG-UK) Consortium | Health Services Laboratories and Alex Alderton, Roberto Amato, Jeffrey Barrett, Sonia Goncalves, Ewan Harrison, David K. Jackson, Ian Johnston, Dominic Kwiatkowski, Cordelia Langford, John Sillitoe on behalf of the Wellcome Sanger Institute COVID-19 Surveillance Team |

| | | | |
|---|---|---|---|
| EPI_ISL_2457104 | Lighthouse Lab in Milton Keynes | Wellcome Sanger Institute for the COVID-19 Genomics UK (COG-UK) Consortium | The Lighthouse Lab in Milton Keynes and Alex Alderton, Roberto Amato, Jeffrey Barrett, Sonia Goncalves, Ewan Harrison, David K. Jackson, Ian Johnston, Dominic Kwiatkowski, Cordelia Langford, John Sillitoe on behalf of the Wellcome Sanger Institute COVID-19 Surveillance Team |
| EPI_ISL_2457105 | Health Services Laboratories | Wellcome Sanger Institute for the COVID-19 Genomics UK (COG-UK) Consortium | Health Services Laboratories and Alex Alderton, Roberto Amato, Jeffrey Barrett, Sonia Goncalves, Ewan Harrison, David K. Jackson, Ian Johnston, Dominic Kwiatkowski, Cordelia Langford, John Sillitoe on behalf of the Wellcome Sanger Institute COVID-19 Surveillance Team |
| EPI_ISL_2457106, EPI_ISL_2457107, EPI_ISL_2457108, EPI_ISL_2457109, EPI_ISL_2457110 | Lighthouse Lab in Milton Keynes | Wellcome Sanger Institute for the COVID-19 Genomics UK (COG-UK) Consortium | The Lighthouse Lab in Milton Keynes and Alex Alderton, Roberto Amato, Jeffrey Barrett, Sonia Goncalves, Ewan Harrison, David K. Jackson, Ian Johnston, Dominic Kwiatkowski, Cordelia Langford, John Sillitoe on behalf of the Wellcome Sanger Institute COVID-19 Surveillance Team |
| EPI_ISL_2457111 | Lighthouse Lab in Alderley Park | Wellcome Sanger Institute for the COVID-19 Genomics UK (COG-UK) Consortium | Jacquelyn Wynn, Mairead Hyland, The Lighthouse Lab in Alderley Park and Alex Alderton, Roberto Amato, Jeffrey Barrett, Sonia Goncalves, Ewan Harrison, David K. Jackson, Ian Johnston, Dominic Kwiatkowski, Cordelia Langford, John Sillitoe on behalf of the Wellcome Sanger Institute COVID-19 Surveillance Team |
| EPI_ISL_2457112 | Lighthouse Lab in Milton Keynes | Wellcome Sanger Institute for the COVID-19 Genomics UK (COG-UK) Consortium | The Lighthouse Lab in Milton Keynes and Alex Alderton, Roberto Amato, Jeffrey Barrett, Sonia Goncalves, Ewan Harrison, David K. Jackson, Ian Johnston, Dominic Kwiatkowski, Cordelia Langford, John Sillitoe on behalf of the Wellcome Sanger Institute COVID-19 Surveillance Team |
| EPI_ISL_2457113 | Lighthouse Lab in Alderley Park | Wellcome Sanger Institute for the COVID-19 Genomics UK (COG-UK) Consortium | Jacquelyn Wynn, Mairead Hyland, The Lighthouse Lab in Alderley Park and Alex Alderton, Roberto Amato, Jeffrey Barrett, Sonia Goncalves, Ewan Harrison, David K. Jackson, Ian Johnston, Dominic Kwiatkowski, Cordelia Langford, John Sillitoe on behalf of the Wellcome Sanger Institute COVID-19 Surveillance Team |
| EPI_ISL_2457114 | Health Services Laboratories | Wellcome Sanger Institute for the COVID-19 Genomics UK (COG-UK) Consortium | Health Services Laboratories and Alex Alderton, Roberto Amato, Jeffrey Barrett, Sonia Goncalves, Ewan Harrison, David K. Jackson, Ian Johnston, Dominic Kwiatkowski, Cordelia Langford, John Sillitoe on behalf of the Wellcome Sanger Institute COVID-19 Surveillance Team |
| EPI_ISL_2457115 | Lighthouse Lab in Milton Keynes | Wellcome Sanger Institute for the COVID-19 Genomics UK (COG-UK) Consortium | The Lighthouse Lab in Milton Keynes and Alex Alderton, Roberto Amato, Jeffrey Barrett, Sonia Goncalves, Ewan Harrison, David K. Jackson, Ian Johnston, Dominic Kwiatkowski, Cordelia Langford, John Sillitoe on behalf of the Wellcome Sanger Institute COVID-19 Surveillance Team |
| EPI_ISL_2457116, EPI_ISL_2457117 | Lighthouse Lab in Alderley Park | Wellcome Sanger Institute for the COVID-19 Genomics UK (COG-UK) Consortium | Jacquelyn Wynn, Mairead Hyland, The Lighthouse Lab in Alderley Park and Alex Alderton, Roberto Amato, Jeffrey Barrett, Sonia Goncalves, Ewan Harrison, David K. Jackson, Ian Johnston, Dominic Kwiatkowski, Cordelia Langford, John Sillitoe on behalf of the Wellcome Sanger Institute COVID-19 Surveillance Team |
| EPI_ISL_2457118 | Lighthouse Lab in Milton Keynes | Wellcome Sanger Institute for the COVID-19 Genomics UK (COG-UK) Consortium | The Lighthouse Lab in Milton Keynes and Alex Alderton, Roberto Amato, Jeffrey Barrett, Sonia Goncalves, Ewan Harrison, David K. Jackson, Ian Johnston, Dominic Kwiatkowski, Cordelia Langford, John Sillitoe on behalf of the Wellcome Sanger Institute COVID-19 Surveillance Team |
| EPI_ISL_2457119, EPI_ISL_2457120, EPI_ISL_2457121 | Lighthouse Lab in Alderley Park | Wellcome Sanger Institute for the COVID-19 Genomics UK (COG-UK) Consortium | Jacquelyn Wynn, Mairead Hyland, The Lighthouse Lab in Alderley Park and Alex Alderton, Roberto Amato, Jeffrey Barrett, Sonia Goncalves, Ewan Harrison, David K. Jackson, Ian Johnston, Dominic Kwiatkowski, Cordelia Langford, John Sillitoe on behalf of the Wellcome Sanger Institute COVID-19 Surveillance Team |
| EPI_ISL_2457122, EPI_ISL_2457123 | Health Services Laboratories | Wellcome Sanger Institute for the COVID-19 Genomics UK (COG-UK) Consortium | Health Services Laboratories and Alex Alderton, Roberto Amato, Jeffrey Barrett, Sonia Goncalves, Ewan Harrison, David K. Jackson, Ian Johnston, Dominic Kwiatkowski, Cordelia Langford, John Sillitoe on behalf of the Wellcome Sanger Institute COVID-19 Surveillance Team |
| EPI_ISL_2457124 | Lighthouse Lab in Milton Keynes | Wellcome Sanger Institute for the COVID-19 Genomics UK (COG-UK) Consortium | The Lighthouse Lab in Milton Keynes and Alex Alderton, Roberto Amato, Jeffrey Barrett, Sonia Goncalves, Ewan Harrison, David K. Jackson, Ian Johnston, Dominic Kwiatkowski, Cordelia Langford, John Sillitoe on behalf of the Wellcome Sanger Institute COVID-19 Surveillance Team |
| EPI_ISL_2457125 | Health Services Laboratories | Wellcome Sanger Institute for the COVID-19 Genomics UK (COG-UK) Consortium | Health Services Laboratories and Alex Alderton, Roberto Amato, Jeffrey Barrett, Sonia Goncalves, Ewan Harrison, David K. Jackson, Ian Johnston, Dominic Kwiatkowski, Cordelia Langford, John Sillitoe on behalf of the Wellcome Sanger Institute COVID-19 Surveillance Team |
| EPI_ISL_2457126 | Lighthouse Lab in Milton Keynes | Wellcome Sanger Institute for the COVID-19 Genomics UK (COG-UK) Consortium | The Lighthouse Lab in Milton Keynes and Alex Alderton, Roberto Amato, Jeffrey Barrett, Sonia Goncalves, Ewan Harrison, David K. Jackson, Ian Johnston, Dominic Kwiatkowski, Cordelia Langford, John Sillitoe on behalf of the Wellcome Sanger Institute COVID-19 Surveillance Team |
| EPI_ISL_2457127 | Health Services Laboratories | Wellcome Sanger Institute for the COVID-19 Genomics UK (COG-UK) Consortium | Health Services Laboratories and Alex Alderton, Roberto Amato, Jeffrey Barrett, Sonia Goncalves, Ewan Harrison, David K. Jackson, Ian Johnston, Dominic Kwiatkowski, Cordelia Langford, John Sillitoe on behalf of the Wellcome Sanger Institute COVID-19 Surveillance Team |
| EPI_ISL_2457128, EPI_ISL_2457129, EPI_ISL_2457130 | Lighthouse Lab in Milton Keynes | Wellcome Sanger Institute for the COVID-19 Genomics UK (COG-UK) Consortium | The Lighthouse Lab in Milton Keynes and Alex Alderton, Roberto Amato, Jeffrey Barrett, Sonia Goncalves, Ewan Harrison, David K. Jackson, Ian Johnston, Dominic Kwiatkowski, Cordelia Langford, John Sillitoe on behalf of the Wellcome Sanger Institute COVID-19 Surveillance Team |
| EPI_ISL_2457131 | Lighthouse Lab in Glasgow | Wellcome Sanger Institute for the COVID-19 Genomics UK (COG-UK) Consortium | Harper VanSteenhouse, Yumi Kasai, David Gray, Carol Clugston, Anna Dominiczak and Alex Alderton, Roberto Amato, Jeffrey Barrett, Sonia Goncalves, Ewan Harrison, David K. Jackson, Ian Johnston, Dominic Kwiatkowski, Cordelia Langford, John Sillitoe on behalf of the Wellcome Sanger Institute COVID-19 Surveillance Team |
| EPI_ISL_2457132 | Lighthouse Lab in Alderley Park | Wellcome Sanger Institute for the COVID-19 Genomics UK (COG-UK) Consortium | Jacquelyn Wynn, Mairead Hyland, The Lighthouse Lab in Alderley Park and Alex Alderton, Roberto Amato, Jeffrey Barrett, Sonia Goncalves, Ewan Harrison, David K. Jackson, Ian Johnston, Dominic Kwiatkowski, Cordelia Langford, John Sillitoe on behalf of the Wellcome Sanger Institute COVID-19 Surveillance Team |
| EPI_ISL_2457133, EPI_ISL_2457134 | Lighthouse Lab in Glasgow | Wellcome Sanger Institute for the COVID-19 Genomics UK (COG-UK) Consortium | Harper VanSteenhouse, Yumi Kasai, David Gray, Carol Clugston, Anna Dominiczak and Alex Alderton, Roberto Amato, Jeffrey Barrett, Sonia Goncalves, Ewan Harrison, David K. Jackson, Ian Johnston, Dominic Kwiatkowski, Cordelia Langford, John Sillitoe on behalf of the Wellcome Sanger Institute COVID-19 Surveillance Team |
| EPI_ISL_2457135, EPI_ISL_2457136, EPI_ISL_2457137, EPI_ISL_2457138, EPI_ISL_2457139, EPI_ISL_2457140 | Lighthouse Lab in Alderley Park | Wellcome Sanger Institute for the COVID-19 Genomics UK (COG-UK) Consortium | Jacquelyn Wynn, Mairead Hyland, The Lighthouse Lab in Alderley Park and Alex Alderton, Roberto Amato, Jeffrey Barrett, Sonia Goncalves, Ewan Harrison, David K. Jackson, Ian Johnston, Dominic Kwiatkowski, Cordelia Langford, John Sillitoe on behalf of the Wellcome Sanger Institute COVID-19 Surveillance Team |
| EPI_ISL_2457141 | Lighthouse Lab in Glasgow | Wellcome Sanger Institute for the COVID-19 Genomics UK (COG-UK) Consortium | Harper VanSteenhouse, Yumi Kasai, David Gray, Carol Clugston, Anna Dominiczak and Alex Alderton, Roberto Amato, Jeffrey Barrett, Sonia Goncalves, Ewan Harrison, David K. Jackson, Ian Johnston, Dominic Kwiatkowski, Cordelia Langford, John Sillitoe on behalf of the Wellcome Sanger Institute COVID-19 Surveillance Team |
| EPI_ISL_2457142, EPI_ISL_2457143, EPI_ISL_2457144 | Lighthouse Lab in Alderley Park | Wellcome Sanger Institute for the COVID-19 Genomics UK (COG-UK) Consortium | Jacquelyn Wynn, Mairead Hyland, The Lighthouse Lab in Alderley Park and Alex Alderton, Roberto Amato, Jeffrey Barrett, Sonia Goncalves, Ewan Harrison, David K. Jackson, Ian Johnston, Dominic Kwiatkowski, Cordelia Langford, John Sillitoe on behalf of the Wellcome Sanger Institute COVID-19 Surveillance Team |
| EPI_ISL_2457145 | Lighthouse Lab in Glasgow | Wellcome Sanger Institute for the COVID-19 Genomics UK (COG-UK) Consortium | Harper VanSteenhouse, Yumi Kasai, David Gray, Carol Clugston, Anna Dominiczak and Alex Alderton, Roberto Amato, Jeffrey Barrett, Sonia Goncalves, Ewan Harrison, David K. Jackson, Ian Johnston, Dominic Kwiatkowski, Cordelia Langford, John Sillitoe on behalf of the Wellcome Sanger Institute COVID-19 Surveillance Team |
| EPI_ISL_2457146, EPI_ISL_2457147, EPI_ISL_2457148 | Lighthouse Lab in Alderley Park | Wellcome Sanger Institute for the COVID-19 Genomics UK (COG-UK) Consortium | Jacquelyn Wynn, Mairead Hyland, The Lighthouse Lab in Alderley Park and Alex Alderton, Roberto Amato, Jeffrey Barrett, Sonia Goncalves, Ewan Harrison, David K. Jackson, Ian Johnston, Dominic Kwiatkowski, Cordelia Langford, John Sillitoe on behalf of the Wellcome Sanger Institute COVID-19 Surveillance Team |
| EPI_ISL_2457149, EPI_ISL_2457150, EPI_ISL_2457151, EPI_ISL_2457152, EPI_ISL_2457153, EPI_ISL_2457154 | Lighthouse Lab in Glasgow | Wellcome Sanger Institute for the COVID-19 Genomics UK (COG-UK) Consortium | Harper VanSteenhouse, Yumi Kasai, David Gray, Carol Clugston, Anna Dominiczak and Alex Alderton, Roberto Amato, Jeffrey Barrett, Sonia Goncalves, Ewan Harrison, David K. Jackson, Ian Johnston, Dominic Kwiatkowski, Cordelia Langford, John Sillitoe on behalf of the Wellcome Sanger Institute COVID-19 Surveillance Team |
| EPI_ISL_2457155, EPI_ISL_2457156, EPI_ISL_2457157 | Lighthouse Lab in Alderley Park | Wellcome Sanger Institute for the COVID-19 Genomics UK (COG-UK) Consortium | Jacquelyn Wynn, Mairead Hyland, The Lighthouse Lab in Alderley Park and Alex Alderton, Roberto Amato, Jeffrey Barrett, Sonia Goncalves, Ewan Harrison, David K. Jackson, Ian Johnston, Dominic Kwiatkowski, Cordelia Langford, John Sillitoe on behalf of the Wellcome Sanger Institute COVID-19 Surveillance Team |
| EPI_ISL_2457158 | Lighthouse Lab in Glasgow | Wellcome Sanger Institute for the COVID-19 Genomics UK (COG-UK) Consortium | Harper VanSteenhouse, Yumi Kasai, David Gray, Carol Clugston, Anna Dominiczak and Alex Alderton, Roberto Amato, Jeffrey Barrett, Sonia Goncalves, Ewan Harrison, David K. Jackson, Ian Johnston, Dominic Kwiatkowski, Cordelia Langford, John Sillitoe on behalf of the Wellcome Sanger Institute COVID-19 Surveillance Team |
| EPI_ISL_2457159 | Lighthouse Lab in Alderley Park | Wellcome Sanger Institute for the COVID-19 Genomics UK (COG-UK) Consortium | Jacquelyn Wynn, Mairead Hyland, The Lighthouse Lab in Alderley Park and Alex Alderton, Roberto Amato, Jeffrey Barrett, Sonia Goncalves, Ewan Harrison, David K. Jackson, Ian Johnston, Dominic Kwiatkowski, Cordelia Langford, John Sillitoe on behalf of the Wellcome Sanger Institute COVID-19 Surveillance Team |
| EPI_ISL_2457160, EPI_ISL_2457161 | Lighthouse Lab in Glasgow | Wellcome Sanger Institute for the COVID-19 Genomics UK (COG-UK) Consortium | Harper VanSteenhouse, Yumi Kasai, David Gray, Carol Clugston, Anna Dominiczak and Alex Alderton, Roberto Amato, Jeffrey Barrett, Sonia Goncalves, Ewan Harrison, David K. Jackson, Ian Johnston, Dominic Kwiatkowski, Cordelia Langford, John Sillitoe on behalf of the Wellcome Sanger Institute COVID-19 Surveillance Team |
| EPI_ISL_2457162, EPI_ISL_2457163, EPI_ISL_2457164 | Lighthouse Lab in Alderley Park | Wellcome Sanger Institute for the COVID-19 Genomics UK (COG-UK) Consortium | Jacquelyn Wynn, Mairead Hyland, The Lighthouse Lab in Alderley Park and Alex Alderton, Roberto Amato, Jeffrey Barrett, Sonia Goncalves, Ewan Harrison, David K. Jackson, Ian Johnston, Dominic Kwiatkowski, Cordelia Langford, John Sillitoe on behalf of the Wellcome Sanger Institute COVID-19 Surveillance Team |
| EPI_ISL_2457165, EPI_ISL_2457166, EPI_ISL_2457167, EPI_ISL_2457168, EPI_ISL_2457169, EPI_ISL_2457171, EPI_ISL_2457172 | Lighthouse Lab in Glasgow | Wellcome Sanger Institute for the COVID-19 Genomics UK (COG-UK) Consortium | Harper VanSteenhouse, Yumi Kasai, David Gray, Carol Clugston, Anna Dominiczak and Alex Alderton, Roberto Amato, Jeffrey Barrett, Sonia Goncalves, Ewan Harrison, David K. Jackson, Ian Johnston, Dominic Kwiatkowski, Cordelia Langford, John Sillitoe on behalf of the Wellcome Sanger Institute COVID-19 Surveillance Team |
| EPI_ISL_2457173 | National Hospital for Tropical Diseases | Oxford University Clinical Research Unit, Hanoi, Vietnam | Pham Ngoc Thach, Nguyen Thi Tam, Van Dinh Trang, Le Van Duyet, Nguyen Thu Trang, Nguyen Thi Hong Thuong, Nguyen Thi Kim Chi, Phan Manh Cuong, Thomas Kesteman, H.Rogier van Doorn on behalf of the OUCRU COVID-19 research group |
| EPI_ISL_2457174 | Lighthouse Lab in Alderley Park | Wellcome Sanger Institute for the COVID-19 Genomics UK (COG-UK) Consortium | Jacquelyn Wynn, Mairead Hyland, The Lighthouse Lab in Alderley Park and Alex Alderton, Roberto Amato, Jeffrey Barrett, Sonia Goncalves, Ewan Harrison, David K. Jackson, Ian Johnston, Dominic Kwiatkowski, Cordelia Langford, John Sillitoe on behalf of the Wellcome Sanger Institute COVID-19 Surveillance Team |
| EPI_ISL_2457175, EPI_ISL_2457176, EPI_ISL_2457177, EPI_ISL_2457178 | Lighthouse Lab in Glasgow | Wellcome Sanger Institute for the COVID-19 Genomics UK (COG-UK) Consortium | Harper VanSteenhouse, Yumi Kasai, David Gray, Carol Clugston, Anna Dominiczak and Alex Alderton, Roberto Amato, Jeffrey Barrett, Sonia Goncalves, Ewan Harrison, David K. Jackson, Ian Johnston, Dominic Kwiatkowski, Cordelia Langford, John Sillitoe on behalf of the Wellcome Sanger Institute COVID-19 Surveillance Team |

| | | | |
|---|---|---|---|
| EPI_ISL_2457179, EPI_ISL_2457180, EPI_ISL_2457181, EPI_ISL_2457182 | Lighthouse Lab in Alderley Park | Wellcome Sanger Institute for the COVID-19 Genomics UK (COG-UK) Consortium | Jacquelyn Wynn, Mairead Hyland, The Lighthouse Lab in Alderley Park and Alex Alderton, Roberto Amato, Jeffrey Barrett, Sonia Goncalves, Ewan Harrison, David K. Jackson, Ian Johnston, Dominic Kwiatkowski, Cordelia Langford, John Sillitoe on behalf of the Wellcome Sanger Institute COVID-19 Surveillance Team |
| EPI_ISL_2457183, EPI_ISL_2457184, EPI_ISL_2457187 | Lighthouse Lab in Glasgow | Wellcome Sanger Institute for the COVID-19 Genomics UK (COG-UK) Consortium | Harper VanSteenhouse, Yumi Kasai, David Gray, Carol Clugston, Anna Dominiczak and Alex Alderton, Roberto Amato, Jeffrey Barrett, Sonia Goncalves, Ewan Harrison, David K. Jackson, Ian Johnston, Dominic Kwiatkowski, Cordelia Langford, John Sillitoe on behalf of the Wellcome Sanger Institute COVID-19 Surveillance Team |
| EPI_ISL_2457188, EPI_ISL_2457189 | Lighthouse Lab in Alderley Park | Wellcome Sanger Institute for the COVID-19 Genomics UK (COG-UK) Consortium | Jacquelyn Wynn, Mairead Hyland, The Lighthouse Lab in Alderley Park and Alex Alderton, Roberto Amato, Jeffrey Barrett, Sonia Goncalves, Ewan Harrison, David K. Jackson, Ian Johnston, Dominic Kwiatkowski, Cordelia Langford, John Sillitoe on behalf of the Wellcome Sanger Institute COVID-19 Surveillance Team |
| EPI_ISL_2457190, EPI_ISL_2457191, EPI_ISL_2457192, EPI_ISL_2457193, EPI_ISL_2457194, EPI_ISL_2457195 | Lighthouse Lab in Glasgow | Wellcome Sanger Institute for the COVID-19 Genomics UK (COG-UK) Consortium | Harper VanSteenhouse, Yumi Kasai, David Gray, Carol Clugston, Anna Dominiczak and Alex Alderton, Roberto Amato, Jeffrey Barrett, Sonia Goncalves, Ewan Harrison, David K. Jackson, Ian Johnston, Dominic Kwiatkowski, Cordelia Langford, John Sillitoe on behalf of the Wellcome Sanger Institute COVID-19 Surveillance Team |
| EPI_ISL_2457196, EPI_ISL_2457197, EPI_ISL_2457198 | Lighthouse Lab in Alderley Park | Wellcome Sanger Institute for the COVID-19 Genomics UK (COG-UK) Consortium | Jacquelyn Wynn, Mairead Hyland, The Lighthouse Lab in Alderley Park and Alex Alderton, Roberto Amato, Jeffrey Barrett, Sonia Goncalves, Ewan Harrison, David K. Jackson, Ian Johnston, Dominic Kwiatkowski, Cordelia Langford, John Sillitoe on behalf of the Wellcome Sanger Institute COVID-19 Surveillance Team |
| EPI_ISL_2457199 | Lighthouse Lab in Glasgow | Wellcome Sanger Institute for the COVID-19 Genomics UK (COG-UK) Consortium | Harper VanSteenhouse, Yumi Kasai, David Gray, Carol Clugston, Anna Dominiczak and Alex Alderton, Roberto Amato, Jeffrey Barrett, Sonia Goncalves, Ewan Harrison, David K. Jackson, Ian Johnston, Dominic Kwiatkowski, Cordelia Langford, John Sillitoe on behalf of the Wellcome Sanger Institute COVID-19 Surveillance Team |
| EPI_ISL_2457200, EPI_ISL_2457201 | Lighthouse Lab in Alderley Park | Wellcome Sanger Institute for the COVID-19 Genomics UK (COG-UK) Consortium | Jacquelyn Wynn, Mairead Hyland, The Lighthouse Lab in Alderley Park and Alex Alderton, Roberto Amato, Jeffrey Barrett, Sonia Goncalves, Ewan Harrison, David K. Jackson, Ian Johnston, Dominic Kwiatkowski, Cordelia Langford, John Sillitoe on behalf of the Wellcome Sanger Institute COVID-19 Surveillance Team |
| EPI_ISL_2457202, EPI_ISL_2457203, EPI_ISL_2457204, EPI_ISL_2457205 | Lighthouse Lab in Glasgow | Wellcome Sanger Institute for the COVID-19 Genomics UK (COG-UK) Consortium | Harper VanSteenhouse, Yumi Kasai, David Gray, Carol Clugston, Anna Dominiczak and Alex Alderton, Roberto Amato, Jeffrey Barrett, Sonia Goncalves, Ewan Harrison, David K. Jackson, Ian Johnston, Dominic Kwiatkowski, Cordelia Langford, John Sillitoe on behalf of the Wellcome Sanger Institute COVID-19 Surveillance Team |
| EPI_ISL_2457206 | Lighthouse Lab in Alderley Park | Wellcome Sanger Institute for the COVID-19 Genomics UK (COG-UK) Consortium | Jacquelyn Wynn, Mairead Hyland, The Lighthouse Lab in Alderley Park and Alex Alderton, Roberto Amato, Jeffrey Barrett, Sonia Goncalves, Ewan Harrison, David K. Jackson, Ian Johnston, Dominic Kwiatkowski, Cordelia Langford, John Sillitoe on behalf of the Wellcome Sanger Institute COVID-19 Surveillance Team |
| EPI_ISL_2457207 | Lighthouse Lab in Glasgow | Wellcome Sanger Institute for the COVID-19 Genomics UK (COG-UK) Consortium | Harper VanSteenhouse, Yumi Kasai, David Gray, Carol Clugston, Anna Dominiczak and Alex Alderton, Roberto Amato, Jeffrey Barrett, Sonia Goncalves, Ewan Harrison, David K. Jackson, Ian Johnston, Dominic Kwiatkowski, Cordelia Langford, John Sillitoe on behalf of the Wellcome Sanger Institute COVID-19 Surveillance Team |
| EPI_ISL_2457208, EPI_ISL_2457209 | Lighthouse Lab in Alderley Park | Wellcome Sanger Institute for the COVID-19 Genomics UK (COG-UK) Consortium | Jacquelyn Wynn, Mairead Hyland, The Lighthouse Lab in Alderley Park and Alex Alderton, Roberto Amato, Jeffrey Barrett, Sonia Goncalves, Ewan Harrison, David K. Jackson, Ian Johnston, Dominic Kwiatkowski, Cordelia Langford, John Sillitoe on behalf of the Wellcome Sanger Institute COVID-19 Surveillance Team |
| EPI_ISL_2457210 | Lighthouse Lab in Glasgow | Wellcome Sanger Institute for the COVID-19 Genomics UK (COG-UK) Consortium | Harper VanSteenhouse, Yumi Kasai, David Gray, Carol Clugston, Anna Dominiczak and Alex Alderton, Roberto Amato, Jeffrey Barrett, Sonia Goncalves, Ewan Harrison, David K. Jackson, Ian Johnston, Dominic Kwiatkowski, Cordelia Langford, John Sillitoe on behalf of the Wellcome Sanger Institute COVID-19 Surveillance Team |
| EPI_ISL_2457211, EPI_ISL_2457212, EPI_ISL_2457213, EPI_ISL_2457214, EPI_ISL_2457215 | Lighthouse Lab in Alderley Park | Wellcome Sanger Institute for the COVID-19 Genomics UK (COG-UK) Consortium | Jacquelyn Wynn, Mairead Hyland, The Lighthouse Lab in Alderley Park and Alex Alderton, Roberto Amato, Jeffrey Barrett, Sonia Goncalves, Ewan Harrison, David K. Jackson, Ian Johnston, Dominic Kwiatkowski, Cordelia Langford, John Sillitoe on behalf of the Wellcome Sanger Institute COVID-19 Surveillance Team |
| EPI_ISL_2457216 | Lighthouse Lab in Glasgow | Wellcome Sanger Institute for the COVID-19 Genomics UK (COG-UK) Consortium | Harper VanSteenhouse, Yumi Kasai, David Gray, Carol Clugston, Anna Dominiczak and Alex Alderton, Roberto Amato, Jeffrey Barrett, Sonia Goncalves, Ewan Harrison, David K. Jackson, Ian Johnston, Dominic Kwiatkowski, Cordelia Langford, John Sillitoe on behalf of the Wellcome Sanger Institute COVID-19 Surveillance Team |
| EPI_ISL_2457217, EPI_ISL_2457218 | Lighthouse Lab in Alderley Park | Wellcome Sanger Institute for the COVID-19 Genomics UK (COG-UK) Consortium | Jacquelyn Wynn, Mairead Hyland, The Lighthouse Lab in Alderley Park and Alex Alderton, Roberto Amato, Jeffrey Barrett, Sonia Goncalves, Ewan Harrison, David K. Jackson, Ian Johnston, Dominic Kwiatkowski, Cordelia Langford, John Sillitoe on behalf of the Wellcome Sanger Institute COVID-19 Surveillance Team |
| EPI_ISL_2457219 | Lighthouse Lab in Glasgow | Wellcome Sanger Institute for the COVID-19 Genomics UK (COG-UK) Consortium | Harper VanSteenhouse, Yumi Kasai, David Gray, Carol Clugston, Anna Dominiczak and Alex Alderton, Roberto Amato, Jeffrey Barrett, Sonia Goncalves, Ewan Harrison, David K. Jackson, Ian Johnston, Dominic Kwiatkowski, Cordelia Langford, John Sillitoe on behalf of the Wellcome Sanger Institute COVID-19 Surveillance Team |
| EPI_ISL_2457220 | Lighthouse Lab in Alderley Park | Wellcome Sanger Institute for the COVID-19 Genomics UK (COG-UK) Consortium | Jacquelyn Wynn, Mairead Hyland, The Lighthouse Lab in Alderley Park and Alex Alderton, Roberto Amato, Jeffrey Barrett, Sonia Goncalves, Ewan Harrison, David K. Jackson, Ian Johnston, Dominic Kwiatkowski, Cordelia Langford, John Sillitoe on behalf of the Wellcome Sanger Institute COVID-19 Surveillance Team |
| EPI_ISL_2457221 | Lighthouse Lab in Glasgow | Wellcome Sanger Institute for the COVID-19 Genomics UK (COG-UK) Consortium | Harper VanSteenhouse, Yumi Kasai, David Gray, Carol Clugston, Anna Dominiczak and Alex Alderton, Roberto Amato, Jeffrey Barrett, Sonia Goncalves, Ewan Harrison, David K. Jackson, Ian Johnston, Dominic Kwiatkowski, Cordelia Langford, John Sillitoe on behalf of the Wellcome Sanger Institute COVID-19 Surveillance Team |
| EPI_ISL_2457223 | Lighthouse Lab in Alderley Park | Wellcome Sanger Institute for the COVID-19 Genomics UK (COG-UK) Consortium | Jacquelyn Wynn, Mairead Hyland, The Lighthouse Lab in Alderley Park and Alex Alderton, Roberto Amato, Jeffrey Barrett, Sonia Goncalves, Ewan Harrison, David K. Jackson, Ian Johnston, Dominic Kwiatkowski, Cordelia Langford, John Sillitoe on behalf of the Wellcome Sanger Institute COVID-19 Surveillance Team |
| EPI_ISL_2457224 | Lighthouse Lab in Glasgow | Wellcome Sanger Institute for the COVID-19 Genomics UK (COG-UK) Consortium | Harper VanSteenhouse, Yumi Kasai, David Gray, Carol Clugston, Anna Dominiczak and Alex Alderton, Roberto Amato, Jeffrey Barrett, Sonia Goncalves, Ewan Harrison, David K. Jackson, Ian Johnston, Dominic Kwiatkowski, Cordelia Langford, John Sillitoe on behalf of the Wellcome Sanger Institute COVID-19 Surveillance Team |
| EPI_ISL_2457225, EPI_ISL_2457226 | Lighthouse Lab in Alderley Park | Wellcome Sanger Institute for the COVID-19 Genomics UK (COG-UK) Consortium | Jacquelyn Wynn, Mairead Hyland, The Lighthouse Lab in Alderley Park and Alex Alderton, Roberto Amato, Jeffrey Barrett, Sonia Goncalves, Ewan Harrison, David K. Jackson, Ian Johnston, Dominic Kwiatkowski, Cordelia Langford, John Sillitoe on behalf of the Wellcome Sanger Institute COVID-19 Surveillance Team |
| EPI_ISL_2457227, EPI_ISL_2457228, EPI_ISL_2457229 | Lighthouse Lab in Glasgow | Wellcome Sanger Institute for the COVID-19 Genomics UK (COG-UK) Consortium | Harper VanSteenhouse, Yumi Kasai, David Gray, Carol Clugston, Anna Dominiczak and Alex Alderton, Roberto Amato, Jeffrey Barrett, Sonia Goncalves, Ewan Harrison, David K. Jackson, Ian Johnston, Dominic Kwiatkowski, Cordelia Langford, John Sillitoe on behalf of the Wellcome Sanger Institute COVID-19 Surveillance Team |
| EPI_ISL_2457230 | Lighthouse Lab in Alderley Park | Wellcome Sanger Institute for the COVID-19 Genomics UK (COG-UK) Consortium | Jacquelyn Wynn, Mairead Hyland, The Lighthouse Lab in Alderley Park and Alex Alderton, Roberto Amato, Jeffrey Barrett, Sonia Goncalves, Ewan Harrison, David K. Jackson, Ian Johnston, Dominic Kwiatkowski, Cordelia Langford, John Sillitoe on behalf of the Wellcome Sanger Institute COVID-19 Surveillance Team |
| EPI_ISL_2457231 | Lighthouse Lab in Glasgow | Wellcome Sanger Institute for the COVID-19 Genomics UK (COG-UK) Consortium | Harper VanSteenhouse, Yumi Kasai, David Gray, Carol Clugston, Anna Dominiczak and Alex Alderton, Roberto Amato, Jeffrey Barrett, Sonia Goncalves, Ewan Harrison, David K. Jackson, Ian Johnston, Dominic Kwiatkowski, Cordelia Langford, John Sillitoe on behalf of the Wellcome Sanger Institute COVID-19 Surveillance Team |
| EPI_ISL_2457232, EPI_ISL_2457233, EPI_ISL_2457234 | Lighthouse Lab in Alderley Park | Wellcome Sanger Institute for the COVID-19 Genomics UK (COG-UK) Consortium | Jacquelyn Wynn, Mairead Hyland, The Lighthouse Lab in Alderley Park and Alex Alderton, Roberto Amato, Jeffrey Barrett, Sonia Goncalves, Ewan Harrison, David K. Jackson, Ian Johnston, Dominic Kwiatkowski, Cordelia Langford, John Sillitoe on behalf of the Wellcome Sanger Institute COVID-19 Surveillance Team |
| EPI_ISL_2457235, EPI_ISL_2457236, EPI_ISL_2457237 | Lighthouse Lab in Glasgow | Wellcome Sanger Institute for the COVID-19 Genomics UK (COG-UK) Consortium | Harper VanSteenhouse, Yumi Kasai, David Gray, Carol Clugston, Anna Dominiczak and Alex Alderton, Roberto Amato, Jeffrey Barrett, Sonia Goncalves, Ewan Harrison, David K. Jackson, Ian Johnston, Dominic Kwiatkowski, Cordelia Langford, John Sillitoe on behalf of the Wellcome Sanger Institute COVID-19 Surveillance Team |
| EPI_ISL_2457238, EPI_ISL_2457239 | Lighthouse Lab in Alderley Park | Wellcome Sanger Institute for the COVID-19 Genomics UK (COG-UK) Consortium | Jacquelyn Wynn, Mairead Hyland, The Lighthouse Lab in Alderley Park and Alex Alderton, Roberto Amato, Jeffrey Barrett, Sonia Goncalves, Ewan Harrison, David K. Jackson, Ian Johnston, Dominic Kwiatkowski, Cordelia Langford, John Sillitoe on behalf of the Wellcome Sanger Institute COVID-19 Surveillance Team |
| EPI_ISL_2457240, EPI_ISL_2457241 | Lighthouse Lab in Glasgow | Wellcome Sanger Institute for the COVID-19 Genomics UK (COG-UK) Consortium | Harper VanSteenhouse, Yumi Kasai, David Gray, Carol Clugston, Anna Dominiczak and Alex Alderton, Roberto Amato, Jeffrey Barrett, Sonia Goncalves, Ewan Harrison, David K. Jackson, Ian Johnston, Dominic Kwiatkowski, Cordelia Langford, John Sillitoe on behalf of the Wellcome Sanger Institute COVID-19 Surveillance Team |
| EPI_ISL_2457242, EPI_ISL_2457243, EPI_ISL_2457245 | Lighthouse Lab in Alderley Park | Wellcome Sanger Institute for the COVID-19 Genomics UK (COG-UK) Consortium | Jacquelyn Wynn, Mairead Hyland, The Lighthouse Lab in Alderley Park and Alex Alderton, Roberto Amato, Jeffrey Barrett, Sonia Goncalves, Ewan Harrison, David K. Jackson, Ian Johnston, Dominic Kwiatkowski, Cordelia Langford, John Sillitoe on behalf of the Wellcome Sanger Institute COVID-19 Surveillance Team |
| EPI_ISL_2457246 | Lighthouse Lab in Glasgow | Wellcome Sanger Institute for the COVID-19 Genomics UK (COG-UK) Consortium | Harper VanSteenhouse, Yumi Kasai, David Gray, Carol Clugston, Anna Dominiczak and Alex Alderton, Roberto Amato, Jeffrey Barrett, Sonia Goncalves, Ewan Harrison, David K. Jackson, Ian Johnston, Dominic Kwiatkowski, Cordelia Langford, John Sillitoe on behalf of the Wellcome Sanger Institute COVID-19 Surveillance Team |
| EPI_ISL_2457247 | Lighthouse Lab in Alderley Park | Wellcome Sanger Institute for the COVID-19 Genomics UK (COG-UK) Consortium | Jacquelyn Wynn, Mairead Hyland, The Lighthouse Lab in Alderley Park and Alex Alderton, Roberto Amato, Jeffrey Barrett, Sonia Goncalves, Ewan Harrison, David K. Jackson, Ian Johnston, Dominic Kwiatkowski, Cordelia Langford, John Sillitoe on behalf of the Wellcome Sanger Institute COVID-19 Surveillance Team |
| EPI_ISL_2457248 | Lighthouse Lab in Glasgow | Wellcome Sanger Institute for the COVID-19 Genomics UK (COG-UK) Consortium | Harper VanSteenhouse, Yumi Kasai, David Gray, Carol Clugston, Anna Dominiczak and Alex Alderton, Roberto Amato, Jeffrey Barrett, Sonia Goncalves, Ewan Harrison, David K. Jackson, Ian Johnston, Dominic Kwiatkowski, Cordelia Langford, John Sillitoe on behalf of the Wellcome Sanger Institute COVID-19 Surveillance Team |
| EPI_ISL_2457249 | Lighthouse Lab in Alderley Park | Wellcome Sanger Institute for the COVID-19 Genomics UK (COG-UK) Consortium | Jacquelyn Wynn, Mairead Hyland, The Lighthouse Lab in Alderley Park and Alex Alderton, Roberto Amato, Jeffrey Barrett, Sonia Goncalves, Ewan Harrison, David K. Jackson, Ian Johnston, Dominic Kwiatkowski, Cordelia Langford, John Sillitoe on behalf of the Wellcome Sanger Institute COVID-19 Surveillance Team |
| EPI_ISL_2457250, EPI_ISL_2457251 | Lighthouse Lab in Glasgow | Wellcome Sanger Institute for the COVID-19 Genomics UK (COG-UK) Consortium | Harper VanSteenhouse, Yumi Kasai, David Gray, Carol Clugston, Anna Dominiczak and Alex Alderton, Roberto Amato, Jeffrey Barrett, Sonia Goncalves, Ewan Harrison, David K. Jackson, Ian Johnston, Dominic Kwiatkowski, Cordelia Langford, John Sillitoe on behalf of the Wellcome Sanger Institute COVID-19 Surveillance Team |
| EPI_ISL_2457252, EPI_ISL_2457253, EPI_ISL_2457254, EPI_ISL_2457255, EPI_ISL_2457256 | Lighthouse Lab in Alderley Park | Wellcome Sanger Institute for the COVID-19 Genomics UK (COG-UK) Consortium | Jacquelyn Wynn, Mairead Hyland, The Lighthouse Lab in Alderley Park and Alex Alderton, Roberto Amato, Jeffrey Barrett, Sonia Goncalves, Ewan Harrison, David K. Jackson, Ian Johnston, Dominic Kwiatkowski, Cordelia Langford, John Sillitoe on behalf of the Wellcome Sanger Institute COVID-19 Surveillance Team |
| EPI_ISL_2457257 | Lighthouse Lab in Glasgow | Wellcome Sanger Institute for the COVID-19 Genomics UK (COG-UK) Consortium | Harper VanSteenhouse, Yumi Kasai, David Gray, Carol Clugston, Anna Dominiczak and Alex Alderton, Roberto Amato, Jeffrey Barrett, Sonia Goncalves, Ewan Harrison, David K. Jackson, Ian Johnston, Dominic Kwiatkowski, Cordelia Langford, John Sillitoe on behalf of the Wellcome Sanger Institute COVID-19 Surveillance Team |
| EPI_ISL_2457258 | Lighthouse Lab in Alderley Park | Wellcome Sanger Institute for the COVID-19 Genomics UK (COG-UK) Consortium | Jacquelyn Wynn, Mairead Hyland, The Lighthouse Lab in Alderley Park and Alex Alderton, Roberto Amato, Jeffrey Barrett, Sonia Goncalves, Ewan Harrison, David K. Jackson, Ian Johnston, Dominic Kwiatkowski, Cordelia Langford, John Sillitoe on behalf of the Wellcome Sanger Institute COVID-19 Surveillance Team |

| | | | |
|---|---|---|---|
| EPI_ISL_2457259 | Lighthouse Lab in Glasgow | Wellcome Sanger Institute for the COVID-19 Genomics UK (COG-UK) Consortium | Harper VanSteenhouse, Yumi Kasai, David Gray, Carol Clugston, Anna Dominiczak and Alex Alderton, Roberto Amato, Jeffrey Barrett, Sonia Goncalves, Ewan Harrison, David K. Jackson, Ian Johnston, Dominic Kwiatkowski, Cordelia Langford, John Sillitoe on behalf of the Wellcome Sanger Institute COVID-19 Surveillance Team |
| EPI_ISL_2457260, EPI_ISL_2457261 | Lighthouse Lab in Alderley Park | Wellcome Sanger Institute for the COVID-19 Genomics UK (COG-UK) Consortium | Jacquelyn Wynn, Mairead Hyland, The Lighthouse Lab in Alderley Park and Alex Alderton, Roberto Amato, Jeffrey Barrett, Sonia Goncalves, Ewan Harrison, David K. Jackson, Ian Johnston, Dominic Kwiatkowski, Cordelia Langford, John Sillitoe on behalf of the Wellcome Sanger Institute COVID-19 Surveillance Team |
| EPI_ISL_2457262 | Lighthouse Lab in Glasgow | Wellcome Sanger Institute for the COVID-19 Genomics UK (COG-UK) Consortium | Harper VanSteenhouse, Yumi Kasai, David Gray, Carol Clugston, Anna Dominiczak and Alex Alderton, Roberto Amato, Jeffrey Barrett, Sonia Goncalves, Ewan Harrison, David K. Jackson, Ian Johnston, Dominic Kwiatkowski, Cordelia Langford, John Sillitoe on behalf of the Wellcome Sanger Institute COVID-19 Surveillance Team |
| EPI_ISL_2457263 | Lighthouse Lab in Alderley Park | Wellcome Sanger Institute for the COVID-19 Genomics UK (COG-UK) Consortium | Jacquelyn Wynn, Mairead Hyland, The Lighthouse Lab in Alderley Park and Alex Alderton, Roberto Amato, Jeffrey Barrett, Sonia Goncalves, Ewan Harrison, David K. Jackson, Ian Johnston, Dominic Kwiatkowski, Cordelia Langford, John Sillitoe on behalf of the Wellcome Sanger Institute COVID-19 Surveillance Team |
| EPI_ISL_2457264 | Lighthouse Lab in Glasgow | Wellcome Sanger Institute for the COVID-19 Genomics UK (COG-UK) Consortium | Harper VanSteenhouse, Yumi Kasai, David Gray, Carol Clugston, Anna Dominiczak and Alex Alderton, Roberto Amato, Jeffrey Barrett, Sonia Goncalves, Ewan Harrison, David K. Jackson, Ian Johnston, Dominic Kwiatkowski, Cordelia Langford, John Sillitoe on behalf of the Wellcome Sanger Institute COVID-19 Surveillance Team |
| EPI_ISL_2457265 | Lighthouse Lab in Alderley Park | Wellcome Sanger Institute for the COVID-19 Genomics UK (COG-UK) Consortium | Jacquelyn Wynn, Mairead Hyland, The Lighthouse Lab in Alderley Park and Alex Alderton, Roberto Amato, Jeffrey Barrett, Sonia Goncalves, Ewan Harrison, David K. Jackson, Ian Johnston, Dominic Kwiatkowski, Cordelia Langford, John Sillitoe on behalf of the Wellcome Sanger Institute COVID-19 Surveillance Team |
| EPI_ISL_2457266 | Lighthouse Lab in Glasgow | Wellcome Sanger Institute for the COVID-19 Genomics UK (COG-UK) Consortium | Harper VanSteenhouse, Yumi Kasai, David Gray, Carol Clugston, Anna Dominiczak and Alex Alderton, Roberto Amato, Jeffrey Barrett, Sonia Goncalves, Ewan Harrison, David K. Jackson, Ian Johnston, Dominic Kwiatkowski, Cordelia Langford, John Sillitoe on behalf of the Wellcome Sanger Institute COVID-19 Surveillance Team |
| EPI_ISL_2457267, EPI_ISL_2457268 | Lighthouse Lab in Alderley Park | Wellcome Sanger Institute for the COVID-19 Genomics UK (COG-UK) Consortium | Jacquelyn Wynn, Mairead Hyland, The Lighthouse Lab in Alderley Park and Alex Alderton, Roberto Amato, Jeffrey Barrett, Sonia Goncalves, Ewan Harrison, David K. Jackson, Ian Johnston, Dominic Kwiatkowski, Cordelia Langford, John Sillitoe on behalf of the Wellcome Sanger Institute COVID-19 Surveillance Team |
| EPI_ISL_2457269 | Lighthouse Lab in Glasgow | Wellcome Sanger Institute for the COVID-19 Genomics UK (COG-UK) Consortium | Harper VanSteenhouse, Yumi Kasai, David Gray, Carol Clugston, Anna Dominiczak and Alex Alderton, Roberto Amato, Jeffrey Barrett, Sonia Goncalves, Ewan Harrison, David K. Jackson, Ian Johnston, Dominic Kwiatkowski, Cordelia Langford, John Sillitoe on behalf of the Wellcome Sanger Institute COVID-19 Surveillance Team |
| EPI_ISL_2457270 | Lighthouse Lab in Alderley Park | Wellcome Sanger Institute for the COVID-19 Genomics UK (COG-UK) Consortium | Jacquelyn Wynn, Mairead Hyland, The Lighthouse Lab in Alderley Park and Alex Alderton, Roberto Amato, Jeffrey Barrett, Sonia Goncalves, Ewan Harrison, David K. Jackson, Ian Johnston, Dominic Kwiatkowski, Cordelia Langford, John Sillitoe on behalf of the Wellcome Sanger Institute COVID-19 Surveillance Team |
| EPI_ISL_2457271, EPI_ISL_2457272, EPI_ISL_2457273 | Lighthouse Lab in Glasgow | Wellcome Sanger Institute for the COVID-19 Genomics UK (COG-UK) Consortium | Harper VanSteenhouse, Yumi Kasai, David Gray, Carol Clugston, Anna Dominiczak and Alex Alderton, Roberto Amato, Jeffrey Barrett, Sonia Goncalves, Ewan Harrison, David K. Jackson, Ian Johnston, Dominic Kwiatkowski, Cordelia Langford, John Sillitoe on behalf of the Wellcome Sanger Institute COVID-19 Surveillance Team |
| EPI_ISL_2457274, EPI_ISL_2457275, EPI_ISL_2457276, EPI_ISL_2457277 | Lighthouse Lab in Alderley Park | Wellcome Sanger Institute for the COVID-19 Genomics UK (COG-UK) Consortium | Jacquelyn Wynn, Mairead Hyland, The Lighthouse Lab in Alderley Park and Alex Alderton, Roberto Amato, Jeffrey Barrett, Sonia Goncalves, Ewan Harrison, David K. Jackson, Ian Johnston, Dominic Kwiatkowski, Cordelia Langford, John Sillitoe on behalf of the Wellcome Sanger Institute COVID-19 Surveillance Team |
| EPI_ISL_2457278 | Lighthouse Lab in Glasgow | Wellcome Sanger Institute for the COVID-19 Genomics UK (COG-UK) Consortium | Harper VanSteenhouse, Yumi Kasai, David Gray, Carol Clugston, Anna Dominiczak and Alex Alderton, Roberto Amato, Jeffrey Barrett, Sonia Goncalves, Ewan Harrison, David K. Jackson, Ian Johnston, Dominic Kwiatkowski, Cordelia Langford, John Sillitoe on behalf of the Wellcome Sanger Institute COVID-19 Surveillance Team |
| EPI_ISL_2457279 | Lighthouse Lab in Alderley Park | Wellcome Sanger Institute for the COVID-19 Genomics UK (COG-UK) Consortium | Jacquelyn Wynn, Mairead Hyland, The Lighthouse Lab in Alderley Park and Alex Alderton, Roberto Amato, Jeffrey Barrett, Sonia Goncalves, Ewan Harrison, David K. Jackson, Ian Johnston, Dominic Kwiatkowski, Cordelia Langford, John Sillitoe on behalf of the Wellcome Sanger Institute COVID-19 Surveillance Team |
| EPI_ISL_2457280, EPI_ISL_2457281 | Lighthouse Lab in Glasgow | Wellcome Sanger Institute for the COVID-19 Genomics UK (COG-UK) Consortium | Harper VanSteenhouse, Yumi Kasai, David Gray, Carol Clugston, Anna Dominiczak and Alex Alderton, Roberto Amato, Jeffrey Barrett, Sonia Goncalves, Ewan Harrison, David K. Jackson, Ian Johnston, Dominic Kwiatkowski, Cordelia Langford, John Sillitoe on behalf of the Wellcome Sanger Institute COVID-19 Surveillance Team |
| EPI_ISL_2457282, EPI_ISL_2457283 | Lighthouse Lab in Alderley Park | Wellcome Sanger Institute for the COVID-19 Genomics UK (COG-UK) Consortium | Jacquelyn Wynn, Mairead Hyland, The Lighthouse Lab in Alderley Park and Alex Alderton, Roberto Amato, Jeffrey Barrett, Sonia Goncalves, Ewan Harrison, David K. Jackson, Ian Johnston, Dominic Kwiatkowski, Cordelia Langford, John Sillitoe on behalf of the Wellcome Sanger Institute COVID-19 Surveillance Team |
| EPI_ISL_2457284 | Lighthouse Lab in Glasgow | Wellcome Sanger Institute for the COVID-19 Genomics UK (COG-UK) Consortium | Harper VanSteenhouse, Yumi Kasai, David Gray, Carol Clugston, Anna Dominiczak and Alex Alderton, Roberto Amato, Jeffrey Barrett, Sonia Goncalves, Ewan Harrison, David K. Jackson, Ian Johnston, Dominic Kwiatkowski, Cordelia Langford, John Sillitoe on behalf of the Wellcome Sanger Institute COVID-19 Surveillance Team |
| EPI_ISL_2457285, EPI_ISL_2457287 | Lighthouse Lab in Alderley Park | Wellcome Sanger Institute for the COVID-19 Genomics UK (COG-UK) Consortium | Jacquelyn Wynn, Mairead Hyland, The Lighthouse Lab in Alderley Park and Alex Alderton, Roberto Amato, Jeffrey Barrett, Sonia Goncalves, Ewan Harrison, David K. Jackson, Ian Johnston, Dominic Kwiatkowski, Cordelia Langford, John Sillitoe on behalf of the Wellcome Sanger Institute COVID-19 Surveillance Team |
| EPI_ISL_2457288 | Lighthouse Lab in Glasgow | Wellcome Sanger Institute for the COVID-19 Genomics UK (COG-UK) Consortium | Harper VanSteenhouse, Yumi Kasai, David Gray, Carol Clugston, Anna Dominiczak and Alex Alderton, Roberto Amato, Jeffrey Barrett, Sonia Goncalves, Ewan Harrison, David K. Jackson, Ian Johnston, Dominic Kwiatkowski, Cordelia Langford, John Sillitoe on behalf of the Wellcome Sanger Institute COVID-19 Surveillance Team |
| EPI_ISL_2457289, EPI_ISL_2457290 | Lighthouse Lab in Alderley Park | Wellcome Sanger Institute for the COVID-19 Genomics UK (COG-UK) Consortium | Jacquelyn Wynn, Mairead Hyland, The Lighthouse Lab in Alderley Park and Alex Alderton, Roberto Amato, Jeffrey Barrett, Sonia Goncalves, Ewan Harrison, David K. Jackson, Ian Johnston, Dominic Kwiatkowski, Cordelia Langford, John Sillitoe on behalf of the Wellcome Sanger Institute COVID-19 Surveillance Team |
| EPI_ISL_2457291, EPI_ISL_2457292 | Lighthouse Lab in Glasgow | Wellcome Sanger Institute for the COVID-19 Genomics UK (COG-UK) Consortium | Harper VanSteenhouse, Yumi Kasai, David Gray, Carol Clugston, Anna Dominiczak and Alex Alderton, Roberto Amato, Jeffrey Barrett, Sonia Goncalves, Ewan Harrison, David K. Jackson, Ian Johnston, Dominic Kwiatkowski, Cordelia Langford, John Sillitoe on behalf of the Wellcome Sanger Institute COVID-19 Surveillance Team |
| EPI_ISL_2457293, EPI_ISL_2457294 | Lighthouse Lab in Alderley Park | Wellcome Sanger Institute for the COVID-19 Genomics UK (COG-UK) Consortium | Jacquelyn Wynn, Mairead Hyland, The Lighthouse Lab in Alderley Park and Alex Alderton, Roberto Amato, Jeffrey Barrett, Sonia Goncalves, Ewan Harrison, David K. Jackson, Ian Johnston, Dominic Kwiatkowski, Cordelia Langford, John Sillitoe on behalf of the Wellcome Sanger Institute COVID-19 Surveillance Team |
| EPI_ISL_2457295, EPI_ISL_2457296, EPI_ISL_2457297, EPI_ISL_2457298, EPI_ISL_2457299 | Lighthouse Lab in Glasgow | Wellcome Sanger Institute for the COVID-19 Genomics UK (COG-UK) Consortium | Harper VanSteenhouse, Yumi Kasai, David Gray, Carol Clugston, Anna Dominiczak and Alex Alderton, Roberto Amato, Jeffrey Barrett, Sonia Goncalves, Ewan Harrison, David K. Jackson, Ian Johnston, Dominic Kwiatkowski, Cordelia Langford, John Sillitoe on behalf of the Wellcome Sanger Institute COVID-19 Surveillance Team |
| EPI_ISL_2457300, EPI_ISL_2457301, EPI_ISL_2457302 | Lighthouse Lab in Alderley Park | Wellcome Sanger Institute for the COVID-19 Genomics UK (COG-UK) Consortium | Jacquelyn Wynn, Mairead Hyland, The Lighthouse Lab in Alderley Park and Alex Alderton, Roberto Amato, Jeffrey Barrett, Sonia Goncalves, Ewan Harrison, David K. Jackson, Ian Johnston, Dominic Kwiatkowski, Cordelia Langford, John Sillitoe on behalf of the Wellcome Sanger Institute COVID-19 Surveillance Team |
| EPI_ISL_2457303 | Lighthouse Lab in Glasgow | Wellcome Sanger Institute for the COVID-19 Genomics UK (COG-UK) Consortium | Harper VanSteenhouse, Yumi Kasai, David Gray, Carol Clugston, Anna Dominiczak and Alex Alderton, Roberto Amato, Jeffrey Barrett, Sonia Goncalves, Ewan Harrison, David K. Jackson, Ian Johnston, Dominic Kwiatkowski, Cordelia Langford, John Sillitoe on behalf of the Wellcome Sanger Institute COVID-19 Surveillance Team |
| EPI_ISL_2457304 | Lighthouse Lab in Alderley Park | Wellcome Sanger Institute for the COVID-19 Genomics UK (COG-UK) Consortium | Jacquelyn Wynn, Mairead Hyland, The Lighthouse Lab in Alderley Park and Alex Alderton, Roberto Amato, Jeffrey Barrett, Sonia Goncalves, Ewan Harrison, David K. Jackson, Ian Johnston, Dominic Kwiatkowski, Cordelia Langford, John Sillitoe on behalf of the Wellcome Sanger Institute COVID-19 Surveillance Team |
| EPI_ISL_2457305 | Lighthouse Lab in Glasgow | Wellcome Sanger Institute for the COVID-19 Genomics UK (COG-UK) Consortium | Harper VanSteenhouse, Yumi Kasai, David Gray, Carol Clugston, Anna Dominiczak and Alex Alderton, Roberto Amato, Jeffrey Barrett, Sonia Goncalves, Ewan Harrison, David K. Jackson, Ian Johnston, Dominic Kwiatkowski, Cordelia Langford, John Sillitoe on behalf of the Wellcome Sanger Institute COVID-19 Surveillance Team |
| EPI_ISL_2457306 | Lighthouse Lab in Alderley Park | Wellcome Sanger Institute for the COVID-19 Genomics UK (COG-UK) Consortium | Jacquelyn Wynn, Mairead Hyland, The Lighthouse Lab in Alderley Park and Alex Alderton, Roberto Amato, Jeffrey Barrett, Sonia Goncalves, Ewan Harrison, David K. Jackson, Ian Johnston, Dominic Kwiatkowski, Cordelia Langford, John Sillitoe on behalf of the Wellcome Sanger Institute COVID-19 Surveillance Team |
| EPI_ISL_2457309, EPI_ISL_2457310 | Lighthouse Lab in Glasgow | Wellcome Sanger Institute for the COVID-19 Genomics UK (COG-UK) Consortium | Harper VanSteenhouse, Yumi Kasai, David Gray, Carol Clugston, Anna Dominiczak and Alex Alderton, Roberto Amato, Jeffrey Barrett, Sonia Goncalves, Ewan Harrison, David K. Jackson, Ian Johnston, Dominic Kwiatkowski, Cordelia Langford, John Sillitoe on behalf of the Wellcome Sanger Institute COVID-19 Surveillance Team |
| EPI_ISL_2457311 | Lighthouse Lab in Alderley Park | Wellcome Sanger Institute for the COVID-19 Genomics UK (COG-UK) Consortium | Jacquelyn Wynn, Mairead Hyland, The Lighthouse Lab in Alderley Park and Alex Alderton, Roberto Amato, Jeffrey Barrett, Sonia Goncalves, Ewan Harrison, David K. Jackson, Ian Johnston, Dominic Kwiatkowski, Cordelia Langford, John Sillitoe on behalf of the Wellcome Sanger Institute COVID-19 Surveillance Team |
| EPI_ISL_2457312 | Lighthouse Lab in Glasgow | Wellcome Sanger Institute for the COVID-19 Genomics UK (COG-UK) Consortium | Harper VanSteenhouse, Yumi Kasai, David Gray, Carol Clugston, Anna Dominiczak and Alex Alderton, Roberto Amato, Jeffrey Barrett, Sonia Goncalves, Ewan Harrison, David K. Jackson, Ian Johnston, Dominic Kwiatkowski, Cordelia Langford, John Sillitoe on behalf of the Wellcome Sanger Institute COVID-19 Surveillance Team |
| EPI_ISL_2457313, EPI_ISL_2457314, EPI_ISL_2457315 | Lighthouse Lab in Alderley Park | Wellcome Sanger Institute for the COVID-19 Genomics UK (COG-UK) Consortium | Jacquelyn Wynn, Mairead Hyland, The Lighthouse Lab in Alderley Park and Alex Alderton, Roberto Amato, Jeffrey Barrett, Sonia Goncalves, Ewan Harrison, David K. Jackson, Ian Johnston, Dominic Kwiatkowski, Cordelia Langford, John Sillitoe on behalf of the Wellcome Sanger Institute COVID-19 Surveillance Team |
| EPI_ISL_2457316 | Lighthouse Lab in Glasgow | Wellcome Sanger Institute for the COVID-19 Genomics UK (COG-UK) Consortium | Harper VanSteenhouse, Yumi Kasai, David Gray, Carol Clugston, Anna Dominiczak and Alex Alderton, Roberto Amato, Jeffrey Barrett, Sonia Goncalves, Ewan Harrison, David K. Jackson, Ian Johnston, Dominic Kwiatkowski, Cordelia Langford, John Sillitoe on behalf of the Wellcome Sanger Institute COVID-19 Surveillance Team |
| EPI_ISL_2457318 | Lighthouse Lab in Alderley Park | Wellcome Sanger Institute for the COVID-19 Genomics UK (COG-UK) Consortium | Jacquelyn Wynn, Mairead Hyland, The Lighthouse Lab in Alderley Park and Alex Alderton, Roberto Amato, Jeffrey Barrett, Sonia Goncalves, Ewan Harrison, David K. Jackson, Ian Johnston, Dominic Kwiatkowski, Cordelia Langford, John Sillitoe on behalf of the Wellcome Sanger Institute COVID-19 Surveillance Team |
| EPI_ISL_2457319, EPI_ISL_2457321 | Lighthouse Lab in Glasgow | Wellcome Sanger Institute for the COVID-19 Genomics UK (COG-UK) Consortium | Harper VanSteenhouse, Yumi Kasai, David Gray, Carol Clugston, Anna Dominiczak and Alex Alderton, Roberto Amato, Jeffrey Barrett, Sonia Goncalves, Ewan Harrison, David K. Jackson, Ian Johnston, Dominic Kwiatkowski, Cordelia Langford, John Sillitoe on behalf of the Wellcome Sanger Institute COVID-19 Surveillance Team |
| EPI_ISL_2457322, EPI_ISL_2457324 | Lighthouse Lab in Alderley Park | Wellcome Sanger Institute for the COVID-19 Genomics UK (COG-UK) Consortium | Jacquelyn Wynn, Mairead Hyland, The Lighthouse Lab in Alderley Park and Alex Alderton, Roberto Amato, Jeffrey Barrett, Sonia Goncalves, Ewan Harrison, David K. Jackson, Ian Johnston, Dominic Kwiatkowski, Cordelia Langford, John Sillitoe on behalf of the Wellcome Sanger Institute COVID-19 Surveillance Team |
| EPI_ISL_2457325 | Lighthouse Lab in Glasgow | Wellcome Sanger Institute for the COVID-19 Genomics UK (COG-UK) Consortium | Harper VanSteenhouse, Yumi Kasai, David Gray, Carol Clugston, Anna Dominiczak and Alex Alderton, Roberto Amato, Jeffrey Barrett, Sonia Goncalves, Ewan Harrison, David K. Jackson, Ian Johnston, Dominic Kwiatkowski, Cordelia Langford, John Sillitoe on behalf of the Wellcome Sanger Institute COVID-19 Surveillance Team |
| EPI_ISL_2457327 | Lighthouse Lab in Alderley Park | Wellcome Sanger Institute for the COVID-19 Genomics UK (COG-UK) Consortium | Jacquelyn Wynn, Mairead Hyland, The Lighthouse Lab in Alderley Park and Alex Alderton, Roberto Amato, Jeffrey Barrett, Sonia Goncalves, Ewan Harrison, David K. Jackson, Ian Johnston, Dominic Kwiatkowski, Cordelia Langford, John Sillitoe on behalf of the Wellcome Sanger Institute COVID-19 Surveillance Team |
| EPI_ISL_2457328, EPI_ISL_2457329, EPI_ISL_2457330 | Lighthouse Lab in Glasgow | Wellcome Sanger Institute for the COVID-19 Genomics UK (COG-UK) Consortium | Harper VanSteenhouse, Yumi Kasai, David Gray, Carol Clugston, Anna Dominiczak and Alex Alderton, Roberto Amato, Jeffrey Barrett, Sonia Goncalves, Ewan Harrison, David K. Jackson, Ian Johnston, Dominic Kwiatkowski, Cordelia Langford, John Sillitoe on behalf of the Wellcome Sanger Institute COVID-19 Surveillance Team |
| EPI_ISL_2457331, EPI_ISL_2457332 | Lighthouse Lab in Alderley Park | Wellcome Sanger Institute for the COVID-19 Genomics UK (COG-UK) Consortium | Jacquelyn Wynn, Mairead Hyland, The Lighthouse Lab in Alderley Park and Alex Alderton, Roberto Amato, Jeffrey Barrett, Sonia Goncalves, Ewan Harrison, David K. Jackson, Ian Johnston, Dominic Kwiatkowski, Cordelia Langford, John Sillitoe on behalf of the Wellcome Sanger Institute COVID-19 Surveillance Team |
| EPI_ISL_2457333 | Lighthouse Lab in Glasgow | Wellcome Sanger Institute for the COVID-19 Genomics UK (COG-UK) Consortium | Harper VanSteenhouse, Yumi Kasai, David Gray, Carol Clugston, Anna Dominiczak and Alex Alderton, Roberto Amato, Jeffrey Barrett, Sonia Goncalves, Ewan Harrison, David K. Jackson, Ian Johnston, Dominic Kwiatkowski, Cordelia Langford, John Sillitoe on behalf of the Wellcome Sanger Institute COVID-19 Surveillance Team |
| EPI_ISL_2457334 | Lighthouse Lab in Alderley Park | Wellcome Sanger Institute for the COVID-19 Genomics UK (COG-UK) | Jacquelyn Wynn, Mairead Hyland, The Lighthouse Lab in Alderley Park and Alex Alderton, Roberto Amato, Jeffrey Barrett, Sonia Goncalves, Ewan Harrison, David K. Jackson, Ian Johnston, Dominic Kwiatkowski, Cordelia Langford, John Sillitoe on |

| | | | |
|---|---|---|---|
| | | Consortium | behalf of the Wellcome Sanger Institute COVID-19 Surveillance Team |
| EPI_ISL_2457335 | Lighthouse Lab in Glasgow | Wellcome Sanger Institute for the COVID-19 Genomics UK (COG-UK) Consortium | Harper VanSteenhouse, Yumi Kasai, David Gray, Carol Clugston, Anna Dominiczak and Alex Alderton, Roberto Amato, Jeffrey Barrett, Sonia Goncalves, Ewan Harrison, David K. Jackson, Ian Johnston, Dominic Kwiatkowski, Cordelia Langford, John Sillitoe on behalf of the Wellcome Sanger Institute COVID-19 Surveillance Team |
| EPI_ISL_2457336, EPI_ISL_2457337, EPI_ISL_2457338 | Lighthouse Lab in Alderley Park | Wellcome Sanger Institute for the COVID-19 Genomics UK (COG-UK) Consortium | Jacquelyn Wynn, Mairead Hyland, The Lighthouse Lab in Alderley Park and Alex Alderton, Roberto Amato, Jeffrey Barrett, Sonia Goncalves, Ewan Harrison, David K. Jackson, Ian Johnston, Dominic Kwiatkowski, Cordelia Langford, John Sillitoe on behalf of the Wellcome Sanger Institute COVID-19 Surveillance Team |
| EPI_ISL_2457339, EPI_ISL_2457340, EPI_ISL_2457341, EPI_ISL_2457342, EPI_ISL_2457343 | Lighthouse Lab in Glasgow | Wellcome Sanger Institute for the COVID-19 Genomics UK (COG-UK) Consortium | Harper VanSteenhouse, Yumi Kasai, David Gray, Carol Clugston, Anna Dominiczak and Alex Alderton, Roberto Amato, Jeffrey Barrett, Sonia Goncalves, Ewan Harrison, David K. Jackson, Ian Johnston, Dominic Kwiatkowski, Cordelia Langford, John Sillitoe on behalf of the Wellcome Sanger Institute COVID-19 Surveillance Team |
| EPI_ISL_2457344, EPI_ISL_2457345 | Lighthouse Lab in Alderley Park | Wellcome Sanger Institute for the COVID-19 Genomics UK (COG-UK) Consortium | Jacquelyn Wynn, Mairead Hyland, The Lighthouse Lab in Alderley Park and Alex Alderton, Roberto Amato, Jeffrey Barrett, Sonia Goncalves, Ewan Harrison, David K. Jackson, Ian Johnston, Dominic Kwiatkowski, Cordelia Langford, John Sillitoe on behalf of the Wellcome Sanger Institute COVID-19 Surveillance Team |
| EPI_ISL_2457346 | Lighthouse Lab in Glasgow | Wellcome Sanger Institute for the COVID-19 Genomics UK (COG-UK) Consortium | Harper VanSteenhouse, Yumi Kasai, David Gray, Carol Clugston, Anna Dominiczak and Alex Alderton, Roberto Amato, Jeffrey Barrett, Sonia Goncalves, Ewan Harrison, David K. Jackson, Ian Johnston, Dominic Kwiatkowski, Cordelia Langford, John Sillitoe on behalf of the Wellcome Sanger Institute COVID-19 Surveillance Team |
| EPI_ISL_2457347 | Lighthouse Lab in Alderley Park | Wellcome Sanger Institute for the COVID-19 Genomics UK (COG-UK) Consortium | Jacquelyn Wynn, Mairead Hyland, The Lighthouse Lab in Alderley Park and Alex Alderton, Roberto Amato, Jeffrey Barrett, Sonia Goncalves, Ewan Harrison, David K. Jackson, Ian Johnston, Dominic Kwiatkowski, Cordelia Langford, John Sillitoe on behalf of the Wellcome Sanger Institute COVID-19 Surveillance Team |
| EPI_ISL_2457348, EPI_ISL_2457349 | Lighthouse Lab in Glasgow | Wellcome Sanger Institute for the COVID-19 Genomics UK (COG-UK) Consortium | Harper VanSteenhouse, Yumi Kasai, David Gray, Carol Clugston, Anna Dominiczak and Alex Alderton, Roberto Amato, Jeffrey Barrett, Sonia Goncalves, Ewan Harrison, David K. Jackson, Ian Johnston, Dominic Kwiatkowski, Cordelia Langford, John Sillitoe on behalf of the Wellcome Sanger Institute COVID-19 Surveillance Team |
| EPI_ISL_2457350 | Lighthouse Lab in Alderley Park | Wellcome Sanger Institute for the COVID-19 Genomics UK (COG-UK) Consortium | Jacquelyn Wynn, Mairead Hyland, The Lighthouse Lab in Alderley Park and Alex Alderton, Roberto Amato, Jeffrey Barrett, Sonia Goncalves, Ewan Harrison, David K. Jackson, Ian Johnston, Dominic Kwiatkowski, Cordelia Langford, John Sillitoe on behalf of the Wellcome Sanger Institute COVID-19 Surveillance Team |
| EPI_ISL_2457351, EPI_ISL_2457352 | Lighthouse Lab in Glasgow | Wellcome Sanger Institute for the COVID-19 Genomics UK (COG-UK) Consortium | Harper VanSteenhouse, Yumi Kasai, David Gray, Carol Clugston, Anna Dominiczak and Alex Alderton, Roberto Amato, Jeffrey Barrett, Sonia Goncalves, Ewan Harrison, David K. Jackson, Ian Johnston, Dominic Kwiatkowski, Cordelia Langford, John Sillitoe on behalf of the Wellcome Sanger Institute COVID-19 Surveillance Team |
| EPI_ISL_2457354, EPI_ISL_2457355, EPI_ISL_2457356, EPI_ISL_2457357, EPI_ISL_2457358 | Lighthouse Lab in Alderley Park | Wellcome Sanger Institute for the COVID-19 Genomics UK (COG-UK) Consortium | Jacquelyn Wynn, Mairead Hyland, The Lighthouse Lab in Alderley Park and Alex Alderton, Roberto Amato, Jeffrey Barrett, Sonia Goncalves, Ewan Harrison, David K. Jackson, Ian Johnston, Dominic Kwiatkowski, Cordelia Langford, John Sillitoe on behalf of the Wellcome Sanger Institute COVID-19 Surveillance Team |
| EPI_ISL_2457359 | Lighthouse Lab in Glasgow | Wellcome Sanger Institute for the COVID-19 Genomics UK (COG-UK) Consortium | Harper VanSteenhouse, Yumi Kasai, David Gray, Carol Clugston, Anna Dominiczak and Alex Alderton, Roberto Amato, Jeffrey Barrett, Sonia Goncalves, Ewan Harrison, David K. Jackson, Ian Johnston, Dominic Kwiatkowski, Cordelia Langford, John Sillitoe on behalf of the Wellcome Sanger Institute COVID-19 Surveillance Team |
| EPI_ISL_2457360 | Lighthouse Lab in Alderley Park | Wellcome Sanger Institute for the COVID-19 Genomics UK (COG-UK) Consortium | Jacquelyn Wynn, Mairead Hyland, The Lighthouse Lab in Alderley Park and Alex Alderton, Roberto Amato, Jeffrey Barrett, Sonia Goncalves, Ewan Harrison, David K. Jackson, Ian Johnston, Dominic Kwiatkowski, Cordelia Langford, John Sillitoe on behalf of the Wellcome Sanger Institute COVID-19 Surveillance Team |
| EPI_ISL_2457361, EPI_ISL_2457362, EPI_ISL_2457363 | Lighthouse Lab in Glasgow | Wellcome Sanger Institute for the COVID-19 Genomics UK (COG-UK) Consortium | Harper VanSteenhouse, Yumi Kasai, David Gray, Carol Clugston, Anna Dominiczak and Alex Alderton, Roberto Amato, Jeffrey Barrett, Sonia Goncalves, Ewan Harrison, David K. Jackson, Ian Johnston, Dominic Kwiatkowski, Cordelia Langford, John Sillitoe on behalf of the Wellcome Sanger Institute COVID-19 Surveillance Team |
| EPI_ISL_2457364, EPI_ISL_2457365 | Lighthouse Lab in Alderley Park | Wellcome Sanger Institute for the COVID-19 Genomics UK (COG-UK) Consortium | Jacquelyn Wynn, Mairead Hyland, The Lighthouse Lab in Alderley Park and Alex Alderton, Roberto Amato, Jeffrey Barrett, Sonia Goncalves, Ewan Harrison, David K. Jackson, Ian Johnston, Dominic Kwiatkowski, Cordelia Langford, John Sillitoe on behalf of the Wellcome Sanger Institute COVID-19 Surveillance Team |
| EPI_ISL_2457366 | Lighthouse Lab in Glasgow | Wellcome Sanger Institute for the COVID-19 Genomics UK (COG-UK) Consortium | Harper VanSteenhouse, Yumi Kasai, David Gray, Carol Clugston, Anna Dominiczak and Alex Alderton, Roberto Amato, Jeffrey Barrett, Sonia Goncalves, Ewan Harrison, David K. Jackson, Ian Johnston, Dominic Kwiatkowski, Cordelia Langford, John Sillitoe on behalf of the Wellcome Sanger Institute COVID-19 Surveillance Team |
| EPI_ISL_2457368 | Lighthouse Lab in Alderley Park | Wellcome Sanger Institute for the COVID-19 Genomics UK (COG-UK) Consortium | Jacquelyn Wynn, Mairead Hyland, The Lighthouse Lab in Alderley Park and Alex Alderton, Roberto Amato, Jeffrey Barrett, Sonia Goncalves, Ewan Harrison, David K. Jackson, Ian Johnston, Dominic Kwiatkowski, Cordelia Langford, John Sillitoe on behalf of the Wellcome Sanger Institute COVID-19 Surveillance Team |
| EPI_ISL_2457369, EPI_ISL_2457370, EPI_ISL_2457371, EPI_ISL_2457372 | Lighthouse Lab in Glasgow | Wellcome Sanger Institute for the COVID-19 Genomics UK (COG-UK) Consortium | Harper VanSteenhouse, Yumi Kasai, David Gray, Carol Clugston, Anna Dominiczak and Alex Alderton, Roberto Amato, Jeffrey Barrett, Sonia Goncalves, Ewan Harrison, David K. Jackson, Ian Johnston, Dominic Kwiatkowski, Cordelia Langford, John Sillitoe on behalf of the Wellcome Sanger Institute COVID-19 Surveillance Team |
| EPI_ISL_2457373 | Lighthouse Lab in Alderley Park | Wellcome Sanger Institute for the COVID-19 Genomics UK (COG-UK) Consortium | Jacquelyn Wynn, Mairead Hyland, The Lighthouse Lab in Alderley Park and Alex Alderton, Roberto Amato, Jeffrey Barrett, Sonia Goncalves, Ewan Harrison, David K. Jackson, Ian Johnston, Dominic Kwiatkowski, Cordelia Langford, John Sillitoe on behalf of the Wellcome Sanger Institute COVID-19 Surveillance Team |
| EPI_ISL_2457374 | Lighthouse Lab in Glasgow | Wellcome Sanger Institute for the COVID-19 Genomics UK (COG-UK) Consortium | Harper VanSteenhouse, Yumi Kasai, David Gray, Carol Clugston, Anna Dominiczak and Alex Alderton, Roberto Amato, Jeffrey Barrett, Sonia Goncalves, Ewan Harrison, David K. Jackson, Ian Johnston, Dominic Kwiatkowski, Cordelia Langford, John Sillitoe on behalf of the Wellcome Sanger Institute COVID-19 Surveillance Team |
| EPI_ISL_2457375 | Lighthouse Lab in Alderley Park | Wellcome Sanger Institute for the COVID-19 Genomics UK (COG-UK) Consortium | Jacquelyn Wynn, Mairead Hyland, The Lighthouse Lab in Alderley Park and Alex Alderton, Roberto Amato, Jeffrey Barrett, Sonia Goncalves, Ewan Harrison, David K. Jackson, Ian Johnston, Dominic Kwiatkowski, Cordelia Langford, John Sillitoe on behalf of the Wellcome Sanger Institute COVID-19 Surveillance Team |
| EPI_ISL_2457376 | Lighthouse Lab in Glasgow | Wellcome Sanger Institute for the COVID-19 Genomics UK (COG-UK) Consortium | Harper VanSteenhouse, Yumi Kasai, David Gray, Carol Clugston, Anna Dominiczak and Alex Alderton, Roberto Amato, Jeffrey Barrett, Sonia Goncalves, Ewan Harrison, David K. Jackson, Ian Johnston, Dominic Kwiatkowski, Cordelia Langford, John Sillitoe on behalf of the Wellcome Sanger Institute COVID-19 Surveillance Team |
| EPI_ISL_2457377, EPI_ISL_2457378, EPI_ISL_2457379, EPI_ISL_2457380 | Lighthouse Lab in Alderley Park | Wellcome Sanger Institute for the COVID-19 Genomics UK (COG-UK) Consortium | Jacquelyn Wynn, Mairead Hyland, The Lighthouse Lab in Alderley Park and Alex Alderton, Roberto Amato, Jeffrey Barrett, Sonia Goncalves, Ewan Harrison, David K. Jackson, Ian Johnston, Dominic Kwiatkowski, Cordelia Langford, John Sillitoe on behalf of the Wellcome Sanger Institute COVID-19 Surveillance Team |
| EPI_ISL_2457381 | Lighthouse Lab in Glasgow | Wellcome Sanger Institute for the COVID-19 Genomics UK (COG-UK) Consortium | Harper VanSteenhouse, Yumi Kasai, David Gray, Carol Clugston, Anna Dominiczak and Alex Alderton, Roberto Amato, Jeffrey Barrett, Sonia Goncalves, Ewan Harrison, David K. Jackson, Ian Johnston, Dominic Kwiatkowski, Cordelia Langford, John Sillitoe on behalf of the Wellcome Sanger Institute COVID-19 Surveillance Team |
| EPI_ISL_2457382, EPI_ISL_2457383, EPI_ISL_2457384 | Lighthouse Lab in Alderley Park | Wellcome Sanger Institute for the COVID-19 Genomics UK (COG-UK) Consortium | Jacquelyn Wynn, Mairead Hyland, The Lighthouse Lab in Alderley Park and Alex Alderton, Roberto Amato, Jeffrey Barrett, Sonia Goncalves, Ewan Harrison, David K. Jackson, Ian Johnston, Dominic Kwiatkowski, Cordelia Langford, John Sillitoe on behalf of the Wellcome Sanger Institute COVID-19 Surveillance Team |
| EPI_ISL_2457385 | Lighthouse Lab in Glasgow | Wellcome Sanger Institute for the COVID-19 Genomics UK (COG-UK) Consortium | Harper VanSteenhouse, Yumi Kasai, David Gray, Carol Clugston, Anna Dominiczak and Alex Alderton, Roberto Amato, Jeffrey Barrett, Sonia Goncalves, Ewan Harrison, David K. Jackson, Ian Johnston, Dominic Kwiatkowski, Cordelia Langford, John Sillitoe on behalf of the Wellcome Sanger Institute COVID-19 Surveillance Team |
| EPI_ISL_2457386, EPI_ISL_2457387, EPI_ISL_2457388 | Lighthouse Lab in Alderley Park | Wellcome Sanger Institute for the COVID-19 Genomics UK (COG-UK) Consortium | Jacquelyn Wynn, Mairead Hyland, The Lighthouse Lab in Alderley Park and Alex Alderton, Roberto Amato, Jeffrey Barrett, Sonia Goncalves, Ewan Harrison, David K. Jackson, Ian Johnston, Dominic Kwiatkowski, Cordelia Langford, John Sillitoe on behalf of the Wellcome Sanger Institute COVID-19 Surveillance Team |
| EPI_ISL_2457389, EPI_ISL_2457390 | Lighthouse Lab in Glasgow | Wellcome Sanger Institute for the COVID-19 Genomics UK (COG-UK) Consortium | Harper VanSteenhouse, Yumi Kasai, David Gray, Carol Clugston, Anna Dominiczak and Alex Alderton, Roberto Amato, Jeffrey Barrett, Sonia Goncalves, Ewan Harrison, David K. Jackson, Ian Johnston, Dominic Kwiatkowski, Cordelia Langford, John Sillitoe on behalf of the Wellcome Sanger Institute COVID-19 Surveillance Team |
| EPI_ISL_2457391, EPI_ISL_2457394, EPI_ISL_2457395 | Lighthouse Lab in Alderley Park | Wellcome Sanger Institute for the COVID-19 Genomics UK (COG-UK) Consortium | Jacquelyn Wynn, Mairead Hyland, The Lighthouse Lab in Alderley Park and Alex Alderton, Roberto Amato, Jeffrey Barrett, Sonia Goncalves, Ewan Harrison, David K. Jackson, Ian Johnston, Dominic Kwiatkowski, Cordelia Langford, John Sillitoe on behalf of the Wellcome Sanger Institute COVID-19 Surveillance Team |
| EPI_ISL_2457396 | Lighthouse Lab in Glasgow | Wellcome Sanger Institute for the COVID-19 Genomics UK (COG-UK) Consortium | Harper VanSteenhouse, Yumi Kasai, David Gray, Carol Clugston, Anna Dominiczak and Alex Alderton, Roberto Amato, Jeffrey Barrett, Sonia Goncalves, Ewan Harrison, David K. Jackson, Ian Johnston, Dominic Kwiatkowski, Cordelia Langford, John Sillitoe on behalf of the Wellcome Sanger Institute COVID-19 Surveillance Team |
| EPI_ISL_2457397 | Lighthouse Lab in Alderley Park | Wellcome Sanger Institute for the COVID-19 Genomics UK (COG-UK) Consortium | Jacquelyn Wynn, Mairead Hyland, The Lighthouse Lab in Alderley Park and Alex Alderton, Roberto Amato, Jeffrey Barrett, Sonia Goncalves, Ewan Harrison, David K. Jackson, Ian Johnston, Dominic Kwiatkowski, Cordelia Langford, John Sillitoe on behalf of the Wellcome Sanger Institute COVID-19 Surveillance Team |
| EPI_ISL_2457398, EPI_ISL_2457399 | Lighthouse Lab in Glasgow | Wellcome Sanger Institute for the COVID-19 Genomics UK (COG-UK) Consortium | Harper VanSteenhouse, Yumi Kasai, David Gray, Carol Clugston, Anna Dominiczak and Alex Alderton, Roberto Amato, Jeffrey Barrett, Sonia Goncalves, Ewan Harrison, David K. Jackson, Ian Johnston, Dominic Kwiatkowski, Cordelia Langford, John Sillitoe on behalf of the Wellcome Sanger Institute COVID-19 Surveillance Team |
| EPI_ISL_2457400, EPI_ISL_2457401 | Lighthouse Lab in Alderley Park | Wellcome Sanger Institute for the COVID-19 Genomics UK (COG-UK) Consortium | Jacquelyn Wynn, Mairead Hyland, The Lighthouse Lab in Alderley Park and Alex Alderton, Roberto Amato, Jeffrey Barrett, Sonia Goncalves, Ewan Harrison, David K. Jackson, Ian Johnston, Dominic Kwiatkowski, Cordelia Langford, John Sillitoe on behalf of the Wellcome Sanger Institute COVID-19 Surveillance Team |
| EPI_ISL_2457402, EPI_ISL_2457403 | Lighthouse Lab in Glasgow | Wellcome Sanger Institute for the COVID-19 Genomics UK (COG-UK) Consortium | Harper VanSteenhouse, Yumi Kasai, David Gray, Carol Clugston, Anna Dominiczak and Alex Alderton, Roberto Amato, Jeffrey Barrett, Sonia Goncalves, Ewan Harrison, David K. Jackson, Ian Johnston, Dominic Kwiatkowski, Cordelia Langford, John Sillitoe on behalf of the Wellcome Sanger Institute COVID-19 Surveillance Team |
| EPI_ISL_2457404, EPI_ISL_2457405, EPI_ISL_2457406 | Lighthouse Lab in Alderley Park | Wellcome Sanger Institute for the COVID-19 Genomics UK (COG-UK) Consortium | Jacquelyn Wynn, Mairead Hyland, The Lighthouse Lab in Alderley Park and Alex Alderton, Roberto Amato, Jeffrey Barrett, Sonia Goncalves, Ewan Harrison, David K. Jackson, Ian Johnston, Dominic Kwiatkowski, Cordelia Langford, John Sillitoe on behalf of the Wellcome Sanger Institute COVID-19 Surveillance Team |
| EPI_ISL_2457408 | Lighthouse Lab in Glasgow | Wellcome Sanger Institute for the COVID-19 Genomics UK (COG-UK) Consortium | Harper VanSteenhouse, Yumi Kasai, David Gray, Carol Clugston, Anna Dominiczak and Alex Alderton, Roberto Amato, Jeffrey Barrett, Sonia Goncalves, Ewan Harrison, David K. Jackson, Ian Johnston, Dominic Kwiatkowski, Cordelia Langford, John Sillitoe on behalf of the Wellcome Sanger Institute COVID-19 Surveillance Team |
| EPI_ISL_2457409, EPI_ISL_2457410 | Lighthouse Lab in Alderley Park | Wellcome Sanger Institute for the COVID-19 Genomics UK (COG-UK) Consortium | Jacquelyn Wynn, Mairead Hyland, The Lighthouse Lab in Alderley Park and Alex Alderton, Roberto Amato, Jeffrey Barrett, Sonia Goncalves, Ewan Harrison, David K. Jackson, Ian Johnston, Dominic Kwiatkowski, Cordelia Langford, John Sillitoe on behalf of the Wellcome Sanger Institute COVID-19 Surveillance Team |
| EPI_ISL_2457411, EPI_ISL_2457412 | Lighthouse Lab in Glasgow | Wellcome Sanger Institute for the COVID-19 Genomics UK (COG-UK) Consortium | Harper VanSteenhouse, Yumi Kasai, David Gray, Carol Clugston, Anna Dominiczak and Alex Alderton, Roberto Amato, Jeffrey Barrett, Sonia Goncalves, Ewan Harrison, David K. Jackson, Ian Johnston, Dominic Kwiatkowski, Cordelia Langford, John Sillitoe on behalf of the Wellcome Sanger Institute COVID-19 Surveillance Team |
| EPI_ISL_2457413, EPI_ISL_2457414, | Lighthouse Lab in Alderley Park | Wellcome Sanger Institute for the COVID-19 Genomics UK (COG-UK) Consortium | Jacquelyn Wynn, Mairead Hyland, The Lighthouse Lab in Alderley Park and Alex Alderton, Roberto Amato, Jeffrey Barrett, Sonia Goncalves, Ewan Harrison, David K. Jackson, Ian Johnston, Dominic Kwiatkowski, Cordelia Langford, John Sillitoe on behalf of the Wellcome Sanger Institute COVID-19 Surveillance Team |

| | | | |
|---|---|---|---|
| EPI_ISL_2457415 | | | |
| EPI_ISL_2457416, EPI_ISL_2457417 | Lighthouse Lab in Glasgow | Wellcome Sanger Institute for the COVID-19 Genomics UK (COG-UK) Consortium | Harper VanSteenhouse, Yumi Kasai, David Gray, Carol Clugston, Anna Dominiczak and Alex Alderton, Roberto Amato, Jeffrey Barrett, Sonia Goncalves, Ewan Harrison, David K. Jackson, Ian Johnston, Dominic Kwiatkowski, Cordelia Langford, John Sillitoe on behalf of the Wellcome Sanger Institute COVID-19 Surveillance Team |
| EPI_ISL_2457418 | Lighthouse Lab in Alderley Park | Wellcome Sanger Institute for the COVID-19 Genomics UK (COG-UK) Consortium | Jacquelyn Wynn, Mairead Hyland, The Lighthouse Lab in Alderley Park and Alex Alderton, Roberto Amato, Jeffrey Barrett, Sonia Goncalves, Ewan Harrison, David K. Jackson, Ian Johnston, Dominic Kwiatkowski, Cordelia Langford, John Sillitoe on behalf of the Wellcome Sanger Institute COVID-19 Surveillance Team |
| EPI_ISL_2457419 | Lighthouse Lab in Glasgow | Wellcome Sanger Institute for the COVID-19 Genomics UK (COG-UK) Consortium | Harper VanSteenhouse, Yumi Kasai, David Gray, Carol Clugston, Anna Dominiczak and Alex Alderton, Roberto Amato, Jeffrey Barrett, Sonia Goncalves, Ewan Harrison, David K. Jackson, Ian Johnston, Dominic Kwiatkowski, Cordelia Langford, John Sillitoe on behalf of the Wellcome Sanger Institute COVID-19 Surveillance Team |
| EPI_ISL_2457421, EPI_ISL_2457422, EPI_ISL_2457423, EPI_ISL_2457424 | Lighthouse Lab in Alderley Park | Wellcome Sanger Institute for the COVID-19 Genomics UK (COG-UK) Consortium | Jacquelyn Wynn, Mairead Hyland, The Lighthouse Lab in Alderley Park and Alex Alderton, Roberto Amato, Jeffrey Barrett, Sonia Goncalves, Ewan Harrison, David K. Jackson, Ian Johnston, Dominic Kwiatkowski, Cordelia Langford, John Sillitoe on behalf of the Wellcome Sanger Institute COVID-19 Surveillance Team |
| EPI_ISL_2457425 | Lighthouse Lab in Glasgow | Wellcome Sanger Institute for the COVID-19 Genomics UK (COG-UK) Consortium | Harper VanSteenhouse, Yumi Kasai, David Gray, Carol Clugston, Anna Dominiczak and Alex Alderton, Roberto Amato, Jeffrey Barrett, Sonia Goncalves, Ewan Harrison, David K. Jackson, Ian Johnston, Dominic Kwiatkowski, Cordelia Langford, John Sillitoe on behalf of the Wellcome Sanger Institute COVID-19 Surveillance Team |
| EPI_ISL_2457426, EPI_ISL_2457428, EPI_ISL_2457429 | Lighthouse Lab in Alderley Park | Wellcome Sanger Institute for the COVID-19 Genomics UK (COG-UK) Consortium | Jacquelyn Wynn, Mairead Hyland, The Lighthouse Lab in Alderley Park and Alex Alderton, Roberto Amato, Jeffrey Barrett, Sonia Goncalves, Ewan Harrison, David K. Jackson, Ian Johnston, Dominic Kwiatkowski, Cordelia Langford, John Sillitoe on behalf of the Wellcome Sanger Institute COVID-19 Surveillance Team |
| EPI_ISL_2457430, EPI_ISL_2457431, EPI_ISL_2457432 | Lighthouse Lab in Glasgow | Wellcome Sanger Institute for the COVID-19 Genomics UK (COG-UK) Consortium | Harper VanSteenhouse, Yumi Kasai, David Gray, Carol Clugston, Anna Dominiczak and Alex Alderton, Roberto Amato, Jeffrey Barrett, Sonia Goncalves, Ewan Harrison, David K. Jackson, Ian Johnston, Dominic Kwiatkowski, Cordelia Langford, John Sillitoe on behalf of the Wellcome Sanger Institute COVID-19 Surveillance Team |
| EPI_ISL_2457434, EPI_ISL_2457436, EPI_ISL_2457437, EPI_ISL_2457438, EPI_ISL_2457439 | Lighthouse Lab in Alderley Park | Wellcome Sanger Institute for the COVID-19 Genomics UK (COG-UK) Consortium | Jacquelyn Wynn, Mairead Hyland, The Lighthouse Lab in Alderley Park and Alex Alderton, Roberto Amato, Jeffrey Barrett, Sonia Goncalves, Ewan Harrison, David K. Jackson, Ian Johnston, Dominic Kwiatkowski, Cordelia Langford, John Sillitoe on behalf of the Wellcome Sanger Institute COVID-19 Surveillance Team |
| EPI_ISL_2457440, EPI_ISL_2457441, EPI_ISL_2457442, EPI_ISL_2457443 | Lighthouse Lab in Glasgow | Wellcome Sanger Institute for the COVID-19 Genomics UK (COG-UK) Consortium | Harper VanSteenhouse, Yumi Kasai, David Gray, Carol Clugston, Anna Dominiczak and Alex Alderton, Roberto Amato, Jeffrey Barrett, Sonia Goncalves, Ewan Harrison, David K. Jackson, Ian Johnston, Dominic Kwiatkowski, Cordelia Langford, John Sillitoe on behalf of the Wellcome Sanger Institute COVID-19 Surveillance Team |
| EPI_ISL_2457444, EPI_ISL_2457445 | Lighthouse Lab in Alderley Park | Wellcome Sanger Institute for the COVID-19 Genomics UK (COG-UK) Consortium | Jacquelyn Wynn, Mairead Hyland, The Lighthouse Lab in Alderley Park and Alex Alderton, Roberto Amato, Jeffrey Barrett, Sonia Goncalves, Ewan Harrison, David K. Jackson, Ian Johnston, Dominic Kwiatkowski, Cordelia Langford, John Sillitoe on behalf of the Wellcome Sanger Institute COVID-19 Surveillance Team |
| EPI_ISL_2457446, EPI_ISL_2457447 | Lighthouse Lab in Glasgow | Wellcome Sanger Institute for the COVID-19 Genomics UK (COG-UK) Consortium | Harper VanSteenhouse, Yumi Kasai, David Gray, Carol Clugston, Anna Dominiczak and Alex Alderton, Roberto Amato, Jeffrey Barrett, Sonia Goncalves, Ewan Harrison, David K. Jackson, Ian Johnston, Dominic Kwiatkowski, Cordelia Langford, John Sillitoe on behalf of the Wellcome Sanger Institute COVID-19 Surveillance Team |
| EPI_ISL_2457448 | Lighthouse Lab in Alderley Park | Wellcome Sanger Institute for the COVID-19 Genomics UK (COG-UK) Consortium | Jacquelyn Wynn, Mairead Hyland, The Lighthouse Lab in Alderley Park and Alex Alderton, Roberto Amato, Jeffrey Barrett, Sonia Goncalves, Ewan Harrison, David K. Jackson, Ian Johnston, Dominic Kwiatkowski, Cordelia Langford, John Sillitoe on behalf of the Wellcome Sanger Institute COVID-19 Surveillance Team |
| EPI_ISL_2457449 | Lighthouse Lab in Glasgow | Wellcome Sanger Institute for the COVID-19 Genomics UK (COG-UK) Consortium | Harper VanSteenhouse, Yumi Kasai, David Gray, Carol Clugston, Anna Dominiczak and Alex Alderton, Roberto Amato, Jeffrey Barrett, Sonia Goncalves, Ewan Harrison, David K. Jackson, Ian Johnston, Dominic Kwiatkowski, Cordelia Langford, John Sillitoe on behalf of the Wellcome Sanger Institute COVID-19 Surveillance Team |
| EPI_ISL_2457450 | Lighthouse Lab in Alderley Park | Wellcome Sanger Institute for the COVID-19 Genomics UK (COG-UK) Consortium | Jacquelyn Wynn, Mairead Hyland, The Lighthouse Lab in Alderley Park and Alex Alderton, Roberto Amato, Jeffrey Barrett, Sonia Goncalves, Ewan Harrison, David K. Jackson, Ian Johnston, Dominic Kwiatkowski, Cordelia Langford, John Sillitoe on behalf of the Wellcome Sanger Institute COVID-19 Surveillance Team |
| EPI_ISL_2457451, EPI_ISL_2457453 | Lighthouse Lab in Glasgow | Wellcome Sanger Institute for the COVID-19 Genomics UK (COG-UK) Consortium | Harper VanSteenhouse, Yumi Kasai, David Gray, Carol Clugston, Anna Dominiczak and Alex Alderton, Roberto Amato, Jeffrey Barrett, Sonia Goncalves, Ewan Harrison, David K. Jackson, Ian Johnston, Dominic Kwiatkowski, Cordelia Langford, John Sillitoe on behalf of the Wellcome Sanger Institute COVID-19 Surveillance Team |
| EPI_ISL_2457454, EPI_ISL_2457455, EPI_ISL_2457456 | Lighthouse Lab in Alderley Park | Wellcome Sanger Institute for the COVID-19 Genomics UK (COG-UK) Consortium | Jacquelyn Wynn, Mairead Hyland, The Lighthouse Lab in Alderley Park and Alex Alderton, Roberto Amato, Jeffrey Barrett, Sonia Goncalves, Ewan Harrison, David K. Jackson, Ian Johnston, Dominic Kwiatkowski, Cordelia Langford, John Sillitoe on behalf of the Wellcome Sanger Institute COVID-19 Surveillance Team |
| EPI_ISL_2457457 | Lighthouse Lab in Glasgow | Wellcome Sanger Institute for the COVID-19 Genomics UK (COG-UK) Consortium | Harper VanSteenhouse, Yumi Kasai, David Gray, Carol Clugston, Anna Dominiczak and Alex Alderton, Roberto Amato, Jeffrey Barrett, Sonia Goncalves, Ewan Harrison, David K. Jackson, Ian Johnston, Dominic Kwiatkowski, Cordelia Langford, John Sillitoe on behalf of the Wellcome Sanger Institute COVID-19 Surveillance Team |
| EPI_ISL_2457458, EPI_ISL_2457459, EPI_ISL_2457460, EPI_ISL_2457461 | Lighthouse Lab in Alderley Park | Wellcome Sanger Institute for the COVID-19 Genomics UK (COG-UK) Consortium | Jacquelyn Wynn, Mairead Hyland, The Lighthouse Lab in Alderley Park and Alex Alderton, Roberto Amato, Jeffrey Barrett, Sonia Goncalves, Ewan Harrison, David K. Jackson, Ian Johnston, Dominic Kwiatkowski, Cordelia Langford, John Sillitoe on behalf of the Wellcome Sanger Institute COVID-19 Surveillance Team |
| EPI_ISL_2457462 | Lighthouse Lab in Glasgow | Wellcome Sanger Institute for the COVID-19 Genomics UK (COG-UK) Consortium | Harper VanSteenhouse, Yumi Kasai, David Gray, Carol Clugston, Anna Dominiczak and Alex Alderton, Roberto Amato, Jeffrey Barrett, Sonia Goncalves, Ewan Harrison, David K. Jackson, Ian Johnston, Dominic Kwiatkowski, Cordelia Langford, John Sillitoe on behalf of the Wellcome Sanger Institute COVID-19 Surveillance Team |
| EPI_ISL_2457463 | Lighthouse Lab in Alderley Park | Wellcome Sanger Institute for the COVID-19 Genomics UK (COG-UK) Consortium | Jacquelyn Wynn, Mairead Hyland, The Lighthouse Lab in Alderley Park and Alex Alderton, Roberto Amato, Jeffrey Barrett, Sonia Goncalves, Ewan Harrison, David K. Jackson, Ian Johnston, Dominic Kwiatkowski, Cordelia Langford, John Sillitoe on behalf of the Wellcome Sanger Institute COVID-19 Surveillance Team |
| EPI_ISL_2457464 | Lighthouse Lab in Glasgow | Wellcome Sanger Institute for the COVID-19 Genomics UK (COG-UK) Consortium | Harper VanSteenhouse, Yumi Kasai, David Gray, Carol Clugston, Anna Dominiczak and Alex Alderton, Roberto Amato, Jeffrey Barrett, Sonia Goncalves, Ewan Harrison, David K. Jackson, Ian Johnston, Dominic Kwiatkowski, Cordelia Langford, John Sillitoe on behalf of the Wellcome Sanger Institute COVID-19 Surveillance Team |
| EPI_ISL_2457465, EPI_ISL_2457466, EPI_ISL_2457467 | Lighthouse Lab in Alderley Park | Wellcome Sanger Institute for the COVID-19 Genomics UK (COG-UK) Consortium | Jacquelyn Wynn, Mairead Hyland, The Lighthouse Lab in Alderley Park and Alex Alderton, Roberto Amato, Jeffrey Barrett, Sonia Goncalves, Ewan Harrison, David K. Jackson, Ian Johnston, Dominic Kwiatkowski, Cordelia Langford, John Sillitoe on behalf of the Wellcome Sanger Institute COVID-19 Surveillance Team |
| EPI_ISL_2457469, EPI_ISL_2457470, EPI_ISL_2457471 | Lighthouse Lab in Glasgow | Wellcome Sanger Institute for the COVID-19 Genomics UK (COG-UK) Consortium | Harper VanSteenhouse, Yumi Kasai, David Gray, Carol Clugston, Anna Dominiczak and Alex Alderton, Roberto Amato, Jeffrey Barrett, Sonia Goncalves, Ewan Harrison, David K. Jackson, Ian Johnston, Dominic Kwiatkowski, Cordelia Langford, John Sillitoe on behalf of the Wellcome Sanger Institute COVID-19 Surveillance Team |
| EPI_ISL_2457472 | Lighthouse Lab in Alderley Park | Wellcome Sanger Institute for the COVID-19 Genomics UK (COG-UK) Consortium | Jacquelyn Wynn, Mairead Hyland, The Lighthouse Lab in Alderley Park and Alex Alderton, Roberto Amato, Jeffrey Barrett, Sonia Goncalves, Ewan Harrison, David K. Jackson, Ian Johnston, Dominic Kwiatkowski, Cordelia Langford, John Sillitoe on behalf of the Wellcome Sanger Institute COVID-19 Surveillance Team |
| EPI_ISL_2457475, EPI_ISL_2457476, EPI_ISL_2457477, EPI_ISL_2457478, EPI_ISL_2457479, EPI_ISL_2457481, EPI_ISL_2457482, EPI_ISL_2457484, EPI_ISL_2457489, EPI_ISL_2457490, EPI_ISL_2457491, EPI_ISL_2457494, EPI_ISL_2457495, EPI_ISL_2457496, EPI_ISL_2457498, EPI_ISL_2457499, EPI_ISL_2457500 | | | |
| see above | Lighthouse Lab in Glasgow | Wellcome Sanger Institute for the COVID-19 Genomics UK (COG-UK) Consortium | Harper VanSteenhouse, Yumi Kasai, David Gray, Carol Clugston, Anna Dominiczak and Alex Alderton, Roberto Amato, Jeffrey Barrett, Sonia Goncalves, Ewan Harrison, David K. Jackson, Ian Johnston, Dominic Kwiatkowski, Cordelia Langford, John Sillitoe on behalf of the Wellcome Sanger Institute COVID-19 Surveillance Team |
| EPI_ISL_2457504 | Lighthouse Lab in Alderley Park | Wellcome Sanger Institute for the COVID-19 Genomics UK (COG-UK) Consortium | Jacquelyn Wynn, Mairead Hyland, The Lighthouse Lab in Alderley Park and Alex Alderton, Roberto Amato, Jeffrey Barrett, Sonia Goncalves, Ewan Harrison, David K. Jackson, Ian Johnston, Dominic Kwiatkowski, Cordelia Langford, John Sillitoe on behalf of the Wellcome Sanger Institute COVID-19 Surveillance Team |
| EPI_ISL_2457505 | Lighthouse Lab in Milton Keynes | Wellcome Sanger Institute for the COVID-19 Genomics UK (COG-UK) Consortium | The Lighthouse Lab in Milton Keynes and Alex Alderton, Roberto Amato, Jeffrey Barrett, Sonia Goncalves, Ewan Harrison, David K. Jackson, Ian Johnston, Dominic Kwiatkowski, Cordelia Langford, John Sillitoe on behalf of the Wellcome Sanger Institute COVID-19 Surveillance Team |
| EPI_ISL_2457507, EPI_ISL_2457508, EPI_ISL_2457509, EPI_ISL_2457510, EPI_ISL_2457511, EPI_ISL_2457513, EPI_ISL_2457514, EPI_ISL_2457516, EPI_ISL_2457517, EPI_ISL_2457518, EPI_ISL_2457519, EPI_ISL_2457520, EPI_ISL_2457521, EPI_ISL_2457522, EPI_ISL_2457524, EPI_ISL_2457525, EPI_ISL_2457526, EPI_ISL_2457527, EPI_ISL_2457530, EPI_ISL_2457531, EPI_ISL_2457532, EPI_ISL_2457533, EPI_ISL_2457534, EPI_ISL_2457535, EPI_ISL_2457536, EPI_ISL_2457537, EPI_ISL_2457541, EPI_ISL_2457542, EPI_ISL_2457543, EPI_ISL_2457544, EPI_ISL_2457545, EPI_ISL_2457546, EPI_ISL_2457548, EPI_ISL_2457549, EPI_ISL_2457550, EPI_ISL_2457551, EPI_ISL_2457552, EPI_ISL_2457553, EPI_ISL_2457554, EPI_ISL_2457555, EPI_ISL_2457556, EPI_ISL_2457557, EPI_ISL_2457558, EPI_ISL_2457559, EPI_ISL_2457560, EPI_ISL_2457561, EPI_ISL_2457562, EPI_ISL_2457563, EPI_ISL_2457564, EPI_ISL_2457567, EPI_ISL_2457568, EPI_ISL_2457569, EPI_ISL_2457571, EPI_ISL_2457572, EPI_ISL_2457574, EPI_ISL_2457575, EPI_ISL_2457576, EPI_ISL_2457577, EPI_ISL_2457578, EPI_ISL_2457579, EPI_ISL_2457580, EPI_ISL_2457581, EPI_ISL_2457584, EPI_ISL_2457585, EPI_ISL_2457587, EPI_ISL_2457588, EPI_ISL_2457590, EPI_ISL_2457591, EPI_ISL_2457592, EPI_ISL_2457593, EPI_ISL_2457594, EPI_ISL_2457595, EPI_ISL_2457597, EPI_ISL_2457598, EPI_ISL_2457599, EPI_ISL_2457600, EPI_ISL_2457601, EPI_ISL_2457602, EPI_ISL_2457603, EPI_ISL_2457604, EPI_ISL_2457605, EPI_ISL_2457606, EPI_ISL_2457607, EPI_ISL_2457608, EPI_ISL_2457609, EPI_ISL_2457611, EPI_ISL_2457612, EPI_ISL_2457613, EPI_ISL_2457614, EPI_ISL_2457616, EPI_ISL_2457617, EPI_ISL_2457619, EPI_ISL_2457621, EPI_ISL_2457622, EPI_ISL_2457623, EPI_ISL_2457624, EPI_ISL_2457625, EPI_ISL_2457626, EPI_ISL_2457627, EPI_ISL_2457628, EPI_ISL_2457629, EPI_ISL_2457630, EPI_ISL_2457632, EPI_ISL_2457634, EPI_ISL_2457636, EPI_ISL_2457639, EPI_ISL_2457641, EPI_ISL_2457642, EPI_ISL_2457643, EPI_ISL_2457645, EPI_ISL_2457646, EPI_ISL_2457647, EPI_ISL_2457648, EPI_ISL_2457649, EPI_ISL_2457650, EPI_ISL_2457651, EPI_ISL_2457652, EPI_ISL_2457653, EPI_ISL_2457654, EPI_ISL_2457655, EPI_ISL_2457656, EPI_ISL_2457657, EPI_ISL_2457658, EPI_ISL_2457660, EPI_ISL_2457664, EPI_ISL_2457665, EPI_ISL_2457666, EPI_ISL_2457668, EPI_ISL_2457669, EPI_ISL_2457670, EPI_ISL_2457671, EPI_ISL_2457673, EPI_ISL_2457674, EPI_ISL_2457676, EPI_ISL_2457679, EPI_ISL_2457680, EPI_ISL_2457681, EPI_ISL_2457684, EPI_ISL_2457686, EPI_ISL_2457690, EPI_ISL_2457693, EPI_ISL_2457695, EPI_ISL_2457698, EPI_ISL_2457699, EPI_ISL_2457700, EPI_ISL_2457701, EPI_ISL_2457702, EPI_ISL_2457703, EPI_ISL_2457705, EPI_ISL_2457707, EPI_ISL_2457709, EPI_ISL_2457710, EPI_ISL_2457713, EPI_ISL_2457714, EPI_ISL_2457716, EPI_ISL_2457721, EPI_ISL_2457726, EPI_ISL_2457727, EPI_ISL_2457730, EPI_ISL_2457731, EPI_ISL_2457735, EPI_ISL_2457738, EPI_ISL_2457741, EPI_ISL_2457743, EPI_ISL_2457744, EPI_ISL_2457745, EPI_ISL_2457748, EPI_ISL_2457750, EPI_ISL_2457752, EPI_ISL_2457753, EPI_ISL_2457755, EPI_ISL_2457756, EPI_ISL_2457761, EPI_ISL_2457763, EPI_ISL_2457767, EPI_ISL_2457768, EPI_ISL_2457770, EPI_ISL_2457771, EPI_ISL_2457774, EPI_ISL_2457775, EPI_ISL_2457776, EPI_ISL_2457777, EPI_ISL_2457783, EPI_ISL_2457784, EPI_ISL_2457785, EPI_ISL_2457787, EPI_ISL_2457788, EPI_ISL_2457789, EPI_ISL_2457790, EPI_ISL_2457791, EPI_ISL_2457792, EPI_ISL_2457793, EPI_ISL_2457794, EPI_ISL_2457797, EPI_ISL_2457798, EPI_ISL_2457799, EPI_ISL_2457800, EPI_ISL_2457801, EPI_ISL_2457802, EPI_ISL_2457806, EPI_ISL_2457808, EPI_ISL_2457809, EPI_ISL_2457811, EPI_ISL_2457812, EPI_ISL_2457822, EPI_ISL_2457823, EPI_ISL_2457825, EPI_ISL_2457828, EPI_ISL_2457829, EPI_ISL_2457831, EPI_ISL_2457832, EPI_ISL_2457834, EPI_ISL_2457836, EPI_ISL_2457837, EPI_ISL_2457838, EPI_ISL_2457840, EPI_ISL_2457844, EPI_ISL_2457849, EPI_ISL_2457851, EPI_ISL_2457854, EPI_ISL_2457855, EPI_ISL_2457856, EPI_ISL_2457859, EPI_ISL_2457861, EPI_ISL_2457862, EPI_ISL_2457864, EPI_ISL_2457865, EPI_ISL_2457866, EPI_ISL_2457868, EPI_ISL_2457871, EPI_ISL_2457874, EPI_ISL_2457877, EPI_ISL_2457880, EPI_ISL_2457882, EPI_ISL_2457883 | | | |
| see above | Lighthouse Lab in Glasgow | Wellcome Sanger Institute for the COVID-19 Genomics UK (COG-UK) Consortium | Harper VanSteenhouse, Yumi Kasai, David Gray, Carol Clugston, Anna Dominiczak and Alex Alderton, Roberto Amato, Jeffrey Barrett, Sonia Goncalves, Ewan Harrison, David K. Jackson, Ian Johnston, Dominic Kwiatkowski, Cordelia Langford, John Sillitoe on behalf of the Wellcome Sanger Institute COVID-19 Surveillance Team |
| EPI_ISL_2457884 | Lighthouse Lab in Alderley Park | Wellcome Sanger Institute for the COVID-19 Genomics UK (COG-UK) Consortium | Jacquelyn Wynn, Mairead Hyland, The Lighthouse Lab in Alderley Park and Alex Alderton, Roberto Amato, Jeffrey Barrett, Sonia Goncalves, Ewan Harrison, David K. Jackson, Ian Johnston, Dominic Kwiatkowski, Cordelia Langford, John Sillitoe on behalf of the Wellcome Sanger Institute COVID-19 Surveillance Team |
| EPI_ISL_2457885 | Lighthouse Laboratory Plymouth | Wellcome Sanger Institute for the COVID-19 Genomics UK (COG-UK) Consortium | Lighthouse Laboratory Plymouth and Alex Alderton, Roberto Amato, Jeffrey Barrett, Sonia Goncalves, Ewan Harrison, David K. Jackson, Ian Johnston, Dominic Kwiatkowski, Cordelia Langford, John Sillitoe on behalf of the Wellcome Sanger Institute COVID-19 Surveillance Team |
| EPI_ISL_2457886 | Lighthouse Lab in Glasgow | Wellcome Sanger Institute for the COVID-19 Genomics UK (COG-UK) Consortium | Harper VanSteenhouse, Yumi Kasai, David Gray, Carol Clugston, Anna Dominiczak and Alex Alderton, Roberto Amato, Jeffrey Barrett, Sonia Goncalves, Ewan Harrison, David K. Jackson, Ian Johnston, Dominic Kwiatkowski, Cordelia Langford, John Sillitoe on behalf of the Wellcome Sanger Institute COVID-19 Surveillance Team |
| EPI_ISL_2457887 | Lighthouse Lab in Milton Keynes | Wellcome Sanger Institute for the COVID-19 Genomics UK (COG-UK) Consortium | The Lighthouse Lab in Milton Keynes and Alex Alderton, Roberto Amato, Jeffrey Barrett, Sonia Goncalves, Ewan Harrison, David K. Jackson, Ian Johnston, Dominic Kwiatkowski, Cordelia Langford, John Sillitoe on behalf of the Wellcome Sanger Institute COVID-19 Surveillance Team |
| EPI_ISL_2457888, EPI_ISL_2457890, EPI_ISL_2457892, EPI_ISL_2457893, EPI_ISL_2457894, EPI_ISL_2457895, EPI_ISL_2457898, EPI_ISL_2457901, EPI_ISL_2457904, EPI_ISL_2457906, EPI_ISL_2457907, EPI_ISL_2457909, EPI_ISL_2457910, EPI_ISL_2457916, EPI_ISL_2457917, EPI_ISL_2457920, EPI_ISL_2457921, EPI_ISL_2457923, EPI_ISL_2457925, EPI_ISL_2457926, EPI_ISL_2457928, EPI_ISL_2457929, EPI_ISL_2457930, EPI_ISL_2457931, EPI_ISL_2457932, EPI_ISL_2457933, EPI_ISL_2457934, EPI_ISL_2457935, EPI_ISL_2457936, EPI_ISL_2457940, EPI_ISL_2457941, EPI_ISL_2457942, EPI_ISL_2457943, EPI_ISL_2457944, EPI_ISL_2457946, EPI_ISL_2457948, EPI_ISL_2457950, EPI_ISL_2457951, EPI_ISL_2457952, EPI_ISL_2457953, EPI_ISL_2457955, EPI_ISL_2457956, EPI_ISL_2457957, EPI_ISL_2457958, EPI_ISL_2457959, EPI_ISL_2457960, EPI_ISL_2457961, EPI_ISL_2457963, EPI_ISL_2457965, EPI_ISL_2457966, EPI_ISL_2457967, EPI_ISL_2457969, EPI_ISL_2457970, EPI_ISL_2457971, EPI_ISL_2457974, EPI_ISL_2457975, EPI_ISL_2457976, EPI_ISL_2457977, EPI_ISL_2457979, EPI_ISL_2457980, EPI_ISL_2457981, EPI_ISL_2457982, EPI_ISL_2457983, EPI_ISL_2457984, EPI_ISL_2457985, EPI_ISL_2457986, EPI_ISL_2457987, EPI_ISL_2457990, EPI_ISL_2457991, EPI_ISL_2457993, EPI_ISL_2457994, EPI_ISL_2457995, EPI_ISL_2457997, EPI_ISL_2457998, EPI_ISL_2457999, EPI_ISL_2458002, EPI_ISL_2458003, EPI_ISL_2458004, EPI_ISL_2458005, EPI_ISL_2458006, EPI_ISL_2458007, EPI_ISL_2458008, EPI_ISL_2458009, EPI_ISL_2458012, EPI_ISL_2458013, EPI_ISL_2458014, EPI_ISL_2458015, EPI_ISL_2458016, | | | |

| | | | |
|---|---|---|---|
| EPI_ISL_2458019, EPI_ISL_2458020, EPI_ISL_2458021, EPI_ISL_2458023, EPI_ISL_2458024, EPI_ISL_2458025, EPI_ISL_2458026, EPI_ISL_2458027, EPI_ISL_2458032, EPI_ISL_2458033, EPI_ISL_2458036, EPI_ISL_2458038, EPI_ISL_2458039, EPI_ISL_2458040, EPI_ISL_2458041, EPI_ISL_2458045, EPI_ISL_2458046, EPI_ISL_2458047, EPI_ISL_2458048, EPI_ISL_2458049, EPI_ISL_2458050, EPI_ISL_2458051, EPI_ISL_2458052, EPI_ISL_2458053, EPI_ISL_2458054, EPI_ISL_2458056 | | | |
| see above | Lighthouse Lab in Glasgow | Wellcome Sanger Institute for the COVID-19 Genomics UK (COG-UK) Consortium | Harper VanSteenhouse, Yumi Kasai, David Gray, Carol Clugston, Anna Dominiczak and Alex Alderton, Roberto Amato, Jeffrey Barrett, Sonia Goncalves, Ewan Harrison, David K. Jackson, Ian Johnston, Dominic Kwiatkowski, Cordelia Langford, John Sillitoe on behalf of the Wellcome Sanger Institute COVID-19 Surveillance Team |
| EPI_ISL_2458057, EPI_ISL_2458058, EPI_ISL_2458059, EPI_ISL_2458060, EPI_ISL_2458061, EPI_ISL_2458062 | National Hospital for Tropical Diseases | Oxford University Clinical Research Unit, Hanoi, Vietnam | Pham Ngoc Thach, Nguyen Thi Tam, Van Dinh Trang, Le Van Duyet, Nguyen Thu Trang, Nguyen Thi Hong Thuong, Nguyen Thi Kim Chi, Phan Manh Cuong, Thomas Kesteman, H.Rogier van Doorn on behalf of the OUCRU COVID-19 research group |
| EPI_ISL_2458122, EPI_ISL_2458125, EPI_ISL_2458127, EPI_ISL_2458129, EPI_ISL_2458130, EPI_ISL_2458140, EPI_ISL_2458144, EPI_ISL_2458145, EPI_ISL_2458146, EPI_ISL_2458151, EPI_ISL_2458152, EPI_ISL_2458155, EPI_ISL_2458156, EPI_ISL_2458160 | | | |
| see above | Hospital Universitari Vall d'Hebron - Vall d'Hebron Institut de Recerca | Hospital Universitari Vall d'Hebron - Vall d'Hebron Institut de Recerca | Cristina Andrés, Maria Piñana, Alejandra González-Sánchez, Damir Garcia-Cehic, Ariadna Rando, Juliana Esperalba, Maria Gema Codina, Carla Castillo, Maria Carmen Martin, Tomàs Pumarola, Josep Quer, Andrés Antón |
| EPI_ISL_2458185 | NYU Langone Health | Departments of Pathology and Medicine, New York University School of Medicine | Adriana Heguy, Dacia Dimartino, Emily Guzman, Christian Marier, Peter Meyn, Sitharam Ramaswami, Gael Westby, Paul Zappile, Yutong Zhang, Paolo Cotzia, Guiqing Wang |
| EPI_ISL_2458246, EPI_ISL_2458250, EPI_ISL_2458252, EPI_ISL_2458261, EPI_ISL_2458262, EPI_ISL_2458272, EPI_ISL_2458273, EPI_ISL_2458277, EPI_ISL_2458285, EPI_ISL_2458287, EPI_ISL_2458288, EPI_ISL_2458290, EPI_ISL_2458293, EPI_ISL_2458303, EPI_ISL_2458312, EPI_ISL_2458313, EPI_ISL_2458314, EPI_ISL_2458318, EPI_ISL_2458323, EPI_ISL_2458329, EPI_ISL_2458466, EPI_ISL_2458467, EPI_ISL_2458509, EPI_ISL_2458515, EPI_ISL_2458516, EPI_ISL_2458517, EPI_ISL_2458518, EPI_ISL_2458519, EPI_ISL_2458520, EPI_ISL_2458521, EPI_ISL_2458522 | | | |
| see above | Utah Public Health Laboratory | Utah Public Health Laboratory | Erin L. Young, Kelly F. Oakeson, Tara Gallagher |
| EPI_ISL_2460052 | National Centre for Disease Control (NCDC) Biotechnology Division, Delhi | NCDC Delhi, Biotechnology Division INSACOG | Uma Sharma, Manoj K Singh, Radhakrishnan V. S, Robin Marwal, Mahesh S Dhar, Kalaiarasan Ponnusamy, Meena Datta, Priyanka Singh, Hemlata Lall, Hema Gogia, Preeti Madan, Sandhya Kabra, Sujeet K Singh, Partha Rakshit |
| EPI_ISL_2460533, EPI_ISL_2460546 | National Centre for Disease Control (NCDC) Biotechnology Division, Delhi | NCDC Delhi, Biotechnology Division INSACOG | Kalaiarasan Ponnusamy, Meena Datta, Priyanka Singh, Uma Sharma, Hemlata Lall, Manoj K Singh, Radhakrishnan V. S, Robin Marwal, Mahesh S Dhar, Hema Gogia, Preeti Madan, Sandhya Kabra, Sujeet K Singh, Partha Rakshit |
| EPI_ISL_2460547 | National Centre for Disease Control (NCDC) Biotechnology Division, Delhi | NCDC Delhi, Biotechnology Division INSACOG | Mahesh S Dhar, Kalaiarasan Ponnusamy, Meena Datta, Priyanka Singh, Uma Sharma, Hemlata Lall, Manoj K Singh, Radhakrishnan V. S, Robin Marwal, Hema Gogia, Preeti Madan, Sandhya Kabra, Sujeet K Singh, Partha Rakshit |
| EPI_ISL_2460550, EPI_ISL_2460605, EPI_ISL_2460627, EPI_ISL_2460633, EPI_ISL_2460681 | National Centre for Disease Control (NCDC) Biotechnology Division, Delhi | NCDC Delhi, Biotechnology Division INSACOG | Kalaiarasan Ponnusamy, Meena Datta, Priyanka Singh, Uma Sharma, Hemlata Lall, Manoj K Singh, Radhakrishnan V. S, Robin Marwal, Mahesh S Dhar, Hema Gogia, Preeti Madan, Sandhya Kabra, Sujeet K Singh, Partha Rakshit |
| EPI_ISL_2460703, EPI_ISL_2460707 | National Centre for Disease Control (NCDC) Biotechnology Division, Delhi | NCDC Delhi, Biotechnology Division INSACOG | Meena Datta, Priyanka Singh, Uma Sharma, Hemlata Lall, Manoj K Singh, Radhakrishnan V. S, Robin Marwal, Mahesh S Dhar, Kalaiarasan Ponnusamy, Hema Gogia, Preeti Madan, Sandhya Kabra, Sujeet K Singh, Partha Rakshit |
| EPI_ISL_2460786, EPI_ISL_2460794, EPI_ISL_2460798, EPI_ISL_2460863, EPI_ISL_2460868, EPI_ISL_2460869, EPI_ISL_2460870, EPI_ISL_2460871, EPI_ISL_2460872, EPI_ISL_2460873, EPI_ISL_2460874 | | | |
| see above | National Centre for Disease Control (NCDC) Biotechnology Division, Delhi | NCDC Delhi, Biotechnology Division INSACOG | Priyanka Singh, Uma Sharma, Hemlata Lall, Manoj K Singh, Radhakrishnan V. S, Robin Marwal, Mahesh S Dhar, Kalaiarasan Ponnusamy, Meena Datta, Hema Gogia, Preeti Madan, Sandhya Kabra, Sujeet K Singh, Partha Rakshit |
| EPI_ISL_2460884, EPI_ISL_2460886, EPI_ISL_2460894, EPI_ISL_2460895, EPI_ISL_2460901, EPI_ISL_2460903, EPI_ISL_2460906, EPI_ISL_2460912, EPI_ISL_2460913, EPI_ISL_2460914, EPI_ISL_2460915, EPI_ISL_2460916, EPI_ISL_2460917, EPI_ISL_2460918, EPI_ISL_2460919, EPI_ISL_2460927, EPI_ISL_2460929, EPI_ISL_2460930, EPI_ISL_2460931, EPI_ISL_2460932, EPI_ISL_2460933, EPI_ISL_2460935, EPI_ISL_2460936, EPI_ISL_2460940, EPI_ISL_2460942, EPI_ISL_2460943, EPI_ISL_2460944, EPI_ISL_2460945, EPI_ISL_2460950, EPI_ISL_2460951, EPI_ISL_2460953, EPI_ISL_2460956, EPI_ISL_2460957, EPI_ISL_2460958, EPI_ISL_2460960, EPI_ISL_2460961, EPI_ISL_2460963, EPI_ISL_2460965, EPI_ISL_2460970, EPI_ISL_2460971, EPI_ISL_2460973, EPI_ISL_2460975 | | | |
| see above | National Centre for Disease Control (NCDC) Biotechnology Division, Delhi | NCDC Delhi, Biotechnology Division INSACOG | Meena Datta, Priyanka Singh, Uma Sharma, Hemlata Lall, Manoj K Singh, Radhakrishnan V. S, Robin Marwal, Mahesh S Dhar, Kalaiarasan Ponnusamy, Hema Gogia, Preeti Madan, Sandhya Kabra, Sujeet K Singh, Partha Rakshit |
| EPI_ISL_2460976, EPI_ISL_2460978, EPI_ISL_2460979, EPI_ISL_2460982, EPI_ISL_2460984, EPI_ISL_2460985, EPI_ISL_2460986, EPI_ISL_2460988, EPI_ISL_2460989, EPI_ISL_2460990, EPI_ISL_2460991, EPI_ISL_2460996, EPI_ISL_2461001, EPI_ISL_2461005, EPI_ISL_2461007, EPI_ISL_2461010, EPI_ISL_2461011, EPI_ISL_2461013, EPI_ISL_2461014, EPI_ISL_2461015, EPI_ISL_2461016, EPI_ISL_2461021, EPI_ISL_2461023, EPI_ISL_2461024, EPI_ISL_2461025, EPI_ISL_2461026, EPI_ISL_2461027, EPI_ISL_2461028, EPI_ISL_2461029, EPI_ISL_2461030, EPI_ISL_2461031, EPI_ISL_2461032, EPI_ISL_2461033, EPI_ISL_2461035, EPI_ISL_2461036, EPI_ISL_2461037, EPI_ISL_2461038, EPI_ISL_2461039, EPI_ISL_2461040, EPI_ISL_2461042, EPI_ISL_2461043, EPI_ISL_2461044, EPI_ISL_2461045, EPI_ISL_2461046, EPI_ISL_2461047, EPI_ISL_2461048, EPI_ISL_2461049, EPI_ISL_2461050, EPI_ISL_2461051, EPI_ISL_2461052, EPI_ISL_2461053, EPI_ISL_2461054, EPI_ISL_2461055, EPI_ISL_2461056, EPI_ISL_2461057, EPI_ISL_2461058, EPI_ISL_2461059, EPI_ISL_2461060, EPI_ISL_2461061, EPI_ISL_2461062, EPI_ISL_2461063, EPI_ISL_2461064, EPI_ISL_2461065, EPI_ISL_2461066, EPI_ISL_2461068, EPI_ISL_2461069, EPI_ISL_2461070, EPI_ISL_2461071, EPI_ISL_2461072, EPI_ISL_2461073, EPI_ISL_2461075, EPI_ISL_2461076, EPI_ISL_2461078, EPI_ISL_2461080, EPI_ISL_2461081, EPI_ISL_2461082, EPI_ISL_2461083, EPI_ISL_2461084, EPI_ISL_2461086, EPI_ISL_2461088, EPI_ISL_2461089, EPI_ISL_2461090, EPI_ISL_2461091, EPI_ISL_2461093, EPI_ISL_2461094, EPI_ISL_2461095, EPI_ISL_2461096, EPI_ISL_2461097, EPI_ISL_2461098, EPI_ISL_2461099, EPI_ISL_2461101, EPI_ISL_2461102, EPI_ISL_2461103, EPI_ISL_2461104, EPI_ISL_2461105, EPI_ISL_2461106, EPI_ISL_2461109, EPI_ISL_2461113, EPI_ISL_2461116, EPI_ISL_2461117, EPI_ISL_2461120, EPI_ISL_2461121, EPI_ISL_2461122, EPI_ISL_2461123, EPI_ISL_2461124, EPI_ISL_2461125, EPI_ISL_2461127, EPI_ISL_2461128, EPI_ISL_2461130 | | | |
| see above | National Centre for Disease Control (NCDC) Biotechnology Division, Delhi | NCDC Delhi, Biotechnology Division INSACOG | Priyanka Singh, Uma Sharma, Hemlata Lall, Manoj K Singh, Radhakrishnan V. S, Robin Marwal, Mahesh S Dhar, Kalaiarasan Ponnusamy, Meena Datta, Hema Gogia, Preeti Madan, Sandhya Kabra, Sujeet K Singh, Partha Rakshit |
| EPI_ISL_2461132 | National Centre for Disease Control (NCDC) Biotechnology Division, Delhi | NCDC Delhi, Biotechnology Division INSACOG | Uma Sharma, Hemlata Lall, Manoj K Singh, Radhakrishnan V. S, Robin Marwal, Mahesh S Dhar, Kalaiarasan Ponnusamy, Meena Datta, Priyanka Singh, Hema Gogia, Preeti Madan, Sandhya Kabra, Sujeet K Singh, Partha Rakshit |
| EPI_ISL_2461135 | National Centre for Disease Control (NCDC) Biotechnology Division, Delhi | NCDC Delhi, Biotechnology Division INSACOG | Hemlata Lall, Manoj K Singh, Radhakrishnan V. S, Robin Marwal, Mahesh S Dhar, Kalaiarasan Ponnusamy, Meena Datta, Priyanka Singh, Uma Sharma, Hema Gogia, Preeti Madan, Sandhya Kabra, Sujeet K Singh, Partha Rakshit |
| EPI_ISL_2461136 | National Centre for Disease Control (NCDC) Biotechnology Division, Delhi | NCDC Delhi, Biotechnology Division INSACOG | Uma Sharma, Hemlata Lall, Manoj K Singh, Radhakrishnan V. S, Robin Marwal, Mahesh S Dhar, Kalaiarasan Ponnusamy, Meena Datta, Priyanka Singh, Hema Gogia, Preeti Madan, Sandhya Kabra, Sujeet K Singh, Partha Rakshit |
| EPI_ISL_2461137 | National Centre for Disease Control (NCDC) Biotechnology Division, Delhi | NCDC Delhi, Biotechnology Division INSACOG | Hemlata Lall, Manoj K Singh, Radhakrishnan V. S, Robin Marwal, Mahesh S Dhar, Kalaiarasan Ponnusamy, Meena Datta, Priyanka Singh, Uma Sharma, Hema Gogia, Preeti Madan, Sandhya Kabra, Sujeet K Singh, Partha Rakshit |
| EPI_ISL_2461139 | National Centre for Disease Control (NCDC) Biotechnology Division, Delhi | NCDC Delhi, Biotechnology Division INSACOG | Uma Sharma, Hemlata Lall, Manoj K Singh, Radhakrishnan V. S, Robin Marwal, Mahesh S Dhar, Kalaiarasan Ponnusamy, Meena Datta, Priyanka Singh, Hema Gogia, Preeti Madan, Sandhya Kabra, Sujeet K Singh, Partha Rakshit |
| EPI_ISL_2461143, EPI_ISL_2461146 | National Centre for Disease Control (NCDC) Biotechnology Division, Delhi | NCDC Delhi, Biotechnology Division INSACOG | Hemlata Lall, Manoj K Singh, Radhakrishnan V. S, Robin Marwal, Mahesh S Dhar, Kalaiarasan Ponnusamy, Meena Datta, Priyanka Singh, Uma Sharma, Hema Gogia, Preeti Madan, Sandhya Kabra, Sujeet K Singh, Partha Rakshit |
| EPI_ISL_2461149 | National Centre for Disease Control (NCDC) Biotechnology Division, Delhi | NCDC Delhi, Biotechnology Division INSACOG | Uma Sharma, Hemlata Lall, Manoj K Singh, Radhakrishnan V. S, Robin Marwal, Mahesh S Dhar, Kalaiarasan Ponnusamy, Meena Datta, Priyanka Singh, Hema Gogia, Preeti Madan, Sandhya Kabra, Sujeet K Singh, Partha Rakshit |
| EPI_ISL_2461150, EPI_ISL_2461151, EPI_ISL_2461152, EPI_ISL_2461153, EPI_ISL_2461154 | National Centre for Disease Control (NCDC) Biotechnology Division, Delhi | NCDC Delhi, Biotechnology Division INSACOG | Hemlata Lall, Manoj K Singh, Radhakrishnan V. S, Robin Marwal, Mahesh S Dhar, Kalaiarasan Ponnusamy, Meena Datta, Priyanka Singh, Uma Sharma, Hema Gogia, Preeti Madan, Sandhya Kabra, Sujeet K Singh, Partha Rakshit |
| EPI_ISL_2461155 | National Centre for Disease Control (NCDC) Biotechnology Division, Delhi | NCDC Delhi, Biotechnology Division INSACOG | Uma Sharma, Hemlata Lall, Manoj K Singh, Radhakrishnan V. S, Robin Marwal, Mahesh S Dhar, Kalaiarasan Ponnusamy, Meena Datta, Priyanka Singh, Hema Gogia, Preeti Madan, Sandhya Kabra, Sujeet K Singh, Partha Rakshit |
| EPI_ISL_2461156 | National Centre for Disease Control (NCDC) Biotechnology Division, Delhi | NCDC Delhi, Biotechnology Division INSACOG | Hemlata Lall, Manoj K Singh, Radhakrishnan V. S, Robin Marwal, Mahesh S Dhar, Kalaiarasan Ponnusamy, Meena Datta, Priyanka Singh, Uma Sharma, Hema Gogia, Preeti Madan, Sandhya Kabra, Sujeet K Singh, Partha Rakshit |
| EPI_ISL_2461158 | National Centre for Disease Control (NCDC) Biotechnology Division, Delhi | NCDC Delhi, Biotechnology Division INSACOG | Uma Sharma, Hemlata Lall, Manoj K Singh, Radhakrishnan V. S, Robin Marwal, Mahesh S Dhar, Kalaiarasan Ponnusamy, Meena Datta, Priyanka Singh, Hema Gogia, Preeti Madan, Sandhya Kabra, Sujeet K Singh, Partha Rakshit |
| EPI_ISL_2461159, EPI_ISL_2461160, EPI_ISL_2461161, EPI_ISL_2461162, EPI_ISL_2461163 | National Centre for Disease Control (NCDC) Biotechnology Division, Delhi | NCDC Delhi, Biotechnology Division INSACOG | Hemlata Lall, Manoj K Singh, Radhakrishnan V. S, Robin Marwal, Mahesh S Dhar, Kalaiarasan Ponnusamy, Meena Datta, Priyanka Singh, Uma Sharma, Hema Gogia, Preeti Madan, Sandhya Kabra, Sujeet K Singh, Partha Rakshit |
| EPI_ISL_2461164 | National Centre for Disease Control (NCDC) Biotechnology Division, Delhi | NCDC Delhi, Biotechnology Division INSACOG | Uma Sharma, Hemlata Lall, Manoj K Singh, Radhakrishnan V. S, Robin Marwal, Mahesh S Dhar, Kalaiarasan Ponnusamy, Meena Datta, Priyanka Singh, Hema Gogia, Preeti Madan, Sandhya Kabra, Sujeet K Singh, Partha Rakshit |
| EPI_ISL_2461165 | National Centre for Disease Control (NCDC) Biotechnology Division, Delhi | NCDC Delhi, Biotechnology Division INSACOG | Hemlata Lall, Manoj K Singh, Radhakrishnan V. S, Robin Marwal, Mahesh S Dhar, Kalaiarasan Ponnusamy, Meena Datta, Priyanka Singh, Uma Sharma, Hema Gogia, Preeti Madan, Sandhya Kabra, Sujeet K Singh, Partha Rakshit |
| EPI_ISL_2461167 | National Centre for Disease Control (NCDC) Biotechnology Division, Delhi | NCDC Delhi, Biotechnology Division INSACOG | Uma Sharma, Hemlata Lall, Manoj K Singh, Radhakrishnan V. S, Robin Marwal, Mahesh S Dhar, Kalaiarasan Ponnusamy, Meena Datta, Priyanka Singh, Hema Gogia, Preeti Madan, Sandhya Kabra, Sujeet K Singh, Partha Rakshit |
| EPI_ISL_2461169 | National Centre for Disease Control (NCDC) Biotechnology Division, Delhi | NCDC Delhi, Biotechnology Division INSACOG | Hemlata Lall, Manoj K Singh, Radhakrishnan V. S, Robin Marwal, Mahesh S Dhar, Kalaiarasan Ponnusamy, Meena Datta, Priyanka Singh, Uma Sharma, Hema Gogia, Preeti Madan, Sandhya Kabra, Sujeet K Singh, Partha Rakshit |
| EPI_ISL_2461170 | National Centre for Disease Control (NCDC) Biotechnology Division, Delhi | NCDC Delhi, Biotechnology Division INSACOG | Uma Sharma, Hemlata Lall, Manoj K Singh, Radhakrishnan V. S, Robin Marwal, Mahesh S Dhar, Kalaiarasan Ponnusamy, Meena Datta, Priyanka Singh, Hema Gogia, Preeti Madan, Sandhya Kabra, Sujeet K Singh, Partha Rakshit |
| EPI_ISL_2461171 | National Centre for Disease Control (NCDC) Biotechnology Division, Delhi | NCDC Delhi, Biotechnology Division INSACOG | Hemlata Lall, Manoj K Singh, Radhakrishnan V. S, Robin Marwal, Mahesh S Dhar, Kalaiarasan Ponnusamy, Meena Datta, Priyanka Singh, Uma Sharma, Hema Gogia, Preeti Madan, Sandhya Kabra, Sujeet K Singh, Partha Rakshit |
| EPI_ISL_2461172, EPI_ISL_2461173 | National Centre for Disease Control (NCDC) Biotechnology Division, Delhi | NCDC Delhi, Biotechnology Division INSACOG | Uma Sharma, Hemlata Lall, Manoj K Singh, Radhakrishnan V. S, Robin Marwal, Mahesh S Dhar, Kalaiarasan Ponnusamy, Meena Datta, Priyanka Singh, Hema Gogia, Preeti Madan, Sandhya Kabra, Sujeet K Singh, Partha Rakshit |
| EPI_ISL_2461174 | National Centre for Disease Control (NCDC) Biotechnology Division, Delhi | NCDC Delhi, Biotechnology Division INSACOG | Hemlata Lall, Manoj K Singh, Radhakrishnan V. S, Robin Marwal, Mahesh S Dhar, Kalaiarasan Ponnusamy, Meena Datta, Priyanka Singh, Uma Sharma, Hema Gogia, Preeti Madan, Sandhya Kabra, Sujeet K Singh, Partha Rakshit |
| EPI_ISL_2461181 | National Centre for Disease Control (NCDC) Biotechnology Division, Delhi | NCDC Delhi, Biotechnology Division INSACOG | Uma Sharma, Hemlata Lall, Manoj K Singh, Radhakrishnan V. S, Robin Marwal, Mahesh S Dhar, Kalaiarasan Ponnusamy, Meena Datta, Priyanka Singh, Hema Gogia, Preeti Madan, Sandhya Kabra, Sujeet K Singh, Partha Rakshit |

| | | | |
|---|---|---|---|
| EPI_ISL_2461184, EPI_ISL_2461185, EPI_ISL_2461186 | National Centre for Disease Control (NCDC) Biotechnology Division, Delhi | NCDC Delhi, Biotechnology Division INSACOG | Hemlata Lall, Manoj K Singh, Radhakrishnan V. S, Robin Marwal, Mahesh S Dhar, Kalaiarasan Ponnusamy, Meena Datta, Priyanka Singh, Uma Sharma, Hema Gogia, Preeti Madan, Sandhya Kabra, Sujeet K Singh, Partha Rakshit |
| EPI_ISL_2461193, EPI_ISL_2461194 | National Centre for Disease Control (NCDC) Biotechnology Division, Delhi | NCDC Delhi, Biotechnology Division INSACOG | Uma Sharma, Hemlata Lall, Manoj K Singh, Radhakrishnan V. S, Robin Marwal, Mahesh S Dhar, Kalaiarasan Ponnusamy, Meena Datta, Priyanka Singh, Hema Gogia, Preeti Madan, Sandhya Kabra, Sujeet K Singh, Partha Rakshit |
| EPI_ISL_2461195, EPI_ISL_2461196 | National Centre for Disease Control (NCDC) Biotechnology Division, Delhi | NCDC Delhi, Biotechnology Division INSACOG | Hemlata Lall, Manoj K Singh, Radhakrishnan V. S, Robin Marwal, Mahesh S Dhar, Kalaiarasan Ponnusamy, Meena Datta, Priyanka Singh, Uma Sharma, Hema Gogia, Preeti Madan, Sandhya Kabra, Sujeet K Singh, Partha Rakshit |
| EPI_ISL_2461202 | Tokyo Metropolitan Institute of Public Health | Tokyo Metropolitan Institute of Public Health | Masaki Hayashi, Takako Yamazaki, Akane Negishi,Hiroshi Koike, Hirofumi Miyake, Takushi Fujiwara, Mami Nagashima, Jun Suzuki, Kenji Sadamasu |
| EPI_ISL_2461203 | National Centre for Disease Control (NCDC) Biotechnology Division, Delhi | NCDC Delhi, Biotechnology Division INSACOG | Manoj K Singh, Radhakrishnan V. S, Robin Marwal, Mahesh S Dhar, Kalaiarasan Ponnusamy, Meena Datta, Priyanka Singh, Uma Sharma, Hemlata Lall, Hema Gogia, Preeti Madan, Sandhya Kabra, Sujeet K Singh, Partha Rakshit |
| EPI_ISL_2461210, EPI_ISL_2461211, EPI_ISL_2461213 | National Centre for Disease Control (NCDC) Biotechnology Division, Delhi | NCDC Delhi, Biotechnology Division INSACOG | Radhakrishnan V. S, Robin Marwal, Mahesh S Dhar, Kalaiarasan Ponnusamy, Meena Datta, Priyanka Singh, Uma Sharma, Hemlata Lall, Manoj K Singh, Hema Gogia, Preeti Madan, Sandhya Kabra, Sujeet K Singh, Partha Rakshit |
| EPI_ISL_2461215, EPI_ISL_2461218 | National Centre for Disease Control (NCDC) Biotechnology Division, Delhi | NCDC Delhi, Biotechnology Division INSACOG | Manoj K Singh, Radhakrishnan V. S, Robin Marwal, Mahesh S Dhar, Kalaiarasan Ponnusamy, Meena Datta, Priyanka Singh, Uma Sharma, Hemlata Lall, Hema Gogia, Preeti Madan, Sandhya Kabra, Sujeet K Singh, Partha Rakshit |
| EPI_ISL_2461219 | National Centre for Disease Control (NCDC) Biotechnology Division, Delhi | NCDC Delhi, Biotechnology Division INSACOG | Radhakrishnan V. S, Robin Marwal, Mahesh S Dhar, Kalaiarasan Ponnusamy, Meena Datta, Priyanka Singh, Uma Sharma, Hemlata Lall, Manoj K Singh, Hema Gogia, Preeti Madan, Sandhya Kabra, Sujeet K Singh, Partha Rakshit |
| EPI_ISL_2461224, EPI_ISL_2461228 | National Centre for Disease Control (NCDC) Biotechnology Division, Delhi | NCDC Delhi, Biotechnology Division INSACOG | Manoj K Singh, Radhakrishnan V. S, Robin Marwal, Mahesh S Dhar, Kalaiarasan Ponnusamy, Meena Datta, Priyanka Singh, Uma Sharma, Hemlata Lall, Hema Gogia, Preeti Madan, Sandhya Kabra, Sujeet K Singh, Partha Rakshit |
| EPI_ISL_2461229, EPI_ISL_2461230, EPI_ISL_2461233, EPI_ISL_2461234, EPI_ISL_2461237, EPI_ISL_2461238, EPI_ISL_2461239, EPI_ISL_2461240, EPI_ISL_2461242 | National Centre for Disease Control (NCDC) Biotechnology Division, Delhi | NCDC Delhi, Biotechnology Division INSACOG | Radhakrishnan V. S, Robin Marwal, Mahesh S Dhar, Kalaiarasan Ponnusamy, Meena Datta, Priyanka Singh, Uma Sharma, Hemlata Lall, Manoj K Singh, Hema Gogia, Preeti Madan, Sandhya Kabra, Sujeet K Singh, Partha Rakshit |
| EPI_ISL_2461243 | National Centre for Disease Control (NCDC) Biotechnology Division, Delhi | NCDC Delhi, Biotechnology Division INSACOG | Manoj K Singh, Radhakrishnan V. S, Robin Marwal, Mahesh S Dhar, Kalaiarasan Ponnusamy, Meena Datta, Priyanka Singh, Uma Sharma, Hemlata Lall, Hema Gogia, Preeti Madan, Sandhya Kabra, Sujeet K Singh, Partha Rakshit |
| EPI_ISL_2461245, EPI_ISL_2461246, EPI_ISL_2461249, EPI_ISL_2461250 | National Centre for Disease Control (NCDC) Biotechnology Division, Delhi | NCDC Delhi, Biotechnology Division INSACOG | Radhakrishnan V. S, Robin Marwal, Mahesh S Dhar, Kalaiarasan Ponnusamy, Meena Datta, Priyanka Singh, Uma Sharma, Hemlata Lall, Manoj K Singh, Hema Gogia, Preeti Madan, Sandhya Kabra, Sujeet K Singh, Partha Rakshit |
| EPI_ISL_2461252 | National Centre for Disease Control (NCDC) Biotechnology Division, Delhi | NCDC Delhi, Biotechnology Division INSACOG | Manoj K Singh, Radhakrishnan V. S, Robin Marwal, Mahesh S Dhar, Kalaiarasan Ponnusamy, Meena Datta, Priyanka Singh, Uma Sharma, Hemlata Lall, Hema Gogia, Preeti Madan, Sandhya Kabra, Sujeet K Singh, Partha Rakshit |
| EPI_ISL_2461253, EPI_ISL_2461254, EPI_ISL_2461255, EPI_ISL_2461256 | National Centre for Disease Control (NCDC) Biotechnology Division, Delhi | NCDC Delhi, Biotechnology Division INSACOG | Radhakrishnan V. S, Robin Marwal, Mahesh S Dhar, Kalaiarasan Ponnusamy, Meena Datta, Priyanka Singh, Uma Sharma, Hemlata Lall, Manoj K Singh, Hema Gogia, Preeti Madan, Sandhya Kabra, Sujeet K Singh, Partha Rakshit |
| EPI_ISL_2461257 | National Centre for Disease Control (NCDC) Biotechnology Division, Delhi | NCDC Delhi, Biotechnology Division INSACOG | Manoj K Singh, Radhakrishnan V. S, Robin Marwal, Mahesh S Dhar, Kalaiarasan Ponnusamy, Meena Datta, Priyanka Singh, Uma Sharma, Hemlata Lall, Hema Gogia, Preeti Madan, Sandhya Kabra, Sujeet K Singh, Partha Rakshit |
| EPI_ISL_2461258, EPI_ISL_2461259, EPI_ISL_2461260 | National Centre for Disease Control (NCDC) Biotechnology Division, Delhi | NCDC Delhi, Biotechnology Division INSACOG | Radhakrishnan V. S, Robin Marwal, Mahesh S Dhar, Kalaiarasan Ponnusamy, Meena Datta, Priyanka Singh, Uma Sharma, Hemlata Lall, Manoj K Singh, Hema Gogia, Preeti Madan, Sandhya Kabra, Sujeet K Singh, Partha Rakshit |
| EPI_ISL_2461261 | National Centre for Disease Control (NCDC) Biotechnology Division, Delhi | NCDC Delhi, Biotechnology Division INSACOG | Manoj K Singh, Radhakrishnan V. S, Robin Marwal, Mahesh S Dhar, Kalaiarasan Ponnusamy, Meena Datta, Priyanka Singh, Uma Sharma, Hemlata Lall, Hema Gogia, Preeti Madan, Sandhya Kabra, Sujeet K Singh, Partha Rakshit |
| EPI_ISL_2461263, EPI_ISL_2461264, EPI_ISL_2461265, EPI_ISL_2461266, EPI_ISL_2461267 | National Centre for Disease Control (NCDC) Biotechnology Division, Delhi | NCDC Delhi, Biotechnology Division INSACOG | Radhakrishnan V. S, Robin Marwal, Mahesh S Dhar, Kalaiarasan Ponnusamy, Meena Datta, Priyanka Singh, Uma Sharma, Hemlata Lall, Manoj K Singh, Hema Gogia, Preeti Madan, Sandhya Kabra, Sujeet K Singh, Partha Rakshit |
| EPI_ISL_2461270 | National Centre for Disease Control (NCDC) Biotechnology Division, Delhi | NCDC Delhi, Biotechnology Division INSACOG | Manoj K Singh, Radhakrishnan V. S, Robin Marwal, Mahesh S Dhar, Kalaiarasan Ponnusamy, Meena Datta, Priyanka Singh, Uma Sharma, Hemlata Lall, Hema Gogia, Preeti Madan, Sandhya Kabra, Sujeet K Singh, Partha Rakshit |
| EPI_ISL_2461271, EPI_ISL_2461272, EPI_ISL_2461273, EPI_ISL_2461274 | National Centre for Disease Control (NCDC) Biotechnology Division, Delhi | NCDC Delhi, Biotechnology Division INSACOG | Radhakrishnan V. S, Robin Marwal, Mahesh S Dhar, Kalaiarasan Ponnusamy, Meena Datta, Priyanka Singh, Uma Sharma, Hemlata Lall, Manoj K Singh, Hema Gogia, Preeti Madan, Sandhya Kabra, Sujeet K Singh, Partha Rakshit |
| EPI_ISL_2461275 | National Centre for Disease Control (NCDC) Biotechnology Division, Delhi | NCDC Delhi, Biotechnology Division INSACOG | Manoj K Singh, Radhakrishnan V. S, Robin Marwal, Mahesh S Dhar, Kalaiarasan Ponnusamy, Meena Datta, Priyanka Singh, Uma Sharma, Hemlata Lall, Hema Gogia, Preeti Madan, Sandhya Kabra, Sujeet K Singh, Partha Rakshit |
| EPI_ISL_2461276, EPI_ISL_2461279, EPI_ISL_2461280 | National Centre for Disease Control (NCDC) Biotechnology Division, Delhi | NCDC Delhi, Biotechnology Division INSACOG | Radhakrishnan V. S, Robin Marwal, Mahesh S Dhar, Kalaiarasan Ponnusamy, Meena Datta, Priyanka Singh, Uma Sharma, Hemlata Lall, Manoj K Singh, Hema Gogia, Preeti Madan, Sandhya Kabra, Sujeet K Singh, Partha Rakshit |
| EPI_ISL_2461282 | National Centre for Disease Control (NCDC) Biotechnology Division, Delhi | NCDC Delhi, Biotechnology Division INSACOG | Manoj K Singh, Radhakrishnan V. S, Robin Marwal, Mahesh S Dhar, Kalaiarasan Ponnusamy, Meena Datta, Priyanka Singh, Uma Sharma, Hemlata Lall, Hema Gogia, Preeti Madan, Sandhya Kabra, Sujeet K Singh, Partha Rakshit |
| EPI_ISL_2461283 | National Centre for Disease Control (NCDC) Biotechnology Division, Delhi | NCDC Delhi, Biotechnology Division INSACOG | Radhakrishnan V. S, Robin Marwal, Mahesh S Dhar, Kalaiarasan Ponnusamy, Meena Datta, Priyanka Singh, Uma Sharma, Hemlata Lall, Manoj K Singh, Hema Gogia, Preeti Madan, Sandhya Kabra, Sujeet K Singh, Partha Rakshit |
| EPI_ISL_2461285 | National Centre for Disease Control (NCDC) Biotechnology Division, Delhi | NCDC Delhi, Biotechnology Division INSACOG | Manoj K Singh, Radhakrishnan V. S, Robin Marwal, Mahesh S Dhar, Kalaiarasan Ponnusamy, Meena Datta, Priyanka Singh, Uma Sharma, Hemlata Lall, Hema Gogia, Preeti Madan, Sandhya Kabra, Sujeet K Singh, Partha Rakshit |
| EPI_ISL_2461286, EPI_ISL_2461287, EPI_ISL_2461316 | National Centre for Disease Control (NCDC) Biotechnology Division, Delhi | NCDC Delhi, Biotechnology Division INSACOG | Radhakrishnan V. S, Robin Marwal, Mahesh S Dhar, Kalaiarasan Ponnusamy, Meena Datta, Priyanka Singh, Uma Sharma, Hemlata Lall, Manoj K Singh, Hema Gogia, Preeti Madan, Sandhya Kabra, Sujeet K Singh, Partha Rakshit |
| EPI_ISL_2461317 | National Centre for Disease Control (NCDC) Biotechnology Division, Delhi | NCDC Delhi, Biotechnology Division INSACOG | Manoj K Singh, Radhakrishnan V. S, Robin Marwal, Mahesh S Dhar, Kalaiarasan Ponnusamy, Meena Datta, Priyanka Singh, Uma Sharma, Hemlata Lall, Hema Gogia, Preeti Madan, Sandhya Kabra, Sujeet K Singh, Partha Rakshit |
| EPI_ISL_2461318, EPI_ISL_2461341 | National Centre for Disease Control (NCDC) Biotechnology Division, Delhi | NCDC Delhi, Biotechnology Division INSACOG | Radhakrishnan V. S, Robin Marwal, Mahesh S Dhar, Kalaiarasan Ponnusamy, Meena Datta, Priyanka Singh, Uma Sharma, Hemlata Lall, Manoj K Singh, Hema Gogia, Preeti Madan, Sandhya Kabra, Sujeet K Singh, Partha Rakshit |
| EPI_ISL_2461359 | National Centre for Disease Control (NCDC) Biotechnology Division, Delhi | NCDC Delhi, Biotechnology Division INSACOG | Manoj K Singh, Radhakrishnan V. S, Robin Marwal, Mahesh S Dhar, Kalaiarasan Ponnusamy, Meena Datta, Priyanka Singh, Uma Sharma, Hemlata Lall, Hema Gogia, Preeti Madan, Sandhya Kabra, Sujeet K Singh, Partha Rakshit |
| EPI_ISL_2461362 | National Centre for Disease Control (NCDC) Biotechnology Division, Delhi | NCDC Delhi, Biotechnology Division INSACOG | Radhakrishnan V. S, Robin Marwal, Mahesh S Dhar, Kalaiarasan Ponnusamy, Meena Datta, Priyanka Singh, Uma Sharma, Hemlata Lall, Manoj K Singh, Hema Gogia, Preeti Madan, Sandhya Kabra, Sujeet K Singh, Partha Rakshit |
| EPI_ISL_2461363, EPI_ISL_2461364 | National Centre for Disease Control (NCDC) Biotechnology Division, Delhi | NCDC Delhi, Biotechnology Division INSACOG | Manoj K Singh, Radhakrishnan V. S, Robin Marwal, Mahesh S Dhar, Kalaiarasan Ponnusamy, Meena Datta, Priyanka Singh, Uma Sharma, Hemlata Lall, Hema Gogia, Preeti Madan, Sandhya Kabra, Sujeet K Singh, Partha Rakshit |
| EPI_ISL_2461365, EPI_ISL_2461366, EPI_ISL_2461367, EPI_ISL_2461368, EPI_ISL_2461369 | National Centre for Disease Control (NCDC) Biotechnology Division, Delhi | NCDC Delhi, Biotechnology Division INSACOG | Radhakrishnan V. S, Robin Marwal, Mahesh S Dhar, Kalaiarasan Ponnusamy, Meena Datta, Priyanka Singh, Uma Sharma, Hemlata Lall, Manoj K Singh, Hema Gogia, Preeti Madan, Sandhya Kabra, Sujeet K Singh, Partha Rakshit |
| EPI_ISL_2461376 | National Centre for Disease Control (NCDC) Biotechnology Division, Delhi | NCDC Delhi, Biotechnology Division INSACOG | Manoj K Singh, Radhakrishnan V. S, Robin Marwal, Mahesh S Dhar, Kalaiarasan Ponnusamy, Meena Datta, Priyanka Singh, Uma Sharma, Hemlata Lall, Hema Gogia, Preeti Madan, Sandhya Kabra, Sujeet K Singh, Partha Rakshit |
| EPI_ISL_2461377, EPI_ISL_2461378, EPI_ISL_2461379, EPI_ISL_2461380, EPI_ISL_2461381 | National Centre for Disease Control (NCDC) Biotechnology Division, Delhi | NCDC Delhi, Biotechnology Division INSACOG | Radhakrishnan V. S, Robin Marwal, Mahesh S Dhar, Kalaiarasan Ponnusamy, Meena Datta, Priyanka Singh, Uma Sharma, Hemlata Lall, Manoj K Singh, Hema Gogia, Preeti Madan, Sandhya Kabra, Sujeet K Singh, Partha Rakshit |

| | | | |
|---|---|---|---|
| EPI_ISL_2461382, EPI_ISL_2461383, EPI_ISL_2461384, EPI_ISL_2461385 | National Centre for Disease Control (NCDC) Biotechnology Division, Delhi | NCDC Delhi, Biotechnology Division INSACOG | Manoj K Singh, Radhakrishnan V. S, Robin Marwal, Mahesh S Dhar, Kalaiarasan Ponnusamy, Meena Datta, Priyanka Singh, Uma Sharma, Hemlata Lall, Hema Gogia, Preeti Madan, Sandhya Kabra, Sujeet K Singh, Partha Rakshit |
| EPI_ISL_2461386, EPI_ISL_2461387, EPI_ISL_2461388, EPI_ISL_2461389, EPI_ISL_2461391, EPI_ISL_2461392, EPI_ISL_2461393, EPI_ISL_2461394 | National Centre for Disease Control (NCDC) Biotechnology Division, Delhi | NCDC Delhi, Biotechnology Division INSACOG | Radhakrishnan V. S, Robin Marwal, Mahesh S Dhar, Kalaiarasan Ponnusamy, Meena Datta, Priyanka Singh, Uma Sharma, Hemlata Lall, Manoj K Singh, Hema Gogia, Preeti Madan, Sandhya Kabra, Sujeet K Singh, Partha Rakshit |
| EPI_ISL_2461395, EPI_ISL_2461396, EPI_ISL_2461397, EPI_ISL_2461398, EPI_ISL_2461399, EPI_ISL_2461400, EPI_ISL_2461401, EPI_ISL_2461402, EPI_ISL_2461403, EPI_ISL_2461404, EPI_ISL_2461405, EPI_ISL_2461406, EPI_ISL_2461407, EPI_ISL_2461408, EPI_ISL_2461409, EPI_ISL_2461410, EPI_ISL_2461411, EPI_ISL_2461412, EPI_ISL_2461414, EPI_ISL_2461415, EPI_ISL_2461416, EPI_ISL_2461417, EPI_ISL_2461418, EPI_ISL_2461419, EPI_ISL_2461420, EPI_ISL_2461421 | | | |
| see above | National Centre for Disease Control (NCDC) Biotechnology Division, Delhi | NCDC Delhi, Biotechnology Division INSACOG | Manoj K Singh, Radhakrishnan V. S, Robin Marwal, Mahesh S Dhar, Kalaiarasan Ponnusamy, Meena Datta, Priyanka Singh, Uma Sharma, Hemlata Lall, Hema Gogia, Preeti Madan, Sandhya Kabra, Sujeet K Singh, Partha Rakshit |
| EPI_ISL_2461422, EPI_ISL_2461423, EPI_ISL_2461424, EPI_ISL_2461425, EPI_ISL_2461426, EPI_ISL_2461427, EPI_ISL_2461428, EPI_ISL_2461429, EPI_ISL_2461430, EPI_ISL_2461431, EPI_ISL_2461432, EPI_ISL_2461433, EPI_ISL_2461434, EPI_ISL_2461435, EPI_ISL_2461436, EPI_ISL_2461437, EPI_ISL_2461438, EPI_ISL_2461439, EPI_ISL_2461440, EPI_ISL_2461441, EPI_ISL_2461442, EPI_ISL_2461443, EPI_ISL_2461444, EPI_ISL_2461445, EPI_ISL_2461446, EPI_ISL_2461447, EPI_ISL_2461448, EPI_ISL_2461449, EPI_ISL_2461450, EPI_ISL_2461451, EPI_ISL_2461452, EPI_ISL_2461453, EPI_ISL_2461454, EPI_ISL_2461455, EPI_ISL_2461456, EPI_ISL_2461457, EPI_ISL_2461458, EPI_ISL_2461459, EPI_ISL_2461460, EPI_ISL_2461461, EPI_ISL_2461462, EPI_ISL_2461463, EPI_ISL_2461464, EPI_ISL_2461465, EPI_ISL_2461466, EPI_ISL_2461467, EPI_ISL_2461468, EPI_ISL_2461469, EPI_ISL_2461470, EPI_ISL_2461471, EPI_ISL_2461472, EPI_ISL_2461473, EPI_ISL_2461474, EPI_ISL_2461475, EPI_ISL_2461496, EPI_ISL_2461501 | | | |
| see above | National Centre for Disease Control (NCDC) Biotechnology Division, Delhi | NCDC Delhi, Biotechnology Division INSACOG | Radhakrishnan V. S, Robin Marwal, Mahesh S Dhar, Kalaiarasan Ponnusamy, Meena Datta, Priyanka Singh, Uma Sharma, Hemlata Lall, Manoj K Singh, Hema Gogia, Preeti Madan, Sandhya Kabra, Sujeet K Singh, Partha Rakshit |
| EPI_ISL_2461505, EPI_ISL_2461506 | Tokyo Metropolitan Institute of Public Health | Tokyo Metropolitan Institute of Public Health | Hirofumi Miyake, Kenshirou Kuroki, Maya Isogai, Masaki Hayashi, Takako Yamazaki, Akane Negishi,Takushi Fujiwara, Mami Nagashima, Jun Suzuki, Kenji Sadamasu |
| EPI_ISL_2461509, EPI_ISL_2461510 | National Centre for Disease Control (NCDC) Biotechnology Division, Delhi | NCDC Delhi, Biotechnology Division INSACOG | Robin Marwal, Mahesh S Dhar, Kalaiarasan Ponnusamy, Meena Datta, Priyanka Singh, Uma Sharma, Hemlata Lall, Manoj K Singh, Radhakrishnan V. S, Hema Gogia, Preeti Madan, Sandhya Kabra, Sujeet K Singh, Partha Rakshit |
| EPI_ISL_2461511 | National Centre for Disease Control (NCDC) Biotechnology Division, Delhi | NCDC Delhi, Biotechnology Division INSACOG | Radhakrishnan V. S, Robin Marwal, Mahesh S Dhar, Kalaiarasan Ponnusamy, Meena Datta, Priyanka Singh, Uma Sharma, Hemlata Lall, Manoj K Singh, Hema Gogia, Preeti Madan, Sandhya Kabra, Sujeet K Singh, Partha Rakshit |
| EPI_ISL_2461513, EPI_ISL_2461514 | National Centre for Disease Control (NCDC) Biotechnology Division, Delhi | NCDC Delhi, Biotechnology Division INSACOG | Mahesh S Dhar, Kalaiarasan Ponnusamy, Meena Datta, Priyanka Singh, Uma Sharma, Hemlata Lall, Manoj K Singh, Radhakrishnan V. S, Robin Marwal, Hema Gogia, Preeti Madan, Sandhya Kabra, Sujeet K Singh, Partha Rakshit |
| EPI_ISL_2461515, EPI_ISL_2461517, EPI_ISL_2461519 | National Centre for Disease Control (NCDC) Biotechnology Division, Delhi | NCDC Delhi, Biotechnology Division INSACOG | Robin Marwal, Mahesh S Dhar, Kalaiarasan Ponnusamy, Meena Datta, Priyanka Singh, Uma Sharma, Hemlata Lall, Manoj K Singh, Radhakrishnan V. S, Hema Gogia, Preeti Madan, Sandhya Kabra, Sujeet K Singh, Partha Rakshit |
| EPI_ISL_2461521 | National Centre for Disease Control (NCDC) Biotechnology Division, Delhi | NCDC Delhi, Biotechnology Division INSACOG | Radhakrishnan V. S, Robin Marwal, Mahesh S Dhar, Kalaiarasan Ponnusamy, Meena Datta, Priyanka Singh, Uma Sharma, Hemlata Lall, Manoj K Singh, Hema Gogia, Preeti Madan, Sandhya Kabra, Sujeet K Singh, Partha Rakshit |
| EPI_ISL_2461522, EPI_ISL_2461524, EPI_ISL_2461526, EPI_ISL_2461527, EPI_ISL_2461528, EPI_ISL_2461530, EPI_ISL_2461531, EPI_ISL_2461532 | National Centre for Disease Control (NCDC) Biotechnology Division, Delhi | NCDC Delhi, Biotechnology Division INSACOG | Mahesh S Dhar, Kalaiarasan Ponnusamy, Meena Datta, Priyanka Singh, Uma Sharma, Hemlata Lall, Manoj K Singh, Radhakrishnan V. S, Robin Marwal, Hema Gogia, Preeti Madan, Sandhya Kabra, Sujeet K Singh, Partha Rakshit |
| EPI_ISL_2461534, EPI_ISL_2461535, EPI_ISL_2461536, EPI_ISL_2461537 | National Centre for Disease Control (NCDC) Biotechnology Division, Delhi | NCDC Delhi, Biotechnology Division INSACOG | Robin Marwal, Mahesh S Dhar, Kalaiarasan Ponnusamy, Meena Datta, Priyanka Singh, Uma Sharma, Hemlata Lall, Manoj K Singh, Radhakrishnan V. S, Hema Gogia, Preeti Madan, Sandhya Kabra, Sujeet K Singh, Partha Rakshit |
| EPI_ISL_2461538 | National Centre for Disease Control (NCDC) Biotechnology Division, Delhi | NCDC Delhi, Biotechnology Division INSACOG | Mahesh S Dhar, Kalaiarasan Ponnusamy, Meena Datta, Priyanka Singh, Uma Sharma, Hemlata Lall, Manoj K Singh, Radhakrishnan V. S, Robin Marwal, Hema Gogia, Preeti Madan, Sandhya Kabra, Sujeet K Singh, Partha Rakshit |
| EPI_ISL_2461539 | National Centre for Disease Control (NCDC) Biotechnology Division, Delhi | NCDC Delhi, Biotechnology Division INSACOG | Robin Marwal, Mahesh S Dhar, Kalaiarasan Ponnusamy, Meena Datta, Priyanka Singh, Uma Sharma, Hemlata Lall, Manoj K Singh, Radhakrishnan V. S, Hema Gogia, Preeti Madan, Sandhya Kabra, Sujeet K Singh, Partha Rakshit |
| EPI_ISL_2461541, EPI_ISL_2461542 | National Centre for Disease Control (NCDC) Biotechnology Division, Delhi | NCDC Delhi, Biotechnology Division INSACOG | Mahesh S Dhar, Kalaiarasan Ponnusamy, Meena Datta, Priyanka Singh, Uma Sharma, Hemlata Lall, Manoj K Singh, Radhakrishnan V. S, Robin Marwal, Hema Gogia, Preeti Madan, Sandhya Kabra, Sujeet K Singh, Partha Rakshit |
| EPI_ISL_2461543 | National Centre for Disease Control (NCDC) Biotechnology Division, Delhi | NCDC Delhi, Biotechnology Division INSACOG | Radhakrishnan V. S, Robin Marwal, Mahesh S Dhar, Kalaiarasan Ponnusamy, Meena Datta, Priyanka Singh, Uma Sharma, Hemlata Lall, Manoj K Singh, Hema Gogia, Preeti Madan, Sandhya Kabra, Sujeet K Singh, Partha Rakshit |
| EPI_ISL_2461544 | National Centre for Disease Control (NCDC) Biotechnology Division, Delhi | NCDC Delhi, Biotechnology Division INSACOG | Robin Marwal, Mahesh S Dhar, Kalaiarasan Ponnusamy, Meena Datta, Priyanka Singh, Uma Sharma, Hemlata Lall, Manoj K Singh, Radhakrishnan V. S, Hema Gogia, Preeti Madan, Sandhya Kabra, Sujeet K Singh, Partha Rakshit |
| EPI_ISL_2461545 | National Centre for Disease Control (NCDC) Biotechnology Division, Delhi | NCDC Delhi, Biotechnology Division INSACOG | Mahesh S Dhar, Kalaiarasan Ponnusamy, Meena Datta, Priyanka Singh, Uma Sharma, Hemlata Lall, Manoj K Singh, Radhakrishnan V. S, Robin Marwal, Hema Gogia, Preeti Madan, Sandhya Kabra, Sujeet K Singh, Partha Rakshit |
| EPI_ISL_2461546 | National Centre for Disease Control (NCDC) Biotechnology Division, Delhi | NCDC Delhi, Biotechnology Division INSACOG | Robin Marwal, Mahesh S Dhar, Kalaiarasan Ponnusamy, Meena Datta, Priyanka Singh, Uma Sharma, Hemlata Lall, Manoj K Singh, Radhakrishnan V. S, Hema Gogia, Preeti Madan, Sandhya Kabra, Sujeet K Singh, Partha Rakshit |
| EPI_ISL_2461547 | National Centre for Disease Control (NCDC) Biotechnology Division, Delhi | NCDC Delhi, Biotechnology Division INSACOG | Radhakrishnan V. S, Robin Marwal, Mahesh S Dhar, Kalaiarasan Ponnusamy, Meena Datta, Priyanka Singh, Uma Sharma, Hemlata Lall, Manoj K Singh, Hema Gogia, Preeti Madan, Sandhya Kabra, Sujeet K Singh, Partha Rakshit |
| EPI_ISL_2461548 | National Centre for Disease Control (NCDC) Biotechnology Division, Delhi | NCDC Delhi, Biotechnology Division INSACOG | Robin Marwal, Mahesh S Dhar, Kalaiarasan Ponnusamy, Meena Datta, Priyanka Singh, Uma Sharma, Hemlata Lall, Manoj K Singh, Radhakrishnan V. S, Hema Gogia, Preeti Madan, Sandhya Kabra, Sujeet K Singh, Partha Rakshit |
| EPI_ISL_2461549 | National Centre for Disease Control (NCDC) Biotechnology Division, Delhi | NCDC Delhi, Biotechnology Division INSACOG | Mahesh S Dhar, Kalaiarasan Ponnusamy, Meena Datta, Priyanka Singh, Uma Sharma, Hemlata Lall, Manoj K Singh, Radhakrishnan V. S, Robin Marwal, Hema Gogia, Preeti Madan, Sandhya Kabra, Sujeet K Singh, Partha Rakshit |
| EPI_ISL_2461550 | National Centre for Disease Control (NCDC) Biotechnology Division, Delhi | NCDC Delhi, Biotechnology Division INSACOG | Robin Marwal, Mahesh S Dhar, Kalaiarasan Ponnusamy, Meena Datta, Priyanka Singh, Uma Sharma, Hemlata Lall, Manoj K Singh, Radhakrishnan V. S, Hema Gogia, Preeti Madan, Sandhya Kabra, Sujeet K Singh, Partha Rakshit |
| EPI_ISL_2461551 | National Centre for Disease Control (NCDC) Biotechnology Division, Delhi | NCDC Delhi, Biotechnology Division INSACOG | Radhakrishnan V. S, Robin Marwal, Mahesh S Dhar, Kalaiarasan Ponnusamy, Meena Datta, Priyanka Singh, Uma Sharma, Hemlata Lall, Manoj K Singh, Hema Gogia, Preeti Madan, Sandhya Kabra, Sujeet K Singh, Partha Rakshit |
| EPI_ISL_2461552 | National Centre for Disease Control (NCDC) Biotechnology Division, Delhi | NCDC Delhi, Biotechnology Division INSACOG | Mahesh S Dhar, Kalaiarasan Ponnusamy, Meena Datta, Priyanka Singh, Uma Sharma, Hemlata Lall, Manoj K Singh, Radhakrishnan V. S, Robin Marwal, Hema Gogia, Preeti Madan, Sandhya Kabra, Sujeet K Singh, Partha Rakshit |
| EPI_ISL_2461554 | National Centre for Disease Control (NCDC) Biotechnology Division, Delhi | NCDC Delhi, Biotechnology Division INSACOG | Radhakrishnan V. S, Robin Marwal, Mahesh S Dhar, Kalaiarasan Ponnusamy, Meena Datta, Priyanka Singh, Uma Sharma, Hemlata Lall, Manoj K Singh, Hema Gogia, Preeti Madan, Sandhya Kabra, Sujeet K Singh, Partha Rakshit |
| EPI_ISL_2461555, EPI_ISL_2461556, EPI_ISL_2461558, EPI_ISL_2461559, EPI_ISL_2461560 | National Centre for Disease Control (NCDC) Biotechnology Division, Delhi | NCDC Delhi, Biotechnology Division INSACOG | Robin Marwal, Mahesh S Dhar, Kalaiarasan Ponnusamy, Meena Datta, Priyanka Singh, Uma Sharma, Hemlata Lall, Manoj K Singh, Radhakrishnan V. S, Hema Gogia, Preeti Madan, Sandhya Kabra, Sujeet K Singh, Partha Rakshit |
| EPI_ISL_2461561, EPI_ISL_2461562, EPI_ISL_2461563 | National Centre for Disease Control (NCDC) Biotechnology Division, Delhi | NCDC Delhi, Biotechnology Division INSACOG | Mahesh S Dhar, Kalaiarasan Ponnusamy, Meena Datta, Priyanka Singh, Uma Sharma, Hemlata Lall, Manoj K Singh, Radhakrishnan V. S, Robin Marwal, Hema Gogia, Preeti Madan, Sandhya Kabra, Sujeet K Singh, Partha Rakshit |
| EPI_ISL_2461564, EPI_ISL_2461565 | National Centre for Disease Control (NCDC) Biotechnology Division, Delhi | NCDC Delhi, Biotechnology Division INSACOG | Robin Marwal, Mahesh S Dhar, Kalaiarasan Ponnusamy, Meena Datta, Priyanka Singh, Uma Sharma, Hemlata Lall, Manoj K Singh, Radhakrishnan V. S, Hema Gogia, Preeti Madan, Sandhya Kabra, Sujeet K Singh, Partha Rakshit |
| EPI_ISL_2461566, EPI_ISL_2461567 | National Centre for Disease Control (NCDC) Biotechnology Division, Delhi | NCDC Delhi, Biotechnology Division INSACOG | Mahesh S Dhar, Kalaiarasan Ponnusamy, Meena Datta, Priyanka Singh, Uma Sharma, Hemlata Lall, Manoj K Singh, Radhakrishnan V. S, Robin Marwal, Hema Gogia, Preeti Madan, Sandhya Kabra, Sujeet K Singh, Partha Rakshit |
| EPI_ISL_2461568, EPI_ISL_2461569 | National Centre for Disease Control (NCDC) Biotechnology Division, Delhi | NCDC Delhi, Biotechnology Division INSACOG | Radhakrishnan V. S, Robin Marwal, Mahesh S Dhar, Kalaiarasan Ponnusamy, Meena Datta, Priyanka Singh, Uma Sharma, Hemlata Lall, Manoj K Singh, Hema Gogia, Preeti Madan, Sandhya Kabra, Sujeet K Singh, Partha Rakshit |
| EPI_ISL_2461570, EPI_ISL_2461571 | National Centre for Disease Control (NCDC) Biotechnology Division, Delhi | NCDC Delhi, Biotechnology Division INSACOG | Robin Marwal, Mahesh S Dhar, Kalaiarasan Ponnusamy, Meena Datta, Priyanka Singh, Uma Sharma, Hemlata Lall, Manoj K Singh, Radhakrishnan V. S, Hema Gogia, Preeti Madan, Sandhya Kabra, Sujeet K Singh, Partha Rakshit |
| EPI_ISL_2461572, EPI_ISL_2461573 | National Centre for Disease Control (NCDC) Biotechnology Division, Delhi | NCDC Delhi, Biotechnology Division INSACOG | Mahesh S Dhar, Kalaiarasan Ponnusamy, Meena Datta, Priyanka Singh, Uma Sharma, Hemlata Lall, Manoj K Singh, Radhakrishnan V. S, Robin Marwal, Hema Gogia, Preeti Madan, Sandhya Kabra, Sujeet K Singh, Partha Rakshit |
| EPI_ISL_2461574, EPI_ISL_2461575 | National Centre for Disease Control (NCDC) Biotechnology Division, Delhi | NCDC Delhi, Biotechnology Division INSACOG | Robin Marwal, Mahesh S Dhar, Kalaiarasan Ponnusamy, Meena Datta, Priyanka Singh, Uma Sharma, Hemlata Lall, Manoj K Singh, Radhakrishnan V. S, Hema Gogia, Preeti Madan, Sandhya Kabra, Sujeet K Singh, Partha Rakshit |

| | | | |
|---|---|---|---|
| EPI_ISL_2461576 | National Centre for Disease Control (NCDC) Biotechnology Division, Delhi | NCDC Delhi, Biotechnology Division INSACOG | Radhakrishnan V. S, Robin Marwal, Mahesh S Dhar, Kalaiarasan Ponnusamy, Meena Datta, Priyanka Singh, Uma Sharma, Hemlata Lall, Manoj K Singh, Hema Gogia, Preeti Madan, Sandhya Kabra, Sujeet K Singh, Partha Rakshit |
| EPI_ISL_2461577 | National Centre for Disease Control (NCDC) Biotechnology Division, Delhi | NCDC Delhi, Biotechnology Division INSACOG | Robin Marwal, Mahesh S Dhar, Kalaiarasan Ponnusamy, Meena Datta, Priyanka Singh, Uma Sharma, Hemlata Lall, Manoj K Singh, Radhakrishnan V. S, Hema Gogia, Preeti Madan, Sandhya Kabra, Sujeet K Singh, Partha Rakshit |
| EPI_ISL_2461578 | National Centre for Disease Control (NCDC) Biotechnology Division, Delhi | NCDC Delhi, Biotechnology Division INSACOG | Mahesh S Dhar, Kalaiarasan Ponnusamy, Meena Datta, Priyanka Singh, Uma Sharma, Hemlata Lall, Manoj K Singh, Radhakrishnan V. S, Robin Marwal, Hema Gogia, Preeti Madan, Sandhya Kabra, Sujeet K Singh, Partha Rakshit |
| EPI_ISL_2461579 | National Centre for Disease Control (NCDC) Biotechnology Division, Delhi | NCDC Delhi, Biotechnology Division INSACOG | Radhakrishnan V. S, Robin Marwal, Mahesh S Dhar, Kalaiarasan Ponnusamy, Meena Datta, Priyanka Singh, Uma Sharma, Hemlata Lall, Manoj K Singh, Hema Gogia, Preeti Madan, Sandhya Kabra, Sujeet K Singh, Partha Rakshit |
| EPI_ISL_2461580 | National Centre for Disease Control (NCDC) Biotechnology Division, Delhi | NCDC Delhi, Biotechnology Division INSACOG | Mahesh S Dhar, Kalaiarasan Ponnusamy, Meena Datta, Priyanka Singh, Uma Sharma, Hemlata Lall, Manoj K Singh, Radhakrishnan V. S, Robin Marwal, Hema Gogia, Preeti Madan, Sandhya Kabra, Sujeet K Singh, Partha Rakshit |
| EPI_ISL_2461581 | National Centre for Disease Control (NCDC) Biotechnology Division, Delhi | NCDC Delhi, Biotechnology Division INSACOG | Robin Marwal, Mahesh S Dhar, Kalaiarasan Ponnusamy, Meena Datta, Priyanka Singh, Uma Sharma, Hemlata Lall, Manoj K Singh, Radhakrishnan V. S, Hema Gogia, Preeti Madan, Sandhya Kabra, Sujeet K Singh, Partha Rakshit |
| EPI_ISL_2461582, EPI_ISL_2461583 | National Centre for Disease Control (NCDC) Biotechnology Division, Delhi | NCDC Delhi, Biotechnology Division INSACOG | Mahesh S Dhar, Kalaiarasan Ponnusamy, Meena Datta, Priyanka Singh, Uma Sharma, Hemlata Lall, Manoj K Singh, Radhakrishnan V. S, Robin Marwal, Hema Gogia, Preeti Madan, Sandhya Kabra, Sujeet K Singh, Partha Rakshit |
| EPI_ISL_2461584 | National Centre for Disease Control (NCDC) Biotechnology Division, Delhi | NCDC Delhi, Biotechnology Division INSACOG | Radhakrishnan V. S, Robin Marwal, Mahesh S Dhar, Kalaiarasan Ponnusamy, Meena Datta, Priyanka Singh, Uma Sharma, Hemlata Lall, Manoj K Singh, Hema Gogia, Preeti Madan, Sandhya Kabra, Sujeet K Singh, Partha Rakshit |
| EPI_ISL_2461585, EPI_ISL_2461586 | National Centre for Disease Control (NCDC) Biotechnology Division, Delhi | NCDC Delhi, Biotechnology Division INSACOG | Robin Marwal, Mahesh S Dhar, Kalaiarasan Ponnusamy, Meena Datta, Priyanka Singh, Uma Sharma, Hemlata Lall, Manoj K Singh, Radhakrishnan V. S, Hema Gogia, Preeti Madan, Sandhya Kabra, Sujeet K Singh, Partha Rakshit |
| EPI_ISL_2461587 | National Centre for Disease Control (NCDC) Biotechnology Division, Delhi | NCDC Delhi, Biotechnology Division INSACOG | Radhakrishnan V. S, Robin Marwal, Mahesh S Dhar, Kalaiarasan Ponnusamy, Meena Datta, Priyanka Singh, Uma Sharma, Hemlata Lall, Manoj K Singh, Hema Gogia, Preeti Madan, Sandhya Kabra, Sujeet K Singh, Partha Rakshit |
| EPI_ISL_2461589, EPI_ISL_2461590, EPI_ISL_2461591, EPI_ISL_2461592, EPI_ISL_2461593, EPI_ISL_2461594 | National Centre for Disease Control (NCDC) Biotechnology Division, Delhi | NCDC Delhi, Biotechnology Division INSACOG | Robin Marwal, Mahesh S Dhar, Kalaiarasan Ponnusamy, Meena Datta, Priyanka Singh, Uma Sharma, Hemlata Lall, Manoj K Singh, Radhakrishnan V. S, Hema Gogia, Preeti Madan, Sandhya Kabra, Sujeet K Singh, Partha Rakshit |
| EPI_ISL_2461595, EPI_ISL_2461596, EPI_ISL_2461597, EPI_ISL_2461598, EPI_ISL_2461599, EPI_ISL_2461600 | National Centre for Disease Control (NCDC) Biotechnology Division, Delhi | NCDC Delhi, Biotechnology Division INSACOG | Mahesh S Dhar, Kalaiarasan Ponnusamy, Meena Datta, Priyanka Singh, Uma Sharma, Hemlata Lall, Manoj K Singh, Radhakrishnan V. S, Robin Marwal, Hema Gogia, Preeti Madan, Sandhya Kabra, Sujeet K Singh, Partha Rakshit |
| EPI_ISL_2461601, EPI_ISL_2461602, EPI_ISL_2461603, EPI_ISL_2461604, EPI_ISL_2461623, EPI_ISL_2461624 | National Centre for Disease Control (NCDC) Biotechnology Division, Delhi | NCDC Delhi, Biotechnology Division INSACOG | Robin Marwal, Mahesh S Dhar, Kalaiarasan Ponnusamy, Meena Datta, Priyanka Singh, Uma Sharma, Hemlata Lall, Manoj K Singh, Radhakrishnan V. S, Hema Gogia, Preeti Madan, Sandhya Kabra, Sujeet K Singh, Partha Rakshit |
| EPI_ISL_2461631 | National Centre for Disease Control (NCDC) Biotechnology Division, Delhi | NCDC Delhi, Biotechnology Division INSACOG | Radhakrishnan V. S, Robin Marwal, Mahesh S Dhar, Kalaiarasan Ponnusamy, Meena Datta, Priyanka Singh, Uma Sharma, Hemlata Lall, Manoj K Singh, Hema Gogia, Preeti Madan, Sandhya Kabra, Sujeet K Singh, Partha Rakshit |
| EPI_ISL_2461635 | National Centre for Disease Control (NCDC) Biotechnology Division, Delhi | NCDC Delhi, Biotechnology Division INSACOG | Mahesh S Dhar, Kalaiarasan Ponnusamy, Meena Datta, Priyanka Singh, Uma Sharma, Hemlata Lall, Manoj K Singh, Radhakrishnan V. S, Robin Marwal, Hema Gogia, Preeti Madan, Sandhya Kabra, Sujeet K Singh, Partha Rakshit |
| EPI_ISL_2461637 | National Centre for Disease Control (NCDC) Biotechnology Division, Delhi | NCDC Delhi, Biotechnology Division INSACOG | Radhakrishnan V. S, Robin Marwal, Mahesh S Dhar, Kalaiarasan Ponnusamy, Meena Datta, Priyanka Singh, Uma Sharma, Hemlata Lall, Manoj K Singh, Hema Gogia, Preeti Madan, Sandhya Kabra, Sujeet K Singh, Partha Rakshit |
| EPI_ISL_2461638 | National Centre for Disease Control (NCDC) Biotechnology Division, Delhi | NCDC Delhi, Biotechnology Division INSACOG | Mahesh S Dhar, Kalaiarasan Ponnusamy, Meena Datta, Priyanka Singh, Uma Sharma, Hemlata Lall, Manoj K Singh, Radhakrishnan V. S, Robin Marwal, Hema Gogia, Preeti Madan, Sandhya Kabra, Sujeet K Singh, Partha Rakshit |
| EPI_ISL_2461640 | National Centre for Disease Control (NCDC) Biotechnology Division, Delhi | NCDC Delhi, Biotechnology Division INSACOG | Radhakrishnan V. S, Robin Marwal, Mahesh S Dhar, Kalaiarasan Ponnusamy, Meena Datta, Priyanka Singh, Uma Sharma, Hemlata Lall, Manoj K Singh, Hema Gogia, Preeti Madan, Sandhya Kabra, Sujeet K Singh, Partha Rakshit |
| EPI_ISL_2461642 | National Centre for Disease Control (NCDC) Biotechnology Division, Delhi | NCDC Delhi, Biotechnology Division INSACOG | Mahesh S Dhar, Kalaiarasan Ponnusamy, Meena Datta, Priyanka Singh, Uma Sharma, Hemlata Lall, Manoj K Singh, Radhakrishnan V. S, Robin Marwal, Hema Gogia, Preeti Madan, Sandhya Kabra, Sujeet K Singh, Partha Rakshit |
| EPI_ISL_2461644, EPI_ISL_2461645 | National Centre for Disease Control (NCDC) Biotechnology Division, Delhi | NCDC Delhi, Biotechnology Division INSACOG | Robin Marwal, Mahesh S Dhar, Kalaiarasan Ponnusamy, Meena Datta, Priyanka Singh, Uma Sharma, Hemlata Lall, Manoj K Singh, Radhakrishnan V. S, Hema Gogia, Preeti Madan, Sandhya Kabra, Sujeet K Singh, Partha Rakshit |
| EPI_ISL_2461647 | National Centre for Disease Control (NCDC) Biotechnology Division, Delhi | NCDC Delhi, Biotechnology Division INSACOG | Radhakrishnan V. S, Robin Marwal, Mahesh S Dhar, Kalaiarasan Ponnusamy, Meena Datta, Priyanka Singh, Uma Sharma, Hemlata Lall, Manoj K Singh, Hema Gogia, Preeti Madan, Sandhya Kabra, Sujeet K Singh, Partha Rakshit |
| EPI_ISL_2461648 | National Centre for Disease Control (NCDC) Biotechnology Division, Delhi | NCDC Delhi, Biotechnology Division INSACOG | Robin Marwal, Mahesh S Dhar, Kalaiarasan Ponnusamy, Meena Datta, Priyanka Singh, Uma Sharma, Hemlata Lall, Manoj K Singh, Radhakrishnan V. S, Hema Gogia, Preeti Madan, Sandhya Kabra, Sujeet K Singh, Partha Rakshit |
| EPI_ISL_2461653 | National Centre for Disease Control (NCDC) Biotechnology Division, Delhi | NCDC Delhi, Biotechnology Division INSACOG | Mahesh S Dhar, Kalaiarasan Ponnusamy, Meena Datta, Priyanka Singh, Uma Sharma, Hemlata Lall, Manoj K Singh, Radhakrishnan V. S, Robin Marwal, Hema Gogia, Preeti Madan, Sandhya Kabra, Sujeet K Singh, Partha Rakshit |
| EPI_ISL_2461661 | National Centre for Disease Control (NCDC) Biotechnology Division, Delhi | NCDC Delhi, Biotechnology Division INSACOG | Robin Marwal, Mahesh S Dhar, Kalaiarasan Ponnusamy, Meena Datta, Priyanka Singh, Uma Sharma, Hemlata Lall, Manoj K Singh, Radhakrishnan V. S, Hema Gogia, Preeti Madan, Sandhya Kabra, Sujeet K Singh, Partha Rakshit |
| EPI_ISL_2461672 | National Centre for Disease Control (NCDC) Biotechnology Division, Delhi | NCDC Delhi, Biotechnology Division INSACOG | Mahesh S Dhar, Kalaiarasan Ponnusamy, Meena Datta, Priyanka Singh, Uma Sharma, Hemlata Lall, Manoj K Singh, Radhakrishnan V. S, Robin Marwal, Hema Gogia, Preeti Madan, Sandhya Kabra, Sujeet K Singh, Partha Rakshit |
| EPI_ISL_2461688 | National Centre for Disease Control (NCDC) Biotechnology Division, Delhi | NCDC Delhi, Biotechnology Division INSACOG | Radhakrishnan V. S, Robin Marwal, Mahesh S Dhar, Kalaiarasan Ponnusamy, Meena Datta, Priyanka Singh, Uma Sharma, Hemlata Lall, Manoj K Singh, Hema Gogia, Preeti Madan, Sandhya Kabra, Sujeet K Singh, Partha Rakshit |
| EPI_ISL_2461690, EPI_ISL_2461692, EPI_ISL_2461693, EPI_ISL_2461694, EPI_ISL_2461696 | National Centre for Disease Control (NCDC) Biotechnology Division, Delhi | NCDC Delhi, Biotechnology Division INSACOG | Robin Marwal, Mahesh S Dhar, Kalaiarasan Ponnusamy, Meena Datta, Priyanka Singh, Uma Sharma, Hemlata Lall, Manoj K Singh, Radhakrishnan V. S, Hema Gogia, Preeti Madan, Sandhya Kabra, Sujeet K Singh, Partha Rakshit |
| EPI_ISL_2461697, EPI_ISL_2461698, EPI_ISL_2461699 | National Centre for Disease Control (NCDC) Biotechnology Division, Delhi | NCDC Delhi, Biotechnology Division INSACOG | Mahesh S Dhar, Kalaiarasan Ponnusamy, Meena Datta, Priyanka Singh, Uma Sharma, Hemlata Lall, Manoj K Singh, Radhakrishnan V. S, Robin Marwal, Hema Gogia, Preeti Madan, Sandhya Kabra, Sujeet K Singh, Partha Rakshit |
| EPI_ISL_2461700 | National Centre for Disease Control (NCDC) Biotechnology Division, Delhi | NCDC Delhi, Biotechnology Division INSACOG | Radhakrishnan V. S, Robin Marwal, Mahesh S Dhar, Kalaiarasan Ponnusamy, Meena Datta, Priyanka Singh, Uma Sharma, Hemlata Lall, Manoj K Singh, Hema Gogia, Preeti Madan, Sandhya Kabra, Sujeet K Singh, Partha Rakshit |
| EPI_ISL_2461701 | National Centre for Disease Control (NCDC) Biotechnology Division, Delhi | NCDC Delhi, Biotechnology Division INSACOG | Robin Marwal, Mahesh S Dhar, Kalaiarasan Ponnusamy, Meena Datta, Priyanka Singh, Uma Sharma, Hemlata Lall, Manoj K Singh, Radhakrishnan V. S, Hema Gogia, Preeti Madan, Sandhya Kabra, Sujeet K Singh, Partha Rakshit |
| EPI_ISL_2461702 | National Centre for Disease Control (NCDC) Biotechnology Division, Delhi | NCDC Delhi, Biotechnology Division INSACOG | Mahesh S Dhar, Kalaiarasan Ponnusamy, Meena Datta, Priyanka Singh, Uma Sharma, Hemlata Lall, Manoj K Singh, Radhakrishnan V. S, Robin Marwal, Hema Gogia, Preeti Madan, Sandhya Kabra, Sujeet K Singh, Partha Rakshit |
| EPI_ISL_2461703, EPI_ISL_2461704, EPI_ISL_2461705, EPI_ISL_2461706, EPI_ISL_2461707, EPI_ISL_2461708, EPI_ISL_2461709, EPI_ISL_2461710, EPI_ISL_2461711, EPI_ISL_2461712, EPI_ISL_2461713, EPI_ISL_2461714, EPI_ISL_2461715, EPI_ISL_2461716, EPI_ISL_2461717, EPI_ISL_2461718, EPI_ISL_2461719, EPI_ISL_2461720, EPI_ISL_2461721, EPI_ISL_2461722 | | | |
| see above | National Centre for Disease Control (NCDC) Biotechnology Division, Delhi | NCDC Delhi, Biotechnology Division INSACOG | Radhakrishnan V. S, Robin Marwal, Mahesh S Dhar, Kalaiarasan Ponnusamy, Meena Datta, Priyanka Singh, Uma Sharma, Hemlata Lall, Manoj K Singh, Hema Gogia, Preeti Madan, Sandhya Kabra, Sujeet K Singh, Partha Rakshit |
| EPI_ISL_2461723, EPI_ISL_2461724, EPI_ISL_2461725, EPI_ISL_2461726, EPI_ISL_2461727, EPI_ISL_2461728, EPI_ISL_2461729, EPI_ISL_2461730, EPI_ISL_2461731, EPI_ISL_2461732, EPI_ISL_2461733, EPI_ISL_2461734, EPI_ISL_2461735, EPI_ISL_2461736, EPI_ISL_2461737, EPI_ISL_2461738, EPI_ISL_2461739, EPI_ISL_2461740, EPI_ISL_2461741, EPI_ISL_2461742, EPI_ISL_2461743, EPI_ISL_2461744, EPI_ISL_2461745, EPI_ISL_2461746, EPI_ISL_2461747, EPI_ISL_2461748, EPI_ISL_2461749, EPI_ISL_2461750 | | | |
| see above | National Centre for Disease Control (NCDC) Biotechnology Division, Delhi | NCDC Delhi, Biotechnology Division INSACOG | Robin Marwal, Mahesh S Dhar, Kalaiarasan Ponnusamy, Meena Datta, Priyanka Singh, Uma Sharma, Hemlata Lall, Manoj K Singh, Radhakrishnan V. S, Hema Gogia, Preeti Madan, Sandhya Kabra, Sujeet K Singh, Partha Rakshit |
| EPI_ISL_2461751, EPI_ISL_2461752, EPI_ISL_2461753, EPI_ISL_2461754, EPI_ISL_2461755, EPI_ISL_2461756, EPI_ISL_2461757, EPI_ISL_2461758, EPI_ISL_2461759 | National Centre for Disease Control (NCDC) Biotechnology Division, Delhi | NCDC Delhi, Biotechnology Division INSACOG | Mahesh S Dhar, Kalaiarasan Ponnusamy, Meena Datta, Priyanka Singh, Uma Sharma, Hemlata Lall, Manoj K Singh, Radhakrishnan V. S, Robin Marwal, Hema Gogia, Preeti Madan, Sandhya Kabra, Sujeet K Singh, Partha Rakshit |

| | | | |
|---|---|---|---|
| EPI_ISL_2461771 | National Centre for Disease Control (NCDC) Biotechnology Division, Delhi | NCDC Delhi, Biotechnology Division INSACOG | Radhakrishnan V. S, Robin Marwal, Mahesh S Dhar, Kalaiarasan Ponnusamy, Meena Datta, Priyanka Singh, Uma Sharma, Hemlata Lall, Manoj K Singh, Hema Gogia, Preeti Madan, Sandhya Kabra, Sujeet K Singh, Partha Rakshit |
| EPI_ISL_2461775, EPI_ISL_2461780, EPI_ISL_2461781 | National Centre for Disease Control (NCDC) Biotechnology Division, Delhi | NCDC Delhi, Biotechnology Division INSACOG | Mahesh S Dhar, Kalaiarasan Ponnusamy, Meena Datta, Priyanka Singh, Uma Sharma, Hemlata Lall, Manoj K Singh, Radhakrishnan V. S, Robin Marwal, Hema Gogia, Preeti Madan, Sandhya Kabra, Sujeet K Singh, Partha Rakshit |
| EPI_ISL_2461784 | National Centre for Disease Control (NCDC) Biotechnology Division, Delhi | NCDC Delhi, Biotechnology Division INSACOG | Kalaiarasan Ponnusamy, Meena Datta, Priyanka Singh, Uma Sharma, Hemlata Lall, Manoj K Singh, Radhakrishnan V. S, Robin Marwal, Mahesh S Dhar, Hema Gogia, Preeti Madan, Sandhya Kabra, Sujeet K Singh, Partha Rakshit |
| EPI_ISL_2461785 | National Centre for Disease Control (NCDC) Biotechnology Division, Delhi | NCDC Delhi, Biotechnology Division INSACOG | Meena Datta, Priyanka Singh, Uma Sharma, Hemlata Lall, Manoj K Singh, Radhakrishnan V. S, Robin Marwal, Mahesh S Dhar, Kalaiarasan Ponnusamy, Hema Gogia, Preeti Madan, Sandhya Kabra, Sujeet K Singh, Partha Rakshit |
| EPI_ISL_2461786, EPI_ISL_2461787, EPI_ISL_2461788, EPI_ISL_2461789, EPI_ISL_2461790 | National Centre for Disease Control (NCDC) Biotechnology Division, Delhi | NCDC Delhi, Biotechnology Division INSACOG | Kalaiarasan Ponnusamy, Meena Datta, Priyanka Singh, Uma Sharma, Hemlata Lall, Manoj K Singh, Radhakrishnan V. S, Robin Marwal, Mahesh S Dhar, Hema Gogia, Preeti Madan, Sandhya Kabra, Sujeet K Singh, Partha Rakshit |
| EPI_ISL_2461791, EPI_ISL_2461793, EPI_ISL_2461795, EPI_ISL_2461796, EPI_ISL_2461798, EPI_ISL_2461801 | National Centre for Disease Control (NCDC) Biotechnology Division, Delhi | NCDC Delhi, Biotechnology Division INSACOG | Meena Datta, Priyanka Singh, Uma Sharma, Hemlata Lall, Manoj K Singh, Radhakrishnan V. S, Robin Marwal, Mahesh S Dhar, Kalaiarasan Ponnusamy, Hema Gogia, Preeti Madan, Sandhya Kabra, Sujeet K Singh, Partha Rakshit |
| EPI_ISL_2461802 | National Centre for Disease Control (NCDC) Biotechnology Division, Delhi | NCDC Delhi, Biotechnology Division INSACOG | Kalaiarasan Ponnusamy, Meena Datta, Priyanka Singh, Uma Sharma, Hemlata Lall, Manoj K Singh, Radhakrishnan V. S, Robin Marwal, Mahesh S Dhar, Hema Gogia, Preeti Madan, Sandhya Kabra, Sujeet K Singh, Partha Rakshit |
| EPI_ISL_2461803 | National Centre for Disease Control (NCDC) Biotechnology Division, Delhi | NCDC Delhi, Biotechnology Division INSACOG | Meena Datta, Priyanka Singh, Uma Sharma, Hemlata Lall, Manoj K Singh, Radhakrishnan V. S, Robin Marwal, Mahesh S Dhar, Kalaiarasan Ponnusamy, Hema Gogia, Preeti Madan, Sandhya Kabra, Sujeet K Singh, Partha Rakshit |
| EPI_ISL_2461804 | National Centre for Disease Control (NCDC) Biotechnology Division, Delhi | NCDC Delhi, Biotechnology Division INSACOG | Kalaiarasan Ponnusamy, Meena Datta, Priyanka Singh, Uma Sharma, Hemlata Lall, Manoj K Singh, Radhakrishnan V. S, Robin Marwal, Mahesh S Dhar, Hema Gogia, Preeti Madan, Sandhya Kabra, Sujeet K Singh, Partha Rakshit |
| EPI_ISL_2461805, EPI_ISL_2461806 | National Centre for Disease Control (NCDC) Biotechnology Division, Delhi | NCDC Delhi, Biotechnology Division INSACOG | Meena Datta, Priyanka Singh, Uma Sharma, Hemlata Lall, Manoj K Singh, Radhakrishnan V. S, Robin Marwal, Mahesh S Dhar, Kalaiarasan Ponnusamy, Hema Gogia, Preeti Madan, Sandhya Kabra, Sujeet K Singh, Partha Rakshit |
| EPI_ISL_2461808, EPI_ISL_2461809 | National Centre for Disease Control (NCDC) Biotechnology Division, Delhi | NCDC Delhi, Biotechnology Division INSACOG | Kalaiarasan Ponnusamy, Meena Datta, Priyanka Singh, Uma Sharma, Hemlata Lall, Manoj K Singh, Radhakrishnan V. S, Robin Marwal, Mahesh S Dhar, Hema Gogia, Preeti Madan, Sandhya Kabra, Sujeet K Singh, Partha Rakshit |
| EPI_ISL_2461810 | National Centre for Disease Control (NCDC) Biotechnology Division, Delhi | NCDC Delhi, Biotechnology Division INSACOG | Meena Datta, Priyanka Singh, Uma Sharma, Hemlata Lall, Manoj K Singh, Radhakrishnan V. S, Robin Marwal, Mahesh S Dhar, Kalaiarasan Ponnusamy, Hema Gogia, Preeti Madan, Sandhya Kabra, Sujeet K Singh, Partha Rakshit |
| EPI_ISL_2461811 | National Centre for Disease Control (NCDC) Biotechnology Division, Delhi | NCDC Delhi, Biotechnology Division INSACOG | Kalaiarasan Ponnusamy, Meena Datta, Priyanka Singh, Uma Sharma, Hemlata Lall, Manoj K Singh, Radhakrishnan V. S, Robin Marwal, Mahesh S Dhar, Hema Gogia, Preeti Madan, Sandhya Kabra, Sujeet K Singh, Partha Rakshit |
| EPI_ISL_2461812 | National Centre for Disease Control (NCDC) Biotechnology Division, Delhi | NCDC Delhi, Biotechnology Division INSACOG | Meena Datta, Priyanka Singh, Uma Sharma, Hemlata Lall, Manoj K Singh, Radhakrishnan V. S, Robin Marwal, Mahesh S Dhar, Kalaiarasan Ponnusamy, Hema Gogia, Preeti Madan, Sandhya Kabra, Sujeet K Singh, Partha Rakshit |
| EPI_ISL_2461814, EPI_ISL_2461815 | National Centre for Disease Control (NCDC) Biotechnology Division, Delhi | NCDC Delhi, Biotechnology Division INSACOG | Kalaiarasan Ponnusamy, Meena Datta, Priyanka Singh, Uma Sharma, Hemlata Lall, Manoj K Singh, Radhakrishnan V. S, Robin Marwal, Mahesh S Dhar, Hema Gogia, Preeti Madan, Sandhya Kabra, Sujeet K Singh, Partha Rakshit |
| EPI_ISL_2461816, EPI_ISL_2461817 | National Centre for Disease Control (NCDC) Biotechnology Division, Delhi | NCDC Delhi, Biotechnology Division INSACOG | Meena Datta, Priyanka Singh, Uma Sharma, Hemlata Lall, Manoj K Singh, Radhakrishnan V. S, Robin Marwal, Mahesh S Dhar, Kalaiarasan Ponnusamy, Hema Gogia, Preeti Madan, Sandhya Kabra, Sujeet K Singh, Partha Rakshit |
| EPI_ISL_2461818 | National Centre for Disease Control (NCDC) Biotechnology Division, Delhi | NCDC Delhi, Biotechnology Division INSACOG | Kalaiarasan Ponnusamy, Meena Datta, Priyanka Singh, Uma Sharma, Hemlata Lall, Manoj K Singh, Radhakrishnan V. S, Robin Marwal, Mahesh S Dhar, Hema Gogia, Preeti Madan, Sandhya Kabra, Sujeet K Singh, Partha Rakshit |
| EPI_ISL_2461819, EPI_ISL_2461820, EPI_ISL_2461821 | National Centre for Disease Control (NCDC) Biotechnology Division, Delhi | NCDC Delhi, Biotechnology Division INSACOG | Meena Datta, Priyanka Singh, Uma Sharma, Hemlata Lall, Manoj K Singh, Radhakrishnan V. S, Robin Marwal, Mahesh S Dhar, Kalaiarasan Ponnusamy, Hema Gogia, Preeti Madan, Sandhya Kabra, Sujeet K Singh, Partha Rakshit |
| EPI_ISL_2461822, EPI_ISL_2461823 | National Centre for Disease Control (NCDC) Biotechnology Division, Delhi | NCDC Delhi, Biotechnology Division INSACOG | Kalaiarasan Ponnusamy, Meena Datta, Priyanka Singh, Uma Sharma, Hemlata Lall, Manoj K Singh, Radhakrishnan V. S, Robin Marwal, Mahesh S Dhar, Hema Gogia, Preeti Madan, Sandhya Kabra, Sujeet K Singh, Partha Rakshit |
| EPI_ISL_2461824 | National Centre for Disease Control (NCDC) Biotechnology Division, Delhi | NCDC Delhi, Biotechnology Division INSACOG | Meena Datta, Priyanka Singh, Uma Sharma, Hemlata Lall, Manoj K Singh, Radhakrishnan V. S, Robin Marwal, Mahesh S Dhar, Kalaiarasan Ponnusamy, Hema Gogia, Preeti Madan, Sandhya Kabra, Sujeet K Singh, Partha Rakshit |
| EPI_ISL_2461825, EPI_ISL_2461826 | National Centre for Disease Control (NCDC) Biotechnology Division, Delhi | NCDC Delhi, Biotechnology Division INSACOG | Kalaiarasan Ponnusamy, Meena Datta, Priyanka Singh, Uma Sharma, Hemlata Lall, Manoj K Singh, Radhakrishnan V. S, Robin Marwal, Mahesh S Dhar, Hema Gogia, Preeti Madan, Sandhya Kabra, Sujeet K Singh, Partha Rakshit |
| EPI_ISL_2461827, EPI_ISL_2461828 | National Centre for Disease Control (NCDC) Biotechnology Division, Delhi | NCDC Delhi, Biotechnology Division INSACOG | Meena Datta, Priyanka Singh, Uma Sharma, Hemlata Lall, Manoj K Singh, Radhakrishnan V. S, Robin Marwal, Mahesh S Dhar, Kalaiarasan Ponnusamy, Hema Gogia, Preeti Madan, Sandhya Kabra, Sujeet K Singh, Partha Rakshit |
| EPI_ISL_2461829 | National Centre for Disease Control (NCDC) Biotechnology Division, Delhi | NCDC Delhi, Biotechnology Division INSACOG | Kalaiarasan Ponnusamy, Meena Datta, Priyanka Singh, Uma Sharma, Hemlata Lall, Manoj K Singh, Radhakrishnan V. S, Robin Marwal, Mahesh S Dhar, Hema Gogia, Preeti Madan, Sandhya Kabra, Sujeet K Singh, Partha Rakshit |
| EPI_ISL_2461830, EPI_ISL_2461831, EPI_ISL_2461832 | National Centre for Disease Control (NCDC) Biotechnology Division, Delhi | NCDC Delhi, Biotechnology Division INSACOG | Meena Datta, Priyanka Singh, Uma Sharma, Hemlata Lall, Manoj K Singh, Radhakrishnan V. S, Robin Marwal, Mahesh S Dhar, Kalaiarasan Ponnusamy, Hema Gogia, Preeti Madan, Sandhya Kabra, Sujeet K Singh, Partha Rakshit |
| EPI_ISL_2461833, EPI_ISL_2461835 | National Centre for Disease Control (NCDC) Biotechnology Division, Delhi | NCDC Delhi, Biotechnology Division INSACOG | Kalaiarasan Ponnusamy, Meena Datta, Priyanka Singh, Uma Sharma, Hemlata Lall, Manoj K Singh, Radhakrishnan V. S, Robin Marwal, Mahesh S Dhar, Hema Gogia, Preeti Madan, Sandhya Kabra, Sujeet K Singh, Partha Rakshit |
| EPI_ISL_2461836 | National Centre for Disease Control (NCDC) Biotechnology Division, Delhi | NCDC Delhi, Biotechnology Division INSACOG | Meena Datta, Priyanka Singh, Uma Sharma, Hemlata Lall, Manoj K Singh, Radhakrishnan V. S, Robin Marwal, Mahesh S Dhar, Kalaiarasan Ponnusamy, Hema Gogia, Preeti Madan, Sandhya Kabra, Sujeet K Singh, Partha Rakshit |
| EPI_ISL_2461922 | National Centre for Disease Control (NCDC) Biotechnology Division, Delhi | NCDC Delhi, Biotechnology Division INSACOG | Kalaiarasan Ponnusamy, Meena Datta, Priyanka Singh, Uma Sharma, Hemlata Lall, Manoj K Singh, Radhakrishnan V. S, Robin Marwal, Mahesh S Dhar, Hema Gogia, Preeti Madan, Sandhya Kabra, Sujeet K Singh, Partha Rakshit |
| EPI_ISL_2461924 | National Centre for Disease Control (NCDC) Biotechnology Division, Delhi | NCDC Delhi, Biotechnology Division INSACOG | Meena Datta, Priyanka Singh, Uma Sharma, Hemlata Lall, Manoj K Singh, Radhakrishnan V. S, Robin Marwal, Mahesh S Dhar, Kalaiarasan Ponnusamy, Hema Gogia, Preeti Madan, Sandhya Kabra, Sujeet K Singh, Partha Rakshit |
| EPI_ISL_2461929 | National Centre for Disease Control (NCDC) Biotechnology Division, Delhi | NCDC Delhi, Biotechnology Division INSACOG | Kalaiarasan Ponnusamy, Meena Datta, Priyanka Singh, Uma Sharma, Hemlata Lall, Manoj K Singh, Radhakrishnan V. S, Robin Marwal, Mahesh S Dhar, Hema Gogia, Preeti Madan, Sandhya Kabra, Sujeet K Singh, Partha Rakshit |
| EPI_ISL_2461930 | National Centre for Disease Control (NCDC) Biotechnology Division, Delhi | NCDC Delhi, Biotechnology Division INSACOG | Meena Datta, Priyanka Singh, Uma Sharma, Hemlata Lall, Manoj K Singh, Radhakrishnan V. S, Robin Marwal, Mahesh S Dhar, Kalaiarasan Ponnusamy, Hema Gogia, Preeti Madan, Sandhya Kabra, Sujeet K Singh, Partha Rakshit |
| EPI_ISL_2461945 | National Centre for Disease Control (NCDC) Biotechnology Division, Delhi | NCDC Delhi, Biotechnology Division INSACOG | Mahesh S Dhar, Kalaiarasan Ponnusamy, Meena Datta, Priyanka Singh, Uma Sharma, Hemlata Lall, Manoj K Singh, Radhakrishnan V. S, Robin Marwal, Hema Gogia, Preeti Madan, Sandhya Kabra, Sujeet K Singh, Partha Rakshit |
| EPI_ISL_2461948 | National Centre for Disease Control (NCDC) Biotechnology Division, Delhi | NCDC Delhi, Biotechnology Division INSACOG | Kalaiarasan Ponnusamy, Meena Datta, Priyanka Singh, Uma Sharma, Hemlata Lall, Manoj K Singh, Radhakrishnan V. S, Robin Marwal, Mahesh S Dhar, Hema Gogia, Preeti Madan, Sandhya Kabra, Sujeet K Singh, Partha Rakshit |
| EPI_ISL_2461949, EPI_ISL_2461954 | National Centre for Disease Control (NCDC) Biotechnology Division, Delhi | NCDC Delhi, Biotechnology Division INSACOG | Mahesh S Dhar, Kalaiarasan Ponnusamy, Meena Datta, Priyanka Singh, Uma Sharma, Hemlata Lall, Manoj K Singh, Radhakrishnan V. S, Robin Marwal, Hema Gogia, Preeti Madan, Sandhya Kabra, Sujeet K Singh, Partha Rakshit |
| EPI_ISL_2461955 | National Centre for Disease Control (NCDC) Biotechnology Division, Delhi | NCDC Delhi, Biotechnology Division INSACOG | Kalaiarasan Ponnusamy, Meena Datta, Priyanka Singh, Uma Sharma, Hemlata Lall, Manoj K Singh, Radhakrishnan V. S, Robin Marwal, Mahesh S Dhar, Hema Gogia, Preeti Madan, Sandhya Kabra, Sujeet K Singh, Partha Rakshit |
| EPI_ISL_2461956, EPI_ISL_2461957 | National Centre for Disease Control (NCDC) Biotechnology Division, Delhi | NCDC Delhi, Biotechnology Division INSACOG | Mahesh S Dhar, Kalaiarasan Ponnusamy, Meena Datta, Priyanka Singh, Uma Sharma, Hemlata Lall, Manoj K Singh, Radhakrishnan V. S, Robin Marwal, Hema Gogia, Preeti Madan, Sandhya Kabra, Sujeet K Singh, Partha Rakshit |
| EPI_ISL_2461958 | National Centre for Disease Control (NCDC) Biotechnology Division, Delhi | NCDC Delhi, Biotechnology Division INSACOG | Kalaiarasan Ponnusamy, Meena Datta, Priyanka Singh, Uma Sharma, Hemlata Lall, Manoj K Singh, Radhakrishnan V. S, Robin Marwal, Mahesh S Dhar, Hema Gogia, Preeti Madan, Sandhya Kabra, Sujeet K Singh, Partha Rakshit |
| EPI_ISL_2461959 | National Centre for Disease Control (NCDC) Biotechnology Division, Delhi | NCDC Delhi, Biotechnology Division INSACOG | Mahesh S Dhar, Kalaiarasan Ponnusamy, Meena Datta, Priyanka Singh, Uma Sharma, Hemlata Lall, Manoj K Singh, Radhakrishnan V. S, Robin Marwal, Hema Gogia, Preeti Madan, Sandhya Kabra, Sujeet K Singh, Partha Rakshit |
| EPI_ISL_2461960 | National Centre for Disease Control (NCDC) Biotechnology Division, Delhi | NCDC Delhi, Biotechnology Division INSACOG | Kalaiarasan Ponnusamy, Meena Datta, Priyanka Singh, Uma Sharma, Hemlata Lall, Manoj K Singh, Radhakrishnan V. S, Robin Marwal, Mahesh S Dhar, Hema Gogia, Preeti Madan, Sandhya Kabra, Sujeet K Singh, Partha Rakshit |
| EPI_ISL_2461961, EPI_ISL_2461962, EPI_ISL_2461963 | National Centre for Disease Control (NCDC) Biotechnology Division, Delhi | NCDC Delhi, Biotechnology Division INSACOG | Mahesh S Dhar, Kalaiarasan Ponnusamy, Meena Datta, Priyanka Singh, Uma Sharma, Hemlata Lall, Manoj K Singh, Radhakrishnan V. S, Robin Marwal, Hema Gogia, Preeti Madan, Sandhya Kabra, Sujeet K Singh, Partha Rakshit |
| EPI_ISL_2461965, EPI_ISL_2461966 | National Centre for Disease Control (NCDC) Biotechnology Division, Delhi | NCDC Delhi, Biotechnology Division INSACOG | Kalaiarasan Ponnusamy, Meena Datta, Priyanka Singh, Uma Sharma, Hemlata Lall, Manoj K Singh, Radhakrishnan V. S, Robin Marwal, Mahesh S Dhar, Hema Gogia, Preeti Madan, Sandhya Kabra, Sujeet K Singh, Partha Rakshit |
| EPI_ISL_2461976, | National Centre for Disease Control (NCDC) Biotechnology | NCDC Delhi, Biotechnology Division INSACOG | Mahesh S Dhar, Kalaiarasan Ponnusamy, Meena Datta, Priyanka Singh, Uma Sharma, Hemlata Lall, Manoj K Singh, Radhakrishnan V. S, Robin Marwal, Hema Gogia, Preeti Madan, Sandhya Kabra, Sujeet K Singh, Partha Rakshit |

| | | | |
|---|---|---|---|
| EPI_ISL_2461979 | Division, Delhi | | |
| EPI_ISL_2461980, EPI_ISL_2461981 | National Centre for Disease Control (NCDC) Biotechnology Division, Delhi | NCDC Delhi, Biotechnology Division INSACOG | Meena Datta, Priyanka Singh, Uma Sharma, Hemlata Lall, Manoj K Singh, Radhakrishnan V. S, Robin Marwal, Mahesh S Dhar, Kalaiarasan Ponnusamy, Hema Gogia, Preeti Madan, Sandhya Kabra, Sujeet K Singh, Partha Rakshit |
| EPI_ISL_2461982 | National Centre for Disease Control (NCDC) Biotechnology Division, Delhi | NCDC Delhi, Biotechnology Division INSACOG | Mahesh S Dhar, Kalaiarasan Ponnusamy, Meena Datta, Priyanka Singh, Uma Sharma, Hemlata Lall, Manoj K Singh, Radhakrishnan V. S, Robin Marwal, Hema Gogia, Preeti Madan, Sandhya Kabra, Sujeet K Singh, Partha Rakshit |
| EPI_ISL_2461984, EPI_ISL_2461985, EPI_ISL_2461986 | National Centre for Disease Control (NCDC) Biotechnology Division, Delhi | NCDC Delhi, Biotechnology Division INSACOG | Kalaiarasan Ponnusamy, Meena Datta, Priyanka Singh, Uma Sharma, Hemlata Lall, Manoj K Singh, Radhakrishnan V. S, Robin Marwal, Mahesh S Dhar, Hema Gogia, Preeti Madan, Sandhya Kabra, Sujeet K Singh, Partha Rakshit |
| EPI_ISL_2461987, EPI_ISL_2461988 | National Centre for Disease Control (NCDC) Biotechnology Division, Delhi | NCDC Delhi, Biotechnology Division INSACOG | Meena Datta, Priyanka Singh, Uma Sharma, Hemlata Lall, Manoj K Singh, Radhakrishnan V. S, Robin Marwal, Mahesh S Dhar, Kalaiarasan Ponnusamy, Hema Gogia, Preeti Madan, Sandhya Kabra, Sujeet K Singh, Partha Rakshit |
| EPI_ISL_2461991, EPI_ISL_2461992, EPI_ISL_2461993, EPI_ISL_2461994, EPI_ISL_2461995, EPI_ISL_2461996, EPI_ISL_2461997, EPI_ISL_2461998, EPI_ISL_2461999, EPI_ISL_2462000, EPI_ISL_2462001, EPI_ISL_2462002, EPI_ISL_2462003, EPI_ISL_2462004, EPI_ISL_2462005, EPI_ISL_2462006, EPI_ISL_2462007, EPI_ISL_2462008, EPI_ISL_2462009, EPI_ISL_2462010, EPI_ISL_2462011, EPI_ISL_2462012, EPI_ISL_2462013, EPI_ISL_2462014, EPI_ISL_2462015 | | | |
| see above | National Centre for Disease Control (NCDC) Biotechnology Division, Delhi | NCDC Delhi, Biotechnology Division INSACOG | Kalaiarasan Ponnusamy, Meena Datta, Priyanka Singh, Uma Sharma, Hemlata Lall, Manoj K Singh, Radhakrishnan V. S, Robin Marwal, Mahesh S Dhar, Hema Gogia, Preeti Madan, Sandhya Kabra, Sujeet K Singh, Partha Rakshit |
| EPI_ISL_2462016, EPI_ISL_2462017, EPI_ISL_2462018, EPI_ISL_2462019, EPI_ISL_2462020, EPI_ISL_2462021, EPI_ISL_2462022, EPI_ISL_2462023, EPI_ISL_2462024, EPI_ISL_2462025, EPI_ISL_2462026, EPI_ISL_2462027 | | | |
| see above | National Centre for Disease Control (NCDC) Biotechnology Division, Delhi | NCDC Delhi, Biotechnology Division INSACOG | Meena Datta, Priyanka Singh, Uma Sharma, Hemlata Lall, Manoj K Singh, Radhakrishnan V. S, Robin Marwal, Mahesh S Dhar, Kalaiarasan Ponnusamy, Hema Gogia, Preeti Madan, Sandhya Kabra, Sujeet K Singh, Partha Rakshit |
| EPI_ISL_2462074 | OHSU Lab Services Molecular Microbiology Lab | Oregon SARS-CoV-2 Genome Sequencing Center | Brendan L. O'Connell, Sonia Acharya, Cierra LaBlanc, Ruth V. Nichols, Alec J. Hirsch, Donna Hansel, Guang Fan, Xuan Qin, Daniel N. Streblow, William B. Messer, Andrew C. Adey, Benjamin N. Bimber, Brian J. O'Roak |
| EPI_ISL_2462342, EPI_ISL_2462343, EPI_ISL_2462344, EPI_ISL_2462345, EPI_ISL_2462346, EPI_ISL_2462348, EPI_ISL_2462352, EPI_ISL_2462359, EPI_ISL_2462364, EPI_ISL_2462365, EPI_ISL_2462367, EPI_ISL_2462368 | | | |
| see above | SA Pathology | SA Pathology | Caitlin Selway, Lex Leong, Chuan Kok Lim, Mark Turra, Ivan Bastian, Geoff Higgins |
| EPI_ISL_2462421 | Sydney South West Pathology Service (SSWPS) - Royal Prince Alfred Hospital - NSW Health Pathology | NSW Health Pathology - Institute of Clinical Pathology and Medical Research; Westmead Hospital; University of Sydney | CIDM-PH et al. |
| EPI_ISL_2462432, EPI_ISL_2462433, EPI_ISL_2462434 | South Eastern Area Laboratory Services (SEALS) | NSW Health Pathology - Institute of Clinical Pathology and Medical Research; Westmead Hospital; University of Sydney | CIDM-PH et al. |
| EPI_ISL_2462532, EPI_ISL_2462539 | Institute for Infectious Diseases | Institute for Infectious Diseases | Stefan Neuenschwander, Christian Baumann, Miguel A Terrazos Miani, Cora Sägesser, Pascal Bittel, Peter Keller, Franziska Suter-Riniker, Stephen L Leib, Alban Ramette |
| EPI_ISL_2462656, EPI_ISL_2462658 | Viollier AG | Department of Biosystems Science and Engineering, ETH Zürich | Christian Beisel, Sarah Nadeau, Chaoran Chen, Ivan Topolsky, Kim Philipp Jablonski, Lara Fuhrmann, Rebecca Denes, Mirjam Feldkamp, Ina Nissen, Natascha Santacroce, Elodie Burcklen, Christiane Beckmann, Maurice Redondo, Olivier Kobel, Christoph Noppen, Niko Beerenwinkel, Tanja Stadler |
| EPI_ISL_2462722 | Viollier AG | Department of Biosystems Science and Engineering, ETH Zürich | Chaoran Chen, Sarah Nadeau, Catharine Aquino, Ivan Topolsky, Kim Philipp Jablonski, Lara Fuhrmann, Daniel Ehrsam, Isabel Stürmer, Andreia Cabral de Gouvea, Maria Domenica Moccia, Simon Grüter, Timothy Sykes, Lennart Opitz, Griffin White, Laura Neff, Doris Popovic, Andrea Patrignani, Jay Tracy, Ralph Schlapbach, Christiane Beckmann, Maurice Redondo, Olivier Kobel, Christoph Noppen, Niko Beerenwinkel, Tanja Stadler |
| EPI_ISL_2462835, EPI_ISL_2462851, EPI_ISL_2462855 | Viollier AG | Department of Biosystems Science and Engineering, ETH Zürich | Christian Beisel, Sarah Nadeau, Chaoran Chen, Ivan Topolsky, Kim Philipp Jablonski, Lara Fuhrmann, Rebecca Denes, Mirjam Feldkamp, Ina Nissen, Natascha Santacroce, Elodie Burcklen, Christiane Beckmann, Maurice Redondo, Olivier Kobel, Christoph Noppen, Niko Beerenwinkel, Tanja Stadler |
| EPI_ISL_2462879 | WHO National Influenza Centre Russian Federation | WHO National Influenza Centre Russian Federation | Andrey Komissarov, Artem Fadeev, Kseniya Komissarova, Oula Mansour, Kirill Varchenko, Mikhail Bakaev, Tamila Musaeva, Maria Timofeeva, Veronika Eder, Maria Pisareva, Nikita Yolshin, Daria Danilenko, Ksenia Safina, Elena Nabieva, Georgii Bazykin, Dmitry Lioznov |
| EPI_ISL_2462881 | City Hospital No 40 | WHO National Influenza Centre Russian Federation | Andrey Komissarov, Artem Fadeev, Kseniya Komissarova, Oula Mansour, Kirill Varchenko, Mikhail Bakaev, Tamila Musaeva, Maria Timofeeva, Veronika Eder, Maria Pisareva, Nikita Yolshin, Daria Danilenko, Olga Shneider, Sergey Scherbak, Ksenia Safina, Elena Nabieva, Georgii Bazykin, Dmitry Lioznov |
| EPI_ISL_2462964 | Labo Analyses Med | National Reference Center for Viruses of Respiratory Infections, Institut Pasteur, Paris | Marion Barbet, Sylvie Behillil, Méline Bizard, Angela Brisebarre, Camille Capel, Vincent Enouf, Louise Lefrançois, Frédéric Lemoine, Christophe Malabat, Corinne Maufrais, Etienne Simon-Lorière, Maud Vanpeene, Sylvie Van der Werf ,Nouredine Sadeg |
| EPI_ISL_2462965 | Hospital | National Reference Center for Viruses of Respiratory Infections, Institut Pasteur, Paris | Marion Barbet, Sylvie Behillil, Méline Bizard, Angela Brisebarre, Camille Capel, Vincent Enouf, Louise Lefrançois, Frédéric Lemoine, Christophe Malabat, Corinne Maufrais, Amaury Vaysse, Etienne Simon-Lorière, Maud Vanpeene, Sylvie Van der Werf ,Nabil Gastli |
| EPI_ISL_2462998 | Jessa | Jessa | Berden et al. on behalf of the Jessa_cmdLab |
| EPI_ISL_2463023, EPI_ISL_2463026, EPI_ISL_2463035, EPI_ISL_2463046, EPI_ISL_2463054, EPI_ISL_2463062, EPI_ISL_2463064, EPI_ISL_2463069, EPI_ISL_2463080, EPI_ISL_2463087, EPI_ISL_2463094, EPI_ISL_2463098, EPI_ISL_2463116, EPI_ISL_2463137, EPI_ISL_2463143, EPI_ISL_2463156, EPI_ISL_2463159, EPI_ISL_2463167, EPI_ISL_2463179, EPI_ISL_2463189, EPI_ISL_2463191, EPI_ISL_2463196, EPI_ISL_2463200, EPI_ISL_2463223, EPI_ISL_2463228, EPI_ISL_2463232, EPI_ISL_2463258, EPI_ISL_2463267, EPI_ISL_2463273, EPI_ISL_2463279 | | | |
| see above | Quest Diagnostics Incorporated | Centers for Disease Control and Prevention Division of Viral Diseases, Pathogen Discovery | Dakota Howard, Dhwani Batra, Peter W. Cook, Kara Moser, Adrian Paskey, Jason Caravas, Benjamin Rambo-Martin, Shatavia Morrison, Christopher Gulvick, Scott Sammons, Yvette Unoarumhi, Darlene Wagner, Matthew Schmerer, S. H. Rosenthal, A. Gerasimova, R. M. Kagan, B. Anderson, M. Hua, Y. Liu, L.E. Bernstein, K.E. Livingston, A. Perez, I. A. Shlyakhter, R. V. Rolando, R. Owen, P. Tanpaiboon, F. Lacbawan, Clinton R. Paden, Duncan MacCannell |
| EPI_ISL_2463312 | Pathcare Laboratory | National Institute for Communicable Diseases of the National Health Laboratory Service | Sisonke Team, Amoako DG, Everatt J, Viana R, Glass A, Bhiman JN |
| EPI_ISL_2463348, EPI_ISL_2463357 | National Laboratory for Health, Environment and Food, OMM, Maribor | NLZOH (National Laboratory for Health, Environment and Food) / CISLD (Clinical Institute of Special Laboratory Diagnostics), University Children's Hospital, University Medical Center Ljubljana | Sandra Janezic, Aleksander Mahnic, Maja Rupnik, Tjasa Žohar Čretnik, Alenka Štorman, Nika Gobec, Aleksander Kocuvan, Kaja Tominc, Maša Jarčič, Mojca Cimerman, Nika Volmajer, Andrej Golle / Jernej Kovač, Barbara Jenko Bizjan, Tine Tesovnik, Robert Šket, Katarina Kozmos, Ana Grom, Maruša Debeljak, Marko Pokorn, Tadej Battelino |
| EPI_ISL_2463709, EPI_ISL_2463710, EPI_ISL_2463713, EPI_ISL_2463714, EPI_ISL_2463715, EPI_ISL_2463716, EPI_ISL_2463718, EPI_ISL_2463719, EPI_ISL_2463720, EPI_ISL_2463721, EPI_ISL_2463722, EPI_ISL_2463726, EPI_ISL_2463727, EPI_ISL_2463728, EPI_ISL_2463729, EPI_ISL_2463732, EPI_ISL_2463733, EPI_ISL_2463734, EPI_ISL_2463736, EPI_ISL_2463738, EPI_ISL_2463739, EPI_ISL_2463740, EPI_ISL_2463741, EPI_ISL_2463742, EPI_ISL_2463743, EPI_ISL_2463744, EPI_ISL_2463745, EPI_ISL_2463746, EPI_ISL_2463747, EPI_ISL_2463748, EPI_ISL_2463749, EPI_ISL_2463750, EPI_ISL_2463751, EPI_ISL_2463752, EPI_ISL_2463753, EPI_ISL_2463754, EPI_ISL_2463755, EPI_ISL_2463756, EPI_ISL_2463757, EPI_ISL_2463758, EPI_ISL_2463759, EPI_ISL_2463760, EPI_ISL_2463762, EPI_ISL_2463763, EPI_ISL_2463765, EPI_ISL_2463766, EPI_ISL_2463767, EPI_ISL_2463768, EPI_ISL_2463769, EPI_ISL_2463770, EPI_ISL_2463771, EPI_ISL_2463772, EPI_ISL_2463773, EPI_ISL_2463774, EPI_ISL_2463775, EPI_ISL_2463776, EPI_ISL_2463777, EPI_ISL_2463778, EPI_ISL_2463779, EPI_ISL_2463780, EPI_ISL_2463781, EPI_ISL_2463782, EPI_ISL_2463783, EPI_ISL_2463784, EPI_ISL_2463785, EPI_ISL_2463786, EPI_ISL_2463787, EPI_ISL_2463788, EPI_ISL_2463789, EPI_ISL_2463790, EPI_ISL_2463791, EPI_ISL_2463792, EPI_ISL_2463793, EPI_ISL_2463794, EPI_ISL_2463795, EPI_ISL_2463796, EPI_ISL_2463797, EPI_ISL_2463798, EPI_ISL_2463799, EPI_ISL_2463801, EPI_ISL_2463802, EPI_ISL_2463803, EPI_ISL_2463804, EPI_ISL_2463805, EPI_ISL_2463806, EPI_ISL_2463807, EPI_ISL_2463808, EPI_ISL_2463809, EPI_ISL_2463810, EPI_ISL_2463811, EPI_ISL_2463812, EPI_ISL_2463813, EPI_ISL_2463814, EPI_ISL_2463815, EPI_ISL_2463816, EPI_ISL_2463817, EPI_ISL_2463818, EPI_ISL_2463819, EPI_ISL_2463820, EPI_ISL_2463821, EPI_ISL_2463822, EPI_ISL_2463823, EPI_ISL_2463825, EPI_ISL_2463826, EPI_ISL_2463828, EPI_ISL_2463830, EPI_ISL_2463831, EPI_ISL_2463832, EPI_ISL_2463833 | | | |
| see above | Dr S raju, Director of public heath and preventive medicine | inStem NCBS - INSACOG | Uma Ramakrishnan Dasaradhi Palakodeti Aswin SaiNarain |
| EPI_ISL_2464181 | Labo Analyses Med | National Reference Center for Viruses of Respiratory Infections, Institut Pasteur, Paris | Marion Barbet, Sylvie Behillil, Méline Bizard, Angela Brisebarre, Camille Capel, Vincent Enouf, Louise Lefrançois, Frédéric Lemoine, Christophe Malabat, Corinne Maufrais, Etienne Simon-Lorière, Maud Vanpeene, Sylvie Van der Werf ,Arditi Cocco |
| EPI_ISL_2464341, EPI_ISL_2464343, EPI_ISL_2464344, EPI_ISL_2464345, EPI_ISL_2464346, EPI_ISL_2464354, EPI_ISL_2464355, EPI_ISL_2464356, EPI_ISL_2464357, EPI_ISL_2464358, EPI_ISL_2464359, EPI_ISL_2464360, EPI_ISL_2464361, EPI_ISL_2464362, EPI_ISL_2464363, EPI_ISL_2464364, EPI_ISL_2464370, EPI_ISL_2464374, EPI_ISL_2464375 | | | |
| see above | Division of Emerging Infectious Diseases, Bureau of Infectious Diseases Diagnosis Control, Korea Disease Control and Prevention Agency | Division of Emerging Infectious Diseases, Bureau of Infectious Diseases Diagnosis Control, Korea Disease Control and Prevention Agency | Ae Kyung Park, Il-Hwan Kim, Heui Man Kim, Jeong-Min Kim, Jeong-Ah Kim, Chae Young Lee, Eun-Jin Kim |
| EPI_ISL_2464419 | Plateau technique Biomer Metz Verriers | Department of Virology, Henri Mondor University Hospital, Assistance Publique Hôpitaux de Paris, Université Paris-Est Créteil, INSERM U955 | Christophe Rodriguez, Slim Fourati, Vanessa Demontant, Guillaume Gricourt, Melissa N'Debi, Alexandre Soulier, Elisabeth Trawinski, Jean-Michel Pawlotsky |
| EPI_ISL_2464428, EPI_ISL_2464429, EPI_ISL_2464430 | Laboratoire Ana-L | Department of Virology, Henri Mondor University Hospital, Assistance Publique Hôpitaux de Paris, Université Paris-Est Créteil, INSERM U955 | Christophe Rodriguez, Slim Fourati, Vanessa Demontant, Guillaume Gricourt, Melissa N'Debi, Alexandre Soulier, Elisabeth Trawinski, Jean-Michel Pawlotsky |
| EPI_ISL_2464464 | Laboratoire LENYS | Department of Virology, Henri Mondor University Hospital, Assistance Publique Hôpitaux de Paris, Université Paris-Est Créteil, INSERM U955 | Christophe Rodriguez, Slim Fourati, Vanessa Demontant, Guillaume Gricourt, Melissa N'Debi, Alexandre Soulier, Elisabeth Trawinski, Jean-Michel Pawlotsky |
| EPI_ISL_2464555, EPI_ISL_2464556, EPI_ISL_2464557, EPI_ISL_2464558, EPI_ISL_2464559 | Singapore General Hospital | Department of Microbiology | Nurdyana Abdul Rahman, Kenneth Xin Long Chan, Chayaporn Suphavilai, Kun Lee Lim, Sui Sin Goh, Kian Sing Chan, Lynette Oon, Kern Rei Chng, James Sim Heng Chiak, Niranjan Nagarajan, Karrie Ko |
| EPI_ISL_2464560, EPI_ISL_2464561, EPI_ISL_2464562, EPI_ISL_2464563, EPI_ISL_2464564 | Singapore General Hospital | Singapore General Hospital | Nurdyana Abdul Rahman, Kenneth Xin Long Chan, Chayaporn Suphavilai, Kun Lee Lim, Sui Sin Goh, Kian Sing Chan, Lynette Oon, Kern Rei Chng, James Sim Heng Chiak, Niranjan Nagarajan, Karrie Ko |
| EPI_ISL_2464723, EPI_ISL_2464727 | Cerballiance Paris IDF est Saint-Denis | Department of Virology, Henri Mondor University Hospital, Assistance Publique Hôpitaux de Paris, Université Paris-Est Créteil, INSERM U955 | Christophe Rodriguez, Slim Fourati, Vanessa Demontant, Guillaume Gricourt, Melissa N'Debi, Alexandre Soulier, Elisabeth Trawinski, Jean-Michel Pawlotsky |
| EPI_ISL_2464982, EPI_ISL_2465025 | CH. ROBERT BALLANGER | Department of Virology, Henri Mondor University Hospital, Assistance Publique Hôpitaux de Paris, Université Paris-Est Créteil, INSERM U955 | Christophe Rodriguez, Slim Fourati, Vanessa Demontant, Guillaume Gricourt, Melissa N'Debi, Alexandre Soulier, Elisabeth Trawinski, Jean-Michel Pawlotsky |
| EPI_ISL_2465122, EPI_ISL_2465190, EPI_ISL_2465199 | Istituto Zooprofilattico Sperimentale del Mezzogiorno | Telethon Institute of Genetics and Medicine (TIGEM) | Antonio Grimaldi Patrizia Annunziata Francesco Panariello Biancamaria Pierri Claudia Tiberio Teresa Giuliano Valentina Bouche Chiara Colantuono Maria Concetta Cuomo Denise Di Concilio Lucio Di Filippo Anna Manfredi Marcello Salvi Antonio Limone Luigi Atripaldi Pellegrino Cerino Andrea Ballabio Davide Cacchiarelli |
| EPI_ISL_2466268 | Laboratorio Central de Saude Publica do Estado do Rio de | Laboratory of Respiratory Viruses and Measles, Oswaldo Cruz | Paola Resende, Luciana Appolinario, Fernando Motta, Anna Carolina Paixao, Ana Carolina Mendonca, Alice Sampaio Rocha, Taina Venas, Elisa Cavalcante Pereira, Renata Serrano Lopes, Andrea Cony Cavalcanti, Marilda Siqueira on behalf of the |

| | | | |
|---|---|---|---|
| | Janeiro (LACEN-RJ) | Institute, FIOCRUZ | Fiocruz COVID-19 Genomic Surveillance Network |
| EPI_ISL_2466276, EPI_ISL_2466277, EPI_ISL_2466280, EPI_ISL_2466281, EPI_ISL_2466282, EPI_ISL_2466283, EPI_ISL_2466284, EPI_ISL_2466285 | Presidio di Brindisi Di Summa - Perrino | Istituto Zooprofilattico Sperimentale della Puglia e della Basilicata | Parisi A., Bianco A., Capozzi L., Del Sambro L., Simone D., Difato L., Pennuzzi G., Santoro A. |
| EPI_ISL_2466286, EPI_ISL_2466287, EPI_ISL_2466288 | Ospedale Vito Fazzi | Istituto Zooprofilattico Sperimentale della Puglia e della Basilicata | Parisi A., Bianco A., Capozzi L., Del Sambro L., Simone D., Difato L., Bruno A. R. |
| EPI_ISL_2466292, EPI_ISL_2466295, EPI_ISL_2466298, EPI_ISL_2466306, EPI_ISL_2466310 | Ospedale Santa Caterina Novella | Istituto Zooprofilattico Sperimentale della Puglia e della Basilicata | Parisi A., Bianco A., Capozzi L., Del Sambro L., Simone D., Difato L., Lobreglio G. |
| EPI_ISL_2466400 | Inst. for Med. Virology, University Hospital Frankfurt, Goethe University Frankfurt | Inst. for Med. Virology, University Hospital Frankfurt, Goethe University Frankfurt | Widera,M., Toptan,T., Wilhelm,A., Pallas,C., Rabenau,H.F., Goetsch,U., Gottschalk,R. and Ciesek,S. |
| EPI_ISL_2466507, EPI_ISL_2466509 | National Institute of Public Health | State Veterinary Institute Prague | Nagy,A;Jirincova,H;Suri,T;Trnka,D;Vecerova,J |
| EPI_ISL_2466570, EPI_ISL_2466573 | Clinical Virology | Clinical Bacteriology | Tim Roloff, Fanny Wegner, Helena MB Seth-Smith, Alfredo Mari, Karoline Leuzinger, Julia Bielicki, Manuel Battegay, Hans Hirsch, Adrian Egli |
| EPI_ISL_2466607 | Felix-Platter Spital | Clinical Bacteriology | Tim Roloff, Fanny Wegner, Helena MB Seth-Smith, Alfredo Mari, Karoline Leuzinger, Julia Bielicki, Manuel Battegay, Hans Hirsch, Adrian Egli |
| EPI_ISL_2466612 | GHOL | Laboratory of genomics and metagenomics | Trestan Pillonel, Damien Jacot, Sébastien Aeby, Gilbert Greub, Claire Bertelli |
| EPI_ISL_2466672, EPI_ISL_2466674, EPI_ISL_2466675, EPI_ISL_2466676, EPI_ISL_2466677, EPI_ISL_2466679 | Communicable Disease Laboratory, Public Health Directorate | Communicable Disease Laboratory, Public Health Directorate | Alwasti,H., AlHujairi,Z., AlAbbas,Z., Marhoon,A., Almoamen,G. |
| EPI_ISL_2466752, EPI_ISL_2466753 | Kansas Health and Environmental Lab | Kansas Health and Environmental Lab | Mike Grose, Katherine Wiggins, Jonathan Barnell, Ben Olsen, and Phil Adam |
| EPI_ISL_2466756, EPI_ISL_2466758, EPI_ISL_2466759, EPI_ISL_2466761, EPI_ISL_2466762 | Wichita State University - Molecular Diagnostics Lab | Kansas Health and Environmental Lab | Mike Grose, Katherine Wiggins, Jonathan Barnell, Ben Olsen, and Phil Adam |
| EPI_ISL_2466773, EPI_ISL_2466790, EPI_ISL_2466795, EPI_ISL_2466826 | Labor Berlin Charité Vivantes GmbH / Institut für Virologie | Charité Universitätsmedizin Berlin, Institut für Virologie/Labor Berlin | Peter Menzel, Christine Stephan, Rolf Schwarzer, Victor M Corman, Barbara Mühlemann, Terry Jones, Christian Drosten |
| EPI_ISL_2466841 | Fimlab Laboratories | Fimlab Laboratories | Minna Paloniemi, Leena Huhti, Sara Lehtinen, Bruno Luukinen, Tapio Seiskari, Mauri Keinänen |
| EPI_ISL_2467383, EPI_ISL_2467384, EPI_ISL_2467385, EPI_ISL_2467411 | Illinois Department of Public Health | Illinois Department of Public Health - Chicago Lab | Vineet K. Dhiman, Ira Heimler, Joel Price |
| EPI_ISL_2467518, EPI_ISL_2467519 | Fimlab Laboratories | Fimlab Laboratories | Minna Paloniemi, Leena Huhti, Sara Lehtinen, Bruno Luukinen, Tapio Seiskari, Mauri Keinänen |
| EPI_ISL_2467812 | Alaska State Virology Laboratory | Alaska State Virology Laboratory | Stephanie DeRonde, Elva House, Jacob Zidek, Lisa Smith, Ph.D., Jack Chen, Ph.D. |
| EPI_ISL_2467871 | Fimlab Laboratories | Fimlab Laboratories | Minna Paloniemi, Leena Huhti, Sara Lehtinen, Bruno Luukinen, Tapio Seiskari, Mauri Keinänen |
| EPI_ISL_2467885 | OK Public Health Laboratory, Oklahoma State DOH | Genomics and Discovery, Respiratory Viruses Branch, Division of Viral Diseases, Centers for Disease Control and Prevention | Ying Tao, Yan Li, Jing Zhang, Anna Kelleher, Brian Lynch, Krista Queen, Anna Uehara, Peter Cook, Han Jia Justin Ng, Rachel Marine, Clinton R. Paden, Dhwani Batra, Haibin Wang, Tara Coalter, Jasmine Padilla, Morgan Davis, Mili Sheth, Sarah Nobles, Mark Burroughs, Justin Lee, Adam Retchless, Suxiang Tong |
| EPI_ISL_2467935, EPI_ISL_2467936 | Fimlab Laboratories | Fimlab Laboratories | Minna Paloniemi, Leena Huhti, Sara Lehtinen, Bruno Luukinen, Tapio Seiskari, Mauri Keinänen |
| EPI_ISL_2467944 | Lighthouse Laboratory Plymouth | Wellcome Sanger Institute for the COVID-19 Genomics UK (COG-UK) Consortium | Lighthouse Laboratory Plymouth and Alex Alderton, Roberto Amato, Jeffrey Barrett, Sonia Goncalves, Ewan Harrison, David K. Jackson, Ian Johnston, Dominic Kwiatkowski, Cordelia Langford, John Sillitoe on behalf of the Wellcome Sanger Institute COVID-19 Surveillance Team |
| EPI_ISL_2467945 | Lighthouse Lab in Alderley Park | Wellcome Sanger Institute for the COVID-19 Genomics UK (COG-UK) Consortium | Jacquelyn Wynn, Mairead Hyland, The Lighthouse Lab in Alderley Park and Alex Alderton, Roberto Amato, Jeffrey Barrett, Sonia Goncalves, Ewan Harrison, David K. Jackson, Ian Johnston, Dominic Kwiatkowski, Cordelia Langford, John Sillitoe on behalf of the Wellcome Sanger Institute COVID-19 Surveillance Team |
| EPI_ISL_2467946 | Lighthouse Lab in Milton Keynes | Wellcome Sanger Institute for the COVID-19 Genomics UK (COG-UK) Consortium | The Lighthouse Lab in Milton Keynes and Alex Alderton, Roberto Amato, Jeffrey Barrett, Sonia Goncalves, Ewan Harrison, David K. Jackson, Ian Johnston, Dominic Kwiatkowski, Cordelia Langford, John Sillitoe on behalf of the Wellcome Sanger Institute COVID-19 Surveillance Team |
| EPI_ISL_2467947 | Lighthouse Lab in Alderley Park | Wellcome Sanger Institute for the COVID-19 Genomics UK (COG-UK) Consortium | Jacquelyn Wynn, Mairead Hyland, The Lighthouse Lab in Alderley Park and Alex Alderton, Roberto Amato, Jeffrey Barrett, Sonia Goncalves, Ewan Harrison, David K. Jackson, Ian Johnston, Dominic Kwiatkowski, Cordelia Langford, John Sillitoe on behalf of the Wellcome Sanger Institute COVID-19 Surveillance Team |
| EPI_ISL_2467948, EPI_ISL_2467949, EPI_ISL_2467950 | Lighthouse Laboratory Plymouth | Wellcome Sanger Institute for the COVID-19 Genomics UK (COG-UK) Consortium | Lighthouse Laboratory Plymouth and Alex Alderton, Roberto Amato, Jeffrey Barrett, Sonia Goncalves, Ewan Harrison, David K. Jackson, Ian Johnston, Dominic Kwiatkowski, Cordelia Langford, John Sillitoe on behalf of the Wellcome Sanger Institute COVID-19 Surveillance Team |
| EPI_ISL_2467951 | Lighthouse Lab in Milton Keynes | Wellcome Sanger Institute for the COVID-19 Genomics UK (COG-UK) Consortium | The Lighthouse Lab in Milton Keynes and Alex Alderton, Roberto Amato, Jeffrey Barrett, Sonia Goncalves, Ewan Harrison, David K. Jackson, Ian Johnston, Dominic Kwiatkowski, Cordelia Langford, John Sillitoe on behalf of the Wellcome Sanger Institute COVID-19 Surveillance Team |
| EPI_ISL_2467952 | Lighthouse Laboratory Plymouth | Wellcome Sanger Institute for the COVID-19 Genomics UK (COG-UK) Consortium | Lighthouse Laboratory Plymouth and Alex Alderton, Roberto Amato, Jeffrey Barrett, Sonia Goncalves, Ewan Harrison, David K. Jackson, Ian Johnston, Dominic Kwiatkowski, Cordelia Langford, John Sillitoe on behalf of the Wellcome Sanger Institute COVID-19 Surveillance Team |
| EPI_ISL_2467953, EPI_ISL_2467954, EPI_ISL_2467955 | Lighthouse Lab in Alderley Park | Wellcome Sanger Institute for the COVID-19 Genomics UK (COG-UK) Consortium | Jacquelyn Wynn, Mairead Hyland, The Lighthouse Lab in Alderley Park and Alex Alderton, Roberto Amato, Jeffrey Barrett, Sonia Goncalves, Ewan Harrison, David K. Jackson, Ian Johnston, Dominic Kwiatkowski, Cordelia Langford, John Sillitoe on behalf of the Wellcome Sanger Institute COVID-19 Surveillance Team |
| EPI_ISL_2467956 | Lighthouse Lab in Milton Keynes | Wellcome Sanger Institute for the COVID-19 Genomics UK (COG-UK) Consortium | The Lighthouse Lab in Milton Keynes and Alex Alderton, Roberto Amato, Jeffrey Barrett, Sonia Goncalves, Ewan Harrison, David K. Jackson, Ian Johnston, Dominic Kwiatkowski, Cordelia Langford, John Sillitoe on behalf of the Wellcome Sanger Institute COVID-19 Surveillance Team |
| EPI_ISL_2467957 | Lighthouse Lab in Alderley Park | Wellcome Sanger Institute for the COVID-19 Genomics UK (COG-UK) Consortium | Jacquelyn Wynn, Mairead Hyland, The Lighthouse Lab in Alderley Park and Alex Alderton, Roberto Amato, Jeffrey Barrett, Sonia Goncalves, Ewan Harrison, David K. Jackson, Ian Johnston, Dominic Kwiatkowski, Cordelia Langford, John Sillitoe on behalf of the Wellcome Sanger Institute COVID-19 Surveillance Team |
| EPI_ISL_2467958 | Lighthouse Laboratory Plymouth | Wellcome Sanger Institute for the COVID-19 Genomics UK (COG-UK) Consortium | Lighthouse Laboratory Plymouth and Alex Alderton, Roberto Amato, Jeffrey Barrett, Sonia Goncalves, Ewan Harrison, David K. Jackson, Ian Johnston, Dominic Kwiatkowski, Cordelia Langford, John Sillitoe on behalf of the Wellcome Sanger Institute COVID-19 Surveillance Team |
| EPI_ISL_2467959, EPI_ISL_2467960 | Lighthouse Lab in Milton Keynes | Wellcome Sanger Institute for the COVID-19 Genomics UK (COG-UK) Consortium | The Lighthouse Lab in Milton Keynes and Alex Alderton, Roberto Amato, Jeffrey Barrett, Sonia Goncalves, Ewan Harrison, David K. Jackson, Ian Johnston, Dominic Kwiatkowski, Cordelia Langford, John Sillitoe on behalf of the Wellcome Sanger Institute COVID-19 Surveillance Team |
| EPI_ISL_2467961 | Lighthouse Lab in Alderley Park | Wellcome Sanger Institute for the COVID-19 Genomics UK (COG-UK) Consortium | Jacquelyn Wynn, Mairead Hyland, The Lighthouse Lab in Alderley Park and Alex Alderton, Roberto Amato, Jeffrey Barrett, Sonia Goncalves, Ewan Harrison, David K. Jackson, Ian Johnston, Dominic Kwiatkowski, Cordelia Langford, John Sillitoe on behalf of the Wellcome Sanger Institute COVID-19 Surveillance Team |
| EPI_ISL_2467962 | Health Services Laboratories | Wellcome Sanger Institute for the COVID-19 Genomics UK (COG-UK) Consortium | Health Services Laboratories and Alex Alderton, Roberto Amato, Jeffrey Barrett, Sonia Goncalves, Ewan Harrison, David K. Jackson, Ian Johnston, Dominic Kwiatkowski, Cordelia Langford, John Sillitoe on behalf of the Wellcome Sanger Institute COVID-19 Surveillance Team |
| EPI_ISL_2467963 | Lighthouse Lab in Alderley Park | Wellcome Sanger Institute for the COVID-19 Genomics UK (COG-UK) Consortium | Jacquelyn Wynn, Mairead Hyland, The Lighthouse Lab in Alderley Park and Alex Alderton, Roberto Amato, Jeffrey Barrett, Sonia Goncalves, Ewan Harrison, David K. Jackson, Ian Johnston, Dominic Kwiatkowski, Cordelia Langford, John Sillitoe on behalf of the Wellcome Sanger Institute COVID-19 Surveillance Team |
| EPI_ISL_2467964 | Lighthouse Laboratory Plymouth | Wellcome Sanger Institute for the COVID-19 Genomics UK (COG-UK) Consortium | Lighthouse Laboratory Plymouth and Alex Alderton, Roberto Amato, Jeffrey Barrett, Sonia Goncalves, Ewan Harrison, David K. Jackson, Ian Johnston, Dominic Kwiatkowski, Cordelia Langford, John Sillitoe on behalf of the Wellcome Sanger Institute COVID-19 Surveillance Team |
| EPI_ISL_2467965, EPI_ISL_2467966, EPI_ISL_2467967 | Lighthouse Lab in Milton Keynes | Wellcome Sanger Institute for the COVID-19 Genomics UK (COG-UK) Consortium | The Lighthouse Lab in Milton Keynes and Alex Alderton, Roberto Amato, Jeffrey Barrett, Sonia Goncalves, Ewan Harrison, David K. Jackson, Ian Johnston, Dominic Kwiatkowski, Cordelia Langford, John Sillitoe on behalf of the Wellcome Sanger Institute COVID-19 Surveillance Team |
| EPI_ISL_2467968 | Lighthouse Lab in Alderley Park | Wellcome Sanger Institute for the COVID-19 Genomics UK (COG-UK) | Jacquelyn Wynn, Mairead Hyland, The Lighthouse Lab in Alderley Park and Alex Alderton, Roberto Amato, Jeffrey Barrett, Sonia Goncalves, Ewan Harrison, David K. Jackson, Ian Johnston, Dominic Kwiatkowski, Cordelia Langford, John Sillitoe on |

| | | | |
|---|---|---|---|
| | | Consortium | behalf of the Wellcome Sanger Institute COVID-19 Surveillance Team |
| EPI_ISL_2467969 | Health Services Laboratories | Wellcome Sanger Institute for the COVID-19 Genomics UK (COG-UK) Consortium | Health Services Laboratories and Alex Alderton, Roberto Amato, Jeffrey Barrett, Sonia Goncalves, Ewan Harrison, David K. Jackson, Ian Johnston, Dominic Kwiatkowski, Cordelia Langford, John Sillitoe on behalf of the Wellcome Sanger Institute COVID-19 Surveillance Team |
| EPI_ISL_2467970 | Lighthouse Lab in Alderley Park | Wellcome Sanger Institute for the COVID-19 Genomics UK (COG-UK) Consortium | Jacquelyn Wynn, Mairead Hyland, The Lighthouse Lab in Alderley Park and Alex Alderton, Roberto Amato, Jeffrey Barrett, Sonia Goncalves, Ewan Harrison, David K. Jackson, Ian Johnston, Dominic Kwiatkowski, Cordelia Langford, John Sillitoe on behalf of the Wellcome Sanger Institute COVID-19 Surveillance Team |
| EPI_ISL_2467972, EPI_ISL_2467974 | Lighthouse Lab in Milton Keynes | Wellcome Sanger Institute for the COVID-19 Genomics UK (COG-UK) Consortium | The Lighthouse Lab in Milton Keynes and Alex Alderton, Roberto Amato, Jeffrey Barrett, Sonia Goncalves, Ewan Harrison, David K. Jackson, Ian Johnston, Dominic Kwiatkowski, Cordelia Langford, John Sillitoe on behalf of the Wellcome Sanger Institute COVID-19 Surveillance Team |
| EPI_ISL_2467975 | Lighthouse Lab in Alderley Park | Wellcome Sanger Institute for the COVID-19 Genomics UK (COG-UK) Consortium | Jacquelyn Wynn, Mairead Hyland, The Lighthouse Lab in Alderley Park and Alex Alderton, Roberto Amato, Jeffrey Barrett, Sonia Goncalves, Ewan Harrison, David K. Jackson, Ian Johnston, Dominic Kwiatkowski, Cordelia Langford, John Sillitoe on behalf of the Wellcome Sanger Institute COVID-19 Surveillance Team |
| EPI_ISL_2467976 | Lighthouse Laboratory Plymouth | Wellcome Sanger Institute for the COVID-19 Genomics UK (COG-UK) Consortium | Lighthouse Laboratory Plymouth and Alex Alderton, Roberto Amato, Jeffrey Barrett, Sonia Goncalves, Ewan Harrison, David K. Jackson, Ian Johnston, Dominic Kwiatkowski, Cordelia Langford, John Sillitoe on behalf of the Wellcome Sanger Institute COVID-19 Surveillance Team |
| EPI_ISL_2467977 | Lighthouse Lab in Milton Keynes | Wellcome Sanger Institute for the COVID-19 Genomics UK (COG-UK) Consortium | The Lighthouse Lab in Milton Keynes and Alex Alderton, Roberto Amato, Jeffrey Barrett, Sonia Goncalves, Ewan Harrison, David K. Jackson, Ian Johnston, Dominic Kwiatkowski, Cordelia Langford, John Sillitoe on behalf of the Wellcome Sanger Institute COVID-19 Surveillance Team |
| EPI_ISL_2467978, EPI_ISL_2467979 | Lighthouse Lab in Alderley Park | Wellcome Sanger Institute for the COVID-19 Genomics UK (COG-UK) Consortium | Jacquelyn Wynn, Mairead Hyland, The Lighthouse Lab in Alderley Park and Alex Alderton, Roberto Amato, Jeffrey Barrett, Sonia Goncalves, Ewan Harrison, David K. Jackson, Ian Johnston, Dominic Kwiatkowski, Cordelia Langford, John Sillitoe on behalf of the Wellcome Sanger Institute COVID-19 Surveillance Team |
| EPI_ISL_2467981 | Lighthouse Lab in Milton Keynes | Wellcome Sanger Institute for the COVID-19 Genomics UK (COG-UK) Consortium | The Lighthouse Lab in Milton Keynes and Alex Alderton, Roberto Amato, Jeffrey Barrett, Sonia Goncalves, Ewan Harrison, David K. Jackson, Ian Johnston, Dominic Kwiatkowski, Cordelia Langford, John Sillitoe on behalf of the Wellcome Sanger Institute COVID-19 Surveillance Team |
| EPI_ISL_2467982 | Lighthouse Lab in Alderley Park | Wellcome Sanger Institute for the COVID-19 Genomics UK (COG-UK) Consortium | Jacquelyn Wynn, Mairead Hyland, The Lighthouse Lab in Alderley Park and Alex Alderton, Roberto Amato, Jeffrey Barrett, Sonia Goncalves, Ewan Harrison, David K. Jackson, Ian Johnston, Dominic Kwiatkowski, Cordelia Langford, John Sillitoe on behalf of the Wellcome Sanger Institute COVID-19 Surveillance Team |
| EPI_ISL_2467983 | Lighthouse Lab in Milton Keynes | Wellcome Sanger Institute for the COVID-19 Genomics UK (COG-UK) Consortium | The Lighthouse Lab in Milton Keynes and Alex Alderton, Roberto Amato, Jeffrey Barrett, Sonia Goncalves, Ewan Harrison, David K. Jackson, Ian Johnston, Dominic Kwiatkowski, Cordelia Langford, John Sillitoe on behalf of the Wellcome Sanger Institute COVID-19 Surveillance Team |
| EPI_ISL_2467984, EPI_ISL_2467985 | Health Services Laboratories | Wellcome Sanger Institute for the COVID-19 Genomics UK (COG-UK) Consortium | Health Services Laboratories and Alex Alderton, Roberto Amato, Jeffrey Barrett, Sonia Goncalves, Ewan Harrison, David K. Jackson, Ian Johnston, Dominic Kwiatkowski, Cordelia Langford, John Sillitoe on behalf of the Wellcome Sanger Institute COVID-19 Surveillance Team |
| EPI_ISL_2467986, EPI_ISL_2467987, EPI_ISL_2467988 | Lighthouse Lab in Alderley Park | Wellcome Sanger Institute for the COVID-19 Genomics UK (COG-UK) Consortium | Jacquelyn Wynn, Mairead Hyland, The Lighthouse Lab in Alderley Park and Alex Alderton, Roberto Amato, Jeffrey Barrett, Sonia Goncalves, Ewan Harrison, David K. Jackson, Ian Johnston, Dominic Kwiatkowski, Cordelia Langford, John Sillitoe on behalf of the Wellcome Sanger Institute COVID-19 Surveillance Team |
| EPI_ISL_2467989 | Health Services Laboratories | Wellcome Sanger Institute for the COVID-19 Genomics UK (COG-UK) Consortium | Health Services Laboratories and Alex Alderton, Roberto Amato, Jeffrey Barrett, Sonia Goncalves, Ewan Harrison, David K. Jackson, Ian Johnston, Dominic Kwiatkowski, Cordelia Langford, John Sillitoe on behalf of the Wellcome Sanger Institute COVID-19 Surveillance Team |
| EPI_ISL_2467990, EPI_ISL_2467991 | Lighthouse Laboratory Plymouth | Wellcome Sanger Institute for the COVID-19 Genomics UK (COG-UK) Consortium | Lighthouse Laboratory Plymouth and Alex Alderton, Roberto Amato, Jeffrey Barrett, Sonia Goncalves, Ewan Harrison, David K. Jackson, Ian Johnston, Dominic Kwiatkowski, Cordelia Langford, John Sillitoe on behalf of the Wellcome Sanger Institute COVID-19 Surveillance Team |
| EPI_ISL_2467992 | Lighthouse Lab in Alderley Park | Wellcome Sanger Institute for the COVID-19 Genomics UK (COG-UK) Consortium | Jacquelyn Wynn, Mairead Hyland, The Lighthouse Lab in Alderley Park and Alex Alderton, Roberto Amato, Jeffrey Barrett, Sonia Goncalves, Ewan Harrison, David K. Jackson, Ian Johnston, Dominic Kwiatkowski, Cordelia Langford, John Sillitoe on behalf of the Wellcome Sanger Institute COVID-19 Surveillance Team |
| EPI_ISL_2467993 | Lighthouse Laboratory Plymouth | Wellcome Sanger Institute for the COVID-19 Genomics UK (COG-UK) Consortium | Lighthouse Laboratory Plymouth and Alex Alderton, Roberto Amato, Jeffrey Barrett, Sonia Goncalves, Ewan Harrison, David K. Jackson, Ian Johnston, Dominic Kwiatkowski, Cordelia Langford, John Sillitoe on behalf of the Wellcome Sanger Institute COVID-19 Surveillance Team |
| EPI_ISL_2467994 | Lighthouse Lab in Alderley Park | Wellcome Sanger Institute for the COVID-19 Genomics UK (COG-UK) Consortium | Jacquelyn Wynn, Mairead Hyland, The Lighthouse Lab in Alderley Park and Alex Alderton, Roberto Amato, Jeffrey Barrett, Sonia Goncalves, Ewan Harrison, David K. Jackson, Ian Johnston, Dominic Kwiatkowski, Cordelia Langford, John Sillitoe on behalf of the Wellcome Sanger Institute COVID-19 Surveillance Team |
| EPI_ISL_2467995 | Lighthouse Lab in Milton Keynes | Wellcome Sanger Institute for the COVID-19 Genomics UK (COG-UK) Consortium | The Lighthouse Lab in Milton Keynes and Alex Alderton, Roberto Amato, Jeffrey Barrett, Sonia Goncalves, Ewan Harrison, David K. Jackson, Ian Johnston, Dominic Kwiatkowski, Cordelia Langford, John Sillitoe on behalf of the Wellcome Sanger Institute COVID-19 Surveillance Team |
| EPI_ISL_2467996 | Lighthouse Laboratory Plymouth | Wellcome Sanger Institute for the COVID-19 Genomics UK (COG-UK) Consortium | Lighthouse Laboratory Plymouth and Alex Alderton, Roberto Amato, Jeffrey Barrett, Sonia Goncalves, Ewan Harrison, David K. Jackson, Ian Johnston, Dominic Kwiatkowski, Cordelia Langford, John Sillitoe on behalf of the Wellcome Sanger Institute COVID-19 Surveillance Team |
| EPI_ISL_2467997, EPI_ISL_2467998, EPI_ISL_2467999 | Lighthouse Lab in Alderley Park | Wellcome Sanger Institute for the COVID-19 Genomics UK (COG-UK) Consortium | Jacquelyn Wynn, Mairead Hyland, The Lighthouse Lab in Alderley Park and Alex Alderton, Roberto Amato, Jeffrey Barrett, Sonia Goncalves, Ewan Harrison, David K. Jackson, Ian Johnston, Dominic Kwiatkowski, Cordelia Langford, John Sillitoe on behalf of the Wellcome Sanger Institute COVID-19 Surveillance Team |
| EPI_ISL_2468001 | Lighthouse Lab in Milton Keynes | Wellcome Sanger Institute for the COVID-19 Genomics UK (COG-UK) Consortium | The Lighthouse Lab in Milton Keynes and Alex Alderton, Roberto Amato, Jeffrey Barrett, Sonia Goncalves, Ewan Harrison, David K. Jackson, Ian Johnston, Dominic Kwiatkowski, Cordelia Langford, John Sillitoe on behalf of the Wellcome Sanger Institute COVID-19 Surveillance Team |
| EPI_ISL_2468002, EPI_ISL_2468003 | Lighthouse Lab in Alderley Park | Wellcome Sanger Institute for the COVID-19 Genomics UK (COG-UK) Consortium | Jacquelyn Wynn, Mairead Hyland, The Lighthouse Lab in Alderley Park and Alex Alderton, Roberto Amato, Jeffrey Barrett, Sonia Goncalves, Ewan Harrison, David K. Jackson, Ian Johnston, Dominic Kwiatkowski, Cordelia Langford, John Sillitoe on behalf of the Wellcome Sanger Institute COVID-19 Surveillance Team |
| EPI_ISL_2468004 | Lighthouse Lab in Milton Keynes | Wellcome Sanger Institute for the COVID-19 Genomics UK (COG-UK) Consortium | The Lighthouse Lab in Milton Keynes and Alex Alderton, Roberto Amato, Jeffrey Barrett, Sonia Goncalves, Ewan Harrison, David K. Jackson, Ian Johnston, Dominic Kwiatkowski, Cordelia Langford, John Sillitoe on behalf of the Wellcome Sanger Institute COVID-19 Surveillance Team |
| EPI_ISL_2468005 | Lighthouse Lab in Alderley Park | Wellcome Sanger Institute for the COVID-19 Genomics UK (COG-UK) Consortium | Jacquelyn Wynn, Mairead Hyland, The Lighthouse Lab in Alderley Park and Alex Alderton, Roberto Amato, Jeffrey Barrett, Sonia Goncalves, Ewan Harrison, David K. Jackson, Ian Johnston, Dominic Kwiatkowski, Cordelia Langford, John Sillitoe on behalf of the Wellcome Sanger Institute COVID-19 Surveillance Team |
| EPI_ISL_2468006 | Lighthouse Lab in Milton Keynes | Wellcome Sanger Institute for the COVID-19 Genomics UK (COG-UK) Consortium | The Lighthouse Lab in Milton Keynes and Alex Alderton, Roberto Amato, Jeffrey Barrett, Sonia Goncalves, Ewan Harrison, David K. Jackson, Ian Johnston, Dominic Kwiatkowski, Cordelia Langford, John Sillitoe on behalf of the Wellcome Sanger Institute COVID-19 Surveillance Team |
| EPI_ISL_2468007 | Lighthouse Lab in Alderley Park | Wellcome Sanger Institute for the COVID-19 Genomics UK (COG-UK) Consortium | Jacquelyn Wynn, Mairead Hyland, The Lighthouse Lab in Alderley Park and Alex Alderton, Roberto Amato, Jeffrey Barrett, Sonia Goncalves, Ewan Harrison, David K. Jackson, Ian Johnston, Dominic Kwiatkowski, Cordelia Langford, John Sillitoe on behalf of the Wellcome Sanger Institute COVID-19 Surveillance Team |
| EPI_ISL_2468008 | Lighthouse Lab in Milton Keynes | Wellcome Sanger Institute for the COVID-19 Genomics UK (COG-UK) Consortium | The Lighthouse Lab in Milton Keynes and Alex Alderton, Roberto Amato, Jeffrey Barrett, Sonia Goncalves, Ewan Harrison, David K. Jackson, Ian Johnston, Dominic Kwiatkowski, Cordelia Langford, John Sillitoe on behalf of the Wellcome Sanger Institute COVID-19 Surveillance Team |
| EPI_ISL_2468009, EPI_ISL_2468010 | Lighthouse Lab in Alderley Park | Wellcome Sanger Institute for the COVID-19 Genomics UK (COG-UK) Consortium | Jacquelyn Wynn, Mairead Hyland, The Lighthouse Lab in Alderley Park and Alex Alderton, Roberto Amato, Jeffrey Barrett, Sonia Goncalves, Ewan Harrison, David K. Jackson, Ian Johnston, Dominic Kwiatkowski, Cordelia Langford, John Sillitoe on behalf of the Wellcome Sanger Institute COVID-19 Surveillance Team |
| EPI_ISL_2468011 | Lighthouse Laboratory Plymouth | Wellcome Sanger Institute for the COVID-19 Genomics UK (COG-UK) Consortium | Lighthouse Laboratory Plymouth and Alex Alderton, Roberto Amato, Jeffrey Barrett, Sonia Goncalves, Ewan Harrison, David K. Jackson, Ian Johnston, Dominic Kwiatkowski, Cordelia Langford, John Sillitoe on behalf of the Wellcome Sanger Institute COVID-19 Surveillance Team |
| EPI_ISL_2468012 | Lighthouse Lab in Alderley Park | Wellcome Sanger Institute for the COVID-19 Genomics UK (COG-UK) Consortium | Jacquelyn Wynn, Mairead Hyland, The Lighthouse Lab in Alderley Park and Alex Alderton, Roberto Amato, Jeffrey Barrett, Sonia Goncalves, Ewan Harrison, David K. Jackson, Ian Johnston, Dominic Kwiatkowski, Cordelia Langford, John Sillitoe on behalf of the Wellcome Sanger Institute COVID-19 Surveillance Team |
| EPI_ISL_2468013 | Lighthouse Laboratory Plymouth | Wellcome Sanger Institute for the COVID-19 Genomics UK (COG-UK) Consortium | Lighthouse Laboratory Plymouth and Alex Alderton, Roberto Amato, Jeffrey Barrett, Sonia Goncalves, Ewan Harrison, David K. Jackson, Ian Johnston, Dominic Kwiatkowski, Cordelia Langford, John Sillitoe on behalf of the Wellcome Sanger Institute COVID-19 Surveillance Team |
| EPI_ISL_2468014 | Health Services Laboratories | Wellcome Sanger Institute for the COVID-19 Genomics UK (COG-UK) Consortium | Health Services Laboratories and Alex Alderton, Roberto Amato, Jeffrey Barrett, Sonia Goncalves, Ewan Harrison, David K. Jackson, Ian Johnston, Dominic Kwiatkowski, Cordelia Langford, John Sillitoe on behalf of the Wellcome Sanger Institute COVID-19 Surveillance Team |
| EPI_ISL_2468015, EPI_ISL_2468016 | Lighthouse Lab in Alderley Park | Wellcome Sanger Institute for the COVID-19 Genomics UK (COG-UK) Consortium | Jacquelyn Wynn, Mairead Hyland, The Lighthouse Lab in Alderley Park and Alex Alderton, Roberto Amato, Jeffrey Barrett, Sonia Goncalves, Ewan Harrison, David K. Jackson, Ian Johnston, Dominic Kwiatkowski, Cordelia Langford, John Sillitoe on behalf of the Wellcome Sanger Institute COVID-19 Surveillance Team |
| EPI_ISL_2468017 | Lighthouse Laboratory Plymouth | Wellcome Sanger Institute for the COVID-19 Genomics UK (COG-UK) Consortium | Lighthouse Laboratory Plymouth and Alex Alderton, Roberto Amato, Jeffrey Barrett, Sonia Goncalves, Ewan Harrison, David K. Jackson, Ian Johnston, Dominic Kwiatkowski, Cordelia Langford, John Sillitoe on behalf of the Wellcome Sanger Institute COVID-19 Surveillance Team |
| EPI_ISL_2468018 | Lighthouse Lab in Milton Keynes | Wellcome Sanger Institute for the COVID-19 Genomics UK (COG-UK) Consortium | The Lighthouse Lab in Milton Keynes and Alex Alderton, Roberto Amato, Jeffrey Barrett, Sonia Goncalves, Ewan Harrison, David K. Jackson, Ian Johnston, Dominic Kwiatkowski, Cordelia Langford, John Sillitoe on behalf of the Wellcome Sanger Institute COVID-19 Surveillance Team |
| EPI_ISL_2468019 | Lighthouse Lab in Alderley Park | Wellcome Sanger Institute for the COVID-19 Genomics UK (COG-UK) Consortium | Jacquelyn Wynn, Mairead Hyland, The Lighthouse Lab in Alderley Park and Alex Alderton, Roberto Amato, Jeffrey Barrett, Sonia Goncalves, Ewan Harrison, David K. Jackson, Ian Johnston, Dominic Kwiatkowski, Cordelia Langford, John Sillitoe on behalf of the Wellcome Sanger Institute COVID-19 Surveillance Team |
| EPI_ISL_2468020 | Lighthouse Laboratory Plymouth | Wellcome Sanger Institute for the COVID-19 Genomics UK (COG-UK) Consortium | Lighthouse Laboratory Plymouth and Alex Alderton, Roberto Amato, Jeffrey Barrett, Sonia Goncalves, Ewan Harrison, David K. Jackson, Ian Johnston, Dominic Kwiatkowski, Cordelia Langford, John Sillitoe on behalf of the Wellcome Sanger Institute COVID-19 Surveillance Team |
| EPI_ISL_2468022, EPI_ISL_2468023, EPI_ISL_2468024 | Lighthouse Lab in Alderley Park | Wellcome Sanger Institute for the COVID-19 Genomics UK (COG-UK) Consortium | Jacquelyn Wynn, Mairead Hyland, The Lighthouse Lab in Alderley Park and Alex Alderton, Roberto Amato, Jeffrey Barrett, Sonia Goncalves, Ewan Harrison, David K. Jackson, Ian Johnston, Dominic Kwiatkowski, Cordelia Langford, John Sillitoe on behalf of the Wellcome Sanger Institute COVID-19 Surveillance Team |
| EPI_ISL_2468025 | Lighthouse Lab in Milton Keynes | Wellcome Sanger Institute for the COVID-19 Genomics UK (COG-UK) Consortium | The Lighthouse Lab in Milton Keynes and Alex Alderton, Roberto Amato, Jeffrey Barrett, Sonia Goncalves, Ewan Harrison, David K. Jackson, Ian Johnston, Dominic Kwiatkowski, Cordelia Langford, John Sillitoe on behalf of the Wellcome Sanger Institute COVID-19 Surveillance Team |
| EPI_ISL_2468026, EPI_ISL_2468027, EPI_ISL_2468028, EPI_ISL_2468029, EPI_ISL_2468030, EPI_ISL_2468031 | Lighthouse Lab in Alderley Park | Wellcome Sanger Institute for the COVID-19 Genomics UK (COG-UK) Consortium | Jacquelyn Wynn, Mairead Hyland, The Lighthouse Lab in Alderley Park and Alex Alderton, Roberto Amato, Jeffrey Barrett, Sonia Goncalves, Ewan Harrison, David K. Jackson, Ian Johnston, Dominic Kwiatkowski, Cordelia Langford, John Sillitoe on behalf of the Wellcome Sanger Institute COVID-19 Surveillance Team |
| EPI_ISL_2468032 | Lighthouse Laboratory Plymouth | Wellcome Sanger Institute for the COVID-19 Genomics UK (COG-UK) Consortium | Lighthouse Laboratory Plymouth and Alex Alderton, Roberto Amato, Jeffrey Barrett, Sonia Goncalves, Ewan Harrison, David K. Jackson, Ian Johnston, Dominic Kwiatkowski, Cordelia Langford, John Sillitoe on behalf of the Wellcome Sanger Institute COVID-19 Surveillance Team |
| EPI_ISL_2468033 | Lighthouse Lab in Alderley Park | Wellcome Sanger Institute for the COVID-19 Genomics UK (COG-UK) Consortium | Jacquelyn Wynn, Mairead Hyland, The Lighthouse Lab in Alderley Park and Alex Alderton, Roberto Amato, Jeffrey Barrett, Sonia Goncalves, Ewan Harrison, David K. Jackson, Ian Johnston, Dominic Kwiatkowski, Cordelia Langford, John Sillitoe on behalf of the Wellcome Sanger Institute COVID-19 Surveillance Team |

| | | | |
|---|---|---|---|
| EPI_ISL_2468034, EPI_ISL_2468035 | Lighthouse Lab in Milton Keynes | Wellcome Sanger Institute for the COVID-19 Genomics UK (COG-UK) Consortium | The Lighthouse Lab in Milton Keynes and Alex Alderton, Roberto Amato, Jeffrey Barrett, Sonia Goncalves, Ewan Harrison, David K. Jackson, Ian Johnston, Dominic Kwiatkowski, Cordelia Langford, John Sillitoe on behalf of the Wellcome Sanger Institute COVID-19 Surveillance Team |
| EPI_ISL_2468036 | Health Services Laboratories | Wellcome Sanger Institute for the COVID-19 Genomics UK (COG-UK) Consortium | Health Services Laboratories and Alex Alderton, Roberto Amato, Jeffrey Barrett, Sonia Goncalves, Ewan Harrison, David K. Jackson, Ian Johnston, Dominic Kwiatkowski, Cordelia Langford, John Sillitoe on behalf of the Wellcome Sanger Institute COVID-19 Surveillance Team |
| EPI_ISL_2468037 | Lighthouse Laboratory Plymouth | Wellcome Sanger Institute for the COVID-19 Genomics UK (COG-UK) Consortium | Lighthouse Laboratory Plymouth and Alex Alderton, Roberto Amato, Jeffrey Barrett, Sonia Goncalves, Ewan Harrison, David K. Jackson, Ian Johnston, Dominic Kwiatkowski, Cordelia Langford, John Sillitoe on behalf of the Wellcome Sanger Institute COVID-19 Surveillance Team |
| EPI_ISL_2468038 | Lighthouse Lab in Milton Keynes | Wellcome Sanger Institute for the COVID-19 Genomics UK (COG-UK) Consortium | The Lighthouse Lab in Milton Keynes and Alex Alderton, Roberto Amato, Jeffrey Barrett, Sonia Goncalves, Ewan Harrison, David K. Jackson, Ian Johnston, Dominic Kwiatkowski, Cordelia Langford, John Sillitoe on behalf of the Wellcome Sanger Institute COVID-19 Surveillance Team |
| EPI_ISL_2468039, EPI_ISL_2468040 | Lighthouse Lab in Alderley Park | Wellcome Sanger Institute for the COVID-19 Genomics UK (COG-UK) Consortium | Jacquelyn Wynn, Mairead Hyland, The Lighthouse Lab in Alderley Park and Alex Alderton, Roberto Amato, Jeffrey Barrett, Sonia Goncalves, Ewan Harrison, David K. Jackson, Ian Johnston, Dominic Kwiatkowski, Cordelia Langford, John Sillitoe on behalf of the Wellcome Sanger Institute COVID-19 Surveillance Team |
| EPI_ISL_2468041 | Lighthouse Laboratory Plymouth | Wellcome Sanger Institute for the COVID-19 Genomics UK (COG-UK) Consortium | Lighthouse Laboratory Plymouth and Alex Alderton, Roberto Amato, Jeffrey Barrett, Sonia Goncalves, Ewan Harrison, David K. Jackson, Ian Johnston, Dominic Kwiatkowski, Cordelia Langford, John Sillitoe on behalf of the Wellcome Sanger Institute COVID-19 Surveillance Team |
| EPI_ISL_2468042, EPI_ISL_2468043 | Lighthouse Lab in Alderley Park | Wellcome Sanger Institute for the COVID-19 Genomics UK (COG-UK) Consortium | Jacquelyn Wynn, Mairead Hyland, The Lighthouse Lab in Alderley Park and Alex Alderton, Roberto Amato, Jeffrey Barrett, Sonia Goncalves, Ewan Harrison, David K. Jackson, Ian Johnston, Dominic Kwiatkowski, Cordelia Langford, John Sillitoe on behalf of the Wellcome Sanger Institute COVID-19 Surveillance Team |
| EPI_ISL_2468044 | Lighthouse Lab in Milton Keynes | Wellcome Sanger Institute for the COVID-19 Genomics UK (COG-UK) Consortium | The Lighthouse Lab in Milton Keynes and Alex Alderton, Roberto Amato, Jeffrey Barrett, Sonia Goncalves, Ewan Harrison, David K. Jackson, Ian Johnston, Dominic Kwiatkowski, Cordelia Langford, John Sillitoe on behalf of the Wellcome Sanger Institute COVID-19 Surveillance Team |
| EPI_ISL_2468045 | Lighthouse Lab in Alderley Park | Wellcome Sanger Institute for the COVID-19 Genomics UK (COG-UK) Consortium | Jacquelyn Wynn, Mairead Hyland, The Lighthouse Lab in Alderley Park and Alex Alderton, Roberto Amato, Jeffrey Barrett, Sonia Goncalves, Ewan Harrison, David K. Jackson, Ian Johnston, Dominic Kwiatkowski, Cordelia Langford, John Sillitoe on behalf of the Wellcome Sanger Institute COVID-19 Surveillance Team |
| EPI_ISL_2468046, EPI_ISL_2468047 | Lighthouse Laboratory Plymouth | Wellcome Sanger Institute for the COVID-19 Genomics UK (COG-UK) Consortium | Lighthouse Laboratory Plymouth and Alex Alderton, Roberto Amato, Jeffrey Barrett, Sonia Goncalves, Ewan Harrison, David K. Jackson, Ian Johnston, Dominic Kwiatkowski, Cordelia Langford, John Sillitoe on behalf of the Wellcome Sanger Institute COVID-19 Surveillance Team |
| EPI_ISL_2468048, EPI_ISL_2468049 | Lighthouse Lab in Alderley Park | Wellcome Sanger Institute for the COVID-19 Genomics UK (COG-UK) Consortium | Jacquelyn Wynn, Mairead Hyland, The Lighthouse Lab in Alderley Park and Alex Alderton, Roberto Amato, Jeffrey Barrett, Sonia Goncalves, Ewan Harrison, David K. Jackson, Ian Johnston, Dominic Kwiatkowski, Cordelia Langford, John Sillitoe on behalf of the Wellcome Sanger Institute COVID-19 Surveillance Team |
| EPI_ISL_2468050 | Lighthouse Lab in Milton Keynes | Wellcome Sanger Institute for the COVID-19 Genomics UK (COG-UK) Consortium | The Lighthouse Lab in Milton Keynes and Alex Alderton, Roberto Amato, Jeffrey Barrett, Sonia Goncalves, Ewan Harrison, David K. Jackson, Ian Johnston, Dominic Kwiatkowski, Cordelia Langford, John Sillitoe on behalf of the Wellcome Sanger Institute COVID-19 Surveillance Team |
| EPI_ISL_2468051 | Lighthouse Laboratory Plymouth | Wellcome Sanger Institute for the COVID-19 Genomics UK (COG-UK) Consortium | Lighthouse Laboratory Plymouth and Alex Alderton, Roberto Amato, Jeffrey Barrett, Sonia Goncalves, Ewan Harrison, David K. Jackson, Ian Johnston, Dominic Kwiatkowski, Cordelia Langford, John Sillitoe on behalf of the Wellcome Sanger Institute COVID-19 Surveillance Team |
| EPI_ISL_2468052, EPI_ISL_2468054 | Lighthouse Lab in Alderley Park | Wellcome Sanger Institute for the COVID-19 Genomics UK (COG-UK) Consortium | Jacquelyn Wynn, Mairead Hyland, The Lighthouse Lab in Alderley Park and Alex Alderton, Roberto Amato, Jeffrey Barrett, Sonia Goncalves, Ewan Harrison, David K. Jackson, Ian Johnston, Dominic Kwiatkowski, Cordelia Langford, John Sillitoe on behalf of the Wellcome Sanger Institute COVID-19 Surveillance Team |
| EPI_ISL_2468055 | Lighthouse Lab in Milton Keynes | Wellcome Sanger Institute for the COVID-19 Genomics UK (COG-UK) Consortium | The Lighthouse Lab in Milton Keynes and Alex Alderton, Roberto Amato, Jeffrey Barrett, Sonia Goncalves, Ewan Harrison, David K. Jackson, Ian Johnston, Dominic Kwiatkowski, Cordelia Langford, John Sillitoe on behalf of the Wellcome Sanger Institute COVID-19 Surveillance Team |
| EPI_ISL_2468056 | Lighthouse Laboratory Plymouth | Wellcome Sanger Institute for the COVID-19 Genomics UK (COG-UK) Consortium | Lighthouse Laboratory Plymouth and Alex Alderton, Roberto Amato, Jeffrey Barrett, Sonia Goncalves, Ewan Harrison, David K. Jackson, Ian Johnston, Dominic Kwiatkowski, Cordelia Langford, John Sillitoe on behalf of the Wellcome Sanger Institute COVID-19 Surveillance Team |
| EPI_ISL_2468057 | Lighthouse Lab in Alderley Park | Wellcome Sanger Institute for the COVID-19 Genomics UK (COG-UK) Consortium | Jacquelyn Wynn, Mairead Hyland, The Lighthouse Lab in Alderley Park and Alex Alderton, Roberto Amato, Jeffrey Barrett, Sonia Goncalves, Ewan Harrison, David K. Jackson, Ian Johnston, Dominic Kwiatkowski, Cordelia Langford, John Sillitoe on behalf of the Wellcome Sanger Institute COVID-19 Surveillance Team |
| EPI_ISL_2468058, EPI_ISL_2468059 | Lighthouse Lab in Milton Keynes | Wellcome Sanger Institute for the COVID-19 Genomics UK (COG-UK) Consortium | The Lighthouse Lab in Milton Keynes and Alex Alderton, Roberto Amato, Jeffrey Barrett, Sonia Goncalves, Ewan Harrison, David K. Jackson, Ian Johnston, Dominic Kwiatkowski, Cordelia Langford, John Sillitoe on behalf of the Wellcome Sanger Institute COVID-19 Surveillance Team |
| EPI_ISL_2468060 | Lighthouse Lab in Alderley Park | Wellcome Sanger Institute for the COVID-19 Genomics UK (COG-UK) Consortium | Jacquelyn Wynn, Mairead Hyland, The Lighthouse Lab in Alderley Park and Alex Alderton, Roberto Amato, Jeffrey Barrett, Sonia Goncalves, Ewan Harrison, David K. Jackson, Ian Johnston, Dominic Kwiatkowski, Cordelia Langford, John Sillitoe on behalf of the Wellcome Sanger Institute COVID-19 Surveillance Team |
| EPI_ISL_2468061 | Lighthouse Lab in Milton Keynes | Wellcome Sanger Institute for the COVID-19 Genomics UK (COG-UK) Consortium | The Lighthouse Lab in Milton Keynes and Alex Alderton, Roberto Amato, Jeffrey Barrett, Sonia Goncalves, Ewan Harrison, David K. Jackson, Ian Johnston, Dominic Kwiatkowski, Cordelia Langford, John Sillitoe on behalf of the Wellcome Sanger Institute COVID-19 Surveillance Team |
| EPI_ISL_2468062 | Lighthouse Lab in Alderley Park | Wellcome Sanger Institute for the COVID-19 Genomics UK (COG-UK) Consortium | Jacquelyn Wynn, Mairead Hyland, The Lighthouse Lab in Alderley Park and Alex Alderton, Roberto Amato, Jeffrey Barrett, Sonia Goncalves, Ewan Harrison, David K. Jackson, Ian Johnston, Dominic Kwiatkowski, Cordelia Langford, John Sillitoe on behalf of the Wellcome Sanger Institute COVID-19 Surveillance Team |
| EPI_ISL_2468063 | Lighthouse Lab in Milton Keynes | Wellcome Sanger Institute for the COVID-19 Genomics UK (COG-UK) Consortium | The Lighthouse Lab in Milton Keynes and Alex Alderton, Roberto Amato, Jeffrey Barrett, Sonia Goncalves, Ewan Harrison, David K. Jackson, Ian Johnston, Dominic Kwiatkowski, Cordelia Langford, John Sillitoe on behalf of the Wellcome Sanger Institute COVID-19 Surveillance Team |
| EPI_ISL_2468064 | Lighthouse Laboratory Plymouth | Wellcome Sanger Institute for the COVID-19 Genomics UK (COG-UK) Consortium | Lighthouse Laboratory Plymouth and Alex Alderton, Roberto Amato, Jeffrey Barrett, Sonia Goncalves, Ewan Harrison, David K. Jackson, Ian Johnston, Dominic Kwiatkowski, Cordelia Langford, John Sillitoe on behalf of the Wellcome Sanger Institute COVID-19 Surveillance Team |
| EPI_ISL_2468065, EPI_ISL_2468066, EPI_ISL_2468067 | Lighthouse Lab in Alderley Park | Wellcome Sanger Institute for the COVID-19 Genomics UK (COG-UK) Consortium | Jacquelyn Wynn, Mairead Hyland, The Lighthouse Lab in Alderley Park and Alex Alderton, Roberto Amato, Jeffrey Barrett, Sonia Goncalves, Ewan Harrison, David K. Jackson, Ian Johnston, Dominic Kwiatkowski, Cordelia Langford, John Sillitoe on behalf of the Wellcome Sanger Institute COVID-19 Surveillance Team |
| EPI_ISL_2468068 | Lighthouse Laboratory Plymouth | Wellcome Sanger Institute for the COVID-19 Genomics UK (COG-UK) Consortium | Lighthouse Laboratory Plymouth and Alex Alderton, Roberto Amato, Jeffrey Barrett, Sonia Goncalves, Ewan Harrison, David K. Jackson, Ian Johnston, Dominic Kwiatkowski, Cordelia Langford, John Sillitoe on behalf of the Wellcome Sanger Institute COVID-19 Surveillance Team |
| EPI_ISL_2468069 | Lighthouse Lab in Alderley Park | Wellcome Sanger Institute for the COVID-19 Genomics UK (COG-UK) Consortium | Jacquelyn Wynn, Mairead Hyland, The Lighthouse Lab in Alderley Park and Alex Alderton, Roberto Amato, Jeffrey Barrett, Sonia Goncalves, Ewan Harrison, David K. Jackson, Ian Johnston, Dominic Kwiatkowski, Cordelia Langford, John Sillitoe on behalf of the Wellcome Sanger Institute COVID-19 Surveillance Team |
| EPI_ISL_2468070 | Lighthouse Lab in Milton Keynes | Wellcome Sanger Institute for the COVID-19 Genomics UK (COG-UK) Consortium | The Lighthouse Lab in Milton Keynes and Alex Alderton, Roberto Amato, Jeffrey Barrett, Sonia Goncalves, Ewan Harrison, David K. Jackson, Ian Johnston, Dominic Kwiatkowski, Cordelia Langford, John Sillitoe on behalf of the Wellcome Sanger Institute COVID-19 Surveillance Team |
| EPI_ISL_2468071 | Lighthouse Lab in Alderley Park | Wellcome Sanger Institute for the COVID-19 Genomics UK (COG-UK) Consortium | Jacquelyn Wynn, Mairead Hyland, The Lighthouse Lab in Alderley Park and Alex Alderton, Roberto Amato, Jeffrey Barrett, Sonia Goncalves, Ewan Harrison, David K. Jackson, Ian Johnston, Dominic Kwiatkowski, Cordelia Langford, John Sillitoe on behalf of the Wellcome Sanger Institute COVID-19 Surveillance Team |
| EPI_ISL_2468072 | Lighthouse Laboratory Plymouth | Wellcome Sanger Institute for the COVID-19 Genomics UK (COG-UK) Consortium | Lighthouse Laboratory Plymouth and Alex Alderton, Roberto Amato, Jeffrey Barrett, Sonia Goncalves, Ewan Harrison, David K. Jackson, Ian Johnston, Dominic Kwiatkowski, Cordelia Langford, John Sillitoe on behalf of the Wellcome Sanger Institute COVID-19 Surveillance Team |
| EPI_ISL_2468073, EPI_ISL_2468074 | Lighthouse Lab in Alderley Park | Wellcome Sanger Institute for the COVID-19 Genomics UK (COG-UK) Consortium | Jacquelyn Wynn, Mairead Hyland, The Lighthouse Lab in Alderley Park and Alex Alderton, Roberto Amato, Jeffrey Barrett, Sonia Goncalves, Ewan Harrison, David K. Jackson, Ian Johnston, Dominic Kwiatkowski, Cordelia Langford, John Sillitoe on behalf of the Wellcome Sanger Institute COVID-19 Surveillance Team |
| EPI_ISL_2468075 | Lighthouse Laboratory Plymouth | Wellcome Sanger Institute for the COVID-19 Genomics UK (COG-UK) Consortium | Lighthouse Laboratory Plymouth and Alex Alderton, Roberto Amato, Jeffrey Barrett, Sonia Goncalves, Ewan Harrison, David K. Jackson, Ian Johnston, Dominic Kwiatkowski, Cordelia Langford, John Sillitoe on behalf of the Wellcome Sanger Institute COVID-19 Surveillance Team |
| EPI_ISL_2468076 | Lighthouse Lab in Milton Keynes | Wellcome Sanger Institute for the COVID-19 Genomics UK (COG-UK) Consortium | The Lighthouse Lab in Milton Keynes and Alex Alderton, Roberto Amato, Jeffrey Barrett, Sonia Goncalves, Ewan Harrison, David K. Jackson, Ian Johnston, Dominic Kwiatkowski, Cordelia Langford, John Sillitoe on behalf of the Wellcome Sanger Institute COVID-19 Surveillance Team |
| EPI_ISL_2468077, EPI_ISL_2468078, EPI_ISL_2468079, EPI_ISL_2468081 | Lighthouse Lab in Alderley Park | Wellcome Sanger Institute for the COVID-19 Genomics UK (COG-UK) Consortium | Jacquelyn Wynn, Mairead Hyland, The Lighthouse Lab in Alderley Park and Alex Alderton, Roberto Amato, Jeffrey Barrett, Sonia Goncalves, Ewan Harrison, David K. Jackson, Ian Johnston, Dominic Kwiatkowski, Cordelia Langford, John Sillitoe on behalf of the Wellcome Sanger Institute COVID-19 Surveillance Team |
| EPI_ISL_2468082 | Lighthouse Lab in Milton Keynes | Wellcome Sanger Institute for the COVID-19 Genomics UK (COG-UK) Consortium | The Lighthouse Lab in Milton Keynes and Alex Alderton, Roberto Amato, Jeffrey Barrett, Sonia Goncalves, Ewan Harrison, David K. Jackson, Ian Johnston, Dominic Kwiatkowski, Cordelia Langford, John Sillitoe on behalf of the Wellcome Sanger Institute COVID-19 Surveillance Team |
| EPI_ISL_2468083, EPI_ISL_2468084 | Lighthouse Lab in Alderley Park | Wellcome Sanger Institute for the COVID-19 Genomics UK (COG-UK) Consortium | Jacquelyn Wynn, Mairead Hyland, The Lighthouse Lab in Alderley Park and Alex Alderton, Roberto Amato, Jeffrey Barrett, Sonia Goncalves, Ewan Harrison, David K. Jackson, Ian Johnston, Dominic Kwiatkowski, Cordelia Langford, John Sillitoe on behalf of the Wellcome Sanger Institute COVID-19 Surveillance Team |
| EPI_ISL_2468085 | Health Services Laboratories | Wellcome Sanger Institute for the COVID-19 Genomics UK (COG-UK) Consortium | Health Services Laboratories and Alex Alderton, Roberto Amato, Jeffrey Barrett, Sonia Goncalves, Ewan Harrison, David K. Jackson, Ian Johnston, Dominic Kwiatkowski, Cordelia Langford, John Sillitoe on behalf of the Wellcome Sanger Institute COVID-19 Surveillance Team |
| EPI_ISL_2468086 | Lighthouse Lab in Milton Keynes | Wellcome Sanger Institute for the COVID-19 Genomics UK (COG-UK) Consortium | The Lighthouse Lab in Milton Keynes and Alex Alderton, Roberto Amato, Jeffrey Barrett, Sonia Goncalves, Ewan Harrison, David K. Jackson, Ian Johnston, Dominic Kwiatkowski, Cordelia Langford, John Sillitoe on behalf of the Wellcome Sanger Institute COVID-19 Surveillance Team |
| EPI_ISL_2468087, EPI_ISL_2468088 | Lighthouse Laboratory Plymouth | Wellcome Sanger Institute for the COVID-19 Genomics UK (COG-UK) Consortium | Lighthouse Laboratory Plymouth and Alex Alderton, Roberto Amato, Jeffrey Barrett, Sonia Goncalves, Ewan Harrison, David K. Jackson, Ian Johnston, Dominic Kwiatkowski, Cordelia Langford, John Sillitoe on behalf of the Wellcome Sanger Institute COVID-19 Surveillance Team |
| EPI_ISL_2468089, EPI_ISL_2468090 | Lighthouse Lab in Alderley Park | Wellcome Sanger Institute for the COVID-19 Genomics UK (COG-UK) Consortium | Jacquelyn Wynn, Mairead Hyland, The Lighthouse Lab in Alderley Park and Alex Alderton, Roberto Amato, Jeffrey Barrett, Sonia Goncalves, Ewan Harrison, David K. Jackson, Ian Johnston, Dominic Kwiatkowski, Cordelia Langford, John Sillitoe on behalf of the Wellcome Sanger Institute COVID-19 Surveillance Team |
| EPI_ISL_2468091 | Lighthouse Lab in Milton Keynes | Wellcome Sanger Institute for the COVID-19 Genomics UK (COG-UK) Consortium | The Lighthouse Lab in Milton Keynes and Alex Alderton, Roberto Amato, Jeffrey Barrett, Sonia Goncalves, Ewan Harrison, David K. Jackson, Ian Johnston, Dominic Kwiatkowski, Cordelia Langford, John Sillitoe on behalf of the Wellcome Sanger Institute COVID-19 Surveillance Team |
| EPI_ISL_2468092 | Lighthouse Lab in Alderley Park | Wellcome Sanger Institute for the COVID-19 Genomics UK (COG-UK) Consortium | Jacquelyn Wynn, Mairead Hyland, The Lighthouse Lab in Alderley Park and Alex Alderton, Roberto Amato, Jeffrey Barrett, Sonia Goncalves, Ewan Harrison, David K. Jackson, Ian Johnston, Dominic Kwiatkowski, Cordelia Langford, John Sillitoe on behalf of the Wellcome Sanger Institute COVID-19 Surveillance Team |
| EPI_ISL_2468093 | Lighthouse Laboratory Plymouth | Wellcome Sanger Institute for the COVID-19 Genomics UK (COG-UK) Consortium | Lighthouse Laboratory Plymouth and Alex Alderton, Roberto Amato, Jeffrey Barrett, Sonia Goncalves, Ewan Harrison, David K. Jackson, Ian Johnston, Dominic Kwiatkowski, Cordelia Langford, John Sillitoe on behalf of the Wellcome Sanger Institute COVID-19 Surveillance Team |
| EPI_ISL_2468095, EPI_ISL_2468096 | Lighthouse Lab in Milton Keynes | Wellcome Sanger Institute for the COVID-19 Genomics UK (COG-UK) Consortium | The Lighthouse Lab in Milton Keynes and Alex Alderton, Roberto Amato, Jeffrey Barrett, Sonia Goncalves, Ewan Harrison, David K. Jackson, Ian Johnston, Dominic Kwiatkowski, Cordelia Langford, John Sillitoe on behalf of the Wellcome Sanger Institute COVID-19 Surveillance Team |
| EPI_ISL_2468097, | Lighthouse Lab in Alderley Park | Wellcome Sanger Institute for the COVID-19 Genomics UK (COG-UK) | Jacquelyn Wynn, Mairead Hyland, The Lighthouse Lab in Alderley Park and Alex Alderton, Roberto Amato, Jeffrey Barrett, Sonia Goncalves, Ewan Harrison, David K. Jackson, Ian Johnston, Dominic Kwiatkowski, Cordelia Langford, John Sillitoe on |

| | | | |
|---|---|---|---|
| EPI_ISL_2468098 | | Consortium | behalf of the Wellcome Sanger Institute COVID-19 Surveillance Team |
| EPI_ISL_2468099 | Lighthouse Lab in Milton Keynes | Wellcome Sanger Institute for the COVID-19 Genomics UK (COG-UK) Consortium | The Lighthouse Lab in Milton Keynes and Alex Alderton, Roberto Amato, Jeffrey Barrett, Sonia Goncalves, Ewan Harrison, David K. Jackson, Ian Johnston, Dominic Kwiatkowski, Cordelia Langford, John Sillitoe on behalf of the Wellcome Sanger Institute COVID-19 Surveillance Team |
| EPI_ISL_2468100 | Lighthouse Lab in Alderley Park | Wellcome Sanger Institute for the COVID-19 Genomics UK (COG-UK) Consortium | Jacquelyn Wynn, Mairead Hyland, The Lighthouse Lab in Alderley Park and Alex Alderton, Roberto Amato, Jeffrey Barrett, Sonia Goncalves, Ewan Harrison, David K. Jackson, Ian Johnston, Dominic Kwiatkowski, Cordelia Langford, John Sillitoe on behalf of the Wellcome Sanger Institute COVID-19 Surveillance Team |
| EPI_ISL_2468101 | Lighthouse Laboratory Plymouth | Wellcome Sanger Institute for the COVID-19 Genomics UK (COG-UK) Consortium | Lighthouse Laboratory Plymouth and Alex Alderton, Roberto Amato, Jeffrey Barrett, Sonia Goncalves, Ewan Harrison, David K. Jackson, Ian Johnston, Dominic Kwiatkowski, Cordelia Langford, John Sillitoe on behalf of the Wellcome Sanger Institute COVID-19 Surveillance Team |
| EPI_ISL_2468102 | Lighthouse Lab in Milton Keynes | Wellcome Sanger Institute for the COVID-19 Genomics UK (COG-UK) Consortium | The Lighthouse Lab in Milton Keynes and Alex Alderton, Roberto Amato, Jeffrey Barrett, Sonia Goncalves, Ewan Harrison, David K. Jackson, Ian Johnston, Dominic Kwiatkowski, Cordelia Langford, John Sillitoe on behalf of the Wellcome Sanger Institute COVID-19 Surveillance Team |
| EPI_ISL_2468103 | Lighthouse Laboratory Plymouth | Wellcome Sanger Institute for the COVID-19 Genomics UK (COG-UK) Consortium | Lighthouse Laboratory Plymouth and Alex Alderton, Roberto Amato, Jeffrey Barrett, Sonia Goncalves, Ewan Harrison, David K. Jackson, Ian Johnston, Dominic Kwiatkowski, Cordelia Langford, John Sillitoe on behalf of the Wellcome Sanger Institute COVID-19 Surveillance Team |
| EPI_ISL_2468104, EPI_ISL_2468105, EPI_ISL_2468106 | Lighthouse Lab in Alderley Park | Wellcome Sanger Institute for the COVID-19 Genomics UK (COG-UK) Consortium | Jacquelyn Wynn, Mairead Hyland, The Lighthouse Lab in Alderley Park and Alex Alderton, Roberto Amato, Jeffrey Barrett, Sonia Goncalves, Ewan Harrison, David K. Jackson, Ian Johnston, Dominic Kwiatkowski, Cordelia Langford, John Sillitoe on behalf of the Wellcome Sanger Institute COVID-19 Surveillance Team |
| EPI_ISL_2468107 | Lighthouse Lab in Milton Keynes | Wellcome Sanger Institute for the COVID-19 Genomics UK (COG-UK) Consortium | The Lighthouse Lab in Milton Keynes and Alex Alderton, Roberto Amato, Jeffrey Barrett, Sonia Goncalves, Ewan Harrison, David K. Jackson, Ian Johnston, Dominic Kwiatkowski, Cordelia Langford, John Sillitoe on behalf of the Wellcome Sanger Institute COVID-19 Surveillance Team |
| EPI_ISL_2468108 | Lighthouse Laboratory Plymouth | Wellcome Sanger Institute for the COVID-19 Genomics UK (COG-UK) Consortium | Lighthouse Laboratory Plymouth and Alex Alderton, Roberto Amato, Jeffrey Barrett, Sonia Goncalves, Ewan Harrison, David K. Jackson, Ian Johnston, Dominic Kwiatkowski, Cordelia Langford, John Sillitoe on behalf of the Wellcome Sanger Institute COVID-19 Surveillance Team |
| EPI_ISL_2468109, EPI_ISL_2468110 | Lighthouse Lab in Alderley Park | Wellcome Sanger Institute for the COVID-19 Genomics UK (COG-UK) Consortium | Jacquelyn Wynn, Mairead Hyland, The Lighthouse Lab in Alderley Park and Alex Alderton, Roberto Amato, Jeffrey Barrett, Sonia Goncalves, Ewan Harrison, David K. Jackson, Ian Johnston, Dominic Kwiatkowski, Cordelia Langford, John Sillitoe on behalf of the Wellcome Sanger Institute COVID-19 Surveillance Team |
| EPI_ISL_2468111 | Lighthouse Lab in Milton Keynes | Wellcome Sanger Institute for the COVID-19 Genomics UK (COG-UK) Consortium | The Lighthouse Lab in Milton Keynes and Alex Alderton, Roberto Amato, Jeffrey Barrett, Sonia Goncalves, Ewan Harrison, David K. Jackson, Ian Johnston, Dominic Kwiatkowski, Cordelia Langford, John Sillitoe on behalf of the Wellcome Sanger Institute COVID-19 Surveillance Team |
| EPI_ISL_2468112, EPI_ISL_2468113, EPI_ISL_2468114 | Lighthouse Lab in Alderley Park | Wellcome Sanger Institute for the COVID-19 Genomics UK (COG-UK) Consortium | Jacquelyn Wynn, Mairead Hyland, The Lighthouse Lab in Alderley Park and Alex Alderton, Roberto Amato, Jeffrey Barrett, Sonia Goncalves, Ewan Harrison, David K. Jackson, Ian Johnston, Dominic Kwiatkowski, Cordelia Langford, John Sillitoe on behalf of the Wellcome Sanger Institute COVID-19 Surveillance Team |
| EPI_ISL_2468115, EPI_ISL_2468116 | Health Services Laboratories | Wellcome Sanger Institute for the COVID-19 Genomics UK (COG-UK) Consortium | Health Services Laboratories and Alex Alderton, Roberto Amato, Jeffrey Barrett, Sonia Goncalves, Ewan Harrison, David K. Jackson, Ian Johnston, Dominic Kwiatkowski, Cordelia Langford, John Sillitoe on behalf of the Wellcome Sanger Institute COVID-19 Surveillance Team |
| EPI_ISL_2468117, EPI_ISL_2468118 | Lighthouse Lab in Alderley Park | Wellcome Sanger Institute for the COVID-19 Genomics UK (COG-UK) Consortium | Jacquelyn Wynn, Mairead Hyland, The Lighthouse Lab in Alderley Park and Alex Alderton, Roberto Amato, Jeffrey Barrett, Sonia Goncalves, Ewan Harrison, David K. Jackson, Ian Johnston, Dominic Kwiatkowski, Cordelia Langford, John Sillitoe on behalf of the Wellcome Sanger Institute COVID-19 Surveillance Team |
| EPI_ISL_2468119 | Health Services Laboratories | Wellcome Sanger Institute for the COVID-19 Genomics UK (COG-UK) Consortium | Health Services Laboratories and Alex Alderton, Roberto Amato, Jeffrey Barrett, Sonia Goncalves, Ewan Harrison, David K. Jackson, Ian Johnston, Dominic Kwiatkowski, Cordelia Langford, John Sillitoe on behalf of the Wellcome Sanger Institute COVID-19 Surveillance Team |
| EPI_ISL_2468120 | Lighthouse Lab in Alderley Park | Wellcome Sanger Institute for the COVID-19 Genomics UK (COG-UK) Consortium | Jacquelyn Wynn, Mairead Hyland, The Lighthouse Lab in Alderley Park and Alex Alderton, Roberto Amato, Jeffrey Barrett, Sonia Goncalves, Ewan Harrison, David K. Jackson, Ian Johnston, Dominic Kwiatkowski, Cordelia Langford, John Sillitoe on behalf of the Wellcome Sanger Institute COVID-19 Surveillance Team |
| EPI_ISL_2468121 | Lighthouse Lab in Milton Keynes | Wellcome Sanger Institute for the COVID-19 Genomics UK (COG-UK) Consortium | The Lighthouse Lab in Milton Keynes and Alex Alderton, Roberto Amato, Jeffrey Barrett, Sonia Goncalves, Ewan Harrison, David K. Jackson, Ian Johnston, Dominic Kwiatkowski, Cordelia Langford, John Sillitoe on behalf of the Wellcome Sanger Institute COVID-19 Surveillance Team |
| EPI_ISL_2468122, EPI_ISL_2468123 | Lighthouse Lab in Alderley Park | Wellcome Sanger Institute for the COVID-19 Genomics UK (COG-UK) Consortium | Jacquelyn Wynn, Mairead Hyland, The Lighthouse Lab in Alderley Park and Alex Alderton, Roberto Amato, Jeffrey Barrett, Sonia Goncalves, Ewan Harrison, David K. Jackson, Ian Johnston, Dominic Kwiatkowski, Cordelia Langford, John Sillitoe on behalf of the Wellcome Sanger Institute COVID-19 Surveillance Team |
| EPI_ISL_2468124 | Lighthouse Lab in Milton Keynes | Wellcome Sanger Institute for the COVID-19 Genomics UK (COG-UK) Consortium | The Lighthouse Lab in Milton Keynes and Alex Alderton, Roberto Amato, Jeffrey Barrett, Sonia Goncalves, Ewan Harrison, David K. Jackson, Ian Johnston, Dominic Kwiatkowski, Cordelia Langford, John Sillitoe on behalf of the Wellcome Sanger Institute COVID-19 Surveillance Team |
| EPI_ISL_2468125, EPI_ISL_2468126, EPI_ISL_2468127 | Lighthouse Lab in Alderley Park | Wellcome Sanger Institute for the COVID-19 Genomics UK (COG-UK) Consortium | Jacquelyn Wynn, Mairead Hyland, The Lighthouse Lab in Alderley Park and Alex Alderton, Roberto Amato, Jeffrey Barrett, Sonia Goncalves, Ewan Harrison, David K. Jackson, Ian Johnston, Dominic Kwiatkowski, Cordelia Langford, John Sillitoe on behalf of the Wellcome Sanger Institute COVID-19 Surveillance Team |
| EPI_ISL_2468128 | Lighthouse Lab in Milton Keynes | Wellcome Sanger Institute for the COVID-19 Genomics UK (COG-UK) Consortium | The Lighthouse Lab in Milton Keynes and Alex Alderton, Roberto Amato, Jeffrey Barrett, Sonia Goncalves, Ewan Harrison, David K. Jackson, Ian Johnston, Dominic Kwiatkowski, Cordelia Langford, John Sillitoe on behalf of the Wellcome Sanger Institute COVID-19 Surveillance Team |
| EPI_ISL_2468129 | Lighthouse Lab in Alderley Park | Wellcome Sanger Institute for the COVID-19 Genomics UK (COG-UK) Consortium | Jacquelyn Wynn, Mairead Hyland, The Lighthouse Lab in Alderley Park and Alex Alderton, Roberto Amato, Jeffrey Barrett, Sonia Goncalves, Ewan Harrison, David K. Jackson, Ian Johnston, Dominic Kwiatkowski, Cordelia Langford, John Sillitoe on behalf of the Wellcome Sanger Institute COVID-19 Surveillance Team |
| EPI_ISL_2468130 | Lighthouse Laboratory Plymouth | Wellcome Sanger Institute for the COVID-19 Genomics UK (COG-UK) Consortium | Lighthouse Laboratory Plymouth and Alex Alderton, Roberto Amato, Jeffrey Barrett, Sonia Goncalves, Ewan Harrison, David K. Jackson, Ian Johnston, Dominic Kwiatkowski, Cordelia Langford, John Sillitoe on behalf of the Wellcome Sanger Institute COVID-19 Surveillance Team |
| EPI_ISL_2468131, EPI_ISL_2468132 | Lighthouse Lab in Alderley Park | Wellcome Sanger Institute for the COVID-19 Genomics UK (COG-UK) Consortium | Jacquelyn Wynn, Mairead Hyland, The Lighthouse Lab in Alderley Park and Alex Alderton, Roberto Amato, Jeffrey Barrett, Sonia Goncalves, Ewan Harrison, David K. Jackson, Ian Johnston, Dominic Kwiatkowski, Cordelia Langford, John Sillitoe on behalf of the Wellcome Sanger Institute COVID-19 Surveillance Team |
| EPI_ISL_2468133 | Lighthouse Lab in Milton Keynes | Wellcome Sanger Institute for the COVID-19 Genomics UK (COG-UK) Consortium | The Lighthouse Lab in Milton Keynes and Alex Alderton, Roberto Amato, Jeffrey Barrett, Sonia Goncalves, Ewan Harrison, David K. Jackson, Ian Johnston, Dominic Kwiatkowski, Cordelia Langford, John Sillitoe on behalf of the Wellcome Sanger Institute COVID-19 Surveillance Team |
| EPI_ISL_2468134 | Lighthouse Laboratory Plymouth | Wellcome Sanger Institute for the COVID-19 Genomics UK (COG-UK) Consortium | Lighthouse Laboratory Plymouth and Alex Alderton, Roberto Amato, Jeffrey Barrett, Sonia Goncalves, Ewan Harrison, David K. Jackson, Ian Johnston, Dominic Kwiatkowski, Cordelia Langford, John Sillitoe on behalf of the Wellcome Sanger Institute COVID-19 Surveillance Team |
| EPI_ISL_2468135, EPI_ISL_2468137 | Lighthouse Lab in Alderley Park | Wellcome Sanger Institute for the COVID-19 Genomics UK (COG-UK) Consortium | Jacquelyn Wynn, Mairead Hyland, The Lighthouse Lab in Alderley Park and Alex Alderton, Roberto Amato, Jeffrey Barrett, Sonia Goncalves, Ewan Harrison, David K. Jackson, Ian Johnston, Dominic Kwiatkowski, Cordelia Langford, John Sillitoe on behalf of the Wellcome Sanger Institute COVID-19 Surveillance Team |
| EPI_ISL_2468138 | Lighthouse Lab in Milton Keynes | Wellcome Sanger Institute for the COVID-19 Genomics UK (COG-UK) Consortium | The Lighthouse Lab in Milton Keynes and Alex Alderton, Roberto Amato, Jeffrey Barrett, Sonia Goncalves, Ewan Harrison, David K. Jackson, Ian Johnston, Dominic Kwiatkowski, Cordelia Langford, John Sillitoe on behalf of the Wellcome Sanger Institute COVID-19 Surveillance Team |
| EPI_ISL_2468139, EPI_ISL_2468140, EPI_ISL_2468141, EPI_ISL_2468142 | Lighthouse Lab in Alderley Park | Wellcome Sanger Institute for the COVID-19 Genomics UK (COG-UK) Consortium | Jacquelyn Wynn, Mairead Hyland, The Lighthouse Lab in Alderley Park and Alex Alderton, Roberto Amato, Jeffrey Barrett, Sonia Goncalves, Ewan Harrison, David K. Jackson, Ian Johnston, Dominic Kwiatkowski, Cordelia Langford, John Sillitoe on behalf of the Wellcome Sanger Institute COVID-19 Surveillance Team |
| EPI_ISL_2468143 | Lighthouse Lab in Milton Keynes | Wellcome Sanger Institute for the COVID-19 Genomics UK (COG-UK) Consortium | The Lighthouse Lab in Milton Keynes and Alex Alderton, Roberto Amato, Jeffrey Barrett, Sonia Goncalves, Ewan Harrison, David K. Jackson, Ian Johnston, Dominic Kwiatkowski, Cordelia Langford, John Sillitoe on behalf of the Wellcome Sanger Institute COVID-19 Surveillance Team |
| EPI_ISL_2468144, EPI_ISL_2468145, EPI_ISL_2468146, EPI_ISL_2468147 | Lighthouse Lab in Alderley Park | Wellcome Sanger Institute for the COVID-19 Genomics UK (COG-UK) Consortium | Jacquelyn Wynn, Mairead Hyland, The Lighthouse Lab in Alderley Park and Alex Alderton, Roberto Amato, Jeffrey Barrett, Sonia Goncalves, Ewan Harrison, David K. Jackson, Ian Johnston, Dominic Kwiatkowski, Cordelia Langford, John Sillitoe on behalf of the Wellcome Sanger Institute COVID-19 Surveillance Team |
| EPI_ISL_2468149, EPI_ISL_2468150, EPI_ISL_2468151 | Lighthouse Lab in Milton Keynes | Wellcome Sanger Institute for the COVID-19 Genomics UK (COG-UK) Consortium | The Lighthouse Lab in Milton Keynes and Alex Alderton, Roberto Amato, Jeffrey Barrett, Sonia Goncalves, Ewan Harrison, David K. Jackson, Ian Johnston, Dominic Kwiatkowski, Cordelia Langford, John Sillitoe on behalf of the Wellcome Sanger Institute COVID-19 Surveillance Team |
| EPI_ISL_2468152, EPI_ISL_2468153 | Lighthouse Lab in Alderley Park | Wellcome Sanger Institute for the COVID-19 Genomics UK (COG-UK) Consortium | Jacquelyn Wynn, Mairead Hyland, The Lighthouse Lab in Alderley Park and Alex Alderton, Roberto Amato, Jeffrey Barrett, Sonia Goncalves, Ewan Harrison, David K. Jackson, Ian Johnston, Dominic Kwiatkowski, Cordelia Langford, John Sillitoe on behalf of the Wellcome Sanger Institute COVID-19 Surveillance Team |
| EPI_ISL_2468154 | Health Services Laboratories | Wellcome Sanger Institute for the COVID-19 Genomics UK (COG-UK) Consortium | Health Services Laboratories and Alex Alderton, Roberto Amato, Jeffrey Barrett, Sonia Goncalves, Ewan Harrison, David K. Jackson, Ian Johnston, Dominic Kwiatkowski, Cordelia Langford, John Sillitoe on behalf of the Wellcome Sanger Institute COVID-19 Surveillance Team |
| EPI_ISL_2468155, EPI_ISL_2468156, EPI_ISL_2468157 | Lighthouse Lab in Alderley Park | Wellcome Sanger Institute for the COVID-19 Genomics UK (COG-UK) Consortium | Jacquelyn Wynn, Mairead Hyland, The Lighthouse Lab in Alderley Park and Alex Alderton, Roberto Amato, Jeffrey Barrett, Sonia Goncalves, Ewan Harrison, David K. Jackson, Ian Johnston, Dominic Kwiatkowski, Cordelia Langford, John Sillitoe on behalf of the Wellcome Sanger Institute COVID-19 Surveillance Team |
| EPI_ISL_2468158 | Health Services Laboratories | Wellcome Sanger Institute for the COVID-19 Genomics UK (COG-UK) Consortium | Health Services Laboratories and Alex Alderton, Roberto Amato, Jeffrey Barrett, Sonia Goncalves, Ewan Harrison, David K. Jackson, Ian Johnston, Dominic Kwiatkowski, Cordelia Langford, John Sillitoe on behalf of the Wellcome Sanger Institute COVID-19 Surveillance Team |
| EPI_ISL_2468159, EPI_ISL_2468160 | Lighthouse Lab in Milton Keynes | Wellcome Sanger Institute for the COVID-19 Genomics UK (COG-UK) Consortium | The Lighthouse Lab in Milton Keynes and Alex Alderton, Roberto Amato, Jeffrey Barrett, Sonia Goncalves, Ewan Harrison, David K. Jackson, Ian Johnston, Dominic Kwiatkowski, Cordelia Langford, John Sillitoe on behalf of the Wellcome Sanger Institute COVID-19 Surveillance Team |
| EPI_ISL_2468161, EPI_ISL_2468162, EPI_ISL_2468163, EPI_ISL_2468164 | Lighthouse Lab in Alderley Park | Wellcome Sanger Institute for the COVID-19 Genomics UK (COG-UK) Consortium | Jacquelyn Wynn, Mairead Hyland, The Lighthouse Lab in Alderley Park and Alex Alderton, Roberto Amato, Jeffrey Barrett, Sonia Goncalves, Ewan Harrison, David K. Jackson, Ian Johnston, Dominic Kwiatkowski, Cordelia Langford, John Sillitoe on behalf of the Wellcome Sanger Institute COVID-19 Surveillance Team |
| EPI_ISL_2468165 | Lighthouse Laboratory Plymouth | Wellcome Sanger Institute for the COVID-19 Genomics UK (COG-UK) Consortium | Lighthouse Laboratory Plymouth and Alex Alderton, Roberto Amato, Jeffrey Barrett, Sonia Goncalves, Ewan Harrison, David K. Jackson, Ian Johnston, Dominic Kwiatkowski, Cordelia Langford, John Sillitoe on behalf of the Wellcome Sanger Institute COVID-19 Surveillance Team |
| EPI_ISL_2468166, EPI_ISL_2468167, EPI_ISL_2468168 | Lighthouse Lab in Alderley Park | Wellcome Sanger Institute for the COVID-19 Genomics UK (COG-UK) Consortium | Jacquelyn Wynn, Mairead Hyland, The Lighthouse Lab in Alderley Park and Alex Alderton, Roberto Amato, Jeffrey Barrett, Sonia Goncalves, Ewan Harrison, David K. Jackson, Ian Johnston, Dominic Kwiatkowski, Cordelia Langford, John Sillitoe on behalf of the Wellcome Sanger Institute COVID-19 Surveillance Team |
| EPI_ISL_2468169 | Lighthouse Lab in Milton Keynes | Wellcome Sanger Institute for the COVID-19 Genomics UK (COG-UK) | The Lighthouse Lab in Milton Keynes and Alex Alderton, Roberto Amato, Jeffrey Barrett, Sonia Goncalves, Ewan Harrison, David K. Jackson, Ian Johnston, Dominic Kwiatkowski, Cordelia Langford, John Sillitoe on behalf of the Wellcome Sanger |

| | | | |
|---|---|---|---|
| | | Consortium | Institute COVID-19 Surveillance Team |
| EPI_ISL_2468171 | Lighthouse Lab in Alderley Park | Wellcome Sanger Institute for the COVID-19 Genomics UK (COG-UK) Consortium | Jacquelyn Wynn, Mairead Hyland, The Lighthouse Lab in Alderley Park and Alex Alderton, Roberto Amato, Jeffrey Barrett, Sonia Goncalves, Ewan Harrison, David K. Jackson, Ian Johnston, Dominic Kwiatkowski, Cordelia Langford, John Sillitoe on behalf of the Wellcome Sanger Institute COVID-19 Surveillance Team |
| EPI_ISL_2468172, EPI_ISL_2468174, EPI_ISL_2468175 | Lighthouse Lab in Milton Keynes | Wellcome Sanger Institute for the COVID-19 Genomics UK (COG-UK) Consortium | The Lighthouse Lab in Milton Keynes and Alex Alderton, Roberto Amato, Jeffrey Barrett, Sonia Goncalves, Ewan Harrison, David K. Jackson, Ian Johnston, Dominic Kwiatkowski, Cordelia Langford, John Sillitoe on behalf of the Wellcome Sanger Institute COVID-19 Surveillance Team |
| EPI_ISL_2468176 | Health Services Laboratories | Wellcome Sanger Institute for the COVID-19 Genomics UK (COG-UK) Consortium | Health Services Laboratories and Alex Alderton, Roberto Amato, Jeffrey Barrett, Sonia Goncalves, Ewan Harrison, David K. Jackson, Ian Johnston, Dominic Kwiatkowski, Cordelia Langford, John Sillitoe on behalf of the Wellcome Sanger Institute COVID-19 Surveillance Team |
| EPI_ISL_2468177, EPI_ISL_2468178, EPI_ISL_2468180, EPI_ISL_2468181, EPI_ISL_2468182 | Lighthouse Lab in Alderley Park | Wellcome Sanger Institute for the COVID-19 Genomics UK (COG-UK) Consortium | Jacquelyn Wynn, Mairead Hyland, The Lighthouse Lab in Alderley Park and Alex Alderton, Roberto Amato, Jeffrey Barrett, Sonia Goncalves, Ewan Harrison, David K. Jackson, Ian Johnston, Dominic Kwiatkowski, Cordelia Langford, John Sillitoe on behalf of the Wellcome Sanger Institute COVID-19 Surveillance Team |
| EPI_ISL_2468183 | Lighthouse Lab in Milton Keynes | Wellcome Sanger Institute for the COVID-19 Genomics UK (COG-UK) Consortium | The Lighthouse Lab in Milton Keynes and Alex Alderton, Roberto Amato, Jeffrey Barrett, Sonia Goncalves, Ewan Harrison, David K. Jackson, Ian Johnston, Dominic Kwiatkowski, Cordelia Langford, John Sillitoe on behalf of the Wellcome Sanger Institute COVID-19 Surveillance Team |
| EPI_ISL_2468184, EPI_ISL_2468185, EPI_ISL_2468186 | Lighthouse Lab in Alderley Park | Wellcome Sanger Institute for the COVID-19 Genomics UK (COG-UK) Consortium | Jacquelyn Wynn, Mairead Hyland, The Lighthouse Lab in Alderley Park and Alex Alderton, Roberto Amato, Jeffrey Barrett, Sonia Goncalves, Ewan Harrison, David K. Jackson, Ian Johnston, Dominic Kwiatkowski, Cordelia Langford, John Sillitoe on behalf of the Wellcome Sanger Institute COVID-19 Surveillance Team |
| EPI_ISL_2468187 | Health Services Laboratories | Wellcome Sanger Institute for the COVID-19 Genomics UK (COG-UK) Consortium | Health Services Laboratories and Alex Alderton, Roberto Amato, Jeffrey Barrett, Sonia Goncalves, Ewan Harrison, David K. Jackson, Ian Johnston, Dominic Kwiatkowski, Cordelia Langford, John Sillitoe on behalf of the Wellcome Sanger Institute COVID-19 Surveillance Team |
| EPI_ISL_2468188 | Lighthouse Lab in Alderley Park | Wellcome Sanger Institute for the COVID-19 Genomics UK (COG-UK) Consortium | Jacquelyn Wynn, Mairead Hyland, The Lighthouse Lab in Alderley Park and Alex Alderton, Roberto Amato, Jeffrey Barrett, Sonia Goncalves, Ewan Harrison, David K. Jackson, Ian Johnston, Dominic Kwiatkowski, Cordelia Langford, John Sillitoe on behalf of the Wellcome Sanger Institute COVID-19 Surveillance Team |
| EPI_ISL_2468189 | Lighthouse Lab in Milton Keynes | Wellcome Sanger Institute for the COVID-19 Genomics UK (COG-UK) Consortium | The Lighthouse Lab in Milton Keynes and Alex Alderton, Roberto Amato, Jeffrey Barrett, Sonia Goncalves, Ewan Harrison, David K. Jackson, Ian Johnston, Dominic Kwiatkowski, Cordelia Langford, John Sillitoe on behalf of the Wellcome Sanger Institute COVID-19 Surveillance Team |
| EPI_ISL_2468190 | Lighthouse Laboratory Plymouth | Wellcome Sanger Institute for the COVID-19 Genomics UK (COG-UK) Consortium | Lighthouse Laboratory Plymouth and Alex Alderton, Roberto Amato, Jeffrey Barrett, Sonia Goncalves, Ewan Harrison, David K. Jackson, Ian Johnston, Dominic Kwiatkowski, Cordelia Langford, John Sillitoe on behalf of the Wellcome Sanger Institute COVID-19 Surveillance Team |
| EPI_ISL_2468191, EPI_ISL_2468193 | Lighthouse Lab in Alderley Park | Wellcome Sanger Institute for the COVID-19 Genomics UK (COG-UK) Consortium | Jacquelyn Wynn, Mairead Hyland, The Lighthouse Lab in Alderley Park and Alex Alderton, Roberto Amato, Jeffrey Barrett, Sonia Goncalves, Ewan Harrison, David K. Jackson, Ian Johnston, Dominic Kwiatkowski, Cordelia Langford, John Sillitoe on behalf of the Wellcome Sanger Institute COVID-19 Surveillance Team |
| EPI_ISL_2468194 | Health Services Laboratories | Wellcome Sanger Institute for the COVID-19 Genomics UK (COG-UK) Consortium | Health Services Laboratories and Alex Alderton, Roberto Amato, Jeffrey Barrett, Sonia Goncalves, Ewan Harrison, David K. Jackson, Ian Johnston, Dominic Kwiatkowski, Cordelia Langford, John Sillitoe on behalf of the Wellcome Sanger Institute COVID-19 Surveillance Team |
| EPI_ISL_2468195 | Lighthouse Lab in Milton Keynes | Wellcome Sanger Institute for the COVID-19 Genomics UK (COG-UK) Consortium | The Lighthouse Lab in Milton Keynes and Alex Alderton, Roberto Amato, Jeffrey Barrett, Sonia Goncalves, Ewan Harrison, David K. Jackson, Ian Johnston, Dominic Kwiatkowski, Cordelia Langford, John Sillitoe on behalf of the Wellcome Sanger Institute COVID-19 Surveillance Team |
| EPI_ISL_2468196, EPI_ISL_2468197, EPI_ISL_2468198 | Lighthouse Lab in Alderley Park | Wellcome Sanger Institute for the COVID-19 Genomics UK (COG-UK) Consortium | Jacquelyn Wynn, Mairead Hyland, The Lighthouse Lab in Alderley Park and Alex Alderton, Roberto Amato, Jeffrey Barrett, Sonia Goncalves, Ewan Harrison, David K. Jackson, Ian Johnston, Dominic Kwiatkowski, Cordelia Langford, John Sillitoe on behalf of the Wellcome Sanger Institute COVID-19 Surveillance Team |
| EPI_ISL_2468199 | Lighthouse Laboratory Plymouth | Wellcome Sanger Institute for the COVID-19 Genomics UK (COG-UK) Consortium | Lighthouse Laboratory Plymouth and Alex Alderton, Roberto Amato, Jeffrey Barrett, Sonia Goncalves, Ewan Harrison, David K. Jackson, Ian Johnston, Dominic Kwiatkowski, Cordelia Langford, John Sillitoe on behalf of the Wellcome Sanger Institute COVID-19 Surveillance Team |
| EPI_ISL_2468200 | Lighthouse Lab in Alderley Park | Wellcome Sanger Institute for the COVID-19 Genomics UK (COG-UK) Consortium | Jacquelyn Wynn, Mairead Hyland, The Lighthouse Lab in Alderley Park and Alex Alderton, Roberto Amato, Jeffrey Barrett, Sonia Goncalves, Ewan Harrison, David K. Jackson, Ian Johnston, Dominic Kwiatkowski, Cordelia Langford, John Sillitoe on behalf of the Wellcome Sanger Institute COVID-19 Surveillance Team |
| EPI_ISL_2468201, EPI_ISL_2468202 | Lighthouse Lab in Milton Keynes | Wellcome Sanger Institute for the COVID-19 Genomics UK (COG-UK) Consortium | The Lighthouse Lab in Milton Keynes and Alex Alderton, Roberto Amato, Jeffrey Barrett, Sonia Goncalves, Ewan Harrison, David K. Jackson, Ian Johnston, Dominic Kwiatkowski, Cordelia Langford, John Sillitoe on behalf of the Wellcome Sanger Institute COVID-19 Surveillance Team |
| EPI_ISL_2468203 | Health Services Laboratories | Wellcome Sanger Institute for the COVID-19 Genomics UK (COG-UK) Consortium | Health Services Laboratories and Alex Alderton, Roberto Amato, Jeffrey Barrett, Sonia Goncalves, Ewan Harrison, David K. Jackson, Ian Johnston, Dominic Kwiatkowski, Cordelia Langford, John Sillitoe on behalf of the Wellcome Sanger Institute COVID-19 Surveillance Team |
| EPI_ISL_2468204, EPI_ISL_2468205 | Lighthouse Lab in Alderley Park | Wellcome Sanger Institute for the COVID-19 Genomics UK (COG-UK) Consortium | Jacquelyn Wynn, Mairead Hyland, The Lighthouse Lab in Alderley Park and Alex Alderton, Roberto Amato, Jeffrey Barrett, Sonia Goncalves, Ewan Harrison, David K. Jackson, Ian Johnston, Dominic Kwiatkowski, Cordelia Langford, John Sillitoe on behalf of the Wellcome Sanger Institute COVID-19 Surveillance Team |
| EPI_ISL_2468206 | Health Services Laboratories | Wellcome Sanger Institute for the COVID-19 Genomics UK (COG-UK) Consortium | Health Services Laboratories and Alex Alderton, Roberto Amato, Jeffrey Barrett, Sonia Goncalves, Ewan Harrison, David K. Jackson, Ian Johnston, Dominic Kwiatkowski, Cordelia Langford, John Sillitoe on behalf of the Wellcome Sanger Institute COVID-19 Surveillance Team |
| EPI_ISL_2468208 | Lighthouse Lab in Milton Keynes | Wellcome Sanger Institute for the COVID-19 Genomics UK (COG-UK) Consortium | The Lighthouse Lab in Milton Keynes and Alex Alderton, Roberto Amato, Jeffrey Barrett, Sonia Goncalves, Ewan Harrison, David K. Jackson, Ian Johnston, Dominic Kwiatkowski, Cordelia Langford, John Sillitoe on behalf of the Wellcome Sanger Institute COVID-19 Surveillance Team |
| EPI_ISL_2468209 | Lighthouse Laboratory Plymouth | Wellcome Sanger Institute for the COVID-19 Genomics UK (COG-UK) Consortium | Lighthouse Laboratory Plymouth and Alex Alderton, Roberto Amato, Jeffrey Barrett, Sonia Goncalves, Ewan Harrison, David K. Jackson, Ian Johnston, Dominic Kwiatkowski, Cordelia Langford, John Sillitoe on behalf of the Wellcome Sanger Institute COVID-19 Surveillance Team |
| EPI_ISL_2468210 | Health Services Laboratories | Wellcome Sanger Institute for the COVID-19 Genomics UK (COG-UK) Consortium | Health Services Laboratories and Alex Alderton, Roberto Amato, Jeffrey Barrett, Sonia Goncalves, Ewan Harrison, David K. Jackson, Ian Johnston, Dominic Kwiatkowski, Cordelia Langford, John Sillitoe on behalf of the Wellcome Sanger Institute COVID-19 Surveillance Team |
| EPI_ISL_2468211 | Lighthouse Lab in Milton Keynes | Wellcome Sanger Institute for the COVID-19 Genomics UK (COG-UK) Consortium | The Lighthouse Lab in Milton Keynes and Alex Alderton, Roberto Amato, Jeffrey Barrett, Sonia Goncalves, Ewan Harrison, David K. Jackson, Ian Johnston, Dominic Kwiatkowski, Cordelia Langford, John Sillitoe on behalf of the Wellcome Sanger Institute COVID-19 Surveillance Team |
| EPI_ISL_2468212, EPI_ISL_2468213, EPI_ISL_2468214 | Lighthouse Lab in Alderley Park | Wellcome Sanger Institute for the COVID-19 Genomics UK (COG-UK) Consortium | Jacquelyn Wynn, Mairead Hyland, The Lighthouse Lab in Alderley Park and Alex Alderton, Roberto Amato, Jeffrey Barrett, Sonia Goncalves, Ewan Harrison, David K. Jackson, Ian Johnston, Dominic Kwiatkowski, Cordelia Langford, John Sillitoe on behalf of the Wellcome Sanger Institute COVID-19 Surveillance Team |
| EPI_ISL_2468215 | Lighthouse Lab in Milton Keynes | Wellcome Sanger Institute for the COVID-19 Genomics UK (COG-UK) Consortium | The Lighthouse Lab in Milton Keynes and Alex Alderton, Roberto Amato, Jeffrey Barrett, Sonia Goncalves, Ewan Harrison, David K. Jackson, Ian Johnston, Dominic Kwiatkowski, Cordelia Langford, John Sillitoe on behalf of the Wellcome Sanger Institute COVID-19 Surveillance Team |
| EPI_ISL_2468216 | Lighthouse Laboratory Plymouth | Wellcome Sanger Institute for the COVID-19 Genomics UK (COG-UK) Consortium | Lighthouse Laboratory Plymouth and Alex Alderton, Roberto Amato, Jeffrey Barrett, Sonia Goncalves, Ewan Harrison, David K. Jackson, Ian Johnston, Dominic Kwiatkowski, Cordelia Langford, John Sillitoe on behalf of the Wellcome Sanger Institute COVID-19 Surveillance Team |
| EPI_ISL_2468217, EPI_ISL_2468218, EPI_ISL_2468219, EPI_ISL_2468220 | Lighthouse Lab in Alderley Park | Wellcome Sanger Institute for the COVID-19 Genomics UK (COG-UK) Consortium | Jacquelyn Wynn, Mairead Hyland, The Lighthouse Lab in Alderley Park and Alex Alderton, Roberto Amato, Jeffrey Barrett, Sonia Goncalves, Ewan Harrison, David K. Jackson, Ian Johnston, Dominic Kwiatkowski, Cordelia Langford, John Sillitoe on behalf of the Wellcome Sanger Institute COVID-19 Surveillance Team |
| EPI_ISL_2468221 | Lighthouse Laboratory Plymouth | Wellcome Sanger Institute for the COVID-19 Genomics UK (COG-UK) Consortium | Lighthouse Laboratory Plymouth and Alex Alderton, Roberto Amato, Jeffrey Barrett, Sonia Goncalves, Ewan Harrison, David K. Jackson, Ian Johnston, Dominic Kwiatkowski, Cordelia Langford, John Sillitoe on behalf of the Wellcome Sanger Institute COVID-19 Surveillance Team |
| EPI_ISL_2468222, EPI_ISL_2468223 | Lighthouse Lab in Milton Keynes | Wellcome Sanger Institute for the COVID-19 Genomics UK (COG-UK) Consortium | The Lighthouse Lab in Milton Keynes and Alex Alderton, Roberto Amato, Jeffrey Barrett, Sonia Goncalves, Ewan Harrison, David K. Jackson, Ian Johnston, Dominic Kwiatkowski, Cordelia Langford, John Sillitoe on behalf of the Wellcome Sanger Institute COVID-19 Surveillance Team |
| EPI_ISL_2468224 | Lighthouse Lab in Alderley Park | Wellcome Sanger Institute for the COVID-19 Genomics UK (COG-UK) Consortium | Jacquelyn Wynn, Mairead Hyland, The Lighthouse Lab in Alderley Park and Alex Alderton, Roberto Amato, Jeffrey Barrett, Sonia Goncalves, Ewan Harrison, David K. Jackson, Ian Johnston, Dominic Kwiatkowski, Cordelia Langford, John Sillitoe on behalf of the Wellcome Sanger Institute COVID-19 Surveillance Team |
| EPI_ISL_2468225, EPI_ISL_2468226 | Lighthouse Lab in Milton Keynes | Wellcome Sanger Institute for the COVID-19 Genomics UK (COG-UK) Consortium | The Lighthouse Lab in Milton Keynes and Alex Alderton, Roberto Amato, Jeffrey Barrett, Sonia Goncalves, Ewan Harrison, David K. Jackson, Ian Johnston, Dominic Kwiatkowski, Cordelia Langford, John Sillitoe on behalf of the Wellcome Sanger Institute COVID-19 Surveillance Team |
| EPI_ISL_2468227, EPI_ISL_2468228 | Lighthouse Lab in Alderley Park | Wellcome Sanger Institute for the COVID-19 Genomics UK (COG-UK) Consortium | Jacquelyn Wynn, Mairead Hyland, The Lighthouse Lab in Alderley Park and Alex Alderton, Roberto Amato, Jeffrey Barrett, Sonia Goncalves, Ewan Harrison, David K. Jackson, Ian Johnston, Dominic Kwiatkowski, Cordelia Langford, John Sillitoe on behalf of the Wellcome Sanger Institute COVID-19 Surveillance Team |
| EPI_ISL_2468229, EPI_ISL_2468230 | Lighthouse Lab in Milton Keynes | Wellcome Sanger Institute for the COVID-19 Genomics UK (COG-UK) Consortium | The Lighthouse Lab in Milton Keynes and Alex Alderton, Roberto Amato, Jeffrey Barrett, Sonia Goncalves, Ewan Harrison, David K. Jackson, Ian Johnston, Dominic Kwiatkowski, Cordelia Langford, John Sillitoe on behalf of the Wellcome Sanger Institute COVID-19 Surveillance Team |
| EPI_ISL_2468231 | Lighthouse Lab in Alderley Park | Wellcome Sanger Institute for the COVID-19 Genomics UK (COG-UK) Consortium | Jacquelyn Wynn, Mairead Hyland, The Lighthouse Lab in Alderley Park and Alex Alderton, Roberto Amato, Jeffrey Barrett, Sonia Goncalves, Ewan Harrison, David K. Jackson, Ian Johnston, Dominic Kwiatkowski, Cordelia Langford, John Sillitoe on behalf of the Wellcome Sanger Institute COVID-19 Surveillance Team |
| EPI_ISL_2468232, EPI_ISL_2468233 | Lighthouse Lab in Milton Keynes | Wellcome Sanger Institute for the COVID-19 Genomics UK (COG-UK) Consortium | The Lighthouse Lab in Milton Keynes and Alex Alderton, Roberto Amato, Jeffrey Barrett, Sonia Goncalves, Ewan Harrison, David K. Jackson, Ian Johnston, Dominic Kwiatkowski, Cordelia Langford, John Sillitoe on behalf of the Wellcome Sanger Institute COVID-19 Surveillance Team |
| EPI_ISL_2468234 | Lighthouse Lab in Alderley Park | Wellcome Sanger Institute for the COVID-19 Genomics UK (COG-UK) Consortium | Jacquelyn Wynn, Mairead Hyland, The Lighthouse Lab in Alderley Park and Alex Alderton, Roberto Amato, Jeffrey Barrett, Sonia Goncalves, Ewan Harrison, David K. Jackson, Ian Johnston, Dominic Kwiatkowski, Cordelia Langford, John Sillitoe on behalf of the Wellcome Sanger Institute COVID-19 Surveillance Team |
| EPI_ISL_2468235, EPI_ISL_2468236 | Lighthouse Laboratory Plymouth | Wellcome Sanger Institute for the COVID-19 Genomics UK (COG-UK) Consortium | Lighthouse Laboratory Plymouth and Alex Alderton, Roberto Amato, Jeffrey Barrett, Sonia Goncalves, Ewan Harrison, David K. Jackson, Ian Johnston, Dominic Kwiatkowski, Cordelia Langford, John Sillitoe on behalf of the Wellcome Sanger Institute COVID-19 Surveillance Team |
| EPI_ISL_2468237 | Lighthouse Lab in Milton Keynes | Wellcome Sanger Institute for the COVID-19 Genomics UK (COG-UK) Consortium | The Lighthouse Lab in Milton Keynes and Alex Alderton, Roberto Amato, Jeffrey Barrett, Sonia Goncalves, Ewan Harrison, David K. Jackson, Ian Johnston, Dominic Kwiatkowski, Cordelia Langford, John Sillitoe on behalf of the Wellcome Sanger Institute COVID-19 Surveillance Team |
| EPI_ISL_2468238, EPI_ISL_2468239, EPI_ISL_2468240 | Lighthouse Lab in Alderley Park | Wellcome Sanger Institute for the COVID-19 Genomics UK (COG-UK) Consortium | Jacquelyn Wynn, Mairead Hyland, The Lighthouse Lab in Alderley Park and Alex Alderton, Roberto Amato, Jeffrey Barrett, Sonia Goncalves, Ewan Harrison, David K. Jackson, Ian Johnston, Dominic Kwiatkowski, Cordelia Langford, John Sillitoe on behalf of the Wellcome Sanger Institute COVID-19 Surveillance Team |
| EPI_ISL_2468241 | Health Services Laboratories | Wellcome Sanger Institute for the COVID-19 Genomics UK (COG-UK) | Health Services Laboratories and Alex Alderton, Roberto Amato, Jeffrey Barrett, Sonia Goncalves, Ewan Harrison, David K. Jackson, Ian Johnston, Dominic Kwiatkowski, Cordelia Langford, John Sillitoe on behalf of the Wellcome Sanger Institute |

| | | | |
|---|---|---|---|
| | | Consortium | COVID-19 Surveillance Team |
| EPI_ISL_2468242 | Lighthouse Lab in Alderley Park | Wellcome Sanger Institute for the COVID-19 Genomics UK (COG-UK) Consortium | Jacquelyn Wynn, Mairead Hyland, The Lighthouse Lab in Alderley Park and Alex Alderton, Roberto Amato, Jeffrey Barrett, Sonia Goncalves, Ewan Harrison, David K. Jackson, Ian Johnston, Dominic Kwiatkowski, Cordelia Langford, John Sillitoe on behalf of the Wellcome Sanger Institute COVID-19 Surveillance Team |
| EPI_ISL_2468243, EPI_ISL_2468244, EPI_ISL_2468245 | Lighthouse Lab in Milton Keynes | Wellcome Sanger Institute for the COVID-19 Genomics UK (COG-UK) Consortium | The Lighthouse Lab in Milton Keynes and Alex Alderton, Roberto Amato, Jeffrey Barrett, Sonia Goncalves, Ewan Harrison, David K. Jackson, Ian Johnston, Dominic Kwiatkowski, Cordelia Langford, John Sillitoe on behalf of the Wellcome Sanger Institute COVID-19 Surveillance Team |
| EPI_ISL_2468246, EPI_ISL_2468247, EPI_ISL_2468248 | Lighthouse Laboratory Plymouth | Wellcome Sanger Institute for the COVID-19 Genomics UK (COG-UK) Consortium | Lighthouse Laboratory Plymouth and Alex Alderton, Roberto Amato, Jeffrey Barrett, Sonia Goncalves, Ewan Harrison, David K. Jackson, Ian Johnston, Dominic Kwiatkowski, Cordelia Langford, John Sillitoe on behalf of the Wellcome Sanger Institute COVID-19 Surveillance Team |
| EPI_ISL_2468249 | Lighthouse Lab in Alderley Park | Wellcome Sanger Institute for the COVID-19 Genomics UK (COG-UK) Consortium | Jacquelyn Wynn, Mairead Hyland, The Lighthouse Lab in Alderley Park and Alex Alderton, Roberto Amato, Jeffrey Barrett, Sonia Goncalves, Ewan Harrison, David K. Jackson, Ian Johnston, Dominic Kwiatkowski, Cordelia Langford, John Sillitoe on behalf of the Wellcome Sanger Institute COVID-19 Surveillance Team |
| EPI_ISL_2468250 | Lighthouse Lab in Milton Keynes | Wellcome Sanger Institute for the COVID-19 Genomics UK (COG-UK) Consortium | The Lighthouse Lab in Milton Keynes and Alex Alderton, Roberto Amato, Jeffrey Barrett, Sonia Goncalves, Ewan Harrison, David K. Jackson, Ian Johnston, Dominic Kwiatkowski, Cordelia Langford, John Sillitoe on behalf of the Wellcome Sanger Institute COVID-19 Surveillance Team |
| EPI_ISL_2468252 | Lighthouse Lab in Alderley Park | Wellcome Sanger Institute for the COVID-19 Genomics UK (COG-UK) Consortium | Jacquelyn Wynn, Mairead Hyland, The Lighthouse Lab in Alderley Park and Alex Alderton, Roberto Amato, Jeffrey Barrett, Sonia Goncalves, Ewan Harrison, David K. Jackson, Ian Johnston, Dominic Kwiatkowski, Cordelia Langford, John Sillitoe on behalf of the Wellcome Sanger Institute COVID-19 Surveillance Team |
| EPI_ISL_2468254 | Lighthouse Laboratory Plymouth | Wellcome Sanger Institute for the COVID-19 Genomics UK (COG-UK) Consortium | Lighthouse Laboratory Plymouth and Alex Alderton, Roberto Amato, Jeffrey Barrett, Sonia Goncalves, Ewan Harrison, David K. Jackson, Ian Johnston, Dominic Kwiatkowski, Cordelia Langford, John Sillitoe on behalf of the Wellcome Sanger Institute COVID-19 Surveillance Team |
| EPI_ISL_2468255, EPI_ISL_2468256, EPI_ISL_2468257 | Lighthouse Lab in Alderley Park | Wellcome Sanger Institute for the COVID-19 Genomics UK (COG-UK) Consortium | Jacquelyn Wynn, Mairead Hyland, The Lighthouse Lab in Alderley Park and Alex Alderton, Roberto Amato, Jeffrey Barrett, Sonia Goncalves, Ewan Harrison, David K. Jackson, Ian Johnston, Dominic Kwiatkowski, Cordelia Langford, John Sillitoe on behalf of the Wellcome Sanger Institute COVID-19 Surveillance Team |
| EPI_ISL_2468258 | Lighthouse Lab in Milton Keynes | Wellcome Sanger Institute for the COVID-19 Genomics UK (COG-UK) Consortium | The Lighthouse Lab in Milton Keynes and Alex Alderton, Roberto Amato, Jeffrey Barrett, Sonia Goncalves, Ewan Harrison, David K. Jackson, Ian Johnston, Dominic Kwiatkowski, Cordelia Langford, John Sillitoe on behalf of the Wellcome Sanger Institute COVID-19 Surveillance Team |
| EPI_ISL_2468259 | Health Services Laboratories | Wellcome Sanger Institute for the COVID-19 Genomics UK (COG-UK) Consortium | Health Services Laboratories and Alex Alderton, Roberto Amato, Jeffrey Barrett, Sonia Goncalves, Ewan Harrison, David K. Jackson, Ian Johnston, Dominic Kwiatkowski, Cordelia Langford, John Sillitoe on behalf of the Wellcome Sanger Institute COVID-19 Surveillance Team |
| EPI_ISL_2468260 | Lighthouse Lab in Milton Keynes | Wellcome Sanger Institute for the COVID-19 Genomics UK (COG-UK) Consortium | The Lighthouse Lab in Milton Keynes and Alex Alderton, Roberto Amato, Jeffrey Barrett, Sonia Goncalves, Ewan Harrison, David K. Jackson, Ian Johnston, Dominic Kwiatkowski, Cordelia Langford, John Sillitoe on behalf of the Wellcome Sanger Institute COVID-19 Surveillance Team |
| EPI_ISL_2468261, EPI_ISL_2468262, EPI_ISL_2468263 | Lighthouse Lab in Alderley Park | Wellcome Sanger Institute for the COVID-19 Genomics UK (COG-UK) Consortium | Jacquelyn Wynn, Mairead Hyland, The Lighthouse Lab in Alderley Park and Alex Alderton, Roberto Amato, Jeffrey Barrett, Sonia Goncalves, Ewan Harrison, David K. Jackson, Ian Johnston, Dominic Kwiatkowski, Cordelia Langford, John Sillitoe on behalf of the Wellcome Sanger Institute COVID-19 Surveillance Team |
| EPI_ISL_2468264 | Health Services Laboratories | Wellcome Sanger Institute for the COVID-19 Genomics UK (COG-UK) Consortium | Health Services Laboratories and Alex Alderton, Roberto Amato, Jeffrey Barrett, Sonia Goncalves, Ewan Harrison, David K. Jackson, Ian Johnston, Dominic Kwiatkowski, Cordelia Langford, John Sillitoe on behalf of the Wellcome Sanger Institute COVID-19 Surveillance Team |
| EPI_ISL_2468265, EPI_ISL_2468266 | Lighthouse Lab in Milton Keynes | Wellcome Sanger Institute for the COVID-19 Genomics UK (COG-UK) Consortium | The Lighthouse Lab in Milton Keynes and Alex Alderton, Roberto Amato, Jeffrey Barrett, Sonia Goncalves, Ewan Harrison, David K. Jackson, Ian Johnston, Dominic Kwiatkowski, Cordelia Langford, John Sillitoe on behalf of the Wellcome Sanger Institute COVID-19 Surveillance Team |
| EPI_ISL_2468267, EPI_ISL_2468268, EPI_ISL_2468269 | Lighthouse Lab in Alderley Park | Wellcome Sanger Institute for the COVID-19 Genomics UK (COG-UK) Consortium | Jacquelyn Wynn, Mairead Hyland, The Lighthouse Lab in Alderley Park and Alex Alderton, Roberto Amato, Jeffrey Barrett, Sonia Goncalves, Ewan Harrison, David K. Jackson, Ian Johnston, Dominic Kwiatkowski, Cordelia Langford, John Sillitoe on behalf of the Wellcome Sanger Institute COVID-19 Surveillance Team |
| EPI_ISL_2468270 | Lighthouse Lab in Milton Keynes | Wellcome Sanger Institute for the COVID-19 Genomics UK (COG-UK) Consortium | The Lighthouse Lab in Milton Keynes and Alex Alderton, Roberto Amato, Jeffrey Barrett, Sonia Goncalves, Ewan Harrison, David K. Jackson, Ian Johnston, Dominic Kwiatkowski, Cordelia Langford, John Sillitoe on behalf of the Wellcome Sanger Institute COVID-19 Surveillance Team |
| EPI_ISL_2468271, EPI_ISL_2468272 | Lighthouse Lab in Alderley Park | Wellcome Sanger Institute for the COVID-19 Genomics UK (COG-UK) Consortium | Jacquelyn Wynn, Mairead Hyland, The Lighthouse Lab in Alderley Park and Alex Alderton, Roberto Amato, Jeffrey Barrett, Sonia Goncalves, Ewan Harrison, David K. Jackson, Ian Johnston, Dominic Kwiatkowski, Cordelia Langford, John Sillitoe on behalf of the Wellcome Sanger Institute COVID-19 Surveillance Team |
| EPI_ISL_2468273 | Lighthouse Lab in Milton Keynes | Wellcome Sanger Institute for the COVID-19 Genomics UK (COG-UK) Consortium | The Lighthouse Lab in Milton Keynes and Alex Alderton, Roberto Amato, Jeffrey Barrett, Sonia Goncalves, Ewan Harrison, David K. Jackson, Ian Johnston, Dominic Kwiatkowski, Cordelia Langford, John Sillitoe on behalf of the Wellcome Sanger Institute COVID-19 Surveillance Team |
| EPI_ISL_2468274 | Lighthouse Lab in Alderley Park | Wellcome Sanger Institute for the COVID-19 Genomics UK (COG-UK) Consortium | Jacquelyn Wynn, Mairead Hyland, The Lighthouse Lab in Alderley Park and Alex Alderton, Roberto Amato, Jeffrey Barrett, Sonia Goncalves, Ewan Harrison, David K. Jackson, Ian Johnston, Dominic Kwiatkowski, Cordelia Langford, John Sillitoe on behalf of the Wellcome Sanger Institute COVID-19 Surveillance Team |
| EPI_ISL_2468275 | Lighthouse Laboratory Plymouth | Wellcome Sanger Institute for the COVID-19 Genomics UK (COG-UK) Consortium | Lighthouse Laboratory Plymouth and Alex Alderton, Roberto Amato, Jeffrey Barrett, Sonia Goncalves, Ewan Harrison, David K. Jackson, Ian Johnston, Dominic Kwiatkowski, Cordelia Langford, John Sillitoe on behalf of the Wellcome Sanger Institute COVID-19 Surveillance Team |
| EPI_ISL_2468277 | Lighthouse Lab in Alderley Park | Wellcome Sanger Institute for the COVID-19 Genomics UK (COG-UK) Consortium | Jacquelyn Wynn, Mairead Hyland, The Lighthouse Lab in Alderley Park and Alex Alderton, Roberto Amato, Jeffrey Barrett, Sonia Goncalves, Ewan Harrison, David K. Jackson, Ian Johnston, Dominic Kwiatkowski, Cordelia Langford, John Sillitoe on behalf of the Wellcome Sanger Institute COVID-19 Surveillance Team |
| EPI_ISL_2468278 | Lighthouse Lab in Milton Keynes | Wellcome Sanger Institute for the COVID-19 Genomics UK (COG-UK) Consortium | The Lighthouse Lab in Milton Keynes and Alex Alderton, Roberto Amato, Jeffrey Barrett, Sonia Goncalves, Ewan Harrison, David K. Jackson, Ian Johnston, Dominic Kwiatkowski, Cordelia Langford, John Sillitoe on behalf of the Wellcome Sanger Institute COVID-19 Surveillance Team |
| EPI_ISL_2468279, EPI_ISL_2468280, EPI_ISL_2468281, EPI_ISL_2468283 | Lighthouse Lab in Alderley Park | Wellcome Sanger Institute for the COVID-19 Genomics UK (COG-UK) Consortium | Jacquelyn Wynn, Mairead Hyland, The Lighthouse Lab in Alderley Park and Alex Alderton, Roberto Amato, Jeffrey Barrett, Sonia Goncalves, Ewan Harrison, David K. Jackson, Ian Johnston, Dominic Kwiatkowski, Cordelia Langford, John Sillitoe on behalf of the Wellcome Sanger Institute COVID-19 Surveillance Team |
| EPI_ISL_2468284 | Lighthouse Lab in Milton Keynes | Wellcome Sanger Institute for the COVID-19 Genomics UK (COG-UK) Consortium | The Lighthouse Lab in Milton Keynes and Alex Alderton, Roberto Amato, Jeffrey Barrett, Sonia Goncalves, Ewan Harrison, David K. Jackson, Ian Johnston, Dominic Kwiatkowski, Cordelia Langford, John Sillitoe on behalf of the Wellcome Sanger Institute COVID-19 Surveillance Team |
| EPI_ISL_2468285 | Lighthouse Laboratory Plymouth | Wellcome Sanger Institute for the COVID-19 Genomics UK (COG-UK) Consortium | Lighthouse Laboratory Plymouth and Alex Alderton, Roberto Amato, Jeffrey Barrett, Sonia Goncalves, Ewan Harrison, David K. Jackson, Ian Johnston, Dominic Kwiatkowski, Cordelia Langford, John Sillitoe on behalf of the Wellcome Sanger Institute COVID-19 Surveillance Team |
| EPI_ISL_2468286 | Lighthouse Lab in Milton Keynes | Wellcome Sanger Institute for the COVID-19 Genomics UK (COG-UK) Consortium | The Lighthouse Lab in Milton Keynes and Alex Alderton, Roberto Amato, Jeffrey Barrett, Sonia Goncalves, Ewan Harrison, David K. Jackson, Ian Johnston, Dominic Kwiatkowski, Cordelia Langford, John Sillitoe on behalf of the Wellcome Sanger Institute COVID-19 Surveillance Team |
| EPI_ISL_2468287 | Lighthouse Lab in Alderley Park | Wellcome Sanger Institute for the COVID-19 Genomics UK (COG-UK) Consortium | Jacquelyn Wynn, Mairead Hyland, The Lighthouse Lab in Alderley Park and Alex Alderton, Roberto Amato, Jeffrey Barrett, Sonia Goncalves, Ewan Harrison, David K. Jackson, Ian Johnston, Dominic Kwiatkowski, Cordelia Langford, John Sillitoe on behalf of the Wellcome Sanger Institute COVID-19 Surveillance Team |
| EPI_ISL_2468288, EPI_ISL_2468289 | Lighthouse Laboratory Plymouth | Wellcome Sanger Institute for the COVID-19 Genomics UK (COG-UK) Consortium | Lighthouse Laboratory Plymouth and Alex Alderton, Roberto Amato, Jeffrey Barrett, Sonia Goncalves, Ewan Harrison, David K. Jackson, Ian Johnston, Dominic Kwiatkowski, Cordelia Langford, John Sillitoe on behalf of the Wellcome Sanger Institute COVID-19 Surveillance Team |
| EPI_ISL_2468291 | Lighthouse Lab in Alderley Park | Wellcome Sanger Institute for the COVID-19 Genomics UK (COG-UK) Consortium | Jacquelyn Wynn, Mairead Hyland, The Lighthouse Lab in Alderley Park and Alex Alderton, Roberto Amato, Jeffrey Barrett, Sonia Goncalves, Ewan Harrison, David K. Jackson, Ian Johnston, Dominic Kwiatkowski, Cordelia Langford, John Sillitoe on behalf of the Wellcome Sanger Institute COVID-19 Surveillance Team |
| EPI_ISL_2468292 | Lighthouse Laboratory Plymouth | Wellcome Sanger Institute for the COVID-19 Genomics UK (COG-UK) Consortium | Lighthouse Laboratory Plymouth and Alex Alderton, Roberto Amato, Jeffrey Barrett, Sonia Goncalves, Ewan Harrison, David K. Jackson, Ian Johnston, Dominic Kwiatkowski, Cordelia Langford, John Sillitoe on behalf of the Wellcome Sanger Institute COVID-19 Surveillance Team |
| EPI_ISL_2468293 | Lighthouse Lab in Milton Keynes | Wellcome Sanger Institute for the COVID-19 Genomics UK (COG-UK) Consortium | The Lighthouse Lab in Milton Keynes and Alex Alderton, Roberto Amato, Jeffrey Barrett, Sonia Goncalves, Ewan Harrison, David K. Jackson, Ian Johnston, Dominic Kwiatkowski, Cordelia Langford, John Sillitoe on behalf of the Wellcome Sanger Institute COVID-19 Surveillance Team |
| EPI_ISL_2468294 | Lighthouse Laboratory Plymouth | Wellcome Sanger Institute for the COVID-19 Genomics UK (COG-UK) Consortium | Lighthouse Laboratory Plymouth and Alex Alderton, Roberto Amato, Jeffrey Barrett, Sonia Goncalves, Ewan Harrison, David K. Jackson, Ian Johnston, Dominic Kwiatkowski, Cordelia Langford, John Sillitoe on behalf of the Wellcome Sanger Institute COVID-19 Surveillance Team |
| EPI_ISL_2468295 | Lighthouse Lab in Alderley Park | Wellcome Sanger Institute for the COVID-19 Genomics UK (COG-UK) Consortium | Jacquelyn Wynn, Mairead Hyland, The Lighthouse Lab in Alderley Park and Alex Alderton, Roberto Amato, Jeffrey Barrett, Sonia Goncalves, Ewan Harrison, David K. Jackson, Ian Johnston, Dominic Kwiatkowski, Cordelia Langford, John Sillitoe on behalf of the Wellcome Sanger Institute COVID-19 Surveillance Team |
| EPI_ISL_2468296, EPI_ISL_2468297 | Health Services Laboratories | Wellcome Sanger Institute for the COVID-19 Genomics UK (COG-UK) Consortium | Health Services Laboratories and Alex Alderton, Roberto Amato, Jeffrey Barrett, Sonia Goncalves, Ewan Harrison, David K. Jackson, Ian Johnston, Dominic Kwiatkowski, Cordelia Langford, John Sillitoe on behalf of the Wellcome Sanger Institute COVID-19 Surveillance Team |
| EPI_ISL_2468298 | Lighthouse Lab in Glasgow | Wellcome Sanger Institute for the COVID-19 Genomics UK (COG-UK) Consortium | Harper VanSteenhouse, Yumi Kasai, David Gray, Carol Clugston, Anna Dominiczak and Alex Alderton, Roberto Amato, Jeffrey Barrett, Sonia Goncalves, Ewan Harrison, David K. Jackson, Ian Johnston, Dominic Kwiatkowski, Cordelia Langford, John Sillitoe on behalf of the Wellcome Sanger Institute COVID-19 Surveillance Team |
| EPI_ISL_2468299 | Lighthouse Lab in Milton Keynes | Wellcome Sanger Institute for the COVID-19 Genomics UK (COG-UK) Consortium | The Lighthouse Lab in Milton Keynes and Alex Alderton, Roberto Amato, Jeffrey Barrett, Sonia Goncalves, Ewan Harrison, David K. Jackson, Ian Johnston, Dominic Kwiatkowski, Cordelia Langford, John Sillitoe on behalf of the Wellcome Sanger Institute COVID-19 Surveillance Team |
| EPI_ISL_2468300, EPI_ISL_2468301 | Lighthouse Lab in Alderley Park | Wellcome Sanger Institute for the COVID-19 Genomics UK (COG-UK) Consortium | Jacquelyn Wynn, Mairead Hyland, The Lighthouse Lab in Alderley Park and Alex Alderton, Roberto Amato, Jeffrey Barrett, Sonia Goncalves, Ewan Harrison, David K. Jackson, Ian Johnston, Dominic Kwiatkowski, Cordelia Langford, John Sillitoe on behalf of the Wellcome Sanger Institute COVID-19 Surveillance Team |
| EPI_ISL_2468302 | Health Services Laboratories | Wellcome Sanger Institute for the COVID-19 Genomics UK (COG-UK) Consortium | Health Services Laboratories and Alex Alderton, Roberto Amato, Jeffrey Barrett, Sonia Goncalves, Ewan Harrison, David K. Jackson, Ian Johnston, Dominic Kwiatkowski, Cordelia Langford, John Sillitoe on behalf of the Wellcome Sanger Institute COVID-19 Surveillance Team |
| EPI_ISL_2468303 | Lighthouse Lab in Alderley Park | Wellcome Sanger Institute for the COVID-19 Genomics UK (COG-UK) Consortium | Jacquelyn Wynn, Mairead Hyland, The Lighthouse Lab in Alderley Park and Alex Alderton, Roberto Amato, Jeffrey Barrett, Sonia Goncalves, Ewan Harrison, David K. Jackson, Ian Johnston, Dominic Kwiatkowski, Cordelia Langford, John Sillitoe on behalf of the Wellcome Sanger Institute COVID-19 Surveillance Team |
| EPI_ISL_2468304 | Lighthouse Laboratory Plymouth | Wellcome Sanger Institute for the COVID-19 Genomics UK (COG-UK) Consortium | Lighthouse Laboratory Plymouth and Alex Alderton, Roberto Amato, Jeffrey Barrett, Sonia Goncalves, Ewan Harrison, David K. Jackson, Ian Johnston, Dominic Kwiatkowski, Cordelia Langford, John Sillitoe on behalf of the Wellcome Sanger Institute COVID-19 Surveillance Team |
| EPI_ISL_2468305 | Lighthouse Lab in Milton Keynes | Wellcome Sanger Institute for the COVID-19 Genomics UK (COG-UK) Consortium | The Lighthouse Lab in Milton Keynes and Alex Alderton, Roberto Amato, Jeffrey Barrett, Sonia Goncalves, Ewan Harrison, David K. Jackson, Ian Johnston, Dominic Kwiatkowski, Cordelia Langford, John Sillitoe on behalf of the Wellcome Sanger Institute COVID-19 Surveillance Team |
| EPI_ISL_2468306 | Health Services Laboratories | Wellcome Sanger Institute for the COVID-19 Genomics UK (COG-UK) Consortium | Health Services Laboratories and Alex Alderton, Roberto Amato, Jeffrey Barrett, Sonia Goncalves, Ewan Harrison, David K. Jackson, Ian Johnston, Dominic Kwiatkowski, Cordelia Langford, John Sillitoe on behalf of the Wellcome Sanger Institute COVID-19 Surveillance Team |

| | | | |
|---|---|---|---|
| EPI_ISL_2468307 | Lighthouse Lab in Glasgow | Wellcome Sanger Institute for the COVID-19 Genomics UK (COG-UK) Consortium | Harper VanSteenhouse, Yumi Kasai, David Gray, Carol Clugston, Anna Dominiczak and Alex Alderton, Roberto Amato, Jeffrey Barrett, Sonia Goncalves, Ewan Harrison, David K. Jackson, Ian Johnston, Dominic Kwiatkowski, Cordelia Langford, John Sillitoe on behalf of the Wellcome Sanger Institute COVID-19 Surveillance Team |
| EPI_ISL_2468308 | Lighthouse Lab in Alderley Park | Wellcome Sanger Institute for the COVID-19 Genomics UK (COG-UK) Consortium | Jacquelyn Wynn, Mairead Hyland, The Lighthouse Lab in Alderley Park and Alex Alderton, Roberto Amato, Jeffrey Barrett, Sonia Goncalves, Ewan Harrison, David K. Jackson, Ian Johnston, Dominic Kwiatkowski, Cordelia Langford, John Sillitoe on behalf of the Wellcome Sanger Institute COVID-19 Surveillance Team |
| EPI_ISL_2468309 | Health Services Laboratories | Wellcome Sanger Institute for the COVID-19 Genomics UK (COG-UK) Consortium | Health Services Laboratories and Alex Alderton, Roberto Amato, Jeffrey Barrett, Sonia Goncalves, Ewan Harrison, David K. Jackson, Ian Johnston, Dominic Kwiatkowski, Cordelia Langford, John Sillitoe on behalf of the Wellcome Sanger Institute COVID-19 Surveillance Team |
| EPI_ISL_2468310 | Lighthouse Lab in Alderley Park | Wellcome Sanger Institute for the COVID-19 Genomics UK (COG-UK) Consortium | Jacquelyn Wynn, Mairead Hyland, The Lighthouse Lab in Alderley Park and Alex Alderton, Roberto Amato, Jeffrey Barrett, Sonia Goncalves, Ewan Harrison, David K. Jackson, Ian Johnston, Dominic Kwiatkowski, Cordelia Langford, John Sillitoe on behalf of the Wellcome Sanger Institute COVID-19 Surveillance Team |
| EPI_ISL_2468311 | Health Services Laboratories | Wellcome Sanger Institute for the COVID-19 Genomics UK (COG-UK) Consortium | Health Services Laboratories and Alex Alderton, Roberto Amato, Jeffrey Barrett, Sonia Goncalves, Ewan Harrison, David K. Jackson, Ian Johnston, Dominic Kwiatkowski, Cordelia Langford, John Sillitoe on behalf of the Wellcome Sanger Institute COVID-19 Surveillance Team |
| EPI_ISL_2468312 | Lighthouse Lab in Alderley Park | Wellcome Sanger Institute for the COVID-19 Genomics UK (COG-UK) Consortium | Jacquelyn Wynn, Mairead Hyland, The Lighthouse Lab in Alderley Park and Alex Alderton, Roberto Amato, Jeffrey Barrett, Sonia Goncalves, Ewan Harrison, David K. Jackson, Ian Johnston, Dominic Kwiatkowski, Cordelia Langford, John Sillitoe on behalf of the Wellcome Sanger Institute COVID-19 Surveillance Team |
| EPI_ISL_2468313 | Health Services Laboratories | Wellcome Sanger Institute for the COVID-19 Genomics UK (COG-UK) Consortium | Health Services Laboratories and Alex Alderton, Roberto Amato, Jeffrey Barrett, Sonia Goncalves, Ewan Harrison, David K. Jackson, Ian Johnston, Dominic Kwiatkowski, Cordelia Langford, John Sillitoe on behalf of the Wellcome Sanger Institute COVID-19 Surveillance Team |
| EPI_ISL_2468314 | Lighthouse Lab in Alderley Park | Wellcome Sanger Institute for the COVID-19 Genomics UK (COG-UK) Consortium | Jacquelyn Wynn, Mairead Hyland, The Lighthouse Lab in Alderley Park and Alex Alderton, Roberto Amato, Jeffrey Barrett, Sonia Goncalves, Ewan Harrison, David K. Jackson, Ian Johnston, Dominic Kwiatkowski, Cordelia Langford, John Sillitoe on behalf of the Wellcome Sanger Institute COVID-19 Surveillance Team |
| EPI_ISL_2468315 | Health Services Laboratories | Wellcome Sanger Institute for the COVID-19 Genomics UK (COG-UK) Consortium | Health Services Laboratories and Alex Alderton, Roberto Amato, Jeffrey Barrett, Sonia Goncalves, Ewan Harrison, David K. Jackson, Ian Johnston, Dominic Kwiatkowski, Cordelia Langford, John Sillitoe on behalf of the Wellcome Sanger Institute COVID-19 Surveillance Team |
| EPI_ISL_2468316 | Lighthouse Lab in Milton Keynes | Wellcome Sanger Institute for the COVID-19 Genomics UK (COG-UK) Consortium | The Lighthouse Lab in Milton Keynes and Alex Alderton, Roberto Amato, Jeffrey Barrett, Sonia Goncalves, Ewan Harrison, David K. Jackson, Ian Johnston, Dominic Kwiatkowski, Cordelia Langford, John Sillitoe on behalf of the Wellcome Sanger Institute COVID-19 Surveillance Team |
| EPI_ISL_2468317 | Health Services Laboratories | Wellcome Sanger Institute for the COVID-19 Genomics UK (COG-UK) Consortium | Health Services Laboratories and Alex Alderton, Roberto Amato, Jeffrey Barrett, Sonia Goncalves, Ewan Harrison, David K. Jackson, Ian Johnston, Dominic Kwiatkowski, Cordelia Langford, John Sillitoe on behalf of the Wellcome Sanger Institute COVID-19 Surveillance Team |
| EPI_ISL_2468318 | Lighthouse Lab in Alderley Park | Wellcome Sanger Institute for the COVID-19 Genomics UK (COG-UK) Consortium | Jacquelyn Wynn, Mairead Hyland, The Lighthouse Lab in Alderley Park and Alex Alderton, Roberto Amato, Jeffrey Barrett, Sonia Goncalves, Ewan Harrison, David K. Jackson, Ian Johnston, Dominic Kwiatkowski, Cordelia Langford, John Sillitoe on behalf of the Wellcome Sanger Institute COVID-19 Surveillance Team |
| EPI_ISL_2468319, EPI_ISL_2468320 | Health Services Laboratories | Wellcome Sanger Institute for the COVID-19 Genomics UK (COG-UK) Consortium | Health Services Laboratories and Alex Alderton, Roberto Amato, Jeffrey Barrett, Sonia Goncalves, Ewan Harrison, David K. Jackson, Ian Johnston, Dominic Kwiatkowski, Cordelia Langford, John Sillitoe on behalf of the Wellcome Sanger Institute COVID-19 Surveillance Team |
| EPI_ISL_2468321 | Lighthouse Lab in Milton Keynes | Wellcome Sanger Institute for the COVID-19 Genomics UK (COG-UK) Consortium | The Lighthouse Lab in Milton Keynes and Alex Alderton, Roberto Amato, Jeffrey Barrett, Sonia Goncalves, Ewan Harrison, David K. Jackson, Ian Johnston, Dominic Kwiatkowski, Cordelia Langford, John Sillitoe on behalf of the Wellcome Sanger Institute COVID-19 Surveillance Team |
| EPI_ISL_2468323 | Lighthouse Lab in Alderley Park | Wellcome Sanger Institute for the COVID-19 Genomics UK (COG-UK) Consortium | Jacquelyn Wynn, Mairead Hyland, The Lighthouse Lab in Alderley Park and Alex Alderton, Roberto Amato, Jeffrey Barrett, Sonia Goncalves, Ewan Harrison, David K. Jackson, Ian Johnston, Dominic Kwiatkowski, Cordelia Langford, John Sillitoe on behalf of the Wellcome Sanger Institute COVID-19 Surveillance Team |
| EPI_ISL_2468324 | Lighthouse Lab in Glasgow | Wellcome Sanger Institute for the COVID-19 Genomics UK (COG-UK) Consortium | Harper VanSteenhouse, Yumi Kasai, David Gray, Carol Clugston, Anna Dominiczak and Alex Alderton, Roberto Amato, Jeffrey Barrett, Sonia Goncalves, Ewan Harrison, David K. Jackson, Ian Johnston, Dominic Kwiatkowski, Cordelia Langford, John Sillitoe on behalf of the Wellcome Sanger Institute COVID-19 Surveillance Team |
| EPI_ISL_2468325, EPI_ISL_2468326, EPI_ISL_2468327, EPI_ISL_2468328 | Lighthouse Lab in Milton Keynes | Wellcome Sanger Institute for the COVID-19 Genomics UK (COG-UK) Consortium | The Lighthouse Lab in Milton Keynes and Alex Alderton, Roberto Amato, Jeffrey Barrett, Sonia Goncalves, Ewan Harrison, David K. Jackson, Ian Johnston, Dominic Kwiatkowski, Cordelia Langford, John Sillitoe on behalf of the Wellcome Sanger Institute COVID-19 Surveillance Team |
| EPI_ISL_2468329 | Lighthouse Lab in Alderley Park | Wellcome Sanger Institute for the COVID-19 Genomics UK (COG-UK) Consortium | Jacquelyn Wynn, Mairead Hyland, The Lighthouse Lab in Alderley Park and Alex Alderton, Roberto Amato, Jeffrey Barrett, Sonia Goncalves, Ewan Harrison, David K. Jackson, Ian Johnston, Dominic Kwiatkowski, Cordelia Langford, John Sillitoe on behalf of the Wellcome Sanger Institute COVID-19 Surveillance Team |
| EPI_ISL_2468330 | Lighthouse Lab in Glasgow | Wellcome Sanger Institute for the COVID-19 Genomics UK (COG-UK) Consortium | Harper VanSteenhouse, Yumi Kasai, David Gray, Carol Clugston, Anna Dominiczak and Alex Alderton, Roberto Amato, Jeffrey Barrett, Sonia Goncalves, Ewan Harrison, David K. Jackson, Ian Johnston, Dominic Kwiatkowski, Cordelia Langford, John Sillitoe on behalf of the Wellcome Sanger Institute COVID-19 Surveillance Team |
| EPI_ISL_2468331 | Health Services Laboratories | Wellcome Sanger Institute for the COVID-19 Genomics UK (COG-UK) Consortium | Health Services Laboratories and Alex Alderton, Roberto Amato, Jeffrey Barrett, Sonia Goncalves, Ewan Harrison, David K. Jackson, Ian Johnston, Dominic Kwiatkowski, Cordelia Langford, John Sillitoe on behalf of the Wellcome Sanger Institute COVID-19 Surveillance Team |
| EPI_ISL_2468332 | Lighthouse Lab in Alderley Park | Wellcome Sanger Institute for the COVID-19 Genomics UK (COG-UK) Consortium | Jacquelyn Wynn, Mairead Hyland, The Lighthouse Lab in Alderley Park and Alex Alderton, Roberto Amato, Jeffrey Barrett, Sonia Goncalves, Ewan Harrison, David K. Jackson, Ian Johnston, Dominic Kwiatkowski, Cordelia Langford, John Sillitoe on behalf of the Wellcome Sanger Institute COVID-19 Surveillance Team |
| EPI_ISL_2468333, EPI_ISL_2468334, EPI_ISL_2468335 | Lighthouse Lab in Milton Keynes | Wellcome Sanger Institute for the COVID-19 Genomics UK (COG-UK) Consortium | The Lighthouse Lab in Milton Keynes and Alex Alderton, Roberto Amato, Jeffrey Barrett, Sonia Goncalves, Ewan Harrison, David K. Jackson, Ian Johnston, Dominic Kwiatkowski, Cordelia Langford, John Sillitoe on behalf of the Wellcome Sanger Institute COVID-19 Surveillance Team |
| EPI_ISL_2468336 | Lighthouse Lab in Alderley Park | Wellcome Sanger Institute for the COVID-19 Genomics UK (COG-UK) Consortium | Jacquelyn Wynn, Mairead Hyland, The Lighthouse Lab in Alderley Park and Alex Alderton, Roberto Amato, Jeffrey Barrett, Sonia Goncalves, Ewan Harrison, David K. Jackson, Ian Johnston, Dominic Kwiatkowski, Cordelia Langford, John Sillitoe on behalf of the Wellcome Sanger Institute COVID-19 Surveillance Team |
| EPI_ISL_2468337 | Health Services Laboratories | Wellcome Sanger Institute for the COVID-19 Genomics UK (COG-UK) Consortium | Health Services Laboratories and Alex Alderton, Roberto Amato, Jeffrey Barrett, Sonia Goncalves, Ewan Harrison, David K. Jackson, Ian Johnston, Dominic Kwiatkowski, Cordelia Langford, John Sillitoe on behalf of the Wellcome Sanger Institute COVID-19 Surveillance Team |
| EPI_ISL_2468338 | Lighthouse Lab in Alderley Park | Wellcome Sanger Institute for the COVID-19 Genomics UK (COG-UK) Consortium | Jacquelyn Wynn, Mairead Hyland, The Lighthouse Lab in Alderley Park and Alex Alderton, Roberto Amato, Jeffrey Barrett, Sonia Goncalves, Ewan Harrison, David K. Jackson, Ian Johnston, Dominic Kwiatkowski, Cordelia Langford, John Sillitoe on behalf of the Wellcome Sanger Institute COVID-19 Surveillance Team |
| EPI_ISL_2468339 | Lighthouse Lab in Milton Keynes | Wellcome Sanger Institute for the COVID-19 Genomics UK (COG-UK) Consortium | The Lighthouse Lab in Milton Keynes and Alex Alderton, Roberto Amato, Jeffrey Barrett, Sonia Goncalves, Ewan Harrison, David K. Jackson, Ian Johnston, Dominic Kwiatkowski, Cordelia Langford, John Sillitoe on behalf of the Wellcome Sanger Institute COVID-19 Surveillance Team |
| EPI_ISL_2468340 | Lighthouse Lab in Alderley Park | Wellcome Sanger Institute for the COVID-19 Genomics UK (COG-UK) Consortium | Jacquelyn Wynn, Mairead Hyland, The Lighthouse Lab in Alderley Park and Alex Alderton, Roberto Amato, Jeffrey Barrett, Sonia Goncalves, Ewan Harrison, David K. Jackson, Ian Johnston, Dominic Kwiatkowski, Cordelia Langford, John Sillitoe on behalf of the Wellcome Sanger Institute COVID-19 Surveillance Team |
| EPI_ISL_2468341 | Lighthouse Lab in Milton Keynes | Wellcome Sanger Institute for the COVID-19 Genomics UK (COG-UK) Consortium | The Lighthouse Lab in Milton Keynes and Alex Alderton, Roberto Amato, Jeffrey Barrett, Sonia Goncalves, Ewan Harrison, David K. Jackson, Ian Johnston, Dominic Kwiatkowski, Cordelia Langford, John Sillitoe on behalf of the Wellcome Sanger Institute COVID-19 Surveillance Team |
| EPI_ISL_2468342 | Health Services Laboratories | Wellcome Sanger Institute for the COVID-19 Genomics UK (COG-UK) Consortium | Health Services Laboratories and Alex Alderton, Roberto Amato, Jeffrey Barrett, Sonia Goncalves, Ewan Harrison, David K. Jackson, Ian Johnston, Dominic Kwiatkowski, Cordelia Langford, John Sillitoe on behalf of the Wellcome Sanger Institute COVID-19 Surveillance Team |
| EPI_ISL_2468343 | Lighthouse Lab in Alderley Park | Wellcome Sanger Institute for the COVID-19 Genomics UK (COG-UK) Consortium | Jacquelyn Wynn, Mairead Hyland, The Lighthouse Lab in Alderley Park and Alex Alderton, Roberto Amato, Jeffrey Barrett, Sonia Goncalves, Ewan Harrison, David K. Jackson, Ian Johnston, Dominic Kwiatkowski, Cordelia Langford, John Sillitoe on behalf of the Wellcome Sanger Institute COVID-19 Surveillance Team |
| EPI_ISL_2468344 | Lighthouse Laboratory Plymouth | Wellcome Sanger Institute for the COVID-19 Genomics UK (COG-UK) Consortium | Lighthouse Laboratory Plymouth and Alex Alderton, Roberto Amato, Jeffrey Barrett, Sonia Goncalves, Ewan Harrison, David K. Jackson, Ian Johnston, Dominic Kwiatkowski, Cordelia Langford, John Sillitoe on behalf of the Wellcome Sanger Institute COVID-19 Surveillance Team |
| EPI_ISL_2468345, EPI_ISL_2468346 | Lighthouse Lab in Milton Keynes | Wellcome Sanger Institute for the COVID-19 Genomics UK (COG-UK) Consortium | The Lighthouse Lab in Milton Keynes and Alex Alderton, Roberto Amato, Jeffrey Barrett, Sonia Goncalves, Ewan Harrison, David K. Jackson, Ian Johnston, Dominic Kwiatkowski, Cordelia Langford, John Sillitoe on behalf of the Wellcome Sanger Institute COVID-19 Surveillance Team |
| EPI_ISL_2468347 | Lighthouse Lab in Alderley Park | Wellcome Sanger Institute for the COVID-19 Genomics UK (COG-UK) Consortium | Jacquelyn Wynn, Mairead Hyland, The Lighthouse Lab in Alderley Park and Alex Alderton, Roberto Amato, Jeffrey Barrett, Sonia Goncalves, Ewan Harrison, David K. Jackson, Ian Johnston, Dominic Kwiatkowski, Cordelia Langford, John Sillitoe on behalf of the Wellcome Sanger Institute COVID-19 Surveillance Team |
| EPI_ISL_2468348 | Health Services Laboratories | Wellcome Sanger Institute for the COVID-19 Genomics UK (COG-UK) Consortium | Health Services Laboratories and Alex Alderton, Roberto Amato, Jeffrey Barrett, Sonia Goncalves, Ewan Harrison, David K. Jackson, Ian Johnston, Dominic Kwiatkowski, Cordelia Langford, John Sillitoe on behalf of the Wellcome Sanger Institute COVID-19 Surveillance Team |
| EPI_ISL_2468349 | Lighthouse Laboratory Plymouth | Wellcome Sanger Institute for the COVID-19 Genomics UK (COG-UK) Consortium | Lighthouse Laboratory Plymouth and Alex Alderton, Roberto Amato, Jeffrey Barrett, Sonia Goncalves, Ewan Harrison, David K. Jackson, Ian Johnston, Dominic Kwiatkowski, Cordelia Langford, John Sillitoe on behalf of the Wellcome Sanger Institute COVID-19 Surveillance Team |
| EPI_ISL_2468350 | Health Services Laboratories | Wellcome Sanger Institute for the COVID-19 Genomics UK (COG-UK) Consortium | Health Services Laboratories and Alex Alderton, Roberto Amato, Jeffrey Barrett, Sonia Goncalves, Ewan Harrison, David K. Jackson, Ian Johnston, Dominic Kwiatkowski, Cordelia Langford, John Sillitoe on behalf of the Wellcome Sanger Institute COVID-19 Surveillance Team |
| EPI_ISL_2468351 | Lighthouse Lab in Alderley Park | Wellcome Sanger Institute for the COVID-19 Genomics UK (COG-UK) Consortium | Jacquelyn Wynn, Mairead Hyland, The Lighthouse Lab in Alderley Park and Alex Alderton, Roberto Amato, Jeffrey Barrett, Sonia Goncalves, Ewan Harrison, David K. Jackson, Ian Johnston, Dominic Kwiatkowski, Cordelia Langford, John Sillitoe on behalf of the Wellcome Sanger Institute COVID-19 Surveillance Team |
| EPI_ISL_2468352 | Lighthouse Laboratory Plymouth | Wellcome Sanger Institute for the COVID-19 Genomics UK (COG-UK) Consortium | Lighthouse Laboratory Plymouth and Alex Alderton, Roberto Amato, Jeffrey Barrett, Sonia Goncalves, Ewan Harrison, David K. Jackson, Ian Johnston, Dominic Kwiatkowski, Cordelia Langford, John Sillitoe on behalf of the Wellcome Sanger Institute COVID-19 Surveillance Team |
| EPI_ISL_2468353 | Lighthouse Lab in Milton Keynes | Wellcome Sanger Institute for the COVID-19 Genomics UK (COG-UK) Consortium | The Lighthouse Lab in Milton Keynes and Alex Alderton, Roberto Amato, Jeffrey Barrett, Sonia Goncalves, Ewan Harrison, David K. Jackson, Ian Johnston, Dominic Kwiatkowski, Cordelia Langford, John Sillitoe on behalf of the Wellcome Sanger Institute COVID-19 Surveillance Team |
| EPI_ISL_2468354 | Lighthouse Lab in Alderley Park | Wellcome Sanger Institute for the COVID-19 Genomics UK (COG-UK) Consortium | Jacquelyn Wynn, Mairead Hyland, The Lighthouse Lab in Alderley Park and Alex Alderton, Roberto Amato, Jeffrey Barrett, Sonia Goncalves, Ewan Harrison, David K. Jackson, Ian Johnston, Dominic Kwiatkowski, Cordelia Langford, John Sillitoe on behalf of the Wellcome Sanger Institute COVID-19 Surveillance Team |
| EPI_ISL_2468355, EPI_ISL_2468356 | Lighthouse Lab in Milton Keynes | Wellcome Sanger Institute for the COVID-19 Genomics UK (COG-UK) Consortium | The Lighthouse Lab in Milton Keynes and Alex Alderton, Roberto Amato, Jeffrey Barrett, Sonia Goncalves, Ewan Harrison, David K. Jackson, Ian Johnston, Dominic Kwiatkowski, Cordelia Langford, John Sillitoe on behalf of the Wellcome Sanger Institute COVID-19 Surveillance Team |
| EPI_ISL_2468357 | Lighthouse Laboratory Plymouth | Wellcome Sanger Institute for the COVID-19 Genomics UK (COG-UK) Consortium | Lighthouse Laboratory Plymouth and Alex Alderton, Roberto Amato, Jeffrey Barrett, Sonia Goncalves, Ewan Harrison, David K. Jackson, Ian Johnston, Dominic Kwiatkowski, Cordelia Langford, John Sillitoe on behalf of the Wellcome Sanger Institute COVID-19 Surveillance Team |
| EPI_ISL_2468358, EPI_ISL_2468359 | Lighthouse Lab in Alderley Park | Wellcome Sanger Institute for the COVID-19 Genomics UK (COG-UK) Consortium | Jacquelyn Wynn, Mairead Hyland, The Lighthouse Lab in Alderley Park and Alex Alderton, Roberto Amato, Jeffrey Barrett, Sonia Goncalves, Ewan Harrison, David K. Jackson, Ian Johnston, Dominic Kwiatkowski, Cordelia Langford, John Sillitoe on behalf of the Wellcome Sanger Institute COVID-19 Surveillance Team |
| EPI_ISL_2468360 | Lighthouse Lab in Milton Keynes | Wellcome Sanger Institute for the COVID-19 Genomics UK (COG-UK) | The Lighthouse Lab in Milton Keynes and Alex Alderton, Roberto Amato, Jeffrey Barrett, Sonia Goncalves, Ewan Harrison, David K. Jackson, Ian Johnston, Dominic Kwiatkowski, Cordelia Langford, John Sillitoe on behalf of the Wellcome Sanger |

| | | | |
|---|---|---|---|
| | | Consortium | Institute COVID-19 Surveillance Team |
| EPI_ISL_2468361 | Health Services Laboratories | Wellcome Sanger Institute for the COVID-19 Genomics UK (COG-UK) Consortium | Health Services Laboratories and Alex Alderton, Roberto Amato, Jeffrey Barrett, Sonia Goncalves, Ewan Harrison, David K. Jackson, Ian Johnston, Dominic Kwiatkowski, Cordelia Langford, John Sillitoe on behalf of the Wellcome Sanger Institute COVID-19 Surveillance Team |
| EPI_ISL_2468362 | Lighthouse Lab in Milton Keynes | Wellcome Sanger Institute for the COVID-19 Genomics UK (COG-UK) Consortium | The Lighthouse Lab in Milton Keynes and Alex Alderton, Roberto Amato, Jeffrey Barrett, Sonia Goncalves, Ewan Harrison, David K. Jackson, Ian Johnston, Dominic Kwiatkowski, Cordelia Langford, John Sillitoe on behalf of the Wellcome Sanger Institute COVID-19 Surveillance Team |
| EPI_ISL_2468363, EPI_ISL_2468364 | Lighthouse Lab in Alderley Park | Wellcome Sanger Institute for the COVID-19 Genomics UK (COG-UK) Consortium | Jacquelyn Wynn, Mairead Hyland, The Lighthouse Lab in Alderley Park and Alex Alderton, Roberto Amato, Jeffrey Barrett, Sonia Goncalves, Ewan Harrison, David K. Jackson, Ian Johnston, Dominic Kwiatkowski, Cordelia Langford, John Sillitoe on behalf of the Wellcome Sanger Institute COVID-19 Surveillance Team |
| EPI_ISL_2468365 | Health Services Laboratories | Wellcome Sanger Institute for the COVID-19 Genomics UK (COG-UK) Consortium | Health Services Laboratories and Alex Alderton, Roberto Amato, Jeffrey Barrett, Sonia Goncalves, Ewan Harrison, David K. Jackson, Ian Johnston, Dominic Kwiatkowski, Cordelia Langford, John Sillitoe on behalf of the Wellcome Sanger Institute COVID-19 Surveillance Team |
| EPI_ISL_2468366 | Lighthouse Lab in Milton Keynes | Wellcome Sanger Institute for the COVID-19 Genomics UK (COG-UK) Consortium | The Lighthouse Lab in Milton Keynes and Alex Alderton, Roberto Amato, Jeffrey Barrett, Sonia Goncalves, Ewan Harrison, David K. Jackson, Ian Johnston, Dominic Kwiatkowski, Cordelia Langford, John Sillitoe on behalf of the Wellcome Sanger Institute COVID-19 Surveillance Team |
| EPI_ISL_2468367 | Lighthouse Laboratory Plymouth | Wellcome Sanger Institute for the COVID-19 Genomics UK (COG-UK) Consortium | Lighthouse Laboratory Plymouth and Alex Alderton, Roberto Amato, Jeffrey Barrett, Sonia Goncalves, Ewan Harrison, David K. Jackson, Ian Johnston, Dominic Kwiatkowski, Cordelia Langford, John Sillitoe on behalf of the Wellcome Sanger Institute COVID-19 Surveillance Team |
| EPI_ISL_2468368 | Lighthouse Lab in Alderley Park | Wellcome Sanger Institute for the COVID-19 Genomics UK (COG-UK) Consortium | Jacquelyn Wynn, Mairead Hyland, The Lighthouse Lab in Alderley Park and Alex Alderton, Roberto Amato, Jeffrey Barrett, Sonia Goncalves, Ewan Harrison, David K. Jackson, Ian Johnston, Dominic Kwiatkowski, Cordelia Langford, John Sillitoe on behalf of the Wellcome Sanger Institute COVID-19 Surveillance Team |
| EPI_ISL_2468369 | Lighthouse Lab in Milton Keynes | Wellcome Sanger Institute for the COVID-19 Genomics UK (COG-UK) Consortium | The Lighthouse Lab in Milton Keynes and Alex Alderton, Roberto Amato, Jeffrey Barrett, Sonia Goncalves, Ewan Harrison, David K. Jackson, Ian Johnston, Dominic Kwiatkowski, Cordelia Langford, John Sillitoe on behalf of the Wellcome Sanger Institute COVID-19 Surveillance Team |
| EPI_ISL_2468370 | Lighthouse Laboratory Plymouth | Wellcome Sanger Institute for the COVID-19 Genomics UK (COG-UK) Consortium | Lighthouse Laboratory Plymouth and Alex Alderton, Roberto Amato, Jeffrey Barrett, Sonia Goncalves, Ewan Harrison, David K. Jackson, Ian Johnston, Dominic Kwiatkowski, Cordelia Langford, John Sillitoe on behalf of the Wellcome Sanger Institute COVID-19 Surveillance Team |
| EPI_ISL_2468371 | Lighthouse Lab in Milton Keynes | Wellcome Sanger Institute for the COVID-19 Genomics UK (COG-UK) Consortium | The Lighthouse Lab in Milton Keynes and Alex Alderton, Roberto Amato, Jeffrey Barrett, Sonia Goncalves, Ewan Harrison, David K. Jackson, Ian Johnston, Dominic Kwiatkowski, Cordelia Langford, John Sillitoe on behalf of the Wellcome Sanger Institute COVID-19 Surveillance Team |
| EPI_ISL_2468373 | Health Services Laboratories | Wellcome Sanger Institute for the COVID-19 Genomics UK (COG-UK) Consortium | Health Services Laboratories and Alex Alderton, Roberto Amato, Jeffrey Barrett, Sonia Goncalves, Ewan Harrison, David K. Jackson, Ian Johnston, Dominic Kwiatkowski, Cordelia Langford, John Sillitoe on behalf of the Wellcome Sanger Institute COVID-19 Surveillance Team |
| EPI_ISL_2468375 | Lighthouse Lab in Alderley Park | Wellcome Sanger Institute for the COVID-19 Genomics UK (COG-UK) Consortium | Jacquelyn Wynn, Mairead Hyland, The Lighthouse Lab in Alderley Park and Alex Alderton, Roberto Amato, Jeffrey Barrett, Sonia Goncalves, Ewan Harrison, David K. Jackson, Ian Johnston, Dominic Kwiatkowski, Cordelia Langford, John Sillitoe on behalf of the Wellcome Sanger Institute COVID-19 Surveillance Team |
| EPI_ISL_2468376 | Lighthouse Lab in Milton Keynes | Wellcome Sanger Institute for the COVID-19 Genomics UK (COG-UK) Consortium | The Lighthouse Lab in Milton Keynes and Alex Alderton, Roberto Amato, Jeffrey Barrett, Sonia Goncalves, Ewan Harrison, David K. Jackson, Ian Johnston, Dominic Kwiatkowski, Cordelia Langford, John Sillitoe on behalf of the Wellcome Sanger Institute COVID-19 Surveillance Team |
| EPI_ISL_2468377, EPI_ISL_2468378 | Lighthouse Lab in Alderley Park | Wellcome Sanger Institute for the COVID-19 Genomics UK (COG-UK) Consortium | Jacquelyn Wynn, Mairead Hyland, The Lighthouse Lab in Alderley Park and Alex Alderton, Roberto Amato, Jeffrey Barrett, Sonia Goncalves, Ewan Harrison, David K. Jackson, Ian Johnston, Dominic Kwiatkowski, Cordelia Langford, John Sillitoe on behalf of the Wellcome Sanger Institute COVID-19 Surveillance Team |
| EPI_ISL_2468379, EPI_ISL_2468380 | Lighthouse Lab in Milton Keynes | Wellcome Sanger Institute for the COVID-19 Genomics UK (COG-UK) Consortium | The Lighthouse Lab in Milton Keynes and Alex Alderton, Roberto Amato, Jeffrey Barrett, Sonia Goncalves, Ewan Harrison, David K. Jackson, Ian Johnston, Dominic Kwiatkowski, Cordelia Langford, John Sillitoe on behalf of the Wellcome Sanger Institute COVID-19 Surveillance Team |
| EPI_ISL_2468381, EPI_ISL_2468382 | Lighthouse Lab in Alderley Park | Wellcome Sanger Institute for the COVID-19 Genomics UK (COG-UK) Consortium | Jacquelyn Wynn, Mairead Hyland, The Lighthouse Lab in Alderley Park and Alex Alderton, Roberto Amato, Jeffrey Barrett, Sonia Goncalves, Ewan Harrison, David K. Jackson, Ian Johnston, Dominic Kwiatkowski, Cordelia Langford, John Sillitoe on behalf of the Wellcome Sanger Institute COVID-19 Surveillance Team |
| EPI_ISL_2468383 | Health Services Laboratories | Wellcome Sanger Institute for the COVID-19 Genomics UK (COG-UK) Consortium | Health Services Laboratories and Alex Alderton, Roberto Amato, Jeffrey Barrett, Sonia Goncalves, Ewan Harrison, David K. Jackson, Ian Johnston, Dominic Kwiatkowski, Cordelia Langford, John Sillitoe on behalf of the Wellcome Sanger Institute COVID-19 Surveillance Team |
| EPI_ISL_2468384 | Lighthouse Lab in Alderley Park | Wellcome Sanger Institute for the COVID-19 Genomics UK (COG-UK) Consortium | Jacquelyn Wynn, Mairead Hyland, The Lighthouse Lab in Alderley Park and Alex Alderton, Roberto Amato, Jeffrey Barrett, Sonia Goncalves, Ewan Harrison, David K. Jackson, Ian Johnston, Dominic Kwiatkowski, Cordelia Langford, John Sillitoe on behalf of the Wellcome Sanger Institute COVID-19 Surveillance Team |
| EPI_ISL_2468385, EPI_ISL_2468386 | Health Services Laboratories | Wellcome Sanger Institute for the COVID-19 Genomics UK (COG-UK) Consortium | Health Services Laboratories and Alex Alderton, Roberto Amato, Jeffrey Barrett, Sonia Goncalves, Ewan Harrison, David K. Jackson, Ian Johnston, Dominic Kwiatkowski, Cordelia Langford, John Sillitoe on behalf of the Wellcome Sanger Institute COVID-19 Surveillance Team |
| EPI_ISL_2468387 | Lighthouse Lab in Milton Keynes | Wellcome Sanger Institute for the COVID-19 Genomics UK (COG-UK) Consortium | The Lighthouse Lab in Milton Keynes and Alex Alderton, Roberto Amato, Jeffrey Barrett, Sonia Goncalves, Ewan Harrison, David K. Jackson, Ian Johnston, Dominic Kwiatkowski, Cordelia Langford, John Sillitoe on behalf of the Wellcome Sanger Institute COVID-19 Surveillance Team |
| EPI_ISL_2468388, EPI_ISL_2468389 | Lighthouse Lab in Alderley Park | Wellcome Sanger Institute for the COVID-19 Genomics UK (COG-UK) Consortium | Jacquelyn Wynn, Mairead Hyland, The Lighthouse Lab in Alderley Park and Alex Alderton, Roberto Amato, Jeffrey Barrett, Sonia Goncalves, Ewan Harrison, David K. Jackson, Ian Johnston, Dominic Kwiatkowski, Cordelia Langford, John Sillitoe on behalf of the Wellcome Sanger Institute COVID-19 Surveillance Team |
| EPI_ISL_2468390, EPI_ISL_2468391 | Lighthouse Lab in Milton Keynes | Wellcome Sanger Institute for the COVID-19 Genomics UK (COG-UK) Consortium | The Lighthouse Lab in Milton Keynes and Alex Alderton, Roberto Amato, Jeffrey Barrett, Sonia Goncalves, Ewan Harrison, David K. Jackson, Ian Johnston, Dominic Kwiatkowski, Cordelia Langford, John Sillitoe on behalf of the Wellcome Sanger Institute COVID-19 Surveillance Team |
| EPI_ISL_2468392 | Lighthouse Lab in Alderley Park | Wellcome Sanger Institute for the COVID-19 Genomics UK (COG-UK) Consortium | Jacquelyn Wynn, Mairead Hyland, The Lighthouse Lab in Alderley Park and Alex Alderton, Roberto Amato, Jeffrey Barrett, Sonia Goncalves, Ewan Harrison, David K. Jackson, Ian Johnston, Dominic Kwiatkowski, Cordelia Langford, John Sillitoe on behalf of the Wellcome Sanger Institute COVID-19 Surveillance Team |
| EPI_ISL_2468393 | Lighthouse Laboratory Plymouth | Wellcome Sanger Institute for the COVID-19 Genomics UK (COG-UK) Consortium | Lighthouse Laboratory Plymouth and Alex Alderton, Roberto Amato, Jeffrey Barrett, Sonia Goncalves, Ewan Harrison, David K. Jackson, Ian Johnston, Dominic Kwiatkowski, Cordelia Langford, John Sillitoe on behalf of the Wellcome Sanger Institute COVID-19 Surveillance Team |
| EPI_ISL_2468394 | Lighthouse Lab in Milton Keynes | Wellcome Sanger Institute for the COVID-19 Genomics UK (COG-UK) Consortium | The Lighthouse Lab in Milton Keynes and Alex Alderton, Roberto Amato, Jeffrey Barrett, Sonia Goncalves, Ewan Harrison, David K. Jackson, Ian Johnston, Dominic Kwiatkowski, Cordelia Langford, John Sillitoe on behalf of the Wellcome Sanger Institute COVID-19 Surveillance Team |
| EPI_ISL_2468395 | Lighthouse Laboratory Plymouth | Wellcome Sanger Institute for the COVID-19 Genomics UK (COG-UK) Consortium | Lighthouse Laboratory Plymouth and Alex Alderton, Roberto Amato, Jeffrey Barrett, Sonia Goncalves, Ewan Harrison, David K. Jackson, Ian Johnston, Dominic Kwiatkowski, Cordelia Langford, John Sillitoe on behalf of the Wellcome Sanger Institute COVID-19 Surveillance Team |
| EPI_ISL_2468396 | Lighthouse Lab in Alderley Park | Wellcome Sanger Institute for the COVID-19 Genomics UK (COG-UK) Consortium | Jacquelyn Wynn, Mairead Hyland, The Lighthouse Lab in Alderley Park and Alex Alderton, Roberto Amato, Jeffrey Barrett, Sonia Goncalves, Ewan Harrison, David K. Jackson, Ian Johnston, Dominic Kwiatkowski, Cordelia Langford, John Sillitoe on behalf of the Wellcome Sanger Institute COVID-19 Surveillance Team |
| EPI_ISL_2468397 | Lighthouse Laboratory Plymouth | Wellcome Sanger Institute for the COVID-19 Genomics UK (COG-UK) Consortium | Lighthouse Laboratory Plymouth and Alex Alderton, Roberto Amato, Jeffrey Barrett, Sonia Goncalves, Ewan Harrison, David K. Jackson, Ian Johnston, Dominic Kwiatkowski, Cordelia Langford, John Sillitoe on behalf of the Wellcome Sanger Institute COVID-19 Surveillance Team |
| EPI_ISL_2468398 | Lighthouse Lab in Alderley Park | Wellcome Sanger Institute for the COVID-19 Genomics UK (COG-UK) Consortium | Jacquelyn Wynn, Mairead Hyland, The Lighthouse Lab in Alderley Park and Alex Alderton, Roberto Amato, Jeffrey Barrett, Sonia Goncalves, Ewan Harrison, David K. Jackson, Ian Johnston, Dominic Kwiatkowski, Cordelia Langford, John Sillitoe on behalf of the Wellcome Sanger Institute COVID-19 Surveillance Team |
| EPI_ISL_2468399 | Health Services Laboratories | Wellcome Sanger Institute for the COVID-19 Genomics UK (COG-UK) Consortium | Health Services Laboratories and Alex Alderton, Roberto Amato, Jeffrey Barrett, Sonia Goncalves, Ewan Harrison, David K. Jackson, Ian Johnston, Dominic Kwiatkowski, Cordelia Langford, John Sillitoe on behalf of the Wellcome Sanger Institute COVID-19 Surveillance Team |
| EPI_ISL_2468400 | Lighthouse Lab in Glasgow | Wellcome Sanger Institute for the COVID-19 Genomics UK (COG-UK) Consortium | Harper VanSteenhouse, Yumi Kasai, David Gray, Carol Clugston, Anna Dominiczak and Alex Alderton, Roberto Amato, Jeffrey Barrett, Sonia Goncalves, Ewan Harrison, David K. Jackson, Ian Johnston, Dominic Kwiatkowski, Cordelia Langford, John Sillitoe on behalf of the Wellcome Sanger Institute COVID-19 Surveillance Team |
| EPI_ISL_2468401 | Health Services Laboratories | Wellcome Sanger Institute for the COVID-19 Genomics UK (COG-UK) Consortium | Health Services Laboratories and Alex Alderton, Roberto Amato, Jeffrey Barrett, Sonia Goncalves, Ewan Harrison, David K. Jackson, Ian Johnston, Dominic Kwiatkowski, Cordelia Langford, John Sillitoe on behalf of the Wellcome Sanger Institute COVID-19 Surveillance Team |
| EPI_ISL_2468402 | Lighthouse Lab in Alderley Park | Wellcome Sanger Institute for the COVID-19 Genomics UK (COG-UK) Consortium | Jacquelyn Wynn, Mairead Hyland, The Lighthouse Lab in Alderley Park and Alex Alderton, Roberto Amato, Jeffrey Barrett, Sonia Goncalves, Ewan Harrison, David K. Jackson, Ian Johnston, Dominic Kwiatkowski, Cordelia Langford, John Sillitoe on behalf of the Wellcome Sanger Institute COVID-19 Surveillance Team |
| EPI_ISL_2468403 | Lighthouse Lab in Milton Keynes | Wellcome Sanger Institute for the COVID-19 Genomics UK (COG-UK) Consortium | The Lighthouse Lab in Milton Keynes and Alex Alderton, Roberto Amato, Jeffrey Barrett, Sonia Goncalves, Ewan Harrison, David K. Jackson, Ian Johnston, Dominic Kwiatkowski, Cordelia Langford, John Sillitoe on behalf of the Wellcome Sanger Institute COVID-19 Surveillance Team |
| EPI_ISL_2468404, EPI_ISL_2468405 | Lighthouse Lab in Alderley Park | Wellcome Sanger Institute for the COVID-19 Genomics UK (COG-UK) Consortium | Jacquelyn Wynn, Mairead Hyland, The Lighthouse Lab in Alderley Park and Alex Alderton, Roberto Amato, Jeffrey Barrett, Sonia Goncalves, Ewan Harrison, David K. Jackson, Ian Johnston, Dominic Kwiatkowski, Cordelia Langford, John Sillitoe on behalf of the Wellcome Sanger Institute COVID-19 Surveillance Team |
| EPI_ISL_2468406 | Lighthouse Lab in Glasgow | Wellcome Sanger Institute for the COVID-19 Genomics UK (COG-UK) Consortium | Harper VanSteenhouse, Yumi Kasai, David Gray, Carol Clugston, Anna Dominiczak and Alex Alderton, Roberto Amato, Jeffrey Barrett, Sonia Goncalves, Ewan Harrison, David K. Jackson, Ian Johnston, Dominic Kwiatkowski, Cordelia Langford, John Sillitoe on behalf of the Wellcome Sanger Institute COVID-19 Surveillance Team |
| EPI_ISL_2468408 | Lighthouse Lab in Alderley Park | Wellcome Sanger Institute for the COVID-19 Genomics UK (COG-UK) Consortium | Jacquelyn Wynn, Mairead Hyland, The Lighthouse Lab in Alderley Park and Alex Alderton, Roberto Amato, Jeffrey Barrett, Sonia Goncalves, Ewan Harrison, David K. Jackson, Ian Johnston, Dominic Kwiatkowski, Cordelia Langford, John Sillitoe on behalf of the Wellcome Sanger Institute COVID-19 Surveillance Team |
| EPI_ISL_2468409, EPI_ISL_2468410, EPI_ISL_2468411, EPI_ISL_2468412 | Lighthouse Lab in Milton Keynes | Wellcome Sanger Institute for the COVID-19 Genomics UK (COG-UK) Consortium | The Lighthouse Lab in Milton Keynes and Alex Alderton, Roberto Amato, Jeffrey Barrett, Sonia Goncalves, Ewan Harrison, David K. Jackson, Ian Johnston, Dominic Kwiatkowski, Cordelia Langford, John Sillitoe on behalf of the Wellcome Sanger Institute COVID-19 Surveillance Team |
| EPI_ISL_2468413 | Lighthouse Laboratory Plymouth | Wellcome Sanger Institute for the COVID-19 Genomics UK (COG-UK) Consortium | Lighthouse Laboratory Plymouth and Alex Alderton, Roberto Amato, Jeffrey Barrett, Sonia Goncalves, Ewan Harrison, David K. Jackson, Ian Johnston, Dominic Kwiatkowski, Cordelia Langford, John Sillitoe on behalf of the Wellcome Sanger Institute COVID-19 Surveillance Team |
| EPI_ISL_2468414 | Lighthouse Lab in Milton Keynes | Wellcome Sanger Institute for the COVID-19 Genomics UK (COG-UK) Consortium | The Lighthouse Lab in Milton Keynes and Alex Alderton, Roberto Amato, Jeffrey Barrett, Sonia Goncalves, Ewan Harrison, David K. Jackson, Ian Johnston, Dominic Kwiatkowski, Cordelia Langford, John Sillitoe on behalf of the Wellcome Sanger Institute COVID-19 Surveillance Team |
| EPI_ISL_2468415 | Health Services Laboratories | Wellcome Sanger Institute for the COVID-19 Genomics UK (COG-UK) Consortium | Health Services Laboratories and Alex Alderton, Roberto Amato, Jeffrey Barrett, Sonia Goncalves, Ewan Harrison, David K. Jackson, Ian Johnston, Dominic Kwiatkowski, Cordelia Langford, John Sillitoe on behalf of the Wellcome Sanger Institute COVID-19 Surveillance Team |
| EPI_ISL_2468416 | Lighthouse Lab in Milton Keynes | Wellcome Sanger Institute for the COVID-19 Genomics UK (COG-UK) Consortium | The Lighthouse Lab in Milton Keynes and Alex Alderton, Roberto Amato, Jeffrey Barrett, Sonia Goncalves, Ewan Harrison, David K. Jackson, Ian Johnston, Dominic Kwiatkowski, Cordelia Langford, John Sillitoe on behalf of the Wellcome Sanger Institute COVID-19 Surveillance Team |
| EPI_ISL_2468417 | Lighthouse Lab in Alderley Park | Wellcome Sanger Institute for the COVID-19 Genomics UK (COG-UK) Consortium | Jacquelyn Wynn, Mairead Hyland, The Lighthouse Lab in Alderley Park and Alex Alderton, Roberto Amato, Jeffrey Barrett, Sonia Goncalves, Ewan Harrison, David K. Jackson, Ian Johnston, Dominic Kwiatkowski, Cordelia Langford, John Sillitoe on behalf of the Wellcome Sanger Institute COVID-19 Surveillance Team |
| EPI_ISL_2468418, | Lighthouse Lab in Milton Keynes | Wellcome Sanger Institute for the COVID-19 Genomics UK (COG-UK) | The Lighthouse Lab in Milton Keynes and Alex Alderton, Roberto Amato, Jeffrey Barrett, Sonia Goncalves, Ewan Harrison, David K. Jackson, Ian Johnston, Dominic Kwiatkowski, Cordelia Langford, John Sillitoe on behalf of the Wellcome Sanger |

| | | | |
|---|---|---|---|
| EPI_ISL_2468419, EPI_ISL_2468420 | | Consortium | Institute COVID-19 Surveillance Team |
| EPI_ISL_2468421 | Health Services Laboratories | Wellcome Sanger Institute for the COVID-19 Genomics UK (COG-UK) Consortium | Health Services Laboratories and Alex Alderton, Roberto Amato, Jeffrey Barrett, Sonia Goncalves, Ewan Harrison, David K. Jackson, Ian Johnston, Dominic Kwiatkowski, Cordelia Langford, John Sillitoe on behalf of the Wellcome Sanger Institute COVID-19 Surveillance Team |
| EPI_ISL_2468422 | Lighthouse Lab in Alderley Park | Wellcome Sanger Institute for the COVID-19 Genomics UK (COG-UK) Consortium | Jacquelyn Wynn, Mairead Hyland, The Lighthouse Lab in Alderley Park and Alex Alderton, Roberto Amato, Jeffrey Barrett, Sonia Goncalves, Ewan Harrison, David K. Jackson, Ian Johnston, Dominic Kwiatkowski, Cordelia Langford, John Sillitoe on behalf of the Wellcome Sanger Institute COVID-19 Surveillance Team |
| EPI_ISL_2468423 | Health Services Laboratories | Wellcome Sanger Institute for the COVID-19 Genomics UK (COG-UK) Consortium | Health Services Laboratories and Alex Alderton, Roberto Amato, Jeffrey Barrett, Sonia Goncalves, Ewan Harrison, David K. Jackson, Ian Johnston, Dominic Kwiatkowski, Cordelia Langford, John Sillitoe on behalf of the Wellcome Sanger Institute COVID-19 Surveillance Team |
| EPI_ISL_2468424 | Lighthouse Lab in Milton Keynes | Wellcome Sanger Institute for the COVID-19 Genomics UK (COG-UK) Consortium | The Lighthouse Lab in Milton Keynes and Alex Alderton, Roberto Amato, Jeffrey Barrett, Sonia Goncalves, Ewan Harrison, David K. Jackson, Ian Johnston, Dominic Kwiatkowski, Cordelia Langford, John Sillitoe on behalf of the Wellcome Sanger Institute COVID-19 Surveillance Team |
| EPI_ISL_2468425 | Lighthouse Lab in Glasgow | Wellcome Sanger Institute for the COVID-19 Genomics UK (COG-UK) Consortium | Harper VanSteenhouse, Yumi Kasai, David Gray, Carol Clugston, Anna Dominiczak and Alex Alderton, Roberto Amato, Jeffrey Barrett, Sonia Goncalves, Ewan Harrison, David K. Jackson, Ian Johnston, Dominic Kwiatkowski, Cordelia Langford, John Sillitoe on behalf of the Wellcome Sanger Institute COVID-19 Surveillance Team |
| EPI_ISL_2468426 | Health Services Laboratories | Wellcome Sanger Institute for the COVID-19 Genomics UK (COG-UK) Consortium | Health Services Laboratories and Alex Alderton, Roberto Amato, Jeffrey Barrett, Sonia Goncalves, Ewan Harrison, David K. Jackson, Ian Johnston, Dominic Kwiatkowski, Cordelia Langford, John Sillitoe on behalf of the Wellcome Sanger Institute COVID-19 Surveillance Team |
| EPI_ISL_2468428, EPI_ISL_2468429 | Lighthouse Lab in Milton Keynes | Wellcome Sanger Institute for the COVID-19 Genomics UK (COG-UK) Consortium | The Lighthouse Lab in Milton Keynes and Alex Alderton, Roberto Amato, Jeffrey Barrett, Sonia Goncalves, Ewan Harrison, David K. Jackson, Ian Johnston, Dominic Kwiatkowski, Cordelia Langford, John Sillitoe on behalf of the Wellcome Sanger Institute COVID-19 Surveillance Team |
| EPI_ISL_2468430, EPI_ISL_2468431 | Health Services Laboratories | Wellcome Sanger Institute for the COVID-19 Genomics UK (COG-UK) Consortium | Health Services Laboratories and Alex Alderton, Roberto Amato, Jeffrey Barrett, Sonia Goncalves, Ewan Harrison, David K. Jackson, Ian Johnston, Dominic Kwiatkowski, Cordelia Langford, John Sillitoe on behalf of the Wellcome Sanger Institute COVID-19 Surveillance Team |
| EPI_ISL_2468433 | Lighthouse Lab in Milton Keynes | Wellcome Sanger Institute for the COVID-19 Genomics UK (COG-UK) Consortium | The Lighthouse Lab in Milton Keynes and Alex Alderton, Roberto Amato, Jeffrey Barrett, Sonia Goncalves, Ewan Harrison, David K. Jackson, Ian Johnston, Dominic Kwiatkowski, Cordelia Langford, John Sillitoe on behalf of the Wellcome Sanger Institute COVID-19 Surveillance Team |
| EPI_ISL_2468434 | Health Services Laboratories | Wellcome Sanger Institute for the COVID-19 Genomics UK (COG-UK) Consortium | Health Services Laboratories and Alex Alderton, Roberto Amato, Jeffrey Barrett, Sonia Goncalves, Ewan Harrison, David K. Jackson, Ian Johnston, Dominic Kwiatkowski, Cordelia Langford, John Sillitoe on behalf of the Wellcome Sanger Institute COVID-19 Surveillance Team |
| EPI_ISL_2468435, EPI_ISL_2468436, EPI_ISL_2468437 | Lighthouse Lab in Milton Keynes | Wellcome Sanger Institute for the COVID-19 Genomics UK (COG-UK) Consortium | The Lighthouse Lab in Milton Keynes and Alex Alderton, Roberto Amato, Jeffrey Barrett, Sonia Goncalves, Ewan Harrison, David K. Jackson, Ian Johnston, Dominic Kwiatkowski, Cordelia Langford, John Sillitoe on behalf of the Wellcome Sanger Institute COVID-19 Surveillance Team |
| EPI_ISL_2468438, EPI_ISL_2468439 | Health Services Laboratories | Wellcome Sanger Institute for the COVID-19 Genomics UK (COG-UK) Consortium | Health Services Laboratories and Alex Alderton, Roberto Amato, Jeffrey Barrett, Sonia Goncalves, Ewan Harrison, David K. Jackson, Ian Johnston, Dominic Kwiatkowski, Cordelia Langford, John Sillitoe on behalf of the Wellcome Sanger Institute COVID-19 Surveillance Team |
| EPI_ISL_2468440 | Lighthouse Lab in Alderley Park | Wellcome Sanger Institute for the COVID-19 Genomics UK (COG-UK) Consortium | Jacquelyn Wynn, Mairead Hyland, The Lighthouse Lab in Alderley Park and Alex Alderton, Roberto Amato, Jeffrey Barrett, Sonia Goncalves, Ewan Harrison, David K. Jackson, Ian Johnston, Dominic Kwiatkowski, Cordelia Langford, John Sillitoe on behalf of the Wellcome Sanger Institute COVID-19 Surveillance Team |
| EPI_ISL_2468441, EPI_ISL_2468442 | Lighthouse Lab in Milton Keynes | Wellcome Sanger Institute for the COVID-19 Genomics UK (COG-UK) Consortium | The Lighthouse Lab in Milton Keynes and Alex Alderton, Roberto Amato, Jeffrey Barrett, Sonia Goncalves, Ewan Harrison, David K. Jackson, Ian Johnston, Dominic Kwiatkowski, Cordelia Langford, John Sillitoe on behalf of the Wellcome Sanger Institute COVID-19 Surveillance Team |
| EPI_ISL_2468444 | Lighthouse Lab in Glasgow | Wellcome Sanger Institute for the COVID-19 Genomics UK (COG-UK) Consortium | Harper VanSteenhouse, Yumi Kasai, David Gray, Carol Clugston, Anna Dominiczak and Alex Alderton, Roberto Amato, Jeffrey Barrett, Sonia Goncalves, Ewan Harrison, David K. Jackson, Ian Johnston, Dominic Kwiatkowski, Cordelia Langford, John Sillitoe on behalf of the Wellcome Sanger Institute COVID-19 Surveillance Team |
| EPI_ISL_2468445 | Health Services Laboratories | Wellcome Sanger Institute for the COVID-19 Genomics UK (COG-UK) Consortium | Health Services Laboratories and Alex Alderton, Roberto Amato, Jeffrey Barrett, Sonia Goncalves, Ewan Harrison, David K. Jackson, Ian Johnston, Dominic Kwiatkowski, Cordelia Langford, John Sillitoe on behalf of the Wellcome Sanger Institute COVID-19 Surveillance Team |
| EPI_ISL_2468446 | Lighthouse Laboratory Plymouth | Wellcome Sanger Institute for the COVID-19 Genomics UK (COG-UK) Consortium | Lighthouse Laboratory Plymouth and Alex Alderton, Roberto Amato, Jeffrey Barrett, Sonia Goncalves, Ewan Harrison, David K. Jackson, Ian Johnston, Dominic Kwiatkowski, Cordelia Langford, John Sillitoe on behalf of the Wellcome Sanger Institute COVID-19 Surveillance Team |
| EPI_ISL_2468447 | Lighthouse Lab in Alderley Park | Wellcome Sanger Institute for the COVID-19 Genomics UK (COG-UK) Consortium | Jacquelyn Wynn, Mairead Hyland, The Lighthouse Lab in Alderley Park and Alex Alderton, Roberto Amato, Jeffrey Barrett, Sonia Goncalves, Ewan Harrison, David K. Jackson, Ian Johnston, Dominic Kwiatkowski, Cordelia Langford, John Sillitoe on behalf of the Wellcome Sanger Institute COVID-19 Surveillance Team |
| EPI_ISL_2468448 | Lighthouse Lab in Milton Keynes | Wellcome Sanger Institute for the COVID-19 Genomics UK (COG-UK) Consortium | The Lighthouse Lab in Milton Keynes and Alex Alderton, Roberto Amato, Jeffrey Barrett, Sonia Goncalves, Ewan Harrison, David K. Jackson, Ian Johnston, Dominic Kwiatkowski, Cordelia Langford, John Sillitoe on behalf of the Wellcome Sanger Institute COVID-19 Surveillance Team |
| EPI_ISL_2468449 | Lighthouse Lab in Glasgow | Wellcome Sanger Institute for the COVID-19 Genomics UK (COG-UK) Consortium | Harper VanSteenhouse, Yumi Kasai, David Gray, Carol Clugston, Anna Dominiczak and Alex Alderton, Roberto Amato, Jeffrey Barrett, Sonia Goncalves, Ewan Harrison, David K. Jackson, Ian Johnston, Dominic Kwiatkowski, Cordelia Langford, John Sillitoe on behalf of the Wellcome Sanger Institute COVID-19 Surveillance Team |
| EPI_ISL_2468450 | Health Services Laboratories | Wellcome Sanger Institute for the COVID-19 Genomics UK (COG-UK) Consortium | Health Services Laboratories and Alex Alderton, Roberto Amato, Jeffrey Barrett, Sonia Goncalves, Ewan Harrison, David K. Jackson, Ian Johnston, Dominic Kwiatkowski, Cordelia Langford, John Sillitoe on behalf of the Wellcome Sanger Institute COVID-19 Surveillance Team |
| EPI_ISL_2468451 | Lighthouse Lab in Alderley Park | Wellcome Sanger Institute for the COVID-19 Genomics UK (COG-UK) Consortium | Jacquelyn Wynn, Mairead Hyland, The Lighthouse Lab in Alderley Park and Alex Alderton, Roberto Amato, Jeffrey Barrett, Sonia Goncalves, Ewan Harrison, David K. Jackson, Ian Johnston, Dominic Kwiatkowski, Cordelia Langford, John Sillitoe on behalf of the Wellcome Sanger Institute COVID-19 Surveillance Team |
| EPI_ISL_2468452 | Lighthouse Lab in Milton Keynes | Wellcome Sanger Institute for the COVID-19 Genomics UK (COG-UK) Consortium | The Lighthouse Lab in Milton Keynes and Alex Alderton, Roberto Amato, Jeffrey Barrett, Sonia Goncalves, Ewan Harrison, David K. Jackson, Ian Johnston, Dominic Kwiatkowski, Cordelia Langford, John Sillitoe on behalf of the Wellcome Sanger Institute COVID-19 Surveillance Team |
| EPI_ISL_2468453 | Lighthouse Lab in Alderley Park | Wellcome Sanger Institute for the COVID-19 Genomics UK (COG-UK) Consortium | Jacquelyn Wynn, Mairead Hyland, The Lighthouse Lab in Alderley Park and Alex Alderton, Roberto Amato, Jeffrey Barrett, Sonia Goncalves, Ewan Harrison, David K. Jackson, Ian Johnston, Dominic Kwiatkowski, Cordelia Langford, John Sillitoe on behalf of the Wellcome Sanger Institute COVID-19 Surveillance Team |
| EPI_ISL_2468454, EPI_ISL_2468455 | Lighthouse Lab in Milton Keynes | Wellcome Sanger Institute for the COVID-19 Genomics UK (COG-UK) Consortium | The Lighthouse Lab in Milton Keynes and Alex Alderton, Roberto Amato, Jeffrey Barrett, Sonia Goncalves, Ewan Harrison, David K. Jackson, Ian Johnston, Dominic Kwiatkowski, Cordelia Langford, John Sillitoe on behalf of the Wellcome Sanger Institute COVID-19 Surveillance Team |
| EPI_ISL_2468457 | Lighthouse Lab in Alderley Park | Wellcome Sanger Institute for the COVID-19 Genomics UK (COG-UK) Consortium | Jacquelyn Wynn, Mairead Hyland, The Lighthouse Lab in Alderley Park and Alex Alderton, Roberto Amato, Jeffrey Barrett, Sonia Goncalves, Ewan Harrison, David K. Jackson, Ian Johnston, Dominic Kwiatkowski, Cordelia Langford, John Sillitoe on behalf of the Wellcome Sanger Institute COVID-19 Surveillance Team |
| EPI_ISL_2468458 | Lighthouse Lab in Milton Keynes | Wellcome Sanger Institute for the COVID-19 Genomics UK (COG-UK) Consortium | The Lighthouse Lab in Milton Keynes and Alex Alderton, Roberto Amato, Jeffrey Barrett, Sonia Goncalves, Ewan Harrison, David K. Jackson, Ian Johnston, Dominic Kwiatkowski, Cordelia Langford, John Sillitoe on behalf of the Wellcome Sanger Institute COVID-19 Surveillance Team |
| EPI_ISL_2468459 | Health Services Laboratories | Wellcome Sanger Institute for the COVID-19 Genomics UK (COG-UK) Consortium | Health Services Laboratories and Alex Alderton, Roberto Amato, Jeffrey Barrett, Sonia Goncalves, Ewan Harrison, David K. Jackson, Ian Johnston, Dominic Kwiatkowski, Cordelia Langford, John Sillitoe on behalf of the Wellcome Sanger Institute COVID-19 Surveillance Team |
| EPI_ISL_2468460 | Lighthouse Lab in Alderley Park | Wellcome Sanger Institute for the COVID-19 Genomics UK (COG-UK) Consortium | Jacquelyn Wynn, Mairead Hyland, The Lighthouse Lab in Alderley Park and Alex Alderton, Roberto Amato, Jeffrey Barrett, Sonia Goncalves, Ewan Harrison, David K. Jackson, Ian Johnston, Dominic Kwiatkowski, Cordelia Langford, John Sillitoe on behalf of the Wellcome Sanger Institute COVID-19 Surveillance Team |
| EPI_ISL_2468461, EPI_ISL_2468462 | Lighthouse Lab in Milton Keynes | Wellcome Sanger Institute for the COVID-19 Genomics UK (COG-UK) Consortium | The Lighthouse Lab in Milton Keynes and Alex Alderton, Roberto Amato, Jeffrey Barrett, Sonia Goncalves, Ewan Harrison, David K. Jackson, Ian Johnston, Dominic Kwiatkowski, Cordelia Langford, John Sillitoe on behalf of the Wellcome Sanger Institute COVID-19 Surveillance Team |
| EPI_ISL_2468463 | Health Services Laboratories | Wellcome Sanger Institute for the COVID-19 Genomics UK (COG-UK) Consortium | Health Services Laboratories and Alex Alderton, Roberto Amato, Jeffrey Barrett, Sonia Goncalves, Ewan Harrison, David K. Jackson, Ian Johnston, Dominic Kwiatkowski, Cordelia Langford, John Sillitoe on behalf of the Wellcome Sanger Institute COVID-19 Surveillance Team |
| EPI_ISL_2468464 | Lighthouse Laboratory Plymouth | Wellcome Sanger Institute for the COVID-19 Genomics UK (COG-UK) Consortium | Lighthouse Laboratory Plymouth and Alex Alderton, Roberto Amato, Jeffrey Barrett, Sonia Goncalves, Ewan Harrison, David K. Jackson, Ian Johnston, Dominic Kwiatkowski, Cordelia Langford, John Sillitoe on behalf of the Wellcome Sanger Institute COVID-19 Surveillance Team |
| EPI_ISL_2468465 | Lighthouse Lab in Alderley Park | Wellcome Sanger Institute for the COVID-19 Genomics UK (COG-UK) Consortium | Jacquelyn Wynn, Mairead Hyland, The Lighthouse Lab in Alderley Park and Alex Alderton, Roberto Amato, Jeffrey Barrett, Sonia Goncalves, Ewan Harrison, David K. Jackson, Ian Johnston, Dominic Kwiatkowski, Cordelia Langford, John Sillitoe on behalf of the Wellcome Sanger Institute COVID-19 Surveillance Team |
| EPI_ISL_2468466 | Lighthouse Laboratory Plymouth | Wellcome Sanger Institute for the COVID-19 Genomics UK (COG-UK) Consortium | Lighthouse Laboratory Plymouth and Alex Alderton, Roberto Amato, Jeffrey Barrett, Sonia Goncalves, Ewan Harrison, David K. Jackson, Ian Johnston, Dominic Kwiatkowski, Cordelia Langford, John Sillitoe on behalf of the Wellcome Sanger Institute COVID-19 Surveillance Team |
| EPI_ISL_2468469, EPI_ISL_2468470, EPI_ISL_2468471, EPI_ISL_2468472 | Lighthouse Lab in Alderley Park | Wellcome Sanger Institute for the COVID-19 Genomics UK (COG-UK) Consortium | Jacquelyn Wynn, Mairead Hyland, The Lighthouse Lab in Alderley Park and Alex Alderton, Roberto Amato, Jeffrey Barrett, Sonia Goncalves, Ewan Harrison, David K. Jackson, Ian Johnston, Dominic Kwiatkowski, Cordelia Langford, John Sillitoe on behalf of the Wellcome Sanger Institute COVID-19 Surveillance Team |
| EPI_ISL_2468473 | Health Services Laboratories | Wellcome Sanger Institute for the COVID-19 Genomics UK (COG-UK) Consortium | Health Services Laboratories and Alex Alderton, Roberto Amato, Jeffrey Barrett, Sonia Goncalves, Ewan Harrison, David K. Jackson, Ian Johnston, Dominic Kwiatkowski, Cordelia Langford, John Sillitoe on behalf of the Wellcome Sanger Institute COVID-19 Surveillance Team |
| EPI_ISL_2468474, EPI_ISL_2468475 | Lighthouse Lab in Alderley Park | Wellcome Sanger Institute for the COVID-19 Genomics UK (COG-UK) Consortium | Jacquelyn Wynn, Mairead Hyland, The Lighthouse Lab in Alderley Park and Alex Alderton, Roberto Amato, Jeffrey Barrett, Sonia Goncalves, Ewan Harrison, David K. Jackson, Ian Johnston, Dominic Kwiatkowski, Cordelia Langford, John Sillitoe on behalf of the Wellcome Sanger Institute COVID-19 Surveillance Team |
| EPI_ISL_2468476 | Health Services Laboratories | Wellcome Sanger Institute for the COVID-19 Genomics UK (COG-UK) Consortium | Health Services Laboratories and Alex Alderton, Roberto Amato, Jeffrey Barrett, Sonia Goncalves, Ewan Harrison, David K. Jackson, Ian Johnston, Dominic Kwiatkowski, Cordelia Langford, John Sillitoe on behalf of the Wellcome Sanger Institute COVID-19 Surveillance Team |
| EPI_ISL_2468477 | Lighthouse Laboratory Plymouth | Wellcome Sanger Institute for the COVID-19 Genomics UK (COG-UK) Consortium | Lighthouse Laboratory Plymouth and Alex Alderton, Roberto Amato, Jeffrey Barrett, Sonia Goncalves, Ewan Harrison, David K. Jackson, Ian Johnston, Dominic Kwiatkowski, Cordelia Langford, John Sillitoe on behalf of the Wellcome Sanger Institute COVID-19 Surveillance Team |
| EPI_ISL_2468478 | Lighthouse Lab in Alderley Park | Wellcome Sanger Institute for the COVID-19 Genomics UK (COG-UK) Consortium | Jacquelyn Wynn, Mairead Hyland, The Lighthouse Lab in Alderley Park and Alex Alderton, Roberto Amato, Jeffrey Barrett, Sonia Goncalves, Ewan Harrison, David K. Jackson, Ian Johnston, Dominic Kwiatkowski, Cordelia Langford, John Sillitoe on behalf of the Wellcome Sanger Institute COVID-19 Surveillance Team |
| EPI_ISL_2468479, EPI_ISL_2468480, EPI_ISL_2468481, EPI_ISL_2468482 | Lighthouse Lab in Milton Keynes | Wellcome Sanger Institute for the COVID-19 Genomics UK (COG-UK) Consortium | The Lighthouse Lab in Milton Keynes and Alex Alderton, Roberto Amato, Jeffrey Barrett, Sonia Goncalves, Ewan Harrison, David K. Jackson, Ian Johnston, Dominic Kwiatkowski, Cordelia Langford, John Sillitoe on behalf of the Wellcome Sanger Institute COVID-19 Surveillance Team |
| EPI_ISL_2468483 | Lighthouse Lab in Alderley Park | Wellcome Sanger Institute for the COVID-19 Genomics UK (COG-UK) Consortium | Jacquelyn Wynn, Mairead Hyland, The Lighthouse Lab in Alderley Park and Alex Alderton, Roberto Amato, Jeffrey Barrett, Sonia Goncalves, Ewan Harrison, David K. Jackson, Ian Johnston, Dominic Kwiatkowski, Cordelia Langford, John Sillitoe on behalf of the Wellcome Sanger Institute COVID-19 Surveillance Team |

| | | | |
|---|---|---|---|
| EPI_ISL_2468484 | Health Services Laboratories | Wellcome Sanger Institute for the COVID-19 Genomics UK (COG-UK) Consortium | Health Services Laboratories and Alex Alderton, Roberto Amato, Jeffrey Barrett, Sonia Goncalves, Ewan Harrison, David K. Jackson, Ian Johnston, Dominic Kwiatkowski, Cordelia Langford, John Sillitoe on behalf of the Wellcome Sanger Institute COVID-19 Surveillance Team |
| EPI_ISL_2468485 | Lighthouse Lab in Milton Keynes | Wellcome Sanger Institute for the COVID-19 Genomics UK (COG-UK) Consortium | The Lighthouse Lab in Milton Keynes and Alex Alderton, Roberto Amato, Jeffrey Barrett, Sonia Goncalves, Ewan Harrison, David K. Jackson, Ian Johnston, Dominic Kwiatkowski, Cordelia Langford, John Sillitoe on behalf of the Wellcome Sanger Institute COVID-19 Surveillance Team |
| EPI_ISL_2468486 | Health Services Laboratories | Wellcome Sanger Institute for the COVID-19 Genomics UK (COG-UK) Consortium | Health Services Laboratories and Alex Alderton, Roberto Amato, Jeffrey Barrett, Sonia Goncalves, Ewan Harrison, David K. Jackson, Ian Johnston, Dominic Kwiatkowski, Cordelia Langford, John Sillitoe on behalf of the Wellcome Sanger Institute COVID-19 Surveillance Team |
| EPI_ISL_2468487 | Lighthouse Lab in Alderley Park | Wellcome Sanger Institute for the COVID-19 Genomics UK (COG-UK) Consortium | Jacquelyn Wynn, Mairead Hyland, The Lighthouse Lab in Alderley Park and Alex Alderton, Roberto Amato, Jeffrey Barrett, Sonia Goncalves, Ewan Harrison, David K. Jackson, Ian Johnston, Dominic Kwiatkowski, Cordelia Langford, John Sillitoe on behalf of the Wellcome Sanger Institute COVID-19 Surveillance Team |
| EPI_ISL_2468488 | Lighthouse Lab in Milton Keynes | Wellcome Sanger Institute for the COVID-19 Genomics UK (COG-UK) Consortium | The Lighthouse Lab in Milton Keynes and Alex Alderton, Roberto Amato, Jeffrey Barrett, Sonia Goncalves, Ewan Harrison, David K. Jackson, Ian Johnston, Dominic Kwiatkowski, Cordelia Langford, John Sillitoe on behalf of the Wellcome Sanger Institute COVID-19 Surveillance Team |
| EPI_ISL_2468489 | Health Services Laboratories | Wellcome Sanger Institute for the COVID-19 Genomics UK (COG-UK) Consortium | Health Services Laboratories and Alex Alderton, Roberto Amato, Jeffrey Barrett, Sonia Goncalves, Ewan Harrison, David K. Jackson, Ian Johnston, Dominic Kwiatkowski, Cordelia Langford, John Sillitoe on behalf of the Wellcome Sanger Institute COVID-19 Surveillance Team |
| EPI_ISL_2468490, EPI_ISL_2468491 | Lighthouse Lab in Alderley Park | Wellcome Sanger Institute for the COVID-19 Genomics UK (COG-UK) Consortium | Jacquelyn Wynn, Mairead Hyland, The Lighthouse Lab in Alderley Park and Alex Alderton, Roberto Amato, Jeffrey Barrett, Sonia Goncalves, Ewan Harrison, David K. Jackson, Ian Johnston, Dominic Kwiatkowski, Cordelia Langford, John Sillitoe on behalf of the Wellcome Sanger Institute COVID-19 Surveillance Team |
| EPI_ISL_2468493 | Health Services Laboratories | Wellcome Sanger Institute for the COVID-19 Genomics UK (COG-UK) Consortium | Health Services Laboratories and Alex Alderton, Roberto Amato, Jeffrey Barrett, Sonia Goncalves, Ewan Harrison, David K. Jackson, Ian Johnston, Dominic Kwiatkowski, Cordelia Langford, John Sillitoe on behalf of the Wellcome Sanger Institute COVID-19 Surveillance Team |
| EPI_ISL_2468494 | Lighthouse Lab in Glasgow | Wellcome Sanger Institute for the COVID-19 Genomics UK (COG-UK) Consortium | Harper VanSteenhouse, Yumi Kasai, David Gray, Carol Clugston, Anna Dominiczak and Alex Alderton, Roberto Amato, Jeffrey Barrett, Sonia Goncalves, Ewan Harrison, David K. Jackson, Ian Johnston, Dominic Kwiatkowski, Cordelia Langford, John Sillitoe on behalf of the Wellcome Sanger Institute COVID-19 Surveillance Team |
| EPI_ISL_2468495, EPI_ISL_2468496, EPI_ISL_2468497 | Lighthouse Lab in Alderley Park | Wellcome Sanger Institute for the COVID-19 Genomics UK (COG-UK) Consortium | Jacquelyn Wynn, Mairead Hyland, The Lighthouse Lab in Alderley Park and Alex Alderton, Roberto Amato, Jeffrey Barrett, Sonia Goncalves, Ewan Harrison, David K. Jackson, Ian Johnston, Dominic Kwiatkowski, Cordelia Langford, John Sillitoe on behalf of the Wellcome Sanger Institute COVID-19 Surveillance Team |
| EPI_ISL_2468498 | Lighthouse Laboratory Plymouth | Wellcome Sanger Institute for the COVID-19 Genomics UK (COG-UK) Consortium | Lighthouse Laboratory Plymouth and Alex Alderton, Roberto Amato, Jeffrey Barrett, Sonia Goncalves, Ewan Harrison, David K. Jackson, Ian Johnston, Dominic Kwiatkowski, Cordelia Langford, John Sillitoe on behalf of the Wellcome Sanger Institute COVID-19 Surveillance Team |
| EPI_ISL_2468499 | Health Services Laboratories | Wellcome Sanger Institute for the COVID-19 Genomics UK (COG-UK) Consortium | Health Services Laboratories and Alex Alderton, Roberto Amato, Jeffrey Barrett, Sonia Goncalves, Ewan Harrison, David K. Jackson, Ian Johnston, Dominic Kwiatkowski, Cordelia Langford, John Sillitoe on behalf of the Wellcome Sanger Institute COVID-19 Surveillance Team |
| EPI_ISL_2468500 | Lighthouse Lab in Milton Keynes | Wellcome Sanger Institute for the COVID-19 Genomics UK (COG-UK) Consortium | The Lighthouse Lab in Milton Keynes and Alex Alderton, Roberto Amato, Jeffrey Barrett, Sonia Goncalves, Ewan Harrison, David K. Jackson, Ian Johnston, Dominic Kwiatkowski, Cordelia Langford, John Sillitoe on behalf of the Wellcome Sanger Institute COVID-19 Surveillance Team |
| EPI_ISL_2468501 | Health Services Laboratories | Wellcome Sanger Institute for the COVID-19 Genomics UK (COG-UK) Consortium | Health Services Laboratories and Alex Alderton, Roberto Amato, Jeffrey Barrett, Sonia Goncalves, Ewan Harrison, David K. Jackson, Ian Johnston, Dominic Kwiatkowski, Cordelia Langford, John Sillitoe on behalf of the Wellcome Sanger Institute COVID-19 Surveillance Team |
| EPI_ISL_2468502, EPI_ISL_2468503 | Lighthouse Lab in Milton Keynes | Wellcome Sanger Institute for the COVID-19 Genomics UK (COG-UK) Consortium | The Lighthouse Lab in Milton Keynes and Alex Alderton, Roberto Amato, Jeffrey Barrett, Sonia Goncalves, Ewan Harrison, David K. Jackson, Ian Johnston, Dominic Kwiatkowski, Cordelia Langford, John Sillitoe on behalf of the Wellcome Sanger Institute COVID-19 Surveillance Team |
| EPI_ISL_2468504 | Lighthouse Lab in Alderley Park | Wellcome Sanger Institute for the COVID-19 Genomics UK (COG-UK) Consortium | Jacquelyn Wynn, Mairead Hyland, The Lighthouse Lab in Alderley Park and Alex Alderton, Roberto Amato, Jeffrey Barrett, Sonia Goncalves, Ewan Harrison, David K. Jackson, Ian Johnston, Dominic Kwiatkowski, Cordelia Langford, John Sillitoe on behalf of the Wellcome Sanger Institute COVID-19 Surveillance Team |
| EPI_ISL_2468505 | Lighthouse Lab in Milton Keynes | Wellcome Sanger Institute for the COVID-19 Genomics UK (COG-UK) Consortium | The Lighthouse Lab in Milton Keynes and Alex Alderton, Roberto Amato, Jeffrey Barrett, Sonia Goncalves, Ewan Harrison, David K. Jackson, Ian Johnston, Dominic Kwiatkowski, Cordelia Langford, John Sillitoe on behalf of the Wellcome Sanger Institute COVID-19 Surveillance Team |
| EPI_ISL_2468506 | Lighthouse Lab in Alderley Park | Wellcome Sanger Institute for the COVID-19 Genomics UK (COG-UK) Consortium | Jacquelyn Wynn, Mairead Hyland, The Lighthouse Lab in Alderley Park and Alex Alderton, Roberto Amato, Jeffrey Barrett, Sonia Goncalves, Ewan Harrison, David K. Jackson, Ian Johnston, Dominic Kwiatkowski, Cordelia Langford, John Sillitoe on behalf of the Wellcome Sanger Institute COVID-19 Surveillance Team |
| EPI_ISL_2468507 | Lighthouse Lab in Milton Keynes | Wellcome Sanger Institute for the COVID-19 Genomics UK (COG-UK) Consortium | The Lighthouse Lab in Milton Keynes and Alex Alderton, Roberto Amato, Jeffrey Barrett, Sonia Goncalves, Ewan Harrison, David K. Jackson, Ian Johnston, Dominic Kwiatkowski, Cordelia Langford, John Sillitoe on behalf of the Wellcome Sanger Institute COVID-19 Surveillance Team |
| EPI_ISL_2468508 | Lighthouse Lab in Glasgow | Wellcome Sanger Institute for the COVID-19 Genomics UK (COG-UK) Consortium | Harper VanSteenhouse, Yumi Kasai, David Gray, Carol Clugston, Anna Dominiczak and Alex Alderton, Roberto Amato, Jeffrey Barrett, Sonia Goncalves, Ewan Harrison, David K. Jackson, Ian Johnston, Dominic Kwiatkowski, Cordelia Langford, John Sillitoe on behalf of the Wellcome Sanger Institute COVID-19 Surveillance Team |
| EPI_ISL_2468509, EPI_ISL_2468510 | Lighthouse Lab in Milton Keynes | Wellcome Sanger Institute for the COVID-19 Genomics UK (COG-UK) Consortium | The Lighthouse Lab in Milton Keynes and Alex Alderton, Roberto Amato, Jeffrey Barrett, Sonia Goncalves, Ewan Harrison, David K. Jackson, Ian Johnston, Dominic Kwiatkowski, Cordelia Langford, John Sillitoe on behalf of the Wellcome Sanger Institute COVID-19 Surveillance Team |
| EPI_ISL_2468511 | Lighthouse Lab in Glasgow | Wellcome Sanger Institute for the COVID-19 Genomics UK (COG-UK) Consortium | Harper VanSteenhouse, Yumi Kasai, David Gray, Carol Clugston, Anna Dominiczak and Alex Alderton, Roberto Amato, Jeffrey Barrett, Sonia Goncalves, Ewan Harrison, David K. Jackson, Ian Johnston, Dominic Kwiatkowski, Cordelia Langford, John Sillitoe on behalf of the Wellcome Sanger Institute COVID-19 Surveillance Team |
| EPI_ISL_2468513 | Lighthouse Lab in Alderley Park | Wellcome Sanger Institute for the COVID-19 Genomics UK (COG-UK) Consortium | Jacquelyn Wynn, Mairead Hyland, The Lighthouse Lab in Alderley Park and Alex Alderton, Roberto Amato, Jeffrey Barrett, Sonia Goncalves, Ewan Harrison, David K. Jackson, Ian Johnston, Dominic Kwiatkowski, Cordelia Langford, John Sillitoe on behalf of the Wellcome Sanger Institute COVID-19 Surveillance Team |
| EPI_ISL_2468515 | Lighthouse Lab in Milton Keynes | Wellcome Sanger Institute for the COVID-19 Genomics UK (COG-UK) Consortium | The Lighthouse Lab in Milton Keynes and Alex Alderton, Roberto Amato, Jeffrey Barrett, Sonia Goncalves, Ewan Harrison, David K. Jackson, Ian Johnston, Dominic Kwiatkowski, Cordelia Langford, John Sillitoe on behalf of the Wellcome Sanger Institute COVID-19 Surveillance Team |
| EPI_ISL_2468516 | Lighthouse Lab in Alderley Park | Wellcome Sanger Institute for the COVID-19 Genomics UK (COG-UK) Consortium | Jacquelyn Wynn, Mairead Hyland, The Lighthouse Lab in Alderley Park and Alex Alderton, Roberto Amato, Jeffrey Barrett, Sonia Goncalves, Ewan Harrison, David K. Jackson, Ian Johnston, Dominic Kwiatkowski, Cordelia Langford, John Sillitoe on behalf of the Wellcome Sanger Institute COVID-19 Surveillance Team |
| EPI_ISL_2468517, EPI_ISL_2468518 | Lighthouse Lab in Milton Keynes | Wellcome Sanger Institute for the COVID-19 Genomics UK (COG-UK) Consortium | The Lighthouse Lab in Milton Keynes and Alex Alderton, Roberto Amato, Jeffrey Barrett, Sonia Goncalves, Ewan Harrison, David K. Jackson, Ian Johnston, Dominic Kwiatkowski, Cordelia Langford, John Sillitoe on behalf of the Wellcome Sanger Institute COVID-19 Surveillance Team |
| EPI_ISL_2468519 | Lighthouse Lab in Alderley Park | Wellcome Sanger Institute for the COVID-19 Genomics UK (COG-UK) Consortium | Jacquelyn Wynn, Mairead Hyland, The Lighthouse Lab in Alderley Park and Alex Alderton, Roberto Amato, Jeffrey Barrett, Sonia Goncalves, Ewan Harrison, David K. Jackson, Ian Johnston, Dominic Kwiatkowski, Cordelia Langford, John Sillitoe on behalf of the Wellcome Sanger Institute COVID-19 Surveillance Team |
| EPI_ISL_2468520 | Health Services Laboratories | Wellcome Sanger Institute for the COVID-19 Genomics UK (COG-UK) Consortium | Health Services Laboratories and Alex Alderton, Roberto Amato, Jeffrey Barrett, Sonia Goncalves, Ewan Harrison, David K. Jackson, Ian Johnston, Dominic Kwiatkowski, Cordelia Langford, John Sillitoe on behalf of the Wellcome Sanger Institute COVID-19 Surveillance Team |
| EPI_ISL_2468521, EPI_ISL_2468522 | Lighthouse Lab in Alderley Park | Wellcome Sanger Institute for the COVID-19 Genomics UK (COG-UK) Consortium | Jacquelyn Wynn, Mairead Hyland, The Lighthouse Lab in Alderley Park and Alex Alderton, Roberto Amato, Jeffrey Barrett, Sonia Goncalves, Ewan Harrison, David K. Jackson, Ian Johnston, Dominic Kwiatkowski, Cordelia Langford, John Sillitoe on behalf of the Wellcome Sanger Institute COVID-19 Surveillance Team |
| EPI_ISL_2468523, EPI_ISL_2468524 | Lighthouse Laboratory Plymouth | Wellcome Sanger Institute for the COVID-19 Genomics UK (COG-UK) Consortium | Lighthouse Laboratory Plymouth and Alex Alderton, Roberto Amato, Jeffrey Barrett, Sonia Goncalves, Ewan Harrison, David K. Jackson, Ian Johnston, Dominic Kwiatkowski, Cordelia Langford, John Sillitoe on behalf of the Wellcome Sanger Institute COVID-19 Surveillance Team |
| EPI_ISL_2468525 | Lighthouse Lab in Alderley Park | Wellcome Sanger Institute for the COVID-19 Genomics UK (COG-UK) Consortium | Jacquelyn Wynn, Mairead Hyland, The Lighthouse Lab in Alderley Park and Alex Alderton, Roberto Amato, Jeffrey Barrett, Sonia Goncalves, Ewan Harrison, David K. Jackson, Ian Johnston, Dominic Kwiatkowski, Cordelia Langford, John Sillitoe on behalf of the Wellcome Sanger Institute COVID-19 Surveillance Team |
| EPI_ISL_2468526 | Health Services Laboratories | Wellcome Sanger Institute for the COVID-19 Genomics UK (COG-UK) Consortium | Health Services Laboratories and Alex Alderton, Roberto Amato, Jeffrey Barrett, Sonia Goncalves, Ewan Harrison, David K. Jackson, Ian Johnston, Dominic Kwiatkowski, Cordelia Langford, John Sillitoe on behalf of the Wellcome Sanger Institute COVID-19 Surveillance Team |
| EPI_ISL_2468527, EPI_ISL_2468528 | Lighthouse Lab in Alderley Park | Wellcome Sanger Institute for the COVID-19 Genomics UK (COG-UK) Consortium | Jacquelyn Wynn, Mairead Hyland, The Lighthouse Lab in Alderley Park and Alex Alderton, Roberto Amato, Jeffrey Barrett, Sonia Goncalves, Ewan Harrison, David K. Jackson, Ian Johnston, Dominic Kwiatkowski, Cordelia Langford, John Sillitoe on behalf of the Wellcome Sanger Institute COVID-19 Surveillance Team |
| EPI_ISL_2468529 | Lighthouse Laboratory Plymouth | Wellcome Sanger Institute for the COVID-19 Genomics UK (COG-UK) Consortium | Lighthouse Laboratory Plymouth and Alex Alderton, Roberto Amato, Jeffrey Barrett, Sonia Goncalves, Ewan Harrison, David K. Jackson, Ian Johnston, Dominic Kwiatkowski, Cordelia Langford, John Sillitoe on behalf of the Wellcome Sanger Institute COVID-19 Surveillance Team |
| EPI_ISL_2468530, EPI_ISL_2468531, EPI_ISL_2468532 | Lighthouse Lab in Alderley Park | Wellcome Sanger Institute for the COVID-19 Genomics UK (COG-UK) Consortium | Jacquelyn Wynn, Mairead Hyland, The Lighthouse Lab in Alderley Park and Alex Alderton, Roberto Amato, Jeffrey Barrett, Sonia Goncalves, Ewan Harrison, David K. Jackson, Ian Johnston, Dominic Kwiatkowski, Cordelia Langford, John Sillitoe on behalf of the Wellcome Sanger Institute COVID-19 Surveillance Team |
| EPI_ISL_2468533, EPI_ISL_2468534 | Lighthouse Lab in Milton Keynes | Wellcome Sanger Institute for the COVID-19 Genomics UK (COG-UK) Consortium | The Lighthouse Lab in Milton Keynes and Alex Alderton, Roberto Amato, Jeffrey Barrett, Sonia Goncalves, Ewan Harrison, David K. Jackson, Ian Johnston, Dominic Kwiatkowski, Cordelia Langford, John Sillitoe on behalf of the Wellcome Sanger Institute COVID-19 Surveillance Team |
| EPI_ISL_2468535 | Lighthouse Laboratory Plymouth | Wellcome Sanger Institute for the COVID-19 Genomics UK (COG-UK) Consortium | Lighthouse Laboratory Plymouth and Alex Alderton, Roberto Amato, Jeffrey Barrett, Sonia Goncalves, Ewan Harrison, David K. Jackson, Ian Johnston, Dominic Kwiatkowski, Cordelia Langford, John Sillitoe on behalf of the Wellcome Sanger Institute COVID-19 Surveillance Team |
| EPI_ISL_2468536, EPI_ISL_2468537, EPI_ISL_2468538 | Health Services Laboratories | Wellcome Sanger Institute for the COVID-19 Genomics UK (COG-UK) Consortium | Health Services Laboratories and Alex Alderton, Roberto Amato, Jeffrey Barrett, Sonia Goncalves, Ewan Harrison, David K. Jackson, Ian Johnston, Dominic Kwiatkowski, Cordelia Langford, John Sillitoe on behalf of the Wellcome Sanger Institute COVID-19 Surveillance Team |
| EPI_ISL_2468539 | Lighthouse Lab in Glasgow | Wellcome Sanger Institute for the COVID-19 Genomics UK (COG-UK) Consortium | Harper VanSteenhouse, Yumi Kasai, David Gray, Carol Clugston, Anna Dominiczak and Alex Alderton, Roberto Amato, Jeffrey Barrett, Sonia Goncalves, Ewan Harrison, David K. Jackson, Ian Johnston, Dominic Kwiatkowski, Cordelia Langford, John Sillitoe on behalf of the Wellcome Sanger Institute COVID-19 Surveillance Team |
| EPI_ISL_2468540 | Lighthouse Lab in Alderley Park | Wellcome Sanger Institute for the COVID-19 Genomics UK (COG-UK) Consortium | Jacquelyn Wynn, Mairead Hyland, The Lighthouse Lab in Alderley Park and Alex Alderton, Roberto Amato, Jeffrey Barrett, Sonia Goncalves, Ewan Harrison, David K. Jackson, Ian Johnston, Dominic Kwiatkowski, Cordelia Langford, John Sillitoe on behalf of the Wellcome Sanger Institute COVID-19 Surveillance Team |
| EPI_ISL_2468541 | Lighthouse Lab in Milton Keynes | Wellcome Sanger Institute for the COVID-19 Genomics UK (COG-UK) Consortium | The Lighthouse Lab in Milton Keynes and Alex Alderton, Roberto Amato, Jeffrey Barrett, Sonia Goncalves, Ewan Harrison, David K. Jackson, Ian Johnston, Dominic Kwiatkowski, Cordelia Langford, John Sillitoe on behalf of the Wellcome Sanger Institute COVID-19 Surveillance Team |
| EPI_ISL_2468542 | Lighthouse Lab in Alderley Park | Wellcome Sanger Institute for the COVID-19 Genomics UK (COG-UK) Consortium | Jacquelyn Wynn, Mairead Hyland, The Lighthouse Lab in Alderley Park and Alex Alderton, Roberto Amato, Jeffrey Barrett, Sonia Goncalves, Ewan Harrison, David K. Jackson, Ian Johnston, Dominic Kwiatkowski, Cordelia Langford, John Sillitoe on behalf of the Wellcome Sanger Institute COVID-19 Surveillance Team |
| EPI_ISL_2468543, EPI_ISL_2468544 | Lighthouse Lab in Milton Keynes | Wellcome Sanger Institute for the COVID-19 Genomics UK (COG-UK) Consortium | The Lighthouse Lab in Milton Keynes and Alex Alderton, Roberto Amato, Jeffrey Barrett, Sonia Goncalves, Ewan Harrison, David K. Jackson, Ian Johnston, Dominic Kwiatkowski, Cordelia Langford, John Sillitoe on behalf of the Wellcome Sanger Institute COVID-19 Surveillance Team |
| EPI_ISL_2468545 | Lighthouse Laboratory Plymouth | Wellcome Sanger Institute for the COVID-19 Genomics UK (COG-UK) Consortium | Lighthouse Laboratory Plymouth and Alex Alderton, Roberto Amato, Jeffrey Barrett, Sonia Goncalves, Ewan Harrison, David K. Jackson, Ian Johnston, Dominic Kwiatkowski, Cordelia Langford, John Sillitoe on behalf of the Wellcome Sanger Institute COVID-19 Surveillance Team |

| | | | |
|---|---|---|---|
| EPI_ISL_2468547 | Health Services Laboratories | Wellcome Sanger Institute for the COVID-19 Genomics UK (COG-UK) Consortium | Health Services Laboratories and Alex Alderton, Roberto Amato, Jeffrey Barrett, Sonia Goncalves, Ewan Harrison, David K. Jackson, Ian Johnston, Dominic Kwiatkowski, Cordelia Langford, John Sillitoe on behalf of the Wellcome Sanger Institute COVID-19 Surveillance Team |
| EPI_ISL_2468548 | Lighthouse Lab in Milton Keynes | Wellcome Sanger Institute for the COVID-19 Genomics UK (COG-UK) Consortium | The Lighthouse Lab in Milton Keynes and Alex Alderton, Roberto Amato, Jeffrey Barrett, Sonia Goncalves, Ewan Harrison, David K. Jackson, Ian Johnston, Dominic Kwiatkowski, Cordelia Langford, John Sillitoe on behalf of the Wellcome Sanger Institute COVID-19 Surveillance Team |
| EPI_ISL_2468549 | Health Services Laboratories | Wellcome Sanger Institute for the COVID-19 Genomics UK (COG-UK) Consortium | Health Services Laboratories and Alex Alderton, Roberto Amato, Jeffrey Barrett, Sonia Goncalves, Ewan Harrison, David K. Jackson, Ian Johnston, Dominic Kwiatkowski, Cordelia Langford, John Sillitoe on behalf of the Wellcome Sanger Institute COVID-19 Surveillance Team |
| EPI_ISL_2468551, EPI_ISL_2468552, EPI_ISL_2468553 | Lighthouse Lab in Alderley Park | Wellcome Sanger Institute for the COVID-19 Genomics UK (COG-UK) Consortium | Jacquelyn Wynn, Mairead Hyland, The Lighthouse Lab in Alderley Park and Alex Alderton, Roberto Amato, Jeffrey Barrett, Sonia Goncalves, Ewan Harrison, David K. Jackson, Ian Johnston, Dominic Kwiatkowski, Cordelia Langford, John Sillitoe on behalf of the Wellcome Sanger Institute COVID-19 Surveillance Team |
| EPI_ISL_2468554, EPI_ISL_2468555 | Health Services Laboratories | Wellcome Sanger Institute for the COVID-19 Genomics UK (COG-UK) Consortium | Health Services Laboratories and Alex Alderton, Roberto Amato, Jeffrey Barrett, Sonia Goncalves, Ewan Harrison, David K. Jackson, Ian Johnston, Dominic Kwiatkowski, Cordelia Langford, John Sillitoe on behalf of the Wellcome Sanger Institute COVID-19 Surveillance Team |
| EPI_ISL_2468556 | Lighthouse Laboratory Plymouth | Wellcome Sanger Institute for the COVID-19 Genomics UK (COG-UK) Consortium | Lighthouse Laboratory Plymouth and Alex Alderton, Roberto Amato, Jeffrey Barrett, Sonia Goncalves, Ewan Harrison, David K. Jackson, Ian Johnston, Dominic Kwiatkowski, Cordelia Langford, John Sillitoe on behalf of the Wellcome Sanger Institute COVID-19 Surveillance Team |
| EPI_ISL_2468558 | Lighthouse Lab in Alderley Park | Wellcome Sanger Institute for the COVID-19 Genomics UK (COG-UK) Consortium | Jacquelyn Wynn, Mairead Hyland, The Lighthouse Lab in Alderley Park and Alex Alderton, Roberto Amato, Jeffrey Barrett, Sonia Goncalves, Ewan Harrison, David K. Jackson, Ian Johnston, Dominic Kwiatkowski, Cordelia Langford, John Sillitoe on behalf of the Wellcome Sanger Institute COVID-19 Surveillance Team |
| EPI_ISL_2468559 | Lighthouse Lab in Milton Keynes | Wellcome Sanger Institute for the COVID-19 Genomics UK (COG-UK) Consortium | The Lighthouse Lab in Milton Keynes and Alex Alderton, Roberto Amato, Jeffrey Barrett, Sonia Goncalves, Ewan Harrison, David K. Jackson, Ian Johnston, Dominic Kwiatkowski, Cordelia Langford, John Sillitoe on behalf of the Wellcome Sanger Institute COVID-19 Surveillance Team |
| EPI_ISL_2468560 | Health Services Laboratories | Wellcome Sanger Institute for the COVID-19 Genomics UK (COG-UK) Consortium | Health Services Laboratories and Alex Alderton, Roberto Amato, Jeffrey Barrett, Sonia Goncalves, Ewan Harrison, David K. Jackson, Ian Johnston, Dominic Kwiatkowski, Cordelia Langford, John Sillitoe on behalf of the Wellcome Sanger Institute COVID-19 Surveillance Team |
| EPI_ISL_2468561 | Lighthouse Laboratory Plymouth | Wellcome Sanger Institute for the COVID-19 Genomics UK (COG-UK) Consortium | Lighthouse Laboratory Plymouth and Alex Alderton, Roberto Amato, Jeffrey Barrett, Sonia Goncalves, Ewan Harrison, David K. Jackson, Ian Johnston, Dominic Kwiatkowski, Cordelia Langford, John Sillitoe on behalf of the Wellcome Sanger Institute COVID-19 Surveillance Team |
| EPI_ISL_2468562 | Health Services Laboratories | Wellcome Sanger Institute for the COVID-19 Genomics UK (COG-UK) Consortium | Health Services Laboratories and Alex Alderton, Roberto Amato, Jeffrey Barrett, Sonia Goncalves, Ewan Harrison, David K. Jackson, Ian Johnston, Dominic Kwiatkowski, Cordelia Langford, John Sillitoe on behalf of the Wellcome Sanger Institute COVID-19 Surveillance Team |
| EPI_ISL_2468563, EPI_ISL_2468564 | Lighthouse Lab in Milton Keynes | Wellcome Sanger Institute for the COVID-19 Genomics UK (COG-UK) Consortium | The Lighthouse Lab in Milton Keynes and Alex Alderton, Roberto Amato, Jeffrey Barrett, Sonia Goncalves, Ewan Harrison, David K. Jackson, Ian Johnston, Dominic Kwiatkowski, Cordelia Langford, John Sillitoe on behalf of the Wellcome Sanger Institute COVID-19 Surveillance Team |
| EPI_ISL_2468565 | Lighthouse Laboratory Plymouth | Wellcome Sanger Institute for the COVID-19 Genomics UK (COG-UK) Consortium | Lighthouse Laboratory Plymouth and Alex Alderton, Roberto Amato, Jeffrey Barrett, Sonia Goncalves, Ewan Harrison, David K. Jackson, Ian Johnston, Dominic Kwiatkowski, Cordelia Langford, John Sillitoe on behalf of the Wellcome Sanger Institute COVID-19 Surveillance Team |
| EPI_ISL_2468567 | Health Services Laboratories | Wellcome Sanger Institute for the COVID-19 Genomics UK (COG-UK) Consortium | Health Services Laboratories and Alex Alderton, Roberto Amato, Jeffrey Barrett, Sonia Goncalves, Ewan Harrison, David K. Jackson, Ian Johnston, Dominic Kwiatkowski, Cordelia Langford, John Sillitoe on behalf of the Wellcome Sanger Institute COVID-19 Surveillance Team |
| EPI_ISL_2468568 | Lighthouse Lab in Alderley Park | Wellcome Sanger Institute for the COVID-19 Genomics UK (COG-UK) Consortium | Jacquelyn Wynn, Mairead Hyland, The Lighthouse Lab in Alderley Park and Alex Alderton, Roberto Amato, Jeffrey Barrett, Sonia Goncalves, Ewan Harrison, David K. Jackson, Ian Johnston, Dominic Kwiatkowski, Cordelia Langford, John Sillitoe on behalf of the Wellcome Sanger Institute COVID-19 Surveillance Team |
| EPI_ISL_2468569 | Lighthouse Lab in Milton Keynes | Wellcome Sanger Institute for the COVID-19 Genomics UK (COG-UK) Consortium | The Lighthouse Lab in Milton Keynes and Alex Alderton, Roberto Amato, Jeffrey Barrett, Sonia Goncalves, Ewan Harrison, David K. Jackson, Ian Johnston, Dominic Kwiatkowski, Cordelia Langford, John Sillitoe on behalf of the Wellcome Sanger Institute COVID-19 Surveillance Team |
| EPI_ISL_2468570 | Lighthouse Lab in Alderley Park | Wellcome Sanger Institute for the COVID-19 Genomics UK (COG-UK) Consortium | Jacquelyn Wynn, Mairead Hyland, The Lighthouse Lab in Alderley Park and Alex Alderton, Roberto Amato, Jeffrey Barrett, Sonia Goncalves, Ewan Harrison, David K. Jackson, Ian Johnston, Dominic Kwiatkowski, Cordelia Langford, John Sillitoe on behalf of the Wellcome Sanger Institute COVID-19 Surveillance Team |
| EPI_ISL_2468571, EPI_ISL_2468572 | Lighthouse Lab in Milton Keynes | Wellcome Sanger Institute for the COVID-19 Genomics UK (COG-UK) Consortium | The Lighthouse Lab in Milton Keynes and Alex Alderton, Roberto Amato, Jeffrey Barrett, Sonia Goncalves, Ewan Harrison, David K. Jackson, Ian Johnston, Dominic Kwiatkowski, Cordelia Langford, John Sillitoe on behalf of the Wellcome Sanger Institute COVID-19 Surveillance Team |
| EPI_ISL_2468573 | Lighthouse Laboratory Plymouth | Wellcome Sanger Institute for the COVID-19 Genomics UK (COG-UK) Consortium | Lighthouse Laboratory Plymouth and Alex Alderton, Roberto Amato, Jeffrey Barrett, Sonia Goncalves, Ewan Harrison, David K. Jackson, Ian Johnston, Dominic Kwiatkowski, Cordelia Langford, John Sillitoe on behalf of the Wellcome Sanger Institute COVID-19 Surveillance Team |
| EPI_ISL_2468574 | Lighthouse Lab in Alderley Park | Wellcome Sanger Institute for the COVID-19 Genomics UK (COG-UK) Consortium | Jacquelyn Wynn, Mairead Hyland, The Lighthouse Lab in Alderley Park and Alex Alderton, Roberto Amato, Jeffrey Barrett, Sonia Goncalves, Ewan Harrison, David K. Jackson, Ian Johnston, Dominic Kwiatkowski, Cordelia Langford, John Sillitoe on behalf of the Wellcome Sanger Institute COVID-19 Surveillance Team |
| EPI_ISL_2468575, EPI_ISL_2468576, EPI_ISL_2468577 | Health Services Laboratories | Wellcome Sanger Institute for the COVID-19 Genomics UK (COG-UK) Consortium | Health Services Laboratories and Alex Alderton, Roberto Amato, Jeffrey Barrett, Sonia Goncalves, Ewan Harrison, David K. Jackson, Ian Johnston, Dominic Kwiatkowski, Cordelia Langford, John Sillitoe on behalf of the Wellcome Sanger Institute COVID-19 Surveillance Team |
| EPI_ISL_2468578 | Lighthouse Lab in Milton Keynes | Wellcome Sanger Institute for the COVID-19 Genomics UK (COG-UK) Consortium | The Lighthouse Lab in Milton Keynes and Alex Alderton, Roberto Amato, Jeffrey Barrett, Sonia Goncalves, Ewan Harrison, David K. Jackson, Ian Johnston, Dominic Kwiatkowski, Cordelia Langford, John Sillitoe on behalf of the Wellcome Sanger Institute COVID-19 Surveillance Team |
| EPI_ISL_2468579 | Health Services Laboratories | Wellcome Sanger Institute for the COVID-19 Genomics UK (COG-UK) Consortium | Health Services Laboratories and Alex Alderton, Roberto Amato, Jeffrey Barrett, Sonia Goncalves, Ewan Harrison, David K. Jackson, Ian Johnston, Dominic Kwiatkowski, Cordelia Langford, John Sillitoe on behalf of the Wellcome Sanger Institute COVID-19 Surveillance Team |
| EPI_ISL_2468580 | Lighthouse Lab in Milton Keynes | Wellcome Sanger Institute for the COVID-19 Genomics UK (COG-UK) Consortium | The Lighthouse Lab in Milton Keynes and Alex Alderton, Roberto Amato, Jeffrey Barrett, Sonia Goncalves, Ewan Harrison, David K. Jackson, Ian Johnston, Dominic Kwiatkowski, Cordelia Langford, John Sillitoe on behalf of the Wellcome Sanger Institute COVID-19 Surveillance Team |
| EPI_ISL_2468581 | Lighthouse Lab in Glasgow | Wellcome Sanger Institute for the COVID-19 Genomics UK (COG-UK) Consortium | Harper VanSteenhouse, Yumi Kasai, David Gray, Carol Clugston, Anna Dominiczak and Alex Alderton, Roberto Amato, Jeffrey Barrett, Sonia Goncalves, Ewan Harrison, David K. Jackson, Ian Johnston, Dominic Kwiatkowski, Cordelia Langford, John Sillitoe on behalf of the Wellcome Sanger Institute COVID-19 Surveillance Team |
| EPI_ISL_2468582 | Lighthouse Lab in Alderley Park | Wellcome Sanger Institute for the COVID-19 Genomics UK (COG-UK) Consortium | Jacquelyn Wynn, Mairead Hyland, The Lighthouse Lab in Alderley Park and Alex Alderton, Roberto Amato, Jeffrey Barrett, Sonia Goncalves, Ewan Harrison, David K. Jackson, Ian Johnston, Dominic Kwiatkowski, Cordelia Langford, John Sillitoe on behalf of the Wellcome Sanger Institute COVID-19 Surveillance Team |
| EPI_ISL_2468583 | Lighthouse Lab in Milton Keynes | Wellcome Sanger Institute for the COVID-19 Genomics UK (COG-UK) Consortium | The Lighthouse Lab in Milton Keynes and Alex Alderton, Roberto Amato, Jeffrey Barrett, Sonia Goncalves, Ewan Harrison, David K. Jackson, Ian Johnston, Dominic Kwiatkowski, Cordelia Langford, John Sillitoe on behalf of the Wellcome Sanger Institute COVID-19 Surveillance Team |
| EPI_ISL_2468585, EPI_ISL_2468586 | Lighthouse Lab in Alderley Park | Wellcome Sanger Institute for the COVID-19 Genomics UK (COG-UK) Consortium | Jacquelyn Wynn, Mairead Hyland, The Lighthouse Lab in Alderley Park and Alex Alderton, Roberto Amato, Jeffrey Barrett, Sonia Goncalves, Ewan Harrison, David K. Jackson, Ian Johnston, Dominic Kwiatkowski, Cordelia Langford, John Sillitoe on behalf of the Wellcome Sanger Institute COVID-19 Surveillance Team |
| EPI_ISL_2468587, EPI_ISL_2468588 | Lighthouse Lab in Milton Keynes | Wellcome Sanger Institute for the COVID-19 Genomics UK (COG-UK) Consortium | The Lighthouse Lab in Milton Keynes and Alex Alderton, Roberto Amato, Jeffrey Barrett, Sonia Goncalves, Ewan Harrison, David K. Jackson, Ian Johnston, Dominic Kwiatkowski, Cordelia Langford, John Sillitoe on behalf of the Wellcome Sanger Institute COVID-19 Surveillance Team |
| EPI_ISL_2468589, EPI_ISL_2468592 | Lighthouse Lab in Alderley Park | Wellcome Sanger Institute for the COVID-19 Genomics UK (COG-UK) Consortium | Jacquelyn Wynn, Mairead Hyland, The Lighthouse Lab in Alderley Park and Alex Alderton, Roberto Amato, Jeffrey Barrett, Sonia Goncalves, Ewan Harrison, David K. Jackson, Ian Johnston, Dominic Kwiatkowski, Cordelia Langford, John Sillitoe on behalf of the Wellcome Sanger Institute COVID-19 Surveillance Team |
| EPI_ISL_2468593 | Lighthouse Lab in Milton Keynes | Wellcome Sanger Institute for the COVID-19 Genomics UK (COG-UK) Consortium | The Lighthouse Lab in Milton Keynes and Alex Alderton, Roberto Amato, Jeffrey Barrett, Sonia Goncalves, Ewan Harrison, David K. Jackson, Ian Johnston, Dominic Kwiatkowski, Cordelia Langford, John Sillitoe on behalf of the Wellcome Sanger Institute COVID-19 Surveillance Team |
| EPI_ISL_2468594 | Lighthouse Lab in Alderley Park | Wellcome Sanger Institute for the COVID-19 Genomics UK (COG-UK) Consortium | Jacquelyn Wynn, Mairead Hyland, The Lighthouse Lab in Alderley Park and Alex Alderton, Roberto Amato, Jeffrey Barrett, Sonia Goncalves, Ewan Harrison, David K. Jackson, Ian Johnston, Dominic Kwiatkowski, Cordelia Langford, John Sillitoe on behalf of the Wellcome Sanger Institute COVID-19 Surveillance Team |
| EPI_ISL_2468595 | Lighthouse Lab in Milton Keynes | Wellcome Sanger Institute for the COVID-19 Genomics UK (COG-UK) Consortium | The Lighthouse Lab in Milton Keynes and Alex Alderton, Roberto Amato, Jeffrey Barrett, Sonia Goncalves, Ewan Harrison, David K. Jackson, Ian Johnston, Dominic Kwiatkowski, Cordelia Langford, John Sillitoe on behalf of the Wellcome Sanger Institute COVID-19 Surveillance Team |
| EPI_ISL_2468596 | Lighthouse Lab in Alderley Park | Wellcome Sanger Institute for the COVID-19 Genomics UK (COG-UK) Consortium | Jacquelyn Wynn, Mairead Hyland, The Lighthouse Lab in Alderley Park and Alex Alderton, Roberto Amato, Jeffrey Barrett, Sonia Goncalves, Ewan Harrison, David K. Jackson, Ian Johnston, Dominic Kwiatkowski, Cordelia Langford, John Sillitoe on behalf of the Wellcome Sanger Institute COVID-19 Surveillance Team |
| EPI_ISL_2468597 | Lighthouse Lab in Milton Keynes | Wellcome Sanger Institute for the COVID-19 Genomics UK (COG-UK) Consortium | The Lighthouse Lab in Milton Keynes and Alex Alderton, Roberto Amato, Jeffrey Barrett, Sonia Goncalves, Ewan Harrison, David K. Jackson, Ian Johnston, Dominic Kwiatkowski, Cordelia Langford, John Sillitoe on behalf of the Wellcome Sanger Institute COVID-19 Surveillance Team |
| EPI_ISL_2468598, EPI_ISL_2468599, EPI_ISL_2468600 | Lighthouse Lab in Alderley Park | Wellcome Sanger Institute for the COVID-19 Genomics UK (COG-UK) Consortium | Jacquelyn Wynn, Mairead Hyland, The Lighthouse Lab in Alderley Park and Alex Alderton, Roberto Amato, Jeffrey Barrett, Sonia Goncalves, Ewan Harrison, David K. Jackson, Ian Johnston, Dominic Kwiatkowski, Cordelia Langford, John Sillitoe on behalf of the Wellcome Sanger Institute COVID-19 Surveillance Team |
| EPI_ISL_2468601 | Lighthouse Lab in Milton Keynes | Wellcome Sanger Institute for the COVID-19 Genomics UK (COG-UK) Consortium | The Lighthouse Lab in Milton Keynes and Alex Alderton, Roberto Amato, Jeffrey Barrett, Sonia Goncalves, Ewan Harrison, David K. Jackson, Ian Johnston, Dominic Kwiatkowski, Cordelia Langford, John Sillitoe on behalf of the Wellcome Sanger Institute COVID-19 Surveillance Team |
| EPI_ISL_2468602, EPI_ISL_2468603, EPI_ISL_2468604, EPI_ISL_2468606, EPI_ISL_2468607, EPI_ISL_2468608, EPI_ISL_2468609, EPI_ISL_2468610, EPI_ISL_2468611 | Lighthouse Lab in Alderley Park | Wellcome Sanger Institute for the COVID-19 Genomics UK (COG-UK) Consortium | Jacquelyn Wynn, Mairead Hyland, The Lighthouse Lab in Alderley Park and Alex Alderton, Roberto Amato, Jeffrey Barrett, Sonia Goncalves, Ewan Harrison, David K. Jackson, Ian Johnston, Dominic Kwiatkowski, Cordelia Langford, John Sillitoe on behalf of the Wellcome Sanger Institute COVID-19 Surveillance Team |
| EPI_ISL_2468612 | Lighthouse Lab in Milton Keynes | Wellcome Sanger Institute for the COVID-19 Genomics UK (COG-UK) Consortium | The Lighthouse Lab in Milton Keynes and Alex Alderton, Roberto Amato, Jeffrey Barrett, Sonia Goncalves, Ewan Harrison, David K. Jackson, Ian Johnston, Dominic Kwiatkowski, Cordelia Langford, John Sillitoe on behalf of the Wellcome Sanger Institute COVID-19 Surveillance Team |
| EPI_ISL_2468613, EPI_ISL_2468614 | Lighthouse Lab in Alderley Park | Wellcome Sanger Institute for the COVID-19 Genomics UK (COG-UK) Consortium | Jacquelyn Wynn, Mairead Hyland, The Lighthouse Lab in Alderley Park and Alex Alderton, Roberto Amato, Jeffrey Barrett, Sonia Goncalves, Ewan Harrison, David K. Jackson, Ian Johnston, Dominic Kwiatkowski, Cordelia Langford, John Sillitoe on behalf of the Wellcome Sanger Institute COVID-19 Surveillance Team |
| EPI_ISL_2468615 | Lighthouse Lab in Milton Keynes | Wellcome Sanger Institute for the COVID-19 Genomics UK (COG-UK) Consortium | The Lighthouse Lab in Milton Keynes and Alex Alderton, Roberto Amato, Jeffrey Barrett, Sonia Goncalves, Ewan Harrison, David K. Jackson, Ian Johnston, Dominic Kwiatkowski, Cordelia Langford, John Sillitoe on behalf of the Wellcome Sanger Institute COVID-19 Surveillance Team |

| | | | |
|---|---|---|---|
| EPI_ISL_2468616, EPI_ISL_2468617, EPI_ISL_2468618, EPI_ISL_2468619 | Lighthouse Lab in Alderley Park | Wellcome Sanger Institute for the COVID-19 Genomics UK (COG-UK) Consortium | Jacquelyn Wynn, Mairead Hyland, The Lighthouse Lab in Alderley Park and Alex Alderton, Roberto Amato, Jeffrey Barrett, Sonia Goncalves, Ewan Harrison, David K. Jackson, Ian Johnston, Dominic Kwiatkowski, Cordelia Langford, John Sillitoe on behalf of the Wellcome Sanger Institute COVID-19 Surveillance Team |
| EPI_ISL_2468620 | Lighthouse Lab in Milton Keynes | Wellcome Sanger Institute for the COVID-19 Genomics UK (COG-UK) Consortium | The Lighthouse Lab in Milton Keynes and Alex Alderton, Roberto Amato, Jeffrey Barrett, Sonia Goncalves, Ewan Harrison, David K. Jackson, Ian Johnston, Dominic Kwiatkowski, Cordelia Langford, John Sillitoe on behalf of the Wellcome Sanger Institute COVID-19 Surveillance Team |
| EPI_ISL_2468622 | Lighthouse Lab in Alderley Park | Wellcome Sanger Institute for the COVID-19 Genomics UK (COG-UK) Consortium | Jacquelyn Wynn, Mairead Hyland, The Lighthouse Lab in Alderley Park and Alex Alderton, Roberto Amato, Jeffrey Barrett, Sonia Goncalves, Ewan Harrison, David K. Jackson, Ian Johnston, Dominic Kwiatkowski, Cordelia Langford, John Sillitoe on behalf of the Wellcome Sanger Institute COVID-19 Surveillance Team |
| EPI_ISL_2468623 | Lighthouse Lab in Milton Keynes | Wellcome Sanger Institute for the COVID-19 Genomics UK (COG-UK) Consortium | The Lighthouse Lab in Milton Keynes and Alex Alderton, Roberto Amato, Jeffrey Barrett, Sonia Goncalves, Ewan Harrison, David K. Jackson, Ian Johnston, Dominic Kwiatkowski, Cordelia Langford, John Sillitoe on behalf of the Wellcome Sanger Institute COVID-19 Surveillance Team |
| EPI_ISL_2468624, EPI_ISL_2468625 | Lighthouse Lab in Alderley Park | Wellcome Sanger Institute for the COVID-19 Genomics UK (COG-UK) Consortium | Jacquelyn Wynn, Mairead Hyland, The Lighthouse Lab in Alderley Park and Alex Alderton, Roberto Amato, Jeffrey Barrett, Sonia Goncalves, Ewan Harrison, David K. Jackson, Ian Johnston, Dominic Kwiatkowski, Cordelia Langford, John Sillitoe on behalf of the Wellcome Sanger Institute COVID-19 Surveillance Team |
| EPI_ISL_2468627 | Lighthouse Lab in Milton Keynes | Wellcome Sanger Institute for the COVID-19 Genomics UK (COG-UK) Consortium | The Lighthouse Lab in Milton Keynes and Alex Alderton, Roberto Amato, Jeffrey Barrett, Sonia Goncalves, Ewan Harrison, David K. Jackson, Ian Johnston, Dominic Kwiatkowski, Cordelia Langford, John Sillitoe on behalf of the Wellcome Sanger Institute COVID-19 Surveillance Team |
| EPI_ISL_2468628, EPI_ISL_2468629, EPI_ISL_2468630, EPI_ISL_2468632, EPI_ISL_2468633, EPI_ISL_2468634, EPI_ISL_2468635 | Lighthouse Lab in Alderley Park | Wellcome Sanger Institute for the COVID-19 Genomics UK (COG-UK) Consortium | Jacquelyn Wynn, Mairead Hyland, The Lighthouse Lab in Alderley Park and Alex Alderton, Roberto Amato, Jeffrey Barrett, Sonia Goncalves, Ewan Harrison, David K. Jackson, Ian Johnston, Dominic Kwiatkowski, Cordelia Langford, John Sillitoe on behalf of the Wellcome Sanger Institute COVID-19 Surveillance Team |
| EPI_ISL_2468636 | Lighthouse Lab in Milton Keynes | Wellcome Sanger Institute for the COVID-19 Genomics UK (COG-UK) Consortium | The Lighthouse Lab in Milton Keynes and Alex Alderton, Roberto Amato, Jeffrey Barrett, Sonia Goncalves, Ewan Harrison, David K. Jackson, Ian Johnston, Dominic Kwiatkowski, Cordelia Langford, John Sillitoe on behalf of the Wellcome Sanger Institute COVID-19 Surveillance Team |
| EPI_ISL_2468637 | Lighthouse Lab in Alderley Park | Wellcome Sanger Institute for the COVID-19 Genomics UK (COG-UK) Consortium | Jacquelyn Wynn, Mairead Hyland, The Lighthouse Lab in Alderley Park and Alex Alderton, Roberto Amato, Jeffrey Barrett, Sonia Goncalves, Ewan Harrison, David K. Jackson, Ian Johnston, Dominic Kwiatkowski, Cordelia Langford, John Sillitoe on behalf of the Wellcome Sanger Institute COVID-19 Surveillance Team |
| EPI_ISL_2468638 | Lighthouse Lab in Milton Keynes | Wellcome Sanger Institute for the COVID-19 Genomics UK (COG-UK) Consortium | The Lighthouse Lab in Milton Keynes and Alex Alderton, Roberto Amato, Jeffrey Barrett, Sonia Goncalves, Ewan Harrison, David K. Jackson, Ian Johnston, Dominic Kwiatkowski, Cordelia Langford, John Sillitoe on behalf of the Wellcome Sanger Institute COVID-19 Surveillance Team |
| EPI_ISL_2468639, EPI_ISL_2468641 | Lighthouse Lab in Alderley Park | Wellcome Sanger Institute for the COVID-19 Genomics UK (COG-UK) Consortium | Jacquelyn Wynn, Mairead Hyland, The Lighthouse Lab in Alderley Park and Alex Alderton, Roberto Amato, Jeffrey Barrett, Sonia Goncalves, Ewan Harrison, David K. Jackson, Ian Johnston, Dominic Kwiatkowski, Cordelia Langford, John Sillitoe on behalf of the Wellcome Sanger Institute COVID-19 Surveillance Team |
| EPI_ISL_2468642 | Lighthouse Lab in Milton Keynes | Wellcome Sanger Institute for the COVID-19 Genomics UK (COG-UK) Consortium | The Lighthouse Lab in Milton Keynes and Alex Alderton, Roberto Amato, Jeffrey Barrett, Sonia Goncalves, Ewan Harrison, David K. Jackson, Ian Johnston, Dominic Kwiatkowski, Cordelia Langford, John Sillitoe on behalf of the Wellcome Sanger Institute COVID-19 Surveillance Team |
| EPI_ISL_2468643 | Lighthouse Lab in Alderley Park | Wellcome Sanger Institute for the COVID-19 Genomics UK (COG-UK) Consortium | Jacquelyn Wynn, Mairead Hyland, The Lighthouse Lab in Alderley Park and Alex Alderton, Roberto Amato, Jeffrey Barrett, Sonia Goncalves, Ewan Harrison, David K. Jackson, Ian Johnston, Dominic Kwiatkowski, Cordelia Langford, John Sillitoe on behalf of the Wellcome Sanger Institute COVID-19 Surveillance Team |
| EPI_ISL_2468644 | Lighthouse Lab in Milton Keynes | Wellcome Sanger Institute for the COVID-19 Genomics UK (COG-UK) Consortium | The Lighthouse Lab in Milton Keynes and Alex Alderton, Roberto Amato, Jeffrey Barrett, Sonia Goncalves, Ewan Harrison, David K. Jackson, Ian Johnston, Dominic Kwiatkowski, Cordelia Langford, John Sillitoe on behalf of the Wellcome Sanger Institute COVID-19 Surveillance Team |
| EPI_ISL_2468645, EPI_ISL_2468646 | Lighthouse Lab in Alderley Park | Wellcome Sanger Institute for the COVID-19 Genomics UK (COG-UK) Consortium | Jacquelyn Wynn, Mairead Hyland, The Lighthouse Lab in Alderley Park and Alex Alderton, Roberto Amato, Jeffrey Barrett, Sonia Goncalves, Ewan Harrison, David K. Jackson, Ian Johnston, Dominic Kwiatkowski, Cordelia Langford, John Sillitoe on behalf of the Wellcome Sanger Institute COVID-19 Surveillance Team |
| EPI_ISL_2468647, EPI_ISL_2468648 | Lighthouse Lab in Milton Keynes | Wellcome Sanger Institute for the COVID-19 Genomics UK (COG-UK) Consortium | The Lighthouse Lab in Milton Keynes and Alex Alderton, Roberto Amato, Jeffrey Barrett, Sonia Goncalves, Ewan Harrison, David K. Jackson, Ian Johnston, Dominic Kwiatkowski, Cordelia Langford, John Sillitoe on behalf of the Wellcome Sanger Institute COVID-19 Surveillance Team |
| EPI_ISL_2468649, EPI_ISL_2468650, EPI_ISL_2468651, EPI_ISL_2468652, EPI_ISL_2468653, EPI_ISL_2468654, EPI_ISL_2468655, EPI_ISL_2468658 | Lighthouse Lab in Alderley Park | Wellcome Sanger Institute for the COVID-19 Genomics UK (COG-UK) Consortium | Jacquelyn Wynn, Mairead Hyland, The Lighthouse Lab in Alderley Park and Alex Alderton, Roberto Amato, Jeffrey Barrett, Sonia Goncalves, Ewan Harrison, David K. Jackson, Ian Johnston, Dominic Kwiatkowski, Cordelia Langford, John Sillitoe on behalf of the Wellcome Sanger Institute COVID-19 Surveillance Team |
| EPI_ISL_2468659 | Lighthouse Lab in Milton Keynes | Wellcome Sanger Institute for the COVID-19 Genomics UK (COG-UK) Consortium | The Lighthouse Lab in Milton Keynes and Alex Alderton, Roberto Amato, Jeffrey Barrett, Sonia Goncalves, Ewan Harrison, David K. Jackson, Ian Johnston, Dominic Kwiatkowski, Cordelia Langford, John Sillitoe on behalf of the Wellcome Sanger Institute COVID-19 Surveillance Team |
| EPI_ISL_2468660 | Lighthouse Lab in Alderley Park | Wellcome Sanger Institute for the COVID-19 Genomics UK (COG-UK) Consortium | Jacquelyn Wynn, Mairead Hyland, The Lighthouse Lab in Alderley Park and Alex Alderton, Roberto Amato, Jeffrey Barrett, Sonia Goncalves, Ewan Harrison, David K. Jackson, Ian Johnston, Dominic Kwiatkowski, Cordelia Langford, John Sillitoe on behalf of the Wellcome Sanger Institute COVID-19 Surveillance Team |
| EPI_ISL_2468661 | Lighthouse Lab in Milton Keynes | Wellcome Sanger Institute for the COVID-19 Genomics UK (COG-UK) Consortium | The Lighthouse Lab in Milton Keynes and Alex Alderton, Roberto Amato, Jeffrey Barrett, Sonia Goncalves, Ewan Harrison, David K. Jackson, Ian Johnston, Dominic Kwiatkowski, Cordelia Langford, John Sillitoe on behalf of the Wellcome Sanger Institute COVID-19 Surveillance Team |
| EPI_ISL_2468662, EPI_ISL_2468663, EPI_ISL_2468664, EPI_ISL_2468665 | Lighthouse Lab in Alderley Park | Wellcome Sanger Institute for the COVID-19 Genomics UK (COG-UK) Consortium | Jacquelyn Wynn, Mairead Hyland, The Lighthouse Lab in Alderley Park and Alex Alderton, Roberto Amato, Jeffrey Barrett, Sonia Goncalves, Ewan Harrison, David K. Jackson, Ian Johnston, Dominic Kwiatkowski, Cordelia Langford, John Sillitoe on behalf of the Wellcome Sanger Institute COVID-19 Surveillance Team |
| EPI_ISL_2468666 | Lighthouse Lab in Milton Keynes | Wellcome Sanger Institute for the COVID-19 Genomics UK (COG-UK) Consortium | The Lighthouse Lab in Milton Keynes and Alex Alderton, Roberto Amato, Jeffrey Barrett, Sonia Goncalves, Ewan Harrison, David K. Jackson, Ian Johnston, Dominic Kwiatkowski, Cordelia Langford, John Sillitoe on behalf of the Wellcome Sanger Institute COVID-19 Surveillance Team |
| EPI_ISL_2468667, EPI_ISL_2468668 | Lighthouse Lab in Alderley Park | Wellcome Sanger Institute for the COVID-19 Genomics UK (COG-UK) Consortium | Jacquelyn Wynn, Mairead Hyland, The Lighthouse Lab in Alderley Park and Alex Alderton, Roberto Amato, Jeffrey Barrett, Sonia Goncalves, Ewan Harrison, David K. Jackson, Ian Johnston, Dominic Kwiatkowski, Cordelia Langford, John Sillitoe on behalf of the Wellcome Sanger Institute COVID-19 Surveillance Team |
| EPI_ISL_2468669 | Lighthouse Lab in Milton Keynes | Wellcome Sanger Institute for the COVID-19 Genomics UK (COG-UK) Consortium | The Lighthouse Lab in Milton Keynes and Alex Alderton, Roberto Amato, Jeffrey Barrett, Sonia Goncalves, Ewan Harrison, David K. Jackson, Ian Johnston, Dominic Kwiatkowski, Cordelia Langford, John Sillitoe on behalf of the Wellcome Sanger Institute COVID-19 Surveillance Team |
| EPI_ISL_2468670, EPI_ISL_2468671, EPI_ISL_2468672 | Lighthouse Lab in Alderley Park | Wellcome Sanger Institute for the COVID-19 Genomics UK (COG-UK) Consortium | Jacquelyn Wynn, Mairead Hyland, The Lighthouse Lab in Alderley Park and Alex Alderton, Roberto Amato, Jeffrey Barrett, Sonia Goncalves, Ewan Harrison, David K. Jackson, Ian Johnston, Dominic Kwiatkowski, Cordelia Langford, John Sillitoe on behalf of the Wellcome Sanger Institute COVID-19 Surveillance Team |
| EPI_ISL_2468673, EPI_ISL_2468674 | Lighthouse Lab in Milton Keynes | Wellcome Sanger Institute for the COVID-19 Genomics UK (COG-UK) Consortium | The Lighthouse Lab in Milton Keynes and Alex Alderton, Roberto Amato, Jeffrey Barrett, Sonia Goncalves, Ewan Harrison, David K. Jackson, Ian Johnston, Dominic Kwiatkowski, Cordelia Langford, John Sillitoe on behalf of the Wellcome Sanger Institute COVID-19 Surveillance Team |
| EPI_ISL_2468676, EPI_ISL_2468677, EPI_ISL_2468679, EPI_ISL_2468680, EPI_ISL_2468681 | Lighthouse Lab in Alderley Park | Wellcome Sanger Institute for the COVID-19 Genomics UK (COG-UK) Consortium | Jacquelyn Wynn, Mairead Hyland, The Lighthouse Lab in Alderley Park and Alex Alderton, Roberto Amato, Jeffrey Barrett, Sonia Goncalves, Ewan Harrison, David K. Jackson, Ian Johnston, Dominic Kwiatkowski, Cordelia Langford, John Sillitoe on behalf of the Wellcome Sanger Institute COVID-19 Surveillance Team |
| EPI_ISL_2468682 | Lighthouse Lab in Milton Keynes | Wellcome Sanger Institute for the COVID-19 Genomics UK (COG-UK) Consortium | The Lighthouse Lab in Milton Keynes and Alex Alderton, Roberto Amato, Jeffrey Barrett, Sonia Goncalves, Ewan Harrison, David K. Jackson, Ian Johnston, Dominic Kwiatkowski, Cordelia Langford, John Sillitoe on behalf of the Wellcome Sanger Institute COVID-19 Surveillance Team |
| EPI_ISL_2468683, EPI_ISL_2468684, EPI_ISL_2468685 | Lighthouse Lab in Alderley Park | Wellcome Sanger Institute for the COVID-19 Genomics UK (COG-UK) Consortium | Jacquelyn Wynn, Mairead Hyland, The Lighthouse Lab in Alderley Park and Alex Alderton, Roberto Amato, Jeffrey Barrett, Sonia Goncalves, Ewan Harrison, David K. Jackson, Ian Johnston, Dominic Kwiatkowski, Cordelia Langford, John Sillitoe on behalf of the Wellcome Sanger Institute COVID-19 Surveillance Team |
| EPI_ISL_2468686 | Lighthouse Lab in Milton Keynes | Wellcome Sanger Institute for the COVID-19 Genomics UK (COG-UK) Consortium | The Lighthouse Lab in Milton Keynes and Alex Alderton, Roberto Amato, Jeffrey Barrett, Sonia Goncalves, Ewan Harrison, David K. Jackson, Ian Johnston, Dominic Kwiatkowski, Cordelia Langford, John Sillitoe on behalf of the Wellcome Sanger Institute COVID-19 Surveillance Team |
| EPI_ISL_2468687 | Lighthouse Lab in Alderley Park | Wellcome Sanger Institute for the COVID-19 Genomics UK (COG-UK) Consortium | Jacquelyn Wynn, Mairead Hyland, The Lighthouse Lab in Alderley Park and Alex Alderton, Roberto Amato, Jeffrey Barrett, Sonia Goncalves, Ewan Harrison, David K. Jackson, Ian Johnston, Dominic Kwiatkowski, Cordelia Langford, John Sillitoe on behalf of the Wellcome Sanger Institute COVID-19 Surveillance Team |
| EPI_ISL_2468688, EPI_ISL_2468689 | Lighthouse Lab in Milton Keynes | Wellcome Sanger Institute for the COVID-19 Genomics UK (COG-UK) Consortium | The Lighthouse Lab in Milton Keynes and Alex Alderton, Roberto Amato, Jeffrey Barrett, Sonia Goncalves, Ewan Harrison, David K. Jackson, Ian Johnston, Dominic Kwiatkowski, Cordelia Langford, John Sillitoe on behalf of the Wellcome Sanger Institute COVID-19 Surveillance Team |
| EPI_ISL_2468690 | Lighthouse Lab in Alderley Park | Wellcome Sanger Institute for the COVID-19 Genomics UK (COG-UK) Consortium | Jacquelyn Wynn, Mairead Hyland, The Lighthouse Lab in Alderley Park and Alex Alderton, Roberto Amato, Jeffrey Barrett, Sonia Goncalves, Ewan Harrison, David K. Jackson, Ian Johnston, Dominic Kwiatkowski, Cordelia Langford, John Sillitoe on behalf of the Wellcome Sanger Institute COVID-19 Surveillance Team |
| EPI_ISL_2468691 | Lighthouse Lab in Milton Keynes | Wellcome Sanger Institute for the COVID-19 Genomics UK (COG-UK) Consortium | The Lighthouse Lab in Milton Keynes and Alex Alderton, Roberto Amato, Jeffrey Barrett, Sonia Goncalves, Ewan Harrison, David K. Jackson, Ian Johnston, Dominic Kwiatkowski, Cordelia Langford, John Sillitoe on behalf of the Wellcome Sanger Institute COVID-19 Surveillance Team |
| EPI_ISL_2468692, EPI_ISL_2468693, EPI_ISL_2468694 | Lighthouse Lab in Alderley Park | Wellcome Sanger Institute for the COVID-19 Genomics UK (COG-UK) Consortium | Jacquelyn Wynn, Mairead Hyland, The Lighthouse Lab in Alderley Park and Alex Alderton, Roberto Amato, Jeffrey Barrett, Sonia Goncalves, Ewan Harrison, David K. Jackson, Ian Johnston, Dominic Kwiatkowski, Cordelia Langford, John Sillitoe on behalf of the Wellcome Sanger Institute COVID-19 Surveillance Team |
| EPI_ISL_2468695 | Lighthouse Lab in Milton Keynes | Wellcome Sanger Institute for the COVID-19 Genomics UK (COG-UK) Consortium | The Lighthouse Lab in Milton Keynes and Alex Alderton, Roberto Amato, Jeffrey Barrett, Sonia Goncalves, Ewan Harrison, David K. Jackson, Ian Johnston, Dominic Kwiatkowski, Cordelia Langford, John Sillitoe on behalf of the Wellcome Sanger Institute COVID-19 Surveillance Team |

EPI_ISL_2468696, EPI_ISL_2468697, EPI_ISL_2468698, EPI_ISL_2468699, EPI_ISL_2468700, EPI_ISL_2468701, EPI_ISL_2468702, EPI_ISL_2468703, EPI_ISL_2468704, EPI_ISL_2468705, EPI_ISL_2468706, EPI_ISL_2468707

| | | | |
|---|---|---|---|
| see above | Lighthouse Lab in Alderley Park | Wellcome Sanger Institute for the COVID-19 Genomics UK (COG-UK) Consortium | Jacquelyn Wynn, Mairead Hyland, The Lighthouse Lab in Alderley Park and Alex Alderton, Roberto Amato, Jeffrey Barrett, Sonia Goncalves, Ewan Harrison, David K. Jackson, Ian Johnston, Dominic Kwiatkowski, Cordelia Langford, John Sillitoe on behalf of the Wellcome Sanger Institute COVID-19 Surveillance Team |
| EPI_ISL_2468709 | Lighthouse Lab in Milton Keynes | Wellcome Sanger Institute for the COVID-19 Genomics UK (COG-UK) Consortium | The Lighthouse Lab in Milton Keynes and Alex Alderton, Roberto Amato, Jeffrey Barrett, Sonia Goncalves, Ewan Harrison, David K. Jackson, Ian Johnston, Dominic Kwiatkowski, Cordelia Langford, John Sillitoe on behalf of the Wellcome Sanger Institute COVID-19 Surveillance Team |
| EPI_ISL_2468710, EPI_ISL_2468711, EPI_ISL_2468713, EPI_ISL_2468714 | Lighthouse Lab in Alderley Park | Wellcome Sanger Institute for the COVID-19 Genomics UK (COG-UK) Consortium | Jacquelyn Wynn, Mairead Hyland, The Lighthouse Lab in Alderley Park and Alex Alderton, Roberto Amato, Jeffrey Barrett, Sonia Goncalves, Ewan Harrison, David K. Jackson, Ian Johnston, Dominic Kwiatkowski, Cordelia Langford, John Sillitoe on behalf of the Wellcome Sanger Institute COVID-19 Surveillance Team |
| EPI_ISL_2468715 | Lighthouse Lab in Milton Keynes | Wellcome Sanger Institute for the COVID-19 Genomics UK (COG-UK) Consortium | The Lighthouse Lab in Milton Keynes and Alex Alderton, Roberto Amato, Jeffrey Barrett, Sonia Goncalves, Ewan Harrison, David K. Jackson, Ian Johnston, Dominic Kwiatkowski, Cordelia Langford, John Sillitoe on behalf of the Wellcome Sanger Institute COVID-19 Surveillance Team |
| EPI_ISL_2468717, EPI_ISL_2468718, EPI_ISL_2468719, EPI_ISL_2468720, EPI_ISL_2468721 | Lighthouse Lab in Alderley Park | Wellcome Sanger Institute for the COVID-19 Genomics UK (COG-UK) Consortium | Jacquelyn Wynn, Mairead Hyland, The Lighthouse Lab in Alderley Park and Alex Alderton, Roberto Amato, Jeffrey Barrett, Sonia Goncalves, Ewan Harrison, David K. Jackson, Ian Johnston, Dominic Kwiatkowski, Cordelia Langford, John Sillitoe on behalf of the Wellcome Sanger Institute COVID-19 Surveillance Team |
| EPI_ISL_2468722 | Lighthouse Lab in Milton Keynes | Wellcome Sanger Institute for the COVID-19 Genomics UK (COG-UK) Consortium | The Lighthouse Lab in Milton Keynes and Alex Alderton, Roberto Amato, Jeffrey Barrett, Sonia Goncalves, Ewan Harrison, David K. Jackson, Ian Johnston, Dominic Kwiatkowski, Cordelia Langford, John Sillitoe on behalf of the Wellcome Sanger Institute COVID-19 Surveillance Team |
| EPI_ISL_2468723 | Lighthouse Lab in Alderley Park | Wellcome Sanger Institute for the COVID-19 Genomics UK (COG-UK) Consortium | Jacquelyn Wynn, Mairead Hyland, The Lighthouse Lab in Alderley Park and Alex Alderton, Roberto Amato, Jeffrey Barrett, Sonia Goncalves, Ewan Harrison, David K. Jackson, Ian Johnston, Dominic Kwiatkowski, Cordelia Langford, John Sillitoe on behalf of the Wellcome Sanger Institute COVID-19 Surveillance Team |
| EPI_ISL_2468724, EPI_ISL_2468725 | Lighthouse Lab in Milton Keynes | Wellcome Sanger Institute for the COVID-19 Genomics UK (COG-UK) Consortium | The Lighthouse Lab in Milton Keynes and Alex Alderton, Roberto Amato, Jeffrey Barrett, Sonia Goncalves, Ewan Harrison, David K. Jackson, Ian Johnston, Dominic Kwiatkowski, Cordelia Langford, John Sillitoe on behalf of the Wellcome Sanger Institute COVID-19 Surveillance Team |
| EPI_ISL_2468726, EPI_ISL_2468727, EPI_ISL_2468728, EPI_ISL_2468729, EPI_ISL_2468730, EPI_ISL_2468733 | Lighthouse Lab in Alderley Park | Wellcome Sanger Institute for the COVID-19 Genomics UK (COG-UK) Consortium | Jacquelyn Wynn, Mairead Hyland, The Lighthouse Lab in Alderley Park and Alex Alderton, Roberto Amato, Jeffrey Barrett, Sonia Goncalves, Ewan Harrison, David K. Jackson, Ian Johnston, Dominic Kwiatkowski, Cordelia Langford, John Sillitoe on behalf of the Wellcome Sanger Institute COVID-19 Surveillance Team |
| EPI_ISL_2468734 | Lighthouse Lab in Milton Keynes | Wellcome Sanger Institute for the COVID-19 Genomics UK (COG-UK) Consortium | The Lighthouse Lab in Milton Keynes and Alex Alderton, Roberto Amato, Jeffrey Barrett, Sonia Goncalves, Ewan Harrison, David K. Jackson, Ian Johnston, Dominic Kwiatkowski, Cordelia Langford, John Sillitoe on behalf of the Wellcome Sanger Institute COVID-19 Surveillance Team |
| EPI_ISL_2468735 | Lighthouse Lab in Alderley Park | Wellcome Sanger Institute for the COVID-19 Genomics UK (COG-UK) Consortium | Jacquelyn Wynn, Mairead Hyland, The Lighthouse Lab in Alderley Park and Alex Alderton, Roberto Amato, Jeffrey Barrett, Sonia Goncalves, Ewan Harrison, David K. Jackson, Ian Johnston, Dominic Kwiatkowski, Cordelia Langford, John Sillitoe on behalf of the Wellcome Sanger Institute COVID-19 Surveillance Team |
| EPI_ISL_2468736 | Lighthouse Lab in Milton Keynes | Wellcome Sanger Institute for the COVID-19 Genomics UK (COG-UK) Consortium | The Lighthouse Lab in Milton Keynes and Alex Alderton, Roberto Amato, Jeffrey Barrett, Sonia Goncalves, Ewan Harrison, David K. Jackson, Ian Johnston, Dominic Kwiatkowski, Cordelia Langford, John Sillitoe on behalf of the Wellcome Sanger Institute COVID-19 Surveillance Team |
| EPI_ISL_2468737, EPI_ISL_2468738, EPI_ISL_2468739, EPI_ISL_2468740, EPI_ISL_2468741, EPI_ISL_2468742, EPI_ISL_2468743 | Lighthouse Lab in Alderley Park | Wellcome Sanger Institute for the COVID-19 Genomics UK (COG-UK) Consortium | Jacquelyn Wynn, Mairead Hyland, The Lighthouse Lab in Alderley Park and Alex Alderton, Roberto Amato, Jeffrey Barrett, Sonia Goncalves, Ewan Harrison, David K. Jackson, Ian Johnston, Dominic Kwiatkowski, Cordelia Langford, John Sillitoe on behalf of the Wellcome Sanger Institute COVID-19 Surveillance Team |
| EPI_ISL_2468744 | Lighthouse Lab in Milton Keynes | Wellcome Sanger Institute for the COVID-19 Genomics UK (COG-UK) Consortium | The Lighthouse Lab in Milton Keynes and Alex Alderton, Roberto Amato, Jeffrey Barrett, Sonia Goncalves, Ewan Harrison, David K. Jackson, Ian Johnston, Dominic Kwiatkowski, Cordelia Langford, John Sillitoe on behalf of the Wellcome Sanger Institute COVID-19 Surveillance Team |
| EPI_ISL_2468745, EPI_ISL_2468746, EPI_ISL_2468747, EPI_ISL_2468748, EPI_ISL_2468749 | Lighthouse Lab in Alderley Park | Wellcome Sanger Institute for the COVID-19 Genomics UK (COG-UK) Consortium | Jacquelyn Wynn, Mairead Hyland, The Lighthouse Lab in Alderley Park and Alex Alderton, Roberto Amato, Jeffrey Barrett, Sonia Goncalves, Ewan Harrison, David K. Jackson, Ian Johnston, Dominic Kwiatkowski, Cordelia Langford, John Sillitoe on behalf of the Wellcome Sanger Institute COVID-19 Surveillance Team |
| EPI_ISL_2468750 | Lighthouse Lab in Milton Keynes | Wellcome Sanger Institute for the COVID-19 Genomics UK (COG-UK) Consortium | The Lighthouse Lab in Milton Keynes and Alex Alderton, Roberto Amato, Jeffrey Barrett, Sonia Goncalves, Ewan Harrison, David K. Jackson, Ian Johnston, Dominic Kwiatkowski, Cordelia Langford, John Sillitoe on behalf of the Wellcome Sanger Institute COVID-19 Surveillance Team |
| EPI_ISL_2468751, EPI_ISL_2468752, EPI_ISL_2468753, EPI_ISL_2468754 | Lighthouse Lab in Alderley Park | Wellcome Sanger Institute for the COVID-19 Genomics UK (COG-UK) Consortium | Jacquelyn Wynn, Mairead Hyland, The Lighthouse Lab in Alderley Park and Alex Alderton, Roberto Amato, Jeffrey Barrett, Sonia Goncalves, Ewan Harrison, David K. Jackson, Ian Johnston, Dominic Kwiatkowski, Cordelia Langford, John Sillitoe on behalf of the Wellcome Sanger Institute COVID-19 Surveillance Team |
| EPI_ISL_2468755 | Lighthouse Lab in Milton Keynes | Wellcome Sanger Institute for the COVID-19 Genomics UK (COG-UK) Consortium | The Lighthouse Lab in Milton Keynes and Alex Alderton, Roberto Amato, Jeffrey Barrett, Sonia Goncalves, Ewan Harrison, David K. Jackson, Ian Johnston, Dominic Kwiatkowski, Cordelia Langford, John Sillitoe on behalf of the Wellcome Sanger Institute COVID-19 Surveillance Team |
| EPI_ISL_2468756, EPI_ISL_2468757, EPI_ISL_2468758 | Lighthouse Lab in Alderley Park | Wellcome Sanger Institute for the COVID-19 Genomics UK (COG-UK) Consortium | Jacquelyn Wynn, Mairead Hyland, The Lighthouse Lab in Alderley Park and Alex Alderton, Roberto Amato, Jeffrey Barrett, Sonia Goncalves, Ewan Harrison, David K. Jackson, Ian Johnston, Dominic Kwiatkowski, Cordelia Langford, John Sillitoe on behalf of the Wellcome Sanger Institute COVID-19 Surveillance Team |
| EPI_ISL_2468759 | Lighthouse Lab in Milton Keynes | Wellcome Sanger Institute for the COVID-19 Genomics UK (COG-UK) Consortium | The Lighthouse Lab in Milton Keynes and Alex Alderton, Roberto Amato, Jeffrey Barrett, Sonia Goncalves, Ewan Harrison, David K. Jackson, Ian Johnston, Dominic Kwiatkowski, Cordelia Langford, John Sillitoe on behalf of the Wellcome Sanger Institute COVID-19 Surveillance Team |
| EPI_ISL_2468760, EPI_ISL_2468761, EPI_ISL_2468762 | Lighthouse Lab in Alderley Park | Wellcome Sanger Institute for the COVID-19 Genomics UK (COG-UK) Consortium | Jacquelyn Wynn, Mairead Hyland, The Lighthouse Lab in Alderley Park and Alex Alderton, Roberto Amato, Jeffrey Barrett, Sonia Goncalves, Ewan Harrison, David K. Jackson, Ian Johnston, Dominic Kwiatkowski, Cordelia Langford, John Sillitoe on behalf of the Wellcome Sanger Institute COVID-19 Surveillance Team |
| EPI_ISL_2468763 | Lighthouse Lab in Milton Keynes | Wellcome Sanger Institute for the COVID-19 Genomics UK (COG-UK) Consortium | The Lighthouse Lab in Milton Keynes and Alex Alderton, Roberto Amato, Jeffrey Barrett, Sonia Goncalves, Ewan Harrison, David K. Jackson, Ian Johnston, Dominic Kwiatkowski, Cordelia Langford, John Sillitoe on behalf of the Wellcome Sanger Institute COVID-19 Surveillance Team |
| EPI_ISL_2468764, EPI_ISL_2468766, EPI_ISL_2468767, EPI_ISL_2468768, EPI_ISL_2468769, EPI_ISL_2468770, EPI_ISL_2468771, EPI_ISL_2468772, EPI_ISL_2468774, EPI_ISL_2468775, EPI_ISL_2468776 | | | |
| see above | Lighthouse Lab in Alderley Park | Wellcome Sanger Institute for the COVID-19 Genomics UK (COG-UK) Consortium | Jacquelyn Wynn, Mairead Hyland, The Lighthouse Lab in Alderley Park and Alex Alderton, Roberto Amato, Jeffrey Barrett, Sonia Goncalves, Ewan Harrison, David K. Jackson, Ian Johnston, Dominic Kwiatkowski, Cordelia Langford, John Sillitoe on behalf of the Wellcome Sanger Institute COVID-19 Surveillance Team |
| EPI_ISL_2468777 | Lighthouse Lab in Milton Keynes | Wellcome Sanger Institute for the COVID-19 Genomics UK (COG-UK) Consortium | The Lighthouse Lab in Milton Keynes and Alex Alderton, Roberto Amato, Jeffrey Barrett, Sonia Goncalves, Ewan Harrison, David K. Jackson, Ian Johnston, Dominic Kwiatkowski, Cordelia Langford, John Sillitoe on behalf of the Wellcome Sanger Institute COVID-19 Surveillance Team |
| EPI_ISL_2468778, EPI_ISL_2468779, EPI_ISL_2468780, EPI_ISL_2468781, EPI_ISL_2468782 | Lighthouse Lab in Alderley Park | Wellcome Sanger Institute for the COVID-19 Genomics UK (COG-UK) Consortium | Jacquelyn Wynn, Mairead Hyland, The Lighthouse Lab in Alderley Park and Alex Alderton, Roberto Amato, Jeffrey Barrett, Sonia Goncalves, Ewan Harrison, David K. Jackson, Ian Johnston, Dominic Kwiatkowski, Cordelia Langford, John Sillitoe on behalf of the Wellcome Sanger Institute COVID-19 Surveillance Team |
| EPI_ISL_2468784 | Lighthouse Lab in Milton Keynes | Wellcome Sanger Institute for the COVID-19 Genomics UK (COG-UK) Consortium | The Lighthouse Lab in Milton Keynes and Alex Alderton, Roberto Amato, Jeffrey Barrett, Sonia Goncalves, Ewan Harrison, David K. Jackson, Ian Johnston, Dominic Kwiatkowski, Cordelia Langford, John Sillitoe on behalf of the Wellcome Sanger Institute COVID-19 Surveillance Team |
| EPI_ISL_2468785 | Lighthouse Lab in Alderley Park | Wellcome Sanger Institute for the COVID-19 Genomics UK (COG-UK) Consortium | Jacquelyn Wynn, Mairead Hyland, The Lighthouse Lab in Alderley Park and Alex Alderton, Roberto Amato, Jeffrey Barrett, Sonia Goncalves, Ewan Harrison, David K. Jackson, Ian Johnston, Dominic Kwiatkowski, Cordelia Langford, John Sillitoe on behalf of the Wellcome Sanger Institute COVID-19 Surveillance Team |
| EPI_ISL_2468786 | Lighthouse Lab in Milton Keynes | Wellcome Sanger Institute for the COVID-19 Genomics UK (COG-UK) Consortium | The Lighthouse Lab in Milton Keynes and Alex Alderton, Roberto Amato, Jeffrey Barrett, Sonia Goncalves, Ewan Harrison, David K. Jackson, Ian Johnston, Dominic Kwiatkowski, Cordelia Langford, John Sillitoe on behalf of the Wellcome Sanger Institute COVID-19 Surveillance Team |
| EPI_ISL_2468787, EPI_ISL_2468788, EPI_ISL_2468789 | Lighthouse Lab in Alderley Park | Wellcome Sanger Institute for the COVID-19 Genomics UK (COG-UK) Consortium | Jacquelyn Wynn, Mairead Hyland, The Lighthouse Lab in Alderley Park and Alex Alderton, Roberto Amato, Jeffrey Barrett, Sonia Goncalves, Ewan Harrison, David K. Jackson, Ian Johnston, Dominic Kwiatkowski, Cordelia Langford, John Sillitoe on behalf of the Wellcome Sanger Institute COVID-19 Surveillance Team |
| EPI_ISL_2468790, EPI_ISL_2468791, EPI_ISL_2468792 | Lighthouse Lab in Milton Keynes | Wellcome Sanger Institute for the COVID-19 Genomics UK (COG-UK) Consortium | The Lighthouse Lab in Milton Keynes and Alex Alderton, Roberto Amato, Jeffrey Barrett, Sonia Goncalves, Ewan Harrison, David K. Jackson, Ian Johnston, Dominic Kwiatkowski, Cordelia Langford, John Sillitoe on behalf of the Wellcome Sanger Institute COVID-19 Surveillance Team |
| EPI_ISL_2468793 | Lighthouse Lab in Alderley Park | Wellcome Sanger Institute for the COVID-19 Genomics UK (COG-UK) Consortium | Jacquelyn Wynn, Mairead Hyland, The Lighthouse Lab in Alderley Park and Alex Alderton, Roberto Amato, Jeffrey Barrett, Sonia Goncalves, Ewan Harrison, David K. Jackson, Ian Johnston, Dominic Kwiatkowski, Cordelia Langford, John Sillitoe on behalf of the Wellcome Sanger Institute COVID-19 Surveillance Team |
| EPI_ISL_2468794 | Lighthouse Lab in Milton Keynes | Wellcome Sanger Institute for the COVID-19 Genomics UK (COG-UK) Consortium | The Lighthouse Lab in Milton Keynes and Alex Alderton, Roberto Amato, Jeffrey Barrett, Sonia Goncalves, Ewan Harrison, David K. Jackson, Ian Johnston, Dominic Kwiatkowski, Cordelia Langford, John Sillitoe on behalf of the Wellcome Sanger Institute COVID-19 Surveillance Team |
| EPI_ISL_2468795, EPI_ISL_2468796, EPI_ISL_2468797, EPI_ISL_2468798, EPI_ISL_2468799, EPI_ISL_2468800, EPI_ISL_2468801, EPI_ISL_2468802, EPI_ISL_2468803, EPI_ISL_2468804, EPI_ISL_2468805, EPI_ISL_2468806, EPI_ISL_2468807, EPI_ISL_2468808 | | | |
| see above | Lighthouse Lab in Alderley Park | Wellcome Sanger Institute for the COVID-19 Genomics UK (COG-UK) Consortium | Jacquelyn Wynn, Mairead Hyland, The Lighthouse Lab in Alderley Park and Alex Alderton, Roberto Amato, Jeffrey Barrett, Sonia Goncalves, Ewan Harrison, David K. Jackson, Ian Johnston, Dominic Kwiatkowski, Cordelia Langford, John Sillitoe on behalf of the Wellcome Sanger Institute COVID-19 Surveillance Team |

| | | | |
|---|---|---|---|
| EPI_ISL_2468809 | Lighthouse Lab in Milton Keynes | Wellcome Sanger Institute for the COVID-19 Genomics UK (COG-UK) Consortium | The Lighthouse Lab in Milton Keynes and Alex Alderton, Roberto Amato, Jeffrey Barrett, Sonia Goncalves, Ewan Harrison, David K. Jackson, Ian Johnston, Dominic Kwiatkowski, Cordelia Langford, John Sillitoe on behalf of the Wellcome Sanger Institute COVID-19 Surveillance Team |
| EPI_ISL_2468810, EPI_ISL_2468811, EPI_ISL_2468812 | Lighthouse Lab in Alderley Park | Wellcome Sanger Institute for the COVID-19 Genomics UK (COG-UK) Consortium | Jacquelyn Wynn, Mairead Hyland, The Lighthouse Lab in Alderley Park and Alex Alderton, Roberto Amato, Jeffrey Barrett, Sonia Goncalves, Ewan Harrison, David K. Jackson, Ian Johnston, Dominic Kwiatkowski, Cordelia Langford, John Sillitoe on behalf of the Wellcome Sanger Institute COVID-19 Surveillance Team |
| EPI_ISL_2468813 | Lighthouse Lab in Milton Keynes | Wellcome Sanger Institute for the COVID-19 Genomics UK (COG-UK) Consortium | The Lighthouse Lab in Milton Keynes and Alex Alderton, Roberto Amato, Jeffrey Barrett, Sonia Goncalves, Ewan Harrison, David K. Jackson, Ian Johnston, Dominic Kwiatkowski, Cordelia Langford, John Sillitoe on behalf of the Wellcome Sanger Institute COVID-19 Surveillance Team |
| EPI_ISL_2468815 | Lighthouse Lab in Alderley Park | Wellcome Sanger Institute for the COVID-19 Genomics UK (COG-UK) Consortium | Jacquelyn Wynn, Mairead Hyland, The Lighthouse Lab in Alderley Park and Alex Alderton, Roberto Amato, Jeffrey Barrett, Sonia Goncalves, Ewan Harrison, David K. Jackson, Ian Johnston, Dominic Kwiatkowski, Cordelia Langford, John Sillitoe on behalf of the Wellcome Sanger Institute COVID-19 Surveillance Team |
| EPI_ISL_2468816 | Lighthouse Lab in Milton Keynes | Wellcome Sanger Institute for the COVID-19 Genomics UK (COG-UK) Consortium | The Lighthouse Lab in Milton Keynes and Alex Alderton, Roberto Amato, Jeffrey Barrett, Sonia Goncalves, Ewan Harrison, David K. Jackson, Ian Johnston, Dominic Kwiatkowski, Cordelia Langford, John Sillitoe on behalf of the Wellcome Sanger Institute COVID-19 Surveillance Team |
| EPI_ISL_2468817, EPI_ISL_2468818, EPI_ISL_2468819, EPI_ISL_2468820, EPI_ISL_2468821, EPI_ISL_2468822, EPI_ISL_2468823, EPI_ISL_2468824, EPI_ISL_2468825 | Lighthouse Lab in Alderley Park | Wellcome Sanger Institute for the COVID-19 Genomics UK (COG-UK) Consortium | Jacquelyn Wynn, Mairead Hyland, The Lighthouse Lab in Alderley Park and Alex Alderton, Roberto Amato, Jeffrey Barrett, Sonia Goncalves, Ewan Harrison, David K. Jackson, Ian Johnston, Dominic Kwiatkowski, Cordelia Langford, John Sillitoe on behalf of the Wellcome Sanger Institute COVID-19 Surveillance Team |
| EPI_ISL_2468826 | Lighthouse Lab in Milton Keynes | Wellcome Sanger Institute for the COVID-19 Genomics UK (COG-UK) Consortium | The Lighthouse Lab in Milton Keynes and Alex Alderton, Roberto Amato, Jeffrey Barrett, Sonia Goncalves, Ewan Harrison, David K. Jackson, Ian Johnston, Dominic Kwiatkowski, Cordelia Langford, John Sillitoe on behalf of the Wellcome Sanger Institute COVID-19 Surveillance Team |
| EPI_ISL_2468827, EPI_ISL_2468828, EPI_ISL_2468829 | Lighthouse Lab in Alderley Park | Wellcome Sanger Institute for the COVID-19 Genomics UK (COG-UK) Consortium | Jacquelyn Wynn, Mairead Hyland, The Lighthouse Lab in Alderley Park and Alex Alderton, Roberto Amato, Jeffrey Barrett, Sonia Goncalves, Ewan Harrison, David K. Jackson, Ian Johnston, Dominic Kwiatkowski, Cordelia Langford, John Sillitoe on behalf of the Wellcome Sanger Institute COVID-19 Surveillance Team |
| EPI_ISL_2468830 | Lighthouse Lab in Milton Keynes | Wellcome Sanger Institute for the COVID-19 Genomics UK (COG-UK) Consortium | The Lighthouse Lab in Milton Keynes and Alex Alderton, Roberto Amato, Jeffrey Barrett, Sonia Goncalves, Ewan Harrison, David K. Jackson, Ian Johnston, Dominic Kwiatkowski, Cordelia Langford, John Sillitoe on behalf of the Wellcome Sanger Institute COVID-19 Surveillance Team |
| EPI_ISL_2468831, EPI_ISL_2468832, EPI_ISL_2468833, EPI_ISL_2468834, EPI_ISL_2468835 | Lighthouse Lab in Alderley Park | Wellcome Sanger Institute for the COVID-19 Genomics UK (COG-UK) Consortium | Jacquelyn Wynn, Mairead Hyland, The Lighthouse Lab in Alderley Park and Alex Alderton, Roberto Amato, Jeffrey Barrett, Sonia Goncalves, Ewan Harrison, David K. Jackson, Ian Johnston, Dominic Kwiatkowski, Cordelia Langford, John Sillitoe on behalf of the Wellcome Sanger Institute COVID-19 Surveillance Team |
| EPI_ISL_2468836 | Lighthouse Lab in Milton Keynes | Wellcome Sanger Institute for the COVID-19 Genomics UK (COG-UK) Consortium | The Lighthouse Lab in Milton Keynes and Alex Alderton, Roberto Amato, Jeffrey Barrett, Sonia Goncalves, Ewan Harrison, David K. Jackson, Ian Johnston, Dominic Kwiatkowski, Cordelia Langford, John Sillitoe on behalf of the Wellcome Sanger Institute COVID-19 Surveillance Team |
| EPI_ISL_2468837 | Lighthouse Lab in Alderley Park | Wellcome Sanger Institute for the COVID-19 Genomics UK (COG-UK) Consortium | Jacquelyn Wynn, Mairead Hyland, The Lighthouse Lab in Alderley Park and Alex Alderton, Roberto Amato, Jeffrey Barrett, Sonia Goncalves, Ewan Harrison, David K. Jackson, Ian Johnston, Dominic Kwiatkowski, Cordelia Langford, John Sillitoe on behalf of the Wellcome Sanger Institute COVID-19 Surveillance Team |
| EPI_ISL_2468838 | Lighthouse Lab in Milton Keynes | Wellcome Sanger Institute for the COVID-19 Genomics UK (COG-UK) Consortium | The Lighthouse Lab in Milton Keynes and Alex Alderton, Roberto Amato, Jeffrey Barrett, Sonia Goncalves, Ewan Harrison, David K. Jackson, Ian Johnston, Dominic Kwiatkowski, Cordelia Langford, John Sillitoe on behalf of the Wellcome Sanger Institute COVID-19 Surveillance Team |
| EPI_ISL_2468839 | Lighthouse Lab in Alderley Park | Wellcome Sanger Institute for the COVID-19 Genomics UK (COG-UK) Consortium | Jacquelyn Wynn, Mairead Hyland, The Lighthouse Lab in Alderley Park and Alex Alderton, Roberto Amato, Jeffrey Barrett, Sonia Goncalves, Ewan Harrison, David K. Jackson, Ian Johnston, Dominic Kwiatkowski, Cordelia Langford, John Sillitoe on behalf of the Wellcome Sanger Institute COVID-19 Surveillance Team |
| EPI_ISL_2468840 | Lighthouse Lab in Milton Keynes | Wellcome Sanger Institute for the COVID-19 Genomics UK (COG-UK) Consortium | The Lighthouse Lab in Milton Keynes and Alex Alderton, Roberto Amato, Jeffrey Barrett, Sonia Goncalves, Ewan Harrison, David K. Jackson, Ian Johnston, Dominic Kwiatkowski, Cordelia Langford, John Sillitoe on behalf of the Wellcome Sanger Institute COVID-19 Surveillance Team |
| EPI_ISL_2468842, EPI_ISL_2468843, EPI_ISL_2468844, EPI_ISL_2468845, EPI_ISL_2468846, EPI_ISL_2468847, EPI_ISL_2468848, EPI_ISL_2468849, EPI_ISL_2468850, EPI_ISL_2468851, EPI_ISL_2468852, EPI_ISL_2468853, EPI_ISL_2468854, EPI_ISL_2468855 | | | |
| see above | Lighthouse Lab in Alderley Park | Wellcome Sanger Institute for the COVID-19 Genomics UK (COG-UK) Consortium | Jacquelyn Wynn, Mairead Hyland, The Lighthouse Lab in Alderley Park and Alex Alderton, Roberto Amato, Jeffrey Barrett, Sonia Goncalves, Ewan Harrison, David K. Jackson, Ian Johnston, Dominic Kwiatkowski, Cordelia Langford, John Sillitoe on behalf of the Wellcome Sanger Institute COVID-19 Surveillance Team |
| EPI_ISL_2468856 | Lighthouse Lab in Milton Keynes | Wellcome Sanger Institute for the COVID-19 Genomics UK (COG-UK) Consortium | The Lighthouse Lab in Milton Keynes and Alex Alderton, Roberto Amato, Jeffrey Barrett, Sonia Goncalves, Ewan Harrison, David K. Jackson, Ian Johnston, Dominic Kwiatkowski, Cordelia Langford, John Sillitoe on behalf of the Wellcome Sanger Institute COVID-19 Surveillance Team |
| EPI_ISL_2468857 | Lighthouse Lab in Alderley Park | Wellcome Sanger Institute for the COVID-19 Genomics UK (COG-UK) Consortium | Jacquelyn Wynn, Mairead Hyland, The Lighthouse Lab in Alderley Park and Alex Alderton, Roberto Amato, Jeffrey Barrett, Sonia Goncalves, Ewan Harrison, David K. Jackson, Ian Johnston, Dominic Kwiatkowski, Cordelia Langford, John Sillitoe on behalf of the Wellcome Sanger Institute COVID-19 Surveillance Team |
| EPI_ISL_2468858 | Lighthouse Lab in Milton Keynes | Wellcome Sanger Institute for the COVID-19 Genomics UK (COG-UK) Consortium | The Lighthouse Lab in Milton Keynes and Alex Alderton, Roberto Amato, Jeffrey Barrett, Sonia Goncalves, Ewan Harrison, David K. Jackson, Ian Johnston, Dominic Kwiatkowski, Cordelia Langford, John Sillitoe on behalf of the Wellcome Sanger Institute COVID-19 Surveillance Team |
| EPI_ISL_2468859, EPI_ISL_2468860, EPI_ISL_2468861, EPI_ISL_2468862, EPI_ISL_2468863, EPI_ISL_2468864, EPI_ISL_2468865 | Lighthouse Lab in Alderley Park | Wellcome Sanger Institute for the COVID-19 Genomics UK (COG-UK) Consortium | Jacquelyn Wynn, Mairead Hyland, The Lighthouse Lab in Alderley Park and Alex Alderton, Roberto Amato, Jeffrey Barrett, Sonia Goncalves, Ewan Harrison, David K. Jackson, Ian Johnston, Dominic Kwiatkowski, Cordelia Langford, John Sillitoe on behalf of the Wellcome Sanger Institute COVID-19 Surveillance Team |
| EPI_ISL_2468866 | Lighthouse Lab in Milton Keynes | Wellcome Sanger Institute for the COVID-19 Genomics UK (COG-UK) Consortium | The Lighthouse Lab in Milton Keynes and Alex Alderton, Roberto Amato, Jeffrey Barrett, Sonia Goncalves, Ewan Harrison, David K. Jackson, Ian Johnston, Dominic Kwiatkowski, Cordelia Langford, John Sillitoe on behalf of the Wellcome Sanger Institute COVID-19 Surveillance Team |
| EPI_ISL_2468867 | Lighthouse Lab in Alderley Park | Wellcome Sanger Institute for the COVID-19 Genomics UK (COG-UK) Consortium | Jacquelyn Wynn, Mairead Hyland, The Lighthouse Lab in Alderley Park and Alex Alderton, Roberto Amato, Jeffrey Barrett, Sonia Goncalves, Ewan Harrison, David K. Jackson, Ian Johnston, Dominic Kwiatkowski, Cordelia Langford, John Sillitoe on behalf of the Wellcome Sanger Institute COVID-19 Surveillance Team |
| EPI_ISL_2468868 | Lighthouse Lab in Milton Keynes | Wellcome Sanger Institute for the COVID-19 Genomics UK (COG-UK) Consortium | The Lighthouse Lab in Milton Keynes and Alex Alderton, Roberto Amato, Jeffrey Barrett, Sonia Goncalves, Ewan Harrison, David K. Jackson, Ian Johnston, Dominic Kwiatkowski, Cordelia Langford, John Sillitoe on behalf of the Wellcome Sanger Institute COVID-19 Surveillance Team |
| EPI_ISL_2468869, EPI_ISL_2468870, EPI_ISL_2468871, EPI_ISL_2468872, EPI_ISL_2468874 | Lighthouse Lab in Alderley Park | Wellcome Sanger Institute for the COVID-19 Genomics UK (COG-UK) Consortium | Jacquelyn Wynn, Mairead Hyland, The Lighthouse Lab in Alderley Park and Alex Alderton, Roberto Amato, Jeffrey Barrett, Sonia Goncalves, Ewan Harrison, David K. Jackson, Ian Johnston, Dominic Kwiatkowski, Cordelia Langford, John Sillitoe on behalf of the Wellcome Sanger Institute COVID-19 Surveillance Team |
| EPI_ISL_2468875, EPI_ISL_2468876 | Lighthouse Lab in Milton Keynes | Wellcome Sanger Institute for the COVID-19 Genomics UK (COG-UK) Consortium | The Lighthouse Lab in Milton Keynes and Alex Alderton, Roberto Amato, Jeffrey Barrett, Sonia Goncalves, Ewan Harrison, David K. Jackson, Ian Johnston, Dominic Kwiatkowski, Cordelia Langford, John Sillitoe on behalf of the Wellcome Sanger Institute COVID-19 Surveillance Team |
| EPI_ISL_2468877, EPI_ISL_2468878, EPI_ISL_2468879, EPI_ISL_2468880 | Lighthouse Lab in Alderley Park | Wellcome Sanger Institute for the COVID-19 Genomics UK (COG-UK) Consortium | Jacquelyn Wynn, Mairead Hyland, The Lighthouse Lab in Alderley Park and Alex Alderton, Roberto Amato, Jeffrey Barrett, Sonia Goncalves, Ewan Harrison, David K. Jackson, Ian Johnston, Dominic Kwiatkowski, Cordelia Langford, John Sillitoe on behalf of the Wellcome Sanger Institute COVID-19 Surveillance Team |
| EPI_ISL_2468881 | Lighthouse Lab in Milton Keynes | Wellcome Sanger Institute for the COVID-19 Genomics UK (COG-UK) Consortium | The Lighthouse Lab in Milton Keynes and Alex Alderton, Roberto Amato, Jeffrey Barrett, Sonia Goncalves, Ewan Harrison, David K. Jackson, Ian Johnston, Dominic Kwiatkowski, Cordelia Langford, John Sillitoe on behalf of the Wellcome Sanger Institute COVID-19 Surveillance Team |
| EPI_ISL_2468883, EPI_ISL_2468884 | Lighthouse Lab in Alderley Park | Wellcome Sanger Institute for the COVID-19 Genomics UK (COG-UK) Consortium | Jacquelyn Wynn, Mairead Hyland, The Lighthouse Lab in Alderley Park and Alex Alderton, Roberto Amato, Jeffrey Barrett, Sonia Goncalves, Ewan Harrison, David K. Jackson, Ian Johnston, Dominic Kwiatkowski, Cordelia Langford, John Sillitoe on behalf of the Wellcome Sanger Institute COVID-19 Surveillance Team |
| EPI_ISL_2468885 | Lighthouse Lab in Milton Keynes | Wellcome Sanger Institute for the COVID-19 Genomics UK (COG-UK) Consortium | The Lighthouse Lab in Milton Keynes and Alex Alderton, Roberto Amato, Jeffrey Barrett, Sonia Goncalves, Ewan Harrison, David K. Jackson, Ian Johnston, Dominic Kwiatkowski, Cordelia Langford, John Sillitoe on behalf of the Wellcome Sanger Institute COVID-19 Surveillance Team |
| EPI_ISL_2468886 | Lighthouse Lab in Alderley Park | Wellcome Sanger Institute for the COVID-19 Genomics UK (COG-UK) Consortium | Jacquelyn Wynn, Mairead Hyland, The Lighthouse Lab in Alderley Park and Alex Alderton, Roberto Amato, Jeffrey Barrett, Sonia Goncalves, Ewan Harrison, David K. Jackson, Ian Johnston, Dominic Kwiatkowski, Cordelia Langford, John Sillitoe on behalf of the Wellcome Sanger Institute COVID-19 Surveillance Team |
| EPI_ISL_2468887 | Lighthouse Lab in Milton Keynes | Wellcome Sanger Institute for the COVID-19 Genomics UK (COG-UK) Consortium | The Lighthouse Lab in Milton Keynes and Alex Alderton, Roberto Amato, Jeffrey Barrett, Sonia Goncalves, Ewan Harrison, David K. Jackson, Ian Johnston, Dominic Kwiatkowski, Cordelia Langford, John Sillitoe on behalf of the Wellcome Sanger Institute COVID-19 Surveillance Team |
| EPI_ISL_2468889 | Lighthouse Lab in Alderley Park | Wellcome Sanger Institute for the COVID-19 Genomics UK (COG-UK) Consortium | Jacquelyn Wynn, Mairead Hyland, The Lighthouse Lab in Alderley Park and Alex Alderton, Roberto Amato, Jeffrey Barrett, Sonia Goncalves, Ewan Harrison, David K. Jackson, Ian Johnston, Dominic Kwiatkowski, Cordelia Langford, John Sillitoe on behalf of the Wellcome Sanger Institute COVID-19 Surveillance Team |
| EPI_ISL_2468890 | Lighthouse Lab in Milton Keynes | Wellcome Sanger Institute for the COVID-19 Genomics UK (COG-UK) Consortium | The Lighthouse Lab in Milton Keynes and Alex Alderton, Roberto Amato, Jeffrey Barrett, Sonia Goncalves, Ewan Harrison, David K. Jackson, Ian Johnston, Dominic Kwiatkowski, Cordelia Langford, John Sillitoe on behalf of the Wellcome Sanger Institute COVID-19 Surveillance Team |
| EPI_ISL_2468891, EPI_ISL_2468892, EPI_ISL_2468893 | Lighthouse Lab in Alderley Park | Wellcome Sanger Institute for the COVID-19 Genomics UK (COG-UK) Consortium | Jacquelyn Wynn, Mairead Hyland, The Lighthouse Lab in Alderley Park and Alex Alderton, Roberto Amato, Jeffrey Barrett, Sonia Goncalves, Ewan Harrison, David K. Jackson, Ian Johnston, Dominic Kwiatkowski, Cordelia Langford, John Sillitoe on behalf of the Wellcome Sanger Institute COVID-19 Surveillance Team |
| EPI_ISL_2468894, EPI_ISL_2468895 | Lighthouse Lab in Milton Keynes | Wellcome Sanger Institute for the COVID-19 Genomics UK (COG-UK) Consortium | The Lighthouse Lab in Milton Keynes and Alex Alderton, Roberto Amato, Jeffrey Barrett, Sonia Goncalves, Ewan Harrison, David K. Jackson, Ian Johnston, Dominic Kwiatkowski, Cordelia Langford, John Sillitoe on behalf of the Wellcome Sanger Institute COVID-19 Surveillance Team |

| | | | |
|---|---|---|---|
| EPI_ISL_2468896, EPI_ISL_2468897, EPI_ISL_2468898 | Lighthouse Lab in Alderley Park | Wellcome Sanger Institute for the COVID-19 Genomics UK (COG-UK) Consortium | Jacquelyn Wynn, Mairead Hyland, The Lighthouse Lab in Alderley Park and Alex Alderton, Roberto Amato, Jeffrey Barrett, Sonia Goncalves, Ewan Harrison, David K. Jackson, Ian Johnston, Dominic Kwiatkowski, Cordelia Langford, John Sillitoe on behalf of the Wellcome Sanger Institute COVID-19 Surveillance Team |
| EPI_ISL_2468899 | Lighthouse Lab in Milton Keynes | Wellcome Sanger Institute for the COVID-19 Genomics UK (COG-UK) Consortium | The Lighthouse Lab in Milton Keynes and Alex Alderton, Roberto Amato, Jeffrey Barrett, Sonia Goncalves, Ewan Harrison, David K. Jackson, Ian Johnston, Dominic Kwiatkowski, Cordelia Langford, John Sillitoe on behalf of the Wellcome Sanger Institute COVID-19 Surveillance Team |
| EPI_ISL_2468900, EPI_ISL_2468901, EPI_ISL_2468902, EPI_ISL_2468903, EPI_ISL_2468904, EPI_ISL_2468905, EPI_ISL_2468906, EPI_ISL_2468907, EPI_ISL_2468908, EPI_ISL_2468909 | Lighthouse Lab in Alderley Park | Wellcome Sanger Institute for the COVID-19 Genomics UK (COG-UK) Consortium | Jacquelyn Wynn, Mairead Hyland, The Lighthouse Lab in Alderley Park and Alex Alderton, Roberto Amato, Jeffrey Barrett, Sonia Goncalves, Ewan Harrison, David K. Jackson, Ian Johnston, Dominic Kwiatkowski, Cordelia Langford, John Sillitoe on behalf of the Wellcome Sanger Institute COVID-19 Surveillance Team |
| EPI_ISL_2468910 | Lighthouse Lab in Milton Keynes | Wellcome Sanger Institute for the COVID-19 Genomics UK (COG-UK) Consortium | The Lighthouse Lab in Milton Keynes and Alex Alderton, Roberto Amato, Jeffrey Barrett, Sonia Goncalves, Ewan Harrison, David K. Jackson, Ian Johnston, Dominic Kwiatkowski, Cordelia Langford, John Sillitoe on behalf of the Wellcome Sanger Institute COVID-19 Surveillance Team |
| EPI_ISL_2468911, EPI_ISL_2468912, EPI_ISL_2468913, EPI_ISL_2468914, EPI_ISL_2468915 | Lighthouse Lab in Alderley Park | Wellcome Sanger Institute for the COVID-19 Genomics UK (COG-UK) Consortium | Jacquelyn Wynn, Mairead Hyland, The Lighthouse Lab in Alderley Park and Alex Alderton, Roberto Amato, Jeffrey Barrett, Sonia Goncalves, Ewan Harrison, David K. Jackson, Ian Johnston, Dominic Kwiatkowski, Cordelia Langford, John Sillitoe on behalf of the Wellcome Sanger Institute COVID-19 Surveillance Team |
| EPI_ISL_2468916 | Lighthouse Lab in Milton Keynes | Wellcome Sanger Institute for the COVID-19 Genomics UK (COG-UK) Consortium | The Lighthouse Lab in Milton Keynes and Alex Alderton, Roberto Amato, Jeffrey Barrett, Sonia Goncalves, Ewan Harrison, David K. Jackson, Ian Johnston, Dominic Kwiatkowski, Cordelia Langford, John Sillitoe on behalf of the Wellcome Sanger Institute COVID-19 Surveillance Team |
| EPI_ISL_2468917 | Lighthouse Lab in Alderley Park | Wellcome Sanger Institute for the COVID-19 Genomics UK (COG-UK) Consortium | Jacquelyn Wynn, Mairead Hyland, The Lighthouse Lab in Alderley Park and Alex Alderton, Roberto Amato, Jeffrey Barrett, Sonia Goncalves, Ewan Harrison, David K. Jackson, Ian Johnston, Dominic Kwiatkowski, Cordelia Langford, John Sillitoe on behalf of the Wellcome Sanger Institute COVID-19 Surveillance Team |
| EPI_ISL_2468918, EPI_ISL_2468919, EPI_ISL_2468920, EPI_ISL_2468922 | Lighthouse Lab in Milton Keynes | Wellcome Sanger Institute for the COVID-19 Genomics UK (COG-UK) Consortium | The Lighthouse Lab in Milton Keynes and Alex Alderton, Roberto Amato, Jeffrey Barrett, Sonia Goncalves, Ewan Harrison, David K. Jackson, Ian Johnston, Dominic Kwiatkowski, Cordelia Langford, John Sillitoe on behalf of the Wellcome Sanger Institute COVID-19 Surveillance Team |
| EPI_ISL_2468923, EPI_ISL_2468925, EPI_ISL_2468926, EPI_ISL_2468927, EPI_ISL_2468928, EPI_ISL_2468929 | Lighthouse Lab in Alderley Park | Wellcome Sanger Institute for the COVID-19 Genomics UK (COG-UK) Consortium | Jacquelyn Wynn, Mairead Hyland, The Lighthouse Lab in Alderley Park and Alex Alderton, Roberto Amato, Jeffrey Barrett, Sonia Goncalves, Ewan Harrison, David K. Jackson, Ian Johnston, Dominic Kwiatkowski, Cordelia Langford, John Sillitoe on behalf of the Wellcome Sanger Institute COVID-19 Surveillance Team |
| EPI_ISL_2468930, EPI_ISL_2468931 | Lighthouse Lab in Milton Keynes | Wellcome Sanger Institute for the COVID-19 Genomics UK (COG-UK) Consortium | The Lighthouse Lab in Milton Keynes and Alex Alderton, Roberto Amato, Jeffrey Barrett, Sonia Goncalves, Ewan Harrison, David K. Jackson, Ian Johnston, Dominic Kwiatkowski, Cordelia Langford, John Sillitoe on behalf of the Wellcome Sanger Institute COVID-19 Surveillance Team |
| EPI_ISL_2468932 | Lighthouse Lab in Alderley Park | Wellcome Sanger Institute for the COVID-19 Genomics UK (COG-UK) Consortium | Jacquelyn Wynn, Mairead Hyland, The Lighthouse Lab in Alderley Park and Alex Alderton, Roberto Amato, Jeffrey Barrett, Sonia Goncalves, Ewan Harrison, David K. Jackson, Ian Johnston, Dominic Kwiatkowski, Cordelia Langford, John Sillitoe on behalf of the Wellcome Sanger Institute COVID-19 Surveillance Team |
| EPI_ISL_2468933 | Lighthouse Lab in Milton Keynes | Wellcome Sanger Institute for the COVID-19 Genomics UK (COG-UK) Consortium | The Lighthouse Lab in Milton Keynes and Alex Alderton, Roberto Amato, Jeffrey Barrett, Sonia Goncalves, Ewan Harrison, David K. Jackson, Ian Johnston, Dominic Kwiatkowski, Cordelia Langford, John Sillitoe on behalf of the Wellcome Sanger Institute COVID-19 Surveillance Team |
| EPI_ISL_2468934 | Lighthouse Lab in Alderley Park | Wellcome Sanger Institute for the COVID-19 Genomics UK (COG-UK) Consortium | Jacquelyn Wynn, Mairead Hyland, The Lighthouse Lab in Alderley Park and Alex Alderton, Roberto Amato, Jeffrey Barrett, Sonia Goncalves, Ewan Harrison, David K. Jackson, Ian Johnston, Dominic Kwiatkowski, Cordelia Langford, John Sillitoe on behalf of the Wellcome Sanger Institute COVID-19 Surveillance Team |
| EPI_ISL_2468935 | Lighthouse Lab in Milton Keynes | Wellcome Sanger Institute for the COVID-19 Genomics UK (COG-UK) Consortium | The Lighthouse Lab in Milton Keynes and Alex Alderton, Roberto Amato, Jeffrey Barrett, Sonia Goncalves, Ewan Harrison, David K. Jackson, Ian Johnston, Dominic Kwiatkowski, Cordelia Langford, John Sillitoe on behalf of the Wellcome Sanger Institute COVID-19 Surveillance Team |
| EPI_ISL_2468936, EPI_ISL_2468937, EPI_ISL_2468938, EPI_ISL_2468939, EPI_ISL_2468940, EPI_ISL_2468941, EPI_ISL_2468942, EPI_ISL_2468943, EPI_ISL_2468945, EPI_ISL_2468946, EPI_ISL_2468947 | | | |
| see above | Lighthouse Lab in Alderley Park | Wellcome Sanger Institute for the COVID-19 Genomics UK (COG-UK) Consortium | Jacquelyn Wynn, Mairead Hyland, The Lighthouse Lab in Alderley Park and Alex Alderton, Roberto Amato, Jeffrey Barrett, Sonia Goncalves, Ewan Harrison, David K. Jackson, Ian Johnston, Dominic Kwiatkowski, Cordelia Langford, John Sillitoe on behalf of the Wellcome Sanger Institute COVID-19 Surveillance Team |
| EPI_ISL_2468948 | Lighthouse Lab in Glasgow | Wellcome Sanger Institute for the COVID-19 Genomics UK (COG-UK) Consortium | Harper VanSteenhouse, Yumi Kasai, David Gray, Carol Clugston, Anna Dominiczak and Alex Alderton, Roberto Amato, Jeffrey Barrett, Sonia Goncalves, Ewan Harrison, David K. Jackson, Ian Johnston, Dominic Kwiatkowski, Cordelia Langford, John Sillitoe on behalf of the Wellcome Sanger Institute COVID-19 Surveillance Team |
| EPI_ISL_2468950, EPI_ISL_2468951, EPI_ISL_2468952, EPI_ISL_2468953, EPI_ISL_2468954, EPI_ISL_2468955, EPI_ISL_2468957, EPI_ISL_2468958, EPI_ISL_2468959, EPI_ISL_2468960 | Lighthouse Lab in Alderley Park | Wellcome Sanger Institute for the COVID-19 Genomics UK (COG-UK) Consortium | Jacquelyn Wynn, Mairead Hyland, The Lighthouse Lab in Alderley Park and Alex Alderton, Roberto Amato, Jeffrey Barrett, Sonia Goncalves, Ewan Harrison, David K. Jackson, Ian Johnston, Dominic Kwiatkowski, Cordelia Langford, John Sillitoe on behalf of the Wellcome Sanger Institute COVID-19 Surveillance Team |
| EPI_ISL_2468961 | Lighthouse Lab in Glasgow | Wellcome Sanger Institute for the COVID-19 Genomics UK (COG-UK) Consortium | Harper VanSteenhouse, Yumi Kasai, David Gray, Carol Clugston, Anna Dominiczak and Alex Alderton, Roberto Amato, Jeffrey Barrett, Sonia Goncalves, Ewan Harrison, David K. Jackson, Ian Johnston, Dominic Kwiatkowski, Cordelia Langford, John Sillitoe on behalf of the Wellcome Sanger Institute COVID-19 Surveillance Team |
| EPI_ISL_2468962, EPI_ISL_2468963, EPI_ISL_2468964, EPI_ISL_2468965, EPI_ISL_2468966, EPI_ISL_2468967, EPI_ISL_2468968, EPI_ISL_2468969, EPI_ISL_2468970, EPI_ISL_2468971, EPI_ISL_2468972, EPI_ISL_2468973, EPI_ISL_2468974, EPI_ISL_2468975, EPI_ISL_2468976, EPI_ISL_2468978, EPI_ISL_2468979, EPI_ISL_2468980, EPI_ISL_2468981, EPI_ISL_2468982, EPI_ISL_2468983 | | | |
| see above | Lighthouse Lab in Alderley Park | Wellcome Sanger Institute for the COVID-19 Genomics UK (COG-UK) Consortium | Jacquelyn Wynn, Mairead Hyland, The Lighthouse Lab in Alderley Park and Alex Alderton, Roberto Amato, Jeffrey Barrett, Sonia Goncalves, Ewan Harrison, David K. Jackson, Ian Johnston, Dominic Kwiatkowski, Cordelia Langford, John Sillitoe on behalf of the Wellcome Sanger Institute COVID-19 Surveillance Team |
| EPI_ISL_2468984 | Lighthouse Lab in Glasgow | Wellcome Sanger Institute for the COVID-19 Genomics UK (COG-UK) Consortium | Harper VanSteenhouse, Yumi Kasai, David Gray, Carol Clugston, Anna Dominiczak and Alex Alderton, Roberto Amato, Jeffrey Barrett, Sonia Goncalves, Ewan Harrison, David K. Jackson, Ian Johnston, Dominic Kwiatkowski, Cordelia Langford, John Sillitoe on behalf of the Wellcome Sanger Institute COVID-19 Surveillance Team |
| EPI_ISL_2468985, EPI_ISL_2468986, EPI_ISL_2468987, EPI_ISL_2468988, EPI_ISL_2468989, EPI_ISL_2468990, EPI_ISL_2468991, EPI_ISL_2468992, EPI_ISL_2468993, EPI_ISL_2468994, EPI_ISL_2468995, EPI_ISL_2468997, EPI_ISL_2468998 | | | |
| see above | Lighthouse Lab in Alderley Park | Wellcome Sanger Institute for the COVID-19 Genomics UK (COG-UK) Consortium | Jacquelyn Wynn, Mairead Hyland, The Lighthouse Lab in Alderley Park and Alex Alderton, Roberto Amato, Jeffrey Barrett, Sonia Goncalves, Ewan Harrison, David K. Jackson, Ian Johnston, Dominic Kwiatkowski, Cordelia Langford, John Sillitoe on behalf of the Wellcome Sanger Institute COVID-19 Surveillance Team |
| EPI_ISL_2468999 | Lighthouse Lab in Glasgow | Wellcome Sanger Institute for the COVID-19 Genomics UK (COG-UK) Consortium | Harper VanSteenhouse, Yumi Kasai, David Gray, Carol Clugston, Anna Dominiczak and Alex Alderton, Roberto Amato, Jeffrey Barrett, Sonia Goncalves, Ewan Harrison, David K. Jackson, Ian Johnston, Dominic Kwiatkowski, Cordelia Langford, John Sillitoe on behalf of the Wellcome Sanger Institute COVID-19 Surveillance Team |
| EPI_ISL_2469000, EPI_ISL_2469001, EPI_ISL_2469002, EPI_ISL_2469003, EPI_ISL_2469004, EPI_ISL_2469005, EPI_ISL_2469006, EPI_ISL_2469007, EPI_ISL_2469008, EPI_ISL_2469009, EPI_ISL_2469010, EPI_ISL_2469011, EPI_ISL_2469012, EPI_ISL_2469013, EPI_ISL_2469014, EPI_ISL_2469015, EPI_ISL_2469016, EPI_ISL_2469017, EPI_ISL_2469018, EPI_ISL_2469019, EPI_ISL_2469020, EPI_ISL_2469021, EPI_ISL_2469022, EPI_ISL_2469023, EPI_ISL_2469024, EPI_ISL_2469025, EPI_ISL_2469026, EPI_ISL_2469027, EPI_ISL_2469028, EPI_ISL_2469029, EPI_ISL_2469030, EPI_ISL_2469031, EPI_ISL_2469033, EPI_ISL_2469034, EPI_ISL_2469035, EPI_ISL_2469036, EPI_ISL_2469037, EPI_ISL_2469038, EPI_ISL_2469040, EPI_ISL_2469041, EPI_ISL_2469042, EPI_ISL_2469043, EPI_ISL_2469044, EPI_ISL_2469045, EPI_ISL_2469046, EPI_ISL_2469047, EPI_ISL_2469048, EPI_ISL_2469049, EPI_ISL_2469050, EPI_ISL_2469051, EPI_ISL_2469052, EPI_ISL_2469053, EPI_ISL_2469054, EPI_ISL_2469055, EPI_ISL_2469056, EPI_ISL_2469058, EPI_ISL_2469059, EPI_ISL_2469060, EPI_ISL_2469061, EPI_ISL_2469062, EPI_ISL_2469063, EPI_ISL_2469064, EPI_ISL_2469065, EPI_ISL_2469066, EPI_ISL_2469067, EPI_ISL_2469069, EPI_ISL_2469070, EPI_ISL_2469071, EPI_ISL_2469073, EPI_ISL_2469074, EPI_ISL_2469075, EPI_ISL_2469076, EPI_ISL_2469077, EPI_ISL_2469078, EPI_ISL_2469079, EPI_ISL_2469080, EPI_ISL_2469081, EPI_ISL_2469082, EPI_ISL_2469083, EPI_ISL_2469084, EPI_ISL_2469085, EPI_ISL_2469086, EPI_ISL_2469087, EPI_ISL_2469088, EPI_ISL_2469089, EPI_ISL_2469090, EPI_ISL_2469091, EPI_ISL_2469093, EPI_ISL_2469094, EPI_ISL_2469095, EPI_ISL_2469096, EPI_ISL_2469097, EPI_ISL_2469098, EPI_ISL_2469099, EPI_ISL_2469100, EPI_ISL_2469101, EPI_ISL_2469102, EPI_ISL_2469103, EPI_ISL_2469104, EPI_ISL_2469105, EPI_ISL_2469106, EPI_ISL_2469107, EPI_ISL_2469108, EPI_ISL_2469109, EPI_ISL_2469110, EPI_ISL_2469111, EPI_ISL_2469112, EPI_ISL_2469113, EPI_ISL_2469114, EPI_ISL_2469115, EPI_ISL_2469117, EPI_ISL_2469118, EPI_ISL_2469119, EPI_ISL_2469120, EPI_ISL_2469121, EPI_ISL_2469122, EPI_ISL_2469123, EPI_ISL_2469124, EPI_ISL_2469125, EPI_ISL_2469126, EPI_ISL_2469127, EPI_ISL_2469128, EPI_ISL_2469129, EPI_ISL_2469130, EPI_ISL_2469131, EPI_ISL_2469132, EPI_ISL_2469133, EPI_ISL_2469135, EPI_ISL_2469136, EPI_ISL_2469137, EPI_ISL_2469139, EPI_ISL_2469140, EPI_ISL_2469141, EPI_ISL_2469142, EPI_ISL_2469143, EPI_ISL_2469144, EPI_ISL_2469145, EPI_ISL_2469146, EPI_ISL_2469147, EPI_ISL_2469148, EPI_ISL_2469150, EPI_ISL_2469151, EPI_ISL_2469152, EPI_ISL_2469153, EPI_ISL_2469154, EPI_ISL_2469155, EPI_ISL_2469156, EPI_ISL_2469157, EPI_ISL_2469158, EPI_ISL_2469159, EPI_ISL_2469160, EPI_ISL_2469161, EPI_ISL_2469162, EPI_ISL_2469163, EPI_ISL_2469164, EPI_ISL_2469165, EPI_ISL_2469166, EPI_ISL_2469167, EPI_ISL_2469168, EPI_ISL_2469169, EPI_ISL_2469170, EPI_ISL_2469171, EPI_ISL_2469172, EPI_ISL_2469173, EPI_ISL_2469174, EPI_ISL_2469175, EPI_ISL_2469176, EPI_ISL_2469178, EPI_ISL_2469179, EPI_ISL_2469180, EPI_ISL_2469181, EPI_ISL_2469182, EPI_ISL_2469183, EPI_ISL_2469184, EPI_ISL_2469185, EPI_ISL_2469186, EPI_ISL_2469187, EPI_ISL_2469188 | | | |
| see above | Lighthouse Lab in Alderley Park | Wellcome Sanger Institute for the COVID-19 Genomics UK (COG-UK) Consortium | Jacquelyn Wynn, Mairead Hyland, The Lighthouse Lab in Alderley Park and Alex Alderton, Roberto Amato, Jeffrey Barrett, Sonia Goncalves, Ewan Harrison, David K. Jackson, Ian Johnston, Dominic Kwiatkowski, Cordelia Langford, John Sillitoe on behalf of the Wellcome Sanger Institute COVID-19 Surveillance Team |
| EPI_ISL_2469189 | Lighthouse Lab in Glasgow | Wellcome Sanger Institute for the COVID-19 Genomics UK (COG-UK) Consortium | Harper VanSteenhouse, Yumi Kasai, David Gray, Carol Clugston, Anna Dominiczak and Alex Alderton, Roberto Amato, Jeffrey Barrett, Sonia Goncalves, Ewan Harrison, David K. Jackson, Ian Johnston, Dominic Kwiatkowski, Cordelia Langford, John Sillitoe on behalf of the Wellcome Sanger Institute COVID-19 Surveillance Team |
| EPI_ISL_2469190, EPI_ISL_2469191, EPI_ISL_2469192, EPI_ISL_2469193, EPI_ISL_2469194, EPI_ISL_2469195, EPI_ISL_2469196, EPI_ISL_2469197, EPI_ISL_2469198, EPI_ISL_2469199, EPI_ISL_2469200, EPI_ISL_2469201, EPI_ISL_2469202, EPI_ISL_2469203, EPI_ISL_2469204, EPI_ISL_2469205, EPI_ISL_2469206, EPI_ISL_2469207, EPI_ISL_2469208, EPI_ISL_2469209, EPI_ISL_2469210, EPI_ISL_2469211 | | | |
| see above | Lighthouse Lab in Alderley Park | Wellcome Sanger Institute for the COVID-19 Genomics UK (COG-UK) Consortium | Jacquelyn Wynn, Mairead Hyland, The Lighthouse Lab in Alderley Park and Alex Alderton, Roberto Amato, Jeffrey Barrett, Sonia Goncalves, Ewan Harrison, David K. Jackson, Ian Johnston, Dominic Kwiatkowski, Cordelia Langford, John Sillitoe on behalf of the Wellcome Sanger Institute COVID-19 Surveillance Team |
| EPI_ISL_2469212 | Lighthouse Lab in Glasgow | Wellcome Sanger Institute for the COVID-19 Genomics UK (COG-UK) Consortium | Harper VanSteenhouse, Yumi Kasai, David Gray, Carol Clugston, Anna Dominiczak and Alex Alderton, Roberto Amato, Jeffrey Barrett, Sonia Goncalves, Ewan Harrison, David K. Jackson, Ian Johnston, Dominic Kwiatkowski, Cordelia Langford, John Sillitoe on behalf of the Wellcome Sanger Institute COVID-19 Surveillance Team |
| EPI_ISL_2469213, EPI_ISL_2469214, EPI_ISL_2469215, EPI_ISL_2469216, EPI_ISL_2469217, | Lighthouse Lab in Alderley Park | Wellcome Sanger Institute for the COVID-19 Genomics UK (COG-UK) Consortium | Jacquelyn Wynn, Mairead Hyland, The Lighthouse Lab in Alderley Park and Alex Alderton, Roberto Amato, Jeffrey Barrett, Sonia Goncalves, Ewan Harrison, David K. Jackson, Ian Johnston, Dominic Kwiatkowski, Cordelia Langford, John Sillitoe on behalf of the Wellcome Sanger Institute COVID-19 Surveillance Team |

| | | | |
|---|---|---|---|
| EPI_ISL_2469218, EPI_ISL_2469220, EPI_ISL_2469221 | | | |
| EPI_ISL_2469222 | Lighthouse Lab in Glasgow | Wellcome Sanger Institute for the COVID-19 Genomics UK (COG-UK) Consortium | Harper VanSteenhouse, Yumi Kasai, David Gray, Carol Clugston, Anna Dominiczak and Alex Alderton, Roberto Amato, Jeffrey Barrett, Sonia Goncalves, Ewan Harrison, David K. Jackson, Ian Johnston, Dominic Kwiatkowski, Cordelia Langford, John Sillitoe on behalf of the Wellcome Sanger Institute COVID-19 Surveillance Team |
| EPI_ISL_2469223, EPI_ISL_2469224, EPI_ISL_2469225, EPI_ISL_2469226 | Lighthouse Lab in Alderley Park | Wellcome Sanger Institute for the COVID-19 Genomics UK (COG-UK) Consortium | Jacquelyn Wynn, Mairead Hyland, The Lighthouse Lab in Alderley Park and Alex Alderton, Roberto Amato, Jeffrey Barrett, Sonia Goncalves, Ewan Harrison, David K. Jackson, Ian Johnston, Dominic Kwiatkowski, Cordelia Langford, John Sillitoe on behalf of the Wellcome Sanger Institute COVID-19 Surveillance Team |
| EPI_ISL_2469227, EPI_ISL_2469228 | Health Services Laboratories | Wellcome Sanger Institute for the COVID-19 Genomics UK (COG-UK) Consortium | Health Services Laboratories and Alex Alderton, Roberto Amato, Jeffrey Barrett, Sonia Goncalves, Ewan Harrison, David K. Jackson, Ian Johnston, Dominic Kwiatkowski, Cordelia Langford, John Sillitoe on behalf of the Wellcome Sanger Institute COVID-19 Surveillance Team |
| EPI_ISL_2469229 | Lighthouse Lab in Alderley Park | Wellcome Sanger Institute for the COVID-19 Genomics UK (COG-UK) Consortium | Jacquelyn Wynn, Mairead Hyland, The Lighthouse Lab in Alderley Park and Alex Alderton, Roberto Amato, Jeffrey Barrett, Sonia Goncalves, Ewan Harrison, David K. Jackson, Ian Johnston, Dominic Kwiatkowski, Cordelia Langford, John Sillitoe on behalf of the Wellcome Sanger Institute COVID-19 Surveillance Team |
| EPI_ISL_2469230 | Health Services Laboratories | Wellcome Sanger Institute for the COVID-19 Genomics UK (COG-UK) Consortium | Health Services Laboratories and Alex Alderton, Roberto Amato, Jeffrey Barrett, Sonia Goncalves, Ewan Harrison, David K. Jackson, Ian Johnston, Dominic Kwiatkowski, Cordelia Langford, John Sillitoe on behalf of the Wellcome Sanger Institute COVID-19 Surveillance Team |
| EPI_ISL_2469231 | Lighthouse Lab in Milton Keynes | Wellcome Sanger Institute for the COVID-19 Genomics UK (COG-UK) Consortium | The Lighthouse Lab in Milton Keynes and Alex Alderton, Roberto Amato, Jeffrey Barrett, Sonia Goncalves, Ewan Harrison, David K. Jackson, Ian Johnston, Dominic Kwiatkowski, Cordelia Langford, John Sillitoe on behalf of the Wellcome Sanger Institute COVID-19 Surveillance Team |
| EPI_ISL_2469232 | Lighthouse Lab in Alderley Park | Wellcome Sanger Institute for the COVID-19 Genomics UK (COG-UK) Consortium | Jacquelyn Wynn, Mairead Hyland, The Lighthouse Lab in Alderley Park and Alex Alderton, Roberto Amato, Jeffrey Barrett, Sonia Goncalves, Ewan Harrison, David K. Jackson, Ian Johnston, Dominic Kwiatkowski, Cordelia Langford, John Sillitoe on behalf of the Wellcome Sanger Institute COVID-19 Surveillance Team |
| EPI_ISL_2469233, EPI_ISL_2469234, EPI_ISL_2469235, EPI_ISL_2469236, EPI_ISL_2469237 | Health Services Laboratories | Wellcome Sanger Institute for the COVID-19 Genomics UK (COG-UK) Consortium | Health Services Laboratories and Alex Alderton, Roberto Amato, Jeffrey Barrett, Sonia Goncalves, Ewan Harrison, David K. Jackson, Ian Johnston, Dominic Kwiatkowski, Cordelia Langford, John Sillitoe on behalf of the Wellcome Sanger Institute COVID-19 Surveillance Team |
| EPI_ISL_2469238 | Lighthouse Lab in Alderley Park | Wellcome Sanger Institute for the COVID-19 Genomics UK (COG-UK) Consortium | Jacquelyn Wynn, Mairead Hyland, The Lighthouse Lab in Alderley Park and Alex Alderton, Roberto Amato, Jeffrey Barrett, Sonia Goncalves, Ewan Harrison, David K. Jackson, Ian Johnston, Dominic Kwiatkowski, Cordelia Langford, John Sillitoe on behalf of the Wellcome Sanger Institute COVID-19 Surveillance Team |
| EPI_ISL_2469239, EPI_ISL_2469240, EPI_ISL_2469241 | Health Services Laboratories | Wellcome Sanger Institute for the COVID-19 Genomics UK (COG-UK) Consortium | Health Services Laboratories and Alex Alderton, Roberto Amato, Jeffrey Barrett, Sonia Goncalves, Ewan Harrison, David K. Jackson, Ian Johnston, Dominic Kwiatkowski, Cordelia Langford, John Sillitoe on behalf of the Wellcome Sanger Institute COVID-19 Surveillance Team |
| EPI_ISL_2469243, EPI_ISL_2469244 | Lighthouse Lab in Alderley Park | Wellcome Sanger Institute for the COVID-19 Genomics UK (COG-UK) Consortium | Jacquelyn Wynn, Mairead Hyland, The Lighthouse Lab in Alderley Park and Alex Alderton, Roberto Amato, Jeffrey Barrett, Sonia Goncalves, Ewan Harrison, David K. Jackson, Ian Johnston, Dominic Kwiatkowski, Cordelia Langford, John Sillitoe on behalf of the Wellcome Sanger Institute COVID-19 Surveillance Team |
| EPI_ISL_2469245, EPI_ISL_2469246 | Health Services Laboratories | Wellcome Sanger Institute for the COVID-19 Genomics UK (COG-UK) Consortium | Health Services Laboratories and Alex Alderton, Roberto Amato, Jeffrey Barrett, Sonia Goncalves, Ewan Harrison, David K. Jackson, Ian Johnston, Dominic Kwiatkowski, Cordelia Langford, John Sillitoe on behalf of the Wellcome Sanger Institute COVID-19 Surveillance Team |
| EPI_ISL_2469247 | Lighthouse Lab in Alderley Park | Wellcome Sanger Institute for the COVID-19 Genomics UK (COG-UK) Consortium | Jacquelyn Wynn, Mairead Hyland, The Lighthouse Lab in Alderley Park and Alex Alderton, Roberto Amato, Jeffrey Barrett, Sonia Goncalves, Ewan Harrison, David K. Jackson, Ian Johnston, Dominic Kwiatkowski, Cordelia Langford, John Sillitoe on behalf of the Wellcome Sanger Institute COVID-19 Surveillance Team |
| EPI_ISL_2469248 | Health Services Laboratories | Wellcome Sanger Institute for the COVID-19 Genomics UK (COG-UK) Consortium | Health Services Laboratories and Alex Alderton, Roberto Amato, Jeffrey Barrett, Sonia Goncalves, Ewan Harrison, David K. Jackson, Ian Johnston, Dominic Kwiatkowski, Cordelia Langford, John Sillitoe on behalf of the Wellcome Sanger Institute COVID-19 Surveillance Team |
| EPI_ISL_2469250 | Lighthouse Lab in Alderley Park | Wellcome Sanger Institute for the COVID-19 Genomics UK (COG-UK) Consortium | Jacquelyn Wynn, Mairead Hyland, The Lighthouse Lab in Alderley Park and Alex Alderton, Roberto Amato, Jeffrey Barrett, Sonia Goncalves, Ewan Harrison, David K. Jackson, Ian Johnston, Dominic Kwiatkowski, Cordelia Langford, John Sillitoe on behalf of the Wellcome Sanger Institute COVID-19 Surveillance Team |
| EPI_ISL_2469251, EPI_ISL_2469252 | Health Services Laboratories | Wellcome Sanger Institute for the COVID-19 Genomics UK (COG-UK) Consortium | Health Services Laboratories and Alex Alderton, Roberto Amato, Jeffrey Barrett, Sonia Goncalves, Ewan Harrison, David K. Jackson, Ian Johnston, Dominic Kwiatkowski, Cordelia Langford, John Sillitoe on behalf of the Wellcome Sanger Institute COVID-19 Surveillance Team |
| EPI_ISL_2469253 | Lighthouse Lab in Milton Keynes | Wellcome Sanger Institute for the COVID-19 Genomics UK (COG-UK) Consortium | The Lighthouse Lab in Milton Keynes and Alex Alderton, Roberto Amato, Jeffrey Barrett, Sonia Goncalves, Ewan Harrison, David K. Jackson, Ian Johnston, Dominic Kwiatkowski, Cordelia Langford, John Sillitoe on behalf of the Wellcome Sanger Institute COVID-19 Surveillance Team |
| EPI_ISL_2469254, EPI_ISL_2469255, EPI_ISL_2469256, EPI_ISL_2469257 | Lighthouse Lab in Alderley Park | Wellcome Sanger Institute for the COVID-19 Genomics UK (COG-UK) Consortium | Jacquelyn Wynn, Mairead Hyland, The Lighthouse Lab in Alderley Park and Alex Alderton, Roberto Amato, Jeffrey Barrett, Sonia Goncalves, Ewan Harrison, David K. Jackson, Ian Johnston, Dominic Kwiatkowski, Cordelia Langford, John Sillitoe on behalf of the Wellcome Sanger Institute COVID-19 Surveillance Team |
| EPI_ISL_2469258 | Health Services Laboratories | Wellcome Sanger Institute for the COVID-19 Genomics UK (COG-UK) Consortium | Health Services Laboratories and Alex Alderton, Roberto Amato, Jeffrey Barrett, Sonia Goncalves, Ewan Harrison, David K. Jackson, Ian Johnston, Dominic Kwiatkowski, Cordelia Langford, John Sillitoe on behalf of the Wellcome Sanger Institute COVID-19 Surveillance Team |
| EPI_ISL_2469259, EPI_ISL_2469260 | Lighthouse Lab in Milton Keynes | Wellcome Sanger Institute for the COVID-19 Genomics UK (COG-UK) Consortium | The Lighthouse Lab in Milton Keynes and Alex Alderton, Roberto Amato, Jeffrey Barrett, Sonia Goncalves, Ewan Harrison, David K. Jackson, Ian Johnston, Dominic Kwiatkowski, Cordelia Langford, John Sillitoe on behalf of the Wellcome Sanger Institute COVID-19 Surveillance Team |
| EPI_ISL_2469262, EPI_ISL_2469263 | Health Services Laboratories | Wellcome Sanger Institute for the COVID-19 Genomics UK (COG-UK) Consortium | Health Services Laboratories and Alex Alderton, Roberto Amato, Jeffrey Barrett, Sonia Goncalves, Ewan Harrison, David K. Jackson, Ian Johnston, Dominic Kwiatkowski, Cordelia Langford, John Sillitoe on behalf of the Wellcome Sanger Institute COVID-19 Surveillance Team |
| EPI_ISL_2469264 | Lighthouse Lab in Alderley Park | Wellcome Sanger Institute for the COVID-19 Genomics UK (COG-UK) Consortium | Jacquelyn Wynn, Mairead Hyland, The Lighthouse Lab in Alderley Park and Alex Alderton, Roberto Amato, Jeffrey Barrett, Sonia Goncalves, Ewan Harrison, David K. Jackson, Ian Johnston, Dominic Kwiatkowski, Cordelia Langford, John Sillitoe on behalf of the Wellcome Sanger Institute COVID-19 Surveillance Team |
| EPI_ISL_2469265, EPI_ISL_2469267 | Health Services Laboratories | Wellcome Sanger Institute for the COVID-19 Genomics UK (COG-UK) Consortium | Health Services Laboratories and Alex Alderton, Roberto Amato, Jeffrey Barrett, Sonia Goncalves, Ewan Harrison, David K. Jackson, Ian Johnston, Dominic Kwiatkowski, Cordelia Langford, John Sillitoe on behalf of the Wellcome Sanger Institute COVID-19 Surveillance Team |
| EPI_ISL_2469268 | Lighthouse Lab in Glasgow | Wellcome Sanger Institute for the COVID-19 Genomics UK (COG-UK) Consortium | Harper VanSteenhouse, Yumi Kasai, David Gray, Carol Clugston, Anna Dominiczak and Alex Alderton, Roberto Amato, Jeffrey Barrett, Sonia Goncalves, Ewan Harrison, David K. Jackson, Ian Johnston, Dominic Kwiatkowski, Cordelia Langford, John Sillitoe on behalf of the Wellcome Sanger Institute COVID-19 Surveillance Team |
| EPI_ISL_2469269, EPI_ISL_2469270, EPI_ISL_2469271 | Lighthouse Lab in Alderley Park | Wellcome Sanger Institute for the COVID-19 Genomics UK (COG-UK) Consortium | Jacquelyn Wynn, Mairead Hyland, The Lighthouse Lab in Alderley Park and Alex Alderton, Roberto Amato, Jeffrey Barrett, Sonia Goncalves, Ewan Harrison, David K. Jackson, Ian Johnston, Dominic Kwiatkowski, Cordelia Langford, John Sillitoe on behalf of the Wellcome Sanger Institute COVID-19 Surveillance Team |
| EPI_ISL_2469273 | Health Services Laboratories | Wellcome Sanger Institute for the COVID-19 Genomics UK (COG-UK) Consortium | Health Services Laboratories and Alex Alderton, Roberto Amato, Jeffrey Barrett, Sonia Goncalves, Ewan Harrison, David K. Jackson, Ian Johnston, Dominic Kwiatkowski, Cordelia Langford, John Sillitoe on behalf of the Wellcome Sanger Institute COVID-19 Surveillance Team |
| EPI_ISL_2469275 | Lighthouse Lab in Milton Keynes | Wellcome Sanger Institute for the COVID-19 Genomics UK (COG-UK) Consortium | The Lighthouse Lab in Milton Keynes and Alex Alderton, Roberto Amato, Jeffrey Barrett, Sonia Goncalves, Ewan Harrison, David K. Jackson, Ian Johnston, Dominic Kwiatkowski, Cordelia Langford, John Sillitoe on behalf of the Wellcome Sanger Institute COVID-19 Surveillance Team |
| EPI_ISL_2469276, EPI_ISL_2469277, EPI_ISL_2469278 | Health Services Laboratories | Wellcome Sanger Institute for the COVID-19 Genomics UK (COG-UK) Consortium | Health Services Laboratories and Alex Alderton, Roberto Amato, Jeffrey Barrett, Sonia Goncalves, Ewan Harrison, David K. Jackson, Ian Johnston, Dominic Kwiatkowski, Cordelia Langford, John Sillitoe on behalf of the Wellcome Sanger Institute COVID-19 Surveillance Team |
| EPI_ISL_2469280 | Lighthouse Lab in Alderley Park | Wellcome Sanger Institute for the COVID-19 Genomics UK (COG-UK) Consortium | Jacquelyn Wynn, Mairead Hyland, The Lighthouse Lab in Alderley Park and Alex Alderton, Roberto Amato, Jeffrey Barrett, Sonia Goncalves, Ewan Harrison, David K. Jackson, Ian Johnston, Dominic Kwiatkowski, Cordelia Langford, John Sillitoe on behalf of the Wellcome Sanger Institute COVID-19 Surveillance Team |
| EPI_ISL_2469281, EPI_ISL_2469282 | Lighthouse Lab in Milton Keynes | Wellcome Sanger Institute for the COVID-19 Genomics UK (COG-UK) Consortium | The Lighthouse Lab in Milton Keynes and Alex Alderton, Roberto Amato, Jeffrey Barrett, Sonia Goncalves, Ewan Harrison, David K. Jackson, Ian Johnston, Dominic Kwiatkowski, Cordelia Langford, John Sillitoe on behalf of the Wellcome Sanger Institute COVID-19 Surveillance Team |
| EPI_ISL_2469283 | Lighthouse Lab in Alderley Park | Wellcome Sanger Institute for the COVID-19 Genomics UK (COG-UK) Consortium | Jacquelyn Wynn, Mairead Hyland, The Lighthouse Lab in Alderley Park and Alex Alderton, Roberto Amato, Jeffrey Barrett, Sonia Goncalves, Ewan Harrison, David K. Jackson, Ian Johnston, Dominic Kwiatkowski, Cordelia Langford, John Sillitoe on behalf of the Wellcome Sanger Institute COVID-19 Surveillance Team |
| EPI_ISL_2469286, EPI_ISL_2469287 | Health Services Laboratories | Wellcome Sanger Institute for the COVID-19 Genomics UK (COG-UK) Consortium | Health Services Laboratories and Alex Alderton, Roberto Amato, Jeffrey Barrett, Sonia Goncalves, Ewan Harrison, David K. Jackson, Ian Johnston, Dominic Kwiatkowski, Cordelia Langford, John Sillitoe on behalf of the Wellcome Sanger Institute COVID-19 Surveillance Team |
| EPI_ISL_2469288 | Lighthouse Lab in Alderley Park | Wellcome Sanger Institute for the COVID-19 Genomics UK (COG-UK) Consortium | Jacquelyn Wynn, Mairead Hyland, The Lighthouse Lab in Alderley Park and Alex Alderton, Roberto Amato, Jeffrey Barrett, Sonia Goncalves, Ewan Harrison, David K. Jackson, Ian Johnston, Dominic Kwiatkowski, Cordelia Langford, John Sillitoe on behalf of the Wellcome Sanger Institute COVID-19 Surveillance Team |
| EPI_ISL_2469289 | Health Services Laboratories | Wellcome Sanger Institute for the COVID-19 Genomics UK (COG-UK) Consortium | Health Services Laboratories and Alex Alderton, Roberto Amato, Jeffrey Barrett, Sonia Goncalves, Ewan Harrison, David K. Jackson, Ian Johnston, Dominic Kwiatkowski, Cordelia Langford, John Sillitoe on behalf of the Wellcome Sanger Institute COVID-19 Surveillance Team |
| EPI_ISL_2469290 | Lighthouse Lab in Alderley Park | Wellcome Sanger Institute for the COVID-19 Genomics UK (COG-UK) Consortium | Jacquelyn Wynn, Mairead Hyland, The Lighthouse Lab in Alderley Park and Alex Alderton, Roberto Amato, Jeffrey Barrett, Sonia Goncalves, Ewan Harrison, David K. Jackson, Ian Johnston, Dominic Kwiatkowski, Cordelia Langford, John Sillitoe on behalf of the Wellcome Sanger Institute COVID-19 Surveillance Team |
| EPI_ISL_2469291 | Lighthouse Lab in Milton Keynes | Wellcome Sanger Institute for the COVID-19 Genomics UK (COG-UK) Consortium | The Lighthouse Lab in Milton Keynes and Alex Alderton, Roberto Amato, Jeffrey Barrett, Sonia Goncalves, Ewan Harrison, David K. Jackson, Ian Johnston, Dominic Kwiatkowski, Cordelia Langford, John Sillitoe on behalf of the Wellcome Sanger Institute COVID-19 Surveillance Team |
| EPI_ISL_2469292 | Lighthouse Lab in Alderley Park | Wellcome Sanger Institute for the COVID-19 Genomics UK (COG-UK) Consortium | Jacquelyn Wynn, Mairead Hyland, The Lighthouse Lab in Alderley Park and Alex Alderton, Roberto Amato, Jeffrey Barrett, Sonia Goncalves, Ewan Harrison, David K. Jackson, Ian Johnston, Dominic Kwiatkowski, Cordelia Langford, John Sillitoe on behalf of the Wellcome Sanger Institute COVID-19 Surveillance Team |
| EPI_ISL_2469293 | Health Services Laboratories | Wellcome Sanger Institute for the COVID-19 Genomics UK (COG-UK) Consortium | Health Services Laboratories and Alex Alderton, Roberto Amato, Jeffrey Barrett, Sonia Goncalves, Ewan Harrison, David K. Jackson, Ian Johnston, Dominic Kwiatkowski, Cordelia Langford, John Sillitoe on behalf of the Wellcome Sanger Institute COVID-19 Surveillance Team |
| EPI_ISL_2469294 | Lighthouse Lab in Alderley Park | Wellcome Sanger Institute for the COVID-19 Genomics UK (COG-UK) Consortium | Jacquelyn Wynn, Mairead Hyland, The Lighthouse Lab in Alderley Park and Alex Alderton, Roberto Amato, Jeffrey Barrett, Sonia Goncalves, Ewan Harrison, David K. Jackson, Ian Johnston, Dominic Kwiatkowski, Cordelia Langford, John Sillitoe on behalf of the Wellcome Sanger Institute COVID-19 Surveillance Team |
| EPI_ISL_2469295 | Health Services Laboratories | Wellcome Sanger Institute for the COVID-19 Genomics UK (COG-UK) Consortium | Health Services Laboratories and Alex Alderton, Roberto Amato, Jeffrey Barrett, Sonia Goncalves, Ewan Harrison, David K. Jackson, Ian Johnston, Dominic Kwiatkowski, Cordelia Langford, John Sillitoe on behalf of the Wellcome Sanger Institute COVID-19 Surveillance Team |

| | | | |
|---|---|---|---|
| EPI_ISL_2469296 | Lighthouse Lab in Alderley Park | Wellcome Sanger Institute for the COVID-19 Genomics UK (COG-UK) Consortium | Jacquelyn Wynn, Mairead Hyland, The Lighthouse Lab in Alderley Park and Alex Alderton, Roberto Amato, Jeffrey Barrett, Sonia Goncalves, Ewan Harrison, David K. Jackson, Ian Johnston, Dominic Kwiatkowski, Cordelia Langford, John Sillitoe on behalf of the Wellcome Sanger Institute COVID-19 Surveillance Team |
| EPI_ISL_2469297 | Lighthouse Lab in Milton Keynes | Wellcome Sanger Institute for the COVID-19 Genomics UK (COG-UK) Consortium | The Lighthouse Lab in Milton Keynes and Alex Alderton, Roberto Amato, Jeffrey Barrett, Sonia Goncalves, Ewan Harrison, David K. Jackson, Ian Johnston, Dominic Kwiatkowski, Cordelia Langford, John Sillitoe on behalf of the Wellcome Sanger Institute COVID-19 Surveillance Team |
| EPI_ISL_2469298 | Health Services Laboratories | Wellcome Sanger Institute for the COVID-19 Genomics UK (COG-UK) Consortium | Health Services Laboratories and Alex Alderton, Roberto Amato, Jeffrey Barrett, Sonia Goncalves, Ewan Harrison, David K. Jackson, Ian Johnston, Dominic Kwiatkowski, Cordelia Langford, John Sillitoe on behalf of the Wellcome Sanger Institute COVID-19 Surveillance Team |
| EPI_ISL_2469299 | Lighthouse Lab in Milton Keynes | Wellcome Sanger Institute for the COVID-19 Genomics UK (COG-UK) Consortium | The Lighthouse Lab in Milton Keynes and Alex Alderton, Roberto Amato, Jeffrey Barrett, Sonia Goncalves, Ewan Harrison, David K. Jackson, Ian Johnston, Dominic Kwiatkowski, Cordelia Langford, John Sillitoe on behalf of the Wellcome Sanger Institute COVID-19 Surveillance Team |
| EPI_ISL_2469300 | Health Services Laboratories | Wellcome Sanger Institute for the COVID-19 Genomics UK (COG-UK) Consortium | Health Services Laboratories and Alex Alderton, Roberto Amato, Jeffrey Barrett, Sonia Goncalves, Ewan Harrison, David K. Jackson, Ian Johnston, Dominic Kwiatkowski, Cordelia Langford, John Sillitoe on behalf of the Wellcome Sanger Institute COVID-19 Surveillance Team |
| EPI_ISL_2469302 | Lighthouse Lab in Milton Keynes | Wellcome Sanger Institute for the COVID-19 Genomics UK (COG-UK) Consortium | The Lighthouse Lab in Milton Keynes and Alex Alderton, Roberto Amato, Jeffrey Barrett, Sonia Goncalves, Ewan Harrison, David K. Jackson, Ian Johnston, Dominic Kwiatkowski, Cordelia Langford, John Sillitoe on behalf of the Wellcome Sanger Institute COVID-19 Surveillance Team |
| EPI_ISL_2469303, EPI_ISL_2469304, EPI_ISL_2469305 | Health Services Laboratories | Wellcome Sanger Institute for the COVID-19 Genomics UK (COG-UK) Consortium | Health Services Laboratories and Alex Alderton, Roberto Amato, Jeffrey Barrett, Sonia Goncalves, Ewan Harrison, David K. Jackson, Ian Johnston, Dominic Kwiatkowski, Cordelia Langford, John Sillitoe on behalf of the Wellcome Sanger Institute COVID-19 Surveillance Team |
| EPI_ISL_2469306 | Lighthouse Lab in Alderley Park | Wellcome Sanger Institute for the COVID-19 Genomics UK (COG-UK) Consortium | Jacquelyn Wynn, Mairead Hyland, The Lighthouse Lab in Alderley Park and Alex Alderton, Roberto Amato, Jeffrey Barrett, Sonia Goncalves, Ewan Harrison, David K. Jackson, Ian Johnston, Dominic Kwiatkowski, Cordelia Langford, John Sillitoe on behalf of the Wellcome Sanger Institute COVID-19 Surveillance Team |
| EPI_ISL_2469309 | Health Services Laboratories | Wellcome Sanger Institute for the COVID-19 Genomics UK (COG-UK) Consortium | Health Services Laboratories and Alex Alderton, Roberto Amato, Jeffrey Barrett, Sonia Goncalves, Ewan Harrison, David K. Jackson, Ian Johnston, Dominic Kwiatkowski, Cordelia Langford, John Sillitoe on behalf of the Wellcome Sanger Institute COVID-19 Surveillance Team |
| EPI_ISL_2469310, EPI_ISL_2469311, EPI_ISL_2469312, EPI_ISL_2469313 | Lighthouse Lab in Alderley Park | Wellcome Sanger Institute for the COVID-19 Genomics UK (COG-UK) Consortium | Jacquelyn Wynn, Mairead Hyland, The Lighthouse Lab in Alderley Park and Alex Alderton, Roberto Amato, Jeffrey Barrett, Sonia Goncalves, Ewan Harrison, David K. Jackson, Ian Johnston, Dominic Kwiatkowski, Cordelia Langford, John Sillitoe on behalf of the Wellcome Sanger Institute COVID-19 Surveillance Team |
| EPI_ISL_2469314 | Lighthouse Lab in Milton Keynes | Wellcome Sanger Institute for the COVID-19 Genomics UK (COG-UK) Consortium | The Lighthouse Lab in Milton Keynes and Alex Alderton, Roberto Amato, Jeffrey Barrett, Sonia Goncalves, Ewan Harrison, David K. Jackson, Ian Johnston, Dominic Kwiatkowski, Cordelia Langford, John Sillitoe on behalf of the Wellcome Sanger Institute COVID-19 Surveillance Team |
| EPI_ISL_2469315 | Health Services Laboratories | Wellcome Sanger Institute for the COVID-19 Genomics UK (COG-UK) Consortium | Health Services Laboratories and Alex Alderton, Roberto Amato, Jeffrey Barrett, Sonia Goncalves, Ewan Harrison, David K. Jackson, Ian Johnston, Dominic Kwiatkowski, Cordelia Langford, John Sillitoe on behalf of the Wellcome Sanger Institute COVID-19 Surveillance Team |
| EPI_ISL_2469316 | Lighthouse Lab in Milton Keynes | Wellcome Sanger Institute for the COVID-19 Genomics UK (COG-UK) Consortium | The Lighthouse Lab in Milton Keynes and Alex Alderton, Roberto Amato, Jeffrey Barrett, Sonia Goncalves, Ewan Harrison, David K. Jackson, Ian Johnston, Dominic Kwiatkowski, Cordelia Langford, John Sillitoe on behalf of the Wellcome Sanger Institute COVID-19 Surveillance Team |
| EPI_ISL_2469317 | Health Services Laboratories | Wellcome Sanger Institute for the COVID-19 Genomics UK (COG-UK) Consortium | Health Services Laboratories and Alex Alderton, Roberto Amato, Jeffrey Barrett, Sonia Goncalves, Ewan Harrison, David K. Jackson, Ian Johnston, Dominic Kwiatkowski, Cordelia Langford, John Sillitoe on behalf of the Wellcome Sanger Institute COVID-19 Surveillance Team |
| EPI_ISL_2469318 | Lighthouse Lab in Glasgow | Wellcome Sanger Institute for the COVID-19 Genomics UK (COG-UK) Consortium | Harper VanSteenhouse, Yumi Kasai, David Gray, Carol Clugston, Anna Dominiczak and Alex Alderton, Roberto Amato, Jeffrey Barrett, Sonia Goncalves, Ewan Harrison, David K. Jackson, Ian Johnston, Dominic Kwiatkowski, Cordelia Langford, John Sillitoe on behalf of the Wellcome Sanger Institute COVID-19 Surveillance Team |
| EPI_ISL_2469319 | Lighthouse Lab in Alderley Park | Wellcome Sanger Institute for the COVID-19 Genomics UK (COG-UK) Consortium | Jacquelyn Wynn, Mairead Hyland, The Lighthouse Lab in Alderley Park and Alex Alderton, Roberto Amato, Jeffrey Barrett, Sonia Goncalves, Ewan Harrison, David K. Jackson, Ian Johnston, Dominic Kwiatkowski, Cordelia Langford, John Sillitoe on behalf of the Wellcome Sanger Institute COVID-19 Surveillance Team |
| EPI_ISL_2469320, EPI_ISL_2469321, EPI_ISL_2469322 | Health Services Laboratories | Wellcome Sanger Institute for the COVID-19 Genomics UK (COG-UK) Consortium | Health Services Laboratories and Alex Alderton, Roberto Amato, Jeffrey Barrett, Sonia Goncalves, Ewan Harrison, David K. Jackson, Ian Johnston, Dominic Kwiatkowski, Cordelia Langford, John Sillitoe on behalf of the Wellcome Sanger Institute COVID-19 Surveillance Team |
| EPI_ISL_2469323 | Lighthouse Lab in Milton Keynes | Wellcome Sanger Institute for the COVID-19 Genomics UK (COG-UK) Consortium | The Lighthouse Lab in Milton Keynes and Alex Alderton, Roberto Amato, Jeffrey Barrett, Sonia Goncalves, Ewan Harrison, David K. Jackson, Ian Johnston, Dominic Kwiatkowski, Cordelia Langford, John Sillitoe on behalf of the Wellcome Sanger Institute COVID-19 Surveillance Team |
| EPI_ISL_2469324, EPI_ISL_2469325 | Lighthouse Lab in Alderley Park | Wellcome Sanger Institute for the COVID-19 Genomics UK (COG-UK) Consortium | Jacquelyn Wynn, Mairead Hyland, The Lighthouse Lab in Alderley Park and Alex Alderton, Roberto Amato, Jeffrey Barrett, Sonia Goncalves, Ewan Harrison, David K. Jackson, Ian Johnston, Dominic Kwiatkowski, Cordelia Langford, John Sillitoe on behalf of the Wellcome Sanger Institute COVID-19 Surveillance Team |
| EPI_ISL_2469326, EPI_ISL_2469327 | Lighthouse Lab in Milton Keynes | Wellcome Sanger Institute for the COVID-19 Genomics UK (COG-UK) Consortium | The Lighthouse Lab in Milton Keynes and Alex Alderton, Roberto Amato, Jeffrey Barrett, Sonia Goncalves, Ewan Harrison, David K. Jackson, Ian Johnston, Dominic Kwiatkowski, Cordelia Langford, John Sillitoe on behalf of the Wellcome Sanger Institute COVID-19 Surveillance Team |
| EPI_ISL_2469328, EPI_ISL_2469329 | Lighthouse Lab in Alderley Park | Wellcome Sanger Institute for the COVID-19 Genomics UK (COG-UK) Consortium | Jacquelyn Wynn, Mairead Hyland, The Lighthouse Lab in Alderley Park and Alex Alderton, Roberto Amato, Jeffrey Barrett, Sonia Goncalves, Ewan Harrison, David K. Jackson, Ian Johnston, Dominic Kwiatkowski, Cordelia Langford, John Sillitoe on behalf of the Wellcome Sanger Institute COVID-19 Surveillance Team |
| EPI_ISL_2469330 | Health Services Laboratories | Wellcome Sanger Institute for the COVID-19 Genomics UK (COG-UK) Consortium | Health Services Laboratories and Alex Alderton, Roberto Amato, Jeffrey Barrett, Sonia Goncalves, Ewan Harrison, David K. Jackson, Ian Johnston, Dominic Kwiatkowski, Cordelia Langford, John Sillitoe on behalf of the Wellcome Sanger Institute COVID-19 Surveillance Team |
| EPI_ISL_2469331, EPI_ISL_2469332 | Lighthouse Lab in Milton Keynes | Wellcome Sanger Institute for the COVID-19 Genomics UK (COG-UK) Consortium | The Lighthouse Lab in Milton Keynes and Alex Alderton, Roberto Amato, Jeffrey Barrett, Sonia Goncalves, Ewan Harrison, David K. Jackson, Ian Johnston, Dominic Kwiatkowski, Cordelia Langford, John Sillitoe on behalf of the Wellcome Sanger Institute COVID-19 Surveillance Team |
| EPI_ISL_2469333, EPI_ISL_2469334, EPI_ISL_2469335 | Lighthouse Lab in Alderley Park | Wellcome Sanger Institute for the COVID-19 Genomics UK (COG-UK) Consortium | Jacquelyn Wynn, Mairead Hyland, The Lighthouse Lab in Alderley Park and Alex Alderton, Roberto Amato, Jeffrey Barrett, Sonia Goncalves, Ewan Harrison, David K. Jackson, Ian Johnston, Dominic Kwiatkowski, Cordelia Langford, John Sillitoe on behalf of the Wellcome Sanger Institute COVID-19 Surveillance Team |
| EPI_ISL_2469336 | Health Services Laboratories | Wellcome Sanger Institute for the COVID-19 Genomics UK (COG-UK) Consortium | Health Services Laboratories and Alex Alderton, Roberto Amato, Jeffrey Barrett, Sonia Goncalves, Ewan Harrison, David K. Jackson, Ian Johnston, Dominic Kwiatkowski, Cordelia Langford, John Sillitoe on behalf of the Wellcome Sanger Institute COVID-19 Surveillance Team |
| EPI_ISL_2469337 | Lighthouse Lab in Milton Keynes | Wellcome Sanger Institute for the COVID-19 Genomics UK (COG-UK) Consortium | The Lighthouse Lab in Milton Keynes and Alex Alderton, Roberto Amato, Jeffrey Barrett, Sonia Goncalves, Ewan Harrison, David K. Jackson, Ian Johnston, Dominic Kwiatkowski, Cordelia Langford, John Sillitoe on behalf of the Wellcome Sanger Institute COVID-19 Surveillance Team |
| EPI_ISL_2469338 | Health Services Laboratories | Wellcome Sanger Institute for the COVID-19 Genomics UK (COG-UK) Consortium | Health Services Laboratories and Alex Alderton, Roberto Amato, Jeffrey Barrett, Sonia Goncalves, Ewan Harrison, David K. Jackson, Ian Johnston, Dominic Kwiatkowski, Cordelia Langford, John Sillitoe on behalf of the Wellcome Sanger Institute COVID-19 Surveillance Team |
| EPI_ISL_2469339 | Lighthouse Lab in Milton Keynes | Wellcome Sanger Institute for the COVID-19 Genomics UK (COG-UK) Consortium | The Lighthouse Lab in Milton Keynes and Alex Alderton, Roberto Amato, Jeffrey Barrett, Sonia Goncalves, Ewan Harrison, David K. Jackson, Ian Johnston, Dominic Kwiatkowski, Cordelia Langford, John Sillitoe on behalf of the Wellcome Sanger Institute COVID-19 Surveillance Team |
| EPI_ISL_2469340 | Health Services Laboratories | Wellcome Sanger Institute for the COVID-19 Genomics UK (COG-UK) Consortium | Health Services Laboratories and Alex Alderton, Roberto Amato, Jeffrey Barrett, Sonia Goncalves, Ewan Harrison, David K. Jackson, Ian Johnston, Dominic Kwiatkowski, Cordelia Langford, John Sillitoe on behalf of the Wellcome Sanger Institute COVID-19 Surveillance Team |
| EPI_ISL_2469341 | Lighthouse Lab in Milton Keynes | Wellcome Sanger Institute for the COVID-19 Genomics UK (COG-UK) Consortium | The Lighthouse Lab in Milton Keynes and Alex Alderton, Roberto Amato, Jeffrey Barrett, Sonia Goncalves, Ewan Harrison, David K. Jackson, Ian Johnston, Dominic Kwiatkowski, Cordelia Langford, John Sillitoe on behalf of the Wellcome Sanger Institute COVID-19 Surveillance Team |
| EPI_ISL_2469342, EPI_ISL_2469343 | Lighthouse Lab in Alderley Park | Wellcome Sanger Institute for the COVID-19 Genomics UK (COG-UK) Consortium | Jacquelyn Wynn, Mairead Hyland, The Lighthouse Lab in Alderley Park and Alex Alderton, Roberto Amato, Jeffrey Barrett, Sonia Goncalves, Ewan Harrison, David K. Jackson, Ian Johnston, Dominic Kwiatkowski, Cordelia Langford, John Sillitoe on behalf of the Wellcome Sanger Institute COVID-19 Surveillance Team |
| EPI_ISL_2469345 | Health Services Laboratories | Wellcome Sanger Institute for the COVID-19 Genomics UK (COG-UK) Consortium | Health Services Laboratories and Alex Alderton, Roberto Amato, Jeffrey Barrett, Sonia Goncalves, Ewan Harrison, David K. Jackson, Ian Johnston, Dominic Kwiatkowski, Cordelia Langford, John Sillitoe on behalf of the Wellcome Sanger Institute COVID-19 Surveillance Team |
| EPI_ISL_2469346 | Lighthouse Lab in Alderley Park | Wellcome Sanger Institute for the COVID-19 Genomics UK (COG-UK) Consortium | Jacquelyn Wynn, Mairead Hyland, The Lighthouse Lab in Alderley Park and Alex Alderton, Roberto Amato, Jeffrey Barrett, Sonia Goncalves, Ewan Harrison, David K. Jackson, Ian Johnston, Dominic Kwiatkowski, Cordelia Langford, John Sillitoe on behalf of the Wellcome Sanger Institute COVID-19 Surveillance Team |
| EPI_ISL_2469347 | Health Services Laboratories | Wellcome Sanger Institute for the COVID-19 Genomics UK (COG-UK) Consortium | Health Services Laboratories and Alex Alderton, Roberto Amato, Jeffrey Barrett, Sonia Goncalves, Ewan Harrison, David K. Jackson, Ian Johnston, Dominic Kwiatkowski, Cordelia Langford, John Sillitoe on behalf of the Wellcome Sanger Institute COVID-19 Surveillance Team |
| EPI_ISL_2469348 | Lighthouse Lab in Alderley Park | Wellcome Sanger Institute for the COVID-19 Genomics UK (COG-UK) Consortium | Jacquelyn Wynn, Mairead Hyland, The Lighthouse Lab in Alderley Park and Alex Alderton, Roberto Amato, Jeffrey Barrett, Sonia Goncalves, Ewan Harrison, David K. Jackson, Ian Johnston, Dominic Kwiatkowski, Cordelia Langford, John Sillitoe on behalf of the Wellcome Sanger Institute COVID-19 Surveillance Team |
| EPI_ISL_2469349 | Health Services Laboratories | Wellcome Sanger Institute for the COVID-19 Genomics UK (COG-UK) Consortium | Health Services Laboratories and Alex Alderton, Roberto Amato, Jeffrey Barrett, Sonia Goncalves, Ewan Harrison, David K. Jackson, Ian Johnston, Dominic Kwiatkowski, Cordelia Langford, John Sillitoe on behalf of the Wellcome Sanger Institute COVID-19 Surveillance Team |
| EPI_ISL_2469350, EPI_ISL_2469351, EPI_ISL_2469352 | Lighthouse Lab in Alderley Park | Wellcome Sanger Institute for the COVID-19 Genomics UK (COG-UK) Consortium | Jacquelyn Wynn, Mairead Hyland, The Lighthouse Lab in Alderley Park and Alex Alderton, Roberto Amato, Jeffrey Barrett, Sonia Goncalves, Ewan Harrison, David K. Jackson, Ian Johnston, Dominic Kwiatkowski, Cordelia Langford, John Sillitoe on behalf of the Wellcome Sanger Institute COVID-19 Surveillance Team |
| EPI_ISL_2469353, EPI_ISL_2469354 | Health Services Laboratories | Wellcome Sanger Institute for the COVID-19 Genomics UK (COG-UK) Consortium | Health Services Laboratories and Alex Alderton, Roberto Amato, Jeffrey Barrett, Sonia Goncalves, Ewan Harrison, David K. Jackson, Ian Johnston, Dominic Kwiatkowski, Cordelia Langford, John Sillitoe on behalf of the Wellcome Sanger Institute COVID-19 Surveillance Team |
| EPI_ISL_2469355, EPI_ISL_2469357 | Lighthouse Lab in Milton Keynes | Wellcome Sanger Institute for the COVID-19 Genomics UK (COG-UK) Consortium | The Lighthouse Lab in Milton Keynes and Alex Alderton, Roberto Amato, Jeffrey Barrett, Sonia Goncalves, Ewan Harrison, David K. Jackson, Ian Johnston, Dominic Kwiatkowski, Cordelia Langford, John Sillitoe on behalf of the Wellcome Sanger Institute COVID-19 Surveillance Team |
| EPI_ISL_2469358, EPI_ISL_2469359 | Lighthouse Lab in Alderley Park | Wellcome Sanger Institute for the COVID-19 Genomics UK (COG-UK) Consortium | Jacquelyn Wynn, Mairead Hyland, The Lighthouse Lab in Alderley Park and Alex Alderton, Roberto Amato, Jeffrey Barrett, Sonia Goncalves, Ewan Harrison, David K. Jackson, Ian Johnston, Dominic Kwiatkowski, Cordelia Langford, John Sillitoe on behalf of the Wellcome Sanger Institute COVID-19 Surveillance Team |
| EPI_ISL_2469360 | Lighthouse Lab in Milton Keynes | Wellcome Sanger Institute for the COVID-19 Genomics UK (COG-UK) Consortium | The Lighthouse Lab in Milton Keynes and Alex Alderton, Roberto Amato, Jeffrey Barrett, Sonia Goncalves, Ewan Harrison, David K. Jackson, Ian Johnston, Dominic Kwiatkowski, Cordelia Langford, John Sillitoe on behalf of the Wellcome Sanger Institute COVID-19 Surveillance Team |
| EPI_ISL_2469361 | Lighthouse Lab in Alderley Park | Wellcome Sanger Institute for the COVID-19 Genomics UK (COG-UK) Consortium | Jacquelyn Wynn, Mairead Hyland, The Lighthouse Lab in Alderley Park and Alex Alderton, Roberto Amato, Jeffrey Barrett, Sonia Goncalves, Ewan Harrison, David K. Jackson, Ian Johnston, Dominic Kwiatkowski, Cordelia Langford, John Sillitoe on behalf of the Wellcome Sanger Institute COVID-19 Surveillance Team |
| EPI_ISL_2469362, | Health Services Laboratories | Wellcome Sanger Institute for the COVID-19 Genomics UK (COG-UK) | Health Services Laboratories and Alex Alderton, Roberto Amato, Jeffrey Barrett, Sonia Goncalves, Ewan Harrison, David K. Jackson, Ian Johnston, Dominic Kwiatkowski, Cordelia Langford, John Sillitoe on behalf of the Wellcome Sanger Institute |

| | | | |
|---|---|---|---|
| EPI_ISL_2469363, EPI_ISL_2469364, EPI_ISL_2469365 | | Consortium | COVID-19 Surveillance Team |
| EPI_ISL_2469366 | Lighthouse Lab in Milton Keynes | Wellcome Sanger Institute for the COVID-19 Genomics UK (COG-UK) Consortium | The Lighthouse Lab in Milton Keynes and Alex Alderton, Roberto Amato, Jeffrey Barrett, Sonia Goncalves, Ewan Harrison, David K. Jackson, Ian Johnston, Dominic Kwiatkowski, Cordelia Langford, John Sillitoe on behalf of the Wellcome Sanger Institute COVID-19 Surveillance Team |
| EPI_ISL_2469367, EPI_ISL_2469368 | Lighthouse Lab in Alderley Park | Wellcome Sanger Institute for the COVID-19 Genomics UK (COG-UK) Consortium | Jacquelyn Wynn, Mairead Hyland, The Lighthouse Lab in Alderley Park and Alex Alderton, Roberto Amato, Jeffrey Barrett, Sonia Goncalves, Ewan Harrison, David K. Jackson, Ian Johnston, Dominic Kwiatkowski, Cordelia Langford, John Sillitoe on behalf of the Wellcome Sanger Institute COVID-19 Surveillance Team |
| EPI_ISL_2469369, EPI_ISL_2469370 | Health Services Laboratories | Wellcome Sanger Institute for the COVID-19 Genomics UK (COG-UK) Consortium | Health Services Laboratories and Alex Alderton, Roberto Amato, Jeffrey Barrett, Sonia Goncalves, Ewan Harrison, David K. Jackson, Ian Johnston, Dominic Kwiatkowski, Cordelia Langford, John Sillitoe on behalf of the Wellcome Sanger Institute COVID-19 Surveillance Team |
| EPI_ISL_2469372 | Lighthouse Lab in Milton Keynes | Wellcome Sanger Institute for the COVID-19 Genomics UK (COG-UK) Consortium | The Lighthouse Lab in Milton Keynes and Alex Alderton, Roberto Amato, Jeffrey Barrett, Sonia Goncalves, Ewan Harrison, David K. Jackson, Ian Johnston, Dominic Kwiatkowski, Cordelia Langford, John Sillitoe on behalf of the Wellcome Sanger Institute COVID-19 Surveillance Team |
| EPI_ISL_2469373 | Health Services Laboratories | Wellcome Sanger Institute for the COVID-19 Genomics UK (COG-UK) Consortium | Health Services Laboratories and Alex Alderton, Roberto Amato, Jeffrey Barrett, Sonia Goncalves, Ewan Harrison, David K. Jackson, Ian Johnston, Dominic Kwiatkowski, Cordelia Langford, John Sillitoe on behalf of the Wellcome Sanger Institute COVID-19 Surveillance Team |
| EPI_ISL_2469374 | Lighthouse Lab in Milton Keynes | Wellcome Sanger Institute for the COVID-19 Genomics UK (COG-UK) Consortium | The Lighthouse Lab in Milton Keynes and Alex Alderton, Roberto Amato, Jeffrey Barrett, Sonia Goncalves, Ewan Harrison, David K. Jackson, Ian Johnston, Dominic Kwiatkowski, Cordelia Langford, John Sillitoe on behalf of the Wellcome Sanger Institute COVID-19 Surveillance Team |
| EPI_ISL_2469375, EPI_ISL_2469377, EPI_ISL_2469378 | Lighthouse Lab in Alderley Park | Wellcome Sanger Institute for the COVID-19 Genomics UK (COG-UK) Consortium | Jacquelyn Wynn, Mairead Hyland, The Lighthouse Lab in Alderley Park and Alex Alderton, Roberto Amato, Jeffrey Barrett, Sonia Goncalves, Ewan Harrison, David K. Jackson, Ian Johnston, Dominic Kwiatkowski, Cordelia Langford, John Sillitoe on behalf of the Wellcome Sanger Institute COVID-19 Surveillance Team |
| EPI_ISL_2469380, EPI_ISL_2469381 | Health Services Laboratories | Wellcome Sanger Institute for the COVID-19 Genomics UK (COG-UK) Consortium | Health Services Laboratories and Alex Alderton, Roberto Amato, Jeffrey Barrett, Sonia Goncalves, Ewan Harrison, David K. Jackson, Ian Johnston, Dominic Kwiatkowski, Cordelia Langford, John Sillitoe on behalf of the Wellcome Sanger Institute COVID-19 Surveillance Team |
| EPI_ISL_2469382 | Lighthouse Lab in Glasgow | Wellcome Sanger Institute for the COVID-19 Genomics UK (COG-UK) Consortium | Harper VanSteenhouse, Yumi Kasai, David Gray, Carol Clugston, Anna Dominiczak and Alex Alderton, Roberto Amato, Jeffrey Barrett, Sonia Goncalves, Ewan Harrison, David K. Jackson, Ian Johnston, Dominic Kwiatkowski, Cordelia Langford, John Sillitoe on behalf of the Wellcome Sanger Institute COVID-19 Surveillance Team |
| EPI_ISL_2469383, EPI_ISL_2469384 | Lighthouse Lab in Alderley Park | Wellcome Sanger Institute for the COVID-19 Genomics UK (COG-UK) Consortium | Jacquelyn Wynn, Mairead Hyland, The Lighthouse Lab in Alderley Park and Alex Alderton, Roberto Amato, Jeffrey Barrett, Sonia Goncalves, Ewan Harrison, David K. Jackson, Ian Johnston, Dominic Kwiatkowski, Cordelia Langford, John Sillitoe on behalf of the Wellcome Sanger Institute COVID-19 Surveillance Team |
| EPI_ISL_2469385, EPI_ISL_2469386 | Health Services Laboratories | Wellcome Sanger Institute for the COVID-19 Genomics UK (COG-UK) Consortium | Health Services Laboratories and Alex Alderton, Roberto Amato, Jeffrey Barrett, Sonia Goncalves, Ewan Harrison, David K. Jackson, Ian Johnston, Dominic Kwiatkowski, Cordelia Langford, John Sillitoe on behalf of the Wellcome Sanger Institute COVID-19 Surveillance Team |
| EPI_ISL_2469387 | Lighthouse Lab in Alderley Park | Wellcome Sanger Institute for the COVID-19 Genomics UK (COG-UK) Consortium | Jacquelyn Wynn, Mairead Hyland, The Lighthouse Lab in Alderley Park and Alex Alderton, Roberto Amato, Jeffrey Barrett, Sonia Goncalves, Ewan Harrison, David K. Jackson, Ian Johnston, Dominic Kwiatkowski, Cordelia Langford, John Sillitoe on behalf of the Wellcome Sanger Institute COVID-19 Surveillance Team |
| EPI_ISL_2469388 | Lighthouse Lab in Milton Keynes | Wellcome Sanger Institute for the COVID-19 Genomics UK (COG-UK) Consortium | The Lighthouse Lab in Milton Keynes and Alex Alderton, Roberto Amato, Jeffrey Barrett, Sonia Goncalves, Ewan Harrison, David K. Jackson, Ian Johnston, Dominic Kwiatkowski, Cordelia Langford, John Sillitoe on behalf of the Wellcome Sanger Institute COVID-19 Surveillance Team |
| EPI_ISL_2469389, EPI_ISL_2469390 | Health Services Laboratories | Wellcome Sanger Institute for the COVID-19 Genomics UK (COG-UK) Consortium | Health Services Laboratories and Alex Alderton, Roberto Amato, Jeffrey Barrett, Sonia Goncalves, Ewan Harrison, David K. Jackson, Ian Johnston, Dominic Kwiatkowski, Cordelia Langford, John Sillitoe on behalf of the Wellcome Sanger Institute COVID-19 Surveillance Team |
| EPI_ISL_2469391 | Lighthouse Lab in Alderley Park | Wellcome Sanger Institute for the COVID-19 Genomics UK (COG-UK) Consortium | Jacquelyn Wynn, Mairead Hyland, The Lighthouse Lab in Alderley Park and Alex Alderton, Roberto Amato, Jeffrey Barrett, Sonia Goncalves, Ewan Harrison, David K. Jackson, Ian Johnston, Dominic Kwiatkowski, Cordelia Langford, John Sillitoe on behalf of the Wellcome Sanger Institute COVID-19 Surveillance Team |
| EPI_ISL_2469392 | Health Services Laboratories | Wellcome Sanger Institute for the COVID-19 Genomics UK (COG-UK) Consortium | Health Services Laboratories and Alex Alderton, Roberto Amato, Jeffrey Barrett, Sonia Goncalves, Ewan Harrison, David K. Jackson, Ian Johnston, Dominic Kwiatkowski, Cordelia Langford, John Sillitoe on behalf of the Wellcome Sanger Institute COVID-19 Surveillance Team |
| EPI_ISL_2469393, EPI_ISL_2469395 | Lighthouse Lab in Alderley Park | Wellcome Sanger Institute for the COVID-19 Genomics UK (COG-UK) Consortium | Jacquelyn Wynn, Mairead Hyland, The Lighthouse Lab in Alderley Park and Alex Alderton, Roberto Amato, Jeffrey Barrett, Sonia Goncalves, Ewan Harrison, David K. Jackson, Ian Johnston, Dominic Kwiatkowski, Cordelia Langford, John Sillitoe on behalf of the Wellcome Sanger Institute COVID-19 Surveillance Team |
| EPI_ISL_2469396 | Lighthouse Lab in Milton Keynes | Wellcome Sanger Institute for the COVID-19 Genomics UK (COG-UK) Consortium | The Lighthouse Lab in Milton Keynes and Alex Alderton, Roberto Amato, Jeffrey Barrett, Sonia Goncalves, Ewan Harrison, David K. Jackson, Ian Johnston, Dominic Kwiatkowski, Cordelia Langford, John Sillitoe on behalf of the Wellcome Sanger Institute COVID-19 Surveillance Team |
| EPI_ISL_2469397 | Health Services Laboratories | Wellcome Sanger Institute for the COVID-19 Genomics UK (COG-UK) Consortium | Health Services Laboratories and Alex Alderton, Roberto Amato, Jeffrey Barrett, Sonia Goncalves, Ewan Harrison, David K. Jackson, Ian Johnston, Dominic Kwiatkowski, Cordelia Langford, John Sillitoe on behalf of the Wellcome Sanger Institute COVID-19 Surveillance Team |
| EPI_ISL_2469398 | Lighthouse Lab in Alderley Park | Wellcome Sanger Institute for the COVID-19 Genomics UK (COG-UK) Consortium | Jacquelyn Wynn, Mairead Hyland, The Lighthouse Lab in Alderley Park and Alex Alderton, Roberto Amato, Jeffrey Barrett, Sonia Goncalves, Ewan Harrison, David K. Jackson, Ian Johnston, Dominic Kwiatkowski, Cordelia Langford, John Sillitoe on behalf of the Wellcome Sanger Institute COVID-19 Surveillance Team |
| EPI_ISL_2469399, EPI_ISL_2469400 | Health Services Laboratories | Wellcome Sanger Institute for the COVID-19 Genomics UK (COG-UK) Consortium | Health Services Laboratories and Alex Alderton, Roberto Amato, Jeffrey Barrett, Sonia Goncalves, Ewan Harrison, David K. Jackson, Ian Johnston, Dominic Kwiatkowski, Cordelia Langford, John Sillitoe on behalf of the Wellcome Sanger Institute COVID-19 Surveillance Team |
| EPI_ISL_2469401 | Lighthouse Lab in Alderley Park | Wellcome Sanger Institute for the COVID-19 Genomics UK (COG-UK) Consortium | Jacquelyn Wynn, Mairead Hyland, The Lighthouse Lab in Alderley Park and Alex Alderton, Roberto Amato, Jeffrey Barrett, Sonia Goncalves, Ewan Harrison, David K. Jackson, Ian Johnston, Dominic Kwiatkowski, Cordelia Langford, John Sillitoe on behalf of the Wellcome Sanger Institute COVID-19 Surveillance Team |
| EPI_ISL_2469402 | Health Services Laboratories | Wellcome Sanger Institute for the COVID-19 Genomics UK (COG-UK) Consortium | Health Services Laboratories and Alex Alderton, Roberto Amato, Jeffrey Barrett, Sonia Goncalves, Ewan Harrison, David K. Jackson, Ian Johnston, Dominic Kwiatkowski, Cordelia Langford, John Sillitoe on behalf of the Wellcome Sanger Institute COVID-19 Surveillance Team |
| EPI_ISL_2469403 | Lighthouse Lab in Milton Keynes | Wellcome Sanger Institute for the COVID-19 Genomics UK (COG-UK) Consortium | The Lighthouse Lab in Milton Keynes and Alex Alderton, Roberto Amato, Jeffrey Barrett, Sonia Goncalves, Ewan Harrison, David K. Jackson, Ian Johnston, Dominic Kwiatkowski, Cordelia Langford, John Sillitoe on behalf of the Wellcome Sanger Institute COVID-19 Surveillance Team |
| EPI_ISL_2469404, EPI_ISL_2469405, EPI_ISL_2469406 | Health Services Laboratories | Wellcome Sanger Institute for the COVID-19 Genomics UK (COG-UK) Consortium | Health Services Laboratories and Alex Alderton, Roberto Amato, Jeffrey Barrett, Sonia Goncalves, Ewan Harrison, David K. Jackson, Ian Johnston, Dominic Kwiatkowski, Cordelia Langford, John Sillitoe on behalf of the Wellcome Sanger Institute COVID-19 Surveillance Team |
| EPI_ISL_2469408, EPI_ISL_2469409 | Lighthouse Lab in Alderley Park | Wellcome Sanger Institute for the COVID-19 Genomics UK (COG-UK) Consortium | Jacquelyn Wynn, Mairead Hyland, The Lighthouse Lab in Alderley Park and Alex Alderton, Roberto Amato, Jeffrey Barrett, Sonia Goncalves, Ewan Harrison, David K. Jackson, Ian Johnston, Dominic Kwiatkowski, Cordelia Langford, John Sillitoe on behalf of the Wellcome Sanger Institute COVID-19 Surveillance Team |
| EPI_ISL_2469412 | Lighthouse Lab in Milton Keynes | Wellcome Sanger Institute for the COVID-19 Genomics UK (COG-UK) Consortium | The Lighthouse Lab in Milton Keynes and Alex Alderton, Roberto Amato, Jeffrey Barrett, Sonia Goncalves, Ewan Harrison, David K. Jackson, Ian Johnston, Dominic Kwiatkowski, Cordelia Langford, John Sillitoe on behalf of the Wellcome Sanger Institute COVID-19 Surveillance Team |
| EPI_ISL_2469413 | Lighthouse Lab in Alderley Park | Wellcome Sanger Institute for the COVID-19 Genomics UK (COG-UK) Consortium | Jacquelyn Wynn, Mairead Hyland, The Lighthouse Lab in Alderley Park and Alex Alderton, Roberto Amato, Jeffrey Barrett, Sonia Goncalves, Ewan Harrison, David K. Jackson, Ian Johnston, Dominic Kwiatkowski, Cordelia Langford, John Sillitoe on behalf of the Wellcome Sanger Institute COVID-19 Surveillance Team |
| EPI_ISL_2469414 | Health Services Laboratories | Wellcome Sanger Institute for the COVID-19 Genomics UK (COG-UK) Consortium | Health Services Laboratories and Alex Alderton, Roberto Amato, Jeffrey Barrett, Sonia Goncalves, Ewan Harrison, David K. Jackson, Ian Johnston, Dominic Kwiatkowski, Cordelia Langford, John Sillitoe on behalf of the Wellcome Sanger Institute COVID-19 Surveillance Team |
| EPI_ISL_2469415, EPI_ISL_2469416 | Lighthouse Lab in Milton Keynes | Wellcome Sanger Institute for the COVID-19 Genomics UK (COG-UK) Consortium | The Lighthouse Lab in Milton Keynes and Alex Alderton, Roberto Amato, Jeffrey Barrett, Sonia Goncalves, Ewan Harrison, David K. Jackson, Ian Johnston, Dominic Kwiatkowski, Cordelia Langford, John Sillitoe on behalf of the Wellcome Sanger Institute COVID-19 Surveillance Team |
| EPI_ISL_2469417, EPI_ISL_2469418 | Health Services Laboratories | Wellcome Sanger Institute for the COVID-19 Genomics UK (COG-UK) Consortium | Health Services Laboratories and Alex Alderton, Roberto Amato, Jeffrey Barrett, Sonia Goncalves, Ewan Harrison, David K. Jackson, Ian Johnston, Dominic Kwiatkowski, Cordelia Langford, John Sillitoe on behalf of the Wellcome Sanger Institute COVID-19 Surveillance Team |
| EPI_ISL_2469419, EPI_ISL_2469420, EPI_ISL_2469421 | Lighthouse Lab in Alderley Park | Wellcome Sanger Institute for the COVID-19 Genomics UK (COG-UK) Consortium | Jacquelyn Wynn, Mairead Hyland, The Lighthouse Lab in Alderley Park and Alex Alderton, Roberto Amato, Jeffrey Barrett, Sonia Goncalves, Ewan Harrison, David K. Jackson, Ian Johnston, Dominic Kwiatkowski, Cordelia Langford, John Sillitoe on behalf of the Wellcome Sanger Institute COVID-19 Surveillance Team |
| EPI_ISL_2469422 | Lighthouse Lab in Milton Keynes | Wellcome Sanger Institute for the COVID-19 Genomics UK (COG-UK) Consortium | The Lighthouse Lab in Milton Keynes and Alex Alderton, Roberto Amato, Jeffrey Barrett, Sonia Goncalves, Ewan Harrison, David K. Jackson, Ian Johnston, Dominic Kwiatkowski, Cordelia Langford, John Sillitoe on behalf of the Wellcome Sanger Institute COVID-19 Surveillance Team |
| EPI_ISL_2469423 | Lighthouse Lab in Alderley Park | Wellcome Sanger Institute for the COVID-19 Genomics UK (COG-UK) Consortium | Jacquelyn Wynn, Mairead Hyland, The Lighthouse Lab in Alderley Park and Alex Alderton, Roberto Amato, Jeffrey Barrett, Sonia Goncalves, Ewan Harrison, David K. Jackson, Ian Johnston, Dominic Kwiatkowski, Cordelia Langford, John Sillitoe on behalf of the Wellcome Sanger Institute COVID-19 Surveillance Team |
| EPI_ISL_2469424 | Lighthouse Lab in Milton Keynes | Wellcome Sanger Institute for the COVID-19 Genomics UK (COG-UK) Consortium | The Lighthouse Lab in Milton Keynes and Alex Alderton, Roberto Amato, Jeffrey Barrett, Sonia Goncalves, Ewan Harrison, David K. Jackson, Ian Johnston, Dominic Kwiatkowski, Cordelia Langford, John Sillitoe on behalf of the Wellcome Sanger Institute COVID-19 Surveillance Team |
| EPI_ISL_2469425 | Lighthouse Lab in Alderley Park | Wellcome Sanger Institute for the COVID-19 Genomics UK (COG-UK) Consortium | Jacquelyn Wynn, Mairead Hyland, The Lighthouse Lab in Alderley Park and Alex Alderton, Roberto Amato, Jeffrey Barrett, Sonia Goncalves, Ewan Harrison, David K. Jackson, Ian Johnston, Dominic Kwiatkowski, Cordelia Langford, John Sillitoe on behalf of the Wellcome Sanger Institute COVID-19 Surveillance Team |
| EPI_ISL_2469426 | Lighthouse Lab in Milton Keynes | Wellcome Sanger Institute for the COVID-19 Genomics UK (COG-UK) Consortium | The Lighthouse Lab in Milton Keynes and Alex Alderton, Roberto Amato, Jeffrey Barrett, Sonia Goncalves, Ewan Harrison, David K. Jackson, Ian Johnston, Dominic Kwiatkowski, Cordelia Langford, John Sillitoe on behalf of the Wellcome Sanger Institute COVID-19 Surveillance Team |
| EPI_ISL_2469428 | Health Services Laboratories | Wellcome Sanger Institute for the COVID-19 Genomics UK (COG-UK) Consortium | Health Services Laboratories and Alex Alderton, Roberto Amato, Jeffrey Barrett, Sonia Goncalves, Ewan Harrison, David K. Jackson, Ian Johnston, Dominic Kwiatkowski, Cordelia Langford, John Sillitoe on behalf of the Wellcome Sanger Institute COVID-19 Surveillance Team |
| EPI_ISL_2469429 | Lighthouse Lab in Alderley Park | Wellcome Sanger Institute for the COVID-19 Genomics UK (COG-UK) Consortium | Jacquelyn Wynn, Mairead Hyland, The Lighthouse Lab in Alderley Park and Alex Alderton, Roberto Amato, Jeffrey Barrett, Sonia Goncalves, Ewan Harrison, David K. Jackson, Ian Johnston, Dominic Kwiatkowski, Cordelia Langford, John Sillitoe on behalf of the Wellcome Sanger Institute COVID-19 Surveillance Team |
| EPI_ISL_2469430 | Health Services Laboratories | Wellcome Sanger Institute for the COVID-19 Genomics UK (COG-UK) Consortium | Health Services Laboratories and Alex Alderton, Roberto Amato, Jeffrey Barrett, Sonia Goncalves, Ewan Harrison, David K. Jackson, Ian Johnston, Dominic Kwiatkowski, Cordelia Langford, John Sillitoe on behalf of the Wellcome Sanger Institute COVID-19 Surveillance Team |
| EPI_ISL_2469431 | Lighthouse Lab in Milton Keynes | Wellcome Sanger Institute for the COVID-19 Genomics UK (COG-UK) Consortium | The Lighthouse Lab in Milton Keynes and Alex Alderton, Roberto Amato, Jeffrey Barrett, Sonia Goncalves, Ewan Harrison, David K. Jackson, Ian Johnston, Dominic Kwiatkowski, Cordelia Langford, John Sillitoe on behalf of the Wellcome Sanger Institute COVID-19 Surveillance Team |
| EPI_ISL_2469432 | Health Services Laboratories | Wellcome Sanger Institute for the COVID-19 Genomics UK (COG-UK) Consortium | Health Services Laboratories and Alex Alderton, Roberto Amato, Jeffrey Barrett, Sonia Goncalves, Ewan Harrison, David K. Jackson, Ian Johnston, Dominic Kwiatkowski, Cordelia Langford, John Sillitoe on behalf of the Wellcome Sanger Institute COVID-19 Surveillance Team |
| EPI_ISL_2469433 | Lighthouse Lab in Alderley Park | Wellcome Sanger Institute for the COVID-19 Genomics UK (COG-UK) | Jacquelyn Wynn, Mairead Hyland, The Lighthouse Lab in Alderley Park and Alex Alderton, Roberto Amato, Jeffrey Barrett, Sonia Goncalves, Ewan Harrison, David K. Jackson, Ian Johnston, Dominic Kwiatkowski, Cordelia Langford, John Sillitoe on |

| | | | |
|---|---|---|---|
| | | Consortium | behalf of the Wellcome Sanger Institute COVID-19 Surveillance Team |
| EPI_ISL_2469434 | Lighthouse Lab in Glasgow | Wellcome Sanger Institute for the COVID-19 Genomics UK (COG-UK) Consortium | Harper VanSteenhouse, Yumi Kasai, David Gray, Carol Clugston, Anna Dominiczak and Alex Alderton, Roberto Amato, Jeffrey Barrett, Sonia Goncalves, Ewan Harrison, David K. Jackson, Ian Johnston, Dominic Kwiatkowski, Cordelia Langford, John Sillitoe on behalf of the Wellcome Sanger Institute COVID-19 Surveillance Team |
| EPI_ISL_2469435 | Lighthouse Lab in Milton Keynes | Wellcome Sanger Institute for the COVID-19 Genomics UK (COG-UK) Consortium | The Lighthouse Lab in Milton Keynes and Alex Alderton, Roberto Amato, Jeffrey Barrett, Sonia Goncalves, Ewan Harrison, David K. Jackson, Ian Johnston, Dominic Kwiatkowski, Cordelia Langford, John Sillitoe on behalf of the Wellcome Sanger Institute COVID-19 Surveillance Team |
| EPI_ISL_2469436 | Health Services Laboratories | Wellcome Sanger Institute for the COVID-19 Genomics UK (COG-UK) Consortium | Health Services Laboratories and Alex Alderton, Roberto Amato, Jeffrey Barrett, Sonia Goncalves, Ewan Harrison, David K. Jackson, Ian Johnston, Dominic Kwiatkowski, Cordelia Langford, John Sillitoe on behalf of the Wellcome Sanger Institute COVID-19 Surveillance Team |
| EPI_ISL_2469437, EPI_ISL_2469438 | Lighthouse Lab in Alderley Park | Wellcome Sanger Institute for the COVID-19 Genomics UK (COG-UK) Consortium | Jacquelyn Wynn, Mairead Hyland, The Lighthouse Lab in Alderley Park and Alex Alderton, Roberto Amato, Jeffrey Barrett, Sonia Goncalves, Ewan Harrison, David K. Jackson, Ian Johnston, Dominic Kwiatkowski, Cordelia Langford, John Sillitoe on behalf of the Wellcome Sanger Institute COVID-19 Surveillance Team |
| EPI_ISL_2469439, EPI_ISL_2469440 | Lighthouse Lab in Milton Keynes | Wellcome Sanger Institute for the COVID-19 Genomics UK (COG-UK) Consortium | The Lighthouse Lab in Milton Keynes and Alex Alderton, Roberto Amato, Jeffrey Barrett, Sonia Goncalves, Ewan Harrison, David K. Jackson, Ian Johnston, Dominic Kwiatkowski, Cordelia Langford, John Sillitoe on behalf of the Wellcome Sanger Institute COVID-19 Surveillance Team |
| EPI_ISL_2469441 | Lighthouse Lab in Alderley Park | Wellcome Sanger Institute for the COVID-19 Genomics UK (COG-UK) Consortium | Jacquelyn Wynn, Mairead Hyland, The Lighthouse Lab in Alderley Park and Alex Alderton, Roberto Amato, Jeffrey Barrett, Sonia Goncalves, Ewan Harrison, David K. Jackson, Ian Johnston, Dominic Kwiatkowski, Cordelia Langford, John Sillitoe on behalf of the Wellcome Sanger Institute COVID-19 Surveillance Team |
| EPI_ISL_2469442 | Lighthouse Lab in Milton Keynes | Wellcome Sanger Institute for the COVID-19 Genomics UK (COG-UK) Consortium | The Lighthouse Lab in Milton Keynes and Alex Alderton, Roberto Amato, Jeffrey Barrett, Sonia Goncalves, Ewan Harrison, David K. Jackson, Ian Johnston, Dominic Kwiatkowski, Cordelia Langford, John Sillitoe on behalf of the Wellcome Sanger Institute COVID-19 Surveillance Team |
| EPI_ISL_2469443 | Lighthouse Lab in Alderley Park | Wellcome Sanger Institute for the COVID-19 Genomics UK (COG-UK) Consortium | Jacquelyn Wynn, Mairead Hyland, The Lighthouse Lab in Alderley Park and Alex Alderton, Roberto Amato, Jeffrey Barrett, Sonia Goncalves, Ewan Harrison, David K. Jackson, Ian Johnston, Dominic Kwiatkowski, Cordelia Langford, John Sillitoe on behalf of the Wellcome Sanger Institute COVID-19 Surveillance Team |
| EPI_ISL_2469444 | Lighthouse Lab in Milton Keynes | Wellcome Sanger Institute for the COVID-19 Genomics UK (COG-UK) Consortium | The Lighthouse Lab in Milton Keynes and Alex Alderton, Roberto Amato, Jeffrey Barrett, Sonia Goncalves, Ewan Harrison, David K. Jackson, Ian Johnston, Dominic Kwiatkowski, Cordelia Langford, John Sillitoe on behalf of the Wellcome Sanger Institute COVID-19 Surveillance Team |
| EPI_ISL_2469445 | Lighthouse Lab in Alderley Park | Wellcome Sanger Institute for the COVID-19 Genomics UK (COG-UK) Consortium | Jacquelyn Wynn, Mairead Hyland, The Lighthouse Lab in Alderley Park and Alex Alderton, Roberto Amato, Jeffrey Barrett, Sonia Goncalves, Ewan Harrison, David K. Jackson, Ian Johnston, Dominic Kwiatkowski, Cordelia Langford, John Sillitoe on behalf of the Wellcome Sanger Institute COVID-19 Surveillance Team |
| EPI_ISL_2469446 | Health Services Laboratories | Wellcome Sanger Institute for the COVID-19 Genomics UK (COG-UK) Consortium | Health Services Laboratories and Alex Alderton, Roberto Amato, Jeffrey Barrett, Sonia Goncalves, Ewan Harrison, David K. Jackson, Ian Johnston, Dominic Kwiatkowski, Cordelia Langford, John Sillitoe on behalf of the Wellcome Sanger Institute COVID-19 Surveillance Team |
| EPI_ISL_2469448, EPI_ISL_2469449 | Lighthouse Lab in Alderley Park | Wellcome Sanger Institute for the COVID-19 Genomics UK (COG-UK) Consortium | Jacquelyn Wynn, Mairead Hyland, The Lighthouse Lab in Alderley Park and Alex Alderton, Roberto Amato, Jeffrey Barrett, Sonia Goncalves, Ewan Harrison, David K. Jackson, Ian Johnston, Dominic Kwiatkowski, Cordelia Langford, John Sillitoe on behalf of the Wellcome Sanger Institute COVID-19 Surveillance Team |
| EPI_ISL_2469450 | Health Services Laboratories | Wellcome Sanger Institute for the COVID-19 Genomics UK (COG-UK) Consortium | Health Services Laboratories and Alex Alderton, Roberto Amato, Jeffrey Barrett, Sonia Goncalves, Ewan Harrison, David K. Jackson, Ian Johnston, Dominic Kwiatkowski, Cordelia Langford, John Sillitoe on behalf of the Wellcome Sanger Institute COVID-19 Surveillance Team |
| EPI_ISL_2469451 | Lighthouse Lab in Alderley Park | Wellcome Sanger Institute for the COVID-19 Genomics UK (COG-UK) Consortium | Jacquelyn Wynn, Mairead Hyland, The Lighthouse Lab in Alderley Park and Alex Alderton, Roberto Amato, Jeffrey Barrett, Sonia Goncalves, Ewan Harrison, David K. Jackson, Ian Johnston, Dominic Kwiatkowski, Cordelia Langford, John Sillitoe on behalf of the Wellcome Sanger Institute COVID-19 Surveillance Team |
| EPI_ISL_2469452, EPI_ISL_2469453, EPI_ISL_2469454, EPI_ISL_2469455, EPI_ISL_2469456, EPI_ISL_2469457, EPI_ISL_2469458, EPI_ISL_2469459, EPI_ISL_2469460, EPI_ISL_2469461, EPI_ISL_2469462, EPI_ISL_2469463, EPI_ISL_2469464 | | | |
| see above | Lighthouse Lab in Milton Keynes | Wellcome Sanger Institute for the COVID-19 Genomics UK (COG-UK) Consortium | The Lighthouse Lab in Milton Keynes and Alex Alderton, Roberto Amato, Jeffrey Barrett, Sonia Goncalves, Ewan Harrison, David K. Jackson, Ian Johnston, Dominic Kwiatkowski, Cordelia Langford, John Sillitoe on behalf of the Wellcome Sanger Institute COVID-19 Surveillance Team |
| EPI_ISL_2469465 | Lighthouse Lab in Alderley Park | Wellcome Sanger Institute for the COVID-19 Genomics UK (COG-UK) Consortium | Jacquelyn Wynn, Mairead Hyland, The Lighthouse Lab in Alderley Park and Alex Alderton, Roberto Amato, Jeffrey Barrett, Sonia Goncalves, Ewan Harrison, David K. Jackson, Ian Johnston, Dominic Kwiatkowski, Cordelia Langford, John Sillitoe on behalf of the Wellcome Sanger Institute COVID-19 Surveillance Team |
| EPI_ISL_2469466 | Health Services Laboratories | Wellcome Sanger Institute for the COVID-19 Genomics UK (COG-UK) Consortium | Health Services Laboratories and Alex Alderton, Roberto Amato, Jeffrey Barrett, Sonia Goncalves, Ewan Harrison, David K. Jackson, Ian Johnston, Dominic Kwiatkowski, Cordelia Langford, John Sillitoe on behalf of the Wellcome Sanger Institute COVID-19 Surveillance Team |
| EPI_ISL_2469467, EPI_ISL_2469468 | Lighthouse Lab in Alderley Park | Wellcome Sanger Institute for the COVID-19 Genomics UK (COG-UK) Consortium | Jacquelyn Wynn, Mairead Hyland, The Lighthouse Lab in Alderley Park and Alex Alderton, Roberto Amato, Jeffrey Barrett, Sonia Goncalves, Ewan Harrison, David K. Jackson, Ian Johnston, Dominic Kwiatkowski, Cordelia Langford, John Sillitoe on behalf of the Wellcome Sanger Institute COVID-19 Surveillance Team |
| EPI_ISL_2469490, EPI_ISL_2469531, EPI_ISL_2469558 | SYNLAB MVZ Leverkusen | Robert Koch Institute | unknown |
| EPI_ISL_2469628 | SYNLAB MVZ Weiden | Robert Koch Institute | unknown |
| EPI_ISL_2469654, EPI_ISL_2469687 | SYNLAB MVZ Trier | Robert Koch Institute | unknown |
| EPI_ISL_2469754, EPI_ISL_2469755, EPI_ISL_2469756, EPI_ISL_2469757, EPI_ISL_2469758, EPI_ISL_2469759, EPI_ISL_2469760, EPI_ISL_2469761 | Med. Labor Prof. Schenk Dr. Ansorge & Kollegen | Robert Koch Institute | unknown |
| EPI_ISL_2469762, EPI_ISL_2469771, EPI_ISL_2469774, EPI_ISL_2469777, EPI_ISL_2469779 | Corona-Testzentrum ifp Institut für Produktqualität GmbH | Robert Koch Institute | unknown |
| EPI_ISL_2469837, EPI_ISL_2469839 | MVZ Dr. Eberhard & Partner Dortmund | Robert Koch Institute | unknown |
| EPI_ISL_2469850 | Medizinisches Labor Wahl Lüdenscheid | Robert Koch Institute | unknown |
| EPI_ISL_2469851, EPI_ISL_2469852 | MVZ Dr. Eberhard & Partner Dortmund | Robert Koch Institute | unknown |
| EPI_ISL_2469859 | Labor Lübeck bzw. Laborärztliche Gemeinschaftspraxis Lübeck | Robert Koch Institute | unknown |
| EPI_ISL_2469860 | Medizinisches Labor Wahl Lüdenscheid | Robert Koch Institute | unknown |
| EPI_ISL_2469871, EPI_ISL_2469876 | MVZ Dr. Eberhard & Partner Dortmund | Robert Koch Institute | unknown |
| EPI_ISL_2469878 | Medizinisches Labor Wahl Lüdenscheid | Robert Koch Institute | unknown |
| EPI_ISL_2469879, EPI_ISL_2469887, EPI_ISL_2469888, EPI_ISL_2469893, EPI_ISL_2469895, EPI_ISL_2469900, EPI_ISL_2469913, EPI_ISL_2469916 | MVZ Dr. Eberhard & Partner Dortmund | Robert Koch Institute | unknown |
| EPI_ISL_2469933 | Bioscientia Labor Wermsdorf | Robert Koch Institute | unknown |
| EPI_ISL_2469969 | Sonic - Dr. Staber & Kollegen GmbH München | Robert Koch Institute | unknown |
| EPI_ISL_2469990 | Labor 28 MVZ GmbH | Robert Koch Institute | unknown |
| EPI_ISL_2469993 | Sonic - Dr. Staber & Kollegen GmbH München | Robert Koch Institute | unknown |
| EPI_ISL_2469998 | Sonic - Bioscientia - MVZ Labor Saar GmbH | Robert Koch Institute | unknown |
| EPI_ISL_2470007 | Bioscientia Labor Wermsdorf | Robert Koch Institute | unknown |
| EPI_ISL_2470015, EPI_ISL_2470023 | Labor 28 MVZ GmbH | Robert Koch Institute | unknown |
| EPI_ISL_2470031 | LabKom - Labor Augsburg MVZ GmbH | Robert Koch Institute | unknown |
| EPI_ISL_2470032 | Bioscientia Labor Wermsdorf | Robert Koch Institute | unknown |
| EPI_ISL_2470043 | Sonic - Dr. Staber & Kollegen GmbH München | Robert Koch Institute | unknown |
| EPI_ISL_2470057 | Bioscientia Labor Wermsdorf | Robert Koch Institute | unknown |
| EPI_ISL_2470121, EPI_ISL_2470151 | Labor 28 MVZ GmbH | Robert Koch Institute | unknown |
| EPI_ISL_2470203 | Universitätsklinikum Heidelberg | Robert Koch Institute | unknown |

| | | | |
|---|---|---|---|
| EPI_ISL_2470207 | Bioscientia Labor Wermsdorf | Robert Koch Institute | unknown |
| EPI_ISL_2470211, EPI_ISL_2470223 | Bioscientia MVZ Labor Karlsruhe GmbH | Robert Koch Institute | unknown |
| EPI_ISL_2470251 | Sonic - MVZ Medizinisches Labor Bremen GmbH | Robert Koch Institute | unknown |
| EPI_ISL_2470257 | Sonic - Bioscientia - MVZ Labor Saar GmbH | Robert Koch Institute | unknown |
| EPI_ISL_2470310 | Bioscientia Labor Wermsdorf | Robert Koch Institute | unknown |
| EPI_ISL_2470320, EPI_ISL_2470321 | Bioscientia MVZ Labor Karlsruhe GmbH | Robert Koch Institute | unknown |
| EPI_ISL_2470349 | Sonic - MVZ Medizinisches Labor Bremen GmbH | Robert Koch Institute | unknown |
| EPI_ISL_2470354 | Bioscientia MVZ Labor Karlsruhe GmbH | Robert Koch Institute | unknown |
| EPI_ISL_2470359 | Sonic - Bioscientia - MVZ Labor Saar GmbH | Robert Koch Institute | unknown |
| EPI_ISL_2470373, EPI_ISL_2470396 | Bioscientia MVZ Labor Karlsruhe GmbH | Robert Koch Institute | unknown |
| EPI_ISL_2470404 | Sonic - Labor Staber MVZ Kassel | Robert Koch Institute | unknown |
| EPI_ISL_2470413, EPI_ISL_2470457, EPI_ISL_2470477, EPI_ISL_2470478 | Bioscientia Labor Wermsdorf | Robert Koch Institute | unknown |
| EPI_ISL_2470670, EPI_ISL_2470684 | Limbach - MVZ Humangenetik Ulm | Robert Koch Institute | unknown |
| EPI_ISL_2470825, EPI_ISL_2470855 | Medizinische Laboratorien Düsseldorf | Robert Koch Institute | unknown |
| EPI_ISL_2471024 | Labormedizin Darmstadt | Robert Koch Institute | unknown |
| EPI_ISL_2471045, EPI_ISL_2471054, EPI_ISL_2471055 | Labor Dr. Heidrich & Kollegen MVZ GmbH Hamburg | Robert Koch Institute | unknown |
| EPI_ISL_2471113 | MVZ Labor Krone GbR | Robert Koch Institute | unknown |
| EPI_ISL_2471310, EPI_ISL_2471456 | CENTOGENE Frankfurt Laboratory: Niederlassung Industriepark Höchst | Robert Koch Institute | unknown |
| EPI_ISL_2471464 | MVZ Labor Dr. Limbach & Kollegen GbR | Robert Koch Institute | unknown |
| EPI_ISL_2471533, EPI_ISL_2471536 | MVZ Labor Dr. Fenner und Kollegen (Standort Hamburg) | Robert Koch Institute | unknown |
| EPI_ISL_2471560, EPI_ISL_2471593 | MVZ für Laboratoriumsmedizin und Mikrobiologie Koblenz-Mittelrhein (Labor Koblenz) | Robert Koch Institute | unknown |
| EPI_ISL_2471609, EPI_ISL_2471610, EPI_ISL_2471617, EPI_ISL_2471619, EPI_ISL_2471620, EPI_ISL_2471622, EPI_ISL_2471627, EPI_ISL_2471633, EPI_ISL_2471638, EPI_ISL_2471640, EPI_ISL_2471646, EPI_ISL_2471652, EPI_ISL_2471659, EPI_ISL_2471660, EPI_ISL_2471661, EPI_ISL_2471662, EPI_ISL_2471665, EPI_ISL_2471667, EPI_ISL_2471669, EPI_ISL_2471671, EPI_ISL_2471672, EPI_ISL_2471673, EPI_ISL_2471677, EPI_ISL_2471683, EPI_ISL_2471684, EPI_ISL_2471685, EPI_ISL_2471686, EPI_ISL_2471687, EPI_ISL_2471688, EPI_ISL_2471689, EPI_ISL_2471690 | | | |
| see above | Eurofins LifeCodexx GmbH | Robert Koch Institute | unknown |
| EPI_ISL_2471693 | MVZ für Laboratoriumsmedizin und Mikrobiologie Würzburg | Robert Koch Institute | unknown |
| EPI_ISL_2471702 | Procomcure Biotech Germany GmbH | Robert Koch Institute | unknown |
| EPI_ISL_2471716 | MVZ Dr. Eberhard & Partner Dortmund | Robert Koch Institute | unknown |
| EPI_ISL_2471745 | MVZ Medizinisches Labor Hannover GmbH | Robert Koch Institute | unknown |
| EPI_ISL_2471761, EPI_ISL_2471770 | Medizinisch-Diagnostisches Labor Kempten allgäulab | Robert Koch Institute | unknown |
| EPI_ISL_2471799 | Labor Mönchengladbach MVZ Dr. Stein + Kollegen GbR | Robert Koch Institute | unknown |
| EPI_ISL_2471822 | Carl-Thiem-Klinikum Cottbus - Zentrum für Laboratoriumsmedizin Mikrobiologie und Krankenhaushygiene | Robert Koch Institute | unknown |
| EPI_ISL_2471872 | Städtisches Klinikum Dresden Institut für Klinische Chemie und Laboratoriumsmedizin | Robert Koch Institute | unknown |
| EPI_ISL_2471884 | Uniklinikum Carl Gustav Carus an der TU Dresden; Institut für Virologie | Robert Koch Institute | unknown |
| EPI_ISL_2471905 | MVZ Labor Dr. Limbach & Kollegen GbR | Robert Koch Institute | unknown |
| EPI_ISL_2471915 | Eurofins MVZ Labor Gelsenkirchen | Robert Koch Institute | unknown |
| EPI_ISL_2471943, EPI_ISL_2471954, EPI_ISL_2471955, EPI_ISL_2471959, EPI_ISL_2471962, EPI_ISL_2471966, EPI_ISL_2471969, EPI_ISL_2471970 | MDI Limbach Berlin GmbH; MVZ Labor Berlin | Robert Koch Institute | unknown |
| EPI_ISL_2471989, EPI_ISL_2471996, EPI_ISL_2472000, EPI_ISL_2472005, EPI_ISL_2472012 | IMD - Labor Oderland | Robert Koch Institute | unknown |
| EPI_ISL_2472053 | Eurofins LifeCodexx GmbH | Robert Koch Institute | unknown |
| EPI_ISL_2472061 | SYNLAB MVZ Trier | Robert Koch Institute | unknown |
| EPI_ISL_2472091, EPI_ISL_2472093 | SYNLAB MVZ Weiden | Robert Koch Institute | unknown |
| EPI_ISL_2472107, EPI_ISL_2472160 | SYNLAB MVZ Leverkusen | Robert Koch Institute | unknown |
| EPI_ISL_2472167 | SYNLAB MVZ Leinfelden-Echterdingen | Robert Koch Institute | unknown |
| EPI_ISL_2472174 | SYNLAB MVZ Leverkusen | Robert Koch Institute | unknown |
| EPI_ISL_2472177 | SYNLAB MVZ Weiden | Robert Koch Institute | unknown |
| EPI_ISL_2472206 | SYNLAB MVZ Leinfelden-Echterdingen | Robert Koch Institute | unknown |
| EPI_ISL_2472217 | SYNLAB MVZ Trier | Robert Koch Institute | unknown |
| EPI_ISL_2472223 | SYNLAB MVZ Leverkusen | Robert Koch Institute | unknown |
| EPI_ISL_2472226 | SYNLAB MVZ Weiden | Robert Koch Institute | unknown |
| EPI_ISL_2472227, EPI_ISL_2472231 | SYNLAB MVZ Trier | Robert Koch Institute | unknown |
| EPI_ISL_2472247 | SYNLAB MVZ Weiden | Robert Koch Institute | unknown |
| EPI_ISL_2472249 | SYNLAB MVZ Trier | Robert Koch Institute | unknown |
| EPI_ISL_2472251 | SYNLAB MVZ Leinfelden-Echterdingen | Robert Koch Institute | unknown |
| EPI_ISL_2472267 | SYNLAB MVZ Weiden | Robert Koch Institute | unknown |
| EPI_ISL_2472283 | SYNLAB MVZ Heidelberg | Robert Koch Institute | unknown |
| EPI_ISL_2472355 | SYNLAB MVZ Leverkusen | Robert Koch Institute | unknown |

| | | | |
|---|---|---|---|
| EPI_ISL_2472374 | MDI Limbach Berlin GmbH; MVZ Labor Berlin | Robert Koch Institute | unknown |
| EPI_ISL_2472407 | MVZ für Laboratoriumsmedizin und Mikrobiologie Koblenz-Mittelrhein (Labor Koblenz) | Robert Koch Institute | unknown |
| EPI_ISL_2472421, EPI_ISL_2472454, EPI_ISL_2472458 | MVZ Labor Dr. Limbach & Kollegen GbR | Robert Koch Institute | unknown |
| EPI_ISL_2472575 | Bayerisches Landesamt für Gesundheit und Lebensmittelsicherheit (LGL) | Robert Koch Institute | unknown |
| EPI_ISL_2472664, EPI_ISL_2472667 | Eurofins LifeCodexx GmbH | Robert Koch Institute | unknown |
| EPI_ISL_2472708, EPI_ISL_2472727, EPI_ISL_2472728, EPI_ISL_2472748, EPI_ISL_2472751 | MDI Limbach Berlin GmbH; MVZ Labor Berlin | Robert Koch Institute | unknown |
| EPI_ISL_2473331, EPI_ISL_2473442, EPI_ISL_2473455, EPI_ISL_2473473, EPI_ISL_2473476, EPI_ISL_2473486, EPI_ISL_2473487, EPI_ISL_2473498, EPI_ISL_2473512, EPI_ISL_2473513, EPI_ISL_2473517, EPI_ISL_2473520, EPI_ISL_2473525, EPI_ISL_2473528, EPI_ISL_2473529, EPI_ISL_2473547, EPI_ISL_2473550, EPI_ISL_2473551, EPI_ISL_2473627 | | | |
| see above | State Testing Facility | Altius Institute for Biomedical Research | Daniel Bates, Rebecca Bruders, Michael Buckley, Mark Frerker, Amanda Gale, Clem Green, Muhammad Halimun, Kneshay Harper, Matt Hartman, Alex Isner, Audra Johnson, Jessica Kunder, Lauren Mitchell, Jemma Nelson, Alex Nguyen, Sofia Olsson, Sadie Patraw, Tobias Ragoczy, Joshua Richards, Jean Robinston, Jacob Rodriguez, John Stamatoyannopoulos, Eric Thorland, Julia Wald |
| EPI_ISL_2473913 | Reditus Laboratories | Reditus Laboratories | Joshua J. Geltz, Ph.D., Robert M. Sgambelluri, Ph.D., Rex Dyer, Ph.D., Cassy Philips, M.S., Alexa Eichelberger, M.S. |
| EPI_ISL_2476259, EPI_ISL_2476260, EPI_ISL_2476261, EPI_ISL_2476262, EPI_ISL_2476263, EPI_ISL_2476311, EPI_ISL_2476312, EPI_ISL_2476313, EPI_ISL_2476314, EPI_ISL_2476315, EPI_ISL_2476316, EPI_ISL_2476317, EPI_ISL_2476318, EPI_ISL_2476319, EPI_ISL_2476320, EPI_ISL_2476321, EPI_ISL_2476322, EPI_ISL_2476323, EPI_ISL_2476324, EPI_ISL_2476325, EPI_ISL_2476326, EPI_ISL_2476327, EPI_ISL_2476328, EPI_ISL_2476329, EPI_ISL_2476330, EPI_ISL_2476331, EPI_ISL_2476332 | | | |
| see above | Dutch COVID-19 response team | National Institute for Public Health and the Environment (RIVM) | Adam Meijer, Harry Vennema, Dirk Eggink, Jeroen Cremer, Sharon van den Brink, Bas van der Veer, AnneMarie van den Brandt, Lisa Wijsman, Kim Freriks, Ryanne Jaarsma, Euníce Then, Lynn Aarts, Sanne Bos, Melissa van Tuil, Linda van de Nes, Florian Zwagemaker, Dennis Schmitz, Annelies Kroneman, Karim Hajji, Chantal Reusken, on behalf of the national COVID-19 response team |
| EPI_ISL_2476346 | LBM AX BIO Ocean | LBM GenBio | A.ZRHIDRI; G.SEVRIN; S.KEMENY;G.EGEA; T.DURET. |
| EPI_ISL_2476408 | Department of Pathology, National Institute of Infectious Diseases | Department of Pathology, National Institute of Infectious Diseases | Harutaka Katano, Shun Iida, Yuichiro Hirata, Tadaki Suzuki |
| EPI_ISL_2477869, EPI_ISL_2478413, EPI_ISL_2478415, EPI_ISL_2478457, EPI_ISL_2478515, EPI_ISL_2478641, EPI_ISL_2478642, EPI_ISL_2478963, EPI_ISL_2478988, EPI_ISL_2479000, EPI_ISL_2479023, EPI_ISL_2479127, EPI_ISL_2479454, EPI_ISL_2479472, EPI_ISL_2479575, EPI_ISL_2479576, EPI_ISL_2479581, EPI_ISL_2479594, EPI_ISL_2479693, EPI_ISL_2479857, EPI_ISL_2479874 | | | |
| see above | Edmonton Provincial Lab | Public Health Agency of Canada (PHAC) National Microbiology Laboratory | Buss, E, Croxen M, Deo A, Dieu P, Gill K, Ferrato C, Khan F, Koleva P, Li V, Lloyd C, Lynch T, Ma R, Murphy S, Pabbaraju K, Shokoples S, Tipples G, Thayer J, Whitehouse M, Wong A, Yu C, Zelyas N |
| EPI_ISL_2479934, EPI_ISL_2479935, EPI_ISL_2479936, EPI_ISL_2479937, EPI_ISL_2479938, EPI_ISL_2479939, EPI_ISL_2479940, EPI_ISL_2479941, EPI_ISL_2479942, EPI_ISL_2479943, EPI_ISL_2479944, EPI_ISL_2479945, EPI_ISL_2479946, EPI_ISL_2479947 | | | |
| see above | National Influenza Center, National Institute of Hygiene and Epidemiology (NIHE) | National Influenza Center, National Institute of Hygiene and Epidemiology (NIHE) | Le Quynh Mai, Taichiro Takemura, Nguyen Le Khanh Hang, Hoang Vu Mai Phuong, Ung Thi Hong Trang, Le Thi Thanh, Nguyen Vu Son, Vuong Duc Cuong, Pham Thi Hien, Tran Thu Huong, Nguyen Phuong Anh, Nguyen Co Thach, Tran Thi Hien, Vũ Hiếu, Nguyen Thi Kieu Anh, Dang Duc Anh |
| EPI_ISL_2479951, EPI_ISL_2479952, EPI_ISL_2479953, EPI_ISL_2479954, EPI_ISL_2479955, EPI_ISL_2479956, EPI_ISL_2479957, EPI_ISL_2479958, EPI_ISL_2479959, EPI_ISL_2479960, EPI_ISL_2479961, EPI_ISL_2479962, EPI_ISL_2479963, EPI_ISL_2479964, EPI_ISL_2479965, EPI_ISL_2479966, EPI_ISL_2479967, EPI_ISL_2479969, EPI_ISL_2479970, EPI_ISL_2479971, EPI_ISL_2479972, EPI_ISL_2479973, EPI_ISL_2479974, EPI_ISL_2479975, EPI_ISL_2479976, EPI_ISL_2479977, EPI_ISL_2479978, EPI_ISL_2479979, EPI_ISL_2479980, EPI_ISL_2479981, EPI_ISL_2479982, EPI_ISL_2479983, EPI_ISL_2479984, EPI_ISL_2479985 | | | |
| see above | National Public Health Laboartory | CSIR-Institute of Genomics and Integrative Biology | Runa Jha, Jyoti Acharya, Lokbandhu Chaudhary, Suni Dangol, Priya Jha |
| EPI_ISL_2480001, EPI_ISL_2480006, EPI_ISL_2480007, EPI_ISL_2480008, EPI_ISL_2480010, EPI_ISL_2480011, EPI_ISL_2480012, EPI_ISL_2480013, EPI_ISL_2480069, EPI_ISL_2480070, EPI_ISL_2480071, EPI_ISL_2480072, EPI_ISL_2480088, EPI_ISL_2480089, EPI_ISL_2480092, EPI_ISL_2480103, EPI_ISL_2480112, EPI_ISL_2480114, EPI_ISL_2480118, EPI_ISL_2480119, EPI_ISL_2480120, EPI_ISL_2480123, EPI_ISL_2480129, EPI_ISL_2480131, EPI_ISL_2480132, EPI_ISL_2480133, EPI_ISL_2480136, EPI_ISL_2480138, EPI_ISL_2480140, EPI_ISL_2480145, EPI_ISL_2480161, EPI_ISL_2480166 | | | |
| see above | Fulgent Genetics | Centers for Disease Control and Prevention Division of Viral Diseases, Pathogen Discovery | Dakota Howard, Dhwani Batra, Peter W. Cook, Kara Moser, Adrian Paskey, Jason Caravas, Benjamin Rambo-Martin, Shatavia Morrison, Christopher Gulvick, Scott Sammons, Yvette Unoarumhi, Darlene Wagner, Matthew Schmerer, Harry Gao, Mickey Li, John Gao, Joseph Fierro, Benafsh Sapra, Becky Tsai, Yan Meng, Doreen Ng, James Xie, Clinton R. Paden, Duncan MacCannell |
| EPI_ISL_2480235, EPI_ISL_2480242, EPI_ISL_2480295, EPI_ISL_2480300, EPI_ISL_2480369, EPI_ISL_2480419, EPI_ISL_2480438, EPI_ISL_2480454, EPI_ISL_2480470, EPI_ISL_2480510, EPI_ISL_2480511, EPI_ISL_2480537, EPI_ISL_2480540, EPI_ISL_2480541, EPI_ISL_2480545, EPI_ISL_2480549, EPI_ISL_2480550, EPI_ISL_2480606, EPI_ISL_2480708, EPI_ISL_2480711, EPI_ISL_2480712, EPI_ISL_2480740, EPI_ISL_2480786, EPI_ISL_2480809, EPI_ISL_2480824, EPI_ISL_2480890, EPI_ISL_2480952, EPI_ISL_2480967, EPI_ISL_2480969, EPI_ISL_2480973, EPI_ISL_2480975, EPI_ISL_2480977, EPI_ISL_2480978, EPI_ISL_2481023, EPI_ISL_2481057, EPI_ISL_2481087, EPI_ISL_2481101, EPI_ISL_2481152, EPI_ISL_2481165, EPI_ISL_2481166, EPI_ISL_2481167, EPI_ISL_2481168, EPI_ISL_2481169, EPI_ISL_2481174, EPI_ISL_2481175, EPI_ISL_2481201, EPI_ISL_2481207, EPI_ISL_2481232, EPI_ISL_2481235, EPI_ISL_2481273, EPI_ISL_2481288, EPI_ISL_2481383 | | | |
| see above | Laboratory Corporation of America | Centers for Disease Control and Prevention Division of Viral Diseases, Pathogen Discovery | Dakota Howard, Dhwani Batra, Peter W. Cook, Kara Moser, Adrian Paskey, Jason Caravas, Benjamin Rambo-Martin, Shatavia Morrison, Christopher Gulvick, Scott Sammons, Yvette Unoarumhi, Darlene Wagner, Matthew Schmerer, Minoo Agarwal, Eyad Almasri, Debbie Boles, Ayla Burns, Nuthawin Charoensri, Oren Cohen, Susan Countryman, Mary Ann Cristobal, Bobbi Croy, Suzanne Dale, Hrushikesh Deshmukh, Amanda Douglas, Vincent Drouillon, Marcia Eisenberg, Howard Engler, Rama Ghatti, Prashant Gupta, Susan Hicks, Jake Humphrey, Lax Iyer, Lisa Pfefferle, Manoj Jain, Matthew Robinson, Mohan Kolli, Brian Krueger, Tim Kuphal, Stanley Letovsky, Michael Levandoski, Craig Lukasik, Jonathan Meltzer, Brian Norvell, Mindy Nye, Scott Parker, Christos Petropoulos, John Pruitt, Steven Ragan, Scott Ryan, Mike Sapeta, Jana Schroth, Suresh Babu Selvaraju, Goran Stevovic, Amanda Suchanek, Andrea Throop, Lyndon Tilson, Thomas Urban, Joe Voshell, Kimberly Wagner, Jonathan Williams, Mary Williamson, Qian Zeng, Tricia Zwiefelhofer, Clinton R. Paden, Duncan MacCannell |
| EPI_ISL_2481390 | Centre for Dengue Research and AICBU, Department of Immunology and Molecular Medicine | Centre for Dengue Research and AICBU, Department of Immunology and Molecular Medicine | Chandima Jeewandara, Deshni Jayathilaka, Dinuka Ariyaratne, Deshan Madhusanka, Diyanath Ranasinghe, Laksiri Gomes, Gathsaurie Neelika Malavige |
| EPI_ISL_2481401, EPI_ISL_2481420, EPI_ISL_2481443, EPI_ISL_2481456, EPI_ISL_2481466, EPI_ISL_2481479, EPI_ISL_2481480, EPI_ISL_2481494, EPI_ISL_2481525, EPI_ISL_2481537, EPI_ISL_2481592, EPI_ISL_2481595, EPI_ISL_2481596, EPI_ISL_2481600, EPI_ISL_2481622, EPI_ISL_2481645, EPI_ISL_2481654, EPI_ISL_2481657, EPI_ISL_2481667, EPI_ISL_2481668, EPI_ISL_2481669, EPI_ISL_2481672, EPI_ISL_2481679, EPI_ISL_2481689, EPI_ISL_2481692, EPI_ISL_2481708, EPI_ISL_2481709, EPI_ISL_2481742, EPI_ISL_2481765, EPI_ISL_2481770, EPI_ISL_2481772, EPI_ISL_2481788, EPI_ISL_2481789, EPI_ISL_2481790, EPI_ISL_2481798, EPI_ISL_2481800, EPI_ISL_2481804, EPI_ISL_2481827, EPI_ISL_2481851, EPI_ISL_2481883, EPI_ISL_2481915, EPI_ISL_2481932, EPI_ISL_2481933, EPI_ISL_2481952, EPI_ISL_2481956, EPI_ISL_2481957, EPI_ISL_2482018, EPI_ISL_2482065, EPI_ISL_2482066, EPI_ISL_2482068, EPI_ISL_2482093, EPI_ISL_2482105, EPI_ISL_2482106, EPI_ISL_2482112, EPI_ISL_2482113, EPI_ISL_2482114, EPI_ISL_2482119, EPI_ISL_2482153, EPI_ISL_2482155, EPI_ISL_2482162, EPI_ISL_2482168, EPI_ISL_2482169, EPI_ISL_2482172, EPI_ISL_2482173, EPI_ISL_2482174, EPI_ISL_2482190, EPI_ISL_2482201, EPI_ISL_2482202, EPI_ISL_2482205, EPI_ISL_2482218, EPI_ISL_2482229, EPI_ISL_2482232, EPI_ISL_2482235, EPI_ISL_2482236, EPI_ISL_2482273, EPI_ISL_2482276, EPI_ISL_2482278 | | | |
| see above | Laboratory Corporation of America | Centers for Disease Control and Prevention Division of Viral Diseases, Pathogen Discovery | Dakota Howard, Dhwani Batra, Peter W. Cook, Kara Moser, Adrian Paskey, Jason Caravas, Benjamin Rambo-Martin, Shatavia Morrison, Christopher Gulvick, Scott Sammons, Yvette Unoarumhi, Darlene Wagner, Matthew Schmerer, Minoo Agarwal, Eyad Almasri, Debbie Boles, Ayla Burns, Nuthawin Charoensri, Oren Cohen, Susan Countryman, Mary Ann Cristobal, Bobbi Croy, Suzanne Dale, Hrushikesh Deshmukh, Amanda Douglas, Vincent Drouillon, Marcia Eisenberg, Howard Engler, Rama Ghatti, Prashant Gupta, Susan Hicks, Jake Humphrey, Lax Iyer, Lisa Pfefferle, Manoj Jain, Matthew Robinson, Mohan Kolli, Brian Krueger, Tim Kuphal, Stanley Letovsky, Michael Levandoski, Craig Lukasik, Jonathan Meltzer, Brian Norvell, Mindy Nye, Scott Parker, Christos Petropoulos, John Pruitt, Steven Ragan, Scott Ryan, Mike Sapeta, Jana Schroth, Suresh Babu Selvaraju, Goran Stevovic, Amanda Suchanek, Andrea Throop, Lyndon Tilson, Thomas Urban, Joe Voshell, Kimberly Wagner, Jonathan Williams, Mary Williamson, Qian Zeng, Tricia Zwiefelhofer, Clinton R. Paden, Duncan MacCannell |
| EPI_ISL_2482434, EPI_ISL_2482436, EPI_ISL_2482442, EPI_ISL_2482443, EPI_ISL_2482445, EPI_ISL_2482448, EPI_ISL_2482454, EPI_ISL_2482455, EPI_ISL_2482456, EPI_ISL_2482457 | Microbiological Diagnostic Unit - Public Health Laboratory (MDU-PHL) | MDU-PHL | Seemann T., Sait, M.L., Sherry, N.L. |
| EPI_ISL_2482761 | DYOMEDEA-LABORATOIRE DE LA SAUVEGARDE | CNR Virus des Infections Respiratoires - France SUD | Antonin Bal, Gregory Destras, Gwendolyne Burfin, Hadrien Regue, Quentin Semanas, Martine Valette, Bruno Lina, Laurence Josset |
| EPI_ISL_2482846 | LBM UNIBIO VALENTIN | CNR Virus des Infections Respiratoires - France SUD | Antonin Bal, Gregory Destras, Gwendolyne Burfin, Hadrien Regue, Quentin Semanas, Martine Valette, Bruno Lina, Laurence Josset |
| EPI_ISL_2482885 | CENTRE HOSPITALIER DU HAUT BUGEY | CNR Virus des Infections Respiratoires - France SUD | Antonin Bal, Gregory Destras, Gwendolyne Burfin, Hadrien Regue, Quentin Semanas, Martine Valette, Bruno Lina, Laurence Josset |
| EPI_ISL_2483024 | CNR Virus des Infections Respiratoires - France SUD | CNR Virus des Infections Respiratoires - France SUD | Antonin Bal, Gregory Destras, Gwendolyne Burfin, Hadrien Regue, Quentin Semanas, Martine Valette, Bruno Lina, Laurence Josset |
| EPI_ISL_2483086 | Akershus University Hospital, Department for Microbiology and Infectious Disease Control | Norwegian Institute of Public Health, Department of Virology | Kathrine Stene-Johansen, Kamilla Heddeland Instefjord, Hilde Elshaug, Garcia Llorente Ignacio, Jon Bråte, Engebretsen Serina Beate,Pedersen Benedikte Nevjen, Line Victoria Moen, Debech Nadia, Atiya R Ali,Marie Paulsen Madsen, Rasmus Riis Kopperud, Hilde Vollan, Karoline Bragstad, Olav Hungnes |
| EPI_ISL_2483123 | Unilabs Laboratory Medicine | Norwegian Institute of Public Health, Department of Virology | Kathrine Stene-Johansen, Kamilla Heddeland Instefjord, Hilde Elshaug, Garcia Llorente Ignacio, Jon Bråte, Engebretsen Serina Beate,Pedersen Benedikte Nevjen, Line Victoria Moen, Debech Nadia, Atiya R Ali,Marie Paulsen Madsen, Rasmus Riis Kopperud, Hilde Vollan, Karoline Bragstad, Olav Hungnes |
| EPI_ISL_2483493, EPI_ISL_2483494, EPI_ISL_2483495, EPI_ISL_2483499, EPI_ISL_2483509 | Dutch COVID-19 response team | Erasmus Medical Center | Bas Oude Munnink, Reina Sikkema, David Nieuwenhuijse, Irina Chestakova, Anne van der Linden, Marjan Boter, Emmanuelle Munger, Corine GeurtsvanKessel, Annemiek van der Eijk, Richard Molenkamp, Marion Koopmans, on behalf of the Dutch national COVID-19 respo |
| EPI_ISL_2483779 | Massachusetts State Public Health Laboratory | Massachusetts State Public Health Laboratory | Andrew Lang, Timelia Fink, Glen Gallagher, Sandra Smole |
| EPI_ISL_2483841, EPI_ISL_2483846 | National Platform bis UMONS/Jolimont | National Platform bis UMONS/Jolimont | François Dufrasne, Guillaume Bayon-Vicente, Florian Juszczak, Gautier Detry, Ruddy Wattiez |
| EPI_ISL_2483893, EPI_ISL_2483899, EPI_ISL_2483900, EPI_ISL_2483903, EPI_ISL_2483905, EPI_ISL_2483909, EPI_ISL_2483911, EPI_ISL_2483921, EPI_ISL_2483927, EPI_ISL_2483931 | SYNLAB | GIGA Medical Genomics | Keith Durkin, Maria Artesi, Sébastien Bontems, Raphaël Boreux, Bouchra Boujemla, Nathalie Renotte, Cécile Meex, Pierrette Melin, Marie-Pierre Hayette, Vincent Bours |

| | | | |
|---|---|---|---|
| EPI_ISL_2483940 | University of Liège COVID-19 testing center | GIGA Medical Genomics | Keith Durkin, Maria Artesi, Bouchra Boujemla, Nathalie Renotte, Cécile Meex, Sébastien Bontems, Fabrice Bureau, Laurent Gillet, Wouter Coppieters, Marie-Pierre Hayette, Vincent Bours |
| EPI_ISL_2483955, EPI_ISL_2483960, EPI_ISL_2483961, EPI_ISL_2483967, EPI_ISL_2484022 | Hospital General Universitario Gregorio Marañón | Hospital General Universitario Gregorio Marañón | Sergio Buenestado Serrano, Pedro Sola Campoy, Laura Pérez-Lago, Cristina Rodriguez-Grande, Marta Herranz Martin, Victor Manuel de la Cueva, Julia Suárez, Pilar Catalán, Patricia Muñoz, Darío García de Viedma |
| EPI_ISL_2484043, EPI_ISL_2484048, EPI_ISL_2484073, EPI_ISL_2484082, EPI_ISL_2484090 | Arizona State University | Arizona State University | Peter T. Skidmore, LaRinda A. Holland, Matthew F. Smith, Rabia Maqsood, Nicholas J. Mellor, Joy M. Blain, Valerie Harris, Joshua LaBaer, Vel Murugan, Efrem S. Lim |
| EPI_ISL_2484620, EPI_ISL_2484621, EPI_ISL_2484622, EPI_ISL_2484623, EPI_ISL_2484624, EPI_ISL_2484625, EPI_ISL_2484626, EPI_ISL_2484627, EPI_ISL_2484628, EPI_ISL_2484629, EPI_ISL_2484630, EPI_ISL_2484631, EPI_ISL_2484632, EPI_ISL_2484633, EPI_ISL_2484634, EPI_ISL_2484635, EPI_ISL_2484636, EPI_ISL_2484637, EPI_ISL_2484638, EPI_ISL_2484639, EPI_ISL_2484640, EPI_ISL_2484641, EPI_ISL_2484642, EPI_ISL_2484643, EPI_ISL_2484644 | | | |
| see above | NL-Dr. Leonard A. Miller Centre for Health Services | National Microbiology Laboratory (NML) | Anna Majer, Shari Tyson, Grace Seo, Philip Mabon, Elsie Grudeski, Rhiannon Huzarewich, Russell Mandes, Anneliese Landgraff, Jennifer Tanner, Natalie Knox, Morag Graham, Gary Van Domselaar, Robert Needle, Yang Yu, Adel Malek, Laura Gilbert, George Zahariadis, Nathalie Bastien, Yan Li, Timothy Booth, Darian Hole, Madison Chapel, Kirsten Biggar, Kerri Smith, CanCOGeN's metadata curation team, Public Health Agency of Canada CanCOGeN team |
| EPI_ISL_2484683 | Arkansas Department of Health | Center for Global Health, University of New Mexico Health Sciences Center | Daryl Domman, Kurt Schwalm, Valerie Morley, Atul Kothari, Joshua L. Kennedy, Darrell Dinwiddie |
| EPI_ISL_2484686, EPI_ISL_2484692 | Baptist Health Medical Center | Center for Global Health, University of New Mexico Health Sciences Center | Daryl Domman, Kurt Schwalm, Valerie Morley, Amanda Novack, Dirk Haselow, Joshua L. Kennedy, Darrell Dinwiddie |
| EPI_ISL_2484694 | Arkansas Children's Hospital | Center for Global Health, University of New Mexico Health Sciences Center | Daryl Domman, Kurt Schwalm, Valerie Morley, Amanda Novack, Dirk Haselow, Joshua L. Kennedy, Darrell Dinwiddie |
| EPI_ISL_2484748, EPI_ISL_2484751 | Arkansas Department of Health | Center for Global Health, University of New Mexico Health Sciences Center | Daryl Domman, Kurt Schwalm, Valerie Morley, Atul Kothari, Joshua L. Kennedy, Darrell Dinwiddie |
| EPI_ISL_2484752 | Arkansas Children's Hospital | Center for Global Health, University of New Mexico Health Sciences Center | Daryl Domman, Kurt Schwalm, Valerie Morley, Amanda Novack, Dirk Haselow, Joshua L. Kennedy, Darrell Dinwiddie |
| EPI_ISL_2484753 | Baptist Health Medical Center | Center for Global Health, University of New Mexico Health Sciences Center | Daryl Domman, Kurt Schwalm, Valerie Morley, Amanda Novack, Dirk Haselow, Joshua L. Kennedy, Darrell Dinwiddie |
| EPI_ISL_2484754, EPI_ISL_2484755 | Arkansas Children's Hospital | Center for Global Health, University of New Mexico Health Sciences Center | Daryl Domman, Kurt Schwalm, Valerie Morley, Amanda Novack, Dirk Haselow, Joshua L. Kennedy, Darrell Dinwiddie |
| EPI_ISL_2484895 | Amdavad Municipal Corporation, Ahmedabad | Gujarat Biotechnology Research Centre | Manish Kumar,Twinkle Soni,Sonal Sharma ,Nitin Savaliya,Dinesh Kumar,Ramesh Pandit,Dalip Singh Rathore,Hiren Mandalia,Chaitanya Joshi,Madhvi Joshi |
| EPI_ISL_2484897 | Amdavad Municipal Corporation, Ahmedabad | Gujarat Biotechnology Research Centre | Sonal Sharma ,Nitin Savaliya,Dinesh Kumar,Ramesh Pandit,Dalip Singh Rathore,Twinkle Soni,Hiren Mandalia,Manish Kumar,Chaitanya Joshi,Madhvi Joshi |
| EPI_ISL_2484898 | Amdavad Municipal Corporation, Ahmedabad | Gujarat Biotechnology Research Centre | Nitin Savaliya,Dinesh Kumar,Ramesh Pandit,Dalip Singh Rathore,Twinkle Soni,Sonal Sharma ,Hiren Mandalia,Manish Kumar,Chaitanya Joshi,Madhvi Joshi |
| EPI_ISL_2484934 | Massachusetts State Public Health Laboratory | Massachusetts State Public Health Laboratory | Andrew Lang, Timelia Fink, Glen Gallagher, Sandra Smole |
| EPI_ISL_2484977, EPI_ISL_2485157 | Austrian Agency for Health and Food Safety (AGES) | Bergthaler laboratory, CeMM Research Center for Molecular Medicine of the Austrian Academy of Sciences | Lukas Endler, Anna Schedl, Fabian Amman, Petr Triska, Thomas Penz, Benedikt Agerer, Maelle Le Moing, Michael Schuster, Bekir Erguner, Jan Laine, Martin Senekowitsch, Christoph Bock, Andreas Bergthaler |
| EPI_ISL_2485257 | Lighthouse Laboratory Plymouth | Wellcome Sanger Institute for the COVID-19 Genomics UK (COG-UK) Consortium | Lighthouse Laboratory Plymouth and Alex Alderton, Roberto Amato, Jeffrey Barrett, Sonia Goncalves, Ewan Harrison, David K. Jackson, Ian Johnston, Dominic Kwiatkowski, Cordelia Langford, John Sillitoe on behalf of the Wellcome Sanger Institute COVID-19 Surveillance Team |
| EPI_ISL_2485258 | Lighthouse Lab in Milton Keynes | Wellcome Sanger Institute for the COVID-19 Genomics UK (COG-UK) Consortium | The Lighthouse Lab in Milton Keynes and Alex Alderton, Roberto Amato, Jeffrey Barrett, Sonia Goncalves, Ewan Harrison, David K. Jackson, Ian Johnston, Dominic Kwiatkowski, Cordelia Langford, John Sillitoe on behalf of the Wellcome Sanger Institute COVID-19 Surveillance Team |
| EPI_ISL_2485259 | Lighthouse Lab in Alderley Park | Wellcome Sanger Institute for the COVID-19 Genomics UK (COG-UK) Consortium | Jacquelyn Wynn, Mairead Hyland, The Lighthouse Lab in Alderley Park and Alex Alderton, Roberto Amato, Jeffrey Barrett, Sonia Goncalves, Ewan Harrison, David K. Jackson, Ian Johnston, Dominic Kwiatkowski, Cordelia Langford, John Sillitoe on behalf of the Wellcome Sanger Institute COVID-19 Surveillance Team |
| EPI_ISL_2485260, EPI_ISL_2485261 | Lighthouse Lab in Milton Keynes | Wellcome Sanger Institute for the COVID-19 Genomics UK (COG-UK) Consortium | The Lighthouse Lab in Milton Keynes and Alex Alderton, Roberto Amato, Jeffrey Barrett, Sonia Goncalves, Ewan Harrison, David K. Jackson, Ian Johnston, Dominic Kwiatkowski, Cordelia Langford, John Sillitoe on behalf of the Wellcome Sanger Institute COVID-19 Surveillance Team |
| EPI_ISL_2485262 | Lighthouse Lab in Alderley Park | Wellcome Sanger Institute for the COVID-19 Genomics UK (COG-UK) Consortium | Jacquelyn Wynn, Mairead Hyland, The Lighthouse Lab in Alderley Park and Alex Alderton, Roberto Amato, Jeffrey Barrett, Sonia Goncalves, Ewan Harrison, David K. Jackson, Ian Johnston, Dominic Kwiatkowski, Cordelia Langford, John Sillitoe on behalf of the Wellcome Sanger Institute COVID-19 Surveillance Team |
| EPI_ISL_2485263 | Lighthouse Laboratory Plymouth | Wellcome Sanger Institute for the COVID-19 Genomics UK (COG-UK) Consortium | Lighthouse Laboratory Plymouth and Alex Alderton, Roberto Amato, Jeffrey Barrett, Sonia Goncalves, Ewan Harrison, David K. Jackson, Ian Johnston, Dominic Kwiatkowski, Cordelia Langford, John Sillitoe on behalf of the Wellcome Sanger Institute COVID-19 Surveillance Team |
| EPI_ISL_2485265, EPI_ISL_2485266, EPI_ISL_2485267, EPI_ISL_2485268, EPI_ISL_2485269, EPI_ISL_2485270, EPI_ISL_2485272 | Lighthouse Lab in Alderley Park | Wellcome Sanger Institute for the COVID-19 Genomics UK (COG-UK) Consortium | Jacquelyn Wynn, Mairead Hyland, The Lighthouse Lab in Alderley Park and Alex Alderton, Roberto Amato, Jeffrey Barrett, Sonia Goncalves, Ewan Harrison, David K. Jackson, Ian Johnston, Dominic Kwiatkowski, Cordelia Langford, John Sillitoe on behalf of the Wellcome Sanger Institute COVID-19 Surveillance Team |
| EPI_ISL_2485273 | Lighthouse Laboratory Plymouth | Wellcome Sanger Institute for the COVID-19 Genomics UK (COG-UK) Consortium | Lighthouse Laboratory Plymouth and Alex Alderton, Roberto Amato, Jeffrey Barrett, Sonia Goncalves, Ewan Harrison, David K. Jackson, Ian Johnston, Dominic Kwiatkowski, Cordelia Langford, John Sillitoe on behalf of the Wellcome Sanger Institute COVID-19 Surveillance Team |
| EPI_ISL_2485274, EPI_ISL_2485275 | Lighthouse Lab in Alderley Park | Wellcome Sanger Institute for the COVID-19 Genomics UK (COG-UK) Consortium | Jacquelyn Wynn, Mairead Hyland, The Lighthouse Lab in Alderley Park and Alex Alderton, Roberto Amato, Jeffrey Barrett, Sonia Goncalves, Ewan Harrison, David K. Jackson, Ian Johnston, Dominic Kwiatkowski, Cordelia Langford, John Sillitoe on behalf of the Wellcome Sanger Institute COVID-19 Surveillance Team |
| EPI_ISL_2485276 | Lighthouse Lab in Milton Keynes | Wellcome Sanger Institute for the COVID-19 Genomics UK (COG-UK) Consortium | The Lighthouse Lab in Milton Keynes and Alex Alderton, Roberto Amato, Jeffrey Barrett, Sonia Goncalves, Ewan Harrison, David K. Jackson, Ian Johnston, Dominic Kwiatkowski, Cordelia Langford, John Sillitoe on behalf of the Wellcome Sanger Institute COVID-19 Surveillance Team |
| EPI_ISL_2485277 | Lighthouse Lab in Alderley Park | Wellcome Sanger Institute for the COVID-19 Genomics UK (COG-UK) Consortium | Jacquelyn Wynn, Mairead Hyland, The Lighthouse Lab in Alderley Park and Alex Alderton, Roberto Amato, Jeffrey Barrett, Sonia Goncalves, Ewan Harrison, David K. Jackson, Ian Johnston, Dominic Kwiatkowski, Cordelia Langford, John Sillitoe on behalf of the Wellcome Sanger Institute COVID-19 Surveillance Team |
| EPI_ISL_2485278, EPI_ISL_2485279 | Lighthouse Lab in Milton Keynes | Wellcome Sanger Institute for the COVID-19 Genomics UK (COG-UK) Consortium | The Lighthouse Lab in Milton Keynes and Alex Alderton, Roberto Amato, Jeffrey Barrett, Sonia Goncalves, Ewan Harrison, David K. Jackson, Ian Johnston, Dominic Kwiatkowski, Cordelia Langford, John Sillitoe on behalf of the Wellcome Sanger Institute COVID-19 Surveillance Team |
| EPI_ISL_2485280, EPI_ISL_2485281 | Lighthouse Lab in Alderley Park | Wellcome Sanger Institute for the COVID-19 Genomics UK (COG-UK) Consortium | Jacquelyn Wynn, Mairead Hyland, The Lighthouse Lab in Alderley Park and Alex Alderton, Roberto Amato, Jeffrey Barrett, Sonia Goncalves, Ewan Harrison, David K. Jackson, Ian Johnston, Dominic Kwiatkowski, Cordelia Langford, John Sillitoe on behalf of the Wellcome Sanger Institute COVID-19 Surveillance Team |
| EPI_ISL_2485282 | Lighthouse Lab in Milton Keynes | Wellcome Sanger Institute for the COVID-19 Genomics UK (COG-UK) Consortium | The Lighthouse Lab in Milton Keynes and Alex Alderton, Roberto Amato, Jeffrey Barrett, Sonia Goncalves, Ewan Harrison, David K. Jackson, Ian Johnston, Dominic Kwiatkowski, Cordelia Langford, John Sillitoe on behalf of the Wellcome Sanger Institute COVID-19 Surveillance Team |
| EPI_ISL_2485283, EPI_ISL_2485284 | Lighthouse Lab in Alderley Park | Wellcome Sanger Institute for the COVID-19 Genomics UK (COG-UK) Consortium | Jacquelyn Wynn, Mairead Hyland, The Lighthouse Lab in Alderley Park and Alex Alderton, Roberto Amato, Jeffrey Barrett, Sonia Goncalves, Ewan Harrison, David K. Jackson, Ian Johnston, Dominic Kwiatkowski, Cordelia Langford, John Sillitoe on behalf of the Wellcome Sanger Institute COVID-19 Surveillance Team |
| EPI_ISL_2485286 | Lighthouse Lab in Milton Keynes | Wellcome Sanger Institute for the COVID-19 Genomics UK (COG-UK) Consortium | The Lighthouse Lab in Milton Keynes and Alex Alderton, Roberto Amato, Jeffrey Barrett, Sonia Goncalves, Ewan Harrison, David K. Jackson, Ian Johnston, Dominic Kwiatkowski, Cordelia Langford, John Sillitoe on behalf of the Wellcome Sanger Institute COVID-19 Surveillance Team |
| EPI_ISL_2485287 | Lighthouse Lab in Alderley Park | Wellcome Sanger Institute for the COVID-19 Genomics UK (COG-UK) Consortium | Jacquelyn Wynn, Mairead Hyland, The Lighthouse Lab in Alderley Park and Alex Alderton, Roberto Amato, Jeffrey Barrett, Sonia Goncalves, Ewan Harrison, David K. Jackson, Ian Johnston, Dominic Kwiatkowski, Cordelia Langford, John Sillitoe on behalf of the Wellcome Sanger Institute COVID-19 Surveillance Team |
| EPI_ISL_2485288, EPI_ISL_2485289 | Lighthouse Laboratory Plymouth | Wellcome Sanger Institute for the COVID-19 Genomics UK (COG-UK) Consortium | Lighthouse Laboratory Plymouth and Alex Alderton, Roberto Amato, Jeffrey Barrett, Sonia Goncalves, Ewan Harrison, David K. Jackson, Ian Johnston, Dominic Kwiatkowski, Cordelia Langford, John Sillitoe on behalf of the Wellcome Sanger Institute COVID-19 Surveillance Team |
| EPI_ISL_2485290, EPI_ISL_2485291, EPI_ISL_2485292 | Lighthouse Lab in Milton Keynes | Wellcome Sanger Institute for the COVID-19 Genomics UK (COG-UK) Consortium | The Lighthouse Lab in Milton Keynes and Alex Alderton, Roberto Amato, Jeffrey Barrett, Sonia Goncalves, Ewan Harrison, David K. Jackson, Ian Johnston, Dominic Kwiatkowski, Cordelia Langford, John Sillitoe on behalf of the Wellcome Sanger Institute COVID-19 Surveillance Team |
| EPI_ISL_2485293 | Lighthouse Lab in Alderley Park | Wellcome Sanger Institute for the COVID-19 Genomics UK (COG-UK) Consortium | Jacquelyn Wynn, Mairead Hyland, The Lighthouse Lab in Alderley Park and Alex Alderton, Roberto Amato, Jeffrey Barrett, Sonia Goncalves, Ewan Harrison, David K. Jackson, Ian Johnston, Dominic Kwiatkowski, Cordelia Langford, John Sillitoe on behalf of the Wellcome Sanger Institute COVID-19 Surveillance Team |
| EPI_ISL_2485294 | Lighthouse Laboratory Plymouth | Wellcome Sanger Institute for the COVID-19 Genomics UK (COG-UK) Consortium | Lighthouse Laboratory Plymouth and Alex Alderton, Roberto Amato, Jeffrey Barrett, Sonia Goncalves, Ewan Harrison, David K. Jackson, Ian Johnston, Dominic Kwiatkowski, Cordelia Langford, John Sillitoe on behalf of the Wellcome Sanger Institute COVID-19 Surveillance Team |
| EPI_ISL_2485295, EPI_ISL_2485296, EPI_ISL_2485298 | Lighthouse Lab in Alderley Park | Wellcome Sanger Institute for the COVID-19 Genomics UK (COG-UK) Consortium | Jacquelyn Wynn, Mairead Hyland, The Lighthouse Lab in Alderley Park and Alex Alderton, Roberto Amato, Jeffrey Barrett, Sonia Goncalves, Ewan Harrison, David K. Jackson, Ian Johnston, Dominic Kwiatkowski, Cordelia Langford, John Sillitoe on behalf of the Wellcome Sanger Institute COVID-19 Surveillance Team |
| EPI_ISL_2485299, EPI_ISL_2485300 | Lighthouse Lab in Milton Keynes | Wellcome Sanger Institute for the COVID-19 Genomics UK (COG-UK) Consortium | The Lighthouse Lab in Milton Keynes and Alex Alderton, Roberto Amato, Jeffrey Barrett, Sonia Goncalves, Ewan Harrison, David K. Jackson, Ian Johnston, Dominic Kwiatkowski, Cordelia Langford, John Sillitoe on behalf of the Wellcome Sanger Institute COVID-19 Surveillance Team |
| EPI_ISL_2485302, EPI_ISL_2485303 | Lighthouse Lab in Alderley Park | Wellcome Sanger Institute for the COVID-19 Genomics UK (COG-UK) Consortium | Jacquelyn Wynn, Mairead Hyland, The Lighthouse Lab in Alderley Park and Alex Alderton, Roberto Amato, Jeffrey Barrett, Sonia Goncalves, Ewan Harrison, David K. Jackson, Ian Johnston, Dominic Kwiatkowski, Cordelia Langford, John Sillitoe on behalf of the Wellcome Sanger Institute COVID-19 Surveillance Team |
| EPI_ISL_2485304 | Lighthouse Lab in Milton Keynes | Wellcome Sanger Institute for the COVID-19 Genomics UK (COG-UK) | The Lighthouse Lab in Milton Keynes and Alex Alderton, Roberto Amato, Jeffrey Barrett, Sonia Goncalves, Ewan Harrison, David K. Jackson, Ian Johnston, Dominic Kwiatkowski, Cordelia Langford, John Sillitoe on behalf of the Wellcome Sanger |

| | | | |
|---|---|---|---|
| | | Consortium | Institute COVID-19 Surveillance Team |
| EPI_ISL_2485305 | Lighthouse Lab in Alderley Park | Wellcome Sanger Institute for the COVID-19 Genomics UK (COG-UK) Consortium | Jacquelyn Wynn, Mairead Hyland, The Lighthouse Lab in Alderley Park and Alex Alderton, Roberto Amato, Jeffrey Barrett, Sonia Goncalves, Ewan Harrison, David K. Jackson, Ian Johnston, Dominic Kwiatkowski, Cordelia Langford, John Sillitoe on behalf of the Wellcome Sanger Institute COVID-19 Surveillance Team |
| EPI_ISL_2485306 | Lighthouse Lab in Milton Keynes | Wellcome Sanger Institute for the COVID-19 Genomics UK (COG-UK) Consortium | The Lighthouse Lab in Milton Keynes and Alex Alderton, Roberto Amato, Jeffrey Barrett, Sonia Goncalves, Ewan Harrison, David K. Jackson, Ian Johnston, Dominic Kwiatkowski, Cordelia Langford, John Sillitoe on behalf of the Wellcome Sanger Institute COVID-19 Surveillance Team |
| EPI_ISL_2485307 | Lighthouse Lab in Alderley Park | Wellcome Sanger Institute for the COVID-19 Genomics UK (COG-UK) Consortium | Jacquelyn Wynn, Mairead Hyland, The Lighthouse Lab in Alderley Park and Alex Alderton, Roberto Amato, Jeffrey Barrett, Sonia Goncalves, Ewan Harrison, David K. Jackson, Ian Johnston, Dominic Kwiatkowski, Cordelia Langford, John Sillitoe on behalf of the Wellcome Sanger Institute COVID-19 Surveillance Team |
| EPI_ISL_2485308 | Lighthouse Lab in Milton Keynes | Wellcome Sanger Institute for the COVID-19 Genomics UK (COG-UK) Consortium | The Lighthouse Lab in Milton Keynes and Alex Alderton, Roberto Amato, Jeffrey Barrett, Sonia Goncalves, Ewan Harrison, David K. Jackson, Ian Johnston, Dominic Kwiatkowski, Cordelia Langford, John Sillitoe on behalf of the Wellcome Sanger Institute COVID-19 Surveillance Team |
| EPI_ISL_2485309, EPI_ISL_2485310, EPI_ISL_2485311 | Lighthouse Lab in Alderley Park | Wellcome Sanger Institute for the COVID-19 Genomics UK (COG-UK) Consortium | Jacquelyn Wynn, Mairead Hyland, The Lighthouse Lab in Alderley Park and Alex Alderton, Roberto Amato, Jeffrey Barrett, Sonia Goncalves, Ewan Harrison, David K. Jackson, Ian Johnston, Dominic Kwiatkowski, Cordelia Langford, John Sillitoe on behalf of the Wellcome Sanger Institute COVID-19 Surveillance Team |
| EPI_ISL_2485312 | Lighthouse Lab in Milton Keynes | Wellcome Sanger Institute for the COVID-19 Genomics UK (COG-UK) Consortium | The Lighthouse Lab in Milton Keynes and Alex Alderton, Roberto Amato, Jeffrey Barrett, Sonia Goncalves, Ewan Harrison, David K. Jackson, Ian Johnston, Dominic Kwiatkowski, Cordelia Langford, John Sillitoe on behalf of the Wellcome Sanger Institute COVID-19 Surveillance Team |
| EPI_ISL_2485313, EPI_ISL_2485314, EPI_ISL_2485315 | Lighthouse Lab in Alderley Park | Wellcome Sanger Institute for the COVID-19 Genomics UK (COG-UK) Consortium | Jacquelyn Wynn, Mairead Hyland, The Lighthouse Lab in Alderley Park and Alex Alderton, Roberto Amato, Jeffrey Barrett, Sonia Goncalves, Ewan Harrison, David K. Jackson, Ian Johnston, Dominic Kwiatkowski, Cordelia Langford, John Sillitoe on behalf of the Wellcome Sanger Institute COVID-19 Surveillance Team |
| EPI_ISL_2485316 | Lighthouse Lab in Milton Keynes | Wellcome Sanger Institute for the COVID-19 Genomics UK (COG-UK) Consortium | The Lighthouse Lab in Milton Keynes and Alex Alderton, Roberto Amato, Jeffrey Barrett, Sonia Goncalves, Ewan Harrison, David K. Jackson, Ian Johnston, Dominic Kwiatkowski, Cordelia Langford, John Sillitoe on behalf of the Wellcome Sanger Institute COVID-19 Surveillance Team |
| EPI_ISL_2485317 | Lighthouse Laboratory Plymouth | Wellcome Sanger Institute for the COVID-19 Genomics UK (COG-UK) Consortium | Lighthouse Laboratory Plymouth and Alex Alderton, Roberto Amato, Jeffrey Barrett, Sonia Goncalves, Ewan Harrison, David K. Jackson, Ian Johnston, Dominic Kwiatkowski, Cordelia Langford, John Sillitoe on behalf of the Wellcome Sanger Institute COVID-19 Surveillance Team |
| EPI_ISL_2485319, EPI_ISL_2485320 | Lighthouse Lab in Alderley Park | Wellcome Sanger Institute for the COVID-19 Genomics UK (COG-UK) Consortium | Jacquelyn Wynn, Mairead Hyland, The Lighthouse Lab in Alderley Park and Alex Alderton, Roberto Amato, Jeffrey Barrett, Sonia Goncalves, Ewan Harrison, David K. Jackson, Ian Johnston, Dominic Kwiatkowski, Cordelia Langford, John Sillitoe on behalf of the Wellcome Sanger Institute COVID-19 Surveillance Team |
| EPI_ISL_2485321 | Lighthouse Laboratory Plymouth | Wellcome Sanger Institute for the COVID-19 Genomics UK (COG-UK) Consortium | Lighthouse Laboratory Plymouth and Alex Alderton, Roberto Amato, Jeffrey Barrett, Sonia Goncalves, Ewan Harrison, David K. Jackson, Ian Johnston, Dominic Kwiatkowski, Cordelia Langford, John Sillitoe on behalf of the Wellcome Sanger Institute COVID-19 Surveillance Team |
| EPI_ISL_2485322, EPI_ISL_2485323, EPI_ISL_2485324 | Lighthouse Lab in Alderley Park | Wellcome Sanger Institute for the COVID-19 Genomics UK (COG-UK) Consortium | Jacquelyn Wynn, Mairead Hyland, The Lighthouse Lab in Alderley Park and Alex Alderton, Roberto Amato, Jeffrey Barrett, Sonia Goncalves, Ewan Harrison, David K. Jackson, Ian Johnston, Dominic Kwiatkowski, Cordelia Langford, John Sillitoe on behalf of the Wellcome Sanger Institute COVID-19 Surveillance Team |
| EPI_ISL_2485325 | Lighthouse Lab in Milton Keynes | Wellcome Sanger Institute for the COVID-19 Genomics UK (COG-UK) Consortium | The Lighthouse Lab in Milton Keynes and Alex Alderton, Roberto Amato, Jeffrey Barrett, Sonia Goncalves, Ewan Harrison, David K. Jackson, Ian Johnston, Dominic Kwiatkowski, Cordelia Langford, John Sillitoe on behalf of the Wellcome Sanger Institute COVID-19 Surveillance Team |
| EPI_ISL_2485326, EPI_ISL_2485327 | Lighthouse Lab in Alderley Park | Wellcome Sanger Institute for the COVID-19 Genomics UK (COG-UK) Consortium | Jacquelyn Wynn, Mairead Hyland, The Lighthouse Lab in Alderley Park and Alex Alderton, Roberto Amato, Jeffrey Barrett, Sonia Goncalves, Ewan Harrison, David K. Jackson, Ian Johnston, Dominic Kwiatkowski, Cordelia Langford, John Sillitoe on behalf of the Wellcome Sanger Institute COVID-19 Surveillance Team |
| EPI_ISL_2485328 | Health Services Laboratories | Wellcome Sanger Institute for the COVID-19 Genomics UK (COG-UK) Consortium | Health Services Laboratories and Alex Alderton, Roberto Amato, Jeffrey Barrett, Sonia Goncalves, Ewan Harrison, David K. Jackson, Ian Johnston, Dominic Kwiatkowski, Cordelia Langford, John Sillitoe on behalf of the Wellcome Sanger Institute COVID-19 Surveillance Team |
| EPI_ISL_2485329 | Lighthouse Lab in Alderley Park | Wellcome Sanger Institute for the COVID-19 Genomics UK (COG-UK) Consortium | Jacquelyn Wynn, Mairead Hyland, The Lighthouse Lab in Alderley Park and Alex Alderton, Roberto Amato, Jeffrey Barrett, Sonia Goncalves, Ewan Harrison, David K. Jackson, Ian Johnston, Dominic Kwiatkowski, Cordelia Langford, John Sillitoe on behalf of the Wellcome Sanger Institute COVID-19 Surveillance Team |
| EPI_ISL_2485330 | Lighthouse Laboratory Plymouth | Wellcome Sanger Institute for the COVID-19 Genomics UK (COG-UK) Consortium | Lighthouse Laboratory Plymouth and Alex Alderton, Roberto Amato, Jeffrey Barrett, Sonia Goncalves, Ewan Harrison, David K. Jackson, Ian Johnston, Dominic Kwiatkowski, Cordelia Langford, John Sillitoe on behalf of the Wellcome Sanger Institute COVID-19 Surveillance Team |
| EPI_ISL_2485331 | Lighthouse Lab in Alderley Park | Wellcome Sanger Institute for the COVID-19 Genomics UK (COG-UK) Consortium | Jacquelyn Wynn, Mairead Hyland, The Lighthouse Lab in Alderley Park and Alex Alderton, Roberto Amato, Jeffrey Barrett, Sonia Goncalves, Ewan Harrison, David K. Jackson, Ian Johnston, Dominic Kwiatkowski, Cordelia Langford, John Sillitoe on behalf of the Wellcome Sanger Institute COVID-19 Surveillance Team |
| EPI_ISL_2485332 | Lighthouse Lab in Milton Keynes | Wellcome Sanger Institute for the COVID-19 Genomics UK (COG-UK) Consortium | The Lighthouse Lab in Milton Keynes and Alex Alderton, Roberto Amato, Jeffrey Barrett, Sonia Goncalves, Ewan Harrison, David K. Jackson, Ian Johnston, Dominic Kwiatkowski, Cordelia Langford, John Sillitoe on behalf of the Wellcome Sanger Institute COVID-19 Surveillance Team |
| EPI_ISL_2485333 | Lighthouse Laboratory Plymouth | Wellcome Sanger Institute for the COVID-19 Genomics UK (COG-UK) Consortium | Lighthouse Laboratory Plymouth and Alex Alderton, Roberto Amato, Jeffrey Barrett, Sonia Goncalves, Ewan Harrison, David K. Jackson, Ian Johnston, Dominic Kwiatkowski, Cordelia Langford, John Sillitoe on behalf of the Wellcome Sanger Institute COVID-19 Surveillance Team |
| EPI_ISL_2485334 | Lighthouse Lab in Alderley Park | Wellcome Sanger Institute for the COVID-19 Genomics UK (COG-UK) Consortium | Jacquelyn Wynn, Mairead Hyland, The Lighthouse Lab in Alderley Park and Alex Alderton, Roberto Amato, Jeffrey Barrett, Sonia Goncalves, Ewan Harrison, David K. Jackson, Ian Johnston, Dominic Kwiatkowski, Cordelia Langford, John Sillitoe on behalf of the Wellcome Sanger Institute COVID-19 Surveillance Team |
| EPI_ISL_2485335, EPI_ISL_2485336, EPI_ISL_2485337 | Lighthouse Lab in Milton Keynes | Wellcome Sanger Institute for the COVID-19 Genomics UK (COG-UK) Consortium | The Lighthouse Lab in Milton Keynes and Alex Alderton, Roberto Amato, Jeffrey Barrett, Sonia Goncalves, Ewan Harrison, David K. Jackson, Ian Johnston, Dominic Kwiatkowski, Cordelia Langford, John Sillitoe on behalf of the Wellcome Sanger Institute COVID-19 Surveillance Team |
| EPI_ISL_2485338, EPI_ISL_2485339, EPI_ISL_2485340, EPI_ISL_2485341, EPI_ISL_2485342 | Lighthouse Lab in Alderley Park | Wellcome Sanger Institute for the COVID-19 Genomics UK (COG-UK) Consortium | Jacquelyn Wynn, Mairead Hyland, The Lighthouse Lab in Alderley Park and Alex Alderton, Roberto Amato, Jeffrey Barrett, Sonia Goncalves, Ewan Harrison, David K. Jackson, Ian Johnston, Dominic Kwiatkowski, Cordelia Langford, John Sillitoe on behalf of the Wellcome Sanger Institute COVID-19 Surveillance Team |
| EPI_ISL_2485343 | Lighthouse Lab in Milton Keynes | Wellcome Sanger Institute for the COVID-19 Genomics UK (COG-UK) Consortium | The Lighthouse Lab in Milton Keynes and Alex Alderton, Roberto Amato, Jeffrey Barrett, Sonia Goncalves, Ewan Harrison, David K. Jackson, Ian Johnston, Dominic Kwiatkowski, Cordelia Langford, John Sillitoe on behalf of the Wellcome Sanger Institute COVID-19 Surveillance Team |
| EPI_ISL_2485344 | Lighthouse Lab in Alderley Park | Wellcome Sanger Institute for the COVID-19 Genomics UK (COG-UK) Consortium | Jacquelyn Wynn, Mairead Hyland, The Lighthouse Lab in Alderley Park and Alex Alderton, Roberto Amato, Jeffrey Barrett, Sonia Goncalves, Ewan Harrison, David K. Jackson, Ian Johnston, Dominic Kwiatkowski, Cordelia Langford, John Sillitoe on behalf of the Wellcome Sanger Institute COVID-19 Surveillance Team |
| EPI_ISL_2485345 | Health Services Laboratories | Wellcome Sanger Institute for the COVID-19 Genomics UK (COG-UK) Consortium | Health Services Laboratories and Alex Alderton, Roberto Amato, Jeffrey Barrett, Sonia Goncalves, Ewan Harrison, David K. Jackson, Ian Johnston, Dominic Kwiatkowski, Cordelia Langford, John Sillitoe on behalf of the Wellcome Sanger Institute COVID-19 Surveillance Team |
| EPI_ISL_2485346 | Lighthouse Laboratory Plymouth | Wellcome Sanger Institute for the COVID-19 Genomics UK (COG-UK) Consortium | Lighthouse Laboratory Plymouth and Alex Alderton, Roberto Amato, Jeffrey Barrett, Sonia Goncalves, Ewan Harrison, David K. Jackson, Ian Johnston, Dominic Kwiatkowski, Cordelia Langford, John Sillitoe on behalf of the Wellcome Sanger Institute COVID-19 Surveillance Team |
| EPI_ISL_2485348 | Lighthouse Lab in Milton Keynes | Wellcome Sanger Institute for the COVID-19 Genomics UK (COG-UK) Consortium | The Lighthouse Lab in Milton Keynes and Alex Alderton, Roberto Amato, Jeffrey Barrett, Sonia Goncalves, Ewan Harrison, David K. Jackson, Ian Johnston, Dominic Kwiatkowski, Cordelia Langford, John Sillitoe on behalf of the Wellcome Sanger Institute COVID-19 Surveillance Team |
| EPI_ISL_2485349 | Lighthouse Laboratory Plymouth | Wellcome Sanger Institute for the COVID-19 Genomics UK (COG-UK) Consortium | Lighthouse Laboratory Plymouth and Alex Alderton, Roberto Amato, Jeffrey Barrett, Sonia Goncalves, Ewan Harrison, David K. Jackson, Ian Johnston, Dominic Kwiatkowski, Cordelia Langford, John Sillitoe on behalf of the Wellcome Sanger Institute COVID-19 Surveillance Team |
| EPI_ISL_2485350 | Lighthouse Lab in Alderley Park | Wellcome Sanger Institute for the COVID-19 Genomics UK (COG-UK) Consortium | Jacquelyn Wynn, Mairead Hyland, The Lighthouse Lab in Alderley Park and Alex Alderton, Roberto Amato, Jeffrey Barrett, Sonia Goncalves, Ewan Harrison, David K. Jackson, Ian Johnston, Dominic Kwiatkowski, Cordelia Langford, John Sillitoe on behalf of the Wellcome Sanger Institute COVID-19 Surveillance Team |
| EPI_ISL_2485352 | Lighthouse Laboratory Plymouth | Wellcome Sanger Institute for the COVID-19 Genomics UK (COG-UK) Consortium | Lighthouse Laboratory Plymouth and Alex Alderton, Roberto Amato, Jeffrey Barrett, Sonia Goncalves, Ewan Harrison, David K. Jackson, Ian Johnston, Dominic Kwiatkowski, Cordelia Langford, John Sillitoe on behalf of the Wellcome Sanger Institute COVID-19 Surveillance Team |
| EPI_ISL_2485353 | Lighthouse Lab in Milton Keynes | Wellcome Sanger Institute for the COVID-19 Genomics UK (COG-UK) Consortium | The Lighthouse Lab in Milton Keynes and Alex Alderton, Roberto Amato, Jeffrey Barrett, Sonia Goncalves, Ewan Harrison, David K. Jackson, Ian Johnston, Dominic Kwiatkowski, Cordelia Langford, John Sillitoe on behalf of the Wellcome Sanger Institute COVID-19 Surveillance Team |
| EPI_ISL_2485354 | Lighthouse Lab in Alderley Park | Wellcome Sanger Institute for the COVID-19 Genomics UK (COG-UK) Consortium | Jacquelyn Wynn, Mairead Hyland, The Lighthouse Lab in Alderley Park and Alex Alderton, Roberto Amato, Jeffrey Barrett, Sonia Goncalves, Ewan Harrison, David K. Jackson, Ian Johnston, Dominic Kwiatkowski, Cordelia Langford, John Sillitoe on behalf of the Wellcome Sanger Institute COVID-19 Surveillance Team |
| EPI_ISL_2485355 | Lighthouse Lab in Milton Keynes | Wellcome Sanger Institute for the COVID-19 Genomics UK (COG-UK) Consortium | The Lighthouse Lab in Milton Keynes and Alex Alderton, Roberto Amato, Jeffrey Barrett, Sonia Goncalves, Ewan Harrison, David K. Jackson, Ian Johnston, Dominic Kwiatkowski, Cordelia Langford, John Sillitoe on behalf of the Wellcome Sanger Institute COVID-19 Surveillance Team |
| EPI_ISL_2485356, EPI_ISL_2485357, EPI_ISL_2485358, EPI_ISL_2485359, EPI_ISL_2485360 | Lighthouse Lab in Alderley Park | Wellcome Sanger Institute for the COVID-19 Genomics UK (COG-UK) Consortium | Jacquelyn Wynn, Mairead Hyland, The Lighthouse Lab in Alderley Park and Alex Alderton, Roberto Amato, Jeffrey Barrett, Sonia Goncalves, Ewan Harrison, David K. Jackson, Ian Johnston, Dominic Kwiatkowski, Cordelia Langford, John Sillitoe on behalf of the Wellcome Sanger Institute COVID-19 Surveillance Team |
| EPI_ISL_2485361 | Lighthouse Lab in Milton Keynes | Wellcome Sanger Institute for the COVID-19 Genomics UK (COG-UK) Consortium | The Lighthouse Lab in Milton Keynes and Alex Alderton, Roberto Amato, Jeffrey Barrett, Sonia Goncalves, Ewan Harrison, David K. Jackson, Ian Johnston, Dominic Kwiatkowski, Cordelia Langford, John Sillitoe on behalf of the Wellcome Sanger Institute COVID-19 Surveillance Team |
| EPI_ISL_2485362, EPI_ISL_2485363 | Lighthouse Lab in Alderley Park | Wellcome Sanger Institute for the COVID-19 Genomics UK (COG-UK) Consortium | Jacquelyn Wynn, Mairead Hyland, The Lighthouse Lab in Alderley Park and Alex Alderton, Roberto Amato, Jeffrey Barrett, Sonia Goncalves, Ewan Harrison, David K. Jackson, Ian Johnston, Dominic Kwiatkowski, Cordelia Langford, John Sillitoe on behalf of the Wellcome Sanger Institute COVID-19 Surveillance Team |
| EPI_ISL_2485364 | Lighthouse Lab in Milton Keynes | Wellcome Sanger Institute for the COVID-19 Genomics UK (COG-UK) Consortium | The Lighthouse Lab in Milton Keynes and Alex Alderton, Roberto Amato, Jeffrey Barrett, Sonia Goncalves, Ewan Harrison, David K. Jackson, Ian Johnston, Dominic Kwiatkowski, Cordelia Langford, John Sillitoe on behalf of the Wellcome Sanger Institute COVID-19 Surveillance Team |
| EPI_ISL_2485365, EPI_ISL_2485366 | Lighthouse Lab in Alderley Park | Wellcome Sanger Institute for the COVID-19 Genomics UK (COG-UK) Consortium | Jacquelyn Wynn, Mairead Hyland, The Lighthouse Lab in Alderley Park and Alex Alderton, Roberto Amato, Jeffrey Barrett, Sonia Goncalves, Ewan Harrison, David K. Jackson, Ian Johnston, Dominic Kwiatkowski, Cordelia Langford, John Sillitoe on behalf of the Wellcome Sanger Institute COVID-19 Surveillance Team |
| EPI_ISL_2485367 | Health Services Laboratories | Wellcome Sanger Institute for the COVID-19 Genomics UK (COG-UK) Consortium | Health Services Laboratories and Alex Alderton, Roberto Amato, Jeffrey Barrett, Sonia Goncalves, Ewan Harrison, David K. Jackson, Ian Johnston, Dominic Kwiatkowski, Cordelia Langford, John Sillitoe on behalf of the Wellcome Sanger Institute COVID-19 Surveillance Team |
| EPI_ISL_2485368, | Lighthouse Lab in Alderley Park | Wellcome Sanger Institute for the COVID-19 Genomics UK (COG-UK) | Jacquelyn Wynn, Mairead Hyland, The Lighthouse Lab in Alderley Park and Alex Alderton, Roberto Amato, Jeffrey Barrett, Sonia Goncalves, Ewan Harrison, David K. Jackson, Ian Johnston, Dominic Kwiatkowski, Cordelia Langford, John Sillitoe on |

| | | | |
|---|---|---|---|
| EPI_ISL_2485369 | | Consortium | behalf of the Wellcome Sanger Institute COVID-19 Surveillance Team |
| EPI_ISL_2485370 | Lighthouse Lab in Milton Keynes | Wellcome Sanger Institute for the COVID-19 Genomics UK (COG-UK) Consortium | The Lighthouse Lab in Milton Keynes and Alex Alderton, Roberto Amato, Jeffrey Barrett, Sonia Goncalves, Ewan Harrison, David K. Jackson, Ian Johnston, Dominic Kwiatkowski, Cordelia Langford, John Sillitoe on behalf of the Wellcome Sanger Institute COVID-19 Surveillance Team |
| EPI_ISL_2485371 | Health Services Laboratories | Wellcome Sanger Institute for the COVID-19 Genomics UK (COG-UK) Consortium | Health Services Laboratories and Alex Alderton, Roberto Amato, Jeffrey Barrett, Sonia Goncalves, Ewan Harrison, David K. Jackson, Ian Johnston, Dominic Kwiatkowski, Cordelia Langford, John Sillitoe on behalf of the Wellcome Sanger Institute COVID-19 Surveillance Team |
| EPI_ISL_2485372 | Lighthouse Lab in Milton Keynes | Wellcome Sanger Institute for the COVID-19 Genomics UK (COG-UK) Consortium | The Lighthouse Lab in Milton Keynes and Alex Alderton, Roberto Amato, Jeffrey Barrett, Sonia Goncalves, Ewan Harrison, David K. Jackson, Ian Johnston, Dominic Kwiatkowski, Cordelia Langford, John Sillitoe on behalf of the Wellcome Sanger Institute COVID-19 Surveillance Team |
| EPI_ISL_2485373 | Lighthouse Lab in Alderley Park | Wellcome Sanger Institute for the COVID-19 Genomics UK (COG-UK) Consortium | Jacquelyn Wynn, Mairead Hyland, The Lighthouse Lab in Alderley Park and Alex Alderton, Roberto Amato, Jeffrey Barrett, Sonia Goncalves, Ewan Harrison, David K. Jackson, Ian Johnston, Dominic Kwiatkowski, Cordelia Langford, John Sillitoe on behalf of the Wellcome Sanger Institute COVID-19 Surveillance Team |
| EPI_ISL_2485374 | Lighthouse Lab in Milton Keynes | Wellcome Sanger Institute for the COVID-19 Genomics UK (COG-UK) Consortium | The Lighthouse Lab in Milton Keynes and Alex Alderton, Roberto Amato, Jeffrey Barrett, Sonia Goncalves, Ewan Harrison, David K. Jackson, Ian Johnston, Dominic Kwiatkowski, Cordelia Langford, John Sillitoe on behalf of the Wellcome Sanger Institute COVID-19 Surveillance Team |
| EPI_ISL_2485375, EPI_ISL_2485376, EPI_ISL_2485377, EPI_ISL_2485378 | Lighthouse Lab in Alderley Park | Wellcome Sanger Institute for the COVID-19 Genomics UK (COG-UK) Consortium | Jacquelyn Wynn, Mairead Hyland, The Lighthouse Lab in Alderley Park and Alex Alderton, Roberto Amato, Jeffrey Barrett, Sonia Goncalves, Ewan Harrison, David K. Jackson, Ian Johnston, Dominic Kwiatkowski, Cordelia Langford, John Sillitoe on behalf of the Wellcome Sanger Institute COVID-19 Surveillance Team |
| EPI_ISL_2485379 | Health Services Laboratories | Wellcome Sanger Institute for the COVID-19 Genomics UK (COG-UK) Consortium | Health Services Laboratories and Alex Alderton, Roberto Amato, Jeffrey Barrett, Sonia Goncalves, Ewan Harrison, David K. Jackson, Ian Johnston, Dominic Kwiatkowski, Cordelia Langford, John Sillitoe on behalf of the Wellcome Sanger Institute COVID-19 Surveillance Team |
| EPI_ISL_2485380, EPI_ISL_2485381 | Lighthouse Lab in Milton Keynes | Wellcome Sanger Institute for the COVID-19 Genomics UK (COG-UK) Consortium | The Lighthouse Lab in Milton Keynes and Alex Alderton, Roberto Amato, Jeffrey Barrett, Sonia Goncalves, Ewan Harrison, David K. Jackson, Ian Johnston, Dominic Kwiatkowski, Cordelia Langford, John Sillitoe on behalf of the Wellcome Sanger Institute COVID-19 Surveillance Team |
| EPI_ISL_2485382 | Lighthouse Laboratory Plymouth | Wellcome Sanger Institute for the COVID-19 Genomics UK (COG-UK) Consortium | Lighthouse Laboratory Plymouth and Alex Alderton, Roberto Amato, Jeffrey Barrett, Sonia Goncalves, Ewan Harrison, David K. Jackson, Ian Johnston, Dominic Kwiatkowski, Cordelia Langford, John Sillitoe on behalf of the Wellcome Sanger Institute COVID-19 Surveillance Team |
| EPI_ISL_2485383, EPI_ISL_2485384 | Lighthouse Lab in Milton Keynes | Wellcome Sanger Institute for the COVID-19 Genomics UK (COG-UK) Consortium | The Lighthouse Lab in Milton Keynes and Alex Alderton, Roberto Amato, Jeffrey Barrett, Sonia Goncalves, Ewan Harrison, David K. Jackson, Ian Johnston, Dominic Kwiatkowski, Cordelia Langford, John Sillitoe on behalf of the Wellcome Sanger Institute COVID-19 Surveillance Team |
| EPI_ISL_2485386 | Lighthouse Lab in Alderley Park | Wellcome Sanger Institute for the COVID-19 Genomics UK (COG-UK) Consortium | Jacquelyn Wynn, Mairead Hyland, The Lighthouse Lab in Alderley Park and Alex Alderton, Roberto Amato, Jeffrey Barrett, Sonia Goncalves, Ewan Harrison, David K. Jackson, Ian Johnston, Dominic Kwiatkowski, Cordelia Langford, John Sillitoe on behalf of the Wellcome Sanger Institute COVID-19 Surveillance Team |
| EPI_ISL_2485387 | Lighthouse Laboratory Plymouth | Wellcome Sanger Institute for the COVID-19 Genomics UK (COG-UK) Consortium | Lighthouse Laboratory Plymouth and Alex Alderton, Roberto Amato, Jeffrey Barrett, Sonia Goncalves, Ewan Harrison, David K. Jackson, Ian Johnston, Dominic Kwiatkowski, Cordelia Langford, John Sillitoe on behalf of the Wellcome Sanger Institute COVID-19 Surveillance Team |
| EPI_ISL_2485388, EPI_ISL_2485389, EPI_ISL_2485390, EPI_ISL_2485391 | Lighthouse Lab in Alderley Park | Wellcome Sanger Institute for the COVID-19 Genomics UK (COG-UK) Consortium | Jacquelyn Wynn, Mairead Hyland, The Lighthouse Lab in Alderley Park and Alex Alderton, Roberto Amato, Jeffrey Barrett, Sonia Goncalves, Ewan Harrison, David K. Jackson, Ian Johnston, Dominic Kwiatkowski, Cordelia Langford, John Sillitoe on behalf of the Wellcome Sanger Institute COVID-19 Surveillance Team |
| EPI_ISL_2485392, EPI_ISL_2485393 | Health Services Laboratories | Wellcome Sanger Institute for the COVID-19 Genomics UK (COG-UK) Consortium | Health Services Laboratories and Alex Alderton, Roberto Amato, Jeffrey Barrett, Sonia Goncalves, Ewan Harrison, David K. Jackson, Ian Johnston, Dominic Kwiatkowski, Cordelia Langford, John Sillitoe on behalf of the Wellcome Sanger Institute COVID-19 Surveillance Team |
| EPI_ISL_2485394, EPI_ISL_2485395, EPI_ISL_2485397, EPI_ISL_2485398, EPI_ISL_2485399 | Lighthouse Lab in Alderley Park | Wellcome Sanger Institute for the COVID-19 Genomics UK (COG-UK) Consortium | Jacquelyn Wynn, Mairead Hyland, The Lighthouse Lab in Alderley Park and Alex Alderton, Roberto Amato, Jeffrey Barrett, Sonia Goncalves, Ewan Harrison, David K. Jackson, Ian Johnston, Dominic Kwiatkowski, Cordelia Langford, John Sillitoe on behalf of the Wellcome Sanger Institute COVID-19 Surveillance Team |
| EPI_ISL_2485400, EPI_ISL_2485401 | Lighthouse Laboratory Plymouth | Wellcome Sanger Institute for the COVID-19 Genomics UK (COG-UK) Consortium | Lighthouse Laboratory Plymouth and Alex Alderton, Roberto Amato, Jeffrey Barrett, Sonia Goncalves, Ewan Harrison, David K. Jackson, Ian Johnston, Dominic Kwiatkowski, Cordelia Langford, John Sillitoe on behalf of the Wellcome Sanger Institute COVID-19 Surveillance Team |
| EPI_ISL_2485402 | Lighthouse Lab in Alderley Park | Wellcome Sanger Institute for the COVID-19 Genomics UK (COG-UK) Consortium | Jacquelyn Wynn, Mairead Hyland, The Lighthouse Lab in Alderley Park and Alex Alderton, Roberto Amato, Jeffrey Barrett, Sonia Goncalves, Ewan Harrison, David K. Jackson, Ian Johnston, Dominic Kwiatkowski, Cordelia Langford, John Sillitoe on behalf of the Wellcome Sanger Institute COVID-19 Surveillance Team |
| EPI_ISL_2485403 | Health Services Laboratories | Wellcome Sanger Institute for the COVID-19 Genomics UK (COG-UK) Consortium | Health Services Laboratories and Alex Alderton, Roberto Amato, Jeffrey Barrett, Sonia Goncalves, Ewan Harrison, David K. Jackson, Ian Johnston, Dominic Kwiatkowski, Cordelia Langford, John Sillitoe on behalf of the Wellcome Sanger Institute COVID-19 Surveillance Team |
| EPI_ISL_2485404, EPI_ISL_2485405 | Lighthouse Lab in Alderley Park | Wellcome Sanger Institute for the COVID-19 Genomics UK (COG-UK) Consortium | Jacquelyn Wynn, Mairead Hyland, The Lighthouse Lab in Alderley Park and Alex Alderton, Roberto Amato, Jeffrey Barrett, Sonia Goncalves, Ewan Harrison, David K. Jackson, Ian Johnston, Dominic Kwiatkowski, Cordelia Langford, John Sillitoe on behalf of the Wellcome Sanger Institute COVID-19 Surveillance Team |
| EPI_ISL_2485406 | Lighthouse Laboratory Plymouth | Wellcome Sanger Institute for the COVID-19 Genomics UK (COG-UK) Consortium | Lighthouse Laboratory Plymouth and Alex Alderton, Roberto Amato, Jeffrey Barrett, Sonia Goncalves, Ewan Harrison, David K. Jackson, Ian Johnston, Dominic Kwiatkowski, Cordelia Langford, John Sillitoe on behalf of the Wellcome Sanger Institute COVID-19 Surveillance Team |
| EPI_ISL_2485407, EPI_ISL_2485408, EPI_ISL_2485409 | Lighthouse Lab in Alderley Park | Wellcome Sanger Institute for the COVID-19 Genomics UK (COG-UK) Consortium | Jacquelyn Wynn, Mairead Hyland, The Lighthouse Lab in Alderley Park and Alex Alderton, Roberto Amato, Jeffrey Barrett, Sonia Goncalves, Ewan Harrison, David K. Jackson, Ian Johnston, Dominic Kwiatkowski, Cordelia Langford, John Sillitoe on behalf of the Wellcome Sanger Institute COVID-19 Surveillance Team |
| EPI_ISL_2485410 | Lighthouse Lab in Milton Keynes | Wellcome Sanger Institute for the COVID-19 Genomics UK (COG-UK) Consortium | The Lighthouse Lab in Milton Keynes and Alex Alderton, Roberto Amato, Jeffrey Barrett, Sonia Goncalves, Ewan Harrison, David K. Jackson, Ian Johnston, Dominic Kwiatkowski, Cordelia Langford, John Sillitoe on behalf of the Wellcome Sanger Institute COVID-19 Surveillance Team |
| EPI_ISL_2485411, EPI_ISL_2485412 | Lighthouse Lab in Alderley Park | Wellcome Sanger Institute for the COVID-19 Genomics UK (COG-UK) Consortium | Jacquelyn Wynn, Mairead Hyland, The Lighthouse Lab in Alderley Park and Alex Alderton, Roberto Amato, Jeffrey Barrett, Sonia Goncalves, Ewan Harrison, David K. Jackson, Ian Johnston, Dominic Kwiatkowski, Cordelia Langford, John Sillitoe on behalf of the Wellcome Sanger Institute COVID-19 Surveillance Team |
| EPI_ISL_2485413, EPI_ISL_2485414 | Lighthouse Lab in Milton Keynes | Wellcome Sanger Institute for the COVID-19 Genomics UK (COG-UK) Consortium | The Lighthouse Lab in Milton Keynes and Alex Alderton, Roberto Amato, Jeffrey Barrett, Sonia Goncalves, Ewan Harrison, David K. Jackson, Ian Johnston, Dominic Kwiatkowski, Cordelia Langford, John Sillitoe on behalf of the Wellcome Sanger Institute COVID-19 Surveillance Team |
| EPI_ISL_2485415 | Lighthouse Laboratory Plymouth | Wellcome Sanger Institute for the COVID-19 Genomics UK (COG-UK) Consortium | Lighthouse Laboratory Plymouth and Alex Alderton, Roberto Amato, Jeffrey Barrett, Sonia Goncalves, Ewan Harrison, David K. Jackson, Ian Johnston, Dominic Kwiatkowski, Cordelia Langford, John Sillitoe on behalf of the Wellcome Sanger Institute COVID-19 Surveillance Team |
| EPI_ISL_2485416 | Lighthouse Lab in Milton Keynes | Wellcome Sanger Institute for the COVID-19 Genomics UK (COG-UK) Consortium | The Lighthouse Lab in Milton Keynes and Alex Alderton, Roberto Amato, Jeffrey Barrett, Sonia Goncalves, Ewan Harrison, David K. Jackson, Ian Johnston, Dominic Kwiatkowski, Cordelia Langford, John Sillitoe on behalf of the Wellcome Sanger Institute COVID-19 Surveillance Team |
| EPI_ISL_2485417 | Lighthouse Lab in Alderley Park | Wellcome Sanger Institute for the COVID-19 Genomics UK (COG-UK) Consortium | Jacquelyn Wynn, Mairead Hyland, The Lighthouse Lab in Alderley Park and Alex Alderton, Roberto Amato, Jeffrey Barrett, Sonia Goncalves, Ewan Harrison, David K. Jackson, Ian Johnston, Dominic Kwiatkowski, Cordelia Langford, John Sillitoe on behalf of the Wellcome Sanger Institute COVID-19 Surveillance Team |
| EPI_ISL_2485418 | Lighthouse Laboratory Plymouth | Wellcome Sanger Institute for the COVID-19 Genomics UK (COG-UK) Consortium | Lighthouse Laboratory Plymouth and Alex Alderton, Roberto Amato, Jeffrey Barrett, Sonia Goncalves, Ewan Harrison, David K. Jackson, Ian Johnston, Dominic Kwiatkowski, Cordelia Langford, John Sillitoe on behalf of the Wellcome Sanger Institute COVID-19 Surveillance Team |
| EPI_ISL_2485419, EPI_ISL_2485420 | Lighthouse Lab in Alderley Park | Wellcome Sanger Institute for the COVID-19 Genomics UK (COG-UK) Consortium | Jacquelyn Wynn, Mairead Hyland, The Lighthouse Lab in Alderley Park and Alex Alderton, Roberto Amato, Jeffrey Barrett, Sonia Goncalves, Ewan Harrison, David K. Jackson, Ian Johnston, Dominic Kwiatkowski, Cordelia Langford, John Sillitoe on behalf of the Wellcome Sanger Institute COVID-19 Surveillance Team |
| EPI_ISL_2485421 | Lighthouse Lab in Milton Keynes | Wellcome Sanger Institute for the COVID-19 Genomics UK (COG-UK) Consortium | The Lighthouse Lab in Milton Keynes and Alex Alderton, Roberto Amato, Jeffrey Barrett, Sonia Goncalves, Ewan Harrison, David K. Jackson, Ian Johnston, Dominic Kwiatkowski, Cordelia Langford, John Sillitoe on behalf of the Wellcome Sanger Institute COVID-19 Surveillance Team |
| EPI_ISL_2485422 | Lighthouse Laboratory Plymouth | Wellcome Sanger Institute for the COVID-19 Genomics UK (COG-UK) Consortium | Lighthouse Laboratory Plymouth and Alex Alderton, Roberto Amato, Jeffrey Barrett, Sonia Goncalves, Ewan Harrison, David K. Jackson, Ian Johnston, Dominic Kwiatkowski, Cordelia Langford, John Sillitoe on behalf of the Wellcome Sanger Institute COVID-19 Surveillance Team |
| EPI_ISL_2485423, EPI_ISL_2485424, EPI_ISL_2485425, EPI_ISL_2485426, EPI_ISL_2485427, EPI_ISL_2485428, EPI_ISL_2485429, EPI_ISL_2485430 | Lighthouse Lab in Alderley Park | Wellcome Sanger Institute for the COVID-19 Genomics UK (COG-UK) Consortium | Jacquelyn Wynn, Mairead Hyland, The Lighthouse Lab in Alderley Park and Alex Alderton, Roberto Amato, Jeffrey Barrett, Sonia Goncalves, Ewan Harrison, David K. Jackson, Ian Johnston, Dominic Kwiatkowski, Cordelia Langford, John Sillitoe on behalf of the Wellcome Sanger Institute COVID-19 Surveillance Team |
| EPI_ISL_2485431 | Lighthouse Lab in Milton Keynes | Wellcome Sanger Institute for the COVID-19 Genomics UK (COG-UK) Consortium | The Lighthouse Lab in Milton Keynes and Alex Alderton, Roberto Amato, Jeffrey Barrett, Sonia Goncalves, Ewan Harrison, David K. Jackson, Ian Johnston, Dominic Kwiatkowski, Cordelia Langford, John Sillitoe on behalf of the Wellcome Sanger Institute COVID-19 Surveillance Team |
| EPI_ISL_2485433 | Lighthouse Lab in Alderley Park | Wellcome Sanger Institute for the COVID-19 Genomics UK (COG-UK) Consortium | Jacquelyn Wynn, Mairead Hyland, The Lighthouse Lab in Alderley Park and Alex Alderton, Roberto Amato, Jeffrey Barrett, Sonia Goncalves, Ewan Harrison, David K. Jackson, Ian Johnston, Dominic Kwiatkowski, Cordelia Langford, John Sillitoe on behalf of the Wellcome Sanger Institute COVID-19 Surveillance Team |
| EPI_ISL_2485435 | Lighthouse Lab in Milton Keynes | Wellcome Sanger Institute for the COVID-19 Genomics UK (COG-UK) Consortium | The Lighthouse Lab in Milton Keynes and Alex Alderton, Roberto Amato, Jeffrey Barrett, Sonia Goncalves, Ewan Harrison, David K. Jackson, Ian Johnston, Dominic Kwiatkowski, Cordelia Langford, John Sillitoe on behalf of the Wellcome Sanger Institute COVID-19 Surveillance Team |
| EPI_ISL_2485436, EPI_ISL_2485437, EPI_ISL_2485438, EPI_ISL_2485439 | Lighthouse Lab in Alderley Park | Wellcome Sanger Institute for the COVID-19 Genomics UK (COG-UK) Consortium | Jacquelyn Wynn, Mairead Hyland, The Lighthouse Lab in Alderley Park and Alex Alderton, Roberto Amato, Jeffrey Barrett, Sonia Goncalves, Ewan Harrison, David K. Jackson, Ian Johnston, Dominic Kwiatkowski, Cordelia Langford, John Sillitoe on behalf of the Wellcome Sanger Institute COVID-19 Surveillance Team |
| EPI_ISL_2485440 | Lighthouse Lab in Milton Keynes | Wellcome Sanger Institute for the COVID-19 Genomics UK (COG-UK) Consortium | The Lighthouse Lab in Milton Keynes and Alex Alderton, Roberto Amato, Jeffrey Barrett, Sonia Goncalves, Ewan Harrison, David K. Jackson, Ian Johnston, Dominic Kwiatkowski, Cordelia Langford, John Sillitoe on behalf of the Wellcome Sanger Institute COVID-19 Surveillance Team |
| EPI_ISL_2485441 | Lighthouse Lab in Alderley Park | Wellcome Sanger Institute for the COVID-19 Genomics UK (COG-UK) Consortium | Jacquelyn Wynn, Mairead Hyland, The Lighthouse Lab in Alderley Park and Alex Alderton, Roberto Amato, Jeffrey Barrett, Sonia Goncalves, Ewan Harrison, David K. Jackson, Ian Johnston, Dominic Kwiatkowski, Cordelia Langford, John Sillitoe on behalf of the Wellcome Sanger Institute COVID-19 Surveillance Team |

| | | | |
|---|---|---|---|
| EPI_ISL_2485442, EPI_ISL_2485443, EPI_ISL_2485444 | Lighthouse Lab in Milton Keynes | Wellcome Sanger Institute for the COVID-19 Genomics UK (COG-UK) Consortium | The Lighthouse Lab in Milton Keynes and Alex Alderton, Roberto Amato, Jeffrey Barrett, Sonia Goncalves, Ewan Harrison, David K. Jackson, Ian Johnston, Dominic Kwiatkowski, Cordelia Langford, John Sillitoe on behalf of the Wellcome Sanger Institute COVID-19 Surveillance Team |
| EPI_ISL_2485445 | Health Services Laboratories | Wellcome Sanger Institute for the COVID-19 Genomics UK (COG-UK) Consortium | Health Services Laboratories and Alex Alderton, Roberto Amato, Jeffrey Barrett, Sonia Goncalves, Ewan Harrison, David K. Jackson, Ian Johnston, Dominic Kwiatkowski, Cordelia Langford, John Sillitoe on behalf of the Wellcome Sanger Institute COVID-19 Surveillance Team |
| EPI_ISL_2485446, EPI_ISL_2485447, EPI_ISL_2485448, EPI_ISL_2485449 | Lighthouse Lab in Alderley Park | Wellcome Sanger Institute for the COVID-19 Genomics UK (COG-UK) Consortium | Jacquelyn Wynn, Mairead Hyland, The Lighthouse Lab in Alderley Park and Alex Alderton, Roberto Amato, Jeffrey Barrett, Sonia Goncalves, Ewan Harrison, David K. Jackson, Ian Johnston, Dominic Kwiatkowski, Cordelia Langford, John Sillitoe on behalf of the Wellcome Sanger Institute COVID-19 Surveillance Team |
| EPI_ISL_2485450, EPI_ISL_2485451 | Lighthouse Lab in Milton Keynes | Wellcome Sanger Institute for the COVID-19 Genomics UK (COG-UK) Consortium | The Lighthouse Lab in Milton Keynes and Alex Alderton, Roberto Amato, Jeffrey Barrett, Sonia Goncalves, Ewan Harrison, David K. Jackson, Ian Johnston, Dominic Kwiatkowski, Cordelia Langford, John Sillitoe on behalf of the Wellcome Sanger Institute COVID-19 Surveillance Team |
| EPI_ISL_2485452 | Lighthouse Lab in Alderley Park | Wellcome Sanger Institute for the COVID-19 Genomics UK (COG-UK) Consortium | Jacquelyn Wynn, Mairead Hyland, The Lighthouse Lab in Alderley Park and Alex Alderton, Roberto Amato, Jeffrey Barrett, Sonia Goncalves, Ewan Harrison, David K. Jackson, Ian Johnston, Dominic Kwiatkowski, Cordelia Langford, John Sillitoe on behalf of the Wellcome Sanger Institute COVID-19 Surveillance Team |
| EPI_ISL_2485453 | Lighthouse Lab in Milton Keynes | Wellcome Sanger Institute for the COVID-19 Genomics UK (COG-UK) Consortium | The Lighthouse Lab in Milton Keynes and Alex Alderton, Roberto Amato, Jeffrey Barrett, Sonia Goncalves, Ewan Harrison, David K. Jackson, Ian Johnston, Dominic Kwiatkowski, Cordelia Langford, John Sillitoe on behalf of the Wellcome Sanger Institute COVID-19 Surveillance Team |
| EPI_ISL_2485454, EPI_ISL_2485455 | Lighthouse Lab in Alderley Park | Wellcome Sanger Institute for the COVID-19 Genomics UK (COG-UK) Consortium | Jacquelyn Wynn, Mairead Hyland, The Lighthouse Lab in Alderley Park and Alex Alderton, Roberto Amato, Jeffrey Barrett, Sonia Goncalves, Ewan Harrison, David K. Jackson, Ian Johnston, Dominic Kwiatkowski, Cordelia Langford, John Sillitoe on behalf of the Wellcome Sanger Institute COVID-19 Surveillance Team |
| EPI_ISL_2485456, EPI_ISL_2485457 | Lighthouse Lab in Milton Keynes | Wellcome Sanger Institute for the COVID-19 Genomics UK (COG-UK) Consortium | The Lighthouse Lab in Milton Keynes and Alex Alderton, Roberto Amato, Jeffrey Barrett, Sonia Goncalves, Ewan Harrison, David K. Jackson, Ian Johnston, Dominic Kwiatkowski, Cordelia Langford, John Sillitoe on behalf of the Wellcome Sanger Institute COVID-19 Surveillance Team |
| EPI_ISL_2485458, EPI_ISL_2485459, EPI_ISL_2485460 | Lighthouse Lab in Alderley Park | Wellcome Sanger Institute for the COVID-19 Genomics UK (COG-UK) Consortium | Jacquelyn Wynn, Mairead Hyland, The Lighthouse Lab in Alderley Park and Alex Alderton, Roberto Amato, Jeffrey Barrett, Sonia Goncalves, Ewan Harrison, David K. Jackson, Ian Johnston, Dominic Kwiatkowski, Cordelia Langford, John Sillitoe on behalf of the Wellcome Sanger Institute COVID-19 Surveillance Team |
| EPI_ISL_2485462 | Lighthouse Lab in Milton Keynes | Wellcome Sanger Institute for the COVID-19 Genomics UK (COG-UK) Consortium | The Lighthouse Lab in Milton Keynes and Alex Alderton, Roberto Amato, Jeffrey Barrett, Sonia Goncalves, Ewan Harrison, David K. Jackson, Ian Johnston, Dominic Kwiatkowski, Cordelia Langford, John Sillitoe on behalf of the Wellcome Sanger Institute COVID-19 Surveillance Team |
| EPI_ISL_2485463, EPI_ISL_2485464 | Lighthouse Lab in Alderley Park | Wellcome Sanger Institute for the COVID-19 Genomics UK (COG-UK) Consortium | Jacquelyn Wynn, Mairead Hyland, The Lighthouse Lab in Alderley Park and Alex Alderton, Roberto Amato, Jeffrey Barrett, Sonia Goncalves, Ewan Harrison, David K. Jackson, Ian Johnston, Dominic Kwiatkowski, Cordelia Langford, John Sillitoe on behalf of the Wellcome Sanger Institute COVID-19 Surveillance Team |
| EPI_ISL_2485465 | Lighthouse Lab in Milton Keynes | Wellcome Sanger Institute for the COVID-19 Genomics UK (COG-UK) Consortium | The Lighthouse Lab in Milton Keynes and Alex Alderton, Roberto Amato, Jeffrey Barrett, Sonia Goncalves, Ewan Harrison, David K. Jackson, Ian Johnston, Dominic Kwiatkowski, Cordelia Langford, John Sillitoe on behalf of the Wellcome Sanger Institute COVID-19 Surveillance Team |
| EPI_ISL_2485466 | Lighthouse Lab in Alderley Park | Wellcome Sanger Institute for the COVID-19 Genomics UK (COG-UK) Consortium | Jacquelyn Wynn, Mairead Hyland, The Lighthouse Lab in Alderley Park and Alex Alderton, Roberto Amato, Jeffrey Barrett, Sonia Goncalves, Ewan Harrison, David K. Jackson, Ian Johnston, Dominic Kwiatkowski, Cordelia Langford, John Sillitoe on behalf of the Wellcome Sanger Institute COVID-19 Surveillance Team |
| EPI_ISL_2485468 | Lighthouse Lab in Milton Keynes | Wellcome Sanger Institute for the COVID-19 Genomics UK (COG-UK) Consortium | The Lighthouse Lab in Milton Keynes and Alex Alderton, Roberto Amato, Jeffrey Barrett, Sonia Goncalves, Ewan Harrison, David K. Jackson, Ian Johnston, Dominic Kwiatkowski, Cordelia Langford, John Sillitoe on behalf of the Wellcome Sanger Institute COVID-19 Surveillance Team |
| EPI_ISL_2485469, EPI_ISL_2485470 | Lighthouse Lab in Alderley Park | Wellcome Sanger Institute for the COVID-19 Genomics UK (COG-UK) Consortium | Jacquelyn Wynn, Mairead Hyland, The Lighthouse Lab in Alderley Park and Alex Alderton, Roberto Amato, Jeffrey Barrett, Sonia Goncalves, Ewan Harrison, David K. Jackson, Ian Johnston, Dominic Kwiatkowski, Cordelia Langford, John Sillitoe on behalf of the Wellcome Sanger Institute COVID-19 Surveillance Team |
| EPI_ISL_2485471 | Health Services Laboratories | Wellcome Sanger Institute for the COVID-19 Genomics UK (COG-UK) Consortium | Health Services Laboratories and Alex Alderton, Roberto Amato, Jeffrey Barrett, Sonia Goncalves, Ewan Harrison, David K. Jackson, Ian Johnston, Dominic Kwiatkowski, Cordelia Langford, John Sillitoe on behalf of the Wellcome Sanger Institute COVID-19 Surveillance Team |
| EPI_ISL_2485472 | Lighthouse Lab in Milton Keynes | Wellcome Sanger Institute for the COVID-19 Genomics UK (COG-UK) Consortium | The Lighthouse Lab in Milton Keynes and Alex Alderton, Roberto Amato, Jeffrey Barrett, Sonia Goncalves, Ewan Harrison, David K. Jackson, Ian Johnston, Dominic Kwiatkowski, Cordelia Langford, John Sillitoe on behalf of the Wellcome Sanger Institute COVID-19 Surveillance Team |
| EPI_ISL_2485473 | Health Services Laboratories | Wellcome Sanger Institute for the COVID-19 Genomics UK (COG-UK) Consortium | Health Services Laboratories and Alex Alderton, Roberto Amato, Jeffrey Barrett, Sonia Goncalves, Ewan Harrison, David K. Jackson, Ian Johnston, Dominic Kwiatkowski, Cordelia Langford, John Sillitoe on behalf of the Wellcome Sanger Institute COVID-19 Surveillance Team |
| EPI_ISL_2485475 | Lighthouse Laboratory Plymouth | Wellcome Sanger Institute for the COVID-19 Genomics UK (COG-UK) Consortium | Lighthouse Laboratory Plymouth and Alex Alderton, Roberto Amato, Jeffrey Barrett, Sonia Goncalves, Ewan Harrison, David K. Jackson, Ian Johnston, Dominic Kwiatkowski, Cordelia Langford, John Sillitoe on behalf of the Wellcome Sanger Institute COVID-19 Surveillance Team |
| EPI_ISL_2485476, EPI_ISL_2485477 | Lighthouse Lab in Milton Keynes | Wellcome Sanger Institute for the COVID-19 Genomics UK (COG-UK) Consortium | The Lighthouse Lab in Milton Keynes and Alex Alderton, Roberto Amato, Jeffrey Barrett, Sonia Goncalves, Ewan Harrison, David K. Jackson, Ian Johnston, Dominic Kwiatkowski, Cordelia Langford, John Sillitoe on behalf of the Wellcome Sanger Institute COVID-19 Surveillance Team |
| EPI_ISL_2485478 | Lighthouse Lab in Alderley Park | Wellcome Sanger Institute for the COVID-19 Genomics UK (COG-UK) Consortium | Jacquelyn Wynn, Mairead Hyland, The Lighthouse Lab in Alderley Park and Alex Alderton, Roberto Amato, Jeffrey Barrett, Sonia Goncalves, Ewan Harrison, David K. Jackson, Ian Johnston, Dominic Kwiatkowski, Cordelia Langford, John Sillitoe on behalf of the Wellcome Sanger Institute COVID-19 Surveillance Team |
| EPI_ISL_2485479 | Lighthouse Laboratory Plymouth | Wellcome Sanger Institute for the COVID-19 Genomics UK (COG-UK) Consortium | Lighthouse Laboratory Plymouth and Alex Alderton, Roberto Amato, Jeffrey Barrett, Sonia Goncalves, Ewan Harrison, David K. Jackson, Ian Johnston, Dominic Kwiatkowski, Cordelia Langford, John Sillitoe on behalf of the Wellcome Sanger Institute COVID-19 Surveillance Team |
| EPI_ISL_2485480, EPI_ISL_2485481 | Lighthouse Lab in Alderley Park | Wellcome Sanger Institute for the COVID-19 Genomics UK (COG-UK) Consortium | Jacquelyn Wynn, Mairead Hyland, The Lighthouse Lab in Alderley Park and Alex Alderton, Roberto Amato, Jeffrey Barrett, Sonia Goncalves, Ewan Harrison, David K. Jackson, Ian Johnston, Dominic Kwiatkowski, Cordelia Langford, John Sillitoe on behalf of the Wellcome Sanger Institute COVID-19 Surveillance Team |
| EPI_ISL_2485482, EPI_ISL_2485483 | Lighthouse Laboratory Plymouth | Wellcome Sanger Institute for the COVID-19 Genomics UK (COG-UK) Consortium | Lighthouse Laboratory Plymouth and Alex Alderton, Roberto Amato, Jeffrey Barrett, Sonia Goncalves, Ewan Harrison, David K. Jackson, Ian Johnston, Dominic Kwiatkowski, Cordelia Langford, John Sillitoe on behalf of the Wellcome Sanger Institute COVID-19 Surveillance Team |
| EPI_ISL_2485484 | Lighthouse Lab in Alderley Park | Wellcome Sanger Institute for the COVID-19 Genomics UK (COG-UK) Consortium | Jacquelyn Wynn, Mairead Hyland, The Lighthouse Lab in Alderley Park and Alex Alderton, Roberto Amato, Jeffrey Barrett, Sonia Goncalves, Ewan Harrison, David K. Jackson, Ian Johnston, Dominic Kwiatkowski, Cordelia Langford, John Sillitoe on behalf of the Wellcome Sanger Institute COVID-19 Surveillance Team |
| EPI_ISL_2485485 | Lighthouse Lab in Milton Keynes | Wellcome Sanger Institute for the COVID-19 Genomics UK (COG-UK) Consortium | The Lighthouse Lab in Milton Keynes and Alex Alderton, Roberto Amato, Jeffrey Barrett, Sonia Goncalves, Ewan Harrison, David K. Jackson, Ian Johnston, Dominic Kwiatkowski, Cordelia Langford, John Sillitoe on behalf of the Wellcome Sanger Institute COVID-19 Surveillance Team |
| EPI_ISL_2485486 | Health Services Laboratories | Wellcome Sanger Institute for the COVID-19 Genomics UK (COG-UK) Consortium | Health Services Laboratories and Alex Alderton, Roberto Amato, Jeffrey Barrett, Sonia Goncalves, Ewan Harrison, David K. Jackson, Ian Johnston, Dominic Kwiatkowski, Cordelia Langford, John Sillitoe on behalf of the Wellcome Sanger Institute COVID-19 Surveillance Team |
| EPI_ISL_2485487 | Lighthouse Laboratory Plymouth | Wellcome Sanger Institute for the COVID-19 Genomics UK (COG-UK) Consortium | Lighthouse Laboratory Plymouth and Alex Alderton, Roberto Amato, Jeffrey Barrett, Sonia Goncalves, Ewan Harrison, David K. Jackson, Ian Johnston, Dominic Kwiatkowski, Cordelia Langford, John Sillitoe on behalf of the Wellcome Sanger Institute COVID-19 Surveillance Team |
| EPI_ISL_2485488, EPI_ISL_2485489 | Lighthouse Lab in Alderley Park | Wellcome Sanger Institute for the COVID-19 Genomics UK (COG-UK) Consortium | Jacquelyn Wynn, Mairead Hyland, The Lighthouse Lab in Alderley Park and Alex Alderton, Roberto Amato, Jeffrey Barrett, Sonia Goncalves, Ewan Harrison, David K. Jackson, Ian Johnston, Dominic Kwiatkowski, Cordelia Langford, John Sillitoe on behalf of the Wellcome Sanger Institute COVID-19 Surveillance Team |
| EPI_ISL_2485490 | Lighthouse Lab in Milton Keynes | Wellcome Sanger Institute for the COVID-19 Genomics UK (COG-UK) Consortium | The Lighthouse Lab in Milton Keynes and Alex Alderton, Roberto Amato, Jeffrey Barrett, Sonia Goncalves, Ewan Harrison, David K. Jackson, Ian Johnston, Dominic Kwiatkowski, Cordelia Langford, John Sillitoe on behalf of the Wellcome Sanger Institute COVID-19 Surveillance Team |
| EPI_ISL_2485491 | Lighthouse Lab in Alderley Park | Wellcome Sanger Institute for the COVID-19 Genomics UK (COG-UK) Consortium | Jacquelyn Wynn, Mairead Hyland, The Lighthouse Lab in Alderley Park and Alex Alderton, Roberto Amato, Jeffrey Barrett, Sonia Goncalves, Ewan Harrison, David K. Jackson, Ian Johnston, Dominic Kwiatkowski, Cordelia Langford, John Sillitoe on behalf of the Wellcome Sanger Institute COVID-19 Surveillance Team |
| EPI_ISL_2485492 | Health Services Laboratories | Wellcome Sanger Institute for the COVID-19 Genomics UK (COG-UK) Consortium | Health Services Laboratories and Alex Alderton, Roberto Amato, Jeffrey Barrett, Sonia Goncalves, Ewan Harrison, David K. Jackson, Ian Johnston, Dominic Kwiatkowski, Cordelia Langford, John Sillitoe on behalf of the Wellcome Sanger Institute COVID-19 Surveillance Team |
| EPI_ISL_2485493 | Lighthouse Lab in Alderley Park | Wellcome Sanger Institute for the COVID-19 Genomics UK (COG-UK) Consortium | Jacquelyn Wynn, Mairead Hyland, The Lighthouse Lab in Alderley Park and Alex Alderton, Roberto Amato, Jeffrey Barrett, Sonia Goncalves, Ewan Harrison, David K. Jackson, Ian Johnston, Dominic Kwiatkowski, Cordelia Langford, John Sillitoe on behalf of the Wellcome Sanger Institute COVID-19 Surveillance Team |
| EPI_ISL_2485494 | Lighthouse Lab in Milton Keynes | Wellcome Sanger Institute for the COVID-19 Genomics UK (COG-UK) Consortium | The Lighthouse Lab in Milton Keynes and Alex Alderton, Roberto Amato, Jeffrey Barrett, Sonia Goncalves, Ewan Harrison, David K. Jackson, Ian Johnston, Dominic Kwiatkowski, Cordelia Langford, John Sillitoe on behalf of the Wellcome Sanger Institute COVID-19 Surveillance Team |
| EPI_ISL_2485495, EPI_ISL_2485496 | Lighthouse Lab in Alderley Park | Wellcome Sanger Institute for the COVID-19 Genomics UK (COG-UK) Consortium | Jacquelyn Wynn, Mairead Hyland, The Lighthouse Lab in Alderley Park and Alex Alderton, Roberto Amato, Jeffrey Barrett, Sonia Goncalves, Ewan Harrison, David K. Jackson, Ian Johnston, Dominic Kwiatkowski, Cordelia Langford, John Sillitoe on behalf of the Wellcome Sanger Institute COVID-19 Surveillance Team |
| EPI_ISL_2485497 | Health Services Laboratories | Wellcome Sanger Institute for the COVID-19 Genomics UK (COG-UK) Consortium | Health Services Laboratories and Alex Alderton, Roberto Amato, Jeffrey Barrett, Sonia Goncalves, Ewan Harrison, David K. Jackson, Ian Johnston, Dominic Kwiatkowski, Cordelia Langford, John Sillitoe on behalf of the Wellcome Sanger Institute COVID-19 Surveillance Team |
| EPI_ISL_2485498, EPI_ISL_2485500, EPI_ISL_2485501, EPI_ISL_2485502, EPI_ISL_2485503 | Lighthouse Lab in Alderley Park | Wellcome Sanger Institute for the COVID-19 Genomics UK (COG-UK) Consortium | Jacquelyn Wynn, Mairead Hyland, The Lighthouse Lab in Alderley Park and Alex Alderton, Roberto Amato, Jeffrey Barrett, Sonia Goncalves, Ewan Harrison, David K. Jackson, Ian Johnston, Dominic Kwiatkowski, Cordelia Langford, John Sillitoe on behalf of the Wellcome Sanger Institute COVID-19 Surveillance Team |
| EPI_ISL_2485504, EPI_ISL_2485505 | Lighthouse Lab in Milton Keynes | Wellcome Sanger Institute for the COVID-19 Genomics UK (COG-UK) Consortium | The Lighthouse Lab in Milton Keynes and Alex Alderton, Roberto Amato, Jeffrey Barrett, Sonia Goncalves, Ewan Harrison, David K. Jackson, Ian Johnston, Dominic Kwiatkowski, Cordelia Langford, John Sillitoe on behalf of the Wellcome Sanger Institute COVID-19 Surveillance Team |
| EPI_ISL_2485507 | Health Services Laboratories | Wellcome Sanger Institute for the COVID-19 Genomics UK (COG-UK) Consortium | Health Services Laboratories and Alex Alderton, Roberto Amato, Jeffrey Barrett, Sonia Goncalves, Ewan Harrison, David K. Jackson, Ian Johnston, Dominic Kwiatkowski, Cordelia Langford, John Sillitoe on behalf of the Wellcome Sanger Institute COVID-19 Surveillance Team |
| EPI_ISL_2485508 | Lighthouse Lab in Alderley Park | Wellcome Sanger Institute for the COVID-19 Genomics UK (COG-UK) Consortium | Jacquelyn Wynn, Mairead Hyland, The Lighthouse Lab in Alderley Park and Alex Alderton, Roberto Amato, Jeffrey Barrett, Sonia Goncalves, Ewan Harrison, David K. Jackson, Ian Johnston, Dominic Kwiatkowski, Cordelia Langford, John Sillitoe on behalf of the Wellcome Sanger Institute COVID-19 Surveillance Team |
| EPI_ISL_2485509 | Health Services Laboratories | Wellcome Sanger Institute for the COVID-19 Genomics UK (COG-UK) Consortium | Health Services Laboratories and Alex Alderton, Roberto Amato, Jeffrey Barrett, Sonia Goncalves, Ewan Harrison, David K. Jackson, Ian Johnston, Dominic Kwiatkowski, Cordelia Langford, John Sillitoe on behalf of the Wellcome Sanger Institute COVID-19 Surveillance Team |
| EPI_ISL_2485510, EPI_ISL_2485511 | Lighthouse Lab in Alderley Park | Wellcome Sanger Institute for the COVID-19 Genomics UK (COG-UK) Consortium | Jacquelyn Wynn, Mairead Hyland, The Lighthouse Lab in Alderley Park and Alex Alderton, Roberto Amato, Jeffrey Barrett, Sonia Goncalves, Ewan Harrison, David K. Jackson, Ian Johnston, Dominic Kwiatkowski, Cordelia Langford, John Sillitoe on behalf of the Wellcome Sanger Institute COVID-19 Surveillance Team |

| | | | |
|---|---|---|---|
| EPI_ISL_2485512 | Lighthouse Laboratory Plymouth | Wellcome Sanger Institute for the COVID-19 Genomics UK (COG-UK) Consortium | Lighthouse Laboratory Plymouth and Alex Alderton, Roberto Amato, Jeffrey Barrett, Sonia Goncalves, Ewan Harrison, David K. Jackson, Ian Johnston, Dominic Kwiatkowski, Cordelia Langford, John Sillitoe on behalf of the Wellcome Sanger Institute COVID-19 Surveillance Team |
| EPI_ISL_2485513 | Lighthouse Lab in Milton Keynes | Wellcome Sanger Institute for the COVID-19 Genomics UK (COG-UK) Consortium | The Lighthouse Lab in Milton Keynes and Alex Alderton, Roberto Amato, Jeffrey Barrett, Sonia Goncalves, Ewan Harrison, David K. Jackson, Ian Johnston, Dominic Kwiatkowski, Cordelia Langford, John Sillitoe on behalf of the Wellcome Sanger Institute COVID-19 Surveillance Team |
| EPI_ISL_2485515, EPI_ISL_2485516 | Lighthouse Lab in Alderley Park | Wellcome Sanger Institute for the COVID-19 Genomics UK (COG-UK) Consortium | Jacquelyn Wynn, Mairead Hyland, The Lighthouse Lab in Alderley Park and Alex Alderton, Roberto Amato, Jeffrey Barrett, Sonia Goncalves, Ewan Harrison, David K. Jackson, Ian Johnston, Dominic Kwiatkowski, Cordelia Langford, John Sillitoe on behalf of the Wellcome Sanger Institute COVID-19 Surveillance Team |
| EPI_ISL_2485517 | Health Services Laboratories | Wellcome Sanger Institute for the COVID-19 Genomics UK (COG-UK) Consortium | Health Services Laboratories and Alex Alderton, Roberto Amato, Jeffrey Barrett, Sonia Goncalves, Ewan Harrison, David K. Jackson, Ian Johnston, Dominic Kwiatkowski, Cordelia Langford, John Sillitoe on behalf of the Wellcome Sanger Institute COVID-19 Surveillance Team |
| EPI_ISL_2485518, EPI_ISL_2485519, EPI_ISL_2485520 | Lighthouse Lab in Alderley Park | Wellcome Sanger Institute for the COVID-19 Genomics UK (COG-UK) Consortium | Jacquelyn Wynn, Mairead Hyland, The Lighthouse Lab in Alderley Park and Alex Alderton, Roberto Amato, Jeffrey Barrett, Sonia Goncalves, Ewan Harrison, David K. Jackson, Ian Johnston, Dominic Kwiatkowski, Cordelia Langford, John Sillitoe on behalf of the Wellcome Sanger Institute COVID-19 Surveillance Team |
| EPI_ISL_2485521 | Health Services Laboratories | Wellcome Sanger Institute for the COVID-19 Genomics UK (COG-UK) Consortium | Health Services Laboratories and Alex Alderton, Roberto Amato, Jeffrey Barrett, Sonia Goncalves, Ewan Harrison, David K. Jackson, Ian Johnston, Dominic Kwiatkowski, Cordelia Langford, John Sillitoe on behalf of the Wellcome Sanger Institute COVID-19 Surveillance Team |
| EPI_ISL_2485522 | Lighthouse Lab in Milton Keynes | Wellcome Sanger Institute for the COVID-19 Genomics UK (COG-UK) Consortium | The Lighthouse Lab in Milton Keynes and Alex Alderton, Roberto Amato, Jeffrey Barrett, Sonia Goncalves, Ewan Harrison, David K. Jackson, Ian Johnston, Dominic Kwiatkowski, Cordelia Langford, John Sillitoe on behalf of the Wellcome Sanger Institute COVID-19 Surveillance Team |
| EPI_ISL_2485523, EPI_ISL_2485524 | Lighthouse Lab in Alderley Park | Wellcome Sanger Institute for the COVID-19 Genomics UK (COG-UK) Consortium | Jacquelyn Wynn, Mairead Hyland, The Lighthouse Lab in Alderley Park and Alex Alderton, Roberto Amato, Jeffrey Barrett, Sonia Goncalves, Ewan Harrison, David K. Jackson, Ian Johnston, Dominic Kwiatkowski, Cordelia Langford, John Sillitoe on behalf of the Wellcome Sanger Institute COVID-19 Surveillance Team |
| EPI_ISL_2485525 | Lighthouse Lab in Milton Keynes | Wellcome Sanger Institute for the COVID-19 Genomics UK (COG-UK) Consortium | The Lighthouse Lab in Milton Keynes and Alex Alderton, Roberto Amato, Jeffrey Barrett, Sonia Goncalves, Ewan Harrison, David K. Jackson, Ian Johnston, Dominic Kwiatkowski, Cordelia Langford, John Sillitoe on behalf of the Wellcome Sanger Institute COVID-19 Surveillance Team |
| EPI_ISL_2485526, EPI_ISL_2485527 | Lighthouse Lab in Alderley Park | Wellcome Sanger Institute for the COVID-19 Genomics UK (COG-UK) Consortium | Jacquelyn Wynn, Mairead Hyland, The Lighthouse Lab in Alderley Park and Alex Alderton, Roberto Amato, Jeffrey Barrett, Sonia Goncalves, Ewan Harrison, David K. Jackson, Ian Johnston, Dominic Kwiatkowski, Cordelia Langford, John Sillitoe on behalf of the Wellcome Sanger Institute COVID-19 Surveillance Team |
| EPI_ISL_2485528, EPI_ISL_2485529 | Lighthouse Lab in Milton Keynes | Wellcome Sanger Institute for the COVID-19 Genomics UK (COG-UK) Consortium | The Lighthouse Lab in Milton Keynes and Alex Alderton, Roberto Amato, Jeffrey Barrett, Sonia Goncalves, Ewan Harrison, David K. Jackson, Ian Johnston, Dominic Kwiatkowski, Cordelia Langford, John Sillitoe on behalf of the Wellcome Sanger Institute COVID-19 Surveillance Team |
| EPI_ISL_2485530 | Lighthouse Lab in Alderley Park | Wellcome Sanger Institute for the COVID-19 Genomics UK (COG-UK) Consortium | Jacquelyn Wynn, Mairead Hyland, The Lighthouse Lab in Alderley Park and Alex Alderton, Roberto Amato, Jeffrey Barrett, Sonia Goncalves, Ewan Harrison, David K. Jackson, Ian Johnston, Dominic Kwiatkowski, Cordelia Langford, John Sillitoe on behalf of the Wellcome Sanger Institute COVID-19 Surveillance Team |
| EPI_ISL_2485531, EPI_ISL_2485532 | Lighthouse Lab in Milton Keynes | Wellcome Sanger Institute for the COVID-19 Genomics UK (COG-UK) Consortium | The Lighthouse Lab in Milton Keynes and Alex Alderton, Roberto Amato, Jeffrey Barrett, Sonia Goncalves, Ewan Harrison, David K. Jackson, Ian Johnston, Dominic Kwiatkowski, Cordelia Langford, John Sillitoe on behalf of the Wellcome Sanger Institute COVID-19 Surveillance Team |
| EPI_ISL_2485533, EPI_ISL_2485534, EPI_ISL_2485535, EPI_ISL_2485536 | Lighthouse Lab in Alderley Park | Wellcome Sanger Institute for the COVID-19 Genomics UK (COG-UK) Consortium | Jacquelyn Wynn, Mairead Hyland, The Lighthouse Lab in Alderley Park and Alex Alderton, Roberto Amato, Jeffrey Barrett, Sonia Goncalves, Ewan Harrison, David K. Jackson, Ian Johnston, Dominic Kwiatkowski, Cordelia Langford, John Sillitoe on behalf of the Wellcome Sanger Institute COVID-19 Surveillance Team |
| EPI_ISL_2485537 | Lighthouse Lab in Milton Keynes | Wellcome Sanger Institute for the COVID-19 Genomics UK (COG-UK) Consortium | The Lighthouse Lab in Milton Keynes and Alex Alderton, Roberto Amato, Jeffrey Barrett, Sonia Goncalves, Ewan Harrison, David K. Jackson, Ian Johnston, Dominic Kwiatkowski, Cordelia Langford, John Sillitoe on behalf of the Wellcome Sanger Institute COVID-19 Surveillance Team |
| EPI_ISL_2485538 | Lighthouse Lab in Alderley Park | Wellcome Sanger Institute for the COVID-19 Genomics UK (COG-UK) Consortium | Jacquelyn Wynn, Mairead Hyland, The Lighthouse Lab in Alderley Park and Alex Alderton, Roberto Amato, Jeffrey Barrett, Sonia Goncalves, Ewan Harrison, David K. Jackson, Ian Johnston, Dominic Kwiatkowski, Cordelia Langford, John Sillitoe on behalf of the Wellcome Sanger Institute COVID-19 Surveillance Team |
| EPI_ISL_2485539, EPI_ISL_2485540, EPI_ISL_2485541 | Lighthouse Lab in Milton Keynes | Wellcome Sanger Institute for the COVID-19 Genomics UK (COG-UK) Consortium | The Lighthouse Lab in Milton Keynes and Alex Alderton, Roberto Amato, Jeffrey Barrett, Sonia Goncalves, Ewan Harrison, David K. Jackson, Ian Johnston, Dominic Kwiatkowski, Cordelia Langford, John Sillitoe on behalf of the Wellcome Sanger Institute COVID-19 Surveillance Team |
| EPI_ISL_2485543 | Lighthouse Lab in Alderley Park | Wellcome Sanger Institute for the COVID-19 Genomics UK (COG-UK) Consortium | Jacquelyn Wynn, Mairead Hyland, The Lighthouse Lab in Alderley Park and Alex Alderton, Roberto Amato, Jeffrey Barrett, Sonia Goncalves, Ewan Harrison, David K. Jackson, Ian Johnston, Dominic Kwiatkowski, Cordelia Langford, John Sillitoe on behalf of the Wellcome Sanger Institute COVID-19 Surveillance Team |
| EPI_ISL_2485544 | Lighthouse Lab in Milton Keynes | Wellcome Sanger Institute for the COVID-19 Genomics UK (COG-UK) Consortium | The Lighthouse Lab in Milton Keynes and Alex Alderton, Roberto Amato, Jeffrey Barrett, Sonia Goncalves, Ewan Harrison, David K. Jackson, Ian Johnston, Dominic Kwiatkowski, Cordelia Langford, John Sillitoe on behalf of the Wellcome Sanger Institute COVID-19 Surveillance Team |
| EPI_ISL_2485545 | Lighthouse Lab in Alderley Park | Wellcome Sanger Institute for the COVID-19 Genomics UK (COG-UK) Consortium | Jacquelyn Wynn, Mairead Hyland, The Lighthouse Lab in Alderley Park and Alex Alderton, Roberto Amato, Jeffrey Barrett, Sonia Goncalves, Ewan Harrison, David K. Jackson, Ian Johnston, Dominic Kwiatkowski, Cordelia Langford, John Sillitoe on behalf of the Wellcome Sanger Institute COVID-19 Surveillance Team |
| EPI_ISL_2485546 | Lighthouse Lab in Milton Keynes | Wellcome Sanger Institute for the COVID-19 Genomics UK (COG-UK) Consortium | The Lighthouse Lab in Milton Keynes and Alex Alderton, Roberto Amato, Jeffrey Barrett, Sonia Goncalves, Ewan Harrison, David K. Jackson, Ian Johnston, Dominic Kwiatkowski, Cordelia Langford, John Sillitoe on behalf of the Wellcome Sanger Institute COVID-19 Surveillance Team |
| EPI_ISL_2485547, EPI_ISL_2485548, EPI_ISL_2485549 | Lighthouse Lab in Alderley Park | Wellcome Sanger Institute for the COVID-19 Genomics UK (COG-UK) Consortium | Jacquelyn Wynn, Mairead Hyland, The Lighthouse Lab in Alderley Park and Alex Alderton, Roberto Amato, Jeffrey Barrett, Sonia Goncalves, Ewan Harrison, David K. Jackson, Ian Johnston, Dominic Kwiatkowski, Cordelia Langford, John Sillitoe on behalf of the Wellcome Sanger Institute COVID-19 Surveillance Team |
| EPI_ISL_2485550 | Lighthouse Lab in Milton Keynes | Wellcome Sanger Institute for the COVID-19 Genomics UK (COG-UK) Consortium | The Lighthouse Lab in Milton Keynes and Alex Alderton, Roberto Amato, Jeffrey Barrett, Sonia Goncalves, Ewan Harrison, David K. Jackson, Ian Johnston, Dominic Kwiatkowski, Cordelia Langford, John Sillitoe on behalf of the Wellcome Sanger Institute COVID-19 Surveillance Team |
| EPI_ISL_2485551, EPI_ISL_2485552, EPI_ISL_2485553 | Lighthouse Lab in Alderley Park | Wellcome Sanger Institute for the COVID-19 Genomics UK (COG-UK) Consortium | Jacquelyn Wynn, Mairead Hyland, The Lighthouse Lab in Alderley Park and Alex Alderton, Roberto Amato, Jeffrey Barrett, Sonia Goncalves, Ewan Harrison, David K. Jackson, Ian Johnston, Dominic Kwiatkowski, Cordelia Langford, John Sillitoe on behalf of the Wellcome Sanger Institute COVID-19 Surveillance Team |
| EPI_ISL_2485554 | Lighthouse Lab in Milton Keynes | Wellcome Sanger Institute for the COVID-19 Genomics UK (COG-UK) Consortium | The Lighthouse Lab in Milton Keynes and Alex Alderton, Roberto Amato, Jeffrey Barrett, Sonia Goncalves, Ewan Harrison, David K. Jackson, Ian Johnston, Dominic Kwiatkowski, Cordelia Langford, John Sillitoe on behalf of the Wellcome Sanger Institute COVID-19 Surveillance Team |
| EPI_ISL_2485555, EPI_ISL_2485556, EPI_ISL_2485557, EPI_ISL_2485558 | Lighthouse Lab in Alderley Park | Wellcome Sanger Institute for the COVID-19 Genomics UK (COG-UK) Consortium | Jacquelyn Wynn, Mairead Hyland, The Lighthouse Lab in Alderley Park and Alex Alderton, Roberto Amato, Jeffrey Barrett, Sonia Goncalves, Ewan Harrison, David K. Jackson, Ian Johnston, Dominic Kwiatkowski, Cordelia Langford, John Sillitoe on behalf of the Wellcome Sanger Institute COVID-19 Surveillance Team |
| EPI_ISL_2485559 | Lighthouse Lab in Glasgow | Wellcome Sanger Institute for the COVID-19 Genomics UK (COG-UK) Consortium | Harper VanSteenhouse, Yumi Kasai, David Gray, Carol Clugston, Anna Dominiczak and Alex Alderton, Roberto Amato, Jeffrey Barrett, Sonia Goncalves, Ewan Harrison, David K. Jackson, Ian Johnston, Dominic Kwiatkowski, Cordelia Langford, John Sillitoe on behalf of the Wellcome Sanger Institute COVID-19 Surveillance Team |
| EPI_ISL_2485560 | Lighthouse Lab in Milton Keynes | Wellcome Sanger Institute for the COVID-19 Genomics UK (COG-UK) Consortium | The Lighthouse Lab in Milton Keynes and Alex Alderton, Roberto Amato, Jeffrey Barrett, Sonia Goncalves, Ewan Harrison, David K. Jackson, Ian Johnston, Dominic Kwiatkowski, Cordelia Langford, John Sillitoe on behalf of the Wellcome Sanger Institute COVID-19 Surveillance Team |
| EPI_ISL_2485561, EPI_ISL_2485562, EPI_ISL_2485563, EPI_ISL_2485565, EPI_ISL_2485566 | Lighthouse Lab in Alderley Park | Wellcome Sanger Institute for the COVID-19 Genomics UK (COG-UK) Consortium | Jacquelyn Wynn, Mairead Hyland, The Lighthouse Lab in Alderley Park and Alex Alderton, Roberto Amato, Jeffrey Barrett, Sonia Goncalves, Ewan Harrison, David K. Jackson, Ian Johnston, Dominic Kwiatkowski, Cordelia Langford, John Sillitoe on behalf of the Wellcome Sanger Institute COVID-19 Surveillance Team |
| EPI_ISL_2485567 | Lighthouse Lab in Milton Keynes | Wellcome Sanger Institute for the COVID-19 Genomics UK (COG-UK) Consortium | The Lighthouse Lab in Milton Keynes and Alex Alderton, Roberto Amato, Jeffrey Barrett, Sonia Goncalves, Ewan Harrison, David K. Jackson, Ian Johnston, Dominic Kwiatkowski, Cordelia Langford, John Sillitoe on behalf of the Wellcome Sanger Institute COVID-19 Surveillance Team |
| EPI_ISL_2485568, EPI_ISL_2485570 | Lighthouse Lab in Alderley Park | Wellcome Sanger Institute for the COVID-19 Genomics UK (COG-UK) Consortium | Jacquelyn Wynn, Mairead Hyland, The Lighthouse Lab in Alderley Park and Alex Alderton, Roberto Amato, Jeffrey Barrett, Sonia Goncalves, Ewan Harrison, David K. Jackson, Ian Johnston, Dominic Kwiatkowski, Cordelia Langford, John Sillitoe on behalf of the Wellcome Sanger Institute COVID-19 Surveillance Team |
| EPI_ISL_2485571 | Lighthouse Lab in Glasgow | Wellcome Sanger Institute for the COVID-19 Genomics UK (COG-UK) Consortium | Harper VanSteenhouse, Yumi Kasai, David Gray, Carol Clugston, Anna Dominiczak and Alex Alderton, Roberto Amato, Jeffrey Barrett, Sonia Goncalves, Ewan Harrison, David K. Jackson, Ian Johnston, Dominic Kwiatkowski, Cordelia Langford, John Sillitoe on behalf of the Wellcome Sanger Institute COVID-19 Surveillance Team |
| EPI_ISL_2485572, EPI_ISL_2485573, EPI_ISL_2485574, EPI_ISL_2485575, EPI_ISL_2485576, EPI_ISL_2485577, EPI_ISL_2485578 | Lighthouse Lab in Alderley Park | Wellcome Sanger Institute for the COVID-19 Genomics UK (COG-UK) Consortium | Jacquelyn Wynn, Mairead Hyland, The Lighthouse Lab in Alderley Park and Alex Alderton, Roberto Amato, Jeffrey Barrett, Sonia Goncalves, Ewan Harrison, David K. Jackson, Ian Johnston, Dominic Kwiatkowski, Cordelia Langford, John Sillitoe on behalf of the Wellcome Sanger Institute COVID-19 Surveillance Team |
| EPI_ISL_2485579, EPI_ISL_2485580, EPI_ISL_2485581, EPI_ISL_2485582 | Lighthouse Lab in Milton Keynes | Wellcome Sanger Institute for the COVID-19 Genomics UK (COG-UK) Consortium | The Lighthouse Lab in Milton Keynes and Alex Alderton, Roberto Amato, Jeffrey Barrett, Sonia Goncalves, Ewan Harrison, David K. Jackson, Ian Johnston, Dominic Kwiatkowski, Cordelia Langford, John Sillitoe on behalf of the Wellcome Sanger Institute COVID-19 Surveillance Team |
| EPI_ISL_2485583 | Lighthouse Lab in Alderley Park | Wellcome Sanger Institute for the COVID-19 Genomics UK (COG-UK) Consortium | Jacquelyn Wynn, Mairead Hyland, The Lighthouse Lab in Alderley Park and Alex Alderton, Roberto Amato, Jeffrey Barrett, Sonia Goncalves, Ewan Harrison, David K. Jackson, Ian Johnston, Dominic Kwiatkowski, Cordelia Langford, John Sillitoe on behalf of the Wellcome Sanger Institute COVID-19 Surveillance Team |
| EPI_ISL_2485584 | Lighthouse Lab in Milton Keynes | Wellcome Sanger Institute for the COVID-19 Genomics UK (COG-UK) Consortium | The Lighthouse Lab in Milton Keynes and Alex Alderton, Roberto Amato, Jeffrey Barrett, Sonia Goncalves, Ewan Harrison, David K. Jackson, Ian Johnston, Dominic Kwiatkowski, Cordelia Langford, John Sillitoe on behalf of the Wellcome Sanger Institute COVID-19 Surveillance Team |
| EPI_ISL_2485585, EPI_ISL_2485587 | Lighthouse Lab in Alderley Park | Wellcome Sanger Institute for the COVID-19 Genomics UK (COG-UK) Consortium | Jacquelyn Wynn, Mairead Hyland, The Lighthouse Lab in Alderley Park and Alex Alderton, Roberto Amato, Jeffrey Barrett, Sonia Goncalves, Ewan Harrison, David K. Jackson, Ian Johnston, Dominic Kwiatkowski, Cordelia Langford, John Sillitoe on behalf of the Wellcome Sanger Institute COVID-19 Surveillance Team |
| EPI_ISL_2485588, EPI_ISL_2485589, | Lighthouse Lab in Milton Keynes | Wellcome Sanger Institute for the COVID-19 Genomics UK (COG-UK) Consortium | The Lighthouse Lab in Milton Keynes and Alex Alderton, Roberto Amato, Jeffrey Barrett, Sonia Goncalves, Ewan Harrison, David K. Jackson, Ian Johnston, Dominic Kwiatkowski, Cordelia Langford, John Sillitoe on behalf of the Wellcome Sanger Institute COVID-19 Surveillance Team |

| | | | |
|---|---|---|---|
| EPI_ISL_2485590 | | | |
| EPI_ISL_2485591 | Lighthouse Lab in Alderley Park | Wellcome Sanger Institute for the COVID-19 Genomics UK (COG-UK) Consortium | Jacquelyn Wynn, Mairead Hyland, The Lighthouse Lab in Alderley Park and Alex Alderton, Roberto Amato, Jeffrey Barrett, Sonia Goncalves, Ewan Harrison, David K. Jackson, Ian Johnston, Dominic Kwiatkowski, Cordelia Langford, John Sillitoe on behalf of the Wellcome Sanger Institute COVID-19 Surveillance Team |
| EPI_ISL_2485592 | Lighthouse Lab in Milton Keynes | Wellcome Sanger Institute for the COVID-19 Genomics UK (COG-UK) Consortium | The Lighthouse Lab in Milton Keynes and Alex Alderton, Roberto Amato, Jeffrey Barrett, Sonia Goncalves, Ewan Harrison, David K. Jackson, Ian Johnston, Dominic Kwiatkowski, Cordelia Langford, John Sillitoe on behalf of the Wellcome Sanger Institute COVID-19 Surveillance Team |
| EPI_ISL_2485593 | Lighthouse Lab in Alderley Park | Wellcome Sanger Institute for the COVID-19 Genomics UK (COG-UK) Consortium | Jacquelyn Wynn, Mairead Hyland, The Lighthouse Lab in Alderley Park and Alex Alderton, Roberto Amato, Jeffrey Barrett, Sonia Goncalves, Ewan Harrison, David K. Jackson, Ian Johnston, Dominic Kwiatkowski, Cordelia Langford, John Sillitoe on behalf of the Wellcome Sanger Institute COVID-19 Surveillance Team |
| EPI_ISL_2485595 | Lighthouse Lab in Glasgow | Wellcome Sanger Institute for the COVID-19 Genomics UK (COG-UK) Consortium | Harper VanSteenhouse, Yumi Kasai, David Gray, Carol Clugston, Anna Dominiczak and Alex Alderton, Roberto Amato, Jeffrey Barrett, Sonia Goncalves, Ewan Harrison, David K. Jackson, Ian Johnston, Dominic Kwiatkowski, Cordelia Langford, John Sillitoe on behalf of the Wellcome Sanger Institute COVID-19 Surveillance Team |
| EPI_ISL_2485596 | Lighthouse Lab in Milton Keynes | Wellcome Sanger Institute for the COVID-19 Genomics UK (COG-UK) Consortium | The Lighthouse Lab in Milton Keynes and Alex Alderton, Roberto Amato, Jeffrey Barrett, Sonia Goncalves, Ewan Harrison, David K. Jackson, Ian Johnston, Dominic Kwiatkowski, Cordelia Langford, John Sillitoe on behalf of the Wellcome Sanger Institute COVID-19 Surveillance Team |
| EPI_ISL_2485599 | Lighthouse Lab in Alderley Park | Wellcome Sanger Institute for the COVID-19 Genomics UK (COG-UK) Consortium | Jacquelyn Wynn, Mairead Hyland, The Lighthouse Lab in Alderley Park and Alex Alderton, Roberto Amato, Jeffrey Barrett, Sonia Goncalves, Ewan Harrison, David K. Jackson, Ian Johnston, Dominic Kwiatkowski, Cordelia Langford, John Sillitoe on behalf of the Wellcome Sanger Institute COVID-19 Surveillance Team |
| EPI_ISL_2485600 | Lighthouse Lab in Milton Keynes | Wellcome Sanger Institute for the COVID-19 Genomics UK (COG-UK) Consortium | The Lighthouse Lab in Milton Keynes and Alex Alderton, Roberto Amato, Jeffrey Barrett, Sonia Goncalves, Ewan Harrison, David K. Jackson, Ian Johnston, Dominic Kwiatkowski, Cordelia Langford, John Sillitoe on behalf of the Wellcome Sanger Institute COVID-19 Surveillance Team |
| EPI_ISL_2485601 | Lighthouse Lab in Alderley Park | Wellcome Sanger Institute for the COVID-19 Genomics UK (COG-UK) Consortium | Jacquelyn Wynn, Mairead Hyland, The Lighthouse Lab in Alderley Park and Alex Alderton, Roberto Amato, Jeffrey Barrett, Sonia Goncalves, Ewan Harrison, David K. Jackson, Ian Johnston, Dominic Kwiatkowski, Cordelia Langford, John Sillitoe on behalf of the Wellcome Sanger Institute COVID-19 Surveillance Team |
| EPI_ISL_2485602 | Lighthouse Lab in Milton Keynes | Wellcome Sanger Institute for the COVID-19 Genomics UK (COG-UK) Consortium | The Lighthouse Lab in Milton Keynes and Alex Alderton, Roberto Amato, Jeffrey Barrett, Sonia Goncalves, Ewan Harrison, David K. Jackson, Ian Johnston, Dominic Kwiatkowski, Cordelia Langford, John Sillitoe on behalf of the Wellcome Sanger Institute COVID-19 Surveillance Team |
| EPI_ISL_2485603, EPI_ISL_2485604 | Lighthouse Lab in Alderley Park | Wellcome Sanger Institute for the COVID-19 Genomics UK (COG-UK) Consortium | Jacquelyn Wynn, Mairead Hyland, The Lighthouse Lab in Alderley Park and Alex Alderton, Roberto Amato, Jeffrey Barrett, Sonia Goncalves, Ewan Harrison, David K. Jackson, Ian Johnston, Dominic Kwiatkowski, Cordelia Langford, John Sillitoe on behalf of the Wellcome Sanger Institute COVID-19 Surveillance Team |
| EPI_ISL_2485605, EPI_ISL_2485606 | Lighthouse Lab in Milton Keynes | Wellcome Sanger Institute for the COVID-19 Genomics UK (COG-UK) Consortium | The Lighthouse Lab in Milton Keynes and Alex Alderton, Roberto Amato, Jeffrey Barrett, Sonia Goncalves, Ewan Harrison, David K. Jackson, Ian Johnston, Dominic Kwiatkowski, Cordelia Langford, John Sillitoe on behalf of the Wellcome Sanger Institute COVID-19 Surveillance Team |
| EPI_ISL_2485607 | Lighthouse Lab in Alderley Park | Wellcome Sanger Institute for the COVID-19 Genomics UK (COG-UK) Consortium | Jacquelyn Wynn, Mairead Hyland, The Lighthouse Lab in Alderley Park and Alex Alderton, Roberto Amato, Jeffrey Barrett, Sonia Goncalves, Ewan Harrison, David K. Jackson, Ian Johnston, Dominic Kwiatkowski, Cordelia Langford, John Sillitoe on behalf of the Wellcome Sanger Institute COVID-19 Surveillance Team |
| EPI_ISL_2485608 | Lighthouse Lab in Milton Keynes | Wellcome Sanger Institute for the COVID-19 Genomics UK (COG-UK) Consortium | The Lighthouse Lab in Milton Keynes and Alex Alderton, Roberto Amato, Jeffrey Barrett, Sonia Goncalves, Ewan Harrison, David K. Jackson, Ian Johnston, Dominic Kwiatkowski, Cordelia Langford, John Sillitoe on behalf of the Wellcome Sanger Institute COVID-19 Surveillance Team |
| EPI_ISL_2485609 | Lighthouse Lab in Alderley Park | Wellcome Sanger Institute for the COVID-19 Genomics UK (COG-UK) Consortium | Jacquelyn Wynn, Mairead Hyland, The Lighthouse Lab in Alderley Park and Alex Alderton, Roberto Amato, Jeffrey Barrett, Sonia Goncalves, Ewan Harrison, David K. Jackson, Ian Johnston, Dominic Kwiatkowski, Cordelia Langford, John Sillitoe on behalf of the Wellcome Sanger Institute COVID-19 Surveillance Team |
| EPI_ISL_2485610 | Lighthouse Lab in Milton Keynes | Wellcome Sanger Institute for the COVID-19 Genomics UK (COG-UK) Consortium | The Lighthouse Lab in Milton Keynes and Alex Alderton, Roberto Amato, Jeffrey Barrett, Sonia Goncalves, Ewan Harrison, David K. Jackson, Ian Johnston, Dominic Kwiatkowski, Cordelia Langford, John Sillitoe on behalf of the Wellcome Sanger Institute COVID-19 Surveillance Team |
| EPI_ISL_2485611 | Lighthouse Lab in Alderley Park | Wellcome Sanger Institute for the COVID-19 Genomics UK (COG-UK) Consortium | Jacquelyn Wynn, Mairead Hyland, The Lighthouse Lab in Alderley Park and Alex Alderton, Roberto Amato, Jeffrey Barrett, Sonia Goncalves, Ewan Harrison, David K. Jackson, Ian Johnston, Dominic Kwiatkowski, Cordelia Langford, John Sillitoe on behalf of the Wellcome Sanger Institute COVID-19 Surveillance Team |
| EPI_ISL_2485612 | Lighthouse Lab in Milton Keynes | Wellcome Sanger Institute for the COVID-19 Genomics UK (COG-UK) Consortium | The Lighthouse Lab in Milton Keynes and Alex Alderton, Roberto Amato, Jeffrey Barrett, Sonia Goncalves, Ewan Harrison, David K. Jackson, Ian Johnston, Dominic Kwiatkowski, Cordelia Langford, John Sillitoe on behalf of the Wellcome Sanger Institute COVID-19 Surveillance Team |
| EPI_ISL_2485613, EPI_ISL_2485614, EPI_ISL_2485615 | Lighthouse Lab in Alderley Park | Wellcome Sanger Institute for the COVID-19 Genomics UK (COG-UK) Consortium | Jacquelyn Wynn, Mairead Hyland, The Lighthouse Lab in Alderley Park and Alex Alderton, Roberto Amato, Jeffrey Barrett, Sonia Goncalves, Ewan Harrison, David K. Jackson, Ian Johnston, Dominic Kwiatkowski, Cordelia Langford, John Sillitoe on behalf of the Wellcome Sanger Institute COVID-19 Surveillance Team |
| EPI_ISL_2485616 | Lighthouse Lab in Milton Keynes | Wellcome Sanger Institute for the COVID-19 Genomics UK (COG-UK) Consortium | The Lighthouse Lab in Milton Keynes and Alex Alderton, Roberto Amato, Jeffrey Barrett, Sonia Goncalves, Ewan Harrison, David K. Jackson, Ian Johnston, Dominic Kwiatkowski, Cordelia Langford, John Sillitoe on behalf of the Wellcome Sanger Institute COVID-19 Surveillance Team |
| EPI_ISL_2485617 | Lighthouse Lab in Alderley Park | Wellcome Sanger Institute for the COVID-19 Genomics UK (COG-UK) Consortium | Jacquelyn Wynn, Mairead Hyland, The Lighthouse Lab in Alderley Park and Alex Alderton, Roberto Amato, Jeffrey Barrett, Sonia Goncalves, Ewan Harrison, David K. Jackson, Ian Johnston, Dominic Kwiatkowski, Cordelia Langford, John Sillitoe on behalf of the Wellcome Sanger Institute COVID-19 Surveillance Team |
| EPI_ISL_2485618, EPI_ISL_2485619 | Lighthouse Lab in Milton Keynes | Wellcome Sanger Institute for the COVID-19 Genomics UK (COG-UK) Consortium | The Lighthouse Lab in Milton Keynes and Alex Alderton, Roberto Amato, Jeffrey Barrett, Sonia Goncalves, Ewan Harrison, David K. Jackson, Ian Johnston, Dominic Kwiatkowski, Cordelia Langford, John Sillitoe on behalf of the Wellcome Sanger Institute COVID-19 Surveillance Team |
| EPI_ISL_2485620, EPI_ISL_2485621 | Lighthouse Lab in Alderley Park | Wellcome Sanger Institute for the COVID-19 Genomics UK (COG-UK) Consortium | Jacquelyn Wynn, Mairead Hyland, The Lighthouse Lab in Alderley Park and Alex Alderton, Roberto Amato, Jeffrey Barrett, Sonia Goncalves, Ewan Harrison, David K. Jackson, Ian Johnston, Dominic Kwiatkowski, Cordelia Langford, John Sillitoe on behalf of the Wellcome Sanger Institute COVID-19 Surveillance Team |
| EPI_ISL_2485622 | Lighthouse Lab in Glasgow | Wellcome Sanger Institute for the COVID-19 Genomics UK (COG-UK) Consortium | Harper VanSteenhouse, Yumi Kasai, David Gray, Carol Clugston, Anna Dominiczak and Alex Alderton, Roberto Amato, Jeffrey Barrett, Sonia Goncalves, Ewan Harrison, David K. Jackson, Ian Johnston, Dominic Kwiatkowski, Cordelia Langford, John Sillitoe on behalf of the Wellcome Sanger Institute COVID-19 Surveillance Team |
| EPI_ISL_2485623 | Lighthouse Lab in Milton Keynes | Wellcome Sanger Institute for the COVID-19 Genomics UK (COG-UK) Consortium | The Lighthouse Lab in Milton Keynes and Alex Alderton, Roberto Amato, Jeffrey Barrett, Sonia Goncalves, Ewan Harrison, David K. Jackson, Ian Johnston, Dominic Kwiatkowski, Cordelia Langford, John Sillitoe on behalf of the Wellcome Sanger Institute COVID-19 Surveillance Team |
| EPI_ISL_2485624, EPI_ISL_2485625, EPI_ISL_2485626, EPI_ISL_2485627 | Lighthouse Lab in Alderley Park | Wellcome Sanger Institute for the COVID-19 Genomics UK (COG-UK) Consortium | Jacquelyn Wynn, Mairead Hyland, The Lighthouse Lab in Alderley Park and Alex Alderton, Roberto Amato, Jeffrey Barrett, Sonia Goncalves, Ewan Harrison, David K. Jackson, Ian Johnston, Dominic Kwiatkowski, Cordelia Langford, John Sillitoe on behalf of the Wellcome Sanger Institute COVID-19 Surveillance Team |
| EPI_ISL_2485628 | Lighthouse Lab in Milton Keynes | Wellcome Sanger Institute for the COVID-19 Genomics UK (COG-UK) Consortium | The Lighthouse Lab in Milton Keynes and Alex Alderton, Roberto Amato, Jeffrey Barrett, Sonia Goncalves, Ewan Harrison, David K. Jackson, Ian Johnston, Dominic Kwiatkowski, Cordelia Langford, John Sillitoe on behalf of the Wellcome Sanger Institute COVID-19 Surveillance Team |
| EPI_ISL_2485629, EPI_ISL_2485630, EPI_ISL_2485631, EPI_ISL_2485632 | Lighthouse Lab in Alderley Park | Wellcome Sanger Institute for the COVID-19 Genomics UK (COG-UK) Consortium | Jacquelyn Wynn, Mairead Hyland, The Lighthouse Lab in Alderley Park and Alex Alderton, Roberto Amato, Jeffrey Barrett, Sonia Goncalves, Ewan Harrison, David K. Jackson, Ian Johnston, Dominic Kwiatkowski, Cordelia Langford, John Sillitoe on behalf of the Wellcome Sanger Institute COVID-19 Surveillance Team |
| EPI_ISL_2485633 | Lighthouse Lab in Milton Keynes | Wellcome Sanger Institute for the COVID-19 Genomics UK (COG-UK) Consortium | The Lighthouse Lab in Milton Keynes and Alex Alderton, Roberto Amato, Jeffrey Barrett, Sonia Goncalves, Ewan Harrison, David K. Jackson, Ian Johnston, Dominic Kwiatkowski, Cordelia Langford, John Sillitoe on behalf of the Wellcome Sanger Institute COVID-19 Surveillance Team |
| EPI_ISL_2485634, EPI_ISL_2485635, EPI_ISL_2485636, EPI_ISL_2485637, EPI_ISL_2485638, EPI_ISL_2485639, EPI_ISL_2485640 | Lighthouse Lab in Alderley Park | Wellcome Sanger Institute for the COVID-19 Genomics UK (COG-UK) Consortium | Jacquelyn Wynn, Mairead Hyland, The Lighthouse Lab in Alderley Park and Alex Alderton, Roberto Amato, Jeffrey Barrett, Sonia Goncalves, Ewan Harrison, David K. Jackson, Ian Johnston, Dominic Kwiatkowski, Cordelia Langford, John Sillitoe on behalf of the Wellcome Sanger Institute COVID-19 Surveillance Team |
| EPI_ISL_2485641, EPI_ISL_2485642, EPI_ISL_2485643 | Lighthouse Lab in Milton Keynes | Wellcome Sanger Institute for the COVID-19 Genomics UK (COG-UK) Consortium | The Lighthouse Lab in Milton Keynes and Alex Alderton, Roberto Amato, Jeffrey Barrett, Sonia Goncalves, Ewan Harrison, David K. Jackson, Ian Johnston, Dominic Kwiatkowski, Cordelia Langford, John Sillitoe on behalf of the Wellcome Sanger Institute COVID-19 Surveillance Team |
| EPI_ISL_2485644, EPI_ISL_2485645 | Lighthouse Lab in Alderley Park | Wellcome Sanger Institute for the COVID-19 Genomics UK (COG-UK) Consortium | Jacquelyn Wynn, Mairead Hyland, The Lighthouse Lab in Alderley Park and Alex Alderton, Roberto Amato, Jeffrey Barrett, Sonia Goncalves, Ewan Harrison, David K. Jackson, Ian Johnston, Dominic Kwiatkowski, Cordelia Langford, John Sillitoe on behalf of the Wellcome Sanger Institute COVID-19 Surveillance Team |
| EPI_ISL_2485646, EPI_ISL_2485647, EPI_ISL_2485648 | Lighthouse Lab in Milton Keynes | Wellcome Sanger Institute for the COVID-19 Genomics UK (COG-UK) Consortium | The Lighthouse Lab in Milton Keynes and Alex Alderton, Roberto Amato, Jeffrey Barrett, Sonia Goncalves, Ewan Harrison, David K. Jackson, Ian Johnston, Dominic Kwiatkowski, Cordelia Langford, John Sillitoe on behalf of the Wellcome Sanger Institute COVID-19 Surveillance Team |
| EPI_ISL_2485649, EPI_ISL_2485650 | Lighthouse Lab in Alderley Park | Wellcome Sanger Institute for the COVID-19 Genomics UK (COG-UK) Consortium | Jacquelyn Wynn, Mairead Hyland, The Lighthouse Lab in Alderley Park and Alex Alderton, Roberto Amato, Jeffrey Barrett, Sonia Goncalves, Ewan Harrison, David K. Jackson, Ian Johnston, Dominic Kwiatkowski, Cordelia Langford, John Sillitoe on behalf of the Wellcome Sanger Institute COVID-19 Surveillance Team |
| EPI_ISL_2485651, EPI_ISL_2485652 | Lighthouse Lab in Milton Keynes | Wellcome Sanger Institute for the COVID-19 Genomics UK (COG-UK) Consortium | The Lighthouse Lab in Milton Keynes and Alex Alderton, Roberto Amato, Jeffrey Barrett, Sonia Goncalves, Ewan Harrison, David K. Jackson, Ian Johnston, Dominic Kwiatkowski, Cordelia Langford, John Sillitoe on behalf of the Wellcome Sanger Institute COVID-19 Surveillance Team |
| EPI_ISL_2485653 | Lighthouse Lab in Alderley Park | Wellcome Sanger Institute for the COVID-19 Genomics UK (COG-UK) Consortium | Jacquelyn Wynn, Mairead Hyland, The Lighthouse Lab in Alderley Park and Alex Alderton, Roberto Amato, Jeffrey Barrett, Sonia Goncalves, Ewan Harrison, David K. Jackson, Ian Johnston, Dominic Kwiatkowski, Cordelia Langford, John Sillitoe on behalf of the Wellcome Sanger Institute COVID-19 Surveillance Team |
| EPI_ISL_2485654 | Health Services Laboratories | Wellcome Sanger Institute for the COVID-19 Genomics UK (COG-UK) Consortium | Health Services Laboratories and Alex Alderton, Roberto Amato, Jeffrey Barrett, Sonia Goncalves, Ewan Harrison, David K. Jackson, Ian Johnston, Dominic Kwiatkowski, Cordelia Langford, John Sillitoe on behalf of the Wellcome Sanger Institute COVID-19 Surveillance Team |
| EPI_ISL_2485655, EPI_ISL_2485656, EPI_ISL_2485657, EPI_ISL_2485658, EPI_ISL_2485659 | Lighthouse Lab in Alderley Park | Wellcome Sanger Institute for the COVID-19 Genomics UK (COG-UK) Consortium | Jacquelyn Wynn, Mairead Hyland, The Lighthouse Lab in Alderley Park and Alex Alderton, Roberto Amato, Jeffrey Barrett, Sonia Goncalves, Ewan Harrison, David K. Jackson, Ian Johnston, Dominic Kwiatkowski, Cordelia Langford, John Sillitoe on behalf of the Wellcome Sanger Institute COVID-19 Surveillance Team |
| EPI_ISL_2485660 | Lighthouse Lab in Glasgow | Wellcome Sanger Institute for the COVID-19 Genomics UK (COG-UK) Consortium | Harper VanSteenhouse, Yumi Kasai, David Gray, Carol Clugston, Anna Dominiczak and Alex Alderton, Roberto Amato, Jeffrey Barrett, Sonia Goncalves, Ewan Harrison, David K. Jackson, Ian Johnston, Dominic Kwiatkowski, Cordelia Langford, John Sillitoe on behalf of the Wellcome Sanger Institute COVID-19 Surveillance Team |
| EPI_ISL_2485661, | Lighthouse Lab in Alderley Park | Wellcome Sanger Institute for the COVID-19 Genomics UK (COG-UK) | Jacquelyn Wynn, Mairead Hyland, The Lighthouse Lab in Alderley Park and Alex Alderton, Roberto Amato, Jeffrey Barrett, Sonia Goncalves, Ewan Harrison, David K. Jackson, Ian Johnston, Dominic Kwiatkowski, Cordelia Langford, John Sillitoe on |

| | | | |
|---|---|---|---|
| EPI_ISL_2485662, EPI_ISL_2485663 | | Consortium | behalf of the Wellcome Sanger Institute COVID-19 Surveillance Team |
| EPI_ISL_2485664, EPI_ISL_2485665, EPI_ISL_2485666, EPI_ISL_2485667 | Lighthouse Lab in Milton Keynes | Wellcome Sanger Institute for the COVID-19 Genomics UK (COG-UK) Consortium | The Lighthouse Lab in Milton Keynes and Alex Alderton, Roberto Amato, Jeffrey Barrett, Sonia Goncalves, Ewan Harrison, David K. Jackson, Ian Johnston, Dominic Kwiatkowski, Cordelia Langford, John Sillitoe on behalf of the Wellcome Sanger Institute COVID-19 Surveillance Team |
| EPI_ISL_2485668, EPI_ISL_2485669, EPI_ISL_2485670, EPI_ISL_2485671 | Lighthouse Lab in Alderley Park | Wellcome Sanger Institute for the COVID-19 Genomics UK (COG-UK) Consortium | Jacquelyn Wynn, Mairead Hyland, The Lighthouse Lab in Alderley Park and Alex Alderton, Roberto Amato, Jeffrey Barrett, Sonia Goncalves, Ewan Harrison, David K. Jackson, Ian Johnston, Dominic Kwiatkowski, Cordelia Langford, John Sillitoe on behalf of the Wellcome Sanger Institute COVID-19 Surveillance Team |
| EPI_ISL_2485672, EPI_ISL_2485673 | Lighthouse Lab in Milton Keynes | Wellcome Sanger Institute for the COVID-19 Genomics UK (COG-UK) Consortium | The Lighthouse Lab in Milton Keynes and Alex Alderton, Roberto Amato, Jeffrey Barrett, Sonia Goncalves, Ewan Harrison, David K. Jackson, Ian Johnston, Dominic Kwiatkowski, Cordelia Langford, John Sillitoe on behalf of the Wellcome Sanger Institute COVID-19 Surveillance Team |
| EPI_ISL_2485674, EPI_ISL_2485675 | Lighthouse Lab in Alderley Park | Wellcome Sanger Institute for the COVID-19 Genomics UK (COG-UK) Consortium | Jacquelyn Wynn, Mairead Hyland, The Lighthouse Lab in Alderley Park and Alex Alderton, Roberto Amato, Jeffrey Barrett, Sonia Goncalves, Ewan Harrison, David K. Jackson, Ian Johnston, Dominic Kwiatkowski, Cordelia Langford, John Sillitoe on behalf of the Wellcome Sanger Institute COVID-19 Surveillance Team |
| EPI_ISL_2485676 | Lighthouse Lab in Glasgow | Wellcome Sanger Institute for the COVID-19 Genomics UK (COG-UK) Consortium | Harper VanSteenhouse, Yumi Kasai, David Gray, Carol Clugston, Anna Dominiczak and Alex Alderton, Roberto Amato, Jeffrey Barrett, Sonia Goncalves, Ewan Harrison, David K. Jackson, Ian Johnston, Dominic Kwiatkowski, Cordelia Langford, John Sillitoe on behalf of the Wellcome Sanger Institute COVID-19 Surveillance Team |
| EPI_ISL_2485677 | Lighthouse Lab in Milton Keynes | Wellcome Sanger Institute for the COVID-19 Genomics UK (COG-UK) Consortium | The Lighthouse Lab in Milton Keynes and Alex Alderton, Roberto Amato, Jeffrey Barrett, Sonia Goncalves, Ewan Harrison, David K. Jackson, Ian Johnston, Dominic Kwiatkowski, Cordelia Langford, John Sillitoe on behalf of the Wellcome Sanger Institute COVID-19 Surveillance Team |
| EPI_ISL_2485678 | Lighthouse Lab in Alderley Park | Wellcome Sanger Institute for the COVID-19 Genomics UK (COG-UK) Consortium | Jacquelyn Wynn, Mairead Hyland, The Lighthouse Lab in Alderley Park and Alex Alderton, Roberto Amato, Jeffrey Barrett, Sonia Goncalves, Ewan Harrison, David K. Jackson, Ian Johnston, Dominic Kwiatkowski, Cordelia Langford, John Sillitoe on behalf of the Wellcome Sanger Institute COVID-19 Surveillance Team |
| EPI_ISL_2485679 | Lighthouse Lab in Glasgow | Wellcome Sanger Institute for the COVID-19 Genomics UK (COG-UK) Consortium | Harper VanSteenhouse, Yumi Kasai, David Gray, Carol Clugston, Anna Dominiczak and Alex Alderton, Roberto Amato, Jeffrey Barrett, Sonia Goncalves, Ewan Harrison, David K. Jackson, Ian Johnston, Dominic Kwiatkowski, Cordelia Langford, John Sillitoe on behalf of the Wellcome Sanger Institute COVID-19 Surveillance Team |
| EPI_ISL_2485680 | Lighthouse Lab in Milton Keynes | Wellcome Sanger Institute for the COVID-19 Genomics UK (COG-UK) Consortium | The Lighthouse Lab in Milton Keynes and Alex Alderton, Roberto Amato, Jeffrey Barrett, Sonia Goncalves, Ewan Harrison, David K. Jackson, Ian Johnston, Dominic Kwiatkowski, Cordelia Langford, John Sillitoe on behalf of the Wellcome Sanger Institute COVID-19 Surveillance Team |
| EPI_ISL_2485681 | Health Services Laboratories | Wellcome Sanger Institute for the COVID-19 Genomics UK (COG-UK) Consortium | Health Services Laboratories and Alex Alderton, Roberto Amato, Jeffrey Barrett, Sonia Goncalves, Ewan Harrison, David K. Jackson, Ian Johnston, Dominic Kwiatkowski, Cordelia Langford, John Sillitoe on behalf of the Wellcome Sanger Institute COVID-19 Surveillance Team |
| EPI_ISL_2485682, EPI_ISL_2485683 | Lighthouse Lab in Alderley Park | Wellcome Sanger Institute for the COVID-19 Genomics UK (COG-UK) Consortium | Jacquelyn Wynn, Mairead Hyland, The Lighthouse Lab in Alderley Park and Alex Alderton, Roberto Amato, Jeffrey Barrett, Sonia Goncalves, Ewan Harrison, David K. Jackson, Ian Johnston, Dominic Kwiatkowski, Cordelia Langford, John Sillitoe on behalf of the Wellcome Sanger Institute COVID-19 Surveillance Team |
| EPI_ISL_2485684 | Lighthouse Lab in Glasgow | Wellcome Sanger Institute for the COVID-19 Genomics UK (COG-UK) Consortium | Harper VanSteenhouse, Yumi Kasai, David Gray, Carol Clugston, Anna Dominiczak and Alex Alderton, Roberto Amato, Jeffrey Barrett, Sonia Goncalves, Ewan Harrison, David K. Jackson, Ian Johnston, Dominic Kwiatkowski, Cordelia Langford, John Sillitoe on behalf of the Wellcome Sanger Institute COVID-19 Surveillance Team |
| EPI_ISL_2485685 | Lighthouse Lab in Milton Keynes | Wellcome Sanger Institute for the COVID-19 Genomics UK (COG-UK) Consortium | The Lighthouse Lab in Milton Keynes and Alex Alderton, Roberto Amato, Jeffrey Barrett, Sonia Goncalves, Ewan Harrison, David K. Jackson, Ian Johnston, Dominic Kwiatkowski, Cordelia Langford, John Sillitoe on behalf of the Wellcome Sanger Institute COVID-19 Surveillance Team |
| EPI_ISL_2485687 | Lighthouse Lab in Alderley Park | Wellcome Sanger Institute for the COVID-19 Genomics UK (COG-UK) Consortium | Jacquelyn Wynn, Mairead Hyland, The Lighthouse Lab in Alderley Park and Alex Alderton, Roberto Amato, Jeffrey Barrett, Sonia Goncalves, Ewan Harrison, David K. Jackson, Ian Johnston, Dominic Kwiatkowski, Cordelia Langford, John Sillitoe on behalf of the Wellcome Sanger Institute COVID-19 Surveillance Team |
| EPI_ISL_2485688 | Lighthouse Lab in Glasgow | Wellcome Sanger Institute for the COVID-19 Genomics UK (COG-UK) Consortium | Harper VanSteenhouse, Yumi Kasai, David Gray, Carol Clugston, Anna Dominiczak and Alex Alderton, Roberto Amato, Jeffrey Barrett, Sonia Goncalves, Ewan Harrison, David K. Jackson, Ian Johnston, Dominic Kwiatkowski, Cordelia Langford, John Sillitoe on behalf of the Wellcome Sanger Institute COVID-19 Surveillance Team |
| EPI_ISL_2485689, EPI_ISL_2485690 | Lighthouse Lab in Milton Keynes | Wellcome Sanger Institute for the COVID-19 Genomics UK (COG-UK) Consortium | The Lighthouse Lab in Milton Keynes and Alex Alderton, Roberto Amato, Jeffrey Barrett, Sonia Goncalves, Ewan Harrison, David K. Jackson, Ian Johnston, Dominic Kwiatkowski, Cordelia Langford, John Sillitoe on behalf of the Wellcome Sanger Institute COVID-19 Surveillance Team |
| EPI_ISL_2485691 | Lighthouse Lab in Alderley Park | Wellcome Sanger Institute for the COVID-19 Genomics UK (COG-UK) Consortium | Jacquelyn Wynn, Mairead Hyland, The Lighthouse Lab in Alderley Park and Alex Alderton, Roberto Amato, Jeffrey Barrett, Sonia Goncalves, Ewan Harrison, David K. Jackson, Ian Johnston, Dominic Kwiatkowski, Cordelia Langford, John Sillitoe on behalf of the Wellcome Sanger Institute COVID-19 Surveillance Team |
| EPI_ISL_2485692, EPI_ISL_2485693, EPI_ISL_2485694 | Lighthouse Lab in Milton Keynes | Wellcome Sanger Institute for the COVID-19 Genomics UK (COG-UK) Consortium | The Lighthouse Lab in Milton Keynes and Alex Alderton, Roberto Amato, Jeffrey Barrett, Sonia Goncalves, Ewan Harrison, David K. Jackson, Ian Johnston, Dominic Kwiatkowski, Cordelia Langford, John Sillitoe on behalf of the Wellcome Sanger Institute COVID-19 Surveillance Team |
| EPI_ISL_2485695, EPI_ISL_2485696 | Lighthouse Lab in Alderley Park | Wellcome Sanger Institute for the COVID-19 Genomics UK (COG-UK) Consortium | Jacquelyn Wynn, Mairead Hyland, The Lighthouse Lab in Alderley Park and Alex Alderton, Roberto Amato, Jeffrey Barrett, Sonia Goncalves, Ewan Harrison, David K. Jackson, Ian Johnston, Dominic Kwiatkowski, Cordelia Langford, John Sillitoe on behalf of the Wellcome Sanger Institute COVID-19 Surveillance Team |
| EPI_ISL_2485698 | Health Services Laboratories | Wellcome Sanger Institute for the COVID-19 Genomics UK (COG-UK) Consortium | Health Services Laboratories and Alex Alderton, Roberto Amato, Jeffrey Barrett, Sonia Goncalves, Ewan Harrison, David K. Jackson, Ian Johnston, Dominic Kwiatkowski, Cordelia Langford, John Sillitoe on behalf of the Wellcome Sanger Institute COVID-19 Surveillance Team |
| EPI_ISL_2485699, EPI_ISL_2485700, EPI_ISL_2485702 | Lighthouse Lab in Alderley Park | Wellcome Sanger Institute for the COVID-19 Genomics UK (COG-UK) Consortium | Jacquelyn Wynn, Mairead Hyland, The Lighthouse Lab in Alderley Park and Alex Alderton, Roberto Amato, Jeffrey Barrett, Sonia Goncalves, Ewan Harrison, David K. Jackson, Ian Johnston, Dominic Kwiatkowski, Cordelia Langford, John Sillitoe on behalf of the Wellcome Sanger Institute COVID-19 Surveillance Team |
| EPI_ISL_2485703, EPI_ISL_2485704 | Lighthouse Lab in Milton Keynes | Wellcome Sanger Institute for the COVID-19 Genomics UK (COG-UK) Consortium | The Lighthouse Lab in Milton Keynes and Alex Alderton, Roberto Amato, Jeffrey Barrett, Sonia Goncalves, Ewan Harrison, David K. Jackson, Ian Johnston, Dominic Kwiatkowski, Cordelia Langford, John Sillitoe on behalf of the Wellcome Sanger Institute COVID-19 Surveillance Team |
| EPI_ISL_2485705 | Lighthouse Lab in Alderley Park | Wellcome Sanger Institute for the COVID-19 Genomics UK (COG-UK) Consortium | Jacquelyn Wynn, Mairead Hyland, The Lighthouse Lab in Alderley Park and Alex Alderton, Roberto Amato, Jeffrey Barrett, Sonia Goncalves, Ewan Harrison, David K. Jackson, Ian Johnston, Dominic Kwiatkowski, Cordelia Langford, John Sillitoe on behalf of the Wellcome Sanger Institute COVID-19 Surveillance Team |
| EPI_ISL_2485706, EPI_ISL_2485707, EPI_ISL_2485708 | Lighthouse Lab in Milton Keynes | Wellcome Sanger Institute for the COVID-19 Genomics UK (COG-UK) Consortium | The Lighthouse Lab in Milton Keynes and Alex Alderton, Roberto Amato, Jeffrey Barrett, Sonia Goncalves, Ewan Harrison, David K. Jackson, Ian Johnston, Dominic Kwiatkowski, Cordelia Langford, John Sillitoe on behalf of the Wellcome Sanger Institute COVID-19 Surveillance Team |
| EPI_ISL_2485709 | Lighthouse Lab in Alderley Park | Wellcome Sanger Institute for the COVID-19 Genomics UK (COG-UK) Consortium | Jacquelyn Wynn, Mairead Hyland, The Lighthouse Lab in Alderley Park and Alex Alderton, Roberto Amato, Jeffrey Barrett, Sonia Goncalves, Ewan Harrison, David K. Jackson, Ian Johnston, Dominic Kwiatkowski, Cordelia Langford, John Sillitoe on behalf of the Wellcome Sanger Institute COVID-19 Surveillance Team |
| EPI_ISL_2485710, EPI_ISL_2485711, EPI_ISL_2485712, EPI_ISL_2485714 | Lighthouse Lab in Milton Keynes | Wellcome Sanger Institute for the COVID-19 Genomics UK (COG-UK) Consortium | The Lighthouse Lab in Milton Keynes and Alex Alderton, Roberto Amato, Jeffrey Barrett, Sonia Goncalves, Ewan Harrison, David K. Jackson, Ian Johnston, Dominic Kwiatkowski, Cordelia Langford, John Sillitoe on behalf of the Wellcome Sanger Institute COVID-19 Surveillance Team |
| EPI_ISL_2485716 | Lighthouse Lab in Glasgow | Wellcome Sanger Institute for the COVID-19 Genomics UK (COG-UK) Consortium | Harper VanSteenhouse, Yumi Kasai, David Gray, Carol Clugston, Anna Dominiczak and Alex Alderton, Roberto Amato, Jeffrey Barrett, Sonia Goncalves, Ewan Harrison, David K. Jackson, Ian Johnston, Dominic Kwiatkowski, Cordelia Langford, John Sillitoe on behalf of the Wellcome Sanger Institute COVID-19 Surveillance Team |
| EPI_ISL_2485717 | Lighthouse Lab in Milton Keynes | Wellcome Sanger Institute for the COVID-19 Genomics UK (COG-UK) Consortium | The Lighthouse Lab in Milton Keynes and Alex Alderton, Roberto Amato, Jeffrey Barrett, Sonia Goncalves, Ewan Harrison, David K. Jackson, Ian Johnston, Dominic Kwiatkowski, Cordelia Langford, John Sillitoe on behalf of the Wellcome Sanger Institute COVID-19 Surveillance Team |
| EPI_ISL_2485718, EPI_ISL_2485720, EPI_ISL_2485721, EPI_ISL_2485722 | Lighthouse Lab in Alderley Park | Wellcome Sanger Institute for the COVID-19 Genomics UK (COG-UK) Consortium | Jacquelyn Wynn, Mairead Hyland, The Lighthouse Lab in Alderley Park and Alex Alderton, Roberto Amato, Jeffrey Barrett, Sonia Goncalves, Ewan Harrison, David K. Jackson, Ian Johnston, Dominic Kwiatkowski, Cordelia Langford, John Sillitoe on behalf of the Wellcome Sanger Institute COVID-19 Surveillance Team |
| EPI_ISL_2485723 | Lighthouse Lab in Milton Keynes | Wellcome Sanger Institute for the COVID-19 Genomics UK (COG-UK) Consortium | The Lighthouse Lab in Milton Keynes and Alex Alderton, Roberto Amato, Jeffrey Barrett, Sonia Goncalves, Ewan Harrison, David K. Jackson, Ian Johnston, Dominic Kwiatkowski, Cordelia Langford, John Sillitoe on behalf of the Wellcome Sanger Institute COVID-19 Surveillance Team |
| EPI_ISL_2485724 | Lighthouse Lab in Alderley Park | Wellcome Sanger Institute for the COVID-19 Genomics UK (COG-UK) Consortium | Jacquelyn Wynn, Mairead Hyland, The Lighthouse Lab in Alderley Park and Alex Alderton, Roberto Amato, Jeffrey Barrett, Sonia Goncalves, Ewan Harrison, David K. Jackson, Ian Johnston, Dominic Kwiatkowski, Cordelia Langford, John Sillitoe on behalf of the Wellcome Sanger Institute COVID-19 Surveillance Team |
| EPI_ISL_2485725, EPI_ISL_2485726, EPI_ISL_2485727 | Lighthouse Lab in Milton Keynes | Wellcome Sanger Institute for the COVID-19 Genomics UK (COG-UK) Consortium | The Lighthouse Lab in Milton Keynes and Alex Alderton, Roberto Amato, Jeffrey Barrett, Sonia Goncalves, Ewan Harrison, David K. Jackson, Ian Johnston, Dominic Kwiatkowski, Cordelia Langford, John Sillitoe on behalf of the Wellcome Sanger Institute COVID-19 Surveillance Team |
| EPI_ISL_2485728 | Lighthouse Lab in Alderley Park | Wellcome Sanger Institute for the COVID-19 Genomics UK (COG-UK) Consortium | Jacquelyn Wynn, Mairead Hyland, The Lighthouse Lab in Alderley Park and Alex Alderton, Roberto Amato, Jeffrey Barrett, Sonia Goncalves, Ewan Harrison, David K. Jackson, Ian Johnston, Dominic Kwiatkowski, Cordelia Langford, John Sillitoe on behalf of the Wellcome Sanger Institute COVID-19 Surveillance Team |
| EPI_ISL_2485729 | Lighthouse Lab in Milton Keynes | Wellcome Sanger Institute for the COVID-19 Genomics UK (COG-UK) Consortium | The Lighthouse Lab in Milton Keynes and Alex Alderton, Roberto Amato, Jeffrey Barrett, Sonia Goncalves, Ewan Harrison, David K. Jackson, Ian Johnston, Dominic Kwiatkowski, Cordelia Langford, John Sillitoe on behalf of the Wellcome Sanger Institute COVID-19 Surveillance Team |
| EPI_ISL_2485731 | Lighthouse Lab in Alderley Park | Wellcome Sanger Institute for the COVID-19 Genomics UK (COG-UK) Consortium | Jacquelyn Wynn, Mairead Hyland, The Lighthouse Lab in Alderley Park and Alex Alderton, Roberto Amato, Jeffrey Barrett, Sonia Goncalves, Ewan Harrison, David K. Jackson, Ian Johnston, Dominic Kwiatkowski, Cordelia Langford, John Sillitoe on behalf of the Wellcome Sanger Institute COVID-19 Surveillance Team |
| EPI_ISL_2485732 | Lighthouse Lab in Milton Keynes | Wellcome Sanger Institute for the COVID-19 Genomics UK (COG-UK) Consortium | The Lighthouse Lab in Milton Keynes and Alex Alderton, Roberto Amato, Jeffrey Barrett, Sonia Goncalves, Ewan Harrison, David K. Jackson, Ian Johnston, Dominic Kwiatkowski, Cordelia Langford, John Sillitoe on behalf of the Wellcome Sanger Institute COVID-19 Surveillance Team |
| EPI_ISL_2485733, EPI_ISL_2485734, EPI_ISL_2485736, EPI_ISL_2485737, EPI_ISL_2485738 | Lighthouse Lab in Alderley Park | Wellcome Sanger Institute for the COVID-19 Genomics UK (COG-UK) Consortium | Jacquelyn Wynn, Mairead Hyland, The Lighthouse Lab in Alderley Park and Alex Alderton, Roberto Amato, Jeffrey Barrett, Sonia Goncalves, Ewan Harrison, David K. Jackson, Ian Johnston, Dominic Kwiatkowski, Cordelia Langford, John Sillitoe on behalf of the Wellcome Sanger Institute COVID-19 Surveillance Team |
| EPI_ISL_2485739, EPI_ISL_2485742, | Lighthouse Lab in Milton Keynes | Wellcome Sanger Institute for the COVID-19 Genomics UK (COG-UK) Consortium | The Lighthouse Lab in Milton Keynes and Alex Alderton, Roberto Amato, Jeffrey Barrett, Sonia Goncalves, Ewan Harrison, David K. Jackson, Ian Johnston, Dominic Kwiatkowski, Cordelia Langford, John Sillitoe on behalf of the Wellcome Sanger Institute COVID-19 Surveillance Team |

| | | | |
|---|---|---|---|
| EPI_ISL_2485743 | | | |
| EPI_ISL_2485744 | Lighthouse Lab in Glasgow | Wellcome Sanger Institute for the COVID-19 Genomics UK (COG-UK) Consortium | Harper VanSteenhouse, Yumi Kasai, David Gray, Carol Clugston, Anna Dominiczak and Alex Alderton, Roberto Amato, Jeffrey Barrett, Sonia Goncalves, Ewan Harrison, David K. Jackson, Ian Johnston, Dominic Kwiatkowski, Cordelia Langford, John Sillitoe on behalf of the Wellcome Sanger Institute COVID-19 Surveillance Team |
| EPI_ISL_2485745, EPI_ISL_2485746 | Lighthouse Lab in Milton Keynes | Wellcome Sanger Institute for the COVID-19 Genomics UK (COG-UK) Consortium | The Lighthouse Lab in Milton Keynes and Alex Alderton, Roberto Amato, Jeffrey Barrett, Sonia Goncalves, Ewan Harrison, David K. Jackson, Ian Johnston, Dominic Kwiatkowski, Cordelia Langford, John Sillitoe on behalf of the Wellcome Sanger Institute COVID-19 Surveillance Team |
| EPI_ISL_2485747 | Lighthouse Lab in Alderley Park | Wellcome Sanger Institute for the COVID-19 Genomics UK (COG-UK) Consortium | Jacquelyn Wynn, Mairead Hyland, The Lighthouse Lab in Alderley Park and Alex Alderton, Roberto Amato, Jeffrey Barrett, Sonia Goncalves, Ewan Harrison, David K. Jackson, Ian Johnston, Dominic Kwiatkowski, Cordelia Langford, John Sillitoe on behalf of the Wellcome Sanger Institute COVID-19 Surveillance Team |
| EPI_ISL_2485750, EPI_ISL_2485751, EPI_ISL_2485752 | Lighthouse Lab in Milton Keynes | Wellcome Sanger Institute for the COVID-19 Genomics UK (COG-UK) Consortium | The Lighthouse Lab in Milton Keynes and Alex Alderton, Roberto Amato, Jeffrey Barrett, Sonia Goncalves, Ewan Harrison, David K. Jackson, Ian Johnston, Dominic Kwiatkowski, Cordelia Langford, John Sillitoe on behalf of the Wellcome Sanger Institute COVID-19 Surveillance Team |
| EPI_ISL_2485753, EPI_ISL_2485754 | Lighthouse Lab in Alderley Park | Wellcome Sanger Institute for the COVID-19 Genomics UK (COG-UK) Consortium | Jacquelyn Wynn, Mairead Hyland, The Lighthouse Lab in Alderley Park and Alex Alderton, Roberto Amato, Jeffrey Barrett, Sonia Goncalves, Ewan Harrison, David K. Jackson, Ian Johnston, Dominic Kwiatkowski, Cordelia Langford, John Sillitoe on behalf of the Wellcome Sanger Institute COVID-19 Surveillance Team |
| EPI_ISL_2485755, EPI_ISL_2485756, EPI_ISL_2485758 | Lighthouse Lab in Milton Keynes | Wellcome Sanger Institute for the COVID-19 Genomics UK (COG-UK) Consortium | The Lighthouse Lab in Milton Keynes and Alex Alderton, Roberto Amato, Jeffrey Barrett, Sonia Goncalves, Ewan Harrison, David K. Jackson, Ian Johnston, Dominic Kwiatkowski, Cordelia Langford, John Sillitoe on behalf of the Wellcome Sanger Institute COVID-19 Surveillance Team |
| EPI_ISL_2485759 | Lighthouse Lab in Alderley Park | Wellcome Sanger Institute for the COVID-19 Genomics UK (COG-UK) Consortium | Jacquelyn Wynn, Mairead Hyland, The Lighthouse Lab in Alderley Park and Alex Alderton, Roberto Amato, Jeffrey Barrett, Sonia Goncalves, Ewan Harrison, David K. Jackson, Ian Johnston, Dominic Kwiatkowski, Cordelia Langford, John Sillitoe on behalf of the Wellcome Sanger Institute COVID-19 Surveillance Team |
| EPI_ISL_2485760 | Lighthouse Lab in Milton Keynes | Wellcome Sanger Institute for the COVID-19 Genomics UK (COG-UK) Consortium | The Lighthouse Lab in Milton Keynes and Alex Alderton, Roberto Amato, Jeffrey Barrett, Sonia Goncalves, Ewan Harrison, David K. Jackson, Ian Johnston, Dominic Kwiatkowski, Cordelia Langford, John Sillitoe on behalf of the Wellcome Sanger Institute COVID-19 Surveillance Team |
| EPI_ISL_2485761 | Lighthouse Lab in Alderley Park | Wellcome Sanger Institute for the COVID-19 Genomics UK (COG-UK) Consortium | Jacquelyn Wynn, Mairead Hyland, The Lighthouse Lab in Alderley Park and Alex Alderton, Roberto Amato, Jeffrey Barrett, Sonia Goncalves, Ewan Harrison, David K. Jackson, Ian Johnston, Dominic Kwiatkowski, Cordelia Langford, John Sillitoe on behalf of the Wellcome Sanger Institute COVID-19 Surveillance Team |
| EPI_ISL_2485763 | Lighthouse Lab in Glasgow | Wellcome Sanger Institute for the COVID-19 Genomics UK (COG-UK) Consortium | Harper VanSteenhouse, Yumi Kasai, David Gray, Carol Clugston, Anna Dominiczak and Alex Alderton, Roberto Amato, Jeffrey Barrett, Sonia Goncalves, Ewan Harrison, David K. Jackson, Ian Johnston, Dominic Kwiatkowski, Cordelia Langford, John Sillitoe on behalf of the Wellcome Sanger Institute COVID-19 Surveillance Team |
| EPI_ISL_2485764 | Lighthouse Lab in Milton Keynes | Wellcome Sanger Institute for the COVID-19 Genomics UK (COG-UK) Consortium | The Lighthouse Lab in Milton Keynes and Alex Alderton, Roberto Amato, Jeffrey Barrett, Sonia Goncalves, Ewan Harrison, David K. Jackson, Ian Johnston, Dominic Kwiatkowski, Cordelia Langford, John Sillitoe on behalf of the Wellcome Sanger Institute COVID-19 Surveillance Team |
| EPI_ISL_2485766 | Lighthouse Lab in Alderley Park | Wellcome Sanger Institute for the COVID-19 Genomics UK (COG-UK) Consortium | Jacquelyn Wynn, Mairead Hyland, The Lighthouse Lab in Alderley Park and Alex Alderton, Roberto Amato, Jeffrey Barrett, Sonia Goncalves, Ewan Harrison, David K. Jackson, Ian Johnston, Dominic Kwiatkowski, Cordelia Langford, John Sillitoe on behalf of the Wellcome Sanger Institute COVID-19 Surveillance Team |
| EPI_ISL_2485767, EPI_ISL_2485768 | Lighthouse Lab in Milton Keynes | Wellcome Sanger Institute for the COVID-19 Genomics UK (COG-UK) Consortium | The Lighthouse Lab in Milton Keynes and Alex Alderton, Roberto Amato, Jeffrey Barrett, Sonia Goncalves, Ewan Harrison, David K. Jackson, Ian Johnston, Dominic Kwiatkowski, Cordelia Langford, John Sillitoe on behalf of the Wellcome Sanger Institute COVID-19 Surveillance Team |
| EPI_ISL_2485769, EPI_ISL_2485770 | Lighthouse Lab in Alderley Park | Wellcome Sanger Institute for the COVID-19 Genomics UK (COG-UK) Consortium | Jacquelyn Wynn, Mairead Hyland, The Lighthouse Lab in Alderley Park and Alex Alderton, Roberto Amato, Jeffrey Barrett, Sonia Goncalves, Ewan Harrison, David K. Jackson, Ian Johnston, Dominic Kwiatkowski, Cordelia Langford, John Sillitoe on behalf of the Wellcome Sanger Institute COVID-19 Surveillance Team |
| EPI_ISL_2485771 | Lighthouse Lab in Milton Keynes | Wellcome Sanger Institute for the COVID-19 Genomics UK (COG-UK) Consortium | The Lighthouse Lab in Milton Keynes and Alex Alderton, Roberto Amato, Jeffrey Barrett, Sonia Goncalves, Ewan Harrison, David K. Jackson, Ian Johnston, Dominic Kwiatkowski, Cordelia Langford, John Sillitoe on behalf of the Wellcome Sanger Institute COVID-19 Surveillance Team |
| EPI_ISL_2485772 | Lighthouse Lab in Alderley Park | Wellcome Sanger Institute for the COVID-19 Genomics UK (COG-UK) Consortium | Jacquelyn Wynn, Mairead Hyland, The Lighthouse Lab in Alderley Park and Alex Alderton, Roberto Amato, Jeffrey Barrett, Sonia Goncalves, Ewan Harrison, David K. Jackson, Ian Johnston, Dominic Kwiatkowski, Cordelia Langford, John Sillitoe on behalf of the Wellcome Sanger Institute COVID-19 Surveillance Team |
| EPI_ISL_2485773 | Lighthouse Lab in Milton Keynes | Wellcome Sanger Institute for the COVID-19 Genomics UK (COG-UK) Consortium | The Lighthouse Lab in Milton Keynes and Alex Alderton, Roberto Amato, Jeffrey Barrett, Sonia Goncalves, Ewan Harrison, David K. Jackson, Ian Johnston, Dominic Kwiatkowski, Cordelia Langford, John Sillitoe on behalf of the Wellcome Sanger Institute COVID-19 Surveillance Team |
| EPI_ISL_2485774, EPI_ISL_2485775, EPI_ISL_2485776 | Lighthouse Lab in Alderley Park | Wellcome Sanger Institute for the COVID-19 Genomics UK (COG-UK) Consortium | Jacquelyn Wynn, Mairead Hyland, The Lighthouse Lab in Alderley Park and Alex Alderton, Roberto Amato, Jeffrey Barrett, Sonia Goncalves, Ewan Harrison, David K. Jackson, Ian Johnston, Dominic Kwiatkowski, Cordelia Langford, John Sillitoe on behalf of the Wellcome Sanger Institute COVID-19 Surveillance Team |
| EPI_ISL_2485777, EPI_ISL_2485779 | Lighthouse Lab in Milton Keynes | Wellcome Sanger Institute for the COVID-19 Genomics UK (COG-UK) Consortium | The Lighthouse Lab in Milton Keynes and Alex Alderton, Roberto Amato, Jeffrey Barrett, Sonia Goncalves, Ewan Harrison, David K. Jackson, Ian Johnston, Dominic Kwiatkowski, Cordelia Langford, John Sillitoe on behalf of the Wellcome Sanger Institute COVID-19 Surveillance Team |
| EPI_ISL_2485780, EPI_ISL_2485781, EPI_ISL_2485782, EPI_ISL_2485783 | Lighthouse Lab in Alderley Park | Wellcome Sanger Institute for the COVID-19 Genomics UK (COG-UK) Consortium | Jacquelyn Wynn, Mairead Hyland, The Lighthouse Lab in Alderley Park and Alex Alderton, Roberto Amato, Jeffrey Barrett, Sonia Goncalves, Ewan Harrison, David K. Jackson, Ian Johnston, Dominic Kwiatkowski, Cordelia Langford, John Sillitoe on behalf of the Wellcome Sanger Institute COVID-19 Surveillance Team |
| EPI_ISL_2485785 | Lighthouse Lab in Milton Keynes | Wellcome Sanger Institute for the COVID-19 Genomics UK (COG-UK) Consortium | The Lighthouse Lab in Milton Keynes and Alex Alderton, Roberto Amato, Jeffrey Barrett, Sonia Goncalves, Ewan Harrison, David K. Jackson, Ian Johnston, Dominic Kwiatkowski, Cordelia Langford, John Sillitoe on behalf of the Wellcome Sanger Institute COVID-19 Surveillance Team |
| EPI_ISL_2485786, EPI_ISL_2485787, EPI_ISL_2485789, EPI_ISL_2485790 | Lighthouse Lab in Alderley Park | Wellcome Sanger Institute for the COVID-19 Genomics UK (COG-UK) Consortium | Jacquelyn Wynn, Mairead Hyland, The Lighthouse Lab in Alderley Park and Alex Alderton, Roberto Amato, Jeffrey Barrett, Sonia Goncalves, Ewan Harrison, David K. Jackson, Ian Johnston, Dominic Kwiatkowski, Cordelia Langford, John Sillitoe on behalf of the Wellcome Sanger Institute COVID-19 Surveillance Team |
| EPI_ISL_2485791, EPI_ISL_2485792 | Lighthouse Lab in Milton Keynes | Wellcome Sanger Institute for the COVID-19 Genomics UK (COG-UK) Consortium | The Lighthouse Lab in Milton Keynes and Alex Alderton, Roberto Amato, Jeffrey Barrett, Sonia Goncalves, Ewan Harrison, David K. Jackson, Ian Johnston, Dominic Kwiatkowski, Cordelia Langford, John Sillitoe on behalf of the Wellcome Sanger Institute COVID-19 Surveillance Team |
| EPI_ISL_2485793 | Lighthouse Lab in Alderley Park | Wellcome Sanger Institute for the COVID-19 Genomics UK (COG-UK) Consortium | Jacquelyn Wynn, Mairead Hyland, The Lighthouse Lab in Alderley Park and Alex Alderton, Roberto Amato, Jeffrey Barrett, Sonia Goncalves, Ewan Harrison, David K. Jackson, Ian Johnston, Dominic Kwiatkowski, Cordelia Langford, John Sillitoe on behalf of the Wellcome Sanger Institute COVID-19 Surveillance Team |
| EPI_ISL_2485794, EPI_ISL_2485795, EPI_ISL_2485796 | Lighthouse Lab in Milton Keynes | Wellcome Sanger Institute for the COVID-19 Genomics UK (COG-UK) Consortium | The Lighthouse Lab in Milton Keynes and Alex Alderton, Roberto Amato, Jeffrey Barrett, Sonia Goncalves, Ewan Harrison, David K. Jackson, Ian Johnston, Dominic Kwiatkowski, Cordelia Langford, John Sillitoe on behalf of the Wellcome Sanger Institute COVID-19 Surveillance Team |
| EPI_ISL_2485797, EPI_ISL_2485798, EPI_ISL_2485799 | Lighthouse Lab in Alderley Park | Wellcome Sanger Institute for the COVID-19 Genomics UK (COG-UK) Consortium | Jacquelyn Wynn, Mairead Hyland, The Lighthouse Lab in Alderley Park and Alex Alderton, Roberto Amato, Jeffrey Barrett, Sonia Goncalves, Ewan Harrison, David K. Jackson, Ian Johnston, Dominic Kwiatkowski, Cordelia Langford, John Sillitoe on behalf of the Wellcome Sanger Institute COVID-19 Surveillance Team |
| EPI_ISL_2485800, EPI_ISL_2485801 | Lighthouse Lab in Milton Keynes | Wellcome Sanger Institute for the COVID-19 Genomics UK (COG-UK) Consortium | The Lighthouse Lab in Milton Keynes and Alex Alderton, Roberto Amato, Jeffrey Barrett, Sonia Goncalves, Ewan Harrison, David K. Jackson, Ian Johnston, Dominic Kwiatkowski, Cordelia Langford, John Sillitoe on behalf of the Wellcome Sanger Institute COVID-19 Surveillance Team |
| EPI_ISL_2485802, EPI_ISL_2485803 | Lighthouse Lab in Alderley Park | Wellcome Sanger Institute for the COVID-19 Genomics UK (COG-UK) Consortium | Jacquelyn Wynn, Mairead Hyland, The Lighthouse Lab in Alderley Park and Alex Alderton, Roberto Amato, Jeffrey Barrett, Sonia Goncalves, Ewan Harrison, David K. Jackson, Ian Johnston, Dominic Kwiatkowski, Cordelia Langford, John Sillitoe on behalf of the Wellcome Sanger Institute COVID-19 Surveillance Team |
| EPI_ISL_2485804 | Lighthouse Lab in Milton Keynes | Wellcome Sanger Institute for the COVID-19 Genomics UK (COG-UK) Consortium | The Lighthouse Lab in Milton Keynes and Alex Alderton, Roberto Amato, Jeffrey Barrett, Sonia Goncalves, Ewan Harrison, David K. Jackson, Ian Johnston, Dominic Kwiatkowski, Cordelia Langford, John Sillitoe on behalf of the Wellcome Sanger Institute COVID-19 Surveillance Team |
| EPI_ISL_2485805 | Lighthouse Lab in Alderley Park | Wellcome Sanger Institute for the COVID-19 Genomics UK (COG-UK) Consortium | Jacquelyn Wynn, Mairead Hyland, The Lighthouse Lab in Alderley Park and Alex Alderton, Roberto Amato, Jeffrey Barrett, Sonia Goncalves, Ewan Harrison, David K. Jackson, Ian Johnston, Dominic Kwiatkowski, Cordelia Langford, John Sillitoe on behalf of the Wellcome Sanger Institute COVID-19 Surveillance Team |
| EPI_ISL_2485806, EPI_ISL_2485807 | Lighthouse Lab in Milton Keynes | Wellcome Sanger Institute for the COVID-19 Genomics UK (COG-UK) Consortium | The Lighthouse Lab in Milton Keynes and Alex Alderton, Roberto Amato, Jeffrey Barrett, Sonia Goncalves, Ewan Harrison, David K. Jackson, Ian Johnston, Dominic Kwiatkowski, Cordelia Langford, John Sillitoe on behalf of the Wellcome Sanger Institute COVID-19 Surveillance Team |
| EPI_ISL_2485808 | Lighthouse Lab in Alderley Park | Wellcome Sanger Institute for the COVID-19 Genomics UK (COG-UK) Consortium | Jacquelyn Wynn, Mairead Hyland, The Lighthouse Lab in Alderley Park and Alex Alderton, Roberto Amato, Jeffrey Barrett, Sonia Goncalves, Ewan Harrison, David K. Jackson, Ian Johnston, Dominic Kwiatkowski, Cordelia Langford, John Sillitoe on behalf of the Wellcome Sanger Institute COVID-19 Surveillance Team |
| EPI_ISL_2485809 | Lighthouse Lab in Milton Keynes | Wellcome Sanger Institute for the COVID-19 Genomics UK (COG-UK) Consortium | The Lighthouse Lab in Milton Keynes and Alex Alderton, Roberto Amato, Jeffrey Barrett, Sonia Goncalves, Ewan Harrison, David K. Jackson, Ian Johnston, Dominic Kwiatkowski, Cordelia Langford, John Sillitoe on behalf of the Wellcome Sanger Institute COVID-19 Surveillance Team |
| EPI_ISL_2485810 | Lighthouse Lab in Alderley Park | Wellcome Sanger Institute for the COVID-19 Genomics UK (COG-UK) Consortium | Jacquelyn Wynn, Mairead Hyland, The Lighthouse Lab in Alderley Park and Alex Alderton, Roberto Amato, Jeffrey Barrett, Sonia Goncalves, Ewan Harrison, David K. Jackson, Ian Johnston, Dominic Kwiatkowski, Cordelia Langford, John Sillitoe on behalf of the Wellcome Sanger Institute COVID-19 Surveillance Team |
| EPI_ISL_2485811 | Lighthouse Lab in Glasgow | Wellcome Sanger Institute for the COVID-19 Genomics UK (COG-UK) Consortium | Harper VanSteenhouse, Yumi Kasai, David Gray, Carol Clugston, Anna Dominiczak and Alex Alderton, Roberto Amato, Jeffrey Barrett, Sonia Goncalves, Ewan Harrison, David K. Jackson, Ian Johnston, Dominic Kwiatkowski, Cordelia Langford, John Sillitoe on behalf of the Wellcome Sanger Institute COVID-19 Surveillance Team |
| EPI_ISL_2485812 | Lighthouse Lab in Milton Keynes | Wellcome Sanger Institute for the COVID-19 Genomics UK (COG-UK) Consortium | The Lighthouse Lab in Milton Keynes and Alex Alderton, Roberto Amato, Jeffrey Barrett, Sonia Goncalves, Ewan Harrison, David K. Jackson, Ian Johnston, Dominic Kwiatkowski, Cordelia Langford, John Sillitoe on behalf of the Wellcome Sanger Institute COVID-19 Surveillance Team |
| EPI_ISL_2485813 | Lighthouse Lab in Alderley Park | Wellcome Sanger Institute for the COVID-19 Genomics UK (COG-UK) Consortium | Jacquelyn Wynn, Mairead Hyland, The Lighthouse Lab in Alderley Park and Alex Alderton, Roberto Amato, Jeffrey Barrett, Sonia Goncalves, Ewan Harrison, David K. Jackson, Ian Johnston, Dominic Kwiatkowski, Cordelia Langford, John Sillitoe on behalf of the Wellcome Sanger Institute COVID-19 Surveillance Team |
| EPI_ISL_2485814, EPI_ISL_2485815 | Lighthouse Lab in Milton Keynes | Wellcome Sanger Institute for the COVID-19 Genomics UK (COG-UK) Consortium | The Lighthouse Lab in Milton Keynes and Alex Alderton, Roberto Amato, Jeffrey Barrett, Sonia Goncalves, Ewan Harrison, David K. Jackson, Ian Johnston, Dominic Kwiatkowski, Cordelia Langford, John Sillitoe on behalf of the Wellcome Sanger Institute COVID-19 Surveillance Team |
| EPI_ISL_2485816, EPI_ISL_2485817, EPI_ISL_2485818 | Lighthouse Lab in Alderley Park | Wellcome Sanger Institute for the COVID-19 Genomics UK (COG-UK) Consortium | Jacquelyn Wynn, Mairead Hyland, The Lighthouse Lab in Alderley Park and Alex Alderton, Roberto Amato, Jeffrey Barrett, Sonia Goncalves, Ewan Harrison, David K. Jackson, Ian Johnston, Dominic Kwiatkowski, Cordelia Langford, John Sillitoe on behalf of the Wellcome Sanger Institute COVID-19 Surveillance Team |
| EPI_ISL_2485819, EPI_ISL_2485821 | Lighthouse Lab in Milton Keynes | Wellcome Sanger Institute for the COVID-19 Genomics UK (COG-UK) Consortium | The Lighthouse Lab in Milton Keynes and Alex Alderton, Roberto Amato, Jeffrey Barrett, Sonia Goncalves, Ewan Harrison, David K. Jackson, Ian Johnston, Dominic Kwiatkowski, Cordelia Langford, John Sillitoe on behalf of the Wellcome Sanger Institute COVID-19 Surveillance Team |
| EPI_ISL_2485823, | Lighthouse Lab in Alderley Park | Wellcome Sanger Institute for the COVID-19 Genomics UK (COG-UK) | Jacquelyn Wynn, Mairead Hyland, The Lighthouse Lab in Alderley Park and Alex Alderton, Roberto Amato, Jeffrey Barrett, Sonia Goncalves, Ewan Harrison, David K. Jackson, Ian Johnston, Dominic Kwiatkowski, Cordelia Langford, John Sillitoe on |

| | | | |
|---|---|---|---|
| EPI_ISL_2485824 | | Consortium | behalf of the Wellcome Sanger Institute COVID-19 Surveillance Team |
| EPI_ISL_2485825 | Lighthouse Lab in Milton Keynes | Wellcome Sanger Institute for the COVID-19 Genomics UK (COG-UK) Consortium | The Lighthouse Lab in Milton Keynes and Alex Alderton, Roberto Amato, Jeffrey Barrett, Sonia Goncalves, Ewan Harrison, David K. Jackson, Ian Johnston, Dominic Kwiatkowski, Cordelia Langford, John Sillitoe on behalf of the Wellcome Sanger Institute COVID-19 Surveillance Team |
| EPI_ISL_2485826, EPI_ISL_2485827, EPI_ISL_2485828 | Lighthouse Lab in Alderley Park | Wellcome Sanger Institute for the COVID-19 Genomics UK (COG-UK) Consortium | Jacquelyn Wynn, Mairead Hyland, The Lighthouse Lab in Alderley Park and Alex Alderton, Roberto Amato, Jeffrey Barrett, Sonia Goncalves, Ewan Harrison, David K. Jackson, Ian Johnston, Dominic Kwiatkowski, Cordelia Langford, John Sillitoe on behalf of the Wellcome Sanger Institute COVID-19 Surveillance Team |
| EPI_ISL_2485830 | Lighthouse Lab in Milton Keynes | Wellcome Sanger Institute for the COVID-19 Genomics UK (COG-UK) Consortium | The Lighthouse Lab in Milton Keynes and Alex Alderton, Roberto Amato, Jeffrey Barrett, Sonia Goncalves, Ewan Harrison, David K. Jackson, Ian Johnston, Dominic Kwiatkowski, Cordelia Langford, John Sillitoe on behalf of the Wellcome Sanger Institute COVID-19 Surveillance Team |
| EPI_ISL_2485831, EPI_ISL_2485832, EPI_ISL_2485833, EPI_ISL_2485834 | Lighthouse Lab in Alderley Park | Wellcome Sanger Institute for the COVID-19 Genomics UK (COG-UK) Consortium | Jacquelyn Wynn, Mairead Hyland, The Lighthouse Lab in Alderley Park and Alex Alderton, Roberto Amato, Jeffrey Barrett, Sonia Goncalves, Ewan Harrison, David K. Jackson, Ian Johnston, Dominic Kwiatkowski, Cordelia Langford, John Sillitoe on behalf of the Wellcome Sanger Institute COVID-19 Surveillance Team |
| EPI_ISL_2485835 | Lighthouse Lab in Milton Keynes | Wellcome Sanger Institute for the COVID-19 Genomics UK (COG-UK) Consortium | The Lighthouse Lab in Milton Keynes and Alex Alderton, Roberto Amato, Jeffrey Barrett, Sonia Goncalves, Ewan Harrison, David K. Jackson, Ian Johnston, Dominic Kwiatkowski, Cordelia Langford, John Sillitoe on behalf of the Wellcome Sanger Institute COVID-19 Surveillance Team |
| EPI_ISL_2485836, EPI_ISL_2485837, EPI_ISL_2485838 | Lighthouse Lab in Alderley Park | Wellcome Sanger Institute for the COVID-19 Genomics UK (COG-UK) Consortium | Jacquelyn Wynn, Mairead Hyland, The Lighthouse Lab in Alderley Park and Alex Alderton, Roberto Amato, Jeffrey Barrett, Sonia Goncalves, Ewan Harrison, David K. Jackson, Ian Johnston, Dominic Kwiatkowski, Cordelia Langford, John Sillitoe on behalf of the Wellcome Sanger Institute COVID-19 Surveillance Team |
| EPI_ISL_2485839 | Lighthouse Lab in Milton Keynes | Wellcome Sanger Institute for the COVID-19 Genomics UK (COG-UK) Consortium | The Lighthouse Lab in Milton Keynes and Alex Alderton, Roberto Amato, Jeffrey Barrett, Sonia Goncalves, Ewan Harrison, David K. Jackson, Ian Johnston, Dominic Kwiatkowski, Cordelia Langford, John Sillitoe on behalf of the Wellcome Sanger Institute COVID-19 Surveillance Team |
| EPI_ISL_2485840, EPI_ISL_2485841, EPI_ISL_2485842, EPI_ISL_2485843 | Lighthouse Lab in Alderley Park | Wellcome Sanger Institute for the COVID-19 Genomics UK (COG-UK) Consortium | Jacquelyn Wynn, Mairead Hyland, The Lighthouse Lab in Alderley Park and Alex Alderton, Roberto Amato, Jeffrey Barrett, Sonia Goncalves, Ewan Harrison, David K. Jackson, Ian Johnston, Dominic Kwiatkowski, Cordelia Langford, John Sillitoe on behalf of the Wellcome Sanger Institute COVID-19 Surveillance Team |
| EPI_ISL_2485844 | Lighthouse Lab in Milton Keynes | Wellcome Sanger Institute for the COVID-19 Genomics UK (COG-UK) Consortium | The Lighthouse Lab in Milton Keynes and Alex Alderton, Roberto Amato, Jeffrey Barrett, Sonia Goncalves, Ewan Harrison, David K. Jackson, Ian Johnston, Dominic Kwiatkowski, Cordelia Langford, John Sillitoe on behalf of the Wellcome Sanger Institute COVID-19 Surveillance Team |
| EPI_ISL_2485845 | Lighthouse Lab in Glasgow | Wellcome Sanger Institute for the COVID-19 Genomics UK (COG-UK) Consortium | Harper VanSteenhouse, Yumi Kasai, David Gray, Carol Clugston, Anna Dominiczak and Alex Alderton, Roberto Amato, Jeffrey Barrett, Sonia Goncalves, Ewan Harrison, David K. Jackson, Ian Johnston, Dominic Kwiatkowski, Cordelia Langford, John Sillitoe on behalf of the Wellcome Sanger Institute COVID-19 Surveillance Team |
| EPI_ISL_2485846 | Lighthouse Lab in Milton Keynes | Wellcome Sanger Institute for the COVID-19 Genomics UK (COG-UK) Consortium | The Lighthouse Lab in Milton Keynes and Alex Alderton, Roberto Amato, Jeffrey Barrett, Sonia Goncalves, Ewan Harrison, David K. Jackson, Ian Johnston, Dominic Kwiatkowski, Cordelia Langford, John Sillitoe on behalf of the Wellcome Sanger Institute COVID-19 Surveillance Team |
| EPI_ISL_2485847, EPI_ISL_2485848 | Lighthouse Lab in Alderley Park | Wellcome Sanger Institute for the COVID-19 Genomics UK (COG-UK) Consortium | Jacquelyn Wynn, Mairead Hyland, The Lighthouse Lab in Alderley Park and Alex Alderton, Roberto Amato, Jeffrey Barrett, Sonia Goncalves, Ewan Harrison, David K. Jackson, Ian Johnston, Dominic Kwiatkowski, Cordelia Langford, John Sillitoe on behalf of the Wellcome Sanger Institute COVID-19 Surveillance Team |
| EPI_ISL_2485849 | Lighthouse Lab in Glasgow | Wellcome Sanger Institute for the COVID-19 Genomics UK (COG-UK) Consortium | Harper VanSteenhouse, Yumi Kasai, David Gray, Carol Clugston, Anna Dominiczak and Alex Alderton, Roberto Amato, Jeffrey Barrett, Sonia Goncalves, Ewan Harrison, David K. Jackson, Ian Johnston, Dominic Kwiatkowski, Cordelia Langford, John Sillitoe on behalf of the Wellcome Sanger Institute COVID-19 Surveillance Team |
| EPI_ISL_2485850 | Lighthouse Lab in Milton Keynes | Wellcome Sanger Institute for the COVID-19 Genomics UK (COG-UK) Consortium | The Lighthouse Lab in Milton Keynes and Alex Alderton, Roberto Amato, Jeffrey Barrett, Sonia Goncalves, Ewan Harrison, David K. Jackson, Ian Johnston, Dominic Kwiatkowski, Cordelia Langford, John Sillitoe on behalf of the Wellcome Sanger Institute COVID-19 Surveillance Team |
| EPI_ISL_2485851 | Lighthouse Lab in Glasgow | Wellcome Sanger Institute for the COVID-19 Genomics UK (COG-UK) Consortium | Harper VanSteenhouse, Yumi Kasai, David Gray, Carol Clugston, Anna Dominiczak and Alex Alderton, Roberto Amato, Jeffrey Barrett, Sonia Goncalves, Ewan Harrison, David K. Jackson, Ian Johnston, Dominic Kwiatkowski, Cordelia Langford, John Sillitoe on behalf of the Wellcome Sanger Institute COVID-19 Surveillance Team |
| EPI_ISL_2485852, EPI_ISL_2485853 | Lighthouse Lab in Alderley Park | Wellcome Sanger Institute for the COVID-19 Genomics UK (COG-UK) Consortium | Jacquelyn Wynn, Mairead Hyland, The Lighthouse Lab in Alderley Park and Alex Alderton, Roberto Amato, Jeffrey Barrett, Sonia Goncalves, Ewan Harrison, David K. Jackson, Ian Johnston, Dominic Kwiatkowski, Cordelia Langford, John Sillitoe on behalf of the Wellcome Sanger Institute COVID-19 Surveillance Team |
| EPI_ISL_2485854 | Lighthouse Lab in Glasgow | Wellcome Sanger Institute for the COVID-19 Genomics UK (COG-UK) Consortium | Harper VanSteenhouse, Yumi Kasai, David Gray, Carol Clugston, Anna Dominiczak and Alex Alderton, Roberto Amato, Jeffrey Barrett, Sonia Goncalves, Ewan Harrison, David K. Jackson, Ian Johnston, Dominic Kwiatkowski, Cordelia Langford, John Sillitoe on behalf of the Wellcome Sanger Institute COVID-19 Surveillance Team |
| EPI_ISL_2485855 | Lighthouse Lab in Milton Keynes | Wellcome Sanger Institute for the COVID-19 Genomics UK (COG-UK) Consortium | The Lighthouse Lab in Milton Keynes and Alex Alderton, Roberto Amato, Jeffrey Barrett, Sonia Goncalves, Ewan Harrison, David K. Jackson, Ian Johnston, Dominic Kwiatkowski, Cordelia Langford, John Sillitoe on behalf of the Wellcome Sanger Institute COVID-19 Surveillance Team |
| EPI_ISL_2485857 | Lighthouse Lab in Alderley Park | Wellcome Sanger Institute for the COVID-19 Genomics UK (COG-UK) Consortium | Jacquelyn Wynn, Mairead Hyland, The Lighthouse Lab in Alderley Park and Alex Alderton, Roberto Amato, Jeffrey Barrett, Sonia Goncalves, Ewan Harrison, David K. Jackson, Ian Johnston, Dominic Kwiatkowski, Cordelia Langford, John Sillitoe on behalf of the Wellcome Sanger Institute COVID-19 Surveillance Team |
| EPI_ISL_2485858 | Lighthouse Lab in Milton Keynes | Wellcome Sanger Institute for the COVID-19 Genomics UK (COG-UK) Consortium | The Lighthouse Lab in Milton Keynes and Alex Alderton, Roberto Amato, Jeffrey Barrett, Sonia Goncalves, Ewan Harrison, David K. Jackson, Ian Johnston, Dominic Kwiatkowski, Cordelia Langford, John Sillitoe on behalf of the Wellcome Sanger Institute COVID-19 Surveillance Team |
| EPI_ISL_2485859, EPI_ISL_2485860 | Lighthouse Lab in Glasgow | Wellcome Sanger Institute for the COVID-19 Genomics UK (COG-UK) Consortium | Harper VanSteenhouse, Yumi Kasai, David Gray, Carol Clugston, Anna Dominiczak and Alex Alderton, Roberto Amato, Jeffrey Barrett, Sonia Goncalves, Ewan Harrison, David K. Jackson, Ian Johnston, Dominic Kwiatkowski, Cordelia Langford, John Sillitoe on behalf of the Wellcome Sanger Institute COVID-19 Surveillance Team |
| EPI_ISL_2485861 | Lighthouse Lab in Alderley Park | Wellcome Sanger Institute for the COVID-19 Genomics UK (COG-UK) Consortium | Jacquelyn Wynn, Mairead Hyland, The Lighthouse Lab in Alderley Park and Alex Alderton, Roberto Amato, Jeffrey Barrett, Sonia Goncalves, Ewan Harrison, David K. Jackson, Ian Johnston, Dominic Kwiatkowski, Cordelia Langford, John Sillitoe on behalf of the Wellcome Sanger Institute COVID-19 Surveillance Team |
| EPI_ISL_2485862, EPI_ISL_2485863 | Lighthouse Lab in Glasgow | Wellcome Sanger Institute for the COVID-19 Genomics UK (COG-UK) Consortium | Harper VanSteenhouse, Yumi Kasai, David Gray, Carol Clugston, Anna Dominiczak and Alex Alderton, Roberto Amato, Jeffrey Barrett, Sonia Goncalves, Ewan Harrison, David K. Jackson, Ian Johnston, Dominic Kwiatkowski, Cordelia Langford, John Sillitoe on behalf of the Wellcome Sanger Institute COVID-19 Surveillance Team |
| EPI_ISL_2485864 | Lighthouse Lab in Milton Keynes | Wellcome Sanger Institute for the COVID-19 Genomics UK (COG-UK) Consortium | The Lighthouse Lab in Milton Keynes and Alex Alderton, Roberto Amato, Jeffrey Barrett, Sonia Goncalves, Ewan Harrison, David K. Jackson, Ian Johnston, Dominic Kwiatkowski, Cordelia Langford, John Sillitoe on behalf of the Wellcome Sanger Institute COVID-19 Surveillance Team |
| EPI_ISL_2485865 | Lighthouse Lab in Glasgow | Wellcome Sanger Institute for the COVID-19 Genomics UK (COG-UK) Consortium | Harper VanSteenhouse, Yumi Kasai, David Gray, Carol Clugston, Anna Dominiczak and Alex Alderton, Roberto Amato, Jeffrey Barrett, Sonia Goncalves, Ewan Harrison, David K. Jackson, Ian Johnston, Dominic Kwiatkowski, Cordelia Langford, John Sillitoe on behalf of the Wellcome Sanger Institute COVID-19 Surveillance Team |
| EPI_ISL_2485866 | Lighthouse Lab in Alderley Park | Wellcome Sanger Institute for the COVID-19 Genomics UK (COG-UK) Consortium | Jacquelyn Wynn, Mairead Hyland, The Lighthouse Lab in Alderley Park and Alex Alderton, Roberto Amato, Jeffrey Barrett, Sonia Goncalves, Ewan Harrison, David K. Jackson, Ian Johnston, Dominic Kwiatkowski, Cordelia Langford, John Sillitoe on behalf of the Wellcome Sanger Institute COVID-19 Surveillance Team |
| EPI_ISL_2485868 | Lighthouse Lab in Milton Keynes | Wellcome Sanger Institute for the COVID-19 Genomics UK (COG-UK) Consortium | The Lighthouse Lab in Milton Keynes and Alex Alderton, Roberto Amato, Jeffrey Barrett, Sonia Goncalves, Ewan Harrison, David K. Jackson, Ian Johnston, Dominic Kwiatkowski, Cordelia Langford, John Sillitoe on behalf of the Wellcome Sanger Institute COVID-19 Surveillance Team |
| EPI_ISL_2485870 | Lighthouse Lab in Alderley Park | Wellcome Sanger Institute for the COVID-19 Genomics UK (COG-UK) Consortium | Jacquelyn Wynn, Mairead Hyland, The Lighthouse Lab in Alderley Park and Alex Alderton, Roberto Amato, Jeffrey Barrett, Sonia Goncalves, Ewan Harrison, David K. Jackson, Ian Johnston, Dominic Kwiatkowski, Cordelia Langford, John Sillitoe on behalf of the Wellcome Sanger Institute COVID-19 Surveillance Team |
| EPI_ISL_2485871, EPI_ISL_2485873, EPI_ISL_2485874 | Lighthouse Lab in Glasgow | Wellcome Sanger Institute for the COVID-19 Genomics UK (COG-UK) Consortium | Harper VanSteenhouse, Yumi Kasai, David Gray, Carol Clugston, Anna Dominiczak and Alex Alderton, Roberto Amato, Jeffrey Barrett, Sonia Goncalves, Ewan Harrison, David K. Jackson, Ian Johnston, Dominic Kwiatkowski, Cordelia Langford, John Sillitoe on behalf of the Wellcome Sanger Institute COVID-19 Surveillance Team |
| EPI_ISL_2485875 | Lighthouse Lab in Milton Keynes | Wellcome Sanger Institute for the COVID-19 Genomics UK (COG-UK) Consortium | The Lighthouse Lab in Milton Keynes and Alex Alderton, Roberto Amato, Jeffrey Barrett, Sonia Goncalves, Ewan Harrison, David K. Jackson, Ian Johnston, Dominic Kwiatkowski, Cordelia Langford, John Sillitoe on behalf of the Wellcome Sanger Institute COVID-19 Surveillance Team |
| EPI_ISL_2485876 | Lighthouse Lab in Alderley Park | Wellcome Sanger Institute for the COVID-19 Genomics UK (COG-UK) Consortium | Jacquelyn Wynn, Mairead Hyland, The Lighthouse Lab in Alderley Park and Alex Alderton, Roberto Amato, Jeffrey Barrett, Sonia Goncalves, Ewan Harrison, David K. Jackson, Ian Johnston, Dominic Kwiatkowski, Cordelia Langford, John Sillitoe on behalf of the Wellcome Sanger Institute COVID-19 Surveillance Team |
| EPI_ISL_2485877 | Lighthouse Lab in Milton Keynes | Wellcome Sanger Institute for the COVID-19 Genomics UK (COG-UK) Consortium | The Lighthouse Lab in Milton Keynes and Alex Alderton, Roberto Amato, Jeffrey Barrett, Sonia Goncalves, Ewan Harrison, David K. Jackson, Ian Johnston, Dominic Kwiatkowski, Cordelia Langford, John Sillitoe on behalf of the Wellcome Sanger Institute COVID-19 Surveillance Team |
| EPI_ISL_2485879 | Lighthouse Lab in Glasgow | Wellcome Sanger Institute for the COVID-19 Genomics UK (COG-UK) Consortium | Harper VanSteenhouse, Yumi Kasai, David Gray, Carol Clugston, Anna Dominiczak and Alex Alderton, Roberto Amato, Jeffrey Barrett, Sonia Goncalves, Ewan Harrison, David K. Jackson, Ian Johnston, Dominic Kwiatkowski, Cordelia Langford, John Sillitoe on behalf of the Wellcome Sanger Institute COVID-19 Surveillance Team |
| EPI_ISL_2485880, EPI_ISL_2485881 | Lighthouse Lab in Alderley Park | Wellcome Sanger Institute for the COVID-19 Genomics UK (COG-UK) Consortium | Jacquelyn Wynn, Mairead Hyland, The Lighthouse Lab in Alderley Park and Alex Alderton, Roberto Amato, Jeffrey Barrett, Sonia Goncalves, Ewan Harrison, David K. Jackson, Ian Johnston, Dominic Kwiatkowski, Cordelia Langford, John Sillitoe on behalf of the Wellcome Sanger Institute COVID-19 Surveillance Team |
| EPI_ISL_2485882 | Lighthouse Lab in Glasgow | Wellcome Sanger Institute for the COVID-19 Genomics UK (COG-UK) Consortium | Harper VanSteenhouse, Yumi Kasai, David Gray, Carol Clugston, Anna Dominiczak and Alex Alderton, Roberto Amato, Jeffrey Barrett, Sonia Goncalves, Ewan Harrison, David K. Jackson, Ian Johnston, Dominic Kwiatkowski, Cordelia Langford, John Sillitoe on behalf of the Wellcome Sanger Institute COVID-19 Surveillance Team |
| EPI_ISL_2485883 | Lighthouse Lab in Milton Keynes | Wellcome Sanger Institute for the COVID-19 Genomics UK (COG-UK) Consortium | The Lighthouse Lab in Milton Keynes and Alex Alderton, Roberto Amato, Jeffrey Barrett, Sonia Goncalves, Ewan Harrison, David K. Jackson, Ian Johnston, Dominic Kwiatkowski, Cordelia Langford, John Sillitoe on behalf of the Wellcome Sanger Institute COVID-19 Surveillance Team |
| EPI_ISL_2485884, EPI_ISL_2485885, EPI_ISL_2485886, EPI_ISL_2485887, EPI_ISL_2485888 | Lighthouse Lab in Glasgow | Wellcome Sanger Institute for the COVID-19 Genomics UK (COG-UK) Consortium | Harper VanSteenhouse, Yumi Kasai, David Gray, Carol Clugston, Anna Dominiczak and Alex Alderton, Roberto Amato, Jeffrey Barrett, Sonia Goncalves, Ewan Harrison, David K. Jackson, Ian Johnston, Dominic Kwiatkowski, Cordelia Langford, John Sillitoe on behalf of the Wellcome Sanger Institute COVID-19 Surveillance Team |
| EPI_ISL_2485890 | Lighthouse Lab in Milton Keynes | Wellcome Sanger Institute for the COVID-19 Genomics UK (COG-UK) Consortium | The Lighthouse Lab in Milton Keynes and Alex Alderton, Roberto Amato, Jeffrey Barrett, Sonia Goncalves, Ewan Harrison, David K. Jackson, Ian Johnston, Dominic Kwiatkowski, Cordelia Langford, John Sillitoe on behalf of the Wellcome Sanger Institute COVID-19 Surveillance Team |
| EPI_ISL_2485891 | Lighthouse Lab in Glasgow | Wellcome Sanger Institute for the COVID-19 Genomics UK (COG-UK) Consortium | Harper VanSteenhouse, Yumi Kasai, David Gray, Carol Clugston, Anna Dominiczak and Alex Alderton, Roberto Amato, Jeffrey Barrett, Sonia Goncalves, Ewan Harrison, David K. Jackson, Ian Johnston, Dominic Kwiatkowski, Cordelia Langford, John Sillitoe on behalf of the Wellcome Sanger Institute COVID-19 Surveillance Team |
| EPI_ISL_2485892, EPI_ISL_2485893, EPI_ISL_2485894 | Lighthouse Lab in Milton Keynes | Wellcome Sanger Institute for the COVID-19 Genomics UK (COG-UK) Consortium | The Lighthouse Lab in Milton Keynes and Alex Alderton, Roberto Amato, Jeffrey Barrett, Sonia Goncalves, Ewan Harrison, David K. Jackson, Ian Johnston, Dominic Kwiatkowski, Cordelia Langford, John Sillitoe on behalf of the Wellcome Sanger Institute COVID-19 Surveillance Team |

| | | | |
|---|---|---|---|
| EPI_ISL_2485895 | Lighthouse Lab in Glasgow | Wellcome Sanger Institute for the COVID-19 Genomics UK (COG-UK) Consortium | Harper VanSteenhouse, Yumi Kasai, David Gray, Carol Clugston, Anna Dominiczak and Alex Alderton, Roberto Amato, Jeffrey Barrett, Sonia Goncalves, Ewan Harrison, David K. Jackson, Ian Johnston, Dominic Kwiatkowski, Cordelia Langford, John Sillitoe on behalf of the Wellcome Sanger Institute COVID-19 Surveillance Team |
| EPI_ISL_2485896 | Lighthouse Lab in Alderley Park | Wellcome Sanger Institute for the COVID-19 Genomics UK (COG-UK) Consortium | Jacquelyn Wynn, Mairead Hyland, The Lighthouse Lab in Alderley Park and Alex Alderton, Roberto Amato, Jeffrey Barrett, Sonia Goncalves, Ewan Harrison, David K. Jackson, Ian Johnston, Dominic Kwiatkowski, Cordelia Langford, John Sillitoe on behalf of the Wellcome Sanger Institute COVID-19 Surveillance Team |
| EPI_ISL_2485897 | Lighthouse Lab in Glasgow | Wellcome Sanger Institute for the COVID-19 Genomics UK (COG-UK) Consortium | Harper VanSteenhouse, Yumi Kasai, David Gray, Carol Clugston, Anna Dominiczak and Alex Alderton, Roberto Amato, Jeffrey Barrett, Sonia Goncalves, Ewan Harrison, David K. Jackson, Ian Johnston, Dominic Kwiatkowski, Cordelia Langford, John Sillitoe on behalf of the Wellcome Sanger Institute COVID-19 Surveillance Team |
| EPI_ISL_2485898 | Lighthouse Lab in Milton Keynes | Wellcome Sanger Institute for the COVID-19 Genomics UK (COG-UK) Consortium | The Lighthouse Lab in Milton Keynes and Alex Alderton, Roberto Amato, Jeffrey Barrett, Sonia Goncalves, Ewan Harrison, David K. Jackson, Ian Johnston, Dominic Kwiatkowski, Cordelia Langford, John Sillitoe on behalf of the Wellcome Sanger Institute COVID-19 Surveillance Team |
| EPI_ISL_2485899, EPI_ISL_2485900 | Lighthouse Lab in Alderley Park | Wellcome Sanger Institute for the COVID-19 Genomics UK (COG-UK) Consortium | Jacquelyn Wynn, Mairead Hyland, The Lighthouse Lab in Alderley Park and Alex Alderton, Roberto Amato, Jeffrey Barrett, Sonia Goncalves, Ewan Harrison, David K. Jackson, Ian Johnston, Dominic Kwiatkowski, Cordelia Langford, John Sillitoe on behalf of the Wellcome Sanger Institute COVID-19 Surveillance Team |
| EPI_ISL_2485901 | Lighthouse Lab in Milton Keynes | Wellcome Sanger Institute for the COVID-19 Genomics UK (COG-UK) Consortium | The Lighthouse Lab in Milton Keynes and Alex Alderton, Roberto Amato, Jeffrey Barrett, Sonia Goncalves, Ewan Harrison, David K. Jackson, Ian Johnston, Dominic Kwiatkowski, Cordelia Langford, John Sillitoe on behalf of the Wellcome Sanger Institute COVID-19 Surveillance Team |
| EPI_ISL_2485903, EPI_ISL_2485905, EPI_ISL_2485906, EPI_ISL_2485907 | Lighthouse Lab in Glasgow | Wellcome Sanger Institute for the COVID-19 Genomics UK (COG-UK) Consortium | Harper VanSteenhouse, Yumi Kasai, David Gray, Carol Clugston, Anna Dominiczak and Alex Alderton, Roberto Amato, Jeffrey Barrett, Sonia Goncalves, Ewan Harrison, David K. Jackson, Ian Johnston, Dominic Kwiatkowski, Cordelia Langford, John Sillitoe on behalf of the Wellcome Sanger Institute COVID-19 Surveillance Team |
| EPI_ISL_2485908 | Lighthouse Lab in Milton Keynes | Wellcome Sanger Institute for the COVID-19 Genomics UK (COG-UK) Consortium | The Lighthouse Lab in Milton Keynes and Alex Alderton, Roberto Amato, Jeffrey Barrett, Sonia Goncalves, Ewan Harrison, David K. Jackson, Ian Johnston, Dominic Kwiatkowski, Cordelia Langford, John Sillitoe on behalf of the Wellcome Sanger Institute COVID-19 Surveillance Team |
| EPI_ISL_2485909 | Lighthouse Lab in Glasgow | Wellcome Sanger Institute for the COVID-19 Genomics UK (COG-UK) Consortium | Harper VanSteenhouse, Yumi Kasai, David Gray, Carol Clugston, Anna Dominiczak and Alex Alderton, Roberto Amato, Jeffrey Barrett, Sonia Goncalves, Ewan Harrison, David K. Jackson, Ian Johnston, Dominic Kwiatkowski, Cordelia Langford, John Sillitoe on behalf of the Wellcome Sanger Institute COVID-19 Surveillance Team |
| EPI_ISL_2485910 | Lighthouse Lab in Alderley Park | Wellcome Sanger Institute for the COVID-19 Genomics UK (COG-UK) Consortium | Jacquelyn Wynn, Mairead Hyland, The Lighthouse Lab in Alderley Park and Alex Alderton, Roberto Amato, Jeffrey Barrett, Sonia Goncalves, Ewan Harrison, David K. Jackson, Ian Johnston, Dominic Kwiatkowski, Cordelia Langford, John Sillitoe on behalf of the Wellcome Sanger Institute COVID-19 Surveillance Team |
| EPI_ISL_2485911 | Lighthouse Lab in Glasgow | Wellcome Sanger Institute for the COVID-19 Genomics UK (COG-UK) Consortium | Harper VanSteenhouse, Yumi Kasai, David Gray, Carol Clugston, Anna Dominiczak and Alex Alderton, Roberto Amato, Jeffrey Barrett, Sonia Goncalves, Ewan Harrison, David K. Jackson, Ian Johnston, Dominic Kwiatkowski, Cordelia Langford, John Sillitoe on behalf of the Wellcome Sanger Institute COVID-19 Surveillance Team |
| EPI_ISL_2485913 | Lighthouse Lab in Milton Keynes | Wellcome Sanger Institute for the COVID-19 Genomics UK (COG-UK) Consortium | The Lighthouse Lab in Milton Keynes and Alex Alderton, Roberto Amato, Jeffrey Barrett, Sonia Goncalves, Ewan Harrison, David K. Jackson, Ian Johnston, Dominic Kwiatkowski, Cordelia Langford, John Sillitoe on behalf of the Wellcome Sanger Institute COVID-19 Surveillance Team |
| EPI_ISL_2485914, EPI_ISL_2485915 | Lighthouse Lab in Glasgow | Wellcome Sanger Institute for the COVID-19 Genomics UK (COG-UK) Consortium | Harper VanSteenhouse, Yumi Kasai, David Gray, Carol Clugston, Anna Dominiczak and Alex Alderton, Roberto Amato, Jeffrey Barrett, Sonia Goncalves, Ewan Harrison, David K. Jackson, Ian Johnston, Dominic Kwiatkowski, Cordelia Langford, John Sillitoe on behalf of the Wellcome Sanger Institute COVID-19 Surveillance Team |
| EPI_ISL_2485916 | Lighthouse Lab in Alderley Park | Wellcome Sanger Institute for the COVID-19 Genomics UK (COG-UK) Consortium | Jacquelyn Wynn, Mairead Hyland, The Lighthouse Lab in Alderley Park and Alex Alderton, Roberto Amato, Jeffrey Barrett, Sonia Goncalves, Ewan Harrison, David K. Jackson, Ian Johnston, Dominic Kwiatkowski, Cordelia Langford, John Sillitoe on behalf of the Wellcome Sanger Institute COVID-19 Surveillance Team |
| EPI_ISL_2485917 | Lighthouse Lab in Glasgow | Wellcome Sanger Institute for the COVID-19 Genomics UK (COG-UK) Consortium | Harper VanSteenhouse, Yumi Kasai, David Gray, Carol Clugston, Anna Dominiczak and Alex Alderton, Roberto Amato, Jeffrey Barrett, Sonia Goncalves, Ewan Harrison, David K. Jackson, Ian Johnston, Dominic Kwiatkowski, Cordelia Langford, John Sillitoe on behalf of the Wellcome Sanger Institute COVID-19 Surveillance Team |
| EPI_ISL_2485918 | Lighthouse Lab in Milton Keynes | Wellcome Sanger Institute for the COVID-19 Genomics UK (COG-UK) Consortium | The Lighthouse Lab in Milton Keynes and Alex Alderton, Roberto Amato, Jeffrey Barrett, Sonia Goncalves, Ewan Harrison, David K. Jackson, Ian Johnston, Dominic Kwiatkowski, Cordelia Langford, John Sillitoe on behalf of the Wellcome Sanger Institute COVID-19 Surveillance Team |
| EPI_ISL_2485919, EPI_ISL_2485920, EPI_ISL_2485921 | Lighthouse Lab in Glasgow | Wellcome Sanger Institute for the COVID-19 Genomics UK (COG-UK) Consortium | Harper VanSteenhouse, Yumi Kasai, David Gray, Carol Clugston, Anna Dominiczak and Alex Alderton, Roberto Amato, Jeffrey Barrett, Sonia Goncalves, Ewan Harrison, David K. Jackson, Ian Johnston, Dominic Kwiatkowski, Cordelia Langford, John Sillitoe on behalf of the Wellcome Sanger Institute COVID-19 Surveillance Team |
| EPI_ISL_2485922, EPI_ISL_2485923 | Lighthouse Lab in Alderley Park | Wellcome Sanger Institute for the COVID-19 Genomics UK (COG-UK) Consortium | Jacquelyn Wynn, Mairead Hyland, The Lighthouse Lab in Alderley Park and Alex Alderton, Roberto Amato, Jeffrey Barrett, Sonia Goncalves, Ewan Harrison, David K. Jackson, Ian Johnston, Dominic Kwiatkowski, Cordelia Langford, John Sillitoe on behalf of the Wellcome Sanger Institute COVID-19 Surveillance Team |
| EPI_ISL_2485924, EPI_ISL_2485925, EPI_ISL_2485926 | Lighthouse Lab in Glasgow | Wellcome Sanger Institute for the COVID-19 Genomics UK (COG-UK) Consortium | Harper VanSteenhouse, Yumi Kasai, David Gray, Carol Clugston, Anna Dominiczak and Alex Alderton, Roberto Amato, Jeffrey Barrett, Sonia Goncalves, Ewan Harrison, David K. Jackson, Ian Johnston, Dominic Kwiatkowski, Cordelia Langford, John Sillitoe on behalf of the Wellcome Sanger Institute COVID-19 Surveillance Team |
| EPI_ISL_2485927 | Lighthouse Lab in Milton Keynes | Wellcome Sanger Institute for the COVID-19 Genomics UK (COG-UK) Consortium | The Lighthouse Lab in Milton Keynes and Alex Alderton, Roberto Amato, Jeffrey Barrett, Sonia Goncalves, Ewan Harrison, David K. Jackson, Ian Johnston, Dominic Kwiatkowski, Cordelia Langford, John Sillitoe on behalf of the Wellcome Sanger Institute COVID-19 Surveillance Team |
| EPI_ISL_2485928, EPI_ISL_2485929 | Lighthouse Lab in Glasgow | Wellcome Sanger Institute for the COVID-19 Genomics UK (COG-UK) Consortium | Harper VanSteenhouse, Yumi Kasai, David Gray, Carol Clugston, Anna Dominiczak and Alex Alderton, Roberto Amato, Jeffrey Barrett, Sonia Goncalves, Ewan Harrison, David K. Jackson, Ian Johnston, Dominic Kwiatkowski, Cordelia Langford, John Sillitoe on behalf of the Wellcome Sanger Institute COVID-19 Surveillance Team |
| EPI_ISL_2485930 | Lighthouse Lab in Milton Keynes | Wellcome Sanger Institute for the COVID-19 Genomics UK (COG-UK) Consortium | The Lighthouse Lab in Milton Keynes and Alex Alderton, Roberto Amato, Jeffrey Barrett, Sonia Goncalves, Ewan Harrison, David K. Jackson, Ian Johnston, Dominic Kwiatkowski, Cordelia Langford, John Sillitoe on behalf of the Wellcome Sanger Institute COVID-19 Surveillance Team |
| EPI_ISL_2485931 | Lighthouse Lab in Glasgow | Wellcome Sanger Institute for the COVID-19 Genomics UK (COG-UK) Consortium | Harper VanSteenhouse, Yumi Kasai, David Gray, Carol Clugston, Anna Dominiczak and Alex Alderton, Roberto Amato, Jeffrey Barrett, Sonia Goncalves, Ewan Harrison, David K. Jackson, Ian Johnston, Dominic Kwiatkowski, Cordelia Langford, John Sillitoe on behalf of the Wellcome Sanger Institute COVID-19 Surveillance Team |
| EPI_ISL_2485932 | Lighthouse Lab in Milton Keynes | Wellcome Sanger Institute for the COVID-19 Genomics UK (COG-UK) Consortium | The Lighthouse Lab in Milton Keynes and Alex Alderton, Roberto Amato, Jeffrey Barrett, Sonia Goncalves, Ewan Harrison, David K. Jackson, Ian Johnston, Dominic Kwiatkowski, Cordelia Langford, John Sillitoe on behalf of the Wellcome Sanger Institute COVID-19 Surveillance Team |
| EPI_ISL_2485933 | Lighthouse Lab in Glasgow | Wellcome Sanger Institute for the COVID-19 Genomics UK (COG-UK) Consortium | Harper VanSteenhouse, Yumi Kasai, David Gray, Carol Clugston, Anna Dominiczak and Alex Alderton, Roberto Amato, Jeffrey Barrett, Sonia Goncalves, Ewan Harrison, David K. Jackson, Ian Johnston, Dominic Kwiatkowski, Cordelia Langford, John Sillitoe on behalf of the Wellcome Sanger Institute COVID-19 Surveillance Team |
| EPI_ISL_2485934 | Lighthouse Lab in Alderley Park | Wellcome Sanger Institute for the COVID-19 Genomics UK (COG-UK) Consortium | Jacquelyn Wynn, Mairead Hyland, The Lighthouse Lab in Alderley Park and Alex Alderton, Roberto Amato, Jeffrey Barrett, Sonia Goncalves, Ewan Harrison, David K. Jackson, Ian Johnston, Dominic Kwiatkowski, Cordelia Langford, John Sillitoe on behalf of the Wellcome Sanger Institute COVID-19 Surveillance Team |
| EPI_ISL_2485935, EPI_ISL_2485936, EPI_ISL_2485938, EPI_ISL_2485939 | Lighthouse Lab in Glasgow | Wellcome Sanger Institute for the COVID-19 Genomics UK (COG-UK) Consortium | Harper VanSteenhouse, Yumi Kasai, David Gray, Carol Clugston, Anna Dominiczak and Alex Alderton, Roberto Amato, Jeffrey Barrett, Sonia Goncalves, Ewan Harrison, David K. Jackson, Ian Johnston, Dominic Kwiatkowski, Cordelia Langford, John Sillitoe on behalf of the Wellcome Sanger Institute COVID-19 Surveillance Team |
| EPI_ISL_2485941, EPI_ISL_2485942 | Lighthouse Lab in Alderley Park | Wellcome Sanger Institute for the COVID-19 Genomics UK (COG-UK) Consortium | Jacquelyn Wynn, Mairead Hyland, The Lighthouse Lab in Alderley Park and Alex Alderton, Roberto Amato, Jeffrey Barrett, Sonia Goncalves, Ewan Harrison, David K. Jackson, Ian Johnston, Dominic Kwiatkowski, Cordelia Langford, John Sillitoe on behalf of the Wellcome Sanger Institute COVID-19 Surveillance Team |
| EPI_ISL_2485944 | Lighthouse Lab in Glasgow | Wellcome Sanger Institute for the COVID-19 Genomics UK (COG-UK) Consortium | Harper VanSteenhouse, Yumi Kasai, David Gray, Carol Clugston, Anna Dominiczak and Alex Alderton, Roberto Amato, Jeffrey Barrett, Sonia Goncalves, Ewan Harrison, David K. Jackson, Ian Johnston, Dominic Kwiatkowski, Cordelia Langford, John Sillitoe on behalf of the Wellcome Sanger Institute COVID-19 Surveillance Team |
| EPI_ISL_2485946 | Lighthouse Lab in Alderley Park | Wellcome Sanger Institute for the COVID-19 Genomics UK (COG-UK) Consortium | Jacquelyn Wynn, Mairead Hyland, The Lighthouse Lab in Alderley Park and Alex Alderton, Roberto Amato, Jeffrey Barrett, Sonia Goncalves, Ewan Harrison, David K. Jackson, Ian Johnston, Dominic Kwiatkowski, Cordelia Langford, John Sillitoe on behalf of the Wellcome Sanger Institute COVID-19 Surveillance Team |
| EPI_ISL_2485947, EPI_ISL_2485948, EPI_ISL_2485949 | Lighthouse Lab in Glasgow | Wellcome Sanger Institute for the COVID-19 Genomics UK (COG-UK) Consortium | Harper VanSteenhouse, Yumi Kasai, David Gray, Carol Clugston, Anna Dominiczak and Alex Alderton, Roberto Amato, Jeffrey Barrett, Sonia Goncalves, Ewan Harrison, David K. Jackson, Ian Johnston, Dominic Kwiatkowski, Cordelia Langford, John Sillitoe on behalf of the Wellcome Sanger Institute COVID-19 Surveillance Team |
| EPI_ISL_2485950 | Lighthouse Lab in Milton Keynes | Wellcome Sanger Institute for the COVID-19 Genomics UK (COG-UK) Consortium | The Lighthouse Lab in Milton Keynes and Alex Alderton, Roberto Amato, Jeffrey Barrett, Sonia Goncalves, Ewan Harrison, David K. Jackson, Ian Johnston, Dominic Kwiatkowski, Cordelia Langford, John Sillitoe on behalf of the Wellcome Sanger Institute COVID-19 Surveillance Team |
| EPI_ISL_2485951, EPI_ISL_2485952 | Lighthouse Lab in Glasgow | Wellcome Sanger Institute for the COVID-19 Genomics UK (COG-UK) Consortium | Harper VanSteenhouse, Yumi Kasai, David Gray, Carol Clugston, Anna Dominiczak and Alex Alderton, Roberto Amato, Jeffrey Barrett, Sonia Goncalves, Ewan Harrison, David K. Jackson, Ian Johnston, Dominic Kwiatkowski, Cordelia Langford, John Sillitoe on behalf of the Wellcome Sanger Institute COVID-19 Surveillance Team |
| EPI_ISL_2485953, EPI_ISL_2485954, EPI_ISL_2485955, EPI_ISL_2485956, EPI_ISL_2485957 | Lighthouse Lab in Alderley Park | Wellcome Sanger Institute for the COVID-19 Genomics UK (COG-UK) Consortium | Jacquelyn Wynn, Mairead Hyland, The Lighthouse Lab in Alderley Park and Alex Alderton, Roberto Amato, Jeffrey Barrett, Sonia Goncalves, Ewan Harrison, David K. Jackson, Ian Johnston, Dominic Kwiatkowski, Cordelia Langford, John Sillitoe on behalf of the Wellcome Sanger Institute COVID-19 Surveillance Team |
| EPI_ISL_2485958, EPI_ISL_2485959, EPI_ISL_2485960, EPI_ISL_2485961 | Lighthouse Lab in Glasgow | Wellcome Sanger Institute for the COVID-19 Genomics UK (COG-UK) Consortium | Harper VanSteenhouse, Yumi Kasai, David Gray, Carol Clugston, Anna Dominiczak and Alex Alderton, Roberto Amato, Jeffrey Barrett, Sonia Goncalves, Ewan Harrison, David K. Jackson, Ian Johnston, Dominic Kwiatkowski, Cordelia Langford, John Sillitoe on behalf of the Wellcome Sanger Institute COVID-19 Surveillance Team |
| EPI_ISL_2485962 | Lighthouse Lab in Milton Keynes | Wellcome Sanger Institute for the COVID-19 Genomics UK (COG-UK) Consortium | The Lighthouse Lab in Milton Keynes and Alex Alderton, Roberto Amato, Jeffrey Barrett, Sonia Goncalves, Ewan Harrison, David K. Jackson, Ian Johnston, Dominic Kwiatkowski, Cordelia Langford, John Sillitoe on behalf of the Wellcome Sanger Institute COVID-19 Surveillance Team |
| EPI_ISL_2485963 | Lighthouse Lab in Glasgow | Wellcome Sanger Institute for the COVID-19 Genomics UK (COG-UK) Consortium | Harper VanSteenhouse, Yumi Kasai, David Gray, Carol Clugston, Anna Dominiczak and Alex Alderton, Roberto Amato, Jeffrey Barrett, Sonia Goncalves, Ewan Harrison, David K. Jackson, Ian Johnston, Dominic Kwiatkowski, Cordelia Langford, John Sillitoe on behalf of the Wellcome Sanger Institute COVID-19 Surveillance Team |
| EPI_ISL_2485965 | Lighthouse Lab in Milton Keynes | Wellcome Sanger Institute for the COVID-19 Genomics UK (COG-UK) Consortium | The Lighthouse Lab in Milton Keynes and Alex Alderton, Roberto Amato, Jeffrey Barrett, Sonia Goncalves, Ewan Harrison, David K. Jackson, Ian Johnston, Dominic Kwiatkowski, Cordelia Langford, John Sillitoe on behalf of the Wellcome Sanger Institute COVID-19 Surveillance Team |
| EPI_ISL_2485966, EPI_ISL_2485967, EPI_ISL_2485968, EPI_ISL_2485969 | Lighthouse Lab in Glasgow | Wellcome Sanger Institute for the COVID-19 Genomics UK (COG-UK) Consortium | Harper VanSteenhouse, Yumi Kasai, David Gray, Carol Clugston, Anna Dominiczak and Alex Alderton, Roberto Amato, Jeffrey Barrett, Sonia Goncalves, Ewan Harrison, David K. Jackson, Ian Johnston, Dominic Kwiatkowski, Cordelia Langford, John Sillitoe on behalf of the Wellcome Sanger Institute COVID-19 Surveillance Team |
| EPI_ISL_2485970 | Lighthouse Lab in Alderley Park | Wellcome Sanger Institute for the COVID-19 Genomics UK (COG-UK) | Jacquelyn Wynn, Mairead Hyland, The Lighthouse Lab in Alderley Park and Alex Alderton, Roberto Amato, Jeffrey Barrett, Sonia Goncalves, Ewan Harrison, David K. Jackson, Ian Johnston, Dominic Kwiatkowski, Cordelia Langford, John Sillitoe on |

| | | | |
|---|---|---|---|
| | | Consortium | behalf of the Wellcome Sanger Institute COVID-19 Surveillance Team |
| EPI_ISL_2485971, EPI_ISL_2485972, EPI_ISL_2485973, EPI_ISL_2485974, EPI_ISL_2485975 | Lighthouse Lab in Glasgow | Wellcome Sanger Institute for the COVID-19 Genomics UK (COG-UK) Consortium | Harper VanSteenhouse, Yumi Kasai, David Gray, Carol Clugston, Anna Dominiczak and Alex Alderton, Roberto Amato, Jeffrey Barrett, Sonia Goncalves, Ewan Harrison, David K. Jackson, Ian Johnston, Dominic Kwiatkowski, Cordelia Langford, John Sillitoe on behalf of the Wellcome Sanger Institute COVID-19 Surveillance Team |
| EPI_ISL_2485976 | Lighthouse Lab in Alderley Park | Wellcome Sanger Institute for the COVID-19 Genomics UK (COG-UK) Consortium | Jacquelyn Wynn, Mairead Hyland, The Lighthouse Lab in Alderley Park and Alex Alderton, Roberto Amato, Jeffrey Barrett, Sonia Goncalves, Ewan Harrison, David K. Jackson, Ian Johnston, Dominic Kwiatkowski, Cordelia Langford, John Sillitoe on behalf of the Wellcome Sanger Institute COVID-19 Surveillance Team |
| EPI_ISL_2485977 | Lighthouse Lab in Milton Keynes | Wellcome Sanger Institute for the COVID-19 Genomics UK (COG-UK) Consortium | The Lighthouse Lab in Milton Keynes and Alex Alderton, Roberto Amato, Jeffrey Barrett, Sonia Goncalves, Ewan Harrison, David K. Jackson, Ian Johnston, Dominic Kwiatkowski, Cordelia Langford, John Sillitoe on behalf of the Wellcome Sanger Institute COVID-19 Surveillance Team |
| EPI_ISL_2485978 | Lighthouse Lab in Alderley Park | Wellcome Sanger Institute for the COVID-19 Genomics UK (COG-UK) Consortium | Jacquelyn Wynn, Mairead Hyland, The Lighthouse Lab in Alderley Park and Alex Alderton, Roberto Amato, Jeffrey Barrett, Sonia Goncalves, Ewan Harrison, David K. Jackson, Ian Johnston, Dominic Kwiatkowski, Cordelia Langford, John Sillitoe on behalf of the Wellcome Sanger Institute COVID-19 Surveillance Team |
| EPI_ISL_2485979 | Lighthouse Lab in Milton Keynes | Wellcome Sanger Institute for the COVID-19 Genomics UK (COG-UK) Consortium | The Lighthouse Lab in Milton Keynes and Alex Alderton, Roberto Amato, Jeffrey Barrett, Sonia Goncalves, Ewan Harrison, David K. Jackson, Ian Johnston, Dominic Kwiatkowski, Cordelia Langford, John Sillitoe on behalf of the Wellcome Sanger Institute COVID-19 Surveillance Team |
| EPI_ISL_2485981, EPI_ISL_2485982 | Lighthouse Lab in Glasgow | Wellcome Sanger Institute for the COVID-19 Genomics UK (COG-UK) Consortium | Harper VanSteenhouse, Yumi Kasai, David Gray, Carol Clugston, Anna Dominiczak and Alex Alderton, Roberto Amato, Jeffrey Barrett, Sonia Goncalves, Ewan Harrison, David K. Jackson, Ian Johnston, Dominic Kwiatkowski, Cordelia Langford, John Sillitoe on behalf of the Wellcome Sanger Institute COVID-19 Surveillance Team |
| EPI_ISL_2485983 | Lighthouse Lab in Alderley Park | Wellcome Sanger Institute for the COVID-19 Genomics UK (COG-UK) Consortium | Jacquelyn Wynn, Mairead Hyland, The Lighthouse Lab in Alderley Park and Alex Alderton, Roberto Amato, Jeffrey Barrett, Sonia Goncalves, Ewan Harrison, David K. Jackson, Ian Johnston, Dominic Kwiatkowski, Cordelia Langford, John Sillitoe on behalf of the Wellcome Sanger Institute COVID-19 Surveillance Team |
| EPI_ISL_2485984, EPI_ISL_2485985 | Lighthouse Lab in Glasgow | Wellcome Sanger Institute for the COVID-19 Genomics UK (COG-UK) Consortium | Harper VanSteenhouse, Yumi Kasai, David Gray, Carol Clugston, Anna Dominiczak and Alex Alderton, Roberto Amato, Jeffrey Barrett, Sonia Goncalves, Ewan Harrison, David K. Jackson, Ian Johnston, Dominic Kwiatkowski, Cordelia Langford, John Sillitoe on behalf of the Wellcome Sanger Institute COVID-19 Surveillance Team |
| EPI_ISL_2485986 | Lighthouse Lab in Milton Keynes | Wellcome Sanger Institute for the COVID-19 Genomics UK (COG-UK) Consortium | The Lighthouse Lab in Milton Keynes and Alex Alderton, Roberto Amato, Jeffrey Barrett, Sonia Goncalves, Ewan Harrison, David K. Jackson, Ian Johnston, Dominic Kwiatkowski, Cordelia Langford, John Sillitoe on behalf of the Wellcome Sanger Institute COVID-19 Surveillance Team |
| EPI_ISL_2485988, EPI_ISL_2485989, EPI_ISL_2485990 | Lighthouse Lab in Glasgow | Wellcome Sanger Institute for the COVID-19 Genomics UK (COG-UK) Consortium | Harper VanSteenhouse, Yumi Kasai, David Gray, Carol Clugston, Anna Dominiczak and Alex Alderton, Roberto Amato, Jeffrey Barrett, Sonia Goncalves, Ewan Harrison, David K. Jackson, Ian Johnston, Dominic Kwiatkowski, Cordelia Langford, John Sillitoe on behalf of the Wellcome Sanger Institute COVID-19 Surveillance Team |
| EPI_ISL_2485991 | Lighthouse Lab in Alderley Park | Wellcome Sanger Institute for the COVID-19 Genomics UK (COG-UK) Consortium | Jacquelyn Wynn, Mairead Hyland, The Lighthouse Lab in Alderley Park and Alex Alderton, Roberto Amato, Jeffrey Barrett, Sonia Goncalves, Ewan Harrison, David K. Jackson, Ian Johnston, Dominic Kwiatkowski, Cordelia Langford, John Sillitoe on behalf of the Wellcome Sanger Institute COVID-19 Surveillance Team |
| EPI_ISL_2485992, EPI_ISL_2485993, EPI_ISL_2485994 | Lighthouse Lab in Glasgow | Wellcome Sanger Institute for the COVID-19 Genomics UK (COG-UK) Consortium | Harper VanSteenhouse, Yumi Kasai, David Gray, Carol Clugston, Anna Dominiczak and Alex Alderton, Roberto Amato, Jeffrey Barrett, Sonia Goncalves, Ewan Harrison, David K. Jackson, Ian Johnston, Dominic Kwiatkowski, Cordelia Langford, John Sillitoe on behalf of the Wellcome Sanger Institute COVID-19 Surveillance Team |
| EPI_ISL_2485995 | Lighthouse Lab in Milton Keynes | Wellcome Sanger Institute for the COVID-19 Genomics UK (COG-UK) Consortium | The Lighthouse Lab in Milton Keynes and Alex Alderton, Roberto Amato, Jeffrey Barrett, Sonia Goncalves, Ewan Harrison, David K. Jackson, Ian Johnston, Dominic Kwiatkowski, Cordelia Langford, John Sillitoe on behalf of the Wellcome Sanger Institute COVID-19 Surveillance Team |
| EPI_ISL_2485996 | Lighthouse Lab in Glasgow | Wellcome Sanger Institute for the COVID-19 Genomics UK (COG-UK) Consortium | Harper VanSteenhouse, Yumi Kasai, David Gray, Carol Clugston, Anna Dominiczak and Alex Alderton, Roberto Amato, Jeffrey Barrett, Sonia Goncalves, Ewan Harrison, David K. Jackson, Ian Johnston, Dominic Kwiatkowski, Cordelia Langford, John Sillitoe on behalf of the Wellcome Sanger Institute COVID-19 Surveillance Team |
| EPI_ISL_2485997, EPI_ISL_2485998 | Lighthouse Lab in Alderley Park | Wellcome Sanger Institute for the COVID-19 Genomics UK (COG-UK) Consortium | Jacquelyn Wynn, Mairead Hyland, The Lighthouse Lab in Alderley Park and Alex Alderton, Roberto Amato, Jeffrey Barrett, Sonia Goncalves, Ewan Harrison, David K. Jackson, Ian Johnston, Dominic Kwiatkowski, Cordelia Langford, John Sillitoe on behalf of the Wellcome Sanger Institute COVID-19 Surveillance Team |
| EPI_ISL_2486000 | Lighthouse Lab in Milton Keynes | Wellcome Sanger Institute for the COVID-19 Genomics UK (COG-UK) Consortium | The Lighthouse Lab in Milton Keynes and Alex Alderton, Roberto Amato, Jeffrey Barrett, Sonia Goncalves, Ewan Harrison, David K. Jackson, Ian Johnston, Dominic Kwiatkowski, Cordelia Langford, John Sillitoe on behalf of the Wellcome Sanger Institute COVID-19 Surveillance Team |
| EPI_ISL_2486001 | Lighthouse Lab in Glasgow | Wellcome Sanger Institute for the COVID-19 Genomics UK (COG-UK) Consortium | Harper VanSteenhouse, Yumi Kasai, David Gray, Carol Clugston, Anna Dominiczak and Alex Alderton, Roberto Amato, Jeffrey Barrett, Sonia Goncalves, Ewan Harrison, David K. Jackson, Ian Johnston, Dominic Kwiatkowski, Cordelia Langford, John Sillitoe on behalf of the Wellcome Sanger Institute COVID-19 Surveillance Team |
| EPI_ISL_2486002, EPI_ISL_2486003 | Lighthouse Lab in Alderley Park | Wellcome Sanger Institute for the COVID-19 Genomics UK (COG-UK) Consortium | Jacquelyn Wynn, Mairead Hyland, The Lighthouse Lab in Alderley Park and Alex Alderton, Roberto Amato, Jeffrey Barrett, Sonia Goncalves, Ewan Harrison, David K. Jackson, Ian Johnston, Dominic Kwiatkowski, Cordelia Langford, John Sillitoe on behalf of the Wellcome Sanger Institute COVID-19 Surveillance Team |
| EPI_ISL_2486004 | Lighthouse Lab in Glasgow | Wellcome Sanger Institute for the COVID-19 Genomics UK (COG-UK) Consortium | Harper VanSteenhouse, Yumi Kasai, David Gray, Carol Clugston, Anna Dominiczak and Alex Alderton, Roberto Amato, Jeffrey Barrett, Sonia Goncalves, Ewan Harrison, David K. Jackson, Ian Johnston, Dominic Kwiatkowski, Cordelia Langford, John Sillitoe on behalf of the Wellcome Sanger Institute COVID-19 Surveillance Team |
| EPI_ISL_2486005 | Lighthouse Lab in Milton Keynes | Wellcome Sanger Institute for the COVID-19 Genomics UK (COG-UK) Consortium | The Lighthouse Lab in Milton Keynes and Alex Alderton, Roberto Amato, Jeffrey Barrett, Sonia Goncalves, Ewan Harrison, David K. Jackson, Ian Johnston, Dominic Kwiatkowski, Cordelia Langford, John Sillitoe on behalf of the Wellcome Sanger Institute COVID-19 Surveillance Team |
| EPI_ISL_2486007 | Lighthouse Lab in Alderley Park | Wellcome Sanger Institute for the COVID-19 Genomics UK (COG-UK) Consortium | Jacquelyn Wynn, Mairead Hyland, The Lighthouse Lab in Alderley Park and Alex Alderton, Roberto Amato, Jeffrey Barrett, Sonia Goncalves, Ewan Harrison, David K. Jackson, Ian Johnston, Dominic Kwiatkowski, Cordelia Langford, John Sillitoe on behalf of the Wellcome Sanger Institute COVID-19 Surveillance Team |
| EPI_ISL_2486008, EPI_ISL_2486010 | Lighthouse Lab in Glasgow | Wellcome Sanger Institute for the COVID-19 Genomics UK (COG-UK) Consortium | Harper VanSteenhouse, Yumi Kasai, David Gray, Carol Clugston, Anna Dominiczak and Alex Alderton, Roberto Amato, Jeffrey Barrett, Sonia Goncalves, Ewan Harrison, David K. Jackson, Ian Johnston, Dominic Kwiatkowski, Cordelia Langford, John Sillitoe on behalf of the Wellcome Sanger Institute COVID-19 Surveillance Team |
| EPI_ISL_2486011 | Lighthouse Lab in Alderley Park | Wellcome Sanger Institute for the COVID-19 Genomics UK (COG-UK) Consortium | Jacquelyn Wynn, Mairead Hyland, The Lighthouse Lab in Alderley Park and Alex Alderton, Roberto Amato, Jeffrey Barrett, Sonia Goncalves, Ewan Harrison, David K. Jackson, Ian Johnston, Dominic Kwiatkowski, Cordelia Langford, John Sillitoe on behalf of the Wellcome Sanger Institute COVID-19 Surveillance Team |
| EPI_ISL_2486012 | Lighthouse Lab in Glasgow | Wellcome Sanger Institute for the COVID-19 Genomics UK (COG-UK) Consortium | Harper VanSteenhouse, Yumi Kasai, David Gray, Carol Clugston, Anna Dominiczak and Alex Alderton, Roberto Amato, Jeffrey Barrett, Sonia Goncalves, Ewan Harrison, David K. Jackson, Ian Johnston, Dominic Kwiatkowski, Cordelia Langford, John Sillitoe on behalf of the Wellcome Sanger Institute COVID-19 Surveillance Team |
| EPI_ISL_2486013 | Lighthouse Lab in Alderley Park | Wellcome Sanger Institute for the COVID-19 Genomics UK (COG-UK) Consortium | Jacquelyn Wynn, Mairead Hyland, The Lighthouse Lab in Alderley Park and Alex Alderton, Roberto Amato, Jeffrey Barrett, Sonia Goncalves, Ewan Harrison, David K. Jackson, Ian Johnston, Dominic Kwiatkowski, Cordelia Langford, John Sillitoe on behalf of the Wellcome Sanger Institute COVID-19 Surveillance Team |
| EPI_ISL_2486014 | Lighthouse Lab in Milton Keynes | Wellcome Sanger Institute for the COVID-19 Genomics UK (COG-UK) Consortium | The Lighthouse Lab in Milton Keynes and Alex Alderton, Roberto Amato, Jeffrey Barrett, Sonia Goncalves, Ewan Harrison, David K. Jackson, Ian Johnston, Dominic Kwiatkowski, Cordelia Langford, John Sillitoe on behalf of the Wellcome Sanger Institute COVID-19 Surveillance Team |
| EPI_ISL_2486015 | Lighthouse Lab in Glasgow | Wellcome Sanger Institute for the COVID-19 Genomics UK (COG-UK) Consortium | Harper VanSteenhouse, Yumi Kasai, David Gray, Carol Clugston, Anna Dominiczak and Alex Alderton, Roberto Amato, Jeffrey Barrett, Sonia Goncalves, Ewan Harrison, David K. Jackson, Ian Johnston, Dominic Kwiatkowski, Cordelia Langford, John Sillitoe on behalf of the Wellcome Sanger Institute COVID-19 Surveillance Team |
| EPI_ISL_2486016 | Lighthouse Lab in Milton Keynes | Wellcome Sanger Institute for the COVID-19 Genomics UK (COG-UK) Consortium | The Lighthouse Lab in Milton Keynes and Alex Alderton, Roberto Amato, Jeffrey Barrett, Sonia Goncalves, Ewan Harrison, David K. Jackson, Ian Johnston, Dominic Kwiatkowski, Cordelia Langford, John Sillitoe on behalf of the Wellcome Sanger Institute COVID-19 Surveillance Team |
| EPI_ISL_2486018, EPI_ISL_2486019, EPI_ISL_2486020, EPI_ISL_2486022 | Lighthouse Lab in Glasgow | Wellcome Sanger Institute for the COVID-19 Genomics UK (COG-UK) Consortium | Harper VanSteenhouse, Yumi Kasai, David Gray, Carol Clugston, Anna Dominiczak and Alex Alderton, Roberto Amato, Jeffrey Barrett, Sonia Goncalves, Ewan Harrison, David K. Jackson, Ian Johnston, Dominic Kwiatkowski, Cordelia Langford, John Sillitoe on behalf of the Wellcome Sanger Institute COVID-19 Surveillance Team |
| EPI_ISL_2486023, EPI_ISL_2486024 | Lighthouse Lab in Milton Keynes | Wellcome Sanger Institute for the COVID-19 Genomics UK (COG-UK) Consortium | The Lighthouse Lab in Milton Keynes and Alex Alderton, Roberto Amato, Jeffrey Barrett, Sonia Goncalves, Ewan Harrison, David K. Jackson, Ian Johnston, Dominic Kwiatkowski, Cordelia Langford, John Sillitoe on behalf of the Wellcome Sanger Institute COVID-19 Surveillance Team |
| EPI_ISL_2486025, EPI_ISL_2486026, EPI_ISL_2486027 | Lighthouse Lab in Glasgow | Wellcome Sanger Institute for the COVID-19 Genomics UK (COG-UK) Consortium | Harper VanSteenhouse, Yumi Kasai, David Gray, Carol Clugston, Anna Dominiczak and Alex Alderton, Roberto Amato, Jeffrey Barrett, Sonia Goncalves, Ewan Harrison, David K. Jackson, Ian Johnston, Dominic Kwiatkowski, Cordelia Langford, John Sillitoe on behalf of the Wellcome Sanger Institute COVID-19 Surveillance Team |
| EPI_ISL_2486028, EPI_ISL_2486029 | Lighthouse Lab in Alderley Park | Wellcome Sanger Institute for the COVID-19 Genomics UK (COG-UK) Consortium | Jacquelyn Wynn, Mairead Hyland, The Lighthouse Lab in Alderley Park and Alex Alderton, Roberto Amato, Jeffrey Barrett, Sonia Goncalves, Ewan Harrison, David K. Jackson, Ian Johnston, Dominic Kwiatkowski, Cordelia Langford, John Sillitoe on behalf of the Wellcome Sanger Institute COVID-19 Surveillance Team |
| EPI_ISL_2486030, EPI_ISL_2486031, EPI_ISL_2486032, EPI_ISL_2486033 | Lighthouse Lab in Milton Keynes | Wellcome Sanger Institute for the COVID-19 Genomics UK (COG-UK) Consortium | The Lighthouse Lab in Milton Keynes and Alex Alderton, Roberto Amato, Jeffrey Barrett, Sonia Goncalves, Ewan Harrison, David K. Jackson, Ian Johnston, Dominic Kwiatkowski, Cordelia Langford, John Sillitoe on behalf of the Wellcome Sanger Institute COVID-19 Surveillance Team |
| EPI_ISL_2486034, EPI_ISL_2486035 | Lighthouse Lab in Alderley Park | Wellcome Sanger Institute for the COVID-19 Genomics UK (COG-UK) Consortium | Jacquelyn Wynn, Mairead Hyland, The Lighthouse Lab in Alderley Park and Alex Alderton, Roberto Amato, Jeffrey Barrett, Sonia Goncalves, Ewan Harrison, David K. Jackson, Ian Johnston, Dominic Kwiatkowski, Cordelia Langford, John Sillitoe on behalf of the Wellcome Sanger Institute COVID-19 Surveillance Team |
| EPI_ISL_2486036 | Lighthouse Lab in Milton Keynes | Wellcome Sanger Institute for the COVID-19 Genomics UK (COG-UK) Consortium | The Lighthouse Lab in Milton Keynes and Alex Alderton, Roberto Amato, Jeffrey Barrett, Sonia Goncalves, Ewan Harrison, David K. Jackson, Ian Johnston, Dominic Kwiatkowski, Cordelia Langford, John Sillitoe on behalf of the Wellcome Sanger Institute COVID-19 Surveillance Team |
| EPI_ISL_2486037 | Lighthouse Lab in Alderley Park | Wellcome Sanger Institute for the COVID-19 Genomics UK (COG-UK) Consortium | Jacquelyn Wynn, Mairead Hyland, The Lighthouse Lab in Alderley Park and Alex Alderton, Roberto Amato, Jeffrey Barrett, Sonia Goncalves, Ewan Harrison, David K. Jackson, Ian Johnston, Dominic Kwiatkowski, Cordelia Langford, John Sillitoe on behalf of the Wellcome Sanger Institute COVID-19 Surveillance Team |
| EPI_ISL_2486038 | Lighthouse Lab in Milton Keynes | Wellcome Sanger Institute for the COVID-19 Genomics UK (COG-UK) Consortium | The Lighthouse Lab in Milton Keynes and Alex Alderton, Roberto Amato, Jeffrey Barrett, Sonia Goncalves, Ewan Harrison, David K. Jackson, Ian Johnston, Dominic Kwiatkowski, Cordelia Langford, John Sillitoe on behalf of the Wellcome Sanger Institute COVID-19 Surveillance Team |
| EPI_ISL_2486039 | Lighthouse Lab in Alderley Park | Wellcome Sanger Institute for the COVID-19 Genomics UK (COG-UK) Consortium | Jacquelyn Wynn, Mairead Hyland, The Lighthouse Lab in Alderley Park and Alex Alderton, Roberto Amato, Jeffrey Barrett, Sonia Goncalves, Ewan Harrison, David K. Jackson, Ian Johnston, Dominic Kwiatkowski, Cordelia Langford, John Sillitoe on behalf of the Wellcome Sanger Institute COVID-19 Surveillance Team |
| EPI_ISL_2486040, EPI_ISL_2486041 | Lighthouse Lab in Glasgow | Wellcome Sanger Institute for the COVID-19 Genomics UK (COG-UK) Consortium | Harper VanSteenhouse, Yumi Kasai, David Gray, Carol Clugston, Anna Dominiczak and Alex Alderton, Roberto Amato, Jeffrey Barrett, Sonia Goncalves, Ewan Harrison, David K. Jackson, Ian Johnston, Dominic Kwiatkowski, Cordelia Langford, John Sillitoe on behalf of the Wellcome Sanger Institute COVID-19 Surveillance Team |
| EPI_ISL_2486042 | Lighthouse Lab in Milton Keynes | Wellcome Sanger Institute for the COVID-19 Genomics UK (COG-UK) Consortium | The Lighthouse Lab in Milton Keynes and Alex Alderton, Roberto Amato, Jeffrey Barrett, Sonia Goncalves, Ewan Harrison, David K. Jackson, Ian Johnston, Dominic Kwiatkowski, Cordelia Langford, John Sillitoe on behalf of the Wellcome Sanger Institute COVID-19 Surveillance Team |
| EPI_ISL_2486043 | Lighthouse Lab in Glasgow | Wellcome Sanger Institute for the COVID-19 Genomics UK (COG-UK) | Harper VanSteenhouse, Yumi Kasai, David Gray, Carol Clugston, Anna Dominiczak and Alex Alderton, Roberto Amato, Jeffrey Barrett, Sonia Goncalves, Ewan Harrison, David K. Jackson, Ian Johnston, Dominic Kwiatkowski, Cordelia Langford, |

| | | | |
|---|---|---|---|
| | | Consortium | John Sillitoe on behalf of the Wellcome Sanger Institute COVID-19 Surveillance Team |
| EPI_ISL_2486044 | Lighthouse Lab in Alderley Park | Wellcome Sanger Institute for the COVID-19 Genomics UK (COG-UK) Consortium | Jacquelyn Wynn, Mairead Hyland, The Lighthouse Lab in Alderley Park and Alex Alderton, Roberto Amato, Jeffrey Barrett, Sonia Goncalves, Ewan Harrison, David K. Jackson, Ian Johnston, Dominic Kwiatkowski, Cordelia Langford, John Sillitoe on behalf of the Wellcome Sanger Institute COVID-19 Surveillance Team |
| EPI_ISL_2486045 | Lighthouse Lab in Milton Keynes | Wellcome Sanger Institute for the COVID-19 Genomics UK (COG-UK) Consortium | The Lighthouse Lab in Milton Keynes and Alex Alderton, Roberto Amato, Jeffrey Barrett, Sonia Goncalves, Ewan Harrison, David K. Jackson, Ian Johnston, Dominic Kwiatkowski, Cordelia Langford, John Sillitoe on behalf of the Wellcome Sanger Institute COVID-19 Surveillance Team |
| EPI_ISL_2486046 | Lighthouse Lab in Glasgow | Wellcome Sanger Institute for the COVID-19 Genomics UK (COG-UK) Consortium | Harper VanSteenhouse, Yumi Kasai, David Gray, Carol Clugston, Anna Dominiczak and Alex Alderton, Roberto Amato, Jeffrey Barrett, Sonia Goncalves, Ewan Harrison, David K. Jackson, Ian Johnston, Dominic Kwiatkowski, Cordelia Langford, John Sillitoe on behalf of the Wellcome Sanger Institute COVID-19 Surveillance Team |
| EPI_ISL_2486047 | Lighthouse Lab in Alderley Park | Wellcome Sanger Institute for the COVID-19 Genomics UK (COG-UK) Consortium | Jacquelyn Wynn, Mairead Hyland, The Lighthouse Lab in Alderley Park and Alex Alderton, Roberto Amato, Jeffrey Barrett, Sonia Goncalves, Ewan Harrison, David K. Jackson, Ian Johnston, Dominic Kwiatkowski, Cordelia Langford, John Sillitoe on behalf of the Wellcome Sanger Institute COVID-19 Surveillance Team |
| EPI_ISL_2486048, EPI_ISL_2486049 | Lighthouse Lab in Glasgow | Wellcome Sanger Institute for the COVID-19 Genomics UK (COG-UK) Consortium | Harper VanSteenhouse, Yumi Kasai, David Gray, Carol Clugston, Anna Dominiczak and Alex Alderton, Roberto Amato, Jeffrey Barrett, Sonia Goncalves, Ewan Harrison, David K. Jackson, Ian Johnston, Dominic Kwiatkowski, Cordelia Langford, John Sillitoe on behalf of the Wellcome Sanger Institute COVID-19 Surveillance Team |
| EPI_ISL_2486050 | Lighthouse Lab in Alderley Park | Wellcome Sanger Institute for the COVID-19 Genomics UK (COG-UK) Consortium | Jacquelyn Wynn, Mairead Hyland, The Lighthouse Lab in Alderley Park and Alex Alderton, Roberto Amato, Jeffrey Barrett, Sonia Goncalves, Ewan Harrison, David K. Jackson, Ian Johnston, Dominic Kwiatkowski, Cordelia Langford, John Sillitoe on behalf of the Wellcome Sanger Institute COVID-19 Surveillance Team |
| EPI_ISL_2486051 | Lighthouse Lab in Milton Keynes | Wellcome Sanger Institute for the COVID-19 Genomics UK (COG-UK) Consortium | The Lighthouse Lab in Milton Keynes and Alex Alderton, Roberto Amato, Jeffrey Barrett, Sonia Goncalves, Ewan Harrison, David K. Jackson, Ian Johnston, Dominic Kwiatkowski, Cordelia Langford, John Sillitoe on behalf of the Wellcome Sanger Institute COVID-19 Surveillance Team |
| EPI_ISL_2486052, EPI_ISL_2486053 | Lighthouse Lab in Glasgow | Wellcome Sanger Institute for the COVID-19 Genomics UK (COG-UK) Consortium | Harper VanSteenhouse, Yumi Kasai, David Gray, Carol Clugston, Anna Dominiczak and Alex Alderton, Roberto Amato, Jeffrey Barrett, Sonia Goncalves, Ewan Harrison, David K. Jackson, Ian Johnston, Dominic Kwiatkowski, Cordelia Langford, John Sillitoe on behalf of the Wellcome Sanger Institute COVID-19 Surveillance Team |
| EPI_ISL_2486054, EPI_ISL_2486055 | Lighthouse Lab in Alderley Park | Wellcome Sanger Institute for the COVID-19 Genomics UK (COG-UK) Consortium | Jacquelyn Wynn, Mairead Hyland, The Lighthouse Lab in Alderley Park and Alex Alderton, Roberto Amato, Jeffrey Barrett, Sonia Goncalves, Ewan Harrison, David K. Jackson, Ian Johnston, Dominic Kwiatkowski, Cordelia Langford, John Sillitoe on behalf of the Wellcome Sanger Institute COVID-19 Surveillance Team |
| EPI_ISL_2486056 | Lighthouse Lab in Milton Keynes | Wellcome Sanger Institute for the COVID-19 Genomics UK (COG-UK) Consortium | The Lighthouse Lab in Milton Keynes and Alex Alderton, Roberto Amato, Jeffrey Barrett, Sonia Goncalves, Ewan Harrison, David K. Jackson, Ian Johnston, Dominic Kwiatkowski, Cordelia Langford, John Sillitoe on behalf of the Wellcome Sanger Institute COVID-19 Surveillance Team |
| EPI_ISL_2486057 | Lighthouse Lab in Glasgow | Wellcome Sanger Institute for the COVID-19 Genomics UK (COG-UK) Consortium | Harper VanSteenhouse, Yumi Kasai, David Gray, Carol Clugston, Anna Dominiczak and Alex Alderton, Roberto Amato, Jeffrey Barrett, Sonia Goncalves, Ewan Harrison, David K. Jackson, Ian Johnston, Dominic Kwiatkowski, Cordelia Langford, John Sillitoe on behalf of the Wellcome Sanger Institute COVID-19 Surveillance Team |
| EPI_ISL_2486058 | Lighthouse Lab in Milton Keynes | Wellcome Sanger Institute for the COVID-19 Genomics UK (COG-UK) Consortium | The Lighthouse Lab in Milton Keynes and Alex Alderton, Roberto Amato, Jeffrey Barrett, Sonia Goncalves, Ewan Harrison, David K. Jackson, Ian Johnston, Dominic Kwiatkowski, Cordelia Langford, John Sillitoe on behalf of the Wellcome Sanger Institute COVID-19 Surveillance Team |
| EPI_ISL_2486059 | Lighthouse Lab in Glasgow | Wellcome Sanger Institute for the COVID-19 Genomics UK (COG-UK) Consortium | Harper VanSteenhouse, Yumi Kasai, David Gray, Carol Clugston, Anna Dominiczak and Alex Alderton, Roberto Amato, Jeffrey Barrett, Sonia Goncalves, Ewan Harrison, David K. Jackson, Ian Johnston, Dominic Kwiatkowski, Cordelia Langford, John Sillitoe on behalf of the Wellcome Sanger Institute COVID-19 Surveillance Team |
| EPI_ISL_2486060 | Lighthouse Lab in Alderley Park | Wellcome Sanger Institute for the COVID-19 Genomics UK (COG-UK) Consortium | Jacquelyn Wynn, Mairead Hyland, The Lighthouse Lab in Alderley Park and Alex Alderton, Roberto Amato, Jeffrey Barrett, Sonia Goncalves, Ewan Harrison, David K. Jackson, Ian Johnston, Dominic Kwiatkowski, Cordelia Langford, John Sillitoe on behalf of the Wellcome Sanger Institute COVID-19 Surveillance Team |
| EPI_ISL_2486061, EPI_ISL_2486063 | Lighthouse Lab in Glasgow | Wellcome Sanger Institute for the COVID-19 Genomics UK (COG-UK) Consortium | Harper VanSteenhouse, Yumi Kasai, David Gray, Carol Clugston, Anna Dominiczak and Alex Alderton, Roberto Amato, Jeffrey Barrett, Sonia Goncalves, Ewan Harrison, David K. Jackson, Ian Johnston, Dominic Kwiatkowski, Cordelia Langford, John Sillitoe on behalf of the Wellcome Sanger Institute COVID-19 Surveillance Team |
| EPI_ISL_2486065 | Lighthouse Lab in Alderley Park | Wellcome Sanger Institute for the COVID-19 Genomics UK (COG-UK) Consortium | Jacquelyn Wynn, Mairead Hyland, The Lighthouse Lab in Alderley Park and Alex Alderton, Roberto Amato, Jeffrey Barrett, Sonia Goncalves, Ewan Harrison, David K. Jackson, Ian Johnston, Dominic Kwiatkowski, Cordelia Langford, John Sillitoe on behalf of the Wellcome Sanger Institute COVID-19 Surveillance Team |
| EPI_ISL_2486066, EPI_ISL_2486067 | Lighthouse Lab in Glasgow | Wellcome Sanger Institute for the COVID-19 Genomics UK (COG-UK) Consortium | Harper VanSteenhouse, Yumi Kasai, David Gray, Carol Clugston, Anna Dominiczak and Alex Alderton, Roberto Amato, Jeffrey Barrett, Sonia Goncalves, Ewan Harrison, David K. Jackson, Ian Johnston, Dominic Kwiatkowski, Cordelia Langford, John Sillitoe on behalf of the Wellcome Sanger Institute COVID-19 Surveillance Team |
| EPI_ISL_2486068 | Lighthouse Lab in Alderley Park | Wellcome Sanger Institute for the COVID-19 Genomics UK (COG-UK) Consortium | Jacquelyn Wynn, Mairead Hyland, The Lighthouse Lab in Alderley Park and Alex Alderton, Roberto Amato, Jeffrey Barrett, Sonia Goncalves, Ewan Harrison, David K. Jackson, Ian Johnston, Dominic Kwiatkowski, Cordelia Langford, John Sillitoe on behalf of the Wellcome Sanger Institute COVID-19 Surveillance Team |
| EPI_ISL_2486069, EPI_ISL_2486070 | Lighthouse Lab in Glasgow | Wellcome Sanger Institute for the COVID-19 Genomics UK (COG-UK) Consortium | Harper VanSteenhouse, Yumi Kasai, David Gray, Carol Clugston, Anna Dominiczak and Alex Alderton, Roberto Amato, Jeffrey Barrett, Sonia Goncalves, Ewan Harrison, David K. Jackson, Ian Johnston, Dominic Kwiatkowski, Cordelia Langford, John Sillitoe on behalf of the Wellcome Sanger Institute COVID-19 Surveillance Team |
| EPI_ISL_2486071 | Lighthouse Lab in Milton Keynes | Wellcome Sanger Institute for the COVID-19 Genomics UK (COG-UK) Consortium | The Lighthouse Lab in Milton Keynes and Alex Alderton, Roberto Amato, Jeffrey Barrett, Sonia Goncalves, Ewan Harrison, David K. Jackson, Ian Johnston, Dominic Kwiatkowski, Cordelia Langford, John Sillitoe on behalf of the Wellcome Sanger Institute COVID-19 Surveillance Team |
| EPI_ISL_2486072, EPI_ISL_2486073, EPI_ISL_2486075 | Lighthouse Lab in Glasgow | Wellcome Sanger Institute for the COVID-19 Genomics UK (COG-UK) Consortium | Harper VanSteenhouse, Yumi Kasai, David Gray, Carol Clugston, Anna Dominiczak and Alex Alderton, Roberto Amato, Jeffrey Barrett, Sonia Goncalves, Ewan Harrison, David K. Jackson, Ian Johnston, Dominic Kwiatkowski, Cordelia Langford, John Sillitoe on behalf of the Wellcome Sanger Institute COVID-19 Surveillance Team |
| EPI_ISL_2486076 | Lighthouse Lab in Milton Keynes | Wellcome Sanger Institute for the COVID-19 Genomics UK (COG-UK) Consortium | The Lighthouse Lab in Milton Keynes and Alex Alderton, Roberto Amato, Jeffrey Barrett, Sonia Goncalves, Ewan Harrison, David K. Jackson, Ian Johnston, Dominic Kwiatkowski, Cordelia Langford, John Sillitoe on behalf of the Wellcome Sanger Institute COVID-19 Surveillance Team |
| EPI_ISL_2486077 | Lighthouse Lab in Alderley Park | Wellcome Sanger Institute for the COVID-19 Genomics UK (COG-UK) Consortium | Jacquelyn Wynn, Mairead Hyland, The Lighthouse Lab in Alderley Park and Alex Alderton, Roberto Amato, Jeffrey Barrett, Sonia Goncalves, Ewan Harrison, David K. Jackson, Ian Johnston, Dominic Kwiatkowski, Cordelia Langford, John Sillitoe on behalf of the Wellcome Sanger Institute COVID-19 Surveillance Team |
| EPI_ISL_2486078 | Lighthouse Lab in Milton Keynes | Wellcome Sanger Institute for the COVID-19 Genomics UK (COG-UK) Consortium | The Lighthouse Lab in Milton Keynes and Alex Alderton, Roberto Amato, Jeffrey Barrett, Sonia Goncalves, Ewan Harrison, David K. Jackson, Ian Johnston, Dominic Kwiatkowski, Cordelia Langford, John Sillitoe on behalf of the Wellcome Sanger Institute COVID-19 Surveillance Team |
| EPI_ISL_2486079 | Lighthouse Lab in Glasgow | Wellcome Sanger Institute for the COVID-19 Genomics UK (COG-UK) Consortium | Harper VanSteenhouse, Yumi Kasai, David Gray, Carol Clugston, Anna Dominiczak and Alex Alderton, Roberto Amato, Jeffrey Barrett, Sonia Goncalves, Ewan Harrison, David K. Jackson, Ian Johnston, Dominic Kwiatkowski, Cordelia Langford, John Sillitoe on behalf of the Wellcome Sanger Institute COVID-19 Surveillance Team |
| EPI_ISL_2486080 | Lighthouse Lab in Alderley Park | Wellcome Sanger Institute for the COVID-19 Genomics UK (COG-UK) Consortium | Jacquelyn Wynn, Mairead Hyland, The Lighthouse Lab in Alderley Park and Alex Alderton, Roberto Amato, Jeffrey Barrett, Sonia Goncalves, Ewan Harrison, David K. Jackson, Ian Johnston, Dominic Kwiatkowski, Cordelia Langford, John Sillitoe on behalf of the Wellcome Sanger Institute COVID-19 Surveillance Team |
| EPI_ISL_2486082 | Lighthouse Lab in Glasgow | Wellcome Sanger Institute for the COVID-19 Genomics UK (COG-UK) Consortium | Harper VanSteenhouse, Yumi Kasai, David Gray, Carol Clugston, Anna Dominiczak and Alex Alderton, Roberto Amato, Jeffrey Barrett, Sonia Goncalves, Ewan Harrison, David K. Jackson, Ian Johnston, Dominic Kwiatkowski, Cordelia Langford, John Sillitoe on behalf of the Wellcome Sanger Institute COVID-19 Surveillance Team |
| EPI_ISL_2486083 | Lighthouse Lab in Milton Keynes | Wellcome Sanger Institute for the COVID-19 Genomics UK (COG-UK) Consortium | The Lighthouse Lab in Milton Keynes and Alex Alderton, Roberto Amato, Jeffrey Barrett, Sonia Goncalves, Ewan Harrison, David K. Jackson, Ian Johnston, Dominic Kwiatkowski, Cordelia Langford, John Sillitoe on behalf of the Wellcome Sanger Institute COVID-19 Surveillance Team |
| EPI_ISL_2486084, EPI_ISL_2486085 | Lighthouse Lab in Alderley Park | Wellcome Sanger Institute for the COVID-19 Genomics UK (COG-UK) Consortium | Jacquelyn Wynn, Mairead Hyland, The Lighthouse Lab in Alderley Park and Alex Alderton, Roberto Amato, Jeffrey Barrett, Sonia Goncalves, Ewan Harrison, David K. Jackson, Ian Johnston, Dominic Kwiatkowski, Cordelia Langford, John Sillitoe on behalf of the Wellcome Sanger Institute COVID-19 Surveillance Team |
| EPI_ISL_2486087 | Lighthouse Lab in Milton Keynes | Wellcome Sanger Institute for the COVID-19 Genomics UK (COG-UK) Consortium | The Lighthouse Lab in Milton Keynes and Alex Alderton, Roberto Amato, Jeffrey Barrett, Sonia Goncalves, Ewan Harrison, David K. Jackson, Ian Johnston, Dominic Kwiatkowski, Cordelia Langford, John Sillitoe on behalf of the Wellcome Sanger Institute COVID-19 Surveillance Team |
| EPI_ISL_2486089, EPI_ISL_2486090, EPI_ISL_2486091, EPI_ISL_2486092 | Lighthouse Lab in Alderley Park | Wellcome Sanger Institute for the COVID-19 Genomics UK (COG-UK) Consortium | Jacquelyn Wynn, Mairead Hyland, The Lighthouse Lab in Alderley Park and Alex Alderton, Roberto Amato, Jeffrey Barrett, Sonia Goncalves, Ewan Harrison, David K. Jackson, Ian Johnston, Dominic Kwiatkowski, Cordelia Langford, John Sillitoe on behalf of the Wellcome Sanger Institute COVID-19 Surveillance Team |
| EPI_ISL_2486093 | Lighthouse Lab in Milton Keynes | Wellcome Sanger Institute for the COVID-19 Genomics UK (COG-UK) Consortium | The Lighthouse Lab in Milton Keynes and Alex Alderton, Roberto Amato, Jeffrey Barrett, Sonia Goncalves, Ewan Harrison, David K. Jackson, Ian Johnston, Dominic Kwiatkowski, Cordelia Langford, John Sillitoe on behalf of the Wellcome Sanger Institute COVID-19 Surveillance Team |
| EPI_ISL_2486094 | Lighthouse Lab in Glasgow | Wellcome Sanger Institute for the COVID-19 Genomics UK (COG-UK) Consortium | Harper VanSteenhouse, Yumi Kasai, David Gray, Carol Clugston, Anna Dominiczak and Alex Alderton, Roberto Amato, Jeffrey Barrett, Sonia Goncalves, Ewan Harrison, David K. Jackson, Ian Johnston, Dominic Kwiatkowski, Cordelia Langford, John Sillitoe on behalf of the Wellcome Sanger Institute COVID-19 Surveillance Team |
| EPI_ISL_2486095, EPI_ISL_2486096, EPI_ISL_2486097, EPI_ISL_2486098, EPI_ISL_2486099, EPI_ISL_2486100, EPI_ISL_2486101, EPI_ISL_2486102 | Lighthouse Lab in Alderley Park | Wellcome Sanger Institute for the COVID-19 Genomics UK (COG-UK) Consortium | Jacquelyn Wynn, Mairead Hyland, The Lighthouse Lab in Alderley Park and Alex Alderton, Roberto Amato, Jeffrey Barrett, Sonia Goncalves, Ewan Harrison, David K. Jackson, Ian Johnston, Dominic Kwiatkowski, Cordelia Langford, John Sillitoe on behalf of the Wellcome Sanger Institute COVID-19 Surveillance Team |
| EPI_ISL_2486103 | Lighthouse Lab in Glasgow | Wellcome Sanger Institute for the COVID-19 Genomics UK (COG-UK) Consortium | Harper VanSteenhouse, Yumi Kasai, David Gray, Carol Clugston, Anna Dominiczak and Alex Alderton, Roberto Amato, Jeffrey Barrett, Sonia Goncalves, Ewan Harrison, David K. Jackson, Ian Johnston, Dominic Kwiatkowski, Cordelia Langford, John Sillitoe on behalf of the Wellcome Sanger Institute COVID-19 Surveillance Team |
| EPI_ISL_2486105 | Lighthouse Lab in Alderley Park | Wellcome Sanger Institute for the COVID-19 Genomics UK (COG-UK) Consortium | Jacquelyn Wynn, Mairead Hyland, The Lighthouse Lab in Alderley Park and Alex Alderton, Roberto Amato, Jeffrey Barrett, Sonia Goncalves, Ewan Harrison, David K. Jackson, Ian Johnston, Dominic Kwiatkowski, Cordelia Langford, John Sillitoe on behalf of the Wellcome Sanger Institute COVID-19 Surveillance Team |
| EPI_ISL_2486107 | Lighthouse Lab in Milton Keynes | Wellcome Sanger Institute for the COVID-19 Genomics UK (COG-UK) Consortium | The Lighthouse Lab in Milton Keynes and Alex Alderton, Roberto Amato, Jeffrey Barrett, Sonia Goncalves, Ewan Harrison, David K. Jackson, Ian Johnston, Dominic Kwiatkowski, Cordelia Langford, John Sillitoe on behalf of the Wellcome Sanger Institute COVID-19 Surveillance Team |
| EPI_ISL_2486108 | Lighthouse Lab in Glasgow | Wellcome Sanger Institute for the COVID-19 Genomics UK (COG-UK) Consortium | Harper VanSteenhouse, Yumi Kasai, David Gray, Carol Clugston, Anna Dominiczak and Alex Alderton, Roberto Amato, Jeffrey Barrett, Sonia Goncalves, Ewan Harrison, David K. Jackson, Ian Johnston, Dominic Kwiatkowski, Cordelia Langford, John Sillitoe on behalf of the Wellcome Sanger Institute COVID-19 Surveillance Team |
| EPI_ISL_2486109 | Lighthouse Lab in Milton Keynes | Wellcome Sanger Institute for the COVID-19 Genomics UK (COG-UK) Consortium | The Lighthouse Lab in Milton Keynes and Alex Alderton, Roberto Amato, Jeffrey Barrett, Sonia Goncalves, Ewan Harrison, David K. Jackson, Ian Johnston, Dominic Kwiatkowski, Cordelia Langford, John Sillitoe on behalf of the Wellcome Sanger Institute COVID-19 Surveillance Team |
| EPI_ISL_2486110 | Lighthouse Lab in Alderley Park | Wellcome Sanger Institute for the COVID-19 Genomics UK (COG-UK) Consortium | Jacquelyn Wynn, Mairead Hyland, The Lighthouse Lab in Alderley Park and Alex Alderton, Roberto Amato, Jeffrey Barrett, Sonia Goncalves, Ewan Harrison, David K. Jackson, Ian Johnston, Dominic Kwiatkowski, Cordelia Langford, John Sillitoe on behalf of the Wellcome Sanger Institute COVID-19 Surveillance Team |
| EPI_ISL_2486111, EPI_ISL_2486112 | Lighthouse Lab in Glasgow | Wellcome Sanger Institute for the COVID-19 Genomics UK (COG-UK) Consortium | Harper VanSteenhouse, Yumi Kasai, David Gray, Carol Clugston, Anna Dominiczak and Alex Alderton, Roberto Amato, Jeffrey Barrett, Sonia Goncalves, Ewan Harrison, David K. Jackson, Ian Johnston, Dominic Kwiatkowski, Cordelia Langford, John Sillitoe on behalf of the Wellcome Sanger Institute COVID-19 Surveillance Team |

| | | | |
|---|---|---|---|
| EPI_ISL_2486113 | Lighthouse Lab in Milton Keynes | Wellcome Sanger Institute for the COVID-19 Genomics UK (COG-UK) Consortium | The Lighthouse Lab in Milton Keynes and Alex Alderton, Roberto Amato, Jeffrey Barrett, Sonia Goncalves, Ewan Harrison, David K. Jackson, Ian Johnston, Dominic Kwiatkowski, Cordelia Langford, John Sillitoe on behalf of the Wellcome Sanger Institute COVID-19 Surveillance Team |
| EPI_ISL_2486114, EPI_ISL_2486115, EPI_ISL_2486116, EPI_ISL_2486117, EPI_ISL_2486118 | Lighthouse Lab in Glasgow | Wellcome Sanger Institute for the COVID-19 Genomics UK (COG-UK) Consortium | Harper VanSteenhouse, Yumi Kasai, David Gray, Carol Clugston, Anna Dominiczak and Alex Alderton, Roberto Amato, Jeffrey Barrett, Sonia Goncalves, Ewan Harrison, David K. Jackson, Ian Johnston, Dominic Kwiatkowski, Cordelia Langford, John Sillitoe on behalf of the Wellcome Sanger Institute COVID-19 Surveillance Team |
| EPI_ISL_2486119 | Lighthouse Lab in Milton Keynes | Wellcome Sanger Institute for the COVID-19 Genomics UK (COG-UK) Consortium | The Lighthouse Lab in Milton Keynes and Alex Alderton, Roberto Amato, Jeffrey Barrett, Sonia Goncalves, Ewan Harrison, David K. Jackson, Ian Johnston, Dominic Kwiatkowski, Cordelia Langford, John Sillitoe on behalf of the Wellcome Sanger Institute COVID-19 Surveillance Team |
| EPI_ISL_2486120, EPI_ISL_2486121 | Lighthouse Lab in Alderley Park | Wellcome Sanger Institute for the COVID-19 Genomics UK (COG-UK) Consortium | Jacquelyn Wynn, Mairead Hyland, The Lighthouse Lab in Alderley Park and Alex Alderton, Roberto Amato, Jeffrey Barrett, Sonia Goncalves, Ewan Harrison, David K. Jackson, Ian Johnston, Dominic Kwiatkowski, Cordelia Langford, John Sillitoe on behalf of the Wellcome Sanger Institute COVID-19 Surveillance Team |
| EPI_ISL_2486122, EPI_ISL_2486123 | Lighthouse Lab in Glasgow | Wellcome Sanger Institute for the COVID-19 Genomics UK (COG-UK) Consortium | Harper VanSteenhouse, Yumi Kasai, David Gray, Carol Clugston, Anna Dominiczak and Alex Alderton, Roberto Amato, Jeffrey Barrett, Sonia Goncalves, Ewan Harrison, David K. Jackson, Ian Johnston, Dominic Kwiatkowski, Cordelia Langford, John Sillitoe on behalf of the Wellcome Sanger Institute COVID-19 Surveillance Team |
| EPI_ISL_2486124 | Lighthouse Lab in Milton Keynes | Wellcome Sanger Institute for the COVID-19 Genomics UK (COG-UK) Consortium | The Lighthouse Lab in Milton Keynes and Alex Alderton, Roberto Amato, Jeffrey Barrett, Sonia Goncalves, Ewan Harrison, David K. Jackson, Ian Johnston, Dominic Kwiatkowski, Cordelia Langford, John Sillitoe on behalf of the Wellcome Sanger Institute COVID-19 Surveillance Team |
| EPI_ISL_2486125 | Lighthouse Lab in Glasgow | Wellcome Sanger Institute for the COVID-19 Genomics UK (COG-UK) Consortium | Harper VanSteenhouse, Yumi Kasai, David Gray, Carol Clugston, Anna Dominiczak and Alex Alderton, Roberto Amato, Jeffrey Barrett, Sonia Goncalves, Ewan Harrison, David K. Jackson, Ian Johnston, Dominic Kwiatkowski, Cordelia Langford, John Sillitoe on behalf of the Wellcome Sanger Institute COVID-19 Surveillance Team |
| EPI_ISL_2486126 | Lighthouse Lab in Alderley Park | Wellcome Sanger Institute for the COVID-19 Genomics UK (COG-UK) Consortium | Jacquelyn Wynn, Mairead Hyland, The Lighthouse Lab in Alderley Park and Alex Alderton, Roberto Amato, Jeffrey Barrett, Sonia Goncalves, Ewan Harrison, David K. Jackson, Ian Johnston, Dominic Kwiatkowski, Cordelia Langford, John Sillitoe on behalf of the Wellcome Sanger Institute COVID-19 Surveillance Team |
| EPI_ISL_2486127, EPI_ISL_2486128 | Lighthouse Lab in Milton Keynes | Wellcome Sanger Institute for the COVID-19 Genomics UK (COG-UK) Consortium | The Lighthouse Lab in Milton Keynes and Alex Alderton, Roberto Amato, Jeffrey Barrett, Sonia Goncalves, Ewan Harrison, David K. Jackson, Ian Johnston, Dominic Kwiatkowski, Cordelia Langford, John Sillitoe on behalf of the Wellcome Sanger Institute COVID-19 Surveillance Team |
| EPI_ISL_2486129 | Lighthouse Lab in Glasgow | Wellcome Sanger Institute for the COVID-19 Genomics UK (COG-UK) Consortium | Harper VanSteenhouse, Yumi Kasai, David Gray, Carol Clugston, Anna Dominiczak and Alex Alderton, Roberto Amato, Jeffrey Barrett, Sonia Goncalves, Ewan Harrison, David K. Jackson, Ian Johnston, Dominic Kwiatkowski, Cordelia Langford, John Sillitoe on behalf of the Wellcome Sanger Institute COVID-19 Surveillance Team |
| EPI_ISL_2486130 | Lighthouse Lab in Alderley Park | Wellcome Sanger Institute for the COVID-19 Genomics UK (COG-UK) Consortium | Jacquelyn Wynn, Mairead Hyland, The Lighthouse Lab in Alderley Park and Alex Alderton, Roberto Amato, Jeffrey Barrett, Sonia Goncalves, Ewan Harrison, David K. Jackson, Ian Johnston, Dominic Kwiatkowski, Cordelia Langford, John Sillitoe on behalf of the Wellcome Sanger Institute COVID-19 Surveillance Team |
| EPI_ISL_2486131 | Lighthouse Lab in Glasgow | Wellcome Sanger Institute for the COVID-19 Genomics UK (COG-UK) Consortium | Harper VanSteenhouse, Yumi Kasai, David Gray, Carol Clugston, Anna Dominiczak and Alex Alderton, Roberto Amato, Jeffrey Barrett, Sonia Goncalves, Ewan Harrison, David K. Jackson, Ian Johnston, Dominic Kwiatkowski, Cordelia Langford, John Sillitoe on behalf of the Wellcome Sanger Institute COVID-19 Surveillance Team |
| EPI_ISL_2486132 | Lighthouse Lab in Milton Keynes | Wellcome Sanger Institute for the COVID-19 Genomics UK (COG-UK) Consortium | The Lighthouse Lab in Milton Keynes and Alex Alderton, Roberto Amato, Jeffrey Barrett, Sonia Goncalves, Ewan Harrison, David K. Jackson, Ian Johnston, Dominic Kwiatkowski, Cordelia Langford, John Sillitoe on behalf of the Wellcome Sanger Institute COVID-19 Surveillance Team |
| EPI_ISL_2486133, EPI_ISL_2486134 | Lighthouse Lab in Glasgow | Wellcome Sanger Institute for the COVID-19 Genomics UK (COG-UK) Consortium | Harper VanSteenhouse, Yumi Kasai, David Gray, Carol Clugston, Anna Dominiczak and Alex Alderton, Roberto Amato, Jeffrey Barrett, Sonia Goncalves, Ewan Harrison, David K. Jackson, Ian Johnston, Dominic Kwiatkowski, Cordelia Langford, John Sillitoe on behalf of the Wellcome Sanger Institute COVID-19 Surveillance Team |
| EPI_ISL_2486135, EPI_ISL_2486136, EPI_ISL_2486137 | Lighthouse Lab in Milton Keynes | Wellcome Sanger Institute for the COVID-19 Genomics UK (COG-UK) Consortium | The Lighthouse Lab in Milton Keynes and Alex Alderton, Roberto Amato, Jeffrey Barrett, Sonia Goncalves, Ewan Harrison, David K. Jackson, Ian Johnston, Dominic Kwiatkowski, Cordelia Langford, John Sillitoe on behalf of the Wellcome Sanger Institute COVID-19 Surveillance Team |
| EPI_ISL_2486138, EPI_ISL_2486139 | Lighthouse Lab in Glasgow | Wellcome Sanger Institute for the COVID-19 Genomics UK (COG-UK) Consortium | Harper VanSteenhouse, Yumi Kasai, David Gray, Carol Clugston, Anna Dominiczak and Alex Alderton, Roberto Amato, Jeffrey Barrett, Sonia Goncalves, Ewan Harrison, David K. Jackson, Ian Johnston, Dominic Kwiatkowski, Cordelia Langford, John Sillitoe on behalf of the Wellcome Sanger Institute COVID-19 Surveillance Team |
| EPI_ISL_2486140 | Lighthouse Lab in Alderley Park | Wellcome Sanger Institute for the COVID-19 Genomics UK (COG-UK) Consortium | Jacquelyn Wynn, Mairead Hyland, The Lighthouse Lab in Alderley Park and Alex Alderton, Roberto Amato, Jeffrey Barrett, Sonia Goncalves, Ewan Harrison, David K. Jackson, Ian Johnston, Dominic Kwiatkowski, Cordelia Langford, John Sillitoe on behalf of the Wellcome Sanger Institute COVID-19 Surveillance Team |
| EPI_ISL_2486142 | Lighthouse Lab in Milton Keynes | Wellcome Sanger Institute for the COVID-19 Genomics UK (COG-UK) Consortium | The Lighthouse Lab in Milton Keynes and Alex Alderton, Roberto Amato, Jeffrey Barrett, Sonia Goncalves, Ewan Harrison, David K. Jackson, Ian Johnston, Dominic Kwiatkowski, Cordelia Langford, John Sillitoe on behalf of the Wellcome Sanger Institute COVID-19 Surveillance Team |
| EPI_ISL_2486143 | Lighthouse Lab in Glasgow | Wellcome Sanger Institute for the COVID-19 Genomics UK (COG-UK) Consortium | Harper VanSteenhouse, Yumi Kasai, David Gray, Carol Clugston, Anna Dominiczak and Alex Alderton, Roberto Amato, Jeffrey Barrett, Sonia Goncalves, Ewan Harrison, David K. Jackson, Ian Johnston, Dominic Kwiatkowski, Cordelia Langford, John Sillitoe on behalf of the Wellcome Sanger Institute COVID-19 Surveillance Team |
| EPI_ISL_2486144, EPI_ISL_2486146 | Lighthouse Lab in Alderley Park | Wellcome Sanger Institute for the COVID-19 Genomics UK (COG-UK) Consortium | Jacquelyn Wynn, Mairead Hyland, The Lighthouse Lab in Alderley Park and Alex Alderton, Roberto Amato, Jeffrey Barrett, Sonia Goncalves, Ewan Harrison, David K. Jackson, Ian Johnston, Dominic Kwiatkowski, Cordelia Langford, John Sillitoe on behalf of the Wellcome Sanger Institute COVID-19 Surveillance Team |
| EPI_ISL_2486147 | Lighthouse Lab in Milton Keynes | Wellcome Sanger Institute for the COVID-19 Genomics UK (COG-UK) Consortium | The Lighthouse Lab in Milton Keynes and Alex Alderton, Roberto Amato, Jeffrey Barrett, Sonia Goncalves, Ewan Harrison, David K. Jackson, Ian Johnston, Dominic Kwiatkowski, Cordelia Langford, John Sillitoe on behalf of the Wellcome Sanger Institute COVID-19 Surveillance Team |
| EPI_ISL_2486148, EPI_ISL_2486149 | Lighthouse Lab in Glasgow | Wellcome Sanger Institute for the COVID-19 Genomics UK (COG-UK) Consortium | Harper VanSteenhouse, Yumi Kasai, David Gray, Carol Clugston, Anna Dominiczak and Alex Alderton, Roberto Amato, Jeffrey Barrett, Sonia Goncalves, Ewan Harrison, David K. Jackson, Ian Johnston, Dominic Kwiatkowski, Cordelia Langford, John Sillitoe on behalf of the Wellcome Sanger Institute COVID-19 Surveillance Team |
| EPI_ISL_2486150 | Lighthouse Lab in Alderley Park | Wellcome Sanger Institute for the COVID-19 Genomics UK (COG-UK) Consortium | Jacquelyn Wynn, Mairead Hyland, The Lighthouse Lab in Alderley Park and Alex Alderton, Roberto Amato, Jeffrey Barrett, Sonia Goncalves, Ewan Harrison, David K. Jackson, Ian Johnston, Dominic Kwiatkowski, Cordelia Langford, John Sillitoe on behalf of the Wellcome Sanger Institute COVID-19 Surveillance Team |
| EPI_ISL_2486152, EPI_ISL_2486153 | Lighthouse Lab in Glasgow | Wellcome Sanger Institute for the COVID-19 Genomics UK (COG-UK) Consortium | Harper VanSteenhouse, Yumi Kasai, David Gray, Carol Clugston, Anna Dominiczak and Alex Alderton, Roberto Amato, Jeffrey Barrett, Sonia Goncalves, Ewan Harrison, David K. Jackson, Ian Johnston, Dominic Kwiatkowski, Cordelia Langford, John Sillitoe on behalf of the Wellcome Sanger Institute COVID-19 Surveillance Team |
| EPI_ISL_2486154 | Lighthouse Lab in Alderley Park | Wellcome Sanger Institute for the COVID-19 Genomics UK (COG-UK) Consortium | Jacquelyn Wynn, Mairead Hyland, The Lighthouse Lab in Alderley Park and Alex Alderton, Roberto Amato, Jeffrey Barrett, Sonia Goncalves, Ewan Harrison, David K. Jackson, Ian Johnston, Dominic Kwiatkowski, Cordelia Langford, John Sillitoe on behalf of the Wellcome Sanger Institute COVID-19 Surveillance Team |
| EPI_ISL_2486155, EPI_ISL_2486156 | Lighthouse Lab in Glasgow | Wellcome Sanger Institute for the COVID-19 Genomics UK (COG-UK) Consortium | Harper VanSteenhouse, Yumi Kasai, David Gray, Carol Clugston, Anna Dominiczak and Alex Alderton, Roberto Amato, Jeffrey Barrett, Sonia Goncalves, Ewan Harrison, David K. Jackson, Ian Johnston, Dominic Kwiatkowski, Cordelia Langford, John Sillitoe on behalf of the Wellcome Sanger Institute COVID-19 Surveillance Team |
| EPI_ISL_2486157 | Lighthouse Lab in Alderley Park | Wellcome Sanger Institute for the COVID-19 Genomics UK (COG-UK) Consortium | Jacquelyn Wynn, Mairead Hyland, The Lighthouse Lab in Alderley Park and Alex Alderton, Roberto Amato, Jeffrey Barrett, Sonia Goncalves, Ewan Harrison, David K. Jackson, Ian Johnston, Dominic Kwiatkowski, Cordelia Langford, John Sillitoe on behalf of the Wellcome Sanger Institute COVID-19 Surveillance Team |
| EPI_ISL_2486158 | Lighthouse Lab in Glasgow | Wellcome Sanger Institute for the COVID-19 Genomics UK (COG-UK) Consortium | Harper VanSteenhouse, Yumi Kasai, David Gray, Carol Clugston, Anna Dominiczak and Alex Alderton, Roberto Amato, Jeffrey Barrett, Sonia Goncalves, Ewan Harrison, David K. Jackson, Ian Johnston, Dominic Kwiatkowski, Cordelia Langford, John Sillitoe on behalf of the Wellcome Sanger Institute COVID-19 Surveillance Team |
| EPI_ISL_2486159 | Lighthouse Lab in Milton Keynes | Wellcome Sanger Institute for the COVID-19 Genomics UK (COG-UK) Consortium | The Lighthouse Lab in Milton Keynes and Alex Alderton, Roberto Amato, Jeffrey Barrett, Sonia Goncalves, Ewan Harrison, David K. Jackson, Ian Johnston, Dominic Kwiatkowski, Cordelia Langford, John Sillitoe on behalf of the Wellcome Sanger Institute COVID-19 Surveillance Team |
| EPI_ISL_2486160 | Lighthouse Lab in Glasgow | Wellcome Sanger Institute for the COVID-19 Genomics UK (COG-UK) Consortium | Harper VanSteenhouse, Yumi Kasai, David Gray, Carol Clugston, Anna Dominiczak and Alex Alderton, Roberto Amato, Jeffrey Barrett, Sonia Goncalves, Ewan Harrison, David K. Jackson, Ian Johnston, Dominic Kwiatkowski, Cordelia Langford, John Sillitoe on behalf of the Wellcome Sanger Institute COVID-19 Surveillance Team |
| EPI_ISL_2486161, EPI_ISL_2486162, EPI_ISL_2486163 | Lighthouse Lab in Alderley Park | Wellcome Sanger Institute for the COVID-19 Genomics UK (COG-UK) Consortium | Jacquelyn Wynn, Mairead Hyland, The Lighthouse Lab in Alderley Park and Alex Alderton, Roberto Amato, Jeffrey Barrett, Sonia Goncalves, Ewan Harrison, David K. Jackson, Ian Johnston, Dominic Kwiatkowski, Cordelia Langford, John Sillitoe on behalf of the Wellcome Sanger Institute COVID-19 Surveillance Team |
| EPI_ISL_2486165, EPI_ISL_2486166, EPI_ISL_2486167, EPI_ISL_2486169 | Lighthouse Lab in Glasgow | Wellcome Sanger Institute for the COVID-19 Genomics UK (COG-UK) Consortium | Harper VanSteenhouse, Yumi Kasai, David Gray, Carol Clugston, Anna Dominiczak and Alex Alderton, Roberto Amato, Jeffrey Barrett, Sonia Goncalves, Ewan Harrison, David K. Jackson, Ian Johnston, Dominic Kwiatkowski, Cordelia Langford, John Sillitoe on behalf of the Wellcome Sanger Institute COVID-19 Surveillance Team |
| EPI_ISL_2486170 | Lighthouse Lab in Milton Keynes | Wellcome Sanger Institute for the COVID-19 Genomics UK (COG-UK) Consortium | The Lighthouse Lab in Milton Keynes and Alex Alderton, Roberto Amato, Jeffrey Barrett, Sonia Goncalves, Ewan Harrison, David K. Jackson, Ian Johnston, Dominic Kwiatkowski, Cordelia Langford, John Sillitoe on behalf of the Wellcome Sanger Institute COVID-19 Surveillance Team |
| EPI_ISL_2486171 | Lighthouse Lab in Alderley Park | Wellcome Sanger Institute for the COVID-19 Genomics UK (COG-UK) Consortium | Jacquelyn Wynn, Mairead Hyland, The Lighthouse Lab in Alderley Park and Alex Alderton, Roberto Amato, Jeffrey Barrett, Sonia Goncalves, Ewan Harrison, David K. Jackson, Ian Johnston, Dominic Kwiatkowski, Cordelia Langford, John Sillitoe on behalf of the Wellcome Sanger Institute COVID-19 Surveillance Team |
| EPI_ISL_2486173 | Lighthouse Lab in Milton Keynes | Wellcome Sanger Institute for the COVID-19 Genomics UK (COG-UK) Consortium | The Lighthouse Lab in Milton Keynes and Alex Alderton, Roberto Amato, Jeffrey Barrett, Sonia Goncalves, Ewan Harrison, David K. Jackson, Ian Johnston, Dominic Kwiatkowski, Cordelia Langford, John Sillitoe on behalf of the Wellcome Sanger Institute COVID-19 Surveillance Team |
| EPI_ISL_2486174 | Lighthouse Lab in Alderley Park | Wellcome Sanger Institute for the COVID-19 Genomics UK (COG-UK) Consortium | Jacquelyn Wynn, Mairead Hyland, The Lighthouse Lab in Alderley Park and Alex Alderton, Roberto Amato, Jeffrey Barrett, Sonia Goncalves, Ewan Harrison, David K. Jackson, Ian Johnston, Dominic Kwiatkowski, Cordelia Langford, John Sillitoe on behalf of the Wellcome Sanger Institute COVID-19 Surveillance Team |
| EPI_ISL_2486175 | Lighthouse Lab in Glasgow | Wellcome Sanger Institute for the COVID-19 Genomics UK (COG-UK) Consortium | Harper VanSteenhouse, Yumi Kasai, David Gray, Carol Clugston, Anna Dominiczak and Alex Alderton, Roberto Amato, Jeffrey Barrett, Sonia Goncalves, Ewan Harrison, David K. Jackson, Ian Johnston, Dominic Kwiatkowski, Cordelia Langford, John Sillitoe on behalf of the Wellcome Sanger Institute COVID-19 Surveillance Team |
| EPI_ISL_2486176 | Lighthouse Lab in Milton Keynes | Wellcome Sanger Institute for the COVID-19 Genomics UK (COG-UK) Consortium | The Lighthouse Lab in Milton Keynes and Alex Alderton, Roberto Amato, Jeffrey Barrett, Sonia Goncalves, Ewan Harrison, David K. Jackson, Ian Johnston, Dominic Kwiatkowski, Cordelia Langford, John Sillitoe on behalf of the Wellcome Sanger Institute COVID-19 Surveillance Team |
| EPI_ISL_2486177, EPI_ISL_2486178, EPI_ISL_2486179 | Lighthouse Lab in Glasgow | Wellcome Sanger Institute for the COVID-19 Genomics UK (COG-UK) Consortium | Harper VanSteenhouse, Yumi Kasai, David Gray, Carol Clugston, Anna Dominiczak and Alex Alderton, Roberto Amato, Jeffrey Barrett, Sonia Goncalves, Ewan Harrison, David K. Jackson, Ian Johnston, Dominic Kwiatkowski, Cordelia Langford, John Sillitoe on behalf of the Wellcome Sanger Institute COVID-19 Surveillance Team |
| EPI_ISL_2486180 | Lighthouse Lab in Alderley Park | Wellcome Sanger Institute for the COVID-19 Genomics UK (COG-UK) Consortium | Jacquelyn Wynn, Mairead Hyland, The Lighthouse Lab in Alderley Park and Alex Alderton, Roberto Amato, Jeffrey Barrett, Sonia Goncalves, Ewan Harrison, David K. Jackson, Ian Johnston, Dominic Kwiatkowski, Cordelia Langford, John Sillitoe on behalf of the Wellcome Sanger Institute COVID-19 Surveillance Team |
| EPI_ISL_2486181 | Lighthouse Lab in Milton Keynes | Wellcome Sanger Institute for the COVID-19 Genomics UK (COG-UK) Consortium | The Lighthouse Lab in Milton Keynes and Alex Alderton, Roberto Amato, Jeffrey Barrett, Sonia Goncalves, Ewan Harrison, David K. Jackson, Ian Johnston, Dominic Kwiatkowski, Cordelia Langford, John Sillitoe on behalf of the Wellcome Sanger Institute COVID-19 Surveillance Team |
| EPI_ISL_2486182 | Lighthouse Lab in Glasgow | Wellcome Sanger Institute for the COVID-19 Genomics UK (COG-UK) Consortium | Harper VanSteenhouse, Yumi Kasai, David Gray, Carol Clugston, Anna Dominiczak and Alex Alderton, Roberto Amato, Jeffrey Barrett, Sonia Goncalves, Ewan Harrison, David K. Jackson, Ian Johnston, Dominic Kwiatkowski, Cordelia Langford, John Sillitoe on behalf of the Wellcome Sanger Institute COVID-19 Surveillance Team |

| | | | |
|---|---|---|---|
| EPI_ISL_2486183 | Lighthouse Lab in Alderley Park | Wellcome Sanger Institute for the COVID-19 Genomics UK (COG-UK) Consortium | Jacquelyn Wynn, Mairead Hyland, The Lighthouse Lab in Alderley Park and Alex Alderton, Roberto Amato, Jeffrey Barrett, Sonia Goncalves, Ewan Harrison, David K. Jackson, Ian Johnston, Dominic Kwiatkowski, Cordelia Langford, John Sillitoe on behalf of the Wellcome Sanger Institute COVID-19 Surveillance Team |
| EPI_ISL_2486184 | Lighthouse Lab in Glasgow | Wellcome Sanger Institute for the COVID-19 Genomics UK (COG-UK) Consortium | Harper VanSteenhouse, Yumi Kasai, David Gray, Carol Clugston, Anna Dominiczak and Alex Alderton, Roberto Amato, Jeffrey Barrett, Sonia Goncalves, Ewan Harrison, David K. Jackson, Ian Johnston, Dominic Kwiatkowski, Cordelia Langford, John Sillitoe on behalf of the Wellcome Sanger Institute COVID-19 Surveillance Team |
| EPI_ISL_2486185, EPI_ISL_2486186 | Lighthouse Lab in Milton Keynes | Wellcome Sanger Institute for the COVID-19 Genomics UK (COG-UK) Consortium | The Lighthouse Lab in Milton Keynes and Alex Alderton, Roberto Amato, Jeffrey Barrett, Sonia Goncalves, Ewan Harrison, David K. Jackson, Ian Johnston, Dominic Kwiatkowski, Cordelia Langford, John Sillitoe on behalf of the Wellcome Sanger Institute COVID-19 Surveillance Team |
| EPI_ISL_2486187, EPI_ISL_2486190 | Lighthouse Lab in Glasgow | Wellcome Sanger Institute for the COVID-19 Genomics UK (COG-UK) Consortium | Harper VanSteenhouse, Yumi Kasai, David Gray, Carol Clugston, Anna Dominiczak and Alex Alderton, Roberto Amato, Jeffrey Barrett, Sonia Goncalves, Ewan Harrison, David K. Jackson, Ian Johnston, Dominic Kwiatkowski, Cordelia Langford, John Sillitoe on behalf of the Wellcome Sanger Institute COVID-19 Surveillance Team |
| EPI_ISL_2486191 | Lighthouse Lab in Milton Keynes | Wellcome Sanger Institute for the COVID-19 Genomics UK (COG-UK) Consortium | The Lighthouse Lab in Milton Keynes and Alex Alderton, Roberto Amato, Jeffrey Barrett, Sonia Goncalves, Ewan Harrison, David K. Jackson, Ian Johnston, Dominic Kwiatkowski, Cordelia Langford, John Sillitoe on behalf of the Wellcome Sanger Institute COVID-19 Surveillance Team |
| EPI_ISL_2486192, EPI_ISL_2486193, EPI_ISL_2486195, EPI_ISL_2486196, EPI_ISL_2486197, EPI_ISL_2486198, EPI_ISL_2486199, EPI_ISL_2486200, EPI_ISL_2486201, EPI_ISL_2486202, EPI_ISL_2486203, EPI_ISL_2486204, EPI_ISL_2486205, EPI_ISL_2486206, EPI_ISL_2486207, EPI_ISL_2486208, EPI_ISL_2486209, EPI_ISL_2486210, EPI_ISL_2486211, EPI_ISL_2486212, EPI_ISL_2486213, EPI_ISL_2486214, EPI_ISL_2486215, EPI_ISL_2486216, EPI_ISL_2486219, EPI_ISL_2486220, EPI_ISL_2486221, EPI_ISL_2486222, EPI_ISL_2486223, EPI_ISL_2486224, EPI_ISL_2486225, EPI_ISL_2486226, EPI_ISL_2486227, EPI_ISL_2486228, EPI_ISL_2486229, EPI_ISL_2486230, EPI_ISL_2486231, EPI_ISL_2486233, EPI_ISL_2486235, EPI_ISL_2486236, EPI_ISL_2486237, EPI_ISL_2486238, EPI_ISL_2486240, EPI_ISL_2486241, EPI_ISL_2486242, EPI_ISL_2486243, EPI_ISL_2486244, EPI_ISL_2486246, EPI_ISL_2486247, EPI_ISL_2486248, EPI_ISL_2486249, EPI_ISL_2486250, EPI_ISL_2486251, EPI_ISL_2486252, EPI_ISL_2486253, EPI_ISL_2486256, EPI_ISL_2486257, EPI_ISL_2486259, EPI_ISL_2486260, EPI_ISL_2486261, EPI_ISL_2486262, EPI_ISL_2486263, EPI_ISL_2486264, EPI_ISL_2486265, EPI_ISL_2486267, EPI_ISL_2486268, EPI_ISL_2486270, EPI_ISL_2486271, EPI_ISL_2486272, EPI_ISL_2486273, EPI_ISL_2486274, EPI_ISL_2486275, EPI_ISL_2486276, EPI_ISL_2486277, EPI_ISL_2486279, EPI_ISL_2486280, EPI_ISL_2486281, EPI_ISL_2486282, EPI_ISL_2486283, EPI_ISL_2486284, EPI_ISL_2486285, EPI_ISL_2486286, EPI_ISL_2486287, EPI_ISL_2486288, EPI_ISL_2486290, EPI_ISL_2486291, EPI_ISL_2486292, EPI_ISL_2486293, EPI_ISL_2486294, EPI_ISL_2486295, EPI_ISL_2486296, EPI_ISL_2486297, EPI_ISL_2486299, EPI_ISL_2486300, EPI_ISL_2486301, EPI_ISL_2486302, EPI_ISL_2486303, EPI_ISL_2486304, EPI_ISL_2486305, EPI_ISL_2486306, EPI_ISL_2486308, EPI_ISL_2486310, EPI_ISL_2486311, EPI_ISL_2486312, EPI_ISL_2486313, EPI_ISL_2486316, EPI_ISL_2486317, EPI_ISL_2486318, EPI_ISL_2486321, EPI_ISL_2486322, EPI_ISL_2486323, EPI_ISL_2486324, EPI_ISL_2486325, EPI_ISL_2486326, EPI_ISL_2486327, EPI_ISL_2486328, EPI_ISL_2486329, EPI_ISL_2486331, EPI_ISL_2486332, EPI_ISL_2486335, EPI_ISL_2486336, EPI_ISL_2486337, EPI_ISL_2486338, EPI_ISL_2486339, EPI_ISL_2486340, EPI_ISL_2486341, EPI_ISL_2486342, EPI_ISL_2486344, EPI_ISL_2486345, EPI_ISL_2486346, EPI_ISL_2486347, EPI_ISL_2486348, EPI_ISL_2486349, EPI_ISL_2486351, EPI_ISL_2486354, EPI_ISL_2486355, EPI_ISL_2486357, EPI_ISL_2486359, EPI_ISL_2486360, EPI_ISL_2486361, EPI_ISL_2486362, EPI_ISL_2486363, EPI_ISL_2486365, EPI_ISL_2486367, EPI_ISL_2486368, EPI_ISL_2486369, EPI_ISL_2486370, EPI_ISL_2486373, EPI_ISL_2486374, EPI_ISL_2486376, EPI_ISL_2486377, EPI_ISL_2486378, EPI_ISL_2486379, EPI_ISL_2486380, EPI_ISL_2486381, EPI_ISL_2486382, EPI_ISL_2486383, EPI_ISL_2486384, EPI_ISL_2486385, EPI_ISL_2486386, EPI_ISL_2486387, EPI_ISL_2486388, EPI_ISL_2486389, EPI_ISL_2486390, EPI_ISL_2486392, EPI_ISL_2486393, EPI_ISL_2486395, EPI_ISL_2486398, EPI_ISL_2486399, EPI_ISL_2486400, EPI_ISL_2486401, EPI_ISL_2486403, EPI_ISL_2486404, EPI_ISL_2486406, EPI_ISL_2486408, EPI_ISL_2486410, EPI_ISL_2486411, EPI_ISL_2486413, EPI_ISL_2486414, EPI_ISL_2486416, EPI_ISL_2486417, EPI_ISL_2486418, EPI_ISL_2486419, EPI_ISL_2486420, EPI_ISL_2486421, EPI_ISL_2486422, EPI_ISL_2486423, EPI_ISL_2486424, EPI_ISL_2486425, EPI_ISL_2486426, EPI_ISL_2486427, EPI_ISL_2486428, EPI_ISL_2486429, EPI_ISL_2486430, EPI_ISL_2486431, EPI_ISL_2486432, EPI_ISL_2486433, EPI_ISL_2486434, EPI_ISL_2486435, EPI_ISL_2486436, EPI_ISL_2486437, EPI_ISL_2486438, EPI_ISL_2486440, EPI_ISL_2486441, EPI_ISL_2486442, EPI_ISL_2486444, EPI_ISL_2486446, EPI_ISL_2486447, EPI_ISL_2486449, EPI_ISL_2486450, EPI_ISL_2486451, EPI_ISL_2486453, EPI_ISL_2486454, EPI_ISL_2486455, EPI_ISL_2486456, EPI_ISL_2486457, EPI_ISL_2486458, EPI_ISL_2486459, EPI_ISL_2486461, EPI_ISL_2486462, EPI_ISL_2486463, EPI_ISL_2486464, EPI_ISL_2486465, EPI_ISL_2486466, EPI_ISL_2486467, EPI_ISL_2486468, EPI_ISL_2486469, EPI_ISL_2486470, EPI_ISL_2486471, EPI_ISL_2486472, EPI_ISL_2486473, EPI_ISL_2486474, EPI_ISL_2486475, EPI_ISL_2486476, EPI_ISL_2486478, EPI_ISL_2486479, EPI_ISL_2486480, EPI_ISL_2486481, EPI_ISL_2486482, EPI_ISL_2486483, EPI_ISL_2486484, EPI_ISL_2486486, EPI_ISL_2486487, EPI_ISL_2486488, EPI_ISL_2486489, EPI_ISL_2486490, EPI_ISL_2486492, EPI_ISL_2486494, EPI_ISL_2486495, EPI_ISL_2486496, EPI_ISL_2486497, EPI_ISL_2486498, EPI_ISL_2486500, EPI_ISL_2486502, EPI_ISL_2486503, EPI_ISL_2486505, EPI_ISL_2486506, EPI_ISL_2486508, EPI_ISL_2486510, EPI_ISL_2486512, EPI_ISL_2486513, EPI_ISL_2486514, EPI_ISL_2486515, EPI_ISL_2486516, EPI_ISL_2486517, EPI_ISL_2486518, EPI_ISL_2486521, EPI_ISL_2486522, EPI_ISL_2486523, EPI_ISL_2486524, EPI_ISL_2486525, EPI_ISL_2486526, EPI_ISL_2486527, EPI_ISL_2486529 | | | |
| see above | Lighthouse Lab in Glasgow | Wellcome Sanger Institute for the COVID-19 Genomics UK (COG-UK) Consortium | Harper VanSteenhouse, Yumi Kasai, David Gray, Carol Clugston, Anna Dominiczak and Alex Alderton, Roberto Amato, Jeffrey Barrett, Sonia Goncalves, Ewan Harrison, David K. Jackson, Ian Johnston, Dominic Kwiatkowski, Cordelia Langford, John Sillitoe on behalf of the Wellcome Sanger Institute COVID-19 Surveillance Team |
| EPI_ISL_2486530, EPI_ISL_2486531, EPI_ISL_2486532, EPI_ISL_2486533, EPI_ISL_2486534, EPI_ISL_2486536, EPI_ISL_2486537, EPI_ISL_2486538, EPI_ISL_2486539, EPI_ISL_2486540, EPI_ISL_2486541, EPI_ISL_2486543, EPI_ISL_2486544, EPI_ISL_2486545, EPI_ISL_2486546, EPI_ISL_2486547, EPI_ISL_2486548, EPI_ISL_2486549, EPI_ISL_2486550, EPI_ISL_2486551, EPI_ISL_2486552, EPI_ISL_2486553, EPI_ISL_2486554, EPI_ISL_2486555, EPI_ISL_2486556, EPI_ISL_2486557, EPI_ISL_2486558, EPI_ISL_2486559, EPI_ISL_2486560, EPI_ISL_2486561, EPI_ISL_2486562, EPI_ISL_2486563, EPI_ISL_2486564, EPI_ISL_2486565, EPI_ISL_2486566, EPI_ISL_2486567, EPI_ISL_2486568, EPI_ISL_2486569, EPI_ISL_2486570, EPI_ISL_2486571, EPI_ISL_2486573, EPI_ISL_2486574, EPI_ISL_2486575, EPI_ISL_2486576, EPI_ISL_2486577, EPI_ISL_2486578, EPI_ISL_2486579, EPI_ISL_2486580, EPI_ISL_2486581, EPI_ISL_2486582, EPI_ISL_2486583, EPI_ISL_2486584, EPI_ISL_2486585, EPI_ISL_2486586, EPI_ISL_2486587, EPI_ISL_2486588, EPI_ISL_2486589, EPI_ISL_2486590, EPI_ISL_2486591, EPI_ISL_2486592, EPI_ISL_2486593, EPI_ISL_2486594, EPI_ISL_2486595, EPI_ISL_2486596, EPI_ISL_2486597, EPI_ISL_2486598, EPI_ISL_2486599, EPI_ISL_2486600, EPI_ISL_2486601, EPI_ISL_2486602, EPI_ISL_2486603, EPI_ISL_2486604, EPI_ISL_2486605, EPI_ISL_2486606, EPI_ISL_2486607, EPI_ISL_2486608, EPI_ISL_2486609, EPI_ISL_2486610, EPI_ISL_2486611, EPI_ISL_2486612, EPI_ISL_2486614, EPI_ISL_2486615, EPI_ISL_2486616, EPI_ISL_2486617, EPI_ISL_2486618, EPI_ISL_2486619, EPI_ISL_2486620, EPI_ISL_2486621, EPI_ISL_2486622, EPI_ISL_2486623, EPI_ISL_2486624, EPI_ISL_2486625, EPI_ISL_2486626, EPI_ISL_2486627, EPI_ISL_2486628, EPI_ISL_2486629, EPI_ISL_2486630, EPI_ISL_2486631, EPI_ISL_2486632, EPI_ISL_2486633, EPI_ISL_2486634, EPI_ISL_2486635, EPI_ISL_2486636, EPI_ISL_2486637, EPI_ISL_2486638, EPI_ISL_2486639, EPI_ISL_2486640, EPI_ISL_2486641, EPI_ISL_2486642, EPI_ISL_2486643, EPI_ISL_2486644, EPI_ISL_2486645, EPI_ISL_2486646, EPI_ISL_2486647, EPI_ISL_2486648, EPI_ISL_2486649, EPI_ISL_2486650, EPI_ISL_2486651, EPI_ISL_2486652, EPI_ISL_2486653, EPI_ISL_2486654, EPI_ISL_2486655, EPI_ISL_2486656, EPI_ISL_2486657, EPI_ISL_2486658, EPI_ISL_2486659, EPI_ISL_2486660, EPI_ISL_2486661, EPI_ISL_2486662, EPI_ISL_2486663, EPI_ISL_2486664, EPI_ISL_2486665, EPI_ISL_2486666, EPI_ISL_2486668, EPI_ISL_2486669, EPI_ISL_2486670, EPI_ISL_2486671, EPI_ISL_2486673, EPI_ISL_2486674, EPI_ISL_2486675, EPI_ISL_2486677, EPI_ISL_2486678, EPI_ISL_2486679, EPI_ISL_2486680, EPI_ISL_2486682, EPI_ISL_2486683, EPI_ISL_2486684, EPI_ISL_2486685, EPI_ISL_2486686, EPI_ISL_2486687, EPI_ISL_2486688, EPI_ISL_2486689, EPI_ISL_2486691, EPI_ISL_2486692, EPI_ISL_2486693, EPI_ISL_2486694, EPI_ISL_2486695, EPI_ISL_2486697, EPI_ISL_2486698, EPI_ISL_2486699, EPI_ISL_2486700, EPI_ISL_2486701, EPI_ISL_2486702, EPI_ISL_2486703, EPI_ISL_2486704, EPI_ISL_2486705, EPI_ISL_2486706, EPI_ISL_2486707, EPI_ISL_2486708, EPI_ISL_2486710, EPI_ISL_2486711, EPI_ISL_2486712, EPI_ISL_2486713, EPI_ISL_2486714, EPI_ISL_2486715, EPI_ISL_2486716, EPI_ISL_2486717, EPI_ISL_2486718, EPI_ISL_2486719, EPI_ISL_2486720, EPI_ISL_2486721, EPI_ISL_2486722, EPI_ISL_2486723, EPI_ISL_2486724, EPI_ISL_2486725, EPI_ISL_2486726, EPI_ISL_2486727, EPI_ISL_2486728, EPI_ISL_2486729, EPI_ISL_2486730, EPI_ISL_2486731, EPI_ISL_2486732, EPI_ISL_2486733, EPI_ISL_2486734, EPI_ISL_2486735, EPI_ISL_2486736, EPI_ISL_2486737, EPI_ISL_2486738, EPI_ISL_2486739, EPI_ISL_2486740, EPI_ISL_2486741, EPI_ISL_2486742, EPI_ISL_2486743, EPI_ISL_2486744, EPI_ISL_2486745, EPI_ISL_2486747, EPI_ISL_2486749, EPI_ISL_2486750, EPI_ISL_2486751, EPI_ISL_2486752, EPI_ISL_2486753, EPI_ISL_2486754, EPI_ISL_2486755, EPI_ISL_2486756, EPI_ISL_2486757, EPI_ISL_2486758, EPI_ISL_2486759, EPI_ISL_2486760, EPI_ISL_2486761, EPI_ISL_2486762, EPI_ISL_2486763, EPI_ISL_2486764, EPI_ISL_2486765, EPI_ISL_2486766, EPI_ISL_2486767, EPI_ISL_2486768, EPI_ISL_2486769, EPI_ISL_2486770, EPI_ISL_2486771, EPI_ISL_2486772, EPI_ISL_2486773, EPI_ISL_2486774, EPI_ISL_2486775, EPI_ISL_2486776, EPI_ISL_2486777, EPI_ISL_2486778, EPI_ISL_2486779, EPI_ISL_2486780, EPI_ISL_2486781, EPI_ISL_2486782, EPI_ISL_2486784, EPI_ISL_2486785, EPI_ISL_2486786, EPI_ISL_2486787, EPI_ISL_2486788, EPI_ISL_2486789, EPI_ISL_2486790, EPI_ISL_2486791, EPI_ISL_2486792, EPI_ISL_2486793, EPI_ISL_2486794, EPI_ISL_2486795, EPI_ISL_2486797, EPI_ISL_2486798, EPI_ISL_2486799, EPI_ISL_2486800, EPI_ISL_2486801, EPI_ISL_2486802, EPI_ISL_2486803, EPI_ISL_2486804, EPI_ISL_2486805, EPI_ISL_2486806, EPI_ISL_2486807, EPI_ISL_2486808, EPI_ISL_2486809, EPI_ISL_2486810, EPI_ISL_2486811, EPI_ISL_2486812, EPI_ISL_2486813, EPI_ISL_2486814, EPI_ISL_2486815, EPI_ISL_2486816, EPI_ISL_2486817, EPI_ISL_2486818, EPI_ISL_2486819, EPI_ISL_2486820, EPI_ISL_2486821, EPI_ISL_2486822, EPI_ISL_2486823, EPI_ISL_2486824, EPI_ISL_2486825, EPI_ISL_2486826, EPI_ISL_2486827, EPI_ISL_2486828, EPI_ISL_2486830, EPI_ISL_2486831, EPI_ISL_2486832, EPI_ISL_2486833, EPI_ISL_2486834, EPI_ISL_2486835, EPI_ISL_2486836, EPI_ISL_2486837, EPI_ISL_2486838, EPI_ISL_2486839, EPI_ISL_2486840, EPI_ISL_2486841, EPI_ISL_2486842, EPI_ISL_2486843, EPI_ISL_2486844, EPI_ISL_2486845, EPI_ISL_2486846, EPI_ISL_2486847, EPI_ISL_2486848, EPI_ISL_2486849, EPI_ISL_2486850, EPI_ISL_2486852, EPI_ISL_2486853, EPI_ISL_2486854, EPI_ISL_2486855, EPI_ISL_2486856, EPI_ISL_2486857, EPI_ISL_2486858, EPI_ISL_2486859, EPI_ISL_2486860, EPI_ISL_2486861, EPI_ISL_2486862, EPI_ISL_2486863, EPI_ISL_2486864, EPI_ISL_2486865, EPI_ISL_2486866, EPI_ISL_2486867, EPI_ISL_2486868, EPI_ISL_2486869, EPI_ISL_2486871, EPI_ISL_2486872, EPI_ISL_2486873, EPI_ISL_2486874, EPI_ISL_2486875, EPI_ISL_2486876, EPI_ISL_2486877, EPI_ISL_2486878, EPI_ISL_2486879, EPI_ISL_2486880, EPI_ISL_2486881, EPI_ISL_2486882, EPI_ISL_2486883, EPI_ISL_2486884, EPI_ISL_2486885, EPI_ISL_2486886, EPI_ISL_2486887, EPI_ISL_2486888, EPI_ISL_2486889, EPI_ISL_2486890, EPI_ISL_2486891, EPI_ISL_2486892, EPI_ISL_2486893, EPI_ISL_2486894 | | | |
| see above | Lighthouse Lab in Alderley Park | Wellcome Sanger Institute for the COVID-19 Genomics UK (COG-UK) Consortium | Jacquelyn Wynn, Mairead Hyland, The Lighthouse Lab in Alderley Park and Alex Alderton, Roberto Amato, Jeffrey Barrett, Sonia Goncalves, Ewan Harrison, David K. Jackson, Ian Johnston, Dominic Kwiatkowski, Cordelia Langford, John Sillitoe on behalf of the Wellcome Sanger Institute COVID-19 Surveillance Team |
| EPI_ISL_2486895 | Health Services Laboratories | Wellcome Sanger Institute for the COVID-19 Genomics UK (COG-UK) Consortium | Health Services Laboratories and Alex Alderton, Roberto Amato, Jeffrey Barrett, Sonia Goncalves, Ewan Harrison, David K. Jackson, Ian Johnston, Dominic Kwiatkowski, Cordelia Langford, John Sillitoe on behalf of the Wellcome Sanger Institute COVID-19 Surveillance Team |
| EPI_ISL_2486896 | Lighthouse Lab in Milton Keynes | Wellcome Sanger Institute for the COVID-19 Genomics UK (COG-UK) Consortium | The Lighthouse Lab in Milton Keynes and Alex Alderton, Roberto Amato, Jeffrey Barrett, Sonia Goncalves, Ewan Harrison, David K. Jackson, Ian Johnston, Dominic Kwiatkowski, Cordelia Langford, John Sillitoe on behalf of the Wellcome Sanger Institute COVID-19 Surveillance Team |
| EPI_ISL_2486897 | Lighthouse Lab in Alderley Park | Wellcome Sanger Institute for the COVID-19 Genomics UK (COG-UK) Consortium | Jacquelyn Wynn, Mairead Hyland, The Lighthouse Lab in Alderley Park and Alex Alderton, Roberto Amato, Jeffrey Barrett, Sonia Goncalves, Ewan Harrison, David K. Jackson, Ian Johnston, Dominic Kwiatkowski, Cordelia Langford, John Sillitoe on behalf of the Wellcome Sanger Institute COVID-19 Surveillance Team |
| EPI_ISL_2486898, EPI_ISL_2486899, EPI_ISL_2486900 | Lighthouse Lab in Milton Keynes | Wellcome Sanger Institute for the COVID-19 Genomics UK (COG-UK) Consortium | The Lighthouse Lab in Milton Keynes and Alex Alderton, Roberto Amato, Jeffrey Barrett, Sonia Goncalves, Ewan Harrison, David K. Jackson, Ian Johnston, Dominic Kwiatkowski, Cordelia Langford, John Sillitoe on behalf of the Wellcome Sanger Institute COVID-19 Surveillance Team |
| EPI_ISL_2486902, EPI_ISL_2486903 | Lighthouse Lab in Alderley Park | Wellcome Sanger Institute for the COVID-19 Genomics UK (COG-UK) Consortium | Jacquelyn Wynn, Mairead Hyland, The Lighthouse Lab in Alderley Park and Alex Alderton, Roberto Amato, Jeffrey Barrett, Sonia Goncalves, Ewan Harrison, David K. Jackson, Ian Johnston, Dominic Kwiatkowski, Cordelia Langford, John Sillitoe on behalf of the Wellcome Sanger Institute COVID-19 Surveillance Team |
| EPI_ISL_2486904 | Lighthouse Lab in Milton Keynes | Wellcome Sanger Institute for the COVID-19 Genomics UK (COG-UK) Consortium | The Lighthouse Lab in Milton Keynes and Alex Alderton, Roberto Amato, Jeffrey Barrett, Sonia Goncalves, Ewan Harrison, David K. Jackson, Ian Johnston, Dominic Kwiatkowski, Cordelia Langford, John Sillitoe on behalf of the Wellcome Sanger Institute COVID-19 Surveillance Team |
| EPI_ISL_2486905, EPI_ISL_2486906, EPI_ISL_2486907 | Lighthouse Lab in Alderley Park | Wellcome Sanger Institute for the COVID-19 Genomics UK (COG-UK) Consortium | Jacquelyn Wynn, Mairead Hyland, The Lighthouse Lab in Alderley Park and Alex Alderton, Roberto Amato, Jeffrey Barrett, Sonia Goncalves, Ewan Harrison, David K. Jackson, Ian Johnston, Dominic Kwiatkowski, Cordelia Langford, John Sillitoe on behalf of the Wellcome Sanger Institute COVID-19 Surveillance Team |
| EPI_ISL_2486908 | Lighthouse Lab in Milton Keynes | Wellcome Sanger Institute for the COVID-19 Genomics UK (COG-UK) Consortium | The Lighthouse Lab in Milton Keynes and Alex Alderton, Roberto Amato, Jeffrey Barrett, Sonia Goncalves, Ewan Harrison, David K. Jackson, Ian Johnston, Dominic Kwiatkowski, Cordelia Langford, John Sillitoe on behalf of the Wellcome Sanger Institute COVID-19 Surveillance Team |
| EPI_ISL_2486909, EPI_ISL_2486910, EPI_ISL_2486911, EPI_ISL_2486912, EPI_ISL_2486913, EPI_ISL_2486914, EPI_ISL_2486915 | Lighthouse Lab in Alderley Park | Wellcome Sanger Institute for the COVID-19 Genomics UK (COG-UK) Consortium | Jacquelyn Wynn, Mairead Hyland, The Lighthouse Lab in Alderley Park and Alex Alderton, Roberto Amato, Jeffrey Barrett, Sonia Goncalves, Ewan Harrison, David K. Jackson, Ian Johnston, Dominic Kwiatkowski, Cordelia Langford, John Sillitoe on behalf of the Wellcome Sanger Institute COVID-19 Surveillance Team |
| EPI_ISL_2486916 | Health Services Laboratories | Wellcome Sanger Institute for the COVID-19 Genomics UK (COG-UK) Consortium | Health Services Laboratories and Alex Alderton, Roberto Amato, Jeffrey Barrett, Sonia Goncalves, Ewan Harrison, David K. Jackson, Ian Johnston, Dominic Kwiatkowski, Cordelia Langford, John Sillitoe on behalf of the Wellcome Sanger Institute COVID-19 Surveillance Team |
| EPI_ISL_2486917 | Lighthouse Lab in Alderley Park | Wellcome Sanger Institute for the COVID-19 Genomics UK (COG-UK) Consortium | Jacquelyn Wynn, Mairead Hyland, The Lighthouse Lab in Alderley Park and Alex Alderton, Roberto Amato, Jeffrey Barrett, Sonia Goncalves, Ewan Harrison, David K. Jackson, Ian Johnston, Dominic Kwiatkowski, Cordelia Langford, John Sillitoe on behalf of the Wellcome Sanger Institute COVID-19 Surveillance Team |
| EPI_ISL_2486918 | Health Services Laboratories | Wellcome Sanger Institute for the COVID-19 Genomics UK (COG-UK) Consortium | Health Services Laboratories and Alex Alderton, Roberto Amato, Jeffrey Barrett, Sonia Goncalves, Ewan Harrison, David K. Jackson, Ian Johnston, Dominic Kwiatkowski, Cordelia Langford, John Sillitoe on behalf of the Wellcome Sanger Institute COVID-19 Surveillance Team |
| EPI_ISL_2486919, EPI_ISL_2486920 | Lighthouse Lab in Alderley Park | Wellcome Sanger Institute for the COVID-19 Genomics UK (COG-UK) Consortium | Jacquelyn Wynn, Mairead Hyland, The Lighthouse Lab in Alderley Park and Alex Alderton, Roberto Amato, Jeffrey Barrett, Sonia Goncalves, Ewan Harrison, David K. Jackson, Ian Johnston, Dominic Kwiatkowski, Cordelia Langford, John Sillitoe on behalf of the Wellcome Sanger Institute COVID-19 Surveillance Team |
| EPI_ISL_2486921 | Lighthouse Lab in Milton Keynes | Wellcome Sanger Institute for the COVID-19 Genomics UK (COG-UK) Consortium | The Lighthouse Lab in Milton Keynes and Alex Alderton, Roberto Amato, Jeffrey Barrett, Sonia Goncalves, Ewan Harrison, David K. Jackson, Ian Johnston, Dominic Kwiatkowski, Cordelia Langford, John Sillitoe on behalf of the Wellcome Sanger Institute COVID-19 Surveillance Team |
| EPI_ISL_2486923 | Lighthouse Lab in Alderley Park | Wellcome Sanger Institute for the COVID-19 Genomics UK (COG-UK) Consortium | Jacquelyn Wynn, Mairead Hyland, The Lighthouse Lab in Alderley Park and Alex Alderton, Roberto Amato, Jeffrey Barrett, Sonia Goncalves, Ewan Harrison, David K. Jackson, Ian Johnston, Dominic Kwiatkowski, Cordelia Langford, John Sillitoe on behalf of the Wellcome Sanger Institute COVID-19 Surveillance Team |
| EPI_ISL_2486924, EPI_ISL_2486925 | Lighthouse Lab in Milton Keynes | Wellcome Sanger Institute for the COVID-19 Genomics UK (COG-UK) Consortium | The Lighthouse Lab in Milton Keynes and Alex Alderton, Roberto Amato, Jeffrey Barrett, Sonia Goncalves, Ewan Harrison, David K. Jackson, Ian Johnston, Dominic Kwiatkowski, Cordelia Langford, John Sillitoe on behalf of the Wellcome Sanger Institute COVID-19 Surveillance Team |
| EPI_ISL_2486926, EPI_ISL_2486927 | Lighthouse Lab in Alderley Park | Wellcome Sanger Institute for the COVID-19 Genomics UK (COG-UK) Consortium | Jacquelyn Wynn, Mairead Hyland, The Lighthouse Lab in Alderley Park and Alex Alderton, Roberto Amato, Jeffrey Barrett, Sonia Goncalves, Ewan Harrison, David K. Jackson, Ian Johnston, Dominic Kwiatkowski, Cordelia Langford, John Sillitoe on behalf of the Wellcome Sanger Institute COVID-19 Surveillance Team |
| EPI_ISL_2486930, EPI_ISL_2486931, EPI_ISL_2486932, EPI_ISL_2486933 | Lighthouse Lab in Milton Keynes | Wellcome Sanger Institute for the COVID-19 Genomics UK (COG-UK) Consortium | The Lighthouse Lab in Milton Keynes and Alex Alderton, Roberto Amato, Jeffrey Barrett, Sonia Goncalves, Ewan Harrison, David K. Jackson, Ian Johnston, Dominic Kwiatkowski, Cordelia Langford, John Sillitoe on behalf of the Wellcome Sanger Institute COVID-19 Surveillance Team |
| EPI_ISL_2486934 | Health Services Laboratories | Wellcome Sanger Institute for the COVID-19 Genomics UK (COG-UK) Consortium | Health Services Laboratories and Alex Alderton, Roberto Amato, Jeffrey Barrett, Sonia Goncalves, Ewan Harrison, David K. Jackson, Ian Johnston, Dominic Kwiatkowski, Cordelia Langford, John Sillitoe on behalf of the Wellcome Sanger Institute COVID-19 Surveillance Team |
| EPI_ISL_2486935, EPI_ISL_2486936 | Lighthouse Lab in Alderley Park | Wellcome Sanger Institute for the COVID-19 Genomics UK (COG-UK) Consortium | Jacquelyn Wynn, Mairead Hyland, The Lighthouse Lab in Alderley Park and Alex Alderton, Roberto Amato, Jeffrey Barrett, Sonia Goncalves, Ewan Harrison, David K. Jackson, Ian Johnston, Dominic Kwiatkowski, Cordelia Langford, John Sillitoe on behalf of the Wellcome Sanger Institute COVID-19 Surveillance Team |
| EPI_ISL_2486938 | Lighthouse Lab in Milton Keynes | Wellcome Sanger Institute for the COVID-19 Genomics UK (COG-UK) Consortium | The Lighthouse Lab in Milton Keynes and Alex Alderton, Roberto Amato, Jeffrey Barrett, Sonia Goncalves, Ewan Harrison, David K. Jackson, Ian Johnston, Dominic Kwiatkowski, Cordelia Langford, John Sillitoe on behalf of the Wellcome Sanger Institute COVID-19 Surveillance Team |
| EPI_ISL_2486939, EPI_ISL_2486940 | Lighthouse Lab in Alderley Park | Wellcome Sanger Institute for the COVID-19 Genomics UK (COG-UK) Consortium | Jacquelyn Wynn, Mairead Hyland, The Lighthouse Lab in Alderley Park and Alex Alderton, Roberto Amato, Jeffrey Barrett, Sonia Goncalves, Ewan Harrison, David K. Jackson, Ian Johnston, Dominic Kwiatkowski, Cordelia Langford, John Sillitoe on behalf of the Wellcome Sanger Institute COVID-19 Surveillance Team |

| | | | |
|---|---|---|---|
| EPI_ISL_2486941, EPI_ISL_2486942 | Lighthouse Lab in Milton Keynes | Wellcome Sanger Institute for the COVID-19 Genomics UK (COG-UK) Consortium | The Lighthouse Lab in Milton Keynes and Alex Alderton, Roberto Amato, Jeffrey Barrett, Sonia Goncalves, Ewan Harrison, David K. Jackson, Ian Johnston, Dominic Kwiatkowski, Cordelia Langford, John Sillitoe on behalf of the Wellcome Sanger Institute COVID-19 Surveillance Team |
| EPI_ISL_2486943 | Lighthouse Lab in Alderley Park | Wellcome Sanger Institute for the COVID-19 Genomics UK (COG-UK) Consortium | Jacquelyn Wynn, Mairead Hyland, The Lighthouse Lab in Alderley Park and Alex Alderton, Roberto Amato, Jeffrey Barrett, Sonia Goncalves, Ewan Harrison, David K. Jackson, Ian Johnston, Dominic Kwiatkowski, Cordelia Langford, John Sillitoe on behalf of the Wellcome Sanger Institute COVID-19 Surveillance Team |
| EPI_ISL_2486944 | Health Services Laboratories | Wellcome Sanger Institute for the COVID-19 Genomics UK (COG-UK) Consortium | Health Services Laboratories and Alex Alderton, Roberto Amato, Jeffrey Barrett, Sonia Goncalves, Ewan Harrison, David K. Jackson, Ian Johnston, Dominic Kwiatkowski, Cordelia Langford, John Sillitoe on behalf of the Wellcome Sanger Institute COVID-19 Surveillance Team |
| EPI_ISL_2486945, EPI_ISL_2486947 | Lighthouse Lab in Milton Keynes | Wellcome Sanger Institute for the COVID-19 Genomics UK (COG-UK) Consortium | The Lighthouse Lab in Milton Keynes and Alex Alderton, Roberto Amato, Jeffrey Barrett, Sonia Goncalves, Ewan Harrison, David K. Jackson, Ian Johnston, Dominic Kwiatkowski, Cordelia Langford, John Sillitoe on behalf of the Wellcome Sanger Institute COVID-19 Surveillance Team |
| EPI_ISL_2486948 | Lighthouse Lab in Alderley Park | Wellcome Sanger Institute for the COVID-19 Genomics UK (COG-UK) Consortium | Jacquelyn Wynn, Mairead Hyland, The Lighthouse Lab in Alderley Park and Alex Alderton, Roberto Amato, Jeffrey Barrett, Sonia Goncalves, Ewan Harrison, David K. Jackson, Ian Johnston, Dominic Kwiatkowski, Cordelia Langford, John Sillitoe on behalf of the Wellcome Sanger Institute COVID-19 Surveillance Team |
| EPI_ISL_2486949, EPI_ISL_2486950 | Lighthouse Lab in Milton Keynes | Wellcome Sanger Institute for the COVID-19 Genomics UK (COG-UK) Consortium | The Lighthouse Lab in Milton Keynes and Alex Alderton, Roberto Amato, Jeffrey Barrett, Sonia Goncalves, Ewan Harrison, David K. Jackson, Ian Johnston, Dominic Kwiatkowski, Cordelia Langford, John Sillitoe on behalf of the Wellcome Sanger Institute COVID-19 Surveillance Team |
| EPI_ISL_2486951 | Health Services Laboratories | Wellcome Sanger Institute for the COVID-19 Genomics UK (COG-UK) Consortium | Health Services Laboratories and Alex Alderton, Roberto Amato, Jeffrey Barrett, Sonia Goncalves, Ewan Harrison, David K. Jackson, Ian Johnston, Dominic Kwiatkowski, Cordelia Langford, John Sillitoe on behalf of the Wellcome Sanger Institute COVID-19 Surveillance Team |
| EPI_ISL_2486952, EPI_ISL_2486953 | Lighthouse Lab in Milton Keynes | Wellcome Sanger Institute for the COVID-19 Genomics UK (COG-UK) Consortium | The Lighthouse Lab in Milton Keynes and Alex Alderton, Roberto Amato, Jeffrey Barrett, Sonia Goncalves, Ewan Harrison, David K. Jackson, Ian Johnston, Dominic Kwiatkowski, Cordelia Langford, John Sillitoe on behalf of the Wellcome Sanger Institute COVID-19 Surveillance Team |
| EPI_ISL_2486954, EPI_ISL_2486955 | Lighthouse Lab in Alderley Park | Wellcome Sanger Institute for the COVID-19 Genomics UK (COG-UK) Consortium | Jacquelyn Wynn, Mairead Hyland, The Lighthouse Lab in Alderley Park and Alex Alderton, Roberto Amato, Jeffrey Barrett, Sonia Goncalves, Ewan Harrison, David K. Jackson, Ian Johnston, Dominic Kwiatkowski, Cordelia Langford, John Sillitoe on behalf of the Wellcome Sanger Institute COVID-19 Surveillance Team |
| EPI_ISL_2486956 | Lighthouse Lab in Milton Keynes | Wellcome Sanger Institute for the COVID-19 Genomics UK (COG-UK) Consortium | The Lighthouse Lab in Milton Keynes and Alex Alderton, Roberto Amato, Jeffrey Barrett, Sonia Goncalves, Ewan Harrison, David K. Jackson, Ian Johnston, Dominic Kwiatkowski, Cordelia Langford, John Sillitoe on behalf of the Wellcome Sanger Institute COVID-19 Surveillance Team |
| EPI_ISL_2486957, EPI_ISL_2486958, EPI_ISL_2486959, EPI_ISL_2486960 | Lighthouse Lab in Alderley Park | Wellcome Sanger Institute for the COVID-19 Genomics UK (COG-UK) Consortium | Jacquelyn Wynn, Mairead Hyland, The Lighthouse Lab in Alderley Park and Alex Alderton, Roberto Amato, Jeffrey Barrett, Sonia Goncalves, Ewan Harrison, David K. Jackson, Ian Johnston, Dominic Kwiatkowski, Cordelia Langford, John Sillitoe on behalf of the Wellcome Sanger Institute COVID-19 Surveillance Team |
| EPI_ISL_2486961 | Lighthouse Lab in Milton Keynes | Wellcome Sanger Institute for the COVID-19 Genomics UK (COG-UK) Consortium | The Lighthouse Lab in Milton Keynes and Alex Alderton, Roberto Amato, Jeffrey Barrett, Sonia Goncalves, Ewan Harrison, David K. Jackson, Ian Johnston, Dominic Kwiatkowski, Cordelia Langford, John Sillitoe on behalf of the Wellcome Sanger Institute COVID-19 Surveillance Team |
| EPI_ISL_2486962 | Lighthouse Lab in Alderley Park | Wellcome Sanger Institute for the COVID-19 Genomics UK (COG-UK) Consortium | Jacquelyn Wynn, Mairead Hyland, The Lighthouse Lab in Alderley Park and Alex Alderton, Roberto Amato, Jeffrey Barrett, Sonia Goncalves, Ewan Harrison, David K. Jackson, Ian Johnston, Dominic Kwiatkowski, Cordelia Langford, John Sillitoe on behalf of the Wellcome Sanger Institute COVID-19 Surveillance Team |
| EPI_ISL_2486963 | Lighthouse Lab in Milton Keynes | Wellcome Sanger Institute for the COVID-19 Genomics UK (COG-UK) Consortium | The Lighthouse Lab in Milton Keynes and Alex Alderton, Roberto Amato, Jeffrey Barrett, Sonia Goncalves, Ewan Harrison, David K. Jackson, Ian Johnston, Dominic Kwiatkowski, Cordelia Langford, John Sillitoe on behalf of the Wellcome Sanger Institute COVID-19 Surveillance Team |
| EPI_ISL_2486964, EPI_ISL_2486965, EPI_ISL_2486966, EPI_ISL_2486967, EPI_ISL_2486968 | Lighthouse Lab in Alderley Park | Wellcome Sanger Institute for the COVID-19 Genomics UK (COG-UK) Consortium | Jacquelyn Wynn, Mairead Hyland, The Lighthouse Lab in Alderley Park and Alex Alderton, Roberto Amato, Jeffrey Barrett, Sonia Goncalves, Ewan Harrison, David K. Jackson, Ian Johnston, Dominic Kwiatkowski, Cordelia Langford, John Sillitoe on behalf of the Wellcome Sanger Institute COVID-19 Surveillance Team |
| EPI_ISL_2486969 | Lighthouse Lab in Milton Keynes | Wellcome Sanger Institute for the COVID-19 Genomics UK (COG-UK) Consortium | The Lighthouse Lab in Milton Keynes and Alex Alderton, Roberto Amato, Jeffrey Barrett, Sonia Goncalves, Ewan Harrison, David K. Jackson, Ian Johnston, Dominic Kwiatkowski, Cordelia Langford, John Sillitoe on behalf of the Wellcome Sanger Institute COVID-19 Surveillance Team |
| EPI_ISL_2486970 | Lighthouse Lab in Alderley Park | Wellcome Sanger Institute for the COVID-19 Genomics UK (COG-UK) Consortium | Jacquelyn Wynn, Mairead Hyland, The Lighthouse Lab in Alderley Park and Alex Alderton, Roberto Amato, Jeffrey Barrett, Sonia Goncalves, Ewan Harrison, David K. Jackson, Ian Johnston, Dominic Kwiatkowski, Cordelia Langford, John Sillitoe on behalf of the Wellcome Sanger Institute COVID-19 Surveillance Team |
| EPI_ISL_2486971 | Lighthouse Lab in Milton Keynes | Wellcome Sanger Institute for the COVID-19 Genomics UK (COG-UK) Consortium | The Lighthouse Lab in Milton Keynes and Alex Alderton, Roberto Amato, Jeffrey Barrett, Sonia Goncalves, Ewan Harrison, David K. Jackson, Ian Johnston, Dominic Kwiatkowski, Cordelia Langford, John Sillitoe on behalf of the Wellcome Sanger Institute COVID-19 Surveillance Team |
| EPI_ISL_2486972, EPI_ISL_2486973, EPI_ISL_2486974, EPI_ISL_2486975 | Lighthouse Lab in Alderley Park | Wellcome Sanger Institute for the COVID-19 Genomics UK (COG-UK) Consortium | Jacquelyn Wynn, Mairead Hyland, The Lighthouse Lab in Alderley Park and Alex Alderton, Roberto Amato, Jeffrey Barrett, Sonia Goncalves, Ewan Harrison, David K. Jackson, Ian Johnston, Dominic Kwiatkowski, Cordelia Langford, John Sillitoe on behalf of the Wellcome Sanger Institute COVID-19 Surveillance Team |
| EPI_ISL_2486976, EPI_ISL_2486977, EPI_ISL_2486978, EPI_ISL_2486979 | Lighthouse Lab in Milton Keynes | Wellcome Sanger Institute for the COVID-19 Genomics UK (COG-UK) Consortium | The Lighthouse Lab in Milton Keynes and Alex Alderton, Roberto Amato, Jeffrey Barrett, Sonia Goncalves, Ewan Harrison, David K. Jackson, Ian Johnston, Dominic Kwiatkowski, Cordelia Langford, John Sillitoe on behalf of the Wellcome Sanger Institute COVID-19 Surveillance Team |
| EPI_ISL_2486980 | Lighthouse Lab in Alderley Park | Wellcome Sanger Institute for the COVID-19 Genomics UK (COG-UK) Consortium | Jacquelyn Wynn, Mairead Hyland, The Lighthouse Lab in Alderley Park and Alex Alderton, Roberto Amato, Jeffrey Barrett, Sonia Goncalves, Ewan Harrison, David K. Jackson, Ian Johnston, Dominic Kwiatkowski, Cordelia Langford, John Sillitoe on behalf of the Wellcome Sanger Institute COVID-19 Surveillance Team |
| EPI_ISL_2486981, EPI_ISL_2486982, EPI_ISL_2486983, EPI_ISL_2486984 | Lighthouse Lab in Milton Keynes | Wellcome Sanger Institute for the COVID-19 Genomics UK (COG-UK) Consortium | The Lighthouse Lab in Milton Keynes and Alex Alderton, Roberto Amato, Jeffrey Barrett, Sonia Goncalves, Ewan Harrison, David K. Jackson, Ian Johnston, Dominic Kwiatkowski, Cordelia Langford, John Sillitoe on behalf of the Wellcome Sanger Institute COVID-19 Surveillance Team |
| EPI_ISL_2486985, EPI_ISL_2486986 | Lighthouse Lab in Alderley Park | Wellcome Sanger Institute for the COVID-19 Genomics UK (COG-UK) Consortium | Jacquelyn Wynn, Mairead Hyland, The Lighthouse Lab in Alderley Park and Alex Alderton, Roberto Amato, Jeffrey Barrett, Sonia Goncalves, Ewan Harrison, David K. Jackson, Ian Johnston, Dominic Kwiatkowski, Cordelia Langford, John Sillitoe on behalf of the Wellcome Sanger Institute COVID-19 Surveillance Team |
| EPI_ISL_2486987 | Lighthouse Lab in Milton Keynes | Wellcome Sanger Institute for the COVID-19 Genomics UK (COG-UK) Consortium | The Lighthouse Lab in Milton Keynes and Alex Alderton, Roberto Amato, Jeffrey Barrett, Sonia Goncalves, Ewan Harrison, David K. Jackson, Ian Johnston, Dominic Kwiatkowski, Cordelia Langford, John Sillitoe on behalf of the Wellcome Sanger Institute COVID-19 Surveillance Team |
| EPI_ISL_2486988 | Lighthouse Lab in Alderley Park | Wellcome Sanger Institute for the COVID-19 Genomics UK (COG-UK) Consortium | Jacquelyn Wynn, Mairead Hyland, The Lighthouse Lab in Alderley Park and Alex Alderton, Roberto Amato, Jeffrey Barrett, Sonia Goncalves, Ewan Harrison, David K. Jackson, Ian Johnston, Dominic Kwiatkowski, Cordelia Langford, John Sillitoe on behalf of the Wellcome Sanger Institute COVID-19 Surveillance Team |
| EPI_ISL_2486989 | Lighthouse Lab in Milton Keynes | Wellcome Sanger Institute for the COVID-19 Genomics UK (COG-UK) Consortium | The Lighthouse Lab in Milton Keynes and Alex Alderton, Roberto Amato, Jeffrey Barrett, Sonia Goncalves, Ewan Harrison, David K. Jackson, Ian Johnston, Dominic Kwiatkowski, Cordelia Langford, John Sillitoe on behalf of the Wellcome Sanger Institute COVID-19 Surveillance Team |
| EPI_ISL_2486991, EPI_ISL_2486992, EPI_ISL_2486993, EPI_ISL_2486994 | Lighthouse Lab in Alderley Park | Wellcome Sanger Institute for the COVID-19 Genomics UK (COG-UK) Consortium | Jacquelyn Wynn, Mairead Hyland, The Lighthouse Lab in Alderley Park and Alex Alderton, Roberto Amato, Jeffrey Barrett, Sonia Goncalves, Ewan Harrison, David K. Jackson, Ian Johnston, Dominic Kwiatkowski, Cordelia Langford, John Sillitoe on behalf of the Wellcome Sanger Institute COVID-19 Surveillance Team |
| EPI_ISL_2486995, EPI_ISL_2486996 | Lighthouse Lab in Milton Keynes | Wellcome Sanger Institute for the COVID-19 Genomics UK (COG-UK) Consortium | The Lighthouse Lab in Milton Keynes and Alex Alderton, Roberto Amato, Jeffrey Barrett, Sonia Goncalves, Ewan Harrison, David K. Jackson, Ian Johnston, Dominic Kwiatkowski, Cordelia Langford, John Sillitoe on behalf of the Wellcome Sanger Institute COVID-19 Surveillance Team |
| EPI_ISL_2486997, EPI_ISL_2486998, EPI_ISL_2486999, EPI_ISL_2487000 | Lighthouse Lab in Alderley Park | Wellcome Sanger Institute for the COVID-19 Genomics UK (COG-UK) Consortium | Jacquelyn Wynn, Mairead Hyland, The Lighthouse Lab in Alderley Park and Alex Alderton, Roberto Amato, Jeffrey Barrett, Sonia Goncalves, Ewan Harrison, David K. Jackson, Ian Johnston, Dominic Kwiatkowski, Cordelia Langford, John Sillitoe on behalf of the Wellcome Sanger Institute COVID-19 Surveillance Team |
| EPI_ISL_2487001 | Lighthouse Lab in Milton Keynes | Wellcome Sanger Institute for the COVID-19 Genomics UK (COG-UK) Consortium | The Lighthouse Lab in Milton Keynes and Alex Alderton, Roberto Amato, Jeffrey Barrett, Sonia Goncalves, Ewan Harrison, David K. Jackson, Ian Johnston, Dominic Kwiatkowski, Cordelia Langford, John Sillitoe on behalf of the Wellcome Sanger Institute COVID-19 Surveillance Team |
| EPI_ISL_2487002, EPI_ISL_2487003, EPI_ISL_2487004 | Lighthouse Lab in Alderley Park | Wellcome Sanger Institute for the COVID-19 Genomics UK (COG-UK) Consortium | Jacquelyn Wynn, Mairead Hyland, The Lighthouse Lab in Alderley Park and Alex Alderton, Roberto Amato, Jeffrey Barrett, Sonia Goncalves, Ewan Harrison, David K. Jackson, Ian Johnston, Dominic Kwiatkowski, Cordelia Langford, John Sillitoe on behalf of the Wellcome Sanger Institute COVID-19 Surveillance Team |
| EPI_ISL_2487005, EPI_ISL_2487006 | Lighthouse Lab in Milton Keynes | Wellcome Sanger Institute for the COVID-19 Genomics UK (COG-UK) Consortium | The Lighthouse Lab in Milton Keynes and Alex Alderton, Roberto Amato, Jeffrey Barrett, Sonia Goncalves, Ewan Harrison, David K. Jackson, Ian Johnston, Dominic Kwiatkowski, Cordelia Langford, John Sillitoe on behalf of the Wellcome Sanger Institute COVID-19 Surveillance Team |
| EPI_ISL_2487007, EPI_ISL_2487008, EPI_ISL_2487009, EPI_ISL_2487010 | Lighthouse Lab in Alderley Park | Wellcome Sanger Institute for the COVID-19 Genomics UK (COG-UK) Consortium | Jacquelyn Wynn, Mairead Hyland, The Lighthouse Lab in Alderley Park and Alex Alderton, Roberto Amato, Jeffrey Barrett, Sonia Goncalves, Ewan Harrison, David K. Jackson, Ian Johnston, Dominic Kwiatkowski, Cordelia Langford, John Sillitoe on behalf of the Wellcome Sanger Institute COVID-19 Surveillance Team |
| EPI_ISL_2487011 | Lighthouse Lab in Milton Keynes | Wellcome Sanger Institute for the COVID-19 Genomics UK (COG-UK) Consortium | The Lighthouse Lab in Milton Keynes and Alex Alderton, Roberto Amato, Jeffrey Barrett, Sonia Goncalves, Ewan Harrison, David K. Jackson, Ian Johnston, Dominic Kwiatkowski, Cordelia Langford, John Sillitoe on behalf of the Wellcome Sanger Institute COVID-19 Surveillance Team |
| EPI_ISL_2487012 | Lighthouse Lab in Alderley Park | Wellcome Sanger Institute for the COVID-19 Genomics UK (COG-UK) Consortium | Jacquelyn Wynn, Mairead Hyland, The Lighthouse Lab in Alderley Park and Alex Alderton, Roberto Amato, Jeffrey Barrett, Sonia Goncalves, Ewan Harrison, David K. Jackson, Ian Johnston, Dominic Kwiatkowski, Cordelia Langford, John Sillitoe on behalf of the Wellcome Sanger Institute COVID-19 Surveillance Team |
| EPI_ISL_2487013, EPI_ISL_2487014, EPI_ISL_2487015 | Lighthouse Lab in Milton Keynes | Wellcome Sanger Institute for the COVID-19 Genomics UK (COG-UK) Consortium | The Lighthouse Lab in Milton Keynes and Alex Alderton, Roberto Amato, Jeffrey Barrett, Sonia Goncalves, Ewan Harrison, David K. Jackson, Ian Johnston, Dominic Kwiatkowski, Cordelia Langford, John Sillitoe on behalf of the Wellcome Sanger Institute COVID-19 Surveillance Team |
| EPI_ISL_2487016 | Lighthouse Lab in Alderley Park | Wellcome Sanger Institute for the COVID-19 Genomics UK (COG-UK) Consortium | Jacquelyn Wynn, Mairead Hyland, The Lighthouse Lab in Alderley Park and Alex Alderton, Roberto Amato, Jeffrey Barrett, Sonia Goncalves, Ewan Harrison, David K. Jackson, Ian Johnston, Dominic Kwiatkowski, Cordelia Langford, John Sillitoe on behalf of the Wellcome Sanger Institute COVID-19 Surveillance Team |

| | | | |
|---|---|---|---|
| EPI_ISL_2487017 | Lighthouse Lab in Milton Keynes | Wellcome Sanger Institute for the COVID-19 Genomics UK (COG-UK) Consortium | The Lighthouse Lab in Milton Keynes and Alex Alderton, Roberto Amato, Jeffrey Barrett, Sonia Goncalves, Ewan Harrison, David K. Jackson, Ian Johnston, Dominic Kwiatkowski, Cordelia Langford, John Sillitoe on behalf of the Wellcome Sanger Institute COVID-19 Surveillance Team |
| EPI_ISL_2487018, EPI_ISL_2487019 | Lighthouse Lab in Alderley Park | Wellcome Sanger Institute for the COVID-19 Genomics UK (COG-UK) Consortium | Jacquelyn Wynn, Mairead Hyland, The Lighthouse Lab in Alderley Park and Alex Alderton, Roberto Amato, Jeffrey Barrett, Sonia Goncalves, Ewan Harrison, David K. Jackson, Ian Johnston, Dominic Kwiatkowski, Cordelia Langford, John Sillitoe on behalf of the Wellcome Sanger Institute COVID-19 Surveillance Team |
| EPI_ISL_2487020 | Lighthouse Lab in Milton Keynes | Wellcome Sanger Institute for the COVID-19 Genomics UK (COG-UK) Consortium | The Lighthouse Lab in Milton Keynes and Alex Alderton, Roberto Amato, Jeffrey Barrett, Sonia Goncalves, Ewan Harrison, David K. Jackson, Ian Johnston, Dominic Kwiatkowski, Cordelia Langford, John Sillitoe on behalf of the Wellcome Sanger Institute COVID-19 Surveillance Team |
| EPI_ISL_2487023, EPI_ISL_2487024 | Lighthouse Lab in Alderley Park | Wellcome Sanger Institute for the COVID-19 Genomics UK (COG-UK) Consortium | Jacquelyn Wynn, Mairead Hyland, The Lighthouse Lab in Alderley Park and Alex Alderton, Roberto Amato, Jeffrey Barrett, Sonia Goncalves, Ewan Harrison, David K. Jackson, Ian Johnston, Dominic Kwiatkowski, Cordelia Langford, John Sillitoe on behalf of the Wellcome Sanger Institute COVID-19 Surveillance Team |
| EPI_ISL_2487026 | Lighthouse Lab in Milton Keynes | Wellcome Sanger Institute for the COVID-19 Genomics UK (COG-UK) Consortium | The Lighthouse Lab in Milton Keynes and Alex Alderton, Roberto Amato, Jeffrey Barrett, Sonia Goncalves, Ewan Harrison, David K. Jackson, Ian Johnston, Dominic Kwiatkowski, Cordelia Langford, John Sillitoe on behalf of the Wellcome Sanger Institute COVID-19 Surveillance Team |
| EPI_ISL_2487027, EPI_ISL_2487028, EPI_ISL_2487029 | Lighthouse Lab in Alderley Park | Wellcome Sanger Institute for the COVID-19 Genomics UK (COG-UK) Consortium | Jacquelyn Wynn, Mairead Hyland, The Lighthouse Lab in Alderley Park and Alex Alderton, Roberto Amato, Jeffrey Barrett, Sonia Goncalves, Ewan Harrison, David K. Jackson, Ian Johnston, Dominic Kwiatkowski, Cordelia Langford, John Sillitoe on behalf of the Wellcome Sanger Institute COVID-19 Surveillance Team |
| EPI_ISL_2487030 | Lighthouse Lab in Milton Keynes | Wellcome Sanger Institute for the COVID-19 Genomics UK (COG-UK) Consortium | The Lighthouse Lab in Milton Keynes and Alex Alderton, Roberto Amato, Jeffrey Barrett, Sonia Goncalves, Ewan Harrison, David K. Jackson, Ian Johnston, Dominic Kwiatkowski, Cordelia Langford, John Sillitoe on behalf of the Wellcome Sanger Institute COVID-19 Surveillance Team |
| EPI_ISL_2487032, EPI_ISL_2487033 | Lighthouse Lab in Alderley Park | Wellcome Sanger Institute for the COVID-19 Genomics UK (COG-UK) Consortium | Jacquelyn Wynn, Mairead Hyland, The Lighthouse Lab in Alderley Park and Alex Alderton, Roberto Amato, Jeffrey Barrett, Sonia Goncalves, Ewan Harrison, David K. Jackson, Ian Johnston, Dominic Kwiatkowski, Cordelia Langford, John Sillitoe on behalf of the Wellcome Sanger Institute COVID-19 Surveillance Team |
| EPI_ISL_2487035 | Health Services Laboratories | Wellcome Sanger Institute for the COVID-19 Genomics UK (COG-UK) Consortium | Health Services Laboratories and Alex Alderton, Roberto Amato, Jeffrey Barrett, Sonia Goncalves, Ewan Harrison, David K. Jackson, Ian Johnston, Dominic Kwiatkowski, Cordelia Langford, John Sillitoe on behalf of the Wellcome Sanger Institute COVID-19 Surveillance Team |
| EPI_ISL_2487036 | Lighthouse Lab in Milton Keynes | Wellcome Sanger Institute for the COVID-19 Genomics UK (COG-UK) Consortium | The Lighthouse Lab in Milton Keynes and Alex Alderton, Roberto Amato, Jeffrey Barrett, Sonia Goncalves, Ewan Harrison, David K. Jackson, Ian Johnston, Dominic Kwiatkowski, Cordelia Langford, John Sillitoe on behalf of the Wellcome Sanger Institute COVID-19 Surveillance Team |
| EPI_ISL_2487037 | Lighthouse Lab in Alderley Park | Wellcome Sanger Institute for the COVID-19 Genomics UK (COG-UK) Consortium | Jacquelyn Wynn, Mairead Hyland, The Lighthouse Lab in Alderley Park and Alex Alderton, Roberto Amato, Jeffrey Barrett, Sonia Goncalves, Ewan Harrison, David K. Jackson, Ian Johnston, Dominic Kwiatkowski, Cordelia Langford, John Sillitoe on behalf of the Wellcome Sanger Institute COVID-19 Surveillance Team |
| EPI_ISL_2487038 | Lighthouse Lab in Milton Keynes | Wellcome Sanger Institute for the COVID-19 Genomics UK (COG-UK) Consortium | The Lighthouse Lab in Milton Keynes and Alex Alderton, Roberto Amato, Jeffrey Barrett, Sonia Goncalves, Ewan Harrison, David K. Jackson, Ian Johnston, Dominic Kwiatkowski, Cordelia Langford, John Sillitoe on behalf of the Wellcome Sanger Institute COVID-19 Surveillance Team |
| EPI_ISL_2487039, EPI_ISL_2487040 | Lighthouse Lab in Alderley Park | Wellcome Sanger Institute for the COVID-19 Genomics UK (COG-UK) Consortium | Jacquelyn Wynn, Mairead Hyland, The Lighthouse Lab in Alderley Park and Alex Alderton, Roberto Amato, Jeffrey Barrett, Sonia Goncalves, Ewan Harrison, David K. Jackson, Ian Johnston, Dominic Kwiatkowski, Cordelia Langford, John Sillitoe on behalf of the Wellcome Sanger Institute COVID-19 Surveillance Team |
| EPI_ISL_2487041, EPI_ISL_2487043, EPI_ISL_2487044 | Lighthouse Lab in Milton Keynes | Wellcome Sanger Institute for the COVID-19 Genomics UK (COG-UK) Consortium | The Lighthouse Lab in Milton Keynes and Alex Alderton, Roberto Amato, Jeffrey Barrett, Sonia Goncalves, Ewan Harrison, David K. Jackson, Ian Johnston, Dominic Kwiatkowski, Cordelia Langford, John Sillitoe on behalf of the Wellcome Sanger Institute COVID-19 Surveillance Team |
| EPI_ISL_2487045, EPI_ISL_2487046, EPI_ISL_2487047 | Lighthouse Lab in Alderley Park | Wellcome Sanger Institute for the COVID-19 Genomics UK (COG-UK) Consortium | Jacquelyn Wynn, Mairead Hyland, The Lighthouse Lab in Alderley Park and Alex Alderton, Roberto Amato, Jeffrey Barrett, Sonia Goncalves, Ewan Harrison, David K. Jackson, Ian Johnston, Dominic Kwiatkowski, Cordelia Langford, John Sillitoe on behalf of the Wellcome Sanger Institute COVID-19 Surveillance Team |
| EPI_ISL_2487048 | Lighthouse Lab in Milton Keynes | Wellcome Sanger Institute for the COVID-19 Genomics UK (COG-UK) Consortium | The Lighthouse Lab in Milton Keynes and Alex Alderton, Roberto Amato, Jeffrey Barrett, Sonia Goncalves, Ewan Harrison, David K. Jackson, Ian Johnston, Dominic Kwiatkowski, Cordelia Langford, John Sillitoe on behalf of the Wellcome Sanger Institute COVID-19 Surveillance Team |
| EPI_ISL_2487049 | Lighthouse Lab in Alderley Park | Wellcome Sanger Institute for the COVID-19 Genomics UK (COG-UK) Consortium | Jacquelyn Wynn, Mairead Hyland, The Lighthouse Lab in Alderley Park and Alex Alderton, Roberto Amato, Jeffrey Barrett, Sonia Goncalves, Ewan Harrison, David K. Jackson, Ian Johnston, Dominic Kwiatkowski, Cordelia Langford, John Sillitoe on behalf of the Wellcome Sanger Institute COVID-19 Surveillance Team |
| EPI_ISL_2487050 | Lighthouse Lab in Milton Keynes | Wellcome Sanger Institute for the COVID-19 Genomics UK (COG-UK) Consortium | The Lighthouse Lab in Milton Keynes and Alex Alderton, Roberto Amato, Jeffrey Barrett, Sonia Goncalves, Ewan Harrison, David K. Jackson, Ian Johnston, Dominic Kwiatkowski, Cordelia Langford, John Sillitoe on behalf of the Wellcome Sanger Institute COVID-19 Surveillance Team |
| EPI_ISL_2487051, EPI_ISL_2487053, EPI_ISL_2487054 | Lighthouse Lab in Alderley Park | Wellcome Sanger Institute for the COVID-19 Genomics UK (COG-UK) Consortium | Jacquelyn Wynn, Mairead Hyland, The Lighthouse Lab in Alderley Park and Alex Alderton, Roberto Amato, Jeffrey Barrett, Sonia Goncalves, Ewan Harrison, David K. Jackson, Ian Johnston, Dominic Kwiatkowski, Cordelia Langford, John Sillitoe on behalf of the Wellcome Sanger Institute COVID-19 Surveillance Team |
| EPI_ISL_2487055 | Lighthouse Lab in Milton Keynes | Wellcome Sanger Institute for the COVID-19 Genomics UK (COG-UK) Consortium | The Lighthouse Lab in Milton Keynes and Alex Alderton, Roberto Amato, Jeffrey Barrett, Sonia Goncalves, Ewan Harrison, David K. Jackson, Ian Johnston, Dominic Kwiatkowski, Cordelia Langford, John Sillitoe on behalf of the Wellcome Sanger Institute COVID-19 Surveillance Team |
| EPI_ISL_2487056 | Lighthouse Lab in Alderley Park | Wellcome Sanger Institute for the COVID-19 Genomics UK (COG-UK) Consortium | Jacquelyn Wynn, Mairead Hyland, The Lighthouse Lab in Alderley Park and Alex Alderton, Roberto Amato, Jeffrey Barrett, Sonia Goncalves, Ewan Harrison, David K. Jackson, Ian Johnston, Dominic Kwiatkowski, Cordelia Langford, John Sillitoe on behalf of the Wellcome Sanger Institute COVID-19 Surveillance Team |
| EPI_ISL_2487057, EPI_ISL_2487058 | Lighthouse Lab in Milton Keynes | Wellcome Sanger Institute for the COVID-19 Genomics UK (COG-UK) Consortium | The Lighthouse Lab in Milton Keynes and Alex Alderton, Roberto Amato, Jeffrey Barrett, Sonia Goncalves, Ewan Harrison, David K. Jackson, Ian Johnston, Dominic Kwiatkowski, Cordelia Langford, John Sillitoe on behalf of the Wellcome Sanger Institute COVID-19 Surveillance Team |
| EPI_ISL_2487059, EPI_ISL_2487060 | Lighthouse Lab in Alderley Park | Wellcome Sanger Institute for the COVID-19 Genomics UK (COG-UK) Consortium | Jacquelyn Wynn, Mairead Hyland, The Lighthouse Lab in Alderley Park and Alex Alderton, Roberto Amato, Jeffrey Barrett, Sonia Goncalves, Ewan Harrison, David K. Jackson, Ian Johnston, Dominic Kwiatkowski, Cordelia Langford, John Sillitoe on behalf of the Wellcome Sanger Institute COVID-19 Surveillance Team |
| EPI_ISL_2487061, EPI_ISL_2487062 | Lighthouse Lab in Milton Keynes | Wellcome Sanger Institute for the COVID-19 Genomics UK (COG-UK) Consortium | The Lighthouse Lab in Milton Keynes and Alex Alderton, Roberto Amato, Jeffrey Barrett, Sonia Goncalves, Ewan Harrison, David K. Jackson, Ian Johnston, Dominic Kwiatkowski, Cordelia Langford, John Sillitoe on behalf of the Wellcome Sanger Institute COVID-19 Surveillance Team |
| EPI_ISL_2487063, EPI_ISL_2487064 | Lighthouse Lab in Alderley Park | Wellcome Sanger Institute for the COVID-19 Genomics UK (COG-UK) Consortium | Jacquelyn Wynn, Mairead Hyland, The Lighthouse Lab in Alderley Park and Alex Alderton, Roberto Amato, Jeffrey Barrett, Sonia Goncalves, Ewan Harrison, David K. Jackson, Ian Johnston, Dominic Kwiatkowski, Cordelia Langford, John Sillitoe on behalf of the Wellcome Sanger Institute COVID-19 Surveillance Team |
| EPI_ISL_2487065 | Lighthouse Lab in Milton Keynes | Wellcome Sanger Institute for the COVID-19 Genomics UK (COG-UK) Consortium | The Lighthouse Lab in Milton Keynes and Alex Alderton, Roberto Amato, Jeffrey Barrett, Sonia Goncalves, Ewan Harrison, David K. Jackson, Ian Johnston, Dominic Kwiatkowski, Cordelia Langford, John Sillitoe on behalf of the Wellcome Sanger Institute COVID-19 Surveillance Team |
| EPI_ISL_2487066 | Lighthouse Lab in Alderley Park | Wellcome Sanger Institute for the COVID-19 Genomics UK (COG-UK) Consortium | Jacquelyn Wynn, Mairead Hyland, The Lighthouse Lab in Alderley Park and Alex Alderton, Roberto Amato, Jeffrey Barrett, Sonia Goncalves, Ewan Harrison, David K. Jackson, Ian Johnston, Dominic Kwiatkowski, Cordelia Langford, John Sillitoe on behalf of the Wellcome Sanger Institute COVID-19 Surveillance Team |
| EPI_ISL_2487067 | Lighthouse Lab in Milton Keynes | Wellcome Sanger Institute for the COVID-19 Genomics UK (COG-UK) Consortium | The Lighthouse Lab in Milton Keynes and Alex Alderton, Roberto Amato, Jeffrey Barrett, Sonia Goncalves, Ewan Harrison, David K. Jackson, Ian Johnston, Dominic Kwiatkowski, Cordelia Langford, John Sillitoe on behalf of the Wellcome Sanger Institute COVID-19 Surveillance Team |
| EPI_ISL_2487068, EPI_ISL_2487069 | Lighthouse Lab in Alderley Park | Wellcome Sanger Institute for the COVID-19 Genomics UK (COG-UK) Consortium | Jacquelyn Wynn, Mairead Hyland, The Lighthouse Lab in Alderley Park and Alex Alderton, Roberto Amato, Jeffrey Barrett, Sonia Goncalves, Ewan Harrison, David K. Jackson, Ian Johnston, Dominic Kwiatkowski, Cordelia Langford, John Sillitoe on behalf of the Wellcome Sanger Institute COVID-19 Surveillance Team |
| EPI_ISL_2487070, EPI_ISL_2487071 | Lighthouse Lab in Milton Keynes | Wellcome Sanger Institute for the COVID-19 Genomics UK (COG-UK) Consortium | The Lighthouse Lab in Milton Keynes and Alex Alderton, Roberto Amato, Jeffrey Barrett, Sonia Goncalves, Ewan Harrison, David K. Jackson, Ian Johnston, Dominic Kwiatkowski, Cordelia Langford, John Sillitoe on behalf of the Wellcome Sanger Institute COVID-19 Surveillance Team |
| EPI_ISL_2487072 | Lighthouse Lab in Alderley Park | Wellcome Sanger Institute for the COVID-19 Genomics UK (COG-UK) Consortium | Jacquelyn Wynn, Mairead Hyland, The Lighthouse Lab in Alderley Park and Alex Alderton, Roberto Amato, Jeffrey Barrett, Sonia Goncalves, Ewan Harrison, David K. Jackson, Ian Johnston, Dominic Kwiatkowski, Cordelia Langford, John Sillitoe on behalf of the Wellcome Sanger Institute COVID-19 Surveillance Team |
| EPI_ISL_2487073 | Lighthouse Lab in Milton Keynes | Wellcome Sanger Institute for the COVID-19 Genomics UK (COG-UK) Consortium | The Lighthouse Lab in Milton Keynes and Alex Alderton, Roberto Amato, Jeffrey Barrett, Sonia Goncalves, Ewan Harrison, David K. Jackson, Ian Johnston, Dominic Kwiatkowski, Cordelia Langford, John Sillitoe on behalf of the Wellcome Sanger Institute COVID-19 Surveillance Team |
| EPI_ISL_2487074, EPI_ISL_2487075 | Lighthouse Lab in Alderley Park | Wellcome Sanger Institute for the COVID-19 Genomics UK (COG-UK) Consortium | Jacquelyn Wynn, Mairead Hyland, The Lighthouse Lab in Alderley Park and Alex Alderton, Roberto Amato, Jeffrey Barrett, Sonia Goncalves, Ewan Harrison, David K. Jackson, Ian Johnston, Dominic Kwiatkowski, Cordelia Langford, John Sillitoe on behalf of the Wellcome Sanger Institute COVID-19 Surveillance Team |
| EPI_ISL_2487077 | Lighthouse Lab in Milton Keynes | Wellcome Sanger Institute for the COVID-19 Genomics UK (COG-UK) Consortium | The Lighthouse Lab in Milton Keynes and Alex Alderton, Roberto Amato, Jeffrey Barrett, Sonia Goncalves, Ewan Harrison, David K. Jackson, Ian Johnston, Dominic Kwiatkowski, Cordelia Langford, John Sillitoe on behalf of the Wellcome Sanger Institute COVID-19 Surveillance Team |
| EPI_ISL_2487078 | Lighthouse Lab in Alderley Park | Wellcome Sanger Institute for the COVID-19 Genomics UK (COG-UK) Consortium | Jacquelyn Wynn, Mairead Hyland, The Lighthouse Lab in Alderley Park and Alex Alderton, Roberto Amato, Jeffrey Barrett, Sonia Goncalves, Ewan Harrison, David K. Jackson, Ian Johnston, Dominic Kwiatkowski, Cordelia Langford, John Sillitoe on behalf of the Wellcome Sanger Institute COVID-19 Surveillance Team |
| EPI_ISL_2487079, EPI_ISL_2487080 | Lighthouse Lab in Milton Keynes | Wellcome Sanger Institute for the COVID-19 Genomics UK (COG-UK) Consortium | The Lighthouse Lab in Milton Keynes and Alex Alderton, Roberto Amato, Jeffrey Barrett, Sonia Goncalves, Ewan Harrison, David K. Jackson, Ian Johnston, Dominic Kwiatkowski, Cordelia Langford, John Sillitoe on behalf of the Wellcome Sanger Institute COVID-19 Surveillance Team |
| EPI_ISL_2487081 | Lighthouse Lab in Alderley Park | Wellcome Sanger Institute for the COVID-19 Genomics UK (COG-UK) Consortium | Jacquelyn Wynn, Mairead Hyland, The Lighthouse Lab in Alderley Park and Alex Alderton, Roberto Amato, Jeffrey Barrett, Sonia Goncalves, Ewan Harrison, David K. Jackson, Ian Johnston, Dominic Kwiatkowski, Cordelia Langford, John Sillitoe on behalf of the Wellcome Sanger Institute COVID-19 Surveillance Team |
| EPI_ISL_2487082 | Lighthouse Lab in Milton Keynes | Wellcome Sanger Institute for the COVID-19 Genomics UK (COG-UK) Consortium | The Lighthouse Lab in Milton Keynes and Alex Alderton, Roberto Amato, Jeffrey Barrett, Sonia Goncalves, Ewan Harrison, David K. Jackson, Ian Johnston, Dominic Kwiatkowski, Cordelia Langford, John Sillitoe on behalf of the Wellcome Sanger Institute COVID-19 Surveillance Team |
| EPI_ISL_2487084, EPI_ISL_2487085, EPI_ISL_2487086, EPI_ISL_2487087 | Lighthouse Lab in Alderley Park | Wellcome Sanger Institute for the COVID-19 Genomics UK (COG-UK) Consortium | Jacquelyn Wynn, Mairead Hyland, The Lighthouse Lab in Alderley Park and Alex Alderton, Roberto Amato, Jeffrey Barrett, Sonia Goncalves, Ewan Harrison, David K. Jackson, Ian Johnston, Dominic Kwiatkowski, Cordelia Langford, John Sillitoe on behalf of the Wellcome Sanger Institute COVID-19 Surveillance Team |
| EPI_ISL_2487088 | Health Services Laboratories | Wellcome Sanger Institute for the COVID-19 Genomics UK (COG-UK) Consortium | Health Services Laboratories and Alex Alderton, Roberto Amato, Jeffrey Barrett, Sonia Goncalves, Ewan Harrison, David K. Jackson, Ian Johnston, Dominic Kwiatkowski, Cordelia Langford, John Sillitoe on behalf of the Wellcome Sanger Institute COVID-19 Surveillance Team |
| EPI_ISL_2487089, EPI_ISL_2487090, EPI_ISL_2487092 | Lighthouse Lab in Milton Keynes | Wellcome Sanger Institute for the COVID-19 Genomics UK (COG-UK) Consortium | The Lighthouse Lab in Milton Keynes and Alex Alderton, Roberto Amato, Jeffrey Barrett, Sonia Goncalves, Ewan Harrison, David K. Jackson, Ian Johnston, Dominic Kwiatkowski, Cordelia Langford, John Sillitoe on behalf of the Wellcome Sanger Institute COVID-19 Surveillance Team |
| EPI_ISL_2487093, EPI_ISL_2487094, EPI_ISL_2487095 | Lighthouse Lab in Alderley Park | Wellcome Sanger Institute for the COVID-19 Genomics UK (COG-UK) Consortium | Jacquelyn Wynn, Mairead Hyland, The Lighthouse Lab in Alderley Park and Alex Alderton, Roberto Amato, Jeffrey Barrett, Sonia Goncalves, Ewan Harrison, David K. Jackson, Ian Johnston, Dominic Kwiatkowski, Cordelia Langford, John Sillitoe on behalf of the Wellcome Sanger Institute COVID-19 Surveillance Team |

| | | | |
|---|---|---|---|
| EPI_ISL_2487096 | Health Services Laboratories | Wellcome Sanger Institute for the COVID-19 Genomics UK (COG-UK) Consortium | Health Services Laboratories and Alex Alderton, Roberto Amato, Jeffrey Barrett, Sonia Goncalves, Ewan Harrison, David K. Jackson, Ian Johnston, Dominic Kwiatkowski, Cordelia Langford, John Sillitoe on behalf of the Wellcome Sanger Institute COVID-19 Surveillance Team |
| EPI_ISL_2487097 | Lighthouse Lab in Milton Keynes | Wellcome Sanger Institute for the COVID-19 Genomics UK (COG-UK) Consortium | The Lighthouse Lab in Milton Keynes and Alex Alderton, Roberto Amato, Jeffrey Barrett, Sonia Goncalves, Ewan Harrison, David K. Jackson, Ian Johnston, Dominic Kwiatkowski, Cordelia Langford, John Sillitoe on behalf of the Wellcome Sanger Institute COVID-19 Surveillance Team |
| EPI_ISL_2487098, EPI_ISL_2487099, EPI_ISL_2487100, EPI_ISL_2487101, EPI_ISL_2487102, EPI_ISL_2487103, EPI_ISL_2487104, EPI_ISL_2487105 | Lighthouse Lab in Alderley Park | Wellcome Sanger Institute for the COVID-19 Genomics UK (COG-UK) Consortium | Jacquelyn Wynn, Mairead Hyland, The Lighthouse Lab in Alderley Park and Alex Alderton, Roberto Amato, Jeffrey Barrett, Sonia Goncalves, Ewan Harrison, David K. Jackson, Ian Johnston, Dominic Kwiatkowski, Cordelia Langford, John Sillitoe on behalf of the Wellcome Sanger Institute COVID-19 Surveillance Team |
| EPI_ISL_2487106 | Lighthouse Lab in Milton Keynes | Wellcome Sanger Institute for the COVID-19 Genomics UK (COG-UK) Consortium | The Lighthouse Lab in Milton Keynes and Alex Alderton, Roberto Amato, Jeffrey Barrett, Sonia Goncalves, Ewan Harrison, David K. Jackson, Ian Johnston, Dominic Kwiatkowski, Cordelia Langford, John Sillitoe on behalf of the Wellcome Sanger Institute COVID-19 Surveillance Team |
| EPI_ISL_2487108, EPI_ISL_2487109, EPI_ISL_2487110 | Lighthouse Lab in Alderley Park | Wellcome Sanger Institute for the COVID-19 Genomics UK (COG-UK) Consortium | Jacquelyn Wynn, Mairead Hyland, The Lighthouse Lab in Alderley Park and Alex Alderton, Roberto Amato, Jeffrey Barrett, Sonia Goncalves, Ewan Harrison, David K. Jackson, Ian Johnston, Dominic Kwiatkowski, Cordelia Langford, John Sillitoe on behalf of the Wellcome Sanger Institute COVID-19 Surveillance Team |
| EPI_ISL_2487113 | Lighthouse Lab in Milton Keynes | Wellcome Sanger Institute for the COVID-19 Genomics UK (COG-UK) Consortium | The Lighthouse Lab in Milton Keynes and Alex Alderton, Roberto Amato, Jeffrey Barrett, Sonia Goncalves, Ewan Harrison, David K. Jackson, Ian Johnston, Dominic Kwiatkowski, Cordelia Langford, John Sillitoe on behalf of the Wellcome Sanger Institute COVID-19 Surveillance Team |
| EPI_ISL_2487115 | Lighthouse Lab in Alderley Park | Wellcome Sanger Institute for the COVID-19 Genomics UK (COG-UK) Consortium | Jacquelyn Wynn, Mairead Hyland, The Lighthouse Lab in Alderley Park and Alex Alderton, Roberto Amato, Jeffrey Barrett, Sonia Goncalves, Ewan Harrison, David K. Jackson, Ian Johnston, Dominic Kwiatkowski, Cordelia Langford, John Sillitoe on behalf of the Wellcome Sanger Institute COVID-19 Surveillance Team |
| EPI_ISL_2487116 | Health Services Laboratories | Wellcome Sanger Institute for the COVID-19 Genomics UK (COG-UK) Consortium | Health Services Laboratories and Alex Alderton, Roberto Amato, Jeffrey Barrett, Sonia Goncalves, Ewan Harrison, David K. Jackson, Ian Johnston, Dominic Kwiatkowski, Cordelia Langford, John Sillitoe on behalf of the Wellcome Sanger Institute COVID-19 Surveillance Team |
| EPI_ISL_2487119, EPI_ISL_2487120 | Lighthouse Lab in Alderley Park | Wellcome Sanger Institute for the COVID-19 Genomics UK (COG-UK) Consortium | Jacquelyn Wynn, Mairead Hyland, The Lighthouse Lab in Alderley Park and Alex Alderton, Roberto Amato, Jeffrey Barrett, Sonia Goncalves, Ewan Harrison, David K. Jackson, Ian Johnston, Dominic Kwiatkowski, Cordelia Langford, John Sillitoe on behalf of the Wellcome Sanger Institute COVID-19 Surveillance Team |
| EPI_ISL_2487121 | Lighthouse Lab in Milton Keynes | Wellcome Sanger Institute for the COVID-19 Genomics UK (COG-UK) Consortium | The Lighthouse Lab in Milton Keynes and Alex Alderton, Roberto Amato, Jeffrey Barrett, Sonia Goncalves, Ewan Harrison, David K. Jackson, Ian Johnston, Dominic Kwiatkowski, Cordelia Langford, John Sillitoe on behalf of the Wellcome Sanger Institute COVID-19 Surveillance Team |
| EPI_ISL_2487122, EPI_ISL_2487123 | Lighthouse Lab in Alderley Park | Wellcome Sanger Institute for the COVID-19 Genomics UK (COG-UK) Consortium | Jacquelyn Wynn, Mairead Hyland, The Lighthouse Lab in Alderley Park and Alex Alderton, Roberto Amato, Jeffrey Barrett, Sonia Goncalves, Ewan Harrison, David K. Jackson, Ian Johnston, Dominic Kwiatkowski, Cordelia Langford, John Sillitoe on behalf of the Wellcome Sanger Institute COVID-19 Surveillance Team |
| EPI_ISL_2487124 | Lighthouse Lab in Milton Keynes | Wellcome Sanger Institute for the COVID-19 Genomics UK (COG-UK) Consortium | The Lighthouse Lab in Milton Keynes and Alex Alderton, Roberto Amato, Jeffrey Barrett, Sonia Goncalves, Ewan Harrison, David K. Jackson, Ian Johnston, Dominic Kwiatkowski, Cordelia Langford, John Sillitoe on behalf of the Wellcome Sanger Institute COVID-19 Surveillance Team |
| EPI_ISL_2487125 | Lighthouse Lab in Alderley Park | Wellcome Sanger Institute for the COVID-19 Genomics UK (COG-UK) Consortium | Jacquelyn Wynn, Mairead Hyland, The Lighthouse Lab in Alderley Park and Alex Alderton, Roberto Amato, Jeffrey Barrett, Sonia Goncalves, Ewan Harrison, David K. Jackson, Ian Johnston, Dominic Kwiatkowski, Cordelia Langford, John Sillitoe on behalf of the Wellcome Sanger Institute COVID-19 Surveillance Team |
| EPI_ISL_2487126 | Lighthouse Lab in Milton Keynes | Wellcome Sanger Institute for the COVID-19 Genomics UK (COG-UK) Consortium | The Lighthouse Lab in Milton Keynes and Alex Alderton, Roberto Amato, Jeffrey Barrett, Sonia Goncalves, Ewan Harrison, David K. Jackson, Ian Johnston, Dominic Kwiatkowski, Cordelia Langford, John Sillitoe on behalf of the Wellcome Sanger Institute COVID-19 Surveillance Team |
| EPI_ISL_2487127 | Lighthouse Lab in Alderley Park | Wellcome Sanger Institute for the COVID-19 Genomics UK (COG-UK) Consortium | Jacquelyn Wynn, Mairead Hyland, The Lighthouse Lab in Alderley Park and Alex Alderton, Roberto Amato, Jeffrey Barrett, Sonia Goncalves, Ewan Harrison, David K. Jackson, Ian Johnston, Dominic Kwiatkowski, Cordelia Langford, John Sillitoe on behalf of the Wellcome Sanger Institute COVID-19 Surveillance Team |
| EPI_ISL_2487129, EPI_ISL_2487130 | Lighthouse Lab in Milton Keynes | Wellcome Sanger Institute for the COVID-19 Genomics UK (COG-UK) Consortium | The Lighthouse Lab in Milton Keynes and Alex Alderton, Roberto Amato, Jeffrey Barrett, Sonia Goncalves, Ewan Harrison, David K. Jackson, Ian Johnston, Dominic Kwiatkowski, Cordelia Langford, John Sillitoe on behalf of the Wellcome Sanger Institute COVID-19 Surveillance Team |
| EPI_ISL_2487131, EPI_ISL_2487132, EPI_ISL_2487133, EPI_ISL_2487134, EPI_ISL_2487135 | Lighthouse Lab in Alderley Park | Wellcome Sanger Institute for the COVID-19 Genomics UK (COG-UK) Consortium | Jacquelyn Wynn, Mairead Hyland, The Lighthouse Lab in Alderley Park and Alex Alderton, Roberto Amato, Jeffrey Barrett, Sonia Goncalves, Ewan Harrison, David K. Jackson, Ian Johnston, Dominic Kwiatkowski, Cordelia Langford, John Sillitoe on behalf of the Wellcome Sanger Institute COVID-19 Surveillance Team |
| EPI_ISL_2487137 | Lighthouse Lab in Milton Keynes | Wellcome Sanger Institute for the COVID-19 Genomics UK (COG-UK) Consortium | The Lighthouse Lab in Milton Keynes and Alex Alderton, Roberto Amato, Jeffrey Barrett, Sonia Goncalves, Ewan Harrison, David K. Jackson, Ian Johnston, Dominic Kwiatkowski, Cordelia Langford, John Sillitoe on behalf of the Wellcome Sanger Institute COVID-19 Surveillance Team |
| EPI_ISL_2487138, EPI_ISL_2487139, EPI_ISL_2487140 | Lighthouse Lab in Alderley Park | Wellcome Sanger Institute for the COVID-19 Genomics UK (COG-UK) Consortium | Jacquelyn Wynn, Mairead Hyland, The Lighthouse Lab in Alderley Park and Alex Alderton, Roberto Amato, Jeffrey Barrett, Sonia Goncalves, Ewan Harrison, David K. Jackson, Ian Johnston, Dominic Kwiatkowski, Cordelia Langford, John Sillitoe on behalf of the Wellcome Sanger Institute COVID-19 Surveillance Team |
| EPI_ISL_2487141 | Lighthouse Lab in Milton Keynes | Wellcome Sanger Institute for the COVID-19 Genomics UK (COG-UK) Consortium | The Lighthouse Lab in Milton Keynes and Alex Alderton, Roberto Amato, Jeffrey Barrett, Sonia Goncalves, Ewan Harrison, David K. Jackson, Ian Johnston, Dominic Kwiatkowski, Cordelia Langford, John Sillitoe on behalf of the Wellcome Sanger Institute COVID-19 Surveillance Team |
| EPI_ISL_2487142 | Lighthouse Lab in Alderley Park | Wellcome Sanger Institute for the COVID-19 Genomics UK (COG-UK) Consortium | Jacquelyn Wynn, Mairead Hyland, The Lighthouse Lab in Alderley Park and Alex Alderton, Roberto Amato, Jeffrey Barrett, Sonia Goncalves, Ewan Harrison, David K. Jackson, Ian Johnston, Dominic Kwiatkowski, Cordelia Langford, John Sillitoe on behalf of the Wellcome Sanger Institute COVID-19 Surveillance Team |
| EPI_ISL_2487144, EPI_ISL_2487145 | Lighthouse Lab in Milton Keynes | Wellcome Sanger Institute for the COVID-19 Genomics UK (COG-UK) Consortium | The Lighthouse Lab in Milton Keynes and Alex Alderton, Roberto Amato, Jeffrey Barrett, Sonia Goncalves, Ewan Harrison, David K. Jackson, Ian Johnston, Dominic Kwiatkowski, Cordelia Langford, John Sillitoe on behalf of the Wellcome Sanger Institute COVID-19 Surveillance Team |
| EPI_ISL_2487146 | Lighthouse Lab in Alderley Park | Wellcome Sanger Institute for the COVID-19 Genomics UK (COG-UK) Consortium | Jacquelyn Wynn, Mairead Hyland, The Lighthouse Lab in Alderley Park and Alex Alderton, Roberto Amato, Jeffrey Barrett, Sonia Goncalves, Ewan Harrison, David K. Jackson, Ian Johnston, Dominic Kwiatkowski, Cordelia Langford, John Sillitoe on behalf of the Wellcome Sanger Institute COVID-19 Surveillance Team |
| EPI_ISL_2487147 | Lighthouse Lab in Milton Keynes | Wellcome Sanger Institute for the COVID-19 Genomics UK (COG-UK) Consortium | The Lighthouse Lab in Milton Keynes and Alex Alderton, Roberto Amato, Jeffrey Barrett, Sonia Goncalves, Ewan Harrison, David K. Jackson, Ian Johnston, Dominic Kwiatkowski, Cordelia Langford, John Sillitoe on behalf of the Wellcome Sanger Institute COVID-19 Surveillance Team |
| EPI_ISL_2487148 | Health Services Laboratories | Wellcome Sanger Institute for the COVID-19 Genomics UK (COG-UK) Consortium | Health Services Laboratories and Alex Alderton, Roberto Amato, Jeffrey Barrett, Sonia Goncalves, Ewan Harrison, David K. Jackson, Ian Johnston, Dominic Kwiatkowski, Cordelia Langford, John Sillitoe on behalf of the Wellcome Sanger Institute COVID-19 Surveillance Team |
| EPI_ISL_2487149 | Lighthouse Lab in Alderley Park | Wellcome Sanger Institute for the COVID-19 Genomics UK (COG-UK) Consortium | Jacquelyn Wynn, Mairead Hyland, The Lighthouse Lab in Alderley Park and Alex Alderton, Roberto Amato, Jeffrey Barrett, Sonia Goncalves, Ewan Harrison, David K. Jackson, Ian Johnston, Dominic Kwiatkowski, Cordelia Langford, John Sillitoe on behalf of the Wellcome Sanger Institute COVID-19 Surveillance Team |
| EPI_ISL_2487150, EPI_ISL_2487151 | Lighthouse Lab in Milton Keynes | Wellcome Sanger Institute for the COVID-19 Genomics UK (COG-UK) Consortium | The Lighthouse Lab in Milton Keynes and Alex Alderton, Roberto Amato, Jeffrey Barrett, Sonia Goncalves, Ewan Harrison, David K. Jackson, Ian Johnston, Dominic Kwiatkowski, Cordelia Langford, John Sillitoe on behalf of the Wellcome Sanger Institute COVID-19 Surveillance Team |
| EPI_ISL_2487152 | Lighthouse Lab in Alderley Park | Wellcome Sanger Institute for the COVID-19 Genomics UK (COG-UK) Consortium | Jacquelyn Wynn, Mairead Hyland, The Lighthouse Lab in Alderley Park and Alex Alderton, Roberto Amato, Jeffrey Barrett, Sonia Goncalves, Ewan Harrison, David K. Jackson, Ian Johnston, Dominic Kwiatkowski, Cordelia Langford, John Sillitoe on behalf of the Wellcome Sanger Institute COVID-19 Surveillance Team |
| EPI_ISL_2487153 | Lighthouse Lab in Milton Keynes | Wellcome Sanger Institute for the COVID-19 Genomics UK (COG-UK) Consortium | The Lighthouse Lab in Milton Keynes and Alex Alderton, Roberto Amato, Jeffrey Barrett, Sonia Goncalves, Ewan Harrison, David K. Jackson, Ian Johnston, Dominic Kwiatkowski, Cordelia Langford, John Sillitoe on behalf of the Wellcome Sanger Institute COVID-19 Surveillance Team |
| EPI_ISL_2487154, EPI_ISL_2487155, EPI_ISL_2487156, EPI_ISL_2487157 | Lighthouse Lab in Alderley Park | Wellcome Sanger Institute for the COVID-19 Genomics UK (COG-UK) Consortium | Jacquelyn Wynn, Mairead Hyland, The Lighthouse Lab in Alderley Park and Alex Alderton, Roberto Amato, Jeffrey Barrett, Sonia Goncalves, Ewan Harrison, David K. Jackson, Ian Johnston, Dominic Kwiatkowski, Cordelia Langford, John Sillitoe on behalf of the Wellcome Sanger Institute COVID-19 Surveillance Team |
| EPI_ISL_2487158, EPI_ISL_2487159 | Lighthouse Lab in Milton Keynes | Wellcome Sanger Institute for the COVID-19 Genomics UK (COG-UK) Consortium | The Lighthouse Lab in Milton Keynes and Alex Alderton, Roberto Amato, Jeffrey Barrett, Sonia Goncalves, Ewan Harrison, David K. Jackson, Ian Johnston, Dominic Kwiatkowski, Cordelia Langford, John Sillitoe on behalf of the Wellcome Sanger Institute COVID-19 Surveillance Team |
| EPI_ISL_2487160 | Lighthouse Lab in Alderley Park | Wellcome Sanger Institute for the COVID-19 Genomics UK (COG-UK) Consortium | Jacquelyn Wynn, Mairead Hyland, The Lighthouse Lab in Alderley Park and Alex Alderton, Roberto Amato, Jeffrey Barrett, Sonia Goncalves, Ewan Harrison, David K. Jackson, Ian Johnston, Dominic Kwiatkowski, Cordelia Langford, John Sillitoe on behalf of the Wellcome Sanger Institute COVID-19 Surveillance Team |
| EPI_ISL_2487161 | Lighthouse Lab in Milton Keynes | Wellcome Sanger Institute for the COVID-19 Genomics UK (COG-UK) Consortium | The Lighthouse Lab in Milton Keynes and Alex Alderton, Roberto Amato, Jeffrey Barrett, Sonia Goncalves, Ewan Harrison, David K. Jackson, Ian Johnston, Dominic Kwiatkowski, Cordelia Langford, John Sillitoe on behalf of the Wellcome Sanger Institute COVID-19 Surveillance Team |
| EPI_ISL_2487162, EPI_ISL_2487163 | Lighthouse Lab in Alderley Park | Wellcome Sanger Institute for the COVID-19 Genomics UK (COG-UK) Consortium | Jacquelyn Wynn, Mairead Hyland, The Lighthouse Lab in Alderley Park and Alex Alderton, Roberto Amato, Jeffrey Barrett, Sonia Goncalves, Ewan Harrison, David K. Jackson, Ian Johnston, Dominic Kwiatkowski, Cordelia Langford, John Sillitoe on behalf of the Wellcome Sanger Institute COVID-19 Surveillance Team |
| EPI_ISL_2487164 | Lighthouse Lab in Milton Keynes | Wellcome Sanger Institute for the COVID-19 Genomics UK (COG-UK) Consortium | The Lighthouse Lab in Milton Keynes and Alex Alderton, Roberto Amato, Jeffrey Barrett, Sonia Goncalves, Ewan Harrison, David K. Jackson, Ian Johnston, Dominic Kwiatkowski, Cordelia Langford, John Sillitoe on behalf of the Wellcome Sanger Institute COVID-19 Surveillance Team |
| EPI_ISL_2487165, EPI_ISL_2487166, EPI_ISL_2487167, EPI_ISL_2487168 | Lighthouse Lab in Alderley Park | Wellcome Sanger Institute for the COVID-19 Genomics UK (COG-UK) Consortium | Jacquelyn Wynn, Mairead Hyland, The Lighthouse Lab in Alderley Park and Alex Alderton, Roberto Amato, Jeffrey Barrett, Sonia Goncalves, Ewan Harrison, David K. Jackson, Ian Johnston, Dominic Kwiatkowski, Cordelia Langford, John Sillitoe on behalf of the Wellcome Sanger Institute COVID-19 Surveillance Team |
| EPI_ISL_2487169, EPI_ISL_2487170 | Lighthouse Lab in Milton Keynes | Wellcome Sanger Institute for the COVID-19 Genomics UK (COG-UK) Consortium | The Lighthouse Lab in Milton Keynes and Alex Alderton, Roberto Amato, Jeffrey Barrett, Sonia Goncalves, Ewan Harrison, David K. Jackson, Ian Johnston, Dominic Kwiatkowski, Cordelia Langford, John Sillitoe on behalf of the Wellcome Sanger Institute COVID-19 Surveillance Team |
| EPI_ISL_2487172 | Health Services Laboratories | Wellcome Sanger Institute for the COVID-19 Genomics UK (COG-UK) Consortium | Health Services Laboratories and Alex Alderton, Roberto Amato, Jeffrey Barrett, Sonia Goncalves, Ewan Harrison, David K. Jackson, Ian Johnston, Dominic Kwiatkowski, Cordelia Langford, John Sillitoe on behalf of the Wellcome Sanger Institute COVID-19 Surveillance Team |
| EPI_ISL_2487174 | Lighthouse Lab in Milton Keynes | Wellcome Sanger Institute for the COVID-19 Genomics UK (COG-UK) Consortium | The Lighthouse Lab in Milton Keynes and Alex Alderton, Roberto Amato, Jeffrey Barrett, Sonia Goncalves, Ewan Harrison, David K. Jackson, Ian Johnston, Dominic Kwiatkowski, Cordelia Langford, John Sillitoe on behalf of the Wellcome Sanger Institute COVID-19 Surveillance Team |

| | | | |
|---|---|---|---|
| EPI_ISL_2487175 | Health Services Laboratories | Wellcome Sanger Institute for the COVID-19 Genomics UK (COG-UK) Consortium | Health Services Laboratories and Alex Alderton, Roberto Amato, Jeffrey Barrett, Sonia Goncalves, Ewan Harrison, David K. Jackson, Ian Johnston, Dominic Kwiatkowski, Cordelia Langford, John Sillitoe on behalf of the Wellcome Sanger Institute COVID-19 Surveillance Team |
| EPI_ISL_2487176 | Lighthouse Lab in Alderley Park | Wellcome Sanger Institute for the COVID-19 Genomics UK (COG-UK) Consortium | Jacquelyn Wynn, Mairead Hyland, The Lighthouse Lab in Alderley Park and Alex Alderton, Roberto Amato, Jeffrey Barrett, Sonia Goncalves, Ewan Harrison, David K. Jackson, Ian Johnston, Dominic Kwiatkowski, Cordelia Langford, John Sillitoe on behalf of the Wellcome Sanger Institute COVID-19 Surveillance Team |
| EPI_ISL_2487177 | Lighthouse Lab in Milton Keynes | Wellcome Sanger Institute for the COVID-19 Genomics UK (COG-UK) Consortium | The Lighthouse Lab in Milton Keynes and Alex Alderton, Roberto Amato, Jeffrey Barrett, Sonia Goncalves, Ewan Harrison, David K. Jackson, Ian Johnston, Dominic Kwiatkowski, Cordelia Langford, John Sillitoe on behalf of the Wellcome Sanger Institute COVID-19 Surveillance Team |
| EPI_ISL_2487178, EPI_ISL_2487179, EPI_ISL_2487180 | Lighthouse Lab in Alderley Park | Wellcome Sanger Institute for the COVID-19 Genomics UK (COG-UK) Consortium | Jacquelyn Wynn, Mairead Hyland, The Lighthouse Lab in Alderley Park and Alex Alderton, Roberto Amato, Jeffrey Barrett, Sonia Goncalves, Ewan Harrison, David K. Jackson, Ian Johnston, Dominic Kwiatkowski, Cordelia Langford, John Sillitoe on behalf of the Wellcome Sanger Institute COVID-19 Surveillance Team |
| EPI_ISL_2487181 | Lighthouse Lab in Milton Keynes | Wellcome Sanger Institute for the COVID-19 Genomics UK (COG-UK) Consortium | The Lighthouse Lab in Milton Keynes and Alex Alderton, Roberto Amato, Jeffrey Barrett, Sonia Goncalves, Ewan Harrison, David K. Jackson, Ian Johnston, Dominic Kwiatkowski, Cordelia Langford, John Sillitoe on behalf of the Wellcome Sanger Institute COVID-19 Surveillance Team |
| EPI_ISL_2487182 | Lighthouse Lab in Alderley Park | Wellcome Sanger Institute for the COVID-19 Genomics UK (COG-UK) Consortium | Jacquelyn Wynn, Mairead Hyland, The Lighthouse Lab in Alderley Park and Alex Alderton, Roberto Amato, Jeffrey Barrett, Sonia Goncalves, Ewan Harrison, David K. Jackson, Ian Johnston, Dominic Kwiatkowski, Cordelia Langford, John Sillitoe on behalf of the Wellcome Sanger Institute COVID-19 Surveillance Team |
| EPI_ISL_2487183 | Lighthouse Lab in Milton Keynes | Wellcome Sanger Institute for the COVID-19 Genomics UK (COG-UK) Consortium | The Lighthouse Lab in Milton Keynes and Alex Alderton, Roberto Amato, Jeffrey Barrett, Sonia Goncalves, Ewan Harrison, David K. Jackson, Ian Johnston, Dominic Kwiatkowski, Cordelia Langford, John Sillitoe on behalf of the Wellcome Sanger Institute COVID-19 Surveillance Team |
| EPI_ISL_2487184, EPI_ISL_2487185 | Lighthouse Lab in Alderley Park | Wellcome Sanger Institute for the COVID-19 Genomics UK (COG-UK) Consortium | Jacquelyn Wynn, Mairead Hyland, The Lighthouse Lab in Alderley Park and Alex Alderton, Roberto Amato, Jeffrey Barrett, Sonia Goncalves, Ewan Harrison, David K. Jackson, Ian Johnston, Dominic Kwiatkowski, Cordelia Langford, John Sillitoe on behalf of the Wellcome Sanger Institute COVID-19 Surveillance Team |
| EPI_ISL_2487187, EPI_ISL_2487188, EPI_ISL_2487189 | Lighthouse Lab in Milton Keynes | Wellcome Sanger Institute for the COVID-19 Genomics UK (COG-UK) Consortium | The Lighthouse Lab in Milton Keynes and Alex Alderton, Roberto Amato, Jeffrey Barrett, Sonia Goncalves, Ewan Harrison, David K. Jackson, Ian Johnston, Dominic Kwiatkowski, Cordelia Langford, John Sillitoe on behalf of the Wellcome Sanger Institute COVID-19 Surveillance Team |
| EPI_ISL_2487190, EPI_ISL_2487191, EPI_ISL_2487193 | Lighthouse Lab in Alderley Park | Wellcome Sanger Institute for the COVID-19 Genomics UK (COG-UK) Consortium | Jacquelyn Wynn, Mairead Hyland, The Lighthouse Lab in Alderley Park and Alex Alderton, Roberto Amato, Jeffrey Barrett, Sonia Goncalves, Ewan Harrison, David K. Jackson, Ian Johnston, Dominic Kwiatkowski, Cordelia Langford, John Sillitoe on behalf of the Wellcome Sanger Institute COVID-19 Surveillance Team |
| EPI_ISL_2487194, EPI_ISL_2487195, EPI_ISL_2487196 | Lighthouse Lab in Milton Keynes | Wellcome Sanger Institute for the COVID-19 Genomics UK (COG-UK) Consortium | The Lighthouse Lab in Milton Keynes and Alex Alderton, Roberto Amato, Jeffrey Barrett, Sonia Goncalves, Ewan Harrison, David K. Jackson, Ian Johnston, Dominic Kwiatkowski, Cordelia Langford, John Sillitoe on behalf of the Wellcome Sanger Institute COVID-19 Surveillance Team |
| EPI_ISL_2487197, EPI_ISL_2487198, EPI_ISL_2487199 | Lighthouse Lab in Alderley Park | Wellcome Sanger Institute for the COVID-19 Genomics UK (COG-UK) Consortium | Jacquelyn Wynn, Mairead Hyland, The Lighthouse Lab in Alderley Park and Alex Alderton, Roberto Amato, Jeffrey Barrett, Sonia Goncalves, Ewan Harrison, David K. Jackson, Ian Johnston, Dominic Kwiatkowski, Cordelia Langford, John Sillitoe on behalf of the Wellcome Sanger Institute COVID-19 Surveillance Team |
| EPI_ISL_2487200, EPI_ISL_2487201, EPI_ISL_2487203 | Lighthouse Lab in Milton Keynes | Wellcome Sanger Institute for the COVID-19 Genomics UK (COG-UK) Consortium | The Lighthouse Lab in Milton Keynes and Alex Alderton, Roberto Amato, Jeffrey Barrett, Sonia Goncalves, Ewan Harrison, David K. Jackson, Ian Johnston, Dominic Kwiatkowski, Cordelia Langford, John Sillitoe on behalf of the Wellcome Sanger Institute COVID-19 Surveillance Team |
| EPI_ISL_2487204, EPI_ISL_2487205 | Lighthouse Lab in Alderley Park | Wellcome Sanger Institute for the COVID-19 Genomics UK (COG-UK) Consortium | Jacquelyn Wynn, Mairead Hyland, The Lighthouse Lab in Alderley Park and Alex Alderton, Roberto Amato, Jeffrey Barrett, Sonia Goncalves, Ewan Harrison, David K. Jackson, Ian Johnston, Dominic Kwiatkowski, Cordelia Langford, John Sillitoe on behalf of the Wellcome Sanger Institute COVID-19 Surveillance Team |
| EPI_ISL_2487206 | Lighthouse Lab in Milton Keynes | Wellcome Sanger Institute for the COVID-19 Genomics UK (COG-UK) Consortium | The Lighthouse Lab in Milton Keynes and Alex Alderton, Roberto Amato, Jeffrey Barrett, Sonia Goncalves, Ewan Harrison, David K. Jackson, Ian Johnston, Dominic Kwiatkowski, Cordelia Langford, John Sillitoe on behalf of the Wellcome Sanger Institute COVID-19 Surveillance Team |
| EPI_ISL_2487207 | Lighthouse Lab in Glasgow | Wellcome Sanger Institute for the COVID-19 Genomics UK (COG-UK) Consortium | Harper VanSteenhouse, Yumi Kasai, David Gray, Carol Clugston, Anna Dominiczak and Alex Alderton, Roberto Amato, Jeffrey Barrett, Sonia Goncalves, Ewan Harrison, David K. Jackson, Ian Johnston, Dominic Kwiatkowski, Cordelia Langford, John Sillitoe on behalf of the Wellcome Sanger Institute COVID-19 Surveillance Team |
| EPI_ISL_2487208 | Lighthouse Lab in Alderley Park | Wellcome Sanger Institute for the COVID-19 Genomics UK (COG-UK) Consortium | Jacquelyn Wynn, Mairead Hyland, The Lighthouse Lab in Alderley Park and Alex Alderton, Roberto Amato, Jeffrey Barrett, Sonia Goncalves, Ewan Harrison, David K. Jackson, Ian Johnston, Dominic Kwiatkowski, Cordelia Langford, John Sillitoe on behalf of the Wellcome Sanger Institute COVID-19 Surveillance Team |
| EPI_ISL_2487209 | Health Services Laboratories | Wellcome Sanger Institute for the COVID-19 Genomics UK (COG-UK) Consortium | Health Services Laboratories and Alex Alderton, Roberto Amato, Jeffrey Barrett, Sonia Goncalves, Ewan Harrison, David K. Jackson, Ian Johnston, Dominic Kwiatkowski, Cordelia Langford, John Sillitoe on behalf of the Wellcome Sanger Institute COVID-19 Surveillance Team |
| EPI_ISL_2487210 | Lighthouse Lab in Milton Keynes | Wellcome Sanger Institute for the COVID-19 Genomics UK (COG-UK) Consortium | The Lighthouse Lab in Milton Keynes and Alex Alderton, Roberto Amato, Jeffrey Barrett, Sonia Goncalves, Ewan Harrison, David K. Jackson, Ian Johnston, Dominic Kwiatkowski, Cordelia Langford, John Sillitoe on behalf of the Wellcome Sanger Institute COVID-19 Surveillance Team |
| EPI_ISL_2487211 | Lighthouse Lab in Alderley Park | Wellcome Sanger Institute for the COVID-19 Genomics UK (COG-UK) Consortium | Jacquelyn Wynn, Mairead Hyland, The Lighthouse Lab in Alderley Park and Alex Alderton, Roberto Amato, Jeffrey Barrett, Sonia Goncalves, Ewan Harrison, David K. Jackson, Ian Johnston, Dominic Kwiatkowski, Cordelia Langford, John Sillitoe on behalf of the Wellcome Sanger Institute COVID-19 Surveillance Team |
| EPI_ISL_2487212 | Lighthouse Lab in Milton Keynes | Wellcome Sanger Institute for the COVID-19 Genomics UK (COG-UK) Consortium | The Lighthouse Lab in Milton Keynes and Alex Alderton, Roberto Amato, Jeffrey Barrett, Sonia Goncalves, Ewan Harrison, David K. Jackson, Ian Johnston, Dominic Kwiatkowski, Cordelia Langford, John Sillitoe on behalf of the Wellcome Sanger Institute COVID-19 Surveillance Team |
| EPI_ISL_2487213 | Health Services Laboratories | Wellcome Sanger Institute for the COVID-19 Genomics UK (COG-UK) Consortium | Health Services Laboratories and Alex Alderton, Roberto Amato, Jeffrey Barrett, Sonia Goncalves, Ewan Harrison, David K. Jackson, Ian Johnston, Dominic Kwiatkowski, Cordelia Langford, John Sillitoe on behalf of the Wellcome Sanger Institute COVID-19 Surveillance Team |
| EPI_ISL_2487214 | Lighthouse Lab in Alderley Park | Wellcome Sanger Institute for the COVID-19 Genomics UK (COG-UK) Consortium | Jacquelyn Wynn, Mairead Hyland, The Lighthouse Lab in Alderley Park and Alex Alderton, Roberto Amato, Jeffrey Barrett, Sonia Goncalves, Ewan Harrison, David K. Jackson, Ian Johnston, Dominic Kwiatkowski, Cordelia Langford, John Sillitoe on behalf of the Wellcome Sanger Institute COVID-19 Surveillance Team |
| EPI_ISL_2487215, EPI_ISL_2487216 | Lighthouse Lab in Milton Keynes | Wellcome Sanger Institute for the COVID-19 Genomics UK (COG-UK) Consortium | The Lighthouse Lab in Milton Keynes and Alex Alderton, Roberto Amato, Jeffrey Barrett, Sonia Goncalves, Ewan Harrison, David K. Jackson, Ian Johnston, Dominic Kwiatkowski, Cordelia Langford, John Sillitoe on behalf of the Wellcome Sanger Institute COVID-19 Surveillance Team |
| EPI_ISL_2487217 | Lighthouse Lab in Alderley Park | Wellcome Sanger Institute for the COVID-19 Genomics UK (COG-UK) Consortium | Jacquelyn Wynn, Mairead Hyland, The Lighthouse Lab in Alderley Park and Alex Alderton, Roberto Amato, Jeffrey Barrett, Sonia Goncalves, Ewan Harrison, David K. Jackson, Ian Johnston, Dominic Kwiatkowski, Cordelia Langford, John Sillitoe on behalf of the Wellcome Sanger Institute COVID-19 Surveillance Team |
| EPI_ISL_2487218, EPI_ISL_2487220, EPI_ISL_2487221, EPI_ISL_2487222, EPI_ISL_2487223, EPI_ISL_2487224, EPI_ISL_2487225, EPI_ISL_2487226, EPI_ISL_2487227, EPI_ISL_2487228, EPI_ISL_2487229, EPI_ISL_2487230, EPI_ISL_2487231, EPI_ISL_2487234, EPI_ISL_2487235, EPI_ISL_2487236, EPI_ISL_2487237, EPI_ISL_2487238, EPI_ISL_2487240, EPI_ISL_2487241, EPI_ISL_2487242, EPI_ISL_2487243, EPI_ISL_2487244, EPI_ISL_2487245, EPI_ISL_2487246, EPI_ISL_2487248, EPI_ISL_2487249, EPI_ISL_2487250, EPI_ISL_2487251, EPI_ISL_2487252, EPI_ISL_2487253, EPI_ISL_2487254, EPI_ISL_2487255, EPI_ISL_2487256, EPI_ISL_2487257, EPI_ISL_2487258, EPI_ISL_2487259, EPI_ISL_2487261, EPI_ISL_2487263, EPI_ISL_2487264, EPI_ISL_2487265, EPI_ISL_2487266, EPI_ISL_2487267, EPI_ISL_2487268, EPI_ISL_2487269, EPI_ISL_2487270, EPI_ISL_2487271, EPI_ISL_2487272, EPI_ISL_2487273, EPI_ISL_2487274, EPI_ISL_2487275, EPI_ISL_2487276, EPI_ISL_2487277, EPI_ISL_2487279, EPI_ISL_2487280 | | | |
| see above | Lighthouse Lab in Glasgow | Wellcome Sanger Institute for the COVID-19 Genomics UK (COG-UK) Consortium | Harper VanSteenhouse, Yumi Kasai, David Gray, Carol Clugston, Anna Dominiczak and Alex Alderton, Roberto Amato, Jeffrey Barrett, Sonia Goncalves, Ewan Harrison, David K. Jackson, Ian Johnston, Dominic Kwiatkowski, Cordelia Langford, John Sillitoe on behalf of the Wellcome Sanger Institute COVID-19 Surveillance Team |
| EPI_ISL_2487282 | Lighthouse Laboratory Plymouth | Wellcome Sanger Institute for the COVID-19 Genomics UK (COG-UK) Consortium | Lighthouse Laboratory Plymouth and Alex Alderton, Roberto Amato, Jeffrey Barrett, Sonia Goncalves, Ewan Harrison, David K. Jackson, Ian Johnston, Dominic Kwiatkowski, Cordelia Langford, John Sillitoe on behalf of the Wellcome Sanger Institute COVID-19 Surveillance Team |
| EPI_ISL_2487284, EPI_ISL_2487285, EPI_ISL_2487286, EPI_ISL_2487287, EPI_ISL_2487289, EPI_ISL_2487290, EPI_ISL_2487291, EPI_ISL_2487292, EPI_ISL_2487293, EPI_ISL_2487294, EPI_ISL_2487295, EPI_ISL_2487296, EPI_ISL_2487297, EPI_ISL_2487298, EPI_ISL_2487299, EPI_ISL_2487300, EPI_ISL_2487301, EPI_ISL_2487302, EPI_ISL_2487303, EPI_ISL_2487306, EPI_ISL_2487308, EPI_ISL_2487312, EPI_ISL_2487313, EPI_ISL_2487315, EPI_ISL_2487316, EPI_ISL_2487318, EPI_ISL_2487320, EPI_ISL_2487321, EPI_ISL_2487322, EPI_ISL_2487324, EPI_ISL_2487325, EPI_ISL_2487326, EPI_ISL_2487327, EPI_ISL_2487328, EPI_ISL_2487329, EPI_ISL_2487331, EPI_ISL_2487332, EPI_ISL_2487333, EPI_ISL_2487337, EPI_ISL_2487338, EPI_ISL_2487339, EPI_ISL_2487341, EPI_ISL_2487343, EPI_ISL_2487344, EPI_ISL_2487345, EPI_ISL_2487346, EPI_ISL_2487347, EPI_ISL_2487348, EPI_ISL_2487349, EPI_ISL_2487350, EPI_ISL_2487351, EPI_ISL_2487352, EPI_ISL_2487353, EPI_ISL_2487354, EPI_ISL_2487355, EPI_ISL_2487356, EPI_ISL_2487357, EPI_ISL_2487358, EPI_ISL_2487359, EPI_ISL_2487360, EPI_ISL_2487361, EPI_ISL_2487362, EPI_ISL_2487363, EPI_ISL_2487365, EPI_ISL_2487366, EPI_ISL_2487368, EPI_ISL_2487369, EPI_ISL_2487370, EPI_ISL_2487372, EPI_ISL_2487375, EPI_ISL_2487377, EPI_ISL_2487378, EPI_ISL_2487380, EPI_ISL_2487381, EPI_ISL_2487382, EPI_ISL_2487383, EPI_ISL_2487385, EPI_ISL_2487386, EPI_ISL_2487387, EPI_ISL_2487388, EPI_ISL_2487389, EPI_ISL_2487390, EPI_ISL_2487391, EPI_ISL_2487392, EPI_ISL_2487393, EPI_ISL_2487394, EPI_ISL_2487395, EPI_ISL_2487396, EPI_ISL_2487398, EPI_ISL_2487401, EPI_ISL_2487403, EPI_ISL_2487404, EPI_ISL_2487406, EPI_ISL_2487407, EPI_ISL_2487408, EPI_ISL_2487409, EPI_ISL_2487411, EPI_ISL_2487412, EPI_ISL_2487413, EPI_ISL_2487414, EPI_ISL_2487415, EPI_ISL_2487419, EPI_ISL_2487420, EPI_ISL_2487421, EPI_ISL_2487422, EPI_ISL_2487423 | | | |
| see above | Lighthouse Lab in Glasgow | Wellcome Sanger Institute for the COVID-19 Genomics UK (COG-UK) Consortium | Harper VanSteenhouse, Yumi Kasai, David Gray, Carol Clugston, Anna Dominiczak and Alex Alderton, Roberto Amato, Jeffrey Barrett, Sonia Goncalves, Ewan Harrison, David K. Jackson, Ian Johnston, Dominic Kwiatkowski, Cordelia Langford, John Sillitoe on behalf of the Wellcome Sanger Institute COVID-19 Surveillance Team |
| EPI_ISL_2487425 | Lighthouse Laboratory Plymouth | Wellcome Sanger Institute for the COVID-19 Genomics UK (COG-UK) Consortium | Lighthouse Laboratory Plymouth and Alex Alderton, Roberto Amato, Jeffrey Barrett, Sonia Goncalves, Ewan Harrison, David K. Jackson, Ian Johnston, Dominic Kwiatkowski, Cordelia Langford, John Sillitoe on behalf of the Wellcome Sanger Institute COVID-19 Surveillance Team |
| EPI_ISL_2487427, EPI_ISL_2487428, EPI_ISL_2487429, EPI_ISL_2487430, EPI_ISL_2487431, EPI_ISL_2487433, EPI_ISL_2487434, EPI_ISL_2487435, EPI_ISL_2487436, EPI_ISL_2487437, EPI_ISL_2487438, EPI_ISL_2487439, EPI_ISL_2487441, EPI_ISL_2487442, EPI_ISL_2487443, EPI_ISL_2487444, EPI_ISL_2487445, EPI_ISL_2487446 | | | |
| see above | Lighthouse Lab in Glasgow | Wellcome Sanger Institute for the COVID-19 Genomics UK (COG-UK) Consortium | Harper VanSteenhouse, Yumi Kasai, David Gray, Carol Clugston, Anna Dominiczak and Alex Alderton, Roberto Amato, Jeffrey Barrett, Sonia Goncalves, Ewan Harrison, David K. Jackson, Ian Johnston, Dominic Kwiatkowski, Cordelia Langford, John Sillitoe on behalf of the Wellcome Sanger Institute COVID-19 Surveillance Team |
| EPI_ISL_2487447 | Lighthouse Laboratory Plymouth | Wellcome Sanger Institute for the COVID-19 Genomics UK (COG-UK) Consortium | Lighthouse Laboratory Plymouth and Alex Alderton, Roberto Amato, Jeffrey Barrett, Sonia Goncalves, Ewan Harrison, David K. Jackson, Ian Johnston, Dominic Kwiatkowski, Cordelia Langford, John Sillitoe on behalf of the Wellcome Sanger Institute COVID-19 Surveillance Team |
| EPI_ISL_2487448, EPI_ISL_2487449, EPI_ISL_2487450, EPI_ISL_2487451, EPI_ISL_2487452, EPI_ISL_2487453, EPI_ISL_2487455, EPI_ISL_2487457, EPI_ISL_2487458, EPI_ISL_2487459, EPI_ISL_2487460, EPI_ISL_2487461 | | | |
| see above | Lighthouse Lab in Glasgow | Wellcome Sanger Institute for the COVID-19 Genomics UK (COG-UK) Consortium | Harper VanSteenhouse, Yumi Kasai, David Gray, Carol Clugston, Anna Dominiczak and Alex Alderton, Roberto Amato, Jeffrey Barrett, Sonia Goncalves, Ewan Harrison, David K. Jackson, Ian Johnston, Dominic Kwiatkowski, Cordelia Langford, John Sillitoe on behalf of the Wellcome Sanger Institute COVID-19 Surveillance Team |
| EPI_ISL_2487462 | Lighthouse Laboratory Plymouth | Wellcome Sanger Institute for the COVID-19 Genomics UK (COG-UK) Consortium | Lighthouse Laboratory Plymouth and Alex Alderton, Roberto Amato, Jeffrey Barrett, Sonia Goncalves, Ewan Harrison, David K. Jackson, Ian Johnston, Dominic Kwiatkowski, Cordelia Langford, John Sillitoe on behalf of the Wellcome Sanger Institute COVID-19 Surveillance Team |
| EPI_ISL_2487463, EPI_ISL_2487464, EPI_ISL_2487465, EPI_ISL_2487466, EPI_ISL_2487468, EPI_ISL_2487471, EPI_ISL_2487472, EPI_ISL_2487473 | Lighthouse Lab in Glasgow | Wellcome Sanger Institute for the COVID-19 Genomics UK (COG-UK) Consortium | Harper VanSteenhouse, Yumi Kasai, David Gray, Carol Clugston, Anna Dominiczak and Alex Alderton, Roberto Amato, Jeffrey Barrett, Sonia Goncalves, Ewan Harrison, David K. Jackson, Ian Johnston, Dominic Kwiatkowski, Cordelia Langford, John Sillitoe on behalf of the Wellcome Sanger Institute COVID-19 Surveillance Team |
| EPI_ISL_2487474 | Lighthouse Laboratory Plymouth | Wellcome Sanger Institute for the COVID-19 Genomics UK (COG-UK) Consortium | Lighthouse Laboratory Plymouth and Alex Alderton, Roberto Amato, Jeffrey Barrett, Sonia Goncalves, Ewan Harrison, David K. Jackson, Ian Johnston, Dominic Kwiatkowski, Cordelia Langford, John Sillitoe on behalf of the Wellcome Sanger Institute COVID-19 Surveillance Team |
| EPI_ISL_2487475, EPI_ISL_2487476, EPI_ISL_2487477, EPI_ISL_2487478, EPI_ISL_2487479, EPI_ISL_2487480, EPI_ISL_2487481, EPI_ISL_2487484, EPI_ISL_2487485, EPI_ISL_2487486, EPI_ISL_2487487, EPI_ISL_2487489, EPI_ISL_2487490, EPI_ISL_2487491, EPI_ISL_2487492, EPI_ISL_2487493, EPI_ISL_2487494, EPI_ISL_2487495, EPI_ISL_2487496, EPI_ISL_2487497, EPI_ISL_2487498, EPI_ISL_2487499, | | | |

| | | | |
|---|---|---|---|
| EPI_ISL_2487500, EPI_ISL_2487502, EPI_ISL_2487503, EPI_ISL_2487504, EPI_ISL_2487505, EPI_ISL_2487506, EPI_ISL_2487509, EPI_ISL_2487511, EPI_ISL_2487512, EPI_ISL_2487513, EPI_ISL_2487514, EPI_ISL_2487515, EPI_ISL_2487516, EPI_ISL_2487517, EPI_ISL_2487519, EPI_ISL_2487520, EPI_ISL_2487521, EPI_ISL_2487522, EPI_ISL_2487523, EPI_ISL_2487524, EPI_ISL_2487526 | | | |
| see above | Lighthouse Lab in Glasgow | Wellcome Sanger Institute for the COVID-19 Genomics UK (COG-UK) Consortium | Harper VanSteenhouse, Yumi Kasai, David Gray, Carol Clugston, Anna Dominiczak and Alex Alderton, Roberto Amato, Jeffrey Barrett, Sonia Goncalves, Ewan Harrison, David K. Jackson, Ian Johnston, Dominic Kwiatkowski, Cordelia Langford, John Sillitoe on behalf of the Wellcome Sanger Institute COVID-19 Surveillance Team |
| EPI_ISL_2487527 | Lighthouse Laboratory Plymouth | Wellcome Sanger Institute for the COVID-19 Genomics UK (COG-UK) Consortium | Lighthouse Laboratory Plymouth and Alex Alderton, Roberto Amato, Jeffrey Barrett, Sonia Goncalves, Ewan Harrison, David K. Jackson, Ian Johnston, Dominic Kwiatkowski, Cordelia Langford, John Sillitoe on behalf of the Wellcome Sanger Institute COVID-19 Surveillance Team |
| EPI_ISL_2487528, EPI_ISL_2487529, EPI_ISL_2487530, EPI_ISL_2487531, EPI_ISL_2487532, EPI_ISL_2487535, EPI_ISL_2487536, EPI_ISL_2487537, EPI_ISL_2487539, EPI_ISL_2487540, EPI_ISL_2487541, EPI_ISL_2487542, EPI_ISL_2487543, EPI_ISL_2487544, EPI_ISL_2487545, EPI_ISL_2487546, EPI_ISL_2487547, EPI_ISL_2487548, EPI_ISL_2487550, EPI_ISL_2487552, EPI_ISL_2487553, EPI_ISL_2487554, EPI_ISL_2487555, EPI_ISL_2487556 | | | |
| see above | Lighthouse Lab in Glasgow | Wellcome Sanger Institute for the COVID-19 Genomics UK (COG-UK) Consortium | Harper VanSteenhouse, Yumi Kasai, David Gray, Carol Clugston, Anna Dominiczak and Alex Alderton, Roberto Amato, Jeffrey Barrett, Sonia Goncalves, Ewan Harrison, David K. Jackson, Ian Johnston, Dominic Kwiatkowski, Cordelia Langford, John Sillitoe on behalf of the Wellcome Sanger Institute COVID-19 Surveillance Team |
| EPI_ISL_2487558 | Lighthouse Laboratory Plymouth | Wellcome Sanger Institute for the COVID-19 Genomics UK (COG-UK) Consortium | Lighthouse Laboratory Plymouth and Alex Alderton, Roberto Amato, Jeffrey Barrett, Sonia Goncalves, Ewan Harrison, David K. Jackson, Ian Johnston, Dominic Kwiatkowski, Cordelia Langford, John Sillitoe on behalf of the Wellcome Sanger Institute COVID-19 Surveillance Team |
| EPI_ISL_2487559, EPI_ISL_2487560, EPI_ISL_2487562, EPI_ISL_2487563, EPI_ISL_2487564, EPI_ISL_2487565, EPI_ISL_2487567, EPI_ISL_2487568, EPI_ISL_2487569, EPI_ISL_2487571, EPI_ISL_2487572, EPI_ISL_2487573, EPI_ISL_2487574, EPI_ISL_2487576, EPI_ISL_2487577, EPI_ISL_2487578, EPI_ISL_2487579, EPI_ISL_2487580, EPI_ISL_2487581, EPI_ISL_2487583, EPI_ISL_2487584 | | | |
| see above | Lighthouse Lab in Glasgow | Wellcome Sanger Institute for the COVID-19 Genomics UK (COG-UK) Consortium | Harper VanSteenhouse, Yumi Kasai, David Gray, Carol Clugston, Anna Dominiczak and Alex Alderton, Roberto Amato, Jeffrey Barrett, Sonia Goncalves, Ewan Harrison, David K. Jackson, Ian Johnston, Dominic Kwiatkowski, Cordelia Langford, John Sillitoe on behalf of the Wellcome Sanger Institute COVID-19 Surveillance Team |
| EPI_ISL_2487585, EPI_ISL_2487586 | Lighthouse Lab in Alderley Park | Wellcome Sanger Institute for the COVID-19 Genomics UK (COG-UK) Consortium | Jacquelyn Wynn, Mairead Hyland, The Lighthouse Lab in Alderley Park and Alex Alderton, Roberto Amato, Jeffrey Barrett, Sonia Goncalves, Ewan Harrison, David K. Jackson, Ian Johnston, Dominic Kwiatkowski, Cordelia Langford, John Sillitoe on behalf of the Wellcome Sanger Institute COVID-19 Surveillance Team |
| EPI_ISL_2487588, EPI_ISL_2487589, EPI_ISL_2487590 | Lighthouse Lab in Glasgow | Wellcome Sanger Institute for the COVID-19 Genomics UK (COG-UK) Consortium | Harper VanSteenhouse, Yumi Kasai, David Gray, Carol Clugston, Anna Dominiczak and Alex Alderton, Roberto Amato, Jeffrey Barrett, Sonia Goncalves, Ewan Harrison, David K. Jackson, Ian Johnston, Dominic Kwiatkowski, Cordelia Langford, John Sillitoe on behalf of the Wellcome Sanger Institute COVID-19 Surveillance Team |
| EPI_ISL_2487591 | Lighthouse Lab in Alderley Park | Wellcome Sanger Institute for the COVID-19 Genomics UK (COG-UK) Consortium | Jacquelyn Wynn, Mairead Hyland, The Lighthouse Lab in Alderley Park and Alex Alderton, Roberto Amato, Jeffrey Barrett, Sonia Goncalves, Ewan Harrison, David K. Jackson, Ian Johnston, Dominic Kwiatkowski, Cordelia Langford, John Sillitoe on behalf of the Wellcome Sanger Institute COVID-19 Surveillance Team |
| EPI_ISL_2487592, EPI_ISL_2487593, EPI_ISL_2487595 | Lighthouse Lab in Glasgow | Wellcome Sanger Institute for the COVID-19 Genomics UK (COG-UK) Consortium | Harper VanSteenhouse, Yumi Kasai, David Gray, Carol Clugston, Anna Dominiczak and Alex Alderton, Roberto Amato, Jeffrey Barrett, Sonia Goncalves, Ewan Harrison, David K. Jackson, Ian Johnston, Dominic Kwiatkowski, Cordelia Langford, John Sillitoe on behalf of the Wellcome Sanger Institute COVID-19 Surveillance Team |
| EPI_ISL_2487596 | Lighthouse Lab in Alderley Park | Wellcome Sanger Institute for the COVID-19 Genomics UK (COG-UK) Consortium | Jacquelyn Wynn, Mairead Hyland, The Lighthouse Lab in Alderley Park and Alex Alderton, Roberto Amato, Jeffrey Barrett, Sonia Goncalves, Ewan Harrison, David K. Jackson, Ian Johnston, Dominic Kwiatkowski, Cordelia Langford, John Sillitoe on behalf of the Wellcome Sanger Institute COVID-19 Surveillance Team |
| EPI_ISL_2487597, EPI_ISL_2487598, EPI_ISL_2487599, EPI_ISL_2487600 | Lighthouse Lab in Glasgow | Wellcome Sanger Institute for the COVID-19 Genomics UK (COG-UK) Consortium | Harper VanSteenhouse, Yumi Kasai, David Gray, Carol Clugston, Anna Dominiczak and Alex Alderton, Roberto Amato, Jeffrey Barrett, Sonia Goncalves, Ewan Harrison, David K. Jackson, Ian Johnston, Dominic Kwiatkowski, Cordelia Langford, John Sillitoe on behalf of the Wellcome Sanger Institute COVID-19 Surveillance Team |
| EPI_ISL_2487601 | Lighthouse Lab in Alderley Park | Wellcome Sanger Institute for the COVID-19 Genomics UK (COG-UK) Consortium | Jacquelyn Wynn, Mairead Hyland, The Lighthouse Lab in Alderley Park and Alex Alderton, Roberto Amato, Jeffrey Barrett, Sonia Goncalves, Ewan Harrison, David K. Jackson, Ian Johnston, Dominic Kwiatkowski, Cordelia Langford, John Sillitoe on behalf of the Wellcome Sanger Institute COVID-19 Surveillance Team |
| EPI_ISL_2487602, EPI_ISL_2487603, EPI_ISL_2487604, EPI_ISL_2487605, EPI_ISL_2487607, EPI_ISL_2487608, EPI_ISL_2487610 | Lighthouse Lab in Glasgow | Wellcome Sanger Institute for the COVID-19 Genomics UK (COG-UK) Consortium | Harper VanSteenhouse, Yumi Kasai, David Gray, Carol Clugston, Anna Dominiczak and Alex Alderton, Roberto Amato, Jeffrey Barrett, Sonia Goncalves, Ewan Harrison, David K. Jackson, Ian Johnston, Dominic Kwiatkowski, Cordelia Langford, John Sillitoe on behalf of the Wellcome Sanger Institute COVID-19 Surveillance Team |
| EPI_ISL_2487611 | Lighthouse Lab in Alderley Park | Wellcome Sanger Institute for the COVID-19 Genomics UK (COG-UK) Consortium | Jacquelyn Wynn, Mairead Hyland, The Lighthouse Lab in Alderley Park and Alex Alderton, Roberto Amato, Jeffrey Barrett, Sonia Goncalves, Ewan Harrison, David K. Jackson, Ian Johnston, Dominic Kwiatkowski, Cordelia Langford, John Sillitoe on behalf of the Wellcome Sanger Institute COVID-19 Surveillance Team |
| EPI_ISL_2487612, EPI_ISL_2487613, EPI_ISL_2487614, EPI_ISL_2487616, EPI_ISL_2487617, EPI_ISL_2487619, EPI_ISL_2487620, EPI_ISL_2487622 | Lighthouse Lab in Glasgow | Wellcome Sanger Institute for the COVID-19 Genomics UK (COG-UK) Consortium | Harper VanSteenhouse, Yumi Kasai, David Gray, Carol Clugston, Anna Dominiczak and Alex Alderton, Roberto Amato, Jeffrey Barrett, Sonia Goncalves, Ewan Harrison, David K. Jackson, Ian Johnston, Dominic Kwiatkowski, Cordelia Langford, John Sillitoe on behalf of the Wellcome Sanger Institute COVID-19 Surveillance Team |
| EPI_ISL_2487623 | Lighthouse Lab in Alderley Park | Wellcome Sanger Institute for the COVID-19 Genomics UK (COG-UK) Consortium | Jacquelyn Wynn, Mairead Hyland, The Lighthouse Lab in Alderley Park and Alex Alderton, Roberto Amato, Jeffrey Barrett, Sonia Goncalves, Ewan Harrison, David K. Jackson, Ian Johnston, Dominic Kwiatkowski, Cordelia Langford, John Sillitoe on behalf of the Wellcome Sanger Institute COVID-19 Surveillance Team |
| EPI_ISL_2487625, EPI_ISL_2487627, EPI_ISL_2487628, EPI_ISL_2487629 | Lighthouse Lab in Glasgow | Wellcome Sanger Institute for the COVID-19 Genomics UK (COG-UK) Consortium | Harper VanSteenhouse, Yumi Kasai, David Gray, Carol Clugston, Anna Dominiczak and Alex Alderton, Roberto Amato, Jeffrey Barrett, Sonia Goncalves, Ewan Harrison, David K. Jackson, Ian Johnston, Dominic Kwiatkowski, Cordelia Langford, John Sillitoe on behalf of the Wellcome Sanger Institute COVID-19 Surveillance Team |
| EPI_ISL_2487630, EPI_ISL_2487631 | Lighthouse Lab in Alderley Park | Wellcome Sanger Institute for the COVID-19 Genomics UK (COG-UK) Consortium | Jacquelyn Wynn, Mairead Hyland, The Lighthouse Lab in Alderley Park and Alex Alderton, Roberto Amato, Jeffrey Barrett, Sonia Goncalves, Ewan Harrison, David K. Jackson, Ian Johnston, Dominic Kwiatkowski, Cordelia Langford, John Sillitoe on behalf of the Wellcome Sanger Institute COVID-19 Surveillance Team |
| EPI_ISL_2487632, EPI_ISL_2487633, EPI_ISL_2487634 | Lighthouse Lab in Glasgow | Wellcome Sanger Institute for the COVID-19 Genomics UK (COG-UK) Consortium | Harper VanSteenhouse, Yumi Kasai, David Gray, Carol Clugston, Anna Dominiczak and Alex Alderton, Roberto Amato, Jeffrey Barrett, Sonia Goncalves, Ewan Harrison, David K. Jackson, Ian Johnston, Dominic Kwiatkowski, Cordelia Langford, John Sillitoe on behalf of the Wellcome Sanger Institute COVID-19 Surveillance Team |
| EPI_ISL_2487635 | Lighthouse Lab in Alderley Park | Wellcome Sanger Institute for the COVID-19 Genomics UK (COG-UK) Consortium | Jacquelyn Wynn, Mairead Hyland, The Lighthouse Lab in Alderley Park and Alex Alderton, Roberto Amato, Jeffrey Barrett, Sonia Goncalves, Ewan Harrison, David K. Jackson, Ian Johnston, Dominic Kwiatkowski, Cordelia Langford, John Sillitoe on behalf of the Wellcome Sanger Institute COVID-19 Surveillance Team |
| EPI_ISL_2487636, EPI_ISL_2487637, EPI_ISL_2487638, EPI_ISL_2487639 | Lighthouse Lab in Glasgow | Wellcome Sanger Institute for the COVID-19 Genomics UK (COG-UK) Consortium | Harper VanSteenhouse, Yumi Kasai, David Gray, Carol Clugston, Anna Dominiczak and Alex Alderton, Roberto Amato, Jeffrey Barrett, Sonia Goncalves, Ewan Harrison, David K. Jackson, Ian Johnston, Dominic Kwiatkowski, Cordelia Langford, John Sillitoe on behalf of the Wellcome Sanger Institute COVID-19 Surveillance Team |
| EPI_ISL_2487640 | Lighthouse Lab in Alderley Park | Wellcome Sanger Institute for the COVID-19 Genomics UK (COG-UK) Consortium | Jacquelyn Wynn, Mairead Hyland, The Lighthouse Lab in Alderley Park and Alex Alderton, Roberto Amato, Jeffrey Barrett, Sonia Goncalves, Ewan Harrison, David K. Jackson, Ian Johnston, Dominic Kwiatkowski, Cordelia Langford, John Sillitoe on behalf of the Wellcome Sanger Institute COVID-19 Surveillance Team |
| EPI_ISL_2487641, EPI_ISL_2487642, EPI_ISL_2487643, EPI_ISL_2487644, EPI_ISL_2487645, EPI_ISL_2487646, EPI_ISL_2487648 | Lighthouse Lab in Glasgow | Wellcome Sanger Institute for the COVID-19 Genomics UK (COG-UK) Consortium | Harper VanSteenhouse, Yumi Kasai, David Gray, Carol Clugston, Anna Dominiczak and Alex Alderton, Roberto Amato, Jeffrey Barrett, Sonia Goncalves, Ewan Harrison, David K. Jackson, Ian Johnston, Dominic Kwiatkowski, Cordelia Langford, John Sillitoe on behalf of the Wellcome Sanger Institute COVID-19 Surveillance Team |
| EPI_ISL_2487649, EPI_ISL_2487650 | Lighthouse Lab in Alderley Park | Wellcome Sanger Institute for the COVID-19 Genomics UK (COG-UK) Consortium | Jacquelyn Wynn, Mairead Hyland, The Lighthouse Lab in Alderley Park and Alex Alderton, Roberto Amato, Jeffrey Barrett, Sonia Goncalves, Ewan Harrison, David K. Jackson, Ian Johnston, Dominic Kwiatkowski, Cordelia Langford, John Sillitoe on behalf of the Wellcome Sanger Institute COVID-19 Surveillance Team |
| EPI_ISL_2487651, EPI_ISL_2487652, EPI_ISL_2487653, EPI_ISL_2487655, EPI_ISL_2487656, EPI_ISL_2487658, EPI_ISL_2487659, EPI_ISL_2487660 | Lighthouse Lab in Glasgow | Wellcome Sanger Institute for the COVID-19 Genomics UK (COG-UK) Consortium | Harper VanSteenhouse, Yumi Kasai, David Gray, Carol Clugston, Anna Dominiczak and Alex Alderton, Roberto Amato, Jeffrey Barrett, Sonia Goncalves, Ewan Harrison, David K. Jackson, Ian Johnston, Dominic Kwiatkowski, Cordelia Langford, John Sillitoe on behalf of the Wellcome Sanger Institute COVID-19 Surveillance Team |
| EPI_ISL_2487662 | Lighthouse Lab in Alderley Park | Wellcome Sanger Institute for the COVID-19 Genomics UK (COG-UK) Consortium | Jacquelyn Wynn, Mairead Hyland, The Lighthouse Lab in Alderley Park and Alex Alderton, Roberto Amato, Jeffrey Barrett, Sonia Goncalves, Ewan Harrison, David K. Jackson, Ian Johnston, Dominic Kwiatkowski, Cordelia Langford, John Sillitoe on behalf of the Wellcome Sanger Institute COVID-19 Surveillance Team |
| EPI_ISL_2487663 | Lighthouse Lab in Glasgow | Wellcome Sanger Institute for the COVID-19 Genomics UK (COG-UK) Consortium | Harper VanSteenhouse, Yumi Kasai, David Gray, Carol Clugston, Anna Dominiczak and Alex Alderton, Roberto Amato, Jeffrey Barrett, Sonia Goncalves, Ewan Harrison, David K. Jackson, Ian Johnston, Dominic Kwiatkowski, Cordelia Langford, John Sillitoe on behalf of the Wellcome Sanger Institute COVID-19 Surveillance Team |
| EPI_ISL_2487664 | Lighthouse Lab in Alderley Park | Wellcome Sanger Institute for the COVID-19 Genomics UK (COG-UK) Consortium | Jacquelyn Wynn, Mairead Hyland, The Lighthouse Lab in Alderley Park and Alex Alderton, Roberto Amato, Jeffrey Barrett, Sonia Goncalves, Ewan Harrison, David K. Jackson, Ian Johnston, Dominic Kwiatkowski, Cordelia Langford, John Sillitoe on behalf of the Wellcome Sanger Institute COVID-19 Surveillance Team |
| EPI_ISL_2487665, EPI_ISL_2487666, EPI_ISL_2487668 | Lighthouse Lab in Glasgow | Wellcome Sanger Institute for the COVID-19 Genomics UK (COG-UK) Consortium | Harper VanSteenhouse, Yumi Kasai, David Gray, Carol Clugston, Anna Dominiczak and Alex Alderton, Roberto Amato, Jeffrey Barrett, Sonia Goncalves, Ewan Harrison, David K. Jackson, Ian Johnston, Dominic Kwiatkowski, Cordelia Langford, John Sillitoe on behalf of the Wellcome Sanger Institute COVID-19 Surveillance Team |
| EPI_ISL_2487670 | Lighthouse Lab in Alderley Park | Wellcome Sanger Institute for the COVID-19 Genomics UK (COG-UK) Consortium | Jacquelyn Wynn, Mairead Hyland, The Lighthouse Lab in Alderley Park and Alex Alderton, Roberto Amato, Jeffrey Barrett, Sonia Goncalves, Ewan Harrison, David K. Jackson, Ian Johnston, Dominic Kwiatkowski, Cordelia Langford, John Sillitoe on behalf of the Wellcome Sanger Institute COVID-19 Surveillance Team |

| | | | |
|---|---|---|---|
| EPI_ISL_2487672, EPI_ISL_2487673, EPI_ISL_2487676, EPI_ISL_2487679, EPI_ISL_2487680, EPI_ISL_2487681, EPI_ISL_2487682, EPI_ISL_2487683, EPI_ISL_2487684, EPI_ISL_2487685, EPI_ISL_2487686 | | | |
| see above | Lighthouse Lab in Glasgow | Wellcome Sanger Institute for the COVID-19 Genomics UK (COG-UK) Consortium | Harper VanSteenhouse, Yumi Kasai, David Gray, Carol Clugston, Anna Dominiczak and Alex Alderton, Roberto Amato, Jeffrey Barrett, Sonia Goncalves, Ewan Harrison, David K. Jackson, Ian Johnston, Dominic Kwiatkowski, Cordelia Langford, John Sillitoe on behalf of the Wellcome Sanger Institute COVID-19 Surveillance Team |
| EPI_ISL_2487687 | Lighthouse Lab in Alderley Park | Wellcome Sanger Institute for the COVID-19 Genomics UK (COG-UK) Consortium | Jacquelyn Wynn, Mairead Hyland, The Lighthouse Lab in Alderley Park and Alex Alderton, Roberto Amato, Jeffrey Barrett, Sonia Goncalves, Ewan Harrison, David K. Jackson, Ian Johnston, Dominic Kwiatkowski, Cordelia Langford, John Sillitoe on behalf of the Wellcome Sanger Institute COVID-19 Surveillance Team |
| EPI_ISL_2487688 | Lighthouse Lab in Glasgow | Wellcome Sanger Institute for the COVID-19 Genomics UK (COG-UK) Consortium | Harper VanSteenhouse, Yumi Kasai, David Gray, Carol Clugston, Anna Dominiczak and Alex Alderton, Roberto Amato, Jeffrey Barrett, Sonia Goncalves, Ewan Harrison, David K. Jackson, Ian Johnston, Dominic Kwiatkowski, Cordelia Langford, John Sillitoe on behalf of the Wellcome Sanger Institute COVID-19 Surveillance Team |
| EPI_ISL_2487689 | Lighthouse Lab in Alderley Park | Wellcome Sanger Institute for the COVID-19 Genomics UK (COG-UK) Consortium | Jacquelyn Wynn, Mairead Hyland, The Lighthouse Lab in Alderley Park and Alex Alderton, Roberto Amato, Jeffrey Barrett, Sonia Goncalves, Ewan Harrison, David K. Jackson, Ian Johnston, Dominic Kwiatkowski, Cordelia Langford, John Sillitoe on behalf of the Wellcome Sanger Institute COVID-19 Surveillance Team |
| EPI_ISL_2487690, EPI_ISL_2487692 | Lighthouse Lab in Glasgow | Wellcome Sanger Institute for the COVID-19 Genomics UK (COG-UK) Consortium | Harper VanSteenhouse, Yumi Kasai, David Gray, Carol Clugston, Anna Dominiczak and Alex Alderton, Roberto Amato, Jeffrey Barrett, Sonia Goncalves, Ewan Harrison, David K. Jackson, Ian Johnston, Dominic Kwiatkowski, Cordelia Langford, John Sillitoe on behalf of the Wellcome Sanger Institute COVID-19 Surveillance Team |
| EPI_ISL_2487693 | Lighthouse Lab in Alderley Park | Wellcome Sanger Institute for the COVID-19 Genomics UK (COG-UK) Consortium | Jacquelyn Wynn, Mairead Hyland, The Lighthouse Lab in Alderley Park and Alex Alderton, Roberto Amato, Jeffrey Barrett, Sonia Goncalves, Ewan Harrison, David K. Jackson, Ian Johnston, Dominic Kwiatkowski, Cordelia Langford, John Sillitoe on behalf of the Wellcome Sanger Institute COVID-19 Surveillance Team |
| EPI_ISL_2487694, EPI_ISL_2487695, EPI_ISL_2487696, EPI_ISL_2487697, EPI_ISL_2487698, EPI_ISL_2487699 | Lighthouse Lab in Glasgow | Wellcome Sanger Institute for the COVID-19 Genomics UK (COG-UK) Consortium | Harper VanSteenhouse, Yumi Kasai, David Gray, Carol Clugston, Anna Dominiczak and Alex Alderton, Roberto Amato, Jeffrey Barrett, Sonia Goncalves, Ewan Harrison, David K. Jackson, Ian Johnston, Dominic Kwiatkowski, Cordelia Langford, John Sillitoe on behalf of the Wellcome Sanger Institute COVID-19 Surveillance Team |
| EPI_ISL_2487700 | Lighthouse Lab in Alderley Park | Wellcome Sanger Institute for the COVID-19 Genomics UK (COG-UK) Consortium | Jacquelyn Wynn, Mairead Hyland, The Lighthouse Lab in Alderley Park and Alex Alderton, Roberto Amato, Jeffrey Barrett, Sonia Goncalves, Ewan Harrison, David K. Jackson, Ian Johnston, Dominic Kwiatkowski, Cordelia Langford, John Sillitoe on behalf of the Wellcome Sanger Institute COVID-19 Surveillance Team |
| EPI_ISL_2487701, EPI_ISL_2487702, EPI_ISL_2487704, EPI_ISL_2487705 | Lighthouse Lab in Glasgow | Wellcome Sanger Institute for the COVID-19 Genomics UK (COG-UK) Consortium | Harper VanSteenhouse, Yumi Kasai, David Gray, Carol Clugston, Anna Dominiczak and Alex Alderton, Roberto Amato, Jeffrey Barrett, Sonia Goncalves, Ewan Harrison, David K. Jackson, Ian Johnston, Dominic Kwiatkowski, Cordelia Langford, John Sillitoe on behalf of the Wellcome Sanger Institute COVID-19 Surveillance Team |
| EPI_ISL_2487706 | Lighthouse Lab in Alderley Park | Wellcome Sanger Institute for the COVID-19 Genomics UK (COG-UK) Consortium | Jacquelyn Wynn, Mairead Hyland, The Lighthouse Lab in Alderley Park and Alex Alderton, Roberto Amato, Jeffrey Barrett, Sonia Goncalves, Ewan Harrison, David K. Jackson, Ian Johnston, Dominic Kwiatkowski, Cordelia Langford, John Sillitoe on behalf of the Wellcome Sanger Institute COVID-19 Surveillance Team |
| EPI_ISL_2487707, EPI_ISL_2487708, EPI_ISL_2487709, EPI_ISL_2487710, EPI_ISL_2487711 | Lighthouse Lab in Glasgow | Wellcome Sanger Institute for the COVID-19 Genomics UK (COG-UK) Consortium | Harper VanSteenhouse, Yumi Kasai, David Gray, Carol Clugston, Anna Dominiczak and Alex Alderton, Roberto Amato, Jeffrey Barrett, Sonia Goncalves, Ewan Harrison, David K. Jackson, Ian Johnston, Dominic Kwiatkowski, Cordelia Langford, John Sillitoe on behalf of the Wellcome Sanger Institute COVID-19 Surveillance Team |
| EPI_ISL_2487712 | Lighthouse Lab in Alderley Park | Wellcome Sanger Institute for the COVID-19 Genomics UK (COG-UK) Consortium | Jacquelyn Wynn, Mairead Hyland, The Lighthouse Lab in Alderley Park and Alex Alderton, Roberto Amato, Jeffrey Barrett, Sonia Goncalves, Ewan Harrison, David K. Jackson, Ian Johnston, Dominic Kwiatkowski, Cordelia Langford, John Sillitoe on behalf of the Wellcome Sanger Institute COVID-19 Surveillance Team |
| EPI_ISL_2487713, EPI_ISL_2487714, EPI_ISL_2487715 | Lighthouse Lab in Glasgow | Wellcome Sanger Institute for the COVID-19 Genomics UK (COG-UK) Consortium | Harper VanSteenhouse, Yumi Kasai, David Gray, Carol Clugston, Anna Dominiczak and Alex Alderton, Roberto Amato, Jeffrey Barrett, Sonia Goncalves, Ewan Harrison, David K. Jackson, Ian Johnston, Dominic Kwiatkowski, Cordelia Langford, John Sillitoe on behalf of the Wellcome Sanger Institute COVID-19 Surveillance Team |
| EPI_ISL_2487716 | Lighthouse Lab in Alderley Park | Wellcome Sanger Institute for the COVID-19 Genomics UK (COG-UK) Consortium | Jacquelyn Wynn, Mairead Hyland, The Lighthouse Lab in Alderley Park and Alex Alderton, Roberto Amato, Jeffrey Barrett, Sonia Goncalves, Ewan Harrison, David K. Jackson, Ian Johnston, Dominic Kwiatkowski, Cordelia Langford, John Sillitoe on behalf of the Wellcome Sanger Institute COVID-19 Surveillance Team |
| EPI_ISL_2487717, EPI_ISL_2487718, EPI_ISL_2487721, EPI_ISL_2487722, EPI_ISL_2487723 | Lighthouse Lab in Glasgow | Wellcome Sanger Institute for the COVID-19 Genomics UK (COG-UK) Consortium | Harper VanSteenhouse, Yumi Kasai, David Gray, Carol Clugston, Anna Dominiczak and Alex Alderton, Roberto Amato, Jeffrey Barrett, Sonia Goncalves, Ewan Harrison, David K. Jackson, Ian Johnston, Dominic Kwiatkowski, Cordelia Langford, John Sillitoe on behalf of the Wellcome Sanger Institute COVID-19 Surveillance Team |
| EPI_ISL_2487724, EPI_ISL_2487725, EPI_ISL_2487726 | Lighthouse Lab in Alderley Park | Wellcome Sanger Institute for the COVID-19 Genomics UK (COG-UK) Consortium | Jacquelyn Wynn, Mairead Hyland, The Lighthouse Lab in Alderley Park and Alex Alderton, Roberto Amato, Jeffrey Barrett, Sonia Goncalves, Ewan Harrison, David K. Jackson, Ian Johnston, Dominic Kwiatkowski, Cordelia Langford, John Sillitoe on behalf of the Wellcome Sanger Institute COVID-19 Surveillance Team |
| EPI_ISL_2487727, EPI_ISL_2487728 | Lighthouse Lab in Glasgow | Wellcome Sanger Institute for the COVID-19 Genomics UK (COG-UK) Consortium | Harper VanSteenhouse, Yumi Kasai, David Gray, Carol Clugston, Anna Dominiczak and Alex Alderton, Roberto Amato, Jeffrey Barrett, Sonia Goncalves, Ewan Harrison, David K. Jackson, Ian Johnston, Dominic Kwiatkowski, Cordelia Langford, John Sillitoe on behalf of the Wellcome Sanger Institute COVID-19 Surveillance Team |
| EPI_ISL_2487729 | Lighthouse Lab in Alderley Park | Wellcome Sanger Institute for the COVID-19 Genomics UK (COG-UK) Consortium | Jacquelyn Wynn, Mairead Hyland, The Lighthouse Lab in Alderley Park and Alex Alderton, Roberto Amato, Jeffrey Barrett, Sonia Goncalves, Ewan Harrison, David K. Jackson, Ian Johnston, Dominic Kwiatkowski, Cordelia Langford, John Sillitoe on behalf of the Wellcome Sanger Institute COVID-19 Surveillance Team |
| EPI_ISL_2487730, EPI_ISL_2487732, EPI_ISL_2487733, EPI_ISL_2487734, EPI_ISL_2487737, EPI_ISL_2487738, EPI_ISL_2487739, EPI_ISL_2487740, EPI_ISL_2487741, EPI_ISL_2487742, EPI_ISL_2487744, EPI_ISL_2487746, EPI_ISL_2487747, EPI_ISL_2487748, EPI_ISL_2487749, EPI_ISL_2487752, EPI_ISL_2487754, EPI_ISL_2487756, EPI_ISL_2487757, EPI_ISL_2487758, EPI_ISL_2487759, EPI_ISL_2487761, EPI_ISL_2487762, EPI_ISL_2487763, EPI_ISL_2487764, EPI_ISL_2487765, EPI_ISL_2487766, EPI_ISL_2487767, EPI_ISL_2487768 | | | |
| see above | Lighthouse Lab in Glasgow | Wellcome Sanger Institute for the COVID-19 Genomics UK (COG-UK) Consortium | Harper VanSteenhouse, Yumi Kasai, David Gray, Carol Clugston, Anna Dominiczak and Alex Alderton, Roberto Amato, Jeffrey Barrett, Sonia Goncalves, Ewan Harrison, David K. Jackson, Ian Johnston, Dominic Kwiatkowski, Cordelia Langford, John Sillitoe on behalf of the Wellcome Sanger Institute COVID-19 Surveillance Team |
| EPI_ISL_2487769 | Lighthouse Lab in Alderley Park | Wellcome Sanger Institute for the COVID-19 Genomics UK (COG-UK) Consortium | Jacquelyn Wynn, Mairead Hyland, The Lighthouse Lab in Alderley Park and Alex Alderton, Roberto Amato, Jeffrey Barrett, Sonia Goncalves, Ewan Harrison, David K. Jackson, Ian Johnston, Dominic Kwiatkowski, Cordelia Langford, John Sillitoe on behalf of the Wellcome Sanger Institute COVID-19 Surveillance Team |
| EPI_ISL_2487770, EPI_ISL_2487773, EPI_ISL_2487774, EPI_ISL_2487776, EPI_ISL_2487777, EPI_ISL_2487778, EPI_ISL_2487780, EPI_ISL_2487781, EPI_ISL_2487782 | Lighthouse Lab in Glasgow | Wellcome Sanger Institute for the COVID-19 Genomics UK (COG-UK) Consortium | Harper VanSteenhouse, Yumi Kasai, David Gray, Carol Clugston, Anna Dominiczak and Alex Alderton, Roberto Amato, Jeffrey Barrett, Sonia Goncalves, Ewan Harrison, David K. Jackson, Ian Johnston, Dominic Kwiatkowski, Cordelia Langford, John Sillitoe on behalf of the Wellcome Sanger Institute COVID-19 Surveillance Team |
| EPI_ISL_2487783 | Lighthouse Lab in Alderley Park | Wellcome Sanger Institute for the COVID-19 Genomics UK (COG-UK) Consortium | Jacquelyn Wynn, Mairead Hyland, The Lighthouse Lab in Alderley Park and Alex Alderton, Roberto Amato, Jeffrey Barrett, Sonia Goncalves, Ewan Harrison, David K. Jackson, Ian Johnston, Dominic Kwiatkowski, Cordelia Langford, John Sillitoe on behalf of the Wellcome Sanger Institute COVID-19 Surveillance Team |
| EPI_ISL_2487784, EPI_ISL_2487785, EPI_ISL_2487786, EPI_ISL_2487788, EPI_ISL_2487790, EPI_ISL_2487792, EPI_ISL_2487794, EPI_ISL_2487795, EPI_ISL_2487797, EPI_ISL_2487799, EPI_ISL_2487800, EPI_ISL_2487801, EPI_ISL_2487802, EPI_ISL_2487804 | | | |
| see above | Lighthouse Lab in Glasgow | Wellcome Sanger Institute for the COVID-19 Genomics UK (COG-UK) Consortium | Harper VanSteenhouse, Yumi Kasai, David Gray, Carol Clugston, Anna Dominiczak and Alex Alderton, Roberto Amato, Jeffrey Barrett, Sonia Goncalves, Ewan Harrison, David K. Jackson, Ian Johnston, Dominic Kwiatkowski, Cordelia Langford, John Sillitoe on behalf of the Wellcome Sanger Institute COVID-19 Surveillance Team |
| EPI_ISL_2487805 | Lighthouse Lab in Alderley Park | Wellcome Sanger Institute for the COVID-19 Genomics UK (COG-UK) Consortium | Jacquelyn Wynn, Mairead Hyland, The Lighthouse Lab in Alderley Park and Alex Alderton, Roberto Amato, Jeffrey Barrett, Sonia Goncalves, Ewan Harrison, David K. Jackson, Ian Johnston, Dominic Kwiatkowski, Cordelia Langford, John Sillitoe on behalf of the Wellcome Sanger Institute COVID-19 Surveillance Team |
| EPI_ISL_2487808 | Lighthouse Lab in Glasgow | Wellcome Sanger Institute for the COVID-19 Genomics UK (COG-UK) Consortium | Harper VanSteenhouse, Yumi Kasai, David Gray, Carol Clugston, Anna Dominiczak and Alex Alderton, Roberto Amato, Jeffrey Barrett, Sonia Goncalves, Ewan Harrison, David K. Jackson, Ian Johnston, Dominic Kwiatkowski, Cordelia Langford, John Sillitoe on behalf of the Wellcome Sanger Institute COVID-19 Surveillance Team |
| EPI_ISL_2487809 | Lighthouse Lab in Alderley Park | Wellcome Sanger Institute for the COVID-19 Genomics UK (COG-UK) Consortium | Jacquelyn Wynn, Mairead Hyland, The Lighthouse Lab in Alderley Park and Alex Alderton, Roberto Amato, Jeffrey Barrett, Sonia Goncalves, Ewan Harrison, David K. Jackson, Ian Johnston, Dominic Kwiatkowski, Cordelia Langford, John Sillitoe on behalf of the Wellcome Sanger Institute COVID-19 Surveillance Team |
| EPI_ISL_2487810, EPI_ISL_2487811, EPI_ISL_2487812 | Lighthouse Lab in Glasgow | Wellcome Sanger Institute for the COVID-19 Genomics UK (COG-UK) Consortium | Harper VanSteenhouse, Yumi Kasai, David Gray, Carol Clugston, Anna Dominiczak and Alex Alderton, Roberto Amato, Jeffrey Barrett, Sonia Goncalves, Ewan Harrison, David K. Jackson, Ian Johnston, Dominic Kwiatkowski, Cordelia Langford, John Sillitoe on behalf of the Wellcome Sanger Institute COVID-19 Surveillance Team |
| EPI_ISL_2487813, EPI_ISL_2487814 | Lighthouse Lab in Alderley Park | Wellcome Sanger Institute for the COVID-19 Genomics UK (COG-UK) Consortium | Jacquelyn Wynn, Mairead Hyland, The Lighthouse Lab in Alderley Park and Alex Alderton, Roberto Amato, Jeffrey Barrett, Sonia Goncalves, Ewan Harrison, David K. Jackson, Ian Johnston, Dominic Kwiatkowski, Cordelia Langford, John Sillitoe on behalf of the Wellcome Sanger Institute COVID-19 Surveillance Team |
| EPI_ISL_2487815, EPI_ISL_2487816, EPI_ISL_2487817 | Lighthouse Lab in Glasgow | Wellcome Sanger Institute for the COVID-19 Genomics UK (COG-UK) Consortium | Harper VanSteenhouse, Yumi Kasai, David Gray, Carol Clugston, Anna Dominiczak and Alex Alderton, Roberto Amato, Jeffrey Barrett, Sonia Goncalves, Ewan Harrison, David K. Jackson, Ian Johnston, Dominic Kwiatkowski, Cordelia Langford, John Sillitoe on behalf of the Wellcome Sanger Institute COVID-19 Surveillance Team |
| EPI_ISL_2487818 | Lighthouse Lab in Alderley Park | Wellcome Sanger Institute for the COVID-19 Genomics UK (COG-UK) Consortium | Jacquelyn Wynn, Mairead Hyland, The Lighthouse Lab in Alderley Park and Alex Alderton, Roberto Amato, Jeffrey Barrett, Sonia Goncalves, Ewan Harrison, David K. Jackson, Ian Johnston, Dominic Kwiatkowski, Cordelia Langford, John Sillitoe on behalf of the Wellcome Sanger Institute COVID-19 Surveillance Team |
| EPI_ISL_2487819, EPI_ISL_2487820, EPI_ISL_2487821, EPI_ISL_2487822 | Lighthouse Lab in Glasgow | Wellcome Sanger Institute for the COVID-19 Genomics UK (COG-UK) Consortium | Harper VanSteenhouse, Yumi Kasai, David Gray, Carol Clugston, Anna Dominiczak and Alex Alderton, Roberto Amato, Jeffrey Barrett, Sonia Goncalves, Ewan Harrison, David K. Jackson, Ian Johnston, Dominic Kwiatkowski, Cordelia Langford, John Sillitoe on behalf of the Wellcome Sanger Institute COVID-19 Surveillance Team |
| EPI_ISL_2487823 | Lighthouse Lab in Alderley Park | Wellcome Sanger Institute for the COVID-19 Genomics UK (COG-UK) Consortium | Jacquelyn Wynn, Mairead Hyland, The Lighthouse Lab in Alderley Park and Alex Alderton, Roberto Amato, Jeffrey Barrett, Sonia Goncalves, Ewan Harrison, David K. Jackson, Ian Johnston, Dominic Kwiatkowski, Cordelia Langford, John Sillitoe on behalf of the Wellcome Sanger Institute COVID-19 Surveillance Team |
| EPI_ISL_2487824, | Lighthouse Lab in Glasgow | Wellcome Sanger Institute for the COVID-19 Genomics UK (COG-UK) | Harper VanSteenhouse, Yumi Kasai, David Gray, Carol Clugston, Anna Dominiczak and Alex Alderton, Roberto Amato, Jeffrey Barrett, Sonia Goncalves, Ewan Harrison, David K. Jackson, Ian Johnston, Dominic Kwiatkowski, Cordelia Langford, |

| | | | |
|---|---|---|---|
| EPI_ISL_2487825, EPI_ISL_2487826 | | Consortium | John Sillitoe on behalf of the Wellcome Sanger Institute COVID-19 Surveillance Team |
| EPI_ISL_2487828 | Lighthouse Lab in Alderley Park | Wellcome Sanger Institute for the COVID-19 Genomics UK (COG-UK) Consortium | Jacquelyn Wynn, Mairead Hyland, The Lighthouse Lab in Alderley Park and Alex Alderton, Roberto Amato, Jeffrey Barrett, Sonia Goncalves, Ewan Harrison, David K. Jackson, Ian Johnston, Dominic Kwiatkowski, Cordelia Langford, John Sillitoe on behalf of the Wellcome Sanger Institute COVID-19 Surveillance Team |
| EPI_ISL_2487829, EPI_ISL_2487830 | Lighthouse Lab in Glasgow | Wellcome Sanger Institute for the COVID-19 Genomics UK (COG-UK) Consortium | Harper VanSteenhouse, Yumi Kasai, David Gray, Carol Clugston, Anna Dominiczak and Alex Alderton, Roberto Amato, Jeffrey Barrett, Sonia Goncalves, Ewan Harrison, David K. Jackson, Ian Johnston, Dominic Kwiatkowski, Cordelia Langford, John Sillitoe on behalf of the Wellcome Sanger Institute COVID-19 Surveillance Team |
| EPI_ISL_2487831 | Lighthouse Lab in Alderley Park | Wellcome Sanger Institute for the COVID-19 Genomics UK (COG-UK) Consortium | Jacquelyn Wynn, Mairead Hyland, The Lighthouse Lab in Alderley Park and Alex Alderton, Roberto Amato, Jeffrey Barrett, Sonia Goncalves, Ewan Harrison, David K. Jackson, Ian Johnston, Dominic Kwiatkowski, Cordelia Langford, John Sillitoe on behalf of the Wellcome Sanger Institute COVID-19 Surveillance Team |
| EPI_ISL_2487832, EPI_ISL_2487833, EPI_ISL_2487835 | Lighthouse Lab in Glasgow | Wellcome Sanger Institute for the COVID-19 Genomics UK (COG-UK) Consortium | Harper VanSteenhouse, Yumi Kasai, David Gray, Carol Clugston, Anna Dominiczak and Alex Alderton, Roberto Amato, Jeffrey Barrett, Sonia Goncalves, Ewan Harrison, David K. Jackson, Ian Johnston, Dominic Kwiatkowski, Cordelia Langford, John Sillitoe on behalf of the Wellcome Sanger Institute COVID-19 Surveillance Team |
| EPI_ISL_2487836 | Lighthouse Lab in Alderley Park | Wellcome Sanger Institute for the COVID-19 Genomics UK (COG-UK) Consortium | Jacquelyn Wynn, Mairead Hyland, The Lighthouse Lab in Alderley Park and Alex Alderton, Roberto Amato, Jeffrey Barrett, Sonia Goncalves, Ewan Harrison, David K. Jackson, Ian Johnston, Dominic Kwiatkowski, Cordelia Langford, John Sillitoe on behalf of the Wellcome Sanger Institute COVID-19 Surveillance Team |
| EPI_ISL_2487837, EPI_ISL_2487838, EPI_ISL_2487839, EPI_ISL_2487840, EPI_ISL_2487841, EPI_ISL_2487842, EPI_ISL_2487843, EPI_ISL_2487844, EPI_ISL_2487845, EPI_ISL_2487846, EPI_ISL_2487847 | | | |
| see above | Lighthouse Lab in Glasgow | Wellcome Sanger Institute for the COVID-19 Genomics UK (COG-UK) Consortium | Harper VanSteenhouse, Yumi Kasai, David Gray, Carol Clugston, Anna Dominiczak and Alex Alderton, Roberto Amato, Jeffrey Barrett, Sonia Goncalves, Ewan Harrison, David K. Jackson, Ian Johnston, Dominic Kwiatkowski, Cordelia Langford, John Sillitoe on behalf of the Wellcome Sanger Institute COVID-19 Surveillance Team |
| EPI_ISL_2487848 | Lighthouse Lab in Alderley Park | Wellcome Sanger Institute for the COVID-19 Genomics UK (COG-UK) Consortium | Jacquelyn Wynn, Mairead Hyland, The Lighthouse Lab in Alderley Park and Alex Alderton, Roberto Amato, Jeffrey Barrett, Sonia Goncalves, Ewan Harrison, David K. Jackson, Ian Johnston, Dominic Kwiatkowski, Cordelia Langford, John Sillitoe on behalf of the Wellcome Sanger Institute COVID-19 Surveillance Team |
| EPI_ISL_2487849, EPI_ISL_2487850, EPI_ISL_2487852, EPI_ISL_2487853, EPI_ISL_2487854, EPI_ISL_2487856, EPI_ISL_2487857 | Lighthouse Lab in Glasgow | Wellcome Sanger Institute for the COVID-19 Genomics UK (COG-UK) Consortium | Harper VanSteenhouse, Yumi Kasai, David Gray, Carol Clugston, Anna Dominiczak and Alex Alderton, Roberto Amato, Jeffrey Barrett, Sonia Goncalves, Ewan Harrison, David K. Jackson, Ian Johnston, Dominic Kwiatkowski, Cordelia Langford, John Sillitoe on behalf of the Wellcome Sanger Institute COVID-19 Surveillance Team |
| EPI_ISL_2487858 | Lighthouse Lab in Alderley Park | Wellcome Sanger Institute for the COVID-19 Genomics UK (COG-UK) Consortium | Jacquelyn Wynn, Mairead Hyland, The Lighthouse Lab in Alderley Park and Alex Alderton, Roberto Amato, Jeffrey Barrett, Sonia Goncalves, Ewan Harrison, David K. Jackson, Ian Johnston, Dominic Kwiatkowski, Cordelia Langford, John Sillitoe on behalf of the Wellcome Sanger Institute COVID-19 Surveillance Team |
| EPI_ISL_2487860 | Lighthouse Lab in Glasgow | Wellcome Sanger Institute for the COVID-19 Genomics UK (COG-UK) Consortium | Harper VanSteenhouse, Yumi Kasai, David Gray, Carol Clugston, Anna Dominiczak and Alex Alderton, Roberto Amato, Jeffrey Barrett, Sonia Goncalves, Ewan Harrison, David K. Jackson, Ian Johnston, Dominic Kwiatkowski, Cordelia Langford, John Sillitoe on behalf of the Wellcome Sanger Institute COVID-19 Surveillance Team |
| EPI_ISL_2487861 | Lighthouse Lab in Alderley Park | Wellcome Sanger Institute for the COVID-19 Genomics UK (COG-UK) Consortium | Jacquelyn Wynn, Mairead Hyland, The Lighthouse Lab in Alderley Park and Alex Alderton, Roberto Amato, Jeffrey Barrett, Sonia Goncalves, Ewan Harrison, David K. Jackson, Ian Johnston, Dominic Kwiatkowski, Cordelia Langford, John Sillitoe on behalf of the Wellcome Sanger Institute COVID-19 Surveillance Team |
| EPI_ISL_2487862, EPI_ISL_2487863, EPI_ISL_2487864, EPI_ISL_2487865, EPI_ISL_2487866 | Lighthouse Lab in Glasgow | Wellcome Sanger Institute for the COVID-19 Genomics UK (COG-UK) Consortium | Harper VanSteenhouse, Yumi Kasai, David Gray, Carol Clugston, Anna Dominiczak and Alex Alderton, Roberto Amato, Jeffrey Barrett, Sonia Goncalves, Ewan Harrison, David K. Jackson, Ian Johnston, Dominic Kwiatkowski, Cordelia Langford, John Sillitoe on behalf of the Wellcome Sanger Institute COVID-19 Surveillance Team |
| EPI_ISL_2487867 | Lighthouse Lab in Alderley Park | Wellcome Sanger Institute for the COVID-19 Genomics UK (COG-UK) Consortium | Jacquelyn Wynn, Mairead Hyland, The Lighthouse Lab in Alderley Park and Alex Alderton, Roberto Amato, Jeffrey Barrett, Sonia Goncalves, Ewan Harrison, David K. Jackson, Ian Johnston, Dominic Kwiatkowski, Cordelia Langford, John Sillitoe on behalf of the Wellcome Sanger Institute COVID-19 Surveillance Team |
| EPI_ISL_2487868, EPI_ISL_2487869, EPI_ISL_2487870, EPI_ISL_2487872, EPI_ISL_2487873, EPI_ISL_2487874, EPI_ISL_2487875, EPI_ISL_2487876, EPI_ISL_2487877, EPI_ISL_2487878, EPI_ISL_2487879, EPI_ISL_2487880 | | | |
| see above | Lighthouse Lab in Glasgow | Wellcome Sanger Institute for the COVID-19 Genomics UK (COG-UK) Consortium | Harper VanSteenhouse, Yumi Kasai, David Gray, Carol Clugston, Anna Dominiczak and Alex Alderton, Roberto Amato, Jeffrey Barrett, Sonia Goncalves, Ewan Harrison, David K. Jackson, Ian Johnston, Dominic Kwiatkowski, Cordelia Langford, John Sillitoe on behalf of the Wellcome Sanger Institute COVID-19 Surveillance Team |
| EPI_ISL_2487881 | Lighthouse Lab in Alderley Park | Wellcome Sanger Institute for the COVID-19 Genomics UK (COG-UK) Consortium | Jacquelyn Wynn, Mairead Hyland, The Lighthouse Lab in Alderley Park and Alex Alderton, Roberto Amato, Jeffrey Barrett, Sonia Goncalves, Ewan Harrison, David K. Jackson, Ian Johnston, Dominic Kwiatkowski, Cordelia Langford, John Sillitoe on behalf of the Wellcome Sanger Institute COVID-19 Surveillance Team |
| EPI_ISL_2487882, EPI_ISL_2487883 | Lighthouse Lab in Glasgow | Wellcome Sanger Institute for the COVID-19 Genomics UK (COG-UK) Consortium | Harper VanSteenhouse, Yumi Kasai, David Gray, Carol Clugston, Anna Dominiczak and Alex Alderton, Roberto Amato, Jeffrey Barrett, Sonia Goncalves, Ewan Harrison, David K. Jackson, Ian Johnston, Dominic Kwiatkowski, Cordelia Langford, John Sillitoe on behalf of the Wellcome Sanger Institute COVID-19 Surveillance Team |
| EPI_ISL_2487884 | Lighthouse Lab in Alderley Park | Wellcome Sanger Institute for the COVID-19 Genomics UK (COG-UK) Consortium | Jacquelyn Wynn, Mairead Hyland, The Lighthouse Lab in Alderley Park and Alex Alderton, Roberto Amato, Jeffrey Barrett, Sonia Goncalves, Ewan Harrison, David K. Jackson, Ian Johnston, Dominic Kwiatkowski, Cordelia Langford, John Sillitoe on behalf of the Wellcome Sanger Institute COVID-19 Surveillance Team |
| EPI_ISL_2487885, EPI_ISL_2487886, EPI_ISL_2487887, EPI_ISL_2487889, EPI_ISL_2487891, EPI_ISL_2487894 | Lighthouse Lab in Glasgow | Wellcome Sanger Institute for the COVID-19 Genomics UK (COG-UK) Consortium | Harper VanSteenhouse, Yumi Kasai, David Gray, Carol Clugston, Anna Dominiczak and Alex Alderton, Roberto Amato, Jeffrey Barrett, Sonia Goncalves, Ewan Harrison, David K. Jackson, Ian Johnston, Dominic Kwiatkowski, Cordelia Langford, John Sillitoe on behalf of the Wellcome Sanger Institute COVID-19 Surveillance Team |
| EPI_ISL_2487895, EPI_ISL_2487896, EPI_ISL_2487897, EPI_ISL_2487898 | Lighthouse Lab in Alderley Park | Wellcome Sanger Institute for the COVID-19 Genomics UK (COG-UK) Consortium | Jacquelyn Wynn, Mairead Hyland, The Lighthouse Lab in Alderley Park and Alex Alderton, Roberto Amato, Jeffrey Barrett, Sonia Goncalves, Ewan Harrison, David K. Jackson, Ian Johnston, Dominic Kwiatkowski, Cordelia Langford, John Sillitoe on behalf of the Wellcome Sanger Institute COVID-19 Surveillance Team |
| EPI_ISL_2487899, EPI_ISL_2487900 | Lighthouse Lab in Glasgow | Wellcome Sanger Institute for the COVID-19 Genomics UK (COG-UK) Consortium | Harper VanSteenhouse, Yumi Kasai, David Gray, Carol Clugston, Anna Dominiczak and Alex Alderton, Roberto Amato, Jeffrey Barrett, Sonia Goncalves, Ewan Harrison, David K. Jackson, Ian Johnston, Dominic Kwiatkowski, Cordelia Langford, John Sillitoe on behalf of the Wellcome Sanger Institute COVID-19 Surveillance Team |
| EPI_ISL_2487902 | Lighthouse Lab in Alderley Park | Wellcome Sanger Institute for the COVID-19 Genomics UK (COG-UK) Consortium | Jacquelyn Wynn, Mairead Hyland, The Lighthouse Lab in Alderley Park and Alex Alderton, Roberto Amato, Jeffrey Barrett, Sonia Goncalves, Ewan Harrison, David K. Jackson, Ian Johnston, Dominic Kwiatkowski, Cordelia Langford, John Sillitoe on behalf of the Wellcome Sanger Institute COVID-19 Surveillance Team |
| EPI_ISL_2487903, EPI_ISL_2487904, EPI_ISL_2487905, EPI_ISL_2487906, EPI_ISL_2487907, EPI_ISL_2487908, EPI_ISL_2487909, EPI_ISL_2487910, EPI_ISL_2487911 | Lighthouse Lab in Glasgow | Wellcome Sanger Institute for the COVID-19 Genomics UK (COG-UK) Consortium | Harper VanSteenhouse, Yumi Kasai, David Gray, Carol Clugston, Anna Dominiczak and Alex Alderton, Roberto Amato, Jeffrey Barrett, Sonia Goncalves, Ewan Harrison, David K. Jackson, Ian Johnston, Dominic Kwiatkowski, Cordelia Langford, John Sillitoe on behalf of the Wellcome Sanger Institute COVID-19 Surveillance Team |
| EPI_ISL_2487913, EPI_ISL_2487914 | Lighthouse Lab in Alderley Park | Wellcome Sanger Institute for the COVID-19 Genomics UK (COG-UK) Consortium | Jacquelyn Wynn, Mairead Hyland, The Lighthouse Lab in Alderley Park and Alex Alderton, Roberto Amato, Jeffrey Barrett, Sonia Goncalves, Ewan Harrison, David K. Jackson, Ian Johnston, Dominic Kwiatkowski, Cordelia Langford, John Sillitoe on behalf of the Wellcome Sanger Institute COVID-19 Surveillance Team |
| EPI_ISL_2487915, EPI_ISL_2487916 | Lighthouse Lab in Glasgow | Wellcome Sanger Institute for the COVID-19 Genomics UK (COG-UK) Consortium | Harper VanSteenhouse, Yumi Kasai, David Gray, Carol Clugston, Anna Dominiczak and Alex Alderton, Roberto Amato, Jeffrey Barrett, Sonia Goncalves, Ewan Harrison, David K. Jackson, Ian Johnston, Dominic Kwiatkowski, Cordelia Langford, John Sillitoe on behalf of the Wellcome Sanger Institute COVID-19 Surveillance Team |
| EPI_ISL_2487917 | Lighthouse Lab in Alderley Park | Wellcome Sanger Institute for the COVID-19 Genomics UK (COG-UK) Consortium | Jacquelyn Wynn, Mairead Hyland, The Lighthouse Lab in Alderley Park and Alex Alderton, Roberto Amato, Jeffrey Barrett, Sonia Goncalves, Ewan Harrison, David K. Jackson, Ian Johnston, Dominic Kwiatkowski, Cordelia Langford, John Sillitoe on behalf of the Wellcome Sanger Institute COVID-19 Surveillance Team |
| EPI_ISL_2487918, EPI_ISL_2487920, EPI_ISL_2487921, EPI_ISL_2487922, EPI_ISL_2487923, EPI_ISL_2487924, EPI_ISL_2487925, EPI_ISL_2487926, EPI_ISL_2487927, EPI_ISL_2487928 | Lighthouse Lab in Glasgow | Wellcome Sanger Institute for the COVID-19 Genomics UK (COG-UK) Consortium | Harper VanSteenhouse, Yumi Kasai, David Gray, Carol Clugston, Anna Dominiczak and Alex Alderton, Roberto Amato, Jeffrey Barrett, Sonia Goncalves, Ewan Harrison, David K. Jackson, Ian Johnston, Dominic Kwiatkowski, Cordelia Langford, John Sillitoe on behalf of the Wellcome Sanger Institute COVID-19 Surveillance Team |
| EPI_ISL_2487929 | Lighthouse Lab in Alderley Park | Wellcome Sanger Institute for the COVID-19 Genomics UK (COG-UK) Consortium | Jacquelyn Wynn, Mairead Hyland, The Lighthouse Lab in Alderley Park and Alex Alderton, Roberto Amato, Jeffrey Barrett, Sonia Goncalves, Ewan Harrison, David K. Jackson, Ian Johnston, Dominic Kwiatkowski, Cordelia Langford, John Sillitoe on behalf of the Wellcome Sanger Institute COVID-19 Surveillance Team |
| EPI_ISL_2487930, EPI_ISL_2487931, EPI_ISL_2487933, EPI_ISL_2487934, EPI_ISL_2487935, EPI_ISL_2487936, EPI_ISL_2487938, EPI_ISL_2487939, EPI_ISL_2487940, EPI_ISL_2487941, EPI_ISL_2487942, EPI_ISL_2487943, EPI_ISL_2487944, EPI_ISL_2487945, EPI_ISL_2487946, EPI_ISL_2487947, EPI_ISL_2487948 | | | |
| see above | Lighthouse Lab in Glasgow | Wellcome Sanger Institute for the COVID-19 Genomics UK (COG-UK) Consortium | Harper VanSteenhouse, Yumi Kasai, David Gray, Carol Clugston, Anna Dominiczak and Alex Alderton, Roberto Amato, Jeffrey Barrett, Sonia Goncalves, Ewan Harrison, David K. Jackson, Ian Johnston, Dominic Kwiatkowski, Cordelia Langford, John Sillitoe on behalf of the Wellcome Sanger Institute COVID-19 Surveillance Team |
| EPI_ISL_2487949 | Health Services Laboratories | Wellcome Sanger Institute for the COVID-19 Genomics UK (COG-UK) Consortium | Health Services Laboratories and Alex Alderton, Roberto Amato, Jeffrey Barrett, Sonia Goncalves, Ewan Harrison, David K. Jackson, Ian Johnston, Dominic Kwiatkowski, Cordelia Langford, John Sillitoe on behalf of the Wellcome Sanger Institute COVID-19 Surveillance Team |
| EPI_ISL_2487950, EPI_ISL_2487951, EPI_ISL_2487952, EPI_ISL_2487953, EPI_ISL_2487954, EPI_ISL_2487955, EPI_ISL_2487956, EPI_ISL_2487957, EPI_ISL_2487958, EPI_ISL_2487959, EPI_ISL_2487962, EPI_ISL_2487963, EPI_ISL_2487964, EPI_ISL_2487965, EPI_ISL_2487966, EPI_ISL_2487968, EPI_ISL_2487969, EPI_ISL_2487970, EPI_ISL_2487971, EPI_ISL_2487972, EPI_ISL_2487975, EPI_ISL_2487979, EPI_ISL_2487981, EPI_ISL_2487982, EPI_ISL_2487984, EPI_ISL_2487985, EPI_ISL_2487986 | | | |

| | | | |
|---|---|---|---|
| see above | Lighthouse Lab in Glasgow | Wellcome Sanger Institute for the COVID-19 Genomics UK (COG-UK) Consortium | Harper VanSteenhouse, Yumi Kasai, David Gray, Carol Clugston, Anna Dominiczak and Alex Alderton, Roberto Amato, Jeffrey Barrett, Sonia Goncalves, Ewan Harrison, David K. Jackson, Ian Johnston, Dominic Kwiatkowski, Cordelia Langford, John Sillitoe on behalf of the Wellcome Sanger Institute COVID-19 Surveillance Team |
| EPI_ISL_2487989 | Lighthouse Laboratory Plymouth | Wellcome Sanger Institute for the COVID-19 Genomics UK (COG-UK) Consortium | Lighthouse Laboratory Plymouth and Alex Alderton, Roberto Amato, Jeffrey Barrett, Sonia Goncalves, Ewan Harrison, David K. Jackson, Ian Johnston, Dominic Kwiatkowski, Cordelia Langford, John Sillitoe on behalf of the Wellcome Sanger Institute COVID-19 Surveillance Team |
| EPI_ISL_2487990, EPI_ISL_2487991, EPI_ISL_2487992, EPI_ISL_2487993, EPI_ISL_2487994, EPI_ISL_2487995, EPI_ISL_2487997, EPI_ISL_2487998, EPI_ISL_2487999, EPI_ISL_2488000, EPI_ISL_2488001, EPI_ISL_2488002, EPI_ISL_2488004, EPI_ISL_2488006, EPI_ISL_2488007, EPI_ISL_2488010, EPI_ISL_2488011, EPI_ISL_2488012, EPI_ISL_2488013, EPI_ISL_2488014, EPI_ISL_2488015, EPI_ISL_2488016, EPI_ISL_2488017, EPI_ISL_2488018, EPI_ISL_2488019, EPI_ISL_2488020, EPI_ISL_2488022, EPI_ISL_2488023, EPI_ISL_2488025, EPI_ISL_2488026, EPI_ISL_2488027, EPI_ISL_2488028, EPI_ISL_2488029, EPI_ISL_2488030, EPI_ISL_2488031, EPI_ISL_2488034, EPI_ISL_2488035, EPI_ISL_2488036, EPI_ISL_2488037, EPI_ISL_2488038, EPI_ISL_2488039, EPI_ISL_2488040, EPI_ISL_2488041, EPI_ISL_2488042, EPI_ISL_2488043, EPI_ISL_2488044, EPI_ISL_2488045, EPI_ISL_2488046, EPI_ISL_2488047, EPI_ISL_2488048, EPI_ISL_2488049, EPI_ISL_2488051, EPI_ISL_2488052, EPI_ISL_2488053, EPI_ISL_2488054, EPI_ISL_2488055, EPI_ISL_2488056, EPI_ISL_2488057, EPI_ISL_2488058, EPI_ISL_2488059, EPI_ISL_2488060, EPI_ISL_2488061, EPI_ISL_2488062, EPI_ISL_2488063, EPI_ISL_2488064, EPI_ISL_2488065, EPI_ISL_2488066, EPI_ISL_2488067, EPI_ISL_2488068, EPI_ISL_2488072, EPI_ISL_2488073, EPI_ISL_2488074, EPI_ISL_2488075, EPI_ISL_2488077, EPI_ISL_2488078, EPI_ISL_2488079 | | | |
| see above | Lighthouse Lab in Glasgow | Wellcome Sanger Institute for the COVID-19 Genomics UK (COG-UK) Consortium | Harper VanSteenhouse, Yumi Kasai, David Gray, Carol Clugston, Anna Dominiczak and Alex Alderton, Roberto Amato, Jeffrey Barrett, Sonia Goncalves, Ewan Harrison, David K. Jackson, Ian Johnston, Dominic Kwiatkowski, Cordelia Langford, John Sillitoe on behalf of the Wellcome Sanger Institute COVID-19 Surveillance Team |
| EPI_ISL_2488080, EPI_ISL_2488081 | Lighthouse Lab in Alderley Park | Wellcome Sanger Institute for the COVID-19 Genomics UK (COG-UK) Consortium | Jacquelyn Wynn, Mairead Hyland, The Lighthouse Lab in Alderley Park and Alex Alderton, Roberto Amato, Jeffrey Barrett, Sonia Goncalves, Ewan Harrison, David K. Jackson, Ian Johnston, Dominic Kwiatkowski, Cordelia Langford, John Sillitoe on behalf of the Wellcome Sanger Institute COVID-19 Surveillance Team |
| EPI_ISL_2488082, EPI_ISL_2488084, EPI_ISL_2488085, EPI_ISL_2488087, EPI_ISL_2488088, EPI_ISL_2488089, EPI_ISL_2488090, EPI_ISL_2488093 | Lighthouse Lab in Glasgow | Wellcome Sanger Institute for the COVID-19 Genomics UK (COG-UK) Consortium | Harper VanSteenhouse, Yumi Kasai, David Gray, Carol Clugston, Anna Dominiczak and Alex Alderton, Roberto Amato, Jeffrey Barrett, Sonia Goncalves, Ewan Harrison, David K. Jackson, Ian Johnston, Dominic Kwiatkowski, Cordelia Langford, John Sillitoe on behalf of the Wellcome Sanger Institute COVID-19 Surveillance Team |
| EPI_ISL_2488094 | Lighthouse Lab in Alderley Park | Wellcome Sanger Institute for the COVID-19 Genomics UK (COG-UK) Consortium | Jacquelyn Wynn, Mairead Hyland, The Lighthouse Lab in Alderley Park and Alex Alderton, Roberto Amato, Jeffrey Barrett, Sonia Goncalves, Ewan Harrison, David K. Jackson, Ian Johnston, Dominic Kwiatkowski, Cordelia Langford, John Sillitoe on behalf of the Wellcome Sanger Institute COVID-19 Surveillance Team |
| EPI_ISL_2488096, EPI_ISL_2488097, EPI_ISL_2488098, EPI_ISL_2488100, EPI_ISL_2488101, EPI_ISL_2488102, EPI_ISL_2488103, EPI_ISL_2488104, EPI_ISL_2488105, EPI_ISL_2488106, EPI_ISL_2488107, EPI_ISL_2488108, EPI_ISL_2488109, EPI_ISL_2488112, EPI_ISL_2488113, EPI_ISL_2488115, EPI_ISL_2488117, EPI_ISL_2488118, EPI_ISL_2488121, EPI_ISL_2488122, EPI_ISL_2488123, EPI_ISL_2488124, EPI_ISL_2488126, EPI_ISL_2488127, EPI_ISL_2488128, EPI_ISL_2488131, EPI_ISL_2488132, EPI_ISL_2488134, EPI_ISL_2488135, EPI_ISL_2488136, EPI_ISL_2488137, EPI_ISL_2488138, EPI_ISL_2488139, EPI_ISL_2488140, EPI_ISL_2488141, EPI_ISL_2488143, EPI_ISL_2488144, EPI_ISL_2488145, EPI_ISL_2488146, EPI_ISL_2488147, EPI_ISL_2488148, EPI_ISL_2488150, EPI_ISL_2488151, EPI_ISL_2488152, EPI_ISL_2488154, EPI_ISL_2488155, EPI_ISL_2488156, EPI_ISL_2488157, EPI_ISL_2488158, EPI_ISL_2488159, EPI_ISL_2488160, EPI_ISL_2488161, EPI_ISL_2488162, EPI_ISL_2488163, EPI_ISL_2488165, EPI_ISL_2488166, EPI_ISL_2488167, EPI_ISL_2488168, EPI_ISL_2488169, EPI_ISL_2488171, EPI_ISL_2488173, EPI_ISL_2488175, EPI_ISL_2488176, EPI_ISL_2488177, EPI_ISL_2488178, EPI_ISL_2488181, EPI_ISL_2488182, EPI_ISL_2488184, EPI_ISL_2488187, EPI_ISL_2488188, EPI_ISL_2488189, EPI_ISL_2488192, EPI_ISL_2488193, EPI_ISL_2488194, EPI_ISL_2488195, EPI_ISL_2488196, EPI_ISL_2488199, EPI_ISL_2488200, EPI_ISL_2488201, EPI_ISL_2488203, EPI_ISL_2488204, EPI_ISL_2488205, EPI_ISL_2488206, EPI_ISL_2488207, EPI_ISL_2488209, EPI_ISL_2488210, EPI_ISL_2488211, EPI_ISL_2488215, EPI_ISL_2488216, EPI_ISL_2488218, EPI_ISL_2488219, EPI_ISL_2488220, EPI_ISL_2488221, EPI_ISL_2488222, EPI_ISL_2488223, EPI_ISL_2488224, EPI_ISL_2488225, EPI_ISL_2488226, EPI_ISL_2488227, EPI_ISL_2488228, EPI_ISL_2488232, EPI_ISL_2488234, EPI_ISL_2488235 | | | |
| see above | Lighthouse Lab in Glasgow | Wellcome Sanger Institute for the COVID-19 Genomics UK (COG-UK) Consortium | Harper VanSteenhouse, Yumi Kasai, David Gray, Carol Clugston, Anna Dominiczak and Alex Alderton, Roberto Amato, Jeffrey Barrett, Sonia Goncalves, Ewan Harrison, David K. Jackson, Ian Johnston, Dominic Kwiatkowski, Cordelia Langford, John Sillitoe on behalf of the Wellcome Sanger Institute COVID-19 Surveillance Team |
| EPI_ISL_2488236, EPI_ISL_2488237 | Lighthouse Laboratory Plymouth | Wellcome Sanger Institute for the COVID-19 Genomics UK (COG-UK) Consortium | Lighthouse Laboratory Plymouth and Alex Alderton, Roberto Amato, Jeffrey Barrett, Sonia Goncalves, Ewan Harrison, David K. Jackson, Ian Johnston, Dominic Kwiatkowski, Cordelia Langford, John Sillitoe on behalf of the Wellcome Sanger Institute COVID-19 Surveillance Team |
| EPI_ISL_2488238 | Lighthouse Lab in Alderley Park | Wellcome Sanger Institute for the COVID-19 Genomics UK (COG-UK) Consortium | Jacquelyn Wynn, Mairead Hyland, The Lighthouse Lab in Alderley Park and Alex Alderton, Roberto Amato, Jeffrey Barrett, Sonia Goncalves, Ewan Harrison, David K. Jackson, Ian Johnston, Dominic Kwiatkowski, Cordelia Langford, John Sillitoe on behalf of the Wellcome Sanger Institute COVID-19 Surveillance Team |
| EPI_ISL_2488239 | Lighthouse Lab in Milton Keynes | Wellcome Sanger Institute for the COVID-19 Genomics UK (COG-UK) Consortium | The Lighthouse Lab in Milton Keynes and Alex Alderton, Roberto Amato, Jeffrey Barrett, Sonia Goncalves, Ewan Harrison, David K. Jackson, Ian Johnston, Dominic Kwiatkowski, Cordelia Langford, John Sillitoe on behalf of the Wellcome Sanger Institute COVID-19 Surveillance Team |
| EPI_ISL_2488240, EPI_ISL_2488241 | Lighthouse Lab in Alderley Park | Wellcome Sanger Institute for the COVID-19 Genomics UK (COG-UK) Consortium | Jacquelyn Wynn, Mairead Hyland, The Lighthouse Lab in Alderley Park and Alex Alderton, Roberto Amato, Jeffrey Barrett, Sonia Goncalves, Ewan Harrison, David K. Jackson, Ian Johnston, Dominic Kwiatkowski, Cordelia Langford, John Sillitoe on behalf of the Wellcome Sanger Institute COVID-19 Surveillance Team |
| EPI_ISL_2488242 | Lighthouse Lab in Milton Keynes | Wellcome Sanger Institute for the COVID-19 Genomics UK (COG-UK) Consortium | The Lighthouse Lab in Milton Keynes and Alex Alderton, Roberto Amato, Jeffrey Barrett, Sonia Goncalves, Ewan Harrison, David K. Jackson, Ian Johnston, Dominic Kwiatkowski, Cordelia Langford, John Sillitoe on behalf of the Wellcome Sanger Institute COVID-19 Surveillance Team |
| EPI_ISL_2488243, EPI_ISL_2488244, EPI_ISL_2488245, EPI_ISL_2488246, EPI_ISL_2488247, EPI_ISL_2488248, EPI_ISL_2488249, EPI_ISL_2488250, EPI_ISL_2488251, EPI_ISL_2488252, EPI_ISL_2488253, EPI_ISL_2488255, EPI_ISL_2488256, EPI_ISL_2488257, EPI_ISL_2488258, EPI_ISL_2488259, EPI_ISL_2488260, EPI_ISL_2488261, EPI_ISL_2488262, EPI_ISL_2488263, EPI_ISL_2488264, EPI_ISL_2488265, EPI_ISL_2488266, EPI_ISL_2488267, EPI_ISL_2488268, EPI_ISL_2488269, EPI_ISL_2488270, EPI_ISL_2488271, EPI_ISL_2488275, EPI_ISL_2488276, EPI_ISL_2488277, EPI_ISL_2488278, EPI_ISL_2488280, EPI_ISL_2488281 | | | |
| see above | Lighthouse Lab in Glasgow | Wellcome Sanger Institute for the COVID-19 Genomics UK (COG-UK) Consortium | Harper VanSteenhouse, Yumi Kasai, David Gray, Carol Clugston, Anna Dominiczak and Alex Alderton, Roberto Amato, Jeffrey Barrett, Sonia Goncalves, Ewan Harrison, David K. Jackson, Ian Johnston, Dominic Kwiatkowski, Cordelia Langford, John Sillitoe on behalf of the Wellcome Sanger Institute COVID-19 Surveillance Team |
| EPI_ISL_2488282 | Lighthouse Lab in Milton Keynes | Wellcome Sanger Institute for the COVID-19 Genomics UK (COG-UK) Consortium | The Lighthouse Lab in Milton Keynes and Alex Alderton, Roberto Amato, Jeffrey Barrett, Sonia Goncalves, Ewan Harrison, David K. Jackson, Ian Johnston, Dominic Kwiatkowski, Cordelia Langford, John Sillitoe on behalf of the Wellcome Sanger Institute COVID-19 Surveillance Team |
| EPI_ISL_2488283, EPI_ISL_2488284, EPI_ISL_2488286, EPI_ISL_2488288, EPI_ISL_2488289, EPI_ISL_2488290, EPI_ISL_2488292, EPI_ISL_2488293, EPI_ISL_2488294, EPI_ISL_2488295, EPI_ISL_2488296, EPI_ISL_2488297, EPI_ISL_2488298, EPI_ISL_2488299, EPI_ISL_2488301, EPI_ISL_2488303, EPI_ISL_2488304, EPI_ISL_2488305, EPI_ISL_2488307, EPI_ISL_2488308, EPI_ISL_2488309, EPI_ISL_2488311, EPI_ISL_2488312 | | | |
| see above | Lighthouse Lab in Glasgow | Wellcome Sanger Institute for the COVID-19 Genomics UK (COG-UK) Consortium | Harper VanSteenhouse, Yumi Kasai, David Gray, Carol Clugston, Anna Dominiczak and Alex Alderton, Roberto Amato, Jeffrey Barrett, Sonia Goncalves, Ewan Harrison, David K. Jackson, Ian Johnston, Dominic Kwiatkowski, Cordelia Langford, John Sillitoe on behalf of the Wellcome Sanger Institute COVID-19 Surveillance Team |
| EPI_ISL_2488315 | Lighthouse Lab in Milton Keynes | Wellcome Sanger Institute for the COVID-19 Genomics UK (COG-UK) Consortium | The Lighthouse Lab in Milton Keynes and Alex Alderton, Roberto Amato, Jeffrey Barrett, Sonia Goncalves, Ewan Harrison, David K. Jackson, Ian Johnston, Dominic Kwiatkowski, Cordelia Langford, John Sillitoe on behalf of the Wellcome Sanger Institute COVID-19 Surveillance Team |
| EPI_ISL_2488316, EPI_ISL_2488317, EPI_ISL_2488318, EPI_ISL_2488319, EPI_ISL_2488320, EPI_ISL_2488321, EPI_ISL_2488322, EPI_ISL_2488323, EPI_ISL_2488324, EPI_ISL_2488325, EPI_ISL_2488326, EPI_ISL_2488329, EPI_ISL_2488330, EPI_ISL_2488332, EPI_ISL_2488333, EPI_ISL_2488334, EPI_ISL_2488336, EPI_ISL_2488337, EPI_ISL_2488338, EPI_ISL_2488339, EPI_ISL_2488340, EPI_ISL_2488341, EPI_ISL_2488343, EPI_ISL_2488345, EPI_ISL_2488346, EPI_ISL_2488347, EPI_ISL_2488348, EPI_ISL_2488350, EPI_ISL_2488351, EPI_ISL_2488352, EPI_ISL_2488353, EPI_ISL_2488354, EPI_ISL_2488355, EPI_ISL_2488356, EPI_ISL_2488357, EPI_ISL_2488358, EPI_ISL_2488359, EPI_ISL_2488360, EPI_ISL_2488361, EPI_ISL_2488365, EPI_ISL_2488366, EPI_ISL_2488367, EPI_ISL_2488368, EPI_ISL_2488369, EPI_ISL_2488370, EPI_ISL_2488372, EPI_ISL_2488373, EPI_ISL_2488374, EPI_ISL_2488376, EPI_ISL_2488378, EPI_ISL_2488380, EPI_ISL_2488381, EPI_ISL_2488382, EPI_ISL_2488383, EPI_ISL_2488384, EPI_ISL_2488385, EPI_ISL_2488387, EPI_ISL_2488388, EPI_ISL_2488389, EPI_ISL_2488390, EPI_ISL_2488391, EPI_ISL_2488392, EPI_ISL_2488395, EPI_ISL_2488398, EPI_ISL_2488400, EPI_ISL_2488402, EPI_ISL_2488403, EPI_ISL_2488404, EPI_ISL_2488406, EPI_ISL_2488407, EPI_ISL_2488408, EPI_ISL_2488409, EPI_ISL_2488410, EPI_ISL_2488411, EPI_ISL_2488412, EPI_ISL_2488415, EPI_ISL_2488416, EPI_ISL_2488418, EPI_ISL_2488419, EPI_ISL_2488420, EPI_ISL_2488421, EPI_ISL_2488422, EPI_ISL_2488423, EPI_ISL_2488424, EPI_ISL_2488426, EPI_ISL_2488427, EPI_ISL_2488428, EPI_ISL_2488430, EPI_ISL_2488432, EPI_ISL_2488434, EPI_ISL_2488435, EPI_ISL_2488436, EPI_ISL_2488437 | | | |
| see above | Lighthouse Lab in Glasgow | Wellcome Sanger Institute for the COVID-19 Genomics UK (COG-UK) Consortium | Harper VanSteenhouse, Yumi Kasai, David Gray, Carol Clugston, Anna Dominiczak and Alex Alderton, Roberto Amato, Jeffrey Barrett, Sonia Goncalves, Ewan Harrison, David K. Jackson, Ian Johnston, Dominic Kwiatkowski, Cordelia Langford, John Sillitoe on behalf of the Wellcome Sanger Institute COVID-19 Surveillance Team |
| EPI_ISL_2488438 | Lighthouse Lab in Alderley Park | Wellcome Sanger Institute for the COVID-19 Genomics UK (COG-UK) Consortium | Jacquelyn Wynn, Mairead Hyland, The Lighthouse Lab in Alderley Park and Alex Alderton, Roberto Amato, Jeffrey Barrett, Sonia Goncalves, Ewan Harrison, David K. Jackson, Ian Johnston, Dominic Kwiatkowski, Cordelia Langford, John Sillitoe on behalf of the Wellcome Sanger Institute COVID-19 Surveillance Team |
| EPI_ISL_2488439, EPI_ISL_2488440, EPI_ISL_2488441, EPI_ISL_2488442, EPI_ISL_2488444, EPI_ISL_2488446, EPI_ISL_2488447, EPI_ISL_2488448, EPI_ISL_2488449, EPI_ISL_2488451, EPI_ISL_2488452, EPI_ISL_2488454, EPI_ISL_2488455, EPI_ISL_2488456, EPI_ISL_2488457, EPI_ISL_2488458, EPI_ISL_2488459, EPI_ISL_2488460, EPI_ISL_2488462, EPI_ISL_2488463, EPI_ISL_2488465, EPI_ISL_2488466, EPI_ISL_2488467, EPI_ISL_2488468, EPI_ISL_2488469, EPI_ISL_2488471, EPI_ISL_2488472, EPI_ISL_2488473, EPI_ISL_2488475, EPI_ISL_2488476, EPI_ISL_2488477, EPI_ISL_2488478, EPI_ISL_2488479, EPI_ISL_2488481, EPI_ISL_2488482, EPI_ISL_2488485, EPI_ISL_2488486, EPI_ISL_2488487, EPI_ISL_2488489, EPI_ISL_2488490, EPI_ISL_2488491, EPI_ISL_2488494, EPI_ISL_2488496, EPI_ISL_2488497, EPI_ISL_2488499, EPI_ISL_2488500, EPI_ISL_2488502, EPI_ISL_2488503, EPI_ISL_2488504 | | | |
| see above | Lighthouse Lab in Glasgow | Wellcome Sanger Institute for the COVID-19 Genomics UK (COG-UK) Consortium | Harper VanSteenhouse, Yumi Kasai, David Gray, Carol Clugston, Anna Dominiczak and Alex Alderton, Roberto Amato, Jeffrey Barrett, Sonia Goncalves, Ewan Harrison, David K. Jackson, Ian Johnston, Dominic Kwiatkowski, Cordelia Langford, John Sillitoe on behalf of the Wellcome Sanger Institute COVID-19 Surveillance Team |
| EPI_ISL_2488505 | Lighthouse Lab in Alderley Park | Wellcome Sanger Institute for the COVID-19 Genomics UK (COG-UK) Consortium | Jacquelyn Wynn, Mairead Hyland, The Lighthouse Lab in Alderley Park and Alex Alderton, Roberto Amato, Jeffrey Barrett, Sonia Goncalves, Ewan Harrison, David K. Jackson, Ian Johnston, Dominic Kwiatkowski, Cordelia Langford, John Sillitoe on behalf of the Wellcome Sanger Institute COVID-19 Surveillance Team |
| EPI_ISL_2488510, EPI_ISL_2488512, EPI_ISL_2488514, EPI_ISL_2488515, EPI_ISL_2488516, EPI_ISL_2488518, EPI_ISL_2488519, EPI_ISL_2488520, EPI_ISL_2488522, EPI_ISL_2488525, EPI_ISL_2488526, EPI_ISL_2488527, EPI_ISL_2488528, EPI_ISL_2488529, EPI_ISL_2488530, EPI_ISL_2488531, EPI_ISL_2488533, EPI_ISL_2488534, EPI_ISL_2488535, EPI_ISL_2488536, EPI_ISL_2488542, EPI_ISL_2488543, EPI_ISL_2488546, EPI_ISL_2488547, EPI_ISL_2488548, EPI_ISL_2488549, EPI_ISL_2488550, EPI_ISL_2488551, EPI_ISL_2488556, EPI_ISL_2488557, EPI_ISL_2488558, EPI_ISL_2488559, EPI_ISL_2488560, EPI_ISL_2488561, EPI_ISL_2488562, EPI_ISL_2488563, EPI_ISL_2488565, EPI_ISL_2488566, EPI_ISL_2488568, EPI_ISL_2488569, EPI_ISL_2488570, EPI_ISL_2488572, EPI_ISL_2488573, EPI_ISL_2488574, EPI_ISL_2488575, EPI_ISL_2488576, EPI_ISL_2488577, EPI_ISL_2488578, EPI_ISL_2488579, EPI_ISL_2488580, EPI_ISL_2488582, EPI_ISL_2488583, EPI_ISL_2488585, EPI_ISL_2488586, EPI_ISL_2488587, EPI_ISL_2488590, EPI_ISL_2488591, EPI_ISL_2488593, EPI_ISL_2488594, EPI_ISL_2488595, EPI_ISL_2488596, EPI_ISL_2488597, EPI_ISL_2488598, EPI_ISL_2488599, EPI_ISL_2488600, EPI_ISL_2488604, EPI_ISL_2488605, EPI_ISL_2488606, EPI_ISL_2488607, EPI_ISL_2488608, EPI_ISL_2488609, EPI_ISL_2488610, EPI_ISL_2488611, EPI_ISL_2488612, EPI_ISL_2488614, EPI_ISL_2488615, EPI_ISL_2488618, EPI_ISL_2488620, EPI_ISL_2488623, EPI_ISL_2488624, EPI_ISL_2488626, EPI_ISL_2488629, EPI_ISL_2488630, EPI_ISL_2488631, EPI_ISL_2488632, EPI_ISL_2488633, EPI_ISL_2488634, EPI_ISL_2488635, EPI_ISL_2488636, EPI_ISL_2488637, EPI_ISL_2488638, EPI_ISL_2488639, EPI_ISL_2488640, EPI_ISL_2488641, EPI_ISL_2488643, EPI_ISL_2488644, EPI_ISL_2488645, EPI_ISL_2488648, EPI_ISL_2488649, EPI_ISL_2488650, EPI_ISL_2488651, EPI_ISL_2488652, EPI_ISL_2488653, EPI_ISL_2488654, EPI_ISL_2488656, EPI_ISL_2488657, EPI_ISL_2488658, EPI_ISL_2488659, EPI_ISL_2488660, EPI_ISL_2488661, EPI_ISL_2488662, EPI_ISL_2488664, EPI_ISL_2488665, EPI_ISL_2488666, EPI_ISL_2488667, EPI_ISL_2488668, EPI_ISL_2488669, EPI_ISL_2488670, EPI_ISL_2488672, EPI_ISL_2488673, EPI_ISL_2488674, EPI_ISL_2488675, EPI_ISL_2488676, EPI_ISL_2488678, EPI_ISL_2488679, EPI_ISL_2488680, EPI_ISL_2488681, EPI_ISL_2488683, EPI_ISL_2488685, EPI_ISL_2488687, EPI_ISL_2488688, EPI_ISL_2488689, EPI_ISL_2488690, EPI_ISL_2488691, EPI_ISL_2488692, EPI_ISL_2488694, EPI_ISL_2488695, EPI_ISL_2488696, EPI_ISL_2488697, EPI_ISL_2488698, EPI_ISL_2488701, EPI_ISL_2488703, EPI_ISL_2488704, EPI_ISL_2488705, EPI_ISL_2488706, EPI_ISL_2488707 | | | |
| see above | Lighthouse Lab in Glasgow | Wellcome Sanger Institute for the COVID-19 Genomics UK (COG-UK) Consortium | Harper VanSteenhouse, Yumi Kasai, David Gray, Carol Clugston, Anna Dominiczak and Alex Alderton, Roberto Amato, Jeffrey Barrett, Sonia Goncalves, Ewan Harrison, David K. Jackson, Ian Johnston, Dominic Kwiatkowski, Cordelia Langford, John Sillitoe on behalf of the Wellcome Sanger Institute COVID-19 Surveillance Team |
| EPI_ISL_2488863, EPI_ISL_2488871, EPI_ISL_2488893, EPI_ISL_2488919 | Mako Medical | Centers for Disease Control and Prevention Division of Viral Diseases, Pathogen Discovery | Dakota Howard, Dhwani Batra, Peter W. Cook, Kara Moser, Adrian Paskey, Jason Caravas, Benjamin Rambo-Martin, Shatavia Morrison, Christopher Gulvick, Scott Sammons, Yvette Unoarumhi, Darlene Wagner, Matthew Schmerer,Matthew Tugwell, Lauren Moon, Clinton R. Paden, Duncan MacCannell |
| EPI_ISL_2489077, EPI_ISL_2489096, EPI_ISL_2489098, EPI_ISL_2489099, EPI_ISL_2489119, EPI_ISL_2489134, EPI_ISL_2489170, EPI_ISL_2489185, EPI_ISL_2489210, EPI_ISL_2489211, EPI_ISL_2489219, EPI_ISL_2489222, EPI_ISL_2489248, EPI_ISL_2489263, EPI_ISL_2489266, EPI_ISL_2489269, EPI_ISL_2489290, EPI_ISL_2489298, EPI_ISL_2489327, EPI_ISL_2489362, EPI_ISL_2489375, EPI_ISL_2489377, EPI_ISL_2489378, EPI_ISL_2489416, EPI_ISL_2489445, EPI_ISL_2489480, EPI_ISL_2489482, EPI_ISL_2489498, EPI_ISL_2489499, EPI_ISL_2489518, EPI_ISL_2489553, EPI_ISL_2489594, EPI_ISL_2489602, EPI_ISL_2489621, EPI_ISL_2489646, EPI_ISL_2489665, EPI_ISL_2489675, EPI_ISL_2489676, EPI_ISL_2489684, EPI_ISL_2489685, EPI_ISL_2489686, EPI_ISL_2489687, EPI_ISL_2489688, EPI_ISL_2489701, EPI_ISL_2489707, EPI_ISL_2489711, EPI_ISL_2489735, EPI_ISL_2489750, EPI_ISL_2489761 | | | |
| see above | Aegis Sciences Corporation | Centers for Disease Control and Prevention Division of Viral Diseases, Pathogen Discovery | Dakota Howard, Dhwani Batra, Peter W. Cook, Kara Moser, Adrian Paskey, Jason Caravas, Benjamin Rambo-Martin, Shatavia Morrison, Christopher Gulvick, Scott Sammons, Yvette Unoarumhi, Darlene Wagner, Matthew Schmerer, Cyndi Clark, Patrick Campbell, Rob Case, Vikramsinha Ghorpade, Holly Houdeshell, Ola Kvalvaag, Dillon Nall, Ethan Sanders, Alec Vest, Shaun Westlund, Matthew Hardison, Clinton R. Paden, Duncan MacCannell |
| EPI_ISL_2489992 | Seattle Flu Study | Seattle Flu Study | Deborah A. Nickerson, Chris D. Frazar, Jover Lee, Benjamin Pelle, Erica Ryke, Matthew Richardson, Amanda Adler, Elisabeth Brandstetter, Peter D. Han, Kairsten Fay, Misja Ilcisin, Kirsten Lacombe, Thomas R. Sibley, Melissa Truong, Caitlin R. Wolf, Karen Cowgill, Stephanie Schrag, Jeff Duchin, Michael Boeckh, Janet A. Englund, Michael Famulare, Barry R. Lutz, Mark J. Rieder, Lea M. Starita, Matthew Thompson, Helen Y. Chu, Trevor Bedford, Jay Shendure |
| EPI_ISL_2490014 | Seattle Flu Study | Seattle Flu Study | Deborah A. Nickerson, Chris D. Frazar, Jover Lee, Benjamin Pelle, Erica Ryke, Matthew Richardson, Amanda Adler, Elisabeth Brandstetter, Peter D. Han, Kairsten Fay, Misja Ilcisin, Kirsten Lacombe, Thomas R. Sibley, Melissa Truong, Caitlin R. Wolf, Michael Boeckh, Janet A. Englund, Michael Famulare, Barry R. Lutz, Mark J. Rieder, Lea M. Starita, Matthew Thompson, Jay Shendure, Trevor Bedford, Helen Y. Chu |
| EPI_ISL_2490043, EPI_ISL_2490044 | Seattle Flu Study | Seattle Flu Study | Deborah A. Nickerson, Chris D. Frazar, Jover Lee, Benjamin Pelle, Erica Ryke, Matthew Richardson, Amanda Adler, Elisabeth Brandstetter, Peter D. Han, Kairsten Fay, Misja Ilcisin, Kirsten Lacombe, Thomas R. Sibley, Melissa Truong, Caitlin R. Wolf, Karen Cowgill, Stephanie Schrag, Jeff Duchin, Michael Boeckh, Janet A. Englund, Michael Famulare, Barry R. Lutz, Mark J. Rieder, Lea M. Starita, Matthew Thompson, Helen Y. Chu, Trevor Bedford, Jay Shendure |
| EPI_ISL_2490307, EPI_ISL_2490308, EPI_ISL_2490309, | Basurto University Hospital: Clinical Microbiology Laboratory | Biocruces Bizkaia | Mikel Urrutikoetxea-Gutiérrez, Mª Carmen Nieto Toboso, Estibaliz Ugalde Zarraga, Mikel Gallego Rodrigo, Ana de la Hoz, José Luis Díaz de Tuesta del Arco |

| | | | |
|---|---|---|---|
| EPI_ISL_2490534 | | | |
| EPI_ISL_2490654, EPI_ISL_2490655, EPI_ISL_2490656, EPI_ISL_2490657, EPI_ISL_2490658, EPI_ISL_2490659, EPI_ISL_2490660 | University Hospitals of Geneva, Laboratory of Virology | HUG, Laboratory of Virology and the Health2030 Genome Center | Samuel Cordey, Ana Rita Goncalves, Laurent Kaiser, Lorenzo Cerutti, Henri Pegeot, Melyssa Elies, Deborah Penet, Keith Harshman, Ioannis Xenarios, Emmanouil Dermitzakis |
| EPI_ISL_2490702, EPI_ISL_2490717, EPI_ISL_2490718, EPI_ISL_2490719, EPI_ISL_2490720, EPI_ISL_2490721 | National Virus Reference Laboratory | National Virus Reference Laboratory | Zoe Yandle, Charlene Bennett, Gabriel Gonzalez, Michael Carr, Jonathan Dean, Cillian F De Gascun |
| EPI_ISL_2490968, EPI_ISL_2490997 | TGen North | TGen North | "Jolene Bowers, Heather Centner, Chris French, Hayley Yaglom, Ashlyn Pfeiffer, Darrin Lemmer, Dave Engelthaler, The Arizona COVID Genomics Union (ACGU)" |
| EPI_ISL_2491119, EPI_ISL_2491120, EPI_ISL_2491121, EPI_ISL_2491123, EPI_ISL_2491124, EPI_ISL_2491125, EPI_ISL_2491127 | National Virus Reference Laboratory | National Virus Reference Laboratory | Zoe Yandle, Charlene Bennett, Gabriel Gonzalez, Michael Carr, Jonathan Dean, Cillian F De Gascun |
| EPI_ISL_2491164, EPI_ISL_2491189, EPI_ISL_2491209 | TGen North | TGen North | "Jolene Bowers, Heather Centner, Chris French, Hayley Yaglom, Ashlyn Pfeiffer, Darrin Lemmer, Dave Engelthaler, The Arizona COVID Genomics Union (ACGU)" |
| EPI_ISL_2491217 | Jessa | Jessa | Berden et al. on behalf of the Jessa_cmdLab |
| EPI_ISL_2491418, EPI_ISL_2491419, EPI_ISL_2491420, EPI_ISL_2491421, EPI_ISL_2491422, EPI_ISL_2491423, EPI_ISL_2491424, EPI_ISL_2491567, EPI_ISL_2491568 | Furst Medical Laboratory | Norwegian Institute of Public Health, Department of Virology | Kathrine Stene-Johansen, Kamilla Heddeland Instefjord, Hilde Elshaug, Garcia Llorente Ignacio, Jon Bråte, Engebretsen Serina Beate,Pedersen Benedikte Nevjen, Line Victoria Moen, Debech Nadia, Atiya R Ali,Marie Paulsen Madsen, Rasmus Riis Kopperud, Hilde Vollan, Karoline Bragstad, Olav Hungnes |
| EPI_ISL_2491569 | Medical Microbiology Unit, Department for Laboratory Medicine, Drammen Hospital, Vestre Viken Health Trust, | Norwegian Institute of Public Health, Department of Virology | Kathrine Stene-Johansen, Kamilla Heddeland Instefjord, Hilde Elshaug, Garcia Llorente Ignacio, Jon Bråte, Engebretsen Serina Beate,Pedersen Benedikte Nevjen, Line Victoria Moen, Debech Nadia, Atiya R Ali,Marie Paulsen Madsen, Rasmus Riis Kopperud, Hilde Vollan, Karoline Bragstad, Olav Hungnes |
| EPI_ISL_2491570 | Akershus University Hospital, Department for Microbiology and Infectious Disease Control | Norwegian Institute of Public Health, Department of Virology | Kathrine Stene-Johansen, Kamilla Heddeland Instefjord, Hilde Elshaug, Garcia Llorente Ignacio, Jon Bråte, Engebretsen Serina Beate,Pedersen Benedikte Nevjen, Line Victoria Moen, Debech Nadia, Atiya R Ali,Marie Paulsen Madsen, Rasmus Riis Kopperud, Hilde Vollan, Karoline Bragstad, Olav Hungnes |
| EPI_ISL_2492325 | Institute of Microbiology and Immunology, Faculty of Medicine, University of Ljubljana | Institute of Microbiology and Immunology, Faculty of Medicine, University of Ljubljana | Alen Suljič, Samo Zakotnik, Tomaž Mark Zorec, Matic Brvar, Doroteja Vlaj, Andraž Celar, Dominika Šturm, Patricija Pozvek, Špela Pleh, Miša Korva, Mario Poljak, Tatjana Avšič – Županc |
| EPI_ISL_2492429, EPI_ISL_2492430, EPI_ISL_2492431, EPI_ISL_2492432, EPI_ISL_2492433, EPI_ISL_2492434, EPI_ISL_2492435, EPI_ISL_2492436 | Cliniques universitaires Saint-Luc | UCLouvain/IREC/MBLG | Jean Ruelle, Benoit Kabamba Mukadi |
| EPI_ISL_2492437, EPI_ISL_2492438, EPI_ISL_2492439 | Clinique Saint-Pierre Ottignies | UCLouvain/IREC/MBLG | Jean Ruelle, Benoit Kabamba Mukadi |
| EPI_ISL_2492945, EPI_ISL_2493032, EPI_ISL_2493068, EPI_ISL_2493069 | National HIV Reference Laboratory, Ministry of Health, Public Health Institute of Malawi | CERI, Centre for Epidemic Response and Innvoation, Stellenbosch University and KRISP, KZN Research Innovation and Sequencing Platform, UKZN. | Mvula B, Chilima B, Chiwaula M, Mwangomba W, Panja L, Kasambara W, Auld A, Kim L, Kampira E, Kaba M, Wadonda N, Maida A, Giandhari J, Pillay S, Naidoo Y, Lessells R, Emmanuel SJ, Tegally H, Wilkinson E, de Oliveira T |
| EPI_ISL_2493114, EPI_ISL_2493122, EPI_ISL_2493132, EPI_ISL_2493136 | Hospital Universitari Vall d'Hebron - Vall d'Hebron Institut de Recerca | Hospital Universitari Vall d'Hebron - Vall d'Hebron Institut de Recerca | Cristina Andrés, Maria Piñana, Alejandra González-Sánchez, Damir Garcia-Cehic, Ariadna Rando, Juliana Esperalba, Maria Gema Codina, Carla Castillo, Maria Carmen Martin, Tomàs Pumarola, Josep Quer, Andrés Antón |
| EPI_ISL_2494782, EPI_ISL_2494899, EPI_ISL_2494937, EPI_ISL_2494966, EPI_ISL_2495038, EPI_ISL_2495039, EPI_ISL_2495042, EPI_ISL_2495043 | National HIV Reference Laboratory, Ministry of Health, Public Health Institute of Malawi | KRISP, KZN Research Innovation and Sequencing Platform | Mvula B, Chilima B, Chiwaula M, Mwangomba W, Panja L, Kasambara W, Auld A, Kim L, Kampira E, Kaba M, Wadonda N, Maida A, Giandhari J, Pillay S, Naidoo Y, Lessells R, Emmanuel SJ, Tegally H, Wilkinson E, de Oliveira T |
| EPI_ISL_2495292 | Diagnostyka Sp. z o.o. | 1. National Institute of Public Health - National Institute of Hygiene; 2. Eurofins Genomics Europe Sequencing GmbH | Wołkowicz Tomasz, Zacharczuk Katarzyna, Sadkowska-Todys Małgorzata, Gierczyńki Rafał, Eurofins Genomics Europe Sequencing Team, ECDC COVID-19 WGS support team |
| EPI_ISL_2495622, EPI_ISL_2495623, EPI_ISL_2495624, EPI_ISL_2495625, EPI_ISL_2495626, EPI_ISL_2495627, EPI_ISL_2495628, EPI_ISL_2495629, EPI_ISL_2495630, EPI_ISL_2495631, EPI_ISL_2495632, EPI_ISL_2495633, EPI_ISL_2495634, EPI_ISL_2495635, EPI_ISL_2495636, EPI_ISL_2495637, EPI_ISL_2495638, EPI_ISL_2495639, EPI_ISL_2495640, EPI_ISL_2495641, EPI_ISL_2495642, EPI_ISL_2495643, EPI_ISL_2495644, EPI_ISL_2495645, EPI_ISL_2495646, EPI_ISL_2495647, EPI_ISL_2495648, EPI_ISL_2495649, EPI_ISL_2495650, EPI_ISL_2495651, EPI_ISL_2495652, EPI_ISL_2495653, EPI_ISL_2495654, EPI_ISL_2495655, EPI_ISL_2495656, EPI_ISL_2495657, EPI_ISL_2495658, EPI_ISL_2495659, EPI_ISL_2495660, EPI_ISL_2495661, EPI_ISL_2495662, EPI_ISL_2495663, EPI_ISL_2495664, EPI_ISL_2495665, EPI_ISL_2495666 | | | |
| see above | MB-Cadham Provincial laboratory | National Microbiology Laboratory (NML) | Anna Majer, Shari Tyson, Grace Seo, Philip Mabon, Elsie Grudeski, Rhiannon Huzarewich, Russell Mandes, Anneliese Landgraff, Jennifer Tanner, Natalie Knox, Morag Graham, Gary Van Domselaar, Paul Van Caeseele, Jared Bullard, David Alexander, Kerry Dust, Nathalie Bastien, Yan Li, Timothy Booth, Darian Hole, Madison Chapel, Kirsten Biggar, CanCOGeN's metadata curation team, Public Health Agency of Canada CanCOGeN team |
| EPI_ISL_2496777, EPI_ISL_2496787 | Santa Clara County Public Health Laboratory | Chan-Zuckerberg Biohub | CZB Cliahub Consortium |
| EPI_ISL_2496919, EPI_ISL_2496920, EPI_ISL_2496924 | Contra Costa County Public Health Lab | Chan-Zuckerberg Biohub | CZB Cliahub Consortium |
| EPI_ISL_2496931 | Santa Clara County Public Health Laboratory | Chan-Zuckerberg Biohub | CZB Cliahub Consortium |
| EPI_ISL_2496932, EPI_ISL_2496933 | Contra Costa County Public Health Lab | Chan-Zuckerberg Biohub | CZB Cliahub Consortium |
| EPI_ISL_2496934 | CA DPH Viral and Rickettsial Disease Laboratory | Chan-Zuckerberg Biohub | CZB Cliahub Consortium |
| EPI_ISL_2496935, EPI_ISL_2496936, EPI_ISL_2496938, EPI_ISL_2496939 | Contra Costa County Public Health Lab | Chan-Zuckerberg Biohub | CZB Cliahub Consortium |
| EPI_ISL_2496940 | CA DPH Viral and Rickettsial Disease Laboratory | Chan-Zuckerberg Biohub | CZB Cliahub Consortium |
| EPI_ISL_2496941 | Santa Clara County Public Health Laboratory | Chan-Zuckerberg Biohub | CZB Cliahub Consortium |
| EPI_ISL_2496942 | Contra Costa County Public Health Lab | Chan-Zuckerberg Biohub | CZB Cliahub Consortium |
| EPI_ISL_2497217, EPI_ISL_2497234, EPI_ISL_2497240, EPI_ISL_2497244, EPI_ISL_2497252, EPI_ISL_2497257, EPI_ISL_2497273, EPI_ISL_2497282, EPI_ISL_2497283, EPI_ISL_2497284, EPI_ISL_2497300, EPI_ISL_2497304, EPI_ISL_2497336, EPI_ISL_2497355, EPI_ISL_2497379, EPI_ISL_2497390, EPI_ISL_2497409, EPI_ISL_2497415 | | | |
| see above | UW Virology Lab | UW Virology Lab | Pavitra Roychoudhury, Hong Xie, Lasata Shrestha, Tien V. Nguyen, Shah Mohamed Bakhash, Michelle Lin, Noah R. Baker, Ricardo Perez, Sean Ellis, Nathan Breit, Robert J. Livingston, Meei-Li Huang, Keith R Jerome, Patrick Mathias, Alexander Greninger |
| EPI_ISL_2497544, EPI_ISL_2497548 | CA DPH Viral and Rickettsial Disease Laboratory | Chan-Zuckerberg Biohub | CZB Cliahub Consortium |

| | | | |
|---|---|---|---|
| EPI_ISL_2497675, EPI_ISL_2497677, EPI_ISL_2497678, EPI_ISL_2497679, EPI_ISL_2497681 | Contra Costa County Public Health Lab | Chan-Zuckerberg Biohub | CZB Cliahub Consortium |
| EPI_ISL_2497893, EPI_ISL_2497894 | University of Mississippi Medical Center, Department of Pathology | University of Mississippi Medical Center, Molecular and Genomics Core Facility | Ashley C. Johnson, Ithiel J. Frame, Krishna K. Ayyalasomayajula, Michael R. Garrett, D. Ashley Robinson |
| EPI_ISL_2497909, EPI_ISL_2497910, EPI_ISL_2497914, EPI_ISL_2497915, EPI_ISL_2497921 | Wichita State University - Molecular Diagnostics Lab | Kansas Health and Environmental Lab | Mike Grose, Katherine Wiggins, Jonathan Barnell, Ben Olsen, and Phil Adam |
| EPI_ISL_2497925 | UW Virology Lab | UW Virology Lab | Pavitra Roychoudhury, Hong Xie, Lasata Shrestha, Tien V. Nguyen, Shah Mohamed Bakhash, Michelle Lin, Noah R. Baker, Ricardo Perez, Sean Ellis, Nathan Breit, Robert J. Livingston, Meei-Li Huang, Keith R Jerome, Patrick Mathias, Alexander Greninger |
| EPI_ISL_2499949, EPI_ISL_2499951, EPI_ISL_2499952, EPI_ISL_2499953 | Kansas Health and Environmental Lab | Kansas Health and Environmental Lab | Mike Grose, Katherine Wiggins, Jonathan Barnell, Ben Olsen, and Phil Adam |
| EPI_ISL_2499955 | Wichita State University - Molecular Diagnostics Lab | Kansas Health and Environmental Lab | Mike Grose, Katherine Wiggins, Jonathan Barnell, Ben Olsen, and Phil Adam |
| EPI_ISL_2499962, EPI_ISL_2499963, EPI_ISL_2499964 | Genomik Solidaritas Indonesia Laboratorium | Genomik Solidaritas Indonesia Laboratorium | Vania Gavrila Wikasa, Carissa Sintca Wijaya, Alfin Mohammad Abdillah, Dhahlia Agustina Cahyono, Normastuti Adhini Tantyo, Annisa Muthiah Sukirman, Gracia Felias Enos Korompis, Pratiwi Sudarmono, Budi Wiweko, Meutia Ayuputeri Kumaheri, Anindya Pradipta Susanto, Louisa Markus, Ariel Pradipta |
| EPI_ISL_2500000, EPI_ISL_2500006, EPI_ISL_2500007, EPI_ISL_2500011, EPI_ISL_2500013 | University of Liège COVID-19 testing center | GIGA Medical Genomics | Keith Durkin, Maria Artesi, Bouchra Boujemla, Nathalie Renotte, Cécile Meex, Sébastien Bontems, Fabrice Bureau, Laurent Gillet, Wouter Coppieters, Marie-Pierre Hayette, Vincent Bours |
| EPI_ISL_2500033 | Santa Clara County Public Health Laboratory | Chan-Zuckerberg Biohub | CZB Cliahub Consortium |
| EPI_ISL_2500069 | Humboldt County Public Health Laboratory | Chan-Zuckerberg Biohub | CZB Cliahub Consortium |
| EPI_ISL_2500090 | CA DPH Viral and Rickettsial Disease Laboratory | Chan-Zuckerberg Biohub | CZB Cliahub Consortium |
| EPI_ISL_2500136 | Santa Clara County Public Health Laboratory | Chan-Zuckerberg Biohub | CZB Cliahub Consortium |
| EPI_ISL_2500168, EPI_ISL_2500194 | CA DPH Viral and Rickettsial Disease Laboratory | Chan-Zuckerberg Biohub | CZB Cliahub Consortium |
| EPI_ISL_2500217 | Genomik Solidaritas Indonesia Laboratorium | Genomik Solidaritas Indonesia Laboratorium | Vania Gavrila Wikasa, Carissa Sintca Wijaya, Alfin Mohammad Abdillah, Dhahlia Agustina Cahyono, Normastuti Adhini Tantyo, Annisa Muthiah Sukirman, Gracia Felias Enos Korompis, Pratiwi Sudarmono, Budi Wiweko, Meutia Ayuputeri Kumaheri, Anindya Pradipta Susanto, Louisa Markus, Ariel Pradipta |
| EPI_ISL_2500365, EPI_ISL_2500366, EPI_ISL_2500367, EPI_ISL_2500369, EPI_ISL_2500370 | A. Krumbholz, Labor Dr. Krause und Kollegen MVZ GmbH, Kiel | Charité Universitätsmedizin Berlin, Institut für Virologie | Victor M Corman, Barbara Mühlemann, Jörn Beheim-Schwarzbach, Talitha Veith, Julia Schneider, Tobias Bleicker, Julia Tesch, Terry Jones, Christian Drosten |
| EPI_ISL_2500466 | Genomik Solidaritas Indonesia Laboratorium | Genomik Solidaritas Indonesia Laboratorium | Vania Gavrila Wikasa, Carissa Sintca Wijaya, Alfin Mohammad Abdillah, Dhahlia Agustina Cahyono, Normastuti Adhini Tantyo, Annisa Muthiah Sukirman, Gracia Felias Enos Korompis, Pratiwi Sudarmono, Budi Wiweko, Meutia Ayuputeri Kumaheri, Anindya Pradipta Susanto, Louisa Markus, Ariel Pradipta |
| EPI_ISL_2500470, EPI_ISL_2500471, EPI_ISL_2500489 | Genomik Solidaritas Indonesia Laboratorium | Genomik Solidaritas Indonesia Laboratorium | Carissa Sintca Wijaya, Vania Gavrila Wikasa, Normastuti Adhini Tantyo, Alfin Mohammad Abdillah, Dhahlia Agustina Cahyono, Gracia Felias Enos Korompis, Annisa Muthiah Sukirman, Pratiwi Sudarmono, Budi Wiweko, Meutia Ayuputeri Kumaheri, Anindya Pradipta Susanto, Louisa Markus, Ariel Pradipta |
| EPI_ISL_2501018 | Gundersen Clinical Microbiology Laboratory | Kabara Cancer Research Institute | Craig S. Richmond, Paraic A. Kenny |
| EPI_ISL_2501034, EPI_ISL_2501035 | Tboung Khmum Rapid Response Team | Virology Unit, Institut Pasteur du Cambodge | Cecile Troupin, Leakhena Pum, Jurre Y Siegers, Ly Sovann, Kraing Sidonn, Yi Sengdoeurn, Chin Savuth, Chau Darapheak, Veasna Duong, Erik A Karlsson |
| EPI_ISL_2501060 | Virology Unit, Institut Pasteur du Cambodge | Virology Unit, Institut Pasteur du Cambodge | Cecile Troupin, Leakhena Pum, Jurre Y Siegers, Ly Sovann, Kraing Sidonn, Yi Sengdoeurn, Chin Savuth, Chau Darapheak, Veasna Duong, Erik A Karlsson |
| EPI_ISL_2501110 | Maple Grove Hospital | Minnesota Department of Health, Public Health Laboratory | Alexandra Lorentz, Jacob Garfin, Matt Plumb, and Xiong Wang |
| EPI_ISL_2501311, EPI_ISL_2501369, EPI_ISL_2501370, EPI_ISL_2501371, EPI_ISL_2501372, EPI_ISL_2501373, EPI_ISL_2501374 | Microbiology Department. Complexo Hospitalario Universitario de Vigo | Microbiology Department. Complexo Hospitalario Universitario de Vigo | Alfaya N, Alonso I, Alvarez M, Cabrera JJ, Carballo R, Cores O, Cortizo S, del-Campo V, Martinez L, Mediero G, Perez S, Potel C, Regueiro B, Rey S, Vasallo FJ |
| EPI_ISL_2501399, EPI_ISL_2501401, EPI_ISL_2501500, EPI_ISL_2501501, EPI_ISL_2501502, EPI_ISL_2501503, EPI_ISL_2501504, EPI_ISL_2501505, EPI_ISL_2501509, EPI_ISL_2501510, EPI_ISL_2501511, EPI_ISL_2501512, EPI_ISL_2501513, EPI_ISL_2501514, EPI_ISL_2501515, EPI_ISL_2501516 | | | |
| see above | UW Virology Lab | UW Virology Lab | Pavitra Roychoudhury, Hong Xie, Lasata Shrestha, Tien V. Nguyen, Shah Mohamed Bakhash, Michelle Lin, Noah R. Baker, Ricardo Perez, Sean Ellis, Nathan Breit, Robert J. Livingston, Meei-Li Huang, Keith R Jerome, Patrick Mathias, Alexander Greninger |
| EPI_ISL_2501668, EPI_ISL_2501721, EPI_ISL_2501722 | Yale Clinical Virology Lab | Grubaugh Lab - Yale School of Public Health | Joseph Fauver, Mallery Breban, Isabel Ott, Tara Alpert, Mary Petrone, Anderson Brito, Chantal Vogels, Annie Watkins, Chaney Kalinich, Jessica Rothman, Marie L. Landry, Nathan Grubaugh |
| EPI_ISL_2501744, EPI_ISL_2501747 | TXDSHS | TXDSHS | Rashmi Tuladhar, Bonnie Oh, Jenny Zhang, Maliha Rahman, Mayela Pedrueza, Anita Pokharel, Karen Bobier, Lorraine Rodriguez, Myong Koag, Chun Wang, Rachel Lee, Grace Kubin |
| EPI_ISL_2501755, EPI_ISL_2501756, EPI_ISL_2501834, EPI_ISL_2501842 | Maryland Genomics, Institute for Genome Sciences, University of Maryland School of Medicine | Maryland Genomics, Institute for Genome Sciences, University of Maryland School of Medicine | Tallon, Luke J; Sadzewicz, Lisa D; Humphrys, Mike; Ott, Sandra; Roussey, Holly; Santana-Cruz, Ivette; Vavikolanu, Kranthi; Lim, Jonathan;Hazen, Tracy;Fraser, Claire M; Ravel, Jacques |
| EPI_ISL_2501860 | Laboratorio analisi, IRCCS Policlinico San Donato | Laboratory of Clinical Microbiology, Virology and Bioemergencies, ASST Fatebenefratelli Sacco - Sacco Hospital | Valeria Micheli, Alessandro Mancon, Alberto Rizzo, Fiorenza Bracchitta, Maria Rita Gismondo |
| EPI_ISL_2501866 | Unità Operativa Complessa Analisi Chimico Cliniche - ASST Brianza | Laboratory of Clinical Microbiology, Virology and Bioemergencies, ASST Fatebenefratelli Sacco - Sacco Hospital | Valeria Micheli, Alessandro Mancon, Alberto Rizzo, Fiorenza Bracchitta, Maria Rita Gismondo |
| EPI_ISL_2501867, EPI_ISL_2501878, EPI_ISL_2501882, EPI_ISL_2501884, EPI_ISL_2501885, EPI_ISL_2501887, EPI_ISL_2501888, EPI_ISL_2501889, EPI_ISL_2501892, EPI_ISL_2501893, EPI_ISL_2501894, EPI_ISL_2501925 | | | |
| see above | Missouri State Public Health Laboratory | Missouri State Public Health Laboratory | Matthew Sinn, Joshua Barry, Ashley New |
| EPI_ISL_2502028, EPI_ISL_2502029 | National Centre for Disease Control (NCDC) Biotechnology Division, Delhi | NCDC Delhi, Biotechnology Division INSACOG | Priyanka Singh, Uma Sharma, Hemlata Lall, Manoj K Singh, Radhakrishnan V. S, Robin Marwal, Mahesh S Dhar, Kalaiarasan Ponnusamy, Meena Datta, Hema Gogia, Preeti Madan, Sandhya Kabra, Sujeet K Singh, Partha Rakshit |
| EPI_ISL_2502030 | National Centre for Disease Control (NCDC) Biotechnology Division, Delhi | NCDC Delhi, Biotechnology Division INSACOG | Uma Sharma, Hemlata Lall, Manoj K Singh, Radhakrishnan V. S, Robin Marwal, Mahesh S Dhar, Kalaiarasan Ponnusamy, Meena Datta, Priyanka Singh, Hema Gogia, Preeti Madan, Sandhya Kabra, Sujeet K Singh, Partha Rakshit |
| EPI_ISL_2502031 | National Centre for Disease Control (NCDC) Biotechnology Division, Delhi | NCDC Delhi, Biotechnology Division INSACOG | Priyanka Singh, Uma Sharma, Hemlata Lall, Manoj K Singh, Radhakrishnan V. S, Robin Marwal, Mahesh S Dhar, Kalaiarasan Ponnusamy, Meena Datta, Hema Gogia, Preeti Madan, Sandhya Kabra, Sujeet K Singh, Partha Rakshit |
| EPI_ISL_2502036 | National Centre for Disease Control (NCDC) Biotechnology Division, Delhi | NCDC Delhi, Biotechnology Division INSACOG | Uma Sharma, Hemlata Lall, Manoj K Singh, Radhakrishnan V. S, Robin Marwal, Mahesh S Dhar, Kalaiarasan Ponnusamy, Meena Datta, Priyanka Singh, Hema Gogia, Preeti Madan, Sandhya Kabra, Sujeet K Singh, Partha Rakshit |
| EPI_ISL_2502059, EPI_ISL_2502060, EPI_ISL_2502065, EPI_ISL_2502071 | National Centre for Disease Control (NCDC) Biotechnology Division, Delhi | NCDC Delhi, Biotechnology Division INSACOG | Priyanka Singh, Uma Sharma, Hemlata Lall, Manoj K Singh, Radhakrishnan V. S, Robin Marwal, Mahesh S Dhar, Kalaiarasan Ponnusamy, Meena Datta, Hema Gogia, Preeti Madan, Sandhya Kabra, Sujeet K Singh, Partha Rakshit |
| EPI_ISL_2502085 | National Centre for Disease Control (NCDC) Biotechnology Division, Delhi | NCDC Delhi, Biotechnology Division INSACOG | Uma Sharma, Hemlata Lall, Manoj K Singh, Radhakrishnan V. S, Robin Marwal, Mahesh S Dhar, Kalaiarasan Ponnusamy, Meena Datta, Priyanka Singh, Hema Gogia, Preeti Madan, Sandhya Kabra, Sujeet K Singh, Partha Rakshit |
| EPI_ISL_2502088, EPI_ISL_2502095, EPI_ISL_2502098, EPI_ISL_2502106 | National Centre for Disease Control (NCDC) Biotechnology Division, Delhi | NCDC Delhi, Biotechnology Division INSACOG | Priyanka Singh, Uma Sharma, Hemlata Lall, Manoj K Singh, Radhakrishnan V. S, Robin Marwal, Mahesh S Dhar, Kalaiarasan Ponnusamy, Meena Datta, Hema Gogia, Preeti Madan, Sandhya Kabra, Sujeet K Singh, Partha Rakshit |

| | | | |
|---|---|---|---|
| EPI_ISL_2502109, EPI_ISL_2502110 | National Centre for Disease Control (NCDC) Biotechnology Division, Delhi | NCDC Delhi, Biotechnology Division INSACOG | Uma Sharma, Hemlata Lall, Manoj K Singh, Radhakrishnan V. S, Robin Marwal, Mahesh S Dhar, Kalaiarasan Ponnusamy, Meena Datta, Priyanka Singh, Hema Gogia, Preeti Madan, Sandhya Kabra, Sujeet K Singh, Partha Rakshit |
| EPI_ISL_2502120 | National Centre for Disease Control (NCDC) Biotechnology Division, Delhi | NCDC Delhi, Biotechnology Division INSACOG | Priyanka Singh, Uma Sharma, Hemlata Lall, Manoj K Singh, Radhakrishnan V. S, Robin Marwal, Mahesh S Dhar, Kalaiarasan Ponnusamy, Meena Datta, Hema Gogia, Preeti Madan, Sandhya Kabra, Sujeet K Singh, Partha Rakshit |
| EPI_ISL_2502121, EPI_ISL_2502126 | National Centre for Disease Control (NCDC) Biotechnology Division, Delhi | NCDC Delhi, Biotechnology Division INSACOG | Uma Sharma, Hemlata Lall, Manoj K Singh, Radhakrishnan V. S, Robin Marwal, Mahesh S Dhar, Kalaiarasan Ponnusamy, Meena Datta, Priyanka Singh, Hema Gogia, Preeti Madan, Sandhya Kabra, Sujeet K Singh, Partha Rakshit |
| EPI_ISL_2502129 | National Centre for Disease Control (NCDC) Biotechnology Division, Delhi | NCDC Delhi, Biotechnology Division INSACOG | Priyanka Singh, Uma Sharma, Hemlata Lall, Manoj K Singh, Radhakrishnan V. S, Robin Marwal, Mahesh S Dhar, Kalaiarasan Ponnusamy, Meena Datta, Hema Gogia, Preeti Madan, Sandhya Kabra, Sujeet K Singh, Partha Rakshit |
| EPI_ISL_2502171 | National Centre for Disease Control (NCDC) Biotechnology Division, Delhi | NCDC Delhi, Biotechnology Division INSACOG | Uma Sharma, Hemlata Lall, Manoj K Singh, Radhakrishnan V. S, Robin Marwal, Mahesh S Dhar, Kalaiarasan Ponnusamy, Meena Datta, Priyanka Singh, Hema Gogia, Preeti Madan, Sandhya Kabra, Sujeet K Singh, Partha Rakshit |
| EPI_ISL_2502172 | National Centre for Disease Control (NCDC) Biotechnology Division, Delhi | NCDC Delhi, Biotechnology Division INSACOG | Priyanka Singh, Uma Sharma, Hemlata Lall, Manoj K Singh, Radhakrishnan V. S, Robin Marwal, Mahesh S Dhar, Kalaiarasan Ponnusamy, Meena Datta, Hema Gogia, Preeti Madan, Sandhya Kabra, Sujeet K Singh, Partha Rakshit |
| EPI_ISL_2502174 | National Centre for Disease Control (NCDC) Biotechnology Division, Delhi | NCDC Delhi, Biotechnology Division INSACOG | Uma Sharma, Hemlata Lall, Manoj K Singh, Radhakrishnan V. S, Robin Marwal, Mahesh S Dhar, Kalaiarasan Ponnusamy, Meena Datta, Priyanka Singh, Hema Gogia, Preeti Madan, Sandhya Kabra, Sujeet K Singh, Partha Rakshit |
| EPI_ISL_2502178 | National Centre for Disease Control (NCDC) Biotechnology Division, Delhi | NCDC Delhi, Biotechnology Division INSACOG | Priyanka Singh, Uma Sharma, Hemlata Lall, Manoj K Singh, Radhakrishnan V. S, Robin Marwal, Mahesh S Dhar, Kalaiarasan Ponnusamy, Meena Datta, Hema Gogia, Preeti Madan, Sandhya Kabra, Sujeet K Singh, Partha Rakshit |
| EPI_ISL_2502180, EPI_ISL_2502182 | National Centre for Disease Control (NCDC) Biotechnology Division, Delhi | NCDC Delhi, Biotechnology Division INSACOG | Uma Sharma, Hemlata Lall, Manoj K Singh, Radhakrishnan V. S, Robin Marwal, Mahesh S Dhar, Kalaiarasan Ponnusamy, Meena Datta, Priyanka Singh, Hema Gogia, Preeti Madan, Sandhya Kabra, Sujeet K Singh, Partha Rakshit |
| EPI_ISL_2502185 | National Centre for Disease Control (NCDC) Biotechnology Division, Delhi | NCDC Delhi, Biotechnology Division INSACOG | Priyanka Singh, Uma Sharma, Hemlata Lall, Manoj K Singh, Radhakrishnan V. S, Robin Marwal, Mahesh S Dhar, Kalaiarasan Ponnusamy, Meena Datta, Hema Gogia, Preeti Madan, Sandhya Kabra, Sujeet K Singh, Partha Rakshit |
| EPI_ISL_2502188, EPI_ISL_2502192 | National Centre for Disease Control (NCDC) Biotechnology Division, Delhi | NCDC Delhi, Biotechnology Division INSACOG | Uma Sharma, Hemlata Lall, Manoj K Singh, Radhakrishnan V. S, Robin Marwal, Mahesh S Dhar, Kalaiarasan Ponnusamy, Meena Datta, Priyanka Singh, Hema Gogia, Preeti Madan, Sandhya Kabra, Sujeet K Singh, Partha Rakshit |
| EPI_ISL_2502196, EPI_ISL_2502212 | National Centre for Disease Control (NCDC) Biotechnology Division, Delhi | NCDC Delhi, Biotechnology Division INSACOG | Priyanka Singh, Uma Sharma, Hemlata Lall, Manoj K Singh, Radhakrishnan V. S, Robin Marwal, Mahesh S Dhar, Kalaiarasan Ponnusamy, Meena Datta, Hema Gogia, Preeti Madan, Sandhya Kabra, Sujeet K Singh, Partha Rakshit |
| EPI_ISL_2502214, EPI_ISL_2502215 | National Centre for Disease Control (NCDC) Biotechnology Division, Delhi | NCDC Delhi, Biotechnology Division INSACOG | Uma Sharma, Hemlata Lall, Manoj K Singh, Radhakrishnan V. S, Robin Marwal, Mahesh S Dhar, Kalaiarasan Ponnusamy, Meena Datta, Priyanka Singh, Hema Gogia, Preeti Madan, Sandhya Kabra, Sujeet K Singh, Partha Rakshit |
| EPI_ISL_2502217, EPI_ISL_2502218 | National Centre for Disease Control (NCDC) Biotechnology Division, Delhi | NCDC Delhi, Biotechnology Division INSACOG | Priyanka Singh, Uma Sharma, Hemlata Lall, Manoj K Singh, Radhakrishnan V. S, Robin Marwal, Mahesh S Dhar, Kalaiarasan Ponnusamy, Meena Datta, Hema Gogia, Preeti Madan, Sandhya Kabra, Sujeet K Singh, Partha Rakshit |
| EPI_ISL_2502222, EPI_ISL_2502223 | National Centre for Disease Control (NCDC) Biotechnology Division, Delhi | NCDC Delhi, Biotechnology Division INSACOG | Uma Sharma, Hemlata Lall, Manoj K Singh, Radhakrishnan V. S, Robin Marwal, Mahesh S Dhar, Kalaiarasan Ponnusamy, Meena Datta, Priyanka Singh, Hema Gogia, Preeti Madan, Sandhya Kabra, Sujeet K Singh, Partha Rakshit |
| EPI_ISL_2502231, EPI_ISL_2502232, EPI_ISL_2502237, EPI_ISL_2502247, EPI_ISL_2502248 | National Centre for Disease Control (NCDC) Biotechnology Division, Delhi | NCDC Delhi, Biotechnology Division INSACOG | Priyanka Singh, Uma Sharma, Hemlata Lall, Manoj K Singh, Radhakrishnan V. S, Robin Marwal, Mahesh S Dhar, Kalaiarasan Ponnusamy, Meena Datta, Hema Gogia, Preeti Madan, Sandhya Kabra, Sujeet K Singh, Partha Rakshit |
| EPI_ISL_2502266, EPI_ISL_2502268, EPI_ISL_2502269, EPI_ISL_2502270, EPI_ISL_2502272 | National Centre for Disease Control (NCDC) Biotechnology Division, Delhi | NCDC Delhi, Biotechnology Division INSACOG | Uma Sharma, Hemlata Lall, Manoj K Singh, Radhakrishnan V. S, Robin Marwal, Mahesh S Dhar, Kalaiarasan Ponnusamy, Meena Datta, Priyanka Singh, Hema Gogia, Preeti Madan, Sandhya Kabra, Sujeet K Singh, Partha Rakshit |
| EPI_ISL_2502273 | National Centre for Disease Control (NCDC) Biotechnology Division, Delhi | NCDC Delhi, Biotechnology Division INSACOG | Priyanka Singh, Uma Sharma, Hemlata Lall, Manoj K Singh, Radhakrishnan V. S, Robin Marwal, Mahesh S Dhar, Kalaiarasan Ponnusamy, Meena Datta, Hema Gogia, Preeti Madan, Sandhya Kabra, Sujeet K Singh, Partha Rakshit |
| EPI_ISL_2502274 | National Centre for Disease Control (NCDC) Biotechnology Division, Delhi | NCDC Delhi, Biotechnology Division INSACOG | Uma Sharma, Hemlata Lall, Manoj K Singh, Radhakrishnan V. S, Robin Marwal, Mahesh S Dhar, Kalaiarasan Ponnusamy, Meena Datta, Priyanka Singh, Hema Gogia, Preeti Madan, Sandhya Kabra, Sujeet K Singh, Partha Rakshit |
| EPI_ISL_2502275 | National Centre for Disease Control (NCDC) Biotechnology Division, Delhi | NCDC Delhi, Biotechnology Division INSACOG | Priyanka Singh, Uma Sharma, Hemlata Lall, Manoj K Singh, Radhakrishnan V. S, Robin Marwal, Mahesh S Dhar, Kalaiarasan Ponnusamy, Meena Datta, Hema Gogia, Preeti Madan, Sandhya Kabra, Sujeet K Singh, Partha Rakshit |
| EPI_ISL_2502278, EPI_ISL_2502279, EPI_ISL_2502289, EPI_ISL_2502291 | National Centre for Disease Control (NCDC) Biotechnology Division, Delhi | NCDC Delhi, Biotechnology Division INSACOG | Uma Sharma, Hemlata Lall, Manoj K Singh, Radhakrishnan V. S, Robin Marwal, Mahesh S Dhar, Kalaiarasan Ponnusamy, Meena Datta, Priyanka Singh, Hema Gogia, Preeti Madan, Sandhya Kabra, Sujeet K Singh, Partha Rakshit |
| EPI_ISL_2502296 | National Centre for Disease Control (NCDC) Biotechnology Division, Delhi | NCDC Delhi, Biotechnology Division INSACOG | Priyanka Singh, Uma Sharma, Hemlata Lall, Manoj K Singh, Radhakrishnan V. S, Robin Marwal, Mahesh S Dhar, Kalaiarasan Ponnusamy, Meena Datta, Hema Gogia, Preeti Madan, Sandhya Kabra, Sujeet K Singh, Partha Rakshit |
| EPI_ISL_2502297, EPI_ISL_2502298 | National Centre for Disease Control (NCDC) Biotechnology Division, Delhi | NCDC Delhi, Biotechnology Division INSACOG | Uma Sharma, Hemlata Lall, Manoj K Singh, Radhakrishnan V. S, Robin Marwal, Mahesh S Dhar, Kalaiarasan Ponnusamy, Meena Datta, Priyanka Singh, Hema Gogia, Preeti Madan, Sandhya Kabra, Sujeet K Singh, Partha Rakshit |
| EPI_ISL_2502299 | National Centre for Disease Control (NCDC) Biotechnology Division, Delhi | NCDC Delhi, Biotechnology Division INSACOG | Priyanka Singh, Uma Sharma, Hemlata Lall, Manoj K Singh, Radhakrishnan V. S, Robin Marwal, Mahesh S Dhar, Kalaiarasan Ponnusamy, Meena Datta, Hema Gogia, Preeti Madan, Sandhya Kabra, Sujeet K Singh, Partha Rakshit |
| EPI_ISL_2502300, EPI_ISL_2502301, EPI_ISL_2502304 | National Centre for Disease Control (NCDC) Biotechnology Division, Delhi | NCDC Delhi, Biotechnology Division INSACOG | Uma Sharma, Hemlata Lall, Manoj K Singh, Radhakrishnan V. S, Robin Marwal, Mahesh S Dhar, Kalaiarasan Ponnusamy, Meena Datta, Priyanka Singh, Hema Gogia, Preeti Madan, Sandhya Kabra, Sujeet K Singh, Partha Rakshit |
| EPI_ISL_2502308, EPI_ISL_2502309, EPI_ISL_2502311, EPI_ISL_2502312, EPI_ISL_2502319, EPI_ISL_2502321, EPI_ISL_2502322, EPI_ISL_2502323, EPI_ISL_2502326, EPI_ISL_2502327, EPI_ISL_2502329, EPI_ISL_2502330, EPI_ISL_2502331, EPI_ISL_2502332, EPI_ISL_2502333, EPI_ISL_2502334, EPI_ISL_2502335, EPI_ISL_2502336, EPI_ISL_2502337, EPI_ISL_2502338, EPI_ISL_2502339, EPI_ISL_2502340, EPI_ISL_2502342, EPI_ISL_2502343, EPI_ISL_2502344, EPI_ISL_2502345, EPI_ISL_2502346, EPI_ISL_2502347, EPI_ISL_2502350, EPI_ISL_2502351, EPI_ISL_2502352, EPI_ISL_2502353, EPI_ISL_2502355, EPI_ISL_2502356, EPI_ISL_2502357, EPI_ISL_2502358, EPI_ISL_2502359, EPI_ISL_2502361, EPI_ISL_2502363, EPI_ISL_2502364, EPI_ISL_2502365, EPI_ISL_2502366, EPI_ISL_2502367, EPI_ISL_2502368, EPI_ISL_2502369, EPI_ISL_2502370, EPI_ISL_2502371, EPI_ISL_2502372, EPI_ISL_2502373, EPI_ISL_2502374 | | | |
| see above | INSACOG-Assam | National Institute of Biomedical Genomics – INSACOG | Arindam Maitra, Ajanta Sharma, Nidhan Kumar Biswas, Lahari Saikia, Sreedhar Chinnaswamy, Kailash Chamuah, Saumitra Das |
| EPI_ISL_2502615 | RSUD dr Loekmono Hadi | Genetics Working Group (Pokja Genetik) Faculty of Medicine, Public Health and Nursing Universitas Gadjah Mada (FK-KMK UGM); Disease Investigation Center Wates Ministry of Agriculture Indonesia; Department of Microbiology FK-KMK UGM; Laboratorium Diagnostik Yayasan Tahija World Mosquito Program (WMP) Yogyakarta Center for Tropical Medicine FK-KMK UGM; Integrated Research Center FK-KMK UGM; Department of Computer Science and Electronics FMIPA UGM; RSUP Dr. Sardjito | Gunadi, Hendra Wibawa, Marcellus, Mohamad S. Hakim, Edwin W. Daniwijaya, Ludhang P. Rizki, Endah Supriyati, Eggi Arguni, Titik Nuryastuti, Tri Wibawa, Dwi AA Nugrahaningsih, Afiahayati, Siswanto,Kristy Iskandar,Nungki Anggorowati, Pramesti G Dewi, Dwi Indaryati,Indarto Sulistiyono, Susan Simanjaya, Dyah A Puspitarani |
| EPI_ISL_2502616 | RSUD dr. LOEKMONO HADI | Genetics Working Group (Pokja Genetik) Faculty of Medicine, Public Health and Nursing Universitas Gadjah Mada (FK-KMK UGM); Disease Investigation Center Wates Ministry of Agriculture Indonesia; Department of Microbiology FK-KMK UGM; Laboratorium Diagnostik Yayasan Tahija World Mosquito Program (WMP) Yogyakarta Center for Tropical Medicine FK-KMK UGM; Integrated Research Center FK-KMK UGM; Department of Computer Science and Electronics FMIPA UGM; RSUP Dr. Sardjito | Gunadi, Hendra Wibawa, Marcellus, Mohamad S. Hakim, Edwin W. Daniwijaya, Ludhang P. Rizki, Endah Supriyati, Eggi Arguni, Titik Nuryastuti, Tri Wibawa, Dwi AA Nugrahaningsih, Afiahayati, Siswanto,Kristy Iskandar,Nungki Anggorowati, Pramesti G Dewi, Dwi Indaryati,Indarto Sulistiyono, Irene Tania, Hana F Hanifin |
| EPI_ISL_2502627 | RSUD dr. LOEKMONO HADI | Genetics Working Group (Pokja Genetik) Faculty of Medicine, Public Health and Nursing Universitas Gadjah Mada (FK-KMK UGM); Disease Investigation Center Wates Ministry of Agriculture Indonesia; Department of Microbiology FK-KMK UGM; Laboratorium Diagnostik Yayasan Tahija World Mosquito Program (WMP) Yogyakarta Center for Tropical Medicine FK-KMK UGM; Integrated Research Center FK-KMK UGM; Department of Computer Science and Electronics FMIPA UGM; RSUP Dr. Sardjito | Gunadi, Hendra Wibawa, Marcellus, Mohamad S. Hakim, Edwin W. Daniwijaya, Ludhang P. Rizki, Endah Supriyati, Eggi Arguni, Titik Nuryastuti, Tri Wibawa, Dwi AA Nugrahaningsih, Afiahayati, Siswanto,Kristy Iskandar,Nungki Anggorowati, Pramesti G Dewi, Dwi Indaryati,Indarto Sulistiyono, Cita S Amalia, Alvina A Setiawan |
| EPI_ISL_2502628 | RSUD dr. LOEKMONO HADI | Genetics Working Group (Pokja Genetik) Faculty of Medicine, Public Health and Nursing Universitas Gadjah Mada (FK-KMK UGM); Disease Investigation Center Wates Ministry of Agriculture Indonesia; Department of Microbiology FK-KMK UGM; Laboratorium Diagnostik Yayasan Tahija World Mosquito Program (WMP) Yogyakarta Center for Tropical Medicine FK-KMK UGM; Integrated Research Center FK-KMK UGM; Department of Computer Science and Electronics FMIPA UGM; RSUP Dr. Sardjito | Gunadi, Hendra Wibawa, Marcellus, Mohamad S. Hakim, Edwin W. Daniwijaya, Ludhang P. Rizki, Endah Supriyati, Eggi Arguni, Titik Nuryastuti, Tri Wibawa, Dwi AA Nugrahaningsih, Afiahayati, Siswanto,Kristy Iskandar,Nungki Anggorowati, Pramesti G Dewi, Dwi Indaryati,Indarto Sulistiyono, Khanza a Vujira, Alvina A Setiawan |
| EPI_ISL_2502629 | RSUD dr. LOEKMONO HADI | Genetics Working Group (Pokja Genetik) Faculty of Medicine, Public Health and Nursing Universitas Gadjah Mada (FK-KMK UGM); | Gunadi, Hendra Wibawa, Marcellus, Mohamad S. Hakim, Edwin W. Daniwijaya, Ludhang P. Rizki, Endah Supriyati, Eggi Arguni, Titik Nuryastuti, Tri Wibawa, Dwi AA Nugrahaningsih, Afiahayati, Siswanto,Kristy Iskandar,Nungki Anggorowati, Pramesti G Dewi, Dwi Indaryati,Indarto Sulistiyono, Susan Simanjaya, Dyah A Puspitarani |

| | | | |
|---|---|---|---|
| | | Disease Investigation Center Wates Ministry of Agriculture Indonesia; Department of Microbiology FK-KMK UGM; Laboratorium Diagnostik Yayasan Tahija World Mosquito Program (WMP) Yogyakarta Center for Tropical Medicine FK-KMK UGM; Integrated Research Center FK-KMK UGM; Department of Computer Science and Electronics FMIPA UGM; RSUP Dr. Sardjito | |
| EPI_ISL_2502667 | RSUD dr. LOEKMONO HADI | Genetics Working Group (Pokja Genetik) Faculty of Medicine, Public Health and Nursing Universitas Gadjah Mada (FK-KMK UGM); Disease Investigation Center Wates Ministry of Agriculture Indonesia; Department of Microbiology FK-KMK UGM; Laboratorium Diagnostik Yayasan Tahija World Mosquito Program (WMP) Yogyakarta Center for Tropical Medicine FK-KMK UGM; Integrated Research Center FK-KMK UGM; Department of Computer Science and Electronics FMIPA UGM; RSUP Dr. Sardjito | Gunadi, Hendra Wibawa, Marcellus, Mohamad S. Hakim, Edwin W. Daniwijaya, Ludhang P. Rizki, Endah Supriyati, Eggi Arguni, Titik Nuryastuti, Tri Wibawa, Dwi AA Nugrahaningsih, Afiahayati, Siswanto,Kristy Iskandar,Nungki Anggorowati, Pramesti G Dewi, Dwi Indaryati,Indarto Sulistiyono, Irene Tania, Hana F Hanifin |
| EPI_ISL_2502687 | RSUD dr. LOEKMONO HADI | Genetics Working Group (Pokja Genetik) Faculty of Medicine, Public Health and Nursing Universitas Gadjah Mada (FK-KMK UGM); Disease Investigation Center Wates Ministry of Agriculture Indonesia; Department of Microbiology FK-KMK UGM; Laboratorium Diagnostik Yayasan Tahija World Mosquito Program (WMP) Yogyakarta Center for Tropical Medicine FK-KMK UGM; Integrated Research Center FK-KMK UGM; Department of Computer Science and Electronics FMIPA UGM; RSUP Dr. Sardjito | Gunadi, Hendra Wibawa, Marcellus, Mohamad S. Hakim, Edwin W. Daniwijaya, Ludhang P. Rizki, Endah Supriyati, Eggi Arguni, Titik Nuryastuti, Tri Wibawa, Dwi AA Nugrahaningsih, Afiahayati, Siswanto,Kristy Iskandar,Nungki Anggorowati, Pramesti G Dewi, Dwi Indaryati,Indarto Sulistiyono, Cita S Amalia, Alvina A Setiawan |
| EPI_ISL_2502717 | RSUD dr. LOEKMONO HADI | Genetics Working Group (Pokja Genetik) Faculty of Medicine, Public Health and Nursing Universitas Gadjah Mada (FK-KMK UGM); Disease Investigation Center Wates Ministry of Agriculture Indonesia; Department of Microbiology FK-KMK UGM; Laboratorium Diagnostik Yayasan Tahija World Mosquito Program (WMP) Yogyakarta Center for Tropical Medicine FK-KMK UGM; Integrated Research Center FK-KMK UGM; Department of Computer Science and Electronics FMIPA UGM; RSUP Dr. Sardjito | Gunadi, Hendra Wibawa, Marcellus, Mohamad S. Hakim, Edwin W. Daniwijaya, Ludhang P. Rizki, Endah Supriyati, Eggi Arguni, Titik Nuryastuti, Tri Wibawa, Dwi AA Nugrahaningsih, Afiahayati, Siswanto,Kristy Iskandar,Nungki Anggorowati, Pramesti G Dewi, Dwi Indaryati,Indarto Sulistiyono, Khanza a Vujira, Alvina A Setiawan |
| EPI_ISL_2502763 | RSUD dr. LOEKMONO HADI | Genetics Working Group (Pokja Genetik) Faculty of Medicine, Public Health and Nursing Universitas Gadjah Mada (FK-KMK UGM); Disease Investigation Center Wates Ministry of Agriculture Indonesia; Department of Microbiology FK-KMK UGM; Laboratorium Diagnostik Yayasan Tahija World Mosquito Program (WMP) Yogyakarta Center for Tropical Medicine FK-KMK UGM; Integrated Research Center FK-KMK UGM; Department of Computer Science and Electronics FMIPA UGM; RSUP Dr. Sardjito | Gunadi, Hendra Wibawa, Marcellus, Mohamad S. Hakim, Edwin W. Daniwijaya, Ludhang P. Rizki, Endah Supriyati, Eggi Arguni, Titik Nuryastuti, Tri Wibawa, Dwi AA Nugrahaningsih, Afiahayati, Siswanto,Kristy Iskandar,Nungki Anggorowati, Pramesti G Dewi, Dwi Indaryati,Indarto Sulistiyono, Susan Simanjaya, Dyah A Puspitarani |
| EPI_ISL_2502764 | RSUD dr. LOEKMONO HADI | Genetics Working Group (Pokja Genetik) Faculty of Medicine, Public Health and Nursing Universitas Gadjah Mada (FK-KMK UGM); Disease Investigation Center Wates Ministry of Agriculture Indonesia; Department of Microbiology FK-KMK UGM; Laboratorium Diagnostik Yayasan Tahija World Mosquito Program (WMP) Yogyakarta Center for Tropical Medicine FK-KMK UGM; Integrated Research Center FK-KMK UGM; Department of Computer Science and Electronics FMIPA UGM; RSUP Dr. Sardjito | Gunadi, Hendra Wibawa, Marcellus, Mohamad S. Hakim, Edwin W. Daniwijaya, Ludhang P. Rizki, Endah Supriyati, Eggi Arguni, Titik Nuryastuti, Tri Wibawa, Dwi AA Nugrahaningsih, Afiahayati, Siswanto,Kristy Iskandar,Nungki Anggorowati, Pramesti G Dewi, Dwi Indaryati,Indarto Sulistiyono, Irene Tania, Hana F Hanifin |
| EPI_ISL_2502765 | RSUD dr. LOEKMONO HADI | Genetics Working Group (Pokja Genetik) Faculty of Medicine, Public Health and Nursing Universitas Gadjah Mada (FK-KMK UGM); Disease Investigation Center Wates Ministry of Agriculture Indonesia; Department of Microbiology FK-KMK UGM; Laboratorium Diagnostik Yayasan Tahija World Mosquito Program (WMP) Yogyakarta Center for Tropical Medicine FK-KMK UGM; Integrated Research Center FK-KMK UGM; Department of Computer Science and Electronics FMIPA UGM; RSUP Dr. Sardjito | Gunadi, Hendra Wibawa, Marcellus, Mohamad S. Hakim, Edwin W. Daniwijaya, Ludhang P. Rizki, Endah Supriyati, Eggi Arguni, Titik Nuryastuti, Tri Wibawa, Dwi AA Nugrahaningsih, Afiahayati, Siswanto,Kristy Iskandar,Nungki Anggorowati, Pramesti G Dewi, Dwi Indaryati,Indarto Sulistiyono, Cita S Amalia, Alvina A Setiawan |
| EPI_ISL_2502898, EPI_ISL_2502925, EPI_ISL_2502986, EPI_ISL_2503005, EPI_ISL_2503017, EPI_ISL_2503018, EPI_ISL_2503049, EPI_ISL_2503087, EPI_ISL_2503106 | UW Virology Lab | UW Virology Lab | Pavitra Roychoudhury, Hong Xie, Lasata Shrestha, Shah Mohamed Bakhash, Tien V. Nguyen, Noah R. Baker, Sean Ellis, Meei-Li Huang, Keith R Jerome, Alexander Greninger |
| EPI_ISL_2503151, EPI_ISL_2503156, EPI_ISL_2503162, EPI_ISL_2503165, EPI_ISL_2503169, EPI_ISL_2503188, EPI_ISL_2503189, EPI_ISL_2503190, EPI_ISL_2503193, EPI_ISL_2503194, EPI_ISL_2503195, EPI_ISL_2503204, EPI_ISL_2503207, EPI_ISL_2503226, EPI_ISL_2503235, EPI_ISL_2503236, EPI_ISL_2503241, EPI_ISL_2503246, EPI_ISL_2503253, EPI_ISL_2503257, EPI_ISL_2503261, EPI_ISL_2503262, EPI_ISL_2503263, EPI_ISL_2503270, EPI_ISL_2503274, EPI_ISL_2503277, EPI_ISL_2503287, EPI_ISL_2503289, EPI_ISL_2503290, EPI_ISL_2503291, EPI_ISL_2503294, EPI_ISL_2503295, EPI_ISL_2503300, EPI_ISL_2503301, EPI_ISL_2503304, EPI_ISL_2503307, EPI_ISL_2503308, EPI_ISL_2503312, EPI_ISL_2503314, EPI_ISL_2503315, EPI_ISL_2503320, EPI_ISL_2503321, EPI_ISL_2503329, EPI_ISL_2503330, EPI_ISL_2503335, EPI_ISL_2503339, EPI_ISL_2503341, EPI_ISL_2503342, EPI_ISL_2503343, EPI_ISL_2503346, EPI_ISL_2503348, EPI_ISL_2503349, EPI_ISL_2503350, EPI_ISL_2503351, EPI_ISL_2503353, EPI_ISL_2503359, EPI_ISL_2503362, EPI_ISL_2503363, EPI_ISL_2503367, EPI_ISL_2503399, EPI_ISL_2503401, EPI_ISL_2503402, EPI_ISL_2503409, EPI_ISL_2503415, EPI_ISL_2503429, EPI_ISL_2503474, EPI_ISL_2503480, EPI_ISL_2503516, EPI_ISL_2503517, EPI_ISL_2503519, EPI_ISL_2503520, EPI_ISL_2503521, EPI_ISL_2503522, EPI_ISL_2503523, EPI_ISL_2503524, EPI_ISL_2503525, EPI_ISL_2503526, EPI_ISL_2503527, EPI_ISL_2503528, EPI_ISL_2503529, EPI_ISL_2503530, EPI_ISL_2503532, EPI_ISL_2503535, EPI_ISL_2503538, EPI_ISL_2503547, EPI_ISL_2503572, EPI_ISL_2503577, EPI_ISL_2503578, EPI_ISL_2503579, EPI_ISL_2503580, EPI_ISL_2503581, EPI_ISL_2503584, EPI_ISL_2503585, EPI_ISL_2503586, EPI_ISL_2503590, EPI_ISL_2503591, EPI_ISL_2503592, EPI_ISL_2503593, EPI_ISL_2503595, EPI_ISL_2503596, EPI_ISL_2503597, EPI_ISL_2503599, EPI_ISL_2503603, EPI_ISL_2503604, EPI_ISL_2503606, EPI_ISL_2503608, EPI_ISL_2503609, EPI_ISL_2503614, EPI_ISL_2503615, EPI_ISL_2503616, EPI_ISL_2503618, EPI_ISL_2503619, EPI_ISL_2503620, EPI_ISL_2503622, EPI_ISL_2503623, EPI_ISL_2503624, EPI_ISL_2503627, EPI_ISL_2503628, EPI_ISL_2503629, EPI_ISL_2503632, EPI_ISL_2503635, EPI_ISL_2503636, EPI_ISL_2503637, EPI_ISL_2503638, EPI_ISL_2503640, EPI_ISL_2503641, EPI_ISL_2503653, EPI_ISL_2503654, EPI_ISL_2503656, EPI_ISL_2503657, EPI_ISL_2503658, EPI_ISL_2503659, EPI_ISL_2503662, EPI_ISL_2503664, EPI_ISL_2503665, EPI_ISL_2503667, EPI_ISL_2503672, EPI_ISL_2503673, EPI_ISL_2503674, EPI_ISL_2503675, EPI_ISL_2503677, EPI_ISL_2503684, EPI_ISL_2503685, EPI_ISL_2503686, EPI_ISL_2503688, EPI_ISL_2503692, EPI_ISL_2503694, EPI_ISL_2503701, EPI_ISL_2503704, EPI_ISL_2503708, EPI_ISL_2503709, EPI_ISL_2503710, EPI_ISL_2503711, EPI_ISL_2503712, EPI_ISL_2503715, EPI_ISL_2503716, EPI_ISL_2503717, EPI_ISL_2503719, EPI_ISL_2503720, EPI_ISL_2503721, EPI_ISL_2503722, EPI_ISL_2503723, EPI_ISL_2503724, EPI_ISL_2503725, EPI_ISL_2503732, EPI_ISL_2503733, EPI_ISL_2503735, EPI_ISL_2503739, EPI_ISL_2503741, EPI_ISL_2503742, EPI_ISL_2503743, EPI_ISL_2503744, EPI_ISL_2503747, EPI_ISL_2503749, EPI_ISL_2503750, EPI_ISL_2503751, EPI_ISL_2503752, EPI_ISL_2503753, EPI_ISL_2503754, EPI_ISL_2503757, EPI_ISL_2503758, EPI_ISL_2503759, EPI_ISL_2503762, EPI_ISL_2503763, EPI_ISL_2503764, EPI_ISL_2503765, EPI_ISL_2503767, EPI_ISL_2503769, EPI_ISL_2503771, EPI_ISL_2503775, EPI_ISL_2503776, EPI_ISL_2503779, EPI_ISL_2503790, EPI_ISL_2503791, EPI_ISL_2503792, EPI_ISL_2503793, EPI_ISL_2503794, EPI_ISL_2503797, EPI_ISL_2503800, EPI_ISL_2503802, EPI_ISL_2503803, EPI_ISL_2503805, EPI_ISL_2503806, EPI_ISL_2503809, EPI_ISL_2503810, EPI_ISL_2503811, EPI_ISL_2503812, EPI_ISL_2503816, EPI_ISL_2503818, EPI_ISL_2503819, EPI_ISL_2503822 | | | |
| see above | INSACOG-WB | National Institute of Biomedical Genomics – INSACOG | Arindam Maitra, Bhaswati Bandyopadhyay, Nidhan Kumar Biswas, Tamal Ghosh, Sreedhar Chinnaswamy, Ajay Chakraborti, Saumitra Das |
| EPI_ISL_2503826, EPI_ISL_2503827, EPI_ISL_2503828, EPI_ISL_2503831, EPI_ISL_2503832, EPI_ISL_2503833, EPI_ISL_2503834, EPI_ISL_2503835, EPI_ISL_2503836 | INSACOG-ARUNACHAL PRADESH | National Institute of Biomedical Genomics – INSACOG | Arindam Maitra, Mika Umpo, Nidhan Kumar Biswas, Lobsang Jampa, Sreedhar Chinnaswamy, Yompe Kamki, Saumitra Das |
| EPI_ISL_2503908, EPI_ISL_2503914 | Diagnostyka Sp. z o.o. | 1. Academic Center for Pathomorphological and Genetic-Molecular Diagnostics ltd, Bialystok, Poland 2. National Institute of Public Health - National Institute of Hygiene, Warsaw, Poland | Radosław Charkiewicz, Jacek Nikliński, Przemysław Biecek. Joanna Reszeć, Piotr Majewski, Anetta Sulewska, Piotr Karabowicz, Joanna Kiśluk, Janusz Dzięcioł, Konrad Raczkowski, Katarzyna Zacharczuk, Magdalena Nowakowska, Małgorzata Sadkowska-Todys, Tomasz Wołkowicz |
| EPI_ISL_2503925 | Vitomed Sp. z o.o. | 1. Academic Center for Pathomorphological and Genetic-Molecular Diagnostics ltd, Bialystok, Poland 2. National Institute of Public Health - National Institute of Hygiene, Warsaw, Poland | Radosław Charkiewicz, Jacek Nikliński, Przemysław Biecek. Joanna Reszeć, Piotr Majewski, Anetta Sulewska, Piotr Karabowicz, Joanna Kiśluk, Janusz Dzięcioł, Konrad Raczkowski, Katarzyna Zacharczuk, Magdalena Nowakowska, Małgorzata Sadkowska-Todys, Tomasz Wołkowicz |
| EPI_ISL_2503955, EPI_ISL_2503960, EPI_ISL_2503966 | Diagnostyka | 1. Academic Center for Pathomorphological and Genetic-Molecular Diagnostics ltd, Bialystok, Poland 2. National Institute of Public Health - National Institute of Hygiene, Warsaw, Poland | Radosław Charkiewicz, Jacek Nikliński, Przemysław Biecek. Joanna Reszeć, Piotr Majewski, Anetta Sulewska, Piotr Karabowicz, Joanna Kiśluk, Janusz Dzięcioł, Konrad Raczkowski, Katarzyna Zacharczuk, Magdalena Nowakowska, Małgorzata Sadkowska-Todys, Tomasz Wołkowicz |
| EPI_ISL_2504008 | Alaska State Virology Laboratory | Alaska State Virology Laboratory | Stephanie DeRonde, Elva House, Jacob Zidek, Lisa Smith, Ph.D., Jack Chen, Ph.D. |
| EPI_ISL_2504015, EPI_ISL_2504025, EPI_ISL_2504039, EPI_ISL_2504043 | Labor Berlin Charité Vivantes GmbH / Institut für Virologie | Charité Universitätsmedizin Berlin, Institut für Virologie/Labor Berlin | Peter Menzel, Christine Stephan, Rolf Schwarzer, Victor M Corman, Barbara Mühlemann, Terry Jones, Christian Drosten |
| EPI_ISL_2504055, EPI_ISL_2504057, EPI_ISL_2504058, EPI_ISL_2504060, EPI_ISL_2504061, EPI_ISL_2504063 | A. Krumbholz, Labor Dr. Krause und Kollegen MVZ GmbH, Kiel | Charité Universitätsmedizin Berlin, Institut für Virologie/Labor Berlin | Victor M Corman, Andi Krumbholz, Joern Beheim-Schwarzbach, Barbara Muehlemann, Julia Schneider, Talitha Veith, Tobias Bleicker, Julia Tesch, Peter Menzel, Christine Stephan, Rolf Schwarzer, Terry Jones, Christian Drosten |

| | | | |
|---|---|---|---|
| EPI_ISL_2504284 | National Centre for Disease Control (NCDC) Biotechnology Division, Delhi | NCDC Delhi, Biotechnology Division INSACOG | Priyanka Singh, Uma Sharma, Manoj K Singh, Radhakrishnan V. S, Robin Marwal, Mahesh S Dhar, Kalaiarasan Ponnusamy, Meena Datta, Hemlata Lall, Hema Gogia, Preeti Madan, Sandhya Kabra, Sujeet K Singh, Partha Rakshit |
| EPI_ISL_2504861 | National Centre for Disease Control (NCDC) Biotechnology Division, Delhi | NCDC Delhi, Biotechnology Division INSACOG | Radhakrishnan V. S, Robin Marwal, Mahesh S Dhar, Kalaiarasan Ponnusamy, Meena Datta, Priyanka Singh, Uma Sharma, Hemlata Lall, Manoj K Singh, Hema Gogia, Preeti Madan, Sandhya Kabra, Sujeet K Singh, Partha Rakshit |
| EPI_ISL_2506039 | RSUD dr. LOEKMONO HADI | Genetics Working Group (Pokja Genetik) Faculty of Medicine, Public Health and Nursing Universitas Gadjah Mada (FK-KMK UGM); Disease Investigation Center Wates Ministry of Agriculture Indonesia; Department of Microbiology FK-KMK UGM; Laboratorium Diagnostik Yayasan Tahija World Mosquito Program (WMP) Yogyakarta Center for Tropical Medicine FK-KMK UGM; Integrated Research Center FK-KMK UGM; Department of Computer Science and Electronics FMIPA UGM; RSUP Dr. Sardjito | Gunadi, Hendra Wibawa, Marcellus, Mohamad S. Hakim, Edwin W. Daniwijaya, Ludhang P. Rizki, Endah Supriyati, Eggi Arguni, Titik Nuryastuti, Tri Wibawa, Dwi AA Nugrahaningsih, Afiahayati, Siswanto,Kristy Iskandar,Nungki Anggorowati, Pramesti G Dewi, Dwi Indaryati,Indarto Sulistiyono, Khanza a Vujira, Alvina A Setiawan |
| EPI_ISL_2506040 | RSUD dr. LOEKMONO HADI | Genetics Working Group (Pokja Genetik) Faculty of Medicine, Public Health and Nursing Universitas Gadjah Mada (FK-KMK UGM); Disease Investigation Center Wates Ministry of Agriculture Indonesia; Department of Microbiology FK-KMK UGM; Laboratorium Diagnostik Yayasan Tahija World Mosquito Program (WMP) Yogyakarta Center for Tropical Medicine FK-KMK UGM; Integrated Research Center FK-KMK UGM; Department of Computer Science and Electronics FMIPA UGM; RSUP Dr. Sardjito | Gunadi, Hendra Wibawa, Marcellus, Mohamad S. Hakim, Edwin W. Daniwijaya, Ludhang P. Rizki, Endah Supriyati, Eggi Arguni, Titik Nuryastuti, Tri Wibawa, Dwi AA Nugrahaningsih, Afiahayati, Siswanto,Kristy Iskandar,Nungki Anggorowati, Pramesti G Dewi, Dwi Indaryati,Indarto Sulistiyono, Susan Simanjaya, Dyah A Puspitarani |
| EPI_ISL_2506041 | RSUD dr. LOEKMONO HADI | Genetics Working Group (Pokja Genetik) Faculty of Medicine, Public Health and Nursing Universitas Gadjah Mada (FK-KMK UGM); Disease Investigation Center Wates Ministry of Agriculture Indonesia; Department of Microbiology FK-KMK UGM; Laboratorium Diagnostik Yayasan Tahija World Mosquito Program (WMP) Yogyakarta Center for Tropical Medicine FK-KMK UGM; Integrated Research Center FK-KMK UGM; Department of Computer Science and Electronics FMIPA UGM; RSUP Dr. Sardjito | Gunadi, Hendra Wibawa, Marcellus, Mohamad S. Hakim, Edwin W. Daniwijaya, Ludhang P. Rizki, Endah Supriyati, Eggi Arguni, Titik Nuryastuti, Tri Wibawa, Dwi AA Nugrahaningsih, Afiahayati, Siswanto,Kristy Iskandar,Nungki Anggorowati, Pramesti G Dewi, Dwi Indaryati,Indarto Sulistiyono, Irene Tania, Hana F Hanifin |
| EPI_ISL_2507084, EPI_ISL_2507085, EPI_ISL_2507086, EPI_ISL_2507090, EPI_ISL_2507091, EPI_ISL_2507092, EPI_ISL_2507093, EPI_ISL_2507094, EPI_ISL_2507095, EPI_ISL_2507096 | Queensland Health Forensic and Scientific Services | Queensland Health Forensic and Scientific Services | Son Nguyen |
| EPI_ISL_2507131, EPI_ISL_2507150, EPI_ISL_2507152, EPI_ISL_2507163, EPI_ISL_2507197, EPI_ISL_2507199, EPI_ISL_2507231, EPI_ISL_2507266, EPI_ISL_2507272, EPI_ISL_2507286, EPI_ISL_2507303, EPI_ISL_2507308, EPI_ISL_2507310, EPI_ISL_2507317, EPI_ISL_2507326, EPI_ISL_2507342, EPI_ISL_2507343, EPI_ISL_2507344, EPI_ISL_2507345, EPI_ISL_2507346, EPI_ISL_2507349, EPI_ISL_2507351, EPI_ISL_2507356 | | | |
| see above | Curative Labs | Curative Labs | Mikhail Hanewich-Hollatz, J.Cesar Ignacio-Espinoza, Elias L. Salfati, Nabjot Sandhu, George Way, Zhiyi Xie, Eugenia Khorosheva, Sophia Quasem, Janet Chen, Vladimir Slepnev |
| EPI_ISL_2508228 | Vita Laboratoriot Oy | Expert Microbiology, National Institute for Health and Welfare | Soile Blomqvist, Jani Halkilahti, Kirsi Liitsola, Haider al-Hello, Päivi Laurila, Erika Lindh, Teemu Smura, Ravi Kant, Phuoc Truong, Olli Vapalahti, Sari Hannula, Pekka Ellonen, Niina Ikonen, Carita Savolainen-Kopra |
| EPI_ISL_2508364, EPI_ISL_2508367 | DeRisi Lab, University of California, San Francisco | DeRisi Lab, University of California, San Francisco | Sabrina Mann, Anthea Mitchell, Jamin Liu, Sara Sunshine, Matthew Laurie, Genay Pilarowski, Amy Kistler, Manu Vanaerschot, Patrick Ayscue, Lucy Li, Eric Chow, IDseq Team, James Peng, Carina Marquez, Luis Rubio, Gabriel Chamie, Diane Jones, Jon Jacobo, Susana Rojas, Susy Rojas, Valerie Tulier-Laiwa, Douglas Black, Jackie Martinez, Maya Petersen, Diane Havlir, and Joseph DeRisi |
| EPI_ISL_2508429, EPI_ISL_2508454, EPI_ISL_2508544 | Microbiology Division, SC DHEC | Microbiology Division, SC DHEC | Freeman,J., Flores,H. |
| EPI_ISL_2508619, EPI_ISL_2508620, EPI_ISL_2508621, EPI_ISL_2508622, EPI_ISL_2508623, EPI_ISL_2508624, EPI_ISL_2508625, EPI_ISL_2508626, EPI_ISL_2508627, EPI_ISL_2508628, EPI_ISL_2508629, EPI_ISL_2508630, EPI_ISL_2508631, EPI_ISL_2508632, EPI_ISL_2508633, EPI_ISL_2508634, EPI_ISL_2508635, EPI_ISL_2508636, EPI_ISL_2508637, EPI_ISL_2508638, EPI_ISL_2508639, EPI_ISL_2508640, EPI_ISL_2508641, EPI_ISL_2508642, EPI_ISL_2508643, EPI_ISL_2508644, EPI_ISL_2508645, EPI_ISL_2508646, EPI_ISL_2508647, EPI_ISL_2508648, EPI_ISL_2508649, EPI_ISL_2508650, EPI_ISL_2508651, EPI_ISL_2508652, EPI_ISL_2508653, EPI_ISL_2508654, EPI_ISL_2508655, EPI_ISL_2508656, EPI_ISL_2508657, EPI_ISL_2508658, EPI_ISL_2508659, EPI_ISL_2508660, EPI_ISL_2508661, EPI_ISL_2508662, EPI_ISL_2508663, EPI_ISL_2508664, EPI_ISL_2508665, EPI_ISL_2508666, EPI_ISL_2508667, EPI_ISL_2508668, EPI_ISL_2508669, EPI_ISL_2508670, EPI_ISL_2508671, EPI_ISL_2508672, EPI_ISL_2508673, EPI_ISL_2508674, EPI_ISL_2508675, EPI_ISL_2508676, EPI_ISL_2508677, EPI_ISL_2508678, EPI_ISL_2508679, EPI_ISL_2508680, EPI_ISL_2508681, EPI_ISL_2508682, EPI_ISL_2508683, EPI_ISL_2508684, EPI_ISL_2508685, EPI_ISL_2508686, EPI_ISL_2508687, EPI_ISL_2508688, EPI_ISL_2508689, EPI_ISL_2508690, EPI_ISL_2508691, EPI_ISL_2508692, EPI_ISL_2508693, EPI_ISL_2508694, EPI_ISL_2508695, EPI_ISL_2508696, EPI_ISL_2508697, EPI_ISL_2508698, EPI_ISL_2508699, EPI_ISL_2508700, EPI_ISL_2508701, EPI_ISL_2508702, EPI_ISL_2508703, EPI_ISL_2508704, EPI_ISL_2508705, EPI_ISL_2508706, EPI_ISL_2508707, EPI_ISL_2508708, EPI_ISL_2508709, EPI_ISL_2508710, EPI_ISL_2508711, EPI_ISL_2508717, EPI_ISL_2508718, EPI_ISL_2508719, EPI_ISL_2508728, EPI_ISL_2508729, EPI_ISL_2508730, EPI_ISL_2508731, EPI_ISL_2508732, EPI_ISL_2508733, EPI_ISL_2508734, EPI_ISL_2508735, EPI_ISL_2508736, EPI_ISL_2508737, EPI_ISL_2508738, EPI_ISL_2508739, EPI_ISL_2508740, EPI_ISL_2508741, EPI_ISL_2508742, EPI_ISL_2508743, EPI_ISL_2508744, EPI_ISL_2508745, EPI_ISL_2508746, EPI_ISL_2508747, EPI_ISL_2508748, EPI_ISL_2508749, EPI_ISL_2508750, EPI_ISL_2508751, EPI_ISL_2508752, EPI_ISL_2508753, EPI_ISL_2508754, EPI_ISL_2508755, EPI_ISL_2508756, EPI_ISL_2508757, EPI_ISL_2508758, EPI_ISL_2508759, EPI_ISL_2508760, EPI_ISL_2508761, EPI_ISL_2508762, EPI_ISL_2508763, EPI_ISL_2508764, EPI_ISL_2508765, EPI_ISL_2508766, EPI_ISL_2508767, EPI_ISL_2508768, EPI_ISL_2508769, EPI_ISL_2508770, EPI_ISL_2508771, EPI_ISL_2508772, EPI_ISL_2508773, EPI_ISL_2508774, EPI_ISL_2508775, EPI_ISL_2508776, EPI_ISL_2508777, EPI_ISL_2508778, EPI_ISL_2508779, EPI_ISL_2508780, EPI_ISL_2508781, EPI_ISL_2508782, EPI_ISL_2508783, EPI_ISL_2508784, EPI_ISL_2508785, EPI_ISL_2508786, EPI_ISL_2508787, EPI_ISL_2508788, EPI_ISL_2508789, EPI_ISL_2508790, EPI_ISL_2508791, EPI_ISL_2508792, EPI_ISL_2508793, EPI_ISL_2508794, EPI_ISL_2508795, EPI_ISL_2508796, EPI_ISL_2508797, EPI_ISL_2508798, EPI_ISL_2508799, EPI_ISL_2508800, EPI_ISL_2508810, EPI_ISL_2508811, EPI_ISL_2508812, EPI_ISL_2508813, EPI_ISL_2508814, EPI_ISL_2508815, EPI_ISL_2508816, EPI_ISL_2508817, EPI_ISL_2508818, EPI_ISL_2508819, EPI_ISL_2508820, EPI_ISL_2508821, EPI_ISL_2508822, EPI_ISL_2508823, EPI_ISL_2508824, EPI_ISL_2508825, EPI_ISL_2508826, EPI_ISL_2508827, EPI_ISL_2508828, EPI_ISL_2508829, EPI_ISL_2508830, EPI_ISL_2508831, EPI_ISL_2508832, EPI_ISL_2508833, EPI_ISL_2508834, EPI_ISL_2508835, EPI_ISL_2508836, EPI_ISL_2508837, EPI_ISL_2508838, EPI_ISL_2508839, EPI_ISL_2508840, EPI_ISL_2508841, EPI_ISL_2508842, EPI_ISL_2508843, EPI_ISL_2508844, EPI_ISL_2508845, EPI_ISL_2508846, EPI_ISL_2508847, EPI_ISL_2508848, EPI_ISL_2508849, EPI_ISL_2508850, EPI_ISL_2508851, EPI_ISL_2508852, EPI_ISL_2508853, EPI_ISL_2508854, EPI_ISL_2508855, EPI_ISL_2508856, EPI_ISL_2508857, EPI_ISL_2508858, EPI_ISL_2508859, EPI_ISL_2508860, EPI_ISL_2508861, EPI_ISL_2508862, EPI_ISL_2508863, EPI_ISL_2508864, EPI_ISL_2508865, EPI_ISL_2508866, EPI_ISL_2508867, EPI_ISL_2508868, EPI_ISL_2508869, EPI_ISL_2508870, EPI_ISL_2508871, EPI_ISL_2508872, EPI_ISL_2508873, EPI_ISL_2508874, EPI_ISL_2508875, EPI_ISL_2508876, EPI_ISL_2508877, EPI_ISL_2508878, EPI_ISL_2508879, EPI_ISL_2508880, EPI_ISL_2508881, EPI_ISL_2508882, EPI_ISL_2508883, EPI_ISL_2508884, EPI_ISL_2508885, EPI_ISL_2508886, EPI_ISL_2508887, EPI_ISL_2508891, EPI_ISL_2508892, EPI_ISL_2508893, EPI_ISL_2508894, EPI_ISL_2508895, EPI_ISL_2508896, EPI_ISL_2508897, EPI_ISL_2508898, EPI_ISL_2508899, EPI_ISL_2508900, EPI_ISL_2508901, EPI_ISL_2508902, EPI_ISL_2508903, EPI_ISL_2508904, EPI_ISL_2508905, EPI_ISL_2508906, EPI_ISL_2508907, EPI_ISL_2508908, EPI_ISL_2508909, EPI_ISL_2508910, EPI_ISL_2508911, EPI_ISL_2508912, EPI_ISL_2508913, EPI_ISL_2508914, EPI_ISL_2508915, EPI_ISL_2508916, EPI_ISL_2508917, EPI_ISL_2508918, EPI_ISL_2508919, EPI_ISL_2508920, EPI_ISL_2508921, EPI_ISL_2508922, EPI_ISL_2508923, EPI_ISL_2508924, EPI_ISL_2508925, EPI_ISL_2508926, EPI_ISL_2508927, EPI_ISL_2508928, EPI_ISL_2508929, EPI_ISL_2508930, EPI_ISL_2508931, EPI_ISL_2508932, EPI_ISL_2508933, EPI_ISL_2508934, EPI_ISL_2508935, EPI_ISL_2508936, EPI_ISL_2508937, EPI_ISL_2508938, EPI_ISL_2508939, EPI_ISL_2508940, EPI_ISL_2508941, EPI_ISL_2508942, EPI_ISL_2508943, EPI_ISL_2508944, EPI_ISL_2508945, EPI_ISL_2508946, EPI_ISL_2508947, EPI_ISL_2508948, EPI_ISL_2508949, EPI_ISL_2508950, EPI_ISL_2508951, EPI_ISL_2508952, EPI_ISL_2508953, EPI_ISL_2508954, EPI_ISL_2508955, EPI_ISL_2508956, EPI_ISL_2508957, EPI_ISL_2508958, EPI_ISL_2508959, EPI_ISL_2508960, EPI_ISL_2508961, EPI_ISL_2508962, EPI_ISL_2508963, EPI_ISL_2508964, EPI_ISL_2508965, EPI_ISL_2508966, EPI_ISL_2508967, EPI_ISL_2508968, EPI_ISL_2508969, EPI_ISL_2508970, EPI_ISL_2508971, EPI_ISL_2508972, EPI_ISL_2508973, EPI_ISL_2508974, EPI_ISL_2508975, EPI_ISL_2508976, EPI_ISL_2508977, EPI_ISL_2508978, EPI_ISL_2508979, EPI_ISL_2508980, EPI_ISL_2508981, EPI_ISL_2508982, EPI_ISL_2508983, EPI_ISL_2508984, EPI_ISL_2508985, EPI_ISL_2508986, EPI_ISL_2508987, EPI_ISL_2508988, EPI_ISL_2508989, EPI_ISL_2508990, EPI_ISL_2508991, EPI_ISL_2508992, EPI_ISL_2508993, EPI_ISL_2508994, EPI_ISL_2508995, EPI_ISL_2508996, EPI_ISL_2508997, EPI_ISL_2508998, EPI_ISL_2508999, EPI_ISL_2509000, EPI_ISL_2509001, EPI_ISL_2509002, EPI_ISL_2509003, EPI_ISL_2509004, EPI_ISL_2509005, EPI_ISL_2509006, EPI_ISL_2509007, EPI_ISL_2509008, EPI_ISL_2509009, EPI_ISL_2509010, EPI_ISL_2509011, EPI_ISL_2509012, EPI_ISL_2509013, EPI_ISL_2509014, EPI_ISL_2509015, EPI_ISL_2509016, EPI_ISL_2509017, EPI_ISL_2509018, EPI_ISL_2509019, EPI_ISL_2509020, EPI_ISL_2509021, EPI_ISL_2509022, EPI_ISL_2509023, EPI_ISL_2509036, EPI_ISL_2509037, EPI_ISL_2509038, EPI_ISL_2509039, EPI_ISL_2509040, EPI_ISL_2509041, EPI_ISL_2509042, EPI_ISL_2509043, EPI_ISL_2509044, EPI_ISL_2509045, EPI_ISL_2509046, EPI_ISL_2509047, EPI_ISL_2509048, EPI_ISL_2509049, EPI_ISL_2509050, EPI_ISL_2509051, EPI_ISL_2509052, EPI_ISL_2509053, EPI_ISL_2509054, EPI_ISL_2509055, EPI_ISL_2509056, EPI_ISL_2509057, EPI_ISL_2509058, EPI_ISL_2509059, EPI_ISL_2509060, EPI_ISL_2509061, EPI_ISL_2509062, EPI_ISL_2509063, EPI_ISL_2509064, EPI_ISL_2509065, EPI_ISL_2509066, EPI_ISL_2509067, EPI_ISL_2509068, EPI_ISL_2509069, EPI_ISL_2509070, EPI_ISL_2509071, EPI_ISL_2509072, EPI_ISL_2509073, EPI_ISL_2509074, EPI_ISL_2509075, EPI_ISL_2509076, EPI_ISL_2509077, EPI_ISL_2509078, EPI_ISL_2509079, EPI_ISL_2509080, EPI_ISL_2509081, EPI_ISL_2509082, EPI_ISL_2509083, EPI_ISL_2509084, EPI_ISL_2509085, EPI_ISL_2509086, EPI_ISL_2509087, EPI_ISL_2509088, EPI_ISL_2509089, EPI_ISL_2509090, EPI_ISL_2509091, EPI_ISL_2509092, EPI_ISL_2509093, EPI_ISL_2509094, EPI_ISL_2509095, EPI_ISL_2509096, EPI_ISL_2509097, EPI_ISL_2509098, EPI_ISL_2509099, EPI_ISL_2509100, EPI_ISL_2509101, EPI_ISL_2509102, EPI_ISL_2509103, EPI_ISL_2509104, EPI_ISL_2509105, EPI_ISL_2509106, EPI_ISL_2509107, EPI_ISL_2509108, EPI_ISL_2509109, EPI_ISL_2509110 | | | |
| see above | National Public Health Laboratory, National Centre for Infectious Diseases | National Public Health Laboratory, National Centre for Infectious Diseases | Tze Minn Mak, Zhenyang Zhou, Royce Ang, Lin Cui, Raymond Tzer Pin Lin |
| EPI_ISL_2509187, EPI_ISL_2509191, EPI_ISL_2509193, EPI_ISL_2509196, EPI_ISL_2509202, EPI_ISL_2509203, EPI_ISL_2509214, EPI_ISL_2509215, EPI_ISL_2509217, EPI_ISL_2509227, EPI_ISL_2509228, EPI_ISL_2509234, EPI_ISL_2509237, EPI_ISL_2509238, EPI_ISL_2509240, EPI_ISL_2509242, EPI_ISL_2509249, EPI_ISL_2509252, EPI_ISL_2509257, EPI_ISL_2509260, EPI_ISL_2509261, EPI_ISL_2509267, EPI_ISL_2509271 | | | |
| see above | Microbiology Department, Laboratori Clínic Metropolitana Nord. Hospital Universitari Germans Trias i Pujol | Can Ruti SARS-CoV-2 Sequencing Hub (HUGTiP/IrsiCaixa/IGTP) | Marc Noguera-Julian, Pilar Armengol, Ignacio Blanco, Antoni E Bordoy, Francesc Catala-Moll, Pere-Joan Cardona, Maria Casadellà, Cristina Casañ, Gemma Clara, Bonaventura Clotet, Cristina Esteban, Montserrat Giménez, Mercedes Guerrero, Anna Not, Roger Paredes, Mariona Parera, Verónica Saludes, Alba Sánchez, and Elisa Martró on behalf of the Can Ruti SARS-CoV-2 Sequencing Hub. |
| EPI_ISL_2509934 | N.H.L Municipal Medical College, Ahmedabad | Gujarat Biotechnology Research Centre | Dinesh Kumar,Twinkle Soni,Sonal Sharma ,Zuber Saiyed,Ramesh Pandit,Janvi Raval,Zarna Patel,Nitin Savaliya,Jayshri Pethani,Umang Mishra,Nitesh Shah,Chaitanya Joshi,Madhvi Joshi |
| EPI_ISL_2509935 | N.H.L Municipal Medical College, Ahmedabad | Gujarat Biotechnology Research Centre | Twinkle Soni,Sonal Sharma ,Zuber Saiyed,Ramesh Pandit,Janvi Raval,Zarna Patel,Nitin Savaliya,Dinesh Kumar,Jayshri Pethani,Umang Mishra,Nitesh Shah,Chaitanya Joshi,Madhvi Joshi |
| EPI_ISL_2509936 | N.H.L Municipal Medical College, Ahmedabad | Gujarat Biotechnology Research Centre | Sonal Sharma ,Zuber Saiyed,Ramesh Pandit,Janvi Raval,Zarna Patel,Nitin Savaliya,Dinesh Kumar,Twinkle Soni,Jayshri Pethani,Umang Mishra,Nitesh Shah,Chaitanya Joshi,Madhvi Joshi |
| EPI_ISL_2509937 | N.H.L Municipal Medical College, Ahmedabad | Gujarat Biotechnology Research Centre | Zuber Saiyed,Ramesh Pandit,Janvi Raval,Zarna Patel,Nitin Savaliya,Dinesh Kumar,Twinkle Soni,Sonal Sharma ,Jayshri Pethani,Umang Mishra,Nitesh Shah,Chaitanya Joshi,Madhvi Joshi |
| EPI_ISL_2509938 | BJ Medical College and Civil Hospital, Ahmedabad | Gujarat Biotechnology Research Centre | Ramesh Pandit,Janvi Raval,Zarna Patel,Nitin Savaliya,Dinesh Kumar,Twinkle Soni,Sonal Sharma ,Zuber Saiyed,Pranay Shah,Umang Mishra,Nitesh Shah,Chaitanya Joshi,Madhvi Joshi |
| EPI_ISL_2509939 | N.H.L Municipal Medical College, Ahmedabad | Gujarat Biotechnology Research Centre | Janvi Raval,Zarna Patel,Nitin Savaliya,Dinesh Kumar,Twinkle Soni,Sonal Sharma ,Zuber Saiyed,Ramesh Pandit,Jayshri Pethani,Umang Mishra,Nitesh Shah,Chaitanya Joshi,Madhvi Joshi |
| EPI_ISL_2509940 | N.H.L Municipal Medical College, Ahmedabad | Gujarat Biotechnology Research Centre | Zarna Patel,Nitin Savaliya,Dinesh Kumar,Twinkle Soni,Sonal Sharma ,Zuber Saiyed,Ramesh Pandit,Janvi Raval,Jayshri Pethani,Umang Mishra,Nitesh Shah,Chaitanya Joshi,Madhvi Joshi |
| EPI_ISL_2509941 | N.H.L Municipal Medical College, Ahmedabad | Gujarat Biotechnology Research Centre | Nitin Savaliya,Dinesh Kumar,Twinkle Soni,Sonal Sharma ,Zuber Saiyed,Ramesh Pandit,Janvi Raval,Zarna Patel,Jayshri Pethani,Umang Mishra,Nitesh Shah,Chaitanya Joshi,Madhvi Joshi |
| EPI_ISL_2509942 | N.H.L Municipal Medical College, Ahmedabad | Gujarat Biotechnology Research Centre | Dinesh Kumar,Twinkle Soni,Sonal Sharma ,Zuber Saiyed,Ramesh Pandit,Janvi Raval,Zarna Patel,Nitin Savaliya,Jayshri Pethani,Umang Mishra,Nitesh Shah,Chaitanya Joshi,Madhvi Joshi |
| EPI_ISL_2509943 | N.H.L Municipal Medical College, Ahmedabad | Gujarat Biotechnology Research Centre | Twinkle Soni,Sonal Sharma ,Zuber Saiyed,Ramesh Pandit,Janvi Raval,Zarna Patel,Nitin Savaliya,Dinesh Kumar,Jayshri Pethani,Umang Mishra,Nitesh Shah,Chaitanya Joshi,Madhvi Joshi |
| EPI_ISL_2509944 | N.H.L Municipal Medical College, Ahmedabad | Gujarat Biotechnology Research Centre | Sonal Sharma ,Zuber Saiyed,Ramesh Pandit,Janvi Raval,Zarna Patel,Nitin Savaliya,Dinesh Kumar,Twinkle Soni,Jayshri Pethani,Umang Mishra,Nitesh Shah,Chaitanya Joshi,Madhvi Joshi |
| EPI_ISL_2509945 | N.H.L Municipal Medical College, Ahmedabad | Gujarat Biotechnology Research Centre | Zuber Saiyed,Ramesh Pandit,Janvi Raval,Zarna Patel,Nitin Savaliya,Dinesh Kumar,Twinkle Soni,Sonal Sharma ,Jayshri Pethani,Umang Mishra,Nitesh Shah,Chaitanya Joshi,Madhvi Joshi |
| EPI_ISL_2509946 | N.H.L Municipal Medical College, Ahmedabad | Gujarat Biotechnology Research Centre | Ramesh Pandit,Janvi Raval,Zarna Patel,Nitin Savaliya,Dinesh Kumar,Twinkle Soni,Sonal Sharma ,Zuber Saiyed,Jayshri Pethani,Umang Mishra,Nitesh Shah,Chaitanya Joshi,Madhvi Joshi |
| EPI_ISL_2509949 | N.H.L Municipal Medical College, Ahmedabad | Gujarat Biotechnology Research Centre | Nitin Savaliya,Dinesh Kumar,Twinkle Soni,Sonal Sharma ,Zuber Saiyed,Ramesh Pandit,Janvi Raval,Zarna Patel,Jayshri Pethani,Umang Mishra,Nitesh Shah,Chaitanya Joshi,Madhvi Joshi |
| EPI_ISL_2509950 | N.H.L Municipal Medical College, Ahmedabad | Gujarat Biotechnology Research Centre | Dinesh Kumar,Twinkle Soni,Sonal Sharma ,Zuber Saiyed,Ramesh Pandit,Janvi Raval,Zarna Patel,Nitin Savaliya,Jayshri Pethani,Umang Mishra,Nitesh Shah,Chaitanya Joshi,Madhvi Joshi |
| EPI_ISL_2509951 | N.H.L Municipal Medical College, Ahmedabad | Gujarat Biotechnology Research Centre | Twinkle Soni,Sonal Sharma ,Zuber Saiyed,Ramesh Pandit,Janvi Raval,Zarna Patel,Nitin Savaliya,Dinesh Kumar,Jayshri Pethani,Umang Mishra,Nitesh Shah,Chaitanya Joshi,Madhvi Joshi |
| EPI_ISL_2509952 | N.H.L Municipal Medical College, Ahmedabad | Gujarat Biotechnology Research Centre | Sonal Sharma ,Zuber Saiyed,Ramesh Pandit,Janvi Raval,Zarna Patel,Nitin Savaliya,Dinesh Kumar,Twinkle Soni,Jayshri Pethani,Umang Mishra,Nitesh Shah,Chaitanya Joshi,Madhvi Joshi |
| EPI_ISL_2509953 | N.H.L Municipal Medical College, Ahmedabad | Gujarat Biotechnology Research Centre | Zuber Saiyed,Ramesh Pandit,Janvi Raval,Zarna Patel,Nitin Savaliya,Dinesh Kumar,Twinkle Soni,Sonal Sharma ,Jayshri Pethani,Umang Mishra,Nitesh Shah,Chaitanya Joshi,Madhvi Joshi |
| EPI_ISL_2509954 | N.H.L Municipal Medical College, Ahmedabad | Gujarat Biotechnology Research Centre | Ramesh Pandit,Janvi Raval,Zarna Patel,Nitin Savaliya,Dinesh Kumar,Twinkle Soni,Sonal Sharma ,Zuber Saiyed,Jayshri Pethani,Umang Mishra,Nitesh Shah,Chaitanya Joshi,Madhvi Joshi |
| EPI_ISL_2509955 | N.H.L Municipal Medical College, Ahmedabad | Gujarat Biotechnology Research Centre | Janvi Raval,Zarna Patel,Nitin Savaliya,Dinesh Kumar,Twinkle Soni,Sonal Sharma ,Zuber Saiyed,Ramesh Pandit,Jayshri Pethani,Umang Mishra,Nitesh Shah,Chaitanya Joshi,Madhvi Joshi |

| | | | |
|---|---|---|---|
| EPI_ISL_2509957 | N.H.L Municipal Medical College, Ahmedabad | Gujarat Biotechnology Research Centre | Nitin Savaliya,Dinesh Kumar,Twinkle Soni,Sonal Sharma ,Zuber Saiyed,Ramesh Pandit,Janvi Raval,Zarna Patel,Jayshri Pethani,Umang Mishra,Nitesh Shah,Chaitanya Joshi,Madhvi Joshi |
| EPI_ISL_2509958 | N.H.L Municipal Medical College, Ahmedabad | Gujarat Biotechnology Research Centre | Dinesh Kumar,Twinkle Soni,Sonal Sharma ,Zuber Saiyed,Ramesh Pandit,Janvi Raval,Zarna Patel,Nitin Savaliya,Jayshri Pethani,Umang Mishra,Nitesh Shah,Chaitanya Joshi,Madhvi Joshi |
| EPI_ISL_2509959 | GMERS Government Medical College (GMC), Gotri | Gujarat Biotechnology Research Centre | Twinkle Soni,Sonal Sharma ,Zuber Saiyed,Ramesh Pandit,Janvi Raval,Zarna Patel,Nitin Savaliya,Dinesh Kumar,Bithika Duttaroy,Umang Mishra,Nitesh Shah,Chaitanya Joshi,Madhvi Joshi |
| EPI_ISL_2510580 | Servicio de Microbiología Clínica (Complejo Hospitalario de Navarra, Pamplona) | Centro de Secuenciación NASERTIC | Carmen Ezpeleta Baquedano, Ana Navascués, Ana Miqueleiz |
| EPI_ISL_2510682 | Cliniques universitaires Saint-Luc | UCLouvain/IREC/MBLG | Jean Ruelle, Benoit Kabamba Mukadi |
| EPI_ISL_2510687 | Surabaya Health Office | Institute of Tropical Disease, Universitas Airlangga | Jezzy R Dewantari, Rima R Prasetya, Krisnoadi Rahardjo, Aldise M Nastri, Febria Rachmanita, Gatot Soegiarto, Laksmi Wulandari, Resti Yudhawati, Yasuko Mori, Soetjipto, Kazufumi Shimizu, Maria I Lusida |
| EPI_ISL_2510689 | King Chulalongkorn Memorial Hospital | Thai Red Cross Emerging Infectious Diseases Clinical Center and Faculty of Medicine, Chulalongkorn University | Opass Putcharoen, Stefan Fernandez, Leilani Paitoonpong, Anthony R. Jones, Gompol Suwanpimolkul, Chonticha Klungthong, Watsamon Jantarabenjakul, Piyawan Chinnawirotpisan, Torvorapanit Pattama, Khajohn Joonlasak, Wudtichai Manasatienkij, Supaporn Wacharapluesadee |
| EPI_ISL_2510859 | Spital MŠnnedorf AG | Institute of Medical Virology | Daniel Ehrsam, Isabel StŸrmer, Catharine Aquino, Joel Wirz, Weihong Qi, Hubert Rehrauer, Verena Kufner, Gabriela Ziltener, Maryam Zaheri, Stefan Schmutz, Annette AudigŽ, Maria GrŸnberg, Kevin Steiner, Jon Huder, Cyril Shah, Riccarda Capaul, Guido Bloemberg, JŸrg Bšni, Michael Huber, Alexandra Trkola |
| EPI_ISL_2510872, EPI_ISL_2510873 | Lighthouse Lab in Milton Keynes | Wellcome Sanger Institute for the COVID-19 Genomics UK (COG-UK) Consortium | The Lighthouse Lab in Milton Keynes and Alex Alderton, Roberto Amato, Jeffrey Barrett, Sonia Goncalves, Ewan Harrison, David K. Jackson, Ian Johnston, Dominic Kwiatkowski, Cordelia Langford, John Sillitoe on behalf of the Wellcome Sanger Institute COVID-19 Surveillance Team |
| EPI_ISL_2510875 | Lighthouse Lab in Glasgow | Wellcome Sanger Institute for the COVID-19 Genomics UK (COG-UK) Consortium | Harper VanSteenhouse, Yumi Kasai, David Gray, Carol Clugston, Anna Dominiczak and Alex Alderton, Roberto Amato, Jeffrey Barrett, Sonia Goncalves, Ewan Harrison, David K. Jackson, Ian Johnston, Dominic Kwiatkowski, Cordelia Langford, John Sillitoe on behalf of the Wellcome Sanger Institute COVID-19 Surveillance Team |
| EPI_ISL_2510876, EPI_ISL_2510878, EPI_ISL_2510879, EPI_ISL_2510881 | Lighthouse Lab in Milton Keynes | Wellcome Sanger Institute for the COVID-19 Genomics UK (COG-UK) Consortium | The Lighthouse Lab in Milton Keynes and Alex Alderton, Roberto Amato, Jeffrey Barrett, Sonia Goncalves, Ewan Harrison, David K. Jackson, Ian Johnston, Dominic Kwiatkowski, Cordelia Langford, John Sillitoe on behalf of the Wellcome Sanger Institute COVID-19 Surveillance Team |
| EPI_ISL_2510882 | Lighthouse Lab in Glasgow | Wellcome Sanger Institute for the COVID-19 Genomics UK (COG-UK) Consortium | Harper VanSteenhouse, Yumi Kasai, David Gray, Carol Clugston, Anna Dominiczak and Alex Alderton, Roberto Amato, Jeffrey Barrett, Sonia Goncalves, Ewan Harrison, David K. Jackson, Ian Johnston, Dominic Kwiatkowski, Cordelia Langford, John Sillitoe on behalf of the Wellcome Sanger Institute COVID-19 Surveillance Team |
| EPI_ISL_2510884 | Lighthouse Lab in Milton Keynes | Wellcome Sanger Institute for the COVID-19 Genomics UK (COG-UK) Consortium | The Lighthouse Lab in Milton Keynes and Alex Alderton, Roberto Amato, Jeffrey Barrett, Sonia Goncalves, Ewan Harrison, David K. Jackson, Ian Johnston, Dominic Kwiatkowski, Cordelia Langford, John Sillitoe on behalf of the Wellcome Sanger Institute COVID-19 Surveillance Team |
| EPI_ISL_2510886 | Lighthouse Lab in Glasgow | Wellcome Sanger Institute for the COVID-19 Genomics UK (COG-UK) Consortium | Harper VanSteenhouse, Yumi Kasai, David Gray, Carol Clugston, Anna Dominiczak and Alex Alderton, Roberto Amato, Jeffrey Barrett, Sonia Goncalves, Ewan Harrison, David K. Jackson, Ian Johnston, Dominic Kwiatkowski, Cordelia Langford, John Sillitoe on behalf of the Wellcome Sanger Institute COVID-19 Surveillance Team |
| EPI_ISL_2510887, EPI_ISL_2510889, EPI_ISL_2510890, EPI_ISL_2510892 | Lighthouse Lab in Milton Keynes | Wellcome Sanger Institute for the COVID-19 Genomics UK (COG-UK) Consortium | The Lighthouse Lab in Milton Keynes and Alex Alderton, Roberto Amato, Jeffrey Barrett, Sonia Goncalves, Ewan Harrison, David K. Jackson, Ian Johnston, Dominic Kwiatkowski, Cordelia Langford, John Sillitoe on behalf of the Wellcome Sanger Institute COVID-19 Surveillance Team |
| EPI_ISL_2510893, EPI_ISL_2510895 | Lighthouse Lab in Glasgow | Wellcome Sanger Institute for the COVID-19 Genomics UK (COG-UK) Consortium | Harper VanSteenhouse, Yumi Kasai, David Gray, Carol Clugston, Anna Dominiczak and Alex Alderton, Roberto Amato, Jeffrey Barrett, Sonia Goncalves, Ewan Harrison, David K. Jackson, Ian Johnston, Dominic Kwiatkowski, Cordelia Langford, John Sillitoe on behalf of the Wellcome Sanger Institute COVID-19 Surveillance Team |
| EPI_ISL_2510896, EPI_ISL_2510898, EPI_ISL_2510899, EPI_ISL_2510901 | Lighthouse Lab in Milton Keynes | Wellcome Sanger Institute for the COVID-19 Genomics UK (COG-UK) Consortium | The Lighthouse Lab in Milton Keynes and Alex Alderton, Roberto Amato, Jeffrey Barrett, Sonia Goncalves, Ewan Harrison, David K. Jackson, Ian Johnston, Dominic Kwiatkowski, Cordelia Langford, John Sillitoe on behalf of the Wellcome Sanger Institute COVID-19 Surveillance Team |
| EPI_ISL_2510902, EPI_ISL_2510904 | Lighthouse Lab in Glasgow | Wellcome Sanger Institute for the COVID-19 Genomics UK (COG-UK) Consortium | Harper VanSteenhouse, Yumi Kasai, David Gray, Carol Clugston, Anna Dominiczak and Alex Alderton, Roberto Amato, Jeffrey Barrett, Sonia Goncalves, Ewan Harrison, David K. Jackson, Ian Johnston, Dominic Kwiatkowski, Cordelia Langford, John Sillitoe on behalf of the Wellcome Sanger Institute COVID-19 Surveillance Team |
| EPI_ISL_2510906, EPI_ISL_2510907, EPI_ISL_2510909 | Lighthouse Lab in Milton Keynes | Wellcome Sanger Institute for the COVID-19 Genomics UK (COG-UK) Consortium | The Lighthouse Lab in Milton Keynes and Alex Alderton, Roberto Amato, Jeffrey Barrett, Sonia Goncalves, Ewan Harrison, David K. Jackson, Ian Johnston, Dominic Kwiatkowski, Cordelia Langford, John Sillitoe on behalf of the Wellcome Sanger Institute COVID-19 Surveillance Team |
| EPI_ISL_2510910 | Lighthouse Lab in Glasgow | Wellcome Sanger Institute for the COVID-19 Genomics UK (COG-UK) Consortium | Harper VanSteenhouse, Yumi Kasai, David Gray, Carol Clugston, Anna Dominiczak and Alex Alderton, Roberto Amato, Jeffrey Barrett, Sonia Goncalves, Ewan Harrison, David K. Jackson, Ian Johnston, Dominic Kwiatkowski, Cordelia Langford, John Sillitoe on behalf of the Wellcome Sanger Institute COVID-19 Surveillance Team |
| EPI_ISL_2510913, EPI_ISL_2510914, EPI_ISL_2510916, EPI_ISL_2510917, EPI_ISL_2510919, EPI_ISL_2510920, EPI_ISL_2510922, EPI_ISL_2510923, EPI_ISL_2510925, EPI_ISL_2510927, EPI_ISL_2510929 | | | |
| see above | Lighthouse Lab in Milton Keynes | Wellcome Sanger Institute for the COVID-19 Genomics UK (COG-UK) Consortium | The Lighthouse Lab in Milton Keynes and Alex Alderton, Roberto Amato, Jeffrey Barrett, Sonia Goncalves, Ewan Harrison, David K. Jackson, Ian Johnston, Dominic Kwiatkowski, Cordelia Langford, John Sillitoe on behalf of the Wellcome Sanger Institute COVID-19 Surveillance Team |
| EPI_ISL_2510930, EPI_ISL_2510932, EPI_ISL_2510933 | Lighthouse Lab in Glasgow | Wellcome Sanger Institute for the COVID-19 Genomics UK (COG-UK) Consortium | Harper VanSteenhouse, Yumi Kasai, David Gray, Carol Clugston, Anna Dominiczak and Alex Alderton, Roberto Amato, Jeffrey Barrett, Sonia Goncalves, Ewan Harrison, David K. Jackson, Ian Johnston, Dominic Kwiatkowski, Cordelia Langford, John Sillitoe on behalf of the Wellcome Sanger Institute COVID-19 Surveillance Team |
| EPI_ISL_2510935, EPI_ISL_2510938, EPI_ISL_2510940, EPI_ISL_2510941, EPI_ISL_2510942, EPI_ISL_2510944, EPI_ISL_2510946, EPI_ISL_2510947, EPI_ISL_2510949, EPI_ISL_2510950, EPI_ISL_2510952 | | | |
| see above | Lighthouse Lab in Milton Keynes | Wellcome Sanger Institute for the COVID-19 Genomics UK (COG-UK) Consortium | The Lighthouse Lab in Milton Keynes and Alex Alderton, Roberto Amato, Jeffrey Barrett, Sonia Goncalves, Ewan Harrison, David K. Jackson, Ian Johnston, Dominic Kwiatkowski, Cordelia Langford, John Sillitoe on behalf of the Wellcome Sanger Institute COVID-19 Surveillance Team |
| EPI_ISL_2510953, EPI_ISL_2510956 | Lighthouse Lab in Glasgow | Wellcome Sanger Institute for the COVID-19 Genomics UK (COG-UK) Consortium | Harper VanSteenhouse, Yumi Kasai, David Gray, Carol Clugston, Anna Dominiczak and Alex Alderton, Roberto Amato, Jeffrey Barrett, Sonia Goncalves, Ewan Harrison, David K. Jackson, Ian Johnston, Dominic Kwiatkowski, Cordelia Langford, John Sillitoe on behalf of the Wellcome Sanger Institute COVID-19 Surveillance Team |
| EPI_ISL_2510959, EPI_ISL_2510961 | Lighthouse Lab in Milton Keynes | Wellcome Sanger Institute for the COVID-19 Genomics UK (COG-UK) Consortium | The Lighthouse Lab in Milton Keynes and Alex Alderton, Roberto Amato, Jeffrey Barrett, Sonia Goncalves, Ewan Harrison, David K. Jackson, Ian Johnston, Dominic Kwiatkowski, Cordelia Langford, John Sillitoe on behalf of the Wellcome Sanger Institute COVID-19 Surveillance Team |
| EPI_ISL_2510962 | Lighthouse Lab in Glasgow | Wellcome Sanger Institute for the COVID-19 Genomics UK (COG-UK) Consortium | Harper VanSteenhouse, Yumi Kasai, David Gray, Carol Clugston, Anna Dominiczak and Alex Alderton, Roberto Amato, Jeffrey Barrett, Sonia Goncalves, Ewan Harrison, David K. Jackson, Ian Johnston, Dominic Kwiatkowski, Cordelia Langford, John Sillitoe on behalf of the Wellcome Sanger Institute COVID-19 Surveillance Team |
| EPI_ISL_2510963 | Lighthouse Lab in Milton Keynes | Wellcome Sanger Institute for the COVID-19 Genomics UK (COG-UK) Consortium | The Lighthouse Lab in Milton Keynes and Alex Alderton, Roberto Amato, Jeffrey Barrett, Sonia Goncalves, Ewan Harrison, David K. Jackson, Ian Johnston, Dominic Kwiatkowski, Cordelia Langford, John Sillitoe on behalf of the Wellcome Sanger Institute COVID-19 Surveillance Team |
| EPI_ISL_2510965 | Lighthouse Lab in Glasgow | Wellcome Sanger Institute for the COVID-19 Genomics UK (COG-UK) Consortium | Harper VanSteenhouse, Yumi Kasai, David Gray, Carol Clugston, Anna Dominiczak and Alex Alderton, Roberto Amato, Jeffrey Barrett, Sonia Goncalves, Ewan Harrison, David K. Jackson, Ian Johnston, Dominic Kwiatkowski, Cordelia Langford, John Sillitoe on behalf of the Wellcome Sanger Institute COVID-19 Surveillance Team |
| EPI_ISL_2510966, EPI_ISL_2510968, EPI_ISL_2510969, EPI_ISL_2510971, EPI_ISL_2510972, EPI_ISL_2510974, EPI_ISL_2510975, EPI_ISL_2510977, EPI_ISL_2510978, EPI_ISL_2510980, EPI_ISL_2510981, EPI_ISL_2510983, EPI_ISL_2510984, EPI_ISL_2510986, EPI_ISL_2510987, EPI_ISL_2510989, EPI_ISL_2510993, EPI_ISL_2510994, EPI_ISL_2510995, EPI_ISL_2510998, EPI_ISL_2511000, EPI_ISL_2511001, EPI_ISL_2511003, EPI_ISL_2511006, EPI_ISL_2511007, EPI_ISL_2511009, EPI_ISL_2511010 | | | |
| see above | Lighthouse Lab in Milton Keynes | Wellcome Sanger Institute for the COVID-19 Genomics UK (COG-UK) Consortium | The Lighthouse Lab in Milton Keynes and Alex Alderton, Roberto Amato, Jeffrey Barrett, Sonia Goncalves, Ewan Harrison, David K. Jackson, Ian Johnston, Dominic Kwiatkowski, Cordelia Langford, John Sillitoe on behalf of the Wellcome Sanger Institute COVID-19 Surveillance Team |
| EPI_ISL_2511011 | Lighthouse Lab in Glasgow | Wellcome Sanger Institute for the COVID-19 Genomics UK (COG-UK) Consortium | Harper VanSteenhouse, Yumi Kasai, David Gray, Carol Clugston, Anna Dominiczak and Alex Alderton, Roberto Amato, Jeffrey Barrett, Sonia Goncalves, Ewan Harrison, David K. Jackson, Ian Johnston, Dominic Kwiatkowski, Cordelia Langford, John Sillitoe on behalf of the Wellcome Sanger Institute COVID-19 Surveillance Team |
| EPI_ISL_2511013 | Lighthouse Lab in Milton Keynes | Wellcome Sanger Institute for the COVID-19 Genomics UK (COG-UK) Consortium | The Lighthouse Lab in Milton Keynes and Alex Alderton, Roberto Amato, Jeffrey Barrett, Sonia Goncalves, Ewan Harrison, David K. Jackson, Ian Johnston, Dominic Kwiatkowski, Cordelia Langford, John Sillitoe on behalf of the Wellcome Sanger Institute COVID-19 Surveillance Team |
| EPI_ISL_2511014 | Lighthouse Lab in Glasgow | Wellcome Sanger Institute for the COVID-19 Genomics UK (COG-UK) Consortium | Harper VanSteenhouse, Yumi Kasai, David Gray, Carol Clugston, Anna Dominiczak and Alex Alderton, Roberto Amato, Jeffrey Barrett, Sonia Goncalves, Ewan Harrison, David K. Jackson, Ian Johnston, Dominic Kwiatkowski, Cordelia Langford, John Sillitoe on behalf of the Wellcome Sanger Institute COVID-19 Surveillance Team |
| EPI_ISL_2511016, EPI_ISL_2511018 | Lighthouse Lab in Milton Keynes | Wellcome Sanger Institute for the COVID-19 Genomics UK (COG-UK) Consortium | The Lighthouse Lab in Milton Keynes and Alex Alderton, Roberto Amato, Jeffrey Barrett, Sonia Goncalves, Ewan Harrison, David K. Jackson, Ian Johnston, Dominic Kwiatkowski, Cordelia Langford, John Sillitoe on behalf of the Wellcome Sanger Institute COVID-19 Surveillance Team |
| EPI_ISL_2511019 | Lighthouse Lab in Glasgow | Wellcome Sanger Institute for the COVID-19 Genomics UK (COG-UK) Consortium | Harper VanSteenhouse, Yumi Kasai, David Gray, Carol Clugston, Anna Dominiczak and Alex Alderton, Roberto Amato, Jeffrey Barrett, Sonia Goncalves, Ewan Harrison, David K. Jackson, Ian Johnston, Dominic Kwiatkowski, Cordelia Langford, John Sillitoe on behalf of the Wellcome Sanger Institute COVID-19 Surveillance Team |
| EPI_ISL_2511020 | Lighthouse Lab in Milton Keynes | Wellcome Sanger Institute for the COVID-19 Genomics UK (COG-UK) Consortium | The Lighthouse Lab in Milton Keynes and Alex Alderton, Roberto Amato, Jeffrey Barrett, Sonia Goncalves, Ewan Harrison, David K. Jackson, Ian Johnston, Dominic Kwiatkowski, Cordelia Langford, John Sillitoe on behalf of the Wellcome Sanger Institute COVID-19 Surveillance Team |
| EPI_ISL_2511021 | Lighthouse Lab in Glasgow | Wellcome Sanger Institute for the COVID-19 Genomics UK (COG-UK) Consortium | Harper VanSteenhouse, Yumi Kasai, David Gray, Carol Clugston, Anna Dominiczak and Alex Alderton, Roberto Amato, Jeffrey Barrett, Sonia Goncalves, Ewan Harrison, David K. Jackson, Ian Johnston, Dominic Kwiatkowski, Cordelia Langford, John Sillitoe on behalf of the Wellcome Sanger Institute COVID-19 Surveillance Team |
| EPI_ISL_2511022, EPI_ISL_2511023, EPI_ISL_2511024, EPI_ISL_2511025, EPI_ISL_2511026, EPI_ISL_2511027 | Lighthouse Lab in Milton Keynes | Wellcome Sanger Institute for the COVID-19 Genomics UK (COG-UK) Consortium | The Lighthouse Lab in Milton Keynes and Alex Alderton, Roberto Amato, Jeffrey Barrett, Sonia Goncalves, Ewan Harrison, David K. Jackson, Ian Johnston, Dominic Kwiatkowski, Cordelia Langford, John Sillitoe on behalf of the Wellcome Sanger Institute COVID-19 Surveillance Team |
| EPI_ISL_2511028 | Lighthouse Lab in Glasgow | Wellcome Sanger Institute for the COVID-19 Genomics UK (COG-UK) Consortium | Harper VanSteenhouse, Yumi Kasai, David Gray, Carol Clugston, Anna Dominiczak and Alex Alderton, Roberto Amato, Jeffrey Barrett, Sonia Goncalves, Ewan Harrison, David K. Jackson, Ian Johnston, Dominic Kwiatkowski, Cordelia Langford, John Sillitoe on behalf of the Wellcome Sanger Institute COVID-19 Surveillance Team |
| EPI_ISL_2511029, EPI_ISL_2511030, EPI_ISL_2511032, EPI_ISL_2511033, EPI_ISL_2511034, | Lighthouse Lab in Milton Keynes | Wellcome Sanger Institute for the COVID-19 Genomics UK (COG-UK) Consortium | The Lighthouse Lab in Milton Keynes and Alex Alderton, Roberto Amato, Jeffrey Barrett, Sonia Goncalves, Ewan Harrison, David K. Jackson, Ian Johnston, Dominic Kwiatkowski, Cordelia Langford, John Sillitoe on behalf of the Wellcome Sanger Institute COVID-19 Surveillance Team |

| | | | |
|---|---|---|---|
| EPI_ISL_2511035, EPI_ISL_2511036 | | | |
| EPI_ISL_2511037 | Lighthouse Lab in Glasgow | Wellcome Sanger Institute for the COVID-19 Genomics UK (COG-UK) Consortium | Harper VanSteenhouse, Yumi Kasai, David Gray, Carol Clugston, Anna Dominiczak and Alex Alderton, Roberto Amato, Jeffrey Barrett, Sonia Goncalves, Ewan Harrison, David K. Jackson, Ian Johnston, Dominic Kwiatkowski, Cordelia Langford, John Sillitoe on behalf of the Wellcome Sanger Institute COVID-19 Surveillance Team |
| EPI_ISL_2511038, EPI_ISL_2511039, EPI_ISL_2511040, EPI_ISL_2511042, EPI_ISL_2511043, EPI_ISL_2511044, EPI_ISL_2511046, EPI_ISL_2511047, EPI_ISL_2511048, EPI_ISL_2511049, EPI_ISL_2511050 | | | |
| see above | Lighthouse Lab in Milton Keynes | Wellcome Sanger Institute for the COVID-19 Genomics UK (COG-UK) Consortium | The Lighthouse Lab in Milton Keynes and Alex Alderton, Roberto Amato, Jeffrey Barrett, Sonia Goncalves, Ewan Harrison, David K. Jackson, Ian Johnston, Dominic Kwiatkowski, Cordelia Langford, John Sillitoe on behalf of the Wellcome Sanger Institute COVID-19 Surveillance Team |
| EPI_ISL_2511051 | Lighthouse Lab in Glasgow | Wellcome Sanger Institute for the COVID-19 Genomics UK (COG-UK) Consortium | Harper VanSteenhouse, Yumi Kasai, David Gray, Carol Clugston, Anna Dominiczak and Alex Alderton, Roberto Amato, Jeffrey Barrett, Sonia Goncalves, Ewan Harrison, David K. Jackson, Ian Johnston, Dominic Kwiatkowski, Cordelia Langford, John Sillitoe on behalf of the Wellcome Sanger Institute COVID-19 Surveillance Team |
| EPI_ISL_2511053, EPI_ISL_2511055, EPI_ISL_2511056, EPI_ISL_2511057, EPI_ISL_2511058, EPI_ISL_2511059, EPI_ISL_2511060 | Lighthouse Lab in Milton Keynes | Wellcome Sanger Institute for the COVID-19 Genomics UK (COG-UK) Consortium | The Lighthouse Lab in Milton Keynes and Alex Alderton, Roberto Amato, Jeffrey Barrett, Sonia Goncalves, Ewan Harrison, David K. Jackson, Ian Johnston, Dominic Kwiatkowski, Cordelia Langford, John Sillitoe on behalf of the Wellcome Sanger Institute COVID-19 Surveillance Team |
| EPI_ISL_2511062 | Lighthouse Lab in Glasgow | Wellcome Sanger Institute for the COVID-19 Genomics UK (COG-UK) Consortium | Harper VanSteenhouse, Yumi Kasai, David Gray, Carol Clugston, Anna Dominiczak and Alex Alderton, Roberto Amato, Jeffrey Barrett, Sonia Goncalves, Ewan Harrison, David K. Jackson, Ian Johnston, Dominic Kwiatkowski, Cordelia Langford, John Sillitoe on behalf of the Wellcome Sanger Institute COVID-19 Surveillance Team |
| EPI_ISL_2511064, EPI_ISL_2511065, EPI_ISL_2511067, EPI_ISL_2511068, EPI_ISL_2511069 | Lighthouse Lab in Milton Keynes | Wellcome Sanger Institute for the COVID-19 Genomics UK (COG-UK) Consortium | The Lighthouse Lab in Milton Keynes and Alex Alderton, Roberto Amato, Jeffrey Barrett, Sonia Goncalves, Ewan Harrison, David K. Jackson, Ian Johnston, Dominic Kwiatkowski, Cordelia Langford, John Sillitoe on behalf of the Wellcome Sanger Institute COVID-19 Surveillance Team |
| EPI_ISL_2511070 | Lighthouse Lab in Glasgow | Wellcome Sanger Institute for the COVID-19 Genomics UK (COG-UK) Consortium | Harper VanSteenhouse, Yumi Kasai, David Gray, Carol Clugston, Anna Dominiczak and Alex Alderton, Roberto Amato, Jeffrey Barrett, Sonia Goncalves, Ewan Harrison, David K. Jackson, Ian Johnston, Dominic Kwiatkowski, Cordelia Langford, John Sillitoe on behalf of the Wellcome Sanger Institute COVID-19 Surveillance Team |
| EPI_ISL_2511073, EPI_ISL_2511074, EPI_ISL_2511075, EPI_ISL_2511076, EPI_ISL_2511077, EPI_ISL_2511078, EPI_ISL_2511079 | Lighthouse Lab in Milton Keynes | Wellcome Sanger Institute for the COVID-19 Genomics UK (COG-UK) Consortium | The Lighthouse Lab in Milton Keynes and Alex Alderton, Roberto Amato, Jeffrey Barrett, Sonia Goncalves, Ewan Harrison, David K. Jackson, Ian Johnston, Dominic Kwiatkowski, Cordelia Langford, John Sillitoe on behalf of the Wellcome Sanger Institute COVID-19 Surveillance Team |
| EPI_ISL_2511080 | Lighthouse Lab in Glasgow | Wellcome Sanger Institute for the COVID-19 Genomics UK (COG-UK) Consortium | Harper VanSteenhouse, Yumi Kasai, David Gray, Carol Clugston, Anna Dominiczak and Alex Alderton, Roberto Amato, Jeffrey Barrett, Sonia Goncalves, Ewan Harrison, David K. Jackson, Ian Johnston, Dominic Kwiatkowski, Cordelia Langford, John Sillitoe on behalf of the Wellcome Sanger Institute COVID-19 Surveillance Team |
| EPI_ISL_2511082, EPI_ISL_2511083, EPI_ISL_2511085, EPI_ISL_2511086 | Lighthouse Lab in Milton Keynes | Wellcome Sanger Institute for the COVID-19 Genomics UK (COG-UK) Consortium | The Lighthouse Lab in Milton Keynes and Alex Alderton, Roberto Amato, Jeffrey Barrett, Sonia Goncalves, Ewan Harrison, David K. Jackson, Ian Johnston, Dominic Kwiatkowski, Cordelia Langford, John Sillitoe on behalf of the Wellcome Sanger Institute COVID-19 Surveillance Team |
| EPI_ISL_2511087 | Lighthouse Lab in Glasgow | Wellcome Sanger Institute for the COVID-19 Genomics UK (COG-UK) Consortium | Harper VanSteenhouse, Yumi Kasai, David Gray, Carol Clugston, Anna Dominiczak and Alex Alderton, Roberto Amato, Jeffrey Barrett, Sonia Goncalves, Ewan Harrison, David K. Jackson, Ian Johnston, Dominic Kwiatkowski, Cordelia Langford, John Sillitoe on behalf of the Wellcome Sanger Institute COVID-19 Surveillance Team |
| EPI_ISL_2511089, EPI_ISL_2511090 | Lighthouse Lab in Milton Keynes | Wellcome Sanger Institute for the COVID-19 Genomics UK (COG-UK) Consortium | The Lighthouse Lab in Milton Keynes and Alex Alderton, Roberto Amato, Jeffrey Barrett, Sonia Goncalves, Ewan Harrison, David K. Jackson, Ian Johnston, Dominic Kwiatkowski, Cordelia Langford, John Sillitoe on behalf of the Wellcome Sanger Institute COVID-19 Surveillance Team |
| EPI_ISL_2511093, EPI_ISL_2511094 | Lighthouse Lab in Glasgow | Wellcome Sanger Institute for the COVID-19 Genomics UK (COG-UK) Consortium | Harper VanSteenhouse, Yumi Kasai, David Gray, Carol Clugston, Anna Dominiczak and Alex Alderton, Roberto Amato, Jeffrey Barrett, Sonia Goncalves, Ewan Harrison, David K. Jackson, Ian Johnston, Dominic Kwiatkowski, Cordelia Langford, John Sillitoe on behalf of the Wellcome Sanger Institute COVID-19 Surveillance Team |
| EPI_ISL_2511095 | Lighthouse Lab in Milton Keynes | Wellcome Sanger Institute for the COVID-19 Genomics UK (COG-UK) Consortium | The Lighthouse Lab in Milton Keynes and Alex Alderton, Roberto Amato, Jeffrey Barrett, Sonia Goncalves, Ewan Harrison, David K. Jackson, Ian Johnston, Dominic Kwiatkowski, Cordelia Langford, John Sillitoe on behalf of the Wellcome Sanger Institute COVID-19 Surveillance Team |
| EPI_ISL_2511096 | Lighthouse Lab in Glasgow | Wellcome Sanger Institute for the COVID-19 Genomics UK (COG-UK) Consortium | Harper VanSteenhouse, Yumi Kasai, David Gray, Carol Clugston, Anna Dominiczak and Alex Alderton, Roberto Amato, Jeffrey Barrett, Sonia Goncalves, Ewan Harrison, David K. Jackson, Ian Johnston, Dominic Kwiatkowski, Cordelia Langford, John Sillitoe on behalf of the Wellcome Sanger Institute COVID-19 Surveillance Team |
| EPI_ISL_2511097, EPI_ISL_2511098 | Health Services Laboratories | Wellcome Sanger Institute for the COVID-19 Genomics UK (COG-UK) Consortium | Health Services Laboratories and Alex Alderton, Roberto Amato, Jeffrey Barrett, Sonia Goncalves, Ewan Harrison, David K. Jackson, Ian Johnston, Dominic Kwiatkowski, Cordelia Langford, John Sillitoe on behalf of the Wellcome Sanger Institute COVID-19 Surveillance Team |
| EPI_ISL_2511099, EPI_ISL_2511100, EPI_ISL_2511101 | Lighthouse Lab in Glasgow | Wellcome Sanger Institute for the COVID-19 Genomics UK (COG-UK) Consortium | Harper VanSteenhouse, Yumi Kasai, David Gray, Carol Clugston, Anna Dominiczak and Alex Alderton, Roberto Amato, Jeffrey Barrett, Sonia Goncalves, Ewan Harrison, David K. Jackson, Ian Johnston, Dominic Kwiatkowski, Cordelia Langford, John Sillitoe on behalf of the Wellcome Sanger Institute COVID-19 Surveillance Team |
| EPI_ISL_2511102 | Health Services Laboratories | Wellcome Sanger Institute for the COVID-19 Genomics UK (COG-UK) Consortium | Health Services Laboratories and Alex Alderton, Roberto Amato, Jeffrey Barrett, Sonia Goncalves, Ewan Harrison, David K. Jackson, Ian Johnston, Dominic Kwiatkowski, Cordelia Langford, John Sillitoe on behalf of the Wellcome Sanger Institute COVID-19 Surveillance Team |
| EPI_ISL_2511103, EPI_ISL_2511104, EPI_ISL_2511105, EPI_ISL_2511106 | Lighthouse Lab in Glasgow | Wellcome Sanger Institute for the COVID-19 Genomics UK (COG-UK) Consortium | Harper VanSteenhouse, Yumi Kasai, David Gray, Carol Clugston, Anna Dominiczak and Alex Alderton, Roberto Amato, Jeffrey Barrett, Sonia Goncalves, Ewan Harrison, David K. Jackson, Ian Johnston, Dominic Kwiatkowski, Cordelia Langford, John Sillitoe on behalf of the Wellcome Sanger Institute COVID-19 Surveillance Team |
| EPI_ISL_2511107 | Health Services Laboratories | Wellcome Sanger Institute for the COVID-19 Genomics UK (COG-UK) Consortium | Health Services Laboratories and Alex Alderton, Roberto Amato, Jeffrey Barrett, Sonia Goncalves, Ewan Harrison, David K. Jackson, Ian Johnston, Dominic Kwiatkowski, Cordelia Langford, John Sillitoe on behalf of the Wellcome Sanger Institute COVID-19 Surveillance Team |
| EPI_ISL_2511108, EPI_ISL_2511109 | Lighthouse Lab in Glasgow | Wellcome Sanger Institute for the COVID-19 Genomics UK (COG-UK) Consortium | Harper VanSteenhouse, Yumi Kasai, David Gray, Carol Clugston, Anna Dominiczak and Alex Alderton, Roberto Amato, Jeffrey Barrett, Sonia Goncalves, Ewan Harrison, David K. Jackson, Ian Johnston, Dominic Kwiatkowski, Cordelia Langford, John Sillitoe on behalf of the Wellcome Sanger Institute COVID-19 Surveillance Team |
| EPI_ISL_2511110 | Health Services Laboratories | Wellcome Sanger Institute for the COVID-19 Genomics UK (COG-UK) Consortium | Health Services Laboratories and Alex Alderton, Roberto Amato, Jeffrey Barrett, Sonia Goncalves, Ewan Harrison, David K. Jackson, Ian Johnston, Dominic Kwiatkowski, Cordelia Langford, John Sillitoe on behalf of the Wellcome Sanger Institute COVID-19 Surveillance Team |
| EPI_ISL_2511111 | Lighthouse Lab in Glasgow | Wellcome Sanger Institute for the COVID-19 Genomics UK (COG-UK) Consortium | Harper VanSteenhouse, Yumi Kasai, David Gray, Carol Clugston, Anna Dominiczak and Alex Alderton, Roberto Amato, Jeffrey Barrett, Sonia Goncalves, Ewan Harrison, David K. Jackson, Ian Johnston, Dominic Kwiatkowski, Cordelia Langford, John Sillitoe on behalf of the Wellcome Sanger Institute COVID-19 Surveillance Team |
| EPI_ISL_2511112 | Health Services Laboratories | Wellcome Sanger Institute for the COVID-19 Genomics UK (COG-UK) Consortium | Health Services Laboratories and Alex Alderton, Roberto Amato, Jeffrey Barrett, Sonia Goncalves, Ewan Harrison, David K. Jackson, Ian Johnston, Dominic Kwiatkowski, Cordelia Langford, John Sillitoe on behalf of the Wellcome Sanger Institute COVID-19 Surveillance Team |
| EPI_ISL_2511113 | Lighthouse Lab in Glasgow | Wellcome Sanger Institute for the COVID-19 Genomics UK (COG-UK) Consortium | Harper VanSteenhouse, Yumi Kasai, David Gray, Carol Clugston, Anna Dominiczak and Alex Alderton, Roberto Amato, Jeffrey Barrett, Sonia Goncalves, Ewan Harrison, David K. Jackson, Ian Johnston, Dominic Kwiatkowski, Cordelia Langford, John Sillitoe on behalf of the Wellcome Sanger Institute COVID-19 Surveillance Team |
| EPI_ISL_2511114, EPI_ISL_2511115, EPI_ISL_2511116, EPI_ISL_2511117, EPI_ISL_2511118 | Health Services Laboratories | Wellcome Sanger Institute for the COVID-19 Genomics UK (COG-UK) Consortium | Health Services Laboratories and Alex Alderton, Roberto Amato, Jeffrey Barrett, Sonia Goncalves, Ewan Harrison, David K. Jackson, Ian Johnston, Dominic Kwiatkowski, Cordelia Langford, John Sillitoe on behalf of the Wellcome Sanger Institute COVID-19 Surveillance Team |
| EPI_ISL_2511119, EPI_ISL_2511120, EPI_ISL_2511121, EPI_ISL_2511122, EPI_ISL_2511123, EPI_ISL_2511124, EPI_ISL_2511125, EPI_ISL_2511126, EPI_ISL_2511127 | Lighthouse Lab in Glasgow | Wellcome Sanger Institute for the COVID-19 Genomics UK (COG-UK) Consortium | Harper VanSteenhouse, Yumi Kasai, David Gray, Carol Clugston, Anna Dominiczak and Alex Alderton, Roberto Amato, Jeffrey Barrett, Sonia Goncalves, Ewan Harrison, David K. Jackson, Ian Johnston, Dominic Kwiatkowski, Cordelia Langford, John Sillitoe on behalf of the Wellcome Sanger Institute COVID-19 Surveillance Team |
| EPI_ISL_2511128, EPI_ISL_2511129 | Health Services Laboratories | Wellcome Sanger Institute for the COVID-19 Genomics UK (COG-UK) Consortium | Health Services Laboratories and Alex Alderton, Roberto Amato, Jeffrey Barrett, Sonia Goncalves, Ewan Harrison, David K. Jackson, Ian Johnston, Dominic Kwiatkowski, Cordelia Langford, John Sillitoe on behalf of the Wellcome Sanger Institute COVID-19 Surveillance Team |
| EPI_ISL_2511130 | Lighthouse Lab in Glasgow | Wellcome Sanger Institute for the COVID-19 Genomics UK (COG-UK) Consortium | Harper VanSteenhouse, Yumi Kasai, David Gray, Carol Clugston, Anna Dominiczak and Alex Alderton, Roberto Amato, Jeffrey Barrett, Sonia Goncalves, Ewan Harrison, David K. Jackson, Ian Johnston, Dominic Kwiatkowski, Cordelia Langford, John Sillitoe on behalf of the Wellcome Sanger Institute COVID-19 Surveillance Team |
| EPI_ISL_2511132 | Health Services Laboratories | Wellcome Sanger Institute for the COVID-19 Genomics UK (COG-UK) Consortium | Health Services Laboratories and Alex Alderton, Roberto Amato, Jeffrey Barrett, Sonia Goncalves, Ewan Harrison, David K. Jackson, Ian Johnston, Dominic Kwiatkowski, Cordelia Langford, John Sillitoe on behalf of the Wellcome Sanger Institute COVID-19 Surveillance Team |
| EPI_ISL_2511133 | Lighthouse Lab in Glasgow | Wellcome Sanger Institute for the COVID-19 Genomics UK (COG-UK) Consortium | Harper VanSteenhouse, Yumi Kasai, David Gray, Carol Clugston, Anna Dominiczak and Alex Alderton, Roberto Amato, Jeffrey Barrett, Sonia Goncalves, Ewan Harrison, David K. Jackson, Ian Johnston, Dominic Kwiatkowski, Cordelia Langford, John Sillitoe on behalf of the Wellcome Sanger Institute COVID-19 Surveillance Team |
| EPI_ISL_2511134 | Lighthouse Lab in Milton Keynes | Wellcome Sanger Institute for the COVID-19 Genomics UK (COG-UK) Consortium | The Lighthouse Lab in Milton Keynes and Alex Alderton, Roberto Amato, Jeffrey Barrett, Sonia Goncalves, Ewan Harrison, David K. Jackson, Ian Johnston, Dominic Kwiatkowski, Cordelia Langford, John Sillitoe on behalf of the Wellcome Sanger Institute COVID-19 Surveillance Team |

| | | | |
|---|---|---|---|
| EPI_ISL_2511135, EPI_ISL_2511136, EPI_ISL_2511137, EPI_ISL_2511138, EPI_ISL_2511139, EPI_ISL_2511140 | Lighthouse Lab in Glasgow | Wellcome Sanger Institute for the COVID-19 Genomics UK (COG-UK) Consortium | Harper VanSteenhouse, Yumi Kasai, David Gray, Carol Clugston, Anna Dominiczak and Alex Alderton, Roberto Amato, Jeffrey Barrett, Sonia Goncalves, Ewan Harrison, David K. Jackson, Ian Johnston, Dominic Kwiatkowski, Cordelia Langford, John Sillitoe on behalf of the Wellcome Sanger Institute COVID-19 Surveillance Team |
| EPI_ISL_2511141 | Health Services Laboratories | Wellcome Sanger Institute for the COVID-19 Genomics UK (COG-UK) Consortium | Health Services Laboratories and Alex Alderton, Roberto Amato, Jeffrey Barrett, Sonia Goncalves, Ewan Harrison, David K. Jackson, Ian Johnston, Dominic Kwiatkowski, Cordelia Langford, John Sillitoe on behalf of the Wellcome Sanger Institute COVID-19 Surveillance Team |
| EPI_ISL_2511142, EPI_ISL_2511143 | Lighthouse Lab in Glasgow | Wellcome Sanger Institute for the COVID-19 Genomics UK (COG-UK) Consortium | Harper VanSteenhouse, Yumi Kasai, David Gray, Carol Clugston, Anna Dominiczak and Alex Alderton, Roberto Amato, Jeffrey Barrett, Sonia Goncalves, Ewan Harrison, David K. Jackson, Ian Johnston, Dominic Kwiatkowski, Cordelia Langford, John Sillitoe on behalf of the Wellcome Sanger Institute COVID-19 Surveillance Team |
| EPI_ISL_2511144 | Health Services Laboratories | Wellcome Sanger Institute for the COVID-19 Genomics UK (COG-UK) Consortium | Health Services Laboratories and Alex Alderton, Roberto Amato, Jeffrey Barrett, Sonia Goncalves, Ewan Harrison, David K. Jackson, Ian Johnston, Dominic Kwiatkowski, Cordelia Langford, John Sillitoe on behalf of the Wellcome Sanger Institute COVID-19 Surveillance Team |
| EPI_ISL_2511145 | Lighthouse Lab in Glasgow | Wellcome Sanger Institute for the COVID-19 Genomics UK (COG-UK) Consortium | Harper VanSteenhouse, Yumi Kasai, David Gray, Carol Clugston, Anna Dominiczak and Alex Alderton, Roberto Amato, Jeffrey Barrett, Sonia Goncalves, Ewan Harrison, David K. Jackson, Ian Johnston, Dominic Kwiatkowski, Cordelia Langford, John Sillitoe on behalf of the Wellcome Sanger Institute COVID-19 Surveillance Team |
| EPI_ISL_2511146 | Health Services Laboratories | Wellcome Sanger Institute for the COVID-19 Genomics UK (COG-UK) Consortium | Health Services Laboratories and Alex Alderton, Roberto Amato, Jeffrey Barrett, Sonia Goncalves, Ewan Harrison, David K. Jackson, Ian Johnston, Dominic Kwiatkowski, Cordelia Langford, John Sillitoe on behalf of the Wellcome Sanger Institute COVID-19 Surveillance Team |
| EPI_ISL_2511147, EPI_ISL_2511148 | Lighthouse Lab in Glasgow | Wellcome Sanger Institute for the COVID-19 Genomics UK (COG-UK) Consortium | Harper VanSteenhouse, Yumi Kasai, David Gray, Carol Clugston, Anna Dominiczak and Alex Alderton, Roberto Amato, Jeffrey Barrett, Sonia Goncalves, Ewan Harrison, David K. Jackson, Ian Johnston, Dominic Kwiatkowski, Cordelia Langford, John Sillitoe on behalf of the Wellcome Sanger Institute COVID-19 Surveillance Team |
| EPI_ISL_2511149 | Health Services Laboratories | Wellcome Sanger Institute for the COVID-19 Genomics UK (COG-UK) Consortium | Health Services Laboratories and Alex Alderton, Roberto Amato, Jeffrey Barrett, Sonia Goncalves, Ewan Harrison, David K. Jackson, Ian Johnston, Dominic Kwiatkowski, Cordelia Langford, John Sillitoe on behalf of the Wellcome Sanger Institute COVID-19 Surveillance Team |
| EPI_ISL_2511150, EPI_ISL_2511151, EPI_ISL_2511152, EPI_ISL_2511153 | Lighthouse Lab in Glasgow | Wellcome Sanger Institute for the COVID-19 Genomics UK (COG-UK) Consortium | Harper VanSteenhouse, Yumi Kasai, David Gray, Carol Clugston, Anna Dominiczak and Alex Alderton, Roberto Amato, Jeffrey Barrett, Sonia Goncalves, Ewan Harrison, David K. Jackson, Ian Johnston, Dominic Kwiatkowski, Cordelia Langford, John Sillitoe on behalf of the Wellcome Sanger Institute COVID-19 Surveillance Team |
| EPI_ISL_2511154 | Health Services Laboratories | Wellcome Sanger Institute for the COVID-19 Genomics UK (COG-UK) Consortium | Health Services Laboratories and Alex Alderton, Roberto Amato, Jeffrey Barrett, Sonia Goncalves, Ewan Harrison, David K. Jackson, Ian Johnston, Dominic Kwiatkowski, Cordelia Langford, John Sillitoe on behalf of the Wellcome Sanger Institute COVID-19 Surveillance Team |
| EPI_ISL_2511155, EPI_ISL_2511156, EPI_ISL_2511157, EPI_ISL_2511158 | Lighthouse Lab in Glasgow | Wellcome Sanger Institute for the COVID-19 Genomics UK (COG-UK) Consortium | Harper VanSteenhouse, Yumi Kasai, David Gray, Carol Clugston, Anna Dominiczak and Alex Alderton, Roberto Amato, Jeffrey Barrett, Sonia Goncalves, Ewan Harrison, David K. Jackson, Ian Johnston, Dominic Kwiatkowski, Cordelia Langford, John Sillitoe on behalf of the Wellcome Sanger Institute COVID-19 Surveillance Team |
| EPI_ISL_2511159, EPI_ISL_2511160, EPI_ISL_2511162, EPI_ISL_2511163 | Health Services Laboratories | Wellcome Sanger Institute for the COVID-19 Genomics UK (COG-UK) Consortium | Health Services Laboratories and Alex Alderton, Roberto Amato, Jeffrey Barrett, Sonia Goncalves, Ewan Harrison, David K. Jackson, Ian Johnston, Dominic Kwiatkowski, Cordelia Langford, John Sillitoe on behalf of the Wellcome Sanger Institute COVID-19 Surveillance Team |
| EPI_ISL_2511164, EPI_ISL_2511165 | Lighthouse Lab in Glasgow | Wellcome Sanger Institute for the COVID-19 Genomics UK (COG-UK) Consortium | Harper VanSteenhouse, Yumi Kasai, David Gray, Carol Clugston, Anna Dominiczak and Alex Alderton, Roberto Amato, Jeffrey Barrett, Sonia Goncalves, Ewan Harrison, David K. Jackson, Ian Johnston, Dominic Kwiatkowski, Cordelia Langford, John Sillitoe on behalf of the Wellcome Sanger Institute COVID-19 Surveillance Team |
| EPI_ISL_2511167 | Health Services Laboratories | Wellcome Sanger Institute for the COVID-19 Genomics UK (COG-UK) Consortium | Health Services Laboratories and Alex Alderton, Roberto Amato, Jeffrey Barrett, Sonia Goncalves, Ewan Harrison, David K. Jackson, Ian Johnston, Dominic Kwiatkowski, Cordelia Langford, John Sillitoe on behalf of the Wellcome Sanger Institute COVID-19 Surveillance Team |
| EPI_ISL_2511168 | Lighthouse Lab in Glasgow | Wellcome Sanger Institute for the COVID-19 Genomics UK (COG-UK) Consortium | Harper VanSteenhouse, Yumi Kasai, David Gray, Carol Clugston, Anna Dominiczak and Alex Alderton, Roberto Amato, Jeffrey Barrett, Sonia Goncalves, Ewan Harrison, David K. Jackson, Ian Johnston, Dominic Kwiatkowski, Cordelia Langford, John Sillitoe on behalf of the Wellcome Sanger Institute COVID-19 Surveillance Team |
| EPI_ISL_2511169 | Health Services Laboratories | Wellcome Sanger Institute for the COVID-19 Genomics UK (COG-UK) Consortium | Health Services Laboratories and Alex Alderton, Roberto Amato, Jeffrey Barrett, Sonia Goncalves, Ewan Harrison, David K. Jackson, Ian Johnston, Dominic Kwiatkowski, Cordelia Langford, John Sillitoe on behalf of the Wellcome Sanger Institute COVID-19 Surveillance Team |
| EPI_ISL_2511170 | Lighthouse Lab in Glasgow | Wellcome Sanger Institute for the COVID-19 Genomics UK (COG-UK) Consortium | Harper VanSteenhouse, Yumi Kasai, David Gray, Carol Clugston, Anna Dominiczak and Alex Alderton, Roberto Amato, Jeffrey Barrett, Sonia Goncalves, Ewan Harrison, David K. Jackson, Ian Johnston, Dominic Kwiatkowski, Cordelia Langford, John Sillitoe on behalf of the Wellcome Sanger Institute COVID-19 Surveillance Team |
| EPI_ISL_2511171 | Health Services Laboratories | Wellcome Sanger Institute for the COVID-19 Genomics UK (COG-UK) Consortium | Health Services Laboratories and Alex Alderton, Roberto Amato, Jeffrey Barrett, Sonia Goncalves, Ewan Harrison, David K. Jackson, Ian Johnston, Dominic Kwiatkowski, Cordelia Langford, John Sillitoe on behalf of the Wellcome Sanger Institute COVID-19 Surveillance Team |
| EPI_ISL_2511172, EPI_ISL_2511173 | Lighthouse Lab in Glasgow | Wellcome Sanger Institute for the COVID-19 Genomics UK (COG-UK) Consortium | Harper VanSteenhouse, Yumi Kasai, David Gray, Carol Clugston, Anna Dominiczak and Alex Alderton, Roberto Amato, Jeffrey Barrett, Sonia Goncalves, Ewan Harrison, David K. Jackson, Ian Johnston, Dominic Kwiatkowski, Cordelia Langford, John Sillitoe on behalf of the Wellcome Sanger Institute COVID-19 Surveillance Team |
| EPI_ISL_2511174 | Health Services Laboratories | Wellcome Sanger Institute for the COVID-19 Genomics UK (COG-UK) Consortium | Health Services Laboratories and Alex Alderton, Roberto Amato, Jeffrey Barrett, Sonia Goncalves, Ewan Harrison, David K. Jackson, Ian Johnston, Dominic Kwiatkowski, Cordelia Langford, John Sillitoe on behalf of the Wellcome Sanger Institute COVID-19 Surveillance Team |
| EPI_ISL_2511175, EPI_ISL_2511176 | Lighthouse Lab in Glasgow | Wellcome Sanger Institute for the COVID-19 Genomics UK (COG-UK) Consortium | Harper VanSteenhouse, Yumi Kasai, David Gray, Carol Clugston, Anna Dominiczak and Alex Alderton, Roberto Amato, Jeffrey Barrett, Sonia Goncalves, Ewan Harrison, David K. Jackson, Ian Johnston, Dominic Kwiatkowski, Cordelia Langford, John Sillitoe on behalf of the Wellcome Sanger Institute COVID-19 Surveillance Team |
| EPI_ISL_2511177 | Health Services Laboratories | Wellcome Sanger Institute for the COVID-19 Genomics UK (COG-UK) Consortium | Health Services Laboratories and Alex Alderton, Roberto Amato, Jeffrey Barrett, Sonia Goncalves, Ewan Harrison, David K. Jackson, Ian Johnston, Dominic Kwiatkowski, Cordelia Langford, John Sillitoe on behalf of the Wellcome Sanger Institute COVID-19 Surveillance Team |
| EPI_ISL_2511178 | Lighthouse Lab in Glasgow | Wellcome Sanger Institute for the COVID-19 Genomics UK (COG-UK) Consortium | Harper VanSteenhouse, Yumi Kasai, David Gray, Carol Clugston, Anna Dominiczak and Alex Alderton, Roberto Amato, Jeffrey Barrett, Sonia Goncalves, Ewan Harrison, David K. Jackson, Ian Johnston, Dominic Kwiatkowski, Cordelia Langford, John Sillitoe on behalf of the Wellcome Sanger Institute COVID-19 Surveillance Team |
| EPI_ISL_2511179, EPI_ISL_2511180, EPI_ISL_2511182 | Health Services Laboratories | Wellcome Sanger Institute for the COVID-19 Genomics UK (COG-UK) Consortium | Health Services Laboratories and Alex Alderton, Roberto Amato, Jeffrey Barrett, Sonia Goncalves, Ewan Harrison, David K. Jackson, Ian Johnston, Dominic Kwiatkowski, Cordelia Langford, John Sillitoe on behalf of the Wellcome Sanger Institute COVID-19 Surveillance Team |
| EPI_ISL_2511183, EPI_ISL_2511184, EPI_ISL_2511185, EPI_ISL_2511186, EPI_ISL_2511188 | Lighthouse Lab in Glasgow | Wellcome Sanger Institute for the COVID-19 Genomics UK (COG-UK) Consortium | Harper VanSteenhouse, Yumi Kasai, David Gray, Carol Clugston, Anna Dominiczak and Alex Alderton, Roberto Amato, Jeffrey Barrett, Sonia Goncalves, Ewan Harrison, David K. Jackson, Ian Johnston, Dominic Kwiatkowski, Cordelia Langford, John Sillitoe on behalf of the Wellcome Sanger Institute COVID-19 Surveillance Team |
| EPI_ISL_2511189, EPI_ISL_2511190, EPI_ISL_2511191, EPI_ISL_2511192, EPI_ISL_2511193, EPI_ISL_2511194 | Health Services Laboratories | Wellcome Sanger Institute for the COVID-19 Genomics UK (COG-UK) Consortium | Health Services Laboratories and Alex Alderton, Roberto Amato, Jeffrey Barrett, Sonia Goncalves, Ewan Harrison, David K. Jackson, Ian Johnston, Dominic Kwiatkowski, Cordelia Langford, John Sillitoe on behalf of the Wellcome Sanger Institute COVID-19 Surveillance Team |
| EPI_ISL_2511195, EPI_ISL_2511196, EPI_ISL_2511197, EPI_ISL_2511198, EPI_ISL_2511199, EPI_ISL_2511200 | Lighthouse Lab in Glasgow | Wellcome Sanger Institute for the COVID-19 Genomics UK (COG-UK) Consortium | Harper VanSteenhouse, Yumi Kasai, David Gray, Carol Clugston, Anna Dominiczak and Alex Alderton, Roberto Amato, Jeffrey Barrett, Sonia Goncalves, Ewan Harrison, David K. Jackson, Ian Johnston, Dominic Kwiatkowski, Cordelia Langford, John Sillitoe on behalf of the Wellcome Sanger Institute COVID-19 Surveillance Team |
| EPI_ISL_2511201 | Health Services Laboratories | Wellcome Sanger Institute for the COVID-19 Genomics UK (COG-UK) Consortium | Health Services Laboratories and Alex Alderton, Roberto Amato, Jeffrey Barrett, Sonia Goncalves, Ewan Harrison, David K. Jackson, Ian Johnston, Dominic Kwiatkowski, Cordelia Langford, John Sillitoe on behalf of the Wellcome Sanger Institute COVID-19 Surveillance Team |
| EPI_ISL_2511202, EPI_ISL_2511203, EPI_ISL_2511204 | Lighthouse Lab in Glasgow | Wellcome Sanger Institute for the COVID-19 Genomics UK (COG-UK) Consortium | Harper VanSteenhouse, Yumi Kasai, David Gray, Carol Clugston, Anna Dominiczak and Alex Alderton, Roberto Amato, Jeffrey Barrett, Sonia Goncalves, Ewan Harrison, David K. Jackson, Ian Johnston, Dominic Kwiatkowski, Cordelia Langford, John Sillitoe on behalf of the Wellcome Sanger Institute COVID-19 Surveillance Team |
| EPI_ISL_2511205 | Health Services Laboratories | Wellcome Sanger Institute for the COVID-19 Genomics UK (COG-UK) Consortium | Health Services Laboratories and Alex Alderton, Roberto Amato, Jeffrey Barrett, Sonia Goncalves, Ewan Harrison, David K. Jackson, Ian Johnston, Dominic Kwiatkowski, Cordelia Langford, John Sillitoe on behalf of the Wellcome Sanger Institute COVID-19 Surveillance Team |
| EPI_ISL_2511206, EPI_ISL_2511207, EPI_ISL_2511210, EPI_ISL_2511211 | Lighthouse Lab in Glasgow | Wellcome Sanger Institute for the COVID-19 Genomics UK (COG-UK) Consortium | Harper VanSteenhouse, Yumi Kasai, David Gray, Carol Clugston, Anna Dominiczak and Alex Alderton, Roberto Amato, Jeffrey Barrett, Sonia Goncalves, Ewan Harrison, David K. Jackson, Ian Johnston, Dominic Kwiatkowski, Cordelia Langford, John Sillitoe on behalf of the Wellcome Sanger Institute COVID-19 Surveillance Team |
| EPI_ISL_2511212, EPI_ISL_2511213 | Health Services Laboratories | Wellcome Sanger Institute for the COVID-19 Genomics UK (COG-UK) Consortium | Health Services Laboratories and Alex Alderton, Roberto Amato, Jeffrey Barrett, Sonia Goncalves, Ewan Harrison, David K. Jackson, Ian Johnston, Dominic Kwiatkowski, Cordelia Langford, John Sillitoe on behalf of the Wellcome Sanger Institute COVID-19 Surveillance Team |
| EPI_ISL_2511214, EPI_ISL_2511215, EPI_ISL_2511216, EPI_ISL_2511217 | Lighthouse Lab in Glasgow | Wellcome Sanger Institute for the COVID-19 Genomics UK (COG-UK) Consortium | Harper VanSteenhouse, Yumi Kasai, David Gray, Carol Clugston, Anna Dominiczak and Alex Alderton, Roberto Amato, Jeffrey Barrett, Sonia Goncalves, Ewan Harrison, David K. Jackson, Ian Johnston, Dominic Kwiatkowski, Cordelia Langford, John Sillitoe on behalf of the Wellcome Sanger Institute COVID-19 Surveillance Team |
| EPI_ISL_2511218 | Health Services Laboratories | Wellcome Sanger Institute for the COVID-19 Genomics UK (COG-UK) Consortium | Health Services Laboratories and Alex Alderton, Roberto Amato, Jeffrey Barrett, Sonia Goncalves, Ewan Harrison, David K. Jackson, Ian Johnston, Dominic Kwiatkowski, Cordelia Langford, John Sillitoe on behalf of the Wellcome Sanger Institute COVID-19 Surveillance Team |

| | | | |
|---|---|---|---|
| EPI_ISL_2511219 | Lighthouse Lab in Glasgow | Wellcome Sanger Institute for the COVID-19 Genomics UK (COG-UK) Consortium | Harper VanSteenhouse, Yumi Kasai, David Gray, Carol Clugston, Anna Dominiczak and Alex Alderton, Roberto Amato, Jeffrey Barrett, Sonia Goncalves, Ewan Harrison, David K. Jackson, Ian Johnston, Dominic Kwiatkowski, Cordelia Langford, John Sillitoe on behalf of the Wellcome Sanger Institute COVID-19 Surveillance Team |
| EPI_ISL_2511220, EPI_ISL_2511222 | Health Services Laboratories | Wellcome Sanger Institute for the COVID-19 Genomics UK (COG-UK) Consortium | Health Services Laboratories and Alex Alderton, Roberto Amato, Jeffrey Barrett, Sonia Goncalves, Ewan Harrison, David K. Jackson, Ian Johnston, Dominic Kwiatkowski, Cordelia Langford, John Sillitoe on behalf of the Wellcome Sanger Institute COVID-19 Surveillance Team |
| EPI_ISL_2511223, EPI_ISL_2511224, EPI_ISL_2511225, EPI_ISL_2511226, EPI_ISL_2511227, EPI_ISL_2511228, EPI_ISL_2511229, EPI_ISL_2511231, EPI_ISL_2511233, EPI_ISL_2511235 | Lighthouse Lab in Glasgow | Wellcome Sanger Institute for the COVID-19 Genomics UK (COG-UK) Consortium | Harper VanSteenhouse, Yumi Kasai, David Gray, Carol Clugston, Anna Dominiczak and Alex Alderton, Roberto Amato, Jeffrey Barrett, Sonia Goncalves, Ewan Harrison, David K. Jackson, Ian Johnston, Dominic Kwiatkowski, Cordelia Langford, John Sillitoe on behalf of the Wellcome Sanger Institute COVID-19 Surveillance Team |
| EPI_ISL_2511236 | Health Services Laboratories | Wellcome Sanger Institute for the COVID-19 Genomics UK (COG-UK) Consortium | Health Services Laboratories and Alex Alderton, Roberto Amato, Jeffrey Barrett, Sonia Goncalves, Ewan Harrison, David K. Jackson, Ian Johnston, Dominic Kwiatkowski, Cordelia Langford, John Sillitoe on behalf of the Wellcome Sanger Institute COVID-19 Surveillance Team |
| EPI_ISL_2511237, EPI_ISL_2511238, EPI_ISL_2511239, EPI_ISL_2511240 | Lighthouse Lab in Glasgow | Wellcome Sanger Institute for the COVID-19 Genomics UK (COG-UK) Consortium | Harper VanSteenhouse, Yumi Kasai, David Gray, Carol Clugston, Anna Dominiczak and Alex Alderton, Roberto Amato, Jeffrey Barrett, Sonia Goncalves, Ewan Harrison, David K. Jackson, Ian Johnston, Dominic Kwiatkowski, Cordelia Langford, John Sillitoe on behalf of the Wellcome Sanger Institute COVID-19 Surveillance Team |
| EPI_ISL_2511241, EPI_ISL_2511242, EPI_ISL_2511243 | Health Services Laboratories | Wellcome Sanger Institute for the COVID-19 Genomics UK (COG-UK) Consortium | Health Services Laboratories and Alex Alderton, Roberto Amato, Jeffrey Barrett, Sonia Goncalves, Ewan Harrison, David K. Jackson, Ian Johnston, Dominic Kwiatkowski, Cordelia Langford, John Sillitoe on behalf of the Wellcome Sanger Institute COVID-19 Surveillance Team |
| EPI_ISL_2511244 | Lighthouse Lab in Glasgow | Wellcome Sanger Institute for the COVID-19 Genomics UK (COG-UK) Consortium | Harper VanSteenhouse, Yumi Kasai, David Gray, Carol Clugston, Anna Dominiczak and Alex Alderton, Roberto Amato, Jeffrey Barrett, Sonia Goncalves, Ewan Harrison, David K. Jackson, Ian Johnston, Dominic Kwiatkowski, Cordelia Langford, John Sillitoe on behalf of the Wellcome Sanger Institute COVID-19 Surveillance Team |
| EPI_ISL_2511245, EPI_ISL_2511246 | Health Services Laboratories | Wellcome Sanger Institute for the COVID-19 Genomics UK (COG-UK) Consortium | Health Services Laboratories and Alex Alderton, Roberto Amato, Jeffrey Barrett, Sonia Goncalves, Ewan Harrison, David K. Jackson, Ian Johnston, Dominic Kwiatkowski, Cordelia Langford, John Sillitoe on behalf of the Wellcome Sanger Institute COVID-19 Surveillance Team |
| EPI_ISL_2511248 | Lighthouse Lab in Glasgow | Wellcome Sanger Institute for the COVID-19 Genomics UK (COG-UK) Consortium | Harper VanSteenhouse, Yumi Kasai, David Gray, Carol Clugston, Anna Dominiczak and Alex Alderton, Roberto Amato, Jeffrey Barrett, Sonia Goncalves, Ewan Harrison, David K. Jackson, Ian Johnston, Dominic Kwiatkowski, Cordelia Langford, John Sillitoe on behalf of the Wellcome Sanger Institute COVID-19 Surveillance Team |
| EPI_ISL_2511249 | Health Services Laboratories | Wellcome Sanger Institute for the COVID-19 Genomics UK (COG-UK) Consortium | Health Services Laboratories and Alex Alderton, Roberto Amato, Jeffrey Barrett, Sonia Goncalves, Ewan Harrison, David K. Jackson, Ian Johnston, Dominic Kwiatkowski, Cordelia Langford, John Sillitoe on behalf of the Wellcome Sanger Institute COVID-19 Surveillance Team |
| EPI_ISL_2511251, EPI_ISL_2511252, EPI_ISL_2511254, EPI_ISL_2511255 | Lighthouse Lab in Glasgow | Wellcome Sanger Institute for the COVID-19 Genomics UK (COG-UK) Consortium | Harper VanSteenhouse, Yumi Kasai, David Gray, Carol Clugston, Anna Dominiczak and Alex Alderton, Roberto Amato, Jeffrey Barrett, Sonia Goncalves, Ewan Harrison, David K. Jackson, Ian Johnston, Dominic Kwiatkowski, Cordelia Langford, John Sillitoe on behalf of the Wellcome Sanger Institute COVID-19 Surveillance Team |
| EPI_ISL_2511256 | Health Services Laboratories | Wellcome Sanger Institute for the COVID-19 Genomics UK (COG-UK) Consortium | Health Services Laboratories and Alex Alderton, Roberto Amato, Jeffrey Barrett, Sonia Goncalves, Ewan Harrison, David K. Jackson, Ian Johnston, Dominic Kwiatkowski, Cordelia Langford, John Sillitoe on behalf of the Wellcome Sanger Institute COVID-19 Surveillance Team |
| EPI_ISL_2511257, EPI_ISL_2511258, EPI_ISL_2511259 | Lighthouse Lab in Glasgow | Wellcome Sanger Institute for the COVID-19 Genomics UK (COG-UK) Consortium | Harper VanSteenhouse, Yumi Kasai, David Gray, Carol Clugston, Anna Dominiczak and Alex Alderton, Roberto Amato, Jeffrey Barrett, Sonia Goncalves, Ewan Harrison, David K. Jackson, Ian Johnston, Dominic Kwiatkowski, Cordelia Langford, John Sillitoe on behalf of the Wellcome Sanger Institute COVID-19 Surveillance Team |
| EPI_ISL_2511260 | Health Services Laboratories | Wellcome Sanger Institute for the COVID-19 Genomics UK (COG-UK) Consortium | Health Services Laboratories and Alex Alderton, Roberto Amato, Jeffrey Barrett, Sonia Goncalves, Ewan Harrison, David K. Jackson, Ian Johnston, Dominic Kwiatkowski, Cordelia Langford, John Sillitoe on behalf of the Wellcome Sanger Institute COVID-19 Surveillance Team |
| EPI_ISL_2511261, EPI_ISL_2511262, EPI_ISL_2511263, EPI_ISL_2511264, EPI_ISL_2511265, EPI_ISL_2511266, EPI_ISL_2511268 | Lighthouse Lab in Glasgow | Wellcome Sanger Institute for the COVID-19 Genomics UK (COG-UK) Consortium | Harper VanSteenhouse, Yumi Kasai, David Gray, Carol Clugston, Anna Dominiczak and Alex Alderton, Roberto Amato, Jeffrey Barrett, Sonia Goncalves, Ewan Harrison, David K. Jackson, Ian Johnston, Dominic Kwiatkowski, Cordelia Langford, John Sillitoe on behalf of the Wellcome Sanger Institute COVID-19 Surveillance Team |
| EPI_ISL_2511269 | Health Services Laboratories | Wellcome Sanger Institute for the COVID-19 Genomics UK (COG-UK) Consortium | Health Services Laboratories and Alex Alderton, Roberto Amato, Jeffrey Barrett, Sonia Goncalves, Ewan Harrison, David K. Jackson, Ian Johnston, Dominic Kwiatkowski, Cordelia Langford, John Sillitoe on behalf of the Wellcome Sanger Institute COVID-19 Surveillance Team |
| EPI_ISL_2511271, EPI_ISL_2511273 | Lighthouse Lab in Glasgow | Wellcome Sanger Institute for the COVID-19 Genomics UK (COG-UK) Consortium | Harper VanSteenhouse, Yumi Kasai, David Gray, Carol Clugston, Anna Dominiczak and Alex Alderton, Roberto Amato, Jeffrey Barrett, Sonia Goncalves, Ewan Harrison, David K. Jackson, Ian Johnston, Dominic Kwiatkowski, Cordelia Langford, John Sillitoe on behalf of the Wellcome Sanger Institute COVID-19 Surveillance Team |
| EPI_ISL_2511274, EPI_ISL_2511275 | Health Services Laboratories | Wellcome Sanger Institute for the COVID-19 Genomics UK (COG-UK) Consortium | Health Services Laboratories and Alex Alderton, Roberto Amato, Jeffrey Barrett, Sonia Goncalves, Ewan Harrison, David K. Jackson, Ian Johnston, Dominic Kwiatkowski, Cordelia Langford, John Sillitoe on behalf of the Wellcome Sanger Institute COVID-19 Surveillance Team |
| EPI_ISL_2511276, EPI_ISL_2511277, EPI_ISL_2511278, EPI_ISL_2511279 | Lighthouse Lab in Glasgow | Wellcome Sanger Institute for the COVID-19 Genomics UK (COG-UK) Consortium | Harper VanSteenhouse, Yumi Kasai, David Gray, Carol Clugston, Anna Dominiczak and Alex Alderton, Roberto Amato, Jeffrey Barrett, Sonia Goncalves, Ewan Harrison, David K. Jackson, Ian Johnston, Dominic Kwiatkowski, Cordelia Langford, John Sillitoe on behalf of the Wellcome Sanger Institute COVID-19 Surveillance Team |
| EPI_ISL_2511280, EPI_ISL_2511281, EPI_ISL_2511282 | Health Services Laboratories | Wellcome Sanger Institute for the COVID-19 Genomics UK (COG-UK) Consortium | Health Services Laboratories and Alex Alderton, Roberto Amato, Jeffrey Barrett, Sonia Goncalves, Ewan Harrison, David K. Jackson, Ian Johnston, Dominic Kwiatkowski, Cordelia Langford, John Sillitoe on behalf of the Wellcome Sanger Institute COVID-19 Surveillance Team |
| EPI_ISL_2511283, EPI_ISL_2511284, EPI_ISL_2511285, EPI_ISL_2511286, EPI_ISL_2511287, EPI_ISL_2511288, EPI_ISL_2511289 | Lighthouse Lab in Glasgow | Wellcome Sanger Institute for the COVID-19 Genomics UK (COG-UK) Consortium | Harper VanSteenhouse, Yumi Kasai, David Gray, Carol Clugston, Anna Dominiczak and Alex Alderton, Roberto Amato, Jeffrey Barrett, Sonia Goncalves, Ewan Harrison, David K. Jackson, Ian Johnston, Dominic Kwiatkowski, Cordelia Langford, John Sillitoe on behalf of the Wellcome Sanger Institute COVID-19 Surveillance Team |
| EPI_ISL_2511290 | Health Services Laboratories | Wellcome Sanger Institute for the COVID-19 Genomics UK (COG-UK) Consortium | Health Services Laboratories and Alex Alderton, Roberto Amato, Jeffrey Barrett, Sonia Goncalves, Ewan Harrison, David K. Jackson, Ian Johnston, Dominic Kwiatkowski, Cordelia Langford, John Sillitoe on behalf of the Wellcome Sanger Institute COVID-19 Surveillance Team |
| EPI_ISL_2511292, EPI_ISL_2511293 | Lighthouse Lab in Glasgow | Wellcome Sanger Institute for the COVID-19 Genomics UK (COG-UK) Consortium | Harper VanSteenhouse, Yumi Kasai, David Gray, Carol Clugston, Anna Dominiczak and Alex Alderton, Roberto Amato, Jeffrey Barrett, Sonia Goncalves, Ewan Harrison, David K. Jackson, Ian Johnston, Dominic Kwiatkowski, Cordelia Langford, John Sillitoe on behalf of the Wellcome Sanger Institute COVID-19 Surveillance Team |
| EPI_ISL_2511294, EPI_ISL_2511295, EPI_ISL_2511296, EPI_ISL_2511297 | Health Services Laboratories | Wellcome Sanger Institute for the COVID-19 Genomics UK (COG-UK) Consortium | Health Services Laboratories and Alex Alderton, Roberto Amato, Jeffrey Barrett, Sonia Goncalves, Ewan Harrison, David K. Jackson, Ian Johnston, Dominic Kwiatkowski, Cordelia Langford, John Sillitoe on behalf of the Wellcome Sanger Institute COVID-19 Surveillance Team |
| EPI_ISL_2511298, EPI_ISL_2511299, EPI_ISL_2511300, EPI_ISL_2511301, EPI_ISL_2511302, EPI_ISL_2511303, EPI_ISL_2511304, EPI_ISL_2511305 | Lighthouse Lab in Glasgow | Wellcome Sanger Institute for the COVID-19 Genomics UK (COG-UK) Consortium | Harper VanSteenhouse, Yumi Kasai, David Gray, Carol Clugston, Anna Dominiczak and Alex Alderton, Roberto Amato, Jeffrey Barrett, Sonia Goncalves, Ewan Harrison, David K. Jackson, Ian Johnston, Dominic Kwiatkowski, Cordelia Langford, John Sillitoe on behalf of the Wellcome Sanger Institute COVID-19 Surveillance Team |
| EPI_ISL_2511306 | Health Services Laboratories | Wellcome Sanger Institute for the COVID-19 Genomics UK (COG-UK) Consortium | Health Services Laboratories and Alex Alderton, Roberto Amato, Jeffrey Barrett, Sonia Goncalves, Ewan Harrison, David K. Jackson, Ian Johnston, Dominic Kwiatkowski, Cordelia Langford, John Sillitoe on behalf of the Wellcome Sanger Institute COVID-19 Surveillance Team |
| EPI_ISL_2511307 | Lighthouse Lab in Glasgow | Wellcome Sanger Institute for the COVID-19 Genomics UK (COG-UK) Consortium | Harper VanSteenhouse, Yumi Kasai, David Gray, Carol Clugston, Anna Dominiczak and Alex Alderton, Roberto Amato, Jeffrey Barrett, Sonia Goncalves, Ewan Harrison, David K. Jackson, Ian Johnston, Dominic Kwiatkowski, Cordelia Langford, John Sillitoe on behalf of the Wellcome Sanger Institute COVID-19 Surveillance Team |
| EPI_ISL_2511308 | Health Services Laboratories | Wellcome Sanger Institute for the COVID-19 Genomics UK (COG-UK) Consortium | Health Services Laboratories and Alex Alderton, Roberto Amato, Jeffrey Barrett, Sonia Goncalves, Ewan Harrison, David K. Jackson, Ian Johnston, Dominic Kwiatkowski, Cordelia Langford, John Sillitoe on behalf of the Wellcome Sanger Institute COVID-19 Surveillance Team |
| EPI_ISL_2511309 | Lighthouse Lab in Glasgow | Wellcome Sanger Institute for the COVID-19 Genomics UK (COG-UK) Consortium | Harper VanSteenhouse, Yumi Kasai, David Gray, Carol Clugston, Anna Dominiczak and Alex Alderton, Roberto Amato, Jeffrey Barrett, Sonia Goncalves, Ewan Harrison, David K. Jackson, Ian Johnston, Dominic Kwiatkowski, Cordelia Langford, John Sillitoe on behalf of the Wellcome Sanger Institute COVID-19 Surveillance Team |
| EPI_ISL_2511310, EPI_ISL_2511311 | Health Services Laboratories | Wellcome Sanger Institute for the COVID-19 Genomics UK (COG-UK) Consortium | Health Services Laboratories and Alex Alderton, Roberto Amato, Jeffrey Barrett, Sonia Goncalves, Ewan Harrison, David K. Jackson, Ian Johnston, Dominic Kwiatkowski, Cordelia Langford, John Sillitoe on behalf of the Wellcome Sanger Institute COVID-19 Surveillance Team |
| EPI_ISL_2511312, EPI_ISL_2511313, | Lighthouse Lab in Glasgow | Wellcome Sanger Institute for the COVID-19 Genomics UK (COG-UK) Consortium | Harper VanSteenhouse, Yumi Kasai, David Gray, Carol Clugston, Anna Dominiczak and Alex Alderton, Roberto Amato, Jeffrey Barrett, Sonia Goncalves, Ewan Harrison, David K. Jackson, Ian Johnston, Dominic Kwiatkowski, Cordelia Langford, John Sillitoe on behalf of the Wellcome Sanger Institute COVID-19 Surveillance Team |

| | | | |
|---|---|---|---|
| EPI_ISL_2511314, EPI_ISL_2511316, EPI_ISL_2511318 | | | |
| EPI_ISL_2511319 | Health Services Laboratories | Wellcome Sanger Institute for the COVID-19 Genomics UK (COG-UK) Consortium | Health Services Laboratories and Alex Alderton, Roberto Amato, Jeffrey Barrett, Sonia Goncalves, Ewan Harrison, David K. Jackson, Ian Johnston, Dominic Kwiatkowski, Cordelia Langford, John Sillitoe on behalf of the Wellcome Sanger Institute COVID-19 Surveillance Team |
| EPI_ISL_2511320 | Lighthouse Lab in Glasgow | Wellcome Sanger Institute for the COVID-19 Genomics UK (COG-UK) Consortium | Harper VanSteenhouse, Yumi Kasai, David Gray, Carol Clugston, Anna Dominiczak and Alex Alderton, Roberto Amato, Jeffrey Barrett, Sonia Goncalves, Ewan Harrison, David K. Jackson, Ian Johnston, Dominic Kwiatkowski, Cordelia Langford, John Sillitoe on behalf of the Wellcome Sanger Institute COVID-19 Surveillance Team |
| EPI_ISL_2511321 | Health Services Laboratories | Wellcome Sanger Institute for the COVID-19 Genomics UK (COG-UK) Consortium | Health Services Laboratories and Alex Alderton, Roberto Amato, Jeffrey Barrett, Sonia Goncalves, Ewan Harrison, David K. Jackson, Ian Johnston, Dominic Kwiatkowski, Cordelia Langford, John Sillitoe on behalf of the Wellcome Sanger Institute COVID-19 Surveillance Team |
| EPI_ISL_2511322 | Lighthouse Lab in Glasgow | Wellcome Sanger Institute for the COVID-19 Genomics UK (COG-UK) Consortium | Harper VanSteenhouse, Yumi Kasai, David Gray, Carol Clugston, Anna Dominiczak and Alex Alderton, Roberto Amato, Jeffrey Barrett, Sonia Goncalves, Ewan Harrison, David K. Jackson, Ian Johnston, Dominic Kwiatkowski, Cordelia Langford, John Sillitoe on behalf of the Wellcome Sanger Institute COVID-19 Surveillance Team |
| EPI_ISL_2511324 | Health Services Laboratories | Wellcome Sanger Institute for the COVID-19 Genomics UK (COG-UK) Consortium | Health Services Laboratories and Alex Alderton, Roberto Amato, Jeffrey Barrett, Sonia Goncalves, Ewan Harrison, David K. Jackson, Ian Johnston, Dominic Kwiatkowski, Cordelia Langford, John Sillitoe on behalf of the Wellcome Sanger Institute COVID-19 Surveillance Team |
| EPI_ISL_2511325 | Lighthouse Lab in Glasgow | Wellcome Sanger Institute for the COVID-19 Genomics UK (COG-UK) Consortium | Harper VanSteenhouse, Yumi Kasai, David Gray, Carol Clugston, Anna Dominiczak and Alex Alderton, Roberto Amato, Jeffrey Barrett, Sonia Goncalves, Ewan Harrison, David K. Jackson, Ian Johnston, Dominic Kwiatkowski, Cordelia Langford, John Sillitoe on behalf of the Wellcome Sanger Institute COVID-19 Surveillance Team |
| EPI_ISL_2511326 | Health Services Laboratories | Wellcome Sanger Institute for the COVID-19 Genomics UK (COG-UK) Consortium | Health Services Laboratories and Alex Alderton, Roberto Amato, Jeffrey Barrett, Sonia Goncalves, Ewan Harrison, David K. Jackson, Ian Johnston, Dominic Kwiatkowski, Cordelia Langford, John Sillitoe on behalf of the Wellcome Sanger Institute COVID-19 Surveillance Team |
| EPI_ISL_2511327 | Lighthouse Lab in Milton Keynes | Wellcome Sanger Institute for the COVID-19 Genomics UK (COG-UK) Consortium | The Lighthouse Lab in Milton Keynes and Alex Alderton, Roberto Amato, Jeffrey Barrett, Sonia Goncalves, Ewan Harrison, David K. Jackson, Ian Johnston, Dominic Kwiatkowski, Cordelia Langford, John Sillitoe on behalf of the Wellcome Sanger Institute COVID-19 Surveillance Team |
| EPI_ISL_2511328 | Lighthouse Lab in Glasgow | Wellcome Sanger Institute for the COVID-19 Genomics UK (COG-UK) Consortium | Harper VanSteenhouse, Yumi Kasai, David Gray, Carol Clugston, Anna Dominiczak and Alex Alderton, Roberto Amato, Jeffrey Barrett, Sonia Goncalves, Ewan Harrison, David K. Jackson, Ian Johnston, Dominic Kwiatkowski, Cordelia Langford, John Sillitoe on behalf of the Wellcome Sanger Institute COVID-19 Surveillance Team |
| EPI_ISL_2511329, EPI_ISL_2511330 | Health Services Laboratories | Wellcome Sanger Institute for the COVID-19 Genomics UK (COG-UK) Consortium | Health Services Laboratories and Alex Alderton, Roberto Amato, Jeffrey Barrett, Sonia Goncalves, Ewan Harrison, David K. Jackson, Ian Johnston, Dominic Kwiatkowski, Cordelia Langford, John Sillitoe on behalf of the Wellcome Sanger Institute COVID-19 Surveillance Team |
| EPI_ISL_2511331 | Lighthouse Lab in Glasgow | Wellcome Sanger Institute for the COVID-19 Genomics UK (COG-UK) Consortium | Harper VanSteenhouse, Yumi Kasai, David Gray, Carol Clugston, Anna Dominiczak and Alex Alderton, Roberto Amato, Jeffrey Barrett, Sonia Goncalves, Ewan Harrison, David K. Jackson, Ian Johnston, Dominic Kwiatkowski, Cordelia Langford, John Sillitoe on behalf of the Wellcome Sanger Institute COVID-19 Surveillance Team |
| EPI_ISL_2511332 | Health Services Laboratories | Wellcome Sanger Institute for the COVID-19 Genomics UK (COG-UK) Consortium | Health Services Laboratories and Alex Alderton, Roberto Amato, Jeffrey Barrett, Sonia Goncalves, Ewan Harrison, David K. Jackson, Ian Johnston, Dominic Kwiatkowski, Cordelia Langford, John Sillitoe on behalf of the Wellcome Sanger Institute COVID-19 Surveillance Team |
| EPI_ISL_2511333 | Lighthouse Lab in Glasgow | Wellcome Sanger Institute for the COVID-19 Genomics UK (COG-UK) Consortium | Harper VanSteenhouse, Yumi Kasai, David Gray, Carol Clugston, Anna Dominiczak and Alex Alderton, Roberto Amato, Jeffrey Barrett, Sonia Goncalves, Ewan Harrison, David K. Jackson, Ian Johnston, Dominic Kwiatkowski, Cordelia Langford, John Sillitoe on behalf of the Wellcome Sanger Institute COVID-19 Surveillance Team |
| EPI_ISL_2511334, EPI_ISL_2511336 | Health Services Laboratories | Wellcome Sanger Institute for the COVID-19 Genomics UK (COG-UK) Consortium | Health Services Laboratories and Alex Alderton, Roberto Amato, Jeffrey Barrett, Sonia Goncalves, Ewan Harrison, David K. Jackson, Ian Johnston, Dominic Kwiatkowski, Cordelia Langford, John Sillitoe on behalf of the Wellcome Sanger Institute COVID-19 Surveillance Team |
| EPI_ISL_2511337 | Lighthouse Lab in Glasgow | Wellcome Sanger Institute for the COVID-19 Genomics UK (COG-UK) Consortium | Harper VanSteenhouse, Yumi Kasai, David Gray, Carol Clugston, Anna Dominiczak and Alex Alderton, Roberto Amato, Jeffrey Barrett, Sonia Goncalves, Ewan Harrison, David K. Jackson, Ian Johnston, Dominic Kwiatkowski, Cordelia Langford, John Sillitoe on behalf of the Wellcome Sanger Institute COVID-19 Surveillance Team |
| EPI_ISL_2511338 | Health Services Laboratories | Wellcome Sanger Institute for the COVID-19 Genomics UK (COG-UK) Consortium | Health Services Laboratories and Alex Alderton, Roberto Amato, Jeffrey Barrett, Sonia Goncalves, Ewan Harrison, David K. Jackson, Ian Johnston, Dominic Kwiatkowski, Cordelia Langford, John Sillitoe on behalf of the Wellcome Sanger Institute COVID-19 Surveillance Team |
| EPI_ISL_2511339 | Lighthouse Lab in Glasgow | Wellcome Sanger Institute for the COVID-19 Genomics UK (COG-UK) Consortium | Harper VanSteenhouse, Yumi Kasai, David Gray, Carol Clugston, Anna Dominiczak and Alex Alderton, Roberto Amato, Jeffrey Barrett, Sonia Goncalves, Ewan Harrison, David K. Jackson, Ian Johnston, Dominic Kwiatkowski, Cordelia Langford, John Sillitoe on behalf of the Wellcome Sanger Institute COVID-19 Surveillance Team |
| EPI_ISL_2511340, EPI_ISL_2511341 | Health Services Laboratories | Wellcome Sanger Institute for the COVID-19 Genomics UK (COG-UK) Consortium | Health Services Laboratories and Alex Alderton, Roberto Amato, Jeffrey Barrett, Sonia Goncalves, Ewan Harrison, David K. Jackson, Ian Johnston, Dominic Kwiatkowski, Cordelia Langford, John Sillitoe on behalf of the Wellcome Sanger Institute COVID-19 Surveillance Team |
| EPI_ISL_2511342 | Lighthouse Lab in Glasgow | Wellcome Sanger Institute for the COVID-19 Genomics UK (COG-UK) Consortium | Harper VanSteenhouse, Yumi Kasai, David Gray, Carol Clugston, Anna Dominiczak and Alex Alderton, Roberto Amato, Jeffrey Barrett, Sonia Goncalves, Ewan Harrison, David K. Jackson, Ian Johnston, Dominic Kwiatkowski, Cordelia Langford, John Sillitoe on behalf of the Wellcome Sanger Institute COVID-19 Surveillance Team |
| EPI_ISL_2511343, EPI_ISL_2511344 | Health Services Laboratories | Wellcome Sanger Institute for the COVID-19 Genomics UK (COG-UK) Consortium | Health Services Laboratories and Alex Alderton, Roberto Amato, Jeffrey Barrett, Sonia Goncalves, Ewan Harrison, David K. Jackson, Ian Johnston, Dominic Kwiatkowski, Cordelia Langford, John Sillitoe on behalf of the Wellcome Sanger Institute COVID-19 Surveillance Team |
| EPI_ISL_2511345, EPI_ISL_2511346, EPI_ISL_2511347, EPI_ISL_2511348 | Lighthouse Lab in Glasgow | Wellcome Sanger Institute for the COVID-19 Genomics UK (COG-UK) Consortium | Harper VanSteenhouse, Yumi Kasai, David Gray, Carol Clugston, Anna Dominiczak and Alex Alderton, Roberto Amato, Jeffrey Barrett, Sonia Goncalves, Ewan Harrison, David K. Jackson, Ian Johnston, Dominic Kwiatkowski, Cordelia Langford, John Sillitoe on behalf of the Wellcome Sanger Institute COVID-19 Surveillance Team |
| EPI_ISL_2511349 | Health Services Laboratories | Wellcome Sanger Institute for the COVID-19 Genomics UK (COG-UK) Consortium | Health Services Laboratories and Alex Alderton, Roberto Amato, Jeffrey Barrett, Sonia Goncalves, Ewan Harrison, David K. Jackson, Ian Johnston, Dominic Kwiatkowski, Cordelia Langford, John Sillitoe on behalf of the Wellcome Sanger Institute COVID-19 Surveillance Team |
| EPI_ISL_2511350 | Lighthouse Lab in Glasgow | Wellcome Sanger Institute for the COVID-19 Genomics UK (COG-UK) Consortium | Harper VanSteenhouse, Yumi Kasai, David Gray, Carol Clugston, Anna Dominiczak and Alex Alderton, Roberto Amato, Jeffrey Barrett, Sonia Goncalves, Ewan Harrison, David K. Jackson, Ian Johnston, Dominic Kwiatkowski, Cordelia Langford, John Sillitoe on behalf of the Wellcome Sanger Institute COVID-19 Surveillance Team |
| EPI_ISL_2511351 | Health Services Laboratories | Wellcome Sanger Institute for the COVID-19 Genomics UK (COG-UK) Consortium | Health Services Laboratories and Alex Alderton, Roberto Amato, Jeffrey Barrett, Sonia Goncalves, Ewan Harrison, David K. Jackson, Ian Johnston, Dominic Kwiatkowski, Cordelia Langford, John Sillitoe on behalf of the Wellcome Sanger Institute COVID-19 Surveillance Team |
| EPI_ISL_2511352, EPI_ISL_2511353, EPI_ISL_2511354, EPI_ISL_2511355, EPI_ISL_2511357, EPI_ISL_2511358 | Lighthouse Lab in Milton Keynes | Wellcome Sanger Institute for the COVID-19 Genomics UK (COG-UK) Consortium | The Lighthouse Lab in Milton Keynes and Alex Alderton, Roberto Amato, Jeffrey Barrett, Sonia Goncalves, Ewan Harrison, David K. Jackson, Ian Johnston, Dominic Kwiatkowski, Cordelia Langford, John Sillitoe on behalf of the Wellcome Sanger Institute COVID-19 Surveillance Team |
| EPI_ISL_2511359 | Lighthouse Lab in Glasgow | Wellcome Sanger Institute for the COVID-19 Genomics UK (COG-UK) Consortium | Harper VanSteenhouse, Yumi Kasai, David Gray, Carol Clugston, Anna Dominiczak and Alex Alderton, Roberto Amato, Jeffrey Barrett, Sonia Goncalves, Ewan Harrison, David K. Jackson, Ian Johnston, Dominic Kwiatkowski, Cordelia Langford, John Sillitoe on behalf of the Wellcome Sanger Institute COVID-19 Surveillance Team |
| EPI_ISL_2511360, EPI_ISL_2511361, EPI_ISL_2511362 | Lighthouse Lab in Alderley Park | Wellcome Sanger Institute for the COVID-19 Genomics UK (COG-UK) Consortium | Jacquelyn Wynn, Mairead Hyland, The Lighthouse Lab in Alderley Park and Alex Alderton, Roberto Amato, Jeffrey Barrett, Sonia Goncalves, Ewan Harrison, David K. Jackson, Ian Johnston, Dominic Kwiatkowski, Cordelia Langford, John Sillitoe on behalf of the Wellcome Sanger Institute COVID-19 Surveillance Team |
| EPI_ISL_2511363, EPI_ISL_2511364, EPI_ISL_2511365 | Lighthouse Lab in Milton Keynes | Wellcome Sanger Institute for the COVID-19 Genomics UK (COG-UK) Consortium | The Lighthouse Lab in Milton Keynes and Alex Alderton, Roberto Amato, Jeffrey Barrett, Sonia Goncalves, Ewan Harrison, David K. Jackson, Ian Johnston, Dominic Kwiatkowski, Cordelia Langford, John Sillitoe on behalf of the Wellcome Sanger Institute COVID-19 Surveillance Team |
| EPI_ISL_2511366, EPI_ISL_2511367 | Lighthouse Lab in Alderley Park | Wellcome Sanger Institute for the COVID-19 Genomics UK (COG-UK) Consortium | Jacquelyn Wynn, Mairead Hyland, The Lighthouse Lab in Alderley Park and Alex Alderton, Roberto Amato, Jeffrey Barrett, Sonia Goncalves, Ewan Harrison, David K. Jackson, Ian Johnston, Dominic Kwiatkowski, Cordelia Langford, John Sillitoe on behalf of the Wellcome Sanger Institute COVID-19 Surveillance Team |
| EPI_ISL_2511368, EPI_ISL_2511369 | Lighthouse Lab in Milton Keynes | Wellcome Sanger Institute for the COVID-19 Genomics UK (COG-UK) Consortium | The Lighthouse Lab in Milton Keynes and Alex Alderton, Roberto Amato, Jeffrey Barrett, Sonia Goncalves, Ewan Harrison, David K. Jackson, Ian Johnston, Dominic Kwiatkowski, Cordelia Langford, John Sillitoe on behalf of the Wellcome Sanger Institute COVID-19 Surveillance Team |
| EPI_ISL_2511370 | Lighthouse Lab in Alderley Park | Wellcome Sanger Institute for the COVID-19 Genomics UK (COG-UK) Consortium | Jacquelyn Wynn, Mairead Hyland, The Lighthouse Lab in Alderley Park and Alex Alderton, Roberto Amato, Jeffrey Barrett, Sonia Goncalves, Ewan Harrison, David K. Jackson, Ian Johnston, Dominic Kwiatkowski, Cordelia Langford, John Sillitoe on behalf of the Wellcome Sanger Institute COVID-19 Surveillance Team |
| EPI_ISL_2511371, EPI_ISL_2511372 | Lighthouse Lab in Milton Keynes | Wellcome Sanger Institute for the COVID-19 Genomics UK (COG-UK) Consortium | The Lighthouse Lab in Milton Keynes and Alex Alderton, Roberto Amato, Jeffrey Barrett, Sonia Goncalves, Ewan Harrison, David K. Jackson, Ian Johnston, Dominic Kwiatkowski, Cordelia Langford, John Sillitoe on behalf of the Wellcome Sanger Institute COVID-19 Surveillance Team |
| EPI_ISL_2511373 | Lighthouse Lab in Alderley Park | Wellcome Sanger Institute for the COVID-19 Genomics UK (COG-UK) Consortium | Jacquelyn Wynn, Mairead Hyland, The Lighthouse Lab in Alderley Park and Alex Alderton, Roberto Amato, Jeffrey Barrett, Sonia Goncalves, Ewan Harrison, David K. Jackson, Ian Johnston, Dominic Kwiatkowski, Cordelia Langford, John Sillitoe on behalf of the Wellcome Sanger Institute COVID-19 Surveillance Team |
| EPI_ISL_2511374 | Lighthouse Laboratory Plymouth | Wellcome Sanger Institute for the COVID-19 Genomics UK (COG-UK) Consortium | Lighthouse Laboratory Plymouth and Alex Alderton, Roberto Amato, Jeffrey Barrett, Sonia Goncalves, Ewan Harrison, David K. Jackson, Ian Johnston, Dominic Kwiatkowski, Cordelia Langford, John Sillitoe on behalf of the Wellcome Sanger Institute COVID-19 Surveillance Team |
| EPI_ISL_2511375 | Lighthouse Lab in Milton Keynes | Wellcome Sanger Institute for the COVID-19 Genomics UK (COG-UK) Consortium | The Lighthouse Lab in Milton Keynes and Alex Alderton, Roberto Amato, Jeffrey Barrett, Sonia Goncalves, Ewan Harrison, David K. Jackson, Ian Johnston, Dominic Kwiatkowski, Cordelia Langford, John Sillitoe on behalf of the Wellcome Sanger Institute COVID-19 Surveillance Team |
| EPI_ISL_2511376, EPI_ISL_2511377, EPI_ISL_2511378, EPI_ISL_2511379 | Lighthouse Lab in Alderley Park | Wellcome Sanger Institute for the COVID-19 Genomics UK (COG-UK) Consortium | Jacquelyn Wynn, Mairead Hyland, The Lighthouse Lab in Alderley Park and Alex Alderton, Roberto Amato, Jeffrey Barrett, Sonia Goncalves, Ewan Harrison, David K. Jackson, Ian Johnston, Dominic Kwiatkowski, Cordelia Langford, John Sillitoe on behalf of the Wellcome Sanger Institute COVID-19 Surveillance Team |
| EPI_ISL_2511380, EPI_ISL_2511381, EPI_ISL_2511382 | Lighthouse Lab in Milton Keynes | Wellcome Sanger Institute for the COVID-19 Genomics UK (COG-UK) Consortium | The Lighthouse Lab in Milton Keynes and Alex Alderton, Roberto Amato, Jeffrey Barrett, Sonia Goncalves, Ewan Harrison, David K. Jackson, Ian Johnston, Dominic Kwiatkowski, Cordelia Langford, John Sillitoe on behalf of the Wellcome Sanger Institute COVID-19 Surveillance Team |
| EPI_ISL_2511383 | Lighthouse Lab in Alderley Park | Wellcome Sanger Institute for the COVID-19 Genomics UK (COG-UK) Consortium | Jacquelyn Wynn, Mairead Hyland, The Lighthouse Lab in Alderley Park and Alex Alderton, Roberto Amato, Jeffrey Barrett, Sonia Goncalves, Ewan Harrison, David K. Jackson, Ian Johnston, Dominic Kwiatkowski, Cordelia Langford, John Sillitoe on behalf of the Wellcome Sanger Institute COVID-19 Surveillance Team |
| EPI_ISL_2511384 | Lighthouse Laboratory Plymouth | Wellcome Sanger Institute for the COVID-19 Genomics UK (COG-UK) Consortium | Lighthouse Laboratory Plymouth and Alex Alderton, Roberto Amato, Jeffrey Barrett, Sonia Goncalves, Ewan Harrison, David K. Jackson, Ian Johnston, Dominic Kwiatkowski, Cordelia Langford, John Sillitoe on behalf of the Wellcome Sanger Institute COVID-19 Surveillance Team |
| EPI_ISL_2511385, | Lighthouse Lab in Alderley Park | Wellcome Sanger Institute for the COVID-19 Genomics UK (COG-UK) | Jacquelyn Wynn, Mairead Hyland, The Lighthouse Lab in Alderley Park and Alex Alderton, Roberto Amato, Jeffrey Barrett, Sonia Goncalves, Ewan Harrison, David K. Jackson, Ian Johnston, Dominic Kwiatkowski, Cordelia Langford, John Sillitoe on |

| | | | |
|---|---|---|---|
| EPI_ISL_2511386 | | Consortium | behalf of the Wellcome Sanger Institute COVID-19 Surveillance Team |
| EPI_ISL_2511387, EPI_ISL_2511388 | Lighthouse Lab in Milton Keynes | Wellcome Sanger Institute for the COVID-19 Genomics UK (COG-UK) Consortium | The Lighthouse Lab in Milton Keynes and Alex Alderton, Roberto Amato, Jeffrey Barrett, Sonia Goncalves, Ewan Harrison, David K. Jackson, Ian Johnston, Dominic Kwiatkowski, Cordelia Langford, John Sillitoe on behalf of the Wellcome Sanger Institute COVID-19 Surveillance Team |
| EPI_ISL_2511389 | Lighthouse Lab in Alderley Park | Wellcome Sanger Institute for the COVID-19 Genomics UK (COG-UK) Consortium | Jacquelyn Wynn, Mairead Hyland, The Lighthouse Lab in Alderley Park and Alex Alderton, Roberto Amato, Jeffrey Barrett, Sonia Goncalves, Ewan Harrison, David K. Jackson, Ian Johnston, Dominic Kwiatkowski, Cordelia Langford, John Sillitoe on behalf of the Wellcome Sanger Institute COVID-19 Surveillance Team |
| EPI_ISL_2511390 | Lighthouse Laboratory Plymouth | Wellcome Sanger Institute for the COVID-19 Genomics UK (COG-UK) Consortium | Lighthouse Laboratory Plymouth and Alex Alderton, Roberto Amato, Jeffrey Barrett, Sonia Goncalves, Ewan Harrison, David K. Jackson, Ian Johnston, Dominic Kwiatkowski, Cordelia Langford, John Sillitoe on behalf of the Wellcome Sanger Institute COVID-19 Surveillance Team |
| EPI_ISL_2511391 | Lighthouse Lab in Alderley Park | Wellcome Sanger Institute for the COVID-19 Genomics UK (COG-UK) Consortium | Jacquelyn Wynn, Mairead Hyland, The Lighthouse Lab in Alderley Park and Alex Alderton, Roberto Amato, Jeffrey Barrett, Sonia Goncalves, Ewan Harrison, David K. Jackson, Ian Johnston, Dominic Kwiatkowski, Cordelia Langford, John Sillitoe on behalf of the Wellcome Sanger Institute COVID-19 Surveillance Team |
| EPI_ISL_2511392 | Lighthouse Lab in Milton Keynes | Wellcome Sanger Institute for the COVID-19 Genomics UK (COG-UK) Consortium | The Lighthouse Lab in Milton Keynes and Alex Alderton, Roberto Amato, Jeffrey Barrett, Sonia Goncalves, Ewan Harrison, David K. Jackson, Ian Johnston, Dominic Kwiatkowski, Cordelia Langford, John Sillitoe on behalf of the Wellcome Sanger Institute COVID-19 Surveillance Team |
| EPI_ISL_2511393, EPI_ISL_2511394, EPI_ISL_2511395 | Lighthouse Lab in Alderley Park | Wellcome Sanger Institute for the COVID-19 Genomics UK (COG-UK) Consortium | Jacquelyn Wynn, Mairead Hyland, The Lighthouse Lab in Alderley Park and Alex Alderton, Roberto Amato, Jeffrey Barrett, Sonia Goncalves, Ewan Harrison, David K. Jackson, Ian Johnston, Dominic Kwiatkowski, Cordelia Langford, John Sillitoe on behalf of the Wellcome Sanger Institute COVID-19 Surveillance Team |
| EPI_ISL_2511396 | Lighthouse Lab in Milton Keynes | Wellcome Sanger Institute for the COVID-19 Genomics UK (COG-UK) Consortium | The Lighthouse Lab in Milton Keynes and Alex Alderton, Roberto Amato, Jeffrey Barrett, Sonia Goncalves, Ewan Harrison, David K. Jackson, Ian Johnston, Dominic Kwiatkowski, Cordelia Langford, John Sillitoe on behalf of the Wellcome Sanger Institute COVID-19 Surveillance Team |
| EPI_ISL_2511397 | Lighthouse Lab in Alderley Park | Wellcome Sanger Institute for the COVID-19 Genomics UK (COG-UK) Consortium | Jacquelyn Wynn, Mairead Hyland, The Lighthouse Lab in Alderley Park and Alex Alderton, Roberto Amato, Jeffrey Barrett, Sonia Goncalves, Ewan Harrison, David K. Jackson, Ian Johnston, Dominic Kwiatkowski, Cordelia Langford, John Sillitoe on behalf of the Wellcome Sanger Institute COVID-19 Surveillance Team |
| EPI_ISL_2511398, EPI_ISL_2511399 | Lighthouse Lab in Milton Keynes | Wellcome Sanger Institute for the COVID-19 Genomics UK (COG-UK) Consortium | The Lighthouse Lab in Milton Keynes and Alex Alderton, Roberto Amato, Jeffrey Barrett, Sonia Goncalves, Ewan Harrison, David K. Jackson, Ian Johnston, Dominic Kwiatkowski, Cordelia Langford, John Sillitoe on behalf of the Wellcome Sanger Institute COVID-19 Surveillance Team |
| EPI_ISL_2511400 | Lighthouse Laboratory Plymouth | Wellcome Sanger Institute for the COVID-19 Genomics UK (COG-UK) Consortium | Lighthouse Laboratory Plymouth and Alex Alderton, Roberto Amato, Jeffrey Barrett, Sonia Goncalves, Ewan Harrison, David K. Jackson, Ian Johnston, Dominic Kwiatkowski, Cordelia Langford, John Sillitoe on behalf of the Wellcome Sanger Institute COVID-19 Surveillance Team |
| EPI_ISL_2511401, EPI_ISL_2511402 | Lighthouse Lab in Milton Keynes | Wellcome Sanger Institute for the COVID-19 Genomics UK (COG-UK) Consortium | The Lighthouse Lab in Milton Keynes and Alex Alderton, Roberto Amato, Jeffrey Barrett, Sonia Goncalves, Ewan Harrison, David K. Jackson, Ian Johnston, Dominic Kwiatkowski, Cordelia Langford, John Sillitoe on behalf of the Wellcome Sanger Institute COVID-19 Surveillance Team |
| EPI_ISL_2511403 | Lighthouse Laboratory Plymouth | Wellcome Sanger Institute for the COVID-19 Genomics UK (COG-UK) Consortium | Lighthouse Laboratory Plymouth and Alex Alderton, Roberto Amato, Jeffrey Barrett, Sonia Goncalves, Ewan Harrison, David K. Jackson, Ian Johnston, Dominic Kwiatkowski, Cordelia Langford, John Sillitoe on behalf of the Wellcome Sanger Institute COVID-19 Surveillance Team |
| EPI_ISL_2511404 | Lighthouse Lab in Glasgow | Wellcome Sanger Institute for the COVID-19 Genomics UK (COG-UK) Consortium | Harper VanSteenhouse, Yumi Kasai, David Gray, Carol Clugston, Anna Dominiczak and Alex Alderton, Roberto Amato, Jeffrey Barrett, Sonia Goncalves, Ewan Harrison, David K. Jackson, Ian Johnston, Dominic Kwiatkowski, Cordelia Langford, John Sillitoe on behalf of the Wellcome Sanger Institute COVID-19 Surveillance Team |
| EPI_ISL_2511405 | Lighthouse Lab in Alderley Park | Wellcome Sanger Institute for the COVID-19 Genomics UK (COG-UK) Consortium | Jacquelyn Wynn, Mairead Hyland, The Lighthouse Lab in Alderley Park and Alex Alderton, Roberto Amato, Jeffrey Barrett, Sonia Goncalves, Ewan Harrison, David K. Jackson, Ian Johnston, Dominic Kwiatkowski, Cordelia Langford, John Sillitoe on behalf of the Wellcome Sanger Institute COVID-19 Surveillance Team |
| EPI_ISL_2511407 | Lighthouse Laboratory Plymouth | Wellcome Sanger Institute for the COVID-19 Genomics UK (COG-UK) Consortium | Lighthouse Laboratory Plymouth and Alex Alderton, Roberto Amato, Jeffrey Barrett, Sonia Goncalves, Ewan Harrison, David K. Jackson, Ian Johnston, Dominic Kwiatkowski, Cordelia Langford, John Sillitoe on behalf of the Wellcome Sanger Institute COVID-19 Surveillance Team |
| EPI_ISL_2511408, EPI_ISL_2511409, EPI_ISL_2511410 | Lighthouse Lab in Alderley Park | Wellcome Sanger Institute for the COVID-19 Genomics UK (COG-UK) Consortium | Jacquelyn Wynn, Mairead Hyland, The Lighthouse Lab in Alderley Park and Alex Alderton, Roberto Amato, Jeffrey Barrett, Sonia Goncalves, Ewan Harrison, David K. Jackson, Ian Johnston, Dominic Kwiatkowski, Cordelia Langford, John Sillitoe on behalf of the Wellcome Sanger Institute COVID-19 Surveillance Team |
| EPI_ISL_2511411 | Lighthouse Laboratory Plymouth | Wellcome Sanger Institute for the COVID-19 Genomics UK (COG-UK) Consortium | Lighthouse Laboratory Plymouth and Alex Alderton, Roberto Amato, Jeffrey Barrett, Sonia Goncalves, Ewan Harrison, David K. Jackson, Ian Johnston, Dominic Kwiatkowski, Cordelia Langford, John Sillitoe on behalf of the Wellcome Sanger Institute COVID-19 Surveillance Team |
| EPI_ISL_2511412, EPI_ISL_2511413 | Lighthouse Lab in Milton Keynes | Wellcome Sanger Institute for the COVID-19 Genomics UK (COG-UK) Consortium | The Lighthouse Lab in Milton Keynes and Alex Alderton, Roberto Amato, Jeffrey Barrett, Sonia Goncalves, Ewan Harrison, David K. Jackson, Ian Johnston, Dominic Kwiatkowski, Cordelia Langford, John Sillitoe on behalf of the Wellcome Sanger Institute COVID-19 Surveillance Team |
| EPI_ISL_2511414 | Lighthouse Laboratory Plymouth | Wellcome Sanger Institute for the COVID-19 Genomics UK (COG-UK) Consortium | Lighthouse Laboratory Plymouth and Alex Alderton, Roberto Amato, Jeffrey Barrett, Sonia Goncalves, Ewan Harrison, David K. Jackson, Ian Johnston, Dominic Kwiatkowski, Cordelia Langford, John Sillitoe on behalf of the Wellcome Sanger Institute COVID-19 Surveillance Team |
| EPI_ISL_2511415, EPI_ISL_2511416, EPI_ISL_2511417, EPI_ISL_2511419 | Lighthouse Lab in Alderley Park | Wellcome Sanger Institute for the COVID-19 Genomics UK (COG-UK) Consortium | Jacquelyn Wynn, Mairead Hyland, The Lighthouse Lab in Alderley Park and Alex Alderton, Roberto Amato, Jeffrey Barrett, Sonia Goncalves, Ewan Harrison, David K. Jackson, Ian Johnston, Dominic Kwiatkowski, Cordelia Langford, John Sillitoe on behalf of the Wellcome Sanger Institute COVID-19 Surveillance Team |
| EPI_ISL_2511420 | Lighthouse Lab in Milton Keynes | Wellcome Sanger Institute for the COVID-19 Genomics UK (COG-UK) Consortium | The Lighthouse Lab in Milton Keynes and Alex Alderton, Roberto Amato, Jeffrey Barrett, Sonia Goncalves, Ewan Harrison, David K. Jackson, Ian Johnston, Dominic Kwiatkowski, Cordelia Langford, John Sillitoe on behalf of the Wellcome Sanger Institute COVID-19 Surveillance Team |
| EPI_ISL_2511421 | Lighthouse Lab in Alderley Park | Wellcome Sanger Institute for the COVID-19 Genomics UK (COG-UK) Consortium | Jacquelyn Wynn, Mairead Hyland, The Lighthouse Lab in Alderley Park and Alex Alderton, Roberto Amato, Jeffrey Barrett, Sonia Goncalves, Ewan Harrison, David K. Jackson, Ian Johnston, Dominic Kwiatkowski, Cordelia Langford, John Sillitoe on behalf of the Wellcome Sanger Institute COVID-19 Surveillance Team |
| EPI_ISL_2511422 | Lighthouse Lab in Milton Keynes | Wellcome Sanger Institute for the COVID-19 Genomics UK (COG-UK) Consortium | The Lighthouse Lab in Milton Keynes and Alex Alderton, Roberto Amato, Jeffrey Barrett, Sonia Goncalves, Ewan Harrison, David K. Jackson, Ian Johnston, Dominic Kwiatkowski, Cordelia Langford, John Sillitoe on behalf of the Wellcome Sanger Institute COVID-19 Surveillance Team |
| EPI_ISL_2511423, EPI_ISL_2511424, EPI_ISL_2511425 | Lighthouse Lab in Alderley Park | Wellcome Sanger Institute for the COVID-19 Genomics UK (COG-UK) Consortium | Jacquelyn Wynn, Mairead Hyland, The Lighthouse Lab in Alderley Park and Alex Alderton, Roberto Amato, Jeffrey Barrett, Sonia Goncalves, Ewan Harrison, David K. Jackson, Ian Johnston, Dominic Kwiatkowski, Cordelia Langford, John Sillitoe on behalf of the Wellcome Sanger Institute COVID-19 Surveillance Team |
| EPI_ISL_2511426 | Lighthouse Laboratory Plymouth | Wellcome Sanger Institute for the COVID-19 Genomics UK (COG-UK) Consortium | Lighthouse Laboratory Plymouth and Alex Alderton, Roberto Amato, Jeffrey Barrett, Sonia Goncalves, Ewan Harrison, David K. Jackson, Ian Johnston, Dominic Kwiatkowski, Cordelia Langford, John Sillitoe on behalf of the Wellcome Sanger Institute COVID-19 Surveillance Team |
| EPI_ISL_2511427 | Lighthouse Lab in Alderley Park | Wellcome Sanger Institute for the COVID-19 Genomics UK (COG-UK) Consortium | Jacquelyn Wynn, Mairead Hyland, The Lighthouse Lab in Alderley Park and Alex Alderton, Roberto Amato, Jeffrey Barrett, Sonia Goncalves, Ewan Harrison, David K. Jackson, Ian Johnston, Dominic Kwiatkowski, Cordelia Langford, John Sillitoe on behalf of the Wellcome Sanger Institute COVID-19 Surveillance Team |
| EPI_ISL_2511428, EPI_ISL_2511429 | Lighthouse Lab in Milton Keynes | Wellcome Sanger Institute for the COVID-19 Genomics UK (COG-UK) Consortium | The Lighthouse Lab in Milton Keynes and Alex Alderton, Roberto Amato, Jeffrey Barrett, Sonia Goncalves, Ewan Harrison, David K. Jackson, Ian Johnston, Dominic Kwiatkowski, Cordelia Langford, John Sillitoe on behalf of the Wellcome Sanger Institute COVID-19 Surveillance Team |
| EPI_ISL_2511430 | Lighthouse Lab in Alderley Park | Wellcome Sanger Institute for the COVID-19 Genomics UK (COG-UK) Consortium | Jacquelyn Wynn, Mairead Hyland, The Lighthouse Lab in Alderley Park and Alex Alderton, Roberto Amato, Jeffrey Barrett, Sonia Goncalves, Ewan Harrison, David K. Jackson, Ian Johnston, Dominic Kwiatkowski, Cordelia Langford, John Sillitoe on behalf of the Wellcome Sanger Institute COVID-19 Surveillance Team |
| EPI_ISL_2511431, EPI_ISL_2511432 | Lighthouse Lab in Milton Keynes | Wellcome Sanger Institute for the COVID-19 Genomics UK (COG-UK) Consortium | The Lighthouse Lab in Milton Keynes and Alex Alderton, Roberto Amato, Jeffrey Barrett, Sonia Goncalves, Ewan Harrison, David K. Jackson, Ian Johnston, Dominic Kwiatkowski, Cordelia Langford, John Sillitoe on behalf of the Wellcome Sanger Institute COVID-19 Surveillance Team |
| EPI_ISL_2511433 | Lighthouse Laboratory Plymouth | Wellcome Sanger Institute for the COVID-19 Genomics UK (COG-UK) Consortium | Lighthouse Laboratory Plymouth and Alex Alderton, Roberto Amato, Jeffrey Barrett, Sonia Goncalves, Ewan Harrison, David K. Jackson, Ian Johnston, Dominic Kwiatkowski, Cordelia Langford, John Sillitoe on behalf of the Wellcome Sanger Institute COVID-19 Surveillance Team |
| EPI_ISL_2511434 | Lighthouse Lab in Alderley Park | Wellcome Sanger Institute for the COVID-19 Genomics UK (COG-UK) Consortium | Jacquelyn Wynn, Mairead Hyland, The Lighthouse Lab in Alderley Park and Alex Alderton, Roberto Amato, Jeffrey Barrett, Sonia Goncalves, Ewan Harrison, David K. Jackson, Ian Johnston, Dominic Kwiatkowski, Cordelia Langford, John Sillitoe on behalf of the Wellcome Sanger Institute COVID-19 Surveillance Team |
| EPI_ISL_2511435 | Lighthouse Lab in Milton Keynes | Wellcome Sanger Institute for the COVID-19 Genomics UK (COG-UK) Consortium | The Lighthouse Lab in Milton Keynes and Alex Alderton, Roberto Amato, Jeffrey Barrett, Sonia Goncalves, Ewan Harrison, David K. Jackson, Ian Johnston, Dominic Kwiatkowski, Cordelia Langford, John Sillitoe on behalf of the Wellcome Sanger Institute COVID-19 Surveillance Team |
| EPI_ISL_2511436 | Lighthouse Laboratory Plymouth | Wellcome Sanger Institute for the COVID-19 Genomics UK (COG-UK) Consortium | Lighthouse Laboratory Plymouth and Alex Alderton, Roberto Amato, Jeffrey Barrett, Sonia Goncalves, Ewan Harrison, David K. Jackson, Ian Johnston, Dominic Kwiatkowski, Cordelia Langford, John Sillitoe on behalf of the Wellcome Sanger Institute COVID-19 Surveillance Team |
| EPI_ISL_2511437 | Lighthouse Lab in Alderley Park | Wellcome Sanger Institute for the COVID-19 Genomics UK (COG-UK) Consortium | Jacquelyn Wynn, Mairead Hyland, The Lighthouse Lab in Alderley Park and Alex Alderton, Roberto Amato, Jeffrey Barrett, Sonia Goncalves, Ewan Harrison, David K. Jackson, Ian Johnston, Dominic Kwiatkowski, Cordelia Langford, John Sillitoe on behalf of the Wellcome Sanger Institute COVID-19 Surveillance Team |
| EPI_ISL_2511438 | Lighthouse Lab in Milton Keynes | Wellcome Sanger Institute for the COVID-19 Genomics UK (COG-UK) Consortium | The Lighthouse Lab in Milton Keynes and Alex Alderton, Roberto Amato, Jeffrey Barrett, Sonia Goncalves, Ewan Harrison, David K. Jackson, Ian Johnston, Dominic Kwiatkowski, Cordelia Langford, John Sillitoe on behalf of the Wellcome Sanger Institute COVID-19 Surveillance Team |
| EPI_ISL_2511439 | Lighthouse Lab in Glasgow | Wellcome Sanger Institute for the COVID-19 Genomics UK (COG-UK) Consortium | Harper VanSteenhouse, Yumi Kasai, David Gray, Carol Clugston, Anna Dominiczak and Alex Alderton, Roberto Amato, Jeffrey Barrett, Sonia Goncalves, Ewan Harrison, David K. Jackson, Ian Johnston, Dominic Kwiatkowski, Cordelia Langford, John Sillitoe on behalf of the Wellcome Sanger Institute COVID-19 Surveillance Team |
| EPI_ISL_2511440 | Lighthouse Lab in Milton Keynes | Wellcome Sanger Institute for the COVID-19 Genomics UK (COG-UK) Consortium | The Lighthouse Lab in Milton Keynes and Alex Alderton, Roberto Amato, Jeffrey Barrett, Sonia Goncalves, Ewan Harrison, David K. Jackson, Ian Johnston, Dominic Kwiatkowski, Cordelia Langford, John Sillitoe on behalf of the Wellcome Sanger Institute COVID-19 Surveillance Team |
| EPI_ISL_2511441 | Lighthouse Lab in Glasgow | Wellcome Sanger Institute for the COVID-19 Genomics UK (COG-UK) Consortium | Harper VanSteenhouse, Yumi Kasai, David Gray, Carol Clugston, Anna Dominiczak and Alex Alderton, Roberto Amato, Jeffrey Barrett, Sonia Goncalves, Ewan Harrison, David K. Jackson, Ian Johnston, Dominic Kwiatkowski, Cordelia Langford, John Sillitoe on behalf of the Wellcome Sanger Institute COVID-19 Surveillance Team |
| EPI_ISL_2511442, EPI_ISL_2511443 | Lighthouse Lab in Alderley Park | Wellcome Sanger Institute for the COVID-19 Genomics UK (COG-UK) Consortium | Jacquelyn Wynn, Mairead Hyland, The Lighthouse Lab in Alderley Park and Alex Alderton, Roberto Amato, Jeffrey Barrett, Sonia Goncalves, Ewan Harrison, David K. Jackson, Ian Johnston, Dominic Kwiatkowski, Cordelia Langford, John Sillitoe on behalf of the Wellcome Sanger Institute COVID-19 Surveillance Team |
| EPI_ISL_2511444, EPI_ISL_2511445 | Lighthouse Lab in Milton Keynes | Wellcome Sanger Institute for the COVID-19 Genomics UK (COG-UK) Consortium | The Lighthouse Lab in Milton Keynes and Alex Alderton, Roberto Amato, Jeffrey Barrett, Sonia Goncalves, Ewan Harrison, David K. Jackson, Ian Johnston, Dominic Kwiatkowski, Cordelia Langford, John Sillitoe on behalf of the Wellcome Sanger Institute COVID-19 Surveillance Team |
| EPI_ISL_2511446 | Lighthouse Lab in Alderley Park | Wellcome Sanger Institute for the COVID-19 Genomics UK (COG-UK) Consortium | Jacquelyn Wynn, Mairead Hyland, The Lighthouse Lab in Alderley Park and Alex Alderton, Roberto Amato, Jeffrey Barrett, Sonia Goncalves, Ewan Harrison, David K. Jackson, Ian Johnston, Dominic Kwiatkowski, Cordelia Langford, John Sillitoe on behalf of the Wellcome Sanger Institute COVID-19 Surveillance Team |
| EPI_ISL_2511448 | Lighthouse Lab in Milton Keynes | Wellcome Sanger Institute for the COVID-19 Genomics UK (COG-UK) Consortium | The Lighthouse Lab in Milton Keynes and Alex Alderton, Roberto Amato, Jeffrey Barrett, Sonia Goncalves, Ewan Harrison, David K. Jackson, Ian Johnston, Dominic Kwiatkowski, Cordelia Langford, John Sillitoe on behalf of the Wellcome Sanger Institute COVID-19 Surveillance Team |
| EPI_ISL_2511449, | Lighthouse Lab in Alderley Park | Wellcome Sanger Institute for the COVID-19 Genomics UK (COG-UK) | Jacquelyn Wynn, Mairead Hyland, The Lighthouse Lab in Alderley Park and Alex Alderton, Roberto Amato, Jeffrey Barrett, Sonia Goncalves, Ewan Harrison, David K. Jackson, Ian Johnston, Dominic Kwiatkowski, Cordelia Langford, John Sillitoe on |

| | | | |
|---|---|---|---|
| EPI_ISL_2511450, EPI_ISL_2511451 | | Consortium | behalf of the Wellcome Sanger Institute COVID-19 Surveillance Team |
| EPI_ISL_2511452 | Lighthouse Lab in Milton Keynes | Wellcome Sanger Institute for the COVID-19 Genomics UK (COG-UK) Consortium | The Lighthouse Lab in Milton Keynes and Alex Alderton, Roberto Amato, Jeffrey Barrett, Sonia Goncalves, Ewan Harrison, David K. Jackson, Ian Johnston, Dominic Kwiatkowski, Cordelia Langford, John Sillitoe on behalf of the Wellcome Sanger Institute COVID-19 Surveillance Team |
| EPI_ISL_2511453 | Lighthouse Lab in Alderley Park | Wellcome Sanger Institute for the COVID-19 Genomics UK (COG-UK) Consortium | Jacquelyn Wynn, Mairead Hyland, The Lighthouse Lab in Alderley Park and Alex Alderton, Roberto Amato, Jeffrey Barrett, Sonia Goncalves, Ewan Harrison, David K. Jackson, Ian Johnston, Dominic Kwiatkowski, Cordelia Langford, John Sillitoe on behalf of the Wellcome Sanger Institute COVID-19 Surveillance Team |
| EPI_ISL_2511454, EPI_ISL_2511457, EPI_ISL_2511458 | Lighthouse Lab in Milton Keynes | Wellcome Sanger Institute for the COVID-19 Genomics UK (COG-UK) Consortium | The Lighthouse Lab in Milton Keynes and Alex Alderton, Roberto Amato, Jeffrey Barrett, Sonia Goncalves, Ewan Harrison, David K. Jackson, Ian Johnston, Dominic Kwiatkowski, Cordelia Langford, John Sillitoe on behalf of the Wellcome Sanger Institute COVID-19 Surveillance Team |
| EPI_ISL_2511459 | Lighthouse Laboratory Plymouth | Wellcome Sanger Institute for the COVID-19 Genomics UK (COG-UK) Consortium | Lighthouse Laboratory Plymouth and Alex Alderton, Roberto Amato, Jeffrey Barrett, Sonia Goncalves, Ewan Harrison, David K. Jackson, Ian Johnston, Dominic Kwiatkowski, Cordelia Langford, John Sillitoe on behalf of the Wellcome Sanger Institute COVID-19 Surveillance Team |
| EPI_ISL_2511460 | Lighthouse Lab in Milton Keynes | Wellcome Sanger Institute for the COVID-19 Genomics UK (COG-UK) Consortium | The Lighthouse Lab in Milton Keynes and Alex Alderton, Roberto Amato, Jeffrey Barrett, Sonia Goncalves, Ewan Harrison, David K. Jackson, Ian Johnston, Dominic Kwiatkowski, Cordelia Langford, John Sillitoe on behalf of the Wellcome Sanger Institute COVID-19 Surveillance Team |
| EPI_ISL_2511461, EPI_ISL_2511462 | Lighthouse Laboratory Plymouth | Wellcome Sanger Institute for the COVID-19 Genomics UK (COG-UK) Consortium | Lighthouse Laboratory Plymouth and Alex Alderton, Roberto Amato, Jeffrey Barrett, Sonia Goncalves, Ewan Harrison, David K. Jackson, Ian Johnston, Dominic Kwiatkowski, Cordelia Langford, John Sillitoe on behalf of the Wellcome Sanger Institute COVID-19 Surveillance Team |
| EPI_ISL_2511464, EPI_ISL_2511466 | Lighthouse Lab in Milton Keynes | Wellcome Sanger Institute for the COVID-19 Genomics UK (COG-UK) Consortium | The Lighthouse Lab in Milton Keynes and Alex Alderton, Roberto Amato, Jeffrey Barrett, Sonia Goncalves, Ewan Harrison, David K. Jackson, Ian Johnston, Dominic Kwiatkowski, Cordelia Langford, John Sillitoe on behalf of the Wellcome Sanger Institute COVID-19 Surveillance Team |
| EPI_ISL_2511468 | Lighthouse Lab in Alderley Park | Wellcome Sanger Institute for the COVID-19 Genomics UK (COG-UK) Consortium | Jacquelyn Wynn, Mairead Hyland, The Lighthouse Lab in Alderley Park and Alex Alderton, Roberto Amato, Jeffrey Barrett, Sonia Goncalves, Ewan Harrison, David K. Jackson, Ian Johnston, Dominic Kwiatkowski, Cordelia Langford, John Sillitoe on behalf of the Wellcome Sanger Institute COVID-19 Surveillance Team |
| EPI_ISL_2511469 | Lighthouse Laboratory Plymouth | Wellcome Sanger Institute for the COVID-19 Genomics UK (COG-UK) Consortium | Lighthouse Laboratory Plymouth and Alex Alderton, Roberto Amato, Jeffrey Barrett, Sonia Goncalves, Ewan Harrison, David K. Jackson, Ian Johnston, Dominic Kwiatkowski, Cordelia Langford, John Sillitoe on behalf of the Wellcome Sanger Institute COVID-19 Surveillance Team |
| EPI_ISL_2511470 | Lighthouse Lab in Milton Keynes | Wellcome Sanger Institute for the COVID-19 Genomics UK (COG-UK) Consortium | The Lighthouse Lab in Milton Keynes and Alex Alderton, Roberto Amato, Jeffrey Barrett, Sonia Goncalves, Ewan Harrison, David K. Jackson, Ian Johnston, Dominic Kwiatkowski, Cordelia Langford, John Sillitoe on behalf of the Wellcome Sanger Institute COVID-19 Surveillance Team |
| EPI_ISL_2511471 | Lighthouse Laboratory Plymouth | Wellcome Sanger Institute for the COVID-19 Genomics UK (COG-UK) Consortium | Lighthouse Laboratory Plymouth and Alex Alderton, Roberto Amato, Jeffrey Barrett, Sonia Goncalves, Ewan Harrison, David K. Jackson, Ian Johnston, Dominic Kwiatkowski, Cordelia Langford, John Sillitoe on behalf of the Wellcome Sanger Institute COVID-19 Surveillance Team |
| EPI_ISL_2511472, EPI_ISL_2511474, EPI_ISL_2511475, EPI_ISL_2511476, EPI_ISL_2511477, EPI_ISL_2511478 | Lighthouse Lab in Milton Keynes | Wellcome Sanger Institute for the COVID-19 Genomics UK (COG-UK) Consortium | The Lighthouse Lab in Milton Keynes and Alex Alderton, Roberto Amato, Jeffrey Barrett, Sonia Goncalves, Ewan Harrison, David K. Jackson, Ian Johnston, Dominic Kwiatkowski, Cordelia Langford, John Sillitoe on behalf of the Wellcome Sanger Institute COVID-19 Surveillance Team |
| EPI_ISL_2511479 | Lighthouse Lab in Alderley Park | Wellcome Sanger Institute for the COVID-19 Genomics UK (COG-UK) Consortium | Jacquelyn Wynn, Mairead Hyland, The Lighthouse Lab in Alderley Park and Alex Alderton, Roberto Amato, Jeffrey Barrett, Sonia Goncalves, Ewan Harrison, David K. Jackson, Ian Johnston, Dominic Kwiatkowski, Cordelia Langford, John Sillitoe on behalf of the Wellcome Sanger Institute COVID-19 Surveillance Team |
| EPI_ISL_2511480 | Lighthouse Laboratory Plymouth | Wellcome Sanger Institute for the COVID-19 Genomics UK (COG-UK) Consortium | Lighthouse Laboratory Plymouth and Alex Alderton, Roberto Amato, Jeffrey Barrett, Sonia Goncalves, Ewan Harrison, David K. Jackson, Ian Johnston, Dominic Kwiatkowski, Cordelia Langford, John Sillitoe on behalf of the Wellcome Sanger Institute COVID-19 Surveillance Team |
| EPI_ISL_2511481, EPI_ISL_2511482 | Lighthouse Lab in Alderley Park | Wellcome Sanger Institute for the COVID-19 Genomics UK (COG-UK) Consortium | Jacquelyn Wynn, Mairead Hyland, The Lighthouse Lab in Alderley Park and Alex Alderton, Roberto Amato, Jeffrey Barrett, Sonia Goncalves, Ewan Harrison, David K. Jackson, Ian Johnston, Dominic Kwiatkowski, Cordelia Langford, John Sillitoe on behalf of the Wellcome Sanger Institute COVID-19 Surveillance Team |
| EPI_ISL_2511483 | Lighthouse Laboratory Plymouth | Wellcome Sanger Institute for the COVID-19 Genomics UK (COG-UK) Consortium | Lighthouse Laboratory Plymouth and Alex Alderton, Roberto Amato, Jeffrey Barrett, Sonia Goncalves, Ewan Harrison, David K. Jackson, Ian Johnston, Dominic Kwiatkowski, Cordelia Langford, John Sillitoe on behalf of the Wellcome Sanger Institute COVID-19 Surveillance Team |
| EPI_ISL_2511484 | Lighthouse Lab in Alderley Park | Wellcome Sanger Institute for the COVID-19 Genomics UK (COG-UK) Consortium | Jacquelyn Wynn, Mairead Hyland, The Lighthouse Lab in Alderley Park and Alex Alderton, Roberto Amato, Jeffrey Barrett, Sonia Goncalves, Ewan Harrison, David K. Jackson, Ian Johnston, Dominic Kwiatkowski, Cordelia Langford, John Sillitoe on behalf of the Wellcome Sanger Institute COVID-19 Surveillance Team |
| EPI_ISL_2511485, EPI_ISL_2511486, EPI_ISL_2511487, EPI_ISL_2511488 | Lighthouse Lab in Milton Keynes | Wellcome Sanger Institute for the COVID-19 Genomics UK (COG-UK) Consortium | The Lighthouse Lab in Milton Keynes and Alex Alderton, Roberto Amato, Jeffrey Barrett, Sonia Goncalves, Ewan Harrison, David K. Jackson, Ian Johnston, Dominic Kwiatkowski, Cordelia Langford, John Sillitoe on behalf of the Wellcome Sanger Institute COVID-19 Surveillance Team |
| EPI_ISL_2511489 | Lighthouse Lab in Glasgow | Wellcome Sanger Institute for the COVID-19 Genomics UK (COG-UK) Consortium | Harper VanSteenhouse, Yumi Kasai, David Gray, Carol Clugston, Anna Dominiczak and Alex Alderton, Roberto Amato, Jeffrey Barrett, Sonia Goncalves, Ewan Harrison, David K. Jackson, Ian Johnston, Dominic Kwiatkowski, Cordelia Langford, John Sillitoe on behalf of the Wellcome Sanger Institute COVID-19 Surveillance Team |
| EPI_ISL_2511490, EPI_ISL_2511491 | Lighthouse Lab in Milton Keynes | Wellcome Sanger Institute for the COVID-19 Genomics UK (COG-UK) Consortium | The Lighthouse Lab in Milton Keynes and Alex Alderton, Roberto Amato, Jeffrey Barrett, Sonia Goncalves, Ewan Harrison, David K. Jackson, Ian Johnston, Dominic Kwiatkowski, Cordelia Langford, John Sillitoe on behalf of the Wellcome Sanger Institute COVID-19 Surveillance Team |
| EPI_ISL_2511492 | Lighthouse Lab in Glasgow | Wellcome Sanger Institute for the COVID-19 Genomics UK (COG-UK) Consortium | Harper VanSteenhouse, Yumi Kasai, David Gray, Carol Clugston, Anna Dominiczak and Alex Alderton, Roberto Amato, Jeffrey Barrett, Sonia Goncalves, Ewan Harrison, David K. Jackson, Ian Johnston, Dominic Kwiatkowski, Cordelia Langford, John Sillitoe on behalf of the Wellcome Sanger Institute COVID-19 Surveillance Team |
| EPI_ISL_2511493 | Lighthouse Lab in Alderley Park | Wellcome Sanger Institute for the COVID-19 Genomics UK (COG-UK) Consortium | Jacquelyn Wynn, Mairead Hyland, The Lighthouse Lab in Alderley Park and Alex Alderton, Roberto Amato, Jeffrey Barrett, Sonia Goncalves, Ewan Harrison, David K. Jackson, Ian Johnston, Dominic Kwiatkowski, Cordelia Langford, John Sillitoe on behalf of the Wellcome Sanger Institute COVID-19 Surveillance Team |
| EPI_ISL_2511494, EPI_ISL_2511495, EPI_ISL_2511496 | Lighthouse Lab in Milton Keynes | Wellcome Sanger Institute for the COVID-19 Genomics UK (COG-UK) Consortium | The Lighthouse Lab in Milton Keynes and Alex Alderton, Roberto Amato, Jeffrey Barrett, Sonia Goncalves, Ewan Harrison, David K. Jackson, Ian Johnston, Dominic Kwiatkowski, Cordelia Langford, John Sillitoe on behalf of the Wellcome Sanger Institute COVID-19 Surveillance Team |
| EPI_ISL_2511497 | Lighthouse Lab in Alderley Park | Wellcome Sanger Institute for the COVID-19 Genomics UK (COG-UK) Consortium | Jacquelyn Wynn, Mairead Hyland, The Lighthouse Lab in Alderley Park and Alex Alderton, Roberto Amato, Jeffrey Barrett, Sonia Goncalves, Ewan Harrison, David K. Jackson, Ian Johnston, Dominic Kwiatkowski, Cordelia Langford, John Sillitoe on behalf of the Wellcome Sanger Institute COVID-19 Surveillance Team |
| EPI_ISL_2511498 | Lighthouse Lab in Milton Keynes | Wellcome Sanger Institute for the COVID-19 Genomics UK (COG-UK) Consortium | The Lighthouse Lab in Milton Keynes and Alex Alderton, Roberto Amato, Jeffrey Barrett, Sonia Goncalves, Ewan Harrison, David K. Jackson, Ian Johnston, Dominic Kwiatkowski, Cordelia Langford, John Sillitoe on behalf of the Wellcome Sanger Institute COVID-19 Surveillance Team |
| EPI_ISL_2511500, EPI_ISL_2511501 | Lighthouse Laboratory Plymouth | Wellcome Sanger Institute for the COVID-19 Genomics UK (COG-UK) Consortium | Lighthouse Laboratory Plymouth and Alex Alderton, Roberto Amato, Jeffrey Barrett, Sonia Goncalves, Ewan Harrison, David K. Jackson, Ian Johnston, Dominic Kwiatkowski, Cordelia Langford, John Sillitoe on behalf of the Wellcome Sanger Institute COVID-19 Surveillance Team |
| EPI_ISL_2511502 | Lighthouse Lab in Glasgow | Wellcome Sanger Institute for the COVID-19 Genomics UK (COG-UK) Consortium | Harper VanSteenhouse, Yumi Kasai, David Gray, Carol Clugston, Anna Dominiczak and Alex Alderton, Roberto Amato, Jeffrey Barrett, Sonia Goncalves, Ewan Harrison, David K. Jackson, Ian Johnston, Dominic Kwiatkowski, Cordelia Langford, John Sillitoe on behalf of the Wellcome Sanger Institute COVID-19 Surveillance Team |
| EPI_ISL_2511503 | Lighthouse Lab in Milton Keynes | Wellcome Sanger Institute for the COVID-19 Genomics UK (COG-UK) Consortium | The Lighthouse Lab in Milton Keynes and Alex Alderton, Roberto Amato, Jeffrey Barrett, Sonia Goncalves, Ewan Harrison, David K. Jackson, Ian Johnston, Dominic Kwiatkowski, Cordelia Langford, John Sillitoe on behalf of the Wellcome Sanger Institute COVID-19 Surveillance Team |
| EPI_ISL_2511504 | Lighthouse Lab in Glasgow | Wellcome Sanger Institute for the COVID-19 Genomics UK (COG-UK) Consortium | Harper VanSteenhouse, Yumi Kasai, David Gray, Carol Clugston, Anna Dominiczak and Alex Alderton, Roberto Amato, Jeffrey Barrett, Sonia Goncalves, Ewan Harrison, David K. Jackson, Ian Johnston, Dominic Kwiatkowski, Cordelia Langford, John Sillitoe on behalf of the Wellcome Sanger Institute COVID-19 Surveillance Team |
| EPI_ISL_2511505 | Lighthouse Lab in Alderley Park | Wellcome Sanger Institute for the COVID-19 Genomics UK (COG-UK) Consortium | Jacquelyn Wynn, Mairead Hyland, The Lighthouse Lab in Alderley Park and Alex Alderton, Roberto Amato, Jeffrey Barrett, Sonia Goncalves, Ewan Harrison, David K. Jackson, Ian Johnston, Dominic Kwiatkowski, Cordelia Langford, John Sillitoe on behalf of the Wellcome Sanger Institute COVID-19 Surveillance Team |
| EPI_ISL_2511507, EPI_ISL_2511508, EPI_ISL_2511509 | Lighthouse Lab in Milton Keynes | Wellcome Sanger Institute for the COVID-19 Genomics UK (COG-UK) Consortium | The Lighthouse Lab in Milton Keynes and Alex Alderton, Roberto Amato, Jeffrey Barrett, Sonia Goncalves, Ewan Harrison, David K. Jackson, Ian Johnston, Dominic Kwiatkowski, Cordelia Langford, John Sillitoe on behalf of the Wellcome Sanger Institute COVID-19 Surveillance Team |
| EPI_ISL_2511510, EPI_ISL_2511511 | Lighthouse Lab in Alderley Park | Wellcome Sanger Institute for the COVID-19 Genomics UK (COG-UK) Consortium | Jacquelyn Wynn, Mairead Hyland, The Lighthouse Lab in Alderley Park and Alex Alderton, Roberto Amato, Jeffrey Barrett, Sonia Goncalves, Ewan Harrison, David K. Jackson, Ian Johnston, Dominic Kwiatkowski, Cordelia Langford, John Sillitoe on behalf of the Wellcome Sanger Institute COVID-19 Surveillance Team |
| EPI_ISL_2511512, EPI_ISL_2511513 | Lighthouse Lab in Milton Keynes | Wellcome Sanger Institute for the COVID-19 Genomics UK (COG-UK) Consortium | The Lighthouse Lab in Milton Keynes and Alex Alderton, Roberto Amato, Jeffrey Barrett, Sonia Goncalves, Ewan Harrison, David K. Jackson, Ian Johnston, Dominic Kwiatkowski, Cordelia Langford, John Sillitoe on behalf of the Wellcome Sanger Institute COVID-19 Surveillance Team |
| EPI_ISL_2511514 | Lighthouse Laboratory Plymouth | Wellcome Sanger Institute for the COVID-19 Genomics UK (COG-UK) Consortium | Lighthouse Laboratory Plymouth and Alex Alderton, Roberto Amato, Jeffrey Barrett, Sonia Goncalves, Ewan Harrison, David K. Jackson, Ian Johnston, Dominic Kwiatkowski, Cordelia Langford, John Sillitoe on behalf of the Wellcome Sanger Institute COVID-19 Surveillance Team |
| EPI_ISL_2511515 | Lighthouse Lab in Milton Keynes | Wellcome Sanger Institute for the COVID-19 Genomics UK (COG-UK) Consortium | The Lighthouse Lab in Milton Keynes and Alex Alderton, Roberto Amato, Jeffrey Barrett, Sonia Goncalves, Ewan Harrison, David K. Jackson, Ian Johnston, Dominic Kwiatkowski, Cordelia Langford, John Sillitoe on behalf of the Wellcome Sanger Institute COVID-19 Surveillance Team |
| EPI_ISL_2511516 | Lighthouse Laboratory Plymouth | Wellcome Sanger Institute for the COVID-19 Genomics UK (COG-UK) Consortium | Lighthouse Laboratory Plymouth and Alex Alderton, Roberto Amato, Jeffrey Barrett, Sonia Goncalves, Ewan Harrison, David K. Jackson, Ian Johnston, Dominic Kwiatkowski, Cordelia Langford, John Sillitoe on behalf of the Wellcome Sanger Institute COVID-19 Surveillance Team |
| EPI_ISL_2511517 | Lighthouse Lab in Milton Keynes | Wellcome Sanger Institute for the COVID-19 Genomics UK (COG-UK) Consortium | The Lighthouse Lab in Milton Keynes and Alex Alderton, Roberto Amato, Jeffrey Barrett, Sonia Goncalves, Ewan Harrison, David K. Jackson, Ian Johnston, Dominic Kwiatkowski, Cordelia Langford, John Sillitoe on behalf of the Wellcome Sanger Institute COVID-19 Surveillance Team |
| EPI_ISL_2511518 | Lighthouse Lab in Alderley Park | Wellcome Sanger Institute for the COVID-19 Genomics UK (COG-UK) Consortium | Jacquelyn Wynn, Mairead Hyland, The Lighthouse Lab in Alderley Park and Alex Alderton, Roberto Amato, Jeffrey Barrett, Sonia Goncalves, Ewan Harrison, David K. Jackson, Ian Johnston, Dominic Kwiatkowski, Cordelia Langford, John Sillitoe on behalf of the Wellcome Sanger Institute COVID-19 Surveillance Team |
| EPI_ISL_2511519, EPI_ISL_2511520, EPI_ISL_2511521 | Lighthouse Lab in Milton Keynes | Wellcome Sanger Institute for the COVID-19 Genomics UK (COG-UK) Consortium | The Lighthouse Lab in Milton Keynes and Alex Alderton, Roberto Amato, Jeffrey Barrett, Sonia Goncalves, Ewan Harrison, David K. Jackson, Ian Johnston, Dominic Kwiatkowski, Cordelia Langford, John Sillitoe on behalf of the Wellcome Sanger Institute COVID-19 Surveillance Team |
| EPI_ISL_2511522 | Lighthouse Laboratory Plymouth | Wellcome Sanger Institute for the COVID-19 Genomics UK (COG-UK) Consortium | Lighthouse Laboratory Plymouth and Alex Alderton, Roberto Amato, Jeffrey Barrett, Sonia Goncalves, Ewan Harrison, David K. Jackson, Ian Johnston, Dominic Kwiatkowski, Cordelia Langford, John Sillitoe on behalf of the Wellcome Sanger Institute COVID-19 Surveillance Team |

| | | | |
|---|---|---|---|
| EPI_ISL_2511523, EPI_ISL_2511524, EPI_ISL_2511525, EPI_ISL_2511526, EPI_ISL_2511527, EPI_ISL_2511529 | Lighthouse Lab in Alderley Park | Wellcome Sanger Institute for the COVID-19 Genomics UK (COG-UK) Consortium | Jacquelyn Wynn, Mairead Hyland, The Lighthouse Lab in Alderley Park and Alex Alderton, Roberto Amato, Jeffrey Barrett, Sonia Goncalves, Ewan Harrison, David K. Jackson, Ian Johnston, Dominic Kwiatkowski, Cordelia Langford, John Sillitoe on behalf of the Wellcome Sanger Institute COVID-19 Surveillance Team |
| EPI_ISL_2511530, EPI_ISL_2511531 | Lighthouse Laboratory Plymouth | Wellcome Sanger Institute for the COVID-19 Genomics UK (COG-UK) Consortium | Lighthouse Laboratory Plymouth and Alex Alderton, Roberto Amato, Jeffrey Barrett, Sonia Goncalves, Ewan Harrison, David K. Jackson, Ian Johnston, Dominic Kwiatkowski, Cordelia Langford, John Sillitoe on behalf of the Wellcome Sanger Institute COVID-19 Surveillance Team |
| EPI_ISL_2511532 | Lighthouse Lab in Alderley Park | Wellcome Sanger Institute for the COVID-19 Genomics UK (COG-UK) Consortium | Jacquelyn Wynn, Mairead Hyland, The Lighthouse Lab in Alderley Park and Alex Alderton, Roberto Amato, Jeffrey Barrett, Sonia Goncalves, Ewan Harrison, David K. Jackson, Ian Johnston, Dominic Kwiatkowski, Cordelia Langford, John Sillitoe on behalf of the Wellcome Sanger Institute COVID-19 Surveillance Team |
| EPI_ISL_2511533, EPI_ISL_2511534, EPI_ISL_2511536 | Lighthouse Laboratory Plymouth | Wellcome Sanger Institute for the COVID-19 Genomics UK (COG-UK) Consortium | Lighthouse Laboratory Plymouth and Alex Alderton, Roberto Amato, Jeffrey Barrett, Sonia Goncalves, Ewan Harrison, David K. Jackson, Ian Johnston, Dominic Kwiatkowski, Cordelia Langford, John Sillitoe on behalf of the Wellcome Sanger Institute COVID-19 Surveillance Team |
| EPI_ISL_2511538 | Lighthouse Lab in Milton Keynes | Wellcome Sanger Institute for the COVID-19 Genomics UK (COG-UK) Consortium | The Lighthouse Lab in Milton Keynes and Alex Alderton, Roberto Amato, Jeffrey Barrett, Sonia Goncalves, Ewan Harrison, David K. Jackson, Ian Johnston, Dominic Kwiatkowski, Cordelia Langford, John Sillitoe on behalf of the Wellcome Sanger Institute COVID-19 Surveillance Team |
| EPI_ISL_2511539, EPI_ISL_2511541 | Lighthouse Lab in Alderley Park | Wellcome Sanger Institute for the COVID-19 Genomics UK (COG-UK) Consortium | Jacquelyn Wynn, Mairead Hyland, The Lighthouse Lab in Alderley Park and Alex Alderton, Roberto Amato, Jeffrey Barrett, Sonia Goncalves, Ewan Harrison, David K. Jackson, Ian Johnston, Dominic Kwiatkowski, Cordelia Langford, John Sillitoe on behalf of the Wellcome Sanger Institute COVID-19 Surveillance Team |
| EPI_ISL_2511542 | Lighthouse Lab in Milton Keynes | Wellcome Sanger Institute for the COVID-19 Genomics UK (COG-UK) Consortium | The Lighthouse Lab in Milton Keynes and Alex Alderton, Roberto Amato, Jeffrey Barrett, Sonia Goncalves, Ewan Harrison, David K. Jackson, Ian Johnston, Dominic Kwiatkowski, Cordelia Langford, John Sillitoe on behalf of the Wellcome Sanger Institute COVID-19 Surveillance Team |
| EPI_ISL_2511543 | Lighthouse Laboratory Plymouth | Wellcome Sanger Institute for the COVID-19 Genomics UK (COG-UK) Consortium | Lighthouse Laboratory Plymouth and Alex Alderton, Roberto Amato, Jeffrey Barrett, Sonia Goncalves, Ewan Harrison, David K. Jackson, Ian Johnston, Dominic Kwiatkowski, Cordelia Langford, John Sillitoe on behalf of the Wellcome Sanger Institute COVID-19 Surveillance Team |
| EPI_ISL_2511544 | Lighthouse Lab in Milton Keynes | Wellcome Sanger Institute for the COVID-19 Genomics UK (COG-UK) Consortium | The Lighthouse Lab in Milton Keynes and Alex Alderton, Roberto Amato, Jeffrey Barrett, Sonia Goncalves, Ewan Harrison, David K. Jackson, Ian Johnston, Dominic Kwiatkowski, Cordelia Langford, John Sillitoe on behalf of the Wellcome Sanger Institute COVID-19 Surveillance Team |
| EPI_ISL_2511545 | Lighthouse Lab in Alderley Park | Wellcome Sanger Institute for the COVID-19 Genomics UK (COG-UK) Consortium | Jacquelyn Wynn, Mairead Hyland, The Lighthouse Lab in Alderley Park and Alex Alderton, Roberto Amato, Jeffrey Barrett, Sonia Goncalves, Ewan Harrison, David K. Jackson, Ian Johnston, Dominic Kwiatkowski, Cordelia Langford, John Sillitoe on behalf of the Wellcome Sanger Institute COVID-19 Surveillance Team |
| EPI_ISL_2511546, EPI_ISL_2511547, EPI_ISL_2511548 | Lighthouse Laboratory Plymouth | Wellcome Sanger Institute for the COVID-19 Genomics UK (COG-UK) Consortium | Lighthouse Laboratory Plymouth and Alex Alderton, Roberto Amato, Jeffrey Barrett, Sonia Goncalves, Ewan Harrison, David K. Jackson, Ian Johnston, Dominic Kwiatkowski, Cordelia Langford, John Sillitoe on behalf of the Wellcome Sanger Institute COVID-19 Surveillance Team |
| EPI_ISL_2511549, EPI_ISL_2511550 | Lighthouse Lab in Milton Keynes | Wellcome Sanger Institute for the COVID-19 Genomics UK (COG-UK) Consortium | The Lighthouse Lab in Milton Keynes and Alex Alderton, Roberto Amato, Jeffrey Barrett, Sonia Goncalves, Ewan Harrison, David K. Jackson, Ian Johnston, Dominic Kwiatkowski, Cordelia Langford, John Sillitoe on behalf of the Wellcome Sanger Institute COVID-19 Surveillance Team |
| EPI_ISL_2511551 | Lighthouse Lab in Alderley Park | Wellcome Sanger Institute for the COVID-19 Genomics UK (COG-UK) Consortium | Jacquelyn Wynn, Mairead Hyland, The Lighthouse Lab in Alderley Park and Alex Alderton, Roberto Amato, Jeffrey Barrett, Sonia Goncalves, Ewan Harrison, David K. Jackson, Ian Johnston, Dominic Kwiatkowski, Cordelia Langford, John Sillitoe on behalf of the Wellcome Sanger Institute COVID-19 Surveillance Team |
| EPI_ISL_2511552, EPI_ISL_2511553, EPI_ISL_2511554 | Lighthouse Laboratory Plymouth | Wellcome Sanger Institute for the COVID-19 Genomics UK (COG-UK) Consortium | Lighthouse Laboratory Plymouth and Alex Alderton, Roberto Amato, Jeffrey Barrett, Sonia Goncalves, Ewan Harrison, David K. Jackson, Ian Johnston, Dominic Kwiatkowski, Cordelia Langford, John Sillitoe on behalf of the Wellcome Sanger Institute COVID-19 Surveillance Team |
| EPI_ISL_2511555, EPI_ISL_2511556, EPI_ISL_2511557 | Lighthouse Lab in Milton Keynes | Wellcome Sanger Institute for the COVID-19 Genomics UK (COG-UK) Consortium | The Lighthouse Lab in Milton Keynes and Alex Alderton, Roberto Amato, Jeffrey Barrett, Sonia Goncalves, Ewan Harrison, David K. Jackson, Ian Johnston, Dominic Kwiatkowski, Cordelia Langford, John Sillitoe on behalf of the Wellcome Sanger Institute COVID-19 Surveillance Team |
| EPI_ISL_2511558 | Lighthouse Lab in Alderley Park | Wellcome Sanger Institute for the COVID-19 Genomics UK (COG-UK) Consortium | Jacquelyn Wynn, Mairead Hyland, The Lighthouse Lab in Alderley Park and Alex Alderton, Roberto Amato, Jeffrey Barrett, Sonia Goncalves, Ewan Harrison, David K. Jackson, Ian Johnston, Dominic Kwiatkowski, Cordelia Langford, John Sillitoe on behalf of the Wellcome Sanger Institute COVID-19 Surveillance Team |
| EPI_ISL_2511559, EPI_ISL_2511560 | Lighthouse Lab in Milton Keynes | Wellcome Sanger Institute for the COVID-19 Genomics UK (COG-UK) Consortium | The Lighthouse Lab in Milton Keynes and Alex Alderton, Roberto Amato, Jeffrey Barrett, Sonia Goncalves, Ewan Harrison, David K. Jackson, Ian Johnston, Dominic Kwiatkowski, Cordelia Langford, John Sillitoe on behalf of the Wellcome Sanger Institute COVID-19 Surveillance Team |
| EPI_ISL_2511561, EPI_ISL_2511562, EPI_ISL_2511563 | Lighthouse Lab in Alderley Park | Wellcome Sanger Institute for the COVID-19 Genomics UK (COG-UK) Consortium | Jacquelyn Wynn, Mairead Hyland, The Lighthouse Lab in Alderley Park and Alex Alderton, Roberto Amato, Jeffrey Barrett, Sonia Goncalves, Ewan Harrison, David K. Jackson, Ian Johnston, Dominic Kwiatkowski, Cordelia Langford, John Sillitoe on behalf of the Wellcome Sanger Institute COVID-19 Surveillance Team |
| EPI_ISL_2511564 | Lighthouse Laboratory Plymouth | Wellcome Sanger Institute for the COVID-19 Genomics UK (COG-UK) Consortium | Lighthouse Laboratory Plymouth and Alex Alderton, Roberto Amato, Jeffrey Barrett, Sonia Goncalves, Ewan Harrison, David K. Jackson, Ian Johnston, Dominic Kwiatkowski, Cordelia Langford, John Sillitoe on behalf of the Wellcome Sanger Institute COVID-19 Surveillance Team |
| EPI_ISL_2511565, EPI_ISL_2511566, EPI_ISL_2511567 | Lighthouse Lab in Alderley Park | Wellcome Sanger Institute for the COVID-19 Genomics UK (COG-UK) Consortium | Jacquelyn Wynn, Mairead Hyland, The Lighthouse Lab in Alderley Park and Alex Alderton, Roberto Amato, Jeffrey Barrett, Sonia Goncalves, Ewan Harrison, David K. Jackson, Ian Johnston, Dominic Kwiatkowski, Cordelia Langford, John Sillitoe on behalf of the Wellcome Sanger Institute COVID-19 Surveillance Team |
| EPI_ISL_2511568 | Lighthouse Lab in Milton Keynes | Wellcome Sanger Institute for the COVID-19 Genomics UK (COG-UK) Consortium | The Lighthouse Lab in Milton Keynes and Alex Alderton, Roberto Amato, Jeffrey Barrett, Sonia Goncalves, Ewan Harrison, David K. Jackson, Ian Johnston, Dominic Kwiatkowski, Cordelia Langford, John Sillitoe on behalf of the Wellcome Sanger Institute COVID-19 Surveillance Team |
| EPI_ISL_2511569 | Lighthouse Laboratory Plymouth | Wellcome Sanger Institute for the COVID-19 Genomics UK (COG-UK) Consortium | Lighthouse Laboratory Plymouth and Alex Alderton, Roberto Amato, Jeffrey Barrett, Sonia Goncalves, Ewan Harrison, David K. Jackson, Ian Johnston, Dominic Kwiatkowski, Cordelia Langford, John Sillitoe on behalf of the Wellcome Sanger Institute COVID-19 Surveillance Team |
| EPI_ISL_2511570 | Lighthouse Lab in Alderley Park | Wellcome Sanger Institute for the COVID-19 Genomics UK (COG-UK) Consortium | Jacquelyn Wynn, Mairead Hyland, The Lighthouse Lab in Alderley Park and Alex Alderton, Roberto Amato, Jeffrey Barrett, Sonia Goncalves, Ewan Harrison, David K. Jackson, Ian Johnston, Dominic Kwiatkowski, Cordelia Langford, John Sillitoe on behalf of the Wellcome Sanger Institute COVID-19 Surveillance Team |
| EPI_ISL_2511571 | Lighthouse Lab in Glasgow | Wellcome Sanger Institute for the COVID-19 Genomics UK (COG-UK) Consortium | Harper VanSteenhouse, Yumi Kasai, David Gray, Carol Clugston, Anna Dominiczak and Alex Alderton, Roberto Amato, Jeffrey Barrett, Sonia Goncalves, Ewan Harrison, David K. Jackson, Ian Johnston, Dominic Kwiatkowski, Cordelia Langford, John Sillitoe on behalf of the Wellcome Sanger Institute COVID-19 Surveillance Team |
| EPI_ISL_2511572 | Lighthouse Lab in Milton Keynes | Wellcome Sanger Institute for the COVID-19 Genomics UK (COG-UK) Consortium | The Lighthouse Lab in Milton Keynes and Alex Alderton, Roberto Amato, Jeffrey Barrett, Sonia Goncalves, Ewan Harrison, David K. Jackson, Ian Johnston, Dominic Kwiatkowski, Cordelia Langford, John Sillitoe on behalf of the Wellcome Sanger Institute COVID-19 Surveillance Team |
| EPI_ISL_2511573 | Lighthouse Lab in Glasgow | Wellcome Sanger Institute for the COVID-19 Genomics UK (COG-UK) Consortium | Harper VanSteenhouse, Yumi Kasai, David Gray, Carol Clugston, Anna Dominiczak and Alex Alderton, Roberto Amato, Jeffrey Barrett, Sonia Goncalves, Ewan Harrison, David K. Jackson, Ian Johnston, Dominic Kwiatkowski, Cordelia Langford, John Sillitoe on behalf of the Wellcome Sanger Institute COVID-19 Surveillance Team |
| EPI_ISL_2511574, EPI_ISL_2511575 | Lighthouse Lab in Alderley Park | Wellcome Sanger Institute for the COVID-19 Genomics UK (COG-UK) Consortium | Jacquelyn Wynn, Mairead Hyland, The Lighthouse Lab in Alderley Park and Alex Alderton, Roberto Amato, Jeffrey Barrett, Sonia Goncalves, Ewan Harrison, David K. Jackson, Ian Johnston, Dominic Kwiatkowski, Cordelia Langford, John Sillitoe on behalf of the Wellcome Sanger Institute COVID-19 Surveillance Team |
| EPI_ISL_2511577, EPI_ISL_2511578, EPI_ISL_2511579 | Lighthouse Lab in Milton Keynes | Wellcome Sanger Institute for the COVID-19 Genomics UK (COG-UK) Consortium | The Lighthouse Lab in Milton Keynes and Alex Alderton, Roberto Amato, Jeffrey Barrett, Sonia Goncalves, Ewan Harrison, David K. Jackson, Ian Johnston, Dominic Kwiatkowski, Cordelia Langford, John Sillitoe on behalf of the Wellcome Sanger Institute COVID-19 Surveillance Team |
| EPI_ISL_2511580 | Lighthouse Lab in Alderley Park | Wellcome Sanger Institute for the COVID-19 Genomics UK (COG-UK) Consortium | Jacquelyn Wynn, Mairead Hyland, The Lighthouse Lab in Alderley Park and Alex Alderton, Roberto Amato, Jeffrey Barrett, Sonia Goncalves, Ewan Harrison, David K. Jackson, Ian Johnston, Dominic Kwiatkowski, Cordelia Langford, John Sillitoe on behalf of the Wellcome Sanger Institute COVID-19 Surveillance Team |
| EPI_ISL_2511581 | Lighthouse Laboratory Plymouth | Wellcome Sanger Institute for the COVID-19 Genomics UK (COG-UK) Consortium | Lighthouse Laboratory Plymouth and Alex Alderton, Roberto Amato, Jeffrey Barrett, Sonia Goncalves, Ewan Harrison, David K. Jackson, Ian Johnston, Dominic Kwiatkowski, Cordelia Langford, John Sillitoe on behalf of the Wellcome Sanger Institute COVID-19 Surveillance Team |
| EPI_ISL_2511582 | Lighthouse Lab in Alderley Park | Wellcome Sanger Institute for the COVID-19 Genomics UK (COG-UK) Consortium | Jacquelyn Wynn, Mairead Hyland, The Lighthouse Lab in Alderley Park and Alex Alderton, Roberto Amato, Jeffrey Barrett, Sonia Goncalves, Ewan Harrison, David K. Jackson, Ian Johnston, Dominic Kwiatkowski, Cordelia Langford, John Sillitoe on behalf of the Wellcome Sanger Institute COVID-19 Surveillance Team |
| EPI_ISL_2511583 | Lighthouse Laboratory Plymouth | Wellcome Sanger Institute for the COVID-19 Genomics UK (COG-UK) Consortium | Lighthouse Laboratory Plymouth and Alex Alderton, Roberto Amato, Jeffrey Barrett, Sonia Goncalves, Ewan Harrison, David K. Jackson, Ian Johnston, Dominic Kwiatkowski, Cordelia Langford, John Sillitoe on behalf of the Wellcome Sanger Institute COVID-19 Surveillance Team |
| EPI_ISL_2511584 | Lighthouse Lab in Milton Keynes | Wellcome Sanger Institute for the COVID-19 Genomics UK (COG-UK) Consortium | The Lighthouse Lab in Milton Keynes and Alex Alderton, Roberto Amato, Jeffrey Barrett, Sonia Goncalves, Ewan Harrison, David K. Jackson, Ian Johnston, Dominic Kwiatkowski, Cordelia Langford, John Sillitoe on behalf of the Wellcome Sanger Institute COVID-19 Surveillance Team |
| EPI_ISL_2511585, EPI_ISL_2511587 | Lighthouse Lab in Alderley Park | Wellcome Sanger Institute for the COVID-19 Genomics UK (COG-UK) Consortium | Jacquelyn Wynn, Mairead Hyland, The Lighthouse Lab in Alderley Park and Alex Alderton, Roberto Amato, Jeffrey Barrett, Sonia Goncalves, Ewan Harrison, David K. Jackson, Ian Johnston, Dominic Kwiatkowski, Cordelia Langford, John Sillitoe on behalf of the Wellcome Sanger Institute COVID-19 Surveillance Team |
| EPI_ISL_2511588, EPI_ISL_2511589 | Lighthouse Lab in Milton Keynes | Wellcome Sanger Institute for the COVID-19 Genomics UK (COG-UK) Consortium | The Lighthouse Lab in Milton Keynes and Alex Alderton, Roberto Amato, Jeffrey Barrett, Sonia Goncalves, Ewan Harrison, David K. Jackson, Ian Johnston, Dominic Kwiatkowski, Cordelia Langford, John Sillitoe on behalf of the Wellcome Sanger Institute COVID-19 Surveillance Team |
| EPI_ISL_2511590, EPI_ISL_2511591, EPI_ISL_2511592 | Lighthouse Laboratory Plymouth | Wellcome Sanger Institute for the COVID-19 Genomics UK (COG-UK) Consortium | Lighthouse Laboratory Plymouth and Alex Alderton, Roberto Amato, Jeffrey Barrett, Sonia Goncalves, Ewan Harrison, David K. Jackson, Ian Johnston, Dominic Kwiatkowski, Cordelia Langford, John Sillitoe on behalf of the Wellcome Sanger Institute COVID-19 Surveillance Team |
| EPI_ISL_2511593 | Lighthouse Lab in Alderley Park | Wellcome Sanger Institute for the COVID-19 Genomics UK (COG-UK) Consortium | Jacquelyn Wynn, Mairead Hyland, The Lighthouse Lab in Alderley Park and Alex Alderton, Roberto Amato, Jeffrey Barrett, Sonia Goncalves, Ewan Harrison, David K. Jackson, Ian Johnston, Dominic Kwiatkowski, Cordelia Langford, John Sillitoe on behalf of the Wellcome Sanger Institute COVID-19 Surveillance Team |
| EPI_ISL_2511594 | Lighthouse Lab in Milton Keynes | Wellcome Sanger Institute for the COVID-19 Genomics UK (COG-UK) Consortium | The Lighthouse Lab in Milton Keynes and Alex Alderton, Roberto Amato, Jeffrey Barrett, Sonia Goncalves, Ewan Harrison, David K. Jackson, Ian Johnston, Dominic Kwiatkowski, Cordelia Langford, John Sillitoe on behalf of the Wellcome Sanger Institute COVID-19 Surveillance Team |
| EPI_ISL_2511595 | Lighthouse Lab in Alderley Park | Wellcome Sanger Institute for the COVID-19 Genomics UK (COG-UK) Consortium | Jacquelyn Wynn, Mairead Hyland, The Lighthouse Lab in Alderley Park and Alex Alderton, Roberto Amato, Jeffrey Barrett, Sonia Goncalves, Ewan Harrison, David K. Jackson, Ian Johnston, Dominic Kwiatkowski, Cordelia Langford, John Sillitoe on behalf of the Wellcome Sanger Institute COVID-19 Surveillance Team |
| EPI_ISL_2511596 | Lighthouse Lab in Glasgow | Wellcome Sanger Institute for the COVID-19 Genomics UK (COG-UK) Consortium | Harper VanSteenhouse, Yumi Kasai, David Gray, Carol Clugston, Anna Dominiczak and Alex Alderton, Roberto Amato, Jeffrey Barrett, Sonia Goncalves, Ewan Harrison, David K. Jackson, Ian Johnston, Dominic Kwiatkowski, Cordelia Langford, John Sillitoe on behalf of the Wellcome Sanger Institute COVID-19 Surveillance Team |
| EPI_ISL_2511598, | Lighthouse Lab in Alderley Park | Wellcome Sanger Institute for the COVID-19 Genomics UK (COG-UK) | Jacquelyn Wynn, Mairead Hyland, The Lighthouse Lab in Alderley Park and Alex Alderton, Roberto Amato, Jeffrey Barrett, Sonia Goncalves, Ewan Harrison, David K. Jackson, Ian Johnston, Dominic Kwiatkowski, Cordelia Langford, John Sillitoe on |

| | | | |
|---|---|---|---|
| EPI_ISL_2511599 | | Consortium | behalf of the Wellcome Sanger Institute COVID-19 Surveillance Team |
| EPI_ISL_2511600 | Lighthouse Lab in Milton Keynes | Wellcome Sanger Institute for the COVID-19 Genomics UK (COG-UK) Consortium | The Lighthouse Lab in Milton Keynes and Alex Alderton, Roberto Amato, Jeffrey Barrett, Sonia Goncalves, Ewan Harrison, David K. Jackson, Ian Johnston, Dominic Kwiatkowski, Cordelia Langford, John Sillitoe on behalf of the Wellcome Sanger Institute COVID-19 Surveillance Team |
| EPI_ISL_2511602 | Lighthouse Lab in Alderley Park | Wellcome Sanger Institute for the COVID-19 Genomics UK (COG-UK) Consortium | Jacquelyn Wynn, Mairead Hyland, The Lighthouse Lab in Alderley Park and Alex Alderton, Roberto Amato, Jeffrey Barrett, Sonia Goncalves, Ewan Harrison, David K. Jackson, Ian Johnston, Dominic Kwiatkowski, Cordelia Langford, John Sillitoe on behalf of the Wellcome Sanger Institute COVID-19 Surveillance Team |
| EPI_ISL_2511603 | Lighthouse Lab in Milton Keynes | Wellcome Sanger Institute for the COVID-19 Genomics UK (COG-UK) Consortium | The Lighthouse Lab in Milton Keynes and Alex Alderton, Roberto Amato, Jeffrey Barrett, Sonia Goncalves, Ewan Harrison, David K. Jackson, Ian Johnston, Dominic Kwiatkowski, Cordelia Langford, John Sillitoe on behalf of the Wellcome Sanger Institute COVID-19 Surveillance Team |
| EPI_ISL_2511604 | Lighthouse Laboratory Plymouth | Wellcome Sanger Institute for the COVID-19 Genomics UK (COG-UK) Consortium | Lighthouse Laboratory Plymouth and Alex Alderton, Roberto Amato, Jeffrey Barrett, Sonia Goncalves, Ewan Harrison, David K. Jackson, Ian Johnston, Dominic Kwiatkowski, Cordelia Langford, John Sillitoe on behalf of the Wellcome Sanger Institute COVID-19 Surveillance Team |
| EPI_ISL_2511605 | Lighthouse Lab in Milton Keynes | Wellcome Sanger Institute for the COVID-19 Genomics UK (COG-UK) Consortium | The Lighthouse Lab in Milton Keynes and Alex Alderton, Roberto Amato, Jeffrey Barrett, Sonia Goncalves, Ewan Harrison, David K. Jackson, Ian Johnston, Dominic Kwiatkowski, Cordelia Langford, John Sillitoe on behalf of the Wellcome Sanger Institute COVID-19 Surveillance Team |
| EPI_ISL_2511606 | Lighthouse Lab in Alderley Park | Wellcome Sanger Institute for the COVID-19 Genomics UK (COG-UK) Consortium | Jacquelyn Wynn, Mairead Hyland, The Lighthouse Lab in Alderley Park and Alex Alderton, Roberto Amato, Jeffrey Barrett, Sonia Goncalves, Ewan Harrison, David K. Jackson, Ian Johnston, Dominic Kwiatkowski, Cordelia Langford, John Sillitoe on behalf of the Wellcome Sanger Institute COVID-19 Surveillance Team |
| EPI_ISL_2511607, EPI_ISL_2511608 | Lighthouse Lab in Milton Keynes | Wellcome Sanger Institute for the COVID-19 Genomics UK (COG-UK) Consortium | The Lighthouse Lab in Milton Keynes and Alex Alderton, Roberto Amato, Jeffrey Barrett, Sonia Goncalves, Ewan Harrison, David K. Jackson, Ian Johnston, Dominic Kwiatkowski, Cordelia Langford, John Sillitoe on behalf of the Wellcome Sanger Institute COVID-19 Surveillance Team |
| EPI_ISL_2511609 | Lighthouse Laboratory Plymouth | Wellcome Sanger Institute for the COVID-19 Genomics UK (COG-UK) Consortium | Lighthouse Laboratory Plymouth and Alex Alderton, Roberto Amato, Jeffrey Barrett, Sonia Goncalves, Ewan Harrison, David K. Jackson, Ian Johnston, Dominic Kwiatkowski, Cordelia Langford, John Sillitoe on behalf of the Wellcome Sanger Institute COVID-19 Surveillance Team |
| EPI_ISL_2511610 | Lighthouse Lab in Alderley Park | Wellcome Sanger Institute for the COVID-19 Genomics UK (COG-UK) Consortium | Jacquelyn Wynn, Mairead Hyland, The Lighthouse Lab in Alderley Park and Alex Alderton, Roberto Amato, Jeffrey Barrett, Sonia Goncalves, Ewan Harrison, David K. Jackson, Ian Johnston, Dominic Kwiatkowski, Cordelia Langford, John Sillitoe on behalf of the Wellcome Sanger Institute COVID-19 Surveillance Team |
| EPI_ISL_2511611 | Lighthouse Lab in Milton Keynes | Wellcome Sanger Institute for the COVID-19 Genomics UK (COG-UK) Consortium | The Lighthouse Lab in Milton Keynes and Alex Alderton, Roberto Amato, Jeffrey Barrett, Sonia Goncalves, Ewan Harrison, David K. Jackson, Ian Johnston, Dominic Kwiatkowski, Cordelia Langford, John Sillitoe on behalf of the Wellcome Sanger Institute COVID-19 Surveillance Team |
| EPI_ISL_2511612 | Lighthouse Lab in Alderley Park | Wellcome Sanger Institute for the COVID-19 Genomics UK (COG-UK) Consortium | Jacquelyn Wynn, Mairead Hyland, The Lighthouse Lab in Alderley Park and Alex Alderton, Roberto Amato, Jeffrey Barrett, Sonia Goncalves, Ewan Harrison, David K. Jackson, Ian Johnston, Dominic Kwiatkowski, Cordelia Langford, John Sillitoe on behalf of the Wellcome Sanger Institute COVID-19 Surveillance Team |
| EPI_ISL_2511613 | Lighthouse Laboratory Plymouth | Wellcome Sanger Institute for the COVID-19 Genomics UK (COG-UK) Consortium | Lighthouse Laboratory Plymouth and Alex Alderton, Roberto Amato, Jeffrey Barrett, Sonia Goncalves, Ewan Harrison, David K. Jackson, Ian Johnston, Dominic Kwiatkowski, Cordelia Langford, John Sillitoe on behalf of the Wellcome Sanger Institute COVID-19 Surveillance Team |
| EPI_ISL_2511614, EPI_ISL_2511615, EPI_ISL_2511616 | Lighthouse Lab in Alderley Park | Wellcome Sanger Institute for the COVID-19 Genomics UK (COG-UK) Consortium | Jacquelyn Wynn, Mairead Hyland, The Lighthouse Lab in Alderley Park and Alex Alderton, Roberto Amato, Jeffrey Barrett, Sonia Goncalves, Ewan Harrison, David K. Jackson, Ian Johnston, Dominic Kwiatkowski, Cordelia Langford, John Sillitoe on behalf of the Wellcome Sanger Institute COVID-19 Surveillance Team |
| EPI_ISL_2511617 | Lighthouse Lab in Milton Keynes | Wellcome Sanger Institute for the COVID-19 Genomics UK (COG-UK) Consortium | The Lighthouse Lab in Milton Keynes and Alex Alderton, Roberto Amato, Jeffrey Barrett, Sonia Goncalves, Ewan Harrison, David K. Jackson, Ian Johnston, Dominic Kwiatkowski, Cordelia Langford, John Sillitoe on behalf of the Wellcome Sanger Institute COVID-19 Surveillance Team |
| EPI_ISL_2511618 | Lighthouse Lab in Alderley Park | Wellcome Sanger Institute for the COVID-19 Genomics UK (COG-UK) Consortium | Jacquelyn Wynn, Mairead Hyland, The Lighthouse Lab in Alderley Park and Alex Alderton, Roberto Amato, Jeffrey Barrett, Sonia Goncalves, Ewan Harrison, David K. Jackson, Ian Johnston, Dominic Kwiatkowski, Cordelia Langford, John Sillitoe on behalf of the Wellcome Sanger Institute COVID-19 Surveillance Team |
| EPI_ISL_2511619, EPI_ISL_2511620 | Lighthouse Lab in Milton Keynes | Wellcome Sanger Institute for the COVID-19 Genomics UK (COG-UK) Consortium | The Lighthouse Lab in Milton Keynes and Alex Alderton, Roberto Amato, Jeffrey Barrett, Sonia Goncalves, Ewan Harrison, David K. Jackson, Ian Johnston, Dominic Kwiatkowski, Cordelia Langford, John Sillitoe on behalf of the Wellcome Sanger Institute COVID-19 Surveillance Team |
| EPI_ISL_2511621 | Lighthouse Laboratory Plymouth | Wellcome Sanger Institute for the COVID-19 Genomics UK (COG-UK) Consortium | Lighthouse Laboratory Plymouth and Alex Alderton, Roberto Amato, Jeffrey Barrett, Sonia Goncalves, Ewan Harrison, David K. Jackson, Ian Johnston, Dominic Kwiatkowski, Cordelia Langford, John Sillitoe on behalf of the Wellcome Sanger Institute COVID-19 Surveillance Team |
| EPI_ISL_2511622, EPI_ISL_2511624, EPI_ISL_2511625 | Lighthouse Lab in Milton Keynes | Wellcome Sanger Institute for the COVID-19 Genomics UK (COG-UK) Consortium | The Lighthouse Lab in Milton Keynes and Alex Alderton, Roberto Amato, Jeffrey Barrett, Sonia Goncalves, Ewan Harrison, David K. Jackson, Ian Johnston, Dominic Kwiatkowski, Cordelia Langford, John Sillitoe on behalf of the Wellcome Sanger Institute COVID-19 Surveillance Team |
| EPI_ISL_2511626 | Lighthouse Lab in Alderley Park | Wellcome Sanger Institute for the COVID-19 Genomics UK (COG-UK) Consortium | Jacquelyn Wynn, Mairead Hyland, The Lighthouse Lab in Alderley Park and Alex Alderton, Roberto Amato, Jeffrey Barrett, Sonia Goncalves, Ewan Harrison, David K. Jackson, Ian Johnston, Dominic Kwiatkowski, Cordelia Langford, John Sillitoe on behalf of the Wellcome Sanger Institute COVID-19 Surveillance Team |
| EPI_ISL_2511627 | Lighthouse Lab in Milton Keynes | Wellcome Sanger Institute for the COVID-19 Genomics UK (COG-UK) Consortium | The Lighthouse Lab in Milton Keynes and Alex Alderton, Roberto Amato, Jeffrey Barrett, Sonia Goncalves, Ewan Harrison, David K. Jackson, Ian Johnston, Dominic Kwiatkowski, Cordelia Langford, John Sillitoe on behalf of the Wellcome Sanger Institute COVID-19 Surveillance Team |
| EPI_ISL_2511628, EPI_ISL_2511629 | Lighthouse Lab in Alderley Park | Wellcome Sanger Institute for the COVID-19 Genomics UK (COG-UK) Consortium | Jacquelyn Wynn, Mairead Hyland, The Lighthouse Lab in Alderley Park and Alex Alderton, Roberto Amato, Jeffrey Barrett, Sonia Goncalves, Ewan Harrison, David K. Jackson, Ian Johnston, Dominic Kwiatkowski, Cordelia Langford, John Sillitoe on behalf of the Wellcome Sanger Institute COVID-19 Surveillance Team |
| EPI_ISL_2511630 | Lighthouse Lab in Milton Keynes | Wellcome Sanger Institute for the COVID-19 Genomics UK (COG-UK) Consortium | The Lighthouse Lab in Milton Keynes and Alex Alderton, Roberto Amato, Jeffrey Barrett, Sonia Goncalves, Ewan Harrison, David K. Jackson, Ian Johnston, Dominic Kwiatkowski, Cordelia Langford, John Sillitoe on behalf of the Wellcome Sanger Institute COVID-19 Surveillance Team |
| EPI_ISL_2511631 | Lighthouse Lab in Alderley Park | Wellcome Sanger Institute for the COVID-19 Genomics UK (COG-UK) Consortium | Jacquelyn Wynn, Mairead Hyland, The Lighthouse Lab in Alderley Park and Alex Alderton, Roberto Amato, Jeffrey Barrett, Sonia Goncalves, Ewan Harrison, David K. Jackson, Ian Johnston, Dominic Kwiatkowski, Cordelia Langford, John Sillitoe on behalf of the Wellcome Sanger Institute COVID-19 Surveillance Team |
| EPI_ISL_2511632 | Lighthouse Laboratory Plymouth | Wellcome Sanger Institute for the COVID-19 Genomics UK (COG-UK) Consortium | Lighthouse Laboratory Plymouth and Alex Alderton, Roberto Amato, Jeffrey Barrett, Sonia Goncalves, Ewan Harrison, David K. Jackson, Ian Johnston, Dominic Kwiatkowski, Cordelia Langford, John Sillitoe on behalf of the Wellcome Sanger Institute COVID-19 Surveillance Team |
| EPI_ISL_2511633 | Lighthouse Lab in Milton Keynes | Wellcome Sanger Institute for the COVID-19 Genomics UK (COG-UK) Consortium | The Lighthouse Lab in Milton Keynes and Alex Alderton, Roberto Amato, Jeffrey Barrett, Sonia Goncalves, Ewan Harrison, David K. Jackson, Ian Johnston, Dominic Kwiatkowski, Cordelia Langford, John Sillitoe on behalf of the Wellcome Sanger Institute COVID-19 Surveillance Team |
| EPI_ISL_2511634 | Lighthouse Lab in Alderley Park | Wellcome Sanger Institute for the COVID-19 Genomics UK (COG-UK) Consortium | Jacquelyn Wynn, Mairead Hyland, The Lighthouse Lab in Alderley Park and Alex Alderton, Roberto Amato, Jeffrey Barrett, Sonia Goncalves, Ewan Harrison, David K. Jackson, Ian Johnston, Dominic Kwiatkowski, Cordelia Langford, John Sillitoe on behalf of the Wellcome Sanger Institute COVID-19 Surveillance Team |
| EPI_ISL_2511635 | Lighthouse Laboratory Plymouth | Wellcome Sanger Institute for the COVID-19 Genomics UK (COG-UK) Consortium | Lighthouse Laboratory Plymouth and Alex Alderton, Roberto Amato, Jeffrey Barrett, Sonia Goncalves, Ewan Harrison, David K. Jackson, Ian Johnston, Dominic Kwiatkowski, Cordelia Langford, John Sillitoe on behalf of the Wellcome Sanger Institute COVID-19 Surveillance Team |
| EPI_ISL_2511636, EPI_ISL_2511637 | Lighthouse Lab in Milton Keynes | Wellcome Sanger Institute for the COVID-19 Genomics UK (COG-UK) Consortium | The Lighthouse Lab in Milton Keynes and Alex Alderton, Roberto Amato, Jeffrey Barrett, Sonia Goncalves, Ewan Harrison, David K. Jackson, Ian Johnston, Dominic Kwiatkowski, Cordelia Langford, John Sillitoe on behalf of the Wellcome Sanger Institute COVID-19 Surveillance Team |
| EPI_ISL_2511638 | Lighthouse Lab in Alderley Park | Wellcome Sanger Institute for the COVID-19 Genomics UK (COG-UK) Consortium | Jacquelyn Wynn, Mairead Hyland, The Lighthouse Lab in Alderley Park and Alex Alderton, Roberto Amato, Jeffrey Barrett, Sonia Goncalves, Ewan Harrison, David K. Jackson, Ian Johnston, Dominic Kwiatkowski, Cordelia Langford, John Sillitoe on behalf of the Wellcome Sanger Institute COVID-19 Surveillance Team |
| EPI_ISL_2511639 | Lighthouse Laboratory Plymouth | Wellcome Sanger Institute for the COVID-19 Genomics UK (COG-UK) Consortium | Lighthouse Laboratory Plymouth and Alex Alderton, Roberto Amato, Jeffrey Barrett, Sonia Goncalves, Ewan Harrison, David K. Jackson, Ian Johnston, Dominic Kwiatkowski, Cordelia Langford, John Sillitoe on behalf of the Wellcome Sanger Institute COVID-19 Surveillance Team |
| EPI_ISL_2511640, EPI_ISL_2511641 | Lighthouse Lab in Milton Keynes | Wellcome Sanger Institute for the COVID-19 Genomics UK (COG-UK) Consortium | The Lighthouse Lab in Milton Keynes and Alex Alderton, Roberto Amato, Jeffrey Barrett, Sonia Goncalves, Ewan Harrison, David K. Jackson, Ian Johnston, Dominic Kwiatkowski, Cordelia Langford, John Sillitoe on behalf of the Wellcome Sanger Institute COVID-19 Surveillance Team |
| EPI_ISL_2511642 | Lighthouse Laboratory Plymouth | Wellcome Sanger Institute for the COVID-19 Genomics UK (COG-UK) Consortium | Lighthouse Laboratory Plymouth and Alex Alderton, Roberto Amato, Jeffrey Barrett, Sonia Goncalves, Ewan Harrison, David K. Jackson, Ian Johnston, Dominic Kwiatkowski, Cordelia Langford, John Sillitoe on behalf of the Wellcome Sanger Institute COVID-19 Surveillance Team |
| EPI_ISL_2511643, EPI_ISL_2511644, EPI_ISL_2511645 | Lighthouse Lab in Glasgow | Wellcome Sanger Institute for the COVID-19 Genomics UK (COG-UK) Consortium | Harper VanSteenhouse, Yumi Kasai, David Gray, Carol Clugston, Anna Dominiczak and Alex Alderton, Roberto Amato, Jeffrey Barrett, Sonia Goncalves, Ewan Harrison, David K. Jackson, Ian Johnston, Dominic Kwiatkowski, Cordelia Langford, John Sillitoe on behalf of the Wellcome Sanger Institute COVID-19 Surveillance Team |
| EPI_ISL_2511646 | Lighthouse Lab in Alderley Park | Wellcome Sanger Institute for the COVID-19 Genomics UK (COG-UK) Consortium | Jacquelyn Wynn, Mairead Hyland, The Lighthouse Lab in Alderley Park and Alex Alderton, Roberto Amato, Jeffrey Barrett, Sonia Goncalves, Ewan Harrison, David K. Jackson, Ian Johnston, Dominic Kwiatkowski, Cordelia Langford, John Sillitoe on behalf of the Wellcome Sanger Institute COVID-19 Surveillance Team |
| EPI_ISL_2511647 | Lighthouse Lab in Milton Keynes | Wellcome Sanger Institute for the COVID-19 Genomics UK (COG-UK) Consortium | The Lighthouse Lab in Milton Keynes and Alex Alderton, Roberto Amato, Jeffrey Barrett, Sonia Goncalves, Ewan Harrison, David K. Jackson, Ian Johnston, Dominic Kwiatkowski, Cordelia Langford, John Sillitoe on behalf of the Wellcome Sanger Institute COVID-19 Surveillance Team |
| EPI_ISL_2511648 | Lighthouse Lab in Alderley Park | Wellcome Sanger Institute for the COVID-19 Genomics UK (COG-UK) Consortium | Jacquelyn Wynn, Mairead Hyland, The Lighthouse Lab in Alderley Park and Alex Alderton, Roberto Amato, Jeffrey Barrett, Sonia Goncalves, Ewan Harrison, David K. Jackson, Ian Johnston, Dominic Kwiatkowski, Cordelia Langford, John Sillitoe on behalf of the Wellcome Sanger Institute COVID-19 Surveillance Team |
| EPI_ISL_2511649 | Lighthouse Lab in Milton Keynes | Wellcome Sanger Institute for the COVID-19 Genomics UK (COG-UK) Consortium | The Lighthouse Lab in Milton Keynes and Alex Alderton, Roberto Amato, Jeffrey Barrett, Sonia Goncalves, Ewan Harrison, David K. Jackson, Ian Johnston, Dominic Kwiatkowski, Cordelia Langford, John Sillitoe on behalf of the Wellcome Sanger Institute COVID-19 Surveillance Team |
| EPI_ISL_2511650, EPI_ISL_2511651, EPI_ISL_2511652, EPI_ISL_2511653 | Lighthouse Lab in Alderley Park | Wellcome Sanger Institute for the COVID-19 Genomics UK (COG-UK) Consortium | Jacquelyn Wynn, Mairead Hyland, The Lighthouse Lab in Alderley Park and Alex Alderton, Roberto Amato, Jeffrey Barrett, Sonia Goncalves, Ewan Harrison, David K. Jackson, Ian Johnston, Dominic Kwiatkowski, Cordelia Langford, John Sillitoe on behalf of the Wellcome Sanger Institute COVID-19 Surveillance Team |
| EPI_ISL_2511654, EPI_ISL_2511655, EPI_ISL_2511656 | Lighthouse Lab in Milton Keynes | Wellcome Sanger Institute for the COVID-19 Genomics UK (COG-UK) Consortium | The Lighthouse Lab in Milton Keynes and Alex Alderton, Roberto Amato, Jeffrey Barrett, Sonia Goncalves, Ewan Harrison, David K. Jackson, Ian Johnston, Dominic Kwiatkowski, Cordelia Langford, John Sillitoe on behalf of the Wellcome Sanger Institute COVID-19 Surveillance Team |
| EPI_ISL_2511657, EPI_ISL_2511658, EPI_ISL_2511659 | Lighthouse Lab in Alderley Park | Wellcome Sanger Institute for the COVID-19 Genomics UK (COG-UK) Consortium | Jacquelyn Wynn, Mairead Hyland, The Lighthouse Lab in Alderley Park and Alex Alderton, Roberto Amato, Jeffrey Barrett, Sonia Goncalves, Ewan Harrison, David K. Jackson, Ian Johnston, Dominic Kwiatkowski, Cordelia Langford, John Sillitoe on behalf of the Wellcome Sanger Institute COVID-19 Surveillance Team |
| EPI_ISL_2511660, EPI_ISL_2511661, EPI_ISL_2511662, EPI_ISL_2511663, | Lighthouse Lab in Milton Keynes | Wellcome Sanger Institute for the COVID-19 Genomics UK (COG-UK) Consortium | The Lighthouse Lab in Milton Keynes and Alex Alderton, Roberto Amato, Jeffrey Barrett, Sonia Goncalves, Ewan Harrison, David K. Jackson, Ian Johnston, Dominic Kwiatkowski, Cordelia Langford, John Sillitoe on behalf of the Wellcome Sanger Institute COVID-19 Surveillance Team |

| | | | |
|---|---|---|---|
| EPI_ISL_2511664 | | | |
| EPI_ISL_2511665 | Lighthouse Lab in Alderley Park | Wellcome Sanger Institute for the COVID-19 Genomics UK (COG-UK) Consortium | Jacquelyn Wynn, Mairead Hyland, The Lighthouse Lab in Alderley Park and Alex Alderton, Roberto Amato, Jeffrey Barrett, Sonia Goncalves, Ewan Harrison, David K. Jackson, Ian Johnston, Dominic Kwiatkowski, Cordelia Langford, John Sillitoe on behalf of the Wellcome Sanger Institute COVID-19 Surveillance Team |
| EPI_ISL_2511667, EPI_ISL_2511668, EPI_ISL_2511669, EPI_ISL_2511670, EPI_ISL_2511671, EPI_ISL_2511672, EPI_ISL_2511673, EPI_ISL_2511674, EPI_ISL_2511675, EPI_ISL_2511676 | Lighthouse Lab in Milton Keynes | Wellcome Sanger Institute for the COVID-19 Genomics UK (COG-UK) Consortium | The Lighthouse Lab in Milton Keynes and Alex Alderton, Roberto Amato, Jeffrey Barrett, Sonia Goncalves, Ewan Harrison, David K. Jackson, Ian Johnston, Dominic Kwiatkowski, Cordelia Langford, John Sillitoe on behalf of the Wellcome Sanger Institute COVID-19 Surveillance Team |
| EPI_ISL_2511677 | Lighthouse Lab in Alderley Park | Wellcome Sanger Institute for the COVID-19 Genomics UK (COG-UK) Consortium | Jacquelyn Wynn, Mairead Hyland, The Lighthouse Lab in Alderley Park and Alex Alderton, Roberto Amato, Jeffrey Barrett, Sonia Goncalves, Ewan Harrison, David K. Jackson, Ian Johnston, Dominic Kwiatkowski, Cordelia Langford, John Sillitoe on behalf of the Wellcome Sanger Institute COVID-19 Surveillance Team |
| EPI_ISL_2511678 | Lighthouse Lab in Milton Keynes | Wellcome Sanger Institute for the COVID-19 Genomics UK (COG-UK) Consortium | The Lighthouse Lab in Milton Keynes and Alex Alderton, Roberto Amato, Jeffrey Barrett, Sonia Goncalves, Ewan Harrison, David K. Jackson, Ian Johnston, Dominic Kwiatkowski, Cordelia Langford, John Sillitoe on behalf of the Wellcome Sanger Institute COVID-19 Surveillance Team |
| EPI_ISL_2511679 | Lighthouse Lab in Glasgow | Wellcome Sanger Institute for the COVID-19 Genomics UK (COG-UK) Consortium | Harper VanSteenhouse, Yumi Kasai, David Gray, Carol Clugston, Anna Dominiczak and Alex Alderton, Roberto Amato, Jeffrey Barrett, Sonia Goncalves, Ewan Harrison, David K. Jackson, Ian Johnston, Dominic Kwiatkowski, Cordelia Langford, John Sillitoe on behalf of the Wellcome Sanger Institute COVID-19 Surveillance Team |
| EPI_ISL_2511680, EPI_ISL_2511681, EPI_ISL_2511682 | Lighthouse Lab in Milton Keynes | Wellcome Sanger Institute for the COVID-19 Genomics UK (COG-UK) Consortium | The Lighthouse Lab in Milton Keynes and Alex Alderton, Roberto Amato, Jeffrey Barrett, Sonia Goncalves, Ewan Harrison, David K. Jackson, Ian Johnston, Dominic Kwiatkowski, Cordelia Langford, John Sillitoe on behalf of the Wellcome Sanger Institute COVID-19 Surveillance Team |
| EPI_ISL_2511683 | Lighthouse Lab in Glasgow | Wellcome Sanger Institute for the COVID-19 Genomics UK (COG-UK) Consortium | Harper VanSteenhouse, Yumi Kasai, David Gray, Carol Clugston, Anna Dominiczak and Alex Alderton, Roberto Amato, Jeffrey Barrett, Sonia Goncalves, Ewan Harrison, David K. Jackson, Ian Johnston, Dominic Kwiatkowski, Cordelia Langford, John Sillitoe on behalf of the Wellcome Sanger Institute COVID-19 Surveillance Team |
| EPI_ISL_2511684, EPI_ISL_2511685, EPI_ISL_2511687 | Lighthouse Lab in Alderley Park | Wellcome Sanger Institute for the COVID-19 Genomics UK (COG-UK) Consortium | Jacquelyn Wynn, Mairead Hyland, The Lighthouse Lab in Alderley Park and Alex Alderton, Roberto Amato, Jeffrey Barrett, Sonia Goncalves, Ewan Harrison, David K. Jackson, Ian Johnston, Dominic Kwiatkowski, Cordelia Langford, John Sillitoe on behalf of the Wellcome Sanger Institute COVID-19 Surveillance Team |
| EPI_ISL_2511688 | Lighthouse Lab in Milton Keynes | Wellcome Sanger Institute for the COVID-19 Genomics UK (COG-UK) Consortium | The Lighthouse Lab in Milton Keynes and Alex Alderton, Roberto Amato, Jeffrey Barrett, Sonia Goncalves, Ewan Harrison, David K. Jackson, Ian Johnston, Dominic Kwiatkowski, Cordelia Langford, John Sillitoe on behalf of the Wellcome Sanger Institute COVID-19 Surveillance Team |
| EPI_ISL_2511689 | Lighthouse Lab in Alderley Park | Wellcome Sanger Institute for the COVID-19 Genomics UK (COG-UK) Consortium | Jacquelyn Wynn, Mairead Hyland, The Lighthouse Lab in Alderley Park and Alex Alderton, Roberto Amato, Jeffrey Barrett, Sonia Goncalves, Ewan Harrison, David K. Jackson, Ian Johnston, Dominic Kwiatkowski, Cordelia Langford, John Sillitoe on behalf of the Wellcome Sanger Institute COVID-19 Surveillance Team |
| EPI_ISL_2511690 | Lighthouse Lab in Glasgow | Wellcome Sanger Institute for the COVID-19 Genomics UK (COG-UK) Consortium | Harper VanSteenhouse, Yumi Kasai, David Gray, Carol Clugston, Anna Dominiczak and Alex Alderton, Roberto Amato, Jeffrey Barrett, Sonia Goncalves, Ewan Harrison, David K. Jackson, Ian Johnston, Dominic Kwiatkowski, Cordelia Langford, John Sillitoe on behalf of the Wellcome Sanger Institute COVID-19 Surveillance Team |
| EPI_ISL_2511691, EPI_ISL_2511692, EPI_ISL_2511693 | Lighthouse Lab in Milton Keynes | Wellcome Sanger Institute for the COVID-19 Genomics UK (COG-UK) Consortium | The Lighthouse Lab in Milton Keynes and Alex Alderton, Roberto Amato, Jeffrey Barrett, Sonia Goncalves, Ewan Harrison, David K. Jackson, Ian Johnston, Dominic Kwiatkowski, Cordelia Langford, John Sillitoe on behalf of the Wellcome Sanger Institute COVID-19 Surveillance Team |
| EPI_ISL_2511694 | Lighthouse Lab in Alderley Park | Wellcome Sanger Institute for the COVID-19 Genomics UK (COG-UK) Consortium | Jacquelyn Wynn, Mairead Hyland, The Lighthouse Lab in Alderley Park and Alex Alderton, Roberto Amato, Jeffrey Barrett, Sonia Goncalves, Ewan Harrison, David K. Jackson, Ian Johnston, Dominic Kwiatkowski, Cordelia Langford, John Sillitoe on behalf of the Wellcome Sanger Institute COVID-19 Surveillance Team |
| EPI_ISL_2511695 | Lighthouse Lab in Milton Keynes | Wellcome Sanger Institute for the COVID-19 Genomics UK (COG-UK) Consortium | The Lighthouse Lab in Milton Keynes and Alex Alderton, Roberto Amato, Jeffrey Barrett, Sonia Goncalves, Ewan Harrison, David K. Jackson, Ian Johnston, Dominic Kwiatkowski, Cordelia Langford, John Sillitoe on behalf of the Wellcome Sanger Institute COVID-19 Surveillance Team |
| EPI_ISL_2511696, EPI_ISL_2511697 | Lighthouse Lab in Alderley Park | Wellcome Sanger Institute for the COVID-19 Genomics UK (COG-UK) Consortium | Jacquelyn Wynn, Mairead Hyland, The Lighthouse Lab in Alderley Park and Alex Alderton, Roberto Amato, Jeffrey Barrett, Sonia Goncalves, Ewan Harrison, David K. Jackson, Ian Johnston, Dominic Kwiatkowski, Cordelia Langford, John Sillitoe on behalf of the Wellcome Sanger Institute COVID-19 Surveillance Team |
| EPI_ISL_2511698 | Lighthouse Lab in Milton Keynes | Wellcome Sanger Institute for the COVID-19 Genomics UK (COG-UK) Consortium | The Lighthouse Lab in Milton Keynes and Alex Alderton, Roberto Amato, Jeffrey Barrett, Sonia Goncalves, Ewan Harrison, David K. Jackson, Ian Johnston, Dominic Kwiatkowski, Cordelia Langford, John Sillitoe on behalf of the Wellcome Sanger Institute COVID-19 Surveillance Team |
| EPI_ISL_2511699 | Lighthouse Lab in Alderley Park | Wellcome Sanger Institute for the COVID-19 Genomics UK (COG-UK) Consortium | Jacquelyn Wynn, Mairead Hyland, The Lighthouse Lab in Alderley Park and Alex Alderton, Roberto Amato, Jeffrey Barrett, Sonia Goncalves, Ewan Harrison, David K. Jackson, Ian Johnston, Dominic Kwiatkowski, Cordelia Langford, John Sillitoe on behalf of the Wellcome Sanger Institute COVID-19 Surveillance Team |
| EPI_ISL_2511700 | Lighthouse Lab in Milton Keynes | Wellcome Sanger Institute for the COVID-19 Genomics UK (COG-UK) Consortium | The Lighthouse Lab in Milton Keynes and Alex Alderton, Roberto Amato, Jeffrey Barrett, Sonia Goncalves, Ewan Harrison, David K. Jackson, Ian Johnston, Dominic Kwiatkowski, Cordelia Langford, John Sillitoe on behalf of the Wellcome Sanger Institute COVID-19 Surveillance Team |
| EPI_ISL_2511701 | Lighthouse Lab in Alderley Park | Wellcome Sanger Institute for the COVID-19 Genomics UK (COG-UK) Consortium | Jacquelyn Wynn, Mairead Hyland, The Lighthouse Lab in Alderley Park and Alex Alderton, Roberto Amato, Jeffrey Barrett, Sonia Goncalves, Ewan Harrison, David K. Jackson, Ian Johnston, Dominic Kwiatkowski, Cordelia Langford, John Sillitoe on behalf of the Wellcome Sanger Institute COVID-19 Surveillance Team |
| EPI_ISL_2511703 | Lighthouse Lab in Milton Keynes | Wellcome Sanger Institute for the COVID-19 Genomics UK (COG-UK) Consortium | The Lighthouse Lab in Milton Keynes and Alex Alderton, Roberto Amato, Jeffrey Barrett, Sonia Goncalves, Ewan Harrison, David K. Jackson, Ian Johnston, Dominic Kwiatkowski, Cordelia Langford, John Sillitoe on behalf of the Wellcome Sanger Institute COVID-19 Surveillance Team |
| EPI_ISL_2511704 | Lighthouse Lab in Glasgow | Wellcome Sanger Institute for the COVID-19 Genomics UK (COG-UK) Consortium | Harper VanSteenhouse, Yumi Kasai, David Gray, Carol Clugston, Anna Dominiczak and Alex Alderton, Roberto Amato, Jeffrey Barrett, Sonia Goncalves, Ewan Harrison, David K. Jackson, Ian Johnston, Dominic Kwiatkowski, Cordelia Langford, John Sillitoe on behalf of the Wellcome Sanger Institute COVID-19 Surveillance Team |
| EPI_ISL_2511705 | Lighthouse Lab in Milton Keynes | Wellcome Sanger Institute for the COVID-19 Genomics UK (COG-UK) Consortium | The Lighthouse Lab in Milton Keynes and Alex Alderton, Roberto Amato, Jeffrey Barrett, Sonia Goncalves, Ewan Harrison, David K. Jackson, Ian Johnston, Dominic Kwiatkowski, Cordelia Langford, John Sillitoe on behalf of the Wellcome Sanger Institute COVID-19 Surveillance Team |
| EPI_ISL_2511706, EPI_ISL_2511707 | Lighthouse Lab in Glasgow | Wellcome Sanger Institute for the COVID-19 Genomics UK (COG-UK) Consortium | Harper VanSteenhouse, Yumi Kasai, David Gray, Carol Clugston, Anna Dominiczak and Alex Alderton, Roberto Amato, Jeffrey Barrett, Sonia Goncalves, Ewan Harrison, David K. Jackson, Ian Johnston, Dominic Kwiatkowski, Cordelia Langford, John Sillitoe on behalf of the Wellcome Sanger Institute COVID-19 Surveillance Team |
| EPI_ISL_2511708, EPI_ISL_2511709 | Lighthouse Lab in Alderley Park | Wellcome Sanger Institute for the COVID-19 Genomics UK (COG-UK) Consortium | Jacquelyn Wynn, Mairead Hyland, The Lighthouse Lab in Alderley Park and Alex Alderton, Roberto Amato, Jeffrey Barrett, Sonia Goncalves, Ewan Harrison, David K. Jackson, Ian Johnston, Dominic Kwiatkowski, Cordelia Langford, John Sillitoe on behalf of the Wellcome Sanger Institute COVID-19 Surveillance Team |
| EPI_ISL_2511710 | Lighthouse Lab in Milton Keynes | Wellcome Sanger Institute for the COVID-19 Genomics UK (COG-UK) Consortium | The Lighthouse Lab in Milton Keynes and Alex Alderton, Roberto Amato, Jeffrey Barrett, Sonia Goncalves, Ewan Harrison, David K. Jackson, Ian Johnston, Dominic Kwiatkowski, Cordelia Langford, John Sillitoe on behalf of the Wellcome Sanger Institute COVID-19 Surveillance Team |
| EPI_ISL_2511711, EPI_ISL_2511713 | Lighthouse Lab in Alderley Park | Wellcome Sanger Institute for the COVID-19 Genomics UK (COG-UK) Consortium | Jacquelyn Wynn, Mairead Hyland, The Lighthouse Lab in Alderley Park and Alex Alderton, Roberto Amato, Jeffrey Barrett, Sonia Goncalves, Ewan Harrison, David K. Jackson, Ian Johnston, Dominic Kwiatkowski, Cordelia Langford, John Sillitoe on behalf of the Wellcome Sanger Institute COVID-19 Surveillance Team |
| EPI_ISL_2511714 | Lighthouse Lab in Glasgow | Wellcome Sanger Institute for the COVID-19 Genomics UK (COG-UK) Consortium | Harper VanSteenhouse, Yumi Kasai, David Gray, Carol Clugston, Anna Dominiczak and Alex Alderton, Roberto Amato, Jeffrey Barrett, Sonia Goncalves, Ewan Harrison, David K. Jackson, Ian Johnston, Dominic Kwiatkowski, Cordelia Langford, John Sillitoe on behalf of the Wellcome Sanger Institute COVID-19 Surveillance Team |
| EPI_ISL_2511715, EPI_ISL_2511716 | Lighthouse Lab in Milton Keynes | Wellcome Sanger Institute for the COVID-19 Genomics UK (COG-UK) Consortium | The Lighthouse Lab in Milton Keynes and Alex Alderton, Roberto Amato, Jeffrey Barrett, Sonia Goncalves, Ewan Harrison, David K. Jackson, Ian Johnston, Dominic Kwiatkowski, Cordelia Langford, John Sillitoe on behalf of the Wellcome Sanger Institute COVID-19 Surveillance Team |
| EPI_ISL_2511717, EPI_ISL_2511718 | Lighthouse Lab in Alderley Park | Wellcome Sanger Institute for the COVID-19 Genomics UK (COG-UK) Consortium | Jacquelyn Wynn, Mairead Hyland, The Lighthouse Lab in Alderley Park and Alex Alderton, Roberto Amato, Jeffrey Barrett, Sonia Goncalves, Ewan Harrison, David K. Jackson, Ian Johnston, Dominic Kwiatkowski, Cordelia Langford, John Sillitoe on behalf of the Wellcome Sanger Institute COVID-19 Surveillance Team |
| EPI_ISL_2511719 | Lighthouse Lab in Milton Keynes | Wellcome Sanger Institute for the COVID-19 Genomics UK (COG-UK) Consortium | The Lighthouse Lab in Milton Keynes and Alex Alderton, Roberto Amato, Jeffrey Barrett, Sonia Goncalves, Ewan Harrison, David K. Jackson, Ian Johnston, Dominic Kwiatkowski, Cordelia Langford, John Sillitoe on behalf of the Wellcome Sanger Institute COVID-19 Surveillance Team |
| EPI_ISL_2511721 | Lighthouse Lab in Alderley Park | Wellcome Sanger Institute for the COVID-19 Genomics UK (COG-UK) Consortium | Jacquelyn Wynn, Mairead Hyland, The Lighthouse Lab in Alderley Park and Alex Alderton, Roberto Amato, Jeffrey Barrett, Sonia Goncalves, Ewan Harrison, David K. Jackson, Ian Johnston, Dominic Kwiatkowski, Cordelia Langford, John Sillitoe on behalf of the Wellcome Sanger Institute COVID-19 Surveillance Team |
| EPI_ISL_2511722 | Lighthouse Lab in Milton Keynes | Wellcome Sanger Institute for the COVID-19 Genomics UK (COG-UK) Consortium | The Lighthouse Lab in Milton Keynes and Alex Alderton, Roberto Amato, Jeffrey Barrett, Sonia Goncalves, Ewan Harrison, David K. Jackson, Ian Johnston, Dominic Kwiatkowski, Cordelia Langford, John Sillitoe on behalf of the Wellcome Sanger Institute COVID-19 Surveillance Team |
| EPI_ISL_2511723 | Lighthouse Lab in Glasgow | Wellcome Sanger Institute for the COVID-19 Genomics UK (COG-UK) Consortium | Harper VanSteenhouse, Yumi Kasai, David Gray, Carol Clugston, Anna Dominiczak and Alex Alderton, Roberto Amato, Jeffrey Barrett, Sonia Goncalves, Ewan Harrison, David K. Jackson, Ian Johnston, Dominic Kwiatkowski, Cordelia Langford, John Sillitoe on behalf of the Wellcome Sanger Institute COVID-19 Surveillance Team |
| EPI_ISL_2511724, EPI_ISL_2511725 | Lighthouse Lab in Milton Keynes | Wellcome Sanger Institute for the COVID-19 Genomics UK (COG-UK) Consortium | The Lighthouse Lab in Milton Keynes and Alex Alderton, Roberto Amato, Jeffrey Barrett, Sonia Goncalves, Ewan Harrison, David K. Jackson, Ian Johnston, Dominic Kwiatkowski, Cordelia Langford, John Sillitoe on behalf of the Wellcome Sanger Institute COVID-19 Surveillance Team |
| EPI_ISL_2511726 | Lighthouse Lab in Alderley Park | Wellcome Sanger Institute for the COVID-19 Genomics UK (COG-UK) Consortium | Jacquelyn Wynn, Mairead Hyland, The Lighthouse Lab in Alderley Park and Alex Alderton, Roberto Amato, Jeffrey Barrett, Sonia Goncalves, Ewan Harrison, David K. Jackson, Ian Johnston, Dominic Kwiatkowski, Cordelia Langford, John Sillitoe on behalf of the Wellcome Sanger Institute COVID-19 Surveillance Team |
| EPI_ISL_2511727 | Lighthouse Lab in Milton Keynes | Wellcome Sanger Institute for the COVID-19 Genomics UK (COG-UK) Consortium | The Lighthouse Lab in Milton Keynes and Alex Alderton, Roberto Amato, Jeffrey Barrett, Sonia Goncalves, Ewan Harrison, David K. Jackson, Ian Johnston, Dominic Kwiatkowski, Cordelia Langford, John Sillitoe on behalf of the Wellcome Sanger Institute COVID-19 Surveillance Team |
| EPI_ISL_2511728, EPI_ISL_2511729 | Lighthouse Lab in Glasgow | Wellcome Sanger Institute for the COVID-19 Genomics UK (COG-UK) Consortium | Harper VanSteenhouse, Yumi Kasai, David Gray, Carol Clugston, Anna Dominiczak and Alex Alderton, Roberto Amato, Jeffrey Barrett, Sonia Goncalves, Ewan Harrison, David K. Jackson, Ian Johnston, Dominic Kwiatkowski, Cordelia Langford, John Sillitoe on behalf of the Wellcome Sanger Institute COVID-19 Surveillance Team |
| EPI_ISL_2511730 | Lighthouse Lab in Milton Keynes | Wellcome Sanger Institute for the COVID-19 Genomics UK (COG-UK) Consortium | The Lighthouse Lab in Milton Keynes and Alex Alderton, Roberto Amato, Jeffrey Barrett, Sonia Goncalves, Ewan Harrison, David K. Jackson, Ian Johnston, Dominic Kwiatkowski, Cordelia Langford, John Sillitoe on behalf of the Wellcome Sanger Institute COVID-19 Surveillance Team |
| EPI_ISL_2511731, EPI_ISL_2511732 | Lighthouse Lab in Alderley Park | Wellcome Sanger Institute for the COVID-19 Genomics UK (COG-UK) Consortium | Jacquelyn Wynn, Mairead Hyland, The Lighthouse Lab in Alderley Park and Alex Alderton, Roberto Amato, Jeffrey Barrett, Sonia Goncalves, Ewan Harrison, David K. Jackson, Ian Johnston, Dominic Kwiatkowski, Cordelia Langford, John Sillitoe on behalf of the Wellcome Sanger Institute COVID-19 Surveillance Team |
| EPI_ISL_2511733, EPI_ISL_2511734 | Lighthouse Lab in Milton Keynes | Wellcome Sanger Institute for the COVID-19 Genomics UK (COG-UK) Consortium | The Lighthouse Lab in Milton Keynes and Alex Alderton, Roberto Amato, Jeffrey Barrett, Sonia Goncalves, Ewan Harrison, David K. Jackson, Ian Johnston, Dominic Kwiatkowski, Cordelia Langford, John Sillitoe on behalf of the Wellcome Sanger Institute COVID-19 Surveillance Team |

| | | | |
|---|---|---|---|
| EPI_ISL_2511735 | Lighthouse Lab in Alderley Park | Wellcome Sanger Institute for the COVID-19 Genomics UK (COG-UK) Consortium | Jacquelyn Wynn, Mairead Hyland, The Lighthouse Lab in Alderley Park and Alex Alderton, Roberto Amato, Jeffrey Barrett, Sonia Goncalves, Ewan Harrison, David K. Jackson, Ian Johnston, Dominic Kwiatkowski, Cordelia Langford, John Sillitoe on behalf of the Wellcome Sanger Institute COVID-19 Surveillance Team |
| EPI_ISL_2511736, EPI_ISL_2511737, EPI_ISL_2511738, EPI_ISL_2511739, EPI_ISL_2511740, EPI_ISL_2511741 | Lighthouse Lab in Milton Keynes | Wellcome Sanger Institute for the COVID-19 Genomics UK (COG-UK) Consortium | The Lighthouse Lab in Milton Keynes and Alex Alderton, Roberto Amato, Jeffrey Barrett, Sonia Goncalves, Ewan Harrison, David K. Jackson, Ian Johnston, Dominic Kwiatkowski, Cordelia Langford, John Sillitoe on behalf of the Wellcome Sanger Institute COVID-19 Surveillance Team |
| EPI_ISL_2511742 | Lighthouse Lab in Alderley Park | Wellcome Sanger Institute for the COVID-19 Genomics UK (COG-UK) Consortium | Jacquelyn Wynn, Mairead Hyland, The Lighthouse Lab in Alderley Park and Alex Alderton, Roberto Amato, Jeffrey Barrett, Sonia Goncalves, Ewan Harrison, David K. Jackson, Ian Johnston, Dominic Kwiatkowski, Cordelia Langford, John Sillitoe on behalf of the Wellcome Sanger Institute COVID-19 Surveillance Team |
| EPI_ISL_2511743 | Lighthouse Lab in Glasgow | Wellcome Sanger Institute for the COVID-19 Genomics UK (COG-UK) Consortium | Harper VanSteenhouse, Yumi Kasai, David Gray, Carol Clugston, Anna Dominiczak and Alex Alderton, Roberto Amato, Jeffrey Barrett, Sonia Goncalves, Ewan Harrison, David K. Jackson, Ian Johnston, Dominic Kwiatkowski, Cordelia Langford, John Sillitoe on behalf of the Wellcome Sanger Institute COVID-19 Surveillance Team |
| EPI_ISL_2511744 | Lighthouse Lab in Alderley Park | Wellcome Sanger Institute for the COVID-19 Genomics UK (COG-UK) Consortium | Jacquelyn Wynn, Mairead Hyland, The Lighthouse Lab in Alderley Park and Alex Alderton, Roberto Amato, Jeffrey Barrett, Sonia Goncalves, Ewan Harrison, David K. Jackson, Ian Johnston, Dominic Kwiatkowski, Cordelia Langford, John Sillitoe on behalf of the Wellcome Sanger Institute COVID-19 Surveillance Team |
| EPI_ISL_2511745 | Lighthouse Lab in Milton Keynes | Wellcome Sanger Institute for the COVID-19 Genomics UK (COG-UK) Consortium | The Lighthouse Lab in Milton Keynes and Alex Alderton, Roberto Amato, Jeffrey Barrett, Sonia Goncalves, Ewan Harrison, David K. Jackson, Ian Johnston, Dominic Kwiatkowski, Cordelia Langford, John Sillitoe on behalf of the Wellcome Sanger Institute COVID-19 Surveillance Team |
| EPI_ISL_2511746 | Lighthouse Lab in Alderley Park | Wellcome Sanger Institute for the COVID-19 Genomics UK (COG-UK) Consortium | Jacquelyn Wynn, Mairead Hyland, The Lighthouse Lab in Alderley Park and Alex Alderton, Roberto Amato, Jeffrey Barrett, Sonia Goncalves, Ewan Harrison, David K. Jackson, Ian Johnston, Dominic Kwiatkowski, Cordelia Langford, John Sillitoe on behalf of the Wellcome Sanger Institute COVID-19 Surveillance Team |
| EPI_ISL_2511747, EPI_ISL_2511748, EPI_ISL_2511749 | Lighthouse Lab in Milton Keynes | Wellcome Sanger Institute for the COVID-19 Genomics UK (COG-UK) Consortium | The Lighthouse Lab in Milton Keynes and Alex Alderton, Roberto Amato, Jeffrey Barrett, Sonia Goncalves, Ewan Harrison, David K. Jackson, Ian Johnston, Dominic Kwiatkowski, Cordelia Langford, John Sillitoe on behalf of the Wellcome Sanger Institute COVID-19 Surveillance Team |
| EPI_ISL_2511750 | Lighthouse Lab in Glasgow | Wellcome Sanger Institute for the COVID-19 Genomics UK (COG-UK) Consortium | Harper VanSteenhouse, Yumi Kasai, David Gray, Carol Clugston, Anna Dominiczak and Alex Alderton, Roberto Amato, Jeffrey Barrett, Sonia Goncalves, Ewan Harrison, David K. Jackson, Ian Johnston, Dominic Kwiatkowski, Cordelia Langford, John Sillitoe on behalf of the Wellcome Sanger Institute COVID-19 Surveillance Team |
| EPI_ISL_2511751, EPI_ISL_2511752, EPI_ISL_2511753 | Lighthouse Lab in Milton Keynes | Wellcome Sanger Institute for the COVID-19 Genomics UK (COG-UK) Consortium | The Lighthouse Lab in Milton Keynes and Alex Alderton, Roberto Amato, Jeffrey Barrett, Sonia Goncalves, Ewan Harrison, David K. Jackson, Ian Johnston, Dominic Kwiatkowski, Cordelia Langford, John Sillitoe on behalf of the Wellcome Sanger Institute COVID-19 Surveillance Team |
| EPI_ISL_2511754 | Lighthouse Lab in Alderley Park | Wellcome Sanger Institute for the COVID-19 Genomics UK (COG-UK) Consortium | Jacquelyn Wynn, Mairead Hyland, The Lighthouse Lab in Alderley Park and Alex Alderton, Roberto Amato, Jeffrey Barrett, Sonia Goncalves, Ewan Harrison, David K. Jackson, Ian Johnston, Dominic Kwiatkowski, Cordelia Langford, John Sillitoe on behalf of the Wellcome Sanger Institute COVID-19 Surveillance Team |
| EPI_ISL_2511755 | Lighthouse Lab in Glasgow | Wellcome Sanger Institute for the COVID-19 Genomics UK (COG-UK) Consortium | Harper VanSteenhouse, Yumi Kasai, David Gray, Carol Clugston, Anna Dominiczak and Alex Alderton, Roberto Amato, Jeffrey Barrett, Sonia Goncalves, Ewan Harrison, David K. Jackson, Ian Johnston, Dominic Kwiatkowski, Cordelia Langford, John Sillitoe on behalf of the Wellcome Sanger Institute COVID-19 Surveillance Team |
| EPI_ISL_2511756 | Lighthouse Lab in Milton Keynes | Wellcome Sanger Institute for the COVID-19 Genomics UK (COG-UK) Consortium | The Lighthouse Lab in Milton Keynes and Alex Alderton, Roberto Amato, Jeffrey Barrett, Sonia Goncalves, Ewan Harrison, David K. Jackson, Ian Johnston, Dominic Kwiatkowski, Cordelia Langford, John Sillitoe on behalf of the Wellcome Sanger Institute COVID-19 Surveillance Team |
| EPI_ISL_2511757 | Lighthouse Lab in Alderley Park | Wellcome Sanger Institute for the COVID-19 Genomics UK (COG-UK) Consortium | Jacquelyn Wynn, Mairead Hyland, The Lighthouse Lab in Alderley Park and Alex Alderton, Roberto Amato, Jeffrey Barrett, Sonia Goncalves, Ewan Harrison, David K. Jackson, Ian Johnston, Dominic Kwiatkowski, Cordelia Langford, John Sillitoe on behalf of the Wellcome Sanger Institute COVID-19 Surveillance Team |
| EPI_ISL_2511758, EPI_ISL_2511759, EPI_ISL_2511760 | Lighthouse Lab in Milton Keynes | Wellcome Sanger Institute for the COVID-19 Genomics UK (COG-UK) Consortium | The Lighthouse Lab in Milton Keynes and Alex Alderton, Roberto Amato, Jeffrey Barrett, Sonia Goncalves, Ewan Harrison, David K. Jackson, Ian Johnston, Dominic Kwiatkowski, Cordelia Langford, John Sillitoe on behalf of the Wellcome Sanger Institute COVID-19 Surveillance Team |
| EPI_ISL_2511761, EPI_ISL_2511762 | Lighthouse Lab in Alderley Park | Wellcome Sanger Institute for the COVID-19 Genomics UK (COG-UK) Consortium | Jacquelyn Wynn, Mairead Hyland, The Lighthouse Lab in Alderley Park and Alex Alderton, Roberto Amato, Jeffrey Barrett, Sonia Goncalves, Ewan Harrison, David K. Jackson, Ian Johnston, Dominic Kwiatkowski, Cordelia Langford, John Sillitoe on behalf of the Wellcome Sanger Institute COVID-19 Surveillance Team |
| EPI_ISL_2511763 | Lighthouse Lab in Glasgow | Wellcome Sanger Institute for the COVID-19 Genomics UK (COG-UK) Consortium | Harper VanSteenhouse, Yumi Kasai, David Gray, Carol Clugston, Anna Dominiczak and Alex Alderton, Roberto Amato, Jeffrey Barrett, Sonia Goncalves, Ewan Harrison, David K. Jackson, Ian Johnston, Dominic Kwiatkowski, Cordelia Langford, John Sillitoe on behalf of the Wellcome Sanger Institute COVID-19 Surveillance Team |
| EPI_ISL_2511764, EPI_ISL_2511765 | Lighthouse Lab in Alderley Park | Wellcome Sanger Institute for the COVID-19 Genomics UK (COG-UK) Consortium | Jacquelyn Wynn, Mairead Hyland, The Lighthouse Lab in Alderley Park and Alex Alderton, Roberto Amato, Jeffrey Barrett, Sonia Goncalves, Ewan Harrison, David K. Jackson, Ian Johnston, Dominic Kwiatkowski, Cordelia Langford, John Sillitoe on behalf of the Wellcome Sanger Institute COVID-19 Surveillance Team |
| EPI_ISL_2511766, EPI_ISL_2511767, EPI_ISL_2511768, EPI_ISL_2511770, EPI_ISL_2511771, EPI_ISL_2511772 | Lighthouse Lab in Milton Keynes | Wellcome Sanger Institute for the COVID-19 Genomics UK (COG-UK) Consortium | The Lighthouse Lab in Milton Keynes and Alex Alderton, Roberto Amato, Jeffrey Barrett, Sonia Goncalves, Ewan Harrison, David K. Jackson, Ian Johnston, Dominic Kwiatkowski, Cordelia Langford, John Sillitoe on behalf of the Wellcome Sanger Institute COVID-19 Surveillance Team |
| EPI_ISL_2511773, EPI_ISL_2511776 | Lighthouse Lab in Alderley Park | Wellcome Sanger Institute for the COVID-19 Genomics UK (COG-UK) Consortium | Jacquelyn Wynn, Mairead Hyland, The Lighthouse Lab in Alderley Park and Alex Alderton, Roberto Amato, Jeffrey Barrett, Sonia Goncalves, Ewan Harrison, David K. Jackson, Ian Johnston, Dominic Kwiatkowski, Cordelia Langford, John Sillitoe on behalf of the Wellcome Sanger Institute COVID-19 Surveillance Team |
| EPI_ISL_2511777 | Lighthouse Lab in Milton Keynes | Wellcome Sanger Institute for the COVID-19 Genomics UK (COG-UK) Consortium | The Lighthouse Lab in Milton Keynes and Alex Alderton, Roberto Amato, Jeffrey Barrett, Sonia Goncalves, Ewan Harrison, David K. Jackson, Ian Johnston, Dominic Kwiatkowski, Cordelia Langford, John Sillitoe on behalf of the Wellcome Sanger Institute COVID-19 Surveillance Team |
| EPI_ISL_2511778 | Lighthouse Lab in Glasgow | Wellcome Sanger Institute for the COVID-19 Genomics UK (COG-UK) Consortium | Harper VanSteenhouse, Yumi Kasai, David Gray, Carol Clugston, Anna Dominiczak and Alex Alderton, Roberto Amato, Jeffrey Barrett, Sonia Goncalves, Ewan Harrison, David K. Jackson, Ian Johnston, Dominic Kwiatkowski, Cordelia Langford, John Sillitoe on behalf of the Wellcome Sanger Institute COVID-19 Surveillance Team |
| EPI_ISL_2511779 | Lighthouse Lab in Alderley Park | Wellcome Sanger Institute for the COVID-19 Genomics UK (COG-UK) Consortium | Jacquelyn Wynn, Mairead Hyland, The Lighthouse Lab in Alderley Park and Alex Alderton, Roberto Amato, Jeffrey Barrett, Sonia Goncalves, Ewan Harrison, David K. Jackson, Ian Johnston, Dominic Kwiatkowski, Cordelia Langford, John Sillitoe on behalf of the Wellcome Sanger Institute COVID-19 Surveillance Team |
| EPI_ISL_2511780 | Lighthouse Lab in Milton Keynes | Wellcome Sanger Institute for the COVID-19 Genomics UK (COG-UK) Consortium | The Lighthouse Lab in Milton Keynes and Alex Alderton, Roberto Amato, Jeffrey Barrett, Sonia Goncalves, Ewan Harrison, David K. Jackson, Ian Johnston, Dominic Kwiatkowski, Cordelia Langford, John Sillitoe on behalf of the Wellcome Sanger Institute COVID-19 Surveillance Team |
| EPI_ISL_2511781 | Lighthouse Lab in Alderley Park | Wellcome Sanger Institute for the COVID-19 Genomics UK (COG-UK) Consortium | Jacquelyn Wynn, Mairead Hyland, The Lighthouse Lab in Alderley Park and Alex Alderton, Roberto Amato, Jeffrey Barrett, Sonia Goncalves, Ewan Harrison, David K. Jackson, Ian Johnston, Dominic Kwiatkowski, Cordelia Langford, John Sillitoe on behalf of the Wellcome Sanger Institute COVID-19 Surveillance Team |
| EPI_ISL_2511783 | Lighthouse Lab in Milton Keynes | Wellcome Sanger Institute for the COVID-19 Genomics UK (COG-UK) Consortium | The Lighthouse Lab in Milton Keynes and Alex Alderton, Roberto Amato, Jeffrey Barrett, Sonia Goncalves, Ewan Harrison, David K. Jackson, Ian Johnston, Dominic Kwiatkowski, Cordelia Langford, John Sillitoe on behalf of the Wellcome Sanger Institute COVID-19 Surveillance Team |
| EPI_ISL_2511784 | Lighthouse Lab in Alderley Park | Wellcome Sanger Institute for the COVID-19 Genomics UK (COG-UK) Consortium | Jacquelyn Wynn, Mairead Hyland, The Lighthouse Lab in Alderley Park and Alex Alderton, Roberto Amato, Jeffrey Barrett, Sonia Goncalves, Ewan Harrison, David K. Jackson, Ian Johnston, Dominic Kwiatkowski, Cordelia Langford, John Sillitoe on behalf of the Wellcome Sanger Institute COVID-19 Surveillance Team |
| EPI_ISL_2511785, EPI_ISL_2511786 | Lighthouse Lab in Milton Keynes | Wellcome Sanger Institute for the COVID-19 Genomics UK (COG-UK) Consortium | The Lighthouse Lab in Milton Keynes and Alex Alderton, Roberto Amato, Jeffrey Barrett, Sonia Goncalves, Ewan Harrison, David K. Jackson, Ian Johnston, Dominic Kwiatkowski, Cordelia Langford, John Sillitoe on behalf of the Wellcome Sanger Institute COVID-19 Surveillance Team |
| EPI_ISL_2511787 | Lighthouse Lab in Alderley Park | Wellcome Sanger Institute for the COVID-19 Genomics UK (COG-UK) Consortium | Jacquelyn Wynn, Mairead Hyland, The Lighthouse Lab in Alderley Park and Alex Alderton, Roberto Amato, Jeffrey Barrett, Sonia Goncalves, Ewan Harrison, David K. Jackson, Ian Johnston, Dominic Kwiatkowski, Cordelia Langford, John Sillitoe on behalf of the Wellcome Sanger Institute COVID-19 Surveillance Team |
| EPI_ISL_2511788, EPI_ISL_2511789 | Lighthouse Lab in Milton Keynes | Wellcome Sanger Institute for the COVID-19 Genomics UK (COG-UK) Consortium | The Lighthouse Lab in Milton Keynes and Alex Alderton, Roberto Amato, Jeffrey Barrett, Sonia Goncalves, Ewan Harrison, David K. Jackson, Ian Johnston, Dominic Kwiatkowski, Cordelia Langford, John Sillitoe on behalf of the Wellcome Sanger Institute COVID-19 Surveillance Team |
| EPI_ISL_2511790 | Lighthouse Lab in Alderley Park | Wellcome Sanger Institute for the COVID-19 Genomics UK (COG-UK) Consortium | Jacquelyn Wynn, Mairead Hyland, The Lighthouse Lab in Alderley Park and Alex Alderton, Roberto Amato, Jeffrey Barrett, Sonia Goncalves, Ewan Harrison, David K. Jackson, Ian Johnston, Dominic Kwiatkowski, Cordelia Langford, John Sillitoe on behalf of the Wellcome Sanger Institute COVID-19 Surveillance Team |
| EPI_ISL_2511791, EPI_ISL_2511792 | Lighthouse Lab in Milton Keynes | Wellcome Sanger Institute for the COVID-19 Genomics UK (COG-UK) Consortium | The Lighthouse Lab in Milton Keynes and Alex Alderton, Roberto Amato, Jeffrey Barrett, Sonia Goncalves, Ewan Harrison, David K. Jackson, Ian Johnston, Dominic Kwiatkowski, Cordelia Langford, John Sillitoe on behalf of the Wellcome Sanger Institute COVID-19 Surveillance Team |
| EPI_ISL_2511793 | Lighthouse Lab in Alderley Park | Wellcome Sanger Institute for the COVID-19 Genomics UK (COG-UK) Consortium | Jacquelyn Wynn, Mairead Hyland, The Lighthouse Lab in Alderley Park and Alex Alderton, Roberto Amato, Jeffrey Barrett, Sonia Goncalves, Ewan Harrison, David K. Jackson, Ian Johnston, Dominic Kwiatkowski, Cordelia Langford, John Sillitoe on behalf of the Wellcome Sanger Institute COVID-19 Surveillance Team |
| EPI_ISL_2511794, EPI_ISL_2511795, EPI_ISL_2511797, EPI_ISL_2511798 | Lighthouse Lab in Milton Keynes | Wellcome Sanger Institute for the COVID-19 Genomics UK (COG-UK) Consortium | The Lighthouse Lab in Milton Keynes and Alex Alderton, Roberto Amato, Jeffrey Barrett, Sonia Goncalves, Ewan Harrison, David K. Jackson, Ian Johnston, Dominic Kwiatkowski, Cordelia Langford, John Sillitoe on behalf of the Wellcome Sanger Institute COVID-19 Surveillance Team |
| EPI_ISL_2511799 | Lighthouse Lab in Alderley Park | Wellcome Sanger Institute for the COVID-19 Genomics UK (COG-UK) Consortium | Jacquelyn Wynn, Mairead Hyland, The Lighthouse Lab in Alderley Park and Alex Alderton, Roberto Amato, Jeffrey Barrett, Sonia Goncalves, Ewan Harrison, David K. Jackson, Ian Johnston, Dominic Kwiatkowski, Cordelia Langford, John Sillitoe on behalf of the Wellcome Sanger Institute COVID-19 Surveillance Team |
| EPI_ISL_2511800, EPI_ISL_2511802, EPI_ISL_2511803 | Lighthouse Lab in Milton Keynes | Wellcome Sanger Institute for the COVID-19 Genomics UK (COG-UK) Consortium | The Lighthouse Lab in Milton Keynes and Alex Alderton, Roberto Amato, Jeffrey Barrett, Sonia Goncalves, Ewan Harrison, David K. Jackson, Ian Johnston, Dominic Kwiatkowski, Cordelia Langford, John Sillitoe on behalf of the Wellcome Sanger Institute COVID-19 Surveillance Team |
| EPI_ISL_2511804 | Lighthouse Lab in Alderley Park | Wellcome Sanger Institute for the COVID-19 Genomics UK (COG-UK) Consortium | Jacquelyn Wynn, Mairead Hyland, The Lighthouse Lab in Alderley Park and Alex Alderton, Roberto Amato, Jeffrey Barrett, Sonia Goncalves, Ewan Harrison, David K. Jackson, Ian Johnston, Dominic Kwiatkowski, Cordelia Langford, John Sillitoe on behalf of the Wellcome Sanger Institute COVID-19 Surveillance Team |
| EPI_ISL_2511805, EPI_ISL_2511806, EPI_ISL_2511808, EPI_ISL_2511809 | Lighthouse Lab in Milton Keynes | Wellcome Sanger Institute for the COVID-19 Genomics UK (COG-UK) Consortium | The Lighthouse Lab in Milton Keynes and Alex Alderton, Roberto Amato, Jeffrey Barrett, Sonia Goncalves, Ewan Harrison, David K. Jackson, Ian Johnston, Dominic Kwiatkowski, Cordelia Langford, John Sillitoe on behalf of the Wellcome Sanger Institute COVID-19 Surveillance Team |
| EPI_ISL_2511810 | Lighthouse Lab in Glasgow | Wellcome Sanger Institute for the COVID-19 Genomics UK (COG-UK) | Harper VanSteenhouse, Yumi Kasai, David Gray, Carol Clugston, Anna Dominiczak and Alex Alderton, Roberto Amato, Jeffrey Barrett, Sonia Goncalves, Ewan Harrison, David K. Jackson, Ian Johnston, Dominic Kwiatkowski, Cordelia Langford, |

| | | | |
|---|---|---|---|
| | | Consortium | John Sillitoe on behalf of the Wellcome Sanger Institute COVID-19 Surveillance Team |
| EPI_ISL_2511811 | Lighthouse Lab in Alderley Park | Wellcome Sanger Institute for the COVID-19 Genomics UK (COG-UK) Consortium | Jacquelyn Wynn, Mairead Hyland, The Lighthouse Lab in Alderley Park and Alex Alderton, Roberto Amato, Jeffrey Barrett, Sonia Goncalves, Ewan Harrison, David K. Jackson, Ian Johnston, Dominic Kwiatkowski, Cordelia Langford, John Sillitoe on behalf of the Wellcome Sanger Institute COVID-19 Surveillance Team |
| EPI_ISL_2511812 | Lighthouse Lab in Milton Keynes | Wellcome Sanger Institute for the COVID-19 Genomics UK (COG-UK) Consortium | The Lighthouse Lab in Milton Keynes and Alex Alderton, Roberto Amato, Jeffrey Barrett, Sonia Goncalves, Ewan Harrison, David K. Jackson, Ian Johnston, Dominic Kwiatkowski, Cordelia Langford, John Sillitoe on behalf of the Wellcome Sanger Institute COVID-19 Surveillance Team |
| EPI_ISL_2511813, EPI_ISL_2511814 | Lighthouse Lab in Alderley Park | Wellcome Sanger Institute for the COVID-19 Genomics UK (COG-UK) Consortium | Jacquelyn Wynn, Mairead Hyland, The Lighthouse Lab in Alderley Park and Alex Alderton, Roberto Amato, Jeffrey Barrett, Sonia Goncalves, Ewan Harrison, David K. Jackson, Ian Johnston, Dominic Kwiatkowski, Cordelia Langford, John Sillitoe on behalf of the Wellcome Sanger Institute COVID-19 Surveillance Team |
| EPI_ISL_2511815 | Lighthouse Lab in Milton Keynes | Wellcome Sanger Institute for the COVID-19 Genomics UK (COG-UK) Consortium | The Lighthouse Lab in Milton Keynes and Alex Alderton, Roberto Amato, Jeffrey Barrett, Sonia Goncalves, Ewan Harrison, David K. Jackson, Ian Johnston, Dominic Kwiatkowski, Cordelia Langford, John Sillitoe on behalf of the Wellcome Sanger Institute COVID-19 Surveillance Team |
| EPI_ISL_2511816 | Lighthouse Lab in Alderley Park | Wellcome Sanger Institute for the COVID-19 Genomics UK (COG-UK) Consortium | Jacquelyn Wynn, Mairead Hyland, The Lighthouse Lab in Alderley Park and Alex Alderton, Roberto Amato, Jeffrey Barrett, Sonia Goncalves, Ewan Harrison, David K. Jackson, Ian Johnston, Dominic Kwiatkowski, Cordelia Langford, John Sillitoe on behalf of the Wellcome Sanger Institute COVID-19 Surveillance Team |
| EPI_ISL_2511817, EPI_ISL_2511818 | Lighthouse Lab in Milton Keynes | Wellcome Sanger Institute for the COVID-19 Genomics UK (COG-UK) Consortium | The Lighthouse Lab in Milton Keynes and Alex Alderton, Roberto Amato, Jeffrey Barrett, Sonia Goncalves, Ewan Harrison, David K. Jackson, Ian Johnston, Dominic Kwiatkowski, Cordelia Langford, John Sillitoe on behalf of the Wellcome Sanger Institute COVID-19 Surveillance Team |
| EPI_ISL_2511819, EPI_ISL_2511820 | Lighthouse Lab in Alderley Park | Wellcome Sanger Institute for the COVID-19 Genomics UK (COG-UK) Consortium | Jacquelyn Wynn, Mairead Hyland, The Lighthouse Lab in Alderley Park and Alex Alderton, Roberto Amato, Jeffrey Barrett, Sonia Goncalves, Ewan Harrison, David K. Jackson, Ian Johnston, Dominic Kwiatkowski, Cordelia Langford, John Sillitoe on behalf of the Wellcome Sanger Institute COVID-19 Surveillance Team |
| EPI_ISL_2511821 | Lighthouse Lab in Glasgow | Wellcome Sanger Institute for the COVID-19 Genomics UK (COG-UK) Consortium | Harper VanSteenhouse, Yumi Kasai, David Gray, Carol Clugston, Anna Dominiczak and Alex Alderton, Roberto Amato, Jeffrey Barrett, Sonia Goncalves, Ewan Harrison, David K. Jackson, Ian Johnston, Dominic Kwiatkowski, Cordelia Langford, John Sillitoe on behalf of the Wellcome Sanger Institute COVID-19 Surveillance Team |
| EPI_ISL_2511822, EPI_ISL_2511823 | Lighthouse Lab in Milton Keynes | Wellcome Sanger Institute for the COVID-19 Genomics UK (COG-UK) Consortium | The Lighthouse Lab in Milton Keynes and Alex Alderton, Roberto Amato, Jeffrey Barrett, Sonia Goncalves, Ewan Harrison, David K. Jackson, Ian Johnston, Dominic Kwiatkowski, Cordelia Langford, John Sillitoe on behalf of the Wellcome Sanger Institute COVID-19 Surveillance Team |
| EPI_ISL_2511824, EPI_ISL_2511825 | Lighthouse Lab in Alderley Park | Wellcome Sanger Institute for the COVID-19 Genomics UK (COG-UK) Consortium | Jacquelyn Wynn, Mairead Hyland, The Lighthouse Lab in Alderley Park and Alex Alderton, Roberto Amato, Jeffrey Barrett, Sonia Goncalves, Ewan Harrison, David K. Jackson, Ian Johnston, Dominic Kwiatkowski, Cordelia Langford, John Sillitoe on behalf of the Wellcome Sanger Institute COVID-19 Surveillance Team |
| EPI_ISL_2511826, EPI_ISL_2511827 | Lighthouse Lab in Milton Keynes | Wellcome Sanger Institute for the COVID-19 Genomics UK (COG-UK) Consortium | The Lighthouse Lab in Milton Keynes and Alex Alderton, Roberto Amato, Jeffrey Barrett, Sonia Goncalves, Ewan Harrison, David K. Jackson, Ian Johnston, Dominic Kwiatkowski, Cordelia Langford, John Sillitoe on behalf of the Wellcome Sanger Institute COVID-19 Surveillance Team |
| EPI_ISL_2511828 | Lighthouse Lab in Alderley Park | Wellcome Sanger Institute for the COVID-19 Genomics UK (COG-UK) Consortium | Jacquelyn Wynn, Mairead Hyland, The Lighthouse Lab in Alderley Park and Alex Alderton, Roberto Amato, Jeffrey Barrett, Sonia Goncalves, Ewan Harrison, David K. Jackson, Ian Johnston, Dominic Kwiatkowski, Cordelia Langford, John Sillitoe on behalf of the Wellcome Sanger Institute COVID-19 Surveillance Team |
| EPI_ISL_2511829 | Lighthouse Lab in Glasgow | Wellcome Sanger Institute for the COVID-19 Genomics UK (COG-UK) Consortium | Harper VanSteenhouse, Yumi Kasai, David Gray, Carol Clugston, Anna Dominiczak and Alex Alderton, Roberto Amato, Jeffrey Barrett, Sonia Goncalves, Ewan Harrison, David K. Jackson, Ian Johnston, Dominic Kwiatkowski, Cordelia Langford, John Sillitoe on behalf of the Wellcome Sanger Institute COVID-19 Surveillance Team |
| EPI_ISL_2511830, EPI_ISL_2511831 | Lighthouse Lab in Milton Keynes | Wellcome Sanger Institute for the COVID-19 Genomics UK (COG-UK) Consortium | The Lighthouse Lab in Milton Keynes and Alex Alderton, Roberto Amato, Jeffrey Barrett, Sonia Goncalves, Ewan Harrison, David K. Jackson, Ian Johnston, Dominic Kwiatkowski, Cordelia Langford, John Sillitoe on behalf of the Wellcome Sanger Institute COVID-19 Surveillance Team |
| EPI_ISL_2511832, EPI_ISL_2511833, EPI_ISL_2511834 | Lighthouse Lab in Alderley Park | Wellcome Sanger Institute for the COVID-19 Genomics UK (COG-UK) Consortium | Jacquelyn Wynn, Mairead Hyland, The Lighthouse Lab in Alderley Park and Alex Alderton, Roberto Amato, Jeffrey Barrett, Sonia Goncalves, Ewan Harrison, David K. Jackson, Ian Johnston, Dominic Kwiatkowski, Cordelia Langford, John Sillitoe on behalf of the Wellcome Sanger Institute COVID-19 Surveillance Team |
| EPI_ISL_2511835 | Lighthouse Lab in Milton Keynes | Wellcome Sanger Institute for the COVID-19 Genomics UK (COG-UK) Consortium | The Lighthouse Lab in Milton Keynes and Alex Alderton, Roberto Amato, Jeffrey Barrett, Sonia Goncalves, Ewan Harrison, David K. Jackson, Ian Johnston, Dominic Kwiatkowski, Cordelia Langford, John Sillitoe on behalf of the Wellcome Sanger Institute COVID-19 Surveillance Team |
| EPI_ISL_2511836 | Lighthouse Lab in Glasgow | Wellcome Sanger Institute for the COVID-19 Genomics UK (COG-UK) Consortium | Harper VanSteenhouse, Yumi Kasai, David Gray, Carol Clugston, Anna Dominiczak and Alex Alderton, Roberto Amato, Jeffrey Barrett, Sonia Goncalves, Ewan Harrison, David K. Jackson, Ian Johnston, Dominic Kwiatkowski, Cordelia Langford, John Sillitoe on behalf of the Wellcome Sanger Institute COVID-19 Surveillance Team |
| EPI_ISL_2511837 | Lighthouse Lab in Milton Keynes | Wellcome Sanger Institute for the COVID-19 Genomics UK (COG-UK) Consortium | The Lighthouse Lab in Milton Keynes and Alex Alderton, Roberto Amato, Jeffrey Barrett, Sonia Goncalves, Ewan Harrison, David K. Jackson, Ian Johnston, Dominic Kwiatkowski, Cordelia Langford, John Sillitoe on behalf of the Wellcome Sanger Institute COVID-19 Surveillance Team |
| EPI_ISL_2511838 | Lighthouse Lab in Glasgow | Wellcome Sanger Institute for the COVID-19 Genomics UK (COG-UK) Consortium | Harper VanSteenhouse, Yumi Kasai, David Gray, Carol Clugston, Anna Dominiczak and Alex Alderton, Roberto Amato, Jeffrey Barrett, Sonia Goncalves, Ewan Harrison, David K. Jackson, Ian Johnston, Dominic Kwiatkowski, Cordelia Langford, John Sillitoe on behalf of the Wellcome Sanger Institute COVID-19 Surveillance Team |
| EPI_ISL_2511839, EPI_ISL_2511840, EPI_ISL_2511841 | Lighthouse Lab in Milton Keynes | Wellcome Sanger Institute for the COVID-19 Genomics UK (COG-UK) Consortium | The Lighthouse Lab in Milton Keynes and Alex Alderton, Roberto Amato, Jeffrey Barrett, Sonia Goncalves, Ewan Harrison, David K. Jackson, Ian Johnston, Dominic Kwiatkowski, Cordelia Langford, John Sillitoe on behalf of the Wellcome Sanger Institute COVID-19 Surveillance Team |
| EPI_ISL_2511842 | Lighthouse Lab in Alderley Park | Wellcome Sanger Institute for the COVID-19 Genomics UK (COG-UK) Consortium | Jacquelyn Wynn, Mairead Hyland, The Lighthouse Lab in Alderley Park and Alex Alderton, Roberto Amato, Jeffrey Barrett, Sonia Goncalves, Ewan Harrison, David K. Jackson, Ian Johnston, Dominic Kwiatkowski, Cordelia Langford, John Sillitoe on behalf of the Wellcome Sanger Institute COVID-19 Surveillance Team |
| EPI_ISL_2511843 | Lighthouse Lab in Milton Keynes | Wellcome Sanger Institute for the COVID-19 Genomics UK (COG-UK) Consortium | The Lighthouse Lab in Milton Keynes and Alex Alderton, Roberto Amato, Jeffrey Barrett, Sonia Goncalves, Ewan Harrison, David K. Jackson, Ian Johnston, Dominic Kwiatkowski, Cordelia Langford, John Sillitoe on behalf of the Wellcome Sanger Institute COVID-19 Surveillance Team |
| EPI_ISL_2511844 | Lighthouse Lab in Glasgow | Wellcome Sanger Institute for the COVID-19 Genomics UK (COG-UK) Consortium | Harper VanSteenhouse, Yumi Kasai, David Gray, Carol Clugston, Anna Dominiczak and Alex Alderton, Roberto Amato, Jeffrey Barrett, Sonia Goncalves, Ewan Harrison, David K. Jackson, Ian Johnston, Dominic Kwiatkowski, Cordelia Langford, John Sillitoe on behalf of the Wellcome Sanger Institute COVID-19 Surveillance Team |
| EPI_ISL_2511845 | Lighthouse Lab in Milton Keynes | Wellcome Sanger Institute for the COVID-19 Genomics UK (COG-UK) Consortium | The Lighthouse Lab in Milton Keynes and Alex Alderton, Roberto Amato, Jeffrey Barrett, Sonia Goncalves, Ewan Harrison, David K. Jackson, Ian Johnston, Dominic Kwiatkowski, Cordelia Langford, John Sillitoe on behalf of the Wellcome Sanger Institute COVID-19 Surveillance Team |
| EPI_ISL_2511846 | Lighthouse Lab in Glasgow | Wellcome Sanger Institute for the COVID-19 Genomics UK (COG-UK) Consortium | Harper VanSteenhouse, Yumi Kasai, David Gray, Carol Clugston, Anna Dominiczak and Alex Alderton, Roberto Amato, Jeffrey Barrett, Sonia Goncalves, Ewan Harrison, David K. Jackson, Ian Johnston, Dominic Kwiatkowski, Cordelia Langford, John Sillitoe on behalf of the Wellcome Sanger Institute COVID-19 Surveillance Team |
| EPI_ISL_2511847, EPI_ISL_2511848 | Lighthouse Lab in Alderley Park | Wellcome Sanger Institute for the COVID-19 Genomics UK (COG-UK) Consortium | Jacquelyn Wynn, Mairead Hyland, The Lighthouse Lab in Alderley Park and Alex Alderton, Roberto Amato, Jeffrey Barrett, Sonia Goncalves, Ewan Harrison, David K. Jackson, Ian Johnston, Dominic Kwiatkowski, Cordelia Langford, John Sillitoe on behalf of the Wellcome Sanger Institute COVID-19 Surveillance Team |
| EPI_ISL_2511850 | Lighthouse Lab in Milton Keynes | Wellcome Sanger Institute for the COVID-19 Genomics UK (COG-UK) Consortium | The Lighthouse Lab in Milton Keynes and Alex Alderton, Roberto Amato, Jeffrey Barrett, Sonia Goncalves, Ewan Harrison, David K. Jackson, Ian Johnston, Dominic Kwiatkowski, Cordelia Langford, John Sillitoe on behalf of the Wellcome Sanger Institute COVID-19 Surveillance Team |
| EPI_ISL_2511851 | Lighthouse Lab in Glasgow | Wellcome Sanger Institute for the COVID-19 Genomics UK (COG-UK) Consortium | Harper VanSteenhouse, Yumi Kasai, David Gray, Carol Clugston, Anna Dominiczak and Alex Alderton, Roberto Amato, Jeffrey Barrett, Sonia Goncalves, Ewan Harrison, David K. Jackson, Ian Johnston, Dominic Kwiatkowski, Cordelia Langford, John Sillitoe on behalf of the Wellcome Sanger Institute COVID-19 Surveillance Team |
| EPI_ISL_2511852, EPI_ISL_2511853 | Lighthouse Lab in Alderley Park | Wellcome Sanger Institute for the COVID-19 Genomics UK (COG-UK) Consortium | Jacquelyn Wynn, Mairead Hyland, The Lighthouse Lab in Alderley Park and Alex Alderton, Roberto Amato, Jeffrey Barrett, Sonia Goncalves, Ewan Harrison, David K. Jackson, Ian Johnston, Dominic Kwiatkowski, Cordelia Langford, John Sillitoe on behalf of the Wellcome Sanger Institute COVID-19 Surveillance Team |
| EPI_ISL_2511854, EPI_ISL_2511856, EPI_ISL_2511858 | Lighthouse Lab in Glasgow | Wellcome Sanger Institute for the COVID-19 Genomics UK (COG-UK) Consortium | Harper VanSteenhouse, Yumi Kasai, David Gray, Carol Clugston, Anna Dominiczak and Alex Alderton, Roberto Amato, Jeffrey Barrett, Sonia Goncalves, Ewan Harrison, David K. Jackson, Ian Johnston, Dominic Kwiatkowski, Cordelia Langford, John Sillitoe on behalf of the Wellcome Sanger Institute COVID-19 Surveillance Team |
| EPI_ISL_2511859, EPI_ISL_2511860, EPI_ISL_2511861 | Lighthouse Lab in Milton Keynes | Wellcome Sanger Institute for the COVID-19 Genomics UK (COG-UK) Consortium | The Lighthouse Lab in Milton Keynes and Alex Alderton, Roberto Amato, Jeffrey Barrett, Sonia Goncalves, Ewan Harrison, David K. Jackson, Ian Johnston, Dominic Kwiatkowski, Cordelia Langford, John Sillitoe on behalf of the Wellcome Sanger Institute COVID-19 Surveillance Team |
| EPI_ISL_2511862, EPI_ISL_2511863, EPI_ISL_2511864 | Lighthouse Lab in Alderley Park | Wellcome Sanger Institute for the COVID-19 Genomics UK (COG-UK) Consortium | Jacquelyn Wynn, Mairead Hyland, The Lighthouse Lab in Alderley Park and Alex Alderton, Roberto Amato, Jeffrey Barrett, Sonia Goncalves, Ewan Harrison, David K. Jackson, Ian Johnston, Dominic Kwiatkowski, Cordelia Langford, John Sillitoe on behalf of the Wellcome Sanger Institute COVID-19 Surveillance Team |
| EPI_ISL_2511865, EPI_ISL_2511866 | Lighthouse Lab in Milton Keynes | Wellcome Sanger Institute for the COVID-19 Genomics UK (COG-UK) Consortium | The Lighthouse Lab in Milton Keynes and Alex Alderton, Roberto Amato, Jeffrey Barrett, Sonia Goncalves, Ewan Harrison, David K. Jackson, Ian Johnston, Dominic Kwiatkowski, Cordelia Langford, John Sillitoe on behalf of the Wellcome Sanger Institute COVID-19 Surveillance Team |
| EPI_ISL_2511867 | Lighthouse Lab in Alderley Park | Wellcome Sanger Institute for the COVID-19 Genomics UK (COG-UK) Consortium | Jacquelyn Wynn, Mairead Hyland, The Lighthouse Lab in Alderley Park and Alex Alderton, Roberto Amato, Jeffrey Barrett, Sonia Goncalves, Ewan Harrison, David K. Jackson, Ian Johnston, Dominic Kwiatkowski, Cordelia Langford, John Sillitoe on behalf of the Wellcome Sanger Institute COVID-19 Surveillance Team |
| EPI_ISL_2511868 | Lighthouse Lab in Milton Keynes | Wellcome Sanger Institute for the COVID-19 Genomics UK (COG-UK) Consortium | The Lighthouse Lab in Milton Keynes and Alex Alderton, Roberto Amato, Jeffrey Barrett, Sonia Goncalves, Ewan Harrison, David K. Jackson, Ian Johnston, Dominic Kwiatkowski, Cordelia Langford, John Sillitoe on behalf of the Wellcome Sanger Institute COVID-19 Surveillance Team |
| EPI_ISL_2511870, EPI_ISL_2511871 | Lighthouse Lab in Alderley Park | Wellcome Sanger Institute for the COVID-19 Genomics UK (COG-UK) Consortium | Jacquelyn Wynn, Mairead Hyland, The Lighthouse Lab in Alderley Park and Alex Alderton, Roberto Amato, Jeffrey Barrett, Sonia Goncalves, Ewan Harrison, David K. Jackson, Ian Johnston, Dominic Kwiatkowski, Cordelia Langford, John Sillitoe on behalf of the Wellcome Sanger Institute COVID-19 Surveillance Team |
| EPI_ISL_2511872 | Lighthouse Lab in Milton Keynes | Wellcome Sanger Institute for the COVID-19 Genomics UK (COG-UK) Consortium | The Lighthouse Lab in Milton Keynes and Alex Alderton, Roberto Amato, Jeffrey Barrett, Sonia Goncalves, Ewan Harrison, David K. Jackson, Ian Johnston, Dominic Kwiatkowski, Cordelia Langford, John Sillitoe on behalf of the Wellcome Sanger Institute COVID-19 Surveillance Team |
| EPI_ISL_2511873, EPI_ISL_2511874, EPI_ISL_2511875 | Lighthouse Lab in Alderley Park | Wellcome Sanger Institute for the COVID-19 Genomics UK (COG-UK) Consortium | Jacquelyn Wynn, Mairead Hyland, The Lighthouse Lab in Alderley Park and Alex Alderton, Roberto Amato, Jeffrey Barrett, Sonia Goncalves, Ewan Harrison, David K. Jackson, Ian Johnston, Dominic Kwiatkowski, Cordelia Langford, John Sillitoe on behalf of the Wellcome Sanger Institute COVID-19 Surveillance Team |
| EPI_ISL_2511876, EPI_ISL_2511877, EPI_ISL_2511878, EPI_ISL_2511879 | Lighthouse Lab in Milton Keynes | Wellcome Sanger Institute for the COVID-19 Genomics UK (COG-UK) Consortium | The Lighthouse Lab in Milton Keynes and Alex Alderton, Roberto Amato, Jeffrey Barrett, Sonia Goncalves, Ewan Harrison, David K. Jackson, Ian Johnston, Dominic Kwiatkowski, Cordelia Langford, John Sillitoe on behalf of the Wellcome Sanger Institute COVID-19 Surveillance Team |
| EPI_ISL_2511880 | Lighthouse Lab in Alderley Park | Wellcome Sanger Institute for the COVID-19 Genomics UK (COG-UK) Consortium | Jacquelyn Wynn, Mairead Hyland, The Lighthouse Lab in Alderley Park and Alex Alderton, Roberto Amato, Jeffrey Barrett, Sonia Goncalves, Ewan Harrison, David K. Jackson, Ian Johnston, Dominic Kwiatkowski, Cordelia Langford, John Sillitoe on behalf of the Wellcome Sanger Institute COVID-19 Surveillance Team |
| EPI_ISL_2511881, EPI_ISL_2511882, EPI_ISL_2511883, | Lighthouse Lab in Milton Keynes | Wellcome Sanger Institute for the COVID-19 Genomics UK (COG-UK) Consortium | The Lighthouse Lab in Milton Keynes and Alex Alderton, Roberto Amato, Jeffrey Barrett, Sonia Goncalves, Ewan Harrison, David K. Jackson, Ian Johnston, Dominic Kwiatkowski, Cordelia Langford, John Sillitoe on behalf of the Wellcome Sanger Institute COVID-19 Surveillance Team |

| | | | |
|---|---|---|---|
| EPI_ISL_2511884, EPI_ISL_2511885 | | | |
| EPI_ISL_2511886, EPI_ISL_2511887 | Lighthouse Lab in Glasgow | Wellcome Sanger Institute for the COVID-19 Genomics UK (COG-UK) Consortium | Harper VanSteenhouse, Yumi Kasai, David Gray, Carol Clugston, Anna Dominiczak and Alex Alderton, Roberto Amato, Jeffrey Barrett, Sonia Goncalves, Ewan Harrison, David K. Jackson, Ian Johnston, Dominic Kwiatkowski, Cordelia Langford, John Sillitoe on behalf of the Wellcome Sanger Institute COVID-19 Surveillance Team |
| EPI_ISL_2511888 | Lighthouse Lab in Milton Keynes | Wellcome Sanger Institute for the COVID-19 Genomics UK (COG-UK) Consortium | The Lighthouse Lab in Milton Keynes and Alex Alderton, Roberto Amato, Jeffrey Barrett, Sonia Goncalves, Ewan Harrison, David K. Jackson, Ian Johnston, Dominic Kwiatkowski, Cordelia Langford, John Sillitoe on behalf of the Wellcome Sanger Institute COVID-19 Surveillance Team |
| EPI_ISL_2511889 | Lighthouse Lab in Alderley Park | Wellcome Sanger Institute for the COVID-19 Genomics UK (COG-UK) Consortium | Jacquelyn Wynn, Mairead Hyland, The Lighthouse Lab in Alderley Park and Alex Alderton, Roberto Amato, Jeffrey Barrett, Sonia Goncalves, Ewan Harrison, David K. Jackson, Ian Johnston, Dominic Kwiatkowski, Cordelia Langford, John Sillitoe on behalf of the Wellcome Sanger Institute COVID-19 Surveillance Team |
| EPI_ISL_2511890, EPI_ISL_2511891, EPI_ISL_2511892 | Lighthouse Lab in Milton Keynes | Wellcome Sanger Institute for the COVID-19 Genomics UK (COG-UK) Consortium | The Lighthouse Lab in Milton Keynes and Alex Alderton, Roberto Amato, Jeffrey Barrett, Sonia Goncalves, Ewan Harrison, David K. Jackson, Ian Johnston, Dominic Kwiatkowski, Cordelia Langford, John Sillitoe on behalf of the Wellcome Sanger Institute COVID-19 Surveillance Team |
| EPI_ISL_2511893, EPI_ISL_2511894, EPI_ISL_2511895 | Lighthouse Lab in Glasgow | Wellcome Sanger Institute for the COVID-19 Genomics UK (COG-UK) Consortium | Harper VanSteenhouse, Yumi Kasai, David Gray, Carol Clugston, Anna Dominiczak and Alex Alderton, Roberto Amato, Jeffrey Barrett, Sonia Goncalves, Ewan Harrison, David K. Jackson, Ian Johnston, Dominic Kwiatkowski, Cordelia Langford, John Sillitoe on behalf of the Wellcome Sanger Institute COVID-19 Surveillance Team |
| EPI_ISL_2511896 | Lighthouse Lab in Milton Keynes | Wellcome Sanger Institute for the COVID-19 Genomics UK (COG-UK) Consortium | The Lighthouse Lab in Milton Keynes and Alex Alderton, Roberto Amato, Jeffrey Barrett, Sonia Goncalves, Ewan Harrison, David K. Jackson, Ian Johnston, Dominic Kwiatkowski, Cordelia Langford, John Sillitoe on behalf of the Wellcome Sanger Institute COVID-19 Surveillance Team |
| EPI_ISL_2511897, EPI_ISL_2511898 | Lighthouse Lab in Alderley Park | Wellcome Sanger Institute for the COVID-19 Genomics UK (COG-UK) Consortium | Jacquelyn Wynn, Mairead Hyland, The Lighthouse Lab in Alderley Park and Alex Alderton, Roberto Amato, Jeffrey Barrett, Sonia Goncalves, Ewan Harrison, David K. Jackson, Ian Johnston, Dominic Kwiatkowski, Cordelia Langford, John Sillitoe on behalf of the Wellcome Sanger Institute COVID-19 Surveillance Team |
| EPI_ISL_2511899 | Lighthouse Lab in Milton Keynes | Wellcome Sanger Institute for the COVID-19 Genomics UK (COG-UK) Consortium | The Lighthouse Lab in Milton Keynes and Alex Alderton, Roberto Amato, Jeffrey Barrett, Sonia Goncalves, Ewan Harrison, David K. Jackson, Ian Johnston, Dominic Kwiatkowski, Cordelia Langford, John Sillitoe on behalf of the Wellcome Sanger Institute COVID-19 Surveillance Team |
| EPI_ISL_2511900 | Lighthouse Lab in Alderley Park | Wellcome Sanger Institute for the COVID-19 Genomics UK (COG-UK) Consortium | Jacquelyn Wynn, Mairead Hyland, The Lighthouse Lab in Alderley Park and Alex Alderton, Roberto Amato, Jeffrey Barrett, Sonia Goncalves, Ewan Harrison, David K. Jackson, Ian Johnston, Dominic Kwiatkowski, Cordelia Langford, John Sillitoe on behalf of the Wellcome Sanger Institute COVID-19 Surveillance Team |
| EPI_ISL_2511901 | Lighthouse Lab in Milton Keynes | Wellcome Sanger Institute for the COVID-19 Genomics UK (COG-UK) Consortium | The Lighthouse Lab in Milton Keynes and Alex Alderton, Roberto Amato, Jeffrey Barrett, Sonia Goncalves, Ewan Harrison, David K. Jackson, Ian Johnston, Dominic Kwiatkowski, Cordelia Langford, John Sillitoe on behalf of the Wellcome Sanger Institute COVID-19 Surveillance Team |
| EPI_ISL_2511902 | Lighthouse Lab in Alderley Park | Wellcome Sanger Institute for the COVID-19 Genomics UK (COG-UK) Consortium | Jacquelyn Wynn, Mairead Hyland, The Lighthouse Lab in Alderley Park and Alex Alderton, Roberto Amato, Jeffrey Barrett, Sonia Goncalves, Ewan Harrison, David K. Jackson, Ian Johnston, Dominic Kwiatkowski, Cordelia Langford, John Sillitoe on behalf of the Wellcome Sanger Institute COVID-19 Surveillance Team |
| EPI_ISL_2511903 | Lighthouse Lab in Milton Keynes | Wellcome Sanger Institute for the COVID-19 Genomics UK (COG-UK) Consortium | The Lighthouse Lab in Milton Keynes and Alex Alderton, Roberto Amato, Jeffrey Barrett, Sonia Goncalves, Ewan Harrison, David K. Jackson, Ian Johnston, Dominic Kwiatkowski, Cordelia Langford, John Sillitoe on behalf of the Wellcome Sanger Institute COVID-19 Surveillance Team |
| EPI_ISL_2511904, EPI_ISL_2511905 | Lighthouse Lab in Alderley Park | Wellcome Sanger Institute for the COVID-19 Genomics UK (COG-UK) Consortium | Jacquelyn Wynn, Mairead Hyland, The Lighthouse Lab in Alderley Park and Alex Alderton, Roberto Amato, Jeffrey Barrett, Sonia Goncalves, Ewan Harrison, David K. Jackson, Ian Johnston, Dominic Kwiatkowski, Cordelia Langford, John Sillitoe on behalf of the Wellcome Sanger Institute COVID-19 Surveillance Team |
| EPI_ISL_2511906, EPI_ISL_2511907 | Lighthouse Lab in Glasgow | Wellcome Sanger Institute for the COVID-19 Genomics UK (COG-UK) Consortium | Harper VanSteenhouse, Yumi Kasai, David Gray, Carol Clugston, Anna Dominiczak and Alex Alderton, Roberto Amato, Jeffrey Barrett, Sonia Goncalves, Ewan Harrison, David K. Jackson, Ian Johnston, Dominic Kwiatkowski, Cordelia Langford, John Sillitoe on behalf of the Wellcome Sanger Institute COVID-19 Surveillance Team |
| EPI_ISL_2511908 | Lighthouse Lab in Milton Keynes | Wellcome Sanger Institute for the COVID-19 Genomics UK (COG-UK) Consortium | The Lighthouse Lab in Milton Keynes and Alex Alderton, Roberto Amato, Jeffrey Barrett, Sonia Goncalves, Ewan Harrison, David K. Jackson, Ian Johnston, Dominic Kwiatkowski, Cordelia Langford, John Sillitoe on behalf of the Wellcome Sanger Institute COVID-19 Surveillance Team |
| EPI_ISL_2511909 | Lighthouse Lab in Alderley Park | Wellcome Sanger Institute for the COVID-19 Genomics UK (COG-UK) Consortium | Jacquelyn Wynn, Mairead Hyland, The Lighthouse Lab in Alderley Park and Alex Alderton, Roberto Amato, Jeffrey Barrett, Sonia Goncalves, Ewan Harrison, David K. Jackson, Ian Johnston, Dominic Kwiatkowski, Cordelia Langford, John Sillitoe on behalf of the Wellcome Sanger Institute COVID-19 Surveillance Team |
| EPI_ISL_2511910 | Lighthouse Lab in Glasgow | Wellcome Sanger Institute for the COVID-19 Genomics UK (COG-UK) Consortium | Harper VanSteenhouse, Yumi Kasai, David Gray, Carol Clugston, Anna Dominiczak and Alex Alderton, Roberto Amato, Jeffrey Barrett, Sonia Goncalves, Ewan Harrison, David K. Jackson, Ian Johnston, Dominic Kwiatkowski, Cordelia Langford, John Sillitoe on behalf of the Wellcome Sanger Institute COVID-19 Surveillance Team |
| EPI_ISL_2511911, EPI_ISL_2511912, EPI_ISL_2511913, EPI_ISL_2511914 | Lighthouse Lab in Milton Keynes | Wellcome Sanger Institute for the COVID-19 Genomics UK (COG-UK) Consortium | The Lighthouse Lab in Milton Keynes and Alex Alderton, Roberto Amato, Jeffrey Barrett, Sonia Goncalves, Ewan Harrison, David K. Jackson, Ian Johnston, Dominic Kwiatkowski, Cordelia Langford, John Sillitoe on behalf of the Wellcome Sanger Institute COVID-19 Surveillance Team |
| EPI_ISL_2511915 | Lighthouse Lab in Glasgow | Wellcome Sanger Institute for the COVID-19 Genomics UK (COG-UK) Consortium | Harper VanSteenhouse, Yumi Kasai, David Gray, Carol Clugston, Anna Dominiczak and Alex Alderton, Roberto Amato, Jeffrey Barrett, Sonia Goncalves, Ewan Harrison, David K. Jackson, Ian Johnston, Dominic Kwiatkowski, Cordelia Langford, John Sillitoe on behalf of the Wellcome Sanger Institute COVID-19 Surveillance Team |
| EPI_ISL_2511916 | Lighthouse Lab in Alderley Park | Wellcome Sanger Institute for the COVID-19 Genomics UK (COG-UK) Consortium | Jacquelyn Wynn, Mairead Hyland, The Lighthouse Lab in Alderley Park and Alex Alderton, Roberto Amato, Jeffrey Barrett, Sonia Goncalves, Ewan Harrison, David K. Jackson, Ian Johnston, Dominic Kwiatkowski, Cordelia Langford, John Sillitoe on behalf of the Wellcome Sanger Institute COVID-19 Surveillance Team |
| EPI_ISL_2511917, EPI_ISL_2511918, EPI_ISL_2511919 | Lighthouse Lab in Milton Keynes | Wellcome Sanger Institute for the COVID-19 Genomics UK (COG-UK) Consortium | The Lighthouse Lab in Milton Keynes and Alex Alderton, Roberto Amato, Jeffrey Barrett, Sonia Goncalves, Ewan Harrison, David K. Jackson, Ian Johnston, Dominic Kwiatkowski, Cordelia Langford, John Sillitoe on behalf of the Wellcome Sanger Institute COVID-19 Surveillance Team |
| EPI_ISL_2511920 | Lighthouse Lab in Alderley Park | Wellcome Sanger Institute for the COVID-19 Genomics UK (COG-UK) Consortium | Jacquelyn Wynn, Mairead Hyland, The Lighthouse Lab in Alderley Park and Alex Alderton, Roberto Amato, Jeffrey Barrett, Sonia Goncalves, Ewan Harrison, David K. Jackson, Ian Johnston, Dominic Kwiatkowski, Cordelia Langford, John Sillitoe on behalf of the Wellcome Sanger Institute COVID-19 Surveillance Team |
| EPI_ISL_2511921 | Lighthouse Lab in Glasgow | Wellcome Sanger Institute for the COVID-19 Genomics UK (COG-UK) Consortium | Harper VanSteenhouse, Yumi Kasai, David Gray, Carol Clugston, Anna Dominiczak and Alex Alderton, Roberto Amato, Jeffrey Barrett, Sonia Goncalves, Ewan Harrison, David K. Jackson, Ian Johnston, Dominic Kwiatkowski, Cordelia Langford, John Sillitoe on behalf of the Wellcome Sanger Institute COVID-19 Surveillance Team |
| EPI_ISL_2511922 | Lighthouse Lab in Milton Keynes | Wellcome Sanger Institute for the COVID-19 Genomics UK (COG-UK) Consortium | The Lighthouse Lab in Milton Keynes and Alex Alderton, Roberto Amato, Jeffrey Barrett, Sonia Goncalves, Ewan Harrison, David K. Jackson, Ian Johnston, Dominic Kwiatkowski, Cordelia Langford, John Sillitoe on behalf of the Wellcome Sanger Institute COVID-19 Surveillance Team |
| EPI_ISL_2511923 | Lighthouse Lab in Glasgow | Wellcome Sanger Institute for the COVID-19 Genomics UK (COG-UK) Consortium | Harper VanSteenhouse, Yumi Kasai, David Gray, Carol Clugston, Anna Dominiczak and Alex Alderton, Roberto Amato, Jeffrey Barrett, Sonia Goncalves, Ewan Harrison, David K. Jackson, Ian Johnston, Dominic Kwiatkowski, Cordelia Langford, John Sillitoe on behalf of the Wellcome Sanger Institute COVID-19 Surveillance Team |
| EPI_ISL_2511924, EPI_ISL_2511925, EPI_ISL_2511926, EPI_ISL_2511927 | Lighthouse Lab in Milton Keynes | Wellcome Sanger Institute for the COVID-19 Genomics UK (COG-UK) Consortium | The Lighthouse Lab in Milton Keynes and Alex Alderton, Roberto Amato, Jeffrey Barrett, Sonia Goncalves, Ewan Harrison, David K. Jackson, Ian Johnston, Dominic Kwiatkowski, Cordelia Langford, John Sillitoe on behalf of the Wellcome Sanger Institute COVID-19 Surveillance Team |
| EPI_ISL_2511928 | Lighthouse Lab in Alderley Park | Wellcome Sanger Institute for the COVID-19 Genomics UK (COG-UK) Consortium | Jacquelyn Wynn, Mairead Hyland, The Lighthouse Lab in Alderley Park and Alex Alderton, Roberto Amato, Jeffrey Barrett, Sonia Goncalves, Ewan Harrison, David K. Jackson, Ian Johnston, Dominic Kwiatkowski, Cordelia Langford, John Sillitoe on behalf of the Wellcome Sanger Institute COVID-19 Surveillance Team |
| EPI_ISL_2511929, EPI_ISL_2511930, EPI_ISL_2511931, EPI_ISL_2511932, EPI_ISL_2511933 | Lighthouse Lab in Milton Keynes | Wellcome Sanger Institute for the COVID-19 Genomics UK (COG-UK) Consortium | The Lighthouse Lab in Milton Keynes and Alex Alderton, Roberto Amato, Jeffrey Barrett, Sonia Goncalves, Ewan Harrison, David K. Jackson, Ian Johnston, Dominic Kwiatkowski, Cordelia Langford, John Sillitoe on behalf of the Wellcome Sanger Institute COVID-19 Surveillance Team |
| EPI_ISL_2511934, EPI_ISL_2511935, EPI_ISL_2511936 | Lighthouse Lab in Alderley Park | Wellcome Sanger Institute for the COVID-19 Genomics UK (COG-UK) Consortium | Jacquelyn Wynn, Mairead Hyland, The Lighthouse Lab in Alderley Park and Alex Alderton, Roberto Amato, Jeffrey Barrett, Sonia Goncalves, Ewan Harrison, David K. Jackson, Ian Johnston, Dominic Kwiatkowski, Cordelia Langford, John Sillitoe on behalf of the Wellcome Sanger Institute COVID-19 Surveillance Team |
| EPI_ISL_2511937 | Lighthouse Lab in Milton Keynes | Wellcome Sanger Institute for the COVID-19 Genomics UK (COG-UK) Consortium | The Lighthouse Lab in Milton Keynes and Alex Alderton, Roberto Amato, Jeffrey Barrett, Sonia Goncalves, Ewan Harrison, David K. Jackson, Ian Johnston, Dominic Kwiatkowski, Cordelia Langford, John Sillitoe on behalf of the Wellcome Sanger Institute COVID-19 Surveillance Team |
| EPI_ISL_2511938, EPI_ISL_2511939 | Health Services Laboratories | Wellcome Sanger Institute for the COVID-19 Genomics UK (COG-UK) Consortium | Health Services Laboratories and Alex Alderton, Roberto Amato, Jeffrey Barrett, Sonia Goncalves, Ewan Harrison, David K. Jackson, Ian Johnston, Dominic Kwiatkowski, Cordelia Langford, John Sillitoe on behalf of the Wellcome Sanger Institute COVID-19 Surveillance Team |
| EPI_ISL_2511940 | Lighthouse Lab in Alderley Park | Wellcome Sanger Institute for the COVID-19 Genomics UK (COG-UK) Consortium | Jacquelyn Wynn, Mairead Hyland, The Lighthouse Lab in Alderley Park and Alex Alderton, Roberto Amato, Jeffrey Barrett, Sonia Goncalves, Ewan Harrison, David K. Jackson, Ian Johnston, Dominic Kwiatkowski, Cordelia Langford, John Sillitoe on behalf of the Wellcome Sanger Institute COVID-19 Surveillance Team |
| EPI_ISL_2511941, EPI_ISL_2511942 | Health Services Laboratories | Wellcome Sanger Institute for the COVID-19 Genomics UK (COG-UK) Consortium | Health Services Laboratories and Alex Alderton, Roberto Amato, Jeffrey Barrett, Sonia Goncalves, Ewan Harrison, David K. Jackson, Ian Johnston, Dominic Kwiatkowski, Cordelia Langford, John Sillitoe on behalf of the Wellcome Sanger Institute COVID-19 Surveillance Team |
| EPI_ISL_2511944 | Lighthouse Lab in Alderley Park | Wellcome Sanger Institute for the COVID-19 Genomics UK (COG-UK) Consortium | Jacquelyn Wynn, Mairead Hyland, The Lighthouse Lab in Alderley Park and Alex Alderton, Roberto Amato, Jeffrey Barrett, Sonia Goncalves, Ewan Harrison, David K. Jackson, Ian Johnston, Dominic Kwiatkowski, Cordelia Langford, John Sillitoe on behalf of the Wellcome Sanger Institute COVID-19 Surveillance Team |
| EPI_ISL_2511945 | Health Services Laboratories | Wellcome Sanger Institute for the COVID-19 Genomics UK (COG-UK) Consortium | Health Services Laboratories and Alex Alderton, Roberto Amato, Jeffrey Barrett, Sonia Goncalves, Ewan Harrison, David K. Jackson, Ian Johnston, Dominic Kwiatkowski, Cordelia Langford, John Sillitoe on behalf of the Wellcome Sanger Institute COVID-19 Surveillance Team |
| EPI_ISL_2511946 | Lighthouse Lab in Alderley Park | Wellcome Sanger Institute for the COVID-19 Genomics UK (COG-UK) Consortium | Jacquelyn Wynn, Mairead Hyland, The Lighthouse Lab in Alderley Park and Alex Alderton, Roberto Amato, Jeffrey Barrett, Sonia Goncalves, Ewan Harrison, David K. Jackson, Ian Johnston, Dominic Kwiatkowski, Cordelia Langford, John Sillitoe on behalf of the Wellcome Sanger Institute COVID-19 Surveillance Team |
| EPI_ISL_2511947 | Health Services Laboratories | Wellcome Sanger Institute for the COVID-19 Genomics UK (COG-UK) Consortium | Health Services Laboratories and Alex Alderton, Roberto Amato, Jeffrey Barrett, Sonia Goncalves, Ewan Harrison, David K. Jackson, Ian Johnston, Dominic Kwiatkowski, Cordelia Langford, John Sillitoe on behalf of the Wellcome Sanger Institute COVID-19 Surveillance Team |
| EPI_ISL_2511948 | Lighthouse Lab in Alderley Park | Wellcome Sanger Institute for the COVID-19 Genomics UK (COG-UK) Consortium | Jacquelyn Wynn, Mairead Hyland, The Lighthouse Lab in Alderley Park and Alex Alderton, Roberto Amato, Jeffrey Barrett, Sonia Goncalves, Ewan Harrison, David K. Jackson, Ian Johnston, Dominic Kwiatkowski, Cordelia Langford, John Sillitoe on behalf of the Wellcome Sanger Institute COVID-19 Surveillance Team |
| EPI_ISL_2511949, EPI_ISL_2511950, EPI_ISL_2511951, EPI_ISL_2511952, | Health Services Laboratories | Wellcome Sanger Institute for the COVID-19 Genomics UK (COG-UK) Consortium | Health Services Laboratories and Alex Alderton, Roberto Amato, Jeffrey Barrett, Sonia Goncalves, Ewan Harrison, David K. Jackson, Ian Johnston, Dominic Kwiatkowski, Cordelia Langford, John Sillitoe on behalf of the Wellcome Sanger Institute COVID-19 Surveillance Team |

| | | | |
|---|---|---|---|
| EPI_ISL_2511954 | | | |
| EPI_ISL_2511956 | Lighthouse Lab in Alderley Park | Wellcome Sanger Institute for the COVID-19 Genomics UK (COG-UK) Consortium | Jacquelyn Wynn, Mairead Hyland, The Lighthouse Lab in Alderley Park and Alex Alderton, Roberto Amato, Jeffrey Barrett, Sonia Goncalves, Ewan Harrison, David K. Jackson, Ian Johnston, Dominic Kwiatkowski, Cordelia Langford, John Sillitoe on behalf of the Wellcome Sanger Institute COVID-19 Surveillance Team |
| EPI_ISL_2511957, EPI_ISL_2511958, EPI_ISL_2511959, EPI_ISL_2511960 | Health Services Laboratories | Wellcome Sanger Institute for the COVID-19 Genomics UK (COG-UK) Consortium | Health Services Laboratories and Alex Alderton, Roberto Amato, Jeffrey Barrett, Sonia Goncalves, Ewan Harrison, David K. Jackson, Ian Johnston, Dominic Kwiatkowski, Cordelia Langford, John Sillitoe on behalf of the Wellcome Sanger Institute COVID-19 Surveillance Team |
| EPI_ISL_2511961 | Lighthouse Lab in Alderley Park | Wellcome Sanger Institute for the COVID-19 Genomics UK (COG-UK) Consortium | Jacquelyn Wynn, Mairead Hyland, The Lighthouse Lab in Alderley Park and Alex Alderton, Roberto Amato, Jeffrey Barrett, Sonia Goncalves, Ewan Harrison, David K. Jackson, Ian Johnston, Dominic Kwiatkowski, Cordelia Langford, John Sillitoe on behalf of the Wellcome Sanger Institute COVID-19 Surveillance Team |
| EPI_ISL_2511962 | Health Services Laboratories | Wellcome Sanger Institute for the COVID-19 Genomics UK (COG-UK) Consortium | Health Services Laboratories and Alex Alderton, Roberto Amato, Jeffrey Barrett, Sonia Goncalves, Ewan Harrison, David K. Jackson, Ian Johnston, Dominic Kwiatkowski, Cordelia Langford, John Sillitoe on behalf of the Wellcome Sanger Institute COVID-19 Surveillance Team |
| EPI_ISL_2511965 | Lighthouse Lab in Alderley Park | Wellcome Sanger Institute for the COVID-19 Genomics UK (COG-UK) Consortium | Jacquelyn Wynn, Mairead Hyland, The Lighthouse Lab in Alderley Park and Alex Alderton, Roberto Amato, Jeffrey Barrett, Sonia Goncalves, Ewan Harrison, David K. Jackson, Ian Johnston, Dominic Kwiatkowski, Cordelia Langford, John Sillitoe on behalf of the Wellcome Sanger Institute COVID-19 Surveillance Team |
| EPI_ISL_2511966, EPI_ISL_2511967, EPI_ISL_2511968, EPI_ISL_2511969, EPI_ISL_2511970, EPI_ISL_2511971, EPI_ISL_2511972 | Health Services Laboratories | Wellcome Sanger Institute for the COVID-19 Genomics UK (COG-UK) Consortium | Health Services Laboratories and Alex Alderton, Roberto Amato, Jeffrey Barrett, Sonia Goncalves, Ewan Harrison, David K. Jackson, Ian Johnston, Dominic Kwiatkowski, Cordelia Langford, John Sillitoe on behalf of the Wellcome Sanger Institute COVID-19 Surveillance Team |
| EPI_ISL_2511973 | Lighthouse Lab in Alderley Park | Wellcome Sanger Institute for the COVID-19 Genomics UK (COG-UK) Consortium | Jacquelyn Wynn, Mairead Hyland, The Lighthouse Lab in Alderley Park and Alex Alderton, Roberto Amato, Jeffrey Barrett, Sonia Goncalves, Ewan Harrison, David K. Jackson, Ian Johnston, Dominic Kwiatkowski, Cordelia Langford, John Sillitoe on behalf of the Wellcome Sanger Institute COVID-19 Surveillance Team |
| EPI_ISL_2511974, EPI_ISL_2511975 | Health Services Laboratories | Wellcome Sanger Institute for the COVID-19 Genomics UK (COG-UK) Consortium | Health Services Laboratories and Alex Alderton, Roberto Amato, Jeffrey Barrett, Sonia Goncalves, Ewan Harrison, David K. Jackson, Ian Johnston, Dominic Kwiatkowski, Cordelia Langford, John Sillitoe on behalf of the Wellcome Sanger Institute COVID-19 Surveillance Team |
| EPI_ISL_2511976 | Lighthouse Lab in Alderley Park | Wellcome Sanger Institute for the COVID-19 Genomics UK (COG-UK) Consortium | Jacquelyn Wynn, Mairead Hyland, The Lighthouse Lab in Alderley Park and Alex Alderton, Roberto Amato, Jeffrey Barrett, Sonia Goncalves, Ewan Harrison, David K. Jackson, Ian Johnston, Dominic Kwiatkowski, Cordelia Langford, John Sillitoe on behalf of the Wellcome Sanger Institute COVID-19 Surveillance Team |
| EPI_ISL_2511977, EPI_ISL_2511979, EPI_ISL_2511980 | Health Services Laboratories | Wellcome Sanger Institute for the COVID-19 Genomics UK (COG-UK) Consortium | Health Services Laboratories and Alex Alderton, Roberto Amato, Jeffrey Barrett, Sonia Goncalves, Ewan Harrison, David K. Jackson, Ian Johnston, Dominic Kwiatkowski, Cordelia Langford, John Sillitoe on behalf of the Wellcome Sanger Institute COVID-19 Surveillance Team |
| EPI_ISL_2511981, EPI_ISL_2511982, EPI_ISL_2511983 | Lighthouse Lab in Alderley Park | Wellcome Sanger Institute for the COVID-19 Genomics UK (COG-UK) Consortium | Jacquelyn Wynn, Mairead Hyland, The Lighthouse Lab in Alderley Park and Alex Alderton, Roberto Amato, Jeffrey Barrett, Sonia Goncalves, Ewan Harrison, David K. Jackson, Ian Johnston, Dominic Kwiatkowski, Cordelia Langford, John Sillitoe on behalf of the Wellcome Sanger Institute COVID-19 Surveillance Team |
| EPI_ISL_2511984, EPI_ISL_2511985, EPI_ISL_2511986, EPI_ISL_2511988, EPI_ISL_2511990, EPI_ISL_2511991, EPI_ISL_2511992, EPI_ISL_2511993, EPI_ISL_2511994 | Health Services Laboratories | Wellcome Sanger Institute for the COVID-19 Genomics UK (COG-UK) Consortium | Health Services Laboratories and Alex Alderton, Roberto Amato, Jeffrey Barrett, Sonia Goncalves, Ewan Harrison, David K. Jackson, Ian Johnston, Dominic Kwiatkowski, Cordelia Langford, John Sillitoe on behalf of the Wellcome Sanger Institute COVID-19 Surveillance Team |
| EPI_ISL_2511995 | Lighthouse Lab in Alderley Park | Wellcome Sanger Institute for the COVID-19 Genomics UK (COG-UK) Consortium | Jacquelyn Wynn, Mairead Hyland, The Lighthouse Lab in Alderley Park and Alex Alderton, Roberto Amato, Jeffrey Barrett, Sonia Goncalves, Ewan Harrison, David K. Jackson, Ian Johnston, Dominic Kwiatkowski, Cordelia Langford, John Sillitoe on behalf of the Wellcome Sanger Institute COVID-19 Surveillance Team |
| EPI_ISL_2511996, EPI_ISL_2511997, EPI_ISL_2511998, EPI_ISL_2511999, EPI_ISL_2512000, EPI_ISL_2512001, EPI_ISL_2512002, EPI_ISL_2512003, EPI_ISL_2512004, EPI_ISL_2512005 | Health Services Laboratories | Wellcome Sanger Institute for the COVID-19 Genomics UK (COG-UK) Consortium | Health Services Laboratories and Alex Alderton, Roberto Amato, Jeffrey Barrett, Sonia Goncalves, Ewan Harrison, David K. Jackson, Ian Johnston, Dominic Kwiatkowski, Cordelia Langford, John Sillitoe on behalf of the Wellcome Sanger Institute COVID-19 Surveillance Team |
| EPI_ISL_2512006 | Lighthouse Lab in Alderley Park | Wellcome Sanger Institute for the COVID-19 Genomics UK (COG-UK) Consortium | Jacquelyn Wynn, Mairead Hyland, The Lighthouse Lab in Alderley Park and Alex Alderton, Roberto Amato, Jeffrey Barrett, Sonia Goncalves, Ewan Harrison, David K. Jackson, Ian Johnston, Dominic Kwiatkowski, Cordelia Langford, John Sillitoe on behalf of the Wellcome Sanger Institute COVID-19 Surveillance Team |
| EPI_ISL_2512007, EPI_ISL_2512008, EPI_ISL_2512009, EPI_ISL_2512010, EPI_ISL_2512011, EPI_ISL_2512012 | Health Services Laboratories | Wellcome Sanger Institute for the COVID-19 Genomics UK (COG-UK) Consortium | Health Services Laboratories and Alex Alderton, Roberto Amato, Jeffrey Barrett, Sonia Goncalves, Ewan Harrison, David K. Jackson, Ian Johnston, Dominic Kwiatkowski, Cordelia Langford, John Sillitoe on behalf of the Wellcome Sanger Institute COVID-19 Surveillance Team |
| EPI_ISL_2512013, EPI_ISL_2512014 | Lighthouse Lab in Alderley Park | Wellcome Sanger Institute for the COVID-19 Genomics UK (COG-UK) Consortium | Jacquelyn Wynn, Mairead Hyland, The Lighthouse Lab in Alderley Park and Alex Alderton, Roberto Amato, Jeffrey Barrett, Sonia Goncalves, Ewan Harrison, David K. Jackson, Ian Johnston, Dominic Kwiatkowski, Cordelia Langford, John Sillitoe on behalf of the Wellcome Sanger Institute COVID-19 Surveillance Team |
| EPI_ISL_2512015, EPI_ISL_2512016, EPI_ISL_2512018, EPI_ISL_2512019, EPI_ISL_2512020 | Health Services Laboratories | Wellcome Sanger Institute for the COVID-19 Genomics UK (COG-UK) Consortium | Health Services Laboratories and Alex Alderton, Roberto Amato, Jeffrey Barrett, Sonia Goncalves, Ewan Harrison, David K. Jackson, Ian Johnston, Dominic Kwiatkowski, Cordelia Langford, John Sillitoe on behalf of the Wellcome Sanger Institute COVID-19 Surveillance Team |
| EPI_ISL_2512022 | Lighthouse Lab in Alderley Park | Wellcome Sanger Institute for the COVID-19 Genomics UK (COG-UK) Consortium | Jacquelyn Wynn, Mairead Hyland, The Lighthouse Lab in Alderley Park and Alex Alderton, Roberto Amato, Jeffrey Barrett, Sonia Goncalves, Ewan Harrison, David K. Jackson, Ian Johnston, Dominic Kwiatkowski, Cordelia Langford, John Sillitoe on behalf of the Wellcome Sanger Institute COVID-19 Surveillance Team |
| EPI_ISL_2512023, EPI_ISL_2512024, EPI_ISL_2512025, EPI_ISL_2512026, EPI_ISL_2512027, EPI_ISL_2512028, EPI_ISL_2512029, EPI_ISL_2512030, EPI_ISL_2512031, EPI_ISL_2512032 | Health Services Laboratories | Wellcome Sanger Institute for the COVID-19 Genomics UK (COG-UK) Consortium | Health Services Laboratories and Alex Alderton, Roberto Amato, Jeffrey Barrett, Sonia Goncalves, Ewan Harrison, David K. Jackson, Ian Johnston, Dominic Kwiatkowski, Cordelia Langford, John Sillitoe on behalf of the Wellcome Sanger Institute COVID-19 Surveillance Team |
| EPI_ISL_2512033 | Lighthouse Lab in Alderley Park | Wellcome Sanger Institute for the COVID-19 Genomics UK (COG-UK) Consortium | Jacquelyn Wynn, Mairead Hyland, The Lighthouse Lab in Alderley Park and Alex Alderton, Roberto Amato, Jeffrey Barrett, Sonia Goncalves, Ewan Harrison, David K. Jackson, Ian Johnston, Dominic Kwiatkowski, Cordelia Langford, John Sillitoe on behalf of the Wellcome Sanger Institute COVID-19 Surveillance Team |
| EPI_ISL_2512034, EPI_ISL_2512035, EPI_ISL_2512036, EPI_ISL_2512038 | Health Services Laboratories | Wellcome Sanger Institute for the COVID-19 Genomics UK (COG-UK) Consortium | Health Services Laboratories and Alex Alderton, Roberto Amato, Jeffrey Barrett, Sonia Goncalves, Ewan Harrison, David K. Jackson, Ian Johnston, Dominic Kwiatkowski, Cordelia Langford, John Sillitoe on behalf of the Wellcome Sanger Institute COVID-19 Surveillance Team |
| EPI_ISL_2512040 | Lighthouse Lab in Alderley Park | Wellcome Sanger Institute for the COVID-19 Genomics UK (COG-UK) Consortium | Jacquelyn Wynn, Mairead Hyland, The Lighthouse Lab in Alderley Park and Alex Alderton, Roberto Amato, Jeffrey Barrett, Sonia Goncalves, Ewan Harrison, David K. Jackson, Ian Johnston, Dominic Kwiatkowski, Cordelia Langford, John Sillitoe on behalf of the Wellcome Sanger Institute COVID-19 Surveillance Team |
| EPI_ISL_2512041, EPI_ISL_2512042 | Health Services Laboratories | Wellcome Sanger Institute for the COVID-19 Genomics UK (COG-UK) Consortium | Health Services Laboratories and Alex Alderton, Roberto Amato, Jeffrey Barrett, Sonia Goncalves, Ewan Harrison, David K. Jackson, Ian Johnston, Dominic Kwiatkowski, Cordelia Langford, John Sillitoe on behalf of the Wellcome Sanger Institute COVID-19 Surveillance Team |
| EPI_ISL_2512043, EPI_ISL_2512044, EPI_ISL_2512045, EPI_ISL_2512046 | Lighthouse Lab in Alderley Park | Wellcome Sanger Institute for the COVID-19 Genomics UK (COG-UK) Consortium | Jacquelyn Wynn, Mairead Hyland, The Lighthouse Lab in Alderley Park and Alex Alderton, Roberto Amato, Jeffrey Barrett, Sonia Goncalves, Ewan Harrison, David K. Jackson, Ian Johnston, Dominic Kwiatkowski, Cordelia Langford, John Sillitoe on behalf of the Wellcome Sanger Institute COVID-19 Surveillance Team |
| EPI_ISL_2512047 | Health Services Laboratories | Wellcome Sanger Institute for the COVID-19 Genomics UK (COG-UK) Consortium | Health Services Laboratories and Alex Alderton, Roberto Amato, Jeffrey Barrett, Sonia Goncalves, Ewan Harrison, David K. Jackson, Ian Johnston, Dominic Kwiatkowski, Cordelia Langford, John Sillitoe on behalf of the Wellcome Sanger Institute COVID-19 Surveillance Team |
| EPI_ISL_2512048, | Lighthouse Lab in Alderley Park | Wellcome Sanger Institute for the COVID-19 Genomics UK (COG-UK) | Jacquelyn Wynn, Mairead Hyland, The Lighthouse Lab in Alderley Park and Alex Alderton, Roberto Amato, Jeffrey Barrett, Sonia Goncalves, Ewan Harrison, David K. Jackson, Ian Johnston, Dominic Kwiatkowski, Cordelia Langford, John Sillitoe on |

| | | | |
|---|---|---|---|
| EPI_ISL_2512049, EPI_ISL_2512050 | | Consortium | behalf of the Wellcome Sanger Institute COVID-19 Surveillance Team |
| EPI_ISL_2512052, EPI_ISL_2512053, EPI_ISL_2512054 | Health Services Laboratories | Wellcome Sanger Institute for the COVID-19 Genomics UK (COG-UK) Consortium | Health Services Laboratories and Alex Alderton, Roberto Amato, Jeffrey Barrett, Sonia Goncalves, Ewan Harrison, David K. Jackson, Ian Johnston, Dominic Kwiatkowski, Cordelia Langford, John Sillitoe on behalf of the Wellcome Sanger Institute COVID-19 Surveillance Team |
| EPI_ISL_2512056 | Lighthouse Laboratory Plymouth | Wellcome Sanger Institute for the COVID-19 Genomics UK (COG-UK) Consortium | Lighthouse Laboratory Plymouth and Alex Alderton, Roberto Amato, Jeffrey Barrett, Sonia Goncalves, Ewan Harrison, David K. Jackson, Ian Johnston, Dominic Kwiatkowski, Cordelia Langford, John Sillitoe on behalf of the Wellcome Sanger Institute COVID-19 Surveillance Team |
| EPI_ISL_2512057, EPI_ISL_2512058, EPI_ISL_2512059, EPI_ISL_2512060, EPI_ISL_2512061 | Health Services Laboratories | Wellcome Sanger Institute for the COVID-19 Genomics UK (COG-UK) Consortium | Health Services Laboratories and Alex Alderton, Roberto Amato, Jeffrey Barrett, Sonia Goncalves, Ewan Harrison, David K. Jackson, Ian Johnston, Dominic Kwiatkowski, Cordelia Langford, John Sillitoe on behalf of the Wellcome Sanger Institute COVID-19 Surveillance Team |
| EPI_ISL_2512062 | Lighthouse Lab in Alderley Park | Wellcome Sanger Institute for the COVID-19 Genomics UK (COG-UK) Consortium | Jacquelyn Wynn, Mairead Hyland, The Lighthouse Lab in Alderley Park and Alex Alderton, Roberto Amato, Jeffrey Barrett, Sonia Goncalves, Ewan Harrison, David K. Jackson, Ian Johnston, Dominic Kwiatkowski, Cordelia Langford, John Sillitoe on behalf of the Wellcome Sanger Institute COVID-19 Surveillance Team |
| EPI_ISL_2512063 | Health Services Laboratories | Wellcome Sanger Institute for the COVID-19 Genomics UK (COG-UK) Consortium | Health Services Laboratories and Alex Alderton, Roberto Amato, Jeffrey Barrett, Sonia Goncalves, Ewan Harrison, David K. Jackson, Ian Johnston, Dominic Kwiatkowski, Cordelia Langford, John Sillitoe on behalf of the Wellcome Sanger Institute COVID-19 Surveillance Team |
| EPI_ISL_2512064 | Lighthouse Lab in Alderley Park | Wellcome Sanger Institute for the COVID-19 Genomics UK (COG-UK) Consortium | Jacquelyn Wynn, Mairead Hyland, The Lighthouse Lab in Alderley Park and Alex Alderton, Roberto Amato, Jeffrey Barrett, Sonia Goncalves, Ewan Harrison, David K. Jackson, Ian Johnston, Dominic Kwiatkowski, Cordelia Langford, John Sillitoe on behalf of the Wellcome Sanger Institute COVID-19 Surveillance Team |
| EPI_ISL_2512065 | Health Services Laboratories | Wellcome Sanger Institute for the COVID-19 Genomics UK (COG-UK) Consortium | Health Services Laboratories and Alex Alderton, Roberto Amato, Jeffrey Barrett, Sonia Goncalves, Ewan Harrison, David K. Jackson, Ian Johnston, Dominic Kwiatkowski, Cordelia Langford, John Sillitoe on behalf of the Wellcome Sanger Institute COVID-19 Surveillance Team |
| EPI_ISL_2512066 | Lighthouse Lab in Alderley Park | Wellcome Sanger Institute for the COVID-19 Genomics UK (COG-UK) Consortium | Jacquelyn Wynn, Mairead Hyland, The Lighthouse Lab in Alderley Park and Alex Alderton, Roberto Amato, Jeffrey Barrett, Sonia Goncalves, Ewan Harrison, David K. Jackson, Ian Johnston, Dominic Kwiatkowski, Cordelia Langford, John Sillitoe on behalf of the Wellcome Sanger Institute COVID-19 Surveillance Team |
| EPI_ISL_2512067, EPI_ISL_2512068, EPI_ISL_2512069, EPI_ISL_2512070 | Health Services Laboratories | Wellcome Sanger Institute for the COVID-19 Genomics UK (COG-UK) Consortium | Health Services Laboratories and Alex Alderton, Roberto Amato, Jeffrey Barrett, Sonia Goncalves, Ewan Harrison, David K. Jackson, Ian Johnston, Dominic Kwiatkowski, Cordelia Langford, John Sillitoe on behalf of the Wellcome Sanger Institute COVID-19 Surveillance Team |
| EPI_ISL_2512071, EPI_ISL_2512074 | Lighthouse Lab in Alderley Park | Wellcome Sanger Institute for the COVID-19 Genomics UK (COG-UK) Consortium | Jacquelyn Wynn, Mairead Hyland, The Lighthouse Lab in Alderley Park and Alex Alderton, Roberto Amato, Jeffrey Barrett, Sonia Goncalves, Ewan Harrison, David K. Jackson, Ian Johnston, Dominic Kwiatkowski, Cordelia Langford, John Sillitoe on behalf of the Wellcome Sanger Institute COVID-19 Surveillance Team |
| EPI_ISL_2512075, EPI_ISL_2512076, EPI_ISL_2512077 | Health Services Laboratories | Wellcome Sanger Institute for the COVID-19 Genomics UK (COG-UK) Consortium | Health Services Laboratories and Alex Alderton, Roberto Amato, Jeffrey Barrett, Sonia Goncalves, Ewan Harrison, David K. Jackson, Ian Johnston, Dominic Kwiatkowski, Cordelia Langford, John Sillitoe on behalf of the Wellcome Sanger Institute COVID-19 Surveillance Team |
| EPI_ISL_2512078 | Lighthouse Lab in Alderley Park | Wellcome Sanger Institute for the COVID-19 Genomics UK (COG-UK) Consortium | Jacquelyn Wynn, Mairead Hyland, The Lighthouse Lab in Alderley Park and Alex Alderton, Roberto Amato, Jeffrey Barrett, Sonia Goncalves, Ewan Harrison, David K. Jackson, Ian Johnston, Dominic Kwiatkowski, Cordelia Langford, John Sillitoe on behalf of the Wellcome Sanger Institute COVID-19 Surveillance Team |
| EPI_ISL_2512079 | Health Services Laboratories | Wellcome Sanger Institute for the COVID-19 Genomics UK (COG-UK) Consortium | Health Services Laboratories and Alex Alderton, Roberto Amato, Jeffrey Barrett, Sonia Goncalves, Ewan Harrison, David K. Jackson, Ian Johnston, Dominic Kwiatkowski, Cordelia Langford, John Sillitoe on behalf of the Wellcome Sanger Institute COVID-19 Surveillance Team |
| EPI_ISL_2512080, EPI_ISL_2512081, EPI_ISL_2512082 | Lighthouse Lab in Alderley Park | Wellcome Sanger Institute for the COVID-19 Genomics UK (COG-UK) Consortium | Jacquelyn Wynn, Mairead Hyland, The Lighthouse Lab in Alderley Park and Alex Alderton, Roberto Amato, Jeffrey Barrett, Sonia Goncalves, Ewan Harrison, David K. Jackson, Ian Johnston, Dominic Kwiatkowski, Cordelia Langford, John Sillitoe on behalf of the Wellcome Sanger Institute COVID-19 Surveillance Team |
| EPI_ISL_2512083 | Lighthouse Laboratory Plymouth | Wellcome Sanger Institute for the COVID-19 Genomics UK (COG-UK) Consortium | Lighthouse Laboratory Plymouth and Alex Alderton, Roberto Amato, Jeffrey Barrett, Sonia Goncalves, Ewan Harrison, David K. Jackson, Ian Johnston, Dominic Kwiatkowski, Cordelia Langford, John Sillitoe on behalf of the Wellcome Sanger Institute COVID-19 Surveillance Team |
| EPI_ISL_2512084 | Lighthouse Lab in Alderley Park | Wellcome Sanger Institute for the COVID-19 Genomics UK (COG-UK) Consortium | Jacquelyn Wynn, Mairead Hyland, The Lighthouse Lab in Alderley Park and Alex Alderton, Roberto Amato, Jeffrey Barrett, Sonia Goncalves, Ewan Harrison, David K. Jackson, Ian Johnston, Dominic Kwiatkowski, Cordelia Langford, John Sillitoe on behalf of the Wellcome Sanger Institute COVID-19 Surveillance Team |
| EPI_ISL_2512085, EPI_ISL_2512087, EPI_ISL_2512088, EPI_ISL_2512089, EPI_ISL_2512090, EPI_ISL_2512091, EPI_ISL_2512092, EPI_ISL_2512093, EPI_ISL_2512094, EPI_ISL_2512095, EPI_ISL_2512096, EPI_ISL_2512097, EPI_ISL_2512099, EPI_ISL_2512100, EPI_ISL_2512101, EPI_ISL_2512102, EPI_ISL_2512103, EPI_ISL_2512104, EPI_ISL_2512105, EPI_ISL_2512106 | | | |
| see above | Health Services Laboratories | Wellcome Sanger Institute for the COVID-19 Genomics UK (COG-UK) Consortium | Health Services Laboratories and Alex Alderton, Roberto Amato, Jeffrey Barrett, Sonia Goncalves, Ewan Harrison, David K. Jackson, Ian Johnston, Dominic Kwiatkowski, Cordelia Langford, John Sillitoe on behalf of the Wellcome Sanger Institute COVID-19 Surveillance Team |
| EPI_ISL_2512107, EPI_ISL_2512108 | Lighthouse Lab in Alderley Park | Wellcome Sanger Institute for the COVID-19 Genomics UK (COG-UK) Consortium | Jacquelyn Wynn, Mairead Hyland, The Lighthouse Lab in Alderley Park and Alex Alderton, Roberto Amato, Jeffrey Barrett, Sonia Goncalves, Ewan Harrison, David K. Jackson, Ian Johnston, Dominic Kwiatkowski, Cordelia Langford, John Sillitoe on behalf of the Wellcome Sanger Institute COVID-19 Surveillance Team |
| EPI_ISL_2512109, EPI_ISL_2512110 | Health Services Laboratories | Wellcome Sanger Institute for the COVID-19 Genomics UK (COG-UK) Consortium | Health Services Laboratories and Alex Alderton, Roberto Amato, Jeffrey Barrett, Sonia Goncalves, Ewan Harrison, David K. Jackson, Ian Johnston, Dominic Kwiatkowski, Cordelia Langford, John Sillitoe on behalf of the Wellcome Sanger Institute COVID-19 Surveillance Team |
| EPI_ISL_2512111 | Lighthouse Lab in Alderley Park | Wellcome Sanger Institute for the COVID-19 Genomics UK (COG-UK) Consortium | Jacquelyn Wynn, Mairead Hyland, The Lighthouse Lab in Alderley Park and Alex Alderton, Roberto Amato, Jeffrey Barrett, Sonia Goncalves, Ewan Harrison, David K. Jackson, Ian Johnston, Dominic Kwiatkowski, Cordelia Langford, John Sillitoe on behalf of the Wellcome Sanger Institute COVID-19 Surveillance Team |
| EPI_ISL_2512112, EPI_ISL_2512113, EPI_ISL_2512114, EPI_ISL_2512115 | Health Services Laboratories | Wellcome Sanger Institute for the COVID-19 Genomics UK (COG-UK) Consortium | Health Services Laboratories and Alex Alderton, Roberto Amato, Jeffrey Barrett, Sonia Goncalves, Ewan Harrison, David K. Jackson, Ian Johnston, Dominic Kwiatkowski, Cordelia Langford, John Sillitoe on behalf of the Wellcome Sanger Institute COVID-19 Surveillance Team |
| EPI_ISL_2512116 | Lighthouse Lab in Alderley Park | Wellcome Sanger Institute for the COVID-19 Genomics UK (COG-UK) Consortium | Jacquelyn Wynn, Mairead Hyland, The Lighthouse Lab in Alderley Park and Alex Alderton, Roberto Amato, Jeffrey Barrett, Sonia Goncalves, Ewan Harrison, David K. Jackson, Ian Johnston, Dominic Kwiatkowski, Cordelia Langford, John Sillitoe on behalf of the Wellcome Sanger Institute COVID-19 Surveillance Team |
| EPI_ISL_2512117, EPI_ISL_2512118, EPI_ISL_2512119 | Health Services Laboratories | Wellcome Sanger Institute for the COVID-19 Genomics UK (COG-UK) Consortium | Health Services Laboratories and Alex Alderton, Roberto Amato, Jeffrey Barrett, Sonia Goncalves, Ewan Harrison, David K. Jackson, Ian Johnston, Dominic Kwiatkowski, Cordelia Langford, John Sillitoe on behalf of the Wellcome Sanger Institute COVID-19 Surveillance Team |
| EPI_ISL_2512120 | Lighthouse Lab in Alderley Park | Wellcome Sanger Institute for the COVID-19 Genomics UK (COG-UK) Consortium | Jacquelyn Wynn, Mairead Hyland, The Lighthouse Lab in Alderley Park and Alex Alderton, Roberto Amato, Jeffrey Barrett, Sonia Goncalves, Ewan Harrison, David K. Jackson, Ian Johnston, Dominic Kwiatkowski, Cordelia Langford, John Sillitoe on behalf of the Wellcome Sanger Institute COVID-19 Surveillance Team |
| EPI_ISL_2512121, EPI_ISL_2512122, EPI_ISL_2512123, EPI_ISL_2512124 | Health Services Laboratories | Wellcome Sanger Institute for the COVID-19 Genomics UK (COG-UK) Consortium | Health Services Laboratories and Alex Alderton, Roberto Amato, Jeffrey Barrett, Sonia Goncalves, Ewan Harrison, David K. Jackson, Ian Johnston, Dominic Kwiatkowski, Cordelia Langford, John Sillitoe on behalf of the Wellcome Sanger Institute COVID-19 Surveillance Team |
| EPI_ISL_2512125, EPI_ISL_2512126, EPI_ISL_2512127 | Lighthouse Lab in Alderley Park | Wellcome Sanger Institute for the COVID-19 Genomics UK (COG-UK) Consortium | Jacquelyn Wynn, Mairead Hyland, The Lighthouse Lab in Alderley Park and Alex Alderton, Roberto Amato, Jeffrey Barrett, Sonia Goncalves, Ewan Harrison, David K. Jackson, Ian Johnston, Dominic Kwiatkowski, Cordelia Langford, John Sillitoe on behalf of the Wellcome Sanger Institute COVID-19 Surveillance Team |
| EPI_ISL_2512128, EPI_ISL_2512129, EPI_ISL_2512130, EPI_ISL_2512131, EPI_ISL_2512132, EPI_ISL_2512133, EPI_ISL_2512134, EPI_ISL_2512135, EPI_ISL_2512136, EPI_ISL_2512137 | Health Services Laboratories | Wellcome Sanger Institute for the COVID-19 Genomics UK (COG-UK) Consortium | Health Services Laboratories and Alex Alderton, Roberto Amato, Jeffrey Barrett, Sonia Goncalves, Ewan Harrison, David K. Jackson, Ian Johnston, Dominic Kwiatkowski, Cordelia Langford, John Sillitoe on behalf of the Wellcome Sanger Institute COVID-19 Surveillance Team |
| EPI_ISL_2512138 | Lighthouse Lab in Alderley Park | Wellcome Sanger Institute for the COVID-19 Genomics UK (COG-UK) Consortium | Jacquelyn Wynn, Mairead Hyland, The Lighthouse Lab in Alderley Park and Alex Alderton, Roberto Amato, Jeffrey Barrett, Sonia Goncalves, Ewan Harrison, David K. Jackson, Ian Johnston, Dominic Kwiatkowski, Cordelia Langford, John Sillitoe on behalf of the Wellcome Sanger Institute COVID-19 Surveillance Team |
| EPI_ISL_2512139 | Health Services Laboratories | Wellcome Sanger Institute for the COVID-19 Genomics UK (COG-UK) Consortium | Health Services Laboratories and Alex Alderton, Roberto Amato, Jeffrey Barrett, Sonia Goncalves, Ewan Harrison, David K. Jackson, Ian Johnston, Dominic Kwiatkowski, Cordelia Langford, John Sillitoe on behalf of the Wellcome Sanger Institute COVID-19 Surveillance Team |
| EPI_ISL_2512140, EPI_ISL_2512141 | Lighthouse Lab in Alderley Park | Wellcome Sanger Institute for the COVID-19 Genomics UK (COG-UK) Consortium | Jacquelyn Wynn, Mairead Hyland, The Lighthouse Lab in Alderley Park and Alex Alderton, Roberto Amato, Jeffrey Barrett, Sonia Goncalves, Ewan Harrison, David K. Jackson, Ian Johnston, Dominic Kwiatkowski, Cordelia Langford, John Sillitoe on behalf of the Wellcome Sanger Institute COVID-19 Surveillance Team |
| EPI_ISL_2512144, EPI_ISL_2512145, EPI_ISL_2512146, EPI_ISL_2512147, EPI_ISL_2512148, EPI_ISL_2512149, EPI_ISL_2512150, EPI_ISL_2512151, EPI_ISL_2512152, EPI_ISL_2512153, EPI_ISL_2512154, EPI_ISL_2512155, EPI_ISL_2512156, EPI_ISL_2512158, EPI_ISL_2512159 | | | |
| see above | Health Services Laboratories | Wellcome Sanger Institute for the COVID-19 Genomics UK (COG-UK) Consortium | Health Services Laboratories and Alex Alderton, Roberto Amato, Jeffrey Barrett, Sonia Goncalves, Ewan Harrison, David K. Jackson, Ian Johnston, Dominic Kwiatkowski, Cordelia Langford, John Sillitoe on behalf of the Wellcome Sanger Institute COVID-19 Surveillance Team |
| EPI_ISL_2512160 | Lighthouse Lab in Alderley Park | Wellcome Sanger Institute for the COVID-19 Genomics UK (COG-UK) Consortium | Jacquelyn Wynn, Mairead Hyland, The Lighthouse Lab in Alderley Park and Alex Alderton, Roberto Amato, Jeffrey Barrett, Sonia Goncalves, Ewan Harrison, David K. Jackson, Ian Johnston, Dominic Kwiatkowski, Cordelia Langford, John Sillitoe on behalf of the Wellcome Sanger Institute COVID-19 Surveillance Team |
| EPI_ISL_2512161, EPI_ISL_2512162, EPI_ISL_2512163, EPI_ISL_2512164 | Health Services Laboratories | Wellcome Sanger Institute for the COVID-19 Genomics UK (COG-UK) Consortium | Health Services Laboratories and Alex Alderton, Roberto Amato, Jeffrey Barrett, Sonia Goncalves, Ewan Harrison, David K. Jackson, Ian Johnston, Dominic Kwiatkowski, Cordelia Langford, John Sillitoe on behalf of the Wellcome Sanger Institute COVID-19 Surveillance Team |

| | | | |
|---|---|---|---|
| EPI_ISL_2512165 | Lighthouse Lab in Alderley Park | Wellcome Sanger Institute for the COVID-19 Genomics UK (COG-UK) Consortium | Jacquelyn Wynn, Mairead Hyland, The Lighthouse Lab in Alderley Park and Alex Alderton, Roberto Amato, Jeffrey Barrett, Sonia Goncalves, Ewan Harrison, David K. Jackson, Ian Johnston, Dominic Kwiatkowski, Cordelia Langford, John Sillitoe on behalf of the Wellcome Sanger Institute COVID-19 Surveillance Team |
| EPI_ISL_2512166 | Health Services Laboratories | Wellcome Sanger Institute for the COVID-19 Genomics UK (COG-UK) Consortium | Health Services Laboratories and Alex Alderton, Roberto Amato, Jeffrey Barrett, Sonia Goncalves, Ewan Harrison, David K. Jackson, Ian Johnston, Dominic Kwiatkowski, Cordelia Langford, John Sillitoe on behalf of the Wellcome Sanger Institute COVID-19 Surveillance Team |
| EPI_ISL_2512167, EPI_ISL_2512168 | Lighthouse Lab in Alderley Park | Wellcome Sanger Institute for the COVID-19 Genomics UK (COG-UK) Consortium | Jacquelyn Wynn, Mairead Hyland, The Lighthouse Lab in Alderley Park and Alex Alderton, Roberto Amato, Jeffrey Barrett, Sonia Goncalves, Ewan Harrison, David K. Jackson, Ian Johnston, Dominic Kwiatkowski, Cordelia Langford, John Sillitoe on behalf of the Wellcome Sanger Institute COVID-19 Surveillance Team |
| EPI_ISL_2512169, EPI_ISL_2512170, EPI_ISL_2512171 | Health Services Laboratories | Wellcome Sanger Institute for the COVID-19 Genomics UK (COG-UK) Consortium | Health Services Laboratories and Alex Alderton, Roberto Amato, Jeffrey Barrett, Sonia Goncalves, Ewan Harrison, David K. Jackson, Ian Johnston, Dominic Kwiatkowski, Cordelia Langford, John Sillitoe on behalf of the Wellcome Sanger Institute COVID-19 Surveillance Team |
| EPI_ISL_2512172, EPI_ISL_2512173 | Lighthouse Lab in Alderley Park | Wellcome Sanger Institute for the COVID-19 Genomics UK (COG-UK) Consortium | Jacquelyn Wynn, Mairead Hyland, The Lighthouse Lab in Alderley Park and Alex Alderton, Roberto Amato, Jeffrey Barrett, Sonia Goncalves, Ewan Harrison, David K. Jackson, Ian Johnston, Dominic Kwiatkowski, Cordelia Langford, John Sillitoe on behalf of the Wellcome Sanger Institute COVID-19 Surveillance Team |
| EPI_ISL_2512174, EPI_ISL_2512175, EPI_ISL_2512176 | Health Services Laboratories | Wellcome Sanger Institute for the COVID-19 Genomics UK (COG-UK) Consortium | Health Services Laboratories and Alex Alderton, Roberto Amato, Jeffrey Barrett, Sonia Goncalves, Ewan Harrison, David K. Jackson, Ian Johnston, Dominic Kwiatkowski, Cordelia Langford, John Sillitoe on behalf of the Wellcome Sanger Institute COVID-19 Surveillance Team |
| EPI_ISL_2512177 | Lighthouse Lab in Alderley Park | Wellcome Sanger Institute for the COVID-19 Genomics UK (COG-UK) Consortium | Jacquelyn Wynn, Mairead Hyland, The Lighthouse Lab in Alderley Park and Alex Alderton, Roberto Amato, Jeffrey Barrett, Sonia Goncalves, Ewan Harrison, David K. Jackson, Ian Johnston, Dominic Kwiatkowski, Cordelia Langford, John Sillitoe on behalf of the Wellcome Sanger Institute COVID-19 Surveillance Team |
| EPI_ISL_2512178, EPI_ISL_2512179, EPI_ISL_2512180, EPI_ISL_2512181 | Health Services Laboratories | Wellcome Sanger Institute for the COVID-19 Genomics UK (COG-UK) Consortium | Health Services Laboratories and Alex Alderton, Roberto Amato, Jeffrey Barrett, Sonia Goncalves, Ewan Harrison, David K. Jackson, Ian Johnston, Dominic Kwiatkowski, Cordelia Langford, John Sillitoe on behalf of the Wellcome Sanger Institute COVID-19 Surveillance Team |
| EPI_ISL_2512182 | Lighthouse Lab in Alderley Park | Wellcome Sanger Institute for the COVID-19 Genomics UK (COG-UK) Consortium | Jacquelyn Wynn, Mairead Hyland, The Lighthouse Lab in Alderley Park and Alex Alderton, Roberto Amato, Jeffrey Barrett, Sonia Goncalves, Ewan Harrison, David K. Jackson, Ian Johnston, Dominic Kwiatkowski, Cordelia Langford, John Sillitoe on behalf of the Wellcome Sanger Institute COVID-19 Surveillance Team |
| EPI_ISL_2512183, EPI_ISL_2512184, EPI_ISL_2512185, EPI_ISL_2512186 | Health Services Laboratories | Wellcome Sanger Institute for the COVID-19 Genomics UK (COG-UK) Consortium | Health Services Laboratories and Alex Alderton, Roberto Amato, Jeffrey Barrett, Sonia Goncalves, Ewan Harrison, David K. Jackson, Ian Johnston, Dominic Kwiatkowski, Cordelia Langford, John Sillitoe on behalf of the Wellcome Sanger Institute COVID-19 Surveillance Team |
| EPI_ISL_2512187 | Lighthouse Lab in Alderley Park | Wellcome Sanger Institute for the COVID-19 Genomics UK (COG-UK) Consortium | Jacquelyn Wynn, Mairead Hyland, The Lighthouse Lab in Alderley Park and Alex Alderton, Roberto Amato, Jeffrey Barrett, Sonia Goncalves, Ewan Harrison, David K. Jackson, Ian Johnston, Dominic Kwiatkowski, Cordelia Langford, John Sillitoe on behalf of the Wellcome Sanger Institute COVID-19 Surveillance Team |
| EPI_ISL_2512188, EPI_ISL_2512189, EPI_ISL_2512190, EPI_ISL_2512191, EPI_ISL_2512192, EPI_ISL_2512194, EPI_ISL_2512195 | Health Services Laboratories | Wellcome Sanger Institute for the COVID-19 Genomics UK (COG-UK) Consortium | Health Services Laboratories and Alex Alderton, Roberto Amato, Jeffrey Barrett, Sonia Goncalves, Ewan Harrison, David K. Jackson, Ian Johnston, Dominic Kwiatkowski, Cordelia Langford, John Sillitoe on behalf of the Wellcome Sanger Institute COVID-19 Surveillance Team |
| EPI_ISL_2512196 | Lighthouse Lab in Alderley Park | Wellcome Sanger Institute for the COVID-19 Genomics UK (COG-UK) Consortium | Jacquelyn Wynn, Mairead Hyland, The Lighthouse Lab in Alderley Park and Alex Alderton, Roberto Amato, Jeffrey Barrett, Sonia Goncalves, Ewan Harrison, David K. Jackson, Ian Johnston, Dominic Kwiatkowski, Cordelia Langford, John Sillitoe on behalf of the Wellcome Sanger Institute COVID-19 Surveillance Team |
| EPI_ISL_2512197 | Health Services Laboratories | Wellcome Sanger Institute for the COVID-19 Genomics UK (COG-UK) Consortium | Health Services Laboratories and Alex Alderton, Roberto Amato, Jeffrey Barrett, Sonia Goncalves, Ewan Harrison, David K. Jackson, Ian Johnston, Dominic Kwiatkowski, Cordelia Langford, John Sillitoe on behalf of the Wellcome Sanger Institute COVID-19 Surveillance Team |
| EPI_ISL_2512198 | Lighthouse Lab in Alderley Park | Wellcome Sanger Institute for the COVID-19 Genomics UK (COG-UK) Consortium | Jacquelyn Wynn, Mairead Hyland, The Lighthouse Lab in Alderley Park and Alex Alderton, Roberto Amato, Jeffrey Barrett, Sonia Goncalves, Ewan Harrison, David K. Jackson, Ian Johnston, Dominic Kwiatkowski, Cordelia Langford, John Sillitoe on behalf of the Wellcome Sanger Institute COVID-19 Surveillance Team |
| EPI_ISL_2512199 | Health Services Laboratories | Wellcome Sanger Institute for the COVID-19 Genomics UK (COG-UK) Consortium | Health Services Laboratories and Alex Alderton, Roberto Amato, Jeffrey Barrett, Sonia Goncalves, Ewan Harrison, David K. Jackson, Ian Johnston, Dominic Kwiatkowski, Cordelia Langford, John Sillitoe on behalf of the Wellcome Sanger Institute COVID-19 Surveillance Team |
| EPI_ISL_2512200 | Lighthouse Lab in Alderley Park | Wellcome Sanger Institute for the COVID-19 Genomics UK (COG-UK) Consortium | Jacquelyn Wynn, Mairead Hyland, The Lighthouse Lab in Alderley Park and Alex Alderton, Roberto Amato, Jeffrey Barrett, Sonia Goncalves, Ewan Harrison, David K. Jackson, Ian Johnston, Dominic Kwiatkowski, Cordelia Langford, John Sillitoe on behalf of the Wellcome Sanger Institute COVID-19 Surveillance Team |
| EPI_ISL_2512201, EPI_ISL_2512203 | Health Services Laboratories | Wellcome Sanger Institute for the COVID-19 Genomics UK (COG-UK) Consortium | Health Services Laboratories and Alex Alderton, Roberto Amato, Jeffrey Barrett, Sonia Goncalves, Ewan Harrison, David K. Jackson, Ian Johnston, Dominic Kwiatkowski, Cordelia Langford, John Sillitoe on behalf of the Wellcome Sanger Institute COVID-19 Surveillance Team |
| EPI_ISL_2512204 | Lighthouse Lab in Alderley Park | Wellcome Sanger Institute for the COVID-19 Genomics UK (COG-UK) Consortium | Jacquelyn Wynn, Mairead Hyland, The Lighthouse Lab in Alderley Park and Alex Alderton, Roberto Amato, Jeffrey Barrett, Sonia Goncalves, Ewan Harrison, David K. Jackson, Ian Johnston, Dominic Kwiatkowski, Cordelia Langford, John Sillitoe on behalf of the Wellcome Sanger Institute COVID-19 Surveillance Team |
| EPI_ISL_2512205, EPI_ISL_2512206 | Health Services Laboratories | Wellcome Sanger Institute for the COVID-19 Genomics UK (COG-UK) Consortium | Health Services Laboratories and Alex Alderton, Roberto Amato, Jeffrey Barrett, Sonia Goncalves, Ewan Harrison, David K. Jackson, Ian Johnston, Dominic Kwiatkowski, Cordelia Langford, John Sillitoe on behalf of the Wellcome Sanger Institute COVID-19 Surveillance Team |
| EPI_ISL_2512207, EPI_ISL_2512208 | Lighthouse Lab in Alderley Park | Wellcome Sanger Institute for the COVID-19 Genomics UK (COG-UK) Consortium | Jacquelyn Wynn, Mairead Hyland, The Lighthouse Lab in Alderley Park and Alex Alderton, Roberto Amato, Jeffrey Barrett, Sonia Goncalves, Ewan Harrison, David K. Jackson, Ian Johnston, Dominic Kwiatkowski, Cordelia Langford, John Sillitoe on behalf of the Wellcome Sanger Institute COVID-19 Surveillance Team |
| EPI_ISL_2512210, EPI_ISL_2512211, EPI_ISL_2512213, EPI_ISL_2512214, EPI_ISL_2512215, EPI_ISL_2512216 | Health Services Laboratories | Wellcome Sanger Institute for the COVID-19 Genomics UK (COG-UK) Consortium | Health Services Laboratories and Alex Alderton, Roberto Amato, Jeffrey Barrett, Sonia Goncalves, Ewan Harrison, David K. Jackson, Ian Johnston, Dominic Kwiatkowski, Cordelia Langford, John Sillitoe on behalf of the Wellcome Sanger Institute COVID-19 Surveillance Team |
| EPI_ISL_2512217 | Lighthouse Lab in Alderley Park | Wellcome Sanger Institute for the COVID-19 Genomics UK (COG-UK) Consortium | Jacquelyn Wynn, Mairead Hyland, The Lighthouse Lab in Alderley Park and Alex Alderton, Roberto Amato, Jeffrey Barrett, Sonia Goncalves, Ewan Harrison, David K. Jackson, Ian Johnston, Dominic Kwiatkowski, Cordelia Langford, John Sillitoe on behalf of the Wellcome Sanger Institute COVID-19 Surveillance Team |
| EPI_ISL_2512218, EPI_ISL_2512220, EPI_ISL_2512222, EPI_ISL_2512223 | Health Services Laboratories | Wellcome Sanger Institute for the COVID-19 Genomics UK (COG-UK) Consortium | Health Services Laboratories and Alex Alderton, Roberto Amato, Jeffrey Barrett, Sonia Goncalves, Ewan Harrison, David K. Jackson, Ian Johnston, Dominic Kwiatkowski, Cordelia Langford, John Sillitoe on behalf of the Wellcome Sanger Institute COVID-19 Surveillance Team |
| EPI_ISL_2512225 | Lighthouse Lab in Alderley Park | Wellcome Sanger Institute for the COVID-19 Genomics UK (COG-UK) Consortium | Jacquelyn Wynn, Mairead Hyland, The Lighthouse Lab in Alderley Park and Alex Alderton, Roberto Amato, Jeffrey Barrett, Sonia Goncalves, Ewan Harrison, David K. Jackson, Ian Johnston, Dominic Kwiatkowski, Cordelia Langford, John Sillitoe on behalf of the Wellcome Sanger Institute COVID-19 Surveillance Team |
| EPI_ISL_2512226, EPI_ISL_2512227, EPI_ISL_2512228, EPI_ISL_2512229, EPI_ISL_2512230, EPI_ISL_2512231 | Health Services Laboratories | Wellcome Sanger Institute for the COVID-19 Genomics UK (COG-UK) Consortium | Health Services Laboratories and Alex Alderton, Roberto Amato, Jeffrey Barrett, Sonia Goncalves, Ewan Harrison, David K. Jackson, Ian Johnston, Dominic Kwiatkowski, Cordelia Langford, John Sillitoe on behalf of the Wellcome Sanger Institute COVID-19 Surveillance Team |
| EPI_ISL_2512232 | Lighthouse Lab in Alderley Park | Wellcome Sanger Institute for the COVID-19 Genomics UK (COG-UK) Consortium | Jacquelyn Wynn, Mairead Hyland, The Lighthouse Lab in Alderley Park and Alex Alderton, Roberto Amato, Jeffrey Barrett, Sonia Goncalves, Ewan Harrison, David K. Jackson, Ian Johnston, Dominic Kwiatkowski, Cordelia Langford, John Sillitoe on behalf of the Wellcome Sanger Institute COVID-19 Surveillance Team |
| EPI_ISL_2512233, EPI_ISL_2512234 | Health Services Laboratories | Wellcome Sanger Institute for the COVID-19 Genomics UK (COG-UK) Consortium | Health Services Laboratories and Alex Alderton, Roberto Amato, Jeffrey Barrett, Sonia Goncalves, Ewan Harrison, David K. Jackson, Ian Johnston, Dominic Kwiatkowski, Cordelia Langford, John Sillitoe on behalf of the Wellcome Sanger Institute COVID-19 Surveillance Team |
| EPI_ISL_2512235 | Lighthouse Lab in Alderley Park | Wellcome Sanger Institute for the COVID-19 Genomics UK (COG-UK) Consortium | Jacquelyn Wynn, Mairead Hyland, The Lighthouse Lab in Alderley Park and Alex Alderton, Roberto Amato, Jeffrey Barrett, Sonia Goncalves, Ewan Harrison, David K. Jackson, Ian Johnston, Dominic Kwiatkowski, Cordelia Langford, John Sillitoe on behalf of the Wellcome Sanger Institute COVID-19 Surveillance Team |
| EPI_ISL_2512236, EPI_ISL_2512237, EPI_ISL_2512238, EPI_ISL_2512239, EPI_ISL_2512240 | Health Services Laboratories | Wellcome Sanger Institute for the COVID-19 Genomics UK (COG-UK) Consortium | Health Services Laboratories and Alex Alderton, Roberto Amato, Jeffrey Barrett, Sonia Goncalves, Ewan Harrison, David K. Jackson, Ian Johnston, Dominic Kwiatkowski, Cordelia Langford, John Sillitoe on behalf of the Wellcome Sanger Institute COVID-19 Surveillance Team |
| EPI_ISL_2512243, EPI_ISL_2512244 | Lighthouse Lab in Alderley Park | Wellcome Sanger Institute for the COVID-19 Genomics UK (COG-UK) Consortium | Jacquelyn Wynn, Mairead Hyland, The Lighthouse Lab in Alderley Park and Alex Alderton, Roberto Amato, Jeffrey Barrett, Sonia Goncalves, Ewan Harrison, David K. Jackson, Ian Johnston, Dominic Kwiatkowski, Cordelia Langford, John Sillitoe on behalf of the Wellcome Sanger Institute COVID-19 Surveillance Team |
| EPI_ISL_2512245, EPI_ISL_2512246, EPI_ISL_2512247, EPI_ISL_2512248 | Health Services Laboratories | Wellcome Sanger Institute for the COVID-19 Genomics UK (COG-UK) Consortium | Health Services Laboratories and Alex Alderton, Roberto Amato, Jeffrey Barrett, Sonia Goncalves, Ewan Harrison, David K. Jackson, Ian Johnston, Dominic Kwiatkowski, Cordelia Langford, John Sillitoe on behalf of the Wellcome Sanger Institute COVID-19 Surveillance Team |
| EPI_ISL_2512249 | Lighthouse Lab in Alderley Park | Wellcome Sanger Institute for the COVID-19 Genomics UK (COG-UK) Consortium | Jacquelyn Wynn, Mairead Hyland, The Lighthouse Lab in Alderley Park and Alex Alderton, Roberto Amato, Jeffrey Barrett, Sonia Goncalves, Ewan Harrison, David K. Jackson, Ian Johnston, Dominic Kwiatkowski, Cordelia Langford, John Sillitoe on behalf of the Wellcome Sanger Institute COVID-19 Surveillance Team |
| EPI_ISL_2512250, EPI_ISL_2512252, EPI_ISL_2512253 | Health Services Laboratories | Wellcome Sanger Institute for the COVID-19 Genomics UK (COG-UK) Consortium | Health Services Laboratories and Alex Alderton, Roberto Amato, Jeffrey Barrett, Sonia Goncalves, Ewan Harrison, David K. Jackson, Ian Johnston, Dominic Kwiatkowski, Cordelia Langford, John Sillitoe on behalf of the Wellcome Sanger Institute COVID-19 Surveillance Team |

| | | | |
|---|---|---|---|
| EPI_ISL_2512254 | Lighthouse Lab in Alderley Park | Wellcome Sanger Institute for the COVID-19 Genomics UK (COG-UK) Consortium | Jacquelyn Wynn, Mairead Hyland, The Lighthouse Lab in Alderley Park and Alex Alderton, Roberto Amato, Jeffrey Barrett, Sonia Goncalves, Ewan Harrison, David K. Jackson, Ian Johnston, Dominic Kwiatkowski, Cordelia Langford, John Sillitoe on behalf of the Wellcome Sanger Institute COVID-19 Surveillance Team |
| EPI_ISL_2512256, EPI_ISL_2512257, EPI_ISL_2512258, EPI_ISL_2512259, EPI_ISL_2512260, EPI_ISL_2512261 | Health Services Laboratories | Wellcome Sanger Institute for the COVID-19 Genomics UK (COG-UK) Consortium | Health Services Laboratories and Alex Alderton, Roberto Amato, Jeffrey Barrett, Sonia Goncalves, Ewan Harrison, David K. Jackson, Ian Johnston, Dominic Kwiatkowski, Cordelia Langford, John Sillitoe on behalf of the Wellcome Sanger Institute COVID-19 Surveillance Team |
| EPI_ISL_2512262 | Lighthouse Lab in Alderley Park | Wellcome Sanger Institute for the COVID-19 Genomics UK (COG-UK) Consortium | Jacquelyn Wynn, Mairead Hyland, The Lighthouse Lab in Alderley Park and Alex Alderton, Roberto Amato, Jeffrey Barrett, Sonia Goncalves, Ewan Harrison, David K. Jackson, Ian Johnston, Dominic Kwiatkowski, Cordelia Langford, John Sillitoe on behalf of the Wellcome Sanger Institute COVID-19 Surveillance Team |
| EPI_ISL_2512263, EPI_ISL_2512265, EPI_ISL_2512266 | Health Services Laboratories | Wellcome Sanger Institute for the COVID-19 Genomics UK (COG-UK) Consortium | Health Services Laboratories and Alex Alderton, Roberto Amato, Jeffrey Barrett, Sonia Goncalves, Ewan Harrison, David K. Jackson, Ian Johnston, Dominic Kwiatkowski, Cordelia Langford, John Sillitoe on behalf of the Wellcome Sanger Institute COVID-19 Surveillance Team |
| EPI_ISL_2512267 | Lighthouse Lab in Alderley Park | Wellcome Sanger Institute for the COVID-19 Genomics UK (COG-UK) Consortium | Jacquelyn Wynn, Mairead Hyland, The Lighthouse Lab in Alderley Park and Alex Alderton, Roberto Amato, Jeffrey Barrett, Sonia Goncalves, Ewan Harrison, David K. Jackson, Ian Johnston, Dominic Kwiatkowski, Cordelia Langford, John Sillitoe on behalf of the Wellcome Sanger Institute COVID-19 Surveillance Team |
| EPI_ISL_2512268 | Health Services Laboratories | Wellcome Sanger Institute for the COVID-19 Genomics UK (COG-UK) Consortium | Health Services Laboratories and Alex Alderton, Roberto Amato, Jeffrey Barrett, Sonia Goncalves, Ewan Harrison, David K. Jackson, Ian Johnston, Dominic Kwiatkowski, Cordelia Langford, John Sillitoe on behalf of the Wellcome Sanger Institute COVID-19 Surveillance Team |
| EPI_ISL_2512269 | Lighthouse Lab in Alderley Park | Wellcome Sanger Institute for the COVID-19 Genomics UK (COG-UK) Consortium | Jacquelyn Wynn, Mairead Hyland, The Lighthouse Lab in Alderley Park and Alex Alderton, Roberto Amato, Jeffrey Barrett, Sonia Goncalves, Ewan Harrison, David K. Jackson, Ian Johnston, Dominic Kwiatkowski, Cordelia Langford, John Sillitoe on behalf of the Wellcome Sanger Institute COVID-19 Surveillance Team |
| EPI_ISL_2512270, EPI_ISL_2512271, EPI_ISL_2512272, EPI_ISL_2512273, EPI_ISL_2512274, EPI_ISL_2512275, EPI_ISL_2512276 | Health Services Laboratories | Wellcome Sanger Institute for the COVID-19 Genomics UK (COG-UK) Consortium | Health Services Laboratories and Alex Alderton, Roberto Amato, Jeffrey Barrett, Sonia Goncalves, Ewan Harrison, David K. Jackson, Ian Johnston, Dominic Kwiatkowski, Cordelia Langford, John Sillitoe on behalf of the Wellcome Sanger Institute COVID-19 Surveillance Team |
| EPI_ISL_2512277, EPI_ISL_2512279 | Lighthouse Lab in Alderley Park | Wellcome Sanger Institute for the COVID-19 Genomics UK (COG-UK) Consortium | Jacquelyn Wynn, Mairead Hyland, The Lighthouse Lab in Alderley Park and Alex Alderton, Roberto Amato, Jeffrey Barrett, Sonia Goncalves, Ewan Harrison, David K. Jackson, Ian Johnston, Dominic Kwiatkowski, Cordelia Langford, John Sillitoe on behalf of the Wellcome Sanger Institute COVID-19 Surveillance Team |
| EPI_ISL_2512280, EPI_ISL_2512281 | Health Services Laboratories | Wellcome Sanger Institute for the COVID-19 Genomics UK (COG-UK) Consortium | Health Services Laboratories and Alex Alderton, Roberto Amato, Jeffrey Barrett, Sonia Goncalves, Ewan Harrison, David K. Jackson, Ian Johnston, Dominic Kwiatkowski, Cordelia Langford, John Sillitoe on behalf of the Wellcome Sanger Institute COVID-19 Surveillance Team |
| EPI_ISL_2512282 | Lighthouse Lab in Alderley Park | Wellcome Sanger Institute for the COVID-19 Genomics UK (COG-UK) Consortium | Jacquelyn Wynn, Mairead Hyland, The Lighthouse Lab in Alderley Park and Alex Alderton, Roberto Amato, Jeffrey Barrett, Sonia Goncalves, Ewan Harrison, David K. Jackson, Ian Johnston, Dominic Kwiatkowski, Cordelia Langford, John Sillitoe on behalf of the Wellcome Sanger Institute COVID-19 Surveillance Team |
| EPI_ISL_2512283, EPI_ISL_2512284 | Health Services Laboratories | Wellcome Sanger Institute for the COVID-19 Genomics UK (COG-UK) Consortium | Health Services Laboratories and Alex Alderton, Roberto Amato, Jeffrey Barrett, Sonia Goncalves, Ewan Harrison, David K. Jackson, Ian Johnston, Dominic Kwiatkowski, Cordelia Langford, John Sillitoe on behalf of the Wellcome Sanger Institute COVID-19 Surveillance Team |
| EPI_ISL_2512285 | Lighthouse Lab in Alderley Park | Wellcome Sanger Institute for the COVID-19 Genomics UK (COG-UK) Consortium | Jacquelyn Wynn, Mairead Hyland, The Lighthouse Lab in Alderley Park and Alex Alderton, Roberto Amato, Jeffrey Barrett, Sonia Goncalves, Ewan Harrison, David K. Jackson, Ian Johnston, Dominic Kwiatkowski, Cordelia Langford, John Sillitoe on behalf of the Wellcome Sanger Institute COVID-19 Surveillance Team |
| EPI_ISL_2512286, EPI_ISL_2512287, EPI_ISL_2512288, EPI_ISL_2512289, EPI_ISL_2512290, EPI_ISL_2512291, EPI_ISL_2512292, EPI_ISL_2512293 | Health Services Laboratories | Wellcome Sanger Institute for the COVID-19 Genomics UK (COG-UK) Consortium | Health Services Laboratories and Alex Alderton, Roberto Amato, Jeffrey Barrett, Sonia Goncalves, Ewan Harrison, David K. Jackson, Ian Johnston, Dominic Kwiatkowski, Cordelia Langford, John Sillitoe on behalf of the Wellcome Sanger Institute COVID-19 Surveillance Team |
| EPI_ISL_2512294, EPI_ISL_2512295 | Lighthouse Lab in Glasgow | Wellcome Sanger Institute for the COVID-19 Genomics UK (COG-UK) Consortium | Harper VanSteenhouse, Yumi Kasai, David Gray, Carol Clugston, Anna Dominiczak and Alex Alderton, Roberto Amato, Jeffrey Barrett, Sonia Goncalves, Ewan Harrison, David K. Jackson, Ian Johnston, Dominic Kwiatkowski, Cordelia Langford, John Sillitoe on behalf of the Wellcome Sanger Institute COVID-19 Surveillance Team |
| EPI_ISL_2512296 | Health Services Laboratories | Wellcome Sanger Institute for the COVID-19 Genomics UK (COG-UK) Consortium | Health Services Laboratories and Alex Alderton, Roberto Amato, Jeffrey Barrett, Sonia Goncalves, Ewan Harrison, David K. Jackson, Ian Johnston, Dominic Kwiatkowski, Cordelia Langford, John Sillitoe on behalf of the Wellcome Sanger Institute COVID-19 Surveillance Team |
| EPI_ISL_2512297, EPI_ISL_2512298, EPI_ISL_2512299, EPI_ISL_2512300, EPI_ISL_2512301, EPI_ISL_2512302, EPI_ISL_2512303 | Lighthouse Lab in Glasgow | Wellcome Sanger Institute for the COVID-19 Genomics UK (COG-UK) Consortium | Harper VanSteenhouse, Yumi Kasai, David Gray, Carol Clugston, Anna Dominiczak and Alex Alderton, Roberto Amato, Jeffrey Barrett, Sonia Goncalves, Ewan Harrison, David K. Jackson, Ian Johnston, Dominic Kwiatkowski, Cordelia Langford, John Sillitoe on behalf of the Wellcome Sanger Institute COVID-19 Surveillance Team |
| EPI_ISL_2512305, EPI_ISL_2512307 | Health Services Laboratories | Wellcome Sanger Institute for the COVID-19 Genomics UK (COG-UK) Consortium | Health Services Laboratories and Alex Alderton, Roberto Amato, Jeffrey Barrett, Sonia Goncalves, Ewan Harrison, David K. Jackson, Ian Johnston, Dominic Kwiatkowski, Cordelia Langford, John Sillitoe on behalf of the Wellcome Sanger Institute COVID-19 Surveillance Team |
| EPI_ISL_2512308 | Lighthouse Lab in Glasgow | Wellcome Sanger Institute for the COVID-19 Genomics UK (COG-UK) Consortium | Harper VanSteenhouse, Yumi Kasai, David Gray, Carol Clugston, Anna Dominiczak and Alex Alderton, Roberto Amato, Jeffrey Barrett, Sonia Goncalves, Ewan Harrison, David K. Jackson, Ian Johnston, Dominic Kwiatkowski, Cordelia Langford, John Sillitoe on behalf of the Wellcome Sanger Institute COVID-19 Surveillance Team |
| EPI_ISL_2512309 | Health Services Laboratories | Wellcome Sanger Institute for the COVID-19 Genomics UK (COG-UK) Consortium | Health Services Laboratories and Alex Alderton, Roberto Amato, Jeffrey Barrett, Sonia Goncalves, Ewan Harrison, David K. Jackson, Ian Johnston, Dominic Kwiatkowski, Cordelia Langford, John Sillitoe on behalf of the Wellcome Sanger Institute COVID-19 Surveillance Team |
| EPI_ISL_2512310 | Lighthouse Lab in Glasgow | Wellcome Sanger Institute for the COVID-19 Genomics UK (COG-UK) Consortium | Harper VanSteenhouse, Yumi Kasai, David Gray, Carol Clugston, Anna Dominiczak and Alex Alderton, Roberto Amato, Jeffrey Barrett, Sonia Goncalves, Ewan Harrison, David K. Jackson, Ian Johnston, Dominic Kwiatkowski, Cordelia Langford, John Sillitoe on behalf of the Wellcome Sanger Institute COVID-19 Surveillance Team |
| EPI_ISL_2512311 | Health Services Laboratories | Wellcome Sanger Institute for the COVID-19 Genomics UK (COG-UK) Consortium | Health Services Laboratories and Alex Alderton, Roberto Amato, Jeffrey Barrett, Sonia Goncalves, Ewan Harrison, David K. Jackson, Ian Johnston, Dominic Kwiatkowski, Cordelia Langford, John Sillitoe on behalf of the Wellcome Sanger Institute COVID-19 Surveillance Team |
| EPI_ISL_2512312 | Lighthouse Lab in Glasgow | Wellcome Sanger Institute for the COVID-19 Genomics UK (COG-UK) Consortium | Harper VanSteenhouse, Yumi Kasai, David Gray, Carol Clugston, Anna Dominiczak and Alex Alderton, Roberto Amato, Jeffrey Barrett, Sonia Goncalves, Ewan Harrison, David K. Jackson, Ian Johnston, Dominic Kwiatkowski, Cordelia Langford, John Sillitoe on behalf of the Wellcome Sanger Institute COVID-19 Surveillance Team |
| EPI_ISL_2512313 | Health Services Laboratories | Wellcome Sanger Institute for the COVID-19 Genomics UK (COG-UK) Consortium | Health Services Laboratories and Alex Alderton, Roberto Amato, Jeffrey Barrett, Sonia Goncalves, Ewan Harrison, David K. Jackson, Ian Johnston, Dominic Kwiatkowski, Cordelia Langford, John Sillitoe on behalf of the Wellcome Sanger Institute COVID-19 Surveillance Team |
| EPI_ISL_2512314, EPI_ISL_2512315, EPI_ISL_2512317 | Lighthouse Lab in Glasgow | Wellcome Sanger Institute for the COVID-19 Genomics UK (COG-UK) Consortium | Harper VanSteenhouse, Yumi Kasai, David Gray, Carol Clugston, Anna Dominiczak and Alex Alderton, Roberto Amato, Jeffrey Barrett, Sonia Goncalves, Ewan Harrison, David K. Jackson, Ian Johnston, Dominic Kwiatkowski, Cordelia Langford, John Sillitoe on behalf of the Wellcome Sanger Institute COVID-19 Surveillance Team |
| EPI_ISL_2512318 | Health Services Laboratories | Wellcome Sanger Institute for the COVID-19 Genomics UK (COG-UK) Consortium | Health Services Laboratories and Alex Alderton, Roberto Amato, Jeffrey Barrett, Sonia Goncalves, Ewan Harrison, David K. Jackson, Ian Johnston, Dominic Kwiatkowski, Cordelia Langford, John Sillitoe on behalf of the Wellcome Sanger Institute COVID-19 Surveillance Team |
| EPI_ISL_2512319 | Lighthouse Lab in Glasgow | Wellcome Sanger Institute for the COVID-19 Genomics UK (COG-UK) Consortium | Harper VanSteenhouse, Yumi Kasai, David Gray, Carol Clugston, Anna Dominiczak and Alex Alderton, Roberto Amato, Jeffrey Barrett, Sonia Goncalves, Ewan Harrison, David K. Jackson, Ian Johnston, Dominic Kwiatkowski, Cordelia Langford, John Sillitoe on behalf of the Wellcome Sanger Institute COVID-19 Surveillance Team |
| EPI_ISL_2512320 | Health Services Laboratories | Wellcome Sanger Institute for the COVID-19 Genomics UK (COG-UK) Consortium | Health Services Laboratories and Alex Alderton, Roberto Amato, Jeffrey Barrett, Sonia Goncalves, Ewan Harrison, David K. Jackson, Ian Johnston, Dominic Kwiatkowski, Cordelia Langford, John Sillitoe on behalf of the Wellcome Sanger Institute COVID-19 Surveillance Team |
| EPI_ISL_2512321, EPI_ISL_2512322 | Lighthouse Lab in Glasgow | Wellcome Sanger Institute for the COVID-19 Genomics UK (COG-UK) Consortium | Harper VanSteenhouse, Yumi Kasai, David Gray, Carol Clugston, Anna Dominiczak and Alex Alderton, Roberto Amato, Jeffrey Barrett, Sonia Goncalves, Ewan Harrison, David K. Jackson, Ian Johnston, Dominic Kwiatkowski, Cordelia Langford, John Sillitoe on behalf of the Wellcome Sanger Institute COVID-19 Surveillance Team |
| EPI_ISL_2512325 | Health Services Laboratories | Wellcome Sanger Institute for the COVID-19 Genomics UK (COG-UK) Consortium | Health Services Laboratories and Alex Alderton, Roberto Amato, Jeffrey Barrett, Sonia Goncalves, Ewan Harrison, David K. Jackson, Ian Johnston, Dominic Kwiatkowski, Cordelia Langford, John Sillitoe on behalf of the Wellcome Sanger Institute COVID-19 Surveillance Team |
| EPI_ISL_2512327, EPI_ISL_2512328, EPI_ISL_2512329 | Lighthouse Lab in Glasgow | Wellcome Sanger Institute for the COVID-19 Genomics UK (COG-UK) Consortium | Harper VanSteenhouse, Yumi Kasai, David Gray, Carol Clugston, Anna Dominiczak and Alex Alderton, Roberto Amato, Jeffrey Barrett, Sonia Goncalves, Ewan Harrison, David K. Jackson, Ian Johnston, Dominic Kwiatkowski, Cordelia Langford, John Sillitoe on behalf of the Wellcome Sanger Institute COVID-19 Surveillance Team |
| EPI_ISL_2512330, EPI_ISL_2512332 | Health Services Laboratories | Wellcome Sanger Institute for the COVID-19 Genomics UK (COG-UK) Consortium | Health Services Laboratories and Alex Alderton, Roberto Amato, Jeffrey Barrett, Sonia Goncalves, Ewan Harrison, David K. Jackson, Ian Johnston, Dominic Kwiatkowski, Cordelia Langford, John Sillitoe on behalf of the Wellcome Sanger Institute COVID-19 Surveillance Team |
| EPI_ISL_2512333, EPI_ISL_2512335, EPI_ISL_2512336, EPI_ISL_2512337, EPI_ISL_2512338, EPI_ISL_2512339, EPI_ISL_2512340, EPI_ISL_2512341, EPI_ISL_2512342, | Lighthouse Lab in Glasgow | Wellcome Sanger Institute for the COVID-19 Genomics UK (COG-UK) Consortium | Harper VanSteenhouse, Yumi Kasai, David Gray, Carol Clugston, Anna Dominiczak and Alex Alderton, Roberto Amato, Jeffrey Barrett, Sonia Goncalves, Ewan Harrison, David K. Jackson, Ian Johnston, Dominic Kwiatkowski, Cordelia Langford, John Sillitoe on behalf of the Wellcome Sanger Institute COVID-19 Surveillance Team |

| | | | |
|---|---|---|---|
| EPI_ISL_2512343 | | | |
| EPI_ISL_2512344 | Health Services Laboratories | Wellcome Sanger Institute for the COVID-19 Genomics UK (COG-UK) Consortium | Health Services Laboratories and Alex Alderton, Roberto Amato, Jeffrey Barrett, Sonia Goncalves, Ewan Harrison, David K. Jackson, Ian Johnston, Dominic Kwiatkowski, Cordelia Langford, John Sillitoe on behalf of the Wellcome Sanger Institute COVID-19 Surveillance Team |
| EPI_ISL_2512345, EPI_ISL_2512346, EPI_ISL_2512347 | Lighthouse Lab in Glasgow | Wellcome Sanger Institute for the COVID-19 Genomics UK (COG-UK) Consortium | Harper VanSteenhouse, Yumi Kasai, David Gray, Carol Clugston, Anna Dominiczak and Alex Alderton, Roberto Amato, Jeffrey Barrett, Sonia Goncalves, Ewan Harrison, David K. Jackson, Ian Johnston, Dominic Kwiatkowski, Cordelia Langford, John Sillitoe on behalf of the Wellcome Sanger Institute COVID-19 Surveillance Team |
| EPI_ISL_2512348 | Health Services Laboratories | Wellcome Sanger Institute for the COVID-19 Genomics UK (COG-UK) Consortium | Health Services Laboratories and Alex Alderton, Roberto Amato, Jeffrey Barrett, Sonia Goncalves, Ewan Harrison, David K. Jackson, Ian Johnston, Dominic Kwiatkowski, Cordelia Langford, John Sillitoe on behalf of the Wellcome Sanger Institute COVID-19 Surveillance Team |
| EPI_ISL_2512349, EPI_ISL_2512350, EPI_ISL_2512351, EPI_ISL_2512352, EPI_ISL_2512355, EPI_ISL_2512356, EPI_ISL_2512357, EPI_ISL_2512358 | Lighthouse Lab in Glasgow | Wellcome Sanger Institute for the COVID-19 Genomics UK (COG-UK) Consortium | Harper VanSteenhouse, Yumi Kasai, David Gray, Carol Clugston, Anna Dominiczak and Alex Alderton, Roberto Amato, Jeffrey Barrett, Sonia Goncalves, Ewan Harrison, David K. Jackson, Ian Johnston, Dominic Kwiatkowski, Cordelia Langford, John Sillitoe on behalf of the Wellcome Sanger Institute COVID-19 Surveillance Team |
| EPI_ISL_2512359 | Health Services Laboratories | Wellcome Sanger Institute for the COVID-19 Genomics UK (COG-UK) Consortium | Health Services Laboratories and Alex Alderton, Roberto Amato, Jeffrey Barrett, Sonia Goncalves, Ewan Harrison, David K. Jackson, Ian Johnston, Dominic Kwiatkowski, Cordelia Langford, John Sillitoe on behalf of the Wellcome Sanger Institute COVID-19 Surveillance Team |
| EPI_ISL_2512361, EPI_ISL_2512362 | Lighthouse Lab in Glasgow | Wellcome Sanger Institute for the COVID-19 Genomics UK (COG-UK) Consortium | Harper VanSteenhouse, Yumi Kasai, David Gray, Carol Clugston, Anna Dominiczak and Alex Alderton, Roberto Amato, Jeffrey Barrett, Sonia Goncalves, Ewan Harrison, David K. Jackson, Ian Johnston, Dominic Kwiatkowski, Cordelia Langford, John Sillitoe on behalf of the Wellcome Sanger Institute COVID-19 Surveillance Team |
| EPI_ISL_2512363, EPI_ISL_2512364 | Health Services Laboratories | Wellcome Sanger Institute for the COVID-19 Genomics UK (COG-UK) Consortium | Health Services Laboratories and Alex Alderton, Roberto Amato, Jeffrey Barrett, Sonia Goncalves, Ewan Harrison, David K. Jackson, Ian Johnston, Dominic Kwiatkowski, Cordelia Langford, John Sillitoe on behalf of the Wellcome Sanger Institute COVID-19 Surveillance Team |
| EPI_ISL_2512365, EPI_ISL_2512366, EPI_ISL_2512368 | Lighthouse Lab in Glasgow | Wellcome Sanger Institute for the COVID-19 Genomics UK (COG-UK) Consortium | Harper VanSteenhouse, Yumi Kasai, David Gray, Carol Clugston, Anna Dominiczak and Alex Alderton, Roberto Amato, Jeffrey Barrett, Sonia Goncalves, Ewan Harrison, David K. Jackson, Ian Johnston, Dominic Kwiatkowski, Cordelia Langford, John Sillitoe on behalf of the Wellcome Sanger Institute COVID-19 Surveillance Team |
| EPI_ISL_2512369 | Health Services Laboratories | Wellcome Sanger Institute for the COVID-19 Genomics UK (COG-UK) Consortium | Health Services Laboratories and Alex Alderton, Roberto Amato, Jeffrey Barrett, Sonia Goncalves, Ewan Harrison, David K. Jackson, Ian Johnston, Dominic Kwiatkowski, Cordelia Langford, John Sillitoe on behalf of the Wellcome Sanger Institute COVID-19 Surveillance Team |
| EPI_ISL_2512370, EPI_ISL_2512371, EPI_ISL_2512372, EPI_ISL_2512373, EPI_ISL_2512374 | Lighthouse Lab in Glasgow | Wellcome Sanger Institute for the COVID-19 Genomics UK (COG-UK) Consortium | Harper VanSteenhouse, Yumi Kasai, David Gray, Carol Clugston, Anna Dominiczak and Alex Alderton, Roberto Amato, Jeffrey Barrett, Sonia Goncalves, Ewan Harrison, David K. Jackson, Ian Johnston, Dominic Kwiatkowski, Cordelia Langford, John Sillitoe on behalf of the Wellcome Sanger Institute COVID-19 Surveillance Team |
| EPI_ISL_2512375, EPI_ISL_2512376, EPI_ISL_2512377, EPI_ISL_2512378, EPI_ISL_2512379, EPI_ISL_2512380, EPI_ISL_2512382, EPI_ISL_2512383 | Lighthouse Lab in Milton Keynes | Wellcome Sanger Institute for the COVID-19 Genomics UK (COG-UK) Consortium | The Lighthouse Lab in Milton Keynes and Alex Alderton, Roberto Amato, Jeffrey Barrett, Sonia Goncalves, Ewan Harrison, David K. Jackson, Ian Johnston, Dominic Kwiatkowski, Cordelia Langford, John Sillitoe on behalf of the Wellcome Sanger Institute COVID-19 Surveillance Team |
| EPI_ISL_2512384 | Lighthouse Lab in Glasgow | Wellcome Sanger Institute for the COVID-19 Genomics UK (COG-UK) Consortium | Harper VanSteenhouse, Yumi Kasai, David Gray, Carol Clugston, Anna Dominiczak and Alex Alderton, Roberto Amato, Jeffrey Barrett, Sonia Goncalves, Ewan Harrison, David K. Jackson, Ian Johnston, Dominic Kwiatkowski, Cordelia Langford, John Sillitoe on behalf of the Wellcome Sanger Institute COVID-19 Surveillance Team |
| EPI_ISL_2512385, EPI_ISL_2512386, EPI_ISL_2512387, EPI_ISL_2512388 | Lighthouse Lab in Milton Keynes | Wellcome Sanger Institute for the COVID-19 Genomics UK (COG-UK) Consortium | The Lighthouse Lab in Milton Keynes and Alex Alderton, Roberto Amato, Jeffrey Barrett, Sonia Goncalves, Ewan Harrison, David K. Jackson, Ian Johnston, Dominic Kwiatkowski, Cordelia Langford, John Sillitoe on behalf of the Wellcome Sanger Institute COVID-19 Surveillance Team |
| EPI_ISL_2512389 | Lighthouse Lab in Glasgow | Wellcome Sanger Institute for the COVID-19 Genomics UK (COG-UK) Consortium | Harper VanSteenhouse, Yumi Kasai, David Gray, Carol Clugston, Anna Dominiczak and Alex Alderton, Roberto Amato, Jeffrey Barrett, Sonia Goncalves, Ewan Harrison, David K. Jackson, Ian Johnston, Dominic Kwiatkowski, Cordelia Langford, John Sillitoe on behalf of the Wellcome Sanger Institute COVID-19 Surveillance Team |
| EPI_ISL_2512390, EPI_ISL_2512391, EPI_ISL_2512393, EPI_ISL_2512394 | Lighthouse Lab in Milton Keynes | Wellcome Sanger Institute for the COVID-19 Genomics UK (COG-UK) Consortium | The Lighthouse Lab in Milton Keynes and Alex Alderton, Roberto Amato, Jeffrey Barrett, Sonia Goncalves, Ewan Harrison, David K. Jackson, Ian Johnston, Dominic Kwiatkowski, Cordelia Langford, John Sillitoe on behalf of the Wellcome Sanger Institute COVID-19 Surveillance Team |
| EPI_ISL_2512395 | Lighthouse Lab in Glasgow | Wellcome Sanger Institute for the COVID-19 Genomics UK (COG-UK) Consortium | Harper VanSteenhouse, Yumi Kasai, David Gray, Carol Clugston, Anna Dominiczak and Alex Alderton, Roberto Amato, Jeffrey Barrett, Sonia Goncalves, Ewan Harrison, David K. Jackson, Ian Johnston, Dominic Kwiatkowski, Cordelia Langford, John Sillitoe on behalf of the Wellcome Sanger Institute COVID-19 Surveillance Team |
| EPI_ISL_2512396 | Lighthouse Lab in Milton Keynes | Wellcome Sanger Institute for the COVID-19 Genomics UK (COG-UK) Consortium | The Lighthouse Lab in Milton Keynes and Alex Alderton, Roberto Amato, Jeffrey Barrett, Sonia Goncalves, Ewan Harrison, David K. Jackson, Ian Johnston, Dominic Kwiatkowski, Cordelia Langford, John Sillitoe on behalf of the Wellcome Sanger Institute COVID-19 Surveillance Team |
| EPI_ISL_2512397 | Lighthouse Lab in Glasgow | Wellcome Sanger Institute for the COVID-19 Genomics UK (COG-UK) Consortium | Harper VanSteenhouse, Yumi Kasai, David Gray, Carol Clugston, Anna Dominiczak and Alex Alderton, Roberto Amato, Jeffrey Barrett, Sonia Goncalves, Ewan Harrison, David K. Jackson, Ian Johnston, Dominic Kwiatkowski, Cordelia Langford, John Sillitoe on behalf of the Wellcome Sanger Institute COVID-19 Surveillance Team |
| EPI_ISL_2512398, EPI_ISL_2512399, EPI_ISL_2512400, EPI_ISL_2512401, EPI_ISL_2512402, EPI_ISL_2512403, EPI_ISL_2512404, EPI_ISL_2512405, EPI_ISL_2512406, EPI_ISL_2512407, EPI_ISL_2512408, EPI_ISL_2512409, EPI_ISL_2512410, EPI_ISL_2512411, EPI_ISL_2512412, EPI_ISL_2512413, EPI_ISL_2512414, EPI_ISL_2512416, EPI_ISL_2512417, EPI_ISL_2512418, EPI_ISL_2512419, EPI_ISL_2512420, EPI_ISL_2512423, EPI_ISL_2512424 | | | |
| see above | Lighthouse Lab in Milton Keynes | Wellcome Sanger Institute for the COVID-19 Genomics UK (COG-UK) Consortium | The Lighthouse Lab in Milton Keynes and Alex Alderton, Roberto Amato, Jeffrey Barrett, Sonia Goncalves, Ewan Harrison, David K. Jackson, Ian Johnston, Dominic Kwiatkowski, Cordelia Langford, John Sillitoe on behalf of the Wellcome Sanger Institute COVID-19 Surveillance Team |
| EPI_ISL_2512425 | Lighthouse Lab in Glasgow | Wellcome Sanger Institute for the COVID-19 Genomics UK (COG-UK) Consortium | Harper VanSteenhouse, Yumi Kasai, David Gray, Carol Clugston, Anna Dominiczak and Alex Alderton, Roberto Amato, Jeffrey Barrett, Sonia Goncalves, Ewan Harrison, David K. Jackson, Ian Johnston, Dominic Kwiatkowski, Cordelia Langford, John Sillitoe on behalf of the Wellcome Sanger Institute COVID-19 Surveillance Team |
| EPI_ISL_2512426, EPI_ISL_2512427, EPI_ISL_2512428, EPI_ISL_2512429, EPI_ISL_2512430, EPI_ISL_2512432, EPI_ISL_2512433, EPI_ISL_2512434 | Lighthouse Lab in Milton Keynes | Wellcome Sanger Institute for the COVID-19 Genomics UK (COG-UK) Consortium | The Lighthouse Lab in Milton Keynes and Alex Alderton, Roberto Amato, Jeffrey Barrett, Sonia Goncalves, Ewan Harrison, David K. Jackson, Ian Johnston, Dominic Kwiatkowski, Cordelia Langford, John Sillitoe on behalf of the Wellcome Sanger Institute COVID-19 Surveillance Team |
| EPI_ISL_2512435, EPI_ISL_2512436 | Lighthouse Lab in Glasgow | Wellcome Sanger Institute for the COVID-19 Genomics UK (COG-UK) Consortium | Harper VanSteenhouse, Yumi Kasai, David Gray, Carol Clugston, Anna Dominiczak and Alex Alderton, Roberto Amato, Jeffrey Barrett, Sonia Goncalves, Ewan Harrison, David K. Jackson, Ian Johnston, Dominic Kwiatkowski, Cordelia Langford, John Sillitoe on behalf of the Wellcome Sanger Institute COVID-19 Surveillance Team |
| EPI_ISL_2512437, EPI_ISL_2512438 | Lighthouse Lab in Milton Keynes | Wellcome Sanger Institute for the COVID-19 Genomics UK (COG-UK) Consortium | The Lighthouse Lab in Milton Keynes and Alex Alderton, Roberto Amato, Jeffrey Barrett, Sonia Goncalves, Ewan Harrison, David K. Jackson, Ian Johnston, Dominic Kwiatkowski, Cordelia Langford, John Sillitoe on behalf of the Wellcome Sanger Institute COVID-19 Surveillance Team |
| EPI_ISL_2512439 | Lighthouse Lab in Glasgow | Wellcome Sanger Institute for the COVID-19 Genomics UK (COG-UK) Consortium | Harper VanSteenhouse, Yumi Kasai, David Gray, Carol Clugston, Anna Dominiczak and Alex Alderton, Roberto Amato, Jeffrey Barrett, Sonia Goncalves, Ewan Harrison, David K. Jackson, Ian Johnston, Dominic Kwiatkowski, Cordelia Langford, John Sillitoe on behalf of the Wellcome Sanger Institute COVID-19 Surveillance Team |
| EPI_ISL_2512440, EPI_ISL_2512441, EPI_ISL_2512443 | Lighthouse Lab in Milton Keynes | Wellcome Sanger Institute for the COVID-19 Genomics UK (COG-UK) Consortium | The Lighthouse Lab in Milton Keynes and Alex Alderton, Roberto Amato, Jeffrey Barrett, Sonia Goncalves, Ewan Harrison, David K. Jackson, Ian Johnston, Dominic Kwiatkowski, Cordelia Langford, John Sillitoe on behalf of the Wellcome Sanger Institute COVID-19 Surveillance Team |
| EPI_ISL_2512444 | Lighthouse Lab in Glasgow | Wellcome Sanger Institute for the COVID-19 Genomics UK (COG-UK) Consortium | Harper VanSteenhouse, Yumi Kasai, David Gray, Carol Clugston, Anna Dominiczak and Alex Alderton, Roberto Amato, Jeffrey Barrett, Sonia Goncalves, Ewan Harrison, David K. Jackson, Ian Johnston, Dominic Kwiatkowski, Cordelia Langford, John Sillitoe on behalf of the Wellcome Sanger Institute COVID-19 Surveillance Team |
| EPI_ISL_2512445, EPI_ISL_2512446, EPI_ISL_2512447, EPI_ISL_2512448, EPI_ISL_2512449, EPI_ISL_2512450, EPI_ISL_2512452 | Lighthouse Lab in Milton Keynes | Wellcome Sanger Institute for the COVID-19 Genomics UK (COG-UK) Consortium | The Lighthouse Lab in Milton Keynes and Alex Alderton, Roberto Amato, Jeffrey Barrett, Sonia Goncalves, Ewan Harrison, David K. Jackson, Ian Johnston, Dominic Kwiatkowski, Cordelia Langford, John Sillitoe on behalf of the Wellcome Sanger Institute COVID-19 Surveillance Team |
| EPI_ISL_2512453 | Lighthouse Lab in Glasgow | Wellcome Sanger Institute for the COVID-19 Genomics UK (COG-UK) Consortium | Harper VanSteenhouse, Yumi Kasai, David Gray, Carol Clugston, Anna Dominiczak and Alex Alderton, Roberto Amato, Jeffrey Barrett, Sonia Goncalves, Ewan Harrison, David K. Jackson, Ian Johnston, Dominic Kwiatkowski, Cordelia Langford, John Sillitoe on behalf of the Wellcome Sanger Institute COVID-19 Surveillance Team |
| EPI_ISL_2512454, EPI_ISL_2512455 | Lighthouse Lab in Milton Keynes | Wellcome Sanger Institute for the COVID-19 Genomics UK (COG-UK) Consortium | The Lighthouse Lab in Milton Keynes and Alex Alderton, Roberto Amato, Jeffrey Barrett, Sonia Goncalves, Ewan Harrison, David K. Jackson, Ian Johnston, Dominic Kwiatkowski, Cordelia Langford, John Sillitoe on behalf of the Wellcome Sanger Institute COVID-19 Surveillance Team |
| EPI_ISL_2512456 | Lighthouse Lab in Glasgow | Wellcome Sanger Institute for the COVID-19 Genomics UK (COG-UK) Consortium | Harper VanSteenhouse, Yumi Kasai, David Gray, Carol Clugston, Anna Dominiczak and Alex Alderton, Roberto Amato, Jeffrey Barrett, Sonia Goncalves, Ewan Harrison, David K. Jackson, Ian Johnston, Dominic Kwiatkowski, Cordelia Langford, John Sillitoe on behalf of the Wellcome Sanger Institute COVID-19 Surveillance Team |

| | | | |
|---|---|---|---|
| EPI_ISL_2512458, EPI_ISL_2512459, EPI_ISL_2512460, EPI_ISL_2512461, EPI_ISL_2512462 | Lighthouse Lab in Milton Keynes | Wellcome Sanger Institute for the COVID-19 Genomics UK (COG-UK) Consortium | The Lighthouse Lab in Milton Keynes and Alex Alderton, Roberto Amato, Jeffrey Barrett, Sonia Goncalves, Ewan Harrison, David K. Jackson, Ian Johnston, Dominic Kwiatkowski, Cordelia Langford, John Sillitoe on behalf of the Wellcome Sanger Institute COVID-19 Surveillance Team |
| EPI_ISL_2512463 | Lighthouse Lab in Glasgow | Wellcome Sanger Institute for the COVID-19 Genomics UK (COG-UK) Consortium | Harper VanSteenhouse, Yumi Kasai, David Gray, Carol Clugston, Anna Dominiczak and Alex Alderton, Roberto Amato, Jeffrey Barrett, Sonia Goncalves, Ewan Harrison, David K. Jackson, Ian Johnston, Dominic Kwiatkowski, Cordelia Langford, John Sillitoe on behalf of the Wellcome Sanger Institute COVID-19 Surveillance Team |
| EPI_ISL_2512464, EPI_ISL_2512465, EPI_ISL_2512466, EPI_ISL_2512467 | Lighthouse Lab in Milton Keynes | Wellcome Sanger Institute for the COVID-19 Genomics UK (COG-UK) Consortium | The Lighthouse Lab in Milton Keynes and Alex Alderton, Roberto Amato, Jeffrey Barrett, Sonia Goncalves, Ewan Harrison, David K. Jackson, Ian Johnston, Dominic Kwiatkowski, Cordelia Langford, John Sillitoe on behalf of the Wellcome Sanger Institute COVID-19 Surveillance Team |
| EPI_ISL_2512469, EPI_ISL_2512470 | Lighthouse Lab in Glasgow | Wellcome Sanger Institute for the COVID-19 Genomics UK (COG-UK) Consortium | Harper VanSteenhouse, Yumi Kasai, David Gray, Carol Clugston, Anna Dominiczak and Alex Alderton, Roberto Amato, Jeffrey Barrett, Sonia Goncalves, Ewan Harrison, David K. Jackson, Ian Johnston, Dominic Kwiatkowski, Cordelia Langford, John Sillitoe on behalf of the Wellcome Sanger Institute COVID-19 Surveillance Team |
| EPI_ISL_2512471, EPI_ISL_2512472, EPI_ISL_2512473, EPI_ISL_2512474, EPI_ISL_2512475, EPI_ISL_2512477, EPI_ISL_2512479 | Lighthouse Lab in Milton Keynes | Wellcome Sanger Institute for the COVID-19 Genomics UK (COG-UK) Consortium | The Lighthouse Lab in Milton Keynes and Alex Alderton, Roberto Amato, Jeffrey Barrett, Sonia Goncalves, Ewan Harrison, David K. Jackson, Ian Johnston, Dominic Kwiatkowski, Cordelia Langford, John Sillitoe on behalf of the Wellcome Sanger Institute COVID-19 Surveillance Team |
| EPI_ISL_2512480 | Lighthouse Lab in Glasgow | Wellcome Sanger Institute for the COVID-19 Genomics UK (COG-UK) Consortium | Harper VanSteenhouse, Yumi Kasai, David Gray, Carol Clugston, Anna Dominiczak and Alex Alderton, Roberto Amato, Jeffrey Barrett, Sonia Goncalves, Ewan Harrison, David K. Jackson, Ian Johnston, Dominic Kwiatkowski, Cordelia Langford, John Sillitoe on behalf of the Wellcome Sanger Institute COVID-19 Surveillance Team |
| EPI_ISL_2512481, EPI_ISL_2512482, EPI_ISL_2512483, EPI_ISL_2512484, EPI_ISL_2512485, EPI_ISL_2512486, EPI_ISL_2512487, EPI_ISL_2512488, EPI_ISL_2512489, EPI_ISL_2512490, EPI_ISL_2512491, EPI_ISL_2512493, EPI_ISL_2512494, EPI_ISL_2512495, EPI_ISL_2512496 | | | |
| see above | Lighthouse Lab in Milton Keynes | Wellcome Sanger Institute for the COVID-19 Genomics UK (COG-UK) Consortium | The Lighthouse Lab in Milton Keynes and Alex Alderton, Roberto Amato, Jeffrey Barrett, Sonia Goncalves, Ewan Harrison, David K. Jackson, Ian Johnston, Dominic Kwiatkowski, Cordelia Langford, John Sillitoe on behalf of the Wellcome Sanger Institute COVID-19 Surveillance Team |
| EPI_ISL_2512497 | Lighthouse Lab in Glasgow | Wellcome Sanger Institute for the COVID-19 Genomics UK (COG-UK) Consortium | Harper VanSteenhouse, Yumi Kasai, David Gray, Carol Clugston, Anna Dominiczak and Alex Alderton, Roberto Amato, Jeffrey Barrett, Sonia Goncalves, Ewan Harrison, David K. Jackson, Ian Johnston, Dominic Kwiatkowski, Cordelia Langford, John Sillitoe on behalf of the Wellcome Sanger Institute COVID-19 Surveillance Team |
| EPI_ISL_2512498, EPI_ISL_2512499, EPI_ISL_2512500, EPI_ISL_2512501, EPI_ISL_2512502, EPI_ISL_2512503, EPI_ISL_2512504, EPI_ISL_2512505 | Lighthouse Lab in Milton Keynes | Wellcome Sanger Institute for the COVID-19 Genomics UK (COG-UK) Consortium | The Lighthouse Lab in Milton Keynes and Alex Alderton, Roberto Amato, Jeffrey Barrett, Sonia Goncalves, Ewan Harrison, David K. Jackson, Ian Johnston, Dominic Kwiatkowski, Cordelia Langford, John Sillitoe on behalf of the Wellcome Sanger Institute COVID-19 Surveillance Team |
| EPI_ISL_2512506, EPI_ISL_2512507 | Lighthouse Lab in Glasgow | Wellcome Sanger Institute for the COVID-19 Genomics UK (COG-UK) Consortium | Harper VanSteenhouse, Yumi Kasai, David Gray, Carol Clugston, Anna Dominiczak and Alex Alderton, Roberto Amato, Jeffrey Barrett, Sonia Goncalves, Ewan Harrison, David K. Jackson, Ian Johnston, Dominic Kwiatkowski, Cordelia Langford, John Sillitoe on behalf of the Wellcome Sanger Institute COVID-19 Surveillance Team |
| EPI_ISL_2512508, EPI_ISL_2512509, EPI_ISL_2512510, EPI_ISL_2512511, EPI_ISL_2512512, EPI_ISL_2512513, EPI_ISL_2512515, EPI_ISL_2512516, EPI_ISL_2512517, EPI_ISL_2512518, EPI_ISL_2512519 | | | |
| see above | Lighthouse Lab in Milton Keynes | Wellcome Sanger Institute for the COVID-19 Genomics UK (COG-UK) Consortium | The Lighthouse Lab in Milton Keynes and Alex Alderton, Roberto Amato, Jeffrey Barrett, Sonia Goncalves, Ewan Harrison, David K. Jackson, Ian Johnston, Dominic Kwiatkowski, Cordelia Langford, John Sillitoe on behalf of the Wellcome Sanger Institute COVID-19 Surveillance Team |
| EPI_ISL_2512520 | Lighthouse Lab in Glasgow | Wellcome Sanger Institute for the COVID-19 Genomics UK (COG-UK) Consortium | Harper VanSteenhouse, Yumi Kasai, David Gray, Carol Clugston, Anna Dominiczak and Alex Alderton, Roberto Amato, Jeffrey Barrett, Sonia Goncalves, Ewan Harrison, David K. Jackson, Ian Johnston, Dominic Kwiatkowski, Cordelia Langford, John Sillitoe on behalf of the Wellcome Sanger Institute COVID-19 Surveillance Team |
| EPI_ISL_2512521, EPI_ISL_2512522, EPI_ISL_2512524 | Lighthouse Lab in Milton Keynes | Wellcome Sanger Institute for the COVID-19 Genomics UK (COG-UK) Consortium | The Lighthouse Lab in Milton Keynes and Alex Alderton, Roberto Amato, Jeffrey Barrett, Sonia Goncalves, Ewan Harrison, David K. Jackson, Ian Johnston, Dominic Kwiatkowski, Cordelia Langford, John Sillitoe on behalf of the Wellcome Sanger Institute COVID-19 Surveillance Team |
| EPI_ISL_2512525 | Lighthouse Lab in Glasgow | Wellcome Sanger Institute for the COVID-19 Genomics UK (COG-UK) Consortium | Harper VanSteenhouse, Yumi Kasai, David Gray, Carol Clugston, Anna Dominiczak and Alex Alderton, Roberto Amato, Jeffrey Barrett, Sonia Goncalves, Ewan Harrison, David K. Jackson, Ian Johnston, Dominic Kwiatkowski, Cordelia Langford, John Sillitoe on behalf of the Wellcome Sanger Institute COVID-19 Surveillance Team |
| EPI_ISL_2512527 | Lighthouse Lab in Milton Keynes | Wellcome Sanger Institute for the COVID-19 Genomics UK (COG-UK) Consortium | The Lighthouse Lab in Milton Keynes and Alex Alderton, Roberto Amato, Jeffrey Barrett, Sonia Goncalves, Ewan Harrison, David K. Jackson, Ian Johnston, Dominic Kwiatkowski, Cordelia Langford, John Sillitoe on behalf of the Wellcome Sanger Institute COVID-19 Surveillance Team |
| EPI_ISL_2512528 | Lighthouse Lab in Glasgow | Wellcome Sanger Institute for the COVID-19 Genomics UK (COG-UK) Consortium | Harper VanSteenhouse, Yumi Kasai, David Gray, Carol Clugston, Anna Dominiczak and Alex Alderton, Roberto Amato, Jeffrey Barrett, Sonia Goncalves, Ewan Harrison, David K. Jackson, Ian Johnston, Dominic Kwiatkowski, Cordelia Langford, John Sillitoe on behalf of the Wellcome Sanger Institute COVID-19 Surveillance Team |
| EPI_ISL_2512529, EPI_ISL_2512530, EPI_ISL_2512531 | Lighthouse Lab in Milton Keynes | Wellcome Sanger Institute for the COVID-19 Genomics UK (COG-UK) Consortium | The Lighthouse Lab in Milton Keynes and Alex Alderton, Roberto Amato, Jeffrey Barrett, Sonia Goncalves, Ewan Harrison, David K. Jackson, Ian Johnston, Dominic Kwiatkowski, Cordelia Langford, John Sillitoe on behalf of the Wellcome Sanger Institute COVID-19 Surveillance Team |
| EPI_ISL_2512532 | Lighthouse Lab in Glasgow | Wellcome Sanger Institute for the COVID-19 Genomics UK (COG-UK) Consortium | Harper VanSteenhouse, Yumi Kasai, David Gray, Carol Clugston, Anna Dominiczak and Alex Alderton, Roberto Amato, Jeffrey Barrett, Sonia Goncalves, Ewan Harrison, David K. Jackson, Ian Johnston, Dominic Kwiatkowski, Cordelia Langford, John Sillitoe on behalf of the Wellcome Sanger Institute COVID-19 Surveillance Team |
| EPI_ISL_2512533, EPI_ISL_2512534, EPI_ISL_2512536, EPI_ISL_2512539, EPI_ISL_2512540, EPI_ISL_2512541, EPI_ISL_2512542, EPI_ISL_2512543, EPI_ISL_2512544, EPI_ISL_2512545, EPI_ISL_2512546, EPI_ISL_2512548, EPI_ISL_2512549, EPI_ISL_2512550, EPI_ISL_2512553, EPI_ISL_2512554, EPI_ISL_2512555, EPI_ISL_2512556, EPI_ISL_2512557, EPI_ISL_2512558, EPI_ISL_2512559, EPI_ISL_2512560, EPI_ISL_2512561, EPI_ISL_2512562, EPI_ISL_2512564, EPI_ISL_2512565, EPI_ISL_2512566, EPI_ISL_2512568, EPI_ISL_2512569, EPI_ISL_2512570, EPI_ISL_2512571, EPI_ISL_2512572, EPI_ISL_2512573, EPI_ISL_2512574 | | | |
| see above | Lighthouse Lab in Milton Keynes | Wellcome Sanger Institute for the COVID-19 Genomics UK (COG-UK) Consortium | The Lighthouse Lab in Milton Keynes and Alex Alderton, Roberto Amato, Jeffrey Barrett, Sonia Goncalves, Ewan Harrison, David K. Jackson, Ian Johnston, Dominic Kwiatkowski, Cordelia Langford, John Sillitoe on behalf of the Wellcome Sanger Institute COVID-19 Surveillance Team |
| EPI_ISL_2512575 | Lighthouse Lab in Glasgow | Wellcome Sanger Institute for the COVID-19 Genomics UK (COG-UK) Consortium | Harper VanSteenhouse, Yumi Kasai, David Gray, Carol Clugston, Anna Dominiczak and Alex Alderton, Roberto Amato, Jeffrey Barrett, Sonia Goncalves, Ewan Harrison, David K. Jackson, Ian Johnston, Dominic Kwiatkowski, Cordelia Langford, John Sillitoe on behalf of the Wellcome Sanger Institute COVID-19 Surveillance Team |
| EPI_ISL_2512576, EPI_ISL_2512577, EPI_ISL_2512578, EPI_ISL_2512579, EPI_ISL_2512580, EPI_ISL_2512581, EPI_ISL_2512582, EPI_ISL_2512585, EPI_ISL_2512586, EPI_ISL_2512587, EPI_ISL_2512588 | | | |
| see above | Lighthouse Lab in Milton Keynes | Wellcome Sanger Institute for the COVID-19 Genomics UK (COG-UK) Consortium | The Lighthouse Lab in Milton Keynes and Alex Alderton, Roberto Amato, Jeffrey Barrett, Sonia Goncalves, Ewan Harrison, David K. Jackson, Ian Johnston, Dominic Kwiatkowski, Cordelia Langford, John Sillitoe on behalf of the Wellcome Sanger Institute COVID-19 Surveillance Team |
| EPI_ISL_2512589 | Lighthouse Lab in Glasgow | Wellcome Sanger Institute for the COVID-19 Genomics UK (COG-UK) Consortium | Harper VanSteenhouse, Yumi Kasai, David Gray, Carol Clugston, Anna Dominiczak and Alex Alderton, Roberto Amato, Jeffrey Barrett, Sonia Goncalves, Ewan Harrison, David K. Jackson, Ian Johnston, Dominic Kwiatkowski, Cordelia Langford, John Sillitoe on behalf of the Wellcome Sanger Institute COVID-19 Surveillance Team |
| EPI_ISL_2512590, EPI_ISL_2512591, EPI_ISL_2512592, EPI_ISL_2512593, EPI_ISL_2512594, EPI_ISL_2512595, EPI_ISL_2512596, EPI_ISL_2512597, EPI_ISL_2512598, EPI_ISL_2512599 | Lighthouse Lab in Milton Keynes | Wellcome Sanger Institute for the COVID-19 Genomics UK (COG-UK) Consortium | The Lighthouse Lab in Milton Keynes and Alex Alderton, Roberto Amato, Jeffrey Barrett, Sonia Goncalves, Ewan Harrison, David K. Jackson, Ian Johnston, Dominic Kwiatkowski, Cordelia Langford, John Sillitoe on behalf of the Wellcome Sanger Institute COVID-19 Surveillance Team |
| EPI_ISL_2512601 | Lighthouse Lab in Glasgow | Wellcome Sanger Institute for the COVID-19 Genomics UK (COG-UK) Consortium | Harper VanSteenhouse, Yumi Kasai, David Gray, Carol Clugston, Anna Dominiczak and Alex Alderton, Roberto Amato, Jeffrey Barrett, Sonia Goncalves, Ewan Harrison, David K. Jackson, Ian Johnston, Dominic Kwiatkowski, Cordelia Langford, John Sillitoe on behalf of the Wellcome Sanger Institute COVID-19 Surveillance Team |
| EPI_ISL_2512602, EPI_ISL_2512603, EPI_ISL_2512604, EPI_ISL_2512605 | Lighthouse Lab in Milton Keynes | Wellcome Sanger Institute for the COVID-19 Genomics UK (COG-UK) Consortium | The Lighthouse Lab in Milton Keynes and Alex Alderton, Roberto Amato, Jeffrey Barrett, Sonia Goncalves, Ewan Harrison, David K. Jackson, Ian Johnston, Dominic Kwiatkowski, Cordelia Langford, John Sillitoe on behalf of the Wellcome Sanger Institute COVID-19 Surveillance Team |
| EPI_ISL_2512606 | Lighthouse Lab in Glasgow | Wellcome Sanger Institute for the COVID-19 Genomics UK (COG-UK) Consortium | Harper VanSteenhouse, Yumi Kasai, David Gray, Carol Clugston, Anna Dominiczak and Alex Alderton, Roberto Amato, Jeffrey Barrett, Sonia Goncalves, Ewan Harrison, David K. Jackson, Ian Johnston, Dominic Kwiatkowski, Cordelia Langford, John Sillitoe on behalf of the Wellcome Sanger Institute COVID-19 Surveillance Team |
| EPI_ISL_2512607, EPI_ISL_2512608, EPI_ISL_2512609, EPI_ISL_2512610, EPI_ISL_2512612, EPI_ISL_2512614, EPI_ISL_2512615, EPI_ISL_2512616, EPI_ISL_2512617, EPI_ISL_2512618, EPI_ISL_2512619, EPI_ISL_2512620, EPI_ISL_2512621, EPI_ISL_2512623, EPI_ISL_2512624, EPI_ISL_2512625, EPI_ISL_2512626, EPI_ISL_2512627, EPI_ISL_2512628, EPI_ISL_2512629, EPI_ISL_2512630, EPI_ISL_2512631, EPI_ISL_2512632, EPI_ISL_2512634, EPI_ISL_2512635, EPI_ISL_2512636, EPI_ISL_2512637, EPI_ISL_2512638, EPI_ISL_2512639, EPI_ISL_2512640, EPI_ISL_2512641, EPI_ISL_2512642, EPI_ISL_2512643, EPI_ISL_2512644, EPI_ISL_2512645 | | | |
| see above | Lighthouse Lab in Milton Keynes | Wellcome Sanger Institute for the COVID-19 Genomics UK (COG-UK) Consortium | The Lighthouse Lab in Milton Keynes and Alex Alderton, Roberto Amato, Jeffrey Barrett, Sonia Goncalves, Ewan Harrison, David K. Jackson, Ian Johnston, Dominic Kwiatkowski, Cordelia Langford, John Sillitoe on behalf of the Wellcome Sanger Institute COVID-19 Surveillance Team |
| EPI_ISL_2512646 | Lighthouse Lab in Glasgow | Wellcome Sanger Institute for the COVID-19 Genomics UK (COG-UK) Consortium | Harper VanSteenhouse, Yumi Kasai, David Gray, Carol Clugston, Anna Dominiczak and Alex Alderton, Roberto Amato, Jeffrey Barrett, Sonia Goncalves, Ewan Harrison, David K. Jackson, Ian Johnston, Dominic Kwiatkowski, Cordelia Langford, John Sillitoe on behalf of the Wellcome Sanger Institute COVID-19 Surveillance Team |
| EPI_ISL_2512647, EPI_ISL_2512648, EPI_ISL_2512649, EPI_ISL_2512650, EPI_ISL_2512651, EPI_ISL_2512652, EPI_ISL_2512653, EPI_ISL_2512654, EPI_ISL_2512655, EPI_ISL_2512656, EPI_ISL_2512657, EPI_ISL_2512658, EPI_ISL_2512659, EPI_ISL_2512660, EPI_ISL_2512661, EPI_ISL_2512662, EPI_ISL_2512663, EPI_ISL_2512664, EPI_ISL_2512665 | | | |
| see above | Lighthouse Lab in Milton Keynes | Wellcome Sanger Institute for the COVID-19 Genomics UK (COG-UK) Consortium | The Lighthouse Lab in Milton Keynes and Alex Alderton, Roberto Amato, Jeffrey Barrett, Sonia Goncalves, Ewan Harrison, David K. Jackson, Ian Johnston, Dominic Kwiatkowski, Cordelia Langford, John Sillitoe on behalf of the Wellcome Sanger Institute COVID-19 Surveillance Team |

| | | | |
|---|---|---|---|
| EPI_ISL_2512666 | Lighthouse Lab in Glasgow | Wellcome Sanger Institute for the COVID-19 Genomics UK (COG-UK) Consortium | Harper VanSteenhouse, Yumi Kasai, David Gray, Carol Clugston, Anna Dominiczak and Alex Alderton, Roberto Amato, Jeffrey Barrett, Sonia Goncalves, Ewan Harrison, David K. Jackson, Ian Johnston, Dominic Kwiatkowski, Cordelia Langford, John Sillitoe on behalf of the Wellcome Sanger Institute COVID-19 Surveillance Team |
| EPI_ISL_2512667, EPI_ISL_2512668, EPI_ISL_2512669, EPI_ISL_2512670, EPI_ISL_2512671, EPI_ISL_2512673, EPI_ISL_2512675, EPI_ISL_2512676, EPI_ISL_2512678, EPI_ISL_2512679, EPI_ISL_2512680, EPI_ISL_2512681, EPI_ISL_2512682, EPI_ISL_2512683 | | | |
| see above | Lighthouse Lab in Milton Keynes | Wellcome Sanger Institute for the COVID-19 Genomics UK (COG-UK) Consortium | The Lighthouse Lab in Milton Keynes and Alex Alderton, Roberto Amato, Jeffrey Barrett, Sonia Goncalves, Ewan Harrison, David K. Jackson, Ian Johnston, Dominic Kwiatkowski, Cordelia Langford, John Sillitoe on behalf of the Wellcome Sanger Institute COVID-19 Surveillance Team |
| EPI_ISL_2512684 | Lighthouse Lab in Glasgow | Wellcome Sanger Institute for the COVID-19 Genomics UK (COG-UK) Consortium | Harper VanSteenhouse, Yumi Kasai, David Gray, Carol Clugston, Anna Dominiczak and Alex Alderton, Roberto Amato, Jeffrey Barrett, Sonia Goncalves, Ewan Harrison, David K. Jackson, Ian Johnston, Dominic Kwiatkowski, Cordelia Langford, John Sillitoe on behalf of the Wellcome Sanger Institute COVID-19 Surveillance Team |
| EPI_ISL_2512685, EPI_ISL_2512686 | Lighthouse Lab in Milton Keynes | Wellcome Sanger Institute for the COVID-19 Genomics UK (COG-UK) Consortium | The Lighthouse Lab in Milton Keynes and Alex Alderton, Roberto Amato, Jeffrey Barrett, Sonia Goncalves, Ewan Harrison, David K. Jackson, Ian Johnston, Dominic Kwiatkowski, Cordelia Langford, John Sillitoe on behalf of the Wellcome Sanger Institute COVID-19 Surveillance Team |
| EPI_ISL_2512687, EPI_ISL_2512688, EPI_ISL_2512689 | Lighthouse Lab in Glasgow | Wellcome Sanger Institute for the COVID-19 Genomics UK (COG-UK) Consortium | Harper VanSteenhouse, Yumi Kasai, David Gray, Carol Clugston, Anna Dominiczak and Alex Alderton, Roberto Amato, Jeffrey Barrett, Sonia Goncalves, Ewan Harrison, David K. Jackson, Ian Johnston, Dominic Kwiatkowski, Cordelia Langford, John Sillitoe on behalf of the Wellcome Sanger Institute COVID-19 Surveillance Team |
| EPI_ISL_2512690 | Lighthouse Lab in Milton Keynes | Wellcome Sanger Institute for the COVID-19 Genomics UK (COG-UK) Consortium | The Lighthouse Lab in Milton Keynes and Alex Alderton, Roberto Amato, Jeffrey Barrett, Sonia Goncalves, Ewan Harrison, David K. Jackson, Ian Johnston, Dominic Kwiatkowski, Cordelia Langford, John Sillitoe on behalf of the Wellcome Sanger Institute COVID-19 Surveillance Team |
| EPI_ISL_2512691, EPI_ISL_2512692 | Lighthouse Lab in Glasgow | Wellcome Sanger Institute for the COVID-19 Genomics UK (COG-UK) Consortium | Harper VanSteenhouse, Yumi Kasai, David Gray, Carol Clugston, Anna Dominiczak and Alex Alderton, Roberto Amato, Jeffrey Barrett, Sonia Goncalves, Ewan Harrison, David K. Jackson, Ian Johnston, Dominic Kwiatkowski, Cordelia Langford, John Sillitoe on behalf of the Wellcome Sanger Institute COVID-19 Surveillance Team |
| EPI_ISL_2512693 | Lighthouse Lab in Milton Keynes | Wellcome Sanger Institute for the COVID-19 Genomics UK (COG-UK) Consortium | The Lighthouse Lab in Milton Keynes and Alex Alderton, Roberto Amato, Jeffrey Barrett, Sonia Goncalves, Ewan Harrison, David K. Jackson, Ian Johnston, Dominic Kwiatkowski, Cordelia Langford, John Sillitoe on behalf of the Wellcome Sanger Institute COVID-19 Surveillance Team |
| EPI_ISL_2512694 | Lighthouse Lab in Glasgow | Wellcome Sanger Institute for the COVID-19 Genomics UK (COG-UK) Consortium | Harper VanSteenhouse, Yumi Kasai, David Gray, Carol Clugston, Anna Dominiczak and Alex Alderton, Roberto Amato, Jeffrey Barrett, Sonia Goncalves, Ewan Harrison, David K. Jackson, Ian Johnston, Dominic Kwiatkowski, Cordelia Langford, John Sillitoe on behalf of the Wellcome Sanger Institute COVID-19 Surveillance Team |
| EPI_ISL_2512695, EPI_ISL_2512696, EPI_ISL_2512697 | Lighthouse Lab in Milton Keynes | Wellcome Sanger Institute for the COVID-19 Genomics UK (COG-UK) Consortium | The Lighthouse Lab in Milton Keynes and Alex Alderton, Roberto Amato, Jeffrey Barrett, Sonia Goncalves, Ewan Harrison, David K. Jackson, Ian Johnston, Dominic Kwiatkowski, Cordelia Langford, John Sillitoe on behalf of the Wellcome Sanger Institute COVID-19 Surveillance Team |
| EPI_ISL_2512698, EPI_ISL_2512699 | Lighthouse Lab in Glasgow | Wellcome Sanger Institute for the COVID-19 Genomics UK (COG-UK) Consortium | Harper VanSteenhouse, Yumi Kasai, David Gray, Carol Clugston, Anna Dominiczak and Alex Alderton, Roberto Amato, Jeffrey Barrett, Sonia Goncalves, Ewan Harrison, David K. Jackson, Ian Johnston, Dominic Kwiatkowski, Cordelia Langford, John Sillitoe on behalf of the Wellcome Sanger Institute COVID-19 Surveillance Team |
| EPI_ISL_2512700, EPI_ISL_2512701, EPI_ISL_2512702, EPI_ISL_2512703, EPI_ISL_2512704 | Lighthouse Lab in Milton Keynes | Wellcome Sanger Institute for the COVID-19 Genomics UK (COG-UK) Consortium | The Lighthouse Lab in Milton Keynes and Alex Alderton, Roberto Amato, Jeffrey Barrett, Sonia Goncalves, Ewan Harrison, David K. Jackson, Ian Johnston, Dominic Kwiatkowski, Cordelia Langford, John Sillitoe on behalf of the Wellcome Sanger Institute COVID-19 Surveillance Team |
| EPI_ISL_2512705 | Lighthouse Lab in Glasgow | Wellcome Sanger Institute for the COVID-19 Genomics UK (COG-UK) Consortium | Harper VanSteenhouse, Yumi Kasai, David Gray, Carol Clugston, Anna Dominiczak and Alex Alderton, Roberto Amato, Jeffrey Barrett, Sonia Goncalves, Ewan Harrison, David K. Jackson, Ian Johnston, Dominic Kwiatkowski, Cordelia Langford, John Sillitoe on behalf of the Wellcome Sanger Institute COVID-19 Surveillance Team |
| EPI_ISL_2512706 | Lighthouse Lab in Milton Keynes | Wellcome Sanger Institute for the COVID-19 Genomics UK (COG-UK) Consortium | The Lighthouse Lab in Milton Keynes and Alex Alderton, Roberto Amato, Jeffrey Barrett, Sonia Goncalves, Ewan Harrison, David K. Jackson, Ian Johnston, Dominic Kwiatkowski, Cordelia Langford, John Sillitoe on behalf of the Wellcome Sanger Institute COVID-19 Surveillance Team |
| EPI_ISL_2512707 | Lighthouse Lab in Glasgow | Wellcome Sanger Institute for the COVID-19 Genomics UK (COG-UK) Consortium | Harper VanSteenhouse, Yumi Kasai, David Gray, Carol Clugston, Anna Dominiczak and Alex Alderton, Roberto Amato, Jeffrey Barrett, Sonia Goncalves, Ewan Harrison, David K. Jackson, Ian Johnston, Dominic Kwiatkowski, Cordelia Langford, John Sillitoe on behalf of the Wellcome Sanger Institute COVID-19 Surveillance Team |
| EPI_ISL_2512708, EPI_ISL_2512709 | Lighthouse Lab in Milton Keynes | Wellcome Sanger Institute for the COVID-19 Genomics UK (COG-UK) Consortium | The Lighthouse Lab in Milton Keynes and Alex Alderton, Roberto Amato, Jeffrey Barrett, Sonia Goncalves, Ewan Harrison, David K. Jackson, Ian Johnston, Dominic Kwiatkowski, Cordelia Langford, John Sillitoe on behalf of the Wellcome Sanger Institute COVID-19 Surveillance Team |
| EPI_ISL_2512710, EPI_ISL_2512711 | Lighthouse Lab in Glasgow | Wellcome Sanger Institute for the COVID-19 Genomics UK (COG-UK) Consortium | Harper VanSteenhouse, Yumi Kasai, David Gray, Carol Clugston, Anna Dominiczak and Alex Alderton, Roberto Amato, Jeffrey Barrett, Sonia Goncalves, Ewan Harrison, David K. Jackson, Ian Johnston, Dominic Kwiatkowski, Cordelia Langford, John Sillitoe on behalf of the Wellcome Sanger Institute COVID-19 Surveillance Team |
| EPI_ISL_2512712 | Lighthouse Lab in Milton Keynes | Wellcome Sanger Institute for the COVID-19 Genomics UK (COG-UK) Consortium | The Lighthouse Lab in Milton Keynes and Alex Alderton, Roberto Amato, Jeffrey Barrett, Sonia Goncalves, Ewan Harrison, David K. Jackson, Ian Johnston, Dominic Kwiatkowski, Cordelia Langford, John Sillitoe on behalf of the Wellcome Sanger Institute COVID-19 Surveillance Team |
| EPI_ISL_2512713, EPI_ISL_2512714 | Lighthouse Lab in Glasgow | Wellcome Sanger Institute for the COVID-19 Genomics UK (COG-UK) Consortium | Harper VanSteenhouse, Yumi Kasai, David Gray, Carol Clugston, Anna Dominiczak and Alex Alderton, Roberto Amato, Jeffrey Barrett, Sonia Goncalves, Ewan Harrison, David K. Jackson, Ian Johnston, Dominic Kwiatkowski, Cordelia Langford, John Sillitoe on behalf of the Wellcome Sanger Institute COVID-19 Surveillance Team |
| EPI_ISL_2512715, EPI_ISL_2512716, EPI_ISL_2512717, EPI_ISL_2512719, EPI_ISL_2512720 | Lighthouse Lab in Milton Keynes | Wellcome Sanger Institute for the COVID-19 Genomics UK (COG-UK) Consortium | The Lighthouse Lab in Milton Keynes and Alex Alderton, Roberto Amato, Jeffrey Barrett, Sonia Goncalves, Ewan Harrison, David K. Jackson, Ian Johnston, Dominic Kwiatkowski, Cordelia Langford, John Sillitoe on behalf of the Wellcome Sanger Institute COVID-19 Surveillance Team |
| EPI_ISL_2512721 | Lighthouse Lab in Glasgow | Wellcome Sanger Institute for the COVID-19 Genomics UK (COG-UK) Consortium | Harper VanSteenhouse, Yumi Kasai, David Gray, Carol Clugston, Anna Dominiczak and Alex Alderton, Roberto Amato, Jeffrey Barrett, Sonia Goncalves, Ewan Harrison, David K. Jackson, Ian Johnston, Dominic Kwiatkowski, Cordelia Langford, John Sillitoe on behalf of the Wellcome Sanger Institute COVID-19 Surveillance Team |
| EPI_ISL_2512722, EPI_ISL_2512723, EPI_ISL_2512724, EPI_ISL_2512725, EPI_ISL_2512726, EPI_ISL_2512727, EPI_ISL_2512729 | Lighthouse Lab in Milton Keynes | Wellcome Sanger Institute for the COVID-19 Genomics UK (COG-UK) Consortium | The Lighthouse Lab in Milton Keynes and Alex Alderton, Roberto Amato, Jeffrey Barrett, Sonia Goncalves, Ewan Harrison, David K. Jackson, Ian Johnston, Dominic Kwiatkowski, Cordelia Langford, John Sillitoe on behalf of the Wellcome Sanger Institute COVID-19 Surveillance Team |
| EPI_ISL_2512730 | Lighthouse Lab in Glasgow | Wellcome Sanger Institute for the COVID-19 Genomics UK (COG-UK) Consortium | Harper VanSteenhouse, Yumi Kasai, David Gray, Carol Clugston, Anna Dominiczak and Alex Alderton, Roberto Amato, Jeffrey Barrett, Sonia Goncalves, Ewan Harrison, David K. Jackson, Ian Johnston, Dominic Kwiatkowski, Cordelia Langford, John Sillitoe on behalf of the Wellcome Sanger Institute COVID-19 Surveillance Team |
| EPI_ISL_2512731 | Lighthouse Lab in Milton Keynes | Wellcome Sanger Institute for the COVID-19 Genomics UK (COG-UK) Consortium | The Lighthouse Lab in Milton Keynes and Alex Alderton, Roberto Amato, Jeffrey Barrett, Sonia Goncalves, Ewan Harrison, David K. Jackson, Ian Johnston, Dominic Kwiatkowski, Cordelia Langford, John Sillitoe on behalf of the Wellcome Sanger Institute COVID-19 Surveillance Team |
| EPI_ISL_2512732 | Lighthouse Lab in Glasgow | Wellcome Sanger Institute for the COVID-19 Genomics UK (COG-UK) Consortium | Harper VanSteenhouse, Yumi Kasai, David Gray, Carol Clugston, Anna Dominiczak and Alex Alderton, Roberto Amato, Jeffrey Barrett, Sonia Goncalves, Ewan Harrison, David K. Jackson, Ian Johnston, Dominic Kwiatkowski, Cordelia Langford, John Sillitoe on behalf of the Wellcome Sanger Institute COVID-19 Surveillance Team |
| EPI_ISL_2512733, EPI_ISL_2512734, EPI_ISL_2512735, EPI_ISL_2512736 | Lighthouse Lab in Milton Keynes | Wellcome Sanger Institute for the COVID-19 Genomics UK (COG-UK) Consortium | The Lighthouse Lab in Milton Keynes and Alex Alderton, Roberto Amato, Jeffrey Barrett, Sonia Goncalves, Ewan Harrison, David K. Jackson, Ian Johnston, Dominic Kwiatkowski, Cordelia Langford, John Sillitoe on behalf of the Wellcome Sanger Institute COVID-19 Surveillance Team |
| EPI_ISL_2512737 | Lighthouse Lab in Glasgow | Wellcome Sanger Institute for the COVID-19 Genomics UK (COG-UK) Consortium | Harper VanSteenhouse, Yumi Kasai, David Gray, Carol Clugston, Anna Dominiczak and Alex Alderton, Roberto Amato, Jeffrey Barrett, Sonia Goncalves, Ewan Harrison, David K. Jackson, Ian Johnston, Dominic Kwiatkowski, Cordelia Langford, John Sillitoe on behalf of the Wellcome Sanger Institute COVID-19 Surveillance Team |
| EPI_ISL_2512738, EPI_ISL_2512739, EPI_ISL_2512740, EPI_ISL_2512741, EPI_ISL_2512742, EPI_ISL_2512743, EPI_ISL_2512744, EPI_ISL_2512745, EPI_ISL_2512746, EPI_ISL_2512747 | Lighthouse Lab in Milton Keynes | Wellcome Sanger Institute for the COVID-19 Genomics UK (COG-UK) Consortium | The Lighthouse Lab in Milton Keynes and Alex Alderton, Roberto Amato, Jeffrey Barrett, Sonia Goncalves, Ewan Harrison, David K. Jackson, Ian Johnston, Dominic Kwiatkowski, Cordelia Langford, John Sillitoe on behalf of the Wellcome Sanger Institute COVID-19 Surveillance Team |
| EPI_ISL_2512748 | Lighthouse Lab in Glasgow | Wellcome Sanger Institute for the COVID-19 Genomics UK (COG-UK) Consortium | Harper VanSteenhouse, Yumi Kasai, David Gray, Carol Clugston, Anna Dominiczak and Alex Alderton, Roberto Amato, Jeffrey Barrett, Sonia Goncalves, Ewan Harrison, David K. Jackson, Ian Johnston, Dominic Kwiatkowski, Cordelia Langford, John Sillitoe on behalf of the Wellcome Sanger Institute COVID-19 Surveillance Team |
| EPI_ISL_2512749, EPI_ISL_2512750, EPI_ISL_2512752, EPI_ISL_2512753 | Lighthouse Lab in Milton Keynes | Wellcome Sanger Institute for the COVID-19 Genomics UK (COG-UK) Consortium | The Lighthouse Lab in Milton Keynes and Alex Alderton, Roberto Amato, Jeffrey Barrett, Sonia Goncalves, Ewan Harrison, David K. Jackson, Ian Johnston, Dominic Kwiatkowski, Cordelia Langford, John Sillitoe on behalf of the Wellcome Sanger Institute COVID-19 Surveillance Team |
| EPI_ISL_2512754, EPI_ISL_2512755 | Lighthouse Lab in Glasgow | Wellcome Sanger Institute for the COVID-19 Genomics UK (COG-UK) Consortium | Harper VanSteenhouse, Yumi Kasai, David Gray, Carol Clugston, Anna Dominiczak and Alex Alderton, Roberto Amato, Jeffrey Barrett, Sonia Goncalves, Ewan Harrison, David K. Jackson, Ian Johnston, Dominic Kwiatkowski, Cordelia Langford, John Sillitoe on behalf of the Wellcome Sanger Institute COVID-19 Surveillance Team |
| EPI_ISL_2512756 | Lighthouse Lab in Milton Keynes | Wellcome Sanger Institute for the COVID-19 Genomics UK (COG-UK) Consortium | The Lighthouse Lab in Milton Keynes and Alex Alderton, Roberto Amato, Jeffrey Barrett, Sonia Goncalves, Ewan Harrison, David K. Jackson, Ian Johnston, Dominic Kwiatkowski, Cordelia Langford, John Sillitoe on behalf of the Wellcome Sanger Institute COVID-19 Surveillance Team |
| EPI_ISL_2512757 | Lighthouse Lab in Glasgow | Wellcome Sanger Institute for the COVID-19 Genomics UK (COG-UK) Consortium | Harper VanSteenhouse, Yumi Kasai, David Gray, Carol Clugston, Anna Dominiczak and Alex Alderton, Roberto Amato, Jeffrey Barrett, Sonia Goncalves, Ewan Harrison, David K. Jackson, Ian Johnston, Dominic Kwiatkowski, Cordelia Langford, John Sillitoe on behalf of the Wellcome Sanger Institute COVID-19 Surveillance Team |
| EPI_ISL_2512759 | Lighthouse Lab in Milton Keynes | Wellcome Sanger Institute for the COVID-19 Genomics UK (COG-UK) Consortium | The Lighthouse Lab in Milton Keynes and Alex Alderton, Roberto Amato, Jeffrey Barrett, Sonia Goncalves, Ewan Harrison, David K. Jackson, Ian Johnston, Dominic Kwiatkowski, Cordelia Langford, John Sillitoe on behalf of the Wellcome Sanger Institute COVID-19 Surveillance Team |
| EPI_ISL_2512760 | Lighthouse Lab in Glasgow | Wellcome Sanger Institute for the COVID-19 Genomics UK (COG-UK) | Harper VanSteenhouse, Yumi Kasai, David Gray, Carol Clugston, Anna Dominiczak and Alex Alderton, Roberto Amato, Jeffrey Barrett, Sonia Goncalves, Ewan Harrison, David K. Jackson, Ian Johnston, Dominic Kwiatkowski, Cordelia Langford, |

| | | | |
|---|---|---|---|
| | | Consortium | John Sillitoe on behalf of the Wellcome Sanger Institute COVID-19 Surveillance Team |
| EPI_ISL_2512761, EPI_ISL_2512762, EPI_ISL_2512763 | Lighthouse Lab in Milton Keynes | Wellcome Sanger Institute for the COVID-19 Genomics UK (COG-UK) Consortium | The Lighthouse Lab in Milton Keynes and Alex Alderton, Roberto Amato, Jeffrey Barrett, Sonia Goncalves, Ewan Harrison, David K. Jackson, Ian Johnston, Dominic Kwiatkowski, Cordelia Langford, John Sillitoe on behalf of the Wellcome Sanger Institute COVID-19 Surveillance Team |
| EPI_ISL_2512764, EPI_ISL_2512766, EPI_ISL_2512767 | Lighthouse Lab in Glasgow | Wellcome Sanger Institute for the COVID-19 Genomics UK (COG-UK) Consortium | Harper VanSteenhouse, Yumi Kasai, David Gray, Carol Clugston, Anna Dominiczak and Alex Alderton, Roberto Amato, Jeffrey Barrett, Sonia Goncalves, Ewan Harrison, David K. Jackson, Ian Johnston, Dominic Kwiatkowski, Cordelia Langford, John Sillitoe on behalf of the Wellcome Sanger Institute COVID-19 Surveillance Team |
| EPI_ISL_2512769 | Lighthouse Lab in Milton Keynes | Wellcome Sanger Institute for the COVID-19 Genomics UK (COG-UK) Consortium | The Lighthouse Lab in Milton Keynes and Alex Alderton, Roberto Amato, Jeffrey Barrett, Sonia Goncalves, Ewan Harrison, David K. Jackson, Ian Johnston, Dominic Kwiatkowski, Cordelia Langford, John Sillitoe on behalf of the Wellcome Sanger Institute COVID-19 Surveillance Team |
| EPI_ISL_2512770 | Lighthouse Lab in Glasgow | Wellcome Sanger Institute for the COVID-19 Genomics UK (COG-UK) Consortium | Harper VanSteenhouse, Yumi Kasai, David Gray, Carol Clugston, Anna Dominiczak and Alex Alderton, Roberto Amato, Jeffrey Barrett, Sonia Goncalves, Ewan Harrison, David K. Jackson, Ian Johnston, Dominic Kwiatkowski, Cordelia Langford, John Sillitoe on behalf of the Wellcome Sanger Institute COVID-19 Surveillance Team |
| EPI_ISL_2512771 | Lighthouse Lab in Milton Keynes | Wellcome Sanger Institute for the COVID-19 Genomics UK (COG-UK) Consortium | The Lighthouse Lab in Milton Keynes and Alex Alderton, Roberto Amato, Jeffrey Barrett, Sonia Goncalves, Ewan Harrison, David K. Jackson, Ian Johnston, Dominic Kwiatkowski, Cordelia Langford, John Sillitoe on behalf of the Wellcome Sanger Institute COVID-19 Surveillance Team |
| EPI_ISL_2512772 | Lighthouse Lab in Glasgow | Wellcome Sanger Institute for the COVID-19 Genomics UK (COG-UK) Consortium | Harper VanSteenhouse, Yumi Kasai, David Gray, Carol Clugston, Anna Dominiczak and Alex Alderton, Roberto Amato, Jeffrey Barrett, Sonia Goncalves, Ewan Harrison, David K. Jackson, Ian Johnston, Dominic Kwiatkowski, Cordelia Langford, John Sillitoe on behalf of the Wellcome Sanger Institute COVID-19 Surveillance Team |
| EPI_ISL_2512774 | Lighthouse Lab in Milton Keynes | Wellcome Sanger Institute for the COVID-19 Genomics UK (COG-UK) Consortium | The Lighthouse Lab in Milton Keynes and Alex Alderton, Roberto Amato, Jeffrey Barrett, Sonia Goncalves, Ewan Harrison, David K. Jackson, Ian Johnston, Dominic Kwiatkowski, Cordelia Langford, John Sillitoe on behalf of the Wellcome Sanger Institute COVID-19 Surveillance Team |
| EPI_ISL_2512775 | Lighthouse Lab in Glasgow | Wellcome Sanger Institute for the COVID-19 Genomics UK (COG-UK) Consortium | Harper VanSteenhouse, Yumi Kasai, David Gray, Carol Clugston, Anna Dominiczak and Alex Alderton, Roberto Amato, Jeffrey Barrett, Sonia Goncalves, Ewan Harrison, David K. Jackson, Ian Johnston, Dominic Kwiatkowski, Cordelia Langford, John Sillitoe on behalf of the Wellcome Sanger Institute COVID-19 Surveillance Team |
| EPI_ISL_2512776, EPI_ISL_2512777 | Lighthouse Lab in Milton Keynes | Wellcome Sanger Institute for the COVID-19 Genomics UK (COG-UK) Consortium | The Lighthouse Lab in Milton Keynes and Alex Alderton, Roberto Amato, Jeffrey Barrett, Sonia Goncalves, Ewan Harrison, David K. Jackson, Ian Johnston, Dominic Kwiatkowski, Cordelia Langford, John Sillitoe on behalf of the Wellcome Sanger Institute COVID-19 Surveillance Team |
| EPI_ISL_2512778 | Lighthouse Lab in Glasgow | Wellcome Sanger Institute for the COVID-19 Genomics UK (COG-UK) Consortium | Harper VanSteenhouse, Yumi Kasai, David Gray, Carol Clugston, Anna Dominiczak and Alex Alderton, Roberto Amato, Jeffrey Barrett, Sonia Goncalves, Ewan Harrison, David K. Jackson, Ian Johnston, Dominic Kwiatkowski, Cordelia Langford, John Sillitoe on behalf of the Wellcome Sanger Institute COVID-19 Surveillance Team |
| EPI_ISL_2512779, EPI_ISL_2512780 | Lighthouse Lab in Milton Keynes | Wellcome Sanger Institute for the COVID-19 Genomics UK (COG-UK) Consortium | The Lighthouse Lab in Milton Keynes and Alex Alderton, Roberto Amato, Jeffrey Barrett, Sonia Goncalves, Ewan Harrison, David K. Jackson, Ian Johnston, Dominic Kwiatkowski, Cordelia Langford, John Sillitoe on behalf of the Wellcome Sanger Institute COVID-19 Surveillance Team |
| EPI_ISL_2512781 | Lighthouse Lab in Glasgow | Wellcome Sanger Institute for the COVID-19 Genomics UK (COG-UK) Consortium | Harper VanSteenhouse, Yumi Kasai, David Gray, Carol Clugston, Anna Dominiczak and Alex Alderton, Roberto Amato, Jeffrey Barrett, Sonia Goncalves, Ewan Harrison, David K. Jackson, Ian Johnston, Dominic Kwiatkowski, Cordelia Langford, John Sillitoe on behalf of the Wellcome Sanger Institute COVID-19 Surveillance Team |
| EPI_ISL_2512782, EPI_ISL_2512783, EPI_ISL_2512785 | Lighthouse Lab in Milton Keynes | Wellcome Sanger Institute for the COVID-19 Genomics UK (COG-UK) Consortium | The Lighthouse Lab in Milton Keynes and Alex Alderton, Roberto Amato, Jeffrey Barrett, Sonia Goncalves, Ewan Harrison, David K. Jackson, Ian Johnston, Dominic Kwiatkowski, Cordelia Langford, John Sillitoe on behalf of the Wellcome Sanger Institute COVID-19 Surveillance Team |
| EPI_ISL_2512786 | Lighthouse Lab in Glasgow | Wellcome Sanger Institute for the COVID-19 Genomics UK (COG-UK) Consortium | Harper VanSteenhouse, Yumi Kasai, David Gray, Carol Clugston, Anna Dominiczak and Alex Alderton, Roberto Amato, Jeffrey Barrett, Sonia Goncalves, Ewan Harrison, David K. Jackson, Ian Johnston, Dominic Kwiatkowski, Cordelia Langford, John Sillitoe on behalf of the Wellcome Sanger Institute COVID-19 Surveillance Team |
| EPI_ISL_2512787, EPI_ISL_2512788 | Lighthouse Lab in Milton Keynes | Wellcome Sanger Institute for the COVID-19 Genomics UK (COG-UK) Consortium | The Lighthouse Lab in Milton Keynes and Alex Alderton, Roberto Amato, Jeffrey Barrett, Sonia Goncalves, Ewan Harrison, David K. Jackson, Ian Johnston, Dominic Kwiatkowski, Cordelia Langford, John Sillitoe on behalf of the Wellcome Sanger Institute COVID-19 Surveillance Team |
| EPI_ISL_2512789 | Lighthouse Lab in Glasgow | Wellcome Sanger Institute for the COVID-19 Genomics UK (COG-UK) Consortium | Harper VanSteenhouse, Yumi Kasai, David Gray, Carol Clugston, Anna Dominiczak and Alex Alderton, Roberto Amato, Jeffrey Barrett, Sonia Goncalves, Ewan Harrison, David K. Jackson, Ian Johnston, Dominic Kwiatkowski, Cordelia Langford, John Sillitoe on behalf of the Wellcome Sanger Institute COVID-19 Surveillance Team |
| EPI_ISL_2512790 | Lighthouse Lab in Milton Keynes | Wellcome Sanger Institute for the COVID-19 Genomics UK (COG-UK) Consortium | The Lighthouse Lab in Milton Keynes and Alex Alderton, Roberto Amato, Jeffrey Barrett, Sonia Goncalves, Ewan Harrison, David K. Jackson, Ian Johnston, Dominic Kwiatkowski, Cordelia Langford, John Sillitoe on behalf of the Wellcome Sanger Institute COVID-19 Surveillance Team |
| EPI_ISL_2512791, EPI_ISL_2512792 | Lighthouse Lab in Glasgow | Wellcome Sanger Institute for the COVID-19 Genomics UK (COG-UK) Consortium | Harper VanSteenhouse, Yumi Kasai, David Gray, Carol Clugston, Anna Dominiczak and Alex Alderton, Roberto Amato, Jeffrey Barrett, Sonia Goncalves, Ewan Harrison, David K. Jackson, Ian Johnston, Dominic Kwiatkowski, Cordelia Langford, John Sillitoe on behalf of the Wellcome Sanger Institute COVID-19 Surveillance Team |
| EPI_ISL_2512793 | Lighthouse Lab in Milton Keynes | Wellcome Sanger Institute for the COVID-19 Genomics UK (COG-UK) Consortium | The Lighthouse Lab in Milton Keynes and Alex Alderton, Roberto Amato, Jeffrey Barrett, Sonia Goncalves, Ewan Harrison, David K. Jackson, Ian Johnston, Dominic Kwiatkowski, Cordelia Langford, John Sillitoe on behalf of the Wellcome Sanger Institute COVID-19 Surveillance Team |
| EPI_ISL_2512794 | Lighthouse Lab in Glasgow | Wellcome Sanger Institute for the COVID-19 Genomics UK (COG-UK) Consortium | Harper VanSteenhouse, Yumi Kasai, David Gray, Carol Clugston, Anna Dominiczak and Alex Alderton, Roberto Amato, Jeffrey Barrett, Sonia Goncalves, Ewan Harrison, David K. Jackson, Ian Johnston, Dominic Kwiatkowski, Cordelia Langford, John Sillitoe on behalf of the Wellcome Sanger Institute COVID-19 Surveillance Team |
| EPI_ISL_2512795, EPI_ISL_2512796, EPI_ISL_2512798, EPI_ISL_2512799, EPI_ISL_2512800, EPI_ISL_2512801 | Lighthouse Lab in Milton Keynes | Wellcome Sanger Institute for the COVID-19 Genomics UK (COG-UK) Consortium | The Lighthouse Lab in Milton Keynes and Alex Alderton, Roberto Amato, Jeffrey Barrett, Sonia Goncalves, Ewan Harrison, David K. Jackson, Ian Johnston, Dominic Kwiatkowski, Cordelia Langford, John Sillitoe on behalf of the Wellcome Sanger Institute COVID-19 Surveillance Team |
| EPI_ISL_2512802 | Lighthouse Lab in Glasgow | Wellcome Sanger Institute for the COVID-19 Genomics UK (COG-UK) Consortium | Harper VanSteenhouse, Yumi Kasai, David Gray, Carol Clugston, Anna Dominiczak and Alex Alderton, Roberto Amato, Jeffrey Barrett, Sonia Goncalves, Ewan Harrison, David K. Jackson, Ian Johnston, Dominic Kwiatkowski, Cordelia Langford, John Sillitoe on behalf of the Wellcome Sanger Institute COVID-19 Surveillance Team |
| EPI_ISL_2512803 | Lighthouse Lab in Milton Keynes | Wellcome Sanger Institute for the COVID-19 Genomics UK (COG-UK) Consortium | The Lighthouse Lab in Milton Keynes and Alex Alderton, Roberto Amato, Jeffrey Barrett, Sonia Goncalves, Ewan Harrison, David K. Jackson, Ian Johnston, Dominic Kwiatkowski, Cordelia Langford, John Sillitoe on behalf of the Wellcome Sanger Institute COVID-19 Surveillance Team |
| EPI_ISL_2512804 | Lighthouse Lab in Glasgow | Wellcome Sanger Institute for the COVID-19 Genomics UK (COG-UK) Consortium | Harper VanSteenhouse, Yumi Kasai, David Gray, Carol Clugston, Anna Dominiczak and Alex Alderton, Roberto Amato, Jeffrey Barrett, Sonia Goncalves, Ewan Harrison, David K. Jackson, Ian Johnston, Dominic Kwiatkowski, Cordelia Langford, John Sillitoe on behalf of the Wellcome Sanger Institute COVID-19 Surveillance Team |
| EPI_ISL_2512805, EPI_ISL_2512806 | Lighthouse Lab in Milton Keynes | Wellcome Sanger Institute for the COVID-19 Genomics UK (COG-UK) Consortium | The Lighthouse Lab in Milton Keynes and Alex Alderton, Roberto Amato, Jeffrey Barrett, Sonia Goncalves, Ewan Harrison, David K. Jackson, Ian Johnston, Dominic Kwiatkowski, Cordelia Langford, John Sillitoe on behalf of the Wellcome Sanger Institute COVID-19 Surveillance Team |
| EPI_ISL_2512807 | Lighthouse Lab in Glasgow | Wellcome Sanger Institute for the COVID-19 Genomics UK (COG-UK) Consortium | Harper VanSteenhouse, Yumi Kasai, David Gray, Carol Clugston, Anna Dominiczak and Alex Alderton, Roberto Amato, Jeffrey Barrett, Sonia Goncalves, Ewan Harrison, David K. Jackson, Ian Johnston, Dominic Kwiatkowski, Cordelia Langford, John Sillitoe on behalf of the Wellcome Sanger Institute COVID-19 Surveillance Team |
| EPI_ISL_2512808 | Lighthouse Lab in Milton Keynes | Wellcome Sanger Institute for the COVID-19 Genomics UK (COG-UK) Consortium | The Lighthouse Lab in Milton Keynes and Alex Alderton, Roberto Amato, Jeffrey Barrett, Sonia Goncalves, Ewan Harrison, David K. Jackson, Ian Johnston, Dominic Kwiatkowski, Cordelia Langford, John Sillitoe on behalf of the Wellcome Sanger Institute COVID-19 Surveillance Team |
| EPI_ISL_2512809, EPI_ISL_2512810 | Lighthouse Lab in Glasgow | Wellcome Sanger Institute for the COVID-19 Genomics UK (COG-UK) Consortium | Harper VanSteenhouse, Yumi Kasai, David Gray, Carol Clugston, Anna Dominiczak and Alex Alderton, Roberto Amato, Jeffrey Barrett, Sonia Goncalves, Ewan Harrison, David K. Jackson, Ian Johnston, Dominic Kwiatkowski, Cordelia Langford, John Sillitoe on behalf of the Wellcome Sanger Institute COVID-19 Surveillance Team |
| EPI_ISL_2512811, EPI_ISL_2512812, EPI_ISL_2512813, EPI_ISL_2512814 | Lighthouse Lab in Milton Keynes | Wellcome Sanger Institute for the COVID-19 Genomics UK (COG-UK) Consortium | The Lighthouse Lab in Milton Keynes and Alex Alderton, Roberto Amato, Jeffrey Barrett, Sonia Goncalves, Ewan Harrison, David K. Jackson, Ian Johnston, Dominic Kwiatkowski, Cordelia Langford, John Sillitoe on behalf of the Wellcome Sanger Institute COVID-19 Surveillance Team |
| EPI_ISL_2512815 | Lighthouse Lab in Glasgow | Wellcome Sanger Institute for the COVID-19 Genomics UK (COG-UK) Consortium | Harper VanSteenhouse, Yumi Kasai, David Gray, Carol Clugston, Anna Dominiczak and Alex Alderton, Roberto Amato, Jeffrey Barrett, Sonia Goncalves, Ewan Harrison, David K. Jackson, Ian Johnston, Dominic Kwiatkowski, Cordelia Langford, John Sillitoe on behalf of the Wellcome Sanger Institute COVID-19 Surveillance Team |
| EPI_ISL_2512816, EPI_ISL_2512817, EPI_ISL_2512818, EPI_ISL_2512819, EPI_ISL_2512820, EPI_ISL_2512822, EPI_ISL_2512823, EPI_ISL_2512824, EPI_ISL_2512825, EPI_ISL_2512826, EPI_ISL_2512827, EPI_ISL_2512828 | | | |
| see above | Lighthouse Lab in Milton Keynes | Wellcome Sanger Institute for the COVID-19 Genomics UK (COG-UK) Consortium | The Lighthouse Lab in Milton Keynes and Alex Alderton, Roberto Amato, Jeffrey Barrett, Sonia Goncalves, Ewan Harrison, David K. Jackson, Ian Johnston, Dominic Kwiatkowski, Cordelia Langford, John Sillitoe on behalf of the Wellcome Sanger Institute COVID-19 Surveillance Team |
| EPI_ISL_2512829 | Lighthouse Lab in Glasgow | Wellcome Sanger Institute for the COVID-19 Genomics UK (COG-UK) Consortium | Harper VanSteenhouse, Yumi Kasai, David Gray, Carol Clugston, Anna Dominiczak and Alex Alderton, Roberto Amato, Jeffrey Barrett, Sonia Goncalves, Ewan Harrison, David K. Jackson, Ian Johnston, Dominic Kwiatkowski, Cordelia Langford, John Sillitoe on behalf of the Wellcome Sanger Institute COVID-19 Surveillance Team |
| EPI_ISL_2512830, EPI_ISL_2512831, EPI_ISL_2512833, EPI_ISL_2512834, EPI_ISL_2512835, EPI_ISL_2512836, EPI_ISL_2512837, EPI_ISL_2512838, EPI_ISL_2512839, EPI_ISL_2512840, EPI_ISL_2512841, EPI_ISL_2512842, EPI_ISL_2512843 | | | |
| see above | Lighthouse Lab in Milton Keynes | Wellcome Sanger Institute for the COVID-19 Genomics UK (COG-UK) Consortium | The Lighthouse Lab in Milton Keynes and Alex Alderton, Roberto Amato, Jeffrey Barrett, Sonia Goncalves, Ewan Harrison, David K. Jackson, Ian Johnston, Dominic Kwiatkowski, Cordelia Langford, John Sillitoe on behalf of the Wellcome Sanger Institute COVID-19 Surveillance Team |
| EPI_ISL_2512844 | Lighthouse Lab in Glasgow | Wellcome Sanger Institute for the COVID-19 Genomics UK (COG-UK) Consortium | Harper VanSteenhouse, Yumi Kasai, David Gray, Carol Clugston, Anna Dominiczak and Alex Alderton, Roberto Amato, Jeffrey Barrett, Sonia Goncalves, Ewan Harrison, David K. Jackson, Ian Johnston, Dominic Kwiatkowski, Cordelia Langford, John Sillitoe on behalf of the Wellcome Sanger Institute COVID-19 Surveillance Team |
| EPI_ISL_2512845, EPI_ISL_2512846, EPI_ISL_2512847, EPI_ISL_2512848, EPI_ISL_2512849, EPI_ISL_2512850 | Lighthouse Lab in Milton Keynes | Wellcome Sanger Institute for the COVID-19 Genomics UK (COG-UK) Consortium | The Lighthouse Lab in Milton Keynes and Alex Alderton, Roberto Amato, Jeffrey Barrett, Sonia Goncalves, Ewan Harrison, David K. Jackson, Ian Johnston, Dominic Kwiatkowski, Cordelia Langford, John Sillitoe on behalf of the Wellcome Sanger Institute COVID-19 Surveillance Team |
| EPI_ISL_2512851 | Lighthouse Lab in Glasgow | Wellcome Sanger Institute for the COVID-19 Genomics UK (COG-UK) Consortium | Harper VanSteenhouse, Yumi Kasai, David Gray, Carol Clugston, Anna Dominiczak and Alex Alderton, Roberto Amato, Jeffrey Barrett, Sonia Goncalves, Ewan Harrison, David K. Jackson, Ian Johnston, Dominic Kwiatkowski, Cordelia Langford, John Sillitoe on behalf of the Wellcome Sanger Institute COVID-19 Surveillance Team |
| EPI_ISL_2512852, EPI_ISL_2512853 | Lighthouse Lab in Milton Keynes | Wellcome Sanger Institute for the COVID-19 Genomics UK (COG-UK) Consortium | The Lighthouse Lab in Milton Keynes and Alex Alderton, Roberto Amato, Jeffrey Barrett, Sonia Goncalves, Ewan Harrison, David K. Jackson, Ian Johnston, Dominic Kwiatkowski, Cordelia Langford, John Sillitoe on behalf of the Wellcome Sanger Institute COVID-19 Surveillance Team |
| EPI_ISL_2512854 | Lighthouse Lab in Glasgow | Wellcome Sanger Institute for the COVID-19 Genomics UK (COG-UK) Consortium | Harper VanSteenhouse, Yumi Kasai, David Gray, Carol Clugston, Anna Dominiczak and Alex Alderton, Roberto Amato, Jeffrey Barrett, Sonia Goncalves, Ewan Harrison, David K. Jackson, Ian Johnston, Dominic Kwiatkowski, Cordelia Langford, John Sillitoe on behalf of the Wellcome Sanger Institute COVID-19 Surveillance Team |

| | | | |
|---|---|---|---|
| EPI_ISL_2512855 | Lighthouse Lab in Milton Keynes | Wellcome Sanger Institute for the COVID-19 Genomics UK (COG-UK) Consortium | The Lighthouse Lab in Milton Keynes and Alex Alderton, Roberto Amato, Jeffrey Barrett, Sonia Goncalves, Ewan Harrison, David K. Jackson, Ian Johnston, Dominic Kwiatkowski, Cordelia Langford, John Sillitoe on behalf of the Wellcome Sanger Institute COVID-19 Surveillance Team |
| EPI_ISL_2512856 | Lighthouse Lab in Glasgow | Wellcome Sanger Institute for the COVID-19 Genomics UK (COG-UK) Consortium | Harper VanSteenhouse, Yumi Kasai, David Gray, Carol Clugston, Anna Dominiczak and Alex Alderton, Roberto Amato, Jeffrey Barrett, Sonia Goncalves, Ewan Harrison, David K. Jackson, Ian Johnston, Dominic Kwiatkowski, Cordelia Langford, John Sillitoe on behalf of the Wellcome Sanger Institute COVID-19 Surveillance Team |
| EPI_ISL_2512858 | Lighthouse Lab in Milton Keynes | Wellcome Sanger Institute for the COVID-19 Genomics UK (COG-UK) Consortium | The Lighthouse Lab in Milton Keynes and Alex Alderton, Roberto Amato, Jeffrey Barrett, Sonia Goncalves, Ewan Harrison, David K. Jackson, Ian Johnston, Dominic Kwiatkowski, Cordelia Langford, John Sillitoe on behalf of the Wellcome Sanger Institute COVID-19 Surveillance Team |
| EPI_ISL_2512859, EPI_ISL_2512861, EPI_ISL_2512862, EPI_ISL_2512863 | Lighthouse Lab in Glasgow | Wellcome Sanger Institute for the COVID-19 Genomics UK (COG-UK) Consortium | Harper VanSteenhouse, Yumi Kasai, David Gray, Carol Clugston, Anna Dominiczak and Alex Alderton, Roberto Amato, Jeffrey Barrett, Sonia Goncalves, Ewan Harrison, David K. Jackson, Ian Johnston, Dominic Kwiatkowski, Cordelia Langford, John Sillitoe on behalf of the Wellcome Sanger Institute COVID-19 Surveillance Team |
| EPI_ISL_2512864 | Lighthouse Lab in Milton Keynes | Wellcome Sanger Institute for the COVID-19 Genomics UK (COG-UK) Consortium | The Lighthouse Lab in Milton Keynes and Alex Alderton, Roberto Amato, Jeffrey Barrett, Sonia Goncalves, Ewan Harrison, David K. Jackson, Ian Johnston, Dominic Kwiatkowski, Cordelia Langford, John Sillitoe on behalf of the Wellcome Sanger Institute COVID-19 Surveillance Team |
| EPI_ISL_2512865 | Lighthouse Lab in Glasgow | Wellcome Sanger Institute for the COVID-19 Genomics UK (COG-UK) Consortium | Harper VanSteenhouse, Yumi Kasai, David Gray, Carol Clugston, Anna Dominiczak and Alex Alderton, Roberto Amato, Jeffrey Barrett, Sonia Goncalves, Ewan Harrison, David K. Jackson, Ian Johnston, Dominic Kwiatkowski, Cordelia Langford, John Sillitoe on behalf of the Wellcome Sanger Institute COVID-19 Surveillance Team |
| EPI_ISL_2512866, EPI_ISL_2512867 | Lighthouse Lab in Milton Keynes | Wellcome Sanger Institute for the COVID-19 Genomics UK (COG-UK) Consortium | The Lighthouse Lab in Milton Keynes and Alex Alderton, Roberto Amato, Jeffrey Barrett, Sonia Goncalves, Ewan Harrison, David K. Jackson, Ian Johnston, Dominic Kwiatkowski, Cordelia Langford, John Sillitoe on behalf of the Wellcome Sanger Institute COVID-19 Surveillance Team |
| EPI_ISL_2512868, EPI_ISL_2512869, EPI_ISL_2512870 | Lighthouse Lab in Glasgow | Wellcome Sanger Institute for the COVID-19 Genomics UK (COG-UK) Consortium | Harper VanSteenhouse, Yumi Kasai, David Gray, Carol Clugston, Anna Dominiczak and Alex Alderton, Roberto Amato, Jeffrey Barrett, Sonia Goncalves, Ewan Harrison, David K. Jackson, Ian Johnston, Dominic Kwiatkowski, Cordelia Langford, John Sillitoe on behalf of the Wellcome Sanger Institute COVID-19 Surveillance Team |
| EPI_ISL_2512871, EPI_ISL_2512872, EPI_ISL_2512873, EPI_ISL_2512874, EPI_ISL_2512875 | Lighthouse Lab in Milton Keynes | Wellcome Sanger Institute for the COVID-19 Genomics UK (COG-UK) Consortium | The Lighthouse Lab in Milton Keynes and Alex Alderton, Roberto Amato, Jeffrey Barrett, Sonia Goncalves, Ewan Harrison, David K. Jackson, Ian Johnston, Dominic Kwiatkowski, Cordelia Langford, John Sillitoe on behalf of the Wellcome Sanger Institute COVID-19 Surveillance Team |
| EPI_ISL_2512876 | Lighthouse Lab in Glasgow | Wellcome Sanger Institute for the COVID-19 Genomics UK (COG-UK) Consortium | Harper VanSteenhouse, Yumi Kasai, David Gray, Carol Clugston, Anna Dominiczak and Alex Alderton, Roberto Amato, Jeffrey Barrett, Sonia Goncalves, Ewan Harrison, David K. Jackson, Ian Johnston, Dominic Kwiatkowski, Cordelia Langford, John Sillitoe on behalf of the Wellcome Sanger Institute COVID-19 Surveillance Team |
| EPI_ISL_2512877, EPI_ISL_2512878, EPI_ISL_2512879, EPI_ISL_2512880, EPI_ISL_2512881, EPI_ISL_2512883 | Lighthouse Lab in Milton Keynes | Wellcome Sanger Institute for the COVID-19 Genomics UK (COG-UK) Consortium | The Lighthouse Lab in Milton Keynes and Alex Alderton, Roberto Amato, Jeffrey Barrett, Sonia Goncalves, Ewan Harrison, David K. Jackson, Ian Johnston, Dominic Kwiatkowski, Cordelia Langford, John Sillitoe on behalf of the Wellcome Sanger Institute COVID-19 Surveillance Team |
| EPI_ISL_2512884, EPI_ISL_2512885 | Lighthouse Lab in Glasgow | Wellcome Sanger Institute for the COVID-19 Genomics UK (COG-UK) Consortium | Harper VanSteenhouse, Yumi Kasai, David Gray, Carol Clugston, Anna Dominiczak and Alex Alderton, Roberto Amato, Jeffrey Barrett, Sonia Goncalves, Ewan Harrison, David K. Jackson, Ian Johnston, Dominic Kwiatkowski, Cordelia Langford, John Sillitoe on behalf of the Wellcome Sanger Institute COVID-19 Surveillance Team |
| EPI_ISL_2512886 | Lighthouse Lab in Milton Keynes | Wellcome Sanger Institute for the COVID-19 Genomics UK (COG-UK) Consortium | The Lighthouse Lab in Milton Keynes and Alex Alderton, Roberto Amato, Jeffrey Barrett, Sonia Goncalves, Ewan Harrison, David K. Jackson, Ian Johnston, Dominic Kwiatkowski, Cordelia Langford, John Sillitoe on behalf of the Wellcome Sanger Institute COVID-19 Surveillance Team |
| EPI_ISL_2512887 | Lighthouse Lab in Glasgow | Wellcome Sanger Institute for the COVID-19 Genomics UK (COG-UK) Consortium | Harper VanSteenhouse, Yumi Kasai, David Gray, Carol Clugston, Anna Dominiczak and Alex Alderton, Roberto Amato, Jeffrey Barrett, Sonia Goncalves, Ewan Harrison, David K. Jackson, Ian Johnston, Dominic Kwiatkowski, Cordelia Langford, John Sillitoe on behalf of the Wellcome Sanger Institute COVID-19 Surveillance Team |
| EPI_ISL_2512888 | Lighthouse Lab in Milton Keynes | Wellcome Sanger Institute for the COVID-19 Genomics UK (COG-UK) Consortium | The Lighthouse Lab in Milton Keynes and Alex Alderton, Roberto Amato, Jeffrey Barrett, Sonia Goncalves, Ewan Harrison, David K. Jackson, Ian Johnston, Dominic Kwiatkowski, Cordelia Langford, John Sillitoe on behalf of the Wellcome Sanger Institute COVID-19 Surveillance Team |
| EPI_ISL_2512889, EPI_ISL_2512890 | Lighthouse Lab in Glasgow | Wellcome Sanger Institute for the COVID-19 Genomics UK (COG-UK) Consortium | Harper VanSteenhouse, Yumi Kasai, David Gray, Carol Clugston, Anna Dominiczak and Alex Alderton, Roberto Amato, Jeffrey Barrett, Sonia Goncalves, Ewan Harrison, David K. Jackson, Ian Johnston, Dominic Kwiatkowski, Cordelia Langford, John Sillitoe on behalf of the Wellcome Sanger Institute COVID-19 Surveillance Team |
| EPI_ISL_2512891, EPI_ISL_2512893, EPI_ISL_2512894, EPI_ISL_2512895, EPI_ISL_2512896 | Lighthouse Lab in Milton Keynes | Wellcome Sanger Institute for the COVID-19 Genomics UK (COG-UK) Consortium | The Lighthouse Lab in Milton Keynes and Alex Alderton, Roberto Amato, Jeffrey Barrett, Sonia Goncalves, Ewan Harrison, David K. Jackson, Ian Johnston, Dominic Kwiatkowski, Cordelia Langford, John Sillitoe on behalf of the Wellcome Sanger Institute COVID-19 Surveillance Team |
| EPI_ISL_2512897, EPI_ISL_2512898 | Lighthouse Lab in Glasgow | Wellcome Sanger Institute for the COVID-19 Genomics UK (COG-UK) Consortium | Harper VanSteenhouse, Yumi Kasai, David Gray, Carol Clugston, Anna Dominiczak and Alex Alderton, Roberto Amato, Jeffrey Barrett, Sonia Goncalves, Ewan Harrison, David K. Jackson, Ian Johnston, Dominic Kwiatkowski, Cordelia Langford, John Sillitoe on behalf of the Wellcome Sanger Institute COVID-19 Surveillance Team |
| EPI_ISL_2512900 | Lighthouse Lab in Milton Keynes | Wellcome Sanger Institute for the COVID-19 Genomics UK (COG-UK) Consortium | The Lighthouse Lab in Milton Keynes and Alex Alderton, Roberto Amato, Jeffrey Barrett, Sonia Goncalves, Ewan Harrison, David K. Jackson, Ian Johnston, Dominic Kwiatkowski, Cordelia Langford, John Sillitoe on behalf of the Wellcome Sanger Institute COVID-19 Surveillance Team |
| EPI_ISL_2512901, EPI_ISL_2512902 | Lighthouse Lab in Glasgow | Wellcome Sanger Institute for the COVID-19 Genomics UK (COG-UK) Consortium | Harper VanSteenhouse, Yumi Kasai, David Gray, Carol Clugston, Anna Dominiczak and Alex Alderton, Roberto Amato, Jeffrey Barrett, Sonia Goncalves, Ewan Harrison, David K. Jackson, Ian Johnston, Dominic Kwiatkowski, Cordelia Langford, John Sillitoe on behalf of the Wellcome Sanger Institute COVID-19 Surveillance Team |
| EPI_ISL_2512903 | Lighthouse Lab in Milton Keynes | Wellcome Sanger Institute for the COVID-19 Genomics UK (COG-UK) Consortium | The Lighthouse Lab in Milton Keynes and Alex Alderton, Roberto Amato, Jeffrey Barrett, Sonia Goncalves, Ewan Harrison, David K. Jackson, Ian Johnston, Dominic Kwiatkowski, Cordelia Langford, John Sillitoe on behalf of the Wellcome Sanger Institute COVID-19 Surveillance Team |
| EPI_ISL_2512904 | Lighthouse Lab in Glasgow | Wellcome Sanger Institute for the COVID-19 Genomics UK (COG-UK) Consortium | Harper VanSteenhouse, Yumi Kasai, David Gray, Carol Clugston, Anna Dominiczak and Alex Alderton, Roberto Amato, Jeffrey Barrett, Sonia Goncalves, Ewan Harrison, David K. Jackson, Ian Johnston, Dominic Kwiatkowski, Cordelia Langford, John Sillitoe on behalf of the Wellcome Sanger Institute COVID-19 Surveillance Team |
| EPI_ISL_2512905, EPI_ISL_2512906 | Lighthouse Lab in Milton Keynes | Wellcome Sanger Institute for the COVID-19 Genomics UK (COG-UK) Consortium | The Lighthouse Lab in Milton Keynes and Alex Alderton, Roberto Amato, Jeffrey Barrett, Sonia Goncalves, Ewan Harrison, David K. Jackson, Ian Johnston, Dominic Kwiatkowski, Cordelia Langford, John Sillitoe on behalf of the Wellcome Sanger Institute COVID-19 Surveillance Team |
| EPI_ISL_2512907, EPI_ISL_2512908 | Lighthouse Lab in Glasgow | Wellcome Sanger Institute for the COVID-19 Genomics UK (COG-UK) Consortium | Harper VanSteenhouse, Yumi Kasai, David Gray, Carol Clugston, Anna Dominiczak and Alex Alderton, Roberto Amato, Jeffrey Barrett, Sonia Goncalves, Ewan Harrison, David K. Jackson, Ian Johnston, Dominic Kwiatkowski, Cordelia Langford, John Sillitoe on behalf of the Wellcome Sanger Institute COVID-19 Surveillance Team |
| EPI_ISL_2512909, EPI_ISL_2512910, EPI_ISL_2512911, EPI_ISL_2512912, EPI_ISL_2512913 | Lighthouse Lab in Milton Keynes | Wellcome Sanger Institute for the COVID-19 Genomics UK (COG-UK) Consortium | The Lighthouse Lab in Milton Keynes and Alex Alderton, Roberto Amato, Jeffrey Barrett, Sonia Goncalves, Ewan Harrison, David K. Jackson, Ian Johnston, Dominic Kwiatkowski, Cordelia Langford, John Sillitoe on behalf of the Wellcome Sanger Institute COVID-19 Surveillance Team |
| EPI_ISL_2512914, EPI_ISL_2512915 | Lighthouse Lab in Glasgow | Wellcome Sanger Institute for the COVID-19 Genomics UK (COG-UK) Consortium | Harper VanSteenhouse, Yumi Kasai, David Gray, Carol Clugston, Anna Dominiczak and Alex Alderton, Roberto Amato, Jeffrey Barrett, Sonia Goncalves, Ewan Harrison, David K. Jackson, Ian Johnston, Dominic Kwiatkowski, Cordelia Langford, John Sillitoe on behalf of the Wellcome Sanger Institute COVID-19 Surveillance Team |
| EPI_ISL_2512916, EPI_ISL_2512917, EPI_ISL_2512918, EPI_ISL_2512920, EPI_ISL_2512921, EPI_ISL_2512922 | Lighthouse Lab in Milton Keynes | Wellcome Sanger Institute for the COVID-19 Genomics UK (COG-UK) Consortium | The Lighthouse Lab in Milton Keynes and Alex Alderton, Roberto Amato, Jeffrey Barrett, Sonia Goncalves, Ewan Harrison, David K. Jackson, Ian Johnston, Dominic Kwiatkowski, Cordelia Langford, John Sillitoe on behalf of the Wellcome Sanger Institute COVID-19 Surveillance Team |
| EPI_ISL_2512924 | Health Services Laboratories | Wellcome Sanger Institute for the COVID-19 Genomics UK (COG-UK) Consortium | Health Services Laboratories and Alex Alderton, Roberto Amato, Jeffrey Barrett, Sonia Goncalves, Ewan Harrison, David K. Jackson, Ian Johnston, Dominic Kwiatkowski, Cordelia Langford, John Sillitoe on behalf of the Wellcome Sanger Institute COVID-19 Surveillance Team |
| EPI_ISL_2512926 | Lighthouse Lab in Alderley Park | Wellcome Sanger Institute for the COVID-19 Genomics UK (COG-UK) Consortium | Jacquelyn Wynn, Mairead Hyland, The Lighthouse Lab in Alderley Park and Alex Alderton, Roberto Amato, Jeffrey Barrett, Sonia Goncalves, Ewan Harrison, David K. Jackson, Ian Johnston, Dominic Kwiatkowski, Cordelia Langford, John Sillitoe on behalf of the Wellcome Sanger Institute COVID-19 Surveillance Team |
| EPI_ISL_2512927 | Lighthouse Lab in Milton Keynes | Wellcome Sanger Institute for the COVID-19 Genomics UK (COG-UK) Consortium | The Lighthouse Lab in Milton Keynes and Alex Alderton, Roberto Amato, Jeffrey Barrett, Sonia Goncalves, Ewan Harrison, David K. Jackson, Ian Johnston, Dominic Kwiatkowski, Cordelia Langford, John Sillitoe on behalf of the Wellcome Sanger Institute COVID-19 Surveillance Team |
| EPI_ISL_2512928 | Lighthouse Lab in Alderley Park | Wellcome Sanger Institute for the COVID-19 Genomics UK (COG-UK) Consortium | Jacquelyn Wynn, Mairead Hyland, The Lighthouse Lab in Alderley Park and Alex Alderton, Roberto Amato, Jeffrey Barrett, Sonia Goncalves, Ewan Harrison, David K. Jackson, Ian Johnston, Dominic Kwiatkowski, Cordelia Langford, John Sillitoe on behalf of the Wellcome Sanger Institute COVID-19 Surveillance Team |
| EPI_ISL_2512929 | Lighthouse Lab in Milton Keynes | Wellcome Sanger Institute for the COVID-19 Genomics UK (COG-UK) Consortium | The Lighthouse Lab in Milton Keynes and Alex Alderton, Roberto Amato, Jeffrey Barrett, Sonia Goncalves, Ewan Harrison, David K. Jackson, Ian Johnston, Dominic Kwiatkowski, Cordelia Langford, John Sillitoe on behalf of the Wellcome Sanger Institute COVID-19 Surveillance Team |
| EPI_ISL_2512930 | Health Services Laboratories | Wellcome Sanger Institute for the COVID-19 Genomics UK (COG-UK) Consortium | Health Services Laboratories and Alex Alderton, Roberto Amato, Jeffrey Barrett, Sonia Goncalves, Ewan Harrison, David K. Jackson, Ian Johnston, Dominic Kwiatkowski, Cordelia Langford, John Sillitoe on behalf of the Wellcome Sanger Institute COVID-19 Surveillance Team |
| EPI_ISL_2512931, EPI_ISL_2512932, EPI_ISL_2512933 | Lighthouse Lab in Milton Keynes | Wellcome Sanger Institute for the COVID-19 Genomics UK (COG-UK) Consortium | The Lighthouse Lab in Milton Keynes and Alex Alderton, Roberto Amato, Jeffrey Barrett, Sonia Goncalves, Ewan Harrison, David K. Jackson, Ian Johnston, Dominic Kwiatkowski, Cordelia Langford, John Sillitoe on behalf of the Wellcome Sanger Institute COVID-19 Surveillance Team |
| EPI_ISL_2512934, EPI_ISL_2512935 | Health Services Laboratories | Wellcome Sanger Institute for the COVID-19 Genomics UK (COG-UK) Consortium | Health Services Laboratories and Alex Alderton, Roberto Amato, Jeffrey Barrett, Sonia Goncalves, Ewan Harrison, David K. Jackson, Ian Johnston, Dominic Kwiatkowski, Cordelia Langford, John Sillitoe on behalf of the Wellcome Sanger Institute COVID-19 Surveillance Team |
| EPI_ISL_2512936 | Lighthouse Lab in Milton Keynes | Wellcome Sanger Institute for the COVID-19 Genomics UK (COG-UK) Consortium | The Lighthouse Lab in Milton Keynes and Alex Alderton, Roberto Amato, Jeffrey Barrett, Sonia Goncalves, Ewan Harrison, David K. Jackson, Ian Johnston, Dominic Kwiatkowski, Cordelia Langford, John Sillitoe on behalf of the Wellcome Sanger Institute COVID-19 Surveillance Team |
| EPI_ISL_2512937, | Lighthouse Lab in Alderley Park | Wellcome Sanger Institute for the COVID-19 Genomics UK (COG-UK) | Jacquelyn Wynn, Mairead Hyland, The Lighthouse Lab in Alderley Park and Alex Alderton, Roberto Amato, Jeffrey Barrett, Sonia Goncalves, Ewan Harrison, David K. Jackson, Ian Johnston, Dominic Kwiatkowski, Cordelia Langford, John Sillitoe on |

| | | | |
|---|---|---|---|
| EPI_ISL_2512938, EPI_ISL_2512939 | | Consortium | behalf of the Wellcome Sanger Institute COVID-19 Surveillance Team |
| EPI_ISL_2512940 | Lighthouse Lab in Milton Keynes | Wellcome Sanger Institute for the COVID-19 Genomics UK (COG-UK) Consortium | The Lighthouse Lab in Milton Keynes and Alex Alderton, Roberto Amato, Jeffrey Barrett, Sonia Goncalves, Ewan Harrison, David K. Jackson, Ian Johnston, Dominic Kwiatkowski, Cordelia Langford, John Sillitoe on behalf of the Wellcome Sanger Institute COVID-19 Surveillance Team |
| EPI_ISL_2512941 | Lighthouse Lab in Alderley Park | Wellcome Sanger Institute for the COVID-19 Genomics UK (COG-UK) Consortium | Jacquelyn Wynn, Mairead Hyland, The Lighthouse Lab in Alderley Park and Alex Alderton, Roberto Amato, Jeffrey Barrett, Sonia Goncalves, Ewan Harrison, David K. Jackson, Ian Johnston, Dominic Kwiatkowski, Cordelia Langford, John Sillitoe on behalf of the Wellcome Sanger Institute COVID-19 Surveillance Team |
| EPI_ISL_2512943, EPI_ISL_2512944, EPI_ISL_2512945, EPI_ISL_2512946 | Health Services Laboratories | Wellcome Sanger Institute for the COVID-19 Genomics UK (COG-UK) Consortium | Health Services Laboratories and Alex Alderton, Roberto Amato, Jeffrey Barrett, Sonia Goncalves, Ewan Harrison, David K. Jackson, Ian Johnston, Dominic Kwiatkowski, Cordelia Langford, John Sillitoe on behalf of the Wellcome Sanger Institute COVID-19 Surveillance Team |
| EPI_ISL_2512947 | Lighthouse Lab in Alderley Park | Wellcome Sanger Institute for the COVID-19 Genomics UK (COG-UK) Consortium | Jacquelyn Wynn, Mairead Hyland, The Lighthouse Lab in Alderley Park and Alex Alderton, Roberto Amato, Jeffrey Barrett, Sonia Goncalves, Ewan Harrison, David K. Jackson, Ian Johnston, Dominic Kwiatkowski, Cordelia Langford, John Sillitoe on behalf of the Wellcome Sanger Institute COVID-19 Surveillance Team |
| EPI_ISL_2512948 | Lighthouse Lab in Milton Keynes | Wellcome Sanger Institute for the COVID-19 Genomics UK (COG-UK) Consortium | The Lighthouse Lab in Milton Keynes and Alex Alderton, Roberto Amato, Jeffrey Barrett, Sonia Goncalves, Ewan Harrison, David K. Jackson, Ian Johnston, Dominic Kwiatkowski, Cordelia Langford, John Sillitoe on behalf of the Wellcome Sanger Institute COVID-19 Surveillance Team |
| EPI_ISL_2512949, EPI_ISL_2512950 | Lighthouse Lab in Alderley Park | Wellcome Sanger Institute for the COVID-19 Genomics UK (COG-UK) Consortium | Jacquelyn Wynn, Mairead Hyland, The Lighthouse Lab in Alderley Park and Alex Alderton, Roberto Amato, Jeffrey Barrett, Sonia Goncalves, Ewan Harrison, David K. Jackson, Ian Johnston, Dominic Kwiatkowski, Cordelia Langford, John Sillitoe on behalf of the Wellcome Sanger Institute COVID-19 Surveillance Team |
| EPI_ISL_2512951 | Health Services Laboratories | Wellcome Sanger Institute for the COVID-19 Genomics UK (COG-UK) Consortium | Health Services Laboratories and Alex Alderton, Roberto Amato, Jeffrey Barrett, Sonia Goncalves, Ewan Harrison, David K. Jackson, Ian Johnston, Dominic Kwiatkowski, Cordelia Langford, John Sillitoe on behalf of the Wellcome Sanger Institute COVID-19 Surveillance Team |
| EPI_ISL_2512952 | Lighthouse Lab in Milton Keynes | Wellcome Sanger Institute for the COVID-19 Genomics UK (COG-UK) Consortium | The Lighthouse Lab in Milton Keynes and Alex Alderton, Roberto Amato, Jeffrey Barrett, Sonia Goncalves, Ewan Harrison, David K. Jackson, Ian Johnston, Dominic Kwiatkowski, Cordelia Langford, John Sillitoe on behalf of the Wellcome Sanger Institute COVID-19 Surveillance Team |
| EPI_ISL_2512953, EPI_ISL_2512954 | Lighthouse Lab in Alderley Park | Wellcome Sanger Institute for the COVID-19 Genomics UK (COG-UK) Consortium | Jacquelyn Wynn, Mairead Hyland, The Lighthouse Lab in Alderley Park and Alex Alderton, Roberto Amato, Jeffrey Barrett, Sonia Goncalves, Ewan Harrison, David K. Jackson, Ian Johnston, Dominic Kwiatkowski, Cordelia Langford, John Sillitoe on behalf of the Wellcome Sanger Institute COVID-19 Surveillance Team |
| EPI_ISL_2512955 | Health Services Laboratories | Wellcome Sanger Institute for the COVID-19 Genomics UK (COG-UK) Consortium | Health Services Laboratories and Alex Alderton, Roberto Amato, Jeffrey Barrett, Sonia Goncalves, Ewan Harrison, David K. Jackson, Ian Johnston, Dominic Kwiatkowski, Cordelia Langford, John Sillitoe on behalf of the Wellcome Sanger Institute COVID-19 Surveillance Team |
| EPI_ISL_2512956 | Lighthouse Lab in Milton Keynes | Wellcome Sanger Institute for the COVID-19 Genomics UK (COG-UK) Consortium | The Lighthouse Lab in Milton Keynes and Alex Alderton, Roberto Amato, Jeffrey Barrett, Sonia Goncalves, Ewan Harrison, David K. Jackson, Ian Johnston, Dominic Kwiatkowski, Cordelia Langford, John Sillitoe on behalf of the Wellcome Sanger Institute COVID-19 Surveillance Team |
| EPI_ISL_2512957, EPI_ISL_2512958, EPI_ISL_2512959, EPI_ISL_2512960 | Lighthouse Lab in Alderley Park | Wellcome Sanger Institute for the COVID-19 Genomics UK (COG-UK) Consortium | Jacquelyn Wynn, Mairead Hyland, The Lighthouse Lab in Alderley Park and Alex Alderton, Roberto Amato, Jeffrey Barrett, Sonia Goncalves, Ewan Harrison, David K. Jackson, Ian Johnston, Dominic Kwiatkowski, Cordelia Langford, John Sillitoe on behalf of the Wellcome Sanger Institute COVID-19 Surveillance Team |
| EPI_ISL_2512961 | Health Services Laboratories | Wellcome Sanger Institute for the COVID-19 Genomics UK (COG-UK) Consortium | Health Services Laboratories and Alex Alderton, Roberto Amato, Jeffrey Barrett, Sonia Goncalves, Ewan Harrison, David K. Jackson, Ian Johnston, Dominic Kwiatkowski, Cordelia Langford, John Sillitoe on behalf of the Wellcome Sanger Institute COVID-19 Surveillance Team |
| EPI_ISL_2512962 | Lighthouse Lab in Milton Keynes | Wellcome Sanger Institute for the COVID-19 Genomics UK (COG-UK) Consortium | The Lighthouse Lab in Milton Keynes and Alex Alderton, Roberto Amato, Jeffrey Barrett, Sonia Goncalves, Ewan Harrison, David K. Jackson, Ian Johnston, Dominic Kwiatkowski, Cordelia Langford, John Sillitoe on behalf of the Wellcome Sanger Institute COVID-19 Surveillance Team |
| EPI_ISL_2512963, EPI_ISL_2512964 | Health Services Laboratories | Wellcome Sanger Institute for the COVID-19 Genomics UK (COG-UK) Consortium | Health Services Laboratories and Alex Alderton, Roberto Amato, Jeffrey Barrett, Sonia Goncalves, Ewan Harrison, David K. Jackson, Ian Johnston, Dominic Kwiatkowski, Cordelia Langford, John Sillitoe on behalf of the Wellcome Sanger Institute COVID-19 Surveillance Team |
| EPI_ISL_2512965, EPI_ISL_2512967, EPI_ISL_2512968 | Lighthouse Lab in Alderley Park | Wellcome Sanger Institute for the COVID-19 Genomics UK (COG-UK) Consortium | Jacquelyn Wynn, Mairead Hyland, The Lighthouse Lab in Alderley Park and Alex Alderton, Roberto Amato, Jeffrey Barrett, Sonia Goncalves, Ewan Harrison, David K. Jackson, Ian Johnston, Dominic Kwiatkowski, Cordelia Langford, John Sillitoe on behalf of the Wellcome Sanger Institute COVID-19 Surveillance Team |
| EPI_ISL_2512969 | Lighthouse Lab in Milton Keynes | Wellcome Sanger Institute for the COVID-19 Genomics UK (COG-UK) Consortium | The Lighthouse Lab in Milton Keynes and Alex Alderton, Roberto Amato, Jeffrey Barrett, Sonia Goncalves, Ewan Harrison, David K. Jackson, Ian Johnston, Dominic Kwiatkowski, Cordelia Langford, John Sillitoe on behalf of the Wellcome Sanger Institute COVID-19 Surveillance Team |
| EPI_ISL_2512970, EPI_ISL_2512971 | Health Services Laboratories | Wellcome Sanger Institute for the COVID-19 Genomics UK (COG-UK) Consortium | Health Services Laboratories and Alex Alderton, Roberto Amato, Jeffrey Barrett, Sonia Goncalves, Ewan Harrison, David K. Jackson, Ian Johnston, Dominic Kwiatkowski, Cordelia Langford, John Sillitoe on behalf of the Wellcome Sanger Institute COVID-19 Surveillance Team |
| EPI_ISL_2512972, EPI_ISL_2512973, EPI_ISL_2512974, EPI_ISL_2512975 | Lighthouse Lab in Alderley Park | Wellcome Sanger Institute for the COVID-19 Genomics UK (COG-UK) Consortium | Jacquelyn Wynn, Mairead Hyland, The Lighthouse Lab in Alderley Park and Alex Alderton, Roberto Amato, Jeffrey Barrett, Sonia Goncalves, Ewan Harrison, David K. Jackson, Ian Johnston, Dominic Kwiatkowski, Cordelia Langford, John Sillitoe on behalf of the Wellcome Sanger Institute COVID-19 Surveillance Team |
| EPI_ISL_2512977, EPI_ISL_2512978 | Health Services Laboratories | Wellcome Sanger Institute for the COVID-19 Genomics UK (COG-UK) Consortium | Health Services Laboratories and Alex Alderton, Roberto Amato, Jeffrey Barrett, Sonia Goncalves, Ewan Harrison, David K. Jackson, Ian Johnston, Dominic Kwiatkowski, Cordelia Langford, John Sillitoe on behalf of the Wellcome Sanger Institute COVID-19 Surveillance Team |
| EPI_ISL_2512979 | Lighthouse Lab in Alderley Park | Wellcome Sanger Institute for the COVID-19 Genomics UK (COG-UK) Consortium | Jacquelyn Wynn, Mairead Hyland, The Lighthouse Lab in Alderley Park and Alex Alderton, Roberto Amato, Jeffrey Barrett, Sonia Goncalves, Ewan Harrison, David K. Jackson, Ian Johnston, Dominic Kwiatkowski, Cordelia Langford, John Sillitoe on behalf of the Wellcome Sanger Institute COVID-19 Surveillance Team |
| EPI_ISL_2512981, EPI_ISL_2512982 | Health Services Laboratories | Wellcome Sanger Institute for the COVID-19 Genomics UK (COG-UK) Consortium | Health Services Laboratories and Alex Alderton, Roberto Amato, Jeffrey Barrett, Sonia Goncalves, Ewan Harrison, David K. Jackson, Ian Johnston, Dominic Kwiatkowski, Cordelia Langford, John Sillitoe on behalf of the Wellcome Sanger Institute COVID-19 Surveillance Team |
| EPI_ISL_2512983 | Lighthouse Lab in Alderley Park | Wellcome Sanger Institute for the COVID-19 Genomics UK (COG-UK) Consortium | Jacquelyn Wynn, Mairead Hyland, The Lighthouse Lab in Alderley Park and Alex Alderton, Roberto Amato, Jeffrey Barrett, Sonia Goncalves, Ewan Harrison, David K. Jackson, Ian Johnston, Dominic Kwiatkowski, Cordelia Langford, John Sillitoe on behalf of the Wellcome Sanger Institute COVID-19 Surveillance Team |
| EPI_ISL_2512984 | Lighthouse Lab in Milton Keynes | Wellcome Sanger Institute for the COVID-19 Genomics UK (COG-UK) Consortium | The Lighthouse Lab in Milton Keynes and Alex Alderton, Roberto Amato, Jeffrey Barrett, Sonia Goncalves, Ewan Harrison, David K. Jackson, Ian Johnston, Dominic Kwiatkowski, Cordelia Langford, John Sillitoe on behalf of the Wellcome Sanger Institute COVID-19 Surveillance Team |
| EPI_ISL_2512985, EPI_ISL_2512986, EPI_ISL_2512987, EPI_ISL_2512988, EPI_ISL_2512989, EPI_ISL_2512991 | Lighthouse Lab in Alderley Park | Wellcome Sanger Institute for the COVID-19 Genomics UK (COG-UK) Consortium | Jacquelyn Wynn, Mairead Hyland, The Lighthouse Lab in Alderley Park and Alex Alderton, Roberto Amato, Jeffrey Barrett, Sonia Goncalves, Ewan Harrison, David K. Jackson, Ian Johnston, Dominic Kwiatkowski, Cordelia Langford, John Sillitoe on behalf of the Wellcome Sanger Institute COVID-19 Surveillance Team |
| EPI_ISL_2512992 | Health Services Laboratories | Wellcome Sanger Institute for the COVID-19 Genomics UK (COG-UK) Consortium | Health Services Laboratories and Alex Alderton, Roberto Amato, Jeffrey Barrett, Sonia Goncalves, Ewan Harrison, David K. Jackson, Ian Johnston, Dominic Kwiatkowski, Cordelia Langford, John Sillitoe on behalf of the Wellcome Sanger Institute COVID-19 Surveillance Team |
| EPI_ISL_2512993, EPI_ISL_2512994, EPI_ISL_2512995, EPI_ISL_2512996 | Lighthouse Lab in Alderley Park | Wellcome Sanger Institute for the COVID-19 Genomics UK (COG-UK) Consortium | Jacquelyn Wynn, Mairead Hyland, The Lighthouse Lab in Alderley Park and Alex Alderton, Roberto Amato, Jeffrey Barrett, Sonia Goncalves, Ewan Harrison, David K. Jackson, Ian Johnston, Dominic Kwiatkowski, Cordelia Langford, John Sillitoe on behalf of the Wellcome Sanger Institute COVID-19 Surveillance Team |
| EPI_ISL_2512997, EPI_ISL_2512998 | Lighthouse Lab in Milton Keynes | Wellcome Sanger Institute for the COVID-19 Genomics UK (COG-UK) Consortium | The Lighthouse Lab in Milton Keynes and Alex Alderton, Roberto Amato, Jeffrey Barrett, Sonia Goncalves, Ewan Harrison, David K. Jackson, Ian Johnston, Dominic Kwiatkowski, Cordelia Langford, John Sillitoe on behalf of the Wellcome Sanger Institute COVID-19 Surveillance Team |
| EPI_ISL_2512999 | Health Services Laboratories | Wellcome Sanger Institute for the COVID-19 Genomics UK (COG-UK) Consortium | Health Services Laboratories and Alex Alderton, Roberto Amato, Jeffrey Barrett, Sonia Goncalves, Ewan Harrison, David K. Jackson, Ian Johnston, Dominic Kwiatkowski, Cordelia Langford, John Sillitoe on behalf of the Wellcome Sanger Institute COVID-19 Surveillance Team |
| EPI_ISL_2513000 | Lighthouse Lab in Milton Keynes | Wellcome Sanger Institute for the COVID-19 Genomics UK (COG-UK) Consortium | The Lighthouse Lab in Milton Keynes and Alex Alderton, Roberto Amato, Jeffrey Barrett, Sonia Goncalves, Ewan Harrison, David K. Jackson, Ian Johnston, Dominic Kwiatkowski, Cordelia Langford, John Sillitoe on behalf of the Wellcome Sanger Institute COVID-19 Surveillance Team |
| EPI_ISL_2513001 | Health Services Laboratories | Wellcome Sanger Institute for the COVID-19 Genomics UK (COG-UK) Consortium | Health Services Laboratories and Alex Alderton, Roberto Amato, Jeffrey Barrett, Sonia Goncalves, Ewan Harrison, David K. Jackson, Ian Johnston, Dominic Kwiatkowski, Cordelia Langford, John Sillitoe on behalf of the Wellcome Sanger Institute COVID-19 Surveillance Team |
| EPI_ISL_2513002, EPI_ISL_2513003, EPI_ISL_2513004 | Lighthouse Lab in Alderley Park | Wellcome Sanger Institute for the COVID-19 Genomics UK (COG-UK) Consortium | Jacquelyn Wynn, Mairead Hyland, The Lighthouse Lab in Alderley Park and Alex Alderton, Roberto Amato, Jeffrey Barrett, Sonia Goncalves, Ewan Harrison, David K. Jackson, Ian Johnston, Dominic Kwiatkowski, Cordelia Langford, John Sillitoe on behalf of the Wellcome Sanger Institute COVID-19 Surveillance Team |
| EPI_ISL_2513005, EPI_ISL_2513006, EPI_ISL_2513007 | Lighthouse Lab in Milton Keynes | Wellcome Sanger Institute for the COVID-19 Genomics UK (COG-UK) Consortium | The Lighthouse Lab in Milton Keynes and Alex Alderton, Roberto Amato, Jeffrey Barrett, Sonia Goncalves, Ewan Harrison, David K. Jackson, Ian Johnston, Dominic Kwiatkowski, Cordelia Langford, John Sillitoe on behalf of the Wellcome Sanger Institute COVID-19 Surveillance Team |
| EPI_ISL_2513008 | Health Services Laboratories | Wellcome Sanger Institute for the COVID-19 Genomics UK (COG-UK) Consortium | Health Services Laboratories and Alex Alderton, Roberto Amato, Jeffrey Barrett, Sonia Goncalves, Ewan Harrison, David K. Jackson, Ian Johnston, Dominic Kwiatkowski, Cordelia Langford, John Sillitoe on behalf of the Wellcome Sanger Institute COVID-19 Surveillance Team |
| EPI_ISL_2513010, EPI_ISL_2513011, EPI_ISL_2513012, EPI_ISL_2513013 | Lighthouse Lab in Alderley Park | Wellcome Sanger Institute for the COVID-19 Genomics UK (COG-UK) Consortium | Jacquelyn Wynn, Mairead Hyland, The Lighthouse Lab in Alderley Park and Alex Alderton, Roberto Amato, Jeffrey Barrett, Sonia Goncalves, Ewan Harrison, David K. Jackson, Ian Johnston, Dominic Kwiatkowski, Cordelia Langford, John Sillitoe on behalf of the Wellcome Sanger Institute COVID-19 Surveillance Team |
| EPI_ISL_2513014 | Lighthouse Lab in Milton Keynes | Wellcome Sanger Institute for the COVID-19 Genomics UK (COG-UK) Consortium | The Lighthouse Lab in Milton Keynes and Alex Alderton, Roberto Amato, Jeffrey Barrett, Sonia Goncalves, Ewan Harrison, David K. Jackson, Ian Johnston, Dominic Kwiatkowski, Cordelia Langford, John Sillitoe on behalf of the Wellcome Sanger Institute COVID-19 Surveillance Team |
| EPI_ISL_2513015, EPI_ISL_2513016, EPI_ISL_2513017 | Lighthouse Lab in Alderley Park | Wellcome Sanger Institute for the COVID-19 Genomics UK (COG-UK) Consortium | Jacquelyn Wynn, Mairead Hyland, The Lighthouse Lab in Alderley Park and Alex Alderton, Roberto Amato, Jeffrey Barrett, Sonia Goncalves, Ewan Harrison, David K. Jackson, Ian Johnston, Dominic Kwiatkowski, Cordelia Langford, John Sillitoe on behalf of the Wellcome Sanger Institute COVID-19 Surveillance Team |

| | | | |
|---|---|---|---|
| EPI_ISL_2513018 | Lighthouse Lab in Milton Keynes | Wellcome Sanger Institute for the COVID-19 Genomics UK (COG-UK) Consortium | The Lighthouse Lab in Milton Keynes and Alex Alderton, Roberto Amato, Jeffrey Barrett, Sonia Goncalves, Ewan Harrison, David K. Jackson, Ian Johnston, Dominic Kwiatkowski, Cordelia Langford, John Sillitoe on behalf of the Wellcome Sanger Institute COVID-19 Surveillance Team |
| EPI_ISL_2513019, EPI_ISL_2513020 | Lighthouse Lab in Alderley Park | Wellcome Sanger Institute for the COVID-19 Genomics UK (COG-UK) Consortium | Jacquelyn Wynn, Mairead Hyland, The Lighthouse Lab in Alderley Park and Alex Alderton, Roberto Amato, Jeffrey Barrett, Sonia Goncalves, Ewan Harrison, David K. Jackson, Ian Johnston, Dominic Kwiatkowski, Cordelia Langford, John Sillitoe on behalf of the Wellcome Sanger Institute COVID-19 Surveillance Team |
| EPI_ISL_2513021, EPI_ISL_2513022 | Lighthouse Lab in Milton Keynes | Wellcome Sanger Institute for the COVID-19 Genomics UK (COG-UK) Consortium | The Lighthouse Lab in Milton Keynes and Alex Alderton, Roberto Amato, Jeffrey Barrett, Sonia Goncalves, Ewan Harrison, David K. Jackson, Ian Johnston, Dominic Kwiatkowski, Cordelia Langford, John Sillitoe on behalf of the Wellcome Sanger Institute COVID-19 Surveillance Team |
| EPI_ISL_2513023 | Health Services Laboratories | Wellcome Sanger Institute for the COVID-19 Genomics UK (COG-UK) Consortium | Health Services Laboratories and Alex Alderton, Roberto Amato, Jeffrey Barrett, Sonia Goncalves, Ewan Harrison, David K. Jackson, Ian Johnston, Dominic Kwiatkowski, Cordelia Langford, John Sillitoe on behalf of the Wellcome Sanger Institute COVID-19 Surveillance Team |
| EPI_ISL_2513024, EPI_ISL_2513025 | Lighthouse Lab in Alderley Park | Wellcome Sanger Institute for the COVID-19 Genomics UK (COG-UK) Consortium | Jacquelyn Wynn, Mairead Hyland, The Lighthouse Lab in Alderley Park and Alex Alderton, Roberto Amato, Jeffrey Barrett, Sonia Goncalves, Ewan Harrison, David K. Jackson, Ian Johnston, Dominic Kwiatkowski, Cordelia Langford, John Sillitoe on behalf of the Wellcome Sanger Institute COVID-19 Surveillance Team |
| EPI_ISL_2513026 | Lighthouse Lab in Milton Keynes | Wellcome Sanger Institute for the COVID-19 Genomics UK (COG-UK) Consortium | The Lighthouse Lab in Milton Keynes and Alex Alderton, Roberto Amato, Jeffrey Barrett, Sonia Goncalves, Ewan Harrison, David K. Jackson, Ian Johnston, Dominic Kwiatkowski, Cordelia Langford, John Sillitoe on behalf of the Wellcome Sanger Institute COVID-19 Surveillance Team |
| EPI_ISL_2513027 | Lighthouse Lab in Alderley Park | Wellcome Sanger Institute for the COVID-19 Genomics UK (COG-UK) Consortium | Jacquelyn Wynn, Mairead Hyland, The Lighthouse Lab in Alderley Park and Alex Alderton, Roberto Amato, Jeffrey Barrett, Sonia Goncalves, Ewan Harrison, David K. Jackson, Ian Johnston, Dominic Kwiatkowski, Cordelia Langford, John Sillitoe on behalf of the Wellcome Sanger Institute COVID-19 Surveillance Team |
| EPI_ISL_2513029 | Health Services Laboratories | Wellcome Sanger Institute for the COVID-19 Genomics UK (COG-UK) Consortium | Health Services Laboratories and Alex Alderton, Roberto Amato, Jeffrey Barrett, Sonia Goncalves, Ewan Harrison, David K. Jackson, Ian Johnston, Dominic Kwiatkowski, Cordelia Langford, John Sillitoe on behalf of the Wellcome Sanger Institute COVID-19 Surveillance Team |
| EPI_ISL_2513030, EPI_ISL_2513031 | Lighthouse Lab in Milton Keynes | Wellcome Sanger Institute for the COVID-19 Genomics UK (COG-UK) Consortium | The Lighthouse Lab in Milton Keynes and Alex Alderton, Roberto Amato, Jeffrey Barrett, Sonia Goncalves, Ewan Harrison, David K. Jackson, Ian Johnston, Dominic Kwiatkowski, Cordelia Langford, John Sillitoe on behalf of the Wellcome Sanger Institute COVID-19 Surveillance Team |
| EPI_ISL_2513033 | Lighthouse Lab in Alderley Park | Wellcome Sanger Institute for the COVID-19 Genomics UK (COG-UK) Consortium | Jacquelyn Wynn, Mairead Hyland, The Lighthouse Lab in Alderley Park and Alex Alderton, Roberto Amato, Jeffrey Barrett, Sonia Goncalves, Ewan Harrison, David K. Jackson, Ian Johnston, Dominic Kwiatkowski, Cordelia Langford, John Sillitoe on behalf of the Wellcome Sanger Institute COVID-19 Surveillance Team |
| EPI_ISL_2513034 | Lighthouse Lab in Milton Keynes | Wellcome Sanger Institute for the COVID-19 Genomics UK (COG-UK) Consortium | The Lighthouse Lab in Milton Keynes and Alex Alderton, Roberto Amato, Jeffrey Barrett, Sonia Goncalves, Ewan Harrison, David K. Jackson, Ian Johnston, Dominic Kwiatkowski, Cordelia Langford, John Sillitoe on behalf of the Wellcome Sanger Institute COVID-19 Surveillance Team |
| EPI_ISL_2513035 | Lighthouse Lab in Alderley Park | Wellcome Sanger Institute for the COVID-19 Genomics UK (COG-UK) Consortium | Jacquelyn Wynn, Mairead Hyland, The Lighthouse Lab in Alderley Park and Alex Alderton, Roberto Amato, Jeffrey Barrett, Sonia Goncalves, Ewan Harrison, David K. Jackson, Ian Johnston, Dominic Kwiatkowski, Cordelia Langford, John Sillitoe on behalf of the Wellcome Sanger Institute COVID-19 Surveillance Team |
| EPI_ISL_2513036 | Lighthouse Lab in Milton Keynes | Wellcome Sanger Institute for the COVID-19 Genomics UK (COG-UK) Consortium | The Lighthouse Lab in Milton Keynes and Alex Alderton, Roberto Amato, Jeffrey Barrett, Sonia Goncalves, Ewan Harrison, David K. Jackson, Ian Johnston, Dominic Kwiatkowski, Cordelia Langford, John Sillitoe on behalf of the Wellcome Sanger Institute COVID-19 Surveillance Team |
| EPI_ISL_2513037 | Health Services Laboratories | Wellcome Sanger Institute for the COVID-19 Genomics UK (COG-UK) Consortium | Health Services Laboratories and Alex Alderton, Roberto Amato, Jeffrey Barrett, Sonia Goncalves, Ewan Harrison, David K. Jackson, Ian Johnston, Dominic Kwiatkowski, Cordelia Langford, John Sillitoe on behalf of the Wellcome Sanger Institute COVID-19 Surveillance Team |
| EPI_ISL_2513038 | Lighthouse Lab in Alderley Park | Wellcome Sanger Institute for the COVID-19 Genomics UK (COG-UK) Consortium | Jacquelyn Wynn, Mairead Hyland, The Lighthouse Lab in Alderley Park and Alex Alderton, Roberto Amato, Jeffrey Barrett, Sonia Goncalves, Ewan Harrison, David K. Jackson, Ian Johnston, Dominic Kwiatkowski, Cordelia Langford, John Sillitoe on behalf of the Wellcome Sanger Institute COVID-19 Surveillance Team |
| EPI_ISL_2513039 | Lighthouse Lab in Milton Keynes | Wellcome Sanger Institute for the COVID-19 Genomics UK (COG-UK) Consortium | The Lighthouse Lab in Milton Keynes and Alex Alderton, Roberto Amato, Jeffrey Barrett, Sonia Goncalves, Ewan Harrison, David K. Jackson, Ian Johnston, Dominic Kwiatkowski, Cordelia Langford, John Sillitoe on behalf of the Wellcome Sanger Institute COVID-19 Surveillance Team |
| EPI_ISL_2513040 | Health Services Laboratories | Wellcome Sanger Institute for the COVID-19 Genomics UK (COG-UK) Consortium | Health Services Laboratories and Alex Alderton, Roberto Amato, Jeffrey Barrett, Sonia Goncalves, Ewan Harrison, David K. Jackson, Ian Johnston, Dominic Kwiatkowski, Cordelia Langford, John Sillitoe on behalf of the Wellcome Sanger Institute COVID-19 Surveillance Team |
| EPI_ISL_2513042 | Lighthouse Lab in Alderley Park | Wellcome Sanger Institute for the COVID-19 Genomics UK (COG-UK) Consortium | Jacquelyn Wynn, Mairead Hyland, The Lighthouse Lab in Alderley Park and Alex Alderton, Roberto Amato, Jeffrey Barrett, Sonia Goncalves, Ewan Harrison, David K. Jackson, Ian Johnston, Dominic Kwiatkowski, Cordelia Langford, John Sillitoe on behalf of the Wellcome Sanger Institute COVID-19 Surveillance Team |
| EPI_ISL_2513043 | Lighthouse Lab in Milton Keynes | Wellcome Sanger Institute for the COVID-19 Genomics UK (COG-UK) Consortium | The Lighthouse Lab in Milton Keynes and Alex Alderton, Roberto Amato, Jeffrey Barrett, Sonia Goncalves, Ewan Harrison, David K. Jackson, Ian Johnston, Dominic Kwiatkowski, Cordelia Langford, John Sillitoe on behalf of the Wellcome Sanger Institute COVID-19 Surveillance Team |
| EPI_ISL_2513044 | Health Services Laboratories | Wellcome Sanger Institute for the COVID-19 Genomics UK (COG-UK) Consortium | Health Services Laboratories and Alex Alderton, Roberto Amato, Jeffrey Barrett, Sonia Goncalves, Ewan Harrison, David K. Jackson, Ian Johnston, Dominic Kwiatkowski, Cordelia Langford, John Sillitoe on behalf of the Wellcome Sanger Institute COVID-19 Surveillance Team |
| EPI_ISL_2513045, EPI_ISL_2513046 | Lighthouse Lab in Alderley Park | Wellcome Sanger Institute for the COVID-19 Genomics UK (COG-UK) Consortium | Jacquelyn Wynn, Mairead Hyland, The Lighthouse Lab in Alderley Park and Alex Alderton, Roberto Amato, Jeffrey Barrett, Sonia Goncalves, Ewan Harrison, David K. Jackson, Ian Johnston, Dominic Kwiatkowski, Cordelia Langford, John Sillitoe on behalf of the Wellcome Sanger Institute COVID-19 Surveillance Team |
| EPI_ISL_2513047 | Lighthouse Lab in Milton Keynes | Wellcome Sanger Institute for the COVID-19 Genomics UK (COG-UK) Consortium | The Lighthouse Lab in Milton Keynes and Alex Alderton, Roberto Amato, Jeffrey Barrett, Sonia Goncalves, Ewan Harrison, David K. Jackson, Ian Johnston, Dominic Kwiatkowski, Cordelia Langford, John Sillitoe on behalf of the Wellcome Sanger Institute COVID-19 Surveillance Team |
| EPI_ISL_2513048 | Lighthouse Lab in Alderley Park | Wellcome Sanger Institute for the COVID-19 Genomics UK (COG-UK) Consortium | Jacquelyn Wynn, Mairead Hyland, The Lighthouse Lab in Alderley Park and Alex Alderton, Roberto Amato, Jeffrey Barrett, Sonia Goncalves, Ewan Harrison, David K. Jackson, Ian Johnston, Dominic Kwiatkowski, Cordelia Langford, John Sillitoe on behalf of the Wellcome Sanger Institute COVID-19 Surveillance Team |
| EPI_ISL_2513050 | Lighthouse Lab in Milton Keynes | Wellcome Sanger Institute for the COVID-19 Genomics UK (COG-UK) Consortium | The Lighthouse Lab in Milton Keynes and Alex Alderton, Roberto Amato, Jeffrey Barrett, Sonia Goncalves, Ewan Harrison, David K. Jackson, Ian Johnston, Dominic Kwiatkowski, Cordelia Langford, John Sillitoe on behalf of the Wellcome Sanger Institute COVID-19 Surveillance Team |
| EPI_ISL_2513051, EPI_ISL_2513052 | Lighthouse Lab in Alderley Park | Wellcome Sanger Institute for the COVID-19 Genomics UK (COG-UK) Consortium | Jacquelyn Wynn, Mairead Hyland, The Lighthouse Lab in Alderley Park and Alex Alderton, Roberto Amato, Jeffrey Barrett, Sonia Goncalves, Ewan Harrison, David K. Jackson, Ian Johnston, Dominic Kwiatkowski, Cordelia Langford, John Sillitoe on behalf of the Wellcome Sanger Institute COVID-19 Surveillance Team |
| EPI_ISL_2513053, EPI_ISL_2513054 | Health Services Laboratories | Wellcome Sanger Institute for the COVID-19 Genomics UK (COG-UK) Consortium | Health Services Laboratories and Alex Alderton, Roberto Amato, Jeffrey Barrett, Sonia Goncalves, Ewan Harrison, David K. Jackson, Ian Johnston, Dominic Kwiatkowski, Cordelia Langford, John Sillitoe on behalf of the Wellcome Sanger Institute COVID-19 Surveillance Team |
| EPI_ISL_2513055 | Lighthouse Lab in Alderley Park | Wellcome Sanger Institute for the COVID-19 Genomics UK (COG-UK) Consortium | Jacquelyn Wynn, Mairead Hyland, The Lighthouse Lab in Alderley Park and Alex Alderton, Roberto Amato, Jeffrey Barrett, Sonia Goncalves, Ewan Harrison, David K. Jackson, Ian Johnston, Dominic Kwiatkowski, Cordelia Langford, John Sillitoe on behalf of the Wellcome Sanger Institute COVID-19 Surveillance Team |
| EPI_ISL_2513056, EPI_ISL_2513057 | Health Services Laboratories | Wellcome Sanger Institute for the COVID-19 Genomics UK (COG-UK) Consortium | Health Services Laboratories and Alex Alderton, Roberto Amato, Jeffrey Barrett, Sonia Goncalves, Ewan Harrison, David K. Jackson, Ian Johnston, Dominic Kwiatkowski, Cordelia Langford, John Sillitoe on behalf of the Wellcome Sanger Institute COVID-19 Surveillance Team |
| EPI_ISL_2513058, EPI_ISL_2513059, EPI_ISL_2513061, EPI_ISL_2513062 | Lighthouse Lab in Alderley Park | Wellcome Sanger Institute for the COVID-19 Genomics UK (COG-UK) Consortium | Jacquelyn Wynn, Mairead Hyland, The Lighthouse Lab in Alderley Park and Alex Alderton, Roberto Amato, Jeffrey Barrett, Sonia Goncalves, Ewan Harrison, David K. Jackson, Ian Johnston, Dominic Kwiatkowski, Cordelia Langford, John Sillitoe on behalf of the Wellcome Sanger Institute COVID-19 Surveillance Team |
| EPI_ISL_2513063 | Lighthouse Lab in Milton Keynes | Wellcome Sanger Institute for the COVID-19 Genomics UK (COG-UK) Consortium | The Lighthouse Lab in Milton Keynes and Alex Alderton, Roberto Amato, Jeffrey Barrett, Sonia Goncalves, Ewan Harrison, David K. Jackson, Ian Johnston, Dominic Kwiatkowski, Cordelia Langford, John Sillitoe on behalf of the Wellcome Sanger Institute COVID-19 Surveillance Team |
| EPI_ISL_2513064, EPI_ISL_2513065, EPI_ISL_2513066, EPI_ISL_2513067, EPI_ISL_2513068, EPI_ISL_2513069, EPI_ISL_2513070 | Lighthouse Lab in Alderley Park | Wellcome Sanger Institute for the COVID-19 Genomics UK (COG-UK) Consortium | Jacquelyn Wynn, Mairead Hyland, The Lighthouse Lab in Alderley Park and Alex Alderton, Roberto Amato, Jeffrey Barrett, Sonia Goncalves, Ewan Harrison, David K. Jackson, Ian Johnston, Dominic Kwiatkowski, Cordelia Langford, John Sillitoe on behalf of the Wellcome Sanger Institute COVID-19 Surveillance Team |
| EPI_ISL_2513071, EPI_ISL_2513072, EPI_ISL_2513073 | Health Services Laboratories | Wellcome Sanger Institute for the COVID-19 Genomics UK (COG-UK) Consortium | Health Services Laboratories and Alex Alderton, Roberto Amato, Jeffrey Barrett, Sonia Goncalves, Ewan Harrison, David K. Jackson, Ian Johnston, Dominic Kwiatkowski, Cordelia Langford, John Sillitoe on behalf of the Wellcome Sanger Institute COVID-19 Surveillance Team |
| EPI_ISL_2513074 | Lighthouse Lab in Alderley Park | Wellcome Sanger Institute for the COVID-19 Genomics UK (COG-UK) Consortium | Jacquelyn Wynn, Mairead Hyland, The Lighthouse Lab in Alderley Park and Alex Alderton, Roberto Amato, Jeffrey Barrett, Sonia Goncalves, Ewan Harrison, David K. Jackson, Ian Johnston, Dominic Kwiatkowski, Cordelia Langford, John Sillitoe on behalf of the Wellcome Sanger Institute COVID-19 Surveillance Team |
| EPI_ISL_2513075 | Lighthouse Lab in Milton Keynes | Wellcome Sanger Institute for the COVID-19 Genomics UK (COG-UK) Consortium | The Lighthouse Lab in Milton Keynes and Alex Alderton, Roberto Amato, Jeffrey Barrett, Sonia Goncalves, Ewan Harrison, David K. Jackson, Ian Johnston, Dominic Kwiatkowski, Cordelia Langford, John Sillitoe on behalf of the Wellcome Sanger Institute COVID-19 Surveillance Team |
| EPI_ISL_2513076, EPI_ISL_2513077, EPI_ISL_2513078 | Lighthouse Lab in Alderley Park | Wellcome Sanger Institute for the COVID-19 Genomics UK (COG-UK) Consortium | Jacquelyn Wynn, Mairead Hyland, The Lighthouse Lab in Alderley Park and Alex Alderton, Roberto Amato, Jeffrey Barrett, Sonia Goncalves, Ewan Harrison, David K. Jackson, Ian Johnston, Dominic Kwiatkowski, Cordelia Langford, John Sillitoe on behalf of the Wellcome Sanger Institute COVID-19 Surveillance Team |
| EPI_ISL_2513079 | Health Services Laboratories | Wellcome Sanger Institute for the COVID-19 Genomics UK (COG-UK) Consortium | Health Services Laboratories and Alex Alderton, Roberto Amato, Jeffrey Barrett, Sonia Goncalves, Ewan Harrison, David K. Jackson, Ian Johnston, Dominic Kwiatkowski, Cordelia Langford, John Sillitoe on behalf of the Wellcome Sanger Institute COVID-19 Surveillance Team |
| EPI_ISL_2513080 | Lighthouse Lab in Milton Keynes | Wellcome Sanger Institute for the COVID-19 Genomics UK (COG-UK) Consortium | The Lighthouse Lab in Milton Keynes and Alex Alderton, Roberto Amato, Jeffrey Barrett, Sonia Goncalves, Ewan Harrison, David K. Jackson, Ian Johnston, Dominic Kwiatkowski, Cordelia Langford, John Sillitoe on behalf of the Wellcome Sanger Institute COVID-19 Surveillance Team |
| EPI_ISL_2513081 | Lighthouse Lab in Alderley Park | Wellcome Sanger Institute for the COVID-19 Genomics UK (COG-UK) Consortium | Jacquelyn Wynn, Mairead Hyland, The Lighthouse Lab in Alderley Park and Alex Alderton, Roberto Amato, Jeffrey Barrett, Sonia Goncalves, Ewan Harrison, David K. Jackson, Ian Johnston, Dominic Kwiatkowski, Cordelia Langford, John Sillitoe on behalf of the Wellcome Sanger Institute COVID-19 Surveillance Team |
| EPI_ISL_2513082 | Health Services Laboratories | Wellcome Sanger Institute for the COVID-19 Genomics UK (COG-UK) Consortium | Health Services Laboratories and Alex Alderton, Roberto Amato, Jeffrey Barrett, Sonia Goncalves, Ewan Harrison, David K. Jackson, Ian Johnston, Dominic Kwiatkowski, Cordelia Langford, John Sillitoe on behalf of the Wellcome Sanger Institute COVID-19 Surveillance Team |
| EPI_ISL_2513083 | Lighthouse Lab in Milton Keynes | Wellcome Sanger Institute for the COVID-19 Genomics UK (COG-UK) Consortium | The Lighthouse Lab in Milton Keynes and Alex Alderton, Roberto Amato, Jeffrey Barrett, Sonia Goncalves, Ewan Harrison, David K. Jackson, Ian Johnston, Dominic Kwiatkowski, Cordelia Langford, John Sillitoe on behalf of the Wellcome Sanger Institute COVID-19 Surveillance Team |
| EPI_ISL_2513084 | Health Services Laboratories | Wellcome Sanger Institute for the COVID-19 Genomics UK (COG-UK) Consortium | Health Services Laboratories and Alex Alderton, Roberto Amato, Jeffrey Barrett, Sonia Goncalves, Ewan Harrison, David K. Jackson, Ian Johnston, Dominic Kwiatkowski, Cordelia Langford, John Sillitoe on behalf of the Wellcome Sanger Institute COVID-19 Surveillance Team |

| | | | |
|---|---|---|---|
| EPI_ISL_2513085, EPI_ISL_2513086, EPI_ISL_2513088 | Lighthouse Lab in Alderley Park | Wellcome Sanger Institute for the COVID-19 Genomics UK (COG-UK) Consortium | Jacquelyn Wynn, Mairead Hyland, The Lighthouse Lab in Alderley Park and Alex Alderton, Roberto Amato, Jeffrey Barrett, Sonia Goncalves, Ewan Harrison, David K. Jackson, Ian Johnston, Dominic Kwiatkowski, Cordelia Langford, John Sillitoe on behalf of the Wellcome Sanger Institute COVID-19 Surveillance Team |
| EPI_ISL_2513089 | Health Services Laboratories | Wellcome Sanger Institute for the COVID-19 Genomics UK (COG-UK) Consortium | Health Services Laboratories and Alex Alderton, Roberto Amato, Jeffrey Barrett, Sonia Goncalves, Ewan Harrison, David K. Jackson, Ian Johnston, Dominic Kwiatkowski, Cordelia Langford, John Sillitoe on behalf of the Wellcome Sanger Institute COVID-19 Surveillance Team |
| EPI_ISL_2513090, EPI_ISL_2513091 | Lighthouse Lab in Milton Keynes | Wellcome Sanger Institute for the COVID-19 Genomics UK (COG-UK) Consortium | The Lighthouse Lab in Milton Keynes and Alex Alderton, Roberto Amato, Jeffrey Barrett, Sonia Goncalves, Ewan Harrison, David K. Jackson, Ian Johnston, Dominic Kwiatkowski, Cordelia Langford, John Sillitoe on behalf of the Wellcome Sanger Institute COVID-19 Surveillance Team |
| EPI_ISL_2513092 | Lighthouse Lab in Alderley Park | Wellcome Sanger Institute for the COVID-19 Genomics UK (COG-UK) Consortium | Jacquelyn Wynn, Mairead Hyland, The Lighthouse Lab in Alderley Park and Alex Alderton, Roberto Amato, Jeffrey Barrett, Sonia Goncalves, Ewan Harrison, David K. Jackson, Ian Johnston, Dominic Kwiatkowski, Cordelia Langford, John Sillitoe on behalf of the Wellcome Sanger Institute COVID-19 Surveillance Team |
| EPI_ISL_2513093, EPI_ISL_2513095 | Lighthouse Lab in Milton Keynes | Wellcome Sanger Institute for the COVID-19 Genomics UK (COG-UK) Consortium | The Lighthouse Lab in Milton Keynes and Alex Alderton, Roberto Amato, Jeffrey Barrett, Sonia Goncalves, Ewan Harrison, David K. Jackson, Ian Johnston, Dominic Kwiatkowski, Cordelia Langford, John Sillitoe on behalf of the Wellcome Sanger Institute COVID-19 Surveillance Team |
| EPI_ISL_2513096 | Lighthouse Lab in Alderley Park | Wellcome Sanger Institute for the COVID-19 Genomics UK (COG-UK) Consortium | Jacquelyn Wynn, Mairead Hyland, The Lighthouse Lab in Alderley Park and Alex Alderton, Roberto Amato, Jeffrey Barrett, Sonia Goncalves, Ewan Harrison, David K. Jackson, Ian Johnston, Dominic Kwiatkowski, Cordelia Langford, John Sillitoe on behalf of the Wellcome Sanger Institute COVID-19 Surveillance Team |
| EPI_ISL_2513097 | Health Services Laboratories | Wellcome Sanger Institute for the COVID-19 Genomics UK (COG-UK) Consortium | Health Services Laboratories and Alex Alderton, Roberto Amato, Jeffrey Barrett, Sonia Goncalves, Ewan Harrison, David K. Jackson, Ian Johnston, Dominic Kwiatkowski, Cordelia Langford, John Sillitoe on behalf of the Wellcome Sanger Institute COVID-19 Surveillance Team |
| EPI_ISL_2513098 | Lighthouse Lab in Alderley Park | Wellcome Sanger Institute for the COVID-19 Genomics UK (COG-UK) Consortium | Jacquelyn Wynn, Mairead Hyland, The Lighthouse Lab in Alderley Park and Alex Alderton, Roberto Amato, Jeffrey Barrett, Sonia Goncalves, Ewan Harrison, David K. Jackson, Ian Johnston, Dominic Kwiatkowski, Cordelia Langford, John Sillitoe on behalf of the Wellcome Sanger Institute COVID-19 Surveillance Team |
| EPI_ISL_2513099 | Health Services Laboratories | Wellcome Sanger Institute for the COVID-19 Genomics UK (COG-UK) Consortium | Health Services Laboratories and Alex Alderton, Roberto Amato, Jeffrey Barrett, Sonia Goncalves, Ewan Harrison, David K. Jackson, Ian Johnston, Dominic Kwiatkowski, Cordelia Langford, John Sillitoe on behalf of the Wellcome Sanger Institute COVID-19 Surveillance Team |
| EPI_ISL_2513100, EPI_ISL_2513102 | Lighthouse Lab in Alderley Park | Wellcome Sanger Institute for the COVID-19 Genomics UK (COG-UK) Consortium | Jacquelyn Wynn, Mairead Hyland, The Lighthouse Lab in Alderley Park and Alex Alderton, Roberto Amato, Jeffrey Barrett, Sonia Goncalves, Ewan Harrison, David K. Jackson, Ian Johnston, Dominic Kwiatkowski, Cordelia Langford, John Sillitoe on behalf of the Wellcome Sanger Institute COVID-19 Surveillance Team |
| EPI_ISL_2513103 | Health Services Laboratories | Wellcome Sanger Institute for the COVID-19 Genomics UK (COG-UK) Consortium | Health Services Laboratories and Alex Alderton, Roberto Amato, Jeffrey Barrett, Sonia Goncalves, Ewan Harrison, David K. Jackson, Ian Johnston, Dominic Kwiatkowski, Cordelia Langford, John Sillitoe on behalf of the Wellcome Sanger Institute COVID-19 Surveillance Team |
| EPI_ISL_2513104 | Lighthouse Lab in Alderley Park | Wellcome Sanger Institute for the COVID-19 Genomics UK (COG-UK) Consortium | Jacquelyn Wynn, Mairead Hyland, The Lighthouse Lab in Alderley Park and Alex Alderton, Roberto Amato, Jeffrey Barrett, Sonia Goncalves, Ewan Harrison, David K. Jackson, Ian Johnston, Dominic Kwiatkowski, Cordelia Langford, John Sillitoe on behalf of the Wellcome Sanger Institute COVID-19 Surveillance Team |
| EPI_ISL_2513105 | Lighthouse Lab in Milton Keynes | Wellcome Sanger Institute for the COVID-19 Genomics UK (COG-UK) Consortium | The Lighthouse Lab in Milton Keynes and Alex Alderton, Roberto Amato, Jeffrey Barrett, Sonia Goncalves, Ewan Harrison, David K. Jackson, Ian Johnston, Dominic Kwiatkowski, Cordelia Langford, John Sillitoe on behalf of the Wellcome Sanger Institute COVID-19 Surveillance Team |
| EPI_ISL_2513106 | Lighthouse Lab in Alderley Park | Wellcome Sanger Institute for the COVID-19 Genomics UK (COG-UK) Consortium | Jacquelyn Wynn, Mairead Hyland, The Lighthouse Lab in Alderley Park and Alex Alderton, Roberto Amato, Jeffrey Barrett, Sonia Goncalves, Ewan Harrison, David K. Jackson, Ian Johnston, Dominic Kwiatkowski, Cordelia Langford, John Sillitoe on behalf of the Wellcome Sanger Institute COVID-19 Surveillance Team |
| EPI_ISL_2513107 | Lighthouse Lab in Milton Keynes | Wellcome Sanger Institute for the COVID-19 Genomics UK (COG-UK) Consortium | The Lighthouse Lab in Milton Keynes and Alex Alderton, Roberto Amato, Jeffrey Barrett, Sonia Goncalves, Ewan Harrison, David K. Jackson, Ian Johnston, Dominic Kwiatkowski, Cordelia Langford, John Sillitoe on behalf of the Wellcome Sanger Institute COVID-19 Surveillance Team |
| EPI_ISL_2513108 | Lighthouse Lab in Alderley Park | Wellcome Sanger Institute for the COVID-19 Genomics UK (COG-UK) Consortium | Jacquelyn Wynn, Mairead Hyland, The Lighthouse Lab in Alderley Park and Alex Alderton, Roberto Amato, Jeffrey Barrett, Sonia Goncalves, Ewan Harrison, David K. Jackson, Ian Johnston, Dominic Kwiatkowski, Cordelia Langford, John Sillitoe on behalf of the Wellcome Sanger Institute COVID-19 Surveillance Team |
| EPI_ISL_2513109 | Lighthouse Lab in Milton Keynes | Wellcome Sanger Institute for the COVID-19 Genomics UK (COG-UK) Consortium | The Lighthouse Lab in Milton Keynes and Alex Alderton, Roberto Amato, Jeffrey Barrett, Sonia Goncalves, Ewan Harrison, David K. Jackson, Ian Johnston, Dominic Kwiatkowski, Cordelia Langford, John Sillitoe on behalf of the Wellcome Sanger Institute COVID-19 Surveillance Team |
| EPI_ISL_2513110, EPI_ISL_2513111, EPI_ISL_2513112, EPI_ISL_2513113 | Lighthouse Lab in Alderley Park | Wellcome Sanger Institute for the COVID-19 Genomics UK (COG-UK) Consortium | Jacquelyn Wynn, Mairead Hyland, The Lighthouse Lab in Alderley Park and Alex Alderton, Roberto Amato, Jeffrey Barrett, Sonia Goncalves, Ewan Harrison, David K. Jackson, Ian Johnston, Dominic Kwiatkowski, Cordelia Langford, John Sillitoe on behalf of the Wellcome Sanger Institute COVID-19 Surveillance Team |
| EPI_ISL_2513114 | Lighthouse Lab in Milton Keynes | Wellcome Sanger Institute for the COVID-19 Genomics UK (COG-UK) Consortium | The Lighthouse Lab in Milton Keynes and Alex Alderton, Roberto Amato, Jeffrey Barrett, Sonia Goncalves, Ewan Harrison, David K. Jackson, Ian Johnston, Dominic Kwiatkowski, Cordelia Langford, John Sillitoe on behalf of the Wellcome Sanger Institute COVID-19 Surveillance Team |
| EPI_ISL_2513117, EPI_ISL_2513118 | Lighthouse Lab in Alderley Park | Wellcome Sanger Institute for the COVID-19 Genomics UK (COG-UK) Consortium | Jacquelyn Wynn, Mairead Hyland, The Lighthouse Lab in Alderley Park and Alex Alderton, Roberto Amato, Jeffrey Barrett, Sonia Goncalves, Ewan Harrison, David K. Jackson, Ian Johnston, Dominic Kwiatkowski, Cordelia Langford, John Sillitoe on behalf of the Wellcome Sanger Institute COVID-19 Surveillance Team |
| EPI_ISL_2513119 | Lighthouse Lab in Milton Keynes | Wellcome Sanger Institute for the COVID-19 Genomics UK (COG-UK) Consortium | The Lighthouse Lab in Milton Keynes and Alex Alderton, Roberto Amato, Jeffrey Barrett, Sonia Goncalves, Ewan Harrison, David K. Jackson, Ian Johnston, Dominic Kwiatkowski, Cordelia Langford, John Sillitoe on behalf of the Wellcome Sanger Institute COVID-19 Surveillance Team |
| EPI_ISL_2513120, EPI_ISL_2513121, EPI_ISL_2513122 | Health Services Laboratories | Wellcome Sanger Institute for the COVID-19 Genomics UK (COG-UK) Consortium | Health Services Laboratories and Alex Alderton, Roberto Amato, Jeffrey Barrett, Sonia Goncalves, Ewan Harrison, David K. Jackson, Ian Johnston, Dominic Kwiatkowski, Cordelia Langford, John Sillitoe on behalf of the Wellcome Sanger Institute COVID-19 Surveillance Team |
| EPI_ISL_2513123 | Lighthouse Lab in Milton Keynes | Wellcome Sanger Institute for the COVID-19 Genomics UK (COG-UK) Consortium | The Lighthouse Lab in Milton Keynes and Alex Alderton, Roberto Amato, Jeffrey Barrett, Sonia Goncalves, Ewan Harrison, David K. Jackson, Ian Johnston, Dominic Kwiatkowski, Cordelia Langford, John Sillitoe on behalf of the Wellcome Sanger Institute COVID-19 Surveillance Team |
| EPI_ISL_2513124 | Health Services Laboratories | Wellcome Sanger Institute for the COVID-19 Genomics UK (COG-UK) Consortium | Health Services Laboratories and Alex Alderton, Roberto Amato, Jeffrey Barrett, Sonia Goncalves, Ewan Harrison, David K. Jackson, Ian Johnston, Dominic Kwiatkowski, Cordelia Langford, John Sillitoe on behalf of the Wellcome Sanger Institute COVID-19 Surveillance Team |
| EPI_ISL_2513125 | Lighthouse Lab in Milton Keynes | Wellcome Sanger Institute for the COVID-19 Genomics UK (COG-UK) Consortium | The Lighthouse Lab in Milton Keynes and Alex Alderton, Roberto Amato, Jeffrey Barrett, Sonia Goncalves, Ewan Harrison, David K. Jackson, Ian Johnston, Dominic Kwiatkowski, Cordelia Langford, John Sillitoe on behalf of the Wellcome Sanger Institute COVID-19 Surveillance Team |
| EPI_ISL_2513126 | Health Services Laboratories | Wellcome Sanger Institute for the COVID-19 Genomics UK (COG-UK) Consortium | Health Services Laboratories and Alex Alderton, Roberto Amato, Jeffrey Barrett, Sonia Goncalves, Ewan Harrison, David K. Jackson, Ian Johnston, Dominic Kwiatkowski, Cordelia Langford, John Sillitoe on behalf of the Wellcome Sanger Institute COVID-19 Surveillance Team |
| EPI_ISL_2513128, EPI_ISL_2513129, EPI_ISL_2513130 | Lighthouse Lab in Alderley Park | Wellcome Sanger Institute for the COVID-19 Genomics UK (COG-UK) Consortium | Jacquelyn Wynn, Mairead Hyland, The Lighthouse Lab in Alderley Park and Alex Alderton, Roberto Amato, Jeffrey Barrett, Sonia Goncalves, Ewan Harrison, David K. Jackson, Ian Johnston, Dominic Kwiatkowski, Cordelia Langford, John Sillitoe on behalf of the Wellcome Sanger Institute COVID-19 Surveillance Team |
| EPI_ISL_2513131 | Health Services Laboratories | Wellcome Sanger Institute for the COVID-19 Genomics UK (COG-UK) Consortium | Health Services Laboratories and Alex Alderton, Roberto Amato, Jeffrey Barrett, Sonia Goncalves, Ewan Harrison, David K. Jackson, Ian Johnston, Dominic Kwiatkowski, Cordelia Langford, John Sillitoe on behalf of the Wellcome Sanger Institute COVID-19 Surveillance Team |
| EPI_ISL_2513132, EPI_ISL_2513134 | Lighthouse Lab in Milton Keynes | Wellcome Sanger Institute for the COVID-19 Genomics UK (COG-UK) Consortium | The Lighthouse Lab in Milton Keynes and Alex Alderton, Roberto Amato, Jeffrey Barrett, Sonia Goncalves, Ewan Harrison, David K. Jackson, Ian Johnston, Dominic Kwiatkowski, Cordelia Langford, John Sillitoe on behalf of the Wellcome Sanger Institute COVID-19 Surveillance Team |
| EPI_ISL_2513135 | Lighthouse Lab in Alderley Park | Wellcome Sanger Institute for the COVID-19 Genomics UK (COG-UK) Consortium | Jacquelyn Wynn, Mairead Hyland, The Lighthouse Lab in Alderley Park and Alex Alderton, Roberto Amato, Jeffrey Barrett, Sonia Goncalves, Ewan Harrison, David K. Jackson, Ian Johnston, Dominic Kwiatkowski, Cordelia Langford, John Sillitoe on behalf of the Wellcome Sanger Institute COVID-19 Surveillance Team |
| EPI_ISL_2513136 | Health Services Laboratories | Wellcome Sanger Institute for the COVID-19 Genomics UK (COG-UK) Consortium | Health Services Laboratories and Alex Alderton, Roberto Amato, Jeffrey Barrett, Sonia Goncalves, Ewan Harrison, David K. Jackson, Ian Johnston, Dominic Kwiatkowski, Cordelia Langford, John Sillitoe on behalf of the Wellcome Sanger Institute COVID-19 Surveillance Team |
| EPI_ISL_2513137 | Lighthouse Lab in Alderley Park | Wellcome Sanger Institute for the COVID-19 Genomics UK (COG-UK) Consortium | Jacquelyn Wynn, Mairead Hyland, The Lighthouse Lab in Alderley Park and Alex Alderton, Roberto Amato, Jeffrey Barrett, Sonia Goncalves, Ewan Harrison, David K. Jackson, Ian Johnston, Dominic Kwiatkowski, Cordelia Langford, John Sillitoe on behalf of the Wellcome Sanger Institute COVID-19 Surveillance Team |
| EPI_ISL_2513138 | Health Services Laboratories | Wellcome Sanger Institute for the COVID-19 Genomics UK (COG-UK) Consortium | Health Services Laboratories and Alex Alderton, Roberto Amato, Jeffrey Barrett, Sonia Goncalves, Ewan Harrison, David K. Jackson, Ian Johnston, Dominic Kwiatkowski, Cordelia Langford, John Sillitoe on behalf of the Wellcome Sanger Institute COVID-19 Surveillance Team |
| EPI_ISL_2513139 | Lighthouse Lab in Alderley Park | Wellcome Sanger Institute for the COVID-19 Genomics UK (COG-UK) Consortium | Jacquelyn Wynn, Mairead Hyland, The Lighthouse Lab in Alderley Park and Alex Alderton, Roberto Amato, Jeffrey Barrett, Sonia Goncalves, Ewan Harrison, David K. Jackson, Ian Johnston, Dominic Kwiatkowski, Cordelia Langford, John Sillitoe on behalf of the Wellcome Sanger Institute COVID-19 Surveillance Team |
| EPI_ISL_2513140 | Health Services Laboratories | Wellcome Sanger Institute for the COVID-19 Genomics UK (COG-UK) Consortium | Health Services Laboratories and Alex Alderton, Roberto Amato, Jeffrey Barrett, Sonia Goncalves, Ewan Harrison, David K. Jackson, Ian Johnston, Dominic Kwiatkowski, Cordelia Langford, John Sillitoe on behalf of the Wellcome Sanger Institute COVID-19 Surveillance Team |
| EPI_ISL_2513141 | Lighthouse Lab in Alderley Park | Wellcome Sanger Institute for the COVID-19 Genomics UK (COG-UK) Consortium | Jacquelyn Wynn, Mairead Hyland, The Lighthouse Lab in Alderley Park and Alex Alderton, Roberto Amato, Jeffrey Barrett, Sonia Goncalves, Ewan Harrison, David K. Jackson, Ian Johnston, Dominic Kwiatkowski, Cordelia Langford, John Sillitoe on behalf of the Wellcome Sanger Institute COVID-19 Surveillance Team |
| EPI_ISL_2513142, EPI_ISL_2513143, EPI_ISL_2513145 | Health Services Laboratories | Wellcome Sanger Institute for the COVID-19 Genomics UK (COG-UK) Consortium | Health Services Laboratories and Alex Alderton, Roberto Amato, Jeffrey Barrett, Sonia Goncalves, Ewan Harrison, David K. Jackson, Ian Johnston, Dominic Kwiatkowski, Cordelia Langford, John Sillitoe on behalf of the Wellcome Sanger Institute COVID-19 Surveillance Team |
| EPI_ISL_2513146 | Lighthouse Lab in Alderley Park | Wellcome Sanger Institute for the COVID-19 Genomics UK (COG-UK) Consortium | Jacquelyn Wynn, Mairead Hyland, The Lighthouse Lab in Alderley Park and Alex Alderton, Roberto Amato, Jeffrey Barrett, Sonia Goncalves, Ewan Harrison, David K. Jackson, Ian Johnston, Dominic Kwiatkowski, Cordelia Langford, John Sillitoe on behalf of the Wellcome Sanger Institute COVID-19 Surveillance Team |
| EPI_ISL_2513147 | Lighthouse Lab in Milton Keynes | Wellcome Sanger Institute for the COVID-19 Genomics UK (COG-UK) Consortium | The Lighthouse Lab in Milton Keynes and Alex Alderton, Roberto Amato, Jeffrey Barrett, Sonia Goncalves, Ewan Harrison, David K. Jackson, Ian Johnston, Dominic Kwiatkowski, Cordelia Langford, John Sillitoe on behalf of the Wellcome Sanger Institute COVID-19 Surveillance Team |
| EPI_ISL_2513149 | Health Services Laboratories | Wellcome Sanger Institute for the COVID-19 Genomics UK (COG-UK) Consortium | Health Services Laboratories and Alex Alderton, Roberto Amato, Jeffrey Barrett, Sonia Goncalves, Ewan Harrison, David K. Jackson, Ian Johnston, Dominic Kwiatkowski, Cordelia Langford, John Sillitoe on behalf of the Wellcome Sanger Institute COVID-19 Surveillance Team |
| EPI_ISL_2513150 | Lighthouse Lab in Alderley Park | Wellcome Sanger Institute for the COVID-19 Genomics UK (COG-UK) Consortium | Jacquelyn Wynn, Mairead Hyland, The Lighthouse Lab in Alderley Park and Alex Alderton, Roberto Amato, Jeffrey Barrett, Sonia Goncalves, Ewan Harrison, David K. Jackson, Ian Johnston, Dominic Kwiatkowski, Cordelia Langford, John Sillitoe on behalf of the Wellcome Sanger Institute COVID-19 Surveillance Team |
| EPI_ISL_2513151 | Health Services Laboratories | Wellcome Sanger Institute for the COVID-19 Genomics UK (COG-UK) Consortium | Health Services Laboratories and Alex Alderton, Roberto Amato, Jeffrey Barrett, Sonia Goncalves, Ewan Harrison, David K. Jackson, Ian Johnston, Dominic Kwiatkowski, Cordelia Langford, John Sillitoe on behalf of the Wellcome Sanger Institute COVID-19 Surveillance Team |
| EPI_ISL_2513152 | Lighthouse Lab in Milton Keynes | Wellcome Sanger Institute for the COVID-19 Genomics UK (COG-UK) Consortium | The Lighthouse Lab in Milton Keynes and Alex Alderton, Roberto Amato, Jeffrey Barrett, Sonia Goncalves, Ewan Harrison, David K. Jackson, Ian Johnston, Dominic Kwiatkowski, Cordelia Langford, John Sillitoe on behalf of the Wellcome Sanger Institute COVID-19 Surveillance Team |

| | | | |
|---|---|---|---|
| EPI_ISL_2513153, EPI_ISL_2513154, EPI_ISL_2513155 | Lighthouse Lab in Alderley Park | Wellcome Sanger Institute for the COVID-19 Genomics UK (COG-UK) Consortium | Jacquelyn Wynn, Mairead Hyland, The Lighthouse Lab in Alderley Park and Alex Alderton, Roberto Amato, Jeffrey Barrett, Sonia Goncalves, Ewan Harrison, David K. Jackson, Ian Johnston, Dominic Kwiatkowski, Cordelia Langford, John Sillitoe on behalf of the Wellcome Sanger Institute COVID-19 Surveillance Team |
| EPI_ISL_2513156, EPI_ISL_2513157 | Lighthouse Lab in Milton Keynes | Wellcome Sanger Institute for the COVID-19 Genomics UK (COG-UK) Consortium | The Lighthouse Lab in Milton Keynes and Alex Alderton, Roberto Amato, Jeffrey Barrett, Sonia Goncalves, Ewan Harrison, David K. Jackson, Ian Johnston, Dominic Kwiatkowski, Cordelia Langford, John Sillitoe on behalf of the Wellcome Sanger Institute COVID-19 Surveillance Team |
| EPI_ISL_2513159, EPI_ISL_2513161, EPI_ISL_2513162, EPI_ISL_2513163 | Lighthouse Lab in Alderley Park | Wellcome Sanger Institute for the COVID-19 Genomics UK (COG-UK) Consortium | Jacquelyn Wynn, Mairead Hyland, The Lighthouse Lab in Alderley Park and Alex Alderton, Roberto Amato, Jeffrey Barrett, Sonia Goncalves, Ewan Harrison, David K. Jackson, Ian Johnston, Dominic Kwiatkowski, Cordelia Langford, John Sillitoe on behalf of the Wellcome Sanger Institute COVID-19 Surveillance Team |
| EPI_ISL_2513164, EPI_ISL_2513165 | Health Services Laboratories | Wellcome Sanger Institute for the COVID-19 Genomics UK (COG-UK) Consortium | Health Services Laboratories and Alex Alderton, Roberto Amato, Jeffrey Barrett, Sonia Goncalves, Ewan Harrison, David K. Jackson, Ian Johnston, Dominic Kwiatkowski, Cordelia Langford, John Sillitoe on behalf of the Wellcome Sanger Institute COVID-19 Surveillance Team |
| EPI_ISL_2513166 | Lighthouse Lab in Alderley Park | Wellcome Sanger Institute for the COVID-19 Genomics UK (COG-UK) Consortium | Jacquelyn Wynn, Mairead Hyland, The Lighthouse Lab in Alderley Park and Alex Alderton, Roberto Amato, Jeffrey Barrett, Sonia Goncalves, Ewan Harrison, David K. Jackson, Ian Johnston, Dominic Kwiatkowski, Cordelia Langford, John Sillitoe on behalf of the Wellcome Sanger Institute COVID-19 Surveillance Team |
| EPI_ISL_2513167 | Lighthouse Lab in Milton Keynes | Wellcome Sanger Institute for the COVID-19 Genomics UK (COG-UK) Consortium | The Lighthouse Lab in Milton Keynes and Alex Alderton, Roberto Amato, Jeffrey Barrett, Sonia Goncalves, Ewan Harrison, David K. Jackson, Ian Johnston, Dominic Kwiatkowski, Cordelia Langford, John Sillitoe on behalf of the Wellcome Sanger Institute COVID-19 Surveillance Team |
| EPI_ISL_2513168 | Lighthouse Lab in Alderley Park | Wellcome Sanger Institute for the COVID-19 Genomics UK (COG-UK) Consortium | Jacquelyn Wynn, Mairead Hyland, The Lighthouse Lab in Alderley Park and Alex Alderton, Roberto Amato, Jeffrey Barrett, Sonia Goncalves, Ewan Harrison, David K. Jackson, Ian Johnston, Dominic Kwiatkowski, Cordelia Langford, John Sillitoe on behalf of the Wellcome Sanger Institute COVID-19 Surveillance Team |
| EPI_ISL_2513169 | Health Services Laboratories | Wellcome Sanger Institute for the COVID-19 Genomics UK (COG-UK) Consortium | Health Services Laboratories and Alex Alderton, Roberto Amato, Jeffrey Barrett, Sonia Goncalves, Ewan Harrison, David K. Jackson, Ian Johnston, Dominic Kwiatkowski, Cordelia Langford, John Sillitoe on behalf of the Wellcome Sanger Institute COVID-19 Surveillance Team |
| EPI_ISL_2513170, EPI_ISL_2513171, EPI_ISL_2513172 | Lighthouse Lab in Alderley Park | Wellcome Sanger Institute for the COVID-19 Genomics UK (COG-UK) Consortium | Jacquelyn Wynn, Mairead Hyland, The Lighthouse Lab in Alderley Park and Alex Alderton, Roberto Amato, Jeffrey Barrett, Sonia Goncalves, Ewan Harrison, David K. Jackson, Ian Johnston, Dominic Kwiatkowski, Cordelia Langford, John Sillitoe on behalf of the Wellcome Sanger Institute COVID-19 Surveillance Team |
| EPI_ISL_2513173 | Health Services Laboratories | Wellcome Sanger Institute for the COVID-19 Genomics UK (COG-UK) Consortium | Health Services Laboratories and Alex Alderton, Roberto Amato, Jeffrey Barrett, Sonia Goncalves, Ewan Harrison, David K. Jackson, Ian Johnston, Dominic Kwiatkowski, Cordelia Langford, John Sillitoe on behalf of the Wellcome Sanger Institute COVID-19 Surveillance Team |
| EPI_ISL_2513174, EPI_ISL_2513175, EPI_ISL_2513177 | Lighthouse Lab in Alderley Park | Wellcome Sanger Institute for the COVID-19 Genomics UK (COG-UK) Consortium | Jacquelyn Wynn, Mairead Hyland, The Lighthouse Lab in Alderley Park and Alex Alderton, Roberto Amato, Jeffrey Barrett, Sonia Goncalves, Ewan Harrison, David K. Jackson, Ian Johnston, Dominic Kwiatkowski, Cordelia Langford, John Sillitoe on behalf of the Wellcome Sanger Institute COVID-19 Surveillance Team |
| EPI_ISL_2513178, EPI_ISL_2513179 | Health Services Laboratories | Wellcome Sanger Institute for the COVID-19 Genomics UK (COG-UK) Consortium | Health Services Laboratories and Alex Alderton, Roberto Amato, Jeffrey Barrett, Sonia Goncalves, Ewan Harrison, David K. Jackson, Ian Johnston, Dominic Kwiatkowski, Cordelia Langford, John Sillitoe on behalf of the Wellcome Sanger Institute COVID-19 Surveillance Team |
| EPI_ISL_2513180, EPI_ISL_2513181, EPI_ISL_2513182, EPI_ISL_2513183, EPI_ISL_2513184 | Lighthouse Lab in Alderley Park | Wellcome Sanger Institute for the COVID-19 Genomics UK (COG-UK) Consortium | Jacquelyn Wynn, Mairead Hyland, The Lighthouse Lab in Alderley Park and Alex Alderton, Roberto Amato, Jeffrey Barrett, Sonia Goncalves, Ewan Harrison, David K. Jackson, Ian Johnston, Dominic Kwiatkowski, Cordelia Langford, John Sillitoe on behalf of the Wellcome Sanger Institute COVID-19 Surveillance Team |
| EPI_ISL_2513185 | Health Services Laboratories | Wellcome Sanger Institute for the COVID-19 Genomics UK (COG-UK) Consortium | Health Services Laboratories and Alex Alderton, Roberto Amato, Jeffrey Barrett, Sonia Goncalves, Ewan Harrison, David K. Jackson, Ian Johnston, Dominic Kwiatkowski, Cordelia Langford, John Sillitoe on behalf of the Wellcome Sanger Institute COVID-19 Surveillance Team |
| EPI_ISL_2513186 | Lighthouse Lab in Milton Keynes | Wellcome Sanger Institute for the COVID-19 Genomics UK (COG-UK) Consortium | The Lighthouse Lab in Milton Keynes and Alex Alderton, Roberto Amato, Jeffrey Barrett, Sonia Goncalves, Ewan Harrison, David K. Jackson, Ian Johnston, Dominic Kwiatkowski, Cordelia Langford, John Sillitoe on behalf of the Wellcome Sanger Institute COVID-19 Surveillance Team |
| EPI_ISL_2513188 | Lighthouse Lab in Alderley Park | Wellcome Sanger Institute for the COVID-19 Genomics UK (COG-UK) Consortium | Jacquelyn Wynn, Mairead Hyland, The Lighthouse Lab in Alderley Park and Alex Alderton, Roberto Amato, Jeffrey Barrett, Sonia Goncalves, Ewan Harrison, David K. Jackson, Ian Johnston, Dominic Kwiatkowski, Cordelia Langford, John Sillitoe on behalf of the Wellcome Sanger Institute COVID-19 Surveillance Team |
| EPI_ISL_2513189 | Health Services Laboratories | Wellcome Sanger Institute for the COVID-19 Genomics UK (COG-UK) Consortium | Health Services Laboratories and Alex Alderton, Roberto Amato, Jeffrey Barrett, Sonia Goncalves, Ewan Harrison, David K. Jackson, Ian Johnston, Dominic Kwiatkowski, Cordelia Langford, John Sillitoe on behalf of the Wellcome Sanger Institute COVID-19 Surveillance Team |
| EPI_ISL_2513190, EPI_ISL_2513191, EPI_ISL_2513192, EPI_ISL_2513193, EPI_ISL_2513194, EPI_ISL_2513195, EPI_ISL_2513196, EPI_ISL_2513197 | Lighthouse Lab in Alderley Park | Wellcome Sanger Institute for the COVID-19 Genomics UK (COG-UK) Consortium | Jacquelyn Wynn, Mairead Hyland, The Lighthouse Lab in Alderley Park and Alex Alderton, Roberto Amato, Jeffrey Barrett, Sonia Goncalves, Ewan Harrison, David K. Jackson, Ian Johnston, Dominic Kwiatkowski, Cordelia Langford, John Sillitoe on behalf of the Wellcome Sanger Institute COVID-19 Surveillance Team |
| EPI_ISL_2513198 | Health Services Laboratories | Wellcome Sanger Institute for the COVID-19 Genomics UK (COG-UK) Consortium | Health Services Laboratories and Alex Alderton, Roberto Amato, Jeffrey Barrett, Sonia Goncalves, Ewan Harrison, David K. Jackson, Ian Johnston, Dominic Kwiatkowski, Cordelia Langford, John Sillitoe on behalf of the Wellcome Sanger Institute COVID-19 Surveillance Team |
| EPI_ISL_2513199, EPI_ISL_2513200 | Lighthouse Lab in Alderley Park | Wellcome Sanger Institute for the COVID-19 Genomics UK (COG-UK) Consortium | Jacquelyn Wynn, Mairead Hyland, The Lighthouse Lab in Alderley Park and Alex Alderton, Roberto Amato, Jeffrey Barrett, Sonia Goncalves, Ewan Harrison, David K. Jackson, Ian Johnston, Dominic Kwiatkowski, Cordelia Langford, John Sillitoe on behalf of the Wellcome Sanger Institute COVID-19 Surveillance Team |
| EPI_ISL_2513201, EPI_ISL_2513202 | Lighthouse Lab in Milton Keynes | Wellcome Sanger Institute for the COVID-19 Genomics UK (COG-UK) Consortium | The Lighthouse Lab in Milton Keynes and Alex Alderton, Roberto Amato, Jeffrey Barrett, Sonia Goncalves, Ewan Harrison, David K. Jackson, Ian Johnston, Dominic Kwiatkowski, Cordelia Langford, John Sillitoe on behalf of the Wellcome Sanger Institute COVID-19 Surveillance Team |
| EPI_ISL_2513203 | Health Services Laboratories | Wellcome Sanger Institute for the COVID-19 Genomics UK (COG-UK) Consortium | Health Services Laboratories and Alex Alderton, Roberto Amato, Jeffrey Barrett, Sonia Goncalves, Ewan Harrison, David K. Jackson, Ian Johnston, Dominic Kwiatkowski, Cordelia Langford, John Sillitoe on behalf of the Wellcome Sanger Institute COVID-19 Surveillance Team |
| EPI_ISL_2513204, EPI_ISL_2513205, EPI_ISL_2513206, EPI_ISL_2513207 | Lighthouse Lab in Alderley Park | Wellcome Sanger Institute for the COVID-19 Genomics UK (COG-UK) Consortium | Jacquelyn Wynn, Mairead Hyland, The Lighthouse Lab in Alderley Park and Alex Alderton, Roberto Amato, Jeffrey Barrett, Sonia Goncalves, Ewan Harrison, David K. Jackson, Ian Johnston, Dominic Kwiatkowski, Cordelia Langford, John Sillitoe on behalf of the Wellcome Sanger Institute COVID-19 Surveillance Team |
| EPI_ISL_2513208 | Health Services Laboratories | Wellcome Sanger Institute for the COVID-19 Genomics UK (COG-UK) Consortium | Health Services Laboratories and Alex Alderton, Roberto Amato, Jeffrey Barrett, Sonia Goncalves, Ewan Harrison, David K. Jackson, Ian Johnston, Dominic Kwiatkowski, Cordelia Langford, John Sillitoe on behalf of the Wellcome Sanger Institute COVID-19 Surveillance Team |
| EPI_ISL_2513210 | Lighthouse Lab in Alderley Park | Wellcome Sanger Institute for the COVID-19 Genomics UK (COG-UK) Consortium | Jacquelyn Wynn, Mairead Hyland, The Lighthouse Lab in Alderley Park and Alex Alderton, Roberto Amato, Jeffrey Barrett, Sonia Goncalves, Ewan Harrison, David K. Jackson, Ian Johnston, Dominic Kwiatkowski, Cordelia Langford, John Sillitoe on behalf of the Wellcome Sanger Institute COVID-19 Surveillance Team |
| EPI_ISL_2513211 | Health Services Laboratories | Wellcome Sanger Institute for the COVID-19 Genomics UK (COG-UK) Consortium | Health Services Laboratories and Alex Alderton, Roberto Amato, Jeffrey Barrett, Sonia Goncalves, Ewan Harrison, David K. Jackson, Ian Johnston, Dominic Kwiatkowski, Cordelia Langford, John Sillitoe on behalf of the Wellcome Sanger Institute COVID-19 Surveillance Team |
| EPI_ISL_2513212 | Lighthouse Lab in Alderley Park | Wellcome Sanger Institute for the COVID-19 Genomics UK (COG-UK) Consortium | Jacquelyn Wynn, Mairead Hyland, The Lighthouse Lab in Alderley Park and Alex Alderton, Roberto Amato, Jeffrey Barrett, Sonia Goncalves, Ewan Harrison, David K. Jackson, Ian Johnston, Dominic Kwiatkowski, Cordelia Langford, John Sillitoe on behalf of the Wellcome Sanger Institute COVID-19 Surveillance Team |
| EPI_ISL_2513214, EPI_ISL_2513216 | Health Services Laboratories | Wellcome Sanger Institute for the COVID-19 Genomics UK (COG-UK) Consortium | Health Services Laboratories and Alex Alderton, Roberto Amato, Jeffrey Barrett, Sonia Goncalves, Ewan Harrison, David K. Jackson, Ian Johnston, Dominic Kwiatkowski, Cordelia Langford, John Sillitoe on behalf of the Wellcome Sanger Institute COVID-19 Surveillance Team |
| EPI_ISL_2513217, EPI_ISL_2513218 | Lighthouse Lab in Alderley Park | Wellcome Sanger Institute for the COVID-19 Genomics UK (COG-UK) Consortium | Jacquelyn Wynn, Mairead Hyland, The Lighthouse Lab in Alderley Park and Alex Alderton, Roberto Amato, Jeffrey Barrett, Sonia Goncalves, Ewan Harrison, David K. Jackson, Ian Johnston, Dominic Kwiatkowski, Cordelia Langford, John Sillitoe on behalf of the Wellcome Sanger Institute COVID-19 Surveillance Team |
| EPI_ISL_2513219, EPI_ISL_2513220 | Health Services Laboratories | Wellcome Sanger Institute for the COVID-19 Genomics UK (COG-UK) Consortium | Health Services Laboratories and Alex Alderton, Roberto Amato, Jeffrey Barrett, Sonia Goncalves, Ewan Harrison, David K. Jackson, Ian Johnston, Dominic Kwiatkowski, Cordelia Langford, John Sillitoe on behalf of the Wellcome Sanger Institute COVID-19 Surveillance Team |
| EPI_ISL_2513221 | Lighthouse Lab in Milton Keynes | Wellcome Sanger Institute for the COVID-19 Genomics UK (COG-UK) Consortium | The Lighthouse Lab in Milton Keynes and Alex Alderton, Roberto Amato, Jeffrey Barrett, Sonia Goncalves, Ewan Harrison, David K. Jackson, Ian Johnston, Dominic Kwiatkowski, Cordelia Langford, John Sillitoe on behalf of the Wellcome Sanger Institute COVID-19 Surveillance Team |
| EPI_ISL_2513222 | Health Services Laboratories | Wellcome Sanger Institute for the COVID-19 Genomics UK (COG-UK) Consortium | Health Services Laboratories and Alex Alderton, Roberto Amato, Jeffrey Barrett, Sonia Goncalves, Ewan Harrison, David K. Jackson, Ian Johnston, Dominic Kwiatkowski, Cordelia Langford, John Sillitoe on behalf of the Wellcome Sanger Institute COVID-19 Surveillance Team |
| EPI_ISL_2513223 | Lighthouse Lab in Alderley Park | Wellcome Sanger Institute for the COVID-19 Genomics UK (COG-UK) Consortium | Jacquelyn Wynn, Mairead Hyland, The Lighthouse Lab in Alderley Park and Alex Alderton, Roberto Amato, Jeffrey Barrett, Sonia Goncalves, Ewan Harrison, David K. Jackson, Ian Johnston, Dominic Kwiatkowski, Cordelia Langford, John Sillitoe on behalf of the Wellcome Sanger Institute COVID-19 Surveillance Team |
| EPI_ISL_2513224 | Health Services Laboratories | Wellcome Sanger Institute for the COVID-19 Genomics UK (COG-UK) Consortium | Health Services Laboratories and Alex Alderton, Roberto Amato, Jeffrey Barrett, Sonia Goncalves, Ewan Harrison, David K. Jackson, Ian Johnston, Dominic Kwiatkowski, Cordelia Langford, John Sillitoe on behalf of the Wellcome Sanger Institute COVID-19 Surveillance Team |
| EPI_ISL_2513225 | Lighthouse Lab in Milton Keynes | Wellcome Sanger Institute for the COVID-19 Genomics UK (COG-UK) Consortium | The Lighthouse Lab in Milton Keynes and Alex Alderton, Roberto Amato, Jeffrey Barrett, Sonia Goncalves, Ewan Harrison, David K. Jackson, Ian Johnston, Dominic Kwiatkowski, Cordelia Langford, John Sillitoe on behalf of the Wellcome Sanger Institute COVID-19 Surveillance Team |
| EPI_ISL_2513226, EPI_ISL_2513227, EPI_ISL_2513228, EPI_ISL_2513229 | Lighthouse Lab in Alderley Park | Wellcome Sanger Institute for the COVID-19 Genomics UK (COG-UK) Consortium | Jacquelyn Wynn, Mairead Hyland, The Lighthouse Lab in Alderley Park and Alex Alderton, Roberto Amato, Jeffrey Barrett, Sonia Goncalves, Ewan Harrison, David K. Jackson, Ian Johnston, Dominic Kwiatkowski, Cordelia Langford, John Sillitoe on behalf of the Wellcome Sanger Institute COVID-19 Surveillance Team |
| EPI_ISL_2513230 | Health Services Laboratories | Wellcome Sanger Institute for the COVID-19 Genomics UK (COG-UK) Consortium | Health Services Laboratories and Alex Alderton, Roberto Amato, Jeffrey Barrett, Sonia Goncalves, Ewan Harrison, David K. Jackson, Ian Johnston, Dominic Kwiatkowski, Cordelia Langford, John Sillitoe on behalf of the Wellcome Sanger Institute COVID-19 Surveillance Team |
| EPI_ISL_2513231, | Lighthouse Lab in Alderley Park | Wellcome Sanger Institute for the COVID-19 Genomics UK (COG-UK) | Jacquelyn Wynn, Mairead Hyland, The Lighthouse Lab in Alderley Park and Alex Alderton, Roberto Amato, Jeffrey Barrett, Sonia Goncalves, Ewan Harrison, David K. Jackson, Ian Johnston, Dominic Kwiatkowski, Cordelia Langford, John Sillitoe on |

| | | | |
|---|---|---|---|
| EPI_ISL_2513232, EPI_ISL_2513233 | | Consortium | behalf of the Wellcome Sanger Institute COVID-19 Surveillance Team |
| EPI_ISL_2513234 | Health Services Laboratories | Wellcome Sanger Institute for the COVID-19 Genomics UK (COG-UK) Consortium | Health Services Laboratories and Alex Alderton, Roberto Amato, Jeffrey Barrett, Sonia Goncalves, Ewan Harrison, David K. Jackson, Ian Johnston, Dominic Kwiatkowski, Cordelia Langford, John Sillitoe on behalf of the Wellcome Sanger Institute COVID-19 Surveillance Team |
| EPI_ISL_2513235, EPI_ISL_2513236, EPI_ISL_2513237, EPI_ISL_2513238 | Lighthouse Lab in Alderley Park | Wellcome Sanger Institute for the COVID-19 Genomics UK (COG-UK) Consortium | Jacquelyn Wynn, Mairead Hyland, The Lighthouse Lab in Alderley Park and Alex Alderton, Roberto Amato, Jeffrey Barrett, Sonia Goncalves, Ewan Harrison, David K. Jackson, Ian Johnston, Dominic Kwiatkowski, Cordelia Langford, John Sillitoe on behalf of the Wellcome Sanger Institute COVID-19 Surveillance Team |
| EPI_ISL_2513239 | Health Services Laboratories | Wellcome Sanger Institute for the COVID-19 Genomics UK (COG-UK) Consortium | Health Services Laboratories and Alex Alderton, Roberto Amato, Jeffrey Barrett, Sonia Goncalves, Ewan Harrison, David K. Jackson, Ian Johnston, Dominic Kwiatkowski, Cordelia Langford, John Sillitoe on behalf of the Wellcome Sanger Institute COVID-19 Surveillance Team |
| EPI_ISL_2513240 | Lighthouse Lab in Alderley Park | Wellcome Sanger Institute for the COVID-19 Genomics UK (COG-UK) Consortium | Jacquelyn Wynn, Mairead Hyland, The Lighthouse Lab in Alderley Park and Alex Alderton, Roberto Amato, Jeffrey Barrett, Sonia Goncalves, Ewan Harrison, David K. Jackson, Ian Johnston, Dominic Kwiatkowski, Cordelia Langford, John Sillitoe on behalf of the Wellcome Sanger Institute COVID-19 Surveillance Team |
| EPI_ISL_2513241 | Lighthouse Lab in Milton Keynes | Wellcome Sanger Institute for the COVID-19 Genomics UK (COG-UK) Consortium | The Lighthouse Lab in Milton Keynes and Alex Alderton, Roberto Amato, Jeffrey Barrett, Sonia Goncalves, Ewan Harrison, David K. Jackson, Ian Johnston, Dominic Kwiatkowski, Cordelia Langford, John Sillitoe on behalf of the Wellcome Sanger Institute COVID-19 Surveillance Team |
| EPI_ISL_2513242, EPI_ISL_2513243, EPI_ISL_2513244, EPI_ISL_2513245 | Lighthouse Lab in Alderley Park | Wellcome Sanger Institute for the COVID-19 Genomics UK (COG-UK) Consortium | Jacquelyn Wynn, Mairead Hyland, The Lighthouse Lab in Alderley Park and Alex Alderton, Roberto Amato, Jeffrey Barrett, Sonia Goncalves, Ewan Harrison, David K. Jackson, Ian Johnston, Dominic Kwiatkowski, Cordelia Langford, John Sillitoe on behalf of the Wellcome Sanger Institute COVID-19 Surveillance Team |
| EPI_ISL_2513246 | Lighthouse Lab in Milton Keynes | Wellcome Sanger Institute for the COVID-19 Genomics UK (COG-UK) Consortium | The Lighthouse Lab in Milton Keynes and Alex Alderton, Roberto Amato, Jeffrey Barrett, Sonia Goncalves, Ewan Harrison, David K. Jackson, Ian Johnston, Dominic Kwiatkowski, Cordelia Langford, John Sillitoe on behalf of the Wellcome Sanger Institute COVID-19 Surveillance Team |
| EPI_ISL_2513247 | Health Services Laboratories | Wellcome Sanger Institute for the COVID-19 Genomics UK (COG-UK) Consortium | Health Services Laboratories and Alex Alderton, Roberto Amato, Jeffrey Barrett, Sonia Goncalves, Ewan Harrison, David K. Jackson, Ian Johnston, Dominic Kwiatkowski, Cordelia Langford, John Sillitoe on behalf of the Wellcome Sanger Institute COVID-19 Surveillance Team |
| EPI_ISL_2513248 | Lighthouse Lab in Milton Keynes | Wellcome Sanger Institute for the COVID-19 Genomics UK (COG-UK) Consortium | The Lighthouse Lab in Milton Keynes and Alex Alderton, Roberto Amato, Jeffrey Barrett, Sonia Goncalves, Ewan Harrison, David K. Jackson, Ian Johnston, Dominic Kwiatkowski, Cordelia Langford, John Sillitoe on behalf of the Wellcome Sanger Institute COVID-19 Surveillance Team |
| EPI_ISL_2513249 | Lighthouse Lab in Alderley Park | Wellcome Sanger Institute for the COVID-19 Genomics UK (COG-UK) Consortium | Jacquelyn Wynn, Mairead Hyland, The Lighthouse Lab in Alderley Park and Alex Alderton, Roberto Amato, Jeffrey Barrett, Sonia Goncalves, Ewan Harrison, David K. Jackson, Ian Johnston, Dominic Kwiatkowski, Cordelia Langford, John Sillitoe on behalf of the Wellcome Sanger Institute COVID-19 Surveillance Team |
| EPI_ISL_2513250 | Lighthouse Lab in Milton Keynes | Wellcome Sanger Institute for the COVID-19 Genomics UK (COG-UK) Consortium | The Lighthouse Lab in Milton Keynes and Alex Alderton, Roberto Amato, Jeffrey Barrett, Sonia Goncalves, Ewan Harrison, David K. Jackson, Ian Johnston, Dominic Kwiatkowski, Cordelia Langford, John Sillitoe on behalf of the Wellcome Sanger Institute COVID-19 Surveillance Team |
| EPI_ISL_2513251, EPI_ISL_2513252 | Lighthouse Lab in Alderley Park | Wellcome Sanger Institute for the COVID-19 Genomics UK (COG-UK) Consortium | Jacquelyn Wynn, Mairead Hyland, The Lighthouse Lab in Alderley Park and Alex Alderton, Roberto Amato, Jeffrey Barrett, Sonia Goncalves, Ewan Harrison, David K. Jackson, Ian Johnston, Dominic Kwiatkowski, Cordelia Langford, John Sillitoe on behalf of the Wellcome Sanger Institute COVID-19 Surveillance Team |
| EPI_ISL_2513253 | Health Services Laboratories | Wellcome Sanger Institute for the COVID-19 Genomics UK (COG-UK) Consortium | Health Services Laboratories and Alex Alderton, Roberto Amato, Jeffrey Barrett, Sonia Goncalves, Ewan Harrison, David K. Jackson, Ian Johnston, Dominic Kwiatkowski, Cordelia Langford, John Sillitoe on behalf of the Wellcome Sanger Institute COVID-19 Surveillance Team |
| EPI_ISL_2513254, EPI_ISL_2513255 | Lighthouse Lab in Alderley Park | Wellcome Sanger Institute for the COVID-19 Genomics UK (COG-UK) Consortium | Jacquelyn Wynn, Mairead Hyland, The Lighthouse Lab in Alderley Park and Alex Alderton, Roberto Amato, Jeffrey Barrett, Sonia Goncalves, Ewan Harrison, David K. Jackson, Ian Johnston, Dominic Kwiatkowski, Cordelia Langford, John Sillitoe on behalf of the Wellcome Sanger Institute COVID-19 Surveillance Team |
| EPI_ISL_2513256 | Health Services Laboratories | Wellcome Sanger Institute for the COVID-19 Genomics UK (COG-UK) Consortium | Health Services Laboratories and Alex Alderton, Roberto Amato, Jeffrey Barrett, Sonia Goncalves, Ewan Harrison, David K. Jackson, Ian Johnston, Dominic Kwiatkowski, Cordelia Langford, John Sillitoe on behalf of the Wellcome Sanger Institute COVID-19 Surveillance Team |
| EPI_ISL_2513257, EPI_ISL_2513258 | Lighthouse Lab in Milton Keynes | Wellcome Sanger Institute for the COVID-19 Genomics UK (COG-UK) Consortium | The Lighthouse Lab in Milton Keynes and Alex Alderton, Roberto Amato, Jeffrey Barrett, Sonia Goncalves, Ewan Harrison, David K. Jackson, Ian Johnston, Dominic Kwiatkowski, Cordelia Langford, John Sillitoe on behalf of the Wellcome Sanger Institute COVID-19 Surveillance Team |
| EPI_ISL_2513259, EPI_ISL_2513261 | Lighthouse Lab in Alderley Park | Wellcome Sanger Institute for the COVID-19 Genomics UK (COG-UK) Consortium | Jacquelyn Wynn, Mairead Hyland, The Lighthouse Lab in Alderley Park and Alex Alderton, Roberto Amato, Jeffrey Barrett, Sonia Goncalves, Ewan Harrison, David K. Jackson, Ian Johnston, Dominic Kwiatkowski, Cordelia Langford, John Sillitoe on behalf of the Wellcome Sanger Institute COVID-19 Surveillance Team |
| EPI_ISL_2513262 | Health Services Laboratories | Wellcome Sanger Institute for the COVID-19 Genomics UK (COG-UK) Consortium | Health Services Laboratories and Alex Alderton, Roberto Amato, Jeffrey Barrett, Sonia Goncalves, Ewan Harrison, David K. Jackson, Ian Johnston, Dominic Kwiatkowski, Cordelia Langford, John Sillitoe on behalf of the Wellcome Sanger Institute COVID-19 Surveillance Team |
| EPI_ISL_2513264 | Lighthouse Lab in Milton Keynes | Wellcome Sanger Institute for the COVID-19 Genomics UK (COG-UK) Consortium | The Lighthouse Lab in Milton Keynes and Alex Alderton, Roberto Amato, Jeffrey Barrett, Sonia Goncalves, Ewan Harrison, David K. Jackson, Ian Johnston, Dominic Kwiatkowski, Cordelia Langford, John Sillitoe on behalf of the Wellcome Sanger Institute COVID-19 Surveillance Team |
| EPI_ISL_2513265, EPI_ISL_2513266 | Lighthouse Lab in Alderley Park | Wellcome Sanger Institute for the COVID-19 Genomics UK (COG-UK) Consortium | Jacquelyn Wynn, Mairead Hyland, The Lighthouse Lab in Alderley Park and Alex Alderton, Roberto Amato, Jeffrey Barrett, Sonia Goncalves, Ewan Harrison, David K. Jackson, Ian Johnston, Dominic Kwiatkowski, Cordelia Langford, John Sillitoe on behalf of the Wellcome Sanger Institute COVID-19 Surveillance Team |
| EPI_ISL_2513268 | Health Services Laboratories | Wellcome Sanger Institute for the COVID-19 Genomics UK (COG-UK) Consortium | Health Services Laboratories and Alex Alderton, Roberto Amato, Jeffrey Barrett, Sonia Goncalves, Ewan Harrison, David K. Jackson, Ian Johnston, Dominic Kwiatkowski, Cordelia Langford, John Sillitoe on behalf of the Wellcome Sanger Institute COVID-19 Surveillance Team |
| EPI_ISL_2513269 | Lighthouse Lab in Alderley Park | Wellcome Sanger Institute for the COVID-19 Genomics UK (COG-UK) Consortium | Jacquelyn Wynn, Mairead Hyland, The Lighthouse Lab in Alderley Park and Alex Alderton, Roberto Amato, Jeffrey Barrett, Sonia Goncalves, Ewan Harrison, David K. Jackson, Ian Johnston, Dominic Kwiatkowski, Cordelia Langford, John Sillitoe on behalf of the Wellcome Sanger Institute COVID-19 Surveillance Team |
| EPI_ISL_2513270, EPI_ISL_2513271 | Health Services Laboratories | Wellcome Sanger Institute for the COVID-19 Genomics UK (COG-UK) Consortium | Health Services Laboratories and Alex Alderton, Roberto Amato, Jeffrey Barrett, Sonia Goncalves, Ewan Harrison, David K. Jackson, Ian Johnston, Dominic Kwiatkowski, Cordelia Langford, John Sillitoe on behalf of the Wellcome Sanger Institute COVID-19 Surveillance Team |
| EPI_ISL_2513272, EPI_ISL_2513273, EPI_ISL_2513274 | Lighthouse Lab in Alderley Park | Wellcome Sanger Institute for the COVID-19 Genomics UK (COG-UK) Consortium | Jacquelyn Wynn, Mairead Hyland, The Lighthouse Lab in Alderley Park and Alex Alderton, Roberto Amato, Jeffrey Barrett, Sonia Goncalves, Ewan Harrison, David K. Jackson, Ian Johnston, Dominic Kwiatkowski, Cordelia Langford, John Sillitoe on behalf of the Wellcome Sanger Institute COVID-19 Surveillance Team |
| EPI_ISL_2513275 | Lighthouse Lab in Milton Keynes | Wellcome Sanger Institute for the COVID-19 Genomics UK (COG-UK) Consortium | The Lighthouse Lab in Milton Keynes and Alex Alderton, Roberto Amato, Jeffrey Barrett, Sonia Goncalves, Ewan Harrison, David K. Jackson, Ian Johnston, Dominic Kwiatkowski, Cordelia Langford, John Sillitoe on behalf of the Wellcome Sanger Institute COVID-19 Surveillance Team |
| EPI_ISL_2513276 | Lighthouse Lab in Alderley Park | Wellcome Sanger Institute for the COVID-19 Genomics UK (COG-UK) Consortium | Jacquelyn Wynn, Mairead Hyland, The Lighthouse Lab in Alderley Park and Alex Alderton, Roberto Amato, Jeffrey Barrett, Sonia Goncalves, Ewan Harrison, David K. Jackson, Ian Johnston, Dominic Kwiatkowski, Cordelia Langford, John Sillitoe on behalf of the Wellcome Sanger Institute COVID-19 Surveillance Team |
| EPI_ISL_2513277 | Lighthouse Lab in Milton Keynes | Wellcome Sanger Institute for the COVID-19 Genomics UK (COG-UK) Consortium | The Lighthouse Lab in Milton Keynes and Alex Alderton, Roberto Amato, Jeffrey Barrett, Sonia Goncalves, Ewan Harrison, David K. Jackson, Ian Johnston, Dominic Kwiatkowski, Cordelia Langford, John Sillitoe on behalf of the Wellcome Sanger Institute COVID-19 Surveillance Team |
| EPI_ISL_2513278 | Lighthouse Lab in Alderley Park | Wellcome Sanger Institute for the COVID-19 Genomics UK (COG-UK) Consortium | Jacquelyn Wynn, Mairead Hyland, The Lighthouse Lab in Alderley Park and Alex Alderton, Roberto Amato, Jeffrey Barrett, Sonia Goncalves, Ewan Harrison, David K. Jackson, Ian Johnston, Dominic Kwiatkowski, Cordelia Langford, John Sillitoe on behalf of the Wellcome Sanger Institute COVID-19 Surveillance Team |
| EPI_ISL_2513279 | Lighthouse Lab in Milton Keynes | Wellcome Sanger Institute for the COVID-19 Genomics UK (COG-UK) Consortium | The Lighthouse Lab in Milton Keynes and Alex Alderton, Roberto Amato, Jeffrey Barrett, Sonia Goncalves, Ewan Harrison, David K. Jackson, Ian Johnston, Dominic Kwiatkowski, Cordelia Langford, John Sillitoe on behalf of the Wellcome Sanger Institute COVID-19 Surveillance Team |
| EPI_ISL_2513280 | Health Services Laboratories | Wellcome Sanger Institute for the COVID-19 Genomics UK (COG-UK) Consortium | Health Services Laboratories and Alex Alderton, Roberto Amato, Jeffrey Barrett, Sonia Goncalves, Ewan Harrison, David K. Jackson, Ian Johnston, Dominic Kwiatkowski, Cordelia Langford, John Sillitoe on behalf of the Wellcome Sanger Institute COVID-19 Surveillance Team |
| EPI_ISL_2513281, EPI_ISL_2513282 | Lighthouse Lab in Alderley Park | Wellcome Sanger Institute for the COVID-19 Genomics UK (COG-UK) Consortium | Jacquelyn Wynn, Mairead Hyland, The Lighthouse Lab in Alderley Park and Alex Alderton, Roberto Amato, Jeffrey Barrett, Sonia Goncalves, Ewan Harrison, David K. Jackson, Ian Johnston, Dominic Kwiatkowski, Cordelia Langford, John Sillitoe on behalf of the Wellcome Sanger Institute COVID-19 Surveillance Team |
| EPI_ISL_2513283 | Lighthouse Lab in Milton Keynes | Wellcome Sanger Institute for the COVID-19 Genomics UK (COG-UK) Consortium | The Lighthouse Lab in Milton Keynes and Alex Alderton, Roberto Amato, Jeffrey Barrett, Sonia Goncalves, Ewan Harrison, David K. Jackson, Ian Johnston, Dominic Kwiatkowski, Cordelia Langford, John Sillitoe on behalf of the Wellcome Sanger Institute COVID-19 Surveillance Team |
| EPI_ISL_2513284, EPI_ISL_2513286, EPI_ISL_2513287, EPI_ISL_2513288 | Lighthouse Lab in Alderley Park | Wellcome Sanger Institute for the COVID-19 Genomics UK (COG-UK) Consortium | Jacquelyn Wynn, Mairead Hyland, The Lighthouse Lab in Alderley Park and Alex Alderton, Roberto Amato, Jeffrey Barrett, Sonia Goncalves, Ewan Harrison, David K. Jackson, Ian Johnston, Dominic Kwiatkowski, Cordelia Langford, John Sillitoe on behalf of the Wellcome Sanger Institute COVID-19 Surveillance Team |
| EPI_ISL_2513289 | Lighthouse Lab in Milton Keynes | Wellcome Sanger Institute for the COVID-19 Genomics UK (COG-UK) Consortium | The Lighthouse Lab in Milton Keynes and Alex Alderton, Roberto Amato, Jeffrey Barrett, Sonia Goncalves, Ewan Harrison, David K. Jackson, Ian Johnston, Dominic Kwiatkowski, Cordelia Langford, John Sillitoe on behalf of the Wellcome Sanger Institute COVID-19 Surveillance Team |
| EPI_ISL_2513291, EPI_ISL_2513292, EPI_ISL_2513293 | Lighthouse Lab in Alderley Park | Wellcome Sanger Institute for the COVID-19 Genomics UK (COG-UK) Consortium | Jacquelyn Wynn, Mairead Hyland, The Lighthouse Lab in Alderley Park and Alex Alderton, Roberto Amato, Jeffrey Barrett, Sonia Goncalves, Ewan Harrison, David K. Jackson, Ian Johnston, Dominic Kwiatkowski, Cordelia Langford, John Sillitoe on behalf of the Wellcome Sanger Institute COVID-19 Surveillance Team |
| EPI_ISL_2513294 | Lighthouse Lab in Glasgow | Wellcome Sanger Institute for the COVID-19 Genomics UK (COG-UK) Consortium | Harper VanSteenhouse, Yumi Kasai, David Gray, Carol Clugston, Anna Dominiczak and Alex Alderton, Roberto Amato, Jeffrey Barrett, Sonia Goncalves, Ewan Harrison, David K. Jackson, Ian Johnston, Dominic Kwiatkowski, Cordelia Langford, John Sillitoe on behalf of the Wellcome Sanger Institute COVID-19 Surveillance Team |
| EPI_ISL_2513295, EPI_ISL_2513296, EPI_ISL_2513297, EPI_ISL_2513299 | Lighthouse Lab in Alderley Park | Wellcome Sanger Institute for the COVID-19 Genomics UK (COG-UK) Consortium | Jacquelyn Wynn, Mairead Hyland, The Lighthouse Lab in Alderley Park and Alex Alderton, Roberto Amato, Jeffrey Barrett, Sonia Goncalves, Ewan Harrison, David K. Jackson, Ian Johnston, Dominic Kwiatkowski, Cordelia Langford, John Sillitoe on behalf of the Wellcome Sanger Institute COVID-19 Surveillance Team |
| EPI_ISL_2513301 | Lighthouse Lab in Glasgow | Wellcome Sanger Institute for the COVID-19 Genomics UK (COG-UK) Consortium | Harper VanSteenhouse, Yumi Kasai, David Gray, Carol Clugston, Anna Dominiczak and Alex Alderton, Roberto Amato, Jeffrey Barrett, Sonia Goncalves, Ewan Harrison, David K. Jackson, Ian Johnston, Dominic Kwiatkowski, Cordelia Langford, John Sillitoe on behalf of the Wellcome Sanger Institute COVID-19 Surveillance Team |
| EPI_ISL_2513302 | Lighthouse Lab in Alderley Park | Wellcome Sanger Institute for the COVID-19 Genomics UK (COG-UK) Consortium | Jacquelyn Wynn, Mairead Hyland, The Lighthouse Lab in Alderley Park and Alex Alderton, Roberto Amato, Jeffrey Barrett, Sonia Goncalves, Ewan Harrison, David K. Jackson, Ian Johnston, Dominic Kwiatkowski, Cordelia Langford, John Sillitoe on behalf of the Wellcome Sanger Institute COVID-19 Surveillance Team |
| EPI_ISL_2513303 | Lighthouse Lab in Glasgow | Wellcome Sanger Institute for the COVID-19 Genomics UK (COG-UK) Consortium | Harper VanSteenhouse, Yumi Kasai, David Gray, Carol Clugston, Anna Dominiczak and Alex Alderton, Roberto Amato, Jeffrey Barrett, Sonia Goncalves, Ewan Harrison, David K. Jackson, Ian Johnston, Dominic Kwiatkowski, Cordelia Langford, John Sillitoe on behalf of the Wellcome Sanger Institute COVID-19 Surveillance Team |

| | | | |
|---|---|---|---|
| EPI_ISL_2513304, EPI_ISL_2513305 | Lighthouse Lab in Alderley Park | Wellcome Sanger Institute for the COVID-19 Genomics UK (COG-UK) Consortium | Jacquelyn Wynn, Mairead Hyland, The Lighthouse Lab in Alderley Park and Alex Alderton, Roberto Amato, Jeffrey Barrett, Sonia Goncalves, Ewan Harrison, David K. Jackson, Ian Johnston, Dominic Kwiatkowski, Cordelia Langford, John Sillitoe on behalf of the Wellcome Sanger Institute COVID-19 Surveillance Team |
| EPI_ISL_2513306 | Lighthouse Lab in Glasgow | Wellcome Sanger Institute for the COVID-19 Genomics UK (COG-UK) Consortium | Harper VanSteenhouse, Yumi Kasai, David Gray, Carol Clugston, Anna Dominiczak and Alex Alderton, Roberto Amato, Jeffrey Barrett, Sonia Goncalves, Ewan Harrison, David K. Jackson, Ian Johnston, Dominic Kwiatkowski, Cordelia Langford, John Sillitoe on behalf of the Wellcome Sanger Institute COVID-19 Surveillance Team |
| EPI_ISL_2513307, EPI_ISL_2513308 | Lighthouse Lab in Alderley Park | Wellcome Sanger Institute for the COVID-19 Genomics UK (COG-UK) Consortium | Jacquelyn Wynn, Mairead Hyland, The Lighthouse Lab in Alderley Park and Alex Alderton, Roberto Amato, Jeffrey Barrett, Sonia Goncalves, Ewan Harrison, David K. Jackson, Ian Johnston, Dominic Kwiatkowski, Cordelia Langford, John Sillitoe on behalf of the Wellcome Sanger Institute COVID-19 Surveillance Team |
| EPI_ISL_2513309 | Lighthouse Lab in Glasgow | Wellcome Sanger Institute for the COVID-19 Genomics UK (COG-UK) Consortium | Harper VanSteenhouse, Yumi Kasai, David Gray, Carol Clugston, Anna Dominiczak and Alex Alderton, Roberto Amato, Jeffrey Barrett, Sonia Goncalves, Ewan Harrison, David K. Jackson, Ian Johnston, Dominic Kwiatkowski, Cordelia Langford, John Sillitoe on behalf of the Wellcome Sanger Institute COVID-19 Surveillance Team |
| EPI_ISL_2513310 | Lighthouse Lab in Alderley Park | Wellcome Sanger Institute for the COVID-19 Genomics UK (COG-UK) Consortium | Jacquelyn Wynn, Mairead Hyland, The Lighthouse Lab in Alderley Park and Alex Alderton, Roberto Amato, Jeffrey Barrett, Sonia Goncalves, Ewan Harrison, David K. Jackson, Ian Johnston, Dominic Kwiatkowski, Cordelia Langford, John Sillitoe on behalf of the Wellcome Sanger Institute COVID-19 Surveillance Team |
| EPI_ISL_2513311 | Lighthouse Lab in Glasgow | Wellcome Sanger Institute for the COVID-19 Genomics UK (COG-UK) Consortium | Harper VanSteenhouse, Yumi Kasai, David Gray, Carol Clugston, Anna Dominiczak and Alex Alderton, Roberto Amato, Jeffrey Barrett, Sonia Goncalves, Ewan Harrison, David K. Jackson, Ian Johnston, Dominic Kwiatkowski, Cordelia Langford, John Sillitoe on behalf of the Wellcome Sanger Institute COVID-19 Surveillance Team |
| EPI_ISL_2513312 | Lighthouse Lab in Alderley Park | Wellcome Sanger Institute for the COVID-19 Genomics UK (COG-UK) Consortium | Jacquelyn Wynn, Mairead Hyland, The Lighthouse Lab in Alderley Park and Alex Alderton, Roberto Amato, Jeffrey Barrett, Sonia Goncalves, Ewan Harrison, David K. Jackson, Ian Johnston, Dominic Kwiatkowski, Cordelia Langford, John Sillitoe on behalf of the Wellcome Sanger Institute COVID-19 Surveillance Team |
| EPI_ISL_2513313, EPI_ISL_2513314 | Lighthouse Lab in Glasgow | Wellcome Sanger Institute for the COVID-19 Genomics UK (COG-UK) Consortium | Harper VanSteenhouse, Yumi Kasai, David Gray, Carol Clugston, Anna Dominiczak and Alex Alderton, Roberto Amato, Jeffrey Barrett, Sonia Goncalves, Ewan Harrison, David K. Jackson, Ian Johnston, Dominic Kwiatkowski, Cordelia Langford, John Sillitoe on behalf of the Wellcome Sanger Institute COVID-19 Surveillance Team |
| EPI_ISL_2513315, EPI_ISL_2513317 | Lighthouse Lab in Alderley Park | Wellcome Sanger Institute for the COVID-19 Genomics UK (COG-UK) Consortium | Jacquelyn Wynn, Mairead Hyland, The Lighthouse Lab in Alderley Park and Alex Alderton, Roberto Amato, Jeffrey Barrett, Sonia Goncalves, Ewan Harrison, David K. Jackson, Ian Johnston, Dominic Kwiatkowski, Cordelia Langford, John Sillitoe on behalf of the Wellcome Sanger Institute COVID-19 Surveillance Team |
| EPI_ISL_2513318, EPI_ISL_2513320 | Lighthouse Lab in Glasgow | Wellcome Sanger Institute for the COVID-19 Genomics UK (COG-UK) Consortium | Harper VanSteenhouse, Yumi Kasai, David Gray, Carol Clugston, Anna Dominiczak and Alex Alderton, Roberto Amato, Jeffrey Barrett, Sonia Goncalves, Ewan Harrison, David K. Jackson, Ian Johnston, Dominic Kwiatkowski, Cordelia Langford, John Sillitoe on behalf of the Wellcome Sanger Institute COVID-19 Surveillance Team |
| EPI_ISL_2513321 | Lighthouse Lab in Alderley Park | Wellcome Sanger Institute for the COVID-19 Genomics UK (COG-UK) Consortium | Jacquelyn Wynn, Mairead Hyland, The Lighthouse Lab in Alderley Park and Alex Alderton, Roberto Amato, Jeffrey Barrett, Sonia Goncalves, Ewan Harrison, David K. Jackson, Ian Johnston, Dominic Kwiatkowski, Cordelia Langford, John Sillitoe on behalf of the Wellcome Sanger Institute COVID-19 Surveillance Team |
| EPI_ISL_2513322 | Lighthouse Lab in Glasgow | Wellcome Sanger Institute for the COVID-19 Genomics UK (COG-UK) Consortium | Harper VanSteenhouse, Yumi Kasai, David Gray, Carol Clugston, Anna Dominiczak and Alex Alderton, Roberto Amato, Jeffrey Barrett, Sonia Goncalves, Ewan Harrison, David K. Jackson, Ian Johnston, Dominic Kwiatkowski, Cordelia Langford, John Sillitoe on behalf of the Wellcome Sanger Institute COVID-19 Surveillance Team |
| EPI_ISL_2513323, EPI_ISL_2513324, EPI_ISL_2513325 | Lighthouse Lab in Alderley Park | Wellcome Sanger Institute for the COVID-19 Genomics UK (COG-UK) Consortium | Jacquelyn Wynn, Mairead Hyland, The Lighthouse Lab in Alderley Park and Alex Alderton, Roberto Amato, Jeffrey Barrett, Sonia Goncalves, Ewan Harrison, David K. Jackson, Ian Johnston, Dominic Kwiatkowski, Cordelia Langford, John Sillitoe on behalf of the Wellcome Sanger Institute COVID-19 Surveillance Team |
| EPI_ISL_2513326 | Lighthouse Lab in Glasgow | Wellcome Sanger Institute for the COVID-19 Genomics UK (COG-UK) Consortium | Harper VanSteenhouse, Yumi Kasai, David Gray, Carol Clugston, Anna Dominiczak and Alex Alderton, Roberto Amato, Jeffrey Barrett, Sonia Goncalves, Ewan Harrison, David K. Jackson, Ian Johnston, Dominic Kwiatkowski, Cordelia Langford, John Sillitoe on behalf of the Wellcome Sanger Institute COVID-19 Surveillance Team |
| EPI_ISL_2513327, EPI_ISL_2513328, EPI_ISL_2513329, EPI_ISL_2513330, EPI_ISL_2513331, EPI_ISL_2513333 | Lighthouse Lab in Alderley Park | Wellcome Sanger Institute for the COVID-19 Genomics UK (COG-UK) Consortium | Jacquelyn Wynn, Mairead Hyland, The Lighthouse Lab in Alderley Park and Alex Alderton, Roberto Amato, Jeffrey Barrett, Sonia Goncalves, Ewan Harrison, David K. Jackson, Ian Johnston, Dominic Kwiatkowski, Cordelia Langford, John Sillitoe on behalf of the Wellcome Sanger Institute COVID-19 Surveillance Team |
| EPI_ISL_2513334 | Lighthouse Lab in Glasgow | Wellcome Sanger Institute for the COVID-19 Genomics UK (COG-UK) Consortium | Harper VanSteenhouse, Yumi Kasai, David Gray, Carol Clugston, Anna Dominiczak and Alex Alderton, Roberto Amato, Jeffrey Barrett, Sonia Goncalves, Ewan Harrison, David K. Jackson, Ian Johnston, Dominic Kwiatkowski, Cordelia Langford, John Sillitoe on behalf of the Wellcome Sanger Institute COVID-19 Surveillance Team |
| EPI_ISL_2513336, EPI_ISL_2513337, EPI_ISL_2513338 | Lighthouse Lab in Alderley Park | Wellcome Sanger Institute for the COVID-19 Genomics UK (COG-UK) Consortium | Jacquelyn Wynn, Mairead Hyland, The Lighthouse Lab in Alderley Park and Alex Alderton, Roberto Amato, Jeffrey Barrett, Sonia Goncalves, Ewan Harrison, David K. Jackson, Ian Johnston, Dominic Kwiatkowski, Cordelia Langford, John Sillitoe on behalf of the Wellcome Sanger Institute COVID-19 Surveillance Team |
| EPI_ISL_2513339 | Lighthouse Lab in Glasgow | Wellcome Sanger Institute for the COVID-19 Genomics UK (COG-UK) Consortium | Harper VanSteenhouse, Yumi Kasai, David Gray, Carol Clugston, Anna Dominiczak and Alex Alderton, Roberto Amato, Jeffrey Barrett, Sonia Goncalves, Ewan Harrison, David K. Jackson, Ian Johnston, Dominic Kwiatkowski, Cordelia Langford, John Sillitoe on behalf of the Wellcome Sanger Institute COVID-19 Surveillance Team |
| EPI_ISL_2513340 | Lighthouse Lab in Alderley Park | Wellcome Sanger Institute for the COVID-19 Genomics UK (COG-UK) Consortium | Jacquelyn Wynn, Mairead Hyland, The Lighthouse Lab in Alderley Park and Alex Alderton, Roberto Amato, Jeffrey Barrett, Sonia Goncalves, Ewan Harrison, David K. Jackson, Ian Johnston, Dominic Kwiatkowski, Cordelia Langford, John Sillitoe on behalf of the Wellcome Sanger Institute COVID-19 Surveillance Team |
| EPI_ISL_2513341, EPI_ISL_2513342 | Lighthouse Lab in Glasgow | Wellcome Sanger Institute for the COVID-19 Genomics UK (COG-UK) Consortium | Harper VanSteenhouse, Yumi Kasai, David Gray, Carol Clugston, Anna Dominiczak and Alex Alderton, Roberto Amato, Jeffrey Barrett, Sonia Goncalves, Ewan Harrison, David K. Jackson, Ian Johnston, Dominic Kwiatkowski, Cordelia Langford, John Sillitoe on behalf of the Wellcome Sanger Institute COVID-19 Surveillance Team |
| EPI_ISL_2513343, EPI_ISL_2513344, EPI_ISL_2513345 | Lighthouse Lab in Alderley Park | Wellcome Sanger Institute for the COVID-19 Genomics UK (COG-UK) Consortium | Jacquelyn Wynn, Mairead Hyland, The Lighthouse Lab in Alderley Park and Alex Alderton, Roberto Amato, Jeffrey Barrett, Sonia Goncalves, Ewan Harrison, David K. Jackson, Ian Johnston, Dominic Kwiatkowski, Cordelia Langford, John Sillitoe on behalf of the Wellcome Sanger Institute COVID-19 Surveillance Team |
| EPI_ISL_2513346 | Lighthouse Lab in Glasgow | Wellcome Sanger Institute for the COVID-19 Genomics UK (COG-UK) Consortium | Harper VanSteenhouse, Yumi Kasai, David Gray, Carol Clugston, Anna Dominiczak and Alex Alderton, Roberto Amato, Jeffrey Barrett, Sonia Goncalves, Ewan Harrison, David K. Jackson, Ian Johnston, Dominic Kwiatkowski, Cordelia Langford, John Sillitoe on behalf of the Wellcome Sanger Institute COVID-19 Surveillance Team |
| EPI_ISL_2513347 | Lighthouse Lab in Alderley Park | Wellcome Sanger Institute for the COVID-19 Genomics UK (COG-UK) Consortium | Jacquelyn Wynn, Mairead Hyland, The Lighthouse Lab in Alderley Park and Alex Alderton, Roberto Amato, Jeffrey Barrett, Sonia Goncalves, Ewan Harrison, David K. Jackson, Ian Johnston, Dominic Kwiatkowski, Cordelia Langford, John Sillitoe on behalf of the Wellcome Sanger Institute COVID-19 Surveillance Team |
| EPI_ISL_2513348 | Lighthouse Lab in Glasgow | Wellcome Sanger Institute for the COVID-19 Genomics UK (COG-UK) Consortium | Harper VanSteenhouse, Yumi Kasai, David Gray, Carol Clugston, Anna Dominiczak and Alex Alderton, Roberto Amato, Jeffrey Barrett, Sonia Goncalves, Ewan Harrison, David K. Jackson, Ian Johnston, Dominic Kwiatkowski, Cordelia Langford, John Sillitoe on behalf of the Wellcome Sanger Institute COVID-19 Surveillance Team |
| EPI_ISL_2513349, EPI_ISL_2513350 | Lighthouse Lab in Alderley Park | Wellcome Sanger Institute for the COVID-19 Genomics UK (COG-UK) Consortium | Jacquelyn Wynn, Mairead Hyland, The Lighthouse Lab in Alderley Park and Alex Alderton, Roberto Amato, Jeffrey Barrett, Sonia Goncalves, Ewan Harrison, David K. Jackson, Ian Johnston, Dominic Kwiatkowski, Cordelia Langford, John Sillitoe on behalf of the Wellcome Sanger Institute COVID-19 Surveillance Team |
| EPI_ISL_2513351 | Lighthouse Lab in Glasgow | Wellcome Sanger Institute for the COVID-19 Genomics UK (COG-UK) Consortium | Harper VanSteenhouse, Yumi Kasai, David Gray, Carol Clugston, Anna Dominiczak and Alex Alderton, Roberto Amato, Jeffrey Barrett, Sonia Goncalves, Ewan Harrison, David K. Jackson, Ian Johnston, Dominic Kwiatkowski, Cordelia Langford, John Sillitoe on behalf of the Wellcome Sanger Institute COVID-19 Surveillance Team |
| EPI_ISL_2513352, EPI_ISL_2513353, EPI_ISL_2513354, EPI_ISL_2513355, EPI_ISL_2513356, EPI_ISL_2513357, EPI_ISL_2513358, EPI_ISL_2513359, EPI_ISL_2513360, EPI_ISL_2513361 | Lighthouse Lab in Alderley Park | Wellcome Sanger Institute for the COVID-19 Genomics UK (COG-UK) Consortium | Jacquelyn Wynn, Mairead Hyland, The Lighthouse Lab in Alderley Park and Alex Alderton, Roberto Amato, Jeffrey Barrett, Sonia Goncalves, Ewan Harrison, David K. Jackson, Ian Johnston, Dominic Kwiatkowski, Cordelia Langford, John Sillitoe on behalf of the Wellcome Sanger Institute COVID-19 Surveillance Team |
| EPI_ISL_2513362 | Lighthouse Lab in Glasgow | Wellcome Sanger Institute for the COVID-19 Genomics UK (COG-UK) Consortium | Harper VanSteenhouse, Yumi Kasai, David Gray, Carol Clugston, Anna Dominiczak and Alex Alderton, Roberto Amato, Jeffrey Barrett, Sonia Goncalves, Ewan Harrison, David K. Jackson, Ian Johnston, Dominic Kwiatkowski, Cordelia Langford, John Sillitoe on behalf of the Wellcome Sanger Institute COVID-19 Surveillance Team |
| EPI_ISL_2513363, EPI_ISL_2513364, EPI_ISL_2513365 | Lighthouse Lab in Alderley Park | Wellcome Sanger Institute for the COVID-19 Genomics UK (COG-UK) Consortium | Jacquelyn Wynn, Mairead Hyland, The Lighthouse Lab in Alderley Park and Alex Alderton, Roberto Amato, Jeffrey Barrett, Sonia Goncalves, Ewan Harrison, David K. Jackson, Ian Johnston, Dominic Kwiatkowski, Cordelia Langford, John Sillitoe on behalf of the Wellcome Sanger Institute COVID-19 Surveillance Team |
| EPI_ISL_2513366 | Lighthouse Lab in Glasgow | Wellcome Sanger Institute for the COVID-19 Genomics UK (COG-UK) Consortium | Harper VanSteenhouse, Yumi Kasai, David Gray, Carol Clugston, Anna Dominiczak and Alex Alderton, Roberto Amato, Jeffrey Barrett, Sonia Goncalves, Ewan Harrison, David K. Jackson, Ian Johnston, Dominic Kwiatkowski, Cordelia Langford, John Sillitoe on behalf of the Wellcome Sanger Institute COVID-19 Surveillance Team |
| EPI_ISL_2513367 | Lighthouse Lab in Alderley Park | Wellcome Sanger Institute for the COVID-19 Genomics UK (COG-UK) Consortium | Jacquelyn Wynn, Mairead Hyland, The Lighthouse Lab in Alderley Park and Alex Alderton, Roberto Amato, Jeffrey Barrett, Sonia Goncalves, Ewan Harrison, David K. Jackson, Ian Johnston, Dominic Kwiatkowski, Cordelia Langford, John Sillitoe on behalf of the Wellcome Sanger Institute COVID-19 Surveillance Team |
| EPI_ISL_2513368 | Lighthouse Lab in Glasgow | Wellcome Sanger Institute for the COVID-19 Genomics UK (COG-UK) Consortium | Harper VanSteenhouse, Yumi Kasai, David Gray, Carol Clugston, Anna Dominiczak and Alex Alderton, Roberto Amato, Jeffrey Barrett, Sonia Goncalves, Ewan Harrison, David K. Jackson, Ian Johnston, Dominic Kwiatkowski, Cordelia Langford, John Sillitoe on behalf of the Wellcome Sanger Institute COVID-19 Surveillance Team |
| EPI_ISL_2513369 | Lighthouse Lab in Alderley Park | Wellcome Sanger Institute for the COVID-19 Genomics UK (COG-UK) Consortium | Jacquelyn Wynn, Mairead Hyland, The Lighthouse Lab in Alderley Park and Alex Alderton, Roberto Amato, Jeffrey Barrett, Sonia Goncalves, Ewan Harrison, David K. Jackson, Ian Johnston, Dominic Kwiatkowski, Cordelia Langford, John Sillitoe on behalf of the Wellcome Sanger Institute COVID-19 Surveillance Team |
| EPI_ISL_2513370 | Lighthouse Lab in Glasgow | Wellcome Sanger Institute for the COVID-19 Genomics UK (COG-UK) Consortium | Harper VanSteenhouse, Yumi Kasai, David Gray, Carol Clugston, Anna Dominiczak and Alex Alderton, Roberto Amato, Jeffrey Barrett, Sonia Goncalves, Ewan Harrison, David K. Jackson, Ian Johnston, Dominic Kwiatkowski, Cordelia Langford, John Sillitoe on behalf of the Wellcome Sanger Institute COVID-19 Surveillance Team |
| EPI_ISL_2513371, EPI_ISL_2513372 | Lighthouse Lab in Alderley Park | Wellcome Sanger Institute for the COVID-19 Genomics UK (COG-UK) Consortium | Jacquelyn Wynn, Mairead Hyland, The Lighthouse Lab in Alderley Park and Alex Alderton, Roberto Amato, Jeffrey Barrett, Sonia Goncalves, Ewan Harrison, David K. Jackson, Ian Johnston, Dominic Kwiatkowski, Cordelia Langford, John Sillitoe on behalf of the Wellcome Sanger Institute COVID-19 Surveillance Team |
| EPI_ISL_2513373 | Lighthouse Lab in Glasgow | Wellcome Sanger Institute for the COVID-19 Genomics UK (COG-UK) Consortium | Harper VanSteenhouse, Yumi Kasai, David Gray, Carol Clugston, Anna Dominiczak and Alex Alderton, Roberto Amato, Jeffrey Barrett, Sonia Goncalves, Ewan Harrison, David K. Jackson, Ian Johnston, Dominic Kwiatkowski, Cordelia Langford, John Sillitoe on behalf of the Wellcome Sanger Institute COVID-19 Surveillance Team |
| EPI_ISL_2513374 | Lighthouse Lab in Alderley Park | Wellcome Sanger Institute for the COVID-19 Genomics UK (COG-UK) Consortium | Jacquelyn Wynn, Mairead Hyland, The Lighthouse Lab in Alderley Park and Alex Alderton, Roberto Amato, Jeffrey Barrett, Sonia Goncalves, Ewan Harrison, David K. Jackson, Ian Johnston, Dominic Kwiatkowski, Cordelia Langford, John Sillitoe on behalf of the Wellcome Sanger Institute COVID-19 Surveillance Team |
| EPI_ISL_2513375, EPI_ISL_2513376, EPI_ISL_2513377 | Lighthouse Lab in Glasgow | Wellcome Sanger Institute for the COVID-19 Genomics UK (COG-UK) Consortium | Harper VanSteenhouse, Yumi Kasai, David Gray, Carol Clugston, Anna Dominiczak and Alex Alderton, Roberto Amato, Jeffrey Barrett, Sonia Goncalves, Ewan Harrison, David K. Jackson, Ian Johnston, Dominic Kwiatkowski, Cordelia Langford, John Sillitoe on behalf of the Wellcome Sanger Institute COVID-19 Surveillance Team |

| | | | |
|---|---|---|---|
| EPI_ISL_2513378 | Lighthouse Lab in Alderley Park | Wellcome Sanger Institute for the COVID-19 Genomics UK (COG-UK) Consortium | Jacquelyn Wynn, Mairead Hyland, The Lighthouse Lab in Alderley Park and Alex Alderton, Roberto Amato, Jeffrey Barrett, Sonia Goncalves, Ewan Harrison, David K. Jackson, Ian Johnston, Dominic Kwiatkowski, Cordelia Langford, John Sillitoe on behalf of the Wellcome Sanger Institute COVID-19 Surveillance Team |
| EPI_ISL_2513379 | Lighthouse Lab in Glasgow | Wellcome Sanger Institute for the COVID-19 Genomics UK (COG-UK) Consortium | Harper VanSteenhouse, Yumi Kasai, David Gray, Carol Clugston, Anna Dominiczak and Alex Alderton, Roberto Amato, Jeffrey Barrett, Sonia Goncalves, Ewan Harrison, David K. Jackson, Ian Johnston, Dominic Kwiatkowski, Cordelia Langford, John Sillitoe on behalf of the Wellcome Sanger Institute COVID-19 Surveillance Team |
| EPI_ISL_2513380, EPI_ISL_2513381, EPI_ISL_2513382, EPI_ISL_2513383, EPI_ISL_2513384, EPI_ISL_2513385, EPI_ISL_2513386, EPI_ISL_2513387, EPI_ISL_2513388, EPI_ISL_2513389 | Lighthouse Lab in Alderley Park | Wellcome Sanger Institute for the COVID-19 Genomics UK (COG-UK) Consortium | Jacquelyn Wynn, Mairead Hyland, The Lighthouse Lab in Alderley Park and Alex Alderton, Roberto Amato, Jeffrey Barrett, Sonia Goncalves, Ewan Harrison, David K. Jackson, Ian Johnston, Dominic Kwiatkowski, Cordelia Langford, John Sillitoe on behalf of the Wellcome Sanger Institute COVID-19 Surveillance Team |
| EPI_ISL_2513390 | Lighthouse Lab in Glasgow | Wellcome Sanger Institute for the COVID-19 Genomics UK (COG-UK) Consortium | Harper VanSteenhouse, Yumi Kasai, David Gray, Carol Clugston, Anna Dominiczak and Alex Alderton, Roberto Amato, Jeffrey Barrett, Sonia Goncalves, Ewan Harrison, David K. Jackson, Ian Johnston, Dominic Kwiatkowski, Cordelia Langford, John Sillitoe on behalf of the Wellcome Sanger Institute COVID-19 Surveillance Team |
| EPI_ISL_2513391, EPI_ISL_2513392, EPI_ISL_2513393 | Lighthouse Lab in Alderley Park | Wellcome Sanger Institute for the COVID-19 Genomics UK (COG-UK) Consortium | Jacquelyn Wynn, Mairead Hyland, The Lighthouse Lab in Alderley Park and Alex Alderton, Roberto Amato, Jeffrey Barrett, Sonia Goncalves, Ewan Harrison, David K. Jackson, Ian Johnston, Dominic Kwiatkowski, Cordelia Langford, John Sillitoe on behalf of the Wellcome Sanger Institute COVID-19 Surveillance Team |
| EPI_ISL_2513394 | Lighthouse Lab in Glasgow | Wellcome Sanger Institute for the COVID-19 Genomics UK (COG-UK) Consortium | Harper VanSteenhouse, Yumi Kasai, David Gray, Carol Clugston, Anna Dominiczak and Alex Alderton, Roberto Amato, Jeffrey Barrett, Sonia Goncalves, Ewan Harrison, David K. Jackson, Ian Johnston, Dominic Kwiatkowski, Cordelia Langford, John Sillitoe on behalf of the Wellcome Sanger Institute COVID-19 Surveillance Team |
| EPI_ISL_2513395 | Lighthouse Lab in Alderley Park | Wellcome Sanger Institute for the COVID-19 Genomics UK (COG-UK) Consortium | Jacquelyn Wynn, Mairead Hyland, The Lighthouse Lab in Alderley Park and Alex Alderton, Roberto Amato, Jeffrey Barrett, Sonia Goncalves, Ewan Harrison, David K. Jackson, Ian Johnston, Dominic Kwiatkowski, Cordelia Langford, John Sillitoe on behalf of the Wellcome Sanger Institute COVID-19 Surveillance Team |
| EPI_ISL_2513396 | Lighthouse Lab in Glasgow | Wellcome Sanger Institute for the COVID-19 Genomics UK (COG-UK) Consortium | Harper VanSteenhouse, Yumi Kasai, David Gray, Carol Clugston, Anna Dominiczak and Alex Alderton, Roberto Amato, Jeffrey Barrett, Sonia Goncalves, Ewan Harrison, David K. Jackson, Ian Johnston, Dominic Kwiatkowski, Cordelia Langford, John Sillitoe on behalf of the Wellcome Sanger Institute COVID-19 Surveillance Team |
| EPI_ISL_2513397, EPI_ISL_2513398, EPI_ISL_2513399, EPI_ISL_2513400 | Lighthouse Lab in Alderley Park | Wellcome Sanger Institute for the COVID-19 Genomics UK (COG-UK) Consortium | Jacquelyn Wynn, Mairead Hyland, The Lighthouse Lab in Alderley Park and Alex Alderton, Roberto Amato, Jeffrey Barrett, Sonia Goncalves, Ewan Harrison, David K. Jackson, Ian Johnston, Dominic Kwiatkowski, Cordelia Langford, John Sillitoe on behalf of the Wellcome Sanger Institute COVID-19 Surveillance Team |
| EPI_ISL_2513401 | Lighthouse Lab in Glasgow | Wellcome Sanger Institute for the COVID-19 Genomics UK (COG-UK) Consortium | Harper VanSteenhouse, Yumi Kasai, David Gray, Carol Clugston, Anna Dominiczak and Alex Alderton, Roberto Amato, Jeffrey Barrett, Sonia Goncalves, Ewan Harrison, David K. Jackson, Ian Johnston, Dominic Kwiatkowski, Cordelia Langford, John Sillitoe on behalf of the Wellcome Sanger Institute COVID-19 Surveillance Team |
| EPI_ISL_2513402, EPI_ISL_2513403 | Lighthouse Lab in Alderley Park | Wellcome Sanger Institute for the COVID-19 Genomics UK (COG-UK) Consortium | Jacquelyn Wynn, Mairead Hyland, The Lighthouse Lab in Alderley Park and Alex Alderton, Roberto Amato, Jeffrey Barrett, Sonia Goncalves, Ewan Harrison, David K. Jackson, Ian Johnston, Dominic Kwiatkowski, Cordelia Langford, John Sillitoe on behalf of the Wellcome Sanger Institute COVID-19 Surveillance Team |
| EPI_ISL_2513404 | Lighthouse Lab in Glasgow | Wellcome Sanger Institute for the COVID-19 Genomics UK (COG-UK) Consortium | Harper VanSteenhouse, Yumi Kasai, David Gray, Carol Clugston, Anna Dominiczak and Alex Alderton, Roberto Amato, Jeffrey Barrett, Sonia Goncalves, Ewan Harrison, David K. Jackson, Ian Johnston, Dominic Kwiatkowski, Cordelia Langford, John Sillitoe on behalf of the Wellcome Sanger Institute COVID-19 Surveillance Team |
| EPI_ISL_2513405 | Lighthouse Lab in Alderley Park | Wellcome Sanger Institute for the COVID-19 Genomics UK (COG-UK) Consortium | Jacquelyn Wynn, Mairead Hyland, The Lighthouse Lab in Alderley Park and Alex Alderton, Roberto Amato, Jeffrey Barrett, Sonia Goncalves, Ewan Harrison, David K. Jackson, Ian Johnston, Dominic Kwiatkowski, Cordelia Langford, John Sillitoe on behalf of the Wellcome Sanger Institute COVID-19 Surveillance Team |
| EPI_ISL_2513406 | Lighthouse Lab in Glasgow | Wellcome Sanger Institute for the COVID-19 Genomics UK (COG-UK) Consortium | Harper VanSteenhouse, Yumi Kasai, David Gray, Carol Clugston, Anna Dominiczak and Alex Alderton, Roberto Amato, Jeffrey Barrett, Sonia Goncalves, Ewan Harrison, David K. Jackson, Ian Johnston, Dominic Kwiatkowski, Cordelia Langford, John Sillitoe on behalf of the Wellcome Sanger Institute COVID-19 Surveillance Team |
| EPI_ISL_2513407, EPI_ISL_2513408, EPI_ISL_2513409 | Lighthouse Lab in Alderley Park | Wellcome Sanger Institute for the COVID-19 Genomics UK (COG-UK) Consortium | Jacquelyn Wynn, Mairead Hyland, The Lighthouse Lab in Alderley Park and Alex Alderton, Roberto Amato, Jeffrey Barrett, Sonia Goncalves, Ewan Harrison, David K. Jackson, Ian Johnston, Dominic Kwiatkowski, Cordelia Langford, John Sillitoe on behalf of the Wellcome Sanger Institute COVID-19 Surveillance Team |
| EPI_ISL_2513411 | Lighthouse Lab in Glasgow | Wellcome Sanger Institute for the COVID-19 Genomics UK (COG-UK) Consortium | Harper VanSteenhouse, Yumi Kasai, David Gray, Carol Clugston, Anna Dominiczak and Alex Alderton, Roberto Amato, Jeffrey Barrett, Sonia Goncalves, Ewan Harrison, David K. Jackson, Ian Johnston, Dominic Kwiatkowski, Cordelia Langford, John Sillitoe on behalf of the Wellcome Sanger Institute COVID-19 Surveillance Team |
| EPI_ISL_2513412 | Lighthouse Lab in Alderley Park | Wellcome Sanger Institute for the COVID-19 Genomics UK (COG-UK) Consortium | Jacquelyn Wynn, Mairead Hyland, The Lighthouse Lab in Alderley Park and Alex Alderton, Roberto Amato, Jeffrey Barrett, Sonia Goncalves, Ewan Harrison, David K. Jackson, Ian Johnston, Dominic Kwiatkowski, Cordelia Langford, John Sillitoe on behalf of the Wellcome Sanger Institute COVID-19 Surveillance Team |
| EPI_ISL_2513413 | Lighthouse Lab in Glasgow | Wellcome Sanger Institute for the COVID-19 Genomics UK (COG-UK) Consortium | Harper VanSteenhouse, Yumi Kasai, David Gray, Carol Clugston, Anna Dominiczak and Alex Alderton, Roberto Amato, Jeffrey Barrett, Sonia Goncalves, Ewan Harrison, David K. Jackson, Ian Johnston, Dominic Kwiatkowski, Cordelia Langford, John Sillitoe on behalf of the Wellcome Sanger Institute COVID-19 Surveillance Team |
| EPI_ISL_2513414, EPI_ISL_2513415, EPI_ISL_2513417, EPI_ISL_2513418, EPI_ISL_2513419, EPI_ISL_2513420 | Lighthouse Lab in Alderley Park | Wellcome Sanger Institute for the COVID-19 Genomics UK (COG-UK) Consortium | Jacquelyn Wynn, Mairead Hyland, The Lighthouse Lab in Alderley Park and Alex Alderton, Roberto Amato, Jeffrey Barrett, Sonia Goncalves, Ewan Harrison, David K. Jackson, Ian Johnston, Dominic Kwiatkowski, Cordelia Langford, John Sillitoe on behalf of the Wellcome Sanger Institute COVID-19 Surveillance Team |
| EPI_ISL_2513421, EPI_ISL_2513422 | Lighthouse Lab in Glasgow | Wellcome Sanger Institute for the COVID-19 Genomics UK (COG-UK) Consortium | Harper VanSteenhouse, Yumi Kasai, David Gray, Carol Clugston, Anna Dominiczak and Alex Alderton, Roberto Amato, Jeffrey Barrett, Sonia Goncalves, Ewan Harrison, David K. Jackson, Ian Johnston, Dominic Kwiatkowski, Cordelia Langford, John Sillitoe on behalf of the Wellcome Sanger Institute COVID-19 Surveillance Team |
| EPI_ISL_2513423, EPI_ISL_2513424, EPI_ISL_2513425, EPI_ISL_2513426, EPI_ISL_2513429, EPI_ISL_2513430, EPI_ISL_2513431, EPI_ISL_2513432, EPI_ISL_2513433 | Lighthouse Lab in Alderley Park | Wellcome Sanger Institute for the COVID-19 Genomics UK (COG-UK) Consortium | Jacquelyn Wynn, Mairead Hyland, The Lighthouse Lab in Alderley Park and Alex Alderton, Roberto Amato, Jeffrey Barrett, Sonia Goncalves, Ewan Harrison, David K. Jackson, Ian Johnston, Dominic Kwiatkowski, Cordelia Langford, John Sillitoe on behalf of the Wellcome Sanger Institute COVID-19 Surveillance Team |
| EPI_ISL_2513434 | Lighthouse Lab in Glasgow | Wellcome Sanger Institute for the COVID-19 Genomics UK (COG-UK) Consortium | Harper VanSteenhouse, Yumi Kasai, David Gray, Carol Clugston, Anna Dominiczak and Alex Alderton, Roberto Amato, Jeffrey Barrett, Sonia Goncalves, Ewan Harrison, David K. Jackson, Ian Johnston, Dominic Kwiatkowski, Cordelia Langford, John Sillitoe on behalf of the Wellcome Sanger Institute COVID-19 Surveillance Team |
| EPI_ISL_2513435, EPI_ISL_2513436, EPI_ISL_2513437, EPI_ISL_2513439, EPI_ISL_2513440, EPI_ISL_2513441 | Lighthouse Lab in Alderley Park | Wellcome Sanger Institute for the COVID-19 Genomics UK (COG-UK) Consortium | Jacquelyn Wynn, Mairead Hyland, The Lighthouse Lab in Alderley Park and Alex Alderton, Roberto Amato, Jeffrey Barrett, Sonia Goncalves, Ewan Harrison, David K. Jackson, Ian Johnston, Dominic Kwiatkowski, Cordelia Langford, John Sillitoe on behalf of the Wellcome Sanger Institute COVID-19 Surveillance Team |
| EPI_ISL_2513442 | Lighthouse Lab in Glasgow | Wellcome Sanger Institute for the COVID-19 Genomics UK (COG-UK) Consortium | Harper VanSteenhouse, Yumi Kasai, David Gray, Carol Clugston, Anna Dominiczak and Alex Alderton, Roberto Amato, Jeffrey Barrett, Sonia Goncalves, Ewan Harrison, David K. Jackson, Ian Johnston, Dominic Kwiatkowski, Cordelia Langford, John Sillitoe on behalf of the Wellcome Sanger Institute COVID-19 Surveillance Team |
| EPI_ISL_2513443, EPI_ISL_2513444, EPI_ISL_2513446 | Lighthouse Lab in Alderley Park | Wellcome Sanger Institute for the COVID-19 Genomics UK (COG-UK) Consortium | Jacquelyn Wynn, Mairead Hyland, The Lighthouse Lab in Alderley Park and Alex Alderton, Roberto Amato, Jeffrey Barrett, Sonia Goncalves, Ewan Harrison, David K. Jackson, Ian Johnston, Dominic Kwiatkowski, Cordelia Langford, John Sillitoe on behalf of the Wellcome Sanger Institute COVID-19 Surveillance Team |
| EPI_ISL_2513447 | Lighthouse Lab in Glasgow | Wellcome Sanger Institute for the COVID-19 Genomics UK (COG-UK) Consortium | Harper VanSteenhouse, Yumi Kasai, David Gray, Carol Clugston, Anna Dominiczak and Alex Alderton, Roberto Amato, Jeffrey Barrett, Sonia Goncalves, Ewan Harrison, David K. Jackson, Ian Johnston, Dominic Kwiatkowski, Cordelia Langford, John Sillitoe on behalf of the Wellcome Sanger Institute COVID-19 Surveillance Team |
| EPI_ISL_2513448 | Lighthouse Lab in Alderley Park | Wellcome Sanger Institute for the COVID-19 Genomics UK (COG-UK) Consortium | Jacquelyn Wynn, Mairead Hyland, The Lighthouse Lab in Alderley Park and Alex Alderton, Roberto Amato, Jeffrey Barrett, Sonia Goncalves, Ewan Harrison, David K. Jackson, Ian Johnston, Dominic Kwiatkowski, Cordelia Langford, John Sillitoe on behalf of the Wellcome Sanger Institute COVID-19 Surveillance Team |
| EPI_ISL_2513450 | Lighthouse Lab in Glasgow | Wellcome Sanger Institute for the COVID-19 Genomics UK (COG-UK) Consortium | Harper VanSteenhouse, Yumi Kasai, David Gray, Carol Clugston, Anna Dominiczak and Alex Alderton, Roberto Amato, Jeffrey Barrett, Sonia Goncalves, Ewan Harrison, David K. Jackson, Ian Johnston, Dominic Kwiatkowski, Cordelia Langford, John Sillitoe on behalf of the Wellcome Sanger Institute COVID-19 Surveillance Team |
| EPI_ISL_2513451, EPI_ISL_2513453, EPI_ISL_2513454 | Lighthouse Lab in Alderley Park | Wellcome Sanger Institute for the COVID-19 Genomics UK (COG-UK) Consortium | Jacquelyn Wynn, Mairead Hyland, The Lighthouse Lab in Alderley Park and Alex Alderton, Roberto Amato, Jeffrey Barrett, Sonia Goncalves, Ewan Harrison, David K. Jackson, Ian Johnston, Dominic Kwiatkowski, Cordelia Langford, John Sillitoe on behalf of the Wellcome Sanger Institute COVID-19 Surveillance Team |
| EPI_ISL_2513455 | Lighthouse Lab in Glasgow | Wellcome Sanger Institute for the COVID-19 Genomics UK (COG-UK) Consortium | Harper VanSteenhouse, Yumi Kasai, David Gray, Carol Clugston, Anna Dominiczak and Alex Alderton, Roberto Amato, Jeffrey Barrett, Sonia Goncalves, Ewan Harrison, David K. Jackson, Ian Johnston, Dominic Kwiatkowski, Cordelia Langford, John Sillitoe on behalf of the Wellcome Sanger Institute COVID-19 Surveillance Team |
| EPI_ISL_2513456 | Lighthouse Lab in Alderley Park | Wellcome Sanger Institute for the COVID-19 Genomics UK (COG-UK) Consortium | Jacquelyn Wynn, Mairead Hyland, The Lighthouse Lab in Alderley Park and Alex Alderton, Roberto Amato, Jeffrey Barrett, Sonia Goncalves, Ewan Harrison, David K. Jackson, Ian Johnston, Dominic Kwiatkowski, Cordelia Langford, John Sillitoe on behalf of the Wellcome Sanger Institute COVID-19 Surveillance Team |
| EPI_ISL_2513457 | Lighthouse Lab in Glasgow | Wellcome Sanger Institute for the COVID-19 Genomics UK (COG-UK) Consortium | Harper VanSteenhouse, Yumi Kasai, David Gray, Carol Clugston, Anna Dominiczak and Alex Alderton, Roberto Amato, Jeffrey Barrett, Sonia Goncalves, Ewan Harrison, David K. Jackson, Ian Johnston, Dominic Kwiatkowski, Cordelia Langford, John Sillitoe on behalf of the Wellcome Sanger Institute COVID-19 Surveillance Team |
| EPI_ISL_2513458, EPI_ISL_2513459, EPI_ISL_2513460, | Lighthouse Lab in Alderley Park | Wellcome Sanger Institute for the COVID-19 Genomics UK (COG-UK) Consortium | Jacquelyn Wynn, Mairead Hyland, The Lighthouse Lab in Alderley Park and Alex Alderton, Roberto Amato, Jeffrey Barrett, Sonia Goncalves, Ewan Harrison, David K. Jackson, Ian Johnston, Dominic Kwiatkowski, Cordelia Langford, John Sillitoe on behalf of the Wellcome Sanger Institute COVID-19 Surveillance Team |

| | | | |
|---|---|---|---|
| EPI_ISL_2513461 | | | |
| EPI_ISL_2513462 | Lighthouse Lab in Glasgow | Wellcome Sanger Institute for the COVID-19 Genomics UK (COG-UK) Consortium | Harper VanSteenhouse, Yumi Kasai, David Gray, Carol Clugston, Anna Dominiczak and Alex Alderton, Roberto Amato, Jeffrey Barrett, Sonia Goncalves, Ewan Harrison, David K. Jackson, Ian Johnston, Dominic Kwiatkowski, Cordelia Langford, John Sillitoe on behalf of the Wellcome Sanger Institute COVID-19 Surveillance Team |
| EPI_ISL_2513463 | Lighthouse Lab in Alderley Park | Wellcome Sanger Institute for the COVID-19 Genomics UK (COG-UK) Consortium | Jacquelyn Wynn, Mairead Hyland, The Lighthouse Lab in Alderley Park and Alex Alderton, Roberto Amato, Jeffrey Barrett, Sonia Goncalves, Ewan Harrison, David K. Jackson, Ian Johnston, Dominic Kwiatkowski, Cordelia Langford, John Sillitoe on behalf of the Wellcome Sanger Institute COVID-19 Surveillance Team |
| EPI_ISL_2513465 | Lighthouse Lab in Glasgow | Wellcome Sanger Institute for the COVID-19 Genomics UK (COG-UK) Consortium | Harper VanSteenhouse, Yumi Kasai, David Gray, Carol Clugston, Anna Dominiczak and Alex Alderton, Roberto Amato, Jeffrey Barrett, Sonia Goncalves, Ewan Harrison, David K. Jackson, Ian Johnston, Dominic Kwiatkowski, Cordelia Langford, John Sillitoe on behalf of the Wellcome Sanger Institute COVID-19 Surveillance Team |
| EPI_ISL_2513466, EPI_ISL_2513467, EPI_ISL_2513468 | Lighthouse Lab in Alderley Park | Wellcome Sanger Institute for the COVID-19 Genomics UK (COG-UK) Consortium | Jacquelyn Wynn, Mairead Hyland, The Lighthouse Lab in Alderley Park and Alex Alderton, Roberto Amato, Jeffrey Barrett, Sonia Goncalves, Ewan Harrison, David K. Jackson, Ian Johnston, Dominic Kwiatkowski, Cordelia Langford, John Sillitoe on behalf of the Wellcome Sanger Institute COVID-19 Surveillance Team |
| EPI_ISL_2513469, EPI_ISL_2513470 | Lighthouse Lab in Glasgow | Wellcome Sanger Institute for the COVID-19 Genomics UK (COG-UK) Consortium | Harper VanSteenhouse, Yumi Kasai, David Gray, Carol Clugston, Anna Dominiczak and Alex Alderton, Roberto Amato, Jeffrey Barrett, Sonia Goncalves, Ewan Harrison, David K. Jackson, Ian Johnston, Dominic Kwiatkowski, Cordelia Langford, John Sillitoe on behalf of the Wellcome Sanger Institute COVID-19 Surveillance Team |
| EPI_ISL_2513471, EPI_ISL_2513472 | Lighthouse Lab in Alderley Park | Wellcome Sanger Institute for the COVID-19 Genomics UK (COG-UK) Consortium | Jacquelyn Wynn, Mairead Hyland, The Lighthouse Lab in Alderley Park and Alex Alderton, Roberto Amato, Jeffrey Barrett, Sonia Goncalves, Ewan Harrison, David K. Jackson, Ian Johnston, Dominic Kwiatkowski, Cordelia Langford, John Sillitoe on behalf of the Wellcome Sanger Institute COVID-19 Surveillance Team |
| EPI_ISL_2513473 | Lighthouse Lab in Glasgow | Wellcome Sanger Institute for the COVID-19 Genomics UK (COG-UK) Consortium | Harper VanSteenhouse, Yumi Kasai, David Gray, Carol Clugston, Anna Dominiczak and Alex Alderton, Roberto Amato, Jeffrey Barrett, Sonia Goncalves, Ewan Harrison, David K. Jackson, Ian Johnston, Dominic Kwiatkowski, Cordelia Langford, John Sillitoe on behalf of the Wellcome Sanger Institute COVID-19 Surveillance Team |
| EPI_ISL_2513474, EPI_ISL_2513475 | Lighthouse Lab in Alderley Park | Wellcome Sanger Institute for the COVID-19 Genomics UK (COG-UK) Consortium | Jacquelyn Wynn, Mairead Hyland, The Lighthouse Lab in Alderley Park and Alex Alderton, Roberto Amato, Jeffrey Barrett, Sonia Goncalves, Ewan Harrison, David K. Jackson, Ian Johnston, Dominic Kwiatkowski, Cordelia Langford, John Sillitoe on behalf of the Wellcome Sanger Institute COVID-19 Surveillance Team |
| EPI_ISL_2513476 | Lighthouse Lab in Glasgow | Wellcome Sanger Institute for the COVID-19 Genomics UK (COG-UK) Consortium | Harper VanSteenhouse, Yumi Kasai, David Gray, Carol Clugston, Anna Dominiczak and Alex Alderton, Roberto Amato, Jeffrey Barrett, Sonia Goncalves, Ewan Harrison, David K. Jackson, Ian Johnston, Dominic Kwiatkowski, Cordelia Langford, John Sillitoe on behalf of the Wellcome Sanger Institute COVID-19 Surveillance Team |
| EPI_ISL_2513477, EPI_ISL_2513479, EPI_ISL_2513480, EPI_ISL_2513481, EPI_ISL_2513484, EPI_ISL_2513486, EPI_ISL_2513487, EPI_ISL_2513488, EPI_ISL_2513489 | Lighthouse Lab in Alderley Park | Wellcome Sanger Institute for the COVID-19 Genomics UK (COG-UK) Consortium | Jacquelyn Wynn, Mairead Hyland, The Lighthouse Lab in Alderley Park and Alex Alderton, Roberto Amato, Jeffrey Barrett, Sonia Goncalves, Ewan Harrison, David K. Jackson, Ian Johnston, Dominic Kwiatkowski, Cordelia Langford, John Sillitoe on behalf of the Wellcome Sanger Institute COVID-19 Surveillance Team |
| EPI_ISL_2513490 | Lighthouse Lab in Glasgow | Wellcome Sanger Institute for the COVID-19 Genomics UK (COG-UK) Consortium | Harper VanSteenhouse, Yumi Kasai, David Gray, Carol Clugston, Anna Dominiczak and Alex Alderton, Roberto Amato, Jeffrey Barrett, Sonia Goncalves, Ewan Harrison, David K. Jackson, Ian Johnston, Dominic Kwiatkowski, Cordelia Langford, John Sillitoe on behalf of the Wellcome Sanger Institute COVID-19 Surveillance Team |
| EPI_ISL_2513491, EPI_ISL_2513493, EPI_ISL_2513494, EPI_ISL_2513495 | Lighthouse Lab in Alderley Park | Wellcome Sanger Institute for the COVID-19 Genomics UK (COG-UK) Consortium | Jacquelyn Wynn, Mairead Hyland, The Lighthouse Lab in Alderley Park and Alex Alderton, Roberto Amato, Jeffrey Barrett, Sonia Goncalves, Ewan Harrison, David K. Jackson, Ian Johnston, Dominic Kwiatkowski, Cordelia Langford, John Sillitoe on behalf of the Wellcome Sanger Institute COVID-19 Surveillance Team |
| EPI_ISL_2513496, EPI_ISL_2513497, EPI_ISL_2513498, EPI_ISL_2513499 | Lighthouse Lab in Glasgow | Wellcome Sanger Institute for the COVID-19 Genomics UK (COG-UK) Consortium | Harper VanSteenhouse, Yumi Kasai, David Gray, Carol Clugston, Anna Dominiczak and Alex Alderton, Roberto Amato, Jeffrey Barrett, Sonia Goncalves, Ewan Harrison, David K. Jackson, Ian Johnston, Dominic Kwiatkowski, Cordelia Langford, John Sillitoe on behalf of the Wellcome Sanger Institute COVID-19 Surveillance Team |
| EPI_ISL_2513501 | Lighthouse Lab in Alderley Park | Wellcome Sanger Institute for the COVID-19 Genomics UK (COG-UK) Consortium | Jacquelyn Wynn, Mairead Hyland, The Lighthouse Lab in Alderley Park and Alex Alderton, Roberto Amato, Jeffrey Barrett, Sonia Goncalves, Ewan Harrison, David K. Jackson, Ian Johnston, Dominic Kwiatkowski, Cordelia Langford, John Sillitoe on behalf of the Wellcome Sanger Institute COVID-19 Surveillance Team |
| EPI_ISL_2513502 | Lighthouse Lab in Glasgow | Wellcome Sanger Institute for the COVID-19 Genomics UK (COG-UK) Consortium | Harper VanSteenhouse, Yumi Kasai, David Gray, Carol Clugston, Anna Dominiczak and Alex Alderton, Roberto Amato, Jeffrey Barrett, Sonia Goncalves, Ewan Harrison, David K. Jackson, Ian Johnston, Dominic Kwiatkowski, Cordelia Langford, John Sillitoe on behalf of the Wellcome Sanger Institute COVID-19 Surveillance Team |
| EPI_ISL_2513504 | Lighthouse Lab in Alderley Park | Wellcome Sanger Institute for the COVID-19 Genomics UK (COG-UK) Consortium | Jacquelyn Wynn, Mairead Hyland, The Lighthouse Lab in Alderley Park and Alex Alderton, Roberto Amato, Jeffrey Barrett, Sonia Goncalves, Ewan Harrison, David K. Jackson, Ian Johnston, Dominic Kwiatkowski, Cordelia Langford, John Sillitoe on behalf of the Wellcome Sanger Institute COVID-19 Surveillance Team |
| EPI_ISL_2513505 | Lighthouse Lab in Glasgow | Wellcome Sanger Institute for the COVID-19 Genomics UK (COG-UK) Consortium | Harper VanSteenhouse, Yumi Kasai, David Gray, Carol Clugston, Anna Dominiczak and Alex Alderton, Roberto Amato, Jeffrey Barrett, Sonia Goncalves, Ewan Harrison, David K. Jackson, Ian Johnston, Dominic Kwiatkowski, Cordelia Langford, John Sillitoe on behalf of the Wellcome Sanger Institute COVID-19 Surveillance Team |
| EPI_ISL_2513506, EPI_ISL_2513507, EPI_ISL_2513508, EPI_ISL_2513509, EPI_ISL_2513511, EPI_ISL_2513512, EPI_ISL_2513513, EPI_ISL_2513514 | Lighthouse Lab in Alderley Park | Wellcome Sanger Institute for the COVID-19 Genomics UK (COG-UK) Consortium | Jacquelyn Wynn, Mairead Hyland, The Lighthouse Lab in Alderley Park and Alex Alderton, Roberto Amato, Jeffrey Barrett, Sonia Goncalves, Ewan Harrison, David K. Jackson, Ian Johnston, Dominic Kwiatkowski, Cordelia Langford, John Sillitoe on behalf of the Wellcome Sanger Institute COVID-19 Surveillance Team |
| EPI_ISL_2513515 | Lighthouse Lab in Glasgow | Wellcome Sanger Institute for the COVID-19 Genomics UK (COG-UK) Consortium | Harper VanSteenhouse, Yumi Kasai, David Gray, Carol Clugston, Anna Dominiczak and Alex Alderton, Roberto Amato, Jeffrey Barrett, Sonia Goncalves, Ewan Harrison, David K. Jackson, Ian Johnston, Dominic Kwiatkowski, Cordelia Langford, John Sillitoe on behalf of the Wellcome Sanger Institute COVID-19 Surveillance Team |
| EPI_ISL_2513516 | Lighthouse Lab in Alderley Park | Wellcome Sanger Institute for the COVID-19 Genomics UK (COG-UK) Consortium | Jacquelyn Wynn, Mairead Hyland, The Lighthouse Lab in Alderley Park and Alex Alderton, Roberto Amato, Jeffrey Barrett, Sonia Goncalves, Ewan Harrison, David K. Jackson, Ian Johnston, Dominic Kwiatkowski, Cordelia Langford, John Sillitoe on behalf of the Wellcome Sanger Institute COVID-19 Surveillance Team |
| EPI_ISL_2513517, EPI_ISL_2513518 | Lighthouse Lab in Glasgow | Wellcome Sanger Institute for the COVID-19 Genomics UK (COG-UK) Consortium | Harper VanSteenhouse, Yumi Kasai, David Gray, Carol Clugston, Anna Dominiczak and Alex Alderton, Roberto Amato, Jeffrey Barrett, Sonia Goncalves, Ewan Harrison, David K. Jackson, Ian Johnston, Dominic Kwiatkowski, Cordelia Langford, John Sillitoe on behalf of the Wellcome Sanger Institute COVID-19 Surveillance Team |
| EPI_ISL_2513519, EPI_ISL_2513520, EPI_ISL_2513522, EPI_ISL_2513523 | Lighthouse Lab in Alderley Park | Wellcome Sanger Institute for the COVID-19 Genomics UK (COG-UK) Consortium | Jacquelyn Wynn, Mairead Hyland, The Lighthouse Lab in Alderley Park and Alex Alderton, Roberto Amato, Jeffrey Barrett, Sonia Goncalves, Ewan Harrison, David K. Jackson, Ian Johnston, Dominic Kwiatkowski, Cordelia Langford, John Sillitoe on behalf of the Wellcome Sanger Institute COVID-19 Surveillance Team |
| EPI_ISL_2513524, EPI_ISL_2513525, EPI_ISL_2513526 | Lighthouse Lab in Glasgow | Wellcome Sanger Institute for the COVID-19 Genomics UK (COG-UK) Consortium | Harper VanSteenhouse, Yumi Kasai, David Gray, Carol Clugston, Anna Dominiczak and Alex Alderton, Roberto Amato, Jeffrey Barrett, Sonia Goncalves, Ewan Harrison, David K. Jackson, Ian Johnston, Dominic Kwiatkowski, Cordelia Langford, John Sillitoe on behalf of the Wellcome Sanger Institute COVID-19 Surveillance Team |
| EPI_ISL_2513527, EPI_ISL_2513528, EPI_ISL_2513529, EPI_ISL_2513530, EPI_ISL_2513531 | Lighthouse Lab in Alderley Park | Wellcome Sanger Institute for the COVID-19 Genomics UK (COG-UK) Consortium | Jacquelyn Wynn, Mairead Hyland, The Lighthouse Lab in Alderley Park and Alex Alderton, Roberto Amato, Jeffrey Barrett, Sonia Goncalves, Ewan Harrison, David K. Jackson, Ian Johnston, Dominic Kwiatkowski, Cordelia Langford, John Sillitoe on behalf of the Wellcome Sanger Institute COVID-19 Surveillance Team |
| EPI_ISL_2513532, EPI_ISL_2513533 | Lighthouse Lab in Glasgow | Wellcome Sanger Institute for the COVID-19 Genomics UK (COG-UK) Consortium | Harper VanSteenhouse, Yumi Kasai, David Gray, Carol Clugston, Anna Dominiczak and Alex Alderton, Roberto Amato, Jeffrey Barrett, Sonia Goncalves, Ewan Harrison, David K. Jackson, Ian Johnston, Dominic Kwiatkowski, Cordelia Langford, John Sillitoe on behalf of the Wellcome Sanger Institute COVID-19 Surveillance Team |
| EPI_ISL_2513534, EPI_ISL_2513535 | Lighthouse Lab in Alderley Park | Wellcome Sanger Institute for the COVID-19 Genomics UK (COG-UK) Consortium | Jacquelyn Wynn, Mairead Hyland, The Lighthouse Lab in Alderley Park and Alex Alderton, Roberto Amato, Jeffrey Barrett, Sonia Goncalves, Ewan Harrison, David K. Jackson, Ian Johnston, Dominic Kwiatkowski, Cordelia Langford, John Sillitoe on behalf of the Wellcome Sanger Institute COVID-19 Surveillance Team |
| EPI_ISL_2513536 | Lighthouse Lab in Glasgow | Wellcome Sanger Institute for the COVID-19 Genomics UK (COG-UK) Consortium | Harper VanSteenhouse, Yumi Kasai, David Gray, Carol Clugston, Anna Dominiczak and Alex Alderton, Roberto Amato, Jeffrey Barrett, Sonia Goncalves, Ewan Harrison, David K. Jackson, Ian Johnston, Dominic Kwiatkowski, Cordelia Langford, John Sillitoe on behalf of the Wellcome Sanger Institute COVID-19 Surveillance Team |
| EPI_ISL_2513537 | Lighthouse Lab in Alderley Park | Wellcome Sanger Institute for the COVID-19 Genomics UK (COG-UK) Consortium | Jacquelyn Wynn, Mairead Hyland, The Lighthouse Lab in Alderley Park and Alex Alderton, Roberto Amato, Jeffrey Barrett, Sonia Goncalves, Ewan Harrison, David K. Jackson, Ian Johnston, Dominic Kwiatkowski, Cordelia Langford, John Sillitoe on behalf of the Wellcome Sanger Institute COVID-19 Surveillance Team |
| EPI_ISL_2513538 | Lighthouse Lab in Glasgow | Wellcome Sanger Institute for the COVID-19 Genomics UK (COG-UK) Consortium | Harper VanSteenhouse, Yumi Kasai, David Gray, Carol Clugston, Anna Dominiczak and Alex Alderton, Roberto Amato, Jeffrey Barrett, Sonia Goncalves, Ewan Harrison, David K. Jackson, Ian Johnston, Dominic Kwiatkowski, Cordelia Langford, John Sillitoe on behalf of the Wellcome Sanger Institute COVID-19 Surveillance Team |
| EPI_ISL_2513539 | Lighthouse Lab in Alderley Park | Wellcome Sanger Institute for the COVID-19 Genomics UK (COG-UK) Consortium | Jacquelyn Wynn, Mairead Hyland, The Lighthouse Lab in Alderley Park and Alex Alderton, Roberto Amato, Jeffrey Barrett, Sonia Goncalves, Ewan Harrison, David K. Jackson, Ian Johnston, Dominic Kwiatkowski, Cordelia Langford, John Sillitoe on behalf of the Wellcome Sanger Institute COVID-19 Surveillance Team |
| EPI_ISL_2513541 | Lighthouse Lab in Glasgow | Wellcome Sanger Institute for the COVID-19 Genomics UK (COG-UK) Consortium | Harper VanSteenhouse, Yumi Kasai, David Gray, Carol Clugston, Anna Dominiczak and Alex Alderton, Roberto Amato, Jeffrey Barrett, Sonia Goncalves, Ewan Harrison, David K. Jackson, Ian Johnston, Dominic Kwiatkowski, Cordelia Langford, John Sillitoe on behalf of the Wellcome Sanger Institute COVID-19 Surveillance Team |
| EPI_ISL_2513543 | Lighthouse Lab in Alderley Park | Wellcome Sanger Institute for the COVID-19 Genomics UK (COG-UK) Consortium | Jacquelyn Wynn, Mairead Hyland, The Lighthouse Lab in Alderley Park and Alex Alderton, Roberto Amato, Jeffrey Barrett, Sonia Goncalves, Ewan Harrison, David K. Jackson, Ian Johnston, Dominic Kwiatkowski, Cordelia Langford, John Sillitoe on behalf of the Wellcome Sanger Institute COVID-19 Surveillance Team |
| EPI_ISL_2513544 | Lighthouse Lab in Glasgow | Wellcome Sanger Institute for the COVID-19 Genomics UK (COG-UK) Consortium | Harper VanSteenhouse, Yumi Kasai, David Gray, Carol Clugston, Anna Dominiczak and Alex Alderton, Roberto Amato, Jeffrey Barrett, Sonia Goncalves, Ewan Harrison, David K. Jackson, Ian Johnston, Dominic Kwiatkowski, Cordelia Langford, John Sillitoe on behalf of the Wellcome Sanger Institute COVID-19 Surveillance Team |
| EPI_ISL_2513545, EPI_ISL_2513546, EPI_ISL_2513547, EPI_ISL_2513548, | Lighthouse Lab in Alderley Park | Wellcome Sanger Institute for the COVID-19 Genomics UK (COG-UK) Consortium | Jacquelyn Wynn, Mairead Hyland, The Lighthouse Lab in Alderley Park and Alex Alderton, Roberto Amato, Jeffrey Barrett, Sonia Goncalves, Ewan Harrison, David K. Jackson, Ian Johnston, Dominic Kwiatkowski, Cordelia Langford, John Sillitoe on behalf of the Wellcome Sanger Institute COVID-19 Surveillance Team |

| | | | |
|---|---|---|---|
| EPI_ISL_2513549 | | | |
| EPI_ISL_2513550 | Lighthouse Lab in Glasgow | Wellcome Sanger Institute for the COVID-19 Genomics UK (COG-UK) Consortium | Harper VanSteenhouse, Yumi Kasai, David Gray, Carol Clugston, Anna Dominiczak and Alex Alderton, Roberto Amato, Jeffrey Barrett, Sonia Goncalves, Ewan Harrison, David K. Jackson, Ian Johnston, Dominic Kwiatkowski, Cordelia Langford, John Sillitoe on behalf of the Wellcome Sanger Institute COVID-19 Surveillance Team |
| EPI_ISL_2513551, EPI_ISL_2513552, EPI_ISL_2513554, EPI_ISL_2513555, EPI_ISL_2513556 | Lighthouse Lab in Alderley Park | Wellcome Sanger Institute for the COVID-19 Genomics UK (COG-UK) Consortium | Jacquelyn Wynn, Mairead Hyland, The Lighthouse Lab in Alderley Park and Alex Alderton, Roberto Amato, Jeffrey Barrett, Sonia Goncalves, Ewan Harrison, David K. Jackson, Ian Johnston, Dominic Kwiatkowski, Cordelia Langford, John Sillitoe on behalf of the Wellcome Sanger Institute COVID-19 Surveillance Team |
| EPI_ISL_2513557, EPI_ISL_2513558 | Lighthouse Lab in Glasgow | Wellcome Sanger Institute for the COVID-19 Genomics UK (COG-UK) Consortium | Harper VanSteenhouse, Yumi Kasai, David Gray, Carol Clugston, Anna Dominiczak and Alex Alderton, Roberto Amato, Jeffrey Barrett, Sonia Goncalves, Ewan Harrison, David K. Jackson, Ian Johnston, Dominic Kwiatkowski, Cordelia Langford, John Sillitoe on behalf of the Wellcome Sanger Institute COVID-19 Surveillance Team |
| EPI_ISL_2513559, EPI_ISL_2513560, EPI_ISL_2513561, EPI_ISL_2513562, EPI_ISL_2513565 | Lighthouse Lab in Alderley Park | Wellcome Sanger Institute for the COVID-19 Genomics UK (COG-UK) Consortium | Jacquelyn Wynn, Mairead Hyland, The Lighthouse Lab in Alderley Park and Alex Alderton, Roberto Amato, Jeffrey Barrett, Sonia Goncalves, Ewan Harrison, David K. Jackson, Ian Johnston, Dominic Kwiatkowski, Cordelia Langford, John Sillitoe on behalf of the Wellcome Sanger Institute COVID-19 Surveillance Team |
| EPI_ISL_2513566, EPI_ISL_2513568 | Lighthouse Lab in Glasgow | Wellcome Sanger Institute for the COVID-19 Genomics UK (COG-UK) Consortium | Harper VanSteenhouse, Yumi Kasai, David Gray, Carol Clugston, Anna Dominiczak and Alex Alderton, Roberto Amato, Jeffrey Barrett, Sonia Goncalves, Ewan Harrison, David K. Jackson, Ian Johnston, Dominic Kwiatkowski, Cordelia Langford, John Sillitoe on behalf of the Wellcome Sanger Institute COVID-19 Surveillance Team |
| EPI_ISL_2513569, EPI_ISL_2513570, EPI_ISL_2513571, EPI_ISL_2513572, EPI_ISL_2513574 | Lighthouse Lab in Alderley Park | Wellcome Sanger Institute for the COVID-19 Genomics UK (COG-UK) Consortium | Jacquelyn Wynn, Mairead Hyland, The Lighthouse Lab in Alderley Park and Alex Alderton, Roberto Amato, Jeffrey Barrett, Sonia Goncalves, Ewan Harrison, David K. Jackson, Ian Johnston, Dominic Kwiatkowski, Cordelia Langford, John Sillitoe on behalf of the Wellcome Sanger Institute COVID-19 Surveillance Team |
| EPI_ISL_2513575 | Lighthouse Lab in Glasgow | Wellcome Sanger Institute for the COVID-19 Genomics UK (COG-UK) Consortium | Harper VanSteenhouse, Yumi Kasai, David Gray, Carol Clugston, Anna Dominiczak and Alex Alderton, Roberto Amato, Jeffrey Barrett, Sonia Goncalves, Ewan Harrison, David K. Jackson, Ian Johnston, Dominic Kwiatkowski, Cordelia Langford, John Sillitoe on behalf of the Wellcome Sanger Institute COVID-19 Surveillance Team |
| EPI_ISL_2513576, EPI_ISL_2513577, EPI_ISL_2513578 | Lighthouse Lab in Alderley Park | Wellcome Sanger Institute for the COVID-19 Genomics UK (COG-UK) Consortium | Jacquelyn Wynn, Mairead Hyland, The Lighthouse Lab in Alderley Park and Alex Alderton, Roberto Amato, Jeffrey Barrett, Sonia Goncalves, Ewan Harrison, David K. Jackson, Ian Johnston, Dominic Kwiatkowski, Cordelia Langford, John Sillitoe on behalf of the Wellcome Sanger Institute COVID-19 Surveillance Team |
| EPI_ISL_2513579 | Lighthouse Lab in Glasgow | Wellcome Sanger Institute for the COVID-19 Genomics UK (COG-UK) Consortium | Harper VanSteenhouse, Yumi Kasai, David Gray, Carol Clugston, Anna Dominiczak and Alex Alderton, Roberto Amato, Jeffrey Barrett, Sonia Goncalves, Ewan Harrison, David K. Jackson, Ian Johnston, Dominic Kwiatkowski, Cordelia Langford, John Sillitoe on behalf of the Wellcome Sanger Institute COVID-19 Surveillance Team |
| EPI_ISL_2513580, EPI_ISL_2513581, EPI_ISL_2513582, EPI_ISL_2513583, EPI_ISL_2513584, EPI_ISL_2513585, EPI_ISL_2513586 | Lighthouse Lab in Alderley Park | Wellcome Sanger Institute for the COVID-19 Genomics UK (COG-UK) Consortium | Jacquelyn Wynn, Mairead Hyland, The Lighthouse Lab in Alderley Park and Alex Alderton, Roberto Amato, Jeffrey Barrett, Sonia Goncalves, Ewan Harrison, David K. Jackson, Ian Johnston, Dominic Kwiatkowski, Cordelia Langford, John Sillitoe on behalf of the Wellcome Sanger Institute COVID-19 Surveillance Team |
| EPI_ISL_2513587, EPI_ISL_2513588 | Lighthouse Lab in Glasgow | Wellcome Sanger Institute for the COVID-19 Genomics UK (COG-UK) Consortium | Harper VanSteenhouse, Yumi Kasai, David Gray, Carol Clugston, Anna Dominiczak and Alex Alderton, Roberto Amato, Jeffrey Barrett, Sonia Goncalves, Ewan Harrison, David K. Jackson, Ian Johnston, Dominic Kwiatkowski, Cordelia Langford, John Sillitoe on behalf of the Wellcome Sanger Institute COVID-19 Surveillance Team |
| EPI_ISL_2513589, EPI_ISL_2513590 | Lighthouse Lab in Alderley Park | Wellcome Sanger Institute for the COVID-19 Genomics UK (COG-UK) Consortium | Jacquelyn Wynn, Mairead Hyland, The Lighthouse Lab in Alderley Park and Alex Alderton, Roberto Amato, Jeffrey Barrett, Sonia Goncalves, Ewan Harrison, David K. Jackson, Ian Johnston, Dominic Kwiatkowski, Cordelia Langford, John Sillitoe on behalf of the Wellcome Sanger Institute COVID-19 Surveillance Team |
| EPI_ISL_2513591 | Lighthouse Lab in Glasgow | Wellcome Sanger Institute for the COVID-19 Genomics UK (COG-UK) Consortium | Harper VanSteenhouse, Yumi Kasai, David Gray, Carol Clugston, Anna Dominiczak and Alex Alderton, Roberto Amato, Jeffrey Barrett, Sonia Goncalves, Ewan Harrison, David K. Jackson, Ian Johnston, Dominic Kwiatkowski, Cordelia Langford, John Sillitoe on behalf of the Wellcome Sanger Institute COVID-19 Surveillance Team |
| EPI_ISL_2513592, EPI_ISL_2513594, EPI_ISL_2513595 | Lighthouse Lab in Alderley Park | Wellcome Sanger Institute for the COVID-19 Genomics UK (COG-UK) Consortium | Jacquelyn Wynn, Mairead Hyland, The Lighthouse Lab in Alderley Park and Alex Alderton, Roberto Amato, Jeffrey Barrett, Sonia Goncalves, Ewan Harrison, David K. Jackson, Ian Johnston, Dominic Kwiatkowski, Cordelia Langford, John Sillitoe on behalf of the Wellcome Sanger Institute COVID-19 Surveillance Team |
| EPI_ISL_2513596 | Lighthouse Lab in Glasgow | Wellcome Sanger Institute for the COVID-19 Genomics UK (COG-UK) Consortium | Harper VanSteenhouse, Yumi Kasai, David Gray, Carol Clugston, Anna Dominiczak and Alex Alderton, Roberto Amato, Jeffrey Barrett, Sonia Goncalves, Ewan Harrison, David K. Jackson, Ian Johnston, Dominic Kwiatkowski, Cordelia Langford, John Sillitoe on behalf of the Wellcome Sanger Institute COVID-19 Surveillance Team |
| EPI_ISL_2513597, EPI_ISL_2513598, EPI_ISL_2513599, EPI_ISL_2513600, EPI_ISL_2513601, EPI_ISL_2513602, EPI_ISL_2513603, EPI_ISL_2513604, EPI_ISL_2513605, EPI_ISL_2513606 | Lighthouse Lab in Alderley Park | Wellcome Sanger Institute for the COVID-19 Genomics UK (COG-UK) Consortium | Jacquelyn Wynn, Mairead Hyland, The Lighthouse Lab in Alderley Park and Alex Alderton, Roberto Amato, Jeffrey Barrett, Sonia Goncalves, Ewan Harrison, David K. Jackson, Ian Johnston, Dominic Kwiatkowski, Cordelia Langford, John Sillitoe on behalf of the Wellcome Sanger Institute COVID-19 Surveillance Team |
| EPI_ISL_2513607 | Lighthouse Lab in Glasgow | Wellcome Sanger Institute for the COVID-19 Genomics UK (COG-UK) Consortium | Harper VanSteenhouse, Yumi Kasai, David Gray, Carol Clugston, Anna Dominiczak and Alex Alderton, Roberto Amato, Jeffrey Barrett, Sonia Goncalves, Ewan Harrison, David K. Jackson, Ian Johnston, Dominic Kwiatkowski, Cordelia Langford, John Sillitoe on behalf of the Wellcome Sanger Institute COVID-19 Surveillance Team |
| EPI_ISL_2513608 | Lighthouse Lab in Alderley Park | Wellcome Sanger Institute for the COVID-19 Genomics UK (COG-UK) Consortium | Jacquelyn Wynn, Mairead Hyland, The Lighthouse Lab in Alderley Park and Alex Alderton, Roberto Amato, Jeffrey Barrett, Sonia Goncalves, Ewan Harrison, David K. Jackson, Ian Johnston, Dominic Kwiatkowski, Cordelia Langford, John Sillitoe on behalf of the Wellcome Sanger Institute COVID-19 Surveillance Team |
| EPI_ISL_2513609 | Lighthouse Lab in Glasgow | Wellcome Sanger Institute for the COVID-19 Genomics UK (COG-UK) Consortium | Harper VanSteenhouse, Yumi Kasai, David Gray, Carol Clugston, Anna Dominiczak and Alex Alderton, Roberto Amato, Jeffrey Barrett, Sonia Goncalves, Ewan Harrison, David K. Jackson, Ian Johnston, Dominic Kwiatkowski, Cordelia Langford, John Sillitoe on behalf of the Wellcome Sanger Institute COVID-19 Surveillance Team |
| EPI_ISL_2513610, EPI_ISL_2513611, EPI_ISL_2513612 | Lighthouse Lab in Alderley Park | Wellcome Sanger Institute for the COVID-19 Genomics UK (COG-UK) Consortium | Jacquelyn Wynn, Mairead Hyland, The Lighthouse Lab in Alderley Park and Alex Alderton, Roberto Amato, Jeffrey Barrett, Sonia Goncalves, Ewan Harrison, David K. Jackson, Ian Johnston, Dominic Kwiatkowski, Cordelia Langford, John Sillitoe on behalf of the Wellcome Sanger Institute COVID-19 Surveillance Team |
| EPI_ISL_2513613, EPI_ISL_2513614 | Lighthouse Lab in Glasgow | Wellcome Sanger Institute for the COVID-19 Genomics UK (COG-UK) Consortium | Harper VanSteenhouse, Yumi Kasai, David Gray, Carol Clugston, Anna Dominiczak and Alex Alderton, Roberto Amato, Jeffrey Barrett, Sonia Goncalves, Ewan Harrison, David K. Jackson, Ian Johnston, Dominic Kwiatkowski, Cordelia Langford, John Sillitoe on behalf of the Wellcome Sanger Institute COVID-19 Surveillance Team |
| EPI_ISL_2513615 | Lighthouse Lab in Alderley Park | Wellcome Sanger Institute for the COVID-19 Genomics UK (COG-UK) Consortium | Jacquelyn Wynn, Mairead Hyland, The Lighthouse Lab in Alderley Park and Alex Alderton, Roberto Amato, Jeffrey Barrett, Sonia Goncalves, Ewan Harrison, David K. Jackson, Ian Johnston, Dominic Kwiatkowski, Cordelia Langford, John Sillitoe on behalf of the Wellcome Sanger Institute COVID-19 Surveillance Team |
| EPI_ISL_2513617 | Lighthouse Lab in Glasgow | Wellcome Sanger Institute for the COVID-19 Genomics UK (COG-UK) Consortium | Harper VanSteenhouse, Yumi Kasai, David Gray, Carol Clugston, Anna Dominiczak and Alex Alderton, Roberto Amato, Jeffrey Barrett, Sonia Goncalves, Ewan Harrison, David K. Jackson, Ian Johnston, Dominic Kwiatkowski, Cordelia Langford, John Sillitoe on behalf of the Wellcome Sanger Institute COVID-19 Surveillance Team |
| EPI_ISL_2513618 | Lighthouse Lab in Alderley Park | Wellcome Sanger Institute for the COVID-19 Genomics UK (COG-UK) Consortium | Jacquelyn Wynn, Mairead Hyland, The Lighthouse Lab in Alderley Park and Alex Alderton, Roberto Amato, Jeffrey Barrett, Sonia Goncalves, Ewan Harrison, David K. Jackson, Ian Johnston, Dominic Kwiatkowski, Cordelia Langford, John Sillitoe on behalf of the Wellcome Sanger Institute COVID-19 Surveillance Team |
| EPI_ISL_2513619, EPI_ISL_2513620 | Lighthouse Lab in Glasgow | Wellcome Sanger Institute for the COVID-19 Genomics UK (COG-UK) Consortium | Harper VanSteenhouse, Yumi Kasai, David Gray, Carol Clugston, Anna Dominiczak and Alex Alderton, Roberto Amato, Jeffrey Barrett, Sonia Goncalves, Ewan Harrison, David K. Jackson, Ian Johnston, Dominic Kwiatkowski, Cordelia Langford, John Sillitoe on behalf of the Wellcome Sanger Institute COVID-19 Surveillance Team |
| EPI_ISL_2513621, EPI_ISL_2513622, EPI_ISL_2513623 | Lighthouse Lab in Alderley Park | Wellcome Sanger Institute for the COVID-19 Genomics UK (COG-UK) Consortium | Jacquelyn Wynn, Mairead Hyland, The Lighthouse Lab in Alderley Park and Alex Alderton, Roberto Amato, Jeffrey Barrett, Sonia Goncalves, Ewan Harrison, David K. Jackson, Ian Johnston, Dominic Kwiatkowski, Cordelia Langford, John Sillitoe on behalf of the Wellcome Sanger Institute COVID-19 Surveillance Team |
| EPI_ISL_2513624 | Lighthouse Lab in Glasgow | Wellcome Sanger Institute for the COVID-19 Genomics UK (COG-UK) Consortium | Harper VanSteenhouse, Yumi Kasai, David Gray, Carol Clugston, Anna Dominiczak and Alex Alderton, Roberto Amato, Jeffrey Barrett, Sonia Goncalves, Ewan Harrison, David K. Jackson, Ian Johnston, Dominic Kwiatkowski, Cordelia Langford, John Sillitoe on behalf of the Wellcome Sanger Institute COVID-19 Surveillance Team |
| EPI_ISL_2513625, EPI_ISL_2513626 | Lighthouse Lab in Alderley Park | Wellcome Sanger Institute for the COVID-19 Genomics UK (COG-UK) Consortium | Jacquelyn Wynn, Mairead Hyland, The Lighthouse Lab in Alderley Park and Alex Alderton, Roberto Amato, Jeffrey Barrett, Sonia Goncalves, Ewan Harrison, David K. Jackson, Ian Johnston, Dominic Kwiatkowski, Cordelia Langford, John Sillitoe on behalf of the Wellcome Sanger Institute COVID-19 Surveillance Team |
| EPI_ISL_2513627 | Lighthouse Lab in Glasgow | Wellcome Sanger Institute for the COVID-19 Genomics UK (COG-UK) Consortium | Harper VanSteenhouse, Yumi Kasai, David Gray, Carol Clugston, Anna Dominiczak and Alex Alderton, Roberto Amato, Jeffrey Barrett, Sonia Goncalves, Ewan Harrison, David K. Jackson, Ian Johnston, Dominic Kwiatkowski, Cordelia Langford, John Sillitoe on behalf of the Wellcome Sanger Institute COVID-19 Surveillance Team |
| EPI_ISL_2513628 | Lighthouse Lab in Alderley Park | Wellcome Sanger Institute for the COVID-19 Genomics UK (COG-UK) Consortium | Jacquelyn Wynn, Mairead Hyland, The Lighthouse Lab in Alderley Park and Alex Alderton, Roberto Amato, Jeffrey Barrett, Sonia Goncalves, Ewan Harrison, David K. Jackson, Ian Johnston, Dominic Kwiatkowski, Cordelia Langford, John Sillitoe on behalf of the Wellcome Sanger Institute COVID-19 Surveillance Team |
| EPI_ISL_2513630 | Lighthouse Lab in Glasgow | Wellcome Sanger Institute for the COVID-19 Genomics UK (COG-UK) Consortium | Harper VanSteenhouse, Yumi Kasai, David Gray, Carol Clugston, Anna Dominiczak and Alex Alderton, Roberto Amato, Jeffrey Barrett, Sonia Goncalves, Ewan Harrison, David K. Jackson, Ian Johnston, Dominic Kwiatkowski, Cordelia Langford, John Sillitoe on behalf of the Wellcome Sanger Institute COVID-19 Surveillance Team |
| EPI_ISL_2513631 | Lighthouse Lab in Alderley Park | Wellcome Sanger Institute for the COVID-19 Genomics UK (COG-UK) Consortium | Jacquelyn Wynn, Mairead Hyland, The Lighthouse Lab in Alderley Park and Alex Alderton, Roberto Amato, Jeffrey Barrett, Sonia Goncalves, Ewan Harrison, David K. Jackson, Ian Johnston, Dominic Kwiatkowski, Cordelia Langford, John Sillitoe on behalf of the Wellcome Sanger Institute COVID-19 Surveillance Team |
| EPI_ISL_2513632, EPI_ISL_2513633, EPI_ISL_2513634 | Lighthouse Lab in Glasgow | Wellcome Sanger Institute for the COVID-19 Genomics UK (COG-UK) Consortium | Harper VanSteenhouse, Yumi Kasai, David Gray, Carol Clugston, Anna Dominiczak and Alex Alderton, Roberto Amato, Jeffrey Barrett, Sonia Goncalves, Ewan Harrison, David K. Jackson, Ian Johnston, Dominic Kwiatkowski, Cordelia Langford, John Sillitoe on behalf of the Wellcome Sanger Institute COVID-19 Surveillance Team |
| EPI_ISL_2513635 | Lighthouse Lab in Alderley Park | Wellcome Sanger Institute for the COVID-19 Genomics UK (COG-UK) | Jacquelyn Wynn, Mairead Hyland, The Lighthouse Lab in Alderley Park and Alex Alderton, Roberto Amato, Jeffrey Barrett, Sonia Goncalves, Ewan Harrison, David K. Jackson, Ian Johnston, Dominic Kwiatkowski, Cordelia Langford, John Sillitoe on |

| | | | |
|---|---|---|---|
| | | Consortium | behalf of the Wellcome Sanger Institute COVID-19 Surveillance Team |
| EPI_ISL_2513636, EPI_ISL_2513637, EPI_ISL_2513638, EPI_ISL_2513639, EPI_ISL_2513642, EPI_ISL_2513643, EPI_ISL_2513644, EPI_ISL_2513645 | Lighthouse Lab in Glasgow | Wellcome Sanger Institute for the COVID-19 Genomics UK (COG-UK) Consortium | Harper VanSteenhouse, Yumi Kasai, David Gray, Carol Clugston, Anna Dominiczak and Alex Alderton, Roberto Amato, Jeffrey Barrett, Sonia Goncalves, Ewan Harrison, David K. Jackson, Ian Johnston, Dominic Kwiatkowski, Cordelia Langford, John Sillitoe on behalf of the Wellcome Sanger Institute COVID-19 Surveillance Team |
| EPI_ISL_2513646, EPI_ISL_2513649 | Lighthouse Lab in Alderley Park | Wellcome Sanger Institute for the COVID-19 Genomics UK (COG-UK) Consortium | Jacquelyn Wynn, Mairead Hyland, The Lighthouse Lab in Alderley Park and Alex Alderton, Roberto Amato, Jeffrey Barrett, Sonia Goncalves, Ewan Harrison, David K. Jackson, Ian Johnston, Dominic Kwiatkowski, Cordelia Langford, John Sillitoe on behalf of the Wellcome Sanger Institute COVID-19 Surveillance Team |
| EPI_ISL_2513650, EPI_ISL_2513651, EPI_ISL_2513652 | Lighthouse Lab in Glasgow | Wellcome Sanger Institute for the COVID-19 Genomics UK (COG-UK) Consortium | Harper VanSteenhouse, Yumi Kasai, David Gray, Carol Clugston, Anna Dominiczak and Alex Alderton, Roberto Amato, Jeffrey Barrett, Sonia Goncalves, Ewan Harrison, David K. Jackson, Ian Johnston, Dominic Kwiatkowski, Cordelia Langford, John Sillitoe on behalf of the Wellcome Sanger Institute COVID-19 Surveillance Team |
| EPI_ISL_2513653 | Lighthouse Lab in Alderley Park | Wellcome Sanger Institute for the COVID-19 Genomics UK (COG-UK) Consortium | Jacquelyn Wynn, Mairead Hyland, The Lighthouse Lab in Alderley Park and Alex Alderton, Roberto Amato, Jeffrey Barrett, Sonia Goncalves, Ewan Harrison, David K. Jackson, Ian Johnston, Dominic Kwiatkowski, Cordelia Langford, John Sillitoe on behalf of the Wellcome Sanger Institute COVID-19 Surveillance Team |
| EPI_ISL_2513654 | Lighthouse Lab in Glasgow | Wellcome Sanger Institute for the COVID-19 Genomics UK (COG-UK) Consortium | Harper VanSteenhouse, Yumi Kasai, David Gray, Carol Clugston, Anna Dominiczak and Alex Alderton, Roberto Amato, Jeffrey Barrett, Sonia Goncalves, Ewan Harrison, David K. Jackson, Ian Johnston, Dominic Kwiatkowski, Cordelia Langford, John Sillitoe on behalf of the Wellcome Sanger Institute COVID-19 Surveillance Team |
| EPI_ISL_2513656 | Lighthouse Lab in Alderley Park | Wellcome Sanger Institute for the COVID-19 Genomics UK (COG-UK) Consortium | Jacquelyn Wynn, Mairead Hyland, The Lighthouse Lab in Alderley Park and Alex Alderton, Roberto Amato, Jeffrey Barrett, Sonia Goncalves, Ewan Harrison, David K. Jackson, Ian Johnston, Dominic Kwiatkowski, Cordelia Langford, John Sillitoe on behalf of the Wellcome Sanger Institute COVID-19 Surveillance Team |
| EPI_ISL_2513657 | Lighthouse Lab in Glasgow | Wellcome Sanger Institute for the COVID-19 Genomics UK (COG-UK) Consortium | Harper VanSteenhouse, Yumi Kasai, David Gray, Carol Clugston, Anna Dominiczak and Alex Alderton, Roberto Amato, Jeffrey Barrett, Sonia Goncalves, Ewan Harrison, David K. Jackson, Ian Johnston, Dominic Kwiatkowski, Cordelia Langford, John Sillitoe on behalf of the Wellcome Sanger Institute COVID-19 Surveillance Team |
| EPI_ISL_2513658, EPI_ISL_2513659 | Lighthouse Lab in Alderley Park | Wellcome Sanger Institute for the COVID-19 Genomics UK (COG-UK) Consortium | Jacquelyn Wynn, Mairead Hyland, The Lighthouse Lab in Alderley Park and Alex Alderton, Roberto Amato, Jeffrey Barrett, Sonia Goncalves, Ewan Harrison, David K. Jackson, Ian Johnston, Dominic Kwiatkowski, Cordelia Langford, John Sillitoe on behalf of the Wellcome Sanger Institute COVID-19 Surveillance Team |
| EPI_ISL_2513661 | Lighthouse Lab in Glasgow | Wellcome Sanger Institute for the COVID-19 Genomics UK (COG-UK) Consortium | Harper VanSteenhouse, Yumi Kasai, David Gray, Carol Clugston, Anna Dominiczak and Alex Alderton, Roberto Amato, Jeffrey Barrett, Sonia Goncalves, Ewan Harrison, David K. Jackson, Ian Johnston, Dominic Kwiatkowski, Cordelia Langford, John Sillitoe on behalf of the Wellcome Sanger Institute COVID-19 Surveillance Team |
| EPI_ISL_2513663, EPI_ISL_2513664 | Lighthouse Lab in Alderley Park | Wellcome Sanger Institute for the COVID-19 Genomics UK (COG-UK) Consortium | Jacquelyn Wynn, Mairead Hyland, The Lighthouse Lab in Alderley Park and Alex Alderton, Roberto Amato, Jeffrey Barrett, Sonia Goncalves, Ewan Harrison, David K. Jackson, Ian Johnston, Dominic Kwiatkowski, Cordelia Langford, John Sillitoe on behalf of the Wellcome Sanger Institute COVID-19 Surveillance Team |
| EPI_ISL_2513665, EPI_ISL_2513666 | Lighthouse Lab in Glasgow | Wellcome Sanger Institute for the COVID-19 Genomics UK (COG-UK) Consortium | Harper VanSteenhouse, Yumi Kasai, David Gray, Carol Clugston, Anna Dominiczak and Alex Alderton, Roberto Amato, Jeffrey Barrett, Sonia Goncalves, Ewan Harrison, David K. Jackson, Ian Johnston, Dominic Kwiatkowski, Cordelia Langford, John Sillitoe on behalf of the Wellcome Sanger Institute COVID-19 Surveillance Team |
| EPI_ISL_2513667, EPI_ISL_2513668 | Lighthouse Lab in Alderley Park | Wellcome Sanger Institute for the COVID-19 Genomics UK (COG-UK) Consortium | Jacquelyn Wynn, Mairead Hyland, The Lighthouse Lab in Alderley Park and Alex Alderton, Roberto Amato, Jeffrey Barrett, Sonia Goncalves, Ewan Harrison, David K. Jackson, Ian Johnston, Dominic Kwiatkowski, Cordelia Langford, John Sillitoe on behalf of the Wellcome Sanger Institute COVID-19 Surveillance Team |
| EPI_ISL_2513669 | Lighthouse Lab in Glasgow | Wellcome Sanger Institute for the COVID-19 Genomics UK (COG-UK) Consortium | Harper VanSteenhouse, Yumi Kasai, David Gray, Carol Clugston, Anna Dominiczak and Alex Alderton, Roberto Amato, Jeffrey Barrett, Sonia Goncalves, Ewan Harrison, David K. Jackson, Ian Johnston, Dominic Kwiatkowski, Cordelia Langford, John Sillitoe on behalf of the Wellcome Sanger Institute COVID-19 Surveillance Team |
| EPI_ISL_2513670, EPI_ISL_2513671 | Lighthouse Lab in Alderley Park | Wellcome Sanger Institute for the COVID-19 Genomics UK (COG-UK) Consortium | Jacquelyn Wynn, Mairead Hyland, The Lighthouse Lab in Alderley Park and Alex Alderton, Roberto Amato, Jeffrey Barrett, Sonia Goncalves, Ewan Harrison, David K. Jackson, Ian Johnston, Dominic Kwiatkowski, Cordelia Langford, John Sillitoe on behalf of the Wellcome Sanger Institute COVID-19 Surveillance Team |
| EPI_ISL_2513673 | Lighthouse Lab in Glasgow | Wellcome Sanger Institute for the COVID-19 Genomics UK (COG-UK) Consortium | Harper VanSteenhouse, Yumi Kasai, David Gray, Carol Clugston, Anna Dominiczak and Alex Alderton, Roberto Amato, Jeffrey Barrett, Sonia Goncalves, Ewan Harrison, David K. Jackson, Ian Johnston, Dominic Kwiatkowski, Cordelia Langford, John Sillitoe on behalf of the Wellcome Sanger Institute COVID-19 Surveillance Team |
| EPI_ISL_2513675 | Lighthouse Lab in Alderley Park | Wellcome Sanger Institute for the COVID-19 Genomics UK (COG-UK) Consortium | Jacquelyn Wynn, Mairead Hyland, The Lighthouse Lab in Alderley Park and Alex Alderton, Roberto Amato, Jeffrey Barrett, Sonia Goncalves, Ewan Harrison, David K. Jackson, Ian Johnston, Dominic Kwiatkowski, Cordelia Langford, John Sillitoe on behalf of the Wellcome Sanger Institute COVID-19 Surveillance Team |
| EPI_ISL_2513676 | Lighthouse Lab in Glasgow | Wellcome Sanger Institute for the COVID-19 Genomics UK (COG-UK) Consortium | Harper VanSteenhouse, Yumi Kasai, David Gray, Carol Clugston, Anna Dominiczak and Alex Alderton, Roberto Amato, Jeffrey Barrett, Sonia Goncalves, Ewan Harrison, David K. Jackson, Ian Johnston, Dominic Kwiatkowski, Cordelia Langford, John Sillitoe on behalf of the Wellcome Sanger Institute COVID-19 Surveillance Team |
| EPI_ISL_2513677 | Lighthouse Lab in Alderley Park | Wellcome Sanger Institute for the COVID-19 Genomics UK (COG-UK) Consortium | Jacquelyn Wynn, Mairead Hyland, The Lighthouse Lab in Alderley Park and Alex Alderton, Roberto Amato, Jeffrey Barrett, Sonia Goncalves, Ewan Harrison, David K. Jackson, Ian Johnston, Dominic Kwiatkowski, Cordelia Langford, John Sillitoe on behalf of the Wellcome Sanger Institute COVID-19 Surveillance Team |
| EPI_ISL_2513678 | Lighthouse Lab in Glasgow | Wellcome Sanger Institute for the COVID-19 Genomics UK (COG-UK) Consortium | Harper VanSteenhouse, Yumi Kasai, David Gray, Carol Clugston, Anna Dominiczak and Alex Alderton, Roberto Amato, Jeffrey Barrett, Sonia Goncalves, Ewan Harrison, David K. Jackson, Ian Johnston, Dominic Kwiatkowski, Cordelia Langford, John Sillitoe on behalf of the Wellcome Sanger Institute COVID-19 Surveillance Team |
| EPI_ISL_2513679 | Lighthouse Lab in Alderley Park | Wellcome Sanger Institute for the COVID-19 Genomics UK (COG-UK) Consortium | Jacquelyn Wynn, Mairead Hyland, The Lighthouse Lab in Alderley Park and Alex Alderton, Roberto Amato, Jeffrey Barrett, Sonia Goncalves, Ewan Harrison, David K. Jackson, Ian Johnston, Dominic Kwiatkowski, Cordelia Langford, John Sillitoe on behalf of the Wellcome Sanger Institute COVID-19 Surveillance Team |
| EPI_ISL_2513680 | Lighthouse Lab in Glasgow | Wellcome Sanger Institute for the COVID-19 Genomics UK (COG-UK) Consortium | Harper VanSteenhouse, Yumi Kasai, David Gray, Carol Clugston, Anna Dominiczak and Alex Alderton, Roberto Amato, Jeffrey Barrett, Sonia Goncalves, Ewan Harrison, David K. Jackson, Ian Johnston, Dominic Kwiatkowski, Cordelia Langford, John Sillitoe on behalf of the Wellcome Sanger Institute COVID-19 Surveillance Team |
| EPI_ISL_2513681 | Lighthouse Lab in Alderley Park | Wellcome Sanger Institute for the COVID-19 Genomics UK (COG-UK) Consortium | Jacquelyn Wynn, Mairead Hyland, The Lighthouse Lab in Alderley Park and Alex Alderton, Roberto Amato, Jeffrey Barrett, Sonia Goncalves, Ewan Harrison, David K. Jackson, Ian Johnston, Dominic Kwiatkowski, Cordelia Langford, John Sillitoe on behalf of the Wellcome Sanger Institute COVID-19 Surveillance Team |
| EPI_ISL_2513682 | Lighthouse Lab in Glasgow | Wellcome Sanger Institute for the COVID-19 Genomics UK (COG-UK) Consortium | Harper VanSteenhouse, Yumi Kasai, David Gray, Carol Clugston, Anna Dominiczak and Alex Alderton, Roberto Amato, Jeffrey Barrett, Sonia Goncalves, Ewan Harrison, David K. Jackson, Ian Johnston, Dominic Kwiatkowski, Cordelia Langford, John Sillitoe on behalf of the Wellcome Sanger Institute COVID-19 Surveillance Team |
| EPI_ISL_2513683 | Lighthouse Lab in Alderley Park | Wellcome Sanger Institute for the COVID-19 Genomics UK (COG-UK) Consortium | Jacquelyn Wynn, Mairead Hyland, The Lighthouse Lab in Alderley Park and Alex Alderton, Roberto Amato, Jeffrey Barrett, Sonia Goncalves, Ewan Harrison, David K. Jackson, Ian Johnston, Dominic Kwiatkowski, Cordelia Langford, John Sillitoe on behalf of the Wellcome Sanger Institute COVID-19 Surveillance Team |
| EPI_ISL_2513684 | Lighthouse Lab in Glasgow | Wellcome Sanger Institute for the COVID-19 Genomics UK (COG-UK) Consortium | Harper VanSteenhouse, Yumi Kasai, David Gray, Carol Clugston, Anna Dominiczak and Alex Alderton, Roberto Amato, Jeffrey Barrett, Sonia Goncalves, Ewan Harrison, David K. Jackson, Ian Johnston, Dominic Kwiatkowski, Cordelia Langford, John Sillitoe on behalf of the Wellcome Sanger Institute COVID-19 Surveillance Team |
| EPI_ISL_2513686, EPI_ISL_2513687, EPI_ISL_2513688 | Lighthouse Lab in Alderley Park | Wellcome Sanger Institute for the COVID-19 Genomics UK (COG-UK) Consortium | Jacquelyn Wynn, Mairead Hyland, The Lighthouse Lab in Alderley Park and Alex Alderton, Roberto Amato, Jeffrey Barrett, Sonia Goncalves, Ewan Harrison, David K. Jackson, Ian Johnston, Dominic Kwiatkowski, Cordelia Langford, John Sillitoe on behalf of the Wellcome Sanger Institute COVID-19 Surveillance Team |
| EPI_ISL_2513689 | Lighthouse Lab in Glasgow | Wellcome Sanger Institute for the COVID-19 Genomics UK (COG-UK) Consortium | Harper VanSteenhouse, Yumi Kasai, David Gray, Carol Clugston, Anna Dominiczak and Alex Alderton, Roberto Amato, Jeffrey Barrett, Sonia Goncalves, Ewan Harrison, David K. Jackson, Ian Johnston, Dominic Kwiatkowski, Cordelia Langford, John Sillitoe on behalf of the Wellcome Sanger Institute COVID-19 Surveillance Team |
| EPI_ISL_2513690 | Lighthouse Lab in Alderley Park | Wellcome Sanger Institute for the COVID-19 Genomics UK (COG-UK) Consortium | Jacquelyn Wynn, Mairead Hyland, The Lighthouse Lab in Alderley Park and Alex Alderton, Roberto Amato, Jeffrey Barrett, Sonia Goncalves, Ewan Harrison, David K. Jackson, Ian Johnston, Dominic Kwiatkowski, Cordelia Langford, John Sillitoe on behalf of the Wellcome Sanger Institute COVID-19 Surveillance Team |
| EPI_ISL_2513692 | Lighthouse Lab in Glasgow | Wellcome Sanger Institute for the COVID-19 Genomics UK (COG-UK) Consortium | Harper VanSteenhouse, Yumi Kasai, David Gray, Carol Clugston, Anna Dominiczak and Alex Alderton, Roberto Amato, Jeffrey Barrett, Sonia Goncalves, Ewan Harrison, David K. Jackson, Ian Johnston, Dominic Kwiatkowski, Cordelia Langford, John Sillitoe on behalf of the Wellcome Sanger Institute COVID-19 Surveillance Team |
| EPI_ISL_2513693 | Lighthouse Lab in Alderley Park | Wellcome Sanger Institute for the COVID-19 Genomics UK (COG-UK) Consortium | Jacquelyn Wynn, Mairead Hyland, The Lighthouse Lab in Alderley Park and Alex Alderton, Roberto Amato, Jeffrey Barrett, Sonia Goncalves, Ewan Harrison, David K. Jackson, Ian Johnston, Dominic Kwiatkowski, Cordelia Langford, John Sillitoe on behalf of the Wellcome Sanger Institute COVID-19 Surveillance Team |
| EPI_ISL_2513694, EPI_ISL_2513695, EPI_ISL_2513698 | Lighthouse Lab in Glasgow | Wellcome Sanger Institute for the COVID-19 Genomics UK (COG-UK) Consortium | Harper VanSteenhouse, Yumi Kasai, David Gray, Carol Clugston, Anna Dominiczak and Alex Alderton, Roberto Amato, Jeffrey Barrett, Sonia Goncalves, Ewan Harrison, David K. Jackson, Ian Johnston, Dominic Kwiatkowski, Cordelia Langford, John Sillitoe on behalf of the Wellcome Sanger Institute COVID-19 Surveillance Team |
| EPI_ISL_2513699, EPI_ISL_2513700 | Lighthouse Lab in Alderley Park | Wellcome Sanger Institute for the COVID-19 Genomics UK (COG-UK) Consortium | Jacquelyn Wynn, Mairead Hyland, The Lighthouse Lab in Alderley Park and Alex Alderton, Roberto Amato, Jeffrey Barrett, Sonia Goncalves, Ewan Harrison, David K. Jackson, Ian Johnston, Dominic Kwiatkowski, Cordelia Langford, John Sillitoe on behalf of the Wellcome Sanger Institute COVID-19 Surveillance Team |
| EPI_ISL_2513702 | Lighthouse Lab in Glasgow | Wellcome Sanger Institute for the COVID-19 Genomics UK (COG-UK) Consortium | Harper VanSteenhouse, Yumi Kasai, David Gray, Carol Clugston, Anna Dominiczak and Alex Alderton, Roberto Amato, Jeffrey Barrett, Sonia Goncalves, Ewan Harrison, David K. Jackson, Ian Johnston, Dominic Kwiatkowski, Cordelia Langford, John Sillitoe on behalf of the Wellcome Sanger Institute COVID-19 Surveillance Team |
| EPI_ISL_2513703, EPI_ISL_2513704, EPI_ISL_2513705 | Lighthouse Lab in Alderley Park | Wellcome Sanger Institute for the COVID-19 Genomics UK (COG-UK) Consortium | Jacquelyn Wynn, Mairead Hyland, The Lighthouse Lab in Alderley Park and Alex Alderton, Roberto Amato, Jeffrey Barrett, Sonia Goncalves, Ewan Harrison, David K. Jackson, Ian Johnston, Dominic Kwiatkowski, Cordelia Langford, John Sillitoe on behalf of the Wellcome Sanger Institute COVID-19 Surveillance Team |
| EPI_ISL_2513706, EPI_ISL_2513708 | Lighthouse Lab in Glasgow | Wellcome Sanger Institute for the COVID-19 Genomics UK (COG-UK) Consortium | Harper VanSteenhouse, Yumi Kasai, David Gray, Carol Clugston, Anna Dominiczak and Alex Alderton, Roberto Amato, Jeffrey Barrett, Sonia Goncalves, Ewan Harrison, David K. Jackson, Ian Johnston, Dominic Kwiatkowski, Cordelia Langford, John Sillitoe on behalf of the Wellcome Sanger Institute COVID-19 Surveillance Team |
| EPI_ISL_2513709 | Lighthouse Lab in Alderley Park | Wellcome Sanger Institute for the COVID-19 Genomics UK (COG-UK) Consortium | Jacquelyn Wynn, Mairead Hyland, The Lighthouse Lab in Alderley Park and Alex Alderton, Roberto Amato, Jeffrey Barrett, Sonia Goncalves, Ewan Harrison, David K. Jackson, Ian Johnston, Dominic Kwiatkowski, Cordelia Langford, John Sillitoe on behalf of the Wellcome Sanger Institute COVID-19 Surveillance Team |
| EPI_ISL_2513710, EPI_ISL_2513711, EPI_ISL_2513712, EPI_ISL_2513713 | Lighthouse Lab in Glasgow | Wellcome Sanger Institute for the COVID-19 Genomics UK (COG-UK) Consortium | Harper VanSteenhouse, Yumi Kasai, David Gray, Carol Clugston, Anna Dominiczak and Alex Alderton, Roberto Amato, Jeffrey Barrett, Sonia Goncalves, Ewan Harrison, David K. Jackson, Ian Johnston, Dominic Kwiatkowski, Cordelia Langford, John Sillitoe on behalf of the Wellcome Sanger Institute COVID-19 Surveillance Team |
| EPI_ISL_2513714 | Lighthouse Lab in Alderley Park | Wellcome Sanger Institute for the COVID-19 Genomics UK (COG-UK) Consortium | Jacquelyn Wynn, Mairead Hyland, The Lighthouse Lab in Alderley Park and Alex Alderton, Roberto Amato, Jeffrey Barrett, Sonia Goncalves, Ewan Harrison, David K. Jackson, Ian Johnston, Dominic Kwiatkowski, Cordelia Langford, John Sillitoe on behalf of the Wellcome Sanger Institute COVID-19 Surveillance Team |
| EPI_ISL_2513715 | Lighthouse Lab in Glasgow | Wellcome Sanger Institute for the COVID-19 Genomics UK (COG-UK) Consortium | Harper VanSteenhouse, Yumi Kasai, David Gray, Carol Clugston, Anna Dominiczak and Alex Alderton, Roberto Amato, Jeffrey Barrett, Sonia Goncalves, Ewan Harrison, David K. Jackson, Ian Johnston, Dominic Kwiatkowski, Cordelia Langford, John Sillitoe on behalf of the Wellcome Sanger Institute COVID-19 Surveillance Team |
| EPI_ISL_2513716 | Lighthouse Lab in Alderley Park | Wellcome Sanger Institute for the COVID-19 Genomics UK (COG-UK) Consortium | Jacquelyn Wynn, Mairead Hyland, The Lighthouse Lab in Alderley Park and Alex Alderton, Roberto Amato, Jeffrey Barrett, Sonia Goncalves, Ewan Harrison, David K. Jackson, Ian Johnston, Dominic Kwiatkowski, Cordelia Langford, John Sillitoe on behalf of the Wellcome Sanger Institute COVID-19 Surveillance Team |

| | | | |
|---|---|---|---|
| EPI_ISL_2513717 | Lighthouse Lab in Glasgow | Wellcome Sanger Institute for the COVID-19 Genomics UK (COG-UK) Consortium | Harper VanSteenhouse, Yumi Kasai, David Gray, Carol Clugston, Anna Dominiczak and Alex Alderton, Roberto Amato, Jeffrey Barrett, Sonia Goncalves, Ewan Harrison, David K. Jackson, Ian Johnston, Dominic Kwiatkowski, Cordelia Langford, John Sillitoe on behalf of the Wellcome Sanger Institute COVID-19 Surveillance Team |
| EPI_ISL_2513718 | Lighthouse Lab in Alderley Park | Wellcome Sanger Institute for the COVID-19 Genomics UK (COG-UK) Consortium | Jacquelyn Wynn, Mairead Hyland, The Lighthouse Lab in Alderley Park and Alex Alderton, Roberto Amato, Jeffrey Barrett, Sonia Goncalves, Ewan Harrison, David K. Jackson, Ian Johnston, Dominic Kwiatkowski, Cordelia Langford, John Sillitoe on behalf of the Wellcome Sanger Institute COVID-19 Surveillance Team |
| EPI_ISL_2513719 | Lighthouse Lab in Glasgow | Wellcome Sanger Institute for the COVID-19 Genomics UK (COG-UK) Consortium | Harper VanSteenhouse, Yumi Kasai, David Gray, Carol Clugston, Anna Dominiczak and Alex Alderton, Roberto Amato, Jeffrey Barrett, Sonia Goncalves, Ewan Harrison, David K. Jackson, Ian Johnston, Dominic Kwiatkowski, Cordelia Langford, John Sillitoe on behalf of the Wellcome Sanger Institute COVID-19 Surveillance Team |
| EPI_ISL_2513720 | Lighthouse Lab in Alderley Park | Wellcome Sanger Institute for the COVID-19 Genomics UK (COG-UK) Consortium | Jacquelyn Wynn, Mairead Hyland, The Lighthouse Lab in Alderley Park and Alex Alderton, Roberto Amato, Jeffrey Barrett, Sonia Goncalves, Ewan Harrison, David K. Jackson, Ian Johnston, Dominic Kwiatkowski, Cordelia Langford, John Sillitoe on behalf of the Wellcome Sanger Institute COVID-19 Surveillance Team |
| EPI_ISL_2513721 | Lighthouse Lab in Glasgow | Wellcome Sanger Institute for the COVID-19 Genomics UK (COG-UK) Consortium | Harper VanSteenhouse, Yumi Kasai, David Gray, Carol Clugston, Anna Dominiczak and Alex Alderton, Roberto Amato, Jeffrey Barrett, Sonia Goncalves, Ewan Harrison, David K. Jackson, Ian Johnston, Dominic Kwiatkowski, Cordelia Langford, John Sillitoe on behalf of the Wellcome Sanger Institute COVID-19 Surveillance Team |
| EPI_ISL_2513722, EPI_ISL_2513723 | Lighthouse Lab in Alderley Park | Wellcome Sanger Institute for the COVID-19 Genomics UK (COG-UK) Consortium | Jacquelyn Wynn, Mairead Hyland, The Lighthouse Lab in Alderley Park and Alex Alderton, Roberto Amato, Jeffrey Barrett, Sonia Goncalves, Ewan Harrison, David K. Jackson, Ian Johnston, Dominic Kwiatkowski, Cordelia Langford, John Sillitoe on behalf of the Wellcome Sanger Institute COVID-19 Surveillance Team |
| EPI_ISL_2513724 | Lighthouse Lab in Glasgow | Wellcome Sanger Institute for the COVID-19 Genomics UK (COG-UK) Consortium | Harper VanSteenhouse, Yumi Kasai, David Gray, Carol Clugston, Anna Dominiczak and Alex Alderton, Roberto Amato, Jeffrey Barrett, Sonia Goncalves, Ewan Harrison, David K. Jackson, Ian Johnston, Dominic Kwiatkowski, Cordelia Langford, John Sillitoe on behalf of the Wellcome Sanger Institute COVID-19 Surveillance Team |
| EPI_ISL_2513726, EPI_ISL_2513727 | Lighthouse Lab in Alderley Park | Wellcome Sanger Institute for the COVID-19 Genomics UK (COG-UK) Consortium | Jacquelyn Wynn, Mairead Hyland, The Lighthouse Lab in Alderley Park and Alex Alderton, Roberto Amato, Jeffrey Barrett, Sonia Goncalves, Ewan Harrison, David K. Jackson, Ian Johnston, Dominic Kwiatkowski, Cordelia Langford, John Sillitoe on behalf of the Wellcome Sanger Institute COVID-19 Surveillance Team |
| EPI_ISL_2513728, EPI_ISL_2513729, EPI_ISL_2513730, EPI_ISL_2513731, EPI_ISL_2513732, EPI_ISL_2513733, EPI_ISL_2513734, EPI_ISL_2513735 | Lighthouse Lab in Glasgow | Wellcome Sanger Institute for the COVID-19 Genomics UK (COG-UK) Consortium | Harper VanSteenhouse, Yumi Kasai, David Gray, Carol Clugston, Anna Dominiczak and Alex Alderton, Roberto Amato, Jeffrey Barrett, Sonia Goncalves, Ewan Harrison, David K. Jackson, Ian Johnston, Dominic Kwiatkowski, Cordelia Langford, John Sillitoe on behalf of the Wellcome Sanger Institute COVID-19 Surveillance Team |
| EPI_ISL_2513738, EPI_ISL_2513739, EPI_ISL_2513740, EPI_ISL_2513741 | Lighthouse Lab in Alderley Park | Wellcome Sanger Institute for the COVID-19 Genomics UK (COG-UK) Consortium | Jacquelyn Wynn, Mairead Hyland, The Lighthouse Lab in Alderley Park and Alex Alderton, Roberto Amato, Jeffrey Barrett, Sonia Goncalves, Ewan Harrison, David K. Jackson, Ian Johnston, Dominic Kwiatkowski, Cordelia Langford, John Sillitoe on behalf of the Wellcome Sanger Institute COVID-19 Surveillance Team |
| EPI_ISL_2513742, EPI_ISL_2513743, EPI_ISL_2513744, EPI_ISL_2513745 | Lighthouse Lab in Glasgow | Wellcome Sanger Institute for the COVID-19 Genomics UK (COG-UK) Consortium | Harper VanSteenhouse, Yumi Kasai, David Gray, Carol Clugston, Anna Dominiczak and Alex Alderton, Roberto Amato, Jeffrey Barrett, Sonia Goncalves, Ewan Harrison, David K. Jackson, Ian Johnston, Dominic Kwiatkowski, Cordelia Langford, John Sillitoe on behalf of the Wellcome Sanger Institute COVID-19 Surveillance Team |
| EPI_ISL_2513747 | Lighthouse Lab in Alderley Park | Wellcome Sanger Institute for the COVID-19 Genomics UK (COG-UK) Consortium | Jacquelyn Wynn, Mairead Hyland, The Lighthouse Lab in Alderley Park and Alex Alderton, Roberto Amato, Jeffrey Barrett, Sonia Goncalves, Ewan Harrison, David K. Jackson, Ian Johnston, Dominic Kwiatkowski, Cordelia Langford, John Sillitoe on behalf of the Wellcome Sanger Institute COVID-19 Surveillance Team |
| EPI_ISL_2513748, EPI_ISL_2513749 | Lighthouse Lab in Glasgow | Wellcome Sanger Institute for the COVID-19 Genomics UK (COG-UK) Consortium | Harper VanSteenhouse, Yumi Kasai, David Gray, Carol Clugston, Anna Dominiczak and Alex Alderton, Roberto Amato, Jeffrey Barrett, Sonia Goncalves, Ewan Harrison, David K. Jackson, Ian Johnston, Dominic Kwiatkowski, Cordelia Langford, John Sillitoe on behalf of the Wellcome Sanger Institute COVID-19 Surveillance Team |
| EPI_ISL_2513750 | Lighthouse Lab in Alderley Park | Wellcome Sanger Institute for the COVID-19 Genomics UK (COG-UK) Consortium | Jacquelyn Wynn, Mairead Hyland, The Lighthouse Lab in Alderley Park and Alex Alderton, Roberto Amato, Jeffrey Barrett, Sonia Goncalves, Ewan Harrison, David K. Jackson, Ian Johnston, Dominic Kwiatkowski, Cordelia Langford, John Sillitoe on behalf of the Wellcome Sanger Institute COVID-19 Surveillance Team |
| EPI_ISL_2513751 | Lighthouse Lab in Glasgow | Wellcome Sanger Institute for the COVID-19 Genomics UK (COG-UK) Consortium | Harper VanSteenhouse, Yumi Kasai, David Gray, Carol Clugston, Anna Dominiczak and Alex Alderton, Roberto Amato, Jeffrey Barrett, Sonia Goncalves, Ewan Harrison, David K. Jackson, Ian Johnston, Dominic Kwiatkowski, Cordelia Langford, John Sillitoe on behalf of the Wellcome Sanger Institute COVID-19 Surveillance Team |
| EPI_ISL_2513754 | Lighthouse Lab in Alderley Park | Wellcome Sanger Institute for the COVID-19 Genomics UK (COG-UK) Consortium | Jacquelyn Wynn, Mairead Hyland, The Lighthouse Lab in Alderley Park and Alex Alderton, Roberto Amato, Jeffrey Barrett, Sonia Goncalves, Ewan Harrison, David K. Jackson, Ian Johnston, Dominic Kwiatkowski, Cordelia Langford, John Sillitoe on behalf of the Wellcome Sanger Institute COVID-19 Surveillance Team |
| EPI_ISL_2513755, EPI_ISL_2513756 | Lighthouse Lab in Glasgow | Wellcome Sanger Institute for the COVID-19 Genomics UK (COG-UK) Consortium | Harper VanSteenhouse, Yumi Kasai, David Gray, Carol Clugston, Anna Dominiczak and Alex Alderton, Roberto Amato, Jeffrey Barrett, Sonia Goncalves, Ewan Harrison, David K. Jackson, Ian Johnston, Dominic Kwiatkowski, Cordelia Langford, John Sillitoe on behalf of the Wellcome Sanger Institute COVID-19 Surveillance Team |
| EPI_ISL_2513757, EPI_ISL_2513758 | Lighthouse Lab in Alderley Park | Wellcome Sanger Institute for the COVID-19 Genomics UK (COG-UK) Consortium | Jacquelyn Wynn, Mairead Hyland, The Lighthouse Lab in Alderley Park and Alex Alderton, Roberto Amato, Jeffrey Barrett, Sonia Goncalves, Ewan Harrison, David K. Jackson, Ian Johnston, Dominic Kwiatkowski, Cordelia Langford, John Sillitoe on behalf of the Wellcome Sanger Institute COVID-19 Surveillance Team |
| EPI_ISL_2513760, EPI_ISL_2513761 | Lighthouse Lab in Glasgow | Wellcome Sanger Institute for the COVID-19 Genomics UK (COG-UK) Consortium | Harper VanSteenhouse, Yumi Kasai, David Gray, Carol Clugston, Anna Dominiczak and Alex Alderton, Roberto Amato, Jeffrey Barrett, Sonia Goncalves, Ewan Harrison, David K. Jackson, Ian Johnston, Dominic Kwiatkowski, Cordelia Langford, John Sillitoe on behalf of the Wellcome Sanger Institute COVID-19 Surveillance Team |
| EPI_ISL_2513762, EPI_ISL_2513764, EPI_ISL_2513766 | Lighthouse Lab in Alderley Park | Wellcome Sanger Institute for the COVID-19 Genomics UK (COG-UK) Consortium | Jacquelyn Wynn, Mairead Hyland, The Lighthouse Lab in Alderley Park and Alex Alderton, Roberto Amato, Jeffrey Barrett, Sonia Goncalves, Ewan Harrison, David K. Jackson, Ian Johnston, Dominic Kwiatkowski, Cordelia Langford, John Sillitoe on behalf of the Wellcome Sanger Institute COVID-19 Surveillance Team |
| EPI_ISL_2513767 | Lighthouse Lab in Glasgow | Wellcome Sanger Institute for the COVID-19 Genomics UK (COG-UK) Consortium | Harper VanSteenhouse, Yumi Kasai, David Gray, Carol Clugston, Anna Dominiczak and Alex Alderton, Roberto Amato, Jeffrey Barrett, Sonia Goncalves, Ewan Harrison, David K. Jackson, Ian Johnston, Dominic Kwiatkowski, Cordelia Langford, John Sillitoe on behalf of the Wellcome Sanger Institute COVID-19 Surveillance Team |
| EPI_ISL_2513768 | Lighthouse Lab in Alderley Park | Wellcome Sanger Institute for the COVID-19 Genomics UK (COG-UK) Consortium | Jacquelyn Wynn, Mairead Hyland, The Lighthouse Lab in Alderley Park and Alex Alderton, Roberto Amato, Jeffrey Barrett, Sonia Goncalves, Ewan Harrison, David K. Jackson, Ian Johnston, Dominic Kwiatkowski, Cordelia Langford, John Sillitoe on behalf of the Wellcome Sanger Institute COVID-19 Surveillance Team |
| EPI_ISL_2513769 | Lighthouse Lab in Glasgow | Wellcome Sanger Institute for the COVID-19 Genomics UK (COG-UK) Consortium | Harper VanSteenhouse, Yumi Kasai, David Gray, Carol Clugston, Anna Dominiczak and Alex Alderton, Roberto Amato, Jeffrey Barrett, Sonia Goncalves, Ewan Harrison, David K. Jackson, Ian Johnston, Dominic Kwiatkowski, Cordelia Langford, John Sillitoe on behalf of the Wellcome Sanger Institute COVID-19 Surveillance Team |
| EPI_ISL_2513770 | Lighthouse Lab in Alderley Park | Wellcome Sanger Institute for the COVID-19 Genomics UK (COG-UK) Consortium | Jacquelyn Wynn, Mairead Hyland, The Lighthouse Lab in Alderley Park and Alex Alderton, Roberto Amato, Jeffrey Barrett, Sonia Goncalves, Ewan Harrison, David K. Jackson, Ian Johnston, Dominic Kwiatkowski, Cordelia Langford, John Sillitoe on behalf of the Wellcome Sanger Institute COVID-19 Surveillance Team |
| EPI_ISL_2513771, EPI_ISL_2513772 | Lighthouse Lab in Glasgow | Wellcome Sanger Institute for the COVID-19 Genomics UK (COG-UK) Consortium | Harper VanSteenhouse, Yumi Kasai, David Gray, Carol Clugston, Anna Dominiczak and Alex Alderton, Roberto Amato, Jeffrey Barrett, Sonia Goncalves, Ewan Harrison, David K. Jackson, Ian Johnston, Dominic Kwiatkowski, Cordelia Langford, John Sillitoe on behalf of the Wellcome Sanger Institute COVID-19 Surveillance Team |
| EPI_ISL_2513774 | Lighthouse Lab in Alderley Park | Wellcome Sanger Institute for the COVID-19 Genomics UK (COG-UK) Consortium | Jacquelyn Wynn, Mairead Hyland, The Lighthouse Lab in Alderley Park and Alex Alderton, Roberto Amato, Jeffrey Barrett, Sonia Goncalves, Ewan Harrison, David K. Jackson, Ian Johnston, Dominic Kwiatkowski, Cordelia Langford, John Sillitoe on behalf of the Wellcome Sanger Institute COVID-19 Surveillance Team |
| EPI_ISL_2513775 | Lighthouse Lab in Glasgow | Wellcome Sanger Institute for the COVID-19 Genomics UK (COG-UK) Consortium | Harper VanSteenhouse, Yumi Kasai, David Gray, Carol Clugston, Anna Dominiczak and Alex Alderton, Roberto Amato, Jeffrey Barrett, Sonia Goncalves, Ewan Harrison, David K. Jackson, Ian Johnston, Dominic Kwiatkowski, Cordelia Langford, John Sillitoe on behalf of the Wellcome Sanger Institute COVID-19 Surveillance Team |
| EPI_ISL_2513776 | Lighthouse Lab in Alderley Park | Wellcome Sanger Institute for the COVID-19 Genomics UK (COG-UK) Consortium | Jacquelyn Wynn, Mairead Hyland, The Lighthouse Lab in Alderley Park and Alex Alderton, Roberto Amato, Jeffrey Barrett, Sonia Goncalves, Ewan Harrison, David K. Jackson, Ian Johnston, Dominic Kwiatkowski, Cordelia Langford, John Sillitoe on behalf of the Wellcome Sanger Institute COVID-19 Surveillance Team |
| EPI_ISL_2513777, EPI_ISL_2513778 | Lighthouse Lab in Glasgow | Wellcome Sanger Institute for the COVID-19 Genomics UK (COG-UK) Consortium | Harper VanSteenhouse, Yumi Kasai, David Gray, Carol Clugston, Anna Dominiczak and Alex Alderton, Roberto Amato, Jeffrey Barrett, Sonia Goncalves, Ewan Harrison, David K. Jackson, Ian Johnston, Dominic Kwiatkowski, Cordelia Langford, John Sillitoe on behalf of the Wellcome Sanger Institute COVID-19 Surveillance Team |
| EPI_ISL_2513779, EPI_ISL_2513780, EPI_ISL_2513783, EPI_ISL_2513784, EPI_ISL_2513785 | Lighthouse Lab in Alderley Park | Wellcome Sanger Institute for the COVID-19 Genomics UK (COG-UK) Consortium | Jacquelyn Wynn, Mairead Hyland, The Lighthouse Lab in Alderley Park and Alex Alderton, Roberto Amato, Jeffrey Barrett, Sonia Goncalves, Ewan Harrison, David K. Jackson, Ian Johnston, Dominic Kwiatkowski, Cordelia Langford, John Sillitoe on behalf of the Wellcome Sanger Institute COVID-19 Surveillance Team |
| EPI_ISL_2513786 | Lighthouse Lab in Glasgow | Wellcome Sanger Institute for the COVID-19 Genomics UK (COG-UK) Consortium | Harper VanSteenhouse, Yumi Kasai, David Gray, Carol Clugston, Anna Dominiczak and Alex Alderton, Roberto Amato, Jeffrey Barrett, Sonia Goncalves, Ewan Harrison, David K. Jackson, Ian Johnston, Dominic Kwiatkowski, Cordelia Langford, John Sillitoe on behalf of the Wellcome Sanger Institute COVID-19 Surveillance Team |
| EPI_ISL_2513787, EPI_ISL_2513788 | Lighthouse Lab in Alderley Park | Wellcome Sanger Institute for the COVID-19 Genomics UK (COG-UK) Consortium | Jacquelyn Wynn, Mairead Hyland, The Lighthouse Lab in Alderley Park and Alex Alderton, Roberto Amato, Jeffrey Barrett, Sonia Goncalves, Ewan Harrison, David K. Jackson, Ian Johnston, Dominic Kwiatkowski, Cordelia Langford, John Sillitoe on behalf of the Wellcome Sanger Institute COVID-19 Surveillance Team |
| EPI_ISL_2513789 | Lighthouse Lab in Glasgow | Wellcome Sanger Institute for the COVID-19 Genomics UK (COG-UK) Consortium | Harper VanSteenhouse, Yumi Kasai, David Gray, Carol Clugston, Anna Dominiczak and Alex Alderton, Roberto Amato, Jeffrey Barrett, Sonia Goncalves, Ewan Harrison, David K. Jackson, Ian Johnston, Dominic Kwiatkowski, Cordelia Langford, John Sillitoe on behalf of the Wellcome Sanger Institute COVID-19 Surveillance Team |
| EPI_ISL_2513790 | Lighthouse Lab in Alderley Park | Wellcome Sanger Institute for the COVID-19 Genomics UK (COG-UK) Consortium | Jacquelyn Wynn, Mairead Hyland, The Lighthouse Lab in Alderley Park and Alex Alderton, Roberto Amato, Jeffrey Barrett, Sonia Goncalves, Ewan Harrison, David K. Jackson, Ian Johnston, Dominic Kwiatkowski, Cordelia Langford, John Sillitoe on behalf of the Wellcome Sanger Institute COVID-19 Surveillance Team |
| EPI_ISL_2513791, EPI_ISL_2513792 | Lighthouse Lab in Glasgow | Wellcome Sanger Institute for the COVID-19 Genomics UK (COG-UK) Consortium | Harper VanSteenhouse, Yumi Kasai, David Gray, Carol Clugston, Anna Dominiczak and Alex Alderton, Roberto Amato, Jeffrey Barrett, Sonia Goncalves, Ewan Harrison, David K. Jackson, Ian Johnston, Dominic Kwiatkowski, Cordelia Langford, John Sillitoe on behalf of the Wellcome Sanger Institute COVID-19 Surveillance Team |
| EPI_ISL_2513793 | Lighthouse Lab in Alderley Park | Wellcome Sanger Institute for the COVID-19 Genomics UK (COG-UK) Consortium | Jacquelyn Wynn, Mairead Hyland, The Lighthouse Lab in Alderley Park and Alex Alderton, Roberto Amato, Jeffrey Barrett, Sonia Goncalves, Ewan Harrison, David K. Jackson, Ian Johnston, Dominic Kwiatkowski, Cordelia Langford, John Sillitoe on behalf of the Wellcome Sanger Institute COVID-19 Surveillance Team |
| EPI_ISL_2513794 | Lighthouse Lab in Glasgow | Wellcome Sanger Institute for the COVID-19 Genomics UK (COG-UK) Consortium | Harper VanSteenhouse, Yumi Kasai, David Gray, Carol Clugston, Anna Dominiczak and Alex Alderton, Roberto Amato, Jeffrey Barrett, Sonia Goncalves, Ewan Harrison, David K. Jackson, Ian Johnston, Dominic Kwiatkowski, Cordelia Langford, John Sillitoe on behalf of the Wellcome Sanger Institute COVID-19 Surveillance Team |
| EPI_ISL_2513795, EPI_ISL_2513796, EPI_ISL_2513798, EPI_ISL_2513799 | Lighthouse Lab in Alderley Park | Wellcome Sanger Institute for the COVID-19 Genomics UK (COG-UK) Consortium | Jacquelyn Wynn, Mairead Hyland, The Lighthouse Lab in Alderley Park and Alex Alderton, Roberto Amato, Jeffrey Barrett, Sonia Goncalves, Ewan Harrison, David K. Jackson, Ian Johnston, Dominic Kwiatkowski, Cordelia Langford, John Sillitoe on behalf of the Wellcome Sanger Institute COVID-19 Surveillance Team |
| EPI_ISL_2513800 | Lighthouse Lab in Glasgow | Wellcome Sanger Institute for the COVID-19 Genomics UK (COG-UK) | Harper VanSteenhouse, Yumi Kasai, David Gray, Carol Clugston, Anna Dominiczak and Alex Alderton, Roberto Amato, Jeffrey Barrett, Sonia Goncalves, Ewan Harrison, David K. Jackson, Ian Johnston, Dominic Kwiatkowski, Cordelia Langford, |

| | | | |
|---|---|---|---|
| | | Consortium | John Sillitoe on behalf of the Wellcome Sanger Institute COVID-19 Surveillance Team |
| EPI_ISL_2513801, EPI_ISL_2513802 | Lighthouse Lab in Alderley Park | Wellcome Sanger Institute for the COVID-19 Genomics UK (COG-UK) Consortium | Jacquelyn Wynn, Mairead Hyland, The Lighthouse Lab in Alderley Park and Alex Alderton, Roberto Amato, Jeffrey Barrett, Sonia Goncalves, Ewan Harrison, David K. Jackson, Ian Johnston, Dominic Kwiatkowski, Cordelia Langford, John Sillitoe on behalf of the Wellcome Sanger Institute COVID-19 Surveillance Team |
| EPI_ISL_2513803, EPI_ISL_2513805 | Health Services Laboratories | Wellcome Sanger Institute for the COVID-19 Genomics UK (COG-UK) Consortium | Health Services Laboratories and Alex Alderton, Roberto Amato, Jeffrey Barrett, Sonia Goncalves, Ewan Harrison, David K. Jackson, Ian Johnston, Dominic Kwiatkowski, Cordelia Langford, John Sillitoe on behalf of the Wellcome Sanger Institute COVID-19 Surveillance Team |
| EPI_ISL_2513806 | Lighthouse Laboratory Plymouth | Wellcome Sanger Institute for the COVID-19 Genomics UK (COG-UK) Consortium | Lighthouse Laboratory Plymouth and Alex Alderton, Roberto Amato, Jeffrey Barrett, Sonia Goncalves, Ewan Harrison, David K. Jackson, Ian Johnston, Dominic Kwiatkowski, Cordelia Langford, John Sillitoe on behalf of the Wellcome Sanger Institute COVID-19 Surveillance Team |
| EPI_ISL_2513808, EPI_ISL_2513809, EPI_ISL_2513810, EPI_ISL_2513811, EPI_ISL_2513812, EPI_ISL_2513814 | Health Services Laboratories | Wellcome Sanger Institute for the COVID-19 Genomics UK (COG-UK) Consortium | Health Services Laboratories and Alex Alderton, Roberto Amato, Jeffrey Barrett, Sonia Goncalves, Ewan Harrison, David K. Jackson, Ian Johnston, Dominic Kwiatkowski, Cordelia Langford, John Sillitoe on behalf of the Wellcome Sanger Institute COVID-19 Surveillance Team |
| EPI_ISL_2513815 | Lighthouse Laboratory Plymouth | Wellcome Sanger Institute for the COVID-19 Genomics UK (COG-UK) Consortium | Lighthouse Laboratory Plymouth and Alex Alderton, Roberto Amato, Jeffrey Barrett, Sonia Goncalves, Ewan Harrison, David K. Jackson, Ian Johnston, Dominic Kwiatkowski, Cordelia Langford, John Sillitoe on behalf of the Wellcome Sanger Institute COVID-19 Surveillance Team |
| EPI_ISL_2513816 | Health Services Laboratories | Wellcome Sanger Institute for the COVID-19 Genomics UK (COG-UK) Consortium | Health Services Laboratories and Alex Alderton, Roberto Amato, Jeffrey Barrett, Sonia Goncalves, Ewan Harrison, David K. Jackson, Ian Johnston, Dominic Kwiatkowski, Cordelia Langford, John Sillitoe on behalf of the Wellcome Sanger Institute COVID-19 Surveillance Team |
| EPI_ISL_2513818, EPI_ISL_2513819 | Lighthouse Laboratory Plymouth | Wellcome Sanger Institute for the COVID-19 Genomics UK (COG-UK) Consortium | Lighthouse Laboratory Plymouth and Alex Alderton, Roberto Amato, Jeffrey Barrett, Sonia Goncalves, Ewan Harrison, David K. Jackson, Ian Johnston, Dominic Kwiatkowski, Cordelia Langford, John Sillitoe on behalf of the Wellcome Sanger Institute COVID-19 Surveillance Team |
| EPI_ISL_2513820 | Health Services Laboratories | Wellcome Sanger Institute for the COVID-19 Genomics UK (COG-UK) Consortium | Health Services Laboratories and Alex Alderton, Roberto Amato, Jeffrey Barrett, Sonia Goncalves, Ewan Harrison, David K. Jackson, Ian Johnston, Dominic Kwiatkowski, Cordelia Langford, John Sillitoe on behalf of the Wellcome Sanger Institute COVID-19 Surveillance Team |
| EPI_ISL_2513821, EPI_ISL_2513823 | Lighthouse Lab in Alderley Park | Wellcome Sanger Institute for the COVID-19 Genomics UK (COG-UK) Consortium | Jacquelyn Wynn, Mairead Hyland, The Lighthouse Lab in Alderley Park and Alex Alderton, Roberto Amato, Jeffrey Barrett, Sonia Goncalves, Ewan Harrison, David K. Jackson, Ian Johnston, Dominic Kwiatkowski, Cordelia Langford, John Sillitoe on behalf of the Wellcome Sanger Institute COVID-19 Surveillance Team |
| EPI_ISL_2513824 | Lighthouse Laboratory Plymouth | Wellcome Sanger Institute for the COVID-19 Genomics UK (COG-UK) Consortium | Lighthouse Laboratory Plymouth and Alex Alderton, Roberto Amato, Jeffrey Barrett, Sonia Goncalves, Ewan Harrison, David K. Jackson, Ian Johnston, Dominic Kwiatkowski, Cordelia Langford, John Sillitoe on behalf of the Wellcome Sanger Institute COVID-19 Surveillance Team |
| EPI_ISL_2513825 | Health Services Laboratories | Wellcome Sanger Institute for the COVID-19 Genomics UK (COG-UK) Consortium | Health Services Laboratories and Alex Alderton, Roberto Amato, Jeffrey Barrett, Sonia Goncalves, Ewan Harrison, David K. Jackson, Ian Johnston, Dominic Kwiatkowski, Cordelia Langford, John Sillitoe on behalf of the Wellcome Sanger Institute COVID-19 Surveillance Team |
| EPI_ISL_2513826, EPI_ISL_2513828, EPI_ISL_2513829, EPI_ISL_2513830 | Lighthouse Lab in Alderley Park | Wellcome Sanger Institute for the COVID-19 Genomics UK (COG-UK) Consortium | Jacquelyn Wynn, Mairead Hyland, The Lighthouse Lab in Alderley Park and Alex Alderton, Roberto Amato, Jeffrey Barrett, Sonia Goncalves, Ewan Harrison, David K. Jackson, Ian Johnston, Dominic Kwiatkowski, Cordelia Langford, John Sillitoe on behalf of the Wellcome Sanger Institute COVID-19 Surveillance Team |
| EPI_ISL_2513832, EPI_ISL_2513833, EPI_ISL_2513834, EPI_ISL_2513835, EPI_ISL_2513837, EPI_ISL_2513838 | Health Services Laboratories | Wellcome Sanger Institute for the COVID-19 Genomics UK (COG-UK) Consortium | Health Services Laboratories and Alex Alderton, Roberto Amato, Jeffrey Barrett, Sonia Goncalves, Ewan Harrison, David K. Jackson, Ian Johnston, Dominic Kwiatkowski, Cordelia Langford, John Sillitoe on behalf of the Wellcome Sanger Institute COVID-19 Surveillance Team |
| EPI_ISL_2513839 | Lighthouse Lab in Alderley Park | Wellcome Sanger Institute for the COVID-19 Genomics UK (COG-UK) Consortium | Jacquelyn Wynn, Mairead Hyland, The Lighthouse Lab in Alderley Park and Alex Alderton, Roberto Amato, Jeffrey Barrett, Sonia Goncalves, Ewan Harrison, David K. Jackson, Ian Johnston, Dominic Kwiatkowski, Cordelia Langford, John Sillitoe on behalf of the Wellcome Sanger Institute COVID-19 Surveillance Team |
| EPI_ISL_2513840 | Health Services Laboratories | Wellcome Sanger Institute for the COVID-19 Genomics UK (COG-UK) Consortium | Health Services Laboratories and Alex Alderton, Roberto Amato, Jeffrey Barrett, Sonia Goncalves, Ewan Harrison, David K. Jackson, Ian Johnston, Dominic Kwiatkowski, Cordelia Langford, John Sillitoe on behalf of the Wellcome Sanger Institute COVID-19 Surveillance Team |
| EPI_ISL_2513841, EPI_ISL_2513842 | Lighthouse Lab in Alderley Park | Wellcome Sanger Institute for the COVID-19 Genomics UK (COG-UK) Consortium | Jacquelyn Wynn, Mairead Hyland, The Lighthouse Lab in Alderley Park and Alex Alderton, Roberto Amato, Jeffrey Barrett, Sonia Goncalves, Ewan Harrison, David K. Jackson, Ian Johnston, Dominic Kwiatkowski, Cordelia Langford, John Sillitoe on behalf of the Wellcome Sanger Institute COVID-19 Surveillance Team |
| EPI_ISL_2513843 | Lighthouse Laboratory Plymouth | Wellcome Sanger Institute for the COVID-19 Genomics UK (COG-UK) Consortium | Lighthouse Laboratory Plymouth and Alex Alderton, Roberto Amato, Jeffrey Barrett, Sonia Goncalves, Ewan Harrison, David K. Jackson, Ian Johnston, Dominic Kwiatkowski, Cordelia Langford, John Sillitoe on behalf of the Wellcome Sanger Institute COVID-19 Surveillance Team |
| EPI_ISL_2513844, EPI_ISL_2513846, EPI_ISL_2513847, EPI_ISL_2513848, EPI_ISL_2513849, EPI_ISL_2513850, EPI_ISL_2513851, EPI_ISL_2513852 | Health Services Laboratories | Wellcome Sanger Institute for the COVID-19 Genomics UK (COG-UK) Consortium | Health Services Laboratories and Alex Alderton, Roberto Amato, Jeffrey Barrett, Sonia Goncalves, Ewan Harrison, David K. Jackson, Ian Johnston, Dominic Kwiatkowski, Cordelia Langford, John Sillitoe on behalf of the Wellcome Sanger Institute COVID-19 Surveillance Team |
| EPI_ISL_2513853 | Lighthouse Laboratory Plymouth | Wellcome Sanger Institute for the COVID-19 Genomics UK (COG-UK) Consortium | Lighthouse Laboratory Plymouth and Alex Alderton, Roberto Amato, Jeffrey Barrett, Sonia Goncalves, Ewan Harrison, David K. Jackson, Ian Johnston, Dominic Kwiatkowski, Cordelia Langford, John Sillitoe on behalf of the Wellcome Sanger Institute COVID-19 Surveillance Team |
| EPI_ISL_2513854, EPI_ISL_2513855, EPI_ISL_2513856, EPI_ISL_2513857, EPI_ISL_2513858, EPI_ISL_2513859, EPI_ISL_2513862 | Health Services Laboratories | Wellcome Sanger Institute for the COVID-19 Genomics UK (COG-UK) Consortium | Health Services Laboratories and Alex Alderton, Roberto Amato, Jeffrey Barrett, Sonia Goncalves, Ewan Harrison, David K. Jackson, Ian Johnston, Dominic Kwiatkowski, Cordelia Langford, John Sillitoe on behalf of the Wellcome Sanger Institute COVID-19 Surveillance Team |
| EPI_ISL_2513863 | Lighthouse Laboratory Plymouth | Wellcome Sanger Institute for the COVID-19 Genomics UK (COG-UK) Consortium | Lighthouse Laboratory Plymouth and Alex Alderton, Roberto Amato, Jeffrey Barrett, Sonia Goncalves, Ewan Harrison, David K. Jackson, Ian Johnston, Dominic Kwiatkowski, Cordelia Langford, John Sillitoe on behalf of the Wellcome Sanger Institute COVID-19 Surveillance Team |
| EPI_ISL_2513864 | Lighthouse Lab in Alderley Park | Wellcome Sanger Institute for the COVID-19 Genomics UK (COG-UK) Consortium | Jacquelyn Wynn, Mairead Hyland, The Lighthouse Lab in Alderley Park and Alex Alderton, Roberto Amato, Jeffrey Barrett, Sonia Goncalves, Ewan Harrison, David K. Jackson, Ian Johnston, Dominic Kwiatkowski, Cordelia Langford, John Sillitoe on behalf of the Wellcome Sanger Institute COVID-19 Surveillance Team |
| EPI_ISL_2513865 | Health Services Laboratories | Wellcome Sanger Institute for the COVID-19 Genomics UK (COG-UK) Consortium | Health Services Laboratories and Alex Alderton, Roberto Amato, Jeffrey Barrett, Sonia Goncalves, Ewan Harrison, David K. Jackson, Ian Johnston, Dominic Kwiatkowski, Cordelia Langford, John Sillitoe on behalf of the Wellcome Sanger Institute COVID-19 Surveillance Team |
| EPI_ISL_2513866, EPI_ISL_2513867 | Lighthouse Laboratory Plymouth | Wellcome Sanger Institute for the COVID-19 Genomics UK (COG-UK) Consortium | Lighthouse Laboratory Plymouth and Alex Alderton, Roberto Amato, Jeffrey Barrett, Sonia Goncalves, Ewan Harrison, David K. Jackson, Ian Johnston, Dominic Kwiatkowski, Cordelia Langford, John Sillitoe on behalf of the Wellcome Sanger Institute COVID-19 Surveillance Team |
| EPI_ISL_2513868 | Health Services Laboratories | Wellcome Sanger Institute for the COVID-19 Genomics UK (COG-UK) Consortium | Health Services Laboratories and Alex Alderton, Roberto Amato, Jeffrey Barrett, Sonia Goncalves, Ewan Harrison, David K. Jackson, Ian Johnston, Dominic Kwiatkowski, Cordelia Langford, John Sillitoe on behalf of the Wellcome Sanger Institute COVID-19 Surveillance Team |
| EPI_ISL_2513869 | Lighthouse Lab in Alderley Park | Wellcome Sanger Institute for the COVID-19 Genomics UK (COG-UK) Consortium | Jacquelyn Wynn, Mairead Hyland, The Lighthouse Lab in Alderley Park and Alex Alderton, Roberto Amato, Jeffrey Barrett, Sonia Goncalves, Ewan Harrison, David K. Jackson, Ian Johnston, Dominic Kwiatkowski, Cordelia Langford, John Sillitoe on behalf of the Wellcome Sanger Institute COVID-19 Surveillance Team |
| EPI_ISL_2513870, EPI_ISL_2513872 | Lighthouse Laboratory Plymouth | Wellcome Sanger Institute for the COVID-19 Genomics UK (COG-UK) Consortium | Lighthouse Laboratory Plymouth and Alex Alderton, Roberto Amato, Jeffrey Barrett, Sonia Goncalves, Ewan Harrison, David K. Jackson, Ian Johnston, Dominic Kwiatkowski, Cordelia Langford, John Sillitoe on behalf of the Wellcome Sanger Institute COVID-19 Surveillance Team |
| EPI_ISL_2513873, EPI_ISL_2513874 | Lighthouse Lab in Alderley Park | Wellcome Sanger Institute for the COVID-19 Genomics UK (COG-UK) Consortium | Jacquelyn Wynn, Mairead Hyland, The Lighthouse Lab in Alderley Park and Alex Alderton, Roberto Amato, Jeffrey Barrett, Sonia Goncalves, Ewan Harrison, David K. Jackson, Ian Johnston, Dominic Kwiatkowski, Cordelia Langford, John Sillitoe on behalf of the Wellcome Sanger Institute COVID-19 Surveillance Team |
| EPI_ISL_2513875 | Health Services Laboratories | Wellcome Sanger Institute for the COVID-19 Genomics UK (COG-UK) Consortium | Health Services Laboratories and Alex Alderton, Roberto Amato, Jeffrey Barrett, Sonia Goncalves, Ewan Harrison, David K. Jackson, Ian Johnston, Dominic Kwiatkowski, Cordelia Langford, John Sillitoe on behalf of the Wellcome Sanger Institute COVID-19 Surveillance Team |
| EPI_ISL_2513876, EPI_ISL_2513877 | Lighthouse Laboratory Plymouth | Wellcome Sanger Institute for the COVID-19 Genomics UK (COG-UK) Consortium | Lighthouse Laboratory Plymouth and Alex Alderton, Roberto Amato, Jeffrey Barrett, Sonia Goncalves, Ewan Harrison, David K. Jackson, Ian Johnston, Dominic Kwiatkowski, Cordelia Langford, John Sillitoe on behalf of the Wellcome Sanger Institute COVID-19 Surveillance Team |
| EPI_ISL_2513878, EPI_ISL_2513879 | Health Services Laboratories | Wellcome Sanger Institute for the COVID-19 Genomics UK (COG-UK) Consortium | Health Services Laboratories and Alex Alderton, Roberto Amato, Jeffrey Barrett, Sonia Goncalves, Ewan Harrison, David K. Jackson, Ian Johnston, Dominic Kwiatkowski, Cordelia Langford, John Sillitoe on behalf of the Wellcome Sanger Institute COVID-19 Surveillance Team |
| EPI_ISL_2513880 | Lighthouse Lab in Alderley Park | Wellcome Sanger Institute for the COVID-19 Genomics UK (COG-UK) Consortium | Jacquelyn Wynn, Mairead Hyland, The Lighthouse Lab in Alderley Park and Alex Alderton, Roberto Amato, Jeffrey Barrett, Sonia Goncalves, Ewan Harrison, David K. Jackson, Ian Johnston, Dominic Kwiatkowski, Cordelia Langford, John Sillitoe on behalf of the Wellcome Sanger Institute COVID-19 Surveillance Team |
| EPI_ISL_2513881, EPI_ISL_2513882 | Health Services Laboratories | Wellcome Sanger Institute for the COVID-19 Genomics UK (COG-UK) Consortium | Health Services Laboratories and Alex Alderton, Roberto Amato, Jeffrey Barrett, Sonia Goncalves, Ewan Harrison, David K. Jackson, Ian Johnston, Dominic Kwiatkowski, Cordelia Langford, John Sillitoe on behalf of the Wellcome Sanger Institute COVID-19 Surveillance Team |
| EPI_ISL_2513883, EPI_ISL_2513884 | Lighthouse Lab in Alderley Park | Wellcome Sanger Institute for the COVID-19 Genomics UK (COG-UK) Consortium | Jacquelyn Wynn, Mairead Hyland, The Lighthouse Lab in Alderley Park and Alex Alderton, Roberto Amato, Jeffrey Barrett, Sonia Goncalves, Ewan Harrison, David K. Jackson, Ian Johnston, Dominic Kwiatkowski, Cordelia Langford, John Sillitoe on behalf of the Wellcome Sanger Institute COVID-19 Surveillance Team |
| EPI_ISL_2513885 | Lighthouse Laboratory Plymouth | Wellcome Sanger Institute for the COVID-19 Genomics UK (COG-UK) Consortium | Lighthouse Laboratory Plymouth and Alex Alderton, Roberto Amato, Jeffrey Barrett, Sonia Goncalves, Ewan Harrison, David K. Jackson, Ian Johnston, Dominic Kwiatkowski, Cordelia Langford, John Sillitoe on behalf of the Wellcome Sanger Institute COVID-19 Surveillance Team |
| EPI_ISL_2513886 | Health Services Laboratories | Wellcome Sanger Institute for the COVID-19 Genomics UK (COG-UK) Consortium | Health Services Laboratories and Alex Alderton, Roberto Amato, Jeffrey Barrett, Sonia Goncalves, Ewan Harrison, David K. Jackson, Ian Johnston, Dominic Kwiatkowski, Cordelia Langford, John Sillitoe on behalf of the Wellcome Sanger Institute COVID-19 Surveillance Team |

| | | | |
|---|---|---|---|
| EPI_ISL_2513887 | Lighthouse Laboratory Plymouth | Wellcome Sanger Institute for the COVID-19 Genomics UK (COG-UK) Consortium | Lighthouse Laboratory Plymouth and Alex Alderton, Roberto Amato, Jeffrey Barrett, Sonia Goncalves, Ewan Harrison, David K. Jackson, Ian Johnston, Dominic Kwiatkowski, Cordelia Langford, John Sillitoe on behalf of the Wellcome Sanger Institute COVID-19 Surveillance Team |
| EPI_ISL_2513888, EPI_ISL_2513889 | Health Services Laboratories | Wellcome Sanger Institute for the COVID-19 Genomics UK (COG-UK) Consortium | Health Services Laboratories and Alex Alderton, Roberto Amato, Jeffrey Barrett, Sonia Goncalves, Ewan Harrison, David K. Jackson, Ian Johnston, Dominic Kwiatkowski, Cordelia Langford, John Sillitoe on behalf of the Wellcome Sanger Institute COVID-19 Surveillance Team |
| EPI_ISL_2513890 | Lighthouse Laboratory Plymouth | Wellcome Sanger Institute for the COVID-19 Genomics UK (COG-UK) Consortium | Lighthouse Laboratory Plymouth and Alex Alderton, Roberto Amato, Jeffrey Barrett, Sonia Goncalves, Ewan Harrison, David K. Jackson, Ian Johnston, Dominic Kwiatkowski, Cordelia Langford, John Sillitoe on behalf of the Wellcome Sanger Institute COVID-19 Surveillance Team |
| EPI_ISL_2513891 | Health Services Laboratories | Wellcome Sanger Institute for the COVID-19 Genomics UK (COG-UK) Consortium | Health Services Laboratories and Alex Alderton, Roberto Amato, Jeffrey Barrett, Sonia Goncalves, Ewan Harrison, David K. Jackson, Ian Johnston, Dominic Kwiatkowski, Cordelia Langford, John Sillitoe on behalf of the Wellcome Sanger Institute COVID-19 Surveillance Team |
| EPI_ISL_2513892 | Lighthouse Lab in Alderley Park | Wellcome Sanger Institute for the COVID-19 Genomics UK (COG-UK) Consortium | Jacquelyn Wynn, Mairead Hyland, The Lighthouse Lab in Alderley Park and Alex Alderton, Roberto Amato, Jeffrey Barrett, Sonia Goncalves, Ewan Harrison, David K. Jackson, Ian Johnston, Dominic Kwiatkowski, Cordelia Langford, John Sillitoe on behalf of the Wellcome Sanger Institute COVID-19 Surveillance Team |
| EPI_ISL_2513893, EPI_ISL_2513894, EPI_ISL_2513895, EPI_ISL_2513896 | Health Services Laboratories | Wellcome Sanger Institute for the COVID-19 Genomics UK (COG-UK) Consortium | Health Services Laboratories and Alex Alderton, Roberto Amato, Jeffrey Barrett, Sonia Goncalves, Ewan Harrison, David K. Jackson, Ian Johnston, Dominic Kwiatkowski, Cordelia Langford, John Sillitoe on behalf of the Wellcome Sanger Institute COVID-19 Surveillance Team |
| EPI_ISL_2513897 | Lighthouse Lab in Alderley Park | Wellcome Sanger Institute for the COVID-19 Genomics UK (COG-UK) Consortium | Jacquelyn Wynn, Mairead Hyland, The Lighthouse Lab in Alderley Park and Alex Alderton, Roberto Amato, Jeffrey Barrett, Sonia Goncalves, Ewan Harrison, David K. Jackson, Ian Johnston, Dominic Kwiatkowski, Cordelia Langford, John Sillitoe on behalf of the Wellcome Sanger Institute COVID-19 Surveillance Team |
| EPI_ISL_2513898, EPI_ISL_2513899 | Health Services Laboratories | Wellcome Sanger Institute for the COVID-19 Genomics UK (COG-UK) Consortium | Health Services Laboratories and Alex Alderton, Roberto Amato, Jeffrey Barrett, Sonia Goncalves, Ewan Harrison, David K. Jackson, Ian Johnston, Dominic Kwiatkowski, Cordelia Langford, John Sillitoe on behalf of the Wellcome Sanger Institute COVID-19 Surveillance Team |
| EPI_ISL_2513900 | Lighthouse Lab in Alderley Park | Wellcome Sanger Institute for the COVID-19 Genomics UK (COG-UK) Consortium | Jacquelyn Wynn, Mairead Hyland, The Lighthouse Lab in Alderley Park and Alex Alderton, Roberto Amato, Jeffrey Barrett, Sonia Goncalves, Ewan Harrison, David K. Jackson, Ian Johnston, Dominic Kwiatkowski, Cordelia Langford, John Sillitoe on behalf of the Wellcome Sanger Institute COVID-19 Surveillance Team |
| EPI_ISL_2513901, EPI_ISL_2513902 | Health Services Laboratories | Wellcome Sanger Institute for the COVID-19 Genomics UK (COG-UK) Consortium | Health Services Laboratories and Alex Alderton, Roberto Amato, Jeffrey Barrett, Sonia Goncalves, Ewan Harrison, David K. Jackson, Ian Johnston, Dominic Kwiatkowski, Cordelia Langford, John Sillitoe on behalf of the Wellcome Sanger Institute COVID-19 Surveillance Team |
| EPI_ISL_2513903, EPI_ISL_2513904, EPI_ISL_2513905 | Lighthouse Lab in Alderley Park | Wellcome Sanger Institute for the COVID-19 Genomics UK (COG-UK) Consortium | Jacquelyn Wynn, Mairead Hyland, The Lighthouse Lab in Alderley Park and Alex Alderton, Roberto Amato, Jeffrey Barrett, Sonia Goncalves, Ewan Harrison, David K. Jackson, Ian Johnston, Dominic Kwiatkowski, Cordelia Langford, John Sillitoe on behalf of the Wellcome Sanger Institute COVID-19 Surveillance Team |
| EPI_ISL_2513906 | Lighthouse Laboratory Plymouth | Wellcome Sanger Institute for the COVID-19 Genomics UK (COG-UK) Consortium | Lighthouse Laboratory Plymouth and Alex Alderton, Roberto Amato, Jeffrey Barrett, Sonia Goncalves, Ewan Harrison, David K. Jackson, Ian Johnston, Dominic Kwiatkowski, Cordelia Langford, John Sillitoe on behalf of the Wellcome Sanger Institute COVID-19 Surveillance Team |
| EPI_ISL_2513907, EPI_ISL_2513908, EPI_ISL_2513909 | Health Services Laboratories | Wellcome Sanger Institute for the COVID-19 Genomics UK (COG-UK) Consortium | Health Services Laboratories and Alex Alderton, Roberto Amato, Jeffrey Barrett, Sonia Goncalves, Ewan Harrison, David K. Jackson, Ian Johnston, Dominic Kwiatkowski, Cordelia Langford, John Sillitoe on behalf of the Wellcome Sanger Institute COVID-19 Surveillance Team |
| EPI_ISL_2513910 | Lighthouse Lab in Glasgow | Wellcome Sanger Institute for the COVID-19 Genomics UK (COG-UK) Consortium | Harper VanSteenhouse, Yumi Kasai, David Gray, Carol Clugston, Anna Dominiczak and Alex Alderton, Roberto Amato, Jeffrey Barrett, Sonia Goncalves, Ewan Harrison, David K. Jackson, Ian Johnston, Dominic Kwiatkowski, Cordelia Langford, John Sillitoe on behalf of the Wellcome Sanger Institute COVID-19 Surveillance Team |
| EPI_ISL_2513912, EPI_ISL_2513913, EPI_ISL_2513914, EPI_ISL_2513915 | Health Services Laboratories | Wellcome Sanger Institute for the COVID-19 Genomics UK (COG-UK) Consortium | Health Services Laboratories and Alex Alderton, Roberto Amato, Jeffrey Barrett, Sonia Goncalves, Ewan Harrison, David K. Jackson, Ian Johnston, Dominic Kwiatkowski, Cordelia Langford, John Sillitoe on behalf of the Wellcome Sanger Institute COVID-19 Surveillance Team |
| EPI_ISL_2513916 | Lighthouse Lab in Alderley Park | Wellcome Sanger Institute for the COVID-19 Genomics UK (COG-UK) Consortium | Jacquelyn Wynn, Mairead Hyland, The Lighthouse Lab in Alderley Park and Alex Alderton, Roberto Amato, Jeffrey Barrett, Sonia Goncalves, Ewan Harrison, David K. Jackson, Ian Johnston, Dominic Kwiatkowski, Cordelia Langford, John Sillitoe on behalf of the Wellcome Sanger Institute COVID-19 Surveillance Team |
| EPI_ISL_2513917 | Lighthouse Laboratory Plymouth | Wellcome Sanger Institute for the COVID-19 Genomics UK (COG-UK) Consortium | Lighthouse Laboratory Plymouth and Alex Alderton, Roberto Amato, Jeffrey Barrett, Sonia Goncalves, Ewan Harrison, David K. Jackson, Ian Johnston, Dominic Kwiatkowski, Cordelia Langford, John Sillitoe on behalf of the Wellcome Sanger Institute COVID-19 Surveillance Team |
| EPI_ISL_2513918 | Lighthouse Lab in Alderley Park | Wellcome Sanger Institute for the COVID-19 Genomics UK (COG-UK) Consortium | Jacquelyn Wynn, Mairead Hyland, The Lighthouse Lab in Alderley Park and Alex Alderton, Roberto Amato, Jeffrey Barrett, Sonia Goncalves, Ewan Harrison, David K. Jackson, Ian Johnston, Dominic Kwiatkowski, Cordelia Langford, John Sillitoe on behalf of the Wellcome Sanger Institute COVID-19 Surveillance Team |
| EPI_ISL_2513919, EPI_ISL_2513920, EPI_ISL_2513921, EPI_ISL_2513922 | Health Services Laboratories | Wellcome Sanger Institute for the COVID-19 Genomics UK (COG-UK) Consortium | Health Services Laboratories and Alex Alderton, Roberto Amato, Jeffrey Barrett, Sonia Goncalves, Ewan Harrison, David K. Jackson, Ian Johnston, Dominic Kwiatkowski, Cordelia Langford, John Sillitoe on behalf of the Wellcome Sanger Institute COVID-19 Surveillance Team |
| EPI_ISL_2513924, EPI_ISL_2513925 | Lighthouse Lab in Alderley Park | Wellcome Sanger Institute for the COVID-19 Genomics UK (COG-UK) Consortium | Jacquelyn Wynn, Mairead Hyland, The Lighthouse Lab in Alderley Park and Alex Alderton, Roberto Amato, Jeffrey Barrett, Sonia Goncalves, Ewan Harrison, David K. Jackson, Ian Johnston, Dominic Kwiatkowski, Cordelia Langford, John Sillitoe on behalf of the Wellcome Sanger Institute COVID-19 Surveillance Team |
| EPI_ISL_2513926, EPI_ISL_2513927 | Health Services Laboratories | Wellcome Sanger Institute for the COVID-19 Genomics UK (COG-UK) Consortium | Health Services Laboratories and Alex Alderton, Roberto Amato, Jeffrey Barrett, Sonia Goncalves, Ewan Harrison, David K. Jackson, Ian Johnston, Dominic Kwiatkowski, Cordelia Langford, John Sillitoe on behalf of the Wellcome Sanger Institute COVID-19 Surveillance Team |
| EPI_ISL_2513928, EPI_ISL_2513929 | Lighthouse Lab in Alderley Park | Wellcome Sanger Institute for the COVID-19 Genomics UK (COG-UK) Consortium | Jacquelyn Wynn, Mairead Hyland, The Lighthouse Lab in Alderley Park and Alex Alderton, Roberto Amato, Jeffrey Barrett, Sonia Goncalves, Ewan Harrison, David K. Jackson, Ian Johnston, Dominic Kwiatkowski, Cordelia Langford, John Sillitoe on behalf of the Wellcome Sanger Institute COVID-19 Surveillance Team |
| EPI_ISL_2513931 | Health Services Laboratories | Wellcome Sanger Institute for the COVID-19 Genomics UK (COG-UK) Consortium | Health Services Laboratories and Alex Alderton, Roberto Amato, Jeffrey Barrett, Sonia Goncalves, Ewan Harrison, David K. Jackson, Ian Johnston, Dominic Kwiatkowski, Cordelia Langford, John Sillitoe on behalf of the Wellcome Sanger Institute COVID-19 Surveillance Team |
| EPI_ISL_2513932 | Lighthouse Lab in Alderley Park | Wellcome Sanger Institute for the COVID-19 Genomics UK (COG-UK) Consortium | Jacquelyn Wynn, Mairead Hyland, The Lighthouse Lab in Alderley Park and Alex Alderton, Roberto Amato, Jeffrey Barrett, Sonia Goncalves, Ewan Harrison, David K. Jackson, Ian Johnston, Dominic Kwiatkowski, Cordelia Langford, John Sillitoe on behalf of the Wellcome Sanger Institute COVID-19 Surveillance Team |
| EPI_ISL_2513933 | Health Services Laboratories | Wellcome Sanger Institute for the COVID-19 Genomics UK (COG-UK) Consortium | Health Services Laboratories and Alex Alderton, Roberto Amato, Jeffrey Barrett, Sonia Goncalves, Ewan Harrison, David K. Jackson, Ian Johnston, Dominic Kwiatkowski, Cordelia Langford, John Sillitoe on behalf of the Wellcome Sanger Institute COVID-19 Surveillance Team |
| EPI_ISL_2513934 | Lighthouse Laboratory Plymouth | Wellcome Sanger Institute for the COVID-19 Genomics UK (COG-UK) Consortium | Lighthouse Laboratory Plymouth and Alex Alderton, Roberto Amato, Jeffrey Barrett, Sonia Goncalves, Ewan Harrison, David K. Jackson, Ian Johnston, Dominic Kwiatkowski, Cordelia Langford, John Sillitoe on behalf of the Wellcome Sanger Institute COVID-19 Surveillance Team |
| EPI_ISL_2513935, EPI_ISL_2513936 | Lighthouse Lab in Alderley Park | Wellcome Sanger Institute for the COVID-19 Genomics UK (COG-UK) Consortium | Jacquelyn Wynn, Mairead Hyland, The Lighthouse Lab in Alderley Park and Alex Alderton, Roberto Amato, Jeffrey Barrett, Sonia Goncalves, Ewan Harrison, David K. Jackson, Ian Johnston, Dominic Kwiatkowski, Cordelia Langford, John Sillitoe on behalf of the Wellcome Sanger Institute COVID-19 Surveillance Team |
| EPI_ISL_2513937, EPI_ISL_2513938 | Health Services Laboratories | Wellcome Sanger Institute for the COVID-19 Genomics UK (COG-UK) Consortium | Health Services Laboratories and Alex Alderton, Roberto Amato, Jeffrey Barrett, Sonia Goncalves, Ewan Harrison, David K. Jackson, Ian Johnston, Dominic Kwiatkowski, Cordelia Langford, John Sillitoe on behalf of the Wellcome Sanger Institute COVID-19 Surveillance Team |
| EPI_ISL_2513939, EPI_ISL_2513940 | Lighthouse Lab in Alderley Park | Wellcome Sanger Institute for the COVID-19 Genomics UK (COG-UK) Consortium | Jacquelyn Wynn, Mairead Hyland, The Lighthouse Lab in Alderley Park and Alex Alderton, Roberto Amato, Jeffrey Barrett, Sonia Goncalves, Ewan Harrison, David K. Jackson, Ian Johnston, Dominic Kwiatkowski, Cordelia Langford, John Sillitoe on behalf of the Wellcome Sanger Institute COVID-19 Surveillance Team |
| EPI_ISL_2513941 | Lighthouse Laboratory Plymouth | Wellcome Sanger Institute for the COVID-19 Genomics UK (COG-UK) Consortium | Lighthouse Laboratory Plymouth and Alex Alderton, Roberto Amato, Jeffrey Barrett, Sonia Goncalves, Ewan Harrison, David K. Jackson, Ian Johnston, Dominic Kwiatkowski, Cordelia Langford, John Sillitoe on behalf of the Wellcome Sanger Institute COVID-19 Surveillance Team |
| EPI_ISL_2513942, EPI_ISL_2513943 | Health Services Laboratories | Wellcome Sanger Institute for the COVID-19 Genomics UK (COG-UK) Consortium | Health Services Laboratories and Alex Alderton, Roberto Amato, Jeffrey Barrett, Sonia Goncalves, Ewan Harrison, David K. Jackson, Ian Johnston, Dominic Kwiatkowski, Cordelia Langford, John Sillitoe on behalf of the Wellcome Sanger Institute COVID-19 Surveillance Team |
| EPI_ISL_2513944 | Lighthouse Laboratory Plymouth | Wellcome Sanger Institute for the COVID-19 Genomics UK (COG-UK) Consortium | Lighthouse Laboratory Plymouth and Alex Alderton, Roberto Amato, Jeffrey Barrett, Sonia Goncalves, Ewan Harrison, David K. Jackson, Ian Johnston, Dominic Kwiatkowski, Cordelia Langford, John Sillitoe on behalf of the Wellcome Sanger Institute COVID-19 Surveillance Team |
| EPI_ISL_2513945, EPI_ISL_2513946 | Lighthouse Lab in Alderley Park | Wellcome Sanger Institute for the COVID-19 Genomics UK (COG-UK) Consortium | Jacquelyn Wynn, Mairead Hyland, The Lighthouse Lab in Alderley Park and Alex Alderton, Roberto Amato, Jeffrey Barrett, Sonia Goncalves, Ewan Harrison, David K. Jackson, Ian Johnston, Dominic Kwiatkowski, Cordelia Langford, John Sillitoe on behalf of the Wellcome Sanger Institute COVID-19 Surveillance Team |
| EPI_ISL_2513947 | Lighthouse Laboratory Plymouth | Wellcome Sanger Institute for the COVID-19 Genomics UK (COG-UK) Consortium | Lighthouse Laboratory Plymouth and Alex Alderton, Roberto Amato, Jeffrey Barrett, Sonia Goncalves, Ewan Harrison, David K. Jackson, Ian Johnston, Dominic Kwiatkowski, Cordelia Langford, John Sillitoe on behalf of the Wellcome Sanger Institute COVID-19 Surveillance Team |
| EPI_ISL_2513948, EPI_ISL_2513949, EPI_ISL_2513950, EPI_ISL_2513951 | Health Services Laboratories | Wellcome Sanger Institute for the COVID-19 Genomics UK (COG-UK) Consortium | Health Services Laboratories and Alex Alderton, Roberto Amato, Jeffrey Barrett, Sonia Goncalves, Ewan Harrison, David K. Jackson, Ian Johnston, Dominic Kwiatkowski, Cordelia Langford, John Sillitoe on behalf of the Wellcome Sanger Institute COVID-19 Surveillance Team |
| EPI_ISL_2513952, EPI_ISL_2513954 | Lighthouse Lab in Alderley Park | Wellcome Sanger Institute for the COVID-19 Genomics UK (COG-UK) Consortium | Jacquelyn Wynn, Mairead Hyland, The Lighthouse Lab in Alderley Park and Alex Alderton, Roberto Amato, Jeffrey Barrett, Sonia Goncalves, Ewan Harrison, David K. Jackson, Ian Johnston, Dominic Kwiatkowski, Cordelia Langford, John Sillitoe on behalf of the Wellcome Sanger Institute COVID-19 Surveillance Team |
| EPI_ISL_2513955, EPI_ISL_2513956 | Health Services Laboratories | Wellcome Sanger Institute for the COVID-19 Genomics UK (COG-UK) Consortium | Health Services Laboratories and Alex Alderton, Roberto Amato, Jeffrey Barrett, Sonia Goncalves, Ewan Harrison, David K. Jackson, Ian Johnston, Dominic Kwiatkowski, Cordelia Langford, John Sillitoe on behalf of the Wellcome Sanger Institute COVID-19 Surveillance Team |
| EPI_ISL_2513957 | Lighthouse Lab in Alderley Park | Wellcome Sanger Institute for the COVID-19 Genomics UK (COG-UK) Consortium | Jacquelyn Wynn, Mairead Hyland, The Lighthouse Lab in Alderley Park and Alex Alderton, Roberto Amato, Jeffrey Barrett, Sonia Goncalves, Ewan Harrison, David K. Jackson, Ian Johnston, Dominic Kwiatkowski, Cordelia Langford, John Sillitoe on behalf of the Wellcome Sanger Institute COVID-19 Surveillance Team |
| EPI_ISL_2513958 | Health Services Laboratories | Wellcome Sanger Institute for the COVID-19 Genomics UK (COG-UK) Consortium | Health Services Laboratories and Alex Alderton, Roberto Amato, Jeffrey Barrett, Sonia Goncalves, Ewan Harrison, David K. Jackson, Ian Johnston, Dominic Kwiatkowski, Cordelia Langford, John Sillitoe on behalf of the Wellcome Sanger Institute COVID-19 Surveillance Team |
| EPI_ISL_2513959, EPI_ISL_2513960, EPI_ISL_2513961, EPI_ISL_2513962, EPI_ISL_2513963 | Lighthouse Lab in Alderley Park | Wellcome Sanger Institute for the COVID-19 Genomics UK (COG-UK) Consortium | Jacquelyn Wynn, Mairead Hyland, The Lighthouse Lab in Alderley Park and Alex Alderton, Roberto Amato, Jeffrey Barrett, Sonia Goncalves, Ewan Harrison, David K. Jackson, Ian Johnston, Dominic Kwiatkowski, Cordelia Langford, John Sillitoe on behalf of the Wellcome Sanger Institute COVID-19 Surveillance Team |

| | | | |
|---|---|---|---|
| EPI_ISL_2513964 | Lighthouse Lab in Glasgow | Wellcome Sanger Institute for the COVID-19 Genomics UK (COG-UK) Consortium | Harper VanSteenhouse, Yumi Kasai, David Gray, Carol Clugston, Anna Dominiczak and Alex Alderton, Roberto Amato, Jeffrey Barrett, Sonia Goncalves, Ewan Harrison, David K. Jackson, Ian Johnston, Dominic Kwiatkowski, Cordelia Langford, John Sillitoe on behalf of the Wellcome Sanger Institute COVID-19 Surveillance Team |
| EPI_ISL_2513965 | Health Services Laboratories | Wellcome Sanger Institute for the COVID-19 Genomics UK (COG-UK) Consortium | Health Services Laboratories and Alex Alderton, Roberto Amato, Jeffrey Barrett, Sonia Goncalves, Ewan Harrison, David K. Jackson, Ian Johnston, Dominic Kwiatkowski, Cordelia Langford, John Sillitoe on behalf of the Wellcome Sanger Institute COVID-19 Surveillance Team |
| EPI_ISL_2513966 | Lighthouse Lab in Alderley Park | Wellcome Sanger Institute for the COVID-19 Genomics UK (COG-UK) Consortium | Jacquelyn Wynn, Mairead Hyland, The Lighthouse Lab in Alderley Park and Alex Alderton, Roberto Amato, Jeffrey Barrett, Sonia Goncalves, Ewan Harrison, David K. Jackson, Ian Johnston, Dominic Kwiatkowski, Cordelia Langford, John Sillitoe on behalf of the Wellcome Sanger Institute COVID-19 Surveillance Team |
| EPI_ISL_2513967, EPI_ISL_2513968 | Health Services Laboratories | Wellcome Sanger Institute for the COVID-19 Genomics UK (COG-UK) Consortium | Health Services Laboratories and Alex Alderton, Roberto Amato, Jeffrey Barrett, Sonia Goncalves, Ewan Harrison, David K. Jackson, Ian Johnston, Dominic Kwiatkowski, Cordelia Langford, John Sillitoe on behalf of the Wellcome Sanger Institute COVID-19 Surveillance Team |
| EPI_ISL_2513969 | Lighthouse Laboratory Plymouth | Wellcome Sanger Institute for the COVID-19 Genomics UK (COG-UK) Consortium | Lighthouse Laboratory Plymouth and Alex Alderton, Roberto Amato, Jeffrey Barrett, Sonia Goncalves, Ewan Harrison, David K. Jackson, Ian Johnston, Dominic Kwiatkowski, Cordelia Langford, John Sillitoe on behalf of the Wellcome Sanger Institute COVID-19 Surveillance Team |
| EPI_ISL_2513970 | Lighthouse Lab in Alderley Park | Wellcome Sanger Institute for the COVID-19 Genomics UK (COG-UK) Consortium | Jacquelyn Wynn, Mairead Hyland, The Lighthouse Lab in Alderley Park and Alex Alderton, Roberto Amato, Jeffrey Barrett, Sonia Goncalves, Ewan Harrison, David K. Jackson, Ian Johnston, Dominic Kwiatkowski, Cordelia Langford, John Sillitoe on behalf of the Wellcome Sanger Institute COVID-19 Surveillance Team |
| EPI_ISL_2513971 | Health Services Laboratories | Wellcome Sanger Institute for the COVID-19 Genomics UK (COG-UK) Consortium | Health Services Laboratories and Alex Alderton, Roberto Amato, Jeffrey Barrett, Sonia Goncalves, Ewan Harrison, David K. Jackson, Ian Johnston, Dominic Kwiatkowski, Cordelia Langford, John Sillitoe on behalf of the Wellcome Sanger Institute COVID-19 Surveillance Team |
| EPI_ISL_2513972, EPI_ISL_2513973, EPI_ISL_2513974 | Lighthouse Lab in Alderley Park | Wellcome Sanger Institute for the COVID-19 Genomics UK (COG-UK) Consortium | Jacquelyn Wynn, Mairead Hyland, The Lighthouse Lab in Alderley Park and Alex Alderton, Roberto Amato, Jeffrey Barrett, Sonia Goncalves, Ewan Harrison, David K. Jackson, Ian Johnston, Dominic Kwiatkowski, Cordelia Langford, John Sillitoe on behalf of the Wellcome Sanger Institute COVID-19 Surveillance Team |
| EPI_ISL_2513975, EPI_ISL_2513976, EPI_ISL_2513977, EPI_ISL_2513978, EPI_ISL_2513979 | Health Services Laboratories | Wellcome Sanger Institute for the COVID-19 Genomics UK (COG-UK) Consortium | Health Services Laboratories and Alex Alderton, Roberto Amato, Jeffrey Barrett, Sonia Goncalves, Ewan Harrison, David K. Jackson, Ian Johnston, Dominic Kwiatkowski, Cordelia Langford, John Sillitoe on behalf of the Wellcome Sanger Institute COVID-19 Surveillance Team |
| EPI_ISL_2513980 | Lighthouse Lab in Alderley Park | Wellcome Sanger Institute for the COVID-19 Genomics UK (COG-UK) Consortium | Jacquelyn Wynn, Mairead Hyland, The Lighthouse Lab in Alderley Park and Alex Alderton, Roberto Amato, Jeffrey Barrett, Sonia Goncalves, Ewan Harrison, David K. Jackson, Ian Johnston, Dominic Kwiatkowski, Cordelia Langford, John Sillitoe on behalf of the Wellcome Sanger Institute COVID-19 Surveillance Team |
| EPI_ISL_2513981, EPI_ISL_2513982 | Health Services Laboratories | Wellcome Sanger Institute for the COVID-19 Genomics UK (COG-UK) Consortium | Health Services Laboratories and Alex Alderton, Roberto Amato, Jeffrey Barrett, Sonia Goncalves, Ewan Harrison, David K. Jackson, Ian Johnston, Dominic Kwiatkowski, Cordelia Langford, John Sillitoe on behalf of the Wellcome Sanger Institute COVID-19 Surveillance Team |
| EPI_ISL_2513983 | Lighthouse Lab in Alderley Park | Wellcome Sanger Institute for the COVID-19 Genomics UK (COG-UK) Consortium | Jacquelyn Wynn, Mairead Hyland, The Lighthouse Lab in Alderley Park and Alex Alderton, Roberto Amato, Jeffrey Barrett, Sonia Goncalves, Ewan Harrison, David K. Jackson, Ian Johnston, Dominic Kwiatkowski, Cordelia Langford, John Sillitoe on behalf of the Wellcome Sanger Institute COVID-19 Surveillance Team |
| EPI_ISL_2513984 | Health Services Laboratories | Wellcome Sanger Institute for the COVID-19 Genomics UK (COG-UK) Consortium | Health Services Laboratories and Alex Alderton, Roberto Amato, Jeffrey Barrett, Sonia Goncalves, Ewan Harrison, David K. Jackson, Ian Johnston, Dominic Kwiatkowski, Cordelia Langford, John Sillitoe on behalf of the Wellcome Sanger Institute COVID-19 Surveillance Team |
| EPI_ISL_2513985 | Lighthouse Lab in Alderley Park | Wellcome Sanger Institute for the COVID-19 Genomics UK (COG-UK) Consortium | Jacquelyn Wynn, Mairead Hyland, The Lighthouse Lab in Alderley Park and Alex Alderton, Roberto Amato, Jeffrey Barrett, Sonia Goncalves, Ewan Harrison, David K. Jackson, Ian Johnston, Dominic Kwiatkowski, Cordelia Langford, John Sillitoe on behalf of the Wellcome Sanger Institute COVID-19 Surveillance Team |
| EPI_ISL_2513987, EPI_ISL_2513989 | Health Services Laboratories | Wellcome Sanger Institute for the COVID-19 Genomics UK (COG-UK) Consortium | Health Services Laboratories and Alex Alderton, Roberto Amato, Jeffrey Barrett, Sonia Goncalves, Ewan Harrison, David K. Jackson, Ian Johnston, Dominic Kwiatkowski, Cordelia Langford, John Sillitoe on behalf of the Wellcome Sanger Institute COVID-19 Surveillance Team |
| EPI_ISL_2513990 | Lighthouse Lab in Alderley Park | Wellcome Sanger Institute for the COVID-19 Genomics UK (COG-UK) Consortium | Jacquelyn Wynn, Mairead Hyland, The Lighthouse Lab in Alderley Park and Alex Alderton, Roberto Amato, Jeffrey Barrett, Sonia Goncalves, Ewan Harrison, David K. Jackson, Ian Johnston, Dominic Kwiatkowski, Cordelia Langford, John Sillitoe on behalf of the Wellcome Sanger Institute COVID-19 Surveillance Team |
| EPI_ISL_2513991 | Lighthouse Laboratory Plymouth | Wellcome Sanger Institute for the COVID-19 Genomics UK (COG-UK) Consortium | Lighthouse Laboratory Plymouth and Alex Alderton, Roberto Amato, Jeffrey Barrett, Sonia Goncalves, Ewan Harrison, David K. Jackson, Ian Johnston, Dominic Kwiatkowski, Cordelia Langford, John Sillitoe on behalf of the Wellcome Sanger Institute COVID-19 Surveillance Team |
| EPI_ISL_2513992 | Health Services Laboratories | Wellcome Sanger Institute for the COVID-19 Genomics UK (COG-UK) Consortium | Health Services Laboratories and Alex Alderton, Roberto Amato, Jeffrey Barrett, Sonia Goncalves, Ewan Harrison, David K. Jackson, Ian Johnston, Dominic Kwiatkowski, Cordelia Langford, John Sillitoe on behalf of the Wellcome Sanger Institute COVID-19 Surveillance Team |
| EPI_ISL_2513994, EPI_ISL_2513995 | Lighthouse Lab in Alderley Park | Wellcome Sanger Institute for the COVID-19 Genomics UK (COG-UK) Consortium | Jacquelyn Wynn, Mairead Hyland, The Lighthouse Lab in Alderley Park and Alex Alderton, Roberto Amato, Jeffrey Barrett, Sonia Goncalves, Ewan Harrison, David K. Jackson, Ian Johnston, Dominic Kwiatkowski, Cordelia Langford, John Sillitoe on behalf of the Wellcome Sanger Institute COVID-19 Surveillance Team |
| EPI_ISL_2513996 | Lighthouse Laboratory Plymouth | Wellcome Sanger Institute for the COVID-19 Genomics UK (COG-UK) Consortium | Lighthouse Laboratory Plymouth and Alex Alderton, Roberto Amato, Jeffrey Barrett, Sonia Goncalves, Ewan Harrison, David K. Jackson, Ian Johnston, Dominic Kwiatkowski, Cordelia Langford, John Sillitoe on behalf of the Wellcome Sanger Institute COVID-19 Surveillance Team |
| EPI_ISL_2513997 | Health Services Laboratories | Wellcome Sanger Institute for the COVID-19 Genomics UK (COG-UK) Consortium | Health Services Laboratories and Alex Alderton, Roberto Amato, Jeffrey Barrett, Sonia Goncalves, Ewan Harrison, David K. Jackson, Ian Johnston, Dominic Kwiatkowski, Cordelia Langford, John Sillitoe on behalf of the Wellcome Sanger Institute COVID-19 Surveillance Team |
| EPI_ISL_2513998 | Lighthouse Lab in Alderley Park | Wellcome Sanger Institute for the COVID-19 Genomics UK (COG-UK) Consortium | Jacquelyn Wynn, Mairead Hyland, The Lighthouse Lab in Alderley Park and Alex Alderton, Roberto Amato, Jeffrey Barrett, Sonia Goncalves, Ewan Harrison, David K. Jackson, Ian Johnston, Dominic Kwiatkowski, Cordelia Langford, John Sillitoe on behalf of the Wellcome Sanger Institute COVID-19 Surveillance Team |
| EPI_ISL_2513999, EPI_ISL_2514000 | Health Services Laboratories | Wellcome Sanger Institute for the COVID-19 Genomics UK (COG-UK) Consortium | Health Services Laboratories and Alex Alderton, Roberto Amato, Jeffrey Barrett, Sonia Goncalves, Ewan Harrison, David K. Jackson, Ian Johnston, Dominic Kwiatkowski, Cordelia Langford, John Sillitoe on behalf of the Wellcome Sanger Institute COVID-19 Surveillance Team |
| EPI_ISL_2514001 | Lighthouse Lab in Alderley Park | Wellcome Sanger Institute for the COVID-19 Genomics UK (COG-UK) Consortium | Jacquelyn Wynn, Mairead Hyland, The Lighthouse Lab in Alderley Park and Alex Alderton, Roberto Amato, Jeffrey Barrett, Sonia Goncalves, Ewan Harrison, David K. Jackson, Ian Johnston, Dominic Kwiatkowski, Cordelia Langford, John Sillitoe on behalf of the Wellcome Sanger Institute COVID-19 Surveillance Team |
| EPI_ISL_2514002, EPI_ISL_2514003, EPI_ISL_2514004, EPI_ISL_2514005 | Health Services Laboratories | Wellcome Sanger Institute for the COVID-19 Genomics UK (COG-UK) Consortium | Health Services Laboratories and Alex Alderton, Roberto Amato, Jeffrey Barrett, Sonia Goncalves, Ewan Harrison, David K. Jackson, Ian Johnston, Dominic Kwiatkowski, Cordelia Langford, John Sillitoe on behalf of the Wellcome Sanger Institute COVID-19 Surveillance Team |
| EPI_ISL_2514006 | Lighthouse Lab in Alderley Park | Wellcome Sanger Institute for the COVID-19 Genomics UK (COG-UK) Consortium | Jacquelyn Wynn, Mairead Hyland, The Lighthouse Lab in Alderley Park and Alex Alderton, Roberto Amato, Jeffrey Barrett, Sonia Goncalves, Ewan Harrison, David K. Jackson, Ian Johnston, Dominic Kwiatkowski, Cordelia Langford, John Sillitoe on behalf of the Wellcome Sanger Institute COVID-19 Surveillance Team |
| EPI_ISL_2514007 | Health Services Laboratories | Wellcome Sanger Institute for the COVID-19 Genomics UK (COG-UK) Consortium | Health Services Laboratories and Alex Alderton, Roberto Amato, Jeffrey Barrett, Sonia Goncalves, Ewan Harrison, David K. Jackson, Ian Johnston, Dominic Kwiatkowski, Cordelia Langford, John Sillitoe on behalf of the Wellcome Sanger Institute COVID-19 Surveillance Team |
| EPI_ISL_2514008 | Lighthouse Lab in Alderley Park | Wellcome Sanger Institute for the COVID-19 Genomics UK (COG-UK) Consortium | Jacquelyn Wynn, Mairead Hyland, The Lighthouse Lab in Alderley Park and Alex Alderton, Roberto Amato, Jeffrey Barrett, Sonia Goncalves, Ewan Harrison, David K. Jackson, Ian Johnston, Dominic Kwiatkowski, Cordelia Langford, John Sillitoe on behalf of the Wellcome Sanger Institute COVID-19 Surveillance Team |
| EPI_ISL_2514009 | Health Services Laboratories | Wellcome Sanger Institute for the COVID-19 Genomics UK (COG-UK) Consortium | Health Services Laboratories and Alex Alderton, Roberto Amato, Jeffrey Barrett, Sonia Goncalves, Ewan Harrison, David K. Jackson, Ian Johnston, Dominic Kwiatkowski, Cordelia Langford, John Sillitoe on behalf of the Wellcome Sanger Institute COVID-19 Surveillance Team |
| EPI_ISL_2514010 | Lighthouse Lab in Alderley Park | Wellcome Sanger Institute for the COVID-19 Genomics UK (COG-UK) Consortium | Jacquelyn Wynn, Mairead Hyland, The Lighthouse Lab in Alderley Park and Alex Alderton, Roberto Amato, Jeffrey Barrett, Sonia Goncalves, Ewan Harrison, David K. Jackson, Ian Johnston, Dominic Kwiatkowski, Cordelia Langford, John Sillitoe on behalf of the Wellcome Sanger Institute COVID-19 Surveillance Team |
| EPI_ISL_2514011 | Health Services Laboratories | Wellcome Sanger Institute for the COVID-19 Genomics UK (COG-UK) Consortium | Health Services Laboratories and Alex Alderton, Roberto Amato, Jeffrey Barrett, Sonia Goncalves, Ewan Harrison, David K. Jackson, Ian Johnston, Dominic Kwiatkowski, Cordelia Langford, John Sillitoe on behalf of the Wellcome Sanger Institute COVID-19 Surveillance Team |
| EPI_ISL_2514012 | Lighthouse Laboratory Plymouth | Wellcome Sanger Institute for the COVID-19 Genomics UK (COG-UK) Consortium | Lighthouse Laboratory Plymouth and Alex Alderton, Roberto Amato, Jeffrey Barrett, Sonia Goncalves, Ewan Harrison, David K. Jackson, Ian Johnston, Dominic Kwiatkowski, Cordelia Langford, John Sillitoe on behalf of the Wellcome Sanger Institute COVID-19 Surveillance Team |
| EPI_ISL_2514014 | Health Services Laboratories | Wellcome Sanger Institute for the COVID-19 Genomics UK (COG-UK) Consortium | Health Services Laboratories and Alex Alderton, Roberto Amato, Jeffrey Barrett, Sonia Goncalves, Ewan Harrison, David K. Jackson, Ian Johnston, Dominic Kwiatkowski, Cordelia Langford, John Sillitoe on behalf of the Wellcome Sanger Institute COVID-19 Surveillance Team |
| EPI_ISL_2514015, EPI_ISL_2514016 | Lighthouse Lab in Alderley Park | Wellcome Sanger Institute for the COVID-19 Genomics UK (COG-UK) Consortium | Jacquelyn Wynn, Mairead Hyland, The Lighthouse Lab in Alderley Park and Alex Alderton, Roberto Amato, Jeffrey Barrett, Sonia Goncalves, Ewan Harrison, David K. Jackson, Ian Johnston, Dominic Kwiatkowski, Cordelia Langford, John Sillitoe on behalf of the Wellcome Sanger Institute COVID-19 Surveillance Team |
| EPI_ISL_2514017 | Lighthouse Laboratory Plymouth | Wellcome Sanger Institute for the COVID-19 Genomics UK (COG-UK) Consortium | Lighthouse Laboratory Plymouth and Alex Alderton, Roberto Amato, Jeffrey Barrett, Sonia Goncalves, Ewan Harrison, David K. Jackson, Ian Johnston, Dominic Kwiatkowski, Cordelia Langford, John Sillitoe on behalf of the Wellcome Sanger Institute COVID-19 Surveillance Team |
| EPI_ISL_2514018 | Lighthouse Lab in Alderley Park | Wellcome Sanger Institute for the COVID-19 Genomics UK (COG-UK) Consortium | Jacquelyn Wynn, Mairead Hyland, The Lighthouse Lab in Alderley Park and Alex Alderton, Roberto Amato, Jeffrey Barrett, Sonia Goncalves, Ewan Harrison, David K. Jackson, Ian Johnston, Dominic Kwiatkowski, Cordelia Langford, John Sillitoe on behalf of the Wellcome Sanger Institute COVID-19 Surveillance Team |
| EPI_ISL_2514019 | Lighthouse Laboratory Plymouth | Wellcome Sanger Institute for the COVID-19 Genomics UK (COG-UK) Consortium | Lighthouse Laboratory Plymouth and Alex Alderton, Roberto Amato, Jeffrey Barrett, Sonia Goncalves, Ewan Harrison, David K. Jackson, Ian Johnston, Dominic Kwiatkowski, Cordelia Langford, John Sillitoe on behalf of the Wellcome Sanger Institute COVID-19 Surveillance Team |
| EPI_ISL_2514021, EPI_ISL_2514022 | Health Services Laboratories | Wellcome Sanger Institute for the COVID-19 Genomics UK (COG-UK) Consortium | Health Services Laboratories and Alex Alderton, Roberto Amato, Jeffrey Barrett, Sonia Goncalves, Ewan Harrison, David K. Jackson, Ian Johnston, Dominic Kwiatkowski, Cordelia Langford, John Sillitoe on behalf of the Wellcome Sanger Institute COVID-19 Surveillance Team |
| EPI_ISL_2514023 | Lighthouse Lab in Alderley Park | Wellcome Sanger Institute for the COVID-19 Genomics UK (COG-UK) Consortium | Jacquelyn Wynn, Mairead Hyland, The Lighthouse Lab in Alderley Park and Alex Alderton, Roberto Amato, Jeffrey Barrett, Sonia Goncalves, Ewan Harrison, David K. Jackson, Ian Johnston, Dominic Kwiatkowski, Cordelia Langford, John Sillitoe on behalf of the Wellcome Sanger Institute COVID-19 Surveillance Team |
| EPI_ISL_2514024, EPI_ISL_2514025, EPI_ISL_2514026 | Health Services Laboratories | Wellcome Sanger Institute for the COVID-19 Genomics UK (COG-UK) Consortium | Health Services Laboratories and Alex Alderton, Roberto Amato, Jeffrey Barrett, Sonia Goncalves, Ewan Harrison, David K. Jackson, Ian Johnston, Dominic Kwiatkowski, Cordelia Langford, John Sillitoe on behalf of the Wellcome Sanger Institute COVID-19 Surveillance Team |
| EPI_ISL_2514027 | Lighthouse Lab in Alderley Park | Wellcome Sanger Institute for the COVID-19 Genomics UK (COG-UK) Consortium | Jacquelyn Wynn, Mairead Hyland, The Lighthouse Lab in Alderley Park and Alex Alderton, Roberto Amato, Jeffrey Barrett, Sonia Goncalves, Ewan Harrison, David K. Jackson, Ian Johnston, Dominic Kwiatkowski, Cordelia Langford, John Sillitoe on behalf of the Wellcome Sanger Institute COVID-19 Surveillance Team |
| EPI_ISL_2514028, EPI_ISL_2514029, | Health Services Laboratories | Wellcome Sanger Institute for the COVID-19 Genomics UK (COG-UK) Consortium | Health Services Laboratories and Alex Alderton, Roberto Amato, Jeffrey Barrett, Sonia Goncalves, Ewan Harrison, David K. Jackson, Ian Johnston, Dominic Kwiatkowski, Cordelia Langford, John Sillitoe on behalf of the Wellcome Sanger Institute COVID-19 Surveillance Team |

| | | | |
|---|---|---|---|
| EPI_ISL_2514031, EPI_ISL_2514033 | | | |
| EPI_ISL_2514034 | Lighthouse Laboratory Plymouth | Wellcome Sanger Institute for the COVID-19 Genomics UK (COG-UK) Consortium | Lighthouse Laboratory Plymouth and Alex Alderton, Roberto Amato, Jeffrey Barrett, Sonia Goncalves, Ewan Harrison, David K. Jackson, Ian Johnston, Dominic Kwiatkowski, Cordelia Langford, John Sillitoe on behalf of the Wellcome Sanger Institute COVID-19 Surveillance Team |
| EPI_ISL_2514035, EPI_ISL_2514036, EPI_ISL_2514037, EPI_ISL_2514038, EPI_ISL_2514039 | Health Services Laboratories | Wellcome Sanger Institute for the COVID-19 Genomics UK (COG-UK) Consortium | Health Services Laboratories and Alex Alderton, Roberto Amato, Jeffrey Barrett, Sonia Goncalves, Ewan Harrison, David K. Jackson, Ian Johnston, Dominic Kwiatkowski, Cordelia Langford, John Sillitoe on behalf of the Wellcome Sanger Institute COVID-19 Surveillance Team |
| EPI_ISL_2514040 | Lighthouse Laboratory Plymouth | Wellcome Sanger Institute for the COVID-19 Genomics UK (COG-UK) Consortium | Lighthouse Laboratory Plymouth and Alex Alderton, Roberto Amato, Jeffrey Barrett, Sonia Goncalves, Ewan Harrison, David K. Jackson, Ian Johnston, Dominic Kwiatkowski, Cordelia Langford, John Sillitoe on behalf of the Wellcome Sanger Institute COVID-19 Surveillance Team |
| EPI_ISL_2514041 | Health Services Laboratories | Wellcome Sanger Institute for the COVID-19 Genomics UK (COG-UK) Consortium | Health Services Laboratories and Alex Alderton, Roberto Amato, Jeffrey Barrett, Sonia Goncalves, Ewan Harrison, David K. Jackson, Ian Johnston, Dominic Kwiatkowski, Cordelia Langford, John Sillitoe on behalf of the Wellcome Sanger Institute COVID-19 Surveillance Team |
| EPI_ISL_2514042 | Lighthouse Lab in Alderley Park | Wellcome Sanger Institute for the COVID-19 Genomics UK (COG-UK) Consortium | Jacquelyn Wynn, Mairead Hyland, The Lighthouse Lab in Alderley Park and Alex Alderton, Roberto Amato, Jeffrey Barrett, Sonia Goncalves, Ewan Harrison, David K. Jackson, Ian Johnston, Dominic Kwiatkowski, Cordelia Langford, John Sillitoe on behalf of the Wellcome Sanger Institute COVID-19 Surveillance Team |
| EPI_ISL_2514043 | Lighthouse Laboratory Plymouth | Wellcome Sanger Institute for the COVID-19 Genomics UK (COG-UK) Consortium | Lighthouse Laboratory Plymouth and Alex Alderton, Roberto Amato, Jeffrey Barrett, Sonia Goncalves, Ewan Harrison, David K. Jackson, Ian Johnston, Dominic Kwiatkowski, Cordelia Langford, John Sillitoe on behalf of the Wellcome Sanger Institute COVID-19 Surveillance Team |
| EPI_ISL_2514045, EPI_ISL_2514046, EPI_ISL_2514047 | Lighthouse Lab in Alderley Park | Wellcome Sanger Institute for the COVID-19 Genomics UK (COG-UK) Consortium | Jacquelyn Wynn, Mairead Hyland, The Lighthouse Lab in Alderley Park and Alex Alderton, Roberto Amato, Jeffrey Barrett, Sonia Goncalves, Ewan Harrison, David K. Jackson, Ian Johnston, Dominic Kwiatkowski, Cordelia Langford, John Sillitoe on behalf of the Wellcome Sanger Institute COVID-19 Surveillance Team |
| EPI_ISL_2514048 | Health Services Laboratories | Wellcome Sanger Institute for the COVID-19 Genomics UK (COG-UK) Consortium | Health Services Laboratories and Alex Alderton, Roberto Amato, Jeffrey Barrett, Sonia Goncalves, Ewan Harrison, David K. Jackson, Ian Johnston, Dominic Kwiatkowski, Cordelia Langford, John Sillitoe on behalf of the Wellcome Sanger Institute COVID-19 Surveillance Team |
| EPI_ISL_2514049 | Lighthouse Lab in Alderley Park | Wellcome Sanger Institute for the COVID-19 Genomics UK (COG-UK) Consortium | Jacquelyn Wynn, Mairead Hyland, The Lighthouse Lab in Alderley Park and Alex Alderton, Roberto Amato, Jeffrey Barrett, Sonia Goncalves, Ewan Harrison, David K. Jackson, Ian Johnston, Dominic Kwiatkowski, Cordelia Langford, John Sillitoe on behalf of the Wellcome Sanger Institute COVID-19 Surveillance Team |
| EPI_ISL_2514050 | Health Services Laboratories | Wellcome Sanger Institute for the COVID-19 Genomics UK (COG-UK) Consortium | Health Services Laboratories and Alex Alderton, Roberto Amato, Jeffrey Barrett, Sonia Goncalves, Ewan Harrison, David K. Jackson, Ian Johnston, Dominic Kwiatkowski, Cordelia Langford, John Sillitoe on behalf of the Wellcome Sanger Institute COVID-19 Surveillance Team |
| EPI_ISL_2514051 | Lighthouse Laboratory Plymouth | Wellcome Sanger Institute for the COVID-19 Genomics UK (COG-UK) Consortium | Lighthouse Laboratory Plymouth and Alex Alderton, Roberto Amato, Jeffrey Barrett, Sonia Goncalves, Ewan Harrison, David K. Jackson, Ian Johnston, Dominic Kwiatkowski, Cordelia Langford, John Sillitoe on behalf of the Wellcome Sanger Institute COVID-19 Surveillance Team |
| EPI_ISL_2514052 | Lighthouse Lab in Alderley Park | Wellcome Sanger Institute for the COVID-19 Genomics UK (COG-UK) Consortium | Jacquelyn Wynn, Mairead Hyland, The Lighthouse Lab in Alderley Park and Alex Alderton, Roberto Amato, Jeffrey Barrett, Sonia Goncalves, Ewan Harrison, David K. Jackson, Ian Johnston, Dominic Kwiatkowski, Cordelia Langford, John Sillitoe on behalf of the Wellcome Sanger Institute COVID-19 Surveillance Team |
| EPI_ISL_2514053 | Health Services Laboratories | Wellcome Sanger Institute for the COVID-19 Genomics UK (COG-UK) Consortium | Health Services Laboratories and Alex Alderton, Roberto Amato, Jeffrey Barrett, Sonia Goncalves, Ewan Harrison, David K. Jackson, Ian Johnston, Dominic Kwiatkowski, Cordelia Langford, John Sillitoe on behalf of the Wellcome Sanger Institute COVID-19 Surveillance Team |
| EPI_ISL_2514054 | Lighthouse Laboratory Plymouth | Wellcome Sanger Institute for the COVID-19 Genomics UK (COG-UK) Consortium | Lighthouse Laboratory Plymouth and Alex Alderton, Roberto Amato, Jeffrey Barrett, Sonia Goncalves, Ewan Harrison, David K. Jackson, Ian Johnston, Dominic Kwiatkowski, Cordelia Langford, John Sillitoe on behalf of the Wellcome Sanger Institute COVID-19 Surveillance Team |
| EPI_ISL_2514055 | Lighthouse Lab in Alderley Park | Wellcome Sanger Institute for the COVID-19 Genomics UK (COG-UK) Consortium | Jacquelyn Wynn, Mairead Hyland, The Lighthouse Lab in Alderley Park and Alex Alderton, Roberto Amato, Jeffrey Barrett, Sonia Goncalves, Ewan Harrison, David K. Jackson, Ian Johnston, Dominic Kwiatkowski, Cordelia Langford, John Sillitoe on behalf of the Wellcome Sanger Institute COVID-19 Surveillance Team |
| EPI_ISL_2514056, EPI_ISL_2514057 | Health Services Laboratories | Wellcome Sanger Institute for the COVID-19 Genomics UK (COG-UK) Consortium | Health Services Laboratories and Alex Alderton, Roberto Amato, Jeffrey Barrett, Sonia Goncalves, Ewan Harrison, David K. Jackson, Ian Johnston, Dominic Kwiatkowski, Cordelia Langford, John Sillitoe on behalf of the Wellcome Sanger Institute COVID-19 Surveillance Team |
| EPI_ISL_2514058, EPI_ISL_2514059 | Lighthouse Lab in Alderley Park | Wellcome Sanger Institute for the COVID-19 Genomics UK (COG-UK) Consortium | Jacquelyn Wynn, Mairead Hyland, The Lighthouse Lab in Alderley Park and Alex Alderton, Roberto Amato, Jeffrey Barrett, Sonia Goncalves, Ewan Harrison, David K. Jackson, Ian Johnston, Dominic Kwiatkowski, Cordelia Langford, John Sillitoe on behalf of the Wellcome Sanger Institute COVID-19 Surveillance Team |
| EPI_ISL_2514060 | Health Services Laboratories | Wellcome Sanger Institute for the COVID-19 Genomics UK (COG-UK) Consortium | Health Services Laboratories and Alex Alderton, Roberto Amato, Jeffrey Barrett, Sonia Goncalves, Ewan Harrison, David K. Jackson, Ian Johnston, Dominic Kwiatkowski, Cordelia Langford, John Sillitoe on behalf of the Wellcome Sanger Institute COVID-19 Surveillance Team |
| EPI_ISL_2514061 | Lighthouse Lab in Alderley Park | Wellcome Sanger Institute for the COVID-19 Genomics UK (COG-UK) Consortium | Jacquelyn Wynn, Mairead Hyland, The Lighthouse Lab in Alderley Park and Alex Alderton, Roberto Amato, Jeffrey Barrett, Sonia Goncalves, Ewan Harrison, David K. Jackson, Ian Johnston, Dominic Kwiatkowski, Cordelia Langford, John Sillitoe on behalf of the Wellcome Sanger Institute COVID-19 Surveillance Team |
| EPI_ISL_2514062 | Health Services Laboratories | Wellcome Sanger Institute for the COVID-19 Genomics UK (COG-UK) Consortium | Health Services Laboratories and Alex Alderton, Roberto Amato, Jeffrey Barrett, Sonia Goncalves, Ewan Harrison, David K. Jackson, Ian Johnston, Dominic Kwiatkowski, Cordelia Langford, John Sillitoe on behalf of the Wellcome Sanger Institute COVID-19 Surveillance Team |
| EPI_ISL_2514064 | Lighthouse Laboratory Plymouth | Wellcome Sanger Institute for the COVID-19 Genomics UK (COG-UK) Consortium | Lighthouse Laboratory Plymouth and Alex Alderton, Roberto Amato, Jeffrey Barrett, Sonia Goncalves, Ewan Harrison, David K. Jackson, Ian Johnston, Dominic Kwiatkowski, Cordelia Langford, John Sillitoe on behalf of the Wellcome Sanger Institute COVID-19 Surveillance Team |
| EPI_ISL_2514065, EPI_ISL_2514066, EPI_ISL_2514067, EPI_ISL_2514068 | Lighthouse Lab in Alderley Park | Wellcome Sanger Institute for the COVID-19 Genomics UK (COG-UK) Consortium | Jacquelyn Wynn, Mairead Hyland, The Lighthouse Lab in Alderley Park and Alex Alderton, Roberto Amato, Jeffrey Barrett, Sonia Goncalves, Ewan Harrison, David K. Jackson, Ian Johnston, Dominic Kwiatkowski, Cordelia Langford, John Sillitoe on behalf of the Wellcome Sanger Institute COVID-19 Surveillance Team |
| EPI_ISL_2514069 | Lighthouse Laboratory Plymouth | Wellcome Sanger Institute for the COVID-19 Genomics UK (COG-UK) Consortium | Lighthouse Laboratory Plymouth and Alex Alderton, Roberto Amato, Jeffrey Barrett, Sonia Goncalves, Ewan Harrison, David K. Jackson, Ian Johnston, Dominic Kwiatkowski, Cordelia Langford, John Sillitoe on behalf of the Wellcome Sanger Institute COVID-19 Surveillance Team |
| EPI_ISL_2514070 | Lighthouse Lab in Alderley Park | Wellcome Sanger Institute for the COVID-19 Genomics UK (COG-UK) Consortium | Jacquelyn Wynn, Mairead Hyland, The Lighthouse Lab in Alderley Park and Alex Alderton, Roberto Amato, Jeffrey Barrett, Sonia Goncalves, Ewan Harrison, David K. Jackson, Ian Johnston, Dominic Kwiatkowski, Cordelia Langford, John Sillitoe on behalf of the Wellcome Sanger Institute COVID-19 Surveillance Team |
| EPI_ISL_2514071 | Health Services Laboratories | Wellcome Sanger Institute for the COVID-19 Genomics UK (COG-UK) Consortium | Health Services Laboratories and Alex Alderton, Roberto Amato, Jeffrey Barrett, Sonia Goncalves, Ewan Harrison, David K. Jackson, Ian Johnston, Dominic Kwiatkowski, Cordelia Langford, John Sillitoe on behalf of the Wellcome Sanger Institute COVID-19 Surveillance Team |
| EPI_ISL_2514073, EPI_ISL_2514074, EPI_ISL_2514075, EPI_ISL_2514076, EPI_ISL_2514077, EPI_ISL_2514078, EPI_ISL_2514079, EPI_ISL_2514080, EPI_ISL_2514081, EPI_ISL_2514082, EPI_ISL_2514083, EPI_ISL_2514084, EPI_ISL_2514085, EPI_ISL_2514086, EPI_ISL_2514087, EPI_ISL_2514088, EPI_ISL_2514089, EPI_ISL_2514090, EPI_ISL_2514091, EPI_ISL_2514096, EPI_ISL_2514097, EPI_ISL_2514098, EPI_ISL_2514099, EPI_ISL_2514100, EPI_ISL_2514101 | | | |
| see above | Lighthouse Lab in Alderley Park | Wellcome Sanger Institute for the COVID-19 Genomics UK (COG-UK) Consortium | Jacquelyn Wynn, Mairead Hyland, The Lighthouse Lab in Alderley Park and Alex Alderton, Roberto Amato, Jeffrey Barrett, Sonia Goncalves, Ewan Harrison, David K. Jackson, Ian Johnston, Dominic Kwiatkowski, Cordelia Langford, John Sillitoe on behalf of the Wellcome Sanger Institute COVID-19 Surveillance Team |
| EPI_ISL_2514102, EPI_ISL_2514103 | Lighthouse Lab in Milton Keynes | Wellcome Sanger Institute for the COVID-19 Genomics UK (COG-UK) Consortium | The Lighthouse Lab in Milton Keynes and Alex Alderton, Roberto Amato, Jeffrey Barrett, Sonia Goncalves, Ewan Harrison, David K. Jackson, Ian Johnston, Dominic Kwiatkowski, Cordelia Langford, John Sillitoe on behalf of the Wellcome Sanger Institute COVID-19 Surveillance Team |
| EPI_ISL_2514104, EPI_ISL_2514105, EPI_ISL_2514106, EPI_ISL_2514108, EPI_ISL_2514109, EPI_ISL_2514110, EPI_ISL_2514111, EPI_ISL_2514112, EPI_ISL_2514113, EPI_ISL_2514114, EPI_ISL_2514115 | | | |
| see above | Lighthouse Lab in Alderley Park | Wellcome Sanger Institute for the COVID-19 Genomics UK (COG-UK) Consortium | Jacquelyn Wynn, Mairead Hyland, The Lighthouse Lab in Alderley Park and Alex Alderton, Roberto Amato, Jeffrey Barrett, Sonia Goncalves, Ewan Harrison, David K. Jackson, Ian Johnston, Dominic Kwiatkowski, Cordelia Langford, John Sillitoe on behalf of the Wellcome Sanger Institute COVID-19 Surveillance Team |
| EPI_ISL_2514116 | Lighthouse Lab in Milton Keynes | Wellcome Sanger Institute for the COVID-19 Genomics UK (COG-UK) Consortium | The Lighthouse Lab in Milton Keynes and Alex Alderton, Roberto Amato, Jeffrey Barrett, Sonia Goncalves, Ewan Harrison, David K. Jackson, Ian Johnston, Dominic Kwiatkowski, Cordelia Langford, John Sillitoe on behalf of the Wellcome Sanger Institute COVID-19 Surveillance Team |
| EPI_ISL_2514117 | Lighthouse Lab in Alderley Park | Wellcome Sanger Institute for the COVID-19 Genomics UK (COG-UK) Consortium | Jacquelyn Wynn, Mairead Hyland, The Lighthouse Lab in Alderley Park and Alex Alderton, Roberto Amato, Jeffrey Barrett, Sonia Goncalves, Ewan Harrison, David K. Jackson, Ian Johnston, Dominic Kwiatkowski, Cordelia Langford, John Sillitoe on behalf of the Wellcome Sanger Institute COVID-19 Surveillance Team |
| EPI_ISL_2514118 | Lighthouse Lab in Milton Keynes | Wellcome Sanger Institute for the COVID-19 Genomics UK (COG-UK) Consortium | The Lighthouse Lab in Milton Keynes and Alex Alderton, Roberto Amato, Jeffrey Barrett, Sonia Goncalves, Ewan Harrison, David K. Jackson, Ian Johnston, Dominic Kwiatkowski, Cordelia Langford, John Sillitoe on behalf of the Wellcome Sanger Institute COVID-19 Surveillance Team |
| EPI_ISL_2514119 | Lighthouse Lab in Alderley Park | Wellcome Sanger Institute for the COVID-19 Genomics UK (COG-UK) Consortium | Jacquelyn Wynn, Mairead Hyland, The Lighthouse Lab in Alderley Park and Alex Alderton, Roberto Amato, Jeffrey Barrett, Sonia Goncalves, Ewan Harrison, David K. Jackson, Ian Johnston, Dominic Kwiatkowski, Cordelia Langford, John Sillitoe on behalf of the Wellcome Sanger Institute COVID-19 Surveillance Team |
| EPI_ISL_2514121 | Lighthouse Lab in Milton Keynes | Wellcome Sanger Institute for the COVID-19 Genomics UK (COG-UK) Consortium | The Lighthouse Lab in Milton Keynes and Alex Alderton, Roberto Amato, Jeffrey Barrett, Sonia Goncalves, Ewan Harrison, David K. Jackson, Ian Johnston, Dominic Kwiatkowski, Cordelia Langford, John Sillitoe on behalf of the Wellcome Sanger Institute COVID-19 Surveillance Team |
| EPI_ISL_2514122, EPI_ISL_2514123, EPI_ISL_2514124 | Lighthouse Lab in Alderley Park | Wellcome Sanger Institute for the COVID-19 Genomics UK (COG-UK) Consortium | Jacquelyn Wynn, Mairead Hyland, The Lighthouse Lab in Alderley Park and Alex Alderton, Roberto Amato, Jeffrey Barrett, Sonia Goncalves, Ewan Harrison, David K. Jackson, Ian Johnston, Dominic Kwiatkowski, Cordelia Langford, John Sillitoe on behalf of the Wellcome Sanger Institute COVID-19 Surveillance Team |
| EPI_ISL_2514125, EPI_ISL_2514126 | Lighthouse Lab in Milton Keynes | Wellcome Sanger Institute for the COVID-19 Genomics UK (COG-UK) Consortium | The Lighthouse Lab in Milton Keynes and Alex Alderton, Roberto Amato, Jeffrey Barrett, Sonia Goncalves, Ewan Harrison, David K. Jackson, Ian Johnston, Dominic Kwiatkowski, Cordelia Langford, John Sillitoe on behalf of the Wellcome Sanger Institute COVID-19 Surveillance Team |
| EPI_ISL_2514127, EPI_ISL_2514128 | Lighthouse Lab in Alderley Park | Wellcome Sanger Institute for the COVID-19 Genomics UK (COG-UK) Consortium | Jacquelyn Wynn, Mairead Hyland, The Lighthouse Lab in Alderley Park and Alex Alderton, Roberto Amato, Jeffrey Barrett, Sonia Goncalves, Ewan Harrison, David K. Jackson, Ian Johnston, Dominic Kwiatkowski, Cordelia Langford, John Sillitoe on behalf of the Wellcome Sanger Institute COVID-19 Surveillance Team |
| EPI_ISL_2514129 | Lighthouse Lab in Milton Keynes | Wellcome Sanger Institute for the COVID-19 Genomics UK (COG-UK) Consortium | The Lighthouse Lab in Milton Keynes and Alex Alderton, Roberto Amato, Jeffrey Barrett, Sonia Goncalves, Ewan Harrison, David K. Jackson, Ian Johnston, Dominic Kwiatkowski, Cordelia Langford, John Sillitoe on behalf of the Wellcome Sanger Institute COVID-19 Surveillance Team |
| EPI_ISL_2514130 | Lighthouse Lab in Alderley Park | Wellcome Sanger Institute for the COVID-19 Genomics UK (COG-UK) Consortium | Jacquelyn Wynn, Mairead Hyland, The Lighthouse Lab in Alderley Park and Alex Alderton, Roberto Amato, Jeffrey Barrett, Sonia Goncalves, Ewan Harrison, David K. Jackson, Ian Johnston, Dominic Kwiatkowski, Cordelia Langford, John Sillitoe on behalf of the Wellcome Sanger Institute COVID-19 Surveillance Team |
| EPI_ISL_2514131 | Lighthouse Lab in Milton Keynes | Wellcome Sanger Institute for the COVID-19 Genomics UK (COG-UK) Consortium | The Lighthouse Lab in Milton Keynes and Alex Alderton, Roberto Amato, Jeffrey Barrett, Sonia Goncalves, Ewan Harrison, David K. Jackson, Ian Johnston, Dominic Kwiatkowski, Cordelia Langford, John Sillitoe on behalf of the Wellcome Sanger Institute COVID-19 Surveillance Team |
| EPI_ISL_2514132, | Lighthouse Lab in Alderley Park | Wellcome Sanger Institute for the COVID-19 Genomics UK (COG-UK) Consortium | Jacquelyn Wynn, Mairead Hyland, The Lighthouse Lab in Alderley Park and Alex Alderton, Roberto Amato, Jeffrey Barrett, Sonia Goncalves, Ewan Harrison, David K. Jackson, Ian Johnston, Dominic Kwiatkowski, Cordelia Langford, John Sillitoe on |

| | | | |
|---|---|---|---|
| EPI_ISL_2514133 | | Consortium | behalf of the Wellcome Sanger Institute COVID-19 Surveillance Team |
| EPI_ISL_2514135, EPI_ISL_2514136 | Lighthouse Lab in Milton Keynes | Wellcome Sanger Institute for the COVID-19 Genomics UK (COG-UK) Consortium | The Lighthouse Lab in Milton Keynes and Alex Alderton, Roberto Amato, Jeffrey Barrett, Sonia Goncalves, Ewan Harrison, David K. Jackson, Ian Johnston, Dominic Kwiatkowski, Cordelia Langford, John Sillitoe on behalf of the Wellcome Sanger Institute COVID-19 Surveillance Team |
| EPI_ISL_2514137, EPI_ISL_2514138, EPI_ISL_2514139, EPI_ISL_2514140, EPI_ISL_2514141, EPI_ISL_2514142, EPI_ISL_2514143 | Lighthouse Lab in Alderley Park | Wellcome Sanger Institute for the COVID-19 Genomics UK (COG-UK) Consortium | Jacquelyn Wynn, Mairead Hyland, The Lighthouse Lab in Alderley Park and Alex Alderton, Roberto Amato, Jeffrey Barrett, Sonia Goncalves, Ewan Harrison, David K. Jackson, Ian Johnston, Dominic Kwiatkowski, Cordelia Langford, John Sillitoe on behalf of the Wellcome Sanger Institute COVID-19 Surveillance Team |
| EPI_ISL_2514144 | Lighthouse Lab in Milton Keynes | Wellcome Sanger Institute for the COVID-19 Genomics UK (COG-UK) Consortium | The Lighthouse Lab in Milton Keynes and Alex Alderton, Roberto Amato, Jeffrey Barrett, Sonia Goncalves, Ewan Harrison, David K. Jackson, Ian Johnston, Dominic Kwiatkowski, Cordelia Langford, John Sillitoe on behalf of the Wellcome Sanger Institute COVID-19 Surveillance Team |
| EPI_ISL_2514145, EPI_ISL_2514146, EPI_ISL_2514147, EPI_ISL_2514148, EPI_ISL_2514149, EPI_ISL_2514150, EPI_ISL_2514151 | Lighthouse Lab in Alderley Park | Wellcome Sanger Institute for the COVID-19 Genomics UK (COG-UK) Consortium | Jacquelyn Wynn, Mairead Hyland, The Lighthouse Lab in Alderley Park and Alex Alderton, Roberto Amato, Jeffrey Barrett, Sonia Goncalves, Ewan Harrison, David K. Jackson, Ian Johnston, Dominic Kwiatkowski, Cordelia Langford, John Sillitoe on behalf of the Wellcome Sanger Institute COVID-19 Surveillance Team |
| EPI_ISL_2514152 | Lighthouse Lab in Milton Keynes | Wellcome Sanger Institute for the COVID-19 Genomics UK (COG-UK) Consortium | The Lighthouse Lab in Milton Keynes and Alex Alderton, Roberto Amato, Jeffrey Barrett, Sonia Goncalves, Ewan Harrison, David K. Jackson, Ian Johnston, Dominic Kwiatkowski, Cordelia Langford, John Sillitoe on behalf of the Wellcome Sanger Institute COVID-19 Surveillance Team |
| EPI_ISL_2514153, EPI_ISL_2514154, EPI_ISL_2514155, EPI_ISL_2514157, EPI_ISL_2514158, EPI_ISL_2514159, EPI_ISL_2514160, EPI_ISL_2514161 | Lighthouse Lab in Alderley Park | Wellcome Sanger Institute for the COVID-19 Genomics UK (COG-UK) Consortium | Jacquelyn Wynn, Mairead Hyland, The Lighthouse Lab in Alderley Park and Alex Alderton, Roberto Amato, Jeffrey Barrett, Sonia Goncalves, Ewan Harrison, David K. Jackson, Ian Johnston, Dominic Kwiatkowski, Cordelia Langford, John Sillitoe on behalf of the Wellcome Sanger Institute COVID-19 Surveillance Team |
| EPI_ISL_2514162 | Lighthouse Lab in Milton Keynes | Wellcome Sanger Institute for the COVID-19 Genomics UK (COG-UK) Consortium | The Lighthouse Lab in Milton Keynes and Alex Alderton, Roberto Amato, Jeffrey Barrett, Sonia Goncalves, Ewan Harrison, David K. Jackson, Ian Johnston, Dominic Kwiatkowski, Cordelia Langford, John Sillitoe on behalf of the Wellcome Sanger Institute COVID-19 Surveillance Team |
| EPI_ISL_2514163, EPI_ISL_2514164, EPI_ISL_2514165, EPI_ISL_2514166, EPI_ISL_2514167 | Lighthouse Lab in Alderley Park | Wellcome Sanger Institute for the COVID-19 Genomics UK (COG-UK) Consortium | Jacquelyn Wynn, Mairead Hyland, The Lighthouse Lab in Alderley Park and Alex Alderton, Roberto Amato, Jeffrey Barrett, Sonia Goncalves, Ewan Harrison, David K. Jackson, Ian Johnston, Dominic Kwiatkowski, Cordelia Langford, John Sillitoe on behalf of the Wellcome Sanger Institute COVID-19 Surveillance Team |
| EPI_ISL_2514168 | Lighthouse Lab in Milton Keynes | Wellcome Sanger Institute for the COVID-19 Genomics UK (COG-UK) Consortium | The Lighthouse Lab in Milton Keynes and Alex Alderton, Roberto Amato, Jeffrey Barrett, Sonia Goncalves, Ewan Harrison, David K. Jackson, Ian Johnston, Dominic Kwiatkowski, Cordelia Langford, John Sillitoe on behalf of the Wellcome Sanger Institute COVID-19 Surveillance Team |
| EPI_ISL_2514170, EPI_ISL_2514172, EPI_ISL_2514173 | Lighthouse Lab in Alderley Park | Wellcome Sanger Institute for the COVID-19 Genomics UK (COG-UK) Consortium | Jacquelyn Wynn, Mairead Hyland, The Lighthouse Lab in Alderley Park and Alex Alderton, Roberto Amato, Jeffrey Barrett, Sonia Goncalves, Ewan Harrison, David K. Jackson, Ian Johnston, Dominic Kwiatkowski, Cordelia Langford, John Sillitoe on behalf of the Wellcome Sanger Institute COVID-19 Surveillance Team |
| EPI_ISL_2514174 | Lighthouse Lab in Milton Keynes | Wellcome Sanger Institute for the COVID-19 Genomics UK (COG-UK) Consortium | The Lighthouse Lab in Milton Keynes and Alex Alderton, Roberto Amato, Jeffrey Barrett, Sonia Goncalves, Ewan Harrison, David K. Jackson, Ian Johnston, Dominic Kwiatkowski, Cordelia Langford, John Sillitoe on behalf of the Wellcome Sanger Institute COVID-19 Surveillance Team |
| EPI_ISL_2514175, EPI_ISL_2514176, EPI_ISL_2514178, EPI_ISL_2514179, EPI_ISL_2514180 | Lighthouse Lab in Alderley Park | Wellcome Sanger Institute for the COVID-19 Genomics UK (COG-UK) Consortium | Jacquelyn Wynn, Mairead Hyland, The Lighthouse Lab in Alderley Park and Alex Alderton, Roberto Amato, Jeffrey Barrett, Sonia Goncalves, Ewan Harrison, David K. Jackson, Ian Johnston, Dominic Kwiatkowski, Cordelia Langford, John Sillitoe on behalf of the Wellcome Sanger Institute COVID-19 Surveillance Team |
| EPI_ISL_2514181 | Lighthouse Lab in Milton Keynes | Wellcome Sanger Institute for the COVID-19 Genomics UK (COG-UK) Consortium | The Lighthouse Lab in Milton Keynes and Alex Alderton, Roberto Amato, Jeffrey Barrett, Sonia Goncalves, Ewan Harrison, David K. Jackson, Ian Johnston, Dominic Kwiatkowski, Cordelia Langford, John Sillitoe on behalf of the Wellcome Sanger Institute COVID-19 Surveillance Team |
| EPI_ISL_2514182 | Lighthouse Lab in Alderley Park | Wellcome Sanger Institute for the COVID-19 Genomics UK (COG-UK) Consortium | Jacquelyn Wynn, Mairead Hyland, The Lighthouse Lab in Alderley Park and Alex Alderton, Roberto Amato, Jeffrey Barrett, Sonia Goncalves, Ewan Harrison, David K. Jackson, Ian Johnston, Dominic Kwiatkowski, Cordelia Langford, John Sillitoe on behalf of the Wellcome Sanger Institute COVID-19 Surveillance Team |
| EPI_ISL_2514183 | Lighthouse Lab in Milton Keynes | Wellcome Sanger Institute for the COVID-19 Genomics UK (COG-UK) Consortium | The Lighthouse Lab in Milton Keynes and Alex Alderton, Roberto Amato, Jeffrey Barrett, Sonia Goncalves, Ewan Harrison, David K. Jackson, Ian Johnston, Dominic Kwiatkowski, Cordelia Langford, John Sillitoe on behalf of the Wellcome Sanger Institute COVID-19 Surveillance Team |
| EPI_ISL_2514184, EPI_ISL_2514186, EPI_ISL_2514187, EPI_ISL_2514188, EPI_ISL_2514189 | Lighthouse Lab in Alderley Park | Wellcome Sanger Institute for the COVID-19 Genomics UK (COG-UK) Consortium | Jacquelyn Wynn, Mairead Hyland, The Lighthouse Lab in Alderley Park and Alex Alderton, Roberto Amato, Jeffrey Barrett, Sonia Goncalves, Ewan Harrison, David K. Jackson, Ian Johnston, Dominic Kwiatkowski, Cordelia Langford, John Sillitoe on behalf of the Wellcome Sanger Institute COVID-19 Surveillance Team |
| EPI_ISL_2514190 | Lighthouse Lab in Milton Keynes | Wellcome Sanger Institute for the COVID-19 Genomics UK (COG-UK) Consortium | The Lighthouse Lab in Milton Keynes and Alex Alderton, Roberto Amato, Jeffrey Barrett, Sonia Goncalves, Ewan Harrison, David K. Jackson, Ian Johnston, Dominic Kwiatkowski, Cordelia Langford, John Sillitoe on behalf of the Wellcome Sanger Institute COVID-19 Surveillance Team |
| EPI_ISL_2514191, EPI_ISL_2514192, EPI_ISL_2514194, EPI_ISL_2514195, EPI_ISL_2514196, EPI_ISL_2514197, EPI_ISL_2514198, EPI_ISL_2514199, EPI_ISL_2514200, EPI_ISL_2514202 | Lighthouse Lab in Alderley Park | Wellcome Sanger Institute for the COVID-19 Genomics UK (COG-UK) Consortium | Jacquelyn Wynn, Mairead Hyland, The Lighthouse Lab in Alderley Park and Alex Alderton, Roberto Amato, Jeffrey Barrett, Sonia Goncalves, Ewan Harrison, David K. Jackson, Ian Johnston, Dominic Kwiatkowski, Cordelia Langford, John Sillitoe on behalf of the Wellcome Sanger Institute COVID-19 Surveillance Team |
| EPI_ISL_2514203 | Lighthouse Lab in Milton Keynes | Wellcome Sanger Institute for the COVID-19 Genomics UK (COG-UK) Consortium | The Lighthouse Lab in Milton Keynes and Alex Alderton, Roberto Amato, Jeffrey Barrett, Sonia Goncalves, Ewan Harrison, David K. Jackson, Ian Johnston, Dominic Kwiatkowski, Cordelia Langford, John Sillitoe on behalf of the Wellcome Sanger Institute COVID-19 Surveillance Team |
| EPI_ISL_2514204, EPI_ISL_2514205, EPI_ISL_2514206, EPI_ISL_2514207, EPI_ISL_2514208, EPI_ISL_2514209, EPI_ISL_2514210 | Lighthouse Lab in Alderley Park | Wellcome Sanger Institute for the COVID-19 Genomics UK (COG-UK) Consortium | Jacquelyn Wynn, Mairead Hyland, The Lighthouse Lab in Alderley Park and Alex Alderton, Roberto Amato, Jeffrey Barrett, Sonia Goncalves, Ewan Harrison, David K. Jackson, Ian Johnston, Dominic Kwiatkowski, Cordelia Langford, John Sillitoe on behalf of the Wellcome Sanger Institute COVID-19 Surveillance Team |
| EPI_ISL_2514211 | Lighthouse Lab in Milton Keynes | Wellcome Sanger Institute for the COVID-19 Genomics UK (COG-UK) Consortium | The Lighthouse Lab in Milton Keynes and Alex Alderton, Roberto Amato, Jeffrey Barrett, Sonia Goncalves, Ewan Harrison, David K. Jackson, Ian Johnston, Dominic Kwiatkowski, Cordelia Langford, John Sillitoe on behalf of the Wellcome Sanger Institute COVID-19 Surveillance Team |
| EPI_ISL_2514212, EPI_ISL_2514213, EPI_ISL_2514214, EPI_ISL_2514215, EPI_ISL_2514216, EPI_ISL_2514217 | Lighthouse Lab in Alderley Park | Wellcome Sanger Institute for the COVID-19 Genomics UK (COG-UK) Consortium | Jacquelyn Wynn, Mairead Hyland, The Lighthouse Lab in Alderley Park and Alex Alderton, Roberto Amato, Jeffrey Barrett, Sonia Goncalves, Ewan Harrison, David K. Jackson, Ian Johnston, Dominic Kwiatkowski, Cordelia Langford, John Sillitoe on behalf of the Wellcome Sanger Institute COVID-19 Surveillance Team |
| EPI_ISL_2514218 | Lighthouse Lab in Milton Keynes | Wellcome Sanger Institute for the COVID-19 Genomics UK (COG-UK) Consortium | The Lighthouse Lab in Milton Keynes and Alex Alderton, Roberto Amato, Jeffrey Barrett, Sonia Goncalves, Ewan Harrison, David K. Jackson, Ian Johnston, Dominic Kwiatkowski, Cordelia Langford, John Sillitoe on behalf of the Wellcome Sanger Institute COVID-19 Surveillance Team |
| EPI_ISL_2514220, EPI_ISL_2514221 | Lighthouse Lab in Alderley Park | Wellcome Sanger Institute for the COVID-19 Genomics UK (COG-UK) Consortium | Jacquelyn Wynn, Mairead Hyland, The Lighthouse Lab in Alderley Park and Alex Alderton, Roberto Amato, Jeffrey Barrett, Sonia Goncalves, Ewan Harrison, David K. Jackson, Ian Johnston, Dominic Kwiatkowski, Cordelia Langford, John Sillitoe on behalf of the Wellcome Sanger Institute COVID-19 Surveillance Team |
| EPI_ISL_2514222 | Lighthouse Lab in Milton Keynes | Wellcome Sanger Institute for the COVID-19 Genomics UK (COG-UK) Consortium | The Lighthouse Lab in Milton Keynes and Alex Alderton, Roberto Amato, Jeffrey Barrett, Sonia Goncalves, Ewan Harrison, David K. Jackson, Ian Johnston, Dominic Kwiatkowski, Cordelia Langford, John Sillitoe on behalf of the Wellcome Sanger Institute COVID-19 Surveillance Team |
| EPI_ISL_2514223, EPI_ISL_2514224, EPI_ISL_2514225, EPI_ISL_2514226, EPI_ISL_2514227, EPI_ISL_2514228, EPI_ISL_2514229, EPI_ISL_2514230, EPI_ISL_2514231, EPI_ISL_2514232, EPI_ISL_2514233 | | | |
| see above | Lighthouse Lab in Alderley Park | Wellcome Sanger Institute for the COVID-19 Genomics UK (COG-UK) Consortium | Jacquelyn Wynn, Mairead Hyland, The Lighthouse Lab in Alderley Park and Alex Alderton, Roberto Amato, Jeffrey Barrett, Sonia Goncalves, Ewan Harrison, David K. Jackson, Ian Johnston, Dominic Kwiatkowski, Cordelia Langford, John Sillitoe on behalf of the Wellcome Sanger Institute COVID-19 Surveillance Team |

| | | | |
|---|---|---|---|
| EPI_ISL_2514234 | Lighthouse Lab in Milton Keynes | Wellcome Sanger Institute for the COVID-19 Genomics UK (COG-UK) Consortium | The Lighthouse Lab in Milton Keynes and Alex Alderton, Roberto Amato, Jeffrey Barrett, Sonia Goncalves, Ewan Harrison, David K. Jackson, Ian Johnston, Dominic Kwiatkowski, Cordelia Langford, John Sillitoe on behalf of the Wellcome Sanger Institute COVID-19 Surveillance Team |
| EPI_ISL_2514235 | Lighthouse Lab in Alderley Park | Wellcome Sanger Institute for the COVID-19 Genomics UK (COG-UK) Consortium | Jacquelyn Wynn, Mairead Hyland, The Lighthouse Lab in Alderley Park and Alex Alderton, Roberto Amato, Jeffrey Barrett, Sonia Goncalves, Ewan Harrison, David K. Jackson, Ian Johnston, Dominic Kwiatkowski, Cordelia Langford, John Sillitoe on behalf of the Wellcome Sanger Institute COVID-19 Surveillance Team |
| EPI_ISL_2514236 | Lighthouse Lab in Milton Keynes | Wellcome Sanger Institute for the COVID-19 Genomics UK (COG-UK) Consortium | The Lighthouse Lab in Milton Keynes and Alex Alderton, Roberto Amato, Jeffrey Barrett, Sonia Goncalves, Ewan Harrison, David K. Jackson, Ian Johnston, Dominic Kwiatkowski, Cordelia Langford, John Sillitoe on behalf of the Wellcome Sanger Institute COVID-19 Surveillance Team |
| EPI_ISL_2514237, EPI_ISL_2514238, EPI_ISL_2514239, EPI_ISL_2514240, EPI_ISL_2514241, EPI_ISL_2514242, EPI_ISL_2514243, EPI_ISL_2514244, EPI_ISL_2514245, EPI_ISL_2514246, EPI_ISL_2514247, EPI_ISL_2514248, EPI_ISL_2514249 | | | |
| see above | Lighthouse Lab in Alderley Park | Wellcome Sanger Institute for the COVID-19 Genomics UK (COG-UK) Consortium | Jacquelyn Wynn, Mairead Hyland, The Lighthouse Lab in Alderley Park and Alex Alderton, Roberto Amato, Jeffrey Barrett, Sonia Goncalves, Ewan Harrison, David K. Jackson, Ian Johnston, Dominic Kwiatkowski, Cordelia Langford, John Sillitoe on behalf of the Wellcome Sanger Institute COVID-19 Surveillance Team |
| EPI_ISL_2514250 | Lighthouse Lab in Milton Keynes | Wellcome Sanger Institute for the COVID-19 Genomics UK (COG-UK) Consortium | The Lighthouse Lab in Milton Keynes and Alex Alderton, Roberto Amato, Jeffrey Barrett, Sonia Goncalves, Ewan Harrison, David K. Jackson, Ian Johnston, Dominic Kwiatkowski, Cordelia Langford, John Sillitoe on behalf of the Wellcome Sanger Institute COVID-19 Surveillance Team |
| EPI_ISL_2514251, EPI_ISL_2514252 | Lighthouse Lab in Alderley Park | Wellcome Sanger Institute for the COVID-19 Genomics UK (COG-UK) Consortium | Jacquelyn Wynn, Mairead Hyland, The Lighthouse Lab in Alderley Park and Alex Alderton, Roberto Amato, Jeffrey Barrett, Sonia Goncalves, Ewan Harrison, David K. Jackson, Ian Johnston, Dominic Kwiatkowski, Cordelia Langford, John Sillitoe on behalf of the Wellcome Sanger Institute COVID-19 Surveillance Team |
| EPI_ISL_2514253, EPI_ISL_2514254 | Lighthouse Lab in Milton Keynes | Wellcome Sanger Institute for the COVID-19 Genomics UK (COG-UK) Consortium | The Lighthouse Lab in Milton Keynes and Alex Alderton, Roberto Amato, Jeffrey Barrett, Sonia Goncalves, Ewan Harrison, David K. Jackson, Ian Johnston, Dominic Kwiatkowski, Cordelia Langford, John Sillitoe on behalf of the Wellcome Sanger Institute COVID-19 Surveillance Team |
| EPI_ISL_2514255, EPI_ISL_2514256, EPI_ISL_2514257, EPI_ISL_2514258, EPI_ISL_2514259, EPI_ISL_2514260, EPI_ISL_2514263, EPI_ISL_2514264, EPI_ISL_2514265, EPI_ISL_2514266, EPI_ISL_2514267 | | | |
| see above | Lighthouse Lab in Alderley Park | Wellcome Sanger Institute for the COVID-19 Genomics UK (COG-UK) Consortium | Jacquelyn Wynn, Mairead Hyland, The Lighthouse Lab in Alderley Park and Alex Alderton, Roberto Amato, Jeffrey Barrett, Sonia Goncalves, Ewan Harrison, David K. Jackson, Ian Johnston, Dominic Kwiatkowski, Cordelia Langford, John Sillitoe on behalf of the Wellcome Sanger Institute COVID-19 Surveillance Team |
| EPI_ISL_2514268 | Lighthouse Lab in Milton Keynes | Wellcome Sanger Institute for the COVID-19 Genomics UK (COG-UK) Consortium | The Lighthouse Lab in Milton Keynes and Alex Alderton, Roberto Amato, Jeffrey Barrett, Sonia Goncalves, Ewan Harrison, David K. Jackson, Ian Johnston, Dominic Kwiatkowski, Cordelia Langford, John Sillitoe on behalf of the Wellcome Sanger Institute COVID-19 Surveillance Team |
| EPI_ISL_2514270, EPI_ISL_2514271, EPI_ISL_2514272, EPI_ISL_2514273, EPI_ISL_2514274, EPI_ISL_2514275, EPI_ISL_2514276, EPI_ISL_2514277, EPI_ISL_2514278, EPI_ISL_2514279, EPI_ISL_2514281, EPI_ISL_2514282, EPI_ISL_2514283 | | | |
| see above | Lighthouse Lab in Alderley Park | Wellcome Sanger Institute for the COVID-19 Genomics UK (COG-UK) Consortium | Jacquelyn Wynn, Mairead Hyland, The Lighthouse Lab in Alderley Park and Alex Alderton, Roberto Amato, Jeffrey Barrett, Sonia Goncalves, Ewan Harrison, David K. Jackson, Ian Johnston, Dominic Kwiatkowski, Cordelia Langford, John Sillitoe on behalf of the Wellcome Sanger Institute COVID-19 Surveillance Team |
| EPI_ISL_2514284 | Lighthouse Lab in Milton Keynes | Wellcome Sanger Institute for the COVID-19 Genomics UK (COG-UK) Consortium | The Lighthouse Lab in Milton Keynes and Alex Alderton, Roberto Amato, Jeffrey Barrett, Sonia Goncalves, Ewan Harrison, David K. Jackson, Ian Johnston, Dominic Kwiatkowski, Cordelia Langford, John Sillitoe on behalf of the Wellcome Sanger Institute COVID-19 Surveillance Team |
| EPI_ISL_2514285, EPI_ISL_2514286, EPI_ISL_2514287, EPI_ISL_2514289, EPI_ISL_2514290 | Lighthouse Lab in Alderley Park | Wellcome Sanger Institute for the COVID-19 Genomics UK (COG-UK) Consortium | Jacquelyn Wynn, Mairead Hyland, The Lighthouse Lab in Alderley Park and Alex Alderton, Roberto Amato, Jeffrey Barrett, Sonia Goncalves, Ewan Harrison, David K. Jackson, Ian Johnston, Dominic Kwiatkowski, Cordelia Langford, John Sillitoe on behalf of the Wellcome Sanger Institute COVID-19 Surveillance Team |
| EPI_ISL_2514291, EPI_ISL_2514292 | Lighthouse Lab in Milton Keynes | Wellcome Sanger Institute for the COVID-19 Genomics UK (COG-UK) Consortium | The Lighthouse Lab in Milton Keynes and Alex Alderton, Roberto Amato, Jeffrey Barrett, Sonia Goncalves, Ewan Harrison, David K. Jackson, Ian Johnston, Dominic Kwiatkowski, Cordelia Langford, John Sillitoe on behalf of the Wellcome Sanger Institute COVID-19 Surveillance Team |
| EPI_ISL_2514293, EPI_ISL_2514294, EPI_ISL_2514295, EPI_ISL_2514296, EPI_ISL_2514297 | Lighthouse Lab in Alderley Park | Wellcome Sanger Institute for the COVID-19 Genomics UK (COG-UK) Consortium | Jacquelyn Wynn, Mairead Hyland, The Lighthouse Lab in Alderley Park and Alex Alderton, Roberto Amato, Jeffrey Barrett, Sonia Goncalves, Ewan Harrison, David K. Jackson, Ian Johnston, Dominic Kwiatkowski, Cordelia Langford, John Sillitoe on behalf of the Wellcome Sanger Institute COVID-19 Surveillance Team |
| EPI_ISL_2514298 | Lighthouse Lab in Milton Keynes | Wellcome Sanger Institute for the COVID-19 Genomics UK (COG-UK) Consortium | The Lighthouse Lab in Milton Keynes and Alex Alderton, Roberto Amato, Jeffrey Barrett, Sonia Goncalves, Ewan Harrison, David K. Jackson, Ian Johnston, Dominic Kwiatkowski, Cordelia Langford, John Sillitoe on behalf of the Wellcome Sanger Institute COVID-19 Surveillance Team |
| EPI_ISL_2514299 | Lighthouse Lab in Alderley Park | Wellcome Sanger Institute for the COVID-19 Genomics UK (COG-UK) Consortium | Jacquelyn Wynn, Mairead Hyland, The Lighthouse Lab in Alderley Park and Alex Alderton, Roberto Amato, Jeffrey Barrett, Sonia Goncalves, Ewan Harrison, David K. Jackson, Ian Johnston, Dominic Kwiatkowski, Cordelia Langford, John Sillitoe on behalf of the Wellcome Sanger Institute COVID-19 Surveillance Team |
| EPI_ISL_2514300 | Lighthouse Lab in Milton Keynes | Wellcome Sanger Institute for the COVID-19 Genomics UK (COG-UK) Consortium | The Lighthouse Lab in Milton Keynes and Alex Alderton, Roberto Amato, Jeffrey Barrett, Sonia Goncalves, Ewan Harrison, David K. Jackson, Ian Johnston, Dominic Kwiatkowski, Cordelia Langford, John Sillitoe on behalf of the Wellcome Sanger Institute COVID-19 Surveillance Team |
| EPI_ISL_2514301, EPI_ISL_2514302, EPI_ISL_2514303 | Lighthouse Lab in Alderley Park | Wellcome Sanger Institute for the COVID-19 Genomics UK (COG-UK) Consortium | Jacquelyn Wynn, Mairead Hyland, The Lighthouse Lab in Alderley Park and Alex Alderton, Roberto Amato, Jeffrey Barrett, Sonia Goncalves, Ewan Harrison, David K. Jackson, Ian Johnston, Dominic Kwiatkowski, Cordelia Langford, John Sillitoe on behalf of the Wellcome Sanger Institute COVID-19 Surveillance Team |
| EPI_ISL_2514304 | Lighthouse Lab in Milton Keynes | Wellcome Sanger Institute for the COVID-19 Genomics UK (COG-UK) Consortium | The Lighthouse Lab in Milton Keynes and Alex Alderton, Roberto Amato, Jeffrey Barrett, Sonia Goncalves, Ewan Harrison, David K. Jackson, Ian Johnston, Dominic Kwiatkowski, Cordelia Langford, John Sillitoe on behalf of the Wellcome Sanger Institute COVID-19 Surveillance Team |
| EPI_ISL_2514305, EPI_ISL_2514306, EPI_ISL_2514307, EPI_ISL_2514308, EPI_ISL_2514309 | Lighthouse Lab in Alderley Park | Wellcome Sanger Institute for the COVID-19 Genomics UK (COG-UK) Consortium | Jacquelyn Wynn, Mairead Hyland, The Lighthouse Lab in Alderley Park and Alex Alderton, Roberto Amato, Jeffrey Barrett, Sonia Goncalves, Ewan Harrison, David K. Jackson, Ian Johnston, Dominic Kwiatkowski, Cordelia Langford, John Sillitoe on behalf of the Wellcome Sanger Institute COVID-19 Surveillance Team |
| EPI_ISL_2514310, EPI_ISL_2514311 | Lighthouse Lab in Milton Keynes | Wellcome Sanger Institute for the COVID-19 Genomics UK (COG-UK) Consortium | The Lighthouse Lab in Milton Keynes and Alex Alderton, Roberto Amato, Jeffrey Barrett, Sonia Goncalves, Ewan Harrison, David K. Jackson, Ian Johnston, Dominic Kwiatkowski, Cordelia Langford, John Sillitoe on behalf of the Wellcome Sanger Institute COVID-19 Surveillance Team |
| EPI_ISL_2514312, EPI_ISL_2514313, EPI_ISL_2514315, EPI_ISL_2514316, EPI_ISL_2514317, EPI_ISL_2514318, EPI_ISL_2514319, EPI_ISL_2514320, EPI_ISL_2514321, EPI_ISL_2514322 | Lighthouse Lab in Alderley Park | Wellcome Sanger Institute for the COVID-19 Genomics UK (COG-UK) Consortium | Jacquelyn Wynn, Mairead Hyland, The Lighthouse Lab in Alderley Park and Alex Alderton, Roberto Amato, Jeffrey Barrett, Sonia Goncalves, Ewan Harrison, David K. Jackson, Ian Johnston, Dominic Kwiatkowski, Cordelia Langford, John Sillitoe on behalf of the Wellcome Sanger Institute COVID-19 Surveillance Team |
| EPI_ISL_2514323 | Lighthouse Lab in Milton Keynes | Wellcome Sanger Institute for the COVID-19 Genomics UK (COG-UK) Consortium | The Lighthouse Lab in Milton Keynes and Alex Alderton, Roberto Amato, Jeffrey Barrett, Sonia Goncalves, Ewan Harrison, David K. Jackson, Ian Johnston, Dominic Kwiatkowski, Cordelia Langford, John Sillitoe on behalf of the Wellcome Sanger Institute COVID-19 Surveillance Team |
| EPI_ISL_2514324, EPI_ISL_2514325, EPI_ISL_2514326 | Lighthouse Lab in Alderley Park | Wellcome Sanger Institute for the COVID-19 Genomics UK (COG-UK) Consortium | Jacquelyn Wynn, Mairead Hyland, The Lighthouse Lab in Alderley Park and Alex Alderton, Roberto Amato, Jeffrey Barrett, Sonia Goncalves, Ewan Harrison, David K. Jackson, Ian Johnston, Dominic Kwiatkowski, Cordelia Langford, John Sillitoe on behalf of the Wellcome Sanger Institute COVID-19 Surveillance Team |
| EPI_ISL_2514327 | Lighthouse Lab in Milton Keynes | Wellcome Sanger Institute for the COVID-19 Genomics UK (COG-UK) Consortium | The Lighthouse Lab in Milton Keynes and Alex Alderton, Roberto Amato, Jeffrey Barrett, Sonia Goncalves, Ewan Harrison, David K. Jackson, Ian Johnston, Dominic Kwiatkowski, Cordelia Langford, John Sillitoe on behalf of the Wellcome Sanger Institute COVID-19 Surveillance Team |
| EPI_ISL_2514328, EPI_ISL_2514329 | Lighthouse Lab in Alderley Park | Wellcome Sanger Institute for the COVID-19 Genomics UK (COG-UK) Consortium | Jacquelyn Wynn, Mairead Hyland, The Lighthouse Lab in Alderley Park and Alex Alderton, Roberto Amato, Jeffrey Barrett, Sonia Goncalves, Ewan Harrison, David K. Jackson, Ian Johnston, Dominic Kwiatkowski, Cordelia Langford, John Sillitoe on behalf of the Wellcome Sanger Institute COVID-19 Surveillance Team |
| EPI_ISL_2514330 | Lighthouse Lab in Milton Keynes | Wellcome Sanger Institute for the COVID-19 Genomics UK (COG-UK) Consortium | The Lighthouse Lab in Milton Keynes and Alex Alderton, Roberto Amato, Jeffrey Barrett, Sonia Goncalves, Ewan Harrison, David K. Jackson, Ian Johnston, Dominic Kwiatkowski, Cordelia Langford, John Sillitoe on behalf of the Wellcome Sanger Institute COVID-19 Surveillance Team |
| EPI_ISL_2514331, EPI_ISL_2514332, EPI_ISL_2514333, EPI_ISL_2514334, EPI_ISL_2514335, EPI_ISL_2514336, EPI_ISL_2514337, EPI_ISL_2514338, EPI_ISL_2514339, EPI_ISL_2514340, EPI_ISL_2514341, EPI_ISL_2514342, EPI_ISL_2514343 | | | |
| see above | Lighthouse Lab in Alderley Park | Wellcome Sanger Institute for the COVID-19 Genomics UK (COG-UK) Consortium | Jacquelyn Wynn, Mairead Hyland, The Lighthouse Lab in Alderley Park and Alex Alderton, Roberto Amato, Jeffrey Barrett, Sonia Goncalves, Ewan Harrison, David K. Jackson, Ian Johnston, Dominic Kwiatkowski, Cordelia Langford, John Sillitoe on behalf of the Wellcome Sanger Institute COVID-19 Surveillance Team |
| EPI_ISL_2514344 | Lighthouse Lab in Milton Keynes | Wellcome Sanger Institute for the COVID-19 Genomics UK (COG-UK) Consortium | The Lighthouse Lab in Milton Keynes and Alex Alderton, Roberto Amato, Jeffrey Barrett, Sonia Goncalves, Ewan Harrison, David K. Jackson, Ian Johnston, Dominic Kwiatkowski, Cordelia Langford, John Sillitoe on behalf of the Wellcome Sanger Institute COVID-19 Surveillance Team |
| EPI_ISL_2514345, EPI_ISL_2514346 | Lighthouse Lab in Alderley Park | Wellcome Sanger Institute for the COVID-19 Genomics UK (COG-UK) Consortium | Jacquelyn Wynn, Mairead Hyland, The Lighthouse Lab in Alderley Park and Alex Alderton, Roberto Amato, Jeffrey Barrett, Sonia Goncalves, Ewan Harrison, David K. Jackson, Ian Johnston, Dominic Kwiatkowski, Cordelia Langford, John Sillitoe on behalf of the Wellcome Sanger Institute COVID-19 Surveillance Team |
| EPI_ISL_2514347 | Lighthouse Lab in Milton Keynes | Wellcome Sanger Institute for the COVID-19 Genomics UK (COG-UK) Consortium | The Lighthouse Lab in Milton Keynes and Alex Alderton, Roberto Amato, Jeffrey Barrett, Sonia Goncalves, Ewan Harrison, David K. Jackson, Ian Johnston, Dominic Kwiatkowski, Cordelia Langford, John Sillitoe on behalf of the Wellcome Sanger Institute COVID-19 Surveillance Team |
| EPI_ISL_2514348, EPI_ISL_2514349, EPI_ISL_2514350, EPI_ISL_2514351, EPI_ISL_2514352, EPI_ISL_2514353 | Lighthouse Lab in Alderley Park | Wellcome Sanger Institute for the COVID-19 Genomics UK (COG-UK) Consortium | Jacquelyn Wynn, Mairead Hyland, The Lighthouse Lab in Alderley Park and Alex Alderton, Roberto Amato, Jeffrey Barrett, Sonia Goncalves, Ewan Harrison, David K. Jackson, Ian Johnston, Dominic Kwiatkowski, Cordelia Langford, John Sillitoe on behalf of the Wellcome Sanger Institute COVID-19 Surveillance Team |
| EPI_ISL_2514354 | Lighthouse Lab in Milton Keynes | Wellcome Sanger Institute for the COVID-19 Genomics UK (COG-UK) Consortium | The Lighthouse Lab in Milton Keynes and Alex Alderton, Roberto Amato, Jeffrey Barrett, Sonia Goncalves, Ewan Harrison, David K. Jackson, Ian Johnston, Dominic Kwiatkowski, Cordelia Langford, John Sillitoe on behalf of the Wellcome Sanger Institute COVID-19 Surveillance Team |
| EPI_ISL_2514355 | Lighthouse Lab in Alderley Park | Wellcome Sanger Institute for the COVID-19 Genomics UK (COG-UK) Consortium | Jacquelyn Wynn, Mairead Hyland, The Lighthouse Lab in Alderley Park and Alex Alderton, Roberto Amato, Jeffrey Barrett, Sonia Goncalves, Ewan Harrison, David K. Jackson, Ian Johnston, Dominic Kwiatkowski, Cordelia Langford, John Sillitoe on |

| | | | |
|---|---|---|---|
| | | Consortium | behalf of the Wellcome Sanger Institute COVID-19 Surveillance Team |
| EPI_ISL_2514356, EPI_ISL_2514357 | Lighthouse Lab in Milton Keynes | Wellcome Sanger Institute for the COVID-19 Genomics UK (COG-UK) Consortium | The Lighthouse Lab in Milton Keynes and Alex Alderton, Roberto Amato, Jeffrey Barrett, Sonia Goncalves, Ewan Harrison, David K. Jackson, Ian Johnston, Dominic Kwiatkowski, Cordelia Langford, John Sillitoe on behalf of the Wellcome Sanger Institute COVID-19 Surveillance Team |
| EPI_ISL_2514358 | Lighthouse Lab in Alderley Park | Wellcome Sanger Institute for the COVID-19 Genomics UK (COG-UK) Consortium | Jacquelyn Wynn, Mairead Hyland, The Lighthouse Lab in Alderley Park and Alex Alderton, Roberto Amato, Jeffrey Barrett, Sonia Goncalves, Ewan Harrison, David K. Jackson, Ian Johnston, Dominic Kwiatkowski, Cordelia Langford, John Sillitoe on behalf of the Wellcome Sanger Institute COVID-19 Surveillance Team |
| EPI_ISL_2514359 | Lighthouse Lab in Milton Keynes | Wellcome Sanger Institute for the COVID-19 Genomics UK (COG-UK) Consortium | The Lighthouse Lab in Milton Keynes and Alex Alderton, Roberto Amato, Jeffrey Barrett, Sonia Goncalves, Ewan Harrison, David K. Jackson, Ian Johnston, Dominic Kwiatkowski, Cordelia Langford, John Sillitoe on behalf of the Wellcome Sanger Institute COVID-19 Surveillance Team |
| EPI_ISL_2514360, EPI_ISL_2514361, EPI_ISL_2514362, EPI_ISL_2514363, EPI_ISL_2514364 | Lighthouse Lab in Alderley Park | Wellcome Sanger Institute for the COVID-19 Genomics UK (COG-UK) Consortium | Jacquelyn Wynn, Mairead Hyland, The Lighthouse Lab in Alderley Park and Alex Alderton, Roberto Amato, Jeffrey Barrett, Sonia Goncalves, Ewan Harrison, David K. Jackson, Ian Johnston, Dominic Kwiatkowski, Cordelia Langford, John Sillitoe on behalf of the Wellcome Sanger Institute COVID-19 Surveillance Team |
| EPI_ISL_2514365 | Lighthouse Lab in Milton Keynes | Wellcome Sanger Institute for the COVID-19 Genomics UK (COG-UK) Consortium | The Lighthouse Lab in Milton Keynes and Alex Alderton, Roberto Amato, Jeffrey Barrett, Sonia Goncalves, Ewan Harrison, David K. Jackson, Ian Johnston, Dominic Kwiatkowski, Cordelia Langford, John Sillitoe on behalf of the Wellcome Sanger Institute COVID-19 Surveillance Team |
| EPI_ISL_2514367, EPI_ISL_2514368, EPI_ISL_2514369 | Lighthouse Lab in Alderley Park | Wellcome Sanger Institute for the COVID-19 Genomics UK (COG-UK) Consortium | Jacquelyn Wynn, Mairead Hyland, The Lighthouse Lab in Alderley Park and Alex Alderton, Roberto Amato, Jeffrey Barrett, Sonia Goncalves, Ewan Harrison, David K. Jackson, Ian Johnston, Dominic Kwiatkowski, Cordelia Langford, John Sillitoe on behalf of the Wellcome Sanger Institute COVID-19 Surveillance Team |
| EPI_ISL_2514370 | Lighthouse Lab in Milton Keynes | Wellcome Sanger Institute for the COVID-19 Genomics UK (COG-UK) Consortium | The Lighthouse Lab in Milton Keynes and Alex Alderton, Roberto Amato, Jeffrey Barrett, Sonia Goncalves, Ewan Harrison, David K. Jackson, Ian Johnston, Dominic Kwiatkowski, Cordelia Langford, John Sillitoe on behalf of the Wellcome Sanger Institute COVID-19 Surveillance Team |
| EPI_ISL_2514372, EPI_ISL_2514373, EPI_ISL_2514374 | Lighthouse Lab in Alderley Park | Wellcome Sanger Institute for the COVID-19 Genomics UK (COG-UK) Consortium | Jacquelyn Wynn, Mairead Hyland, The Lighthouse Lab in Alderley Park and Alex Alderton, Roberto Amato, Jeffrey Barrett, Sonia Goncalves, Ewan Harrison, David K. Jackson, Ian Johnston, Dominic Kwiatkowski, Cordelia Langford, John Sillitoe on behalf of the Wellcome Sanger Institute COVID-19 Surveillance Team |
| EPI_ISL_2514375, EPI_ISL_2514376 | Lighthouse Lab in Milton Keynes | Wellcome Sanger Institute for the COVID-19 Genomics UK (COG-UK) Consortium | The Lighthouse Lab in Milton Keynes and Alex Alderton, Roberto Amato, Jeffrey Barrett, Sonia Goncalves, Ewan Harrison, David K. Jackson, Ian Johnston, Dominic Kwiatkowski, Cordelia Langford, John Sillitoe on behalf of the Wellcome Sanger Institute COVID-19 Surveillance Team |
| EPI_ISL_2514377, EPI_ISL_2514378, EPI_ISL_2514380, EPI_ISL_2514381 | Lighthouse Lab in Alderley Park | Wellcome Sanger Institute for the COVID-19 Genomics UK (COG-UK) Consortium | Jacquelyn Wynn, Mairead Hyland, The Lighthouse Lab in Alderley Park and Alex Alderton, Roberto Amato, Jeffrey Barrett, Sonia Goncalves, Ewan Harrison, David K. Jackson, Ian Johnston, Dominic Kwiatkowski, Cordelia Langford, John Sillitoe on behalf of the Wellcome Sanger Institute COVID-19 Surveillance Team |
| EPI_ISL_2514383 | Lighthouse Lab in Milton Keynes | Wellcome Sanger Institute for the COVID-19 Genomics UK (COG-UK) Consortium | The Lighthouse Lab in Milton Keynes and Alex Alderton, Roberto Amato, Jeffrey Barrett, Sonia Goncalves, Ewan Harrison, David K. Jackson, Ian Johnston, Dominic Kwiatkowski, Cordelia Langford, John Sillitoe on behalf of the Wellcome Sanger Institute COVID-19 Surveillance Team |
| EPI_ISL_2514384 | Lighthouse Lab in Alderley Park | Wellcome Sanger Institute for the COVID-19 Genomics UK (COG-UK) Consortium | Jacquelyn Wynn, Mairead Hyland, The Lighthouse Lab in Alderley Park and Alex Alderton, Roberto Amato, Jeffrey Barrett, Sonia Goncalves, Ewan Harrison, David K. Jackson, Ian Johnston, Dominic Kwiatkowski, Cordelia Langford, John Sillitoe on behalf of the Wellcome Sanger Institute COVID-19 Surveillance Team |
| EPI_ISL_2514386 | Lighthouse Lab in Milton Keynes | Wellcome Sanger Institute for the COVID-19 Genomics UK (COG-UK) Consortium | The Lighthouse Lab in Milton Keynes and Alex Alderton, Roberto Amato, Jeffrey Barrett, Sonia Goncalves, Ewan Harrison, David K. Jackson, Ian Johnston, Dominic Kwiatkowski, Cordelia Langford, John Sillitoe on behalf of the Wellcome Sanger Institute COVID-19 Surveillance Team |
| EPI_ISL_2514387, EPI_ISL_2514388 | Lighthouse Lab in Alderley Park | Wellcome Sanger Institute for the COVID-19 Genomics UK (COG-UK) Consortium | Jacquelyn Wynn, Mairead Hyland, The Lighthouse Lab in Alderley Park and Alex Alderton, Roberto Amato, Jeffrey Barrett, Sonia Goncalves, Ewan Harrison, David K. Jackson, Ian Johnston, Dominic Kwiatkowski, Cordelia Langford, John Sillitoe on behalf of the Wellcome Sanger Institute COVID-19 Surveillance Team |
| EPI_ISL_2514389 | Lighthouse Lab in Milton Keynes | Wellcome Sanger Institute for the COVID-19 Genomics UK (COG-UK) Consortium | The Lighthouse Lab in Milton Keynes and Alex Alderton, Roberto Amato, Jeffrey Barrett, Sonia Goncalves, Ewan Harrison, David K. Jackson, Ian Johnston, Dominic Kwiatkowski, Cordelia Langford, John Sillitoe on behalf of the Wellcome Sanger Institute COVID-19 Surveillance Team |
| EPI_ISL_2514390, EPI_ISL_2514391, EPI_ISL_2514392, EPI_ISL_2514393, EPI_ISL_2514394 | Lighthouse Lab in Alderley Park | Wellcome Sanger Institute for the COVID-19 Genomics UK (COG-UK) Consortium | Jacquelyn Wynn, Mairead Hyland, The Lighthouse Lab in Alderley Park and Alex Alderton, Roberto Amato, Jeffrey Barrett, Sonia Goncalves, Ewan Harrison, David K. Jackson, Ian Johnston, Dominic Kwiatkowski, Cordelia Langford, John Sillitoe on behalf of the Wellcome Sanger Institute COVID-19 Surveillance Team |
| EPI_ISL_2514395, EPI_ISL_2514396 | Lighthouse Lab in Milton Keynes | Wellcome Sanger Institute for the COVID-19 Genomics UK (COG-UK) Consortium | The Lighthouse Lab in Milton Keynes and Alex Alderton, Roberto Amato, Jeffrey Barrett, Sonia Goncalves, Ewan Harrison, David K. Jackson, Ian Johnston, Dominic Kwiatkowski, Cordelia Langford, John Sillitoe on behalf of the Wellcome Sanger Institute COVID-19 Surveillance Team |
| EPI_ISL_2514398, EPI_ISL_2514399, EPI_ISL_2514400 | Lighthouse Lab in Alderley Park | Wellcome Sanger Institute for the COVID-19 Genomics UK (COG-UK) Consortium | Jacquelyn Wynn, Mairead Hyland, The Lighthouse Lab in Alderley Park and Alex Alderton, Roberto Amato, Jeffrey Barrett, Sonia Goncalves, Ewan Harrison, David K. Jackson, Ian Johnston, Dominic Kwiatkowski, Cordelia Langford, John Sillitoe on behalf of the Wellcome Sanger Institute COVID-19 Surveillance Team |
| EPI_ISL_2514401 | Lighthouse Lab in Milton Keynes | Wellcome Sanger Institute for the COVID-19 Genomics UK (COG-UK) Consortium | The Lighthouse Lab in Milton Keynes and Alex Alderton, Roberto Amato, Jeffrey Barrett, Sonia Goncalves, Ewan Harrison, David K. Jackson, Ian Johnston, Dominic Kwiatkowski, Cordelia Langford, John Sillitoe on behalf of the Wellcome Sanger Institute COVID-19 Surveillance Team |
| EPI_ISL_2514402 | Lighthouse Lab in Alderley Park | Wellcome Sanger Institute for the COVID-19 Genomics UK (COG-UK) Consortium | Jacquelyn Wynn, Mairead Hyland, The Lighthouse Lab in Alderley Park and Alex Alderton, Roberto Amato, Jeffrey Barrett, Sonia Goncalves, Ewan Harrison, David K. Jackson, Ian Johnston, Dominic Kwiatkowski, Cordelia Langford, John Sillitoe on behalf of the Wellcome Sanger Institute COVID-19 Surveillance Team |
| EPI_ISL_2514403 | Lighthouse Lab in Milton Keynes | Wellcome Sanger Institute for the COVID-19 Genomics UK (COG-UK) Consortium | The Lighthouse Lab in Milton Keynes and Alex Alderton, Roberto Amato, Jeffrey Barrett, Sonia Goncalves, Ewan Harrison, David K. Jackson, Ian Johnston, Dominic Kwiatkowski, Cordelia Langford, John Sillitoe on behalf of the Wellcome Sanger Institute COVID-19 Surveillance Team |
| EPI_ISL_2514404, EPI_ISL_2514405, EPI_ISL_2514406, EPI_ISL_2514407, EPI_ISL_2514408 | Lighthouse Lab in Alderley Park | Wellcome Sanger Institute for the COVID-19 Genomics UK (COG-UK) Consortium | Jacquelyn Wynn, Mairead Hyland, The Lighthouse Lab in Alderley Park and Alex Alderton, Roberto Amato, Jeffrey Barrett, Sonia Goncalves, Ewan Harrison, David K. Jackson, Ian Johnston, Dominic Kwiatkowski, Cordelia Langford, John Sillitoe on behalf of the Wellcome Sanger Institute COVID-19 Surveillance Team |
| EPI_ISL_2514410 | Lighthouse Lab in Milton Keynes | Wellcome Sanger Institute for the COVID-19 Genomics UK (COG-UK) Consortium | The Lighthouse Lab in Milton Keynes and Alex Alderton, Roberto Amato, Jeffrey Barrett, Sonia Goncalves, Ewan Harrison, David K. Jackson, Ian Johnston, Dominic Kwiatkowski, Cordelia Langford, John Sillitoe on behalf of the Wellcome Sanger Institute COVID-19 Surveillance Team |
| EPI_ISL_2514411 | Lighthouse Lab in Alderley Park | Wellcome Sanger Institute for the COVID-19 Genomics UK (COG-UK) Consortium | Jacquelyn Wynn, Mairead Hyland, The Lighthouse Lab in Alderley Park and Alex Alderton, Roberto Amato, Jeffrey Barrett, Sonia Goncalves, Ewan Harrison, David K. Jackson, Ian Johnston, Dominic Kwiatkowski, Cordelia Langford, John Sillitoe on behalf of the Wellcome Sanger Institute COVID-19 Surveillance Team |
| EPI_ISL_2514412 | Lighthouse Lab in Milton Keynes | Wellcome Sanger Institute for the COVID-19 Genomics UK (COG-UK) Consortium | The Lighthouse Lab in Milton Keynes and Alex Alderton, Roberto Amato, Jeffrey Barrett, Sonia Goncalves, Ewan Harrison, David K. Jackson, Ian Johnston, Dominic Kwiatkowski, Cordelia Langford, John Sillitoe on behalf of the Wellcome Sanger Institute COVID-19 Surveillance Team |
| EPI_ISL_2514413, EPI_ISL_2514414 | Lighthouse Lab in Alderley Park | Wellcome Sanger Institute for the COVID-19 Genomics UK (COG-UK) Consortium | Jacquelyn Wynn, Mairead Hyland, The Lighthouse Lab in Alderley Park and Alex Alderton, Roberto Amato, Jeffrey Barrett, Sonia Goncalves, Ewan Harrison, David K. Jackson, Ian Johnston, Dominic Kwiatkowski, Cordelia Langford, John Sillitoe on behalf of the Wellcome Sanger Institute COVID-19 Surveillance Team |
| EPI_ISL_2514415, EPI_ISL_2514416 | Lighthouse Lab in Milton Keynes | Wellcome Sanger Institute for the COVID-19 Genomics UK (COG-UK) Consortium | The Lighthouse Lab in Milton Keynes and Alex Alderton, Roberto Amato, Jeffrey Barrett, Sonia Goncalves, Ewan Harrison, David K. Jackson, Ian Johnston, Dominic Kwiatkowski, Cordelia Langford, John Sillitoe on behalf of the Wellcome Sanger Institute COVID-19 Surveillance Team |
| EPI_ISL_2514417 | Lighthouse Lab in Alderley Park | Wellcome Sanger Institute for the COVID-19 Genomics UK (COG-UK) Consortium | Jacquelyn Wynn, Mairead Hyland, The Lighthouse Lab in Alderley Park and Alex Alderton, Roberto Amato, Jeffrey Barrett, Sonia Goncalves, Ewan Harrison, David K. Jackson, Ian Johnston, Dominic Kwiatkowski, Cordelia Langford, John Sillitoe on behalf of the Wellcome Sanger Institute COVID-19 Surveillance Team |
| EPI_ISL_2514418, EPI_ISL_2514419 | Lighthouse Lab in Milton Keynes | Wellcome Sanger Institute for the COVID-19 Genomics UK (COG-UK) Consortium | The Lighthouse Lab in Milton Keynes and Alex Alderton, Roberto Amato, Jeffrey Barrett, Sonia Goncalves, Ewan Harrison, David K. Jackson, Ian Johnston, Dominic Kwiatkowski, Cordelia Langford, John Sillitoe on behalf of the Wellcome Sanger Institute COVID-19 Surveillance Team |
| EPI_ISL_2514420 | Lighthouse Lab in Alderley Park | Wellcome Sanger Institute for the COVID-19 Genomics UK (COG-UK) Consortium | Jacquelyn Wynn, Mairead Hyland, The Lighthouse Lab in Alderley Park and Alex Alderton, Roberto Amato, Jeffrey Barrett, Sonia Goncalves, Ewan Harrison, David K. Jackson, Ian Johnston, Dominic Kwiatkowski, Cordelia Langford, John Sillitoe on behalf of the Wellcome Sanger Institute COVID-19 Surveillance Team |
| EPI_ISL_2514421, EPI_ISL_2514422 | Lighthouse Lab in Milton Keynes | Wellcome Sanger Institute for the COVID-19 Genomics UK (COG-UK) Consortium | The Lighthouse Lab in Milton Keynes and Alex Alderton, Roberto Amato, Jeffrey Barrett, Sonia Goncalves, Ewan Harrison, David K. Jackson, Ian Johnston, Dominic Kwiatkowski, Cordelia Langford, John Sillitoe on behalf of the Wellcome Sanger Institute COVID-19 Surveillance Team |
| EPI_ISL_2514423, EPI_ISL_2514424, EPI_ISL_2514425 | Lighthouse Lab in Alderley Park | Wellcome Sanger Institute for the COVID-19 Genomics UK (COG-UK) Consortium | Jacquelyn Wynn, Mairead Hyland, The Lighthouse Lab in Alderley Park and Alex Alderton, Roberto Amato, Jeffrey Barrett, Sonia Goncalves, Ewan Harrison, David K. Jackson, Ian Johnston, Dominic Kwiatkowski, Cordelia Langford, John Sillitoe on behalf of the Wellcome Sanger Institute COVID-19 Surveillance Team |
| EPI_ISL_2514426, EPI_ISL_2514427 | Lighthouse Lab in Milton Keynes | Wellcome Sanger Institute for the COVID-19 Genomics UK (COG-UK) Consortium | The Lighthouse Lab in Milton Keynes and Alex Alderton, Roberto Amato, Jeffrey Barrett, Sonia Goncalves, Ewan Harrison, David K. Jackson, Ian Johnston, Dominic Kwiatkowski, Cordelia Langford, John Sillitoe on behalf of the Wellcome Sanger Institute COVID-19 Surveillance Team |
| EPI_ISL_2514428, EPI_ISL_2514429, EPI_ISL_2514430, EPI_ISL_2514431, EPI_ISL_2514432, EPI_ISL_2514433, EPI_ISL_2514434, EPI_ISL_2514435 | Lighthouse Lab in Alderley Park | Wellcome Sanger Institute for the COVID-19 Genomics UK (COG-UK) Consortium | Jacquelyn Wynn, Mairead Hyland, The Lighthouse Lab in Alderley Park and Alex Alderton, Roberto Amato, Jeffrey Barrett, Sonia Goncalves, Ewan Harrison, David K. Jackson, Ian Johnston, Dominic Kwiatkowski, Cordelia Langford, John Sillitoe on behalf of the Wellcome Sanger Institute COVID-19 Surveillance Team |
| EPI_ISL_2514437 | Lighthouse Lab in Milton Keynes | Wellcome Sanger Institute for the COVID-19 Genomics UK (COG-UK) Consortium | The Lighthouse Lab in Milton Keynes and Alex Alderton, Roberto Amato, Jeffrey Barrett, Sonia Goncalves, Ewan Harrison, David K. Jackson, Ian Johnston, Dominic Kwiatkowski, Cordelia Langford, John Sillitoe on behalf of the Wellcome Sanger Institute COVID-19 Surveillance Team |
| EPI_ISL_2514438 | Lighthouse Lab in Alderley Park | Wellcome Sanger Institute for the COVID-19 Genomics UK (COG-UK) | Jacquelyn Wynn, Mairead Hyland, The Lighthouse Lab in Alderley Park and Alex Alderton, Roberto Amato, Jeffrey Barrett, Sonia Goncalves, Ewan Harrison, David K. Jackson, Ian Johnston, Dominic Kwiatkowski, Cordelia Langford, John Sillitoe on |

| | | | |
|---|---|---|---|
| | | Consortium | behalf of the Wellcome Sanger Institute COVID-19 Surveillance Team |
| EPI_ISL_2514439 | Lighthouse Lab in Milton Keynes | Wellcome Sanger Institute for the COVID-19 Genomics UK (COG-UK) Consortium | The Lighthouse Lab in Milton Keynes and Alex Alderton, Roberto Amato, Jeffrey Barrett, Sonia Goncalves, Ewan Harrison, David K. Jackson, Ian Johnston, Dominic Kwiatkowski, Cordelia Langford, John Sillitoe on behalf of the Wellcome Sanger Institute COVID-19 Surveillance Team |
| EPI_ISL_2514440 | Lighthouse Lab in Alderley Park | Wellcome Sanger Institute for the COVID-19 Genomics UK (COG-UK) Consortium | Jacquelyn Wynn, Mairead Hyland, The Lighthouse Lab in Alderley Park and Alex Alderton, Roberto Amato, Jeffrey Barrett, Sonia Goncalves, Ewan Harrison, David K. Jackson, Ian Johnston, Dominic Kwiatkowski, Cordelia Langford, John Sillitoe on behalf of the Wellcome Sanger Institute COVID-19 Surveillance Team |
| EPI_ISL_2514441, EPI_ISL_2514442 | Lighthouse Lab in Milton Keynes | Wellcome Sanger Institute for the COVID-19 Genomics UK (COG-UK) Consortium | The Lighthouse Lab in Milton Keynes and Alex Alderton, Roberto Amato, Jeffrey Barrett, Sonia Goncalves, Ewan Harrison, David K. Jackson, Ian Johnston, Dominic Kwiatkowski, Cordelia Langford, John Sillitoe on behalf of the Wellcome Sanger Institute COVID-19 Surveillance Team |
| EPI_ISL_2514443, EPI_ISL_2514444, EPI_ISL_2514445 | Lighthouse Lab in Alderley Park | Wellcome Sanger Institute for the COVID-19 Genomics UK (COG-UK) Consortium | Jacquelyn Wynn, Mairead Hyland, The Lighthouse Lab in Alderley Park and Alex Alderton, Roberto Amato, Jeffrey Barrett, Sonia Goncalves, Ewan Harrison, David K. Jackson, Ian Johnston, Dominic Kwiatkowski, Cordelia Langford, John Sillitoe on behalf of the Wellcome Sanger Institute COVID-19 Surveillance Team |
| EPI_ISL_2514446 | Lighthouse Lab in Milton Keynes | Wellcome Sanger Institute for the COVID-19 Genomics UK (COG-UK) Consortium | The Lighthouse Lab in Milton Keynes and Alex Alderton, Roberto Amato, Jeffrey Barrett, Sonia Goncalves, Ewan Harrison, David K. Jackson, Ian Johnston, Dominic Kwiatkowski, Cordelia Langford, John Sillitoe on behalf of the Wellcome Sanger Institute COVID-19 Surveillance Team |
| EPI_ISL_2514447 | Lighthouse Lab in Alderley Park | Wellcome Sanger Institute for the COVID-19 Genomics UK (COG-UK) Consortium | Jacquelyn Wynn, Mairead Hyland, The Lighthouse Lab in Alderley Park and Alex Alderton, Roberto Amato, Jeffrey Barrett, Sonia Goncalves, Ewan Harrison, David K. Jackson, Ian Johnston, Dominic Kwiatkowski, Cordelia Langford, John Sillitoe on behalf of the Wellcome Sanger Institute COVID-19 Surveillance Team |
| EPI_ISL_2514448 | Lighthouse Lab in Milton Keynes | Wellcome Sanger Institute for the COVID-19 Genomics UK (COG-UK) Consortium | The Lighthouse Lab in Milton Keynes and Alex Alderton, Roberto Amato, Jeffrey Barrett, Sonia Goncalves, Ewan Harrison, David K. Jackson, Ian Johnston, Dominic Kwiatkowski, Cordelia Langford, John Sillitoe on behalf of the Wellcome Sanger Institute COVID-19 Surveillance Team |
| EPI_ISL_2514449 | Lighthouse Lab in Alderley Park | Wellcome Sanger Institute for the COVID-19 Genomics UK (COG-UK) Consortium | Jacquelyn Wynn, Mairead Hyland, The Lighthouse Lab in Alderley Park and Alex Alderton, Roberto Amato, Jeffrey Barrett, Sonia Goncalves, Ewan Harrison, David K. Jackson, Ian Johnston, Dominic Kwiatkowski, Cordelia Langford, John Sillitoe on behalf of the Wellcome Sanger Institute COVID-19 Surveillance Team |
| EPI_ISL_2514450, EPI_ISL_2514451, EPI_ISL_2514452, EPI_ISL_2514453 | Lighthouse Lab in Milton Keynes | Wellcome Sanger Institute for the COVID-19 Genomics UK (COG-UK) Consortium | The Lighthouse Lab in Milton Keynes and Alex Alderton, Roberto Amato, Jeffrey Barrett, Sonia Goncalves, Ewan Harrison, David K. Jackson, Ian Johnston, Dominic Kwiatkowski, Cordelia Langford, John Sillitoe on behalf of the Wellcome Sanger Institute COVID-19 Surveillance Team |
| EPI_ISL_2514454 | Lighthouse Lab in Alderley Park | Wellcome Sanger Institute for the COVID-19 Genomics UK (COG-UK) Consortium | Jacquelyn Wynn, Mairead Hyland, The Lighthouse Lab in Alderley Park and Alex Alderton, Roberto Amato, Jeffrey Barrett, Sonia Goncalves, Ewan Harrison, David K. Jackson, Ian Johnston, Dominic Kwiatkowski, Cordelia Langford, John Sillitoe on behalf of the Wellcome Sanger Institute COVID-19 Surveillance Team |
| EPI_ISL_2514455 | Lighthouse Lab in Milton Keynes | Wellcome Sanger Institute for the COVID-19 Genomics UK (COG-UK) Consortium | The Lighthouse Lab in Milton Keynes and Alex Alderton, Roberto Amato, Jeffrey Barrett, Sonia Goncalves, Ewan Harrison, David K. Jackson, Ian Johnston, Dominic Kwiatkowski, Cordelia Langford, John Sillitoe on behalf of the Wellcome Sanger Institute COVID-19 Surveillance Team |
| EPI_ISL_2514456 | Lighthouse Lab in Alderley Park | Wellcome Sanger Institute for the COVID-19 Genomics UK (COG-UK) Consortium | Jacquelyn Wynn, Mairead Hyland, The Lighthouse Lab in Alderley Park and Alex Alderton, Roberto Amato, Jeffrey Barrett, Sonia Goncalves, Ewan Harrison, David K. Jackson, Ian Johnston, Dominic Kwiatkowski, Cordelia Langford, John Sillitoe on behalf of the Wellcome Sanger Institute COVID-19 Surveillance Team |
| EPI_ISL_2514458, EPI_ISL_2514459, EPI_ISL_2514460 | Lighthouse Lab in Milton Keynes | Wellcome Sanger Institute for the COVID-19 Genomics UK (COG-UK) Consortium | The Lighthouse Lab in Milton Keynes and Alex Alderton, Roberto Amato, Jeffrey Barrett, Sonia Goncalves, Ewan Harrison, David K. Jackson, Ian Johnston, Dominic Kwiatkowski, Cordelia Langford, John Sillitoe on behalf of the Wellcome Sanger Institute COVID-19 Surveillance Team |
| EPI_ISL_2514461, EPI_ISL_2514462, EPI_ISL_2514464, EPI_ISL_2514466 | Lighthouse Lab in Alderley Park | Wellcome Sanger Institute for the COVID-19 Genomics UK (COG-UK) Consortium | Jacquelyn Wynn, Mairead Hyland, The Lighthouse Lab in Alderley Park and Alex Alderton, Roberto Amato, Jeffrey Barrett, Sonia Goncalves, Ewan Harrison, David K. Jackson, Ian Johnston, Dominic Kwiatkowski, Cordelia Langford, John Sillitoe on behalf of the Wellcome Sanger Institute COVID-19 Surveillance Team |
| EPI_ISL_2514467 | Lighthouse Lab in Milton Keynes | Wellcome Sanger Institute for the COVID-19 Genomics UK (COG-UK) Consortium | The Lighthouse Lab in Milton Keynes and Alex Alderton, Roberto Amato, Jeffrey Barrett, Sonia Goncalves, Ewan Harrison, David K. Jackson, Ian Johnston, Dominic Kwiatkowski, Cordelia Langford, John Sillitoe on behalf of the Wellcome Sanger Institute COVID-19 Surveillance Team |
| EPI_ISL_2514468 | Lighthouse Lab in Alderley Park | Wellcome Sanger Institute for the COVID-19 Genomics UK (COG-UK) Consortium | Jacquelyn Wynn, Mairead Hyland, The Lighthouse Lab in Alderley Park and Alex Alderton, Roberto Amato, Jeffrey Barrett, Sonia Goncalves, Ewan Harrison, David K. Jackson, Ian Johnston, Dominic Kwiatkowski, Cordelia Langford, John Sillitoe on behalf of the Wellcome Sanger Institute COVID-19 Surveillance Team |
| EPI_ISL_2514469 | Lighthouse Lab in Milton Keynes | Wellcome Sanger Institute for the COVID-19 Genomics UK (COG-UK) Consortium | The Lighthouse Lab in Milton Keynes and Alex Alderton, Roberto Amato, Jeffrey Barrett, Sonia Goncalves, Ewan Harrison, David K. Jackson, Ian Johnston, Dominic Kwiatkowski, Cordelia Langford, John Sillitoe on behalf of the Wellcome Sanger Institute COVID-19 Surveillance Team |
| EPI_ISL_2514470, EPI_ISL_2514471, EPI_ISL_2514472, EPI_ISL_2514473, EPI_ISL_2514474 | Lighthouse Lab in Alderley Park | Wellcome Sanger Institute for the COVID-19 Genomics UK (COG-UK) Consortium | Jacquelyn Wynn, Mairead Hyland, The Lighthouse Lab in Alderley Park and Alex Alderton, Roberto Amato, Jeffrey Barrett, Sonia Goncalves, Ewan Harrison, David K. Jackson, Ian Johnston, Dominic Kwiatkowski, Cordelia Langford, John Sillitoe on behalf of the Wellcome Sanger Institute COVID-19 Surveillance Team |
| EPI_ISL_2514476 | Lighthouse Lab in Milton Keynes | Wellcome Sanger Institute for the COVID-19 Genomics UK (COG-UK) Consortium | The Lighthouse Lab in Milton Keynes and Alex Alderton, Roberto Amato, Jeffrey Barrett, Sonia Goncalves, Ewan Harrison, David K. Jackson, Ian Johnston, Dominic Kwiatkowski, Cordelia Langford, John Sillitoe on behalf of the Wellcome Sanger Institute COVID-19 Surveillance Team |
| EPI_ISL_2514477 | Lighthouse Lab in Alderley Park | Wellcome Sanger Institute for the COVID-19 Genomics UK (COG-UK) Consortium | Jacquelyn Wynn, Mairead Hyland, The Lighthouse Lab in Alderley Park and Alex Alderton, Roberto Amato, Jeffrey Barrett, Sonia Goncalves, Ewan Harrison, David K. Jackson, Ian Johnston, Dominic Kwiatkowski, Cordelia Langford, John Sillitoe on behalf of the Wellcome Sanger Institute COVID-19 Surveillance Team |
| EPI_ISL_2514479 | Lighthouse Lab in Milton Keynes | Wellcome Sanger Institute for the COVID-19 Genomics UK (COG-UK) Consortium | The Lighthouse Lab in Milton Keynes and Alex Alderton, Roberto Amato, Jeffrey Barrett, Sonia Goncalves, Ewan Harrison, David K. Jackson, Ian Johnston, Dominic Kwiatkowski, Cordelia Langford, John Sillitoe on behalf of the Wellcome Sanger Institute COVID-19 Surveillance Team |
| EPI_ISL_2514480, EPI_ISL_2514481, EPI_ISL_2514482 | Lighthouse Lab in Alderley Park | Wellcome Sanger Institute for the COVID-19 Genomics UK (COG-UK) Consortium | Jacquelyn Wynn, Mairead Hyland, The Lighthouse Lab in Alderley Park and Alex Alderton, Roberto Amato, Jeffrey Barrett, Sonia Goncalves, Ewan Harrison, David K. Jackson, Ian Johnston, Dominic Kwiatkowski, Cordelia Langford, John Sillitoe on behalf of the Wellcome Sanger Institute COVID-19 Surveillance Team |
| EPI_ISL_2514483 | Lighthouse Lab in Milton Keynes | Wellcome Sanger Institute for the COVID-19 Genomics UK (COG-UK) Consortium | The Lighthouse Lab in Milton Keynes and Alex Alderton, Roberto Amato, Jeffrey Barrett, Sonia Goncalves, Ewan Harrison, David K. Jackson, Ian Johnston, Dominic Kwiatkowski, Cordelia Langford, John Sillitoe on behalf of the Wellcome Sanger Institute COVID-19 Surveillance Team |
| EPI_ISL_2514484, EPI_ISL_2514485 | Lighthouse Lab in Alderley Park | Wellcome Sanger Institute for the COVID-19 Genomics UK (COG-UK) Consortium | Jacquelyn Wynn, Mairead Hyland, The Lighthouse Lab in Alderley Park and Alex Alderton, Roberto Amato, Jeffrey Barrett, Sonia Goncalves, Ewan Harrison, David K. Jackson, Ian Johnston, Dominic Kwiatkowski, Cordelia Langford, John Sillitoe on behalf of the Wellcome Sanger Institute COVID-19 Surveillance Team |
| EPI_ISL_2514486 | Lighthouse Lab in Milton Keynes | Wellcome Sanger Institute for the COVID-19 Genomics UK (COG-UK) Consortium | The Lighthouse Lab in Milton Keynes and Alex Alderton, Roberto Amato, Jeffrey Barrett, Sonia Goncalves, Ewan Harrison, David K. Jackson, Ian Johnston, Dominic Kwiatkowski, Cordelia Langford, John Sillitoe on behalf of the Wellcome Sanger Institute COVID-19 Surveillance Team |
| EPI_ISL_2514487, EPI_ISL_2514488 | Lighthouse Lab in Alderley Park | Wellcome Sanger Institute for the COVID-19 Genomics UK (COG-UK) Consortium | Jacquelyn Wynn, Mairead Hyland, The Lighthouse Lab in Alderley Park and Alex Alderton, Roberto Amato, Jeffrey Barrett, Sonia Goncalves, Ewan Harrison, David K. Jackson, Ian Johnston, Dominic Kwiatkowski, Cordelia Langford, John Sillitoe on behalf of the Wellcome Sanger Institute COVID-19 Surveillance Team |
| EPI_ISL_2514489 | Lighthouse Lab in Milton Keynes | Wellcome Sanger Institute for the COVID-19 Genomics UK (COG-UK) Consortium | The Lighthouse Lab in Milton Keynes and Alex Alderton, Roberto Amato, Jeffrey Barrett, Sonia Goncalves, Ewan Harrison, David K. Jackson, Ian Johnston, Dominic Kwiatkowski, Cordelia Langford, John Sillitoe on behalf of the Wellcome Sanger Institute COVID-19 Surveillance Team |
| EPI_ISL_2514490, EPI_ISL_2514491 | Lighthouse Lab in Alderley Park | Wellcome Sanger Institute for the COVID-19 Genomics UK (COG-UK) Consortium | Jacquelyn Wynn, Mairead Hyland, The Lighthouse Lab in Alderley Park and Alex Alderton, Roberto Amato, Jeffrey Barrett, Sonia Goncalves, Ewan Harrison, David K. Jackson, Ian Johnston, Dominic Kwiatkowski, Cordelia Langford, John Sillitoe on behalf of the Wellcome Sanger Institute COVID-19 Surveillance Team |
| EPI_ISL_2514492, EPI_ISL_2514494 | Lighthouse Lab in Milton Keynes | Wellcome Sanger Institute for the COVID-19 Genomics UK (COG-UK) Consortium | The Lighthouse Lab in Milton Keynes and Alex Alderton, Roberto Amato, Jeffrey Barrett, Sonia Goncalves, Ewan Harrison, David K. Jackson, Ian Johnston, Dominic Kwiatkowski, Cordelia Langford, John Sillitoe on behalf of the Wellcome Sanger Institute COVID-19 Surveillance Team |
| EPI_ISL_2514495, EPI_ISL_2514496 | Lighthouse Lab in Alderley Park | Wellcome Sanger Institute for the COVID-19 Genomics UK (COG-UK) Consortium | Jacquelyn Wynn, Mairead Hyland, The Lighthouse Lab in Alderley Park and Alex Alderton, Roberto Amato, Jeffrey Barrett, Sonia Goncalves, Ewan Harrison, David K. Jackson, Ian Johnston, Dominic Kwiatkowski, Cordelia Langford, John Sillitoe on behalf of the Wellcome Sanger Institute COVID-19 Surveillance Team |
| EPI_ISL_2514497 | Lighthouse Lab in Milton Keynes | Wellcome Sanger Institute for the COVID-19 Genomics UK (COG-UK) Consortium | The Lighthouse Lab in Milton Keynes and Alex Alderton, Roberto Amato, Jeffrey Barrett, Sonia Goncalves, Ewan Harrison, David K. Jackson, Ian Johnston, Dominic Kwiatkowski, Cordelia Langford, John Sillitoe on behalf of the Wellcome Sanger Institute COVID-19 Surveillance Team |
| EPI_ISL_2514498, EPI_ISL_2514499 | Lighthouse Lab in Alderley Park | Wellcome Sanger Institute for the COVID-19 Genomics UK (COG-UK) Consortium | Jacquelyn Wynn, Mairead Hyland, The Lighthouse Lab in Alderley Park and Alex Alderton, Roberto Amato, Jeffrey Barrett, Sonia Goncalves, Ewan Harrison, David K. Jackson, Ian Johnston, Dominic Kwiatkowski, Cordelia Langford, John Sillitoe on behalf of the Wellcome Sanger Institute COVID-19 Surveillance Team |
| EPI_ISL_2514500 | Lighthouse Lab in Milton Keynes | Wellcome Sanger Institute for the COVID-19 Genomics UK (COG-UK) Consortium | The Lighthouse Lab in Milton Keynes and Alex Alderton, Roberto Amato, Jeffrey Barrett, Sonia Goncalves, Ewan Harrison, David K. Jackson, Ian Johnston, Dominic Kwiatkowski, Cordelia Langford, John Sillitoe on behalf of the Wellcome Sanger Institute COVID-19 Surveillance Team |
| EPI_ISL_2514501 | Lighthouse Lab in Alderley Park | Wellcome Sanger Institute for the COVID-19 Genomics UK (COG-UK) Consortium | Jacquelyn Wynn, Mairead Hyland, The Lighthouse Lab in Alderley Park and Alex Alderton, Roberto Amato, Jeffrey Barrett, Sonia Goncalves, Ewan Harrison, David K. Jackson, Ian Johnston, Dominic Kwiatkowski, Cordelia Langford, John Sillitoe on behalf of the Wellcome Sanger Institute COVID-19 Surveillance Team |
| EPI_ISL_2514502 | Lighthouse Lab in Milton Keynes | Wellcome Sanger Institute for the COVID-19 Genomics UK (COG-UK) Consortium | The Lighthouse Lab in Milton Keynes and Alex Alderton, Roberto Amato, Jeffrey Barrett, Sonia Goncalves, Ewan Harrison, David K. Jackson, Ian Johnston, Dominic Kwiatkowski, Cordelia Langford, John Sillitoe on behalf of the Wellcome Sanger Institute COVID-19 Surveillance Team |
| EPI_ISL_2514503, EPI_ISL_2514504 | Lighthouse Lab in Alderley Park | Wellcome Sanger Institute for the COVID-19 Genomics UK (COG-UK) Consortium | Jacquelyn Wynn, Mairead Hyland, The Lighthouse Lab in Alderley Park and Alex Alderton, Roberto Amato, Jeffrey Barrett, Sonia Goncalves, Ewan Harrison, David K. Jackson, Ian Johnston, Dominic Kwiatkowski, Cordelia Langford, John Sillitoe on behalf of the Wellcome Sanger Institute COVID-19 Surveillance Team |
| EPI_ISL_2514505 | Lighthouse Lab in Milton Keynes | Wellcome Sanger Institute for the COVID-19 Genomics UK (COG-UK) Consortium | The Lighthouse Lab in Milton Keynes and Alex Alderton, Roberto Amato, Jeffrey Barrett, Sonia Goncalves, Ewan Harrison, David K. Jackson, Ian Johnston, Dominic Kwiatkowski, Cordelia Langford, John Sillitoe on behalf of the Wellcome Sanger Institute COVID-19 Surveillance Team |
| EPI_ISL_2514506 | Lighthouse Lab in Alderley Park | Wellcome Sanger Institute for the COVID-19 Genomics UK (COG-UK) Consortium | Jacquelyn Wynn, Mairead Hyland, The Lighthouse Lab in Alderley Park and Alex Alderton, Roberto Amato, Jeffrey Barrett, Sonia Goncalves, Ewan Harrison, David K. Jackson, Ian Johnston, Dominic Kwiatkowski, Cordelia Langford, John Sillitoe on behalf of the Wellcome Sanger Institute COVID-19 Surveillance Team |
| EPI_ISL_2514507 | Lighthouse Lab in Milton Keynes | Wellcome Sanger Institute for the COVID-19 Genomics UK (COG-UK) Consortium | The Lighthouse Lab in Milton Keynes and Alex Alderton, Roberto Amato, Jeffrey Barrett, Sonia Goncalves, Ewan Harrison, David K. Jackson, Ian Johnston, Dominic Kwiatkowski, Cordelia Langford, John Sillitoe on behalf of the Wellcome Sanger Institute COVID-19 Surveillance Team |
| EPI_ISL_2514508 | Lighthouse Lab in Alderley Park | Wellcome Sanger Institute for the COVID-19 Genomics UK (COG-UK) Consortium | Jacquelyn Wynn, Mairead Hyland, The Lighthouse Lab in Alderley Park and Alex Alderton, Roberto Amato, Jeffrey Barrett, Sonia Goncalves, Ewan Harrison, David K. Jackson, Ian Johnston, Dominic Kwiatkowski, Cordelia Langford, John Sillitoe on behalf of the Wellcome Sanger Institute COVID-19 Surveillance Team |
| EPI_ISL_2514510 | Lighthouse Lab in Milton Keynes | Wellcome Sanger Institute for the COVID-19 Genomics UK (COG-UK) | The Lighthouse Lab in Milton Keynes and Alex Alderton, Roberto Amato, Jeffrey Barrett, Sonia Goncalves, Ewan Harrison, David K. Jackson, Ian Johnston, Dominic Kwiatkowski, Cordelia Langford, John Sillitoe on behalf of the Wellcome Sanger |

| | | | |
|---|---|---|---|
| | | Consortium | Institute COVID-19 Surveillance Team |
| EPI_ISL_2514511 | Lighthouse Lab in Alderley Park | Wellcome Sanger Institute for the COVID-19 Genomics UK (COG-UK) Consortium | Jacquelyn Wynn, Mairead Hyland, The Lighthouse Lab in Alderley Park and Alex Alderton, Roberto Amato, Jeffrey Barrett, Sonia Goncalves, Ewan Harrison, David K. Jackson, Ian Johnston, Dominic Kwiatkowski, Cordelia Langford, John Sillitoe on behalf of the Wellcome Sanger Institute COVID-19 Surveillance Team |
| EPI_ISL_2514512 | Lighthouse Lab in Milton Keynes | Wellcome Sanger Institute for the COVID-19 Genomics UK (COG-UK) Consortium | The Lighthouse Lab in Milton Keynes and Alex Alderton, Roberto Amato, Jeffrey Barrett, Sonia Goncalves, Ewan Harrison, David K. Jackson, Ian Johnston, Dominic Kwiatkowski, Cordelia Langford, John Sillitoe on behalf of the Wellcome Sanger Institute COVID-19 Surveillance Team |
| EPI_ISL_2514513, EPI_ISL_2514514 | Lighthouse Lab in Alderley Park | Wellcome Sanger Institute for the COVID-19 Genomics UK (COG-UK) Consortium | Jacquelyn Wynn, Mairead Hyland, The Lighthouse Lab in Alderley Park and Alex Alderton, Roberto Amato, Jeffrey Barrett, Sonia Goncalves, Ewan Harrison, David K. Jackson, Ian Johnston, Dominic Kwiatkowski, Cordelia Langford, John Sillitoe on behalf of the Wellcome Sanger Institute COVID-19 Surveillance Team |
| EPI_ISL_2514515 | Lighthouse Lab in Milton Keynes | Wellcome Sanger Institute for the COVID-19 Genomics UK (COG-UK) Consortium | The Lighthouse Lab in Milton Keynes and Alex Alderton, Roberto Amato, Jeffrey Barrett, Sonia Goncalves, Ewan Harrison, David K. Jackson, Ian Johnston, Dominic Kwiatkowski, Cordelia Langford, John Sillitoe on behalf of the Wellcome Sanger Institute COVID-19 Surveillance Team |
| EPI_ISL_2514516 | Lighthouse Lab in Alderley Park | Wellcome Sanger Institute for the COVID-19 Genomics UK (COG-UK) Consortium | Jacquelyn Wynn, Mairead Hyland, The Lighthouse Lab in Alderley Park and Alex Alderton, Roberto Amato, Jeffrey Barrett, Sonia Goncalves, Ewan Harrison, David K. Jackson, Ian Johnston, Dominic Kwiatkowski, Cordelia Langford, John Sillitoe on behalf of the Wellcome Sanger Institute COVID-19 Surveillance Team |
| EPI_ISL_2514517 | Lighthouse Lab in Milton Keynes | Wellcome Sanger Institute for the COVID-19 Genomics UK (COG-UK) Consortium | The Lighthouse Lab in Milton Keynes and Alex Alderton, Roberto Amato, Jeffrey Barrett, Sonia Goncalves, Ewan Harrison, David K. Jackson, Ian Johnston, Dominic Kwiatkowski, Cordelia Langford, John Sillitoe on behalf of the Wellcome Sanger Institute COVID-19 Surveillance Team |
| EPI_ISL_2514519 | Lighthouse Lab in Alderley Park | Wellcome Sanger Institute for the COVID-19 Genomics UK (COG-UK) Consortium | Jacquelyn Wynn, Mairead Hyland, The Lighthouse Lab in Alderley Park and Alex Alderton, Roberto Amato, Jeffrey Barrett, Sonia Goncalves, Ewan Harrison, David K. Jackson, Ian Johnston, Dominic Kwiatkowski, Cordelia Langford, John Sillitoe on behalf of the Wellcome Sanger Institute COVID-19 Surveillance Team |
| EPI_ISL_2514520, EPI_ISL_2514521 | Lighthouse Lab in Milton Keynes | Wellcome Sanger Institute for the COVID-19 Genomics UK (COG-UK) Consortium | The Lighthouse Lab in Milton Keynes and Alex Alderton, Roberto Amato, Jeffrey Barrett, Sonia Goncalves, Ewan Harrison, David K. Jackson, Ian Johnston, Dominic Kwiatkowski, Cordelia Langford, John Sillitoe on behalf of the Wellcome Sanger Institute COVID-19 Surveillance Team |
| EPI_ISL_2514522 | Lighthouse Lab in Alderley Park | Wellcome Sanger Institute for the COVID-19 Genomics UK (COG-UK) Consortium | Jacquelyn Wynn, Mairead Hyland, The Lighthouse Lab in Alderley Park and Alex Alderton, Roberto Amato, Jeffrey Barrett, Sonia Goncalves, Ewan Harrison, David K. Jackson, Ian Johnston, Dominic Kwiatkowski, Cordelia Langford, John Sillitoe on behalf of the Wellcome Sanger Institute COVID-19 Surveillance Team |
| EPI_ISL_2514523 | Lighthouse Lab in Milton Keynes | Wellcome Sanger Institute for the COVID-19 Genomics UK (COG-UK) Consortium | The Lighthouse Lab in Milton Keynes and Alex Alderton, Roberto Amato, Jeffrey Barrett, Sonia Goncalves, Ewan Harrison, David K. Jackson, Ian Johnston, Dominic Kwiatkowski, Cordelia Langford, John Sillitoe on behalf of the Wellcome Sanger Institute COVID-19 Surveillance Team |
| EPI_ISL_2514524 | Lighthouse Lab in Alderley Park | Wellcome Sanger Institute for the COVID-19 Genomics UK (COG-UK) Consortium | Jacquelyn Wynn, Mairead Hyland, The Lighthouse Lab in Alderley Park and Alex Alderton, Roberto Amato, Jeffrey Barrett, Sonia Goncalves, Ewan Harrison, David K. Jackson, Ian Johnston, Dominic Kwiatkowski, Cordelia Langford, John Sillitoe on behalf of the Wellcome Sanger Institute COVID-19 Surveillance Team |
| EPI_ISL_2514525, EPI_ISL_2514526 | Lighthouse Lab in Milton Keynes | Wellcome Sanger Institute for the COVID-19 Genomics UK (COG-UK) Consortium | The Lighthouse Lab in Milton Keynes and Alex Alderton, Roberto Amato, Jeffrey Barrett, Sonia Goncalves, Ewan Harrison, David K. Jackson, Ian Johnston, Dominic Kwiatkowski, Cordelia Langford, John Sillitoe on behalf of the Wellcome Sanger Institute COVID-19 Surveillance Team |
| EPI_ISL_2514527, EPI_ISL_2514528 | Lighthouse Lab in Alderley Park | Wellcome Sanger Institute for the COVID-19 Genomics UK (COG-UK) Consortium | Jacquelyn Wynn, Mairead Hyland, The Lighthouse Lab in Alderley Park and Alex Alderton, Roberto Amato, Jeffrey Barrett, Sonia Goncalves, Ewan Harrison, David K. Jackson, Ian Johnston, Dominic Kwiatkowski, Cordelia Langford, John Sillitoe on behalf of the Wellcome Sanger Institute COVID-19 Surveillance Team |
| EPI_ISL_2514529, EPI_ISL_2514530, EPI_ISL_2514531, EPI_ISL_2514532 | Lighthouse Lab in Milton Keynes | Wellcome Sanger Institute for the COVID-19 Genomics UK (COG-UK) Consortium | The Lighthouse Lab in Milton Keynes and Alex Alderton, Roberto Amato, Jeffrey Barrett, Sonia Goncalves, Ewan Harrison, David K. Jackson, Ian Johnston, Dominic Kwiatkowski, Cordelia Langford, John Sillitoe on behalf of the Wellcome Sanger Institute COVID-19 Surveillance Team |
| EPI_ISL_2514533, EPI_ISL_2514534, EPI_ISL_2514535 | Lighthouse Lab in Alderley Park | Wellcome Sanger Institute for the COVID-19 Genomics UK (COG-UK) Consortium | Jacquelyn Wynn, Mairead Hyland, The Lighthouse Lab in Alderley Park and Alex Alderton, Roberto Amato, Jeffrey Barrett, Sonia Goncalves, Ewan Harrison, David K. Jackson, Ian Johnston, Dominic Kwiatkowski, Cordelia Langford, John Sillitoe on behalf of the Wellcome Sanger Institute COVID-19 Surveillance Team |
| EPI_ISL_2514536, EPI_ISL_2514537 | Lighthouse Lab in Milton Keynes | Wellcome Sanger Institute for the COVID-19 Genomics UK (COG-UK) Consortium | The Lighthouse Lab in Milton Keynes and Alex Alderton, Roberto Amato, Jeffrey Barrett, Sonia Goncalves, Ewan Harrison, David K. Jackson, Ian Johnston, Dominic Kwiatkowski, Cordelia Langford, John Sillitoe on behalf of the Wellcome Sanger Institute COVID-19 Surveillance Team |
| EPI_ISL_2514538 | Lighthouse Lab in Alderley Park | Wellcome Sanger Institute for the COVID-19 Genomics UK (COG-UK) Consortium | Jacquelyn Wynn, Mairead Hyland, The Lighthouse Lab in Alderley Park and Alex Alderton, Roberto Amato, Jeffrey Barrett, Sonia Goncalves, Ewan Harrison, David K. Jackson, Ian Johnston, Dominic Kwiatkowski, Cordelia Langford, John Sillitoe on behalf of the Wellcome Sanger Institute COVID-19 Surveillance Team |
| EPI_ISL_2514539 | Lighthouse Lab in Milton Keynes | Wellcome Sanger Institute for the COVID-19 Genomics UK (COG-UK) Consortium | The Lighthouse Lab in Milton Keynes and Alex Alderton, Roberto Amato, Jeffrey Barrett, Sonia Goncalves, Ewan Harrison, David K. Jackson, Ian Johnston, Dominic Kwiatkowski, Cordelia Langford, John Sillitoe on behalf of the Wellcome Sanger Institute COVID-19 Surveillance Team |
| EPI_ISL_2514540, EPI_ISL_2514541, EPI_ISL_2514542, EPI_ISL_2514543 | Lighthouse Lab in Alderley Park | Wellcome Sanger Institute for the COVID-19 Genomics UK (COG-UK) Consortium | Jacquelyn Wynn, Mairead Hyland, The Lighthouse Lab in Alderley Park and Alex Alderton, Roberto Amato, Jeffrey Barrett, Sonia Goncalves, Ewan Harrison, David K. Jackson, Ian Johnston, Dominic Kwiatkowski, Cordelia Langford, John Sillitoe on behalf of the Wellcome Sanger Institute COVID-19 Surveillance Team |
| EPI_ISL_2514544 | Lighthouse Lab in Milton Keynes | Wellcome Sanger Institute for the COVID-19 Genomics UK (COG-UK) Consortium | The Lighthouse Lab in Milton Keynes and Alex Alderton, Roberto Amato, Jeffrey Barrett, Sonia Goncalves, Ewan Harrison, David K. Jackson, Ian Johnston, Dominic Kwiatkowski, Cordelia Langford, John Sillitoe on behalf of the Wellcome Sanger Institute COVID-19 Surveillance Team |
| EPI_ISL_2514545, EPI_ISL_2514546, EPI_ISL_2514547 | Lighthouse Lab in Alderley Park | Wellcome Sanger Institute for the COVID-19 Genomics UK (COG-UK) Consortium | Jacquelyn Wynn, Mairead Hyland, The Lighthouse Lab in Alderley Park and Alex Alderton, Roberto Amato, Jeffrey Barrett, Sonia Goncalves, Ewan Harrison, David K. Jackson, Ian Johnston, Dominic Kwiatkowski, Cordelia Langford, John Sillitoe on behalf of the Wellcome Sanger Institute COVID-19 Surveillance Team |
| EPI_ISL_2514548 | Lighthouse Lab in Milton Keynes | Wellcome Sanger Institute for the COVID-19 Genomics UK (COG-UK) Consortium | The Lighthouse Lab in Milton Keynes and Alex Alderton, Roberto Amato, Jeffrey Barrett, Sonia Goncalves, Ewan Harrison, David K. Jackson, Ian Johnston, Dominic Kwiatkowski, Cordelia Langford, John Sillitoe on behalf of the Wellcome Sanger Institute COVID-19 Surveillance Team |
| EPI_ISL_2514549 | Lighthouse Lab in Alderley Park | Wellcome Sanger Institute for the COVID-19 Genomics UK (COG-UK) Consortium | Jacquelyn Wynn, Mairead Hyland, The Lighthouse Lab in Alderley Park and Alex Alderton, Roberto Amato, Jeffrey Barrett, Sonia Goncalves, Ewan Harrison, David K. Jackson, Ian Johnston, Dominic Kwiatkowski, Cordelia Langford, John Sillitoe on behalf of the Wellcome Sanger Institute COVID-19 Surveillance Team |
| EPI_ISL_2514550 | Lighthouse Lab in Milton Keynes | Wellcome Sanger Institute for the COVID-19 Genomics UK (COG-UK) Consortium | The Lighthouse Lab in Milton Keynes and Alex Alderton, Roberto Amato, Jeffrey Barrett, Sonia Goncalves, Ewan Harrison, David K. Jackson, Ian Johnston, Dominic Kwiatkowski, Cordelia Langford, John Sillitoe on behalf of the Wellcome Sanger Institute COVID-19 Surveillance Team |
| EPI_ISL_2514551 | Lighthouse Lab in Alderley Park | Wellcome Sanger Institute for the COVID-19 Genomics UK (COG-UK) Consortium | Jacquelyn Wynn, Mairead Hyland, The Lighthouse Lab in Alderley Park and Alex Alderton, Roberto Amato, Jeffrey Barrett, Sonia Goncalves, Ewan Harrison, David K. Jackson, Ian Johnston, Dominic Kwiatkowski, Cordelia Langford, John Sillitoe on behalf of the Wellcome Sanger Institute COVID-19 Surveillance Team |
| EPI_ISL_2514552, EPI_ISL_2514553, EPI_ISL_2514554 | Lighthouse Lab in Milton Keynes | Wellcome Sanger Institute for the COVID-19 Genomics UK (COG-UK) Consortium | The Lighthouse Lab in Milton Keynes and Alex Alderton, Roberto Amato, Jeffrey Barrett, Sonia Goncalves, Ewan Harrison, David K. Jackson, Ian Johnston, Dominic Kwiatkowski, Cordelia Langford, John Sillitoe on behalf of the Wellcome Sanger Institute COVID-19 Surveillance Team |
| EPI_ISL_2514555 | Lighthouse Lab in Alderley Park | Wellcome Sanger Institute for the COVID-19 Genomics UK (COG-UK) Consortium | Jacquelyn Wynn, Mairead Hyland, The Lighthouse Lab in Alderley Park and Alex Alderton, Roberto Amato, Jeffrey Barrett, Sonia Goncalves, Ewan Harrison, David K. Jackson, Ian Johnston, Dominic Kwiatkowski, Cordelia Langford, John Sillitoe on behalf of the Wellcome Sanger Institute COVID-19 Surveillance Team |
| EPI_ISL_2514556 | Lighthouse Lab in Milton Keynes | Wellcome Sanger Institute for the COVID-19 Genomics UK (COG-UK) Consortium | The Lighthouse Lab in Milton Keynes and Alex Alderton, Roberto Amato, Jeffrey Barrett, Sonia Goncalves, Ewan Harrison, David K. Jackson, Ian Johnston, Dominic Kwiatkowski, Cordelia Langford, John Sillitoe on behalf of the Wellcome Sanger Institute COVID-19 Surveillance Team |
| EPI_ISL_2514557, EPI_ISL_2514558 | Lighthouse Lab in Alderley Park | Wellcome Sanger Institute for the COVID-19 Genomics UK (COG-UK) Consortium | Jacquelyn Wynn, Mairead Hyland, The Lighthouse Lab in Alderley Park and Alex Alderton, Roberto Amato, Jeffrey Barrett, Sonia Goncalves, Ewan Harrison, David K. Jackson, Ian Johnston, Dominic Kwiatkowski, Cordelia Langford, John Sillitoe on behalf of the Wellcome Sanger Institute COVID-19 Surveillance Team |
| EPI_ISL_2514559, EPI_ISL_2514560, EPI_ISL_2514561, EPI_ISL_2514562, EPI_ISL_2514563 | Lighthouse Lab in Milton Keynes | Wellcome Sanger Institute for the COVID-19 Genomics UK (COG-UK) Consortium | The Lighthouse Lab in Milton Keynes and Alex Alderton, Roberto Amato, Jeffrey Barrett, Sonia Goncalves, Ewan Harrison, David K. Jackson, Ian Johnston, Dominic Kwiatkowski, Cordelia Langford, John Sillitoe on behalf of the Wellcome Sanger Institute COVID-19 Surveillance Team |
| EPI_ISL_2514564, EPI_ISL_2514565, EPI_ISL_2514566 | Lighthouse Lab in Alderley Park | Wellcome Sanger Institute for the COVID-19 Genomics UK (COG-UK) Consortium | Jacquelyn Wynn, Mairead Hyland, The Lighthouse Lab in Alderley Park and Alex Alderton, Roberto Amato, Jeffrey Barrett, Sonia Goncalves, Ewan Harrison, David K. Jackson, Ian Johnston, Dominic Kwiatkowski, Cordelia Langford, John Sillitoe on behalf of the Wellcome Sanger Institute COVID-19 Surveillance Team |
| EPI_ISL_2514567 | Lighthouse Lab in Milton Keynes | Wellcome Sanger Institute for the COVID-19 Genomics UK (COG-UK) Consortium | The Lighthouse Lab in Milton Keynes and Alex Alderton, Roberto Amato, Jeffrey Barrett, Sonia Goncalves, Ewan Harrison, David K. Jackson, Ian Johnston, Dominic Kwiatkowski, Cordelia Langford, John Sillitoe on behalf of the Wellcome Sanger Institute COVID-19 Surveillance Team |
| EPI_ISL_2514568, EPI_ISL_2514572, EPI_ISL_2514573 | Lighthouse Lab in Alderley Park | Wellcome Sanger Institute for the COVID-19 Genomics UK (COG-UK) Consortium | Jacquelyn Wynn, Mairead Hyland, The Lighthouse Lab in Alderley Park and Alex Alderton, Roberto Amato, Jeffrey Barrett, Sonia Goncalves, Ewan Harrison, David K. Jackson, Ian Johnston, Dominic Kwiatkowski, Cordelia Langford, John Sillitoe on behalf of the Wellcome Sanger Institute COVID-19 Surveillance Team |
| EPI_ISL_2514574, EPI_ISL_2514575 | Lighthouse Lab in Milton Keynes | Wellcome Sanger Institute for the COVID-19 Genomics UK (COG-UK) Consortium | The Lighthouse Lab in Milton Keynes and Alex Alderton, Roberto Amato, Jeffrey Barrett, Sonia Goncalves, Ewan Harrison, David K. Jackson, Ian Johnston, Dominic Kwiatkowski, Cordelia Langford, John Sillitoe on behalf of the Wellcome Sanger Institute COVID-19 Surveillance Team |
| EPI_ISL_2514576 | Lighthouse Lab in Alderley Park | Wellcome Sanger Institute for the COVID-19 Genomics UK (COG-UK) Consortium | Jacquelyn Wynn, Mairead Hyland, The Lighthouse Lab in Alderley Park and Alex Alderton, Roberto Amato, Jeffrey Barrett, Sonia Goncalves, Ewan Harrison, David K. Jackson, Ian Johnston, Dominic Kwiatkowski, Cordelia Langford, John Sillitoe on behalf of the Wellcome Sanger Institute COVID-19 Surveillance Team |
| EPI_ISL_2514577 | Lighthouse Lab in Milton Keynes | Wellcome Sanger Institute for the COVID-19 Genomics UK (COG-UK) Consortium | The Lighthouse Lab in Milton Keynes and Alex Alderton, Roberto Amato, Jeffrey Barrett, Sonia Goncalves, Ewan Harrison, David K. Jackson, Ian Johnston, Dominic Kwiatkowski, Cordelia Langford, John Sillitoe on behalf of the Wellcome Sanger Institute COVID-19 Surveillance Team |
| EPI_ISL_2514578, EPI_ISL_2514579, EPI_ISL_2514581 | Lighthouse Lab in Alderley Park | Wellcome Sanger Institute for the COVID-19 Genomics UK (COG-UK) Consortium | Jacquelyn Wynn, Mairead Hyland, The Lighthouse Lab in Alderley Park and Alex Alderton, Roberto Amato, Jeffrey Barrett, Sonia Goncalves, Ewan Harrison, David K. Jackson, Ian Johnston, Dominic Kwiatkowski, Cordelia Langford, John Sillitoe on behalf of the Wellcome Sanger Institute COVID-19 Surveillance Team |
| EPI_ISL_2514582 | Lighthouse Lab in Milton Keynes | Wellcome Sanger Institute for the COVID-19 Genomics UK (COG-UK) Consortium | The Lighthouse Lab in Milton Keynes and Alex Alderton, Roberto Amato, Jeffrey Barrett, Sonia Goncalves, Ewan Harrison, David K. Jackson, Ian Johnston, Dominic Kwiatkowski, Cordelia Langford, John Sillitoe on behalf of the Wellcome Sanger Institute COVID-19 Surveillance Team |
| EPI_ISL_2514583, EPI_ISL_2514584, | Lighthouse Lab in Alderley Park | Wellcome Sanger Institute for the COVID-19 Genomics UK (COG-UK) Consortium | Jacquelyn Wynn, Mairead Hyland, The Lighthouse Lab in Alderley Park and Alex Alderton, Roberto Amato, Jeffrey Barrett, Sonia Goncalves, Ewan Harrison, David K. Jackson, Ian Johnston, Dominic Kwiatkowski, Cordelia Langford, John Sillitoe on behalf of the Wellcome Sanger Institute COVID-19 Surveillance Team |

| | | | |
|---|---|---|---|
| EPI_ISL_2514585, EPI_ISL_2514586, EPI_ISL_2514587, EPI_ISL_2514588, EPI_ISL_2514589 | | | |
| EPI_ISL_2514590, EPI_ISL_2514591, EPI_ISL_2514592 | Lighthouse Lab in Milton Keynes | Wellcome Sanger Institute for the COVID-19 Genomics UK (COG-UK) Consortium | The Lighthouse Lab in Milton Keynes and Alex Alderton, Roberto Amato, Jeffrey Barrett, Sonia Goncalves, Ewan Harrison, David K. Jackson, Ian Johnston, Dominic Kwiatkowski, Cordelia Langford, John Sillitoe on behalf of the Wellcome Sanger Institute COVID-19 Surveillance Team |
| EPI_ISL_2514593, EPI_ISL_2514594, EPI_ISL_2514595, EPI_ISL_2514596, EPI_ISL_2514597 | Lighthouse Lab in Alderley Park | Wellcome Sanger Institute for the COVID-19 Genomics UK (COG-UK) Consortium | Jacquelyn Wynn, Mairead Hyland, The Lighthouse Lab in Alderley Park and Alex Alderton, Roberto Amato, Jeffrey Barrett, Sonia Goncalves, Ewan Harrison, David K. Jackson, Ian Johnston, Dominic Kwiatkowski, Cordelia Langford, John Sillitoe on behalf of the Wellcome Sanger Institute COVID-19 Surveillance Team |
| EPI_ISL_2514599 | Lighthouse Lab in Milton Keynes | Wellcome Sanger Institute for the COVID-19 Genomics UK (COG-UK) Consortium | The Lighthouse Lab in Milton Keynes and Alex Alderton, Roberto Amato, Jeffrey Barrett, Sonia Goncalves, Ewan Harrison, David K. Jackson, Ian Johnston, Dominic Kwiatkowski, Cordelia Langford, John Sillitoe on behalf of the Wellcome Sanger Institute COVID-19 Surveillance Team |
| EPI_ISL_2514600 | Lighthouse Lab in Alderley Park | Wellcome Sanger Institute for the COVID-19 Genomics UK (COG-UK) Consortium | Jacquelyn Wynn, Mairead Hyland, The Lighthouse Lab in Alderley Park and Alex Alderton, Roberto Amato, Jeffrey Barrett, Sonia Goncalves, Ewan Harrison, David K. Jackson, Ian Johnston, Dominic Kwiatkowski, Cordelia Langford, John Sillitoe on behalf of the Wellcome Sanger Institute COVID-19 Surveillance Team |
| EPI_ISL_2514601 | Lighthouse Lab in Milton Keynes | Wellcome Sanger Institute for the COVID-19 Genomics UK (COG-UK) Consortium | The Lighthouse Lab in Milton Keynes and Alex Alderton, Roberto Amato, Jeffrey Barrett, Sonia Goncalves, Ewan Harrison, David K. Jackson, Ian Johnston, Dominic Kwiatkowski, Cordelia Langford, John Sillitoe on behalf of the Wellcome Sanger Institute COVID-19 Surveillance Team |
| EPI_ISL_2514602, EPI_ISL_2514603, EPI_ISL_2514604, EPI_ISL_2514605, EPI_ISL_2514607 | Lighthouse Lab in Alderley Park | Wellcome Sanger Institute for the COVID-19 Genomics UK (COG-UK) Consortium | Jacquelyn Wynn, Mairead Hyland, The Lighthouse Lab in Alderley Park and Alex Alderton, Roberto Amato, Jeffrey Barrett, Sonia Goncalves, Ewan Harrison, David K. Jackson, Ian Johnston, Dominic Kwiatkowski, Cordelia Langford, John Sillitoe on behalf of the Wellcome Sanger Institute COVID-19 Surveillance Team |
| EPI_ISL_2514608, EPI_ISL_2514609, EPI_ISL_2514610, EPI_ISL_2514611, EPI_ISL_2514612 | Lighthouse Lab in Milton Keynes | Wellcome Sanger Institute for the COVID-19 Genomics UK (COG-UK) Consortium | The Lighthouse Lab in Milton Keynes and Alex Alderton, Roberto Amato, Jeffrey Barrett, Sonia Goncalves, Ewan Harrison, David K. Jackson, Ian Johnston, Dominic Kwiatkowski, Cordelia Langford, John Sillitoe on behalf of the Wellcome Sanger Institute COVID-19 Surveillance Team |
| EPI_ISL_2514614 | Lighthouse Lab in Alderley Park | Wellcome Sanger Institute for the COVID-19 Genomics UK (COG-UK) Consortium | Jacquelyn Wynn, Mairead Hyland, The Lighthouse Lab in Alderley Park and Alex Alderton, Roberto Amato, Jeffrey Barrett, Sonia Goncalves, Ewan Harrison, David K. Jackson, Ian Johnston, Dominic Kwiatkowski, Cordelia Langford, John Sillitoe on behalf of the Wellcome Sanger Institute COVID-19 Surveillance Team |
| EPI_ISL_2514615, EPI_ISL_2514616, EPI_ISL_2514617 | Lighthouse Lab in Milton Keynes | Wellcome Sanger Institute for the COVID-19 Genomics UK (COG-UK) Consortium | The Lighthouse Lab in Milton Keynes and Alex Alderton, Roberto Amato, Jeffrey Barrett, Sonia Goncalves, Ewan Harrison, David K. Jackson, Ian Johnston, Dominic Kwiatkowski, Cordelia Langford, John Sillitoe on behalf of the Wellcome Sanger Institute COVID-19 Surveillance Team |
| EPI_ISL_2514618, EPI_ISL_2514620, EPI_ISL_2514621 | Lighthouse Lab in Alderley Park | Wellcome Sanger Institute for the COVID-19 Genomics UK (COG-UK) Consortium | Jacquelyn Wynn, Mairead Hyland, The Lighthouse Lab in Alderley Park and Alex Alderton, Roberto Amato, Jeffrey Barrett, Sonia Goncalves, Ewan Harrison, David K. Jackson, Ian Johnston, Dominic Kwiatkowski, Cordelia Langford, John Sillitoe on behalf of the Wellcome Sanger Institute COVID-19 Surveillance Team |
| EPI_ISL_2514622, EPI_ISL_2514623 | Lighthouse Lab in Milton Keynes | Wellcome Sanger Institute for the COVID-19 Genomics UK (COG-UK) Consortium | The Lighthouse Lab in Milton Keynes and Alex Alderton, Roberto Amato, Jeffrey Barrett, Sonia Goncalves, Ewan Harrison, David K. Jackson, Ian Johnston, Dominic Kwiatkowski, Cordelia Langford, John Sillitoe on behalf of the Wellcome Sanger Institute COVID-19 Surveillance Team |
| EPI_ISL_2514624 | Lighthouse Lab in Alderley Park | Wellcome Sanger Institute for the COVID-19 Genomics UK (COG-UK) Consortium | Jacquelyn Wynn, Mairead Hyland, The Lighthouse Lab in Alderley Park and Alex Alderton, Roberto Amato, Jeffrey Barrett, Sonia Goncalves, Ewan Harrison, David K. Jackson, Ian Johnston, Dominic Kwiatkowski, Cordelia Langford, John Sillitoe on behalf of the Wellcome Sanger Institute COVID-19 Surveillance Team |
| EPI_ISL_2514626 | Lighthouse Lab in Milton Keynes | Wellcome Sanger Institute for the COVID-19 Genomics UK (COG-UK) Consortium | The Lighthouse Lab in Milton Keynes and Alex Alderton, Roberto Amato, Jeffrey Barrett, Sonia Goncalves, Ewan Harrison, David K. Jackson, Ian Johnston, Dominic Kwiatkowski, Cordelia Langford, John Sillitoe on behalf of the Wellcome Sanger Institute COVID-19 Surveillance Team |
| EPI_ISL_2514627, EPI_ISL_2514628, EPI_ISL_2514629 | Lighthouse Lab in Alderley Park | Wellcome Sanger Institute for the COVID-19 Genomics UK (COG-UK) Consortium | Jacquelyn Wynn, Mairead Hyland, The Lighthouse Lab in Alderley Park and Alex Alderton, Roberto Amato, Jeffrey Barrett, Sonia Goncalves, Ewan Harrison, David K. Jackson, Ian Johnston, Dominic Kwiatkowski, Cordelia Langford, John Sillitoe on behalf of the Wellcome Sanger Institute COVID-19 Surveillance Team |
| EPI_ISL_2514630, EPI_ISL_2514631 | Lighthouse Lab in Milton Keynes | Wellcome Sanger Institute for the COVID-19 Genomics UK (COG-UK) Consortium | The Lighthouse Lab in Milton Keynes and Alex Alderton, Roberto Amato, Jeffrey Barrett, Sonia Goncalves, Ewan Harrison, David K. Jackson, Ian Johnston, Dominic Kwiatkowski, Cordelia Langford, John Sillitoe on behalf of the Wellcome Sanger Institute COVID-19 Surveillance Team |
| EPI_ISL_2514632 | Lighthouse Lab in Alderley Park | Wellcome Sanger Institute for the COVID-19 Genomics UK (COG-UK) Consortium | Jacquelyn Wynn, Mairead Hyland, The Lighthouse Lab in Alderley Park and Alex Alderton, Roberto Amato, Jeffrey Barrett, Sonia Goncalves, Ewan Harrison, David K. Jackson, Ian Johnston, Dominic Kwiatkowski, Cordelia Langford, John Sillitoe on behalf of the Wellcome Sanger Institute COVID-19 Surveillance Team |
| EPI_ISL_2514633, EPI_ISL_2514634 | Lighthouse Lab in Milton Keynes | Wellcome Sanger Institute for the COVID-19 Genomics UK (COG-UK) Consortium | The Lighthouse Lab in Milton Keynes and Alex Alderton, Roberto Amato, Jeffrey Barrett, Sonia Goncalves, Ewan Harrison, David K. Jackson, Ian Johnston, Dominic Kwiatkowski, Cordelia Langford, John Sillitoe on behalf of the Wellcome Sanger Institute COVID-19 Surveillance Team |
| EPI_ISL_2514635, EPI_ISL_2514637 | Lighthouse Lab in Alderley Park | Wellcome Sanger Institute for the COVID-19 Genomics UK (COG-UK) Consortium | Jacquelyn Wynn, Mairead Hyland, The Lighthouse Lab in Alderley Park and Alex Alderton, Roberto Amato, Jeffrey Barrett, Sonia Goncalves, Ewan Harrison, David K. Jackson, Ian Johnston, Dominic Kwiatkowski, Cordelia Langford, John Sillitoe on behalf of the Wellcome Sanger Institute COVID-19 Surveillance Team |
| EPI_ISL_2514639, EPI_ISL_2514640 | Lighthouse Lab in Milton Keynes | Wellcome Sanger Institute for the COVID-19 Genomics UK (COG-UK) Consortium | The Lighthouse Lab in Milton Keynes and Alex Alderton, Roberto Amato, Jeffrey Barrett, Sonia Goncalves, Ewan Harrison, David K. Jackson, Ian Johnston, Dominic Kwiatkowski, Cordelia Langford, John Sillitoe on behalf of the Wellcome Sanger Institute COVID-19 Surveillance Team |
| EPI_ISL_2514641, EPI_ISL_2514642, EPI_ISL_2514643, EPI_ISL_2514644 | Lighthouse Lab in Alderley Park | Wellcome Sanger Institute for the COVID-19 Genomics UK (COG-UK) Consortium | Jacquelyn Wynn, Mairead Hyland, The Lighthouse Lab in Alderley Park and Alex Alderton, Roberto Amato, Jeffrey Barrett, Sonia Goncalves, Ewan Harrison, David K. Jackson, Ian Johnston, Dominic Kwiatkowski, Cordelia Langford, John Sillitoe on behalf of the Wellcome Sanger Institute COVID-19 Surveillance Team |
| EPI_ISL_2514645, EPI_ISL_2514646 | Lighthouse Lab in Milton Keynes | Wellcome Sanger Institute for the COVID-19 Genomics UK (COG-UK) Consortium | The Lighthouse Lab in Milton Keynes and Alex Alderton, Roberto Amato, Jeffrey Barrett, Sonia Goncalves, Ewan Harrison, David K. Jackson, Ian Johnston, Dominic Kwiatkowski, Cordelia Langford, John Sillitoe on behalf of the Wellcome Sanger Institute COVID-19 Surveillance Team |
| EPI_ISL_2514647, EPI_ISL_2514648, EPI_ISL_2514649, EPI_ISL_2514650, EPI_ISL_2514651, EPI_ISL_2514652, EPI_ISL_2514653 | Lighthouse Lab in Alderley Park | Wellcome Sanger Institute for the COVID-19 Genomics UK (COG-UK) Consortium | Jacquelyn Wynn, Mairead Hyland, The Lighthouse Lab in Alderley Park and Alex Alderton, Roberto Amato, Jeffrey Barrett, Sonia Goncalves, Ewan Harrison, David K. Jackson, Ian Johnston, Dominic Kwiatkowski, Cordelia Langford, John Sillitoe on behalf of the Wellcome Sanger Institute COVID-19 Surveillance Team |
| EPI_ISL_2514654, EPI_ISL_2514655 | Lighthouse Lab in Milton Keynes | Wellcome Sanger Institute for the COVID-19 Genomics UK (COG-UK) Consortium | The Lighthouse Lab in Milton Keynes and Alex Alderton, Roberto Amato, Jeffrey Barrett, Sonia Goncalves, Ewan Harrison, David K. Jackson, Ian Johnston, Dominic Kwiatkowski, Cordelia Langford, John Sillitoe on behalf of the Wellcome Sanger Institute COVID-19 Surveillance Team |
| EPI_ISL_2514656, EPI_ISL_2514657, EPI_ISL_2514658, EPI_ISL_2514659 | Lighthouse Lab in Alderley Park | Wellcome Sanger Institute for the COVID-19 Genomics UK (COG-UK) Consortium | Jacquelyn Wynn, Mairead Hyland, The Lighthouse Lab in Alderley Park and Alex Alderton, Roberto Amato, Jeffrey Barrett, Sonia Goncalves, Ewan Harrison, David K. Jackson, Ian Johnston, Dominic Kwiatkowski, Cordelia Langford, John Sillitoe on behalf of the Wellcome Sanger Institute COVID-19 Surveillance Team |
| EPI_ISL_2514660 | Lighthouse Lab in Milton Keynes | Wellcome Sanger Institute for the COVID-19 Genomics UK (COG-UK) Consortium | The Lighthouse Lab in Milton Keynes and Alex Alderton, Roberto Amato, Jeffrey Barrett, Sonia Goncalves, Ewan Harrison, David K. Jackson, Ian Johnston, Dominic Kwiatkowski, Cordelia Langford, John Sillitoe on behalf of the Wellcome Sanger Institute COVID-19 Surveillance Team |
| EPI_ISL_2514661, EPI_ISL_2514662, EPI_ISL_2514663, EPI_ISL_2514664, EPI_ISL_2514665 | Lighthouse Lab in Alderley Park | Wellcome Sanger Institute for the COVID-19 Genomics UK (COG-UK) Consortium | Jacquelyn Wynn, Mairead Hyland, The Lighthouse Lab in Alderley Park and Alex Alderton, Roberto Amato, Jeffrey Barrett, Sonia Goncalves, Ewan Harrison, David K. Jackson, Ian Johnston, Dominic Kwiatkowski, Cordelia Langford, John Sillitoe on behalf of the Wellcome Sanger Institute COVID-19 Surveillance Team |
| EPI_ISL_2514666, EPI_ISL_2514667, EPI_ISL_2514668, EPI_ISL_2514669 | Lighthouse Lab in Milton Keynes | Wellcome Sanger Institute for the COVID-19 Genomics UK (COG-UK) Consortium | The Lighthouse Lab in Milton Keynes and Alex Alderton, Roberto Amato, Jeffrey Barrett, Sonia Goncalves, Ewan Harrison, David K. Jackson, Ian Johnston, Dominic Kwiatkowski, Cordelia Langford, John Sillitoe on behalf of the Wellcome Sanger Institute COVID-19 Surveillance Team |
| EPI_ISL_2514670 | Lighthouse Lab in Alderley Park | Wellcome Sanger Institute for the COVID-19 Genomics UK (COG-UK) Consortium | Jacquelyn Wynn, Mairead Hyland, The Lighthouse Lab in Alderley Park and Alex Alderton, Roberto Amato, Jeffrey Barrett, Sonia Goncalves, Ewan Harrison, David K. Jackson, Ian Johnston, Dominic Kwiatkowski, Cordelia Langford, John Sillitoe on behalf of the Wellcome Sanger Institute COVID-19 Surveillance Team |
| EPI_ISL_2514671 | Lighthouse Lab in Milton Keynes | Wellcome Sanger Institute for the COVID-19 Genomics UK (COG-UK) Consortium | The Lighthouse Lab in Milton Keynes and Alex Alderton, Roberto Amato, Jeffrey Barrett, Sonia Goncalves, Ewan Harrison, David K. Jackson, Ian Johnston, Dominic Kwiatkowski, Cordelia Langford, John Sillitoe on behalf of the Wellcome Sanger Institute COVID-19 Surveillance Team |
| EPI_ISL_2514672, EPI_ISL_2514673, EPI_ISL_2514675, EPI_ISL_2514676, EPI_ISL_2514677 | Lighthouse Lab in Alderley Park | Wellcome Sanger Institute for the COVID-19 Genomics UK (COG-UK) Consortium | Jacquelyn Wynn, Mairead Hyland, The Lighthouse Lab in Alderley Park and Alex Alderton, Roberto Amato, Jeffrey Barrett, Sonia Goncalves, Ewan Harrison, David K. Jackson, Ian Johnston, Dominic Kwiatkowski, Cordelia Langford, John Sillitoe on behalf of the Wellcome Sanger Institute COVID-19 Surveillance Team |
| EPI_ISL_2514680 | Lighthouse Lab in Milton Keynes | Wellcome Sanger Institute for the COVID-19 Genomics UK (COG-UK) Consortium | The Lighthouse Lab in Milton Keynes and Alex Alderton, Roberto Amato, Jeffrey Barrett, Sonia Goncalves, Ewan Harrison, David K. Jackson, Ian Johnston, Dominic Kwiatkowski, Cordelia Langford, John Sillitoe on behalf of the Wellcome Sanger Institute COVID-19 Surveillance Team |

| | | | |
|---|---|---|---|
| EPI_ISL_2514681, EPI_ISL_2514682 | Lighthouse Lab in Alderley Park | Wellcome Sanger Institute for the COVID-19 Genomics UK (COG-UK) Consortium | Jacquelyn Wynn, Mairead Hyland, The Lighthouse Lab in Alderley Park and Alex Alderton, Roberto Amato, Jeffrey Barrett, Sonia Goncalves, Ewan Harrison, David K. Jackson, Ian Johnston, Dominic Kwiatkowski, Cordelia Langford, John Sillitoe on behalf of the Wellcome Sanger Institute COVID-19 Surveillance Team |
| EPI_ISL_2514684, EPI_ISL_2514685 | Lighthouse Lab in Milton Keynes | Wellcome Sanger Institute for the COVID-19 Genomics UK (COG-UK) Consortium | The Lighthouse Lab in Milton Keynes and Alex Alderton, Roberto Amato, Jeffrey Barrett, Sonia Goncalves, Ewan Harrison, David K. Jackson, Ian Johnston, Dominic Kwiatkowski, Cordelia Langford, John Sillitoe on behalf of the Wellcome Sanger Institute COVID-19 Surveillance Team |
| EPI_ISL_2514686 | Lighthouse Lab in Alderley Park | Wellcome Sanger Institute for the COVID-19 Genomics UK (COG-UK) Consortium | Jacquelyn Wynn, Mairead Hyland, The Lighthouse Lab in Alderley Park and Alex Alderton, Roberto Amato, Jeffrey Barrett, Sonia Goncalves, Ewan Harrison, David K. Jackson, Ian Johnston, Dominic Kwiatkowski, Cordelia Langford, John Sillitoe on behalf of the Wellcome Sanger Institute COVID-19 Surveillance Team |
| EPI_ISL_2514687, EPI_ISL_2514688 | Lighthouse Lab in Milton Keynes | Wellcome Sanger Institute for the COVID-19 Genomics UK (COG-UK) Consortium | The Lighthouse Lab in Milton Keynes and Alex Alderton, Roberto Amato, Jeffrey Barrett, Sonia Goncalves, Ewan Harrison, David K. Jackson, Ian Johnston, Dominic Kwiatkowski, Cordelia Langford, John Sillitoe on behalf of the Wellcome Sanger Institute COVID-19 Surveillance Team |
| EPI_ISL_2514689, EPI_ISL_2514690, EPI_ISL_2514691 | Lighthouse Lab in Alderley Park | Wellcome Sanger Institute for the COVID-19 Genomics UK (COG-UK) Consortium | Jacquelyn Wynn, Mairead Hyland, The Lighthouse Lab in Alderley Park and Alex Alderton, Roberto Amato, Jeffrey Barrett, Sonia Goncalves, Ewan Harrison, David K. Jackson, Ian Johnston, Dominic Kwiatkowski, Cordelia Langford, John Sillitoe on behalf of the Wellcome Sanger Institute COVID-19 Surveillance Team |
| EPI_ISL_2514692, EPI_ISL_2514693, EPI_ISL_2514694 | Lighthouse Lab in Milton Keynes | Wellcome Sanger Institute for the COVID-19 Genomics UK (COG-UK) Consortium | The Lighthouse Lab in Milton Keynes and Alex Alderton, Roberto Amato, Jeffrey Barrett, Sonia Goncalves, Ewan Harrison, David K. Jackson, Ian Johnston, Dominic Kwiatkowski, Cordelia Langford, John Sillitoe on behalf of the Wellcome Sanger Institute COVID-19 Surveillance Team |
| EPI_ISL_2514695, EPI_ISL_2514696 | Lighthouse Lab in Alderley Park | Wellcome Sanger Institute for the COVID-19 Genomics UK (COG-UK) Consortium | Jacquelyn Wynn, Mairead Hyland, The Lighthouse Lab in Alderley Park and Alex Alderton, Roberto Amato, Jeffrey Barrett, Sonia Goncalves, Ewan Harrison, David K. Jackson, Ian Johnston, Dominic Kwiatkowski, Cordelia Langford, John Sillitoe on behalf of the Wellcome Sanger Institute COVID-19 Surveillance Team |
| EPI_ISL_2514697, EPI_ISL_2514698, EPI_ISL_2514699 | Lighthouse Lab in Milton Keynes | Wellcome Sanger Institute for the COVID-19 Genomics UK (COG-UK) Consortium | The Lighthouse Lab in Milton Keynes and Alex Alderton, Roberto Amato, Jeffrey Barrett, Sonia Goncalves, Ewan Harrison, David K. Jackson, Ian Johnston, Dominic Kwiatkowski, Cordelia Langford, John Sillitoe on behalf of the Wellcome Sanger Institute COVID-19 Surveillance Team |
| EPI_ISL_2514700, EPI_ISL_2514701 | Lighthouse Lab in Alderley Park | Wellcome Sanger Institute for the COVID-19 Genomics UK (COG-UK) Consortium | Jacquelyn Wynn, Mairead Hyland, The Lighthouse Lab in Alderley Park and Alex Alderton, Roberto Amato, Jeffrey Barrett, Sonia Goncalves, Ewan Harrison, David K. Jackson, Ian Johnston, Dominic Kwiatkowski, Cordelia Langford, John Sillitoe on behalf of the Wellcome Sanger Institute COVID-19 Surveillance Team |
| EPI_ISL_2514702, EPI_ISL_2514703 | Lighthouse Lab in Milton Keynes | Wellcome Sanger Institute for the COVID-19 Genomics UK (COG-UK) Consortium | The Lighthouse Lab in Milton Keynes and Alex Alderton, Roberto Amato, Jeffrey Barrett, Sonia Goncalves, Ewan Harrison, David K. Jackson, Ian Johnston, Dominic Kwiatkowski, Cordelia Langford, John Sillitoe on behalf of the Wellcome Sanger Institute COVID-19 Surveillance Team |
| EPI_ISL_2514704, EPI_ISL_2514705, EPI_ISL_2514706, EPI_ISL_2514707, EPI_ISL_2514708, EPI_ISL_2514709 | Lighthouse Lab in Alderley Park | Wellcome Sanger Institute for the COVID-19 Genomics UK (COG-UK) Consortium | Jacquelyn Wynn, Mairead Hyland, The Lighthouse Lab in Alderley Park and Alex Alderton, Roberto Amato, Jeffrey Barrett, Sonia Goncalves, Ewan Harrison, David K. Jackson, Ian Johnston, Dominic Kwiatkowski, Cordelia Langford, John Sillitoe on behalf of the Wellcome Sanger Institute COVID-19 Surveillance Team |
| EPI_ISL_2514710, EPI_ISL_2514711, EPI_ISL_2514712 | Lighthouse Lab in Milton Keynes | Wellcome Sanger Institute for the COVID-19 Genomics UK (COG-UK) Consortium | The Lighthouse Lab in Milton Keynes and Alex Alderton, Roberto Amato, Jeffrey Barrett, Sonia Goncalves, Ewan Harrison, David K. Jackson, Ian Johnston, Dominic Kwiatkowski, Cordelia Langford, John Sillitoe on behalf of the Wellcome Sanger Institute COVID-19 Surveillance Team |
| EPI_ISL_2514713, EPI_ISL_2514714 | Lighthouse Lab in Alderley Park | Wellcome Sanger Institute for the COVID-19 Genomics UK (COG-UK) Consortium | Jacquelyn Wynn, Mairead Hyland, The Lighthouse Lab in Alderley Park and Alex Alderton, Roberto Amato, Jeffrey Barrett, Sonia Goncalves, Ewan Harrison, David K. Jackson, Ian Johnston, Dominic Kwiatkowski, Cordelia Langford, John Sillitoe on behalf of the Wellcome Sanger Institute COVID-19 Surveillance Team |
| EPI_ISL_2514715, EPI_ISL_2514716, EPI_ISL_2514719 | Lighthouse Lab in Milton Keynes | Wellcome Sanger Institute for the COVID-19 Genomics UK (COG-UK) Consortium | The Lighthouse Lab in Milton Keynes and Alex Alderton, Roberto Amato, Jeffrey Barrett, Sonia Goncalves, Ewan Harrison, David K. Jackson, Ian Johnston, Dominic Kwiatkowski, Cordelia Langford, John Sillitoe on behalf of the Wellcome Sanger Institute COVID-19 Surveillance Team |
| EPI_ISL_2514720, EPI_ISL_2514721, EPI_ISL_2514722, EPI_ISL_2514723 | Lighthouse Lab in Alderley Park | Wellcome Sanger Institute for the COVID-19 Genomics UK (COG-UK) Consortium | Jacquelyn Wynn, Mairead Hyland, The Lighthouse Lab in Alderley Park and Alex Alderton, Roberto Amato, Jeffrey Barrett, Sonia Goncalves, Ewan Harrison, David K. Jackson, Ian Johnston, Dominic Kwiatkowski, Cordelia Langford, John Sillitoe on behalf of the Wellcome Sanger Institute COVID-19 Surveillance Team |
| EPI_ISL_2514725, EPI_ISL_2514726 | Lighthouse Lab in Milton Keynes | Wellcome Sanger Institute for the COVID-19 Genomics UK (COG-UK) Consortium | The Lighthouse Lab in Milton Keynes and Alex Alderton, Roberto Amato, Jeffrey Barrett, Sonia Goncalves, Ewan Harrison, David K. Jackson, Ian Johnston, Dominic Kwiatkowski, Cordelia Langford, John Sillitoe on behalf of the Wellcome Sanger Institute COVID-19 Surveillance Team |
| EPI_ISL_2514727 | Lighthouse Lab in Alderley Park | Wellcome Sanger Institute for the COVID-19 Genomics UK (COG-UK) Consortium | Jacquelyn Wynn, Mairead Hyland, The Lighthouse Lab in Alderley Park and Alex Alderton, Roberto Amato, Jeffrey Barrett, Sonia Goncalves, Ewan Harrison, David K. Jackson, Ian Johnston, Dominic Kwiatkowski, Cordelia Langford, John Sillitoe on behalf of the Wellcome Sanger Institute COVID-19 Surveillance Team |
| EPI_ISL_2514728 | Lighthouse Lab in Milton Keynes | Wellcome Sanger Institute for the COVID-19 Genomics UK (COG-UK) Consortium | The Lighthouse Lab in Milton Keynes and Alex Alderton, Roberto Amato, Jeffrey Barrett, Sonia Goncalves, Ewan Harrison, David K. Jackson, Ian Johnston, Dominic Kwiatkowski, Cordelia Langford, John Sillitoe on behalf of the Wellcome Sanger Institute COVID-19 Surveillance Team |
| EPI_ISL_2514729 | Lighthouse Lab in Glasgow | Wellcome Sanger Institute for the COVID-19 Genomics UK (COG-UK) Consortium | Harper VanSteenhouse, Yumi Kasai, David Gray, Carol Clugston, Anna Dominiczak and Alex Alderton, Roberto Amato, Jeffrey Barrett, Sonia Goncalves, Ewan Harrison, David K. Jackson, Ian Johnston, Dominic Kwiatkowski, Cordelia Langford, John Sillitoe on behalf of the Wellcome Sanger Institute COVID-19 Surveillance Team |
| EPI_ISL_2514730 | Lighthouse Lab in Milton Keynes | Wellcome Sanger Institute for the COVID-19 Genomics UK (COG-UK) Consortium | The Lighthouse Lab in Milton Keynes and Alex Alderton, Roberto Amato, Jeffrey Barrett, Sonia Goncalves, Ewan Harrison, David K. Jackson, Ian Johnston, Dominic Kwiatkowski, Cordelia Langford, John Sillitoe on behalf of the Wellcome Sanger Institute COVID-19 Surveillance Team |
| EPI_ISL_2514731, EPI_ISL_2514732, EPI_ISL_2514733, EPI_ISL_2514734, EPI_ISL_2514735, EPI_ISL_2514737, EPI_ISL_2514738, EPI_ISL_2514739, EPI_ISL_2514740 | Lighthouse Lab in Glasgow | Wellcome Sanger Institute for the COVID-19 Genomics UK (COG-UK) Consortium | Harper VanSteenhouse, Yumi Kasai, David Gray, Carol Clugston, Anna Dominiczak and Alex Alderton, Roberto Amato, Jeffrey Barrett, Sonia Goncalves, Ewan Harrison, David K. Jackson, Ian Johnston, Dominic Kwiatkowski, Cordelia Langford, John Sillitoe on behalf of the Wellcome Sanger Institute COVID-19 Surveillance Team |
| EPI_ISL_2514741, EPI_ISL_2514742 | Lighthouse Lab in Milton Keynes | Wellcome Sanger Institute for the COVID-19 Genomics UK (COG-UK) Consortium | The Lighthouse Lab in Milton Keynes and Alex Alderton, Roberto Amato, Jeffrey Barrett, Sonia Goncalves, Ewan Harrison, David K. Jackson, Ian Johnston, Dominic Kwiatkowski, Cordelia Langford, John Sillitoe on behalf of the Wellcome Sanger Institute COVID-19 Surveillance Team |
| EPI_ISL_2514743 | Lighthouse Lab in Glasgow | Wellcome Sanger Institute for the COVID-19 Genomics UK (COG-UK) Consortium | Harper VanSteenhouse, Yumi Kasai, David Gray, Carol Clugston, Anna Dominiczak and Alex Alderton, Roberto Amato, Jeffrey Barrett, Sonia Goncalves, Ewan Harrison, David K. Jackson, Ian Johnston, Dominic Kwiatkowski, Cordelia Langford, John Sillitoe on behalf of the Wellcome Sanger Institute COVID-19 Surveillance Team |
| EPI_ISL_2514744 | Lighthouse Lab in Milton Keynes | Wellcome Sanger Institute for the COVID-19 Genomics UK (COG-UK) Consortium | The Lighthouse Lab in Milton Keynes and Alex Alderton, Roberto Amato, Jeffrey Barrett, Sonia Goncalves, Ewan Harrison, David K. Jackson, Ian Johnston, Dominic Kwiatkowski, Cordelia Langford, John Sillitoe on behalf of the Wellcome Sanger Institute COVID-19 Surveillance Team |
| EPI_ISL_2514745 | Lighthouse Lab in Glasgow | Wellcome Sanger Institute for the COVID-19 Genomics UK (COG-UK) Consortium | Harper VanSteenhouse, Yumi Kasai, David Gray, Carol Clugston, Anna Dominiczak and Alex Alderton, Roberto Amato, Jeffrey Barrett, Sonia Goncalves, Ewan Harrison, David K. Jackson, Ian Johnston, Dominic Kwiatkowski, Cordelia Langford, John Sillitoe on behalf of the Wellcome Sanger Institute COVID-19 Surveillance Team |
| EPI_ISL_2514746 | Lighthouse Lab in Milton Keynes | Wellcome Sanger Institute for the COVID-19 Genomics UK (COG-UK) Consortium | The Lighthouse Lab in Milton Keynes and Alex Alderton, Roberto Amato, Jeffrey Barrett, Sonia Goncalves, Ewan Harrison, David K. Jackson, Ian Johnston, Dominic Kwiatkowski, Cordelia Langford, John Sillitoe on behalf of the Wellcome Sanger Institute COVID-19 Surveillance Team |
| EPI_ISL_2514748, EPI_ISL_2514749 | Lighthouse Lab in Glasgow | Wellcome Sanger Institute for the COVID-19 Genomics UK (COG-UK) Consortium | Harper VanSteenhouse, Yumi Kasai, David Gray, Carol Clugston, Anna Dominiczak and Alex Alderton, Roberto Amato, Jeffrey Barrett, Sonia Goncalves, Ewan Harrison, David K. Jackson, Ian Johnston, Dominic Kwiatkowski, Cordelia Langford, John Sillitoe on behalf of the Wellcome Sanger Institute COVID-19 Surveillance Team |
| EPI_ISL_2514750 | Lighthouse Lab in Milton Keynes | Wellcome Sanger Institute for the COVID-19 Genomics UK (COG-UK) Consortium | The Lighthouse Lab in Milton Keynes and Alex Alderton, Roberto Amato, Jeffrey Barrett, Sonia Goncalves, Ewan Harrison, David K. Jackson, Ian Johnston, Dominic Kwiatkowski, Cordelia Langford, John Sillitoe on behalf of the Wellcome Sanger Institute COVID-19 Surveillance Team |
| EPI_ISL_2514752, EPI_ISL_2514753 | Lighthouse Lab in Glasgow | Wellcome Sanger Institute for the COVID-19 Genomics UK (COG-UK) Consortium | Harper VanSteenhouse, Yumi Kasai, David Gray, Carol Clugston, Anna Dominiczak and Alex Alderton, Roberto Amato, Jeffrey Barrett, Sonia Goncalves, Ewan Harrison, David K. Jackson, Ian Johnston, Dominic Kwiatkowski, Cordelia Langford, John Sillitoe on behalf of the Wellcome Sanger Institute COVID-19 Surveillance Team |
| EPI_ISL_2514754, EPI_ISL_2514755, EPI_ISL_2514756, EPI_ISL_2514757 | Lighthouse Lab in Milton Keynes | Wellcome Sanger Institute for the COVID-19 Genomics UK (COG-UK) Consortium | The Lighthouse Lab in Milton Keynes and Alex Alderton, Roberto Amato, Jeffrey Barrett, Sonia Goncalves, Ewan Harrison, David K. Jackson, Ian Johnston, Dominic Kwiatkowski, Cordelia Langford, John Sillitoe on behalf of the Wellcome Sanger Institute COVID-19 Surveillance Team |
| EPI_ISL_2514758, EPI_ISL_2514761 | Lighthouse Lab in Glasgow | Wellcome Sanger Institute for the COVID-19 Genomics UK (COG-UK) Consortium | Harper VanSteenhouse, Yumi Kasai, David Gray, Carol Clugston, Anna Dominiczak and Alex Alderton, Roberto Amato, Jeffrey Barrett, Sonia Goncalves, Ewan Harrison, David K. Jackson, Ian Johnston, Dominic Kwiatkowski, Cordelia Langford, John Sillitoe on behalf of the Wellcome Sanger Institute COVID-19 Surveillance Team |
| EPI_ISL_2514762, EPI_ISL_2514763, EPI_ISL_2514764 | Lighthouse Lab in Milton Keynes | Wellcome Sanger Institute for the COVID-19 Genomics UK (COG-UK) Consortium | The Lighthouse Lab in Milton Keynes and Alex Alderton, Roberto Amato, Jeffrey Barrett, Sonia Goncalves, Ewan Harrison, David K. Jackson, Ian Johnston, Dominic Kwiatkowski, Cordelia Langford, John Sillitoe on behalf of the Wellcome Sanger Institute COVID-19 Surveillance Team |
| EPI_ISL_2514765, EPI_ISL_2514766 | Lighthouse Lab in Glasgow | Wellcome Sanger Institute for the COVID-19 Genomics UK (COG-UK) Consortium | Harper VanSteenhouse, Yumi Kasai, David Gray, Carol Clugston, Anna Dominiczak and Alex Alderton, Roberto Amato, Jeffrey Barrett, Sonia Goncalves, Ewan Harrison, David K. Jackson, Ian Johnston, Dominic Kwiatkowski, Cordelia Langford, John Sillitoe on behalf of the Wellcome Sanger Institute COVID-19 Surveillance Team |
| EPI_ISL_2514768 | Lighthouse Lab in Milton Keynes | Wellcome Sanger Institute for the COVID-19 Genomics UK (COG-UK) Consortium | The Lighthouse Lab in Milton Keynes and Alex Alderton, Roberto Amato, Jeffrey Barrett, Sonia Goncalves, Ewan Harrison, David K. Jackson, Ian Johnston, Dominic Kwiatkowski, Cordelia Langford, John Sillitoe on behalf of the Wellcome Sanger Institute COVID-19 Surveillance Team |
| EPI_ISL_2514769, EPI_ISL_2514770, EPI_ISL_2514771, EPI_ISL_2514772, EPI_ISL_2514775, EPI_ISL_2514776, | Lighthouse Lab in Glasgow | Wellcome Sanger Institute for the COVID-19 Genomics UK (COG-UK) Consortium | Harper VanSteenhouse, Yumi Kasai, David Gray, Carol Clugston, Anna Dominiczak and Alex Alderton, Roberto Amato, Jeffrey Barrett, Sonia Goncalves, Ewan Harrison, David K. Jackson, Ian Johnston, Dominic Kwiatkowski, Cordelia Langford, John Sillitoe on behalf of the Wellcome Sanger Institute COVID-19 Surveillance Team |

| | | | |
|---|---|---|---|
| EPI_ISL_2514777 | | | |
| EPI_ISL_2514778 | Lighthouse Lab in Milton Keynes | Wellcome Sanger Institute for the COVID-19 Genomics UK (COG-UK) Consortium | The Lighthouse Lab in Milton Keynes and Alex Alderton, Roberto Amato, Jeffrey Barrett, Sonia Goncalves, Ewan Harrison, David K. Jackson, Ian Johnston, Dominic Kwiatkowski, Cordelia Langford, John Sillitoe on behalf of the Wellcome Sanger Institute COVID-19 Surveillance Team |
| EPI_ISL_2514779, EPI_ISL_2514780 | Lighthouse Lab in Glasgow | Wellcome Sanger Institute for the COVID-19 Genomics UK (COG-UK) Consortium | Harper VanSteenhouse, Yumi Kasai, David Gray, Carol Clugston, Anna Dominiczak and Alex Alderton, Roberto Amato, Jeffrey Barrett, Sonia Goncalves, Ewan Harrison, David K. Jackson, Ian Johnston, Dominic Kwiatkowski, Cordelia Langford, John Sillitoe on behalf of the Wellcome Sanger Institute COVID-19 Surveillance Team |
| EPI_ISL_2514781 | Lighthouse Lab in Milton Keynes | Wellcome Sanger Institute for the COVID-19 Genomics UK (COG-UK) Consortium | The Lighthouse Lab in Milton Keynes and Alex Alderton, Roberto Amato, Jeffrey Barrett, Sonia Goncalves, Ewan Harrison, David K. Jackson, Ian Johnston, Dominic Kwiatkowski, Cordelia Langford, John Sillitoe on behalf of the Wellcome Sanger Institute COVID-19 Surveillance Team |
| EPI_ISL_2514782 | Lighthouse Lab in Glasgow | Wellcome Sanger Institute for the COVID-19 Genomics UK (COG-UK) Consortium | Harper VanSteenhouse, Yumi Kasai, David Gray, Carol Clugston, Anna Dominiczak and Alex Alderton, Roberto Amato, Jeffrey Barrett, Sonia Goncalves, Ewan Harrison, David K. Jackson, Ian Johnston, Dominic Kwiatkowski, Cordelia Langford, John Sillitoe on behalf of the Wellcome Sanger Institute COVID-19 Surveillance Team |
| EPI_ISL_2514783, EPI_ISL_2514784, EPI_ISL_2514785, EPI_ISL_2514786 | Lighthouse Lab in Milton Keynes | Wellcome Sanger Institute for the COVID-19 Genomics UK (COG-UK) Consortium | The Lighthouse Lab in Milton Keynes and Alex Alderton, Roberto Amato, Jeffrey Barrett, Sonia Goncalves, Ewan Harrison, David K. Jackson, Ian Johnston, Dominic Kwiatkowski, Cordelia Langford, John Sillitoe on behalf of the Wellcome Sanger Institute COVID-19 Surveillance Team |
| EPI_ISL_2514787, EPI_ISL_2514788, EPI_ISL_2514789, EPI_ISL_2514790 | Lighthouse Lab in Glasgow | Wellcome Sanger Institute for the COVID-19 Genomics UK (COG-UK) Consortium | Harper VanSteenhouse, Yumi Kasai, David Gray, Carol Clugston, Anna Dominiczak and Alex Alderton, Roberto Amato, Jeffrey Barrett, Sonia Goncalves, Ewan Harrison, David K. Jackson, Ian Johnston, Dominic Kwiatkowski, Cordelia Langford, John Sillitoe on behalf of the Wellcome Sanger Institute COVID-19 Surveillance Team |
| EPI_ISL_2514792 | Lighthouse Lab in Milton Keynes | Wellcome Sanger Institute for the COVID-19 Genomics UK (COG-UK) Consortium | The Lighthouse Lab in Milton Keynes and Alex Alderton, Roberto Amato, Jeffrey Barrett, Sonia Goncalves, Ewan Harrison, David K. Jackson, Ian Johnston, Dominic Kwiatkowski, Cordelia Langford, John Sillitoe on behalf of the Wellcome Sanger Institute COVID-19 Surveillance Team |
| EPI_ISL_2514793, EPI_ISL_2514794 | Lighthouse Lab in Glasgow | Wellcome Sanger Institute for the COVID-19 Genomics UK (COG-UK) Consortium | Harper VanSteenhouse, Yumi Kasai, David Gray, Carol Clugston, Anna Dominiczak and Alex Alderton, Roberto Amato, Jeffrey Barrett, Sonia Goncalves, Ewan Harrison, David K. Jackson, Ian Johnston, Dominic Kwiatkowski, Cordelia Langford, John Sillitoe on behalf of the Wellcome Sanger Institute COVID-19 Surveillance Team |
| EPI_ISL_2514795, EPI_ISL_2514796, EPI_ISL_2514797, EPI_ISL_2514798 | Lighthouse Lab in Milton Keynes | Wellcome Sanger Institute for the COVID-19 Genomics UK (COG-UK) Consortium | The Lighthouse Lab in Milton Keynes and Alex Alderton, Roberto Amato, Jeffrey Barrett, Sonia Goncalves, Ewan Harrison, David K. Jackson, Ian Johnston, Dominic Kwiatkowski, Cordelia Langford, John Sillitoe on behalf of the Wellcome Sanger Institute COVID-19 Surveillance Team |
| EPI_ISL_2514799 | Lighthouse Lab in Glasgow | Wellcome Sanger Institute for the COVID-19 Genomics UK (COG-UK) Consortium | Harper VanSteenhouse, Yumi Kasai, David Gray, Carol Clugston, Anna Dominiczak and Alex Alderton, Roberto Amato, Jeffrey Barrett, Sonia Goncalves, Ewan Harrison, David K. Jackson, Ian Johnston, Dominic Kwiatkowski, Cordelia Langford, John Sillitoe on behalf of the Wellcome Sanger Institute COVID-19 Surveillance Team |
| EPI_ISL_2514800 | Lighthouse Lab in Milton Keynes | Wellcome Sanger Institute for the COVID-19 Genomics UK (COG-UK) Consortium | The Lighthouse Lab in Milton Keynes and Alex Alderton, Roberto Amato, Jeffrey Barrett, Sonia Goncalves, Ewan Harrison, David K. Jackson, Ian Johnston, Dominic Kwiatkowski, Cordelia Langford, John Sillitoe on behalf of the Wellcome Sanger Institute COVID-19 Surveillance Team |
| EPI_ISL_2514801, EPI_ISL_2514802 | Lighthouse Lab in Glasgow | Wellcome Sanger Institute for the COVID-19 Genomics UK (COG-UK) Consortium | Harper VanSteenhouse, Yumi Kasai, David Gray, Carol Clugston, Anna Dominiczak and Alex Alderton, Roberto Amato, Jeffrey Barrett, Sonia Goncalves, Ewan Harrison, David K. Jackson, Ian Johnston, Dominic Kwiatkowski, Cordelia Langford, John Sillitoe on behalf of the Wellcome Sanger Institute COVID-19 Surveillance Team |
| EPI_ISL_2514803, EPI_ISL_2514804, EPI_ISL_2514805 | Lighthouse Lab in Milton Keynes | Wellcome Sanger Institute for the COVID-19 Genomics UK (COG-UK) Consortium | The Lighthouse Lab in Milton Keynes and Alex Alderton, Roberto Amato, Jeffrey Barrett, Sonia Goncalves, Ewan Harrison, David K. Jackson, Ian Johnston, Dominic Kwiatkowski, Cordelia Langford, John Sillitoe on behalf of the Wellcome Sanger Institute COVID-19 Surveillance Team |
| EPI_ISL_2514806, EPI_ISL_2514807, EPI_ISL_2514808 | Lighthouse Lab in Glasgow | Wellcome Sanger Institute for the COVID-19 Genomics UK (COG-UK) Consortium | Harper VanSteenhouse, Yumi Kasai, David Gray, Carol Clugston, Anna Dominiczak and Alex Alderton, Roberto Amato, Jeffrey Barrett, Sonia Goncalves, Ewan Harrison, David K. Jackson, Ian Johnston, Dominic Kwiatkowski, Cordelia Langford, John Sillitoe on behalf of the Wellcome Sanger Institute COVID-19 Surveillance Team |
| EPI_ISL_2514809, EPI_ISL_2514810 | Lighthouse Lab in Milton Keynes | Wellcome Sanger Institute for the COVID-19 Genomics UK (COG-UK) Consortium | The Lighthouse Lab in Milton Keynes and Alex Alderton, Roberto Amato, Jeffrey Barrett, Sonia Goncalves, Ewan Harrison, David K. Jackson, Ian Johnston, Dominic Kwiatkowski, Cordelia Langford, John Sillitoe on behalf of the Wellcome Sanger Institute COVID-19 Surveillance Team |
| EPI_ISL_2514811 | Lighthouse Lab in Glasgow | Wellcome Sanger Institute for the COVID-19 Genomics UK (COG-UK) Consortium | Harper VanSteenhouse, Yumi Kasai, David Gray, Carol Clugston, Anna Dominiczak and Alex Alderton, Roberto Amato, Jeffrey Barrett, Sonia Goncalves, Ewan Harrison, David K. Jackson, Ian Johnston, Dominic Kwiatkowski, Cordelia Langford, John Sillitoe on behalf of the Wellcome Sanger Institute COVID-19 Surveillance Team |
| EPI_ISL_2514812, EPI_ISL_2514813 | Lighthouse Lab in Milton Keynes | Wellcome Sanger Institute for the COVID-19 Genomics UK (COG-UK) Consortium | The Lighthouse Lab in Milton Keynes and Alex Alderton, Roberto Amato, Jeffrey Barrett, Sonia Goncalves, Ewan Harrison, David K. Jackson, Ian Johnston, Dominic Kwiatkowski, Cordelia Langford, John Sillitoe on behalf of the Wellcome Sanger Institute COVID-19 Surveillance Team |
| EPI_ISL_2514814 | Lighthouse Lab in Glasgow | Wellcome Sanger Institute for the COVID-19 Genomics UK (COG-UK) Consortium | Harper VanSteenhouse, Yumi Kasai, David Gray, Carol Clugston, Anna Dominiczak and Alex Alderton, Roberto Amato, Jeffrey Barrett, Sonia Goncalves, Ewan Harrison, David K. Jackson, Ian Johnston, Dominic Kwiatkowski, Cordelia Langford, John Sillitoe on behalf of the Wellcome Sanger Institute COVID-19 Surveillance Team |
| EPI_ISL_2514815, EPI_ISL_2514816, EPI_ISL_2514817 | Lighthouse Lab in Milton Keynes | Wellcome Sanger Institute for the COVID-19 Genomics UK (COG-UK) Consortium | The Lighthouse Lab in Milton Keynes and Alex Alderton, Roberto Amato, Jeffrey Barrett, Sonia Goncalves, Ewan Harrison, David K. Jackson, Ian Johnston, Dominic Kwiatkowski, Cordelia Langford, John Sillitoe on behalf of the Wellcome Sanger Institute COVID-19 Surveillance Team |
| EPI_ISL_2514818 | Lighthouse Lab in Glasgow | Wellcome Sanger Institute for the COVID-19 Genomics UK (COG-UK) Consortium | Harper VanSteenhouse, Yumi Kasai, David Gray, Carol Clugston, Anna Dominiczak and Alex Alderton, Roberto Amato, Jeffrey Barrett, Sonia Goncalves, Ewan Harrison, David K. Jackson, Ian Johnston, Dominic Kwiatkowski, Cordelia Langford, John Sillitoe on behalf of the Wellcome Sanger Institute COVID-19 Surveillance Team |
| EPI_ISL_2514819 | Lighthouse Lab in Milton Keynes | Wellcome Sanger Institute for the COVID-19 Genomics UK (COG-UK) Consortium | The Lighthouse Lab in Milton Keynes and Alex Alderton, Roberto Amato, Jeffrey Barrett, Sonia Goncalves, Ewan Harrison, David K. Jackson, Ian Johnston, Dominic Kwiatkowski, Cordelia Langford, John Sillitoe on behalf of the Wellcome Sanger Institute COVID-19 Surveillance Team |
| EPI_ISL_2514820, EPI_ISL_2514821, EPI_ISL_2514822, EPI_ISL_2514823, EPI_ISL_2514824, EPI_ISL_2514825, EPI_ISL_2514827, EPI_ISL_2514828 | Lighthouse Lab in Glasgow | Wellcome Sanger Institute for the COVID-19 Genomics UK (COG-UK) Consortium | Harper VanSteenhouse, Yumi Kasai, David Gray, Carol Clugston, Anna Dominiczak and Alex Alderton, Roberto Amato, Jeffrey Barrett, Sonia Goncalves, Ewan Harrison, David K. Jackson, Ian Johnston, Dominic Kwiatkowski, Cordelia Langford, John Sillitoe on behalf of the Wellcome Sanger Institute COVID-19 Surveillance Team |
| EPI_ISL_2514829, EPI_ISL_2514830 | Lighthouse Lab in Milton Keynes | Wellcome Sanger Institute for the COVID-19 Genomics UK (COG-UK) Consortium | The Lighthouse Lab in Milton Keynes and Alex Alderton, Roberto Amato, Jeffrey Barrett, Sonia Goncalves, Ewan Harrison, David K. Jackson, Ian Johnston, Dominic Kwiatkowski, Cordelia Langford, John Sillitoe on behalf of the Wellcome Sanger Institute COVID-19 Surveillance Team |
| EPI_ISL_2514831, EPI_ISL_2514832 | Lighthouse Lab in Glasgow | Wellcome Sanger Institute for the COVID-19 Genomics UK (COG-UK) Consortium | Harper VanSteenhouse, Yumi Kasai, David Gray, Carol Clugston, Anna Dominiczak and Alex Alderton, Roberto Amato, Jeffrey Barrett, Sonia Goncalves, Ewan Harrison, David K. Jackson, Ian Johnston, Dominic Kwiatkowski, Cordelia Langford, John Sillitoe on behalf of the Wellcome Sanger Institute COVID-19 Surveillance Team |
| EPI_ISL_2514833, EPI_ISL_2514834 | Lighthouse Lab in Milton Keynes | Wellcome Sanger Institute for the COVID-19 Genomics UK (COG-UK) Consortium | The Lighthouse Lab in Milton Keynes and Alex Alderton, Roberto Amato, Jeffrey Barrett, Sonia Goncalves, Ewan Harrison, David K. Jackson, Ian Johnston, Dominic Kwiatkowski, Cordelia Langford, John Sillitoe on behalf of the Wellcome Sanger Institute COVID-19 Surveillance Team |
| EPI_ISL_2514835, EPI_ISL_2514836 | Lighthouse Lab in Glasgow | Wellcome Sanger Institute for the COVID-19 Genomics UK (COG-UK) Consortium | Harper VanSteenhouse, Yumi Kasai, David Gray, Carol Clugston, Anna Dominiczak and Alex Alderton, Roberto Amato, Jeffrey Barrett, Sonia Goncalves, Ewan Harrison, David K. Jackson, Ian Johnston, Dominic Kwiatkowski, Cordelia Langford, John Sillitoe on behalf of the Wellcome Sanger Institute COVID-19 Surveillance Team |
| EPI_ISL_2514837 | Lighthouse Lab in Milton Keynes | Wellcome Sanger Institute for the COVID-19 Genomics UK (COG-UK) Consortium | The Lighthouse Lab in Milton Keynes and Alex Alderton, Roberto Amato, Jeffrey Barrett, Sonia Goncalves, Ewan Harrison, David K. Jackson, Ian Johnston, Dominic Kwiatkowski, Cordelia Langford, John Sillitoe on behalf of the Wellcome Sanger Institute COVID-19 Surveillance Team |
| EPI_ISL_2514838, EPI_ISL_2514839, EPI_ISL_2514840, EPI_ISL_2514841 | Lighthouse Lab in Glasgow | Wellcome Sanger Institute for the COVID-19 Genomics UK (COG-UK) Consortium | Harper VanSteenhouse, Yumi Kasai, David Gray, Carol Clugston, Anna Dominiczak and Alex Alderton, Roberto Amato, Jeffrey Barrett, Sonia Goncalves, Ewan Harrison, David K. Jackson, Ian Johnston, Dominic Kwiatkowski, Cordelia Langford, John Sillitoe on behalf of the Wellcome Sanger Institute COVID-19 Surveillance Team |
| EPI_ISL_2514844, EPI_ISL_2514845 | Lighthouse Lab in Milton Keynes | Wellcome Sanger Institute for the COVID-19 Genomics UK (COG-UK) Consortium | The Lighthouse Lab in Milton Keynes and Alex Alderton, Roberto Amato, Jeffrey Barrett, Sonia Goncalves, Ewan Harrison, David K. Jackson, Ian Johnston, Dominic Kwiatkowski, Cordelia Langford, John Sillitoe on behalf of the Wellcome Sanger Institute COVID-19 Surveillance Team |
| EPI_ISL_2514846, EPI_ISL_2514847, EPI_ISL_2514848, EPI_ISL_2514849 | Lighthouse Lab in Glasgow | Wellcome Sanger Institute for the COVID-19 Genomics UK (COG-UK) Consortium | Harper VanSteenhouse, Yumi Kasai, David Gray, Carol Clugston, Anna Dominiczak and Alex Alderton, Roberto Amato, Jeffrey Barrett, Sonia Goncalves, Ewan Harrison, David K. Jackson, Ian Johnston, Dominic Kwiatkowski, Cordelia Langford, John Sillitoe on behalf of the Wellcome Sanger Institute COVID-19 Surveillance Team |
| EPI_ISL_2514850 | Lighthouse Lab in Milton Keynes | Wellcome Sanger Institute for the COVID-19 Genomics UK (COG-UK) Consortium | The Lighthouse Lab in Milton Keynes and Alex Alderton, Roberto Amato, Jeffrey Barrett, Sonia Goncalves, Ewan Harrison, David K. Jackson, Ian Johnston, Dominic Kwiatkowski, Cordelia Langford, John Sillitoe on behalf of the Wellcome Sanger Institute COVID-19 Surveillance Team |
| EPI_ISL_2514851, EPI_ISL_2514852, EPI_ISL_2514853 | Lighthouse Lab in Glasgow | Wellcome Sanger Institute for the COVID-19 Genomics UK (COG-UK) Consortium | Harper VanSteenhouse, Yumi Kasai, David Gray, Carol Clugston, Anna Dominiczak and Alex Alderton, Roberto Amato, Jeffrey Barrett, Sonia Goncalves, Ewan Harrison, David K. Jackson, Ian Johnston, Dominic Kwiatkowski, Cordelia Langford, John Sillitoe on behalf of the Wellcome Sanger Institute COVID-19 Surveillance Team |
| EPI_ISL_2514854 | Lighthouse Lab in Milton Keynes | Wellcome Sanger Institute for the COVID-19 Genomics UK (COG-UK) Consortium | The Lighthouse Lab in Milton Keynes and Alex Alderton, Roberto Amato, Jeffrey Barrett, Sonia Goncalves, Ewan Harrison, David K. Jackson, Ian Johnston, Dominic Kwiatkowski, Cordelia Langford, John Sillitoe on behalf of the Wellcome Sanger Institute COVID-19 Surveillance Team |
| EPI_ISL_2514855, EPI_ISL_2514856 | Lighthouse Lab in Glasgow | Wellcome Sanger Institute for the COVID-19 Genomics UK (COG-UK) Consortium | Harper VanSteenhouse, Yumi Kasai, David Gray, Carol Clugston, Anna Dominiczak and Alex Alderton, Roberto Amato, Jeffrey Barrett, Sonia Goncalves, Ewan Harrison, David K. Jackson, Ian Johnston, Dominic Kwiatkowski, Cordelia Langford, John Sillitoe on behalf of the Wellcome Sanger Institute COVID-19 Surveillance Team |
| EPI_ISL_2514857 | Lighthouse Lab in Milton Keynes | Wellcome Sanger Institute for the COVID-19 Genomics UK (COG-UK) Consortium | The Lighthouse Lab in Milton Keynes and Alex Alderton, Roberto Amato, Jeffrey Barrett, Sonia Goncalves, Ewan Harrison, David K. Jackson, Ian Johnston, Dominic Kwiatkowski, Cordelia Langford, John Sillitoe on behalf of the Wellcome Sanger Institute COVID-19 Surveillance Team |
| EPI_ISL_2514858, EPI_ISL_2514859 | Lighthouse Lab in Glasgow | Wellcome Sanger Institute for the COVID-19 Genomics UK (COG-UK) Consortium | Harper VanSteenhouse, Yumi Kasai, David Gray, Carol Clugston, Anna Dominiczak and Alex Alderton, Roberto Amato, Jeffrey Barrett, Sonia Goncalves, Ewan Harrison, David K. Jackson, Ian Johnston, Dominic Kwiatkowski, Cordelia Langford, John Sillitoe on behalf of the Wellcome Sanger Institute COVID-19 Surveillance Team |
| EPI_ISL_2514860, | Lighthouse Lab in Milton Keynes | Wellcome Sanger Institute for the COVID-19 Genomics UK (COG-UK) | The Lighthouse Lab in Milton Keynes and Alex Alderton, Roberto Amato, Jeffrey Barrett, Sonia Goncalves, Ewan Harrison, David K. Jackson, Ian Johnston, Dominic Kwiatkowski, Cordelia Langford, John Sillitoe on behalf of the Wellcome Sanger |

| | | | |
|---|---|---|---|
| EPI_ISL_2514861 | | Consortium | Institute COVID-19 Surveillance Team |
| EPI_ISL_2514862, EPI_ISL_2514863 | Lighthouse Lab in Glasgow | Wellcome Sanger Institute for the COVID-19 Genomics UK (COG-UK) Consortium | Harper VanSteenhouse, Yumi Kasai, David Gray, Carol Clugston, Anna Dominiczak and Alex Alderton, Roberto Amato, Jeffrey Barrett, Sonia Goncalves, Ewan Harrison, David K. Jackson, Ian Johnston, Dominic Kwiatkowski, Cordelia Langford, John Sillitoe on behalf of the Wellcome Sanger Institute COVID-19 Surveillance Team |
| EPI_ISL_2514864, EPI_ISL_2514866, EPI_ISL_2514867, EPI_ISL_2514868, EPI_ISL_2514869, EPI_ISL_2514871 | Lighthouse Lab in Milton Keynes | Wellcome Sanger Institute for the COVID-19 Genomics UK (COG-UK) Consortium | The Lighthouse Lab in Milton Keynes and Alex Alderton, Roberto Amato, Jeffrey Barrett, Sonia Goncalves, Ewan Harrison, David K. Jackson, Ian Johnston, Dominic Kwiatkowski, Cordelia Langford, John Sillitoe on behalf of the Wellcome Sanger Institute COVID-19 Surveillance Team |
| EPI_ISL_2514872 | Lighthouse Lab in Glasgow | Wellcome Sanger Institute for the COVID-19 Genomics UK (COG-UK) Consortium | Harper VanSteenhouse, Yumi Kasai, David Gray, Carol Clugston, Anna Dominiczak and Alex Alderton, Roberto Amato, Jeffrey Barrett, Sonia Goncalves, Ewan Harrison, David K. Jackson, Ian Johnston, Dominic Kwiatkowski, Cordelia Langford, John Sillitoe on behalf of the Wellcome Sanger Institute COVID-19 Surveillance Team |
| EPI_ISL_2514873, EPI_ISL_2514874 | Lighthouse Lab in Milton Keynes | Wellcome Sanger Institute for the COVID-19 Genomics UK (COG-UK) Consortium | The Lighthouse Lab in Milton Keynes and Alex Alderton, Roberto Amato, Jeffrey Barrett, Sonia Goncalves, Ewan Harrison, David K. Jackson, Ian Johnston, Dominic Kwiatkowski, Cordelia Langford, John Sillitoe on behalf of the Wellcome Sanger Institute COVID-19 Surveillance Team |
| EPI_ISL_2514875 | Lighthouse Lab in Glasgow | Wellcome Sanger Institute for the COVID-19 Genomics UK (COG-UK) Consortium | Harper VanSteenhouse, Yumi Kasai, David Gray, Carol Clugston, Anna Dominiczak and Alex Alderton, Roberto Amato, Jeffrey Barrett, Sonia Goncalves, Ewan Harrison, David K. Jackson, Ian Johnston, Dominic Kwiatkowski, Cordelia Langford, John Sillitoe on behalf of the Wellcome Sanger Institute COVID-19 Surveillance Team |
| EPI_ISL_2514876 | Lighthouse Lab in Milton Keynes | Wellcome Sanger Institute for the COVID-19 Genomics UK (COG-UK) Consortium | The Lighthouse Lab in Milton Keynes and Alex Alderton, Roberto Amato, Jeffrey Barrett, Sonia Goncalves, Ewan Harrison, David K. Jackson, Ian Johnston, Dominic Kwiatkowski, Cordelia Langford, John Sillitoe on behalf of the Wellcome Sanger Institute COVID-19 Surveillance Team |
| EPI_ISL_2514877 | Lighthouse Lab in Glasgow | Wellcome Sanger Institute for the COVID-19 Genomics UK (COG-UK) Consortium | Harper VanSteenhouse, Yumi Kasai, David Gray, Carol Clugston, Anna Dominiczak and Alex Alderton, Roberto Amato, Jeffrey Barrett, Sonia Goncalves, Ewan Harrison, David K. Jackson, Ian Johnston, Dominic Kwiatkowski, Cordelia Langford, John Sillitoe on behalf of the Wellcome Sanger Institute COVID-19 Surveillance Team |
| EPI_ISL_2514881, EPI_ISL_2514882, EPI_ISL_2514883 | Lighthouse Lab in Milton Keynes | Wellcome Sanger Institute for the COVID-19 Genomics UK (COG-UK) Consortium | The Lighthouse Lab in Milton Keynes and Alex Alderton, Roberto Amato, Jeffrey Barrett, Sonia Goncalves, Ewan Harrison, David K. Jackson, Ian Johnston, Dominic Kwiatkowski, Cordelia Langford, John Sillitoe on behalf of the Wellcome Sanger Institute COVID-19 Surveillance Team |
| EPI_ISL_2514884 | Lighthouse Lab in Glasgow | Wellcome Sanger Institute for the COVID-19 Genomics UK (COG-UK) Consortium | Harper VanSteenhouse, Yumi Kasai, David Gray, Carol Clugston, Anna Dominiczak and Alex Alderton, Roberto Amato, Jeffrey Barrett, Sonia Goncalves, Ewan Harrison, David K. Jackson, Ian Johnston, Dominic Kwiatkowski, Cordelia Langford, John Sillitoe on behalf of the Wellcome Sanger Institute COVID-19 Surveillance Team |
| EPI_ISL_2514885 | Lighthouse Lab in Milton Keynes | Wellcome Sanger Institute for the COVID-19 Genomics UK (COG-UK) Consortium | The Lighthouse Lab in Milton Keynes and Alex Alderton, Roberto Amato, Jeffrey Barrett, Sonia Goncalves, Ewan Harrison, David K. Jackson, Ian Johnston, Dominic Kwiatkowski, Cordelia Langford, John Sillitoe on behalf of the Wellcome Sanger Institute COVID-19 Surveillance Team |
| EPI_ISL_2514886, EPI_ISL_2514887, EPI_ISL_2514888, EPI_ISL_2514889, EPI_ISL_2514890 | Lighthouse Lab in Glasgow | Wellcome Sanger Institute for the COVID-19 Genomics UK (COG-UK) Consortium | Harper VanSteenhouse, Yumi Kasai, David Gray, Carol Clugston, Anna Dominiczak and Alex Alderton, Roberto Amato, Jeffrey Barrett, Sonia Goncalves, Ewan Harrison, David K. Jackson, Ian Johnston, Dominic Kwiatkowski, Cordelia Langford, John Sillitoe on behalf of the Wellcome Sanger Institute COVID-19 Surveillance Team |
| EPI_ISL_2514891, EPI_ISL_2514892, EPI_ISL_2514893 | Lighthouse Lab in Milton Keynes | Wellcome Sanger Institute for the COVID-19 Genomics UK (COG-UK) Consortium | The Lighthouse Lab in Milton Keynes and Alex Alderton, Roberto Amato, Jeffrey Barrett, Sonia Goncalves, Ewan Harrison, David K. Jackson, Ian Johnston, Dominic Kwiatkowski, Cordelia Langford, John Sillitoe on behalf of the Wellcome Sanger Institute COVID-19 Surveillance Team |
| EPI_ISL_2514894 | Lighthouse Lab in Glasgow | Wellcome Sanger Institute for the COVID-19 Genomics UK (COG-UK) Consortium | Harper VanSteenhouse, Yumi Kasai, David Gray, Carol Clugston, Anna Dominiczak and Alex Alderton, Roberto Amato, Jeffrey Barrett, Sonia Goncalves, Ewan Harrison, David K. Jackson, Ian Johnston, Dominic Kwiatkowski, Cordelia Langford, John Sillitoe on behalf of the Wellcome Sanger Institute COVID-19 Surveillance Team |
| EPI_ISL_2514895, EPI_ISL_2514896, EPI_ISL_2514897 | Lighthouse Lab in Milton Keynes | Wellcome Sanger Institute for the COVID-19 Genomics UK (COG-UK) Consortium | The Lighthouse Lab in Milton Keynes and Alex Alderton, Roberto Amato, Jeffrey Barrett, Sonia Goncalves, Ewan Harrison, David K. Jackson, Ian Johnston, Dominic Kwiatkowski, Cordelia Langford, John Sillitoe on behalf of the Wellcome Sanger Institute COVID-19 Surveillance Team |
| EPI_ISL_2514898, EPI_ISL_2514899 | Lighthouse Lab in Glasgow | Wellcome Sanger Institute for the COVID-19 Genomics UK (COG-UK) Consortium | Harper VanSteenhouse, Yumi Kasai, David Gray, Carol Clugston, Anna Dominiczak and Alex Alderton, Roberto Amato, Jeffrey Barrett, Sonia Goncalves, Ewan Harrison, David K. Jackson, Ian Johnston, Dominic Kwiatkowski, Cordelia Langford, John Sillitoe on behalf of the Wellcome Sanger Institute COVID-19 Surveillance Team |
| EPI_ISL_2514900, EPI_ISL_2514901 | Lighthouse Lab in Milton Keynes | Wellcome Sanger Institute for the COVID-19 Genomics UK (COG-UK) Consortium | The Lighthouse Lab in Milton Keynes and Alex Alderton, Roberto Amato, Jeffrey Barrett, Sonia Goncalves, Ewan Harrison, David K. Jackson, Ian Johnston, Dominic Kwiatkowski, Cordelia Langford, John Sillitoe on behalf of the Wellcome Sanger Institute COVID-19 Surveillance Team |
| EPI_ISL_2514903, EPI_ISL_2514904 | Lighthouse Lab in Glasgow | Wellcome Sanger Institute for the COVID-19 Genomics UK (COG-UK) Consortium | Harper VanSteenhouse, Yumi Kasai, David Gray, Carol Clugston, Anna Dominiczak and Alex Alderton, Roberto Amato, Jeffrey Barrett, Sonia Goncalves, Ewan Harrison, David K. Jackson, Ian Johnston, Dominic Kwiatkowski, Cordelia Langford, John Sillitoe on behalf of the Wellcome Sanger Institute COVID-19 Surveillance Team |
| EPI_ISL_2514905, EPI_ISL_2514906, EPI_ISL_2514907, EPI_ISL_2514908, EPI_ISL_2514909 | Lighthouse Lab in Milton Keynes | Wellcome Sanger Institute for the COVID-19 Genomics UK (COG-UK) Consortium | The Lighthouse Lab in Milton Keynes and Alex Alderton, Roberto Amato, Jeffrey Barrett, Sonia Goncalves, Ewan Harrison, David K. Jackson, Ian Johnston, Dominic Kwiatkowski, Cordelia Langford, John Sillitoe on behalf of the Wellcome Sanger Institute COVID-19 Surveillance Team |
| EPI_ISL_2514910, EPI_ISL_2514911, EPI_ISL_2514912, EPI_ISL_2514913 | Lighthouse Lab in Glasgow | Wellcome Sanger Institute for the COVID-19 Genomics UK (COG-UK) Consortium | Harper VanSteenhouse, Yumi Kasai, David Gray, Carol Clugston, Anna Dominiczak and Alex Alderton, Roberto Amato, Jeffrey Barrett, Sonia Goncalves, Ewan Harrison, David K. Jackson, Ian Johnston, Dominic Kwiatkowski, Cordelia Langford, John Sillitoe on behalf of the Wellcome Sanger Institute COVID-19 Surveillance Team |
| EPI_ISL_2514914, EPI_ISL_2514915, EPI_ISL_2514916 | Lighthouse Lab in Milton Keynes | Wellcome Sanger Institute for the COVID-19 Genomics UK (COG-UK) Consortium | The Lighthouse Lab in Milton Keynes and Alex Alderton, Roberto Amato, Jeffrey Barrett, Sonia Goncalves, Ewan Harrison, David K. Jackson, Ian Johnston, Dominic Kwiatkowski, Cordelia Langford, John Sillitoe on behalf of the Wellcome Sanger Institute COVID-19 Surveillance Team |
| EPI_ISL_2514917, EPI_ISL_2514918 | Lighthouse Lab in Glasgow | Wellcome Sanger Institute for the COVID-19 Genomics UK (COG-UK) Consortium | Harper VanSteenhouse, Yumi Kasai, David Gray, Carol Clugston, Anna Dominiczak and Alex Alderton, Roberto Amato, Jeffrey Barrett, Sonia Goncalves, Ewan Harrison, David K. Jackson, Ian Johnston, Dominic Kwiatkowski, Cordelia Langford, John Sillitoe on behalf of the Wellcome Sanger Institute COVID-19 Surveillance Team |
| EPI_ISL_2514919, EPI_ISL_2514922 | Lighthouse Lab in Milton Keynes | Wellcome Sanger Institute for the COVID-19 Genomics UK (COG-UK) Consortium | The Lighthouse Lab in Milton Keynes and Alex Alderton, Roberto Amato, Jeffrey Barrett, Sonia Goncalves, Ewan Harrison, David K. Jackson, Ian Johnston, Dominic Kwiatkowski, Cordelia Langford, John Sillitoe on behalf of the Wellcome Sanger Institute COVID-19 Surveillance Team |
| EPI_ISL_2514923, EPI_ISL_2514924 | Lighthouse Lab in Glasgow | Wellcome Sanger Institute for the COVID-19 Genomics UK (COG-UK) Consortium | Harper VanSteenhouse, Yumi Kasai, David Gray, Carol Clugston, Anna Dominiczak and Alex Alderton, Roberto Amato, Jeffrey Barrett, Sonia Goncalves, Ewan Harrison, David K. Jackson, Ian Johnston, Dominic Kwiatkowski, Cordelia Langford, John Sillitoe on behalf of the Wellcome Sanger Institute COVID-19 Surveillance Team |
| EPI_ISL_2514925, EPI_ISL_2514926 | Lighthouse Lab in Milton Keynes | Wellcome Sanger Institute for the COVID-19 Genomics UK (COG-UK) Consortium | The Lighthouse Lab in Milton Keynes and Alex Alderton, Roberto Amato, Jeffrey Barrett, Sonia Goncalves, Ewan Harrison, David K. Jackson, Ian Johnston, Dominic Kwiatkowski, Cordelia Langford, John Sillitoe on behalf of the Wellcome Sanger Institute COVID-19 Surveillance Team |
| EPI_ISL_2514927 | Lighthouse Lab in Glasgow | Wellcome Sanger Institute for the COVID-19 Genomics UK (COG-UK) Consortium | Harper VanSteenhouse, Yumi Kasai, David Gray, Carol Clugston, Anna Dominiczak and Alex Alderton, Roberto Amato, Jeffrey Barrett, Sonia Goncalves, Ewan Harrison, David K. Jackson, Ian Johnston, Dominic Kwiatkowski, Cordelia Langford, John Sillitoe on behalf of the Wellcome Sanger Institute COVID-19 Surveillance Team |
| EPI_ISL_2514928, EPI_ISL_2514929 | Lighthouse Lab in Alderley Park | Wellcome Sanger Institute for the COVID-19 Genomics UK (COG-UK) Consortium | Jacquelyn Wynn, Mairead Hyland, The Lighthouse Lab in Alderley Park and Alex Alderton, Roberto Amato, Jeffrey Barrett, Sonia Goncalves, Ewan Harrison, David K. Jackson, Ian Johnston, Dominic Kwiatkowski, Cordelia Langford, John Sillitoe on behalf of the Wellcome Sanger Institute COVID-19 Surveillance Team |
| EPI_ISL_2514930, EPI_ISL_2514931, EPI_ISL_2514932 | Lighthouse Laboratory Plymouth | Wellcome Sanger Institute for the COVID-19 Genomics UK (COG-UK) Consortium | Lighthouse Laboratory Plymouth and Alex Alderton, Roberto Amato, Jeffrey Barrett, Sonia Goncalves, Ewan Harrison, David K. Jackson, Ian Johnston, Dominic Kwiatkowski, Cordelia Langford, John Sillitoe on behalf of the Wellcome Sanger Institute COVID-19 Surveillance Team |
| EPI_ISL_2514933, EPI_ISL_2514934 | Lighthouse Lab in Glasgow | Wellcome Sanger Institute for the COVID-19 Genomics UK (COG-UK) Consortium | Harper VanSteenhouse, Yumi Kasai, David Gray, Carol Clugston, Anna Dominiczak and Alex Alderton, Roberto Amato, Jeffrey Barrett, Sonia Goncalves, Ewan Harrison, David K. Jackson, Ian Johnston, Dominic Kwiatkowski, Cordelia Langford, John Sillitoe on behalf of the Wellcome Sanger Institute COVID-19 Surveillance Team |
| EPI_ISL_2514935 | Lighthouse Lab in Alderley Park | Wellcome Sanger Institute for the COVID-19 Genomics UK (COG-UK) Consortium | Jacquelyn Wynn, Mairead Hyland, The Lighthouse Lab in Alderley Park and Alex Alderton, Roberto Amato, Jeffrey Barrett, Sonia Goncalves, Ewan Harrison, David K. Jackson, Ian Johnston, Dominic Kwiatkowski, Cordelia Langford, John Sillitoe on behalf of the Wellcome Sanger Institute COVID-19 Surveillance Team |
| EPI_ISL_2514936 | Lighthouse Laboratory Plymouth | Wellcome Sanger Institute for the COVID-19 Genomics UK (COG-UK) Consortium | Lighthouse Laboratory Plymouth and Alex Alderton, Roberto Amato, Jeffrey Barrett, Sonia Goncalves, Ewan Harrison, David K. Jackson, Ian Johnston, Dominic Kwiatkowski, Cordelia Langford, John Sillitoe on behalf of the Wellcome Sanger Institute COVID-19 Surveillance Team |
| EPI_ISL_2514937, EPI_ISL_2514938, EPI_ISL_2514939, EPI_ISL_2514941, EPI_ISL_2514942 | Lighthouse Lab in Alderley Park | Wellcome Sanger Institute for the COVID-19 Genomics UK (COG-UK) Consortium | Jacquelyn Wynn, Mairead Hyland, The Lighthouse Lab in Alderley Park and Alex Alderton, Roberto Amato, Jeffrey Barrett, Sonia Goncalves, Ewan Harrison, David K. Jackson, Ian Johnston, Dominic Kwiatkowski, Cordelia Langford, John Sillitoe on behalf of the Wellcome Sanger Institute COVID-19 Surveillance Team |
| EPI_ISL_2514943, EPI_ISL_2514944 | Lighthouse Laboratory Plymouth | Wellcome Sanger Institute for the COVID-19 Genomics UK (COG-UK) Consortium | Lighthouse Laboratory Plymouth and Alex Alderton, Roberto Amato, Jeffrey Barrett, Sonia Goncalves, Ewan Harrison, David K. Jackson, Ian Johnston, Dominic Kwiatkowski, Cordelia Langford, John Sillitoe on behalf of the Wellcome Sanger Institute COVID-19 Surveillance Team |
| EPI_ISL_2514945, EPI_ISL_2514946, EPI_ISL_2514947 | Lighthouse Lab in Alderley Park | Wellcome Sanger Institute for the COVID-19 Genomics UK (COG-UK) Consortium | Jacquelyn Wynn, Mairead Hyland, The Lighthouse Lab in Alderley Park and Alex Alderton, Roberto Amato, Jeffrey Barrett, Sonia Goncalves, Ewan Harrison, David K. Jackson, Ian Johnston, Dominic Kwiatkowski, Cordelia Langford, John Sillitoe on behalf of the Wellcome Sanger Institute COVID-19 Surveillance Team |
| EPI_ISL_2514948 | Lighthouse Lab in Glasgow | Wellcome Sanger Institute for the COVID-19 Genomics UK (COG-UK) Consortium | Harper VanSteenhouse, Yumi Kasai, David Gray, Carol Clugston, Anna Dominiczak and Alex Alderton, Roberto Amato, Jeffrey Barrett, Sonia Goncalves, Ewan Harrison, David K. Jackson, Ian Johnston, Dominic Kwiatkowski, Cordelia Langford, John Sillitoe on behalf of the Wellcome Sanger Institute COVID-19 Surveillance Team |
| EPI_ISL_2514949 | Lighthouse Laboratory Plymouth | Wellcome Sanger Institute for the COVID-19 Genomics UK (COG-UK) Consortium | Lighthouse Laboratory Plymouth and Alex Alderton, Roberto Amato, Jeffrey Barrett, Sonia Goncalves, Ewan Harrison, David K. Jackson, Ian Johnston, Dominic Kwiatkowski, Cordelia Langford, John Sillitoe on behalf of the Wellcome Sanger Institute COVID-19 Surveillance Team |
| EPI_ISL_2514950, EPI_ISL_2514951 | Lighthouse Lab in Alderley Park | Wellcome Sanger Institute for the COVID-19 Genomics UK (COG-UK) Consortium | Jacquelyn Wynn, Mairead Hyland, The Lighthouse Lab in Alderley Park and Alex Alderton, Roberto Amato, Jeffrey Barrett, Sonia Goncalves, Ewan Harrison, David K. Jackson, Ian Johnston, Dominic Kwiatkowski, Cordelia Langford, John Sillitoe on behalf of the Wellcome Sanger Institute COVID-19 Surveillance Team |

| | | | |
|---|---|---|---|
| EPI_ISL_2514952, EPI_ISL_2514953 | Lighthouse Laboratory Plymouth | Wellcome Sanger Institute for the COVID-19 Genomics UK (COG-UK) Consortium | Lighthouse Laboratory Plymouth and Alex Alderton, Roberto Amato, Jeffrey Barrett, Sonia Goncalves, Ewan Harrison, David K. Jackson, Ian Johnston, Dominic Kwiatkowski, Cordelia Langford, John Sillitoe on behalf of the Wellcome Sanger Institute COVID-19 Surveillance Team |
| EPI_ISL_2514954 | Lighthouse Lab in Alderley Park | Wellcome Sanger Institute for the COVID-19 Genomics UK (COG-UK) Consortium | Jacquelyn Wynn, Mairead Hyland, The Lighthouse Lab in Alderley Park and Alex Alderton, Roberto Amato, Jeffrey Barrett, Sonia Goncalves, Ewan Harrison, David K. Jackson, Ian Johnston, Dominic Kwiatkowski, Cordelia Langford, John Sillitoe on behalf of the Wellcome Sanger Institute COVID-19 Surveillance Team |
| EPI_ISL_2514955 | Lighthouse Lab in Glasgow | Wellcome Sanger Institute for the COVID-19 Genomics UK (COG-UK) Consortium | Harper VanSteenhouse, Yumi Kasai, David Gray, Carol Clugston, Anna Dominiczak and Alex Alderton, Roberto Amato, Jeffrey Barrett, Sonia Goncalves, Ewan Harrison, David K. Jackson, Ian Johnston, Dominic Kwiatkowski, Cordelia Langford, John Sillitoe on behalf of the Wellcome Sanger Institute COVID-19 Surveillance Team |
| EPI_ISL_2514956 | Lighthouse Lab in Alderley Park | Wellcome Sanger Institute for the COVID-19 Genomics UK (COG-UK) Consortium | Jacquelyn Wynn, Mairead Hyland, The Lighthouse Lab in Alderley Park and Alex Alderton, Roberto Amato, Jeffrey Barrett, Sonia Goncalves, Ewan Harrison, David K. Jackson, Ian Johnston, Dominic Kwiatkowski, Cordelia Langford, John Sillitoe on behalf of the Wellcome Sanger Institute COVID-19 Surveillance Team |
| EPI_ISL_2514957 | Lighthouse Laboratory Plymouth | Wellcome Sanger Institute for the COVID-19 Genomics UK (COG-UK) Consortium | Lighthouse Laboratory Plymouth and Alex Alderton, Roberto Amato, Jeffrey Barrett, Sonia Goncalves, Ewan Harrison, David K. Jackson, Ian Johnston, Dominic Kwiatkowski, Cordelia Langford, John Sillitoe on behalf of the Wellcome Sanger Institute COVID-19 Surveillance Team |
| EPI_ISL_2514958, EPI_ISL_2514959, EPI_ISL_2514960, EPI_ISL_2514961, EPI_ISL_2514962 | Lighthouse Lab in Alderley Park | Wellcome Sanger Institute for the COVID-19 Genomics UK (COG-UK) Consortium | Jacquelyn Wynn, Mairead Hyland, The Lighthouse Lab in Alderley Park and Alex Alderton, Roberto Amato, Jeffrey Barrett, Sonia Goncalves, Ewan Harrison, David K. Jackson, Ian Johnston, Dominic Kwiatkowski, Cordelia Langford, John Sillitoe on behalf of the Wellcome Sanger Institute COVID-19 Surveillance Team |
| EPI_ISL_2514963 | Lighthouse Laboratory Plymouth | Wellcome Sanger Institute for the COVID-19 Genomics UK (COG-UK) Consortium | Lighthouse Laboratory Plymouth and Alex Alderton, Roberto Amato, Jeffrey Barrett, Sonia Goncalves, Ewan Harrison, David K. Jackson, Ian Johnston, Dominic Kwiatkowski, Cordelia Langford, John Sillitoe on behalf of the Wellcome Sanger Institute COVID-19 Surveillance Team |
| EPI_ISL_2514964 | Lighthouse Lab in Glasgow | Wellcome Sanger Institute for the COVID-19 Genomics UK (COG-UK) Consortium | Harper VanSteenhouse, Yumi Kasai, David Gray, Carol Clugston, Anna Dominiczak and Alex Alderton, Roberto Amato, Jeffrey Barrett, Sonia Goncalves, Ewan Harrison, David K. Jackson, Ian Johnston, Dominic Kwiatkowski, Cordelia Langford, John Sillitoe on behalf of the Wellcome Sanger Institute COVID-19 Surveillance Team |
| EPI_ISL_2514965, EPI_ISL_2514966, EPI_ISL_2514967 | Lighthouse Lab in Alderley Park | Wellcome Sanger Institute for the COVID-19 Genomics UK (COG-UK) Consortium | Jacquelyn Wynn, Mairead Hyland, The Lighthouse Lab in Alderley Park and Alex Alderton, Roberto Amato, Jeffrey Barrett, Sonia Goncalves, Ewan Harrison, David K. Jackson, Ian Johnston, Dominic Kwiatkowski, Cordelia Langford, John Sillitoe on behalf of the Wellcome Sanger Institute COVID-19 Surveillance Team |
| EPI_ISL_2514968 | Lighthouse Lab in Glasgow | Wellcome Sanger Institute for the COVID-19 Genomics UK (COG-UK) Consortium | Harper VanSteenhouse, Yumi Kasai, David Gray, Carol Clugston, Anna Dominiczak and Alex Alderton, Roberto Amato, Jeffrey Barrett, Sonia Goncalves, Ewan Harrison, David K. Jackson, Ian Johnston, Dominic Kwiatkowski, Cordelia Langford, John Sillitoe on behalf of the Wellcome Sanger Institute COVID-19 Surveillance Team |
| EPI_ISL_2514969 | Lighthouse Lab in Alderley Park | Wellcome Sanger Institute for the COVID-19 Genomics UK (COG-UK) Consortium | Jacquelyn Wynn, Mairead Hyland, The Lighthouse Lab in Alderley Park and Alex Alderton, Roberto Amato, Jeffrey Barrett, Sonia Goncalves, Ewan Harrison, David K. Jackson, Ian Johnston, Dominic Kwiatkowski, Cordelia Langford, John Sillitoe on behalf of the Wellcome Sanger Institute COVID-19 Surveillance Team |
| EPI_ISL_2514970 | Lighthouse Laboratory Plymouth | Wellcome Sanger Institute for the COVID-19 Genomics UK (COG-UK) Consortium | Lighthouse Laboratory Plymouth and Alex Alderton, Roberto Amato, Jeffrey Barrett, Sonia Goncalves, Ewan Harrison, David K. Jackson, Ian Johnston, Dominic Kwiatkowski, Cordelia Langford, John Sillitoe on behalf of the Wellcome Sanger Institute COVID-19 Surveillance Team |
| EPI_ISL_2514971, EPI_ISL_2514972 | Lighthouse Lab in Alderley Park | Wellcome Sanger Institute for the COVID-19 Genomics UK (COG-UK) Consortium | Jacquelyn Wynn, Mairead Hyland, The Lighthouse Lab in Alderley Park and Alex Alderton, Roberto Amato, Jeffrey Barrett, Sonia Goncalves, Ewan Harrison, David K. Jackson, Ian Johnston, Dominic Kwiatkowski, Cordelia Langford, John Sillitoe on behalf of the Wellcome Sanger Institute COVID-19 Surveillance Team |
| EPI_ISL_2514973, EPI_ISL_2514975 | Lighthouse Lab in Glasgow | Wellcome Sanger Institute for the COVID-19 Genomics UK (COG-UK) Consortium | Harper VanSteenhouse, Yumi Kasai, David Gray, Carol Clugston, Anna Dominiczak and Alex Alderton, Roberto Amato, Jeffrey Barrett, Sonia Goncalves, Ewan Harrison, David K. Jackson, Ian Johnston, Dominic Kwiatkowski, Cordelia Langford, John Sillitoe on behalf of the Wellcome Sanger Institute COVID-19 Surveillance Team |
| EPI_ISL_2514976, EPI_ISL_2514977 | Lighthouse Lab in Alderley Park | Wellcome Sanger Institute for the COVID-19 Genomics UK (COG-UK) Consortium | Jacquelyn Wynn, Mairead Hyland, The Lighthouse Lab in Alderley Park and Alex Alderton, Roberto Amato, Jeffrey Barrett, Sonia Goncalves, Ewan Harrison, David K. Jackson, Ian Johnston, Dominic Kwiatkowski, Cordelia Langford, John Sillitoe on behalf of the Wellcome Sanger Institute COVID-19 Surveillance Team |
| EPI_ISL_2514978 | Lighthouse Lab in Glasgow | Wellcome Sanger Institute for the COVID-19 Genomics UK (COG-UK) Consortium | Harper VanSteenhouse, Yumi Kasai, David Gray, Carol Clugston, Anna Dominiczak and Alex Alderton, Roberto Amato, Jeffrey Barrett, Sonia Goncalves, Ewan Harrison, David K. Jackson, Ian Johnston, Dominic Kwiatkowski, Cordelia Langford, John Sillitoe on behalf of the Wellcome Sanger Institute COVID-19 Surveillance Team |
| EPI_ISL_2514979 | Lighthouse Lab in Alderley Park | Wellcome Sanger Institute for the COVID-19 Genomics UK (COG-UK) Consortium | Jacquelyn Wynn, Mairead Hyland, The Lighthouse Lab in Alderley Park and Alex Alderton, Roberto Amato, Jeffrey Barrett, Sonia Goncalves, Ewan Harrison, David K. Jackson, Ian Johnston, Dominic Kwiatkowski, Cordelia Langford, John Sillitoe on behalf of the Wellcome Sanger Institute COVID-19 Surveillance Team |
| EPI_ISL_2514980, EPI_ISL_2514981 | Lighthouse Laboratory Plymouth | Wellcome Sanger Institute for the COVID-19 Genomics UK (COG-UK) Consortium | Lighthouse Laboratory Plymouth and Alex Alderton, Roberto Amato, Jeffrey Barrett, Sonia Goncalves, Ewan Harrison, David K. Jackson, Ian Johnston, Dominic Kwiatkowski, Cordelia Langford, John Sillitoe on behalf of the Wellcome Sanger Institute COVID-19 Surveillance Team |
| EPI_ISL_2514982, EPI_ISL_2514983 | Lighthouse Lab in Glasgow | Wellcome Sanger Institute for the COVID-19 Genomics UK (COG-UK) Consortium | Harper VanSteenhouse, Yumi Kasai, David Gray, Carol Clugston, Anna Dominiczak and Alex Alderton, Roberto Amato, Jeffrey Barrett, Sonia Goncalves, Ewan Harrison, David K. Jackson, Ian Johnston, Dominic Kwiatkowski, Cordelia Langford, John Sillitoe on behalf of the Wellcome Sanger Institute COVID-19 Surveillance Team |
| EPI_ISL_2514984 | Lighthouse Lab in Alderley Park | Wellcome Sanger Institute for the COVID-19 Genomics UK (COG-UK) Consortium | Jacquelyn Wynn, Mairead Hyland, The Lighthouse Lab in Alderley Park and Alex Alderton, Roberto Amato, Jeffrey Barrett, Sonia Goncalves, Ewan Harrison, David K. Jackson, Ian Johnston, Dominic Kwiatkowski, Cordelia Langford, John Sillitoe on behalf of the Wellcome Sanger Institute COVID-19 Surveillance Team |
| EPI_ISL_2514985 | Lighthouse Lab in Glasgow | Wellcome Sanger Institute for the COVID-19 Genomics UK (COG-UK) Consortium | Harper VanSteenhouse, Yumi Kasai, David Gray, Carol Clugston, Anna Dominiczak and Alex Alderton, Roberto Amato, Jeffrey Barrett, Sonia Goncalves, Ewan Harrison, David K. Jackson, Ian Johnston, Dominic Kwiatkowski, Cordelia Langford, John Sillitoe on behalf of the Wellcome Sanger Institute COVID-19 Surveillance Team |
| EPI_ISL_2514986 | Lighthouse Lab in Alderley Park | Wellcome Sanger Institute for the COVID-19 Genomics UK (COG-UK) Consortium | Jacquelyn Wynn, Mairead Hyland, The Lighthouse Lab in Alderley Park and Alex Alderton, Roberto Amato, Jeffrey Barrett, Sonia Goncalves, Ewan Harrison, David K. Jackson, Ian Johnston, Dominic Kwiatkowski, Cordelia Langford, John Sillitoe on behalf of the Wellcome Sanger Institute COVID-19 Surveillance Team |
| EPI_ISL_2514987 | Lighthouse Laboratory Plymouth | Wellcome Sanger Institute for the COVID-19 Genomics UK (COG-UK) Consortium | Lighthouse Laboratory Plymouth and Alex Alderton, Roberto Amato, Jeffrey Barrett, Sonia Goncalves, Ewan Harrison, David K. Jackson, Ian Johnston, Dominic Kwiatkowski, Cordelia Langford, John Sillitoe on behalf of the Wellcome Sanger Institute COVID-19 Surveillance Team |
| EPI_ISL_2514988, EPI_ISL_2514989 | Lighthouse Lab in Glasgow | Wellcome Sanger Institute for the COVID-19 Genomics UK (COG-UK) Consortium | Harper VanSteenhouse, Yumi Kasai, David Gray, Carol Clugston, Anna Dominiczak and Alex Alderton, Roberto Amato, Jeffrey Barrett, Sonia Goncalves, Ewan Harrison, David K. Jackson, Ian Johnston, Dominic Kwiatkowski, Cordelia Langford, John Sillitoe on behalf of the Wellcome Sanger Institute COVID-19 Surveillance Team |
| EPI_ISL_2514990 | Lighthouse Laboratory Plymouth | Wellcome Sanger Institute for the COVID-19 Genomics UK (COG-UK) Consortium | Lighthouse Laboratory Plymouth and Alex Alderton, Roberto Amato, Jeffrey Barrett, Sonia Goncalves, Ewan Harrison, David K. Jackson, Ian Johnston, Dominic Kwiatkowski, Cordelia Langford, John Sillitoe on behalf of the Wellcome Sanger Institute COVID-19 Surveillance Team |
| EPI_ISL_2514992, EPI_ISL_2514993 | Lighthouse Lab in Alderley Park | Wellcome Sanger Institute for the COVID-19 Genomics UK (COG-UK) Consortium | Jacquelyn Wynn, Mairead Hyland, The Lighthouse Lab in Alderley Park and Alex Alderton, Roberto Amato, Jeffrey Barrett, Sonia Goncalves, Ewan Harrison, David K. Jackson, Ian Johnston, Dominic Kwiatkowski, Cordelia Langford, John Sillitoe on behalf of the Wellcome Sanger Institute COVID-19 Surveillance Team |
| EPI_ISL_2514994 | Lighthouse Laboratory Plymouth | Wellcome Sanger Institute for the COVID-19 Genomics UK (COG-UK) Consortium | Lighthouse Laboratory Plymouth and Alex Alderton, Roberto Amato, Jeffrey Barrett, Sonia Goncalves, Ewan Harrison, David K. Jackson, Ian Johnston, Dominic Kwiatkowski, Cordelia Langford, John Sillitoe on behalf of the Wellcome Sanger Institute COVID-19 Surveillance Team |
| EPI_ISL_2514995, EPI_ISL_2514996, EPI_ISL_2514997 | Lighthouse Lab in Alderley Park | Wellcome Sanger Institute for the COVID-19 Genomics UK (COG-UK) Consortium | Jacquelyn Wynn, Mairead Hyland, The Lighthouse Lab in Alderley Park and Alex Alderton, Roberto Amato, Jeffrey Barrett, Sonia Goncalves, Ewan Harrison, David K. Jackson, Ian Johnston, Dominic Kwiatkowski, Cordelia Langford, John Sillitoe on behalf of the Wellcome Sanger Institute COVID-19 Surveillance Team |
| EPI_ISL_2514998 | Lighthouse Lab in Glasgow | Wellcome Sanger Institute for the COVID-19 Genomics UK (COG-UK) Consortium | Harper VanSteenhouse, Yumi Kasai, David Gray, Carol Clugston, Anna Dominiczak and Alex Alderton, Roberto Amato, Jeffrey Barrett, Sonia Goncalves, Ewan Harrison, David K. Jackson, Ian Johnston, Dominic Kwiatkowski, Cordelia Langford, John Sillitoe on behalf of the Wellcome Sanger Institute COVID-19 Surveillance Team |
| EPI_ISL_2514999 | Lighthouse Laboratory Plymouth | Wellcome Sanger Institute for the COVID-19 Genomics UK (COG-UK) Consortium | Lighthouse Laboratory Plymouth and Alex Alderton, Roberto Amato, Jeffrey Barrett, Sonia Goncalves, Ewan Harrison, David K. Jackson, Ian Johnston, Dominic Kwiatkowski, Cordelia Langford, John Sillitoe on behalf of the Wellcome Sanger Institute COVID-19 Surveillance Team |
| EPI_ISL_2515000, EPI_ISL_2515002, EPI_ISL_2515003 | Lighthouse Lab in Alderley Park | Wellcome Sanger Institute for the COVID-19 Genomics UK (COG-UK) Consortium | Jacquelyn Wynn, Mairead Hyland, The Lighthouse Lab in Alderley Park and Alex Alderton, Roberto Amato, Jeffrey Barrett, Sonia Goncalves, Ewan Harrison, David K. Jackson, Ian Johnston, Dominic Kwiatkowski, Cordelia Langford, John Sillitoe on behalf of the Wellcome Sanger Institute COVID-19 Surveillance Team |
| EPI_ISL_2515004 | Lighthouse Lab in Glasgow | Wellcome Sanger Institute for the COVID-19 Genomics UK (COG-UK) Consortium | Harper VanSteenhouse, Yumi Kasai, David Gray, Carol Clugston, Anna Dominiczak and Alex Alderton, Roberto Amato, Jeffrey Barrett, Sonia Goncalves, Ewan Harrison, David K. Jackson, Ian Johnston, Dominic Kwiatkowski, Cordelia Langford, John Sillitoe on behalf of the Wellcome Sanger Institute COVID-19 Surveillance Team |
| EPI_ISL_2515005 | Lighthouse Lab in Alderley Park | Wellcome Sanger Institute for the COVID-19 Genomics UK (COG-UK) Consortium | Jacquelyn Wynn, Mairead Hyland, The Lighthouse Lab in Alderley Park and Alex Alderton, Roberto Amato, Jeffrey Barrett, Sonia Goncalves, Ewan Harrison, David K. Jackson, Ian Johnston, Dominic Kwiatkowski, Cordelia Langford, John Sillitoe on behalf of the Wellcome Sanger Institute COVID-19 Surveillance Team |
| EPI_ISL_2515006, EPI_ISL_2515007, EPI_ISL_2515008 | Lighthouse Laboratory Plymouth | Wellcome Sanger Institute for the COVID-19 Genomics UK (COG-UK) Consortium | Lighthouse Laboratory Plymouth and Alex Alderton, Roberto Amato, Jeffrey Barrett, Sonia Goncalves, Ewan Harrison, David K. Jackson, Ian Johnston, Dominic Kwiatkowski, Cordelia Langford, John Sillitoe on behalf of the Wellcome Sanger Institute COVID-19 Surveillance Team |
| EPI_ISL_2515009 | Lighthouse Lab in Glasgow | Wellcome Sanger Institute for the COVID-19 Genomics UK (COG-UK) Consortium | Harper VanSteenhouse, Yumi Kasai, David Gray, Carol Clugston, Anna Dominiczak and Alex Alderton, Roberto Amato, Jeffrey Barrett, Sonia Goncalves, Ewan Harrison, David K. Jackson, Ian Johnston, Dominic Kwiatkowski, Cordelia Langford, John Sillitoe on behalf of the Wellcome Sanger Institute COVID-19 Surveillance Team |
| EPI_ISL_2515010, EPI_ISL_2515011 | Lighthouse Laboratory Plymouth | Wellcome Sanger Institute for the COVID-19 Genomics UK (COG-UK) Consortium | Lighthouse Laboratory Plymouth and Alex Alderton, Roberto Amato, Jeffrey Barrett, Sonia Goncalves, Ewan Harrison, David K. Jackson, Ian Johnston, Dominic Kwiatkowski, Cordelia Langford, John Sillitoe on behalf of the Wellcome Sanger Institute COVID-19 Surveillance Team |
| EPI_ISL_2515012, EPI_ISL_2515013, EPI_ISL_2515014, EPI_ISL_2515015, EPI_ISL_2515016 | Lighthouse Lab in Alderley Park | Wellcome Sanger Institute for the COVID-19 Genomics UK (COG-UK) Consortium | Jacquelyn Wynn, Mairead Hyland, The Lighthouse Lab in Alderley Park and Alex Alderton, Roberto Amato, Jeffrey Barrett, Sonia Goncalves, Ewan Harrison, David K. Jackson, Ian Johnston, Dominic Kwiatkowski, Cordelia Langford, John Sillitoe on behalf of the Wellcome Sanger Institute COVID-19 Surveillance Team |
| EPI_ISL_2515017, EPI_ISL_2515018 | Lighthouse Laboratory Plymouth | Wellcome Sanger Institute for the COVID-19 Genomics UK (COG-UK) Consortium | Lighthouse Laboratory Plymouth and Alex Alderton, Roberto Amato, Jeffrey Barrett, Sonia Goncalves, Ewan Harrison, David K. Jackson, Ian Johnston, Dominic Kwiatkowski, Cordelia Langford, John Sillitoe on behalf of the Wellcome Sanger Institute COVID-19 Surveillance Team |
| EPI_ISL_2515019 | Lighthouse Lab in Alderley Park | Wellcome Sanger Institute for the COVID-19 Genomics UK (COG-UK) Consortium | Jacquelyn Wynn, Mairead Hyland, The Lighthouse Lab in Alderley Park and Alex Alderton, Roberto Amato, Jeffrey Barrett, Sonia Goncalves, Ewan Harrison, David K. Jackson, Ian Johnston, Dominic Kwiatkowski, Cordelia Langford, John Sillitoe on behalf of the Wellcome Sanger Institute COVID-19 Surveillance Team |
| EPI_ISL_2515020 | Lighthouse Lab in Glasgow | Wellcome Sanger Institute for the COVID-19 Genomics UK (COG-UK) Consortium | Harper VanSteenhouse, Yumi Kasai, David Gray, Carol Clugston, Anna Dominiczak and Alex Alderton, Roberto Amato, Jeffrey Barrett, Sonia Goncalves, Ewan Harrison, David K. Jackson, Ian Johnston, Dominic Kwiatkowski, Cordelia Langford, John Sillitoe on behalf of the Wellcome Sanger Institute COVID-19 Surveillance Team |
| EPI_ISL_2515021, EPI_ISL_2515022 | Lighthouse Laboratory Plymouth | Wellcome Sanger Institute for the COVID-19 Genomics UK (COG-UK) Consortium | Lighthouse Laboratory Plymouth and Alex Alderton, Roberto Amato, Jeffrey Barrett, Sonia Goncalves, Ewan Harrison, David K. Jackson, Ian Johnston, Dominic Kwiatkowski, Cordelia Langford, John Sillitoe on behalf of the Wellcome Sanger Institute COVID-19 Surveillance Team |

| | | | |
|---|---|---|---|
| EPI_ISL_2515023, EPI_ISL_2515024, EPI_ISL_2515025, EPI_ISL_2515026, EPI_ISL_2515027 | Lighthouse Lab in Alderley Park | Wellcome Sanger Institute for the COVID-19 Genomics UK (COG-UK) Consortium | Jacquelyn Wynn, Mairead Hyland, The Lighthouse Lab in Alderley Park and Alex Alderton, Roberto Amato, Jeffrey Barrett, Sonia Goncalves, Ewan Harrison, David K. Jackson, Ian Johnston, Dominic Kwiatkowski, Cordelia Langford, John Sillitoe on behalf of the Wellcome Sanger Institute COVID-19 Surveillance Team |
| EPI_ISL_2515028 | Lighthouse Laboratory Plymouth | Wellcome Sanger Institute for the COVID-19 Genomics UK (COG-UK) Consortium | Lighthouse Laboratory Plymouth and Alex Alderton, Roberto Amato, Jeffrey Barrett, Sonia Goncalves, Ewan Harrison, David K. Jackson, Ian Johnston, Dominic Kwiatkowski, Cordelia Langford, John Sillitoe on behalf of the Wellcome Sanger Institute COVID-19 Surveillance Team |
| EPI_ISL_2515029, EPI_ISL_2515030, EPI_ISL_2515031 | Lighthouse Lab in Alderley Park | Wellcome Sanger Institute for the COVID-19 Genomics UK (COG-UK) Consortium | Jacquelyn Wynn, Mairead Hyland, The Lighthouse Lab in Alderley Park and Alex Alderton, Roberto Amato, Jeffrey Barrett, Sonia Goncalves, Ewan Harrison, David K. Jackson, Ian Johnston, Dominic Kwiatkowski, Cordelia Langford, John Sillitoe on behalf of the Wellcome Sanger Institute COVID-19 Surveillance Team |
| EPI_ISL_2515032 | Lighthouse Laboratory Plymouth | Wellcome Sanger Institute for the COVID-19 Genomics UK (COG-UK) Consortium | Lighthouse Laboratory Plymouth and Alex Alderton, Roberto Amato, Jeffrey Barrett, Sonia Goncalves, Ewan Harrison, David K. Jackson, Ian Johnston, Dominic Kwiatkowski, Cordelia Langford, John Sillitoe on behalf of the Wellcome Sanger Institute COVID-19 Surveillance Team |
| EPI_ISL_2515033, EPI_ISL_2515034, EPI_ISL_2515035 | Lighthouse Lab in Alderley Park | Wellcome Sanger Institute for the COVID-19 Genomics UK (COG-UK) Consortium | Jacquelyn Wynn, Mairead Hyland, The Lighthouse Lab in Alderley Park and Alex Alderton, Roberto Amato, Jeffrey Barrett, Sonia Goncalves, Ewan Harrison, David K. Jackson, Ian Johnston, Dominic Kwiatkowski, Cordelia Langford, John Sillitoe on behalf of the Wellcome Sanger Institute COVID-19 Surveillance Team |
| EPI_ISL_2515036 | Lighthouse Laboratory Plymouth | Wellcome Sanger Institute for the COVID-19 Genomics UK (COG-UK) Consortium | Lighthouse Laboratory Plymouth and Alex Alderton, Roberto Amato, Jeffrey Barrett, Sonia Goncalves, Ewan Harrison, David K. Jackson, Ian Johnston, Dominic Kwiatkowski, Cordelia Langford, John Sillitoe on behalf of the Wellcome Sanger Institute COVID-19 Surveillance Team |
| EPI_ISL_2515037, EPI_ISL_2515040 | Lighthouse Lab in Alderley Park | Wellcome Sanger Institute for the COVID-19 Genomics UK (COG-UK) Consortium | Jacquelyn Wynn, Mairead Hyland, The Lighthouse Lab in Alderley Park and Alex Alderton, Roberto Amato, Jeffrey Barrett, Sonia Goncalves, Ewan Harrison, David K. Jackson, Ian Johnston, Dominic Kwiatkowski, Cordelia Langford, John Sillitoe on behalf of the Wellcome Sanger Institute COVID-19 Surveillance Team |
| EPI_ISL_2515042 | Lighthouse Lab in Glasgow | Wellcome Sanger Institute for the COVID-19 Genomics UK (COG-UK) Consortium | Harper VanSteenhouse, Yumi Kasai, David Gray, Carol Clugston, Anna Dominiczak and Alex Alderton, Roberto Amato, Jeffrey Barrett, Sonia Goncalves, Ewan Harrison, David K. Jackson, Ian Johnston, Dominic Kwiatkowski, Cordelia Langford, John Sillitoe on behalf of the Wellcome Sanger Institute COVID-19 Surveillance Team |
| EPI_ISL_2515043 | Lighthouse Laboratory Plymouth | Wellcome Sanger Institute for the COVID-19 Genomics UK (COG-UK) Consortium | Lighthouse Laboratory Plymouth and Alex Alderton, Roberto Amato, Jeffrey Barrett, Sonia Goncalves, Ewan Harrison, David K. Jackson, Ian Johnston, Dominic Kwiatkowski, Cordelia Langford, John Sillitoe on behalf of the Wellcome Sanger Institute COVID-19 Surveillance Team |
| EPI_ISL_2515044 | Lighthouse Lab in Glasgow | Wellcome Sanger Institute for the COVID-19 Genomics UK (COG-UK) Consortium | Harper VanSteenhouse, Yumi Kasai, David Gray, Carol Clugston, Anna Dominiczak and Alex Alderton, Roberto Amato, Jeffrey Barrett, Sonia Goncalves, Ewan Harrison, David K. Jackson, Ian Johnston, Dominic Kwiatkowski, Cordelia Langford, John Sillitoe on behalf of the Wellcome Sanger Institute COVID-19 Surveillance Team |
| EPI_ISL_2515045 | Lighthouse Lab in Alderley Park | Wellcome Sanger Institute for the COVID-19 Genomics UK (COG-UK) Consortium | Jacquelyn Wynn, Mairead Hyland, The Lighthouse Lab in Alderley Park and Alex Alderton, Roberto Amato, Jeffrey Barrett, Sonia Goncalves, Ewan Harrison, David K. Jackson, Ian Johnston, Dominic Kwiatkowski, Cordelia Langford, John Sillitoe on behalf of the Wellcome Sanger Institute COVID-19 Surveillance Team |
| EPI_ISL_2515046 | Lighthouse Lab in Glasgow | Wellcome Sanger Institute for the COVID-19 Genomics UK (COG-UK) Consortium | Harper VanSteenhouse, Yumi Kasai, David Gray, Carol Clugston, Anna Dominiczak and Alex Alderton, Roberto Amato, Jeffrey Barrett, Sonia Goncalves, Ewan Harrison, David K. Jackson, Ian Johnston, Dominic Kwiatkowski, Cordelia Langford, John Sillitoe on behalf of the Wellcome Sanger Institute COVID-19 Surveillance Team |
| EPI_ISL_2515047 | Lighthouse Lab in Alderley Park | Wellcome Sanger Institute for the COVID-19 Genomics UK (COG-UK) Consortium | Jacquelyn Wynn, Mairead Hyland, The Lighthouse Lab in Alderley Park and Alex Alderton, Roberto Amato, Jeffrey Barrett, Sonia Goncalves, Ewan Harrison, David K. Jackson, Ian Johnston, Dominic Kwiatkowski, Cordelia Langford, John Sillitoe on behalf of the Wellcome Sanger Institute COVID-19 Surveillance Team |
| EPI_ISL_2515049 | Lighthouse Lab in Glasgow | Wellcome Sanger Institute for the COVID-19 Genomics UK (COG-UK) Consortium | Harper VanSteenhouse, Yumi Kasai, David Gray, Carol Clugston, Anna Dominiczak and Alex Alderton, Roberto Amato, Jeffrey Barrett, Sonia Goncalves, Ewan Harrison, David K. Jackson, Ian Johnston, Dominic Kwiatkowski, Cordelia Langford, John Sillitoe on behalf of the Wellcome Sanger Institute COVID-19 Surveillance Team |
| EPI_ISL_2515050 | Lighthouse Laboratory Plymouth | Wellcome Sanger Institute for the COVID-19 Genomics UK (COG-UK) Consortium | Lighthouse Laboratory Plymouth and Alex Alderton, Roberto Amato, Jeffrey Barrett, Sonia Goncalves, Ewan Harrison, David K. Jackson, Ian Johnston, Dominic Kwiatkowski, Cordelia Langford, John Sillitoe on behalf of the Wellcome Sanger Institute COVID-19 Surveillance Team |
| EPI_ISL_2515051 | Lighthouse Lab in Alderley Park | Wellcome Sanger Institute for the COVID-19 Genomics UK (COG-UK) Consortium | Jacquelyn Wynn, Mairead Hyland, The Lighthouse Lab in Alderley Park and Alex Alderton, Roberto Amato, Jeffrey Barrett, Sonia Goncalves, Ewan Harrison, David K. Jackson, Ian Johnston, Dominic Kwiatkowski, Cordelia Langford, John Sillitoe on behalf of the Wellcome Sanger Institute COVID-19 Surveillance Team |
| EPI_ISL_2515052 | Lighthouse Lab in Glasgow | Wellcome Sanger Institute for the COVID-19 Genomics UK (COG-UK) Consortium | Harper VanSteenhouse, Yumi Kasai, David Gray, Carol Clugston, Anna Dominiczak and Alex Alderton, Roberto Amato, Jeffrey Barrett, Sonia Goncalves, Ewan Harrison, David K. Jackson, Ian Johnston, Dominic Kwiatkowski, Cordelia Langford, John Sillitoe on behalf of the Wellcome Sanger Institute COVID-19 Surveillance Team |
| EPI_ISL_2515053, EPI_ISL_2515055, EPI_ISL_2515056 | Lighthouse Laboratory Plymouth | Wellcome Sanger Institute for the COVID-19 Genomics UK (COG-UK) Consortium | Lighthouse Laboratory Plymouth and Alex Alderton, Roberto Amato, Jeffrey Barrett, Sonia Goncalves, Ewan Harrison, David K. Jackson, Ian Johnston, Dominic Kwiatkowski, Cordelia Langford, John Sillitoe on behalf of the Wellcome Sanger Institute COVID-19 Surveillance Team |
| EPI_ISL_2515057, EPI_ISL_2515058 | Lighthouse Lab in Glasgow | Wellcome Sanger Institute for the COVID-19 Genomics UK (COG-UK) Consortium | Harper VanSteenhouse, Yumi Kasai, David Gray, Carol Clugston, Anna Dominiczak and Alex Alderton, Roberto Amato, Jeffrey Barrett, Sonia Goncalves, Ewan Harrison, David K. Jackson, Ian Johnston, Dominic Kwiatkowski, Cordelia Langford, John Sillitoe on behalf of the Wellcome Sanger Institute COVID-19 Surveillance Team |
| EPI_ISL_2515059, EPI_ISL_2515060, EPI_ISL_2515062, EPI_ISL_2515063 | Lighthouse Lab in Alderley Park | Wellcome Sanger Institute for the COVID-19 Genomics UK (COG-UK) Consortium | Jacquelyn Wynn, Mairead Hyland, The Lighthouse Lab in Alderley Park and Alex Alderton, Roberto Amato, Jeffrey Barrett, Sonia Goncalves, Ewan Harrison, David K. Jackson, Ian Johnston, Dominic Kwiatkowski, Cordelia Langford, John Sillitoe on behalf of the Wellcome Sanger Institute COVID-19 Surveillance Team |
| EPI_ISL_2515064 | Lighthouse Laboratory Plymouth | Wellcome Sanger Institute for the COVID-19 Genomics UK (COG-UK) Consortium | Lighthouse Laboratory Plymouth and Alex Alderton, Roberto Amato, Jeffrey Barrett, Sonia Goncalves, Ewan Harrison, David K. Jackson, Ian Johnston, Dominic Kwiatkowski, Cordelia Langford, John Sillitoe on behalf of the Wellcome Sanger Institute COVID-19 Surveillance Team |
| EPI_ISL_2515065 | Lighthouse Lab in Glasgow | Wellcome Sanger Institute for the COVID-19 Genomics UK (COG-UK) Consortium | Harper VanSteenhouse, Yumi Kasai, David Gray, Carol Clugston, Anna Dominiczak and Alex Alderton, Roberto Amato, Jeffrey Barrett, Sonia Goncalves, Ewan Harrison, David K. Jackson, Ian Johnston, Dominic Kwiatkowski, Cordelia Langford, John Sillitoe on behalf of the Wellcome Sanger Institute COVID-19 Surveillance Team |
| EPI_ISL_2515066 | Lighthouse Lab in Alderley Park | Wellcome Sanger Institute for the COVID-19 Genomics UK (COG-UK) Consortium | Jacquelyn Wynn, Mairead Hyland, The Lighthouse Lab in Alderley Park and Alex Alderton, Roberto Amato, Jeffrey Barrett, Sonia Goncalves, Ewan Harrison, David K. Jackson, Ian Johnston, Dominic Kwiatkowski, Cordelia Langford, John Sillitoe on behalf of the Wellcome Sanger Institute COVID-19 Surveillance Team |
| EPI_ISL_2515068, EPI_ISL_2515069, EPI_ISL_2515070 | Lighthouse Laboratory Plymouth | Wellcome Sanger Institute for the COVID-19 Genomics UK (COG-UK) Consortium | Lighthouse Laboratory Plymouth and Alex Alderton, Roberto Amato, Jeffrey Barrett, Sonia Goncalves, Ewan Harrison, David K. Jackson, Ian Johnston, Dominic Kwiatkowski, Cordelia Langford, John Sillitoe on behalf of the Wellcome Sanger Institute COVID-19 Surveillance Team |
| EPI_ISL_2515071 | Lighthouse Lab in Alderley Park | Wellcome Sanger Institute for the COVID-19 Genomics UK (COG-UK) Consortium | Jacquelyn Wynn, Mairead Hyland, The Lighthouse Lab in Alderley Park and Alex Alderton, Roberto Amato, Jeffrey Barrett, Sonia Goncalves, Ewan Harrison, David K. Jackson, Ian Johnston, Dominic Kwiatkowski, Cordelia Langford, John Sillitoe on behalf of the Wellcome Sanger Institute COVID-19 Surveillance Team |
| EPI_ISL_2515072, EPI_ISL_2515073 | Lighthouse Lab in Glasgow | Wellcome Sanger Institute for the COVID-19 Genomics UK (COG-UK) Consortium | Harper VanSteenhouse, Yumi Kasai, David Gray, Carol Clugston, Anna Dominiczak and Alex Alderton, Roberto Amato, Jeffrey Barrett, Sonia Goncalves, Ewan Harrison, David K. Jackson, Ian Johnston, Dominic Kwiatkowski, Cordelia Langford, John Sillitoe on behalf of the Wellcome Sanger Institute COVID-19 Surveillance Team |
| EPI_ISL_2515074, EPI_ISL_2515075, EPI_ISL_2515076 | Lighthouse Lab in Alderley Park | Wellcome Sanger Institute for the COVID-19 Genomics UK (COG-UK) Consortium | Jacquelyn Wynn, Mairead Hyland, The Lighthouse Lab in Alderley Park and Alex Alderton, Roberto Amato, Jeffrey Barrett, Sonia Goncalves, Ewan Harrison, David K. Jackson, Ian Johnston, Dominic Kwiatkowski, Cordelia Langford, John Sillitoe on behalf of the Wellcome Sanger Institute COVID-19 Surveillance Team |
| EPI_ISL_2515077, EPI_ISL_2515078 | Lighthouse Laboratory Plymouth | Wellcome Sanger Institute for the COVID-19 Genomics UK (COG-UK) Consortium | Lighthouse Laboratory Plymouth and Alex Alderton, Roberto Amato, Jeffrey Barrett, Sonia Goncalves, Ewan Harrison, David K. Jackson, Ian Johnston, Dominic Kwiatkowski, Cordelia Langford, John Sillitoe on behalf of the Wellcome Sanger Institute COVID-19 Surveillance Team |
| EPI_ISL_2515079 | Lighthouse Lab in Glasgow | Wellcome Sanger Institute for the COVID-19 Genomics UK (COG-UK) Consortium | Harper VanSteenhouse, Yumi Kasai, David Gray, Carol Clugston, Anna Dominiczak and Alex Alderton, Roberto Amato, Jeffrey Barrett, Sonia Goncalves, Ewan Harrison, David K. Jackson, Ian Johnston, Dominic Kwiatkowski, Cordelia Langford, John Sillitoe on behalf of the Wellcome Sanger Institute COVID-19 Surveillance Team |
| EPI_ISL_2515080 | Lighthouse Laboratory Plymouth | Wellcome Sanger Institute for the COVID-19 Genomics UK (COG-UK) Consortium | Lighthouse Laboratory Plymouth and Alex Alderton, Roberto Amato, Jeffrey Barrett, Sonia Goncalves, Ewan Harrison, David K. Jackson, Ian Johnston, Dominic Kwiatkowski, Cordelia Langford, John Sillitoe on behalf of the Wellcome Sanger Institute COVID-19 Surveillance Team |
| EPI_ISL_2515081 | Lighthouse Lab in Alderley Park | Wellcome Sanger Institute for the COVID-19 Genomics UK (COG-UK) Consortium | Jacquelyn Wynn, Mairead Hyland, The Lighthouse Lab in Alderley Park and Alex Alderton, Roberto Amato, Jeffrey Barrett, Sonia Goncalves, Ewan Harrison, David K. Jackson, Ian Johnston, Dominic Kwiatkowski, Cordelia Langford, John Sillitoe on behalf of the Wellcome Sanger Institute COVID-19 Surveillance Team |
| EPI_ISL_2515082 | Lighthouse Lab in Glasgow | Wellcome Sanger Institute for the COVID-19 Genomics UK (COG-UK) Consortium | Harper VanSteenhouse, Yumi Kasai, David Gray, Carol Clugston, Anna Dominiczak and Alex Alderton, Roberto Amato, Jeffrey Barrett, Sonia Goncalves, Ewan Harrison, David K. Jackson, Ian Johnston, Dominic Kwiatkowski, Cordelia Langford, John Sillitoe on behalf of the Wellcome Sanger Institute COVID-19 Surveillance Team |
| EPI_ISL_2515083 | Lighthouse Laboratory Plymouth | Wellcome Sanger Institute for the COVID-19 Genomics UK (COG-UK) Consortium | Lighthouse Laboratory Plymouth and Alex Alderton, Roberto Amato, Jeffrey Barrett, Sonia Goncalves, Ewan Harrison, David K. Jackson, Ian Johnston, Dominic Kwiatkowski, Cordelia Langford, John Sillitoe on behalf of the Wellcome Sanger Institute COVID-19 Surveillance Team |
| EPI_ISL_2515084, EPI_ISL_2515085 | Lighthouse Lab in Alderley Park | Wellcome Sanger Institute for the COVID-19 Genomics UK (COG-UK) Consortium | Jacquelyn Wynn, Mairead Hyland, The Lighthouse Lab in Alderley Park and Alex Alderton, Roberto Amato, Jeffrey Barrett, Sonia Goncalves, Ewan Harrison, David K. Jackson, Ian Johnston, Dominic Kwiatkowski, Cordelia Langford, John Sillitoe on behalf of the Wellcome Sanger Institute COVID-19 Surveillance Team |
| EPI_ISL_2515086 | Lighthouse Laboratory Plymouth | Wellcome Sanger Institute for the COVID-19 Genomics UK (COG-UK) Consortium | Lighthouse Laboratory Plymouth and Alex Alderton, Roberto Amato, Jeffrey Barrett, Sonia Goncalves, Ewan Harrison, David K. Jackson, Ian Johnston, Dominic Kwiatkowski, Cordelia Langford, John Sillitoe on behalf of the Wellcome Sanger Institute COVID-19 Surveillance Team |
| EPI_ISL_2515087 | Lighthouse Lab in Glasgow | Wellcome Sanger Institute for the COVID-19 Genomics UK (COG-UK) Consortium | Harper VanSteenhouse, Yumi Kasai, David Gray, Carol Clugston, Anna Dominiczak and Alex Alderton, Roberto Amato, Jeffrey Barrett, Sonia Goncalves, Ewan Harrison, David K. Jackson, Ian Johnston, Dominic Kwiatkowski, Cordelia Langford, John Sillitoe on behalf of the Wellcome Sanger Institute COVID-19 Surveillance Team |
| EPI_ISL_2515088 | Lighthouse Lab in Alderley Park | Wellcome Sanger Institute for the COVID-19 Genomics UK (COG-UK) Consortium | Jacquelyn Wynn, Mairead Hyland, The Lighthouse Lab in Alderley Park and Alex Alderton, Roberto Amato, Jeffrey Barrett, Sonia Goncalves, Ewan Harrison, David K. Jackson, Ian Johnston, Dominic Kwiatkowski, Cordelia Langford, John Sillitoe on behalf of the Wellcome Sanger Institute COVID-19 Surveillance Team |
| EPI_ISL_2515089 | Lighthouse Laboratory Plymouth | Wellcome Sanger Institute for the COVID-19 Genomics UK (COG-UK) Consortium | Lighthouse Laboratory Plymouth and Alex Alderton, Roberto Amato, Jeffrey Barrett, Sonia Goncalves, Ewan Harrison, David K. Jackson, Ian Johnston, Dominic Kwiatkowski, Cordelia Langford, John Sillitoe on behalf of the Wellcome Sanger Institute COVID-19 Surveillance Team |
| EPI_ISL_2515090 | Lighthouse Lab in Glasgow | Wellcome Sanger Institute for the COVID-19 Genomics UK (COG-UK) Consortium | Harper VanSteenhouse, Yumi Kasai, David Gray, Carol Clugston, Anna Dominiczak and Alex Alderton, Roberto Amato, Jeffrey Barrett, Sonia Goncalves, Ewan Harrison, David K. Jackson, Ian Johnston, Dominic Kwiatkowski, Cordelia Langford, John Sillitoe on behalf of the Wellcome Sanger Institute COVID-19 Surveillance Team |
| EPI_ISL_2515091 | Lighthouse Lab in Alderley Park | Wellcome Sanger Institute for the COVID-19 Genomics UK (COG-UK) Consortium | Jacquelyn Wynn, Mairead Hyland, The Lighthouse Lab in Alderley Park and Alex Alderton, Roberto Amato, Jeffrey Barrett, Sonia Goncalves, Ewan Harrison, David K. Jackson, Ian Johnston, Dominic Kwiatkowski, Cordelia Langford, John Sillitoe on behalf of the Wellcome Sanger Institute COVID-19 Surveillance Team |
| EPI_ISL_2515092, EPI_ISL_2515093 | Lighthouse Laboratory Plymouth | Wellcome Sanger Institute for the COVID-19 Genomics UK (COG-UK) Consortium | Lighthouse Laboratory Plymouth and Alex Alderton, Roberto Amato, Jeffrey Barrett, Sonia Goncalves, Ewan Harrison, David K. Jackson, Ian Johnston, Dominic Kwiatkowski, Cordelia Langford, John Sillitoe on behalf of the Wellcome Sanger Institute COVID-19 Surveillance Team |

| | | | |
|---|---|---|---|
| EPI_ISL_2515094 | Lighthouse Lab in Alderley Park | Wellcome Sanger Institute for the COVID-19 Genomics UK (COG-UK) Consortium | Jacquelyn Wynn, Mairead Hyland, The Lighthouse Lab in Alderley Park and Alex Alderton, Roberto Amato, Jeffrey Barrett, Sonia Goncalves, Ewan Harrison, David K. Jackson, Ian Johnston, Dominic Kwiatkowski, Cordelia Langford, John Sillitoe on behalf of the Wellcome Sanger Institute COVID-19 Surveillance Team |
| EPI_ISL_2515095 | Lighthouse Laboratory Plymouth | Wellcome Sanger Institute for the COVID-19 Genomics UK (COG-UK) Consortium | Lighthouse Laboratory Plymouth and Alex Alderton, Roberto Amato, Jeffrey Barrett, Sonia Goncalves, Ewan Harrison, David K. Jackson, Ian Johnston, Dominic Kwiatkowski, Cordelia Langford, John Sillitoe on behalf of the Wellcome Sanger Institute COVID-19 Surveillance Team |
| EPI_ISL_2515096 | Lighthouse Lab in Glasgow | Wellcome Sanger Institute for the COVID-19 Genomics UK (COG-UK) Consortium | Harper VanSteenhouse, Yumi Kasai, David Gray, Carol Clugston, Anna Dominiczak and Alex Alderton, Roberto Amato, Jeffrey Barrett, Sonia Goncalves, Ewan Harrison, David K. Jackson, Ian Johnston, Dominic Kwiatkowski, Cordelia Langford, John Sillitoe on behalf of the Wellcome Sanger Institute COVID-19 Surveillance Team |
| EPI_ISL_2515097 | Lighthouse Laboratory Plymouth | Wellcome Sanger Institute for the COVID-19 Genomics UK (COG-UK) Consortium | Lighthouse Laboratory Plymouth and Alex Alderton, Roberto Amato, Jeffrey Barrett, Sonia Goncalves, Ewan Harrison, David K. Jackson, Ian Johnston, Dominic Kwiatkowski, Cordelia Langford, John Sillitoe on behalf of the Wellcome Sanger Institute COVID-19 Surveillance Team |
| EPI_ISL_2515098, EPI_ISL_2515099 | Lighthouse Lab in Glasgow | Wellcome Sanger Institute for the COVID-19 Genomics UK (COG-UK) Consortium | Harper VanSteenhouse, Yumi Kasai, David Gray, Carol Clugston, Anna Dominiczak and Alex Alderton, Roberto Amato, Jeffrey Barrett, Sonia Goncalves, Ewan Harrison, David K. Jackson, Ian Johnston, Dominic Kwiatkowski, Cordelia Langford, John Sillitoe on behalf of the Wellcome Sanger Institute COVID-19 Surveillance Team |
| EPI_ISL_2515100 | Lighthouse Lab in Alderley Park | Wellcome Sanger Institute for the COVID-19 Genomics UK (COG-UK) Consortium | Jacquelyn Wynn, Mairead Hyland, The Lighthouse Lab in Alderley Park and Alex Alderton, Roberto Amato, Jeffrey Barrett, Sonia Goncalves, Ewan Harrison, David K. Jackson, Ian Johnston, Dominic Kwiatkowski, Cordelia Langford, John Sillitoe on behalf of the Wellcome Sanger Institute COVID-19 Surveillance Team |
| EPI_ISL_2515101, EPI_ISL_2515102 | Lighthouse Laboratory Plymouth | Wellcome Sanger Institute for the COVID-19 Genomics UK (COG-UK) Consortium | Lighthouse Laboratory Plymouth and Alex Alderton, Roberto Amato, Jeffrey Barrett, Sonia Goncalves, Ewan Harrison, David K. Jackson, Ian Johnston, Dominic Kwiatkowski, Cordelia Langford, John Sillitoe on behalf of the Wellcome Sanger Institute COVID-19 Surveillance Team |
| EPI_ISL_2515103, EPI_ISL_2515104, EPI_ISL_2515105, EPI_ISL_2515106 | Lighthouse Lab in Alderley Park | Wellcome Sanger Institute for the COVID-19 Genomics UK (COG-UK) Consortium | Jacquelyn Wynn, Mairead Hyland, The Lighthouse Lab in Alderley Park and Alex Alderton, Roberto Amato, Jeffrey Barrett, Sonia Goncalves, Ewan Harrison, David K. Jackson, Ian Johnston, Dominic Kwiatkowski, Cordelia Langford, John Sillitoe on behalf of the Wellcome Sanger Institute COVID-19 Surveillance Team |
| EPI_ISL_2515107 | Lighthouse Lab in Glasgow | Wellcome Sanger Institute for the COVID-19 Genomics UK (COG-UK) Consortium | Harper VanSteenhouse, Yumi Kasai, David Gray, Carol Clugston, Anna Dominiczak and Alex Alderton, Roberto Amato, Jeffrey Barrett, Sonia Goncalves, Ewan Harrison, David K. Jackson, Ian Johnston, Dominic Kwiatkowski, Cordelia Langford, John Sillitoe on behalf of the Wellcome Sanger Institute COVID-19 Surveillance Team |
| EPI_ISL_2515108 | Lighthouse Lab in Alderley Park | Wellcome Sanger Institute for the COVID-19 Genomics UK (COG-UK) Consortium | Jacquelyn Wynn, Mairead Hyland, The Lighthouse Lab in Alderley Park and Alex Alderton, Roberto Amato, Jeffrey Barrett, Sonia Goncalves, Ewan Harrison, David K. Jackson, Ian Johnston, Dominic Kwiatkowski, Cordelia Langford, John Sillitoe on behalf of the Wellcome Sanger Institute COVID-19 Surveillance Team |
| EPI_ISL_2515110 | Lighthouse Laboratory Plymouth | Wellcome Sanger Institute for the COVID-19 Genomics UK (COG-UK) Consortium | Lighthouse Laboratory Plymouth and Alex Alderton, Roberto Amato, Jeffrey Barrett, Sonia Goncalves, Ewan Harrison, David K. Jackson, Ian Johnston, Dominic Kwiatkowski, Cordelia Langford, John Sillitoe on behalf of the Wellcome Sanger Institute COVID-19 Surveillance Team |
| EPI_ISL_2515111, EPI_ISL_2515112 | Lighthouse Lab in Alderley Park | Wellcome Sanger Institute for the COVID-19 Genomics UK (COG-UK) Consortium | Jacquelyn Wynn, Mairead Hyland, The Lighthouse Lab in Alderley Park and Alex Alderton, Roberto Amato, Jeffrey Barrett, Sonia Goncalves, Ewan Harrison, David K. Jackson, Ian Johnston, Dominic Kwiatkowski, Cordelia Langford, John Sillitoe on behalf of the Wellcome Sanger Institute COVID-19 Surveillance Team |
| EPI_ISL_2515113 | Lighthouse Laboratory Plymouth | Wellcome Sanger Institute for the COVID-19 Genomics UK (COG-UK) Consortium | Lighthouse Laboratory Plymouth and Alex Alderton, Roberto Amato, Jeffrey Barrett, Sonia Goncalves, Ewan Harrison, David K. Jackson, Ian Johnston, Dominic Kwiatkowski, Cordelia Langford, John Sillitoe on behalf of the Wellcome Sanger Institute COVID-19 Surveillance Team |
| EPI_ISL_2515114 | Lighthouse Lab in Glasgow | Wellcome Sanger Institute for the COVID-19 Genomics UK (COG-UK) Consortium | Harper VanSteenhouse, Yumi Kasai, David Gray, Carol Clugston, Anna Dominiczak and Alex Alderton, Roberto Amato, Jeffrey Barrett, Sonia Goncalves, Ewan Harrison, David K. Jackson, Ian Johnston, Dominic Kwiatkowski, Cordelia Langford, John Sillitoe on behalf of the Wellcome Sanger Institute COVID-19 Surveillance Team |
| EPI_ISL_2515115, EPI_ISL_2515116 | Lighthouse Laboratory Plymouth | Wellcome Sanger Institute for the COVID-19 Genomics UK (COG-UK) Consortium | Lighthouse Laboratory Plymouth and Alex Alderton, Roberto Amato, Jeffrey Barrett, Sonia Goncalves, Ewan Harrison, David K. Jackson, Ian Johnston, Dominic Kwiatkowski, Cordelia Langford, John Sillitoe on behalf of the Wellcome Sanger Institute COVID-19 Surveillance Team |
| EPI_ISL_2515117, EPI_ISL_2515118, EPI_ISL_2515119, EPI_ISL_2515120, EPI_ISL_2515121, EPI_ISL_2515122, EPI_ISL_2515123, EPI_ISL_2515124 | Lighthouse Lab in Alderley Park | Wellcome Sanger Institute for the COVID-19 Genomics UK (COG-UK) Consortium | Jacquelyn Wynn, Mairead Hyland, The Lighthouse Lab in Alderley Park and Alex Alderton, Roberto Amato, Jeffrey Barrett, Sonia Goncalves, Ewan Harrison, David K. Jackson, Ian Johnston, Dominic Kwiatkowski, Cordelia Langford, John Sillitoe on behalf of the Wellcome Sanger Institute COVID-19 Surveillance Team |
| EPI_ISL_2515125 | Lighthouse Lab in Glasgow | Wellcome Sanger Institute for the COVID-19 Genomics UK (COG-UK) Consortium | Harper VanSteenhouse, Yumi Kasai, David Gray, Carol Clugston, Anna Dominiczak and Alex Alderton, Roberto Amato, Jeffrey Barrett, Sonia Goncalves, Ewan Harrison, David K. Jackson, Ian Johnston, Dominic Kwiatkowski, Cordelia Langford, John Sillitoe on behalf of the Wellcome Sanger Institute COVID-19 Surveillance Team |
| EPI_ISL_2515126, EPI_ISL_2515127, EPI_ISL_2515128, EPI_ISL_2515129, EPI_ISL_2515130, EPI_ISL_2515131, EPI_ISL_2515132, EPI_ISL_2515133, EPI_ISL_2515134, EPI_ISL_2515135, EPI_ISL_2515136, EPI_ISL_2515137, EPI_ISL_2515138, EPI_ISL_2515140, EPI_ISL_2515141, EPI_ISL_2515142, EPI_ISL_2515143, EPI_ISL_2515144 | | | |
| see above | Lighthouse Lab in Alderley Park | Wellcome Sanger Institute for the COVID-19 Genomics UK (COG-UK) Consortium | Jacquelyn Wynn, Mairead Hyland, The Lighthouse Lab in Alderley Park and Alex Alderton, Roberto Amato, Jeffrey Barrett, Sonia Goncalves, Ewan Harrison, David K. Jackson, Ian Johnston, Dominic Kwiatkowski, Cordelia Langford, John Sillitoe on behalf of the Wellcome Sanger Institute COVID-19 Surveillance Team |
| EPI_ISL_2515145 | Lighthouse Lab in Glasgow | Wellcome Sanger Institute for the COVID-19 Genomics UK (COG-UK) Consortium | Harper VanSteenhouse, Yumi Kasai, David Gray, Carol Clugston, Anna Dominiczak and Alex Alderton, Roberto Amato, Jeffrey Barrett, Sonia Goncalves, Ewan Harrison, David K. Jackson, Ian Johnston, Dominic Kwiatkowski, Cordelia Langford, John Sillitoe on behalf of the Wellcome Sanger Institute COVID-19 Surveillance Team |
| EPI_ISL_2515146, EPI_ISL_2515147, EPI_ISL_2515149, EPI_ISL_2515150, EPI_ISL_2515151 | Lighthouse Lab in Alderley Park | Wellcome Sanger Institute for the COVID-19 Genomics UK (COG-UK) Consortium | Jacquelyn Wynn, Mairead Hyland, The Lighthouse Lab in Alderley Park and Alex Alderton, Roberto Amato, Jeffrey Barrett, Sonia Goncalves, Ewan Harrison, David K. Jackson, Ian Johnston, Dominic Kwiatkowski, Cordelia Langford, John Sillitoe on behalf of the Wellcome Sanger Institute COVID-19 Surveillance Team |
| EPI_ISL_2515152 | Lighthouse Lab in Glasgow | Wellcome Sanger Institute for the COVID-19 Genomics UK (COG-UK) Consortium | Harper VanSteenhouse, Yumi Kasai, David Gray, Carol Clugston, Anna Dominiczak and Alex Alderton, Roberto Amato, Jeffrey Barrett, Sonia Goncalves, Ewan Harrison, David K. Jackson, Ian Johnston, Dominic Kwiatkowski, Cordelia Langford, John Sillitoe on behalf of the Wellcome Sanger Institute COVID-19 Surveillance Team |
| EPI_ISL_2515153, EPI_ISL_2515154 | Lighthouse Lab in Alderley Park | Wellcome Sanger Institute for the COVID-19 Genomics UK (COG-UK) Consortium | Jacquelyn Wynn, Mairead Hyland, The Lighthouse Lab in Alderley Park and Alex Alderton, Roberto Amato, Jeffrey Barrett, Sonia Goncalves, Ewan Harrison, David K. Jackson, Ian Johnston, Dominic Kwiatkowski, Cordelia Langford, John Sillitoe on behalf of the Wellcome Sanger Institute COVID-19 Surveillance Team |
| EPI_ISL_2515155 | Lighthouse Lab in Glasgow | Wellcome Sanger Institute for the COVID-19 Genomics UK (COG-UK) Consortium | Harper VanSteenhouse, Yumi Kasai, David Gray, Carol Clugston, Anna Dominiczak and Alex Alderton, Roberto Amato, Jeffrey Barrett, Sonia Goncalves, Ewan Harrison, David K. Jackson, Ian Johnston, Dominic Kwiatkowski, Cordelia Langford, John Sillitoe on behalf of the Wellcome Sanger Institute COVID-19 Surveillance Team |
| EPI_ISL_2515157, EPI_ISL_2515158, EPI_ISL_2515159, EPI_ISL_2515160, EPI_ISL_2515161, EPI_ISL_2515162, EPI_ISL_2515163, EPI_ISL_2515164, EPI_ISL_2515166, EPI_ISL_2515167, EPI_ISL_2515168, EPI_ISL_2515169, EPI_ISL_2515170, EPI_ISL_2515171, EPI_ISL_2515172, EPI_ISL_2515173 | | | |
| see above | Lighthouse Lab in Alderley Park | Wellcome Sanger Institute for the COVID-19 Genomics UK (COG-UK) Consortium | Jacquelyn Wynn, Mairead Hyland, The Lighthouse Lab in Alderley Park and Alex Alderton, Roberto Amato, Jeffrey Barrett, Sonia Goncalves, Ewan Harrison, David K. Jackson, Ian Johnston, Dominic Kwiatkowski, Cordelia Langford, John Sillitoe on behalf of the Wellcome Sanger Institute COVID-19 Surveillance Team |
| EPI_ISL_2515175 | Lighthouse Lab in Glasgow | Wellcome Sanger Institute for the COVID-19 Genomics UK (COG-UK) Consortium | Harper VanSteenhouse, Yumi Kasai, David Gray, Carol Clugston, Anna Dominiczak and Alex Alderton, Roberto Amato, Jeffrey Barrett, Sonia Goncalves, Ewan Harrison, David K. Jackson, Ian Johnston, Dominic Kwiatkowski, Cordelia Langford, John Sillitoe on behalf of the Wellcome Sanger Institute COVID-19 Surveillance Team |
| EPI_ISL_2515177, EPI_ISL_2515179, EPI_ISL_2515180, EPI_ISL_2515181, EPI_ISL_2515182, EPI_ISL_2515183, EPI_ISL_2515184, EPI_ISL_2515185, EPI_ISL_2515186, EPI_ISL_2515188, EPI_ISL_2515189, EPI_ISL_2515190, EPI_ISL_2515191, EPI_ISL_2515192, EPI_ISL_2515193 | | | |
| see above | Lighthouse Lab in Alderley Park | Wellcome Sanger Institute for the COVID-19 Genomics UK (COG-UK) Consortium | Jacquelyn Wynn, Mairead Hyland, The Lighthouse Lab in Alderley Park and Alex Alderton, Roberto Amato, Jeffrey Barrett, Sonia Goncalves, Ewan Harrison, David K. Jackson, Ian Johnston, Dominic Kwiatkowski, Cordelia Langford, John Sillitoe on behalf of the Wellcome Sanger Institute COVID-19 Surveillance Team |
| EPI_ISL_2515194 | Lighthouse Lab in Glasgow | Wellcome Sanger Institute for the COVID-19 Genomics UK (COG-UK) Consortium | Harper VanSteenhouse, Yumi Kasai, David Gray, Carol Clugston, Anna Dominiczak and Alex Alderton, Roberto Amato, Jeffrey Barrett, Sonia Goncalves, Ewan Harrison, David K. Jackson, Ian Johnston, Dominic Kwiatkowski, Cordelia Langford, John Sillitoe on behalf of the Wellcome Sanger Institute COVID-19 Surveillance Team |
| EPI_ISL_2515195, EPI_ISL_2515197, EPI_ISL_2515198, EPI_ISL_2515199, EPI_ISL_2515200, EPI_ISL_2515201, EPI_ISL_2515202 | Lighthouse Lab in Alderley Park | Wellcome Sanger Institute for the COVID-19 Genomics UK (COG-UK) Consortium | Jacquelyn Wynn, Mairead Hyland, The Lighthouse Lab in Alderley Park and Alex Alderton, Roberto Amato, Jeffrey Barrett, Sonia Goncalves, Ewan Harrison, David K. Jackson, Ian Johnston, Dominic Kwiatkowski, Cordelia Langford, John Sillitoe on behalf of the Wellcome Sanger Institute COVID-19 Surveillance Team |
| EPI_ISL_2515203 | Lighthouse Lab in Glasgow | Wellcome Sanger Institute for the COVID-19 Genomics UK (COG-UK) Consortium | Harper VanSteenhouse, Yumi Kasai, David Gray, Carol Clugston, Anna Dominiczak and Alex Alderton, Roberto Amato, Jeffrey Barrett, Sonia Goncalves, Ewan Harrison, David K. Jackson, Ian Johnston, Dominic Kwiatkowski, Cordelia Langford, John Sillitoe on behalf of the Wellcome Sanger Institute COVID-19 Surveillance Team |
| EPI_ISL_2515204, EPI_ISL_2515205, EPI_ISL_2515206, EPI_ISL_2515208, EPI_ISL_2515209, EPI_ISL_2515210, EPI_ISL_2515211, EPI_ISL_2515212, EPI_ISL_2515213, EPI_ISL_2515214, EPI_ISL_2515215, EPI_ISL_2515216, EPI_ISL_2515217, EPI_ISL_2515218, EPI_ISL_2515219, EPI_ISL_2515220, EPI_ISL_2515221, EPI_ISL_2515222, EPI_ISL_2515223, EPI_ISL_2515224, EPI_ISL_2515225, EPI_ISL_2515226, EPI_ISL_2515227, EPI_ISL_2515228, EPI_ISL_2515229, EPI_ISL_2515230 | | | |
| see above | Lighthouse Lab in Alderley Park | Wellcome Sanger Institute for the COVID-19 Genomics UK (COG-UK) Consortium | Jacquelyn Wynn, Mairead Hyland, The Lighthouse Lab in Alderley Park and Alex Alderton, Roberto Amato, Jeffrey Barrett, Sonia Goncalves, Ewan Harrison, David K. Jackson, Ian Johnston, Dominic Kwiatkowski, Cordelia Langford, John Sillitoe on behalf of the Wellcome Sanger Institute COVID-19 Surveillance Team |
| EPI_ISL_2515231 | Lighthouse Lab in Glasgow | Wellcome Sanger Institute for the COVID-19 Genomics UK (COG-UK) Consortium | Harper VanSteenhouse, Yumi Kasai, David Gray, Carol Clugston, Anna Dominiczak and Alex Alderton, Roberto Amato, Jeffrey Barrett, Sonia Goncalves, Ewan Harrison, David K. Jackson, Ian Johnston, Dominic Kwiatkowski, Cordelia Langford, John Sillitoe on behalf of the Wellcome Sanger Institute COVID-19 Surveillance Team |
| EPI_ISL_2515232, EPI_ISL_2515233, EPI_ISL_2515235, EPI_ISL_2515236, EPI_ISL_2515237, EPI_ISL_2515238, EPI_ISL_2515239, EPI_ISL_2515240, EPI_ISL_2515241, EPI_ISL_2515243, EPI_ISL_2515244, EPI_ISL_2515245, EPI_ISL_2515246, EPI_ISL_2515247, EPI_ISL_2515248 | | | |
| see above | Lighthouse Lab in Alderley Park | Wellcome Sanger Institute for the COVID-19 Genomics UK (COG-UK) Consortium | Jacquelyn Wynn, Mairead Hyland, The Lighthouse Lab in Alderley Park and Alex Alderton, Roberto Amato, Jeffrey Barrett, Sonia Goncalves, Ewan Harrison, David K. Jackson, Ian Johnston, Dominic Kwiatkowski, Cordelia Langford, John Sillitoe on behalf of the Wellcome Sanger Institute COVID-19 Surveillance Team |
| EPI_ISL_2515249 | Lighthouse Lab in Glasgow | Wellcome Sanger Institute for the COVID-19 Genomics UK (COG-UK) Consortium | Harper VanSteenhouse, Yumi Kasai, David Gray, Carol Clugston, Anna Dominiczak and Alex Alderton, Roberto Amato, Jeffrey Barrett, Sonia Goncalves, Ewan Harrison, David K. Jackson, Ian Johnston, Dominic Kwiatkowski, Cordelia Langford, John Sillitoe on behalf of the Wellcome Sanger Institute COVID-19 Surveillance Team |
| EPI_ISL_2515250, EPI_ISL_2515251, EPI_ISL_2515252, EPI_ISL_2515253, EPI_ISL_2515254, EPI_ISL_2515255, EPI_ISL_2515256, EPI_ISL_2515257, EPI_ISL_2515258, EPI_ISL_2515259, EPI_ISL_2515260, EPI_ISL_2515261, EPI_ISL_2515262, EPI_ISL_2515263 | | | |
| see above | Lighthouse Lab in Alderley Park | Wellcome Sanger Institute for the COVID-19 Genomics UK (COG-UK) Consortium | Jacquelyn Wynn, Mairead Hyland, The Lighthouse Lab in Alderley Park and Alex Alderton, Roberto Amato, Jeffrey Barrett, Sonia Goncalves, Ewan Harrison, David K. Jackson, Ian Johnston, Dominic Kwiatkowski, Cordelia Langford, John Sillitoe on behalf of the Wellcome Sanger Institute COVID-19 Surveillance Team |
| EPI_ISL_2515265 | Lighthouse Lab in Glasgow | Wellcome Sanger Institute for the COVID-19 Genomics UK (COG-UK) | Harper VanSteenhouse, Yumi Kasai, David Gray, Carol Clugston, Anna Dominiczak and Alex Alderton, Roberto Amato, Jeffrey Barrett, Sonia Goncalves, Ewan Harrison, David K. Jackson, Ian Johnston, Dominic Kwiatkowski, Cordelia Langford, |

| | | | |
|---|---|---|---|
| | | Consortium | John Sillitoe on behalf of the Wellcome Sanger Institute COVID-19 Surveillance Team |
| EPI_ISL_2515266, EPI_ISL_2515267, EPI_ISL_2515268, EPI_ISL_2515269 | Lighthouse Lab in Alderley Park | Wellcome Sanger Institute for the COVID-19 Genomics UK (COG-UK) Consortium | Jacquelyn Wynn, Mairead Hyland, The Lighthouse Lab in Alderley Park and Alex Alderton, Roberto Amato, Jeffrey Barrett, Sonia Goncalves, Ewan Harrison, David K. Jackson, Ian Johnston, Dominic Kwiatkowski, Cordelia Langford, John Sillitoe on behalf of the Wellcome Sanger Institute COVID-19 Surveillance Team |
| EPI_ISL_2515270 | Lighthouse Lab in Glasgow | Wellcome Sanger Institute for the COVID-19 Genomics UK (COG-UK) Consortium | Harper VanSteenhouse, Yumi Kasai, David Gray, Carol Clugston, Anna Dominiczak and Alex Alderton, Roberto Amato, Jeffrey Barrett, Sonia Goncalves, Ewan Harrison, David K. Jackson, Ian Johnston, Dominic Kwiatkowski, Cordelia Langford, John Sillitoe on behalf of the Wellcome Sanger Institute COVID-19 Surveillance Team |
| EPI_ISL_2515271, EPI_ISL_2515272, EPI_ISL_2515273, EPI_ISL_2515274 | Lighthouse Lab in Alderley Park | Wellcome Sanger Institute for the COVID-19 Genomics UK (COG-UK) Consortium | Jacquelyn Wynn, Mairead Hyland, The Lighthouse Lab in Alderley Park and Alex Alderton, Roberto Amato, Jeffrey Barrett, Sonia Goncalves, Ewan Harrison, David K. Jackson, Ian Johnston, Dominic Kwiatkowski, Cordelia Langford, John Sillitoe on behalf of the Wellcome Sanger Institute COVID-19 Surveillance Team |
| EPI_ISL_2515275 | Lighthouse Lab in Glasgow | Wellcome Sanger Institute for the COVID-19 Genomics UK (COG-UK) Consortium | Harper VanSteenhouse, Yumi Kasai, David Gray, Carol Clugston, Anna Dominiczak and Alex Alderton, Roberto Amato, Jeffrey Barrett, Sonia Goncalves, Ewan Harrison, David K. Jackson, Ian Johnston, Dominic Kwiatkowski, Cordelia Langford, John Sillitoe on behalf of the Wellcome Sanger Institute COVID-19 Surveillance Team |
| EPI_ISL_2515276, EPI_ISL_2515277, EPI_ISL_2515278, EPI_ISL_2515279, EPI_ISL_2515280, EPI_ISL_2515281, EPI_ISL_2515282, EPI_ISL_2515283, EPI_ISL_2515284, EPI_ISL_2515285, EPI_ISL_2515286, EPI_ISL_2515287, EPI_ISL_2515288, EPI_ISL_2515289, EPI_ISL_2515290, EPI_ISL_2515291, EPI_ISL_2515292, EPI_ISL_2515293, EPI_ISL_2515294 | | | |
| see above | Lighthouse Lab in Alderley Park | Wellcome Sanger Institute for the COVID-19 Genomics UK (COG-UK) Consortium | Jacquelyn Wynn, Mairead Hyland, The Lighthouse Lab in Alderley Park and Alex Alderton, Roberto Amato, Jeffrey Barrett, Sonia Goncalves, Ewan Harrison, David K. Jackson, Ian Johnston, Dominic Kwiatkowski, Cordelia Langford, John Sillitoe on behalf of the Wellcome Sanger Institute COVID-19 Surveillance Team |
| EPI_ISL_2515295 | Lighthouse Lab in Glasgow | Wellcome Sanger Institute for the COVID-19 Genomics UK (COG-UK) Consortium | Harper VanSteenhouse, Yumi Kasai, David Gray, Carol Clugston, Anna Dominiczak and Alex Alderton, Roberto Amato, Jeffrey Barrett, Sonia Goncalves, Ewan Harrison, David K. Jackson, Ian Johnston, Dominic Kwiatkowski, Cordelia Langford, John Sillitoe on behalf of the Wellcome Sanger Institute COVID-19 Surveillance Team |
| EPI_ISL_2515296, EPI_ISL_2515297, EPI_ISL_2515299, EPI_ISL_2515300, EPI_ISL_2515301, EPI_ISL_2515303, EPI_ISL_2515304, EPI_ISL_2515305, EPI_ISL_2515306, EPI_ISL_2515308, EPI_ISL_2515309, EPI_ISL_2515311, EPI_ISL_2515312, EPI_ISL_2515313, EPI_ISL_2515314, EPI_ISL_2515315, EPI_ISL_2515316, EPI_ISL_2515317, EPI_ISL_2515318, EPI_ISL_2515319, EPI_ISL_2515320, EPI_ISL_2515321, EPI_ISL_2515322, EPI_ISL_2515323 | | | |
| see above | Lighthouse Lab in Alderley Park | Wellcome Sanger Institute for the COVID-19 Genomics UK (COG-UK) Consortium | Jacquelyn Wynn, Mairead Hyland, The Lighthouse Lab in Alderley Park and Alex Alderton, Roberto Amato, Jeffrey Barrett, Sonia Goncalves, Ewan Harrison, David K. Jackson, Ian Johnston, Dominic Kwiatkowski, Cordelia Langford, John Sillitoe on behalf of the Wellcome Sanger Institute COVID-19 Surveillance Team |
| EPI_ISL_2515324 | Lighthouse Lab in Glasgow | Wellcome Sanger Institute for the COVID-19 Genomics UK (COG-UK) Consortium | Harper VanSteenhouse, Yumi Kasai, David Gray, Carol Clugston, Anna Dominiczak and Alex Alderton, Roberto Amato, Jeffrey Barrett, Sonia Goncalves, Ewan Harrison, David K. Jackson, Ian Johnston, Dominic Kwiatkowski, Cordelia Langford, John Sillitoe on behalf of the Wellcome Sanger Institute COVID-19 Surveillance Team |
| EPI_ISL_2515325, EPI_ISL_2515326, EPI_ISL_2515327, EPI_ISL_2515328, EPI_ISL_2515329, EPI_ISL_2515330, EPI_ISL_2515332, EPI_ISL_2515333 | Lighthouse Lab in Alderley Park | Wellcome Sanger Institute for the COVID-19 Genomics UK (COG-UK) Consortium | Jacquelyn Wynn, Mairead Hyland, The Lighthouse Lab in Alderley Park and Alex Alderton, Roberto Amato, Jeffrey Barrett, Sonia Goncalves, Ewan Harrison, David K. Jackson, Ian Johnston, Dominic Kwiatkowski, Cordelia Langford, John Sillitoe on behalf of the Wellcome Sanger Institute COVID-19 Surveillance Team |
| EPI_ISL_2515334 | Lighthouse Lab in Glasgow | Wellcome Sanger Institute for the COVID-19 Genomics UK (COG-UK) Consortium | Harper VanSteenhouse, Yumi Kasai, David Gray, Carol Clugston, Anna Dominiczak and Alex Alderton, Roberto Amato, Jeffrey Barrett, Sonia Goncalves, Ewan Harrison, David K. Jackson, Ian Johnston, Dominic Kwiatkowski, Cordelia Langford, John Sillitoe on behalf of the Wellcome Sanger Institute COVID-19 Surveillance Team |
| EPI_ISL_2515335, EPI_ISL_2515336, EPI_ISL_2515337, EPI_ISL_2515338, EPI_ISL_2515339, EPI_ISL_2515340, EPI_ISL_2515341, EPI_ISL_2515342, EPI_ISL_2515343, EPI_ISL_2515344, EPI_ISL_2515345, EPI_ISL_2515346, EPI_ISL_2515347, EPI_ISL_2515348, EPI_ISL_2515349, EPI_ISL_2515350, EPI_ISL_2515351, EPI_ISL_2515352, EPI_ISL_2515353, EPI_ISL_2515354, EPI_ISL_2515355, EPI_ISL_2515358 | | | |
| see above | Lighthouse Lab in Alderley Park | Wellcome Sanger Institute for the COVID-19 Genomics UK (COG-UK) Consortium | Jacquelyn Wynn, Mairead Hyland, The Lighthouse Lab in Alderley Park and Alex Alderton, Roberto Amato, Jeffrey Barrett, Sonia Goncalves, Ewan Harrison, David K. Jackson, Ian Johnston, Dominic Kwiatkowski, Cordelia Langford, John Sillitoe on behalf of the Wellcome Sanger Institute COVID-19 Surveillance Team |
| EPI_ISL_2515359 | Lighthouse Lab in Glasgow | Wellcome Sanger Institute for the COVID-19 Genomics UK (COG-UK) Consortium | Harper VanSteenhouse, Yumi Kasai, David Gray, Carol Clugston, Anna Dominiczak and Alex Alderton, Roberto Amato, Jeffrey Barrett, Sonia Goncalves, Ewan Harrison, David K. Jackson, Ian Johnston, Dominic Kwiatkowski, Cordelia Langford, John Sillitoe on behalf of the Wellcome Sanger Institute COVID-19 Surveillance Team |
| EPI_ISL_2515360, EPI_ISL_2515361, EPI_ISL_2515362, EPI_ISL_2515363, EPI_ISL_2515364, EPI_ISL_2515365, EPI_ISL_2515366, EPI_ISL_2515367, EPI_ISL_2515368, EPI_ISL_2515369, EPI_ISL_2515371, EPI_ISL_2515372, EPI_ISL_2515373, EPI_ISL_2515375, EPI_ISL_2515376, EPI_ISL_2515377, EPI_ISL_2515378, EPI_ISL_2515379, EPI_ISL_2515380, EPI_ISL_2515382 | | | |
| see above | Lighthouse Lab in Alderley Park | Wellcome Sanger Institute for the COVID-19 Genomics UK (COG-UK) Consortium | Jacquelyn Wynn, Mairead Hyland, The Lighthouse Lab in Alderley Park and Alex Alderton, Roberto Amato, Jeffrey Barrett, Sonia Goncalves, Ewan Harrison, David K. Jackson, Ian Johnston, Dominic Kwiatkowski, Cordelia Langford, John Sillitoe on behalf of the Wellcome Sanger Institute COVID-19 Surveillance Team |
| EPI_ISL_2515383 | Lighthouse Lab in Glasgow | Wellcome Sanger Institute for the COVID-19 Genomics UK (COG-UK) Consortium | Harper VanSteenhouse, Yumi Kasai, David Gray, Carol Clugston, Anna Dominiczak and Alex Alderton, Roberto Amato, Jeffrey Barrett, Sonia Goncalves, Ewan Harrison, David K. Jackson, Ian Johnston, Dominic Kwiatkowski, Cordelia Langford, John Sillitoe on behalf of the Wellcome Sanger Institute COVID-19 Surveillance Team |
| EPI_ISL_2515384, EPI_ISL_2515385, EPI_ISL_2515386, EPI_ISL_2515387, EPI_ISL_2515388, EPI_ISL_2515389, EPI_ISL_2515390, EPI_ISL_2515391, EPI_ISL_2515392, EPI_ISL_2515393, EPI_ISL_2515394, EPI_ISL_2515395, EPI_ISL_2515396, EPI_ISL_2515397 | | | |
| see above | Lighthouse Lab in Alderley Park | Wellcome Sanger Institute for the COVID-19 Genomics UK (COG-UK) Consortium | Jacquelyn Wynn, Mairead Hyland, The Lighthouse Lab in Alderley Park and Alex Alderton, Roberto Amato, Jeffrey Barrett, Sonia Goncalves, Ewan Harrison, David K. Jackson, Ian Johnston, Dominic Kwiatkowski, Cordelia Langford, John Sillitoe on behalf of the Wellcome Sanger Institute COVID-19 Surveillance Team |
| EPI_ISL_2515398 | Lighthouse Lab in Glasgow | Wellcome Sanger Institute for the COVID-19 Genomics UK (COG-UK) Consortium | Harper VanSteenhouse, Yumi Kasai, David Gray, Carol Clugston, Anna Dominiczak and Alex Alderton, Roberto Amato, Jeffrey Barrett, Sonia Goncalves, Ewan Harrison, David K. Jackson, Ian Johnston, Dominic Kwiatkowski, Cordelia Langford, John Sillitoe on behalf of the Wellcome Sanger Institute COVID-19 Surveillance Team |
| EPI_ISL_2515399 | Lighthouse Lab in Alderley Park | Wellcome Sanger Institute for the COVID-19 Genomics UK (COG-UK) Consortium | Jacquelyn Wynn, Mairead Hyland, The Lighthouse Lab in Alderley Park and Alex Alderton, Roberto Amato, Jeffrey Barrett, Sonia Goncalves, Ewan Harrison, David K. Jackson, Ian Johnston, Dominic Kwiatkowski, Cordelia Langford, John Sillitoe on behalf of the Wellcome Sanger Institute COVID-19 Surveillance Team |
| EPI_ISL_2515400 | Lighthouse Lab in Milton Keynes | Wellcome Sanger Institute for the COVID-19 Genomics UK (COG-UK) Consortium | The Lighthouse Lab in Milton Keynes and Alex Alderton, Roberto Amato, Jeffrey Barrett, Sonia Goncalves, Ewan Harrison, David K. Jackson, Ian Johnston, Dominic Kwiatkowski, Cordelia Langford, John Sillitoe on behalf of the Wellcome Sanger Institute COVID-19 Surveillance Team |
| EPI_ISL_2515401 | Lighthouse Lab in Alderley Park | Wellcome Sanger Institute for the COVID-19 Genomics UK (COG-UK) Consortium | Jacquelyn Wynn, Mairead Hyland, The Lighthouse Lab in Alderley Park and Alex Alderton, Roberto Amato, Jeffrey Barrett, Sonia Goncalves, Ewan Harrison, David K. Jackson, Ian Johnston, Dominic Kwiatkowski, Cordelia Langford, John Sillitoe on behalf of the Wellcome Sanger Institute COVID-19 Surveillance Team |
| EPI_ISL_2515402 | Lighthouse Laboratory Plymouth | Wellcome Sanger Institute for the COVID-19 Genomics UK (COG-UK) Consortium | Lighthouse Laboratory Plymouth and Alex Alderton, Roberto Amato, Jeffrey Barrett, Sonia Goncalves, Ewan Harrison, David K. Jackson, Ian Johnston, Dominic Kwiatkowski, Cordelia Langford, John Sillitoe on behalf of the Wellcome Sanger Institute COVID-19 Surveillance Team |
| EPI_ISL_2515403 | Lighthouse Lab in Alderley Park | Wellcome Sanger Institute for the COVID-19 Genomics UK (COG-UK) Consortium | Jacquelyn Wynn, Mairead Hyland, The Lighthouse Lab in Alderley Park and Alex Alderton, Roberto Amato, Jeffrey Barrett, Sonia Goncalves, Ewan Harrison, David K. Jackson, Ian Johnston, Dominic Kwiatkowski, Cordelia Langford, John Sillitoe on behalf of the Wellcome Sanger Institute COVID-19 Surveillance Team |
| EPI_ISL_2515404, EPI_ISL_2515405 | Lighthouse Lab in Milton Keynes | Wellcome Sanger Institute for the COVID-19 Genomics UK (COG-UK) Consortium | The Lighthouse Lab in Milton Keynes and Alex Alderton, Roberto Amato, Jeffrey Barrett, Sonia Goncalves, Ewan Harrison, David K. Jackson, Ian Johnston, Dominic Kwiatkowski, Cordelia Langford, John Sillitoe on behalf of the Wellcome Sanger Institute COVID-19 Surveillance Team |
| EPI_ISL_2515406, EPI_ISL_2515407 | Lighthouse Lab in Alderley Park | Wellcome Sanger Institute for the COVID-19 Genomics UK (COG-UK) Consortium | Jacquelyn Wynn, Mairead Hyland, The Lighthouse Lab in Alderley Park and Alex Alderton, Roberto Amato, Jeffrey Barrett, Sonia Goncalves, Ewan Harrison, David K. Jackson, Ian Johnston, Dominic Kwiatkowski, Cordelia Langford, John Sillitoe on behalf of the Wellcome Sanger Institute COVID-19 Surveillance Team |
| EPI_ISL_2515408 | Lighthouse Lab in Milton Keynes | Wellcome Sanger Institute for the COVID-19 Genomics UK (COG-UK) Consortium | The Lighthouse Lab in Milton Keynes and Alex Alderton, Roberto Amato, Jeffrey Barrett, Sonia Goncalves, Ewan Harrison, David K. Jackson, Ian Johnston, Dominic Kwiatkowski, Cordelia Langford, John Sillitoe on behalf of the Wellcome Sanger Institute COVID-19 Surveillance Team |
| EPI_ISL_2515409 | Lighthouse Lab in Alderley Park | Wellcome Sanger Institute for the COVID-19 Genomics UK (COG-UK) Consortium | Jacquelyn Wynn, Mairead Hyland, The Lighthouse Lab in Alderley Park and Alex Alderton, Roberto Amato, Jeffrey Barrett, Sonia Goncalves, Ewan Harrison, David K. Jackson, Ian Johnston, Dominic Kwiatkowski, Cordelia Langford, John Sillitoe on behalf of the Wellcome Sanger Institute COVID-19 Surveillance Team |
| EPI_ISL_2515410 | Lighthouse Lab in Milton Keynes | Wellcome Sanger Institute for the COVID-19 Genomics UK (COG-UK) Consortium | The Lighthouse Lab in Milton Keynes and Alex Alderton, Roberto Amato, Jeffrey Barrett, Sonia Goncalves, Ewan Harrison, David K. Jackson, Ian Johnston, Dominic Kwiatkowski, Cordelia Langford, John Sillitoe on behalf of the Wellcome Sanger Institute COVID-19 Surveillance Team |
| EPI_ISL_2515411, EPI_ISL_2515412, EPI_ISL_2515414 | Lighthouse Lab in Glasgow | Wellcome Sanger Institute for the COVID-19 Genomics UK (COG-UK) Consortium | Harper VanSteenhouse, Yumi Kasai, David Gray, Carol Clugston, Anna Dominiczak and Alex Alderton, Roberto Amato, Jeffrey Barrett, Sonia Goncalves, Ewan Harrison, David K. Jackson, Ian Johnston, Dominic Kwiatkowski, Cordelia Langford, John Sillitoe on behalf of the Wellcome Sanger Institute COVID-19 Surveillance Team |
| EPI_ISL_2515415, EPI_ISL_2515416 | Lighthouse Lab in Milton Keynes | Wellcome Sanger Institute for the COVID-19 Genomics UK (COG-UK) Consortium | The Lighthouse Lab in Milton Keynes and Alex Alderton, Roberto Amato, Jeffrey Barrett, Sonia Goncalves, Ewan Harrison, David K. Jackson, Ian Johnston, Dominic Kwiatkowski, Cordelia Langford, John Sillitoe on behalf of the Wellcome Sanger Institute COVID-19 Surveillance Team |
| EPI_ISL_2515417 | Lighthouse Lab in Glasgow | Wellcome Sanger Institute for the COVID-19 Genomics UK (COG-UK) Consortium | Harper VanSteenhouse, Yumi Kasai, David Gray, Carol Clugston, Anna Dominiczak and Alex Alderton, Roberto Amato, Jeffrey Barrett, Sonia Goncalves, Ewan Harrison, David K. Jackson, Ian Johnston, Dominic Kwiatkowski, Cordelia Langford, John Sillitoe on behalf of the Wellcome Sanger Institute COVID-19 Surveillance Team |
| EPI_ISL_2515418 | Lighthouse Laboratory Plymouth | Wellcome Sanger Institute for the COVID-19 Genomics UK (COG-UK) Consortium | Lighthouse Laboratory Plymouth and Alex Alderton, Roberto Amato, Jeffrey Barrett, Sonia Goncalves, Ewan Harrison, David K. Jackson, Ian Johnston, Dominic Kwiatkowski, Cordelia Langford, John Sillitoe on behalf of the Wellcome Sanger Institute COVID-19 Surveillance Team |
| EPI_ISL_2515420 | Lighthouse Lab in Milton Keynes | Wellcome Sanger Institute for the COVID-19 Genomics UK (COG-UK) Consortium | The Lighthouse Lab in Milton Keynes and Alex Alderton, Roberto Amato, Jeffrey Barrett, Sonia Goncalves, Ewan Harrison, David K. Jackson, Ian Johnston, Dominic Kwiatkowski, Cordelia Langford, John Sillitoe on behalf of the Wellcome Sanger Institute COVID-19 Surveillance Team |
| EPI_ISL_2515421 | Lighthouse Lab in Alderley Park | Wellcome Sanger Institute for the COVID-19 Genomics UK (COG-UK) Consortium | Jacquelyn Wynn, Mairead Hyland, The Lighthouse Lab in Alderley Park and Alex Alderton, Roberto Amato, Jeffrey Barrett, Sonia Goncalves, Ewan Harrison, David K. Jackson, Ian Johnston, Dominic Kwiatkowski, Cordelia Langford, John Sillitoe on behalf of the Wellcome Sanger Institute COVID-19 Surveillance Team |
| EPI_ISL_2515422, EPI_ISL_2515423 | Lighthouse Lab in Glasgow | Wellcome Sanger Institute for the COVID-19 Genomics UK (COG-UK) Consortium | Harper VanSteenhouse, Yumi Kasai, David Gray, Carol Clugston, Anna Dominiczak and Alex Alderton, Roberto Amato, Jeffrey Barrett, Sonia Goncalves, Ewan Harrison, David K. Jackson, Ian Johnston, Dominic Kwiatkowski, Cordelia Langford, John Sillitoe on behalf of the Wellcome Sanger Institute COVID-19 Surveillance Team |
| EPI_ISL_2515424, EPI_ISL_2515425 | Lighthouse Lab in Alderley Park | Wellcome Sanger Institute for the COVID-19 Genomics UK (COG-UK) Consortium | Jacquelyn Wynn, Mairead Hyland, The Lighthouse Lab in Alderley Park and Alex Alderton, Roberto Amato, Jeffrey Barrett, Sonia Goncalves, Ewan Harrison, David K. Jackson, Ian Johnston, Dominic Kwiatkowski, Cordelia Langford, John Sillitoe on behalf of the Wellcome Sanger Institute COVID-19 Surveillance Team |
| EPI_ISL_2515426 | Lighthouse Lab in Milton Keynes | Wellcome Sanger Institute for the COVID-19 Genomics UK (COG-UK) Consortium | The Lighthouse Lab in Milton Keynes and Alex Alderton, Roberto Amato, Jeffrey Barrett, Sonia Goncalves, Ewan Harrison, David K. Jackson, Ian Johnston, Dominic Kwiatkowski, Cordelia Langford, John Sillitoe on behalf of the Wellcome Sanger Institute COVID-19 Surveillance Team |
| EPI_ISL_2515427, EPI_ISL_2515428, EPI_ISL_2515429 | Lighthouse Lab in Alderley Park | Wellcome Sanger Institute for the COVID-19 Genomics UK (COG-UK) Consortium | Jacquelyn Wynn, Mairead Hyland, The Lighthouse Lab in Alderley Park and Alex Alderton, Roberto Amato, Jeffrey Barrett, Sonia Goncalves, Ewan Harrison, David K. Jackson, Ian Johnston, Dominic Kwiatkowski, Cordelia Langford, John Sillitoe on behalf of the Wellcome Sanger Institute COVID-19 Surveillance Team |

| | | | |
|---|---|---|---|
| EPI_ISL_2515430, EPI_ISL_2515431 | Lighthouse Lab in Milton Keynes | Wellcome Sanger Institute for the COVID-19 Genomics UK (COG-UK) Consortium | The Lighthouse Lab in Milton Keynes and Alex Alderton, Roberto Amato, Jeffrey Barrett, Sonia Goncalves, Ewan Harrison, David K. Jackson, Ian Johnston, Dominic Kwiatkowski, Cordelia Langford, John Sillitoe on behalf of the Wellcome Sanger Institute COVID-19 Surveillance Team |
| EPI_ISL_2515432 | Lighthouse Laboratory Plymouth | Wellcome Sanger Institute for the COVID-19 Genomics UK (COG-UK) Consortium | Lighthouse Laboratory Plymouth and Alex Alderton, Roberto Amato, Jeffrey Barrett, Sonia Goncalves, Ewan Harrison, David K. Jackson, Ian Johnston, Dominic Kwiatkowski, Cordelia Langford, John Sillitoe on behalf of the Wellcome Sanger Institute COVID-19 Surveillance Team |
| EPI_ISL_2515433 | Lighthouse Lab in Milton Keynes | Wellcome Sanger Institute for the COVID-19 Genomics UK (COG-UK) Consortium | The Lighthouse Lab in Milton Keynes and Alex Alderton, Roberto Amato, Jeffrey Barrett, Sonia Goncalves, Ewan Harrison, David K. Jackson, Ian Johnston, Dominic Kwiatkowski, Cordelia Langford, John Sillitoe on behalf of the Wellcome Sanger Institute COVID-19 Surveillance Team |
| EPI_ISL_2515434 | Lighthouse Lab in Glasgow | Wellcome Sanger Institute for the COVID-19 Genomics UK (COG-UK) Consortium | Harper VanSteenhouse, Yumi Kasai, David Gray, Carol Clugston, Anna Dominiczak and Alex Alderton, Roberto Amato, Jeffrey Barrett, Sonia Goncalves, Ewan Harrison, David K. Jackson, Ian Johnston, Dominic Kwiatkowski, Cordelia Langford, John Sillitoe on behalf of the Wellcome Sanger Institute COVID-19 Surveillance Team |
| EPI_ISL_2515435 | Lighthouse Lab in Alderley Park | Wellcome Sanger Institute for the COVID-19 Genomics UK (COG-UK) Consortium | Jacquelyn Wynn, Mairead Hyland, The Lighthouse Lab in Alderley Park and Alex Alderton, Roberto Amato, Jeffrey Barrett, Sonia Goncalves, Ewan Harrison, David K. Jackson, Ian Johnston, Dominic Kwiatkowski, Cordelia Langford, John Sillitoe on behalf of the Wellcome Sanger Institute COVID-19 Surveillance Team |
| EPI_ISL_2515436 | Lighthouse Lab in Glasgow | Wellcome Sanger Institute for the COVID-19 Genomics UK (COG-UK) Consortium | Harper VanSteenhouse, Yumi Kasai, David Gray, Carol Clugston, Anna Dominiczak and Alex Alderton, Roberto Amato, Jeffrey Barrett, Sonia Goncalves, Ewan Harrison, David K. Jackson, Ian Johnston, Dominic Kwiatkowski, Cordelia Langford, John Sillitoe on behalf of the Wellcome Sanger Institute COVID-19 Surveillance Team |
| EPI_ISL_2515437 | Lighthouse Lab in Milton Keynes | Wellcome Sanger Institute for the COVID-19 Genomics UK (COG-UK) Consortium | The Lighthouse Lab in Milton Keynes and Alex Alderton, Roberto Amato, Jeffrey Barrett, Sonia Goncalves, Ewan Harrison, David K. Jackson, Ian Johnston, Dominic Kwiatkowski, Cordelia Langford, John Sillitoe on behalf of the Wellcome Sanger Institute COVID-19 Surveillance Team |
| EPI_ISL_2515438 | Lighthouse Lab in Alderley Park | Wellcome Sanger Institute for the COVID-19 Genomics UK (COG-UK) Consortium | Jacquelyn Wynn, Mairead Hyland, The Lighthouse Lab in Alderley Park and Alex Alderton, Roberto Amato, Jeffrey Barrett, Sonia Goncalves, Ewan Harrison, David K. Jackson, Ian Johnston, Dominic Kwiatkowski, Cordelia Langford, John Sillitoe on behalf of the Wellcome Sanger Institute COVID-19 Surveillance Team |
| EPI_ISL_2515439 | Lighthouse Lab in Milton Keynes | Wellcome Sanger Institute for the COVID-19 Genomics UK (COG-UK) Consortium | The Lighthouse Lab in Milton Keynes and Alex Alderton, Roberto Amato, Jeffrey Barrett, Sonia Goncalves, Ewan Harrison, David K. Jackson, Ian Johnston, Dominic Kwiatkowski, Cordelia Langford, John Sillitoe on behalf of the Wellcome Sanger Institute COVID-19 Surveillance Team |
| EPI_ISL_2515440 | Lighthouse Lab in Alderley Park | Wellcome Sanger Institute for the COVID-19 Genomics UK (COG-UK) Consortium | Jacquelyn Wynn, Mairead Hyland, The Lighthouse Lab in Alderley Park and Alex Alderton, Roberto Amato, Jeffrey Barrett, Sonia Goncalves, Ewan Harrison, David K. Jackson, Ian Johnston, Dominic Kwiatkowski, Cordelia Langford, John Sillitoe on behalf of the Wellcome Sanger Institute COVID-19 Surveillance Team |
| EPI_ISL_2515441 | Lighthouse Lab in Glasgow | Wellcome Sanger Institute for the COVID-19 Genomics UK (COG-UK) Consortium | Harper VanSteenhouse, Yumi Kasai, David Gray, Carol Clugston, Anna Dominiczak and Alex Alderton, Roberto Amato, Jeffrey Barrett, Sonia Goncalves, Ewan Harrison, David K. Jackson, Ian Johnston, Dominic Kwiatkowski, Cordelia Langford, John Sillitoe on behalf of the Wellcome Sanger Institute COVID-19 Surveillance Team |
| EPI_ISL_2515443, EPI_ISL_2515445, EPI_ISL_2515446, EPI_ISL_2515447 | Lighthouse Lab in Alderley Park | Wellcome Sanger Institute for the COVID-19 Genomics UK (COG-UK) Consortium | Jacquelyn Wynn, Mairead Hyland, The Lighthouse Lab in Alderley Park and Alex Alderton, Roberto Amato, Jeffrey Barrett, Sonia Goncalves, Ewan Harrison, David K. Jackson, Ian Johnston, Dominic Kwiatkowski, Cordelia Langford, John Sillitoe on behalf of the Wellcome Sanger Institute COVID-19 Surveillance Team |
| EPI_ISL_2515448 | Lighthouse Laboratory Plymouth | Wellcome Sanger Institute for the COVID-19 Genomics UK (COG-UK) Consortium | Lighthouse Laboratory Plymouth and Alex Alderton, Roberto Amato, Jeffrey Barrett, Sonia Goncalves, Ewan Harrison, David K. Jackson, Ian Johnston, Dominic Kwiatkowski, Cordelia Langford, John Sillitoe on behalf of the Wellcome Sanger Institute COVID-19 Surveillance Team |
| EPI_ISL_2515449, EPI_ISL_2515450 | Lighthouse Lab in Glasgow | Wellcome Sanger Institute for the COVID-19 Genomics UK (COG-UK) Consortium | Harper VanSteenhouse, Yumi Kasai, David Gray, Carol Clugston, Anna Dominiczak and Alex Alderton, Roberto Amato, Jeffrey Barrett, Sonia Goncalves, Ewan Harrison, David K. Jackson, Ian Johnston, Dominic Kwiatkowski, Cordelia Langford, John Sillitoe on behalf of the Wellcome Sanger Institute COVID-19 Surveillance Team |
| EPI_ISL_2515451 | Lighthouse Lab in Milton Keynes | Wellcome Sanger Institute for the COVID-19 Genomics UK (COG-UK) Consortium | The Lighthouse Lab in Milton Keynes and Alex Alderton, Roberto Amato, Jeffrey Barrett, Sonia Goncalves, Ewan Harrison, David K. Jackson, Ian Johnston, Dominic Kwiatkowski, Cordelia Langford, John Sillitoe on behalf of the Wellcome Sanger Institute COVID-19 Surveillance Team |
| EPI_ISL_2515452 | Lighthouse Lab in Alderley Park | Wellcome Sanger Institute for the COVID-19 Genomics UK (COG-UK) Consortium | Jacquelyn Wynn, Mairead Hyland, The Lighthouse Lab in Alderley Park and Alex Alderton, Roberto Amato, Jeffrey Barrett, Sonia Goncalves, Ewan Harrison, David K. Jackson, Ian Johnston, Dominic Kwiatkowski, Cordelia Langford, John Sillitoe on behalf of the Wellcome Sanger Institute COVID-19 Surveillance Team |
| EPI_ISL_2515453 | Lighthouse Lab in Glasgow | Wellcome Sanger Institute for the COVID-19 Genomics UK (COG-UK) Consortium | Harper VanSteenhouse, Yumi Kasai, David Gray, Carol Clugston, Anna Dominiczak and Alex Alderton, Roberto Amato, Jeffrey Barrett, Sonia Goncalves, Ewan Harrison, David K. Jackson, Ian Johnston, Dominic Kwiatkowski, Cordelia Langford, John Sillitoe on behalf of the Wellcome Sanger Institute COVID-19 Surveillance Team |
| EPI_ISL_2515454 | Lighthouse Lab in Alderley Park | Wellcome Sanger Institute for the COVID-19 Genomics UK (COG-UK) Consortium | Jacquelyn Wynn, Mairead Hyland, The Lighthouse Lab in Alderley Park and Alex Alderton, Roberto Amato, Jeffrey Barrett, Sonia Goncalves, Ewan Harrison, David K. Jackson, Ian Johnston, Dominic Kwiatkowski, Cordelia Langford, John Sillitoe on behalf of the Wellcome Sanger Institute COVID-19 Surveillance Team |
| EPI_ISL_2515455, EPI_ISL_2515456 | Lighthouse Lab in Glasgow | Wellcome Sanger Institute for the COVID-19 Genomics UK (COG-UK) Consortium | Harper VanSteenhouse, Yumi Kasai, David Gray, Carol Clugston, Anna Dominiczak and Alex Alderton, Roberto Amato, Jeffrey Barrett, Sonia Goncalves, Ewan Harrison, David K. Jackson, Ian Johnston, Dominic Kwiatkowski, Cordelia Langford, John Sillitoe on behalf of the Wellcome Sanger Institute COVID-19 Surveillance Team |
| EPI_ISL_2515457, EPI_ISL_2515458, EPI_ISL_2515459 | Lighthouse Lab in Milton Keynes | Wellcome Sanger Institute for the COVID-19 Genomics UK (COG-UK) Consortium | The Lighthouse Lab in Milton Keynes and Alex Alderton, Roberto Amato, Jeffrey Barrett, Sonia Goncalves, Ewan Harrison, David K. Jackson, Ian Johnston, Dominic Kwiatkowski, Cordelia Langford, John Sillitoe on behalf of the Wellcome Sanger Institute COVID-19 Surveillance Team |
| EPI_ISL_2515460 | Lighthouse Lab in Glasgow | Wellcome Sanger Institute for the COVID-19 Genomics UK (COG-UK) Consortium | Harper VanSteenhouse, Yumi Kasai, David Gray, Carol Clugston, Anna Dominiczak and Alex Alderton, Roberto Amato, Jeffrey Barrett, Sonia Goncalves, Ewan Harrison, David K. Jackson, Ian Johnston, Dominic Kwiatkowski, Cordelia Langford, John Sillitoe on behalf of the Wellcome Sanger Institute COVID-19 Surveillance Team |
| EPI_ISL_2515462 | Lighthouse Lab in Alderley Park | Wellcome Sanger Institute for the COVID-19 Genomics UK (COG-UK) Consortium | Jacquelyn Wynn, Mairead Hyland, The Lighthouse Lab in Alderley Park and Alex Alderton, Roberto Amato, Jeffrey Barrett, Sonia Goncalves, Ewan Harrison, David K. Jackson, Ian Johnston, Dominic Kwiatkowski, Cordelia Langford, John Sillitoe on behalf of the Wellcome Sanger Institute COVID-19 Surveillance Team |
| EPI_ISL_2515463, EPI_ISL_2515464 | Lighthouse Lab in Glasgow | Wellcome Sanger Institute for the COVID-19 Genomics UK (COG-UK) Consortium | Harper VanSteenhouse, Yumi Kasai, David Gray, Carol Clugston, Anna Dominiczak and Alex Alderton, Roberto Amato, Jeffrey Barrett, Sonia Goncalves, Ewan Harrison, David K. Jackson, Ian Johnston, Dominic Kwiatkowski, Cordelia Langford, John Sillitoe on behalf of the Wellcome Sanger Institute COVID-19 Surveillance Team |
| EPI_ISL_2515465 | Lighthouse Lab in Alderley Park | Wellcome Sanger Institute for the COVID-19 Genomics UK (COG-UK) Consortium | Jacquelyn Wynn, Mairead Hyland, The Lighthouse Lab in Alderley Park and Alex Alderton, Roberto Amato, Jeffrey Barrett, Sonia Goncalves, Ewan Harrison, David K. Jackson, Ian Johnston, Dominic Kwiatkowski, Cordelia Langford, John Sillitoe on behalf of the Wellcome Sanger Institute COVID-19 Surveillance Team |
| EPI_ISL_2515466, EPI_ISL_2515467 | Lighthouse Lab in Milton Keynes | Wellcome Sanger Institute for the COVID-19 Genomics UK (COG-UK) Consortium | The Lighthouse Lab in Milton Keynes and Alex Alderton, Roberto Amato, Jeffrey Barrett, Sonia Goncalves, Ewan Harrison, David K. Jackson, Ian Johnston, Dominic Kwiatkowski, Cordelia Langford, John Sillitoe on behalf of the Wellcome Sanger Institute COVID-19 Surveillance Team |
| EPI_ISL_2515468 | Lighthouse Lab in Alderley Park | Wellcome Sanger Institute for the COVID-19 Genomics UK (COG-UK) Consortium | Jacquelyn Wynn, Mairead Hyland, The Lighthouse Lab in Alderley Park and Alex Alderton, Roberto Amato, Jeffrey Barrett, Sonia Goncalves, Ewan Harrison, David K. Jackson, Ian Johnston, Dominic Kwiatkowski, Cordelia Langford, John Sillitoe on behalf of the Wellcome Sanger Institute COVID-19 Surveillance Team |
| EPI_ISL_2515469 | Lighthouse Laboratory Plymouth | Wellcome Sanger Institute for the COVID-19 Genomics UK (COG-UK) Consortium | Lighthouse Laboratory Plymouth and Alex Alderton, Roberto Amato, Jeffrey Barrett, Sonia Goncalves, Ewan Harrison, David K. Jackson, Ian Johnston, Dominic Kwiatkowski, Cordelia Langford, John Sillitoe on behalf of the Wellcome Sanger Institute COVID-19 Surveillance Team |
| EPI_ISL_2515471 | Lighthouse Lab in Milton Keynes | Wellcome Sanger Institute for the COVID-19 Genomics UK (COG-UK) Consortium | The Lighthouse Lab in Milton Keynes and Alex Alderton, Roberto Amato, Jeffrey Barrett, Sonia Goncalves, Ewan Harrison, David K. Jackson, Ian Johnston, Dominic Kwiatkowski, Cordelia Langford, John Sillitoe on behalf of the Wellcome Sanger Institute COVID-19 Surveillance Team |
| EPI_ISL_2515472 | Lighthouse Laboratory Plymouth | Wellcome Sanger Institute for the COVID-19 Genomics UK (COG-UK) Consortium | Lighthouse Laboratory Plymouth and Alex Alderton, Roberto Amato, Jeffrey Barrett, Sonia Goncalves, Ewan Harrison, David K. Jackson, Ian Johnston, Dominic Kwiatkowski, Cordelia Langford, John Sillitoe on behalf of the Wellcome Sanger Institute COVID-19 Surveillance Team |
| EPI_ISL_2515473 | Lighthouse Lab in Milton Keynes | Wellcome Sanger Institute for the COVID-19 Genomics UK (COG-UK) Consortium | The Lighthouse Lab in Milton Keynes and Alex Alderton, Roberto Amato, Jeffrey Barrett, Sonia Goncalves, Ewan Harrison, David K. Jackson, Ian Johnston, Dominic Kwiatkowski, Cordelia Langford, John Sillitoe on behalf of the Wellcome Sanger Institute COVID-19 Surveillance Team |
| EPI_ISL_2515474 | Lighthouse Laboratory Plymouth | Wellcome Sanger Institute for the COVID-19 Genomics UK (COG-UK) Consortium | Lighthouse Laboratory Plymouth and Alex Alderton, Roberto Amato, Jeffrey Barrett, Sonia Goncalves, Ewan Harrison, David K. Jackson, Ian Johnston, Dominic Kwiatkowski, Cordelia Langford, John Sillitoe on behalf of the Wellcome Sanger Institute COVID-19 Surveillance Team |
| EPI_ISL_2515475 | Lighthouse Lab in Alderley Park | Wellcome Sanger Institute for the COVID-19 Genomics UK (COG-UK) Consortium | Jacquelyn Wynn, Mairead Hyland, The Lighthouse Lab in Alderley Park and Alex Alderton, Roberto Amato, Jeffrey Barrett, Sonia Goncalves, Ewan Harrison, David K. Jackson, Ian Johnston, Dominic Kwiatkowski, Cordelia Langford, John Sillitoe on behalf of the Wellcome Sanger Institute COVID-19 Surveillance Team |
| EPI_ISL_2515476 | Lighthouse Laboratory Plymouth | Wellcome Sanger Institute for the COVID-19 Genomics UK (COG-UK) Consortium | Lighthouse Laboratory Plymouth and Alex Alderton, Roberto Amato, Jeffrey Barrett, Sonia Goncalves, Ewan Harrison, David K. Jackson, Ian Johnston, Dominic Kwiatkowski, Cordelia Langford, John Sillitoe on behalf of the Wellcome Sanger Institute COVID-19 Surveillance Team |
| EPI_ISL_2515477 | Lighthouse Lab in Glasgow | Wellcome Sanger Institute for the COVID-19 Genomics UK (COG-UK) Consortium | Harper VanSteenhouse, Yumi Kasai, David Gray, Carol Clugston, Anna Dominiczak and Alex Alderton, Roberto Amato, Jeffrey Barrett, Sonia Goncalves, Ewan Harrison, David K. Jackson, Ian Johnston, Dominic Kwiatkowski, Cordelia Langford, John Sillitoe on behalf of the Wellcome Sanger Institute COVID-19 Surveillance Team |
| EPI_ISL_2515478 | Lighthouse Lab in Alderley Park | Wellcome Sanger Institute for the COVID-19 Genomics UK (COG-UK) Consortium | Jacquelyn Wynn, Mairead Hyland, The Lighthouse Lab in Alderley Park and Alex Alderton, Roberto Amato, Jeffrey Barrett, Sonia Goncalves, Ewan Harrison, David K. Jackson, Ian Johnston, Dominic Kwiatkowski, Cordelia Langford, John Sillitoe on behalf of the Wellcome Sanger Institute COVID-19 Surveillance Team |
| EPI_ISL_2515479 | Lighthouse Lab in Glasgow | Wellcome Sanger Institute for the COVID-19 Genomics UK (COG-UK) Consortium | Harper VanSteenhouse, Yumi Kasai, David Gray, Carol Clugston, Anna Dominiczak and Alex Alderton, Roberto Amato, Jeffrey Barrett, Sonia Goncalves, Ewan Harrison, David K. Jackson, Ian Johnston, Dominic Kwiatkowski, Cordelia Langford, John Sillitoe on behalf of the Wellcome Sanger Institute COVID-19 Surveillance Team |
| EPI_ISL_2515480, EPI_ISL_2515481 | Lighthouse Lab in Milton Keynes | Wellcome Sanger Institute for the COVID-19 Genomics UK (COG-UK) Consortium | The Lighthouse Lab in Milton Keynes and Alex Alderton, Roberto Amato, Jeffrey Barrett, Sonia Goncalves, Ewan Harrison, David K. Jackson, Ian Johnston, Dominic Kwiatkowski, Cordelia Langford, John Sillitoe on behalf of the Wellcome Sanger Institute COVID-19 Surveillance Team |
| EPI_ISL_2515482, EPI_ISL_2515483, EPI_ISL_2515484 | Lighthouse Lab in Alderley Park | Wellcome Sanger Institute for the COVID-19 Genomics UK (COG-UK) Consortium | Jacquelyn Wynn, Mairead Hyland, The Lighthouse Lab in Alderley Park and Alex Alderton, Roberto Amato, Jeffrey Barrett, Sonia Goncalves, Ewan Harrison, David K. Jackson, Ian Johnston, Dominic Kwiatkowski, Cordelia Langford, John Sillitoe on behalf of the Wellcome Sanger Institute COVID-19 Surveillance Team |
| EPI_ISL_2515485, EPI_ISL_2515486, EPI_ISL_2515489, EPI_ISL_2515490 | Lighthouse Lab in Milton Keynes | Wellcome Sanger Institute for the COVID-19 Genomics UK (COG-UK) Consortium | The Lighthouse Lab in Milton Keynes and Alex Alderton, Roberto Amato, Jeffrey Barrett, Sonia Goncalves, Ewan Harrison, David K. Jackson, Ian Johnston, Dominic Kwiatkowski, Cordelia Langford, John Sillitoe on behalf of the Wellcome Sanger Institute COVID-19 Surveillance Team |
| EPI_ISL_2515491, EPI_ISL_2515492, EPI_ISL_2515493, EPI_ISL_2515494, EPI_ISL_2515495 | Lighthouse Lab in Alderley Park | Wellcome Sanger Institute for the COVID-19 Genomics UK (COG-UK) Consortium | Jacquelyn Wynn, Mairead Hyland, The Lighthouse Lab in Alderley Park and Alex Alderton, Roberto Amato, Jeffrey Barrett, Sonia Goncalves, Ewan Harrison, David K. Jackson, Ian Johnston, Dominic Kwiatkowski, Cordelia Langford, John Sillitoe on behalf of the Wellcome Sanger Institute COVID-19 Surveillance Team |
| EPI_ISL_2515496 | Lighthouse Lab in Glasgow | Wellcome Sanger Institute for the COVID-19 Genomics UK (COG-UK) Consortium | Harper VanSteenhouse, Yumi Kasai, David Gray, Carol Clugston, Anna Dominiczak and Alex Alderton, Roberto Amato, Jeffrey Barrett, Sonia Goncalves, Ewan Harrison, David K. Jackson, Ian Johnston, Dominic Kwiatkowski, Cordelia Langford, John Sillitoe on behalf of the Wellcome Sanger Institute COVID-19 Surveillance Team |
| EPI_ISL_2515498, | Lighthouse Lab in Milton Keynes | Wellcome Sanger Institute for the COVID-19 Genomics UK (COG-UK) | The Lighthouse Lab in Milton Keynes and Alex Alderton, Roberto Amato, Jeffrey Barrett, Sonia Goncalves, Ewan Harrison, David K. Jackson, Ian Johnston, Dominic Kwiatkowski, Cordelia Langford, John Sillitoe on behalf of the Wellcome Sanger |

| | | | |
|---|---|---|---|
| EPI_ISL_2515499 | | Consortium | Institute COVID-19 Surveillance Team |
| EPI_ISL_2515501 | Lighthouse Lab in Alderley Park | Wellcome Sanger Institute for the COVID-19 Genomics UK (COG-UK) Consortium | Jacquelyn Wynn, Mairead Hyland, The Lighthouse Lab in Alderley Park and Alex Alderton, Roberto Amato, Jeffrey Barrett, Sonia Goncalves, Ewan Harrison, David K. Jackson, Ian Johnston, Dominic Kwiatkowski, Cordelia Langford, John Sillitoe on behalf of the Wellcome Sanger Institute COVID-19 Surveillance Team |
| EPI_ISL_2515502, EPI_ISL_2515503, EPI_ISL_2515504 | Lighthouse Lab in Glasgow | Wellcome Sanger Institute for the COVID-19 Genomics UK (COG-UK) Consortium | Harper VanSteenhouse, Yumi Kasai, David Gray, Carol Clugston, Anna Dominiczak and Alex Alderton, Roberto Amato, Jeffrey Barrett, Sonia Goncalves, Ewan Harrison, David K. Jackson, Ian Johnston, Dominic Kwiatkowski, Cordelia Langford, John Sillitoe on behalf of the Wellcome Sanger Institute COVID-19 Surveillance Team |
| EPI_ISL_2515505 | Lighthouse Lab in Milton Keynes | Wellcome Sanger Institute for the COVID-19 Genomics UK (COG-UK) Consortium | The Lighthouse Lab in Milton Keynes and Alex Alderton, Roberto Amato, Jeffrey Barrett, Sonia Goncalves, Ewan Harrison, David K. Jackson, Ian Johnston, Dominic Kwiatkowski, Cordelia Langford, John Sillitoe on behalf of the Wellcome Sanger Institute COVID-19 Surveillance Team |
| EPI_ISL_2515506 | Lighthouse Lab in Glasgow | Wellcome Sanger Institute for the COVID-19 Genomics UK (COG-UK) Consortium | Harper VanSteenhouse, Yumi Kasai, David Gray, Carol Clugston, Anna Dominiczak and Alex Alderton, Roberto Amato, Jeffrey Barrett, Sonia Goncalves, Ewan Harrison, David K. Jackson, Ian Johnston, Dominic Kwiatkowski, Cordelia Langford, John Sillitoe on behalf of the Wellcome Sanger Institute COVID-19 Surveillance Team |
| EPI_ISL_2515507 | Lighthouse Lab in Alderley Park | Wellcome Sanger Institute for the COVID-19 Genomics UK (COG-UK) Consortium | Jacquelyn Wynn, Mairead Hyland, The Lighthouse Lab in Alderley Park and Alex Alderton, Roberto Amato, Jeffrey Barrett, Sonia Goncalves, Ewan Harrison, David K. Jackson, Ian Johnston, Dominic Kwiatkowski, Cordelia Langford, John Sillitoe on behalf of the Wellcome Sanger Institute COVID-19 Surveillance Team |
| EPI_ISL_2515508 | Lighthouse Lab in Milton Keynes | Wellcome Sanger Institute for the COVID-19 Genomics UK (COG-UK) Consortium | The Lighthouse Lab in Milton Keynes and Alex Alderton, Roberto Amato, Jeffrey Barrett, Sonia Goncalves, Ewan Harrison, David K. Jackson, Ian Johnston, Dominic Kwiatkowski, Cordelia Langford, John Sillitoe on behalf of the Wellcome Sanger Institute COVID-19 Surveillance Team |
| EPI_ISL_2515509 | Lighthouse Lab in Alderley Park | Wellcome Sanger Institute for the COVID-19 Genomics UK (COG-UK) Consortium | Jacquelyn Wynn, Mairead Hyland, The Lighthouse Lab in Alderley Park and Alex Alderton, Roberto Amato, Jeffrey Barrett, Sonia Goncalves, Ewan Harrison, David K. Jackson, Ian Johnston, Dominic Kwiatkowski, Cordelia Langford, John Sillitoe on behalf of the Wellcome Sanger Institute COVID-19 Surveillance Team |
| EPI_ISL_2515510 | Lighthouse Lab in Milton Keynes | Wellcome Sanger Institute for the COVID-19 Genomics UK (COG-UK) Consortium | The Lighthouse Lab in Milton Keynes and Alex Alderton, Roberto Amato, Jeffrey Barrett, Sonia Goncalves, Ewan Harrison, David K. Jackson, Ian Johnston, Dominic Kwiatkowski, Cordelia Langford, John Sillitoe on behalf of the Wellcome Sanger Institute COVID-19 Surveillance Team |
| EPI_ISL_2515511 | Lighthouse Laboratory Plymouth | Wellcome Sanger Institute for the COVID-19 Genomics UK (COG-UK) Consortium | Lighthouse Laboratory Plymouth and Alex Alderton, Roberto Amato, Jeffrey Barrett, Sonia Goncalves, Ewan Harrison, David K. Jackson, Ian Johnston, Dominic Kwiatkowski, Cordelia Langford, John Sillitoe on behalf of the Wellcome Sanger Institute COVID-19 Surveillance Team |
| EPI_ISL_2515512 | Lighthouse Lab in Alderley Park | Wellcome Sanger Institute for the COVID-19 Genomics UK (COG-UK) Consortium | Jacquelyn Wynn, Mairead Hyland, The Lighthouse Lab in Alderley Park and Alex Alderton, Roberto Amato, Jeffrey Barrett, Sonia Goncalves, Ewan Harrison, David K. Jackson, Ian Johnston, Dominic Kwiatkowski, Cordelia Langford, John Sillitoe on behalf of the Wellcome Sanger Institute COVID-19 Surveillance Team |
| EPI_ISL_2515513 | Lighthouse Lab in Milton Keynes | Wellcome Sanger Institute for the COVID-19 Genomics UK (COG-UK) Consortium | The Lighthouse Lab in Milton Keynes and Alex Alderton, Roberto Amato, Jeffrey Barrett, Sonia Goncalves, Ewan Harrison, David K. Jackson, Ian Johnston, Dominic Kwiatkowski, Cordelia Langford, John Sillitoe on behalf of the Wellcome Sanger Institute COVID-19 Surveillance Team |
| EPI_ISL_2515514, EPI_ISL_2515515 | Lighthouse Lab in Alderley Park | Wellcome Sanger Institute for the COVID-19 Genomics UK (COG-UK) Consortium | Jacquelyn Wynn, Mairead Hyland, The Lighthouse Lab in Alderley Park and Alex Alderton, Roberto Amato, Jeffrey Barrett, Sonia Goncalves, Ewan Harrison, David K. Jackson, Ian Johnston, Dominic Kwiatkowski, Cordelia Langford, John Sillitoe on behalf of the Wellcome Sanger Institute COVID-19 Surveillance Team |
| EPI_ISL_2515516 | Lighthouse Lab in Glasgow | Wellcome Sanger Institute for the COVID-19 Genomics UK (COG-UK) Consortium | Harper VanSteenhouse, Yumi Kasai, David Gray, Carol Clugston, Anna Dominiczak and Alex Alderton, Roberto Amato, Jeffrey Barrett, Sonia Goncalves, Ewan Harrison, David K. Jackson, Ian Johnston, Dominic Kwiatkowski, Cordelia Langford, John Sillitoe on behalf of the Wellcome Sanger Institute COVID-19 Surveillance Team |
| EPI_ISL_2515517 | Lighthouse Lab in Alderley Park | Wellcome Sanger Institute for the COVID-19 Genomics UK (COG-UK) Consortium | Jacquelyn Wynn, Mairead Hyland, The Lighthouse Lab in Alderley Park and Alex Alderton, Roberto Amato, Jeffrey Barrett, Sonia Goncalves, Ewan Harrison, David K. Jackson, Ian Johnston, Dominic Kwiatkowski, Cordelia Langford, John Sillitoe on behalf of the Wellcome Sanger Institute COVID-19 Surveillance Team |
| EPI_ISL_2515518, EPI_ISL_2515519, EPI_ISL_2515520, EPI_ISL_2515521 | Lighthouse Lab in Milton Keynes | Wellcome Sanger Institute for the COVID-19 Genomics UK (COG-UK) Consortium | The Lighthouse Lab in Milton Keynes and Alex Alderton, Roberto Amato, Jeffrey Barrett, Sonia Goncalves, Ewan Harrison, David K. Jackson, Ian Johnston, Dominic Kwiatkowski, Cordelia Langford, John Sillitoe on behalf of the Wellcome Sanger Institute COVID-19 Surveillance Team |
| EPI_ISL_2515522 | Lighthouse Lab in Alderley Park | Wellcome Sanger Institute for the COVID-19 Genomics UK (COG-UK) Consortium | Jacquelyn Wynn, Mairead Hyland, The Lighthouse Lab in Alderley Park and Alex Alderton, Roberto Amato, Jeffrey Barrett, Sonia Goncalves, Ewan Harrison, David K. Jackson, Ian Johnston, Dominic Kwiatkowski, Cordelia Langford, John Sillitoe on behalf of the Wellcome Sanger Institute COVID-19 Surveillance Team |
| EPI_ISL_2515523 | Lighthouse Lab in Milton Keynes | Wellcome Sanger Institute for the COVID-19 Genomics UK (COG-UK) Consortium | The Lighthouse Lab in Milton Keynes and Alex Alderton, Roberto Amato, Jeffrey Barrett, Sonia Goncalves, Ewan Harrison, David K. Jackson, Ian Johnston, Dominic Kwiatkowski, Cordelia Langford, John Sillitoe on behalf of the Wellcome Sanger Institute COVID-19 Surveillance Team |
| EPI_ISL_2515524 | Lighthouse Lab in Alderley Park | Wellcome Sanger Institute for the COVID-19 Genomics UK (COG-UK) Consortium | Jacquelyn Wynn, Mairead Hyland, The Lighthouse Lab in Alderley Park and Alex Alderton, Roberto Amato, Jeffrey Barrett, Sonia Goncalves, Ewan Harrison, David K. Jackson, Ian Johnston, Dominic Kwiatkowski, Cordelia Langford, John Sillitoe on behalf of the Wellcome Sanger Institute COVID-19 Surveillance Team |
| EPI_ISL_2515525 | Lighthouse Lab in Glasgow | Wellcome Sanger Institute for the COVID-19 Genomics UK (COG-UK) Consortium | Harper VanSteenhouse, Yumi Kasai, David Gray, Carol Clugston, Anna Dominiczak and Alex Alderton, Roberto Amato, Jeffrey Barrett, Sonia Goncalves, Ewan Harrison, David K. Jackson, Ian Johnston, Dominic Kwiatkowski, Cordelia Langford, John Sillitoe on behalf of the Wellcome Sanger Institute COVID-19 Surveillance Team |
| EPI_ISL_2515527 | Lighthouse Lab in Alderley Park | Wellcome Sanger Institute for the COVID-19 Genomics UK (COG-UK) Consortium | Jacquelyn Wynn, Mairead Hyland, The Lighthouse Lab in Alderley Park and Alex Alderton, Roberto Amato, Jeffrey Barrett, Sonia Goncalves, Ewan Harrison, David K. Jackson, Ian Johnston, Dominic Kwiatkowski, Cordelia Langford, John Sillitoe on behalf of the Wellcome Sanger Institute COVID-19 Surveillance Team |
| EPI_ISL_2515528 | Lighthouse Laboratory Plymouth | Wellcome Sanger Institute for the COVID-19 Genomics UK (COG-UK) Consortium | Lighthouse Laboratory Plymouth and Alex Alderton, Roberto Amato, Jeffrey Barrett, Sonia Goncalves, Ewan Harrison, David K. Jackson, Ian Johnston, Dominic Kwiatkowski, Cordelia Langford, John Sillitoe on behalf of the Wellcome Sanger Institute COVID-19 Surveillance Team |
| EPI_ISL_2515529 | Lighthouse Lab in Alderley Park | Wellcome Sanger Institute for the COVID-19 Genomics UK (COG-UK) Consortium | Jacquelyn Wynn, Mairead Hyland, The Lighthouse Lab in Alderley Park and Alex Alderton, Roberto Amato, Jeffrey Barrett, Sonia Goncalves, Ewan Harrison, David K. Jackson, Ian Johnston, Dominic Kwiatkowski, Cordelia Langford, John Sillitoe on behalf of the Wellcome Sanger Institute COVID-19 Surveillance Team |
| EPI_ISL_2515530 | Lighthouse Lab in Glasgow | Wellcome Sanger Institute for the COVID-19 Genomics UK (COG-UK) Consortium | Harper VanSteenhouse, Yumi Kasai, David Gray, Carol Clugston, Anna Dominiczak and Alex Alderton, Roberto Amato, Jeffrey Barrett, Sonia Goncalves, Ewan Harrison, David K. Jackson, Ian Johnston, Dominic Kwiatkowski, Cordelia Langford, John Sillitoe on behalf of the Wellcome Sanger Institute COVID-19 Surveillance Team |
| EPI_ISL_2515531, EPI_ISL_2515532 | Lighthouse Lab in Milton Keynes | Wellcome Sanger Institute for the COVID-19 Genomics UK (COG-UK) Consortium | The Lighthouse Lab in Milton Keynes and Alex Alderton, Roberto Amato, Jeffrey Barrett, Sonia Goncalves, Ewan Harrison, David K. Jackson, Ian Johnston, Dominic Kwiatkowski, Cordelia Langford, John Sillitoe on behalf of the Wellcome Sanger Institute COVID-19 Surveillance Team |
| EPI_ISL_2515533 | Lighthouse Lab in Glasgow | Wellcome Sanger Institute for the COVID-19 Genomics UK (COG-UK) Consortium | Harper VanSteenhouse, Yumi Kasai, David Gray, Carol Clugston, Anna Dominiczak and Alex Alderton, Roberto Amato, Jeffrey Barrett, Sonia Goncalves, Ewan Harrison, David K. Jackson, Ian Johnston, Dominic Kwiatkowski, Cordelia Langford, John Sillitoe on behalf of the Wellcome Sanger Institute COVID-19 Surveillance Team |
| EPI_ISL_2515534 | Lighthouse Lab in Milton Keynes | Wellcome Sanger Institute for the COVID-19 Genomics UK (COG-UK) Consortium | The Lighthouse Lab in Milton Keynes and Alex Alderton, Roberto Amato, Jeffrey Barrett, Sonia Goncalves, Ewan Harrison, David K. Jackson, Ian Johnston, Dominic Kwiatkowski, Cordelia Langford, John Sillitoe on behalf of the Wellcome Sanger Institute COVID-19 Surveillance Team |
| EPI_ISL_2515535 | Lighthouse Laboratory Plymouth | Wellcome Sanger Institute for the COVID-19 Genomics UK (COG-UK) Consortium | Lighthouse Laboratory Plymouth and Alex Alderton, Roberto Amato, Jeffrey Barrett, Sonia Goncalves, Ewan Harrison, David K. Jackson, Ian Johnston, Dominic Kwiatkowski, Cordelia Langford, John Sillitoe on behalf of the Wellcome Sanger Institute COVID-19 Surveillance Team |
| EPI_ISL_2515536, EPI_ISL_2515537 | Lighthouse Lab in Milton Keynes | Wellcome Sanger Institute for the COVID-19 Genomics UK (COG-UK) Consortium | The Lighthouse Lab in Milton Keynes and Alex Alderton, Roberto Amato, Jeffrey Barrett, Sonia Goncalves, Ewan Harrison, David K. Jackson, Ian Johnston, Dominic Kwiatkowski, Cordelia Langford, John Sillitoe on behalf of the Wellcome Sanger Institute COVID-19 Surveillance Team |
| EPI_ISL_2515538 | Lighthouse Lab in Alderley Park | Wellcome Sanger Institute for the COVID-19 Genomics UK (COG-UK) Consortium | Jacquelyn Wynn, Mairead Hyland, The Lighthouse Lab in Alderley Park and Alex Alderton, Roberto Amato, Jeffrey Barrett, Sonia Goncalves, Ewan Harrison, David K. Jackson, Ian Johnston, Dominic Kwiatkowski, Cordelia Langford, John Sillitoe on behalf of the Wellcome Sanger Institute COVID-19 Surveillance Team |
| EPI_ISL_2515539 | Lighthouse Lab in Milton Keynes | Wellcome Sanger Institute for the COVID-19 Genomics UK (COG-UK) Consortium | The Lighthouse Lab in Milton Keynes and Alex Alderton, Roberto Amato, Jeffrey Barrett, Sonia Goncalves, Ewan Harrison, David K. Jackson, Ian Johnston, Dominic Kwiatkowski, Cordelia Langford, John Sillitoe on behalf of the Wellcome Sanger Institute COVID-19 Surveillance Team |
| EPI_ISL_2515540, EPI_ISL_2515541, EPI_ISL_2515542 | Lighthouse Lab in Alderley Park | Wellcome Sanger Institute for the COVID-19 Genomics UK (COG-UK) Consortium | Jacquelyn Wynn, Mairead Hyland, The Lighthouse Lab in Alderley Park and Alex Alderton, Roberto Amato, Jeffrey Barrett, Sonia Goncalves, Ewan Harrison, David K. Jackson, Ian Johnston, Dominic Kwiatkowski, Cordelia Langford, John Sillitoe on behalf of the Wellcome Sanger Institute COVID-19 Surveillance Team |
| EPI_ISL_2515543 | Lighthouse Lab in Milton Keynes | Wellcome Sanger Institute for the COVID-19 Genomics UK (COG-UK) Consortium | The Lighthouse Lab in Milton Keynes and Alex Alderton, Roberto Amato, Jeffrey Barrett, Sonia Goncalves, Ewan Harrison, David K. Jackson, Ian Johnston, Dominic Kwiatkowski, Cordelia Langford, John Sillitoe on behalf of the Wellcome Sanger Institute COVID-19 Surveillance Team |
| EPI_ISL_2515544 | Lighthouse Lab in Alderley Park | Wellcome Sanger Institute for the COVID-19 Genomics UK (COG-UK) Consortium | Jacquelyn Wynn, Mairead Hyland, The Lighthouse Lab in Alderley Park and Alex Alderton, Roberto Amato, Jeffrey Barrett, Sonia Goncalves, Ewan Harrison, David K. Jackson, Ian Johnston, Dominic Kwiatkowski, Cordelia Langford, John Sillitoe on behalf of the Wellcome Sanger Institute COVID-19 Surveillance Team |
| EPI_ISL_2515545 | Lighthouse Laboratory Plymouth | Wellcome Sanger Institute for the COVID-19 Genomics UK (COG-UK) Consortium | Lighthouse Laboratory Plymouth and Alex Alderton, Roberto Amato, Jeffrey Barrett, Sonia Goncalves, Ewan Harrison, David K. Jackson, Ian Johnston, Dominic Kwiatkowski, Cordelia Langford, John Sillitoe on behalf of the Wellcome Sanger Institute COVID-19 Surveillance Team |
| EPI_ISL_2515547 | Lighthouse Lab in Alderley Park | Wellcome Sanger Institute for the COVID-19 Genomics UK (COG-UK) Consortium | Jacquelyn Wynn, Mairead Hyland, The Lighthouse Lab in Alderley Park and Alex Alderton, Roberto Amato, Jeffrey Barrett, Sonia Goncalves, Ewan Harrison, David K. Jackson, Ian Johnston, Dominic Kwiatkowski, Cordelia Langford, John Sillitoe on behalf of the Wellcome Sanger Institute COVID-19 Surveillance Team |
| EPI_ISL_2515548 | Lighthouse Lab in Glasgow | Wellcome Sanger Institute for the COVID-19 Genomics UK (COG-UK) Consortium | Harper VanSteenhouse, Yumi Kasai, David Gray, Carol Clugston, Anna Dominiczak and Alex Alderton, Roberto Amato, Jeffrey Barrett, Sonia Goncalves, Ewan Harrison, David K. Jackson, Ian Johnston, Dominic Kwiatkowski, Cordelia Langford, John Sillitoe on behalf of the Wellcome Sanger Institute COVID-19 Surveillance Team |
| EPI_ISL_2515549, EPI_ISL_2515550, EPI_ISL_2515551 | Lighthouse Lab in Alderley Park | Wellcome Sanger Institute for the COVID-19 Genomics UK (COG-UK) Consortium | Jacquelyn Wynn, Mairead Hyland, The Lighthouse Lab in Alderley Park and Alex Alderton, Roberto Amato, Jeffrey Barrett, Sonia Goncalves, Ewan Harrison, David K. Jackson, Ian Johnston, Dominic Kwiatkowski, Cordelia Langford, John Sillitoe on behalf of the Wellcome Sanger Institute COVID-19 Surveillance Team |
| EPI_ISL_2515552, EPI_ISL_2515553 | Lighthouse Lab in Glasgow | Wellcome Sanger Institute for the COVID-19 Genomics UK (COG-UK) Consortium | Harper VanSteenhouse, Yumi Kasai, David Gray, Carol Clugston, Anna Dominiczak and Alex Alderton, Roberto Amato, Jeffrey Barrett, Sonia Goncalves, Ewan Harrison, David K. Jackson, Ian Johnston, Dominic Kwiatkowski, Cordelia Langford, John Sillitoe on behalf of the Wellcome Sanger Institute COVID-19 Surveillance Team |
| EPI_ISL_2515554, EPI_ISL_2515555, EPI_ISL_2515557, EPI_ISL_2515558 | Lighthouse Lab in Alderley Park | Wellcome Sanger Institute for the COVID-19 Genomics UK (COG-UK) Consortium | Jacquelyn Wynn, Mairead Hyland, The Lighthouse Lab in Alderley Park and Alex Alderton, Roberto Amato, Jeffrey Barrett, Sonia Goncalves, Ewan Harrison, David K. Jackson, Ian Johnston, Dominic Kwiatkowski, Cordelia Langford, John Sillitoe on behalf of the Wellcome Sanger Institute COVID-19 Surveillance Team |
| EPI_ISL_2515559 | Lighthouse Lab in Milton Keynes | Wellcome Sanger Institute for the COVID-19 Genomics UK (COG-UK) Consortium | The Lighthouse Lab in Milton Keynes and Alex Alderton, Roberto Amato, Jeffrey Barrett, Sonia Goncalves, Ewan Harrison, David K. Jackson, Ian Johnston, Dominic Kwiatkowski, Cordelia Langford, John Sillitoe on behalf of the Wellcome Sanger Institute COVID-19 Surveillance Team |
| EPI_ISL_2515560 | Lighthouse Lab in Glasgow | Wellcome Sanger Institute for the COVID-19 Genomics UK (COG-UK) Consortium | Harper VanSteenhouse, Yumi Kasai, David Gray, Carol Clugston, Anna Dominiczak and Alex Alderton, Roberto Amato, Jeffrey Barrett, Sonia Goncalves, Ewan Harrison, David K. Jackson, Ian Johnston, Dominic Kwiatkowski, Cordelia Langford, John Sillitoe on behalf of the Wellcome Sanger Institute COVID-19 Surveillance Team |
| EPI_ISL_2515561 | Lighthouse Lab in Milton Keynes | Wellcome Sanger Institute for the COVID-19 Genomics UK (COG-UK) | The Lighthouse Lab in Milton Keynes and Alex Alderton, Roberto Amato, Jeffrey Barrett, Sonia Goncalves, Ewan Harrison, David K. Jackson, Ian Johnston, Dominic Kwiatkowski, Cordelia Langford, John Sillitoe on behalf of the Wellcome Sanger |

| | | | |
|---|---|---|---|
| | | Consortium | Institute COVID-19 Surveillance Team |
| EPI_ISL_2515562, EPI_ISL_2515563, EPI_ISL_2515564 | Lighthouse Lab in Alderley Park | Wellcome Sanger Institute for the COVID-19 Genomics UK (COG-UK) Consortium | Jacquelyn Wynn, Mairead Hyland, The Lighthouse Lab in Alderley Park and Alex Alderton, Roberto Amato, Jeffrey Barrett, Sonia Goncalves, Ewan Harrison, David K. Jackson, Ian Johnston, Dominic Kwiatkowski, Cordelia Langford, John Sillitoe on behalf of the Wellcome Sanger Institute COVID-19 Surveillance Team |
| EPI_ISL_2515565 | Lighthouse Laboratory Plymouth | Wellcome Sanger Institute for the COVID-19 Genomics UK (COG-UK) Consortium | Lighthouse Laboratory Plymouth and Alex Alderton, Roberto Amato, Jeffrey Barrett, Sonia Goncalves, Ewan Harrison, David K. Jackson, Ian Johnston, Dominic Kwiatkowski, Cordelia Langford, John Sillitoe on behalf of the Wellcome Sanger Institute COVID-19 Surveillance Team |
| EPI_ISL_2515566 | Lighthouse Lab in Alderley Park | Wellcome Sanger Institute for the COVID-19 Genomics UK (COG-UK) Consortium | Jacquelyn Wynn, Mairead Hyland, The Lighthouse Lab in Alderley Park and Alex Alderton, Roberto Amato, Jeffrey Barrett, Sonia Goncalves, Ewan Harrison, David K. Jackson, Ian Johnston, Dominic Kwiatkowski, Cordelia Langford, John Sillitoe on behalf of the Wellcome Sanger Institute COVID-19 Surveillance Team |
| EPI_ISL_2515567, EPI_ISL_2515568 | Lighthouse Lab in Milton Keynes | Wellcome Sanger Institute for the COVID-19 Genomics UK (COG-UK) Consortium | The Lighthouse Lab in Milton Keynes and Alex Alderton, Roberto Amato, Jeffrey Barrett, Sonia Goncalves, Ewan Harrison, David K. Jackson, Ian Johnston, Dominic Kwiatkowski, Cordelia Langford, John Sillitoe on behalf of the Wellcome Sanger Institute COVID-19 Surveillance Team |
| EPI_ISL_2515569 | Lighthouse Lab in Alderley Park | Wellcome Sanger Institute for the COVID-19 Genomics UK (COG-UK) Consortium | Jacquelyn Wynn, Mairead Hyland, The Lighthouse Lab in Alderley Park and Alex Alderton, Roberto Amato, Jeffrey Barrett, Sonia Goncalves, Ewan Harrison, David K. Jackson, Ian Johnston, Dominic Kwiatkowski, Cordelia Langford, John Sillitoe on behalf of the Wellcome Sanger Institute COVID-19 Surveillance Team |
| EPI_ISL_2515570 | Lighthouse Lab in Milton Keynes | Wellcome Sanger Institute for the COVID-19 Genomics UK (COG-UK) Consortium | The Lighthouse Lab in Milton Keynes and Alex Alderton, Roberto Amato, Jeffrey Barrett, Sonia Goncalves, Ewan Harrison, David K. Jackson, Ian Johnston, Dominic Kwiatkowski, Cordelia Langford, John Sillitoe on behalf of the Wellcome Sanger Institute COVID-19 Surveillance Team |
| EPI_ISL_2515571, EPI_ISL_2515572, EPI_ISL_2515573 | Lighthouse Lab in Alderley Park | Wellcome Sanger Institute for the COVID-19 Genomics UK (COG-UK) Consortium | Jacquelyn Wynn, Mairead Hyland, The Lighthouse Lab in Alderley Park and Alex Alderton, Roberto Amato, Jeffrey Barrett, Sonia Goncalves, Ewan Harrison, David K. Jackson, Ian Johnston, Dominic Kwiatkowski, Cordelia Langford, John Sillitoe on behalf of the Wellcome Sanger Institute COVID-19 Surveillance Team |
| EPI_ISL_2515574 | Lighthouse Lab in Glasgow | Wellcome Sanger Institute for the COVID-19 Genomics UK (COG-UK) Consortium | Harper VanSteenhouse, Yumi Kasai, David Gray, Carol Clugston, Anna Dominiczak and Alex Alderton, Roberto Amato, Jeffrey Barrett, Sonia Goncalves, Ewan Harrison, David K. Jackson, Ian Johnston, Dominic Kwiatkowski, Cordelia Langford, John Sillitoe on behalf of the Wellcome Sanger Institute COVID-19 Surveillance Team |
| EPI_ISL_2515575, EPI_ISL_2515576, EPI_ISL_2515578 | Lighthouse Lab in Milton Keynes | Wellcome Sanger Institute for the COVID-19 Genomics UK (COG-UK) Consortium | The Lighthouse Lab in Milton Keynes and Alex Alderton, Roberto Amato, Jeffrey Barrett, Sonia Goncalves, Ewan Harrison, David K. Jackson, Ian Johnston, Dominic Kwiatkowski, Cordelia Langford, John Sillitoe on behalf of the Wellcome Sanger Institute COVID-19 Surveillance Team |
| EPI_ISL_2515579, EPI_ISL_2515580, EPI_ISL_2515581 | Lighthouse Lab in Alderley Park | Wellcome Sanger Institute for the COVID-19 Genomics UK (COG-UK) Consortium | Jacquelyn Wynn, Mairead Hyland, The Lighthouse Lab in Alderley Park and Alex Alderton, Roberto Amato, Jeffrey Barrett, Sonia Goncalves, Ewan Harrison, David K. Jackson, Ian Johnston, Dominic Kwiatkowski, Cordelia Langford, John Sillitoe on behalf of the Wellcome Sanger Institute COVID-19 Surveillance Team |
| EPI_ISL_2515582 | Lighthouse Lab in Milton Keynes | Wellcome Sanger Institute for the COVID-19 Genomics UK (COG-UK) Consortium | The Lighthouse Lab in Milton Keynes and Alex Alderton, Roberto Amato, Jeffrey Barrett, Sonia Goncalves, Ewan Harrison, David K. Jackson, Ian Johnston, Dominic Kwiatkowski, Cordelia Langford, John Sillitoe on behalf of the Wellcome Sanger Institute COVID-19 Surveillance Team |
| EPI_ISL_2515583, EPI_ISL_2515584, EPI_ISL_2515585 | Lighthouse Lab in Alderley Park | Wellcome Sanger Institute for the COVID-19 Genomics UK (COG-UK) Consortium | Jacquelyn Wynn, Mairead Hyland, The Lighthouse Lab in Alderley Park and Alex Alderton, Roberto Amato, Jeffrey Barrett, Sonia Goncalves, Ewan Harrison, David K. Jackson, Ian Johnston, Dominic Kwiatkowski, Cordelia Langford, John Sillitoe on behalf of the Wellcome Sanger Institute COVID-19 Surveillance Team |
| EPI_ISL_2515586 | Lighthouse Lab in Glasgow | Wellcome Sanger Institute for the COVID-19 Genomics UK (COG-UK) Consortium | Harper VanSteenhouse, Yumi Kasai, David Gray, Carol Clugston, Anna Dominiczak and Alex Alderton, Roberto Amato, Jeffrey Barrett, Sonia Goncalves, Ewan Harrison, David K. Jackson, Ian Johnston, Dominic Kwiatkowski, Cordelia Langford, John Sillitoe on behalf of the Wellcome Sanger Institute COVID-19 Surveillance Team |
| EPI_ISL_2515587 | Lighthouse Lab in Milton Keynes | Wellcome Sanger Institute for the COVID-19 Genomics UK (COG-UK) Consortium | The Lighthouse Lab in Milton Keynes and Alex Alderton, Roberto Amato, Jeffrey Barrett, Sonia Goncalves, Ewan Harrison, David K. Jackson, Ian Johnston, Dominic Kwiatkowski, Cordelia Langford, John Sillitoe on behalf of the Wellcome Sanger Institute COVID-19 Surveillance Team |
| EPI_ISL_2515588 | Lighthouse Lab in Alderley Park | Wellcome Sanger Institute for the COVID-19 Genomics UK (COG-UK) Consortium | Jacquelyn Wynn, Mairead Hyland, The Lighthouse Lab in Alderley Park and Alex Alderton, Roberto Amato, Jeffrey Barrett, Sonia Goncalves, Ewan Harrison, David K. Jackson, Ian Johnston, Dominic Kwiatkowski, Cordelia Langford, John Sillitoe on behalf of the Wellcome Sanger Institute COVID-19 Surveillance Team |
| EPI_ISL_2515589 | Lighthouse Lab in Glasgow | Wellcome Sanger Institute for the COVID-19 Genomics UK (COG-UK) Consortium | Harper VanSteenhouse, Yumi Kasai, David Gray, Carol Clugston, Anna Dominiczak and Alex Alderton, Roberto Amato, Jeffrey Barrett, Sonia Goncalves, Ewan Harrison, David K. Jackson, Ian Johnston, Dominic Kwiatkowski, Cordelia Langford, John Sillitoe on behalf of the Wellcome Sanger Institute COVID-19 Surveillance Team |
| EPI_ISL_2515590, EPI_ISL_2515592 | Lighthouse Lab in Milton Keynes | Wellcome Sanger Institute for the COVID-19 Genomics UK (COG-UK) Consortium | The Lighthouse Lab in Milton Keynes and Alex Alderton, Roberto Amato, Jeffrey Barrett, Sonia Goncalves, Ewan Harrison, David K. Jackson, Ian Johnston, Dominic Kwiatkowski, Cordelia Langford, John Sillitoe on behalf of the Wellcome Sanger Institute COVID-19 Surveillance Team |
| EPI_ISL_2515593 | Lighthouse Laboratory Plymouth | Wellcome Sanger Institute for the COVID-19 Genomics UK (COG-UK) Consortium | Lighthouse Laboratory Plymouth and Alex Alderton, Roberto Amato, Jeffrey Barrett, Sonia Goncalves, Ewan Harrison, David K. Jackson, Ian Johnston, Dominic Kwiatkowski, Cordelia Langford, John Sillitoe on behalf of the Wellcome Sanger Institute COVID-19 Surveillance Team |
| EPI_ISL_2515594 | Lighthouse Lab in Glasgow | Wellcome Sanger Institute for the COVID-19 Genomics UK (COG-UK) Consortium | Harper VanSteenhouse, Yumi Kasai, David Gray, Carol Clugston, Anna Dominiczak and Alex Alderton, Roberto Amato, Jeffrey Barrett, Sonia Goncalves, Ewan Harrison, David K. Jackson, Ian Johnston, Dominic Kwiatkowski, Cordelia Langford, John Sillitoe on behalf of the Wellcome Sanger Institute COVID-19 Surveillance Team |
| EPI_ISL_2515595 | Lighthouse Lab in Milton Keynes | Wellcome Sanger Institute for the COVID-19 Genomics UK (COG-UK) Consortium | The Lighthouse Lab in Milton Keynes and Alex Alderton, Roberto Amato, Jeffrey Barrett, Sonia Goncalves, Ewan Harrison, David K. Jackson, Ian Johnston, Dominic Kwiatkowski, Cordelia Langford, John Sillitoe on behalf of the Wellcome Sanger Institute COVID-19 Surveillance Team |
| EPI_ISL_2515596 | Lighthouse Lab in Glasgow | Wellcome Sanger Institute for the COVID-19 Genomics UK (COG-UK) Consortium | Harper VanSteenhouse, Yumi Kasai, David Gray, Carol Clugston, Anna Dominiczak and Alex Alderton, Roberto Amato, Jeffrey Barrett, Sonia Goncalves, Ewan Harrison, David K. Jackson, Ian Johnston, Dominic Kwiatkowski, Cordelia Langford, John Sillitoe on behalf of the Wellcome Sanger Institute COVID-19 Surveillance Team |
| EPI_ISL_2515597 | Lighthouse Lab in Milton Keynes | Wellcome Sanger Institute for the COVID-19 Genomics UK (COG-UK) Consortium | The Lighthouse Lab in Milton Keynes and Alex Alderton, Roberto Amato, Jeffrey Barrett, Sonia Goncalves, Ewan Harrison, David K. Jackson, Ian Johnston, Dominic Kwiatkowski, Cordelia Langford, John Sillitoe on behalf of the Wellcome Sanger Institute COVID-19 Surveillance Team |
| EPI_ISL_2515598, EPI_ISL_2515599 | Lighthouse Lab in Alderley Park | Wellcome Sanger Institute for the COVID-19 Genomics UK (COG-UK) Consortium | Jacquelyn Wynn, Mairead Hyland, The Lighthouse Lab in Alderley Park and Alex Alderton, Roberto Amato, Jeffrey Barrett, Sonia Goncalves, Ewan Harrison, David K. Jackson, Ian Johnston, Dominic Kwiatkowski, Cordelia Langford, John Sillitoe on behalf of the Wellcome Sanger Institute COVID-19 Surveillance Team |
| EPI_ISL_2515600 | Lighthouse Laboratory Plymouth | Wellcome Sanger Institute for the COVID-19 Genomics UK (COG-UK) Consortium | Lighthouse Laboratory Plymouth and Alex Alderton, Roberto Amato, Jeffrey Barrett, Sonia Goncalves, Ewan Harrison, David K. Jackson, Ian Johnston, Dominic Kwiatkowski, Cordelia Langford, John Sillitoe on behalf of the Wellcome Sanger Institute COVID-19 Surveillance Team |
| EPI_ISL_2515601, EPI_ISL_2515602 | Lighthouse Lab in Alderley Park | Wellcome Sanger Institute for the COVID-19 Genomics UK (COG-UK) Consortium | Jacquelyn Wynn, Mairead Hyland, The Lighthouse Lab in Alderley Park and Alex Alderton, Roberto Amato, Jeffrey Barrett, Sonia Goncalves, Ewan Harrison, David K. Jackson, Ian Johnston, Dominic Kwiatkowski, Cordelia Langford, John Sillitoe on behalf of the Wellcome Sanger Institute COVID-19 Surveillance Team |
| EPI_ISL_2515603 | Lighthouse Laboratory Plymouth | Wellcome Sanger Institute for the COVID-19 Genomics UK (COG-UK) Consortium | Lighthouse Laboratory Plymouth and Alex Alderton, Roberto Amato, Jeffrey Barrett, Sonia Goncalves, Ewan Harrison, David K. Jackson, Ian Johnston, Dominic Kwiatkowski, Cordelia Langford, John Sillitoe on behalf of the Wellcome Sanger Institute COVID-19 Surveillance Team |
| EPI_ISL_2515604 | Lighthouse Lab in Milton Keynes | Wellcome Sanger Institute for the COVID-19 Genomics UK (COG-UK) Consortium | The Lighthouse Lab in Milton Keynes and Alex Alderton, Roberto Amato, Jeffrey Barrett, Sonia Goncalves, Ewan Harrison, David K. Jackson, Ian Johnston, Dominic Kwiatkowski, Cordelia Langford, John Sillitoe on behalf of the Wellcome Sanger Institute COVID-19 Surveillance Team |
| EPI_ISL_2515606 | Lighthouse Lab in Alderley Park | Wellcome Sanger Institute for the COVID-19 Genomics UK (COG-UK) Consortium | Jacquelyn Wynn, Mairead Hyland, The Lighthouse Lab in Alderley Park and Alex Alderton, Roberto Amato, Jeffrey Barrett, Sonia Goncalves, Ewan Harrison, David K. Jackson, Ian Johnston, Dominic Kwiatkowski, Cordelia Langford, John Sillitoe on behalf of the Wellcome Sanger Institute COVID-19 Surveillance Team |
| EPI_ISL_2515607 | Lighthouse Laboratory Plymouth | Wellcome Sanger Institute for the COVID-19 Genomics UK (COG-UK) Consortium | Lighthouse Laboratory Plymouth and Alex Alderton, Roberto Amato, Jeffrey Barrett, Sonia Goncalves, Ewan Harrison, David K. Jackson, Ian Johnston, Dominic Kwiatkowski, Cordelia Langford, John Sillitoe on behalf of the Wellcome Sanger Institute COVID-19 Surveillance Team |
| EPI_ISL_2515608, EPI_ISL_2515609 | Lighthouse Lab in Glasgow | Wellcome Sanger Institute for the COVID-19 Genomics UK (COG-UK) Consortium | Harper VanSteenhouse, Yumi Kasai, David Gray, Carol Clugston, Anna Dominiczak and Alex Alderton, Roberto Amato, Jeffrey Barrett, Sonia Goncalves, Ewan Harrison, David K. Jackson, Ian Johnston, Dominic Kwiatkowski, Cordelia Langford, John Sillitoe on behalf of the Wellcome Sanger Institute COVID-19 Surveillance Team |
| EPI_ISL_2515610, EPI_ISL_2515611, EPI_ISL_2515612, EPI_ISL_2515613 | Lighthouse Lab in Milton Keynes | Wellcome Sanger Institute for the COVID-19 Genomics UK (COG-UK) Consortium | The Lighthouse Lab in Milton Keynes and Alex Alderton, Roberto Amato, Jeffrey Barrett, Sonia Goncalves, Ewan Harrison, David K. Jackson, Ian Johnston, Dominic Kwiatkowski, Cordelia Langford, John Sillitoe on behalf of the Wellcome Sanger Institute COVID-19 Surveillance Team |
| EPI_ISL_2515614 | Lighthouse Lab in Glasgow | Wellcome Sanger Institute for the COVID-19 Genomics UK (COG-UK) Consortium | Harper VanSteenhouse, Yumi Kasai, David Gray, Carol Clugston, Anna Dominiczak and Alex Alderton, Roberto Amato, Jeffrey Barrett, Sonia Goncalves, Ewan Harrison, David K. Jackson, Ian Johnston, Dominic Kwiatkowski, Cordelia Langford, John Sillitoe on behalf of the Wellcome Sanger Institute COVID-19 Surveillance Team |
| EPI_ISL_2515615, EPI_ISL_2515616 | Lighthouse Lab in Milton Keynes | Wellcome Sanger Institute for the COVID-19 Genomics UK (COG-UK) Consortium | The Lighthouse Lab in Milton Keynes and Alex Alderton, Roberto Amato, Jeffrey Barrett, Sonia Goncalves, Ewan Harrison, David K. Jackson, Ian Johnston, Dominic Kwiatkowski, Cordelia Langford, John Sillitoe on behalf of the Wellcome Sanger Institute COVID-19 Surveillance Team |
| EPI_ISL_2515617 | Lighthouse Lab in Alderley Park | Wellcome Sanger Institute for the COVID-19 Genomics UK (COG-UK) Consortium | Jacquelyn Wynn, Mairead Hyland, The Lighthouse Lab in Alderley Park and Alex Alderton, Roberto Amato, Jeffrey Barrett, Sonia Goncalves, Ewan Harrison, David K. Jackson, Ian Johnston, Dominic Kwiatkowski, Cordelia Langford, John Sillitoe on behalf of the Wellcome Sanger Institute COVID-19 Surveillance Team |
| EPI_ISL_2515618 | Lighthouse Lab in Glasgow | Wellcome Sanger Institute for the COVID-19 Genomics UK (COG-UK) Consortium | Harper VanSteenhouse, Yumi Kasai, David Gray, Carol Clugston, Anna Dominiczak and Alex Alderton, Roberto Amato, Jeffrey Barrett, Sonia Goncalves, Ewan Harrison, David K. Jackson, Ian Johnston, Dominic Kwiatkowski, Cordelia Langford, John Sillitoe on behalf of the Wellcome Sanger Institute COVID-19 Surveillance Team |
| EPI_ISL_2515619 | Lighthouse Lab in Milton Keynes | Wellcome Sanger Institute for the COVID-19 Genomics UK (COG-UK) Consortium | The Lighthouse Lab in Milton Keynes and Alex Alderton, Roberto Amato, Jeffrey Barrett, Sonia Goncalves, Ewan Harrison, David K. Jackson, Ian Johnston, Dominic Kwiatkowski, Cordelia Langford, John Sillitoe on behalf of the Wellcome Sanger Institute COVID-19 Surveillance Team |
| EPI_ISL_2515620 | Lighthouse Laboratory Plymouth | Wellcome Sanger Institute for the COVID-19 Genomics UK (COG-UK) Consortium | Lighthouse Laboratory Plymouth and Alex Alderton, Roberto Amato, Jeffrey Barrett, Sonia Goncalves, Ewan Harrison, David K. Jackson, Ian Johnston, Dominic Kwiatkowski, Cordelia Langford, John Sillitoe on behalf of the Wellcome Sanger Institute COVID-19 Surveillance Team |
| EPI_ISL_2515621 | Lighthouse Lab in Glasgow | Wellcome Sanger Institute for the COVID-19 Genomics UK (COG-UK) Consortium | Harper VanSteenhouse, Yumi Kasai, David Gray, Carol Clugston, Anna Dominiczak and Alex Alderton, Roberto Amato, Jeffrey Barrett, Sonia Goncalves, Ewan Harrison, David K. Jackson, Ian Johnston, Dominic Kwiatkowski, Cordelia Langford, John Sillitoe on behalf of the Wellcome Sanger Institute COVID-19 Surveillance Team |
| EPI_ISL_2515622, EPI_ISL_2515623 | Lighthouse Lab in Alderley Park | Wellcome Sanger Institute for the COVID-19 Genomics UK (COG-UK) Consortium | Jacquelyn Wynn, Mairead Hyland, The Lighthouse Lab in Alderley Park and Alex Alderton, Roberto Amato, Jeffrey Barrett, Sonia Goncalves, Ewan Harrison, David K. Jackson, Ian Johnston, Dominic Kwiatkowski, Cordelia Langford, John Sillitoe on behalf of the Wellcome Sanger Institute COVID-19 Surveillance Team |
| EPI_ISL_2515624 | Lighthouse Lab in Milton Keynes | Wellcome Sanger Institute for the COVID-19 Genomics UK (COG-UK) Consortium | The Lighthouse Lab in Milton Keynes and Alex Alderton, Roberto Amato, Jeffrey Barrett, Sonia Goncalves, Ewan Harrison, David K. Jackson, Ian Johnston, Dominic Kwiatkowski, Cordelia Langford, John Sillitoe on behalf of the Wellcome Sanger Institute COVID-19 Surveillance Team |
| EPI_ISL_2515625, EPI_ISL_2515626 | Lighthouse Lab in Alderley Park | Wellcome Sanger Institute for the COVID-19 Genomics UK (COG-UK) Consortium | Jacquelyn Wynn, Mairead Hyland, The Lighthouse Lab in Alderley Park and Alex Alderton, Roberto Amato, Jeffrey Barrett, Sonia Goncalves, Ewan Harrison, David K. Jackson, Ian Johnston, Dominic Kwiatkowski, Cordelia Langford, John Sillitoe on behalf of the Wellcome Sanger Institute COVID-19 Surveillance Team |
| EPI_ISL_2515627 | Lighthouse Lab in Glasgow | Wellcome Sanger Institute for the COVID-19 Genomics UK (COG-UK) Consortium | Harper VanSteenhouse, Yumi Kasai, David Gray, Carol Clugston, Anna Dominiczak and Alex Alderton, Roberto Amato, Jeffrey Barrett, Sonia Goncalves, Ewan Harrison, David K. Jackson, Ian Johnston, Dominic Kwiatkowski, Cordelia Langford, John Sillitoe on behalf of the Wellcome Sanger Institute COVID-19 Surveillance Team |

| | | | |
|---|---|---|---|
| EPI_ISL_2515628 | Lighthouse Laboratory Plymouth | Wellcome Sanger Institute for the COVID-19 Genomics UK (COG-UK) Consortium | Lighthouse Laboratory Plymouth and Alex Alderton, Roberto Amato, Jeffrey Barrett, Sonia Goncalves, Ewan Harrison, David K. Jackson, Ian Johnston, Dominic Kwiatkowski, Cordelia Langford, John Sillitoe on behalf of the Wellcome Sanger Institute COVID-19 Surveillance Team |
| EPI_ISL_2515629, EPI_ISL_2515630 | Lighthouse Lab in Alderley Park | Wellcome Sanger Institute for the COVID-19 Genomics UK (COG-UK) Consortium | Jacquelyn Wynn, Mairead Hyland, The Lighthouse Lab in Alderley Park and Alex Alderton, Roberto Amato, Jeffrey Barrett, Sonia Goncalves, Ewan Harrison, David K. Jackson, Ian Johnston, Dominic Kwiatkowski, Cordelia Langford, John Sillitoe on behalf of the Wellcome Sanger Institute COVID-19 Surveillance Team |
| EPI_ISL_2515631 | Lighthouse Lab in Milton Keynes | Wellcome Sanger Institute for the COVID-19 Genomics UK (COG-UK) Consortium | The Lighthouse Lab in Milton Keynes and Alex Alderton, Roberto Amato, Jeffrey Barrett, Sonia Goncalves, Ewan Harrison, David K. Jackson, Ian Johnston, Dominic Kwiatkowski, Cordelia Langford, John Sillitoe on behalf of the Wellcome Sanger Institute COVID-19 Surveillance Team |
| EPI_ISL_2515632 | Lighthouse Lab in Alderley Park | Wellcome Sanger Institute for the COVID-19 Genomics UK (COG-UK) Consortium | Jacquelyn Wynn, Mairead Hyland, The Lighthouse Lab in Alderley Park and Alex Alderton, Roberto Amato, Jeffrey Barrett, Sonia Goncalves, Ewan Harrison, David K. Jackson, Ian Johnston, Dominic Kwiatkowski, Cordelia Langford, John Sillitoe on behalf of the Wellcome Sanger Institute COVID-19 Surveillance Team |
| EPI_ISL_2515633 | Lighthouse Lab in Glasgow | Wellcome Sanger Institute for the COVID-19 Genomics UK (COG-UK) Consortium | Harper VanSteenhouse, Yumi Kasai, David Gray, Carol Clugston, Anna Dominiczak and Alex Alderton, Roberto Amato, Jeffrey Barrett, Sonia Goncalves, Ewan Harrison, David K. Jackson, Ian Johnston, Dominic Kwiatkowski, Cordelia Langford, John Sillitoe on behalf of the Wellcome Sanger Institute COVID-19 Surveillance Team |
| EPI_ISL_2515634, EPI_ISL_2515635 | Lighthouse Lab in Milton Keynes | Wellcome Sanger Institute for the COVID-19 Genomics UK (COG-UK) Consortium | The Lighthouse Lab in Milton Keynes and Alex Alderton, Roberto Amato, Jeffrey Barrett, Sonia Goncalves, Ewan Harrison, David K. Jackson, Ian Johnston, Dominic Kwiatkowski, Cordelia Langford, John Sillitoe on behalf of the Wellcome Sanger Institute COVID-19 Surveillance Team |
| EPI_ISL_2515636 | Lighthouse Lab in Glasgow | Wellcome Sanger Institute for the COVID-19 Genomics UK (COG-UK) Consortium | Harper VanSteenhouse, Yumi Kasai, David Gray, Carol Clugston, Anna Dominiczak and Alex Alderton, Roberto Amato, Jeffrey Barrett, Sonia Goncalves, Ewan Harrison, David K. Jackson, Ian Johnston, Dominic Kwiatkowski, Cordelia Langford, John Sillitoe on behalf of the Wellcome Sanger Institute COVID-19 Surveillance Team |
| EPI_ISL_2515637, EPI_ISL_2515638 | Lighthouse Lab in Milton Keynes | Wellcome Sanger Institute for the COVID-19 Genomics UK (COG-UK) Consortium | The Lighthouse Lab in Milton Keynes and Alex Alderton, Roberto Amato, Jeffrey Barrett, Sonia Goncalves, Ewan Harrison, David K. Jackson, Ian Johnston, Dominic Kwiatkowski, Cordelia Langford, John Sillitoe on behalf of the Wellcome Sanger Institute COVID-19 Surveillance Team |
| EPI_ISL_2515639 | Lighthouse Laboratory Plymouth | Wellcome Sanger Institute for the COVID-19 Genomics UK (COG-UK) Consortium | Lighthouse Laboratory Plymouth and Alex Alderton, Roberto Amato, Jeffrey Barrett, Sonia Goncalves, Ewan Harrison, David K. Jackson, Ian Johnston, Dominic Kwiatkowski, Cordelia Langford, John Sillitoe on behalf of the Wellcome Sanger Institute COVID-19 Surveillance Team |
| EPI_ISL_2515640 | Lighthouse Lab in Alderley Park | Wellcome Sanger Institute for the COVID-19 Genomics UK (COG-UK) Consortium | Jacquelyn Wynn, Mairead Hyland, The Lighthouse Lab in Alderley Park and Alex Alderton, Roberto Amato, Jeffrey Barrett, Sonia Goncalves, Ewan Harrison, David K. Jackson, Ian Johnston, Dominic Kwiatkowski, Cordelia Langford, John Sillitoe on behalf of the Wellcome Sanger Institute COVID-19 Surveillance Team |
| EPI_ISL_2515641, EPI_ISL_2515642, EPI_ISL_2515643 | Lighthouse Lab in Milton Keynes | Wellcome Sanger Institute for the COVID-19 Genomics UK (COG-UK) Consortium | The Lighthouse Lab in Milton Keynes and Alex Alderton, Roberto Amato, Jeffrey Barrett, Sonia Goncalves, Ewan Harrison, David K. Jackson, Ian Johnston, Dominic Kwiatkowski, Cordelia Langford, John Sillitoe on behalf of the Wellcome Sanger Institute COVID-19 Surveillance Team |
| EPI_ISL_2515644 | Health Services Laboratories | Wellcome Sanger Institute for the COVID-19 Genomics UK (COG-UK) Consortium | Health Services Laboratories and Alex Alderton, Roberto Amato, Jeffrey Barrett, Sonia Goncalves, Ewan Harrison, David K. Jackson, Ian Johnston, Dominic Kwiatkowski, Cordelia Langford, John Sillitoe on behalf of the Wellcome Sanger Institute COVID-19 Surveillance Team |
| EPI_ISL_2515646 | Lighthouse Lab in Milton Keynes | Wellcome Sanger Institute for the COVID-19 Genomics UK (COG-UK) Consortium | The Lighthouse Lab in Milton Keynes and Alex Alderton, Roberto Amato, Jeffrey Barrett, Sonia Goncalves, Ewan Harrison, David K. Jackson, Ian Johnston, Dominic Kwiatkowski, Cordelia Langford, John Sillitoe on behalf of the Wellcome Sanger Institute COVID-19 Surveillance Team |
| EPI_ISL_2515648, EPI_ISL_2515649, EPI_ISL_2515650, EPI_ISL_2515651, EPI_ISL_2515652, EPI_ISL_2515653, EPI_ISL_2515654, EPI_ISL_2515655, EPI_ISL_2515656, EPI_ISL_2515657, EPI_ISL_2515658, EPI_ISL_2515659, EPI_ISL_2515660, EPI_ISL_2515661, EPI_ISL_2515662, EPI_ISL_2515663, EPI_ISL_2515664, EPI_ISL_2515665 | | | |
| see above | Lighthouse Lab in Alderley Park | Wellcome Sanger Institute for the COVID-19 Genomics UK (COG-UK) Consortium | Jacquelyn Wynn, Mairead Hyland, The Lighthouse Lab in Alderley Park and Alex Alderton, Roberto Amato, Jeffrey Barrett, Sonia Goncalves, Ewan Harrison, David K. Jackson, Ian Johnston, Dominic Kwiatkowski, Cordelia Langford, John Sillitoe on behalf of the Wellcome Sanger Institute COVID-19 Surveillance Team |
| EPI_ISL_2515666 | Lighthouse Lab in Milton Keynes | Wellcome Sanger Institute for the COVID-19 Genomics UK (COG-UK) Consortium | The Lighthouse Lab in Milton Keynes and Alex Alderton, Roberto Amato, Jeffrey Barrett, Sonia Goncalves, Ewan Harrison, David K. Jackson, Ian Johnston, Dominic Kwiatkowski, Cordelia Langford, John Sillitoe on behalf of the Wellcome Sanger Institute COVID-19 Surveillance Team |
| EPI_ISL_2515667 | Lighthouse Laboratory Plymouth | Wellcome Sanger Institute for the COVID-19 Genomics UK (COG-UK) Consortium | Lighthouse Laboratory Plymouth and Alex Alderton, Roberto Amato, Jeffrey Barrett, Sonia Goncalves, Ewan Harrison, David K. Jackson, Ian Johnston, Dominic Kwiatkowski, Cordelia Langford, John Sillitoe on behalf of the Wellcome Sanger Institute COVID-19 Surveillance Team |
| EPI_ISL_2515668 | Lighthouse Lab in Milton Keynes | Wellcome Sanger Institute for the COVID-19 Genomics UK (COG-UK) Consortium | The Lighthouse Lab in Milton Keynes and Alex Alderton, Roberto Amato, Jeffrey Barrett, Sonia Goncalves, Ewan Harrison, David K. Jackson, Ian Johnston, Dominic Kwiatkowski, Cordelia Langford, John Sillitoe on behalf of the Wellcome Sanger Institute COVID-19 Surveillance Team |
| EPI_ISL_2515669, EPI_ISL_2515670, EPI_ISL_2515671 | Lighthouse Lab in Alderley Park | Wellcome Sanger Institute for the COVID-19 Genomics UK (COG-UK) Consortium | Jacquelyn Wynn, Mairead Hyland, The Lighthouse Lab in Alderley Park and Alex Alderton, Roberto Amato, Jeffrey Barrett, Sonia Goncalves, Ewan Harrison, David K. Jackson, Ian Johnston, Dominic Kwiatkowski, Cordelia Langford, John Sillitoe on behalf of the Wellcome Sanger Institute COVID-19 Surveillance Team |
| EPI_ISL_2515672 | Lighthouse Laboratory Plymouth | Wellcome Sanger Institute for the COVID-19 Genomics UK (COG-UK) Consortium | Lighthouse Laboratory Plymouth and Alex Alderton, Roberto Amato, Jeffrey Barrett, Sonia Goncalves, Ewan Harrison, David K. Jackson, Ian Johnston, Dominic Kwiatkowski, Cordelia Langford, John Sillitoe on behalf of the Wellcome Sanger Institute COVID-19 Surveillance Team |
| EPI_ISL_2515673, EPI_ISL_2515674, EPI_ISL_2515676, EPI_ISL_2515677, EPI_ISL_2515678 | Lighthouse Lab in Alderley Park | Wellcome Sanger Institute for the COVID-19 Genomics UK (COG-UK) Consortium | Jacquelyn Wynn, Mairead Hyland, The Lighthouse Lab in Alderley Park and Alex Alderton, Roberto Amato, Jeffrey Barrett, Sonia Goncalves, Ewan Harrison, David K. Jackson, Ian Johnston, Dominic Kwiatkowski, Cordelia Langford, John Sillitoe on behalf of the Wellcome Sanger Institute COVID-19 Surveillance Team |
| EPI_ISL_2515679 | Lighthouse Laboratory Plymouth | Wellcome Sanger Institute for the COVID-19 Genomics UK (COG-UK) Consortium | Lighthouse Laboratory Plymouth and Alex Alderton, Roberto Amato, Jeffrey Barrett, Sonia Goncalves, Ewan Harrison, David K. Jackson, Ian Johnston, Dominic Kwiatkowski, Cordelia Langford, John Sillitoe on behalf of the Wellcome Sanger Institute COVID-19 Surveillance Team |
| EPI_ISL_2515680, EPI_ISL_2515682, EPI_ISL_2515683, EPI_ISL_2515684 | Lighthouse Lab in Alderley Park | Wellcome Sanger Institute for the COVID-19 Genomics UK (COG-UK) Consortium | Jacquelyn Wynn, Mairead Hyland, The Lighthouse Lab in Alderley Park and Alex Alderton, Roberto Amato, Jeffrey Barrett, Sonia Goncalves, Ewan Harrison, David K. Jackson, Ian Johnston, Dominic Kwiatkowski, Cordelia Langford, John Sillitoe on behalf of the Wellcome Sanger Institute COVID-19 Surveillance Team |
| EPI_ISL_2515685, EPI_ISL_2515686 | Lighthouse Laboratory Plymouth | Wellcome Sanger Institute for the COVID-19 Genomics UK (COG-UK) Consortium | Lighthouse Laboratory Plymouth and Alex Alderton, Roberto Amato, Jeffrey Barrett, Sonia Goncalves, Ewan Harrison, David K. Jackson, Ian Johnston, Dominic Kwiatkowski, Cordelia Langford, John Sillitoe on behalf of the Wellcome Sanger Institute COVID-19 Surveillance Team |
| EPI_ISL_2515687, EPI_ISL_2515688, EPI_ISL_2515689, EPI_ISL_2515690, EPI_ISL_2515691, EPI_ISL_2515692, EPI_ISL_2515693, EPI_ISL_2515694, EPI_ISL_2515696, EPI_ISL_2515697 | Lighthouse Lab in Alderley Park | Wellcome Sanger Institute for the COVID-19 Genomics UK (COG-UK) Consortium | Jacquelyn Wynn, Mairead Hyland, The Lighthouse Lab in Alderley Park and Alex Alderton, Roberto Amato, Jeffrey Barrett, Sonia Goncalves, Ewan Harrison, David K. Jackson, Ian Johnston, Dominic Kwiatkowski, Cordelia Langford, John Sillitoe on behalf of the Wellcome Sanger Institute COVID-19 Surveillance Team |
| EPI_ISL_2515698 | Lighthouse Laboratory Plymouth | Wellcome Sanger Institute for the COVID-19 Genomics UK (COG-UK) Consortium | Lighthouse Laboratory Plymouth and Alex Alderton, Roberto Amato, Jeffrey Barrett, Sonia Goncalves, Ewan Harrison, David K. Jackson, Ian Johnston, Dominic Kwiatkowski, Cordelia Langford, John Sillitoe on behalf of the Wellcome Sanger Institute COVID-19 Surveillance Team |
| EPI_ISL_2515699 | Lighthouse Lab in Alderley Park | Wellcome Sanger Institute for the COVID-19 Genomics UK (COG-UK) Consortium | Jacquelyn Wynn, Mairead Hyland, The Lighthouse Lab in Alderley Park and Alex Alderton, Roberto Amato, Jeffrey Barrett, Sonia Goncalves, Ewan Harrison, David K. Jackson, Ian Johnston, Dominic Kwiatkowski, Cordelia Langford, John Sillitoe on behalf of the Wellcome Sanger Institute COVID-19 Surveillance Team |
| EPI_ISL_2515700 | Lighthouse Lab in Milton Keynes | Wellcome Sanger Institute for the COVID-19 Genomics UK (COG-UK) Consortium | The Lighthouse Lab in Milton Keynes and Alex Alderton, Roberto Amato, Jeffrey Barrett, Sonia Goncalves, Ewan Harrison, David K. Jackson, Ian Johnston, Dominic Kwiatkowski, Cordelia Langford, John Sillitoe on behalf of the Wellcome Sanger Institute COVID-19 Surveillance Team |
| EPI_ISL_2515701 | Lighthouse Laboratory Plymouth | Wellcome Sanger Institute for the COVID-19 Genomics UK (COG-UK) Consortium | Lighthouse Laboratory Plymouth and Alex Alderton, Roberto Amato, Jeffrey Barrett, Sonia Goncalves, Ewan Harrison, David K. Jackson, Ian Johnston, Dominic Kwiatkowski, Cordelia Langford, John Sillitoe on behalf of the Wellcome Sanger Institute COVID-19 Surveillance Team |
| EPI_ISL_2515702, EPI_ISL_2515703 | Lighthouse Lab in Alderley Park | Wellcome Sanger Institute for the COVID-19 Genomics UK (COG-UK) Consortium | Jacquelyn Wynn, Mairead Hyland, The Lighthouse Lab in Alderley Park and Alex Alderton, Roberto Amato, Jeffrey Barrett, Sonia Goncalves, Ewan Harrison, David K. Jackson, Ian Johnston, Dominic Kwiatkowski, Cordelia Langford, John Sillitoe on behalf of the Wellcome Sanger Institute COVID-19 Surveillance Team |
| EPI_ISL_2515704 | Lighthouse Laboratory Plymouth | Wellcome Sanger Institute for the COVID-19 Genomics UK (COG-UK) Consortium | Lighthouse Laboratory Plymouth and Alex Alderton, Roberto Amato, Jeffrey Barrett, Sonia Goncalves, Ewan Harrison, David K. Jackson, Ian Johnston, Dominic Kwiatkowski, Cordelia Langford, John Sillitoe on behalf of the Wellcome Sanger Institute COVID-19 Surveillance Team |
| EPI_ISL_2515705, EPI_ISL_2515706, EPI_ISL_2515707, EPI_ISL_2515708, EPI_ISL_2515709, EPI_ISL_2515710, EPI_ISL_2515711 | Lighthouse Lab in Alderley Park | Wellcome Sanger Institute for the COVID-19 Genomics UK (COG-UK) Consortium | Jacquelyn Wynn, Mairead Hyland, The Lighthouse Lab in Alderley Park and Alex Alderton, Roberto Amato, Jeffrey Barrett, Sonia Goncalves, Ewan Harrison, David K. Jackson, Ian Johnston, Dominic Kwiatkowski, Cordelia Langford, John Sillitoe on behalf of the Wellcome Sanger Institute COVID-19 Surveillance Team |
| EPI_ISL_2515712 | Lighthouse Laboratory Plymouth | Wellcome Sanger Institute for the COVID-19 Genomics UK (COG-UK) Consortium | Lighthouse Laboratory Plymouth and Alex Alderton, Roberto Amato, Jeffrey Barrett, Sonia Goncalves, Ewan Harrison, David K. Jackson, Ian Johnston, Dominic Kwiatkowski, Cordelia Langford, John Sillitoe on behalf of the Wellcome Sanger Institute COVID-19 Surveillance Team |
| EPI_ISL_2515713, EPI_ISL_2515714 | Lighthouse Lab in Alderley Park | Wellcome Sanger Institute for the COVID-19 Genomics UK (COG-UK) Consortium | Jacquelyn Wynn, Mairead Hyland, The Lighthouse Lab in Alderley Park and Alex Alderton, Roberto Amato, Jeffrey Barrett, Sonia Goncalves, Ewan Harrison, David K. Jackson, Ian Johnston, Dominic Kwiatkowski, Cordelia Langford, John Sillitoe on behalf of the Wellcome Sanger Institute COVID-19 Surveillance Team |
| EPI_ISL_2515715, EPI_ISL_2515716 | Lighthouse Laboratory Plymouth | Wellcome Sanger Institute for the COVID-19 Genomics UK (COG-UK) Consortium | Lighthouse Laboratory Plymouth and Alex Alderton, Roberto Amato, Jeffrey Barrett, Sonia Goncalves, Ewan Harrison, David K. Jackson, Ian Johnston, Dominic Kwiatkowski, Cordelia Langford, John Sillitoe on behalf of the Wellcome Sanger Institute COVID-19 Surveillance Team |
| EPI_ISL_2515718 | Lighthouse Lab in Alderley Park | Wellcome Sanger Institute for the COVID-19 Genomics UK (COG-UK) Consortium | Jacquelyn Wynn, Mairead Hyland, The Lighthouse Lab in Alderley Park and Alex Alderton, Roberto Amato, Jeffrey Barrett, Sonia Goncalves, Ewan Harrison, David K. Jackson, Ian Johnston, Dominic Kwiatkowski, Cordelia Langford, John Sillitoe on behalf of the Wellcome Sanger Institute COVID-19 Surveillance Team |
| EPI_ISL_2515719 | Lighthouse Laboratory Plymouth | Wellcome Sanger Institute for the COVID-19 Genomics UK (COG-UK) Consortium | Lighthouse Laboratory Plymouth and Alex Alderton, Roberto Amato, Jeffrey Barrett, Sonia Goncalves, Ewan Harrison, David K. Jackson, Ian Johnston, Dominic Kwiatkowski, Cordelia Langford, John Sillitoe on behalf of the Wellcome Sanger Institute COVID-19 Surveillance Team |

| | | | |
|---|---|---|---|
| EPI_ISL_2515720 | Lighthouse Lab in Alderley Park | Wellcome Sanger Institute for the COVID-19 Genomics UK (COG-UK) Consortium | Jacquelyn Wynn, Mairead Hyland, The Lighthouse Lab in Alderley Park and Alex Alderton, Roberto Amato, Jeffrey Barrett, Sonia Goncalves, Ewan Harrison, David K. Jackson, Ian Johnston, Dominic Kwiatkowski, Cordelia Langford, John Sillitoe on behalf of the Wellcome Sanger Institute COVID-19 Surveillance Team |
| EPI_ISL_2515721 | Lighthouse Laboratory Plymouth | Wellcome Sanger Institute for the COVID-19 Genomics UK (COG-UK) Consortium | Lighthouse Laboratory Plymouth and Alex Alderton, Roberto Amato, Jeffrey Barrett, Sonia Goncalves, Ewan Harrison, David K. Jackson, Ian Johnston, Dominic Kwiatkowski, Cordelia Langford, John Sillitoe on behalf of the Wellcome Sanger Institute COVID-19 Surveillance Team |
| EPI_ISL_2515722 | Lighthouse Lab in Alderley Park | Wellcome Sanger Institute for the COVID-19 Genomics UK (COG-UK) Consortium | Jacquelyn Wynn, Mairead Hyland, The Lighthouse Lab in Alderley Park and Alex Alderton, Roberto Amato, Jeffrey Barrett, Sonia Goncalves, Ewan Harrison, David K. Jackson, Ian Johnston, Dominic Kwiatkowski, Cordelia Langford, John Sillitoe on behalf of the Wellcome Sanger Institute COVID-19 Surveillance Team |
| EPI_ISL_2515724, EPI_ISL_2515725 | Lighthouse Lab in Milton Keynes | Wellcome Sanger Institute for the COVID-19 Genomics UK (COG-UK) Consortium | The Lighthouse Lab in Milton Keynes and Alex Alderton, Roberto Amato, Jeffrey Barrett, Sonia Goncalves, Ewan Harrison, David K. Jackson, Ian Johnston, Dominic Kwiatkowski, Cordelia Langford, John Sillitoe on behalf of the Wellcome Sanger Institute COVID-19 Surveillance Team |
| EPI_ISL_2515726 | Lighthouse Lab in Alderley Park | Wellcome Sanger Institute for the COVID-19 Genomics UK (COG-UK) Consortium | Jacquelyn Wynn, Mairead Hyland, The Lighthouse Lab in Alderley Park and Alex Alderton, Roberto Amato, Jeffrey Barrett, Sonia Goncalves, Ewan Harrison, David K. Jackson, Ian Johnston, Dominic Kwiatkowski, Cordelia Langford, John Sillitoe on behalf of the Wellcome Sanger Institute COVID-19 Surveillance Team |
| EPI_ISL_2515727 | Lighthouse Lab in Milton Keynes | Wellcome Sanger Institute for the COVID-19 Genomics UK (COG-UK) Consortium | The Lighthouse Lab in Milton Keynes and Alex Alderton, Roberto Amato, Jeffrey Barrett, Sonia Goncalves, Ewan Harrison, David K. Jackson, Ian Johnston, Dominic Kwiatkowski, Cordelia Langford, John Sillitoe on behalf of the Wellcome Sanger Institute COVID-19 Surveillance Team |
| EPI_ISL_2515728, EPI_ISL_2515729 | Lighthouse Lab in Alderley Park | Wellcome Sanger Institute for the COVID-19 Genomics UK (COG-UK) Consortium | Jacquelyn Wynn, Mairead Hyland, The Lighthouse Lab in Alderley Park and Alex Alderton, Roberto Amato, Jeffrey Barrett, Sonia Goncalves, Ewan Harrison, David K. Jackson, Ian Johnston, Dominic Kwiatkowski, Cordelia Langford, John Sillitoe on behalf of the Wellcome Sanger Institute COVID-19 Surveillance Team |
| EPI_ISL_2515730 | Lighthouse Laboratory Plymouth | Wellcome Sanger Institute for the COVID-19 Genomics UK (COG-UK) Consortium | Lighthouse Laboratory Plymouth and Alex Alderton, Roberto Amato, Jeffrey Barrett, Sonia Goncalves, Ewan Harrison, David K. Jackson, Ian Johnston, Dominic Kwiatkowski, Cordelia Langford, John Sillitoe on behalf of the Wellcome Sanger Institute COVID-19 Surveillance Team |
| EPI_ISL_2515731 | Lighthouse Lab in Alderley Park | Wellcome Sanger Institute for the COVID-19 Genomics UK (COG-UK) Consortium | Jacquelyn Wynn, Mairead Hyland, The Lighthouse Lab in Alderley Park and Alex Alderton, Roberto Amato, Jeffrey Barrett, Sonia Goncalves, Ewan Harrison, David K. Jackson, Ian Johnston, Dominic Kwiatkowski, Cordelia Langford, John Sillitoe on behalf of the Wellcome Sanger Institute COVID-19 Surveillance Team |
| EPI_ISL_2515732 | Lighthouse Laboratory Plymouth | Wellcome Sanger Institute for the COVID-19 Genomics UK (COG-UK) Consortium | Lighthouse Laboratory Plymouth and Alex Alderton, Roberto Amato, Jeffrey Barrett, Sonia Goncalves, Ewan Harrison, David K. Jackson, Ian Johnston, Dominic Kwiatkowski, Cordelia Langford, John Sillitoe on behalf of the Wellcome Sanger Institute COVID-19 Surveillance Team |
| EPI_ISL_2515733, EPI_ISL_2515734, EPI_ISL_2515735, EPI_ISL_2515737, EPI_ISL_2515738, EPI_ISL_2515739, EPI_ISL_2515740, EPI_ISL_2515741, EPI_ISL_2515742, EPI_ISL_2515743, EPI_ISL_2515744, EPI_ISL_2515745, EPI_ISL_2515746, EPI_ISL_2515747 | | | |
| see above | Lighthouse Lab in Alderley Park | Wellcome Sanger Institute for the COVID-19 Genomics UK (COG-UK) Consortium | Jacquelyn Wynn, Mairead Hyland, The Lighthouse Lab in Alderley Park and Alex Alderton, Roberto Amato, Jeffrey Barrett, Sonia Goncalves, Ewan Harrison, David K. Jackson, Ian Johnston, Dominic Kwiatkowski, Cordelia Langford, John Sillitoe on behalf of the Wellcome Sanger Institute COVID-19 Surveillance Team |
| EPI_ISL_2515748 | Lighthouse Laboratory Plymouth | Wellcome Sanger Institute for the COVID-19 Genomics UK (COG-UK) Consortium | Lighthouse Laboratory Plymouth and Alex Alderton, Roberto Amato, Jeffrey Barrett, Sonia Goncalves, Ewan Harrison, David K. Jackson, Ian Johnston, Dominic Kwiatkowski, Cordelia Langford, John Sillitoe on behalf of the Wellcome Sanger Institute COVID-19 Surveillance Team |
| EPI_ISL_2515749, EPI_ISL_2515750 | Lighthouse Lab in Alderley Park | Wellcome Sanger Institute for the COVID-19 Genomics UK (COG-UK) Consortium | Jacquelyn Wynn, Mairead Hyland, The Lighthouse Lab in Alderley Park and Alex Alderton, Roberto Amato, Jeffrey Barrett, Sonia Goncalves, Ewan Harrison, David K. Jackson, Ian Johnston, Dominic Kwiatkowski, Cordelia Langford, John Sillitoe on behalf of the Wellcome Sanger Institute COVID-19 Surveillance Team |
| EPI_ISL_2515751 | Lighthouse Laboratory Plymouth | Wellcome Sanger Institute for the COVID-19 Genomics UK (COG-UK) Consortium | Lighthouse Laboratory Plymouth and Alex Alderton, Roberto Amato, Jeffrey Barrett, Sonia Goncalves, Ewan Harrison, David K. Jackson, Ian Johnston, Dominic Kwiatkowski, Cordelia Langford, John Sillitoe on behalf of the Wellcome Sanger Institute COVID-19 Surveillance Team |
| EPI_ISL_2515752, EPI_ISL_2515753, EPI_ISL_2515754, EPI_ISL_2515755 | Lighthouse Lab in Alderley Park | Wellcome Sanger Institute for the COVID-19 Genomics UK (COG-UK) Consortium | Jacquelyn Wynn, Mairead Hyland, The Lighthouse Lab in Alderley Park and Alex Alderton, Roberto Amato, Jeffrey Barrett, Sonia Goncalves, Ewan Harrison, David K. Jackson, Ian Johnston, Dominic Kwiatkowski, Cordelia Langford, John Sillitoe on behalf of the Wellcome Sanger Institute COVID-19 Surveillance Team |
| EPI_ISL_2515756, EPI_ISL_2515757 | Lighthouse Laboratory Plymouth | Wellcome Sanger Institute for the COVID-19 Genomics UK (COG-UK) Consortium | Lighthouse Laboratory Plymouth and Alex Alderton, Roberto Amato, Jeffrey Barrett, Sonia Goncalves, Ewan Harrison, David K. Jackson, Ian Johnston, Dominic Kwiatkowski, Cordelia Langford, John Sillitoe on behalf of the Wellcome Sanger Institute COVID-19 Surveillance Team |
| EPI_ISL_2515758, EPI_ISL_2515759, EPI_ISL_2515760, EPI_ISL_2515761, EPI_ISL_2515762 | Lighthouse Lab in Alderley Park | Wellcome Sanger Institute for the COVID-19 Genomics UK (COG-UK) Consortium | Jacquelyn Wynn, Mairead Hyland, The Lighthouse Lab in Alderley Park and Alex Alderton, Roberto Amato, Jeffrey Barrett, Sonia Goncalves, Ewan Harrison, David K. Jackson, Ian Johnston, Dominic Kwiatkowski, Cordelia Langford, John Sillitoe on behalf of the Wellcome Sanger Institute COVID-19 Surveillance Team |
| EPI_ISL_2515763 | Lighthouse Laboratory Plymouth | Wellcome Sanger Institute for the COVID-19 Genomics UK (COG-UK) Consortium | Lighthouse Laboratory Plymouth and Alex Alderton, Roberto Amato, Jeffrey Barrett, Sonia Goncalves, Ewan Harrison, David K. Jackson, Ian Johnston, Dominic Kwiatkowski, Cordelia Langford, John Sillitoe on behalf of the Wellcome Sanger Institute COVID-19 Surveillance Team |
| EPI_ISL_2515764 | Lighthouse Lab in Alderley Park | Wellcome Sanger Institute for the COVID-19 Genomics UK (COG-UK) Consortium | Jacquelyn Wynn, Mairead Hyland, The Lighthouse Lab in Alderley Park and Alex Alderton, Roberto Amato, Jeffrey Barrett, Sonia Goncalves, Ewan Harrison, David K. Jackson, Ian Johnston, Dominic Kwiatkowski, Cordelia Langford, John Sillitoe on behalf of the Wellcome Sanger Institute COVID-19 Surveillance Team |
| EPI_ISL_2515765 | Lighthouse Laboratory Plymouth | Wellcome Sanger Institute for the COVID-19 Genomics UK (COG-UK) Consortium | Lighthouse Laboratory Plymouth and Alex Alderton, Roberto Amato, Jeffrey Barrett, Sonia Goncalves, Ewan Harrison, David K. Jackson, Ian Johnston, Dominic Kwiatkowski, Cordelia Langford, John Sillitoe on behalf of the Wellcome Sanger Institute COVID-19 Surveillance Team |
| EPI_ISL_2515766, EPI_ISL_2515768, EPI_ISL_2515769, EPI_ISL_2515770, EPI_ISL_2515771 | Lighthouse Lab in Alderley Park | Wellcome Sanger Institute for the COVID-19 Genomics UK (COG-UK) Consortium | Jacquelyn Wynn, Mairead Hyland, The Lighthouse Lab in Alderley Park and Alex Alderton, Roberto Amato, Jeffrey Barrett, Sonia Goncalves, Ewan Harrison, David K. Jackson, Ian Johnston, Dominic Kwiatkowski, Cordelia Langford, John Sillitoe on behalf of the Wellcome Sanger Institute COVID-19 Surveillance Team |
| EPI_ISL_2515772, EPI_ISL_2515773 | Lighthouse Laboratory Plymouth | Wellcome Sanger Institute for the COVID-19 Genomics UK (COG-UK) Consortium | Lighthouse Laboratory Plymouth and Alex Alderton, Roberto Amato, Jeffrey Barrett, Sonia Goncalves, Ewan Harrison, David K. Jackson, Ian Johnston, Dominic Kwiatkowski, Cordelia Langford, John Sillitoe on behalf of the Wellcome Sanger Institute COVID-19 Surveillance Team |
| EPI_ISL_2515774, EPI_ISL_2515775, EPI_ISL_2515776, EPI_ISL_2515777 | Lighthouse Lab in Alderley Park | Wellcome Sanger Institute for the COVID-19 Genomics UK (COG-UK) Consortium | Jacquelyn Wynn, Mairead Hyland, The Lighthouse Lab in Alderley Park and Alex Alderton, Roberto Amato, Jeffrey Barrett, Sonia Goncalves, Ewan Harrison, David K. Jackson, Ian Johnston, Dominic Kwiatkowski, Cordelia Langford, John Sillitoe on behalf of the Wellcome Sanger Institute COVID-19 Surveillance Team |
| EPI_ISL_2515778 | Lighthouse Laboratory Plymouth | Wellcome Sanger Institute for the COVID-19 Genomics UK (COG-UK) Consortium | Lighthouse Laboratory Plymouth and Alex Alderton, Roberto Amato, Jeffrey Barrett, Sonia Goncalves, Ewan Harrison, David K. Jackson, Ian Johnston, Dominic Kwiatkowski, Cordelia Langford, John Sillitoe on behalf of the Wellcome Sanger Institute COVID-19 Surveillance Team |
| EPI_ISL_2515779, EPI_ISL_2515780, EPI_ISL_2515781, EPI_ISL_2515782, EPI_ISL_2515783, EPI_ISL_2515784, EPI_ISL_2515785, EPI_ISL_2515786, EPI_ISL_2515787, EPI_ISL_2515788 | Lighthouse Lab in Alderley Park | Wellcome Sanger Institute for the COVID-19 Genomics UK (COG-UK) Consortium | Jacquelyn Wynn, Mairead Hyland, The Lighthouse Lab in Alderley Park and Alex Alderton, Roberto Amato, Jeffrey Barrett, Sonia Goncalves, Ewan Harrison, David K. Jackson, Ian Johnston, Dominic Kwiatkowski, Cordelia Langford, John Sillitoe on behalf of the Wellcome Sanger Institute COVID-19 Surveillance Team |
| EPI_ISL_2515789 | Lighthouse Laboratory Plymouth | Wellcome Sanger Institute for the COVID-19 Genomics UK (COG-UK) Consortium | Lighthouse Laboratory Plymouth and Alex Alderton, Roberto Amato, Jeffrey Barrett, Sonia Goncalves, Ewan Harrison, David K. Jackson, Ian Johnston, Dominic Kwiatkowski, Cordelia Langford, John Sillitoe on behalf of the Wellcome Sanger Institute COVID-19 Surveillance Team |
| EPI_ISL_2515790, EPI_ISL_2515791, EPI_ISL_2515792 | Lighthouse Lab in Alderley Park | Wellcome Sanger Institute for the COVID-19 Genomics UK (COG-UK) Consortium | Jacquelyn Wynn, Mairead Hyland, The Lighthouse Lab in Alderley Park and Alex Alderton, Roberto Amato, Jeffrey Barrett, Sonia Goncalves, Ewan Harrison, David K. Jackson, Ian Johnston, Dominic Kwiatkowski, Cordelia Langford, John Sillitoe on behalf of the Wellcome Sanger Institute COVID-19 Surveillance Team |
| EPI_ISL_2515793, EPI_ISL_2515794 | Lighthouse Laboratory Plymouth | Wellcome Sanger Institute for the COVID-19 Genomics UK (COG-UK) Consortium | Lighthouse Laboratory Plymouth and Alex Alderton, Roberto Amato, Jeffrey Barrett, Sonia Goncalves, Ewan Harrison, David K. Jackson, Ian Johnston, Dominic Kwiatkowski, Cordelia Langford, John Sillitoe on behalf of the Wellcome Sanger Institute COVID-19 Surveillance Team |
| EPI_ISL_2515795, EPI_ISL_2515796 | Lighthouse Lab in Alderley Park | Wellcome Sanger Institute for the COVID-19 Genomics UK (COG-UK) Consortium | Jacquelyn Wynn, Mairead Hyland, The Lighthouse Lab in Alderley Park and Alex Alderton, Roberto Amato, Jeffrey Barrett, Sonia Goncalves, Ewan Harrison, David K. Jackson, Ian Johnston, Dominic Kwiatkowski, Cordelia Langford, John Sillitoe on behalf of the Wellcome Sanger Institute COVID-19 Surveillance Team |
| EPI_ISL_2515797 | Lighthouse Laboratory Plymouth | Wellcome Sanger Institute for the COVID-19 Genomics UK (COG-UK) Consortium | Lighthouse Laboratory Plymouth and Alex Alderton, Roberto Amato, Jeffrey Barrett, Sonia Goncalves, Ewan Harrison, David K. Jackson, Ian Johnston, Dominic Kwiatkowski, Cordelia Langford, John Sillitoe on behalf of the Wellcome Sanger Institute COVID-19 Surveillance Team |
| EPI_ISL_2515798, EPI_ISL_2515799 | Lighthouse Lab in Alderley Park | Wellcome Sanger Institute for the COVID-19 Genomics UK (COG-UK) Consortium | Jacquelyn Wynn, Mairead Hyland, The Lighthouse Lab in Alderley Park and Alex Alderton, Roberto Amato, Jeffrey Barrett, Sonia Goncalves, Ewan Harrison, David K. Jackson, Ian Johnston, Dominic Kwiatkowski, Cordelia Langford, John Sillitoe on behalf of the Wellcome Sanger Institute COVID-19 Surveillance Team |
| EPI_ISL_2515800 | Lighthouse Laboratory Plymouth | Wellcome Sanger Institute for the COVID-19 Genomics UK (COG-UK) Consortium | Lighthouse Laboratory Plymouth and Alex Alderton, Roberto Amato, Jeffrey Barrett, Sonia Goncalves, Ewan Harrison, David K. Jackson, Ian Johnston, Dominic Kwiatkowski, Cordelia Langford, John Sillitoe on behalf of the Wellcome Sanger Institute COVID-19 Surveillance Team |
| EPI_ISL_2515802, EPI_ISL_2515803, EPI_ISL_2515804, EPI_ISL_2515805, EPI_ISL_2515806, EPI_ISL_2515807 | Lighthouse Lab in Alderley Park | Wellcome Sanger Institute for the COVID-19 Genomics UK (COG-UK) Consortium | Jacquelyn Wynn, Mairead Hyland, The Lighthouse Lab in Alderley Park and Alex Alderton, Roberto Amato, Jeffrey Barrett, Sonia Goncalves, Ewan Harrison, David K. Jackson, Ian Johnston, Dominic Kwiatkowski, Cordelia Langford, John Sillitoe on behalf of the Wellcome Sanger Institute COVID-19 Surveillance Team |
| EPI_ISL_2515808 | Lighthouse Laboratory Plymouth | Wellcome Sanger Institute for the COVID-19 Genomics UK (COG-UK) Consortium | Lighthouse Laboratory Plymouth and Alex Alderton, Roberto Amato, Jeffrey Barrett, Sonia Goncalves, Ewan Harrison, David K. Jackson, Ian Johnston, Dominic Kwiatkowski, Cordelia Langford, John Sillitoe on behalf of the Wellcome Sanger Institute COVID-19 Surveillance Team |
| EPI_ISL_2515809, EPI_ISL_2515810, | Lighthouse Lab in Alderley Park | Wellcome Sanger Institute for the COVID-19 Genomics UK (COG-UK) Consortium | Jacquelyn Wynn, Mairead Hyland, The Lighthouse Lab in Alderley Park and Alex Alderton, Roberto Amato, Jeffrey Barrett, Sonia Goncalves, Ewan Harrison, David K. Jackson, Ian Johnston, Dominic Kwiatkowski, Cordelia Langford, John Sillitoe on behalf of the Wellcome Sanger Institute COVID-19 Surveillance Team |

| | | | |
|---|---|---|---|
| EPI_ISL_2515811, EPI_ISL_2515812, EPI_ISL_2515814, EPI_ISL_2515815 | | | |
| EPI_ISL_2515816 | Lighthouse Laboratory Plymouth | Wellcome Sanger Institute for the COVID-19 Genomics UK (COG-UK) Consortium | Lighthouse Laboratory Plymouth and Alex Alderton, Roberto Amato, Jeffrey Barrett, Sonia Goncalves, Ewan Harrison, David K. Jackson, Ian Johnston, Dominic Kwiatkowski, Cordelia Langford, John Sillitoe on behalf of the Wellcome Sanger Institute COVID-19 Surveillance Team |
| EPI_ISL_2515817, EPI_ISL_2515818, EPI_ISL_2515819, EPI_ISL_2515820, EPI_ISL_2515821, EPI_ISL_2515822 | Lighthouse Lab in Alderley Park | Wellcome Sanger Institute for the COVID-19 Genomics UK (COG-UK) Consortium | Jacquelyn Wynn, Mairead Hyland, The Lighthouse Lab in Alderley Park and Alex Alderton, Roberto Amato, Jeffrey Barrett, Sonia Goncalves, Ewan Harrison, David K. Jackson, Ian Johnston, Dominic Kwiatkowski, Cordelia Langford, John Sillitoe on behalf of the Wellcome Sanger Institute COVID-19 Surveillance Team |
| EPI_ISL_2515823 | Lighthouse Laboratory Plymouth | Wellcome Sanger Institute for the COVID-19 Genomics UK (COG-UK) Consortium | Lighthouse Laboratory Plymouth and Alex Alderton, Roberto Amato, Jeffrey Barrett, Sonia Goncalves, Ewan Harrison, David K. Jackson, Ian Johnston, Dominic Kwiatkowski, Cordelia Langford, John Sillitoe on behalf of the Wellcome Sanger Institute COVID-19 Surveillance Team |
| EPI_ISL_2515825 | Lighthouse Lab in Alderley Park | Wellcome Sanger Institute for the COVID-19 Genomics UK (COG-UK) Consortium | Jacquelyn Wynn, Mairead Hyland, The Lighthouse Lab in Alderley Park and Alex Alderton, Roberto Amato, Jeffrey Barrett, Sonia Goncalves, Ewan Harrison, David K. Jackson, Ian Johnston, Dominic Kwiatkowski, Cordelia Langford, John Sillitoe on behalf of the Wellcome Sanger Institute COVID-19 Surveillance Team |
| EPI_ISL_2515826 | Lighthouse Laboratory Plymouth | Wellcome Sanger Institute for the COVID-19 Genomics UK (COG-UK) Consortium | Lighthouse Laboratory Plymouth and Alex Alderton, Roberto Amato, Jeffrey Barrett, Sonia Goncalves, Ewan Harrison, David K. Jackson, Ian Johnston, Dominic Kwiatkowski, Cordelia Langford, John Sillitoe on behalf of the Wellcome Sanger Institute COVID-19 Surveillance Team |
| EPI_ISL_2515827, EPI_ISL_2515828, EPI_ISL_2515829, EPI_ISL_2515830, EPI_ISL_2515831, EPI_ISL_2515832, EPI_ISL_2515833, EPI_ISL_2515834, EPI_ISL_2515835, EPI_ISL_2515836, EPI_ISL_2515837, EPI_ISL_2515838, EPI_ISL_2515839, EPI_ISL_2515840, EPI_ISL_2515841, EPI_ISL_2515842, EPI_ISL_2515843, EPI_ISL_2515845 | | | |
| see above | Lighthouse Lab in Alderley Park | Wellcome Sanger Institute for the COVID-19 Genomics UK (COG-UK) Consortium | Jacquelyn Wynn, Mairead Hyland, The Lighthouse Lab in Alderley Park and Alex Alderton, Roberto Amato, Jeffrey Barrett, Sonia Goncalves, Ewan Harrison, David K. Jackson, Ian Johnston, Dominic Kwiatkowski, Cordelia Langford, John Sillitoe on behalf of the Wellcome Sanger Institute COVID-19 Surveillance Team |
| EPI_ISL_2515846 | Lighthouse Lab in Milton Keynes | Wellcome Sanger Institute for the COVID-19 Genomics UK (COG-UK) Consortium | The Lighthouse Lab in Milton Keynes and Alex Alderton, Roberto Amato, Jeffrey Barrett, Sonia Goncalves, Ewan Harrison, David K. Jackson, Ian Johnston, Dominic Kwiatkowski, Cordelia Langford, John Sillitoe on behalf of the Wellcome Sanger Institute COVID-19 Surveillance Team |
| EPI_ISL_2515847, EPI_ISL_2515848 | Lighthouse Lab in Alderley Park | Wellcome Sanger Institute for the COVID-19 Genomics UK (COG-UK) Consortium | Jacquelyn Wynn, Mairead Hyland, The Lighthouse Lab in Alderley Park and Alex Alderton, Roberto Amato, Jeffrey Barrett, Sonia Goncalves, Ewan Harrison, David K. Jackson, Ian Johnston, Dominic Kwiatkowski, Cordelia Langford, John Sillitoe on behalf of the Wellcome Sanger Institute COVID-19 Surveillance Team |
| EPI_ISL_2515849 | Lighthouse Laboratory Plymouth | Wellcome Sanger Institute for the COVID-19 Genomics UK (COG-UK) Consortium | Lighthouse Laboratory Plymouth and Alex Alderton, Roberto Amato, Jeffrey Barrett, Sonia Goncalves, Ewan Harrison, David K. Jackson, Ian Johnston, Dominic Kwiatkowski, Cordelia Langford, John Sillitoe on behalf of the Wellcome Sanger Institute COVID-19 Surveillance Team |
| EPI_ISL_2515850 | Lighthouse Lab in Alderley Park | Wellcome Sanger Institute for the COVID-19 Genomics UK (COG-UK) Consortium | Jacquelyn Wynn, Mairead Hyland, The Lighthouse Lab in Alderley Park and Alex Alderton, Roberto Amato, Jeffrey Barrett, Sonia Goncalves, Ewan Harrison, David K. Jackson, Ian Johnston, Dominic Kwiatkowski, Cordelia Langford, John Sillitoe on behalf of the Wellcome Sanger Institute COVID-19 Surveillance Team |
| EPI_ISL_2515851 | Lighthouse Laboratory Plymouth | Wellcome Sanger Institute for the COVID-19 Genomics UK (COG-UK) Consortium | Lighthouse Laboratory Plymouth and Alex Alderton, Roberto Amato, Jeffrey Barrett, Sonia Goncalves, Ewan Harrison, David K. Jackson, Ian Johnston, Dominic Kwiatkowski, Cordelia Langford, John Sillitoe on behalf of the Wellcome Sanger Institute COVID-19 Surveillance Team |
| EPI_ISL_2515852, EPI_ISL_2515853, EPI_ISL_2515854, EPI_ISL_2515855, EPI_ISL_2515856, EPI_ISL_2515857, EPI_ISL_2515858, EPI_ISL_2515859 | Lighthouse Lab in Alderley Park | Wellcome Sanger Institute for the COVID-19 Genomics UK (COG-UK) Consortium | Jacquelyn Wynn, Mairead Hyland, The Lighthouse Lab in Alderley Park and Alex Alderton, Roberto Amato, Jeffrey Barrett, Sonia Goncalves, Ewan Harrison, David K. Jackson, Ian Johnston, Dominic Kwiatkowski, Cordelia Langford, John Sillitoe on behalf of the Wellcome Sanger Institute COVID-19 Surveillance Team |
| EPI_ISL_2515860 | Lighthouse Laboratory Plymouth | Wellcome Sanger Institute for the COVID-19 Genomics UK (COG-UK) Consortium | Lighthouse Laboratory Plymouth and Alex Alderton, Roberto Amato, Jeffrey Barrett, Sonia Goncalves, Ewan Harrison, David K. Jackson, Ian Johnston, Dominic Kwiatkowski, Cordelia Langford, John Sillitoe on behalf of the Wellcome Sanger Institute COVID-19 Surveillance Team |
| EPI_ISL_2515861, EPI_ISL_2515862, EPI_ISL_2515863, EPI_ISL_2515864, EPI_ISL_2515865 | Lighthouse Lab in Alderley Park | Wellcome Sanger Institute for the COVID-19 Genomics UK (COG-UK) Consortium | Jacquelyn Wynn, Mairead Hyland, The Lighthouse Lab in Alderley Park and Alex Alderton, Roberto Amato, Jeffrey Barrett, Sonia Goncalves, Ewan Harrison, David K. Jackson, Ian Johnston, Dominic Kwiatkowski, Cordelia Langford, John Sillitoe on behalf of the Wellcome Sanger Institute COVID-19 Surveillance Team |
| EPI_ISL_2515866 | Lighthouse Laboratory Plymouth | Wellcome Sanger Institute for the COVID-19 Genomics UK (COG-UK) Consortium | Lighthouse Laboratory Plymouth and Alex Alderton, Roberto Amato, Jeffrey Barrett, Sonia Goncalves, Ewan Harrison, David K. Jackson, Ian Johnston, Dominic Kwiatkowski, Cordelia Langford, John Sillitoe on behalf of the Wellcome Sanger Institute COVID-19 Surveillance Team |
| EPI_ISL_2515867 | Lighthouse Lab in Alderley Park | Wellcome Sanger Institute for the COVID-19 Genomics UK (COG-UK) Consortium | Jacquelyn Wynn, Mairead Hyland, The Lighthouse Lab in Alderley Park and Alex Alderton, Roberto Amato, Jeffrey Barrett, Sonia Goncalves, Ewan Harrison, David K. Jackson, Ian Johnston, Dominic Kwiatkowski, Cordelia Langford, John Sillitoe on behalf of the Wellcome Sanger Institute COVID-19 Surveillance Team |
| EPI_ISL_2515868 | Lighthouse Laboratory Plymouth | Wellcome Sanger Institute for the COVID-19 Genomics UK (COG-UK) Consortium | Lighthouse Laboratory Plymouth and Alex Alderton, Roberto Amato, Jeffrey Barrett, Sonia Goncalves, Ewan Harrison, David K. Jackson, Ian Johnston, Dominic Kwiatkowski, Cordelia Langford, John Sillitoe on behalf of the Wellcome Sanger Institute COVID-19 Surveillance Team |
| EPI_ISL_2515869, EPI_ISL_2515870, EPI_ISL_2515871 | Lighthouse Lab in Alderley Park | Wellcome Sanger Institute for the COVID-19 Genomics UK (COG-UK) Consortium | Jacquelyn Wynn, Mairead Hyland, The Lighthouse Lab in Alderley Park and Alex Alderton, Roberto Amato, Jeffrey Barrett, Sonia Goncalves, Ewan Harrison, David K. Jackson, Ian Johnston, Dominic Kwiatkowski, Cordelia Langford, John Sillitoe on behalf of the Wellcome Sanger Institute COVID-19 Surveillance Team |
| EPI_ISL_2515872 | Lighthouse Laboratory Plymouth | Wellcome Sanger Institute for the COVID-19 Genomics UK (COG-UK) Consortium | Lighthouse Laboratory Plymouth and Alex Alderton, Roberto Amato, Jeffrey Barrett, Sonia Goncalves, Ewan Harrison, David K. Jackson, Ian Johnston, Dominic Kwiatkowski, Cordelia Langford, John Sillitoe on behalf of the Wellcome Sanger Institute COVID-19 Surveillance Team |
| EPI_ISL_2515873 | Lighthouse Lab in Alderley Park | Wellcome Sanger Institute for the COVID-19 Genomics UK (COG-UK) Consortium | Jacquelyn Wynn, Mairead Hyland, The Lighthouse Lab in Alderley Park and Alex Alderton, Roberto Amato, Jeffrey Barrett, Sonia Goncalves, Ewan Harrison, David K. Jackson, Ian Johnston, Dominic Kwiatkowski, Cordelia Langford, John Sillitoe on behalf of the Wellcome Sanger Institute COVID-19 Surveillance Team |
| EPI_ISL_2515874 | Lighthouse Lab in Milton Keynes | Wellcome Sanger Institute for the COVID-19 Genomics UK (COG-UK) Consortium | The Lighthouse Lab in Milton Keynes and Alex Alderton, Roberto Amato, Jeffrey Barrett, Sonia Goncalves, Ewan Harrison, David K. Jackson, Ian Johnston, Dominic Kwiatkowski, Cordelia Langford, John Sillitoe on behalf of the Wellcome Sanger Institute COVID-19 Surveillance Team |
| EPI_ISL_2515875, EPI_ISL_2515876 | Lighthouse Lab in Alderley Park | Wellcome Sanger Institute for the COVID-19 Genomics UK (COG-UK) Consortium | Jacquelyn Wynn, Mairead Hyland, The Lighthouse Lab in Alderley Park and Alex Alderton, Roberto Amato, Jeffrey Barrett, Sonia Goncalves, Ewan Harrison, David K. Jackson, Ian Johnston, Dominic Kwiatkowski, Cordelia Langford, John Sillitoe on behalf of the Wellcome Sanger Institute COVID-19 Surveillance Team |
| EPI_ISL_2515878 | Lighthouse Laboratory Plymouth | Wellcome Sanger Institute for the COVID-19 Genomics UK (COG-UK) Consortium | Lighthouse Laboratory Plymouth and Alex Alderton, Roberto Amato, Jeffrey Barrett, Sonia Goncalves, Ewan Harrison, David K. Jackson, Ian Johnston, Dominic Kwiatkowski, Cordelia Langford, John Sillitoe on behalf of the Wellcome Sanger Institute COVID-19 Surveillance Team |
| EPI_ISL_2515879 | Lighthouse Lab in Alderley Park | Wellcome Sanger Institute for the COVID-19 Genomics UK (COG-UK) Consortium | Jacquelyn Wynn, Mairead Hyland, The Lighthouse Lab in Alderley Park and Alex Alderton, Roberto Amato, Jeffrey Barrett, Sonia Goncalves, Ewan Harrison, David K. Jackson, Ian Johnston, Dominic Kwiatkowski, Cordelia Langford, John Sillitoe on behalf of the Wellcome Sanger Institute COVID-19 Surveillance Team |
| EPI_ISL_2515880 | Lighthouse Lab in Milton Keynes | Wellcome Sanger Institute for the COVID-19 Genomics UK (COG-UK) Consortium | The Lighthouse Lab in Milton Keynes and Alex Alderton, Roberto Amato, Jeffrey Barrett, Sonia Goncalves, Ewan Harrison, David K. Jackson, Ian Johnston, Dominic Kwiatkowski, Cordelia Langford, John Sillitoe on behalf of the Wellcome Sanger Institute COVID-19 Surveillance Team |
| EPI_ISL_2515881, EPI_ISL_2515882, EPI_ISL_2515883 | Lighthouse Lab in Alderley Park | Wellcome Sanger Institute for the COVID-19 Genomics UK (COG-UK) Consortium | Jacquelyn Wynn, Mairead Hyland, The Lighthouse Lab in Alderley Park and Alex Alderton, Roberto Amato, Jeffrey Barrett, Sonia Goncalves, Ewan Harrison, David K. Jackson, Ian Johnston, Dominic Kwiatkowski, Cordelia Langford, John Sillitoe on behalf of the Wellcome Sanger Institute COVID-19 Surveillance Team |
| EPI_ISL_2515884, EPI_ISL_2515885, EPI_ISL_2515886 | Lighthouse Laboratory Plymouth | Wellcome Sanger Institute for the COVID-19 Genomics UK (COG-UK) Consortium | Lighthouse Laboratory Plymouth and Alex Alderton, Roberto Amato, Jeffrey Barrett, Sonia Goncalves, Ewan Harrison, David K. Jackson, Ian Johnston, Dominic Kwiatkowski, Cordelia Langford, John Sillitoe on behalf of the Wellcome Sanger Institute COVID-19 Surveillance Team |
| EPI_ISL_2515887, EPI_ISL_2515889, EPI_ISL_2515890, EPI_ISL_2515891, EPI_ISL_2515892, EPI_ISL_2515893 | Lighthouse Lab in Alderley Park | Wellcome Sanger Institute for the COVID-19 Genomics UK (COG-UK) Consortium | Jacquelyn Wynn, Mairead Hyland, The Lighthouse Lab in Alderley Park and Alex Alderton, Roberto Amato, Jeffrey Barrett, Sonia Goncalves, Ewan Harrison, David K. Jackson, Ian Johnston, Dominic Kwiatkowski, Cordelia Langford, John Sillitoe on behalf of the Wellcome Sanger Institute COVID-19 Surveillance Team |
| EPI_ISL_2515894 | Lighthouse Laboratory Plymouth | Wellcome Sanger Institute for the COVID-19 Genomics UK (COG-UK) Consortium | Lighthouse Laboratory Plymouth and Alex Alderton, Roberto Amato, Jeffrey Barrett, Sonia Goncalves, Ewan Harrison, David K. Jackson, Ian Johnston, Dominic Kwiatkowski, Cordelia Langford, John Sillitoe on behalf of the Wellcome Sanger Institute COVID-19 Surveillance Team |
| EPI_ISL_2515895 | Lighthouse Lab in Alderley Park | Wellcome Sanger Institute for the COVID-19 Genomics UK (COG-UK) Consortium | Jacquelyn Wynn, Mairead Hyland, The Lighthouse Lab in Alderley Park and Alex Alderton, Roberto Amato, Jeffrey Barrett, Sonia Goncalves, Ewan Harrison, David K. Jackson, Ian Johnston, Dominic Kwiatkowski, Cordelia Langford, John Sillitoe on behalf of the Wellcome Sanger Institute COVID-19 Surveillance Team |
| EPI_ISL_2515896 | Lighthouse Laboratory Plymouth | Wellcome Sanger Institute for the COVID-19 Genomics UK (COG-UK) Consortium | Lighthouse Laboratory Plymouth and Alex Alderton, Roberto Amato, Jeffrey Barrett, Sonia Goncalves, Ewan Harrison, David K. Jackson, Ian Johnston, Dominic Kwiatkowski, Cordelia Langford, John Sillitoe on behalf of the Wellcome Sanger Institute COVID-19 Surveillance Team |
| EPI_ISL_2515897 | Lighthouse Lab in Alderley Park | Wellcome Sanger Institute for the COVID-19 Genomics UK (COG-UK) Consortium | Jacquelyn Wynn, Mairead Hyland, The Lighthouse Lab in Alderley Park and Alex Alderton, Roberto Amato, Jeffrey Barrett, Sonia Goncalves, Ewan Harrison, David K. Jackson, Ian Johnston, Dominic Kwiatkowski, Cordelia Langford, John Sillitoe on behalf of the Wellcome Sanger Institute COVID-19 Surveillance Team |
| EPI_ISL_2515898 | Lighthouse Laboratory Plymouth | Wellcome Sanger Institute for the COVID-19 Genomics UK (COG-UK) Consortium | Lighthouse Laboratory Plymouth and Alex Alderton, Roberto Amato, Jeffrey Barrett, Sonia Goncalves, Ewan Harrison, David K. Jackson, Ian Johnston, Dominic Kwiatkowski, Cordelia Langford, John Sillitoe on behalf of the Wellcome Sanger Institute COVID-19 Surveillance Team |
| EPI_ISL_2515899, EPI_ISL_2515900, EPI_ISL_2515901 | Lighthouse Lab in Alderley Park | Wellcome Sanger Institute for the COVID-19 Genomics UK (COG-UK) Consortium | Jacquelyn Wynn, Mairead Hyland, The Lighthouse Lab in Alderley Park and Alex Alderton, Roberto Amato, Jeffrey Barrett, Sonia Goncalves, Ewan Harrison, David K. Jackson, Ian Johnston, Dominic Kwiatkowski, Cordelia Langford, John Sillitoe on behalf of the Wellcome Sanger Institute COVID-19 Surveillance Team |

| | | | |
|---|---|---|---|
| EPI_ISL_2515902 | Lighthouse Laboratory Plymouth | Wellcome Sanger Institute for the COVID-19 Genomics UK (COG-UK) Consortium | Lighthouse Laboratory Plymouth and Alex Alderton, Roberto Amato, Jeffrey Barrett, Sonia Goncalves, Ewan Harrison, David K. Jackson, Ian Johnston, Dominic Kwiatkowski, Cordelia Langford, John Sillitoe on behalf of the Wellcome Sanger Institute COVID-19 Surveillance Team |
| EPI_ISL_2515903 | Lighthouse Lab in Alderley Park | Wellcome Sanger Institute for the COVID-19 Genomics UK (COG-UK) Consortium | Jacquelyn Wynn, Mairead Hyland, The Lighthouse Lab in Alderley Park and Alex Alderton, Roberto Amato, Jeffrey Barrett, Sonia Goncalves, Ewan Harrison, David K. Jackson, Ian Johnston, Dominic Kwiatkowski, Cordelia Langford, John Sillitoe on behalf of the Wellcome Sanger Institute COVID-19 Surveillance Team |
| EPI_ISL_2515904, EPI_ISL_2515905 | Lighthouse Laboratory Plymouth | Wellcome Sanger Institute for the COVID-19 Genomics UK (COG-UK) Consortium | Lighthouse Laboratory Plymouth and Alex Alderton, Roberto Amato, Jeffrey Barrett, Sonia Goncalves, Ewan Harrison, David K. Jackson, Ian Johnston, Dominic Kwiatkowski, Cordelia Langford, John Sillitoe on behalf of the Wellcome Sanger Institute COVID-19 Surveillance Team |
| EPI_ISL_2515906 | Lighthouse Lab in Alderley Park | Wellcome Sanger Institute for the COVID-19 Genomics UK (COG-UK) Consortium | Jacquelyn Wynn, Mairead Hyland, The Lighthouse Lab in Alderley Park and Alex Alderton, Roberto Amato, Jeffrey Barrett, Sonia Goncalves, Ewan Harrison, David K. Jackson, Ian Johnston, Dominic Kwiatkowski, Cordelia Langford, John Sillitoe on behalf of the Wellcome Sanger Institute COVID-19 Surveillance Team |
| EPI_ISL_2515907 | Lighthouse Laboratory Plymouth | Wellcome Sanger Institute for the COVID-19 Genomics UK (COG-UK) Consortium | Lighthouse Laboratory Plymouth and Alex Alderton, Roberto Amato, Jeffrey Barrett, Sonia Goncalves, Ewan Harrison, David K. Jackson, Ian Johnston, Dominic Kwiatkowski, Cordelia Langford, John Sillitoe on behalf of the Wellcome Sanger Institute COVID-19 Surveillance Team |
| EPI_ISL_2515908 | Lighthouse Lab in Alderley Park | Wellcome Sanger Institute for the COVID-19 Genomics UK (COG-UK) Consortium | Jacquelyn Wynn, Mairead Hyland, The Lighthouse Lab in Alderley Park and Alex Alderton, Roberto Amato, Jeffrey Barrett, Sonia Goncalves, Ewan Harrison, David K. Jackson, Ian Johnston, Dominic Kwiatkowski, Cordelia Langford, John Sillitoe on behalf of the Wellcome Sanger Institute COVID-19 Surveillance Team |
| EPI_ISL_2515909, EPI_ISL_2515910 | Lighthouse Laboratory Plymouth | Wellcome Sanger Institute for the COVID-19 Genomics UK (COG-UK) Consortium | Lighthouse Laboratory Plymouth and Alex Alderton, Roberto Amato, Jeffrey Barrett, Sonia Goncalves, Ewan Harrison, David K. Jackson, Ian Johnston, Dominic Kwiatkowski, Cordelia Langford, John Sillitoe on behalf of the Wellcome Sanger Institute COVID-19 Surveillance Team |
| EPI_ISL_2515911, EPI_ISL_2515912, EPI_ISL_2515913, EPI_ISL_2515914 | Lighthouse Lab in Alderley Park | Wellcome Sanger Institute for the COVID-19 Genomics UK (COG-UK) Consortium | Jacquelyn Wynn, Mairead Hyland, The Lighthouse Lab in Alderley Park and Alex Alderton, Roberto Amato, Jeffrey Barrett, Sonia Goncalves, Ewan Harrison, David K. Jackson, Ian Johnston, Dominic Kwiatkowski, Cordelia Langford, John Sillitoe on behalf of the Wellcome Sanger Institute COVID-19 Surveillance Team |
| EPI_ISL_2515915 | Lighthouse Lab in Milton Keynes | Wellcome Sanger Institute for the COVID-19 Genomics UK (COG-UK) Consortium | The Lighthouse Lab in Milton Keynes and Alex Alderton, Roberto Amato, Jeffrey Barrett, Sonia Goncalves, Ewan Harrison, David K. Jackson, Ian Johnston, Dominic Kwiatkowski, Cordelia Langford, John Sillitoe on behalf of the Wellcome Sanger Institute COVID-19 Surveillance Team |
| EPI_ISL_2515916, EPI_ISL_2515917 | Lighthouse Lab in Alderley Park | Wellcome Sanger Institute for the COVID-19 Genomics UK (COG-UK) Consortium | Jacquelyn Wynn, Mairead Hyland, The Lighthouse Lab in Alderley Park and Alex Alderton, Roberto Amato, Jeffrey Barrett, Sonia Goncalves, Ewan Harrison, David K. Jackson, Ian Johnston, Dominic Kwiatkowski, Cordelia Langford, John Sillitoe on behalf of the Wellcome Sanger Institute COVID-19 Surveillance Team |
| EPI_ISL_2515918 | Lighthouse Laboratory Plymouth | Wellcome Sanger Institute for the COVID-19 Genomics UK (COG-UK) Consortium | Lighthouse Laboratory Plymouth and Alex Alderton, Roberto Amato, Jeffrey Barrett, Sonia Goncalves, Ewan Harrison, David K. Jackson, Ian Johnston, Dominic Kwiatkowski, Cordelia Langford, John Sillitoe on behalf of the Wellcome Sanger Institute COVID-19 Surveillance Team |
| EPI_ISL_2515919, EPI_ISL_2515920, EPI_ISL_2515921, EPI_ISL_2515922, EPI_ISL_2515923, EPI_ISL_2515924, EPI_ISL_2515925, EPI_ISL_2515926, EPI_ISL_2515927, EPI_ISL_2515928, EPI_ISL_2515929, EPI_ISL_2515930, EPI_ISL_2515931, EPI_ISL_2515932 | | | |
| see above | Lighthouse Lab in Alderley Park | Wellcome Sanger Institute for the COVID-19 Genomics UK (COG-UK) Consortium | Jacquelyn Wynn, Mairead Hyland, The Lighthouse Lab in Alderley Park and Alex Alderton, Roberto Amato, Jeffrey Barrett, Sonia Goncalves, Ewan Harrison, David K. Jackson, Ian Johnston, Dominic Kwiatkowski, Cordelia Langford, John Sillitoe on behalf of the Wellcome Sanger Institute COVID-19 Surveillance Team |
| EPI_ISL_2515933 | Lighthouse Laboratory Plymouth | Wellcome Sanger Institute for the COVID-19 Genomics UK (COG-UK) Consortium | Lighthouse Laboratory Plymouth and Alex Alderton, Roberto Amato, Jeffrey Barrett, Sonia Goncalves, Ewan Harrison, David K. Jackson, Ian Johnston, Dominic Kwiatkowski, Cordelia Langford, John Sillitoe on behalf of the Wellcome Sanger Institute COVID-19 Surveillance Team |
| EPI_ISL_2515934, EPI_ISL_2515935, EPI_ISL_2515936 | Lighthouse Lab in Alderley Park | Wellcome Sanger Institute for the COVID-19 Genomics UK (COG-UK) Consortium | Jacquelyn Wynn, Mairead Hyland, The Lighthouse Lab in Alderley Park and Alex Alderton, Roberto Amato, Jeffrey Barrett, Sonia Goncalves, Ewan Harrison, David K. Jackson, Ian Johnston, Dominic Kwiatkowski, Cordelia Langford, John Sillitoe on behalf of the Wellcome Sanger Institute COVID-19 Surveillance Team |
| EPI_ISL_2515937 | Lighthouse Laboratory Plymouth | Wellcome Sanger Institute for the COVID-19 Genomics UK (COG-UK) Consortium | Lighthouse Laboratory Plymouth and Alex Alderton, Roberto Amato, Jeffrey Barrett, Sonia Goncalves, Ewan Harrison, David K. Jackson, Ian Johnston, Dominic Kwiatkowski, Cordelia Langford, John Sillitoe on behalf of the Wellcome Sanger Institute COVID-19 Surveillance Team |
| EPI_ISL_2515938 | Lighthouse Lab in Milton Keynes | Wellcome Sanger Institute for the COVID-19 Genomics UK (COG-UK) Consortium | The Lighthouse Lab in Milton Keynes and Alex Alderton, Roberto Amato, Jeffrey Barrett, Sonia Goncalves, Ewan Harrison, David K. Jackson, Ian Johnston, Dominic Kwiatkowski, Cordelia Langford, John Sillitoe on behalf of the Wellcome Sanger Institute COVID-19 Surveillance Team |
| EPI_ISL_2515939, EPI_ISL_2515940, EPI_ISL_2515941, EPI_ISL_2515942, EPI_ISL_2515943, EPI_ISL_2515944, EPI_ISL_2515945, EPI_ISL_2515946, EPI_ISL_2515947 | Lighthouse Lab in Alderley Park | Wellcome Sanger Institute for the COVID-19 Genomics UK (COG-UK) Consortium | Jacquelyn Wynn, Mairead Hyland, The Lighthouse Lab in Alderley Park and Alex Alderton, Roberto Amato, Jeffrey Barrett, Sonia Goncalves, Ewan Harrison, David K. Jackson, Ian Johnston, Dominic Kwiatkowski, Cordelia Langford, John Sillitoe on behalf of the Wellcome Sanger Institute COVID-19 Surveillance Team |
| EPI_ISL_2515948, EPI_ISL_2515949 | Lighthouse Laboratory Plymouth | Wellcome Sanger Institute for the COVID-19 Genomics UK (COG-UK) Consortium | Lighthouse Laboratory Plymouth and Alex Alderton, Roberto Amato, Jeffrey Barrett, Sonia Goncalves, Ewan Harrison, David K. Jackson, Ian Johnston, Dominic Kwiatkowski, Cordelia Langford, John Sillitoe on behalf of the Wellcome Sanger Institute COVID-19 Surveillance Team |
| EPI_ISL_2515950, EPI_ISL_2515951, EPI_ISL_2515952, EPI_ISL_2515953, EPI_ISL_2515954, EPI_ISL_2515955, EPI_ISL_2515956, EPI_ISL_2515957, EPI_ISL_2515958, EPI_ISL_2515959, EPI_ISL_2515960, EPI_ISL_2515961, EPI_ISL_2515962 | | | |
| see above | Lighthouse Lab in Alderley Park | Wellcome Sanger Institute for the COVID-19 Genomics UK (COG-UK) Consortium | Jacquelyn Wynn, Mairead Hyland, The Lighthouse Lab in Alderley Park and Alex Alderton, Roberto Amato, Jeffrey Barrett, Sonia Goncalves, Ewan Harrison, David K. Jackson, Ian Johnston, Dominic Kwiatkowski, Cordelia Langford, John Sillitoe on behalf of the Wellcome Sanger Institute COVID-19 Surveillance Team |
| EPI_ISL_2515963, EPI_ISL_2515964 | Lighthouse Lab in Milton Keynes | Wellcome Sanger Institute for the COVID-19 Genomics UK (COG-UK) Consortium | The Lighthouse Lab in Milton Keynes and Alex Alderton, Roberto Amato, Jeffrey Barrett, Sonia Goncalves, Ewan Harrison, David K. Jackson, Ian Johnston, Dominic Kwiatkowski, Cordelia Langford, John Sillitoe on behalf of the Wellcome Sanger Institute COVID-19 Surveillance Team |
| EPI_ISL_2515965, EPI_ISL_2515966, EPI_ISL_2515967, EPI_ISL_2515968, EPI_ISL_2515969, EPI_ISL_2515970 | Lighthouse Lab in Alderley Park | Wellcome Sanger Institute for the COVID-19 Genomics UK (COG-UK) Consortium | Jacquelyn Wynn, Mairead Hyland, The Lighthouse Lab in Alderley Park and Alex Alderton, Roberto Amato, Jeffrey Barrett, Sonia Goncalves, Ewan Harrison, David K. Jackson, Ian Johnston, Dominic Kwiatkowski, Cordelia Langford, John Sillitoe on behalf of the Wellcome Sanger Institute COVID-19 Surveillance Team |
| EPI_ISL_2515971 | Lighthouse Laboratory Plymouth | Wellcome Sanger Institute for the COVID-19 Genomics UK (COG-UK) Consortium | Lighthouse Laboratory Plymouth and Alex Alderton, Roberto Amato, Jeffrey Barrett, Sonia Goncalves, Ewan Harrison, David K. Jackson, Ian Johnston, Dominic Kwiatkowski, Cordelia Langford, John Sillitoe on behalf of the Wellcome Sanger Institute COVID-19 Surveillance Team |
| EPI_ISL_2515972 | Lighthouse Lab in Alderley Park | Wellcome Sanger Institute for the COVID-19 Genomics UK (COG-UK) Consortium | Jacquelyn Wynn, Mairead Hyland, The Lighthouse Lab in Alderley Park and Alex Alderton, Roberto Amato, Jeffrey Barrett, Sonia Goncalves, Ewan Harrison, David K. Jackson, Ian Johnston, Dominic Kwiatkowski, Cordelia Langford, John Sillitoe on behalf of the Wellcome Sanger Institute COVID-19 Surveillance Team |
| EPI_ISL_2515973 | Lighthouse Lab in Milton Keynes | Wellcome Sanger Institute for the COVID-19 Genomics UK (COG-UK) Consortium | The Lighthouse Lab in Milton Keynes and Alex Alderton, Roberto Amato, Jeffrey Barrett, Sonia Goncalves, Ewan Harrison, David K. Jackson, Ian Johnston, Dominic Kwiatkowski, Cordelia Langford, John Sillitoe on behalf of the Wellcome Sanger Institute COVID-19 Surveillance Team |
| EPI_ISL_2515974, EPI_ISL_2515975, EPI_ISL_2515976, EPI_ISL_2515977, EPI_ISL_2515979, EPI_ISL_2515980, EPI_ISL_2515981 | Lighthouse Lab in Alderley Park | Wellcome Sanger Institute for the COVID-19 Genomics UK (COG-UK) Consortium | Jacquelyn Wynn, Mairead Hyland, The Lighthouse Lab in Alderley Park and Alex Alderton, Roberto Amato, Jeffrey Barrett, Sonia Goncalves, Ewan Harrison, David K. Jackson, Ian Johnston, Dominic Kwiatkowski, Cordelia Langford, John Sillitoe on behalf of the Wellcome Sanger Institute COVID-19 Surveillance Team |
| EPI_ISL_2515982 | Lighthouse Laboratory Plymouth | Wellcome Sanger Institute for the COVID-19 Genomics UK (COG-UK) Consortium | Lighthouse Laboratory Plymouth and Alex Alderton, Roberto Amato, Jeffrey Barrett, Sonia Goncalves, Ewan Harrison, David K. Jackson, Ian Johnston, Dominic Kwiatkowski, Cordelia Langford, John Sillitoe on behalf of the Wellcome Sanger Institute COVID-19 Surveillance Team |
| EPI_ISL_2515983, EPI_ISL_2515984, EPI_ISL_2515985 | Lighthouse Lab in Alderley Park | Wellcome Sanger Institute for the COVID-19 Genomics UK (COG-UK) Consortium | Jacquelyn Wynn, Mairead Hyland, The Lighthouse Lab in Alderley Park and Alex Alderton, Roberto Amato, Jeffrey Barrett, Sonia Goncalves, Ewan Harrison, David K. Jackson, Ian Johnston, Dominic Kwiatkowski, Cordelia Langford, John Sillitoe on behalf of the Wellcome Sanger Institute COVID-19 Surveillance Team |
| EPI_ISL_2515986 | Lighthouse Laboratory Plymouth | Wellcome Sanger Institute for the COVID-19 Genomics UK (COG-UK) Consortium | Lighthouse Laboratory Plymouth and Alex Alderton, Roberto Amato, Jeffrey Barrett, Sonia Goncalves, Ewan Harrison, David K. Jackson, Ian Johnston, Dominic Kwiatkowski, Cordelia Langford, John Sillitoe on behalf of the Wellcome Sanger Institute COVID-19 Surveillance Team |
| EPI_ISL_2515987 | Lighthouse Lab in Alderley Park | Wellcome Sanger Institute for the COVID-19 Genomics UK (COG-UK) Consortium | Jacquelyn Wynn, Mairead Hyland, The Lighthouse Lab in Alderley Park and Alex Alderton, Roberto Amato, Jeffrey Barrett, Sonia Goncalves, Ewan Harrison, David K. Jackson, Ian Johnston, Dominic Kwiatkowski, Cordelia Langford, John Sillitoe on behalf of the Wellcome Sanger Institute COVID-19 Surveillance Team |
| EPI_ISL_2515988, EPI_ISL_2515989 | Lighthouse Laboratory Plymouth | Wellcome Sanger Institute for the COVID-19 Genomics UK (COG-UK) Consortium | Lighthouse Laboratory Plymouth and Alex Alderton, Roberto Amato, Jeffrey Barrett, Sonia Goncalves, Ewan Harrison, David K. Jackson, Ian Johnston, Dominic Kwiatkowski, Cordelia Langford, John Sillitoe on behalf of the Wellcome Sanger Institute COVID-19 Surveillance Team |
| EPI_ISL_2515990, EPI_ISL_2515991, EPI_ISL_2515992, EPI_ISL_2515993, EPI_ISL_2515994, EPI_ISL_2515995, EPI_ISL_2515996 | Lighthouse Lab in Alderley Park | Wellcome Sanger Institute for the COVID-19 Genomics UK (COG-UK) Consortium | Jacquelyn Wynn, Mairead Hyland, The Lighthouse Lab in Alderley Park and Alex Alderton, Roberto Amato, Jeffrey Barrett, Sonia Goncalves, Ewan Harrison, David K. Jackson, Ian Johnston, Dominic Kwiatkowski, Cordelia Langford, John Sillitoe on behalf of the Wellcome Sanger Institute COVID-19 Surveillance Team |
| EPI_ISL_2515997 | Lighthouse Laboratory Plymouth | Wellcome Sanger Institute for the COVID-19 Genomics UK (COG-UK) Consortium | Lighthouse Laboratory Plymouth and Alex Alderton, Roberto Amato, Jeffrey Barrett, Sonia Goncalves, Ewan Harrison, David K. Jackson, Ian Johnston, Dominic Kwiatkowski, Cordelia Langford, John Sillitoe on behalf of the Wellcome Sanger Institute COVID-19 Surveillance Team |
| EPI_ISL_2515998, EPI_ISL_2515999 | Lighthouse Lab in Alderley Park | Wellcome Sanger Institute for the COVID-19 Genomics UK (COG-UK) Consortium | Jacquelyn Wynn, Mairead Hyland, The Lighthouse Lab in Alderley Park and Alex Alderton, Roberto Amato, Jeffrey Barrett, Sonia Goncalves, Ewan Harrison, David K. Jackson, Ian Johnston, Dominic Kwiatkowski, Cordelia Langford, John Sillitoe on behalf of the Wellcome Sanger Institute COVID-19 Surveillance Team |
| EPI_ISL_2516000 | Lighthouse Lab in Milton Keynes | Wellcome Sanger Institute for the COVID-19 Genomics UK (COG-UK) Consortium | The Lighthouse Lab in Milton Keynes and Alex Alderton, Roberto Amato, Jeffrey Barrett, Sonia Goncalves, Ewan Harrison, David K. Jackson, Ian Johnston, Dominic Kwiatkowski, Cordelia Langford, John Sillitoe on behalf of the Wellcome Sanger Institute COVID-19 Surveillance Team |

| | | | |
|---|---|---|---|
| EPI_ISL_2516001, EPI_ISL_2516002, EPI_ISL_2516003, EPI_ISL_2516004, EPI_ISL_2516005, EPI_ISL_2516006 | Lighthouse Lab in Alderley Park | Wellcome Sanger Institute for the COVID-19 Genomics UK (COG-UK) Consortium | Jacquelyn Wynn, Mairead Hyland, The Lighthouse Lab in Alderley Park and Alex Alderton, Roberto Amato, Jeffrey Barrett, Sonia Goncalves, Ewan Harrison, David K. Jackson, Ian Johnston, Dominic Kwiatkowski, Cordelia Langford, John Sillitoe on behalf of the Wellcome Sanger Institute COVID-19 Surveillance Team |
| EPI_ISL_2516007 | Lighthouse Laboratory Plymouth | Wellcome Sanger Institute for the COVID-19 Genomics UK (COG-UK) Consortium | Lighthouse Laboratory Plymouth and Alex Alderton, Roberto Amato, Jeffrey Barrett, Sonia Goncalves, Ewan Harrison, David K. Jackson, Ian Johnston, Dominic Kwiatkowski, Cordelia Langford, John Sillitoe on behalf of the Wellcome Sanger Institute COVID-19 Surveillance Team |
| EPI_ISL_2516008, EPI_ISL_2516009 | Lighthouse Lab in Alderley Park | Wellcome Sanger Institute for the COVID-19 Genomics UK (COG-UK) Consortium | Jacquelyn Wynn, Mairead Hyland, The Lighthouse Lab in Alderley Park and Alex Alderton, Roberto Amato, Jeffrey Barrett, Sonia Goncalves, Ewan Harrison, David K. Jackson, Ian Johnston, Dominic Kwiatkowski, Cordelia Langford, John Sillitoe on behalf of the Wellcome Sanger Institute COVID-19 Surveillance Team |
| EPI_ISL_2516010, EPI_ISL_2516011 | Health Services Laboratories | Wellcome Sanger Institute for the COVID-19 Genomics UK (COG-UK) Consortium | Health Services Laboratories and Alex Alderton, Roberto Amato, Jeffrey Barrett, Sonia Goncalves, Ewan Harrison, David K. Jackson, Ian Johnston, Dominic Kwiatkowski, Cordelia Langford, John Sillitoe on behalf of the Wellcome Sanger Institute COVID-19 Surveillance Team |
| EPI_ISL_2516012, EPI_ISL_2516013 | Lighthouse Lab in Alderley Park | Wellcome Sanger Institute for the COVID-19 Genomics UK (COG-UK) Consortium | Jacquelyn Wynn, Mairead Hyland, The Lighthouse Lab in Alderley Park and Alex Alderton, Roberto Amato, Jeffrey Barrett, Sonia Goncalves, Ewan Harrison, David K. Jackson, Ian Johnston, Dominic Kwiatkowski, Cordelia Langford, John Sillitoe on behalf of the Wellcome Sanger Institute COVID-19 Surveillance Team |
| EPI_ISL_2516014 | Lighthouse Lab in Glasgow | Wellcome Sanger Institute for the COVID-19 Genomics UK (COG-UK) Consortium | Harper VanSteenhouse, Yumi Kasai, David Gray, Carol Clugston, Anna Dominiczak and Alex Alderton, Roberto Amato, Jeffrey Barrett, Sonia Goncalves, Ewan Harrison, David K. Jackson, Ian Johnston, Dominic Kwiatkowski, Cordelia Langford, John Sillitoe on behalf of the Wellcome Sanger Institute COVID-19 Surveillance Team |
| EPI_ISL_2516015, EPI_ISL_2516016 | Lighthouse Lab in Alderley Park | Wellcome Sanger Institute for the COVID-19 Genomics UK (COG-UK) Consortium | Jacquelyn Wynn, Mairead Hyland, The Lighthouse Lab in Alderley Park and Alex Alderton, Roberto Amato, Jeffrey Barrett, Sonia Goncalves, Ewan Harrison, David K. Jackson, Ian Johnston, Dominic Kwiatkowski, Cordelia Langford, John Sillitoe on behalf of the Wellcome Sanger Institute COVID-19 Surveillance Team |
| EPI_ISL_2516017 | Health Services Laboratories | Wellcome Sanger Institute for the COVID-19 Genomics UK (COG-UK) Consortium | Health Services Laboratories and Alex Alderton, Roberto Amato, Jeffrey Barrett, Sonia Goncalves, Ewan Harrison, David K. Jackson, Ian Johnston, Dominic Kwiatkowski, Cordelia Langford, John Sillitoe on behalf of the Wellcome Sanger Institute COVID-19 Surveillance Team |
| EPI_ISL_2516018 | Lighthouse Lab in Alderley Park | Wellcome Sanger Institute for the COVID-19 Genomics UK (COG-UK) Consortium | Jacquelyn Wynn, Mairead Hyland, The Lighthouse Lab in Alderley Park and Alex Alderton, Roberto Amato, Jeffrey Barrett, Sonia Goncalves, Ewan Harrison, David K. Jackson, Ian Johnston, Dominic Kwiatkowski, Cordelia Langford, John Sillitoe on behalf of the Wellcome Sanger Institute COVID-19 Surveillance Team |
| EPI_ISL_2516021 | Lighthouse Lab in Glasgow | Wellcome Sanger Institute for the COVID-19 Genomics UK (COG-UK) Consortium | Harper VanSteenhouse, Yumi Kasai, David Gray, Carol Clugston, Anna Dominiczak and Alex Alderton, Roberto Amato, Jeffrey Barrett, Sonia Goncalves, Ewan Harrison, David K. Jackson, Ian Johnston, Dominic Kwiatkowski, Cordelia Langford, John Sillitoe on behalf of the Wellcome Sanger Institute COVID-19 Surveillance Team |
| EPI_ISL_2516023 | Lighthouse Lab in Alderley Park | Wellcome Sanger Institute for the COVID-19 Genomics UK (COG-UK) Consortium | Jacquelyn Wynn, Mairead Hyland, The Lighthouse Lab in Alderley Park and Alex Alderton, Roberto Amato, Jeffrey Barrett, Sonia Goncalves, Ewan Harrison, David K. Jackson, Ian Johnston, Dominic Kwiatkowski, Cordelia Langford, John Sillitoe on behalf of the Wellcome Sanger Institute COVID-19 Surveillance Team |
| EPI_ISL_2516024 | Health Services Laboratories | Wellcome Sanger Institute for the COVID-19 Genomics UK (COG-UK) Consortium | Health Services Laboratories and Alex Alderton, Roberto Amato, Jeffrey Barrett, Sonia Goncalves, Ewan Harrison, David K. Jackson, Ian Johnston, Dominic Kwiatkowski, Cordelia Langford, John Sillitoe on behalf of the Wellcome Sanger Institute COVID-19 Surveillance Team |
| EPI_ISL_2516025, EPI_ISL_2516026 | Lighthouse Lab in Alderley Park | Wellcome Sanger Institute for the COVID-19 Genomics UK (COG-UK) Consortium | Jacquelyn Wynn, Mairead Hyland, The Lighthouse Lab in Alderley Park and Alex Alderton, Roberto Amato, Jeffrey Barrett, Sonia Goncalves, Ewan Harrison, David K. Jackson, Ian Johnston, Dominic Kwiatkowski, Cordelia Langford, John Sillitoe on behalf of the Wellcome Sanger Institute COVID-19 Surveillance Team |
| EPI_ISL_2516027 | Health Services Laboratories | Wellcome Sanger Institute for the COVID-19 Genomics UK (COG-UK) Consortium | Health Services Laboratories and Alex Alderton, Roberto Amato, Jeffrey Barrett, Sonia Goncalves, Ewan Harrison, David K. Jackson, Ian Johnston, Dominic Kwiatkowski, Cordelia Langford, John Sillitoe on behalf of the Wellcome Sanger Institute COVID-19 Surveillance Team |
| EPI_ISL_2516029, EPI_ISL_2516030, EPI_ISL_2516031 | Lighthouse Lab in Alderley Park | Wellcome Sanger Institute for the COVID-19 Genomics UK (COG-UK) Consortium | Jacquelyn Wynn, Mairead Hyland, The Lighthouse Lab in Alderley Park and Alex Alderton, Roberto Amato, Jeffrey Barrett, Sonia Goncalves, Ewan Harrison, David K. Jackson, Ian Johnston, Dominic Kwiatkowski, Cordelia Langford, John Sillitoe on behalf of the Wellcome Sanger Institute COVID-19 Surveillance Team |
| EPI_ISL_2516032 | Health Services Laboratories | Wellcome Sanger Institute for the COVID-19 Genomics UK (COG-UK) Consortium | Health Services Laboratories and Alex Alderton, Roberto Amato, Jeffrey Barrett, Sonia Goncalves, Ewan Harrison, David K. Jackson, Ian Johnston, Dominic Kwiatkowski, Cordelia Langford, John Sillitoe on behalf of the Wellcome Sanger Institute COVID-19 Surveillance Team |
| EPI_ISL_2516033, EPI_ISL_2516034 | Lighthouse Lab in Alderley Park | Wellcome Sanger Institute for the COVID-19 Genomics UK (COG-UK) Consortium | Jacquelyn Wynn, Mairead Hyland, The Lighthouse Lab in Alderley Park and Alex Alderton, Roberto Amato, Jeffrey Barrett, Sonia Goncalves, Ewan Harrison, David K. Jackson, Ian Johnston, Dominic Kwiatkowski, Cordelia Langford, John Sillitoe on behalf of the Wellcome Sanger Institute COVID-19 Surveillance Team |
| EPI_ISL_2516035 | Health Services Laboratories | Wellcome Sanger Institute for the COVID-19 Genomics UK (COG-UK) Consortium | Health Services Laboratories and Alex Alderton, Roberto Amato, Jeffrey Barrett, Sonia Goncalves, Ewan Harrison, David K. Jackson, Ian Johnston, Dominic Kwiatkowski, Cordelia Langford, John Sillitoe on behalf of the Wellcome Sanger Institute COVID-19 Surveillance Team |
| EPI_ISL_2516036, EPI_ISL_2516037, EPI_ISL_2516038, EPI_ISL_2516039, EPI_ISL_2516040 | Lighthouse Lab in Alderley Park | Wellcome Sanger Institute for the COVID-19 Genomics UK (COG-UK) Consortium | Jacquelyn Wynn, Mairead Hyland, The Lighthouse Lab in Alderley Park and Alex Alderton, Roberto Amato, Jeffrey Barrett, Sonia Goncalves, Ewan Harrison, David K. Jackson, Ian Johnston, Dominic Kwiatkowski, Cordelia Langford, John Sillitoe on behalf of the Wellcome Sanger Institute COVID-19 Surveillance Team |
| EPI_ISL_2516041 | Lighthouse Lab in Glasgow | Wellcome Sanger Institute for the COVID-19 Genomics UK (COG-UK) Consortium | Harper VanSteenhouse, Yumi Kasai, David Gray, Carol Clugston, Anna Dominiczak and Alex Alderton, Roberto Amato, Jeffrey Barrett, Sonia Goncalves, Ewan Harrison, David K. Jackson, Ian Johnston, Dominic Kwiatkowski, Cordelia Langford, John Sillitoe on behalf of the Wellcome Sanger Institute COVID-19 Surveillance Team |
| EPI_ISL_2516042, EPI_ISL_2516043, EPI_ISL_2516044, EPI_ISL_2516045 | Lighthouse Lab in Alderley Park | Wellcome Sanger Institute for the COVID-19 Genomics UK (COG-UK) Consortium | Jacquelyn Wynn, Mairead Hyland, The Lighthouse Lab in Alderley Park and Alex Alderton, Roberto Amato, Jeffrey Barrett, Sonia Goncalves, Ewan Harrison, David K. Jackson, Ian Johnston, Dominic Kwiatkowski, Cordelia Langford, John Sillitoe on behalf of the Wellcome Sanger Institute COVID-19 Surveillance Team |
| EPI_ISL_2516047 | Health Services Laboratories | Wellcome Sanger Institute for the COVID-19 Genomics UK (COG-UK) Consortium | Health Services Laboratories and Alex Alderton, Roberto Amato, Jeffrey Barrett, Sonia Goncalves, Ewan Harrison, David K. Jackson, Ian Johnston, Dominic Kwiatkowski, Cordelia Langford, John Sillitoe on behalf of the Wellcome Sanger Institute COVID-19 Surveillance Team |
| EPI_ISL_2516048 | Lighthouse Lab in Alderley Park | Wellcome Sanger Institute for the COVID-19 Genomics UK (COG-UK) Consortium | Jacquelyn Wynn, Mairead Hyland, The Lighthouse Lab in Alderley Park and Alex Alderton, Roberto Amato, Jeffrey Barrett, Sonia Goncalves, Ewan Harrison, David K. Jackson, Ian Johnston, Dominic Kwiatkowski, Cordelia Langford, John Sillitoe on behalf of the Wellcome Sanger Institute COVID-19 Surveillance Team |
| EPI_ISL_2516050 | Health Services Laboratories | Wellcome Sanger Institute for the COVID-19 Genomics UK (COG-UK) Consortium | Health Services Laboratories and Alex Alderton, Roberto Amato, Jeffrey Barrett, Sonia Goncalves, Ewan Harrison, David K. Jackson, Ian Johnston, Dominic Kwiatkowski, Cordelia Langford, John Sillitoe on behalf of the Wellcome Sanger Institute COVID-19 Surveillance Team |
| EPI_ISL_2516051 | Lighthouse Lab in Alderley Park | Wellcome Sanger Institute for the COVID-19 Genomics UK (COG-UK) Consortium | Jacquelyn Wynn, Mairead Hyland, The Lighthouse Lab in Alderley Park and Alex Alderton, Roberto Amato, Jeffrey Barrett, Sonia Goncalves, Ewan Harrison, David K. Jackson, Ian Johnston, Dominic Kwiatkowski, Cordelia Langford, John Sillitoe on behalf of the Wellcome Sanger Institute COVID-19 Surveillance Team |
| EPI_ISL_2516052, EPI_ISL_2516053 | Health Services Laboratories | Wellcome Sanger Institute for the COVID-19 Genomics UK (COG-UK) Consortium | Health Services Laboratories and Alex Alderton, Roberto Amato, Jeffrey Barrett, Sonia Goncalves, Ewan Harrison, David K. Jackson, Ian Johnston, Dominic Kwiatkowski, Cordelia Langford, John Sillitoe on behalf of the Wellcome Sanger Institute COVID-19 Surveillance Team |
| EPI_ISL_2516054, EPI_ISL_2516055, EPI_ISL_2516056, EPI_ISL_2516057, EPI_ISL_2516058, EPI_ISL_2516059, EPI_ISL_2516061, EPI_ISL_2516062 | Lighthouse Lab in Alderley Park | Wellcome Sanger Institute for the COVID-19 Genomics UK (COG-UK) Consortium | Jacquelyn Wynn, Mairead Hyland, The Lighthouse Lab in Alderley Park and Alex Alderton, Roberto Amato, Jeffrey Barrett, Sonia Goncalves, Ewan Harrison, David K. Jackson, Ian Johnston, Dominic Kwiatkowski, Cordelia Langford, John Sillitoe on behalf of the Wellcome Sanger Institute COVID-19 Surveillance Team |
| EPI_ISL_2516063 | Health Services Laboratories | Wellcome Sanger Institute for the COVID-19 Genomics UK (COG-UK) Consortium | Health Services Laboratories and Alex Alderton, Roberto Amato, Jeffrey Barrett, Sonia Goncalves, Ewan Harrison, David K. Jackson, Ian Johnston, Dominic Kwiatkowski, Cordelia Langford, John Sillitoe on behalf of the Wellcome Sanger Institute COVID-19 Surveillance Team |
| EPI_ISL_2516064 | Lighthouse Lab in Alderley Park | Wellcome Sanger Institute for the COVID-19 Genomics UK (COG-UK) Consortium | Jacquelyn Wynn, Mairead Hyland, The Lighthouse Lab in Alderley Park and Alex Alderton, Roberto Amato, Jeffrey Barrett, Sonia Goncalves, Ewan Harrison, David K. Jackson, Ian Johnston, Dominic Kwiatkowski, Cordelia Langford, John Sillitoe on behalf of the Wellcome Sanger Institute COVID-19 Surveillance Team |
| EPI_ISL_2516065 | Health Services Laboratories | Wellcome Sanger Institute for the COVID-19 Genomics UK (COG-UK) Consortium | Health Services Laboratories and Alex Alderton, Roberto Amato, Jeffrey Barrett, Sonia Goncalves, Ewan Harrison, David K. Jackson, Ian Johnston, Dominic Kwiatkowski, Cordelia Langford, John Sillitoe on behalf of the Wellcome Sanger Institute COVID-19 Surveillance Team |
| EPI_ISL_2516066 | Lighthouse Lab in Alderley Park | Wellcome Sanger Institute for the COVID-19 Genomics UK (COG-UK) Consortium | Jacquelyn Wynn, Mairead Hyland, The Lighthouse Lab in Alderley Park and Alex Alderton, Roberto Amato, Jeffrey Barrett, Sonia Goncalves, Ewan Harrison, David K. Jackson, Ian Johnston, Dominic Kwiatkowski, Cordelia Langford, John Sillitoe on behalf of the Wellcome Sanger Institute COVID-19 Surveillance Team |
| EPI_ISL_2516067 | Health Services Laboratories | Wellcome Sanger Institute for the COVID-19 Genomics UK (COG-UK) Consortium | Health Services Laboratories and Alex Alderton, Roberto Amato, Jeffrey Barrett, Sonia Goncalves, Ewan Harrison, David K. Jackson, Ian Johnston, Dominic Kwiatkowski, Cordelia Langford, John Sillitoe on behalf of the Wellcome Sanger Institute COVID-19 Surveillance Team |
| EPI_ISL_2516068 | Lighthouse Lab in Alderley Park | Wellcome Sanger Institute for the COVID-19 Genomics UK (COG-UK) Consortium | Jacquelyn Wynn, Mairead Hyland, The Lighthouse Lab in Alderley Park and Alex Alderton, Roberto Amato, Jeffrey Barrett, Sonia Goncalves, Ewan Harrison, David K. Jackson, Ian Johnston, Dominic Kwiatkowski, Cordelia Langford, John Sillitoe on behalf of the Wellcome Sanger Institute COVID-19 Surveillance Team |
| EPI_ISL_2516069 | Health Services Laboratories | Wellcome Sanger Institute for the COVID-19 Genomics UK (COG-UK) Consortium | Health Services Laboratories and Alex Alderton, Roberto Amato, Jeffrey Barrett, Sonia Goncalves, Ewan Harrison, David K. Jackson, Ian Johnston, Dominic Kwiatkowski, Cordelia Langford, John Sillitoe on behalf of the Wellcome Sanger Institute COVID-19 Surveillance Team |
| EPI_ISL_2516070, EPI_ISL_2516072, EPI_ISL_2516073 | Lighthouse Lab in Alderley Park | Wellcome Sanger Institute for the COVID-19 Genomics UK (COG-UK) Consortium | Jacquelyn Wynn, Mairead Hyland, The Lighthouse Lab in Alderley Park and Alex Alderton, Roberto Amato, Jeffrey Barrett, Sonia Goncalves, Ewan Harrison, David K. Jackson, Ian Johnston, Dominic Kwiatkowski, Cordelia Langford, John Sillitoe on behalf of the Wellcome Sanger Institute COVID-19 Surveillance Team |
| EPI_ISL_2516074 | Health Services Laboratories | Wellcome Sanger Institute for the COVID-19 Genomics UK (COG-UK) Consortium | Health Services Laboratories and Alex Alderton, Roberto Amato, Jeffrey Barrett, Sonia Goncalves, Ewan Harrison, David K. Jackson, Ian Johnston, Dominic Kwiatkowski, Cordelia Langford, John Sillitoe on behalf of the Wellcome Sanger Institute COVID-19 Surveillance Team |
| EPI_ISL_2516075, EPI_ISL_2516076, EPI_ISL_2516077, EPI_ISL_2516078, EPI_ISL_2516079, | Lighthouse Lab in Alderley Park | Wellcome Sanger Institute for the COVID-19 Genomics UK (COG-UK) Consortium | Jacquelyn Wynn, Mairead Hyland, The Lighthouse Lab in Alderley Park and Alex Alderton, Roberto Amato, Jeffrey Barrett, Sonia Goncalves, Ewan Harrison, David K. Jackson, Ian Johnston, Dominic Kwiatkowski, Cordelia Langford, John Sillitoe on behalf of the Wellcome Sanger Institute COVID-19 Surveillance Team |

| | | | |
|---|---|---|---|
| EPI_ISL_2516080, EPI_ISL_2516081, EPI_ISL_2516082 | | | |
| EPI_ISL_2516083 | Health Services Laboratories | Wellcome Sanger Institute for the COVID-19 Genomics UK (COG-UK) Consortium | Health Services Laboratories and Alex Alderton, Roberto Amato, Jeffrey Barrett, Sonia Goncalves, Ewan Harrison, David K. Jackson, Ian Johnston, Dominic Kwiatkowski, Cordelia Langford, John Sillitoe on behalf of the Wellcome Sanger Institute COVID-19 Surveillance Team |
| EPI_ISL_2516084, EPI_ISL_2516085 | Lighthouse Lab in Alderley Park | Wellcome Sanger Institute for the COVID-19 Genomics UK (COG-UK) Consortium | Jacquelyn Wynn, Mairead Hyland, The Lighthouse Lab in Alderley Park and Alex Alderton, Roberto Amato, Jeffrey Barrett, Sonia Goncalves, Ewan Harrison, David K. Jackson, Ian Johnston, Dominic Kwiatkowski, Cordelia Langford, John Sillitoe on behalf of the Wellcome Sanger Institute COVID-19 Surveillance Team |
| EPI_ISL_2516086 | Lighthouse Lab in Glasgow | Wellcome Sanger Institute for the COVID-19 Genomics UK (COG-UK) Consortium | Harper VanSteenhouse, Yumi Kasai, David Gray, Carol Clugston, Anna Dominiczak and Alex Alderton, Roberto Amato, Jeffrey Barrett, Sonia Goncalves, Ewan Harrison, David K. Jackson, Ian Johnston, Dominic Kwiatkowski, Cordelia Langford, John Sillitoe on behalf of the Wellcome Sanger Institute COVID-19 Surveillance Team |
| EPI_ISL_2516087, EPI_ISL_2516088, EPI_ISL_2516089 | Lighthouse Lab in Alderley Park | Wellcome Sanger Institute for the COVID-19 Genomics UK (COG-UK) Consortium | Jacquelyn Wynn, Mairead Hyland, The Lighthouse Lab in Alderley Park and Alex Alderton, Roberto Amato, Jeffrey Barrett, Sonia Goncalves, Ewan Harrison, David K. Jackson, Ian Johnston, Dominic Kwiatkowski, Cordelia Langford, John Sillitoe on behalf of the Wellcome Sanger Institute COVID-19 Surveillance Team |
| EPI_ISL_2516090 | Lighthouse Lab in Glasgow | Wellcome Sanger Institute for the COVID-19 Genomics UK (COG-UK) Consortium | Harper VanSteenhouse, Yumi Kasai, David Gray, Carol Clugston, Anna Dominiczak and Alex Alderton, Roberto Amato, Jeffrey Barrett, Sonia Goncalves, Ewan Harrison, David K. Jackson, Ian Johnston, Dominic Kwiatkowski, Cordelia Langford, John Sillitoe on behalf of the Wellcome Sanger Institute COVID-19 Surveillance Team |
| EPI_ISL_2516091 | Health Services Laboratories | Wellcome Sanger Institute for the COVID-19 Genomics UK (COG-UK) Consortium | Health Services Laboratories and Alex Alderton, Roberto Amato, Jeffrey Barrett, Sonia Goncalves, Ewan Harrison, David K. Jackson, Ian Johnston, Dominic Kwiatkowski, Cordelia Langford, John Sillitoe on behalf of the Wellcome Sanger Institute COVID-19 Surveillance Team |
| EPI_ISL_2516092, EPI_ISL_2516093, EPI_ISL_2516094 | Lighthouse Lab in Alderley Park | Wellcome Sanger Institute for the COVID-19 Genomics UK (COG-UK) Consortium | Jacquelyn Wynn, Mairead Hyland, The Lighthouse Lab in Alderley Park and Alex Alderton, Roberto Amato, Jeffrey Barrett, Sonia Goncalves, Ewan Harrison, David K. Jackson, Ian Johnston, Dominic Kwiatkowski, Cordelia Langford, John Sillitoe on behalf of the Wellcome Sanger Institute COVID-19 Surveillance Team |
| EPI_ISL_2516095 | Health Services Laboratories | Wellcome Sanger Institute for the COVID-19 Genomics UK (COG-UK) Consortium | Health Services Laboratories and Alex Alderton, Roberto Amato, Jeffrey Barrett, Sonia Goncalves, Ewan Harrison, David K. Jackson, Ian Johnston, Dominic Kwiatkowski, Cordelia Langford, John Sillitoe on behalf of the Wellcome Sanger Institute COVID-19 Surveillance Team |
| EPI_ISL_2516096, EPI_ISL_2516097, EPI_ISL_2516098 | Lighthouse Lab in Alderley Park | Wellcome Sanger Institute for the COVID-19 Genomics UK (COG-UK) Consortium | Jacquelyn Wynn, Mairead Hyland, The Lighthouse Lab in Alderley Park and Alex Alderton, Roberto Amato, Jeffrey Barrett, Sonia Goncalves, Ewan Harrison, David K. Jackson, Ian Johnston, Dominic Kwiatkowski, Cordelia Langford, John Sillitoe on behalf of the Wellcome Sanger Institute COVID-19 Surveillance Team |
| EPI_ISL_2516099 | Health Services Laboratories | Wellcome Sanger Institute for the COVID-19 Genomics UK (COG-UK) Consortium | Health Services Laboratories and Alex Alderton, Roberto Amato, Jeffrey Barrett, Sonia Goncalves, Ewan Harrison, David K. Jackson, Ian Johnston, Dominic Kwiatkowski, Cordelia Langford, John Sillitoe on behalf of the Wellcome Sanger Institute COVID-19 Surveillance Team |
| EPI_ISL_2516100 | Lighthouse Lab in Alderley Park | Wellcome Sanger Institute for the COVID-19 Genomics UK (COG-UK) Consortium | Jacquelyn Wynn, Mairead Hyland, The Lighthouse Lab in Alderley Park and Alex Alderton, Roberto Amato, Jeffrey Barrett, Sonia Goncalves, Ewan Harrison, David K. Jackson, Ian Johnston, Dominic Kwiatkowski, Cordelia Langford, John Sillitoe on behalf of the Wellcome Sanger Institute COVID-19 Surveillance Team |
| EPI_ISL_2516101, EPI_ISL_2516102 | Health Services Laboratories | Wellcome Sanger Institute for the COVID-19 Genomics UK (COG-UK) Consortium | Health Services Laboratories and Alex Alderton, Roberto Amato, Jeffrey Barrett, Sonia Goncalves, Ewan Harrison, David K. Jackson, Ian Johnston, Dominic Kwiatkowski, Cordelia Langford, John Sillitoe on behalf of the Wellcome Sanger Institute COVID-19 Surveillance Team |
| EPI_ISL_2516104, EPI_ISL_2516105, EPI_ISL_2516106, EPI_ISL_2516107 | Lighthouse Lab in Alderley Park | Wellcome Sanger Institute for the COVID-19 Genomics UK (COG-UK) Consortium | Jacquelyn Wynn, Mairead Hyland, The Lighthouse Lab in Alderley Park and Alex Alderton, Roberto Amato, Jeffrey Barrett, Sonia Goncalves, Ewan Harrison, David K. Jackson, Ian Johnston, Dominic Kwiatkowski, Cordelia Langford, John Sillitoe on behalf of the Wellcome Sanger Institute COVID-19 Surveillance Team |
| EPI_ISL_2516108 | Health Services Laboratories | Wellcome Sanger Institute for the COVID-19 Genomics UK (COG-UK) Consortium | Health Services Laboratories and Alex Alderton, Roberto Amato, Jeffrey Barrett, Sonia Goncalves, Ewan Harrison, David K. Jackson, Ian Johnston, Dominic Kwiatkowski, Cordelia Langford, John Sillitoe on behalf of the Wellcome Sanger Institute COVID-19 Surveillance Team |
| EPI_ISL_2516109 | Lighthouse Lab in Glasgow | Wellcome Sanger Institute for the COVID-19 Genomics UK (COG-UK) Consortium | Harper VanSteenhouse, Yumi Kasai, David Gray, Carol Clugston, Anna Dominiczak and Alex Alderton, Roberto Amato, Jeffrey Barrett, Sonia Goncalves, Ewan Harrison, David K. Jackson, Ian Johnston, Dominic Kwiatkowski, Cordelia Langford, John Sillitoe on behalf of the Wellcome Sanger Institute COVID-19 Surveillance Team |
| EPI_ISL_2516110 | Health Services Laboratories | Wellcome Sanger Institute for the COVID-19 Genomics UK (COG-UK) Consortium | Health Services Laboratories and Alex Alderton, Roberto Amato, Jeffrey Barrett, Sonia Goncalves, Ewan Harrison, David K. Jackson, Ian Johnston, Dominic Kwiatkowski, Cordelia Langford, John Sillitoe on behalf of the Wellcome Sanger Institute COVID-19 Surveillance Team |
| EPI_ISL_2516111 | Lighthouse Lab in Alderley Park | Wellcome Sanger Institute for the COVID-19 Genomics UK (COG-UK) Consortium | Jacquelyn Wynn, Mairead Hyland, The Lighthouse Lab in Alderley Park and Alex Alderton, Roberto Amato, Jeffrey Barrett, Sonia Goncalves, Ewan Harrison, David K. Jackson, Ian Johnston, Dominic Kwiatkowski, Cordelia Langford, John Sillitoe on behalf of the Wellcome Sanger Institute COVID-19 Surveillance Team |
| EPI_ISL_2516112 | Lighthouse Lab in Glasgow | Wellcome Sanger Institute for the COVID-19 Genomics UK (COG-UK) Consortium | Harper VanSteenhouse, Yumi Kasai, David Gray, Carol Clugston, Anna Dominiczak and Alex Alderton, Roberto Amato, Jeffrey Barrett, Sonia Goncalves, Ewan Harrison, David K. Jackson, Ian Johnston, Dominic Kwiatkowski, Cordelia Langford, John Sillitoe on behalf of the Wellcome Sanger Institute COVID-19 Surveillance Team |
| EPI_ISL_2516113, EPI_ISL_2516114 | Lighthouse Lab in Alderley Park | Wellcome Sanger Institute for the COVID-19 Genomics UK (COG-UK) Consortium | Jacquelyn Wynn, Mairead Hyland, The Lighthouse Lab in Alderley Park and Alex Alderton, Roberto Amato, Jeffrey Barrett, Sonia Goncalves, Ewan Harrison, David K. Jackson, Ian Johnston, Dominic Kwiatkowski, Cordelia Langford, John Sillitoe on behalf of the Wellcome Sanger Institute COVID-19 Surveillance Team |
| EPI_ISL_2516115, EPI_ISL_2516116 | Health Services Laboratories | Wellcome Sanger Institute for the COVID-19 Genomics UK (COG-UK) Consortium | Health Services Laboratories and Alex Alderton, Roberto Amato, Jeffrey Barrett, Sonia Goncalves, Ewan Harrison, David K. Jackson, Ian Johnston, Dominic Kwiatkowski, Cordelia Langford, John Sillitoe on behalf of the Wellcome Sanger Institute COVID-19 Surveillance Team |
| EPI_ISL_2516117, EPI_ISL_2516118 | Lighthouse Lab in Alderley Park | Wellcome Sanger Institute for the COVID-19 Genomics UK (COG-UK) Consortium | Jacquelyn Wynn, Mairead Hyland, The Lighthouse Lab in Alderley Park and Alex Alderton, Roberto Amato, Jeffrey Barrett, Sonia Goncalves, Ewan Harrison, David K. Jackson, Ian Johnston, Dominic Kwiatkowski, Cordelia Langford, John Sillitoe on behalf of the Wellcome Sanger Institute COVID-19 Surveillance Team |
| EPI_ISL_2516119 | Health Services Laboratories | Wellcome Sanger Institute for the COVID-19 Genomics UK (COG-UK) Consortium | Health Services Laboratories and Alex Alderton, Roberto Amato, Jeffrey Barrett, Sonia Goncalves, Ewan Harrison, David K. Jackson, Ian Johnston, Dominic Kwiatkowski, Cordelia Langford, John Sillitoe on behalf of the Wellcome Sanger Institute COVID-19 Surveillance Team |
| EPI_ISL_2516120, EPI_ISL_2516121 | Lighthouse Lab in Alderley Park | Wellcome Sanger Institute for the COVID-19 Genomics UK (COG-UK) Consortium | Jacquelyn Wynn, Mairead Hyland, The Lighthouse Lab in Alderley Park and Alex Alderton, Roberto Amato, Jeffrey Barrett, Sonia Goncalves, Ewan Harrison, David K. Jackson, Ian Johnston, Dominic Kwiatkowski, Cordelia Langford, John Sillitoe on behalf of the Wellcome Sanger Institute COVID-19 Surveillance Team |
| EPI_ISL_2516122 | Health Services Laboratories | Wellcome Sanger Institute for the COVID-19 Genomics UK (COG-UK) Consortium | Health Services Laboratories and Alex Alderton, Roberto Amato, Jeffrey Barrett, Sonia Goncalves, Ewan Harrison, David K. Jackson, Ian Johnston, Dominic Kwiatkowski, Cordelia Langford, John Sillitoe on behalf of the Wellcome Sanger Institute COVID-19 Surveillance Team |
| EPI_ISL_2516123, EPI_ISL_2516124 | Lighthouse Lab in Alderley Park | Wellcome Sanger Institute for the COVID-19 Genomics UK (COG-UK) Consortium | Jacquelyn Wynn, Mairead Hyland, The Lighthouse Lab in Alderley Park and Alex Alderton, Roberto Amato, Jeffrey Barrett, Sonia Goncalves, Ewan Harrison, David K. Jackson, Ian Johnston, Dominic Kwiatkowski, Cordelia Langford, John Sillitoe on behalf of the Wellcome Sanger Institute COVID-19 Surveillance Team |
| EPI_ISL_2516125 | Lighthouse Lab in Glasgow | Wellcome Sanger Institute for the COVID-19 Genomics UK (COG-UK) Consortium | Harper VanSteenhouse, Yumi Kasai, David Gray, Carol Clugston, Anna Dominiczak and Alex Alderton, Roberto Amato, Jeffrey Barrett, Sonia Goncalves, Ewan Harrison, David K. Jackson, Ian Johnston, Dominic Kwiatkowski, Cordelia Langford, John Sillitoe on behalf of the Wellcome Sanger Institute COVID-19 Surveillance Team |
| EPI_ISL_2516126 | Health Services Laboratories | Wellcome Sanger Institute for the COVID-19 Genomics UK (COG-UK) Consortium | Health Services Laboratories and Alex Alderton, Roberto Amato, Jeffrey Barrett, Sonia Goncalves, Ewan Harrison, David K. Jackson, Ian Johnston, Dominic Kwiatkowski, Cordelia Langford, John Sillitoe on behalf of the Wellcome Sanger Institute COVID-19 Surveillance Team |
| EPI_ISL_2516127 | Lighthouse Lab in Glasgow | Wellcome Sanger Institute for the COVID-19 Genomics UK (COG-UK) Consortium | Harper VanSteenhouse, Yumi Kasai, David Gray, Carol Clugston, Anna Dominiczak and Alex Alderton, Roberto Amato, Jeffrey Barrett, Sonia Goncalves, Ewan Harrison, David K. Jackson, Ian Johnston, Dominic Kwiatkowski, Cordelia Langford, John Sillitoe on behalf of the Wellcome Sanger Institute COVID-19 Surveillance Team |
| EPI_ISL_2516128, EPI_ISL_2516129, EPI_ISL_2516130 | Lighthouse Lab in Alderley Park | Wellcome Sanger Institute for the COVID-19 Genomics UK (COG-UK) Consortium | Jacquelyn Wynn, Mairead Hyland, The Lighthouse Lab in Alderley Park and Alex Alderton, Roberto Amato, Jeffrey Barrett, Sonia Goncalves, Ewan Harrison, David K. Jackson, Ian Johnston, Dominic Kwiatkowski, Cordelia Langford, John Sillitoe on behalf of the Wellcome Sanger Institute COVID-19 Surveillance Team |
| EPI_ISL_2516131 | Lighthouse Lab in Glasgow | Wellcome Sanger Institute for the COVID-19 Genomics UK (COG-UK) Consortium | Harper VanSteenhouse, Yumi Kasai, David Gray, Carol Clugston, Anna Dominiczak and Alex Alderton, Roberto Amato, Jeffrey Barrett, Sonia Goncalves, Ewan Harrison, David K. Jackson, Ian Johnston, Dominic Kwiatkowski, Cordelia Langford, John Sillitoe on behalf of the Wellcome Sanger Institute COVID-19 Surveillance Team |
| EPI_ISL_2516132, EPI_ISL_2516133, EPI_ISL_2516134, EPI_ISL_2516135 | Lighthouse Lab in Alderley Park | Wellcome Sanger Institute for the COVID-19 Genomics UK (COG-UK) Consortium | Jacquelyn Wynn, Mairead Hyland, The Lighthouse Lab in Alderley Park and Alex Alderton, Roberto Amato, Jeffrey Barrett, Sonia Goncalves, Ewan Harrison, David K. Jackson, Ian Johnston, Dominic Kwiatkowski, Cordelia Langford, John Sillitoe on behalf of the Wellcome Sanger Institute COVID-19 Surveillance Team |
| EPI_ISL_2516136, EPI_ISL_2516138, EPI_ISL_2516139 | Health Services Laboratories | Wellcome Sanger Institute for the COVID-19 Genomics UK (COG-UK) Consortium | Health Services Laboratories and Alex Alderton, Roberto Amato, Jeffrey Barrett, Sonia Goncalves, Ewan Harrison, David K. Jackson, Ian Johnston, Dominic Kwiatkowski, Cordelia Langford, John Sillitoe on behalf of the Wellcome Sanger Institute COVID-19 Surveillance Team |
| EPI_ISL_2516140, EPI_ISL_2516141, EPI_ISL_2516142 | Lighthouse Lab in Alderley Park | Wellcome Sanger Institute for the COVID-19 Genomics UK (COG-UK) Consortium | Jacquelyn Wynn, Mairead Hyland, The Lighthouse Lab in Alderley Park and Alex Alderton, Roberto Amato, Jeffrey Barrett, Sonia Goncalves, Ewan Harrison, David K. Jackson, Ian Johnston, Dominic Kwiatkowski, Cordelia Langford, John Sillitoe on behalf of the Wellcome Sanger Institute COVID-19 Surveillance Team |
| EPI_ISL_2516143 | Lighthouse Lab in Glasgow | Wellcome Sanger Institute for the COVID-19 Genomics UK (COG-UK) Consortium | Harper VanSteenhouse, Yumi Kasai, David Gray, Carol Clugston, Anna Dominiczak and Alex Alderton, Roberto Amato, Jeffrey Barrett, Sonia Goncalves, Ewan Harrison, David K. Jackson, Ian Johnston, Dominic Kwiatkowski, Cordelia Langford, John Sillitoe on behalf of the Wellcome Sanger Institute COVID-19 Surveillance Team |
| EPI_ISL_2516144 | Health Services Laboratories | Wellcome Sanger Institute for the COVID-19 Genomics UK (COG-UK) Consortium | Health Services Laboratories and Alex Alderton, Roberto Amato, Jeffrey Barrett, Sonia Goncalves, Ewan Harrison, David K. Jackson, Ian Johnston, Dominic Kwiatkowski, Cordelia Langford, John Sillitoe on behalf of the Wellcome Sanger Institute COVID-19 Surveillance Team |
| EPI_ISL_2516145 | Lighthouse Lab in Glasgow | Wellcome Sanger Institute for the COVID-19 Genomics UK (COG-UK) Consortium | Harper VanSteenhouse, Yumi Kasai, David Gray, Carol Clugston, Anna Dominiczak and Alex Alderton, Roberto Amato, Jeffrey Barrett, Sonia Goncalves, Ewan Harrison, David K. Jackson, Ian Johnston, Dominic Kwiatkowski, Cordelia Langford, John Sillitoe on behalf of the Wellcome Sanger Institute COVID-19 Surveillance Team |
| EPI_ISL_2516146 | Lighthouse Lab in Alderley Park | Wellcome Sanger Institute for the COVID-19 Genomics UK (COG-UK) Consortium | Jacquelyn Wynn, Mairead Hyland, The Lighthouse Lab in Alderley Park and Alex Alderton, Roberto Amato, Jeffrey Barrett, Sonia Goncalves, Ewan Harrison, David K. Jackson, Ian Johnston, Dominic Kwiatkowski, Cordelia Langford, John Sillitoe on behalf of the Wellcome Sanger Institute COVID-19 Surveillance Team |
| EPI_ISL_2516147 | Health Services Laboratories | Wellcome Sanger Institute for the COVID-19 Genomics UK (COG-UK) Consortium | Health Services Laboratories and Alex Alderton, Roberto Amato, Jeffrey Barrett, Sonia Goncalves, Ewan Harrison, David K. Jackson, Ian Johnston, Dominic Kwiatkowski, Cordelia Langford, John Sillitoe on behalf of the Wellcome Sanger Institute COVID-19 Surveillance Team |
| EPI_ISL_2516148 | Lighthouse Lab in Alderley Park | Wellcome Sanger Institute for the COVID-19 Genomics UK (COG-UK) Consortium | Jacquelyn Wynn, Mairead Hyland, The Lighthouse Lab in Alderley Park and Alex Alderton, Roberto Amato, Jeffrey Barrett, Sonia Goncalves, Ewan Harrison, David K. Jackson, Ian Johnston, Dominic Kwiatkowski, Cordelia Langford, John Sillitoe on behalf of the Wellcome Sanger Institute COVID-19 Surveillance Team |
| EPI_ISL_2516149 | Lighthouse Lab in Glasgow | Wellcome Sanger Institute for the COVID-19 Genomics UK (COG-UK) | Harper VanSteenhouse, Yumi Kasai, David Gray, Carol Clugston, Anna Dominiczak and Alex Alderton, Roberto Amato, Jeffrey Barrett, Sonia Goncalves, Ewan Harrison, David K. Jackson, Ian Johnston, Dominic Kwiatkowski, Cordelia Langford, |

| | | | |
|---|---|---|---|
| | | Consortium | John Sillitoe on behalf of the Wellcome Sanger Institute COVID-19 Surveillance Team |
| EPI_ISL_2516150, EPI_ISL_2516151, EPI_ISL_2516152 | Lighthouse Lab in Alderley Park | Wellcome Sanger Institute for the COVID-19 Genomics UK (COG-UK) Consortium | Jacquelyn Wynn, Mairead Hyland, The Lighthouse Lab in Alderley Park and Alex Alderton, Roberto Amato, Jeffrey Barrett, Sonia Goncalves, Ewan Harrison, David K. Jackson, Ian Johnston, Dominic Kwiatkowski, Cordelia Langford, John Sillitoe on behalf of the Wellcome Sanger Institute COVID-19 Surveillance Team |
| EPI_ISL_2516153 | Health Services Laboratories | Wellcome Sanger Institute for the COVID-19 Genomics UK (COG-UK) Consortium | Health Services Laboratories and Alex Alderton, Roberto Amato, Jeffrey Barrett, Sonia Goncalves, Ewan Harrison, David K. Jackson, Ian Johnston, Dominic Kwiatkowski, Cordelia Langford, John Sillitoe on behalf of the Wellcome Sanger Institute COVID-19 Surveillance Team |
| EPI_ISL_2516154 | Lighthouse Lab in Alderley Park | Wellcome Sanger Institute for the COVID-19 Genomics UK (COG-UK) Consortium | Jacquelyn Wynn, Mairead Hyland, The Lighthouse Lab in Alderley Park and Alex Alderton, Roberto Amato, Jeffrey Barrett, Sonia Goncalves, Ewan Harrison, David K. Jackson, Ian Johnston, Dominic Kwiatkowski, Cordelia Langford, John Sillitoe on behalf of the Wellcome Sanger Institute COVID-19 Surveillance Team |
| EPI_ISL_2516155 | Health Services Laboratories | Wellcome Sanger Institute for the COVID-19 Genomics UK (COG-UK) Consortium | Health Services Laboratories and Alex Alderton, Roberto Amato, Jeffrey Barrett, Sonia Goncalves, Ewan Harrison, David K. Jackson, Ian Johnston, Dominic Kwiatkowski, Cordelia Langford, John Sillitoe on behalf of the Wellcome Sanger Institute COVID-19 Surveillance Team |
| EPI_ISL_2516156, EPI_ISL_2516157 | Lighthouse Lab in Alderley Park | Wellcome Sanger Institute for the COVID-19 Genomics UK (COG-UK) Consortium | Jacquelyn Wynn, Mairead Hyland, The Lighthouse Lab in Alderley Park and Alex Alderton, Roberto Amato, Jeffrey Barrett, Sonia Goncalves, Ewan Harrison, David K. Jackson, Ian Johnston, Dominic Kwiatkowski, Cordelia Langford, John Sillitoe on behalf of the Wellcome Sanger Institute COVID-19 Surveillance Team |
| EPI_ISL_2516158 | Lighthouse Lab in Glasgow | Wellcome Sanger Institute for the COVID-19 Genomics UK (COG-UK) Consortium | Harper VanSteenhouse, Yumi Kasai, David Gray, Carol Clugston, Anna Dominiczak and Alex Alderton, Roberto Amato, Jeffrey Barrett, Sonia Goncalves, Ewan Harrison, David K. Jackson, Ian Johnston, Dominic Kwiatkowski, Cordelia Langford, John Sillitoe on behalf of the Wellcome Sanger Institute COVID-19 Surveillance Team |
| EPI_ISL_2516159 | Lighthouse Lab in Alderley Park | Wellcome Sanger Institute for the COVID-19 Genomics UK (COG-UK) Consortium | Jacquelyn Wynn, Mairead Hyland, The Lighthouse Lab in Alderley Park and Alex Alderton, Roberto Amato, Jeffrey Barrett, Sonia Goncalves, Ewan Harrison, David K. Jackson, Ian Johnston, Dominic Kwiatkowski, Cordelia Langford, John Sillitoe on behalf of the Wellcome Sanger Institute COVID-19 Surveillance Team |
| EPI_ISL_2516160 | Health Services Laboratories | Wellcome Sanger Institute for the COVID-19 Genomics UK (COG-UK) Consortium | Health Services Laboratories and Alex Alderton, Roberto Amato, Jeffrey Barrett, Sonia Goncalves, Ewan Harrison, David K. Jackson, Ian Johnston, Dominic Kwiatkowski, Cordelia Langford, John Sillitoe on behalf of the Wellcome Sanger Institute COVID-19 Surveillance Team |
| EPI_ISL_2516161 | Lighthouse Lab in Alderley Park | Wellcome Sanger Institute for the COVID-19 Genomics UK (COG-UK) Consortium | Jacquelyn Wynn, Mairead Hyland, The Lighthouse Lab in Alderley Park and Alex Alderton, Roberto Amato, Jeffrey Barrett, Sonia Goncalves, Ewan Harrison, David K. Jackson, Ian Johnston, Dominic Kwiatkowski, Cordelia Langford, John Sillitoe on behalf of the Wellcome Sanger Institute COVID-19 Surveillance Team |
| EPI_ISL_2516162, EPI_ISL_2516163 | Health Services Laboratories | Wellcome Sanger Institute for the COVID-19 Genomics UK (COG-UK) Consortium | Health Services Laboratories and Alex Alderton, Roberto Amato, Jeffrey Barrett, Sonia Goncalves, Ewan Harrison, David K. Jackson, Ian Johnston, Dominic Kwiatkowski, Cordelia Langford, John Sillitoe on behalf of the Wellcome Sanger Institute COVID-19 Surveillance Team |
| EPI_ISL_2516164, EPI_ISL_2516165, EPI_ISL_2516166, EPI_ISL_2516167 | Lighthouse Lab in Alderley Park | Wellcome Sanger Institute for the COVID-19 Genomics UK (COG-UK) Consortium | Jacquelyn Wynn, Mairead Hyland, The Lighthouse Lab in Alderley Park and Alex Alderton, Roberto Amato, Jeffrey Barrett, Sonia Goncalves, Ewan Harrison, David K. Jackson, Ian Johnston, Dominic Kwiatkowski, Cordelia Langford, John Sillitoe on behalf of the Wellcome Sanger Institute COVID-19 Surveillance Team |
| EPI_ISL_2516168 | Health Services Laboratories | Wellcome Sanger Institute for the COVID-19 Genomics UK (COG-UK) Consortium | Health Services Laboratories and Alex Alderton, Roberto Amato, Jeffrey Barrett, Sonia Goncalves, Ewan Harrison, David K. Jackson, Ian Johnston, Dominic Kwiatkowski, Cordelia Langford, John Sillitoe on behalf of the Wellcome Sanger Institute COVID-19 Surveillance Team |
| EPI_ISL_2516169 | Lighthouse Lab in Glasgow | Wellcome Sanger Institute for the COVID-19 Genomics UK (COG-UK) Consortium | Harper VanSteenhouse, Yumi Kasai, David Gray, Carol Clugston, Anna Dominiczak and Alex Alderton, Roberto Amato, Jeffrey Barrett, Sonia Goncalves, Ewan Harrison, David K. Jackson, Ian Johnston, Dominic Kwiatkowski, Cordelia Langford, John Sillitoe on behalf of the Wellcome Sanger Institute COVID-19 Surveillance Team |
| EPI_ISL_2516170 | Health Services Laboratories | Wellcome Sanger Institute for the COVID-19 Genomics UK (COG-UK) Consortium | Health Services Laboratories and Alex Alderton, Roberto Amato, Jeffrey Barrett, Sonia Goncalves, Ewan Harrison, David K. Jackson, Ian Johnston, Dominic Kwiatkowski, Cordelia Langford, John Sillitoe on behalf of the Wellcome Sanger Institute COVID-19 Surveillance Team |
| EPI_ISL_2516171, EPI_ISL_2516172, EPI_ISL_2516173, EPI_ISL_2516174, EPI_ISL_2516175 | Lighthouse Lab in Alderley Park | Wellcome Sanger Institute for the COVID-19 Genomics UK (COG-UK) Consortium | Jacquelyn Wynn, Mairead Hyland, The Lighthouse Lab in Alderley Park and Alex Alderton, Roberto Amato, Jeffrey Barrett, Sonia Goncalves, Ewan Harrison, David K. Jackson, Ian Johnston, Dominic Kwiatkowski, Cordelia Langford, John Sillitoe on behalf of the Wellcome Sanger Institute COVID-19 Surveillance Team |
| EPI_ISL_2516177 | Health Services Laboratories | Wellcome Sanger Institute for the COVID-19 Genomics UK (COG-UK) Consortium | Health Services Laboratories and Alex Alderton, Roberto Amato, Jeffrey Barrett, Sonia Goncalves, Ewan Harrison, David K. Jackson, Ian Johnston, Dominic Kwiatkowski, Cordelia Langford, John Sillitoe on behalf of the Wellcome Sanger Institute COVID-19 Surveillance Team |
| EPI_ISL_2516178, EPI_ISL_2516179, EPI_ISL_2516180, EPI_ISL_2516181, EPI_ISL_2516182 | Lighthouse Lab in Alderley Park | Wellcome Sanger Institute for the COVID-19 Genomics UK (COG-UK) Consortium | Jacquelyn Wynn, Mairead Hyland, The Lighthouse Lab in Alderley Park and Alex Alderton, Roberto Amato, Jeffrey Barrett, Sonia Goncalves, Ewan Harrison, David K. Jackson, Ian Johnston, Dominic Kwiatkowski, Cordelia Langford, John Sillitoe on behalf of the Wellcome Sanger Institute COVID-19 Surveillance Team |
| EPI_ISL_2516183 | Lighthouse Lab in Glasgow | Wellcome Sanger Institute for the COVID-19 Genomics UK (COG-UK) Consortium | Harper VanSteenhouse, Yumi Kasai, David Gray, Carol Clugston, Anna Dominiczak and Alex Alderton, Roberto Amato, Jeffrey Barrett, Sonia Goncalves, Ewan Harrison, David K. Jackson, Ian Johnston, Dominic Kwiatkowski, Cordelia Langford, John Sillitoe on behalf of the Wellcome Sanger Institute COVID-19 Surveillance Team |
| EPI_ISL_2516184 | Lighthouse Lab in Alderley Park | Wellcome Sanger Institute for the COVID-19 Genomics UK (COG-UK) Consortium | Jacquelyn Wynn, Mairead Hyland, The Lighthouse Lab in Alderley Park and Alex Alderton, Roberto Amato, Jeffrey Barrett, Sonia Goncalves, Ewan Harrison, David K. Jackson, Ian Johnston, Dominic Kwiatkowski, Cordelia Langford, John Sillitoe on behalf of the Wellcome Sanger Institute COVID-19 Surveillance Team |
| EPI_ISL_2516185 | Health Services Laboratories | Wellcome Sanger Institute for the COVID-19 Genomics UK (COG-UK) Consortium | Health Services Laboratories and Alex Alderton, Roberto Amato, Jeffrey Barrett, Sonia Goncalves, Ewan Harrison, David K. Jackson, Ian Johnston, Dominic Kwiatkowski, Cordelia Langford, John Sillitoe on behalf of the Wellcome Sanger Institute COVID-19 Surveillance Team |
| EPI_ISL_2516186, EPI_ISL_2516187, EPI_ISL_2516188, EPI_ISL_2516189, EPI_ISL_2516190, EPI_ISL_2516191, EPI_ISL_2516192, EPI_ISL_2516193, EPI_ISL_2516194 | Lighthouse Lab in Alderley Park | Wellcome Sanger Institute for the COVID-19 Genomics UK (COG-UK) Consortium | Jacquelyn Wynn, Mairead Hyland, The Lighthouse Lab in Alderley Park and Alex Alderton, Roberto Amato, Jeffrey Barrett, Sonia Goncalves, Ewan Harrison, David K. Jackson, Ian Johnston, Dominic Kwiatkowski, Cordelia Langford, John Sillitoe on behalf of the Wellcome Sanger Institute COVID-19 Surveillance Team |
| EPI_ISL_2516195 | Health Services Laboratories | Wellcome Sanger Institute for the COVID-19 Genomics UK (COG-UK) Consortium | Health Services Laboratories and Alex Alderton, Roberto Amato, Jeffrey Barrett, Sonia Goncalves, Ewan Harrison, David K. Jackson, Ian Johnston, Dominic Kwiatkowski, Cordelia Langford, John Sillitoe on behalf of the Wellcome Sanger Institute COVID-19 Surveillance Team |
| EPI_ISL_2516196, EPI_ISL_2516197, EPI_ISL_2516198 | Lighthouse Lab in Alderley Park | Wellcome Sanger Institute for the COVID-19 Genomics UK (COG-UK) Consortium | Jacquelyn Wynn, Mairead Hyland, The Lighthouse Lab in Alderley Park and Alex Alderton, Roberto Amato, Jeffrey Barrett, Sonia Goncalves, Ewan Harrison, David K. Jackson, Ian Johnston, Dominic Kwiatkowski, Cordelia Langford, John Sillitoe on behalf of the Wellcome Sanger Institute COVID-19 Surveillance Team |
| EPI_ISL_2516199 | Health Services Laboratories | Wellcome Sanger Institute for the COVID-19 Genomics UK (COG-UK) Consortium | Health Services Laboratories and Alex Alderton, Roberto Amato, Jeffrey Barrett, Sonia Goncalves, Ewan Harrison, David K. Jackson, Ian Johnston, Dominic Kwiatkowski, Cordelia Langford, John Sillitoe on behalf of the Wellcome Sanger Institute COVID-19 Surveillance Team |
| EPI_ISL_2516200, EPI_ISL_2516201 | Lighthouse Lab in Alderley Park | Wellcome Sanger Institute for the COVID-19 Genomics UK (COG-UK) Consortium | Jacquelyn Wynn, Mairead Hyland, The Lighthouse Lab in Alderley Park and Alex Alderton, Roberto Amato, Jeffrey Barrett, Sonia Goncalves, Ewan Harrison, David K. Jackson, Ian Johnston, Dominic Kwiatkowski, Cordelia Langford, John Sillitoe on behalf of the Wellcome Sanger Institute COVID-19 Surveillance Team |
| EPI_ISL_2516202, EPI_ISL_2516203, EPI_ISL_2516204 | Health Services Laboratories | Wellcome Sanger Institute for the COVID-19 Genomics UK (COG-UK) Consortium | Health Services Laboratories and Alex Alderton, Roberto Amato, Jeffrey Barrett, Sonia Goncalves, Ewan Harrison, David K. Jackson, Ian Johnston, Dominic Kwiatkowski, Cordelia Langford, John Sillitoe on behalf of the Wellcome Sanger Institute COVID-19 Surveillance Team |
| EPI_ISL_2516205, EPI_ISL_2516206, EPI_ISL_2516207 | Lighthouse Lab in Alderley Park | Wellcome Sanger Institute for the COVID-19 Genomics UK (COG-UK) Consortium | Jacquelyn Wynn, Mairead Hyland, The Lighthouse Lab in Alderley Park and Alex Alderton, Roberto Amato, Jeffrey Barrett, Sonia Goncalves, Ewan Harrison, David K. Jackson, Ian Johnston, Dominic Kwiatkowski, Cordelia Langford, John Sillitoe on behalf of the Wellcome Sanger Institute COVID-19 Surveillance Team |
| EPI_ISL_2516208 | Health Services Laboratories | Wellcome Sanger Institute for the COVID-19 Genomics UK (COG-UK) Consortium | Health Services Laboratories and Alex Alderton, Roberto Amato, Jeffrey Barrett, Sonia Goncalves, Ewan Harrison, David K. Jackson, Ian Johnston, Dominic Kwiatkowski, Cordelia Langford, John Sillitoe on behalf of the Wellcome Sanger Institute COVID-19 Surveillance Team |
| EPI_ISL_2516209, EPI_ISL_2516210 | Lighthouse Lab in Alderley Park | Wellcome Sanger Institute for the COVID-19 Genomics UK (COG-UK) Consortium | Jacquelyn Wynn, Mairead Hyland, The Lighthouse Lab in Alderley Park and Alex Alderton, Roberto Amato, Jeffrey Barrett, Sonia Goncalves, Ewan Harrison, David K. Jackson, Ian Johnston, Dominic Kwiatkowski, Cordelia Langford, John Sillitoe on behalf of the Wellcome Sanger Institute COVID-19 Surveillance Team |
| EPI_ISL_2516212 | Health Services Laboratories | Wellcome Sanger Institute for the COVID-19 Genomics UK (COG-UK) Consortium | Health Services Laboratories and Alex Alderton, Roberto Amato, Jeffrey Barrett, Sonia Goncalves, Ewan Harrison, David K. Jackson, Ian Johnston, Dominic Kwiatkowski, Cordelia Langford, John Sillitoe on behalf of the Wellcome Sanger Institute COVID-19 Surveillance Team |
| EPI_ISL_2516213, EPI_ISL_2516214 | Lighthouse Lab in Alderley Park | Wellcome Sanger Institute for the COVID-19 Genomics UK (COG-UK) Consortium | Jacquelyn Wynn, Mairead Hyland, The Lighthouse Lab in Alderley Park and Alex Alderton, Roberto Amato, Jeffrey Barrett, Sonia Goncalves, Ewan Harrison, David K. Jackson, Ian Johnston, Dominic Kwiatkowski, Cordelia Langford, John Sillitoe on behalf of the Wellcome Sanger Institute COVID-19 Surveillance Team |
| EPI_ISL_2516216, EPI_ISL_2516217 | Health Services Laboratories | Wellcome Sanger Institute for the COVID-19 Genomics UK (COG-UK) Consortium | Health Services Laboratories and Alex Alderton, Roberto Amato, Jeffrey Barrett, Sonia Goncalves, Ewan Harrison, David K. Jackson, Ian Johnston, Dominic Kwiatkowski, Cordelia Langford, John Sillitoe on behalf of the Wellcome Sanger Institute COVID-19 Surveillance Team |
| EPI_ISL_2516218, EPI_ISL_2516219, EPI_ISL_2516220, EPI_ISL_2516221, EPI_ISL_2516223 | Lighthouse Lab in Alderley Park | Wellcome Sanger Institute for the COVID-19 Genomics UK (COG-UK) Consortium | Jacquelyn Wynn, Mairead Hyland, The Lighthouse Lab in Alderley Park and Alex Alderton, Roberto Amato, Jeffrey Barrett, Sonia Goncalves, Ewan Harrison, David K. Jackson, Ian Johnston, Dominic Kwiatkowski, Cordelia Langford, John Sillitoe on behalf of the Wellcome Sanger Institute COVID-19 Surveillance Team |
| EPI_ISL_2516224 | Lighthouse Lab in Glasgow | Wellcome Sanger Institute for the COVID-19 Genomics UK (COG-UK) Consortium | Harper VanSteenhouse, Yumi Kasai, David Gray, Carol Clugston, Anna Dominiczak and Alex Alderton, Roberto Amato, Jeffrey Barrett, Sonia Goncalves, Ewan Harrison, David K. Jackson, Ian Johnston, Dominic Kwiatkowski, Cordelia Langford, John Sillitoe on behalf of the Wellcome Sanger Institute COVID-19 Surveillance Team |
| EPI_ISL_2516225, EPI_ISL_2516227, EPI_ISL_2516228, | Lighthouse Lab in Alderley Park | Wellcome Sanger Institute for the COVID-19 Genomics UK (COG-UK) Consortium | Jacquelyn Wynn, Mairead Hyland, The Lighthouse Lab in Alderley Park and Alex Alderton, Roberto Amato, Jeffrey Barrett, Sonia Goncalves, Ewan Harrison, David K. Jackson, Ian Johnston, Dominic Kwiatkowski, Cordelia Langford, John Sillitoe on behalf of the Wellcome Sanger Institute COVID-19 Surveillance Team |

| | | | |
|---|---|---|---|
| EPI_ISL_2516229 | | | |
| EPI_ISL_2516230 | Health Services Laboratories | Wellcome Sanger Institute for the COVID-19 Genomics UK (COG-UK) Consortium | Health Services Laboratories and Alex Alderton, Roberto Amato, Jeffrey Barrett, Sonia Goncalves, Ewan Harrison, David K. Jackson, Ian Johnston, Dominic Kwiatkowski, Cordelia Langford, John Sillitoe on behalf of the Wellcome Sanger Institute COVID-19 Surveillance Team |
| EPI_ISL_2516231, EPI_ISL_2516232, EPI_ISL_2516233, EPI_ISL_2516234 | Lighthouse Lab in Alderley Park | Wellcome Sanger Institute for the COVID-19 Genomics UK (COG-UK) Consortium | Jacquelyn Wynn, Mairead Hyland, The Lighthouse Lab in Alderley Park and Alex Alderton, Roberto Amato, Jeffrey Barrett, Sonia Goncalves, Ewan Harrison, David K. Jackson, Ian Johnston, Dominic Kwiatkowski, Cordelia Langford, John Sillitoe on behalf of the Wellcome Sanger Institute COVID-19 Surveillance Team |
| EPI_ISL_2516235, EPI_ISL_2516236 | Health Services Laboratories | Wellcome Sanger Institute for the COVID-19 Genomics UK (COG-UK) Consortium | Health Services Laboratories and Alex Alderton, Roberto Amato, Jeffrey Barrett, Sonia Goncalves, Ewan Harrison, David K. Jackson, Ian Johnston, Dominic Kwiatkowski, Cordelia Langford, John Sillitoe on behalf of the Wellcome Sanger Institute COVID-19 Surveillance Team |
| EPI_ISL_2516237, EPI_ISL_2516238, EPI_ISL_2516240 | Lighthouse Lab in Alderley Park | Wellcome Sanger Institute for the COVID-19 Genomics UK (COG-UK) Consortium | Jacquelyn Wynn, Mairead Hyland, The Lighthouse Lab in Alderley Park and Alex Alderton, Roberto Amato, Jeffrey Barrett, Sonia Goncalves, Ewan Harrison, David K. Jackson, Ian Johnston, Dominic Kwiatkowski, Cordelia Langford, John Sillitoe on behalf of the Wellcome Sanger Institute COVID-19 Surveillance Team |
| EPI_ISL_2516241 | Health Services Laboratories | Wellcome Sanger Institute for the COVID-19 Genomics UK (COG-UK) Consortium | Health Services Laboratories and Alex Alderton, Roberto Amato, Jeffrey Barrett, Sonia Goncalves, Ewan Harrison, David K. Jackson, Ian Johnston, Dominic Kwiatkowski, Cordelia Langford, John Sillitoe on behalf of the Wellcome Sanger Institute COVID-19 Surveillance Team |
| EPI_ISL_2516242 | Lighthouse Lab in Alderley Park | Wellcome Sanger Institute for the COVID-19 Genomics UK (COG-UK) Consortium | Jacquelyn Wynn, Mairead Hyland, The Lighthouse Lab in Alderley Park and Alex Alderton, Roberto Amato, Jeffrey Barrett, Sonia Goncalves, Ewan Harrison, David K. Jackson, Ian Johnston, Dominic Kwiatkowski, Cordelia Langford, John Sillitoe on behalf of the Wellcome Sanger Institute COVID-19 Surveillance Team |
| EPI_ISL_2516243 | Lighthouse Lab in Glasgow | Wellcome Sanger Institute for the COVID-19 Genomics UK (COG-UK) Consortium | Harper VanSteenhouse, Yumi Kasai, David Gray, Carol Clugston, Anna Dominiczak and Alex Alderton, Roberto Amato, Jeffrey Barrett, Sonia Goncalves, Ewan Harrison, David K. Jackson, Ian Johnston, Dominic Kwiatkowski, Cordelia Langford, John Sillitoe on behalf of the Wellcome Sanger Institute COVID-19 Surveillance Team |
| EPI_ISL_2516244, EPI_ISL_2516245 | Lighthouse Lab in Alderley Park | Wellcome Sanger Institute for the COVID-19 Genomics UK (COG-UK) Consortium | Jacquelyn Wynn, Mairead Hyland, The Lighthouse Lab in Alderley Park and Alex Alderton, Roberto Amato, Jeffrey Barrett, Sonia Goncalves, Ewan Harrison, David K. Jackson, Ian Johnston, Dominic Kwiatkowski, Cordelia Langford, John Sillitoe on behalf of the Wellcome Sanger Institute COVID-19 Surveillance Team |
| EPI_ISL_2516246 | Lighthouse Lab in Glasgow | Wellcome Sanger Institute for the COVID-19 Genomics UK (COG-UK) Consortium | Harper VanSteenhouse, Yumi Kasai, David Gray, Carol Clugston, Anna Dominiczak and Alex Alderton, Roberto Amato, Jeffrey Barrett, Sonia Goncalves, Ewan Harrison, David K. Jackson, Ian Johnston, Dominic Kwiatkowski, Cordelia Langford, John Sillitoe on behalf of the Wellcome Sanger Institute COVID-19 Surveillance Team |
| EPI_ISL_2516247 | Lighthouse Lab in Alderley Park | Wellcome Sanger Institute for the COVID-19 Genomics UK (COG-UK) Consortium | Jacquelyn Wynn, Mairead Hyland, The Lighthouse Lab in Alderley Park and Alex Alderton, Roberto Amato, Jeffrey Barrett, Sonia Goncalves, Ewan Harrison, David K. Jackson, Ian Johnston, Dominic Kwiatkowski, Cordelia Langford, John Sillitoe on behalf of the Wellcome Sanger Institute COVID-19 Surveillance Team |
| EPI_ISL_2516248 | Health Services Laboratories | Wellcome Sanger Institute for the COVID-19 Genomics UK (COG-UK) Consortium | Health Services Laboratories and Alex Alderton, Roberto Amato, Jeffrey Barrett, Sonia Goncalves, Ewan Harrison, David K. Jackson, Ian Johnston, Dominic Kwiatkowski, Cordelia Langford, John Sillitoe on behalf of the Wellcome Sanger Institute COVID-19 Surveillance Team |
| EPI_ISL_2516249, EPI_ISL_2516250, EPI_ISL_2516251 | Lighthouse Lab in Alderley Park | Wellcome Sanger Institute for the COVID-19 Genomics UK (COG-UK) Consortium | Jacquelyn Wynn, Mairead Hyland, The Lighthouse Lab in Alderley Park and Alex Alderton, Roberto Amato, Jeffrey Barrett, Sonia Goncalves, Ewan Harrison, David K. Jackson, Ian Johnston, Dominic Kwiatkowski, Cordelia Langford, John Sillitoe on behalf of the Wellcome Sanger Institute COVID-19 Surveillance Team |
| EPI_ISL_2516252 | Health Services Laboratories | Wellcome Sanger Institute for the COVID-19 Genomics UK (COG-UK) Consortium | Health Services Laboratories and Alex Alderton, Roberto Amato, Jeffrey Barrett, Sonia Goncalves, Ewan Harrison, David K. Jackson, Ian Johnston, Dominic Kwiatkowski, Cordelia Langford, John Sillitoe on behalf of the Wellcome Sanger Institute COVID-19 Surveillance Team |
| EPI_ISL_2516253 | Lighthouse Lab in Alderley Park | Wellcome Sanger Institute for the COVID-19 Genomics UK (COG-UK) Consortium | Jacquelyn Wynn, Mairead Hyland, The Lighthouse Lab in Alderley Park and Alex Alderton, Roberto Amato, Jeffrey Barrett, Sonia Goncalves, Ewan Harrison, David K. Jackson, Ian Johnston, Dominic Kwiatkowski, Cordelia Langford, John Sillitoe on behalf of the Wellcome Sanger Institute COVID-19 Surveillance Team |
| EPI_ISL_2516254 | Health Services Laboratories | Wellcome Sanger Institute for the COVID-19 Genomics UK (COG-UK) Consortium | Health Services Laboratories and Alex Alderton, Roberto Amato, Jeffrey Barrett, Sonia Goncalves, Ewan Harrison, David K. Jackson, Ian Johnston, Dominic Kwiatkowski, Cordelia Langford, John Sillitoe on behalf of the Wellcome Sanger Institute COVID-19 Surveillance Team |
| EPI_ISL_2516255, EPI_ISL_2516256 | Lighthouse Lab in Alderley Park | Wellcome Sanger Institute for the COVID-19 Genomics UK (COG-UK) Consortium | Jacquelyn Wynn, Mairead Hyland, The Lighthouse Lab in Alderley Park and Alex Alderton, Roberto Amato, Jeffrey Barrett, Sonia Goncalves, Ewan Harrison, David K. Jackson, Ian Johnston, Dominic Kwiatkowski, Cordelia Langford, John Sillitoe on behalf of the Wellcome Sanger Institute COVID-19 Surveillance Team |
| EPI_ISL_2516257 | Health Services Laboratories | Wellcome Sanger Institute for the COVID-19 Genomics UK (COG-UK) Consortium | Health Services Laboratories and Alex Alderton, Roberto Amato, Jeffrey Barrett, Sonia Goncalves, Ewan Harrison, David K. Jackson, Ian Johnston, Dominic Kwiatkowski, Cordelia Langford, John Sillitoe on behalf of the Wellcome Sanger Institute COVID-19 Surveillance Team |
| EPI_ISL_2516258 | Lighthouse Lab in Alderley Park | Wellcome Sanger Institute for the COVID-19 Genomics UK (COG-UK) Consortium | Jacquelyn Wynn, Mairead Hyland, The Lighthouse Lab in Alderley Park and Alex Alderton, Roberto Amato, Jeffrey Barrett, Sonia Goncalves, Ewan Harrison, David K. Jackson, Ian Johnston, Dominic Kwiatkowski, Cordelia Langford, John Sillitoe on behalf of the Wellcome Sanger Institute COVID-19 Surveillance Team |
| EPI_ISL_2516259 | Health Services Laboratories | Wellcome Sanger Institute for the COVID-19 Genomics UK (COG-UK) Consortium | Health Services Laboratories and Alex Alderton, Roberto Amato, Jeffrey Barrett, Sonia Goncalves, Ewan Harrison, David K. Jackson, Ian Johnston, Dominic Kwiatkowski, Cordelia Langford, John Sillitoe on behalf of the Wellcome Sanger Institute COVID-19 Surveillance Team |
| EPI_ISL_2516260 | Lighthouse Lab in Glasgow | Wellcome Sanger Institute for the COVID-19 Genomics UK (COG-UK) Consortium | Harper VanSteenhouse, Yumi Kasai, David Gray, Carol Clugston, Anna Dominiczak and Alex Alderton, Roberto Amato, Jeffrey Barrett, Sonia Goncalves, Ewan Harrison, David K. Jackson, Ian Johnston, Dominic Kwiatkowski, Cordelia Langford, John Sillitoe on behalf of the Wellcome Sanger Institute COVID-19 Surveillance Team |
| EPI_ISL_2516261, EPI_ISL_2516262, EPI_ISL_2516263 | Lighthouse Lab in Alderley Park | Wellcome Sanger Institute for the COVID-19 Genomics UK (COG-UK) Consortium | Jacquelyn Wynn, Mairead Hyland, The Lighthouse Lab in Alderley Park and Alex Alderton, Roberto Amato, Jeffrey Barrett, Sonia Goncalves, Ewan Harrison, David K. Jackson, Ian Johnston, Dominic Kwiatkowski, Cordelia Langford, John Sillitoe on behalf of the Wellcome Sanger Institute COVID-19 Surveillance Team |
| EPI_ISL_2516264 | Health Services Laboratories | Wellcome Sanger Institute for the COVID-19 Genomics UK (COG-UK) Consortium | Health Services Laboratories and Alex Alderton, Roberto Amato, Jeffrey Barrett, Sonia Goncalves, Ewan Harrison, David K. Jackson, Ian Johnston, Dominic Kwiatkowski, Cordelia Langford, John Sillitoe on behalf of the Wellcome Sanger Institute COVID-19 Surveillance Team |
| EPI_ISL_2516265 | Lighthouse Lab in Alderley Park | Wellcome Sanger Institute for the COVID-19 Genomics UK (COG-UK) Consortium | Jacquelyn Wynn, Mairead Hyland, The Lighthouse Lab in Alderley Park and Alex Alderton, Roberto Amato, Jeffrey Barrett, Sonia Goncalves, Ewan Harrison, David K. Jackson, Ian Johnston, Dominic Kwiatkowski, Cordelia Langford, John Sillitoe on behalf of the Wellcome Sanger Institute COVID-19 Surveillance Team |
| EPI_ISL_2516266 | Health Services Laboratories | Wellcome Sanger Institute for the COVID-19 Genomics UK (COG-UK) Consortium | Health Services Laboratories and Alex Alderton, Roberto Amato, Jeffrey Barrett, Sonia Goncalves, Ewan Harrison, David K. Jackson, Ian Johnston, Dominic Kwiatkowski, Cordelia Langford, John Sillitoe on behalf of the Wellcome Sanger Institute COVID-19 Surveillance Team |
| EPI_ISL_2516267, EPI_ISL_2516269 | Lighthouse Lab in Alderley Park | Wellcome Sanger Institute for the COVID-19 Genomics UK (COG-UK) Consortium | Jacquelyn Wynn, Mairead Hyland, The Lighthouse Lab in Alderley Park and Alex Alderton, Roberto Amato, Jeffrey Barrett, Sonia Goncalves, Ewan Harrison, David K. Jackson, Ian Johnston, Dominic Kwiatkowski, Cordelia Langford, John Sillitoe on behalf of the Wellcome Sanger Institute COVID-19 Surveillance Team |
| EPI_ISL_2516270 | Health Services Laboratories | Wellcome Sanger Institute for the COVID-19 Genomics UK (COG-UK) Consortium | Health Services Laboratories and Alex Alderton, Roberto Amato, Jeffrey Barrett, Sonia Goncalves, Ewan Harrison, David K. Jackson, Ian Johnston, Dominic Kwiatkowski, Cordelia Langford, John Sillitoe on behalf of the Wellcome Sanger Institute COVID-19 Surveillance Team |
| EPI_ISL_2516271, EPI_ISL_2516272, EPI_ISL_2516273 | Lighthouse Lab in Alderley Park | Wellcome Sanger Institute for the COVID-19 Genomics UK (COG-UK) Consortium | Jacquelyn Wynn, Mairead Hyland, The Lighthouse Lab in Alderley Park and Alex Alderton, Roberto Amato, Jeffrey Barrett, Sonia Goncalves, Ewan Harrison, David K. Jackson, Ian Johnston, Dominic Kwiatkowski, Cordelia Langford, John Sillitoe on behalf of the Wellcome Sanger Institute COVID-19 Surveillance Team |
| EPI_ISL_2516274 | Health Services Laboratories | Wellcome Sanger Institute for the COVID-19 Genomics UK (COG-UK) Consortium | Health Services Laboratories and Alex Alderton, Roberto Amato, Jeffrey Barrett, Sonia Goncalves, Ewan Harrison, David K. Jackson, Ian Johnston, Dominic Kwiatkowski, Cordelia Langford, John Sillitoe on behalf of the Wellcome Sanger Institute COVID-19 Surveillance Team |
| EPI_ISL_2516275, EPI_ISL_2516276, EPI_ISL_2516277, EPI_ISL_2516278, EPI_ISL_2516279 | Lighthouse Lab in Alderley Park | Wellcome Sanger Institute for the COVID-19 Genomics UK (COG-UK) Consortium | Jacquelyn Wynn, Mairead Hyland, The Lighthouse Lab in Alderley Park and Alex Alderton, Roberto Amato, Jeffrey Barrett, Sonia Goncalves, Ewan Harrison, David K. Jackson, Ian Johnston, Dominic Kwiatkowski, Cordelia Langford, John Sillitoe on behalf of the Wellcome Sanger Institute COVID-19 Surveillance Team |
| EPI_ISL_2516280, EPI_ISL_2516281, EPI_ISL_2516282 | Health Services Laboratories | Wellcome Sanger Institute for the COVID-19 Genomics UK (COG-UK) Consortium | Health Services Laboratories and Alex Alderton, Roberto Amato, Jeffrey Barrett, Sonia Goncalves, Ewan Harrison, David K. Jackson, Ian Johnston, Dominic Kwiatkowski, Cordelia Langford, John Sillitoe on behalf of the Wellcome Sanger Institute COVID-19 Surveillance Team |
| EPI_ISL_2516283 | Lighthouse Lab in Alderley Park | Wellcome Sanger Institute for the COVID-19 Genomics UK (COG-UK) Consortium | Jacquelyn Wynn, Mairead Hyland, The Lighthouse Lab in Alderley Park and Alex Alderton, Roberto Amato, Jeffrey Barrett, Sonia Goncalves, Ewan Harrison, David K. Jackson, Ian Johnston, Dominic Kwiatkowski, Cordelia Langford, John Sillitoe on behalf of the Wellcome Sanger Institute COVID-19 Surveillance Team |
| EPI_ISL_2516284 | Health Services Laboratories | Wellcome Sanger Institute for the COVID-19 Genomics UK (COG-UK) Consortium | Health Services Laboratories and Alex Alderton, Roberto Amato, Jeffrey Barrett, Sonia Goncalves, Ewan Harrison, David K. Jackson, Ian Johnston, Dominic Kwiatkowski, Cordelia Langford, John Sillitoe on behalf of the Wellcome Sanger Institute COVID-19 Surveillance Team |
| EPI_ISL_2516285 | Lighthouse Lab in Alderley Park | Wellcome Sanger Institute for the COVID-19 Genomics UK (COG-UK) Consortium | Jacquelyn Wynn, Mairead Hyland, The Lighthouse Lab in Alderley Park and Alex Alderton, Roberto Amato, Jeffrey Barrett, Sonia Goncalves, Ewan Harrison, David K. Jackson, Ian Johnston, Dominic Kwiatkowski, Cordelia Langford, John Sillitoe on behalf of the Wellcome Sanger Institute COVID-19 Surveillance Team |
| EPI_ISL_2516286 | Lighthouse Lab in Glasgow | Wellcome Sanger Institute for the COVID-19 Genomics UK (COG-UK) Consortium | Harper VanSteenhouse, Yumi Kasai, David Gray, Carol Clugston, Anna Dominiczak and Alex Alderton, Roberto Amato, Jeffrey Barrett, Sonia Goncalves, Ewan Harrison, David K. Jackson, Ian Johnston, Dominic Kwiatkowski, Cordelia Langford, John Sillitoe on behalf of the Wellcome Sanger Institute COVID-19 Surveillance Team |
| EPI_ISL_2516287 | Lighthouse Lab in Alderley Park | Wellcome Sanger Institute for the COVID-19 Genomics UK (COG-UK) Consortium | Jacquelyn Wynn, Mairead Hyland, The Lighthouse Lab in Alderley Park and Alex Alderton, Roberto Amato, Jeffrey Barrett, Sonia Goncalves, Ewan Harrison, David K. Jackson, Ian Johnston, Dominic Kwiatkowski, Cordelia Langford, John Sillitoe on behalf of the Wellcome Sanger Institute COVID-19 Surveillance Team |
| EPI_ISL_2516288 | Lighthouse Lab in Glasgow | Wellcome Sanger Institute for the COVID-19 Genomics UK (COG-UK) Consortium | Harper VanSteenhouse, Yumi Kasai, David Gray, Carol Clugston, Anna Dominiczak and Alex Alderton, Roberto Amato, Jeffrey Barrett, Sonia Goncalves, Ewan Harrison, David K. Jackson, Ian Johnston, Dominic Kwiatkowski, Cordelia Langford, John Sillitoe on behalf of the Wellcome Sanger Institute COVID-19 Surveillance Team |
| EPI_ISL_2516289, EPI_ISL_2516290 | Health Services Laboratories | Wellcome Sanger Institute for the COVID-19 Genomics UK (COG-UK) Consortium | Health Services Laboratories and Alex Alderton, Roberto Amato, Jeffrey Barrett, Sonia Goncalves, Ewan Harrison, David K. Jackson, Ian Johnston, Dominic Kwiatkowski, Cordelia Langford, John Sillitoe on behalf of the Wellcome Sanger Institute COVID-19 Surveillance Team |
| EPI_ISL_2516291 | Lighthouse Lab in Alderley Park | Wellcome Sanger Institute for the COVID-19 Genomics UK (COG-UK) Consortium | Jacquelyn Wynn, Mairead Hyland, The Lighthouse Lab in Alderley Park and Alex Alderton, Roberto Amato, Jeffrey Barrett, Sonia Goncalves, Ewan Harrison, David K. Jackson, Ian Johnston, Dominic Kwiatkowski, Cordelia Langford, John Sillitoe on behalf of the Wellcome Sanger Institute COVID-19 Surveillance Team |
| EPI_ISL_2516292 | Health Services Laboratories | Wellcome Sanger Institute for the COVID-19 Genomics UK (COG-UK) Consortium | Health Services Laboratories and Alex Alderton, Roberto Amato, Jeffrey Barrett, Sonia Goncalves, Ewan Harrison, David K. Jackson, Ian Johnston, Dominic Kwiatkowski, Cordelia Langford, John Sillitoe on behalf of the Wellcome Sanger Institute COVID-19 Surveillance Team |
| EPI_ISL_2516293 | Lighthouse Lab in Glasgow | Wellcome Sanger Institute for the COVID-19 Genomics UK (COG-UK) Consortium | Harper VanSteenhouse, Yumi Kasai, David Gray, Carol Clugston, Anna Dominiczak and Alex Alderton, Roberto Amato, Jeffrey Barrett, Sonia Goncalves, Ewan Harrison, David K. Jackson, Ian Johnston, Dominic Kwiatkowski, Cordelia Langford, John Sillitoe on behalf of the Wellcome Sanger Institute COVID-19 Surveillance Team |
| EPI_ISL_2516294 | Health Services Laboratories | Wellcome Sanger Institute for the COVID-19 Genomics UK (COG-UK) | Health Services Laboratories and Alex Alderton, Roberto Amato, Jeffrey Barrett, Sonia Goncalves, Ewan Harrison, David K. Jackson, Ian Johnston, Dominic Kwiatkowski, Cordelia Langford, John Sillitoe on behalf of the Wellcome Sanger Institute |

| | | | |
|---|---|---|---|
| | | Consortium | COVID-19 Surveillance Team |
| EPI_ISL_2516295 | Lighthouse Lab in Alderley Park | Wellcome Sanger Institute for the COVID-19 Genomics UK (COG-UK) Consortium | Jacquelyn Wynn, Mairead Hyland, The Lighthouse Lab in Alderley Park and Alex Alderton, Roberto Amato, Jeffrey Barrett, Sonia Goncalves, Ewan Harrison, David K. Jackson, Ian Johnston, Dominic Kwiatkowski, Cordelia Langford, John Sillitoe on behalf of the Wellcome Sanger Institute COVID-19 Surveillance Team |
| EPI_ISL_2516296 | Lighthouse Laboratory Plymouth | Wellcome Sanger Institute for the COVID-19 Genomics UK (COG-UK) Consortium | Lighthouse Laboratory Plymouth and Alex Alderton, Roberto Amato, Jeffrey Barrett, Sonia Goncalves, Ewan Harrison, David K. Jackson, Ian Johnston, Dominic Kwiatkowski, Cordelia Langford, John Sillitoe on behalf of the Wellcome Sanger Institute COVID-19 Surveillance Team |
| EPI_ISL_2516297 | Health Services Laboratories | Wellcome Sanger Institute for the COVID-19 Genomics UK (COG-UK) Consortium | Health Services Laboratories and Alex Alderton, Roberto Amato, Jeffrey Barrett, Sonia Goncalves, Ewan Harrison, David K. Jackson, Ian Johnston, Dominic Kwiatkowski, Cordelia Langford, John Sillitoe on behalf of the Wellcome Sanger Institute COVID-19 Surveillance Team |
| EPI_ISL_2516298 | Lighthouse Lab in Alderley Park | Wellcome Sanger Institute for the COVID-19 Genomics UK (COG-UK) Consortium | Jacquelyn Wynn, Mairead Hyland, The Lighthouse Lab in Alderley Park and Alex Alderton, Roberto Amato, Jeffrey Barrett, Sonia Goncalves, Ewan Harrison, David K. Jackson, Ian Johnston, Dominic Kwiatkowski, Cordelia Langford, John Sillitoe on behalf of the Wellcome Sanger Institute COVID-19 Surveillance Team |
| EPI_ISL_2516299 | Health Services Laboratories | Wellcome Sanger Institute for the COVID-19 Genomics UK (COG-UK) Consortium | Health Services Laboratories and Alex Alderton, Roberto Amato, Jeffrey Barrett, Sonia Goncalves, Ewan Harrison, David K. Jackson, Ian Johnston, Dominic Kwiatkowski, Cordelia Langford, John Sillitoe on behalf of the Wellcome Sanger Institute COVID-19 Surveillance Team |
| EPI_ISL_2516300 | Lighthouse Lab in Alderley Park | Wellcome Sanger Institute for the COVID-19 Genomics UK (COG-UK) Consortium | Jacquelyn Wynn, Mairead Hyland, The Lighthouse Lab in Alderley Park and Alex Alderton, Roberto Amato, Jeffrey Barrett, Sonia Goncalves, Ewan Harrison, David K. Jackson, Ian Johnston, Dominic Kwiatkowski, Cordelia Langford, John Sillitoe on behalf of the Wellcome Sanger Institute COVID-19 Surveillance Team |
| EPI_ISL_2516301, EPI_ISL_2516302 | Health Services Laboratories | Wellcome Sanger Institute for the COVID-19 Genomics UK (COG-UK) Consortium | Health Services Laboratories and Alex Alderton, Roberto Amato, Jeffrey Barrett, Sonia Goncalves, Ewan Harrison, David K. Jackson, Ian Johnston, Dominic Kwiatkowski, Cordelia Langford, John Sillitoe on behalf of the Wellcome Sanger Institute COVID-19 Surveillance Team |
| EPI_ISL_2516303 | Lighthouse Lab in Glasgow | Wellcome Sanger Institute for the COVID-19 Genomics UK (COG-UK) Consortium | Harper VanSteenhouse, Yumi Kasai, David Gray, Carol Clugston, Anna Dominiczak and Alex Alderton, Roberto Amato, Jeffrey Barrett, Sonia Goncalves, Ewan Harrison, David K. Jackson, Ian Johnston, Dominic Kwiatkowski, Cordelia Langford, John Sillitoe on behalf of the Wellcome Sanger Institute COVID-19 Surveillance Team |
| EPI_ISL_2516304, EPI_ISL_2516305 | Lighthouse Lab in Alderley Park | Wellcome Sanger Institute for the COVID-19 Genomics UK (COG-UK) Consortium | Jacquelyn Wynn, Mairead Hyland, The Lighthouse Lab in Alderley Park and Alex Alderton, Roberto Amato, Jeffrey Barrett, Sonia Goncalves, Ewan Harrison, David K. Jackson, Ian Johnston, Dominic Kwiatkowski, Cordelia Langford, John Sillitoe on behalf of the Wellcome Sanger Institute COVID-19 Surveillance Team |
| EPI_ISL_2516306 | Health Services Laboratories | Wellcome Sanger Institute for the COVID-19 Genomics UK (COG-UK) Consortium | Health Services Laboratories and Alex Alderton, Roberto Amato, Jeffrey Barrett, Sonia Goncalves, Ewan Harrison, David K. Jackson, Ian Johnston, Dominic Kwiatkowski, Cordelia Langford, John Sillitoe on behalf of the Wellcome Sanger Institute COVID-19 Surveillance Team |
| EPI_ISL_2516307, EPI_ISL_2516308, EPI_ISL_2516309, EPI_ISL_2516311, EPI_ISL_2516312, EPI_ISL_2516313, EPI_ISL_2516314, EPI_ISL_2516315 | Lighthouse Lab in Alderley Park | Wellcome Sanger Institute for the COVID-19 Genomics UK (COG-UK) Consortium | Jacquelyn Wynn, Mairead Hyland, The Lighthouse Lab in Alderley Park and Alex Alderton, Roberto Amato, Jeffrey Barrett, Sonia Goncalves, Ewan Harrison, David K. Jackson, Ian Johnston, Dominic Kwiatkowski, Cordelia Langford, John Sillitoe on behalf of the Wellcome Sanger Institute COVID-19 Surveillance Team |
| EPI_ISL_2516316 | Lighthouse Laboratory Plymouth | Wellcome Sanger Institute for the COVID-19 Genomics UK (COG-UK) Consortium | Lighthouse Laboratory Plymouth and Alex Alderton, Roberto Amato, Jeffrey Barrett, Sonia Goncalves, Ewan Harrison, David K. Jackson, Ian Johnston, Dominic Kwiatkowski, Cordelia Langford, John Sillitoe on behalf of the Wellcome Sanger Institute COVID-19 Surveillance Team |
| EPI_ISL_2516462, EPI_ISL_2516482 | Hospital Universitari Bellvitge | Microbiology Department | Sara Marti, Aida Gonzalez-Diaz, Laura Calatayud, Jordi Niubó, Miguel Fernandez-Huerta, Carmen Ardanuy, Jordi Camara, M Angeles Domínguez |
| EPI_ISL_2516500 | CLILAB | Microbiology Department | Sara Marti, Aida Gonzalez-Diaz, Laura Calatayud, Jordi Niubó, Miguel Fernandez-Huerta, Carmen Ardanuy, Jordi Camara, M Angeles Domínguez |
| EPI_ISL_2516514, EPI_ISL_2516529, EPI_ISL_2516551, EPI_ISL_2516552, EPI_ISL_2516553, EPI_ISL_2516554, EPI_ISL_2516555, EPI_ISL_2516557, EPI_ISL_2516558 | Hospital Universitari Bellvitge | Microbiology Department | Sara Marti, Aida Gonzalez-Diaz, Laura Calatayud, Jordi Niubó, Miguel Fernandez-Huerta, Carmen Ardanuy, Jordi Camara, M Angeles Domínguez |
| EPI_ISL_2516568 | CLILAB | Microbiology Department | Sara Marti, Aida Gonzalez-Diaz, Laura Calatayud, Jordi Niubó, Miguel Fernandez-Huerta, Carmen Ardanuy, Jordi Camara, M Angeles Domínguez |
| EPI_ISL_2516570, EPI_ISL_2516571, EPI_ISL_2516575, EPI_ISL_2516579, EPI_ISL_2516583 | Hospital Universitari Bellvitge | Microbiology Department | Sara Marti, Aida Gonzalez-Diaz, Laura Calatayud, Jordi Niubó, Miguel Fernandez-Huerta, Carmen Ardanuy, Jordi Camara, M Angeles Domínguez |
| EPI_ISL_2516680, EPI_ISL_2516692, EPI_ISL_2516722, EPI_ISL_2516851, EPI_ISL_2516852, EPI_ISL_2516853, EPI_ISL_2516854, EPI_ISL_2516878, EPI_ISL_2516935, EPI_ISL_2516936 | Servicio de Microbiología Clínica (Complejo Hospitalario de Navarra, Pamplona) | Centro de Secuenciación NASERTIC | Carmen Ezpeleta Baquedano, Ana Navascués, Ana Miqueleiz |
| EPI_ISL_2517066, EPI_ISL_2517068, EPI_ISL_2517089, EPI_ISL_2517101, EPI_ISL_2517122, EPI_ISL_2517124, EPI_ISL_2517145, EPI_ISL_2517147 | Azienda Sanitaria dell'Alto Adige Laboratorio Aziendale di Microbiologia e Virologia | Istituto di Genomica Applicata | Elisabetta Pagani, Irene Bianconi, Elisabetta Giacobazzi, Elisa Masi, Stefanie Wieser, Irena Jurman, Vera Vendramin, Gabriele Magris, Eleonora Paparelli, Davide Scaglione, Michele Morgante |
| EPI_ISL_2517231, EPI_ISL_2517232, EPI_ISL_2517233, EPI_ISL_2517235, EPI_ISL_2517236, EPI_ISL_2517237, EPI_ISL_2517240, EPI_ISL_2517241, EPI_ISL_2517242, EPI_ISL_2517244, EPI_ISL_2517245, EPI_ISL_2517246, EPI_ISL_2517248, EPI_ISL_2517249, EPI_ISL_2517250, EPI_ISL_2517251, EPI_ISL_2517252, EPI_ISL_2517253, EPI_ISL_2517255, EPI_ISL_2517256, EPI_ISL_2517258, EPI_ISL_2517260, EPI_ISL_2517261, EPI_ISL_2517263, EPI_ISL_2517264, EPI_ISL_2517265, EPI_ISL_2517266, EPI_ISL_2517267, EPI_ISL_2517268, EPI_ISL_2517269, EPI_ISL_2517271, EPI_ISL_2517272, EPI_ISL_2517275, EPI_ISL_2517276, EPI_ISL_2517277, EPI_ISL_2517278, EPI_ISL_2517279, EPI_ISL_2517282, EPI_ISL_2517283, EPI_ISL_2517286, EPI_ISL_2517287, EPI_ISL_2517289, EPI_ISL_2517290 | | | |
| see above | West of Scotland Specialist Virology Centre, NHSGGC / MRC-University of Glasgow Centre for Virus Research | COVID-19 Genomics UK (COG-UK) Consortium | Ana da Silva Filipe, Natasha Johnson, Kathy Smollett, Daniel Mair, Stephen Carmichael, Alice Broos, Lily Tong, Jenna Nichols, Kyriaki Nomikou; Sarah McDonald; Ioulia Tsatsani, Richard Orton, Joseph Hughes, Sreenu Vattipally, David L Robertson; Alasdair MacLean, Rory Gunson; Sharif Shaaban, Matthew Holden; Rachel Blacow, Guy Mollett, Kathy Li, James Shepherd, Antonia Ho, Emma Thomson |
| EPI_ISL_2517291, EPI_ISL_2517292, EPI_ISL_2517293, EPI_ISL_2517295, EPI_ISL_2517296, EPI_ISL_2517297, EPI_ISL_2517298, EPI_ISL_2517299, EPI_ISL_2517300, EPI_ISL_2517301 | Virology Department, Royal Infirmary of Edinburgh, NHS Lothian / School of Biological Sciences, University of Edinburgh | COVID-19 Genomics UK (COG-UK) Consortium | McHugh M, Dewar R, Cotton S, Gallagher A, Maloney D, Rooke S, O'Toole Á, Scher E, Hill V, McCrone JT, Colquhoun R, Yu X, Jackson B, Rambaut A, Templeton K |
| EPI_ISL_2517305, EPI_ISL_2517306, EPI_ISL_2517307, EPI_ISL_2517308, EPI_ISL_2517309, EPI_ISL_2517312, EPI_ISL_2517313, EPI_ISL_2517314, EPI_ISL_2517316, EPI_ISL_2517317, EPI_ISL_2517318, EPI_ISL_2517319, EPI_ISL_2517320, EPI_ISL_2517321 | | | |
| see above | Barts Health NHS Trust | COVID-19 Genomics UK (COG-UK) Consortium | CUTINO-MOGUEL, Maria-Teresa; HARRINGTON, David; OWOYEMI, Dola; KULASEGARAN-SHYLINI, Raghavendran; BROAD, Claire; KELE, Beatrix |
| EPI_ISL_2517322, EPI_ISL_2517323, EPI_ISL_2517324, EPI_ISL_2517325, EPI_ISL_2517326, EPI_ISL_2517327, EPI_ISL_2517328, EPI_ISL_2517329, EPI_ISL_2517330, EPI_ISL_2517332, EPI_ISL_2517333, EPI_ISL_2517334, EPI_ISL_2517335, EPI_ISL_2517336, EPI_ISL_2517337, EPI_ISL_2517338, EPI_ISL_2517339, EPI_ISL_2517341, EPI_ISL_2517342, EPI_ISL_2517343, EPI_ISL_2517344, EPI_ISL_2517345, EPI_ISL_2517346, EPI_ISL_2517347, EPI_ISL_2517348, EPI_ISL_2517349, EPI_ISL_2517350, EPI_ISL_2517351, EPI_ISL_2517352, EPI_ISL_2517353, EPI_ISL_2517354, EPI_ISL_2517356, EPI_ISL_2517357, EPI_ISL_2517358, EPI_ISL_2517359, EPI_ISL_2517360, EPI_ISL_2517362, EPI_ISL_2517363, EPI_ISL_2517364, EPI_ISL_2517365, EPI_ISL_2517366, EPI_ISL_2517367, EPI_ISL_2517368, EPI_ISL_2517369, EPI_ISL_2517370, EPI_ISL_2517371, EPI_ISL_2517372, EPI_ISL_2517373, EPI_ISL_2517374, EPI_ISL_2517375, EPI_ISL_2517376, EPI_ISL_2517377, EPI_ISL_2517378, EPI_ISL_2517379, EPI_ISL_2517380, EPI_ISL_2517381, EPI_ISL_2517383, EPI_ISL_2517384, EPI_ISL_2517385, EPI_ISL_2517386, EPI_ISL_2517387, EPI_ISL_2517388, EPI_ISL_2517389, EPI_ISL_2517390, EPI_ISL_2517391, EPI_ISL_2517392, EPI_ISL_2517393, EPI_ISL_2517394, EPI_ISL_2517396, EPI_ISL_2517397, EPI_ISL_2517399, EPI_ISL_2517400, EPI_ISL_2517401, EPI_ISL_2517402, EPI_ISL_2517403, EPI_ISL_2517404, EPI_ISL_2517407, EPI_ISL_2517408, EPI_ISL_2517409, EPI_ISL_2517410, EPI_ISL_2517411, EPI_ISL_2517412, EPI_ISL_2517413, EPI_ISL_2517414, EPI_ISL_2517415, EPI_ISL_2517416, EPI_ISL_2517417, EPI_ISL_2517418, EPI_ISL_2517419, EPI_ISL_2517420, EPI_ISL_2517421, EPI_ISL_2517422, EPI_ISL_2517423, EPI_ISL_2517424, EPI_ISL_2517425, EPI_ISL_2517426, EPI_ISL_2517427, EPI_ISL_2517428, EPI_ISL_2517429, EPI_ISL_2517431, EPI_ISL_2517432, EPI_ISL_2517433, EPI_ISL_2517435, EPI_ISL_2517436, EPI_ISL_2517437, EPI_ISL_2517438, EPI_ISL_2517439, EPI_ISL_2517440, EPI_ISL_2517441, EPI_ISL_2517442, EPI_ISL_2517443, EPI_ISL_2517444, EPI_ISL_2517445, EPI_ISL_2517446, EPI_ISL_2517448, EPI_ISL_2517449, EPI_ISL_2517450, EPI_ISL_2517451, EPI_ISL_2517452, EPI_ISL_2517453, EPI_ISL_2517454, EPI_ISL_2517456, EPI_ISL_2517457, EPI_ISL_2517458, EPI_ISL_2517459, EPI_ISL_2517460, EPI_ISL_2517461, EPI_ISL_2517462, EPI_ISL_2517463, EPI_ISL_2517464, EPI_ISL_2517465, EPI_ISL_2517466, EPI_ISL_2517468, EPI_ISL_2517469, EPI_ISL_2517470, EPI_ISL_2517471, EPI_ISL_2517472, EPI_ISL_2517473, EPI_ISL_2517474, EPI_ISL_2517475, EPI_ISL_2517476, EPI_ISL_2517477, EPI_ISL_2517478, EPI_ISL_2517479, EPI_ISL_2517480, EPI_ISL_2517481, EPI_ISL_2517482, EPI_ISL_2517483, EPI_ISL_2517484, EPI_ISL_2517485, EPI_ISL_2517486, EPI_ISL_2517487, EPI_ISL_2517488, EPI_ISL_2517489, EPI_ISL_2517491, EPI_ISL_2517492, EPI_ISL_2517493, EPI_ISL_2517494, EPI_ISL_2517496, EPI_ISL_2517497, EPI_ISL_2517498, EPI_ISL_2517499, EPI_ISL_2517500, EPI_ISL_2517501, EPI_ISL_2517503, EPI_ISL_2517504, EPI_ISL_2517505, EPI_ISL_2517506, EPI_ISL_2517507, EPI_ISL_2517508, EPI_ISL_2517509, EPI_ISL_2517510, EPI_ISL_2517511, EPI_ISL_2517512, EPI_ISL_2517513, EPI_ISL_2517514, EPI_ISL_2517515, EPI_ISL_2517516, EPI_ISL_2517517, EPI_ISL_2517518, EPI_ISL_2517519, EPI_ISL_2517520, EPI_ISL_2517521, EPI_ISL_2517522, EPI_ISL_2517523, EPI_ISL_2517524, EPI_ISL_2517526, EPI_ISL_2517527, EPI_ISL_2517528, EPI_ISL_2517529, EPI_ISL_2517530, EPI_ISL_2517531, EPI_ISL_2517532, EPI_ISL_2517533, EPI_ISL_2517534, EPI_ISL_2517535, EPI_ISL_2517536, EPI_ISL_2517537, EPI_ISL_2517538, EPI_ISL_2517539, EPI_ISL_2517540, EPI_ISL_2517541, EPI_ISL_2517542, EPI_ISL_2517543, EPI_ISL_2517544, EPI_ISL_2517545, EPI_ISL_2517546, EPI_ISL_2517547, EPI_ISL_2517548, EPI_ISL_2517549, EPI_ISL_2517550, EPI_ISL_2517551, EPI_ISL_2517552, EPI_ISL_2517553, EPI_ISL_2517554, EPI_ISL_2517555, EPI_ISL_2517556, EPI_ISL_2517557, EPI_ISL_2517558, EPI_ISL_2517559, EPI_ISL_2517560, EPI_ISL_2517561, EPI_ISL_2517562, EPI_ISL_2517563, EPI_ISL_2517564, EPI_ISL_2517565, EPI_ISL_2517566, EPI_ISL_2517568, EPI_ISL_2517569, EPI_ISL_2517570, EPI_ISL_2517571, EPI_ISL_2517572, EPI_ISL_2517573, EPI_ISL_2517574, EPI_ISL_2517575, EPI_ISL_2517576, EPI_ISL_2517577, EPI_ISL_2517578, EPI_ISL_2517579, EPI_ISL_2517580, EPI_ISL_2517581, EPI_ISL_2517582, EPI_ISL_2517583, EPI_ISL_2517584, EPI_ISL_2517586, EPI_ISL_2517587, EPI_ISL_2517588, EPI_ISL_2517589, EPI_ISL_2517590, EPI_ISL_2517591, EPI_ISL_2517592, EPI_ISL_2517593, EPI_ISL_2517594, EPI_ISL_2517595, EPI_ISL_2517596, EPI_ISL_2517598, EPI_ISL_2517600, EPI_ISL_2517601, EPI_ISL_2517603, EPI_ISL_2517604, EPI_ISL_2517605, EPI_ISL_2517607, EPI_ISL_2517608, EPI_ISL_2517609, EPI_ISL_2517610, EPI_ISL_2517611, EPI_ISL_2517613, EPI_ISL_2517614, EPI_ISL_2517615, EPI_ISL_2517616, EPI_ISL_2517617, EPI_ISL_2517618, EPI_ISL_2517619, EPI_ISL_2517620, EPI_ISL_2517621, EPI_ISL_2517622, EPI_ISL_2517623, EPI_ISL_2517624, EPI_ISL_2517625, EPI_ISL_2517626, EPI_ISL_2517627, EPI_ISL_2517628, EPI_ISL_2517629, EPI_ISL_2517630, EPI_ISL_2517631, EPI_ISL_2517632, EPI_ISL_2517633, EPI_ISL_2517634, EPI_ISL_2517635, EPI_ISL_2517636, EPI_ISL_2517638, EPI_ISL_2517639, EPI_ISL_2517641, EPI_ISL_2517642, EPI_ISL_2517643, EPI_ISL_2517644, EPI_ISL_2517645, EPI_ISL_2517646, EPI_ISL_2517647, EPI_ISL_2517648, EPI_ISL_2517649, EPI_ISL_2517650, EPI_ISL_2517651, EPI_ISL_2517652, EPI_ISL_2517653, EPI_ISL_2517654, EPI_ISL_2517655, EPI_ISL_2517656, EPI_ISL_2517657, EPI_ISL_2517658, EPI_ISL_2517659, EPI_ISL_2517660, EPI_ISL_2517662, EPI_ISL_2517663, EPI_ISL_2517664, EPI_ISL_2517665, EPI_ISL_2517666, EPI_ISL_2517667, EPI_ISL_2517668, EPI_ISL_2517669, EPI_ISL_2517671, EPI_ISL_2517672, EPI_ISL_2517673, EPI_ISL_2517674, EPI_ISL_2517675, EPI_ISL_2517676, EPI_ISL_2517677, EPI_ISL_2517678, EPI_ISL_2517679, EPI_ISL_2517680, EPI_ISL_2517681, EPI_ISL_2517682, EPI_ISL_2517683, EPI_ISL_2517684, EPI_ISL_2517685, EPI_ISL_2517686, EPI_ISL_2517687, EPI_ISL_2517688, EPI_ISL_2517689, EPI_ISL_2517690, EPI_ISL_2517691, EPI_ISL_2517692, EPI_ISL_2517695, EPI_ISL_2517696, EPI_ISL_2517697, EPI_ISL_2517698, EPI_ISL_2517699, EPI_ISL_2517700, EPI_ISL_2517701, EPI_ISL_2517702, EPI_ISL_2517703, EPI_ISL_2517704, | | | |

EPI_ISL_2517705, EPI_ISL_2517706, EPI_ISL_2517707, EPI_ISL_2517708, EPI_ISL_2517709, EPI_ISL_2517710, EPI_ISL_2517711, EPI_ISL_2517712, EPI_ISL_2517713, EPI_ISL_2517714, EPI_ISL_2517715, EPI_ISL_2517716, EPI_ISL_2517717, EPI_ISL_2517718, EPI_ISL_2517719, EPI_ISL_2517720, EPI_ISL_2517721, EPI_ISL_2517722, EPI_ISL_2517723, EPI_ISL_2517725, EPI_ISL_2517728, EPI_ISL_2517729, EPI_ISL_2517730, EPI_ISL_2517731, EPI_ISL_2517732, EPI_ISL_2517733, EPI_ISL_2517734, EPI_ISL_2517735, EPI_ISL_2517736, EPI_ISL_2517737, EPI_ISL_2517738, EPI_ISL_2517739, EPI_ISL_2517740, EPI_ISL_2517741, EPI_ISL_2517742, EPI_ISL_2517743, EPI_ISL_2517744, EPI_ISL_2517745, EPI_ISL_2517746, EPI_ISL_2517747, EPI_ISL_2517748, EPI_ISL_2517749, EPI_ISL_2517750, EPI_ISL_2517751, EPI_ISL_2517752, EPI_ISL_2517753, EPI_ISL_2517754, EPI_ISL_2517755, EPI_ISL_2517756, EPI_ISL_2517757, EPI_ISL_2517758, EPI_ISL_2517759, EPI_ISL_2517760, EPI_ISL_2517761, EPI_ISL_2517762, EPI_ISL_2517763, EPI_ISL_2517764, EPI_ISL_2517766, EPI_ISL_2517767, EPI_ISL_2517768, EPI_ISL_2517769, EPI_ISL_2517771, EPI_ISL_2517774, EPI_ISL_2517775, EPI_ISL_2517776, EPI_ISL_2517777, EPI_ISL_2517778, EPI_ISL_2517779, EPI_ISL_2517780, EPI_ISL_2517781, EPI_ISL_2517782, EPI_ISL_2517783, EPI_ISL_2517784, EPI_ISL_2517785, EPI_ISL_2517786, EPI_ISL_2517787, EPI_ISL_2517789, EPI_ISL_2517790, EPI_ISL_2517791, EPI_ISL_2517792, EPI_ISL_2517793, EPI_ISL_2517794, EPI_ISL_2517795, EPI_ISL_2517797, EPI_ISL_2517798, EPI_ISL_2517799, EPI_ISL_2517800, EPI_ISL_2517801, EPI_ISL_2517802, EPI_ISL_2517803, EPI_ISL_2517804, EPI_ISL_2517805, EPI_ISL_2517806, EPI_ISL_2517807, EPI_ISL_2517808, EPI_ISL_2517809, EPI_ISL_2517810, EPI_ISL_2517811, EPI_ISL_2517812, EPI_ISL_2517813, EPI_ISL_2517814, EPI_ISL_2517815, EPI_ISL_2517816, EPI_ISL_2517817, EPI_ISL_2517818, EPI_ISL_2517819, EPI_ISL_2517820, EPI_ISL_2517821, EPI_ISL_2517822, EPI_ISL_2517823, EPI_ISL_2517824, EPI_ISL_2517825, EPI_ISL_2517826, EPI_ISL_2517827, EPI_ISL_2517828, EPI_ISL_2517829, EPI_ISL_2517830, EPI_ISL_2517831, EPI_ISL_2517832, EPI_ISL_2517833, EPI_ISL_2517834, EPI_ISL_2517835, EPI_ISL_2517836, EPI_ISL_2517837, EPI_ISL_2517838, EPI_ISL_2517839, EPI_ISL_2517840, EPI_ISL_2517841, EPI_ISL_2517842, EPI_ISL_2517844, EPI_ISL_2517845, EPI_ISL_2517846, EPI_ISL_2517847, EPI_ISL_2517848, EPI_ISL_2517849, EPI_ISL_2517850, EPI_ISL_2517851, EPI_ISL_2517852, EPI_ISL_2517853, EPI_ISL_2517854, EPI_ISL_2517855, EPI_ISL_2517856, EPI_ISL_2517857, EPI_ISL_2517858

see above | University College London, Great Ormond Street Hospital for Children NHS Foundation Trust, Imperial College Healthcare NHS Trust | COVID-19 Genomics UK (COG-UK) Consortium | Sergi Castellano, Rachel Williams, Mark Kristiansen, Paola Resende Silva, Sunando Roy, Tony Brooks, Helena Tutill, Paola Niola, Patricia Dyal, Charlotte Williams, Leysa Forrest, Yasmin Panchbhaya, Jacqueline Findlay, Samuel Weeks, Julianne Brown, Kathryn Harris, Paul Randell, James Price, Alison Holmes, Judith Breuer

EPI_ISL_2517859, EPI_ISL_2517861, EPI_ISL_2517862, EPI_ISL_2517867, EPI_ISL_2517870, EPI_ISL_2517873, EPI_ISL_2517879, EPI_ISL_2517880, EPI_ISL_2517882, EPI_ISL_2517886, EPI_ISL_2517888, EPI_ISL_2517889, EPI_ISL_2517893, EPI_ISL_2517894, EPI_ISL_2517896, EPI_ISL_2517898, EPI_ISL_2517900, EPI_ISL_2517901, EPI_ISL_2517907, EPI_ISL_2517908, EPI_ISL_2517912, EPI_ISL_2517913, EPI_ISL_2517915, EPI_ISL_2517921, EPI_ISL_2517922, EPI_ISL_2517923, EPI_ISL_2517924, EPI_ISL_2517926, EPI_ISL_2517927, EPI_ISL_2517928, EPI_ISL_2517930, EPI_ISL_2517931, EPI_ISL_2517933, EPI_ISL_2517934, EPI_ISL_2517937, EPI_ISL_2517942, EPI_ISL_2517949, EPI_ISL_2517950, EPI_ISL_2517951, EPI_ISL_2517955, EPI_ISL_2517958, EPI_ISL_2517961, EPI_ISL_2517963, EPI_ISL_2517970, EPI_ISL_2517971, EPI_ISL_2517976, EPI_ISL_2517979, EPI_ISL_2517980, EPI_ISL_2517981, EPI_ISL_2517982, EPI_ISL_2517985, EPI_ISL_2517986, EPI_ISL_2517987, EPI_ISL_2517988, EPI_ISL_2517989, EPI_ISL_2517990, EPI_ISL_2517991, EPI_ISL_2517992, EPI_ISL_2517994, EPI_ISL_2517995, EPI_ISL_2517996, EPI_ISL_2517998, EPI_ISL_2518000, EPI_ISL_2518003, EPI_ISL_2518004, EPI_ISL_2518005, EPI_ISL_2518006, EPI_ISL_2518010, EPI_ISL_2518011, EPI_ISL_2518012, EPI_ISL_2518013, EPI_ISL_2518019, EPI_ISL_2518020, EPI_ISL_2518023, EPI_ISL_2518028, EPI_ISL_2518029, EPI_ISL_2518030, EPI_ISL_2518032, EPI_ISL_2518036, EPI_ISL_2518037, EPI_ISL_2518039, EPI_ISL_2518042, EPI_ISL_2518043, EPI_ISL_2518045, EPI_ISL_2518049, EPI_ISL_2518051, EPI_ISL_2518052, EPI_ISL_2518054, EPI_ISL_2518055, EPI_ISL_2518058, EPI_ISL_2518060, EPI_ISL_2518061, EPI_ISL_2518062, EPI_ISL_2518063, EPI_ISL_2518066, EPI_ISL_2518068, EPI_ISL_2518069, EPI_ISL_2518071, EPI_ISL_2518072, EPI_ISL_2518073, EPI_ISL_2518077, EPI_ISL_2518078, EPI_ISL_2518080, EPI_ISL_2518082, EPI_ISL_2518085, EPI_ISL_2518086, EPI_ISL_2518088, EPI_ISL_2518090, EPI_ISL_2518094, EPI_ISL_2518095, EPI_ISL_2518098, EPI_ISL_2518100, EPI_ISL_2518101, EPI_ISL_2518103, EPI_ISL_2518104, EPI_ISL_2518105, EPI_ISL_2518106, EPI_ISL_2518107, EPI_ISL_2518108, EPI_ISL_2518110, EPI_ISL_2518114, EPI_ISL_2518115, EPI_ISL_2518116, EPI_ISL_2518117, EPI_ISL_2518121, EPI_ISL_2518122, EPI_ISL_2518123, EPI_ISL_2518125, EPI_ISL_2518128, EPI_ISL_2518134, EPI_ISL_2518135, EPI_ISL_2518136, EPI_ISL_2518139, EPI_ISL_2518141, EPI_ISL_2518144, EPI_ISL_2518148, EPI_ISL_2518149, EPI_ISL_2518152, EPI_ISL_2518154, EPI_ISL_2518155, EPI_ISL_2518156, EPI_ISL_2518159, EPI_ISL_2518160, EPI_ISL_2518161, EPI_ISL_2518164, EPI_ISL_2518166, EPI_ISL_2518167, EPI_ISL_2518168, EPI_ISL_2518170, EPI_ISL_2518172, EPI_ISL_2518174, EPI_ISL_2518177, EPI_ISL_2518178, EPI_ISL_2518183, EPI_ISL_2518184, EPI_ISL_2518185, EPI_ISL_2518186, EPI_ISL_2518190, EPI_ISL_2518194, EPI_ISL_2518197, EPI_ISL_2518200, EPI_ISL_2518201, EPI_ISL_2518202, EPI_ISL_2518203, EPI_ISL_2518205, EPI_ISL_2518207, EPI_ISL_2518211, EPI_ISL_2518213, EPI_ISL_2518215, EPI_ISL_2518224, EPI_ISL_2518226, EPI_ISL_2518231, EPI_ISL_2518232, EPI_ISL_2518234, EPI_ISL_2518235, EPI_ISL_2518236, EPI_ISL_2518237, EPI_ISL_2518242, EPI_ISL_2518244, EPI_ISL_2518245, EPI_ISL_2518247, EPI_ISL_2518248, EPI_ISL_2518251, EPI_ISL_2518252, EPI_ISL_2518255, EPI_ISL_2518258, EPI_ISL_2518259, EPI_ISL_2518260, EPI_ISL_2518261, EPI_ISL_2518262, EPI_ISL_2518265, EPI_ISL_2518267, EPI_ISL_2518272, EPI_ISL_2518273, EPI_ISL_2518274, EPI_ISL_2518275, EPI_ISL_2518279, EPI_ISL_2518280, EPI_ISL_2518283, EPI_ISL_2518286, EPI_ISL_2518289, EPI_ISL_2518293, EPI_ISL_2518296, EPI_ISL_2518298, EPI_ISL_2518300, EPI_ISL_2518302, EPI_ISL_2518303, EPI_ISL_2518307, EPI_ISL_2518313, EPI_ISL_2518314, EPI_ISL_2518316, EPI_ISL_2518317, EPI_ISL_2518318, EPI_ISL_2518319, EPI_ISL_2518320, EPI_ISL_2518321, EPI_ISL_2518322, EPI_ISL_2518323, EPI_ISL_2518325, EPI_ISL_2518328, EPI_ISL_2518331, EPI_ISL_2518332, EPI_ISL_2518333, EPI_ISL_2518335, EPI_ISL_2518336, EPI_ISL_2518337, EPI_ISL_2518339, EPI_ISL_2518340, EPI_ISL_2518341, EPI_ISL_2518344, EPI_ISL_2518345, EPI_ISL_2518347, EPI_ISL_2518348, EPI_ISL_2518349, EPI_ISL_2518350, EPI_ISL_2518353, EPI_ISL_2518355, EPI_ISL_2518356, EPI_ISL_2518357, EPI_ISL_2518358, EPI_ISL_2518359, EPI_ISL_2518360, EPI_ISL_2518361, EPI_ISL_2518362, EPI_ISL_2518363, EPI_ISL_2518365, EPI_ISL_2518366, EPI_ISL_2518368, EPI_ISL_2518370, EPI_ISL_2518371, EPI_ISL_2518372, EPI_ISL_2518373, EPI_ISL_2518376, EPI_ISL_2518377, EPI_ISL_2518379, EPI_ISL_2518380, EPI_ISL_2518382, EPI_ISL_2518384, EPI_ISL_2518386, EPI_ISL_2518387, EPI_ISL_2518388, EPI_ISL_2518394, EPI_ISL_2518395, EPI_ISL_2518396, EPI_ISL_2518397, EPI_ISL_2518401, EPI_ISL_2518402, EPI_ISL_2518403, EPI_ISL_2518404, EPI_ISL_2518405, EPI_ISL_2518407, EPI_ISL_2518412, EPI_ISL_2518414, EPI_ISL_2518415, EPI_ISL_2518416, EPI_ISL_2518417, EPI_ISL_2518419, EPI_ISL_2518421, EPI_ISL_2518423, EPI_ISL_2518424, EPI_ISL_2518425, EPI_ISL_2518429, EPI_ISL_2518431, EPI_ISL_2518434, EPI_ISL_2518435, EPI_ISL_2518436, EPI_ISL_2518437, EPI_ISL_2518440, EPI_ISL_2518441, EPI_ISL_2518444, EPI_ISL_2518445, EPI_ISL_2518446, EPI_ISL_2518447, EPI_ISL_2518448, EPI_ISL_2518449, EPI_ISL_2518451, EPI_ISL_2518453, EPI_ISL_2518454, EPI_ISL_2518455, EPI_ISL_2518456, EPI_ISL_2518457, EPI_ISL_2518464, EPI_ISL_2518465, EPI_ISL_2518466, EPI_ISL_2518467, EPI_ISL_2518468, EPI_ISL_2518470, EPI_ISL_2518471, EPI_ISL_2518473, EPI_ISL_2518475, EPI_ISL_2518476, EPI_ISL_2518480, EPI_ISL_2518481, EPI_ISL_2518482, EPI_ISL_2518485, EPI_ISL_2518487, EPI_ISL_2518488, EPI_ISL_2518490, EPI_ISL_2518491, EPI_ISL_2518493, EPI_ISL_2518494, EPI_ISL_2518495, EPI_ISL_2518497, EPI_ISL_2518498, EPI_ISL_2518499, EPI_ISL_2518500, EPI_ISL_2518505, EPI_ISL_2518506, EPI_ISL_2518507, EPI_ISL_2518510, EPI_ISL_2518511, EPI_ISL_2518513, EPI_ISL_2518515, EPI_ISL_2518517, EPI_ISL_2518518, EPI_ISL_2518520, EPI_ISL_2518522, EPI_ISL_2518524, EPI_ISL_2518525, EPI_ISL_2518526, EPI_ISL_2518527, EPI_ISL_2518529, EPI_ISL_2518531, EPI_ISL_2518532, EPI_ISL_2518535, EPI_ISL_2518536, EPI_ISL_2518538, EPI_ISL_2518541, EPI_ISL_2518542, EPI_ISL_2518543, EPI_ISL_2518547, EPI_ISL_2518549, EPI_ISL_2518551, EPI_ISL_2518552, EPI_ISL_2518553, EPI_ISL_2518557, EPI_ISL_2518559, EPI_ISL_2518564, EPI_ISL_2518567, EPI_ISL_2518570, EPI_ISL_2518571, EPI_ISL_2518572, EPI_ISL_2518579, EPI_ISL_2518580, EPI_ISL_2518581, EPI_ISL_2518582, EPI_ISL_2518583, EPI_ISL_2518584, EPI_ISL_2518585, EPI_ISL_2518586, EPI_ISL_2518588, EPI_ISL_2518589, EPI_ISL_2518593, EPI_ISL_2518594, EPI_ISL_2518597, EPI_ISL_2518598, EPI_ISL_2518600, EPI_ISL_2518602, EPI_ISL_2518606, EPI_ISL_2518607, EPI_ISL_2518610, EPI_ISL_2518612, EPI_ISL_2518613, EPI_ISL_2518614, EPI_ISL_2518615, EPI_ISL_2518617, EPI_ISL_2518618, EPI_ISL_2518621, EPI_ISL_2518623, EPI_ISL_2518624, EPI_ISL_2518625, EPI_ISL_2518628, EPI_ISL_2518630, EPI_ISL_2518631, EPI_ISL_2518632, EPI_ISL_2518634, EPI_ISL_2518635, EPI_ISL_2518637, EPI_ISL_2518638, EPI_ISL_2518639, EPI_ISL_2518640, EPI_ISL_2518641, EPI_ISL_2518642, EPI_ISL_2518645, EPI_ISL_2518647, EPI_ISL_2518649, EPI_ISL_2518653, EPI_ISL_2518655, EPI_ISL_2518656, EPI_ISL_2518657, EPI_ISL_2518659, EPI_ISL_2518662, EPI_ISL_2518663, EPI_ISL_2518664, EPI_ISL_2518666, EPI_ISL_2518667, EPI_ISL_2518672, EPI_ISL_2518673, EPI_ISL_2518674, EPI_ISL_2518675, EPI_ISL_2518676, EPI_ISL_2518677, EPI_ISL_2518679, EPI_ISL_2518680, EPI_ISL_2518681, EPI_ISL_2518683, EPI_ISL_2518685, EPI_ISL_2518686, EPI_ISL_2518688, EPI_ISL_2518689, EPI_ISL_2518692, EPI_ISL_2518693, EPI_ISL_2518694, EPI_ISL_2518698, EPI_ISL_2518699, EPI_ISL_2518702, EPI_ISL_2518705, EPI_ISL_2518706, EPI_ISL_2518707, EPI_ISL_2518708, EPI_ISL_2518713, EPI_ISL_2518715, EPI_ISL_2518716, EPI_ISL_2518717, EPI_ISL_2518718, EPI_ISL_2518719, EPI_ISL_2518720, EPI_ISL_2518721, EPI_ISL_2518723, EPI_ISL_2518724, EPI_ISL_2518725, EPI_ISL_2518727, EPI_ISL_2518728, EPI_ISL_2518729, EPI_ISL_2518730, EPI_ISL_2518732, EPI_ISL_2518734, EPI_ISL_2518735, EPI_ISL_2518736, EPI_ISL_2518737, EPI_ISL_2518738, EPI_ISL_2518739, EPI_ISL_2518740, EPI_ISL_2518742, EPI_ISL_2518743, EPI_ISL_2518744, EPI_ISL_2518745, EPI_ISL_2518746, EPI_ISL_2518747, EPI_ISL_2518748, EPI_ISL_2518749, EPI_ISL_2518751, EPI_ISL_2518752, EPI_ISL_2518753, EPI_ISL_2518754, EPI_ISL_2518755, EPI_ISL_2518756, EPI_ISL_2518757, EPI_ISL_2518758, EPI_ISL_2518759, EPI_ISL_2518760, EPI_ISL_2518761, EPI_ISL_2518762, EPI_ISL_2518764, EPI_ISL_2518765, EPI_ISL_2518767, EPI_ISL_2518768, EPI_ISL_2518769, EPI_ISL_2518770, EPI_ISL_2518771, EPI_ISL_2518772, EPI_ISL_2518773, EPI_ISL_2518775, EPI_ISL_2518777, EPI_ISL_2518781, EPI_ISL_2518782, EPI_ISL_2518783, EPI_ISL_2518784, EPI_ISL_2518785, EPI_ISL_2518786, EPI_ISL_2518787, EPI_ISL_2518788, EPI_ISL_2518789, EPI_ISL_2518790, EPI_ISL_2518791, EPI_ISL_2518792, EPI_ISL_2518794, EPI_ISL_2518795, EPI_ISL_2518796, EPI_ISL_2518797, EPI_ISL_2518798, EPI_ISL_2518799, EPI_ISL_2518800, EPI_ISL_2518802, EPI_ISL_2518804, EPI_ISL_2518805, EPI_ISL_2518808, EPI_ISL_2518810, EPI_ISL_2518812, EPI_ISL_2518813, EPI_ISL_2518816, EPI_ISL_2518818, EPI_ISL_2518819, EPI_ISL_2518820, EPI_ISL_2518821, EPI_ISL_2518824, EPI_ISL_2518825, EPI_ISL_2518826, EPI_ISL_2518827, EPI_ISL_2518828, EPI_ISL_2518829, EPI_ISL_2518831, EPI_ISL_2518832, EPI_ISL_2518835, EPI_ISL_2518836, EPI_ISL_2518838, EPI_ISL_2518839, EPI_ISL_2518843, EPI_ISL_2518844, EPI_ISL_2518845, EPI_ISL_2518849, EPI_ISL_2518850, EPI_ISL_2518851, EPI_ISL_2518854, EPI_ISL_2518855, EPI_ISL_2518857, EPI_ISL_2518858, EPI_ISL_2518860, EPI_ISL_2518865, EPI_ISL_2518866, EPI_ISL_2518867, EPI_ISL_2518868, EPI_ISL_2518869, EPI_ISL_2518870, EPI_ISL_2518871, EPI_ISL_2518872, EPI_ISL_2518875, EPI_ISL_2518876, EPI_ISL_2518877, EPI_ISL_2518878, EPI_ISL_2518879, EPI_ISL_2518880, EPI_ISL_2518881, EPI_ISL_2518882, EPI_ISL_2518883, EPI_ISL_2518884, EPI_ISL_2518886, EPI_ISL_2518888, EPI_ISL_2518889, EPI_ISL_2518890, EPI_ISL_2518891, EPI_ISL_2518893, EPI_ISL_2518894, EPI_ISL_2518896, EPI_ISL_2518897, EPI_ISL_2518898, EPI_ISL_2518899, EPI_ISL_2518900, EPI_ISL_2518901, EPI_ISL_2518902, EPI_ISL_2518904, EPI_ISL_2518905, EPI_ISL_2518906, EPI_ISL_2518908, EPI_ISL_2518910, EPI_ISL_2518911, EPI_ISL_2518913, EPI_ISL_2518914, EPI_ISL_2518915, EPI_ISL_2518916, EPI_ISL_2518917, EPI_ISL_2518918, EPI_ISL_2518919, EPI_ISL_2518920, EPI_ISL_2518921, EPI_ISL_2518922, EPI_ISL_2518923, EPI_ISL_2518924, EPI_ISL_2518927, EPI_ISL_2518928, EPI_ISL_2518929, EPI_ISL_2518930, EPI_ISL_2518931, EPI_ISL_2518932, EPI_ISL_2518933, EPI_ISL_2518934, EPI_ISL_2518936, EPI_ISL_2518937, EPI_ISL_2518938, EPI_ISL_2518939, EPI_ISL_2518941, EPI_ISL_2518942, EPI_ISL_2518943, EPI_ISL_2518944, EPI_ISL_2518945, EPI_ISL_2518946, EPI_ISL_2518947, EPI_ISL_2518948, EPI_ISL_2518950, EPI_ISL_2518951, EPI_ISL_2518952, EPI_ISL_2518953, EPI_ISL_2518954, EPI_ISL_2518956, EPI_ISL_2518957, EPI_ISL_2518958, EPI_ISL_2518960, EPI_ISL_2518961, EPI_ISL_2518962, EPI_ISL_2518964, EPI_ISL_2518965, EPI_ISL_2518966, EPI_ISL_2518967, EPI_ISL_2518968, EPI_ISL_2518970

see above | Northumbria University / South Tees Hospitals NHS Foundation Trust / North Cumbria Integrated Care NHS Foundation Trust / North Tees and Hartlepool NHS Foundation Trust / Newcastle Hospitals NHS Foundation Trust | COVID-19 Genomics UK (COG-UK) Consortium | Darren L Smith,Andrew Nelson,Matthew Bashton,Gregory R Young,Joshua Loh,John Allan,Mohammad A Tariq,Giles S Holt,Gary Black,Wen C Yew,Lynn Dover,Paul Baker,Steve Liggett,Sarah Essex,Jane Greenaway,Debra Padgett,Clive Graham,Garren Scott,Edward Barton,Emma Swindells,Brendan Payne,Jennifer Collins,Yusri Taha,Gary Eltringham

EPI_ISL_2518971, EPI_ISL_2518972, EPI_ISL_2518973, EPI_ISL_2518974, EPI_ISL_2518975, EPI_ISL_2518976, EPI_ISL_2518977, EPI_ISL_2518978, EPI_ISL_2518979, EPI_ISL_2518980, EPI_ISL_2518981, EPI_ISL_2518982, EPI_ISL_2518983, EPI_ISL_2518984, EPI_ISL_2518985, EPI_ISL_2518986, EPI_ISL_2518987, EPI_ISL_2518988, EPI_ISL_2518989, EPI_ISL_2518990, EPI_ISL_2518991, EPI_ISL_2518992, EPI_ISL_2518993, EPI_ISL_2518995, EPI_ISL_2518996, EPI_ISL_2518997, EPI_ISL_2518998, EPI_ISL_2518999, EPI_ISL_2519001, EPI_ISL_2519002, EPI_ISL_2519003, EPI_ISL_2519004, EPI_ISL_2519005, EPI_ISL_2519006, EPI_ISL_2519007, EPI_ISL_2519008, EPI_ISL_2519009, EPI_ISL_2519010, EPI_ISL_2519011, EPI_ISL_2519012, EPI_ISL_2519013, EPI_ISL_2519014, EPI_ISL_2519015, EPI_ISL_2519016, EPI_ISL_2519017, EPI_ISL_2519018, EPI_ISL_2519019, EPI_ISL_2519020, EPI_ISL_2519021, EPI_ISL_2519022, EPI_ISL_2519023, EPI_ISL_2519024, EPI_ISL_2519025, EPI_ISL_2519027, EPI_ISL_2519028, EPI_ISL_2519029, EPI_ISL_2519030, EPI_ISL_2519031, EPI_ISL_2519032, EPI_ISL_2519033, EPI_ISL_2519034, EPI_ISL_2519035, EPI_ISL_2519036, EPI_ISL_2519037, EPI_ISL_2519038, EPI_ISL_2519039, EPI_ISL_2519040, EPI_ISL_2519041, EPI_ISL_2519042, EPI_ISL_2519043, EPI_ISL_2519044, EPI_ISL_2519045, EPI_ISL_2519046, EPI_ISL_2519047, EPI_ISL_2519048, EPI_ISL_2519049, EPI_ISL_2519050, EPI_ISL_2519051, EPI_ISL_2519052, EPI_ISL_2519053, EPI_ISL_2519054, EPI_ISL_2519055, EPI_ISL_2519056, EPI_ISL_2519057, EPI_ISL_2519058, EPI_ISL_2519059, EPI_ISL_2519060, EPI_ISL_2519061, EPI_ISL_2519062, EPI_ISL_2519063, EPI_ISL_2519064, EPI_ISL_2519065, EPI_ISL_2519067, EPI_ISL_2519068, EPI_ISL_2519069, EPI_ISL_2519070, EPI_ISL_2519071, EPI_ISL_2519074, EPI_ISL_2519075, EPI_ISL_2519076, EPI_ISL_2519077, EPI_ISL_2519078, EPI_ISL_2519079, EPI_ISL_2519080, EPI_ISL_2519082, EPI_ISL_2519083, EPI_ISL_2519084, EPI_ISL_2519085, EPI_ISL_2519086, EPI_ISL_2519087, EPI_ISL_2519088, EPI_ISL_2519089, EPI_ISL_2519090, EPI_ISL_2519091, EPI_ISL_2519092, EPI_ISL_2519094, EPI_ISL_2519095, EPI_ISL_2519096, EPI_ISL_2519097, EPI_ISL_2519098, EPI_ISL_2519099, EPI_ISL_2519100, EPI_ISL_2519101, EPI_ISL_2519102, EPI_ISL_2519103, EPI_ISL_2519104, EPI_ISL_2519105, EPI_ISL_2519106, EPI_ISL_2519107, EPI_ISL_2519108, EPI_ISL_2519109, EPI_ISL_2519110, EPI_ISL_2519111, EPI_ISL_2519112, EPI_ISL_2519113, EPI_ISL_2519114, EPI_ISL_2519115, EPI_ISL_2519116, EPI_ISL_2519118, EPI_ISL_2519120, EPI_ISL_2519121, EPI_ISL_2519122, EPI_ISL_2519123, EPI_ISL_2519125, EPI_ISL_2519126, EPI_ISL_2519127, EPI_ISL_2519128, EPI_ISL_2519129, EPI_ISL_2519130, EPI_ISL_2519131, EPI_ISL_2519132, EPI_ISL_2519133, EPI_ISL_2519134, EPI_ISL_2519135, EPI_ISL_2519136, EPI_ISL_2519137, EPI_ISL_2519138, EPI_ISL_2519139, EPI_ISL_2519140, EPI_ISL_2519141, EPI_ISL_2519142, EPI_ISL_2519143, EPI_ISL_2519144, EPI_ISL_2519145, EPI_ISL_2519146, EPI_ISL_2519147, EPI_ISL_2519148, EPI_ISL_2519149, EPI_ISL_2519150, EPI_ISL_2519151, EPI_ISL_2519152, EPI_ISL_2519153, EPI_ISL_2519154, EPI_ISL_2519155, EPI_ISL_2519156, EPI_ISL_2519157, EPI_ISL_2519158, EPI_ISL_2519159, EPI_ISL_2519160, EPI_ISL_2519161, EPI_ISL_2519162, EPI_ISL_2519163, EPI_ISL_2519164, EPI_ISL_2519165, EPI_ISL_2519166, EPI_ISL_2519167, EPI_ISL_2519168, EPI_ISL_2519169, EPI_ISL_2519170, EPI_ISL_2519171, EPI_ISL_2519172, EPI_ISL_2519173, EPI_ISL_2519174, EPI_ISL_2519175, EPI_ISL_2519176, EPI_ISL_2519177, EPI_ISL_2519180, EPI_ISL_2519181, EPI_ISL_2519182, EPI_ISL_2519183, EPI_ISL_2519184, EPI_ISL_2519185, EPI_ISL_2519186, EPI_ISL_2519187, EPI_ISL_2519188, EPI_ISL_2519190, EPI_ISL_2519191, EPI_ISL_2519192, EPI_ISL_2519193, EPI_ISL_2519195, EPI_ISL_2519196, EPI_ISL_2519197, EPI_ISL_2519198, EPI_ISL_2519199, EPI_ISL_2519200, EPI_ISL_2519201, EPI_ISL_2519202, EPI_ISL_2519203, EPI_ISL_2519204, EPI_ISL_2519205, EPI_ISL_2519206, EPI_ISL_2519207, EPI_ISL_2519208, EPI_ISL_2519209, EPI_ISL_2519210, EPI_ISL_2519211, EPI_ISL_2519212, EPI_ISL_2519213, EPI_ISL_2519215, EPI_ISL_2519216, EPI_ISL_2519219, EPI_ISL_2519220, EPI_ISL_2519221, EPI_ISL_2519222, EPI_ISL_2519223, EPI_ISL_2519224, EPI_ISL_2519225, EPI_ISL_2519226, EPI_ISL_2519227, EPI_ISL_2519228, EPI_ISL_2519229, EPI_ISL_2519230, EPI_ISL_2519231, EPI_ISL_2519232, EPI_ISL_2519233, EPI_ISL_2519234, EPI_ISL_2519235, EPI_ISL_2519236, EPI_ISL_2519237, EPI_ISL_2519238, EPI_ISL_2519239, EPI_ISL_2519240, EPI_ISL_2519241, EPI_ISL_2519242, EPI_ISL_2519243, EPI_ISL_2519244, EPI_ISL_2519245, EPI_ISL_2519246, EPI_ISL_2519247, EPI_ISL_2519248, EPI_ISL_2519249, EPI_ISL_2519250, EPI_ISL_2519251, EPI_ISL_2519252, EPI_ISL_2519253, EPI_ISL_2519254, EPI_ISL_2519255, EPI_ISL_2519256, EPI_ISL_2519257, EPI_ISL_2519258, EPI_ISL_2519259, EPI_ISL_2519260, EPI_ISL_2519261, EPI_ISL_2519262, EPI_ISL_2519263, EPI_ISL_2519264, EPI_ISL_2519265, EPI_ISL_2519266, EPI_ISL_2519267, EPI_ISL_2519268, EPI_ISL_2519269, EPI_ISL_2519271, EPI_ISL_2519272, EPI_ISL_2519273, EPI_ISL_2519274, EPI_ISL_2519275, EPI_ISL_2519276, EPI_ISL_2519277, EPI_ISL_2519278, EPI_ISL_2519280, EPI_ISL_2519281, EPI_ISL_2519282, EPI_ISL_2519284, EPI_ISL_2519285, EPI_ISL_2519287, EPI_ISL_2519288, EPI_ISL_2519290, EPI_ISL_2519291, EPI_ISL_2519292, EPI_ISL_2519293, EPI_ISL_2519295, EPI_ISL_2519297, EPI_ISL_2519298, EPI_ISL_2519300, EPI_ISL_2519301, EPI_ISL_2519302, EPI_ISL_2519303, EPI_ISL_2519304, EPI_ISL_2519305, EPI_ISL_2519306, EPI_ISL_2519307, EPI_ISL_2519308, EPI_ISL_2519310, EPI_ISL_2519311, EPI_ISL_2519312, EPI_ISL_2519314, EPI_ISL_2519315, EPI_ISL_2519316, EPI_ISL_2519317, EPI_ISL_2519318, EPI_ISL_2519319, EPI_ISL_2519320, EPI_ISL_2519321, EPI_ISL_2519322, EPI_ISL_2519323, EPI_ISL_2519324, EPI_ISL_2519325, EPI_ISL_2519326, EPI_ISL_2519327, EPI_ISL_2519328, EPI_ISL_2519329, EPI_ISL_2519330, EPI_ISL_2519332, EPI_ISL_2519334, EPI_ISL_2519335, EPI_ISL_2519336, EPI_ISL_2519337, EPI_ISL_2519339, EPI_ISL_2519340, EPI_ISL_2519341, EPI_ISL_2519342, EPI_ISL_2519343, EPI_ISL_2519344, EPI_ISL_2519345, EPI_ISL_2519346, EPI_ISL_2519347, EPI_ISL_2519348, EPI_ISL_2519349, EPI_ISL_2519350, EPI_ISL_2519351, EPI_ISL_2519352, EPI_ISL_2519353, EPI_ISL_2519354, EPI_ISL_2519355, EPI_ISL_2519356, EPI_ISL_2519357, EPI_ISL_2519358, EPI_ISL_2519359, EPI_ISL_2519360, EPI_ISL_2519361, EPI_ISL_2519362, EPI_ISL_2519363, EPI_ISL_2519364, EPI_ISL_2519365, EPI_ISL_2519367, EPI_ISL_2519368, EPI_ISL_2519369, EPI_ISL_2519370, EPI_ISL_2519371, EPI_ISL_2519372, EPI_ISL_2519373, EPI_ISL_2519374, EPI_ISL_2519375, EPI_ISL_2519378, EPI_ISL_2519379, EPI_ISL_2519380, EPI_ISL_2519381, EPI_ISL_2519383, EPI_ISL_2519384, EPI_ISL_2519385, EPI_ISL_2519386, EPI_ISL_2519388, EPI_ISL_2519389, EPI_ISL_2519390, EPI_ISL_2519391, EPI_ISL_2519392, EPI_ISL_2519393, EPI_ISL_2519395, EPI_ISL_2519397, EPI_ISL_2519398, EPI_ISL_2519399, EPI_ISL_2519400, EPI_ISL_2519401, EPI_ISL_2519402, EPI_ISL_2519403, EPI_ISL_2519404, EPI_ISL_2519405, EPI_ISL_2519406, EPI_ISL_2519407, EPI_ISL_2519408, EPI_ISL_2519409, EPI_ISL_2519410, EPI_ISL_2519411, EPI_ISL_2519412, EPI_ISL_2519413, EPI_ISL_2519414, EPI_ISL_2519415, EPI_ISL_2519416, EPI_ISL_2519417, EPI_ISL_2519418, EPI_ISL_2519419, EPI_ISL_2519420, EPI_ISL_2519421, EPI_ISL_2519422, EPI_ISL_2519423, EPI_ISL_2519424, EPI_ISL_2519425, EPI_ISL_2519426, EPI_ISL_2519427, EPI_ISL_2519428, EPI_ISL_2519429, EPI_ISL_2519430, EPI_ISL_2519431, EPI_ISL_2519432, EPI_ISL_2519433, EPI_ISL_2519434, EPI_ISL_2519435, EPI_ISL_2519436, EPI_ISL_2519438, EPI_ISL_2519439, EPI_ISL_2519440, EPI_ISL_2519441, EPI_ISL_2519442, EPI_ISL_2519443, EPI_ISL_2519444, EPI_ISL_2519445, EPI_ISL_2519446, EPI_ISL_2519447, EPI_ISL_2519448, EPI_ISL_2519449, EPI_ISL_2519450, EPI_ISL_2519451, EPI_ISL_2519452, EPI_ISL_2519453, EPI_ISL_2519454, EPI_ISL_2519455, EPI_ISL_2519456, EPI_ISL_2519457, EPI_ISL_2519458, EPI_ISL_2519459, EPI_ISL_2519460, EPI_ISL_2519461, EPI_ISL_2519462, EPI_ISL_2519463, EPI_ISL_2519464, EPI_ISL_2519465, EPI_ISL_2519466, EPI_ISL_2519467, EPI_ISL_2519468, EPI_ISL_2519469, EPI_ISL_2519470, EPI_ISL_2519471, EPI_ISL_2519472, EPI_ISL_2519473, EPI_ISL_2519474, EPI_ISL_2519475, EPI_ISL_2519476, EPI_ISL_2519478, EPI_ISL_2519479, EPI_ISL_2519480, EPI_ISL_2519481, EPI_ISL_2519482, EPI_ISL_2519484, EPI_ISL_2519485, EPI_ISL_2519486, EPI_ISL_2519487, EPI_ISL_2519488, EPI_ISL_2519489, EPI_ISL_2519490, EPI_ISL_2519491, EPI_ISL_2519492, EPI_ISL_2519493, EPI_ISL_2519494, EPI_ISL_2519495, EPI_ISL_2519496, EPI_ISL_2519497, EPI_ISL_2519498, EPI_ISL_2519499, EPI_ISL_2519500, EPI_ISL_2519501, EPI_ISL_2519502, EPI_ISL_2519503, EPI_ISL_2519504, EPI_ISL_2519505, EPI_ISL_2519507, EPI_ISL_2519508, EPI_ISL_2519509, EPI_ISL_2519510, EPI_ISL_2519511, EPI_ISL_2519512, EPI_ISL_2519513, EPI_ISL_2519514, EPI_ISL_2519515, EPI_ISL_2519516, EPI_ISL_2519517, EPI_ISL_2519519, EPI_ISL_2519520, EPI_ISL_2519521, EPI_ISL_2519522, EPI_ISL_2519523, EPI_ISL_2519524, EPI_ISL_2519526, EPI_ISL_2519527, EPI_ISL_2519528, EPI_ISL_2519530, EPI_ISL_2519531, EPI_ISL_2519532, EPI_ISL_2519533, EPI_ISL_2519534, EPI_ISL_2519535, EPI_ISL_2519536, EPI_ISL_2519537, EPI_ISL_2519538, EPI_ISL_2519540, EPI_ISL_2519542, EPI_ISL_2519543, EPI_ISL_2519544, EPI_ISL_2519545, EPI_ISL_2519546, EPI_ISL_2519547, EPI_ISL_2519548, EPI_ISL_2519549, EPI_ISL_2519550, EPI_ISL_2519551, EPI_ISL_2519552, EPI_ISL_2519553, EPI_ISL_2519554, EPI_ISL_2519555, EPI_ISL_2519556, EPI_ISL_2519557, EPI_ISL_2519558, EPI_ISL_2519559, EPI_ISL_2519560, EPI_ISL_2519561, EPI_ISL_2519562, EPI_ISL_2519563, EPI_ISL_2519564, EPI_ISL_2519565, EPI_ISL_2519566, EPI_ISL_2519567, EPI_ISL_2519568, EPI_ISL_2519570, EPI_ISL_2519571, EPI_ISL_2519572, EPI_ISL_2519573, EPI_ISL_2519574, EPI_ISL_2519575, EPI_ISL_2519576, EPI_ISL_2519578, EPI_ISL_2519579, EPI_ISL_2519580, EPI_ISL_2519582, EPI_ISL_2519583, EPI_ISL_2519584, EPI_ISL_2519585, EPI_ISL_2519586, EPI_ISL_2519588, EPI_ISL_2519589, EPI_ISL_2519590, EPI_ISL_2519591, EPI_ISL_2519592, EPI_ISL_2519593, EPI_ISL_2519594, EPI_ISL_2519596, EPI_ISL_2519598, EPI_ISL_2519599, EPI_ISL_2519601, EPI_ISL_2519602, EPI_ISL_2519603, EPI_ISL_2519604, EPI_ISL_2519605, EPI_ISL_2519606, EPI_ISL_2519607, EPI_ISL_2519608, EPI_ISL_2519609, EPI_ISL_2519610, EPI_ISL_2519611, EPI_ISL_2519612, EPI_ISL_2519613, EPI_ISL_2519614, EPI_ISL_2519615, EPI_ISL_2519616, EPI_ISL_2519618, EPI_ISL_2519619, EPI_ISL_2519620, EPI_ISL_2519621, EPI_ISL_2519622, EPI_ISL_2519623, EPI_ISL_2519624, EPI_ISL_2519625, EPI_ISL_2519626, EPI_ISL_2519627, EPI_ISL_2519628, EPI_ISL_2519629, EPI_ISL_2519630, EPI_ISL_2519631, EPI_ISL_2519632, EPI_ISL_2519633, EPI_ISL_2519636, EPI_ISL_2519637, EPI_ISL_2519638, EPI_ISL_2519639, EPI_ISL_2519640, EPI_ISL_2519641, EPI_ISL_2519642, EPI_ISL_2519643, EPI_ISL_2519645, EPI_ISL_2519646, EPI_ISL_2519648, EPI_ISL_2519649, EPI_ISL_2519650, EPI_ISL_2519651, EPI_ISL_2519652, EPI_ISL_2519653, EPI_ISL_2519654, EPI_ISL_2519655, EPI_ISL_2519656, EPI_ISL_2519657, EPI_ISL_2519658, EPI_ISL_2519659, EPI_ISL_2519662, EPI_ISL_2519663, EPI_ISL_2519664, EPI_ISL_2519665, EPI_ISL_2519666, EPI_ISL_2519667, EPI_ISL_2519668, EPI_ISL_2519669, EPI_ISL_2519670, EPI_ISL_2519671, EPI_ISL_2519672, EPI_ISL_2519673, EPI_ISL_2519674, EPI_ISL_2519675, EPI_ISL_2519676, EPI_ISL_2519677, EPI_ISL_2519678, EPI_ISL_2519679, EPI_ISL_2519680, EPI_ISL_2519681, EPI_ISL_2519682, EPI_ISL_2519683, EPI_ISL_2519684, EPI_ISL_2519685, EPI_ISL_2519686, EPI_ISL_2519687, EPI_ISL_2519688, EPI_ISL_2519690, EPI_ISL_2519691, EPI_ISL_2519692, EPI_ISL_2519693, EPI_ISL_2519694, EPI_ISL_2519695, EPI_ISL_2519696, EPI_ISL_2519697, EPI_ISL_2519698, EPI_ISL_2519699, EPI_ISL_2519700, EPI_ISL_2519701, EPI_ISL_2519703, EPI_ISL_2519704, EPI_ISL_2519705, EPI_ISL_2519707, EPI_ISL_2519708, EPI_ISL_2519709, EPI_ISL_2519710, EPI_ISL_2519711, EPI_ISL_2519712, EPI_ISL_2519713, EPI_ISL_2519714, EPI_ISL_2519715, EPI_ISL_2519716, EPI_ISL_2519717, EPI_ISL_2519718, EPI_ISL_2519719, EPI_ISL_2519720, EPI_ISL_2519721, EPI_ISL_2519722, EPI_ISL_2519723, EPI_ISL_2519724, EPI_ISL_2519725, EPI_ISL_2519726, EPI_ISL_2519727, EPI_ISL_2519728, EPI_ISL_2519730, EPI_ISL_2519731, EPI_ISL_2519732, EPI_ISL_2519733, EPI_ISL_2519734, EPI_ISL_2519735, EPI_ISL_2519736, EPI_ISL_2519738, EPI_ISL_2519739, EPI_ISL_2519740, EPI_ISL_2519741, EPI_ISL_2519742, EPI_ISL_2519743, EPI_ISL_2519744, EPI_ISL_2519745, EPI_ISL_2519746, EPI_ISL_2519747, EPI_ISL_2519748, EPI_ISL_2519749, EPI_ISL_2519750, EPI_ISL_2519751, EPI_ISL_2519752, EPI_ISL_2519753, EPI_ISL_2519754, EPI_ISL_2519755, EPI_ISL_2519756, EPI_ISL_2519757, EPI_ISL_2519758, EPI_ISL_2519759, EPI_ISL_2519760, EPI_ISL_2519761, EPI_ISL_2519762, EPI_ISL_2519763, EPI_ISL_2519764, EPI_ISL_2519765, EPI_ISL_2519766, EPI_ISL_2519767, EPI_ISL_2519768, EPI_ISL_2519769, EPI_ISL_2519770, EPI_ISL_2519772, EPI_ISL_2519773, EPI_ISL_2519775, EPI_ISL_2519776, EPI_ISL_2519777, EPI_ISL_2519778, EPI_ISL_2519779, EPI_ISL_2519780, EPI_ISL_2519781, EPI_ISL_2519782, EPI_ISL_2519783, EPI_ISL_2519784, EPI_ISL_2519785, EPI_ISL_2519786, EPI_ISL_2519787, EPI_ISL_2519790, EPI_ISL_2519791, EPI_ISL_2519793, EPI_ISL_2519794, EPI_ISL_2519796, EPI_ISL_2519797, EPI_ISL_2519798, EPI_ISL_2519800, EPI_ISL_2519801, EPI_ISL_2519802, EPI_ISL_2519803, EPI_ISL_2519804, EPI_ISL_2519805, EPI_ISL_2519806, EPI_ISL_2519808, EPI_ISL_2519809, EPI_ISL_2519811, EPI_ISL_2519813, EPI_ISL_2519814, EPI_ISL_2519815, EPI_ISL_2519816, EPI_ISL_2519817, EPI_ISL_2519818, EPI_ISL_2519819, EPI_ISL_2519820, EPI_ISL_2519821, EPI_ISL_2519822, EPI_ISL_2519823, EPI_ISL_2519824, EPI_ISL_2519826, EPI_ISL_2519827, EPI_ISL_2519829, EPI_ISL_2519831, EPI_ISL_2519832, EPI_ISL_2519833, EPI_ISL_2519834, EPI_ISL_2519835, EPI_ISL_2519837, EPI_ISL_2519838, EPI_ISL_2519839, EPI_ISL_2519840, EPI_ISL_2519841, EPI_ISL_2519842, EPI_ISL_2519843, EPI_ISL_2519844, EPI_ISL_2519848, EPI_ISL_2519849, EPI_ISL_2519850, EPI_ISL_2519851, EPI_ISL_2519852, EPI_ISL_2519853, EPI_ISL_2519854, EPI_ISL_2519855, EPI_ISL_2519856, EPI_ISL_2519857, EPI_ISL_2519858, EPI_ISL_2519859, EPI_ISL_2519860, EPI_ISL_2519862, EPI_ISL_2519864, EPI_ISL_2519865, EPI_ISL_2519866, EPI_ISL_2519867, EPI_ISL_2519868, EPI_ISL_2519869, EPI_ISL_2519870, EPI_ISL_2519871, EPI_ISL_2519873, EPI_ISL_2519874, EPI_ISL_2519875, EPI_ISL_2519876, EPI_ISL_2519877, EPI_ISL_2519878, EPI_ISL_2519880, EPI_ISL_2519881, EPI_ISL_2519882, EPI_ISL_2519883, EPI_ISL_2519884, EPI_ISL_2519885, EPI_ISL_2519886, EPI_ISL_2519887, EPI_ISL_2519888, EPI_ISL_2519890, EPI_ISL_2519891, EPI_ISL_2519892, EPI_ISL_2519893, EPI_ISL_2519894, EPI_ISL_2519895, EPI_ISL_2519896, EPI_ISL_2519898, EPI_ISL_2519900, EPI_ISL_2519901, EPI_ISL_2519902, EPI_ISL_2519903, EPI_ISL_2519904, EPI_ISL_2519905, EPI_ISL_2519906, EPI_ISL_2519907, EPI_ISL_2519908, EPI_ISL_2519911, EPI_ISL_2519912, EPI_ISL_2519913, EPI_ISL_2519914, EPI_ISL_2519915, EPI_ISL_2519916, EPI_ISL_2519917, EPI_ISL_2519918, EPI_ISL_2519919, EPI_ISL_2519920, EPI_ISL_2519921, EPI_ISL_2519922, EPI_ISL_2519923, EPI_ISL_2519924, EPI_ISL_2519925, EPI_ISL_2519927, EPI_ISL_2519928, EPI_ISL_2519929, EPI_ISL_2519930, EPI_ISL_2519932, EPI_ISL_2519933, EPI_ISL_2519934, EPI_ISL_2519935, EPI_ISL_2519936, EPI_ISL_2519937, EPI_ISL_2519938, EPI_ISL_2519939, EPI_ISL_2519940, EPI_ISL_2519941, EPI_ISL_2519942, EPI_ISL_2519943, EPI_ISL_2519944, EPI_ISL_2519945, EPI_ISL_2519946, EPI_ISL_2519947, EPI_ISL_2519948, EPI_ISL_2519950, EPI_ISL_2519951, EPI_ISL_2519952, EPI_ISL_2519953, EPI_ISL_2519955, EPI_ISL_2519956, EPI_ISL_2519958, EPI_ISL_2519959, EPI_ISL_2519960, EPI_ISL_2519961, EPI_ISL_2519963, EPI_ISL_2519964, EPI_ISL_2519965, EPI_ISL_2519966, EPI_ISL_2519967, EPI_ISL_2519968, EPI_ISL_2519969, EPI_ISL_2519970, EPI_ISL_2519971, EPI_ISL_2519973, EPI_ISL_2519974, EPI_ISL_2519975, EPI_ISL_2519976, EPI_ISL_2519977, EPI_ISL_2519978, EPI_ISL_2519979, EPI_ISL_2519980, EPI_ISL_2519981, EPI_ISL_2519982, EPI_ISL_2519983, EPI_ISL_2519985, EPI_ISL_2519986, EPI_ISL_2519987, EPI_ISL_2519990, EPI_ISL_2519991, EPI_ISL_2519992, EPI_ISL_2519993, EPI_ISL_2519995, EPI_ISL_2519996, EPI_ISL_2519998, EPI_ISL_2519999, EPI_ISL_2520001, EPI_ISL_2520002, EPI_ISL_2520003, EPI_ISL_2520004, EPI_ISL_2520005, EPI_ISL_2520006, EPI_ISL_2520008, EPI_ISL_2520009, EPI_ISL_2520010, EPI_ISL_2520011, EPI_ISL_2520012, EPI_ISL_2520013, EPI_ISL_2520015, EPI_ISL_2520016, EPI_ISL_2520017, EPI_ISL_2520018, EPI_ISL_2520019, EPI_ISL_2520020, EPI_ISL_2520021, EPI_ISL_2520023, EPI_ISL_2520024, EPI_ISL_2520026, EPI_ISL_2520027, EPI_ISL_2520029, EPI_ISL_2520030, EPI_ISL_2520031, EPI_ISL_2520032, EPI_ISL_2520033, EPI_ISL_2520034, EPI_ISL_2520035, EPI_ISL_2520036, EPI_ISL_2520037, EPI_ISL_2520038, EPI_ISL_2520041, EPI_ISL_2520042, EPI_ISL_2520043, EPI_ISL_2520044, EPI_ISL_2520045, EPI_ISL_2520046, EPI_ISL_2520048, EPI_ISL_2520050, EPI_ISL_2520051, EPI_ISL_2520052, EPI_ISL_2520053, EPI_ISL_2520054, EPI_ISL_2520055, EPI_ISL_2520057, EPI_ISL_2520058, EPI_ISL_2520059, EPI_ISL_2520060, EPI_ISL_2520061, EPI_ISL_2520062, EPI_ISL_2520063, EPI_ISL_2520064, EPI_ISL_2520065, EPI_ISL_2520066, EPI_ISL_2520067, EPI_ISL_2520068, EPI_ISL_2520069, EPI_ISL_2520070, EPI_ISL_2520071, EPI_ISL_2520072, EPI_ISL_2520073, EPI_ISL_2520074, EPI_ISL_2520075, EPI_ISL_2520076, EPI_ISL_2520077, EPI_ISL_2520079, EPI_ISL_2520080, EPI_ISL_2520081, EPI_ISL_2520082, EPI_ISL_2520083, EPI_ISL_2520084, EPI_ISL_2520085, EPI_ISL_2520086, EPI_ISL_2520087, EPI_ISL_2520088, EPI_ISL_2520089, EPI_ISL_2520091, EPI_ISL_2520092, EPI_ISL_2520093, EPI_ISL_2520094, EPI_ISL_2520095, EPI_ISL_2520096, EPI_ISL_2520097, EPI_ISL_2520099, EPI_ISL_2520100, EPI_ISL_2520101, EPI_ISL_2520102, EPI_ISL_2520104, EPI_ISL_2520105, EPI_ISL_2520106, EPI_ISL_2520107, EPI_ISL_2520108, EPI_ISL_2520109, EPI_ISL_2520110, EPI_ISL_2520111, EPI_ISL_2520112, EPI_ISL_2520113, EPI_ISL_2520114, EPI_ISL_2520115, EPI_ISL_2520116, EPI_ISL_2520117, EPI_ISL_2520118, EPI_ISL_2520119, EPI_ISL_2520120, EPI_ISL_2520121, EPI_ISL_2520122, EPI_ISL_2520123, EPI_ISL_2520124, EPI_ISL_2520125, EPI_ISL_2520126, EPI_ISL_2520127, EPI_ISL_2520128, EPI_ISL_2520129, EPI_ISL_2520130, EPI_ISL_2520132, EPI_ISL_2520133, EPI_ISL_2520134, EPI_ISL_2520135, EPI_ISL_2520136, EPI_ISL_2520137, EPI_ISL_2520138, EPI_ISL_2520139, EPI_ISL_2520140, EPI_ISL_2520141, EPI_ISL_2520142, EPI_ISL_2520143, EPI_ISL_2520144, EPI_ISL_2520145, EPI_ISL_2520146, EPI_ISL_2520147, EPI_ISL_2520148, EPI_ISL_2520149, EPI_ISL_2520150, EPI_ISL_2520153, EPI_ISL_2520154, EPI_ISL_2520155, EPI_ISL_2520156, EPI_ISL_2520157, EPI_ISL_2520158, EPI_ISL_2520159, EPI_ISL_2520160, EPI_ISL_2520161, EPI_ISL_2520162, EPI_ISL_2520163, EPI_ISL_2520164, EPI_ISL_2520165, EPI_ISL_2520166, EPI_ISL_2520167, EPI_ISL_2520168, EPI_ISL_2520169, EPI_ISL_2520170, EPI_ISL_2520171, EPI_ISL_2520172, EPI_ISL_2520173, EPI_ISL_2520175, EPI_ISL_2520176, EPI_ISL_2520177, EPI_ISL_2520178, EPI_ISL_2520179, EPI_ISL_2520180, EPI_ISL_2520181, EPI_ISL_2520182, EPI_ISL_2520183, EPI_ISL_2520184, EPI_ISL_2520186, EPI_ISL_2520187, EPI_ISL_2520188, EPI_ISL_2520189, EPI_ISL_2520190, EPI_ISL_2520191, EPI_ISL_2520192, EPI_ISL_2520193, EPI_ISL_2520194, EPI_ISL_2520195, EPI_ISL_2520196, EPI_ISL_2520197, EPI_ISL_2520198, EPI_ISL_2520199, EPI_ISL_2520200, EPI_ISL_2520201, EPI_ISL_2520202, EPI_ISL_2520203, EPI_ISL_2520204, EPI_ISL_2520205, EPI_ISL_2520206, EPI_ISL_2520207, EPI_ISL_2520208, EPI_ISL_2520209, EPI_ISL_2520210, EPI_ISL_2520211, EPI_ISL_2520213, EPI_ISL_2520214, EPI_ISL_2520215, EPI_ISL_2520216, EPI_ISL_2520217, EPI_ISL_2520218, EPI_ISL_2520219, EPI_ISL_2520220, EPI_ISL_2520221, EPI_ISL_2520222, EPI_ISL_2520223, EPI_ISL_2520225, EPI_ISL_2520226, EPI_ISL_2520227, EPI_ISL_2520228, EPI_ISL_2520229, EPI_ISL_2520230, EPI_ISL_2520231, EPI_ISL_2520232, EPI_ISL_2520233, EPI_ISL_2520236, EPI_ISL_2520239, EPI_ISL_2520240, EPI_ISL_2520241, EPI_ISL_2520242, EPI_ISL_2520243, EPI_ISL_2520244, EPI_ISL_2520245, EPI_ISL_2520246, EPI_ISL_2520247, EPI_ISL_2520248, EPI_ISL_2520249, EPI_ISL_2520250, EPI_ISL_2520251, EPI_ISL_2520252, EPI_ISL_2520254, EPI_ISL_2520255, EPI_ISL_2520257, EPI_ISL_2520258, EPI_ISL_2520259, EPI_ISL_2520260, EPI_ISL_2520261, EPI_ISL_2520262, EPI_ISL_2520263, EPI_ISL_2520264, EPI_ISL_2520265, EPI_ISL_2520266, EPI_ISL_2520268, EPI_ISL_2520269, EPI_ISL_2520270, EPI_ISL_2520271, EPI_ISL_2520272, EPI_ISL_2520273, EPI_ISL_2520274, EPI_ISL_2520275, EPI_ISL_2520276, EPI_ISL_2520277, EPI_ISL_2520278, EPI_ISL_2520279, EPI_ISL_2520280, EPI_ISL_2520281, EPI_ISL_2520282, EPI_ISL_2520283, EPI_ISL_2520284, EPI_ISL_2520285, EPI_ISL_2520286, EPI_ISL_2520287, EPI_ISL_2520288, EPI_ISL_2520290, EPI_ISL_2520291, EPI_ISL_2520293, EPI_ISL_2520294, EPI_ISL_2520295, EPI_ISL_2520296, EPI_ISL_2520297, EPI_ISL_2520298, EPI_ISL_2520299, EPI_ISL_2520300, EPI_ISL_2520301, EPI_ISL_2520302, EPI_ISL_2520303, EPI_ISL_2520304, EPI_ISL_2520305, EPI_ISL_2520306, EPI_ISL_2520307, EPI_ISL_2520308, EPI_ISL_2520309, EPI_ISL_2520310, EPI_ISL_2520311, EPI_ISL_2520312, EPI_ISL_2520313, EPI_ISL_2520315, EPI_ISL_2520316, EPI_ISL_2520317, EPI_ISL_2520318, EPI_ISL_2520319, EPI_ISL_2520320, EPI_ISL_2520321, EPI_ISL_2520323, EPI_ISL_2520324, EPI_ISL_2520325, EPI_ISL_2520326, EPI_ISL_2520327, EPI_ISL_2520328, EPI_ISL_2520329, EPI_ISL_2520330, EPI_ISL_2520331, EPI_ISL_2520332, EPI_ISL_2520333, EPI_ISL_2520334, EPI_ISL_2520335, EPI_ISL_2520337, EPI_ISL_2520338, EPI_ISL_2520339, EPI_ISL_2520340, EPI_ISL_2520341, EPI_ISL_2520342, EPI_ISL_2520343, EPI_ISL_2520344, EPI_ISL_2520345, EPI_ISL_2520347, EPI_ISL_2520348, EPI_ISL_2520349, EPI_ISL_2520350, EPI_ISL_2520351, EPI_ISL_2520352, EPI_ISL_2520354, EPI_ISL_2520355, EPI_ISL_2520356, EPI_ISL_2520357, EPI_ISL_2520358, EPI_ISL_2520359, EPI_ISL_2520360, EPI_ISL_2520361, EPI_ISL_2520362, EPI_ISL_2520363, EPI_ISL_2520364, EPI_ISL_2520365, EPI_ISL_2520366, EPI_ISL_2520368, EPI_ISL_2520369, EPI_ISL_2520370, EPI_ISL_2520371, EPI_ISL_2520372, EPI_ISL_2520373, EPI_ISL_2520374, EPI_ISL_2520375, EPI_ISL_2520377, EPI_ISL_2520378, EPI_ISL_2520380, EPI_ISL_2520381, EPI_ISL_2520382

see above | Quadram Institute Bioscience | COVID-19 Genomics UK (COG-UK) Consortium | Dave J. Baker, Gemma L. Kay, Alp Aydin, Thanh Le-Viet, Steven Rudder, Ana P. Tedim, Anastasia Kolyva, Maria Diaz, Leonardo de Oliveira Martins, Nabil-Fareed Alikhan, Lizzie Meadows, Rachael Stanley, Ngozi Elumogo, Muhammed Yasir, Nicholas M. Thomson, Alexander J Trotter, Rachel Gilroy, Samuel Bloomfield, Claire Stuart, Andrew Bell, Reenesh Prakash, Samir Dervisevic, Alison E. Mather, John Wain, Mark Webber, Andrew J. Page, Justin O'Grady

EPI_ISL_2520383, EPI_ISL_2520384, EPI_ISL_2520385, EPI_ISL_2520386, EPI_ISL_2520387, EPI_ISL_2520388, EPI_ISL_2520389, EPI_ISL_2520391, EPI_ISL_2520393, EPI_ISL_2520394, EPI_ISL_2520395, EPI_ISL_2520396, EPI_ISL_2520397, EPI_ISL_2520398, EPI_ISL_2520399, EPI_ISL_2520400, EPI_ISL_2520401, EPI_ISL_2520402, EPI_ISL_2520403, EPI_ISL_2520404, EPI_ISL_2520405, EPI_ISL_2520406, EPI_ISL_2520407, EPI_ISL_2520408, EPI_ISL_2520409, EPI_ISL_2520410, EPI_ISL_2520411, EPI_ISL_2520412, EPI_ISL_2520413, EPI_ISL_2520414, EPI_ISL_2520415, EPI_ISL_2520416, EPI_ISL_2520417, EPI_ISL_2520418, EPI_ISL_2520419, EPI_ISL_2520420, EPI_ISL_2520422, EPI_ISL_2520423, EPI_ISL_2520424, EPI_ISL_2520425, EPI_ISL_2520427, EPI_ISL_2520428, EPI_ISL_2520429, EPI_ISL_2520430, EPI_ISL_2520433, EPI_ISL_2520434, EPI_ISL_2520437, EPI_ISL_2520438, EPI_ISL_2520439, EPI_ISL_2520441, EPI_ISL_2520443, EPI_ISL_2520444, EPI_ISL_2520445, EPI_ISL_2520446, EPI_ISL_2520447, EPI_ISL_2520448, EPI_ISL_2520449, EPI_ISL_2520450, EPI_ISL_2520451, EPI_ISL_2520452, EPI_ISL_2520456, EPI_ISL_2520457, EPI_ISL_2520459, EPI_ISL_2520460, EPI_ISL_2520462, EPI_ISL_2520463, EPI_ISL_2520464, EPI_ISL_2520465, EPI_ISL_2520466, EPI_ISL_2520467, EPI_ISL_2520468, EPI_ISL_2520469, EPI_ISL_2520470, EPI_ISL_2520471, EPI_ISL_2520472, EPI_ISL_2520473, EPI_ISL_2520474, EPI_ISL_2520475, EPI_ISL_2520476, EPI_ISL_2520478, EPI_ISL_2520479, EPI_ISL_2520481, EPI_ISL_2520482, EPI_ISL_2520483, EPI_ISL_2520484,

| | | | |
|---|---|---|---|
| EPI_ISL_2520485, EPI_ISL_2520486, EPI_ISL_2520487, EPI_ISL_2520488, EPI_ISL_2520489, EPI_ISL_2520490, EPI_ISL_2520491, EPI_ISL_2520492, EPI_ISL_2520493, EPI_ISL_2520494, EPI_ISL_2520495, EPI_ISL_2520496, EPI_ISL_2520497, EPI_ISL_2520498, EPI_ISL_2520499, EPI_ISL_2520500, EPI_ISL_2520501, EPI_ISL_2520502, EPI_ISL_2520503, EPI_ISL_2520504, EPI_ISL_2520505, EPI_ISL_2520506, EPI_ISL_2520507, EPI_ISL_2520508, EPI_ISL_2520509, EPI_ISL_2520510, EPI_ISL_2520511, EPI_ISL_2520512, EPI_ISL_2520513, EPI_ISL_2520514, EPI_ISL_2520516, EPI_ISL_2520517, EPI_ISL_2520519, EPI_ISL_2520520, EPI_ISL_2520521, EPI_ISL_2520522, EPI_ISL_2520523, EPI_ISL_2520524, EPI_ISL_2520525, EPI_ISL_2520527, EPI_ISL_2520528, EPI_ISL_2520529, EPI_ISL_2520531, EPI_ISL_2520532, EPI_ISL_2520533 | | | |
| see above | Queens Medical Centre, Clinical Microbiology Department / DeepSeq Nottingham | COVID-19 Genomics UK (COG-UK) Consortium | Gemma Clark, Wendy Smith, Manjinder Khakh, Vicki M Fleming, Michelle M Lister, Hannah Howson-Wells, Jonathan Ball, Timothy Byaruhanga, Jayasree Dey, Emily Park, Jack Hill, Patrick McClure, Joseph Chappell, Theocharis Tsoleridis, Nadine Holmes, Matthew Carlisle, Christopher Moore, Fei Sang, Johnny Debebe, Victoria Wright, Matthew Loose |
| EPI_ISL_2520534, EPI_ISL_2520535, EPI_ISL_2520536, EPI_ISL_2520537, EPI_ISL_2520538, EPI_ISL_2520539, EPI_ISL_2520540, EPI_ISL_2520541, EPI_ISL_2520542, EPI_ISL_2520543, EPI_ISL_2520544, EPI_ISL_2520545, EPI_ISL_2520546, EPI_ISL_2520547, EPI_ISL_2520548, EPI_ISL_2520549, EPI_ISL_2520550, EPI_ISL_2520551, EPI_ISL_2520552, EPI_ISL_2520553, EPI_ISL_2520554, EPI_ISL_2520555, EPI_ISL_2520556, EPI_ISL_2520558, EPI_ISL_2520560, EPI_ISL_2520561, EPI_ISL_2520562, EPI_ISL_2520564, EPI_ISL_2520565, EPI_ISL_2520566, EPI_ISL_2520567, EPI_ISL_2520568, EPI_ISL_2520569, EPI_ISL_2520570, EPI_ISL_2520572, EPI_ISL_2520574, EPI_ISL_2520575, EPI_ISL_2520576, EPI_ISL_2520583, EPI_ISL_2520584, EPI_ISL_2520585, EPI_ISL_2520589, EPI_ISL_2520590, EPI_ISL_2520593, EPI_ISL_2520596, EPI_ISL_2520597, EPI_ISL_2520598, EPI_ISL_2520599, EPI_ISL_2520600, EPI_ISL_2520601, EPI_ISL_2520602, EPI_ISL_2520605, EPI_ISL_2520606, EPI_ISL_2520607, EPI_ISL_2520610, EPI_ISL_2520612, EPI_ISL_2520614, EPI_ISL_2520615, EPI_ISL_2520617, EPI_ISL_2520618, EPI_ISL_2520624, EPI_ISL_2520626, EPI_ISL_2520627, EPI_ISL_2520628, EPI_ISL_2520633, EPI_ISL_2520634, EPI_ISL_2520635, EPI_ISL_2520636, EPI_ISL_2520637, EPI_ISL_2520640, EPI_ISL_2520641, EPI_ISL_2520642, EPI_ISL_2520643, EPI_ISL_2520644, EPI_ISL_2520645, EPI_ISL_2520646, EPI_ISL_2520647, EPI_ISL_2520648, EPI_ISL_2520649, EPI_ISL_2520650, EPI_ISL_2520651, EPI_ISL_2520653, EPI_ISL_2520654, EPI_ISL_2520656, EPI_ISL_2520657, EPI_ISL_2520658, EPI_ISL_2520659, EPI_ISL_2520660, EPI_ISL_2520662, EPI_ISL_2520664, EPI_ISL_2520665, EPI_ISL_2520666, EPI_ISL_2520667, EPI_ISL_2520668, EPI_ISL_2520669, EPI_ISL_2520670, EPI_ISL_2520671, EPI_ISL_2520672, EPI_ISL_2520673, EPI_ISL_2520676, EPI_ISL_2520678, EPI_ISL_2520679, EPI_ISL_2520680, EPI_ISL_2520681, EPI_ISL_2520683, EPI_ISL_2520684, EPI_ISL_2520685, EPI_ISL_2520686, EPI_ISL_2520687, EPI_ISL_2520688, EPI_ISL_2520689, EPI_ISL_2520690, EPI_ISL_2520691, EPI_ISL_2520692, EPI_ISL_2520693, EPI_ISL_2520694, EPI_ISL_2520695, EPI_ISL_2520696, EPI_ISL_2520697, EPI_ISL_2520698, EPI_ISL_2520699, EPI_ISL_2520700, EPI_ISL_2520701, EPI_ISL_2520702, EPI_ISL_2520703, EPI_ISL_2520704, EPI_ISL_2520705, EPI_ISL_2520706, EPI_ISL_2520707, EPI_ISL_2520708, EPI_ISL_2520709, EPI_ISL_2520710, EPI_ISL_2520711, EPI_ISL_2520712, EPI_ISL_2520713, EPI_ISL_2520714, EPI_ISL_2520715, EPI_ISL_2520716, EPI_ISL_2520717, EPI_ISL_2520718, EPI_ISL_2520719, EPI_ISL_2520720, EPI_ISL_2520721, EPI_ISL_2520722, EPI_ISL_2520723, EPI_ISL_2520725, EPI_ISL_2520726, EPI_ISL_2520727, EPI_ISL_2520735, EPI_ISL_2520736, EPI_ISL_2520740, EPI_ISL_2520741, EPI_ISL_2520745, EPI_ISL_2520746, EPI_ISL_2520750, EPI_ISL_2520751, EPI_ISL_2520752, EPI_ISL_2520754, EPI_ISL_2520755, EPI_ISL_2520756, EPI_ISL_2520757, EPI_ISL_2520758, EPI_ISL_2520759, EPI_ISL_2520761, EPI_ISL_2520762, EPI_ISL_2520763, EPI_ISL_2520764, EPI_ISL_2520765, EPI_ISL_2520769, EPI_ISL_2520771, EPI_ISL_2520772, EPI_ISL_2520774, EPI_ISL_2520776, EPI_ISL_2520777, EPI_ISL_2520778, EPI_ISL_2520780, EPI_ISL_2520781, EPI_ISL_2520783, EPI_ISL_2520784, EPI_ISL_2520786, EPI_ISL_2520787, EPI_ISL_2520788, EPI_ISL_2520792, EPI_ISL_2520794, EPI_ISL_2520796, EPI_ISL_2520802, EPI_ISL_2520803, EPI_ISL_2520805, EPI_ISL_2520808, EPI_ISL_2520809, EPI_ISL_2520810, EPI_ISL_2520811 | | | |
| see above | Wales Specialist Virology Centre Sequencing lab: Pathogen Genomics Unit | Public Health Wales Microbiology Cardiff Wales Specialist Virology Centre | Catherine Moore, Johnathan Evans, Laura Gifford, Malorie Perry, Simon Cottrell, Angela Marchbank, Alec Birchley, Alexander Adams, Amy Gaskin, Bree Gatica-Wilcox, Jason Coombes, Joel Southgate, Lauren Gilbert, Lee Graham, Nicole Pacchiarini, Sara Kumziene-Summerhayes, Sarah Taylor, Sophie Jones, Sara Rey, Matthew Bull, Joanne Watkins, Sally Corden, Tom Connor |
| EPI_ISL_2520813, EPI_ISL_2520814, EPI_ISL_2520815, EPI_ISL_2520816, EPI_ISL_2520817, EPI_ISL_2520818, EPI_ISL_2520819, EPI_ISL_2520820, EPI_ISL_2520821, EPI_ISL_2520822, EPI_ISL_2520824, EPI_ISL_2520825, EPI_ISL_2520826, EPI_ISL_2520827, EPI_ISL_2520828, EPI_ISL_2520829, EPI_ISL_2520830, EPI_ISL_2520832, EPI_ISL_2520833, EPI_ISL_2520834, EPI_ISL_2520835, EPI_ISL_2520836, EPI_ISL_2520837, EPI_ISL_2520838, EPI_ISL_2520839, EPI_ISL_2520840, EPI_ISL_2520841, EPI_ISL_2520842, EPI_ISL_2520843, EPI_ISL_2520844, EPI_ISL_2520845, EPI_ISL_2520846, EPI_ISL_2520847, EPI_ISL_2520848, EPI_ISL_2520850, EPI_ISL_2520851, EPI_ISL_2520852, EPI_ISL_2520853, EPI_ISL_2520854, EPI_ISL_2520855, EPI_ISL_2520856, EPI_ISL_2520857, EPI_ISL_2520858, EPI_ISL_2520859, EPI_ISL_2520861, EPI_ISL_2520862, EPI_ISL_2520863, EPI_ISL_2520864, EPI_ISL_2520865, EPI_ISL_2520866, EPI_ISL_2520867, EPI_ISL_2520868, EPI_ISL_2520869, EPI_ISL_2520870, EPI_ISL_2520871, EPI_ISL_2520872, EPI_ISL_2520873, EPI_ISL_2520874, EPI_ISL_2520875, EPI_ISL_2520876, EPI_ISL_2520877, EPI_ISL_2520878, EPI_ISL_2520879, EPI_ISL_2520880, EPI_ISL_2520881, EPI_ISL_2520882, EPI_ISL_2520883, EPI_ISL_2520884, EPI_ISL_2520885, EPI_ISL_2520886, EPI_ISL_2520887, EPI_ISL_2520888, EPI_ISL_2520889, EPI_ISL_2520890, EPI_ISL_2520891, EPI_ISL_2520893, EPI_ISL_2520894, EPI_ISL_2520895, EPI_ISL_2520896, EPI_ISL_2520897, EPI_ISL_2520898, EPI_ISL_2520900, EPI_ISL_2520901, EPI_ISL_2520902, EPI_ISL_2520903, EPI_ISL_2520904, EPI_ISL_2520905, EPI_ISL_2520906, EPI_ISL_2520907, EPI_ISL_2520908, EPI_ISL_2520909, EPI_ISL_2520910, EPI_ISL_2520911, EPI_ISL_2520912, EPI_ISL_2520913, EPI_ISL_2520914, EPI_ISL_2520915, EPI_ISL_2520916, EPI_ISL_2520917, EPI_ISL_2520918, EPI_ISL_2520919, EPI_ISL_2520920, EPI_ISL_2520921, EPI_ISL_2520923, EPI_ISL_2520924, EPI_ISL_2520925, EPI_ISL_2520926, EPI_ISL_2520927 | | | |
| see above | Centre for Enzyme Innovation, University of Portsmouth / Translational Research Laboratory, Portsmouth Hospitals NHS Trust / Hampshire Hospitals NHS Foundation Trust / Bournemouth University / University Hospitals Dorset NHS Foundation Trust / Brighton and Sussex University Hospitals NHS Trust / Dartford and Gravesham NHS Trust / Isle of Wight NHS Trust / Maidstone and Tunbridge Wells NHS Trust / University Hospital Southampton NHS Foundation Trust | COVID-19 Genomics UK (COG-UK) Consortium | Samuel Robson, Angela Beckett, Salman Goudarzi, Christopher Fearn, Kate Cook, Katie Loveson, Sharon Glaysher, Scott Elliott, Kelly Bicknell, Sarah Wyllie, Allyson Lloyd, Robert Impey, Anoop Chauhan, Stephen Kidd, Nathan Moore, Nick Cortes, Claire Thomas, Anna Mantzaouratou, Sarah Buchan, Magdalena Barrow, Andrew Butt, Liz Sheridan, Jonnie Seymour, Dorian Crudgington, Ben Macklin, Mohammed Hassan-Ibrahim, Cassandra Malone, Benjamin Cogger, Kevin Tucker, Samirakhon Raupova, Rachael Jeremiah, Anibolina Castigador, Emily Macnaughton, Karen Withell, Kordo Saeed, Jacqui Prieto, Adhyana Mahanama, Buddhini Samaraweera, Siona Silvieri, Emanuela Pelosi, Eleri Wilson-Davies, Sarah Jeremiah, Helen Wheeler, Matthew Harvey, Thea Sass, Helen Umpleby, Stephen Aplin |
| EPI_ISL_2520928, EPI_ISL_2520929, EPI_ISL_2520930, EPI_ISL_2520933, EPI_ISL_2520934, EPI_ISL_2520935, EPI_ISL_2520936, EPI_ISL_2520938, EPI_ISL_2520939, EPI_ISL_2520940, EPI_ISL_2520941, EPI_ISL_2520943, EPI_ISL_2520944, EPI_ISL_2520945, EPI_ISL_2520946, EPI_ISL_2520948, EPI_ISL_2520949, EPI_ISL_2520951, EPI_ISL_2520952, EPI_ISL_2520953, EPI_ISL_2520955, EPI_ISL_2520956, EPI_ISL_2520958, EPI_ISL_2520959, EPI_ISL_2520960, EPI_ISL_2520962, EPI_ISL_2520963, EPI_ISL_2520964, EPI_ISL_2520966, EPI_ISL_2520967, EPI_ISL_2520968, EPI_ISL_2520970, EPI_ISL_2520971, EPI_ISL_2520973, EPI_ISL_2520974, EPI_ISL_2520980, EPI_ISL_2520981, EPI_ISL_2520983, EPI_ISL_2520984, EPI_ISL_2520985, EPI_ISL_2520986, EPI_ISL_2520987, EPI_ISL_2520988, EPI_ISL_2520989, EPI_ISL_2520991, EPI_ISL_2520992, EPI_ISL_2520993, EPI_ISL_2520994, EPI_ISL_2520995, EPI_ISL_2520996, EPI_ISL_2520997, EPI_ISL_2520998, EPI_ISL_2520999, EPI_ISL_2521002, EPI_ISL_2521003, EPI_ISL_2521004, EPI_ISL_2521005, EPI_ISL_2521006, EPI_ISL_2521007, EPI_ISL_2521008, EPI_ISL_2521009, EPI_ISL_2521010, EPI_ISL_2521011, EPI_ISL_2521012, EPI_ISL_2521015, EPI_ISL_2521016, EPI_ISL_2521017, EPI_ISL_2521018, EPI_ISL_2521019, EPI_ISL_2521023, EPI_ISL_2521025, EPI_ISL_2521026, EPI_ISL_2521028, EPI_ISL_2521031, EPI_ISL_2521032, EPI_ISL_2521033, EPI_ISL_2521034, EPI_ISL_2521035, EPI_ISL_2521036, EPI_ISL_2521037, EPI_ISL_2521038, EPI_ISL_2521039, EPI_ISL_2521040, EPI_ISL_2521043, EPI_ISL_2521047, EPI_ISL_2521048, EPI_ISL_2521050, EPI_ISL_2521051, EPI_ISL_2521052, EPI_ISL_2521053, EPI_ISL_2521054, EPI_ISL_2521055, EPI_ISL_2521056, EPI_ISL_2521057, EPI_ISL_2521058, EPI_ISL_2521061, EPI_ISL_2521063, EPI_ISL_2521064, EPI_ISL_2521065, EPI_ISL_2521066, EPI_ISL_2521067, EPI_ISL_2521069, EPI_ISL_2521071, EPI_ISL_2521072, EPI_ISL_2521073, EPI_ISL_2521075, EPI_ISL_2521077, EPI_ISL_2521078, EPI_ISL_2521079, EPI_ISL_2521080, EPI_ISL_2521081, EPI_ISL_2521084, EPI_ISL_2521085, EPI_ISL_2521086, EPI_ISL_2521087, EPI_ISL_2521089, EPI_ISL_2521091, EPI_ISL_2521092, EPI_ISL_2521094, EPI_ISL_2521095, EPI_ISL_2521096, EPI_ISL_2521097, EPI_ISL_2521098, EPI_ISL_2521099, EPI_ISL_2521100, EPI_ISL_2521101, EPI_ISL_2521102, EPI_ISL_2521103, EPI_ISL_2521104, EPI_ISL_2521105, EPI_ISL_2521106, EPI_ISL_2521107, EPI_ISL_2521108, EPI_ISL_2521109, EPI_ISL_2521110, EPI_ISL_2521111, EPI_ISL_2521112, EPI_ISL_2521113, EPI_ISL_2521114, EPI_ISL_2521115, EPI_ISL_2521116, EPI_ISL_2521119, EPI_ISL_2521120, EPI_ISL_2521121, EPI_ISL_2521122, EPI_ISL_2521123, EPI_ISL_2521124, EPI_ISL_2521125, EPI_ISL_2521126, EPI_ISL_2521127, EPI_ISL_2521129, EPI_ISL_2521130, EPI_ISL_2521131, EPI_ISL_2521132, EPI_ISL_2521133, EPI_ISL_2521135, EPI_ISL_2521136, EPI_ISL_2521137, EPI_ISL_2521138, EPI_ISL_2521140, EPI_ISL_2521141, EPI_ISL_2521142, EPI_ISL_2521143, EPI_ISL_2521144, EPI_ISL_2521145, EPI_ISL_2521148, EPI_ISL_2521149 | | | |
| see above | Virology Department, Sheffield Teaching Hospitals NHS Foundation Trust/Department of Infection, Immunity and Cardiovascular Disease, The Medical School, University of Sheffield | COVID-19 Genomics UK (COG-UK) Consortium | Thushan de Silva, Matthew Parker, Nikki Smith, Adri Angyal, Rebecca Brown, Luke Green, Rachel Tucker, Paul Parsons, Danielle Groves, Katie Johnson, Laura Carrilero, Alex Keeley, Dave Partridge, Matthew Wyles, Benjamin Lindsey, Mehmet Yavuz, Mohammad Raza, Cariad Evans |
| EPI_ISL_2521350, EPI_ISL_2521371 | Valais Hospital, Central Institute | Valais Hospital, Central Institute | Alexis Dumoulin, Lorenzo Cerutti, Henri Pegeot, Melyssa Elies, Deborah Penet, Keith Harshman, Ioannis Xenarios, Emmanouil Dermitzakis |
| EPI_ISL_2521601, EPI_ISL_2521605, EPI_ISL_2521606, EPI_ISL_2521609 | INSACOG-Mizoram | National Institute of Biomedical Genomics – INSACOG | Arindam Maitra, Swagnik Roy, Nidhan Kumar Biswas, Gracy Laldinmawii, Sreedhar Chinnaswamy, N Senthil Kumar, Saumitra Das |
| EPI_ISL_2521698 | Plateforme de testing Namuroise | Plateforme de testing Namuroise | Lesly Nyinkeu Kemamen; Demars Aurore ; Maschietto Céline, Nobis Chloé ; Otto Gaetan ; Denis Olivier ; Degosserie Jonathan ; Mullier François |
| EPI_ISL_2521701 | Libramont | Plateforme de testing Namuroise | Lesly Nyinkeu Kemamen; Demars Aurore ; Maschietto Céline, Nobis Chloé ; Otto Gaetan ; Denis Olivier ; Degosserie Jonathan ; Mullier François |
| EPI_ISL_2521771, EPI_ISL_2521772, EPI_ISL_2521773, EPI_ISL_2521774, EPI_ISL_2521775, EPI_ISL_2521776, EPI_ISL_2521777, EPI_ISL_2521778, EPI_ISL_2521779, EPI_ISL_2521780, EPI_ISL_2521781, EPI_ISL_2521782, EPI_ISL_2521783, EPI_ISL_2521789, EPI_ISL_2521790, EPI_ISL_2521792, EPI_ISL_2521793, EPI_ISL_2521795, EPI_ISL_2521797, EPI_ISL_2521798, EPI_ISL_2521800, EPI_ISL_2521802, EPI_ISL_2521803, EPI_ISL_2521804, EPI_ISL_2521805, EPI_ISL_2521806, EPI_ISL_2521807, EPI_ISL_2521808, EPI_ISL_2521809, EPI_ISL_2521810, EPI_ISL_2521811, EPI_ISL_2521812, EPI_ISL_2521813, EPI_ISL_2521814, EPI_ISL_2521815, EPI_ISL_2521816, EPI_ISL_2521817, EPI_ISL_2521818, EPI_ISL_2521819, EPI_ISL_2521820, EPI_ISL_2521821, EPI_ISL_2521823, EPI_ISL_2521826, EPI_ISL_2521827, EPI_ISL_2521828, EPI_ISL_2521830, EPI_ISL_2521831, EPI_ISL_2521832, EPI_ISL_2521833, EPI_ISL_2521834, EPI_ISL_2521836, EPI_ISL_2521837, EPI_ISL_2521838, EPI_ISL_2521839, EPI_ISL_2521840, EPI_ISL_2521841, EPI_ISL_2521842, EPI_ISL_2521843, EPI_ISL_2521844, EPI_ISL_2521845, EPI_ISL_2521846, EPI_ISL_2521847, EPI_ISL_2521848, EPI_ISL_2521849, EPI_ISL_2521850, EPI_ISL_2521851, EPI_ISL_2521852, EPI_ISL_2521853, EPI_ISL_2521854, EPI_ISL_2521855, EPI_ISL_2521856, EPI_ISL_2521857, EPI_ISL_2521858, EPI_ISL_2521859, EPI_ISL_2521860, EPI_ISL_2521861, EPI_ISL_2521862, EPI_ISL_2521863, EPI_ISL_2521864, EPI_ISL_2521865, EPI_ISL_2521866, EPI_ISL_2521868, EPI_ISL_2521869, EPI_ISL_2521870, EPI_ISL_2521871, EPI_ISL_2521872, EPI_ISL_2521873, EPI_ISL_2521874, EPI_ISL_2521875, EPI_ISL_2521876, EPI_ISL_2521877, EPI_ISL_2521878, EPI_ISL_2521880, EPI_ISL_2521881, EPI_ISL_2521882, EPI_ISL_2521884, EPI_ISL_2521885, EPI_ISL_2521886, EPI_ISL_2521887, EPI_ISL_2521888, EPI_ISL_2521889, EPI_ISL_2521890, EPI_ISL_2521891, EPI_ISL_2521892, EPI_ISL_2521893, EPI_ISL_2521894, EPI_ISL_2521895, EPI_ISL_2521896, EPI_ISL_2521898, EPI_ISL_2521899, EPI_ISL_2521900, EPI_ISL_2521901, EPI_ISL_2521902, EPI_ISL_2521903, EPI_ISL_2521905, EPI_ISL_2521906, EPI_ISL_2521907, EPI_ISL_2521908, EPI_ISL_2521909, EPI_ISL_2521910, EPI_ISL_2521911, EPI_ISL_2521912, EPI_ISL_2521913, EPI_ISL_2521914, EPI_ISL_2521915, EPI_ISL_2521918, EPI_ISL_2521919, EPI_ISL_2521920, EPI_ISL_2521921, EPI_ISL_2521922, EPI_ISL_2521923, EPI_ISL_2521924, EPI_ISL_2521925, EPI_ISL_2521926, EPI_ISL_2521927, EPI_ISL_2521928, EPI_ISL_2521929, EPI_ISL_2521931, EPI_ISL_2521932, EPI_ISL_2521933, EPI_ISL_2521934, EPI_ISL_2521935, EPI_ISL_2521936, EPI_ISL_2521937, EPI_ISL_2521938, EPI_ISL_2521939, EPI_ISL_2521940, EPI_ISL_2521941, EPI_ISL_2521942, EPI_ISL_2521943, EPI_ISL_2521944, EPI_ISL_2521947, EPI_ISL_2521948, EPI_ISL_2521949, EPI_ISL_2521950, EPI_ISL_2521951, EPI_ISL_2521955, EPI_ISL_2521956, EPI_ISL_2521957, EPI_ISL_2521958, EPI_ISL_2521959 | | | |
| see above | Dr S raju, Director of public heath and preventive medicine | inStem NCBS - INSACOG | Uma Ramakrishnan Dasaradhi Palakodeti Aswin SaiNarain |
| EPI_ISL_2521962, EPI_ISL_2521963, EPI_ISL_2521968, EPI_ISL_2521971, EPI_ISL_2521979 | Plateforme de testing Namuroise | Plateforme de testing Namuroise | Nicolas Gilliard; Demars Aurore ; Maschietto Céline ; Nobis Chloé ; Otto Gaetan ; Denis Olivier ; Degosserie Jonathan ; Mullier François |
| EPI_ISL_2522099 | Ospedale S.M. Goretti di Latina - ASL Latina presidio ospedaliero Nord | INMI Lazzaro Spallanzani IRCCS | F Messina, M Rueca, G Bonfiglio, O Butera, CEM Gruber, F Santini, B Bartolini, E Giombini, A Lucci, L Di Biase, S Pignalosa, A Di Caro, MR Capobianchi |
| EPI_ISL_2523389, EPI_ISL_2523395, EPI_ISL_2523406, EPI_ISL_2523409, EPI_ISL_2523420, EPI_ISL_2523422, EPI_ISL_2523428, EPI_ISL_2523430, EPI_ISL_2523433, EPI_ISL_2523436, EPI_ISL_2523438, EPI_ISL_2523442, EPI_ISL_2523444, EPI_ISL_2523451, EPI_ISL_2523454, EPI_ISL_2523460, EPI_ISL_2523463, EPI_ISL_2523466, EPI_ISL_2523469, EPI_ISL_2523473, EPI_ISL_2523480, EPI_ISL_2523482, EPI_ISL_2523485, EPI_ISL_2523488, EPI_ISL_2523491, EPI_ISL_2523494, EPI_ISL_2523497, EPI_ISL_2523500, EPI_ISL_2523503, EPI_ISL_2523507, EPI_ISL_2523511, EPI_ISL_2523513, EPI_ISL_2523516, EPI_ISL_2523518, EPI_ISL_2523522, EPI_ISL_2523525, EPI_ISL_2523531, EPI_ISL_2523534, EPI_ISL_2523536, EPI_ISL_2523538, EPI_ISL_2523543, EPI_ISL_2523546, EPI_ISL_2523549, EPI_ISL_2523552, EPI_ISL_2523559, EPI_ISL_2523562, EPI_ISL_2523565, EPI_ISL_2523568, EPI_ISL_2523571, EPI_ISL_2523575, EPI_ISL_2523578, EPI_ISL_2523584, EPI_ISL_2523587, EPI_ISL_2523594, EPI_ISL_2523596, EPI_ISL_2523599, EPI_ISL_2523602 | | | |
| see above | WHO National Influenza Centre Russian Federation | WHO National Influenza Centre Russian Federation | Andrey Komissarov, Artem Fadeev, Kseniya Komissarova, Oula Mansour, Kirill Varchenko, Mikhail Bakaev, Tamila Musaeva, Maria Timofeeva, Veronika Eder, Maria Pisareva, Nikita Yolshin, Daria Danilenko, Ksenia Safina, Elena Nabieva, Georgii Bazykin, Dmitry Lioznov |
| EPI_ISL_2523614, EPI_ISL_2523620, EPI_ISL_2523627, EPI_ISL_2523633, EPI_ISL_2523639, EPI_ISL_2523641, EPI_ISL_2523664 | HELIX LLC | WHO National Influenza Centre Russian Federation | Andrey Komissarov, Artem Fadeev, Kseniya Komissarova, Oula Mansour, Kirill Varchenko, Mikhail Bakaev, Tamila Musaeva, Maria Timofeeva, Veronika Eder, Maria Pisareva, Nikita Yolshin, Daria Danilenko, Ksenia Safina, Elena Nabieva, Georgii Bazykin, Dmitry Lioznov |
| EPI_ISL_2523669, EPI_ISL_2523670, EPI_ISL_2523672, EPI_ISL_2523675, EPI_ISL_2523678, EPI_ISL_2523682, EPI_ISL_2523684, EPI_ISL_2523687, EPI_ISL_2523690, EPI_ISL_2523694, EPI_ISL_2523700, EPI_ISL_2523702, EPI_ISL_2523708, EPI_ISL_2523711, EPI_ISL_2523714, EPI_ISL_2523718, EPI_ISL_2523722, EPI_ISL_2523724, EPI_ISL_2523726, EPI_ISL_2523729, EPI_ISL_2523732, EPI_ISL_2523738, EPI_ISL_2523741, EPI_ISL_2523744, EPI_ISL_2523747, EPI_ISL_2523755, EPI_ISL_2523762, EPI_ISL_2523765, EPI_ISL_2523768, EPI_ISL_2523774, EPI_ISL_2523777, EPI_ISL_2523780, EPI_ISL_2523783, EPI_ISL_2523786, EPI_ISL_2523789, EPI_ISL_2523796, EPI_ISL_2523799, EPI_ISL_2523803 | | | |
| see above | WHO National Influenza Centre Russian Federation | WHO National Influenza Centre Russian Federation | Andrey Komissarov, Artem Fadeev, Kseniya Komissarova, Oula Mansour, Kirill Varchenko, Mikhail Bakaev, Tamila Musaeva, Maria Timofeeva, Veronika Eder, Maria Pisareva, Nikita Yolshin, Daria Danilenko, Ksenia Safina, Elena Nabieva, Georgii Bazykin, Dmitry Lioznov |
| EPI_ISL_2523806, EPI_ISL_2523808, EPI_ISL_2523811, EPI_ISL_2523821, EPI_ISL_2523824, EPI_ISL_2523830, EPI_ISL_2523833, EPI_ISL_2523836, EPI_ISL_2523839, EPI_ISL_2523846, EPI_ISL_2523849, EPI_ISL_2523853, EPI_ISL_2523856, EPI_ISL_2523859 | | | |
| see above | City Hospital No 40 | WHO National Influenza Centre Russian Federation | Andrey Komissarov, Artem Fadeev, Kseniya Komissarova, Oula Mansour, Kirill Varchenko, Mikhail Bakaev, Tamila Musaeva, Maria Timofeeva, Veronika Eder, Maria Pisareva, Nikita Yolshin, Daria Danilenko, Olga Shneider, Sergey Scherbak, Ksenia Safina, Elena Nabieva, Georgii Bazykin, Dmitry Lioznov |
| EPI_ISL_2524728, EPI_ISL_2524732, EPI_ISL_2524739, EPI_ISL_2524747, EPI_ISL_2524749, EPI_ISL_2524751, EPI_ISL_2524752, EPI_ISL_2524763 | Hospital Universitari Vall d'Hebron - Vall d'Hebron Institut de Recerca | Hospital Universitari Vall d'Hebron - Vall d'Hebron Institut de Recerca | Cristina Andrés, Maria Piñana, Alejandra González-Sánchez, Damir Garcia-Cehic, Ariadna Rando, Juliana Esperalba, Maria Gema Codina, Carla Castillo, Maria Carmen Martin, Tomàs Pumarola, Josep Quer, Andrés Antón |
| EPI_ISL_2524964, EPI_ISL_2525019, EPI_ISL_2525057, EPI_ISL_2525107, EPI_ISL_2525156, EPI_ISL_2525208, EPI_ISL_2525225, | Mako Medical | Centers for Disease Control and Prevention Division of Viral Diseases, Pathogen Discovery | Dakota Howard, Dhwani Batra, Peter W. Cook, Kara Moser, Adrian Paskey, Jason Caravas, Benjamin Rambo-Martin, Shatavia Morrison, Christopher Gulvick, Scott Sammons, Yvette Unoarumhi, Darlene Wagner, Matthew Schmerer,Matthew Tugwell, Lauren Moon, Clinton R. Paden, Duncan MacCannell |

| | | | |
|---|---|---|---|
| EPI_ISL_2525301 | | | |
| EPI_ISL_2525601, EPI_ISL_2525602, EPI_ISL_2525611, EPI_ISL_2525613, EPI_ISL_2525615, EPI_ISL_2525617, EPI_ISL_2525619, EPI_ISL_2525629, EPI_ISL_2525733, EPI_ISL_2525737, EPI_ISL_2525739, EPI_ISL_2525746, EPI_ISL_2525759, EPI_ISL_2525761, EPI_ISL_2525769, EPI_ISL_2525771, EPI_ISL_2525776, EPI_ISL_2525778, EPI_ISL_2525816, EPI_ISL_2525820, EPI_ISL_2525991 | | | |
| see above | Fulgent Genetics | Centers for Disease Control and Prevention Division of Viral Diseases, Pathogen Discovery | Dakota Howard, Dhwani Batra, Peter W. Cook, Kara Moser, Adrian Paskey, Jason Caravas, Benjamin Rambo-Martin, Shatavia Morrison, Christopher Gulvick, Scott Sammons, Yvette Unoarumhi, Darlene Wagner, Matthew Schmerer, Harry Gao, Mickey Li, John Gao, Joseph Fierro, Benafsh Sapra, Becky Tsai, Yan Meng, Doreen Ng, James Xie, Clinton R. Paden, Duncan MacCannell |
| EPI_ISL_2526761, EPI_ISL_2526766, EPI_ISL_2526788, EPI_ISL_2526794, EPI_ISL_2526800, EPI_ISL_2526820, EPI_ISL_2526833, EPI_ISL_2526851, EPI_ISL_2526853, EPI_ISL_2526873, EPI_ISL_2526877, EPI_ISL_2526894, EPI_ISL_2526896, EPI_ISL_2526913, EPI_ISL_2526948, EPI_ISL_2526952, EPI_ISL_2526959, EPI_ISL_2526969, EPI_ISL_2526995, EPI_ISL_2527003, EPI_ISL_2527016, EPI_ISL_2527026, EPI_ISL_2527030, EPI_ISL_2527032 | | | |
| see above | Quest Diagnostics Incorporated | Centers for Disease Control and Prevention Division of Viral Diseases, Pathogen Discovery | Dakota Howard, Dhwani Batra, Peter W. Cook, Kara Moser, Adrian Paskey, Jason Caravas, Benjamin Rambo-Martin, Shatavia Morrison, Christopher Gulvick, Scott Sammons, Yvette Unoarumhi, Darlene Wagner, Matthew Schmerer, S. H. Rosenthal, A. Gerasimova, R. M. Kagan, B. Anderson, M. Hua, Y. Liu, L.E. Bernstein, K.E. Livingston, A. Perez, I. A. Shlyakhter, R. V. Rolando, R. Owen, P. Tanpaiboon, F. Lacbawan, Clinton R. Paden, Duncan MacCannell |
| EPI_ISL_2527563, EPI_ISL_2527590, EPI_ISL_2527597, EPI_ISL_2527599, EPI_ISL_2527603, EPI_ISL_2527610, EPI_ISL_2527612, EPI_ISL_2527616, EPI_ISL_2527620, EPI_ISL_2527622, EPI_ISL_2527624, EPI_ISL_2527629, EPI_ISL_2527638, EPI_ISL_2527652, EPI_ISL_2527654, EPI_ISL_2527658, EPI_ISL_2527667, EPI_ISL_2527672, EPI_ISL_2527685, EPI_ISL_2527687, EPI_ISL_2527688, EPI_ISL_2527693, EPI_ISL_2527695, EPI_ISL_2527697, EPI_ISL_2527700, EPI_ISL_2527708, EPI_ISL_2527710, EPI_ISL_2527727, EPI_ISL_2527755, EPI_ISL_2527764, EPI_ISL_2527768, EPI_ISL_2527770, EPI_ISL_2527772, EPI_ISL_2527774, EPI_ISL_2527775, EPI_ISL_2527777, EPI_ISL_2527779, EPI_ISL_2527784, EPI_ISL_2527786, EPI_ISL_2527788, EPI_ISL_2527790, EPI_ISL_2527792, EPI_ISL_2527796, EPI_ISL_2527803, EPI_ISL_2527809, EPI_ISL_2527813, EPI_ISL_2527820, EPI_ISL_2527825, EPI_ISL_2527833, EPI_ISL_2527835, EPI_ISL_2527837, EPI_ISL_2527839, EPI_ISL_2527841, EPI_ISL_2527843 | | | |
| see above | Fulgent Genetics | Centers for Disease Control and Prevention Division of Viral Diseases, Pathogen Discovery | Dakota Howard, Dhwani Batra, Peter W. Cook, Kara Moser, Adrian Paskey, Jason Caravas, Benjamin Rambo-Martin, Shatavia Morrison, Christopher Gulvick, Scott Sammons, Yvette Unoarumhi, Darlene Wagner, Matthew Schmerer, Harry Gao, Mickey Li, John Gao, Joseph Fierro, Benafsh Sapra, Becky Tsai, Yan Meng, Doreen Ng, James Xie, Clinton R. Paden, Duncan MacCannell |
| EPI_ISL_2527927, EPI_ISL_2527956, EPI_ISL_2527978, EPI_ISL_2528016, EPI_ISL_2528091, EPI_ISL_2528109, EPI_ISL_2528120, EPI_ISL_2528153, EPI_ISL_2528159, EPI_ISL_2528161, EPI_ISL_2528171, EPI_ISL_2528177, EPI_ISL_2528179, EPI_ISL_2528193, EPI_ISL_2528197, EPI_ISL_2528205, EPI_ISL_2528224, EPI_ISL_2528243, EPI_ISL_2528263, EPI_ISL_2528271, EPI_ISL_2528287, EPI_ISL_2528289, EPI_ISL_2528312, EPI_ISL_2528316, EPI_ISL_2528324, EPI_ISL_2528337, EPI_ISL_2528339, EPI_ISL_2528341, EPI_ISL_2528343, EPI_ISL_2528350, EPI_ISL_2528356, EPI_ISL_2528358, EPI_ISL_2528363, EPI_ISL_2528365, EPI_ISL_2528434, EPI_ISL_2528444, EPI_ISL_2528492, EPI_ISL_2528496, EPI_ISL_2528500, EPI_ISL_2528502, EPI_ISL_2528561, EPI_ISL_2528591, EPI_ISL_2528635, EPI_ISL_2528640, EPI_ISL_2528644, EPI_ISL_2528692, EPI_ISL_2528764, EPI_ISL_2528772, EPI_ISL_2528787, EPI_ISL_2528793, EPI_ISL_2528811, EPI_ISL_2528834, EPI_ISL_2528847, EPI_ISL_2528851, EPI_ISL_2528879, EPI_ISL_2528923, EPI_ISL_2528925, EPI_ISL_2528937, EPI_ISL_2528944 | | | |
| see above | Aegis Sciences Corporation | Centers for Disease Control and Prevention Division of Viral Diseases, Pathogen Discovery | Dakota Howard, Dhwani Batra, Peter W. Cook, Kara Moser, Adrian Paskey, Jason Caravas, Benjamin Rambo-Martin, Shatavia Morrison, Christopher Gulvick, Scott Sammons, Yvette Unoarumhi, Darlene Wagner, Matthew Schmerer, Cyndi Clark, Patrick Campbell, Rob Case, Vikramsinha Ghorpade, Holly Houdeshell, Ola Kvalvaag, Dillon Nall, Ethan Sanders, Alec Vest, Shaun Westlund, Matthew Hardison, Clinton R. Paden, Duncan MacCannell |
| EPI_ISL_2529080, EPI_ISL_2529083, EPI_ISL_2529089, EPI_ISL_2529113, EPI_ISL_2529119, EPI_ISL_2529138, EPI_ISL_2529144, EPI_ISL_2529146, EPI_ISL_2529172, EPI_ISL_2529185, EPI_ISL_2529206, EPI_ISL_2529208, EPI_ISL_2529216, EPI_ISL_2529228, EPI_ISL_2529230, EPI_ISL_2529240, EPI_ISL_2529281, EPI_ISL_2529283, EPI_ISL_2529286 | | | |
| see above | Quest Diagnostics Incorporated | Centers for Disease Control and Prevention Division of Viral Diseases, Pathogen Discovery | Dakota Howard, Dhwani Batra, Peter W. Cook, Kara Moser, Adrian Paskey, Jason Caravas, Benjamin Rambo-Martin, Shatavia Morrison, Christopher Gulvick, Scott Sammons, Yvette Unoarumhi, Darlene Wagner, Matthew Schmerer, S. H. Rosenthal, A. Gerasimova, R. M. Kagan, B. Anderson, M. Hua, Y. Liu, L.E. Bernstein, K.E. Livingston, A. Perez, I. A. Shlyakhter, R. V. Rolando, R. Owen, P. Tanpaiboon, F. Lacbawan, Clinton R. Paden, Duncan MacCannell |
| EPI_ISL_2529474, EPI_ISL_2529477 | Virology, Universitätsklinikum des Saarlandes | Epigenetics, Saarland University | Kathrin Kattler, Nastasia Seiwert, Stefan Lohse, Sascha Tierling, Thorsten Pfuhl, Sigrun Smola, Jörn Walter |
| EPI_ISL_2529589 | BCCDC Public Health Laboratory | BCCDC Public Health Laboratory | Prystajecky Natalie, Linda Hoang, Dan Fornika, John Tyson, Shannon Russell, Kim Macdonald, Kimia Kamelian, Ana Pacagnella, Corrinne Ng, Loretta Janz, Robert Azana, Mel Krajden |
| EPI_ISL_2529850, EPI_ISL_2529852, EPI_ISL_2529854, EPI_ISL_2529856, EPI_ISL_2529858, EPI_ISL_2529861 | Baylor College of Medicine/ GCID | Baylor College of Medicine/ GCID | Timothy Farinholt, Qing Meng, Qinchang Meng, Vipin Menon, Xiang Qin, Harsha Doddapaneni, Paige Farinholt, Charu Agrawal, Donna Muzny, Pedro A. Piedra, Richard A. Gibbs, Joseph Pertosino |
| EPI_ISL_2530115, EPI_ISL_2530117, EPI_ISL_2530119, EPI_ISL_2530132, EPI_ISL_2530134, EPI_ISL_2530143, EPI_ISL_2530148, EPI_ISL_2530160, EPI_ISL_2530161, EPI_ISL_2530163, EPI_ISL_2530169, EPI_ISL_2530174, EPI_ISL_2530179, EPI_ISL_2530185, EPI_ISL_2530186, EPI_ISL_2530191, EPI_ISL_2530193, EPI_ISL_2530195, EPI_ISL_2530196, EPI_ISL_2530198, EPI_ISL_2530199, EPI_ISL_2530201, EPI_ISL_2530202, EPI_ISL_2530204, EPI_ISL_2530206, EPI_ISL_2530207, EPI_ISL_2530209, EPI_ISL_2530210, EPI_ISL_2530212, EPI_ISL_2530214, EPI_ISL_2530217, EPI_ISL_2530219, EPI_ISL_2530221, EPI_ISL_2530222, EPI_ISL_2530224, EPI_ISL_2530226, EPI_ISL_2530227, EPI_ISL_2530229, EPI_ISL_2530230, EPI_ISL_2530232, EPI_ISL_2530233, EPI_ISL_2530235, EPI_ISL_2530236, EPI_ISL_2530239, EPI_ISL_2530241, EPI_ISL_2530243, EPI_ISL_2530245, EPI_ISL_2530246, EPI_ISL_2530248, EPI_ISL_2530250, EPI_ISL_2530252, EPI_ISL_2530253, EPI_ISL_2530255, EPI_ISL_2530256, EPI_ISL_2530258, EPI_ISL_2530259, EPI_ISL_2530262, EPI_ISL_2530264, EPI_ISL_2530265, EPI_ISL_2530268, EPI_ISL_2530270, EPI_ISL_2530290, EPI_ISL_2530299, EPI_ISL_2530304, EPI_ISL_2530308, EPI_ISL_2530319, EPI_ISL_2530373, EPI_ISL_2530374, EPI_ISL_2530378, EPI_ISL_2530379, EPI_ISL_2530384, EPI_ISL_2530388, EPI_ISL_2530390, EPI_ISL_2530391, EPI_ISL_2530392, EPI_ISL_2530394 | | | |
| see above | Hematopathology Laboratory, ACTREC, TMC | Hematopathology Laboratory, ACTREC, TMC | Hematopathology Laboratory, ACTREC |
| EPI_ISL_2530862, EPI_ISL_2530880, EPI_ISL_2530909, EPI_ISL_2530922 | BCCDC Public Health Laboratory | BCCDC Public Health Laboratory | Prystajecky Natalie, Linda Hoang, Dan Fornika, John Tyson, Shannon Russell, Kim Macdonald, Kimia Kamelian, Ana Pacagnella, Corrinne Ng, Loretta Janz, Robert Azana, Mel Krajden |
| EPI_ISL_2530938 | Aegis Sciences Corporation | Centers for Disease Control and Prevention Division of Viral Diseases, Pathogen Discovery | Dakota Howard, Dhwani Batra, Peter W. Cook, Kara Moser, Adrian Paskey, Jason Caravas, Benjamin Rambo-Martin, Shatavia Morrison, Christopher Gulvick, Scott Sammons, Yvette Unoarumhi, Darlene Wagner, Matthew Schmerer, Cyndi Clark, Patrick Campbell, Rob Case, Vikramsinha Ghorpade, Holly Houdeshell, Ola Kvalvaag, Dillon Nall, Ethan Sanders, Alec Vest, Shaun Westlund, Matthew Hardison, Clinton R. Paden, Duncan MacCannell |
| EPI_ISL_2530949 | BCCDC Public Health Laboratory | BCCDC Public Health Laboratory | Prystajecky Natalie, Linda Hoang, Dan Fornika, John Tyson, Shannon Russell, Kim Macdonald, Kimia Kamelian, Ana Pacagnella, Corrinne Ng, Loretta Janz, Robert Azana, Mel Krajden |
| EPI_ISL_2530950 | Aegis Sciences Corporation | Centers for Disease Control and Prevention Division of Viral Diseases, Pathogen Discovery | Dakota Howard, Dhwani Batra, Peter W. Cook, Kara Moser, Adrian Paskey, Jason Caravas, Benjamin Rambo-Martin, Shatavia Morrison, Christopher Gulvick, Scott Sammons, Yvette Unoarumhi, Darlene Wagner, Matthew Schmerer, Cyndi Clark, Patrick Campbell, Rob Case, Vikramsinha Ghorpade, Holly Houdeshell, Ola Kvalvaag, Dillon Nall, Ethan Sanders, Alec Vest, Shaun Westlund, Matthew Hardison, Clinton R. Paden, Duncan MacCannell |
| EPI_ISL_2530951, EPI_ISL_2530954 | BCCDC Public Health Laboratory | BCCDC Public Health Laboratory | Prystajecky Natalie, Linda Hoang, Dan Fornika, John Tyson, Shannon Russell, Kim Macdonald, Kimia Kamelian, Ana Pacagnella, Corrinne Ng, Loretta Janz, Robert Azana, Mel Krajden |
| EPI_ISL_2530988 | Aegis Sciences Corporation | Centers for Disease Control and Prevention Division of Viral Diseases, Pathogen Discovery | Dakota Howard, Dhwani Batra, Peter W. Cook, Kara Moser, Adrian Paskey, Jason Caravas, Benjamin Rambo-Martin, Shatavia Morrison, Christopher Gulvick, Scott Sammons, Yvette Unoarumhi, Darlene Wagner, Matthew Schmerer, Cyndi Clark, Patrick Campbell, Rob Case, Vikramsinha Ghorpade, Holly Houdeshell, Ola Kvalvaag, Dillon Nall, Ethan Sanders, Alec Vest, Shaun Westlund, Matthew Hardison, Clinton R. Paden, Duncan MacCannell |
| EPI_ISL_2531020 | BCCDC Public Health Laboratory | BCCDC Public Health Laboratory | Prystajecky Natalie, Linda Hoang, Dan Fornika, John Tyson, Shannon Russell, Kim Macdonald, Kimia Kamelian, Ana Pacagnella, Corrinne Ng, Loretta Janz, Robert Azana, Mel Krajden |
| EPI_ISL_2531021 | Aegis Sciences Corporation | Centers for Disease Control and Prevention Division of Viral Diseases, Pathogen Discovery | Dakota Howard, Dhwani Batra, Peter W. Cook, Kara Moser, Adrian Paskey, Jason Caravas, Benjamin Rambo-Martin, Shatavia Morrison, Christopher Gulvick, Scott Sammons, Yvette Unoarumhi, Darlene Wagner, Matthew Schmerer, Cyndi Clark, Patrick Campbell, Rob Case, Vikramsinha Ghorpade, Holly Houdeshell, Ola Kvalvaag, Dillon Nall, Ethan Sanders, Alec Vest, Shaun Westlund, Matthew Hardison, Clinton R. Paden, Duncan MacCannell |
| EPI_ISL_2531022, EPI_ISL_2531043, EPI_ISL_2531046, EPI_ISL_2531048 | BCCDC Public Health Laboratory | BCCDC Public Health Laboratory | Prystajecky Natalie, Linda Hoang, Dan Fornika, John Tyson, Shannon Russell, Kim Macdonald, Kimia Kamelian, Ana Pacagnella, Corrinne Ng, Loretta Janz, Robert Azana, Mel Krajden |
| EPI_ISL_2531049 | Aegis Sciences Corporation | Centers for Disease Control and Prevention Division of Viral Diseases, Pathogen Discovery | Dakota Howard, Dhwani Batra, Peter W. Cook, Kara Moser, Adrian Paskey, Jason Caravas, Benjamin Rambo-Martin, Shatavia Morrison, Christopher Gulvick, Scott Sammons, Yvette Unoarumhi, Darlene Wagner, Matthew Schmerer, Cyndi Clark, Patrick Campbell, Rob Case, Vikramsinha Ghorpade, Holly Houdeshell, Ola Kvalvaag, Dillon Nall, Ethan Sanders, Alec Vest, Shaun Westlund, Matthew Hardison, Clinton R. Paden, Duncan MacCannell |
| EPI_ISL_2531050 | BCCDC Public Health Laboratory | BCCDC Public Health Laboratory | Prystajecky Natalie, Linda Hoang, Dan Fornika, John Tyson, Shannon Russell, Kim Macdonald, Kimia Kamelian, Ana Pacagnella, Corrinne Ng, Loretta Janz, Robert Azana, Mel Krajden |
| EPI_ISL_2531053 | Aegis Sciences Corporation | Centers for Disease Control and Prevention Division of Viral Diseases, Pathogen Discovery | Dakota Howard, Dhwani Batra, Peter W. Cook, Kara Moser, Adrian Paskey, Jason Caravas, Benjamin Rambo-Martin, Shatavia Morrison, Christopher Gulvick, Scott Sammons, Yvette Unoarumhi, Darlene Wagner, Matthew Schmerer, Cyndi Clark, Patrick Campbell, Rob Case, Vikramsinha Ghorpade, Holly Houdeshell, Ola Kvalvaag, Dillon Nall, Ethan Sanders, Alec Vest, Shaun Westlund, Matthew Hardison, Clinton R. Paden, Duncan MacCannell |
| EPI_ISL_2531058, EPI_ISL_2531067, EPI_ISL_2531077, EPI_ISL_2531079, EPI_ISL_2531082, EPI_ISL_2531084, EPI_ISL_2531086, EPI_ISL_2531090, EPI_ISL_2531092, EPI_ISL_2531094, EPI_ISL_2531096, EPI_ISL_2531097, EPI_ISL_2531098, EPI_ISL_2531099, EPI_ISL_2531100, EPI_ISL_2531104, EPI_ISL_2531105 | | | |
| see above | BCCDC Public Health Laboratory | BCCDC Public Health Laboratory | Prystajecky Natalie, Linda Hoang, Dan Fornika, John Tyson, Shannon Russell, Kim Macdonald, Kimia Kamelian, Ana Pacagnella, Corrinne Ng, Loretta Janz, Robert Azana, Mel Krajden |
| EPI_ISL_2531894, EPI_ISL_2531897, EPI_ISL_2531901 | RIIP | National Reference Center for Viruses of Respiratory Infections, Institut Pasteur, Paris | Marion Barbet, Sylvie Behillil, Méline Bizard, Angela Brisebarre, Camille Capel, Vincent Enouf, Louise Lefrançois, Frédéric Lemoine, Christophe Malabat, Corinne Maufrais, Etienne Simon-Lorière, Maud Vanpeene, Sylvie Van der Werf ,StéPhanie Guyomard-Rabenirina |
| EPI_ISL_2531945, EPI_ISL_2531947, EPI_ISL_2531948, EPI_ISL_2531953 | Labo Analyses Med | National Reference Center for Viruses of Respiratory Infections, Institut Pasteur, Paris | Marion Barbet, Sylvie Behillil, Méline Bizard, Angela Brisebarre, Camille Capel, Vincent Enouf, Louise Lefrançois, Frédéric Lemoine, Christophe Malabat, Corinne Maufrais, Etienne Simon-Lorière, Maud Vanpeene, Sylvie Van der Werf ,Hugues Leroy |
| EPI_ISL_2532472 | Presidio di Brindisi Di Summa - Perrino | Istituto Zooprofilattico Sperimentale della Puglia e della Basilicata | Parisi A., Bianco A., Capozzi L., Del Sambro L., Simone D., Difato L., Pennuzzi G., Santoro A. |
| EPI_ISL_2532758 | Plateforme de testing Namuroise | Plateforme de testing Namuroise | Nicolas Gilliard; Demars Aurore ; Maschietto Céline ; Nobis Chloé ; Otto Gaetan ; Denis Olivier ; Degosserie Jonathan ; Mullier François |
| EPI_ISL_2532779, EPI_ISL_2532780, EPI_ISL_2532781 | Lifebrain Covid Labor GmbH | Dept. of Laboratory Medicine | Harald Esterbauer, Anna Gschaider, Petra Jurkowitsch, Fabian Konig, Sabina Plumer, Robert Strassl, Claudia Weber, Oswald Wagner |
| EPI_ISL_2533167, EPI_ISL_2533276, EPI_ISL_2533277, EPI_ISL_2533314, EPI_ISL_2533316, EPI_ISL_2533384, EPI_ISL_2533393, EPI_ISL_2533395, EPI_ISL_2533414, EPI_ISL_2533449, EPI_ISL_2533453, EPI_ISL_2533454, EPI_ISL_2533459, EPI_ISL_2533606, EPI_ISL_2533618, EPI_ISL_2533635, EPI_ISL_2533637, EPI_ISL_2533638, EPI_ISL_2533639, EPI_ISL_2533690, EPI_ISL_2533721 | | | |
| see above | National Virus Reference Laboratory | National Virus Reference Laboratory | Zoe Yandle, Charlene Bennett, Gabriel Gonzalez, Michael Carr, Jonathan Dean, Cillian F De Gascun |
| EPI_ISL_2533817 | LESP Ciudad de Mexico | Instituto de Diagnostico y Referencia Epidemiologicos (INDRE) | Claudia Wong-Arambula, Abril Rodriguez-Maldonado, Vanessa Rivero-Arredondo, Ariadna Medina-Benitez, Joaquin Quiroz-Mercado, Sergio Rangel-Guerrero, Natividad Cruz-Ortiz, Tatiana Nunez-Garcia, Gisela Barrera-Badillo, Lucia Hernandez-Rivas, Irma Lopez-Martinez, Ernesto Ramirez-Gonzalez. |
| EPI_ISL_2533818 | LESP Baja California Sur | Instituto de Diagnostico y Referencia Epidemiologicos (INDRE) | Claudia Wong-Arambula, Abril Rodriguez-Maldonado, Vanessa Rivero-Arredondo, Ariadna Medina-Benitez, Joaquin Quiroz-Mercado, Sergio Rangel-Guerrero, Natividad Cruz-Ortiz, Tatiana Nunez-Garcia, Gisela Barrera-Badillo, Lucia Hernandez-Rivas, Irma Lopez-Martinez, Ernesto Ramirez-Gonzalez. |
| EPI_ISL_2533819 | LESP Yucatan | Instituto de Diagnostico y Referencia Epidemiologicos (INDRE) | Claudia Wong-Arambula, Abril Rodriguez-Maldonado, Vanessa Rivero-Arredondo, Ariadna Medina-Benitez, Joaquin Quiroz-Mercado, Sergio Rangel-Guerrero, Natividad Cruz-Ortiz, Tatiana Nunez-Garcia, Gisela Barrera-Badillo, Lucia Hernandez-Rivas, Irma Lopez-Martinez, Ernesto Ramirez-Gonzalez. |
| EPI_ISL_2533820 | LESP Quintana Roo | Instituto de Diagnostico y Referencia Epidemiologicos (INDRE) | Claudia Wong-Arambula, Abril Rodriguez-Maldonado, Vanessa Rivero-Arredondo, Ariadna Medina-Benitez, Joaquin Quiroz-Mercado, Sergio Rangel-Guerrero, Natividad Cruz-Ortiz, Tatiana Nunez-Garcia, Gisela Barrera-Badillo, Lucia Hernandez-Rivas, Irma Lopez-Martinez, Ernesto Ramirez-Gonzalez. |
| EPI_ISL_2533821 | LESP Ciudad de México /DIAGNOMOL | Instituto de Diagnostico y Referencia Epidemiologicos (INDRE) | Claudia Wong-Arambula, Abril Rodriguez-Maldonado, Vanessa Rivero-Arredondo, Ariadna Medina-Benitez, Joaquin Quiroz-Mercado, Sergio Rangel-Guerrero, Natividad Cruz-Ortiz, Tatiana Nunez-Garcia, Gisela Barrera-Badillo, Lucia Hernandez-Rivas, Irma Lopez-Martinez, Ernesto Ramirez-Gonzalez. |
| EPI_ISL_2533828 | Laboratoire Virologie Saint Louis APHP | Laboratoire Virologie Saint Louis APHP | Maud Salmona, Marie Laure Chaix, Severine Mercier Delarue, Marie Laure Néré, Linda Feghoul, Jérome Le Goff, Constance Delauguerre, Sophia Achaibou |
| EPI_ISL_2533891, EPI_ISL_2533896, EPI_ISL_2533899, EPI_ISL_2533902 | Montana Public Health Laboratory | Montana Public Health Laboratory | Joy Ritter, Michelle Mozer, Carrie Biskupiak, Deborah Gibson, Michael Dills |

| | | | |
|---|---|---|---|
| EPI_ISL_2533922, EPI_ISL_2533923, EPI_ISL_2533924 | Hospital Sharp | Microbial Genomics Laboratory | Consorcio Mexicano de Vigilancia Genómica (CoViGen-Mex). Authors (in alphabetical order): Julio Elias Alvarado-Yaah, Carlos F. Arias, Santiago Ávila-Ríos, Eduardo Becerril-Vargas, Víctor Hugo Borja-Aburto, Celia Boukadida, Cristóbal Cháidez-Quiróz, Juan Bautista Chale-Dzul, Ricardo Ciria Merce, Andreu Comas-García, Célida Duque Molina, Julissa Enciso-Ibarra, José Antonio Enciso-Moreno, Gloria Elena Espinosa-Ayala, Fernando Fontove-Herrera, Daniel Fregoso-Rueda, Victor Eduardo García-Arias, Alejandra García-Gasca, Bruno Gómez-Gil, Jean Pierre González, Irvin González-López, Concepción Grajales-Muñiz, Ricardo Grande, Rosa María Gutiérrez Rios, Jesús Hernández, Alejandra Hernández-Terán, Alfredo Herrera-Estrella, Carla Ivón Herrera-Najera, Pavel Isa, Verónica Mata-Haro, Daniel Lira Morales, Susana Lopez, Antonio Loza Román, Brenda Irasema Maldonado-Meza, Bernardo Martínez-Miguel, José Arturo Martínez-Orozco, Célida Martínez-Rodríguez, Margarita Matías-Florentino, Fidencio Mejía-Nepomuceno, María Guadalupe de Jesús Mireles-Rivera, Gloria María Molina-Salinas, Hector Montoya-Fuentes, , Mario Mújica-Sánchez, José Esteban Muñoz-Medina, José de Jesús Nuñez-Contreras, Alicia Ocaña-Mondragón, Luis Alberto Ochoa-Carrera, Hector Esteban Paz-Juárez, Marissa Perez-Garcia, Francisco Pulido, Helen Haydee Fernanda Ramírez-Plascencia, Jorge Salas-Hernández, Angel Gustavo Salas-Lais, Alejandro Sánchez-Flores, Clara Esperanza Santacruz-Tinoco, María Guadalupe Santiago-Mauricio, Selene Zárate, Nelly Sélem-Mojica, Blanca Taboada, Gloria Vazquez, Joel Armando Vázquez-Pérez. |
| EPI_ISL_2533925 | Imágenes Diagnósticas de Mazatlán | Microbial Genomics Laboratory | Consorcio Mexicano de Vigilancia Genómica (CoViGen-Mex). Authors (in alphabetical order): Julio Elias Alvarado-Yaah, Carlos F. Arias, Santiago Ávila-Ríos, Eduardo Becerril-Vargas, Víctor Hugo Borja-Aburto, Celia Boukadida, Cristóbal Cháidez-Quiróz, Juan Bautista Chale-Dzul, Ricardo Ciria Merce, Andreu Comas-García, Célida Duque Molina, Julissa Enciso-Ibarra, José Antonio Enciso-Moreno, Gloria Elena Espinosa-Ayala, Fernando Fontove-Herrera, Daniel Fregoso-Rueda, Victor Eduardo García-Arias, Alejandra García-Gasca, Bruno Gómez-Gil, Jean Pierre González, Irvin González-López, Concepción Grajales-Muñiz, Ricardo Grande, Rosa María Gutiérrez Rios, Jesús Hernández, Alejandra Hernández-Terán, Alfredo Herrera-Estrella, Carla Ivón Herrera-Najera, Pavel Isa, Verónica Mata-Haro, Daniel Lira Morales, Susana Lopez, Antonio Loza Román, Brenda Irasema Maldonado-Meza, Bernardo Martínez-Miguel, José Arturo Martínez-Orozco, Célida Martínez-Rodríguez, Margarita Matías-Florentino, Fidencio Mejía-Nepomuceno, María Guadalupe de Jesús Mireles-Rivera, Gloria María Molina-Salinas, Hector Montoya-Fuentes, , Mario Mújica-Sánchez, José Esteban Muñoz-Medina, José de Jesús Nuñez-Contreras, Alicia Ocaña-Mondragón, Luis Alberto Ochoa-Carrera, Hector Esteban Paz-Juárez, Marissa Perez-Garcia, Francisco Pulido, Helen Haydee Fernanda Ramírez-Plascencia, Jorge Salas-Hernández, Angel Gustavo Salas-Lais, Alejandro Sánchez-Flores, Clara Esperanza Santacruz-Tinoco, María Guadalupe Santiago-Mauricio, Selene Zárate, Nelly Sélem-Mojica, Blanca Taboada, Gloria Vazquez, Joel Armando Vázquez-Pérez. |
| EPI_ISL_2533926, EPI_ISL_2533927 | Hospital Sharp | Microbial Genomics Laboratory | Consorcio Mexicano de Vigilancia Genómica (CoViGen-Mex). Authors (in alphabetical order): Julio Elias Alvarado-Yaah, Carlos F. Arias, Santiago Ávila-Ríos, Eduardo Becerril-Vargas, Víctor Hugo Borja-Aburto, Celia Boukadida, Cristóbal Cháidez-Quiróz, Juan Bautista Chale-Dzul, Ricardo Ciria Merce, Andreu Comas-García, Célida Duque Molina, Julissa Enciso-Ibarra, José Antonio Enciso-Moreno, Gloria Elena Espinosa-Ayala, Fernando Fontove-Herrera, Daniel Fregoso-Rueda, Victor Eduardo García-Arias, Alejandra García-Gasca, Bruno Gómez-Gil, Jean Pierre González, Irvin González-López, Concepción Grajales-Muñiz, Ricardo Grande, Rosa María Gutiérrez Rios, Jesús Hernández, Alejandra Hernández-Terán, Alfredo Herrera-Estrella, Carla Ivón Herrera-Najera, Pavel Isa, Verónica Mata-Haro, Daniel Lira Morales, Susana Lopez, Antonio Loza Román, Brenda Irasema Maldonado-Meza, Bernardo Martínez-Miguel, José Arturo Martínez-Orozco, Célida Martínez-Rodríguez, Margarita Matías-Florentino, Fidencio Mejía-Nepomuceno, María Guadalupe de Jesús Mireles-Rivera, Gloria María Molina-Salinas, Hector Montoya-Fuentes, , Mario Mújica-Sánchez, José Esteban Muñoz-Medina, José de Jesús Nuñez-Contreras, Alicia Ocaña-Mondragón, Luis Alberto Ochoa-Carrera, Hector Esteban Paz-Juárez, Marissa Perez-Garcia, Francisco Pulido, Helen Haydee Fernanda Ramírez-Plascencia, Jorge Salas-Hernández, Angel Gustavo Salas-Lais, Alejandro Sánchez-Flores, Clara Esperanza Santacruz-Tinoco, María Guadalupe Santiago-Mauricio, Selene Zárate, Nelly Sélem-Mojica, Blanca Taboada, Gloria Vazquez, Joel Armando Vázquez-Pérez. |
| EPI_ISL_2534078 | RSUD dr. LOEKMONO HADI | Genetics Working Group (Pokja Genetik) Faculty of Medicine, Public Health and Nursing Universitas Gadjah Mada (FK-KMK UGM); Disease Investigation Center Wates Ministry of Agriculture Indonesia; Department of Microbiology FK-KMK UGM; Laboratorium Diagnostik Yayasan Tahija World Mosquito Program (WMP) Yogyakarta Center for Tropical Medicine FK-KMK UGM; Integrated Research Center FK-KMK UGM; Department of Computer Science and Electronics FMIPA UGM; RSUP Dr. Sardjito | Gunadi, Hendra Wibawa, Marcellus, Mohamad S. Hakim, Edwin W. Daniwijaya, Ludhang P. Rizki, Endah Supriyati, Eggi Arguni, Titik Nuryastuti, Tri Wibawa, Dwi AA Nugrahaningsih, Afiahayati, Siswanto,Kristy Iskandar,Nungki Anggorowati, Pramesti G Dewi, Dwi Indaryati,Indarto Sulistiyono, Khanza a Vujira, Alvina A Setiawan |
| EPI_ISL_2534079 | RSUD dr. LOEKMONO HADI | Genetics Working Group (Pokja Genetik) Faculty of Medicine, Public Health and Nursing Universitas Gadjah Mada (FK-KMK UGM); Disease Investigation Center Wates Ministry of Agriculture Indonesia; Department of Microbiology FK-KMK UGM; Laboratorium Diagnostik Yayasan Tahija World Mosquito Program (WMP) Yogyakarta Center for Tropical Medicine FK-KMK UGM; Integrated Research Center FK-KMK UGM; Department of Computer Science and Electronics FMIPA UGM; RSUP Dr. Sardjito | Gunadi, Hendra Wibawa, Marcellus, Mohamad S. Hakim, Edwin W. Daniwijaya, Ludhang P. Rizki, Endah Supriyati, Eggi Arguni, Titik Nuryastuti, Tri Wibawa, Dwi AA Nugrahaningsih, Afiahayati, Siswanto,Kristy Iskandar,Nungki Anggorowati, Pramesti G Dewi, Dwi Indaryati,Indarto Sulistiyono, Susan Simanjaya, Dyah A Puspitarani |
| EPI_ISL_2534082 | RSUD dr. LOEKMONO HADI | Genetics Working Group (Pokja Genetik) Faculty of Medicine, Public Health and Nursing Universitas Gadjah Mada (FK-KMK UGM); Disease Investigation Center Wates Ministry of Agriculture Indonesia; Department of Microbiology FK-KMK UGM; Laboratorium Diagnostik Yayasan Tahija World Mosquito Program (WMP) Yogyakarta Center for Tropical Medicine FK-KMK UGM; Integrated Research Center FK-KMK UGM; Department of Computer Science and Electronics FMIPA UGM; RSUP Dr. Sardjito | Gunadi, Hendra Wibawa, Marcellus, Mohamad S. Hakim, Edwin W. Daniwijaya, Ludhang P. Rizki, Endah Supriyati, Eggi Arguni, Titik Nuryastuti, Tri Wibawa, Dwi AA Nugrahaningsih, Afiahayati, Siswanto,Kristy Iskandar,Nungki Anggorowati, Pramesti G Dewi, Dwi Indaryati,Indarto Sulistiyono, Irene Tania, Hana F Hanifin |
| EPI_ISL_2534236, EPI_ISL_2534237 | Clinical Reference Laboratory | Kansas Health and Environmental Lab | Mike Grose, Katherine Wiggins, Jonathan Barnell, Ben Olsen, and Phil Adam |
| EPI_ISL_2534238, EPI_ISL_2534239, EPI_ISL_2534242, EPI_ISL_2534246, EPI_ISL_2534250 | Kansas Health and Environmental Lab | Kansas Health and Environmental Lab | Mike Grose, Katherine Wiggins, Jonathan Barnell, Ben Olsen, and Phil Adam |
| EPI_ISL_2534251, EPI_ISL_2534252, EPI_ISL_2534253, EPI_ISL_2534254, EPI_ISL_2534255, EPI_ISL_2534257, EPI_ISL_2534258, EPI_ISL_2534259, EPI_ISL_2534260, EPI_ISL_2534263 | Wichita State University - Molecular Diagnostics Lab | Kansas Health and Environmental Lab | Mike Grose, Katherine Wiggins, Jonathan Barnell, Ben Olsen, and Phil Adam |
| EPI_ISL_2534289 | RSUD dr. LOEKMONO HADI | Genetics Working Group (Pokja Genetik) Faculty of Medicine, Public Health and Nursing Universitas Gadjah Mada (FK-KMK UGM); Disease Investigation Center Wates Ministry of Agriculture Indonesia; Department of Microbiology FK-KMK UGM; Laboratorium Diagnostik Yayasan Tahija World Mosquito Program (WMP) Yogyakarta Center for Tropical Medicine FK-KMK UGM; Integrated Research Center FK-KMK UGM; Department of Computer Science and Electronics FMIPA UGM; RSUP Dr. Sardjito | Gunadi, Hendra Wibawa, Marcellus, Mohamad S. Hakim, Edwin W. Daniwijaya, Ludhang P. Rizki, Endah Supriyati, Eggi Arguni, Titik Nuryastuti, Tri Wibawa, Dwi AA Nugrahaningsih, Afiahayati, Siswanto,Kristy Iskandar,Nungki Anggorowati, Pramesti G Dewi, Dwi Indaryati,Indarto Sulistiyono, Cita S Amalia, Alvina A Setiawan |
| EPI_ISL_2534315, EPI_ISL_2534316, EPI_ISL_2534317 | The Ohio State University Applied Microbiology Services Laboratory | The Ohio State University Applied Microbiology Services Laboratory | Seth A. Faith PhD |
| EPI_ISL_2534338 | RSUD dr. LOEKMONO HADI | Genetics Working Group (Pokja Genetik) Faculty of Medicine, Public Health and Nursing Universitas Gadjah Mada (FK-KMK UGM); Disease Investigation Center Wates Ministry of Agriculture Indonesia; Department of Microbiology FK-KMK UGM; Laboratorium Diagnostik Yayasan Tahija World Mosquito Program (WMP) Yogyakarta Center for Tropical Medicine FK-KMK UGM; Integrated Research Center FK-KMK UGM; Department of Computer Science and Electronics FMIPA UGM; RSUP Dr. Sardjito | Gunadi, Hendra Wibawa, Marcellus, Mohamad S. Hakim, Edwin W. Daniwijaya, Ludhang P. Rizki, Endah Supriyati, Eggi Arguni, Titik Nuryastuti, Tri Wibawa, Dwi AA Nugrahaningsih, Afiahayati, Siswanto,Kristy Iskandar,Nungki Anggorowati, Pramesti G Dewi, Dwi Indaryati,Indarto Sulistiyono, Khanza a Vujira, Alvina A Setiawan |
| EPI_ISL_2534339 | RSUD dr. LOEKMONO HADI | Genetics Working Group (Pokja Genetik) Faculty of Medicine, Public Health and Nursing Universitas Gadjah Mada (FK-KMK UGM); Disease Investigation Center Wates Ministry of Agriculture Indonesia; Department of Microbiology FK-KMK UGM; Laboratorium Diagnostik Yayasan Tahija World Mosquito Program (WMP) Yogyakarta Center for Tropical Medicine FK-KMK UGM; Integrated Research Center FK-KMK UGM; Department of Computer Science and Electronics FMIPA UGM; RSUP Dr. Sardjito | Gunadi, Hendra Wibawa, Marcellus, Mohamad S. Hakim, Edwin W. Daniwijaya, Ludhang P. Rizki, Endah Supriyati, Eggi Arguni, Titik Nuryastuti, Tri Wibawa, Dwi AA Nugrahaningsih, Afiahayati, Siswanto,Kristy Iskandar,Nungki Anggorowati, Pramesti G Dewi, Dwi Indaryati,Indarto Sulistiyono, Susan Simanjaya, Dyah A Puspitarani |
| EPI_ISL_2534346 | RSUD dr. LOEKMONO HADI | Genetics Working Group (Pokja Genetik) Faculty of Medicine, Public Health and Nursing Universitas Gadjah Mada (FK-KMK UGM); Disease Investigation Center Wates Ministry of Agriculture Indonesia; Department of Microbiology FK-KMK UGM; Laboratorium Diagnostik Yayasan Tahija World Mosquito Program (WMP) Yogyakarta Center for Tropical Medicine FK-KMK UGM; Integrated Research Center FK-KMK UGM; Department of Computer Science and Electronics FMIPA UGM; RSUP Dr. Sardjito | Gunadi, Hendra Wibawa, Marcellus, Mohamad S. Hakim, Edwin W. Daniwijaya, Ludhang P. Rizki, Endah Supriyati, Eggi Arguni, Titik Nuryastuti, Tri Wibawa, Dwi AA Nugrahaningsih, Afiahayati, Siswanto,Kristy Iskandar,Nungki Anggorowati, Pramesti G Dewi, Dwi Indaryati,Indarto Sulistiyono, Irene Tania, Hana F Hanifin |
| EPI_ISL_2534376 | RSUD dr. LOEKMONO HADI | Genetics Working Group (Pokja Genetik) Faculty of Medicine, Public Health and Nursing Universitas Gadjah Mada (FK-KMK UGM); Disease Investigation Center Wates Ministry of Agriculture Indonesia; Department of Microbiology FK-KMK UGM; Laboratorium Diagnostik Yayasan Tahija World Mosquito Program (WMP) Yogyakarta Center for Tropical Medicine FK-KMK UGM; Integrated Research Center FK-KMK UGM; Department of Computer Science | Gunadi, Hendra Wibawa, Marcellus, Mohamad S. Hakim, Edwin W. Daniwijaya, Ludhang P. Rizki, Endah Supriyati, Eggi Arguni, Titik Nuryastuti, Tri Wibawa, Dwi AA Nugrahaningsih, Afiahayati, Siswanto,Kristy Iskandar,Nungki Anggorowati, Pramesti G Dewi, Dwi Indaryati,Indarto Sulistiyono, Cita S Amalia, Alvina A Setiawan |

| | | | |
|---|---|---|---|
| | | and Electronics FMIPA UGM; RSUP Dr. Sardjito | |
| EPI_ISL_2534377 | RSUD dr. LOEKMONO HADI | Genetics Working Group (Pokja Genetik) Faculty of Medicine, Public Health and Nursing Universitas Gadjah Mada (FK-KMK UGM); Disease Investigation Center Wates Ministry of Agriculture Indonesia; Department of Microbiology FK-KMK UGM; Laboratorium Diagnostik Yayasan Tahija World Mosquito Program (WMP) Yogyakarta Center for Tropical Medicine FK-KMK UGM; Integrated Research Center FK-KMK UGM; Department of Computer Science and Electronics FMIPA UGM; RSUP Dr. Sardjito | Gunadi, Hendra Wibawa, Marcellus, Mohamad S. Hakim, Edwin W. Daniwijaya, Ludhang P. Rizki, Endah Supriyati, Eggi Arguni, Titik Nuryastuti, Tri Wibawa, Dwi AA Nugrahaningsih, Afiahayati, Siswanto,Kristy Iskandar,Nungki Anggorowati, Pramesti G Dewi, Dwi Indaryati,Indarto Sulistiyono, Khanza a Vujira, Alvina A Setiawan |
| EPI_ISL_2534378 | RSUD dr. LOEKMONO HADI | Genetics Working Group (Pokja Genetik) Faculty of Medicine, Public Health and Nursing Universitas Gadjah Mada (FK-KMK UGM); Disease Investigation Center Wates Ministry of Agriculture Indonesia; Department of Microbiology FK-KMK UGM; Laboratorium Diagnostik Yayasan Tahija World Mosquito Program (WMP) Yogyakarta Center for Tropical Medicine FK-KMK UGM; Integrated Research Center FK-KMK UGM; Department of Computer Science and Electronics FMIPA UGM; RSUP Dr. Sardjito | Gunadi, Hendra Wibawa, Marcellus, Mohamad S. Hakim, Edwin W. Daniwijaya, Ludhang P. Rizki, Endah Supriyati, Eggi Arguni, Titik Nuryastuti, Tri Wibawa, Dwi AA Nugrahaningsih, Afiahayati, Siswanto,Kristy Iskandar,Nungki Anggorowati, Pramesti G Dewi, Dwi Indaryati,Indarto Sulistiyono, Irene Tania, Hana F Hanifin |
| EPI_ISL_2534379 | RSUD dr. LOEKMONO HADI | Genetics Working Group (Pokja Genetik) Faculty of Medicine, Public Health and Nursing Universitas Gadjah Mada (FK-KMK UGM); Disease Investigation Center Wates Ministry of Agriculture Indonesia; Department of Microbiology FK-KMK UGM; Laboratorium Diagnostik Yayasan Tahija World Mosquito Program (WMP) Yogyakarta Center for Tropical Medicine FK-KMK UGM; Integrated Research Center FK-KMK UGM; Department of Computer Science and Electronics FMIPA UGM; RSUP Dr. Sardjito | Gunadi, Hendra Wibawa, Marcellus, Mohamad S. Hakim, Edwin W. Daniwijaya, Ludhang P. Rizki, Endah Supriyati, Eggi Arguni, Titik Nuryastuti, Tri Wibawa, Dwi AA Nugrahaningsih, Afiahayati, Siswanto,Kristy Iskandar,Nungki Anggorowati, Pramesti G Dewi, Dwi Indaryati,Indarto Sulistiyono, Cita S Amalia, Alvina A Setiawan |
| EPI_ISL_2534392, EPI_ISL_2534395, EPI_ISL_2534410, EPI_ISL_2534418, EPI_ISL_2534420 | HOSPITAL UNIVERSITARIO SON ESPASES | HOSPITAL UNIVERSITARIO SON ESPASES | Servicio de Microbiología, Hospital Universitario Son Espases |
| EPI_ISL_2534431 | HOSPITAL CAN MISSES | HOSPITAL UNIVERSITARIO SON ESPASES | Servicio de Microbiología, Hospital Universitario Son Espases |
| EPI_ISL_2534438 | HOSPITAL UNIVERSITARIO SON ESPASES | HOSPITAL UNIVERSITARIO SON ESPASES | Servicio de Microbiología, Hospital Universitario Son Espases |
| EPI_ISL_2534440, EPI_ISL_2534452 | HOSPITAL CAN MISSES | HOSPITAL UNIVERSITARIO SON ESPASES | Servicio de Microbiología, Hospital Universitario Son Espases |
| EPI_ISL_2534454 | RSUD dr. LOEKMONO HADI | Genetics Working Group (Pokja Genetik) Faculty of Medicine, Public Health and Nursing Universitas Gadjah Mada (FK-KMK UGM); Disease Investigation Center Wates Ministry of Agriculture Indonesia; Department of Microbiology FK-KMK UGM; Laboratorium Diagnostik Yayasan Tahija World Mosquito Program (WMP) Yogyakarta Center for Tropical Medicine FK-KMK UGM; Integrated Research Center FK-KMK UGM; Department of Computer Science and Electronics FMIPA UGM; RSUP Dr. Sardjito | Gunadi, Hendra Wibawa, Marcellus, Mohamad S. Hakim, Edwin W. Daniwijaya, Ludhang P. Rizki, Endah Supriyati, Eggi Arguni, Titik Nuryastuti, Tri Wibawa, Dwi AA Nugrahaningsih, Afiahayati, Siswanto,Kristy Iskandar,Nungki Anggorowati, Pramesti G Dewi, Dwi Indaryati,Indarto Sulistiyono, Khanza a Vujira, Alvina A Setiawan |
| EPI_ISL_2534455 | RSUD dr. LOEKMONO HADI | Genetics Working Group (Pokja Genetik) Faculty of Medicine, Public Health and Nursing Universitas Gadjah Mada (FK-KMK UGM); Disease Investigation Center Wates Ministry of Agriculture Indonesia; Department of Microbiology FK-KMK UGM; Laboratorium Diagnostik Yayasan Tahija World Mosquito Program (WMP) Yogyakarta Center for Tropical Medicine FK-KMK UGM; Integrated Research Center FK-KMK UGM; Department of Computer Science and Electronics FMIPA UGM; RSUP Dr. Sardjito | Gunadi, Hendra Wibawa, Marcellus, Mohamad S. Hakim, Edwin W. Daniwijaya, Ludhang P. Rizki, Endah Supriyati, Eggi Arguni, Titik Nuryastuti, Tri Wibawa, Dwi AA Nugrahaningsih, Afiahayati, Siswanto,Kristy Iskandar,Nungki Anggorowati, Pramesti G Dewi, Dwi Indaryati,Indarto Sulistiyono, Susan Simanjaya, Dyah A Puspitarani |
| EPI_ISL_2534456 | RSUD dr. LOEKMONO HADI | Genetics Working Group (Pokja Genetik) Faculty of Medicine, Public Health and Nursing Universitas Gadjah Mada (FK-KMK UGM); Disease Investigation Center Wates Ministry of Agriculture Indonesia; Department of Microbiology FK-KMK UGM; Laboratorium Diagnostik Yayasan Tahija World Mosquito Program (WMP) Yogyakarta Center for Tropical Medicine FK-KMK UGM; Integrated Research Center FK-KMK UGM; Department of Computer Science and Electronics FMIPA UGM; RSUP Dr. Sardjito | Gunadi, Hendra Wibawa, Marcellus, Mohamad S. Hakim, Edwin W. Daniwijaya, Ludhang P. Rizki, Endah Supriyati, Eggi Arguni, Titik Nuryastuti, Tri Wibawa, Dwi AA Nugrahaningsih, Afiahayati, Siswanto,Kristy Iskandar,Nungki Anggorowati, Pramesti G Dewi, Dwi Indaryati,Indarto Sulistiyono, Irene Tania, Hana F Hanifin |
| EPI_ISL_2534457 | RSUD dr. LOEKMONO HADI | Genetics Working Group (Pokja Genetik) Faculty of Medicine, Public Health and Nursing Universitas Gadjah Mada (FK-KMK UGM); Disease Investigation Center Wates Ministry of Agriculture Indonesia; Department of Microbiology FK-KMK UGM; Laboratorium Diagnostik Yayasan Tahija World Mosquito Program (WMP) Yogyakarta Center for Tropical Medicine FK-KMK UGM; Integrated Research Center FK-KMK UGM; Department of Computer Science and Electronics FMIPA UGM; RSUP Dr. Sardjito | Gunadi, Hendra Wibawa, Marcellus, Mohamad S. Hakim, Edwin W. Daniwijaya, Ludhang P. Rizki, Endah Supriyati, Eggi Arguni, Titik Nuryastuti, Tri Wibawa, Dwi AA Nugrahaningsih, Afiahayati, Siswanto,Kristy Iskandar,Nungki Anggorowati, Pramesti G Dewi, Dwi Indaryati,Indarto Sulistiyono, Cita S Amalia, Alvina A Setiawan |
| EPI_ISL_2534458 | RSUD dr. LOEKMONO HADI | Genetics Working Group (Pokja Genetik) Faculty of Medicine, Public Health and Nursing Universitas Gadjah Mada (FK-KMK UGM); Disease Investigation Center Wates Ministry of Agriculture Indonesia; Department of Microbiology FK-KMK UGM; Laboratorium Diagnostik Yayasan Tahija World Mosquito Program (WMP) Yogyakarta Center for Tropical Medicine FK-KMK UGM; Integrated Research Center FK-KMK UGM; Department of Computer Science and Electronics FMIPA UGM; RSUP Dr. Sardjito | Gunadi, Hendra Wibawa, Marcellus, Mohamad S. Hakim, Edwin W. Daniwijaya, Ludhang P. Rizki, Endah Supriyati, Eggi Arguni, Titik Nuryastuti, Tri Wibawa, Dwi AA Nugrahaningsih, Afiahayati, Siswanto,Kristy Iskandar,Nungki Anggorowati, Pramesti G Dewi, Dwi Indaryati,Indarto Sulistiyono, Khanza a Vujira, Alvina A Setiawan |
| EPI_ISL_2534468 | CAP TORELLÓ | Banc de Sang i Teixits | Francisco Vidal, Irene Corrales, Nina Borràs, Carlos Hobeich, Natália Comes, Lorena Ramírez, Noemí Gonzalez, Silvia Sauleda, Maria Glòria Soria |
| EPI_ISL_2534472 | CAP MANLLEU | Banc de Sang i Teixits | Francisco Vidal, Irene Corrales, Nina Borràs, Carlos Hobeich, Natália Comes, Lorena Ramírez, Noemí Gonzalez, Silvia Sauleda, Maria Glòria Soria |
| EPI_ISL_2534478, EPI_ISL_2534486 | CAP SARRIÀ | Banc de Sang i Teixits | Francisco Vidal, Irene Corrales, Nina Borràs, Carlos Hobeich, Natália Comes, Lorena Ramírez, Noemí Gonzalez, Silvia Sauleda, Maria Glòria Soria |
| EPI_ISL_2534489 | RSUD dr. LOEKMONO HADI | Genetics Working Group (Pokja Genetik) Faculty of Medicine, Public Health and Nursing Universitas Gadjah Mada (FK-KMK UGM); Disease Investigation Center Wates Ministry of Agriculture Indonesia; Department of Microbiology FK-KMK UGM; Laboratorium Diagnostik Yayasan Tahija World Mosquito Program (WMP) Yogyakarta Center for Tropical Medicine FK-KMK UGM; Integrated Research Center FK-KMK UGM; Department of Computer Science and Electronics FMIPA UGM; RSUP Dr. Sardjito | Gunadi, Hendra Wibawa, Marcellus, Mohamad S. Hakim, Edwin W. Daniwijaya, Ludhang P. Rizki, Endah Supriyati, Eggi Arguni, Titik Nuryastuti, Tri Wibawa, Dwi AA Nugrahaningsih, Afiahayati, Siswanto,Kristy Iskandar,Nungki Anggorowati, Pramesti G Dewi, Dwi Indaryati,Indarto Sulistiyono, Cita S Amalia, Alvina A Setiawan |
| EPI_ISL_2534490 | RSUD dr. LOEKMONO HADI | Genetics Working Group (Pokja Genetik) Faculty of Medicine, Public Health and Nursing Universitas Gadjah Mada (FK-KMK UGM); Disease Investigation Center Wates Ministry of Agriculture Indonesia; Department of Microbiology FK-KMK UGM; Laboratorium Diagnostik Yayasan Tahija World Mosquito Program (WMP) Yogyakarta Center for Tropical Medicine FK-KMK UGM; Integrated Research Center FK-KMK UGM; Department of Computer Science and Electronics FMIPA UGM; RSUP Dr. Sardjito | Gunadi, Hendra Wibawa, Marcellus, Mohamad S. Hakim, Edwin W. Daniwijaya, Ludhang P. Rizki, Endah Supriyati, Eggi Arguni, Titik Nuryastuti, Tri Wibawa, Dwi AA Nugrahaningsih, Afiahayati, Siswanto,Kristy Iskandar,Nungki Anggorowati, Pramesti G Dewi, Dwi Indaryati,Indarto Sulistiyono, Khanza a Vujira, Alvina A Setiawan |
| EPI_ISL_2534518 | RSUD dr. LOEKMONO HADI | Genetics Working Group (Pokja Genetik) Faculty of Medicine, Public Health and Nursing Universitas Gadjah Mada (FK-KMK UGM); Disease Investigation Center Wates Ministry of Agriculture Indonesia; Department of Microbiology FK-KMK UGM; Laboratorium Diagnostik Yayasan Tahija World Mosquito Program (WMP) Yogyakarta Center for Tropical Medicine FK-KMK UGM; Integrated Research Center FK-KMK UGM; Department of Computer Science and Electronics FMIPA UGM; RSUP Dr. Sardjito | Gunadi, Hendra Wibawa, Marcellus, Mohamad S. Hakim, Edwin W. Daniwijaya, Ludhang P. Rizki, Endah Supriyati, Eggi Arguni, Titik Nuryastuti, Tri Wibawa, Dwi AA Nugrahaningsih, Afiahayati, Siswanto,Kristy Iskandar,Nungki Anggorowati, Pramesti G Dewi, Dwi Indaryati,Indarto Sulistiyono, Susan Simanjaya, Dyah A Puspitarani |

| | | | |
|---|---|---|---|
| EPI_ISL_2534535, EPI_ISL_2534543, EPI_ISL_2534547 | Aegis Sciences Corporation | Centers for Disease Control and Prevention Division of Viral Diseases, Pathogen Discovery | Dakota Howard, Dhwani Batra, Peter W. Cook, Kara Moser, Adrian Paskey, Jason Caravas, Benjamin Rambo-Martin, Shatavia Morrison, Christopher Gulvick, Scott Sammons, Yvette Unoarumhi, Darlene Wagner, Matthew Schmerer, Cyndi Clark, Patrick Campbell, Rob Case, Vikramsinha Ghorpade, Holly Houdeshell, Ola Kvalvaag, Dillon Nall, Ethan Sanders, Alec Vest, Shaun Westlund, Matthew Hardison, Clinton R. Paden, Duncan MacCannell |
| EPI_ISL_2534584, EPI_ISL_2534586, EPI_ISL_2534587 | Area of Virology, Serology and Virology Division (SAViD), New South Wales Health Pathology Randwick | Virology Research Laboratory; Area of Virology, Serology and Virology Division (SAViD), New South Wales Health Pathology Randwick | Foster, C.; Au, J.; Ruiz Silva, M.; Deveson, I.; Bull, R.; Van Hal, S.; Rawlinson, W. |
| EPI_ISL_2535265 | Labo Analyses Med | National Reference Center for Viruses of Respiratory Infections, Institut Pasteur, Paris | Marion Barbet, Sylvie Behillil, Méline Bizard, Angela Brisebarre, Camille Capel, Vincent Enouf, Louise Lefrançois, Frédéric Lemoine, Christophe Malabat, Corinne Maufrais, Etienne Simon-Lorière, Maud Vanpeene, Sylvie Van der Werf ,Hugues Leroy |
| EPI_ISL_2535369, EPI_ISL_2535411, EPI_ISL_2535417, EPI_ISL_2535433 | Institute of Virology, Medical Center, University of Freiburg, Freiburg, Germany | Institute of Virology, Clinial Virus Genomics, Medical Center, University of Freiburg, Freiburg, Germany | Jonas Fuchs, Lisa Kern, Lena Jaki, Sandra Reuter, Hajo Grundmann, Marcus Panning |
| EPI_ISL_2535475 | Hospital | National Reference Center for Viruses of Respiratory Infections, Institut Pasteur, Paris | Marion Barbet, Sylvie Behillil, Méline Bizard, Angela Brisebarre, Camille Capel, Vincent Enouf, Louise Lefrançois, Frédéric Lemoine, Christophe Malabat, Corinne Maufrais, Etienne Simon-Lorière, Maud Vanpeene, Sylvie Van der Werf ,LéA Pilorge |
| EPI_ISL_2535862, EPI_ISL_2535879, EPI_ISL_2535995 | Gravity Diagnostics, LLC | Gravity Diagnostics, LLC | Gravity Diagnostics |
| EPI_ISL_2536040 | INSA | Instituto Nacional de Saude (INSA) | Borges et al |
| EPI_ISL_2536047, EPI_ISL_2536048, EPI_ISL_2536049, EPI_ISL_2536050, EPI_ISL_2536051, EPI_ISL_2536052, EPI_ISL_2536053, EPI_ISL_2536054, EPI_ISL_2536055, EPI_ISL_2536056, EPI_ISL_2536058, EPI_ISL_2536059, EPI_ISL_2536060, EPI_ISL_2536061, EPI_ISL_2536062, EPI_ISL_2536063, EPI_ISL_2536064, EPI_ISL_2536065, EPI_ISL_2536066, EPI_ISL_2536067, EPI_ISL_2536068, EPI_ISL_2536069, EPI_ISL_2536070, EPI_ISL_2536072, EPI_ISL_2536073, EPI_ISL_2536074, EPI_ISL_2536075, EPI_ISL_2536076, EPI_ISL_2536077, EPI_ISL_2536078, EPI_ISL_2536079, EPI_ISL_2536080, EPI_ISL_2536081, EPI_ISL_2536082 | | | |
| see above | CHLC | Instituto Nacional de Saude (INSA) | Borges et al |
| EPI_ISL_2536083 | CHLN | Instituto Nacional de Saude (INSA) | Borges et al |
| EPI_ISL_2536089 | Cintramedica | Instituto Nacional de Saude (INSA) | Borges et al |
| EPI_ISL_2536091, EPI_ISL_2536092, EPI_ISL_2536093, EPI_ISL_2536094, EPI_ISL_2536095, EPI_ISL_2536096 | Germano de Sousa | Instituto Nacional de Saude (INSA) | Borges et al |
| EPI_ISL_2536098 | H Forcas Armadas - Polo Lisboa | Instituto Nacional de Saude (INSA) | Borges et al |
| EPI_ISL_2536099 | IMM | Instituto Nacional de Saude (INSA) | Borges et al |
| EPI_ISL_2536100, EPI_ISL_2536101 | INSA | Instituto Nacional de Saude (INSA) | Borges et al |
| EPI_ISL_2536104 | Joaquim Chaves Saude | Instituto Nacional de Saude (INSA) | Borges et al |
| EPI_ISL_2536105, EPI_ISL_2536106, EPI_ISL_2536107 | Labeto - CAB - Leiria | Instituto Nacional de Saude (INSA) | Borges et al |
| EPI_ISL_2536115, EPI_ISL_2536117, EPI_ISL_2536118, EPI_ISL_2536119, EPI_ISL_2536120 | SYNLAB | Instituto Nacional de Saude (INSA) | Borges et al |
| EPI_ISL_2536122, EPI_ISL_2536123, EPI_ISL_2536124, EPI_ISL_2536125, EPI_ISL_2536126, EPI_ISL_2536127, EPI_ISL_2536128, EPI_ISL_2536129 | LAC UCoimbra | Instituto Nacional de Saude (INSA) | Borges et al |
| EPI_ISL_2536130 | Instituto Nacional de Saude (INSA) | Instituto Nacional de Saude (INSA) | Borges et al |
| EPI_ISL_2536199, EPI_ISL_2536207 | Laboratorio HUB -Azienda Ospedaliero Universitaria - AOU – Cagliari | Laboratorio SPOKE Biologia Molecolare -Azienda Ospedaliero Universitaria - AOU – Cagliari | Germano Orrù, Sara Fais, Valentina Medda, Alessandra Scano, Miriam Loddo, Riccardo Cappai, Ferdinando Coghe |
| EPI_ISL_2536217, EPI_ISL_2536218 | SISP Lecce | University of Bari Biomedical Sciences and Human Oncology | Maria Chironna |
| EPI_ISL_2536220 | SISP Lecce | University of Bari Biomedical Sciences and Human Oncology | Chironna M., Sallustio A., Loconsole D., Accogli M. |
| EPI_ISL_2536630, EPI_ISL_2536633 | Overlake Hospital | Seattle Flu Study | Deborah A. Nickerson, Chris D. Frazar, Jover Lee, Benjamin Pelle, Erica Ryke, Matthew Richardson, Amanda Adler, Elisabeth Brandstetter, Peter D. Han, Kairsten Fay, Misja Ilcisin, Kirsten Lacombe, Thomas R. Sibley, Melissa Truong, Caitlin R. Wolf, Romesh Gautom, Geoff Melly, Brian Hiatt, Philip Dykema, Scott Lindquist, Michael Boeckh, Janet A. Englund, Michael Famulare, Barry R. Lutz, Mark J. Rieder, Lea M. Starita, Matthew Thompson, Helen Y. Chu, Jay Shendure, Trevor Bedford |
| EPI_ISL_2536907 | HealthPartners Central Lab | Minnesota Department of Health, Public Health Laboratory | Alexandra Lorentz, Jacob Garfin, Matt Plumb, and Xiong Wang |
| EPI_ISL_2537134, EPI_ISL_2537185, EPI_ISL_2537240, EPI_ISL_2537254, EPI_ISL_2537262, EPI_ISL_2537263, EPI_ISL_2537266 | Gravity Diagnostics, LLC | Gravity Diagnostics, LLC | Gravity Diagnostics |
| EPI_ISL_2537356, EPI_ISL_2537358, EPI_ISL_2537373, EPI_ISL_2537379, EPI_ISL_2537383, EPI_ISL_2537384 | Montana Public Health Laboratory | Montana Public Health Laboratory | Joy Ritter, Michelle Mozer, Carrie Biskupiak, Deborah Gibson, Michael Dills |
| EPI_ISL_2537412, EPI_ISL_2537461 | California Department of Public Health | California Department of Public Health | CDPH IDLB COVIDNet et al |
| EPI_ISL_2537480 | Laboratori de Referencia de Catalunya | Laboratori de Referencia de Catalunya | Ramirez A., Bellosillo B., Padilla E., Vilas A., Hernandez JJ., Canal M. |
| EPI_ISL_2537486, EPI_ISL_2537488, EPI_ISL_2537489, EPI_ISL_2537491, EPI_ISL_2537492 | Balai Besar Penelitian dan Pengembangan dan Reservoir Penyakit, Salatiga | Genomik Solidaritas Indonesia Laboratorium | Carissa Sintca Wijaya, Vania Gavrila Wikasa, Dhahlia Agustina Cahyono, Alfin Mohammad Abdillah, Normastuti Adhini Tantyo, Yudistira Aditya Rahmat, Joko Waluyo, Annisa Muthiah Sukirman, Gracia Felias Enos Korompis, Pratiwi Sudarmono, Budi Wiweko, Louisa Markus, Anuraj Shankar, Meutia Ayuputeri Kumaheri, Anindya Pradipta Susanto, Ariel Pradipta |
| EPI_ISL_2537493 | Lighthouse Lab in Milton Keynes | Wellcome Sanger Institute for the COVID-19 Genomics UK (COG-UK) Consortium | The Lighthouse Lab in Milton Keynes and Alex Alderton, Roberto Amato, Jeffrey Barrett, Sonia Goncalves, Ewan Harrison, David K. Jackson, Ian Johnston, Dominic Kwiatkowski, Cordelia Langford, John Sillitoe on behalf of the Wellcome Sanger Institute COVID-19 Surveillance Team |
| EPI_ISL_2537494 | Health Services Laboratories | Wellcome Sanger Institute for the COVID-19 Genomics UK (COG-UK) Consortium | Health Services Laboratories and Alex Alderton, Roberto Amato, Jeffrey Barrett, Sonia Goncalves, Ewan Harrison, David K. Jackson, Ian Johnston, Dominic Kwiatkowski, Cordelia Langford, John Sillitoe on behalf of the Wellcome Sanger Institute COVID-19 Surveillance Team |
| EPI_ISL_2537495 | Lighthouse Lab in Milton Keynes | Wellcome Sanger Institute for the COVID-19 Genomics UK (COG-UK) Consortium | The Lighthouse Lab in Milton Keynes and Alex Alderton, Roberto Amato, Jeffrey Barrett, Sonia Goncalves, Ewan Harrison, David K. Jackson, Ian Johnston, Dominic Kwiatkowski, Cordelia Langford, John Sillitoe on behalf of the Wellcome Sanger Institute COVID-19 Surveillance Team |
| EPI_ISL_2537496 | Lighthouse Lab in Alderley Park | Wellcome Sanger Institute for the COVID-19 Genomics UK (COG-UK) Consortium | Jacquelyn Wynn, Mairead Hyland, The Lighthouse Lab in Alderley Park and Alex Alderton, Roberto Amato, Jeffrey Barrett, Sonia Goncalves, Ewan Harrison, David K. Jackson, Ian Johnston, Dominic Kwiatkowski, Cordelia Langford, John Sillitoe on behalf of the Wellcome Sanger Institute COVID-19 Surveillance Team |
| EPI_ISL_2537497, EPI_ISL_2537498 | Lighthouse Lab in Milton Keynes | Wellcome Sanger Institute for the COVID-19 Genomics UK (COG-UK) Consortium | The Lighthouse Lab in Milton Keynes and Alex Alderton, Roberto Amato, Jeffrey Barrett, Sonia Goncalves, Ewan Harrison, David K. Jackson, Ian Johnston, Dominic Kwiatkowski, Cordelia Langford, John Sillitoe on behalf of the Wellcome Sanger Institute COVID-19 Surveillance Team |
| EPI_ISL_2537500, EPI_ISL_2537502, EPI_ISL_2537503 | Lighthouse Lab in Alderley Park | Wellcome Sanger Institute for the COVID-19 Genomics UK (COG-UK) Consortium | Jacquelyn Wynn, Mairead Hyland, The Lighthouse Lab in Alderley Park and Alex Alderton, Roberto Amato, Jeffrey Barrett, Sonia Goncalves, Ewan Harrison, David K. Jackson, Ian Johnston, Dominic Kwiatkowski, Cordelia Langford, John Sillitoe on behalf of the Wellcome Sanger Institute COVID-19 Surveillance Team |
| EPI_ISL_2537504 | Lighthouse Lab in Milton Keynes | Wellcome Sanger Institute for the COVID-19 Genomics UK (COG-UK) | The Lighthouse Lab in Milton Keynes and Alex Alderton, Roberto Amato, Jeffrey Barrett, Sonia Goncalves, Ewan Harrison, David K. Jackson, Ian Johnston, Dominic Kwiatkowski, Cordelia Langford, John Sillitoe on behalf of the Wellcome Sanger |

| | | | |
|---|---|---|---|
| | | Consortium | Institute COVID-19 Surveillance Team |
| EPI_ISL_2537505 | Lighthouse Lab in Alderley Park | Wellcome Sanger Institute for the COVID-19 Genomics UK (COG-UK) Consortium | Jacquelyn Wynn, Mairead Hyland, The Lighthouse Lab in Alderley Park and Alex Alderton, Roberto Amato, Jeffrey Barrett, Sonia Goncalves, Ewan Harrison, David K. Jackson, Ian Johnston, Dominic Kwiatkowski, Cordelia Langford, John Sillitoe on behalf of the Wellcome Sanger Institute COVID-19 Surveillance Team |
| EPI_ISL_2537507 | Lighthouse Lab in Milton Keynes | Wellcome Sanger Institute for the COVID-19 Genomics UK (COG-UK) Consortium | The Lighthouse Lab in Milton Keynes and Alex Alderton, Roberto Amato, Jeffrey Barrett, Sonia Goncalves, Ewan Harrison, David K. Jackson, Ian Johnston, Dominic Kwiatkowski, Cordelia Langford, John Sillitoe on behalf of the Wellcome Sanger Institute COVID-19 Surveillance Team |
| EPI_ISL_2537508 | Lighthouse Lab in Alderley Park | Wellcome Sanger Institute for the COVID-19 Genomics UK (COG-UK) Consortium | Jacquelyn Wynn, Mairead Hyland, The Lighthouse Lab in Alderley Park and Alex Alderton, Roberto Amato, Jeffrey Barrett, Sonia Goncalves, Ewan Harrison, David K. Jackson, Ian Johnston, Dominic Kwiatkowski, Cordelia Langford, John Sillitoe on behalf of the Wellcome Sanger Institute COVID-19 Surveillance Team |
| EPI_ISL_2537510 | Health Services Laboratories | Wellcome Sanger Institute for the COVID-19 Genomics UK (COG-UK) Consortium | Health Services Laboratories and Alex Alderton, Roberto Amato, Jeffrey Barrett, Sonia Goncalves, Ewan Harrison, David K. Jackson, Ian Johnston, Dominic Kwiatkowski, Cordelia Langford, John Sillitoe on behalf of the Wellcome Sanger Institute COVID-19 Surveillance Team |
| EPI_ISL_2537511 | Lighthouse Lab in Milton Keynes | Wellcome Sanger Institute for the COVID-19 Genomics UK (COG-UK) Consortium | The Lighthouse Lab in Milton Keynes and Alex Alderton, Roberto Amato, Jeffrey Barrett, Sonia Goncalves, Ewan Harrison, David K. Jackson, Ian Johnston, Dominic Kwiatkowski, Cordelia Langford, John Sillitoe on behalf of the Wellcome Sanger Institute COVID-19 Surveillance Team |
| EPI_ISL_2537512 | Lighthouse Lab in Alderley Park | Wellcome Sanger Institute for the COVID-19 Genomics UK (COG-UK) Consortium | Jacquelyn Wynn, Mairead Hyland, The Lighthouse Lab in Alderley Park and Alex Alderton, Roberto Amato, Jeffrey Barrett, Sonia Goncalves, Ewan Harrison, David K. Jackson, Ian Johnston, Dominic Kwiatkowski, Cordelia Langford, John Sillitoe on behalf of the Wellcome Sanger Institute COVID-19 Surveillance Team |
| EPI_ISL_2537514 | Lighthouse Lab in Milton Keynes | Wellcome Sanger Institute for the COVID-19 Genomics UK (COG-UK) Consortium | The Lighthouse Lab in Milton Keynes and Alex Alderton, Roberto Amato, Jeffrey Barrett, Sonia Goncalves, Ewan Harrison, David K. Jackson, Ian Johnston, Dominic Kwiatkowski, Cordelia Langford, John Sillitoe on behalf of the Wellcome Sanger Institute COVID-19 Surveillance Team |
| EPI_ISL_2537515 | Health Services Laboratories | Wellcome Sanger Institute for the COVID-19 Genomics UK (COG-UK) Consortium | Health Services Laboratories and Alex Alderton, Roberto Amato, Jeffrey Barrett, Sonia Goncalves, Ewan Harrison, David K. Jackson, Ian Johnston, Dominic Kwiatkowski, Cordelia Langford, John Sillitoe on behalf of the Wellcome Sanger Institute COVID-19 Surveillance Team |
| EPI_ISL_2537516, EPI_ISL_2537517, EPI_ISL_2537518 | Lighthouse Lab in Milton Keynes | Wellcome Sanger Institute for the COVID-19 Genomics UK (COG-UK) Consortium | The Lighthouse Lab in Milton Keynes and Alex Alderton, Roberto Amato, Jeffrey Barrett, Sonia Goncalves, Ewan Harrison, David K. Jackson, Ian Johnston, Dominic Kwiatkowski, Cordelia Langford, John Sillitoe on behalf of the Wellcome Sanger Institute COVID-19 Surveillance Team |
| EPI_ISL_2537519 | Lighthouse Lab in Alderley Park | Wellcome Sanger Institute for the COVID-19 Genomics UK (COG-UK) Consortium | Jacquelyn Wynn, Mairead Hyland, The Lighthouse Lab in Alderley Park and Alex Alderton, Roberto Amato, Jeffrey Barrett, Sonia Goncalves, Ewan Harrison, David K. Jackson, Ian Johnston, Dominic Kwiatkowski, Cordelia Langford, John Sillitoe on behalf of the Wellcome Sanger Institute COVID-19 Surveillance Team |
| EPI_ISL_2537520, EPI_ISL_2537521 | Lighthouse Lab in Milton Keynes | Wellcome Sanger Institute for the COVID-19 Genomics UK (COG-UK) Consortium | The Lighthouse Lab in Milton Keynes and Alex Alderton, Roberto Amato, Jeffrey Barrett, Sonia Goncalves, Ewan Harrison, David K. Jackson, Ian Johnston, Dominic Kwiatkowski, Cordelia Langford, John Sillitoe on behalf of the Wellcome Sanger Institute COVID-19 Surveillance Team |
| EPI_ISL_2537522 | Health Services Laboratories | Wellcome Sanger Institute for the COVID-19 Genomics UK (COG-UK) Consortium | Health Services Laboratories and Alex Alderton, Roberto Amato, Jeffrey Barrett, Sonia Goncalves, Ewan Harrison, David K. Jackson, Ian Johnston, Dominic Kwiatkowski, Cordelia Langford, John Sillitoe on behalf of the Wellcome Sanger Institute COVID-19 Surveillance Team |
| EPI_ISL_2537523 | Lighthouse Lab in Alderley Park | Wellcome Sanger Institute for the COVID-19 Genomics UK (COG-UK) Consortium | Jacquelyn Wynn, Mairead Hyland, The Lighthouse Lab in Alderley Park and Alex Alderton, Roberto Amato, Jeffrey Barrett, Sonia Goncalves, Ewan Harrison, David K. Jackson, Ian Johnston, Dominic Kwiatkowski, Cordelia Langford, John Sillitoe on behalf of the Wellcome Sanger Institute COVID-19 Surveillance Team |
| EPI_ISL_2537524, EPI_ISL_2537526 | Lighthouse Lab in Milton Keynes | Wellcome Sanger Institute for the COVID-19 Genomics UK (COG-UK) Consortium | The Lighthouse Lab in Milton Keynes and Alex Alderton, Roberto Amato, Jeffrey Barrett, Sonia Goncalves, Ewan Harrison, David K. Jackson, Ian Johnston, Dominic Kwiatkowski, Cordelia Langford, John Sillitoe on behalf of the Wellcome Sanger Institute COVID-19 Surveillance Team |
| EPI_ISL_2537527 | Lighthouse Lab in Alderley Park | Wellcome Sanger Institute for the COVID-19 Genomics UK (COG-UK) Consortium | Jacquelyn Wynn, Mairead Hyland, The Lighthouse Lab in Alderley Park and Alex Alderton, Roberto Amato, Jeffrey Barrett, Sonia Goncalves, Ewan Harrison, David K. Jackson, Ian Johnston, Dominic Kwiatkowski, Cordelia Langford, John Sillitoe on behalf of the Wellcome Sanger Institute COVID-19 Surveillance Team |
| EPI_ISL_2537528, EPI_ISL_2537529, EPI_ISL_2537530, EPI_ISL_2537531 | Lighthouse Lab in Milton Keynes | Wellcome Sanger Institute for the COVID-19 Genomics UK (COG-UK) Consortium | The Lighthouse Lab in Milton Keynes and Alex Alderton, Roberto Amato, Jeffrey Barrett, Sonia Goncalves, Ewan Harrison, David K. Jackson, Ian Johnston, Dominic Kwiatkowski, Cordelia Langford, John Sillitoe on behalf of the Wellcome Sanger Institute COVID-19 Surveillance Team |
| EPI_ISL_2537532 | Health Services Laboratories | Wellcome Sanger Institute for the COVID-19 Genomics UK (COG-UK) Consortium | Health Services Laboratories and Alex Alderton, Roberto Amato, Jeffrey Barrett, Sonia Goncalves, Ewan Harrison, David K. Jackson, Ian Johnston, Dominic Kwiatkowski, Cordelia Langford, John Sillitoe on behalf of the Wellcome Sanger Institute COVID-19 Surveillance Team |
| EPI_ISL_2537533 | Lighthouse Lab in Alderley Park | Wellcome Sanger Institute for the COVID-19 Genomics UK (COG-UK) Consortium | Jacquelyn Wynn, Mairead Hyland, The Lighthouse Lab in Alderley Park and Alex Alderton, Roberto Amato, Jeffrey Barrett, Sonia Goncalves, Ewan Harrison, David K. Jackson, Ian Johnston, Dominic Kwiatkowski, Cordelia Langford, John Sillitoe on behalf of the Wellcome Sanger Institute COVID-19 Surveillance Team |
| EPI_ISL_2537534, EPI_ISL_2537535 | Lighthouse Lab in Milton Keynes | Wellcome Sanger Institute for the COVID-19 Genomics UK (COG-UK) Consortium | The Lighthouse Lab in Milton Keynes and Alex Alderton, Roberto Amato, Jeffrey Barrett, Sonia Goncalves, Ewan Harrison, David K. Jackson, Ian Johnston, Dominic Kwiatkowski, Cordelia Langford, John Sillitoe on behalf of the Wellcome Sanger Institute COVID-19 Surveillance Team |
| EPI_ISL_2537536 | Lighthouse Lab in Alderley Park | Wellcome Sanger Institute for the COVID-19 Genomics UK (COG-UK) Consortium | Jacquelyn Wynn, Mairead Hyland, The Lighthouse Lab in Alderley Park and Alex Alderton, Roberto Amato, Jeffrey Barrett, Sonia Goncalves, Ewan Harrison, David K. Jackson, Ian Johnston, Dominic Kwiatkowski, Cordelia Langford, John Sillitoe on behalf of the Wellcome Sanger Institute COVID-19 Surveillance Team |
| EPI_ISL_2537537 | Lighthouse Lab in Milton Keynes | Wellcome Sanger Institute for the COVID-19 Genomics UK (COG-UK) Consortium | The Lighthouse Lab in Milton Keynes and Alex Alderton, Roberto Amato, Jeffrey Barrett, Sonia Goncalves, Ewan Harrison, David K. Jackson, Ian Johnston, Dominic Kwiatkowski, Cordelia Langford, John Sillitoe on behalf of the Wellcome Sanger Institute COVID-19 Surveillance Team |
| EPI_ISL_2537538 | Lighthouse Lab in Alderley Park | Wellcome Sanger Institute for the COVID-19 Genomics UK (COG-UK) Consortium | Jacquelyn Wynn, Mairead Hyland, The Lighthouse Lab in Alderley Park and Alex Alderton, Roberto Amato, Jeffrey Barrett, Sonia Goncalves, Ewan Harrison, David K. Jackson, Ian Johnston, Dominic Kwiatkowski, Cordelia Langford, John Sillitoe on behalf of the Wellcome Sanger Institute COVID-19 Surveillance Team |
| EPI_ISL_2537539, EPI_ISL_2537540, EPI_ISL_2537541 | Lighthouse Lab in Milton Keynes | Wellcome Sanger Institute for the COVID-19 Genomics UK (COG-UK) Consortium | The Lighthouse Lab in Milton Keynes and Alex Alderton, Roberto Amato, Jeffrey Barrett, Sonia Goncalves, Ewan Harrison, David K. Jackson, Ian Johnston, Dominic Kwiatkowski, Cordelia Langford, John Sillitoe on behalf of the Wellcome Sanger Institute COVID-19 Surveillance Team |
| EPI_ISL_2537542, EPI_ISL_2537543 | Lighthouse Lab in Alderley Park | Wellcome Sanger Institute for the COVID-19 Genomics UK (COG-UK) Consortium | Jacquelyn Wynn, Mairead Hyland, The Lighthouse Lab in Alderley Park and Alex Alderton, Roberto Amato, Jeffrey Barrett, Sonia Goncalves, Ewan Harrison, David K. Jackson, Ian Johnston, Dominic Kwiatkowski, Cordelia Langford, John Sillitoe on behalf of the Wellcome Sanger Institute COVID-19 Surveillance Team |
| EPI_ISL_2537544, EPI_ISL_2537545, EPI_ISL_2537546, EPI_ISL_2537548 | Lighthouse Lab in Milton Keynes | Wellcome Sanger Institute for the COVID-19 Genomics UK (COG-UK) Consortium | The Lighthouse Lab in Milton Keynes and Alex Alderton, Roberto Amato, Jeffrey Barrett, Sonia Goncalves, Ewan Harrison, David K. Jackson, Ian Johnston, Dominic Kwiatkowski, Cordelia Langford, John Sillitoe on behalf of the Wellcome Sanger Institute COVID-19 Surveillance Team |
| EPI_ISL_2537549, EPI_ISL_2537550 | Lighthouse Lab in Alderley Park | Wellcome Sanger Institute for the COVID-19 Genomics UK (COG-UK) Consortium | Jacquelyn Wynn, Mairead Hyland, The Lighthouse Lab in Alderley Park and Alex Alderton, Roberto Amato, Jeffrey Barrett, Sonia Goncalves, Ewan Harrison, David K. Jackson, Ian Johnston, Dominic Kwiatkowski, Cordelia Langford, John Sillitoe on behalf of the Wellcome Sanger Institute COVID-19 Surveillance Team |
| EPI_ISL_2537551, EPI_ISL_2537552, EPI_ISL_2537554 | Lighthouse Lab in Milton Keynes | Wellcome Sanger Institute for the COVID-19 Genomics UK (COG-UK) Consortium | The Lighthouse Lab in Milton Keynes and Alex Alderton, Roberto Amato, Jeffrey Barrett, Sonia Goncalves, Ewan Harrison, David K. Jackson, Ian Johnston, Dominic Kwiatkowski, Cordelia Langford, John Sillitoe on behalf of the Wellcome Sanger Institute COVID-19 Surveillance Team |
| EPI_ISL_2537555, EPI_ISL_2537556, EPI_ISL_2537557 | Health Services Laboratories | Wellcome Sanger Institute for the COVID-19 Genomics UK (COG-UK) Consortium | Health Services Laboratories and Alex Alderton, Roberto Amato, Jeffrey Barrett, Sonia Goncalves, Ewan Harrison, David K. Jackson, Ian Johnston, Dominic Kwiatkowski, Cordelia Langford, John Sillitoe on behalf of the Wellcome Sanger Institute COVID-19 Surveillance Team |
| EPI_ISL_2537558, EPI_ISL_2537559, EPI_ISL_2537560 | Lighthouse Lab in Alderley Park | Wellcome Sanger Institute for the COVID-19 Genomics UK (COG-UK) Consortium | Jacquelyn Wynn, Mairead Hyland, The Lighthouse Lab in Alderley Park and Alex Alderton, Roberto Amato, Jeffrey Barrett, Sonia Goncalves, Ewan Harrison, David K. Jackson, Ian Johnston, Dominic Kwiatkowski, Cordelia Langford, John Sillitoe on behalf of the Wellcome Sanger Institute COVID-19 Surveillance Team |
| EPI_ISL_2537561 | Health Services Laboratories | Wellcome Sanger Institute for the COVID-19 Genomics UK (COG-UK) Consortium | Health Services Laboratories and Alex Alderton, Roberto Amato, Jeffrey Barrett, Sonia Goncalves, Ewan Harrison, David K. Jackson, Ian Johnston, Dominic Kwiatkowski, Cordelia Langford, John Sillitoe on behalf of the Wellcome Sanger Institute COVID-19 Surveillance Team |
| EPI_ISL_2537562, EPI_ISL_2537563 | Lighthouse Lab in Milton Keynes | Wellcome Sanger Institute for the COVID-19 Genomics UK (COG-UK) Consortium | The Lighthouse Lab in Milton Keynes and Alex Alderton, Roberto Amato, Jeffrey Barrett, Sonia Goncalves, Ewan Harrison, David K. Jackson, Ian Johnston, Dominic Kwiatkowski, Cordelia Langford, John Sillitoe on behalf of the Wellcome Sanger Institute COVID-19 Surveillance Team |
| EPI_ISL_2537564 | Lighthouse Lab in Alderley Park | Wellcome Sanger Institute for the COVID-19 Genomics UK (COG-UK) Consortium | Jacquelyn Wynn, Mairead Hyland, The Lighthouse Lab in Alderley Park and Alex Alderton, Roberto Amato, Jeffrey Barrett, Sonia Goncalves, Ewan Harrison, David K. Jackson, Ian Johnston, Dominic Kwiatkowski, Cordelia Langford, John Sillitoe on behalf of the Wellcome Sanger Institute COVID-19 Surveillance Team |
| EPI_ISL_2537565, EPI_ISL_2537566 | Lighthouse Lab in Milton Keynes | Wellcome Sanger Institute for the COVID-19 Genomics UK (COG-UK) Consortium | The Lighthouse Lab in Milton Keynes and Alex Alderton, Roberto Amato, Jeffrey Barrett, Sonia Goncalves, Ewan Harrison, David K. Jackson, Ian Johnston, Dominic Kwiatkowski, Cordelia Langford, John Sillitoe on behalf of the Wellcome Sanger Institute COVID-19 Surveillance Team |
| EPI_ISL_2537567 | Health Services Laboratories | Wellcome Sanger Institute for the COVID-19 Genomics UK (COG-UK) Consortium | Health Services Laboratories and Alex Alderton, Roberto Amato, Jeffrey Barrett, Sonia Goncalves, Ewan Harrison, David K. Jackson, Ian Johnston, Dominic Kwiatkowski, Cordelia Langford, John Sillitoe on behalf of the Wellcome Sanger Institute COVID-19 Surveillance Team |
| EPI_ISL_2537568 | Lighthouse Lab in Alderley Park | Wellcome Sanger Institute for the COVID-19 Genomics UK (COG-UK) Consortium | Jacquelyn Wynn, Mairead Hyland, The Lighthouse Lab in Alderley Park and Alex Alderton, Roberto Amato, Jeffrey Barrett, Sonia Goncalves, Ewan Harrison, David K. Jackson, Ian Johnston, Dominic Kwiatkowski, Cordelia Langford, John Sillitoe on behalf of the Wellcome Sanger Institute COVID-19 Surveillance Team |
| EPI_ISL_2537569 | Lighthouse Lab in Milton Keynes | Wellcome Sanger Institute for the COVID-19 Genomics UK (COG-UK) Consortium | The Lighthouse Lab in Milton Keynes and Alex Alderton, Roberto Amato, Jeffrey Barrett, Sonia Goncalves, Ewan Harrison, David K. Jackson, Ian Johnston, Dominic Kwiatkowski, Cordelia Langford, John Sillitoe on behalf of the Wellcome Sanger Institute COVID-19 Surveillance Team |
| EPI_ISL_2537570 | Health Services Laboratories | Wellcome Sanger Institute for the COVID-19 Genomics UK (COG-UK) Consortium | Health Services Laboratories and Alex Alderton, Roberto Amato, Jeffrey Barrett, Sonia Goncalves, Ewan Harrison, David K. Jackson, Ian Johnston, Dominic Kwiatkowski, Cordelia Langford, John Sillitoe on behalf of the Wellcome Sanger Institute COVID-19 Surveillance Team |
| EPI_ISL_2537571, EPI_ISL_2537572, EPI_ISL_2537573, EPI_ISL_2537574, EPI_ISL_2537575 | Lighthouse Lab in Milton Keynes | Wellcome Sanger Institute for the COVID-19 Genomics UK (COG-UK) Consortium | The Lighthouse Lab in Milton Keynes and Alex Alderton, Roberto Amato, Jeffrey Barrett, Sonia Goncalves, Ewan Harrison, David K. Jackson, Ian Johnston, Dominic Kwiatkowski, Cordelia Langford, John Sillitoe on behalf of the Wellcome Sanger Institute COVID-19 Surveillance Team |
| EPI_ISL_2537576 | Health Services Laboratories | Wellcome Sanger Institute for the COVID-19 Genomics UK (COG-UK) Consortium | Health Services Laboratories and Alex Alderton, Roberto Amato, Jeffrey Barrett, Sonia Goncalves, Ewan Harrison, David K. Jackson, Ian Johnston, Dominic Kwiatkowski, Cordelia Langford, John Sillitoe on behalf of the Wellcome Sanger Institute COVID-19 Surveillance Team |
| EPI_ISL_2537577 | Lighthouse Lab in Milton Keynes | Wellcome Sanger Institute for the COVID-19 Genomics UK (COG-UK) | The Lighthouse Lab in Milton Keynes and Alex Alderton, Roberto Amato, Jeffrey Barrett, Sonia Goncalves, Ewan Harrison, David K. Jackson, Ian Johnston, Dominic Kwiatkowski, Cordelia Langford, John Sillitoe on behalf of the Wellcome Sanger |

| | | | |
|---|---|---|---|
| | | Consortium | Institute COVID-19 Surveillance Team |
| EPI_ISL_2537578 | Health Services Laboratories | Wellcome Sanger Institute for the COVID-19 Genomics UK (COG-UK) Consortium | Health Services Laboratories and Alex Alderton, Roberto Amato, Jeffrey Barrett, Sonia Goncalves, Ewan Harrison, David K. Jackson, Ian Johnston, Dominic Kwiatkowski, Cordelia Langford, John Sillitoe on behalf of the Wellcome Sanger Institute COVID-19 Surveillance Team |
| EPI_ISL_2537579 | Lighthouse Lab in Alderley Park | Wellcome Sanger Institute for the COVID-19 Genomics UK (COG-UK) Consortium | Jacquelyn Wynn, Mairead Hyland, The Lighthouse Lab in Alderley Park and Alex Alderton, Roberto Amato, Jeffrey Barrett, Sonia Goncalves, Ewan Harrison, David K. Jackson, Ian Johnston, Dominic Kwiatkowski, Cordelia Langford, John Sillitoe on behalf of the Wellcome Sanger Institute COVID-19 Surveillance Team |
| EPI_ISL_2537580, EPI_ISL_2537581, EPI_ISL_2537582 | Lighthouse Lab in Milton Keynes | Wellcome Sanger Institute for the COVID-19 Genomics UK (COG-UK) Consortium | The Lighthouse Lab in Milton Keynes and Alex Alderton, Roberto Amato, Jeffrey Barrett, Sonia Goncalves, Ewan Harrison, David K. Jackson, Ian Johnston, Dominic Kwiatkowski, Cordelia Langford, John Sillitoe on behalf of the Wellcome Sanger Institute COVID-19 Surveillance Team |
| EPI_ISL_2537583 | Lighthouse Lab in Alderley Park | Wellcome Sanger Institute for the COVID-19 Genomics UK (COG-UK) Consortium | Jacquelyn Wynn, Mairead Hyland, The Lighthouse Lab in Alderley Park and Alex Alderton, Roberto Amato, Jeffrey Barrett, Sonia Goncalves, Ewan Harrison, David K. Jackson, Ian Johnston, Dominic Kwiatkowski, Cordelia Langford, John Sillitoe on behalf of the Wellcome Sanger Institute COVID-19 Surveillance Team |
| EPI_ISL_2537584 | Health Services Laboratories | Wellcome Sanger Institute for the COVID-19 Genomics UK (COG-UK) Consortium | Health Services Laboratories and Alex Alderton, Roberto Amato, Jeffrey Barrett, Sonia Goncalves, Ewan Harrison, David K. Jackson, Ian Johnston, Dominic Kwiatkowski, Cordelia Langford, John Sillitoe on behalf of the Wellcome Sanger Institute COVID-19 Surveillance Team |
| EPI_ISL_2537585, EPI_ISL_2537586, EPI_ISL_2537587 | Lighthouse Lab in Milton Keynes | Wellcome Sanger Institute for the COVID-19 Genomics UK (COG-UK) Consortium | The Lighthouse Lab in Milton Keynes and Alex Alderton, Roberto Amato, Jeffrey Barrett, Sonia Goncalves, Ewan Harrison, David K. Jackson, Ian Johnston, Dominic Kwiatkowski, Cordelia Langford, John Sillitoe on behalf of the Wellcome Sanger Institute COVID-19 Surveillance Team |
| EPI_ISL_2537588 | Lighthouse Lab in Alderley Park | Wellcome Sanger Institute for the COVID-19 Genomics UK (COG-UK) Consortium | Jacquelyn Wynn, Mairead Hyland, The Lighthouse Lab in Alderley Park and Alex Alderton, Roberto Amato, Jeffrey Barrett, Sonia Goncalves, Ewan Harrison, David K. Jackson, Ian Johnston, Dominic Kwiatkowski, Cordelia Langford, John Sillitoe on behalf of the Wellcome Sanger Institute COVID-19 Surveillance Team |
| EPI_ISL_2537589 | Health Services Laboratories | Wellcome Sanger Institute for the COVID-19 Genomics UK (COG-UK) Consortium | Health Services Laboratories and Alex Alderton, Roberto Amato, Jeffrey Barrett, Sonia Goncalves, Ewan Harrison, David K. Jackson, Ian Johnston, Dominic Kwiatkowski, Cordelia Langford, John Sillitoe on behalf of the Wellcome Sanger Institute COVID-19 Surveillance Team |
| EPI_ISL_2537590 | Lighthouse Lab in Milton Keynes | Wellcome Sanger Institute for the COVID-19 Genomics UK (COG-UK) Consortium | The Lighthouse Lab in Milton Keynes and Alex Alderton, Roberto Amato, Jeffrey Barrett, Sonia Goncalves, Ewan Harrison, David K. Jackson, Ian Johnston, Dominic Kwiatkowski, Cordelia Langford, John Sillitoe on behalf of the Wellcome Sanger Institute COVID-19 Surveillance Team |
| EPI_ISL_2537591 | Lighthouse Lab in Alderley Park | Wellcome Sanger Institute for the COVID-19 Genomics UK (COG-UK) Consortium | Jacquelyn Wynn, Mairead Hyland, The Lighthouse Lab in Alderley Park and Alex Alderton, Roberto Amato, Jeffrey Barrett, Sonia Goncalves, Ewan Harrison, David K. Jackson, Ian Johnston, Dominic Kwiatkowski, Cordelia Langford, John Sillitoe on behalf of the Wellcome Sanger Institute COVID-19 Surveillance Team |
| EPI_ISL_2537593, EPI_ISL_2537595, EPI_ISL_2537596, EPI_ISL_2537597, EPI_ISL_2537598, EPI_ISL_2537599 | Lighthouse Lab in Milton Keynes | Wellcome Sanger Institute for the COVID-19 Genomics UK (COG-UK) Consortium | The Lighthouse Lab in Milton Keynes and Alex Alderton, Roberto Amato, Jeffrey Barrett, Sonia Goncalves, Ewan Harrison, David K. Jackson, Ian Johnston, Dominic Kwiatkowski, Cordelia Langford, John Sillitoe on behalf of the Wellcome Sanger Institute COVID-19 Surveillance Team |
| EPI_ISL_2537600 | Lighthouse Lab in Alderley Park | Wellcome Sanger Institute for the COVID-19 Genomics UK (COG-UK) Consortium | Jacquelyn Wynn, Mairead Hyland, The Lighthouse Lab in Alderley Park and Alex Alderton, Roberto Amato, Jeffrey Barrett, Sonia Goncalves, Ewan Harrison, David K. Jackson, Ian Johnston, Dominic Kwiatkowski, Cordelia Langford, John Sillitoe on behalf of the Wellcome Sanger Institute COVID-19 Surveillance Team |
| EPI_ISL_2537601 | Lighthouse Lab in Milton Keynes | Wellcome Sanger Institute for the COVID-19 Genomics UK (COG-UK) Consortium | The Lighthouse Lab in Milton Keynes and Alex Alderton, Roberto Amato, Jeffrey Barrett, Sonia Goncalves, Ewan Harrison, David K. Jackson, Ian Johnston, Dominic Kwiatkowski, Cordelia Langford, John Sillitoe on behalf of the Wellcome Sanger Institute COVID-19 Surveillance Team |
| EPI_ISL_2537602 | Lighthouse Lab in Alderley Park | Wellcome Sanger Institute for the COVID-19 Genomics UK (COG-UK) Consortium | Jacquelyn Wynn, Mairead Hyland, The Lighthouse Lab in Alderley Park and Alex Alderton, Roberto Amato, Jeffrey Barrett, Sonia Goncalves, Ewan Harrison, David K. Jackson, Ian Johnston, Dominic Kwiatkowski, Cordelia Langford, John Sillitoe on behalf of the Wellcome Sanger Institute COVID-19 Surveillance Team |
| EPI_ISL_2537603 | Lighthouse Lab in Milton Keynes | Wellcome Sanger Institute for the COVID-19 Genomics UK (COG-UK) Consortium | The Lighthouse Lab in Milton Keynes and Alex Alderton, Roberto Amato, Jeffrey Barrett, Sonia Goncalves, Ewan Harrison, David K. Jackson, Ian Johnston, Dominic Kwiatkowski, Cordelia Langford, John Sillitoe on behalf of the Wellcome Sanger Institute COVID-19 Surveillance Team |
| EPI_ISL_2537604 | Health Services Laboratories | Wellcome Sanger Institute for the COVID-19 Genomics UK (COG-UK) Consortium | Health Services Laboratories and Alex Alderton, Roberto Amato, Jeffrey Barrett, Sonia Goncalves, Ewan Harrison, David K. Jackson, Ian Johnston, Dominic Kwiatkowski, Cordelia Langford, John Sillitoe on behalf of the Wellcome Sanger Institute COVID-19 Surveillance Team |
| EPI_ISL_2537605, EPI_ISL_2537606, EPI_ISL_2537607 | Lighthouse Lab in Alderley Park | Wellcome Sanger Institute for the COVID-19 Genomics UK (COG-UK) Consortium | Jacquelyn Wynn, Mairead Hyland, The Lighthouse Lab in Alderley Park and Alex Alderton, Roberto Amato, Jeffrey Barrett, Sonia Goncalves, Ewan Harrison, David K. Jackson, Ian Johnston, Dominic Kwiatkowski, Cordelia Langford, John Sillitoe on behalf of the Wellcome Sanger Institute COVID-19 Surveillance Team |
| EPI_ISL_2537608 | Lighthouse Lab in Milton Keynes | Wellcome Sanger Institute for the COVID-19 Genomics UK (COG-UK) Consortium | The Lighthouse Lab in Milton Keynes and Alex Alderton, Roberto Amato, Jeffrey Barrett, Sonia Goncalves, Ewan Harrison, David K. Jackson, Ian Johnston, Dominic Kwiatkowski, Cordelia Langford, John Sillitoe on behalf of the Wellcome Sanger Institute COVID-19 Surveillance Team |
| EPI_ISL_2537609 | Lighthouse Lab in Alderley Park | Wellcome Sanger Institute for the COVID-19 Genomics UK (COG-UK) Consortium | Jacquelyn Wynn, Mairead Hyland, The Lighthouse Lab in Alderley Park and Alex Alderton, Roberto Amato, Jeffrey Barrett, Sonia Goncalves, Ewan Harrison, David K. Jackson, Ian Johnston, Dominic Kwiatkowski, Cordelia Langford, John Sillitoe on behalf of the Wellcome Sanger Institute COVID-19 Surveillance Team |
| EPI_ISL_2537610, EPI_ISL_2537611, EPI_ISL_2537612, EPI_ISL_2537613, EPI_ISL_2537614 | Lighthouse Lab in Milton Keynes | Wellcome Sanger Institute for the COVID-19 Genomics UK (COG-UK) Consortium | The Lighthouse Lab in Milton Keynes and Alex Alderton, Roberto Amato, Jeffrey Barrett, Sonia Goncalves, Ewan Harrison, David K. Jackson, Ian Johnston, Dominic Kwiatkowski, Cordelia Langford, John Sillitoe on behalf of the Wellcome Sanger Institute COVID-19 Surveillance Team |
| EPI_ISL_2537615, EPI_ISL_2537616 | Health Services Laboratories | Wellcome Sanger Institute for the COVID-19 Genomics UK (COG-UK) Consortium | Health Services Laboratories and Alex Alderton, Roberto Amato, Jeffrey Barrett, Sonia Goncalves, Ewan Harrison, David K. Jackson, Ian Johnston, Dominic Kwiatkowski, Cordelia Langford, John Sillitoe on behalf of the Wellcome Sanger Institute COVID-19 Surveillance Team |
| EPI_ISL_2537618, EPI_ISL_2537619 | Lighthouse Lab in Milton Keynes | Wellcome Sanger Institute for the COVID-19 Genomics UK (COG-UK) Consortium | The Lighthouse Lab in Milton Keynes and Alex Alderton, Roberto Amato, Jeffrey Barrett, Sonia Goncalves, Ewan Harrison, David K. Jackson, Ian Johnston, Dominic Kwiatkowski, Cordelia Langford, John Sillitoe on behalf of the Wellcome Sanger Institute COVID-19 Surveillance Team |
| EPI_ISL_2537620, EPI_ISL_2537621 | Health Services Laboratories | Wellcome Sanger Institute for the COVID-19 Genomics UK (COG-UK) Consortium | Health Services Laboratories and Alex Alderton, Roberto Amato, Jeffrey Barrett, Sonia Goncalves, Ewan Harrison, David K. Jackson, Ian Johnston, Dominic Kwiatkowski, Cordelia Langford, John Sillitoe on behalf of the Wellcome Sanger Institute COVID-19 Surveillance Team |
| EPI_ISL_2537622 | Lighthouse Lab in Milton Keynes | Wellcome Sanger Institute for the COVID-19 Genomics UK (COG-UK) Consortium | The Lighthouse Lab in Milton Keynes and Alex Alderton, Roberto Amato, Jeffrey Barrett, Sonia Goncalves, Ewan Harrison, David K. Jackson, Ian Johnston, Dominic Kwiatkowski, Cordelia Langford, John Sillitoe on behalf of the Wellcome Sanger Institute COVID-19 Surveillance Team |
| EPI_ISL_2537623 | Lighthouse Lab in Alderley Park | Wellcome Sanger Institute for the COVID-19 Genomics UK (COG-UK) Consortium | Jacquelyn Wynn, Mairead Hyland, The Lighthouse Lab in Alderley Park and Alex Alderton, Roberto Amato, Jeffrey Barrett, Sonia Goncalves, Ewan Harrison, David K. Jackson, Ian Johnston, Dominic Kwiatkowski, Cordelia Langford, John Sillitoe on behalf of the Wellcome Sanger Institute COVID-19 Surveillance Team |
| EPI_ISL_2537624 | Health Services Laboratories | Wellcome Sanger Institute for the COVID-19 Genomics UK (COG-UK) Consortium | Health Services Laboratories and Alex Alderton, Roberto Amato, Jeffrey Barrett, Sonia Goncalves, Ewan Harrison, David K. Jackson, Ian Johnston, Dominic Kwiatkowski, Cordelia Langford, John Sillitoe on behalf of the Wellcome Sanger Institute COVID-19 Surveillance Team |
| EPI_ISL_2537625 | Lighthouse Lab in Alderley Park | Wellcome Sanger Institute for the COVID-19 Genomics UK (COG-UK) Consortium | Jacquelyn Wynn, Mairead Hyland, The Lighthouse Lab in Alderley Park and Alex Alderton, Roberto Amato, Jeffrey Barrett, Sonia Goncalves, Ewan Harrison, David K. Jackson, Ian Johnston, Dominic Kwiatkowski, Cordelia Langford, John Sillitoe on behalf of the Wellcome Sanger Institute COVID-19 Surveillance Team |
| EPI_ISL_2537626, EPI_ISL_2537627 | Lighthouse Lab in Milton Keynes | Wellcome Sanger Institute for the COVID-19 Genomics UK (COG-UK) Consortium | The Lighthouse Lab in Milton Keynes and Alex Alderton, Roberto Amato, Jeffrey Barrett, Sonia Goncalves, Ewan Harrison, David K. Jackson, Ian Johnston, Dominic Kwiatkowski, Cordelia Langford, John Sillitoe on behalf of the Wellcome Sanger Institute COVID-19 Surveillance Team |
| EPI_ISL_2537628, EPI_ISL_2537629 | Lighthouse Lab in Alderley Park | Wellcome Sanger Institute for the COVID-19 Genomics UK (COG-UK) Consortium | Jacquelyn Wynn, Mairead Hyland, The Lighthouse Lab in Alderley Park and Alex Alderton, Roberto Amato, Jeffrey Barrett, Sonia Goncalves, Ewan Harrison, David K. Jackson, Ian Johnston, Dominic Kwiatkowski, Cordelia Langford, John Sillitoe on behalf of the Wellcome Sanger Institute COVID-19 Surveillance Team |
| EPI_ISL_2537631, EPI_ISL_2537632, EPI_ISL_2537633 | Lighthouse Lab in Milton Keynes | Wellcome Sanger Institute for the COVID-19 Genomics UK (COG-UK) Consortium | The Lighthouse Lab in Milton Keynes and Alex Alderton, Roberto Amato, Jeffrey Barrett, Sonia Goncalves, Ewan Harrison, David K. Jackson, Ian Johnston, Dominic Kwiatkowski, Cordelia Langford, John Sillitoe on behalf of the Wellcome Sanger Institute COVID-19 Surveillance Team |
| EPI_ISL_2537634 | Lighthouse Lab in Alderley Park | Wellcome Sanger Institute for the COVID-19 Genomics UK (COG-UK) Consortium | Jacquelyn Wynn, Mairead Hyland, The Lighthouse Lab in Alderley Park and Alex Alderton, Roberto Amato, Jeffrey Barrett, Sonia Goncalves, Ewan Harrison, David K. Jackson, Ian Johnston, Dominic Kwiatkowski, Cordelia Langford, John Sillitoe on behalf of the Wellcome Sanger Institute COVID-19 Surveillance Team |
| EPI_ISL_2537635, EPI_ISL_2537636, EPI_ISL_2537637, EPI_ISL_2537638, EPI_ISL_2537639, EPI_ISL_2537640, EPI_ISL_2537641 | Lighthouse Lab in Milton Keynes | Wellcome Sanger Institute for the COVID-19 Genomics UK (COG-UK) Consortium | The Lighthouse Lab in Milton Keynes and Alex Alderton, Roberto Amato, Jeffrey Barrett, Sonia Goncalves, Ewan Harrison, David K. Jackson, Ian Johnston, Dominic Kwiatkowski, Cordelia Langford, John Sillitoe on behalf of the Wellcome Sanger Institute COVID-19 Surveillance Team |
| EPI_ISL_2537642 | Lighthouse Lab in Alderley Park | Wellcome Sanger Institute for the COVID-19 Genomics UK (COG-UK) Consortium | Jacquelyn Wynn, Mairead Hyland, The Lighthouse Lab in Alderley Park and Alex Alderton, Roberto Amato, Jeffrey Barrett, Sonia Goncalves, Ewan Harrison, David K. Jackson, Ian Johnston, Dominic Kwiatkowski, Cordelia Langford, John Sillitoe on behalf of the Wellcome Sanger Institute COVID-19 Surveillance Team |
| EPI_ISL_2537643 | Lighthouse Lab in Milton Keynes | Wellcome Sanger Institute for the COVID-19 Genomics UK (COG-UK) Consortium | The Lighthouse Lab in Milton Keynes and Alex Alderton, Roberto Amato, Jeffrey Barrett, Sonia Goncalves, Ewan Harrison, David K. Jackson, Ian Johnston, Dominic Kwiatkowski, Cordelia Langford, John Sillitoe on behalf of the Wellcome Sanger Institute COVID-19 Surveillance Team |
| EPI_ISL_2537644, EPI_ISL_2537645, EPI_ISL_2537646 | Lighthouse Lab in Alderley Park | Wellcome Sanger Institute for the COVID-19 Genomics UK (COG-UK) Consortium | Jacquelyn Wynn, Mairead Hyland, The Lighthouse Lab in Alderley Park and Alex Alderton, Roberto Amato, Jeffrey Barrett, Sonia Goncalves, Ewan Harrison, David K. Jackson, Ian Johnston, Dominic Kwiatkowski, Cordelia Langford, John Sillitoe on behalf of the Wellcome Sanger Institute COVID-19 Surveillance Team |
| EPI_ISL_2537647 | Lighthouse Lab in Milton Keynes | Wellcome Sanger Institute for the COVID-19 Genomics UK (COG-UK) Consortium | The Lighthouse Lab in Milton Keynes and Alex Alderton, Roberto Amato, Jeffrey Barrett, Sonia Goncalves, Ewan Harrison, David K. Jackson, Ian Johnston, Dominic Kwiatkowski, Cordelia Langford, John Sillitoe on behalf of the Wellcome Sanger Institute COVID-19 Surveillance Team |
| EPI_ISL_2537648 | Lighthouse Lab in Alderley Park | Wellcome Sanger Institute for the COVID-19 Genomics UK (COG-UK) Consortium | Jacquelyn Wynn, Mairead Hyland, The Lighthouse Lab in Alderley Park and Alex Alderton, Roberto Amato, Jeffrey Barrett, Sonia Goncalves, Ewan Harrison, David K. Jackson, Ian Johnston, Dominic Kwiatkowski, Cordelia Langford, John Sillitoe on behalf of the Wellcome Sanger Institute COVID-19 Surveillance Team |
| EPI_ISL_2537649, | Health Services Laboratories | Wellcome Sanger Institute for the COVID-19 Genomics UK (COG-UK) | Health Services Laboratories and Alex Alderton, Roberto Amato, Jeffrey Barrett, Sonia Goncalves, Ewan Harrison, David K. Jackson, Ian Johnston, Dominic Kwiatkowski, Cordelia Langford, John Sillitoe on behalf of the Wellcome Sanger Institute |

| | | | |
|---|---|---|---|
| EPI_ISL_2537650 | | Consortium | COVID-19 Surveillance Team |
| EPI_ISL_2537652 | Lighthouse Lab in Alderley Park | Wellcome Sanger Institute for the COVID-19 Genomics UK (COG-UK) Consortium | Jacquelyn Wynn, Mairead Hyland, The Lighthouse Lab in Alderley Park and Alex Alderton, Roberto Amato, Jeffrey Barrett, Sonia Goncalves, Ewan Harrison, David K. Jackson, Ian Johnston, Dominic Kwiatkowski, Cordelia Langford, John Sillitoe on behalf of the Wellcome Sanger Institute COVID-19 Surveillance Team |
| EPI_ISL_2537653, EPI_ISL_2537654 | Lighthouse Lab in Milton Keynes | Wellcome Sanger Institute for the COVID-19 Genomics UK (COG-UK) Consortium | The Lighthouse Lab in Milton Keynes and Alex Alderton, Roberto Amato, Jeffrey Barrett, Sonia Goncalves, Ewan Harrison, David K. Jackson, Ian Johnston, Dominic Kwiatkowski, Cordelia Langford, John Sillitoe on behalf of the Wellcome Sanger Institute COVID-19 Surveillance Team |
| EPI_ISL_2537655, EPI_ISL_2537656 | Lighthouse Lab in Alderley Park | Wellcome Sanger Institute for the COVID-19 Genomics UK (COG-UK) Consortium | Jacquelyn Wynn, Mairead Hyland, The Lighthouse Lab in Alderley Park and Alex Alderton, Roberto Amato, Jeffrey Barrett, Sonia Goncalves, Ewan Harrison, David K. Jackson, Ian Johnston, Dominic Kwiatkowski, Cordelia Langford, John Sillitoe on behalf of the Wellcome Sanger Institute COVID-19 Surveillance Team |
| EPI_ISL_2537657, EPI_ISL_2537658 | Lighthouse Lab in Milton Keynes | Wellcome Sanger Institute for the COVID-19 Genomics UK (COG-UK) Consortium | The Lighthouse Lab in Milton Keynes and Alex Alderton, Roberto Amato, Jeffrey Barrett, Sonia Goncalves, Ewan Harrison, David K. Jackson, Ian Johnston, Dominic Kwiatkowski, Cordelia Langford, John Sillitoe on behalf of the Wellcome Sanger Institute COVID-19 Surveillance Team |
| EPI_ISL_2537659 | Health Services Laboratories | Wellcome Sanger Institute for the COVID-19 Genomics UK (COG-UK) Consortium | Health Services Laboratories and Alex Alderton, Roberto Amato, Jeffrey Barrett, Sonia Goncalves, Ewan Harrison, David K. Jackson, Ian Johnston, Dominic Kwiatkowski, Cordelia Langford, John Sillitoe on behalf of the Wellcome Sanger Institute COVID-19 Surveillance Team |
| EPI_ISL_2537660 | Lighthouse Lab in Alderley Park | Wellcome Sanger Institute for the COVID-19 Genomics UK (COG-UK) Consortium | Jacquelyn Wynn, Mairead Hyland, The Lighthouse Lab in Alderley Park and Alex Alderton, Roberto Amato, Jeffrey Barrett, Sonia Goncalves, Ewan Harrison, David K. Jackson, Ian Johnston, Dominic Kwiatkowski, Cordelia Langford, John Sillitoe on behalf of the Wellcome Sanger Institute COVID-19 Surveillance Team |
| EPI_ISL_2537661 | Lighthouse Lab in Milton Keynes | Wellcome Sanger Institute for the COVID-19 Genomics UK (COG-UK) Consortium | The Lighthouse Lab in Milton Keynes and Alex Alderton, Roberto Amato, Jeffrey Barrett, Sonia Goncalves, Ewan Harrison, David K. Jackson, Ian Johnston, Dominic Kwiatkowski, Cordelia Langford, John Sillitoe on behalf of the Wellcome Sanger Institute COVID-19 Surveillance Team |
| EPI_ISL_2537662, EPI_ISL_2537663 | Lighthouse Lab in Alderley Park | Wellcome Sanger Institute for the COVID-19 Genomics UK (COG-UK) Consortium | Jacquelyn Wynn, Mairead Hyland, The Lighthouse Lab in Alderley Park and Alex Alderton, Roberto Amato, Jeffrey Barrett, Sonia Goncalves, Ewan Harrison, David K. Jackson, Ian Johnston, Dominic Kwiatkowski, Cordelia Langford, John Sillitoe on behalf of the Wellcome Sanger Institute COVID-19 Surveillance Team |
| EPI_ISL_2537664, EPI_ISL_2537665, EPI_ISL_2537666, EPI_ISL_2537667, EPI_ISL_2537668, EPI_ISL_2537669, EPI_ISL_2537670 | Lighthouse Lab in Milton Keynes | Wellcome Sanger Institute for the COVID-19 Genomics UK (COG-UK) Consortium | The Lighthouse Lab in Milton Keynes and Alex Alderton, Roberto Amato, Jeffrey Barrett, Sonia Goncalves, Ewan Harrison, David K. Jackson, Ian Johnston, Dominic Kwiatkowski, Cordelia Langford, John Sillitoe on behalf of the Wellcome Sanger Institute COVID-19 Surveillance Team |
| EPI_ISL_2537671 | Lighthouse Lab in Alderley Park | Wellcome Sanger Institute for the COVID-19 Genomics UK (COG-UK) Consortium | Jacquelyn Wynn, Mairead Hyland, The Lighthouse Lab in Alderley Park and Alex Alderton, Roberto Amato, Jeffrey Barrett, Sonia Goncalves, Ewan Harrison, David K. Jackson, Ian Johnston, Dominic Kwiatkowski, Cordelia Langford, John Sillitoe on behalf of the Wellcome Sanger Institute COVID-19 Surveillance Team |
| EPI_ISL_2537672, EPI_ISL_2537673, EPI_ISL_2537674 | Lighthouse Lab in Milton Keynes | Wellcome Sanger Institute for the COVID-19 Genomics UK (COG-UK) Consortium | The Lighthouse Lab in Milton Keynes and Alex Alderton, Roberto Amato, Jeffrey Barrett, Sonia Goncalves, Ewan Harrison, David K. Jackson, Ian Johnston, Dominic Kwiatkowski, Cordelia Langford, John Sillitoe on behalf of the Wellcome Sanger Institute COVID-19 Surveillance Team |
| EPI_ISL_2537675 | Lighthouse Lab in Alderley Park | Wellcome Sanger Institute for the COVID-19 Genomics UK (COG-UK) Consortium | Jacquelyn Wynn, Mairead Hyland, The Lighthouse Lab in Alderley Park and Alex Alderton, Roberto Amato, Jeffrey Barrett, Sonia Goncalves, Ewan Harrison, David K. Jackson, Ian Johnston, Dominic Kwiatkowski, Cordelia Langford, John Sillitoe on behalf of the Wellcome Sanger Institute COVID-19 Surveillance Team |
| EPI_ISL_2537676, EPI_ISL_2537677 | Lighthouse Lab in Milton Keynes | Wellcome Sanger Institute for the COVID-19 Genomics UK (COG-UK) Consortium | The Lighthouse Lab in Milton Keynes and Alex Alderton, Roberto Amato, Jeffrey Barrett, Sonia Goncalves, Ewan Harrison, David K. Jackson, Ian Johnston, Dominic Kwiatkowski, Cordelia Langford, John Sillitoe on behalf of the Wellcome Sanger Institute COVID-19 Surveillance Team |
| EPI_ISL_2537678 | Lighthouse Lab in Alderley Park | Wellcome Sanger Institute for the COVID-19 Genomics UK (COG-UK) Consortium | Jacquelyn Wynn, Mairead Hyland, The Lighthouse Lab in Alderley Park and Alex Alderton, Roberto Amato, Jeffrey Barrett, Sonia Goncalves, Ewan Harrison, David K. Jackson, Ian Johnston, Dominic Kwiatkowski, Cordelia Langford, John Sillitoe on behalf of the Wellcome Sanger Institute COVID-19 Surveillance Team |
| EPI_ISL_2537680, EPI_ISL_2537681 | Lighthouse Lab in Milton Keynes | Wellcome Sanger Institute for the COVID-19 Genomics UK (COG-UK) Consortium | The Lighthouse Lab in Milton Keynes and Alex Alderton, Roberto Amato, Jeffrey Barrett, Sonia Goncalves, Ewan Harrison, David K. Jackson, Ian Johnston, Dominic Kwiatkowski, Cordelia Langford, John Sillitoe on behalf of the Wellcome Sanger Institute COVID-19 Surveillance Team |
| EPI_ISL_2537682 | Lighthouse Lab in Alderley Park | Wellcome Sanger Institute for the COVID-19 Genomics UK (COG-UK) Consortium | Jacquelyn Wynn, Mairead Hyland, The Lighthouse Lab in Alderley Park and Alex Alderton, Roberto Amato, Jeffrey Barrett, Sonia Goncalves, Ewan Harrison, David K. Jackson, Ian Johnston, Dominic Kwiatkowski, Cordelia Langford, John Sillitoe on behalf of the Wellcome Sanger Institute COVID-19 Surveillance Team |
| EPI_ISL_2537683 | Lighthouse Lab in Milton Keynes | Wellcome Sanger Institute for the COVID-19 Genomics UK (COG-UK) Consortium | The Lighthouse Lab in Milton Keynes and Alex Alderton, Roberto Amato, Jeffrey Barrett, Sonia Goncalves, Ewan Harrison, David K. Jackson, Ian Johnston, Dominic Kwiatkowski, Cordelia Langford, John Sillitoe on behalf of the Wellcome Sanger Institute COVID-19 Surveillance Team |
| EPI_ISL_2537684, EPI_ISL_2537685 | Lighthouse Lab in Alderley Park | Wellcome Sanger Institute for the COVID-19 Genomics UK (COG-UK) Consortium | Jacquelyn Wynn, Mairead Hyland, The Lighthouse Lab in Alderley Park and Alex Alderton, Roberto Amato, Jeffrey Barrett, Sonia Goncalves, Ewan Harrison, David K. Jackson, Ian Johnston, Dominic Kwiatkowski, Cordelia Langford, John Sillitoe on behalf of the Wellcome Sanger Institute COVID-19 Surveillance Team |
| EPI_ISL_2537686, EPI_ISL_2537687, EPI_ISL_2537688, EPI_ISL_2537689, EPI_ISL_2537690 | Lighthouse Lab in Milton Keynes | Wellcome Sanger Institute for the COVID-19 Genomics UK (COG-UK) Consortium | The Lighthouse Lab in Milton Keynes and Alex Alderton, Roberto Amato, Jeffrey Barrett, Sonia Goncalves, Ewan Harrison, David K. Jackson, Ian Johnston, Dominic Kwiatkowski, Cordelia Langford, John Sillitoe on behalf of the Wellcome Sanger Institute COVID-19 Surveillance Team |
| EPI_ISL_2537691 | Lighthouse Lab in Alderley Park | Wellcome Sanger Institute for the COVID-19 Genomics UK (COG-UK) Consortium | Jacquelyn Wynn, Mairead Hyland, The Lighthouse Lab in Alderley Park and Alex Alderton, Roberto Amato, Jeffrey Barrett, Sonia Goncalves, Ewan Harrison, David K. Jackson, Ian Johnston, Dominic Kwiatkowski, Cordelia Langford, John Sillitoe on behalf of the Wellcome Sanger Institute COVID-19 Surveillance Team |
| EPI_ISL_2537692, EPI_ISL_2537693 | Lighthouse Lab in Milton Keynes | Wellcome Sanger Institute for the COVID-19 Genomics UK (COG-UK) Consortium | The Lighthouse Lab in Milton Keynes and Alex Alderton, Roberto Amato, Jeffrey Barrett, Sonia Goncalves, Ewan Harrison, David K. Jackson, Ian Johnston, Dominic Kwiatkowski, Cordelia Langford, John Sillitoe on behalf of the Wellcome Sanger Institute COVID-19 Surveillance Team |
| EPI_ISL_2537694 | Health Services Laboratories | Wellcome Sanger Institute for the COVID-19 Genomics UK (COG-UK) Consortium | Health Services Laboratories and Alex Alderton, Roberto Amato, Jeffrey Barrett, Sonia Goncalves, Ewan Harrison, David K. Jackson, Ian Johnston, Dominic Kwiatkowski, Cordelia Langford, John Sillitoe on behalf of the Wellcome Sanger Institute COVID-19 Surveillance Team |
| EPI_ISL_2537695, EPI_ISL_2537696 | Lighthouse Lab in Milton Keynes | Wellcome Sanger Institute for the COVID-19 Genomics UK (COG-UK) Consortium | The Lighthouse Lab in Milton Keynes and Alex Alderton, Roberto Amato, Jeffrey Barrett, Sonia Goncalves, Ewan Harrison, David K. Jackson, Ian Johnston, Dominic Kwiatkowski, Cordelia Langford, John Sillitoe on behalf of the Wellcome Sanger Institute COVID-19 Surveillance Team |
| EPI_ISL_2537697 | Health Services Laboratories | Wellcome Sanger Institute for the COVID-19 Genomics UK (COG-UK) Consortium | Health Services Laboratories and Alex Alderton, Roberto Amato, Jeffrey Barrett, Sonia Goncalves, Ewan Harrison, David K. Jackson, Ian Johnston, Dominic Kwiatkowski, Cordelia Langford, John Sillitoe on behalf of the Wellcome Sanger Institute COVID-19 Surveillance Team |
| EPI_ISL_2537698, EPI_ISL_2537699 | Lighthouse Lab in Alderley Park | Wellcome Sanger Institute for the COVID-19 Genomics UK (COG-UK) Consortium | Jacquelyn Wynn, Mairead Hyland, The Lighthouse Lab in Alderley Park and Alex Alderton, Roberto Amato, Jeffrey Barrett, Sonia Goncalves, Ewan Harrison, David K. Jackson, Ian Johnston, Dominic Kwiatkowski, Cordelia Langford, John Sillitoe on behalf of the Wellcome Sanger Institute COVID-19 Surveillance Team |
| EPI_ISL_2537700, EPI_ISL_2537701, EPI_ISL_2537702 | Lighthouse Lab in Milton Keynes | Wellcome Sanger Institute for the COVID-19 Genomics UK (COG-UK) Consortium | The Lighthouse Lab in Milton Keynes and Alex Alderton, Roberto Amato, Jeffrey Barrett, Sonia Goncalves, Ewan Harrison, David K. Jackson, Ian Johnston, Dominic Kwiatkowski, Cordelia Langford, John Sillitoe on behalf of the Wellcome Sanger Institute COVID-19 Surveillance Team |
| EPI_ISL_2537703, EPI_ISL_2537704 | Health Services Laboratories | Wellcome Sanger Institute for the COVID-19 Genomics UK (COG-UK) Consortium | Health Services Laboratories and Alex Alderton, Roberto Amato, Jeffrey Barrett, Sonia Goncalves, Ewan Harrison, David K. Jackson, Ian Johnston, Dominic Kwiatkowski, Cordelia Langford, John Sillitoe on behalf of the Wellcome Sanger Institute COVID-19 Surveillance Team |
| EPI_ISL_2537705 | Lighthouse Lab in Milton Keynes | Wellcome Sanger Institute for the COVID-19 Genomics UK (COG-UK) Consortium | The Lighthouse Lab in Milton Keynes and Alex Alderton, Roberto Amato, Jeffrey Barrett, Sonia Goncalves, Ewan Harrison, David K. Jackson, Ian Johnston, Dominic Kwiatkowski, Cordelia Langford, John Sillitoe on behalf of the Wellcome Sanger Institute COVID-19 Surveillance Team |
| EPI_ISL_2537707 | Lighthouse Lab in Alderley Park | Wellcome Sanger Institute for the COVID-19 Genomics UK (COG-UK) Consortium | Jacquelyn Wynn, Mairead Hyland, The Lighthouse Lab in Alderley Park and Alex Alderton, Roberto Amato, Jeffrey Barrett, Sonia Goncalves, Ewan Harrison, David K. Jackson, Ian Johnston, Dominic Kwiatkowski, Cordelia Langford, John Sillitoe on behalf of the Wellcome Sanger Institute COVID-19 Surveillance Team |
| EPI_ISL_2537708 | Lighthouse Lab in Milton Keynes | Wellcome Sanger Institute for the COVID-19 Genomics UK (COG-UK) Consortium | The Lighthouse Lab in Milton Keynes and Alex Alderton, Roberto Amato, Jeffrey Barrett, Sonia Goncalves, Ewan Harrison, David K. Jackson, Ian Johnston, Dominic Kwiatkowski, Cordelia Langford, John Sillitoe on behalf of the Wellcome Sanger Institute COVID-19 Surveillance Team |
| EPI_ISL_2537709, EPI_ISL_2537710, EPI_ISL_2537711, EPI_ISL_2537712 | Lighthouse Lab in Alderley Park | Wellcome Sanger Institute for the COVID-19 Genomics UK (COG-UK) Consortium | Jacquelyn Wynn, Mairead Hyland, The Lighthouse Lab in Alderley Park and Alex Alderton, Roberto Amato, Jeffrey Barrett, Sonia Goncalves, Ewan Harrison, David K. Jackson, Ian Johnston, Dominic Kwiatkowski, Cordelia Langford, John Sillitoe on behalf of the Wellcome Sanger Institute COVID-19 Surveillance Team |
| EPI_ISL_2537713, EPI_ISL_2537714 | Lighthouse Lab in Milton Keynes | Wellcome Sanger Institute for the COVID-19 Genomics UK (COG-UK) Consortium | The Lighthouse Lab in Milton Keynes and Alex Alderton, Roberto Amato, Jeffrey Barrett, Sonia Goncalves, Ewan Harrison, David K. Jackson, Ian Johnston, Dominic Kwiatkowski, Cordelia Langford, John Sillitoe on behalf of the Wellcome Sanger Institute COVID-19 Surveillance Team |
| EPI_ISL_2537715, EPI_ISL_2537716 | Lighthouse Lab in Alderley Park | Wellcome Sanger Institute for the COVID-19 Genomics UK (COG-UK) Consortium | Jacquelyn Wynn, Mairead Hyland, The Lighthouse Lab in Alderley Park and Alex Alderton, Roberto Amato, Jeffrey Barrett, Sonia Goncalves, Ewan Harrison, David K. Jackson, Ian Johnston, Dominic Kwiatkowski, Cordelia Langford, John Sillitoe on behalf of the Wellcome Sanger Institute COVID-19 Surveillance Team |
| EPI_ISL_2537717 | Lighthouse Lab in Milton Keynes | Wellcome Sanger Institute for the COVID-19 Genomics UK (COG-UK) Consortium | The Lighthouse Lab in Milton Keynes and Alex Alderton, Roberto Amato, Jeffrey Barrett, Sonia Goncalves, Ewan Harrison, David K. Jackson, Ian Johnston, Dominic Kwiatkowski, Cordelia Langford, John Sillitoe on behalf of the Wellcome Sanger Institute COVID-19 Surveillance Team |
| EPI_ISL_2537718 | Health Services Laboratories | Wellcome Sanger Institute for the COVID-19 Genomics UK (COG-UK) Consortium | Health Services Laboratories and Alex Alderton, Roberto Amato, Jeffrey Barrett, Sonia Goncalves, Ewan Harrison, David K. Jackson, Ian Johnston, Dominic Kwiatkowski, Cordelia Langford, John Sillitoe on behalf of the Wellcome Sanger Institute COVID-19 Surveillance Team |
| EPI_ISL_2537719 | Lighthouse Lab in Alderley Park | Wellcome Sanger Institute for the COVID-19 Genomics UK (COG-UK) Consortium | Jacquelyn Wynn, Mairead Hyland, The Lighthouse Lab in Alderley Park and Alex Alderton, Roberto Amato, Jeffrey Barrett, Sonia Goncalves, Ewan Harrison, David K. Jackson, Ian Johnston, Dominic Kwiatkowski, Cordelia Langford, John Sillitoe on behalf of the Wellcome Sanger Institute COVID-19 Surveillance Team |
| EPI_ISL_2537720 | Lighthouse Lab in Milton Keynes | Wellcome Sanger Institute for the COVID-19 Genomics UK (COG-UK) Consortium | The Lighthouse Lab in Milton Keynes and Alex Alderton, Roberto Amato, Jeffrey Barrett, Sonia Goncalves, Ewan Harrison, David K. Jackson, Ian Johnston, Dominic Kwiatkowski, Cordelia Langford, John Sillitoe on behalf of the Wellcome Sanger Institute COVID-19 Surveillance Team |
| EPI_ISL_2537721, EPI_ISL_2537722, EPI_ISL_2537723 | Lighthouse Lab in Alderley Park | Wellcome Sanger Institute for the COVID-19 Genomics UK (COG-UK) Consortium | Jacquelyn Wynn, Mairead Hyland, The Lighthouse Lab in Alderley Park and Alex Alderton, Roberto Amato, Jeffrey Barrett, Sonia Goncalves, Ewan Harrison, David K. Jackson, Ian Johnston, Dominic Kwiatkowski, Cordelia Langford, John Sillitoe on behalf of the Wellcome Sanger Institute COVID-19 Surveillance Team |
| EPI_ISL_2537724, EPI_ISL_2537725, | Lighthouse Lab in Milton Keynes | Wellcome Sanger Institute for the COVID-19 Genomics UK (COG-UK) Consortium | The Lighthouse Lab in Milton Keynes and Alex Alderton, Roberto Amato, Jeffrey Barrett, Sonia Goncalves, Ewan Harrison, David K. Jackson, Ian Johnston, Dominic Kwiatkowski, Cordelia Langford, John Sillitoe on behalf of the Wellcome Sanger Institute COVID-19 Surveillance Team |

| | | | |
|---|---|---|---|
| EPI_ISL_2537726, EPI_ISL_2537727 | | | |
| EPI_ISL_2537728 | Lighthouse Lab in Alderley Park | Wellcome Sanger Institute for the COVID-19 Genomics UK (COG-UK) Consortium | Jacquelyn Wynn, Mairead Hyland, The Lighthouse Lab in Alderley Park and Alex Alderton, Roberto Amato, Jeffrey Barrett, Sonia Goncalves, Ewan Harrison, David K. Jackson, Ian Johnston, Dominic Kwiatkowski, Cordelia Langford, John Sillitoe on behalf of the Wellcome Sanger Institute COVID-19 Surveillance Team |
| EPI_ISL_2537729 | Lighthouse Lab in Milton Keynes | Wellcome Sanger Institute for the COVID-19 Genomics UK (COG-UK) Consortium | The Lighthouse Lab in Milton Keynes and Alex Alderton, Roberto Amato, Jeffrey Barrett, Sonia Goncalves, Ewan Harrison, David K. Jackson, Ian Johnston, Dominic Kwiatkowski, Cordelia Langford, John Sillitoe on behalf of the Wellcome Sanger Institute COVID-19 Surveillance Team |
| EPI_ISL_2537730, EPI_ISL_2537731 | Lighthouse Lab in Alderley Park | Wellcome Sanger Institute for the COVID-19 Genomics UK (COG-UK) Consortium | Jacquelyn Wynn, Mairead Hyland, The Lighthouse Lab in Alderley Park and Alex Alderton, Roberto Amato, Jeffrey Barrett, Sonia Goncalves, Ewan Harrison, David K. Jackson, Ian Johnston, Dominic Kwiatkowski, Cordelia Langford, John Sillitoe on behalf of the Wellcome Sanger Institute COVID-19 Surveillance Team |
| EPI_ISL_2537732 | Lighthouse Lab in Milton Keynes | Wellcome Sanger Institute for the COVID-19 Genomics UK (COG-UK) Consortium | The Lighthouse Lab in Milton Keynes and Alex Alderton, Roberto Amato, Jeffrey Barrett, Sonia Goncalves, Ewan Harrison, David K. Jackson, Ian Johnston, Dominic Kwiatkowski, Cordelia Langford, John Sillitoe on behalf of the Wellcome Sanger Institute COVID-19 Surveillance Team |
| EPI_ISL_2537733 | Lighthouse Lab in Alderley Park | Wellcome Sanger Institute for the COVID-19 Genomics UK (COG-UK) Consortium | Jacquelyn Wynn, Mairead Hyland, The Lighthouse Lab in Alderley Park and Alex Alderton, Roberto Amato, Jeffrey Barrett, Sonia Goncalves, Ewan Harrison, David K. Jackson, Ian Johnston, Dominic Kwiatkowski, Cordelia Langford, John Sillitoe on behalf of the Wellcome Sanger Institute COVID-19 Surveillance Team |
| EPI_ISL_2537734 | Lighthouse Lab in Milton Keynes | Wellcome Sanger Institute for the COVID-19 Genomics UK (COG-UK) Consortium | The Lighthouse Lab in Milton Keynes and Alex Alderton, Roberto Amato, Jeffrey Barrett, Sonia Goncalves, Ewan Harrison, David K. Jackson, Ian Johnston, Dominic Kwiatkowski, Cordelia Langford, John Sillitoe on behalf of the Wellcome Sanger Institute COVID-19 Surveillance Team |
| EPI_ISL_2537735 | Health Services Laboratories | Wellcome Sanger Institute for the COVID-19 Genomics UK (COG-UK) Consortium | Health Services Laboratories and Alex Alderton, Roberto Amato, Jeffrey Barrett, Sonia Goncalves, Ewan Harrison, David K. Jackson, Ian Johnston, Dominic Kwiatkowski, Cordelia Langford, John Sillitoe on behalf of the Wellcome Sanger Institute COVID-19 Surveillance Team |
| EPI_ISL_2537736 | Lighthouse Lab in Milton Keynes | Wellcome Sanger Institute for the COVID-19 Genomics UK (COG-UK) Consortium | The Lighthouse Lab in Milton Keynes and Alex Alderton, Roberto Amato, Jeffrey Barrett, Sonia Goncalves, Ewan Harrison, David K. Jackson, Ian Johnston, Dominic Kwiatkowski, Cordelia Langford, John Sillitoe on behalf of the Wellcome Sanger Institute COVID-19 Surveillance Team |
| EPI_ISL_2537737, EPI_ISL_2537738 | Lighthouse Lab in Alderley Park | Wellcome Sanger Institute for the COVID-19 Genomics UK (COG-UK) Consortium | Jacquelyn Wynn, Mairead Hyland, The Lighthouse Lab in Alderley Park and Alex Alderton, Roberto Amato, Jeffrey Barrett, Sonia Goncalves, Ewan Harrison, David K. Jackson, Ian Johnston, Dominic Kwiatkowski, Cordelia Langford, John Sillitoe on behalf of the Wellcome Sanger Institute COVID-19 Surveillance Team |
| EPI_ISL_2537739, EPI_ISL_2537740 | Lighthouse Lab in Milton Keynes | Wellcome Sanger Institute for the COVID-19 Genomics UK (COG-UK) Consortium | The Lighthouse Lab in Milton Keynes and Alex Alderton, Roberto Amato, Jeffrey Barrett, Sonia Goncalves, Ewan Harrison, David K. Jackson, Ian Johnston, Dominic Kwiatkowski, Cordelia Langford, John Sillitoe on behalf of the Wellcome Sanger Institute COVID-19 Surveillance Team |
| EPI_ISL_2537741, EPI_ISL_2537742 | Lighthouse Lab in Alderley Park | Wellcome Sanger Institute for the COVID-19 Genomics UK (COG-UK) Consortium | Jacquelyn Wynn, Mairead Hyland, The Lighthouse Lab in Alderley Park and Alex Alderton, Roberto Amato, Jeffrey Barrett, Sonia Goncalves, Ewan Harrison, David K. Jackson, Ian Johnston, Dominic Kwiatkowski, Cordelia Langford, John Sillitoe on behalf of the Wellcome Sanger Institute COVID-19 Surveillance Team |
| EPI_ISL_2537743 | Lighthouse Lab in Milton Keynes | Wellcome Sanger Institute for the COVID-19 Genomics UK (COG-UK) Consortium | The Lighthouse Lab in Milton Keynes and Alex Alderton, Roberto Amato, Jeffrey Barrett, Sonia Goncalves, Ewan Harrison, David K. Jackson, Ian Johnston, Dominic Kwiatkowski, Cordelia Langford, John Sillitoe on behalf of the Wellcome Sanger Institute COVID-19 Surveillance Team |
| EPI_ISL_2537744, EPI_ISL_2537745, EPI_ISL_2537746, EPI_ISL_2537747 | Lighthouse Lab in Alderley Park | Wellcome Sanger Institute for the COVID-19 Genomics UK (COG-UK) Consortium | Jacquelyn Wynn, Mairead Hyland, The Lighthouse Lab in Alderley Park and Alex Alderton, Roberto Amato, Jeffrey Barrett, Sonia Goncalves, Ewan Harrison, David K. Jackson, Ian Johnston, Dominic Kwiatkowski, Cordelia Langford, John Sillitoe on behalf of the Wellcome Sanger Institute COVID-19 Surveillance Team |
| EPI_ISL_2537748 | Lighthouse Lab in Milton Keynes | Wellcome Sanger Institute for the COVID-19 Genomics UK (COG-UK) Consortium | The Lighthouse Lab in Milton Keynes and Alex Alderton, Roberto Amato, Jeffrey Barrett, Sonia Goncalves, Ewan Harrison, David K. Jackson, Ian Johnston, Dominic Kwiatkowski, Cordelia Langford, John Sillitoe on behalf of the Wellcome Sanger Institute COVID-19 Surveillance Team |
| EPI_ISL_2537749, EPI_ISL_2537751, EPI_ISL_2537752 | Lighthouse Lab in Alderley Park | Wellcome Sanger Institute for the COVID-19 Genomics UK (COG-UK) Consortium | Jacquelyn Wynn, Mairead Hyland, The Lighthouse Lab in Alderley Park and Alex Alderton, Roberto Amato, Jeffrey Barrett, Sonia Goncalves, Ewan Harrison, David K. Jackson, Ian Johnston, Dominic Kwiatkowski, Cordelia Langford, John Sillitoe on behalf of the Wellcome Sanger Institute COVID-19 Surveillance Team |
| EPI_ISL_2537753 | Lighthouse Lab in Milton Keynes | Wellcome Sanger Institute for the COVID-19 Genomics UK (COG-UK) Consortium | The Lighthouse Lab in Milton Keynes and Alex Alderton, Roberto Amato, Jeffrey Barrett, Sonia Goncalves, Ewan Harrison, David K. Jackson, Ian Johnston, Dominic Kwiatkowski, Cordelia Langford, John Sillitoe on behalf of the Wellcome Sanger Institute COVID-19 Surveillance Team |
| EPI_ISL_2537755 | Health Services Laboratories | Wellcome Sanger Institute for the COVID-19 Genomics UK (COG-UK) Consortium | Health Services Laboratories and Alex Alderton, Roberto Amato, Jeffrey Barrett, Sonia Goncalves, Ewan Harrison, David K. Jackson, Ian Johnston, Dominic Kwiatkowski, Cordelia Langford, John Sillitoe on behalf of the Wellcome Sanger Institute COVID-19 Surveillance Team |
| EPI_ISL_2537756 | Lighthouse Lab in Alderley Park | Wellcome Sanger Institute for the COVID-19 Genomics UK (COG-UK) Consortium | Jacquelyn Wynn, Mairead Hyland, The Lighthouse Lab in Alderley Park and Alex Alderton, Roberto Amato, Jeffrey Barrett, Sonia Goncalves, Ewan Harrison, David K. Jackson, Ian Johnston, Dominic Kwiatkowski, Cordelia Langford, John Sillitoe on behalf of the Wellcome Sanger Institute COVID-19 Surveillance Team |
| EPI_ISL_2537757, EPI_ISL_2537758, EPI_ISL_2537759 | Lighthouse Lab in Milton Keynes | Wellcome Sanger Institute for the COVID-19 Genomics UK (COG-UK) Consortium | The Lighthouse Lab in Milton Keynes and Alex Alderton, Roberto Amato, Jeffrey Barrett, Sonia Goncalves, Ewan Harrison, David K. Jackson, Ian Johnston, Dominic Kwiatkowski, Cordelia Langford, John Sillitoe on behalf of the Wellcome Sanger Institute COVID-19 Surveillance Team |
| EPI_ISL_2537760 | Health Services Laboratories | Wellcome Sanger Institute for the COVID-19 Genomics UK (COG-UK) Consortium | Health Services Laboratories and Alex Alderton, Roberto Amato, Jeffrey Barrett, Sonia Goncalves, Ewan Harrison, David K. Jackson, Ian Johnston, Dominic Kwiatkowski, Cordelia Langford, John Sillitoe on behalf of the Wellcome Sanger Institute COVID-19 Surveillance Team |
| EPI_ISL_2537761 | Lighthouse Lab in Milton Keynes | Wellcome Sanger Institute for the COVID-19 Genomics UK (COG-UK) Consortium | The Lighthouse Lab in Milton Keynes and Alex Alderton, Roberto Amato, Jeffrey Barrett, Sonia Goncalves, Ewan Harrison, David K. Jackson, Ian Johnston, Dominic Kwiatkowski, Cordelia Langford, John Sillitoe on behalf of the Wellcome Sanger Institute COVID-19 Surveillance Team |
| EPI_ISL_2537762, EPI_ISL_2537763, EPI_ISL_2537764 | Lighthouse Lab in Alderley Park | Wellcome Sanger Institute for the COVID-19 Genomics UK (COG-UK) Consortium | Jacquelyn Wynn, Mairead Hyland, The Lighthouse Lab in Alderley Park and Alex Alderton, Roberto Amato, Jeffrey Barrett, Sonia Goncalves, Ewan Harrison, David K. Jackson, Ian Johnston, Dominic Kwiatkowski, Cordelia Langford, John Sillitoe on behalf of the Wellcome Sanger Institute COVID-19 Surveillance Team |
| EPI_ISL_2537765 | Health Services Laboratories | Wellcome Sanger Institute for the COVID-19 Genomics UK (COG-UK) Consortium | Health Services Laboratories and Alex Alderton, Roberto Amato, Jeffrey Barrett, Sonia Goncalves, Ewan Harrison, David K. Jackson, Ian Johnston, Dominic Kwiatkowski, Cordelia Langford, John Sillitoe on behalf of the Wellcome Sanger Institute COVID-19 Surveillance Team |
| EPI_ISL_2537766 | Lighthouse Lab in Alderley Park | Wellcome Sanger Institute for the COVID-19 Genomics UK (COG-UK) Consortium | Jacquelyn Wynn, Mairead Hyland, The Lighthouse Lab in Alderley Park and Alex Alderton, Roberto Amato, Jeffrey Barrett, Sonia Goncalves, Ewan Harrison, David K. Jackson, Ian Johnston, Dominic Kwiatkowski, Cordelia Langford, John Sillitoe on behalf of the Wellcome Sanger Institute COVID-19 Surveillance Team |
| EPI_ISL_2537767 | Lighthouse Lab in Milton Keynes | Wellcome Sanger Institute for the COVID-19 Genomics UK (COG-UK) Consortium | The Lighthouse Lab in Milton Keynes and Alex Alderton, Roberto Amato, Jeffrey Barrett, Sonia Goncalves, Ewan Harrison, David K. Jackson, Ian Johnston, Dominic Kwiatkowski, Cordelia Langford, John Sillitoe on behalf of the Wellcome Sanger Institute COVID-19 Surveillance Team |
| EPI_ISL_2537768 | Lighthouse Lab in Alderley Park | Wellcome Sanger Institute for the COVID-19 Genomics UK (COG-UK) Consortium | Jacquelyn Wynn, Mairead Hyland, The Lighthouse Lab in Alderley Park and Alex Alderton, Roberto Amato, Jeffrey Barrett, Sonia Goncalves, Ewan Harrison, David K. Jackson, Ian Johnston, Dominic Kwiatkowski, Cordelia Langford, John Sillitoe on behalf of the Wellcome Sanger Institute COVID-19 Surveillance Team |
| EPI_ISL_2537769, EPI_ISL_2537770, EPI_ISL_2537771 | Lighthouse Lab in Milton Keynes | Wellcome Sanger Institute for the COVID-19 Genomics UK (COG-UK) Consortium | The Lighthouse Lab in Milton Keynes and Alex Alderton, Roberto Amato, Jeffrey Barrett, Sonia Goncalves, Ewan Harrison, David K. Jackson, Ian Johnston, Dominic Kwiatkowski, Cordelia Langford, John Sillitoe on behalf of the Wellcome Sanger Institute COVID-19 Surveillance Team |
| EPI_ISL_2537772 | Lighthouse Lab in Alderley Park | Wellcome Sanger Institute for the COVID-19 Genomics UK (COG-UK) Consortium | Jacquelyn Wynn, Mairead Hyland, The Lighthouse Lab in Alderley Park and Alex Alderton, Roberto Amato, Jeffrey Barrett, Sonia Goncalves, Ewan Harrison, David K. Jackson, Ian Johnston, Dominic Kwiatkowski, Cordelia Langford, John Sillitoe on behalf of the Wellcome Sanger Institute COVID-19 Surveillance Team |
| EPI_ISL_2537773, EPI_ISL_2537774 | Lighthouse Lab in Milton Keynes | Wellcome Sanger Institute for the COVID-19 Genomics UK (COG-UK) Consortium | The Lighthouse Lab in Milton Keynes and Alex Alderton, Roberto Amato, Jeffrey Barrett, Sonia Goncalves, Ewan Harrison, David K. Jackson, Ian Johnston, Dominic Kwiatkowski, Cordelia Langford, John Sillitoe on behalf of the Wellcome Sanger Institute COVID-19 Surveillance Team |
| EPI_ISL_2537775 | Lighthouse Lab in Alderley Park | Wellcome Sanger Institute for the COVID-19 Genomics UK (COG-UK) Consortium | Jacquelyn Wynn, Mairead Hyland, The Lighthouse Lab in Alderley Park and Alex Alderton, Roberto Amato, Jeffrey Barrett, Sonia Goncalves, Ewan Harrison, David K. Jackson, Ian Johnston, Dominic Kwiatkowski, Cordelia Langford, John Sillitoe on behalf of the Wellcome Sanger Institute COVID-19 Surveillance Team |
| EPI_ISL_2537776, EPI_ISL_2537777, EPI_ISL_2537778, EPI_ISL_2537779 | Lighthouse Lab in Milton Keynes | Wellcome Sanger Institute for the COVID-19 Genomics UK (COG-UK) Consortium | The Lighthouse Lab in Milton Keynes and Alex Alderton, Roberto Amato, Jeffrey Barrett, Sonia Goncalves, Ewan Harrison, David K. Jackson, Ian Johnston, Dominic Kwiatkowski, Cordelia Langford, John Sillitoe on behalf of the Wellcome Sanger Institute COVID-19 Surveillance Team |
| EPI_ISL_2537780 | Lighthouse Lab in Alderley Park | Wellcome Sanger Institute for the COVID-19 Genomics UK (COG-UK) Consortium | Jacquelyn Wynn, Mairead Hyland, The Lighthouse Lab in Alderley Park and Alex Alderton, Roberto Amato, Jeffrey Barrett, Sonia Goncalves, Ewan Harrison, David K. Jackson, Ian Johnston, Dominic Kwiatkowski, Cordelia Langford, John Sillitoe on behalf of the Wellcome Sanger Institute COVID-19 Surveillance Team |
| EPI_ISL_2537781 | Lighthouse Lab in Milton Keynes | Wellcome Sanger Institute for the COVID-19 Genomics UK (COG-UK) Consortium | The Lighthouse Lab in Milton Keynes and Alex Alderton, Roberto Amato, Jeffrey Barrett, Sonia Goncalves, Ewan Harrison, David K. Jackson, Ian Johnston, Dominic Kwiatkowski, Cordelia Langford, John Sillitoe on behalf of the Wellcome Sanger Institute COVID-19 Surveillance Team |
| EPI_ISL_2537782 | Lighthouse Lab in Alderley Park | Wellcome Sanger Institute for the COVID-19 Genomics UK (COG-UK) Consortium | Jacquelyn Wynn, Mairead Hyland, The Lighthouse Lab in Alderley Park and Alex Alderton, Roberto Amato, Jeffrey Barrett, Sonia Goncalves, Ewan Harrison, David K. Jackson, Ian Johnston, Dominic Kwiatkowski, Cordelia Langford, John Sillitoe on behalf of the Wellcome Sanger Institute COVID-19 Surveillance Team |
| EPI_ISL_2537783, EPI_ISL_2537784 | Health Services Laboratories | Wellcome Sanger Institute for the COVID-19 Genomics UK (COG-UK) Consortium | Health Services Laboratories and Alex Alderton, Roberto Amato, Jeffrey Barrett, Sonia Goncalves, Ewan Harrison, David K. Jackson, Ian Johnston, Dominic Kwiatkowski, Cordelia Langford, John Sillitoe on behalf of the Wellcome Sanger Institute COVID-19 Surveillance Team |
| EPI_ISL_2537786, EPI_ISL_2537787 | Lighthouse Lab in Alderley Park | Wellcome Sanger Institute for the COVID-19 Genomics UK (COG-UK) Consortium | Jacquelyn Wynn, Mairead Hyland, The Lighthouse Lab in Alderley Park and Alex Alderton, Roberto Amato, Jeffrey Barrett, Sonia Goncalves, Ewan Harrison, David K. Jackson, Ian Johnston, Dominic Kwiatkowski, Cordelia Langford, John Sillitoe on behalf of the Wellcome Sanger Institute COVID-19 Surveillance Team |
| EPI_ISL_2537788 | Lighthouse Lab in Milton Keynes | Wellcome Sanger Institute for the COVID-19 Genomics UK (COG-UK) Consortium | The Lighthouse Lab in Milton Keynes and Alex Alderton, Roberto Amato, Jeffrey Barrett, Sonia Goncalves, Ewan Harrison, David K. Jackson, Ian Johnston, Dominic Kwiatkowski, Cordelia Langford, John Sillitoe on behalf of the Wellcome Sanger Institute COVID-19 Surveillance Team |
| EPI_ISL_2537789 | Lighthouse Lab in Alderley Park | Wellcome Sanger Institute for the COVID-19 Genomics UK (COG-UK) Consortium | Jacquelyn Wynn, Mairead Hyland, The Lighthouse Lab in Alderley Park and Alex Alderton, Roberto Amato, Jeffrey Barrett, Sonia Goncalves, Ewan Harrison, David K. Jackson, Ian Johnston, Dominic Kwiatkowski, Cordelia Langford, John Sillitoe on behalf of the Wellcome Sanger Institute COVID-19 Surveillance Team |
| EPI_ISL_2537790, EPI_ISL_2537791 | Lighthouse Lab in Milton Keynes | Wellcome Sanger Institute for the COVID-19 Genomics UK (COG-UK) Consortium | The Lighthouse Lab in Milton Keynes and Alex Alderton, Roberto Amato, Jeffrey Barrett, Sonia Goncalves, Ewan Harrison, David K. Jackson, Ian Johnston, Dominic Kwiatkowski, Cordelia Langford, John Sillitoe on behalf of the Wellcome Sanger Institute COVID-19 Surveillance Team |
| EPI_ISL_2537792, EPI_ISL_2537793 | Lighthouse Lab in Alderley Park | Wellcome Sanger Institute for the COVID-19 Genomics UK (COG-UK) Consortium | Jacquelyn Wynn, Mairead Hyland, The Lighthouse Lab in Alderley Park and Alex Alderton, Roberto Amato, Jeffrey Barrett, Sonia Goncalves, Ewan Harrison, David K. Jackson, Ian Johnston, Dominic Kwiatkowski, Cordelia Langford, John Sillitoe on behalf of the Wellcome Sanger Institute COVID-19 Surveillance Team |
| EPI_ISL_2537794 | Lighthouse Lab in Milton Keynes | Wellcome Sanger Institute for the COVID-19 Genomics UK (COG-UK) Consortium | The Lighthouse Lab in Milton Keynes and Alex Alderton, Roberto Amato, Jeffrey Barrett, Sonia Goncalves, Ewan Harrison, David K. Jackson, Ian Johnston, Dominic Kwiatkowski, Cordelia Langford, John Sillitoe on behalf of the Wellcome Sanger Institute COVID-19 Surveillance Team |

| | | | |
|---|---|---|---|
| EPI_ISL_2537795, EPI_ISL_2537796 | Lighthouse Lab in Alderley Park | Wellcome Sanger Institute for the COVID-19 Genomics UK (COG-UK) Consortium | Jacquelyn Wynn, Mairead Hyland, The Lighthouse Lab in Alderley Park and Alex Alderton, Roberto Amato, Jeffrey Barrett, Sonia Goncalves, Ewan Harrison, David K. Jackson, Ian Johnston, Dominic Kwiatkowski, Cordelia Langford, John Sillitoe on behalf of the Wellcome Sanger Institute COVID-19 Surveillance Team |
| EPI_ISL_2537797 | Lighthouse Lab in Milton Keynes | Wellcome Sanger Institute for the COVID-19 Genomics UK (COG-UK) Consortium | The Lighthouse Lab in Milton Keynes and Alex Alderton, Roberto Amato, Jeffrey Barrett, Sonia Goncalves, Ewan Harrison, David K. Jackson, Ian Johnston, Dominic Kwiatkowski, Cordelia Langford, John Sillitoe on behalf of the Wellcome Sanger Institute COVID-19 Surveillance Team |
| EPI_ISL_2537798, EPI_ISL_2537799 | Health Services Laboratories | Wellcome Sanger Institute for the COVID-19 Genomics UK (COG-UK) Consortium | Health Services Laboratories and Alex Alderton, Roberto Amato, Jeffrey Barrett, Sonia Goncalves, Ewan Harrison, David K. Jackson, Ian Johnston, Dominic Kwiatkowski, Cordelia Langford, John Sillitoe on behalf of the Wellcome Sanger Institute COVID-19 Surveillance Team |
| EPI_ISL_2537800, EPI_ISL_2537801 | Lighthouse Lab in Alderley Park | Wellcome Sanger Institute for the COVID-19 Genomics UK (COG-UK) Consortium | Jacquelyn Wynn, Mairead Hyland, The Lighthouse Lab in Alderley Park and Alex Alderton, Roberto Amato, Jeffrey Barrett, Sonia Goncalves, Ewan Harrison, David K. Jackson, Ian Johnston, Dominic Kwiatkowski, Cordelia Langford, John Sillitoe on behalf of the Wellcome Sanger Institute COVID-19 Surveillance Team |
| EPI_ISL_2537802, EPI_ISL_2537803, EPI_ISL_2537804 | Lighthouse Lab in Milton Keynes | Wellcome Sanger Institute for the COVID-19 Genomics UK (COG-UK) Consortium | The Lighthouse Lab in Milton Keynes and Alex Alderton, Roberto Amato, Jeffrey Barrett, Sonia Goncalves, Ewan Harrison, David K. Jackson, Ian Johnston, Dominic Kwiatkowski, Cordelia Langford, John Sillitoe on behalf of the Wellcome Sanger Institute COVID-19 Surveillance Team |
| EPI_ISL_2537805 | Health Services Laboratories | Wellcome Sanger Institute for the COVID-19 Genomics UK (COG-UK) Consortium | Health Services Laboratories and Alex Alderton, Roberto Amato, Jeffrey Barrett, Sonia Goncalves, Ewan Harrison, David K. Jackson, Ian Johnston, Dominic Kwiatkowski, Cordelia Langford, John Sillitoe on behalf of the Wellcome Sanger Institute COVID-19 Surveillance Team |
| EPI_ISL_2537807 | Lighthouse Lab in Milton Keynes | Wellcome Sanger Institute for the COVID-19 Genomics UK (COG-UK) Consortium | The Lighthouse Lab in Milton Keynes and Alex Alderton, Roberto Amato, Jeffrey Barrett, Sonia Goncalves, Ewan Harrison, David K. Jackson, Ian Johnston, Dominic Kwiatkowski, Cordelia Langford, John Sillitoe on behalf of the Wellcome Sanger Institute COVID-19 Surveillance Team |
| EPI_ISL_2537808, EPI_ISL_2537809, EPI_ISL_2537810, EPI_ISL_2537811, EPI_ISL_2537812, EPI_ISL_2537813, EPI_ISL_2537814, EPI_ISL_2537815, EPI_ISL_2537816, EPI_ISL_2537817, EPI_ISL_2537818, EPI_ISL_2537819, EPI_ISL_2537820, EPI_ISL_2537821, EPI_ISL_2537822, EPI_ISL_2537823, EPI_ISL_2537824, EPI_ISL_2537825, EPI_ISL_2537826, EPI_ISL_2537827, EPI_ISL_2537828, EPI_ISL_2537829, EPI_ISL_2537830, EPI_ISL_2537831, EPI_ISL_2537832, EPI_ISL_2537833, EPI_ISL_2537834, EPI_ISL_2537835, EPI_ISL_2537837, EPI_ISL_2537838, EPI_ISL_2537839, EPI_ISL_2537840, EPI_ISL_2537841, EPI_ISL_2537842, EPI_ISL_2537843, EPI_ISL_2537844, EPI_ISL_2537845, EPI_ISL_2537846, EPI_ISL_2537847, EPI_ISL_2537848, EPI_ISL_2537849, EPI_ISL_2537850, EPI_ISL_2537851, EPI_ISL_2537853, EPI_ISL_2537854, EPI_ISL_2537855, EPI_ISL_2537856, EPI_ISL_2537857, EPI_ISL_2537858, EPI_ISL_2537859, EPI_ISL_2537860, EPI_ISL_2537861, EPI_ISL_2537862, EPI_ISL_2537863, EPI_ISL_2537864, EPI_ISL_2537865, EPI_ISL_2537866, EPI_ISL_2537867, EPI_ISL_2537868, EPI_ISL_2537869, EPI_ISL_2537870, EPI_ISL_2537872, EPI_ISL_2537873, EPI_ISL_2537875, EPI_ISL_2537876, EPI_ISL_2537877, EPI_ISL_2537878, EPI_ISL_2537879, EPI_ISL_2537880, EPI_ISL_2537881, EPI_ISL_2537882, EPI_ISL_2537883, EPI_ISL_2537884, EPI_ISL_2537885, EPI_ISL_2537886, EPI_ISL_2537887, EPI_ISL_2537888, EPI_ISL_2537889, EPI_ISL_2537890, EPI_ISL_2537891, EPI_ISL_2537892, EPI_ISL_2537893, EPI_ISL_2537894, EPI_ISL_2537897, EPI_ISL_2537898, EPI_ISL_2537899, EPI_ISL_2537900, EPI_ISL_2537901, EPI_ISL_2537902, EPI_ISL_2537903, EPI_ISL_2537904, EPI_ISL_2537905, EPI_ISL_2537906, EPI_ISL_2537907, EPI_ISL_2537908, EPI_ISL_2537909, EPI_ISL_2537910, EPI_ISL_2537911, EPI_ISL_2537912, EPI_ISL_2537913, EPI_ISL_2537914, EPI_ISL_2537915, EPI_ISL_2537916, EPI_ISL_2537918, EPI_ISL_2537919, EPI_ISL_2537920, EPI_ISL_2537921, EPI_ISL_2537922, EPI_ISL_2537923, EPI_ISL_2537924, EPI_ISL_2537925, EPI_ISL_2537926, EPI_ISL_2537927, EPI_ISL_2537928, EPI_ISL_2537930, EPI_ISL_2537931, EPI_ISL_2537932, EPI_ISL_2537933, EPI_ISL_2537934, EPI_ISL_2537935, EPI_ISL_2537936, EPI_ISL_2537937, EPI_ISL_2537938, EPI_ISL_2537939, EPI_ISL_2537940, EPI_ISL_2537941, EPI_ISL_2537942, EPI_ISL_2537943, EPI_ISL_2537946, EPI_ISL_2537947, EPI_ISL_2537948, EPI_ISL_2537949, EPI_ISL_2537950, EPI_ISL_2537951, EPI_ISL_2537952, EPI_ISL_2537953, EPI_ISL_2537954, EPI_ISL_2537955, EPI_ISL_2537956, EPI_ISL_2537957, EPI_ISL_2537958, EPI_ISL_2537959, EPI_ISL_2537960, EPI_ISL_2537961, EPI_ISL_2537962, EPI_ISL_2537963, EPI_ISL_2537964, EPI_ISL_2537965, EPI_ISL_2537966, EPI_ISL_2537967, EPI_ISL_2537968, EPI_ISL_2537969, EPI_ISL_2537970, EPI_ISL_2537971, EPI_ISL_2537972, EPI_ISL_2537973, EPI_ISL_2537974, EPI_ISL_2537975, EPI_ISL_2537976, EPI_ISL_2537977, EPI_ISL_2537978, EPI_ISL_2537979, EPI_ISL_2537980, EPI_ISL_2537981, EPI_ISL_2537982, EPI_ISL_2537983, EPI_ISL_2537984, EPI_ISL_2537985, EPI_ISL_2537986, EPI_ISL_2537987, EPI_ISL_2537988, EPI_ISL_2537989, EPI_ISL_2537990, EPI_ISL_2537991, EPI_ISL_2537992, EPI_ISL_2537993, EPI_ISL_2537994, EPI_ISL_2537995, EPI_ISL_2537996, EPI_ISL_2537997, EPI_ISL_2537998, EPI_ISL_2537999, EPI_ISL_2538000, EPI_ISL_2538001, EPI_ISL_2538002, EPI_ISL_2538003, EPI_ISL_2538004, EPI_ISL_2538005, EPI_ISL_2538006, EPI_ISL_2538007, EPI_ISL_2538008, EPI_ISL_2538009, EPI_ISL_2538010, EPI_ISL_2538011, EPI_ISL_2538012, EPI_ISL_2538013, EPI_ISL_2538014, EPI_ISL_2538015, EPI_ISL_2538016, EPI_ISL_2538017, EPI_ISL_2538018, EPI_ISL_2538019, EPI_ISL_2538020, EPI_ISL_2538021, EPI_ISL_2538022, EPI_ISL_2538023, EPI_ISL_2538024, EPI_ISL_2538025, EPI_ISL_2538026, EPI_ISL_2538027, EPI_ISL_2538028, EPI_ISL_2538029, EPI_ISL_2538030, EPI_ISL_2538031, EPI_ISL_2538033, EPI_ISL_2538034, EPI_ISL_2538035, EPI_ISL_2538036, EPI_ISL_2538037, EPI_ISL_2538038, EPI_ISL_2538039, EPI_ISL_2538040, EPI_ISL_2538041, EPI_ISL_2538042, EPI_ISL_2538043, EPI_ISL_2538044, EPI_ISL_2538045, EPI_ISL_2538046, EPI_ISL_2538047, EPI_ISL_2538048, EPI_ISL_2538050, EPI_ISL_2538051, EPI_ISL_2538052, EPI_ISL_2538053, EPI_ISL_2538054, EPI_ISL_2538056, EPI_ISL_2538057, EPI_ISL_2538059, EPI_ISL_2538060, EPI_ISL_2538062, EPI_ISL_2538063, EPI_ISL_2538065, EPI_ISL_2538067, EPI_ISL_2538068, EPI_ISL_2538069, EPI_ISL_2538070, EPI_ISL_2538071, EPI_ISL_2538072, EPI_ISL_2538073, EPI_ISL_2538075, EPI_ISL_2538076, EPI_ISL_2538077, EPI_ISL_2538078, EPI_ISL_2538079, EPI_ISL_2538080, EPI_ISL_2538081, EPI_ISL_2538083, EPI_ISL_2538084, EPI_ISL_2538085, EPI_ISL_2538086, EPI_ISL_2538087, EPI_ISL_2538088, EPI_ISL_2538089, EPI_ISL_2538090, EPI_ISL_2538091, EPI_ISL_2538092, EPI_ISL_2538093, EPI_ISL_2538094, EPI_ISL_2538095, EPI_ISL_2538096, EPI_ISL_2538097, EPI_ISL_2538098, EPI_ISL_2538099, EPI_ISL_2538100, EPI_ISL_2538101, EPI_ISL_2538102, EPI_ISL_2538103, EPI_ISL_2538104, EPI_ISL_2538105, EPI_ISL_2538106, EPI_ISL_2538107, EPI_ISL_2538108, EPI_ISL_2538109, EPI_ISL_2538110, EPI_ISL_2538112, EPI_ISL_2538113, EPI_ISL_2538114, EPI_ISL_2538115, EPI_ISL_2538116, EPI_ISL_2538117, EPI_ISL_2538118, EPI_ISL_2538119, EPI_ISL_2538120, EPI_ISL_2538121, EPI_ISL_2538122, EPI_ISL_2538123, EPI_ISL_2538124, EPI_ISL_2538125 | | | |
| see above | Lighthouse Lab in Alderley Park | Wellcome Sanger Institute for the COVID-19 Genomics UK (COG-UK) Consortium | Jacquelyn Wynn, Mairead Hyland, The Lighthouse Lab in Alderley Park and Alex Alderton, Roberto Amato, Jeffrey Barrett, Sonia Goncalves, Ewan Harrison, David K. Jackson, Ian Johnston, Dominic Kwiatkowski, Cordelia Langford, John Sillitoe on behalf of the Wellcome Sanger Institute COVID-19 Surveillance Team |
| EPI_ISL_2538126 | Balai Besar Penelitian dan Pengembangan dan Reservoir Penyakit, Salatiga | Genomik Solidaritas Indonesia Laboratorium | Carissa Sintca Wijaya, Vania Gavrila Wikasa, Dhahlia Agustina Cahyono, Alfin Mohammad Abdillah, Normastuti Adhini Tantyo, Yudistira Aditya Rahmat, Joko Waluyo, Annisa Muthiah Sukirman, Gracia Felias Enos Korompis, Pratiwi Sudarmono, Budi Wiweko, Louisa Markus, Anuraj Shankar, Meutia Ayuputeri Kumaheri, Anindya Pradipta Susanto, Ariel Pradipta |
| EPI_ISL_2538127, EPI_ISL_2538128, EPI_ISL_2538129, EPI_ISL_2538130, EPI_ISL_2538131, EPI_ISL_2538132, EPI_ISL_2538133, EPI_ISL_2538134, EPI_ISL_2538135, EPI_ISL_2538136, EPI_ISL_2538137, EPI_ISL_2538138, EPI_ISL_2538139, EPI_ISL_2538140, EPI_ISL_2538141, EPI_ISL_2538142, EPI_ISL_2538143, EPI_ISL_2538144, EPI_ISL_2538147, EPI_ISL_2538148, EPI_ISL_2538149, EPI_ISL_2538150, EPI_ISL_2538151, EPI_ISL_2538152, EPI_ISL_2538153, EPI_ISL_2538154, EPI_ISL_2538155, EPI_ISL_2538156, EPI_ISL_2538157, EPI_ISL_2538158, EPI_ISL_2538159, EPI_ISL_2538160, EPI_ISL_2538161, EPI_ISL_2538163, EPI_ISL_2538164, EPI_ISL_2538165, EPI_ISL_2538166, EPI_ISL_2538167, EPI_ISL_2538168, EPI_ISL_2538169, EPI_ISL_2538170, EPI_ISL_2538171, EPI_ISL_2538172, EPI_ISL_2538173, EPI_ISL_2538174, EPI_ISL_2538175, EPI_ISL_2538176, EPI_ISL_2538177, EPI_ISL_2538178, EPI_ISL_2538179, EPI_ISL_2538180, EPI_ISL_2538181, EPI_ISL_2538182, EPI_ISL_2538183, EPI_ISL_2538184, EPI_ISL_2538185, EPI_ISL_2538186, EPI_ISL_2538187, EPI_ISL_2538189, EPI_ISL_2538190, EPI_ISL_2538191, EPI_ISL_2538192, EPI_ISL_2538193, EPI_ISL_2538194, EPI_ISL_2538195, EPI_ISL_2538196, EPI_ISL_2538197, EPI_ISL_2538198, EPI_ISL_2538199, EPI_ISL_2538200, EPI_ISL_2538201, EPI_ISL_2538203, EPI_ISL_2538204, EPI_ISL_2538205, EPI_ISL_2538206, EPI_ISL_2538207, EPI_ISL_2538208, EPI_ISL_2538209, EPI_ISL_2538211, EPI_ISL_2538212, EPI_ISL_2538213, EPI_ISL_2538214, EPI_ISL_2538215, EPI_ISL_2538216, EPI_ISL_2538218, EPI_ISL_2538220, EPI_ISL_2538221, EPI_ISL_2538222, EPI_ISL_2538223, EPI_ISL_2538224, EPI_ISL_2538225, EPI_ISL_2538226, EPI_ISL_2538227, EPI_ISL_2538228, EPI_ISL_2538230, EPI_ISL_2538231, EPI_ISL_2538232, EPI_ISL_2538233, EPI_ISL_2538234, EPI_ISL_2538235, EPI_ISL_2538236, EPI_ISL_2538237, EPI_ISL_2538239, EPI_ISL_2538240, EPI_ISL_2538241, EPI_ISL_2538242, EPI_ISL_2538243, EPI_ISL_2538244, EPI_ISL_2538245, EPI_ISL_2538246, EPI_ISL_2538247, EPI_ISL_2538248, EPI_ISL_2538249, EPI_ISL_2538250, EPI_ISL_2538251, EPI_ISL_2538252, EPI_ISL_2538253, EPI_ISL_2538254, EPI_ISL_2538255, EPI_ISL_2538256, EPI_ISL_2538257, EPI_ISL_2538258, EPI_ISL_2538259, EPI_ISL_2538260, EPI_ISL_2538261, EPI_ISL_2538262, EPI_ISL_2538263, EPI_ISL_2538264, EPI_ISL_2538265, EPI_ISL_2538268, EPI_ISL_2538269, EPI_ISL_2538270, EPI_ISL_2538272, EPI_ISL_2538273, EPI_ISL_2538274, EPI_ISL_2538275, EPI_ISL_2538276, EPI_ISL_2538278, EPI_ISL_2538279, EPI_ISL_2538280, EPI_ISL_2538281, EPI_ISL_2538282, EPI_ISL_2538283, EPI_ISL_2538284, EPI_ISL_2538285, EPI_ISL_2538286, EPI_ISL_2538287, EPI_ISL_2538288, EPI_ISL_2538289, EPI_ISL_2538290, EPI_ISL_2538291, EPI_ISL_2538292, EPI_ISL_2538293, EPI_ISL_2538294, EPI_ISL_2538295, EPI_ISL_2538296, EPI_ISL_2538297, EPI_ISL_2538299, EPI_ISL_2538300, EPI_ISL_2538301, EPI_ISL_2538303, EPI_ISL_2538304, EPI_ISL_2538305, EPI_ISL_2538306, EPI_ISL_2538307, EPI_ISL_2538308, EPI_ISL_2538309, EPI_ISL_2538310, EPI_ISL_2538311, EPI_ISL_2538312, EPI_ISL_2538313, EPI_ISL_2538314, EPI_ISL_2538315, EPI_ISL_2538316, EPI_ISL_2538317, EPI_ISL_2538318, EPI_ISL_2538319, EPI_ISL_2538320, EPI_ISL_2538321, EPI_ISL_2538322, EPI_ISL_2538323, EPI_ISL_2538324, EPI_ISL_2538325, EPI_ISL_2538326, EPI_ISL_2538327, EPI_ISL_2538328, EPI_ISL_2538329, EPI_ISL_2538330, EPI_ISL_2538331, EPI_ISL_2538332, EPI_ISL_2538333, EPI_ISL_2538334, EPI_ISL_2538335, EPI_ISL_2538336, EPI_ISL_2538337, EPI_ISL_2538338, EPI_ISL_2538339, EPI_ISL_2538340, EPI_ISL_2538341, EPI_ISL_2538342, EPI_ISL_2538343, EPI_ISL_2538344, EPI_ISL_2538346, EPI_ISL_2538347, EPI_ISL_2538349, EPI_ISL_2538350, EPI_ISL_2538351, EPI_ISL_2538352, EPI_ISL_2538354, EPI_ISL_2538355, EPI_ISL_2538356, EPI_ISL_2538357, EPI_ISL_2538358, EPI_ISL_2538359, EPI_ISL_2538360, EPI_ISL_2538361, EPI_ISL_2538362, EPI_ISL_2538363, EPI_ISL_2538364, EPI_ISL_2538365, EPI_ISL_2538366, EPI_ISL_2538367, EPI_ISL_2538368, EPI_ISL_2538369, EPI_ISL_2538370, EPI_ISL_2538372, EPI_ISL_2538373, EPI_ISL_2538374, EPI_ISL_2538375, EPI_ISL_2538376, EPI_ISL_2538377, EPI_ISL_2538378, EPI_ISL_2538379, EPI_ISL_2538380, EPI_ISL_2538381, EPI_ISL_2538382, EPI_ISL_2538383, EPI_ISL_2538384, EPI_ISL_2538385, EPI_ISL_2538386, EPI_ISL_2538388, EPI_ISL_2538389, EPI_ISL_2538390, EPI_ISL_2538391, EPI_ISL_2538392, EPI_ISL_2538393, EPI_ISL_2538394, EPI_ISL_2538395, EPI_ISL_2538396, EPI_ISL_2538397, EPI_ISL_2538398, EPI_ISL_2538399, EPI_ISL_2538400, EPI_ISL_2538401, EPI_ISL_2538402, EPI_ISL_2538403, EPI_ISL_2538404, EPI_ISL_2538405, EPI_ISL_2538406, EPI_ISL_2538407, EPI_ISL_2538408, EPI_ISL_2538409, EPI_ISL_2538410, EPI_ISL_2538411, EPI_ISL_2538412, EPI_ISL_2538413, EPI_ISL_2538414, EPI_ISL_2538416, EPI_ISL_2538417, EPI_ISL_2538418, EPI_ISL_2538419, EPI_ISL_2538420, EPI_ISL_2538421, EPI_ISL_2538422, EPI_ISL_2538423, EPI_ISL_2538424, EPI_ISL_2538425, EPI_ISL_2538428, EPI_ISL_2538429, EPI_ISL_2538430, EPI_ISL_2538433, EPI_ISL_2538434, EPI_ISL_2538435, EPI_ISL_2538436, EPI_ISL_2538437, EPI_ISL_2538438, EPI_ISL_2538439, EPI_ISL_2538440, EPI_ISL_2538441, EPI_ISL_2538442, EPI_ISL_2538443, EPI_ISL_2538444, EPI_ISL_2538445, EPI_ISL_2538446, EPI_ISL_2538447, EPI_ISL_2538448, EPI_ISL_2538449, EPI_ISL_2538450, EPI_ISL_2538451, EPI_ISL_2538452, EPI_ISL_2538453, EPI_ISL_2538455, EPI_ISL_2538456, EPI_ISL_2538458, EPI_ISL_2538459, EPI_ISL_2538460, EPI_ISL_2538461, EPI_ISL_2538462, EPI_ISL_2538463, EPI_ISL_2538464, EPI_ISL_2538465, EPI_ISL_2538466, EPI_ISL_2538467, EPI_ISL_2538468, EPI_ISL_2538469, EPI_ISL_2538470, EPI_ISL_2538471, EPI_ISL_2538472, EPI_ISL_2538473, EPI_ISL_2538474, EPI_ISL_2538475, EPI_ISL_2538476, EPI_ISL_2538477, EPI_ISL_2538478, EPI_ISL_2538479, EPI_ISL_2538480, EPI_ISL_2538481, EPI_ISL_2538482, EPI_ISL_2538483, EPI_ISL_2538484, EPI_ISL_2538485, EPI_ISL_2538486, EPI_ISL_2538487, EPI_ISL_2538488, EPI_ISL_2538489, EPI_ISL_2538490, EPI_ISL_2538491, EPI_ISL_2538492, EPI_ISL_2538493, EPI_ISL_2538494, EPI_ISL_2538495, EPI_ISL_2538496, EPI_ISL_2538498, EPI_ISL_2538499, EPI_ISL_2538500, EPI_ISL_2538501, EPI_ISL_2538502, EPI_ISL_2538503, EPI_ISL_2538504, EPI_ISL_2538506, EPI_ISL_2538507, EPI_ISL_2538508, EPI_ISL_2538509, EPI_ISL_2538510, EPI_ISL_2538511, EPI_ISL_2538512, EPI_ISL_2538513, EPI_ISL_2538514, EPI_ISL_2538515, EPI_ISL_2538516, EPI_ISL_2538519, EPI_ISL_2538520, EPI_ISL_2538521, EPI_ISL_2538522, EPI_ISL_2538523, EPI_ISL_2538524, EPI_ISL_2538525, EPI_ISL_2538527, EPI_ISL_2538528, EPI_ISL_2538529, EPI_ISL_2538530, EPI_ISL_2538531, EPI_ISL_2538532, EPI_ISL_2538533 | | | |
| see above | Lighthouse Lab in Alderley Park | Wellcome Sanger Institute for the COVID-19 Genomics UK (COG-UK) Consortium | Jacquelyn Wynn, Mairead Hyland, The Lighthouse Lab in Alderley Park and Alex Alderton, Roberto Amato, Jeffrey Barrett, Sonia Goncalves, Ewan Harrison, David K. Jackson, Ian Johnston, Dominic Kwiatkowski, Cordelia Langford, John Sillitoe on behalf of the Wellcome Sanger Institute COVID-19 Surveillance Team |
| EPI_ISL_2538534, EPI_ISL_2538535 | Lighthouse Lab in Glasgow | Wellcome Sanger Institute for the COVID-19 Genomics UK (COG-UK) Consortium | Harper VanSteenhouse, Yumi Kasai, David Gray, Carol Clugston, Anna Dominiczak and Alex Alderton, Roberto Amato, Jeffrey Barrett, Sonia Goncalves, Ewan Harrison, David K. Jackson, Ian Johnston, Dominic Kwiatkowski, Cordelia Langford, John Sillitoe on behalf of the Wellcome Sanger Institute COVID-19 Surveillance Team |
| EPI_ISL_2538536, EPI_ISL_2538537 | Lighthouse Lab in Alderley Park | Wellcome Sanger Institute for the COVID-19 Genomics UK (COG-UK) Consortium | Jacquelyn Wynn, Mairead Hyland, The Lighthouse Lab in Alderley Park and Alex Alderton, Roberto Amato, Jeffrey Barrett, Sonia Goncalves, Ewan Harrison, David K. Jackson, Ian Johnston, Dominic Kwiatkowski, Cordelia Langford, John Sillitoe on behalf of the Wellcome Sanger Institute COVID-19 Surveillance Team |
| EPI_ISL_2538538 | Lighthouse Lab in Milton Keynes | Wellcome Sanger Institute for the COVID-19 Genomics UK (COG-UK) Consortium | The Lighthouse Lab in Milton Keynes and Alex Alderton, Roberto Amato, Jeffrey Barrett, Sonia Goncalves, Ewan Harrison, David K. Jackson, Ian Johnston, Dominic Kwiatkowski, Cordelia Langford, John Sillitoe on behalf of the Wellcome Sanger Institute COVID-19 Surveillance Team |
| EPI_ISL_2538539 | Lighthouse Lab in Glasgow | Wellcome Sanger Institute for the COVID-19 Genomics UK (COG-UK) Consortium | Harper VanSteenhouse, Yumi Kasai, David Gray, Carol Clugston, Anna Dominiczak and Alex Alderton, Roberto Amato, Jeffrey Barrett, Sonia Goncalves, Ewan Harrison, David K. Jackson, Ian Johnston, Dominic Kwiatkowski, Cordelia Langford, John Sillitoe on behalf of the Wellcome Sanger Institute COVID-19 Surveillance Team |
| EPI_ISL_2538540 | Lighthouse Lab in Milton Keynes | Wellcome Sanger Institute for the COVID-19 Genomics UK (COG-UK) Consortium | The Lighthouse Lab in Milton Keynes and Alex Alderton, Roberto Amato, Jeffrey Barrett, Sonia Goncalves, Ewan Harrison, David K. Jackson, Ian Johnston, Dominic Kwiatkowski, Cordelia Langford, John Sillitoe on behalf of the Wellcome Sanger Institute COVID-19 Surveillance Team |
| EPI_ISL_2538541 | Lighthouse Lab in Alderley Park | Wellcome Sanger Institute for the COVID-19 Genomics UK (COG-UK) Consortium | Jacquelyn Wynn, Mairead Hyland, The Lighthouse Lab in Alderley Park and Alex Alderton, Roberto Amato, Jeffrey Barrett, Sonia Goncalves, Ewan Harrison, David K. Jackson, Ian Johnston, Dominic Kwiatkowski, Cordelia Langford, John Sillitoe on behalf of the Wellcome Sanger Institute COVID-19 Surveillance Team |
| EPI_ISL_2538542 | Lighthouse Lab in Milton Keynes | Wellcome Sanger Institute for the COVID-19 Genomics UK (COG-UK) Consortium | The Lighthouse Lab in Milton Keynes and Alex Alderton, Roberto Amato, Jeffrey Barrett, Sonia Goncalves, Ewan Harrison, David K. Jackson, Ian Johnston, Dominic Kwiatkowski, Cordelia Langford, John Sillitoe on behalf of the Wellcome Sanger Institute COVID-19 Surveillance Team |
| EPI_ISL_2538543, EPI_ISL_2538544, EPI_ISL_2538545, EPI_ISL_2538546, EPI_ISL_2538547, EPI_ISL_2538548, EPI_ISL_2538549, EPI_ISL_2538550, EPI_ISL_2538551, EPI_ISL_2538552, EPI_ISL_2538553, EPI_ISL_2538554, EPI_ISL_2538555 | | | |
| see above | Lighthouse Lab in Alderley Park | Wellcome Sanger Institute for the COVID-19 Genomics UK (COG-UK) Consortium | Jacquelyn Wynn, Mairead Hyland, The Lighthouse Lab in Alderley Park and Alex Alderton, Roberto Amato, Jeffrey Barrett, Sonia Goncalves, Ewan Harrison, David K. Jackson, Ian Johnston, Dominic Kwiatkowski, Cordelia Langford, John Sillitoe on behalf of the Wellcome Sanger Institute COVID-19 Surveillance Team |
| EPI_ISL_2538556 | Lighthouse Lab in Glasgow | Wellcome Sanger Institute for the COVID-19 Genomics UK (COG-UK) Consortium | Harper VanSteenhouse, Yumi Kasai, David Gray, Carol Clugston, Anna Dominiczak and Alex Alderton, Roberto Amato, Jeffrey Barrett, Sonia Goncalves, Ewan Harrison, David K. Jackson, Ian Johnston, Dominic Kwiatkowski, Cordelia Langford, John Sillitoe on behalf of the Wellcome Sanger Institute COVID-19 Surveillance Team |
| EPI_ISL_2538557, EPI_ISL_2538558, EPI_ISL_2538559 | Lighthouse Lab in Alderley Park | Wellcome Sanger Institute for the COVID-19 Genomics UK (COG-UK) Consortium | Jacquelyn Wynn, Mairead Hyland, The Lighthouse Lab in Alderley Park and Alex Alderton, Roberto Amato, Jeffrey Barrett, Sonia Goncalves, Ewan Harrison, David K. Jackson, Ian Johnston, Dominic Kwiatkowski, Cordelia Langford, John Sillitoe on behalf of the Wellcome Sanger Institute COVID-19 Surveillance Team |
| EPI_ISL_2538560, EPI_ISL_2538561 | Lighthouse Lab in Milton Keynes | Wellcome Sanger Institute for the COVID-19 Genomics UK (COG-UK) Consortium | The Lighthouse Lab in Milton Keynes and Alex Alderton, Roberto Amato, Jeffrey Barrett, Sonia Goncalves, Ewan Harrison, David K. Jackson, Ian Johnston, Dominic Kwiatkowski, Cordelia Langford, John Sillitoe on behalf of the Wellcome Sanger Institute COVID-19 Surveillance Team |
| EPI_ISL_2538562 | Lighthouse Lab in Alderley Park | Wellcome Sanger Institute for the COVID-19 Genomics UK (COG-UK) Consortium | Jacquelyn Wynn, Mairead Hyland, The Lighthouse Lab in Alderley Park and Alex Alderton, Roberto Amato, Jeffrey Barrett, Sonia Goncalves, Ewan Harrison, David K. Jackson, Ian Johnston, Dominic Kwiatkowski, Cordelia Langford, John Sillitoe on behalf of the Wellcome Sanger Institute COVID-19 Surveillance Team |
| EPI_ISL_2538563 | Lighthouse Lab in Glasgow | Wellcome Sanger Institute for the COVID-19 Genomics UK (COG-UK) Consortium | Harper VanSteenhouse, Yumi Kasai, David Gray, Carol Clugston, Anna Dominiczak and Alex Alderton, Roberto Amato, Jeffrey Barrett, Sonia Goncalves, Ewan Harrison, David K. Jackson, Ian Johnston, Dominic Kwiatkowski, Cordelia Langford, John Sillitoe on behalf of the Wellcome Sanger Institute COVID-19 Surveillance Team |
| EPI_ISL_2538564, EPI_ISL_2538565, EPI_ISL_2538566 | Lighthouse Lab in Alderley Park | Wellcome Sanger Institute for the COVID-19 Genomics UK (COG-UK) Consortium | Jacquelyn Wynn, Mairead Hyland, The Lighthouse Lab in Alderley Park and Alex Alderton, Roberto Amato, Jeffrey Barrett, Sonia Goncalves, Ewan Harrison, David K. Jackson, Ian Johnston, Dominic Kwiatkowski, Cordelia Langford, John Sillitoe on behalf of the Wellcome Sanger Institute COVID-19 Surveillance Team |
| EPI_ISL_2538567 | Lighthouse Lab in Glasgow | Wellcome Sanger Institute for the COVID-19 Genomics UK (COG-UK) Consortium | Harper VanSteenhouse, Yumi Kasai, David Gray, Carol Clugston, Anna Dominiczak and Alex Alderton, Roberto Amato, Jeffrey Barrett, Sonia Goncalves, Ewan Harrison, David K. Jackson, Ian Johnston, Dominic Kwiatkowski, Cordelia Langford, John Sillitoe on behalf of the Wellcome Sanger Institute COVID-19 Surveillance Team |
| EPI_ISL_2538568 | Lighthouse Lab in Milton Keynes | Wellcome Sanger Institute for the COVID-19 Genomics UK (COG-UK) Consortium | The Lighthouse Lab in Milton Keynes and Alex Alderton, Roberto Amato, Jeffrey Barrett, Sonia Goncalves, Ewan Harrison, David K. Jackson, Ian Johnston, Dominic Kwiatkowski, Cordelia Langford, John Sillitoe on behalf of the Wellcome Sanger Institute COVID-19 Surveillance Team |
| EPI_ISL_2538569 | Lighthouse Lab in Alderley Park | Wellcome Sanger Institute for the COVID-19 Genomics UK (COG-UK) Consortium | Jacquelyn Wynn, Mairead Hyland, The Lighthouse Lab in Alderley Park and Alex Alderton, Roberto Amato, Jeffrey Barrett, Sonia Goncalves, Ewan Harrison, David K. Jackson, Ian Johnston, Dominic Kwiatkowski, Cordelia Langford, John Sillitoe on behalf of the Wellcome Sanger Institute COVID-19 Surveillance Team |
| EPI_ISL_2538570 | Lighthouse Lab in Glasgow | Wellcome Sanger Institute for the COVID-19 Genomics UK (COG-UK) Consortium | Harper VanSteenhouse, Yumi Kasai, David Gray, Carol Clugston, Anna Dominiczak and Alex Alderton, Roberto Amato, Jeffrey Barrett, Sonia Goncalves, Ewan Harrison, David K. Jackson, Ian Johnston, Dominic Kwiatkowski, Cordelia Langford, John Sillitoe on behalf of the Wellcome Sanger Institute COVID-19 Surveillance Team |
| EPI_ISL_2538571, EPI_ISL_2538572, EPI_ISL_2538573, EPI_ISL_2538574, EPI_ISL_2538575, EPI_ISL_2538576 | Lighthouse Lab in Alderley Park | Wellcome Sanger Institute for the COVID-19 Genomics UK (COG-UK) Consortium | Jacquelyn Wynn, Mairead Hyland, The Lighthouse Lab in Alderley Park and Alex Alderton, Roberto Amato, Jeffrey Barrett, Sonia Goncalves, Ewan Harrison, David K. Jackson, Ian Johnston, Dominic Kwiatkowski, Cordelia Langford, John Sillitoe on behalf of the Wellcome Sanger Institute COVID-19 Surveillance Team |

| | | | |
|---|---|---|---|
| EPI_ISL_2538577 | Lighthouse Lab in Glasgow | Wellcome Sanger Institute for the COVID-19 Genomics UK (COG-UK) Consortium | Harper VanSteenhouse, Yumi Kasai, David Gray, Carol Clugston, Anna Dominiczak and Alex Alderton, Roberto Amato, Jeffrey Barrett, Sonia Goncalves, Ewan Harrison, David K. Jackson, Ian Johnston, Dominic Kwiatkowski, Cordelia Langford, John Sillitoe on behalf of the Wellcome Sanger Institute COVID-19 Surveillance Team |
| EPI_ISL_2538578, EPI_ISL_2538579, EPI_ISL_2538580, EPI_ISL_2538581 | Lighthouse Lab in Alderley Park | Wellcome Sanger Institute for the COVID-19 Genomics UK (COG-UK) Consortium | Jacquelyn Wynn, Mairead Hyland, The Lighthouse Lab in Alderley Park and Alex Alderton, Roberto Amato, Jeffrey Barrett, Sonia Goncalves, Ewan Harrison, David K. Jackson, Ian Johnston, Dominic Kwiatkowski, Cordelia Langford, John Sillitoe on behalf of the Wellcome Sanger Institute COVID-19 Surveillance Team |
| EPI_ISL_2538582 | Lighthouse Lab in Milton Keynes | Wellcome Sanger Institute for the COVID-19 Genomics UK (COG-UK) Consortium | The Lighthouse Lab in Milton Keynes and Alex Alderton, Roberto Amato, Jeffrey Barrett, Sonia Goncalves, Ewan Harrison, David K. Jackson, Ian Johnston, Dominic Kwiatkowski, Cordelia Langford, John Sillitoe on behalf of the Wellcome Sanger Institute COVID-19 Surveillance Team |
| EPI_ISL_2538584, EPI_ISL_2538585, EPI_ISL_2538586 | Lighthouse Lab in Alderley Park | Wellcome Sanger Institute for the COVID-19 Genomics UK (COG-UK) Consortium | Jacquelyn Wynn, Mairead Hyland, The Lighthouse Lab in Alderley Park and Alex Alderton, Roberto Amato, Jeffrey Barrett, Sonia Goncalves, Ewan Harrison, David K. Jackson, Ian Johnston, Dominic Kwiatkowski, Cordelia Langford, John Sillitoe on behalf of the Wellcome Sanger Institute COVID-19 Surveillance Team |
| EPI_ISL_2538587 | Lighthouse Lab in Glasgow | Wellcome Sanger Institute for the COVID-19 Genomics UK (COG-UK) Consortium | Harper VanSteenhouse, Yumi Kasai, David Gray, Carol Clugston, Anna Dominiczak and Alex Alderton, Roberto Amato, Jeffrey Barrett, Sonia Goncalves, Ewan Harrison, David K. Jackson, Ian Johnston, Dominic Kwiatkowski, Cordelia Langford, John Sillitoe on behalf of the Wellcome Sanger Institute COVID-19 Surveillance Team |
| EPI_ISL_2538588, EPI_ISL_2538589, EPI_ISL_2538590, EPI_ISL_2538591, EPI_ISL_2538592 | Lighthouse Lab in Alderley Park | Wellcome Sanger Institute for the COVID-19 Genomics UK (COG-UK) Consortium | Jacquelyn Wynn, Mairead Hyland, The Lighthouse Lab in Alderley Park and Alex Alderton, Roberto Amato, Jeffrey Barrett, Sonia Goncalves, Ewan Harrison, David K. Jackson, Ian Johnston, Dominic Kwiatkowski, Cordelia Langford, John Sillitoe on behalf of the Wellcome Sanger Institute COVID-19 Surveillance Team |
| EPI_ISL_2538593, EPI_ISL_2538594 | Lighthouse Lab in Glasgow | Wellcome Sanger Institute for the COVID-19 Genomics UK (COG-UK) Consortium | Harper VanSteenhouse, Yumi Kasai, David Gray, Carol Clugston, Anna Dominiczak and Alex Alderton, Roberto Amato, Jeffrey Barrett, Sonia Goncalves, Ewan Harrison, David K. Jackson, Ian Johnston, Dominic Kwiatkowski, Cordelia Langford, John Sillitoe on behalf of the Wellcome Sanger Institute COVID-19 Surveillance Team |
| EPI_ISL_2538596, EPI_ISL_2538597 | Lighthouse Lab in Alderley Park | Wellcome Sanger Institute for the COVID-19 Genomics UK (COG-UK) Consortium | Jacquelyn Wynn, Mairead Hyland, The Lighthouse Lab in Alderley Park and Alex Alderton, Roberto Amato, Jeffrey Barrett, Sonia Goncalves, Ewan Harrison, David K. Jackson, Ian Johnston, Dominic Kwiatkowski, Cordelia Langford, John Sillitoe on behalf of the Wellcome Sanger Institute COVID-19 Surveillance Team |
| EPI_ISL_2538598 | Lighthouse Lab in Milton Keynes | Wellcome Sanger Institute for the COVID-19 Genomics UK (COG-UK) Consortium | The Lighthouse Lab in Milton Keynes and Alex Alderton, Roberto Amato, Jeffrey Barrett, Sonia Goncalves, Ewan Harrison, David K. Jackson, Ian Johnston, Dominic Kwiatkowski, Cordelia Langford, John Sillitoe on behalf of the Wellcome Sanger Institute COVID-19 Surveillance Team |
| EPI_ISL_2538599, EPI_ISL_2538600 | Lighthouse Lab in Alderley Park | Wellcome Sanger Institute for the COVID-19 Genomics UK (COG-UK) Consortium | Jacquelyn Wynn, Mairead Hyland, The Lighthouse Lab in Alderley Park and Alex Alderton, Roberto Amato, Jeffrey Barrett, Sonia Goncalves, Ewan Harrison, David K. Jackson, Ian Johnston, Dominic Kwiatkowski, Cordelia Langford, John Sillitoe on behalf of the Wellcome Sanger Institute COVID-19 Surveillance Team |
| EPI_ISL_2538601 | Lighthouse Lab in Glasgow | Wellcome Sanger Institute for the COVID-19 Genomics UK (COG-UK) Consortium | Harper VanSteenhouse, Yumi Kasai, David Gray, Carol Clugston, Anna Dominiczak and Alex Alderton, Roberto Amato, Jeffrey Barrett, Sonia Goncalves, Ewan Harrison, David K. Jackson, Ian Johnston, Dominic Kwiatkowski, Cordelia Langford, John Sillitoe on behalf of the Wellcome Sanger Institute COVID-19 Surveillance Team |
| EPI_ISL_2538602, EPI_ISL_2538604, EPI_ISL_2538605, EPI_ISL_2538606 | Lighthouse Lab in Alderley Park | Wellcome Sanger Institute for the COVID-19 Genomics UK (COG-UK) Consortium | Jacquelyn Wynn, Mairead Hyland, The Lighthouse Lab in Alderley Park and Alex Alderton, Roberto Amato, Jeffrey Barrett, Sonia Goncalves, Ewan Harrison, David K. Jackson, Ian Johnston, Dominic Kwiatkowski, Cordelia Langford, John Sillitoe on behalf of the Wellcome Sanger Institute COVID-19 Surveillance Team |
| EPI_ISL_2538607, EPI_ISL_2538608, EPI_ISL_2538609 | Lighthouse Lab in Glasgow | Wellcome Sanger Institute for the COVID-19 Genomics UK (COG-UK) Consortium | Harper VanSteenhouse, Yumi Kasai, David Gray, Carol Clugston, Anna Dominiczak and Alex Alderton, Roberto Amato, Jeffrey Barrett, Sonia Goncalves, Ewan Harrison, David K. Jackson, Ian Johnston, Dominic Kwiatkowski, Cordelia Langford, John Sillitoe on behalf of the Wellcome Sanger Institute COVID-19 Surveillance Team |
| EPI_ISL_2538610 | Lighthouse Lab in Alderley Park | Wellcome Sanger Institute for the COVID-19 Genomics UK (COG-UK) Consortium | Jacquelyn Wynn, Mairead Hyland, The Lighthouse Lab in Alderley Park and Alex Alderton, Roberto Amato, Jeffrey Barrett, Sonia Goncalves, Ewan Harrison, David K. Jackson, Ian Johnston, Dominic Kwiatkowski, Cordelia Langford, John Sillitoe on behalf of the Wellcome Sanger Institute COVID-19 Surveillance Team |
| EPI_ISL_2538611 | Lighthouse Lab in Milton Keynes | Wellcome Sanger Institute for the COVID-19 Genomics UK (COG-UK) Consortium | The Lighthouse Lab in Milton Keynes and Alex Alderton, Roberto Amato, Jeffrey Barrett, Sonia Goncalves, Ewan Harrison, David K. Jackson, Ian Johnston, Dominic Kwiatkowski, Cordelia Langford, John Sillitoe on behalf of the Wellcome Sanger Institute COVID-19 Surveillance Team |
| EPI_ISL_2538612 | Lighthouse Lab in Alderley Park | Wellcome Sanger Institute for the COVID-19 Genomics UK (COG-UK) Consortium | Jacquelyn Wynn, Mairead Hyland, The Lighthouse Lab in Alderley Park and Alex Alderton, Roberto Amato, Jeffrey Barrett, Sonia Goncalves, Ewan Harrison, David K. Jackson, Ian Johnston, Dominic Kwiatkowski, Cordelia Langford, John Sillitoe on behalf of the Wellcome Sanger Institute COVID-19 Surveillance Team |
| EPI_ISL_2538613 | Lighthouse Lab in Glasgow | Wellcome Sanger Institute for the COVID-19 Genomics UK (COG-UK) Consortium | Harper VanSteenhouse, Yumi Kasai, David Gray, Carol Clugston, Anna Dominiczak and Alex Alderton, Roberto Amato, Jeffrey Barrett, Sonia Goncalves, Ewan Harrison, David K. Jackson, Ian Johnston, Dominic Kwiatkowski, Cordelia Langford, John Sillitoe on behalf of the Wellcome Sanger Institute COVID-19 Surveillance Team |
| EPI_ISL_2538614, EPI_ISL_2538615, EPI_ISL_2538616, EPI_ISL_2538617, EPI_ISL_2538618, EPI_ISL_2538619 | Lighthouse Lab in Alderley Park | Wellcome Sanger Institute for the COVID-19 Genomics UK (COG-UK) Consortium | Jacquelyn Wynn, Mairead Hyland, The Lighthouse Lab in Alderley Park and Alex Alderton, Roberto Amato, Jeffrey Barrett, Sonia Goncalves, Ewan Harrison, David K. Jackson, Ian Johnston, Dominic Kwiatkowski, Cordelia Langford, John Sillitoe on behalf of the Wellcome Sanger Institute COVID-19 Surveillance Team |
| EPI_ISL_2538620 | Lighthouse Lab in Milton Keynes | Wellcome Sanger Institute for the COVID-19 Genomics UK (COG-UK) Consortium | The Lighthouse Lab in Milton Keynes and Alex Alderton, Roberto Amato, Jeffrey Barrett, Sonia Goncalves, Ewan Harrison, David K. Jackson, Ian Johnston, Dominic Kwiatkowski, Cordelia Langford, John Sillitoe on behalf of the Wellcome Sanger Institute COVID-19 Surveillance Team |
| EPI_ISL_2538621, EPI_ISL_2538622 | Lighthouse Lab in Alderley Park | Wellcome Sanger Institute for the COVID-19 Genomics UK (COG-UK) Consortium | Jacquelyn Wynn, Mairead Hyland, The Lighthouse Lab in Alderley Park and Alex Alderton, Roberto Amato, Jeffrey Barrett, Sonia Goncalves, Ewan Harrison, David K. Jackson, Ian Johnston, Dominic Kwiatkowski, Cordelia Langford, John Sillitoe on behalf of the Wellcome Sanger Institute COVID-19 Surveillance Team |
| EPI_ISL_2538623 | Lighthouse Lab in Milton Keynes | Wellcome Sanger Institute for the COVID-19 Genomics UK (COG-UK) Consortium | The Lighthouse Lab in Milton Keynes and Alex Alderton, Roberto Amato, Jeffrey Barrett, Sonia Goncalves, Ewan Harrison, David K. Jackson, Ian Johnston, Dominic Kwiatkowski, Cordelia Langford, John Sillitoe on behalf of the Wellcome Sanger Institute COVID-19 Surveillance Team |
| EPI_ISL_2538624, EPI_ISL_2538625 | Lighthouse Lab in Alderley Park | Wellcome Sanger Institute for the COVID-19 Genomics UK (COG-UK) Consortium | Jacquelyn Wynn, Mairead Hyland, The Lighthouse Lab in Alderley Park and Alex Alderton, Roberto Amato, Jeffrey Barrett, Sonia Goncalves, Ewan Harrison, David K. Jackson, Ian Johnston, Dominic Kwiatkowski, Cordelia Langford, John Sillitoe on behalf of the Wellcome Sanger Institute COVID-19 Surveillance Team |
| EPI_ISL_2538626 | Lighthouse Lab in Milton Keynes | Wellcome Sanger Institute for the COVID-19 Genomics UK (COG-UK) Consortium | The Lighthouse Lab in Milton Keynes and Alex Alderton, Roberto Amato, Jeffrey Barrett, Sonia Goncalves, Ewan Harrison, David K. Jackson, Ian Johnston, Dominic Kwiatkowski, Cordelia Langford, John Sillitoe on behalf of the Wellcome Sanger Institute COVID-19 Surveillance Team |
| EPI_ISL_2538627, EPI_ISL_2538628, EPI_ISL_2538629, EPI_ISL_2538630 | Lighthouse Lab in Alderley Park | Wellcome Sanger Institute for the COVID-19 Genomics UK (COG-UK) Consortium | Jacquelyn Wynn, Mairead Hyland, The Lighthouse Lab in Alderley Park and Alex Alderton, Roberto Amato, Jeffrey Barrett, Sonia Goncalves, Ewan Harrison, David K. Jackson, Ian Johnston, Dominic Kwiatkowski, Cordelia Langford, John Sillitoe on behalf of the Wellcome Sanger Institute COVID-19 Surveillance Team |
| EPI_ISL_2538631 | Lighthouse Lab in Glasgow | Wellcome Sanger Institute for the COVID-19 Genomics UK (COG-UK) Consortium | Harper VanSteenhouse, Yumi Kasai, David Gray, Carol Clugston, Anna Dominiczak and Alex Alderton, Roberto Amato, Jeffrey Barrett, Sonia Goncalves, Ewan Harrison, David K. Jackson, Ian Johnston, Dominic Kwiatkowski, Cordelia Langford, John Sillitoe on behalf of the Wellcome Sanger Institute COVID-19 Surveillance Team |
| EPI_ISL_2538632, EPI_ISL_2538633, EPI_ISL_2538634, EPI_ISL_2538635, EPI_ISL_2538636, EPI_ISL_2538637, EPI_ISL_2538639, EPI_ISL_2538640 | Lighthouse Lab in Alderley Park | Wellcome Sanger Institute for the COVID-19 Genomics UK (COG-UK) Consortium | Jacquelyn Wynn, Mairead Hyland, The Lighthouse Lab in Alderley Park and Alex Alderton, Roberto Amato, Jeffrey Barrett, Sonia Goncalves, Ewan Harrison, David K. Jackson, Ian Johnston, Dominic Kwiatkowski, Cordelia Langford, John Sillitoe on behalf of the Wellcome Sanger Institute COVID-19 Surveillance Team |
| EPI_ISL_2538641 | Lighthouse Lab in Glasgow | Wellcome Sanger Institute for the COVID-19 Genomics UK (COG-UK) Consortium | Harper VanSteenhouse, Yumi Kasai, David Gray, Carol Clugston, Anna Dominiczak and Alex Alderton, Roberto Amato, Jeffrey Barrett, Sonia Goncalves, Ewan Harrison, David K. Jackson, Ian Johnston, Dominic Kwiatkowski, Cordelia Langford, John Sillitoe on behalf of the Wellcome Sanger Institute COVID-19 Surveillance Team |
| EPI_ISL_2538642 | Lighthouse Lab in Milton Keynes | Wellcome Sanger Institute for the COVID-19 Genomics UK (COG-UK) Consortium | The Lighthouse Lab in Milton Keynes and Alex Alderton, Roberto Amato, Jeffrey Barrett, Sonia Goncalves, Ewan Harrison, David K. Jackson, Ian Johnston, Dominic Kwiatkowski, Cordelia Langford, John Sillitoe on behalf of the Wellcome Sanger Institute COVID-19 Surveillance Team |

EPI_ISL_2538643, EPI_ISL_2538645, EPI_ISL_2538646, EPI_ISL_2538647, EPI_ISL_2538648, EPI_ISL_2538649, EPI_ISL_2538650, EPI_ISL_2538651, EPI_ISL_2538652, EPI_ISL_2538653, EPI_ISL_2538655, EPI_ISL_2538656, EPI_ISL_2538657, EPI_ISL_2538658, EPI_ISL_2538659, EPI_ISL_2538660, EPI_ISL_2538661, EPI_ISL_2538662, EPI_ISL_2538663, EPI_ISL_2538664, EPI_ISL_2538665, EPI_ISL_2538666, EPI_ISL_2538667

| | | | |
|---|---|---|---|
| see above | Lighthouse Lab in Alderley Park | Wellcome Sanger Institute for the COVID-19 Genomics UK (COG-UK) Consortium | Jacquelyn Wynn, Mairead Hyland, The Lighthouse Lab in Alderley Park and Alex Alderton, Roberto Amato, Jeffrey Barrett, Sonia Goncalves, Ewan Harrison, David K. Jackson, Ian Johnston, Dominic Kwiatkowski, Cordelia Langford, John Sillitoe on behalf of the Wellcome Sanger Institute COVID-19 Surveillance Team |
| EPI_ISL_2538668 | Lighthouse Lab in Glasgow | Wellcome Sanger Institute for the COVID-19 Genomics UK (COG-UK) Consortium | Harper VanSteenhouse, Yumi Kasai, David Gray, Carol Clugston, Anna Dominiczak and Alex Alderton, Roberto Amato, Jeffrey Barrett, Sonia Goncalves, Ewan Harrison, David K. Jackson, Ian Johnston, Dominic Kwiatkowski, Cordelia Langford, John Sillitoe on behalf of the Wellcome Sanger Institute COVID-19 Surveillance Team |
| EPI_ISL_2538669 | Lighthouse Lab in Alderley Park | Wellcome Sanger Institute for the COVID-19 Genomics UK (COG-UK) Consortium | Jacquelyn Wynn, Mairead Hyland, The Lighthouse Lab in Alderley Park and Alex Alderton, Roberto Amato, Jeffrey Barrett, Sonia Goncalves, Ewan Harrison, David K. Jackson, Ian Johnston, Dominic Kwiatkowski, Cordelia Langford, John Sillitoe on behalf of the Wellcome Sanger Institute COVID-19 Surveillance Team |
| EPI_ISL_2538670 | Lighthouse Lab in Glasgow | Wellcome Sanger Institute for the COVID-19 Genomics UK (COG-UK) Consortium | Harper VanSteenhouse, Yumi Kasai, David Gray, Carol Clugston, Anna Dominiczak and Alex Alderton, Roberto Amato, Jeffrey Barrett, Sonia Goncalves, Ewan Harrison, David K. Jackson, Ian Johnston, Dominic Kwiatkowski, Cordelia Langford, John Sillitoe on behalf of the Wellcome Sanger Institute COVID-19 Surveillance Team |
| EPI_ISL_2538671 | Lighthouse Lab in Alderley Park | Wellcome Sanger Institute for the COVID-19 Genomics UK (COG-UK) Consortium | Jacquelyn Wynn, Mairead Hyland, The Lighthouse Lab in Alderley Park and Alex Alderton, Roberto Amato, Jeffrey Barrett, Sonia Goncalves, Ewan Harrison, David K. Jackson, Ian Johnston, Dominic Kwiatkowski, Cordelia Langford, John Sillitoe on behalf of the Wellcome Sanger Institute COVID-19 Surveillance Team |
| EPI_ISL_2538672 | Lighthouse Lab in Milton Keynes | Wellcome Sanger Institute for the COVID-19 Genomics UK (COG-UK) Consortium | The Lighthouse Lab in Milton Keynes and Alex Alderton, Roberto Amato, Jeffrey Barrett, Sonia Goncalves, Ewan Harrison, David K. Jackson, Ian Johnston, Dominic Kwiatkowski, Cordelia Langford, John Sillitoe on behalf of the Wellcome Sanger Institute COVID-19 Surveillance Team |
| EPI_ISL_2538673, EPI_ISL_2538674, EPI_ISL_2538676, | Lighthouse Lab in Alderley Park | Wellcome Sanger Institute for the COVID-19 Genomics UK (COG-UK) Consortium | Jacquelyn Wynn, Mairead Hyland, The Lighthouse Lab in Alderley Park and Alex Alderton, Roberto Amato, Jeffrey Barrett, Sonia Goncalves, Ewan Harrison, David K. Jackson, Ian Johnston, Dominic Kwiatkowski, Cordelia Langford, John Sillitoe on behalf of the Wellcome Sanger Institute COVID-19 Surveillance Team |

| | | | |
|---|---|---|---|
| EPI_ISL_2538677 | | | |
| EPI_ISL_2538678 | Lighthouse Lab in Glasgow | Wellcome Sanger Institute for the COVID-19 Genomics UK (COG-UK) Consortium | Harper VanSteenhouse, Yumi Kasai, David Gray, Carol Clugston, Anna Dominiczak and Alex Alderton, Roberto Amato, Jeffrey Barrett, Sonia Goncalves, Ewan Harrison, David K. Jackson, Ian Johnston, Dominic Kwiatkowski, Cordelia Langford, John Sillitoe on behalf of the Wellcome Sanger Institute COVID-19 Surveillance Team |
| EPI_ISL_2538679, EPI_ISL_2538680, EPI_ISL_2538681, EPI_ISL_2538682, EPI_ISL_2538683, EPI_ISL_2538684, EPI_ISL_2538685, EPI_ISL_2538686, EPI_ISL_2538687, EPI_ISL_2538688, EPI_ISL_2538689, EPI_ISL_2538690, EPI_ISL_2538691, EPI_ISL_2538692 | | | |
| see above | Lighthouse Lab in Alderley Park | Wellcome Sanger Institute for the COVID-19 Genomics UK (COG-UK) Consortium | Jacquelyn Wynn, Mairead Hyland, The Lighthouse Lab in Alderley Park and Alex Alderton, Roberto Amato, Jeffrey Barrett, Sonia Goncalves, Ewan Harrison, David K. Jackson, Ian Johnston, Dominic Kwiatkowski, Cordelia Langford, John Sillitoe on behalf of the Wellcome Sanger Institute COVID-19 Surveillance Team |
| EPI_ISL_2538694 | Lighthouse Lab in Glasgow | Wellcome Sanger Institute for the COVID-19 Genomics UK (COG-UK) Consortium | Harper VanSteenhouse, Yumi Kasai, David Gray, Carol Clugston, Anna Dominiczak and Alex Alderton, Roberto Amato, Jeffrey Barrett, Sonia Goncalves, Ewan Harrison, David K. Jackson, Ian Johnston, Dominic Kwiatkowski, Cordelia Langford, John Sillitoe on behalf of the Wellcome Sanger Institute COVID-19 Surveillance Team |
| EPI_ISL_2538695 | Lighthouse Lab in Alderley Park | Wellcome Sanger Institute for the COVID-19 Genomics UK (COG-UK) Consortium | Jacquelyn Wynn, Mairead Hyland, The Lighthouse Lab in Alderley Park and Alex Alderton, Roberto Amato, Jeffrey Barrett, Sonia Goncalves, Ewan Harrison, David K. Jackson, Ian Johnston, Dominic Kwiatkowski, Cordelia Langford, John Sillitoe on behalf of the Wellcome Sanger Institute COVID-19 Surveillance Team |
| EPI_ISL_2538696 | Lighthouse Lab in Glasgow | Wellcome Sanger Institute for the COVID-19 Genomics UK (COG-UK) Consortium | Harper VanSteenhouse, Yumi Kasai, David Gray, Carol Clugston, Anna Dominiczak and Alex Alderton, Roberto Amato, Jeffrey Barrett, Sonia Goncalves, Ewan Harrison, David K. Jackson, Ian Johnston, Dominic Kwiatkowski, Cordelia Langford, John Sillitoe on behalf of the Wellcome Sanger Institute COVID-19 Surveillance Team |
| EPI_ISL_2538697 | Lighthouse Lab in Alderley Park | Wellcome Sanger Institute for the COVID-19 Genomics UK (COG-UK) Consortium | Jacquelyn Wynn, Mairead Hyland, The Lighthouse Lab in Alderley Park and Alex Alderton, Roberto Amato, Jeffrey Barrett, Sonia Goncalves, Ewan Harrison, David K. Jackson, Ian Johnston, Dominic Kwiatkowski, Cordelia Langford, John Sillitoe on behalf of the Wellcome Sanger Institute COVID-19 Surveillance Team |
| EPI_ISL_2538699, EPI_ISL_2538700 | Lighthouse Lab in Glasgow | Wellcome Sanger Institute for the COVID-19 Genomics UK (COG-UK) Consortium | Harper VanSteenhouse, Yumi Kasai, David Gray, Carol Clugston, Anna Dominiczak and Alex Alderton, Roberto Amato, Jeffrey Barrett, Sonia Goncalves, Ewan Harrison, David K. Jackson, Ian Johnston, Dominic Kwiatkowski, Cordelia Langford, John Sillitoe on behalf of the Wellcome Sanger Institute COVID-19 Surveillance Team |
| EPI_ISL_2538701 | Lighthouse Lab in Alderley Park | Wellcome Sanger Institute for the COVID-19 Genomics UK (COG-UK) Consortium | Jacquelyn Wynn, Mairead Hyland, The Lighthouse Lab in Alderley Park and Alex Alderton, Roberto Amato, Jeffrey Barrett, Sonia Goncalves, Ewan Harrison, David K. Jackson, Ian Johnston, Dominic Kwiatkowski, Cordelia Langford, John Sillitoe on behalf of the Wellcome Sanger Institute COVID-19 Surveillance Team |
| EPI_ISL_2538702 | Lighthouse Lab in Milton Keynes | Wellcome Sanger Institute for the COVID-19 Genomics UK (COG-UK) Consortium | The Lighthouse Lab in Milton Keynes and Alex Alderton, Roberto Amato, Jeffrey Barrett, Sonia Goncalves, Ewan Harrison, David K. Jackson, Ian Johnston, Dominic Kwiatkowski, Cordelia Langford, John Sillitoe on behalf of the Wellcome Sanger Institute COVID-19 Surveillance Team |
| EPI_ISL_2538704, EPI_ISL_2538705, EPI_ISL_2538706, EPI_ISL_2538707 | Lighthouse Lab in Alderley Park | Wellcome Sanger Institute for the COVID-19 Genomics UK (COG-UK) Consortium | Jacquelyn Wynn, Mairead Hyland, The Lighthouse Lab in Alderley Park and Alex Alderton, Roberto Amato, Jeffrey Barrett, Sonia Goncalves, Ewan Harrison, David K. Jackson, Ian Johnston, Dominic Kwiatkowski, Cordelia Langford, John Sillitoe on behalf of the Wellcome Sanger Institute COVID-19 Surveillance Team |
| EPI_ISL_2538708 | Lighthouse Lab in Glasgow | Wellcome Sanger Institute for the COVID-19 Genomics UK (COG-UK) Consortium | Harper VanSteenhouse, Yumi Kasai, David Gray, Carol Clugston, Anna Dominiczak and Alex Alderton, Roberto Amato, Jeffrey Barrett, Sonia Goncalves, Ewan Harrison, David K. Jackson, Ian Johnston, Dominic Kwiatkowski, Cordelia Langford, John Sillitoe on behalf of the Wellcome Sanger Institute COVID-19 Surveillance Team |
| EPI_ISL_2538709 | Lighthouse Lab in Alderley Park | Wellcome Sanger Institute for the COVID-19 Genomics UK (COG-UK) Consortium | Jacquelyn Wynn, Mairead Hyland, The Lighthouse Lab in Alderley Park and Alex Alderton, Roberto Amato, Jeffrey Barrett, Sonia Goncalves, Ewan Harrison, David K. Jackson, Ian Johnston, Dominic Kwiatkowski, Cordelia Langford, John Sillitoe on behalf of the Wellcome Sanger Institute COVID-19 Surveillance Team |
| EPI_ISL_2538711 | Lighthouse Lab in Milton Keynes | Wellcome Sanger Institute for the COVID-19 Genomics UK (COG-UK) Consortium | The Lighthouse Lab in Milton Keynes and Alex Alderton, Roberto Amato, Jeffrey Barrett, Sonia Goncalves, Ewan Harrison, David K. Jackson, Ian Johnston, Dominic Kwiatkowski, Cordelia Langford, John Sillitoe on behalf of the Wellcome Sanger Institute COVID-19 Surveillance Team |
| EPI_ISL_2538712, EPI_ISL_2538713 | Lighthouse Lab in Glasgow | Wellcome Sanger Institute for the COVID-19 Genomics UK (COG-UK) Consortium | Harper VanSteenhouse, Yumi Kasai, David Gray, Carol Clugston, Anna Dominiczak and Alex Alderton, Roberto Amato, Jeffrey Barrett, Sonia Goncalves, Ewan Harrison, David K. Jackson, Ian Johnston, Dominic Kwiatkowski, Cordelia Langford, John Sillitoe on behalf of the Wellcome Sanger Institute COVID-19 Surveillance Team |
| EPI_ISL_2538715, EPI_ISL_2538716 | Lighthouse Lab in Alderley Park | Wellcome Sanger Institute for the COVID-19 Genomics UK (COG-UK) Consortium | Jacquelyn Wynn, Mairead Hyland, The Lighthouse Lab in Alderley Park and Alex Alderton, Roberto Amato, Jeffrey Barrett, Sonia Goncalves, Ewan Harrison, David K. Jackson, Ian Johnston, Dominic Kwiatkowski, Cordelia Langford, John Sillitoe on behalf of the Wellcome Sanger Institute COVID-19 Surveillance Team |
| EPI_ISL_2538717 | Lighthouse Lab in Glasgow | Wellcome Sanger Institute for the COVID-19 Genomics UK (COG-UK) Consortium | Harper VanSteenhouse, Yumi Kasai, David Gray, Carol Clugston, Anna Dominiczak and Alex Alderton, Roberto Amato, Jeffrey Barrett, Sonia Goncalves, Ewan Harrison, David K. Jackson, Ian Johnston, Dominic Kwiatkowski, Cordelia Langford, John Sillitoe on behalf of the Wellcome Sanger Institute COVID-19 Surveillance Team |
| EPI_ISL_2538718 | Lighthouse Lab in Alderley Park | Wellcome Sanger Institute for the COVID-19 Genomics UK (COG-UK) Consortium | Jacquelyn Wynn, Mairead Hyland, The Lighthouse Lab in Alderley Park and Alex Alderton, Roberto Amato, Jeffrey Barrett, Sonia Goncalves, Ewan Harrison, David K. Jackson, Ian Johnston, Dominic Kwiatkowski, Cordelia Langford, John Sillitoe on behalf of the Wellcome Sanger Institute COVID-19 Surveillance Team |
| EPI_ISL_2538719 | Lighthouse Lab in Glasgow | Wellcome Sanger Institute for the COVID-19 Genomics UK (COG-UK) Consortium | Harper VanSteenhouse, Yumi Kasai, David Gray, Carol Clugston, Anna Dominiczak and Alex Alderton, Roberto Amato, Jeffrey Barrett, Sonia Goncalves, Ewan Harrison, David K. Jackson, Ian Johnston, Dominic Kwiatkowski, Cordelia Langford, John Sillitoe on behalf of the Wellcome Sanger Institute COVID-19 Surveillance Team |
| EPI_ISL_2538720, EPI_ISL_2538721, EPI_ISL_2538722 | Lighthouse Lab in Alderley Park | Wellcome Sanger Institute for the COVID-19 Genomics UK (COG-UK) Consortium | Jacquelyn Wynn, Mairead Hyland, The Lighthouse Lab in Alderley Park and Alex Alderton, Roberto Amato, Jeffrey Barrett, Sonia Goncalves, Ewan Harrison, David K. Jackson, Ian Johnston, Dominic Kwiatkowski, Cordelia Langford, John Sillitoe on behalf of the Wellcome Sanger Institute COVID-19 Surveillance Team |
| EPI_ISL_2538723 | Lighthouse Lab in Milton Keynes | Wellcome Sanger Institute for the COVID-19 Genomics UK (COG-UK) Consortium | The Lighthouse Lab in Milton Keynes and Alex Alderton, Roberto Amato, Jeffrey Barrett, Sonia Goncalves, Ewan Harrison, David K. Jackson, Ian Johnston, Dominic Kwiatkowski, Cordelia Langford, John Sillitoe on behalf of the Wellcome Sanger Institute COVID-19 Surveillance Team |
| EPI_ISL_2538724, EPI_ISL_2538725, EPI_ISL_2538726, EPI_ISL_2538728, EPI_ISL_2538729, EPI_ISL_2538730, EPI_ISL_2538731 | Lighthouse Lab in Alderley Park | Wellcome Sanger Institute for the COVID-19 Genomics UK (COG-UK) Consortium | Jacquelyn Wynn, Mairead Hyland, The Lighthouse Lab in Alderley Park and Alex Alderton, Roberto Amato, Jeffrey Barrett, Sonia Goncalves, Ewan Harrison, David K. Jackson, Ian Johnston, Dominic Kwiatkowski, Cordelia Langford, John Sillitoe on behalf of the Wellcome Sanger Institute COVID-19 Surveillance Team |
| EPI_ISL_2538732 | Lighthouse Lab in Milton Keynes | Wellcome Sanger Institute for the COVID-19 Genomics UK (COG-UK) Consortium | The Lighthouse Lab in Milton Keynes and Alex Alderton, Roberto Amato, Jeffrey Barrett, Sonia Goncalves, Ewan Harrison, David K. Jackson, Ian Johnston, Dominic Kwiatkowski, Cordelia Langford, John Sillitoe on behalf of the Wellcome Sanger Institute COVID-19 Surveillance Team |
| EPI_ISL_2538733 | Lighthouse Lab in Glasgow | Wellcome Sanger Institute for the COVID-19 Genomics UK (COG-UK) Consortium | Harper VanSteenhouse, Yumi Kasai, David Gray, Carol Clugston, Anna Dominiczak and Alex Alderton, Roberto Amato, Jeffrey Barrett, Sonia Goncalves, Ewan Harrison, David K. Jackson, Ian Johnston, Dominic Kwiatkowski, Cordelia Langford, John Sillitoe on behalf of the Wellcome Sanger Institute COVID-19 Surveillance Team |
| EPI_ISL_2538734, EPI_ISL_2538735, EPI_ISL_2538736, EPI_ISL_2538737, EPI_ISL_2538738, EPI_ISL_2538739 | Lighthouse Lab in Alderley Park | Wellcome Sanger Institute for the COVID-19 Genomics UK (COG-UK) Consortium | Jacquelyn Wynn, Mairead Hyland, The Lighthouse Lab in Alderley Park and Alex Alderton, Roberto Amato, Jeffrey Barrett, Sonia Goncalves, Ewan Harrison, David K. Jackson, Ian Johnston, Dominic Kwiatkowski, Cordelia Langford, John Sillitoe on behalf of the Wellcome Sanger Institute COVID-19 Surveillance Team |
| EPI_ISL_2538740 | Lighthouse Lab in Glasgow | Wellcome Sanger Institute for the COVID-19 Genomics UK (COG-UK) Consortium | Harper VanSteenhouse, Yumi Kasai, David Gray, Carol Clugston, Anna Dominiczak and Alex Alderton, Roberto Amato, Jeffrey Barrett, Sonia Goncalves, Ewan Harrison, David K. Jackson, Ian Johnston, Dominic Kwiatkowski, Cordelia Langford, John Sillitoe on behalf of the Wellcome Sanger Institute COVID-19 Surveillance Team |
| EPI_ISL_2538741, EPI_ISL_2538742, EPI_ISL_2538743, EPI_ISL_2538744 | Lighthouse Lab in Alderley Park | Wellcome Sanger Institute for the COVID-19 Genomics UK (COG-UK) Consortium | Jacquelyn Wynn, Mairead Hyland, The Lighthouse Lab in Alderley Park and Alex Alderton, Roberto Amato, Jeffrey Barrett, Sonia Goncalves, Ewan Harrison, David K. Jackson, Ian Johnston, Dominic Kwiatkowski, Cordelia Langford, John Sillitoe on behalf of the Wellcome Sanger Institute COVID-19 Surveillance Team |
| EPI_ISL_2538745 | Lighthouse Lab in Milton Keynes | Wellcome Sanger Institute for the COVID-19 Genomics UK (COG-UK) Consortium | The Lighthouse Lab in Milton Keynes and Alex Alderton, Roberto Amato, Jeffrey Barrett, Sonia Goncalves, Ewan Harrison, David K. Jackson, Ian Johnston, Dominic Kwiatkowski, Cordelia Langford, John Sillitoe on behalf of the Wellcome Sanger Institute COVID-19 Surveillance Team |
| EPI_ISL_2538746, EPI_ISL_2538747 | Lighthouse Lab in Alderley Park | Wellcome Sanger Institute for the COVID-19 Genomics UK (COG-UK) Consortium | Jacquelyn Wynn, Mairead Hyland, The Lighthouse Lab in Alderley Park and Alex Alderton, Roberto Amato, Jeffrey Barrett, Sonia Goncalves, Ewan Harrison, David K. Jackson, Ian Johnston, Dominic Kwiatkowski, Cordelia Langford, John Sillitoe on behalf of the Wellcome Sanger Institute COVID-19 Surveillance Team |
| EPI_ISL_2538748 | Lighthouse Lab in Glasgow | Wellcome Sanger Institute for the COVID-19 Genomics UK (COG-UK) Consortium | Harper VanSteenhouse, Yumi Kasai, David Gray, Carol Clugston, Anna Dominiczak and Alex Alderton, Roberto Amato, Jeffrey Barrett, Sonia Goncalves, Ewan Harrison, David K. Jackson, Ian Johnston, Dominic Kwiatkowski, Cordelia Langford, John Sillitoe on behalf of the Wellcome Sanger Institute COVID-19 Surveillance Team |
| EPI_ISL_2538749, EPI_ISL_2538751, EPI_ISL_2538752, EPI_ISL_2538754, EPI_ISL_2538755, EPI_ISL_2538756 | Lighthouse Lab in Alderley Park | Wellcome Sanger Institute for the COVID-19 Genomics UK (COG-UK) Consortium | Jacquelyn Wynn, Mairead Hyland, The Lighthouse Lab in Alderley Park and Alex Alderton, Roberto Amato, Jeffrey Barrett, Sonia Goncalves, Ewan Harrison, David K. Jackson, Ian Johnston, Dominic Kwiatkowski, Cordelia Langford, John Sillitoe on behalf of the Wellcome Sanger Institute COVID-19 Surveillance Team |
| EPI_ISL_2538757, EPI_ISL_2538758 | Lighthouse Lab in Milton Keynes | Wellcome Sanger Institute for the COVID-19 Genomics UK (COG-UK) Consortium | The Lighthouse Lab in Milton Keynes and Alex Alderton, Roberto Amato, Jeffrey Barrett, Sonia Goncalves, Ewan Harrison, David K. Jackson, Ian Johnston, Dominic Kwiatkowski, Cordelia Langford, John Sillitoe on behalf of the Wellcome Sanger Institute COVID-19 Surveillance Team |
| EPI_ISL_2538759, EPI_ISL_2538760, EPI_ISL_2538761 | Lighthouse Lab in Alderley Park | Wellcome Sanger Institute for the COVID-19 Genomics UK (COG-UK) Consortium | Jacquelyn Wynn, Mairead Hyland, The Lighthouse Lab in Alderley Park and Alex Alderton, Roberto Amato, Jeffrey Barrett, Sonia Goncalves, Ewan Harrison, David K. Jackson, Ian Johnston, Dominic Kwiatkowski, Cordelia Langford, John Sillitoe on behalf of the Wellcome Sanger Institute COVID-19 Surveillance Team |
| EPI_ISL_2538762 | Lighthouse Lab in Glasgow | Wellcome Sanger Institute for the COVID-19 Genomics UK (COG-UK) Consortium | Harper VanSteenhouse, Yumi Kasai, David Gray, Carol Clugston, Anna Dominiczak and Alex Alderton, Roberto Amato, Jeffrey Barrett, Sonia Goncalves, Ewan Harrison, David K. Jackson, Ian Johnston, Dominic Kwiatkowski, Cordelia Langford, John Sillitoe on behalf of the Wellcome Sanger Institute COVID-19 Surveillance Team |
| EPI_ISL_2538763 | Lighthouse Lab in Alderley Park | Wellcome Sanger Institute for the COVID-19 Genomics UK (COG-UK) Consortium | Jacquelyn Wynn, Mairead Hyland, The Lighthouse Lab in Alderley Park and Alex Alderton, Roberto Amato, Jeffrey Barrett, Sonia Goncalves, Ewan Harrison, David K. Jackson, Ian Johnston, Dominic Kwiatkowski, Cordelia Langford, John Sillitoe on behalf of the Wellcome Sanger Institute COVID-19 Surveillance Team |
| EPI_ISL_2538764 | Lighthouse Lab in Glasgow | Wellcome Sanger Institute for the COVID-19 Genomics UK (COG-UK) Consortium | Harper VanSteenhouse, Yumi Kasai, David Gray, Carol Clugston, Anna Dominiczak and Alex Alderton, Roberto Amato, Jeffrey Barrett, Sonia Goncalves, Ewan Harrison, David K. Jackson, Ian Johnston, Dominic Kwiatkowski, Cordelia Langford, John Sillitoe on behalf of the Wellcome Sanger Institute COVID-19 Surveillance Team |
| EPI_ISL_2538765, EPI_ISL_2538766, EPI_ISL_2538767 | Lighthouse Lab in Alderley Park | Wellcome Sanger Institute for the COVID-19 Genomics UK (COG-UK) Consortium | Jacquelyn Wynn, Mairead Hyland, The Lighthouse Lab in Alderley Park and Alex Alderton, Roberto Amato, Jeffrey Barrett, Sonia Goncalves, Ewan Harrison, David K. Jackson, Ian Johnston, Dominic Kwiatkowski, Cordelia Langford, John Sillitoe on behalf of the Wellcome Sanger Institute COVID-19 Surveillance Team |

| | | | |
|---|---|---|---|
| EPI_ISL_2538768 | Lighthouse Lab in Milton Keynes | Wellcome Sanger Institute for the COVID-19 Genomics UK (COG-UK) Consortium | The Lighthouse Lab in Milton Keynes and Alex Alderton, Roberto Amato, Jeffrey Barrett, Sonia Goncalves, Ewan Harrison, David K. Jackson, Ian Johnston, Dominic Kwiatkowski, Cordelia Langford, John Sillitoe on behalf of the Wellcome Sanger Institute COVID-19 Surveillance Team |
| EPI_ISL_2538769, EPI_ISL_2538770, EPI_ISL_2538771, EPI_ISL_2538772, EPI_ISL_2538773, EPI_ISL_2538774 | Lighthouse Lab in Alderley Park | Wellcome Sanger Institute for the COVID-19 Genomics UK (COG-UK) Consortium | Jacquelyn Wynn, Mairead Hyland, The Lighthouse Lab in Alderley Park and Alex Alderton, Roberto Amato, Jeffrey Barrett, Sonia Goncalves, Ewan Harrison, David K. Jackson, Ian Johnston, Dominic Kwiatkowski, Cordelia Langford, John Sillitoe on behalf of the Wellcome Sanger Institute COVID-19 Surveillance Team |
| EPI_ISL_2538775 | Lighthouse Lab in Glasgow | Wellcome Sanger Institute for the COVID-19 Genomics UK (COG-UK) Consortium | Harper VanSteenhouse, Yumi Kasai, David Gray, Carol Clugston, Anna Dominiczak and Alex Alderton, Roberto Amato, Jeffrey Barrett, Sonia Goncalves, Ewan Harrison, David K. Jackson, Ian Johnston, Dominic Kwiatkowski, Cordelia Langford, John Sillitoe on behalf of the Wellcome Sanger Institute COVID-19 Surveillance Team |
| EPI_ISL_2538776 | Lighthouse Lab in Alderley Park | Wellcome Sanger Institute for the COVID-19 Genomics UK (COG-UK) Consortium | Jacquelyn Wynn, Mairead Hyland, The Lighthouse Lab in Alderley Park and Alex Alderton, Roberto Amato, Jeffrey Barrett, Sonia Goncalves, Ewan Harrison, David K. Jackson, Ian Johnston, Dominic Kwiatkowski, Cordelia Langford, John Sillitoe on behalf of the Wellcome Sanger Institute COVID-19 Surveillance Team |
| EPI_ISL_2538777, EPI_ISL_2538779 | Lighthouse Lab in Milton Keynes | Wellcome Sanger Institute for the COVID-19 Genomics UK (COG-UK) Consortium | The Lighthouse Lab in Milton Keynes and Alex Alderton, Roberto Amato, Jeffrey Barrett, Sonia Goncalves, Ewan Harrison, David K. Jackson, Ian Johnston, Dominic Kwiatkowski, Cordelia Langford, John Sillitoe on behalf of the Wellcome Sanger Institute COVID-19 Surveillance Team |
| EPI_ISL_2538780, EPI_ISL_2538781, EPI_ISL_2538782 | Lighthouse Lab in Alderley Park | Wellcome Sanger Institute for the COVID-19 Genomics UK (COG-UK) Consortium | Jacquelyn Wynn, Mairead Hyland, The Lighthouse Lab in Alderley Park and Alex Alderton, Roberto Amato, Jeffrey Barrett, Sonia Goncalves, Ewan Harrison, David K. Jackson, Ian Johnston, Dominic Kwiatkowski, Cordelia Langford, John Sillitoe on behalf of the Wellcome Sanger Institute COVID-19 Surveillance Team |
| EPI_ISL_2538783 | Lighthouse Lab in Milton Keynes | Wellcome Sanger Institute for the COVID-19 Genomics UK (COG-UK) Consortium | The Lighthouse Lab in Milton Keynes and Alex Alderton, Roberto Amato, Jeffrey Barrett, Sonia Goncalves, Ewan Harrison, David K. Jackson, Ian Johnston, Dominic Kwiatkowski, Cordelia Langford, John Sillitoe on behalf of the Wellcome Sanger Institute COVID-19 Surveillance Team |
| EPI_ISL_2538784 | Lighthouse Lab in Alderley Park | Wellcome Sanger Institute for the COVID-19 Genomics UK (COG-UK) Consortium | Jacquelyn Wynn, Mairead Hyland, The Lighthouse Lab in Alderley Park and Alex Alderton, Roberto Amato, Jeffrey Barrett, Sonia Goncalves, Ewan Harrison, David K. Jackson, Ian Johnston, Dominic Kwiatkowski, Cordelia Langford, John Sillitoe on behalf of the Wellcome Sanger Institute COVID-19 Surveillance Team |
| EPI_ISL_2538785 | Lighthouse Lab in Glasgow | Wellcome Sanger Institute for the COVID-19 Genomics UK (COG-UK) Consortium | Harper VanSteenhouse, Yumi Kasai, David Gray, Carol Clugston, Anna Dominiczak and Alex Alderton, Roberto Amato, Jeffrey Barrett, Sonia Goncalves, Ewan Harrison, David K. Jackson, Ian Johnston, Dominic Kwiatkowski, Cordelia Langford, John Sillitoe on behalf of the Wellcome Sanger Institute COVID-19 Surveillance Team |
| EPI_ISL_2538786 | Lighthouse Lab in Alderley Park | Wellcome Sanger Institute for the COVID-19 Genomics UK (COG-UK) Consortium | Jacquelyn Wynn, Mairead Hyland, The Lighthouse Lab in Alderley Park and Alex Alderton, Roberto Amato, Jeffrey Barrett, Sonia Goncalves, Ewan Harrison, David K. Jackson, Ian Johnston, Dominic Kwiatkowski, Cordelia Langford, John Sillitoe on behalf of the Wellcome Sanger Institute COVID-19 Surveillance Team |
| EPI_ISL_2538787 | Lighthouse Lab in Glasgow | Wellcome Sanger Institute for the COVID-19 Genomics UK (COG-UK) Consortium | Harper VanSteenhouse, Yumi Kasai, David Gray, Carol Clugston, Anna Dominiczak and Alex Alderton, Roberto Amato, Jeffrey Barrett, Sonia Goncalves, Ewan Harrison, David K. Jackson, Ian Johnston, Dominic Kwiatkowski, Cordelia Langford, John Sillitoe on behalf of the Wellcome Sanger Institute COVID-19 Surveillance Team |
| EPI_ISL_2538788, EPI_ISL_2538789, EPI_ISL_2538790, EPI_ISL_2538791, EPI_ISL_2538792, EPI_ISL_2538793, EPI_ISL_2538794, EPI_ISL_2538795, EPI_ISL_2538796, EPI_ISL_2538797, EPI_ISL_2538798 | | | |
| see above | Lighthouse Lab in Alderley Park | Wellcome Sanger Institute for the COVID-19 Genomics UK (COG-UK) Consortium | Jacquelyn Wynn, Mairead Hyland, The Lighthouse Lab in Alderley Park and Alex Alderton, Roberto Amato, Jeffrey Barrett, Sonia Goncalves, Ewan Harrison, David K. Jackson, Ian Johnston, Dominic Kwiatkowski, Cordelia Langford, John Sillitoe on behalf of the Wellcome Sanger Institute COVID-19 Surveillance Team |
| EPI_ISL_2538799 | Lighthouse Lab in Milton Keynes | Wellcome Sanger Institute for the COVID-19 Genomics UK (COG-UK) Consortium | The Lighthouse Lab in Milton Keynes and Alex Alderton, Roberto Amato, Jeffrey Barrett, Sonia Goncalves, Ewan Harrison, David K. Jackson, Ian Johnston, Dominic Kwiatkowski, Cordelia Langford, John Sillitoe on behalf of the Wellcome Sanger Institute COVID-19 Surveillance Team |
| EPI_ISL_2538800, EPI_ISL_2538801 | Lighthouse Lab in Alderley Park | Wellcome Sanger Institute for the COVID-19 Genomics UK (COG-UK) Consortium | Jacquelyn Wynn, Mairead Hyland, The Lighthouse Lab in Alderley Park and Alex Alderton, Roberto Amato, Jeffrey Barrett, Sonia Goncalves, Ewan Harrison, David K. Jackson, Ian Johnston, Dominic Kwiatkowski, Cordelia Langford, John Sillitoe on behalf of the Wellcome Sanger Institute COVID-19 Surveillance Team |
| EPI_ISL_2538802 | Lighthouse Lab in Milton Keynes | Wellcome Sanger Institute for the COVID-19 Genomics UK (COG-UK) Consortium | The Lighthouse Lab in Milton Keynes and Alex Alderton, Roberto Amato, Jeffrey Barrett, Sonia Goncalves, Ewan Harrison, David K. Jackson, Ian Johnston, Dominic Kwiatkowski, Cordelia Langford, John Sillitoe on behalf of the Wellcome Sanger Institute COVID-19 Surveillance Team |
| EPI_ISL_2538803, EPI_ISL_2538804, EPI_ISL_2538805, EPI_ISL_2538806, EPI_ISL_2538807, EPI_ISL_2538808, EPI_ISL_2538809 | Lighthouse Lab in Alderley Park | Wellcome Sanger Institute for the COVID-19 Genomics UK (COG-UK) Consortium | Jacquelyn Wynn, Mairead Hyland, The Lighthouse Lab in Alderley Park and Alex Alderton, Roberto Amato, Jeffrey Barrett, Sonia Goncalves, Ewan Harrison, David K. Jackson, Ian Johnston, Dominic Kwiatkowski, Cordelia Langford, John Sillitoe on behalf of the Wellcome Sanger Institute COVID-19 Surveillance Team |
| EPI_ISL_2538810, EPI_ISL_2538811 | Lighthouse Lab in Glasgow | Wellcome Sanger Institute for the COVID-19 Genomics UK (COG-UK) Consortium | Harper VanSteenhouse, Yumi Kasai, David Gray, Carol Clugston, Anna Dominiczak and Alex Alderton, Roberto Amato, Jeffrey Barrett, Sonia Goncalves, Ewan Harrison, David K. Jackson, Ian Johnston, Dominic Kwiatkowski, Cordelia Langford, John Sillitoe on behalf of the Wellcome Sanger Institute COVID-19 Surveillance Team |
| EPI_ISL_2538812 | Lighthouse Lab in Alderley Park | Wellcome Sanger Institute for the COVID-19 Genomics UK (COG-UK) Consortium | Jacquelyn Wynn, Mairead Hyland, The Lighthouse Lab in Alderley Park and Alex Alderton, Roberto Amato, Jeffrey Barrett, Sonia Goncalves, Ewan Harrison, David K. Jackson, Ian Johnston, Dominic Kwiatkowski, Cordelia Langford, John Sillitoe on behalf of the Wellcome Sanger Institute COVID-19 Surveillance Team |
| EPI_ISL_2538813 | Lighthouse Lab in Glasgow | Wellcome Sanger Institute for the COVID-19 Genomics UK (COG-UK) Consortium | Harper VanSteenhouse, Yumi Kasai, David Gray, Carol Clugston, Anna Dominiczak and Alex Alderton, Roberto Amato, Jeffrey Barrett, Sonia Goncalves, Ewan Harrison, David K. Jackson, Ian Johnston, Dominic Kwiatkowski, Cordelia Langford, John Sillitoe on behalf of the Wellcome Sanger Institute COVID-19 Surveillance Team |
| EPI_ISL_2538814, EPI_ISL_2538815 | Lighthouse Lab in Alderley Park | Wellcome Sanger Institute for the COVID-19 Genomics UK (COG-UK) Consortium | Jacquelyn Wynn, Mairead Hyland, The Lighthouse Lab in Alderley Park and Alex Alderton, Roberto Amato, Jeffrey Barrett, Sonia Goncalves, Ewan Harrison, David K. Jackson, Ian Johnston, Dominic Kwiatkowski, Cordelia Langford, John Sillitoe on behalf of the Wellcome Sanger Institute COVID-19 Surveillance Team |
| EPI_ISL_2538816 | Lighthouse Lab in Milton Keynes | Wellcome Sanger Institute for the COVID-19 Genomics UK (COG-UK) Consortium | The Lighthouse Lab in Milton Keynes and Alex Alderton, Roberto Amato, Jeffrey Barrett, Sonia Goncalves, Ewan Harrison, David K. Jackson, Ian Johnston, Dominic Kwiatkowski, Cordelia Langford, John Sillitoe on behalf of the Wellcome Sanger Institute COVID-19 Surveillance Team |
| EPI_ISL_2538817 | Lighthouse Lab in Alderley Park | Wellcome Sanger Institute for the COVID-19 Genomics UK (COG-UK) Consortium | Jacquelyn Wynn, Mairead Hyland, The Lighthouse Lab in Alderley Park and Alex Alderton, Roberto Amato, Jeffrey Barrett, Sonia Goncalves, Ewan Harrison, David K. Jackson, Ian Johnston, Dominic Kwiatkowski, Cordelia Langford, John Sillitoe on behalf of the Wellcome Sanger Institute COVID-19 Surveillance Team |
| EPI_ISL_2538818 | Lighthouse Lab in Milton Keynes | Wellcome Sanger Institute for the COVID-19 Genomics UK (COG-UK) Consortium | The Lighthouse Lab in Milton Keynes and Alex Alderton, Roberto Amato, Jeffrey Barrett, Sonia Goncalves, Ewan Harrison, David K. Jackson, Ian Johnston, Dominic Kwiatkowski, Cordelia Langford, John Sillitoe on behalf of the Wellcome Sanger Institute COVID-19 Surveillance Team |
| EPI_ISL_2538819, EPI_ISL_2538820, EPI_ISL_2538821, EPI_ISL_2538822, EPI_ISL_2538823, EPI_ISL_2538824, EPI_ISL_2538825, EPI_ISL_2538826, EPI_ISL_2538827, EPI_ISL_2538828, EPI_ISL_2538829, EPI_ISL_2538831, EPI_ISL_2538832, EPI_ISL_2538833, EPI_ISL_2538834, EPI_ISL_2538835, EPI_ISL_2538836, EPI_ISL_2538837, EPI_ISL_2538838, EPI_ISL_2538839, EPI_ISL_2538840, EPI_ISL_2538841, EPI_ISL_2538842, EPI_ISL_2538843, EPI_ISL_2538844, EPI_ISL_2538845, EPI_ISL_2538846, EPI_ISL_2538847, EPI_ISL_2538848, EPI_ISL_2538849, EPI_ISL_2538850, EPI_ISL_2538851, EPI_ISL_2538853, EPI_ISL_2538854, EPI_ISL_2538857, EPI_ISL_2538858, EPI_ISL_2538859, EPI_ISL_2538860, EPI_ISL_2538861 | | | |
| see above | Lighthouse Lab in Alderley Park | Wellcome Sanger Institute for the COVID-19 Genomics UK (COG-UK) Consortium | Jacquelyn Wynn, Mairead Hyland, The Lighthouse Lab in Alderley Park and Alex Alderton, Roberto Amato, Jeffrey Barrett, Sonia Goncalves, Ewan Harrison, David K. Jackson, Ian Johnston, Dominic Kwiatkowski, Cordelia Langford, John Sillitoe on behalf of the Wellcome Sanger Institute COVID-19 Surveillance Team |
| EPI_ISL_2538862 | Lighthouse Lab in Glasgow | Wellcome Sanger Institute for the COVID-19 Genomics UK (COG-UK) Consortium | Harper VanSteenhouse, Yumi Kasai, David Gray, Carol Clugston, Anna Dominiczak and Alex Alderton, Roberto Amato, Jeffrey Barrett, Sonia Goncalves, Ewan Harrison, David K. Jackson, Ian Johnston, Dominic Kwiatkowski, Cordelia Langford, John Sillitoe on behalf of the Wellcome Sanger Institute COVID-19 Surveillance Team |
| EPI_ISL_2538863, EPI_ISL_2538864, EPI_ISL_2538865, EPI_ISL_2538866, EPI_ISL_2538867, EPI_ISL_2538868, EPI_ISL_2538869, EPI_ISL_2538870, EPI_ISL_2538871, EPI_ISL_2538872, EPI_ISL_2538873, EPI_ISL_2538874, EPI_ISL_2538875, EPI_ISL_2538876, EPI_ISL_2538877, EPI_ISL_2538878, EPI_ISL_2538880, EPI_ISL_2538881, EPI_ISL_2538882, EPI_ISL_2538883, EPI_ISL_2538884, EPI_ISL_2538885, EPI_ISL_2538886, EPI_ISL_2538887, EPI_ISL_2538888, EPI_ISL_2538889, EPI_ISL_2538890, EPI_ISL_2538891, EPI_ISL_2538892, EPI_ISL_2538893, EPI_ISL_2538894, EPI_ISL_2538895, EPI_ISL_2538896, EPI_ISL_2538897, EPI_ISL_2538898, EPI_ISL_2538899, EPI_ISL_2538900, EPI_ISL_2538901, EPI_ISL_2538902, EPI_ISL_2538903, EPI_ISL_2538904, EPI_ISL_2538905, EPI_ISL_2538906, EPI_ISL_2538907, EPI_ISL_2538908, EPI_ISL_2538909, EPI_ISL_2538910, EPI_ISL_2538911, EPI_ISL_2538912, EPI_ISL_2538913, EPI_ISL_2538914 | | | |
| see above | Lighthouse Lab in Alderley Park | Wellcome Sanger Institute for the COVID-19 Genomics UK (COG-UK) Consortium | Jacquelyn Wynn, Mairead Hyland, The Lighthouse Lab in Alderley Park and Alex Alderton, Roberto Amato, Jeffrey Barrett, Sonia Goncalves, Ewan Harrison, David K. Jackson, Ian Johnston, Dominic Kwiatkowski, Cordelia Langford, John Sillitoe on behalf of the Wellcome Sanger Institute COVID-19 Surveillance Team |
| EPI_ISL_2538915 | Lighthouse Lab in Glasgow | Wellcome Sanger Institute for the COVID-19 Genomics UK (COG-UK) Consortium | Harper VanSteenhouse, Yumi Kasai, David Gray, Carol Clugston, Anna Dominiczak and Alex Alderton, Roberto Amato, Jeffrey Barrett, Sonia Goncalves, Ewan Harrison, David K. Jackson, Ian Johnston, Dominic Kwiatkowski, Cordelia Langford, John Sillitoe on behalf of the Wellcome Sanger Institute COVID-19 Surveillance Team |
| EPI_ISL_2538917, EPI_ISL_2538918, EPI_ISL_2538919, EPI_ISL_2538920, EPI_ISL_2538921, EPI_ISL_2538922, EPI_ISL_2538923, EPI_ISL_2538924, EPI_ISL_2538925, EPI_ISL_2538926, EPI_ISL_2538927, EPI_ISL_2538928, EPI_ISL_2538929 | | | |
| see above | Lighthouse Lab in Alderley Park | Wellcome Sanger Institute for the COVID-19 Genomics UK (COG-UK) Consortium | Jacquelyn Wynn, Mairead Hyland, The Lighthouse Lab in Alderley Park and Alex Alderton, Roberto Amato, Jeffrey Barrett, Sonia Goncalves, Ewan Harrison, David K. Jackson, Ian Johnston, Dominic Kwiatkowski, Cordelia Langford, John Sillitoe on behalf of the Wellcome Sanger Institute COVID-19 Surveillance Team |
| EPI_ISL_2538930 | Lighthouse Lab in Glasgow | Wellcome Sanger Institute for the COVID-19 Genomics UK (COG-UK) Consortium | Harper VanSteenhouse, Yumi Kasai, David Gray, Carol Clugston, Anna Dominiczak and Alex Alderton, Roberto Amato, Jeffrey Barrett, Sonia Goncalves, Ewan Harrison, David K. Jackson, Ian Johnston, Dominic Kwiatkowski, Cordelia Langford, John Sillitoe on behalf of the Wellcome Sanger Institute COVID-19 Surveillance Team |
| EPI_ISL_2538931, EPI_ISL_2538932, EPI_ISL_2538933, EPI_ISL_2538934, EPI_ISL_2538936, EPI_ISL_2538937, EPI_ISL_2538938, EPI_ISL_2538939, EPI_ISL_2538940, EPI_ISL_2538941, EPI_ISL_2538942, EPI_ISL_2538943, EPI_ISL_2538944, EPI_ISL_2538945, EPI_ISL_2538946, EPI_ISL_2538947, EPI_ISL_2538948, EPI_ISL_2538949, EPI_ISL_2538950, EPI_ISL_2538952, EPI_ISL_2538953, EPI_ISL_2538954, EPI_ISL_2538956, EPI_ISL_2538957, EPI_ISL_2538958, EPI_ISL_2538959, EPI_ISL_2538960, EPI_ISL_2538961, EPI_ISL_2538962, EPI_ISL_2538963, EPI_ISL_2538964, EPI_ISL_2538965, EPI_ISL_2538966, EPI_ISL_2538967, EPI_ISL_2538968, EPI_ISL_2538969, EPI_ISL_2538970, EPI_ISL_2538972, EPI_ISL_2538973, EPI_ISL_2538974, EPI_ISL_2538975, EPI_ISL_2538976, EPI_ISL_2538977, EPI_ISL_2538978, EPI_ISL_2538979, EPI_ISL_2538981, EPI_ISL_2538982, EPI_ISL_2538983, EPI_ISL_2538984, EPI_ISL_2538985, EPI_ISL_2538986, EPI_ISL_2538987, EPI_ISL_2538988, EPI_ISL_2538989, EPI_ISL_2538990, EPI_ISL_2538991, EPI_ISL_2538992, EPI_ISL_2538993, EPI_ISL_2538994, EPI_ISL_2538995, EPI_ISL_2538996, EPI_ISL_2538997, EPI_ISL_2538998, EPI_ISL_2538999, EPI_ISL_2539000, EPI_ISL_2539001, EPI_ISL_2539002, EPI_ISL_2539003, EPI_ISL_2539004, EPI_ISL_2539005, EPI_ISL_2539006, EPI_ISL_2539007, EPI_ISL_2539008, EPI_ISL_2539009, EPI_ISL_2539010, EPI_ISL_2539011, EPI_ISL_2539012, EPI_ISL_2539013, EPI_ISL_2539015, EPI_ISL_2539016, EPI_ISL_2539017, EPI_ISL_2539018, EPI_ISL_2539019, EPI_ISL_2539020, EPI_ISL_2539021, EPI_ISL_2539022, EPI_ISL_2539023, EPI_ISL_2539025, EPI_ISL_2539026, EPI_ISL_2539027, EPI_ISL_2539028, EPI_ISL_2539029, EPI_ISL_2539030, EPI_ISL_2539032, EPI_ISL_2539033, EPI_ISL_2539034, EPI_ISL_2539035, EPI_ISL_2539036, EPI_ISL_2539037, EPI_ISL_2539038, EPI_ISL_2539039, EPI_ISL_2539040, EPI_ISL_2539041, EPI_ISL_2539042, EPI_ISL_2539043, EPI_ISL_2539044, EPI_ISL_2539045, EPI_ISL_2539046, EPI_ISL_2539047, EPI_ISL_2539048, EPI_ISL_2539050, EPI_ISL_2539051, EPI_ISL_2539052, EPI_ISL_2539053, EPI_ISL_2539054, EPI_ISL_2539055, EPI_ISL_2539056, EPI_ISL_2539057, EPI_ISL_2539058, EPI_ISL_2539059, EPI_ISL_2539060, EPI_ISL_2539061, EPI_ISL_2539062, EPI_ISL_2539063, EPI_ISL_2539064, EPI_ISL_2539065, EPI_ISL_2539066, EPI_ISL_2539067, EPI_ISL_2539068, EPI_ISL_2539069, EPI_ISL_2539070, EPI_ISL_2539071, EPI_ISL_2539072, EPI_ISL_2539073, EPI_ISL_2539074, EPI_ISL_2539075, EPI_ISL_2539076, EPI_ISL_2539077, EPI_ISL_2539078, EPI_ISL_2539079, EPI_ISL_2539080, EPI_ISL_2539081, EPI_ISL_2539082, EPI_ISL_2539083, EPI_ISL_2539084, EPI_ISL_2539085, EPI_ISL_2539086, EPI_ISL_2539087, EPI_ISL_2539088, EPI_ISL_2539089, EPI_ISL_2539090, EPI_ISL_2539091, EPI_ISL_2539092, EPI_ISL_2539093, EPI_ISL_2539094, EPI_ISL_2539095, EPI_ISL_2539096, EPI_ISL_2539097, EPI_ISL_2539098, EPI_ISL_2539099, EPI_ISL_2539100, EPI_ISL_2539101, EPI_ISL_2539102, EPI_ISL_2539103, EPI_ISL_2539104, EPI_ISL_2539105, EPI_ISL_2539106, EPI_ISL_2539107, EPI_ISL_2539108, EPI_ISL_2539109, EPI_ISL_2539111, EPI_ISL_2539112, EPI_ISL_2539113, EPI_ISL_2539114, EPI_ISL_2539115, EPI_ISL_2539116, EPI_ISL_2539117, EPI_ISL_2539118, EPI_ISL_2539119, EPI_ISL_2539120, EPI_ISL_2539121, EPI_ISL_2539122, EPI_ISL_2539124, EPI_ISL_2539125, EPI_ISL_2539126, EPI_ISL_2539127, EPI_ISL_2539128, EPI_ISL_2539129, EPI_ISL_2539130, EPI_ISL_2539131, EPI_ISL_2539132, EPI_ISL_2539133, EPI_ISL_2539134, EPI_ISL_2539135, EPI_ISL_2539136, EPI_ISL_2539137, EPI_ISL_2539138, EPI_ISL_2539139, EPI_ISL_2539140, EPI_ISL_2539141, EPI_ISL_2539142, EPI_ISL_2539143, EPI_ISL_2539144, EPI_ISL_2539145, EPI_ISL_2539146, EPI_ISL_2539147, EPI_ISL_2539148, EPI_ISL_2539149, EPI_ISL_2539150, EPI_ISL_2539151, EPI_ISL_2539153, EPI_ISL_2539154, EPI_ISL_2539155, EPI_ISL_2539156, EPI_ISL_2539157, EPI_ISL_2539158, EPI_ISL_2539160, EPI_ISL_2539161, EPI_ISL_2539162, EPI_ISL_2539163, EPI_ISL_2539164, EPI_ISL_2539165, EPI_ISL_2539166, EPI_ISL_2539167, EPI_ISL_2539168, EPI_ISL_2539169, EPI_ISL_2539170, EPI_ISL_2539171 | | | |
| see above | Lighthouse Lab in Alderley Park | Wellcome Sanger Institute for the COVID-19 Genomics UK (COG-UK) Consortium | Jacquelyn Wynn, Mairead Hyland, The Lighthouse Lab in Alderley Park and Alex Alderton, Roberto Amato, Jeffrey Barrett, Sonia Goncalves, Ewan Harrison, David K. Jackson, Ian Johnston, Dominic Kwiatkowski, Cordelia Langford, John Sillitoe on behalf of the Wellcome Sanger Institute COVID-19 Surveillance Team |
| EPI_ISL_2539172, EPI_ISL_2539173 | Lighthouse Lab in Milton Keynes | Wellcome Sanger Institute for the COVID-19 Genomics UK (COG-UK) Consortium | The Lighthouse Lab in Milton Keynes and Alex Alderton, Roberto Amato, Jeffrey Barrett, Sonia Goncalves, Ewan Harrison, David K. Jackson, Ian Johnston, Dominic Kwiatkowski, Cordelia Langford, John Sillitoe on behalf of the Wellcome Sanger Institute COVID-19 Surveillance Team |
| EPI_ISL_2539174 | Lighthouse Lab in Alderley Park | Wellcome Sanger Institute for the COVID-19 Genomics UK (COG-UK) Consortium | Jacquelyn Wynn, Mairead Hyland, The Lighthouse Lab in Alderley Park and Alex Alderton, Roberto Amato, Jeffrey Barrett, Sonia Goncalves, Ewan Harrison, David K. Jackson, Ian Johnston, Dominic Kwiatkowski, Cordelia Langford, John Sillitoe on behalf of the Wellcome Sanger Institute COVID-19 Surveillance Team |
| EPI_ISL_2539175, EPI_ISL_2539176, | Lighthouse Lab in Milton Keynes | Wellcome Sanger Institute for the COVID-19 Genomics UK (COG-UK) Consortium | The Lighthouse Lab in Milton Keynes and Alex Alderton, Roberto Amato, Jeffrey Barrett, Sonia Goncalves, Ewan Harrison, David K. Jackson, Ian Johnston, Dominic Kwiatkowski, Cordelia Langford, John Sillitoe on behalf of the Wellcome Sanger Institute COVID-19 Surveillance Team |

| | | | |
|---|---|---|---|
| EPI_ISL_2539178, EPI_ISL_2539179, EPI_ISL_2539180 | | | |
| EPI_ISL_2539181 | Lighthouse Lab in Alderley Park | Wellcome Sanger Institute for the COVID-19 Genomics UK (COG-UK) Consortium | Jacquelyn Wynn, Mairead Hyland, The Lighthouse Lab in Alderley Park and Alex Alderton, Roberto Amato, Jeffrey Barrett, Sonia Goncalves, Ewan Harrison, David K. Jackson, Ian Johnston, Dominic Kwiatkowski, Cordelia Langford, John Sillitoe on behalf of the Wellcome Sanger Institute COVID-19 Surveillance Team |
| EPI_ISL_2539182, EPI_ISL_2539183, EPI_ISL_2539184, EPI_ISL_2539185, EPI_ISL_2539186 | Lighthouse Lab in Milton Keynes | Wellcome Sanger Institute for the COVID-19 Genomics UK (COG-UK) Consortium | The Lighthouse Lab in Milton Keynes and Alex Alderton, Roberto Amato, Jeffrey Barrett, Sonia Goncalves, Ewan Harrison, David K. Jackson, Ian Johnston, Dominic Kwiatkowski, Cordelia Langford, John Sillitoe on behalf of the Wellcome Sanger Institute COVID-19 Surveillance Team |
| EPI_ISL_2539187, EPI_ISL_2539188, EPI_ISL_2539190, EPI_ISL_2539191 | Lighthouse Lab in Alderley Park | Wellcome Sanger Institute for the COVID-19 Genomics UK (COG-UK) Consortium | Jacquelyn Wynn, Mairead Hyland, The Lighthouse Lab in Alderley Park and Alex Alderton, Roberto Amato, Jeffrey Barrett, Sonia Goncalves, Ewan Harrison, David K. Jackson, Ian Johnston, Dominic Kwiatkowski, Cordelia Langford, John Sillitoe on behalf of the Wellcome Sanger Institute COVID-19 Surveillance Team |
| EPI_ISL_2539192, EPI_ISL_2539193, EPI_ISL_2539194, EPI_ISL_2539195, EPI_ISL_2539196 | Lighthouse Lab in Milton Keynes | Wellcome Sanger Institute for the COVID-19 Genomics UK (COG-UK) Consortium | The Lighthouse Lab in Milton Keynes and Alex Alderton, Roberto Amato, Jeffrey Barrett, Sonia Goncalves, Ewan Harrison, David K. Jackson, Ian Johnston, Dominic Kwiatkowski, Cordelia Langford, John Sillitoe on behalf of the Wellcome Sanger Institute COVID-19 Surveillance Team |
| EPI_ISL_2539197, EPI_ISL_2539198, EPI_ISL_2539199 | Lighthouse Lab in Alderley Park | Wellcome Sanger Institute for the COVID-19 Genomics UK (COG-UK) Consortium | Jacquelyn Wynn, Mairead Hyland, The Lighthouse Lab in Alderley Park and Alex Alderton, Roberto Amato, Jeffrey Barrett, Sonia Goncalves, Ewan Harrison, David K. Jackson, Ian Johnston, Dominic Kwiatkowski, Cordelia Langford, John Sillitoe on behalf of the Wellcome Sanger Institute COVID-19 Surveillance Team |
| EPI_ISL_2539200 | Lighthouse Lab in Milton Keynes | Wellcome Sanger Institute for the COVID-19 Genomics UK (COG-UK) Consortium | The Lighthouse Lab in Milton Keynes and Alex Alderton, Roberto Amato, Jeffrey Barrett, Sonia Goncalves, Ewan Harrison, David K. Jackson, Ian Johnston, Dominic Kwiatkowski, Cordelia Langford, John Sillitoe on behalf of the Wellcome Sanger Institute COVID-19 Surveillance Team |
| EPI_ISL_2539201, EPI_ISL_2539202, EPI_ISL_2539204, EPI_ISL_2539205, EPI_ISL_2539206 | Lighthouse Lab in Alderley Park | Wellcome Sanger Institute for the COVID-19 Genomics UK (COG-UK) Consortium | Jacquelyn Wynn, Mairead Hyland, The Lighthouse Lab in Alderley Park and Alex Alderton, Roberto Amato, Jeffrey Barrett, Sonia Goncalves, Ewan Harrison, David K. Jackson, Ian Johnston, Dominic Kwiatkowski, Cordelia Langford, John Sillitoe on behalf of the Wellcome Sanger Institute COVID-19 Surveillance Team |
| EPI_ISL_2539207 | Lighthouse Lab in Milton Keynes | Wellcome Sanger Institute for the COVID-19 Genomics UK (COG-UK) Consortium | The Lighthouse Lab in Milton Keynes and Alex Alderton, Roberto Amato, Jeffrey Barrett, Sonia Goncalves, Ewan Harrison, David K. Jackson, Ian Johnston, Dominic Kwiatkowski, Cordelia Langford, John Sillitoe on behalf of the Wellcome Sanger Institute COVID-19 Surveillance Team |
| EPI_ISL_2539208 | Lighthouse Lab in Alderley Park | Wellcome Sanger Institute for the COVID-19 Genomics UK (COG-UK) Consortium | Jacquelyn Wynn, Mairead Hyland, The Lighthouse Lab in Alderley Park and Alex Alderton, Roberto Amato, Jeffrey Barrett, Sonia Goncalves, Ewan Harrison, David K. Jackson, Ian Johnston, Dominic Kwiatkowski, Cordelia Langford, John Sillitoe on behalf of the Wellcome Sanger Institute COVID-19 Surveillance Team |
| EPI_ISL_2539209, EPI_ISL_2539211, EPI_ISL_2539212, EPI_ISL_2539213, EPI_ISL_2539214, EPI_ISL_2539215 | Lighthouse Lab in Milton Keynes | Wellcome Sanger Institute for the COVID-19 Genomics UK (COG-UK) Consortium | The Lighthouse Lab in Milton Keynes and Alex Alderton, Roberto Amato, Jeffrey Barrett, Sonia Goncalves, Ewan Harrison, David K. Jackson, Ian Johnston, Dominic Kwiatkowski, Cordelia Langford, John Sillitoe on behalf of the Wellcome Sanger Institute COVID-19 Surveillance Team |
| EPI_ISL_2539216 | Lighthouse Lab in Alderley Park | Wellcome Sanger Institute for the COVID-19 Genomics UK (COG-UK) Consortium | Jacquelyn Wynn, Mairead Hyland, The Lighthouse Lab in Alderley Park and Alex Alderton, Roberto Amato, Jeffrey Barrett, Sonia Goncalves, Ewan Harrison, David K. Jackson, Ian Johnston, Dominic Kwiatkowski, Cordelia Langford, John Sillitoe on behalf of the Wellcome Sanger Institute COVID-19 Surveillance Team |
| EPI_ISL_2539217 | Lighthouse Lab in Milton Keynes | Wellcome Sanger Institute for the COVID-19 Genomics UK (COG-UK) Consortium | The Lighthouse Lab in Milton Keynes and Alex Alderton, Roberto Amato, Jeffrey Barrett, Sonia Goncalves, Ewan Harrison, David K. Jackson, Ian Johnston, Dominic Kwiatkowski, Cordelia Langford, John Sillitoe on behalf of the Wellcome Sanger Institute COVID-19 Surveillance Team |
| EPI_ISL_2539218, EPI_ISL_2539219, EPI_ISL_2539220 | Lighthouse Lab in Alderley Park | Wellcome Sanger Institute for the COVID-19 Genomics UK (COG-UK) Consortium | Jacquelyn Wynn, Mairead Hyland, The Lighthouse Lab in Alderley Park and Alex Alderton, Roberto Amato, Jeffrey Barrett, Sonia Goncalves, Ewan Harrison, David K. Jackson, Ian Johnston, Dominic Kwiatkowski, Cordelia Langford, John Sillitoe on behalf of the Wellcome Sanger Institute COVID-19 Surveillance Team |
| EPI_ISL_2539221, EPI_ISL_2539222 | Lighthouse Lab in Milton Keynes | Wellcome Sanger Institute for the COVID-19 Genomics UK (COG-UK) Consortium | The Lighthouse Lab in Milton Keynes and Alex Alderton, Roberto Amato, Jeffrey Barrett, Sonia Goncalves, Ewan Harrison, David K. Jackson, Ian Johnston, Dominic Kwiatkowski, Cordelia Langford, John Sillitoe on behalf of the Wellcome Sanger Institute COVID-19 Surveillance Team |
| EPI_ISL_2539224 | Lighthouse Lab in Alderley Park | Wellcome Sanger Institute for the COVID-19 Genomics UK (COG-UK) Consortium | Jacquelyn Wynn, Mairead Hyland, The Lighthouse Lab in Alderley Park and Alex Alderton, Roberto Amato, Jeffrey Barrett, Sonia Goncalves, Ewan Harrison, David K. Jackson, Ian Johnston, Dominic Kwiatkowski, Cordelia Langford, John Sillitoe on behalf of the Wellcome Sanger Institute COVID-19 Surveillance Team |
| EPI_ISL_2539225 | Lighthouse Lab in Glasgow | Wellcome Sanger Institute for the COVID-19 Genomics UK (COG-UK) Consortium | Harper VanSteenhouse, Yumi Kasai, David Gray, Carol Clugston, Anna Dominiczak and Alex Alderton, Roberto Amato, Jeffrey Barrett, Sonia Goncalves, Ewan Harrison, David K. Jackson, Ian Johnston, Dominic Kwiatkowski, Cordelia Langford, John Sillitoe on behalf of the Wellcome Sanger Institute COVID-19 Surveillance Team |
| EPI_ISL_2539226 | Lighthouse Lab in Milton Keynes | Wellcome Sanger Institute for the COVID-19 Genomics UK (COG-UK) Consortium | The Lighthouse Lab in Milton Keynes and Alex Alderton, Roberto Amato, Jeffrey Barrett, Sonia Goncalves, Ewan Harrison, David K. Jackson, Ian Johnston, Dominic Kwiatkowski, Cordelia Langford, John Sillitoe on behalf of the Wellcome Sanger Institute COVID-19 Surveillance Team |
| EPI_ISL_2539227, EPI_ISL_2539230 | Lighthouse Lab in Alderley Park | Wellcome Sanger Institute for the COVID-19 Genomics UK (COG-UK) Consortium | Jacquelyn Wynn, Mairead Hyland, The Lighthouse Lab in Alderley Park and Alex Alderton, Roberto Amato, Jeffrey Barrett, Sonia Goncalves, Ewan Harrison, David K. Jackson, Ian Johnston, Dominic Kwiatkowski, Cordelia Langford, John Sillitoe on behalf of the Wellcome Sanger Institute COVID-19 Surveillance Team |
| EPI_ISL_2539232, EPI_ISL_2539233, EPI_ISL_2539235 | Lighthouse Lab in Milton Keynes | Wellcome Sanger Institute for the COVID-19 Genomics UK (COG-UK) Consortium | The Lighthouse Lab in Milton Keynes and Alex Alderton, Roberto Amato, Jeffrey Barrett, Sonia Goncalves, Ewan Harrison, David K. Jackson, Ian Johnston, Dominic Kwiatkowski, Cordelia Langford, John Sillitoe on behalf of the Wellcome Sanger Institute COVID-19 Surveillance Team |
| EPI_ISL_2539236 | Lighthouse Lab in Alderley Park | Wellcome Sanger Institute for the COVID-19 Genomics UK (COG-UK) Consortium | Jacquelyn Wynn, Mairead Hyland, The Lighthouse Lab in Alderley Park and Alex Alderton, Roberto Amato, Jeffrey Barrett, Sonia Goncalves, Ewan Harrison, David K. Jackson, Ian Johnston, Dominic Kwiatkowski, Cordelia Langford, John Sillitoe on behalf of the Wellcome Sanger Institute COVID-19 Surveillance Team |
| EPI_ISL_2539237 | Lighthouse Lab in Milton Keynes | Wellcome Sanger Institute for the COVID-19 Genomics UK (COG-UK) Consortium | The Lighthouse Lab in Milton Keynes and Alex Alderton, Roberto Amato, Jeffrey Barrett, Sonia Goncalves, Ewan Harrison, David K. Jackson, Ian Johnston, Dominic Kwiatkowski, Cordelia Langford, John Sillitoe on behalf of the Wellcome Sanger Institute COVID-19 Surveillance Team |
| EPI_ISL_2539238 | Lighthouse Lab in Alderley Park | Wellcome Sanger Institute for the COVID-19 Genomics UK (COG-UK) Consortium | Jacquelyn Wynn, Mairead Hyland, The Lighthouse Lab in Alderley Park and Alex Alderton, Roberto Amato, Jeffrey Barrett, Sonia Goncalves, Ewan Harrison, David K. Jackson, Ian Johnston, Dominic Kwiatkowski, Cordelia Langford, John Sillitoe on behalf of the Wellcome Sanger Institute COVID-19 Surveillance Team |
| EPI_ISL_2539240, EPI_ISL_2539241 | Lighthouse Lab in Milton Keynes | Wellcome Sanger Institute for the COVID-19 Genomics UK (COG-UK) Consortium | The Lighthouse Lab in Milton Keynes and Alex Alderton, Roberto Amato, Jeffrey Barrett, Sonia Goncalves, Ewan Harrison, David K. Jackson, Ian Johnston, Dominic Kwiatkowski, Cordelia Langford, John Sillitoe on behalf of the Wellcome Sanger Institute COVID-19 Surveillance Team |
| EPI_ISL_2539242 | Lighthouse Lab in Alderley Park | Wellcome Sanger Institute for the COVID-19 Genomics UK (COG-UK) Consortium | Jacquelyn Wynn, Mairead Hyland, The Lighthouse Lab in Alderley Park and Alex Alderton, Roberto Amato, Jeffrey Barrett, Sonia Goncalves, Ewan Harrison, David K. Jackson, Ian Johnston, Dominic Kwiatkowski, Cordelia Langford, John Sillitoe on behalf of the Wellcome Sanger Institute COVID-19 Surveillance Team |
| EPI_ISL_2539243, EPI_ISL_2539245, EPI_ISL_2539246, EPI_ISL_2539247 | Lighthouse Lab in Milton Keynes | Wellcome Sanger Institute for the COVID-19 Genomics UK (COG-UK) Consortium | The Lighthouse Lab in Milton Keynes and Alex Alderton, Roberto Amato, Jeffrey Barrett, Sonia Goncalves, Ewan Harrison, David K. Jackson, Ian Johnston, Dominic Kwiatkowski, Cordelia Langford, John Sillitoe on behalf of the Wellcome Sanger Institute COVID-19 Surveillance Team |
| EPI_ISL_2539248 | Lighthouse Lab in Alderley Park | Wellcome Sanger Institute for the COVID-19 Genomics UK (COG-UK) Consortium | Jacquelyn Wynn, Mairead Hyland, The Lighthouse Lab in Alderley Park and Alex Alderton, Roberto Amato, Jeffrey Barrett, Sonia Goncalves, Ewan Harrison, David K. Jackson, Ian Johnston, Dominic Kwiatkowski, Cordelia Langford, John Sillitoe on behalf of the Wellcome Sanger Institute COVID-19 Surveillance Team |
| EPI_ISL_2539249, EPI_ISL_2539250, EPI_ISL_2539252, EPI_ISL_2539253 | Lighthouse Lab in Milton Keynes | Wellcome Sanger Institute for the COVID-19 Genomics UK (COG-UK) Consortium | The Lighthouse Lab in Milton Keynes and Alex Alderton, Roberto Amato, Jeffrey Barrett, Sonia Goncalves, Ewan Harrison, David K. Jackson, Ian Johnston, Dominic Kwiatkowski, Cordelia Langford, John Sillitoe on behalf of the Wellcome Sanger Institute COVID-19 Surveillance Team |
| EPI_ISL_2539254, EPI_ISL_2539255, EPI_ISL_2539256 | Lighthouse Lab in Alderley Park | Wellcome Sanger Institute for the COVID-19 Genomics UK (COG-UK) Consortium | Jacquelyn Wynn, Mairead Hyland, The Lighthouse Lab in Alderley Park and Alex Alderton, Roberto Amato, Jeffrey Barrett, Sonia Goncalves, Ewan Harrison, David K. Jackson, Ian Johnston, Dominic Kwiatkowski, Cordelia Langford, John Sillitoe on behalf of the Wellcome Sanger Institute COVID-19 Surveillance Team |
| EPI_ISL_2539257 | Lighthouse Lab in Milton Keynes | Wellcome Sanger Institute for the COVID-19 Genomics UK (COG-UK) Consortium | The Lighthouse Lab in Milton Keynes and Alex Alderton, Roberto Amato, Jeffrey Barrett, Sonia Goncalves, Ewan Harrison, David K. Jackson, Ian Johnston, Dominic Kwiatkowski, Cordelia Langford, John Sillitoe on behalf of the Wellcome Sanger Institute COVID-19 Surveillance Team |
| EPI_ISL_2539258, EPI_ISL_2539259, EPI_ISL_2539260 | Lighthouse Lab in Alderley Park | Wellcome Sanger Institute for the COVID-19 Genomics UK (COG-UK) Consortium | Jacquelyn Wynn, Mairead Hyland, The Lighthouse Lab in Alderley Park and Alex Alderton, Roberto Amato, Jeffrey Barrett, Sonia Goncalves, Ewan Harrison, David K. Jackson, Ian Johnston, Dominic Kwiatkowski, Cordelia Langford, John Sillitoe on behalf of the Wellcome Sanger Institute COVID-19 Surveillance Team |
| EPI_ISL_2539261, EPI_ISL_2539262, EPI_ISL_2539263, EPI_ISL_2539264, EPI_ISL_2539265, EPI_ISL_2539266 | Lighthouse Lab in Milton Keynes | Wellcome Sanger Institute for the COVID-19 Genomics UK (COG-UK) Consortium | The Lighthouse Lab in Milton Keynes and Alex Alderton, Roberto Amato, Jeffrey Barrett, Sonia Goncalves, Ewan Harrison, David K. Jackson, Ian Johnston, Dominic Kwiatkowski, Cordelia Langford, John Sillitoe on behalf of the Wellcome Sanger Institute COVID-19 Surveillance Team |
| EPI_ISL_2539267 | Lighthouse Lab in Alderley Park | Wellcome Sanger Institute for the COVID-19 Genomics UK (COG-UK) Consortium | Jacquelyn Wynn, Mairead Hyland, The Lighthouse Lab in Alderley Park and Alex Alderton, Roberto Amato, Jeffrey Barrett, Sonia Goncalves, Ewan Harrison, David K. Jackson, Ian Johnston, Dominic Kwiatkowski, Cordelia Langford, John Sillitoe on behalf of the Wellcome Sanger Institute COVID-19 Surveillance Team |
| EPI_ISL_2539268 | Lighthouse Lab in Milton Keynes | Wellcome Sanger Institute for the COVID-19 Genomics UK (COG-UK) | The Lighthouse Lab in Milton Keynes and Alex Alderton, Roberto Amato, Jeffrey Barrett, Sonia Goncalves, Ewan Harrison, David K. Jackson, Ian Johnston, Dominic Kwiatkowski, Cordelia Langford, John Sillitoe on behalf of the Wellcome Sanger |

| | | | |
|---|---|---|---|
| | | Consortium | Institute COVID-19 Surveillance Team |
| EPI_ISL_2539270 | Lighthouse Lab in Alderley Park | Wellcome Sanger Institute for the COVID-19 Genomics UK (COG-UK) Consortium | Jacquelyn Wynn, Mairead Hyland, The Lighthouse Lab in Alderley Park and Alex Alderton, Roberto Amato, Jeffrey Barrett, Sonia Goncalves, Ewan Harrison, David K. Jackson, Ian Johnston, Dominic Kwiatkowski, Cordelia Langford, John Sillitoe on behalf of the Wellcome Sanger Institute COVID-19 Surveillance Team |
| EPI_ISL_2539271, EPI_ISL_2539272, EPI_ISL_2539273 | Lighthouse Lab in Milton Keynes | Wellcome Sanger Institute for the COVID-19 Genomics UK (COG-UK) Consortium | The Lighthouse Lab in Milton Keynes and Alex Alderton, Roberto Amato, Jeffrey Barrett, Sonia Goncalves, Ewan Harrison, David K. Jackson, Ian Johnston, Dominic Kwiatkowski, Cordelia Langford, John Sillitoe on behalf of the Wellcome Sanger Institute COVID-19 Surveillance Team |
| EPI_ISL_2539274, EPI_ISL_2539275 | Lighthouse Lab in Alderley Park | Wellcome Sanger Institute for the COVID-19 Genomics UK (COG-UK) Consortium | Jacquelyn Wynn, Mairead Hyland, The Lighthouse Lab in Alderley Park and Alex Alderton, Roberto Amato, Jeffrey Barrett, Sonia Goncalves, Ewan Harrison, David K. Jackson, Ian Johnston, Dominic Kwiatkowski, Cordelia Langford, John Sillitoe on behalf of the Wellcome Sanger Institute COVID-19 Surveillance Team |
| EPI_ISL_2539276, EPI_ISL_2539277, EPI_ISL_2539278, EPI_ISL_2539279, EPI_ISL_2539280, EPI_ISL_2539281, EPI_ISL_2539282 | Lighthouse Lab in Milton Keynes | Wellcome Sanger Institute for the COVID-19 Genomics UK (COG-UK) Consortium | The Lighthouse Lab in Milton Keynes and Alex Alderton, Roberto Amato, Jeffrey Barrett, Sonia Goncalves, Ewan Harrison, David K. Jackson, Ian Johnston, Dominic Kwiatkowski, Cordelia Langford, John Sillitoe on behalf of the Wellcome Sanger Institute COVID-19 Surveillance Team |
| EPI_ISL_2539283 | Lighthouse Lab in Alderley Park | Wellcome Sanger Institute for the COVID-19 Genomics UK (COG-UK) Consortium | Jacquelyn Wynn, Mairead Hyland, The Lighthouse Lab in Alderley Park and Alex Alderton, Roberto Amato, Jeffrey Barrett, Sonia Goncalves, Ewan Harrison, David K. Jackson, Ian Johnston, Dominic Kwiatkowski, Cordelia Langford, John Sillitoe on behalf of the Wellcome Sanger Institute COVID-19 Surveillance Team |
| EPI_ISL_2539285 | Lighthouse Lab in Milton Keynes | Wellcome Sanger Institute for the COVID-19 Genomics UK (COG-UK) Consortium | The Lighthouse Lab in Milton Keynes and Alex Alderton, Roberto Amato, Jeffrey Barrett, Sonia Goncalves, Ewan Harrison, David K. Jackson, Ian Johnston, Dominic Kwiatkowski, Cordelia Langford, John Sillitoe on behalf of the Wellcome Sanger Institute COVID-19 Surveillance Team |
| EPI_ISL_2539286 | Lighthouse Lab in Alderley Park | Wellcome Sanger Institute for the COVID-19 Genomics UK (COG-UK) Consortium | Jacquelyn Wynn, Mairead Hyland, The Lighthouse Lab in Alderley Park and Alex Alderton, Roberto Amato, Jeffrey Barrett, Sonia Goncalves, Ewan Harrison, David K. Jackson, Ian Johnston, Dominic Kwiatkowski, Cordelia Langford, John Sillitoe on behalf of the Wellcome Sanger Institute COVID-19 Surveillance Team |
| EPI_ISL_2539287, EPI_ISL_2539288 | Lighthouse Lab in Milton Keynes | Wellcome Sanger Institute for the COVID-19 Genomics UK (COG-UK) Consortium | The Lighthouse Lab in Milton Keynes and Alex Alderton, Roberto Amato, Jeffrey Barrett, Sonia Goncalves, Ewan Harrison, David K. Jackson, Ian Johnston, Dominic Kwiatkowski, Cordelia Langford, John Sillitoe on behalf of the Wellcome Sanger Institute COVID-19 Surveillance Team |
| EPI_ISL_2539289 | Lighthouse Lab in Alderley Park | Wellcome Sanger Institute for the COVID-19 Genomics UK (COG-UK) Consortium | Jacquelyn Wynn, Mairead Hyland, The Lighthouse Lab in Alderley Park and Alex Alderton, Roberto Amato, Jeffrey Barrett, Sonia Goncalves, Ewan Harrison, David K. Jackson, Ian Johnston, Dominic Kwiatkowski, Cordelia Langford, John Sillitoe on behalf of the Wellcome Sanger Institute COVID-19 Surveillance Team |
| EPI_ISL_2539290 | Lighthouse Lab in Milton Keynes | Wellcome Sanger Institute for the COVID-19 Genomics UK (COG-UK) Consortium | The Lighthouse Lab in Milton Keynes and Alex Alderton, Roberto Amato, Jeffrey Barrett, Sonia Goncalves, Ewan Harrison, David K. Jackson, Ian Johnston, Dominic Kwiatkowski, Cordelia Langford, John Sillitoe on behalf of the Wellcome Sanger Institute COVID-19 Surveillance Team |
| EPI_ISL_2539291, EPI_ISL_2539292 | Lighthouse Lab in Alderley Park | Wellcome Sanger Institute for the COVID-19 Genomics UK (COG-UK) Consortium | Jacquelyn Wynn, Mairead Hyland, The Lighthouse Lab in Alderley Park and Alex Alderton, Roberto Amato, Jeffrey Barrett, Sonia Goncalves, Ewan Harrison, David K. Jackson, Ian Johnston, Dominic Kwiatkowski, Cordelia Langford, John Sillitoe on behalf of the Wellcome Sanger Institute COVID-19 Surveillance Team |
| EPI_ISL_2539293 | Lighthouse Lab in Milton Keynes | Wellcome Sanger Institute for the COVID-19 Genomics UK (COG-UK) Consortium | The Lighthouse Lab in Milton Keynes and Alex Alderton, Roberto Amato, Jeffrey Barrett, Sonia Goncalves, Ewan Harrison, David K. Jackson, Ian Johnston, Dominic Kwiatkowski, Cordelia Langford, John Sillitoe on behalf of the Wellcome Sanger Institute COVID-19 Surveillance Team |
| EPI_ISL_2539294 | Lighthouse Lab in Alderley Park | Wellcome Sanger Institute for the COVID-19 Genomics UK (COG-UK) Consortium | Jacquelyn Wynn, Mairead Hyland, The Lighthouse Lab in Alderley Park and Alex Alderton, Roberto Amato, Jeffrey Barrett, Sonia Goncalves, Ewan Harrison, David K. Jackson, Ian Johnston, Dominic Kwiatkowski, Cordelia Langford, John Sillitoe on behalf of the Wellcome Sanger Institute COVID-19 Surveillance Team |
| EPI_ISL_2539295, EPI_ISL_2539297, EPI_ISL_2539298, EPI_ISL_2539299 | Lighthouse Lab in Milton Keynes | Wellcome Sanger Institute for the COVID-19 Genomics UK (COG-UK) Consortium | The Lighthouse Lab in Milton Keynes and Alex Alderton, Roberto Amato, Jeffrey Barrett, Sonia Goncalves, Ewan Harrison, David K. Jackson, Ian Johnston, Dominic Kwiatkowski, Cordelia Langford, John Sillitoe on behalf of the Wellcome Sanger Institute COVID-19 Surveillance Team |
| EPI_ISL_2539301, EPI_ISL_2539302 | Lighthouse Lab in Alderley Park | Wellcome Sanger Institute for the COVID-19 Genomics UK (COG-UK) Consortium | Jacquelyn Wynn, Mairead Hyland, The Lighthouse Lab in Alderley Park and Alex Alderton, Roberto Amato, Jeffrey Barrett, Sonia Goncalves, Ewan Harrison, David K. Jackson, Ian Johnston, Dominic Kwiatkowski, Cordelia Langford, John Sillitoe on behalf of the Wellcome Sanger Institute COVID-19 Surveillance Team |
| EPI_ISL_2539303 | Lighthouse Lab in Milton Keynes | Wellcome Sanger Institute for the COVID-19 Genomics UK (COG-UK) Consortium | The Lighthouse Lab in Milton Keynes and Alex Alderton, Roberto Amato, Jeffrey Barrett, Sonia Goncalves, Ewan Harrison, David K. Jackson, Ian Johnston, Dominic Kwiatkowski, Cordelia Langford, John Sillitoe on behalf of the Wellcome Sanger Institute COVID-19 Surveillance Team |
| EPI_ISL_2539304, EPI_ISL_2539305, EPI_ISL_2539306 | Lighthouse Lab in Alderley Park | Wellcome Sanger Institute for the COVID-19 Genomics UK (COG-UK) Consortium | Jacquelyn Wynn, Mairead Hyland, The Lighthouse Lab in Alderley Park and Alex Alderton, Roberto Amato, Jeffrey Barrett, Sonia Goncalves, Ewan Harrison, David K. Jackson, Ian Johnston, Dominic Kwiatkowski, Cordelia Langford, John Sillitoe on behalf of the Wellcome Sanger Institute COVID-19 Surveillance Team |
| EPI_ISL_2539307 | Lighthouse Lab in Milton Keynes | Wellcome Sanger Institute for the COVID-19 Genomics UK (COG-UK) Consortium | The Lighthouse Lab in Milton Keynes and Alex Alderton, Roberto Amato, Jeffrey Barrett, Sonia Goncalves, Ewan Harrison, David K. Jackson, Ian Johnston, Dominic Kwiatkowski, Cordelia Langford, John Sillitoe on behalf of the Wellcome Sanger Institute COVID-19 Surveillance Team |
| EPI_ISL_2539308 | Lighthouse Lab in Alderley Park | Wellcome Sanger Institute for the COVID-19 Genomics UK (COG-UK) Consortium | Jacquelyn Wynn, Mairead Hyland, The Lighthouse Lab in Alderley Park and Alex Alderton, Roberto Amato, Jeffrey Barrett, Sonia Goncalves, Ewan Harrison, David K. Jackson, Ian Johnston, Dominic Kwiatkowski, Cordelia Langford, John Sillitoe on behalf of the Wellcome Sanger Institute COVID-19 Surveillance Team |
| EPI_ISL_2539310, EPI_ISL_2539313 | Lighthouse Lab in Milton Keynes | Wellcome Sanger Institute for the COVID-19 Genomics UK (COG-UK) Consortium | The Lighthouse Lab in Milton Keynes and Alex Alderton, Roberto Amato, Jeffrey Barrett, Sonia Goncalves, Ewan Harrison, David K. Jackson, Ian Johnston, Dominic Kwiatkowski, Cordelia Langford, John Sillitoe on behalf of the Wellcome Sanger Institute COVID-19 Surveillance Team |
| EPI_ISL_2539314, EPI_ISL_2539315 | Lighthouse Lab in Alderley Park | Wellcome Sanger Institute for the COVID-19 Genomics UK (COG-UK) Consortium | Jacquelyn Wynn, Mairead Hyland, The Lighthouse Lab in Alderley Park and Alex Alderton, Roberto Amato, Jeffrey Barrett, Sonia Goncalves, Ewan Harrison, David K. Jackson, Ian Johnston, Dominic Kwiatkowski, Cordelia Langford, John Sillitoe on behalf of the Wellcome Sanger Institute COVID-19 Surveillance Team |
| EPI_ISL_2539316 | Lighthouse Lab in Milton Keynes | Wellcome Sanger Institute for the COVID-19 Genomics UK (COG-UK) Consortium | The Lighthouse Lab in Milton Keynes and Alex Alderton, Roberto Amato, Jeffrey Barrett, Sonia Goncalves, Ewan Harrison, David K. Jackson, Ian Johnston, Dominic Kwiatkowski, Cordelia Langford, John Sillitoe on behalf of the Wellcome Sanger Institute COVID-19 Surveillance Team |
| EPI_ISL_2539317, EPI_ISL_2539318 | Lighthouse Lab in Alderley Park | Wellcome Sanger Institute for the COVID-19 Genomics UK (COG-UK) Consortium | Jacquelyn Wynn, Mairead Hyland, The Lighthouse Lab in Alderley Park and Alex Alderton, Roberto Amato, Jeffrey Barrett, Sonia Goncalves, Ewan Harrison, David K. Jackson, Ian Johnston, Dominic Kwiatkowski, Cordelia Langford, John Sillitoe on behalf of the Wellcome Sanger Institute COVID-19 Surveillance Team |
| EPI_ISL_2539319, EPI_ISL_2539321, EPI_ISL_2539323 | Lighthouse Lab in Milton Keynes | Wellcome Sanger Institute for the COVID-19 Genomics UK (COG-UK) Consortium | The Lighthouse Lab in Milton Keynes and Alex Alderton, Roberto Amato, Jeffrey Barrett, Sonia Goncalves, Ewan Harrison, David K. Jackson, Ian Johnston, Dominic Kwiatkowski, Cordelia Langford, John Sillitoe on behalf of the Wellcome Sanger Institute COVID-19 Surveillance Team |
| EPI_ISL_2539324 | Lighthouse Lab in Alderley Park | Wellcome Sanger Institute for the COVID-19 Genomics UK (COG-UK) Consortium | Jacquelyn Wynn, Mairead Hyland, The Lighthouse Lab in Alderley Park and Alex Alderton, Roberto Amato, Jeffrey Barrett, Sonia Goncalves, Ewan Harrison, David K. Jackson, Ian Johnston, Dominic Kwiatkowski, Cordelia Langford, John Sillitoe on behalf of the Wellcome Sanger Institute COVID-19 Surveillance Team |
| EPI_ISL_2539325 | Lighthouse Lab in Milton Keynes | Wellcome Sanger Institute for the COVID-19 Genomics UK (COG-UK) Consortium | The Lighthouse Lab in Milton Keynes and Alex Alderton, Roberto Amato, Jeffrey Barrett, Sonia Goncalves, Ewan Harrison, David K. Jackson, Ian Johnston, Dominic Kwiatkowski, Cordelia Langford, John Sillitoe on behalf of the Wellcome Sanger Institute COVID-19 Surveillance Team |
| EPI_ISL_2539326 | Lighthouse Lab in Alderley Park | Wellcome Sanger Institute for the COVID-19 Genomics UK (COG-UK) Consortium | Jacquelyn Wynn, Mairead Hyland, The Lighthouse Lab in Alderley Park and Alex Alderton, Roberto Amato, Jeffrey Barrett, Sonia Goncalves, Ewan Harrison, David K. Jackson, Ian Johnston, Dominic Kwiatkowski, Cordelia Langford, John Sillitoe on behalf of the Wellcome Sanger Institute COVID-19 Surveillance Team |
| EPI_ISL_2539327 | Lighthouse Lab in Milton Keynes | Wellcome Sanger Institute for the COVID-19 Genomics UK (COG-UK) Consortium | The Lighthouse Lab in Milton Keynes and Alex Alderton, Roberto Amato, Jeffrey Barrett, Sonia Goncalves, Ewan Harrison, David K. Jackson, Ian Johnston, Dominic Kwiatkowski, Cordelia Langford, John Sillitoe on behalf of the Wellcome Sanger Institute COVID-19 Surveillance Team |
| EPI_ISL_2539328 | Lighthouse Lab in Alderley Park | Wellcome Sanger Institute for the COVID-19 Genomics UK (COG-UK) Consortium | Jacquelyn Wynn, Mairead Hyland, The Lighthouse Lab in Alderley Park and Alex Alderton, Roberto Amato, Jeffrey Barrett, Sonia Goncalves, Ewan Harrison, David K. Jackson, Ian Johnston, Dominic Kwiatkowski, Cordelia Langford, John Sillitoe on behalf of the Wellcome Sanger Institute COVID-19 Surveillance Team |
| EPI_ISL_2539330, EPI_ISL_2539331, EPI_ISL_2539332 | Lighthouse Lab in Milton Keynes | Wellcome Sanger Institute for the COVID-19 Genomics UK (COG-UK) Consortium | The Lighthouse Lab in Milton Keynes and Alex Alderton, Roberto Amato, Jeffrey Barrett, Sonia Goncalves, Ewan Harrison, David K. Jackson, Ian Johnston, Dominic Kwiatkowski, Cordelia Langford, John Sillitoe on behalf of the Wellcome Sanger Institute COVID-19 Surveillance Team |
| EPI_ISL_2539333 | Lighthouse Lab in Alderley Park | Wellcome Sanger Institute for the COVID-19 Genomics UK (COG-UK) Consortium | Jacquelyn Wynn, Mairead Hyland, The Lighthouse Lab in Alderley Park and Alex Alderton, Roberto Amato, Jeffrey Barrett, Sonia Goncalves, Ewan Harrison, David K. Jackson, Ian Johnston, Dominic Kwiatkowski, Cordelia Langford, John Sillitoe on behalf of the Wellcome Sanger Institute COVID-19 Surveillance Team |
| EPI_ISL_2539334, EPI_ISL_2539335, EPI_ISL_2539336 | Lighthouse Lab in Milton Keynes | Wellcome Sanger Institute for the COVID-19 Genomics UK (COG-UK) Consortium | The Lighthouse Lab in Milton Keynes and Alex Alderton, Roberto Amato, Jeffrey Barrett, Sonia Goncalves, Ewan Harrison, David K. Jackson, Ian Johnston, Dominic Kwiatkowski, Cordelia Langford, John Sillitoe on behalf of the Wellcome Sanger Institute COVID-19 Surveillance Team |
| EPI_ISL_2539337 | Lighthouse Lab in Alderley Park | Wellcome Sanger Institute for the COVID-19 Genomics UK (COG-UK) Consortium | Jacquelyn Wynn, Mairead Hyland, The Lighthouse Lab in Alderley Park and Alex Alderton, Roberto Amato, Jeffrey Barrett, Sonia Goncalves, Ewan Harrison, David K. Jackson, Ian Johnston, Dominic Kwiatkowski, Cordelia Langford, John Sillitoe on behalf of the Wellcome Sanger Institute COVID-19 Surveillance Team |
| EPI_ISL_2539338, EPI_ISL_2539339, EPI_ISL_2539341, EPI_ISL_2539342, EPI_ISL_2539344, EPI_ISL_2539345, EPI_ISL_2539346, EPI_ISL_2539347 | Lighthouse Lab in Milton Keynes | Wellcome Sanger Institute for the COVID-19 Genomics UK (COG-UK) Consortium | The Lighthouse Lab in Milton Keynes and Alex Alderton, Roberto Amato, Jeffrey Barrett, Sonia Goncalves, Ewan Harrison, David K. Jackson, Ian Johnston, Dominic Kwiatkowski, Cordelia Langford, John Sillitoe on behalf of the Wellcome Sanger Institute COVID-19 Surveillance Team |
| EPI_ISL_2539348, EPI_ISL_2539349, EPI_ISL_2539351 | Lighthouse Lab in Alderley Park | Wellcome Sanger Institute for the COVID-19 Genomics UK (COG-UK) Consortium | Jacquelyn Wynn, Mairead Hyland, The Lighthouse Lab in Alderley Park and Alex Alderton, Roberto Amato, Jeffrey Barrett, Sonia Goncalves, Ewan Harrison, David K. Jackson, Ian Johnston, Dominic Kwiatkowski, Cordelia Langford, John Sillitoe on behalf of the Wellcome Sanger Institute COVID-19 Surveillance Team |
| EPI_ISL_2539352 | Lighthouse Lab in Glasgow | Wellcome Sanger Institute for the COVID-19 Genomics UK (COG-UK) Consortium | Harper VanSteenhouse, Yumi Kasai, David Gray, Carol Clugston, Anna Dominiczak and Alex Alderton, Roberto Amato, Jeffrey Barrett, Sonia Goncalves, Ewan Harrison, David K. Jackson, Ian Johnston, Dominic Kwiatkowski, Cordelia Langford, John Sillitoe on behalf of the Wellcome Sanger Institute COVID-19 Surveillance Team |
| EPI_ISL_2539353 | Lighthouse Lab in Alderley Park | Wellcome Sanger Institute for the COVID-19 Genomics UK (COG-UK) | Jacquelyn Wynn, Mairead Hyland, The Lighthouse Lab in Alderley Park and Alex Alderton, Roberto Amato, Jeffrey Barrett, Sonia Goncalves, Ewan Harrison, David K. Jackson, Ian Johnston, Dominic Kwiatkowski, Cordelia Langford, John Sillitoe on |

| | | | |
|---|---|---|---|
| | | Consortium | behalf of the Wellcome Sanger Institute COVID-19 Surveillance Team |
| EPI_ISL_2539354 | Lighthouse Lab in Milton Keynes | Wellcome Sanger Institute for the COVID-19 Genomics UK (COG-UK) Consortium | The Lighthouse Lab in Milton Keynes and Alex Alderton, Roberto Amato, Jeffrey Barrett, Sonia Goncalves, Ewan Harrison, David K. Jackson, Ian Johnston, Dominic Kwiatkowski, Cordelia Langford, John Sillitoe on behalf of the Wellcome Sanger Institute COVID-19 Surveillance Team |
| EPI_ISL_2539355 | Lighthouse Lab in Alderley Park | Wellcome Sanger Institute for the COVID-19 Genomics UK (COG-UK) Consortium | Jacquelyn Wynn, Mairead Hyland, The Lighthouse Lab in Alderley Park and Alex Alderton, Roberto Amato, Jeffrey Barrett, Sonia Goncalves, Ewan Harrison, David K. Jackson, Ian Johnston, Dominic Kwiatkowski, Cordelia Langford, John Sillitoe on behalf of the Wellcome Sanger Institute COVID-19 Surveillance Team |
| EPI_ISL_2539356, EPI_ISL_2539357, EPI_ISL_2539359, EPI_ISL_2539360, EPI_ISL_2539361 | Lighthouse Lab in Milton Keynes | Wellcome Sanger Institute for the COVID-19 Genomics UK (COG-UK) Consortium | The Lighthouse Lab in Milton Keynes and Alex Alderton, Roberto Amato, Jeffrey Barrett, Sonia Goncalves, Ewan Harrison, David K. Jackson, Ian Johnston, Dominic Kwiatkowski, Cordelia Langford, John Sillitoe on behalf of the Wellcome Sanger Institute COVID-19 Surveillance Team |
| EPI_ISL_2539362 | Lighthouse Lab in Alderley Park | Wellcome Sanger Institute for the COVID-19 Genomics UK (COG-UK) Consortium | Jacquelyn Wynn, Mairead Hyland, The Lighthouse Lab in Alderley Park and Alex Alderton, Roberto Amato, Jeffrey Barrett, Sonia Goncalves, Ewan Harrison, David K. Jackson, Ian Johnston, Dominic Kwiatkowski, Cordelia Langford, John Sillitoe on behalf of the Wellcome Sanger Institute COVID-19 Surveillance Team |
| EPI_ISL_2539363, EPI_ISL_2539364 | Lighthouse Lab in Milton Keynes | Wellcome Sanger Institute for the COVID-19 Genomics UK (COG-UK) Consortium | The Lighthouse Lab in Milton Keynes and Alex Alderton, Roberto Amato, Jeffrey Barrett, Sonia Goncalves, Ewan Harrison, David K. Jackson, Ian Johnston, Dominic Kwiatkowski, Cordelia Langford, John Sillitoe on behalf of the Wellcome Sanger Institute COVID-19 Surveillance Team |
| EPI_ISL_2539365 | Lighthouse Lab in Alderley Park | Wellcome Sanger Institute for the COVID-19 Genomics UK (COG-UK) Consortium | Jacquelyn Wynn, Mairead Hyland, The Lighthouse Lab in Alderley Park and Alex Alderton, Roberto Amato, Jeffrey Barrett, Sonia Goncalves, Ewan Harrison, David K. Jackson, Ian Johnston, Dominic Kwiatkowski, Cordelia Langford, John Sillitoe on behalf of the Wellcome Sanger Institute COVID-19 Surveillance Team |
| EPI_ISL_2539366, EPI_ISL_2539367, EPI_ISL_2539368, EPI_ISL_2539369 | Lighthouse Lab in Milton Keynes | Wellcome Sanger Institute for the COVID-19 Genomics UK (COG-UK) Consortium | The Lighthouse Lab in Milton Keynes and Alex Alderton, Roberto Amato, Jeffrey Barrett, Sonia Goncalves, Ewan Harrison, David K. Jackson, Ian Johnston, Dominic Kwiatkowski, Cordelia Langford, John Sillitoe on behalf of the Wellcome Sanger Institute COVID-19 Surveillance Team |
| EPI_ISL_2539370, EPI_ISL_2539371, EPI_ISL_2539372 | Lighthouse Lab in Alderley Park | Wellcome Sanger Institute for the COVID-19 Genomics UK (COG-UK) Consortium | Jacquelyn Wynn, Mairead Hyland, The Lighthouse Lab in Alderley Park and Alex Alderton, Roberto Amato, Jeffrey Barrett, Sonia Goncalves, Ewan Harrison, David K. Jackson, Ian Johnston, Dominic Kwiatkowski, Cordelia Langford, John Sillitoe on behalf of the Wellcome Sanger Institute COVID-19 Surveillance Team |
| EPI_ISL_2539373, EPI_ISL_2539374, EPI_ISL_2539375 | Lighthouse Lab in Milton Keynes | Wellcome Sanger Institute for the COVID-19 Genomics UK (COG-UK) Consortium | The Lighthouse Lab in Milton Keynes and Alex Alderton, Roberto Amato, Jeffrey Barrett, Sonia Goncalves, Ewan Harrison, David K. Jackson, Ian Johnston, Dominic Kwiatkowski, Cordelia Langford, John Sillitoe on behalf of the Wellcome Sanger Institute COVID-19 Surveillance Team |
| EPI_ISL_2539376 | Lighthouse Lab in Alderley Park | Wellcome Sanger Institute for the COVID-19 Genomics UK (COG-UK) Consortium | Jacquelyn Wynn, Mairead Hyland, The Lighthouse Lab in Alderley Park and Alex Alderton, Roberto Amato, Jeffrey Barrett, Sonia Goncalves, Ewan Harrison, David K. Jackson, Ian Johnston, Dominic Kwiatkowski, Cordelia Langford, John Sillitoe on behalf of the Wellcome Sanger Institute COVID-19 Surveillance Team |
| EPI_ISL_2539377, EPI_ISL_2539378, EPI_ISL_2539379, EPI_ISL_2539380, EPI_ISL_2539381 | Lighthouse Lab in Milton Keynes | Wellcome Sanger Institute for the COVID-19 Genomics UK (COG-UK) Consortium | The Lighthouse Lab in Milton Keynes and Alex Alderton, Roberto Amato, Jeffrey Barrett, Sonia Goncalves, Ewan Harrison, David K. Jackson, Ian Johnston, Dominic Kwiatkowski, Cordelia Langford, John Sillitoe on behalf of the Wellcome Sanger Institute COVID-19 Surveillance Team |
| EPI_ISL_2539382 | Lighthouse Lab in Alderley Park | Wellcome Sanger Institute for the COVID-19 Genomics UK (COG-UK) Consortium | Jacquelyn Wynn, Mairead Hyland, The Lighthouse Lab in Alderley Park and Alex Alderton, Roberto Amato, Jeffrey Barrett, Sonia Goncalves, Ewan Harrison, David K. Jackson, Ian Johnston, Dominic Kwiatkowski, Cordelia Langford, John Sillitoe on behalf of the Wellcome Sanger Institute COVID-19 Surveillance Team |
| EPI_ISL_2539384, EPI_ISL_2539385 | Lighthouse Lab in Milton Keynes | Wellcome Sanger Institute for the COVID-19 Genomics UK (COG-UK) Consortium | The Lighthouse Lab in Milton Keynes and Alex Alderton, Roberto Amato, Jeffrey Barrett, Sonia Goncalves, Ewan Harrison, David K. Jackson, Ian Johnston, Dominic Kwiatkowski, Cordelia Langford, John Sillitoe on behalf of the Wellcome Sanger Institute COVID-19 Surveillance Team |
| EPI_ISL_2539386, EPI_ISL_2539387 | Lighthouse Lab in Alderley Park | Wellcome Sanger Institute for the COVID-19 Genomics UK (COG-UK) Consortium | Jacquelyn Wynn, Mairead Hyland, The Lighthouse Lab in Alderley Park and Alex Alderton, Roberto Amato, Jeffrey Barrett, Sonia Goncalves, Ewan Harrison, David K. Jackson, Ian Johnston, Dominic Kwiatkowski, Cordelia Langford, John Sillitoe on behalf of the Wellcome Sanger Institute COVID-19 Surveillance Team |
| EPI_ISL_2539388 | Lighthouse Lab in Milton Keynes | Wellcome Sanger Institute for the COVID-19 Genomics UK (COG-UK) Consortium | The Lighthouse Lab in Milton Keynes and Alex Alderton, Roberto Amato, Jeffrey Barrett, Sonia Goncalves, Ewan Harrison, David K. Jackson, Ian Johnston, Dominic Kwiatkowski, Cordelia Langford, John Sillitoe on behalf of the Wellcome Sanger Institute COVID-19 Surveillance Team |
| EPI_ISL_2539389, EPI_ISL_2539391 | Lighthouse Lab in Alderley Park | Wellcome Sanger Institute for the COVID-19 Genomics UK (COG-UK) Consortium | Jacquelyn Wynn, Mairead Hyland, The Lighthouse Lab in Alderley Park and Alex Alderton, Roberto Amato, Jeffrey Barrett, Sonia Goncalves, Ewan Harrison, David K. Jackson, Ian Johnston, Dominic Kwiatkowski, Cordelia Langford, John Sillitoe on behalf of the Wellcome Sanger Institute COVID-19 Surveillance Team |
| EPI_ISL_2539392 | Lighthouse Lab in Milton Keynes | Wellcome Sanger Institute for the COVID-19 Genomics UK (COG-UK) Consortium | The Lighthouse Lab in Milton Keynes and Alex Alderton, Roberto Amato, Jeffrey Barrett, Sonia Goncalves, Ewan Harrison, David K. Jackson, Ian Johnston, Dominic Kwiatkowski, Cordelia Langford, John Sillitoe on behalf of the Wellcome Sanger Institute COVID-19 Surveillance Team |
| EPI_ISL_2539393 | Lighthouse Lab in Alderley Park | Wellcome Sanger Institute for the COVID-19 Genomics UK (COG-UK) Consortium | Jacquelyn Wynn, Mairead Hyland, The Lighthouse Lab in Alderley Park and Alex Alderton, Roberto Amato, Jeffrey Barrett, Sonia Goncalves, Ewan Harrison, David K. Jackson, Ian Johnston, Dominic Kwiatkowski, Cordelia Langford, John Sillitoe on behalf of the Wellcome Sanger Institute COVID-19 Surveillance Team |
| EPI_ISL_2539395, EPI_ISL_2539396, EPI_ISL_2539397 | Lighthouse Lab in Milton Keynes | Wellcome Sanger Institute for the COVID-19 Genomics UK (COG-UK) Consortium | The Lighthouse Lab in Milton Keynes and Alex Alderton, Roberto Amato, Jeffrey Barrett, Sonia Goncalves, Ewan Harrison, David K. Jackson, Ian Johnston, Dominic Kwiatkowski, Cordelia Langford, John Sillitoe on behalf of the Wellcome Sanger Institute COVID-19 Surveillance Team |
| EPI_ISL_2539398 | Lighthouse Lab in Alderley Park | Wellcome Sanger Institute for the COVID-19 Genomics UK (COG-UK) Consortium | Jacquelyn Wynn, Mairead Hyland, The Lighthouse Lab in Alderley Park and Alex Alderton, Roberto Amato, Jeffrey Barrett, Sonia Goncalves, Ewan Harrison, David K. Jackson, Ian Johnston, Dominic Kwiatkowski, Cordelia Langford, John Sillitoe on behalf of the Wellcome Sanger Institute COVID-19 Surveillance Team |
| EPI_ISL_2539400 | Lighthouse Lab in Milton Keynes | Wellcome Sanger Institute for the COVID-19 Genomics UK (COG-UK) Consortium | The Lighthouse Lab in Milton Keynes and Alex Alderton, Roberto Amato, Jeffrey Barrett, Sonia Goncalves, Ewan Harrison, David K. Jackson, Ian Johnston, Dominic Kwiatkowski, Cordelia Langford, John Sillitoe on behalf of the Wellcome Sanger Institute COVID-19 Surveillance Team |
| EPI_ISL_2539401, EPI_ISL_2539402 | Lighthouse Lab in Alderley Park | Wellcome Sanger Institute for the COVID-19 Genomics UK (COG-UK) Consortium | Jacquelyn Wynn, Mairead Hyland, The Lighthouse Lab in Alderley Park and Alex Alderton, Roberto Amato, Jeffrey Barrett, Sonia Goncalves, Ewan Harrison, David K. Jackson, Ian Johnston, Dominic Kwiatkowski, Cordelia Langford, John Sillitoe on behalf of the Wellcome Sanger Institute COVID-19 Surveillance Team |
| EPI_ISL_2539403, EPI_ISL_2539405, EPI_ISL_2539406 | Lighthouse Lab in Milton Keynes | Wellcome Sanger Institute for the COVID-19 Genomics UK (COG-UK) Consortium | The Lighthouse Lab in Milton Keynes and Alex Alderton, Roberto Amato, Jeffrey Barrett, Sonia Goncalves, Ewan Harrison, David K. Jackson, Ian Johnston, Dominic Kwiatkowski, Cordelia Langford, John Sillitoe on behalf of the Wellcome Sanger Institute COVID-19 Surveillance Team |
| EPI_ISL_2539407 | Lighthouse Lab in Alderley Park | Wellcome Sanger Institute for the COVID-19 Genomics UK (COG-UK) Consortium | Jacquelyn Wynn, Mairead Hyland, The Lighthouse Lab in Alderley Park and Alex Alderton, Roberto Amato, Jeffrey Barrett, Sonia Goncalves, Ewan Harrison, David K. Jackson, Ian Johnston, Dominic Kwiatkowski, Cordelia Langford, John Sillitoe on behalf of the Wellcome Sanger Institute COVID-19 Surveillance Team |
| EPI_ISL_2539408 | Lighthouse Lab in Milton Keynes | Wellcome Sanger Institute for the COVID-19 Genomics UK (COG-UK) Consortium | The Lighthouse Lab in Milton Keynes and Alex Alderton, Roberto Amato, Jeffrey Barrett, Sonia Goncalves, Ewan Harrison, David K. Jackson, Ian Johnston, Dominic Kwiatkowski, Cordelia Langford, John Sillitoe on behalf of the Wellcome Sanger Institute COVID-19 Surveillance Team |
| EPI_ISL_2539409, EPI_ISL_2539410 | Lighthouse Lab in Alderley Park | Wellcome Sanger Institute for the COVID-19 Genomics UK (COG-UK) Consortium | Jacquelyn Wynn, Mairead Hyland, The Lighthouse Lab in Alderley Park and Alex Alderton, Roberto Amato, Jeffrey Barrett, Sonia Goncalves, Ewan Harrison, David K. Jackson, Ian Johnston, Dominic Kwiatkowski, Cordelia Langford, John Sillitoe on behalf of the Wellcome Sanger Institute COVID-19 Surveillance Team |
| EPI_ISL_2539411 | Lighthouse Lab in Milton Keynes | Wellcome Sanger Institute for the COVID-19 Genomics UK (COG-UK) Consortium | The Lighthouse Lab in Milton Keynes and Alex Alderton, Roberto Amato, Jeffrey Barrett, Sonia Goncalves, Ewan Harrison, David K. Jackson, Ian Johnston, Dominic Kwiatkowski, Cordelia Langford, John Sillitoe on behalf of the Wellcome Sanger Institute COVID-19 Surveillance Team |
| EPI_ISL_2539412, EPI_ISL_2539413, EPI_ISL_2539414 | Lighthouse Lab in Alderley Park | Wellcome Sanger Institute for the COVID-19 Genomics UK (COG-UK) Consortium | Jacquelyn Wynn, Mairead Hyland, The Lighthouse Lab in Alderley Park and Alex Alderton, Roberto Amato, Jeffrey Barrett, Sonia Goncalves, Ewan Harrison, David K. Jackson, Ian Johnston, Dominic Kwiatkowski, Cordelia Langford, John Sillitoe on behalf of the Wellcome Sanger Institute COVID-19 Surveillance Team |
| EPI_ISL_2539415 | Lighthouse Lab in Glasgow | Wellcome Sanger Institute for the COVID-19 Genomics UK (COG-UK) Consortium | Harper VanSteenhouse, Yumi Kasai, David Gray, Carol Clugston, Anna Dominiczak and Alex Alderton, Roberto Amato, Jeffrey Barrett, Sonia Goncalves, Ewan Harrison, David K. Jackson, Ian Johnston, Dominic Kwiatkowski, Cordelia Langford, John Sillitoe on behalf of the Wellcome Sanger Institute COVID-19 Surveillance Team |
| EPI_ISL_2539416 | Lighthouse Lab in Alderley Park | Wellcome Sanger Institute for the COVID-19 Genomics UK (COG-UK) Consortium | Jacquelyn Wynn, Mairead Hyland, The Lighthouse Lab in Alderley Park and Alex Alderton, Roberto Amato, Jeffrey Barrett, Sonia Goncalves, Ewan Harrison, David K. Jackson, Ian Johnston, Dominic Kwiatkowski, Cordelia Langford, John Sillitoe on behalf of the Wellcome Sanger Institute COVID-19 Surveillance Team |
| EPI_ISL_2539417, EPI_ISL_2539418 | Lighthouse Lab in Milton Keynes | Wellcome Sanger Institute for the COVID-19 Genomics UK (COG-UK) Consortium | The Lighthouse Lab in Milton Keynes and Alex Alderton, Roberto Amato, Jeffrey Barrett, Sonia Goncalves, Ewan Harrison, David K. Jackson, Ian Johnston, Dominic Kwiatkowski, Cordelia Langford, John Sillitoe on behalf of the Wellcome Sanger Institute COVID-19 Surveillance Team |
| EPI_ISL_2539419 | Lighthouse Lab in Alderley Park | Wellcome Sanger Institute for the COVID-19 Genomics UK (COG-UK) Consortium | Jacquelyn Wynn, Mairead Hyland, The Lighthouse Lab in Alderley Park and Alex Alderton, Roberto Amato, Jeffrey Barrett, Sonia Goncalves, Ewan Harrison, David K. Jackson, Ian Johnston, Dominic Kwiatkowski, Cordelia Langford, John Sillitoe on behalf of the Wellcome Sanger Institute COVID-19 Surveillance Team |
| EPI_ISL_2539420, EPI_ISL_2539421, EPI_ISL_2539422 | Lighthouse Lab in Milton Keynes | Wellcome Sanger Institute for the COVID-19 Genomics UK (COG-UK) Consortium | The Lighthouse Lab in Milton Keynes and Alex Alderton, Roberto Amato, Jeffrey Barrett, Sonia Goncalves, Ewan Harrison, David K. Jackson, Ian Johnston, Dominic Kwiatkowski, Cordelia Langford, John Sillitoe on behalf of the Wellcome Sanger Institute COVID-19 Surveillance Team |
| EPI_ISL_2539423 | Lighthouse Lab in Alderley Park | Wellcome Sanger Institute for the COVID-19 Genomics UK (COG-UK) Consortium | Jacquelyn Wynn, Mairead Hyland, The Lighthouse Lab in Alderley Park and Alex Alderton, Roberto Amato, Jeffrey Barrett, Sonia Goncalves, Ewan Harrison, David K. Jackson, Ian Johnston, Dominic Kwiatkowski, Cordelia Langford, John Sillitoe on behalf of the Wellcome Sanger Institute COVID-19 Surveillance Team |
| EPI_ISL_2539424 | Lighthouse Lab in Glasgow | Wellcome Sanger Institute for the COVID-19 Genomics UK (COG-UK) Consortium | Harper VanSteenhouse, Yumi Kasai, David Gray, Carol Clugston, Anna Dominiczak and Alex Alderton, Roberto Amato, Jeffrey Barrett, Sonia Goncalves, Ewan Harrison, David K. Jackson, Ian Johnston, Dominic Kwiatkowski, Cordelia Langford, John Sillitoe on behalf of the Wellcome Sanger Institute COVID-19 Surveillance Team |
| EPI_ISL_2539425 | Lighthouse Lab in Alderley Park | Wellcome Sanger Institute for the COVID-19 Genomics UK (COG-UK) Consortium | Jacquelyn Wynn, Mairead Hyland, The Lighthouse Lab in Alderley Park and Alex Alderton, Roberto Amato, Jeffrey Barrett, Sonia Goncalves, Ewan Harrison, David K. Jackson, Ian Johnston, Dominic Kwiatkowski, Cordelia Langford, John Sillitoe on behalf of the Wellcome Sanger Institute COVID-19 Surveillance Team |
| EPI_ISL_2539427 | Lighthouse Lab in Glasgow | Wellcome Sanger Institute for the COVID-19 Genomics UK (COG-UK) Consortium | Harper VanSteenhouse, Yumi Kasai, David Gray, Carol Clugston, Anna Dominiczak and Alex Alderton, Roberto Amato, Jeffrey Barrett, Sonia Goncalves, Ewan Harrison, David K. Jackson, Ian Johnston, Dominic Kwiatkowski, Cordelia Langford, John Sillitoe on behalf of the Wellcome Sanger Institute COVID-19 Surveillance Team |
| EPI_ISL_2539428, EPI_ISL_2539429, EPI_ISL_2539430, EPI_ISL_2539431 | Lighthouse Lab in Milton Keynes | Wellcome Sanger Institute for the COVID-19 Genomics UK (COG-UK) Consortium | The Lighthouse Lab in Milton Keynes and Alex Alderton, Roberto Amato, Jeffrey Barrett, Sonia Goncalves, Ewan Harrison, David K. Jackson, Ian Johnston, Dominic Kwiatkowski, Cordelia Langford, John Sillitoe on behalf of the Wellcome Sanger Institute COVID-19 Surveillance Team |

| | | | |
|---|---|---|---|
| EPI_ISL_2539432, EPI_ISL_2539433, EPI_ISL_2539434 | Lighthouse Lab in Alderley Park | Wellcome Sanger Institute for the COVID-19 Genomics UK (COG-UK) Consortium | Jacquelyn Wynn, Mairead Hyland, The Lighthouse Lab in Alderley Park and Alex Alderton, Roberto Amato, Jeffrey Barrett, Sonia Goncalves, Ewan Harrison, David K. Jackson, Ian Johnston, Dominic Kwiatkowski, Cordelia Langford, John Sillitoe on behalf of the Wellcome Sanger Institute COVID-19 Surveillance Team |
| EPI_ISL_2539435 | Lighthouse Lab in Milton Keynes | Wellcome Sanger Institute for the COVID-19 Genomics UK (COG-UK) Consortium | The Lighthouse Lab in Milton Keynes and Alex Alderton, Roberto Amato, Jeffrey Barrett, Sonia Goncalves, Ewan Harrison, David K. Jackson, Ian Johnston, Dominic Kwiatkowski, Cordelia Langford, John Sillitoe on behalf of the Wellcome Sanger Institute COVID-19 Surveillance Team |
| EPI_ISL_2539436 | Lighthouse Lab in Alderley Park | Wellcome Sanger Institute for the COVID-19 Genomics UK (COG-UK) Consortium | Jacquelyn Wynn, Mairead Hyland, The Lighthouse Lab in Alderley Park and Alex Alderton, Roberto Amato, Jeffrey Barrett, Sonia Goncalves, Ewan Harrison, David K. Jackson, Ian Johnston, Dominic Kwiatkowski, Cordelia Langford, John Sillitoe on behalf of the Wellcome Sanger Institute COVID-19 Surveillance Team |
| EPI_ISL_2539437, EPI_ISL_2539438 | Lighthouse Lab in Milton Keynes | Wellcome Sanger Institute for the COVID-19 Genomics UK (COG-UK) Consortium | The Lighthouse Lab in Milton Keynes and Alex Alderton, Roberto Amato, Jeffrey Barrett, Sonia Goncalves, Ewan Harrison, David K. Jackson, Ian Johnston, Dominic Kwiatkowski, Cordelia Langford, John Sillitoe on behalf of the Wellcome Sanger Institute COVID-19 Surveillance Team |
| EPI_ISL_2539439, EPI_ISL_2539440 | Lighthouse Lab in Glasgow | Wellcome Sanger Institute for the COVID-19 Genomics UK (COG-UK) Consortium | Harper VanSteenhouse, Yumi Kasai, David Gray, Carol Clugston, Anna Dominiczak and Alex Alderton, Roberto Amato, Jeffrey Barrett, Sonia Goncalves, Ewan Harrison, David K. Jackson, Ian Johnston, Dominic Kwiatkowski, Cordelia Langford, John Sillitoe on behalf of the Wellcome Sanger Institute COVID-19 Surveillance Team |
| EPI_ISL_2539441 | Lighthouse Lab in Milton Keynes | Wellcome Sanger Institute for the COVID-19 Genomics UK (COG-UK) Consortium | The Lighthouse Lab in Milton Keynes and Alex Alderton, Roberto Amato, Jeffrey Barrett, Sonia Goncalves, Ewan Harrison, David K. Jackson, Ian Johnston, Dominic Kwiatkowski, Cordelia Langford, John Sillitoe on behalf of the Wellcome Sanger Institute COVID-19 Surveillance Team |
| EPI_ISL_2539442 | Lighthouse Lab in Alderley Park | Wellcome Sanger Institute for the COVID-19 Genomics UK (COG-UK) Consortium | Jacquelyn Wynn, Mairead Hyland, The Lighthouse Lab in Alderley Park and Alex Alderton, Roberto Amato, Jeffrey Barrett, Sonia Goncalves, Ewan Harrison, David K. Jackson, Ian Johnston, Dominic Kwiatkowski, Cordelia Langford, John Sillitoe on behalf of the Wellcome Sanger Institute COVID-19 Surveillance Team |
| EPI_ISL_2539443, EPI_ISL_2539444 | Lighthouse Lab in Milton Keynes | Wellcome Sanger Institute for the COVID-19 Genomics UK (COG-UK) Consortium | The Lighthouse Lab in Milton Keynes and Alex Alderton, Roberto Amato, Jeffrey Barrett, Sonia Goncalves, Ewan Harrison, David K. Jackson, Ian Johnston, Dominic Kwiatkowski, Cordelia Langford, John Sillitoe on behalf of the Wellcome Sanger Institute COVID-19 Surveillance Team |
| EPI_ISL_2539445 | Lighthouse Lab in Alderley Park | Wellcome Sanger Institute for the COVID-19 Genomics UK (COG-UK) Consortium | Jacquelyn Wynn, Mairead Hyland, The Lighthouse Lab in Alderley Park and Alex Alderton, Roberto Amato, Jeffrey Barrett, Sonia Goncalves, Ewan Harrison, David K. Jackson, Ian Johnston, Dominic Kwiatkowski, Cordelia Langford, John Sillitoe on behalf of the Wellcome Sanger Institute COVID-19 Surveillance Team |
| EPI_ISL_2539446, EPI_ISL_2539447 | Lighthouse Lab in Glasgow | Wellcome Sanger Institute for the COVID-19 Genomics UK (COG-UK) Consortium | Harper VanSteenhouse, Yumi Kasai, David Gray, Carol Clugston, Anna Dominiczak and Alex Alderton, Roberto Amato, Jeffrey Barrett, Sonia Goncalves, Ewan Harrison, David K. Jackson, Ian Johnston, Dominic Kwiatkowski, Cordelia Langford, John Sillitoe on behalf of the Wellcome Sanger Institute COVID-19 Surveillance Team |
| EPI_ISL_2539448, EPI_ISL_2539449 | Lighthouse Lab in Milton Keynes | Wellcome Sanger Institute for the COVID-19 Genomics UK (COG-UK) Consortium | The Lighthouse Lab in Milton Keynes and Alex Alderton, Roberto Amato, Jeffrey Barrett, Sonia Goncalves, Ewan Harrison, David K. Jackson, Ian Johnston, Dominic Kwiatkowski, Cordelia Langford, John Sillitoe on behalf of the Wellcome Sanger Institute COVID-19 Surveillance Team |
| EPI_ISL_2539450 | Lighthouse Lab in Alderley Park | Wellcome Sanger Institute for the COVID-19 Genomics UK (COG-UK) Consortium | Jacquelyn Wynn, Mairead Hyland, The Lighthouse Lab in Alderley Park and Alex Alderton, Roberto Amato, Jeffrey Barrett, Sonia Goncalves, Ewan Harrison, David K. Jackson, Ian Johnston, Dominic Kwiatkowski, Cordelia Langford, John Sillitoe on behalf of the Wellcome Sanger Institute COVID-19 Surveillance Team |
| EPI_ISL_2539451 | Lighthouse Lab in Milton Keynes | Wellcome Sanger Institute for the COVID-19 Genomics UK (COG-UK) Consortium | The Lighthouse Lab in Milton Keynes and Alex Alderton, Roberto Amato, Jeffrey Barrett, Sonia Goncalves, Ewan Harrison, David K. Jackson, Ian Johnston, Dominic Kwiatkowski, Cordelia Langford, John Sillitoe on behalf of the Wellcome Sanger Institute COVID-19 Surveillance Team |
| EPI_ISL_2539452 | Lighthouse Lab in Alderley Park | Wellcome Sanger Institute for the COVID-19 Genomics UK (COG-UK) Consortium | Jacquelyn Wynn, Mairead Hyland, The Lighthouse Lab in Alderley Park and Alex Alderton, Roberto Amato, Jeffrey Barrett, Sonia Goncalves, Ewan Harrison, David K. Jackson, Ian Johnston, Dominic Kwiatkowski, Cordelia Langford, John Sillitoe on behalf of the Wellcome Sanger Institute COVID-19 Surveillance Team |
| EPI_ISL_2539453 | Lighthouse Lab in Milton Keynes | Wellcome Sanger Institute for the COVID-19 Genomics UK (COG-UK) Consortium | The Lighthouse Lab in Milton Keynes and Alex Alderton, Roberto Amato, Jeffrey Barrett, Sonia Goncalves, Ewan Harrison, David K. Jackson, Ian Johnston, Dominic Kwiatkowski, Cordelia Langford, John Sillitoe on behalf of the Wellcome Sanger Institute COVID-19 Surveillance Team |
| EPI_ISL_2539454 | Lighthouse Lab in Alderley Park | Wellcome Sanger Institute for the COVID-19 Genomics UK (COG-UK) Consortium | Jacquelyn Wynn, Mairead Hyland, The Lighthouse Lab in Alderley Park and Alex Alderton, Roberto Amato, Jeffrey Barrett, Sonia Goncalves, Ewan Harrison, David K. Jackson, Ian Johnston, Dominic Kwiatkowski, Cordelia Langford, John Sillitoe on behalf of the Wellcome Sanger Institute COVID-19 Surveillance Team |
| EPI_ISL_2539455 | Lighthouse Lab in Glasgow | Wellcome Sanger Institute for the COVID-19 Genomics UK (COG-UK) Consortium | Harper VanSteenhouse, Yumi Kasai, David Gray, Carol Clugston, Anna Dominiczak and Alex Alderton, Roberto Amato, Jeffrey Barrett, Sonia Goncalves, Ewan Harrison, David K. Jackson, Ian Johnston, Dominic Kwiatkowski, Cordelia Langford, John Sillitoe on behalf of the Wellcome Sanger Institute COVID-19 Surveillance Team |
| EPI_ISL_2539456, EPI_ISL_2539457 | Lighthouse Lab in Alderley Park | Wellcome Sanger Institute for the COVID-19 Genomics UK (COG-UK) Consortium | Jacquelyn Wynn, Mairead Hyland, The Lighthouse Lab in Alderley Park and Alex Alderton, Roberto Amato, Jeffrey Barrett, Sonia Goncalves, Ewan Harrison, David K. Jackson, Ian Johnston, Dominic Kwiatkowski, Cordelia Langford, John Sillitoe on behalf of the Wellcome Sanger Institute COVID-19 Surveillance Team |
| EPI_ISL_2539458 | Lighthouse Lab in Glasgow | Wellcome Sanger Institute for the COVID-19 Genomics UK (COG-UK) Consortium | Harper VanSteenhouse, Yumi Kasai, David Gray, Carol Clugston, Anna Dominiczak and Alex Alderton, Roberto Amato, Jeffrey Barrett, Sonia Goncalves, Ewan Harrison, David K. Jackson, Ian Johnston, Dominic Kwiatkowski, Cordelia Langford, John Sillitoe on behalf of the Wellcome Sanger Institute COVID-19 Surveillance Team |
| EPI_ISL_2539459 | Lighthouse Lab in Milton Keynes | Wellcome Sanger Institute for the COVID-19 Genomics UK (COG-UK) Consortium | The Lighthouse Lab in Milton Keynes and Alex Alderton, Roberto Amato, Jeffrey Barrett, Sonia Goncalves, Ewan Harrison, David K. Jackson, Ian Johnston, Dominic Kwiatkowski, Cordelia Langford, John Sillitoe on behalf of the Wellcome Sanger Institute COVID-19 Surveillance Team |
| EPI_ISL_2539460 | Lighthouse Lab in Glasgow | Wellcome Sanger Institute for the COVID-19 Genomics UK (COG-UK) Consortium | Harper VanSteenhouse, Yumi Kasai, David Gray, Carol Clugston, Anna Dominiczak and Alex Alderton, Roberto Amato, Jeffrey Barrett, Sonia Goncalves, Ewan Harrison, David K. Jackson, Ian Johnston, Dominic Kwiatkowski, Cordelia Langford, John Sillitoe on behalf of the Wellcome Sanger Institute COVID-19 Surveillance Team |
| EPI_ISL_2539461 | Lighthouse Lab in Milton Keynes | Wellcome Sanger Institute for the COVID-19 Genomics UK (COG-UK) Consortium | The Lighthouse Lab in Milton Keynes and Alex Alderton, Roberto Amato, Jeffrey Barrett, Sonia Goncalves, Ewan Harrison, David K. Jackson, Ian Johnston, Dominic Kwiatkowski, Cordelia Langford, John Sillitoe on behalf of the Wellcome Sanger Institute COVID-19 Surveillance Team |
| EPI_ISL_2539462 | Lighthouse Lab in Alderley Park | Wellcome Sanger Institute for the COVID-19 Genomics UK (COG-UK) Consortium | Jacquelyn Wynn, Mairead Hyland, The Lighthouse Lab in Alderley Park and Alex Alderton, Roberto Amato, Jeffrey Barrett, Sonia Goncalves, Ewan Harrison, David K. Jackson, Ian Johnston, Dominic Kwiatkowski, Cordelia Langford, John Sillitoe on behalf of the Wellcome Sanger Institute COVID-19 Surveillance Team |
| EPI_ISL_2539463 | Lighthouse Lab in Glasgow | Wellcome Sanger Institute for the COVID-19 Genomics UK (COG-UK) Consortium | Harper VanSteenhouse, Yumi Kasai, David Gray, Carol Clugston, Anna Dominiczak and Alex Alderton, Roberto Amato, Jeffrey Barrett, Sonia Goncalves, Ewan Harrison, David K. Jackson, Ian Johnston, Dominic Kwiatkowski, Cordelia Langford, John Sillitoe on behalf of the Wellcome Sanger Institute COVID-19 Surveillance Team |
| EPI_ISL_2539464, EPI_ISL_2539465, EPI_ISL_2539466, EPI_ISL_2539467, EPI_ISL_2539469, EPI_ISL_2539470, EPI_ISL_2539471, EPI_ISL_2539472 | Lighthouse Lab in Alderley Park | Wellcome Sanger Institute for the COVID-19 Genomics UK (COG-UK) Consortium | Jacquelyn Wynn, Mairead Hyland, The Lighthouse Lab in Alderley Park and Alex Alderton, Roberto Amato, Jeffrey Barrett, Sonia Goncalves, Ewan Harrison, David K. Jackson, Ian Johnston, Dominic Kwiatkowski, Cordelia Langford, John Sillitoe on behalf of the Wellcome Sanger Institute COVID-19 Surveillance Team |
| EPI_ISL_2539473 | Lighthouse Lab in Glasgow | Wellcome Sanger Institute for the COVID-19 Genomics UK (COG-UK) Consortium | Harper VanSteenhouse, Yumi Kasai, David Gray, Carol Clugston, Anna Dominiczak and Alex Alderton, Roberto Amato, Jeffrey Barrett, Sonia Goncalves, Ewan Harrison, David K. Jackson, Ian Johnston, Dominic Kwiatkowski, Cordelia Langford, John Sillitoe on behalf of the Wellcome Sanger Institute COVID-19 Surveillance Team |
| EPI_ISL_2539474 | Lighthouse Lab in Alderley Park | Wellcome Sanger Institute for the COVID-19 Genomics UK (COG-UK) Consortium | Jacquelyn Wynn, Mairead Hyland, The Lighthouse Lab in Alderley Park and Alex Alderton, Roberto Amato, Jeffrey Barrett, Sonia Goncalves, Ewan Harrison, David K. Jackson, Ian Johnston, Dominic Kwiatkowski, Cordelia Langford, John Sillitoe on behalf of the Wellcome Sanger Institute COVID-19 Surveillance Team |
| EPI_ISL_2539475, EPI_ISL_2539476, EPI_ISL_2539478 | Lighthouse Lab in Glasgow | Wellcome Sanger Institute for the COVID-19 Genomics UK (COG-UK) Consortium | Harper VanSteenhouse, Yumi Kasai, David Gray, Carol Clugston, Anna Dominiczak and Alex Alderton, Roberto Amato, Jeffrey Barrett, Sonia Goncalves, Ewan Harrison, David K. Jackson, Ian Johnston, Dominic Kwiatkowski, Cordelia Langford, John Sillitoe on behalf of the Wellcome Sanger Institute COVID-19 Surveillance Team |
| EPI_ISL_2539479, EPI_ISL_2539481 | Lighthouse Lab in Milton Keynes | Wellcome Sanger Institute for the COVID-19 Genomics UK (COG-UK) Consortium | The Lighthouse Lab in Milton Keynes and Alex Alderton, Roberto Amato, Jeffrey Barrett, Sonia Goncalves, Ewan Harrison, David K. Jackson, Ian Johnston, Dominic Kwiatkowski, Cordelia Langford, John Sillitoe on behalf of the Wellcome Sanger Institute COVID-19 Surveillance Team |
| EPI_ISL_2539482 | Lighthouse Lab in Alderley Park | Wellcome Sanger Institute for the COVID-19 Genomics UK (COG-UK) Consortium | Jacquelyn Wynn, Mairead Hyland, The Lighthouse Lab in Alderley Park and Alex Alderton, Roberto Amato, Jeffrey Barrett, Sonia Goncalves, Ewan Harrison, David K. Jackson, Ian Johnston, Dominic Kwiatkowski, Cordelia Langford, John Sillitoe on behalf of the Wellcome Sanger Institute COVID-19 Surveillance Team |
| EPI_ISL_2539483, EPI_ISL_2539484 | Lighthouse Lab in Milton Keynes | Wellcome Sanger Institute for the COVID-19 Genomics UK (COG-UK) Consortium | The Lighthouse Lab in Milton Keynes and Alex Alderton, Roberto Amato, Jeffrey Barrett, Sonia Goncalves, Ewan Harrison, David K. Jackson, Ian Johnston, Dominic Kwiatkowski, Cordelia Langford, John Sillitoe on behalf of the Wellcome Sanger Institute COVID-19 Surveillance Team |
| EPI_ISL_2539485 | Lighthouse Lab in Alderley Park | Wellcome Sanger Institute for the COVID-19 Genomics UK (COG-UK) Consortium | Jacquelyn Wynn, Mairead Hyland, The Lighthouse Lab in Alderley Park and Alex Alderton, Roberto Amato, Jeffrey Barrett, Sonia Goncalves, Ewan Harrison, David K. Jackson, Ian Johnston, Dominic Kwiatkowski, Cordelia Langford, John Sillitoe on behalf of the Wellcome Sanger Institute COVID-19 Surveillance Team |
| EPI_ISL_2539486 | Lighthouse Lab in Milton Keynes | Wellcome Sanger Institute for the COVID-19 Genomics UK (COG-UK) Consortium | The Lighthouse Lab in Milton Keynes and Alex Alderton, Roberto Amato, Jeffrey Barrett, Sonia Goncalves, Ewan Harrison, David K. Jackson, Ian Johnston, Dominic Kwiatkowski, Cordelia Langford, John Sillitoe on behalf of the Wellcome Sanger Institute COVID-19 Surveillance Team |
| EPI_ISL_2539487, EPI_ISL_2539489 | Lighthouse Lab in Alderley Park | Wellcome Sanger Institute for the COVID-19 Genomics UK (COG-UK) Consortium | Jacquelyn Wynn, Mairead Hyland, The Lighthouse Lab in Alderley Park and Alex Alderton, Roberto Amato, Jeffrey Barrett, Sonia Goncalves, Ewan Harrison, David K. Jackson, Ian Johnston, Dominic Kwiatkowski, Cordelia Langford, John Sillitoe on behalf of the Wellcome Sanger Institute COVID-19 Surveillance Team |
| EPI_ISL_2539490, EPI_ISL_2539491, EPI_ISL_2539492, EPI_ISL_2539493 | Lighthouse Lab in Glasgow | Wellcome Sanger Institute for the COVID-19 Genomics UK (COG-UK) Consortium | Harper VanSteenhouse, Yumi Kasai, David Gray, Carol Clugston, Anna Dominiczak and Alex Alderton, Roberto Amato, Jeffrey Barrett, Sonia Goncalves, Ewan Harrison, David K. Jackson, Ian Johnston, Dominic Kwiatkowski, Cordelia Langford, John Sillitoe on behalf of the Wellcome Sanger Institute COVID-19 Surveillance Team |
| EPI_ISL_2539494 | Lighthouse Lab in Alderley Park | Wellcome Sanger Institute for the COVID-19 Genomics UK (COG-UK) Consortium | Jacquelyn Wynn, Mairead Hyland, The Lighthouse Lab in Alderley Park and Alex Alderton, Roberto Amato, Jeffrey Barrett, Sonia Goncalves, Ewan Harrison, David K. Jackson, Ian Johnston, Dominic Kwiatkowski, Cordelia Langford, John Sillitoe on behalf of the Wellcome Sanger Institute COVID-19 Surveillance Team |
| EPI_ISL_2539495, EPI_ISL_2539497 | Lighthouse Lab in Milton Keynes | Wellcome Sanger Institute for the COVID-19 Genomics UK (COG-UK) Consortium | The Lighthouse Lab in Milton Keynes and Alex Alderton, Roberto Amato, Jeffrey Barrett, Sonia Goncalves, Ewan Harrison, David K. Jackson, Ian Johnston, Dominic Kwiatkowski, Cordelia Langford, John Sillitoe on behalf of the Wellcome Sanger Institute COVID-19 Surveillance Team |
| EPI_ISL_2539498 | Lighthouse Lab in Alderley Park | Wellcome Sanger Institute for the COVID-19 Genomics UK (COG-UK) Consortium | Jacquelyn Wynn, Mairead Hyland, The Lighthouse Lab in Alderley Park and Alex Alderton, Roberto Amato, Jeffrey Barrett, Sonia Goncalves, Ewan Harrison, David K. Jackson, Ian Johnston, Dominic Kwiatkowski, Cordelia Langford, John Sillitoe on behalf of the Wellcome Sanger Institute COVID-19 Surveillance Team |
| EPI_ISL_2539499, EPI_ISL_2539500, EPI_ISL_2539501, EPI_ISL_2539502 | Lighthouse Lab in Glasgow | Wellcome Sanger Institute for the COVID-19 Genomics UK (COG-UK) Consortium | Harper VanSteenhouse, Yumi Kasai, David Gray, Carol Clugston, Anna Dominiczak and Alex Alderton, Roberto Amato, Jeffrey Barrett, Sonia Goncalves, Ewan Harrison, David K. Jackson, Ian Johnston, Dominic Kwiatkowski, Cordelia Langford, John Sillitoe on behalf of the Wellcome Sanger Institute COVID-19 Surveillance Team |
| EPI_ISL_2539503, EPI_ISL_2539504 | Lighthouse Lab in Alderley Park | Wellcome Sanger Institute for the COVID-19 Genomics UK (COG-UK) Consortium | Jacquelyn Wynn, Mairead Hyland, The Lighthouse Lab in Alderley Park and Alex Alderton, Roberto Amato, Jeffrey Barrett, Sonia Goncalves, Ewan Harrison, David K. Jackson, Ian Johnston, Dominic Kwiatkowski, Cordelia Langford, John Sillitoe on behalf of the Wellcome Sanger Institute COVID-19 Surveillance Team |
| EPI_ISL_2539505 | Lighthouse Lab in Milton Keynes | Wellcome Sanger Institute for the COVID-19 Genomics UK (COG-UK) | The Lighthouse Lab in Milton Keynes and Alex Alderton, Roberto Amato, Jeffrey Barrett, Sonia Goncalves, Ewan Harrison, David K. Jackson, Ian Johnston, Dominic Kwiatkowski, Cordelia Langford, John Sillitoe on behalf of the Wellcome Sanger |

| | | | |
|---|---|---|---|
| | | Consortium | Institute COVID-19 Surveillance Team |
| EPI_ISL_2539506 | Lighthouse Lab in Glasgow | Wellcome Sanger Institute for the COVID-19 Genomics UK (COG-UK) Consortium | Harper VanSteenhouse, Yumi Kasai, David Gray, Carol Clugston, Anna Dominiczak and Alex Alderton, Roberto Amato, Jeffrey Barrett, Sonia Goncalves, Ewan Harrison, David K. Jackson, Ian Johnston, Dominic Kwiatkowski, Cordelia Langford, John Sillitoe on behalf of the Wellcome Sanger Institute COVID-19 Surveillance Team |
| EPI_ISL_2539507 | Lighthouse Lab in Alderley Park | Wellcome Sanger Institute for the COVID-19 Genomics UK (COG-UK) Consortium | Jacquelyn Wynn, Mairead Hyland, The Lighthouse Lab in Alderley Park and Alex Alderton, Roberto Amato, Jeffrey Barrett, Sonia Goncalves, Ewan Harrison, David K. Jackson, Ian Johnston, Dominic Kwiatkowski, Cordelia Langford, John Sillitoe on behalf of the Wellcome Sanger Institute COVID-19 Surveillance Team |
| EPI_ISL_2539508, EPI_ISL_2539509, EPI_ISL_2539510 | Lighthouse Lab in Milton Keynes | Wellcome Sanger Institute for the COVID-19 Genomics UK (COG-UK) Consortium | The Lighthouse Lab in Milton Keynes and Alex Alderton, Roberto Amato, Jeffrey Barrett, Sonia Goncalves, Ewan Harrison, David K. Jackson, Ian Johnston, Dominic Kwiatkowski, Cordelia Langford, John Sillitoe on behalf of the Wellcome Sanger Institute COVID-19 Surveillance Team |
| EPI_ISL_2539511 | Lighthouse Lab in Alderley Park | Wellcome Sanger Institute for the COVID-19 Genomics UK (COG-UK) Consortium | Jacquelyn Wynn, Mairead Hyland, The Lighthouse Lab in Alderley Park and Alex Alderton, Roberto Amato, Jeffrey Barrett, Sonia Goncalves, Ewan Harrison, David K. Jackson, Ian Johnston, Dominic Kwiatkowski, Cordelia Langford, John Sillitoe on behalf of the Wellcome Sanger Institute COVID-19 Surveillance Team |
| EPI_ISL_2539512, EPI_ISL_2539513 | Lighthouse Lab in Milton Keynes | Wellcome Sanger Institute for the COVID-19 Genomics UK (COG-UK) Consortium | The Lighthouse Lab in Milton Keynes and Alex Alderton, Roberto Amato, Jeffrey Barrett, Sonia Goncalves, Ewan Harrison, David K. Jackson, Ian Johnston, Dominic Kwiatkowski, Cordelia Langford, John Sillitoe on behalf of the Wellcome Sanger Institute COVID-19 Surveillance Team |
| EPI_ISL_2539514 | Lighthouse Lab in Glasgow | Wellcome Sanger Institute for the COVID-19 Genomics UK (COG-UK) Consortium | Harper VanSteenhouse, Yumi Kasai, David Gray, Carol Clugston, Anna Dominiczak and Alex Alderton, Roberto Amato, Jeffrey Barrett, Sonia Goncalves, Ewan Harrison, David K. Jackson, Ian Johnston, Dominic Kwiatkowski, Cordelia Langford, John Sillitoe on behalf of the Wellcome Sanger Institute COVID-19 Surveillance Team |
| EPI_ISL_2539515 | Lighthouse Lab in Alderley Park | Wellcome Sanger Institute for the COVID-19 Genomics UK (COG-UK) Consortium | Jacquelyn Wynn, Mairead Hyland, The Lighthouse Lab in Alderley Park and Alex Alderton, Roberto Amato, Jeffrey Barrett, Sonia Goncalves, Ewan Harrison, David K. Jackson, Ian Johnston, Dominic Kwiatkowski, Cordelia Langford, John Sillitoe on behalf of the Wellcome Sanger Institute COVID-19 Surveillance Team |
| EPI_ISL_2539516 | Lighthouse Lab in Milton Keynes | Wellcome Sanger Institute for the COVID-19 Genomics UK (COG-UK) Consortium | The Lighthouse Lab in Milton Keynes and Alex Alderton, Roberto Amato, Jeffrey Barrett, Sonia Goncalves, Ewan Harrison, David K. Jackson, Ian Johnston, Dominic Kwiatkowski, Cordelia Langford, John Sillitoe on behalf of the Wellcome Sanger Institute COVID-19 Surveillance Team |
| EPI_ISL_2539517 | Lighthouse Lab in Alderley Park | Wellcome Sanger Institute for the COVID-19 Genomics UK (COG-UK) Consortium | Jacquelyn Wynn, Mairead Hyland, The Lighthouse Lab in Alderley Park and Alex Alderton, Roberto Amato, Jeffrey Barrett, Sonia Goncalves, Ewan Harrison, David K. Jackson, Ian Johnston, Dominic Kwiatkowski, Cordelia Langford, John Sillitoe on behalf of the Wellcome Sanger Institute COVID-19 Surveillance Team |
| EPI_ISL_2539518 | Lighthouse Lab in Milton Keynes | Wellcome Sanger Institute for the COVID-19 Genomics UK (COG-UK) Consortium | The Lighthouse Lab in Milton Keynes and Alex Alderton, Roberto Amato, Jeffrey Barrett, Sonia Goncalves, Ewan Harrison, David K. Jackson, Ian Johnston, Dominic Kwiatkowski, Cordelia Langford, John Sillitoe on behalf of the Wellcome Sanger Institute COVID-19 Surveillance Team |
| EPI_ISL_2539519, EPI_ISL_2539520, EPI_ISL_2539521, EPI_ISL_2539522 | Lighthouse Lab in Alderley Park | Wellcome Sanger Institute for the COVID-19 Genomics UK (COG-UK) Consortium | Jacquelyn Wynn, Mairead Hyland, The Lighthouse Lab in Alderley Park and Alex Alderton, Roberto Amato, Jeffrey Barrett, Sonia Goncalves, Ewan Harrison, David K. Jackson, Ian Johnston, Dominic Kwiatkowski, Cordelia Langford, John Sillitoe on behalf of the Wellcome Sanger Institute COVID-19 Surveillance Team |
| EPI_ISL_2539523, EPI_ISL_2539524, EPI_ISL_2539525 | Lighthouse Lab in Milton Keynes | Wellcome Sanger Institute for the COVID-19 Genomics UK (COG-UK) Consortium | The Lighthouse Lab in Milton Keynes and Alex Alderton, Roberto Amato, Jeffrey Barrett, Sonia Goncalves, Ewan Harrison, David K. Jackson, Ian Johnston, Dominic Kwiatkowski, Cordelia Langford, John Sillitoe on behalf of the Wellcome Sanger Institute COVID-19 Surveillance Team |
| EPI_ISL_2539526 | Lighthouse Lab in Glasgow | Wellcome Sanger Institute for the COVID-19 Genomics UK (COG-UK) Consortium | Harper VanSteenhouse, Yumi Kasai, David Gray, Carol Clugston, Anna Dominiczak and Alex Alderton, Roberto Amato, Jeffrey Barrett, Sonia Goncalves, Ewan Harrison, David K. Jackson, Ian Johnston, Dominic Kwiatkowski, Cordelia Langford, John Sillitoe on behalf of the Wellcome Sanger Institute COVID-19 Surveillance Team |
| EPI_ISL_2539527, EPI_ISL_2539528, EPI_ISL_2539529 | Lighthouse Lab in Milton Keynes | Wellcome Sanger Institute for the COVID-19 Genomics UK (COG-UK) Consortium | The Lighthouse Lab in Milton Keynes and Alex Alderton, Roberto Amato, Jeffrey Barrett, Sonia Goncalves, Ewan Harrison, David K. Jackson, Ian Johnston, Dominic Kwiatkowski, Cordelia Langford, John Sillitoe on behalf of the Wellcome Sanger Institute COVID-19 Surveillance Team |
| EPI_ISL_2539530 | Lighthouse Lab in Alderley Park | Wellcome Sanger Institute for the COVID-19 Genomics UK (COG-UK) Consortium | Jacquelyn Wynn, Mairead Hyland, The Lighthouse Lab in Alderley Park and Alex Alderton, Roberto Amato, Jeffrey Barrett, Sonia Goncalves, Ewan Harrison, David K. Jackson, Ian Johnston, Dominic Kwiatkowski, Cordelia Langford, John Sillitoe on behalf of the Wellcome Sanger Institute COVID-19 Surveillance Team |
| EPI_ISL_2539531 | Lighthouse Lab in Milton Keynes | Wellcome Sanger Institute for the COVID-19 Genomics UK (COG-UK) Consortium | The Lighthouse Lab in Milton Keynes and Alex Alderton, Roberto Amato, Jeffrey Barrett, Sonia Goncalves, Ewan Harrison, David K. Jackson, Ian Johnston, Dominic Kwiatkowski, Cordelia Langford, John Sillitoe on behalf of the Wellcome Sanger Institute COVID-19 Surveillance Team |
| EPI_ISL_2539532, EPI_ISL_2539534, EPI_ISL_2539535 | Lighthouse Lab in Alderley Park | Wellcome Sanger Institute for the COVID-19 Genomics UK (COG-UK) Consortium | Jacquelyn Wynn, Mairead Hyland, The Lighthouse Lab in Alderley Park and Alex Alderton, Roberto Amato, Jeffrey Barrett, Sonia Goncalves, Ewan Harrison, David K. Jackson, Ian Johnston, Dominic Kwiatkowski, Cordelia Langford, John Sillitoe on behalf of the Wellcome Sanger Institute COVID-19 Surveillance Team |
| EPI_ISL_2539536, EPI_ISL_2539538 | Lighthouse Lab in Milton Keynes | Wellcome Sanger Institute for the COVID-19 Genomics UK (COG-UK) Consortium | The Lighthouse Lab in Milton Keynes and Alex Alderton, Roberto Amato, Jeffrey Barrett, Sonia Goncalves, Ewan Harrison, David K. Jackson, Ian Johnston, Dominic Kwiatkowski, Cordelia Langford, John Sillitoe on behalf of the Wellcome Sanger Institute COVID-19 Surveillance Team |
| EPI_ISL_2539539 | Lighthouse Lab in Glasgow | Wellcome Sanger Institute for the COVID-19 Genomics UK (COG-UK) Consortium | Harper VanSteenhouse, Yumi Kasai, David Gray, Carol Clugston, Anna Dominiczak and Alex Alderton, Roberto Amato, Jeffrey Barrett, Sonia Goncalves, Ewan Harrison, David K. Jackson, Ian Johnston, Dominic Kwiatkowski, Cordelia Langford, John Sillitoe on behalf of the Wellcome Sanger Institute COVID-19 Surveillance Team |
| EPI_ISL_2539540, EPI_ISL_2539541 | Lighthouse Lab in Milton Keynes | Wellcome Sanger Institute for the COVID-19 Genomics UK (COG-UK) Consortium | The Lighthouse Lab in Milton Keynes and Alex Alderton, Roberto Amato, Jeffrey Barrett, Sonia Goncalves, Ewan Harrison, David K. Jackson, Ian Johnston, Dominic Kwiatkowski, Cordelia Langford, John Sillitoe on behalf of the Wellcome Sanger Institute COVID-19 Surveillance Team |
| EPI_ISL_2539542 | Lighthouse Lab in Glasgow | Wellcome Sanger Institute for the COVID-19 Genomics UK (COG-UK) Consortium | Harper VanSteenhouse, Yumi Kasai, David Gray, Carol Clugston, Anna Dominiczak and Alex Alderton, Roberto Amato, Jeffrey Barrett, Sonia Goncalves, Ewan Harrison, David K. Jackson, Ian Johnston, Dominic Kwiatkowski, Cordelia Langford, John Sillitoe on behalf of the Wellcome Sanger Institute COVID-19 Surveillance Team |
| EPI_ISL_2539543 | Lighthouse Lab in Milton Keynes | Wellcome Sanger Institute for the COVID-19 Genomics UK (COG-UK) Consortium | The Lighthouse Lab in Milton Keynes and Alex Alderton, Roberto Amato, Jeffrey Barrett, Sonia Goncalves, Ewan Harrison, David K. Jackson, Ian Johnston, Dominic Kwiatkowski, Cordelia Langford, John Sillitoe on behalf of the Wellcome Sanger Institute COVID-19 Surveillance Team |
| EPI_ISL_2539544 | Lighthouse Lab in Alderley Park | Wellcome Sanger Institute for the COVID-19 Genomics UK (COG-UK) Consortium | Jacquelyn Wynn, Mairead Hyland, The Lighthouse Lab in Alderley Park and Alex Alderton, Roberto Amato, Jeffrey Barrett, Sonia Goncalves, Ewan Harrison, David K. Jackson, Ian Johnston, Dominic Kwiatkowski, Cordelia Langford, John Sillitoe on behalf of the Wellcome Sanger Institute COVID-19 Surveillance Team |
| EPI_ISL_2539545 | Lighthouse Lab in Milton Keynes | Wellcome Sanger Institute for the COVID-19 Genomics UK (COG-UK) Consortium | The Lighthouse Lab in Milton Keynes and Alex Alderton, Roberto Amato, Jeffrey Barrett, Sonia Goncalves, Ewan Harrison, David K. Jackson, Ian Johnston, Dominic Kwiatkowski, Cordelia Langford, John Sillitoe on behalf of the Wellcome Sanger Institute COVID-19 Surveillance Team |
| EPI_ISL_2539546 | Lighthouse Lab in Glasgow | Wellcome Sanger Institute for the COVID-19 Genomics UK (COG-UK) Consortium | Harper VanSteenhouse, Yumi Kasai, David Gray, Carol Clugston, Anna Dominiczak and Alex Alderton, Roberto Amato, Jeffrey Barrett, Sonia Goncalves, Ewan Harrison, David K. Jackson, Ian Johnston, Dominic Kwiatkowski, Cordelia Langford, John Sillitoe on behalf of the Wellcome Sanger Institute COVID-19 Surveillance Team |
| EPI_ISL_2539547 | Lighthouse Lab in Milton Keynes | Wellcome Sanger Institute for the COVID-19 Genomics UK (COG-UK) Consortium | The Lighthouse Lab in Milton Keynes and Alex Alderton, Roberto Amato, Jeffrey Barrett, Sonia Goncalves, Ewan Harrison, David K. Jackson, Ian Johnston, Dominic Kwiatkowski, Cordelia Langford, John Sillitoe on behalf of the Wellcome Sanger Institute COVID-19 Surveillance Team |
| EPI_ISL_2539548 | Lighthouse Lab in Glasgow | Wellcome Sanger Institute for the COVID-19 Genomics UK (COG-UK) Consortium | Harper VanSteenhouse, Yumi Kasai, David Gray, Carol Clugston, Anna Dominiczak and Alex Alderton, Roberto Amato, Jeffrey Barrett, Sonia Goncalves, Ewan Harrison, David K. Jackson, Ian Johnston, Dominic Kwiatkowski, Cordelia Langford, John Sillitoe on behalf of the Wellcome Sanger Institute COVID-19 Surveillance Team |
| EPI_ISL_2539549 | Lighthouse Lab in Milton Keynes | Wellcome Sanger Institute for the COVID-19 Genomics UK (COG-UK) Consortium | The Lighthouse Lab in Milton Keynes and Alex Alderton, Roberto Amato, Jeffrey Barrett, Sonia Goncalves, Ewan Harrison, David K. Jackson, Ian Johnston, Dominic Kwiatkowski, Cordelia Langford, John Sillitoe on behalf of the Wellcome Sanger Institute COVID-19 Surveillance Team |
| EPI_ISL_2539550, EPI_ISL_2539551 | Lighthouse Lab in Glasgow | Wellcome Sanger Institute for the COVID-19 Genomics UK (COG-UK) Consortium | Harper VanSteenhouse, Yumi Kasai, David Gray, Carol Clugston, Anna Dominiczak and Alex Alderton, Roberto Amato, Jeffrey Barrett, Sonia Goncalves, Ewan Harrison, David K. Jackson, Ian Johnston, Dominic Kwiatkowski, Cordelia Langford, John Sillitoe on behalf of the Wellcome Sanger Institute COVID-19 Surveillance Team |
| EPI_ISL_2539552, EPI_ISL_2539553 | Lighthouse Lab in Alderley Park | Wellcome Sanger Institute for the COVID-19 Genomics UK (COG-UK) Consortium | Jacquelyn Wynn, Mairead Hyland, The Lighthouse Lab in Alderley Park and Alex Alderton, Roberto Amato, Jeffrey Barrett, Sonia Goncalves, Ewan Harrison, David K. Jackson, Ian Johnston, Dominic Kwiatkowski, Cordelia Langford, John Sillitoe on behalf of the Wellcome Sanger Institute COVID-19 Surveillance Team |
| EPI_ISL_2539554 | Lighthouse Lab in Glasgow | Wellcome Sanger Institute for the COVID-19 Genomics UK (COG-UK) Consortium | Harper VanSteenhouse, Yumi Kasai, David Gray, Carol Clugston, Anna Dominiczak and Alex Alderton, Roberto Amato, Jeffrey Barrett, Sonia Goncalves, Ewan Harrison, David K. Jackson, Ian Johnston, Dominic Kwiatkowski, Cordelia Langford, John Sillitoe on behalf of the Wellcome Sanger Institute COVID-19 Surveillance Team |
| EPI_ISL_2539555 | Lighthouse Lab in Milton Keynes | Wellcome Sanger Institute for the COVID-19 Genomics UK (COG-UK) Consortium | The Lighthouse Lab in Milton Keynes and Alex Alderton, Roberto Amato, Jeffrey Barrett, Sonia Goncalves, Ewan Harrison, David K. Jackson, Ian Johnston, Dominic Kwiatkowski, Cordelia Langford, John Sillitoe on behalf of the Wellcome Sanger Institute COVID-19 Surveillance Team |
| EPI_ISL_2539556 | Lighthouse Lab in Glasgow | Wellcome Sanger Institute for the COVID-19 Genomics UK (COG-UK) Consortium | Harper VanSteenhouse, Yumi Kasai, David Gray, Carol Clugston, Anna Dominiczak and Alex Alderton, Roberto Amato, Jeffrey Barrett, Sonia Goncalves, Ewan Harrison, David K. Jackson, Ian Johnston, Dominic Kwiatkowski, Cordelia Langford, John Sillitoe on behalf of the Wellcome Sanger Institute COVID-19 Surveillance Team |
| EPI_ISL_2539557, EPI_ISL_2539558, EPI_ISL_2539560 | Lighthouse Lab in Alderley Park | Wellcome Sanger Institute for the COVID-19 Genomics UK (COG-UK) Consortium | Jacquelyn Wynn, Mairead Hyland, The Lighthouse Lab in Alderley Park and Alex Alderton, Roberto Amato, Jeffrey Barrett, Sonia Goncalves, Ewan Harrison, David K. Jackson, Ian Johnston, Dominic Kwiatkowski, Cordelia Langford, John Sillitoe on behalf of the Wellcome Sanger Institute COVID-19 Surveillance Team |
| EPI_ISL_2539561, EPI_ISL_2539563 | Lighthouse Lab in Milton Keynes | Wellcome Sanger Institute for the COVID-19 Genomics UK (COG-UK) Consortium | The Lighthouse Lab in Milton Keynes and Alex Alderton, Roberto Amato, Jeffrey Barrett, Sonia Goncalves, Ewan Harrison, David K. Jackson, Ian Johnston, Dominic Kwiatkowski, Cordelia Langford, John Sillitoe on behalf of the Wellcome Sanger Institute COVID-19 Surveillance Team |
| EPI_ISL_2539564 | Lighthouse Lab in Glasgow | Wellcome Sanger Institute for the COVID-19 Genomics UK (COG-UK) Consortium | Harper VanSteenhouse, Yumi Kasai, David Gray, Carol Clugston, Anna Dominiczak and Alex Alderton, Roberto Amato, Jeffrey Barrett, Sonia Goncalves, Ewan Harrison, David K. Jackson, Ian Johnston, Dominic Kwiatkowski, Cordelia Langford, John Sillitoe on behalf of the Wellcome Sanger Institute COVID-19 Surveillance Team |
| EPI_ISL_2539565, EPI_ISL_2539566, EPI_ISL_2539567 | Lighthouse Lab in Alderley Park | Wellcome Sanger Institute for the COVID-19 Genomics UK (COG-UK) Consortium | Jacquelyn Wynn, Mairead Hyland, The Lighthouse Lab in Alderley Park and Alex Alderton, Roberto Amato, Jeffrey Barrett, Sonia Goncalves, Ewan Harrison, David K. Jackson, Ian Johnston, Dominic Kwiatkowski, Cordelia Langford, John Sillitoe on behalf of the Wellcome Sanger Institute COVID-19 Surveillance Team |
| EPI_ISL_2539568, EPI_ISL_2539569, EPI_ISL_2539571 | Lighthouse Lab in Glasgow | Wellcome Sanger Institute for the COVID-19 Genomics UK (COG-UK) Consortium | Harper VanSteenhouse, Yumi Kasai, David Gray, Carol Clugston, Anna Dominiczak and Alex Alderton, Roberto Amato, Jeffrey Barrett, Sonia Goncalves, Ewan Harrison, David K. Jackson, Ian Johnston, Dominic Kwiatkowski, Cordelia Langford, John Sillitoe on behalf of the Wellcome Sanger Institute COVID-19 Surveillance Team |
| EPI_ISL_2539572 | Lighthouse Lab in Alderley Park | Wellcome Sanger Institute for the COVID-19 Genomics UK (COG-UK) Consortium | Jacquelyn Wynn, Mairead Hyland, The Lighthouse Lab in Alderley Park and Alex Alderton, Roberto Amato, Jeffrey Barrett, Sonia Goncalves, Ewan Harrison, David K. Jackson, Ian Johnston, Dominic Kwiatkowski, Cordelia Langford, John Sillitoe on behalf of the Wellcome Sanger Institute COVID-19 Surveillance Team |
| EPI_ISL_2539573, EPI_ISL_2539574 | Lighthouse Lab in Milton Keynes | Wellcome Sanger Institute for the COVID-19 Genomics UK (COG-UK) Consortium | The Lighthouse Lab in Milton Keynes and Alex Alderton, Roberto Amato, Jeffrey Barrett, Sonia Goncalves, Ewan Harrison, David K. Jackson, Ian Johnston, Dominic Kwiatkowski, Cordelia Langford, John Sillitoe on behalf of the Wellcome Sanger Institute COVID-19 Surveillance Team |
| EPI_ISL_2539575, EPI_ISL_2539576 | Lighthouse Lab in Glasgow | Wellcome Sanger Institute for the COVID-19 Genomics UK (COG-UK) Consortium | Harper VanSteenhouse, Yumi Kasai, David Gray, Carol Clugston, Anna Dominiczak and Alex Alderton, Roberto Amato, Jeffrey Barrett, Sonia Goncalves, Ewan Harrison, David K. Jackson, Ian Johnston, Dominic Kwiatkowski, Cordelia Langford, John Sillitoe on behalf of the Wellcome Sanger Institute COVID-19 Surveillance Team |

| | | | |
|---|---|---|---|
| EPI_ISL_2539577 | Lighthouse Lab in Alderley Park | Wellcome Sanger Institute for the COVID-19 Genomics UK (COG-UK) Consortium | Jacquelyn Wynn, Mairead Hyland, The Lighthouse Lab in Alderley Park and Alex Alderton, Roberto Amato, Jeffrey Barrett, Sonia Goncalves, Ewan Harrison, David K. Jackson, Ian Johnston, Dominic Kwiatkowski, Cordelia Langford, John Sillitoe on behalf of the Wellcome Sanger Institute COVID-19 Surveillance Team |
| EPI_ISL_2539578 | Lighthouse Lab in Glasgow | Wellcome Sanger Institute for the COVID-19 Genomics UK (COG-UK) Consortium | Harper VanSteenhouse, Yumi Kasai, David Gray, Carol Clugston, Anna Dominiczak and Alex Alderton, Roberto Amato, Jeffrey Barrett, Sonia Goncalves, Ewan Harrison, David K. Jackson, Ian Johnston, Dominic Kwiatkowski, Cordelia Langford, John Sillitoe on behalf of the Wellcome Sanger Institute COVID-19 Surveillance Team |
| EPI_ISL_2539579, EPI_ISL_2539580 | Lighthouse Lab in Milton Keynes | Wellcome Sanger Institute for the COVID-19 Genomics UK (COG-UK) Consortium | The Lighthouse Lab in Milton Keynes and Alex Alderton, Roberto Amato, Jeffrey Barrett, Sonia Goncalves, Ewan Harrison, David K. Jackson, Ian Johnston, Dominic Kwiatkowski, Cordelia Langford, John Sillitoe on behalf of the Wellcome Sanger Institute COVID-19 Surveillance Team |
| EPI_ISL_2539581 | Lighthouse Lab in Glasgow | Wellcome Sanger Institute for the COVID-19 Genomics UK (COG-UK) Consortium | Harper VanSteenhouse, Yumi Kasai, David Gray, Carol Clugston, Anna Dominiczak and Alex Alderton, Roberto Amato, Jeffrey Barrett, Sonia Goncalves, Ewan Harrison, David K. Jackson, Ian Johnston, Dominic Kwiatkowski, Cordelia Langford, John Sillitoe on behalf of the Wellcome Sanger Institute COVID-19 Surveillance Team |
| EPI_ISL_2539582 | Lighthouse Lab in Alderley Park | Wellcome Sanger Institute for the COVID-19 Genomics UK (COG-UK) Consortium | Jacquelyn Wynn, Mairead Hyland, The Lighthouse Lab in Alderley Park and Alex Alderton, Roberto Amato, Jeffrey Barrett, Sonia Goncalves, Ewan Harrison, David K. Jackson, Ian Johnston, Dominic Kwiatkowski, Cordelia Langford, John Sillitoe on behalf of the Wellcome Sanger Institute COVID-19 Surveillance Team |
| EPI_ISL_2539583, EPI_ISL_2539584 | Lighthouse Lab in Milton Keynes | Wellcome Sanger Institute for the COVID-19 Genomics UK (COG-UK) Consortium | The Lighthouse Lab in Milton Keynes and Alex Alderton, Roberto Amato, Jeffrey Barrett, Sonia Goncalves, Ewan Harrison, David K. Jackson, Ian Johnston, Dominic Kwiatkowski, Cordelia Langford, John Sillitoe on behalf of the Wellcome Sanger Institute COVID-19 Surveillance Team |
| EPI_ISL_2539585, EPI_ISL_2539586 | Lighthouse Lab in Glasgow | Wellcome Sanger Institute for the COVID-19 Genomics UK (COG-UK) Consortium | Harper VanSteenhouse, Yumi Kasai, David Gray, Carol Clugston, Anna Dominiczak and Alex Alderton, Roberto Amato, Jeffrey Barrett, Sonia Goncalves, Ewan Harrison, David K. Jackson, Ian Johnston, Dominic Kwiatkowski, Cordelia Langford, John Sillitoe on behalf of the Wellcome Sanger Institute COVID-19 Surveillance Team |
| EPI_ISL_2539587, EPI_ISL_2539588, EPI_ISL_2539589 | Lighthouse Lab in Milton Keynes | Wellcome Sanger Institute for the COVID-19 Genomics UK (COG-UK) Consortium | The Lighthouse Lab in Milton Keynes and Alex Alderton, Roberto Amato, Jeffrey Barrett, Sonia Goncalves, Ewan Harrison, David K. Jackson, Ian Johnston, Dominic Kwiatkowski, Cordelia Langford, John Sillitoe on behalf of the Wellcome Sanger Institute COVID-19 Surveillance Team |
| EPI_ISL_2539590 | Lighthouse Lab in Alderley Park | Wellcome Sanger Institute for the COVID-19 Genomics UK (COG-UK) Consortium | Jacquelyn Wynn, Mairead Hyland, The Lighthouse Lab in Alderley Park and Alex Alderton, Roberto Amato, Jeffrey Barrett, Sonia Goncalves, Ewan Harrison, David K. Jackson, Ian Johnston, Dominic Kwiatkowski, Cordelia Langford, John Sillitoe on behalf of the Wellcome Sanger Institute COVID-19 Surveillance Team |
| EPI_ISL_2539591, EPI_ISL_2539592, EPI_ISL_2539593 | Lighthouse Lab in Milton Keynes | Wellcome Sanger Institute for the COVID-19 Genomics UK (COG-UK) Consortium | The Lighthouse Lab in Milton Keynes and Alex Alderton, Roberto Amato, Jeffrey Barrett, Sonia Goncalves, Ewan Harrison, David K. Jackson, Ian Johnston, Dominic Kwiatkowski, Cordelia Langford, John Sillitoe on behalf of the Wellcome Sanger Institute COVID-19 Surveillance Team |
| EPI_ISL_2539594, EPI_ISL_2539595, EPI_ISL_2539596, EPI_ISL_2539597, EPI_ISL_2539598 | Lighthouse Lab in Alderley Park | Wellcome Sanger Institute for the COVID-19 Genomics UK (COG-UK) Consortium | Jacquelyn Wynn, Mairead Hyland, The Lighthouse Lab in Alderley Park and Alex Alderton, Roberto Amato, Jeffrey Barrett, Sonia Goncalves, Ewan Harrison, David K. Jackson, Ian Johnston, Dominic Kwiatkowski, Cordelia Langford, John Sillitoe on behalf of the Wellcome Sanger Institute COVID-19 Surveillance Team |
| EPI_ISL_2539599, EPI_ISL_2539600 | Lighthouse Lab in Glasgow | Wellcome Sanger Institute for the COVID-19 Genomics UK (COG-UK) Consortium | Harper VanSteenhouse, Yumi Kasai, David Gray, Carol Clugston, Anna Dominiczak and Alex Alderton, Roberto Amato, Jeffrey Barrett, Sonia Goncalves, Ewan Harrison, David K. Jackson, Ian Johnston, Dominic Kwiatkowski, Cordelia Langford, John Sillitoe on behalf of the Wellcome Sanger Institute COVID-19 Surveillance Team |
| EPI_ISL_2539601 | Lighthouse Lab in Alderley Park | Wellcome Sanger Institute for the COVID-19 Genomics UK (COG-UK) Consortium | Jacquelyn Wynn, Mairead Hyland, The Lighthouse Lab in Alderley Park and Alex Alderton, Roberto Amato, Jeffrey Barrett, Sonia Goncalves, Ewan Harrison, David K. Jackson, Ian Johnston, Dominic Kwiatkowski, Cordelia Langford, John Sillitoe on behalf of the Wellcome Sanger Institute COVID-19 Surveillance Team |
| EPI_ISL_2539602 | Lighthouse Lab in Milton Keynes | Wellcome Sanger Institute for the COVID-19 Genomics UK (COG-UK) Consortium | The Lighthouse Lab in Milton Keynes and Alex Alderton, Roberto Amato, Jeffrey Barrett, Sonia Goncalves, Ewan Harrison, David K. Jackson, Ian Johnston, Dominic Kwiatkowski, Cordelia Langford, John Sillitoe on behalf of the Wellcome Sanger Institute COVID-19 Surveillance Team |
| EPI_ISL_2539603 | Lighthouse Lab in Alderley Park | Wellcome Sanger Institute for the COVID-19 Genomics UK (COG-UK) Consortium | Jacquelyn Wynn, Mairead Hyland, The Lighthouse Lab in Alderley Park and Alex Alderton, Roberto Amato, Jeffrey Barrett, Sonia Goncalves, Ewan Harrison, David K. Jackson, Ian Johnston, Dominic Kwiatkowski, Cordelia Langford, John Sillitoe on behalf of the Wellcome Sanger Institute COVID-19 Surveillance Team |
| EPI_ISL_2539604 | Lighthouse Lab in Glasgow | Wellcome Sanger Institute for the COVID-19 Genomics UK (COG-UK) Consortium | Harper VanSteenhouse, Yumi Kasai, David Gray, Carol Clugston, Anna Dominiczak and Alex Alderton, Roberto Amato, Jeffrey Barrett, Sonia Goncalves, Ewan Harrison, David K. Jackson, Ian Johnston, Dominic Kwiatkowski, Cordelia Langford, John Sillitoe on behalf of the Wellcome Sanger Institute COVID-19 Surveillance Team |
| EPI_ISL_2539605, EPI_ISL_2539606 | Lighthouse Lab in Alderley Park | Wellcome Sanger Institute for the COVID-19 Genomics UK (COG-UK) Consortium | Jacquelyn Wynn, Mairead Hyland, The Lighthouse Lab in Alderley Park and Alex Alderton, Roberto Amato, Jeffrey Barrett, Sonia Goncalves, Ewan Harrison, David K. Jackson, Ian Johnston, Dominic Kwiatkowski, Cordelia Langford, John Sillitoe on behalf of the Wellcome Sanger Institute COVID-19 Surveillance Team |
| EPI_ISL_2539607 | Lighthouse Lab in Milton Keynes | Wellcome Sanger Institute for the COVID-19 Genomics UK (COG-UK) Consortium | The Lighthouse Lab in Milton Keynes and Alex Alderton, Roberto Amato, Jeffrey Barrett, Sonia Goncalves, Ewan Harrison, David K. Jackson, Ian Johnston, Dominic Kwiatkowski, Cordelia Langford, John Sillitoe on behalf of the Wellcome Sanger Institute COVID-19 Surveillance Team |
| EPI_ISL_2539608 | Lighthouse Lab in Glasgow | Wellcome Sanger Institute for the COVID-19 Genomics UK (COG-UK) Consortium | Harper VanSteenhouse, Yumi Kasai, David Gray, Carol Clugston, Anna Dominiczak and Alex Alderton, Roberto Amato, Jeffrey Barrett, Sonia Goncalves, Ewan Harrison, David K. Jackson, Ian Johnston, Dominic Kwiatkowski, Cordelia Langford, John Sillitoe on behalf of the Wellcome Sanger Institute COVID-19 Surveillance Team |
| EPI_ISL_2539609, EPI_ISL_2539610 | Lighthouse Lab in Alderley Park | Wellcome Sanger Institute for the COVID-19 Genomics UK (COG-UK) Consortium | Jacquelyn Wynn, Mairead Hyland, The Lighthouse Lab in Alderley Park and Alex Alderton, Roberto Amato, Jeffrey Barrett, Sonia Goncalves, Ewan Harrison, David K. Jackson, Ian Johnston, Dominic Kwiatkowski, Cordelia Langford, John Sillitoe on behalf of the Wellcome Sanger Institute COVID-19 Surveillance Team |
| EPI_ISL_2539611 | Lighthouse Lab in Milton Keynes | Wellcome Sanger Institute for the COVID-19 Genomics UK (COG-UK) Consortium | The Lighthouse Lab in Milton Keynes and Alex Alderton, Roberto Amato, Jeffrey Barrett, Sonia Goncalves, Ewan Harrison, David K. Jackson, Ian Johnston, Dominic Kwiatkowski, Cordelia Langford, John Sillitoe on behalf of the Wellcome Sanger Institute COVID-19 Surveillance Team |
| EPI_ISL_2539612, EPI_ISL_2539613 | Lighthouse Lab in Alderley Park | Wellcome Sanger Institute for the COVID-19 Genomics UK (COG-UK) Consortium | Jacquelyn Wynn, Mairead Hyland, The Lighthouse Lab in Alderley Park and Alex Alderton, Roberto Amato, Jeffrey Barrett, Sonia Goncalves, Ewan Harrison, David K. Jackson, Ian Johnston, Dominic Kwiatkowski, Cordelia Langford, John Sillitoe on behalf of the Wellcome Sanger Institute COVID-19 Surveillance Team |
| EPI_ISL_2539614 | Lighthouse Lab in Glasgow | Wellcome Sanger Institute for the COVID-19 Genomics UK (COG-UK) Consortium | Harper VanSteenhouse, Yumi Kasai, David Gray, Carol Clugston, Anna Dominiczak and Alex Alderton, Roberto Amato, Jeffrey Barrett, Sonia Goncalves, Ewan Harrison, David K. Jackson, Ian Johnston, Dominic Kwiatkowski, Cordelia Langford, John Sillitoe on behalf of the Wellcome Sanger Institute COVID-19 Surveillance Team |
| EPI_ISL_2539615, EPI_ISL_2539616 | Lighthouse Lab in Alderley Park | Wellcome Sanger Institute for the COVID-19 Genomics UK (COG-UK) Consortium | Jacquelyn Wynn, Mairead Hyland, The Lighthouse Lab in Alderley Park and Alex Alderton, Roberto Amato, Jeffrey Barrett, Sonia Goncalves, Ewan Harrison, David K. Jackson, Ian Johnston, Dominic Kwiatkowski, Cordelia Langford, John Sillitoe on behalf of the Wellcome Sanger Institute COVID-19 Surveillance Team |
| EPI_ISL_2539617 | Lighthouse Lab in Glasgow | Wellcome Sanger Institute for the COVID-19 Genomics UK (COG-UK) Consortium | Harper VanSteenhouse, Yumi Kasai, David Gray, Carol Clugston, Anna Dominiczak and Alex Alderton, Roberto Amato, Jeffrey Barrett, Sonia Goncalves, Ewan Harrison, David K. Jackson, Ian Johnston, Dominic Kwiatkowski, Cordelia Langford, John Sillitoe on behalf of the Wellcome Sanger Institute COVID-19 Surveillance Team |
| EPI_ISL_2539618, EPI_ISL_2539619, EPI_ISL_2539620 | Lighthouse Lab in Milton Keynes | Wellcome Sanger Institute for the COVID-19 Genomics UK (COG-UK) Consortium | The Lighthouse Lab in Milton Keynes and Alex Alderton, Roberto Amato, Jeffrey Barrett, Sonia Goncalves, Ewan Harrison, David K. Jackson, Ian Johnston, Dominic Kwiatkowski, Cordelia Langford, John Sillitoe on behalf of the Wellcome Sanger Institute COVID-19 Surveillance Team |
| EPI_ISL_2539621 | Lighthouse Lab in Glasgow | Wellcome Sanger Institute for the COVID-19 Genomics UK (COG-UK) Consortium | Harper VanSteenhouse, Yumi Kasai, David Gray, Carol Clugston, Anna Dominiczak and Alex Alderton, Roberto Amato, Jeffrey Barrett, Sonia Goncalves, Ewan Harrison, David K. Jackson, Ian Johnston, Dominic Kwiatkowski, Cordelia Langford, John Sillitoe on behalf of the Wellcome Sanger Institute COVID-19 Surveillance Team |
| EPI_ISL_2539622, EPI_ISL_2539623 | Lighthouse Lab in Alderley Park | Wellcome Sanger Institute for the COVID-19 Genomics UK (COG-UK) Consortium | Jacquelyn Wynn, Mairead Hyland, The Lighthouse Lab in Alderley Park and Alex Alderton, Roberto Amato, Jeffrey Barrett, Sonia Goncalves, Ewan Harrison, David K. Jackson, Ian Johnston, Dominic Kwiatkowski, Cordelia Langford, John Sillitoe on behalf of the Wellcome Sanger Institute COVID-19 Surveillance Team |
| EPI_ISL_2539624 | Lighthouse Lab in Milton Keynes | Wellcome Sanger Institute for the COVID-19 Genomics UK (COG-UK) Consortium | The Lighthouse Lab in Milton Keynes and Alex Alderton, Roberto Amato, Jeffrey Barrett, Sonia Goncalves, Ewan Harrison, David K. Jackson, Ian Johnston, Dominic Kwiatkowski, Cordelia Langford, John Sillitoe on behalf of the Wellcome Sanger Institute COVID-19 Surveillance Team |
| EPI_ISL_2539625, EPI_ISL_2539626 | Lighthouse Lab in Alderley Park | Wellcome Sanger Institute for the COVID-19 Genomics UK (COG-UK) Consortium | Jacquelyn Wynn, Mairead Hyland, The Lighthouse Lab in Alderley Park and Alex Alderton, Roberto Amato, Jeffrey Barrett, Sonia Goncalves, Ewan Harrison, David K. Jackson, Ian Johnston, Dominic Kwiatkowski, Cordelia Langford, John Sillitoe on behalf of the Wellcome Sanger Institute COVID-19 Surveillance Team |
| EPI_ISL_2539627 | Lighthouse Lab in Glasgow | Wellcome Sanger Institute for the COVID-19 Genomics UK (COG-UK) Consortium | Harper VanSteenhouse, Yumi Kasai, David Gray, Carol Clugston, Anna Dominiczak and Alex Alderton, Roberto Amato, Jeffrey Barrett, Sonia Goncalves, Ewan Harrison, David K. Jackson, Ian Johnston, Dominic Kwiatkowski, Cordelia Langford, John Sillitoe on behalf of the Wellcome Sanger Institute COVID-19 Surveillance Team |
| EPI_ISL_2539628 | Lighthouse Lab in Alderley Park | Wellcome Sanger Institute for the COVID-19 Genomics UK (COG-UK) Consortium | Jacquelyn Wynn, Mairead Hyland, The Lighthouse Lab in Alderley Park and Alex Alderton, Roberto Amato, Jeffrey Barrett, Sonia Goncalves, Ewan Harrison, David K. Jackson, Ian Johnston, Dominic Kwiatkowski, Cordelia Langford, John Sillitoe on behalf of the Wellcome Sanger Institute COVID-19 Surveillance Team |
| EPI_ISL_2539629 | Lighthouse Lab in Glasgow | Wellcome Sanger Institute for the COVID-19 Genomics UK (COG-UK) Consortium | Harper VanSteenhouse, Yumi Kasai, David Gray, Carol Clugston, Anna Dominiczak and Alex Alderton, Roberto Amato, Jeffrey Barrett, Sonia Goncalves, Ewan Harrison, David K. Jackson, Ian Johnston, Dominic Kwiatkowski, Cordelia Langford, John Sillitoe on behalf of the Wellcome Sanger Institute COVID-19 Surveillance Team |
| EPI_ISL_2539630 | Lighthouse Lab in Milton Keynes | Wellcome Sanger Institute for the COVID-19 Genomics UK (COG-UK) Consortium | The Lighthouse Lab in Milton Keynes and Alex Alderton, Roberto Amato, Jeffrey Barrett, Sonia Goncalves, Ewan Harrison, David K. Jackson, Ian Johnston, Dominic Kwiatkowski, Cordelia Langford, John Sillitoe on behalf of the Wellcome Sanger Institute COVID-19 Surveillance Team |
| EPI_ISL_2539631 | Lighthouse Lab in Glasgow | Wellcome Sanger Institute for the COVID-19 Genomics UK (COG-UK) Consortium | Harper VanSteenhouse, Yumi Kasai, David Gray, Carol Clugston, Anna Dominiczak and Alex Alderton, Roberto Amato, Jeffrey Barrett, Sonia Goncalves, Ewan Harrison, David K. Jackson, Ian Johnston, Dominic Kwiatkowski, Cordelia Langford, John Sillitoe on behalf of the Wellcome Sanger Institute COVID-19 Surveillance Team |
| EPI_ISL_2539632 | Lighthouse Lab in Alderley Park | Wellcome Sanger Institute for the COVID-19 Genomics UK (COG-UK) Consortium | Jacquelyn Wynn, Mairead Hyland, The Lighthouse Lab in Alderley Park and Alex Alderton, Roberto Amato, Jeffrey Barrett, Sonia Goncalves, Ewan Harrison, David K. Jackson, Ian Johnston, Dominic Kwiatkowski, Cordelia Langford, John Sillitoe on behalf of the Wellcome Sanger Institute COVID-19 Surveillance Team |
| EPI_ISL_2539633, EPI_ISL_2539634 | Lighthouse Lab in Glasgow | Wellcome Sanger Institute for the COVID-19 Genomics UK (COG-UK) Consortium | Harper VanSteenhouse, Yumi Kasai, David Gray, Carol Clugston, Anna Dominiczak and Alex Alderton, Roberto Amato, Jeffrey Barrett, Sonia Goncalves, Ewan Harrison, David K. Jackson, Ian Johnston, Dominic Kwiatkowski, Cordelia Langford, John Sillitoe on behalf of the Wellcome Sanger Institute COVID-19 Surveillance Team |
| EPI_ISL_2539635 | Lighthouse Lab in Alderley Park | Wellcome Sanger Institute for the COVID-19 Genomics UK (COG-UK) Consortium | Jacquelyn Wynn, Mairead Hyland, The Lighthouse Lab in Alderley Park and Alex Alderton, Roberto Amato, Jeffrey Barrett, Sonia Goncalves, Ewan Harrison, David K. Jackson, Ian Johnston, Dominic Kwiatkowski, Cordelia Langford, John Sillitoe on behalf of the Wellcome Sanger Institute COVID-19 Surveillance Team |
| EPI_ISL_2539636 | Lighthouse Lab in Milton Keynes | Wellcome Sanger Institute for the COVID-19 Genomics UK (COG-UK) Consortium | The Lighthouse Lab in Milton Keynes and Alex Alderton, Roberto Amato, Jeffrey Barrett, Sonia Goncalves, Ewan Harrison, David K. Jackson, Ian Johnston, Dominic Kwiatkowski, Cordelia Langford, John Sillitoe on behalf of the Wellcome Sanger Institute COVID-19 Surveillance Team |
| EPI_ISL_2539637 | Lighthouse Lab in Alderley Park | Wellcome Sanger Institute for the COVID-19 Genomics UK (COG-UK) Consortium | Jacquelyn Wynn, Mairead Hyland, The Lighthouse Lab in Alderley Park and Alex Alderton, Roberto Amato, Jeffrey Barrett, Sonia Goncalves, Ewan Harrison, David K. Jackson, Ian Johnston, Dominic Kwiatkowski, Cordelia Langford, John Sillitoe on behalf of the Wellcome Sanger Institute COVID-19 Surveillance Team |
| EPI_ISL_2539638 | Lighthouse Lab in Milton Keynes | Wellcome Sanger Institute for the COVID-19 Genomics UK (COG-UK) Consortium | The Lighthouse Lab in Milton Keynes and Alex Alderton, Roberto Amato, Jeffrey Barrett, Sonia Goncalves, Ewan Harrison, David K. Jackson, Ian Johnston, Dominic Kwiatkowski, Cordelia Langford, John Sillitoe on behalf of the Wellcome Sanger Institute COVID-19 Surveillance Team |
| EPI_ISL_2539639 | Lighthouse Lab in Glasgow | Wellcome Sanger Institute for the COVID-19 Genomics UK (COG-UK) Consortium | Harper VanSteenhouse, Yumi Kasai, David Gray, Carol Clugston, Anna Dominiczak and Alex Alderton, Roberto Amato, Jeffrey Barrett, Sonia Goncalves, Ewan Harrison, David K. Jackson, Ian Johnston, Dominic Kwiatkowski, Cordelia Langford, John Sillitoe on behalf of the Wellcome Sanger Institute COVID-19 Surveillance Team |
| EPI_ISL_2539640 | Lighthouse Lab in Milton Keynes | Wellcome Sanger Institute for the COVID-19 Genomics UK (COG-UK) | The Lighthouse Lab in Milton Keynes and Alex Alderton, Roberto Amato, Jeffrey Barrett, Sonia Goncalves, Ewan Harrison, David K. Jackson, Ian Johnston, Dominic Kwiatkowski, Cordelia Langford, John Sillitoe on behalf of the Wellcome Sanger |

| | | | |
|---|---|---|---|
| | | Consortium | Institute COVID-19 Surveillance Team |
| EPI_ISL_2539641 | Lighthouse Lab in Glasgow | Wellcome Sanger Institute for the COVID-19 Genomics UK (COG-UK) Consortium | Harper VanSteenhouse, Yumi Kasai, David Gray, Carol Clugston, Anna Dominiczak and Alex Alderton, Roberto Amato, Jeffrey Barrett, Sonia Goncalves, Ewan Harrison, David K. Jackson, Ian Johnston, Dominic Kwiatkowski, Cordelia Langford, John Sillitoe on behalf of the Wellcome Sanger Institute COVID-19 Surveillance Team |
| EPI_ISL_2539642 | Lighthouse Lab in Milton Keynes | Wellcome Sanger Institute for the COVID-19 Genomics UK (COG-UK) Consortium | The Lighthouse Lab in Milton Keynes and Alex Alderton, Roberto Amato, Jeffrey Barrett, Sonia Goncalves, Ewan Harrison, David K. Jackson, Ian Johnston, Dominic Kwiatkowski, Cordelia Langford, John Sillitoe on behalf of the Wellcome Sanger Institute COVID-19 Surveillance Team |
| EPI_ISL_2539643, EPI_ISL_2539644 | Lighthouse Lab in Glasgow | Wellcome Sanger Institute for the COVID-19 Genomics UK (COG-UK) Consortium | Harper VanSteenhouse, Yumi Kasai, David Gray, Carol Clugston, Anna Dominiczak and Alex Alderton, Roberto Amato, Jeffrey Barrett, Sonia Goncalves, Ewan Harrison, David K. Jackson, Ian Johnston, Dominic Kwiatkowski, Cordelia Langford, John Sillitoe on behalf of the Wellcome Sanger Institute COVID-19 Surveillance Team |
| EPI_ISL_2539646, EPI_ISL_2539647, EPI_ISL_2539648, EPI_ISL_2539649, EPI_ISL_2539650 | Lighthouse Lab in Milton Keynes | Wellcome Sanger Institute for the COVID-19 Genomics UK (COG-UK) Consortium | The Lighthouse Lab in Milton Keynes and Alex Alderton, Roberto Amato, Jeffrey Barrett, Sonia Goncalves, Ewan Harrison, David K. Jackson, Ian Johnston, Dominic Kwiatkowski, Cordelia Langford, John Sillitoe on behalf of the Wellcome Sanger Institute COVID-19 Surveillance Team |
| EPI_ISL_2539651 | Lighthouse Lab in Glasgow | Wellcome Sanger Institute for the COVID-19 Genomics UK (COG-UK) Consortium | Harper VanSteenhouse, Yumi Kasai, David Gray, Carol Clugston, Anna Dominiczak and Alex Alderton, Roberto Amato, Jeffrey Barrett, Sonia Goncalves, Ewan Harrison, David K. Jackson, Ian Johnston, Dominic Kwiatkowski, Cordelia Langford, John Sillitoe on behalf of the Wellcome Sanger Institute COVID-19 Surveillance Team |
| EPI_ISL_2539652 | Lighthouse Lab in Milton Keynes | Wellcome Sanger Institute for the COVID-19 Genomics UK (COG-UK) Consortium | The Lighthouse Lab in Milton Keynes and Alex Alderton, Roberto Amato, Jeffrey Barrett, Sonia Goncalves, Ewan Harrison, David K. Jackson, Ian Johnston, Dominic Kwiatkowski, Cordelia Langford, John Sillitoe on behalf of the Wellcome Sanger Institute COVID-19 Surveillance Team |
| EPI_ISL_2539653, EPI_ISL_2539654, EPI_ISL_2539655 | Lighthouse Lab in Alderley Park | Wellcome Sanger Institute for the COVID-19 Genomics UK (COG-UK) Consortium | Jacquelyn Wynn, Mairead Hyland, The Lighthouse Lab in Alderley Park and Alex Alderton, Roberto Amato, Jeffrey Barrett, Sonia Goncalves, Ewan Harrison, David K. Jackson, Ian Johnston, Dominic Kwiatkowski, Cordelia Langford, John Sillitoe on behalf of the Wellcome Sanger Institute COVID-19 Surveillance Team |
| EPI_ISL_2539656 | Lighthouse Lab in Glasgow | Wellcome Sanger Institute for the COVID-19 Genomics UK (COG-UK) Consortium | Harper VanSteenhouse, Yumi Kasai, David Gray, Carol Clugston, Anna Dominiczak and Alex Alderton, Roberto Amato, Jeffrey Barrett, Sonia Goncalves, Ewan Harrison, David K. Jackson, Ian Johnston, Dominic Kwiatkowski, Cordelia Langford, John Sillitoe on behalf of the Wellcome Sanger Institute COVID-19 Surveillance Team |
| EPI_ISL_2539657 | Lighthouse Lab in Alderley Park | Wellcome Sanger Institute for the COVID-19 Genomics UK (COG-UK) Consortium | Jacquelyn Wynn, Mairead Hyland, The Lighthouse Lab in Alderley Park and Alex Alderton, Roberto Amato, Jeffrey Barrett, Sonia Goncalves, Ewan Harrison, David K. Jackson, Ian Johnston, Dominic Kwiatkowski, Cordelia Langford, John Sillitoe on behalf of the Wellcome Sanger Institute COVID-19 Surveillance Team |
| EPI_ISL_2539658, EPI_ISL_2539659 | Lighthouse Lab in Glasgow | Wellcome Sanger Institute for the COVID-19 Genomics UK (COG-UK) Consortium | Harper VanSteenhouse, Yumi Kasai, David Gray, Carol Clugston, Anna Dominiczak and Alex Alderton, Roberto Amato, Jeffrey Barrett, Sonia Goncalves, Ewan Harrison, David K. Jackson, Ian Johnston, Dominic Kwiatkowski, Cordelia Langford, John Sillitoe on behalf of the Wellcome Sanger Institute COVID-19 Surveillance Team |
| EPI_ISL_2539660 | Lighthouse Lab in Alderley Park | Wellcome Sanger Institute for the COVID-19 Genomics UK (COG-UK) Consortium | Jacquelyn Wynn, Mairead Hyland, The Lighthouse Lab in Alderley Park and Alex Alderton, Roberto Amato, Jeffrey Barrett, Sonia Goncalves, Ewan Harrison, David K. Jackson, Ian Johnston, Dominic Kwiatkowski, Cordelia Langford, John Sillitoe on behalf of the Wellcome Sanger Institute COVID-19 Surveillance Team |
| EPI_ISL_2539661 | Lighthouse Lab in Milton Keynes | Wellcome Sanger Institute for the COVID-19 Genomics UK (COG-UK) Consortium | The Lighthouse Lab in Milton Keynes and Alex Alderton, Roberto Amato, Jeffrey Barrett, Sonia Goncalves, Ewan Harrison, David K. Jackson, Ian Johnston, Dominic Kwiatkowski, Cordelia Langford, John Sillitoe on behalf of the Wellcome Sanger Institute COVID-19 Surveillance Team |
| EPI_ISL_2539662 | Lighthouse Lab in Alderley Park | Wellcome Sanger Institute for the COVID-19 Genomics UK (COG-UK) Consortium | Jacquelyn Wynn, Mairead Hyland, The Lighthouse Lab in Alderley Park and Alex Alderton, Roberto Amato, Jeffrey Barrett, Sonia Goncalves, Ewan Harrison, David K. Jackson, Ian Johnston, Dominic Kwiatkowski, Cordelia Langford, John Sillitoe on behalf of the Wellcome Sanger Institute COVID-19 Surveillance Team |
| EPI_ISL_2539663, EPI_ISL_2539664, EPI_ISL_2539665 | Lighthouse Lab in Milton Keynes | Wellcome Sanger Institute for the COVID-19 Genomics UK (COG-UK) Consortium | The Lighthouse Lab in Milton Keynes and Alex Alderton, Roberto Amato, Jeffrey Barrett, Sonia Goncalves, Ewan Harrison, David K. Jackson, Ian Johnston, Dominic Kwiatkowski, Cordelia Langford, John Sillitoe on behalf of the Wellcome Sanger Institute COVID-19 Surveillance Team |
| EPI_ISL_2539666 | Lighthouse Lab in Alderley Park | Wellcome Sanger Institute for the COVID-19 Genomics UK (COG-UK) Consortium | Jacquelyn Wynn, Mairead Hyland, The Lighthouse Lab in Alderley Park and Alex Alderton, Roberto Amato, Jeffrey Barrett, Sonia Goncalves, Ewan Harrison, David K. Jackson, Ian Johnston, Dominic Kwiatkowski, Cordelia Langford, John Sillitoe on behalf of the Wellcome Sanger Institute COVID-19 Surveillance Team |
| EPI_ISL_2539668 | Lighthouse Lab in Glasgow | Wellcome Sanger Institute for the COVID-19 Genomics UK (COG-UK) Consortium | Harper VanSteenhouse, Yumi Kasai, David Gray, Carol Clugston, Anna Dominiczak and Alex Alderton, Roberto Amato, Jeffrey Barrett, Sonia Goncalves, Ewan Harrison, David K. Jackson, Ian Johnston, Dominic Kwiatkowski, Cordelia Langford, John Sillitoe on behalf of the Wellcome Sanger Institute COVID-19 Surveillance Team |
| EPI_ISL_2539669, EPI_ISL_2539670, EPI_ISL_2539671 | Lighthouse Lab in Alderley Park | Wellcome Sanger Institute for the COVID-19 Genomics UK (COG-UK) Consortium | Jacquelyn Wynn, Mairead Hyland, The Lighthouse Lab in Alderley Park and Alex Alderton, Roberto Amato, Jeffrey Barrett, Sonia Goncalves, Ewan Harrison, David K. Jackson, Ian Johnston, Dominic Kwiatkowski, Cordelia Langford, John Sillitoe on behalf of the Wellcome Sanger Institute COVID-19 Surveillance Team |
| EPI_ISL_2539672 | Lighthouse Lab in Milton Keynes | Wellcome Sanger Institute for the COVID-19 Genomics UK (COG-UK) Consortium | The Lighthouse Lab in Milton Keynes and Alex Alderton, Roberto Amato, Jeffrey Barrett, Sonia Goncalves, Ewan Harrison, David K. Jackson, Ian Johnston, Dominic Kwiatkowski, Cordelia Langford, John Sillitoe on behalf of the Wellcome Sanger Institute COVID-19 Surveillance Team |
| EPI_ISL_2539673, EPI_ISL_2539674 | Lighthouse Lab in Glasgow | Wellcome Sanger Institute for the COVID-19 Genomics UK (COG-UK) Consortium | Harper VanSteenhouse, Yumi Kasai, David Gray, Carol Clugston, Anna Dominiczak and Alex Alderton, Roberto Amato, Jeffrey Barrett, Sonia Goncalves, Ewan Harrison, David K. Jackson, Ian Johnston, Dominic Kwiatkowski, Cordelia Langford, John Sillitoe on behalf of the Wellcome Sanger Institute COVID-19 Surveillance Team |
| EPI_ISL_2539675 | Lighthouse Lab in Alderley Park | Wellcome Sanger Institute for the COVID-19 Genomics UK (COG-UK) Consortium | Jacquelyn Wynn, Mairead Hyland, The Lighthouse Lab in Alderley Park and Alex Alderton, Roberto Amato, Jeffrey Barrett, Sonia Goncalves, Ewan Harrison, David K. Jackson, Ian Johnston, Dominic Kwiatkowski, Cordelia Langford, John Sillitoe on behalf of the Wellcome Sanger Institute COVID-19 Surveillance Team |
| EPI_ISL_2539676 | Lighthouse Lab in Glasgow | Wellcome Sanger Institute for the COVID-19 Genomics UK (COG-UK) Consortium | Harper VanSteenhouse, Yumi Kasai, David Gray, Carol Clugston, Anna Dominiczak and Alex Alderton, Roberto Amato, Jeffrey Barrett, Sonia Goncalves, Ewan Harrison, David K. Jackson, Ian Johnston, Dominic Kwiatkowski, Cordelia Langford, John Sillitoe on behalf of the Wellcome Sanger Institute COVID-19 Surveillance Team |
| EPI_ISL_2539677, EPI_ISL_2539678 | Lighthouse Lab in Milton Keynes | Wellcome Sanger Institute for the COVID-19 Genomics UK (COG-UK) Consortium | The Lighthouse Lab in Milton Keynes and Alex Alderton, Roberto Amato, Jeffrey Barrett, Sonia Goncalves, Ewan Harrison, David K. Jackson, Ian Johnston, Dominic Kwiatkowski, Cordelia Langford, John Sillitoe on behalf of the Wellcome Sanger Institute COVID-19 Surveillance Team |
| EPI_ISL_2539680, EPI_ISL_2539681 | Lighthouse Lab in Glasgow | Wellcome Sanger Institute for the COVID-19 Genomics UK (COG-UK) Consortium | Harper VanSteenhouse, Yumi Kasai, David Gray, Carol Clugston, Anna Dominiczak and Alex Alderton, Roberto Amato, Jeffrey Barrett, Sonia Goncalves, Ewan Harrison, David K. Jackson, Ian Johnston, Dominic Kwiatkowski, Cordelia Langford, John Sillitoe on behalf of the Wellcome Sanger Institute COVID-19 Surveillance Team |
| EPI_ISL_2539682, EPI_ISL_2539683, EPI_ISL_2539684 | Lighthouse Lab in Alderley Park | Wellcome Sanger Institute for the COVID-19 Genomics UK (COG-UK) Consortium | Jacquelyn Wynn, Mairead Hyland, The Lighthouse Lab in Alderley Park and Alex Alderton, Roberto Amato, Jeffrey Barrett, Sonia Goncalves, Ewan Harrison, David K. Jackson, Ian Johnston, Dominic Kwiatkowski, Cordelia Langford, John Sillitoe on behalf of the Wellcome Sanger Institute COVID-19 Surveillance Team |
| EPI_ISL_2539685 | Lighthouse Lab in Glasgow | Wellcome Sanger Institute for the COVID-19 Genomics UK (COG-UK) Consortium | Harper VanSteenhouse, Yumi Kasai, David Gray, Carol Clugston, Anna Dominiczak and Alex Alderton, Roberto Amato, Jeffrey Barrett, Sonia Goncalves, Ewan Harrison, David K. Jackson, Ian Johnston, Dominic Kwiatkowski, Cordelia Langford, John Sillitoe on behalf of the Wellcome Sanger Institute COVID-19 Surveillance Team |
| EPI_ISL_2539687, EPI_ISL_2539688, EPI_ISL_2539689 | Lighthouse Lab in Alderley Park | Wellcome Sanger Institute for the COVID-19 Genomics UK (COG-UK) Consortium | Jacquelyn Wynn, Mairead Hyland, The Lighthouse Lab in Alderley Park and Alex Alderton, Roberto Amato, Jeffrey Barrett, Sonia Goncalves, Ewan Harrison, David K. Jackson, Ian Johnston, Dominic Kwiatkowski, Cordelia Langford, John Sillitoe on behalf of the Wellcome Sanger Institute COVID-19 Surveillance Team |
| EPI_ISL_2539690 | Lighthouse Lab in Milton Keynes | Wellcome Sanger Institute for the COVID-19 Genomics UK (COG-UK) Consortium | The Lighthouse Lab in Milton Keynes and Alex Alderton, Roberto Amato, Jeffrey Barrett, Sonia Goncalves, Ewan Harrison, David K. Jackson, Ian Johnston, Dominic Kwiatkowski, Cordelia Langford, John Sillitoe on behalf of the Wellcome Sanger Institute COVID-19 Surveillance Team |
| EPI_ISL_2539691 | Lighthouse Lab in Glasgow | Wellcome Sanger Institute for the COVID-19 Genomics UK (COG-UK) Consortium | Harper VanSteenhouse, Yumi Kasai, David Gray, Carol Clugston, Anna Dominiczak and Alex Alderton, Roberto Amato, Jeffrey Barrett, Sonia Goncalves, Ewan Harrison, David K. Jackson, Ian Johnston, Dominic Kwiatkowski, Cordelia Langford, John Sillitoe on behalf of the Wellcome Sanger Institute COVID-19 Surveillance Team |
| EPI_ISL_2539692, EPI_ISL_2539693 | Lighthouse Lab in Milton Keynes | Wellcome Sanger Institute for the COVID-19 Genomics UK (COG-UK) Consortium | The Lighthouse Lab in Milton Keynes and Alex Alderton, Roberto Amato, Jeffrey Barrett, Sonia Goncalves, Ewan Harrison, David K. Jackson, Ian Johnston, Dominic Kwiatkowski, Cordelia Langford, John Sillitoe on behalf of the Wellcome Sanger Institute COVID-19 Surveillance Team |
| EPI_ISL_2539694 | Lighthouse Lab in Alderley Park | Wellcome Sanger Institute for the COVID-19 Genomics UK (COG-UK) Consortium | Jacquelyn Wynn, Mairead Hyland, The Lighthouse Lab in Alderley Park and Alex Alderton, Roberto Amato, Jeffrey Barrett, Sonia Goncalves, Ewan Harrison, David K. Jackson, Ian Johnston, Dominic Kwiatkowski, Cordelia Langford, John Sillitoe on behalf of the Wellcome Sanger Institute COVID-19 Surveillance Team |
| EPI_ISL_2539695 | Lighthouse Lab in Milton Keynes | Wellcome Sanger Institute for the COVID-19 Genomics UK (COG-UK) Consortium | The Lighthouse Lab in Milton Keynes and Alex Alderton, Roberto Amato, Jeffrey Barrett, Sonia Goncalves, Ewan Harrison, David K. Jackson, Ian Johnston, Dominic Kwiatkowski, Cordelia Langford, John Sillitoe on behalf of the Wellcome Sanger Institute COVID-19 Surveillance Team |
| EPI_ISL_2539696 | Lighthouse Lab in Alderley Park | Wellcome Sanger Institute for the COVID-19 Genomics UK (COG-UK) Consortium | Jacquelyn Wynn, Mairead Hyland, The Lighthouse Lab in Alderley Park and Alex Alderton, Roberto Amato, Jeffrey Barrett, Sonia Goncalves, Ewan Harrison, David K. Jackson, Ian Johnston, Dominic Kwiatkowski, Cordelia Langford, John Sillitoe on behalf of the Wellcome Sanger Institute COVID-19 Surveillance Team |
| EPI_ISL_2539697 | Lighthouse Lab in Milton Keynes | Wellcome Sanger Institute for the COVID-19 Genomics UK (COG-UK) Consortium | The Lighthouse Lab in Milton Keynes and Alex Alderton, Roberto Amato, Jeffrey Barrett, Sonia Goncalves, Ewan Harrison, David K. Jackson, Ian Johnston, Dominic Kwiatkowski, Cordelia Langford, John Sillitoe on behalf of the Wellcome Sanger Institute COVID-19 Surveillance Team |
| EPI_ISL_2539698, EPI_ISL_2539699 | Lighthouse Lab in Alderley Park | Wellcome Sanger Institute for the COVID-19 Genomics UK (COG-UK) Consortium | Jacquelyn Wynn, Mairead Hyland, The Lighthouse Lab in Alderley Park and Alex Alderton, Roberto Amato, Jeffrey Barrett, Sonia Goncalves, Ewan Harrison, David K. Jackson, Ian Johnston, Dominic Kwiatkowski, Cordelia Langford, John Sillitoe on behalf of the Wellcome Sanger Institute COVID-19 Surveillance Team |
| EPI_ISL_2539700 | Lighthouse Lab in Glasgow | Wellcome Sanger Institute for the COVID-19 Genomics UK (COG-UK) Consortium | Harper VanSteenhouse, Yumi Kasai, David Gray, Carol Clugston, Anna Dominiczak and Alex Alderton, Roberto Amato, Jeffrey Barrett, Sonia Goncalves, Ewan Harrison, David K. Jackson, Ian Johnston, Dominic Kwiatkowski, Cordelia Langford, John Sillitoe on behalf of the Wellcome Sanger Institute COVID-19 Surveillance Team |
| EPI_ISL_2539702 | Lighthouse Lab in Milton Keynes | Wellcome Sanger Institute for the COVID-19 Genomics UK (COG-UK) Consortium | The Lighthouse Lab in Milton Keynes and Alex Alderton, Roberto Amato, Jeffrey Barrett, Sonia Goncalves, Ewan Harrison, David K. Jackson, Ian Johnston, Dominic Kwiatkowski, Cordelia Langford, John Sillitoe on behalf of the Wellcome Sanger Institute COVID-19 Surveillance Team |
| EPI_ISL_2539703, EPI_ISL_2539704 | Lighthouse Lab in Glasgow | Wellcome Sanger Institute for the COVID-19 Genomics UK (COG-UK) Consortium | Harper VanSteenhouse, Yumi Kasai, David Gray, Carol Clugston, Anna Dominiczak and Alex Alderton, Roberto Amato, Jeffrey Barrett, Sonia Goncalves, Ewan Harrison, David K. Jackson, Ian Johnston, Dominic Kwiatkowski, Cordelia Langford, John Sillitoe on behalf of the Wellcome Sanger Institute COVID-19 Surveillance Team |
| EPI_ISL_2539705 | Lighthouse Lab in Milton Keynes | Wellcome Sanger Institute for the COVID-19 Genomics UK (COG-UK) Consortium | The Lighthouse Lab in Milton Keynes and Alex Alderton, Roberto Amato, Jeffrey Barrett, Sonia Goncalves, Ewan Harrison, David K. Jackson, Ian Johnston, Dominic Kwiatkowski, Cordelia Langford, John Sillitoe on behalf of the Wellcome Sanger Institute COVID-19 Surveillance Team |
| EPI_ISL_2539706, EPI_ISL_2539707, EPI_ISL_2539708 | Lighthouse Lab in Alderley Park | Wellcome Sanger Institute for the COVID-19 Genomics UK (COG-UK) Consortium | Jacquelyn Wynn, Mairead Hyland, The Lighthouse Lab in Alderley Park and Alex Alderton, Roberto Amato, Jeffrey Barrett, Sonia Goncalves, Ewan Harrison, David K. Jackson, Ian Johnston, Dominic Kwiatkowski, Cordelia Langford, John Sillitoe on behalf of the Wellcome Sanger Institute COVID-19 Surveillance Team |
| EPI_ISL_2539709 | Lighthouse Lab in Glasgow | Wellcome Sanger Institute for the COVID-19 Genomics UK (COG-UK) Consortium | Harper VanSteenhouse, Yumi Kasai, David Gray, Carol Clugston, Anna Dominiczak and Alex Alderton, Roberto Amato, Jeffrey Barrett, Sonia Goncalves, Ewan Harrison, David K. Jackson, Ian Johnston, Dominic Kwiatkowski, Cordelia Langford, John Sillitoe on behalf of the Wellcome Sanger Institute COVID-19 Surveillance Team |
| EPI_ISL_2539710 | Lighthouse Lab in Alderley Park | Wellcome Sanger Institute for the COVID-19 Genomics UK (COG-UK) Consortium | Jacquelyn Wynn, Mairead Hyland, The Lighthouse Lab in Alderley Park and Alex Alderton, Roberto Amato, Jeffrey Barrett, Sonia Goncalves, Ewan Harrison, David K. Jackson, Ian Johnston, Dominic Kwiatkowski, Cordelia Langford, John Sillitoe on behalf of the Wellcome Sanger Institute COVID-19 Surveillance Team |

| | | | |
|---|---|---|---|
| EPI_ISL_2539711, EPI_ISL_2539712 | Lighthouse Lab in Milton Keynes | Wellcome Sanger Institute for the COVID-19 Genomics UK (COG-UK) Consortium | The Lighthouse Lab in Milton Keynes and Alex Alderton, Roberto Amato, Jeffrey Barrett, Sonia Goncalves, Ewan Harrison, David K. Jackson, Ian Johnston, Dominic Kwiatkowski, Cordelia Langford, John Sillitoe on behalf of the Wellcome Sanger Institute COVID-19 Surveillance Team |
| EPI_ISL_2539713 | Lighthouse Lab in Alderley Park | Wellcome Sanger Institute for the COVID-19 Genomics UK (COG-UK) Consortium | Jacquelyn Wynn, Mairead Hyland, The Lighthouse Lab in Alderley Park and Alex Alderton, Roberto Amato, Jeffrey Barrett, Sonia Goncalves, Ewan Harrison, David K. Jackson, Ian Johnston, Dominic Kwiatkowski, Cordelia Langford, John Sillitoe on behalf of the Wellcome Sanger Institute COVID-19 Surveillance Team |
| EPI_ISL_2539714, EPI_ISL_2539715 | Lighthouse Lab in Milton Keynes | Wellcome Sanger Institute for the COVID-19 Genomics UK (COG-UK) Consortium | The Lighthouse Lab in Milton Keynes and Alex Alderton, Roberto Amato, Jeffrey Barrett, Sonia Goncalves, Ewan Harrison, David K. Jackson, Ian Johnston, Dominic Kwiatkowski, Cordelia Langford, John Sillitoe on behalf of the Wellcome Sanger Institute COVID-19 Surveillance Team |
| EPI_ISL_2539716 | Lighthouse Lab in Glasgow | Wellcome Sanger Institute for the COVID-19 Genomics UK (COG-UK) Consortium | Harper VanSteenhouse, Yumi Kasai, David Gray, Carol Clugston, Anna Dominiczak and Alex Alderton, Roberto Amato, Jeffrey Barrett, Sonia Goncalves, Ewan Harrison, David K. Jackson, Ian Johnston, Dominic Kwiatkowski, Cordelia Langford, John Sillitoe on behalf of the Wellcome Sanger Institute COVID-19 Surveillance Team |
| EPI_ISL_2539717 | Lighthouse Lab in Alderley Park | Wellcome Sanger Institute for the COVID-19 Genomics UK (COG-UK) Consortium | Jacquelyn Wynn, Mairead Hyland, The Lighthouse Lab in Alderley Park and Alex Alderton, Roberto Amato, Jeffrey Barrett, Sonia Goncalves, Ewan Harrison, David K. Jackson, Ian Johnston, Dominic Kwiatkowski, Cordelia Langford, John Sillitoe on behalf of the Wellcome Sanger Institute COVID-19 Surveillance Team |
| EPI_ISL_2539718 | Lighthouse Lab in Glasgow | Wellcome Sanger Institute for the COVID-19 Genomics UK (COG-UK) Consortium | Harper VanSteenhouse, Yumi Kasai, David Gray, Carol Clugston, Anna Dominiczak and Alex Alderton, Roberto Amato, Jeffrey Barrett, Sonia Goncalves, Ewan Harrison, David K. Jackson, Ian Johnston, Dominic Kwiatkowski, Cordelia Langford, John Sillitoe on behalf of the Wellcome Sanger Institute COVID-19 Surveillance Team |
| EPI_ISL_2539719, EPI_ISL_2539720 | Lighthouse Lab in Alderley Park | Wellcome Sanger Institute for the COVID-19 Genomics UK (COG-UK) Consortium | Jacquelyn Wynn, Mairead Hyland, The Lighthouse Lab in Alderley Park and Alex Alderton, Roberto Amato, Jeffrey Barrett, Sonia Goncalves, Ewan Harrison, David K. Jackson, Ian Johnston, Dominic Kwiatkowski, Cordelia Langford, John Sillitoe on behalf of the Wellcome Sanger Institute COVID-19 Surveillance Team |
| EPI_ISL_2539721 | Lighthouse Lab in Milton Keynes | Wellcome Sanger Institute for the COVID-19 Genomics UK (COG-UK) Consortium | The Lighthouse Lab in Milton Keynes and Alex Alderton, Roberto Amato, Jeffrey Barrett, Sonia Goncalves, Ewan Harrison, David K. Jackson, Ian Johnston, Dominic Kwiatkowski, Cordelia Langford, John Sillitoe on behalf of the Wellcome Sanger Institute COVID-19 Surveillance Team |
| EPI_ISL_2539722 | Lighthouse Lab in Alderley Park | Wellcome Sanger Institute for the COVID-19 Genomics UK (COG-UK) Consortium | Jacquelyn Wynn, Mairead Hyland, The Lighthouse Lab in Alderley Park and Alex Alderton, Roberto Amato, Jeffrey Barrett, Sonia Goncalves, Ewan Harrison, David K. Jackson, Ian Johnston, Dominic Kwiatkowski, Cordelia Langford, John Sillitoe on behalf of the Wellcome Sanger Institute COVID-19 Surveillance Team |
| EPI_ISL_2539723 | Lighthouse Lab in Milton Keynes | Wellcome Sanger Institute for the COVID-19 Genomics UK (COG-UK) Consortium | The Lighthouse Lab in Milton Keynes and Alex Alderton, Roberto Amato, Jeffrey Barrett, Sonia Goncalves, Ewan Harrison, David K. Jackson, Ian Johnston, Dominic Kwiatkowski, Cordelia Langford, John Sillitoe on behalf of the Wellcome Sanger Institute COVID-19 Surveillance Team |
| EPI_ISL_2539724 | Lighthouse Lab in Alderley Park | Wellcome Sanger Institute for the COVID-19 Genomics UK (COG-UK) Consortium | Jacquelyn Wynn, Mairead Hyland, The Lighthouse Lab in Alderley Park and Alex Alderton, Roberto Amato, Jeffrey Barrett, Sonia Goncalves, Ewan Harrison, David K. Jackson, Ian Johnston, Dominic Kwiatkowski, Cordelia Langford, John Sillitoe on behalf of the Wellcome Sanger Institute COVID-19 Surveillance Team |
| EPI_ISL_2539725 | Lighthouse Lab in Milton Keynes | Wellcome Sanger Institute for the COVID-19 Genomics UK (COG-UK) Consortium | The Lighthouse Lab in Milton Keynes and Alex Alderton, Roberto Amato, Jeffrey Barrett, Sonia Goncalves, Ewan Harrison, David K. Jackson, Ian Johnston, Dominic Kwiatkowski, Cordelia Langford, John Sillitoe on behalf of the Wellcome Sanger Institute COVID-19 Surveillance Team |
| EPI_ISL_2539726 | Lighthouse Lab in Glasgow | Wellcome Sanger Institute for the COVID-19 Genomics UK (COG-UK) Consortium | Harper VanSteenhouse, Yumi Kasai, David Gray, Carol Clugston, Anna Dominiczak and Alex Alderton, Roberto Amato, Jeffrey Barrett, Sonia Goncalves, Ewan Harrison, David K. Jackson, Ian Johnston, Dominic Kwiatkowski, Cordelia Langford, John Sillitoe on behalf of the Wellcome Sanger Institute COVID-19 Surveillance Team |
| EPI_ISL_2539727, EPI_ISL_2539728 | Lighthouse Lab in Milton Keynes | Wellcome Sanger Institute for the COVID-19 Genomics UK (COG-UK) Consortium | The Lighthouse Lab in Milton Keynes and Alex Alderton, Roberto Amato, Jeffrey Barrett, Sonia Goncalves, Ewan Harrison, David K. Jackson, Ian Johnston, Dominic Kwiatkowski, Cordelia Langford, John Sillitoe on behalf of the Wellcome Sanger Institute COVID-19 Surveillance Team |
| EPI_ISL_2539729 | Lighthouse Lab in Glasgow | Wellcome Sanger Institute for the COVID-19 Genomics UK (COG-UK) Consortium | Harper VanSteenhouse, Yumi Kasai, David Gray, Carol Clugston, Anna Dominiczak and Alex Alderton, Roberto Amato, Jeffrey Barrett, Sonia Goncalves, Ewan Harrison, David K. Jackson, Ian Johnston, Dominic Kwiatkowski, Cordelia Langford, John Sillitoe on behalf of the Wellcome Sanger Institute COVID-19 Surveillance Team |
| EPI_ISL_2539730 | Lighthouse Lab in Alderley Park | Wellcome Sanger Institute for the COVID-19 Genomics UK (COG-UK) Consortium | Jacquelyn Wynn, Mairead Hyland, The Lighthouse Lab in Alderley Park and Alex Alderton, Roberto Amato, Jeffrey Barrett, Sonia Goncalves, Ewan Harrison, David K. Jackson, Ian Johnston, Dominic Kwiatkowski, Cordelia Langford, John Sillitoe on behalf of the Wellcome Sanger Institute COVID-19 Surveillance Team |
| EPI_ISL_2539731 | Lighthouse Lab in Glasgow | Wellcome Sanger Institute for the COVID-19 Genomics UK (COG-UK) Consortium | Harper VanSteenhouse, Yumi Kasai, David Gray, Carol Clugston, Anna Dominiczak and Alex Alderton, Roberto Amato, Jeffrey Barrett, Sonia Goncalves, Ewan Harrison, David K. Jackson, Ian Johnston, Dominic Kwiatkowski, Cordelia Langford, John Sillitoe on behalf of the Wellcome Sanger Institute COVID-19 Surveillance Team |
| EPI_ISL_2539732, EPI_ISL_2539733 | Lighthouse Lab in Milton Keynes | Wellcome Sanger Institute for the COVID-19 Genomics UK (COG-UK) Consortium | The Lighthouse Lab in Milton Keynes and Alex Alderton, Roberto Amato, Jeffrey Barrett, Sonia Goncalves, Ewan Harrison, David K. Jackson, Ian Johnston, Dominic Kwiatkowski, Cordelia Langford, John Sillitoe on behalf of the Wellcome Sanger Institute COVID-19 Surveillance Team |
| EPI_ISL_2539734 | Lighthouse Lab in Glasgow | Wellcome Sanger Institute for the COVID-19 Genomics UK (COG-UK) Consortium | Harper VanSteenhouse, Yumi Kasai, David Gray, Carol Clugston, Anna Dominiczak and Alex Alderton, Roberto Amato, Jeffrey Barrett, Sonia Goncalves, Ewan Harrison, David K. Jackson, Ian Johnston, Dominic Kwiatkowski, Cordelia Langford, John Sillitoe on behalf of the Wellcome Sanger Institute COVID-19 Surveillance Team |
| EPI_ISL_2539736, EPI_ISL_2539737, EPI_ISL_2539738, EPI_ISL_2539739 | Lighthouse Lab in Milton Keynes | Wellcome Sanger Institute for the COVID-19 Genomics UK (COG-UK) Consortium | The Lighthouse Lab in Milton Keynes and Alex Alderton, Roberto Amato, Jeffrey Barrett, Sonia Goncalves, Ewan Harrison, David K. Jackson, Ian Johnston, Dominic Kwiatkowski, Cordelia Langford, John Sillitoe on behalf of the Wellcome Sanger Institute COVID-19 Surveillance Team |
| EPI_ISL_2539740, EPI_ISL_2539741 | Lighthouse Laboratory Plymouth | Wellcome Sanger Institute for the COVID-19 Genomics UK (COG-UK) Consortium | Lighthouse Laboratory Plymouth and Alex Alderton, Roberto Amato, Jeffrey Barrett, Sonia Goncalves, Ewan Harrison, David K. Jackson, Ian Johnston, Dominic Kwiatkowski, Cordelia Langford, John Sillitoe on behalf of the Wellcome Sanger Institute COVID-19 Surveillance Team |
| EPI_ISL_2539742, EPI_ISL_2539743 | Lighthouse Lab in Milton Keynes | Wellcome Sanger Institute for the COVID-19 Genomics UK (COG-UK) Consortium | The Lighthouse Lab in Milton Keynes and Alex Alderton, Roberto Amato, Jeffrey Barrett, Sonia Goncalves, Ewan Harrison, David K. Jackson, Ian Johnston, Dominic Kwiatkowski, Cordelia Langford, John Sillitoe on behalf of the Wellcome Sanger Institute COVID-19 Surveillance Team |
| EPI_ISL_2539744 | Lighthouse Laboratory Plymouth | Wellcome Sanger Institute for the COVID-19 Genomics UK (COG-UK) Consortium | Lighthouse Laboratory Plymouth and Alex Alderton, Roberto Amato, Jeffrey Barrett, Sonia Goncalves, Ewan Harrison, David K. Jackson, Ian Johnston, Dominic Kwiatkowski, Cordelia Langford, John Sillitoe on behalf of the Wellcome Sanger Institute COVID-19 Surveillance Team |
| EPI_ISL_2539745 | Lighthouse Lab in Milton Keynes | Wellcome Sanger Institute for the COVID-19 Genomics UK (COG-UK) Consortium | The Lighthouse Lab in Milton Keynes and Alex Alderton, Roberto Amato, Jeffrey Barrett, Sonia Goncalves, Ewan Harrison, David K. Jackson, Ian Johnston, Dominic Kwiatkowski, Cordelia Langford, John Sillitoe on behalf of the Wellcome Sanger Institute COVID-19 Surveillance Team |
| EPI_ISL_2539746 | Health Services Laboratories | Wellcome Sanger Institute for the COVID-19 Genomics UK (COG-UK) Consortium | Health Services Laboratories and Alex Alderton, Roberto Amato, Jeffrey Barrett, Sonia Goncalves, Ewan Harrison, David K. Jackson, Ian Johnston, Dominic Kwiatkowski, Cordelia Langford, John Sillitoe on behalf of the Wellcome Sanger Institute COVID-19 Surveillance Team |
| EPI_ISL_2539747 | Lighthouse Laboratory Plymouth | Wellcome Sanger Institute for the COVID-19 Genomics UK (COG-UK) Consortium | Lighthouse Laboratory Plymouth and Alex Alderton, Roberto Amato, Jeffrey Barrett, Sonia Goncalves, Ewan Harrison, David K. Jackson, Ian Johnston, Dominic Kwiatkowski, Cordelia Langford, John Sillitoe on behalf of the Wellcome Sanger Institute COVID-19 Surveillance Team |
| EPI_ISL_2539749 | Lighthouse Lab in Milton Keynes | Wellcome Sanger Institute for the COVID-19 Genomics UK (COG-UK) Consortium | The Lighthouse Lab in Milton Keynes and Alex Alderton, Roberto Amato, Jeffrey Barrett, Sonia Goncalves, Ewan Harrison, David K. Jackson, Ian Johnston, Dominic Kwiatkowski, Cordelia Langford, John Sillitoe on behalf of the Wellcome Sanger Institute COVID-19 Surveillance Team |
| EPI_ISL_2539750, EPI_ISL_2539751 | Lighthouse Laboratory Plymouth | Wellcome Sanger Institute for the COVID-19 Genomics UK (COG-UK) Consortium | Lighthouse Laboratory Plymouth and Alex Alderton, Roberto Amato, Jeffrey Barrett, Sonia Goncalves, Ewan Harrison, David K. Jackson, Ian Johnston, Dominic Kwiatkowski, Cordelia Langford, John Sillitoe on behalf of the Wellcome Sanger Institute COVID-19 Surveillance Team |
| EPI_ISL_2539752 | Lighthouse Lab in Milton Keynes | Wellcome Sanger Institute for the COVID-19 Genomics UK (COG-UK) Consortium | The Lighthouse Lab in Milton Keynes and Alex Alderton, Roberto Amato, Jeffrey Barrett, Sonia Goncalves, Ewan Harrison, David K. Jackson, Ian Johnston, Dominic Kwiatkowski, Cordelia Langford, John Sillitoe on behalf of the Wellcome Sanger Institute COVID-19 Surveillance Team |
| EPI_ISL_2539753, EPI_ISL_2539755, EPI_ISL_2539756 | Lighthouse Laboratory Plymouth | Wellcome Sanger Institute for the COVID-19 Genomics UK (COG-UK) Consortium | Lighthouse Laboratory Plymouth and Alex Alderton, Roberto Amato, Jeffrey Barrett, Sonia Goncalves, Ewan Harrison, David K. Jackson, Ian Johnston, Dominic Kwiatkowski, Cordelia Langford, John Sillitoe on behalf of the Wellcome Sanger Institute COVID-19 Surveillance Team |
| EPI_ISL_2539757, EPI_ISL_2539758, EPI_ISL_2539759, EPI_ISL_2539760 | Lighthouse Lab in Milton Keynes | Wellcome Sanger Institute for the COVID-19 Genomics UK (COG-UK) Consortium | The Lighthouse Lab in Milton Keynes and Alex Alderton, Roberto Amato, Jeffrey Barrett, Sonia Goncalves, Ewan Harrison, David K. Jackson, Ian Johnston, Dominic Kwiatkowski, Cordelia Langford, John Sillitoe on behalf of the Wellcome Sanger Institute COVID-19 Surveillance Team |
| EPI_ISL_2539761 | Health Services Laboratories | Wellcome Sanger Institute for the COVID-19 Genomics UK (COG-UK) Consortium | Health Services Laboratories and Alex Alderton, Roberto Amato, Jeffrey Barrett, Sonia Goncalves, Ewan Harrison, David K. Jackson, Ian Johnston, Dominic Kwiatkowski, Cordelia Langford, John Sillitoe on behalf of the Wellcome Sanger Institute COVID-19 Surveillance Team |
| EPI_ISL_2539762 | Lighthouse Lab in Milton Keynes | Wellcome Sanger Institute for the COVID-19 Genomics UK (COG-UK) Consortium | The Lighthouse Lab in Milton Keynes and Alex Alderton, Roberto Amato, Jeffrey Barrett, Sonia Goncalves, Ewan Harrison, David K. Jackson, Ian Johnston, Dominic Kwiatkowski, Cordelia Langford, John Sillitoe on behalf of the Wellcome Sanger Institute COVID-19 Surveillance Team |
| EPI_ISL_2539763 | Health Services Laboratories | Wellcome Sanger Institute for the COVID-19 Genomics UK (COG-UK) Consortium | Health Services Laboratories and Alex Alderton, Roberto Amato, Jeffrey Barrett, Sonia Goncalves, Ewan Harrison, David K. Jackson, Ian Johnston, Dominic Kwiatkowski, Cordelia Langford, John Sillitoe on behalf of the Wellcome Sanger Institute COVID-19 Surveillance Team |
| EPI_ISL_2539764, EPI_ISL_2539765 | Lighthouse Laboratory Plymouth | Wellcome Sanger Institute for the COVID-19 Genomics UK (COG-UK) Consortium | Lighthouse Laboratory Plymouth and Alex Alderton, Roberto Amato, Jeffrey Barrett, Sonia Goncalves, Ewan Harrison, David K. Jackson, Ian Johnston, Dominic Kwiatkowski, Cordelia Langford, John Sillitoe on behalf of the Wellcome Sanger Institute COVID-19 Surveillance Team |
| EPI_ISL_2539766, EPI_ISL_2539767, EPI_ISL_2539768, EPI_ISL_2539769, EPI_ISL_2539770 | Lighthouse Lab in Milton Keynes | Wellcome Sanger Institute for the COVID-19 Genomics UK (COG-UK) Consortium | The Lighthouse Lab in Milton Keynes and Alex Alderton, Roberto Amato, Jeffrey Barrett, Sonia Goncalves, Ewan Harrison, David K. Jackson, Ian Johnston, Dominic Kwiatkowski, Cordelia Langford, John Sillitoe on behalf of the Wellcome Sanger Institute COVID-19 Surveillance Team |
| EPI_ISL_2539771 | Health Services Laboratories | Wellcome Sanger Institute for the COVID-19 Genomics UK (COG-UK) Consortium | Health Services Laboratories and Alex Alderton, Roberto Amato, Jeffrey Barrett, Sonia Goncalves, Ewan Harrison, David K. Jackson, Ian Johnston, Dominic Kwiatkowski, Cordelia Langford, John Sillitoe on behalf of the Wellcome Sanger Institute COVID-19 Surveillance Team |
| EPI_ISL_2539772, EPI_ISL_2539773, EPI_ISL_2539774 | Lighthouse Lab in Milton Keynes | Wellcome Sanger Institute for the COVID-19 Genomics UK (COG-UK) Consortium | The Lighthouse Lab in Milton Keynes and Alex Alderton, Roberto Amato, Jeffrey Barrett, Sonia Goncalves, Ewan Harrison, David K. Jackson, Ian Johnston, Dominic Kwiatkowski, Cordelia Langford, John Sillitoe on behalf of the Wellcome Sanger Institute COVID-19 Surveillance Team |
| EPI_ISL_2539775 | Lighthouse Laboratory Plymouth | Wellcome Sanger Institute for the COVID-19 Genomics UK (COG-UK) Consortium | Lighthouse Laboratory Plymouth and Alex Alderton, Roberto Amato, Jeffrey Barrett, Sonia Goncalves, Ewan Harrison, David K. Jackson, Ian Johnston, Dominic Kwiatkowski, Cordelia Langford, John Sillitoe on behalf of the Wellcome Sanger Institute COVID-19 Surveillance Team |
| EPI_ISL_2539776, EPI_ISL_2539777, EPI_ISL_2539778 | Lighthouse Lab in Milton Keynes | Wellcome Sanger Institute for the COVID-19 Genomics UK (COG-UK) Consortium | The Lighthouse Lab in Milton Keynes and Alex Alderton, Roberto Amato, Jeffrey Barrett, Sonia Goncalves, Ewan Harrison, David K. Jackson, Ian Johnston, Dominic Kwiatkowski, Cordelia Langford, John Sillitoe on behalf of the Wellcome Sanger Institute COVID-19 Surveillance Team |
| EPI_ISL_2539779 | Lighthouse Laboratory Plymouth | Wellcome Sanger Institute for the COVID-19 Genomics UK (COG-UK) | Lighthouse Laboratory Plymouth and Alex Alderton, Roberto Amato, Jeffrey Barrett, Sonia Goncalves, Ewan Harrison, David K. Jackson, Ian Johnston, Dominic Kwiatkowski, Cordelia Langford, John Sillitoe on behalf of the Wellcome Sanger |

| | | | |
|---|---|---|---|
| | | Consortium | Institute COVID-19 Surveillance Team |
| EPI_ISL_2539780 | Lighthouse Lab in Milton Keynes | Wellcome Sanger Institute for the COVID-19 Genomics UK (COG-UK) Consortium | The Lighthouse Lab in Milton Keynes and Alex Alderton, Roberto Amato, Jeffrey Barrett, Sonia Goncalves, Ewan Harrison, David K. Jackson, Ian Johnston, Dominic Kwiatkowski, Cordelia Langford, John Sillitoe on behalf of the Wellcome Sanger Institute COVID-19 Surveillance Team |
| EPI_ISL_2539781, EPI_ISL_2539782, EPI_ISL_2539783, EPI_ISL_2539784, EPI_ISL_2539785, EPI_ISL_2539786 | Lighthouse Laboratory Plymouth | Wellcome Sanger Institute for the COVID-19 Genomics UK (COG-UK) Consortium | Lighthouse Laboratory Plymouth and Alex Alderton, Roberto Amato, Jeffrey Barrett, Sonia Goncalves, Ewan Harrison, David K. Jackson, Ian Johnston, Dominic Kwiatkowski, Cordelia Langford, John Sillitoe on behalf of the Wellcome Sanger Institute COVID-19 Surveillance Team |
| EPI_ISL_2539787, EPI_ISL_2539788, EPI_ISL_2539789 | Lighthouse Lab in Milton Keynes | Wellcome Sanger Institute for the COVID-19 Genomics UK (COG-UK) Consortium | The Lighthouse Lab in Milton Keynes and Alex Alderton, Roberto Amato, Jeffrey Barrett, Sonia Goncalves, Ewan Harrison, David K. Jackson, Ian Johnston, Dominic Kwiatkowski, Cordelia Langford, John Sillitoe on behalf of the Wellcome Sanger Institute COVID-19 Surveillance Team |
| EPI_ISL_2539790 | Health Services Laboratories | Wellcome Sanger Institute for the COVID-19 Genomics UK (COG-UK) Consortium | Health Services Laboratories and Alex Alderton, Roberto Amato, Jeffrey Barrett, Sonia Goncalves, Ewan Harrison, David K. Jackson, Ian Johnston, Dominic Kwiatkowski, Cordelia Langford, John Sillitoe on behalf of the Wellcome Sanger Institute COVID-19 Surveillance Team |
| EPI_ISL_2539791 | Lighthouse Lab in Milton Keynes | Wellcome Sanger Institute for the COVID-19 Genomics UK (COG-UK) Consortium | The Lighthouse Lab in Milton Keynes and Alex Alderton, Roberto Amato, Jeffrey Barrett, Sonia Goncalves, Ewan Harrison, David K. Jackson, Ian Johnston, Dominic Kwiatkowski, Cordelia Langford, John Sillitoe on behalf of the Wellcome Sanger Institute COVID-19 Surveillance Team |
| EPI_ISL_2539792 | Lighthouse Laboratory Plymouth | Wellcome Sanger Institute for the COVID-19 Genomics UK (COG-UK) Consortium | Lighthouse Laboratory Plymouth and Alex Alderton, Roberto Amato, Jeffrey Barrett, Sonia Goncalves, Ewan Harrison, David K. Jackson, Ian Johnston, Dominic Kwiatkowski, Cordelia Langford, John Sillitoe on behalf of the Wellcome Sanger Institute COVID-19 Surveillance Team |
| EPI_ISL_2539793 | Lighthouse Lab in Milton Keynes | Wellcome Sanger Institute for the COVID-19 Genomics UK (COG-UK) Consortium | The Lighthouse Lab in Milton Keynes and Alex Alderton, Roberto Amato, Jeffrey Barrett, Sonia Goncalves, Ewan Harrison, David K. Jackson, Ian Johnston, Dominic Kwiatkowski, Cordelia Langford, John Sillitoe on behalf of the Wellcome Sanger Institute COVID-19 Surveillance Team |
| EPI_ISL_2539796 | Lighthouse Laboratory Plymouth | Wellcome Sanger Institute for the COVID-19 Genomics UK (COG-UK) Consortium | Lighthouse Laboratory Plymouth and Alex Alderton, Roberto Amato, Jeffrey Barrett, Sonia Goncalves, Ewan Harrison, David K. Jackson, Ian Johnston, Dominic Kwiatkowski, Cordelia Langford, John Sillitoe on behalf of the Wellcome Sanger Institute COVID-19 Surveillance Team |
| EPI_ISL_2539797, EPI_ISL_2539798, EPI_ISL_2539799 | Lighthouse Lab in Milton Keynes | Wellcome Sanger Institute for the COVID-19 Genomics UK (COG-UK) Consortium | The Lighthouse Lab in Milton Keynes and Alex Alderton, Roberto Amato, Jeffrey Barrett, Sonia Goncalves, Ewan Harrison, David K. Jackson, Ian Johnston, Dominic Kwiatkowski, Cordelia Langford, John Sillitoe on behalf of the Wellcome Sanger Institute COVID-19 Surveillance Team |
| EPI_ISL_2539800, EPI_ISL_2539802 | Lighthouse Laboratory Plymouth | Wellcome Sanger Institute for the COVID-19 Genomics UK (COG-UK) Consortium | Lighthouse Laboratory Plymouth and Alex Alderton, Roberto Amato, Jeffrey Barrett, Sonia Goncalves, Ewan Harrison, David K. Jackson, Ian Johnston, Dominic Kwiatkowski, Cordelia Langford, John Sillitoe on behalf of the Wellcome Sanger Institute COVID-19 Surveillance Team |
| EPI_ISL_2539803, EPI_ISL_2539804, EPI_ISL_2539805 | Lighthouse Lab in Milton Keynes | Wellcome Sanger Institute for the COVID-19 Genomics UK (COG-UK) Consortium | The Lighthouse Lab in Milton Keynes and Alex Alderton, Roberto Amato, Jeffrey Barrett, Sonia Goncalves, Ewan Harrison, David K. Jackson, Ian Johnston, Dominic Kwiatkowski, Cordelia Langford, John Sillitoe on behalf of the Wellcome Sanger Institute COVID-19 Surveillance Team |
| EPI_ISL_2539806 | Health Services Laboratories | Wellcome Sanger Institute for the COVID-19 Genomics UK (COG-UK) Consortium | Health Services Laboratories and Alex Alderton, Roberto Amato, Jeffrey Barrett, Sonia Goncalves, Ewan Harrison, David K. Jackson, Ian Johnston, Dominic Kwiatkowski, Cordelia Langford, John Sillitoe on behalf of the Wellcome Sanger Institute COVID-19 Surveillance Team |
| EPI_ISL_2539807, EPI_ISL_2539808, EPI_ISL_2539809, EPI_ISL_2539810 | Lighthouse Lab in Milton Keynes | Wellcome Sanger Institute for the COVID-19 Genomics UK (COG-UK) Consortium | The Lighthouse Lab in Milton Keynes and Alex Alderton, Roberto Amato, Jeffrey Barrett, Sonia Goncalves, Ewan Harrison, David K. Jackson, Ian Johnston, Dominic Kwiatkowski, Cordelia Langford, John Sillitoe on behalf of the Wellcome Sanger Institute COVID-19 Surveillance Team |
| EPI_ISL_2539811 | Lighthouse Laboratory Plymouth | Wellcome Sanger Institute for the COVID-19 Genomics UK (COG-UK) Consortium | Lighthouse Laboratory Plymouth and Alex Alderton, Roberto Amato, Jeffrey Barrett, Sonia Goncalves, Ewan Harrison, David K. Jackson, Ian Johnston, Dominic Kwiatkowski, Cordelia Langford, John Sillitoe on behalf of the Wellcome Sanger Institute COVID-19 Surveillance Team |
| EPI_ISL_2539812 | Health Services Laboratories | Wellcome Sanger Institute for the COVID-19 Genomics UK (COG-UK) Consortium | Health Services Laboratories and Alex Alderton, Roberto Amato, Jeffrey Barrett, Sonia Goncalves, Ewan Harrison, David K. Jackson, Ian Johnston, Dominic Kwiatkowski, Cordelia Langford, John Sillitoe on behalf of the Wellcome Sanger Institute COVID-19 Surveillance Team |
| EPI_ISL_2539813, EPI_ISL_2539814 | Lighthouse Laboratory Plymouth | Wellcome Sanger Institute for the COVID-19 Genomics UK (COG-UK) Consortium | Lighthouse Laboratory Plymouth and Alex Alderton, Roberto Amato, Jeffrey Barrett, Sonia Goncalves, Ewan Harrison, David K. Jackson, Ian Johnston, Dominic Kwiatkowski, Cordelia Langford, John Sillitoe on behalf of the Wellcome Sanger Institute COVID-19 Surveillance Team |
| EPI_ISL_2539815 | Lighthouse Lab in Milton Keynes | Wellcome Sanger Institute for the COVID-19 Genomics UK (COG-UK) Consortium | The Lighthouse Lab in Milton Keynes and Alex Alderton, Roberto Amato, Jeffrey Barrett, Sonia Goncalves, Ewan Harrison, David K. Jackson, Ian Johnston, Dominic Kwiatkowski, Cordelia Langford, John Sillitoe on behalf of the Wellcome Sanger Institute COVID-19 Surveillance Team |
| EPI_ISL_2539816 | Lighthouse Laboratory Plymouth | Wellcome Sanger Institute for the COVID-19 Genomics UK (COG-UK) Consortium | Lighthouse Laboratory Plymouth and Alex Alderton, Roberto Amato, Jeffrey Barrett, Sonia Goncalves, Ewan Harrison, David K. Jackson, Ian Johnston, Dominic Kwiatkowski, Cordelia Langford, John Sillitoe on behalf of the Wellcome Sanger Institute COVID-19 Surveillance Team |
| EPI_ISL_2539817 | Health Services Laboratories | Wellcome Sanger Institute for the COVID-19 Genomics UK (COG-UK) Consortium | Health Services Laboratories and Alex Alderton, Roberto Amato, Jeffrey Barrett, Sonia Goncalves, Ewan Harrison, David K. Jackson, Ian Johnston, Dominic Kwiatkowski, Cordelia Langford, John Sillitoe on behalf of the Wellcome Sanger Institute COVID-19 Surveillance Team |
| EPI_ISL_2539818 | Lighthouse Laboratory Plymouth | Wellcome Sanger Institute for the COVID-19 Genomics UK (COG-UK) Consortium | Lighthouse Laboratory Plymouth and Alex Alderton, Roberto Amato, Jeffrey Barrett, Sonia Goncalves, Ewan Harrison, David K. Jackson, Ian Johnston, Dominic Kwiatkowski, Cordelia Langford, John Sillitoe on behalf of the Wellcome Sanger Institute COVID-19 Surveillance Team |
| EPI_ISL_2539819, EPI_ISL_2539820, EPI_ISL_2539821 | Lighthouse Lab in Milton Keynes | Wellcome Sanger Institute for the COVID-19 Genomics UK (COG-UK) Consortium | The Lighthouse Lab in Milton Keynes and Alex Alderton, Roberto Amato, Jeffrey Barrett, Sonia Goncalves, Ewan Harrison, David K. Jackson, Ian Johnston, Dominic Kwiatkowski, Cordelia Langford, John Sillitoe on behalf of the Wellcome Sanger Institute COVID-19 Surveillance Team |
| EPI_ISL_2539822, EPI_ISL_2539823 | Lighthouse Laboratory Plymouth | Wellcome Sanger Institute for the COVID-19 Genomics UK (COG-UK) Consortium | Lighthouse Laboratory Plymouth and Alex Alderton, Roberto Amato, Jeffrey Barrett, Sonia Goncalves, Ewan Harrison, David K. Jackson, Ian Johnston, Dominic Kwiatkowski, Cordelia Langford, John Sillitoe on behalf of the Wellcome Sanger Institute COVID-19 Surveillance Team |
| EPI_ISL_2539824, EPI_ISL_2539825, EPI_ISL_2539826, EPI_ISL_2539827 | Lighthouse Lab in Milton Keynes | Wellcome Sanger Institute for the COVID-19 Genomics UK (COG-UK) Consortium | The Lighthouse Lab in Milton Keynes and Alex Alderton, Roberto Amato, Jeffrey Barrett, Sonia Goncalves, Ewan Harrison, David K. Jackson, Ian Johnston, Dominic Kwiatkowski, Cordelia Langford, John Sillitoe on behalf of the Wellcome Sanger Institute COVID-19 Surveillance Team |
| EPI_ISL_2539828 | Lighthouse Laboratory Plymouth | Wellcome Sanger Institute for the COVID-19 Genomics UK (COG-UK) Consortium | Lighthouse Laboratory Plymouth and Alex Alderton, Roberto Amato, Jeffrey Barrett, Sonia Goncalves, Ewan Harrison, David K. Jackson, Ian Johnston, Dominic Kwiatkowski, Cordelia Langford, John Sillitoe on behalf of the Wellcome Sanger Institute COVID-19 Surveillance Team |
| EPI_ISL_2539829, EPI_ISL_2539830, EPI_ISL_2539831, EPI_ISL_2539832 | Lighthouse Lab in Milton Keynes | Wellcome Sanger Institute for the COVID-19 Genomics UK (COG-UK) Consortium | The Lighthouse Lab in Milton Keynes and Alex Alderton, Roberto Amato, Jeffrey Barrett, Sonia Goncalves, Ewan Harrison, David K. Jackson, Ian Johnston, Dominic Kwiatkowski, Cordelia Langford, John Sillitoe on behalf of the Wellcome Sanger Institute COVID-19 Surveillance Team |
| EPI_ISL_2539833, EPI_ISL_2539834 | Lighthouse Laboratory Plymouth | Wellcome Sanger Institute for the COVID-19 Genomics UK (COG-UK) Consortium | Lighthouse Laboratory Plymouth and Alex Alderton, Roberto Amato, Jeffrey Barrett, Sonia Goncalves, Ewan Harrison, David K. Jackson, Ian Johnston, Dominic Kwiatkowski, Cordelia Langford, John Sillitoe on behalf of the Wellcome Sanger Institute COVID-19 Surveillance Team |
| EPI_ISL_2539835, EPI_ISL_2539836 | Lighthouse Lab in Milton Keynes | Wellcome Sanger Institute for the COVID-19 Genomics UK (COG-UK) Consortium | The Lighthouse Lab in Milton Keynes and Alex Alderton, Roberto Amato, Jeffrey Barrett, Sonia Goncalves, Ewan Harrison, David K. Jackson, Ian Johnston, Dominic Kwiatkowski, Cordelia Langford, John Sillitoe on behalf of the Wellcome Sanger Institute COVID-19 Surveillance Team |
| EPI_ISL_2539837 | Lighthouse Laboratory Plymouth | Wellcome Sanger Institute for the COVID-19 Genomics UK (COG-UK) Consortium | Lighthouse Laboratory Plymouth and Alex Alderton, Roberto Amato, Jeffrey Barrett, Sonia Goncalves, Ewan Harrison, David K. Jackson, Ian Johnston, Dominic Kwiatkowski, Cordelia Langford, John Sillitoe on behalf of the Wellcome Sanger Institute COVID-19 Surveillance Team |
| EPI_ISL_2539838 | Lighthouse Lab in Milton Keynes | Wellcome Sanger Institute for the COVID-19 Genomics UK (COG-UK) Consortium | The Lighthouse Lab in Milton Keynes and Alex Alderton, Roberto Amato, Jeffrey Barrett, Sonia Goncalves, Ewan Harrison, David K. Jackson, Ian Johnston, Dominic Kwiatkowski, Cordelia Langford, John Sillitoe on behalf of the Wellcome Sanger Institute COVID-19 Surveillance Team |
| EPI_ISL_2539839 | Health Services Laboratories | Wellcome Sanger Institute for the COVID-19 Genomics UK (COG-UK) Consortium | Health Services Laboratories and Alex Alderton, Roberto Amato, Jeffrey Barrett, Sonia Goncalves, Ewan Harrison, David K. Jackson, Ian Johnston, Dominic Kwiatkowski, Cordelia Langford, John Sillitoe on behalf of the Wellcome Sanger Institute COVID-19 Surveillance Team |
| EPI_ISL_2539840 | Lighthouse Lab in Milton Keynes | Wellcome Sanger Institute for the COVID-19 Genomics UK (COG-UK) Consortium | The Lighthouse Lab in Milton Keynes and Alex Alderton, Roberto Amato, Jeffrey Barrett, Sonia Goncalves, Ewan Harrison, David K. Jackson, Ian Johnston, Dominic Kwiatkowski, Cordelia Langford, John Sillitoe on behalf of the Wellcome Sanger Institute COVID-19 Surveillance Team |
| EPI_ISL_2539841 | Health Services Laboratories | Wellcome Sanger Institute for the COVID-19 Genomics UK (COG-UK) Consortium | Health Services Laboratories and Alex Alderton, Roberto Amato, Jeffrey Barrett, Sonia Goncalves, Ewan Harrison, David K. Jackson, Ian Johnston, Dominic Kwiatkowski, Cordelia Langford, John Sillitoe on behalf of the Wellcome Sanger Institute COVID-19 Surveillance Team |
| EPI_ISL_2539842 | Lighthouse Laboratory Plymouth | Wellcome Sanger Institute for the COVID-19 Genomics UK (COG-UK) Consortium | Lighthouse Laboratory Plymouth and Alex Alderton, Roberto Amato, Jeffrey Barrett, Sonia Goncalves, Ewan Harrison, David K. Jackson, Ian Johnston, Dominic Kwiatkowski, Cordelia Langford, John Sillitoe on behalf of the Wellcome Sanger Institute COVID-19 Surveillance Team |
| EPI_ISL_2539843 | Lighthouse Lab in Milton Keynes | Wellcome Sanger Institute for the COVID-19 Genomics UK (COG-UK) Consortium | The Lighthouse Lab in Milton Keynes and Alex Alderton, Roberto Amato, Jeffrey Barrett, Sonia Goncalves, Ewan Harrison, David K. Jackson, Ian Johnston, Dominic Kwiatkowski, Cordelia Langford, John Sillitoe on behalf of the Wellcome Sanger Institute COVID-19 Surveillance Team |
| EPI_ISL_2539844 | Lighthouse Laboratory Plymouth | Wellcome Sanger Institute for the COVID-19 Genomics UK (COG-UK) Consortium | Lighthouse Laboratory Plymouth and Alex Alderton, Roberto Amato, Jeffrey Barrett, Sonia Goncalves, Ewan Harrison, David K. Jackson, Ian Johnston, Dominic Kwiatkowski, Cordelia Langford, John Sillitoe on behalf of the Wellcome Sanger Institute COVID-19 Surveillance Team |
| EPI_ISL_2539845, EPI_ISL_2539846, EPI_ISL_2539847, EPI_ISL_2539848, EPI_ISL_2539849, EPI_ISL_2539850, EPI_ISL_2539851 | Lighthouse Lab in Milton Keynes | Wellcome Sanger Institute for the COVID-19 Genomics UK (COG-UK) Consortium | The Lighthouse Lab in Milton Keynes and Alex Alderton, Roberto Amato, Jeffrey Barrett, Sonia Goncalves, Ewan Harrison, David K. Jackson, Ian Johnston, Dominic Kwiatkowski, Cordelia Langford, John Sillitoe on behalf of the Wellcome Sanger Institute COVID-19 Surveillance Team |
| EPI_ISL_2539852, EPI_ISL_2539853 | Health Services Laboratories | Wellcome Sanger Institute for the COVID-19 Genomics UK (COG-UK) Consortium | Health Services Laboratories and Alex Alderton, Roberto Amato, Jeffrey Barrett, Sonia Goncalves, Ewan Harrison, David K. Jackson, Ian Johnston, Dominic Kwiatkowski, Cordelia Langford, John Sillitoe on behalf of the Wellcome Sanger Institute COVID-19 Surveillance Team |

| | | | |
|---|---|---|---|
| EPI_ISL_2539854 | Lighthouse Laboratory Plymouth | Wellcome Sanger Institute for the COVID-19 Genomics UK (COG-UK) Consortium | Lighthouse Laboratory Plymouth and Alex Alderton, Roberto Amato, Jeffrey Barrett, Sonia Goncalves, Ewan Harrison, David K. Jackson, Ian Johnston, Dominic Kwiatkowski, Cordelia Langford, John Sillitoe on behalf of the Wellcome Sanger Institute COVID-19 Surveillance Team |
| EPI_ISL_2539855, EPI_ISL_2539856, EPI_ISL_2539857 | Lighthouse Lab in Milton Keynes | Wellcome Sanger Institute for the COVID-19 Genomics UK (COG-UK) Consortium | The Lighthouse Lab in Milton Keynes and Alex Alderton, Roberto Amato, Jeffrey Barrett, Sonia Goncalves, Ewan Harrison, David K. Jackson, Ian Johnston, Dominic Kwiatkowski, Cordelia Langford, John Sillitoe on behalf of the Wellcome Sanger Institute COVID-19 Surveillance Team |
| EPI_ISL_2539858, EPI_ISL_2539859 | Lighthouse Laboratory Plymouth | Wellcome Sanger Institute for the COVID-19 Genomics UK (COG-UK) Consortium | Lighthouse Laboratory Plymouth and Alex Alderton, Roberto Amato, Jeffrey Barrett, Sonia Goncalves, Ewan Harrison, David K. Jackson, Ian Johnston, Dominic Kwiatkowski, Cordelia Langford, John Sillitoe on behalf of the Wellcome Sanger Institute COVID-19 Surveillance Team |
| EPI_ISL_2539860 | Lighthouse Lab in Milton Keynes | Wellcome Sanger Institute for the COVID-19 Genomics UK (COG-UK) Consortium | The Lighthouse Lab in Milton Keynes and Alex Alderton, Roberto Amato, Jeffrey Barrett, Sonia Goncalves, Ewan Harrison, David K. Jackson, Ian Johnston, Dominic Kwiatkowski, Cordelia Langford, John Sillitoe on behalf of the Wellcome Sanger Institute COVID-19 Surveillance Team |
| EPI_ISL_2539861 | Health Services Laboratories | Wellcome Sanger Institute for the COVID-19 Genomics UK (COG-UK) Consortium | Health Services Laboratories and Alex Alderton, Roberto Amato, Jeffrey Barrett, Sonia Goncalves, Ewan Harrison, David K. Jackson, Ian Johnston, Dominic Kwiatkowski, Cordelia Langford, John Sillitoe on behalf of the Wellcome Sanger Institute COVID-19 Surveillance Team |
| EPI_ISL_2539862 | Lighthouse Laboratory Plymouth | Wellcome Sanger Institute for the COVID-19 Genomics UK (COG-UK) Consortium | Lighthouse Laboratory Plymouth and Alex Alderton, Roberto Amato, Jeffrey Barrett, Sonia Goncalves, Ewan Harrison, David K. Jackson, Ian Johnston, Dominic Kwiatkowski, Cordelia Langford, John Sillitoe on behalf of the Wellcome Sanger Institute COVID-19 Surveillance Team |
| EPI_ISL_2539863, EPI_ISL_2539864 | Health Services Laboratories | Wellcome Sanger Institute for the COVID-19 Genomics UK (COG-UK) Consortium | Health Services Laboratories and Alex Alderton, Roberto Amato, Jeffrey Barrett, Sonia Goncalves, Ewan Harrison, David K. Jackson, Ian Johnston, Dominic Kwiatkowski, Cordelia Langford, John Sillitoe on behalf of the Wellcome Sanger Institute COVID-19 Surveillance Team |
| EPI_ISL_2539865, EPI_ISL_2539866, EPI_ISL_2539867, EPI_ISL_2539868, EPI_ISL_2539869, EPI_ISL_2539870, EPI_ISL_2539871, EPI_ISL_2539872 | Lighthouse Lab in Milton Keynes | Wellcome Sanger Institute for the COVID-19 Genomics UK (COG-UK) Consortium | The Lighthouse Lab in Milton Keynes and Alex Alderton, Roberto Amato, Jeffrey Barrett, Sonia Goncalves, Ewan Harrison, David K. Jackson, Ian Johnston, Dominic Kwiatkowski, Cordelia Langford, John Sillitoe on behalf of the Wellcome Sanger Institute COVID-19 Surveillance Team |
| EPI_ISL_2539873, EPI_ISL_2539874 | Lighthouse Laboratory Plymouth | Wellcome Sanger Institute for the COVID-19 Genomics UK (COG-UK) Consortium | Lighthouse Laboratory Plymouth and Alex Alderton, Roberto Amato, Jeffrey Barrett, Sonia Goncalves, Ewan Harrison, David K. Jackson, Ian Johnston, Dominic Kwiatkowski, Cordelia Langford, John Sillitoe on behalf of the Wellcome Sanger Institute COVID-19 Surveillance Team |
| EPI_ISL_2539875, EPI_ISL_2539876 | Lighthouse Lab in Milton Keynes | Wellcome Sanger Institute for the COVID-19 Genomics UK (COG-UK) Consortium | The Lighthouse Lab in Milton Keynes and Alex Alderton, Roberto Amato, Jeffrey Barrett, Sonia Goncalves, Ewan Harrison, David K. Jackson, Ian Johnston, Dominic Kwiatkowski, Cordelia Langford, John Sillitoe on behalf of the Wellcome Sanger Institute COVID-19 Surveillance Team |
| EPI_ISL_2539878 | Lighthouse Laboratory Plymouth | Wellcome Sanger Institute for the COVID-19 Genomics UK (COG-UK) Consortium | Lighthouse Laboratory Plymouth and Alex Alderton, Roberto Amato, Jeffrey Barrett, Sonia Goncalves, Ewan Harrison, David K. Jackson, Ian Johnston, Dominic Kwiatkowski, Cordelia Langford, John Sillitoe on behalf of the Wellcome Sanger Institute COVID-19 Surveillance Team |
| EPI_ISL_2539879 | Health Services Laboratories | Wellcome Sanger Institute for the COVID-19 Genomics UK (COG-UK) Consortium | Health Services Laboratories and Alex Alderton, Roberto Amato, Jeffrey Barrett, Sonia Goncalves, Ewan Harrison, David K. Jackson, Ian Johnston, Dominic Kwiatkowski, Cordelia Langford, John Sillitoe on behalf of the Wellcome Sanger Institute COVID-19 Surveillance Team |
| EPI_ISL_2539880, EPI_ISL_2539881 | Lighthouse Lab in Milton Keynes | Wellcome Sanger Institute for the COVID-19 Genomics UK (COG-UK) Consortium | The Lighthouse Lab in Milton Keynes and Alex Alderton, Roberto Amato, Jeffrey Barrett, Sonia Goncalves, Ewan Harrison, David K. Jackson, Ian Johnston, Dominic Kwiatkowski, Cordelia Langford, John Sillitoe on behalf of the Wellcome Sanger Institute COVID-19 Surveillance Team |
| EPI_ISL_2539882, EPI_ISL_2539883, EPI_ISL_2539884 | Lighthouse Laboratory Plymouth | Wellcome Sanger Institute for the COVID-19 Genomics UK (COG-UK) Consortium | Lighthouse Laboratory Plymouth and Alex Alderton, Roberto Amato, Jeffrey Barrett, Sonia Goncalves, Ewan Harrison, David K. Jackson, Ian Johnston, Dominic Kwiatkowski, Cordelia Langford, John Sillitoe on behalf of the Wellcome Sanger Institute COVID-19 Surveillance Team |
| EPI_ISL_2539885 | Lighthouse Lab in Milton Keynes | Wellcome Sanger Institute for the COVID-19 Genomics UK (COG-UK) Consortium | The Lighthouse Lab in Milton Keynes and Alex Alderton, Roberto Amato, Jeffrey Barrett, Sonia Goncalves, Ewan Harrison, David K. Jackson, Ian Johnston, Dominic Kwiatkowski, Cordelia Langford, John Sillitoe on behalf of the Wellcome Sanger Institute COVID-19 Surveillance Team |
| EPI_ISL_2539886 | Health Services Laboratories | Wellcome Sanger Institute for the COVID-19 Genomics UK (COG-UK) Consortium | Health Services Laboratories and Alex Alderton, Roberto Amato, Jeffrey Barrett, Sonia Goncalves, Ewan Harrison, David K. Jackson, Ian Johnston, Dominic Kwiatkowski, Cordelia Langford, John Sillitoe on behalf of the Wellcome Sanger Institute COVID-19 Surveillance Team |
| EPI_ISL_2539887 | Lighthouse Lab in Milton Keynes | Wellcome Sanger Institute for the COVID-19 Genomics UK (COG-UK) Consortium | The Lighthouse Lab in Milton Keynes and Alex Alderton, Roberto Amato, Jeffrey Barrett, Sonia Goncalves, Ewan Harrison, David K. Jackson, Ian Johnston, Dominic Kwiatkowski, Cordelia Langford, John Sillitoe on behalf of the Wellcome Sanger Institute COVID-19 Surveillance Team |
| EPI_ISL_2539888 | Lighthouse Laboratory Plymouth | Wellcome Sanger Institute for the COVID-19 Genomics UK (COG-UK) Consortium | Lighthouse Laboratory Plymouth and Alex Alderton, Roberto Amato, Jeffrey Barrett, Sonia Goncalves, Ewan Harrison, David K. Jackson, Ian Johnston, Dominic Kwiatkowski, Cordelia Langford, John Sillitoe on behalf of the Wellcome Sanger Institute COVID-19 Surveillance Team |
| EPI_ISL_2539889 | Lighthouse Lab in Milton Keynes | Wellcome Sanger Institute for the COVID-19 Genomics UK (COG-UK) Consortium | The Lighthouse Lab in Milton Keynes and Alex Alderton, Roberto Amato, Jeffrey Barrett, Sonia Goncalves, Ewan Harrison, David K. Jackson, Ian Johnston, Dominic Kwiatkowski, Cordelia Langford, John Sillitoe on behalf of the Wellcome Sanger Institute COVID-19 Surveillance Team |
| EPI_ISL_2539890 | Health Services Laboratories | Wellcome Sanger Institute for the COVID-19 Genomics UK (COG-UK) Consortium | Health Services Laboratories and Alex Alderton, Roberto Amato, Jeffrey Barrett, Sonia Goncalves, Ewan Harrison, David K. Jackson, Ian Johnston, Dominic Kwiatkowski, Cordelia Langford, John Sillitoe on behalf of the Wellcome Sanger Institute COVID-19 Surveillance Team |
| EPI_ISL_2539891, EPI_ISL_2539892 | Lighthouse Lab in Milton Keynes | Wellcome Sanger Institute for the COVID-19 Genomics UK (COG-UK) Consortium | The Lighthouse Lab in Milton Keynes and Alex Alderton, Roberto Amato, Jeffrey Barrett, Sonia Goncalves, Ewan Harrison, David K. Jackson, Ian Johnston, Dominic Kwiatkowski, Cordelia Langford, John Sillitoe on behalf of the Wellcome Sanger Institute COVID-19 Surveillance Team |
| EPI_ISL_2539893 | Lighthouse Laboratory Plymouth | Wellcome Sanger Institute for the COVID-19 Genomics UK (COG-UK) Consortium | Lighthouse Laboratory Plymouth and Alex Alderton, Roberto Amato, Jeffrey Barrett, Sonia Goncalves, Ewan Harrison, David K. Jackson, Ian Johnston, Dominic Kwiatkowski, Cordelia Langford, John Sillitoe on behalf of the Wellcome Sanger Institute COVID-19 Surveillance Team |
| EPI_ISL_2539894 | Lighthouse Lab in Milton Keynes | Wellcome Sanger Institute for the COVID-19 Genomics UK (COG-UK) Consortium | The Lighthouse Lab in Milton Keynes and Alex Alderton, Roberto Amato, Jeffrey Barrett, Sonia Goncalves, Ewan Harrison, David K. Jackson, Ian Johnston, Dominic Kwiatkowski, Cordelia Langford, John Sillitoe on behalf of the Wellcome Sanger Institute COVID-19 Surveillance Team |
| EPI_ISL_2539895 | Health Services Laboratories | Wellcome Sanger Institute for the COVID-19 Genomics UK (COG-UK) Consortium | Health Services Laboratories and Alex Alderton, Roberto Amato, Jeffrey Barrett, Sonia Goncalves, Ewan Harrison, David K. Jackson, Ian Johnston, Dominic Kwiatkowski, Cordelia Langford, John Sillitoe on behalf of the Wellcome Sanger Institute COVID-19 Surveillance Team |
| EPI_ISL_2539896 | Lighthouse Laboratory Plymouth | Wellcome Sanger Institute for the COVID-19 Genomics UK (COG-UK) Consortium | Lighthouse Laboratory Plymouth and Alex Alderton, Roberto Amato, Jeffrey Barrett, Sonia Goncalves, Ewan Harrison, David K. Jackson, Ian Johnston, Dominic Kwiatkowski, Cordelia Langford, John Sillitoe on behalf of the Wellcome Sanger Institute COVID-19 Surveillance Team |
| EPI_ISL_2539897 | Lighthouse Lab in Milton Keynes | Wellcome Sanger Institute for the COVID-19 Genomics UK (COG-UK) Consortium | The Lighthouse Lab in Milton Keynes and Alex Alderton, Roberto Amato, Jeffrey Barrett, Sonia Goncalves, Ewan Harrison, David K. Jackson, Ian Johnston, Dominic Kwiatkowski, Cordelia Langford, John Sillitoe on behalf of the Wellcome Sanger Institute COVID-19 Surveillance Team |
| EPI_ISL_2539898 | Health Services Laboratories | Wellcome Sanger Institute for the COVID-19 Genomics UK (COG-UK) Consortium | Health Services Laboratories and Alex Alderton, Roberto Amato, Jeffrey Barrett, Sonia Goncalves, Ewan Harrison, David K. Jackson, Ian Johnston, Dominic Kwiatkowski, Cordelia Langford, John Sillitoe on behalf of the Wellcome Sanger Institute COVID-19 Surveillance Team |
| EPI_ISL_2539899 | Lighthouse Lab in Milton Keynes | Wellcome Sanger Institute for the COVID-19 Genomics UK (COG-UK) Consortium | The Lighthouse Lab in Milton Keynes and Alex Alderton, Roberto Amato, Jeffrey Barrett, Sonia Goncalves, Ewan Harrison, David K. Jackson, Ian Johnston, Dominic Kwiatkowski, Cordelia Langford, John Sillitoe on behalf of the Wellcome Sanger Institute COVID-19 Surveillance Team |
| EPI_ISL_2539900 | Health Services Laboratories | Wellcome Sanger Institute for the COVID-19 Genomics UK (COG-UK) Consortium | Health Services Laboratories and Alex Alderton, Roberto Amato, Jeffrey Barrett, Sonia Goncalves, Ewan Harrison, David K. Jackson, Ian Johnston, Dominic Kwiatkowski, Cordelia Langford, John Sillitoe on behalf of the Wellcome Sanger Institute COVID-19 Surveillance Team |
| EPI_ISL_2539901 | Lighthouse Laboratory Plymouth | Wellcome Sanger Institute for the COVID-19 Genomics UK (COG-UK) Consortium | Lighthouse Laboratory Plymouth and Alex Alderton, Roberto Amato, Jeffrey Barrett, Sonia Goncalves, Ewan Harrison, David K. Jackson, Ian Johnston, Dominic Kwiatkowski, Cordelia Langford, John Sillitoe on behalf of the Wellcome Sanger Institute COVID-19 Surveillance Team |
| EPI_ISL_2539902 | Health Services Laboratories | Wellcome Sanger Institute for the COVID-19 Genomics UK (COG-UK) Consortium | Health Services Laboratories and Alex Alderton, Roberto Amato, Jeffrey Barrett, Sonia Goncalves, Ewan Harrison, David K. Jackson, Ian Johnston, Dominic Kwiatkowski, Cordelia Langford, John Sillitoe on behalf of the Wellcome Sanger Institute COVID-19 Surveillance Team |
| EPI_ISL_2539903 | Lighthouse Lab in Milton Keynes | Wellcome Sanger Institute for the COVID-19 Genomics UK (COG-UK) Consortium | The Lighthouse Lab in Milton Keynes and Alex Alderton, Roberto Amato, Jeffrey Barrett, Sonia Goncalves, Ewan Harrison, David K. Jackson, Ian Johnston, Dominic Kwiatkowski, Cordelia Langford, John Sillitoe on behalf of the Wellcome Sanger Institute COVID-19 Surveillance Team |
| EPI_ISL_2539905 | Lighthouse Laboratory Plymouth | Wellcome Sanger Institute for the COVID-19 Genomics UK (COG-UK) Consortium | Lighthouse Laboratory Plymouth and Alex Alderton, Roberto Amato, Jeffrey Barrett, Sonia Goncalves, Ewan Harrison, David K. Jackson, Ian Johnston, Dominic Kwiatkowski, Cordelia Langford, John Sillitoe on behalf of the Wellcome Sanger Institute COVID-19 Surveillance Team |
| EPI_ISL_2539906 | Lighthouse Lab in Milton Keynes | Wellcome Sanger Institute for the COVID-19 Genomics UK (COG-UK) Consortium | The Lighthouse Lab in Milton Keynes and Alex Alderton, Roberto Amato, Jeffrey Barrett, Sonia Goncalves, Ewan Harrison, David K. Jackson, Ian Johnston, Dominic Kwiatkowski, Cordelia Langford, John Sillitoe on behalf of the Wellcome Sanger Institute COVID-19 Surveillance Team |
| EPI_ISL_2539907 | Health Services Laboratories | Wellcome Sanger Institute for the COVID-19 Genomics UK (COG-UK) Consortium | Health Services Laboratories and Alex Alderton, Roberto Amato, Jeffrey Barrett, Sonia Goncalves, Ewan Harrison, David K. Jackson, Ian Johnston, Dominic Kwiatkowski, Cordelia Langford, John Sillitoe on behalf of the Wellcome Sanger Institute COVID-19 Surveillance Team |
| EPI_ISL_2539908, EPI_ISL_2539911 | Lighthouse Laboratory Plymouth | Wellcome Sanger Institute for the COVID-19 Genomics UK (COG-UK) Consortium | Lighthouse Laboratory Plymouth and Alex Alderton, Roberto Amato, Jeffrey Barrett, Sonia Goncalves, Ewan Harrison, David K. Jackson, Ian Johnston, Dominic Kwiatkowski, Cordelia Langford, John Sillitoe on behalf of the Wellcome Sanger Institute COVID-19 Surveillance Team |
| EPI_ISL_2539912, EPI_ISL_2539913 | Lighthouse Lab in Milton Keynes | Wellcome Sanger Institute for the COVID-19 Genomics UK (COG-UK) Consortium | The Lighthouse Lab in Milton Keynes and Alex Alderton, Roberto Amato, Jeffrey Barrett, Sonia Goncalves, Ewan Harrison, David K. Jackson, Ian Johnston, Dominic Kwiatkowski, Cordelia Langford, John Sillitoe on behalf of the Wellcome Sanger Institute COVID-19 Surveillance Team |
| EPI_ISL_2539914 | Health Services Laboratories | Wellcome Sanger Institute for the COVID-19 Genomics UK (COG-UK) Consortium | Health Services Laboratories and Alex Alderton, Roberto Amato, Jeffrey Barrett, Sonia Goncalves, Ewan Harrison, David K. Jackson, Ian Johnston, Dominic Kwiatkowski, Cordelia Langford, John Sillitoe on behalf of the Wellcome Sanger Institute COVID-19 Surveillance Team |
| EPI_ISL_2539915, EPI_ISL_2539916, EPI_ISL_2539918, EPI_ISL_2539919, EPI_ISL_2539920 | Lighthouse Lab in Milton Keynes | Wellcome Sanger Institute for the COVID-19 Genomics UK (COG-UK) Consortium | The Lighthouse Lab in Milton Keynes and Alex Alderton, Roberto Amato, Jeffrey Barrett, Sonia Goncalves, Ewan Harrison, David K. Jackson, Ian Johnston, Dominic Kwiatkowski, Cordelia Langford, John Sillitoe on behalf of the Wellcome Sanger Institute COVID-19 Surveillance Team |
| EPI_ISL_2539921 | Health Services Laboratories | Wellcome Sanger Institute for the COVID-19 Genomics UK (COG-UK) Consortium | Health Services Laboratories and Alex Alderton, Roberto Amato, Jeffrey Barrett, Sonia Goncalves, Ewan Harrison, David K. Jackson, Ian Johnston, Dominic Kwiatkowski, Cordelia Langford, John Sillitoe on behalf of the Wellcome Sanger Institute COVID-19 Surveillance Team |
| EPI_ISL_2539922, | Lighthouse Lab in Milton Keynes | Wellcome Sanger Institute for the COVID-19 Genomics UK (COG-UK) | The Lighthouse Lab in Milton Keynes and Alex Alderton, Roberto Amato, Jeffrey Barrett, Sonia Goncalves, Ewan Harrison, David K. Jackson, Ian Johnston, Dominic Kwiatkowski, Cordelia Langford, John Sillitoe on behalf of the Wellcome Sanger |

| | | | |
|---|---|---|---|
| EPI_ISL_2539923 | | Consortium | Institute COVID-19 Surveillance Team |
| EPI_ISL_2539924 | Health Services Laboratories | Wellcome Sanger Institute for the COVID-19 Genomics UK (COG-UK) Consortium | Health Services Laboratories and Alex Alderton, Roberto Amato, Jeffrey Barrett, Sonia Goncalves, Ewan Harrison, David K. Jackson, Ian Johnston, Dominic Kwiatkowski, Cordelia Langford, John Sillitoe on behalf of the Wellcome Sanger Institute COVID-19 Surveillance Team |
| EPI_ISL_2539925, EPI_ISL_2539926 | Lighthouse Lab in Milton Keynes | Wellcome Sanger Institute for the COVID-19 Genomics UK (COG-UK) Consortium | The Lighthouse Lab in Milton Keynes and Alex Alderton, Roberto Amato, Jeffrey Barrett, Sonia Goncalves, Ewan Harrison, David K. Jackson, Ian Johnston, Dominic Kwiatkowski, Cordelia Langford, John Sillitoe on behalf of the Wellcome Sanger Institute COVID-19 Surveillance Team |
| EPI_ISL_2539927 | Health Services Laboratories | Wellcome Sanger Institute for the COVID-19 Genomics UK (COG-UK) Consortium | Health Services Laboratories and Alex Alderton, Roberto Amato, Jeffrey Barrett, Sonia Goncalves, Ewan Harrison, David K. Jackson, Ian Johnston, Dominic Kwiatkowski, Cordelia Langford, John Sillitoe on behalf of the Wellcome Sanger Institute COVID-19 Surveillance Team |
| EPI_ISL_2539928 | Lighthouse Lab in Milton Keynes | Wellcome Sanger Institute for the COVID-19 Genomics UK (COG-UK) Consortium | The Lighthouse Lab in Milton Keynes and Alex Alderton, Roberto Amato, Jeffrey Barrett, Sonia Goncalves, Ewan Harrison, David K. Jackson, Ian Johnston, Dominic Kwiatkowski, Cordelia Langford, John Sillitoe on behalf of the Wellcome Sanger Institute COVID-19 Surveillance Team |
| EPI_ISL_2539929 | Lighthouse Laboratory Plymouth | Wellcome Sanger Institute for the COVID-19 Genomics UK (COG-UK) Consortium | Lighthouse Laboratory Plymouth and Alex Alderton, Roberto Amato, Jeffrey Barrett, Sonia Goncalves, Ewan Harrison, David K. Jackson, Ian Johnston, Dominic Kwiatkowski, Cordelia Langford, John Sillitoe on behalf of the Wellcome Sanger Institute COVID-19 Surveillance Team |
| EPI_ISL_2539930 | Lighthouse Lab in Milton Keynes | Wellcome Sanger Institute for the COVID-19 Genomics UK (COG-UK) Consortium | The Lighthouse Lab in Milton Keynes and Alex Alderton, Roberto Amato, Jeffrey Barrett, Sonia Goncalves, Ewan Harrison, David K. Jackson, Ian Johnston, Dominic Kwiatkowski, Cordelia Langford, John Sillitoe on behalf of the Wellcome Sanger Institute COVID-19 Surveillance Team |
| EPI_ISL_2539931 | Health Services Laboratories | Wellcome Sanger Institute for the COVID-19 Genomics UK (COG-UK) Consortium | Health Services Laboratories and Alex Alderton, Roberto Amato, Jeffrey Barrett, Sonia Goncalves, Ewan Harrison, David K. Jackson, Ian Johnston, Dominic Kwiatkowski, Cordelia Langford, John Sillitoe on behalf of the Wellcome Sanger Institute COVID-19 Surveillance Team |
| EPI_ISL_2539932 | Lighthouse Laboratory Plymouth | Wellcome Sanger Institute for the COVID-19 Genomics UK (COG-UK) Consortium | Lighthouse Laboratory Plymouth and Alex Alderton, Roberto Amato, Jeffrey Barrett, Sonia Goncalves, Ewan Harrison, David K. Jackson, Ian Johnston, Dominic Kwiatkowski, Cordelia Langford, John Sillitoe on behalf of the Wellcome Sanger Institute COVID-19 Surveillance Team |
| EPI_ISL_2539933, EPI_ISL_2539934 | Lighthouse Lab in Milton Keynes | Wellcome Sanger Institute for the COVID-19 Genomics UK (COG-UK) Consortium | The Lighthouse Lab in Milton Keynes and Alex Alderton, Roberto Amato, Jeffrey Barrett, Sonia Goncalves, Ewan Harrison, David K. Jackson, Ian Johnston, Dominic Kwiatkowski, Cordelia Langford, John Sillitoe on behalf of the Wellcome Sanger Institute COVID-19 Surveillance Team |
| EPI_ISL_2539935 | Lighthouse Laboratory Plymouth | Wellcome Sanger Institute for the COVID-19 Genomics UK (COG-UK) Consortium | Lighthouse Laboratory Plymouth and Alex Alderton, Roberto Amato, Jeffrey Barrett, Sonia Goncalves, Ewan Harrison, David K. Jackson, Ian Johnston, Dominic Kwiatkowski, Cordelia Langford, John Sillitoe on behalf of the Wellcome Sanger Institute COVID-19 Surveillance Team |
| EPI_ISL_2539936 | Lighthouse Lab in Milton Keynes | Wellcome Sanger Institute for the COVID-19 Genomics UK (COG-UK) Consortium | The Lighthouse Lab in Milton Keynes and Alex Alderton, Roberto Amato, Jeffrey Barrett, Sonia Goncalves, Ewan Harrison, David K. Jackson, Ian Johnston, Dominic Kwiatkowski, Cordelia Langford, John Sillitoe on behalf of the Wellcome Sanger Institute COVID-19 Surveillance Team |
| EPI_ISL_2539937 | Lighthouse Laboratory Plymouth | Wellcome Sanger Institute for the COVID-19 Genomics UK (COG-UK) Consortium | Lighthouse Laboratory Plymouth and Alex Alderton, Roberto Amato, Jeffrey Barrett, Sonia Goncalves, Ewan Harrison, David K. Jackson, Ian Johnston, Dominic Kwiatkowski, Cordelia Langford, John Sillitoe on behalf of the Wellcome Sanger Institute COVID-19 Surveillance Team |
| EPI_ISL_2539938 | Lighthouse Lab in Milton Keynes | Wellcome Sanger Institute for the COVID-19 Genomics UK (COG-UK) Consortium | The Lighthouse Lab in Milton Keynes and Alex Alderton, Roberto Amato, Jeffrey Barrett, Sonia Goncalves, Ewan Harrison, David K. Jackson, Ian Johnston, Dominic Kwiatkowski, Cordelia Langford, John Sillitoe on behalf of the Wellcome Sanger Institute COVID-19 Surveillance Team |
| EPI_ISL_2539939 | Health Services Laboratories | Wellcome Sanger Institute for the COVID-19 Genomics UK (COG-UK) Consortium | Health Services Laboratories and Alex Alderton, Roberto Amato, Jeffrey Barrett, Sonia Goncalves, Ewan Harrison, David K. Jackson, Ian Johnston, Dominic Kwiatkowski, Cordelia Langford, John Sillitoe on behalf of the Wellcome Sanger Institute COVID-19 Surveillance Team |
| EPI_ISL_2539940 | Lighthouse Laboratory Plymouth | Wellcome Sanger Institute for the COVID-19 Genomics UK (COG-UK) Consortium | Lighthouse Laboratory Plymouth and Alex Alderton, Roberto Amato, Jeffrey Barrett, Sonia Goncalves, Ewan Harrison, David K. Jackson, Ian Johnston, Dominic Kwiatkowski, Cordelia Langford, John Sillitoe on behalf of the Wellcome Sanger Institute COVID-19 Surveillance Team |
| EPI_ISL_2539941 | Health Services Laboratories | Wellcome Sanger Institute for the COVID-19 Genomics UK (COG-UK) Consortium | Health Services Laboratories and Alex Alderton, Roberto Amato, Jeffrey Barrett, Sonia Goncalves, Ewan Harrison, David K. Jackson, Ian Johnston, Dominic Kwiatkowski, Cordelia Langford, John Sillitoe on behalf of the Wellcome Sanger Institute COVID-19 Surveillance Team |
| EPI_ISL_2539942 | Lighthouse Lab in Milton Keynes | Wellcome Sanger Institute for the COVID-19 Genomics UK (COG-UK) Consortium | The Lighthouse Lab in Milton Keynes and Alex Alderton, Roberto Amato, Jeffrey Barrett, Sonia Goncalves, Ewan Harrison, David K. Jackson, Ian Johnston, Dominic Kwiatkowski, Cordelia Langford, John Sillitoe on behalf of the Wellcome Sanger Institute COVID-19 Surveillance Team |
| EPI_ISL_2539943 | Health Services Laboratories | Wellcome Sanger Institute for the COVID-19 Genomics UK (COG-UK) Consortium | Health Services Laboratories and Alex Alderton, Roberto Amato, Jeffrey Barrett, Sonia Goncalves, Ewan Harrison, David K. Jackson, Ian Johnston, Dominic Kwiatkowski, Cordelia Langford, John Sillitoe on behalf of the Wellcome Sanger Institute COVID-19 Surveillance Team |
| EPI_ISL_2539944 | Lighthouse Laboratory Plymouth | Wellcome Sanger Institute for the COVID-19 Genomics UK (COG-UK) Consortium | Lighthouse Laboratory Plymouth and Alex Alderton, Roberto Amato, Jeffrey Barrett, Sonia Goncalves, Ewan Harrison, David K. Jackson, Ian Johnston, Dominic Kwiatkowski, Cordelia Langford, John Sillitoe on behalf of the Wellcome Sanger Institute COVID-19 Surveillance Team |
| EPI_ISL_2539945, EPI_ISL_2539946, EPI_ISL_2539947 | Lighthouse Lab in Milton Keynes | Wellcome Sanger Institute for the COVID-19 Genomics UK (COG-UK) Consortium | The Lighthouse Lab in Milton Keynes and Alex Alderton, Roberto Amato, Jeffrey Barrett, Sonia Goncalves, Ewan Harrison, David K. Jackson, Ian Johnston, Dominic Kwiatkowski, Cordelia Langford, John Sillitoe on behalf of the Wellcome Sanger Institute COVID-19 Surveillance Team |
| EPI_ISL_2539948 | Lighthouse Laboratory Plymouth | Wellcome Sanger Institute for the COVID-19 Genomics UK (COG-UK) Consortium | Lighthouse Laboratory Plymouth and Alex Alderton, Roberto Amato, Jeffrey Barrett, Sonia Goncalves, Ewan Harrison, David K. Jackson, Ian Johnston, Dominic Kwiatkowski, Cordelia Langford, John Sillitoe on behalf of the Wellcome Sanger Institute COVID-19 Surveillance Team |
| EPI_ISL_2539950 | Health Services Laboratories | Wellcome Sanger Institute for the COVID-19 Genomics UK (COG-UK) Consortium | Health Services Laboratories and Alex Alderton, Roberto Amato, Jeffrey Barrett, Sonia Goncalves, Ewan Harrison, David K. Jackson, Ian Johnston, Dominic Kwiatkowski, Cordelia Langford, John Sillitoe on behalf of the Wellcome Sanger Institute COVID-19 Surveillance Team |
| EPI_ISL_2539951, EPI_ISL_2539952 | Lighthouse Laboratory Plymouth | Wellcome Sanger Institute for the COVID-19 Genomics UK (COG-UK) Consortium | Lighthouse Laboratory Plymouth and Alex Alderton, Roberto Amato, Jeffrey Barrett, Sonia Goncalves, Ewan Harrison, David K. Jackson, Ian Johnston, Dominic Kwiatkowski, Cordelia Langford, John Sillitoe on behalf of the Wellcome Sanger Institute COVID-19 Surveillance Team |
| EPI_ISL_2539953, EPI_ISL_2539954, EPI_ISL_2539955, EPI_ISL_2539956 | Lighthouse Lab in Milton Keynes | Wellcome Sanger Institute for the COVID-19 Genomics UK (COG-UK) Consortium | The Lighthouse Lab in Milton Keynes and Alex Alderton, Roberto Amato, Jeffrey Barrett, Sonia Goncalves, Ewan Harrison, David K. Jackson, Ian Johnston, Dominic Kwiatkowski, Cordelia Langford, John Sillitoe on behalf of the Wellcome Sanger Institute COVID-19 Surveillance Team |
| EPI_ISL_2539957 | Lighthouse Laboratory Plymouth | Wellcome Sanger Institute for the COVID-19 Genomics UK (COG-UK) Consortium | Lighthouse Laboratory Plymouth and Alex Alderton, Roberto Amato, Jeffrey Barrett, Sonia Goncalves, Ewan Harrison, David K. Jackson, Ian Johnston, Dominic Kwiatkowski, Cordelia Langford, John Sillitoe on behalf of the Wellcome Sanger Institute COVID-19 Surveillance Team |
| EPI_ISL_2539958, EPI_ISL_2539959, EPI_ISL_2539960, EPI_ISL_2539961 | Lighthouse Lab in Milton Keynes | Wellcome Sanger Institute for the COVID-19 Genomics UK (COG-UK) Consortium | The Lighthouse Lab in Milton Keynes and Alex Alderton, Roberto Amato, Jeffrey Barrett, Sonia Goncalves, Ewan Harrison, David K. Jackson, Ian Johnston, Dominic Kwiatkowski, Cordelia Langford, John Sillitoe on behalf of the Wellcome Sanger Institute COVID-19 Surveillance Team |
| EPI_ISL_2539962 | Lighthouse Laboratory Plymouth | Wellcome Sanger Institute for the COVID-19 Genomics UK (COG-UK) Consortium | Lighthouse Laboratory Plymouth and Alex Alderton, Roberto Amato, Jeffrey Barrett, Sonia Goncalves, Ewan Harrison, David K. Jackson, Ian Johnston, Dominic Kwiatkowski, Cordelia Langford, John Sillitoe on behalf of the Wellcome Sanger Institute COVID-19 Surveillance Team |
| EPI_ISL_2539963 | Health Services Laboratories | Wellcome Sanger Institute for the COVID-19 Genomics UK (COG-UK) Consortium | Health Services Laboratories and Alex Alderton, Roberto Amato, Jeffrey Barrett, Sonia Goncalves, Ewan Harrison, David K. Jackson, Ian Johnston, Dominic Kwiatkowski, Cordelia Langford, John Sillitoe on behalf of the Wellcome Sanger Institute COVID-19 Surveillance Team |
| EPI_ISL_2539964 | Lighthouse Lab in Milton Keynes | Wellcome Sanger Institute for the COVID-19 Genomics UK (COG-UK) Consortium | The Lighthouse Lab in Milton Keynes and Alex Alderton, Roberto Amato, Jeffrey Barrett, Sonia Goncalves, Ewan Harrison, David K. Jackson, Ian Johnston, Dominic Kwiatkowski, Cordelia Langford, John Sillitoe on behalf of the Wellcome Sanger Institute COVID-19 Surveillance Team |
| EPI_ISL_2539965 | Lighthouse Laboratory Plymouth | Wellcome Sanger Institute for the COVID-19 Genomics UK (COG-UK) Consortium | Lighthouse Laboratory Plymouth and Alex Alderton, Roberto Amato, Jeffrey Barrett, Sonia Goncalves, Ewan Harrison, David K. Jackson, Ian Johnston, Dominic Kwiatkowski, Cordelia Langford, John Sillitoe on behalf of the Wellcome Sanger Institute COVID-19 Surveillance Team |
| EPI_ISL_2539966 | Lighthouse Lab in Milton Keynes | Wellcome Sanger Institute for the COVID-19 Genomics UK (COG-UK) Consortium | The Lighthouse Lab in Milton Keynes and Alex Alderton, Roberto Amato, Jeffrey Barrett, Sonia Goncalves, Ewan Harrison, David K. Jackson, Ian Johnston, Dominic Kwiatkowski, Cordelia Langford, John Sillitoe on behalf of the Wellcome Sanger Institute COVID-19 Surveillance Team |
| EPI_ISL_2539967 | Lighthouse Laboratory Plymouth | Wellcome Sanger Institute for the COVID-19 Genomics UK (COG-UK) Consortium | Lighthouse Laboratory Plymouth and Alex Alderton, Roberto Amato, Jeffrey Barrett, Sonia Goncalves, Ewan Harrison, David K. Jackson, Ian Johnston, Dominic Kwiatkowski, Cordelia Langford, John Sillitoe on behalf of the Wellcome Sanger Institute COVID-19 Surveillance Team |
| EPI_ISL_2539968 | Lighthouse Lab in Milton Keynes | Wellcome Sanger Institute for the COVID-19 Genomics UK (COG-UK) Consortium | The Lighthouse Lab in Milton Keynes and Alex Alderton, Roberto Amato, Jeffrey Barrett, Sonia Goncalves, Ewan Harrison, David K. Jackson, Ian Johnston, Dominic Kwiatkowski, Cordelia Langford, John Sillitoe on behalf of the Wellcome Sanger Institute COVID-19 Surveillance Team |
| EPI_ISL_2539969 | Health Services Laboratories | Wellcome Sanger Institute for the COVID-19 Genomics UK (COG-UK) Consortium | Health Services Laboratories and Alex Alderton, Roberto Amato, Jeffrey Barrett, Sonia Goncalves, Ewan Harrison, David K. Jackson, Ian Johnston, Dominic Kwiatkowski, Cordelia Langford, John Sillitoe on behalf of the Wellcome Sanger Institute COVID-19 Surveillance Team |
| EPI_ISL_2539970 | Lighthouse Lab in Milton Keynes | Wellcome Sanger Institute for the COVID-19 Genomics UK (COG-UK) Consortium | The Lighthouse Lab in Milton Keynes and Alex Alderton, Roberto Amato, Jeffrey Barrett, Sonia Goncalves, Ewan Harrison, David K. Jackson, Ian Johnston, Dominic Kwiatkowski, Cordelia Langford, John Sillitoe on behalf of the Wellcome Sanger Institute COVID-19 Surveillance Team |
| EPI_ISL_2539971 | Health Services Laboratories | Wellcome Sanger Institute for the COVID-19 Genomics UK (COG-UK) Consortium | Health Services Laboratories and Alex Alderton, Roberto Amato, Jeffrey Barrett, Sonia Goncalves, Ewan Harrison, David K. Jackson, Ian Johnston, Dominic Kwiatkowski, Cordelia Langford, John Sillitoe on behalf of the Wellcome Sanger Institute COVID-19 Surveillance Team |
| EPI_ISL_2539972, EPI_ISL_2539973 | Lighthouse Lab in Milton Keynes | Wellcome Sanger Institute for the COVID-19 Genomics UK (COG-UK) Consortium | The Lighthouse Lab in Milton Keynes and Alex Alderton, Roberto Amato, Jeffrey Barrett, Sonia Goncalves, Ewan Harrison, David K. Jackson, Ian Johnston, Dominic Kwiatkowski, Cordelia Langford, John Sillitoe on behalf of the Wellcome Sanger Institute COVID-19 Surveillance Team |
| EPI_ISL_2539974 | Lighthouse Laboratory Plymouth | Wellcome Sanger Institute for the COVID-19 Genomics UK (COG-UK) Consortium | Lighthouse Laboratory Plymouth and Alex Alderton, Roberto Amato, Jeffrey Barrett, Sonia Goncalves, Ewan Harrison, David K. Jackson, Ian Johnston, Dominic Kwiatkowski, Cordelia Langford, John Sillitoe on behalf of the Wellcome Sanger Institute COVID-19 Surveillance Team |
| EPI_ISL_2539975 | Lighthouse Lab in Milton Keynes | Wellcome Sanger Institute for the COVID-19 Genomics UK (COG-UK) Consortium | The Lighthouse Lab in Milton Keynes and Alex Alderton, Roberto Amato, Jeffrey Barrett, Sonia Goncalves, Ewan Harrison, David K. Jackson, Ian Johnston, Dominic Kwiatkowski, Cordelia Langford, John Sillitoe on behalf of the Wellcome Sanger Institute COVID-19 Surveillance Team |
| EPI_ISL_2539977 | Health Services Laboratories | Wellcome Sanger Institute for the COVID-19 Genomics UK (COG-UK) Consortium | Health Services Laboratories and Alex Alderton, Roberto Amato, Jeffrey Barrett, Sonia Goncalves, Ewan Harrison, David K. Jackson, Ian Johnston, Dominic Kwiatkowski, Cordelia Langford, John Sillitoe on behalf of the Wellcome Sanger Institute COVID-19 Surveillance Team |
| EPI_ISL_2539978 | Lighthouse Lab in Milton Keynes | Wellcome Sanger Institute for the COVID-19 Genomics UK (COG-UK) Consortium | The Lighthouse Lab in Milton Keynes and Alex Alderton, Roberto Amato, Jeffrey Barrett, Sonia Goncalves, Ewan Harrison, David K. Jackson, Ian Johnston, Dominic Kwiatkowski, Cordelia Langford, John Sillitoe on behalf of the Wellcome Sanger Institute COVID-19 Surveillance Team |
| EPI_ISL_2539979 | Health Services Laboratories | Wellcome Sanger Institute for the COVID-19 Genomics UK (COG-UK) Consortium | Health Services Laboratories and Alex Alderton, Roberto Amato, Jeffrey Barrett, Sonia Goncalves, Ewan Harrison, David K. Jackson, Ian Johnston, Dominic Kwiatkowski, Cordelia Langford, John Sillitoe on behalf of the Wellcome Sanger Institute COVID-19 Surveillance Team |
| EPI_ISL_2539980, | Lighthouse Lab in Milton Keynes | Wellcome Sanger Institute for the COVID-19 Genomics UK (COG-UK) | The Lighthouse Lab in Milton Keynes and Alex Alderton, Roberto Amato, Jeffrey Barrett, Sonia Goncalves, Ewan Harrison, David K. Jackson, Ian Johnston, Dominic Kwiatkowski, Cordelia Langford, John Sillitoe on behalf of the Wellcome Sanger |

| | | | |
|---|---|---|---|
| EPI_ISL_2539981 | | Consortium | Institute COVID-19 Surveillance Team |
| EPI_ISL_2539982 | Lighthouse Laboratory Plymouth | Wellcome Sanger Institute for the COVID-19 Genomics UK (COG-UK) Consortium | Lighthouse Laboratory Plymouth and Alex Alderton, Roberto Amato, Jeffrey Barrett, Sonia Goncalves, Ewan Harrison, David K. Jackson, Ian Johnston, Dominic Kwiatkowski, Cordelia Langford, John Sillitoe on behalf of the Wellcome Sanger Institute COVID-19 Surveillance Team |
| EPI_ISL_2539983, EPI_ISL_2539984, EPI_ISL_2539985 | Lighthouse Lab in Milton Keynes | Wellcome Sanger Institute for the COVID-19 Genomics UK (COG-UK) Consortium | The Lighthouse Lab in Milton Keynes and Alex Alderton, Roberto Amato, Jeffrey Barrett, Sonia Goncalves, Ewan Harrison, David K. Jackson, Ian Johnston, Dominic Kwiatkowski, Cordelia Langford, John Sillitoe on behalf of the Wellcome Sanger Institute COVID-19 Surveillance Team |
| EPI_ISL_2539986 | Lighthouse Laboratory Plymouth | Wellcome Sanger Institute for the COVID-19 Genomics UK (COG-UK) Consortium | Lighthouse Laboratory Plymouth and Alex Alderton, Roberto Amato, Jeffrey Barrett, Sonia Goncalves, Ewan Harrison, David K. Jackson, Ian Johnston, Dominic Kwiatkowski, Cordelia Langford, John Sillitoe on behalf of the Wellcome Sanger Institute COVID-19 Surveillance Team |
| EPI_ISL_2539987 | Lighthouse Lab in Milton Keynes | Wellcome Sanger Institute for the COVID-19 Genomics UK (COG-UK) Consortium | The Lighthouse Lab in Milton Keynes and Alex Alderton, Roberto Amato, Jeffrey Barrett, Sonia Goncalves, Ewan Harrison, David K. Jackson, Ian Johnston, Dominic Kwiatkowski, Cordelia Langford, John Sillitoe on behalf of the Wellcome Sanger Institute COVID-19 Surveillance Team |
| EPI_ISL_2539988 | Health Services Laboratories | Wellcome Sanger Institute for the COVID-19 Genomics UK (COG-UK) Consortium | Health Services Laboratories and Alex Alderton, Roberto Amato, Jeffrey Barrett, Sonia Goncalves, Ewan Harrison, David K. Jackson, Ian Johnston, Dominic Kwiatkowski, Cordelia Langford, John Sillitoe on behalf of the Wellcome Sanger Institute COVID-19 Surveillance Team |
| EPI_ISL_2539989 | Lighthouse Laboratory Plymouth | Wellcome Sanger Institute for the COVID-19 Genomics UK (COG-UK) Consortium | Lighthouse Laboratory Plymouth and Alex Alderton, Roberto Amato, Jeffrey Barrett, Sonia Goncalves, Ewan Harrison, David K. Jackson, Ian Johnston, Dominic Kwiatkowski, Cordelia Langford, John Sillitoe on behalf of the Wellcome Sanger Institute COVID-19 Surveillance Team |
| EPI_ISL_2539990 | Health Services Laboratories | Wellcome Sanger Institute for the COVID-19 Genomics UK (COG-UK) Consortium | Health Services Laboratories and Alex Alderton, Roberto Amato, Jeffrey Barrett, Sonia Goncalves, Ewan Harrison, David K. Jackson, Ian Johnston, Dominic Kwiatkowski, Cordelia Langford, John Sillitoe on behalf of the Wellcome Sanger Institute COVID-19 Surveillance Team |
| EPI_ISL_2539991 | Lighthouse Laboratory Plymouth | Wellcome Sanger Institute for the COVID-19 Genomics UK (COG-UK) Consortium | Lighthouse Laboratory Plymouth and Alex Alderton, Roberto Amato, Jeffrey Barrett, Sonia Goncalves, Ewan Harrison, David K. Jackson, Ian Johnston, Dominic Kwiatkowski, Cordelia Langford, John Sillitoe on behalf of the Wellcome Sanger Institute COVID-19 Surveillance Team |
| EPI_ISL_2539992, EPI_ISL_2539993, EPI_ISL_2539994 | Lighthouse Lab in Milton Keynes | Wellcome Sanger Institute for the COVID-19 Genomics UK (COG-UK) Consortium | The Lighthouse Lab in Milton Keynes and Alex Alderton, Roberto Amato, Jeffrey Barrett, Sonia Goncalves, Ewan Harrison, David K. Jackson, Ian Johnston, Dominic Kwiatkowski, Cordelia Langford, John Sillitoe on behalf of the Wellcome Sanger Institute COVID-19 Surveillance Team |
| EPI_ISL_2539995 | Health Services Laboratories | Wellcome Sanger Institute for the COVID-19 Genomics UK (COG-UK) Consortium | Health Services Laboratories and Alex Alderton, Roberto Amato, Jeffrey Barrett, Sonia Goncalves, Ewan Harrison, David K. Jackson, Ian Johnston, Dominic Kwiatkowski, Cordelia Langford, John Sillitoe on behalf of the Wellcome Sanger Institute COVID-19 Surveillance Team |
| EPI_ISL_2539996, EPI_ISL_2539997, EPI_ISL_2539998, EPI_ISL_2539999, EPI_ISL_2540000, EPI_ISL_2540001 | Lighthouse Lab in Milton Keynes | Wellcome Sanger Institute for the COVID-19 Genomics UK (COG-UK) Consortium | The Lighthouse Lab in Milton Keynes and Alex Alderton, Roberto Amato, Jeffrey Barrett, Sonia Goncalves, Ewan Harrison, David K. Jackson, Ian Johnston, Dominic Kwiatkowski, Cordelia Langford, John Sillitoe on behalf of the Wellcome Sanger Institute COVID-19 Surveillance Team |
| EPI_ISL_2540003, EPI_ISL_2540004 | Lighthouse Laboratory Plymouth | Wellcome Sanger Institute for the COVID-19 Genomics UK (COG-UK) Consortium | Lighthouse Laboratory Plymouth and Alex Alderton, Roberto Amato, Jeffrey Barrett, Sonia Goncalves, Ewan Harrison, David K. Jackson, Ian Johnston, Dominic Kwiatkowski, Cordelia Langford, John Sillitoe on behalf of the Wellcome Sanger Institute COVID-19 Surveillance Team |
| EPI_ISL_2540005 | Health Services Laboratories | Wellcome Sanger Institute for the COVID-19 Genomics UK (COG-UK) Consortium | Health Services Laboratories and Alex Alderton, Roberto Amato, Jeffrey Barrett, Sonia Goncalves, Ewan Harrison, David K. Jackson, Ian Johnston, Dominic Kwiatkowski, Cordelia Langford, John Sillitoe on behalf of the Wellcome Sanger Institute COVID-19 Surveillance Team |
| EPI_ISL_2540006, EPI_ISL_2540007, EPI_ISL_2540008, EPI_ISL_2540009, EPI_ISL_2540010, EPI_ISL_2540011, EPI_ISL_2540012, EPI_ISL_2540013, EPI_ISL_2540014 | Lighthouse Lab in Milton Keynes | Wellcome Sanger Institute for the COVID-19 Genomics UK (COG-UK) Consortium | The Lighthouse Lab in Milton Keynes and Alex Alderton, Roberto Amato, Jeffrey Barrett, Sonia Goncalves, Ewan Harrison, David K. Jackson, Ian Johnston, Dominic Kwiatkowski, Cordelia Langford, John Sillitoe on behalf of the Wellcome Sanger Institute COVID-19 Surveillance Team |
| EPI_ISL_2540015 | Lighthouse Laboratory Plymouth | Wellcome Sanger Institute for the COVID-19 Genomics UK (COG-UK) Consortium | Lighthouse Laboratory Plymouth and Alex Alderton, Roberto Amato, Jeffrey Barrett, Sonia Goncalves, Ewan Harrison, David K. Jackson, Ian Johnston, Dominic Kwiatkowski, Cordelia Langford, John Sillitoe on behalf of the Wellcome Sanger Institute COVID-19 Surveillance Team |
| EPI_ISL_2540016 | Health Services Laboratories | Wellcome Sanger Institute for the COVID-19 Genomics UK (COG-UK) Consortium | Health Services Laboratories and Alex Alderton, Roberto Amato, Jeffrey Barrett, Sonia Goncalves, Ewan Harrison, David K. Jackson, Ian Johnston, Dominic Kwiatkowski, Cordelia Langford, John Sillitoe on behalf of the Wellcome Sanger Institute COVID-19 Surveillance Team |
| EPI_ISL_2540017, EPI_ISL_2540019, EPI_ISL_2540020, EPI_ISL_2540021, EPI_ISL_2540022, EPI_ISL_2540023 | Lighthouse Lab in Milton Keynes | Wellcome Sanger Institute for the COVID-19 Genomics UK (COG-UK) Consortium | The Lighthouse Lab in Milton Keynes and Alex Alderton, Roberto Amato, Jeffrey Barrett, Sonia Goncalves, Ewan Harrison, David K. Jackson, Ian Johnston, Dominic Kwiatkowski, Cordelia Langford, John Sillitoe on behalf of the Wellcome Sanger Institute COVID-19 Surveillance Team |
| EPI_ISL_2540024 | Lighthouse Laboratory Plymouth | Wellcome Sanger Institute for the COVID-19 Genomics UK (COG-UK) Consortium | Lighthouse Laboratory Plymouth and Alex Alderton, Roberto Amato, Jeffrey Barrett, Sonia Goncalves, Ewan Harrison, David K. Jackson, Ian Johnston, Dominic Kwiatkowski, Cordelia Langford, John Sillitoe on behalf of the Wellcome Sanger Institute COVID-19 Surveillance Team |
| EPI_ISL_2540025, EPI_ISL_2540027, EPI_ISL_2540028 | Lighthouse Lab in Milton Keynes | Wellcome Sanger Institute for the COVID-19 Genomics UK (COG-UK) Consortium | The Lighthouse Lab in Milton Keynes and Alex Alderton, Roberto Amato, Jeffrey Barrett, Sonia Goncalves, Ewan Harrison, David K. Jackson, Ian Johnston, Dominic Kwiatkowski, Cordelia Langford, John Sillitoe on behalf of the Wellcome Sanger Institute COVID-19 Surveillance Team |
| EPI_ISL_2540030, EPI_ISL_2540031 | Lighthouse Laboratory Plymouth | Wellcome Sanger Institute for the COVID-19 Genomics UK (COG-UK) Consortium | Lighthouse Laboratory Plymouth and Alex Alderton, Roberto Amato, Jeffrey Barrett, Sonia Goncalves, Ewan Harrison, David K. Jackson, Ian Johnston, Dominic Kwiatkowski, Cordelia Langford, John Sillitoe on behalf of the Wellcome Sanger Institute COVID-19 Surveillance Team |
| EPI_ISL_2540032, EPI_ISL_2540033, EPI_ISL_2540034, EPI_ISL_2540035, EPI_ISL_2540036, EPI_ISL_2540037, EPI_ISL_2540038, EPI_ISL_2540039, EPI_ISL_2540042, EPI_ISL_2540043, EPI_ISL_2540044, EPI_ISL_2540045, EPI_ISL_2540046, EPI_ISL_2540047, EPI_ISL_2540048, EPI_ISL_2540049, EPI_ISL_2540050, EPI_ISL_2540051, EPI_ISL_2540052, EPI_ISL_2540053, EPI_ISL_2540055, EPI_ISL_2540056, EPI_ISL_2540057, EPI_ISL_2540059, EPI_ISL_2540060, EPI_ISL_2540061, EPI_ISL_2540062, EPI_ISL_2540063, EPI_ISL_2540064, EPI_ISL_2540065, EPI_ISL_2540066, EPI_ISL_2540067, EPI_ISL_2540068, EPI_ISL_2540069, EPI_ISL_2540070, EPI_ISL_2540071, EPI_ISL_2540072, EPI_ISL_2540073, EPI_ISL_2540074, EPI_ISL_2540075, EPI_ISL_2540076, EPI_ISL_2540077, EPI_ISL_2540078, EPI_ISL_2540079, EPI_ISL_2540080, EPI_ISL_2540081, EPI_ISL_2540082, EPI_ISL_2540083, EPI_ISL_2540085, EPI_ISL_2540086, EPI_ISL_2540088, EPI_ISL_2540091, EPI_ISL_2540092, EPI_ISL_2540093, EPI_ISL_2540094, EPI_ISL_2540095, EPI_ISL_2540097, EPI_ISL_2540098, EPI_ISL_2540099, EPI_ISL_2540100, EPI_ISL_2540101, EPI_ISL_2540102, EPI_ISL_2540103, EPI_ISL_2540104, EPI_ISL_2540105, EPI_ISL_2540106, EPI_ISL_2540107, EPI_ISL_2540108, EPI_ISL_2540110, EPI_ISL_2540111, EPI_ISL_2540112, EPI_ISL_2540113, EPI_ISL_2540114, EPI_ISL_2540115, EPI_ISL_2540116, EPI_ISL_2540117, EPI_ISL_2540119, EPI_ISL_2540120, EPI_ISL_2540121, EPI_ISL_2540122, EPI_ISL_2540124, EPI_ISL_2540125, EPI_ISL_2540127, EPI_ISL_2540128, EPI_ISL_2540129, EPI_ISL_2540130, EPI_ISL_2540131, EPI_ISL_2540132, EPI_ISL_2540133, EPI_ISL_2540134, EPI_ISL_2540135, EPI_ISL_2540136, EPI_ISL_2540137, EPI_ISL_2540139, EPI_ISL_2540140, EPI_ISL_2540141, EPI_ISL_2540142, EPI_ISL_2540143, EPI_ISL_2540144, EPI_ISL_2540146, EPI_ISL_2540147, EPI_ISL_2540148, EPI_ISL_2540149, EPI_ISL_2540150, EPI_ISL_2540151, EPI_ISL_2540152, EPI_ISL_2540153, EPI_ISL_2540154, EPI_ISL_2540155, EPI_ISL_2540156, EPI_ISL_2540157, EPI_ISL_2540158, EPI_ISL_2540159, EPI_ISL_2540161, EPI_ISL_2540162, EPI_ISL_2540163, EPI_ISL_2540164, EPI_ISL_2540166, EPI_ISL_2540167, EPI_ISL_2540168, EPI_ISL_2540171, EPI_ISL_2540173, EPI_ISL_2540174, EPI_ISL_2540175, EPI_ISL_2540176, EPI_ISL_2540177, EPI_ISL_2540178, EPI_ISL_2540179, EPI_ISL_2540180, EPI_ISL_2540181, EPI_ISL_2540183, EPI_ISL_2540184, EPI_ISL_2540185, EPI_ISL_2540186, EPI_ISL_2540187, EPI_ISL_2540188, EPI_ISL_2540189, EPI_ISL_2540190, EPI_ISL_2540191, EPI_ISL_2540192, EPI_ISL_2540193, EPI_ISL_2540194, EPI_ISL_2540195, EPI_ISL_2540196, EPI_ISL_2540197, EPI_ISL_2540198, EPI_ISL_2540199, EPI_ISL_2540200, EPI_ISL_2540202, EPI_ISL_2540203, EPI_ISL_2540204, EPI_ISL_2540205, EPI_ISL_2540206, EPI_ISL_2540207, EPI_ISL_2540208, EPI_ISL_2540209, EPI_ISL_2540210, EPI_ISL_2540211, EPI_ISL_2540212, EPI_ISL_2540213, EPI_ISL_2540214, EPI_ISL_2540215, EPI_ISL_2540216, EPI_ISL_2540217, EPI_ISL_2540218, EPI_ISL_2540219, EPI_ISL_2540221, EPI_ISL_2540222, EPI_ISL_2540224, EPI_ISL_2540225, EPI_ISL_2540227, EPI_ISL_2540228, EPI_ISL_2540229, EPI_ISL_2540231, EPI_ISL_2540232, EPI_ISL_2540233, EPI_ISL_2540234, EPI_ISL_2540235, EPI_ISL_2540236, EPI_ISL_2540237, EPI_ISL_2540238, EPI_ISL_2540239, EPI_ISL_2540240, EPI_ISL_2540241, EPI_ISL_2540242, EPI_ISL_2540243, EPI_ISL_2540244, EPI_ISL_2540245, EPI_ISL_2540247, EPI_ISL_2540248, EPI_ISL_2540251, EPI_ISL_2540252, EPI_ISL_2540253, EPI_ISL_2540254, EPI_ISL_2540255, EPI_ISL_2540256, EPI_ISL_2540257, EPI_ISL_2540258, EPI_ISL_2540259, EPI_ISL_2540260, EPI_ISL_2540261, EPI_ISL_2540262, EPI_ISL_2540263, EPI_ISL_2540264, EPI_ISL_2540265, EPI_ISL_2540266, EPI_ISL_2540267, EPI_ISL_2540268, EPI_ISL_2540269, EPI_ISL_2540270, EPI_ISL_2540272, EPI_ISL_2540273, EPI_ISL_2540274, EPI_ISL_2540276, EPI_ISL_2540277, EPI_ISL_2540278, EPI_ISL_2540279, EPI_ISL_2540280, EPI_ISL_2540281, EPI_ISL_2540283, EPI_ISL_2540284, EPI_ISL_2540285, EPI_ISL_2540286, EPI_ISL_2540287, EPI_ISL_2540288, EPI_ISL_2540289, EPI_ISL_2540290, EPI_ISL_2540291, EPI_ISL_2540293, EPI_ISL_2540294, EPI_ISL_2540295, EPI_ISL_2540297, EPI_ISL_2540301, EPI_ISL_2540302, EPI_ISL_2540303, EPI_ISL_2540304, EPI_ISL_2540305, EPI_ISL_2540306, EPI_ISL_2540307, EPI_ISL_2540308, EPI_ISL_2540309, EPI_ISL_2540310, EPI_ISL_2540312, EPI_ISL_2540313, EPI_ISL_2540314, EPI_ISL_2540315, EPI_ISL_2540316, EPI_ISL_2540317, EPI_ISL_2540318, EPI_ISL_2540319, EPI_ISL_2540321, EPI_ISL_2540322, EPI_ISL_2540323, EPI_ISL_2540324, EPI_ISL_2540325, EPI_ISL_2540326, EPI_ISL_2540327, EPI_ISL_2540328, EPI_ISL_2540329, EPI_ISL_2540330, EPI_ISL_2540331, EPI_ISL_2540332, EPI_ISL_2540333, EPI_ISL_2540334, EPI_ISL_2540335, EPI_ISL_2540336, EPI_ISL_2540337, EPI_ISL_2540338, EPI_ISL_2540339, EPI_ISL_2540340, EPI_ISL_2540341, EPI_ISL_2540342, EPI_ISL_2540343, EPI_ISL_2540344, EPI_ISL_2540345, EPI_ISL_2540346, EPI_ISL_2540347, EPI_ISL_2540348, EPI_ISL_2540349, EPI_ISL_2540350, EPI_ISL_2540351, EPI_ISL_2540352, EPI_ISL_2540353, EPI_ISL_2540354, EPI_ISL_2540355, EPI_ISL_2540357, EPI_ISL_2540358, EPI_ISL_2540359, EPI_ISL_2540360, EPI_ISL_2540361, EPI_ISL_2540362, EPI_ISL_2540363, EPI_ISL_2540364, EPI_ISL_2540365, EPI_ISL_2540366, EPI_ISL_2540368, EPI_ISL_2540369, EPI_ISL_2540370, EPI_ISL_2540371, EPI_ISL_2540372 | | | |
| see above | Lighthouse Lab in Milton Keynes | Wellcome Sanger Institute for the COVID-19 Genomics UK (COG-UK) Consortium | The Lighthouse Lab in Milton Keynes and Alex Alderton, Roberto Amato, Jeffrey Barrett, Sonia Goncalves, Ewan Harrison, David K. Jackson, Ian Johnston, Dominic Kwiatkowski, Cordelia Langford, John Sillitoe on behalf of the Wellcome Sanger Institute COVID-19 Surveillance Team |
| EPI_ISL_2540373 | Lighthouse Lab in Alderley Park | Wellcome Sanger Institute for the COVID-19 Genomics UK (COG-UK) Consortium | Jacquelyn Wynn, Mairead Hyland, The Lighthouse Lab in Alderley Park and Alex Alderton, Roberto Amato, Jeffrey Barrett, Sonia Goncalves, Ewan Harrison, David K. Jackson, Ian Johnston, Dominic Kwiatkowski, Cordelia Langford, John Sillitoe on behalf of the Wellcome Sanger Institute COVID-19 Surveillance Team |
| EPI_ISL_2540374, EPI_ISL_2540375 | Lighthouse Lab in Glasgow | Wellcome Sanger Institute for the COVID-19 Genomics UK (COG-UK) Consortium | Harper VanSteenhouse, Yumi Kasai, David Gray, Carol Clugston, Anna Dominiczak and Alex Alderton, Roberto Amato, Jeffrey Barrett, Sonia Goncalves, Ewan Harrison, David K. Jackson, Ian Johnston, Dominic Kwiatkowski, Cordelia Langford, John Sillitoe on behalf of the Wellcome Sanger Institute COVID-19 Surveillance Team |
| EPI_ISL_2540376, EPI_ISL_2540377, EPI_ISL_2540378 | Lighthouse Lab in Alderley Park | Wellcome Sanger Institute for the COVID-19 Genomics UK (COG-UK) Consortium | Jacquelyn Wynn, Mairead Hyland, The Lighthouse Lab in Alderley Park and Alex Alderton, Roberto Amato, Jeffrey Barrett, Sonia Goncalves, Ewan Harrison, David K. Jackson, Ian Johnston, Dominic Kwiatkowski, Cordelia Langford, John Sillitoe on behalf of the Wellcome Sanger Institute COVID-19 Surveillance Team |
| EPI_ISL_2540380 | Lighthouse Lab in Glasgow | Wellcome Sanger Institute for the COVID-19 Genomics UK (COG-UK) Consortium | Harper VanSteenhouse, Yumi Kasai, David Gray, Carol Clugston, Anna Dominiczak and Alex Alderton, Roberto Amato, Jeffrey Barrett, Sonia Goncalves, Ewan Harrison, David K. Jackson, Ian Johnston, Dominic Kwiatkowski, Cordelia Langford, John Sillitoe on behalf of the Wellcome Sanger Institute COVID-19 Surveillance Team |
| EPI_ISL_2540381 | Lighthouse Lab in Alderley Park | Wellcome Sanger Institute for the COVID-19 Genomics UK (COG-UK) Consortium | Jacquelyn Wynn, Mairead Hyland, The Lighthouse Lab in Alderley Park and Alex Alderton, Roberto Amato, Jeffrey Barrett, Sonia Goncalves, Ewan Harrison, David K. Jackson, Ian Johnston, Dominic Kwiatkowski, Cordelia Langford, John Sillitoe on behalf of the Wellcome Sanger Institute COVID-19 Surveillance Team |
| EPI_ISL_2540382 | Lighthouse Lab in Glasgow | Wellcome Sanger Institute for the COVID-19 Genomics UK (COG-UK) Consortium | Harper VanSteenhouse, Yumi Kasai, David Gray, Carol Clugston, Anna Dominiczak and Alex Alderton, Roberto Amato, Jeffrey Barrett, Sonia Goncalves, Ewan Harrison, David K. Jackson, Ian Johnston, Dominic Kwiatkowski, Cordelia Langford, John Sillitoe on behalf of the Wellcome Sanger Institute COVID-19 Surveillance Team |
| EPI_ISL_2540383, EPI_ISL_2540385, EPI_ISL_2540386, EPI_ISL_2540387 | Lighthouse Lab in Alderley Park | Wellcome Sanger Institute for the COVID-19 Genomics UK (COG-UK) Consortium | Jacquelyn Wynn, Mairead Hyland, The Lighthouse Lab in Alderley Park and Alex Alderton, Roberto Amato, Jeffrey Barrett, Sonia Goncalves, Ewan Harrison, David K. Jackson, Ian Johnston, Dominic Kwiatkowski, Cordelia Langford, John Sillitoe on behalf of the Wellcome Sanger Institute COVID-19 Surveillance Team |
| EPI_ISL_2540389 | Lighthouse Lab in Glasgow | Wellcome Sanger Institute for the COVID-19 Genomics UK (COG-UK) Consortium | Harper VanSteenhouse, Yumi Kasai, David Gray, Carol Clugston, Anna Dominiczak and Alex Alderton, Roberto Amato, Jeffrey Barrett, Sonia Goncalves, Ewan Harrison, David K. Jackson, Ian Johnston, Dominic Kwiatkowski, Cordelia Langford, John Sillitoe on behalf of the Wellcome Sanger Institute COVID-19 Surveillance Team |
| EPI_ISL_2540390, EPI_ISL_2540391, | Lighthouse Lab in Alderley Park | Wellcome Sanger Institute for the COVID-19 Genomics UK (COG-UK) Consortium | Jacquelyn Wynn, Mairead Hyland, The Lighthouse Lab in Alderley Park and Alex Alderton, Roberto Amato, Jeffrey Barrett, Sonia Goncalves, Ewan Harrison, David K. Jackson, Ian Johnston, Dominic Kwiatkowski, Cordelia Langford, John Sillitoe on behalf of the Wellcome Sanger Institute COVID-19 Surveillance Team |

| | | | |
|---|---|---|---|
| EPI_ISL_2540392 | | | |
| EPI_ISL_2540393 | Lighthouse Lab in Glasgow | Wellcome Sanger Institute for the COVID-19 Genomics UK (COG-UK) Consortium | Harper VanSteenhouse, Yumi Kasai, David Gray, Carol Clugston, Anna Dominiczak and Alex Alderton, Roberto Amato, Jeffrey Barrett, Sonia Goncalves, Ewan Harrison, David K. Jackson, Ian Johnston, Dominic Kwiatkowski, Cordelia Langford, John Sillitoe on behalf of the Wellcome Sanger Institute COVID-19 Surveillance Team |
| EPI_ISL_2540394, EPI_ISL_2540395, EPI_ISL_2540396, EPI_ISL_2540397 | Lighthouse Lab in Alderley Park | Wellcome Sanger Institute for the COVID-19 Genomics UK (COG-UK) Consortium | Jacquelyn Wynn, Mairead Hyland, The Lighthouse Lab in Alderley Park and Alex Alderton, Roberto Amato, Jeffrey Barrett, Sonia Goncalves, Ewan Harrison, David K. Jackson, Ian Johnston, Dominic Kwiatkowski, Cordelia Langford, John Sillitoe on behalf of the Wellcome Sanger Institute COVID-19 Surveillance Team |
| EPI_ISL_2540398 | Lighthouse Lab in Glasgow | Wellcome Sanger Institute for the COVID-19 Genomics UK (COG-UK) Consortium | Harper VanSteenhouse, Yumi Kasai, David Gray, Carol Clugston, Anna Dominiczak and Alex Alderton, Roberto Amato, Jeffrey Barrett, Sonia Goncalves, Ewan Harrison, David K. Jackson, Ian Johnston, Dominic Kwiatkowski, Cordelia Langford, John Sillitoe on behalf of the Wellcome Sanger Institute COVID-19 Surveillance Team |
| EPI_ISL_2540399 | Lighthouse Lab in Alderley Park | Wellcome Sanger Institute for the COVID-19 Genomics UK (COG-UK) Consortium | Jacquelyn Wynn, Mairead Hyland, The Lighthouse Lab in Alderley Park and Alex Alderton, Roberto Amato, Jeffrey Barrett, Sonia Goncalves, Ewan Harrison, David K. Jackson, Ian Johnston, Dominic Kwiatkowski, Cordelia Langford, John Sillitoe on behalf of the Wellcome Sanger Institute COVID-19 Surveillance Team |
| EPI_ISL_2540400 | Lighthouse Lab in Glasgow | Wellcome Sanger Institute for the COVID-19 Genomics UK (COG-UK) Consortium | Harper VanSteenhouse, Yumi Kasai, David Gray, Carol Clugston, Anna Dominiczak and Alex Alderton, Roberto Amato, Jeffrey Barrett, Sonia Goncalves, Ewan Harrison, David K. Jackson, Ian Johnston, Dominic Kwiatkowski, Cordelia Langford, John Sillitoe on behalf of the Wellcome Sanger Institute COVID-19 Surveillance Team |
| EPI_ISL_2540401 | Lighthouse Lab in Alderley Park | Wellcome Sanger Institute for the COVID-19 Genomics UK (COG-UK) Consortium | Jacquelyn Wynn, Mairead Hyland, The Lighthouse Lab in Alderley Park and Alex Alderton, Roberto Amato, Jeffrey Barrett, Sonia Goncalves, Ewan Harrison, David K. Jackson, Ian Johnston, Dominic Kwiatkowski, Cordelia Langford, John Sillitoe on behalf of the Wellcome Sanger Institute COVID-19 Surveillance Team |
| EPI_ISL_2540402 | Lighthouse Lab in Glasgow | Wellcome Sanger Institute for the COVID-19 Genomics UK (COG-UK) Consortium | Harper VanSteenhouse, Yumi Kasai, David Gray, Carol Clugston, Anna Dominiczak and Alex Alderton, Roberto Amato, Jeffrey Barrett, Sonia Goncalves, Ewan Harrison, David K. Jackson, Ian Johnston, Dominic Kwiatkowski, Cordelia Langford, John Sillitoe on behalf of the Wellcome Sanger Institute COVID-19 Surveillance Team |
| EPI_ISL_2540403, EPI_ISL_2540404 | Lighthouse Lab in Alderley Park | Wellcome Sanger Institute for the COVID-19 Genomics UK (COG-UK) Consortium | Jacquelyn Wynn, Mairead Hyland, The Lighthouse Lab in Alderley Park and Alex Alderton, Roberto Amato, Jeffrey Barrett, Sonia Goncalves, Ewan Harrison, David K. Jackson, Ian Johnston, Dominic Kwiatkowski, Cordelia Langford, John Sillitoe on behalf of the Wellcome Sanger Institute COVID-19 Surveillance Team |
| EPI_ISL_2540405, EPI_ISL_2540406 | Lighthouse Lab in Glasgow | Wellcome Sanger Institute for the COVID-19 Genomics UK (COG-UK) Consortium | Harper VanSteenhouse, Yumi Kasai, David Gray, Carol Clugston, Anna Dominiczak and Alex Alderton, Roberto Amato, Jeffrey Barrett, Sonia Goncalves, Ewan Harrison, David K. Jackson, Ian Johnston, Dominic Kwiatkowski, Cordelia Langford, John Sillitoe on behalf of the Wellcome Sanger Institute COVID-19 Surveillance Team |
| EPI_ISL_2540407, EPI_ISL_2540408, EPI_ISL_2540410, EPI_ISL_2540411, EPI_ISL_2540412, EPI_ISL_2540413, EPI_ISL_2540414 | Lighthouse Lab in Alderley Park | Wellcome Sanger Institute for the COVID-19 Genomics UK (COG-UK) Consortium | Jacquelyn Wynn, Mairead Hyland, The Lighthouse Lab in Alderley Park and Alex Alderton, Roberto Amato, Jeffrey Barrett, Sonia Goncalves, Ewan Harrison, David K. Jackson, Ian Johnston, Dominic Kwiatkowski, Cordelia Langford, John Sillitoe on behalf of the Wellcome Sanger Institute COVID-19 Surveillance Team |
| EPI_ISL_2540415, EPI_ISL_2540416 | Lighthouse Lab in Glasgow | Wellcome Sanger Institute for the COVID-19 Genomics UK (COG-UK) Consortium | Harper VanSteenhouse, Yumi Kasai, David Gray, Carol Clugston, Anna Dominiczak and Alex Alderton, Roberto Amato, Jeffrey Barrett, Sonia Goncalves, Ewan Harrison, David K. Jackson, Ian Johnston, Dominic Kwiatkowski, Cordelia Langford, John Sillitoe on behalf of the Wellcome Sanger Institute COVID-19 Surveillance Team |
| EPI_ISL_2540417, EPI_ISL_2540418, EPI_ISL_2540419, EPI_ISL_2540420 | Lighthouse Lab in Alderley Park | Wellcome Sanger Institute for the COVID-19 Genomics UK (COG-UK) Consortium | Jacquelyn Wynn, Mairead Hyland, The Lighthouse Lab in Alderley Park and Alex Alderton, Roberto Amato, Jeffrey Barrett, Sonia Goncalves, Ewan Harrison, David K. Jackson, Ian Johnston, Dominic Kwiatkowski, Cordelia Langford, John Sillitoe on behalf of the Wellcome Sanger Institute COVID-19 Surveillance Team |
| EPI_ISL_2540421 | Lighthouse Lab in Glasgow | Wellcome Sanger Institute for the COVID-19 Genomics UK (COG-UK) Consortium | Harper VanSteenhouse, Yumi Kasai, David Gray, Carol Clugston, Anna Dominiczak and Alex Alderton, Roberto Amato, Jeffrey Barrett, Sonia Goncalves, Ewan Harrison, David K. Jackson, Ian Johnston, Dominic Kwiatkowski, Cordelia Langford, John Sillitoe on behalf of the Wellcome Sanger Institute COVID-19 Surveillance Team |
| EPI_ISL_2540422, EPI_ISL_2540423, EPI_ISL_2540424, EPI_ISL_2540425, EPI_ISL_2540426, EPI_ISL_2540427, EPI_ISL_2540428, EPI_ISL_2540429, EPI_ISL_2540430, EPI_ISL_2540431, EPI_ISL_2540432, EPI_ISL_2540433, EPI_ISL_2540434 | | | |
| see above | Lighthouse Lab in Alderley Park | Wellcome Sanger Institute for the COVID-19 Genomics UK (COG-UK) Consortium | Jacquelyn Wynn, Mairead Hyland, The Lighthouse Lab in Alderley Park and Alex Alderton, Roberto Amato, Jeffrey Barrett, Sonia Goncalves, Ewan Harrison, David K. Jackson, Ian Johnston, Dominic Kwiatkowski, Cordelia Langford, John Sillitoe on behalf of the Wellcome Sanger Institute COVID-19 Surveillance Team |
| EPI_ISL_2540435 | Lighthouse Lab in Glasgow | Wellcome Sanger Institute for the COVID-19 Genomics UK (COG-UK) Consortium | Harper VanSteenhouse, Yumi Kasai, David Gray, Carol Clugston, Anna Dominiczak and Alex Alderton, Roberto Amato, Jeffrey Barrett, Sonia Goncalves, Ewan Harrison, David K. Jackson, Ian Johnston, Dominic Kwiatkowski, Cordelia Langford, John Sillitoe on behalf of the Wellcome Sanger Institute COVID-19 Surveillance Team |
| EPI_ISL_2540436, EPI_ISL_2540437 | Lighthouse Lab in Alderley Park | Wellcome Sanger Institute for the COVID-19 Genomics UK (COG-UK) Consortium | Jacquelyn Wynn, Mairead Hyland, The Lighthouse Lab in Alderley Park and Alex Alderton, Roberto Amato, Jeffrey Barrett, Sonia Goncalves, Ewan Harrison, David K. Jackson, Ian Johnston, Dominic Kwiatkowski, Cordelia Langford, John Sillitoe on behalf of the Wellcome Sanger Institute COVID-19 Surveillance Team |
| EPI_ISL_2540438 | Lighthouse Lab in Glasgow | Wellcome Sanger Institute for the COVID-19 Genomics UK (COG-UK) Consortium | Harper VanSteenhouse, Yumi Kasai, David Gray, Carol Clugston, Anna Dominiczak and Alex Alderton, Roberto Amato, Jeffrey Barrett, Sonia Goncalves, Ewan Harrison, David K. Jackson, Ian Johnston, Dominic Kwiatkowski, Cordelia Langford, John Sillitoe on behalf of the Wellcome Sanger Institute COVID-19 Surveillance Team |
| EPI_ISL_2540439 | Lighthouse Lab in Alderley Park | Wellcome Sanger Institute for the COVID-19 Genomics UK (COG-UK) Consortium | Jacquelyn Wynn, Mairead Hyland, The Lighthouse Lab in Alderley Park and Alex Alderton, Roberto Amato, Jeffrey Barrett, Sonia Goncalves, Ewan Harrison, David K. Jackson, Ian Johnston, Dominic Kwiatkowski, Cordelia Langford, John Sillitoe on behalf of the Wellcome Sanger Institute COVID-19 Surveillance Team |
| EPI_ISL_2540440 | Lighthouse Lab in Glasgow | Wellcome Sanger Institute for the COVID-19 Genomics UK (COG-UK) Consortium | Harper VanSteenhouse, Yumi Kasai, David Gray, Carol Clugston, Anna Dominiczak and Alex Alderton, Roberto Amato, Jeffrey Barrett, Sonia Goncalves, Ewan Harrison, David K. Jackson, Ian Johnston, Dominic Kwiatkowski, Cordelia Langford, John Sillitoe on behalf of the Wellcome Sanger Institute COVID-19 Surveillance Team |
| EPI_ISL_2540441 | Lighthouse Lab in Alderley Park | Wellcome Sanger Institute for the COVID-19 Genomics UK (COG-UK) Consortium | Jacquelyn Wynn, Mairead Hyland, The Lighthouse Lab in Alderley Park and Alex Alderton, Roberto Amato, Jeffrey Barrett, Sonia Goncalves, Ewan Harrison, David K. Jackson, Ian Johnston, Dominic Kwiatkowski, Cordelia Langford, John Sillitoe on behalf of the Wellcome Sanger Institute COVID-19 Surveillance Team |
| EPI_ISL_2540442 | Lighthouse Lab in Glasgow | Wellcome Sanger Institute for the COVID-19 Genomics UK (COG-UK) Consortium | Harper VanSteenhouse, Yumi Kasai, David Gray, Carol Clugston, Anna Dominiczak and Alex Alderton, Roberto Amato, Jeffrey Barrett, Sonia Goncalves, Ewan Harrison, David K. Jackson, Ian Johnston, Dominic Kwiatkowski, Cordelia Langford, John Sillitoe on behalf of the Wellcome Sanger Institute COVID-19 Surveillance Team |
| EPI_ISL_2540443, EPI_ISL_2540444, EPI_ISL_2540445 | Lighthouse Lab in Alderley Park | Wellcome Sanger Institute for the COVID-19 Genomics UK (COG-UK) Consortium | Jacquelyn Wynn, Mairead Hyland, The Lighthouse Lab in Alderley Park and Alex Alderton, Roberto Amato, Jeffrey Barrett, Sonia Goncalves, Ewan Harrison, David K. Jackson, Ian Johnston, Dominic Kwiatkowski, Cordelia Langford, John Sillitoe on behalf of the Wellcome Sanger Institute COVID-19 Surveillance Team |
| EPI_ISL_2540446 | Lighthouse Lab in Glasgow | Wellcome Sanger Institute for the COVID-19 Genomics UK (COG-UK) Consortium | Harper VanSteenhouse, Yumi Kasai, David Gray, Carol Clugston, Anna Dominiczak and Alex Alderton, Roberto Amato, Jeffrey Barrett, Sonia Goncalves, Ewan Harrison, David K. Jackson, Ian Johnston, Dominic Kwiatkowski, Cordelia Langford, John Sillitoe on behalf of the Wellcome Sanger Institute COVID-19 Surveillance Team |
| EPI_ISL_2540447 | Lighthouse Lab in Alderley Park | Wellcome Sanger Institute for the COVID-19 Genomics UK (COG-UK) Consortium | Jacquelyn Wynn, Mairead Hyland, The Lighthouse Lab in Alderley Park and Alex Alderton, Roberto Amato, Jeffrey Barrett, Sonia Goncalves, Ewan Harrison, David K. Jackson, Ian Johnston, Dominic Kwiatkowski, Cordelia Langford, John Sillitoe on behalf of the Wellcome Sanger Institute COVID-19 Surveillance Team |
| EPI_ISL_2540448 | Lighthouse Lab in Glasgow | Wellcome Sanger Institute for the COVID-19 Genomics UK (COG-UK) Consortium | Harper VanSteenhouse, Yumi Kasai, David Gray, Carol Clugston, Anna Dominiczak and Alex Alderton, Roberto Amato, Jeffrey Barrett, Sonia Goncalves, Ewan Harrison, David K. Jackson, Ian Johnston, Dominic Kwiatkowski, Cordelia Langford, John Sillitoe on behalf of the Wellcome Sanger Institute COVID-19 Surveillance Team |
| EPI_ISL_2540449, EPI_ISL_2540450, EPI_ISL_2540451, EPI_ISL_2540452, EPI_ISL_2540453, EPI_ISL_2540454 | Lighthouse Lab in Alderley Park | Wellcome Sanger Institute for the COVID-19 Genomics UK (COG-UK) Consortium | Jacquelyn Wynn, Mairead Hyland, The Lighthouse Lab in Alderley Park and Alex Alderton, Roberto Amato, Jeffrey Barrett, Sonia Goncalves, Ewan Harrison, David K. Jackson, Ian Johnston, Dominic Kwiatkowski, Cordelia Langford, John Sillitoe on behalf of the Wellcome Sanger Institute COVID-19 Surveillance Team |
| EPI_ISL_2540455 | Lighthouse Lab in Glasgow | Wellcome Sanger Institute for the COVID-19 Genomics UK (COG-UK) Consortium | Harper VanSteenhouse, Yumi Kasai, David Gray, Carol Clugston, Anna Dominiczak and Alex Alderton, Roberto Amato, Jeffrey Barrett, Sonia Goncalves, Ewan Harrison, David K. Jackson, Ian Johnston, Dominic Kwiatkowski, Cordelia Langford, John Sillitoe on behalf of the Wellcome Sanger Institute COVID-19 Surveillance Team |
| EPI_ISL_2540456, EPI_ISL_2540458 | Lighthouse Lab in Alderley Park | Wellcome Sanger Institute for the COVID-19 Genomics UK (COG-UK) Consortium | Jacquelyn Wynn, Mairead Hyland, The Lighthouse Lab in Alderley Park and Alex Alderton, Roberto Amato, Jeffrey Barrett, Sonia Goncalves, Ewan Harrison, David K. Jackson, Ian Johnston, Dominic Kwiatkowski, Cordelia Langford, John Sillitoe on behalf of the Wellcome Sanger Institute COVID-19 Surveillance Team |
| EPI_ISL_2540459 | Lighthouse Lab in Glasgow | Wellcome Sanger Institute for the COVID-19 Genomics UK (COG-UK) Consortium | Harper VanSteenhouse, Yumi Kasai, David Gray, Carol Clugston, Anna Dominiczak and Alex Alderton, Roberto Amato, Jeffrey Barrett, Sonia Goncalves, Ewan Harrison, David K. Jackson, Ian Johnston, Dominic Kwiatkowski, Cordelia Langford, John Sillitoe on behalf of the Wellcome Sanger Institute COVID-19 Surveillance Team |
| EPI_ISL_2540460, EPI_ISL_2540461, EPI_ISL_2540462, EPI_ISL_2540463, EPI_ISL_2540464 | Lighthouse Lab in Alderley Park | Wellcome Sanger Institute for the COVID-19 Genomics UK (COG-UK) Consortium | Jacquelyn Wynn, Mairead Hyland, The Lighthouse Lab in Alderley Park and Alex Alderton, Roberto Amato, Jeffrey Barrett, Sonia Goncalves, Ewan Harrison, David K. Jackson, Ian Johnston, Dominic Kwiatkowski, Cordelia Langford, John Sillitoe on behalf of the Wellcome Sanger Institute COVID-19 Surveillance Team |
| EPI_ISL_2540465, EPI_ISL_2540466 | Lighthouse Lab in Glasgow | Wellcome Sanger Institute for the COVID-19 Genomics UK (COG-UK) Consortium | Harper VanSteenhouse, Yumi Kasai, David Gray, Carol Clugston, Anna Dominiczak and Alex Alderton, Roberto Amato, Jeffrey Barrett, Sonia Goncalves, Ewan Harrison, David K. Jackson, Ian Johnston, Dominic Kwiatkowski, Cordelia Langford, John Sillitoe on behalf of the Wellcome Sanger Institute COVID-19 Surveillance Team |
| EPI_ISL_2540467, EPI_ISL_2540468, EPI_ISL_2540469, EPI_ISL_2540470 | Lighthouse Lab in Alderley Park | Wellcome Sanger Institute for the COVID-19 Genomics UK (COG-UK) Consortium | Jacquelyn Wynn, Mairead Hyland, The Lighthouse Lab in Alderley Park and Alex Alderton, Roberto Amato, Jeffrey Barrett, Sonia Goncalves, Ewan Harrison, David K. Jackson, Ian Johnston, Dominic Kwiatkowski, Cordelia Langford, John Sillitoe on behalf of the Wellcome Sanger Institute COVID-19 Surveillance Team |
| EPI_ISL_2540471 | Lighthouse Lab in Glasgow | Wellcome Sanger Institute for the COVID-19 Genomics UK (COG-UK) Consortium | Harper VanSteenhouse, Yumi Kasai, David Gray, Carol Clugston, Anna Dominiczak and Alex Alderton, Roberto Amato, Jeffrey Barrett, Sonia Goncalves, Ewan Harrison, David K. Jackson, Ian Johnston, Dominic Kwiatkowski, Cordelia Langford, John Sillitoe on behalf of the Wellcome Sanger Institute COVID-19 Surveillance Team |
| EPI_ISL_2540472 | Lighthouse Lab in Alderley Park | Wellcome Sanger Institute for the COVID-19 Genomics UK (COG-UK) Consortium | Jacquelyn Wynn, Mairead Hyland, The Lighthouse Lab in Alderley Park and Alex Alderton, Roberto Amato, Jeffrey Barrett, Sonia Goncalves, Ewan Harrison, David K. Jackson, Ian Johnston, Dominic Kwiatkowski, Cordelia Langford, John Sillitoe on behalf of the Wellcome Sanger Institute COVID-19 Surveillance Team |
| EPI_ISL_2540473 | Lighthouse Lab in Glasgow | Wellcome Sanger Institute for the COVID-19 Genomics UK (COG-UK) Consortium | Harper VanSteenhouse, Yumi Kasai, David Gray, Carol Clugston, Anna Dominiczak and Alex Alderton, Roberto Amato, Jeffrey Barrett, Sonia Goncalves, Ewan Harrison, David K. Jackson, Ian Johnston, Dominic Kwiatkowski, Cordelia Langford, John Sillitoe on behalf of the Wellcome Sanger Institute COVID-19 Surveillance Team |
| EPI_ISL_2540475 | Lighthouse Lab in Alderley Park | Wellcome Sanger Institute for the COVID-19 Genomics UK (COG-UK) Consortium | Jacquelyn Wynn, Mairead Hyland, The Lighthouse Lab in Alderley Park and Alex Alderton, Roberto Amato, Jeffrey Barrett, Sonia Goncalves, Ewan Harrison, David K. Jackson, Ian Johnston, Dominic Kwiatkowski, Cordelia Langford, John Sillitoe on behalf of the Wellcome Sanger Institute COVID-19 Surveillance Team |

| | | | |
|---|---|---|---|
| EPI_ISL_2540476 | Lighthouse Lab in Glasgow | Wellcome Sanger Institute for the COVID-19 Genomics UK (COG-UK) Consortium | Harper VanSteenhouse, Yumi Kasai, David Gray, Carol Clugston, Anna Dominiczak and Alex Alderton, Roberto Amato, Jeffrey Barrett, Sonia Goncalves, Ewan Harrison, David K. Jackson, Ian Johnston, Dominic Kwiatkowski, Cordelia Langford, John Sillitoe on behalf of the Wellcome Sanger Institute COVID-19 Surveillance Team |
| EPI_ISL_2540477, EPI_ISL_2540478, EPI_ISL_2540479, EPI_ISL_2540480, EPI_ISL_2540481, EPI_ISL_2540482, EPI_ISL_2540483, EPI_ISL_2540484 | Lighthouse Lab in Alderley Park | Wellcome Sanger Institute for the COVID-19 Genomics UK (COG-UK) Consortium | Jacquelyn Wynn, Mairead Hyland, The Lighthouse Lab in Alderley Park and Alex Alderton, Roberto Amato, Jeffrey Barrett, Sonia Goncalves, Ewan Harrison, David K. Jackson, Ian Johnston, Dominic Kwiatkowski, Cordelia Langford, John Sillitoe on behalf of the Wellcome Sanger Institute COVID-19 Surveillance Team |
| EPI_ISL_2540485, EPI_ISL_2540486 | Lighthouse Lab in Glasgow | Wellcome Sanger Institute for the COVID-19 Genomics UK (COG-UK) Consortium | Harper VanSteenhouse, Yumi Kasai, David Gray, Carol Clugston, Anna Dominiczak and Alex Alderton, Roberto Amato, Jeffrey Barrett, Sonia Goncalves, Ewan Harrison, David K. Jackson, Ian Johnston, Dominic Kwiatkowski, Cordelia Langford, John Sillitoe on behalf of the Wellcome Sanger Institute COVID-19 Surveillance Team |
| EPI_ISL_2540488, EPI_ISL_2540489, EPI_ISL_2540490 | Lighthouse Lab in Alderley Park | Wellcome Sanger Institute for the COVID-19 Genomics UK (COG-UK) Consortium | Jacquelyn Wynn, Mairead Hyland, The Lighthouse Lab in Alderley Park and Alex Alderton, Roberto Amato, Jeffrey Barrett, Sonia Goncalves, Ewan Harrison, David K. Jackson, Ian Johnston, Dominic Kwiatkowski, Cordelia Langford, John Sillitoe on behalf of the Wellcome Sanger Institute COVID-19 Surveillance Team |
| EPI_ISL_2540491 | Lighthouse Lab in Glasgow | Wellcome Sanger Institute for the COVID-19 Genomics UK (COG-UK) Consortium | Harper VanSteenhouse, Yumi Kasai, David Gray, Carol Clugston, Anna Dominiczak and Alex Alderton, Roberto Amato, Jeffrey Barrett, Sonia Goncalves, Ewan Harrison, David K. Jackson, Ian Johnston, Dominic Kwiatkowski, Cordelia Langford, John Sillitoe on behalf of the Wellcome Sanger Institute COVID-19 Surveillance Team |
| EPI_ISL_2540492, EPI_ISL_2540493, EPI_ISL_2540494, EPI_ISL_2540496, EPI_ISL_2540497, EPI_ISL_2540498, EPI_ISL_2540499, EPI_ISL_2540500, EPI_ISL_2540501, EPI_ISL_2540502 | Lighthouse Lab in Alderley Park | Wellcome Sanger Institute for the COVID-19 Genomics UK (COG-UK) Consortium | Jacquelyn Wynn, Mairead Hyland, The Lighthouse Lab in Alderley Park and Alex Alderton, Roberto Amato, Jeffrey Barrett, Sonia Goncalves, Ewan Harrison, David K. Jackson, Ian Johnston, Dominic Kwiatkowski, Cordelia Langford, John Sillitoe on behalf of the Wellcome Sanger Institute COVID-19 Surveillance Team |
| EPI_ISL_2540503, EPI_ISL_2540504 | Lighthouse Lab in Glasgow | Wellcome Sanger Institute for the COVID-19 Genomics UK (COG-UK) Consortium | Harper VanSteenhouse, Yumi Kasai, David Gray, Carol Clugston, Anna Dominiczak and Alex Alderton, Roberto Amato, Jeffrey Barrett, Sonia Goncalves, Ewan Harrison, David K. Jackson, Ian Johnston, Dominic Kwiatkowski, Cordelia Langford, John Sillitoe on behalf of the Wellcome Sanger Institute COVID-19 Surveillance Team |
| EPI_ISL_2540505, EPI_ISL_2540506 | Lighthouse Lab in Alderley Park | Wellcome Sanger Institute for the COVID-19 Genomics UK (COG-UK) Consortium | Jacquelyn Wynn, Mairead Hyland, The Lighthouse Lab in Alderley Park and Alex Alderton, Roberto Amato, Jeffrey Barrett, Sonia Goncalves, Ewan Harrison, David K. Jackson, Ian Johnston, Dominic Kwiatkowski, Cordelia Langford, John Sillitoe on behalf of the Wellcome Sanger Institute COVID-19 Surveillance Team |
| EPI_ISL_2540507 | Lighthouse Lab in Glasgow | Wellcome Sanger Institute for the COVID-19 Genomics UK (COG-UK) Consortium | Harper VanSteenhouse, Yumi Kasai, David Gray, Carol Clugston, Anna Dominiczak and Alex Alderton, Roberto Amato, Jeffrey Barrett, Sonia Goncalves, Ewan Harrison, David K. Jackson, Ian Johnston, Dominic Kwiatkowski, Cordelia Langford, John Sillitoe on behalf of the Wellcome Sanger Institute COVID-19 Surveillance Team |
| EPI_ISL_2540508, EPI_ISL_2540509, EPI_ISL_2540510 | Lighthouse Lab in Alderley Park | Wellcome Sanger Institute for the COVID-19 Genomics UK (COG-UK) Consortium | Jacquelyn Wynn, Mairead Hyland, The Lighthouse Lab in Alderley Park and Alex Alderton, Roberto Amato, Jeffrey Barrett, Sonia Goncalves, Ewan Harrison, David K. Jackson, Ian Johnston, Dominic Kwiatkowski, Cordelia Langford, John Sillitoe on behalf of the Wellcome Sanger Institute COVID-19 Surveillance Team |
| EPI_ISL_2540511, EPI_ISL_2540512 | Lighthouse Lab in Glasgow | Wellcome Sanger Institute for the COVID-19 Genomics UK (COG-UK) Consortium | Harper VanSteenhouse, Yumi Kasai, David Gray, Carol Clugston, Anna Dominiczak and Alex Alderton, Roberto Amato, Jeffrey Barrett, Sonia Goncalves, Ewan Harrison, David K. Jackson, Ian Johnston, Dominic Kwiatkowski, Cordelia Langford, John Sillitoe on behalf of the Wellcome Sanger Institute COVID-19 Surveillance Team |
| EPI_ISL_2540513, EPI_ISL_2540514 | Lighthouse Lab in Alderley Park | Wellcome Sanger Institute for the COVID-19 Genomics UK (COG-UK) Consortium | Jacquelyn Wynn, Mairead Hyland, The Lighthouse Lab in Alderley Park and Alex Alderton, Roberto Amato, Jeffrey Barrett, Sonia Goncalves, Ewan Harrison, David K. Jackson, Ian Johnston, Dominic Kwiatkowski, Cordelia Langford, John Sillitoe on behalf of the Wellcome Sanger Institute COVID-19 Surveillance Team |
| EPI_ISL_2540515 | Lighthouse Lab in Glasgow | Wellcome Sanger Institute for the COVID-19 Genomics UK (COG-UK) Consortium | Harper VanSteenhouse, Yumi Kasai, David Gray, Carol Clugston, Anna Dominiczak and Alex Alderton, Roberto Amato, Jeffrey Barrett, Sonia Goncalves, Ewan Harrison, David K. Jackson, Ian Johnston, Dominic Kwiatkowski, Cordelia Langford, John Sillitoe on behalf of the Wellcome Sanger Institute COVID-19 Surveillance Team |
| EPI_ISL_2540516, EPI_ISL_2540517 | Lighthouse Lab in Alderley Park | Wellcome Sanger Institute for the COVID-19 Genomics UK (COG-UK) Consortium | Jacquelyn Wynn, Mairead Hyland, The Lighthouse Lab in Alderley Park and Alex Alderton, Roberto Amato, Jeffrey Barrett, Sonia Goncalves, Ewan Harrison, David K. Jackson, Ian Johnston, Dominic Kwiatkowski, Cordelia Langford, John Sillitoe on behalf of the Wellcome Sanger Institute COVID-19 Surveillance Team |
| EPI_ISL_2540518, EPI_ISL_2540519 | Lighthouse Lab in Glasgow | Wellcome Sanger Institute for the COVID-19 Genomics UK (COG-UK) Consortium | Harper VanSteenhouse, Yumi Kasai, David Gray, Carol Clugston, Anna Dominiczak and Alex Alderton, Roberto Amato, Jeffrey Barrett, Sonia Goncalves, Ewan Harrison, David K. Jackson, Ian Johnston, Dominic Kwiatkowski, Cordelia Langford, John Sillitoe on behalf of the Wellcome Sanger Institute COVID-19 Surveillance Team |
| EPI_ISL_2540520 | Lighthouse Lab in Alderley Park | Wellcome Sanger Institute for the COVID-19 Genomics UK (COG-UK) Consortium | Jacquelyn Wynn, Mairead Hyland, The Lighthouse Lab in Alderley Park and Alex Alderton, Roberto Amato, Jeffrey Barrett, Sonia Goncalves, Ewan Harrison, David K. Jackson, Ian Johnston, Dominic Kwiatkowski, Cordelia Langford, John Sillitoe on behalf of the Wellcome Sanger Institute COVID-19 Surveillance Team |
| EPI_ISL_2540521 | Lighthouse Lab in Glasgow | Wellcome Sanger Institute for the COVID-19 Genomics UK (COG-UK) Consortium | Harper VanSteenhouse, Yumi Kasai, David Gray, Carol Clugston, Anna Dominiczak and Alex Alderton, Roberto Amato, Jeffrey Barrett, Sonia Goncalves, Ewan Harrison, David K. Jackson, Ian Johnston, Dominic Kwiatkowski, Cordelia Langford, |
| EPI_ISL_2540522, EPI_ISL_2540523, EPI_ISL_2540524, EPI_ISL_2540525, EPI_ISL_2540526, EPI_ISL_2540527, EPI_ISL_2540528, EPI_ISL_2540529, EPI_ISL_2540530, EPI_ISL_2540531, EPI_ISL_2540532, EPI_ISL_2540533, EPI_ISL_2540534, EPI_ISL_2540535, EPI_ISL_2540536, EPI_ISL_2540537, EPI_ISL_2540538, EPI_ISL_2540539, EPI_ISL_2540540, EPI_ISL_2540541 | | | |
| see above | Lighthouse Lab in Alderley Park | Wellcome Sanger Institute for the COVID-19 Genomics UK (COG-UK) Consortium | Jacquelyn Wynn, Mairead Hyland, The Lighthouse Lab in Alderley Park and Alex Alderton, Roberto Amato, Jeffrey Barrett, Sonia Goncalves, Ewan Harrison, David K. Jackson, Ian Johnston, Dominic Kwiatkowski, Cordelia Langford, John Sillitoe on behalf of the Wellcome Sanger Institute COVID-19 Surveillance Team |
| EPI_ISL_2540542 | Lighthouse Lab in Glasgow | Wellcome Sanger Institute for the COVID-19 Genomics UK (COG-UK) Consortium | Harper VanSteenhouse, Yumi Kasai, David Gray, Carol Clugston, Anna Dominiczak and Alex Alderton, Roberto Amato, Jeffrey Barrett, Sonia Goncalves, Ewan Harrison, David K. Jackson, Ian Johnston, Dominic Kwiatkowski, Cordelia Langford, John Sillitoe on behalf of the Wellcome Sanger Institute COVID-19 Surveillance Team |
| EPI_ISL_2540543 | Lighthouse Lab in Alderley Park | Wellcome Sanger Institute for the COVID-19 Genomics UK (COG-UK) Consortium | Jacquelyn Wynn, Mairead Hyland, The Lighthouse Lab in Alderley Park and Alex Alderton, Roberto Amato, Jeffrey Barrett, Sonia Goncalves, Ewan Harrison, David K. Jackson, Ian Johnston, Dominic Kwiatkowski, Cordelia Langford, John Sillitoe on behalf of the Wellcome Sanger Institute COVID-19 Surveillance Team |
| EPI_ISL_2540544 | Lighthouse Lab in Glasgow | Wellcome Sanger Institute for the COVID-19 Genomics UK (COG-UK) Consortium | Harper VanSteenhouse, Yumi Kasai, David Gray, Carol Clugston, Anna Dominiczak and Alex Alderton, Roberto Amato, Jeffrey Barrett, Sonia Goncalves, Ewan Harrison, David K. Jackson, Ian Johnston, Dominic Kwiatkowski, Cordelia Langford, John Sillitoe on behalf of the Wellcome Sanger Institute COVID-19 Surveillance Team |
| EPI_ISL_2540546, EPI_ISL_2540547, EPI_ISL_2540548 | Lighthouse Lab in Alderley Park | Wellcome Sanger Institute for the COVID-19 Genomics UK (COG-UK) Consortium | Jacquelyn Wynn, Mairead Hyland, The Lighthouse Lab in Alderley Park and Alex Alderton, Roberto Amato, Jeffrey Barrett, Sonia Goncalves, Ewan Harrison, David K. Jackson, Ian Johnston, Dominic Kwiatkowski, Cordelia Langford, John Sillitoe on behalf of the Wellcome Sanger Institute COVID-19 Surveillance Team |
| EPI_ISL_2540549 | Lighthouse Lab in Glasgow | Wellcome Sanger Institute for the COVID-19 Genomics UK (COG-UK) Consortium | Harper VanSteenhouse, Yumi Kasai, David Gray, Carol Clugston, Anna Dominiczak and Alex Alderton, Roberto Amato, Jeffrey Barrett, Sonia Goncalves, Ewan Harrison, David K. Jackson, Ian Johnston, Dominic Kwiatkowski, Cordelia Langford, John Sillitoe on behalf of the Wellcome Sanger Institute COVID-19 Surveillance Team |
| EPI_ISL_2540550, EPI_ISL_2540551, EPI_ISL_2540552, EPI_ISL_2540553, EPI_ISL_2540554, EPI_ISL_2540555, EPI_ISL_2540556, EPI_ISL_2540557 | Lighthouse Lab in Alderley Park | Wellcome Sanger Institute for the COVID-19 Genomics UK (COG-UK) Consortium | Jacquelyn Wynn, Mairead Hyland, The Lighthouse Lab in Alderley Park and Alex Alderton, Roberto Amato, Jeffrey Barrett, Sonia Goncalves, Ewan Harrison, David K. Jackson, Ian Johnston, Dominic Kwiatkowski, Cordelia Langford, John Sillitoe on behalf of the Wellcome Sanger Institute COVID-19 Surveillance Team |
| EPI_ISL_2540558 | Lighthouse Lab in Glasgow | Wellcome Sanger Institute for the COVID-19 Genomics UK (COG-UK) Consortium | Harper VanSteenhouse, Yumi Kasai, David Gray, Carol Clugston, Anna Dominiczak and Alex Alderton, Roberto Amato, Jeffrey Barrett, Sonia Goncalves, Ewan Harrison, David K. Jackson, Ian Johnston, Dominic Kwiatkowski, Cordelia Langford, John Sillitoe on behalf of the Wellcome Sanger Institute COVID-19 Surveillance Team |
| EPI_ISL_2540559, EPI_ISL_2540560, EPI_ISL_2540561, EPI_ISL_2540562, EPI_ISL_2540563, EPI_ISL_2540564, EPI_ISL_2540565, EPI_ISL_2540566, EPI_ISL_2540567 | Lighthouse Lab in Alderley Park | Wellcome Sanger Institute for the COVID-19 Genomics UK (COG-UK) Consortium | Jacquelyn Wynn, Mairead Hyland, The Lighthouse Lab in Alderley Park and Alex Alderton, Roberto Amato, Jeffrey Barrett, Sonia Goncalves, Ewan Harrison, David K. Jackson, Ian Johnston, Dominic Kwiatkowski, Cordelia Langford, John Sillitoe on behalf of the Wellcome Sanger Institute COVID-19 Surveillance Team |
| EPI_ISL_2540568 | Lighthouse Lab in Glasgow | Wellcome Sanger Institute for the COVID-19 Genomics UK (COG-UK) Consortium | Harper VanSteenhouse, Yumi Kasai, David Gray, Carol Clugston, Anna Dominiczak and Alex Alderton, Roberto Amato, Jeffrey Barrett, Sonia Goncalves, Ewan Harrison, David K. Jackson, Ian Johnston, Dominic Kwiatkowski, Cordelia Langford, John Sillitoe on behalf of the Wellcome Sanger Institute COVID-19 Surveillance Team |
| EPI_ISL_2540569, EPI_ISL_2540570, EPI_ISL_2540571, EPI_ISL_2540572 | Lighthouse Lab in Alderley Park | Wellcome Sanger Institute for the COVID-19 Genomics UK (COG-UK) Consortium | Jacquelyn Wynn, Mairead Hyland, The Lighthouse Lab in Alderley Park and Alex Alderton, Roberto Amato, Jeffrey Barrett, Sonia Goncalves, Ewan Harrison, David K. Jackson, Ian Johnston, Dominic Kwiatkowski, Cordelia Langford, John Sillitoe on behalf of the Wellcome Sanger Institute COVID-19 Surveillance Team |
| EPI_ISL_2540573 | Lighthouse Lab in Glasgow | Wellcome Sanger Institute for the COVID-19 Genomics UK (COG-UK) Consortium | Harper VanSteenhouse, Yumi Kasai, David Gray, Carol Clugston, Anna Dominiczak and Alex Alderton, Roberto Amato, Jeffrey Barrett, Sonia Goncalves, Ewan Harrison, David K. Jackson, Ian Johnston, Dominic Kwiatkowski, Cordelia Langford, John Sillitoe on behalf of the Wellcome Sanger Institute COVID-19 Surveillance Team |
| EPI_ISL_2540574 | Lighthouse Lab in Alderley Park | Wellcome Sanger Institute for the COVID-19 Genomics UK (COG-UK) Consortium | Jacquelyn Wynn, Mairead Hyland, The Lighthouse Lab in Alderley Park and Alex Alderton, Roberto Amato, Jeffrey Barrett, Sonia Goncalves, Ewan Harrison, David K. Jackson, Ian Johnston, Dominic Kwiatkowski, Cordelia Langford, John Sillitoe on behalf of the Wellcome Sanger Institute COVID-19 Surveillance Team |
| EPI_ISL_2540575 | Lighthouse Lab in Glasgow | Wellcome Sanger Institute for the COVID-19 Genomics UK (COG-UK) Consortium | Harper VanSteenhouse, Yumi Kasai, David Gray, Carol Clugston, Anna Dominiczak and Alex Alderton, Roberto Amato, Jeffrey Barrett, Sonia Goncalves, Ewan Harrison, David K. Jackson, Ian Johnston, Dominic Kwiatkowski, Cordelia Langford, John Sillitoe on behalf of the Wellcome Sanger Institute COVID-19 Surveillance Team |

| | | | |
|---|---|---|---|
| EPI_ISL_2540576, EPI_ISL_2540577 | Lighthouse Lab in Alderley Park | Wellcome Sanger Institute for the COVID-19 Genomics UK (COG-UK) Consortium | Jacquelyn Wynn, Mairead Hyland, The Lighthouse Lab in Alderley Park and Alex Alderton, Roberto Amato, Jeffrey Barrett, Sonia Goncalves, Ewan Harrison, David K. Jackson, Ian Johnston, Dominic Kwiatkowski, Cordelia Langford, John Sillitoe on behalf of the Wellcome Sanger Institute COVID-19 Surveillance Team |
| EPI_ISL_2540578 | Lighthouse Lab in Glasgow | Wellcome Sanger Institute for the COVID-19 Genomics UK (COG-UK) Consortium | Harper VanSteenhouse, Yumi Kasai, David Gray, Carol Clugston, Anna Dominiczak and Alex Alderton, Roberto Amato, Jeffrey Barrett, Sonia Goncalves, Ewan Harrison, David K. Jackson, Ian Johnston, Dominic Kwiatkowski, Cordelia Langford, John Sillitoe on behalf of the Wellcome Sanger Institute COVID-19 Surveillance Team |
| EPI_ISL_2540579, EPI_ISL_2540580, EPI_ISL_2540581, EPI_ISL_2540582, EPI_ISL_2540583, EPI_ISL_2540584 | Lighthouse Lab in Alderley Park | Wellcome Sanger Institute for the COVID-19 Genomics UK (COG-UK) Consortium | Jacquelyn Wynn, Mairead Hyland, The Lighthouse Lab in Alderley Park and Alex Alderton, Roberto Amato, Jeffrey Barrett, Sonia Goncalves, Ewan Harrison, David K. Jackson, Ian Johnston, Dominic Kwiatkowski, Cordelia Langford, John Sillitoe on behalf of the Wellcome Sanger Institute COVID-19 Surveillance Team |
| EPI_ISL_2540585 | Lighthouse Lab in Glasgow | Wellcome Sanger Institute for the COVID-19 Genomics UK (COG-UK) Consortium | Harper VanSteenhouse, Yumi Kasai, David Gray, Carol Clugston, Anna Dominiczak and Alex Alderton, Roberto Amato, Jeffrey Barrett, Sonia Goncalves, Ewan Harrison, David K. Jackson, Ian Johnston, Dominic Kwiatkowski, Cordelia Langford, John Sillitoe on behalf of the Wellcome Sanger Institute COVID-19 Surveillance Team |
| EPI_ISL_2540587, EPI_ISL_2540588, EPI_ISL_2540590 | Lighthouse Lab in Alderley Park | Wellcome Sanger Institute for the COVID-19 Genomics UK (COG-UK) Consortium | Jacquelyn Wynn, Mairead Hyland, The Lighthouse Lab in Alderley Park and Alex Alderton, Roberto Amato, Jeffrey Barrett, Sonia Goncalves, Ewan Harrison, David K. Jackson, Ian Johnston, Dominic Kwiatkowski, Cordelia Langford, John Sillitoe on behalf of the Wellcome Sanger Institute COVID-19 Surveillance Team |
| EPI_ISL_2540591 | Lighthouse Laboratory Plymouth | Wellcome Sanger Institute for the COVID-19 Genomics UK (COG-UK) Consortium | Lighthouse Laboratory Plymouth and Alex Alderton, Roberto Amato, Jeffrey Barrett, Sonia Goncalves, Ewan Harrison, David K. Jackson, Ian Johnston, Dominic Kwiatkowski, Cordelia Langford, John Sillitoe on behalf of the Wellcome Sanger Institute COVID-19 Surveillance Team |
| EPI_ISL_2540592, EPI_ISL_2540593 | Lighthouse Lab in Alderley Park | Wellcome Sanger Institute for the COVID-19 Genomics UK (COG-UK) Consortium | Jacquelyn Wynn, Mairead Hyland, The Lighthouse Lab in Alderley Park and Alex Alderton, Roberto Amato, Jeffrey Barrett, Sonia Goncalves, Ewan Harrison, David K. Jackson, Ian Johnston, Dominic Kwiatkowski, Cordelia Langford, John Sillitoe on behalf of the Wellcome Sanger Institute COVID-19 Surveillance Team |
| EPI_ISL_2540594 | Lighthouse Lab in Milton Keynes | Wellcome Sanger Institute for the COVID-19 Genomics UK (COG-UK) Consortium | The Lighthouse Lab in Milton Keynes and Alex Alderton, Roberto Amato, Jeffrey Barrett, Sonia Goncalves, Ewan Harrison, David K. Jackson, Ian Johnston, Dominic Kwiatkowski, Cordelia Langford, John Sillitoe on behalf of the Wellcome Sanger Institute COVID-19 Surveillance Team |
| EPI_ISL_2540595, EPI_ISL_2540596, EPI_ISL_2540597, EPI_ISL_2540598 | Lighthouse Lab in Alderley Park | Wellcome Sanger Institute for the COVID-19 Genomics UK (COG-UK) Consortium | Jacquelyn Wynn, Mairead Hyland, The Lighthouse Lab in Alderley Park and Alex Alderton, Roberto Amato, Jeffrey Barrett, Sonia Goncalves, Ewan Harrison, David K. Jackson, Ian Johnston, Dominic Kwiatkowski, Cordelia Langford, John Sillitoe on behalf of the Wellcome Sanger Institute COVID-19 Surveillance Team |
| EPI_ISL_2540599, EPI_ISL_2540600, EPI_ISL_2540601 | Lighthouse Lab in Milton Keynes | Wellcome Sanger Institute for the COVID-19 Genomics UK (COG-UK) Consortium | The Lighthouse Lab in Milton Keynes and Alex Alderton, Roberto Amato, Jeffrey Barrett, Sonia Goncalves, Ewan Harrison, David K. Jackson, Ian Johnston, Dominic Kwiatkowski, Cordelia Langford, John Sillitoe on behalf of the Wellcome Sanger Institute COVID-19 Surveillance Team |
| EPI_ISL_2540602 | Lighthouse Lab in Alderley Park | Wellcome Sanger Institute for the COVID-19 Genomics UK (COG-UK) Consortium | Jacquelyn Wynn, Mairead Hyland, The Lighthouse Lab in Alderley Park and Alex Alderton, Roberto Amato, Jeffrey Barrett, Sonia Goncalves, Ewan Harrison, David K. Jackson, Ian Johnston, Dominic Kwiatkowski, Cordelia Langford, John Sillitoe on behalf of the Wellcome Sanger Institute COVID-19 Surveillance Team |
| EPI_ISL_2540603 | Lighthouse Lab in Milton Keynes | Wellcome Sanger Institute for the COVID-19 Genomics UK (COG-UK) Consortium | The Lighthouse Lab in Milton Keynes and Alex Alderton, Roberto Amato, Jeffrey Barrett, Sonia Goncalves, Ewan Harrison, David K. Jackson, Ian Johnston, Dominic Kwiatkowski, Cordelia Langford, John Sillitoe on behalf of the Wellcome Sanger Institute COVID-19 Surveillance Team |
| EPI_ISL_2540604 | Lighthouse Laboratory Plymouth | Wellcome Sanger Institute for the COVID-19 Genomics UK (COG-UK) Consortium | Lighthouse Laboratory Plymouth and Alex Alderton, Roberto Amato, Jeffrey Barrett, Sonia Goncalves, Ewan Harrison, David K. Jackson, Ian Johnston, Dominic Kwiatkowski, Cordelia Langford, John Sillitoe on behalf of the Wellcome Sanger Institute COVID-19 Surveillance Team |
| EPI_ISL_2540605 | Lighthouse Lab in Alderley Park | Wellcome Sanger Institute for the COVID-19 Genomics UK (COG-UK) Consortium | Jacquelyn Wynn, Mairead Hyland, The Lighthouse Lab in Alderley Park and Alex Alderton, Roberto Amato, Jeffrey Barrett, Sonia Goncalves, Ewan Harrison, David K. Jackson, Ian Johnston, Dominic Kwiatkowski, Cordelia Langford, John Sillitoe on behalf of the Wellcome Sanger Institute COVID-19 Surveillance Team |
| EPI_ISL_2540606 | Lighthouse Lab in Milton Keynes | Wellcome Sanger Institute for the COVID-19 Genomics UK (COG-UK) Consortium | The Lighthouse Lab in Milton Keynes and Alex Alderton, Roberto Amato, Jeffrey Barrett, Sonia Goncalves, Ewan Harrison, David K. Jackson, Ian Johnston, Dominic Kwiatkowski, Cordelia Langford, John Sillitoe on behalf of the Wellcome Sanger Institute COVID-19 Surveillance Team |
| EPI_ISL_2540607 | Lighthouse Laboratory Plymouth | Wellcome Sanger Institute for the COVID-19 Genomics UK (COG-UK) Consortium | Lighthouse Laboratory Plymouth and Alex Alderton, Roberto Amato, Jeffrey Barrett, Sonia Goncalves, Ewan Harrison, David K. Jackson, Ian Johnston, Dominic Kwiatkowski, Cordelia Langford, John Sillitoe on behalf of the Wellcome Sanger Institute COVID-19 Surveillance Team |
| EPI_ISL_2540608, EPI_ISL_2540609 | Lighthouse Lab in Milton Keynes | Wellcome Sanger Institute for the COVID-19 Genomics UK (COG-UK) Consortium | The Lighthouse Lab in Milton Keynes and Alex Alderton, Roberto Amato, Jeffrey Barrett, Sonia Goncalves, Ewan Harrison, David K. Jackson, Ian Johnston, Dominic Kwiatkowski, Cordelia Langford, John Sillitoe on behalf of the Wellcome Sanger Institute COVID-19 Surveillance Team |
| EPI_ISL_2540610 | Lighthouse Lab in Alderley Park | Wellcome Sanger Institute for the COVID-19 Genomics UK (COG-UK) Consortium | Jacquelyn Wynn, Mairead Hyland, The Lighthouse Lab in Alderley Park and Alex Alderton, Roberto Amato, Jeffrey Barrett, Sonia Goncalves, Ewan Harrison, David K. Jackson, Ian Johnston, Dominic Kwiatkowski, Cordelia Langford, John Sillitoe on behalf of the Wellcome Sanger Institute COVID-19 Surveillance Team |
| EPI_ISL_2540611, EPI_ISL_2540612, EPI_ISL_2540613, EPI_ISL_2540614 | Lighthouse Lab in Milton Keynes | Wellcome Sanger Institute for the COVID-19 Genomics UK (COG-UK) Consortium | The Lighthouse Lab in Milton Keynes and Alex Alderton, Roberto Amato, Jeffrey Barrett, Sonia Goncalves, Ewan Harrison, David K. Jackson, Ian Johnston, Dominic Kwiatkowski, Cordelia Langford, John Sillitoe on behalf of the Wellcome Sanger Institute COVID-19 Surveillance Team |
| EPI_ISL_2540615 | Lighthouse Laboratory Plymouth | Wellcome Sanger Institute for the COVID-19 Genomics UK (COG-UK) Consortium | Lighthouse Laboratory Plymouth and Alex Alderton, Roberto Amato, Jeffrey Barrett, Sonia Goncalves, Ewan Harrison, David K. Jackson, Ian Johnston, Dominic Kwiatkowski, Cordelia Langford, John Sillitoe on behalf of the Wellcome Sanger Institute COVID-19 Surveillance Team |
| EPI_ISL_2540616, EPI_ISL_2540617, EPI_ISL_2540618, EPI_ISL_2540619 | Lighthouse Lab in Milton Keynes | Wellcome Sanger Institute for the COVID-19 Genomics UK (COG-UK) Consortium | The Lighthouse Lab in Milton Keynes and Alex Alderton, Roberto Amato, Jeffrey Barrett, Sonia Goncalves, Ewan Harrison, David K. Jackson, Ian Johnston, Dominic Kwiatkowski, Cordelia Langford, John Sillitoe on behalf of the Wellcome Sanger Institute COVID-19 Surveillance Team |
| EPI_ISL_2540620, EPI_ISL_2540622 | Lighthouse Laboratory Plymouth | Wellcome Sanger Institute for the COVID-19 Genomics UK (COG-UK) Consortium | Lighthouse Laboratory Plymouth and Alex Alderton, Roberto Amato, Jeffrey Barrett, Sonia Goncalves, Ewan Harrison, David K. Jackson, Ian Johnston, Dominic Kwiatkowski, Cordelia Langford, John Sillitoe on behalf of the Wellcome Sanger Institute COVID-19 Surveillance Team |
| EPI_ISL_2540623 | Lighthouse Lab in Milton Keynes | Wellcome Sanger Institute for the COVID-19 Genomics UK (COG-UK) Consortium | The Lighthouse Lab in Milton Keynes and Alex Alderton, Roberto Amato, Jeffrey Barrett, Sonia Goncalves, Ewan Harrison, David K. Jackson, Ian Johnston, Dominic Kwiatkowski, Cordelia Langford, John Sillitoe on behalf of the Wellcome Sanger Institute COVID-19 Surveillance Team |
| EPI_ISL_2540624 | Lighthouse Lab in Alderley Park | Wellcome Sanger Institute for the COVID-19 Genomics UK (COG-UK) Consortium | Jacquelyn Wynn, Mairead Hyland, The Lighthouse Lab in Alderley Park and Alex Alderton, Roberto Amato, Jeffrey Barrett, Sonia Goncalves, Ewan Harrison, David K. Jackson, Ian Johnston, Dominic Kwiatkowski, Cordelia Langford, John Sillitoe on behalf of the Wellcome Sanger Institute COVID-19 Surveillance Team |
| EPI_ISL_2540625, EPI_ISL_2540626 | Lighthouse Lab in Milton Keynes | Wellcome Sanger Institute for the COVID-19 Genomics UK (COG-UK) Consortium | The Lighthouse Lab in Milton Keynes and Alex Alderton, Roberto Amato, Jeffrey Barrett, Sonia Goncalves, Ewan Harrison, David K. Jackson, Ian Johnston, Dominic Kwiatkowski, Cordelia Langford, John Sillitoe on behalf of the Wellcome Sanger Institute COVID-19 Surveillance Team |
| EPI_ISL_2540627 | Lighthouse Lab in Alderley Park | Wellcome Sanger Institute for the COVID-19 Genomics UK (COG-UK) Consortium | Jacquelyn Wynn, Mairead Hyland, The Lighthouse Lab in Alderley Park and Alex Alderton, Roberto Amato, Jeffrey Barrett, Sonia Goncalves, Ewan Harrison, David K. Jackson, Ian Johnston, Dominic Kwiatkowski, Cordelia Langford, John Sillitoe on behalf of the Wellcome Sanger Institute COVID-19 Surveillance Team |
| EPI_ISL_2540628 | Lighthouse Lab in Milton Keynes | Wellcome Sanger Institute for the COVID-19 Genomics UK (COG-UK) Consortium | The Lighthouse Lab in Milton Keynes and Alex Alderton, Roberto Amato, Jeffrey Barrett, Sonia Goncalves, Ewan Harrison, David K. Jackson, Ian Johnston, Dominic Kwiatkowski, Cordelia Langford, John Sillitoe on behalf of the Wellcome Sanger Institute COVID-19 Surveillance Team |
| EPI_ISL_2540629 | Lighthouse Laboratory Plymouth | Wellcome Sanger Institute for the COVID-19 Genomics UK (COG-UK) Consortium | Lighthouse Laboratory Plymouth and Alex Alderton, Roberto Amato, Jeffrey Barrett, Sonia Goncalves, Ewan Harrison, David K. Jackson, Ian Johnston, Dominic Kwiatkowski, Cordelia Langford, John Sillitoe on behalf of the Wellcome Sanger Institute COVID-19 Surveillance Team |
| EPI_ISL_2540630, EPI_ISL_2540631, EPI_ISL_2540633, EPI_ISL_2540634, EPI_ISL_2540635 | Lighthouse Lab in Milton Keynes | Wellcome Sanger Institute for the COVID-19 Genomics UK (COG-UK) Consortium | The Lighthouse Lab in Milton Keynes and Alex Alderton, Roberto Amato, Jeffrey Barrett, Sonia Goncalves, Ewan Harrison, David K. Jackson, Ian Johnston, Dominic Kwiatkowski, Cordelia Langford, John Sillitoe on behalf of the Wellcome Sanger Institute COVID-19 Surveillance Team |
| EPI_ISL_2540636, EPI_ISL_2540637 | Lighthouse Lab in Alderley Park | Wellcome Sanger Institute for the COVID-19 Genomics UK (COG-UK) Consortium | Jacquelyn Wynn, Mairead Hyland, The Lighthouse Lab in Alderley Park and Alex Alderton, Roberto Amato, Jeffrey Barrett, Sonia Goncalves, Ewan Harrison, David K. Jackson, Ian Johnston, Dominic Kwiatkowski, Cordelia Langford, John Sillitoe on behalf of the Wellcome Sanger Institute COVID-19 Surveillance Team |
| EPI_ISL_2540638 | Lighthouse Laboratory Plymouth | Wellcome Sanger Institute for the COVID-19 Genomics UK (COG-UK) Consortium | Lighthouse Laboratory Plymouth and Alex Alderton, Roberto Amato, Jeffrey Barrett, Sonia Goncalves, Ewan Harrison, David K. Jackson, Ian Johnston, Dominic Kwiatkowski, Cordelia Langford, John Sillitoe on behalf of the Wellcome Sanger Institute COVID-19 Surveillance Team |
| EPI_ISL_2540639 | Lighthouse Lab in Alderley Park | Wellcome Sanger Institute for the COVID-19 Genomics UK (COG-UK) Consortium | Jacquelyn Wynn, Mairead Hyland, The Lighthouse Lab in Alderley Park and Alex Alderton, Roberto Amato, Jeffrey Barrett, Sonia Goncalves, Ewan Harrison, David K. Jackson, Ian Johnston, Dominic Kwiatkowski, Cordelia Langford, John Sillitoe on behalf of the Wellcome Sanger Institute COVID-19 Surveillance Team |
| EPI_ISL_2540640 | Lighthouse Lab in Milton Keynes | Wellcome Sanger Institute for the COVID-19 Genomics UK (COG-UK) Consortium | The Lighthouse Lab in Milton Keynes and Alex Alderton, Roberto Amato, Jeffrey Barrett, Sonia Goncalves, Ewan Harrison, David K. Jackson, Ian Johnston, Dominic Kwiatkowski, Cordelia Langford, John Sillitoe on behalf of the Wellcome Sanger Institute COVID-19 Surveillance Team |
| EPI_ISL_2540641 | Lighthouse Laboratory Plymouth | Wellcome Sanger Institute for the COVID-19 Genomics UK (COG-UK) Consortium | Lighthouse Laboratory Plymouth and Alex Alderton, Roberto Amato, Jeffrey Barrett, Sonia Goncalves, Ewan Harrison, David K. Jackson, Ian Johnston, Dominic Kwiatkowski, Cordelia Langford, John Sillitoe on behalf of the Wellcome Sanger Institute COVID-19 Surveillance Team |
| EPI_ISL_2540642 | Lighthouse Lab in Milton Keynes | Wellcome Sanger Institute for the COVID-19 Genomics UK (COG-UK) Consortium | The Lighthouse Lab in Milton Keynes and Alex Alderton, Roberto Amato, Jeffrey Barrett, Sonia Goncalves, Ewan Harrison, David K. Jackson, Ian Johnston, Dominic Kwiatkowski, Cordelia Langford, John Sillitoe on behalf of the Wellcome Sanger Institute COVID-19 Surveillance Team |
| EPI_ISL_2540643, EPI_ISL_2540644 | Lighthouse Lab in Alderley Park | Wellcome Sanger Institute for the COVID-19 Genomics UK (COG-UK) Consortium | Jacquelyn Wynn, Mairead Hyland, The Lighthouse Lab in Alderley Park and Alex Alderton, Roberto Amato, Jeffrey Barrett, Sonia Goncalves, Ewan Harrison, David K. Jackson, Ian Johnston, Dominic Kwiatkowski, Cordelia Langford, John Sillitoe on behalf of the Wellcome Sanger Institute COVID-19 Surveillance Team |
| EPI_ISL_2540645 | Lighthouse Lab in Milton Keynes | Wellcome Sanger Institute for the COVID-19 Genomics UK (COG-UK) Consortium | The Lighthouse Lab in Milton Keynes and Alex Alderton, Roberto Amato, Jeffrey Barrett, Sonia Goncalves, Ewan Harrison, David K. Jackson, Ian Johnston, Dominic Kwiatkowski, Cordelia Langford, John Sillitoe on behalf of the Wellcome Sanger Institute COVID-19 Surveillance Team |
| EPI_ISL_2540646, EPI_ISL_2540647, EPI_ISL_2540648, EPI_ISL_2540649 | Lighthouse Lab in Alderley Park | Wellcome Sanger Institute for the COVID-19 Genomics UK (COG-UK) Consortium | Jacquelyn Wynn, Mairead Hyland, The Lighthouse Lab in Alderley Park and Alex Alderton, Roberto Amato, Jeffrey Barrett, Sonia Goncalves, Ewan Harrison, David K. Jackson, Ian Johnston, Dominic Kwiatkowski, Cordelia Langford, John Sillitoe on behalf of the Wellcome Sanger Institute COVID-19 Surveillance Team |

| | | | |
|---|---|---|---|
| EPI_ISL_2540650 | Lighthouse Lab in Milton Keynes | Wellcome Sanger Institute for the COVID-19 Genomics UK (COG-UK) Consortium | The Lighthouse Lab in Milton Keynes and Alex Alderton, Roberto Amato, Jeffrey Barrett, Sonia Goncalves, Ewan Harrison, David K. Jackson, Ian Johnston, Dominic Kwiatkowski, Cordelia Langford, John Sillitoe on behalf of the Wellcome Sanger Institute COVID-19 Surveillance Team |
| EPI_ISL_2540651 | Lighthouse Laboratory Plymouth | Wellcome Sanger Institute for the COVID-19 Genomics UK (COG-UK) Consortium | Lighthouse Laboratory Plymouth and Alex Alderton, Roberto Amato, Jeffrey Barrett, Sonia Goncalves, Ewan Harrison, David K. Jackson, Ian Johnston, Dominic Kwiatkowski, Cordelia Langford, John Sillitoe on behalf of the Wellcome Sanger Institute COVID-19 Surveillance Team |
| EPI_ISL_2540652, EPI_ISL_2540653 | Lighthouse Lab in Milton Keynes | Wellcome Sanger Institute for the COVID-19 Genomics UK (COG-UK) Consortium | The Lighthouse Lab in Milton Keynes and Alex Alderton, Roberto Amato, Jeffrey Barrett, Sonia Goncalves, Ewan Harrison, David K. Jackson, Ian Johnston, Dominic Kwiatkowski, Cordelia Langford, John Sillitoe on behalf of the Wellcome Sanger Institute COVID-19 Surveillance Team |
| EPI_ISL_2540654 | Lighthouse Lab in Alderley Park | Wellcome Sanger Institute for the COVID-19 Genomics UK (COG-UK) Consortium | Jacquelyn Wynn, Mairead Hyland, The Lighthouse Lab in Alderley Park and Alex Alderton, Roberto Amato, Jeffrey Barrett, Sonia Goncalves, Ewan Harrison, David K. Jackson, Ian Johnston, Dominic Kwiatkowski, Cordelia Langford, John Sillitoe on behalf of the Wellcome Sanger Institute COVID-19 Surveillance Team |
| EPI_ISL_2540655, EPI_ISL_2540656, EPI_ISL_2540657 | Lighthouse Lab in Milton Keynes | Wellcome Sanger Institute for the COVID-19 Genomics UK (COG-UK) Consortium | The Lighthouse Lab in Milton Keynes and Alex Alderton, Roberto Amato, Jeffrey Barrett, Sonia Goncalves, Ewan Harrison, David K. Jackson, Ian Johnston, Dominic Kwiatkowski, Cordelia Langford, John Sillitoe on behalf of the Wellcome Sanger Institute COVID-19 Surveillance Team |
| EPI_ISL_2540658 | Lighthouse Lab in Alderley Park | Wellcome Sanger Institute for the COVID-19 Genomics UK (COG-UK) Consortium | Jacquelyn Wynn, Mairead Hyland, The Lighthouse Lab in Alderley Park and Alex Alderton, Roberto Amato, Jeffrey Barrett, Sonia Goncalves, Ewan Harrison, David K. Jackson, Ian Johnston, Dominic Kwiatkowski, Cordelia Langford, John Sillitoe on behalf of the Wellcome Sanger Institute COVID-19 Surveillance Team |
| EPI_ISL_2540659 | Lighthouse Lab in Milton Keynes | Wellcome Sanger Institute for the COVID-19 Genomics UK (COG-UK) Consortium | The Lighthouse Lab in Milton Keynes and Alex Alderton, Roberto Amato, Jeffrey Barrett, Sonia Goncalves, Ewan Harrison, David K. Jackson, Ian Johnston, Dominic Kwiatkowski, Cordelia Langford, John Sillitoe on behalf of the Wellcome Sanger Institute COVID-19 Surveillance Team |
| EPI_ISL_2540660 | Lighthouse Laboratory Plymouth | Wellcome Sanger Institute for the COVID-19 Genomics UK (COG-UK) Consortium | Lighthouse Laboratory Plymouth and Alex Alderton, Roberto Amato, Jeffrey Barrett, Sonia Goncalves, Ewan Harrison, David K. Jackson, Ian Johnston, Dominic Kwiatkowski, Cordelia Langford, John Sillitoe on behalf of the Wellcome Sanger Institute COVID-19 Surveillance Team |
| EPI_ISL_2540661 | Lighthouse Lab in Milton Keynes | Wellcome Sanger Institute for the COVID-19 Genomics UK (COG-UK) Consortium | The Lighthouse Lab in Milton Keynes and Alex Alderton, Roberto Amato, Jeffrey Barrett, Sonia Goncalves, Ewan Harrison, David K. Jackson, Ian Johnston, Dominic Kwiatkowski, Cordelia Langford, John Sillitoe on behalf of the Wellcome Sanger Institute COVID-19 Surveillance Team |
| EPI_ISL_2540662 | Lighthouse Lab in Alderley Park | Wellcome Sanger Institute for the COVID-19 Genomics UK (COG-UK) Consortium | Jacquelyn Wynn, Mairead Hyland, The Lighthouse Lab in Alderley Park and Alex Alderton, Roberto Amato, Jeffrey Barrett, Sonia Goncalves, Ewan Harrison, David K. Jackson, Ian Johnston, Dominic Kwiatkowski, Cordelia Langford, John Sillitoe on behalf of the Wellcome Sanger Institute COVID-19 Surveillance Team |
| EPI_ISL_2540663, EPI_ISL_2540664 | Lighthouse Laboratory Plymouth | Wellcome Sanger Institute for the COVID-19 Genomics UK (COG-UK) Consortium | Lighthouse Laboratory Plymouth and Alex Alderton, Roberto Amato, Jeffrey Barrett, Sonia Goncalves, Ewan Harrison, David K. Jackson, Ian Johnston, Dominic Kwiatkowski, Cordelia Langford, John Sillitoe on behalf of the Wellcome Sanger Institute COVID-19 Surveillance Team |
| EPI_ISL_2540665, EPI_ISL_2540666, EPI_ISL_2540667, EPI_ISL_2540668 | Lighthouse Lab in Milton Keynes | Wellcome Sanger Institute for the COVID-19 Genomics UK (COG-UK) Consortium | The Lighthouse Lab in Milton Keynes and Alex Alderton, Roberto Amato, Jeffrey Barrett, Sonia Goncalves, Ewan Harrison, David K. Jackson, Ian Johnston, Dominic Kwiatkowski, Cordelia Langford, John Sillitoe on behalf of the Wellcome Sanger Institute COVID-19 Surveillance Team |
| EPI_ISL_2540669, EPI_ISL_2540670 | Lighthouse Laboratory Plymouth | Wellcome Sanger Institute for the COVID-19 Genomics UK (COG-UK) Consortium | Lighthouse Laboratory Plymouth and Alex Alderton, Roberto Amato, Jeffrey Barrett, Sonia Goncalves, Ewan Harrison, David K. Jackson, Ian Johnston, Dominic Kwiatkowski, Cordelia Langford, John Sillitoe on behalf of the Wellcome Sanger Institute COVID-19 Surveillance Team |
| EPI_ISL_2540672, EPI_ISL_2540673 | Lighthouse Lab in Alderley Park | Wellcome Sanger Institute for the COVID-19 Genomics UK (COG-UK) Consortium | Jacquelyn Wynn, Mairead Hyland, The Lighthouse Lab in Alderley Park and Alex Alderton, Roberto Amato, Jeffrey Barrett, Sonia Goncalves, Ewan Harrison, David K. Jackson, Ian Johnston, Dominic Kwiatkowski, Cordelia Langford, John Sillitoe on behalf of the Wellcome Sanger Institute COVID-19 Surveillance Team |
| EPI_ISL_2540674 | Lighthouse Laboratory Plymouth | Wellcome Sanger Institute for the COVID-19 Genomics UK (COG-UK) Consortium | Lighthouse Laboratory Plymouth and Alex Alderton, Roberto Amato, Jeffrey Barrett, Sonia Goncalves, Ewan Harrison, David K. Jackson, Ian Johnston, Dominic Kwiatkowski, Cordelia Langford, John Sillitoe on behalf of the Wellcome Sanger Institute COVID-19 Surveillance Team |
| EPI_ISL_2540676 | Lighthouse Lab in Milton Keynes | Wellcome Sanger Institute for the COVID-19 Genomics UK (COG-UK) Consortium | The Lighthouse Lab in Milton Keynes and Alex Alderton, Roberto Amato, Jeffrey Barrett, Sonia Goncalves, Ewan Harrison, David K. Jackson, Ian Johnston, Dominic Kwiatkowski, Cordelia Langford, John Sillitoe on behalf of the Wellcome Sanger Institute COVID-19 Surveillance Team |
| EPI_ISL_2540677 | Lighthouse Laboratory Plymouth | Wellcome Sanger Institute for the COVID-19 Genomics UK (COG-UK) Consortium | Lighthouse Laboratory Plymouth and Alex Alderton, Roberto Amato, Jeffrey Barrett, Sonia Goncalves, Ewan Harrison, David K. Jackson, Ian Johnston, Dominic Kwiatkowski, Cordelia Langford, John Sillitoe on behalf of the Wellcome Sanger Institute COVID-19 Surveillance Team |
| EPI_ISL_2540678 | Lighthouse Lab in Milton Keynes | Wellcome Sanger Institute for the COVID-19 Genomics UK (COG-UK) Consortium | The Lighthouse Lab in Milton Keynes and Alex Alderton, Roberto Amato, Jeffrey Barrett, Sonia Goncalves, Ewan Harrison, David K. Jackson, Ian Johnston, Dominic Kwiatkowski, Cordelia Langford, John Sillitoe on behalf of the Wellcome Sanger Institute COVID-19 Surveillance Team |
| EPI_ISL_2540679 | Lighthouse Lab in Alderley Park | Wellcome Sanger Institute for the COVID-19 Genomics UK (COG-UK) Consortium | Jacquelyn Wynn, Mairead Hyland, The Lighthouse Lab in Alderley Park and Alex Alderton, Roberto Amato, Jeffrey Barrett, Sonia Goncalves, Ewan Harrison, David K. Jackson, Ian Johnston, Dominic Kwiatkowski, Cordelia Langford, John Sillitoe on behalf of the Wellcome Sanger Institute COVID-19 Surveillance Team |
| EPI_ISL_2540681, EPI_ISL_2540682 | Lighthouse Lab in Milton Keynes | Wellcome Sanger Institute for the COVID-19 Genomics UK (COG-UK) Consortium | The Lighthouse Lab in Milton Keynes and Alex Alderton, Roberto Amato, Jeffrey Barrett, Sonia Goncalves, Ewan Harrison, David K. Jackson, Ian Johnston, Dominic Kwiatkowski, Cordelia Langford, John Sillitoe on behalf of the Wellcome Sanger Institute COVID-19 Surveillance Team |
| EPI_ISL_2540683 | Lighthouse Laboratory Plymouth | Wellcome Sanger Institute for the COVID-19 Genomics UK (COG-UK) Consortium | Lighthouse Laboratory Plymouth and Alex Alderton, Roberto Amato, Jeffrey Barrett, Sonia Goncalves, Ewan Harrison, David K. Jackson, Ian Johnston, Dominic Kwiatkowski, Cordelia Langford, John Sillitoe on behalf of the Wellcome Sanger Institute COVID-19 Surveillance Team |
| EPI_ISL_2540684, EPI_ISL_2540685, EPI_ISL_2540686, EPI_ISL_2540687, EPI_ISL_2540688 | Lighthouse Lab in Milton Keynes | Wellcome Sanger Institute for the COVID-19 Genomics UK (COG-UK) Consortium | The Lighthouse Lab in Milton Keynes and Alex Alderton, Roberto Amato, Jeffrey Barrett, Sonia Goncalves, Ewan Harrison, David K. Jackson, Ian Johnston, Dominic Kwiatkowski, Cordelia Langford, John Sillitoe on behalf of the Wellcome Sanger Institute COVID-19 Surveillance Team |
| EPI_ISL_2540689 | Lighthouse Lab in Alderley Park | Wellcome Sanger Institute for the COVID-19 Genomics UK (COG-UK) Consortium | Jacquelyn Wynn, Mairead Hyland, The Lighthouse Lab in Alderley Park and Alex Alderton, Roberto Amato, Jeffrey Barrett, Sonia Goncalves, Ewan Harrison, David K. Jackson, Ian Johnston, Dominic Kwiatkowski, Cordelia Langford, John Sillitoe on behalf of the Wellcome Sanger Institute COVID-19 Surveillance Team |
| EPI_ISL_2540690, EPI_ISL_2540691 | Lighthouse Laboratory Plymouth | Wellcome Sanger Institute for the COVID-19 Genomics UK (COG-UK) Consortium | Lighthouse Laboratory Plymouth and Alex Alderton, Roberto Amato, Jeffrey Barrett, Sonia Goncalves, Ewan Harrison, David K. Jackson, Ian Johnston, Dominic Kwiatkowski, Cordelia Langford, John Sillitoe on behalf of the Wellcome Sanger Institute COVID-19 Surveillance Team |
| EPI_ISL_2540692 | Lighthouse Lab in Milton Keynes | Wellcome Sanger Institute for the COVID-19 Genomics UK (COG-UK) Consortium | The Lighthouse Lab in Milton Keynes and Alex Alderton, Roberto Amato, Jeffrey Barrett, Sonia Goncalves, Ewan Harrison, David K. Jackson, Ian Johnston, Dominic Kwiatkowski, Cordelia Langford, John Sillitoe on behalf of the Wellcome Sanger Institute COVID-19 Surveillance Team |
| EPI_ISL_2540693 | Lighthouse Lab in Alderley Park | Wellcome Sanger Institute for the COVID-19 Genomics UK (COG-UK) Consortium | Jacquelyn Wynn, Mairead Hyland, The Lighthouse Lab in Alderley Park and Alex Alderton, Roberto Amato, Jeffrey Barrett, Sonia Goncalves, Ewan Harrison, David K. Jackson, Ian Johnston, Dominic Kwiatkowski, Cordelia Langford, John Sillitoe on behalf of the Wellcome Sanger Institute COVID-19 Surveillance Team |
| EPI_ISL_2540694 | Lighthouse Lab in Milton Keynes | Wellcome Sanger Institute for the COVID-19 Genomics UK (COG-UK) Consortium | The Lighthouse Lab in Milton Keynes and Alex Alderton, Roberto Amato, Jeffrey Barrett, Sonia Goncalves, Ewan Harrison, David K. Jackson, Ian Johnston, Dominic Kwiatkowski, Cordelia Langford, John Sillitoe on behalf of the Wellcome Sanger Institute COVID-19 Surveillance Team |
| EPI_ISL_2540695, EPI_ISL_2540696 | Lighthouse Lab in Alderley Park | Wellcome Sanger Institute for the COVID-19 Genomics UK (COG-UK) Consortium | Jacquelyn Wynn, Mairead Hyland, The Lighthouse Lab in Alderley Park and Alex Alderton, Roberto Amato, Jeffrey Barrett, Sonia Goncalves, Ewan Harrison, David K. Jackson, Ian Johnston, Dominic Kwiatkowski, Cordelia Langford, John Sillitoe on behalf of the Wellcome Sanger Institute COVID-19 Surveillance Team |
| EPI_ISL_2540697, EPI_ISL_2540698 | Lighthouse Lab in Milton Keynes | Wellcome Sanger Institute for the COVID-19 Genomics UK (COG-UK) Consortium | The Lighthouse Lab in Milton Keynes and Alex Alderton, Roberto Amato, Jeffrey Barrett, Sonia Goncalves, Ewan Harrison, David K. Jackson, Ian Johnston, Dominic Kwiatkowski, Cordelia Langford, John Sillitoe on behalf of the Wellcome Sanger Institute COVID-19 Surveillance Team |
| EPI_ISL_2540699 | Lighthouse Laboratory Plymouth | Wellcome Sanger Institute for the COVID-19 Genomics UK (COG-UK) Consortium | Lighthouse Laboratory Plymouth and Alex Alderton, Roberto Amato, Jeffrey Barrett, Sonia Goncalves, Ewan Harrison, David K. Jackson, Ian Johnston, Dominic Kwiatkowski, Cordelia Langford, John Sillitoe on behalf of the Wellcome Sanger Institute COVID-19 Surveillance Team |
| EPI_ISL_2540701 | Lighthouse Lab in Alderley Park | Wellcome Sanger Institute for the COVID-19 Genomics UK (COG-UK) Consortium | Jacquelyn Wynn, Mairead Hyland, The Lighthouse Lab in Alderley Park and Alex Alderton, Roberto Amato, Jeffrey Barrett, Sonia Goncalves, Ewan Harrison, David K. Jackson, Ian Johnston, Dominic Kwiatkowski, Cordelia Langford, John Sillitoe on behalf of the Wellcome Sanger Institute COVID-19 Surveillance Team |
| EPI_ISL_2540702 | Lighthouse Lab in Milton Keynes | Wellcome Sanger Institute for the COVID-19 Genomics UK (COG-UK) Consortium | The Lighthouse Lab in Milton Keynes and Alex Alderton, Roberto Amato, Jeffrey Barrett, Sonia Goncalves, Ewan Harrison, David K. Jackson, Ian Johnston, Dominic Kwiatkowski, Cordelia Langford, John Sillitoe on behalf of the Wellcome Sanger Institute COVID-19 Surveillance Team |
| EPI_ISL_2540703 | Lighthouse Lab in Alderley Park | Wellcome Sanger Institute for the COVID-19 Genomics UK (COG-UK) Consortium | Jacquelyn Wynn, Mairead Hyland, The Lighthouse Lab in Alderley Park and Alex Alderton, Roberto Amato, Jeffrey Barrett, Sonia Goncalves, Ewan Harrison, David K. Jackson, Ian Johnston, Dominic Kwiatkowski, Cordelia Langford, John Sillitoe on behalf of the Wellcome Sanger Institute COVID-19 Surveillance Team |
| EPI_ISL_2540705 | Lighthouse Lab in Milton Keynes | Wellcome Sanger Institute for the COVID-19 Genomics UK (COG-UK) Consortium | The Lighthouse Lab in Milton Keynes and Alex Alderton, Roberto Amato, Jeffrey Barrett, Sonia Goncalves, Ewan Harrison, David K. Jackson, Ian Johnston, Dominic Kwiatkowski, Cordelia Langford, John Sillitoe on behalf of the Wellcome Sanger Institute COVID-19 Surveillance Team |
| EPI_ISL_2540706 | Lighthouse Laboratory Plymouth | Wellcome Sanger Institute for the COVID-19 Genomics UK (COG-UK) Consortium | Lighthouse Laboratory Plymouth and Alex Alderton, Roberto Amato, Jeffrey Barrett, Sonia Goncalves, Ewan Harrison, David K. Jackson, Ian Johnston, Dominic Kwiatkowski, Cordelia Langford, John Sillitoe on behalf of the Wellcome Sanger Institute COVID-19 Surveillance Team |
| EPI_ISL_2540707, EPI_ISL_2540708 | Lighthouse Lab in Milton Keynes | Wellcome Sanger Institute for the COVID-19 Genomics UK (COG-UK) Consortium | The Lighthouse Lab in Milton Keynes and Alex Alderton, Roberto Amato, Jeffrey Barrett, Sonia Goncalves, Ewan Harrison, David K. Jackson, Ian Johnston, Dominic Kwiatkowski, Cordelia Langford, John Sillitoe on behalf of the Wellcome Sanger Institute COVID-19 Surveillance Team |
| EPI_ISL_2540709 | Lighthouse Lab in Alderley Park | Wellcome Sanger Institute for the COVID-19 Genomics UK (COG-UK) Consortium | Jacquelyn Wynn, Mairead Hyland, The Lighthouse Lab in Alderley Park and Alex Alderton, Roberto Amato, Jeffrey Barrett, Sonia Goncalves, Ewan Harrison, David K. Jackson, Ian Johnston, Dominic Kwiatkowski, Cordelia Langford, John Sillitoe on behalf of the Wellcome Sanger Institute COVID-19 Surveillance Team |
| EPI_ISL_2540710, EPI_ISL_2540711, EPI_ISL_2540712, EPI_ISL_2540713, EPI_ISL_2540714 | Lighthouse Lab in Milton Keynes | Wellcome Sanger Institute for the COVID-19 Genomics UK (COG-UK) Consortium | The Lighthouse Lab in Milton Keynes and Alex Alderton, Roberto Amato, Jeffrey Barrett, Sonia Goncalves, Ewan Harrison, David K. Jackson, Ian Johnston, Dominic Kwiatkowski, Cordelia Langford, John Sillitoe on behalf of the Wellcome Sanger Institute COVID-19 Surveillance Team |
| EPI_ISL_2540715 | Lighthouse Lab in Alderley Park | Wellcome Sanger Institute for the COVID-19 Genomics UK (COG-UK) Consortium | Jacquelyn Wynn, Mairead Hyland, The Lighthouse Lab in Alderley Park and Alex Alderton, Roberto Amato, Jeffrey Barrett, Sonia Goncalves, Ewan Harrison, David K. Jackson, Ian Johnston, Dominic Kwiatkowski, Cordelia Langford, John Sillitoe on behalf of the Wellcome Sanger Institute COVID-19 Surveillance Team |
| EPI_ISL_2540717 | Lighthouse Laboratory Plymouth | Wellcome Sanger Institute for the COVID-19 Genomics UK (COG-UK) Consortium | Lighthouse Laboratory Plymouth and Alex Alderton, Roberto Amato, Jeffrey Barrett, Sonia Goncalves, Ewan Harrison, David K. Jackson, Ian Johnston, Dominic Kwiatkowski, Cordelia Langford, John Sillitoe on behalf of the Wellcome Sanger Institute COVID-19 Surveillance Team |
| EPI_ISL_2540718 | Lighthouse Lab in Milton Keynes | Wellcome Sanger Institute for the COVID-19 Genomics UK (COG-UK) Consortium | The Lighthouse Lab in Milton Keynes and Alex Alderton, Roberto Amato, Jeffrey Barrett, Sonia Goncalves, Ewan Harrison, David K. Jackson, Ian Johnston, Dominic Kwiatkowski, Cordelia Langford, John Sillitoe on behalf of the Wellcome Sanger Institute COVID-19 Surveillance Team |

| | | | |
|---|---|---|---|
| EPI_ISL_2540719 | Lighthouse Lab in Alderley Park | Wellcome Sanger Institute for the COVID-19 Genomics UK (COG-UK) Consortium | Jacquelyn Wynn, Mairead Hyland, The Lighthouse Lab in Alderley Park and Alex Alderton, Roberto Amato, Jeffrey Barrett, Sonia Goncalves, Ewan Harrison, David K. Jackson, Ian Johnston, Dominic Kwiatkowski, Cordelia Langford, John Sillitoe on behalf of the Wellcome Sanger Institute COVID-19 Surveillance Team |
| EPI_ISL_2540720, EPI_ISL_2540721 | Lighthouse Laboratory Plymouth | Wellcome Sanger Institute for the COVID-19 Genomics UK (COG-UK) Consortium | Lighthouse Laboratory Plymouth and Alex Alderton, Roberto Amato, Jeffrey Barrett, Sonia Goncalves, Ewan Harrison, David K. Jackson, Ian Johnston, Dominic Kwiatkowski, Cordelia Langford, John Sillitoe on behalf of the Wellcome Sanger Institute COVID-19 Surveillance Team |
| EPI_ISL_2540722 | Lighthouse Lab in Alderley Park | Wellcome Sanger Institute for the COVID-19 Genomics UK (COG-UK) Consortium | Jacquelyn Wynn, Mairead Hyland, The Lighthouse Lab in Alderley Park and Alex Alderton, Roberto Amato, Jeffrey Barrett, Sonia Goncalves, Ewan Harrison, David K. Jackson, Ian Johnston, Dominic Kwiatkowski, Cordelia Langford, John Sillitoe on behalf of the Wellcome Sanger Institute COVID-19 Surveillance Team |
| EPI_ISL_2540723 | Lighthouse Laboratory Plymouth | Wellcome Sanger Institute for the COVID-19 Genomics UK (COG-UK) Consortium | Lighthouse Laboratory Plymouth and Alex Alderton, Roberto Amato, Jeffrey Barrett, Sonia Goncalves, Ewan Harrison, David K. Jackson, Ian Johnston, Dominic Kwiatkowski, Cordelia Langford, John Sillitoe on behalf of the Wellcome Sanger Institute COVID-19 Surveillance Team |
| EPI_ISL_2540724, EPI_ISL_2540725, EPI_ISL_2540726 | Lighthouse Lab in Milton Keynes | Wellcome Sanger Institute for the COVID-19 Genomics UK (COG-UK) Consortium | The Lighthouse Lab in Milton Keynes and Alex Alderton, Roberto Amato, Jeffrey Barrett, Sonia Goncalves, Ewan Harrison, David K. Jackson, Ian Johnston, Dominic Kwiatkowski, Cordelia Langford, John Sillitoe on behalf of the Wellcome Sanger Institute COVID-19 Surveillance Team |
| EPI_ISL_2540727 | Lighthouse Lab in Alderley Park | Wellcome Sanger Institute for the COVID-19 Genomics UK (COG-UK) Consortium | Jacquelyn Wynn, Mairead Hyland, The Lighthouse Lab in Alderley Park and Alex Alderton, Roberto Amato, Jeffrey Barrett, Sonia Goncalves, Ewan Harrison, David K. Jackson, Ian Johnston, Dominic Kwiatkowski, Cordelia Langford, John Sillitoe on behalf of the Wellcome Sanger Institute COVID-19 Surveillance Team |
| EPI_ISL_2540728, EPI_ISL_2540729 | Lighthouse Lab in Milton Keynes | Wellcome Sanger Institute for the COVID-19 Genomics UK (COG-UK) Consortium | The Lighthouse Lab in Milton Keynes and Alex Alderton, Roberto Amato, Jeffrey Barrett, Sonia Goncalves, Ewan Harrison, David K. Jackson, Ian Johnston, Dominic Kwiatkowski, Cordelia Langford, John Sillitoe on behalf of the Wellcome Sanger Institute COVID-19 Surveillance Team |
| EPI_ISL_2540730 | Lighthouse Lab in Alderley Park | Wellcome Sanger Institute for the COVID-19 Genomics UK (COG-UK) Consortium | Jacquelyn Wynn, Mairead Hyland, The Lighthouse Lab in Alderley Park and Alex Alderton, Roberto Amato, Jeffrey Barrett, Sonia Goncalves, Ewan Harrison, David K. Jackson, Ian Johnston, Dominic Kwiatkowski, Cordelia Langford, John Sillitoe on behalf of the Wellcome Sanger Institute COVID-19 Surveillance Team |
| EPI_ISL_2540731, EPI_ISL_2540732 | Lighthouse Lab in Milton Keynes | Wellcome Sanger Institute for the COVID-19 Genomics UK (COG-UK) Consortium | The Lighthouse Lab in Milton Keynes and Alex Alderton, Roberto Amato, Jeffrey Barrett, Sonia Goncalves, Ewan Harrison, David K. Jackson, Ian Johnston, Dominic Kwiatkowski, Cordelia Langford, John Sillitoe on behalf of the Wellcome Sanger Institute COVID-19 Surveillance Team |
| EPI_ISL_2540733 | Lighthouse Laboratory Plymouth | Wellcome Sanger Institute for the COVID-19 Genomics UK (COG-UK) Consortium | Lighthouse Laboratory Plymouth and Alex Alderton, Roberto Amato, Jeffrey Barrett, Sonia Goncalves, Ewan Harrison, David K. Jackson, Ian Johnston, Dominic Kwiatkowski, Cordelia Langford, John Sillitoe on behalf of the Wellcome Sanger Institute COVID-19 Surveillance Team |
| EPI_ISL_2540734 | Lighthouse Lab in Milton Keynes | Wellcome Sanger Institute for the COVID-19 Genomics UK (COG-UK) Consortium | The Lighthouse Lab in Milton Keynes and Alex Alderton, Roberto Amato, Jeffrey Barrett, Sonia Goncalves, Ewan Harrison, David K. Jackson, Ian Johnston, Dominic Kwiatkowski, Cordelia Langford, John Sillitoe on behalf of the Wellcome Sanger Institute COVID-19 Surveillance Team |
| EPI_ISL_2540735 | Lighthouse Laboratory Plymouth | Wellcome Sanger Institute for the COVID-19 Genomics UK (COG-UK) Consortium | Lighthouse Laboratory Plymouth and Alex Alderton, Roberto Amato, Jeffrey Barrett, Sonia Goncalves, Ewan Harrison, David K. Jackson, Ian Johnston, Dominic Kwiatkowski, Cordelia Langford, John Sillitoe on behalf of the Wellcome Sanger Institute COVID-19 Surveillance Team |
| EPI_ISL_2540736, EPI_ISL_2540737, EPI_ISL_2540738 | Lighthouse Lab in Milton Keynes | Wellcome Sanger Institute for the COVID-19 Genomics UK (COG-UK) Consortium | The Lighthouse Lab in Milton Keynes and Alex Alderton, Roberto Amato, Jeffrey Barrett, Sonia Goncalves, Ewan Harrison, David K. Jackson, Ian Johnston, Dominic Kwiatkowski, Cordelia Langford, John Sillitoe on behalf of the Wellcome Sanger Institute COVID-19 Surveillance Team |
| EPI_ISL_2540739, EPI_ISL_2540740, EPI_ISL_2540741, EPI_ISL_2540742 | Lighthouse Lab in Alderley Park | Wellcome Sanger Institute for the COVID-19 Genomics UK (COG-UK) Consortium | Jacquelyn Wynn, Mairead Hyland, The Lighthouse Lab in Alderley Park and Alex Alderton, Roberto Amato, Jeffrey Barrett, Sonia Goncalves, Ewan Harrison, David K. Jackson, Ian Johnston, Dominic Kwiatkowski, Cordelia Langford, John Sillitoe on behalf of the Wellcome Sanger Institute COVID-19 Surveillance Team |
| EPI_ISL_2540744 | Lighthouse Laboratory Plymouth | Wellcome Sanger Institute for the COVID-19 Genomics UK (COG-UK) Consortium | Lighthouse Laboratory Plymouth and Alex Alderton, Roberto Amato, Jeffrey Barrett, Sonia Goncalves, Ewan Harrison, David K. Jackson, Ian Johnston, Dominic Kwiatkowski, Cordelia Langford, John Sillitoe on behalf of the Wellcome Sanger Institute COVID-19 Surveillance Team |
| EPI_ISL_2540745, EPI_ISL_2540747 | Lighthouse Lab in Alderley Park | Wellcome Sanger Institute for the COVID-19 Genomics UK (COG-UK) Consortium | Jacquelyn Wynn, Mairead Hyland, The Lighthouse Lab in Alderley Park and Alex Alderton, Roberto Amato, Jeffrey Barrett, Sonia Goncalves, Ewan Harrison, David K. Jackson, Ian Johnston, Dominic Kwiatkowski, Cordelia Langford, John Sillitoe on behalf of the Wellcome Sanger Institute COVID-19 Surveillance Team |
| EPI_ISL_2540748 | Lighthouse Laboratory Plymouth | Wellcome Sanger Institute for the COVID-19 Genomics UK (COG-UK) Consortium | Lighthouse Laboratory Plymouth and Alex Alderton, Roberto Amato, Jeffrey Barrett, Sonia Goncalves, Ewan Harrison, David K. Jackson, Ian Johnston, Dominic Kwiatkowski, Cordelia Langford, John Sillitoe on behalf of the Wellcome Sanger Institute COVID-19 Surveillance Team |
| EPI_ISL_2540749 | Lighthouse Lab in Milton Keynes | Wellcome Sanger Institute for the COVID-19 Genomics UK (COG-UK) Consortium | The Lighthouse Lab in Milton Keynes and Alex Alderton, Roberto Amato, Jeffrey Barrett, Sonia Goncalves, Ewan Harrison, David K. Jackson, Ian Johnston, Dominic Kwiatkowski, Cordelia Langford, John Sillitoe on behalf of the Wellcome Sanger Institute COVID-19 Surveillance Team |
| EPI_ISL_2540750, EPI_ISL_2540751 | Lighthouse Laboratory Plymouth | Wellcome Sanger Institute for the COVID-19 Genomics UK (COG-UK) Consortium | Lighthouse Laboratory Plymouth and Alex Alderton, Roberto Amato, Jeffrey Barrett, Sonia Goncalves, Ewan Harrison, David K. Jackson, Ian Johnston, Dominic Kwiatkowski, Cordelia Langford, John Sillitoe on behalf of the Wellcome Sanger Institute COVID-19 Surveillance Team |
| EPI_ISL_2540752 | Lighthouse Lab in Milton Keynes | Wellcome Sanger Institute for the COVID-19 Genomics UK (COG-UK) Consortium | The Lighthouse Lab in Milton Keynes and Alex Alderton, Roberto Amato, Jeffrey Barrett, Sonia Goncalves, Ewan Harrison, David K. Jackson, Ian Johnston, Dominic Kwiatkowski, Cordelia Langford, John Sillitoe on behalf of the Wellcome Sanger Institute COVID-19 Surveillance Team |
| EPI_ISL_2540753 | Lighthouse Laboratory Plymouth | Wellcome Sanger Institute for the COVID-19 Genomics UK (COG-UK) Consortium | Lighthouse Laboratory Plymouth and Alex Alderton, Roberto Amato, Jeffrey Barrett, Sonia Goncalves, Ewan Harrison, David K. Jackson, Ian Johnston, Dominic Kwiatkowski, Cordelia Langford, John Sillitoe on behalf of the Wellcome Sanger Institute COVID-19 Surveillance Team |
| EPI_ISL_2540754, EPI_ISL_2540755, EPI_ISL_2540756, EPI_ISL_2540757 | Lighthouse Lab in Milton Keynes | Wellcome Sanger Institute for the COVID-19 Genomics UK (COG-UK) Consortium | The Lighthouse Lab in Milton Keynes and Alex Alderton, Roberto Amato, Jeffrey Barrett, Sonia Goncalves, Ewan Harrison, David K. Jackson, Ian Johnston, Dominic Kwiatkowski, Cordelia Langford, John Sillitoe on behalf of the Wellcome Sanger Institute COVID-19 Surveillance Team |
| EPI_ISL_2540758 | Lighthouse Lab in Alderley Park | Wellcome Sanger Institute for the COVID-19 Genomics UK (COG-UK) Consortium | Jacquelyn Wynn, Mairead Hyland, The Lighthouse Lab in Alderley Park and Alex Alderton, Roberto Amato, Jeffrey Barrett, Sonia Goncalves, Ewan Harrison, David K. Jackson, Ian Johnston, Dominic Kwiatkowski, Cordelia Langford, John Sillitoe on behalf of the Wellcome Sanger Institute COVID-19 Surveillance Team |
| EPI_ISL_2540759 | Lighthouse Laboratory Plymouth | Wellcome Sanger Institute for the COVID-19 Genomics UK (COG-UK) Consortium | Lighthouse Laboratory Plymouth and Alex Alderton, Roberto Amato, Jeffrey Barrett, Sonia Goncalves, Ewan Harrison, David K. Jackson, Ian Johnston, Dominic Kwiatkowski, Cordelia Langford, John Sillitoe on behalf of the Wellcome Sanger Institute COVID-19 Surveillance Team |
| EPI_ISL_2540760 | Lighthouse Lab in Milton Keynes | Wellcome Sanger Institute for the COVID-19 Genomics UK (COG-UK) Consortium | The Lighthouse Lab in Milton Keynes and Alex Alderton, Roberto Amato, Jeffrey Barrett, Sonia Goncalves, Ewan Harrison, David K. Jackson, Ian Johnston, Dominic Kwiatkowski, Cordelia Langford, John Sillitoe on behalf of the Wellcome Sanger Institute COVID-19 Surveillance Team |
| EPI_ISL_2540761 | Lighthouse Lab in Alderley Park | Wellcome Sanger Institute for the COVID-19 Genomics UK (COG-UK) Consortium | Jacquelyn Wynn, Mairead Hyland, The Lighthouse Lab in Alderley Park and Alex Alderton, Roberto Amato, Jeffrey Barrett, Sonia Goncalves, Ewan Harrison, David K. Jackson, Ian Johnston, Dominic Kwiatkowski, Cordelia Langford, John Sillitoe on behalf of the Wellcome Sanger Institute COVID-19 Surveillance Team |
| EPI_ISL_2540762 | Lighthouse Laboratory Plymouth | Wellcome Sanger Institute for the COVID-19 Genomics UK (COG-UK) Consortium | Lighthouse Laboratory Plymouth and Alex Alderton, Roberto Amato, Jeffrey Barrett, Sonia Goncalves, Ewan Harrison, David K. Jackson, Ian Johnston, Dominic Kwiatkowski, Cordelia Langford, John Sillitoe on behalf of the Wellcome Sanger Institute COVID-19 Surveillance Team |
| EPI_ISL_2540763 | Lighthouse Lab in Alderley Park | Wellcome Sanger Institute for the COVID-19 Genomics UK (COG-UK) Consortium | Jacquelyn Wynn, Mairead Hyland, The Lighthouse Lab in Alderley Park and Alex Alderton, Roberto Amato, Jeffrey Barrett, Sonia Goncalves, Ewan Harrison, David K. Jackson, Ian Johnston, Dominic Kwiatkowski, Cordelia Langford, John Sillitoe on behalf of the Wellcome Sanger Institute COVID-19 Surveillance Team |
| EPI_ISL_2540764, EPI_ISL_2540765, EPI_ISL_2540766 | Lighthouse Lab in Milton Keynes | Wellcome Sanger Institute for the COVID-19 Genomics UK (COG-UK) Consortium | The Lighthouse Lab in Milton Keynes and Alex Alderton, Roberto Amato, Jeffrey Barrett, Sonia Goncalves, Ewan Harrison, David K. Jackson, Ian Johnston, Dominic Kwiatkowski, Cordelia Langford, John Sillitoe on behalf of the Wellcome Sanger Institute COVID-19 Surveillance Team |
| EPI_ISL_2540767, EPI_ISL_2540768, EPI_ISL_2540769 | Lighthouse Lab in Alderley Park | Wellcome Sanger Institute for the COVID-19 Genomics UK (COG-UK) Consortium | Jacquelyn Wynn, Mairead Hyland, The Lighthouse Lab in Alderley Park and Alex Alderton, Roberto Amato, Jeffrey Barrett, Sonia Goncalves, Ewan Harrison, David K. Jackson, Ian Johnston, Dominic Kwiatkowski, Cordelia Langford, John Sillitoe on behalf of the Wellcome Sanger Institute COVID-19 Surveillance Team |
| EPI_ISL_2540770 | Lighthouse Laboratory Plymouth | Wellcome Sanger Institute for the COVID-19 Genomics UK (COG-UK) Consortium | Lighthouse Laboratory Plymouth and Alex Alderton, Roberto Amato, Jeffrey Barrett, Sonia Goncalves, Ewan Harrison, David K. Jackson, Ian Johnston, Dominic Kwiatkowski, Cordelia Langford, John Sillitoe on behalf of the Wellcome Sanger Institute COVID-19 Surveillance Team |
| EPI_ISL_2540771 | Lighthouse Lab in Milton Keynes | Wellcome Sanger Institute for the COVID-19 Genomics UK (COG-UK) Consortium | The Lighthouse Lab in Milton Keynes and Alex Alderton, Roberto Amato, Jeffrey Barrett, Sonia Goncalves, Ewan Harrison, David K. Jackson, Ian Johnston, Dominic Kwiatkowski, Cordelia Langford, John Sillitoe on behalf of the Wellcome Sanger Institute COVID-19 Surveillance Team |
| EPI_ISL_2540772, EPI_ISL_2540773, EPI_ISL_2540774 | Lighthouse Laboratory Plymouth | Wellcome Sanger Institute for the COVID-19 Genomics UK (COG-UK) Consortium | Lighthouse Laboratory Plymouth and Alex Alderton, Roberto Amato, Jeffrey Barrett, Sonia Goncalves, Ewan Harrison, David K. Jackson, Ian Johnston, Dominic Kwiatkowski, Cordelia Langford, John Sillitoe on behalf of the Wellcome Sanger Institute COVID-19 Surveillance Team |
| EPI_ISL_2540775 | Lighthouse Lab in Milton Keynes | Wellcome Sanger Institute for the COVID-19 Genomics UK (COG-UK) Consortium | The Lighthouse Lab in Milton Keynes and Alex Alderton, Roberto Amato, Jeffrey Barrett, Sonia Goncalves, Ewan Harrison, David K. Jackson, Ian Johnston, Dominic Kwiatkowski, Cordelia Langford, John Sillitoe on behalf of the Wellcome Sanger Institute COVID-19 Surveillance Team |
| EPI_ISL_2540776, EPI_ISL_2540779 | Lighthouse Laboratory Plymouth | Wellcome Sanger Institute for the COVID-19 Genomics UK (COG-UK) Consortium | Lighthouse Laboratory Plymouth and Alex Alderton, Roberto Amato, Jeffrey Barrett, Sonia Goncalves, Ewan Harrison, David K. Jackson, Ian Johnston, Dominic Kwiatkowski, Cordelia Langford, John Sillitoe on behalf of the Wellcome Sanger Institute COVID-19 Surveillance Team |
| EPI_ISL_2540781 | Lighthouse Lab in Milton Keynes | Wellcome Sanger Institute for the COVID-19 Genomics UK (COG-UK) Consortium | The Lighthouse Lab in Milton Keynes and Alex Alderton, Roberto Amato, Jeffrey Barrett, Sonia Goncalves, Ewan Harrison, David K. Jackson, Ian Johnston, Dominic Kwiatkowski, Cordelia Langford, John Sillitoe on behalf of the Wellcome Sanger Institute COVID-19 Surveillance Team |
| EPI_ISL_2540782 | Lighthouse Lab in Alderley Park | Wellcome Sanger Institute for the COVID-19 Genomics UK (COG-UK) Consortium | Jacquelyn Wynn, Mairead Hyland, The Lighthouse Lab in Alderley Park and Alex Alderton, Roberto Amato, Jeffrey Barrett, Sonia Goncalves, Ewan Harrison, David K. Jackson, Ian Johnston, Dominic Kwiatkowski, Cordelia Langford, John Sillitoe on behalf of the Wellcome Sanger Institute COVID-19 Surveillance Team |
| EPI_ISL_2540783, EPI_ISL_2540784, EPI_ISL_2540786, EPI_ISL_2540789 | Lighthouse Lab in Milton Keynes | Wellcome Sanger Institute for the COVID-19 Genomics UK (COG-UK) Consortium | The Lighthouse Lab in Milton Keynes and Alex Alderton, Roberto Amato, Jeffrey Barrett, Sonia Goncalves, Ewan Harrison, David K. Jackson, Ian Johnston, Dominic Kwiatkowski, Cordelia Langford, John Sillitoe on behalf of the Wellcome Sanger Institute COVID-19 Surveillance Team |
| EPI_ISL_2540790 | Lighthouse Lab in Alderley Park | Wellcome Sanger Institute for the COVID-19 Genomics UK (COG-UK) Consortium | Jacquelyn Wynn, Mairead Hyland, The Lighthouse Lab in Alderley Park and Alex Alderton, Roberto Amato, Jeffrey Barrett, Sonia Goncalves, Ewan Harrison, David K. Jackson, Ian Johnston, Dominic Kwiatkowski, Cordelia Langford, John Sillitoe on behalf of the Wellcome Sanger Institute COVID-19 Surveillance Team |
| EPI_ISL_2540792 | Lighthouse Lab in Milton Keynes | Wellcome Sanger Institute for the COVID-19 Genomics UK (COG-UK) Consortium | The Lighthouse Lab in Milton Keynes and Alex Alderton, Roberto Amato, Jeffrey Barrett, Sonia Goncalves, Ewan Harrison, David K. Jackson, Ian Johnston, Dominic Kwiatkowski, Cordelia Langford, John Sillitoe on behalf of the Wellcome Sanger Institute COVID-19 Surveillance Team |
| EPI_ISL_2540793 | Lighthouse Laboratory Plymouth | Wellcome Sanger Institute for the COVID-19 Genomics UK (COG-UK) | Lighthouse Laboratory Plymouth and Alex Alderton, Roberto Amato, Jeffrey Barrett, Sonia Goncalves, Ewan Harrison, David K. Jackson, Ian Johnston, Dominic Kwiatkowski, Cordelia Langford, John Sillitoe on behalf of the Wellcome Sanger |

| | | | |
|---|---|---|---|
| | | Consortium | Institute COVID-19 Surveillance Team |
| EPI_ISL_2540794 | Lighthouse Lab in Alderley Park | Wellcome Sanger Institute for the COVID-19 Genomics UK (COG-UK) Consortium | Jacquelyn Wynn, Mairead Hyland, The Lighthouse Lab in Alderley Park and Alex Alderton, Roberto Amato, Jeffrey Barrett, Sonia Goncalves, Ewan Harrison, David K. Jackson, Ian Johnston, Dominic Kwiatkowski, Cordelia Langford, John Sillitoe on behalf of the Wellcome Sanger Institute COVID-19 Surveillance Team |
| EPI_ISL_2540795 | Lighthouse Lab in Milton Keynes | Wellcome Sanger Institute for the COVID-19 Genomics UK (COG-UK) Consortium | The Lighthouse Lab in Milton Keynes and Alex Alderton, Roberto Amato, Jeffrey Barrett, Sonia Goncalves, Ewan Harrison, David K. Jackson, Ian Johnston, Dominic Kwiatkowski, Cordelia Langford, John Sillitoe on behalf of the Wellcome Sanger Institute COVID-19 Surveillance Team |
| EPI_ISL_2540796 | Lighthouse Laboratory Plymouth | Wellcome Sanger Institute for the COVID-19 Genomics UK (COG-UK) Consortium | Lighthouse Laboratory Plymouth and Alex Alderton, Roberto Amato, Jeffrey Barrett, Sonia Goncalves, Ewan Harrison, David K. Jackson, Ian Johnston, Dominic Kwiatkowski, Cordelia Langford, John Sillitoe on behalf of the Wellcome Sanger Institute COVID-19 Surveillance Team |
| EPI_ISL_2540797 | Lighthouse Lab in Milton Keynes | Wellcome Sanger Institute for the COVID-19 Genomics UK (COG-UK) Consortium | The Lighthouse Lab in Milton Keynes and Alex Alderton, Roberto Amato, Jeffrey Barrett, Sonia Goncalves, Ewan Harrison, David K. Jackson, Ian Johnston, Dominic Kwiatkowski, Cordelia Langford, John Sillitoe on behalf of the Wellcome Sanger Institute COVID-19 Surveillance Team |
| EPI_ISL_2540798, EPI_ISL_2540799, EPI_ISL_2540800 | Lighthouse Lab in Alderley Park | Wellcome Sanger Institute for the COVID-19 Genomics UK (COG-UK) Consortium | Jacquelyn Wynn, Mairead Hyland, The Lighthouse Lab in Alderley Park and Alex Alderton, Roberto Amato, Jeffrey Barrett, Sonia Goncalves, Ewan Harrison, David K. Jackson, Ian Johnston, Dominic Kwiatkowski, Cordelia Langford, John Sillitoe on behalf of the Wellcome Sanger Institute COVID-19 Surveillance Team |
| EPI_ISL_2540801 | Lighthouse Lab in Milton Keynes | Wellcome Sanger Institute for the COVID-19 Genomics UK (COG-UK) Consortium | The Lighthouse Lab in Milton Keynes and Alex Alderton, Roberto Amato, Jeffrey Barrett, Sonia Goncalves, Ewan Harrison, David K. Jackson, Ian Johnston, Dominic Kwiatkowski, Cordelia Langford, John Sillitoe on behalf of the Wellcome Sanger Institute COVID-19 Surveillance Team |
| EPI_ISL_2540802, EPI_ISL_2540803, EPI_ISL_2540804, EPI_ISL_2540805, EPI_ISL_2540806 | Lighthouse Lab in Alderley Park | Wellcome Sanger Institute for the COVID-19 Genomics UK (COG-UK) Consortium | Jacquelyn Wynn, Mairead Hyland, The Lighthouse Lab in Alderley Park and Alex Alderton, Roberto Amato, Jeffrey Barrett, Sonia Goncalves, Ewan Harrison, David K. Jackson, Ian Johnston, Dominic Kwiatkowski, Cordelia Langford, John Sillitoe on behalf of the Wellcome Sanger Institute COVID-19 Surveillance Team |
| EPI_ISL_2540807 | Lighthouse Laboratory Plymouth | Wellcome Sanger Institute for the COVID-19 Genomics UK (COG-UK) Consortium | Lighthouse Laboratory Plymouth and Alex Alderton, Roberto Amato, Jeffrey Barrett, Sonia Goncalves, Ewan Harrison, David K. Jackson, Ian Johnston, Dominic Kwiatkowski, Cordelia Langford, John Sillitoe on behalf of the Wellcome Sanger Institute COVID-19 Surveillance Team |
| EPI_ISL_2540808 | Lighthouse Lab in Milton Keynes | Wellcome Sanger Institute for the COVID-19 Genomics UK (COG-UK) Consortium | The Lighthouse Lab in Milton Keynes and Alex Alderton, Roberto Amato, Jeffrey Barrett, Sonia Goncalves, Ewan Harrison, David K. Jackson, Ian Johnston, Dominic Kwiatkowski, Cordelia Langford, John Sillitoe on behalf of the Wellcome Sanger Institute COVID-19 Surveillance Team |
| EPI_ISL_2540809 | Lighthouse Lab in Alderley Park | Wellcome Sanger Institute for the COVID-19 Genomics UK (COG-UK) Consortium | Jacquelyn Wynn, Mairead Hyland, The Lighthouse Lab in Alderley Park and Alex Alderton, Roberto Amato, Jeffrey Barrett, Sonia Goncalves, Ewan Harrison, David K. Jackson, Ian Johnston, Dominic Kwiatkowski, Cordelia Langford, John Sillitoe on behalf of the Wellcome Sanger Institute COVID-19 Surveillance Team |
| EPI_ISL_2540810, EPI_ISL_2540811, EPI_ISL_2540812, EPI_ISL_2540813, EPI_ISL_2540814, EPI_ISL_2540815 | Lighthouse Lab in Milton Keynes | Wellcome Sanger Institute for the COVID-19 Genomics UK (COG-UK) Consortium | The Lighthouse Lab in Milton Keynes and Alex Alderton, Roberto Amato, Jeffrey Barrett, Sonia Goncalves, Ewan Harrison, David K. Jackson, Ian Johnston, Dominic Kwiatkowski, Cordelia Langford, John Sillitoe on behalf of the Wellcome Sanger Institute COVID-19 Surveillance Team |
| EPI_ISL_2540816 | Lighthouse Lab in Alderley Park | Wellcome Sanger Institute for the COVID-19 Genomics UK (COG-UK) Consortium | Jacquelyn Wynn, Mairead Hyland, The Lighthouse Lab in Alderley Park and Alex Alderton, Roberto Amato, Jeffrey Barrett, Sonia Goncalves, Ewan Harrison, David K. Jackson, Ian Johnston, Dominic Kwiatkowski, Cordelia Langford, John Sillitoe on behalf of the Wellcome Sanger Institute COVID-19 Surveillance Team |
| EPI_ISL_2540817, EPI_ISL_2540818, EPI_ISL_2540819 | Lighthouse Lab in Milton Keynes | Wellcome Sanger Institute for the COVID-19 Genomics UK (COG-UK) Consortium | The Lighthouse Lab in Milton Keynes and Alex Alderton, Roberto Amato, Jeffrey Barrett, Sonia Goncalves, Ewan Harrison, David K. Jackson, Ian Johnston, Dominic Kwiatkowski, Cordelia Langford, John Sillitoe on behalf of the Wellcome Sanger Institute COVID-19 Surveillance Team |
| EPI_ISL_2540820, EPI_ISL_2540821 | Lighthouse Laboratory Plymouth | Wellcome Sanger Institute for the COVID-19 Genomics UK (COG-UK) Consortium | Lighthouse Laboratory Plymouth and Alex Alderton, Roberto Amato, Jeffrey Barrett, Sonia Goncalves, Ewan Harrison, David K. Jackson, Ian Johnston, Dominic Kwiatkowski, Cordelia Langford, John Sillitoe on behalf of the Wellcome Sanger Institute COVID-19 Surveillance Team |
| EPI_ISL_2540822, EPI_ISL_2540823, EPI_ISL_2540824 | Lighthouse Lab in Milton Keynes | Wellcome Sanger Institute for the COVID-19 Genomics UK (COG-UK) Consortium | The Lighthouse Lab in Milton Keynes and Alex Alderton, Roberto Amato, Jeffrey Barrett, Sonia Goncalves, Ewan Harrison, David K. Jackson, Ian Johnston, Dominic Kwiatkowski, Cordelia Langford, John Sillitoe on behalf of the Wellcome Sanger Institute COVID-19 Surveillance Team |
| EPI_ISL_2540825, EPI_ISL_2540826, EPI_ISL_2540827, EPI_ISL_2540829 | Lighthouse Laboratory Plymouth | Wellcome Sanger Institute for the COVID-19 Genomics UK (COG-UK) Consortium | Lighthouse Laboratory Plymouth and Alex Alderton, Roberto Amato, Jeffrey Barrett, Sonia Goncalves, Ewan Harrison, David K. Jackson, Ian Johnston, Dominic Kwiatkowski, Cordelia Langford, John Sillitoe on behalf of the Wellcome Sanger Institute COVID-19 Surveillance Team |
| EPI_ISL_2540830 | Lighthouse Lab in Milton Keynes | Wellcome Sanger Institute for the COVID-19 Genomics UK (COG-UK) Consortium | The Lighthouse Lab in Milton Keynes and Alex Alderton, Roberto Amato, Jeffrey Barrett, Sonia Goncalves, Ewan Harrison, David K. Jackson, Ian Johnston, Dominic Kwiatkowski, Cordelia Langford, John Sillitoe on behalf of the Wellcome Sanger Institute COVID-19 Surveillance Team |
| EPI_ISL_2540831, EPI_ISL_2540832 | Lighthouse Lab in Alderley Park | Wellcome Sanger Institute for the COVID-19 Genomics UK (COG-UK) Consortium | Jacquelyn Wynn, Mairead Hyland, The Lighthouse Lab in Alderley Park and Alex Alderton, Roberto Amato, Jeffrey Barrett, Sonia Goncalves, Ewan Harrison, David K. Jackson, Ian Johnston, Dominic Kwiatkowski, Cordelia Langford, John Sillitoe on behalf of the Wellcome Sanger Institute COVID-19 Surveillance Team |
| EPI_ISL_2540833, EPI_ISL_2540834, EPI_ISL_2540835 | Lighthouse Lab in Milton Keynes | Wellcome Sanger Institute for the COVID-19 Genomics UK (COG-UK) Consortium | The Lighthouse Lab in Milton Keynes and Alex Alderton, Roberto Amato, Jeffrey Barrett, Sonia Goncalves, Ewan Harrison, David K. Jackson, Ian Johnston, Dominic Kwiatkowski, Cordelia Langford, John Sillitoe on behalf of the Wellcome Sanger Institute COVID-19 Surveillance Team |
| EPI_ISL_2540836 | Lighthouse Lab in Alderley Park | Wellcome Sanger Institute for the COVID-19 Genomics UK (COG-UK) Consortium | Jacquelyn Wynn, Mairead Hyland, The Lighthouse Lab in Alderley Park and Alex Alderton, Roberto Amato, Jeffrey Barrett, Sonia Goncalves, Ewan Harrison, David K. Jackson, Ian Johnston, Dominic Kwiatkowski, Cordelia Langford, John Sillitoe on behalf of the Wellcome Sanger Institute COVID-19 Surveillance Team |
| EPI_ISL_2540838 | Lighthouse Lab in Milton Keynes | Wellcome Sanger Institute for the COVID-19 Genomics UK (COG-UK) Consortium | The Lighthouse Lab in Milton Keynes and Alex Alderton, Roberto Amato, Jeffrey Barrett, Sonia Goncalves, Ewan Harrison, David K. Jackson, Ian Johnston, Dominic Kwiatkowski, Cordelia Langford, John Sillitoe on behalf of the Wellcome Sanger Institute COVID-19 Surveillance Team |
| EPI_ISL_2540839 | Lighthouse Laboratory Plymouth | Wellcome Sanger Institute for the COVID-19 Genomics UK (COG-UK) Consortium | Lighthouse Laboratory Plymouth and Alex Alderton, Roberto Amato, Jeffrey Barrett, Sonia Goncalves, Ewan Harrison, David K. Jackson, Ian Johnston, Dominic Kwiatkowski, Cordelia Langford, John Sillitoe on behalf of the Wellcome Sanger Institute COVID-19 Surveillance Team |
| EPI_ISL_2540840, EPI_ISL_2540841 | Lighthouse Lab in Alderley Park | Wellcome Sanger Institute for the COVID-19 Genomics UK (COG-UK) Consortium | Jacquelyn Wynn, Mairead Hyland, The Lighthouse Lab in Alderley Park and Alex Alderton, Roberto Amato, Jeffrey Barrett, Sonia Goncalves, Ewan Harrison, David K. Jackson, Ian Johnston, Dominic Kwiatkowski, Cordelia Langford, John Sillitoe on behalf of the Wellcome Sanger Institute COVID-19 Surveillance Team |
| EPI_ISL_2540842, EPI_ISL_2540843 | Lighthouse Laboratory Plymouth | Wellcome Sanger Institute for the COVID-19 Genomics UK (COG-UK) Consortium | Lighthouse Laboratory Plymouth and Alex Alderton, Roberto Amato, Jeffrey Barrett, Sonia Goncalves, Ewan Harrison, David K. Jackson, Ian Johnston, Dominic Kwiatkowski, Cordelia Langford, John Sillitoe on behalf of the Wellcome Sanger Institute COVID-19 Surveillance Team |
| EPI_ISL_2540844, EPI_ISL_2540845 | Lighthouse Lab in Milton Keynes | Wellcome Sanger Institute for the COVID-19 Genomics UK (COG-UK) Consortium | The Lighthouse Lab in Milton Keynes and Alex Alderton, Roberto Amato, Jeffrey Barrett, Sonia Goncalves, Ewan Harrison, David K. Jackson, Ian Johnston, Dominic Kwiatkowski, Cordelia Langford, John Sillitoe on behalf of the Wellcome Sanger Institute COVID-19 Surveillance Team |
| EPI_ISL_2540846 | Lighthouse Lab in Alderley Park | Wellcome Sanger Institute for the COVID-19 Genomics UK (COG-UK) Consortium | Jacquelyn Wynn, Mairead Hyland, The Lighthouse Lab in Alderley Park and Alex Alderton, Roberto Amato, Jeffrey Barrett, Sonia Goncalves, Ewan Harrison, David K. Jackson, Ian Johnston, Dominic Kwiatkowski, Cordelia Langford, John Sillitoe on behalf of the Wellcome Sanger Institute COVID-19 Surveillance Team |
| EPI_ISL_2540847, EPI_ISL_2540848, EPI_ISL_2540850 | Lighthouse Lab in Milton Keynes | Wellcome Sanger Institute for the COVID-19 Genomics UK (COG-UK) Consortium | The Lighthouse Lab in Milton Keynes and Alex Alderton, Roberto Amato, Jeffrey Barrett, Sonia Goncalves, Ewan Harrison, David K. Jackson, Ian Johnston, Dominic Kwiatkowski, Cordelia Langford, John Sillitoe on behalf of the Wellcome Sanger Institute COVID-19 Surveillance Team |
| EPI_ISL_2540851 | Lighthouse Laboratory Plymouth | Wellcome Sanger Institute for the COVID-19 Genomics UK (COG-UK) Consortium | Lighthouse Laboratory Plymouth and Alex Alderton, Roberto Amato, Jeffrey Barrett, Sonia Goncalves, Ewan Harrison, David K. Jackson, Ian Johnston, Dominic Kwiatkowski, Cordelia Langford, John Sillitoe on behalf of the Wellcome Sanger Institute COVID-19 Surveillance Team |
| EPI_ISL_2540852 | Lighthouse Lab in Milton Keynes | Wellcome Sanger Institute for the COVID-19 Genomics UK (COG-UK) Consortium | The Lighthouse Lab in Milton Keynes and Alex Alderton, Roberto Amato, Jeffrey Barrett, Sonia Goncalves, Ewan Harrison, David K. Jackson, Ian Johnston, Dominic Kwiatkowski, Cordelia Langford, John Sillitoe on behalf of the Wellcome Sanger Institute COVID-19 Surveillance Team |
| EPI_ISL_2540853 | Lighthouse Lab in Alderley Park | Wellcome Sanger Institute for the COVID-19 Genomics UK (COG-UK) Consortium | Jacquelyn Wynn, Mairead Hyland, The Lighthouse Lab in Alderley Park and Alex Alderton, Roberto Amato, Jeffrey Barrett, Sonia Goncalves, Ewan Harrison, David K. Jackson, Ian Johnston, Dominic Kwiatkowski, Cordelia Langford, John Sillitoe on behalf of the Wellcome Sanger Institute COVID-19 Surveillance Team |
| EPI_ISL_2540854, EPI_ISL_2540855 | Lighthouse Laboratory Plymouth | Wellcome Sanger Institute for the COVID-19 Genomics UK (COG-UK) Consortium | Lighthouse Laboratory Plymouth and Alex Alderton, Roberto Amato, Jeffrey Barrett, Sonia Goncalves, Ewan Harrison, David K. Jackson, Ian Johnston, Dominic Kwiatkowski, Cordelia Langford, John Sillitoe on behalf of the Wellcome Sanger Institute COVID-19 Surveillance Team |
| EPI_ISL_2540856, EPI_ISL_2540857, EPI_ISL_2540858, EPI_ISL_2540859 | Lighthouse Lab in Milton Keynes | Wellcome Sanger Institute for the COVID-19 Genomics UK (COG-UK) Consortium | The Lighthouse Lab in Milton Keynes and Alex Alderton, Roberto Amato, Jeffrey Barrett, Sonia Goncalves, Ewan Harrison, David K. Jackson, Ian Johnston, Dominic Kwiatkowski, Cordelia Langford, John Sillitoe on behalf of the Wellcome Sanger Institute COVID-19 Surveillance Team |
| EPI_ISL_2540860 | Lighthouse Laboratory Plymouth | Wellcome Sanger Institute for the COVID-19 Genomics UK (COG-UK) Consortium | Lighthouse Laboratory Plymouth and Alex Alderton, Roberto Amato, Jeffrey Barrett, Sonia Goncalves, Ewan Harrison, David K. Jackson, Ian Johnston, Dominic Kwiatkowski, Cordelia Langford, John Sillitoe on behalf of the Wellcome Sanger Institute COVID-19 Surveillance Team |
| EPI_ISL_2540861 | Lighthouse Lab in Milton Keynes | Wellcome Sanger Institute for the COVID-19 Genomics UK (COG-UK) Consortium | The Lighthouse Lab in Milton Keynes and Alex Alderton, Roberto Amato, Jeffrey Barrett, Sonia Goncalves, Ewan Harrison, David K. Jackson, Ian Johnston, Dominic Kwiatkowski, Cordelia Langford, John Sillitoe on behalf of the Wellcome Sanger Institute COVID-19 Surveillance Team |
| EPI_ISL_2540862 | Lighthouse Laboratory Plymouth | Wellcome Sanger Institute for the COVID-19 Genomics UK (COG-UK) Consortium | Lighthouse Laboratory Plymouth and Alex Alderton, Roberto Amato, Jeffrey Barrett, Sonia Goncalves, Ewan Harrison, David K. Jackson, Ian Johnston, Dominic Kwiatkowski, Cordelia Langford, John Sillitoe on behalf of the Wellcome Sanger Institute COVID-19 Surveillance Team |
| EPI_ISL_2540863 | Lighthouse Lab in Alderley Park | Wellcome Sanger Institute for the COVID-19 Genomics UK (COG-UK) Consortium | Jacquelyn Wynn, Mairead Hyland, The Lighthouse Lab in Alderley Park and Alex Alderton, Roberto Amato, Jeffrey Barrett, Sonia Goncalves, Ewan Harrison, David K. Jackson, Ian Johnston, Dominic Kwiatkowski, Cordelia Langford, John Sillitoe on behalf of the Wellcome Sanger Institute COVID-19 Surveillance Team |
| EPI_ISL_2540864, EPI_ISL_2540865, EPI_ISL_2540866 | Lighthouse Lab in Milton Keynes | Wellcome Sanger Institute for the COVID-19 Genomics UK (COG-UK) Consortium | The Lighthouse Lab in Milton Keynes and Alex Alderton, Roberto Amato, Jeffrey Barrett, Sonia Goncalves, Ewan Harrison, David K. Jackson, Ian Johnston, Dominic Kwiatkowski, Cordelia Langford, John Sillitoe on behalf of the Wellcome Sanger Institute COVID-19 Surveillance Team |
| EPI_ISL_2540867, | Lighthouse Lab in Alderley Park | Wellcome Sanger Institute for the COVID-19 Genomics UK (COG-UK) | Jacquelyn Wynn, Mairead Hyland, The Lighthouse Lab in Alderley Park and Alex Alderton, Roberto Amato, Jeffrey Barrett, Sonia Goncalves, Ewan Harrison, David K. Jackson, Ian Johnston, Dominic Kwiatkowski, Cordelia Langford, John Sillitoe on |

| | | | |
|---|---|---|---|
| EPI_ISL_2540868 | | Consortium | behalf of the Wellcome Sanger Institute COVID-19 Surveillance Team |
| EPI_ISL_2540869 | Lighthouse Laboratory Plymouth | Wellcome Sanger Institute for the COVID-19 Genomics UK (COG-UK) Consortium | Lighthouse Laboratory Plymouth and Alex Alderton, Roberto Amato, Jeffrey Barrett, Sonia Goncalves, Ewan Harrison, David K. Jackson, Ian Johnston, Dominic Kwiatkowski, Cordelia Langford, John Sillitoe on behalf of the Wellcome Sanger Institute COVID-19 Surveillance Team |
| EPI_ISL_2540871 | Lighthouse Lab in Milton Keynes | Wellcome Sanger Institute for the COVID-19 Genomics UK (COG-UK) Consortium | The Lighthouse Lab in Milton Keynes and Alex Alderton, Roberto Amato, Jeffrey Barrett, Sonia Goncalves, Ewan Harrison, David K. Jackson, Ian Johnston, Dominic Kwiatkowski, Cordelia Langford, John Sillitoe on behalf of the Wellcome Sanger Institute COVID-19 Surveillance Team |
| EPI_ISL_2540872 | Lighthouse Laboratory Plymouth | Wellcome Sanger Institute for the COVID-19 Genomics UK (COG-UK) Consortium | Lighthouse Laboratory Plymouth and Alex Alderton, Roberto Amato, Jeffrey Barrett, Sonia Goncalves, Ewan Harrison, David K. Jackson, Ian Johnston, Dominic Kwiatkowski, Cordelia Langford, John Sillitoe on behalf of the Wellcome Sanger Institute COVID-19 Surveillance Team |
| EPI_ISL_2540873, EPI_ISL_2540874 | Lighthouse Lab in Milton Keynes | Wellcome Sanger Institute for the COVID-19 Genomics UK (COG-UK) Consortium | The Lighthouse Lab in Milton Keynes and Alex Alderton, Roberto Amato, Jeffrey Barrett, Sonia Goncalves, Ewan Harrison, David K. Jackson, Ian Johnston, Dominic Kwiatkowski, Cordelia Langford, John Sillitoe on behalf of the Wellcome Sanger Institute COVID-19 Surveillance Team |
| EPI_ISL_2540875, EPI_ISL_2540876 | Lighthouse Laboratory Plymouth | Wellcome Sanger Institute for the COVID-19 Genomics UK (COG-UK) Consortium | Lighthouse Laboratory Plymouth and Alex Alderton, Roberto Amato, Jeffrey Barrett, Sonia Goncalves, Ewan Harrison, David K. Jackson, Ian Johnston, Dominic Kwiatkowski, Cordelia Langford, John Sillitoe on behalf of the Wellcome Sanger Institute COVID-19 Surveillance Team |
| EPI_ISL_2540877, EPI_ISL_2540878 | Lighthouse Lab in Milton Keynes | Wellcome Sanger Institute for the COVID-19 Genomics UK (COG-UK) Consortium | The Lighthouse Lab in Milton Keynes and Alex Alderton, Roberto Amato, Jeffrey Barrett, Sonia Goncalves, Ewan Harrison, David K. Jackson, Ian Johnston, Dominic Kwiatkowski, Cordelia Langford, John Sillitoe on behalf of the Wellcome Sanger Institute COVID-19 Surveillance Team |
| EPI_ISL_2540879 | Lighthouse Laboratory Plymouth | Wellcome Sanger Institute for the COVID-19 Genomics UK (COG-UK) Consortium | Lighthouse Laboratory Plymouth and Alex Alderton, Roberto Amato, Jeffrey Barrett, Sonia Goncalves, Ewan Harrison, David K. Jackson, Ian Johnston, Dominic Kwiatkowski, Cordelia Langford, John Sillitoe on behalf of the Wellcome Sanger Institute COVID-19 Surveillance Team |
| EPI_ISL_2540880 | Lighthouse Lab in Milton Keynes | Wellcome Sanger Institute for the COVID-19 Genomics UK (COG-UK) Consortium | The Lighthouse Lab in Milton Keynes and Alex Alderton, Roberto Amato, Jeffrey Barrett, Sonia Goncalves, Ewan Harrison, David K. Jackson, Ian Johnston, Dominic Kwiatkowski, Cordelia Langford, John Sillitoe on behalf of the Wellcome Sanger Institute COVID-19 Surveillance Team |
| EPI_ISL_2540881, EPI_ISL_2540882, EPI_ISL_2540884 | Lighthouse Laboratory Plymouth | Wellcome Sanger Institute for the COVID-19 Genomics UK (COG-UK) Consortium | Lighthouse Laboratory Plymouth and Alex Alderton, Roberto Amato, Jeffrey Barrett, Sonia Goncalves, Ewan Harrison, David K. Jackson, Ian Johnston, Dominic Kwiatkowski, Cordelia Langford, John Sillitoe on behalf of the Wellcome Sanger Institute COVID-19 Surveillance Team |
| EPI_ISL_2540885, EPI_ISL_2540886, EPI_ISL_2540887 | Lighthouse Lab in Milton Keynes | Wellcome Sanger Institute for the COVID-19 Genomics UK (COG-UK) Consortium | The Lighthouse Lab in Milton Keynes and Alex Alderton, Roberto Amato, Jeffrey Barrett, Sonia Goncalves, Ewan Harrison, David K. Jackson, Ian Johnston, Dominic Kwiatkowski, Cordelia Langford, John Sillitoe on behalf of the Wellcome Sanger Institute COVID-19 Surveillance Team |
| EPI_ISL_2540889 | Lighthouse Laboratory Plymouth | Wellcome Sanger Institute for the COVID-19 Genomics UK (COG-UK) Consortium | Lighthouse Laboratory Plymouth and Alex Alderton, Roberto Amato, Jeffrey Barrett, Sonia Goncalves, Ewan Harrison, David K. Jackson, Ian Johnston, Dominic Kwiatkowski, Cordelia Langford, John Sillitoe on behalf of the Wellcome Sanger Institute COVID-19 Surveillance Team |
| EPI_ISL_2540890, EPI_ISL_2540891 | Lighthouse Lab in Milton Keynes | Wellcome Sanger Institute for the COVID-19 Genomics UK (COG-UK) Consortium | The Lighthouse Lab in Milton Keynes and Alex Alderton, Roberto Amato, Jeffrey Barrett, Sonia Goncalves, Ewan Harrison, David K. Jackson, Ian Johnston, Dominic Kwiatkowski, Cordelia Langford, John Sillitoe on behalf of the Wellcome Sanger Institute COVID-19 Surveillance Team |
| EPI_ISL_2540892, EPI_ISL_2540893 | Lighthouse Lab in Alderley Park | Wellcome Sanger Institute for the COVID-19 Genomics UK (COG-UK) Consortium | Jacquelyn Wynn, Mairead Hyland, The Lighthouse Lab in Alderley Park and Alex Alderton, Roberto Amato, Jeffrey Barrett, Sonia Goncalves, Ewan Harrison, David K. Jackson, Ian Johnston, Dominic Kwiatkowski, Cordelia Langford, John Sillitoe on behalf of the Wellcome Sanger Institute COVID-19 Surveillance Team |
| EPI_ISL_2540894 | Lighthouse Laboratory Plymouth | Wellcome Sanger Institute for the COVID-19 Genomics UK (COG-UK) Consortium | Lighthouse Laboratory Plymouth and Alex Alderton, Roberto Amato, Jeffrey Barrett, Sonia Goncalves, Ewan Harrison, David K. Jackson, Ian Johnston, Dominic Kwiatkowski, Cordelia Langford, John Sillitoe on behalf of the Wellcome Sanger Institute COVID-19 Surveillance Team |
| EPI_ISL_2540895 | Lighthouse Lab in Milton Keynes | Wellcome Sanger Institute for the COVID-19 Genomics UK (COG-UK) Consortium | The Lighthouse Lab in Milton Keynes and Alex Alderton, Roberto Amato, Jeffrey Barrett, Sonia Goncalves, Ewan Harrison, David K. Jackson, Ian Johnston, Dominic Kwiatkowski, Cordelia Langford, John Sillitoe on behalf of the Wellcome Sanger Institute COVID-19 Surveillance Team |
| EPI_ISL_2540896, EPI_ISL_2540897, EPI_ISL_2540898 | Lighthouse Laboratory Plymouth | Wellcome Sanger Institute for the COVID-19 Genomics UK (COG-UK) Consortium | Lighthouse Laboratory Plymouth and Alex Alderton, Roberto Amato, Jeffrey Barrett, Sonia Goncalves, Ewan Harrison, David K. Jackson, Ian Johnston, Dominic Kwiatkowski, Cordelia Langford, John Sillitoe on behalf of the Wellcome Sanger Institute COVID-19 Surveillance Team |
| EPI_ISL_2540899 | Lighthouse Lab in Milton Keynes | Wellcome Sanger Institute for the COVID-19 Genomics UK (COG-UK) Consortium | The Lighthouse Lab in Milton Keynes and Alex Alderton, Roberto Amato, Jeffrey Barrett, Sonia Goncalves, Ewan Harrison, David K. Jackson, Ian Johnston, Dominic Kwiatkowski, Cordelia Langford, John Sillitoe on behalf of the Wellcome Sanger Institute COVID-19 Surveillance Team |
| EPI_ISL_2540900 | Lighthouse Lab in Alderley Park | Wellcome Sanger Institute for the COVID-19 Genomics UK (COG-UK) Consortium | Jacquelyn Wynn, Mairead Hyland, The Lighthouse Lab in Alderley Park and Alex Alderton, Roberto Amato, Jeffrey Barrett, Sonia Goncalves, Ewan Harrison, David K. Jackson, Ian Johnston, Dominic Kwiatkowski, Cordelia Langford, John Sillitoe on behalf of the Wellcome Sanger Institute COVID-19 Surveillance Team |
| EPI_ISL_2540901 | Lighthouse Lab in Milton Keynes | Wellcome Sanger Institute for the COVID-19 Genomics UK (COG-UK) Consortium | The Lighthouse Lab in Milton Keynes and Alex Alderton, Roberto Amato, Jeffrey Barrett, Sonia Goncalves, Ewan Harrison, David K. Jackson, Ian Johnston, Dominic Kwiatkowski, Cordelia Langford, John Sillitoe on behalf of the Wellcome Sanger Institute COVID-19 Surveillance Team |
| EPI_ISL_2540902, EPI_ISL_2540903 | Lighthouse Laboratory Plymouth | Wellcome Sanger Institute for the COVID-19 Genomics UK (COG-UK) Consortium | Lighthouse Laboratory Plymouth and Alex Alderton, Roberto Amato, Jeffrey Barrett, Sonia Goncalves, Ewan Harrison, David K. Jackson, Ian Johnston, Dominic Kwiatkowski, Cordelia Langford, John Sillitoe on behalf of the Wellcome Sanger Institute COVID-19 Surveillance Team |
| EPI_ISL_2540904 | Lighthouse Lab in Alderley Park | Wellcome Sanger Institute for the COVID-19 Genomics UK (COG-UK) Consortium | Jacquelyn Wynn, Mairead Hyland, The Lighthouse Lab in Alderley Park and Alex Alderton, Roberto Amato, Jeffrey Barrett, Sonia Goncalves, Ewan Harrison, David K. Jackson, Ian Johnston, Dominic Kwiatkowski, Cordelia Langford, John Sillitoe on behalf of the Wellcome Sanger Institute COVID-19 Surveillance Team |
| EPI_ISL_2540905 | Lighthouse Lab in Milton Keynes | Wellcome Sanger Institute for the COVID-19 Genomics UK (COG-UK) Consortium | The Lighthouse Lab in Milton Keynes and Alex Alderton, Roberto Amato, Jeffrey Barrett, Sonia Goncalves, Ewan Harrison, David K. Jackson, Ian Johnston, Dominic Kwiatkowski, Cordelia Langford, John Sillitoe on behalf of the Wellcome Sanger Institute COVID-19 Surveillance Team |
| EPI_ISL_2540906 | Lighthouse Laboratory Plymouth | Wellcome Sanger Institute for the COVID-19 Genomics UK (COG-UK) Consortium | Lighthouse Laboratory Plymouth and Alex Alderton, Roberto Amato, Jeffrey Barrett, Sonia Goncalves, Ewan Harrison, David K. Jackson, Ian Johnston, Dominic Kwiatkowski, Cordelia Langford, John Sillitoe on behalf of the Wellcome Sanger Institute COVID-19 Surveillance Team |
| EPI_ISL_2540907, EPI_ISL_2540908 | Lighthouse Lab in Milton Keynes | Wellcome Sanger Institute for the COVID-19 Genomics UK (COG-UK) Consortium | The Lighthouse Lab in Milton Keynes and Alex Alderton, Roberto Amato, Jeffrey Barrett, Sonia Goncalves, Ewan Harrison, David K. Jackson, Ian Johnston, Dominic Kwiatkowski, Cordelia Langford, John Sillitoe on behalf of the Wellcome Sanger Institute COVID-19 Surveillance Team |
| EPI_ISL_2540909 | Lighthouse Lab in Alderley Park | Wellcome Sanger Institute for the COVID-19 Genomics UK (COG-UK) Consortium | Jacquelyn Wynn, Mairead Hyland, The Lighthouse Lab in Alderley Park and Alex Alderton, Roberto Amato, Jeffrey Barrett, Sonia Goncalves, Ewan Harrison, David K. Jackson, Ian Johnston, Dominic Kwiatkowski, Cordelia Langford, John Sillitoe on behalf of the Wellcome Sanger Institute COVID-19 Surveillance Team |
| EPI_ISL_2540910, EPI_ISL_2540911, EPI_ISL_2540912, EPI_ISL_2540914, EPI_ISL_2540915 | Lighthouse Lab in Milton Keynes | Wellcome Sanger Institute for the COVID-19 Genomics UK (COG-UK) Consortium | The Lighthouse Lab in Milton Keynes and Alex Alderton, Roberto Amato, Jeffrey Barrett, Sonia Goncalves, Ewan Harrison, David K. Jackson, Ian Johnston, Dominic Kwiatkowski, Cordelia Langford, John Sillitoe on behalf of the Wellcome Sanger Institute COVID-19 Surveillance Team |
| EPI_ISL_2540916 | Lighthouse Laboratory Plymouth | Wellcome Sanger Institute for the COVID-19 Genomics UK (COG-UK) Consortium | Lighthouse Laboratory Plymouth and Alex Alderton, Roberto Amato, Jeffrey Barrett, Sonia Goncalves, Ewan Harrison, David K. Jackson, Ian Johnston, Dominic Kwiatkowski, Cordelia Langford, John Sillitoe on behalf of the Wellcome Sanger Institute COVID-19 Surveillance Team |
| EPI_ISL_2540917 | Lighthouse Lab in Alderley Park | Wellcome Sanger Institute for the COVID-19 Genomics UK (COG-UK) Consortium | Jacquelyn Wynn, Mairead Hyland, The Lighthouse Lab in Alderley Park and Alex Alderton, Roberto Amato, Jeffrey Barrett, Sonia Goncalves, Ewan Harrison, David K. Jackson, Ian Johnston, Dominic Kwiatkowski, Cordelia Langford, John Sillitoe on behalf of the Wellcome Sanger Institute COVID-19 Surveillance Team |
| EPI_ISL_2540918 | Lighthouse Lab in Milton Keynes | Wellcome Sanger Institute for the COVID-19 Genomics UK (COG-UK) Consortium | The Lighthouse Lab in Milton Keynes and Alex Alderton, Roberto Amato, Jeffrey Barrett, Sonia Goncalves, Ewan Harrison, David K. Jackson, Ian Johnston, Dominic Kwiatkowski, Cordelia Langford, John Sillitoe on behalf of the Wellcome Sanger Institute COVID-19 Surveillance Team |
| EPI_ISL_2540920 | Lighthouse Laboratory Plymouth | Wellcome Sanger Institute for the COVID-19 Genomics UK (COG-UK) Consortium | Lighthouse Laboratory Plymouth and Alex Alderton, Roberto Amato, Jeffrey Barrett, Sonia Goncalves, Ewan Harrison, David K. Jackson, Ian Johnston, Dominic Kwiatkowski, Cordelia Langford, John Sillitoe on behalf of the Wellcome Sanger Institute COVID-19 Surveillance Team |
| EPI_ISL_2540921 | Lighthouse Lab in Alderley Park | Wellcome Sanger Institute for the COVID-19 Genomics UK (COG-UK) Consortium | Jacquelyn Wynn, Mairead Hyland, The Lighthouse Lab in Alderley Park and Alex Alderton, Roberto Amato, Jeffrey Barrett, Sonia Goncalves, Ewan Harrison, David K. Jackson, Ian Johnston, Dominic Kwiatkowski, Cordelia Langford, John Sillitoe on behalf of the Wellcome Sanger Institute COVID-19 Surveillance Team |
| EPI_ISL_2540922 | Lighthouse Lab in Milton Keynes | Wellcome Sanger Institute for the COVID-19 Genomics UK (COG-UK) Consortium | The Lighthouse Lab in Milton Keynes and Alex Alderton, Roberto Amato, Jeffrey Barrett, Sonia Goncalves, Ewan Harrison, David K. Jackson, Ian Johnston, Dominic Kwiatkowski, Cordelia Langford, John Sillitoe on behalf of the Wellcome Sanger Institute COVID-19 Surveillance Team |
| EPI_ISL_2540923 | Lighthouse Lab in Alderley Park | Wellcome Sanger Institute for the COVID-19 Genomics UK (COG-UK) Consortium | Jacquelyn Wynn, Mairead Hyland, The Lighthouse Lab in Alderley Park and Alex Alderton, Roberto Amato, Jeffrey Barrett, Sonia Goncalves, Ewan Harrison, David K. Jackson, Ian Johnston, Dominic Kwiatkowski, Cordelia Langford, John Sillitoe on behalf of the Wellcome Sanger Institute COVID-19 Surveillance Team |
| EPI_ISL_2540924 | Lighthouse Laboratory Plymouth | Wellcome Sanger Institute for the COVID-19 Genomics UK (COG-UK) Consortium | Lighthouse Laboratory Plymouth and Alex Alderton, Roberto Amato, Jeffrey Barrett, Sonia Goncalves, Ewan Harrison, David K. Jackson, Ian Johnston, Dominic Kwiatkowski, Cordelia Langford, John Sillitoe on behalf of the Wellcome Sanger Institute COVID-19 Surveillance Team |
| EPI_ISL_2540925 | Lighthouse Lab in Milton Keynes | Wellcome Sanger Institute for the COVID-19 Genomics UK (COG-UK) Consortium | The Lighthouse Lab in Milton Keynes and Alex Alderton, Roberto Amato, Jeffrey Barrett, Sonia Goncalves, Ewan Harrison, David K. Jackson, Ian Johnston, Dominic Kwiatkowski, Cordelia Langford, John Sillitoe on behalf of the Wellcome Sanger Institute COVID-19 Surveillance Team |
| EPI_ISL_2540926 | Lighthouse Laboratory Plymouth | Wellcome Sanger Institute for the COVID-19 Genomics UK (COG-UK) Consortium | Lighthouse Laboratory Plymouth and Alex Alderton, Roberto Amato, Jeffrey Barrett, Sonia Goncalves, Ewan Harrison, David K. Jackson, Ian Johnston, Dominic Kwiatkowski, Cordelia Langford, John Sillitoe on behalf of the Wellcome Sanger Institute COVID-19 Surveillance Team |
| EPI_ISL_2540927, EPI_ISL_2540928 | Lighthouse Lab in Milton Keynes | Wellcome Sanger Institute for the COVID-19 Genomics UK (COG-UK) Consortium | The Lighthouse Lab in Milton Keynes and Alex Alderton, Roberto Amato, Jeffrey Barrett, Sonia Goncalves, Ewan Harrison, David K. Jackson, Ian Johnston, Dominic Kwiatkowski, Cordelia Langford, John Sillitoe on behalf of the Wellcome Sanger Institute COVID-19 Surveillance Team |
| EPI_ISL_2540929, EPI_ISL_2540930 | Lighthouse Lab in Alderley Park | Wellcome Sanger Institute for the COVID-19 Genomics UK (COG-UK) Consortium | Jacquelyn Wynn, Mairead Hyland, The Lighthouse Lab in Alderley Park and Alex Alderton, Roberto Amato, Jeffrey Barrett, Sonia Goncalves, Ewan Harrison, David K. Jackson, Ian Johnston, Dominic Kwiatkowski, Cordelia Langford, John Sillitoe on behalf of the Wellcome Sanger Institute COVID-19 Surveillance Team |
| EPI_ISL_2540932 | Lighthouse Lab in Milton Keynes | Wellcome Sanger Institute for the COVID-19 Genomics UK (COG-UK) Consortium | The Lighthouse Lab in Milton Keynes and Alex Alderton, Roberto Amato, Jeffrey Barrett, Sonia Goncalves, Ewan Harrison, David K. Jackson, Ian Johnston, Dominic Kwiatkowski, Cordelia Langford, John Sillitoe on behalf of the Wellcome Sanger Institute COVID-19 Surveillance Team |
| EPI_ISL_2540933, EPI_ISL_2540934, EPI_ISL_2540935 | Lighthouse Laboratory Plymouth | Wellcome Sanger Institute for the COVID-19 Genomics UK (COG-UK) Consortium | Lighthouse Laboratory Plymouth and Alex Alderton, Roberto Amato, Jeffrey Barrett, Sonia Goncalves, Ewan Harrison, David K. Jackson, Ian Johnston, Dominic Kwiatkowski, Cordelia Langford, John Sillitoe on behalf of the Wellcome Sanger Institute COVID-19 Surveillance Team |
| EPI_ISL_2540936, EPI_ISL_2540937 | Lighthouse Lab in Alderley Park | Wellcome Sanger Institute for the COVID-19 Genomics UK (COG-UK) Consortium | Jacquelyn Wynn, Mairead Hyland, The Lighthouse Lab in Alderley Park and Alex Alderton, Roberto Amato, Jeffrey Barrett, Sonia Goncalves, Ewan Harrison, David K. Jackson, Ian Johnston, Dominic Kwiatkowski, Cordelia Langford, John Sillitoe on behalf of the Wellcome Sanger Institute COVID-19 Surveillance Team |
| EPI_ISL_2540938 | Lighthouse Lab in Milton Keynes | Wellcome Sanger Institute for the COVID-19 Genomics UK (COG-UK) | The Lighthouse Lab in Milton Keynes and Alex Alderton, Roberto Amato, Jeffrey Barrett, Sonia Goncalves, Ewan Harrison, David K. Jackson, Ian Johnston, Dominic Kwiatkowski, Cordelia Langford, John Sillitoe on behalf of the Wellcome Sanger |

| | | | |
|---|---|---|---|
| | | Consortium | Institute COVID-19 Surveillance Team |
| EPI_ISL_2540939 | Lighthouse Laboratory Plymouth | Wellcome Sanger Institute for the COVID-19 Genomics UK (COG-UK) Consortium | Lighthouse Laboratory Plymouth and Alex Alderton, Roberto Amato, Jeffrey Barrett, Sonia Goncalves, Ewan Harrison, David K. Jackson, Ian Johnston, Dominic Kwiatkowski, Cordelia Langford, John Sillitoe on behalf of the Wellcome Sanger Institute COVID-19 Surveillance Team |
| EPI_ISL_2540940, EPI_ISL_2540941, EPI_ISL_2540942, EPI_ISL_2540943 | Lighthouse Lab in Milton Keynes | Wellcome Sanger Institute for the COVID-19 Genomics UK (COG-UK) Consortium | The Lighthouse Lab in Milton Keynes and Alex Alderton, Roberto Amato, Jeffrey Barrett, Sonia Goncalves, Ewan Harrison, David K. Jackson, Ian Johnston, Dominic Kwiatkowski, Cordelia Langford, John Sillitoe on behalf of the Wellcome Sanger Institute COVID-19 Surveillance Team |
| EPI_ISL_2540944 | Lighthouse Laboratory Plymouth | Wellcome Sanger Institute for the COVID-19 Genomics UK (COG-UK) Consortium | Lighthouse Laboratory Plymouth and Alex Alderton, Roberto Amato, Jeffrey Barrett, Sonia Goncalves, Ewan Harrison, David K. Jackson, Ian Johnston, Dominic Kwiatkowski, Cordelia Langford, John Sillitoe on behalf of the Wellcome Sanger Institute COVID-19 Surveillance Team |
| EPI_ISL_2540946, EPI_ISL_2540947, EPI_ISL_2540948, EPI_ISL_2540949, EPI_ISL_2540950 | Lighthouse Lab in Milton Keynes | Wellcome Sanger Institute for the COVID-19 Genomics UK (COG-UK) Consortium | The Lighthouse Lab in Milton Keynes and Alex Alderton, Roberto Amato, Jeffrey Barrett, Sonia Goncalves, Ewan Harrison, David K. Jackson, Ian Johnston, Dominic Kwiatkowski, Cordelia Langford, John Sillitoe on behalf of the Wellcome Sanger Institute COVID-19 Surveillance Team |
| EPI_ISL_2540951 | Health Services Laboratories | Wellcome Sanger Institute for the COVID-19 Genomics UK (COG-UK) Consortium | Health Services Laboratories and Alex Alderton, Roberto Amato, Jeffrey Barrett, Sonia Goncalves, Ewan Harrison, David K. Jackson, Ian Johnston, Dominic Kwiatkowski, Cordelia Langford, John Sillitoe on behalf of the Wellcome Sanger Institute COVID-19 Surveillance Team |
| EPI_ISL_2540953 | Lighthouse Lab in Milton Keynes | Wellcome Sanger Institute for the COVID-19 Genomics UK (COG-UK) Consortium | The Lighthouse Lab in Milton Keynes and Alex Alderton, Roberto Amato, Jeffrey Barrett, Sonia Goncalves, Ewan Harrison, David K. Jackson, Ian Johnston, Dominic Kwiatkowski, Cordelia Langford, John Sillitoe on behalf of the Wellcome Sanger Institute COVID-19 Surveillance Team |
| EPI_ISL_2540954 | Health Services Laboratories | Wellcome Sanger Institute for the COVID-19 Genomics UK (COG-UK) Consortium | Health Services Laboratories and Alex Alderton, Roberto Amato, Jeffrey Barrett, Sonia Goncalves, Ewan Harrison, David K. Jackson, Ian Johnston, Dominic Kwiatkowski, Cordelia Langford, John Sillitoe on behalf of the Wellcome Sanger Institute COVID-19 Surveillance Team |
| EPI_ISL_2540955, EPI_ISL_2540956, EPI_ISL_2540957, EPI_ISL_2540958, EPI_ISL_2540959, EPI_ISL_2540960, EPI_ISL_2540961, EPI_ISL_2540962, EPI_ISL_2540963, EPI_ISL_2540964, EPI_ISL_2540966, EPI_ISL_2540967, EPI_ISL_2540968, EPI_ISL_2540969, EPI_ISL_2540970 | | | |
| see above | Lighthouse Lab in Milton Keynes | Wellcome Sanger Institute for the COVID-19 Genomics UK (COG-UK) Consortium | The Lighthouse Lab in Milton Keynes and Alex Alderton, Roberto Amato, Jeffrey Barrett, Sonia Goncalves, Ewan Harrison, David K. Jackson, Ian Johnston, Dominic Kwiatkowski, Cordelia Langford, John Sillitoe on behalf of the Wellcome Sanger Institute COVID-19 Surveillance Team |
| EPI_ISL_2540971 | Health Services Laboratories | Wellcome Sanger Institute for the COVID-19 Genomics UK (COG-UK) Consortium | Health Services Laboratories and Alex Alderton, Roberto Amato, Jeffrey Barrett, Sonia Goncalves, Ewan Harrison, David K. Jackson, Ian Johnston, Dominic Kwiatkowski, Cordelia Langford, John Sillitoe on behalf of the Wellcome Sanger Institute COVID-19 Surveillance Team |
| EPI_ISL_2540972, EPI_ISL_2540973 | Lighthouse Lab in Milton Keynes | Wellcome Sanger Institute for the COVID-19 Genomics UK (COG-UK) Consortium | The Lighthouse Lab in Milton Keynes and Alex Alderton, Roberto Amato, Jeffrey Barrett, Sonia Goncalves, Ewan Harrison, David K. Jackson, Ian Johnston, Dominic Kwiatkowski, Cordelia Langford, John Sillitoe on behalf of the Wellcome Sanger Institute COVID-19 Surveillance Team |
| EPI_ISL_2540974 | Health Services Laboratories | Wellcome Sanger Institute for the COVID-19 Genomics UK (COG-UK) Consortium | Health Services Laboratories and Alex Alderton, Roberto Amato, Jeffrey Barrett, Sonia Goncalves, Ewan Harrison, David K. Jackson, Ian Johnston, Dominic Kwiatkowski, Cordelia Langford, John Sillitoe on behalf of the Wellcome Sanger Institute COVID-19 Surveillance Team |
| EPI_ISL_2540975, EPI_ISL_2540976 | Lighthouse Lab in Milton Keynes | Wellcome Sanger Institute for the COVID-19 Genomics UK (COG-UK) Consortium | The Lighthouse Lab in Milton Keynes and Alex Alderton, Roberto Amato, Jeffrey Barrett, Sonia Goncalves, Ewan Harrison, David K. Jackson, Ian Johnston, Dominic Kwiatkowski, Cordelia Langford, John Sillitoe on behalf of the Wellcome Sanger Institute COVID-19 Surveillance Team |
| EPI_ISL_2540977 | Health Services Laboratories | Wellcome Sanger Institute for the COVID-19 Genomics UK (COG-UK) Consortium | Health Services Laboratories and Alex Alderton, Roberto Amato, Jeffrey Barrett, Sonia Goncalves, Ewan Harrison, David K. Jackson, Ian Johnston, Dominic Kwiatkowski, Cordelia Langford, John Sillitoe on behalf of the Wellcome Sanger Institute COVID-19 Surveillance Team |
| EPI_ISL_2540978 | Lighthouse Lab in Glasgow | Wellcome Sanger Institute for the COVID-19 Genomics UK (COG-UK) Consortium | Harper VanSteenhouse, Yumi Kasai, David Gray, Carol Clugston, Anna Dominiczak and Alex Alderton, Roberto Amato, Jeffrey Barrett, Sonia Goncalves, Ewan Harrison, David K. Jackson, Ian Johnston, Dominic Kwiatkowski, Cordelia Langford, John Sillitoe on behalf of the Wellcome Sanger Institute COVID-19 Surveillance Team |
| EPI_ISL_2540979 | Health Services Laboratories | Wellcome Sanger Institute for the COVID-19 Genomics UK (COG-UK) Consortium | Health Services Laboratories and Alex Alderton, Roberto Amato, Jeffrey Barrett, Sonia Goncalves, Ewan Harrison, David K. Jackson, Ian Johnston, Dominic Kwiatkowski, Cordelia Langford, John Sillitoe on behalf of the Wellcome Sanger Institute COVID-19 Surveillance Team |
| EPI_ISL_2540980, EPI_ISL_2540981, EPI_ISL_2540982, EPI_ISL_2540983 | Lighthouse Lab in Milton Keynes | Wellcome Sanger Institute for the COVID-19 Genomics UK (COG-UK) Consortium | The Lighthouse Lab in Milton Keynes and Alex Alderton, Roberto Amato, Jeffrey Barrett, Sonia Goncalves, Ewan Harrison, David K. Jackson, Ian Johnston, Dominic Kwiatkowski, Cordelia Langford, John Sillitoe on behalf of the Wellcome Sanger Institute COVID-19 Surveillance Team |
| EPI_ISL_2540984 | Health Services Laboratories | Wellcome Sanger Institute for the COVID-19 Genomics UK (COG-UK) Consortium | Health Services Laboratories and Alex Alderton, Roberto Amato, Jeffrey Barrett, Sonia Goncalves, Ewan Harrison, David K. Jackson, Ian Johnston, Dominic Kwiatkowski, Cordelia Langford, John Sillitoe on behalf of the Wellcome Sanger Institute COVID-19 Surveillance Team |
| EPI_ISL_2540985, EPI_ISL_2540986, EPI_ISL_2540987, EPI_ISL_2540988, EPI_ISL_2540989, EPI_ISL_2540990, EPI_ISL_2540991 | Lighthouse Lab in Milton Keynes | Wellcome Sanger Institute for the COVID-19 Genomics UK (COG-UK) Consortium | The Lighthouse Lab in Milton Keynes and Alex Alderton, Roberto Amato, Jeffrey Barrett, Sonia Goncalves, Ewan Harrison, David K. Jackson, Ian Johnston, Dominic Kwiatkowski, Cordelia Langford, John Sillitoe on behalf of the Wellcome Sanger Institute COVID-19 Surveillance Team |
| EPI_ISL_2540992 | Health Services Laboratories | Wellcome Sanger Institute for the COVID-19 Genomics UK (COG-UK) Consortium | Health Services Laboratories and Alex Alderton, Roberto Amato, Jeffrey Barrett, Sonia Goncalves, Ewan Harrison, David K. Jackson, Ian Johnston, Dominic Kwiatkowski, Cordelia Langford, John Sillitoe on behalf of the Wellcome Sanger Institute COVID-19 Surveillance Team |
| EPI_ISL_2540993, EPI_ISL_2540995, EPI_ISL_2540996 | Lighthouse Lab in Milton Keynes | Wellcome Sanger Institute for the COVID-19 Genomics UK (COG-UK) Consortium | The Lighthouse Lab in Milton Keynes and Alex Alderton, Roberto Amato, Jeffrey Barrett, Sonia Goncalves, Ewan Harrison, David K. Jackson, Ian Johnston, Dominic Kwiatkowski, Cordelia Langford, John Sillitoe on behalf of the Wellcome Sanger Institute COVID-19 Surveillance Team |
| EPI_ISL_2540997 | Health Services Laboratories | Wellcome Sanger Institute for the COVID-19 Genomics UK (COG-UK) Consortium | Health Services Laboratories and Alex Alderton, Roberto Amato, Jeffrey Barrett, Sonia Goncalves, Ewan Harrison, David K. Jackson, Ian Johnston, Dominic Kwiatkowski, Cordelia Langford, John Sillitoe on behalf of the Wellcome Sanger Institute COVID-19 Surveillance Team |
| EPI_ISL_2540998 | Lighthouse Lab in Milton Keynes | Wellcome Sanger Institute for the COVID-19 Genomics UK (COG-UK) Consortium | The Lighthouse Lab in Milton Keynes and Alex Alderton, Roberto Amato, Jeffrey Barrett, Sonia Goncalves, Ewan Harrison, David K. Jackson, Ian Johnston, Dominic Kwiatkowski, Cordelia Langford, John Sillitoe on behalf of the Wellcome Sanger Institute COVID-19 Surveillance Team |
| EPI_ISL_2540999 | Lighthouse Lab in Glasgow | Wellcome Sanger Institute for the COVID-19 Genomics UK (COG-UK) Consortium | Harper VanSteenhouse, Yumi Kasai, David Gray, Carol Clugston, Anna Dominiczak and Alex Alderton, Roberto Amato, Jeffrey Barrett, Sonia Goncalves, Ewan Harrison, David K. Jackson, Ian Johnston, Dominic Kwiatkowski, Cordelia Langford, John Sillitoe on behalf of the Wellcome Sanger Institute COVID-19 Surveillance Team |
| EPI_ISL_2541000 | Health Services Laboratories | Wellcome Sanger Institute for the COVID-19 Genomics UK (COG-UK) Consortium | Health Services Laboratories and Alex Alderton, Roberto Amato, Jeffrey Barrett, Sonia Goncalves, Ewan Harrison, David K. Jackson, Ian Johnston, Dominic Kwiatkowski, Cordelia Langford, John Sillitoe on behalf of the Wellcome Sanger Institute COVID-19 Surveillance Team |
| EPI_ISL_2541001, EPI_ISL_2541002 | Lighthouse Lab in Milton Keynes | Wellcome Sanger Institute for the COVID-19 Genomics UK (COG-UK) Consortium | The Lighthouse Lab in Milton Keynes and Alex Alderton, Roberto Amato, Jeffrey Barrett, Sonia Goncalves, Ewan Harrison, David K. Jackson, Ian Johnston, Dominic Kwiatkowski, Cordelia Langford, John Sillitoe on behalf of the Wellcome Sanger Institute COVID-19 Surveillance Team |
| EPI_ISL_2541003 | Health Services Laboratories | Wellcome Sanger Institute for the COVID-19 Genomics UK (COG-UK) Consortium | Health Services Laboratories and Alex Alderton, Roberto Amato, Jeffrey Barrett, Sonia Goncalves, Ewan Harrison, David K. Jackson, Ian Johnston, Dominic Kwiatkowski, Cordelia Langford, John Sillitoe on behalf of the Wellcome Sanger Institute COVID-19 Surveillance Team |
| EPI_ISL_2541004 | Lighthouse Lab in Milton Keynes | Wellcome Sanger Institute for the COVID-19 Genomics UK (COG-UK) Consortium | The Lighthouse Lab in Milton Keynes and Alex Alderton, Roberto Amato, Jeffrey Barrett, Sonia Goncalves, Ewan Harrison, David K. Jackson, Ian Johnston, Dominic Kwiatkowski, Cordelia Langford, John Sillitoe on behalf of the Wellcome Sanger Institute COVID-19 Surveillance Team |
| EPI_ISL_2541005 | Health Services Laboratories | Wellcome Sanger Institute for the COVID-19 Genomics UK (COG-UK) Consortium | Health Services Laboratories and Alex Alderton, Roberto Amato, Jeffrey Barrett, Sonia Goncalves, Ewan Harrison, David K. Jackson, Ian Johnston, Dominic Kwiatkowski, Cordelia Langford, John Sillitoe on behalf of the Wellcome Sanger Institute COVID-19 Surveillance Team |
| EPI_ISL_2541006, EPI_ISL_2541007, EPI_ISL_2541008 | Lighthouse Lab in Milton Keynes | Wellcome Sanger Institute for the COVID-19 Genomics UK (COG-UK) Consortium | The Lighthouse Lab in Milton Keynes and Alex Alderton, Roberto Amato, Jeffrey Barrett, Sonia Goncalves, Ewan Harrison, David K. Jackson, Ian Johnston, Dominic Kwiatkowski, Cordelia Langford, John Sillitoe on behalf of the Wellcome Sanger Institute COVID-19 Surveillance Team |
| EPI_ISL_2541009 | Health Services Laboratories | Wellcome Sanger Institute for the COVID-19 Genomics UK (COG-UK) Consortium | Health Services Laboratories and Alex Alderton, Roberto Amato, Jeffrey Barrett, Sonia Goncalves, Ewan Harrison, David K. Jackson, Ian Johnston, Dominic Kwiatkowski, Cordelia Langford, John Sillitoe on behalf of the Wellcome Sanger Institute COVID-19 Surveillance Team |
| EPI_ISL_2541010, EPI_ISL_2541011, EPI_ISL_2541012, EPI_ISL_2541013 | Lighthouse Lab in Milton Keynes | Wellcome Sanger Institute for the COVID-19 Genomics UK (COG-UK) Consortium | The Lighthouse Lab in Milton Keynes and Alex Alderton, Roberto Amato, Jeffrey Barrett, Sonia Goncalves, Ewan Harrison, David K. Jackson, Ian Johnston, Dominic Kwiatkowski, Cordelia Langford, John Sillitoe on behalf of the Wellcome Sanger Institute COVID-19 Surveillance Team |
| EPI_ISL_2541014 | Health Services Laboratories | Wellcome Sanger Institute for the COVID-19 Genomics UK (COG-UK) Consortium | Health Services Laboratories and Alex Alderton, Roberto Amato, Jeffrey Barrett, Sonia Goncalves, Ewan Harrison, David K. Jackson, Ian Johnston, Dominic Kwiatkowski, Cordelia Langford, John Sillitoe on behalf of the Wellcome Sanger Institute COVID-19 Surveillance Team |
| EPI_ISL_2541015, EPI_ISL_2541016, EPI_ISL_2541017, EPI_ISL_2541018, EPI_ISL_2541019, EPI_ISL_2541020, EPI_ISL_2541021 | Lighthouse Lab in Milton Keynes | Wellcome Sanger Institute for the COVID-19 Genomics UK (COG-UK) Consortium | The Lighthouse Lab in Milton Keynes and Alex Alderton, Roberto Amato, Jeffrey Barrett, Sonia Goncalves, Ewan Harrison, David K. Jackson, Ian Johnston, Dominic Kwiatkowski, Cordelia Langford, John Sillitoe on behalf of the Wellcome Sanger Institute COVID-19 Surveillance Team |
| EPI_ISL_2541022 | Health Services Laboratories | Wellcome Sanger Institute for the COVID-19 Genomics UK (COG-UK) Consortium | Health Services Laboratories and Alex Alderton, Roberto Amato, Jeffrey Barrett, Sonia Goncalves, Ewan Harrison, David K. Jackson, Ian Johnston, Dominic Kwiatkowski, Cordelia Langford, John Sillitoe on behalf of the Wellcome Sanger Institute COVID-19 Surveillance Team |
| EPI_ISL_2541023, EPI_ISL_2541024, EPI_ISL_2541025, | Lighthouse Lab in Milton Keynes | Wellcome Sanger Institute for the COVID-19 Genomics UK (COG-UK) Consortium | The Lighthouse Lab in Milton Keynes and Alex Alderton, Roberto Amato, Jeffrey Barrett, Sonia Goncalves, Ewan Harrison, David K. Jackson, Ian Johnston, Dominic Kwiatkowski, Cordelia Langford, John Sillitoe on behalf of the Wellcome Sanger Institute COVID-19 Surveillance Team |

| | | | |
|---|---|---|---|
| EPI_ISL_2541026 | | | |
| EPI_ISL_2541027, EPI_ISL_2541028 | Health Services Laboratories | Wellcome Sanger Institute for the COVID-19 Genomics UK (COG-UK) Consortium | Health Services Laboratories and Alex Alderton, Roberto Amato, Jeffrey Barrett, Sonia Goncalves, Ewan Harrison, David K. Jackson, Ian Johnston, Dominic Kwiatkowski, Cordelia Langford, John Sillitoe on behalf of the Wellcome Sanger Institute COVID-19 Surveillance Team |
| EPI_ISL_2541029, EPI_ISL_2541030 | Lighthouse Lab in Milton Keynes | Wellcome Sanger Institute for the COVID-19 Genomics UK (COG-UK) Consortium | The Lighthouse Lab in Milton Keynes and Alex Alderton, Roberto Amato, Jeffrey Barrett, Sonia Goncalves, Ewan Harrison, David K. Jackson, Ian Johnston, Dominic Kwiatkowski, Cordelia Langford, John Sillitoe on behalf of the Wellcome Sanger Institute COVID-19 Surveillance Team |
| EPI_ISL_2541031 | Lighthouse Lab in Glasgow | Wellcome Sanger Institute for the COVID-19 Genomics UK (COG-UK) Consortium | Harper VanSteenhouse, Yumi Kasai, David Gray, Carol Clugston, Anna Dominiczak and Alex Alderton, Roberto Amato, Jeffrey Barrett, Sonia Goncalves, Ewan Harrison, David K. Jackson, Ian Johnston, Dominic Kwiatkowski, Cordelia Langford, John Sillitoe on behalf of the Wellcome Sanger Institute COVID-19 Surveillance Team |
| EPI_ISL_2541032, EPI_ISL_2541033 | Lighthouse Lab in Milton Keynes | Wellcome Sanger Institute for the COVID-19 Genomics UK (COG-UK) Consortium | The Lighthouse Lab in Milton Keynes and Alex Alderton, Roberto Amato, Jeffrey Barrett, Sonia Goncalves, Ewan Harrison, David K. Jackson, Ian Johnston, Dominic Kwiatkowski, Cordelia Langford, John Sillitoe on behalf of the Wellcome Sanger Institute COVID-19 Surveillance Team |
| EPI_ISL_2541035 | Health Services Laboratories | Wellcome Sanger Institute for the COVID-19 Genomics UK (COG-UK) Consortium | Health Services Laboratories and Alex Alderton, Roberto Amato, Jeffrey Barrett, Sonia Goncalves, Ewan Harrison, David K. Jackson, Ian Johnston, Dominic Kwiatkowski, Cordelia Langford, John Sillitoe on behalf of the Wellcome Sanger Institute COVID-19 Surveillance Team |
| EPI_ISL_2541037, EPI_ISL_2541038, EPI_ISL_2541039, EPI_ISL_2541041, EPI_ISL_2541042, EPI_ISL_2541043, EPI_ISL_2541045, EPI_ISL_2541046, EPI_ISL_2541047, EPI_ISL_2541048 | Lighthouse Lab in Milton Keynes | Wellcome Sanger Institute for the COVID-19 Genomics UK (COG-UK) Consortium | The Lighthouse Lab in Milton Keynes and Alex Alderton, Roberto Amato, Jeffrey Barrett, Sonia Goncalves, Ewan Harrison, David K. Jackson, Ian Johnston, Dominic Kwiatkowski, Cordelia Langford, John Sillitoe on behalf of the Wellcome Sanger Institute COVID-19 Surveillance Team |
| EPI_ISL_2541050 | Lighthouse Lab in Glasgow | Wellcome Sanger Institute for the COVID-19 Genomics UK (COG-UK) Consortium | Harper VanSteenhouse, Yumi Kasai, David Gray, Carol Clugston, Anna Dominiczak and Alex Alderton, Roberto Amato, Jeffrey Barrett, Sonia Goncalves, Ewan Harrison, David K. Jackson, Ian Johnston, Dominic Kwiatkowski, Cordelia Langford, John Sillitoe on behalf of the Wellcome Sanger Institute COVID-19 Surveillance Team |
| EPI_ISL_2541052, EPI_ISL_2541053, EPI_ISL_2541054 | Lighthouse Lab in Milton Keynes | Wellcome Sanger Institute for the COVID-19 Genomics UK (COG-UK) Consortium | The Lighthouse Lab in Milton Keynes and Alex Alderton, Roberto Amato, Jeffrey Barrett, Sonia Goncalves, Ewan Harrison, David K. Jackson, Ian Johnston, Dominic Kwiatkowski, Cordelia Langford, John Sillitoe on behalf of the Wellcome Sanger Institute COVID-19 Surveillance Team |
| EPI_ISL_2541055 | Health Services Laboratories | Wellcome Sanger Institute for the COVID-19 Genomics UK (COG-UK) Consortium | Health Services Laboratories and Alex Alderton, Roberto Amato, Jeffrey Barrett, Sonia Goncalves, Ewan Harrison, David K. Jackson, Ian Johnston, Dominic Kwiatkowski, Cordelia Langford, John Sillitoe on behalf of the Wellcome Sanger Institute COVID-19 Surveillance Team |
| EPI_ISL_2541056, EPI_ISL_2541057, EPI_ISL_2541058, EPI_ISL_2541059 | Lighthouse Lab in Milton Keynes | Wellcome Sanger Institute for the COVID-19 Genomics UK (COG-UK) Consortium | The Lighthouse Lab in Milton Keynes and Alex Alderton, Roberto Amato, Jeffrey Barrett, Sonia Goncalves, Ewan Harrison, David K. Jackson, Ian Johnston, Dominic Kwiatkowski, Cordelia Langford, John Sillitoe on behalf of the Wellcome Sanger Institute COVID-19 Surveillance Team |
| EPI_ISL_2541060 | Health Services Laboratories | Wellcome Sanger Institute for the COVID-19 Genomics UK (COG-UK) Consortium | Health Services Laboratories and Alex Alderton, Roberto Amato, Jeffrey Barrett, Sonia Goncalves, Ewan Harrison, David K. Jackson, Ian Johnston, Dominic Kwiatkowski, Cordelia Langford, John Sillitoe on behalf of the Wellcome Sanger Institute COVID-19 Surveillance Team |
| EPI_ISL_2541062, EPI_ISL_2541063, EPI_ISL_2541064, EPI_ISL_2541065, EPI_ISL_2541066, EPI_ISL_2541067, EPI_ISL_2541068, EPI_ISL_2541069, EPI_ISL_2541070, EPI_ISL_2541071 | Lighthouse Lab in Milton Keynes | Wellcome Sanger Institute for the COVID-19 Genomics UK (COG-UK) Consortium | The Lighthouse Lab in Milton Keynes and Alex Alderton, Roberto Amato, Jeffrey Barrett, Sonia Goncalves, Ewan Harrison, David K. Jackson, Ian Johnston, Dominic Kwiatkowski, Cordelia Langford, John Sillitoe on behalf of the Wellcome Sanger Institute COVID-19 Surveillance Team |
| EPI_ISL_2541072 | Health Services Laboratories | Wellcome Sanger Institute for the COVID-19 Genomics UK (COG-UK) Consortium | Health Services Laboratories and Alex Alderton, Roberto Amato, Jeffrey Barrett, Sonia Goncalves, Ewan Harrison, David K. Jackson, Ian Johnston, Dominic Kwiatkowski, Cordelia Langford, John Sillitoe on behalf of the Wellcome Sanger Institute COVID-19 Surveillance Team |
| EPI_ISL_2541073 | Lighthouse Lab in Milton Keynes | Wellcome Sanger Institute for the COVID-19 Genomics UK (COG-UK) Consortium | The Lighthouse Lab in Milton Keynes and Alex Alderton, Roberto Amato, Jeffrey Barrett, Sonia Goncalves, Ewan Harrison, David K. Jackson, Ian Johnston, Dominic Kwiatkowski, Cordelia Langford, John Sillitoe on behalf of the Wellcome Sanger Institute COVID-19 Surveillance Team |
| EPI_ISL_2541074 | Health Services Laboratories | Wellcome Sanger Institute for the COVID-19 Genomics UK (COG-UK) Consortium | Health Services Laboratories and Alex Alderton, Roberto Amato, Jeffrey Barrett, Sonia Goncalves, Ewan Harrison, David K. Jackson, Ian Johnston, Dominic Kwiatkowski, Cordelia Langford, John Sillitoe on behalf of the Wellcome Sanger Institute COVID-19 Surveillance Team |
| EPI_ISL_2541075 | Lighthouse Lab in Glasgow | Wellcome Sanger Institute for the COVID-19 Genomics UK (COG-UK) Consortium | Harper VanSteenhouse, Yumi Kasai, David Gray, Carol Clugston, Anna Dominiczak and Alex Alderton, Roberto Amato, Jeffrey Barrett, Sonia Goncalves, Ewan Harrison, David K. Jackson, Ian Johnston, Dominic Kwiatkowski, Cordelia Langford, John Sillitoe on behalf of the Wellcome Sanger Institute COVID-19 Surveillance Team |
| EPI_ISL_2541076, EPI_ISL_2541077, EPI_ISL_2541078, EPI_ISL_2541079 | Health Services Laboratories | Wellcome Sanger Institute for the COVID-19 Genomics UK (COG-UK) Consortium | Health Services Laboratories and Alex Alderton, Roberto Amato, Jeffrey Barrett, Sonia Goncalves, Ewan Harrison, David K. Jackson, Ian Johnston, Dominic Kwiatkowski, Cordelia Langford, John Sillitoe on behalf of the Wellcome Sanger Institute COVID-19 Surveillance Team |
| EPI_ISL_2541080, EPI_ISL_2541081, EPI_ISL_2541082, EPI_ISL_2541083 | Lighthouse Lab in Milton Keynes | Wellcome Sanger Institute for the COVID-19 Genomics UK (COG-UK) Consortium | The Lighthouse Lab in Milton Keynes and Alex Alderton, Roberto Amato, Jeffrey Barrett, Sonia Goncalves, Ewan Harrison, David K. Jackson, Ian Johnston, Dominic Kwiatkowski, Cordelia Langford, John Sillitoe on behalf of the Wellcome Sanger Institute COVID-19 Surveillance Team |
| EPI_ISL_2541084 | Health Services Laboratories | Wellcome Sanger Institute for the COVID-19 Genomics UK (COG-UK) Consortium | Health Services Laboratories and Alex Alderton, Roberto Amato, Jeffrey Barrett, Sonia Goncalves, Ewan Harrison, David K. Jackson, Ian Johnston, Dominic Kwiatkowski, Cordelia Langford, John Sillitoe on behalf of the Wellcome Sanger Institute COVID-19 Surveillance Team |
| EPI_ISL_2541085, EPI_ISL_2541086, EPI_ISL_2541087, EPI_ISL_2541088, EPI_ISL_2541089, EPI_ISL_2541090, EPI_ISL_2541091, EPI_ISL_2541092, EPI_ISL_2541093 | Lighthouse Lab in Milton Keynes | Wellcome Sanger Institute for the COVID-19 Genomics UK (COG-UK) Consortium | The Lighthouse Lab in Milton Keynes and Alex Alderton, Roberto Amato, Jeffrey Barrett, Sonia Goncalves, Ewan Harrison, David K. Jackson, Ian Johnston, Dominic Kwiatkowski, Cordelia Langford, John Sillitoe on behalf of the Wellcome Sanger Institute COVID-19 Surveillance Team |
| EPI_ISL_2541094 | Health Services Laboratories | Wellcome Sanger Institute for the COVID-19 Genomics UK (COG-UK) Consortium | Health Services Laboratories and Alex Alderton, Roberto Amato, Jeffrey Barrett, Sonia Goncalves, Ewan Harrison, David K. Jackson, Ian Johnston, Dominic Kwiatkowski, Cordelia Langford, John Sillitoe on behalf of the Wellcome Sanger Institute COVID-19 Surveillance Team |
| EPI_ISL_2541095, EPI_ISL_2541096, EPI_ISL_2541097, EPI_ISL_2541098, EPI_ISL_2541099, EPI_ISL_2541100, EPI_ISL_2541101, EPI_ISL_2541102 | Lighthouse Lab in Milton Keynes | Wellcome Sanger Institute for the COVID-19 Genomics UK (COG-UK) Consortium | The Lighthouse Lab in Milton Keynes and Alex Alderton, Roberto Amato, Jeffrey Barrett, Sonia Goncalves, Ewan Harrison, David K. Jackson, Ian Johnston, Dominic Kwiatkowski, Cordelia Langford, John Sillitoe on behalf of the Wellcome Sanger Institute COVID-19 Surveillance Team |
| EPI_ISL_2541103 | Health Services Laboratories | Wellcome Sanger Institute for the COVID-19 Genomics UK (COG-UK) Consortium | Health Services Laboratories and Alex Alderton, Roberto Amato, Jeffrey Barrett, Sonia Goncalves, Ewan Harrison, David K. Jackson, Ian Johnston, Dominic Kwiatkowski, Cordelia Langford, John Sillitoe on behalf of the Wellcome Sanger Institute COVID-19 Surveillance Team |
| EPI_ISL_2541104, EPI_ISL_2541105, EPI_ISL_2541106 | Lighthouse Lab in Milton Keynes | Wellcome Sanger Institute for the COVID-19 Genomics UK (COG-UK) Consortium | The Lighthouse Lab in Milton Keynes and Alex Alderton, Roberto Amato, Jeffrey Barrett, Sonia Goncalves, Ewan Harrison, David K. Jackson, Ian Johnston, Dominic Kwiatkowski, Cordelia Langford, John Sillitoe on behalf of the Wellcome Sanger Institute COVID-19 Surveillance Team |
| EPI_ISL_2541107 | Health Services Laboratories | Wellcome Sanger Institute for the COVID-19 Genomics UK (COG-UK) Consortium | Health Services Laboratories and Alex Alderton, Roberto Amato, Jeffrey Barrett, Sonia Goncalves, Ewan Harrison, David K. Jackson, Ian Johnston, Dominic Kwiatkowski, Cordelia Langford, John Sillitoe on behalf of the Wellcome Sanger Institute COVID-19 Surveillance Team |
| EPI_ISL_2541108, EPI_ISL_2541109, EPI_ISL_2541110, EPI_ISL_2541111, EPI_ISL_2541113, EPI_ISL_2541114, EPI_ISL_2541115 | Lighthouse Lab in Milton Keynes | Wellcome Sanger Institute for the COVID-19 Genomics UK (COG-UK) Consortium | The Lighthouse Lab in Milton Keynes and Alex Alderton, Roberto Amato, Jeffrey Barrett, Sonia Goncalves, Ewan Harrison, David K. Jackson, Ian Johnston, Dominic Kwiatkowski, Cordelia Langford, John Sillitoe on behalf of the Wellcome Sanger Institute COVID-19 Surveillance Team |
| EPI_ISL_2541116 | Health Services Laboratories | Wellcome Sanger Institute for the COVID-19 Genomics UK (COG-UK) Consortium | Health Services Laboratories and Alex Alderton, Roberto Amato, Jeffrey Barrett, Sonia Goncalves, Ewan Harrison, David K. Jackson, Ian Johnston, Dominic Kwiatkowski, Cordelia Langford, John Sillitoe on behalf of the Wellcome Sanger Institute COVID-19 Surveillance Team |
| EPI_ISL_2541117, EPI_ISL_2541118, EPI_ISL_2541119, EPI_ISL_2541122, EPI_ISL_2541123, EPI_ISL_2541124, EPI_ISL_2541125, EPI_ISL_2541126, EPI_ISL_2541127, EPI_ISL_2541128, EPI_ISL_2541133, EPI_ISL_2541134, EPI_ISL_2541135, EPI_ISL_2541136, EPI_ISL_2541137, EPI_ISL_2541138, EPI_ISL_2541139, EPI_ISL_2541140, EPI_ISL_2541141 | | | |

| | | | |
|---|---|---|---|
| see above | Lighthouse Lab in Milton Keynes | Wellcome Sanger Institute for the COVID-19 Genomics UK (COG-UK) Consortium | The Lighthouse Lab in Milton Keynes and Alex Alderton, Roberto Amato, Jeffrey Barrett, Sonia Goncalves, Ewan Harrison, David K. Jackson, Ian Johnston, Dominic Kwiatkowski, Cordelia Langford, John Sillitoe on behalf of the Wellcome Sanger Institute COVID-19 Surveillance Team |
| EPI_ISL_2541142 | Health Services Laboratories | Wellcome Sanger Institute for the COVID-19 Genomics UK (COG-UK) Consortium | Health Services Laboratories and Alex Alderton, Roberto Amato, Jeffrey Barrett, Sonia Goncalves, Ewan Harrison, David K. Jackson, Ian Johnston, Dominic Kwiatkowski, Cordelia Langford, John Sillitoe on behalf of the Wellcome Sanger Institute COVID-19 Surveillance Team |
| EPI_ISL_2541144, EPI_ISL_2541145, EPI_ISL_2541146 | Lighthouse Lab in Milton Keynes | Wellcome Sanger Institute for the COVID-19 Genomics UK (COG-UK) Consortium | The Lighthouse Lab in Milton Keynes and Alex Alderton, Roberto Amato, Jeffrey Barrett, Sonia Goncalves, Ewan Harrison, David K. Jackson, Ian Johnston, Dominic Kwiatkowski, Cordelia Langford, John Sillitoe on behalf of the Wellcome Sanger Institute COVID-19 Surveillance Team |
| EPI_ISL_2541147 | Health Services Laboratories | Wellcome Sanger Institute for the COVID-19 Genomics UK (COG-UK) Consortium | Health Services Laboratories and Alex Alderton, Roberto Amato, Jeffrey Barrett, Sonia Goncalves, Ewan Harrison, David K. Jackson, Ian Johnston, Dominic Kwiatkowski, Cordelia Langford, John Sillitoe on behalf of the Wellcome Sanger Institute COVID-19 Surveillance Team |
| EPI_ISL_2541148, EPI_ISL_2541149, EPI_ISL_2541151, EPI_ISL_2541152, EPI_ISL_2541153, EPI_ISL_2541154, EPI_ISL_2541155, EPI_ISL_2541156 | Lighthouse Lab in Milton Keynes | Wellcome Sanger Institute for the COVID-19 Genomics UK (COG-UK) Consortium | The Lighthouse Lab in Milton Keynes and Alex Alderton, Roberto Amato, Jeffrey Barrett, Sonia Goncalves, Ewan Harrison, David K. Jackson, Ian Johnston, Dominic Kwiatkowski, Cordelia Langford, John Sillitoe on behalf of the Wellcome Sanger Institute COVID-19 Surveillance Team |
| EPI_ISL_2541157 | Health Services Laboratories | Wellcome Sanger Institute for the COVID-19 Genomics UK (COG-UK) Consortium | Health Services Laboratories and Alex Alderton, Roberto Amato, Jeffrey Barrett, Sonia Goncalves, Ewan Harrison, David K. Jackson, Ian Johnston, Dominic Kwiatkowski, Cordelia Langford, John Sillitoe on behalf of the Wellcome Sanger Institute COVID-19 Surveillance Team |
| EPI_ISL_2541158, EPI_ISL_2541159, EPI_ISL_2541160, EPI_ISL_2541161, EPI_ISL_2541162 | Lighthouse Lab in Milton Keynes | Wellcome Sanger Institute for the COVID-19 Genomics UK (COG-UK) Consortium | The Lighthouse Lab in Milton Keynes and Alex Alderton, Roberto Amato, Jeffrey Barrett, Sonia Goncalves, Ewan Harrison, David K. Jackson, Ian Johnston, Dominic Kwiatkowski, Cordelia Langford, John Sillitoe on behalf of the Wellcome Sanger Institute COVID-19 Surveillance Team |
| EPI_ISL_2541163 | Health Services Laboratories | Wellcome Sanger Institute for the COVID-19 Genomics UK (COG-UK) Consortium | Health Services Laboratories and Alex Alderton, Roberto Amato, Jeffrey Barrett, Sonia Goncalves, Ewan Harrison, David K. Jackson, Ian Johnston, Dominic Kwiatkowski, Cordelia Langford, John Sillitoe on behalf of the Wellcome Sanger Institute COVID-19 Surveillance Team |
| EPI_ISL_2541164, EPI_ISL_2541165, EPI_ISL_2541166, EPI_ISL_2541167, EPI_ISL_2541168 | Lighthouse Lab in Milton Keynes | Wellcome Sanger Institute for the COVID-19 Genomics UK (COG-UK) Consortium | The Lighthouse Lab in Milton Keynes and Alex Alderton, Roberto Amato, Jeffrey Barrett, Sonia Goncalves, Ewan Harrison, David K. Jackson, Ian Johnston, Dominic Kwiatkowski, Cordelia Langford, John Sillitoe on behalf of the Wellcome Sanger Institute COVID-19 Surveillance Team |
| EPI_ISL_2541170 | Health Services Laboratories | Wellcome Sanger Institute for the COVID-19 Genomics UK (COG-UK) Consortium | Health Services Laboratories and Alex Alderton, Roberto Amato, Jeffrey Barrett, Sonia Goncalves, Ewan Harrison, David K. Jackson, Ian Johnston, Dominic Kwiatkowski, Cordelia Langford, John Sillitoe on behalf of the Wellcome Sanger Institute COVID-19 Surveillance Team |
| EPI_ISL_2541171, EPI_ISL_2541172, EPI_ISL_2541173, EPI_ISL_2541174, EPI_ISL_2541175, EPI_ISL_2541176 | Lighthouse Lab in Milton Keynes | Wellcome Sanger Institute for the COVID-19 Genomics UK (COG-UK) Consortium | The Lighthouse Lab in Milton Keynes and Alex Alderton, Roberto Amato, Jeffrey Barrett, Sonia Goncalves, Ewan Harrison, David K. Jackson, Ian Johnston, Dominic Kwiatkowski, Cordelia Langford, John Sillitoe on behalf of the Wellcome Sanger Institute COVID-19 Surveillance Team |
| EPI_ISL_2541177 | Health Services Laboratories | Wellcome Sanger Institute for the COVID-19 Genomics UK (COG-UK) Consortium | Health Services Laboratories and Alex Alderton, Roberto Amato, Jeffrey Barrett, Sonia Goncalves, Ewan Harrison, David K. Jackson, Ian Johnston, Dominic Kwiatkowski, Cordelia Langford, John Sillitoe on behalf of the Wellcome Sanger Institute COVID-19 Surveillance Team |
| EPI_ISL_2541178, EPI_ISL_2541179, EPI_ISL_2541180 | Lighthouse Lab in Milton Keynes | Wellcome Sanger Institute for the COVID-19 Genomics UK (COG-UK) Consortium | The Lighthouse Lab in Milton Keynes and Alex Alderton, Roberto Amato, Jeffrey Barrett, Sonia Goncalves, Ewan Harrison, David K. Jackson, Ian Johnston, Dominic Kwiatkowski, Cordelia Langford, John Sillitoe on behalf of the Wellcome Sanger Institute COVID-19 Surveillance Team |
| EPI_ISL_2541181 | Health Services Laboratories | Wellcome Sanger Institute for the COVID-19 Genomics UK (COG-UK) Consortium | Health Services Laboratories and Alex Alderton, Roberto Amato, Jeffrey Barrett, Sonia Goncalves, Ewan Harrison, David K. Jackson, Ian Johnston, Dominic Kwiatkowski, Cordelia Langford, John Sillitoe on behalf of the Wellcome Sanger Institute COVID-19 Surveillance Team |
| EPI_ISL_2541182, EPI_ISL_2541183 | Lighthouse Lab in Milton Keynes | Wellcome Sanger Institute for the COVID-19 Genomics UK (COG-UK) Consortium | The Lighthouse Lab in Milton Keynes and Alex Alderton, Roberto Amato, Jeffrey Barrett, Sonia Goncalves, Ewan Harrison, David K. Jackson, Ian Johnston, Dominic Kwiatkowski, Cordelia Langford, John Sillitoe on behalf of the Wellcome Sanger Institute COVID-19 Surveillance Team |
| EPI_ISL_2541184, EPI_ISL_2541185 | Health Services Laboratories | Wellcome Sanger Institute for the COVID-19 Genomics UK (COG-UK) Consortium | Health Services Laboratories and Alex Alderton, Roberto Amato, Jeffrey Barrett, Sonia Goncalves, Ewan Harrison, David K. Jackson, Ian Johnston, Dominic Kwiatkowski, Cordelia Langford, John Sillitoe on behalf of the Wellcome Sanger Institute COVID-19 Surveillance Team |
| EPI_ISL_2541187, EPI_ISL_2541188, EPI_ISL_2541189, EPI_ISL_2541190, EPI_ISL_2541191, EPI_ISL_2541192, EPI_ISL_2541193 | Lighthouse Lab in Milton Keynes | Wellcome Sanger Institute for the COVID-19 Genomics UK (COG-UK) Consortium | The Lighthouse Lab in Milton Keynes and Alex Alderton, Roberto Amato, Jeffrey Barrett, Sonia Goncalves, Ewan Harrison, David K. Jackson, Ian Johnston, Dominic Kwiatkowski, Cordelia Langford, John Sillitoe on behalf of the Wellcome Sanger Institute COVID-19 Surveillance Team |
| EPI_ISL_2541194 | Health Services Laboratories | Wellcome Sanger Institute for the COVID-19 Genomics UK (COG-UK) Consortium | Health Services Laboratories and Alex Alderton, Roberto Amato, Jeffrey Barrett, Sonia Goncalves, Ewan Harrison, David K. Jackson, Ian Johnston, Dominic Kwiatkowski, Cordelia Langford, John Sillitoe on behalf of the Wellcome Sanger Institute COVID-19 Surveillance Team |
| EPI_ISL_2541195, EPI_ISL_2541196, EPI_ISL_2541197, EPI_ISL_2541198, EPI_ISL_2541200 | Lighthouse Lab in Milton Keynes | Wellcome Sanger Institute for the COVID-19 Genomics UK (COG-UK) Consortium | The Lighthouse Lab in Milton Keynes and Alex Alderton, Roberto Amato, Jeffrey Barrett, Sonia Goncalves, Ewan Harrison, David K. Jackson, Ian Johnston, Dominic Kwiatkowski, Cordelia Langford, John Sillitoe on behalf of the Wellcome Sanger Institute COVID-19 Surveillance Team |
| EPI_ISL_2541201 | Health Services Laboratories | Wellcome Sanger Institute for the COVID-19 Genomics UK (COG-UK) Consortium | Health Services Laboratories and Alex Alderton, Roberto Amato, Jeffrey Barrett, Sonia Goncalves, Ewan Harrison, David K. Jackson, Ian Johnston, Dominic Kwiatkowski, Cordelia Langford, John Sillitoe on behalf of the Wellcome Sanger Institute COVID-19 Surveillance Team |
| EPI_ISL_2541202, EPI_ISL_2541203, EPI_ISL_2541205, EPI_ISL_2541206, EPI_ISL_2541207, EPI_ISL_2541208, EPI_ISL_2541210, EPI_ISL_2541211 | Lighthouse Lab in Milton Keynes | Wellcome Sanger Institute for the COVID-19 Genomics UK (COG-UK) Consortium | The Lighthouse Lab in Milton Keynes and Alex Alderton, Roberto Amato, Jeffrey Barrett, Sonia Goncalves, Ewan Harrison, David K. Jackson, Ian Johnston, Dominic Kwiatkowski, Cordelia Langford, John Sillitoe on behalf of the Wellcome Sanger Institute COVID-19 Surveillance Team |
| EPI_ISL_2541212 | Health Services Laboratories | Wellcome Sanger Institute for the COVID-19 Genomics UK (COG-UK) Consortium | Health Services Laboratories and Alex Alderton, Roberto Amato, Jeffrey Barrett, Sonia Goncalves, Ewan Harrison, David K. Jackson, Ian Johnston, Dominic Kwiatkowski, Cordelia Langford, John Sillitoe on behalf of the Wellcome Sanger Institute COVID-19 Surveillance Team |
| EPI_ISL_2541213, EPI_ISL_2541214, EPI_ISL_2541215, EPI_ISL_2541216, EPI_ISL_2541217, EPI_ISL_2541218, EPI_ISL_2541219, EPI_ISL_2541220, EPI_ISL_2541222, EPI_ISL_2541223 | Lighthouse Lab in Milton Keynes | Wellcome Sanger Institute for the COVID-19 Genomics UK (COG-UK) Consortium | The Lighthouse Lab in Milton Keynes and Alex Alderton, Roberto Amato, Jeffrey Barrett, Sonia Goncalves, Ewan Harrison, David K. Jackson, Ian Johnston, Dominic Kwiatkowski, Cordelia Langford, John Sillitoe on behalf of the Wellcome Sanger Institute COVID-19 Surveillance Team |
| EPI_ISL_2541225 | Health Services Laboratories | Wellcome Sanger Institute for the COVID-19 Genomics UK (COG-UK) Consortium | Health Services Laboratories and Alex Alderton, Roberto Amato, Jeffrey Barrett, Sonia Goncalves, Ewan Harrison, David K. Jackson, Ian Johnston, Dominic Kwiatkowski, Cordelia Langford, John Sillitoe on behalf of the Wellcome Sanger Institute COVID-19 Surveillance Team |
| EPI_ISL_2541226 | Lighthouse Lab in Milton Keynes | Wellcome Sanger Institute for the COVID-19 Genomics UK (COG-UK) Consortium | The Lighthouse Lab in Milton Keynes and Alex Alderton, Roberto Amato, Jeffrey Barrett, Sonia Goncalves, Ewan Harrison, David K. Jackson, Ian Johnston, Dominic Kwiatkowski, Cordelia Langford, John Sillitoe on behalf of the Wellcome Sanger Institute COVID-19 Surveillance Team |
| EPI_ISL_2541227 | Lighthouse Laboratory Plymouth | Wellcome Sanger Institute for the COVID-19 Genomics UK (COG-UK) Consortium | Lighthouse Laboratory Plymouth and Alex Alderton, Roberto Amato, Jeffrey Barrett, Sonia Goncalves, Ewan Harrison, David K. Jackson, Ian Johnston, Dominic Kwiatkowski, Cordelia Langford, John Sillitoe on behalf of the Wellcome Sanger Institute COVID-19 Surveillance Team |
| EPI_ISL_2541228, EPI_ISL_2541229 | Health Services Laboratories | Wellcome Sanger Institute for the COVID-19 Genomics UK (COG-UK) Consortium | Health Services Laboratories and Alex Alderton, Roberto Amato, Jeffrey Barrett, Sonia Goncalves, Ewan Harrison, David K. Jackson, Ian Johnston, Dominic Kwiatkowski, Cordelia Langford, John Sillitoe on behalf of the Wellcome Sanger Institute COVID-19 Surveillance Team |
| EPI_ISL_2541230 | Lighthouse Lab in Milton Keynes | Wellcome Sanger Institute for the COVID-19 Genomics UK (COG-UK) Consortium | The Lighthouse Lab in Milton Keynes and Alex Alderton, Roberto Amato, Jeffrey Barrett, Sonia Goncalves, Ewan Harrison, David K. Jackson, Ian Johnston, Dominic Kwiatkowski, Cordelia Langford, John Sillitoe on behalf of the Wellcome Sanger Institute COVID-19 Surveillance Team |
| EPI_ISL_2541231 | Health Services Laboratories | Wellcome Sanger Institute for the COVID-19 Genomics UK (COG-UK) | Health Services Laboratories and Alex Alderton, Roberto Amato, Jeffrey Barrett, Sonia Goncalves, Ewan Harrison, David K. Jackson, Ian Johnston, Dominic Kwiatkowski, Cordelia Langford, John Sillitoe on behalf of the Wellcome Sanger Institute |

| | | | |
|---|---|---|---|
| | | Consortium | COVID-19 Surveillance Team |
| EPI_ISL_2541232 | Lighthouse Lab in Milton Keynes | Wellcome Sanger Institute for the COVID-19 Genomics UK (COG-UK) Consortium | The Lighthouse Lab in Milton Keynes and Alex Alderton, Roberto Amato, Jeffrey Barrett, Sonia Goncalves, Ewan Harrison, David K. Jackson, Ian Johnston, Dominic Kwiatkowski, Cordelia Langford, John Sillitoe on behalf of the Wellcome Sanger Institute COVID-19 Surveillance Team |
| EPI_ISL_2541233, EPI_ISL_2541234, EPI_ISL_2541235 | Lighthouse Laboratory Plymouth | Wellcome Sanger Institute for the COVID-19 Genomics UK (COG-UK) Consortium | Lighthouse Laboratory Plymouth and Alex Alderton, Roberto Amato, Jeffrey Barrett, Sonia Goncalves, Ewan Harrison, David K. Jackson, Ian Johnston, Dominic Kwiatkowski, Cordelia Langford, John Sillitoe on behalf of the Wellcome Sanger Institute COVID-19 Surveillance Team |
| EPI_ISL_2541236, EPI_ISL_2541237, EPI_ISL_2541239, EPI_ISL_2541240 | Lighthouse Lab in Milton Keynes | Wellcome Sanger Institute for the COVID-19 Genomics UK (COG-UK) Consortium | The Lighthouse Lab in Milton Keynes and Alex Alderton, Roberto Amato, Jeffrey Barrett, Sonia Goncalves, Ewan Harrison, David K. Jackson, Ian Johnston, Dominic Kwiatkowski, Cordelia Langford, John Sillitoe on behalf of the Wellcome Sanger Institute COVID-19 Surveillance Team |
| EPI_ISL_2541241 | Health Services Laboratories | Wellcome Sanger Institute for the COVID-19 Genomics UK (COG-UK) Consortium | Health Services Laboratories and Alex Alderton, Roberto Amato, Jeffrey Barrett, Sonia Goncalves, Ewan Harrison, David K. Jackson, Ian Johnston, Dominic Kwiatkowski, Cordelia Langford, John Sillitoe on behalf of the Wellcome Sanger Institute COVID-19 Surveillance Team |
| EPI_ISL_2541242 | Lighthouse Lab in Milton Keynes | Wellcome Sanger Institute for the COVID-19 Genomics UK (COG-UK) Consortium | The Lighthouse Lab in Milton Keynes and Alex Alderton, Roberto Amato, Jeffrey Barrett, Sonia Goncalves, Ewan Harrison, David K. Jackson, Ian Johnston, Dominic Kwiatkowski, Cordelia Langford, John Sillitoe on behalf of the Wellcome Sanger Institute COVID-19 Surveillance Team |
| EPI_ISL_2541243 | Lighthouse Laboratory Plymouth | Wellcome Sanger Institute for the COVID-19 Genomics UK (COG-UK) Consortium | Lighthouse Laboratory Plymouth and Alex Alderton, Roberto Amato, Jeffrey Barrett, Sonia Goncalves, Ewan Harrison, David K. Jackson, Ian Johnston, Dominic Kwiatkowski, Cordelia Langford, John Sillitoe on behalf of the Wellcome Sanger Institute COVID-19 Surveillance Team |
| EPI_ISL_2541244 | Health Services Laboratories | Wellcome Sanger Institute for the COVID-19 Genomics UK (COG-UK) Consortium | Health Services Laboratories and Alex Alderton, Roberto Amato, Jeffrey Barrett, Sonia Goncalves, Ewan Harrison, David K. Jackson, Ian Johnston, Dominic Kwiatkowski, Cordelia Langford, John Sillitoe on behalf of the Wellcome Sanger Institute COVID-19 Surveillance Team |
| EPI_ISL_2541245, EPI_ISL_2541246, EPI_ISL_2541247, EPI_ISL_2541248 | Lighthouse Lab in Milton Keynes | Wellcome Sanger Institute for the COVID-19 Genomics UK (COG-UK) Consortium | The Lighthouse Lab in Milton Keynes and Alex Alderton, Roberto Amato, Jeffrey Barrett, Sonia Goncalves, Ewan Harrison, David K. Jackson, Ian Johnston, Dominic Kwiatkowski, Cordelia Langford, John Sillitoe on behalf of the Wellcome Sanger Institute COVID-19 Surveillance Team |
| EPI_ISL_2541249, EPI_ISL_2541250 | Lighthouse Laboratory Plymouth | Wellcome Sanger Institute for the COVID-19 Genomics UK (COG-UK) Consortium | Lighthouse Laboratory Plymouth and Alex Alderton, Roberto Amato, Jeffrey Barrett, Sonia Goncalves, Ewan Harrison, David K. Jackson, Ian Johnston, Dominic Kwiatkowski, Cordelia Langford, John Sillitoe on behalf of the Wellcome Sanger Institute COVID-19 Surveillance Team |
| EPI_ISL_2541251 | Health Services Laboratories | Wellcome Sanger Institute for the COVID-19 Genomics UK (COG-UK) Consortium | Health Services Laboratories and Alex Alderton, Roberto Amato, Jeffrey Barrett, Sonia Goncalves, Ewan Harrison, David K. Jackson, Ian Johnston, Dominic Kwiatkowski, Cordelia Langford, John Sillitoe on behalf of the Wellcome Sanger Institute COVID-19 Surveillance Team |
| EPI_ISL_2541252, EPI_ISL_2541253, EPI_ISL_2541254, EPI_ISL_2541255, EPI_ISL_2541256, EPI_ISL_2541257 | Lighthouse Lab in Milton Keynes | Wellcome Sanger Institute for the COVID-19 Genomics UK (COG-UK) Consortium | The Lighthouse Lab in Milton Keynes and Alex Alderton, Roberto Amato, Jeffrey Barrett, Sonia Goncalves, Ewan Harrison, David K. Jackson, Ian Johnston, Dominic Kwiatkowski, Cordelia Langford, John Sillitoe on behalf of the Wellcome Sanger Institute COVID-19 Surveillance Team |
| EPI_ISL_2541258 | Health Services Laboratories | Wellcome Sanger Institute for the COVID-19 Genomics UK (COG-UK) Consortium | Health Services Laboratories and Alex Alderton, Roberto Amato, Jeffrey Barrett, Sonia Goncalves, Ewan Harrison, David K. Jackson, Ian Johnston, Dominic Kwiatkowski, Cordelia Langford, John Sillitoe on behalf of the Wellcome Sanger Institute COVID-19 Surveillance Team |
| EPI_ISL_2541259, EPI_ISL_2541260, EPI_ISL_2541261, EPI_ISL_2541262, EPI_ISL_2541264, EPI_ISL_2541265, EPI_ISL_2541266, EPI_ISL_2541267, EPI_ISL_2541268, EPI_ISL_2541269, EPI_ISL_2541270, EPI_ISL_2541271, EPI_ISL_2541272, EPI_ISL_2541273, EPI_ISL_2541274, EPI_ISL_2541275, EPI_ISL_2541278, EPI_ISL_2541279, EPI_ISL_2541281, EPI_ISL_2541282, EPI_ISL_2541283, EPI_ISL_2541284, EPI_ISL_2541285, EPI_ISL_2541286 | | | |
| see above | Lighthouse Lab in Milton Keynes | Wellcome Sanger Institute for the COVID-19 Genomics UK (COG-UK) Consortium | The Lighthouse Lab in Milton Keynes and Alex Alderton, Roberto Amato, Jeffrey Barrett, Sonia Goncalves, Ewan Harrison, David K. Jackson, Ian Johnston, Dominic Kwiatkowski, Cordelia Langford, John Sillitoe on behalf of the Wellcome Sanger Institute COVID-19 Surveillance Team |
| EPI_ISL_2541287 | Health Services Laboratories | Wellcome Sanger Institute for the COVID-19 Genomics UK (COG-UK) Consortium | Health Services Laboratories and Alex Alderton, Roberto Amato, Jeffrey Barrett, Sonia Goncalves, Ewan Harrison, David K. Jackson, Ian Johnston, Dominic Kwiatkowski, Cordelia Langford, John Sillitoe on behalf of the Wellcome Sanger Institute COVID-19 Surveillance Team |
| EPI_ISL_2541289, EPI_ISL_2541290 | Lighthouse Lab in Milton Keynes | Wellcome Sanger Institute for the COVID-19 Genomics UK (COG-UK) Consortium | The Lighthouse Lab in Milton Keynes and Alex Alderton, Roberto Amato, Jeffrey Barrett, Sonia Goncalves, Ewan Harrison, David K. Jackson, Ian Johnston, Dominic Kwiatkowski, Cordelia Langford, John Sillitoe on behalf of the Wellcome Sanger Institute COVID-19 Surveillance Team |
| EPI_ISL_2541291 | Lighthouse Laboratory Plymouth | Wellcome Sanger Institute for the COVID-19 Genomics UK (COG-UK) Consortium | Lighthouse Laboratory Plymouth and Alex Alderton, Roberto Amato, Jeffrey Barrett, Sonia Goncalves, Ewan Harrison, David K. Jackson, Ian Johnston, Dominic Kwiatkowski, Cordelia Langford, John Sillitoe on behalf of the Wellcome Sanger Institute COVID-19 Surveillance Team |
| EPI_ISL_2541292, EPI_ISL_2541293, EPI_ISL_2541294, EPI_ISL_2541295, EPI_ISL_2541296, EPI_ISL_2541297 | Lighthouse Lab in Milton Keynes | Wellcome Sanger Institute for the COVID-19 Genomics UK (COG-UK) Consortium | The Lighthouse Lab in Milton Keynes and Alex Alderton, Roberto Amato, Jeffrey Barrett, Sonia Goncalves, Ewan Harrison, David K. Jackson, Ian Johnston, Dominic Kwiatkowski, Cordelia Langford, John Sillitoe on behalf of the Wellcome Sanger Institute COVID-19 Surveillance Team |
| EPI_ISL_2541298 | Health Services Laboratories | Wellcome Sanger Institute for the COVID-19 Genomics UK (COG-UK) Consortium | Health Services Laboratories and Alex Alderton, Roberto Amato, Jeffrey Barrett, Sonia Goncalves, Ewan Harrison, David K. Jackson, Ian Johnston, Dominic Kwiatkowski, Cordelia Langford, John Sillitoe on behalf of the Wellcome Sanger Institute COVID-19 Surveillance Team |
| EPI_ISL_2541299 | Lighthouse Lab in Milton Keynes | Wellcome Sanger Institute for the COVID-19 Genomics UK (COG-UK) Consortium | The Lighthouse Lab in Milton Keynes and Alex Alderton, Roberto Amato, Jeffrey Barrett, Sonia Goncalves, Ewan Harrison, David K. Jackson, Ian Johnston, Dominic Kwiatkowski, Cordelia Langford, John Sillitoe on behalf of the Wellcome Sanger Institute COVID-19 Surveillance Team |
| EPI_ISL_2541300 | Health Services Laboratories | Wellcome Sanger Institute for the COVID-19 Genomics UK (COG-UK) Consortium | Health Services Laboratories and Alex Alderton, Roberto Amato, Jeffrey Barrett, Sonia Goncalves, Ewan Harrison, David K. Jackson, Ian Johnston, Dominic Kwiatkowski, Cordelia Langford, John Sillitoe on behalf of the Wellcome Sanger Institute COVID-19 Surveillance Team |
| EPI_ISL_2541301, EPI_ISL_2541302, EPI_ISL_2541303, EPI_ISL_2541304 | Lighthouse Lab in Milton Keynes | Wellcome Sanger Institute for the COVID-19 Genomics UK (COG-UK) Consortium | The Lighthouse Lab in Milton Keynes and Alex Alderton, Roberto Amato, Jeffrey Barrett, Sonia Goncalves, Ewan Harrison, David K. Jackson, Ian Johnston, Dominic Kwiatkowski, Cordelia Langford, John Sillitoe on behalf of the Wellcome Sanger Institute COVID-19 Surveillance Team |
| EPI_ISL_2541305 | Lighthouse Laboratory Plymouth | Wellcome Sanger Institute for the COVID-19 Genomics UK (COG-UK) Consortium | Lighthouse Laboratory Plymouth and Alex Alderton, Roberto Amato, Jeffrey Barrett, Sonia Goncalves, Ewan Harrison, David K. Jackson, Ian Johnston, Dominic Kwiatkowski, Cordelia Langford, John Sillitoe on behalf of the Wellcome Sanger Institute COVID-19 Surveillance Team |
| EPI_ISL_2541306 | Lighthouse Lab in Milton Keynes | Wellcome Sanger Institute for the COVID-19 Genomics UK (COG-UK) Consortium | The Lighthouse Lab in Milton Keynes and Alex Alderton, Roberto Amato, Jeffrey Barrett, Sonia Goncalves, Ewan Harrison, David K. Jackson, Ian Johnston, Dominic Kwiatkowski, Cordelia Langford, John Sillitoe on behalf of the Wellcome Sanger Institute COVID-19 Surveillance Team |
| EPI_ISL_2541307 | Lighthouse Laboratory Plymouth | Wellcome Sanger Institute for the COVID-19 Genomics UK (COG-UK) Consortium | Lighthouse Laboratory Plymouth and Alex Alderton, Roberto Amato, Jeffrey Barrett, Sonia Goncalves, Ewan Harrison, David K. Jackson, Ian Johnston, Dominic Kwiatkowski, Cordelia Langford, John Sillitoe on behalf of the Wellcome Sanger Institute COVID-19 Surveillance Team |
| EPI_ISL_2541308, EPI_ISL_2541309, EPI_ISL_2541310 | Lighthouse Lab in Milton Keynes | Wellcome Sanger Institute for the COVID-19 Genomics UK (COG-UK) Consortium | The Lighthouse Lab in Milton Keynes and Alex Alderton, Roberto Amato, Jeffrey Barrett, Sonia Goncalves, Ewan Harrison, David K. Jackson, Ian Johnston, Dominic Kwiatkowski, Cordelia Langford, John Sillitoe on behalf of the Wellcome Sanger Institute COVID-19 Surveillance Team |
| EPI_ISL_2541311 | Lighthouse Laboratory Plymouth | Wellcome Sanger Institute for the COVID-19 Genomics UK (COG-UK) Consortium | Lighthouse Laboratory Plymouth and Alex Alderton, Roberto Amato, Jeffrey Barrett, Sonia Goncalves, Ewan Harrison, David K. Jackson, Ian Johnston, Dominic Kwiatkowski, Cordelia Langford, John Sillitoe on behalf of the Wellcome Sanger Institute COVID-19 Surveillance Team |
| EPI_ISL_2541312 | Health Services Laboratories | Wellcome Sanger Institute for the COVID-19 Genomics UK (COG-UK) Consortium | Health Services Laboratories and Alex Alderton, Roberto Amato, Jeffrey Barrett, Sonia Goncalves, Ewan Harrison, David K. Jackson, Ian Johnston, Dominic Kwiatkowski, Cordelia Langford, John Sillitoe on behalf of the Wellcome Sanger Institute COVID-19 Surveillance Team |
| EPI_ISL_2541313, EPI_ISL_2541314 | Lighthouse Lab in Milton Keynes | Wellcome Sanger Institute for the COVID-19 Genomics UK (COG-UK) Consortium | The Lighthouse Lab in Milton Keynes and Alex Alderton, Roberto Amato, Jeffrey Barrett, Sonia Goncalves, Ewan Harrison, David K. Jackson, Ian Johnston, Dominic Kwiatkowski, Cordelia Langford, John Sillitoe on behalf of the Wellcome Sanger Institute COVID-19 Surveillance Team |
| EPI_ISL_2541315, EPI_ISL_2541316 | Lighthouse Laboratory Plymouth | Wellcome Sanger Institute for the COVID-19 Genomics UK (COG-UK) Consortium | Lighthouse Laboratory Plymouth and Alex Alderton, Roberto Amato, Jeffrey Barrett, Sonia Goncalves, Ewan Harrison, David K. Jackson, Ian Johnston, Dominic Kwiatkowski, Cordelia Langford, John Sillitoe on behalf of the Wellcome Sanger Institute COVID-19 Surveillance Team |
| EPI_ISL_2541317 | Health Services Laboratories | Wellcome Sanger Institute for the COVID-19 Genomics UK (COG-UK) Consortium | Health Services Laboratories and Alex Alderton, Roberto Amato, Jeffrey Barrett, Sonia Goncalves, Ewan Harrison, David K. Jackson, Ian Johnston, Dominic Kwiatkowski, Cordelia Langford, John Sillitoe on behalf of the Wellcome Sanger Institute COVID-19 Surveillance Team |
| EPI_ISL_2541318 | Lighthouse Lab in Milton Keynes | Wellcome Sanger Institute for the COVID-19 Genomics UK (COG-UK) Consortium | The Lighthouse Lab in Milton Keynes and Alex Alderton, Roberto Amato, Jeffrey Barrett, Sonia Goncalves, Ewan Harrison, David K. Jackson, Ian Johnston, Dominic Kwiatkowski, Cordelia Langford, John Sillitoe on behalf of the Wellcome Sanger Institute COVID-19 Surveillance Team |
| EPI_ISL_2541319 | Health Services Laboratories | Wellcome Sanger Institute for the COVID-19 Genomics UK (COG-UK) Consortium | Health Services Laboratories and Alex Alderton, Roberto Amato, Jeffrey Barrett, Sonia Goncalves, Ewan Harrison, David K. Jackson, Ian Johnston, Dominic Kwiatkowski, Cordelia Langford, John Sillitoe on behalf of the Wellcome Sanger Institute COVID-19 Surveillance Team |
| EPI_ISL_2541320 | Lighthouse Lab in Milton Keynes | Wellcome Sanger Institute for the COVID-19 Genomics UK (COG-UK) Consortium | The Lighthouse Lab in Milton Keynes and Alex Alderton, Roberto Amato, Jeffrey Barrett, Sonia Goncalves, Ewan Harrison, David K. Jackson, Ian Johnston, Dominic Kwiatkowski, Cordelia Langford, John Sillitoe on behalf of the Wellcome Sanger Institute COVID-19 Surveillance Team |
| EPI_ISL_2541321 | Health Services Laboratories | Wellcome Sanger Institute for the COVID-19 Genomics UK (COG-UK) Consortium | Health Services Laboratories and Alex Alderton, Roberto Amato, Jeffrey Barrett, Sonia Goncalves, Ewan Harrison, David K. Jackson, Ian Johnston, Dominic Kwiatkowski, Cordelia Langford, John Sillitoe on behalf of the Wellcome Sanger Institute COVID-19 Surveillance Team |
| EPI_ISL_2541322, EPI_ISL_2541323, EPI_ISL_2541324, EPI_ISL_2541325, EPI_ISL_2541326, EPI_ISL_2541327, EPI_ISL_2541328, | Lighthouse Lab in Milton Keynes | Wellcome Sanger Institute for the COVID-19 Genomics UK (COG-UK) Consortium | The Lighthouse Lab in Milton Keynes and Alex Alderton, Roberto Amato, Jeffrey Barrett, Sonia Goncalves, Ewan Harrison, David K. Jackson, Ian Johnston, Dominic Kwiatkowski, Cordelia Langford, John Sillitoe on behalf of the Wellcome Sanger Institute COVID-19 Surveillance Team |

| | | | |
|---|---|---|---|
| EPI_ISL_2541329, EPI_ISL_2541330, EPI_ISL_2541331 | | | |
| EPI_ISL_2541332 | Lighthouse Laboratory Plymouth | Wellcome Sanger Institute for the COVID-19 Genomics UK (COG-UK) Consortium | Lighthouse Laboratory Plymouth and Alex Alderton, Roberto Amato, Jeffrey Barrett, Sonia Goncalves, Ewan Harrison, David K. Jackson, Ian Johnston, Dominic Kwiatkowski, Cordelia Langford, John Sillitoe on behalf of the Wellcome Sanger Institute COVID-19 Surveillance Team |
| EPI_ISL_2541333, EPI_ISL_2541334, EPI_ISL_2541335, EPI_ISL_2541336 | Lighthouse Lab in Milton Keynes | Wellcome Sanger Institute for the COVID-19 Genomics UK (COG-UK) Consortium | The Lighthouse Lab in Milton Keynes and Alex Alderton, Roberto Amato, Jeffrey Barrett, Sonia Goncalves, Ewan Harrison, David K. Jackson, Ian Johnston, Dominic Kwiatkowski, Cordelia Langford, John Sillitoe on behalf of the Wellcome Sanger Institute COVID-19 Surveillance Team |
| EPI_ISL_2541337 | Lighthouse Laboratory Plymouth | Wellcome Sanger Institute for the COVID-19 Genomics UK (COG-UK) Consortium | Lighthouse Laboratory Plymouth and Alex Alderton, Roberto Amato, Jeffrey Barrett, Sonia Goncalves, Ewan Harrison, David K. Jackson, Ian Johnston, Dominic Kwiatkowski, Cordelia Langford, John Sillitoe on behalf of the Wellcome Sanger Institute COVID-19 Surveillance Team |
| EPI_ISL_2541338 | Lighthouse Lab in Milton Keynes | Wellcome Sanger Institute for the COVID-19 Genomics UK (COG-UK) Consortium | The Lighthouse Lab in Milton Keynes and Alex Alderton, Roberto Amato, Jeffrey Barrett, Sonia Goncalves, Ewan Harrison, David K. Jackson, Ian Johnston, Dominic Kwiatkowski, Cordelia Langford, John Sillitoe on behalf of the Wellcome Sanger Institute COVID-19 Surveillance Team |
| EPI_ISL_2541339, EPI_ISL_2541340 | Lighthouse Laboratory Plymouth | Wellcome Sanger Institute for the COVID-19 Genomics UK (COG-UK) Consortium | Lighthouse Laboratory Plymouth and Alex Alderton, Roberto Amato, Jeffrey Barrett, Sonia Goncalves, Ewan Harrison, David K. Jackson, Ian Johnston, Dominic Kwiatkowski, Cordelia Langford, John Sillitoe on behalf of the Wellcome Sanger Institute COVID-19 Surveillance Team |
| EPI_ISL_2541341, EPI_ISL_2541343, EPI_ISL_2541344, EPI_ISL_2541345, EPI_ISL_2541349, EPI_ISL_2541350, EPI_ISL_2541353 | Lighthouse Lab in Milton Keynes | Wellcome Sanger Institute for the COVID-19 Genomics UK (COG-UK) Consortium | The Lighthouse Lab in Milton Keynes and Alex Alderton, Roberto Amato, Jeffrey Barrett, Sonia Goncalves, Ewan Harrison, David K. Jackson, Ian Johnston, Dominic Kwiatkowski, Cordelia Langford, John Sillitoe on behalf of the Wellcome Sanger Institute COVID-19 Surveillance Team |
| EPI_ISL_2541354 | Lighthouse Laboratory Plymouth | Wellcome Sanger Institute for the COVID-19 Genomics UK (COG-UK) Consortium | Lighthouse Laboratory Plymouth and Alex Alderton, Roberto Amato, Jeffrey Barrett, Sonia Goncalves, Ewan Harrison, David K. Jackson, Ian Johnston, Dominic Kwiatkowski, Cordelia Langford, John Sillitoe on behalf of the Wellcome Sanger Institute COVID-19 Surveillance Team |
| EPI_ISL_2541356, EPI_ISL_2541357, EPI_ISL_2541360, EPI_ISL_2541362 | Lighthouse Lab in Milton Keynes | Wellcome Sanger Institute for the COVID-19 Genomics UK (COG-UK) Consortium | The Lighthouse Lab in Milton Keynes and Alex Alderton, Roberto Amato, Jeffrey Barrett, Sonia Goncalves, Ewan Harrison, David K. Jackson, Ian Johnston, Dominic Kwiatkowski, Cordelia Langford, John Sillitoe on behalf of the Wellcome Sanger Institute COVID-19 Surveillance Team |
| EPI_ISL_2541364 | Health Services Laboratories | Wellcome Sanger Institute for the COVID-19 Genomics UK (COG-UK) Consortium | Health Services Laboratories and Alex Alderton, Roberto Amato, Jeffrey Barrett, Sonia Goncalves, Ewan Harrison, David K. Jackson, Ian Johnston, Dominic Kwiatkowski, Cordelia Langford, John Sillitoe on behalf of the Wellcome Sanger Institute COVID-19 Surveillance Team |
| EPI_ISL_2541365, EPI_ISL_2541366, EPI_ISL_2541368, EPI_ISL_2541369, EPI_ISL_2541371, EPI_ISL_2541372, EPI_ISL_2541374, EPI_ISL_2541375, EPI_ISL_2541377, EPI_ISL_2541378 | Lighthouse Lab in Milton Keynes | Wellcome Sanger Institute for the COVID-19 Genomics UK (COG-UK) Consortium | The Lighthouse Lab in Milton Keynes and Alex Alderton, Roberto Amato, Jeffrey Barrett, Sonia Goncalves, Ewan Harrison, David K. Jackson, Ian Johnston, Dominic Kwiatkowski, Cordelia Langford, John Sillitoe on behalf of the Wellcome Sanger Institute COVID-19 Surveillance Team |
| EPI_ISL_2541380 | Health Services Laboratories | Wellcome Sanger Institute for the COVID-19 Genomics UK (COG-UK) Consortium | Health Services Laboratories and Alex Alderton, Roberto Amato, Jeffrey Barrett, Sonia Goncalves, Ewan Harrison, David K. Jackson, Ian Johnston, Dominic Kwiatkowski, Cordelia Langford, John Sillitoe on behalf of the Wellcome Sanger Institute COVID-19 Surveillance Team |
| EPI_ISL_2541382, EPI_ISL_2541383 | Lighthouse Lab in Milton Keynes | Wellcome Sanger Institute for the COVID-19 Genomics UK (COG-UK) Consortium | The Lighthouse Lab in Milton Keynes and Alex Alderton, Roberto Amato, Jeffrey Barrett, Sonia Goncalves, Ewan Harrison, David K. Jackson, Ian Johnston, Dominic Kwiatkowski, Cordelia Langford, John Sillitoe on behalf of the Wellcome Sanger Institute COVID-19 Surveillance Team |
| EPI_ISL_2541386 | Lighthouse Laboratory Plymouth | Wellcome Sanger Institute for the COVID-19 Genomics UK (COG-UK) Consortium | Lighthouse Laboratory Plymouth and Alex Alderton, Roberto Amato, Jeffrey Barrett, Sonia Goncalves, Ewan Harrison, David K. Jackson, Ian Johnston, Dominic Kwiatkowski, Cordelia Langford, John Sillitoe on behalf of the Wellcome Sanger Institute COVID-19 Surveillance Team |
| EPI_ISL_2541388, EPI_ISL_2541389 | Lighthouse Lab in Milton Keynes | Wellcome Sanger Institute for the COVID-19 Genomics UK (COG-UK) Consortium | The Lighthouse Lab in Milton Keynes and Alex Alderton, Roberto Amato, Jeffrey Barrett, Sonia Goncalves, Ewan Harrison, David K. Jackson, Ian Johnston, Dominic Kwiatkowski, Cordelia Langford, John Sillitoe on behalf of the Wellcome Sanger Institute COVID-19 Surveillance Team |
| EPI_ISL_2541391 | Lighthouse Laboratory Plymouth | Wellcome Sanger Institute for the COVID-19 Genomics UK (COG-UK) Consortium | Lighthouse Laboratory Plymouth and Alex Alderton, Roberto Amato, Jeffrey Barrett, Sonia Goncalves, Ewan Harrison, David K. Jackson, Ian Johnston, Dominic Kwiatkowski, Cordelia Langford, John Sillitoe on behalf of the Wellcome Sanger Institute COVID-19 Surveillance Team |
| EPI_ISL_2541392, EPI_ISL_2541395, EPI_ISL_2541398, EPI_ISL_2541399, EPI_ISL_2541401, EPI_ISL_2541402, EPI_ISL_2541404, EPI_ISL_2541405, EPI_ISL_2541407 | Lighthouse Lab in Milton Keynes | Wellcome Sanger Institute for the COVID-19 Genomics UK (COG-UK) Consortium | The Lighthouse Lab in Milton Keynes and Alex Alderton, Roberto Amato, Jeffrey Barrett, Sonia Goncalves, Ewan Harrison, David K. Jackson, Ian Johnston, Dominic Kwiatkowski, Cordelia Langford, John Sillitoe on behalf of the Wellcome Sanger Institute COVID-19 Surveillance Team |
| EPI_ISL_2541408, EPI_ISL_2541410 | Health Services Laboratories | Wellcome Sanger Institute for the COVID-19 Genomics UK (COG-UK) Consortium | Health Services Laboratories and Alex Alderton, Roberto Amato, Jeffrey Barrett, Sonia Goncalves, Ewan Harrison, David K. Jackson, Ian Johnston, Dominic Kwiatkowski, Cordelia Langford, John Sillitoe on behalf of the Wellcome Sanger Institute COVID-19 Surveillance Team |
| EPI_ISL_2541411, EPI_ISL_2541413 | Lighthouse Lab in Milton Keynes | Wellcome Sanger Institute for the COVID-19 Genomics UK (COG-UK) Consortium | The Lighthouse Lab in Milton Keynes and Alex Alderton, Roberto Amato, Jeffrey Barrett, Sonia Goncalves, Ewan Harrison, David K. Jackson, Ian Johnston, Dominic Kwiatkowski, Cordelia Langford, John Sillitoe on behalf of the Wellcome Sanger Institute COVID-19 Surveillance Team |
| EPI_ISL_2541414 | Health Services Laboratories | Wellcome Sanger Institute for the COVID-19 Genomics UK (COG-UK) Consortium | Health Services Laboratories and Alex Alderton, Roberto Amato, Jeffrey Barrett, Sonia Goncalves, Ewan Harrison, David K. Jackson, Ian Johnston, Dominic Kwiatkowski, Cordelia Langford, John Sillitoe on behalf of the Wellcome Sanger Institute COVID-19 Surveillance Team |
| EPI_ISL_2541416, EPI_ISL_2541419, EPI_ISL_2541421, EPI_ISL_2541422 | Lighthouse Lab in Milton Keynes | Wellcome Sanger Institute for the COVID-19 Genomics UK (COG-UK) Consortium | The Lighthouse Lab in Milton Keynes and Alex Alderton, Roberto Amato, Jeffrey Barrett, Sonia Goncalves, Ewan Harrison, David K. Jackson, Ian Johnston, Dominic Kwiatkowski, Cordelia Langford, John Sillitoe on behalf of the Wellcome Sanger Institute COVID-19 Surveillance Team |
| EPI_ISL_2541424 | Lighthouse Laboratory Plymouth | Wellcome Sanger Institute for the COVID-19 Genomics UK (COG-UK) Consortium | Lighthouse Laboratory Plymouth and Alex Alderton, Roberto Amato, Jeffrey Barrett, Sonia Goncalves, Ewan Harrison, David K. Jackson, Ian Johnston, Dominic Kwiatkowski, Cordelia Langford, John Sillitoe on behalf of the Wellcome Sanger Institute COVID-19 Surveillance Team |
| EPI_ISL_2541425, EPI_ISL_2541427, EPI_ISL_2541429, EPI_ISL_2541430, EPI_ISL_2541432, EPI_ISL_2541435, EPI_ISL_2541436, EPI_ISL_2541438, EPI_ISL_2541440, EPI_ISL_2541441, EPI_ISL_2541443, EPI_ISL_2541444, EPI_ISL_2541446, EPI_ISL_2541447, EPI_ISL_2541449, EPI_ISL_2541450 | | | |
| see above | Lighthouse Lab in Milton Keynes | Wellcome Sanger Institute for the COVID-19 Genomics UK (COG-UK) Consortium | The Lighthouse Lab in Milton Keynes and Alex Alderton, Roberto Amato, Jeffrey Barrett, Sonia Goncalves, Ewan Harrison, David K. Jackson, Ian Johnston, Dominic Kwiatkowski, Cordelia Langford, John Sillitoe on behalf of the Wellcome Sanger Institute COVID-19 Surveillance Team |
| EPI_ISL_2541452 | Health Services Laboratories | Wellcome Sanger Institute for the COVID-19 Genomics UK (COG-UK) Consortium | Health Services Laboratories and Alex Alderton, Roberto Amato, Jeffrey Barrett, Sonia Goncalves, Ewan Harrison, David K. Jackson, Ian Johnston, Dominic Kwiatkowski, Cordelia Langford, John Sillitoe on behalf of the Wellcome Sanger Institute COVID-19 Surveillance Team |
| EPI_ISL_2541454, EPI_ISL_2541455, EPI_ISL_2541457 | Lighthouse Lab in Milton Keynes | Wellcome Sanger Institute for the COVID-19 Genomics UK (COG-UK) Consortium | The Lighthouse Lab in Milton Keynes and Alex Alderton, Roberto Amato, Jeffrey Barrett, Sonia Goncalves, Ewan Harrison, David K. Jackson, Ian Johnston, Dominic Kwiatkowski, Cordelia Langford, John Sillitoe on behalf of the Wellcome Sanger Institute COVID-19 Surveillance Team |
| EPI_ISL_2541459 | Health Services Laboratories | Wellcome Sanger Institute for the COVID-19 Genomics UK (COG-UK) Consortium | Health Services Laboratories and Alex Alderton, Roberto Amato, Jeffrey Barrett, Sonia Goncalves, Ewan Harrison, David K. Jackson, Ian Johnston, Dominic Kwiatkowski, Cordelia Langford, John Sillitoe on behalf of the Wellcome Sanger Institute COVID-19 Surveillance Team |
| EPI_ISL_2541460, EPI_ISL_2541462, EPI_ISL_2541464 | Lighthouse Lab in Milton Keynes | Wellcome Sanger Institute for the COVID-19 Genomics UK (COG-UK) Consortium | The Lighthouse Lab in Milton Keynes and Alex Alderton, Roberto Amato, Jeffrey Barrett, Sonia Goncalves, Ewan Harrison, David K. Jackson, Ian Johnston, Dominic Kwiatkowski, Cordelia Langford, John Sillitoe on behalf of the Wellcome Sanger Institute COVID-19 Surveillance Team |
| EPI_ISL_2541466 | Lighthouse Laboratory Plymouth | Wellcome Sanger Institute for the COVID-19 Genomics UK (COG-UK) Consortium | Lighthouse Laboratory Plymouth and Alex Alderton, Roberto Amato, Jeffrey Barrett, Sonia Goncalves, Ewan Harrison, David K. Jackson, Ian Johnston, Dominic Kwiatkowski, Cordelia Langford, John Sillitoe on behalf of the Wellcome Sanger Institute COVID-19 Surveillance Team |
| EPI_ISL_2541467, EPI_ISL_2541469 | Lighthouse Lab in Milton Keynes | Wellcome Sanger Institute for the COVID-19 Genomics UK (COG-UK) Consortium | The Lighthouse Lab in Milton Keynes and Alex Alderton, Roberto Amato, Jeffrey Barrett, Sonia Goncalves, Ewan Harrison, David K. Jackson, Ian Johnston, Dominic Kwiatkowski, Cordelia Langford, John Sillitoe on behalf of the Wellcome Sanger Institute COVID-19 Surveillance Team |
| EPI_ISL_2541471 | Health Services Laboratories | Wellcome Sanger Institute for the COVID-19 Genomics UK (COG-UK) Consortium | Health Services Laboratories and Alex Alderton, Roberto Amato, Jeffrey Barrett, Sonia Goncalves, Ewan Harrison, David K. Jackson, Ian Johnston, Dominic Kwiatkowski, Cordelia Langford, John Sillitoe on behalf of the Wellcome Sanger Institute COVID-19 Surveillance Team |
| EPI_ISL_2541472, EPI_ISL_2541474, EPI_ISL_2541475 | Lighthouse Lab in Milton Keynes | Wellcome Sanger Institute for the COVID-19 Genomics UK (COG-UK) Consortium | The Lighthouse Lab in Milton Keynes and Alex Alderton, Roberto Amato, Jeffrey Barrett, Sonia Goncalves, Ewan Harrison, David K. Jackson, Ian Johnston, Dominic Kwiatkowski, Cordelia Langford, John Sillitoe on behalf of the Wellcome Sanger Institute COVID-19 Surveillance Team |
| EPI_ISL_2541476 | Health Services Laboratories | Wellcome Sanger Institute for the COVID-19 Genomics UK (COG-UK) Consortium | Health Services Laboratories and Alex Alderton, Roberto Amato, Jeffrey Barrett, Sonia Goncalves, Ewan Harrison, David K. Jackson, Ian Johnston, Dominic Kwiatkowski, Cordelia Langford, John Sillitoe on behalf of the Wellcome Sanger Institute COVID-19 Surveillance Team |
| EPI_ISL_2541477 | Lighthouse Laboratory Plymouth | Wellcome Sanger Institute for the COVID-19 Genomics UK (COG-UK) | Lighthouse Laboratory Plymouth and Alex Alderton, Roberto Amato, Jeffrey Barrett, Sonia Goncalves, Ewan Harrison, David K. Jackson, Ian Johnston, Dominic Kwiatkowski, Cordelia Langford, John Sillitoe on behalf of the Wellcome Sanger |

| | | | |
|---|---|---|---|
| | | Consortium | Institute COVID-19 Surveillance Team |
| EPI_ISL_2541478, EPI_ISL_2541479 | Lighthouse Lab in Milton Keynes | Wellcome Sanger Institute for the COVID-19 Genomics UK (COG-UK) Consortium | The Lighthouse Lab in Milton Keynes and Alex Alderton, Roberto Amato, Jeffrey Barrett, Sonia Goncalves, Ewan Harrison, David K. Jackson, Ian Johnston, Dominic Kwiatkowski, Cordelia Langford, John Sillitoe on behalf of the Wellcome Sanger Institute COVID-19 Surveillance Team |
| EPI_ISL_2541480 | Lighthouse Laboratory Plymouth | Wellcome Sanger Institute for the COVID-19 Genomics UK (COG-UK) Consortium | Lighthouse Laboratory Plymouth and Alex Alderton, Roberto Amato, Jeffrey Barrett, Sonia Goncalves, Ewan Harrison, David K. Jackson, Ian Johnston, Dominic Kwiatkowski, Cordelia Langford, John Sillitoe on behalf of the Wellcome Sanger Institute COVID-19 Surveillance Team |
| EPI_ISL_2541481, EPI_ISL_2541483, EPI_ISL_2541484, EPI_ISL_2541485 | Lighthouse Lab in Milton Keynes | Wellcome Sanger Institute for the COVID-19 Genomics UK (COG-UK) Consortium | The Lighthouse Lab in Milton Keynes and Alex Alderton, Roberto Amato, Jeffrey Barrett, Sonia Goncalves, Ewan Harrison, David K. Jackson, Ian Johnston, Dominic Kwiatkowski, Cordelia Langford, John Sillitoe on behalf of the Wellcome Sanger Institute COVID-19 Surveillance Team |
| EPI_ISL_2541486 | Lighthouse Laboratory Plymouth | Wellcome Sanger Institute for the COVID-19 Genomics UK (COG-UK) Consortium | Lighthouse Laboratory Plymouth and Alex Alderton, Roberto Amato, Jeffrey Barrett, Sonia Goncalves, Ewan Harrison, David K. Jackson, Ian Johnston, Dominic Kwiatkowski, Cordelia Langford, John Sillitoe on behalf of the Wellcome Sanger Institute COVID-19 Surveillance Team |
| EPI_ISL_2541487, EPI_ISL_2541488, EPI_ISL_2541489, EPI_ISL_2541490, EPI_ISL_2541491, EPI_ISL_2541492, EPI_ISL_2541493, EPI_ISL_2541494 | Lighthouse Lab in Milton Keynes | Wellcome Sanger Institute for the COVID-19 Genomics UK (COG-UK) Consortium | The Lighthouse Lab in Milton Keynes and Alex Alderton, Roberto Amato, Jeffrey Barrett, Sonia Goncalves, Ewan Harrison, David K. Jackson, Ian Johnston, Dominic Kwiatkowski, Cordelia Langford, John Sillitoe on behalf of the Wellcome Sanger Institute COVID-19 Surveillance Team |
| EPI_ISL_2541495 | Lighthouse Laboratory Plymouth | Wellcome Sanger Institute for the COVID-19 Genomics UK (COG-UK) Consortium | Lighthouse Laboratory Plymouth and Alex Alderton, Roberto Amato, Jeffrey Barrett, Sonia Goncalves, Ewan Harrison, David K. Jackson, Ian Johnston, Dominic Kwiatkowski, Cordelia Langford, John Sillitoe on behalf of the Wellcome Sanger Institute COVID-19 Surveillance Team |
| EPI_ISL_2541496, EPI_ISL_2541497, EPI_ISL_2541498, EPI_ISL_2541500, EPI_ISL_2541501 | Lighthouse Lab in Milton Keynes | Wellcome Sanger Institute for the COVID-19 Genomics UK (COG-UK) Consortium | The Lighthouse Lab in Milton Keynes and Alex Alderton, Roberto Amato, Jeffrey Barrett, Sonia Goncalves, Ewan Harrison, David K. Jackson, Ian Johnston, Dominic Kwiatkowski, Cordelia Langford, John Sillitoe on behalf of the Wellcome Sanger Institute COVID-19 Surveillance Team |
| EPI_ISL_2541502, EPI_ISL_2541503 | Lighthouse Laboratory Plymouth | Wellcome Sanger Institute for the COVID-19 Genomics UK (COG-UK) Consortium | Lighthouse Laboratory Plymouth and Alex Alderton, Roberto Amato, Jeffrey Barrett, Sonia Goncalves, Ewan Harrison, David K. Jackson, Ian Johnston, Dominic Kwiatkowski, Cordelia Langford, John Sillitoe on behalf of the Wellcome Sanger Institute COVID-19 Surveillance Team |
| EPI_ISL_2541504, EPI_ISL_2541505, EPI_ISL_2541506 | Lighthouse Lab in Milton Keynes | Wellcome Sanger Institute for the COVID-19 Genomics UK (COG-UK) Consortium | The Lighthouse Lab in Milton Keynes and Alex Alderton, Roberto Amato, Jeffrey Barrett, Sonia Goncalves, Ewan Harrison, David K. Jackson, Ian Johnston, Dominic Kwiatkowski, Cordelia Langford, John Sillitoe on behalf of the Wellcome Sanger Institute COVID-19 Surveillance Team |
| EPI_ISL_2541507 | Lighthouse Laboratory Plymouth | Wellcome Sanger Institute for the COVID-19 Genomics UK (COG-UK) Consortium | Lighthouse Laboratory Plymouth and Alex Alderton, Roberto Amato, Jeffrey Barrett, Sonia Goncalves, Ewan Harrison, David K. Jackson, Ian Johnston, Dominic Kwiatkowski, Cordelia Langford, John Sillitoe on behalf of the Wellcome Sanger Institute COVID-19 Surveillance Team |
| EPI_ISL_2541508, EPI_ISL_2541509 | Lighthouse Lab in Milton Keynes | Wellcome Sanger Institute for the COVID-19 Genomics UK (COG-UK) Consortium | The Lighthouse Lab in Milton Keynes and Alex Alderton, Roberto Amato, Jeffrey Barrett, Sonia Goncalves, Ewan Harrison, David K. Jackson, Ian Johnston, Dominic Kwiatkowski, Cordelia Langford, John Sillitoe on behalf of the Wellcome Sanger Institute COVID-19 Surveillance Team |
| EPI_ISL_2541510, EPI_ISL_2541511 | Health Services Laboratories | Wellcome Sanger Institute for the COVID-19 Genomics UK (COG-UK) Consortium | Health Services Laboratories and Alex Alderton, Roberto Amato, Jeffrey Barrett, Sonia Goncalves, Ewan Harrison, David K. Jackson, Ian Johnston, Dominic Kwiatkowski, Cordelia Langford, John Sillitoe on behalf of the Wellcome Sanger Institute COVID-19 Surveillance Team |
| EPI_ISL_2541512 | Lighthouse Laboratory Plymouth | Wellcome Sanger Institute for the COVID-19 Genomics UK (COG-UK) Consortium | Lighthouse Laboratory Plymouth and Alex Alderton, Roberto Amato, Jeffrey Barrett, Sonia Goncalves, Ewan Harrison, David K. Jackson, Ian Johnston, Dominic Kwiatkowski, Cordelia Langford, John Sillitoe on behalf of the Wellcome Sanger Institute COVID-19 Surveillance Team |
| EPI_ISL_2541513 | Lighthouse Lab in Milton Keynes | Wellcome Sanger Institute for the COVID-19 Genomics UK (COG-UK) Consortium | The Lighthouse Lab in Milton Keynes and Alex Alderton, Roberto Amato, Jeffrey Barrett, Sonia Goncalves, Ewan Harrison, David K. Jackson, Ian Johnston, Dominic Kwiatkowski, Cordelia Langford, John Sillitoe on behalf of the Wellcome Sanger Institute COVID-19 Surveillance Team |
| EPI_ISL_2541514 | Lighthouse Laboratory Plymouth | Wellcome Sanger Institute for the COVID-19 Genomics UK (COG-UK) Consortium | Lighthouse Laboratory Plymouth and Alex Alderton, Roberto Amato, Jeffrey Barrett, Sonia Goncalves, Ewan Harrison, David K. Jackson, Ian Johnston, Dominic Kwiatkowski, Cordelia Langford, John Sillitoe on behalf of the Wellcome Sanger Institute COVID-19 Surveillance Team |
| EPI_ISL_2541515, EPI_ISL_2541516, EPI_ISL_2541517, EPI_ISL_2541518, EPI_ISL_2541519, EPI_ISL_2541520, EPI_ISL_2541521 | Lighthouse Lab in Milton Keynes | Wellcome Sanger Institute for the COVID-19 Genomics UK (COG-UK) Consortium | The Lighthouse Lab in Milton Keynes and Alex Alderton, Roberto Amato, Jeffrey Barrett, Sonia Goncalves, Ewan Harrison, David K. Jackson, Ian Johnston, Dominic Kwiatkowski, Cordelia Langford, John Sillitoe on behalf of the Wellcome Sanger Institute COVID-19 Surveillance Team |
| EPI_ISL_2541522, EPI_ISL_2541523 | Health Services Laboratories | Wellcome Sanger Institute for the COVID-19 Genomics UK (COG-UK) Consortium | Health Services Laboratories and Alex Alderton, Roberto Amato, Jeffrey Barrett, Sonia Goncalves, Ewan Harrison, David K. Jackson, Ian Johnston, Dominic Kwiatkowski, Cordelia Langford, John Sillitoe on behalf of the Wellcome Sanger Institute COVID-19 Surveillance Team |
| EPI_ISL_2541525, EPI_ISL_2541527 | Lighthouse Lab in Milton Keynes | Wellcome Sanger Institute for the COVID-19 Genomics UK (COG-UK) Consortium | The Lighthouse Lab in Milton Keynes and Alex Alderton, Roberto Amato, Jeffrey Barrett, Sonia Goncalves, Ewan Harrison, David K. Jackson, Ian Johnston, Dominic Kwiatkowski, Cordelia Langford, John Sillitoe on behalf of the Wellcome Sanger Institute COVID-19 Surveillance Team |
| EPI_ISL_2541528 | Health Services Laboratories | Wellcome Sanger Institute for the COVID-19 Genomics UK (COG-UK) Consortium | Health Services Laboratories and Alex Alderton, Roberto Amato, Jeffrey Barrett, Sonia Goncalves, Ewan Harrison, David K. Jackson, Ian Johnston, Dominic Kwiatkowski, Cordelia Langford, John Sillitoe on behalf of the Wellcome Sanger Institute COVID-19 Surveillance Team |
| EPI_ISL_2541529, EPI_ISL_2541530, EPI_ISL_2541531, EPI_ISL_2541532, EPI_ISL_2541533, EPI_ISL_2541534, EPI_ISL_2541535, EPI_ISL_2541536, EPI_ISL_2541537, EPI_ISL_2541538 | Lighthouse Lab in Milton Keynes | Wellcome Sanger Institute for the COVID-19 Genomics UK (COG-UK) Consortium | The Lighthouse Lab in Milton Keynes and Alex Alderton, Roberto Amato, Jeffrey Barrett, Sonia Goncalves, Ewan Harrison, David K. Jackson, Ian Johnston, Dominic Kwiatkowski, Cordelia Langford, John Sillitoe on behalf of the Wellcome Sanger Institute COVID-19 Surveillance Team |
| EPI_ISL_2541539 | Lighthouse Laboratory Plymouth | Wellcome Sanger Institute for the COVID-19 Genomics UK (COG-UK) Consortium | Lighthouse Laboratory Plymouth and Alex Alderton, Roberto Amato, Jeffrey Barrett, Sonia Goncalves, Ewan Harrison, David K. Jackson, Ian Johnston, Dominic Kwiatkowski, Cordelia Langford, John Sillitoe on behalf of the Wellcome Sanger Institute COVID-19 Surveillance Team |
| EPI_ISL_2541540, EPI_ISL_2541541, EPI_ISL_2541542, EPI_ISL_2541543 | Lighthouse Lab in Milton Keynes | Wellcome Sanger Institute for the COVID-19 Genomics UK (COG-UK) Consortium | The Lighthouse Lab in Milton Keynes and Alex Alderton, Roberto Amato, Jeffrey Barrett, Sonia Goncalves, Ewan Harrison, David K. Jackson, Ian Johnston, Dominic Kwiatkowski, Cordelia Langford, John Sillitoe on behalf of the Wellcome Sanger Institute COVID-19 Surveillance Team |
| EPI_ISL_2541544 | Health Services Laboratories | Wellcome Sanger Institute for the COVID-19 Genomics UK (COG-UK) Consortium | Health Services Laboratories and Alex Alderton, Roberto Amato, Jeffrey Barrett, Sonia Goncalves, Ewan Harrison, David K. Jackson, Ian Johnston, Dominic Kwiatkowski, Cordelia Langford, John Sillitoe on behalf of the Wellcome Sanger Institute COVID-19 Surveillance Team |
| EPI_ISL_2541545, EPI_ISL_2541546, EPI_ISL_2541547, EPI_ISL_2541548, EPI_ISL_2541549 | Lighthouse Lab in Milton Keynes | Wellcome Sanger Institute for the COVID-19 Genomics UK (COG-UK) Consortium | The Lighthouse Lab in Milton Keynes and Alex Alderton, Roberto Amato, Jeffrey Barrett, Sonia Goncalves, Ewan Harrison, David K. Jackson, Ian Johnston, Dominic Kwiatkowski, Cordelia Langford, John Sillitoe on behalf of the Wellcome Sanger Institute COVID-19 Surveillance Team |
| EPI_ISL_2541550 | Lighthouse Laboratory Plymouth | Wellcome Sanger Institute for the COVID-19 Genomics UK (COG-UK) Consortium | Lighthouse Laboratory Plymouth and Alex Alderton, Roberto Amato, Jeffrey Barrett, Sonia Goncalves, Ewan Harrison, David K. Jackson, Ian Johnston, Dominic Kwiatkowski, Cordelia Langford, John Sillitoe on behalf of the Wellcome Sanger Institute COVID-19 Surveillance Team |
| EPI_ISL_2541551 | Health Services Laboratories | Wellcome Sanger Institute for the COVID-19 Genomics UK (COG-UK) Consortium | Health Services Laboratories and Alex Alderton, Roberto Amato, Jeffrey Barrett, Sonia Goncalves, Ewan Harrison, David K. Jackson, Ian Johnston, Dominic Kwiatkowski, Cordelia Langford, John Sillitoe on behalf of the Wellcome Sanger Institute COVID-19 Surveillance Team |
| EPI_ISL_2541552, EPI_ISL_2541553, EPI_ISL_2541554, EPI_ISL_2541555, EPI_ISL_2541556 | Lighthouse Lab in Milton Keynes | Wellcome Sanger Institute for the COVID-19 Genomics UK (COG-UK) Consortium | The Lighthouse Lab in Milton Keynes and Alex Alderton, Roberto Amato, Jeffrey Barrett, Sonia Goncalves, Ewan Harrison, David K. Jackson, Ian Johnston, Dominic Kwiatkowski, Cordelia Langford, John Sillitoe on behalf of the Wellcome Sanger Institute COVID-19 Surveillance Team |
| EPI_ISL_2541557 | Lighthouse Laboratory Plymouth | Wellcome Sanger Institute for the COVID-19 Genomics UK (COG-UK) Consortium | Lighthouse Laboratory Plymouth and Alex Alderton, Roberto Amato, Jeffrey Barrett, Sonia Goncalves, Ewan Harrison, David K. Jackson, Ian Johnston, Dominic Kwiatkowski, Cordelia Langford, John Sillitoe on behalf of the Wellcome Sanger Institute COVID-19 Surveillance Team |
| EPI_ISL_2541559, EPI_ISL_2541560, EPI_ISL_2541561, EPI_ISL_2541562 | Lighthouse Lab in Milton Keynes | Wellcome Sanger Institute for the COVID-19 Genomics UK (COG-UK) Consortium | The Lighthouse Lab in Milton Keynes and Alex Alderton, Roberto Amato, Jeffrey Barrett, Sonia Goncalves, Ewan Harrison, David K. Jackson, Ian Johnston, Dominic Kwiatkowski, Cordelia Langford, John Sillitoe on behalf of the Wellcome Sanger Institute COVID-19 Surveillance Team |
| EPI_ISL_2541563 | Lighthouse Laboratory Plymouth | Wellcome Sanger Institute for the COVID-19 Genomics UK (COG-UK) Consortium | Lighthouse Laboratory Plymouth and Alex Alderton, Roberto Amato, Jeffrey Barrett, Sonia Goncalves, Ewan Harrison, David K. Jackson, Ian Johnston, Dominic Kwiatkowski, Cordelia Langford, John Sillitoe on behalf of the Wellcome Sanger Institute COVID-19 Surveillance Team |

| | | | |
|---|---|---|---|
| EPI_ISL_2541564, EPI_ISL_2541565, EPI_ISL_2541566, EPI_ISL_2541567 | Lighthouse Lab in Milton Keynes | Wellcome Sanger Institute for the COVID-19 Genomics UK (COG-UK) Consortium | The Lighthouse Lab in Milton Keynes and Alex Alderton, Roberto Amato, Jeffrey Barrett, Sonia Goncalves, Ewan Harrison, David K. Jackson, Ian Johnston, Dominic Kwiatkowski, Cordelia Langford, John Sillitoe on behalf of the Wellcome Sanger Institute COVID-19 Surveillance Team |
| EPI_ISL_2541568 | Lighthouse Laboratory Plymouth | Wellcome Sanger Institute for the COVID-19 Genomics UK (COG-UK) Consortium | Lighthouse Laboratory Plymouth and Alex Alderton, Roberto Amato, Jeffrey Barrett, Sonia Goncalves, Ewan Harrison, David K. Jackson, Ian Johnston, Dominic Kwiatkowski, Cordelia Langford, John Sillitoe on behalf of the Wellcome Sanger Institute COVID-19 Surveillance Team |
| EPI_ISL_2541569, EPI_ISL_2541570, EPI_ISL_2541571 | Lighthouse Lab in Milton Keynes | Wellcome Sanger Institute for the COVID-19 Genomics UK (COG-UK) Consortium | The Lighthouse Lab in Milton Keynes and Alex Alderton, Roberto Amato, Jeffrey Barrett, Sonia Goncalves, Ewan Harrison, David K. Jackson, Ian Johnston, Dominic Kwiatkowski, Cordelia Langford, John Sillitoe on behalf of the Wellcome Sanger Institute COVID-19 Surveillance Team |
| EPI_ISL_2541572 | Lighthouse Laboratory Plymouth | Wellcome Sanger Institute for the COVID-19 Genomics UK (COG-UK) Consortium | Lighthouse Laboratory Plymouth and Alex Alderton, Roberto Amato, Jeffrey Barrett, Sonia Goncalves, Ewan Harrison, David K. Jackson, Ian Johnston, Dominic Kwiatkowski, Cordelia Langford, John Sillitoe on behalf of the Wellcome Sanger Institute COVID-19 Surveillance Team |
| EPI_ISL_2541573, EPI_ISL_2541574 | Lighthouse Lab in Milton Keynes | Wellcome Sanger Institute for the COVID-19 Genomics UK (COG-UK) Consortium | The Lighthouse Lab in Milton Keynes and Alex Alderton, Roberto Amato, Jeffrey Barrett, Sonia Goncalves, Ewan Harrison, David K. Jackson, Ian Johnston, Dominic Kwiatkowski, Cordelia Langford, John Sillitoe on behalf of the Wellcome Sanger Institute COVID-19 Surveillance Team |
| EPI_ISL_2541575 | Lighthouse Laboratory Plymouth | Wellcome Sanger Institute for the COVID-19 Genomics UK (COG-UK) Consortium | Lighthouse Laboratory Plymouth and Alex Alderton, Roberto Amato, Jeffrey Barrett, Sonia Goncalves, Ewan Harrison, David K. Jackson, Ian Johnston, Dominic Kwiatkowski, Cordelia Langford, John Sillitoe on behalf of the Wellcome Sanger Institute COVID-19 Surveillance Team |
| EPI_ISL_2541576, EPI_ISL_2541577, EPI_ISL_2541578, EPI_ISL_2541579, EPI_ISL_2541580 | Lighthouse Lab in Milton Keynes | Wellcome Sanger Institute for the COVID-19 Genomics UK (COG-UK) Consortium | The Lighthouse Lab in Milton Keynes and Alex Alderton, Roberto Amato, Jeffrey Barrett, Sonia Goncalves, Ewan Harrison, David K. Jackson, Ian Johnston, Dominic Kwiatkowski, Cordelia Langford, John Sillitoe on behalf of the Wellcome Sanger Institute COVID-19 Surveillance Team |
| EPI_ISL_2541581, EPI_ISL_2541582 | Health Services Laboratories | Wellcome Sanger Institute for the COVID-19 Genomics UK (COG-UK) Consortium | Health Services Laboratories and Alex Alderton, Roberto Amato, Jeffrey Barrett, Sonia Goncalves, Ewan Harrison, David K. Jackson, Ian Johnston, Dominic Kwiatkowski, Cordelia Langford, John Sillitoe on behalf of the Wellcome Sanger Institute COVID-19 Surveillance Team |
| EPI_ISL_2541583, EPI_ISL_2541584 | Lighthouse Lab in Milton Keynes | Wellcome Sanger Institute for the COVID-19 Genomics UK (COG-UK) Consortium | The Lighthouse Lab in Milton Keynes and Alex Alderton, Roberto Amato, Jeffrey Barrett, Sonia Goncalves, Ewan Harrison, David K. Jackson, Ian Johnston, Dominic Kwiatkowski, Cordelia Langford, John Sillitoe on behalf of the Wellcome Sanger Institute COVID-19 Surveillance Team |
| EPI_ISL_2541586 | Lighthouse Laboratory Plymouth | Wellcome Sanger Institute for the COVID-19 Genomics UK (COG-UK) Consortium | Lighthouse Laboratory Plymouth and Alex Alderton, Roberto Amato, Jeffrey Barrett, Sonia Goncalves, Ewan Harrison, David K. Jackson, Ian Johnston, Dominic Kwiatkowski, Cordelia Langford, John Sillitoe on behalf of the Wellcome Sanger Institute COVID-19 Surveillance Team |
| EPI_ISL_2541587, EPI_ISL_2541588 | Lighthouse Lab in Milton Keynes | Wellcome Sanger Institute for the COVID-19 Genomics UK (COG-UK) Consortium | The Lighthouse Lab in Milton Keynes and Alex Alderton, Roberto Amato, Jeffrey Barrett, Sonia Goncalves, Ewan Harrison, David K. Jackson, Ian Johnston, Dominic Kwiatkowski, Cordelia Langford, John Sillitoe on behalf of the Wellcome Sanger Institute COVID-19 Surveillance Team |
| EPI_ISL_2541589, EPI_ISL_2541590 | Lighthouse Laboratory Plymouth | Wellcome Sanger Institute for the COVID-19 Genomics UK (COG-UK) Consortium | Lighthouse Laboratory Plymouth and Alex Alderton, Roberto Amato, Jeffrey Barrett, Sonia Goncalves, Ewan Harrison, David K. Jackson, Ian Johnston, Dominic Kwiatkowski, Cordelia Langford, John Sillitoe on behalf of the Wellcome Sanger Institute COVID-19 Surveillance Team |
| EPI_ISL_2541591 | Lighthouse Lab in Milton Keynes | Wellcome Sanger Institute for the COVID-19 Genomics UK (COG-UK) Consortium | The Lighthouse Lab in Milton Keynes and Alex Alderton, Roberto Amato, Jeffrey Barrett, Sonia Goncalves, Ewan Harrison, David K. Jackson, Ian Johnston, Dominic Kwiatkowski, Cordelia Langford, John Sillitoe on behalf of the Wellcome Sanger Institute COVID-19 Surveillance Team |
| EPI_ISL_2541592 | Lighthouse Laboratory Plymouth | Wellcome Sanger Institute for the COVID-19 Genomics UK (COG-UK) Consortium | Lighthouse Laboratory Plymouth and Alex Alderton, Roberto Amato, Jeffrey Barrett, Sonia Goncalves, Ewan Harrison, David K. Jackson, Ian Johnston, Dominic Kwiatkowski, Cordelia Langford, John Sillitoe on behalf of the Wellcome Sanger Institute COVID-19 Surveillance Team |
| EPI_ISL_2541593, EPI_ISL_2541594, EPI_ISL_2541596, EPI_ISL_2541598, EPI_ISL_2541599, EPI_ISL_2541600, EPI_ISL_2541601 | Lighthouse Lab in Milton Keynes | Wellcome Sanger Institute for the COVID-19 Genomics UK (COG-UK) Consortium | The Lighthouse Lab in Milton Keynes and Alex Alderton, Roberto Amato, Jeffrey Barrett, Sonia Goncalves, Ewan Harrison, David K. Jackson, Ian Johnston, Dominic Kwiatkowski, Cordelia Langford, John Sillitoe on behalf of the Wellcome Sanger Institute COVID-19 Surveillance Team |
| EPI_ISL_2541602 | Health Services Laboratories | Wellcome Sanger Institute for the COVID-19 Genomics UK (COG-UK) Consortium | Health Services Laboratories and Alex Alderton, Roberto Amato, Jeffrey Barrett, Sonia Goncalves, Ewan Harrison, David K. Jackson, Ian Johnston, Dominic Kwiatkowski, Cordelia Langford, John Sillitoe on behalf of the Wellcome Sanger Institute COVID-19 Surveillance Team |
| EPI_ISL_2541603 | Lighthouse Lab in Milton Keynes | Wellcome Sanger Institute for the COVID-19 Genomics UK (COG-UK) Consortium | The Lighthouse Lab in Milton Keynes and Alex Alderton, Roberto Amato, Jeffrey Barrett, Sonia Goncalves, Ewan Harrison, David K. Jackson, Ian Johnston, Dominic Kwiatkowski, Cordelia Langford, John Sillitoe on behalf of the Wellcome Sanger Institute COVID-19 Surveillance Team |
| EPI_ISL_2541604, EPI_ISL_2541605 | Lighthouse Laboratory Plymouth | Wellcome Sanger Institute for the COVID-19 Genomics UK (COG-UK) Consortium | Lighthouse Laboratory Plymouth and Alex Alderton, Roberto Amato, Jeffrey Barrett, Sonia Goncalves, Ewan Harrison, David K. Jackson, Ian Johnston, Dominic Kwiatkowski, Cordelia Langford, John Sillitoe on behalf of the Wellcome Sanger Institute COVID-19 Surveillance Team |
| EPI_ISL_2541606, EPI_ISL_2541608 | Lighthouse Lab in Milton Keynes | Wellcome Sanger Institute for the COVID-19 Genomics UK (COG-UK) Consortium | The Lighthouse Lab in Milton Keynes and Alex Alderton, Roberto Amato, Jeffrey Barrett, Sonia Goncalves, Ewan Harrison, David K. Jackson, Ian Johnston, Dominic Kwiatkowski, Cordelia Langford, John Sillitoe on behalf of the Wellcome Sanger Institute COVID-19 Surveillance Team |
| EPI_ISL_2541609 | Health Services Laboratories | Wellcome Sanger Institute for the COVID-19 Genomics UK (COG-UK) Consortium | Health Services Laboratories and Alex Alderton, Roberto Amato, Jeffrey Barrett, Sonia Goncalves, Ewan Harrison, David K. Jackson, Ian Johnston, Dominic Kwiatkowski, Cordelia Langford, John Sillitoe on behalf of the Wellcome Sanger Institute COVID-19 Surveillance Team |
| EPI_ISL_2541610 | Lighthouse Laboratory Plymouth | Wellcome Sanger Institute for the COVID-19 Genomics UK (COG-UK) Consortium | Lighthouse Laboratory Plymouth and Alex Alderton, Roberto Amato, Jeffrey Barrett, Sonia Goncalves, Ewan Harrison, David K. Jackson, Ian Johnston, Dominic Kwiatkowski, Cordelia Langford, John Sillitoe on behalf of the Wellcome Sanger Institute COVID-19 Surveillance Team |
| EPI_ISL_2541611, EPI_ISL_2541612 | Health Services Laboratories | Wellcome Sanger Institute for the COVID-19 Genomics UK (COG-UK) Consortium | Health Services Laboratories and Alex Alderton, Roberto Amato, Jeffrey Barrett, Sonia Goncalves, Ewan Harrison, David K. Jackson, Ian Johnston, Dominic Kwiatkowski, Cordelia Langford, John Sillitoe on behalf of the Wellcome Sanger Institute COVID-19 Surveillance Team |
| EPI_ISL_2541613 | Lighthouse Lab in Milton Keynes | Wellcome Sanger Institute for the COVID-19 Genomics UK (COG-UK) Consortium | The Lighthouse Lab in Milton Keynes and Alex Alderton, Roberto Amato, Jeffrey Barrett, Sonia Goncalves, Ewan Harrison, David K. Jackson, Ian Johnston, Dominic Kwiatkowski, Cordelia Langford, John Sillitoe on behalf of the Wellcome Sanger Institute COVID-19 Surveillance Team |
| EPI_ISL_2541614, EPI_ISL_2541615 | Lighthouse Laboratory Plymouth | Wellcome Sanger Institute for the COVID-19 Genomics UK (COG-UK) Consortium | Lighthouse Laboratory Plymouth and Alex Alderton, Roberto Amato, Jeffrey Barrett, Sonia Goncalves, Ewan Harrison, David K. Jackson, Ian Johnston, Dominic Kwiatkowski, Cordelia Langford, John Sillitoe on behalf of the Wellcome Sanger Institute COVID-19 Surveillance Team |
| EPI_ISL_2541616 | Health Services Laboratories | Wellcome Sanger Institute for the COVID-19 Genomics UK (COG-UK) Consortium | Health Services Laboratories and Alex Alderton, Roberto Amato, Jeffrey Barrett, Sonia Goncalves, Ewan Harrison, David K. Jackson, Ian Johnston, Dominic Kwiatkowski, Cordelia Langford, John Sillitoe on behalf of the Wellcome Sanger Institute COVID-19 Surveillance Team |
| EPI_ISL_2541617, EPI_ISL_2541618 | Lighthouse Lab in Milton Keynes | Wellcome Sanger Institute for the COVID-19 Genomics UK (COG-UK) Consortium | The Lighthouse Lab in Milton Keynes and Alex Alderton, Roberto Amato, Jeffrey Barrett, Sonia Goncalves, Ewan Harrison, David K. Jackson, Ian Johnston, Dominic Kwiatkowski, Cordelia Langford, John Sillitoe on behalf of the Wellcome Sanger Institute COVID-19 Surveillance Team |
| EPI_ISL_2541620 | Lighthouse Laboratory Plymouth | Wellcome Sanger Institute for the COVID-19 Genomics UK (COG-UK) Consortium | Lighthouse Laboratory Plymouth and Alex Alderton, Roberto Amato, Jeffrey Barrett, Sonia Goncalves, Ewan Harrison, David K. Jackson, Ian Johnston, Dominic Kwiatkowski, Cordelia Langford, John Sillitoe on behalf of the Wellcome Sanger Institute COVID-19 Surveillance Team |
| EPI_ISL_2541621 | Lighthouse Lab in Milton Keynes | Wellcome Sanger Institute for the COVID-19 Genomics UK (COG-UK) Consortium | The Lighthouse Lab in Milton Keynes and Alex Alderton, Roberto Amato, Jeffrey Barrett, Sonia Goncalves, Ewan Harrison, David K. Jackson, Ian Johnston, Dominic Kwiatkowski, Cordelia Langford, John Sillitoe on behalf of the Wellcome Sanger Institute COVID-19 Surveillance Team |
| EPI_ISL_2541622 | Health Services Laboratories | Wellcome Sanger Institute for the COVID-19 Genomics UK (COG-UK) Consortium | Health Services Laboratories and Alex Alderton, Roberto Amato, Jeffrey Barrett, Sonia Goncalves, Ewan Harrison, David K. Jackson, Ian Johnston, Dominic Kwiatkowski, Cordelia Langford, John Sillitoe on behalf of the Wellcome Sanger Institute COVID-19 Surveillance Team |
| EPI_ISL_2541623, EPI_ISL_2541624, EPI_ISL_2541625, EPI_ISL_2541626, EPI_ISL_2541627, EPI_ISL_2541628 | Lighthouse Lab in Milton Keynes | Wellcome Sanger Institute for the COVID-19 Genomics UK (COG-UK) Consortium | The Lighthouse Lab in Milton Keynes and Alex Alderton, Roberto Amato, Jeffrey Barrett, Sonia Goncalves, Ewan Harrison, David K. Jackson, Ian Johnston, Dominic Kwiatkowski, Cordelia Langford, John Sillitoe on behalf of the Wellcome Sanger Institute COVID-19 Surveillance Team |
| EPI_ISL_2541630 | Health Services Laboratories | Wellcome Sanger Institute for the COVID-19 Genomics UK (COG-UK) Consortium | Health Services Laboratories and Alex Alderton, Roberto Amato, Jeffrey Barrett, Sonia Goncalves, Ewan Harrison, David K. Jackson, Ian Johnston, Dominic Kwiatkowski, Cordelia Langford, John Sillitoe on behalf of the Wellcome Sanger Institute COVID-19 Surveillance Team |
| EPI_ISL_2541631 | Lighthouse Lab in Milton Keynes | Wellcome Sanger Institute for the COVID-19 Genomics UK (COG-UK) Consortium | The Lighthouse Lab in Milton Keynes and Alex Alderton, Roberto Amato, Jeffrey Barrett, Sonia Goncalves, Ewan Harrison, David K. Jackson, Ian Johnston, Dominic Kwiatkowski, Cordelia Langford, John Sillitoe on behalf of the Wellcome Sanger Institute COVID-19 Surveillance Team |
| EPI_ISL_2541633 | Lighthouse Laboratory Plymouth | Wellcome Sanger Institute for the COVID-19 Genomics UK (COG-UK) Consortium | Lighthouse Laboratory Plymouth and Alex Alderton, Roberto Amato, Jeffrey Barrett, Sonia Goncalves, Ewan Harrison, David K. Jackson, Ian Johnston, Dominic Kwiatkowski, Cordelia Langford, John Sillitoe on behalf of the Wellcome Sanger Institute COVID-19 Surveillance Team |
| EPI_ISL_2541634 | Lighthouse Lab in Milton Keynes | Wellcome Sanger Institute for the COVID-19 Genomics UK (COG-UK) Consortium | The Lighthouse Lab in Milton Keynes and Alex Alderton, Roberto Amato, Jeffrey Barrett, Sonia Goncalves, Ewan Harrison, David K. Jackson, Ian Johnston, Dominic Kwiatkowski, Cordelia Langford, John Sillitoe on behalf of the Wellcome Sanger Institute COVID-19 Surveillance Team |
| EPI_ISL_2541635 | Health Services Laboratories | Wellcome Sanger Institute for the COVID-19 Genomics UK (COG-UK) Consortium | Health Services Laboratories and Alex Alderton, Roberto Amato, Jeffrey Barrett, Sonia Goncalves, Ewan Harrison, David K. Jackson, Ian Johnston, Dominic Kwiatkowski, Cordelia Langford, John Sillitoe on behalf of the Wellcome Sanger Institute COVID-19 Surveillance Team |

EPI_ISL_2541636, EPI_ISL_2541637, EPI_ISL_2541638, EPI_ISL_2541639, EPI_ISL_2541640, EPI_ISL_2541641, EPI_ISL_2541642, EPI_ISL_2541643, EPI_ISL_2541644, EPI_ISL_2541645, EPI_ISL_2541646, EPI_ISL_2541647, EPI_ISL_2541648, EPI_ISL_2541649, EPI_ISL_2541650, EPI_ISL_2541651, EPI_ISL_2541652, EPI_ISL_2541653, EPI_ISL_2541654, EPI_ISL_2541655, EPI_ISL_2541657, EPI_ISL_2541658, EPI_ISL_2541659, EPI_ISL_2541660, EPI_ISL_2541661, EPI_ISL_2541662, EPI_ISL_2541663, EPI_ISL_2541664, EPI_ISL_2541665, EPI_ISL_2541666, EPI_ISL_2541668, EPI_ISL_2541669, EPI_ISL_2541670, EPI_ISL_2541671, EPI_ISL_2541672, EPI_ISL_2541673, EPI_ISL_2541674, EPI_ISL_2541675, EPI_ISL_2541676, EPI_ISL_2541677, EPI_ISL_2541678, EPI_ISL_2541679, EPI_ISL_2541680, EPI_ISL_2541681, EPI_ISL_2541682, EPI_ISL_2541683, EPI_ISL_2541684, EPI_ISL_2541685, EPI_ISL_2541686, EPI_ISL_2541687, EPI_ISL_2541688, EPI_ISL_2541689, EPI_ISL_2541690, EPI_ISL_2541691, EPI_ISL_2541692, EPI_ISL_2541693, EPI_ISL_2541695, EPI_ISL_2541696, EPI_ISL_2541697, EPI_ISL_2541698, EPI_ISL_2541699, EPI_ISL_2541700, EPI_ISL_2541701, EPI_ISL_2541702, EPI_ISL_2541703, EPI_ISL_2541704, EPI_ISL_2541705, EPI_ISL_2541706, EPI_ISL_2541707, EPI_ISL_2541708, EPI_ISL_2541709, EPI_ISL_2541710, EPI_ISL_2541711, EPI_ISL_2541712, EPI_ISL_2541713, EPI_ISL_2541714, EPI_ISL_2541715, EPI_ISL_2541716, EPI_ISL_2541717, EPI_ISL_2541718, EPI_ISL_2541719, EPI_ISL_2541720, EPI_ISL_2541721, EPI_ISL_2541722, EPI_ISL_2541723, EPI_ISL_2541724, EPI_ISL_2541725, EPI_ISL_2541726, EPI_ISL_2541727, EPI_ISL_2541728, EPI_ISL_2541729, EPI_ISL_2541731, EPI_ISL_2541732, EPI_ISL_2541733, EPI_ISL_2541734, EPI_ISL_2541735, EPI_ISL_2541736, EPI_ISL_2541737, EPI_ISL_2541738, EPI_ISL_2541739, EPI_ISL_2541740, EPI_ISL_2541741, EPI_ISL_2541742, EPI_ISL_2541743, EPI_ISL_2541744, EPI_ISL_2541745, EPI_ISL_2541746, EPI_ISL_2541747, EPI_ISL_2541748, EPI_ISL_2541749, EPI_ISL_2541750, EPI_ISL_2541751, EPI_ISL_2541752, EPI_ISL_2541753, EPI_ISL_2541754, EPI_ISL_2541755, EPI_ISL_2541756, EPI_ISL_2541757, EPI_ISL_2541758, EPI_ISL_2541759, EPI_ISL_2541760, EPI_ISL_2541761, EPI_ISL_2541762, EPI_ISL_2541763, EPI_ISL_2541764, EPI_ISL_2541766, EPI_ISL_2541767, EPI_ISL_2541768, EPI_ISL_2541769, EPI_ISL_2541771, EPI_ISL_2541772, EPI_ISL_2541773, EPI_ISL_2541774, EPI_ISL_2541775, EPI_ISL_2541776, EPI_ISL_2541777, EPI_ISL_2541778, EPI_ISL_2541779, EPI_ISL_2541780, EPI_ISL_2541781, EPI_ISL_2541782, EPI_ISL_2541783, EPI_ISL_2541784, EPI_ISL_2541785, EPI_ISL_2541787, EPI_ISL_2541788, EPI_ISL_2541789, EPI_ISL_2541790, EPI_ISL_2541791, EPI_ISL_2541792, EPI_ISL_2541793, EPI_ISL_2541794, EPI_ISL_2541795, EPI_ISL_2541796, EPI_ISL_2541797, EPI_ISL_2541798, EPI_ISL_2541799, EPI_ISL_2541800, EPI_ISL_2541801, EPI_ISL_2541802, EPI_ISL_2541803, EPI_ISL_2541804, EPI_ISL_2541805, EPI_ISL_2541806, EPI_ISL_2541807, EPI_ISL_2541808, EPI_ISL_2541809, EPI_ISL_2541810, EPI_ISL_2541811, EPI_ISL_2541812, EPI_ISL_2541813, EPI_ISL_2541815, EPI_ISL_2541816, EPI_ISL_2541817, EPI_ISL_2541818, EPI_ISL_2541819, EPI_ISL_2541820, EPI_ISL_2541821, EPI_ISL_2541822, EPI_ISL_2541823, EPI_ISL_2541824, EPI_ISL_2541825, EPI_ISL_2541828, EPI_ISL_2541829, EPI_ISL_2541830, EPI_ISL_2541831, EPI_ISL_2541832, EPI_ISL_2541833, EPI_ISL_2541834, EPI_ISL_2541835, EPI_ISL_2541837, EPI_ISL_2541838, EPI_ISL_2541839, EPI_ISL_2541840, EPI_ISL_2541841, EPI_ISL_2541842, EPI_ISL_2541843, EPI_ISL_2541844, EPI_ISL_2541845, EPI_ISL_2541846, EPI_ISL_2541847, EPI_ISL_2541848, EPI_ISL_2541849, EPI_ISL_2541851, EPI_ISL_2541852, EPI_ISL_2541853, EPI_ISL_2541854, EPI_ISL_2541855, EPI_ISL_2541856, EPI_ISL_2541857, EPI_ISL_2541858, EPI_ISL_2541859, EPI_ISL_2541860, EPI_ISL_2541861, EPI_ISL_2541863, EPI_ISL_2541864, EPI_ISL_2541865, EPI_ISL_2541866, EPI_ISL_2541867, EPI_ISL_2541868,

| | | | |
|---|---|---|---|
| EPI_ISL_2541869, EPI_ISL_2541870, EPI_ISL_2541871, EPI_ISL_2541872, EPI_ISL_2541873, EPI_ISL_2541874, EPI_ISL_2541875, EPI_ISL_2541876, EPI_ISL_2541877, EPI_ISL_2541878, EPI_ISL_2541879, EPI_ISL_2541880, EPI_ISL_2541881, EPI_ISL_2541882, EPI_ISL_2541883, EPI_ISL_2541884, EPI_ISL_2541885, EPI_ISL_2541886, EPI_ISL_2541887, EPI_ISL_2541888, EPI_ISL_2541889, EPI_ISL_2541890, EPI_ISL_2541891, EPI_ISL_2541892, EPI_ISL_2541893, EPI_ISL_2541894, EPI_ISL_2541895, EPI_ISL_2541896, EPI_ISL_2541897, EPI_ISL_2541898, EPI_ISL_2541899, EPI_ISL_2541900, EPI_ISL_2541901, EPI_ISL_2541902, EPI_ISL_2541903, EPI_ISL_2541904, EPI_ISL_2541905, EPI_ISL_2541906, EPI_ISL_2541907, EPI_ISL_2541908, EPI_ISL_2541909, EPI_ISL_2541910, EPI_ISL_2541911, EPI_ISL_2541912, EPI_ISL_2541913, EPI_ISL_2541914, EPI_ISL_2541915, EPI_ISL_2541916, EPI_ISL_2541917, EPI_ISL_2541918, EPI_ISL_2541919, EPI_ISL_2541920, EPI_ISL_2541921, EPI_ISL_2541922, EPI_ISL_2541923, EPI_ISL_2541924, EPI_ISL_2541925, EPI_ISL_2541926, EPI_ISL_2541927, EPI_ISL_2541928, EPI_ISL_2541929, EPI_ISL_2541932, EPI_ISL_2541933, EPI_ISL_2541934, EPI_ISL_2541935, EPI_ISL_2541936, EPI_ISL_2541937, EPI_ISL_2541938, EPI_ISL_2541939, EPI_ISL_2541940, EPI_ISL_2541941, EPI_ISL_2541942, EPI_ISL_2541943, EPI_ISL_2541944, EPI_ISL_2541945, EPI_ISL_2541946, EPI_ISL_2541947, EPI_ISL_2541948, EPI_ISL_2541949, EPI_ISL_2541950, EPI_ISL_2541951, EPI_ISL_2541952, EPI_ISL_2541953, EPI_ISL_2541955, EPI_ISL_2541956, EPI_ISL_2541957, EPI_ISL_2541958, EPI_ISL_2541959, EPI_ISL_2541960, EPI_ISL_2541961, EPI_ISL_2541962, EPI_ISL_2541963, EPI_ISL_2541964, EPI_ISL_2541965, EPI_ISL_2541966, EPI_ISL_2541967, EPI_ISL_2541969, EPI_ISL_2541970, EPI_ISL_2541971, EPI_ISL_2541972, EPI_ISL_2541973, EPI_ISL_2541974, EPI_ISL_2541975, EPI_ISL_2541976, EPI_ISL_2541977, EPI_ISL_2541978, EPI_ISL_2541979, EPI_ISL_2541980, EPI_ISL_2541981, EPI_ISL_2541982, EPI_ISL_2541983, EPI_ISL_2541984, EPI_ISL_2541985, EPI_ISL_2541986, EPI_ISL_2541987 | | | |
| see above | Lighthouse Lab in Alderley Park | Wellcome Sanger Institute for the COVID-19 Genomics UK (COG-UK) Consortium | Jacquelyn Wynn, Mairead Hyland, The Lighthouse Lab in Alderley Park and Alex Alderton, Roberto Amato, Jeffrey Barrett, Sonia Goncalves, Ewan Harrison, David K. Jackson, Ian Johnston, Dominic Kwiatkowski, Cordelia Langford, John Sillitoe on behalf of the Wellcome Sanger Institute COVID-19 Surveillance Team |
| EPI_ISL_2541989, EPI_ISL_2541990, EPI_ISL_2541991, EPI_ISL_2541993, EPI_ISL_2541995, EPI_ISL_2541996, EPI_ISL_2541997, EPI_ISL_2541998, EPI_ISL_2542000, EPI_ISL_2542001, EPI_ISL_2542002, EPI_ISL_2542003, EPI_ISL_2542005, EPI_ISL_2542007, EPI_ISL_2542009, EPI_ISL_2542010, EPI_ISL_2542011, EPI_ISL_2542012, EPI_ISL_2542014, EPI_ISL_2542015, EPI_ISL_2542016, EPI_ISL_2542017, EPI_ISL_2542019, EPI_ISL_2542020 | | | |
| see above | Lighthouse Lab in Glasgow | Wellcome Sanger Institute for the COVID-19 Genomics UK (COG-UK) Consortium | Harper VanSteenhouse, Yumi Kasai, David Gray, Carol Clugston, Anna Dominiczak and Alex Alderton, Roberto Amato, Jeffrey Barrett, Sonia Goncalves, Ewan Harrison, David K. Jackson, Ian Johnston, Dominic Kwiatkowski, Cordelia Langford, John Sillitoe on behalf of the Wellcome Sanger Institute COVID-19 Surveillance Team |
| EPI_ISL_2542023 | Lighthouse Laboratory Plymouth | Wellcome Sanger Institute for the COVID-19 Genomics UK (COG-UK) Consortium | Lighthouse Laboratory Plymouth and Alex Alderton, Roberto Amato, Jeffrey Barrett, Sonia Goncalves, Ewan Harrison, David K. Jackson, Ian Johnston, Dominic Kwiatkowski, Cordelia Langford, John Sillitoe on behalf of the Wellcome Sanger Institute COVID-19 Surveillance Team |
| EPI_ISL_2542024, EPI_ISL_2542025, EPI_ISL_2542026, EPI_ISL_2542027 | Lighthouse Lab in Glasgow | Wellcome Sanger Institute for the COVID-19 Genomics UK (COG-UK) Consortium | Harper VanSteenhouse, Yumi Kasai, David Gray, Carol Clugston, Anna Dominiczak and Alex Alderton, Roberto Amato, Jeffrey Barrett, Sonia Goncalves, Ewan Harrison, David K. Jackson, Ian Johnston, Dominic Kwiatkowski, Cordelia Langford, John Sillitoe on behalf of the Wellcome Sanger Institute COVID-19 Surveillance Team |
| EPI_ISL_2542028 | Lighthouse Laboratory Plymouth | Wellcome Sanger Institute for the COVID-19 Genomics UK (COG-UK) Consortium | Lighthouse Laboratory Plymouth and Alex Alderton, Roberto Amato, Jeffrey Barrett, Sonia Goncalves, Ewan Harrison, David K. Jackson, Ian Johnston, Dominic Kwiatkowski, Cordelia Langford, John Sillitoe on behalf of the Wellcome Sanger Institute COVID-19 Surveillance Team |
| EPI_ISL_2542029, EPI_ISL_2542030, EPI_ISL_2542031, EPI_ISL_2542032, EPI_ISL_2542033, EPI_ISL_2542034, EPI_ISL_2542035, EPI_ISL_2542036, EPI_ISL_2542037, EPI_ISL_2542038, EPI_ISL_2542039, EPI_ISL_2542040, EPI_ISL_2542041, EPI_ISL_2542043, EPI_ISL_2542044, EPI_ISL_2542046, EPI_ISL_2542047, EPI_ISL_2542048, EPI_ISL_2542049, EPI_ISL_2542050, EPI_ISL_2542051, EPI_ISL_2542052, EPI_ISL_2542053, EPI_ISL_2542054, EPI_ISL_2542055, EPI_ISL_2542056, EPI_ISL_2542057, EPI_ISL_2542059, EPI_ISL_2542060, EPI_ISL_2542065, EPI_ISL_2542066, EPI_ISL_2542067, EPI_ISL_2542068, EPI_ISL_2542069, EPI_ISL_2542070, EPI_ISL_2542071, EPI_ISL_2542072, EPI_ISL_2542076, EPI_ISL_2542078, EPI_ISL_2542083, EPI_ISL_2542084, EPI_ISL_2542087, EPI_ISL_2542090, EPI_ISL_2542091, EPI_ISL_2542092, EPI_ISL_2542093, EPI_ISL_2542094, EPI_ISL_2542095, EPI_ISL_2542096, EPI_ISL_2542098, EPI_ISL_2542099, EPI_ISL_2542100, EPI_ISL_2542101, EPI_ISL_2542103 | | | |
| see above | Lighthouse Lab in Glasgow | Wellcome Sanger Institute for the COVID-19 Genomics UK (COG-UK) Consortium | Harper VanSteenhouse, Yumi Kasai, David Gray, Carol Clugston, Anna Dominiczak and Alex Alderton, Roberto Amato, Jeffrey Barrett, Sonia Goncalves, Ewan Harrison, David K. Jackson, Ian Johnston, Dominic Kwiatkowski, Cordelia Langford, John Sillitoe on behalf of the Wellcome Sanger Institute COVID-19 Surveillance Team |
| EPI_ISL_2542104, EPI_ISL_2542105 | Health Services Laboratories | Wellcome Sanger Institute for the COVID-19 Genomics UK (COG-UK) Consortium | Health Services Laboratories and Alex Alderton, Roberto Amato, Jeffrey Barrett, Sonia Goncalves, Ewan Harrison, David K. Jackson, Ian Johnston, Dominic Kwiatkowski, Cordelia Langford, John Sillitoe on behalf of the Wellcome Sanger Institute COVID-19 Surveillance Team |
| EPI_ISL_2542106, EPI_ISL_2542107, EPI_ISL_2542109, EPI_ISL_2542110, EPI_ISL_2542112, EPI_ISL_2542113, EPI_ISL_2542114, EPI_ISL_2542115, EPI_ISL_2542116, EPI_ISL_2542117, EPI_ISL_2542118, EPI_ISL_2542119, EPI_ISL_2542122, EPI_ISL_2542124, EPI_ISL_2542125, EPI_ISL_2542126, EPI_ISL_2542128, EPI_ISL_2542129, EPI_ISL_2542130, EPI_ISL_2542131, EPI_ISL_2542132, EPI_ISL_2542133, EPI_ISL_2542134, EPI_ISL_2542135, EPI_ISL_2542136, EPI_ISL_2542137, EPI_ISL_2542138, EPI_ISL_2542140, EPI_ISL_2542142, EPI_ISL_2542143, EPI_ISL_2542144, EPI_ISL_2542145, EPI_ISL_2542146, EPI_ISL_2542147, EPI_ISL_2542148, EPI_ISL_2542149, EPI_ISL_2542151, EPI_ISL_2542152, EPI_ISL_2542153, EPI_ISL_2542154, EPI_ISL_2542155, EPI_ISL_2542156, EPI_ISL_2542157, EPI_ISL_2542158, EPI_ISL_2542159, EPI_ISL_2542160, EPI_ISL_2542161, EPI_ISL_2542162, EPI_ISL_2542163, EPI_ISL_2542165, EPI_ISL_2542168, EPI_ISL_2542169, EPI_ISL_2542170, EPI_ISL_2542172, EPI_ISL_2542173, EPI_ISL_2542174, EPI_ISL_2542176, EPI_ISL_2542177, EPI_ISL_2542179, EPI_ISL_2542180, EPI_ISL_2542181, EPI_ISL_2542182, EPI_ISL_2542183, EPI_ISL_2542186, EPI_ISL_2542187, EPI_ISL_2542188, EPI_ISL_2542189, EPI_ISL_2542190, EPI_ISL_2542191, EPI_ISL_2542193, EPI_ISL_2542194, EPI_ISL_2542199, EPI_ISL_2542201, EPI_ISL_2542203, EPI_ISL_2542204, EPI_ISL_2542207, EPI_ISL_2542208, EPI_ISL_2542209, EPI_ISL_2542210, EPI_ISL_2542211, EPI_ISL_2542213, EPI_ISL_2542214, EPI_ISL_2542215, EPI_ISL_2542217, EPI_ISL_2542218, EPI_ISL_2542219, EPI_ISL_2542220, EPI_ISL_2542221, EPI_ISL_2542222, EPI_ISL_2542224, EPI_ISL_2542226, EPI_ISL_2542227, EPI_ISL_2542228, EPI_ISL_2542229, EPI_ISL_2542230, EPI_ISL_2542231, EPI_ISL_2542232, EPI_ISL_2542233, EPI_ISL_2542234, EPI_ISL_2542235, EPI_ISL_2542236, EPI_ISL_2542237, EPI_ISL_2542238, EPI_ISL_2542240, EPI_ISL_2542242, EPI_ISL_2542243, EPI_ISL_2542244, EPI_ISL_2542245, EPI_ISL_2542246, EPI_ISL_2542247, EPI_ISL_2542249, EPI_ISL_2542250, EPI_ISL_2542255, EPI_ISL_2542256, EPI_ISL_2542257, EPI_ISL_2542258, EPI_ISL_2542262, EPI_ISL_2542263, EPI_ISL_2542266, EPI_ISL_2542267, EPI_ISL_2542269, EPI_ISL_2542270, EPI_ISL_2542271, EPI_ISL_2542272, EPI_ISL_2542273, EPI_ISL_2542274, EPI_ISL_2542277, EPI_ISL_2542278, EPI_ISL_2542280, EPI_ISL_2542281, EPI_ISL_2542283, EPI_ISL_2542285, EPI_ISL_2542286, EPI_ISL_2542287, EPI_ISL_2542288, EPI_ISL_2542289, EPI_ISL_2542290, EPI_ISL_2542293, EPI_ISL_2542294, EPI_ISL_2542296, EPI_ISL_2542297, EPI_ISL_2542298, EPI_ISL_2542299, EPI_ISL_2542300, EPI_ISL_2542301, EPI_ISL_2542302, EPI_ISL_2542303, EPI_ISL_2542304, EPI_ISL_2542305, EPI_ISL_2542307, EPI_ISL_2542309, EPI_ISL_2542310, EPI_ISL_2542312, EPI_ISL_2542313, EPI_ISL_2542314, EPI_ISL_2542315, EPI_ISL_2542316, EPI_ISL_2542319, EPI_ISL_2542320, EPI_ISL_2542321, EPI_ISL_2542322, EPI_ISL_2542323, EPI_ISL_2542324, EPI_ISL_2542325, EPI_ISL_2542327, EPI_ISL_2542329, EPI_ISL_2542330, EPI_ISL_2542333, EPI_ISL_2542334, EPI_ISL_2542335, EPI_ISL_2542336, EPI_ISL_2542337, EPI_ISL_2542340, EPI_ISL_2542341, EPI_ISL_2542342, EPI_ISL_2542343, EPI_ISL_2542344, EPI_ISL_2542346, EPI_ISL_2542347, EPI_ISL_2542348, EPI_ISL_2542349, EPI_ISL_2542350, EPI_ISL_2542351, EPI_ISL_2542352, EPI_ISL_2542353, EPI_ISL_2542354, EPI_ISL_2542355, EPI_ISL_2542356, EPI_ISL_2542357, EPI_ISL_2542358, EPI_ISL_2542359, EPI_ISL_2542361, EPI_ISL_2542362, EPI_ISL_2542363, EPI_ISL_2542364, EPI_ISL_2542367, EPI_ISL_2542368, EPI_ISL_2542369, EPI_ISL_2542370, EPI_ISL_2542372, EPI_ISL_2542373, EPI_ISL_2542374, EPI_ISL_2542375, EPI_ISL_2542376, EPI_ISL_2542377, EPI_ISL_2542378, EPI_ISL_2542379, EPI_ISL_2542380, EPI_ISL_2542381, EPI_ISL_2542382, EPI_ISL_2542383, EPI_ISL_2542384, EPI_ISL_2542385, EPI_ISL_2542386, EPI_ISL_2542387, EPI_ISL_2542388, EPI_ISL_2542389, EPI_ISL_2542390, EPI_ISL_2542391, EPI_ISL_2542393, EPI_ISL_2542394, EPI_ISL_2542395, EPI_ISL_2542396, EPI_ISL_2542397, EPI_ISL_2542398, EPI_ISL_2542400, EPI_ISL_2542401, EPI_ISL_2542403, EPI_ISL_2542405, EPI_ISL_2542406, EPI_ISL_2542407, EPI_ISL_2542409, EPI_ISL_2542410, EPI_ISL_2542412, EPI_ISL_2542413, EPI_ISL_2542414, EPI_ISL_2542415, EPI_ISL_2542416, EPI_ISL_2542418, EPI_ISL_2542419, EPI_ISL_2542421, EPI_ISL_2542422, EPI_ISL_2542423, EPI_ISL_2542424, EPI_ISL_2542425, EPI_ISL_2542426, EPI_ISL_2542427, EPI_ISL_2542428, EPI_ISL_2542430, EPI_ISL_2542431, EPI_ISL_2542433, EPI_ISL_2542434, EPI_ISL_2542435, EPI_ISL_2542436, EPI_ISL_2542438, EPI_ISL_2542439, EPI_ISL_2542441, EPI_ISL_2542442, EPI_ISL_2542443, EPI_ISL_2542444, EPI_ISL_2542445, EPI_ISL_2542446, EPI_ISL_2542447, EPI_ISL_2542448, EPI_ISL_2542449, EPI_ISL_2542450, EPI_ISL_2542451, EPI_ISL_2542452, EPI_ISL_2542454, EPI_ISL_2542455, EPI_ISL_2542456, EPI_ISL_2542457, EPI_ISL_2542459, EPI_ISL_2542460, EPI_ISL_2542461, EPI_ISL_2542462, EPI_ISL_2542463, EPI_ISL_2542464, EPI_ISL_2542465, EPI_ISL_2542468, EPI_ISL_2542469, EPI_ISL_2542470, EPI_ISL_2542471, EPI_ISL_2542472, EPI_ISL_2542473, EPI_ISL_2542474, EPI_ISL_2542476, EPI_ISL_2542479, EPI_ISL_2542482, EPI_ISL_2542483, EPI_ISL_2542484, EPI_ISL_2542485, EPI_ISL_2542486, EPI_ISL_2542488, EPI_ISL_2542489, EPI_ISL_2542490, EPI_ISL_2542491, EPI_ISL_2542492, EPI_ISL_2542493, EPI_ISL_2542494, EPI_ISL_2542495, EPI_ISL_2542496, EPI_ISL_2542497, EPI_ISL_2542498, EPI_ISL_2542499, EPI_ISL_2542500, EPI_ISL_2542502, EPI_ISL_2542503, EPI_ISL_2542504, EPI_ISL_2542505, EPI_ISL_2542508, EPI_ISL_2542510, EPI_ISL_2542511, EPI_ISL_2542512, EPI_ISL_2542513, EPI_ISL_2542514, EPI_ISL_2542516, EPI_ISL_2542517, EPI_ISL_2542518, EPI_ISL_2542519, EPI_ISL_2542520, EPI_ISL_2542522, EPI_ISL_2542523, EPI_ISL_2542524, EPI_ISL_2542525, EPI_ISL_2542528, EPI_ISL_2542529, EPI_ISL_2542530, EPI_ISL_2542533, EPI_ISL_2542534, EPI_ISL_2542535, EPI_ISL_2542536, EPI_ISL_2542537, EPI_ISL_2542538, EPI_ISL_2542539, EPI_ISL_2542540, EPI_ISL_2542541, EPI_ISL_2542542, EPI_ISL_2542546, EPI_ISL_2542547, EPI_ISL_2542548, EPI_ISL_2542549, EPI_ISL_2542550, EPI_ISL_2542552, EPI_ISL_2542553, EPI_ISL_2542555, EPI_ISL_2542556, EPI_ISL_2542557, EPI_ISL_2542558, EPI_ISL_2542561, EPI_ISL_2542562, EPI_ISL_2542565, EPI_ISL_2542567, EPI_ISL_2542568, EPI_ISL_2542569, EPI_ISL_2542570, EPI_ISL_2542571, EPI_ISL_2542573, EPI_ISL_2542575, EPI_ISL_2542576, EPI_ISL_2542577, EPI_ISL_2542578, EPI_ISL_2542581, EPI_ISL_2542582, EPI_ISL_2542584, EPI_ISL_2542585, EPI_ISL_2542589, EPI_ISL_2542590, EPI_ISL_2542591, EPI_ISL_2542593, EPI_ISL_2542594, EPI_ISL_2542597, EPI_ISL_2542599, EPI_ISL_2542600, EPI_ISL_2542601, EPI_ISL_2542603, EPI_ISL_2542604, EPI_ISL_2542606, EPI_ISL_2542607, EPI_ISL_2542609, EPI_ISL_2542610, EPI_ISL_2542612, EPI_ISL_2542615, EPI_ISL_2542616, EPI_ISL_2542617, EPI_ISL_2542618, EPI_ISL_2542620, EPI_ISL_2542621, EPI_ISL_2542623, EPI_ISL_2542625, EPI_ISL_2542626, EPI_ISL_2542627, EPI_ISL_2542628, EPI_ISL_2542629, EPI_ISL_2542630, EPI_ISL_2542631, EPI_ISL_2542632, EPI_ISL_2542634, EPI_ISL_2542635, EPI_ISL_2542637, EPI_ISL_2542639, EPI_ISL_2542643, EPI_ISL_2542644, EPI_ISL_2542645, EPI_ISL_2542646, EPI_ISL_2542647, EPI_ISL_2542648, EPI_ISL_2542649, EPI_ISL_2542650, EPI_ISL_2542651, EPI_ISL_2542653, EPI_ISL_2542655, EPI_ISL_2542656, EPI_ISL_2542657, EPI_ISL_2542659, EPI_ISL_2542660, EPI_ISL_2542662, EPI_ISL_2542664, EPI_ISL_2542665, EPI_ISL_2542666, EPI_ISL_2542669, EPI_ISL_2542670, EPI_ISL_2542671, EPI_ISL_2542672, EPI_ISL_2542673, EPI_ISL_2542674, EPI_ISL_2542676, EPI_ISL_2542678, EPI_ISL_2542682, EPI_ISL_2542683, EPI_ISL_2542685, EPI_ISL_2542687, EPI_ISL_2542689, EPI_ISL_2542691, EPI_ISL_2542694, EPI_ISL_2542695, EPI_ISL_2542696, EPI_ISL_2542697, EPI_ISL_2542698, EPI_ISL_2542699, EPI_ISL_2542700, EPI_ISL_2542702, EPI_ISL_2542703, EPI_ISL_2542704, EPI_ISL_2542706, EPI_ISL_2542708, EPI_ISL_2542709, EPI_ISL_2542711, EPI_ISL_2542712, EPI_ISL_2542714, EPI_ISL_2542716, EPI_ISL_2542718, EPI_ISL_2542719, EPI_ISL_2542722, EPI_ISL_2542723, EPI_ISL_2542724, EPI_ISL_2542726, EPI_ISL_2542727, EPI_ISL_2542728, EPI_ISL_2542730, EPI_ISL_2542731, EPI_ISL_2542733, EPI_ISL_2542734, EPI_ISL_2542735, EPI_ISL_2542736, EPI_ISL_2542737, EPI_ISL_2542738, EPI_ISL_2542740, EPI_ISL_2542741, EPI_ISL_2542742, EPI_ISL_2542744, EPI_ISL_2542745, EPI_ISL_2542746, EPI_ISL_2542747, EPI_ISL_2542749, EPI_ISL_2542750, EPI_ISL_2542751, EPI_ISL_2542752, EPI_ISL_2542754, EPI_ISL_2542755, EPI_ISL_2542756, EPI_ISL_2542758, EPI_ISL_2542759, EPI_ISL_2542760, EPI_ISL_2542761, EPI_ISL_2542762, EPI_ISL_2542763, EPI_ISL_2542764, EPI_ISL_2542766, EPI_ISL_2542769, EPI_ISL_2542772, EPI_ISL_2542773, EPI_ISL_2542774, EPI_ISL_2542775, EPI_ISL_2542776, EPI_ISL_2542778, EPI_ISL_2542779, EPI_ISL_2542780, EPI_ISL_2542782, EPI_ISL_2542783, EPI_ISL_2542784 | | | |
| see above | Lighthouse Lab in Glasgow | Wellcome Sanger Institute for the COVID-19 Genomics UK (COG-UK) Consortium | Harper VanSteenhouse, Yumi Kasai, David Gray, Carol Clugston, Anna Dominiczak and Alex Alderton, Roberto Amato, Jeffrey Barrett, Sonia Goncalves, Ewan Harrison, David K. Jackson, Ian Johnston, Dominic Kwiatkowski, Cordelia Langford, John Sillitoe on behalf of the Wellcome Sanger Institute COVID-19 Surveillance Team |
| EPI_ISL_2542786, EPI_ISL_2542787, EPI_ISL_2542788 | Lighthouse Lab in Alderley Park | Wellcome Sanger Institute for the COVID-19 Genomics UK (COG-UK) Consortium | Jacquelyn Wynn, Mairead Hyland, The Lighthouse Lab in Alderley Park and Alex Alderton, Roberto Amato, Jeffrey Barrett, Sonia Goncalves, Ewan Harrison, David K. Jackson, Ian Johnston, Dominic Kwiatkowski, Cordelia Langford, John Sillitoe on behalf of the Wellcome Sanger Institute COVID-19 Surveillance Team |
| EPI_ISL_2542789, EPI_ISL_2542790, EPI_ISL_2542791, EPI_ISL_2542792, EPI_ISL_2542794, EPI_ISL_2542795, EPI_ISL_2542796, EPI_ISL_2542797, EPI_ISL_2542798, EPI_ISL_2542799, EPI_ISL_2542800, EPI_ISL_2542802, EPI_ISL_2542803, EPI_ISL_2542804, EPI_ISL_2542805, EPI_ISL_2542807, EPI_ISL_2542809, EPI_ISL_2542812, EPI_ISL_2542813, EPI_ISL_2542814, EPI_ISL_2542815, EPI_ISL_2542817, EPI_ISL_2542818, EPI_ISL_2542819, EPI_ISL_2542820, EPI_ISL_2542821, EPI_ISL_2542822, EPI_ISL_2542823, EPI_ISL_2542826, EPI_ISL_2542828, EPI_ISL_2542829, EPI_ISL_2542831, EPI_ISL_2542832, EPI_ISL_2542834, EPI_ISL_2542835, EPI_ISL_2542837, EPI_ISL_2542838, EPI_ISL_2542839, EPI_ISL_2542840, EPI_ISL_2542841, EPI_ISL_2542842, EPI_ISL_2542843, EPI_ISL_2542844, EPI_ISL_2542845, EPI_ISL_2542846, EPI_ISL_2542847, EPI_ISL_2542849, EPI_ISL_2542850, EPI_ISL_2542852, EPI_ISL_2542853, EPI_ISL_2542854, EPI_ISL_2542857, EPI_ISL_2542858, EPI_ISL_2542859, EPI_ISL_2542860, EPI_ISL_2542861, EPI_ISL_2542863, EPI_ISL_2542864, EPI_ISL_2542865, EPI_ISL_2542867, EPI_ISL_2542868, EPI_ISL_2542869, EPI_ISL_2542871, EPI_ISL_2542872, EPI_ISL_2542873, EPI_ISL_2542874, EPI_ISL_2542875, EPI_ISL_2542876, EPI_ISL_2542879, EPI_ISL_2542880, EPI_ISL_2542881, EPI_ISL_2542882, EPI_ISL_2542884, EPI_ISL_2542885, EPI_ISL_2542887, EPI_ISL_2542888, EPI_ISL_2542889, EPI_ISL_2542890, EPI_ISL_2542892, EPI_ISL_2542893, EPI_ISL_2542895, EPI_ISL_2542897, EPI_ISL_2542899, EPI_ISL_2542901, EPI_ISL_2542903, EPI_ISL_2542904, EPI_ISL_2542905, EPI_ISL_2542907, EPI_ISL_2542908, EPI_ISL_2542911, EPI_ISL_2542912, EPI_ISL_2542914, EPI_ISL_2542918, EPI_ISL_2542920, EPI_ISL_2542922, EPI_ISL_2542924, EPI_ISL_2542925, EPI_ISL_2542926, EPI_ISL_2542928, EPI_ISL_2542930, EPI_ISL_2542931, EPI_ISL_2542932, EPI_ISL_2542933, EPI_ISL_2542935, EPI_ISL_2542936, EPI_ISL_2542938, EPI_ISL_2542939, EPI_ISL_2542940, EPI_ISL_2542941, EPI_ISL_2542942, EPI_ISL_2542943, EPI_ISL_2542944, EPI_ISL_2542946, EPI_ISL_2542948, EPI_ISL_2542950, EPI_ISL_2542952, EPI_ISL_2542953, EPI_ISL_2542954, EPI_ISL_2542955, EPI_ISL_2542957, EPI_ISL_2542958, EPI_ISL_2542959, EPI_ISL_2542960, EPI_ISL_2542961, EPI_ISL_2542962, EPI_ISL_2542964, EPI_ISL_2542965, EPI_ISL_2542966, EPI_ISL_2542967, EPI_ISL_2542968, EPI_ISL_2542970, EPI_ISL_2542971, EPI_ISL_2542972, EPI_ISL_2542974, EPI_ISL_2542975, EPI_ISL_2542976, EPI_ISL_2542978, EPI_ISL_2542979, EPI_ISL_2542980, EPI_ISL_2542982, EPI_ISL_2542985, EPI_ISL_2542986, EPI_ISL_2542987, EPI_ISL_2542988, EPI_ISL_2542990, EPI_ISL_2542991, EPI_ISL_2542992, EPI_ISL_2542993, EPI_ISL_2542994, EPI_ISL_2542995, EPI_ISL_2542997, EPI_ISL_2542998, EPI_ISL_2542999, EPI_ISL_2543000, EPI_ISL_2543001, EPI_ISL_2543002, EPI_ISL_2543003, EPI_ISL_2543004, EPI_ISL_2543007, EPI_ISL_2543008, EPI_ISL_2543009, EPI_ISL_2543011, EPI_ISL_2543012, EPI_ISL_2543013, EPI_ISL_2543015, EPI_ISL_2543016, EPI_ISL_2543017, EPI_ISL_2543018, EPI_ISL_2543019, EPI_ISL_2543020, EPI_ISL_2543021, EPI_ISL_2543022, EPI_ISL_2543023, EPI_ISL_2543024, EPI_ISL_2543025, EPI_ISL_2543026, EPI_ISL_2543027, EPI_ISL_2543029, EPI_ISL_2543030, EPI_ISL_2543031, EPI_ISL_2543033, EPI_ISL_2543035, EPI_ISL_2543036, EPI_ISL_2543037, EPI_ISL_2543039, EPI_ISL_2543040, EPI_ISL_2543041, EPI_ISL_2543042, EPI_ISL_2543044, EPI_ISL_2543045 | | | |
| see above | Lighthouse Lab in Glasgow | Wellcome Sanger Institute for the COVID-19 Genomics UK (COG-UK) Consortium | Harper VanSteenhouse, Yumi Kasai, David Gray, Carol Clugston, Anna Dominiczak and Alex Alderton, Roberto Amato, Jeffrey Barrett, Sonia Goncalves, Ewan Harrison, David K. Jackson, Ian Johnston, Dominic Kwiatkowski, Cordelia Langford, John Sillitoe on behalf of the Wellcome Sanger Institute COVID-19 Surveillance Team |
| EPI_ISL_2543048 | Lighthouse Lab in Alderley Park | Wellcome Sanger Institute for the COVID-19 Genomics UK (COG-UK) Consortium | Jacquelyn Wynn, Mairead Hyland, The Lighthouse Lab in Alderley Park and Alex Alderton, Roberto Amato, Jeffrey Barrett, Sonia Goncalves, Ewan Harrison, David K. Jackson, Ian Johnston, Dominic Kwiatkowski, Cordelia Langford, John Sillitoe on behalf of the Wellcome Sanger Institute COVID-19 Surveillance Team |
| EPI_ISL_2543050, EPI_ISL_2543051, EPI_ISL_2543053, EPI_ISL_2543054, EPI_ISL_2543055, EPI_ISL_2543056, EPI_ISL_2543057, EPI_ISL_2543061, EPI_ISL_2543062, EPI_ISL_2543063, EPI_ISL_2543064, EPI_ISL_2543066, EPI_ISL_2543067, EPI_ISL_2543068, EPI_ISL_2543069, EPI_ISL_2543071, EPI_ISL_2543072, EPI_ISL_2543073, EPI_ISL_2543074, EPI_ISL_2543075, EPI_ISL_2543076, EPI_ISL_2543077, EPI_ISL_2543078, EPI_ISL_2543079, EPI_ISL_2543080, EPI_ISL_2543081, EPI_ISL_2543083, EPI_ISL_2543084, EPI_ISL_2543085, EPI_ISL_2543086, EPI_ISL_2543087, EPI_ISL_2543088, EPI_ISL_2543093, EPI_ISL_2543095, EPI_ISL_2543096, EPI_ISL_2543097, EPI_ISL_2543098, EPI_ISL_2543099, EPI_ISL_2543100, EPI_ISL_2543101, EPI_ISL_2543102, EPI_ISL_2543103, EPI_ISL_2543104, EPI_ISL_2543107, EPI_ISL_2543108, EPI_ISL_2543111, EPI_ISL_2543112, EPI_ISL_2543113, EPI_ISL_2543114, EPI_ISL_2543116, EPI_ISL_2543117, EPI_ISL_2543118, EPI_ISL_2543119, EPI_ISL_2543121, EPI_ISL_2543122, EPI_ISL_2543123, EPI_ISL_2543124, EPI_ISL_2543125, EPI_ISL_2543126, EPI_ISL_2543127, EPI_ISL_2543128, EPI_ISL_2543129, EPI_ISL_2543130, EPI_ISL_2543132, EPI_ISL_2543133, EPI_ISL_2543134, EPI_ISL_2543135, EPI_ISL_2543136, EPI_ISL_2543137, EPI_ISL_2543138, EPI_ISL_2543139, EPI_ISL_2543140, EPI_ISL_2543141, EPI_ISL_2543142, EPI_ISL_2543144, EPI_ISL_2543145, EPI_ISL_2543146, EPI_ISL_2543147 | | | |
| see above | Lighthouse Lab in Glasgow | Wellcome Sanger Institute for the COVID-19 Genomics UK (COG-UK) Consortium | Harper VanSteenhouse, Yumi Kasai, David Gray, Carol Clugston, Anna Dominiczak and Alex Alderton, Roberto Amato, Jeffrey Barrett, Sonia Goncalves, Ewan Harrison, David K. Jackson, Ian Johnston, Dominic Kwiatkowski, Cordelia Langford, John Sillitoe on behalf of the Wellcome Sanger Institute COVID-19 Surveillance Team |
| EPI_ISL_2543148 | Lighthouse Lab in Alderley Park | Wellcome Sanger Institute for the COVID-19 Genomics UK (COG-UK) Consortium | Jacquelyn Wynn, Mairead Hyland, The Lighthouse Lab in Alderley Park and Alex Alderton, Roberto Amato, Jeffrey Barrett, Sonia Goncalves, Ewan Harrison, David K. Jackson, Ian Johnston, Dominic Kwiatkowski, Cordelia Langford, John Sillitoe on behalf of the Wellcome Sanger Institute COVID-19 Surveillance Team |
| EPI_ISL_2543150, EPI_ISL_2543151, EPI_ISL_2543152, EPI_ISL_2543153, EPI_ISL_2543156, EPI_ISL_2543157, EPI_ISL_2543158, EPI_ISL_2543159, EPI_ISL_2543160, EPI_ISL_2543161, EPI_ISL_2543162, EPI_ISL_2543163, EPI_ISL_2543164, EPI_ISL_2543166, EPI_ISL_2543167, EPI_ISL_2543169, EPI_ISL_2543171, EPI_ISL_2543172, EPI_ISL_2543173, EPI_ISL_2543174, EPI_ISL_2543176, EPI_ISL_2543177, EPI_ISL_2543178, EPI_ISL_2543179, EPI_ISL_2543180, EPI_ISL_2543181, EPI_ISL_2543182, EPI_ISL_2543185, EPI_ISL_2543186, EPI_ISL_2543187, EPI_ISL_2543189, EPI_ISL_2543190, EPI_ISL_2543192, EPI_ISL_2543193, EPI_ISL_2543194, EPI_ISL_2543195, EPI_ISL_2543196, EPI_ISL_2543197, EPI_ISL_2543198, EPI_ISL_2543199, EPI_ISL_2543200, EPI_ISL_2543202, EPI_ISL_2543203, EPI_ISL_2543204, EPI_ISL_2543205, EPI_ISL_2543206, EPI_ISL_2543207, EPI_ISL_2543209, EPI_ISL_2543211, EPI_ISL_2543213, EPI_ISL_2543214, EPI_ISL_2543215, EPI_ISL_2543216, EPI_ISL_2543217, EPI_ISL_2543218, EPI_ISL_2543219, EPI_ISL_2543220, EPI_ISL_2543221 | | | |
| see above | Lighthouse Lab in Glasgow | Wellcome Sanger Institute for the COVID-19 Genomics UK (COG-UK) Consortium | Harper VanSteenhouse, Yumi Kasai, David Gray, Carol Clugston, Anna Dominiczak and Alex Alderton, Roberto Amato, Jeffrey Barrett, Sonia Goncalves, Ewan Harrison, David K. Jackson, Ian Johnston, Dominic Kwiatkowski, Cordelia Langford, John Sillitoe on behalf of the Wellcome Sanger Institute COVID-19 Surveillance Team |
| EPI_ISL_2543222 | Lighthouse Lab in Alderley Park | Wellcome Sanger Institute for the COVID-19 Genomics UK (COG-UK) Consortium | Jacquelyn Wynn, Mairead Hyland, The Lighthouse Lab in Alderley Park and Alex Alderton, Roberto Amato, Jeffrey Barrett, Sonia Goncalves, Ewan Harrison, David K. Jackson, Ian Johnston, Dominic Kwiatkowski, Cordelia Langford, John Sillitoe on behalf of the Wellcome Sanger Institute COVID-19 Surveillance Team |
| EPI_ISL_2543223, EPI_ISL_2543224, EPI_ISL_2543225, EPI_ISL_2543226, EPI_ISL_2543230, EPI_ISL_2543231, EPI_ISL_2543232, EPI_ISL_2543235, EPI_ISL_2543237, EPI_ISL_2543238, EPI_ISL_2543239, EPI_ISL_2543240, EPI_ISL_2543241, EPI_ISL_2543242, EPI_ISL_2543243, EPI_ISL_2543245, EPI_ISL_2543246, EPI_ISL_2543248, EPI_ISL_2543250, EPI_ISL_2543251, EPI_ISL_2543254, EPI_ISL_2543255, EPI_ISL_2543256, EPI_ISL_2543257, EPI_ISL_2543258, EPI_ISL_2543259, EPI_ISL_2543260, EPI_ISL_2543261, EPI_ISL_2543262, EPI_ISL_2543263, EPI_ISL_2543264, EPI_ISL_2543265, EPI_ISL_2543266, EPI_ISL_2543268, EPI_ISL_2543269, EPI_ISL_2543270, EPI_ISL_2543271, EPI_ISL_2543272, EPI_ISL_2543273, EPI_ISL_2543274, EPI_ISL_2543275, EPI_ISL_2543276, EPI_ISL_2543277, EPI_ISL_2543279, EPI_ISL_2543280, EPI_ISL_2543282, EPI_ISL_2543283, EPI_ISL_2543284, EPI_ISL_2543285, EPI_ISL_2543286, EPI_ISL_2543287, EPI_ISL_2543288, EPI_ISL_2543289, EPI_ISL_2543290, EPI_ISL_2543291, EPI_ISL_2543293, EPI_ISL_2543294, EPI_ISL_2543295, EPI_ISL_2543297, EPI_ISL_2543299, EPI_ISL_2543301, EPI_ISL_2543302, EPI_ISL_2543303, EPI_ISL_2543304, EPI_ISL_2543306, EPI_ISL_2543309, EPI_ISL_2543310, EPI_ISL_2543314, EPI_ISL_2543316, EPI_ISL_2543317, EPI_ISL_2543319, EPI_ISL_2543321, EPI_ISL_2543322, EPI_ISL_2543325, EPI_ISL_2543326, EPI_ISL_2543327, EPI_ISL_2543328, EPI_ISL_2543329, EPI_ISL_2543330, EPI_ISL_2543331 | | | |
| see above | Lighthouse Lab in Glasgow | Wellcome Sanger Institute for the COVID-19 Genomics UK (COG-UK) Consortium | Harper VanSteenhouse, Yumi Kasai, David Gray, Carol Clugston, Anna Dominiczak and Alex Alderton, Roberto Amato, Jeffrey Barrett, Sonia Goncalves, Ewan Harrison, David K. Jackson, Ian Johnston, Dominic Kwiatkowski, Cordelia Langford, John Sillitoe on behalf of the Wellcome Sanger Institute COVID-19 Surveillance Team |
| EPI_ISL_2543334, EPI_ISL_2543335 | Lighthouse Laboratory Plymouth | Wellcome Sanger Institute for the COVID-19 Genomics UK (COG-UK) Consortium | Lighthouse Laboratory Plymouth and Alex Alderton, Roberto Amato, Jeffrey Barrett, Sonia Goncalves, Ewan Harrison, David K. Jackson, Ian Johnston, Dominic Kwiatkowski, Cordelia Langford, John Sillitoe on behalf of the Wellcome Sanger Institute COVID-19 Surveillance Team |
| EPI_ISL_2543336, EPI_ISL_2543337, EPI_ISL_2543338, EPI_ISL_2543339, EPI_ISL_2543341, EPI_ISL_2543342, EPI_ISL_2543343, EPI_ISL_2543345, EPI_ISL_2543346, EPI_ISL_2543347, EPI_ISL_2543348, EPI_ISL_2543349, EPI_ISL_2543350, EPI_ISL_2543351, EPI_ISL_2543352, EPI_ISL_2543353, EPI_ISL_2543354, EPI_ISL_2543357, EPI_ISL_2543358, EPI_ISL_2543359, EPI_ISL_2543360, EPI_ISL_2543362, EPI_ISL_2543363, EPI_ISL_2543364, EPI_ISL_2543365, EPI_ISL_2543368, EPI_ISL_2543369, EPI_ISL_2543371, EPI_ISL_2543373, EPI_ISL_2543374, EPI_ISL_2543375, EPI_ISL_2543376, EPI_ISL_2543377, EPI_ISL_2543379, EPI_ISL_2543380, EPI_ISL_2543381, EPI_ISL_2543384, EPI_ISL_2543385, EPI_ISL_2543387, EPI_ISL_2543388, EPI_ISL_2543389, EPI_ISL_2543391, EPI_ISL_2543392, EPI_ISL_2543394, EPI_ISL_2543395, EPI_ISL_2543396, EPI_ISL_2543397, EPI_ISL_2543399, EPI_ISL_2543401, EPI_ISL_2543402, EPI_ISL_2543403, EPI_ISL_2543404, EPI_ISL_2543405, EPI_ISL_2543406, EPI_ISL_2543408, EPI_ISL_2543409, EPI_ISL_2543410, EPI_ISL_2543412, EPI_ISL_2543413, EPI_ISL_2543414, EPI_ISL_2543415, EPI_ISL_2543416, EPI_ISL_2543417, EPI_ISL_2543418, EPI_ISL_2543419, EPI_ISL_2543421, EPI_ISL_2543422, EPI_ISL_2543423, EPI_ISL_2543424, EPI_ISL_2543425, EPI_ISL_2543426, EPI_ISL_2543427, EPI_ISL_2543428, EPI_ISL_2543432 | | | |
| see above | Lighthouse Lab in Glasgow | Wellcome Sanger Institute for the COVID-19 Genomics UK (COG-UK) Consortium | Harper VanSteenhouse, Yumi Kasai, David Gray, Carol Clugston, Anna Dominiczak and Alex Alderton, Roberto Amato, Jeffrey Barrett, Sonia Goncalves, Ewan Harrison, David K. Jackson, Ian Johnston, Dominic Kwiatkowski, Cordelia Langford, John Sillitoe on behalf of the Wellcome Sanger Institute COVID-19 Surveillance Team |
| EPI_ISL_2543614 | Balai Besar Penelitian dan Pengembangan dan Reservoir | Genomik Solidaritas Indonesia Laboratorium | Carissa Sintca Wijaya, Vania Gavrila Wikasa, Dhahlia Agustina Cahyono, Alfin Mohammad Abdillah, Normastuti Adhini Tantyo, Yudistira Aditya Rahmat, Joko Waluyo, Annisa Muthiah Sukirman, Gracia Felias Enos Korompis, Pratiwi Sudarmono, |

| | | | |
|---|---|---|---|
| | Penyakit, Salatiga | | Budi Wiweko, Louisa Markus, Anuraj Shankar, Meutia Ayuputeri Kumaheri, Anindya Pradipta Susanto, Ariel Pradipta |
| EPI_ISL_2543667 | Provincial Public Health Reference Laboratory (PPHRL) P&SHD | Provincial Public Health Reference Laboratory (PPHRL) P&SHD | ANDLEEB HANIF |
| EPI_ISL_2543668 | Balai Besar Penelitian dan Pengembangan dan Reservoir Penyakit, Salatiga | Genomik Solidaritas Indonesia Laboratorium | Carissa Sintca Wijaya, Vania Gavrila Wikasa, Dhahlia Agustina Cahyono, Alfin Mohammad Abdillah, Normastuti Adhini Tantyo, Yudistira Aditya Rahmat, Joko Waluyo, Annisa Muthiah Sukirman, Gracia Felias Enos Korompis, Pratiwi Sudarmono, Budi Wiweko, Louisa Markus, Anuraj Shankar, Meutia Ayuputeri Kumaheri, Anindya Pradipta Susanto, Ariel Pradipta |
| EPI_ISL_2543732 | UM im. Karola Marcinkowskiego w Poznaniu Laboratorium UCA_Covid-19 | 1. National Institute of Public Health - National Institute of Hygiene, Warsaw, Poland 2. Biobank Lab, University of Lodz 3. Laboratory of Respiratory Viruses, Teaching and Clinical Center of the Medical University of Lodz | Dominik Strapagiel, Marta Sobalska-Kwapis, Klaudyna Królikowska, Jakub Lach, Marcin Słomka, Tomasz Płoszaj, Magdalena Traczyk-Borszyńska, Izabela Dróżdż, Maciej Borowiec, Katarzyna Zacharczuk, Magdalena Nowakowska, Małgorzata Sadkowska-Todys, Tomasz Wołkowicz |
| EPI_ISL_2543750 | Zakład Mikrobiologii Klinicznej WSZ im dr R. Ostrzyckiego w Koninie | 1. National Institute of Public Health - National Institute of Hygiene, Warsaw, Poland 2. Biobank Lab, University of Lodz 3. Laboratory of Respiratory Viruses, Teaching and Clinical Center of the Medical University of Lodz | Dominik Strapagiel, Marta Sobalska-Kwapis, Klaudyna Królikowska, Jakub Lach, Marcin Słomka, Tomasz Płoszaj, Magdalena Traczyk-Borszyńska, Izabela Dróżdż, Maciej Borowiec, Katarzyna Zacharczuk, Magdalena Nowakowska, Małgorzata Sadkowska-Todys, Tomasz Wołkowicz |
| EPI_ISL_2543891 | N.H.L Municipal Medical College, Ahmedabad | Gujarat Biotechnology Research Centre | Sonal Sharma ,Zuber Saiyed,Ramesh Pandit,Janvi Raval,Zarna Patel,Nitin Savaliya,Dinesh Kumar,Twinkle Soni,Jayshri Pethani,Umang Mishra,Nitesh Shah,Chaitanya Joshi,Madhvi Joshi |
| EPI_ISL_2543892 | N.H.L Municipal Medical College, Ahmedabad | Gujarat Biotechnology Research Centre | Zuber Saiyed,Ramesh Pandit,Janvi Raval,Zarna Patel,Nitin Savaliya,Dinesh Kumar,Twinkle Soni,Sonal Sharma ,Jayshri Pethani,Umang Mishra,Nitesh Shah,Chaitanya Joshi,Madhvi Joshi |
| EPI_ISL_2543893 | N.H.L Municipal Medical College, Ahmedabad | Gujarat Biotechnology Research Centre | Ramesh Pandit,Janvi Raval,Zarna Patel,Nitin Savaliya,Dinesh Kumar,Twinkle Soni,Sonal Sharma ,Zuber Saiyed,Jayshri Pethani,Umang Mishra,Nitesh Shah,Chaitanya Joshi,Madhvi Joshi |
| EPI_ISL_2543894 | N.H.L Municipal Medical College, Ahmedabad | Gujarat Biotechnology Research Centre | Janvi Raval,Zarna Patel,Nitin Savaliya,Dinesh Kumar,Twinkle Soni,Sonal Sharma ,Zuber Saiyed,Ramesh Pandit,Jayshri Pethani,Umang Mishra,Nitesh Shah,Chaitanya Joshi,Madhvi Joshi |
| EPI_ISL_2543895 | N.H.L Municipal Medical College, Ahmedabad | Gujarat Biotechnology Research Centre | Zarna Patel,Nitin Savaliya,Dinesh Kumar,Twinkle Soni,Sonal Sharma ,Zuber Saiyed,Ramesh Pandit,Janvi Raval,Jayshri Pethani,Umang Mishra,Nitesh Shah,Chaitanya Joshi,Madhvi Joshi |
| EPI_ISL_2543896 | N.H.L Municipal Medical College, Ahmedabad | Gujarat Biotechnology Research Centre | Nitin Savaliya,Dinesh Kumar,Twinkle Soni,Sonal Sharma ,Zuber Saiyed,Ramesh Pandit,Janvi Raval,Zarna Patel,Jayshri Pethani,Umang Mishra,Nitesh Shah,Chaitanya Joshi,Madhvi Joshi |
| EPI_ISL_2543897 | N.H.L Municipal Medical College, Ahmedabad | Gujarat Biotechnology Research Centre | Dinesh Kumar,Twinkle Soni,Sonal Sharma ,Zuber Saiyed,Ramesh Pandit,Janvi Raval,Zarna Patel,Nitin Savaliya,Jayshri Pethani,Umang Mishra,Nitesh Shah,Chaitanya Joshi,Madhvi Joshi |
| EPI_ISL_2543898 | N.H.L Municipal Medical College, Ahmedabad | Gujarat Biotechnology Research Centre | Twinkle Soni,Sonal Sharma ,Zuber Saiyed,Ramesh Pandit,Janvi Raval,Zarna Patel,Nitin Savaliya,Dinesh Kumar,Jayshri Pethani,Umang Mishra,Nitesh Shah,Chaitanya Joshi,Madhvi Joshi |
| EPI_ISL_2543899 | Sterling Hospital, Memnagar, Ahmedabad | Gujarat Biotechnology Research Centre | Atulbhai Patel, Sonal Sharma ,Zuber Saiyed,Ramesh Pandit,Janvi Raval,Zarna Patel,Nitin Savaliya,Dinesh Kumar,Twinkle Soni,Umang Mishra,Nitesh Shah,Chaitanya Joshi,Madhvi Joshi |
| EPI_ISL_2543900 | N.H.L Municipal Medical College, Ahmedabad | Gujarat Biotechnology Research Centre | Zuber Saiyed,Ramesh Pandit,Janvi Raval,Zarna Patel,Nitin Savaliya,Dinesh Kumar,Twinkle Soni,Sonal Sharma ,Jayshri Pethani,Umang Mishra,Nitesh Shah,Chaitanya Joshi,Madhvi Joshi |
| EPI_ISL_2543901 | N.H.L Municipal Medical College, Ahmedabad | Gujarat Biotechnology Research Centre | Ramesh Pandit,Janvi Raval,Zarna Patel,Nitin Savaliya,Dinesh Kumar,Twinkle Soni,Sonal Sharma ,Zuber Saiyed,Jayshri Pethani,Umang Mishra,Nitesh Shah,Chaitanya Joshi,Madhvi Joshi |
| EPI_ISL_2543902 | N.H.L Municipal Medical College, Ahmedabad | Gujarat Biotechnology Research Centre | Janvi Raval,Zarna Patel,Nitin Savaliya,Dinesh Kumar,Twinkle Soni,Sonal Sharma ,Zuber Saiyed,Ramesh Pandit,Jayshri Pethani,Umang Mishra,Nitesh Shah,Chaitanya Joshi,Madhvi Joshi |
| EPI_ISL_2543903 | N.H.L Municipal Medical College, Ahmedabad | Gujarat Biotechnology Research Centre | Zarna Patel,Nitin Savaliya,Dinesh Kumar,Twinkle Soni,Sonal Sharma ,Zuber Saiyed,Ramesh Pandit,Janvi Raval,Jayshri Pethani,Umang Mishra,Nitesh Shah,Chaitanya Joshi,Madhvi Joshi |
| EPI_ISL_2543904 | N.H.L Municipal Medical College, Ahmedabad | Gujarat Biotechnology Research Centre | Nitin Savaliya,Dinesh Kumar,Twinkle Soni,Sonal Sharma ,Zuber Saiyed,Ramesh Pandit,Janvi Raval,Zarna Patel,Jayshri Pethani,Umang Mishra,Nitesh Shah,Chaitanya Joshi,Madhvi Joshi |
| EPI_ISL_2543905 | N.H.L Municipal Medical College, Ahmedabad | Gujarat Biotechnology Research Centre | Dinesh Kumar,Twinkle Soni,Sonal Sharma ,Zuber Saiyed,Ramesh Pandit,Janvi Raval,Zarna Patel,Nitin Savaliya,Jayshri Pethani,Umang Mishra,Nitesh Shah,Chaitanya Joshi,Madhvi Joshi |
| EPI_ISL_2543906 | N.H.L Municipal Medical College, Ahmedabad | Gujarat Biotechnology Research Centre | Twinkle Soni,Sonal Sharma ,Zuber Saiyed,Ramesh Pandit,Janvi Raval,Zarna Patel,Nitin Savaliya,Dinesh Kumar,Jayshri Pethani,Umang Mishra,Nitesh Shah,Chaitanya Joshi,Madhvi Joshi |
| EPI_ISL_2543907 | N.H.L Municipal Medical College, Ahmedabad | Gujarat Biotechnology Research Centre | Sonal Sharma ,Zuber Saiyed,Ramesh Pandit,Janvi Raval,Zarna Patel,Nitin Savaliya,Dinesh Kumar,Twinkle Soni,Jayshri Pethani,Umang Mishra,Nitesh Shah,Chaitanya Joshi,Madhvi Joshi |
| EPI_ISL_2543908 | N.H.L Municipal Medical College, Ahmedabad | Gujarat Biotechnology Research Centre | Zuber Saiyed,Ramesh Pandit,Janvi Raval,Zarna Patel,Nitin Savaliya,Dinesh Kumar,Twinkle Soni,Sonal Sharma ,Jayshri Pethani,Umang Mishra,Nitesh Shah,Chaitanya Joshi,Madhvi Joshi |
| EPI_ISL_2543909 | N.H.L Municipal Medical College, Ahmedabad | Gujarat Biotechnology Research Centre | Ramesh Pandit,Janvi Raval,Zarna Patel,Nitin Savaliya,Dinesh Kumar,Twinkle Soni,Sonal Sharma ,Zuber Saiyed,Jayshri Pethani,Umang Mishra,Nitesh Shah,Chaitanya Joshi,Madhvi Joshi |
| EPI_ISL_2543910 | N.H.L Municipal Medical College, Ahmedabad | Gujarat Biotechnology Research Centre | Janvi Raval,Zarna Patel,Nitin Savaliya,Dinesh Kumar,Twinkle Soni,Sonal Sharma ,Zuber Saiyed,Ramesh Pandit,Jayshri Pethani,Umang Mishra,Nitesh Shah,Chaitanya Joshi,Madhvi Joshi |
| EPI_ISL_2543911 | N.H.L Municipal Medical College, Ahmedabad | Gujarat Biotechnology Research Centre | Zarna Patel,Nitin Savaliya,Dinesh Kumar,Twinkle Soni,Sonal Sharma ,Zuber Saiyed,Ramesh Pandit,Janvi Raval,Jayshri Pethani,Umang Mishra,Nitesh Shah,Chaitanya Joshi,Madhvi Joshi |
| EPI_ISL_2543912 | N.H.L Municipal Medical College, Ahmedabad | Gujarat Biotechnology Research Centre | Nitin Savaliya,Dinesh Kumar,Twinkle Soni,Sonal Sharma ,Zuber Saiyed,Ramesh Pandit,Janvi Raval,Zarna Patel,Jayshri Pethani,Umang Mishra,Nitesh Shah,Chaitanya Joshi,Madhvi Joshi |
| EPI_ISL_2543913 | N.H.L Municipal Medical College, Ahmedabad | Gujarat Biotechnology Research Centre | Dinesh Kumar,Twinkle Soni,Sonal Sharma ,Zuber Saiyed,Ramesh Pandit,Janvi Raval,Zarna Patel,Nitin Savaliya,Jayshri Pethani,Umang Mishra,Nitesh Shah,Chaitanya Joshi,Madhvi Joshi |
| EPI_ISL_2543914 | N.H.L Municipal Medical College, Ahmedabad | Gujarat Biotechnology Research Centre | Twinkle Soni,Sonal Sharma ,Zuber Saiyed,Ramesh Pandit,Janvi Raval,Zarna Patel,Nitin Savaliya,Dinesh Kumar,Jayshri Pethani,Umang Mishra,Nitesh Shah,Chaitanya Joshi,Madhvi Joshi |
| EPI_ISL_2543915 | N.H.L Municipal Medical College, Ahmedabad | Gujarat Biotechnology Research Centre | Sonal Sharma ,Zuber Saiyed,Ramesh Pandit,Janvi Raval,Zarna Patel,Nitin Savaliya,Dinesh Kumar,Twinkle Soni,Jayshri Pethani,Umang Mishra,Nitesh Shah,Chaitanya Joshi,Madhvi Joshi |
| EPI_ISL_2544220, EPI_ISL_2544231, EPI_ISL_2544246 | Laboratorio di Microbiologia | Laboratorio di Microbiologia | Martinetti Lucchini Gladys, Valeria Spina |
| EPI_ISL_2544314, EPI_ISL_2544325, EPI_ISL_2544326, EPI_ISL_2544358, EPI_ISL_2544374, EPI_ISL_2544422 | Viollier AG | Department of Biosystems Science and Engineering, ETH Zürich | Chaoran Chen, Sarah Nadeau, Catharine Aquino, Ivan Topolsky, Kim Philipp Jablonski, Lara Fuhrmann, Daniel Ehrsam, Isabel Stürmer, Andreia Cabral de Gouvea, Maria Domenica Moccia, Simon Grüter, Timothy Sykes, Lennart Opitz, Griffin White, Laura Neff, Doris Popovic, Andrea Patrignani, Jay Tracy, Ralph Schlapbach, Christiane Beckmann, Maurice Redondo, Olivier Kobel, Christoph Noppen, Niko Beerenwinkel, Tanja Stadler |
| EPI_ISL_2544529 | Basurto University Hospital: Clinical Microbiology Laboratory | Biocruces Bizkaia | Mikel Urrutikoetxea-Gutiérrez, Mª Carmen Nieto Toboso, Estibaliz Ugalde Zarraga, Mikel Gallego Rodrigo, Ana de la Hoz, José Luis Díaz de Tuesta del Arco |
| EPI_ISL_2544534, EPI_ISL_2544544, EPI_ISL_2544564, EPI_ISL_2544579, EPI_ISL_2544581, EPI_ISL_2544594, EPI_ISL_2544596, EPI_ISL_2544610, EPI_ISL_2544611, EPI_ISL_2544612, EPI_ISL_2544614, EPI_ISL_2544642, EPI_ISL_2544644, EPI_ISL_2544649, EPI_ISL_2544650, EPI_ISL_2544653, EPI_ISL_2544657, EPI_ISL_2544663, EPI_ISL_2544665, EPI_ISL_2544666, EPI_ISL_2544669, EPI_ISL_2544670, EPI_ISL_2544675, EPI_ISL_2544676 | | | |
| see above | IN State Department of Health Laboratory Services | IN State Department of Health Laboratory Services | Cassandra Campion, Jamie Yeadon, Brian Pope, Lixia Liu, Kyle Brownlee, Melissa Hindenlang, Mark Glazier |
| EPI_ISL_2544686, EPI_ISL_2544687, EPI_ISL_2544688, EPI_ISL_2544690 | Kanagawa Prefectural Institute of Public Health | Kanagawa Prefectural Institute of Public Health | Takayuki Hishiki, Rieko Suzuki, Takako Sano, Makiko Kondo, Sumi Watanabe, Itsumi Toyokura, Junichi Sakuragi, Tomohiko Takasaki, Tsuyoshi Sekizuka, Kentaro Itokawa, Rina Tanaka, Masanori Hashino, Makoto Kuroda |
| EPI_ISL_2544782, EPI_ISL_2544786, EPI_ISL_2544788, EPI_ISL_2544797 | MD PHL | MD PHL | Maryland Department of Health Laboratories Administration |
| EPI_ISL_2544800, EPI_ISL_2544801 | Balai Besar Penelitian dan Pengembangan dan Reservoir Penyakit, Salatiga | Genomik Solidaritas Indonesia Laboratorium | Carissa Sintca Wijaya, Vania Gavrila Wikasa, Dhahlia Agustina Cahyono, Alfin Mohammad Abdillah, Normastuti Adhini Tantyo, Yudistira Aditya Rahmat, Joko Waluyo, Annisa Muthiah Sukirman, Gracia Felias Enos Korompis, Pratiwi Sudarmono, Budi Wiweko, Louisa Markus, Anuraj Shankar, Meutia Ayuputeri Kumaheri, Anindya Pradipta Susanto, Ariel Pradipta |
| EPI_ISL_2544804 | LBM ALPHABIO, Marseille | LBM ALPHABIO, Marseille | Vincent GARCIA |
| EPI_ISL_2544812 | GMERS Government Medical College (GMC), Gandhinagar | Gujarat Biotechnology Research Centre | Zuber Saiyed,Ramesh Pandit,Janvi Raval,Zarna Patel,Nitin Savaliya,Dinesh Kumar,Twinkle Soni,Sonal Sharma ,Gaurishankar Shrimali,Umang Mishra,Nitesh Shah,Chaitanya Joshi,Madhvi Joshi |
| EPI_ISL_2544813 | GMERS Government Medical College (GMC), Gandhinagar | Gujarat Biotechnology Research Centre | Ramesh Pandit,Janvi Raval,Zarna Patel,Nitin Savaliya,Dinesh Kumar,Twinkle Soni,Sonal Sharma ,Zuber Saiyed,Gaurishankar Shrimali,Umang Mishra,Nitesh Shah,Chaitanya Joshi,Madhvi Joshi |
| EPI_ISL_2544814 | GMERS Government Medical College (GMC), Gandhinagar | Gujarat Biotechnology Research Centre | Janvi Raval,Zarna Patel,Nitin Savaliya,Dinesh Kumar,Twinkle Soni,Sonal Sharma ,Zuber Saiyed,Ramesh Pandit,Gaurishankar Shrimali,Umang Mishra,Nitesh Shah,Chaitanya Joshi,Madhvi Joshi |
| EPI_ISL_2544815 | GMERS Government Medical College (GMC), Gandhinagar | Gujarat Biotechnology Research Centre | Zarna Patel,Nitin Savaliya,Dinesh Kumar,Twinkle Soni,Sonal Sharma ,Zuber Saiyed,Ramesh Pandit,Janvi Raval,Gaurishankar Shrimali,Umang Mishra,Nitesh Shah,Chaitanya Joshi,Madhvi Joshi |
| EPI_ISL_2544816 | GMERS Government Medical College (GMC), Gandhinagar | Gujarat Biotechnology Research Centre | Nitin Savaliya,Dinesh Kumar,Twinkle Soni,Sonal Sharma ,Zuber Saiyed,Ramesh Pandit,Janvi Raval,Zarna Patel,Gaurishankar Shrimali,Umang Mishra,Nitesh Shah,Chaitanya Joshi,Madhvi Joshi |
| EPI_ISL_2544817 | GMERS Government Medical College (GMC), Gandhinagar | Gujarat Biotechnology Research Centre | Dinesh Kumar,Twinkle Soni,Sonal Sharma ,Zuber Saiyed,Ramesh Pandit,Janvi Raval,Zarna Patel,Nitin Savaliya,Gaurishankar Shrimali,Umang Mishra,Nitesh Shah,Chaitanya Joshi,Madhvi Joshi |
| EPI_ISL_2544818 | GMERS Government Medical College (GMC), Gandhinagar | Gujarat Biotechnology Research Centre | Twinkle Soni,Sonal Sharma ,Zuber Saiyed,Ramesh Pandit,Janvi Raval,Zarna Patel,Nitin Savaliya,Dinesh Kumar,Gaurishankar Shrimali,Umang Mishra,Nitesh Shah,Chaitanya Joshi,Madhvi Joshi |
| EPI_ISL_2544819 | GMERS Government Medical College (GMC), Gandhinagar | Gujarat Biotechnology Research Centre | Sonal Sharma ,Zuber Saiyed,Ramesh Pandit,Janvi Raval,Zarna Patel,Nitin Savaliya,Dinesh Kumar,Twinkle Soni,Gaurishankar Shrimali,Umang Mishra,Nitesh Shah,Chaitanya Joshi,Madhvi Joshi |
| EPI_ISL_2544820 | GMERS Government Medical College (GMC), Gandhinagar | Gujarat Biotechnology Research Centre | Zuber Saiyed,Ramesh Pandit,Janvi Raval,Zarna Patel,Nitin Savaliya,Dinesh Kumar,Twinkle Soni,Sonal Sharma ,Gaurishankar Shrimali,Umang Mishra,Nitesh Shah,Chaitanya Joshi,Madhvi Joshi |
| EPI_ISL_2544821 | GMERS Government Medical College (GMC), Gandhinagar | Gujarat Biotechnology Research Centre | Ramesh Pandit,Janvi Raval,Zarna Patel,Nitin Savaliya,Dinesh Kumar,Twinkle Soni,Sonal Sharma ,Zuber Saiyed,Gaurishankar Shrimali,Umang Mishra,Nitesh Shah,Chaitanya Joshi,Madhvi Joshi |
| EPI_ISL_2544822 | N.H.L Municipal Medical College, Ahmedabad | Gujarat Biotechnology Research Centre | Janvi Raval,Zarna Patel,Nitin Savaliya,Dinesh Kumar,Twinkle Soni,Sonal Sharma ,Zuber Saiyed,Ramesh Pandit,Jayshri Pethani,Umang Mishra,Nitesh Shah,Chaitanya Joshi,Madhvi Joshi |
| EPI_ISL_2544823 | GMERS Government Medical College (GMC), Gotri | Gujarat Biotechnology Research Centre | Zarna Patel,Nitin Savaliya,Dinesh Kumar,Twinkle Soni,Sonal Sharma ,Zuber Saiyed,Ramesh Pandit,Janvi Raval,Bithika Duttaroy,Umang Mishra,Nitesh Shah,Chaitanya Joshi,Madhvi Joshi |
| EPI_ISL_2544824 | GMERS Government Medical College (GMC), Gotri | Gujarat Biotechnology Research Centre | Nitin Savaliya,Dinesh Kumar,Twinkle Soni,Sonal Sharma ,Zuber Saiyed,Ramesh Pandit,Janvi Raval,Zarna Patel,Bithika Duttaroy,Umang Mishra,Nitesh Shah,Chaitanya Joshi,Madhvi Joshi |
| EPI_ISL_2544825 | GMERS Government Medical College (GMC), Gotri | Gujarat Biotechnology Research Centre | Dinesh Kumar,Twinkle Soni,Sonal Sharma ,Zuber Saiyed,Ramesh Pandit,Janvi Raval,Zarna Patel,Nitin Savaliya,Bithika Duttaroy,Umang Mishra,Nitesh Shah,Chaitanya Joshi,Madhvi Joshi |
| EPI_ISL_2544826 | GMERS Government Medical College (GMC), Gotri | Gujarat Biotechnology Research Centre | Twinkle Soni,Sonal Sharma ,Zuber Saiyed,Ramesh Pandit,Janvi Raval,Zarna Patel,Nitin Savaliya,Dinesh Kumar,Bithika Duttaroy,Umang Mishra,Nitesh Shah,Chaitanya Joshi,Madhvi Joshi |
| EPI_ISL_2544827 | GMERS Government Medical College (GMC), Gotri | Gujarat Biotechnology Research Centre | Sonal Sharma ,Zuber Saiyed,Ramesh Pandit,Janvi Raval,Zarna Patel,Nitin Savaliya,Dinesh Kumar,Twinkle Soni,Bithika Duttaroy,Umang Mishra,Nitesh Shah,Chaitanya Joshi,Madhvi Joshi |
| EPI_ISL_2544828 | GMERS Government Medical College (GMC), Gotri | Gujarat Biotechnology Research Centre | Zuber Saiyed,Ramesh Pandit,Janvi Raval,Zarna Patel,Nitin Savaliya,Dinesh Kumar,Twinkle Soni,Sonal Sharma ,Bithika Duttaroy,Umang Mishra,Nitesh Shah,Chaitanya Joshi,Madhvi Joshi |
| EPI_ISL_2544829 | GMERS Government Medical College (GMC), Gotri | Gujarat Biotechnology Research Centre | Ramesh Pandit,Janvi Raval,Zarna Patel,Nitin Savaliya,Dinesh Kumar,Twinkle Soni,Sonal Sharma ,Zuber Saiyed,Bithika Duttaroy,Umang Mishra,Nitesh Shah,Chaitanya Joshi,Madhvi Joshi |
| EPI_ISL_2544830 | GMERS Government Medical College (GMC), Gotri | Gujarat Biotechnology Research Centre | Janvi Raval,Zarna Patel,Nitin Savaliya,Dinesh Kumar,Twinkle Soni,Sonal Sharma ,Zuber Saiyed,Ramesh Pandit,Bithika Duttaroy,Umang Mishra,Nitesh Shah,Chaitanya Joshi,Madhvi Joshi |
| EPI_ISL_2544831 | GMERS Government Medical College (GMC), Gotri | Gujarat Biotechnology Research Centre | Zarna Patel,Nitin Savaliya,Dinesh Kumar,Twinkle Soni,Sonal Sharma ,Zuber Saiyed,Ramesh Pandit,Janvi Raval,Bithika Duttaroy,Umang Mishra,Nitesh Shah,Chaitanya Joshi,Madhvi Joshi |
| EPI_ISL_2544832 | GMERS Government Medical College (GMC), Gotri | Gujarat Biotechnology Research Centre | Nitin Savaliya,Dinesh Kumar,Twinkle Soni,Sonal Sharma ,Zuber Saiyed,Ramesh Pandit,Janvi Raval,Zarna Patel,Bithika Duttaroy,Umang Mishra,Nitesh Shah,Chaitanya Joshi,Madhvi Joshi |
| EPI_ISL_2544833 | GMERS Government Medical College (GMC), Gotri | Gujarat Biotechnology Research Centre | Dinesh Kumar,Twinkle Soni,Sonal Sharma ,Zuber Saiyed,Ramesh Pandit,Janvi Raval,Zarna Patel,Nitin Savaliya,Bithika Duttaroy,Umang Mishra,Nitesh Shah,Chaitanya Joshi,Madhvi Joshi |

| | | | |
|---|---|---|---|
| EPI_ISL_2544834 | GMERS Government Medical College (GMC), Gotri | Gujarat Biotechnology Research Centre | Twinkle Soni,Sonal Sharma ,Zuber Saiyed,Ramesh Pandit,Janvi Raval,Zarna Patel,Nitin Savaliya,Dinesh Kumar,Bithika Duttaroy,Umang Mishra,Nitesh Shah,Chaitanya Joshi,Madhvi Joshi |
| EPI_ISL_2544835 | GMERS Government Medical College (GMC), Gotri | Gujarat Biotechnology Research Centre | Sonal Sharma ,Zuber Saiyed,Ramesh Pandit,Janvi Raval,Zarna Patel,Nitin Savaliya,Dinesh Kumar,Twinkle Soni,Bithika Duttaroy,Umang Mishra,Nitesh Shah,Chaitanya Joshi,Madhvi Joshi |
| EPI_ISL_2544836 | GMERS Government Medical College (GMC), Gotri | Gujarat Biotechnology Research Centre | Zuber Saiyed,Ramesh Pandit,Janvi Raval,Zarna Patel,Nitin Savaliya,Dinesh Kumar,Twinkle Soni,Sonal Sharma ,Bithika Duttaroy,Umang Mishra,Nitesh Shah,Chaitanya Joshi,Madhvi Joshi |
| EPI_ISL_2544905, EPI_ISL_2544907 | OLVZ Aalst | OLVZ Aalst | Anne Vankeerberghen |
| EPI_ISL_2544913, EPI_ISL_2544914, EPI_ISL_2544927, EPI_ISL_2544935, EPI_ISL_2544943, EPI_ISL_2544957, EPI_ISL_2544961, EPI_ISL_2544962, EPI_ISL_2544965, EPI_ISL_2544973, EPI_ISL_2544975, EPI_ISL_2544977, EPI_ISL_2544981, EPI_ISL_2545034, EPI_ISL_2545046, EPI_ISL_2545048, EPI_ISL_2545049, EPI_ISL_2545069, EPI_ISL_2545083, EPI_ISL_2545086, EPI_ISL_2545088, EPI_ISL_2545101, EPI_ISL_2545102, EPI_ISL_2545106, EPI_ISL_2545114, EPI_ISL_2545115, EPI_ISL_2545120, EPI_ISL_2545132, EPI_ISL_2545133, EPI_ISL_2545137, EPI_ISL_2545145, EPI_ISL_2545179, EPI_ISL_2545184 | | | |
| see above | UW Virology Lab | UW Virology Lab | Pavitra Roychoudhury, Hong Xie, Lasata Shrestha, Shah Mohamed Bakhash, Tien V. Nguyen, Noah R. Baker, Sean Ellis, Meei-Li Huang, Keith R Jerome, Alexander Greninger |
| EPI_ISL_2545211 | KEMPEGOWDA INTERNATIONAL AIRPORT | INSACOG-KA, NIMHANS | Chitra Pattabiraman, Pramada Prasad, Anson Kunjumon George, Ananthapadmanabha Kotambail, Darshan Sreenivas, Chetan G K, Gautham Arunachal Udupi, Anita S Desai, V Ravi |
| EPI_ISL_2545226, EPI_ISL_2545227, EPI_ISL_2545234, EPI_ISL_2545236, EPI_ISL_2545240, EPI_ISL_2545243, EPI_ISL_2545246, EPI_ISL_2545247, EPI_ISL_2545251, EPI_ISL_2545254, EPI_ISL_2545257, EPI_ISL_2545262, EPI_ISL_2545264, EPI_ISL_2545265, EPI_ISL_2545267, EPI_ISL_2545270, EPI_ISL_2545276, EPI_ISL_2545277, EPI_ISL_2545290, EPI_ISL_2545292, EPI_ISL_2545295, EPI_ISL_2545296, EPI_ISL_2545297, EPI_ISL_2545299, EPI_ISL_2545302, EPI_ISL_2545303 | | | |
| see above | Shamir Medical Center (Asaf Harofe) | Shamir Medical Center (Asaf Harofe) | Nir Rainy,Chen Weiner,Reut Sorek Abramovich,Yevgeni Yegorov,Anna Vishnevsky,Patricia Benveniste-Lekovitz,Netta Zuckerman,Abu Hamad Ramzia,Adina Bar Chaim |
| EPI_ISL_2545435, EPI_ISL_2545447, EPI_ISL_2545452, EPI_ISL_2545453, EPI_ISL_2545454, EPI_ISL_2545461, EPI_ISL_2545470, EPI_ISL_2545473 | National Platform bis UMONS/Jolimont | National Platform bis UMONS/Jolimont | François Dufrasne, Guillaume Bayon-Vicente, Florian Juszczak, Gautier Detry, Ruddy Wattiez |
| EPI_ISL_2545595, EPI_ISL_2545603, EPI_ISL_2545604, EPI_ISL_2545607, EPI_ISL_2545608, EPI_ISL_2545612 | Platform BIS UZA/UAntwerpen | Labo Klinische Biologie, UZA | Jasmine Coppens, Marie Le Mercier, Basil Britto Xavier, Christine Lammens, Veerle Matheeussen, Herman Goossens |
| EPI_ISL_2545627, EPI_ISL_2545629 | Algemeen Medisch Labo | Labo Klinische Biologie, UZA | Jasmine Coppens, Marie Le Mercier, Basil Britto Xavier, Christine Lammens, Veerle Matheeussen, Herman Goossens |
| EPI_ISL_2545634, EPI_ISL_2545635, EPI_ISL_2545642 | Platform BIS UZA/UAntwerpen | Labo Klinische Biologie, UZA | Jasmine Coppens, Marie Le Mercier, Basil Britto Xavier, Christine Lammens, Veerle Matheeussen, Herman Goossens |
| EPI_ISL_2545652, EPI_ISL_2545663 | LHUB-ULB | Labo Klinische Biologie, UZA | Jasmine Coppens, Marie Le Mercier, Basil Britto Xavier, Christine Lammens, Veerle Matheeussen, Herman Goossens |
| EPI_ISL_2545667 | Tripler Army Medical Center | Tripler Army Medical Center | Rachael L. Downham, Jonathan A. D'Ambrozio, Kendra N. Carter, Lindsey L. O'Neal, Zakariae Bram, and Catherine F. Uyehara |
| EPI_ISL_2545990, EPI_ISL_2545991, EPI_ISL_2545992, EPI_ISL_2545993, EPI_ISL_2545995, EPI_ISL_2546013, EPI_ISL_2546023, EPI_ISL_2546034, EPI_ISL_2546053, EPI_ISL_2546063, EPI_ISL_2546068, EPI_ISL_2546078, EPI_ISL_2546085, EPI_ISL_2546089, EPI_ISL_2546099, EPI_ISL_2546102, EPI_ISL_2546104, EPI_ISL_2546106, EPI_ISL_2546107, EPI_ISL_2546122, EPI_ISL_2546136, EPI_ISL_2546145, EPI_ISL_2546154, EPI_ISL_2546156, EPI_ISL_2546157, EPI_ISL_2546158, EPI_ISL_2546163, EPI_ISL_2546164, EPI_ISL_2546187, EPI_ISL_2546228, EPI_ISL_2546230, EPI_ISL_2546233, EPI_ISL_2546275, EPI_ISL_2546276, EPI_ISL_2546278, EPI_ISL_2546279, EPI_ISL_2546281, EPI_ISL_2546289, EPI_ISL_2546291, EPI_ISL_2546294, EPI_ISL_2546295, EPI_ISL_2546297, EPI_ISL_2546298, EPI_ISL_2546304, EPI_ISL_2546308, EPI_ISL_2546321, EPI_ISL_2546327, EPI_ISL_2546330, EPI_ISL_2546336, EPI_ISL_2546337, EPI_ISL_2546349, EPI_ISL_2546350, EPI_ISL_2546356, EPI_ISL_2546357, EPI_ISL_2546359, EPI_ISL_2546360, EPI_ISL_2546362, EPI_ISL_2546364, EPI_ISL_2546365, EPI_ISL_2546366, EPI_ISL_2546367, EPI_ISL_2546368, EPI_ISL_2546374, EPI_ISL_2546375, EPI_ISL_2546381, EPI_ISL_2546387, EPI_ISL_2546388, EPI_ISL_2546389, EPI_ISL_2546396, EPI_ISL_2546397, EPI_ISL_2546401, EPI_ISL_2546403, EPI_ISL_2546411, EPI_ISL_2546414, EPI_ISL_2546416, EPI_ISL_2546417, EPI_ISL_2546418, EPI_ISL_2546419, EPI_ISL_2546422, EPI_ISL_2546425, EPI_ISL_2546431, EPI_ISL_2546432, EPI_ISL_2546434, EPI_ISL_2546435, EPI_ISL_2546437, EPI_ISL_2546438, EPI_ISL_2546443, EPI_ISL_2546444, EPI_ISL_2546447, EPI_ISL_2546487, EPI_ISL_2546489, EPI_ISL_2546490, EPI_ISL_2546493, EPI_ISL_2546495, EPI_ISL_2546496, EPI_ISL_2546497, EPI_ISL_2546499, EPI_ISL_2546500, EPI_ISL_2546501, EPI_ISL_2546502, EPI_ISL_2546504, EPI_ISL_2546505, EPI_ISL_2546506, EPI_ISL_2546508, EPI_ISL_2546509, EPI_ISL_2546510, EPI_ISL_2546511, EPI_ISL_2546512, EPI_ISL_2546514, EPI_ISL_2546515, EPI_ISL_2546516, EPI_ISL_2546518, EPI_ISL_2546520, EPI_ISL_2546522, EPI_ISL_2546523, EPI_ISL_2546524, EPI_ISL_2546526, EPI_ISL_2546527, EPI_ISL_2546528, EPI_ISL_2546529, EPI_ISL_2546530, EPI_ISL_2546532, EPI_ISL_2546534, EPI_ISL_2546536, EPI_ISL_2546537, EPI_ISL_2546539, EPI_ISL_2546540, EPI_ISL_2546541, EPI_ISL_2546542, EPI_ISL_2546543, EPI_ISL_2546544, EPI_ISL_2546547, EPI_ISL_2546548, EPI_ISL_2546549, EPI_ISL_2546550, EPI_ISL_2546552, EPI_ISL_2546553, EPI_ISL_2546555, EPI_ISL_2546558, EPI_ISL_2546559, EPI_ISL_2546562, EPI_ISL_2546563, EPI_ISL_2546564, EPI_ISL_2546565, EPI_ISL_2546566, EPI_ISL_2546567, EPI_ISL_2546568, EPI_ISL_2546569, EPI_ISL_2546570, EPI_ISL_2546572, EPI_ISL_2546576, EPI_ISL_2546577, EPI_ISL_2546578, EPI_ISL_2546580, EPI_ISL_2546581, EPI_ISL_2546582, EPI_ISL_2546583, EPI_ISL_2546584, EPI_ISL_2546585, EPI_ISL_2546586, EPI_ISL_2546587, EPI_ISL_2546589, EPI_ISL_2546591, EPI_ISL_2546592, EPI_ISL_2546593, EPI_ISL_2546594, EPI_ISL_2546595, EPI_ISL_2546596, EPI_ISL_2546597, EPI_ISL_2546598, EPI_ISL_2546599, EPI_ISL_2546600, EPI_ISL_2546601, EPI_ISL_2546602, EPI_ISL_2546603, EPI_ISL_2546604, EPI_ISL_2546605, EPI_ISL_2546606, EPI_ISL_2546607, EPI_ISL_2546609, EPI_ISL_2546610, EPI_ISL_2546611, EPI_ISL_2546612, EPI_ISL_2546613, EPI_ISL_2546614, EPI_ISL_2546615, EPI_ISL_2546616, EPI_ISL_2546617, EPI_ISL_2546619, EPI_ISL_2546620, EPI_ISL_2546622, EPI_ISL_2546623, EPI_ISL_2546624, EPI_ISL_2546625, EPI_ISL_2546627, EPI_ISL_2546628, EPI_ISL_2546630, EPI_ISL_2546631, EPI_ISL_2546632, EPI_ISL_2546633, EPI_ISL_2546634, EPI_ISL_2546635, EPI_ISL_2546636, EPI_ISL_2546638, EPI_ISL_2546639, EPI_ISL_2546640, EPI_ISL_2546641, EPI_ISL_2546642, EPI_ISL_2546644, EPI_ISL_2546645, EPI_ISL_2546646, EPI_ISL_2546647, EPI_ISL_2546648, EPI_ISL_2546649, EPI_ISL_2546650, EPI_ISL_2546652, EPI_ISL_2546653, EPI_ISL_2546654, EPI_ISL_2546655, EPI_ISL_2546656, EPI_ISL_2546657, EPI_ISL_2546658, EPI_ISL_2546659, EPI_ISL_2546660, EPI_ISL_2546661, EPI_ISL_2546682, EPI_ISL_2546683, EPI_ISL_2546684, EPI_ISL_2546685, EPI_ISL_2546686, EPI_ISL_2546687, EPI_ISL_2546689, EPI_ISL_2546690, EPI_ISL_2546691, EPI_ISL_2546692, EPI_ISL_2546693, EPI_ISL_2546694, EPI_ISL_2546695, EPI_ISL_2546696, EPI_ISL_2546697, EPI_ISL_2546698, EPI_ISL_2546699, EPI_ISL_2546700, EPI_ISL_2546702, EPI_ISL_2546703, EPI_ISL_2546704, EPI_ISL_2546705, EPI_ISL_2546707, EPI_ISL_2546708, EPI_ISL_2546709, EPI_ISL_2546710, EPI_ISL_2546711, EPI_ISL_2546712, EPI_ISL_2546713, EPI_ISL_2546715, EPI_ISL_2546716, EPI_ISL_2546717, EPI_ISL_2546718, EPI_ISL_2546719, EPI_ISL_2546720, EPI_ISL_2546721, EPI_ISL_2546722, EPI_ISL_2546724, EPI_ISL_2546725, EPI_ISL_2546726, EPI_ISL_2546727, EPI_ISL_2546728, EPI_ISL_2546729, EPI_ISL_2546730, EPI_ISL_2546731, EPI_ISL_2546732, EPI_ISL_2546733, EPI_ISL_2546734, EPI_ISL_2546735, EPI_ISL_2546736, EPI_ISL_2546737, EPI_ISL_2546738, EPI_ISL_2546739, EPI_ISL_2546740, EPI_ISL_2546741, EPI_ISL_2546742, EPI_ISL_2546743, EPI_ISL_2546744, EPI_ISL_2546746, EPI_ISL_2546747, EPI_ISL_2546748, EPI_ISL_2546749, EPI_ISL_2546750, EPI_ISL_2546751, EPI_ISL_2546752, EPI_ISL_2546753, EPI_ISL_2546754, EPI_ISL_2546755, EPI_ISL_2546756, EPI_ISL_2546757, EPI_ISL_2546758, EPI_ISL_2546759, EPI_ISL_2546760, EPI_ISL_2546761, EPI_ISL_2546762, EPI_ISL_2546763, EPI_ISL_2546764, EPI_ISL_2546765, EPI_ISL_2546766, EPI_ISL_2546767, EPI_ISL_2546768, EPI_ISL_2546769, EPI_ISL_2546770, EPI_ISL_2546771, EPI_ISL_2546772, EPI_ISL_2546773, EPI_ISL_2546774, EPI_ISL_2546775, EPI_ISL_2546776, EPI_ISL_2546777, EPI_ISL_2546778, EPI_ISL_2546779, EPI_ISL_2546780, EPI_ISL_2546781, EPI_ISL_2546782, EPI_ISL_2546783, EPI_ISL_2546784, EPI_ISL_2546785, EPI_ISL_2546786, EPI_ISL_2546787, EPI_ISL_2546788, EPI_ISL_2546789, EPI_ISL_2546790, EPI_ISL_2546791, EPI_ISL_2546792, EPI_ISL_2546793, EPI_ISL_2546794, EPI_ISL_2546795, EPI_ISL_2546796, EPI_ISL_2546797, EPI_ISL_2546798, EPI_ISL_2546799, EPI_ISL_2546800, EPI_ISL_2546801, EPI_ISL_2546802, EPI_ISL_2546803, EPI_ISL_2546804, EPI_ISL_2546805, EPI_ISL_2546806, EPI_ISL_2546807, EPI_ISL_2546808, EPI_ISL_2546809, EPI_ISL_2546811, EPI_ISL_2546812, EPI_ISL_2546813, EPI_ISL_2546814, EPI_ISL_2546815, EPI_ISL_2546816, EPI_ISL_2546817, EPI_ISL_2546818, EPI_ISL_2546819, EPI_ISL_2546820, EPI_ISL_2546821, EPI_ISL_2546822, EPI_ISL_2546823, EPI_ISL_2546824, EPI_ISL_2546825, EPI_ISL_2546826, EPI_ISL_2546827, EPI_ISL_2546828, EPI_ISL_2546829, EPI_ISL_2546830, EPI_ISL_2546831, EPI_ISL_2546832, EPI_ISL_2546833, EPI_ISL_2546834, EPI_ISL_2546835, EPI_ISL_2546836, EPI_ISL_2546838, EPI_ISL_2546839, EPI_ISL_2546840, EPI_ISL_2546841, EPI_ISL_2546842, EPI_ISL_2546843, EPI_ISL_2546844, EPI_ISL_2546845, EPI_ISL_2546846, EPI_ISL_2546847, EPI_ISL_2546848, EPI_ISL_2546849, EPI_ISL_2546850, EPI_ISL_2546851, EPI_ISL_2546852, EPI_ISL_2546853, EPI_ISL_2546854, EPI_ISL_2546855, EPI_ISL_2546856, EPI_ISL_2546857, EPI_ISL_2546858, EPI_ISL_2546859, EPI_ISL_2546860, EPI_ISL_2546861, EPI_ISL_2546863, EPI_ISL_2546864, EPI_ISL_2546865, EPI_ISL_2546866, EPI_ISL_2546867, EPI_ISL_2546868, EPI_ISL_2546869, EPI_ISL_2546870, EPI_ISL_2546871, EPI_ISL_2546872, EPI_ISL_2546873, EPI_ISL_2546874, EPI_ISL_2546875, EPI_ISL_2546877, EPI_ISL_2546878, EPI_ISL_2546879, EPI_ISL_2546880, EPI_ISL_2546881, EPI_ISL_2546882, EPI_ISL_2546883, EPI_ISL_2546884, EPI_ISL_2546885, EPI_ISL_2546886, EPI_ISL_2546887, EPI_ISL_2546888, EPI_ISL_2546889, EPI_ISL_2546890, EPI_ISL_2546891, EPI_ISL_2546892, EPI_ISL_2546893, EPI_ISL_2546894, EPI_ISL_2546895, EPI_ISL_2546896, EPI_ISL_2546897, EPI_ISL_2546898, EPI_ISL_2546899, EPI_ISL_2546900, EPI_ISL_2547069, EPI_ISL_2547070, EPI_ISL_2547071, EPI_ISL_2547072, EPI_ISL_2547073, EPI_ISL_2547074, EPI_ISL_2547075, EPI_ISL_2547076, EPI_ISL_2547077, EPI_ISL_2547078, EPI_ISL_2547079, EPI_ISL_2547080, EPI_ISL_2547081, EPI_ISL_2547082, EPI_ISL_2547083, EPI_ISL_2547084, EPI_ISL_2547085, EPI_ISL_2547086, EPI_ISL_2547087, EPI_ISL_2547088, EPI_ISL_2547089, EPI_ISL_2547090, EPI_ISL_2547091, EPI_ISL_2547092, EPI_ISL_2547093, EPI_ISL_2547094, EPI_ISL_2547095, EPI_ISL_2547096, EPI_ISL_2547097, EPI_ISL_2547098, EPI_ISL_2547099, EPI_ISL_2547100, EPI_ISL_2547101, EPI_ISL_2547102, EPI_ISL_2547103, EPI_ISL_2547104, EPI_ISL_2547105, EPI_ISL_2547106, EPI_ISL_2547107, EPI_ISL_2547108, EPI_ISL_2547109, EPI_ISL_2547110, EPI_ISL_2547111, EPI_ISL_2547112, EPI_ISL_2547113, EPI_ISL_2547114, EPI_ISL_2547115, EPI_ISL_2547116, EPI_ISL_2547117, EPI_ISL_2547118, EPI_ISL_2547119, EPI_ISL_2547120, EPI_ISL_2547122, EPI_ISL_2547123, EPI_ISL_2547124, EPI_ISL_2547125, EPI_ISL_2547126, EPI_ISL_2547127, EPI_ISL_2547128, EPI_ISL_2547129, EPI_ISL_2547130, EPI_ISL_2547131, EPI_ISL_2547132, EPI_ISL_2547133, EPI_ISL_2547134, EPI_ISL_2547135, EPI_ISL_2547136, EPI_ISL_2547137, EPI_ISL_2547138, EPI_ISL_2547139, EPI_ISL_2547140, EPI_ISL_2547141, EPI_ISL_2547142, EPI_ISL_2547143, EPI_ISL_2547144, EPI_ISL_2547145, EPI_ISL_2547146, EPI_ISL_2547147, EPI_ISL_2547148, EPI_ISL_2547149, EPI_ISL_2547150, EPI_ISL_2547151, EPI_ISL_2547152, EPI_ISL_2547153, EPI_ISL_2547154, EPI_ISL_2547155, EPI_ISL_2547156, EPI_ISL_2547157, EPI_ISL_2547158, EPI_ISL_2547159, EPI_ISL_2547160, EPI_ISL_2547161, EPI_ISL_2547162, EPI_ISL_2547163, EPI_ISL_2547164, EPI_ISL_2547165, EPI_ISL_2547166, EPI_ISL_2547167, EPI_ISL_2547168, EPI_ISL_2547169, EPI_ISL_2547170, EPI_ISL_2547171, EPI_ISL_2547172, EPI_ISL_2547173, EPI_ISL_2547174, EPI_ISL_2547175, EPI_ISL_2547176, EPI_ISL_2547177, EPI_ISL_2547178, EPI_ISL_2547179, EPI_ISL_2547180, EPI_ISL_2547181, EPI_ISL_2547182, EPI_ISL_2547183, EPI_ISL_2547184, EPI_ISL_2547185, EPI_ISL_2547186, EPI_ISL_2547187, EPI_ISL_2547188, EPI_ISL_2547189, EPI_ISL_2547190, EPI_ISL_2547191, EPI_ISL_2547192, EPI_ISL_2547193, EPI_ISL_2547194, EPI_ISL_2547195, EPI_ISL_2547196, EPI_ISL_2547197, EPI_ISL_2547198, EPI_ISL_2547199, EPI_ISL_2547200, EPI_ISL_2547201, EPI_ISL_2547202, EPI_ISL_2547203, EPI_ISL_2547204, EPI_ISL_2547205, EPI_ISL_2547206, EPI_ISL_2547207, EPI_ISL_2547208, EPI_ISL_2547209, EPI_ISL_2547210, EPI_ISL_2547211, EPI_ISL_2547212, EPI_ISL_2547213, EPI_ISL_2547214, EPI_ISL_2547215, EPI_ISL_2547216, EPI_ISL_2547217, EPI_ISL_2547218, EPI_ISL_2547219, EPI_ISL_2547220, EPI_ISL_2547221, EPI_ISL_2547222, EPI_ISL_2547223, EPI_ISL_2547224, EPI_ISL_2547225, EPI_ISL_2547226, EPI_ISL_2547227, EPI_ISL_2547228, EPI_ISL_2547229, EPI_ISL_2547230, EPI_ISL_2547231, EPI_ISL_2547232, EPI_ISL_2547233, EPI_ISL_2547234, EPI_ISL_2547235, EPI_ISL_2547236, EPI_ISL_2547237, EPI_ISL_2547238, EPI_ISL_2547239, EPI_ISL_2547240, EPI_ISL_2547241, EPI_ISL_2547242, EPI_ISL_2547243, EPI_ISL_2547244, EPI_ISL_2547245, EPI_ISL_2547246, EPI_ISL_2547247, EPI_ISL_2547248, EPI_ISL_2547249, EPI_ISL_2547250, EPI_ISL_2547251, EPI_ISL_2547252, EPI_ISL_2547253, EPI_ISL_2547254, EPI_ISL_2547255, EPI_ISL_2547256, EPI_ISL_2547257, EPI_ISL_2547258, EPI_ISL_2547259, EPI_ISL_2547260, EPI_ISL_2547261, EPI_ISL_2547262, EPI_ISL_2547263, EPI_ISL_2547264, EPI_ISL_2547265, EPI_ISL_2547266, EPI_ISL_2547267, EPI_ISL_2547268, EPI_ISL_2547269, EPI_ISL_2547270, EPI_ISL_2547271, EPI_ISL_2547272, EPI_ISL_2547273, EPI_ISL_2547274, EPI_ISL_2547275, EPI_ISL_2547276, EPI_ISL_2547277, EPI_ISL_2547278, EPI_ISL_2547279, EPI_ISL_2547280, EPI_ISL_2547281, EPI_ISL_2547282, EPI_ISL_2547283, EPI_ISL_2547284 | | | |
| see above | ICMR-National Institute of Virology - INSACOG | NIV Influenza | Dr. Varsha Potdar and NIC Team |
| EPI_ISL_2547439, EPI_ISL_2547440, EPI_ISL_2547441, EPI_ISL_2547444, EPI_ISL_2547445, EPI_ISL_2547447, EPI_ISL_2547448, EPI_ISL_2547450, EPI_ISL_2547451, EPI_ISL_2547459, EPI_ISL_2547460, EPI_ISL_2547461, EPI_ISL_2547464, EPI_ISL_2547465, EPI_ISL_2547466, EPI_ISL_2547467, EPI_ISL_2547468, EPI_ISL_2547469, EPI_ISL_2547470, EPI_ISL_2547471, EPI_ISL_2547474, EPI_ISL_2547485, EPI_ISL_2547486, EPI_ISL_2547490, EPI_ISL_2547491, EPI_ISL_2547493, EPI_ISL_2547494, EPI_ISL_2547495, EPI_ISL_2547496, EPI_ISL_2547497, EPI_ISL_2547498, EPI_ISL_2547499, EPI_ISL_2547501, EPI_ISL_2547502, EPI_ISL_2547503, EPI_ISL_2547505, EPI_ISL_2547506, EPI_ISL_2547507, EPI_ISL_2547509, EPI_ISL_2547511, EPI_ISL_2547512, EPI_ISL_2547513, EPI_ISL_2547515, EPI_ISL_2547516, EPI_ISL_2547517, EPI_ISL_2547518, EPI_ISL_2547519, EPI_ISL_2547520, EPI_ISL_2547521 | | | |
| see above | INSACOG-MEGHALAYA | National Institute of Biomedical Genomics – INSACOG | Arindam Maitra, R. Lyngdoh, Nidhan Kumar Biswas, E Shadap, Sreedhar Chinnaswamy, E Marak, Saumitra Das |
| EPI_ISL_2547554, EPI_ISL_2547569, EPI_ISL_2547576, EPI_ISL_2547596, EPI_ISL_2547598, EPI_ISL_2547601, EPI_ISL_2547602, EPI_ISL_2547603, EPI_ISL_2547605, EPI_ISL_2547606, EPI_ISL_2547607, EPI_ISL_2547612, EPI_ISL_2547614, EPI_ISL_2547617, EPI_ISL_2547622, EPI_ISL_2547631, EPI_ISL_2547632, EPI_ISL_2547635, EPI_ISL_2547642, EPI_ISL_2547643, EPI_ISL_2547668, EPI_ISL_2547678, EPI_ISL_2547679, EPI_ISL_2547682, EPI_ISL_2547684, EPI_ISL_2547688, EPI_ISL_2547694, EPI_ISL_2547709, EPI_ISL_2547758, EPI_ISL_2547788, EPI_ISL_2547791, EPI_ISL_2547797, EPI_ISL_2547798, EPI_ISL_2547799, EPI_ISL_2547810, EPI_ISL_2547816, EPI_ISL_2547820, EPI_ISL_2547825, EPI_ISL_2547830, EPI_ISL_2547833, EPI_ISL_2547836, EPI_ISL_2547837, EPI_ISL_2547838, EPI_ISL_2547840, EPI_ISL_2547843, EPI_ISL_2547850, EPI_ISL_2547851, EPI_ISL_2547856, EPI_ISL_2547858, EPI_ISL_2547861, EPI_ISL_2547863, EPI_ISL_2547865, EPI_ISL_2547867, EPI_ISL_2547869, EPI_ISL_2547870, EPI_ISL_2547880, EPI_ISL_2547882, EPI_ISL_2547886, EPI_ISL_2547891, EPI_ISL_2547895, EPI_ISL_2547897, EPI_ISL_2547903, EPI_ISL_2547904, EPI_ISL_2547905, EPI_ISL_2547906, EPI_ISL_2547907, EPI_ISL_2547908, EPI_ISL_2547948, EPI_ISL_2547952, EPI_ISL_2547956, EPI_ISL_2547957, EPI_ISL_2547985, EPI_ISL_2547991, EPI_ISL_2548001, EPI_ISL_2548031, EPI_ISL_2548038, EPI_ISL_2548046, EPI_ISL_2548047, EPI_ISL_2548056, EPI_ISL_2548058, EPI_ISL_2548079, EPI_ISL_2548083, EPI_ISL_2548087, EPI_ISL_2548109, EPI_ISL_2548115, EPI_ISL_2548125, EPI_ISL_2548128, EPI_ISL_2548129, EPI_ISL_2548134, EPI_ISL_2548135, EPI_ISL_2548137, EPI_ISL_2548141, EPI_ISL_2548152, EPI_ISL_2548156, EPI_ISL_2548164, EPI_ISL_2548202, EPI_ISL_2548248, EPI_ISL_2548266, EPI_ISL_2548281, EPI_ISL_2548289, EPI_ISL_2548300, EPI_ISL_2548305, EPI_ISL_2548332, EPI_ISL_2548333, EPI_ISL_2548341, EPI_ISL_2548342, EPI_ISL_2548349, EPI_ISL_2548361 | | | |
| see above | Public Health Ontario Laboratory | Public Health Ontario Laboratory | Vanessa G Allen, Philip Banh, Yao Chen, Richard de Borja, Alireza Eshaghi, Nahuel Fittipaldi, Christine Frantz, Jonathan B Gubbay, Jennifer L Guthrie, Lawrence Heisler, Esha Joshi, Michael Laszloffy, Aimin Li, Michael CY Li, Dean Maxwell, Sandeep Nagra, Samir N Patel, Jared Simpson, Karthikeyan Sivaraman, Ashleigh Sullivan, Yogi Sundaravadanam, Sarah Teatero, Andre Villegas, Matthew Watson, Sandra Zittermann |
| EPI_ISL_2548370 | National Platform bis UMONS/Jolimont | National Platform bis UMONS/Jolimont | François Dufrasne, Guillaume Bayon-Vicente, Florian Juszczak, Gautier Detry, Ruddy Wattiez |
| EPI_ISL_2548371, EPI_ISL_2548372, EPI_ISL_2548373, EPI_ISL_2548375, EPI_ISL_2548376, EPI_ISL_2548377, EPI_ISL_2548378, EPI_ISL_2548380, EPI_ISL_2548381, EPI_ISL_2548382, EPI_ISL_2548383, EPI_ISL_2548384, EPI_ISL_2548385, EPI_ISL_2548386, EPI_ISL_2548387, EPI_ISL_2548388, EPI_ISL_2548389, EPI_ISL_2548390, EPI_ISL_2548391, EPI_ISL_2548392, EPI_ISL_2548393, EPI_ISL_2548394, EPI_ISL_2548395, EPI_ISL_2548396, EPI_ISL_2548397, EPI_ISL_2548398, EPI_ISL_2548399, EPI_ISL_2548400, EPI_ISL_2548401, EPI_ISL_2548402, EPI_ISL_2548403, EPI_ISL_2548404, EPI_ISL_2548405, EPI_ISL_2548406, EPI_ISL_2548407, EPI_ISL_2548408, EPI_ISL_2548409, EPI_ISL_2548410, EPI_ISL_2548411, EPI_ISL_2548412, EPI_ISL_2548413, EPI_ISL_2548414, EPI_ISL_2548415, EPI_ISL_2548416, EPI_ISL_2548417, EPI_ISL_2548418, EPI_ISL_2548419, EPI_ISL_2548420, EPI_ISL_2548421, EPI_ISL_2548423, EPI_ISL_2548424, EPI_ISL_2548425, EPI_ISL_2548426, EPI_ISL_2548427, EPI_ISL_2548428, EPI_ISL_2548429, EPI_ISL_2548430, EPI_ISL_2548431, EPI_ISL_2548432, EPI_ISL_2548433, EPI_ISL_2548434, EPI_ISL_2548435, EPI_ISL_2548436, EPI_ISL_2548437, EPI_ISL_2548438, EPI_ISL_2548439, EPI_ISL_2548440, EPI_ISL_2548441, EPI_ISL_2548442, EPI_ISL_2548443, EPI_ISL_2548445, EPI_ISL_2548446, EPI_ISL_2548447, EPI_ISL_2548448, EPI_ISL_2548449, EPI_ISL_2548450, EPI_ISL_2548451, EPI_ISL_2548452, EPI_ISL_2548453, EPI_ISL_2548454, EPI_ISL_2548455, EPI_ISL_2548456, EPI_ISL_2548457, EPI_ISL_2548458, EPI_ISL_2548459, EPI_ISL_2548460, EPI_ISL_2548461, EPI_ISL_2548462, EPI_ISL_2548463, EPI_ISL_2548464, EPI_ISL_2548465, EPI_ISL_2548466, EPI_ISL_2548467, EPI_ISL_2548468, EPI_ISL_2548469, EPI_ISL_2548470, EPI_ISL_2548471, EPI_ISL_2548472, EPI_ISL_2548473, EPI_ISL_2548474, EPI_ISL_2548475, EPI_ISL_2548476, EPI_ISL_2548477, EPI_ISL_2548478, EPI_ISL_2548479, EPI_ISL_2548480, EPI_ISL_2548481, EPI_ISL_2548482, EPI_ISL_2548483, EPI_ISL_2548484, EPI_ISL_2548485, EPI_ISL_2548486, EPI_ISL_2548487, EPI_ISL_2548488, EPI_ISL_2548489, EPI_ISL_2548490, EPI_ISL_2548491, EPI_ISL_2548492, EPI_ISL_2548493, EPI_ISL_2548494, EPI_ISL_2548495, EPI_ISL_2548496, EPI_ISL_2548497, EPI_ISL_2548498, EPI_ISL_2548499, EPI_ISL_2548500, EPI_ISL_2548503, EPI_ISL_2548504, EPI_ISL_2548505, EPI_ISL_2548506, EPI_ISL_2548508, EPI_ISL_2548509, EPI_ISL_2548510, EPI_ISL_2548511, EPI_ISL_2548512, EPI_ISL_2548513, EPI_ISL_2548514, EPI_ISL_2548515, EPI_ISL_2548516, EPI_ISL_2548517, EPI_ISL_2548518, EPI_ISL_2548519, EPI_ISL_2548520, EPI_ISL_2548521, EPI_ISL_2548522, EPI_ISL_2548523, EPI_ISL_2548524, EPI_ISL_2548525, EPI_ISL_2548526, EPI_ISL_2548528, EPI_ISL_2548532, EPI_ISL_2548533, EPI_ISL_2548534, EPI_ISL_2548536, EPI_ISL_2548537, EPI_ISL_2548538 | | | |
| see above | INSACOG-WB | National Institute of Biomedical Genomics – INSACOG | Arindam Maitra, Bhaswati Bandyopadhyay, Nidhan Kumar Biswas, Tamal Ghosh, Sreedhar Chinnaswamy, Ajay Chakraborti, Saumitra Das |
| EPI_ISL_2548539 | National Platform bis UMONS/Jolimont | National Platform bis UMONS/Jolimont | François Dufrasne, Guillaume Bayon-Vicente, Florian Juszczak, Gautier Detry, Ruddy Wattiez |
| EPI_ISL_2548540, EPI_ISL_2548541, EPI_ISL_2548542, EPI_ISL_2548543, EPI_ISL_2548544, EPI_ISL_2548545, EPI_ISL_2548546, EPI_ISL_2548547, EPI_ISL_2548548, EPI_ISL_2548550, EPI_ISL_2548551, EPI_ISL_2548552, EPI_ISL_2548553, EPI_ISL_2548554, EPI_ISL_2548555, EPI_ISL_2548556, EPI_ISL_2548557, EPI_ISL_2548558, EPI_ISL_2548559, EPI_ISL_2548560, EPI_ISL_2548561, EPI_ISL_2548562, EPI_ISL_2548563, EPI_ISL_2548564, EPI_ISL_2548565, EPI_ISL_2548566, EPI_ISL_2548567, EPI_ISL_2548568, EPI_ISL_2548570, EPI_ISL_2548571, EPI_ISL_2548572, EPI_ISL_2548573, EPI_ISL_2548574, EPI_ISL_2548575, EPI_ISL_2548576, EPI_ISL_2548578, EPI_ISL_2548579, EPI_ISL_2548580, EPI_ISL_2548581, EPI_ISL_2548582, EPI_ISL_2548583, EPI_ISL_2548584, EPI_ISL_2548586, EPI_ISL_2548587, EPI_ISL_2548588, EPI_ISL_2548589, EPI_ISL_2548590, EPI_ISL_2548591, EPI_ISL_2548592, EPI_ISL_2548593, EPI_ISL_2548594, EPI_ISL_2548595, EPI_ISL_2548596, EPI_ISL_2548597, EPI_ISL_2548598, EPI_ISL_2548600, EPI_ISL_2548601, EPI_ISL_2548602, EPI_ISL_2548603, EPI_ISL_2548604, EPI_ISL_2548605, EPI_ISL_2548606, EPI_ISL_2548607, EPI_ISL_2548608, EPI_ISL_2548610, EPI_ISL_2548612, EPI_ISL_2548613, EPI_ISL_2548614, EPI_ISL_2548615, EPI_ISL_2548616, EPI_ISL_2548618, EPI_ISL_2548619, EPI_ISL_2548620, EPI_ISL_2548622, EPI_ISL_2548623, EPI_ISL_2548627, EPI_ISL_2548630, EPI_ISL_2548631, EPI_ISL_2548632, EPI_ISL_2548633, EPI_ISL_2548635, EPI_ISL_2548636, EPI_ISL_2548641, EPI_ISL_2548646, EPI_ISL_2548647, EPI_ISL_2548649, EPI_ISL_2548650, EPI_ISL_2548653, EPI_ISL_2548655, EPI_ISL_2548656, EPI_ISL_2548657, EPI_ISL_2548659, EPI_ISL_2548661, EPI_ISL_2548664, EPI_ISL_2548670, EPI_ISL_2548671, EPI_ISL_2548672, EPI_ISL_2548673, EPI_ISL_2548675, EPI_ISL_2548676, EPI_ISL_2548677, EPI_ISL_2548678, EPI_ISL_2548684, EPI_ISL_2548690, EPI_ISL_2548694, EPI_ISL_2548698, EPI_ISL_2548699, EPI_ISL_2548700, EPI_ISL_2548701, EPI_ISL_2548706, EPI_ISL_2548708, EPI_ISL_2548710, EPI_ISL_2548711, EPI_ISL_2548717, EPI_ISL_2548724, EPI_ISL_2548725, EPI_ISL_2548728, EPI_ISL_2548731, EPI_ISL_2548732, EPI_ISL_2548733, EPI_ISL_2548736, EPI_ISL_2548737, EPI_ISL_2548739, EPI_ISL_2548740, EPI_ISL_2548741, EPI_ISL_2548742, EPI_ISL_2548743, EPI_ISL_2548744, EPI_ISL_2548745, EPI_ISL_2548747, EPI_ISL_2548748, EPI_ISL_2548749, EPI_ISL_2548751, EPI_ISL_2548752, EPI_ISL_2548753, EPI_ISL_2548758, EPI_ISL_2548759, EPI_ISL_2548760, EPI_ISL_2548761, EPI_ISL_2548762, EPI_ISL_2548764, EPI_ISL_2548765, EPI_ISL_2548766, EPI_ISL_2548768, EPI_ISL_2548769, EPI_ISL_2548773, EPI_ISL_2548774, EPI_ISL_2548776, EPI_ISL_2548779, EPI_ISL_2548780, EPI_ISL_2548782, EPI_ISL_2548786, EPI_ISL_2548789, EPI_ISL_2548790, EPI_ISL_2548791, EPI_ISL_2548792, EPI_ISL_2548794, EPI_ISL_2548800, EPI_ISL_2548801, EPI_ISL_2548802, EPI_ISL_2548803, EPI_ISL_2548806, EPI_ISL_2548808, EPI_ISL_2548809, EPI_ISL_2548810, EPI_ISL_2548811, EPI_ISL_2548813, EPI_ISL_2548815, EPI_ISL_2548817, EPI_ISL_2548818, EPI_ISL_2548819, EPI_ISL_2548820, EPI_ISL_2548821, EPI_ISL_2548823, EPI_ISL_2548824, EPI_ISL_2548827, EPI_ISL_2548828, EPI_ISL_2548832, EPI_ISL_2548833, EPI_ISL_2548834, EPI_ISL_2548836, EPI_ISL_2548837, EPI_ISL_2548838, EPI_ISL_2548839, EPI_ISL_2548840, EPI_ISL_2548841, EPI_ISL_2548842, EPI_ISL_2548843, EPI_ISL_2548844, EPI_ISL_2548845, EPI_ISL_2548846, EPI_ISL_2548848, EPI_ISL_2548849, EPI_ISL_2548850, EPI_ISL_2548851, EPI_ISL_2548852, EPI_ISL_2548853, EPI_ISL_2548854, EPI_ISL_2548855, EPI_ISL_2548857, EPI_ISL_2548858, EPI_ISL_2548859, EPI_ISL_2548860, EPI_ISL_2548861, EPI_ISL_2548862, EPI_ISL_2548863, EPI_ISL_2548864, EPI_ISL_2548865, EPI_ISL_2548866, EPI_ISL_2548867, EPI_ISL_2548868, EPI_ISL_2548870, EPI_ISL_2548871, EPI_ISL_2548873, EPI_ISL_2548874, EPI_ISL_2548875, EPI_ISL_2548876, EPI_ISL_2548877, EPI_ISL_2548878, EPI_ISL_2548880, EPI_ISL_2548881, EPI_ISL_2548882, EPI_ISL_2548883, EPI_ISL_2548884, EPI_ISL_2548885, EPI_ISL_2548886, EPI_ISL_2548887, EPI_ISL_2548888, EPI_ISL_2548889, EPI_ISL_2548890, EPI_ISL_2548891, EPI_ISL_2548892, EPI_ISL_2548893, EPI_ISL_2548894, EPI_ISL_2548895, EPI_ISL_2548897, EPI_ISL_2548898, EPI_ISL_2548900, EPI_ISL_2548901, EPI_ISL_2548902, EPI_ISL_2548903, EPI_ISL_2548904, EPI_ISL_2548905, EPI_ISL_2548906, EPI_ISL_2548907, EPI_ISL_2548908, EPI_ISL_2548909, EPI_ISL_2548910, EPI_ISL_2548911, EPI_ISL_2548912, EPI_ISL_2548913, EPI_ISL_2548914, EPI_ISL_2548915, EPI_ISL_2548918, EPI_ISL_2548919, EPI_ISL_2548920, EPI_ISL_2548921, EPI_ISL_2548922, EPI_ISL_2548923, EPI_ISL_2548924, EPI_ISL_2548925, EPI_ISL_2548926, EPI_ISL_2548927, EPI_ISL_2548928, | | | |

| | | | |
|---|---|---|---|
| EPI_ISL_2548929, EPI_ISL_2548930, EPI_ISL_2548931, EPI_ISL_2548932, EPI_ISL_2548933, EPI_ISL_2548934, EPI_ISL_2548935, EPI_ISL_2548936, EPI_ISL_2548937, EPI_ISL_2548938, EPI_ISL_2548939, EPI_ISL_2548940, EPI_ISL_2548942, EPI_ISL_2548943, EPI_ISL_2548944, EPI_ISL_2548945, EPI_ISL_2548946, EPI_ISL_2548947, EPI_ISL_2548948, EPI_ISL_2548949, EPI_ISL_2548950, EPI_ISL_2548951, EPI_ISL_2548952, EPI_ISL_2548953, EPI_ISL_2548954, EPI_ISL_2548955, EPI_ISL_2548956, EPI_ISL_2548957, EPI_ISL_2548958, EPI_ISL_2548959, EPI_ISL_2548960, EPI_ISL_2548961, EPI_ISL_2548962, EPI_ISL_2548963, EPI_ISL_2548964, EPI_ISL_2548965, EPI_ISL_2548966, EPI_ISL_2548967, EPI_ISL_2548969, EPI_ISL_2548970, EPI_ISL_2548971, EPI_ISL_2548972, EPI_ISL_2548973, EPI_ISL_2548974, EPI_ISL_2548975, EPI_ISL_2548976, EPI_ISL_2548977, EPI_ISL_2548978, EPI_ISL_2548979, EPI_ISL_2548980, EPI_ISL_2548981, EPI_ISL_2548982, EPI_ISL_2548983, EPI_ISL_2548984, EPI_ISL_2548985, EPI_ISL_2548986, EPI_ISL_2548989, EPI_ISL_2548990, EPI_ISL_2548991, EPI_ISL_2548992, EPI_ISL_2548993, EPI_ISL_2548994, EPI_ISL_2548995, EPI_ISL_2548996, EPI_ISL_2548997, EPI_ISL_2548998, EPI_ISL_2548999, EPI_ISL_2549000, EPI_ISL_2549003, EPI_ISL_2549004, EPI_ISL_2549005, EPI_ISL_2549006, EPI_ISL_2549007, EPI_ISL_2549008, EPI_ISL_2549009, EPI_ISL_2549010, EPI_ISL_2549012, EPI_ISL_2549013, EPI_ISL_2549014, EPI_ISL_2549015, EPI_ISL_2549016, EPI_ISL_2549018, EPI_ISL_2549019, EPI_ISL_2549021, EPI_ISL_2549022, EPI_ISL_2549023, EPI_ISL_2549024, EPI_ISL_2549025, EPI_ISL_2549026, EPI_ISL_2549027, EPI_ISL_2549028, EPI_ISL_2549029, EPI_ISL_2549030, EPI_ISL_2549031, EPI_ISL_2549032, EPI_ISL_2549033, EPI_ISL_2549034, EPI_ISL_2549036, EPI_ISL_2549037, EPI_ISL_2549038, EPI_ISL_2549039, EPI_ISL_2549041, EPI_ISL_2549042, EPI_ISL_2549048, EPI_ISL_2549054, EPI_ISL_2549055, EPI_ISL_2549056, EPI_ISL_2549057, EPI_ISL_2549058, EPI_ISL_2549059, EPI_ISL_2549060, EPI_ISL_2549061, EPI_ISL_2549062, EPI_ISL_2549063, EPI_ISL_2549064, EPI_ISL_2549065, EPI_ISL_2549066, EPI_ISL_2549067, EPI_ISL_2549068, EPI_ISL_2549069, EPI_ISL_2549070, EPI_ISL_2549072 | | | |
| see above | INSACOG-WB | National Institute of Biomedical Genomics – INSACOG | Arindam Maitra, Bhaswati Bandyopadhyay, Nidhan Kumar Biswas, Tamal Ghosh, Sreedhar Chinnaswamy, Ajay Chakraborti, Saumitra Das |
| EPI_ISL_2549073, EPI_ISL_2549074, EPI_ISL_2549075, EPI_ISL_2549076 | Furst Medical Laboratory | Norwegian Institute of Public Health, Department of Virology | Kathrine Stene-Johansen, Kamilla Heddeland Instefjord, Hilde Elshaug, Garcia Llorente Ignacio, Jon Bråte, Engebretsen Serina Beate,Pedersen Benedikte Nevjen, Line Victoria Moen, Debech Nadia, Atiya R Ali,Marie Paulsen Madsen, Rasmus Riis Kopperud, Hilde Vollan, Karoline Bragstad, Olav Hungnes |
| EPI_ISL_2549095 | Akershus University Hospital, Department for Microbiology and Infectious Disease Control | Norwegian Institute of Public Health, Department of Virology | Kathrine Stene-Johansen, Kamilla Heddeland Instefjord, Hilde Elshaug, Garcia Llorente Ignacio, Jon Bråte, Engebretsen Serina Beate,Pedersen Benedikte Nevjen, Line Victoria Moen, Debech Nadia, Atiya R Ali,Marie Paulsen Madsen, Rasmus Riis Kopperud, Hilde Vollan, Karoline Bragstad, Olav Hungnes |
| EPI_ISL_2549096, EPI_ISL_2549098 | Furst Medical Laboratory | Norwegian Institute of Public Health, Department of Virology | Kathrine Stene-Johansen, Kamilla Heddeland Instefjord, Hilde Elshaug, Garcia Llorente Ignacio, Jon Bråte, Engebretsen Serina Beate,Pedersen Benedikte Nevjen, Line Victoria Moen, Debech Nadia, Atiya R Ali,Marie Paulsen Madsen, Rasmus Riis Kopperud, Hilde Vollan, Karoline Bragstad, Olav Hungnes |
| EPI_ISL_2549196 | National Platform bis UMONS/Jolimont | National Platform bis UMONS/Jolimont | François Dufrasne, Guillaume Bayon-Vicente, Florian Juszczak, Gautier Detry, Ruddy Wattiez |
| EPI_ISL_2549198, EPI_ISL_2549199 | Virology Department, Central Health Laboratory | The Francis Crick Institute | Ramuth M, Manraj SS, Sonoo J, Bahadoor BS, Crawford M, Patel H, Harvey R, Goldstone R, Daniels RS, Nicod J |
| EPI_ISL_2549200, EPI_ISL_2549201, EPI_ISL_2549202, EPI_ISL_2549203, EPI_ISL_2549204, EPI_ISL_2549205, EPI_ISL_2549206, EPI_ISL_2549207, EPI_ISL_2549208, EPI_ISL_2549209, EPI_ISL_2549210, EPI_ISL_2549211, EPI_ISL_2549212, EPI_ISL_2549213, EPI_ISL_2549214, EPI_ISL_2549215, EPI_ISL_2549216, EPI_ISL_2549217, EPI_ISL_2549218, EPI_ISL_2549219, EPI_ISL_2549220, EPI_ISL_2549221, EPI_ISL_2549222, EPI_ISL_2549224, EPI_ISL_2549225, EPI_ISL_2549226, EPI_ISL_2549227, EPI_ISL_2549228, EPI_ISL_2549229, EPI_ISL_2549230, EPI_ISL_2549231, EPI_ISL_2549232, EPI_ISL_2549233, EPI_ISL_2549234, EPI_ISL_2549235, EPI_ISL_2549236, EPI_ISL_2549237, EPI_ISL_2549238, EPI_ISL_2549239, EPI_ISL_2549240, EPI_ISL_2549241, EPI_ISL_2549242, EPI_ISL_2549243, EPI_ISL_2549245, EPI_ISL_2549246, EPI_ISL_2549247, EPI_ISL_2549248, EPI_ISL_2549249, EPI_ISL_2549250, EPI_ISL_2549251, EPI_ISL_2549252, EPI_ISL_2549254, EPI_ISL_2549255, EPI_ISL_2549257, EPI_ISL_2549259, EPI_ISL_2549260, EPI_ISL_2549261, EPI_ISL_2549263, EPI_ISL_2549265, EPI_ISL_2549266, EPI_ISL_2549267, EPI_ISL_2549268, EPI_ISL_2549270, EPI_ISL_2549271, EPI_ISL_2549272, EPI_ISL_2549273, EPI_ISL_2549274, EPI_ISL_2549275, EPI_ISL_2549276, EPI_ISL_2549277, EPI_ISL_2549278, EPI_ISL_2549279, EPI_ISL_2549280, EPI_ISL_2549281, EPI_ISL_2549282, EPI_ISL_2549283, EPI_ISL_2549285 | | | |
| see above | Indian Council of Medical Research-National Institute of Virology, Microbial Containment Complex | Indian Council of Medical Research-National Institute of Virology, Microbial Containment Complex | Pragya D. Yadav |
| EPI_ISL_2549327, EPI_ISL_2549331, EPI_ISL_2549332, EPI_ISL_2549334, EPI_ISL_2549336, EPI_ISL_2549340, EPI_ISL_2549342, EPI_ISL_2549344, EPI_ISL_2549345, EPI_ISL_2549352, EPI_ISL_2549356, EPI_ISL_2549365, EPI_ISL_2549374, EPI_ISL_2549376, EPI_ISL_2549379, EPI_ISL_2549387, EPI_ISL_2549390, EPI_ISL_2549392, EPI_ISL_2549394, EPI_ISL_2549400, EPI_ISL_2549407, EPI_ISL_2549411, EPI_ISL_2549414, EPI_ISL_2549422, EPI_ISL_2549423, EPI_ISL_2549428, EPI_ISL_2549437, EPI_ISL_2549441, EPI_ISL_2549444, EPI_ISL_2549445, EPI_ISL_2549448, EPI_ISL_2549460, EPI_ISL_2549465, EPI_ISL_2549466, EPI_ISL_2549473, EPI_ISL_2549485, EPI_ISL_2549488, EPI_ISL_2549492, EPI_ISL_2549498, EPI_ISL_2549518, EPI_ISL_2549522, EPI_ISL_2549526, EPI_ISL_2549529, EPI_ISL_2549530, EPI_ISL_2549552 | | | |
| see above | UW Virology Lab | UW Virology Lab | Pavitra Roychoudhury, Hong Xie, Lasata Shrestha, Tien V. Nguyen, Shah Mohamed Bakhash, Michelle Lin, Noah R. Baker, Ricardo Perez, Sean Ellis, Nathan Breit, Robert J. Livingston, Meei-Li Huang, Keith R Jerome, Patrick Mathias, Alexander Greninger |
| EPI_ISL_2549562 | Gribbles Pathology | Institute for Medical Research, Infectious Disease Research Centre, National Institutes of Health, Ministry of Health Malaysia | Suppiah J, Kamel K, Mohd Zawawi Z, Azizan MA, Ramly N, Robert F, Thayan R |
| EPI_ISL_2549576 | Hospital Tuanku Fauziah | Institute for Medical Research, Infectious Disease Research Centre, National Institutes of Health, Ministry of Health Malaysia | Suppiah J, Kamel K, Mohd Zawawi Z, Azizan MA, Ramly N, Robert F, Thayan R |
| EPI_ISL_2549741 | Labuan Hospital | Institute for Medical Research, Infectious Disease Research Centre, National Institutes of Health, Ministry of Health Malaysia | Suppiah J, Kamel K, Mohd Zawawi Z, Azizan MA, Ramly N, Robert F, Thayan R |
| EPI_ISL_2549853, EPI_ISL_2549906, EPI_ISL_2549910, EPI_ISL_2549911, EPI_ISL_2549912, EPI_ISL_2549929, EPI_ISL_2550012, EPI_ISL_2550019, EPI_ISL_2550021, EPI_ISL_2550026, EPI_ISL_2550031, EPI_ISL_2550048, EPI_ISL_2550054, EPI_ISL_2550108, EPI_ISL_2550134, EPI_ISL_2550143, EPI_ISL_2550152, EPI_ISL_2550153, EPI_ISL_2550154, EPI_ISL_2550155, EPI_ISL_2550157, EPI_ISL_2550264, EPI_ISL_2550265, EPI_ISL_2550312, EPI_ISL_2550313, EPI_ISL_2550314, EPI_ISL_2550315, EPI_ISL_2550318, EPI_ISL_2550319, EPI_ISL_2550320, EPI_ISL_2550373, EPI_ISL_2550381, EPI_ISL_2550382, EPI_ISL_2550385, EPI_ISL_2550388, EPI_ISL_2550398, EPI_ISL_2550405, EPI_ISL_2550407, EPI_ISL_2550408, EPI_ISL_2550441, EPI_ISL_2550442, EPI_ISL_2550448, EPI_ISL_2550449, EPI_ISL_2550463, EPI_ISL_2550464, EPI_ISL_2550468, EPI_ISL_2550475, EPI_ISL_2550505 | | | |
| see above | Colorado Department of Public Health and Environment | Colorado Department of Public Health and Environment | Laura Bankers, Molly C. Hetherington-Rauth, Diana Ir, Alexandria Rossheim, Shannon R. Matzinger, Sarah Elizabeth Totten, Emily A. Travanty |
| EPI_ISL_2550605 | NB-Hôpital Georges L. Dumont | National Microbiology Laboratory (NML) | Anna Majer, Shari Tyson, Grace Seo, Philip Mabon, Elsie Grudeski, Rhiannon Huzarewich, Russell Mandes, Anneliese Landgraff, Jennifer Tanner, Natalie Knox, Morag Graham, Gary Van Domselaar, Richard Garceau, Guillaume Desnoyers, Nathalie Bastien, Yan Li, Timothy Booth, Darian Hole, Madison Chapel, Kirsten Biggar, CanCOGeN's metadata curation team, Public Health Agency of Canada CanCOGeN team |
| EPI_ISL_2550614, EPI_ISL_2550616, EPI_ISL_2550640, EPI_ISL_2550649, EPI_ISL_2550655, EPI_ISL_2550678, EPI_ISL_2550687, EPI_ISL_2550688, EPI_ISL_2550689, EPI_ISL_2550690, EPI_ISL_2550695, EPI_ISL_2550701, EPI_ISL_2550704, EPI_ISL_2550705 | | | |
| see above | Azienda Sanitaria dell'Alto Adige Laboratorio Aziendale di Microbiologia e Virologia | Istituto di Genomica Applicata | Elisabetta Pagani, Irene Bianconi, Elisabetta Giacobazzi, Elisa Masi, Stefanie Wieser, Irena Jurman, Vera Vendramin, Gabriele Magris, Eleonora Paparelli, Davide Scaglione, Michele Morgante |
| EPI_ISL_2550714 | Johor Bahru Public Health Laboratory | Institute for Medical Research, Infectious Disease Research Centre, National Institutes of Health, Ministry of Health Malaysia | Suppiah J, Kamel K, Mohd Zawawi Z, Azizan MA, Ramly N, Robert F, Thayan R |
| EPI_ISL_2550727 | Laboratorio analisi, IRCCS Policlinico San Donato | Laboratory of Clinical Microbiology, Virology and Bioemergencies, ASST Fatebenefratelli Sacco - Sacco Hospital | Valeria Micheli, Alessandro Mancon, Alberto Rizzo, Fiorenza Bracchitta, Maria Rita Gismondo |
| EPI_ISL_2550728 | Unità Operativa Laboratorio di Analisi Chimico Cliniche e Microbiologiche, IRCCS Istituto Ortopedico Galeazzi | Laboratory of Clinical Microbiology, Virology and Bioemergencies, ASST Fatebenefratelli Sacco - Sacco Hospital | Valeria Micheli, Alessandro Mancon, Alberto Rizzo, Fiorenza Bracchitta, Maria Rita Gismondo |
| EPI_ISL_2550731 | Johor Bahru Public Health Laboratory | Institute for Medical Research, Infectious Disease Research Centre, National Institutes of Health, Ministry of Health Malaysia | Suppiah J, Kamel K, Mohd Zawawi Z, Azizan MA, Ramly N, Robert F, Thayan R |
| EPI_ISL_2550733 | Hospital Labuan | Institute for Medical Research, Infectious Disease Research Centre, National Institutes of Health, Ministry of Health Malaysia | Suppiah J, Kamel K, Mohd Zawawi Z, Azizan MA, Ramly N, Robert F, Thayan R |
| EPI_ISL_2550746, EPI_ISL_2550750 | Salud Digna | Instituto Nacional de Medicina Genomica | Hidalgo-Miranda A, Cedro-Tanda A, Mendoza-Vargas A, Reyes-Grajeda JP, Abraham Campos-Romero, Moreno-Camacho José Luis, Rodriguez-Gallegos Jorge, Luna-Ruiz Marco, Gonzalez-Barrera D, Rangel-DeLeon D, Munguia-Garza P, Ramirez-Vega O, Escobar-Arrazola, M, Herrera-Montalvo LA. |
| EPI_ISL_2550758, EPI_ISL_2550759 | Port Dickson Hospital | Institute for Medical Research, Infectious Disease Research Centre, National Institutes of Health, Ministry of Health Malaysia | Suppiah J, Kamel K, Mohd Zawawi Z, Azizan MA, Ramly N, Robert F, Thayan R |
| EPI_ISL_2550778 | Maryland Public Health Laboratory (MD PHL) | Maryland Public Health Laboratory (MD PHL) | Maryland Department of Health Laboratories Administration |
| EPI_ISL_2551170 | Wyoming Public Health Laboratory | Wyoming Public Health Laboratory | Jim Mildenberger, Taylor Fearing, Channing Weber, Ashley Norberg, Chayse Rowley, Marley Goetz, Brian Dominguez, Elliot Thomasson, Sam Britz, Cari Sloma, Robert Petit, and Rob Christensen |
| EPI_ISL_2551183 | Department of Microbiology, Tochigi Prefectural Institute of Public Health and Environmental Science | Genome Analysis Center, Kamma Memorial Hospital | Fuminori Mizukoshi, Yoshiko Hitomi, Hanako Yazawa, Jun Ishii, Satoko Soma, Kaori Watanabe, Takuya Yazawa, Michiaki Masuda, Hiroshi Kamma |
| EPI_ISL_2551452 | Providence Regional Medical Center Everett | Providence St. Joseph Health Molecular Genomics Laboratory | Alexa K Dowdell, Brian D Piening, Fred L Robinson, Carlo B Bifulco, Mary Campbell, Rogan Rattray |
| EPI_ISL_2551519, EPI_ISL_2551520 | Department of Microbiology, Tochigi Prefectural Institute of Public Health and Environmental Science | Genome Analysis Center, Kamma Memorial Hospital | Fuminori Mizukoshi, Yoshiko Hitomi, Hanako Yazawa, Jun Ishii, Satoko Soma, Kaori Watanabe, Takuya Yazawa, Michiaki Masuda, Hiroshi Kamma |
| EPI_ISL_2551582 | Tochigi Prefectural Institute of Environment and Health | Genome Analysis Center, Kamma Memorial Hospital | Fuminori Mizukoshi, Yoshiko Hitomi, Hanako Yazawa, Jun Ishii, Satoko Soma, Kaori Watanabe, Takuya Yazawa, Michiaki Masuda, Hiroshi Kamma |
| EPI_ISL_2551583, EPI_ISL_2551584, EPI_ISL_2551585, EPI_ISL_2551586 | Lighthouse Lab in Alderley Park | Wellcome Sanger Institute for the COVID-19 Genomics UK (COG-UK) Consortium | Jacquelyn Wynn, Mairead Hyland, The Lighthouse Lab in Alderley Park and Alex Alderton, Roberto Amato, Jeffrey Barrett, Sonia Goncalves, Ewan Harrison, David K. Jackson, Ian Johnston, Dominic Kwiatkowski, Cordelia Langford, John Sillitoe on behalf of the Wellcome Sanger Institute COVID-19 Surveillance Team |
| EPI_ISL_2551587 | Lighthouse Lab in Milton Keynes | Wellcome Sanger Institute for the COVID-19 Genomics UK (COG-UK) Consortium | The Lighthouse Lab in Milton Keynes and Alex Alderton, Roberto Amato, Jeffrey Barrett, Sonia Goncalves, Ewan Harrison, David K. Jackson, Ian Johnston, Dominic Kwiatkowski, Cordelia Langford, John Sillitoe on behalf of the Wellcome Sanger Institute COVID-19 Surveillance Team |
| EPI_ISL_2551588, EPI_ISL_2551589, EPI_ISL_2551590, EPI_ISL_2551591, EPI_ISL_2551592, EPI_ISL_2551593, EPI_ISL_2551594, EPI_ISL_2551595, EPI_ISL_2551596, EPI_ISL_2551597, EPI_ISL_2551598, EPI_ISL_2551599, EPI_ISL_2551600, EPI_ISL_2551602, EPI_ISL_2551603 | | | |
| see above | Lighthouse Lab in Alderley Park | Wellcome Sanger Institute for the COVID-19 Genomics UK (COG-UK) Consortium | Jacquelyn Wynn, Mairead Hyland, The Lighthouse Lab in Alderley Park and Alex Alderton, Roberto Amato, Jeffrey Barrett, Sonia Goncalves, Ewan Harrison, David K. Jackson, Ian Johnston, Dominic Kwiatkowski, Cordelia Langford, John Sillitoe on behalf of the Wellcome Sanger Institute COVID-19 Surveillance Team |
| EPI_ISL_2551604 | Lighthouse Lab in Milton Keynes | Wellcome Sanger Institute for the COVID-19 Genomics UK (COG-UK) Consortium | The Lighthouse Lab in Milton Keynes and Alex Alderton, Roberto Amato, Jeffrey Barrett, Sonia Goncalves, Ewan Harrison, David K. Jackson, Ian Johnston, Dominic Kwiatkowski, Cordelia Langford, John Sillitoe on behalf of the Wellcome Sanger Institute COVID-19 Surveillance Team |
| EPI_ISL_2551605, EPI_ISL_2551606, EPI_ISL_2551607, EPI_ISL_2551608, EPI_ISL_2551610, EPI_ISL_2551611, EPI_ISL_2551612, EPI_ISL_2551613 | Lighthouse Lab in Alderley Park | Wellcome Sanger Institute for the COVID-19 Genomics UK (COG-UK) Consortium | Jacquelyn Wynn, Mairead Hyland, The Lighthouse Lab in Alderley Park and Alex Alderton, Roberto Amato, Jeffrey Barrett, Sonia Goncalves, Ewan Harrison, David K. Jackson, Ian Johnston, Dominic Kwiatkowski, Cordelia Langford, John Sillitoe on behalf of the Wellcome Sanger Institute COVID-19 Surveillance Team |
| EPI_ISL_2551614, EPI_ISL_2551615 | Lighthouse Lab in Milton Keynes | Wellcome Sanger Institute for the COVID-19 Genomics UK (COG-UK) Consortium | The Lighthouse Lab in Milton Keynes and Alex Alderton, Roberto Amato, Jeffrey Barrett, Sonia Goncalves, Ewan Harrison, David K. Jackson, Ian Johnston, Dominic Kwiatkowski, Cordelia Langford, John Sillitoe on behalf of the Wellcome Sanger Institute COVID-19 Surveillance Team |
| EPI_ISL_2551616, EPI_ISL_2551617, EPI_ISL_2551618 | Lighthouse Lab in Alderley Park | Wellcome Sanger Institute for the COVID-19 Genomics UK (COG-UK) Consortium | Jacquelyn Wynn, Mairead Hyland, The Lighthouse Lab in Alderley Park and Alex Alderton, Roberto Amato, Jeffrey Barrett, Sonia Goncalves, Ewan Harrison, David K. Jackson, Ian Johnston, Dominic Kwiatkowski, Cordelia Langford, John Sillitoe on behalf of the Wellcome Sanger Institute COVID-19 Surveillance Team |
| EPI_ISL_2551619 | Lighthouse Lab in Milton Keynes | Wellcome Sanger Institute for the COVID-19 Genomics UK (COG-UK) Consortium | The Lighthouse Lab in Milton Keynes and Alex Alderton, Roberto Amato, Jeffrey Barrett, Sonia Goncalves, Ewan Harrison, David K. Jackson, Ian Johnston, Dominic Kwiatkowski, Cordelia Langford, John Sillitoe on behalf of the Wellcome Sanger Institute COVID-19 Surveillance Team |

| | | | |
|---|---|---|---|
| EPI_ISL_2551620 | Lighthouse Lab in Alderley Park | Wellcome Sanger Institute for the COVID-19 Genomics UK (COG-UK) Consortium | Jacquelyn Wynn, Mairead Hyland, The Lighthouse Lab in Alderley Park and Alex Alderton, Roberto Amato, Jeffrey Barrett, Sonia Goncalves, Ewan Harrison, David K. Jackson, Ian Johnston, Dominic Kwiatkowski, Cordelia Langford, John Sillitoe on behalf of the Wellcome Sanger Institute COVID-19 Surveillance Team |
| EPI_ISL_2551621 | Lighthouse Lab in Milton Keynes | Wellcome Sanger Institute for the COVID-19 Genomics UK (COG-UK) Consortium | The Lighthouse Lab in Milton Keynes and Alex Alderton, Roberto Amato, Jeffrey Barrett, Sonia Goncalves, Ewan Harrison, David K. Jackson, Ian Johnston, Dominic Kwiatkowski, Cordelia Langford, John Sillitoe on behalf of the Wellcome Sanger Institute COVID-19 Surveillance Team |
| EPI_ISL_2551622, EPI_ISL_2551623 | Lighthouse Lab in Alderley Park | Wellcome Sanger Institute for the COVID-19 Genomics UK (COG-UK) Consortium | Jacquelyn Wynn, Mairead Hyland, The Lighthouse Lab in Alderley Park and Alex Alderton, Roberto Amato, Jeffrey Barrett, Sonia Goncalves, Ewan Harrison, David K. Jackson, Ian Johnston, Dominic Kwiatkowski, Cordelia Langford, John Sillitoe on behalf of the Wellcome Sanger Institute COVID-19 Surveillance Team |
| EPI_ISL_2551624 | Lighthouse Lab in Milton Keynes | Wellcome Sanger Institute for the COVID-19 Genomics UK (COG-UK) Consortium | The Lighthouse Lab in Milton Keynes and Alex Alderton, Roberto Amato, Jeffrey Barrett, Sonia Goncalves, Ewan Harrison, David K. Jackson, Ian Johnston, Dominic Kwiatkowski, Cordelia Langford, John Sillitoe on behalf of the Wellcome Sanger Institute COVID-19 Surveillance Team |
| EPI_ISL_2551625, EPI_ISL_2551626, EPI_ISL_2551627 | Lighthouse Lab in Alderley Park | Wellcome Sanger Institute for the COVID-19 Genomics UK (COG-UK) Consortium | Jacquelyn Wynn, Mairead Hyland, The Lighthouse Lab in Alderley Park and Alex Alderton, Roberto Amato, Jeffrey Barrett, Sonia Goncalves, Ewan Harrison, David K. Jackson, Ian Johnston, Dominic Kwiatkowski, Cordelia Langford, John Sillitoe on behalf of the Wellcome Sanger Institute COVID-19 Surveillance Team |
| EPI_ISL_2551628, EPI_ISL_2551630 | Lighthouse Lab in Milton Keynes | Wellcome Sanger Institute for the COVID-19 Genomics UK (COG-UK) Consortium | The Lighthouse Lab in Milton Keynes and Alex Alderton, Roberto Amato, Jeffrey Barrett, Sonia Goncalves, Ewan Harrison, David K. Jackson, Ian Johnston, Dominic Kwiatkowski, Cordelia Langford, John Sillitoe on behalf of the Wellcome Sanger Institute COVID-19 Surveillance Team |
| EPI_ISL_2551631 | Lighthouse Lab in Alderley Park | Wellcome Sanger Institute for the COVID-19 Genomics UK (COG-UK) Consortium | Jacquelyn Wynn, Mairead Hyland, The Lighthouse Lab in Alderley Park and Alex Alderton, Roberto Amato, Jeffrey Barrett, Sonia Goncalves, Ewan Harrison, David K. Jackson, Ian Johnston, Dominic Kwiatkowski, Cordelia Langford, John Sillitoe on behalf of the Wellcome Sanger Institute COVID-19 Surveillance Team |
| EPI_ISL_2551632 | Lighthouse Lab in Milton Keynes | Wellcome Sanger Institute for the COVID-19 Genomics UK (COG-UK) Consortium | The Lighthouse Lab in Milton Keynes and Alex Alderton, Roberto Amato, Jeffrey Barrett, Sonia Goncalves, Ewan Harrison, David K. Jackson, Ian Johnston, Dominic Kwiatkowski, Cordelia Langford, John Sillitoe on behalf of the Wellcome Sanger Institute COVID-19 Surveillance Team |
| EPI_ISL_2551633, EPI_ISL_2551634, EPI_ISL_2551635, EPI_ISL_2551636, EPI_ISL_2551637, EPI_ISL_2551638, EPI_ISL_2551640, EPI_ISL_2551641, EPI_ISL_2551642, EPI_ISL_2551643, EPI_ISL_2551644, EPI_ISL_2551645, EPI_ISL_2551646, EPI_ISL_2551647 | | | |
| see above | Lighthouse Lab in Alderley Park | Wellcome Sanger Institute for the COVID-19 Genomics UK (COG-UK) Consortium | Jacquelyn Wynn, Mairead Hyland, The Lighthouse Lab in Alderley Park and Alex Alderton, Roberto Amato, Jeffrey Barrett, Sonia Goncalves, Ewan Harrison, David K. Jackson, Ian Johnston, Dominic Kwiatkowski, Cordelia Langford, John Sillitoe on behalf of the Wellcome Sanger Institute COVID-19 Surveillance Team |
| EPI_ISL_2551648 | Lighthouse Lab in Milton Keynes | Wellcome Sanger Institute for the COVID-19 Genomics UK (COG-UK) Consortium | The Lighthouse Lab in Milton Keynes and Alex Alderton, Roberto Amato, Jeffrey Barrett, Sonia Goncalves, Ewan Harrison, David K. Jackson, Ian Johnston, Dominic Kwiatkowski, Cordelia Langford, John Sillitoe on behalf of the Wellcome Sanger Institute COVID-19 Surveillance Team |
| EPI_ISL_2551649 | Lighthouse Lab in Alderley Park | Wellcome Sanger Institute for the COVID-19 Genomics UK (COG-UK) Consortium | Jacquelyn Wynn, Mairead Hyland, The Lighthouse Lab in Alderley Park and Alex Alderton, Roberto Amato, Jeffrey Barrett, Sonia Goncalves, Ewan Harrison, David K. Jackson, Ian Johnston, Dominic Kwiatkowski, Cordelia Langford, John Sillitoe on behalf of the Wellcome Sanger Institute COVID-19 Surveillance Team |
| EPI_ISL_2551650 | Lighthouse Lab in Milton Keynes | Wellcome Sanger Institute for the COVID-19 Genomics UK (COG-UK) Consortium | The Lighthouse Lab in Milton Keynes and Alex Alderton, Roberto Amato, Jeffrey Barrett, Sonia Goncalves, Ewan Harrison, David K. Jackson, Ian Johnston, Dominic Kwiatkowski, Cordelia Langford, John Sillitoe on behalf of the Wellcome Sanger Institute COVID-19 Surveillance Team |
| EPI_ISL_2551651, EPI_ISL_2551652 | Lighthouse Lab in Alderley Park | Wellcome Sanger Institute for the COVID-19 Genomics UK (COG-UK) Consortium | Jacquelyn Wynn, Mairead Hyland, The Lighthouse Lab in Alderley Park and Alex Alderton, Roberto Amato, Jeffrey Barrett, Sonia Goncalves, Ewan Harrison, David K. Jackson, Ian Johnston, Dominic Kwiatkowski, Cordelia Langford, John Sillitoe on behalf of the Wellcome Sanger Institute COVID-19 Surveillance Team |
| EPI_ISL_2551653 | Lighthouse Lab in Milton Keynes | Wellcome Sanger Institute for the COVID-19 Genomics UK (COG-UK) Consortium | The Lighthouse Lab in Milton Keynes and Alex Alderton, Roberto Amato, Jeffrey Barrett, Sonia Goncalves, Ewan Harrison, David K. Jackson, Ian Johnston, Dominic Kwiatkowski, Cordelia Langford, John Sillitoe on behalf of the Wellcome Sanger Institute COVID-19 Surveillance Team |
| EPI_ISL_2551654, EPI_ISL_2551655, EPI_ISL_2551658, EPI_ISL_2551659, EPI_ISL_2551660, EPI_ISL_2551661, EPI_ISL_2551662, EPI_ISL_2551663 | Lighthouse Lab in Alderley Park | Wellcome Sanger Institute for the COVID-19 Genomics UK (COG-UK) Consortium | Jacquelyn Wynn, Mairead Hyland, The Lighthouse Lab in Alderley Park and Alex Alderton, Roberto Amato, Jeffrey Barrett, Sonia Goncalves, Ewan Harrison, David K. Jackson, Ian Johnston, Dominic Kwiatkowski, Cordelia Langford, John Sillitoe on behalf of the Wellcome Sanger Institute COVID-19 Surveillance Team |
| EPI_ISL_2551664, EPI_ISL_2551665, EPI_ISL_2551666, EPI_ISL_2551667, EPI_ISL_2551668 | Lighthouse Lab in Milton Keynes | Wellcome Sanger Institute for the COVID-19 Genomics UK (COG-UK) Consortium | The Lighthouse Lab in Milton Keynes and Alex Alderton, Roberto Amato, Jeffrey Barrett, Sonia Goncalves, Ewan Harrison, David K. Jackson, Ian Johnston, Dominic Kwiatkowski, Cordelia Langford, John Sillitoe on behalf of the Wellcome Sanger Institute COVID-19 Surveillance Team |
| EPI_ISL_2551669, EPI_ISL_2551670 | Lighthouse Lab in Alderley Park | Wellcome Sanger Institute for the COVID-19 Genomics UK (COG-UK) Consortium | Jacquelyn Wynn, Mairead Hyland, The Lighthouse Lab in Alderley Park and Alex Alderton, Roberto Amato, Jeffrey Barrett, Sonia Goncalves, Ewan Harrison, David K. Jackson, Ian Johnston, Dominic Kwiatkowski, Cordelia Langford, John Sillitoe on behalf of the Wellcome Sanger Institute COVID-19 Surveillance Team |
| EPI_ISL_2551671 | Lighthouse Lab in Milton Keynes | Wellcome Sanger Institute for the COVID-19 Genomics UK (COG-UK) Consortium | The Lighthouse Lab in Milton Keynes and Alex Alderton, Roberto Amato, Jeffrey Barrett, Sonia Goncalves, Ewan Harrison, David K. Jackson, Ian Johnston, Dominic Kwiatkowski, Cordelia Langford, John Sillitoe on behalf of the Wellcome Sanger Institute COVID-19 Surveillance Team |
| EPI_ISL_2551672, EPI_ISL_2551673, EPI_ISL_2551674, EPI_ISL_2551675 | Lighthouse Lab in Alderley Park | Wellcome Sanger Institute for the COVID-19 Genomics UK (COG-UK) Consortium | Jacquelyn Wynn, Mairead Hyland, The Lighthouse Lab in Alderley Park and Alex Alderton, Roberto Amato, Jeffrey Barrett, Sonia Goncalves, Ewan Harrison, David K. Jackson, Ian Johnston, Dominic Kwiatkowski, Cordelia Langford, John Sillitoe on behalf of the Wellcome Sanger Institute COVID-19 Surveillance Team |
| EPI_ISL_2551676 | Lighthouse Lab in Milton Keynes | Wellcome Sanger Institute for the COVID-19 Genomics UK (COG-UK) Consortium | The Lighthouse Lab in Milton Keynes and Alex Alderton, Roberto Amato, Jeffrey Barrett, Sonia Goncalves, Ewan Harrison, David K. Jackson, Ian Johnston, Dominic Kwiatkowski, Cordelia Langford, John Sillitoe on behalf of the Wellcome Sanger Institute COVID-19 Surveillance Team |
| EPI_ISL_2551677, EPI_ISL_2551678, EPI_ISL_2551679, EPI_ISL_2551680, EPI_ISL_2551681 | Lighthouse Lab in Alderley Park | Wellcome Sanger Institute for the COVID-19 Genomics UK (COG-UK) Consortium | Jacquelyn Wynn, Mairead Hyland, The Lighthouse Lab in Alderley Park and Alex Alderton, Roberto Amato, Jeffrey Barrett, Sonia Goncalves, Ewan Harrison, David K. Jackson, Ian Johnston, Dominic Kwiatkowski, Cordelia Langford, John Sillitoe on behalf of the Wellcome Sanger Institute COVID-19 Surveillance Team |
| EPI_ISL_2551682, EPI_ISL_2551683 | Lighthouse Lab in Milton Keynes | Wellcome Sanger Institute for the COVID-19 Genomics UK (COG-UK) Consortium | The Lighthouse Lab in Milton Keynes and Alex Alderton, Roberto Amato, Jeffrey Barrett, Sonia Goncalves, Ewan Harrison, David K. Jackson, Ian Johnston, Dominic Kwiatkowski, Cordelia Langford, John Sillitoe on behalf of the Wellcome Sanger Institute COVID-19 Surveillance Team |
| EPI_ISL_2551684, EPI_ISL_2551686, EPI_ISL_2551687, EPI_ISL_2551688, EPI_ISL_2551689, EPI_ISL_2551690, EPI_ISL_2551691, EPI_ISL_2551692, EPI_ISL_2551693 | Lighthouse Lab in Alderley Park | Wellcome Sanger Institute for the COVID-19 Genomics UK (COG-UK) Consortium | Jacquelyn Wynn, Mairead Hyland, The Lighthouse Lab in Alderley Park and Alex Alderton, Roberto Amato, Jeffrey Barrett, Sonia Goncalves, Ewan Harrison, David K. Jackson, Ian Johnston, Dominic Kwiatkowski, Cordelia Langford, John Sillitoe on behalf of the Wellcome Sanger Institute COVID-19 Surveillance Team |
| EPI_ISL_2551694 | Lighthouse Lab in Milton Keynes | Wellcome Sanger Institute for the COVID-19 Genomics UK (COG-UK) Consortium | The Lighthouse Lab in Milton Keynes and Alex Alderton, Roberto Amato, Jeffrey Barrett, Sonia Goncalves, Ewan Harrison, David K. Jackson, Ian Johnston, Dominic Kwiatkowski, Cordelia Langford, John Sillitoe on behalf of the Wellcome Sanger Institute COVID-19 Surveillance Team |
| EPI_ISL_2551696, EPI_ISL_2551697 | Lighthouse Lab in Alderley Park | Wellcome Sanger Institute for the COVID-19 Genomics UK (COG-UK) Consortium | Jacquelyn Wynn, Mairead Hyland, The Lighthouse Lab in Alderley Park and Alex Alderton, Roberto Amato, Jeffrey Barrett, Sonia Goncalves, Ewan Harrison, David K. Jackson, Ian Johnston, Dominic Kwiatkowski, Cordelia Langford, John Sillitoe on behalf of the Wellcome Sanger Institute COVID-19 Surveillance Team |
| EPI_ISL_2551698, EPI_ISL_2551699, EPI_ISL_2551700, EPI_ISL_2551701 | Lighthouse Lab in Milton Keynes | Wellcome Sanger Institute for the COVID-19 Genomics UK (COG-UK) Consortium | The Lighthouse Lab in Milton Keynes and Alex Alderton, Roberto Amato, Jeffrey Barrett, Sonia Goncalves, Ewan Harrison, David K. Jackson, Ian Johnston, Dominic Kwiatkowski, Cordelia Langford, John Sillitoe on behalf of the Wellcome Sanger Institute COVID-19 Surveillance Team |
| EPI_ISL_2551702, EPI_ISL_2551703, EPI_ISL_2551704 | Lighthouse Lab in Alderley Park | Wellcome Sanger Institute for the COVID-19 Genomics UK (COG-UK) Consortium | Jacquelyn Wynn, Mairead Hyland, The Lighthouse Lab in Alderley Park and Alex Alderton, Roberto Amato, Jeffrey Barrett, Sonia Goncalves, Ewan Harrison, David K. Jackson, Ian Johnston, Dominic Kwiatkowski, Cordelia Langford, John Sillitoe on behalf of the Wellcome Sanger Institute COVID-19 Surveillance Team |
| EPI_ISL_2551705 | Lighthouse Lab in Milton Keynes | Wellcome Sanger Institute for the COVID-19 Genomics UK (COG-UK) Consortium | The Lighthouse Lab in Milton Keynes and Alex Alderton, Roberto Amato, Jeffrey Barrett, Sonia Goncalves, Ewan Harrison, David K. Jackson, Ian Johnston, Dominic Kwiatkowski, Cordelia Langford, John Sillitoe on behalf of the Wellcome Sanger Institute COVID-19 Surveillance Team |
| EPI_ISL_2551706 | Lighthouse Lab in Alderley Park | Wellcome Sanger Institute for the COVID-19 Genomics UK (COG-UK) Consortium | Jacquelyn Wynn, Mairead Hyland, The Lighthouse Lab in Alderley Park and Alex Alderton, Roberto Amato, Jeffrey Barrett, Sonia Goncalves, Ewan Harrison, David K. Jackson, Ian Johnston, Dominic Kwiatkowski, Cordelia Langford, John Sillitoe on behalf of the Wellcome Sanger Institute COVID-19 Surveillance Team |
| EPI_ISL_2551707 | Lighthouse Lab in Milton Keynes | Wellcome Sanger Institute for the COVID-19 Genomics UK (COG-UK) Consortium | The Lighthouse Lab in Milton Keynes and Alex Alderton, Roberto Amato, Jeffrey Barrett, Sonia Goncalves, Ewan Harrison, David K. Jackson, Ian Johnston, Dominic Kwiatkowski, Cordelia Langford, John Sillitoe on behalf of the Wellcome Sanger Institute COVID-19 Surveillance Team |
| EPI_ISL_2551708, EPI_ISL_2551709, EPI_ISL_2551710 | Lighthouse Lab in Alderley Park | Wellcome Sanger Institute for the COVID-19 Genomics UK (COG-UK) Consortium | Jacquelyn Wynn, Mairead Hyland, The Lighthouse Lab in Alderley Park and Alex Alderton, Roberto Amato, Jeffrey Barrett, Sonia Goncalves, Ewan Harrison, David K. Jackson, Ian Johnston, Dominic Kwiatkowski, Cordelia Langford, John Sillitoe on behalf of the Wellcome Sanger Institute COVID-19 Surveillance Team |
| EPI_ISL_2551711, EPI_ISL_2551712, EPI_ISL_2551713 | Lighthouse Lab in Milton Keynes | Wellcome Sanger Institute for the COVID-19 Genomics UK (COG-UK) Consortium | The Lighthouse Lab in Milton Keynes and Alex Alderton, Roberto Amato, Jeffrey Barrett, Sonia Goncalves, Ewan Harrison, David K. Jackson, Ian Johnston, Dominic Kwiatkowski, Cordelia Langford, John Sillitoe on behalf of the Wellcome Sanger Institute COVID-19 Surveillance Team |
| EPI_ISL_2551714, EPI_ISL_2551715, EPI_ISL_2551716 | Lighthouse Lab in Alderley Park | Wellcome Sanger Institute for the COVID-19 Genomics UK (COG-UK) Consortium | Jacquelyn Wynn, Mairead Hyland, The Lighthouse Lab in Alderley Park and Alex Alderton, Roberto Amato, Jeffrey Barrett, Sonia Goncalves, Ewan Harrison, David K. Jackson, Ian Johnston, Dominic Kwiatkowski, Cordelia Langford, John Sillitoe on behalf of the Wellcome Sanger Institute COVID-19 Surveillance Team |

| | | | |
|---|---|---|---|
| EPI_ISL_2551717 | Lighthouse Lab in Milton Keynes | Wellcome Sanger Institute for the COVID-19 Genomics UK (COG-UK) Consortium | The Lighthouse Lab in Milton Keynes and Alex Alderton, Roberto Amato, Jeffrey Barrett, Sonia Goncalves, Ewan Harrison, David K. Jackson, Ian Johnston, Dominic Kwiatkowski, Cordelia Langford, John Sillitoe on behalf of the Wellcome Sanger Institute COVID-19 Surveillance Team |
| EPI_ISL_2551718, EPI_ISL_2551719 | Lighthouse Lab in Alderley Park | Wellcome Sanger Institute for the COVID-19 Genomics UK (COG-UK) Consortium | Jacquelyn Wynn, Mairead Hyland, The Lighthouse Lab in Alderley Park and Alex Alderton, Roberto Amato, Jeffrey Barrett, Sonia Goncalves, Ewan Harrison, David K. Jackson, Ian Johnston, Dominic Kwiatkowski, Cordelia Langford, John Sillitoe on behalf of the Wellcome Sanger Institute COVID-19 Surveillance Team |
| EPI_ISL_2551720 | Lighthouse Lab in Milton Keynes | Wellcome Sanger Institute for the COVID-19 Genomics UK (COG-UK) Consortium | The Lighthouse Lab in Milton Keynes and Alex Alderton, Roberto Amato, Jeffrey Barrett, Sonia Goncalves, Ewan Harrison, David K. Jackson, Ian Johnston, Dominic Kwiatkowski, Cordelia Langford, John Sillitoe on behalf of the Wellcome Sanger Institute COVID-19 Surveillance Team |
| EPI_ISL_2551721, EPI_ISL_2551723 | Lighthouse Lab in Alderley Park | Wellcome Sanger Institute for the COVID-19 Genomics UK (COG-UK) Consortium | Jacquelyn Wynn, Mairead Hyland, The Lighthouse Lab in Alderley Park and Alex Alderton, Roberto Amato, Jeffrey Barrett, Sonia Goncalves, Ewan Harrison, David K. Jackson, Ian Johnston, Dominic Kwiatkowski, Cordelia Langford, John Sillitoe on behalf of the Wellcome Sanger Institute COVID-19 Surveillance Team |
| EPI_ISL_2551724 | Lighthouse Lab in Milton Keynes | Wellcome Sanger Institute for the COVID-19 Genomics UK (COG-UK) Consortium | The Lighthouse Lab in Milton Keynes and Alex Alderton, Roberto Amato, Jeffrey Barrett, Sonia Goncalves, Ewan Harrison, David K. Jackson, Ian Johnston, Dominic Kwiatkowski, Cordelia Langford, John Sillitoe on behalf of the Wellcome Sanger Institute COVID-19 Surveillance Team |
| EPI_ISL_2551725, EPI_ISL_2551726, EPI_ISL_2551727 | Lighthouse Lab in Alderley Park | Wellcome Sanger Institute for the COVID-19 Genomics UK (COG-UK) Consortium | Jacquelyn Wynn, Mairead Hyland, The Lighthouse Lab in Alderley Park and Alex Alderton, Roberto Amato, Jeffrey Barrett, Sonia Goncalves, Ewan Harrison, David K. Jackson, Ian Johnston, Dominic Kwiatkowski, Cordelia Langford, John Sillitoe on behalf of the Wellcome Sanger Institute COVID-19 Surveillance Team |
| EPI_ISL_2551728 | Lighthouse Lab in Milton Keynes | Wellcome Sanger Institute for the COVID-19 Genomics UK (COG-UK) Consortium | The Lighthouse Lab in Milton Keynes and Alex Alderton, Roberto Amato, Jeffrey Barrett, Sonia Goncalves, Ewan Harrison, David K. Jackson, Ian Johnston, Dominic Kwiatkowski, Cordelia Langford, John Sillitoe on behalf of the Wellcome Sanger Institute COVID-19 Surveillance Team |
| EPI_ISL_2551729, EPI_ISL_2551730, EPI_ISL_2551731 | Lighthouse Lab in Alderley Park | Wellcome Sanger Institute for the COVID-19 Genomics UK (COG-UK) Consortium | Jacquelyn Wynn, Mairead Hyland, The Lighthouse Lab in Alderley Park and Alex Alderton, Roberto Amato, Jeffrey Barrett, Sonia Goncalves, Ewan Harrison, David K. Jackson, Ian Johnston, Dominic Kwiatkowski, Cordelia Langford, John Sillitoe on behalf of the Wellcome Sanger Institute COVID-19 Surveillance Team |
| EPI_ISL_2551732 | Lighthouse Lab in Milton Keynes | Wellcome Sanger Institute for the COVID-19 Genomics UK (COG-UK) Consortium | The Lighthouse Lab in Milton Keynes and Alex Alderton, Roberto Amato, Jeffrey Barrett, Sonia Goncalves, Ewan Harrison, David K. Jackson, Ian Johnston, Dominic Kwiatkowski, Cordelia Langford, John Sillitoe on behalf of the Wellcome Sanger Institute COVID-19 Surveillance Team |
| EPI_ISL_2551733 | Lighthouse Lab in Alderley Park | Wellcome Sanger Institute for the COVID-19 Genomics UK (COG-UK) Consortium | Jacquelyn Wynn, Mairead Hyland, The Lighthouse Lab in Alderley Park and Alex Alderton, Roberto Amato, Jeffrey Barrett, Sonia Goncalves, Ewan Harrison, David K. Jackson, Ian Johnston, Dominic Kwiatkowski, Cordelia Langford, John Sillitoe on behalf of the Wellcome Sanger Institute COVID-19 Surveillance Team |
| EPI_ISL_2551734 | Lighthouse Lab in Milton Keynes | Wellcome Sanger Institute for the COVID-19 Genomics UK (COG-UK) Consortium | The Lighthouse Lab in Milton Keynes and Alex Alderton, Roberto Amato, Jeffrey Barrett, Sonia Goncalves, Ewan Harrison, David K. Jackson, Ian Johnston, Dominic Kwiatkowski, Cordelia Langford, John Sillitoe on behalf of the Wellcome Sanger Institute COVID-19 Surveillance Team |
| EPI_ISL_2551735, EPI_ISL_2551736, EPI_ISL_2551737, EPI_ISL_2551738, EPI_ISL_2551739 | Lighthouse Lab in Alderley Park | Wellcome Sanger Institute for the COVID-19 Genomics UK (COG-UK) Consortium | Jacquelyn Wynn, Mairead Hyland, The Lighthouse Lab in Alderley Park and Alex Alderton, Roberto Amato, Jeffrey Barrett, Sonia Goncalves, Ewan Harrison, David K. Jackson, Ian Johnston, Dominic Kwiatkowski, Cordelia Langford, John Sillitoe on behalf of the Wellcome Sanger Institute COVID-19 Surveillance Team |
| EPI_ISL_2551740 | Lighthouse Lab in Milton Keynes | Wellcome Sanger Institute for the COVID-19 Genomics UK (COG-UK) Consortium | The Lighthouse Lab in Milton Keynes and Alex Alderton, Roberto Amato, Jeffrey Barrett, Sonia Goncalves, Ewan Harrison, David K. Jackson, Ian Johnston, Dominic Kwiatkowski, Cordelia Langford, John Sillitoe on behalf of the Wellcome Sanger Institute COVID-19 Surveillance Team |
| EPI_ISL_2551741, EPI_ISL_2551742, EPI_ISL_2551743, EPI_ISL_2551744, EPI_ISL_2551745 | Lighthouse Lab in Alderley Park | Wellcome Sanger Institute for the COVID-19 Genomics UK (COG-UK) Consortium | Jacquelyn Wynn, Mairead Hyland, The Lighthouse Lab in Alderley Park and Alex Alderton, Roberto Amato, Jeffrey Barrett, Sonia Goncalves, Ewan Harrison, David K. Jackson, Ian Johnston, Dominic Kwiatkowski, Cordelia Langford, John Sillitoe on behalf of the Wellcome Sanger Institute COVID-19 Surveillance Team |
| EPI_ISL_2551746, EPI_ISL_2551747 | Lighthouse Lab in Milton Keynes | Wellcome Sanger Institute for the COVID-19 Genomics UK (COG-UK) Consortium | The Lighthouse Lab in Milton Keynes and Alex Alderton, Roberto Amato, Jeffrey Barrett, Sonia Goncalves, Ewan Harrison, David K. Jackson, Ian Johnston, Dominic Kwiatkowski, Cordelia Langford, John Sillitoe on behalf of the Wellcome Sanger Institute COVID-19 Surveillance Team |
| EPI_ISL_2551748, EPI_ISL_2551749, EPI_ISL_2551750 | Lighthouse Lab in Alderley Park | Wellcome Sanger Institute for the COVID-19 Genomics UK (COG-UK) Consortium | Jacquelyn Wynn, Mairead Hyland, The Lighthouse Lab in Alderley Park and Alex Alderton, Roberto Amato, Jeffrey Barrett, Sonia Goncalves, Ewan Harrison, David K. Jackson, Ian Johnston, Dominic Kwiatkowski, Cordelia Langford, John Sillitoe on behalf of the Wellcome Sanger Institute COVID-19 Surveillance Team |
| EPI_ISL_2551751 | Lighthouse Lab in Milton Keynes | Wellcome Sanger Institute for the COVID-19 Genomics UK (COG-UK) Consortium | The Lighthouse Lab in Milton Keynes and Alex Alderton, Roberto Amato, Jeffrey Barrett, Sonia Goncalves, Ewan Harrison, David K. Jackson, Ian Johnston, Dominic Kwiatkowski, Cordelia Langford, John Sillitoe on behalf of the Wellcome Sanger Institute COVID-19 Surveillance Team |
| EPI_ISL_2551752, EPI_ISL_2551753, EPI_ISL_2551755, EPI_ISL_2551756 | Lighthouse Lab in Alderley Park | Wellcome Sanger Institute for the COVID-19 Genomics UK (COG-UK) Consortium | Jacquelyn Wynn, Mairead Hyland, The Lighthouse Lab in Alderley Park and Alex Alderton, Roberto Amato, Jeffrey Barrett, Sonia Goncalves, Ewan Harrison, David K. Jackson, Ian Johnston, Dominic Kwiatkowski, Cordelia Langford, John Sillitoe on behalf of the Wellcome Sanger Institute COVID-19 Surveillance Team |
| EPI_ISL_2551757, EPI_ISL_2551758 | Lighthouse Lab in Milton Keynes | Wellcome Sanger Institute for the COVID-19 Genomics UK (COG-UK) Consortium | The Lighthouse Lab in Milton Keynes and Alex Alderton, Roberto Amato, Jeffrey Barrett, Sonia Goncalves, Ewan Harrison, David K. Jackson, Ian Johnston, Dominic Kwiatkowski, Cordelia Langford, John Sillitoe on behalf of the Wellcome Sanger Institute COVID-19 Surveillance Team |
| EPI_ISL_2551759, EPI_ISL_2551760, EPI_ISL_2551761, EPI_ISL_2551762, EPI_ISL_2551763 | Lighthouse Lab in Alderley Park | Wellcome Sanger Institute for the COVID-19 Genomics UK (COG-UK) Consortium | Jacquelyn Wynn, Mairead Hyland, The Lighthouse Lab in Alderley Park and Alex Alderton, Roberto Amato, Jeffrey Barrett, Sonia Goncalves, Ewan Harrison, David K. Jackson, Ian Johnston, Dominic Kwiatkowski, Cordelia Langford, John Sillitoe on behalf of the Wellcome Sanger Institute COVID-19 Surveillance Team |
| EPI_ISL_2551764 | Lighthouse Lab in Milton Keynes | Wellcome Sanger Institute for the COVID-19 Genomics UK (COG-UK) Consortium | The Lighthouse Lab in Milton Keynes and Alex Alderton, Roberto Amato, Jeffrey Barrett, Sonia Goncalves, Ewan Harrison, David K. Jackson, Ian Johnston, Dominic Kwiatkowski, Cordelia Langford, John Sillitoe on behalf of the Wellcome Sanger Institute COVID-19 Surveillance Team |
| EPI_ISL_2551765 | Lighthouse Lab in Alderley Park | Wellcome Sanger Institute for the COVID-19 Genomics UK (COG-UK) Consortium | Jacquelyn Wynn, Mairead Hyland, The Lighthouse Lab in Alderley Park and Alex Alderton, Roberto Amato, Jeffrey Barrett, Sonia Goncalves, Ewan Harrison, David K. Jackson, Ian Johnston, Dominic Kwiatkowski, Cordelia Langford, John Sillitoe on behalf of the Wellcome Sanger Institute COVID-19 Surveillance Team |
| EPI_ISL_2551766 | Lighthouse Lab in Milton Keynes | Wellcome Sanger Institute for the COVID-19 Genomics UK (COG-UK) Consortium | The Lighthouse Lab in Milton Keynes and Alex Alderton, Roberto Amato, Jeffrey Barrett, Sonia Goncalves, Ewan Harrison, David K. Jackson, Ian Johnston, Dominic Kwiatkowski, Cordelia Langford, John Sillitoe on behalf of the Wellcome Sanger Institute COVID-19 Surveillance Team |
| EPI_ISL_2551767, EPI_ISL_2551768, EPI_ISL_2551769 | Lighthouse Lab in Alderley Park | Wellcome Sanger Institute for the COVID-19 Genomics UK (COG-UK) Consortium | Jacquelyn Wynn, Mairead Hyland, The Lighthouse Lab in Alderley Park and Alex Alderton, Roberto Amato, Jeffrey Barrett, Sonia Goncalves, Ewan Harrison, David K. Jackson, Ian Johnston, Dominic Kwiatkowski, Cordelia Langford, John Sillitoe on behalf of the Wellcome Sanger Institute COVID-19 Surveillance Team |
| EPI_ISL_2551770 | Lighthouse Lab in Milton Keynes | Wellcome Sanger Institute for the COVID-19 Genomics UK (COG-UK) Consortium | The Lighthouse Lab in Milton Keynes and Alex Alderton, Roberto Amato, Jeffrey Barrett, Sonia Goncalves, Ewan Harrison, David K. Jackson, Ian Johnston, Dominic Kwiatkowski, Cordelia Langford, John Sillitoe on behalf of the Wellcome Sanger Institute COVID-19 Surveillance Team |
| EPI_ISL_2551771, EPI_ISL_2551772, EPI_ISL_2551773, EPI_ISL_2551774 | Lighthouse Lab in Alderley Park | Wellcome Sanger Institute for the COVID-19 Genomics UK (COG-UK) Consortium | Jacquelyn Wynn, Mairead Hyland, The Lighthouse Lab in Alderley Park and Alex Alderton, Roberto Amato, Jeffrey Barrett, Sonia Goncalves, Ewan Harrison, David K. Jackson, Ian Johnston, Dominic Kwiatkowski, Cordelia Langford, John Sillitoe on behalf of the Wellcome Sanger Institute COVID-19 Surveillance Team |
| EPI_ISL_2551775, EPI_ISL_2551776 | Lighthouse Lab in Milton Keynes | Wellcome Sanger Institute for the COVID-19 Genomics UK (COG-UK) Consortium | The Lighthouse Lab in Milton Keynes and Alex Alderton, Roberto Amato, Jeffrey Barrett, Sonia Goncalves, Ewan Harrison, David K. Jackson, Ian Johnston, Dominic Kwiatkowski, Cordelia Langford, John Sillitoe on behalf of the Wellcome Sanger Institute COVID-19 Surveillance Team |
| EPI_ISL_2551777, EPI_ISL_2551778, EPI_ISL_2551779, EPI_ISL_2551780, EPI_ISL_2551781, EPI_ISL_2551782, EPI_ISL_2551783 | Lighthouse Lab in Alderley Park | Wellcome Sanger Institute for the COVID-19 Genomics UK (COG-UK) Consortium | Jacquelyn Wynn, Mairead Hyland, The Lighthouse Lab in Alderley Park and Alex Alderton, Roberto Amato, Jeffrey Barrett, Sonia Goncalves, Ewan Harrison, David K. Jackson, Ian Johnston, Dominic Kwiatkowski, Cordelia Langford, John Sillitoe on behalf of the Wellcome Sanger Institute COVID-19 Surveillance Team |
| EPI_ISL_2551784, EPI_ISL_2551785 | Lighthouse Lab in Milton Keynes | Wellcome Sanger Institute for the COVID-19 Genomics UK (COG-UK) Consortium | The Lighthouse Lab in Milton Keynes and Alex Alderton, Roberto Amato, Jeffrey Barrett, Sonia Goncalves, Ewan Harrison, David K. Jackson, Ian Johnston, Dominic Kwiatkowski, Cordelia Langford, John Sillitoe on behalf of the Wellcome Sanger Institute COVID-19 Surveillance Team |
| EPI_ISL_2551786, EPI_ISL_2551787 | Lighthouse Lab in Alderley Park | Wellcome Sanger Institute for the COVID-19 Genomics UK (COG-UK) Consortium | Jacquelyn Wynn, Mairead Hyland, The Lighthouse Lab in Alderley Park and Alex Alderton, Roberto Amato, Jeffrey Barrett, Sonia Goncalves, Ewan Harrison, David K. Jackson, Ian Johnston, Dominic Kwiatkowski, Cordelia Langford, John Sillitoe on behalf of the Wellcome Sanger Institute COVID-19 Surveillance Team |
| EPI_ISL_2551788 | Lighthouse Lab in Milton Keynes | Wellcome Sanger Institute for the COVID-19 Genomics UK (COG-UK) Consortium | The Lighthouse Lab in Milton Keynes and Alex Alderton, Roberto Amato, Jeffrey Barrett, Sonia Goncalves, Ewan Harrison, David K. Jackson, Ian Johnston, Dominic Kwiatkowski, Cordelia Langford, John Sillitoe on behalf of the Wellcome Sanger Institute COVID-19 Surveillance Team |
| EPI_ISL_2551789, EPI_ISL_2551790, EPI_ISL_2551791, EPI_ISL_2551792, EPI_ISL_2551793, EPI_ISL_2551794, EPI_ISL_2551795 | Lighthouse Lab in Alderley Park | Wellcome Sanger Institute for the COVID-19 Genomics UK (COG-UK) Consortium | Jacquelyn Wynn, Mairead Hyland, The Lighthouse Lab in Alderley Park and Alex Alderton, Roberto Amato, Jeffrey Barrett, Sonia Goncalves, Ewan Harrison, David K. Jackson, Ian Johnston, Dominic Kwiatkowski, Cordelia Langford, John Sillitoe on behalf of the Wellcome Sanger Institute COVID-19 Surveillance Team |
| EPI_ISL_2551796 | Lighthouse Lab in Milton Keynes | Wellcome Sanger Institute for the COVID-19 Genomics UK (COG-UK) Consortium | The Lighthouse Lab in Milton Keynes and Alex Alderton, Roberto Amato, Jeffrey Barrett, Sonia Goncalves, Ewan Harrison, David K. Jackson, Ian Johnston, Dominic Kwiatkowski, Cordelia Langford, John Sillitoe on behalf of the Wellcome Sanger Institute COVID-19 Surveillance Team |
| EPI_ISL_2551797, EPI_ISL_2551798, | Lighthouse Lab in Alderley Park | Wellcome Sanger Institute for the COVID-19 Genomics UK (COG-UK) Consortium | Jacquelyn Wynn, Mairead Hyland, The Lighthouse Lab in Alderley Park and Alex Alderton, Roberto Amato, Jeffrey Barrett, Sonia Goncalves, Ewan Harrison, David K. Jackson, Ian Johnston, Dominic Kwiatkowski, Cordelia Langford, John Sillitoe on behalf of the Wellcome Sanger Institute COVID-19 Surveillance Team |

| | | | |
|---|---|---|---|
| EPI_ISL_2551799, EPI_ISL_2551800, EPI_ISL_2551802, EPI_ISL_2551803, EPI_ISL_2551804 | | | |
| EPI_ISL_2551805, EPI_ISL_2551806 | Lighthouse Lab in Milton Keynes | Wellcome Sanger Institute for the COVID-19 Genomics UK (COG-UK) Consortium | The Lighthouse Lab in Milton Keynes and Alex Alderton, Roberto Amato, Jeffrey Barrett, Sonia Goncalves, Ewan Harrison, David K. Jackson, Ian Johnston, Dominic Kwiatkowski, Cordelia Langford, John Sillitoe on behalf of the Wellcome Sanger Institute COVID-19 Surveillance Team |
| EPI_ISL_2551807 | Lighthouse Lab in Alderley Park | Wellcome Sanger Institute for the COVID-19 Genomics UK (COG-UK) Consortium | Jacquelyn Wynn, Mairead Hyland, The Lighthouse Lab in Alderley Park and Alex Alderton, Roberto Amato, Jeffrey Barrett, Sonia Goncalves, Ewan Harrison, David K. Jackson, Ian Johnston, Dominic Kwiatkowski, Cordelia Langford, John Sillitoe on behalf of the Wellcome Sanger Institute COVID-19 Surveillance Team |
| EPI_ISL_2551808 | Lighthouse Lab in Milton Keynes | Wellcome Sanger Institute for the COVID-19 Genomics UK (COG-UK) Consortium | The Lighthouse Lab in Milton Keynes and Alex Alderton, Roberto Amato, Jeffrey Barrett, Sonia Goncalves, Ewan Harrison, David K. Jackson, Ian Johnston, Dominic Kwiatkowski, Cordelia Langford, John Sillitoe on behalf of the Wellcome Sanger Institute COVID-19 Surveillance Team |
| EPI_ISL_2551809, EPI_ISL_2551810 | Lighthouse Lab in Alderley Park | Wellcome Sanger Institute for the COVID-19 Genomics UK (COG-UK) Consortium | Jacquelyn Wynn, Mairead Hyland, The Lighthouse Lab in Alderley Park and Alex Alderton, Roberto Amato, Jeffrey Barrett, Sonia Goncalves, Ewan Harrison, David K. Jackson, Ian Johnston, Dominic Kwiatkowski, Cordelia Langford, John Sillitoe on behalf of the Wellcome Sanger Institute COVID-19 Surveillance Team |
| EPI_ISL_2551811 | Lighthouse Lab in Milton Keynes | Wellcome Sanger Institute for the COVID-19 Genomics UK (COG-UK) Consortium | The Lighthouse Lab in Milton Keynes and Alex Alderton, Roberto Amato, Jeffrey Barrett, Sonia Goncalves, Ewan Harrison, David K. Jackson, Ian Johnston, Dominic Kwiatkowski, Cordelia Langford, John Sillitoe on behalf of the Wellcome Sanger Institute COVID-19 Surveillance Team |
| EPI_ISL_2551812 | Lighthouse Lab in Alderley Park | Wellcome Sanger Institute for the COVID-19 Genomics UK (COG-UK) Consortium | Jacquelyn Wynn, Mairead Hyland, The Lighthouse Lab in Alderley Park and Alex Alderton, Roberto Amato, Jeffrey Barrett, Sonia Goncalves, Ewan Harrison, David K. Jackson, Ian Johnston, Dominic Kwiatkowski, Cordelia Langford, John Sillitoe on behalf of the Wellcome Sanger Institute COVID-19 Surveillance Team |
| EPI_ISL_2551813 | Lighthouse Lab in Milton Keynes | Wellcome Sanger Institute for the COVID-19 Genomics UK (COG-UK) Consortium | The Lighthouse Lab in Milton Keynes and Alex Alderton, Roberto Amato, Jeffrey Barrett, Sonia Goncalves, Ewan Harrison, David K. Jackson, Ian Johnston, Dominic Kwiatkowski, Cordelia Langford, John Sillitoe on behalf of the Wellcome Sanger Institute COVID-19 Surveillance Team |
| EPI_ISL_2551814, EPI_ISL_2551815, EPI_ISL_2551816, EPI_ISL_2551817 | Lighthouse Lab in Alderley Park | Wellcome Sanger Institute for the COVID-19 Genomics UK (COG-UK) Consortium | Jacquelyn Wynn, Mairead Hyland, The Lighthouse Lab in Alderley Park and Alex Alderton, Roberto Amato, Jeffrey Barrett, Sonia Goncalves, Ewan Harrison, David K. Jackson, Ian Johnston, Dominic Kwiatkowski, Cordelia Langford, John Sillitoe on behalf of the Wellcome Sanger Institute COVID-19 Surveillance Team |
| EPI_ISL_2551818, EPI_ISL_2551819 | Lighthouse Lab in Milton Keynes | Wellcome Sanger Institute for the COVID-19 Genomics UK (COG-UK) Consortium | The Lighthouse Lab in Milton Keynes and Alex Alderton, Roberto Amato, Jeffrey Barrett, Sonia Goncalves, Ewan Harrison, David K. Jackson, Ian Johnston, Dominic Kwiatkowski, Cordelia Langford, John Sillitoe on behalf of the Wellcome Sanger Institute COVID-19 Surveillance Team |
| EPI_ISL_2551820 | Lighthouse Lab in Alderley Park | Wellcome Sanger Institute for the COVID-19 Genomics UK (COG-UK) Consortium | Jacquelyn Wynn, Mairead Hyland, The Lighthouse Lab in Alderley Park and Alex Alderton, Roberto Amato, Jeffrey Barrett, Sonia Goncalves, Ewan Harrison, David K. Jackson, Ian Johnston, Dominic Kwiatkowski, Cordelia Langford, John Sillitoe on behalf of the Wellcome Sanger Institute COVID-19 Surveillance Team |
| EPI_ISL_2551821 | Lighthouse Lab in Milton Keynes | Wellcome Sanger Institute for the COVID-19 Genomics UK (COG-UK) Consortium | The Lighthouse Lab in Milton Keynes and Alex Alderton, Roberto Amato, Jeffrey Barrett, Sonia Goncalves, Ewan Harrison, David K. Jackson, Ian Johnston, Dominic Kwiatkowski, Cordelia Langford, John Sillitoe on behalf of the Wellcome Sanger Institute COVID-19 Surveillance Team |
| EPI_ISL_2551822, EPI_ISL_2551823, EPI_ISL_2551824, EPI_ISL_2551825 | Lighthouse Lab in Alderley Park | Wellcome Sanger Institute for the COVID-19 Genomics UK (COG-UK) Consortium | Jacquelyn Wynn, Mairead Hyland, The Lighthouse Lab in Alderley Park and Alex Alderton, Roberto Amato, Jeffrey Barrett, Sonia Goncalves, Ewan Harrison, David K. Jackson, Ian Johnston, Dominic Kwiatkowski, Cordelia Langford, John Sillitoe on behalf of the Wellcome Sanger Institute COVID-19 Surveillance Team |
| EPI_ISL_2551826 | Lighthouse Lab in Milton Keynes | Wellcome Sanger Institute for the COVID-19 Genomics UK (COG-UK) Consortium | The Lighthouse Lab in Milton Keynes and Alex Alderton, Roberto Amato, Jeffrey Barrett, Sonia Goncalves, Ewan Harrison, David K. Jackson, Ian Johnston, Dominic Kwiatkowski, Cordelia Langford, John Sillitoe on behalf of the Wellcome Sanger Institute COVID-19 Surveillance Team |
| EPI_ISL_2551827 | Lighthouse Lab in Alderley Park | Wellcome Sanger Institute for the COVID-19 Genomics UK (COG-UK) Consortium | Jacquelyn Wynn, Mairead Hyland, The Lighthouse Lab in Alderley Park and Alex Alderton, Roberto Amato, Jeffrey Barrett, Sonia Goncalves, Ewan Harrison, David K. Jackson, Ian Johnston, Dominic Kwiatkowski, Cordelia Langford, John Sillitoe on behalf of the Wellcome Sanger Institute COVID-19 Surveillance Team |
| EPI_ISL_2551828, EPI_ISL_2551829 | Lighthouse Lab in Milton Keynes | Wellcome Sanger Institute for the COVID-19 Genomics UK (COG-UK) Consortium | The Lighthouse Lab in Milton Keynes and Alex Alderton, Roberto Amato, Jeffrey Barrett, Sonia Goncalves, Ewan Harrison, David K. Jackson, Ian Johnston, Dominic Kwiatkowski, Cordelia Langford, John Sillitoe on behalf of the Wellcome Sanger Institute COVID-19 Surveillance Team |
| EPI_ISL_2551830, EPI_ISL_2551831, EPI_ISL_2551832 | Lighthouse Lab in Alderley Park | Wellcome Sanger Institute for the COVID-19 Genomics UK (COG-UK) Consortium | Jacquelyn Wynn, Mairead Hyland, The Lighthouse Lab in Alderley Park and Alex Alderton, Roberto Amato, Jeffrey Barrett, Sonia Goncalves, Ewan Harrison, David K. Jackson, Ian Johnston, Dominic Kwiatkowski, Cordelia Langford, John Sillitoe on behalf of the Wellcome Sanger Institute COVID-19 Surveillance Team |
| EPI_ISL_2551833 | Lighthouse Lab in Milton Keynes | Wellcome Sanger Institute for the COVID-19 Genomics UK (COG-UK) Consortium | The Lighthouse Lab in Milton Keynes and Alex Alderton, Roberto Amato, Jeffrey Barrett, Sonia Goncalves, Ewan Harrison, David K. Jackson, Ian Johnston, Dominic Kwiatkowski, Cordelia Langford, John Sillitoe on behalf of the Wellcome Sanger Institute COVID-19 Surveillance Team |
| EPI_ISL_2551834, EPI_ISL_2551835, EPI_ISL_2551836, EPI_ISL_2551837 | Lighthouse Lab in Alderley Park | Wellcome Sanger Institute for the COVID-19 Genomics UK (COG-UK) Consortium | Jacquelyn Wynn, Mairead Hyland, The Lighthouse Lab in Alderley Park and Alex Alderton, Roberto Amato, Jeffrey Barrett, Sonia Goncalves, Ewan Harrison, David K. Jackson, Ian Johnston, Dominic Kwiatkowski, Cordelia Langford, John Sillitoe on behalf of the Wellcome Sanger Institute COVID-19 Surveillance Team |
| EPI_ISL_2551838 | Lighthouse Lab in Milton Keynes | Wellcome Sanger Institute for the COVID-19 Genomics UK (COG-UK) Consortium | The Lighthouse Lab in Milton Keynes and Alex Alderton, Roberto Amato, Jeffrey Barrett, Sonia Goncalves, Ewan Harrison, David K. Jackson, Ian Johnston, Dominic Kwiatkowski, Cordelia Langford, John Sillitoe on behalf of the Wellcome Sanger Institute COVID-19 Surveillance Team |
| EPI_ISL_2551839, EPI_ISL_2551840 | Lighthouse Lab in Alderley Park | Wellcome Sanger Institute for the COVID-19 Genomics UK (COG-UK) Consortium | Jacquelyn Wynn, Mairead Hyland, The Lighthouse Lab in Alderley Park and Alex Alderton, Roberto Amato, Jeffrey Barrett, Sonia Goncalves, Ewan Harrison, David K. Jackson, Ian Johnston, Dominic Kwiatkowski, Cordelia Langford, John Sillitoe on behalf of the Wellcome Sanger Institute COVID-19 Surveillance Team |
| EPI_ISL_2551841 | Lighthouse Lab in Milton Keynes | Wellcome Sanger Institute for the COVID-19 Genomics UK (COG-UK) Consortium | The Lighthouse Lab in Milton Keynes and Alex Alderton, Roberto Amato, Jeffrey Barrett, Sonia Goncalves, Ewan Harrison, David K. Jackson, Ian Johnston, Dominic Kwiatkowski, Cordelia Langford, John Sillitoe on behalf of the Wellcome Sanger Institute COVID-19 Surveillance Team |
| EPI_ISL_2551842 | Lighthouse Lab in Alderley Park | Wellcome Sanger Institute for the COVID-19 Genomics UK (COG-UK) Consortium | Jacquelyn Wynn, Mairead Hyland, The Lighthouse Lab in Alderley Park and Alex Alderton, Roberto Amato, Jeffrey Barrett, Sonia Goncalves, Ewan Harrison, David K. Jackson, Ian Johnston, Dominic Kwiatkowski, Cordelia Langford, John Sillitoe on behalf of the Wellcome Sanger Institute COVID-19 Surveillance Team |
| EPI_ISL_2551843 | Lighthouse Lab in Milton Keynes | Wellcome Sanger Institute for the COVID-19 Genomics UK (COG-UK) Consortium | The Lighthouse Lab in Milton Keynes and Alex Alderton, Roberto Amato, Jeffrey Barrett, Sonia Goncalves, Ewan Harrison, David K. Jackson, Ian Johnston, Dominic Kwiatkowski, Cordelia Langford, John Sillitoe on behalf of the Wellcome Sanger Institute COVID-19 Surveillance Team |
| EPI_ISL_2551844, EPI_ISL_2551845, EPI_ISL_2551846, EPI_ISL_2551847, EPI_ISL_2551848, EPI_ISL_2551849 | Lighthouse Lab in Alderley Park | Wellcome Sanger Institute for the COVID-19 Genomics UK (COG-UK) Consortium | Jacquelyn Wynn, Mairead Hyland, The Lighthouse Lab in Alderley Park and Alex Alderton, Roberto Amato, Jeffrey Barrett, Sonia Goncalves, Ewan Harrison, David K. Jackson, Ian Johnston, Dominic Kwiatkowski, Cordelia Langford, John Sillitoe on behalf of the Wellcome Sanger Institute COVID-19 Surveillance Team |
| EPI_ISL_2551850 | Lighthouse Lab in Milton Keynes | Wellcome Sanger Institute for the COVID-19 Genomics UK (COG-UK) Consortium | The Lighthouse Lab in Milton Keynes and Alex Alderton, Roberto Amato, Jeffrey Barrett, Sonia Goncalves, Ewan Harrison, David K. Jackson, Ian Johnston, Dominic Kwiatkowski, Cordelia Langford, John Sillitoe on behalf of the Wellcome Sanger Institute COVID-19 Surveillance Team |
| EPI_ISL_2551851, EPI_ISL_2551853, EPI_ISL_2551854, EPI_ISL_2551855, EPI_ISL_2551856, EPI_ISL_2551858, EPI_ISL_2551859 | Lighthouse Lab in Alderley Park | Wellcome Sanger Institute for the COVID-19 Genomics UK (COG-UK) Consortium | Jacquelyn Wynn, Mairead Hyland, The Lighthouse Lab in Alderley Park and Alex Alderton, Roberto Amato, Jeffrey Barrett, Sonia Goncalves, Ewan Harrison, David K. Jackson, Ian Johnston, Dominic Kwiatkowski, Cordelia Langford, John Sillitoe on behalf of the Wellcome Sanger Institute COVID-19 Surveillance Team |
| EPI_ISL_2551860 | Lighthouse Lab in Milton Keynes | Wellcome Sanger Institute for the COVID-19 Genomics UK (COG-UK) Consortium | The Lighthouse Lab in Milton Keynes and Alex Alderton, Roberto Amato, Jeffrey Barrett, Sonia Goncalves, Ewan Harrison, David K. Jackson, Ian Johnston, Dominic Kwiatkowski, Cordelia Langford, John Sillitoe on behalf of the Wellcome Sanger Institute COVID-19 Surveillance Team |
| EPI_ISL_2551861 | Lighthouse Lab in Alderley Park | Wellcome Sanger Institute for the COVID-19 Genomics UK (COG-UK) Consortium | Jacquelyn Wynn, Mairead Hyland, The Lighthouse Lab in Alderley Park and Alex Alderton, Roberto Amato, Jeffrey Barrett, Sonia Goncalves, Ewan Harrison, David K. Jackson, Ian Johnston, Dominic Kwiatkowski, Cordelia Langford, John Sillitoe on behalf of the Wellcome Sanger Institute COVID-19 Surveillance Team |
| EPI_ISL_2551862 | Lighthouse Lab in Milton Keynes | Wellcome Sanger Institute for the COVID-19 Genomics UK (COG-UK) Consortium | The Lighthouse Lab in Milton Keynes and Alex Alderton, Roberto Amato, Jeffrey Barrett, Sonia Goncalves, Ewan Harrison, David K. Jackson, Ian Johnston, Dominic Kwiatkowski, Cordelia Langford, John Sillitoe on behalf of the Wellcome Sanger Institute COVID-19 Surveillance Team |
| EPI_ISL_2551863 | Lighthouse Lab in Alderley Park | Wellcome Sanger Institute for the COVID-19 Genomics UK (COG-UK) Consortium | Jacquelyn Wynn, Mairead Hyland, The Lighthouse Lab in Alderley Park and Alex Alderton, Roberto Amato, Jeffrey Barrett, Sonia Goncalves, Ewan Harrison, David K. Jackson, Ian Johnston, Dominic Kwiatkowski, Cordelia Langford, John Sillitoe on behalf of the Wellcome Sanger Institute COVID-19 Surveillance Team |
| EPI_ISL_2551864 | Lighthouse Lab in Milton Keynes | Wellcome Sanger Institute for the COVID-19 Genomics UK (COG-UK) Consortium | The Lighthouse Lab in Milton Keynes and Alex Alderton, Roberto Amato, Jeffrey Barrett, Sonia Goncalves, Ewan Harrison, David K. Jackson, Ian Johnston, Dominic Kwiatkowski, Cordelia Langford, John Sillitoe on behalf of the Wellcome Sanger Institute COVID-19 Surveillance Team |
| EPI_ISL_2551865, EPI_ISL_2551866, EPI_ISL_2551867 | Lighthouse Lab in Alderley Park | Wellcome Sanger Institute for the COVID-19 Genomics UK (COG-UK) Consortium | Jacquelyn Wynn, Mairead Hyland, The Lighthouse Lab in Alderley Park and Alex Alderton, Roberto Amato, Jeffrey Barrett, Sonia Goncalves, Ewan Harrison, David K. Jackson, Ian Johnston, Dominic Kwiatkowski, Cordelia Langford, John Sillitoe on behalf of the Wellcome Sanger Institute COVID-19 Surveillance Team |
| EPI_ISL_2551868, EPI_ISL_2551869, EPI_ISL_2551870, EPI_ISL_2551871, EPI_ISL_2551872, EPI_ISL_2551873 | Lighthouse Lab in Milton Keynes | Wellcome Sanger Institute for the COVID-19 Genomics UK (COG-UK) Consortium | The Lighthouse Lab in Milton Keynes and Alex Alderton, Roberto Amato, Jeffrey Barrett, Sonia Goncalves, Ewan Harrison, David K. Jackson, Ian Johnston, Dominic Kwiatkowski, Cordelia Langford, John Sillitoe on behalf of the Wellcome Sanger Institute COVID-19 Surveillance Team |

EPI_ISL_2551874, EPI_ISL_2551875, EPI_ISL_2551876, EPI_ISL_2551877, EPI_ISL_2551878, EPI_ISL_2551879, EPI_ISL_2551880, EPI_ISL_2551881, EPI_ISL_2551882, EPI_ISL_2551883, EPI_ISL_2551884, EPI_ISL_2551885, EPI_ISL_2551886, EPI_ISL_2551887, EPI_ISL_2551888, EPI_ISL_2551889, EPI_ISL_2551890

| | | | |
|---|---|---|---|
| see above | Lighthouse Lab in Alderley Park | Wellcome Sanger Institute for the COVID-19 Genomics UK (COG-UK) Consortium | Jacquelyn Wynn, Mairead Hyland, The Lighthouse Lab in Alderley Park and Alex Alderton, Roberto Amato, Jeffrey Barrett, Sonia Goncalves, Ewan Harrison, David K. Jackson, Ian Johnston, Dominic Kwiatkowski, Cordelia Langford, John Sillitoe on behalf of the Wellcome Sanger Institute COVID-19 Surveillance Team |

| | | | |
|---|---|---|---|
| EPI_ISL_2551891 | Lighthouse Lab in Milton Keynes | Wellcome Sanger Institute for the COVID-19 Genomics UK (COG-UK) Consortium | The Lighthouse Lab in Milton Keynes and Alex Alderton, Roberto Amato, Jeffrey Barrett, Sonia Goncalves, Ewan Harrison, David K. Jackson, Ian Johnston, Dominic Kwiatkowski, Cordelia Langford, John Sillitoe on behalf of the Wellcome Sanger Institute COVID-19 Surveillance Team |
| EPI_ISL_2551892 | Lighthouse Lab in Alderley Park | Wellcome Sanger Institute for the COVID-19 Genomics UK (COG-UK) Consortium | Jacquelyn Wynn, Mairead Hyland, The Lighthouse Lab in Alderley Park and Alex Alderton, Roberto Amato, Jeffrey Barrett, Sonia Goncalves, Ewan Harrison, David K. Jackson, Ian Johnston, Dominic Kwiatkowski, Cordelia Langford, John Sillitoe on behalf of the Wellcome Sanger Institute COVID-19 Surveillance Team |
| EPI_ISL_2551894, EPI_ISL_2551895 | Lighthouse Lab in Milton Keynes | Wellcome Sanger Institute for the COVID-19 Genomics UK (COG-UK) Consortium | The Lighthouse Lab in Milton Keynes and Alex Alderton, Roberto Amato, Jeffrey Barrett, Sonia Goncalves, Ewan Harrison, David K. Jackson, Ian Johnston, Dominic Kwiatkowski, Cordelia Langford, John Sillitoe on behalf of the Wellcome Sanger Institute COVID-19 Surveillance Team |
| EPI_ISL_2551897, EPI_ISL_2551898, EPI_ISL_2551899, EPI_ISL_2551900 | Lighthouse Lab in Alderley Park | Wellcome Sanger Institute for the COVID-19 Genomics UK (COG-UK) Consortium | Jacquelyn Wynn, Mairead Hyland, The Lighthouse Lab in Alderley Park and Alex Alderton, Roberto Amato, Jeffrey Barrett, Sonia Goncalves, Ewan Harrison, David K. Jackson, Ian Johnston, Dominic Kwiatkowski, Cordelia Langford, John Sillitoe on behalf of the Wellcome Sanger Institute COVID-19 Surveillance Team |
| EPI_ISL_2551901 | Lighthouse Lab in Milton Keynes | Wellcome Sanger Institute for the COVID-19 Genomics UK (COG-UK) Consortium | The Lighthouse Lab in Milton Keynes and Alex Alderton, Roberto Amato, Jeffrey Barrett, Sonia Goncalves, Ewan Harrison, David K. Jackson, Ian Johnston, Dominic Kwiatkowski, Cordelia Langford, John Sillitoe on behalf of the Wellcome Sanger Institute COVID-19 Surveillance Team |
| EPI_ISL_2551902, EPI_ISL_2551903, EPI_ISL_2551904, EPI_ISL_2551905, EPI_ISL_2551906, EPI_ISL_2551907, EPI_ISL_2551908 | Lighthouse Lab in Alderley Park | Wellcome Sanger Institute for the COVID-19 Genomics UK (COG-UK) Consortium | Jacquelyn Wynn, Mairead Hyland, The Lighthouse Lab in Alderley Park and Alex Alderton, Roberto Amato, Jeffrey Barrett, Sonia Goncalves, Ewan Harrison, David K. Jackson, Ian Johnston, Dominic Kwiatkowski, Cordelia Langford, John Sillitoe on behalf of the Wellcome Sanger Institute COVID-19 Surveillance Team |
| EPI_ISL_2551909 | Lighthouse Lab in Milton Keynes | Wellcome Sanger Institute for the COVID-19 Genomics UK (COG-UK) Consortium | The Lighthouse Lab in Milton Keynes and Alex Alderton, Roberto Amato, Jeffrey Barrett, Sonia Goncalves, Ewan Harrison, David K. Jackson, Ian Johnston, Dominic Kwiatkowski, Cordelia Langford, John Sillitoe on behalf of the Wellcome Sanger Institute COVID-19 Surveillance Team |
| EPI_ISL_2551910, EPI_ISL_2551911, EPI_ISL_2551912, EPI_ISL_2551914, EPI_ISL_2551915, EPI_ISL_2551916, EPI_ISL_2551917, EPI_ISL_2551918 | Lighthouse Lab in Alderley Park | Wellcome Sanger Institute for the COVID-19 Genomics UK (COG-UK) Consortium | Jacquelyn Wynn, Mairead Hyland, The Lighthouse Lab in Alderley Park and Alex Alderton, Roberto Amato, Jeffrey Barrett, Sonia Goncalves, Ewan Harrison, David K. Jackson, Ian Johnston, Dominic Kwiatkowski, Cordelia Langford, John Sillitoe on behalf of the Wellcome Sanger Institute COVID-19 Surveillance Team |
| EPI_ISL_2551919, EPI_ISL_2551921 | Lighthouse Lab in Milton Keynes | Wellcome Sanger Institute for the COVID-19 Genomics UK (COG-UK) Consortium | The Lighthouse Lab in Milton Keynes and Alex Alderton, Roberto Amato, Jeffrey Barrett, Sonia Goncalves, Ewan Harrison, David K. Jackson, Ian Johnston, Dominic Kwiatkowski, Cordelia Langford, John Sillitoe on behalf of the Wellcome Sanger Institute COVID-19 Surveillance Team |
| EPI_ISL_2551923 | Lighthouse Lab in Alderley Park | Wellcome Sanger Institute for the COVID-19 Genomics UK (COG-UK) Consortium | Jacquelyn Wynn, Mairead Hyland, The Lighthouse Lab in Alderley Park and Alex Alderton, Roberto Amato, Jeffrey Barrett, Sonia Goncalves, Ewan Harrison, David K. Jackson, Ian Johnston, Dominic Kwiatkowski, Cordelia Langford, John Sillitoe on behalf of the Wellcome Sanger Institute COVID-19 Surveillance Team |
| EPI_ISL_2551924 | Lighthouse Lab in Milton Keynes | Wellcome Sanger Institute for the COVID-19 Genomics UK (COG-UK) Consortium | The Lighthouse Lab in Milton Keynes and Alex Alderton, Roberto Amato, Jeffrey Barrett, Sonia Goncalves, Ewan Harrison, David K. Jackson, Ian Johnston, Dominic Kwiatkowski, Cordelia Langford, John Sillitoe on behalf of the Wellcome Sanger Institute COVID-19 Surveillance Team |
| EPI_ISL_2551925, EPI_ISL_2551926 | Lighthouse Lab in Alderley Park | Wellcome Sanger Institute for the COVID-19 Genomics UK (COG-UK) Consortium | Jacquelyn Wynn, Mairead Hyland, The Lighthouse Lab in Alderley Park and Alex Alderton, Roberto Amato, Jeffrey Barrett, Sonia Goncalves, Ewan Harrison, David K. Jackson, Ian Johnston, Dominic Kwiatkowski, Cordelia Langford, John Sillitoe on behalf of the Wellcome Sanger Institute COVID-19 Surveillance Team |
| EPI_ISL_2551927, EPI_ISL_2551928, EPI_ISL_2551929, EPI_ISL_2551930, EPI_ISL_2551931, EPI_ISL_2551932, EPI_ISL_2551933, EPI_ISL_2551934, EPI_ISL_2551935, EPI_ISL_2551936, EPI_ISL_2551937, EPI_ISL_2551938, EPI_ISL_2551939, EPI_ISL_2551940, EPI_ISL_2551941, EPI_ISL_2551942, EPI_ISL_2551943, EPI_ISL_2551944, EPI_ISL_2551945, EPI_ISL_2551947, EPI_ISL_2551948, EPI_ISL_2551949, EPI_ISL_2551950, EPI_ISL_2551951, EPI_ISL_2551952, EPI_ISL_2551953, EPI_ISL_2551954, EPI_ISL_2551955, EPI_ISL_2551956, EPI_ISL_2551957, EPI_ISL_2551958, EPI_ISL_2551959, EPI_ISL_2551960, EPI_ISL_2551961, EPI_ISL_2551962, EPI_ISL_2551963, EPI_ISL_2551964, EPI_ISL_2551965, EPI_ISL_2551966, EPI_ISL_2551967, EPI_ISL_2551968, EPI_ISL_2551969, EPI_ISL_2551970, EPI_ISL_2551971, EPI_ISL_2551972, EPI_ISL_2551973, EPI_ISL_2551974, EPI_ISL_2551975, EPI_ISL_2551976, EPI_ISL_2551977, EPI_ISL_2551978, EPI_ISL_2551979, EPI_ISL_2551980, EPI_ISL_2551981, EPI_ISL_2551982, EPI_ISL_2551983, EPI_ISL_2551984, EPI_ISL_2551985, EPI_ISL_2551986, EPI_ISL_2551987, EPI_ISL_2551988, EPI_ISL_2551989, EPI_ISL_2551990, EPI_ISL_2551991, EPI_ISL_2551992, EPI_ISL_2551993, EPI_ISL_2551994, EPI_ISL_2551995, EPI_ISL_2551996, EPI_ISL_2551997, EPI_ISL_2551998, EPI_ISL_2552000, EPI_ISL_2552001, EPI_ISL_2552002, EPI_ISL_2552003, EPI_ISL_2552004, EPI_ISL_2552005, EPI_ISL_2552006, EPI_ISL_2552008, EPI_ISL_2552009, EPI_ISL_2552010, EPI_ISL_2552011, EPI_ISL_2552012, EPI_ISL_2552013, EPI_ISL_2552014, EPI_ISL_2552015, EPI_ISL_2552016, EPI_ISL_2552017 | | | |
| see above | Lighthouse Lab in Glasgow | Wellcome Sanger Institute for the COVID-19 Genomics UK (COG-UK) Consortium | Harper VanSteenhouse, Yumi Kasai, David Gray, Carol Clugston, Anna Dominiczak and Alex Alderton, Roberto Amato, Jeffrey Barrett, Sonia Goncalves, Ewan Harrison, David K. Jackson, Ian Johnston, Dominic Kwiatkowski, Cordelia Langford, John Sillitoe on behalf of the Wellcome Sanger Institute COVID-19 Surveillance Team |
| EPI_ISL_2552018, EPI_ISL_2552019 | Lighthouse Lab in Milton Keynes | Wellcome Sanger Institute for the COVID-19 Genomics UK (COG-UK) Consortium | The Lighthouse Lab in Milton Keynes and Alex Alderton, Roberto Amato, Jeffrey Barrett, Sonia Goncalves, Ewan Harrison, David K. Jackson, Ian Johnston, Dominic Kwiatkowski, Cordelia Langford, John Sillitoe on behalf of the Wellcome Sanger Institute COVID-19 Surveillance Team |
| EPI_ISL_2552020 | Lighthouse Laboratory Plymouth | Wellcome Sanger Institute for the COVID-19 Genomics UK (COG-UK) Consortium | Lighthouse Laboratory Plymouth and Alex Alderton, Roberto Amato, Jeffrey Barrett, Sonia Goncalves, Ewan Harrison, David K. Jackson, Ian Johnston, Dominic Kwiatkowski, Cordelia Langford, John Sillitoe on behalf of the Wellcome Sanger Institute COVID-19 Surveillance Team |
| EPI_ISL_2552021, EPI_ISL_2552022, EPI_ISL_2552023, EPI_ISL_2552024, EPI_ISL_2552025, EPI_ISL_2552026 | Lighthouse Lab in Milton Keynes | Wellcome Sanger Institute for the COVID-19 Genomics UK (COG-UK) Consortium | The Lighthouse Lab in Milton Keynes and Alex Alderton, Roberto Amato, Jeffrey Barrett, Sonia Goncalves, Ewan Harrison, David K. Jackson, Ian Johnston, Dominic Kwiatkowski, Cordelia Langford, John Sillitoe on behalf of the Wellcome Sanger Institute COVID-19 Surveillance Team |
| EPI_ISL_2552027 | Lighthouse Lab in Glasgow | Wellcome Sanger Institute for the COVID-19 Genomics UK (COG-UK) Consortium | Harper VanSteenhouse, Yumi Kasai, David Gray, Carol Clugston, Anna Dominiczak and Alex Alderton, Roberto Amato, Jeffrey Barrett, Sonia Goncalves, Ewan Harrison, David K. Jackson, Ian Johnston, Dominic Kwiatkowski, Cordelia Langford, John Sillitoe on behalf of the Wellcome Sanger Institute COVID-19 Surveillance Team |
| EPI_ISL_2552028, EPI_ISL_2552029, EPI_ISL_2552030 | Lighthouse Lab in Milton Keynes | Wellcome Sanger Institute for the COVID-19 Genomics UK (COG-UK) Consortium | The Lighthouse Lab in Milton Keynes and Alex Alderton, Roberto Amato, Jeffrey Barrett, Sonia Goncalves, Ewan Harrison, David K. Jackson, Ian Johnston, Dominic Kwiatkowski, Cordelia Langford, John Sillitoe on behalf of the Wellcome Sanger Institute COVID-19 Surveillance Team |
| EPI_ISL_2552031, EPI_ISL_2552032 | Lighthouse Lab in Glasgow | Wellcome Sanger Institute for the COVID-19 Genomics UK (COG-UK) Consortium | Harper VanSteenhouse, Yumi Kasai, David Gray, Carol Clugston, Anna Dominiczak and Alex Alderton, Roberto Amato, Jeffrey Barrett, Sonia Goncalves, Ewan Harrison, David K. Jackson, Ian Johnston, Dominic Kwiatkowski, Cordelia Langford, John Sillitoe on behalf of the Wellcome Sanger Institute COVID-19 Surveillance Team |
| EPI_ISL_2552033, EPI_ISL_2552034, EPI_ISL_2552035, EPI_ISL_2552037, EPI_ISL_2552038, EPI_ISL_2552039, EPI_ISL_2552040, EPI_ISL_2552041, EPI_ISL_2552042, EPI_ISL_2552043, EPI_ISL_2552044, EPI_ISL_2552045, EPI_ISL_2552046, EPI_ISL_2552047, EPI_ISL_2552048, EPI_ISL_2552049 | | | |
| see above | Lighthouse Lab in Milton Keynes | Wellcome Sanger Institute for the COVID-19 Genomics UK (COG-UK) Consortium | The Lighthouse Lab in Milton Keynes and Alex Alderton, Roberto Amato, Jeffrey Barrett, Sonia Goncalves, Ewan Harrison, David K. Jackson, Ian Johnston, Dominic Kwiatkowski, Cordelia Langford, John Sillitoe on behalf of the Wellcome Sanger Institute COVID-19 Surveillance Team |
| EPI_ISL_2552051 | Lighthouse Lab in Glasgow | Wellcome Sanger Institute for the COVID-19 Genomics UK (COG-UK) Consortium | Harper VanSteenhouse, Yumi Kasai, David Gray, Carol Clugston, Anna Dominiczak and Alex Alderton, Roberto Amato, Jeffrey Barrett, Sonia Goncalves, Ewan Harrison, David K. Jackson, Ian Johnston, Dominic Kwiatkowski, Cordelia Langford, John Sillitoe on behalf of the Wellcome Sanger Institute COVID-19 Surveillance Team |
| EPI_ISL_2552052 | Lighthouse Laboratory Plymouth | Wellcome Sanger Institute for the COVID-19 Genomics UK (COG-UK) Consortium | Lighthouse Laboratory Plymouth and Alex Alderton, Roberto Amato, Jeffrey Barrett, Sonia Goncalves, Ewan Harrison, David K. Jackson, Ian Johnston, Dominic Kwiatkowski, Cordelia Langford, John Sillitoe on behalf of the Wellcome Sanger Institute COVID-19 Surveillance Team |
| EPI_ISL_2552054, EPI_ISL_2552055, EPI_ISL_2552056, EPI_ISL_2552057 | Lighthouse Lab in Milton Keynes | Wellcome Sanger Institute for the COVID-19 Genomics UK (COG-UK) Consortium | The Lighthouse Lab in Milton Keynes and Alex Alderton, Roberto Amato, Jeffrey Barrett, Sonia Goncalves, Ewan Harrison, David K. Jackson, Ian Johnston, Dominic Kwiatkowski, Cordelia Langford, John Sillitoe on behalf of the Wellcome Sanger Institute COVID-19 Surveillance Team |
| EPI_ISL_2552058 | Lighthouse Lab in Glasgow | Wellcome Sanger Institute for the COVID-19 Genomics UK (COG-UK) Consortium | Harper VanSteenhouse, Yumi Kasai, David Gray, Carol Clugston, Anna Dominiczak and Alex Alderton, Roberto Amato, Jeffrey Barrett, Sonia Goncalves, Ewan Harrison, David K. Jackson, Ian Johnston, Dominic Kwiatkowski, Cordelia Langford, John Sillitoe on behalf of the Wellcome Sanger Institute COVID-19 Surveillance Team |
| EPI_ISL_2552059, EPI_ISL_2552060, EPI_ISL_2552061 | Lighthouse Lab in Milton Keynes | Wellcome Sanger Institute for the COVID-19 Genomics UK (COG-UK) Consortium | The Lighthouse Lab in Milton Keynes and Alex Alderton, Roberto Amato, Jeffrey Barrett, Sonia Goncalves, Ewan Harrison, David K. Jackson, Ian Johnston, Dominic Kwiatkowski, Cordelia Langford, John Sillitoe on behalf of the Wellcome Sanger Institute COVID-19 Surveillance Team |
| EPI_ISL_2552062 | Lighthouse Laboratory Plymouth | Wellcome Sanger Institute for the COVID-19 Genomics UK (COG-UK) Consortium | Lighthouse Laboratory Plymouth and Alex Alderton, Roberto Amato, Jeffrey Barrett, Sonia Goncalves, Ewan Harrison, David K. Jackson, Ian Johnston, Dominic Kwiatkowski, Cordelia Langford, John Sillitoe on behalf of the Wellcome Sanger Institute COVID-19 Surveillance Team |
| EPI_ISL_2552063 | Lighthouse Lab in Milton Keynes | Wellcome Sanger Institute for the COVID-19 Genomics UK (COG-UK) Consortium | The Lighthouse Lab in Milton Keynes and Alex Alderton, Roberto Amato, Jeffrey Barrett, Sonia Goncalves, Ewan Harrison, David K. Jackson, Ian Johnston, Dominic Kwiatkowski, Cordelia Langford, John Sillitoe on behalf of the Wellcome Sanger Institute COVID-19 Surveillance Team |
| EPI_ISL_2552064 | Lighthouse Lab in Glasgow | Wellcome Sanger Institute for the COVID-19 Genomics UK (COG-UK) Consortium | Harper VanSteenhouse, Yumi Kasai, David Gray, Carol Clugston, Anna Dominiczak and Alex Alderton, Roberto Amato, Jeffrey Barrett, Sonia Goncalves, Ewan Harrison, David K. Jackson, Ian Johnston, Dominic Kwiatkowski, Cordelia Langford, John Sillitoe on behalf of the Wellcome Sanger Institute COVID-19 Surveillance Team |
| EPI_ISL_2552065, EPI_ISL_2552066 | Lighthouse Lab in Milton Keynes | Wellcome Sanger Institute for the COVID-19 Genomics UK (COG-UK) Consortium | The Lighthouse Lab in Milton Keynes and Alex Alderton, Roberto Amato, Jeffrey Barrett, Sonia Goncalves, Ewan Harrison, David K. Jackson, Ian Johnston, Dominic Kwiatkowski, Cordelia Langford, John Sillitoe on behalf of the Wellcome Sanger Institute COVID-19 Surveillance Team |
| EPI_ISL_2552067 | Lighthouse Lab in Glasgow | Wellcome Sanger Institute for the COVID-19 Genomics UK (COG-UK) Consortium | Harper VanSteenhouse, Yumi Kasai, David Gray, Carol Clugston, Anna Dominiczak and Alex Alderton, Roberto Amato, Jeffrey Barrett, Sonia Goncalves, Ewan Harrison, David K. Jackson, Ian Johnston, Dominic Kwiatkowski, Cordelia Langford, John Sillitoe on behalf of the Wellcome Sanger Institute COVID-19 Surveillance Team |
| EPI_ISL_2552068 | Lighthouse Lab in Milton Keynes | Wellcome Sanger Institute for the COVID-19 Genomics UK (COG-UK) Consortium | The Lighthouse Lab in Milton Keynes and Alex Alderton, Roberto Amato, Jeffrey Barrett, Sonia Goncalves, Ewan Harrison, David K. Jackson, Ian Johnston, Dominic Kwiatkowski, Cordelia Langford, John Sillitoe on behalf of the Wellcome Sanger Institute COVID-19 Surveillance Team |
| EPI_ISL_2552069 | Lighthouse Lab in Glasgow | Wellcome Sanger Institute for the COVID-19 Genomics UK (COG-UK) Consortium | Harper VanSteenhouse, Yumi Kasai, David Gray, Carol Clugston, Anna Dominiczak and Alex Alderton, Roberto Amato, Jeffrey Barrett, Sonia Goncalves, Ewan Harrison, David K. Jackson, Ian Johnston, Dominic Kwiatkowski, Cordelia Langford, John Sillitoe on behalf of the Wellcome Sanger Institute COVID-19 Surveillance Team |
| EPI_ISL_2552071 | Lighthouse Lab in Milton Keynes | Wellcome Sanger Institute for the COVID-19 Genomics UK (COG-UK) Consortium | The Lighthouse Lab in Milton Keynes and Alex Alderton, Roberto Amato, Jeffrey Barrett, Sonia Goncalves, Ewan Harrison, David K. Jackson, Ian Johnston, Dominic Kwiatkowski, Cordelia Langford, John Sillitoe on behalf of the Wellcome Sanger Institute COVID-19 Surveillance Team |
| EPI_ISL_2552072 | Lighthouse Lab in Glasgow | Wellcome Sanger Institute for the COVID-19 Genomics UK (COG-UK) Consortium | Harper VanSteenhouse, Yumi Kasai, David Gray, Carol Clugston, Anna Dominiczak and Alex Alderton, Roberto Amato, Jeffrey Barrett, Sonia Goncalves, Ewan Harrison, David K. Jackson, Ian Johnston, Dominic Kwiatkowski, Cordelia Langford, John Sillitoe on behalf of the Wellcome Sanger Institute COVID-19 Surveillance Team |

| | | | |
|---|---|---|---|
| EPI_ISL_2552073, EPI_ISL_2552074, EPI_ISL_2552075, EPI_ISL_2552076, EPI_ISL_2552077, EPI_ISL_2552078 | Lighthouse Lab in Milton Keynes | Wellcome Sanger Institute for the COVID-19 Genomics UK (COG-UK) Consortium | The Lighthouse Lab in Milton Keynes and Alex Alderton, Roberto Amato, Jeffrey Barrett, Sonia Goncalves, Ewan Harrison, David K. Jackson, Ian Johnston, Dominic Kwiatkowski, Cordelia Langford, John Sillitoe on behalf of the Wellcome Sanger Institute COVID-19 Surveillance Team |
| EPI_ISL_2552079 | Lighthouse Laboratory Plymouth | Wellcome Sanger Institute for the COVID-19 Genomics UK (COG-UK) Consortium | Lighthouse Laboratory Plymouth and Alex Alderton, Roberto Amato, Jeffrey Barrett, Sonia Goncalves, Ewan Harrison, David K. Jackson, Ian Johnston, Dominic Kwiatkowski, Cordelia Langford, John Sillitoe on behalf of the Wellcome Sanger Institute COVID-19 Surveillance Team |
| EPI_ISL_2552080 | Lighthouse Lab in Milton Keynes | Wellcome Sanger Institute for the COVID-19 Genomics UK (COG-UK) Consortium | The Lighthouse Lab in Milton Keynes and Alex Alderton, Roberto Amato, Jeffrey Barrett, Sonia Goncalves, Ewan Harrison, David K. Jackson, Ian Johnston, Dominic Kwiatkowski, Cordelia Langford, John Sillitoe on behalf of the Wellcome Sanger Institute COVID-19 Surveillance Team |
| EPI_ISL_2552081 | Lighthouse Lab in Glasgow | Wellcome Sanger Institute for the COVID-19 Genomics UK (COG-UK) Consortium | Harper VanSteenhouse, Yumi Kasai, David Gray, Carol Clugston, Anna Dominiczak and Alex Alderton, Roberto Amato, Jeffrey Barrett, Sonia Goncalves, Ewan Harrison, David K. Jackson, Ian Johnston, Dominic Kwiatkowski, Cordelia Langford, John Sillitoe on behalf of the Wellcome Sanger Institute COVID-19 Surveillance Team |
| EPI_ISL_2552082 | Lighthouse Lab in Milton Keynes | Wellcome Sanger Institute for the COVID-19 Genomics UK (COG-UK) Consortium | The Lighthouse Lab in Milton Keynes and Alex Alderton, Roberto Amato, Jeffrey Barrett, Sonia Goncalves, Ewan Harrison, David K. Jackson, Ian Johnston, Dominic Kwiatkowski, Cordelia Langford, John Sillitoe on behalf of the Wellcome Sanger Institute COVID-19 Surveillance Team |
| EPI_ISL_2552083 | Lighthouse Laboratory Plymouth | Wellcome Sanger Institute for the COVID-19 Genomics UK (COG-UK) Consortium | Lighthouse Laboratory Plymouth and Alex Alderton, Roberto Amato, Jeffrey Barrett, Sonia Goncalves, Ewan Harrison, David K. Jackson, Ian Johnston, Dominic Kwiatkowski, Cordelia Langford, John Sillitoe on behalf of the Wellcome Sanger Institute COVID-19 Surveillance Team |
| EPI_ISL_2552084, EPI_ISL_2552085, EPI_ISL_2552086, EPI_ISL_2552087, EPI_ISL_2552088, EPI_ISL_2552089 | Lighthouse Lab in Milton Keynes | Wellcome Sanger Institute for the COVID-19 Genomics UK (COG-UK) Consortium | The Lighthouse Lab in Milton Keynes and Alex Alderton, Roberto Amato, Jeffrey Barrett, Sonia Goncalves, Ewan Harrison, David K. Jackson, Ian Johnston, Dominic Kwiatkowski, Cordelia Langford, John Sillitoe on behalf of the Wellcome Sanger Institute COVID-19 Surveillance Team |
| EPI_ISL_2552090 | Lighthouse Lab in Glasgow | Wellcome Sanger Institute for the COVID-19 Genomics UK (COG-UK) Consortium | Harper VanSteenhouse, Yumi Kasai, David Gray, Carol Clugston, Anna Dominiczak and Alex Alderton, Roberto Amato, Jeffrey Barrett, Sonia Goncalves, Ewan Harrison, David K. Jackson, Ian Johnston, Dominic Kwiatkowski, Cordelia Langford, John Sillitoe on behalf of the Wellcome Sanger Institute COVID-19 Surveillance Team |
| EPI_ISL_2552091, EPI_ISL_2552092, EPI_ISL_2552093 | Lighthouse Lab in Milton Keynes | Wellcome Sanger Institute for the COVID-19 Genomics UK (COG-UK) Consortium | The Lighthouse Lab in Milton Keynes and Alex Alderton, Roberto Amato, Jeffrey Barrett, Sonia Goncalves, Ewan Harrison, David K. Jackson, Ian Johnston, Dominic Kwiatkowski, Cordelia Langford, John Sillitoe on behalf of the Wellcome Sanger Institute COVID-19 Surveillance Team |
| EPI_ISL_2552094, EPI_ISL_2552095 | Lighthouse Lab in Glasgow | Wellcome Sanger Institute for the COVID-19 Genomics UK (COG-UK) Consortium | Harper VanSteenhouse, Yumi Kasai, David Gray, Carol Clugston, Anna Dominiczak and Alex Alderton, Roberto Amato, Jeffrey Barrett, Sonia Goncalves, Ewan Harrison, David K. Jackson, Ian Johnston, Dominic Kwiatkowski, Cordelia Langford, John Sillitoe on behalf of the Wellcome Sanger Institute COVID-19 Surveillance Team |
| EPI_ISL_2552096, EPI_ISL_2552097 | Lighthouse Lab in Milton Keynes | Wellcome Sanger Institute for the COVID-19 Genomics UK (COG-UK) Consortium | The Lighthouse Lab in Milton Keynes and Alex Alderton, Roberto Amato, Jeffrey Barrett, Sonia Goncalves, Ewan Harrison, David K. Jackson, Ian Johnston, Dominic Kwiatkowski, Cordelia Langford, John Sillitoe on behalf of the Wellcome Sanger Institute COVID-19 Surveillance Team |
| EPI_ISL_2552098 | Lighthouse Lab in Glasgow | Wellcome Sanger Institute for the COVID-19 Genomics UK (COG-UK) Consortium | Harper VanSteenhouse, Yumi Kasai, David Gray, Carol Clugston, Anna Dominiczak and Alex Alderton, Roberto Amato, Jeffrey Barrett, Sonia Goncalves, Ewan Harrison, David K. Jackson, Ian Johnston, Dominic Kwiatkowski, Cordelia Langford, John Sillitoe on behalf of the Wellcome Sanger Institute COVID-19 Surveillance Team |
| EPI_ISL_2552099, EPI_ISL_2552100, EPI_ISL_2552101, EPI_ISL_2552102, EPI_ISL_2552103, EPI_ISL_2552104 | Lighthouse Lab in Milton Keynes | Wellcome Sanger Institute for the COVID-19 Genomics UK (COG-UK) Consortium | The Lighthouse Lab in Milton Keynes and Alex Alderton, Roberto Amato, Jeffrey Barrett, Sonia Goncalves, Ewan Harrison, David K. Jackson, Ian Johnston, Dominic Kwiatkowski, Cordelia Langford, John Sillitoe on behalf of the Wellcome Sanger Institute COVID-19 Surveillance Team |
| EPI_ISL_2552105 | Lighthouse Laboratory Plymouth | Wellcome Sanger Institute for the COVID-19 Genomics UK (COG-UK) Consortium | Lighthouse Laboratory Plymouth and Alex Alderton, Roberto Amato, Jeffrey Barrett, Sonia Goncalves, Ewan Harrison, David K. Jackson, Ian Johnston, Dominic Kwiatkowski, Cordelia Langford, John Sillitoe on behalf of the Wellcome Sanger Institute COVID-19 Surveillance Team |
| EPI_ISL_2552106 | Lighthouse Lab in Glasgow | Wellcome Sanger Institute for the COVID-19 Genomics UK (COG-UK) Consortium | Harper VanSteenhouse, Yumi Kasai, David Gray, Carol Clugston, Anna Dominiczak and Alex Alderton, Roberto Amato, Jeffrey Barrett, Sonia Goncalves, Ewan Harrison, David K. Jackson, Ian Johnston, Dominic Kwiatkowski, Cordelia Langford, John Sillitoe on behalf of the Wellcome Sanger Institute COVID-19 Surveillance Team |
| EPI_ISL_2552107, EPI_ISL_2552108, EPI_ISL_2552109, EPI_ISL_2552110 | Lighthouse Lab in Milton Keynes | Wellcome Sanger Institute for the COVID-19 Genomics UK (COG-UK) Consortium | The Lighthouse Lab in Milton Keynes and Alex Alderton, Roberto Amato, Jeffrey Barrett, Sonia Goncalves, Ewan Harrison, David K. Jackson, Ian Johnston, Dominic Kwiatkowski, Cordelia Langford, John Sillitoe on behalf of the Wellcome Sanger Institute COVID-19 Surveillance Team |
| EPI_ISL_2552111, EPI_ISL_2552112 | Lighthouse Lab in Glasgow | Wellcome Sanger Institute for the COVID-19 Genomics UK (COG-UK) Consortium | Harper VanSteenhouse, Yumi Kasai, David Gray, Carol Clugston, Anna Dominiczak and Alex Alderton, Roberto Amato, Jeffrey Barrett, Sonia Goncalves, Ewan Harrison, David K. Jackson, Ian Johnston, Dominic Kwiatkowski, Cordelia Langford, John Sillitoe on behalf of the Wellcome Sanger Institute COVID-19 Surveillance Team |
| EPI_ISL_2552113 | Lighthouse Lab in Milton Keynes | Wellcome Sanger Institute for the COVID-19 Genomics UK (COG-UK) Consortium | The Lighthouse Lab in Milton Keynes and Alex Alderton, Roberto Amato, Jeffrey Barrett, Sonia Goncalves, Ewan Harrison, David K. Jackson, Ian Johnston, Dominic Kwiatkowski, Cordelia Langford, John Sillitoe on behalf of the Wellcome Sanger Institute COVID-19 Surveillance Team |
| EPI_ISL_2552114 | Lighthouse Lab in Glasgow | Wellcome Sanger Institute for the COVID-19 Genomics UK (COG-UK) Consortium | Harper VanSteenhouse, Yumi Kasai, David Gray, Carol Clugston, Anna Dominiczak and Alex Alderton, Roberto Amato, Jeffrey Barrett, Sonia Goncalves, Ewan Harrison, David K. Jackson, Ian Johnston, Dominic Kwiatkowski, Cordelia Langford, John Sillitoe on behalf of the Wellcome Sanger Institute COVID-19 Surveillance Team |
| EPI_ISL_2552115 | Lighthouse Lab in Milton Keynes | Wellcome Sanger Institute for the COVID-19 Genomics UK (COG-UK) Consortium | The Lighthouse Lab in Milton Keynes and Alex Alderton, Roberto Amato, Jeffrey Barrett, Sonia Goncalves, Ewan Harrison, David K. Jackson, Ian Johnston, Dominic Kwiatkowski, Cordelia Langford, John Sillitoe on behalf of the Wellcome Sanger Institute COVID-19 Surveillance Team |
| EPI_ISL_2552116 | Lighthouse Lab in Glasgow | Wellcome Sanger Institute for the COVID-19 Genomics UK (COG-UK) Consortium | Harper VanSteenhouse, Yumi Kasai, David Gray, Carol Clugston, Anna Dominiczak and Alex Alderton, Roberto Amato, Jeffrey Barrett, Sonia Goncalves, Ewan Harrison, David K. Jackson, Ian Johnston, Dominic Kwiatkowski, Cordelia Langford, John Sillitoe on behalf of the Wellcome Sanger Institute COVID-19 Surveillance Team |
| EPI_ISL_2552117, EPI_ISL_2552118, EPI_ISL_2552119, EPI_ISL_2552120, EPI_ISL_2552121, EPI_ISL_2552122 | Lighthouse Lab in Milton Keynes | Wellcome Sanger Institute for the COVID-19 Genomics UK (COG-UK) Consortium | The Lighthouse Lab in Milton Keynes and Alex Alderton, Roberto Amato, Jeffrey Barrett, Sonia Goncalves, Ewan Harrison, David K. Jackson, Ian Johnston, Dominic Kwiatkowski, Cordelia Langford, John Sillitoe on behalf of the Wellcome Sanger Institute COVID-19 Surveillance Team |
| EPI_ISL_2552123, EPI_ISL_2552124, EPI_ISL_2552125 | Lighthouse Lab in Glasgow | Wellcome Sanger Institute for the COVID-19 Genomics UK (COG-UK) Consortium | Harper VanSteenhouse, Yumi Kasai, David Gray, Carol Clugston, Anna Dominiczak and Alex Alderton, Roberto Amato, Jeffrey Barrett, Sonia Goncalves, Ewan Harrison, David K. Jackson, Ian Johnston, Dominic Kwiatkowski, Cordelia Langford, John Sillitoe on behalf of the Wellcome Sanger Institute COVID-19 Surveillance Team |
| EPI_ISL_2552126, EPI_ISL_2552127, EPI_ISL_2552128, EPI_ISL_2552129, EPI_ISL_2552130, EPI_ISL_2552131 | Lighthouse Lab in Milton Keynes | Wellcome Sanger Institute for the COVID-19 Genomics UK (COG-UK) Consortium | The Lighthouse Lab in Milton Keynes and Alex Alderton, Roberto Amato, Jeffrey Barrett, Sonia Goncalves, Ewan Harrison, David K. Jackson, Ian Johnston, Dominic Kwiatkowski, Cordelia Langford, John Sillitoe on behalf of the Wellcome Sanger Institute COVID-19 Surveillance Team |
| EPI_ISL_2552132, EPI_ISL_2552133 | Lighthouse Lab in Glasgow | Wellcome Sanger Institute for the COVID-19 Genomics UK (COG-UK) Consortium | Harper VanSteenhouse, Yumi Kasai, David Gray, Carol Clugston, Anna Dominiczak and Alex Alderton, Roberto Amato, Jeffrey Barrett, Sonia Goncalves, Ewan Harrison, David K. Jackson, Ian Johnston, Dominic Kwiatkowski, Cordelia Langford, John Sillitoe on behalf of the Wellcome Sanger Institute COVID-19 Surveillance Team |
| EPI_ISL_2552134 | Lighthouse Lab in Milton Keynes | Wellcome Sanger Institute for the COVID-19 Genomics UK (COG-UK) Consortium | The Lighthouse Lab in Milton Keynes and Alex Alderton, Roberto Amato, Jeffrey Barrett, Sonia Goncalves, Ewan Harrison, David K. Jackson, Ian Johnston, Dominic Kwiatkowski, Cordelia Langford, John Sillitoe on behalf of the Wellcome Sanger Institute COVID-19 Surveillance Team |
| EPI_ISL_2552135 | Lighthouse Lab in Glasgow | Wellcome Sanger Institute for the COVID-19 Genomics UK (COG-UK) Consortium | Harper VanSteenhouse, Yumi Kasai, David Gray, Carol Clugston, Anna Dominiczak and Alex Alderton, Roberto Amato, Jeffrey Barrett, Sonia Goncalves, Ewan Harrison, David K. Jackson, Ian Johnston, Dominic Kwiatkowski, Cordelia Langford, John Sillitoe on behalf of the Wellcome Sanger Institute COVID-19 Surveillance Team |
| EPI_ISL_2552136, EPI_ISL_2552137, EPI_ISL_2552138 | Lighthouse Lab in Milton Keynes | Wellcome Sanger Institute for the COVID-19 Genomics UK (COG-UK) Consortium | The Lighthouse Lab in Milton Keynes and Alex Alderton, Roberto Amato, Jeffrey Barrett, Sonia Goncalves, Ewan Harrison, David K. Jackson, Ian Johnston, Dominic Kwiatkowski, Cordelia Langford, John Sillitoe on behalf of the Wellcome Sanger Institute COVID-19 Surveillance Team |
| EPI_ISL_2552139, EPI_ISL_2552140 | Lighthouse Lab in Glasgow | Wellcome Sanger Institute for the COVID-19 Genomics UK (COG-UK) Consortium | Harper VanSteenhouse, Yumi Kasai, David Gray, Carol Clugston, Anna Dominiczak and Alex Alderton, Roberto Amato, Jeffrey Barrett, Sonia Goncalves, Ewan Harrison, David K. Jackson, Ian Johnston, Dominic Kwiatkowski, Cordelia Langford, John Sillitoe on behalf of the Wellcome Sanger Institute COVID-19 Surveillance Team |
| EPI_ISL_2552141, EPI_ISL_2552142 | Lighthouse Laboratory Plymouth | Wellcome Sanger Institute for the COVID-19 Genomics UK (COG-UK) Consortium | Lighthouse Laboratory Plymouth and Alex Alderton, Roberto Amato, Jeffrey Barrett, Sonia Goncalves, Ewan Harrison, David K. Jackson, Ian Johnston, Dominic Kwiatkowski, Cordelia Langford, John Sillitoe on behalf of the Wellcome Sanger Institute COVID-19 Surveillance Team |
| EPI_ISL_2552143 | Lighthouse Lab in Glasgow | Wellcome Sanger Institute for the COVID-19 Genomics UK (COG-UK) Consortium | Harper VanSteenhouse, Yumi Kasai, David Gray, Carol Clugston, Anna Dominiczak and Alex Alderton, Roberto Amato, Jeffrey Barrett, Sonia Goncalves, Ewan Harrison, David K. Jackson, Ian Johnston, Dominic Kwiatkowski, Cordelia Langford, John Sillitoe on behalf of the Wellcome Sanger Institute COVID-19 Surveillance Team |
| EPI_ISL_2552144 | Lighthouse Lab in Milton Keynes | Wellcome Sanger Institute for the COVID-19 Genomics UK (COG-UK) Consortium | The Lighthouse Lab in Milton Keynes and Alex Alderton, Roberto Amato, Jeffrey Barrett, Sonia Goncalves, Ewan Harrison, David K. Jackson, Ian Johnston, Dominic Kwiatkowski, Cordelia Langford, John Sillitoe on behalf of the Wellcome Sanger Institute COVID-19 Surveillance Team |
| EPI_ISL_2552145 | Lighthouse Lab in Glasgow | Wellcome Sanger Institute for the COVID-19 Genomics UK (COG-UK) Consortium | Harper VanSteenhouse, Yumi Kasai, David Gray, Carol Clugston, Anna Dominiczak and Alex Alderton, Roberto Amato, Jeffrey Barrett, Sonia Goncalves, Ewan Harrison, David K. Jackson, Ian Johnston, Dominic Kwiatkowski, Cordelia Langford, John Sillitoe on behalf of the Wellcome Sanger Institute COVID-19 Surveillance Team |
| EPI_ISL_2552146, EPI_ISL_2552147, EPI_ISL_2552148 | Lighthouse Lab in Milton Keynes | Wellcome Sanger Institute for the COVID-19 Genomics UK (COG-UK) Consortium | The Lighthouse Lab in Milton Keynes and Alex Alderton, Roberto Amato, Jeffrey Barrett, Sonia Goncalves, Ewan Harrison, David K. Jackson, Ian Johnston, Dominic Kwiatkowski, Cordelia Langford, John Sillitoe on behalf of the Wellcome Sanger Institute COVID-19 Surveillance Team |
| EPI_ISL_2552149 | Lighthouse Lab in Glasgow | Wellcome Sanger Institute for the COVID-19 Genomics UK (COG-UK) | Harper VanSteenhouse, Yumi Kasai, David Gray, Carol Clugston, Anna Dominiczak and Alex Alderton, Roberto Amato, Jeffrey Barrett, Sonia Goncalves, Ewan Harrison, David K. Jackson, Ian Johnston, Dominic Kwiatkowski, Cordelia Langford, |

| | | | |
|---|---|---|---|
| | | Consortium | John Sillitoe on behalf of the Wellcome Sanger Institute COVID-19 Surveillance Team |
| EPI_ISL_2552150, EPI_ISL_2552151, EPI_ISL_2552152, EPI_ISL_2552153 | Lighthouse Lab in Milton Keynes | Wellcome Sanger Institute for the COVID-19 Genomics UK (COG-UK) Consortium | The Lighthouse Lab in Milton Keynes and Alex Alderton, Roberto Amato, Jeffrey Barrett, Sonia Goncalves, Ewan Harrison, David K. Jackson, Ian Johnston, Dominic Kwiatkowski, Cordelia Langford, John Sillitoe on behalf of the Wellcome Sanger Institute COVID-19 Surveillance Team |
| EPI_ISL_2552154 | Lighthouse Lab in Glasgow | Wellcome Sanger Institute for the COVID-19 Genomics UK (COG-UK) Consortium | Harper VanSteenhouse, Yumi Kasai, David Gray, Carol Clugston, Anna Dominiczak and Alex Alderton, Roberto Amato, Jeffrey Barrett, Sonia Goncalves, Ewan Harrison, David K. Jackson, Ian Johnston, Dominic Kwiatkowski, Cordelia Langford, John Sillitoe on behalf of the Wellcome Sanger Institute COVID-19 Surveillance Team |
| EPI_ISL_2552155, EPI_ISL_2552156 | Lighthouse Lab in Milton Keynes | Wellcome Sanger Institute for the COVID-19 Genomics UK (COG-UK) Consortium | The Lighthouse Lab in Milton Keynes and Alex Alderton, Roberto Amato, Jeffrey Barrett, Sonia Goncalves, Ewan Harrison, David K. Jackson, Ian Johnston, Dominic Kwiatkowski, Cordelia Langford, John Sillitoe on behalf of the Wellcome Sanger Institute COVID-19 Surveillance Team |
| EPI_ISL_2552157 | Lighthouse Lab in Glasgow | Wellcome Sanger Institute for the COVID-19 Genomics UK (COG-UK) Consortium | Harper VanSteenhouse, Yumi Kasai, David Gray, Carol Clugston, Anna Dominiczak and Alex Alderton, Roberto Amato, Jeffrey Barrett, Sonia Goncalves, Ewan Harrison, David K. Jackson, Ian Johnston, Dominic Kwiatkowski, Cordelia Langford, John Sillitoe on behalf of the Wellcome Sanger Institute COVID-19 Surveillance Team |
| EPI_ISL_2552158, EPI_ISL_2552162, EPI_ISL_2552163, EPI_ISL_2552164 | Lighthouse Lab in Milton Keynes | Wellcome Sanger Institute for the COVID-19 Genomics UK (COG-UK) Consortium | The Lighthouse Lab in Milton Keynes and Alex Alderton, Roberto Amato, Jeffrey Barrett, Sonia Goncalves, Ewan Harrison, David K. Jackson, Ian Johnston, Dominic Kwiatkowski, Cordelia Langford, John Sillitoe on behalf of the Wellcome Sanger Institute COVID-19 Surveillance Team |
| EPI_ISL_2552166 | Lighthouse Lab in Glasgow | Wellcome Sanger Institute for the COVID-19 Genomics UK (COG-UK) Consortium | Harper VanSteenhouse, Yumi Kasai, David Gray, Carol Clugston, Anna Dominiczak and Alex Alderton, Roberto Amato, Jeffrey Barrett, Sonia Goncalves, Ewan Harrison, David K. Jackson, Ian Johnston, Dominic Kwiatkowski, Cordelia Langford, John Sillitoe on behalf of the Wellcome Sanger Institute COVID-19 Surveillance Team |
| EPI_ISL_2552167 | Lighthouse Lab in Milton Keynes | Wellcome Sanger Institute for the COVID-19 Genomics UK (COG-UK) Consortium | The Lighthouse Lab in Milton Keynes and Alex Alderton, Roberto Amato, Jeffrey Barrett, Sonia Goncalves, Ewan Harrison, David K. Jackson, Ian Johnston, Dominic Kwiatkowski, Cordelia Langford, John Sillitoe on behalf of the Wellcome Sanger Institute COVID-19 Surveillance Team |
| EPI_ISL_2552168 | Lighthouse Lab in Glasgow | Wellcome Sanger Institute for the COVID-19 Genomics UK (COG-UK) Consortium | Harper VanSteenhouse, Yumi Kasai, David Gray, Carol Clugston, Anna Dominiczak and Alex Alderton, Roberto Amato, Jeffrey Barrett, Sonia Goncalves, Ewan Harrison, David K. Jackson, Ian Johnston, Dominic Kwiatkowski, Cordelia Langford, John Sillitoe on behalf of the Wellcome Sanger Institute COVID-19 Surveillance Team |
| EPI_ISL_2552169 | Lighthouse Lab in Milton Keynes | Wellcome Sanger Institute for the COVID-19 Genomics UK (COG-UK) Consortium | The Lighthouse Lab in Milton Keynes and Alex Alderton, Roberto Amato, Jeffrey Barrett, Sonia Goncalves, Ewan Harrison, David K. Jackson, Ian Johnston, Dominic Kwiatkowski, Cordelia Langford, John Sillitoe on behalf of the Wellcome Sanger Institute COVID-19 Surveillance Team |
| EPI_ISL_2552170 | Lighthouse Lab in Glasgow | Wellcome Sanger Institute for the COVID-19 Genomics UK (COG-UK) Consortium | Harper VanSteenhouse, Yumi Kasai, David Gray, Carol Clugston, Anna Dominiczak and Alex Alderton, Roberto Amato, Jeffrey Barrett, Sonia Goncalves, Ewan Harrison, David K. Jackson, Ian Johnston, Dominic Kwiatkowski, Cordelia Langford, John Sillitoe on behalf of the Wellcome Sanger Institute COVID-19 Surveillance Team |
| EPI_ISL_2552171, EPI_ISL_2552172, EPI_ISL_2552173 | Lighthouse Lab in Milton Keynes | Wellcome Sanger Institute for the COVID-19 Genomics UK (COG-UK) Consortium | The Lighthouse Lab in Milton Keynes and Alex Alderton, Roberto Amato, Jeffrey Barrett, Sonia Goncalves, Ewan Harrison, David K. Jackson, Ian Johnston, Dominic Kwiatkowski, Cordelia Langford, John Sillitoe on behalf of the Wellcome Sanger Institute COVID-19 Surveillance Team |
| EPI_ISL_2552174, EPI_ISL_2552175 | Lighthouse Lab in Glasgow | Wellcome Sanger Institute for the COVID-19 Genomics UK (COG-UK) Consortium | Harper VanSteenhouse, Yumi Kasai, David Gray, Carol Clugston, Anna Dominiczak and Alex Alderton, Roberto Amato, Jeffrey Barrett, Sonia Goncalves, Ewan Harrison, David K. Jackson, Ian Johnston, Dominic Kwiatkowski, Cordelia Langford, John Sillitoe on behalf of the Wellcome Sanger Institute COVID-19 Surveillance Team |
| EPI_ISL_2552176, EPI_ISL_2552177 | Lighthouse Lab in Milton Keynes | Wellcome Sanger Institute for the COVID-19 Genomics UK (COG-UK) Consortium | The Lighthouse Lab in Milton Keynes and Alex Alderton, Roberto Amato, Jeffrey Barrett, Sonia Goncalves, Ewan Harrison, David K. Jackson, Ian Johnston, Dominic Kwiatkowski, Cordelia Langford, John Sillitoe on behalf of the Wellcome Sanger Institute COVID-19 Surveillance Team |
| EPI_ISL_2552179 | Lighthouse Lab in Glasgow | Wellcome Sanger Institute for the COVID-19 Genomics UK (COG-UK) Consortium | Harper VanSteenhouse, Yumi Kasai, David Gray, Carol Clugston, Anna Dominiczak and Alex Alderton, Roberto Amato, Jeffrey Barrett, Sonia Goncalves, Ewan Harrison, David K. Jackson, Ian Johnston, Dominic Kwiatkowski, Cordelia Langford, John Sillitoe on behalf of the Wellcome Sanger Institute COVID-19 Surveillance Team |
| EPI_ISL_2552180, EPI_ISL_2552181, EPI_ISL_2552182, EPI_ISL_2552183 | Lighthouse Lab in Milton Keynes | Wellcome Sanger Institute for the COVID-19 Genomics UK (COG-UK) Consortium | The Lighthouse Lab in Milton Keynes and Alex Alderton, Roberto Amato, Jeffrey Barrett, Sonia Goncalves, Ewan Harrison, David K. Jackson, Ian Johnston, Dominic Kwiatkowski, Cordelia Langford, John Sillitoe on behalf of the Wellcome Sanger Institute COVID-19 Surveillance Team |
| EPI_ISL_2552184 | Lighthouse Laboratory Plymouth | Wellcome Sanger Institute for the COVID-19 Genomics UK (COG-UK) Consortium | Lighthouse Laboratory Plymouth and Alex Alderton, Roberto Amato, Jeffrey Barrett, Sonia Goncalves, Ewan Harrison, David K. Jackson, Ian Johnston, Dominic Kwiatkowski, Cordelia Langford, John Sillitoe on behalf of the Wellcome Sanger Institute COVID-19 Surveillance Team |
| EPI_ISL_2552185 | Lighthouse Lab in Glasgow | Wellcome Sanger Institute for the COVID-19 Genomics UK (COG-UK) Consortium | Harper VanSteenhouse, Yumi Kasai, David Gray, Carol Clugston, Anna Dominiczak and Alex Alderton, Roberto Amato, Jeffrey Barrett, Sonia Goncalves, Ewan Harrison, David K. Jackson, Ian Johnston, Dominic Kwiatkowski, Cordelia Langford, John Sillitoe on behalf of the Wellcome Sanger Institute COVID-19 Surveillance Team |
| EPI_ISL_2552186, EPI_ISL_2552187, EPI_ISL_2552188, EPI_ISL_2552189 | Lighthouse Lab in Milton Keynes | Wellcome Sanger Institute for the COVID-19 Genomics UK (COG-UK) Consortium | The Lighthouse Lab in Milton Keynes and Alex Alderton, Roberto Amato, Jeffrey Barrett, Sonia Goncalves, Ewan Harrison, David K. Jackson, Ian Johnston, Dominic Kwiatkowski, Cordelia Langford, John Sillitoe on behalf of the Wellcome Sanger Institute COVID-19 Surveillance Team |
| EPI_ISL_2552190 | Lighthouse Laboratory Plymouth | Wellcome Sanger Institute for the COVID-19 Genomics UK (COG-UK) Consortium | Lighthouse Laboratory Plymouth and Alex Alderton, Roberto Amato, Jeffrey Barrett, Sonia Goncalves, Ewan Harrison, David K. Jackson, Ian Johnston, Dominic Kwiatkowski, Cordelia Langford, John Sillitoe on behalf of the Wellcome Sanger Institute COVID-19 Surveillance Team |
| EPI_ISL_2552191 | Lighthouse Lab in Milton Keynes | Wellcome Sanger Institute for the COVID-19 Genomics UK (COG-UK) Consortium | The Lighthouse Lab in Milton Keynes and Alex Alderton, Roberto Amato, Jeffrey Barrett, Sonia Goncalves, Ewan Harrison, David K. Jackson, Ian Johnston, Dominic Kwiatkowski, Cordelia Langford, John Sillitoe on behalf of the Wellcome Sanger Institute COVID-19 Surveillance Team |
| EPI_ISL_2552192, EPI_ISL_2552193 | Lighthouse Lab in Glasgow | Wellcome Sanger Institute for the COVID-19 Genomics UK (COG-UK) Consortium | Harper VanSteenhouse, Yumi Kasai, David Gray, Carol Clugston, Anna Dominiczak and Alex Alderton, Roberto Amato, Jeffrey Barrett, Sonia Goncalves, Ewan Harrison, David K. Jackson, Ian Johnston, Dominic Kwiatkowski, Cordelia Langford, John Sillitoe on behalf of the Wellcome Sanger Institute COVID-19 Surveillance Team |
| EPI_ISL_2552194, EPI_ISL_2552196, EPI_ISL_2552197, EPI_ISL_2552198, EPI_ISL_2552199, EPI_ISL_2552200, EPI_ISL_2552201, EPI_ISL_2552202, EPI_ISL_2552203, EPI_ISL_2552204, EPI_ISL_2552205, EPI_ISL_2552206 | | | |
| see above | Lighthouse Lab in Milton Keynes | Wellcome Sanger Institute for the COVID-19 Genomics UK (COG-UK) Consortium | The Lighthouse Lab in Milton Keynes and Alex Alderton, Roberto Amato, Jeffrey Barrett, Sonia Goncalves, Ewan Harrison, David K. Jackson, Ian Johnston, Dominic Kwiatkowski, Cordelia Langford, John Sillitoe on behalf of the Wellcome Sanger Institute COVID-19 Surveillance Team |
| EPI_ISL_2552208 | Lighthouse Lab in Glasgow | Wellcome Sanger Institute for the COVID-19 Genomics UK (COG-UK) Consortium | Harper VanSteenhouse, Yumi Kasai, David Gray, Carol Clugston, Anna Dominiczak and Alex Alderton, Roberto Amato, Jeffrey Barrett, Sonia Goncalves, Ewan Harrison, David K. Jackson, Ian Johnston, Dominic Kwiatkowski, Cordelia Langford, John Sillitoe on behalf of the Wellcome Sanger Institute COVID-19 Surveillance Team |
| EPI_ISL_2552209, EPI_ISL_2552210, EPI_ISL_2552211, EPI_ISL_2552212, EPI_ISL_2552214, EPI_ISL_2552215 | Lighthouse Lab in Milton Keynes | Wellcome Sanger Institute for the COVID-19 Genomics UK (COG-UK) Consortium | The Lighthouse Lab in Milton Keynes and Alex Alderton, Roberto Amato, Jeffrey Barrett, Sonia Goncalves, Ewan Harrison, David K. Jackson, Ian Johnston, Dominic Kwiatkowski, Cordelia Langford, John Sillitoe on behalf of the Wellcome Sanger Institute COVID-19 Surveillance Team |
| EPI_ISL_2552216 | Lighthouse Lab in Glasgow | Wellcome Sanger Institute for the COVID-19 Genomics UK (COG-UK) Consortium | Harper VanSteenhouse, Yumi Kasai, David Gray, Carol Clugston, Anna Dominiczak and Alex Alderton, Roberto Amato, Jeffrey Barrett, Sonia Goncalves, Ewan Harrison, David K. Jackson, Ian Johnston, Dominic Kwiatkowski, Cordelia Langford, John Sillitoe on behalf of the Wellcome Sanger Institute COVID-19 Surveillance Team |
| EPI_ISL_2552217, EPI_ISL_2552218, EPI_ISL_2552219, EPI_ISL_2552220 | Lighthouse Lab in Milton Keynes | Wellcome Sanger Institute for the COVID-19 Genomics UK (COG-UK) Consortium | The Lighthouse Lab in Milton Keynes and Alex Alderton, Roberto Amato, Jeffrey Barrett, Sonia Goncalves, Ewan Harrison, David K. Jackson, Ian Johnston, Dominic Kwiatkowski, Cordelia Langford, John Sillitoe on behalf of the Wellcome Sanger Institute COVID-19 Surveillance Team |
| EPI_ISL_2552221 | Lighthouse Lab in Glasgow | Wellcome Sanger Institute for the COVID-19 Genomics UK (COG-UK) Consortium | Harper VanSteenhouse, Yumi Kasai, David Gray, Carol Clugston, Anna Dominiczak and Alex Alderton, Roberto Amato, Jeffrey Barrett, Sonia Goncalves, Ewan Harrison, David K. Jackson, Ian Johnston, Dominic Kwiatkowski, Cordelia Langford, John Sillitoe on behalf of the Wellcome Sanger Institute COVID-19 Surveillance Team |
| EPI_ISL_2552222, EPI_ISL_2552223, EPI_ISL_2552224, EPI_ISL_2552225, EPI_ISL_2552226, EPI_ISL_2552227, EPI_ISL_2552228, EPI_ISL_2552229 | Lighthouse Lab in Milton Keynes | Wellcome Sanger Institute for the COVID-19 Genomics UK (COG-UK) Consortium | The Lighthouse Lab in Milton Keynes and Alex Alderton, Roberto Amato, Jeffrey Barrett, Sonia Goncalves, Ewan Harrison, David K. Jackson, Ian Johnston, Dominic Kwiatkowski, Cordelia Langford, John Sillitoe on behalf of the Wellcome Sanger Institute COVID-19 Surveillance Team |
| EPI_ISL_2552230, EPI_ISL_2552231 | Lighthouse Lab in Glasgow | Wellcome Sanger Institute for the COVID-19 Genomics UK (COG-UK) Consortium | Harper VanSteenhouse, Yumi Kasai, David Gray, Carol Clugston, Anna Dominiczak and Alex Alderton, Roberto Amato, Jeffrey Barrett, Sonia Goncalves, Ewan Harrison, David K. Jackson, Ian Johnston, Dominic Kwiatkowski, Cordelia Langford, John Sillitoe on behalf of the Wellcome Sanger Institute COVID-19 Surveillance Team |
| EPI_ISL_2552232, EPI_ISL_2552233 | Lighthouse Lab in Milton Keynes | Wellcome Sanger Institute for the COVID-19 Genomics UK (COG-UK) Consortium | The Lighthouse Lab in Milton Keynes and Alex Alderton, Roberto Amato, Jeffrey Barrett, Sonia Goncalves, Ewan Harrison, David K. Jackson, Ian Johnston, Dominic Kwiatkowski, Cordelia Langford, John Sillitoe on behalf of the Wellcome Sanger Institute COVID-19 Surveillance Team |
| EPI_ISL_2552234 | Lighthouse Lab in Glasgow | Wellcome Sanger Institute for the COVID-19 Genomics UK (COG-UK) Consortium | Harper VanSteenhouse, Yumi Kasai, David Gray, Carol Clugston, Anna Dominiczak and Alex Alderton, Roberto Amato, Jeffrey Barrett, Sonia Goncalves, Ewan Harrison, David K. Jackson, Ian Johnston, Dominic Kwiatkowski, Cordelia Langford, John Sillitoe on behalf of the Wellcome Sanger Institute COVID-19 Surveillance Team |
| EPI_ISL_2552235 | Lighthouse Laboratory Plymouth | Wellcome Sanger Institute for the COVID-19 Genomics UK (COG-UK) Consortium | Lighthouse Laboratory Plymouth and Alex Alderton, Roberto Amato, Jeffrey Barrett, Sonia Goncalves, Ewan Harrison, David K. Jackson, Ian Johnston, Dominic Kwiatkowski, Cordelia Langford, John Sillitoe on behalf of the Wellcome Sanger Institute COVID-19 Surveillance Team |
| EPI_ISL_2552236, EPI_ISL_2552237 | Lighthouse Lab in Milton Keynes | Wellcome Sanger Institute for the COVID-19 Genomics UK (COG-UK) Consortium | The Lighthouse Lab in Milton Keynes and Alex Alderton, Roberto Amato, Jeffrey Barrett, Sonia Goncalves, Ewan Harrison, David K. Jackson, Ian Johnston, Dominic Kwiatkowski, Cordelia Langford, John Sillitoe on behalf of the Wellcome Sanger Institute COVID-19 Surveillance Team |
| EPI_ISL_2552238 | Lighthouse Lab in Glasgow | Wellcome Sanger Institute for the COVID-19 Genomics UK (COG-UK) Consortium | Harper VanSteenhouse, Yumi Kasai, David Gray, Carol Clugston, Anna Dominiczak and Alex Alderton, Roberto Amato, Jeffrey Barrett, Sonia Goncalves, Ewan Harrison, David K. Jackson, Ian Johnston, Dominic Kwiatkowski, Cordelia Langford, John Sillitoe on behalf of the Wellcome Sanger Institute COVID-19 Surveillance Team |
| EPI_ISL_2552240 | Lighthouse Laboratory Plymouth | Wellcome Sanger Institute for the COVID-19 Genomics UK (COG-UK) Consortium | Lighthouse Laboratory Plymouth and Alex Alderton, Roberto Amato, Jeffrey Barrett, Sonia Goncalves, Ewan Harrison, David K. Jackson, Ian Johnston, Dominic Kwiatkowski, Cordelia Langford, John Sillitoe on behalf of the Wellcome Sanger Institute COVID-19 Surveillance Team |
| EPI_ISL_2552241 | Lighthouse Lab in Milton Keynes | Wellcome Sanger Institute for the COVID-19 Genomics UK (COG-UK) | The Lighthouse Lab in Milton Keynes and Alex Alderton, Roberto Amato, Jeffrey Barrett, Sonia Goncalves, Ewan Harrison, David K. Jackson, Ian Johnston, Dominic Kwiatkowski, Cordelia Langford, John Sillitoe on behalf of the Wellcome Sanger |

| | | | |
|---|---|---|---|
| | | Consortium | Institute COVID-19 Surveillance Team |
| EPI_ISL_2552242 | Lighthouse Lab in Glasgow | Wellcome Sanger Institute for the COVID-19 Genomics UK (COG-UK) Consortium | Harper VanSteenhouse, Yumi Kasai, David Gray, Carol Clugston, Anna Dominiczak and Alex Alderton, Roberto Amato, Jeffrey Barrett, Sonia Goncalves, Ewan Harrison, David K. Jackson, Ian Johnston, Dominic Kwiatkowski, Cordelia Langford, John Sillitoe on behalf of the Wellcome Sanger Institute COVID-19 Surveillance Team |
| EPI_ISL_2552243, EPI_ISL_2552244, EPI_ISL_2552245, EPI_ISL_2552246, EPI_ISL_2552247, EPI_ISL_2552248, EPI_ISL_2552249, EPI_ISL_2552250 | Lighthouse Lab in Milton Keynes | Wellcome Sanger Institute for the COVID-19 Genomics UK (COG-UK) Consortium | The Lighthouse Lab in Milton Keynes and Alex Alderton, Roberto Amato, Jeffrey Barrett, Sonia Goncalves, Ewan Harrison, David K. Jackson, Ian Johnston, Dominic Kwiatkowski, Cordelia Langford, John Sillitoe on behalf of the Wellcome Sanger Institute COVID-19 Surveillance Team |
| EPI_ISL_2552252 | Lighthouse Lab in Glasgow | Wellcome Sanger Institute for the COVID-19 Genomics UK (COG-UK) Consortium | Harper VanSteenhouse, Yumi Kasai, David Gray, Carol Clugston, Anna Dominiczak and Alex Alderton, Roberto Amato, Jeffrey Barrett, Sonia Goncalves, Ewan Harrison, David K. Jackson, Ian Johnston, Dominic Kwiatkowski, Cordelia Langford, John Sillitoe on behalf of the Wellcome Sanger Institute COVID-19 Surveillance Team |
| EPI_ISL_2552253, EPI_ISL_2552254 | Lighthouse Lab in Milton Keynes | Wellcome Sanger Institute for the COVID-19 Genomics UK (COG-UK) Consortium | The Lighthouse Lab in Milton Keynes and Alex Alderton, Roberto Amato, Jeffrey Barrett, Sonia Goncalves, Ewan Harrison, David K. Jackson, Ian Johnston, Dominic Kwiatkowski, Cordelia Langford, John Sillitoe on behalf of the Wellcome Sanger Institute COVID-19 Surveillance Team |
| EPI_ISL_2552255, EPI_ISL_2552256 | Lighthouse Lab in Glasgow | Wellcome Sanger Institute for the COVID-19 Genomics UK (COG-UK) Consortium | Harper VanSteenhouse, Yumi Kasai, David Gray, Carol Clugston, Anna Dominiczak and Alex Alderton, Roberto Amato, Jeffrey Barrett, Sonia Goncalves, Ewan Harrison, David K. Jackson, Ian Johnston, Dominic Kwiatkowski, Cordelia Langford, John Sillitoe on behalf of the Wellcome Sanger Institute COVID-19 Surveillance Team |
| EPI_ISL_2552257, EPI_ISL_2552258, EPI_ISL_2552259 | Lighthouse Lab in Milton Keynes | Wellcome Sanger Institute for the COVID-19 Genomics UK (COG-UK) Consortium | The Lighthouse Lab in Milton Keynes and Alex Alderton, Roberto Amato, Jeffrey Barrett, Sonia Goncalves, Ewan Harrison, David K. Jackson, Ian Johnston, Dominic Kwiatkowski, Cordelia Langford, John Sillitoe on behalf of the Wellcome Sanger Institute COVID-19 Surveillance Team |
| EPI_ISL_2552260, EPI_ISL_2552261, EPI_ISL_2552262, EPI_ISL_2552263, EPI_ISL_2552264, EPI_ISL_2552265, EPI_ISL_2552266, EPI_ISL_2552267, EPI_ISL_2552269, EPI_ISL_2552270, EPI_ISL_2552271, EPI_ISL_2552273, EPI_ISL_2552274, EPI_ISL_2552275, EPI_ISL_2552276, EPI_ISL_2552277, EPI_ISL_2552278, EPI_ISL_2552280, EPI_ISL_2552281, EPI_ISL_2552282, EPI_ISL_2552283, EPI_ISL_2552284, EPI_ISL_2552285, EPI_ISL_2552286, EPI_ISL_2552287, EPI_ISL_2552288, EPI_ISL_2552289, EPI_ISL_2552290, EPI_ISL_2552291, EPI_ISL_2552292, EPI_ISL_2552294, EPI_ISL_2552295, EPI_ISL_2552296, EPI_ISL_2552297, EPI_ISL_2552298, EPI_ISL_2552299, EPI_ISL_2552300, EPI_ISL_2552301, EPI_ISL_2552303, EPI_ISL_2552304, EPI_ISL_2552305, EPI_ISL_2552306, EPI_ISL_2552307, EPI_ISL_2552308, EPI_ISL_2552309, EPI_ISL_2552310, EPI_ISL_2552311, EPI_ISL_2552312, EPI_ISL_2552314, EPI_ISL_2552315, EPI_ISL_2552316, EPI_ISL_2552317, EPI_ISL_2552318, EPI_ISL_2552319, EPI_ISL_2552320, EPI_ISL_2552321, EPI_ISL_2552322, EPI_ISL_2552323, EPI_ISL_2552324, EPI_ISL_2552325, EPI_ISL_2552326, EPI_ISL_2552327, EPI_ISL_2552328, EPI_ISL_2552329, EPI_ISL_2552330, EPI_ISL_2552331, EPI_ISL_2552332, EPI_ISL_2552333, EPI_ISL_2552334, EPI_ISL_2552335, EPI_ISL_2552336, EPI_ISL_2552337, EPI_ISL_2552338, EPI_ISL_2552339, EPI_ISL_2552340, EPI_ISL_2552341, EPI_ISL_2552342, EPI_ISL_2552343, EPI_ISL_2552344, EPI_ISL_2552345, EPI_ISL_2552346, EPI_ISL_2552347, EPI_ISL_2552348, EPI_ISL_2552349, EPI_ISL_2552350, EPI_ISL_2552351, EPI_ISL_2552352, EPI_ISL_2552353, EPI_ISL_2552354, EPI_ISL_2552355, EPI_ISL_2552356, EPI_ISL_2552357, EPI_ISL_2552358, EPI_ISL_2552359, EPI_ISL_2552360, EPI_ISL_2552361, EPI_ISL_2552362, EPI_ISL_2552363, EPI_ISL_2552364, EPI_ISL_2552365, EPI_ISL_2552366, EPI_ISL_2552368, EPI_ISL_2552369, EPI_ISL_2552370, EPI_ISL_2552371, EPI_ISL_2552372, EPI_ISL_2552373, EPI_ISL_2552374, EPI_ISL_2552375, EPI_ISL_2552376, EPI_ISL_2552377, EPI_ISL_2552378, EPI_ISL_2552379, EPI_ISL_2552380, EPI_ISL_2552381, EPI_ISL_2552382, EPI_ISL_2552383, EPI_ISL_2552384, EPI_ISL_2552385, EPI_ISL_2552386, EPI_ISL_2552387, EPI_ISL_2552388, EPI_ISL_2552389, EPI_ISL_2552391, EPI_ISL_2552392, EPI_ISL_2552393, EPI_ISL_2552394, EPI_ISL_2552395, EPI_ISL_2552396, EPI_ISL_2552397, EPI_ISL_2552398, EPI_ISL_2552399, EPI_ISL_2552400, EPI_ISL_2552401, EPI_ISL_2552402, EPI_ISL_2552403, EPI_ISL_2552404, EPI_ISL_2552405, EPI_ISL_2552406, EPI_ISL_2552407, EPI_ISL_2552408, EPI_ISL_2552409, EPI_ISL_2552410, EPI_ISL_2552411, EPI_ISL_2552412, EPI_ISL_2552413, EPI_ISL_2552414, EPI_ISL_2552415, EPI_ISL_2552416, EPI_ISL_2552417, EPI_ISL_2552419, EPI_ISL_2552420, EPI_ISL_2552421, EPI_ISL_2552422, EPI_ISL_2552423, EPI_ISL_2552424, EPI_ISL_2552425, EPI_ISL_2552427, EPI_ISL_2552428, EPI_ISL_2552429, EPI_ISL_2552430, EPI_ISL_2552431, EPI_ISL_2552432, EPI_ISL_2552433, EPI_ISL_2552434, EPI_ISL_2552435, EPI_ISL_2552436, EPI_ISL_2552437, EPI_ISL_2552438, EPI_ISL_2552439, EPI_ISL_2552440, EPI_ISL_2552441, EPI_ISL_2552442, EPI_ISL_2552443, EPI_ISL_2552444, EPI_ISL_2552445, EPI_ISL_2552446, EPI_ISL_2552447, EPI_ISL_2552448, EPI_ISL_2552449, EPI_ISL_2552450, EPI_ISL_2552451, EPI_ISL_2552452, EPI_ISL_2552453, EPI_ISL_2552454, EPI_ISL_2552455, EPI_ISL_2552456, EPI_ISL_2552457, EPI_ISL_2552458, EPI_ISL_2552459, EPI_ISL_2552460, EPI_ISL_2552461, EPI_ISL_2552462, EPI_ISL_2552463, EPI_ISL_2552464, EPI_ISL_2552465, EPI_ISL_2552466, EPI_ISL_2552467, EPI_ISL_2552468, EPI_ISL_2552469, EPI_ISL_2552470, EPI_ISL_2552471, EPI_ISL_2552473, EPI_ISL_2552475, EPI_ISL_2552476, EPI_ISL_2552477, EPI_ISL_2552478, EPI_ISL_2552479, EPI_ISL_2552480, EPI_ISL_2552481, EPI_ISL_2552482, EPI_ISL_2552483, EPI_ISL_2552485, EPI_ISL_2552486, EPI_ISL_2552487, EPI_ISL_2552489, EPI_ISL_2552490, EPI_ISL_2552491, EPI_ISL_2552492, EPI_ISL_2552493, EPI_ISL_2552494, EPI_ISL_2552495, EPI_ISL_2552496, EPI_ISL_2552497, EPI_ISL_2552498, EPI_ISL_2552499, EPI_ISL_2552500, EPI_ISL_2552501, EPI_ISL_2552502, EPI_ISL_2552503, EPI_ISL_2552504, EPI_ISL_2552505, EPI_ISL_2552506, EPI_ISL_2552507, EPI_ISL_2552508, EPI_ISL_2552509, EPI_ISL_2552510, EPI_ISL_2552511, EPI_ISL_2552512, EPI_ISL_2552513, EPI_ISL_2552514, EPI_ISL_2552515, EPI_ISL_2552516, EPI_ISL_2552517, EPI_ISL_2552518, EPI_ISL_2552519, EPI_ISL_2552520, EPI_ISL_2552522, EPI_ISL_2552523, EPI_ISL_2552524, EPI_ISL_2552525, EPI_ISL_2552526, EPI_ISL_2552527, EPI_ISL_2552529, EPI_ISL_2552530, EPI_ISL_2552531, EPI_ISL_2552532, EPI_ISL_2552533, EPI_ISL_2552535, EPI_ISL_2552536, EPI_ISL_2552537, EPI_ISL_2552538, EPI_ISL_2552540, EPI_ISL_2552541, EPI_ISL_2552542, EPI_ISL_2552543, EPI_ISL_2552544, EPI_ISL_2552547, EPI_ISL_2552548, EPI_ISL_2552549, EPI_ISL_2552550, EPI_ISL_2552551, EPI_ISL_2552552, EPI_ISL_2552553, EPI_ISL_2552554, EPI_ISL_2552555, EPI_ISL_2552556, EPI_ISL_2552557, EPI_ISL_2552558, EPI_ISL_2552559, EPI_ISL_2552560, EPI_ISL_2552561, EPI_ISL_2552562, EPI_ISL_2552563, EPI_ISL_2552564, EPI_ISL_2552565, EPI_ISL_2552566, EPI_ISL_2552567, EPI_ISL_2552568, EPI_ISL_2552569, EPI_ISL_2552570, EPI_ISL_2552571, EPI_ISL_2552572, EPI_ISL_2552573, EPI_ISL_2552574, EPI_ISL_2552575, EPI_ISL_2552576, EPI_ISL_2552577, EPI_ISL_2552578, EPI_ISL_2552579, EPI_ISL_2552580, EPI_ISL_2552581, EPI_ISL_2552582, EPI_ISL_2552583, EPI_ISL_2552584, EPI_ISL_2552585, EPI_ISL_2552586, EPI_ISL_2552587, EPI_ISL_2552588, EPI_ISL_2552589, EPI_ISL_2552590, EPI_ISL_2552591, EPI_ISL_2552592, EPI_ISL_2552593, EPI_ISL_2552594, EPI_ISL_2552595, EPI_ISL_2552596, EPI_ISL_2552597, EPI_ISL_2552598, EPI_ISL_2552599, EPI_ISL_2552600, EPI_ISL_2552601, EPI_ISL_2552602, EPI_ISL_2552603, EPI_ISL_2552604, EPI_ISL_2552605, EPI_ISL_2552606 | | | |
| see above | Lighthouse Lab in Alderley Park | Wellcome Sanger Institute for the COVID-19 Genomics UK (COG-UK) Consortium | Jacquelyn Wynn, Mairead Hyland, The Lighthouse Lab in Alderley Park and Alex Alderton, Roberto Amato, Jeffrey Barrett, Sonia Goncalves, Ewan Harrison, David K. Jackson, Ian Johnston, Dominic Kwiatkowski, Cordelia Langford, John Sillitoe on behalf of the Wellcome Sanger Institute COVID-19 Surveillance Team |
| EPI_ISL_2552607 | Lighthouse Lab in Glasgow | Wellcome Sanger Institute for the COVID-19 Genomics UK (COG-UK) Consortium | Harper VanSteenhouse, Yumi Kasai, David Gray, Carol Clugston, Anna Dominiczak and Alex Alderton, Roberto Amato, Jeffrey Barrett, Sonia Goncalves, Ewan Harrison, David K. Jackson, Ian Johnston, Dominic Kwiatkowski, Cordelia Langford, John Sillitoe on behalf of the Wellcome Sanger Institute COVID-19 Surveillance Team |
| EPI_ISL_2552608 | Lighthouse Lab in Milton Keynes | Wellcome Sanger Institute for the COVID-19 Genomics UK (COG-UK) Consortium | The Lighthouse Lab in Milton Keynes and Alex Alderton, Roberto Amato, Jeffrey Barrett, Sonia Goncalves, Ewan Harrison, David K. Jackson, Ian Johnston, Dominic Kwiatkowski, Cordelia Langford, John Sillitoe on behalf of the Wellcome Sanger Institute COVID-19 Surveillance Team |
| EPI_ISL_2552609 | Lighthouse Lab in Alderley Park | Wellcome Sanger Institute for the COVID-19 Genomics UK (COG-UK) Consortium | Jacquelyn Wynn, Mairead Hyland, The Lighthouse Lab in Alderley Park and Alex Alderton, Roberto Amato, Jeffrey Barrett, Sonia Goncalves, Ewan Harrison, David K. Jackson, Ian Johnston, Dominic Kwiatkowski, Cordelia Langford, John Sillitoe on behalf of the Wellcome Sanger Institute COVID-19 Surveillance Team |
| EPI_ISL_2552610 | Lighthouse Lab in Milton Keynes | Wellcome Sanger Institute for the COVID-19 Genomics UK (COG-UK) Consortium | The Lighthouse Lab in Milton Keynes and Alex Alderton, Roberto Amato, Jeffrey Barrett, Sonia Goncalves, Ewan Harrison, David K. Jackson, Ian Johnston, Dominic Kwiatkowski, Cordelia Langford, John Sillitoe on behalf of the Wellcome Sanger Institute COVID-19 Surveillance Team |
| EPI_ISL_2552611 | Lighthouse Lab in Alderley Park | Wellcome Sanger Institute for the COVID-19 Genomics UK (COG-UK) Consortium | Jacquelyn Wynn, Mairead Hyland, The Lighthouse Lab in Alderley Park and Alex Alderton, Roberto Amato, Jeffrey Barrett, Sonia Goncalves, Ewan Harrison, David K. Jackson, Ian Johnston, Dominic Kwiatkowski, Cordelia Langford, John Sillitoe on behalf of the Wellcome Sanger Institute COVID-19 Surveillance Team |
| EPI_ISL_2552612, EPI_ISL_2552613, EPI_ISL_2552614 | Health Services Laboratories | Wellcome Sanger Institute for the COVID-19 Genomics UK (COG-UK) Consortium | Health Services Laboratories and Alex Alderton, Roberto Amato, Jeffrey Barrett, Sonia Goncalves, Ewan Harrison, David K. Jackson, Ian Johnston, Dominic Kwiatkowski, Cordelia Langford, John Sillitoe on behalf of the Wellcome Sanger Institute COVID-19 Surveillance Team |
| EPI_ISL_2552615 | Lighthouse Lab in Milton Keynes | Wellcome Sanger Institute for the COVID-19 Genomics UK (COG-UK) Consortium | The Lighthouse Lab in Milton Keynes and Alex Alderton, Roberto Amato, Jeffrey Barrett, Sonia Goncalves, Ewan Harrison, David K. Jackson, Ian Johnston, Dominic Kwiatkowski, Cordelia Langford, John Sillitoe on behalf of the Wellcome Sanger Institute COVID-19 Surveillance Team |
| EPI_ISL_2552616 | Lighthouse Lab in Alderley Park | Wellcome Sanger Institute for the COVID-19 Genomics UK (COG-UK) Consortium | Jacquelyn Wynn, Mairead Hyland, The Lighthouse Lab in Alderley Park and Alex Alderton, Roberto Amato, Jeffrey Barrett, Sonia Goncalves, Ewan Harrison, David K. Jackson, Ian Johnston, Dominic Kwiatkowski, Cordelia Langford, John Sillitoe on behalf of the Wellcome Sanger Institute COVID-19 Surveillance Team |
| EPI_ISL_2552617, EPI_ISL_2552618 | Lighthouse Lab in Milton Keynes | Wellcome Sanger Institute for the COVID-19 Genomics UK (COG-UK) Consortium | The Lighthouse Lab in Milton Keynes and Alex Alderton, Roberto Amato, Jeffrey Barrett, Sonia Goncalves, Ewan Harrison, David K. Jackson, Ian Johnston, Dominic Kwiatkowski, Cordelia Langford, John Sillitoe on behalf of the Wellcome Sanger Institute COVID-19 Surveillance Team |
| EPI_ISL_2552619 | Health Services Laboratories | Wellcome Sanger Institute for the COVID-19 Genomics UK (COG-UK) Consortium | Health Services Laboratories and Alex Alderton, Roberto Amato, Jeffrey Barrett, Sonia Goncalves, Ewan Harrison, David K. Jackson, Ian Johnston, Dominic Kwiatkowski, Cordelia Langford, John Sillitoe on behalf of the Wellcome Sanger Institute COVID-19 Surveillance Team |
| EPI_ISL_2552620, EPI_ISL_2552621 | Lighthouse Lab in Milton Keynes | Wellcome Sanger Institute for the COVID-19 Genomics UK (COG-UK) Consortium | The Lighthouse Lab in Milton Keynes and Alex Alderton, Roberto Amato, Jeffrey Barrett, Sonia Goncalves, Ewan Harrison, David K. Jackson, Ian Johnston, Dominic Kwiatkowski, Cordelia Langford, John Sillitoe on behalf of the Wellcome Sanger Institute COVID-19 Surveillance Team |
| EPI_ISL_2552622, EPI_ISL_2552623 | Lighthouse Lab in Alderley Park | Wellcome Sanger Institute for the COVID-19 Genomics UK (COG-UK) Consortium | Jacquelyn Wynn, Mairead Hyland, The Lighthouse Lab in Alderley Park and Alex Alderton, Roberto Amato, Jeffrey Barrett, Sonia Goncalves, Ewan Harrison, David K. Jackson, Ian Johnston, Dominic Kwiatkowski, Cordelia Langford, John Sillitoe on behalf of the Wellcome Sanger Institute COVID-19 Surveillance Team |
| EPI_ISL_2552624 | Health Services Laboratories | Wellcome Sanger Institute for the COVID-19 Genomics UK (COG-UK) Consortium | Health Services Laboratories and Alex Alderton, Roberto Amato, Jeffrey Barrett, Sonia Goncalves, Ewan Harrison, David K. Jackson, Ian Johnston, Dominic Kwiatkowski, Cordelia Langford, John Sillitoe on behalf of the Wellcome Sanger Institute COVID-19 Surveillance Team |
| EPI_ISL_2552625 | Lighthouse Lab in Milton Keynes | Wellcome Sanger Institute for the COVID-19 Genomics UK (COG-UK) Consortium | The Lighthouse Lab in Milton Keynes and Alex Alderton, Roberto Amato, Jeffrey Barrett, Sonia Goncalves, Ewan Harrison, David K. Jackson, Ian Johnston, Dominic Kwiatkowski, Cordelia Langford, John Sillitoe on behalf of the Wellcome Sanger Institute COVID-19 Surveillance Team |
| EPI_ISL_2552626, EPI_ISL_2552627 | Lighthouse Lab in Alderley Park | Wellcome Sanger Institute for the COVID-19 Genomics UK (COG-UK) Consortium | Jacquelyn Wynn, Mairead Hyland, The Lighthouse Lab in Alderley Park and Alex Alderton, Roberto Amato, Jeffrey Barrett, Sonia Goncalves, Ewan Harrison, David K. Jackson, Ian Johnston, Dominic Kwiatkowski, Cordelia Langford, John Sillitoe on behalf of the Wellcome Sanger Institute COVID-19 Surveillance Team |
| EPI_ISL_2552628 | Lighthouse Lab in Milton Keynes | Wellcome Sanger Institute for the COVID-19 Genomics UK (COG-UK) Consortium | The Lighthouse Lab in Milton Keynes and Alex Alderton, Roberto Amato, Jeffrey Barrett, Sonia Goncalves, Ewan Harrison, David K. Jackson, Ian Johnston, Dominic Kwiatkowski, Cordelia Langford, John Sillitoe on behalf of the Wellcome Sanger Institute COVID-19 Surveillance Team |
| EPI_ISL_2552630, EPI_ISL_2552631 | Lighthouse Lab in Glasgow | Wellcome Sanger Institute for the COVID-19 Genomics UK (COG-UK) Consortium | Harper VanSteenhouse, Yumi Kasai, David Gray, Carol Clugston, Anna Dominiczak and Alex Alderton, Roberto Amato, Jeffrey Barrett, Sonia Goncalves, Ewan Harrison, David K. Jackson, Ian Johnston, Dominic Kwiatkowski, Cordelia Langford, John Sillitoe on behalf of the Wellcome Sanger Institute COVID-19 Surveillance Team |
| EPI_ISL_2552632 | Lighthouse Lab in Alderley Park | Wellcome Sanger Institute for the COVID-19 Genomics UK (COG-UK) Consortium | Jacquelyn Wynn, Mairead Hyland, The Lighthouse Lab in Alderley Park and Alex Alderton, Roberto Amato, Jeffrey Barrett, Sonia Goncalves, Ewan Harrison, David K. Jackson, Ian Johnston, Dominic Kwiatkowski, Cordelia Langford, John Sillitoe on behalf of the Wellcome Sanger Institute COVID-19 Surveillance Team |
| EPI_ISL_2552633, EPI_ISL_2552634 | Health Services Laboratories | Wellcome Sanger Institute for the COVID-19 Genomics UK (COG-UK) Consortium | Health Services Laboratories and Alex Alderton, Roberto Amato, Jeffrey Barrett, Sonia Goncalves, Ewan Harrison, David K. Jackson, Ian Johnston, Dominic Kwiatkowski, Cordelia Langford, John Sillitoe on behalf of the Wellcome Sanger Institute COVID-19 Surveillance Team |
| EPI_ISL_2552635 | Lighthouse Lab in Alderley Park | Wellcome Sanger Institute for the COVID-19 Genomics UK (COG-UK) Consortium | Jacquelyn Wynn, Mairead Hyland, The Lighthouse Lab in Alderley Park and Alex Alderton, Roberto Amato, Jeffrey Barrett, Sonia Goncalves, Ewan Harrison, David K. Jackson, Ian Johnston, Dominic Kwiatkowski, Cordelia Langford, John Sillitoe on behalf of the Wellcome Sanger Institute COVID-19 Surveillance Team |
| EPI_ISL_2552636 | Health Services Laboratories | Wellcome Sanger Institute for the COVID-19 Genomics UK (COG-UK) Consortium | Health Services Laboratories and Alex Alderton, Roberto Amato, Jeffrey Barrett, Sonia Goncalves, Ewan Harrison, David K. Jackson, Ian Johnston, Dominic Kwiatkowski, Cordelia Langford, John Sillitoe on behalf of the Wellcome Sanger Institute COVID-19 Surveillance Team |
| EPI_ISL_2552638 | Lighthouse Lab in Glasgow | Wellcome Sanger Institute for the COVID-19 Genomics UK (COG-UK) Consortium | Harper VanSteenhouse, Yumi Kasai, David Gray, Carol Clugston, Anna Dominiczak and Alex Alderton, Roberto Amato, Jeffrey Barrett, Sonia Goncalves, Ewan Harrison, David K. Jackson, Ian Johnston, Dominic Kwiatkowski, Cordelia Langford, John Sillitoe on behalf of the Wellcome Sanger Institute COVID-19 Surveillance Team |
| EPI_ISL_2552639 | Lighthouse Lab in Alderley Park | Wellcome Sanger Institute for the COVID-19 Genomics UK (COG-UK) Consortium | Jacquelyn Wynn, Mairead Hyland, The Lighthouse Lab in Alderley Park and Alex Alderton, Roberto Amato, Jeffrey Barrett, Sonia Goncalves, Ewan Harrison, David K. Jackson, Ian Johnston, Dominic Kwiatkowski, Cordelia Langford, John Sillitoe on behalf of the Wellcome Sanger Institute COVID-19 Surveillance Team |
| EPI_ISL_2552640 | Health Services Laboratories | Wellcome Sanger Institute for the COVID-19 Genomics UK (COG-UK) Consortium | Health Services Laboratories and Alex Alderton, Roberto Amato, Jeffrey Barrett, Sonia Goncalves, Ewan Harrison, David K. Jackson, Ian Johnston, Dominic Kwiatkowski, Cordelia Langford, John Sillitoe on behalf of the Wellcome Sanger Institute COVID-19 Surveillance Team |
| EPI_ISL_2552641 | Lighthouse Lab in Glasgow | Wellcome Sanger Institute for the COVID-19 Genomics UK (COG-UK) Consortium | Harper VanSteenhouse, Yumi Kasai, David Gray, Carol Clugston, Anna Dominiczak and Alex Alderton, Roberto Amato, Jeffrey Barrett, Sonia Goncalves, Ewan Harrison, David K. Jackson, Ian Johnston, Dominic Kwiatkowski, Cordelia Langford, John Sillitoe on behalf of the Wellcome Sanger Institute COVID-19 Surveillance Team |
| EPI_ISL_2552642 | Lighthouse Lab in Alderley Park | Wellcome Sanger Institute for the COVID-19 Genomics UK (COG-UK) Consortium | Jacquelyn Wynn, Mairead Hyland, The Lighthouse Lab in Alderley Park and Alex Alderton, Roberto Amato, Jeffrey Barrett, Sonia Goncalves, Ewan Harrison, David K. Jackson, Ian Johnston, Dominic Kwiatkowski, Cordelia Langford, John Sillitoe on behalf of the Wellcome Sanger Institute COVID-19 Surveillance Team |
| EPI_ISL_2552643, EPI_ISL_2552644, EPI_ISL_2552645 | Lighthouse Lab in Glasgow | Wellcome Sanger Institute for the COVID-19 Genomics UK (COG-UK) Consortium | Harper VanSteenhouse, Yumi Kasai, David Gray, Carol Clugston, Anna Dominiczak and Alex Alderton, Roberto Amato, Jeffrey Barrett, Sonia Goncalves, Ewan Harrison, David K. Jackson, Ian Johnston, Dominic Kwiatkowski, Cordelia Langford, John Sillitoe on behalf of the Wellcome Sanger Institute COVID-19 Surveillance Team |
| EPI_ISL_2552646, | Health Services Laboratories | Wellcome Sanger Institute for the COVID-19 Genomics UK (COG-UK) | Health Services Laboratories and Alex Alderton, Roberto Amato, Jeffrey Barrett, Sonia Goncalves, Ewan Harrison, David K. Jackson, Ian Johnston, Dominic Kwiatkowski, Cordelia Langford, John Sillitoe on behalf of the Wellcome Sanger Institute |

| | | | |
|---|---|---|---|
| EPI_ISL_2552648 | | Consortium | COVID-19 Surveillance Team |
| EPI_ISL_2552649 | Lighthouse Lab in Alderley Park | Wellcome Sanger Institute for the COVID-19 Genomics UK (COG-UK) Consortium | Jacquelyn Wynn, Mairead Hyland, The Lighthouse Lab in Alderley Park and Alex Alderton, Roberto Amato, Jeffrey Barrett, Sonia Goncalves, Ewan Harrison, David K. Jackson, Ian Johnston, Dominic Kwiatkowski, Cordelia Langford, John Sillitoe on behalf of the Wellcome Sanger Institute COVID-19 Surveillance Team |
| EPI_ISL_2552651 | Health Services Laboratories | Wellcome Sanger Institute for the COVID-19 Genomics UK (COG-UK) Consortium | Health Services Laboratories and Alex Alderton, Roberto Amato, Jeffrey Barrett, Sonia Goncalves, Ewan Harrison, David K. Jackson, Ian Johnston, Dominic Kwiatkowski, Cordelia Langford, John Sillitoe on behalf of the Wellcome Sanger Institute COVID-19 Surveillance Team |
| EPI_ISL_2552652 | Lighthouse Lab in Glasgow | Wellcome Sanger Institute for the COVID-19 Genomics UK (COG-UK) Consortium | Harper VanSteenhouse, Yumi Kasai, David Gray, Carol Clugston, Anna Dominiczak and Alex Alderton, Roberto Amato, Jeffrey Barrett, Sonia Goncalves, Ewan Harrison, David K. Jackson, Ian Johnston, Dominic Kwiatkowski, Cordelia Langford, John Sillitoe on behalf of the Wellcome Sanger Institute COVID-19 Surveillance Team |
| EPI_ISL_2552653 | Health Services Laboratories | Wellcome Sanger Institute for the COVID-19 Genomics UK (COG-UK) Consortium | Health Services Laboratories and Alex Alderton, Roberto Amato, Jeffrey Barrett, Sonia Goncalves, Ewan Harrison, David K. Jackson, Ian Johnston, Dominic Kwiatkowski, Cordelia Langford, John Sillitoe on behalf of the Wellcome Sanger Institute COVID-19 Surveillance Team |
| EPI_ISL_2552654 | Lighthouse Lab in Glasgow | Wellcome Sanger Institute for the COVID-19 Genomics UK (COG-UK) Consortium | Harper VanSteenhouse, Yumi Kasai, David Gray, Carol Clugston, Anna Dominiczak and Alex Alderton, Roberto Amato, Jeffrey Barrett, Sonia Goncalves, Ewan Harrison, David K. Jackson, Ian Johnston, Dominic Kwiatkowski, Cordelia Langford, John Sillitoe on behalf of the Wellcome Sanger Institute COVID-19 Surveillance Team |
| EPI_ISL_2552655 | Lighthouse Lab in Alderley Park | Wellcome Sanger Institute for the COVID-19 Genomics UK (COG-UK) Consortium | Jacquelyn Wynn, Mairead Hyland, The Lighthouse Lab in Alderley Park and Alex Alderton, Roberto Amato, Jeffrey Barrett, Sonia Goncalves, Ewan Harrison, David K. Jackson, Ian Johnston, Dominic Kwiatkowski, Cordelia Langford, John Sillitoe on behalf of the Wellcome Sanger Institute COVID-19 Surveillance Team |
| EPI_ISL_2552656, EPI_ISL_2552657 | Lighthouse Lab in Milton Keynes | Wellcome Sanger Institute for the COVID-19 Genomics UK (COG-UK) Consortium | The Lighthouse Lab in Milton Keynes and Alex Alderton, Roberto Amato, Jeffrey Barrett, Sonia Goncalves, Ewan Harrison, David K. Jackson, Ian Johnston, Dominic Kwiatkowski, Cordelia Langford, John Sillitoe on behalf of the Wellcome Sanger Institute COVID-19 Surveillance Team |
| EPI_ISL_2552658 | Lighthouse Lab in Alderley Park | Wellcome Sanger Institute for the COVID-19 Genomics UK (COG-UK) Consortium | Jacquelyn Wynn, Mairead Hyland, The Lighthouse Lab in Alderley Park and Alex Alderton, Roberto Amato, Jeffrey Barrett, Sonia Goncalves, Ewan Harrison, David K. Jackson, Ian Johnston, Dominic Kwiatkowski, Cordelia Langford, John Sillitoe on behalf of the Wellcome Sanger Institute COVID-19 Surveillance Team |
| EPI_ISL_2552659 | Lighthouse Lab in Glasgow | Wellcome Sanger Institute for the COVID-19 Genomics UK (COG-UK) Consortium | Harper VanSteenhouse, Yumi Kasai, David Gray, Carol Clugston, Anna Dominiczak and Alex Alderton, Roberto Amato, Jeffrey Barrett, Sonia Goncalves, Ewan Harrison, David K. Jackson, Ian Johnston, Dominic Kwiatkowski, Cordelia Langford, John Sillitoe on behalf of the Wellcome Sanger Institute COVID-19 Surveillance Team |
| EPI_ISL_2552660, EPI_ISL_2552661 | Lighthouse Lab in Alderley Park | Wellcome Sanger Institute for the COVID-19 Genomics UK (COG-UK) Consortium | Jacquelyn Wynn, Mairead Hyland, The Lighthouse Lab in Alderley Park and Alex Alderton, Roberto Amato, Jeffrey Barrett, Sonia Goncalves, Ewan Harrison, David K. Jackson, Ian Johnston, Dominic Kwiatkowski, Cordelia Langford, John Sillitoe on behalf of the Wellcome Sanger Institute COVID-19 Surveillance Team |
| EPI_ISL_2552662 | Health Services Laboratories | Wellcome Sanger Institute for the COVID-19 Genomics UK (COG-UK) Consortium | Health Services Laboratories and Alex Alderton, Roberto Amato, Jeffrey Barrett, Sonia Goncalves, Ewan Harrison, David K. Jackson, Ian Johnston, Dominic Kwiatkowski, Cordelia Langford, John Sillitoe on behalf of the Wellcome Sanger Institute COVID-19 Surveillance Team |
| EPI_ISL_2552663 | Lighthouse Lab in Alderley Park | Wellcome Sanger Institute for the COVID-19 Genomics UK (COG-UK) Consortium | Jacquelyn Wynn, Mairead Hyland, The Lighthouse Lab in Alderley Park and Alex Alderton, Roberto Amato, Jeffrey Barrett, Sonia Goncalves, Ewan Harrison, David K. Jackson, Ian Johnston, Dominic Kwiatkowski, Cordelia Langford, John Sillitoe on behalf of the Wellcome Sanger Institute COVID-19 Surveillance Team |
| EPI_ISL_2552664 | Health Services Laboratories | Wellcome Sanger Institute for the COVID-19 Genomics UK (COG-UK) Consortium | Health Services Laboratories and Alex Alderton, Roberto Amato, Jeffrey Barrett, Sonia Goncalves, Ewan Harrison, David K. Jackson, Ian Johnston, Dominic Kwiatkowski, Cordelia Langford, John Sillitoe on behalf of the Wellcome Sanger Institute COVID-19 Surveillance Team |
| EPI_ISL_2552665 | Lighthouse Lab in Glasgow | Wellcome Sanger Institute for the COVID-19 Genomics UK (COG-UK) Consortium | Harper VanSteenhouse, Yumi Kasai, David Gray, Carol Clugston, Anna Dominiczak and Alex Alderton, Roberto Amato, Jeffrey Barrett, Sonia Goncalves, Ewan Harrison, David K. Jackson, Ian Johnston, Dominic Kwiatkowski, Cordelia Langford, John Sillitoe on behalf of the Wellcome Sanger Institute COVID-19 Surveillance Team |
| EPI_ISL_2552666 | Health Services Laboratories | Wellcome Sanger Institute for the COVID-19 Genomics UK (COG-UK) Consortium | Health Services Laboratories and Alex Alderton, Roberto Amato, Jeffrey Barrett, Sonia Goncalves, Ewan Harrison, David K. Jackson, Ian Johnston, Dominic Kwiatkowski, Cordelia Langford, John Sillitoe on behalf of the Wellcome Sanger Institute COVID-19 Surveillance Team |
| EPI_ISL_2552667 | Lighthouse Lab in Milton Keynes | Wellcome Sanger Institute for the COVID-19 Genomics UK (COG-UK) Consortium | The Lighthouse Lab in Milton Keynes and Alex Alderton, Roberto Amato, Jeffrey Barrett, Sonia Goncalves, Ewan Harrison, David K. Jackson, Ian Johnston, Dominic Kwiatkowski, Cordelia Langford, John Sillitoe on behalf of the Wellcome Sanger Institute COVID-19 Surveillance Team |
| EPI_ISL_2552668 | Health Services Laboratories | Wellcome Sanger Institute for the COVID-19 Genomics UK (COG-UK) Consortium | Health Services Laboratories and Alex Alderton, Roberto Amato, Jeffrey Barrett, Sonia Goncalves, Ewan Harrison, David K. Jackson, Ian Johnston, Dominic Kwiatkowski, Cordelia Langford, John Sillitoe on behalf of the Wellcome Sanger Institute COVID-19 Surveillance Team |
| EPI_ISL_2552669 | Lighthouse Lab in Milton Keynes | Wellcome Sanger Institute for the COVID-19 Genomics UK (COG-UK) Consortium | The Lighthouse Lab in Milton Keynes and Alex Alderton, Roberto Amato, Jeffrey Barrett, Sonia Goncalves, Ewan Harrison, David K. Jackson, Ian Johnston, Dominic Kwiatkowski, Cordelia Langford, John Sillitoe on behalf of the Wellcome Sanger Institute COVID-19 Surveillance Team |
| EPI_ISL_2552670 | Health Services Laboratories | Wellcome Sanger Institute for the COVID-19 Genomics UK (COG-UK) Consortium | Health Services Laboratories and Alex Alderton, Roberto Amato, Jeffrey Barrett, Sonia Goncalves, Ewan Harrison, David K. Jackson, Ian Johnston, Dominic Kwiatkowski, Cordelia Langford, John Sillitoe on behalf of the Wellcome Sanger Institute COVID-19 Surveillance Team |
| EPI_ISL_2552671, EPI_ISL_2552673 | Lighthouse Lab in Glasgow | Wellcome Sanger Institute for the COVID-19 Genomics UK (COG-UK) Consortium | Harper VanSteenhouse, Yumi Kasai, David Gray, Carol Clugston, Anna Dominiczak and Alex Alderton, Roberto Amato, Jeffrey Barrett, Sonia Goncalves, Ewan Harrison, David K. Jackson, Ian Johnston, Dominic Kwiatkowski, Cordelia Langford, John Sillitoe on behalf of the Wellcome Sanger Institute COVID-19 Surveillance Team |
| EPI_ISL_2552674, EPI_ISL_2552675 | Health Services Laboratories | Wellcome Sanger Institute for the COVID-19 Genomics UK (COG-UK) Consortium | Health Services Laboratories and Alex Alderton, Roberto Amato, Jeffrey Barrett, Sonia Goncalves, Ewan Harrison, David K. Jackson, Ian Johnston, Dominic Kwiatkowski, Cordelia Langford, John Sillitoe on behalf of the Wellcome Sanger Institute COVID-19 Surveillance Team |
| EPI_ISL_2552676 | Lighthouse Lab in Milton Keynes | Wellcome Sanger Institute for the COVID-19 Genomics UK (COG-UK) Consortium | The Lighthouse Lab in Milton Keynes and Alex Alderton, Roberto Amato, Jeffrey Barrett, Sonia Goncalves, Ewan Harrison, David K. Jackson, Ian Johnston, Dominic Kwiatkowski, Cordelia Langford, John Sillitoe on behalf of the Wellcome Sanger Institute COVID-19 Surveillance Team |
| EPI_ISL_2552677 | Health Services Laboratories | Wellcome Sanger Institute for the COVID-19 Genomics UK (COG-UK) Consortium | Health Services Laboratories and Alex Alderton, Roberto Amato, Jeffrey Barrett, Sonia Goncalves, Ewan Harrison, David K. Jackson, Ian Johnston, Dominic Kwiatkowski, Cordelia Langford, John Sillitoe on behalf of the Wellcome Sanger Institute COVID-19 Surveillance Team |
| EPI_ISL_2552678, EPI_ISL_2552679 | Lighthouse Lab in Milton Keynes | Wellcome Sanger Institute for the COVID-19 Genomics UK (COG-UK) Consortium | The Lighthouse Lab in Milton Keynes and Alex Alderton, Roberto Amato, Jeffrey Barrett, Sonia Goncalves, Ewan Harrison, David K. Jackson, Ian Johnston, Dominic Kwiatkowski, Cordelia Langford, John Sillitoe on behalf of the Wellcome Sanger Institute COVID-19 Surveillance Team |
| EPI_ISL_2552680, EPI_ISL_2552681 | Lighthouse Lab in Alderley Park | Wellcome Sanger Institute for the COVID-19 Genomics UK (COG-UK) Consortium | Jacquelyn Wynn, Mairead Hyland, The Lighthouse Lab in Alderley Park and Alex Alderton, Roberto Amato, Jeffrey Barrett, Sonia Goncalves, Ewan Harrison, David K. Jackson, Ian Johnston, Dominic Kwiatkowski, Cordelia Langford, John Sillitoe on behalf of the Wellcome Sanger Institute COVID-19 Surveillance Team |
| EPI_ISL_2552682, EPI_ISL_2552683 | Lighthouse Lab in Milton Keynes | Wellcome Sanger Institute for the COVID-19 Genomics UK (COG-UK) Consortium | The Lighthouse Lab in Milton Keynes and Alex Alderton, Roberto Amato, Jeffrey Barrett, Sonia Goncalves, Ewan Harrison, David K. Jackson, Ian Johnston, Dominic Kwiatkowski, Cordelia Langford, John Sillitoe on behalf of the Wellcome Sanger Institute COVID-19 Surveillance Team |
| EPI_ISL_2552684 | Lighthouse Lab in Alderley Park | Wellcome Sanger Institute for the COVID-19 Genomics UK (COG-UK) Consortium | Jacquelyn Wynn, Mairead Hyland, The Lighthouse Lab in Alderley Park and Alex Alderton, Roberto Amato, Jeffrey Barrett, Sonia Goncalves, Ewan Harrison, David K. Jackson, Ian Johnston, Dominic Kwiatkowski, Cordelia Langford, John Sillitoe on behalf of the Wellcome Sanger Institute COVID-19 Surveillance Team |
| EPI_ISL_2552685 | Health Services Laboratories | Wellcome Sanger Institute for the COVID-19 Genomics UK (COG-UK) Consortium | Health Services Laboratories and Alex Alderton, Roberto Amato, Jeffrey Barrett, Sonia Goncalves, Ewan Harrison, David K. Jackson, Ian Johnston, Dominic Kwiatkowski, Cordelia Langford, John Sillitoe on behalf of the Wellcome Sanger Institute COVID-19 Surveillance Team |
| EPI_ISL_2552686, EPI_ISL_2552687 | Lighthouse Lab in Glasgow | Wellcome Sanger Institute for the COVID-19 Genomics UK (COG-UK) Consortium | Harper VanSteenhouse, Yumi Kasai, David Gray, Carol Clugston, Anna Dominiczak and Alex Alderton, Roberto Amato, Jeffrey Barrett, Sonia Goncalves, Ewan Harrison, David K. Jackson, Ian Johnston, Dominic Kwiatkowski, Cordelia Langford, John Sillitoe on behalf of the Wellcome Sanger Institute COVID-19 Surveillance Team |
| EPI_ISL_2552689, EPI_ISL_2552690, EPI_ISL_2552691, EPI_ISL_2552692 | Lighthouse Lab in Alderley Park | Wellcome Sanger Institute for the COVID-19 Genomics UK (COG-UK) Consortium | Jacquelyn Wynn, Mairead Hyland, The Lighthouse Lab in Alderley Park and Alex Alderton, Roberto Amato, Jeffrey Barrett, Sonia Goncalves, Ewan Harrison, David K. Jackson, Ian Johnston, Dominic Kwiatkowski, Cordelia Langford, John Sillitoe on behalf of the Wellcome Sanger Institute COVID-19 Surveillance Team |
| EPI_ISL_2552693 | Lighthouse Lab in Glasgow | Wellcome Sanger Institute for the COVID-19 Genomics UK (COG-UK) Consortium | Harper VanSteenhouse, Yumi Kasai, David Gray, Carol Clugston, Anna Dominiczak and Alex Alderton, Roberto Amato, Jeffrey Barrett, Sonia Goncalves, Ewan Harrison, David K. Jackson, Ian Johnston, Dominic Kwiatkowski, Cordelia Langford, John Sillitoe on behalf of the Wellcome Sanger Institute COVID-19 Surveillance Team |
| EPI_ISL_2552694 | Lighthouse Lab in Milton Keynes | Wellcome Sanger Institute for the COVID-19 Genomics UK (COG-UK) Consortium | The Lighthouse Lab in Milton Keynes and Alex Alderton, Roberto Amato, Jeffrey Barrett, Sonia Goncalves, Ewan Harrison, David K. Jackson, Ian Johnston, Dominic Kwiatkowski, Cordelia Langford, John Sillitoe on behalf of the Wellcome Sanger Institute COVID-19 Surveillance Team |
| EPI_ISL_2552695 | Lighthouse Lab in Alderley Park | Wellcome Sanger Institute for the COVID-19 Genomics UK (COG-UK) Consortium | Jacquelyn Wynn, Mairead Hyland, The Lighthouse Lab in Alderley Park and Alex Alderton, Roberto Amato, Jeffrey Barrett, Sonia Goncalves, Ewan Harrison, David K. Jackson, Ian Johnston, Dominic Kwiatkowski, Cordelia Langford, John Sillitoe on behalf of the Wellcome Sanger Institute COVID-19 Surveillance Team |
| EPI_ISL_2552696, EPI_ISL_2552698, EPI_ISL_2552699, EPI_ISL_2552700 | Health Services Laboratories | Wellcome Sanger Institute for the COVID-19 Genomics UK (COG-UK) Consortium | Health Services Laboratories and Alex Alderton, Roberto Amato, Jeffrey Barrett, Sonia Goncalves, Ewan Harrison, David K. Jackson, Ian Johnston, Dominic Kwiatkowski, Cordelia Langford, John Sillitoe on behalf of the Wellcome Sanger Institute COVID-19 Surveillance Team |
| EPI_ISL_2552701 | Lighthouse Lab in Milton Keynes | Wellcome Sanger Institute for the COVID-19 Genomics UK (COG-UK) Consortium | The Lighthouse Lab in Milton Keynes and Alex Alderton, Roberto Amato, Jeffrey Barrett, Sonia Goncalves, Ewan Harrison, David K. Jackson, Ian Johnston, Dominic Kwiatkowski, Cordelia Langford, John Sillitoe on behalf of the Wellcome Sanger Institute COVID-19 Surveillance Team |
| EPI_ISL_2552702 | Lighthouse Lab in Glasgow | Wellcome Sanger Institute for the COVID-19 Genomics UK (COG-UK) Consortium | Harper VanSteenhouse, Yumi Kasai, David Gray, Carol Clugston, Anna Dominiczak and Alex Alderton, Roberto Amato, Jeffrey Barrett, Sonia Goncalves, Ewan Harrison, David K. Jackson, Ian Johnston, Dominic Kwiatkowski, Cordelia Langford, John Sillitoe on behalf of the Wellcome Sanger Institute COVID-19 Surveillance Team |
| EPI_ISL_2552703 | Lighthouse Lab in Alderley Park | Wellcome Sanger Institute for the COVID-19 Genomics UK (COG-UK) Consortium | Jacquelyn Wynn, Mairead Hyland, The Lighthouse Lab in Alderley Park and Alex Alderton, Roberto Amato, Jeffrey Barrett, Sonia Goncalves, Ewan Harrison, David K. Jackson, Ian Johnston, Dominic Kwiatkowski, Cordelia Langford, John Sillitoe on behalf of the Wellcome Sanger Institute COVID-19 Surveillance Team |
| EPI_ISL_2552704, EPI_ISL_2552705 | Lighthouse Lab in Milton Keynes | Wellcome Sanger Institute for the COVID-19 Genomics UK (COG-UK) Consortium | The Lighthouse Lab in Milton Keynes and Alex Alderton, Roberto Amato, Jeffrey Barrett, Sonia Goncalves, Ewan Harrison, David K. Jackson, Ian Johnston, Dominic Kwiatkowski, Cordelia Langford, John Sillitoe on behalf of the Wellcome Sanger Institute COVID-19 Surveillance Team |
| EPI_ISL_2552706 | Lighthouse Lab in Alderley Park | Wellcome Sanger Institute for the COVID-19 Genomics UK (COG-UK) Consortium | Jacquelyn Wynn, Mairead Hyland, The Lighthouse Lab in Alderley Park and Alex Alderton, Roberto Amato, Jeffrey Barrett, Sonia Goncalves, Ewan Harrison, David K. Jackson, Ian Johnston, Dominic Kwiatkowski, Cordelia Langford, John Sillitoe on behalf of the Wellcome Sanger Institute COVID-19 Surveillance Team |
| EPI_ISL_2552707, EPI_ISL_2552708 | Lighthouse Lab in Glasgow | Wellcome Sanger Institute for the COVID-19 Genomics UK (COG-UK) Consortium | Harper VanSteenhouse, Yumi Kasai, David Gray, Carol Clugston, Anna Dominiczak and Alex Alderton, Roberto Amato, Jeffrey Barrett, Sonia Goncalves, Ewan Harrison, David K. Jackson, Ian Johnston, Dominic Kwiatkowski, Cordelia Langford, John Sillitoe on behalf of the Wellcome Sanger Institute COVID-19 Surveillance Team |
| EPI_ISL_2552709 | Health Services Laboratories | Wellcome Sanger Institute for the COVID-19 Genomics UK (COG-UK) Consortium | Health Services Laboratories and Alex Alderton, Roberto Amato, Jeffrey Barrett, Sonia Goncalves, Ewan Harrison, David K. Jackson, Ian Johnston, Dominic Kwiatkowski, Cordelia Langford, John Sillitoe on behalf of the Wellcome Sanger Institute COVID-19 Surveillance Team |
| EPI_ISL_2552710 | Lighthouse Lab in Alderley Park | Wellcome Sanger Institute for the COVID-19 Genomics UK (COG-UK) Consortium | Jacquelyn Wynn, Mairead Hyland, The Lighthouse Lab in Alderley Park and Alex Alderton, Roberto Amato, Jeffrey Barrett, Sonia Goncalves, Ewan Harrison, David K. Jackson, Ian Johnston, Dominic Kwiatkowski, Cordelia Langford, John Sillitoe on behalf of the Wellcome Sanger Institute COVID-19 Surveillance Team |
| EPI_ISL_2552711 | Lighthouse Lab in Milton Keynes | Wellcome Sanger Institute for the COVID-19 Genomics UK (COG-UK) Consortium | The Lighthouse Lab in Milton Keynes and Alex Alderton, Roberto Amato, Jeffrey Barrett, Sonia Goncalves, Ewan Harrison, David K. Jackson, Ian Johnston, Dominic Kwiatkowski, Cordelia Langford, John Sillitoe on behalf of the Wellcome Sanger Institute COVID-19 Surveillance Team |

| | | | |
|---|---|---|---|
| EPI_ISL_2552712 | Health Services Laboratories | Wellcome Sanger Institute for the COVID-19 Genomics UK (COG-UK) Consortium | Health Services Laboratories and Alex Alderton, Roberto Amato, Jeffrey Barrett, Sonia Goncalves, Ewan Harrison, David K. Jackson, Ian Johnston, Dominic Kwiatkowski, Cordelia Langford, John Sillitoe on behalf of the Wellcome Sanger Institute COVID-19 Surveillance Team |
| EPI_ISL_2552713 | Lighthouse Lab in Alderley Park | Wellcome Sanger Institute for the COVID-19 Genomics UK (COG-UK) Consortium | Jacquelyn Wynn, Mairead Hyland, The Lighthouse Lab in Alderley Park and Alex Alderton, Roberto Amato, Jeffrey Barrett, Sonia Goncalves, Ewan Harrison, David K. Jackson, Ian Johnston, Dominic Kwiatkowski, Cordelia Langford, John Sillitoe on behalf of the Wellcome Sanger Institute COVID-19 Surveillance Team |
| EPI_ISL_2552714 | Lighthouse Lab in Glasgow | Wellcome Sanger Institute for the COVID-19 Genomics UK (COG-UK) Consortium | Harper VanSteenhouse, Yumi Kasai, David Gray, Carol Clugston, Anna Dominiczak and Alex Alderton, Roberto Amato, Jeffrey Barrett, Sonia Goncalves, Ewan Harrison, David K. Jackson, Ian Johnston, Dominic Kwiatkowski, Cordelia Langford, John Sillitoe on behalf of the Wellcome Sanger Institute COVID-19 Surveillance Team |
| EPI_ISL_2552715 | Lighthouse Lab in Alderley Park | Wellcome Sanger Institute for the COVID-19 Genomics UK (COG-UK) Consortium | Jacquelyn Wynn, Mairead Hyland, The Lighthouse Lab in Alderley Park and Alex Alderton, Roberto Amato, Jeffrey Barrett, Sonia Goncalves, Ewan Harrison, David K. Jackson, Ian Johnston, Dominic Kwiatkowski, Cordelia Langford, John Sillitoe on behalf of the Wellcome Sanger Institute COVID-19 Surveillance Team |
| EPI_ISL_2552716 | Lighthouse Lab in Milton Keynes | Wellcome Sanger Institute for the COVID-19 Genomics UK (COG-UK) Consortium | The Lighthouse Lab in Milton Keynes and Alex Alderton, Roberto Amato, Jeffrey Barrett, Sonia Goncalves, Ewan Harrison, David K. Jackson, Ian Johnston, Dominic Kwiatkowski, Cordelia Langford, John Sillitoe on behalf of the Wellcome Sanger Institute COVID-19 Surveillance Team |
| EPI_ISL_2552717, EPI_ISL_2552719, EPI_ISL_2552720 | Health Services Laboratories | Wellcome Sanger Institute for the COVID-19 Genomics UK (COG-UK) Consortium | Health Services Laboratories and Alex Alderton, Roberto Amato, Jeffrey Barrett, Sonia Goncalves, Ewan Harrison, David K. Jackson, Ian Johnston, Dominic Kwiatkowski, Cordelia Langford, John Sillitoe on behalf of the Wellcome Sanger Institute COVID-19 Surveillance Team |
| EPI_ISL_2552721 | Lighthouse Lab in Glasgow | Wellcome Sanger Institute for the COVID-19 Genomics UK (COG-UK) Consortium | Harper VanSteenhouse, Yumi Kasai, David Gray, Carol Clugston, Anna Dominiczak and Alex Alderton, Roberto Amato, Jeffrey Barrett, Sonia Goncalves, Ewan Harrison, David K. Jackson, Ian Johnston, Dominic Kwiatkowski, Cordelia Langford, John Sillitoe on behalf of the Wellcome Sanger Institute COVID-19 Surveillance Team |
| EPI_ISL_2552722 | Lighthouse Lab in Milton Keynes | Wellcome Sanger Institute for the COVID-19 Genomics UK (COG-UK) Consortium | The Lighthouse Lab in Milton Keynes and Alex Alderton, Roberto Amato, Jeffrey Barrett, Sonia Goncalves, Ewan Harrison, David K. Jackson, Ian Johnston, Dominic Kwiatkowski, Cordelia Langford, John Sillitoe on behalf of the Wellcome Sanger Institute COVID-19 Surveillance Team |
| EPI_ISL_2552723 | Health Services Laboratories | Wellcome Sanger Institute for the COVID-19 Genomics UK (COG-UK) Consortium | Health Services Laboratories and Alex Alderton, Roberto Amato, Jeffrey Barrett, Sonia Goncalves, Ewan Harrison, David K. Jackson, Ian Johnston, Dominic Kwiatkowski, Cordelia Langford, John Sillitoe on behalf of the Wellcome Sanger Institute COVID-19 Surveillance Team |
| EPI_ISL_2552724 | Lighthouse Lab in Glasgow | Wellcome Sanger Institute for the COVID-19 Genomics UK (COG-UK) Consortium | Harper VanSteenhouse, Yumi Kasai, David Gray, Carol Clugston, Anna Dominiczak and Alex Alderton, Roberto Amato, Jeffrey Barrett, Sonia Goncalves, Ewan Harrison, David K. Jackson, Ian Johnston, Dominic Kwiatkowski, Cordelia Langford, John Sillitoe on behalf of the Wellcome Sanger Institute COVID-19 Surveillance Team |
| EPI_ISL_2552725 | Health Services Laboratories | Wellcome Sanger Institute for the COVID-19 Genomics UK (COG-UK) Consortium | Health Services Laboratories and Alex Alderton, Roberto Amato, Jeffrey Barrett, Sonia Goncalves, Ewan Harrison, David K. Jackson, Ian Johnston, Dominic Kwiatkowski, Cordelia Langford, John Sillitoe on behalf of the Wellcome Sanger Institute COVID-19 Surveillance Team |
| EPI_ISL_2552726 | Lighthouse Lab in Glasgow | Wellcome Sanger Institute for the COVID-19 Genomics UK (COG-UK) Consortium | Harper VanSteenhouse, Yumi Kasai, David Gray, Carol Clugston, Anna Dominiczak and Alex Alderton, Roberto Amato, Jeffrey Barrett, Sonia Goncalves, Ewan Harrison, David K. Jackson, Ian Johnston, Dominic Kwiatkowski, Cordelia Langford, John Sillitoe on behalf of the Wellcome Sanger Institute COVID-19 Surveillance Team |
| EPI_ISL_2552727, EPI_ISL_2552728 | Lighthouse Lab in Milton Keynes | Wellcome Sanger Institute for the COVID-19 Genomics UK (COG-UK) Consortium | The Lighthouse Lab in Milton Keynes and Alex Alderton, Roberto Amato, Jeffrey Barrett, Sonia Goncalves, Ewan Harrison, David K. Jackson, Ian Johnston, Dominic Kwiatkowski, Cordelia Langford, John Sillitoe on behalf of the Wellcome Sanger Institute COVID-19 Surveillance Team |
| EPI_ISL_2552729 | Lighthouse Lab in Glasgow | Wellcome Sanger Institute for the COVID-19 Genomics UK (COG-UK) Consortium | Harper VanSteenhouse, Yumi Kasai, David Gray, Carol Clugston, Anna Dominiczak and Alex Alderton, Roberto Amato, Jeffrey Barrett, Sonia Goncalves, Ewan Harrison, David K. Jackson, Ian Johnston, Dominic Kwiatkowski, Cordelia Langford, John Sillitoe on behalf of the Wellcome Sanger Institute COVID-19 Surveillance Team |
| EPI_ISL_2552730, EPI_ISL_2552731 | Health Services Laboratories | Wellcome Sanger Institute for the COVID-19 Genomics UK (COG-UK) Consortium | Health Services Laboratories and Alex Alderton, Roberto Amato, Jeffrey Barrett, Sonia Goncalves, Ewan Harrison, David K. Jackson, Ian Johnston, Dominic Kwiatkowski, Cordelia Langford, John Sillitoe on behalf of the Wellcome Sanger Institute COVID-19 Surveillance Team |
| EPI_ISL_2552732 | Lighthouse Lab in Glasgow | Wellcome Sanger Institute for the COVID-19 Genomics UK (COG-UK) Consortium | Harper VanSteenhouse, Yumi Kasai, David Gray, Carol Clugston, Anna Dominiczak and Alex Alderton, Roberto Amato, Jeffrey Barrett, Sonia Goncalves, Ewan Harrison, David K. Jackson, Ian Johnston, Dominic Kwiatkowski, Cordelia Langford, John Sillitoe on behalf of the Wellcome Sanger Institute COVID-19 Surveillance Team |
| EPI_ISL_2552733 | Lighthouse Lab in Milton Keynes | Wellcome Sanger Institute for the COVID-19 Genomics UK (COG-UK) Consortium | The Lighthouse Lab in Milton Keynes and Alex Alderton, Roberto Amato, Jeffrey Barrett, Sonia Goncalves, Ewan Harrison, David K. Jackson, Ian Johnston, Dominic Kwiatkowski, Cordelia Langford, John Sillitoe on behalf of the Wellcome Sanger Institute COVID-19 Surveillance Team |
| EPI_ISL_2552734 | Health Services Laboratories | Wellcome Sanger Institute for the COVID-19 Genomics UK (COG-UK) Consortium | Health Services Laboratories and Alex Alderton, Roberto Amato, Jeffrey Barrett, Sonia Goncalves, Ewan Harrison, David K. Jackson, Ian Johnston, Dominic Kwiatkowski, Cordelia Langford, John Sillitoe on behalf of the Wellcome Sanger Institute COVID-19 Surveillance Team |
| EPI_ISL_2552735, EPI_ISL_2552736 | Lighthouse Lab in Glasgow | Wellcome Sanger Institute for the COVID-19 Genomics UK (COG-UK) Consortium | Harper VanSteenhouse, Yumi Kasai, David Gray, Carol Clugston, Anna Dominiczak and Alex Alderton, Roberto Amato, Jeffrey Barrett, Sonia Goncalves, Ewan Harrison, David K. Jackson, Ian Johnston, Dominic Kwiatkowski, Cordelia Langford, John Sillitoe on behalf of the Wellcome Sanger Institute COVID-19 Surveillance Team |
| EPI_ISL_2552738 | Lighthouse Lab in Alderley Park | Wellcome Sanger Institute for the COVID-19 Genomics UK (COG-UK) Consortium | Jacquelyn Wynn, Mairead Hyland, The Lighthouse Lab in Alderley Park and Alex Alderton, Roberto Amato, Jeffrey Barrett, Sonia Goncalves, Ewan Harrison, David K. Jackson, Ian Johnston, Dominic Kwiatkowski, Cordelia Langford, John Sillitoe on behalf of the Wellcome Sanger Institute COVID-19 Surveillance Team |
| EPI_ISL_2552739, EPI_ISL_2552740 | Lighthouse Lab in Milton Keynes | Wellcome Sanger Institute for the COVID-19 Genomics UK (COG-UK) Consortium | The Lighthouse Lab in Milton Keynes and Alex Alderton, Roberto Amato, Jeffrey Barrett, Sonia Goncalves, Ewan Harrison, David K. Jackson, Ian Johnston, Dominic Kwiatkowski, Cordelia Langford, John Sillitoe on behalf of the Wellcome Sanger Institute COVID-19 Surveillance Team |
| EPI_ISL_2552741, EPI_ISL_2552742 | Lighthouse Lab in Alderley Park | Wellcome Sanger Institute for the COVID-19 Genomics UK (COG-UK) Consortium | Jacquelyn Wynn, Mairead Hyland, The Lighthouse Lab in Alderley Park and Alex Alderton, Roberto Amato, Jeffrey Barrett, Sonia Goncalves, Ewan Harrison, David K. Jackson, Ian Johnston, Dominic Kwiatkowski, Cordelia Langford, John Sillitoe on behalf of the Wellcome Sanger Institute COVID-19 Surveillance Team |
| EPI_ISL_2552743 | Lighthouse Lab in Glasgow | Wellcome Sanger Institute for the COVID-19 Genomics UK (COG-UK) Consortium | Harper VanSteenhouse, Yumi Kasai, David Gray, Carol Clugston, Anna Dominiczak and Alex Alderton, Roberto Amato, Jeffrey Barrett, Sonia Goncalves, Ewan Harrison, David K. Jackson, Ian Johnston, Dominic Kwiatkowski, Cordelia Langford, John Sillitoe on behalf of the Wellcome Sanger Institute COVID-19 Surveillance Team |
| EPI_ISL_2552744, EPI_ISL_2552745, EPI_ISL_2552746 | Health Services Laboratories | Wellcome Sanger Institute for the COVID-19 Genomics UK (COG-UK) Consortium | Health Services Laboratories and Alex Alderton, Roberto Amato, Jeffrey Barrett, Sonia Goncalves, Ewan Harrison, David K. Jackson, Ian Johnston, Dominic Kwiatkowski, Cordelia Langford, John Sillitoe on behalf of the Wellcome Sanger Institute COVID-19 Surveillance Team |
| EPI_ISL_2552747 | Lighthouse Lab in Glasgow | Wellcome Sanger Institute for the COVID-19 Genomics UK (COG-UK) Consortium | Harper VanSteenhouse, Yumi Kasai, David Gray, Carol Clugston, Anna Dominiczak and Alex Alderton, Roberto Amato, Jeffrey Barrett, Sonia Goncalves, Ewan Harrison, David K. Jackson, Ian Johnston, Dominic Kwiatkowski, Cordelia Langford, John Sillitoe on behalf of the Wellcome Sanger Institute COVID-19 Surveillance Team |
| EPI_ISL_2552748, EPI_ISL_2552749 | Health Services Laboratories | Wellcome Sanger Institute for the COVID-19 Genomics UK (COG-UK) Consortium | Health Services Laboratories and Alex Alderton, Roberto Amato, Jeffrey Barrett, Sonia Goncalves, Ewan Harrison, David K. Jackson, Ian Johnston, Dominic Kwiatkowski, Cordelia Langford, John Sillitoe on behalf of the Wellcome Sanger Institute COVID-19 Surveillance Team |
| EPI_ISL_2552750 | Lighthouse Lab in Alderley Park | Wellcome Sanger Institute for the COVID-19 Genomics UK (COG-UK) Consortium | Jacquelyn Wynn, Mairead Hyland, The Lighthouse Lab in Alderley Park and Alex Alderton, Roberto Amato, Jeffrey Barrett, Sonia Goncalves, Ewan Harrison, David K. Jackson, Ian Johnston, Dominic Kwiatkowski, Cordelia Langford, John Sillitoe on behalf of the Wellcome Sanger Institute COVID-19 Surveillance Team |
| EPI_ISL_2552751 | Health Services Laboratories | Wellcome Sanger Institute for the COVID-19 Genomics UK (COG-UK) Consortium | Health Services Laboratories and Alex Alderton, Roberto Amato, Jeffrey Barrett, Sonia Goncalves, Ewan Harrison, David K. Jackson, Ian Johnston, Dominic Kwiatkowski, Cordelia Langford, John Sillitoe on behalf of the Wellcome Sanger Institute COVID-19 Surveillance Team |
| EPI_ISL_2552752, EPI_ISL_2552753 | Lighthouse Lab in Milton Keynes | Wellcome Sanger Institute for the COVID-19 Genomics UK (COG-UK) Consortium | The Lighthouse Lab in Milton Keynes and Alex Alderton, Roberto Amato, Jeffrey Barrett, Sonia Goncalves, Ewan Harrison, David K. Jackson, Ian Johnston, Dominic Kwiatkowski, Cordelia Langford, John Sillitoe on behalf of the Wellcome Sanger Institute COVID-19 Surveillance Team |
| EPI_ISL_2552754 | Lighthouse Lab in Alderley Park | Wellcome Sanger Institute for the COVID-19 Genomics UK (COG-UK) Consortium | Jacquelyn Wynn, Mairead Hyland, The Lighthouse Lab in Alderley Park and Alex Alderton, Roberto Amato, Jeffrey Barrett, Sonia Goncalves, Ewan Harrison, David K. Jackson, Ian Johnston, Dominic Kwiatkowski, Cordelia Langford, John Sillitoe on behalf of the Wellcome Sanger Institute COVID-19 Surveillance Team |
| EPI_ISL_2552755 | Lighthouse Lab in Milton Keynes | Wellcome Sanger Institute for the COVID-19 Genomics UK (COG-UK) Consortium | The Lighthouse Lab in Milton Keynes and Alex Alderton, Roberto Amato, Jeffrey Barrett, Sonia Goncalves, Ewan Harrison, David K. Jackson, Ian Johnston, Dominic Kwiatkowski, Cordelia Langford, John Sillitoe on behalf of the Wellcome Sanger Institute COVID-19 Surveillance Team |
| EPI_ISL_2552756 | Health Services Laboratories | Wellcome Sanger Institute for the COVID-19 Genomics UK (COG-UK) Consortium | Health Services Laboratories and Alex Alderton, Roberto Amato, Jeffrey Barrett, Sonia Goncalves, Ewan Harrison, David K. Jackson, Ian Johnston, Dominic Kwiatkowski, Cordelia Langford, John Sillitoe on behalf of the Wellcome Sanger Institute COVID-19 Surveillance Team |
| EPI_ISL_2552757, EPI_ISL_2552758 | Lighthouse Lab in Milton Keynes | Wellcome Sanger Institute for the COVID-19 Genomics UK (COG-UK) Consortium | The Lighthouse Lab in Milton Keynes and Alex Alderton, Roberto Amato, Jeffrey Barrett, Sonia Goncalves, Ewan Harrison, David K. Jackson, Ian Johnston, Dominic Kwiatkowski, Cordelia Langford, John Sillitoe on behalf of the Wellcome Sanger Institute COVID-19 Surveillance Team |
| EPI_ISL_2552759 | Lighthouse Lab in Alderley Park | Wellcome Sanger Institute for the COVID-19 Genomics UK (COG-UK) Consortium | Jacquelyn Wynn, Mairead Hyland, The Lighthouse Lab in Alderley Park and Alex Alderton, Roberto Amato, Jeffrey Barrett, Sonia Goncalves, Ewan Harrison, David K. Jackson, Ian Johnston, Dominic Kwiatkowski, Cordelia Langford, John Sillitoe on behalf of the Wellcome Sanger Institute COVID-19 Surveillance Team |
| EPI_ISL_2552760 | Lighthouse Lab in Milton Keynes | Wellcome Sanger Institute for the COVID-19 Genomics UK (COG-UK) Consortium | The Lighthouse Lab in Milton Keynes and Alex Alderton, Roberto Amato, Jeffrey Barrett, Sonia Goncalves, Ewan Harrison, David K. Jackson, Ian Johnston, Dominic Kwiatkowski, Cordelia Langford, John Sillitoe on behalf of the Wellcome Sanger Institute COVID-19 Surveillance Team |
| EPI_ISL_2552761, EPI_ISL_2552762 | Lighthouse Lab in Glasgow | Wellcome Sanger Institute for the COVID-19 Genomics UK (COG-UK) Consortium | Harper VanSteenhouse, Yumi Kasai, David Gray, Carol Clugston, Anna Dominiczak and Alex Alderton, Roberto Amato, Jeffrey Barrett, Sonia Goncalves, Ewan Harrison, David K. Jackson, Ian Johnston, Dominic Kwiatkowski, Cordelia Langford, John Sillitoe on behalf of the Wellcome Sanger Institute COVID-19 Surveillance Team |
| EPI_ISL_2552763 | Lighthouse Lab in Alderley Park | Wellcome Sanger Institute for the COVID-19 Genomics UK (COG-UK) Consortium | Jacquelyn Wynn, Mairead Hyland, The Lighthouse Lab in Alderley Park and Alex Alderton, Roberto Amato, Jeffrey Barrett, Sonia Goncalves, Ewan Harrison, David K. Jackson, Ian Johnston, Dominic Kwiatkowski, Cordelia Langford, John Sillitoe on behalf of the Wellcome Sanger Institute COVID-19 Surveillance Team |
| EPI_ISL_2552764 | Lighthouse Lab in Milton Keynes | Wellcome Sanger Institute for the COVID-19 Genomics UK (COG-UK) Consortium | The Lighthouse Lab in Milton Keynes and Alex Alderton, Roberto Amato, Jeffrey Barrett, Sonia Goncalves, Ewan Harrison, David K. Jackson, Ian Johnston, Dominic Kwiatkowski, Cordelia Langford, John Sillitoe on behalf of the Wellcome Sanger Institute COVID-19 Surveillance Team |
| EPI_ISL_2552765 | Lighthouse Lab in Alderley Park | Wellcome Sanger Institute for the COVID-19 Genomics UK (COG-UK) Consortium | Jacquelyn Wynn, Mairead Hyland, The Lighthouse Lab in Alderley Park and Alex Alderton, Roberto Amato, Jeffrey Barrett, Sonia Goncalves, Ewan Harrison, David K. Jackson, Ian Johnston, Dominic Kwiatkowski, Cordelia Langford, John Sillitoe on behalf of the Wellcome Sanger Institute COVID-19 Surveillance Team |
| EPI_ISL_2552766 | Lighthouse Lab in Glasgow | Wellcome Sanger Institute for the COVID-19 Genomics UK (COG-UK) Consortium | Harper VanSteenhouse, Yumi Kasai, David Gray, Carol Clugston, Anna Dominiczak and Alex Alderton, Roberto Amato, Jeffrey Barrett, Sonia Goncalves, Ewan Harrison, David K. Jackson, Ian Johnston, Dominic Kwiatkowski, Cordelia Langford, John Sillitoe on behalf of the Wellcome Sanger Institute COVID-19 Surveillance Team |
| EPI_ISL_2552767, EPI_ISL_2552768 | Health Services Laboratories | Wellcome Sanger Institute for the COVID-19 Genomics UK (COG-UK) Consortium | Health Services Laboratories and Alex Alderton, Roberto Amato, Jeffrey Barrett, Sonia Goncalves, Ewan Harrison, David K. Jackson, Ian Johnston, Dominic Kwiatkowski, Cordelia Langford, John Sillitoe on behalf of the Wellcome Sanger Institute COVID-19 Surveillance Team |
| EPI_ISL_2552769 | Lighthouse Lab in Milton Keynes | Wellcome Sanger Institute for the COVID-19 Genomics UK (COG-UK) Consortium | The Lighthouse Lab in Milton Keynes and Alex Alderton, Roberto Amato, Jeffrey Barrett, Sonia Goncalves, Ewan Harrison, David K. Jackson, Ian Johnston, Dominic Kwiatkowski, Cordelia Langford, John Sillitoe on behalf of the Wellcome Sanger Institute COVID-19 Surveillance Team |
| EPI_ISL_2552770 | Lighthouse Lab in Alderley Park | Wellcome Sanger Institute for the COVID-19 Genomics UK (COG-UK) Consortium | Jacquelyn Wynn, Mairead Hyland, The Lighthouse Lab in Alderley Park and Alex Alderton, Roberto Amato, Jeffrey Barrett, Sonia Goncalves, Ewan Harrison, David K. Jackson, Ian Johnston, Dominic Kwiatkowski, Cordelia Langford, John Sillitoe on behalf of the Wellcome Sanger Institute COVID-19 Surveillance Team |
| EPI_ISL_2552771 | Lighthouse Lab in Glasgow | Wellcome Sanger Institute for the COVID-19 Genomics UK (COG-UK) Consortium | Harper VanSteenhouse, Yumi Kasai, David Gray, Carol Clugston, Anna Dominiczak and Alex Alderton, Roberto Amato, Jeffrey Barrett, Sonia Goncalves, Ewan Harrison, David K. Jackson, Ian Johnston, Dominic Kwiatkowski, Cordelia Langford, John Sillitoe on behalf of the Wellcome Sanger Institute COVID-19 Surveillance Team |
| EPI_ISL_2552772, | Health Services Laboratories | Wellcome Sanger Institute for the COVID-19 Genomics UK (COG-UK) | Health Services Laboratories and Alex Alderton, Roberto Amato, Jeffrey Barrett, Sonia Goncalves, Ewan Harrison, David K. Jackson, Ian Johnston, Dominic Kwiatkowski, Cordelia Langford, John Sillitoe on behalf of the Wellcome Sanger Institute |

| | | | |
|---|---|---|---|
| EPI_ISL_2552773 | | Consortium | COVID-19 Surveillance Team |
| EPI_ISL_2552774 | Lighthouse Lab in Glasgow | Wellcome Sanger Institute for the COVID-19 Genomics UK (COG-UK) Consortium | Harper VanSteenhouse, Yumi Kasai, David Gray, Carol Clugston, Anna Dominiczak and Alex Alderton, Roberto Amato, Jeffrey Barrett, Sonia Goncalves, Ewan Harrison, David K. Jackson, Ian Johnston, Dominic Kwiatkowski, Cordelia Langford, John Sillitoe on behalf of the Wellcome Sanger Institute COVID-19 Surveillance Team |
| EPI_ISL_2552775 | Health Services Laboratories | Wellcome Sanger Institute for the COVID-19 Genomics UK (COG-UK) Consortium | Health Services Laboratories and Alex Alderton, Roberto Amato, Jeffrey Barrett, Sonia Goncalves, Ewan Harrison, David K. Jackson, Ian Johnston, Dominic Kwiatkowski, Cordelia Langford, John Sillitoe on behalf of the Wellcome Sanger Institute COVID-19 Surveillance Team |
| EPI_ISL_2552776 | Lighthouse Lab in Milton Keynes | Wellcome Sanger Institute for the COVID-19 Genomics UK (COG-UK) Consortium | The Lighthouse Lab in Milton Keynes and Alex Alderton, Roberto Amato, Jeffrey Barrett, Sonia Goncalves, Ewan Harrison, David K. Jackson, Ian Johnston, Dominic Kwiatkowski, Cordelia Langford, John Sillitoe on behalf of the Wellcome Sanger Institute COVID-19 Surveillance Team |
| EPI_ISL_2552777 | Health Services Laboratories | Wellcome Sanger Institute for the COVID-19 Genomics UK (COG-UK) Consortium | Health Services Laboratories and Alex Alderton, Roberto Amato, Jeffrey Barrett, Sonia Goncalves, Ewan Harrison, David K. Jackson, Ian Johnston, Dominic Kwiatkowski, Cordelia Langford, John Sillitoe on behalf of the Wellcome Sanger Institute COVID-19 Surveillance Team |
| EPI_ISL_2552778 | Lighthouse Lab in Glasgow | Wellcome Sanger Institute for the COVID-19 Genomics UK (COG-UK) Consortium | Harper VanSteenhouse, Yumi Kasai, David Gray, Carol Clugston, Anna Dominiczak and Alex Alderton, Roberto Amato, Jeffrey Barrett, Sonia Goncalves, Ewan Harrison, David K. Jackson, Ian Johnston, Dominic Kwiatkowski, Cordelia Langford, John Sillitoe on behalf of the Wellcome Sanger Institute COVID-19 Surveillance Team |
| EPI_ISL_2552779 | Lighthouse Lab in Alderley Park | Wellcome Sanger Institute for the COVID-19 Genomics UK (COG-UK) Consortium | Jacquelyn Wynn, Mairead Hyland, The Lighthouse Lab in Alderley Park and Alex Alderton, Roberto Amato, Jeffrey Barrett, Sonia Goncalves, Ewan Harrison, David K. Jackson, Ian Johnston, Dominic Kwiatkowski, Cordelia Langford, John Sillitoe on behalf of the Wellcome Sanger Institute COVID-19 Surveillance Team |
| EPI_ISL_2552780 | Lighthouse Lab in Milton Keynes | Wellcome Sanger Institute for the COVID-19 Genomics UK (COG-UK) Consortium | The Lighthouse Lab in Milton Keynes and Alex Alderton, Roberto Amato, Jeffrey Barrett, Sonia Goncalves, Ewan Harrison, David K. Jackson, Ian Johnston, Dominic Kwiatkowski, Cordelia Langford, John Sillitoe on behalf of the Wellcome Sanger Institute COVID-19 Surveillance Team |
| EPI_ISL_2552781, EPI_ISL_2552782 | Health Services Laboratories | Wellcome Sanger Institute for the COVID-19 Genomics UK (COG-UK) Consortium | Health Services Laboratories and Alex Alderton, Roberto Amato, Jeffrey Barrett, Sonia Goncalves, Ewan Harrison, David K. Jackson, Ian Johnston, Dominic Kwiatkowski, Cordelia Langford, John Sillitoe on behalf of the Wellcome Sanger Institute COVID-19 Surveillance Team |
| EPI_ISL_2552785 | Lighthouse Lab in Milton Keynes | Wellcome Sanger Institute for the COVID-19 Genomics UK (COG-UK) Consortium | The Lighthouse Lab in Milton Keynes and Alex Alderton, Roberto Amato, Jeffrey Barrett, Sonia Goncalves, Ewan Harrison, David K. Jackson, Ian Johnston, Dominic Kwiatkowski, Cordelia Langford, John Sillitoe on behalf of the Wellcome Sanger Institute COVID-19 Surveillance Team |
| EPI_ISL_2552786 | Lighthouse Lab in Glasgow | Wellcome Sanger Institute for the COVID-19 Genomics UK (COG-UK) Consortium | Harper VanSteenhouse, Yumi Kasai, David Gray, Carol Clugston, Anna Dominiczak and Alex Alderton, Roberto Amato, Jeffrey Barrett, Sonia Goncalves, Ewan Harrison, David K. Jackson, Ian Johnston, Dominic Kwiatkowski, Cordelia Langford, John Sillitoe on behalf of the Wellcome Sanger Institute COVID-19 Surveillance Team |
| EPI_ISL_2552787 | Lighthouse Lab in Alderley Park | Wellcome Sanger Institute for the COVID-19 Genomics UK (COG-UK) Consortium | Jacquelyn Wynn, Mairead Hyland, The Lighthouse Lab in Alderley Park and Alex Alderton, Roberto Amato, Jeffrey Barrett, Sonia Goncalves, Ewan Harrison, David K. Jackson, Ian Johnston, Dominic Kwiatkowski, Cordelia Langford, John Sillitoe on behalf of the Wellcome Sanger Institute COVID-19 Surveillance Team |
| EPI_ISL_2552788 | Lighthouse Lab in Glasgow | Wellcome Sanger Institute for the COVID-19 Genomics UK (COG-UK) Consortium | Harper VanSteenhouse, Yumi Kasai, David Gray, Carol Clugston, Anna Dominiczak and Alex Alderton, Roberto Amato, Jeffrey Barrett, Sonia Goncalves, Ewan Harrison, David K. Jackson, Ian Johnston, Dominic Kwiatkowski, Cordelia Langford, John Sillitoe on behalf of the Wellcome Sanger Institute COVID-19 Surveillance Team |
| EPI_ISL_2552789 | Lighthouse Lab in Milton Keynes | Wellcome Sanger Institute for the COVID-19 Genomics UK (COG-UK) Consortium | The Lighthouse Lab in Milton Keynes and Alex Alderton, Roberto Amato, Jeffrey Barrett, Sonia Goncalves, Ewan Harrison, David K. Jackson, Ian Johnston, Dominic Kwiatkowski, Cordelia Langford, John Sillitoe on behalf of the Wellcome Sanger Institute COVID-19 Surveillance Team |
| EPI_ISL_2552790 | Health Services Laboratories | Wellcome Sanger Institute for the COVID-19 Genomics UK (COG-UK) Consortium | Health Services Laboratories and Alex Alderton, Roberto Amato, Jeffrey Barrett, Sonia Goncalves, Ewan Harrison, David K. Jackson, Ian Johnston, Dominic Kwiatkowski, Cordelia Langford, John Sillitoe on behalf of the Wellcome Sanger Institute COVID-19 Surveillance Team |
| EPI_ISL_2552791, EPI_ISL_2552792 | Lighthouse Lab in Glasgow | Wellcome Sanger Institute for the COVID-19 Genomics UK (COG-UK) Consortium | Harper VanSteenhouse, Yumi Kasai, David Gray, Carol Clugston, Anna Dominiczak and Alex Alderton, Roberto Amato, Jeffrey Barrett, Sonia Goncalves, Ewan Harrison, David K. Jackson, Ian Johnston, Dominic Kwiatkowski, Cordelia Langford, John Sillitoe on behalf of the Wellcome Sanger Institute COVID-19 Surveillance Team |
| EPI_ISL_2552793 | Lighthouse Lab in Milton Keynes | Wellcome Sanger Institute for the COVID-19 Genomics UK (COG-UK) Consortium | The Lighthouse Lab in Milton Keynes and Alex Alderton, Roberto Amato, Jeffrey Barrett, Sonia Goncalves, Ewan Harrison, David K. Jackson, Ian Johnston, Dominic Kwiatkowski, Cordelia Langford, John Sillitoe on behalf of the Wellcome Sanger Institute COVID-19 Surveillance Team |
| EPI_ISL_2552794, EPI_ISL_2552795 | Health Services Laboratories | Wellcome Sanger Institute for the COVID-19 Genomics UK (COG-UK) Consortium | Health Services Laboratories and Alex Alderton, Roberto Amato, Jeffrey Barrett, Sonia Goncalves, Ewan Harrison, David K. Jackson, Ian Johnston, Dominic Kwiatkowski, Cordelia Langford, John Sillitoe on behalf of the Wellcome Sanger Institute COVID-19 Surveillance Team |
| EPI_ISL_2552796 | Lighthouse Lab in Glasgow | Wellcome Sanger Institute for the COVID-19 Genomics UK (COG-UK) Consortium | Harper VanSteenhouse, Yumi Kasai, David Gray, Carol Clugston, Anna Dominiczak and Alex Alderton, Roberto Amato, Jeffrey Barrett, Sonia Goncalves, Ewan Harrison, David K. Jackson, Ian Johnston, Dominic Kwiatkowski, Cordelia Langford, John Sillitoe on behalf of the Wellcome Sanger Institute COVID-19 Surveillance Team |
| EPI_ISL_2552797, EPI_ISL_2552798, EPI_ISL_2552799, EPI_ISL_2552800 | Lighthouse Lab in Milton Keynes | Wellcome Sanger Institute for the COVID-19 Genomics UK (COG-UK) Consortium | The Lighthouse Lab in Milton Keynes and Alex Alderton, Roberto Amato, Jeffrey Barrett, Sonia Goncalves, Ewan Harrison, David K. Jackson, Ian Johnston, Dominic Kwiatkowski, Cordelia Langford, John Sillitoe on behalf of the Wellcome Sanger Institute COVID-19 Surveillance Team |
| EPI_ISL_2552801 | Health Services Laboratories | Wellcome Sanger Institute for the COVID-19 Genomics UK (COG-UK) Consortium | Health Services Laboratories and Alex Alderton, Roberto Amato, Jeffrey Barrett, Sonia Goncalves, Ewan Harrison, David K. Jackson, Ian Johnston, Dominic Kwiatkowski, Cordelia Langford, John Sillitoe on behalf of the Wellcome Sanger Institute COVID-19 Surveillance Team |
| EPI_ISL_2552802 | Lighthouse Lab in Milton Keynes | Wellcome Sanger Institute for the COVID-19 Genomics UK (COG-UK) Consortium | The Lighthouse Lab in Milton Keynes and Alex Alderton, Roberto Amato, Jeffrey Barrett, Sonia Goncalves, Ewan Harrison, David K. Jackson, Ian Johnston, Dominic Kwiatkowski, Cordelia Langford, John Sillitoe on behalf of the Wellcome Sanger Institute COVID-19 Surveillance Team |
| EPI_ISL_2552803 | Health Services Laboratories | Wellcome Sanger Institute for the COVID-19 Genomics UK (COG-UK) Consortium | Health Services Laboratories and Alex Alderton, Roberto Amato, Jeffrey Barrett, Sonia Goncalves, Ewan Harrison, David K. Jackson, Ian Johnston, Dominic Kwiatkowski, Cordelia Langford, John Sillitoe on behalf of the Wellcome Sanger Institute COVID-19 Surveillance Team |
| EPI_ISL_2552804 | Lighthouse Lab in Glasgow | Wellcome Sanger Institute for the COVID-19 Genomics UK (COG-UK) Consortium | Harper VanSteenhouse, Yumi Kasai, David Gray, Carol Clugston, Anna Dominiczak and Alex Alderton, Roberto Amato, Jeffrey Barrett, Sonia Goncalves, Ewan Harrison, David K. Jackson, Ian Johnston, Dominic Kwiatkowski, Cordelia Langford, John Sillitoe on behalf of the Wellcome Sanger Institute COVID-19 Surveillance Team |
| EPI_ISL_2552805 | Health Services Laboratories | Wellcome Sanger Institute for the COVID-19 Genomics UK (COG-UK) Consortium | Health Services Laboratories and Alex Alderton, Roberto Amato, Jeffrey Barrett, Sonia Goncalves, Ewan Harrison, David K. Jackson, Ian Johnston, Dominic Kwiatkowski, Cordelia Langford, John Sillitoe on behalf of the Wellcome Sanger Institute COVID-19 Surveillance Team |
| EPI_ISL_2552806 | Lighthouse Lab in Glasgow | Wellcome Sanger Institute for the COVID-19 Genomics UK (COG-UK) Consortium | Harper VanSteenhouse, Yumi Kasai, David Gray, Carol Clugston, Anna Dominiczak and Alex Alderton, Roberto Amato, Jeffrey Barrett, Sonia Goncalves, Ewan Harrison, David K. Jackson, Ian Johnston, Dominic Kwiatkowski, Cordelia Langford, John Sillitoe on behalf of the Wellcome Sanger Institute COVID-19 Surveillance Team |
| EPI_ISL_2552807, EPI_ISL_2552810 | Health Services Laboratories | Wellcome Sanger Institute for the COVID-19 Genomics UK (COG-UK) Consortium | Health Services Laboratories and Alex Alderton, Roberto Amato, Jeffrey Barrett, Sonia Goncalves, Ewan Harrison, David K. Jackson, Ian Johnston, Dominic Kwiatkowski, Cordelia Langford, John Sillitoe on behalf of the Wellcome Sanger Institute COVID-19 Surveillance Team |
| EPI_ISL_2552811 | Lighthouse Lab in Milton Keynes | Wellcome Sanger Institute for the COVID-19 Genomics UK (COG-UK) Consortium | The Lighthouse Lab in Milton Keynes and Alex Alderton, Roberto Amato, Jeffrey Barrett, Sonia Goncalves, Ewan Harrison, David K. Jackson, Ian Johnston, Dominic Kwiatkowski, Cordelia Langford, John Sillitoe on behalf of the Wellcome Sanger Institute COVID-19 Surveillance Team |
| EPI_ISL_2552812 | Lighthouse Lab in Alderley Park | Wellcome Sanger Institute for the COVID-19 Genomics UK (COG-UK) Consortium | Jacquelyn Wynn, Mairead Hyland, The Lighthouse Lab in Alderley Park and Alex Alderton, Roberto Amato, Jeffrey Barrett, Sonia Goncalves, Ewan Harrison, David K. Jackson, Ian Johnston, Dominic Kwiatkowski, Cordelia Langford, John Sillitoe on behalf of the Wellcome Sanger Institute COVID-19 Surveillance Team |
| EPI_ISL_2552813 | Lighthouse Lab in Milton Keynes | Wellcome Sanger Institute for the COVID-19 Genomics UK (COG-UK) Consortium | The Lighthouse Lab in Milton Keynes and Alex Alderton, Roberto Amato, Jeffrey Barrett, Sonia Goncalves, Ewan Harrison, David K. Jackson, Ian Johnston, Dominic Kwiatkowski, Cordelia Langford, John Sillitoe on behalf of the Wellcome Sanger Institute COVID-19 Surveillance Team |
| EPI_ISL_2552814 | Health Services Laboratories | Wellcome Sanger Institute for the COVID-19 Genomics UK (COG-UK) Consortium | Health Services Laboratories and Alex Alderton, Roberto Amato, Jeffrey Barrett, Sonia Goncalves, Ewan Harrison, David K. Jackson, Ian Johnston, Dominic Kwiatkowski, Cordelia Langford, John Sillitoe on behalf of the Wellcome Sanger Institute COVID-19 Surveillance Team |
| EPI_ISL_2552815 | Lighthouse Lab in Milton Keynes | Wellcome Sanger Institute for the COVID-19 Genomics UK (COG-UK) Consortium | The Lighthouse Lab in Milton Keynes and Alex Alderton, Roberto Amato, Jeffrey Barrett, Sonia Goncalves, Ewan Harrison, David K. Jackson, Ian Johnston, Dominic Kwiatkowski, Cordelia Langford, John Sillitoe on behalf of the Wellcome Sanger Institute COVID-19 Surveillance Team |
| EPI_ISL_2552816 | Health Services Laboratories | Wellcome Sanger Institute for the COVID-19 Genomics UK (COG-UK) Consortium | Health Services Laboratories and Alex Alderton, Roberto Amato, Jeffrey Barrett, Sonia Goncalves, Ewan Harrison, David K. Jackson, Ian Johnston, Dominic Kwiatkowski, Cordelia Langford, John Sillitoe on behalf of the Wellcome Sanger Institute COVID-19 Surveillance Team |
| EPI_ISL_2552817 | Lighthouse Lab in Alderley Park | Wellcome Sanger Institute for the COVID-19 Genomics UK (COG-UK) Consortium | Jacquelyn Wynn, Mairead Hyland, The Lighthouse Lab in Alderley Park and Alex Alderton, Roberto Amato, Jeffrey Barrett, Sonia Goncalves, Ewan Harrison, David K. Jackson, Ian Johnston, Dominic Kwiatkowski, Cordelia Langford, John Sillitoe on behalf of the Wellcome Sanger Institute COVID-19 Surveillance Team |
| EPI_ISL_2552818 | Lighthouse Lab in Glasgow | Wellcome Sanger Institute for the COVID-19 Genomics UK (COG-UK) Consortium | Harper VanSteenhouse, Yumi Kasai, David Gray, Carol Clugston, Anna Dominiczak and Alex Alderton, Roberto Amato, Jeffrey Barrett, Sonia Goncalves, Ewan Harrison, David K. Jackson, Ian Johnston, Dominic Kwiatkowski, Cordelia Langford, John Sillitoe on behalf of the Wellcome Sanger Institute COVID-19 Surveillance Team |
| EPI_ISL_2552819, EPI_ISL_2552820 | Health Services Laboratories | Wellcome Sanger Institute for the COVID-19 Genomics UK (COG-UK) Consortium | Health Services Laboratories and Alex Alderton, Roberto Amato, Jeffrey Barrett, Sonia Goncalves, Ewan Harrison, David K. Jackson, Ian Johnston, Dominic Kwiatkowski, Cordelia Langford, John Sillitoe on behalf of the Wellcome Sanger Institute COVID-19 Surveillance Team |
| EPI_ISL_2552821 | Lighthouse Lab in Glasgow | Wellcome Sanger Institute for the COVID-19 Genomics UK (COG-UK) Consortium | Harper VanSteenhouse, Yumi Kasai, David Gray, Carol Clugston, Anna Dominiczak and Alex Alderton, Roberto Amato, Jeffrey Barrett, Sonia Goncalves, Ewan Harrison, David K. Jackson, Ian Johnston, Dominic Kwiatkowski, Cordelia Langford, John Sillitoe on behalf of the Wellcome Sanger Institute COVID-19 Surveillance Team |
| EPI_ISL_2552823 | Health Services Laboratories | Wellcome Sanger Institute for the COVID-19 Genomics UK (COG-UK) Consortium | Health Services Laboratories and Alex Alderton, Roberto Amato, Jeffrey Barrett, Sonia Goncalves, Ewan Harrison, David K. Jackson, Ian Johnston, Dominic Kwiatkowski, Cordelia Langford, John Sillitoe on behalf of the Wellcome Sanger Institute COVID-19 Surveillance Team |
| EPI_ISL_2552824 | Lighthouse Lab in Alderley Park | Wellcome Sanger Institute for the COVID-19 Genomics UK (COG-UK) Consortium | Jacquelyn Wynn, Mairead Hyland, The Lighthouse Lab in Alderley Park and Alex Alderton, Roberto Amato, Jeffrey Barrett, Sonia Goncalves, Ewan Harrison, David K. Jackson, Ian Johnston, Dominic Kwiatkowski, Cordelia Langford, John Sillitoe on behalf of the Wellcome Sanger Institute COVID-19 Surveillance Team |
| EPI_ISL_2552825 | Lighthouse Lab in Milton Keynes | Wellcome Sanger Institute for the COVID-19 Genomics UK (COG-UK) Consortium | The Lighthouse Lab in Milton Keynes and Alex Alderton, Roberto Amato, Jeffrey Barrett, Sonia Goncalves, Ewan Harrison, David K. Jackson, Ian Johnston, Dominic Kwiatkowski, Cordelia Langford, John Sillitoe on behalf of the Wellcome Sanger Institute COVID-19 Surveillance Team |
| EPI_ISL_2552826, EPI_ISL_2552827, EPI_ISL_2552828, EPI_ISL_2552829 | Lighthouse Lab in Glasgow | Wellcome Sanger Institute for the COVID-19 Genomics UK (COG-UK) Consortium | Harper VanSteenhouse, Yumi Kasai, David Gray, Carol Clugston, Anna Dominiczak and Alex Alderton, Roberto Amato, Jeffrey Barrett, Sonia Goncalves, Ewan Harrison, David K. Jackson, Ian Johnston, Dominic Kwiatkowski, Cordelia Langford, John Sillitoe on behalf of the Wellcome Sanger Institute COVID-19 Surveillance Team |
| EPI_ISL_2552830 | Lighthouse Lab in Milton Keynes | Wellcome Sanger Institute for the COVID-19 Genomics UK (COG-UK) Consortium | The Lighthouse Lab in Milton Keynes and Alex Alderton, Roberto Amato, Jeffrey Barrett, Sonia Goncalves, Ewan Harrison, David K. Jackson, Ian Johnston, Dominic Kwiatkowski, Cordelia Langford, John Sillitoe on behalf of the Wellcome Sanger Institute COVID-19 Surveillance Team |
| EPI_ISL_2552832 | Lighthouse Lab in Glasgow | Wellcome Sanger Institute for the COVID-19 Genomics UK (COG-UK) Consortium | Harper VanSteenhouse, Yumi Kasai, David Gray, Carol Clugston, Anna Dominiczak and Alex Alderton, Roberto Amato, Jeffrey Barrett, Sonia Goncalves, Ewan Harrison, David K. Jackson, Ian Johnston, Dominic Kwiatkowski, Cordelia Langford, John Sillitoe on behalf of the Wellcome Sanger Institute COVID-19 Surveillance Team |
| EPI_ISL_2552833, EPI_ISL_2552834 | Lighthouse Lab in Alderley Park | Wellcome Sanger Institute for the COVID-19 Genomics UK (COG-UK) Consortium | Jacquelyn Wynn, Mairead Hyland, The Lighthouse Lab in Alderley Park and Alex Alderton, Roberto Amato, Jeffrey Barrett, Sonia Goncalves, Ewan Harrison, David K. Jackson, Ian Johnston, Dominic Kwiatkowski, Cordelia Langford, John Sillitoe on behalf of the Wellcome Sanger Institute COVID-19 Surveillance Team |

| | | | |
|---|---|---|---|
| EPI_ISL_2552835 | Lighthouse Lab in Milton Keynes | Wellcome Sanger Institute for the COVID-19 Genomics UK (COG-UK) Consortium | The Lighthouse Lab in Milton Keynes and Alex Alderton, Roberto Amato, Jeffrey Barrett, Sonia Goncalves, Ewan Harrison, David K. Jackson, Ian Johnston, Dominic Kwiatkowski, Cordelia Langford, John Sillitoe on behalf of the Wellcome Sanger Institute COVID-19 Surveillance Team |
| EPI_ISL_2552836, EPI_ISL_2552837 | Lighthouse Lab in Alderley Park | Wellcome Sanger Institute for the COVID-19 Genomics UK (COG-UK) Consortium | Jacquelyn Wynn, Mairead Hyland, The Lighthouse Lab in Alderley Park and Alex Alderton, Roberto Amato, Jeffrey Barrett, Sonia Goncalves, Ewan Harrison, David K. Jackson, Ian Johnston, Dominic Kwiatkowski, Cordelia Langford, John Sillitoe on behalf of the Wellcome Sanger Institute COVID-19 Surveillance Team |
| EPI_ISL_2552838, EPI_ISL_2552839 | Health Services Laboratories | Wellcome Sanger Institute for the COVID-19 Genomics UK (COG-UK) Consortium | Health Services Laboratories and Alex Alderton, Roberto Amato, Jeffrey Barrett, Sonia Goncalves, Ewan Harrison, David K. Jackson, Ian Johnston, Dominic Kwiatkowski, Cordelia Langford, John Sillitoe on behalf of the Wellcome Sanger Institute COVID-19 Surveillance Team |
| EPI_ISL_2552840, EPI_ISL_2552841 | Lighthouse Lab in Alderley Park | Wellcome Sanger Institute for the COVID-19 Genomics UK (COG-UK) Consortium | Jacquelyn Wynn, Mairead Hyland, The Lighthouse Lab in Alderley Park and Alex Alderton, Roberto Amato, Jeffrey Barrett, Sonia Goncalves, Ewan Harrison, David K. Jackson, Ian Johnston, Dominic Kwiatkowski, Cordelia Langford, John Sillitoe on behalf of the Wellcome Sanger Institute COVID-19 Surveillance Team |
| EPI_ISL_2552843 | Lighthouse Lab in Glasgow | Wellcome Sanger Institute for the COVID-19 Genomics UK (COG-UK) Consortium | Harper VanSteenhouse, Yumi Kasai, David Gray, Carol Clugston, Anna Dominiczak and Alex Alderton, Roberto Amato, Jeffrey Barrett, Sonia Goncalves, Ewan Harrison, David K. Jackson, Ian Johnston, Dominic Kwiatkowski, Cordelia Langford, John Sillitoe on behalf of the Wellcome Sanger Institute COVID-19 Surveillance Team |
| EPI_ISL_2552844 | Health Services Laboratories | Wellcome Sanger Institute for the COVID-19 Genomics UK (COG-UK) Consortium | Health Services Laboratories and Alex Alderton, Roberto Amato, Jeffrey Barrett, Sonia Goncalves, Ewan Harrison, David K. Jackson, Ian Johnston, Dominic Kwiatkowski, Cordelia Langford, John Sillitoe on behalf of the Wellcome Sanger Institute COVID-19 Surveillance Team |
| EPI_ISL_2552845 | Lighthouse Lab in Alderley Park | Wellcome Sanger Institute for the COVID-19 Genomics UK (COG-UK) Consortium | Jacquelyn Wynn, Mairead Hyland, The Lighthouse Lab in Alderley Park and Alex Alderton, Roberto Amato, Jeffrey Barrett, Sonia Goncalves, Ewan Harrison, David K. Jackson, Ian Johnston, Dominic Kwiatkowski, Cordelia Langford, John Sillitoe on behalf of the Wellcome Sanger Institute COVID-19 Surveillance Team |
| EPI_ISL_2552846 | Lighthouse Lab in Milton Keynes | Wellcome Sanger Institute for the COVID-19 Genomics UK (COG-UK) Consortium | The Lighthouse Lab in Milton Keynes and Alex Alderton, Roberto Amato, Jeffrey Barrett, Sonia Goncalves, Ewan Harrison, David K. Jackson, Ian Johnston, Dominic Kwiatkowski, Cordelia Langford, John Sillitoe on behalf of the Wellcome Sanger Institute COVID-19 Surveillance Team |
| EPI_ISL_2552847 | Lighthouse Lab in Glasgow | Wellcome Sanger Institute for the COVID-19 Genomics UK (COG-UK) Consortium | Harper VanSteenhouse, Yumi Kasai, David Gray, Carol Clugston, Anna Dominiczak and Alex Alderton, Roberto Amato, Jeffrey Barrett, Sonia Goncalves, Ewan Harrison, David K. Jackson, Ian Johnston, Dominic Kwiatkowski, Cordelia Langford, John Sillitoe on behalf of the Wellcome Sanger Institute COVID-19 Surveillance Team |
| EPI_ISL_2552848, EPI_ISL_2552849 | Health Services Laboratories | Wellcome Sanger Institute for the COVID-19 Genomics UK (COG-UK) Consortium | Health Services Laboratories and Alex Alderton, Roberto Amato, Jeffrey Barrett, Sonia Goncalves, Ewan Harrison, David K. Jackson, Ian Johnston, Dominic Kwiatkowski, Cordelia Langford, John Sillitoe on behalf of the Wellcome Sanger Institute COVID-19 Surveillance Team |
| EPI_ISL_2552850 | Lighthouse Lab in Glasgow | Wellcome Sanger Institute for the COVID-19 Genomics UK (COG-UK) Consortium | Harper VanSteenhouse, Yumi Kasai, David Gray, Carol Clugston, Anna Dominiczak and Alex Alderton, Roberto Amato, Jeffrey Barrett, Sonia Goncalves, Ewan Harrison, David K. Jackson, Ian Johnston, Dominic Kwiatkowski, Cordelia Langford, John Sillitoe on behalf of the Wellcome Sanger Institute COVID-19 Surveillance Team |
| EPI_ISL_2552851 | Lighthouse Lab in Alderley Park | Wellcome Sanger Institute for the COVID-19 Genomics UK (COG-UK) Consortium | Jacquelyn Wynn, Mairead Hyland, The Lighthouse Lab in Alderley Park and Alex Alderton, Roberto Amato, Jeffrey Barrett, Sonia Goncalves, Ewan Harrison, David K. Jackson, Ian Johnston, Dominic Kwiatkowski, Cordelia Langford, John Sillitoe on behalf of the Wellcome Sanger Institute COVID-19 Surveillance Team |
| EPI_ISL_2552852 | Health Services Laboratories | Wellcome Sanger Institute for the COVID-19 Genomics UK (COG-UK) Consortium | Health Services Laboratories and Alex Alderton, Roberto Amato, Jeffrey Barrett, Sonia Goncalves, Ewan Harrison, David K. Jackson, Ian Johnston, Dominic Kwiatkowski, Cordelia Langford, John Sillitoe on behalf of the Wellcome Sanger Institute COVID-19 Surveillance Team |
| EPI_ISL_2552853 | Lighthouse Lab in Glasgow | Wellcome Sanger Institute for the COVID-19 Genomics UK (COG-UK) Consortium | Harper VanSteenhouse, Yumi Kasai, David Gray, Carol Clugston, Anna Dominiczak and Alex Alderton, Roberto Amato, Jeffrey Barrett, Sonia Goncalves, Ewan Harrison, David K. Jackson, Ian Johnston, Dominic Kwiatkowski, Cordelia Langford, John Sillitoe on behalf of the Wellcome Sanger Institute COVID-19 Surveillance Team |
| EPI_ISL_2552854 | Lighthouse Lab in Alderley Park | Wellcome Sanger Institute for the COVID-19 Genomics UK (COG-UK) Consortium | Jacquelyn Wynn, Mairead Hyland, The Lighthouse Lab in Alderley Park and Alex Alderton, Roberto Amato, Jeffrey Barrett, Sonia Goncalves, Ewan Harrison, David K. Jackson, Ian Johnston, Dominic Kwiatkowski, Cordelia Langford, John Sillitoe on behalf of the Wellcome Sanger Institute COVID-19 Surveillance Team |
| EPI_ISL_2552855 | Lighthouse Lab in Glasgow | Wellcome Sanger Institute for the COVID-19 Genomics UK (COG-UK) Consortium | Harper VanSteenhouse, Yumi Kasai, David Gray, Carol Clugston, Anna Dominiczak and Alex Alderton, Roberto Amato, Jeffrey Barrett, Sonia Goncalves, Ewan Harrison, David K. Jackson, Ian Johnston, Dominic Kwiatkowski, Cordelia Langford, John Sillitoe on behalf of the Wellcome Sanger Institute COVID-19 Surveillance Team |
| EPI_ISL_2552856 | Lighthouse Lab in Milton Keynes | Wellcome Sanger Institute for the COVID-19 Genomics UK (COG-UK) Consortium | The Lighthouse Lab in Milton Keynes and Alex Alderton, Roberto Amato, Jeffrey Barrett, Sonia Goncalves, Ewan Harrison, David K. Jackson, Ian Johnston, Dominic Kwiatkowski, Cordelia Langford, John Sillitoe on behalf of the Wellcome Sanger Institute COVID-19 Surveillance Team |
| EPI_ISL_2552857 | Lighthouse Lab in Alderley Park | Wellcome Sanger Institute for the COVID-19 Genomics UK (COG-UK) Consortium | Jacquelyn Wynn, Mairead Hyland, The Lighthouse Lab in Alderley Park and Alex Alderton, Roberto Amato, Jeffrey Barrett, Sonia Goncalves, Ewan Harrison, David K. Jackson, Ian Johnston, Dominic Kwiatkowski, Cordelia Langford, John Sillitoe on behalf of the Wellcome Sanger Institute COVID-19 Surveillance Team |
| EPI_ISL_2552858 | Lighthouse Lab in Glasgow | Wellcome Sanger Institute for the COVID-19 Genomics UK (COG-UK) Consortium | Harper VanSteenhouse, Yumi Kasai, David Gray, Carol Clugston, Anna Dominiczak and Alex Alderton, Roberto Amato, Jeffrey Barrett, Sonia Goncalves, Ewan Harrison, David K. Jackson, Ian Johnston, Dominic Kwiatkowski, Cordelia Langford, John Sillitoe on behalf of the Wellcome Sanger Institute COVID-19 Surveillance Team |
| EPI_ISL_2552859, EPI_ISL_2552860 | Health Services Laboratories | Wellcome Sanger Institute for the COVID-19 Genomics UK (COG-UK) Consortium | Health Services Laboratories and Alex Alderton, Roberto Amato, Jeffrey Barrett, Sonia Goncalves, Ewan Harrison, David K. Jackson, Ian Johnston, Dominic Kwiatkowski, Cordelia Langford, John Sillitoe on behalf of the Wellcome Sanger Institute COVID-19 Surveillance Team |
| EPI_ISL_2552861 | Lighthouse Lab in Alderley Park | Wellcome Sanger Institute for the COVID-19 Genomics UK (COG-UK) Consortium | Jacquelyn Wynn, Mairead Hyland, The Lighthouse Lab in Alderley Park and Alex Alderton, Roberto Amato, Jeffrey Barrett, Sonia Goncalves, Ewan Harrison, David K. Jackson, Ian Johnston, Dominic Kwiatkowski, Cordelia Langford, John Sillitoe on behalf of the Wellcome Sanger Institute COVID-19 Surveillance Team |
| EPI_ISL_2552862, EPI_ISL_2552863 | Lighthouse Lab in Glasgow | Wellcome Sanger Institute for the COVID-19 Genomics UK (COG-UK) Consortium | Harper VanSteenhouse, Yumi Kasai, David Gray, Carol Clugston, Anna Dominiczak and Alex Alderton, Roberto Amato, Jeffrey Barrett, Sonia Goncalves, Ewan Harrison, David K. Jackson, Ian Johnston, Dominic Kwiatkowski, Cordelia Langford, John Sillitoe on behalf of the Wellcome Sanger Institute COVID-19 Surveillance Team |
| EPI_ISL_2552864 | Health Services Laboratories | Wellcome Sanger Institute for the COVID-19 Genomics UK (COG-UK) Consortium | Health Services Laboratories and Alex Alderton, Roberto Amato, Jeffrey Barrett, Sonia Goncalves, Ewan Harrison, David K. Jackson, Ian Johnston, Dominic Kwiatkowski, Cordelia Langford, John Sillitoe on behalf of the Wellcome Sanger Institute COVID-19 Surveillance Team |
| EPI_ISL_2552865, EPI_ISL_2552866 | Lighthouse Lab in Alderley Park | Wellcome Sanger Institute for the COVID-19 Genomics UK (COG-UK) Consortium | Jacquelyn Wynn, Mairead Hyland, The Lighthouse Lab in Alderley Park and Alex Alderton, Roberto Amato, Jeffrey Barrett, Sonia Goncalves, Ewan Harrison, David K. Jackson, Ian Johnston, Dominic Kwiatkowski, Cordelia Langford, John Sillitoe on behalf of the Wellcome Sanger Institute COVID-19 Surveillance Team |
| EPI_ISL_2552867 | Lighthouse Lab in Glasgow | Wellcome Sanger Institute for the COVID-19 Genomics UK (COG-UK) Consortium | Harper VanSteenhouse, Yumi Kasai, David Gray, Carol Clugston, Anna Dominiczak and Alex Alderton, Roberto Amato, Jeffrey Barrett, Sonia Goncalves, Ewan Harrison, David K. Jackson, Ian Johnston, Dominic Kwiatkowski, Cordelia Langford, John Sillitoe on behalf of the Wellcome Sanger Institute COVID-19 Surveillance Team |
| EPI_ISL_2552868 | Lighthouse Lab in Milton Keynes | Wellcome Sanger Institute for the COVID-19 Genomics UK (COG-UK) Consortium | The Lighthouse Lab in Milton Keynes and Alex Alderton, Roberto Amato, Jeffrey Barrett, Sonia Goncalves, Ewan Harrison, David K. Jackson, Ian Johnston, Dominic Kwiatkowski, Cordelia Langford, John Sillitoe on behalf of the Wellcome Sanger Institute COVID-19 Surveillance Team |
| EPI_ISL_2552869 | Lighthouse Lab in Glasgow | Wellcome Sanger Institute for the COVID-19 Genomics UK (COG-UK) Consortium | Harper VanSteenhouse, Yumi Kasai, David Gray, Carol Clugston, Anna Dominiczak and Alex Alderton, Roberto Amato, Jeffrey Barrett, Sonia Goncalves, Ewan Harrison, David K. Jackson, Ian Johnston, Dominic Kwiatkowski, Cordelia Langford, John Sillitoe on behalf of the Wellcome Sanger Institute COVID-19 Surveillance Team |
| EPI_ISL_2552870 | Lighthouse Lab in Alderley Park | Wellcome Sanger Institute for the COVID-19 Genomics UK (COG-UK) Consortium | Jacquelyn Wynn, Mairead Hyland, The Lighthouse Lab in Alderley Park and Alex Alderton, Roberto Amato, Jeffrey Barrett, Sonia Goncalves, Ewan Harrison, David K. Jackson, Ian Johnston, Dominic Kwiatkowski, Cordelia Langford, John Sillitoe on behalf of the Wellcome Sanger Institute COVID-19 Surveillance Team |
| EPI_ISL_2552871 | Health Services Laboratories | Wellcome Sanger Institute for the COVID-19 Genomics UK (COG-UK) Consortium | Health Services Laboratories and Alex Alderton, Roberto Amato, Jeffrey Barrett, Sonia Goncalves, Ewan Harrison, David K. Jackson, Ian Johnston, Dominic Kwiatkowski, Cordelia Langford, John Sillitoe on behalf of the Wellcome Sanger Institute COVID-19 Surveillance Team |
| EPI_ISL_2552872 | Lighthouse Lab in Milton Keynes | Wellcome Sanger Institute for the COVID-19 Genomics UK (COG-UK) Consortium | The Lighthouse Lab in Milton Keynes and Alex Alderton, Roberto Amato, Jeffrey Barrett, Sonia Goncalves, Ewan Harrison, David K. Jackson, Ian Johnston, Dominic Kwiatkowski, Cordelia Langford, John Sillitoe on behalf of the Wellcome Sanger Institute COVID-19 Surveillance Team |
| EPI_ISL_2552873, EPI_ISL_2552874 | Lighthouse Lab in Glasgow | Wellcome Sanger Institute for the COVID-19 Genomics UK (COG-UK) Consortium | Harper VanSteenhouse, Yumi Kasai, David Gray, Carol Clugston, Anna Dominiczak and Alex Alderton, Roberto Amato, Jeffrey Barrett, Sonia Goncalves, Ewan Harrison, David K. Jackson, Ian Johnston, Dominic Kwiatkowski, Cordelia Langford, John Sillitoe on behalf of the Wellcome Sanger Institute COVID-19 Surveillance Team |
| EPI_ISL_2552875 | Health Services Laboratories | Wellcome Sanger Institute for the COVID-19 Genomics UK (COG-UK) Consortium | Health Services Laboratories and Alex Alderton, Roberto Amato, Jeffrey Barrett, Sonia Goncalves, Ewan Harrison, David K. Jackson, Ian Johnston, Dominic Kwiatkowski, Cordelia Langford, John Sillitoe on behalf of the Wellcome Sanger Institute COVID-19 Surveillance Team |
| EPI_ISL_2552876 | Lighthouse Lab in Glasgow | Wellcome Sanger Institute for the COVID-19 Genomics UK (COG-UK) Consortium | Harper VanSteenhouse, Yumi Kasai, David Gray, Carol Clugston, Anna Dominiczak and Alex Alderton, Roberto Amato, Jeffrey Barrett, Sonia Goncalves, Ewan Harrison, David K. Jackson, Ian Johnston, Dominic Kwiatkowski, Cordelia Langford, John Sillitoe on behalf of the Wellcome Sanger Institute COVID-19 Surveillance Team |
| EPI_ISL_2552877 | Health Services Laboratories | Wellcome Sanger Institute for the COVID-19 Genomics UK (COG-UK) Consortium | Health Services Laboratories and Alex Alderton, Roberto Amato, Jeffrey Barrett, Sonia Goncalves, Ewan Harrison, David K. Jackson, Ian Johnston, Dominic Kwiatkowski, Cordelia Langford, John Sillitoe on behalf of the Wellcome Sanger Institute COVID-19 Surveillance Team |
| EPI_ISL_2552878, EPI_ISL_2552879 | Lighthouse Lab in Alderley Park | Wellcome Sanger Institute for the COVID-19 Genomics UK (COG-UK) Consortium | Jacquelyn Wynn, Mairead Hyland, The Lighthouse Lab in Alderley Park and Alex Alderton, Roberto Amato, Jeffrey Barrett, Sonia Goncalves, Ewan Harrison, David K. Jackson, Ian Johnston, Dominic Kwiatkowski, Cordelia Langford, John Sillitoe on behalf of the Wellcome Sanger Institute COVID-19 Surveillance Team |
| EPI_ISL_2552880 | Lighthouse Lab in Milton Keynes | Wellcome Sanger Institute for the COVID-19 Genomics UK (COG-UK) Consortium | The Lighthouse Lab in Milton Keynes and Alex Alderton, Roberto Amato, Jeffrey Barrett, Sonia Goncalves, Ewan Harrison, David K. Jackson, Ian Johnston, Dominic Kwiatkowski, Cordelia Langford, John Sillitoe on behalf of the Wellcome Sanger Institute COVID-19 Surveillance Team |
| EPI_ISL_2552881 | Lighthouse Lab in Alderley Park | Wellcome Sanger Institute for the COVID-19 Genomics UK (COG-UK) Consortium | Jacquelyn Wynn, Mairead Hyland, The Lighthouse Lab in Alderley Park and Alex Alderton, Roberto Amato, Jeffrey Barrett, Sonia Goncalves, Ewan Harrison, David K. Jackson, Ian Johnston, Dominic Kwiatkowski, Cordelia Langford, John Sillitoe on behalf of the Wellcome Sanger Institute COVID-19 Surveillance Team |
| EPI_ISL_2552882 | Health Services Laboratories | Wellcome Sanger Institute for the COVID-19 Genomics UK (COG-UK) Consortium | Health Services Laboratories and Alex Alderton, Roberto Amato, Jeffrey Barrett, Sonia Goncalves, Ewan Harrison, David K. Jackson, Ian Johnston, Dominic Kwiatkowski, Cordelia Langford, John Sillitoe on behalf of the Wellcome Sanger Institute COVID-19 Surveillance Team |
| EPI_ISL_2552884 | Lighthouse Lab in Milton Keynes | Wellcome Sanger Institute for the COVID-19 Genomics UK (COG-UK) Consortium | The Lighthouse Lab in Milton Keynes and Alex Alderton, Roberto Amato, Jeffrey Barrett, Sonia Goncalves, Ewan Harrison, David K. Jackson, Ian Johnston, Dominic Kwiatkowski, Cordelia Langford, John Sillitoe on behalf of the Wellcome Sanger Institute COVID-19 Surveillance Team |
| EPI_ISL_2552885 | Lighthouse Lab in Alderley Park | Wellcome Sanger Institute for the COVID-19 Genomics UK (COG-UK) Consortium | Jacquelyn Wynn, Mairead Hyland, The Lighthouse Lab in Alderley Park and Alex Alderton, Roberto Amato, Jeffrey Barrett, Sonia Goncalves, Ewan Harrison, David K. Jackson, Ian Johnston, Dominic Kwiatkowski, Cordelia Langford, John Sillitoe on behalf of the Wellcome Sanger Institute COVID-19 Surveillance Team |
| EPI_ISL_2552886 | Lighthouse Lab in Milton Keynes | Wellcome Sanger Institute for the COVID-19 Genomics UK (COG-UK) Consortium | The Lighthouse Lab in Milton Keynes and Alex Alderton, Roberto Amato, Jeffrey Barrett, Sonia Goncalves, Ewan Harrison, David K. Jackson, Ian Johnston, Dominic Kwiatkowski, Cordelia Langford, John Sillitoe on behalf of the Wellcome Sanger Institute COVID-19 Surveillance Team |
| EPI_ISL_2552887, EPI_ISL_2552888, EPI_ISL_2552889, EPI_ISL_2552890 | Health Services Laboratories | Wellcome Sanger Institute for the COVID-19 Genomics UK (COG-UK) Consortium | Health Services Laboratories and Alex Alderton, Roberto Amato, Jeffrey Barrett, Sonia Goncalves, Ewan Harrison, David K. Jackson, Ian Johnston, Dominic Kwiatkowski, Cordelia Langford, John Sillitoe on behalf of the Wellcome Sanger Institute COVID-19 Surveillance Team |
| EPI_ISL_2552892, EPI_ISL_2552893 | Lighthouse Lab in Alderley Park | Wellcome Sanger Institute for the COVID-19 Genomics UK (COG-UK) Consortium | Jacquelyn Wynn, Mairead Hyland, The Lighthouse Lab in Alderley Park and Alex Alderton, Roberto Amato, Jeffrey Barrett, Sonia Goncalves, Ewan Harrison, David K. Jackson, Ian Johnston, Dominic Kwiatkowski, Cordelia Langford, John Sillitoe on behalf of the Wellcome Sanger Institute COVID-19 Surveillance Team |
| EPI_ISL_2552894, EPI_ISL_2552895, EPI_ISL_2552896 | Lighthouse Lab in Glasgow | Wellcome Sanger Institute for the COVID-19 Genomics UK (COG-UK) Consortium | Harper VanSteenhouse, Yumi Kasai, David Gray, Carol Clugston, Anna Dominiczak and Alex Alderton, Roberto Amato, Jeffrey Barrett, Sonia Goncalves, Ewan Harrison, David K. Jackson, Ian Johnston, Dominic Kwiatkowski, Cordelia Langford, John Sillitoe on behalf of the Wellcome Sanger Institute COVID-19 Surveillance Team |

| | | | |
|---|---|---|---|
| EPI_ISL_2552897 | Lighthouse Lab in Milton Keynes | Wellcome Sanger Institute for the COVID-19 Genomics UK (COG-UK) Consortium | The Lighthouse Lab in Milton Keynes and Alex Alderton, Roberto Amato, Jeffrey Barrett, Sonia Goncalves, Ewan Harrison, David K. Jackson, Ian Johnston, Dominic Kwiatkowski, Cordelia Langford, John Sillitoe on behalf of the Wellcome Sanger Institute COVID-19 Surveillance Team |
| EPI_ISL_2552898 | Lighthouse Lab in Alderley Park | Wellcome Sanger Institute for the COVID-19 Genomics UK (COG-UK) Consortium | Jacquelyn Wynn, Mairead Hyland, The Lighthouse Lab in Alderley Park and Alex Alderton, Roberto Amato, Jeffrey Barrett, Sonia Goncalves, Ewan Harrison, David K. Jackson, Ian Johnston, Dominic Kwiatkowski, Cordelia Langford, John Sillitoe on behalf of the Wellcome Sanger Institute COVID-19 Surveillance Team |
| EPI_ISL_2552899 | Lighthouse Lab in Milton Keynes | Wellcome Sanger Institute for the COVID-19 Genomics UK (COG-UK) Consortium | The Lighthouse Lab in Milton Keynes and Alex Alderton, Roberto Amato, Jeffrey Barrett, Sonia Goncalves, Ewan Harrison, David K. Jackson, Ian Johnston, Dominic Kwiatkowski, Cordelia Langford, John Sillitoe on behalf of the Wellcome Sanger Institute COVID-19 Surveillance Team |
| EPI_ISL_2552900 | Lighthouse Lab in Glasgow | Wellcome Sanger Institute for the COVID-19 Genomics UK (COG-UK) Consortium | Harper VanSteenhouse, Yumi Kasai, David Gray, Carol Clugston, Anna Dominiczak and Alex Alderton, Roberto Amato, Jeffrey Barrett, Sonia Goncalves, Ewan Harrison, David K. Jackson, Ian Johnston, Dominic Kwiatkowski, Cordelia Langford, John Sillitoe on behalf of the Wellcome Sanger Institute COVID-19 Surveillance Team |
| EPI_ISL_2552901 | Health Services Laboratories | Wellcome Sanger Institute for the COVID-19 Genomics UK (COG-UK) Consortium | Health Services Laboratories and Alex Alderton, Roberto Amato, Jeffrey Barrett, Sonia Goncalves, Ewan Harrison, David K. Jackson, Ian Johnston, Dominic Kwiatkowski, Cordelia Langford, John Sillitoe on behalf of the Wellcome Sanger Institute COVID-19 Surveillance Team |
| EPI_ISL_2552902 | Lighthouse Lab in Glasgow | Wellcome Sanger Institute for the COVID-19 Genomics UK (COG-UK) Consortium | Harper VanSteenhouse, Yumi Kasai, David Gray, Carol Clugston, Anna Dominiczak and Alex Alderton, Roberto Amato, Jeffrey Barrett, Sonia Goncalves, Ewan Harrison, David K. Jackson, Ian Johnston, Dominic Kwiatkowski, Cordelia Langford, John Sillitoe on behalf of the Wellcome Sanger Institute COVID-19 Surveillance Team |
| EPI_ISL_2552903, EPI_ISL_2552904 | Lighthouse Lab in Milton Keynes | Wellcome Sanger Institute for the COVID-19 Genomics UK (COG-UK) Consortium | The Lighthouse Lab in Milton Keynes and Alex Alderton, Roberto Amato, Jeffrey Barrett, Sonia Goncalves, Ewan Harrison, David K. Jackson, Ian Johnston, Dominic Kwiatkowski, Cordelia Langford, John Sillitoe on behalf of the Wellcome Sanger Institute COVID-19 Surveillance Team |
| EPI_ISL_2552905 | Health Services Laboratories | Wellcome Sanger Institute for the COVID-19 Genomics UK (COG-UK) Consortium | Health Services Laboratories and Alex Alderton, Roberto Amato, Jeffrey Barrett, Sonia Goncalves, Ewan Harrison, David K. Jackson, Ian Johnston, Dominic Kwiatkowski, Cordelia Langford, John Sillitoe on behalf of the Wellcome Sanger Institute COVID-19 Surveillance Team |
| EPI_ISL_2552907, EPI_ISL_2552908, EPI_ISL_2552909 | Lighthouse Lab in Milton Keynes | Wellcome Sanger Institute for the COVID-19 Genomics UK (COG-UK) Consortium | The Lighthouse Lab in Milton Keynes and Alex Alderton, Roberto Amato, Jeffrey Barrett, Sonia Goncalves, Ewan Harrison, David K. Jackson, Ian Johnston, Dominic Kwiatkowski, Cordelia Langford, John Sillitoe on behalf of the Wellcome Sanger Institute COVID-19 Surveillance Team |
| EPI_ISL_2552910, EPI_ISL_2552911 | Lighthouse Lab in Glasgow | Wellcome Sanger Institute for the COVID-19 Genomics UK (COG-UK) Consortium | Harper VanSteenhouse, Yumi Kasai, David Gray, Carol Clugston, Anna Dominiczak and Alex Alderton, Roberto Amato, Jeffrey Barrett, Sonia Goncalves, Ewan Harrison, David K. Jackson, Ian Johnston, Dominic Kwiatkowski, Cordelia Langford, John Sillitoe on behalf of the Wellcome Sanger Institute COVID-19 Surveillance Team |
| EPI_ISL_2552912 | Health Services Laboratories | Wellcome Sanger Institute for the COVID-19 Genomics UK (COG-UK) Consortium | Health Services Laboratories and Alex Alderton, Roberto Amato, Jeffrey Barrett, Sonia Goncalves, Ewan Harrison, David K. Jackson, Ian Johnston, Dominic Kwiatkowski, Cordelia Langford, John Sillitoe on behalf of the Wellcome Sanger Institute COVID-19 Surveillance Team |
| EPI_ISL_2552913 | Lighthouse Lab in Glasgow | Wellcome Sanger Institute for the COVID-19 Genomics UK (COG-UK) Consortium | Harper VanSteenhouse, Yumi Kasai, David Gray, Carol Clugston, Anna Dominiczak and Alex Alderton, Roberto Amato, Jeffrey Barrett, Sonia Goncalves, Ewan Harrison, David K. Jackson, Ian Johnston, Dominic Kwiatkowski, Cordelia Langford, John Sillitoe on behalf of the Wellcome Sanger Institute COVID-19 Surveillance Team |
| EPI_ISL_2552914 | Health Services Laboratories | Wellcome Sanger Institute for the COVID-19 Genomics UK (COG-UK) Consortium | Health Services Laboratories and Alex Alderton, Roberto Amato, Jeffrey Barrett, Sonia Goncalves, Ewan Harrison, David K. Jackson, Ian Johnston, Dominic Kwiatkowski, Cordelia Langford, John Sillitoe on behalf of the Wellcome Sanger Institute COVID-19 Surveillance Team |
| EPI_ISL_2552915 | Lighthouse Lab in Glasgow | Wellcome Sanger Institute for the COVID-19 Genomics UK (COG-UK) Consortium | Harper VanSteenhouse, Yumi Kasai, David Gray, Carol Clugston, Anna Dominiczak and Alex Alderton, Roberto Amato, Jeffrey Barrett, Sonia Goncalves, Ewan Harrison, David K. Jackson, Ian Johnston, Dominic Kwiatkowski, Cordelia Langford, John Sillitoe on behalf of the Wellcome Sanger Institute COVID-19 Surveillance Team |
| EPI_ISL_2552916, EPI_ISL_2552917, EPI_ISL_2552918 | Lighthouse Lab in Milton Keynes | Wellcome Sanger Institute for the COVID-19 Genomics UK (COG-UK) Consortium | The Lighthouse Lab in Milton Keynes and Alex Alderton, Roberto Amato, Jeffrey Barrett, Sonia Goncalves, Ewan Harrison, David K. Jackson, Ian Johnston, Dominic Kwiatkowski, Cordelia Langford, John Sillitoe on behalf of the Wellcome Sanger Institute COVID-19 Surveillance Team |
| EPI_ISL_2552919 | Health Services Laboratories | Wellcome Sanger Institute for the COVID-19 Genomics UK (COG-UK) Consortium | Health Services Laboratories and Alex Alderton, Roberto Amato, Jeffrey Barrett, Sonia Goncalves, Ewan Harrison, David K. Jackson, Ian Johnston, Dominic Kwiatkowski, Cordelia Langford, John Sillitoe on behalf of the Wellcome Sanger Institute COVID-19 Surveillance Team |
| EPI_ISL_2552920, EPI_ISL_2552921 | Lighthouse Lab in Glasgow | Wellcome Sanger Institute for the COVID-19 Genomics UK (COG-UK) Consortium | Harper VanSteenhouse, Yumi Kasai, David Gray, Carol Clugston, Anna Dominiczak and Alex Alderton, Roberto Amato, Jeffrey Barrett, Sonia Goncalves, Ewan Harrison, David K. Jackson, Ian Johnston, Dominic Kwiatkowski, Cordelia Langford, John Sillitoe on behalf of the Wellcome Sanger Institute COVID-19 Surveillance Team |
| EPI_ISL_2552922 | Lighthouse Lab in Milton Keynes | Wellcome Sanger Institute for the COVID-19 Genomics UK (COG-UK) Consortium | The Lighthouse Lab in Milton Keynes and Alex Alderton, Roberto Amato, Jeffrey Barrett, Sonia Goncalves, Ewan Harrison, David K. Jackson, Ian Johnston, Dominic Kwiatkowski, Cordelia Langford, John Sillitoe on behalf of the Wellcome Sanger Institute COVID-19 Surveillance Team |
| EPI_ISL_2552923 | Health Services Laboratories | Wellcome Sanger Institute for the COVID-19 Genomics UK (COG-UK) Consortium | Health Services Laboratories and Alex Alderton, Roberto Amato, Jeffrey Barrett, Sonia Goncalves, Ewan Harrison, David K. Jackson, Ian Johnston, Dominic Kwiatkowski, Cordelia Langford, John Sillitoe on behalf of the Wellcome Sanger Institute COVID-19 Surveillance Team |
| EPI_ISL_2552924, EPI_ISL_2552925, EPI_ISL_2552926, EPI_ISL_2552927, EPI_ISL_2552928, EPI_ISL_2552929, EPI_ISL_2552930, EPI_ISL_2552931, EPI_ISL_2552932, EPI_ISL_2552933 | Lighthouse Lab in Alderley Park | Wellcome Sanger Institute for the COVID-19 Genomics UK (COG-UK) Consortium | Jacquelyn Wynn, Mairead Hyland, The Lighthouse Lab in Alderley Park and Alex Alderton, Roberto Amato, Jeffrey Barrett, Sonia Goncalves, Ewan Harrison, David K. Jackson, Ian Johnston, Dominic Kwiatkowski, Cordelia Langford, John Sillitoe on behalf of the Wellcome Sanger Institute COVID-19 Surveillance Team |
| EPI_ISL_2552934, EPI_ISL_2552935 | Lighthouse Lab in Glasgow | Wellcome Sanger Institute for the COVID-19 Genomics UK (COG-UK) Consortium | Harper VanSteenhouse, Yumi Kasai, David Gray, Carol Clugston, Anna Dominiczak and Alex Alderton, Roberto Amato, Jeffrey Barrett, Sonia Goncalves, Ewan Harrison, David K. Jackson, Ian Johnston, Dominic Kwiatkowski, Cordelia Langford, John Sillitoe on behalf of the Wellcome Sanger Institute COVID-19 Surveillance Team |
| EPI_ISL_2552936, EPI_ISL_2552937, EPI_ISL_2552938 | Lighthouse Lab in Alderley Park | Wellcome Sanger Institute for the COVID-19 Genomics UK (COG-UK) Consortium | Jacquelyn Wynn, Mairead Hyland, The Lighthouse Lab in Alderley Park and Alex Alderton, Roberto Amato, Jeffrey Barrett, Sonia Goncalves, Ewan Harrison, David K. Jackson, Ian Johnston, Dominic Kwiatkowski, Cordelia Langford, John Sillitoe on behalf of the Wellcome Sanger Institute COVID-19 Surveillance Team |
| EPI_ISL_2552939, EPI_ISL_2552940 | Lighthouse Lab in Glasgow | Wellcome Sanger Institute for the COVID-19 Genomics UK (COG-UK) Consortium | Harper VanSteenhouse, Yumi Kasai, David Gray, Carol Clugston, Anna Dominiczak and Alex Alderton, Roberto Amato, Jeffrey Barrett, Sonia Goncalves, Ewan Harrison, David K. Jackson, Ian Johnston, Dominic Kwiatkowski, Cordelia Langford, John Sillitoe on behalf of the Wellcome Sanger Institute COVID-19 Surveillance Team |
| EPI_ISL_2552941, EPI_ISL_2552942, EPI_ISL_2552943 | Lighthouse Lab in Alderley Park | Wellcome Sanger Institute for the COVID-19 Genomics UK (COG-UK) Consortium | Jacquelyn Wynn, Mairead Hyland, The Lighthouse Lab in Alderley Park and Alex Alderton, Roberto Amato, Jeffrey Barrett, Sonia Goncalves, Ewan Harrison, David K. Jackson, Ian Johnston, Dominic Kwiatkowski, Cordelia Langford, John Sillitoe on behalf of the Wellcome Sanger Institute COVID-19 Surveillance Team |
| EPI_ISL_2552944 | Lighthouse Lab in Glasgow | Wellcome Sanger Institute for the COVID-19 Genomics UK (COG-UK) Consortium | Harper VanSteenhouse, Yumi Kasai, David Gray, Carol Clugston, Anna Dominiczak and Alex Alderton, Roberto Amato, Jeffrey Barrett, Sonia Goncalves, Ewan Harrison, David K. Jackson, Ian Johnston, Dominic Kwiatkowski, Cordelia Langford, John Sillitoe on behalf of the Wellcome Sanger Institute COVID-19 Surveillance Team |
| EPI_ISL_2552945 | Lighthouse Lab in Alderley Park | Wellcome Sanger Institute for the COVID-19 Genomics UK (COG-UK) Consortium | Jacquelyn Wynn, Mairead Hyland, The Lighthouse Lab in Alderley Park and Alex Alderton, Roberto Amato, Jeffrey Barrett, Sonia Goncalves, Ewan Harrison, David K. Jackson, Ian Johnston, Dominic Kwiatkowski, Cordelia Langford, John Sillitoe on behalf of the Wellcome Sanger Institute COVID-19 Surveillance Team |
| EPI_ISL_2552946, EPI_ISL_2552947 | Lighthouse Lab in Glasgow | Wellcome Sanger Institute for the COVID-19 Genomics UK (COG-UK) Consortium | Harper VanSteenhouse, Yumi Kasai, David Gray, Carol Clugston, Anna Dominiczak and Alex Alderton, Roberto Amato, Jeffrey Barrett, Sonia Goncalves, Ewan Harrison, David K. Jackson, Ian Johnston, Dominic Kwiatkowski, Cordelia Langford, John Sillitoe on behalf of the Wellcome Sanger Institute COVID-19 Surveillance Team |
| EPI_ISL_2552948, EPI_ISL_2552949, EPI_ISL_2552950, EPI_ISL_2552951, EPI_ISL_2552952 | Lighthouse Lab in Alderley Park | Wellcome Sanger Institute for the COVID-19 Genomics UK (COG-UK) Consortium | Jacquelyn Wynn, Mairead Hyland, The Lighthouse Lab in Alderley Park and Alex Alderton, Roberto Amato, Jeffrey Barrett, Sonia Goncalves, Ewan Harrison, David K. Jackson, Ian Johnston, Dominic Kwiatkowski, Cordelia Langford, John Sillitoe on behalf of the Wellcome Sanger Institute COVID-19 Surveillance Team |
| EPI_ISL_2552953 | Lighthouse Lab in Glasgow | Wellcome Sanger Institute for the COVID-19 Genomics UK (COG-UK) Consortium | Harper VanSteenhouse, Yumi Kasai, David Gray, Carol Clugston, Anna Dominiczak and Alex Alderton, Roberto Amato, Jeffrey Barrett, Sonia Goncalves, Ewan Harrison, David K. Jackson, Ian Johnston, Dominic Kwiatkowski, Cordelia Langford, John Sillitoe on behalf of the Wellcome Sanger Institute COVID-19 Surveillance Team |
| EPI_ISL_2552954, EPI_ISL_2552955 | Lighthouse Lab in Alderley Park | Wellcome Sanger Institute for the COVID-19 Genomics UK (COG-UK) Consortium | Jacquelyn Wynn, Mairead Hyland, The Lighthouse Lab in Alderley Park and Alex Alderton, Roberto Amato, Jeffrey Barrett, Sonia Goncalves, Ewan Harrison, David K. Jackson, Ian Johnston, Dominic Kwiatkowski, Cordelia Langford, John Sillitoe on behalf of the Wellcome Sanger Institute COVID-19 Surveillance Team |
| EPI_ISL_2552956, EPI_ISL_2552957 | Lighthouse Lab in Glasgow | Wellcome Sanger Institute for the COVID-19 Genomics UK (COG-UK) Consortium | Harper VanSteenhouse, Yumi Kasai, David Gray, Carol Clugston, Anna Dominiczak and Alex Alderton, Roberto Amato, Jeffrey Barrett, Sonia Goncalves, Ewan Harrison, David K. Jackson, Ian Johnston, Dominic Kwiatkowski, Cordelia Langford, John Sillitoe on behalf of the Wellcome Sanger Institute COVID-19 Surveillance Team |
| EPI_ISL_2552958, EPI_ISL_2552959, EPI_ISL_2552960, EPI_ISL_2552961, EPI_ISL_2552962, EPI_ISL_2552963, EPI_ISL_2552964 | Lighthouse Lab in Alderley Park | Wellcome Sanger Institute for the COVID-19 Genomics UK (COG-UK) Consortium | Jacquelyn Wynn, Mairead Hyland, The Lighthouse Lab in Alderley Park and Alex Alderton, Roberto Amato, Jeffrey Barrett, Sonia Goncalves, Ewan Harrison, David K. Jackson, Ian Johnston, Dominic Kwiatkowski, Cordelia Langford, John Sillitoe on behalf of the Wellcome Sanger Institute COVID-19 Surveillance Team |
| EPI_ISL_2552965 | Lighthouse Lab in Glasgow | Wellcome Sanger Institute for the COVID-19 Genomics UK (COG-UK) Consortium | Harper VanSteenhouse, Yumi Kasai, David Gray, Carol Clugston, Anna Dominiczak and Alex Alderton, Roberto Amato, Jeffrey Barrett, Sonia Goncalves, Ewan Harrison, David K. Jackson, Ian Johnston, Dominic Kwiatkowski, Cordelia Langford, John Sillitoe on behalf of the Wellcome Sanger Institute COVID-19 Surveillance Team |
| EPI_ISL_2552966, EPI_ISL_2552967, EPI_ISL_2552968, EPI_ISL_2552969, EPI_ISL_2552970 | Lighthouse Lab in Alderley Park | Wellcome Sanger Institute for the COVID-19 Genomics UK (COG-UK) Consortium | Jacquelyn Wynn, Mairead Hyland, The Lighthouse Lab in Alderley Park and Alex Alderton, Roberto Amato, Jeffrey Barrett, Sonia Goncalves, Ewan Harrison, David K. Jackson, Ian Johnston, Dominic Kwiatkowski, Cordelia Langford, John Sillitoe on behalf of the Wellcome Sanger Institute COVID-19 Surveillance Team |
| EPI_ISL_2552971 | Lighthouse Lab in Glasgow | Wellcome Sanger Institute for the COVID-19 Genomics UK (COG-UK) Consortium | Harper VanSteenhouse, Yumi Kasai, David Gray, Carol Clugston, Anna Dominiczak and Alex Alderton, Roberto Amato, Jeffrey Barrett, Sonia Goncalves, Ewan Harrison, David K. Jackson, Ian Johnston, Dominic Kwiatkowski, Cordelia Langford, John Sillitoe on behalf of the Wellcome Sanger Institute COVID-19 Surveillance Team |
| EPI_ISL_2552972 | Lighthouse Lab in Alderley Park | Wellcome Sanger Institute for the COVID-19 Genomics UK (COG-UK) | Jacquelyn Wynn, Mairead Hyland, The Lighthouse Lab in Alderley Park and Alex Alderton, Roberto Amato, Jeffrey Barrett, Sonia Goncalves, Ewan Harrison, David K. Jackson, Ian Johnston, Dominic Kwiatkowski, Cordelia Langford, John Sillitoe on |

| | | | |
|---|---|---|---|
| | | Consortium | behalf of the Wellcome Sanger Institute COVID-19 Surveillance Team |
| EPI_ISL_2552973, EPI_ISL_2552974 | Lighthouse Lab in Glasgow | Wellcome Sanger Institute for the COVID-19 Genomics UK (COG-UK) Consortium | Harper VanSteenhouse, Yumi Kasai, David Gray, Carol Clugston, Anna Dominiczak and Alex Alderton, Roberto Amato, Jeffrey Barrett, Sonia Goncalves, Ewan Harrison, David K. Jackson, Ian Johnston, Dominic Kwiatkowski, Cordelia Langford, John Sillitoe on behalf of the Wellcome Sanger Institute COVID-19 Surveillance Team |
| EPI_ISL_2552975, EPI_ISL_2552976, EPI_ISL_2552978 | Lighthouse Lab in Alderley Park | Wellcome Sanger Institute for the COVID-19 Genomics UK (COG-UK) Consortium | Jacquelyn Wynn, Mairead Hyland, The Lighthouse Lab in Alderley Park and Alex Alderton, Roberto Amato, Jeffrey Barrett, Sonia Goncalves, Ewan Harrison, David K. Jackson, Ian Johnston, Dominic Kwiatkowski, Cordelia Langford, John Sillitoe on behalf of the Wellcome Sanger Institute COVID-19 Surveillance Team |
| EPI_ISL_2552979 | Lighthouse Lab in Glasgow | Wellcome Sanger Institute for the COVID-19 Genomics UK (COG-UK) Consortium | Harper VanSteenhouse, Yumi Kasai, David Gray, Carol Clugston, Anna Dominiczak and Alex Alderton, Roberto Amato, Jeffrey Barrett, Sonia Goncalves, Ewan Harrison, David K. Jackson, Ian Johnston, Dominic Kwiatkowski, Cordelia Langford, John Sillitoe on behalf of the Wellcome Sanger Institute COVID-19 Surveillance Team |
| EPI_ISL_2552980, EPI_ISL_2552981 | Lighthouse Lab in Alderley Park | Wellcome Sanger Institute for the COVID-19 Genomics UK (COG-UK) Consortium | Jacquelyn Wynn, Mairead Hyland, The Lighthouse Lab in Alderley Park and Alex Alderton, Roberto Amato, Jeffrey Barrett, Sonia Goncalves, Ewan Harrison, David K. Jackson, Ian Johnston, Dominic Kwiatkowski, Cordelia Langford, John Sillitoe on behalf of the Wellcome Sanger Institute COVID-19 Surveillance Team |
| EPI_ISL_2552982 | Lighthouse Lab in Glasgow | Wellcome Sanger Institute for the COVID-19 Genomics UK (COG-UK) Consortium | Harper VanSteenhouse, Yumi Kasai, David Gray, Carol Clugston, Anna Dominiczak and Alex Alderton, Roberto Amato, Jeffrey Barrett, Sonia Goncalves, Ewan Harrison, David K. Jackson, Ian Johnston, Dominic Kwiatkowski, Cordelia Langford, John Sillitoe on behalf of the Wellcome Sanger Institute COVID-19 Surveillance Team |
| EPI_ISL_2552983, EPI_ISL_2552984, EPI_ISL_2552985, EPI_ISL_2552986, EPI_ISL_2552987, EPI_ISL_2552988, EPI_ISL_2552989, EPI_ISL_2552990, EPI_ISL_2552991, EPI_ISL_2552992 | Lighthouse Lab in Alderley Park | Wellcome Sanger Institute for the COVID-19 Genomics UK (COG-UK) Consortium | Jacquelyn Wynn, Mairead Hyland, The Lighthouse Lab in Alderley Park and Alex Alderton, Roberto Amato, Jeffrey Barrett, Sonia Goncalves, Ewan Harrison, David K. Jackson, Ian Johnston, Dominic Kwiatkowski, Cordelia Langford, John Sillitoe on behalf of the Wellcome Sanger Institute COVID-19 Surveillance Team |
| EPI_ISL_2552993 | Lighthouse Lab in Glasgow | Wellcome Sanger Institute for the COVID-19 Genomics UK (COG-UK) Consortium | Harper VanSteenhouse, Yumi Kasai, David Gray, Carol Clugston, Anna Dominiczak and Alex Alderton, Roberto Amato, Jeffrey Barrett, Sonia Goncalves, Ewan Harrison, David K. Jackson, Ian Johnston, Dominic Kwiatkowski, Cordelia Langford, John Sillitoe on behalf of the Wellcome Sanger Institute COVID-19 Surveillance Team |
| EPI_ISL_2552994, EPI_ISL_2552995, EPI_ISL_2552996, EPI_ISL_2552997, EPI_ISL_2552998, EPI_ISL_2552999, EPI_ISL_2553000, EPI_ISL_2553001, EPI_ISL_2553002, EPI_ISL_2553003, EPI_ISL_2553004, EPI_ISL_2553005, EPI_ISL_2553006, EPI_ISL_2553007 | | | |
| see above | Lighthouse Lab in Alderley Park | Wellcome Sanger Institute for the COVID-19 Genomics UK (COG-UK) Consortium | Jacquelyn Wynn, Mairead Hyland, The Lighthouse Lab in Alderley Park and Alex Alderton, Roberto Amato, Jeffrey Barrett, Sonia Goncalves, Ewan Harrison, David K. Jackson, Ian Johnston, Dominic Kwiatkowski, Cordelia Langford, John Sillitoe on behalf of the Wellcome Sanger Institute COVID-19 Surveillance Team |
| EPI_ISL_2553008, EPI_ISL_2553009, EPI_ISL_2553010 | Lighthouse Lab in Glasgow | Wellcome Sanger Institute for the COVID-19 Genomics UK (COG-UK) Consortium | Harper VanSteenhouse, Yumi Kasai, David Gray, Carol Clugston, Anna Dominiczak and Alex Alderton, Roberto Amato, Jeffrey Barrett, Sonia Goncalves, Ewan Harrison, David K. Jackson, Ian Johnston, Dominic Kwiatkowski, Cordelia Langford, John Sillitoe on behalf of the Wellcome Sanger Institute COVID-19 Surveillance Team |
| EPI_ISL_2553011 | Lighthouse Lab in Alderley Park | Wellcome Sanger Institute for the COVID-19 Genomics UK (COG-UK) Consortium | Jacquelyn Wynn, Mairead Hyland, The Lighthouse Lab in Alderley Park and Alex Alderton, Roberto Amato, Jeffrey Barrett, Sonia Goncalves, Ewan Harrison, David K. Jackson, Ian Johnston, Dominic Kwiatkowski, Cordelia Langford, John Sillitoe on behalf of the Wellcome Sanger Institute COVID-19 Surveillance Team |
| EPI_ISL_2553012, EPI_ISL_2553013 | Lighthouse Lab in Glasgow | Wellcome Sanger Institute for the COVID-19 Genomics UK (COG-UK) Consortium | Harper VanSteenhouse, Yumi Kasai, David Gray, Carol Clugston, Anna Dominiczak and Alex Alderton, Roberto Amato, Jeffrey Barrett, Sonia Goncalves, Ewan Harrison, David K. Jackson, Ian Johnston, Dominic Kwiatkowski, Cordelia Langford, John Sillitoe on behalf of the Wellcome Sanger Institute COVID-19 Surveillance Team |
| EPI_ISL_2553014 | Lighthouse Lab in Alderley Park | Wellcome Sanger Institute for the COVID-19 Genomics UK (COG-UK) Consortium | Jacquelyn Wynn, Mairead Hyland, The Lighthouse Lab in Alderley Park and Alex Alderton, Roberto Amato, Jeffrey Barrett, Sonia Goncalves, Ewan Harrison, David K. Jackson, Ian Johnston, Dominic Kwiatkowski, Cordelia Langford, John Sillitoe on behalf of the Wellcome Sanger Institute COVID-19 Surveillance Team |
| EPI_ISL_2553015 | Lighthouse Lab in Glasgow | Wellcome Sanger Institute for the COVID-19 Genomics UK (COG-UK) Consortium | Harper VanSteenhouse, Yumi Kasai, David Gray, Carol Clugston, Anna Dominiczak and Alex Alderton, Roberto Amato, Jeffrey Barrett, Sonia Goncalves, Ewan Harrison, David K. Jackson, Ian Johnston, Dominic Kwiatkowski, Cordelia Langford, John Sillitoe on behalf of the Wellcome Sanger Institute COVID-19 Surveillance Team |
| EPI_ISL_2553016, EPI_ISL_2553017, EPI_ISL_2553018, EPI_ISL_2553019, EPI_ISL_2553020, EPI_ISL_2553022, EPI_ISL_2553023, EPI_ISL_2553024, EPI_ISL_2553025, EPI_ISL_2553026, EPI_ISL_2553027, EPI_ISL_2553028, EPI_ISL_2553029, EPI_ISL_2553030 | | | |
| see above | Lighthouse Lab in Alderley Park | Wellcome Sanger Institute for the COVID-19 Genomics UK (COG-UK) Consortium | Jacquelyn Wynn, Mairead Hyland, The Lighthouse Lab in Alderley Park and Alex Alderton, Roberto Amato, Jeffrey Barrett, Sonia Goncalves, Ewan Harrison, David K. Jackson, Ian Johnston, Dominic Kwiatkowski, Cordelia Langford, John Sillitoe on behalf of the Wellcome Sanger Institute COVID-19 Surveillance Team |
| EPI_ISL_2553031 | Lighthouse Lab in Glasgow | Wellcome Sanger Institute for the COVID-19 Genomics UK (COG-UK) Consortium | Harper VanSteenhouse, Yumi Kasai, David Gray, Carol Clugston, Anna Dominiczak and Alex Alderton, Roberto Amato, Jeffrey Barrett, Sonia Goncalves, Ewan Harrison, David K. Jackson, Ian Johnston, Dominic Kwiatkowski, Cordelia Langford, John Sillitoe on behalf of the Wellcome Sanger Institute COVID-19 Surveillance Team |
| EPI_ISL_2553032, EPI_ISL_2553033, EPI_ISL_2553034, EPI_ISL_2553035 | Lighthouse Lab in Alderley Park | Wellcome Sanger Institute for the COVID-19 Genomics UK (COG-UK) Consortium | Jacquelyn Wynn, Mairead Hyland, The Lighthouse Lab in Alderley Park and Alex Alderton, Roberto Amato, Jeffrey Barrett, Sonia Goncalves, Ewan Harrison, David K. Jackson, Ian Johnston, Dominic Kwiatkowski, Cordelia Langford, John Sillitoe on behalf of the Wellcome Sanger Institute COVID-19 Surveillance Team |
| EPI_ISL_2553036 | Lighthouse Lab in Glasgow | Wellcome Sanger Institute for the COVID-19 Genomics UK (COG-UK) Consortium | Harper VanSteenhouse, Yumi Kasai, David Gray, Carol Clugston, Anna Dominiczak and Alex Alderton, Roberto Amato, Jeffrey Barrett, Sonia Goncalves, Ewan Harrison, David K. Jackson, Ian Johnston, Dominic Kwiatkowski, Cordelia Langford, John Sillitoe on behalf of the Wellcome Sanger Institute COVID-19 Surveillance Team |
| EPI_ISL_2553037, EPI_ISL_2553038, EPI_ISL_2553039, EPI_ISL_2553040 | Lighthouse Lab in Alderley Park | Wellcome Sanger Institute for the COVID-19 Genomics UK (COG-UK) Consortium | Jacquelyn Wynn, Mairead Hyland, The Lighthouse Lab in Alderley Park and Alex Alderton, Roberto Amato, Jeffrey Barrett, Sonia Goncalves, Ewan Harrison, David K. Jackson, Ian Johnston, Dominic Kwiatkowski, Cordelia Langford, John Sillitoe on behalf of the Wellcome Sanger Institute COVID-19 Surveillance Team |
| EPI_ISL_2553041 | Lighthouse Lab in Glasgow | Wellcome Sanger Institute for the COVID-19 Genomics UK (COG-UK) Consortium | Harper VanSteenhouse, Yumi Kasai, David Gray, Carol Clugston, Anna Dominiczak and Alex Alderton, Roberto Amato, Jeffrey Barrett, Sonia Goncalves, Ewan Harrison, David K. Jackson, Ian Johnston, Dominic Kwiatkowski, Cordelia Langford, John Sillitoe on behalf of the Wellcome Sanger Institute COVID-19 Surveillance Team |
| EPI_ISL_2553042, EPI_ISL_2553043, EPI_ISL_2553044, EPI_ISL_2553045 | Lighthouse Lab in Alderley Park | Wellcome Sanger Institute for the COVID-19 Genomics UK (COG-UK) Consortium | Jacquelyn Wynn, Mairead Hyland, The Lighthouse Lab in Alderley Park and Alex Alderton, Roberto Amato, Jeffrey Barrett, Sonia Goncalves, Ewan Harrison, David K. Jackson, Ian Johnston, Dominic Kwiatkowski, Cordelia Langford, John Sillitoe on behalf of the Wellcome Sanger Institute COVID-19 Surveillance Team |
| EPI_ISL_2553046 | Lighthouse Lab in Glasgow | Wellcome Sanger Institute for the COVID-19 Genomics UK (COG-UK) Consortium | Harper VanSteenhouse, Yumi Kasai, David Gray, Carol Clugston, Anna Dominiczak and Alex Alderton, Roberto Amato, Jeffrey Barrett, Sonia Goncalves, Ewan Harrison, David K. Jackson, Ian Johnston, Dominic Kwiatkowski, Cordelia Langford, John Sillitoe on behalf of the Wellcome Sanger Institute COVID-19 Surveillance Team |
| EPI_ISL_2553047, EPI_ISL_2553048 | Lighthouse Lab in Alderley Park | Wellcome Sanger Institute for the COVID-19 Genomics UK (COG-UK) Consortium | Jacquelyn Wynn, Mairead Hyland, The Lighthouse Lab in Alderley Park and Alex Alderton, Roberto Amato, Jeffrey Barrett, Sonia Goncalves, Ewan Harrison, David K. Jackson, Ian Johnston, Dominic Kwiatkowski, Cordelia Langford, John Sillitoe on behalf of the Wellcome Sanger Institute COVID-19 Surveillance Team |
| EPI_ISL_2553049, EPI_ISL_2553051 | Lighthouse Lab in Glasgow | Wellcome Sanger Institute for the COVID-19 Genomics UK (COG-UK) Consortium | Harper VanSteenhouse, Yumi Kasai, David Gray, Carol Clugston, Anna Dominiczak and Alex Alderton, Roberto Amato, Jeffrey Barrett, Sonia Goncalves, Ewan Harrison, David K. Jackson, Ian Johnston, Dominic Kwiatkowski, Cordelia Langford, John Sillitoe on behalf of the Wellcome Sanger Institute COVID-19 Surveillance Team |
| EPI_ISL_2553052, EPI_ISL_2553053, EPI_ISL_2553054, EPI_ISL_2553055 | Lighthouse Lab in Alderley Park | Wellcome Sanger Institute for the COVID-19 Genomics UK (COG-UK) Consortium | Jacquelyn Wynn, Mairead Hyland, The Lighthouse Lab in Alderley Park and Alex Alderton, Roberto Amato, Jeffrey Barrett, Sonia Goncalves, Ewan Harrison, David K. Jackson, Ian Johnston, Dominic Kwiatkowski, Cordelia Langford, John Sillitoe on behalf of the Wellcome Sanger Institute COVID-19 Surveillance Team |
| EPI_ISL_2553056 | Lighthouse Lab in Glasgow | Wellcome Sanger Institute for the COVID-19 Genomics UK (COG-UK) Consortium | Harper VanSteenhouse, Yumi Kasai, David Gray, Carol Clugston, Anna Dominiczak and Alex Alderton, Roberto Amato, Jeffrey Barrett, Sonia Goncalves, Ewan Harrison, David K. Jackson, Ian Johnston, Dominic Kwiatkowski, Cordelia Langford, John Sillitoe on behalf of the Wellcome Sanger Institute COVID-19 Surveillance Team |
| EPI_ISL_2553057, EPI_ISL_2553059, EPI_ISL_2553060 | Lighthouse Lab in Alderley Park | Wellcome Sanger Institute for the COVID-19 Genomics UK (COG-UK) Consortium | Jacquelyn Wynn, Mairead Hyland, The Lighthouse Lab in Alderley Park and Alex Alderton, Roberto Amato, Jeffrey Barrett, Sonia Goncalves, Ewan Harrison, David K. Jackson, Ian Johnston, Dominic Kwiatkowski, Cordelia Langford, John Sillitoe on behalf of the Wellcome Sanger Institute COVID-19 Surveillance Team |
| EPI_ISL_2553061 | Lighthouse Lab in Glasgow | Wellcome Sanger Institute for the COVID-19 Genomics UK (COG-UK) Consortium | Harper VanSteenhouse, Yumi Kasai, David Gray, Carol Clugston, Anna Dominiczak and Alex Alderton, Roberto Amato, Jeffrey Barrett, Sonia Goncalves, Ewan Harrison, David K. Jackson, Ian Johnston, Dominic Kwiatkowski, Cordelia Langford, John Sillitoe on behalf of the Wellcome Sanger Institute COVID-19 Surveillance Team |
| EPI_ISL_2553062, EPI_ISL_2553064 | Lighthouse Lab in Alderley Park | Wellcome Sanger Institute for the COVID-19 Genomics UK (COG-UK) Consortium | Jacquelyn Wynn, Mairead Hyland, The Lighthouse Lab in Alderley Park and Alex Alderton, Roberto Amato, Jeffrey Barrett, Sonia Goncalves, Ewan Harrison, David K. Jackson, Ian Johnston, Dominic Kwiatkowski, Cordelia Langford, John Sillitoe on behalf of the Wellcome Sanger Institute COVID-19 Surveillance Team |
| EPI_ISL_2553066 | Lighthouse Lab in Glasgow | Wellcome Sanger Institute for the COVID-19 Genomics UK (COG-UK) Consortium | Harper VanSteenhouse, Yumi Kasai, David Gray, Carol Clugston, Anna Dominiczak and Alex Alderton, Roberto Amato, Jeffrey Barrett, Sonia Goncalves, Ewan Harrison, David K. Jackson, Ian Johnston, Dominic Kwiatkowski, Cordelia Langford, John Sillitoe on behalf of the Wellcome Sanger Institute COVID-19 Surveillance Team |
| EPI_ISL_2553067, EPI_ISL_2553068, EPI_ISL_2553069 | Lighthouse Lab in Alderley Park | Wellcome Sanger Institute for the COVID-19 Genomics UK (COG-UK) Consortium | Jacquelyn Wynn, Mairead Hyland, The Lighthouse Lab in Alderley Park and Alex Alderton, Roberto Amato, Jeffrey Barrett, Sonia Goncalves, Ewan Harrison, David K. Jackson, Ian Johnston, Dominic Kwiatkowski, Cordelia Langford, John Sillitoe on behalf of the Wellcome Sanger Institute COVID-19 Surveillance Team |
| EPI_ISL_2553070 | Lighthouse Lab in Glasgow | Wellcome Sanger Institute for the COVID-19 Genomics UK (COG-UK) Consortium | Harper VanSteenhouse, Yumi Kasai, David Gray, Carol Clugston, Anna Dominiczak and Alex Alderton, Roberto Amato, Jeffrey Barrett, Sonia Goncalves, Ewan Harrison, David K. Jackson, Ian Johnston, Dominic Kwiatkowski, Cordelia Langford, John Sillitoe on behalf of the Wellcome Sanger Institute COVID-19 Surveillance Team |
| EPI_ISL_2553071, EPI_ISL_2553072 | Lighthouse Lab in Alderley Park | Wellcome Sanger Institute for the COVID-19 Genomics UK (COG-UK) Consortium | Jacquelyn Wynn, Mairead Hyland, The Lighthouse Lab in Alderley Park and Alex Alderton, Roberto Amato, Jeffrey Barrett, Sonia Goncalves, Ewan Harrison, David K. Jackson, Ian Johnston, Dominic Kwiatkowski, Cordelia Langford, John Sillitoe on behalf of the Wellcome Sanger Institute COVID-19 Surveillance Team |
| EPI_ISL_2553073 | Lighthouse Lab in Glasgow | Wellcome Sanger Institute for the COVID-19 Genomics UK (COG-UK) Consortium | Harper VanSteenhouse, Yumi Kasai, David Gray, Carol Clugston, Anna Dominiczak and Alex Alderton, Roberto Amato, Jeffrey Barrett, Sonia Goncalves, Ewan Harrison, David K. Jackson, Ian Johnston, Dominic Kwiatkowski, Cordelia Langford, John Sillitoe on behalf of the Wellcome Sanger Institute COVID-19 Surveillance Team |
| EPI_ISL_2553074, EPI_ISL_2553075, EPI_ISL_2553076, EPI_ISL_2553077, EPI_ISL_2553078, | Lighthouse Lab in Alderley Park | Wellcome Sanger Institute for the COVID-19 Genomics UK (COG-UK) Consortium | Jacquelyn Wynn, Mairead Hyland, The Lighthouse Lab in Alderley Park and Alex Alderton, Roberto Amato, Jeffrey Barrett, Sonia Goncalves, Ewan Harrison, David K. Jackson, Ian Johnston, Dominic Kwiatkowski, Cordelia Langford, John Sillitoe on behalf of the Wellcome Sanger Institute COVID-19 Surveillance Team |

| | | | |
|---|---|---|---|
| EPI_ISL_2553079, EPI_ISL_2553080, EPI_ISL_2553082, EPI_ISL_2553083, EPI_ISL_2553084 | | | |
| EPI_ISL_2553085 | Lighthouse Lab in Glasgow | Wellcome Sanger Institute for the COVID-19 Genomics UK (COG-UK) Consortium | Harper VanSteenhouse, Yumi Kasai, David Gray, Carol Clugston, Anna Dominiczak and Alex Alderton, Roberto Amato, Jeffrey Barrett, Sonia Goncalves, Ewan Harrison, David K. Jackson, Ian Johnston, Dominic Kwiatkowski, Cordelia Langford, John Sillitoe on behalf of the Wellcome Sanger Institute COVID-19 Surveillance Team |
| EPI_ISL_2553086, EPI_ISL_2553087, EPI_ISL_2553088 | Lighthouse Lab in Alderley Park | Wellcome Sanger Institute for the COVID-19 Genomics UK (COG-UK) Consortium | Jacquelyn Wynn, Mairead Hyland, The Lighthouse Lab in Alderley Park and Alex Alderton, Roberto Amato, Jeffrey Barrett, Sonia Goncalves, Ewan Harrison, David K. Jackson, Ian Johnston, Dominic Kwiatkowski, Cordelia Langford, John Sillitoe on behalf of the Wellcome Sanger Institute COVID-19 Surveillance Team |
| EPI_ISL_2553090 | Lighthouse Lab in Milton Keynes | Wellcome Sanger Institute for the COVID-19 Genomics UK (COG-UK) Consortium | The Lighthouse Lab in Milton Keynes and Alex Alderton, Roberto Amato, Jeffrey Barrett, Sonia Goncalves, Ewan Harrison, David K. Jackson, Ian Johnston, Dominic Kwiatkowski, Cordelia Langford, John Sillitoe on behalf of the Wellcome Sanger Institute COVID-19 Surveillance Team |
| EPI_ISL_2553091, EPI_ISL_2553092, EPI_ISL_2553093 | Lighthouse Lab in Alderley Park | Wellcome Sanger Institute for the COVID-19 Genomics UK (COG-UK) Consortium | Jacquelyn Wynn, Mairead Hyland, The Lighthouse Lab in Alderley Park and Alex Alderton, Roberto Amato, Jeffrey Barrett, Sonia Goncalves, Ewan Harrison, David K. Jackson, Ian Johnston, Dominic Kwiatkowski, Cordelia Langford, John Sillitoe on behalf of the Wellcome Sanger Institute COVID-19 Surveillance Team |
| EPI_ISL_2553094, EPI_ISL_2553095, EPI_ISL_2553096, EPI_ISL_2553097 | Lighthouse Lab in Glasgow | Wellcome Sanger Institute for the COVID-19 Genomics UK (COG-UK) Consortium | Harper VanSteenhouse, Yumi Kasai, David Gray, Carol Clugston, Anna Dominiczak and Alex Alderton, Roberto Amato, Jeffrey Barrett, Sonia Goncalves, Ewan Harrison, David K. Jackson, Ian Johnston, Dominic Kwiatkowski, Cordelia Langford, John Sillitoe on behalf of the Wellcome Sanger Institute COVID-19 Surveillance Team |
| EPI_ISL_2553098 | Lighthouse Lab in Alderley Park | Wellcome Sanger Institute for the COVID-19 Genomics UK (COG-UK) Consortium | Jacquelyn Wynn, Mairead Hyland, The Lighthouse Lab in Alderley Park and Alex Alderton, Roberto Amato, Jeffrey Barrett, Sonia Goncalves, Ewan Harrison, David K. Jackson, Ian Johnston, Dominic Kwiatkowski, Cordelia Langford, John Sillitoe on behalf of the Wellcome Sanger Institute COVID-19 Surveillance Team |
| EPI_ISL_2553099, EPI_ISL_2553100, EPI_ISL_2553101 | Lighthouse Lab in Glasgow | Wellcome Sanger Institute for the COVID-19 Genomics UK (COG-UK) Consortium | Harper VanSteenhouse, Yumi Kasai, David Gray, Carol Clugston, Anna Dominiczak and Alex Alderton, Roberto Amato, Jeffrey Barrett, Sonia Goncalves, Ewan Harrison, David K. Jackson, Ian Johnston, Dominic Kwiatkowski, Cordelia Langford, John Sillitoe on behalf of the Wellcome Sanger Institute COVID-19 Surveillance Team |
| EPI_ISL_2553102 | Lighthouse Lab in Milton Keynes | Wellcome Sanger Institute for the COVID-19 Genomics UK (COG-UK) Consortium | The Lighthouse Lab in Milton Keynes and Alex Alderton, Roberto Amato, Jeffrey Barrett, Sonia Goncalves, Ewan Harrison, David K. Jackson, Ian Johnston, Dominic Kwiatkowski, Cordelia Langford, John Sillitoe on behalf of the Wellcome Sanger Institute COVID-19 Surveillance Team |
| EPI_ISL_2553103, EPI_ISL_2553104, EPI_ISL_2553105, EPI_ISL_2553106, EPI_ISL_2553107 | Lighthouse Lab in Glasgow | Wellcome Sanger Institute for the COVID-19 Genomics UK (COG-UK) Consortium | Harper VanSteenhouse, Yumi Kasai, David Gray, Carol Clugston, Anna Dominiczak and Alex Alderton, Roberto Amato, Jeffrey Barrett, Sonia Goncalves, Ewan Harrison, David K. Jackson, Ian Johnston, Dominic Kwiatkowski, Cordelia Langford, John Sillitoe on behalf of the Wellcome Sanger Institute COVID-19 Surveillance Team |
| EPI_ISL_2553108, EPI_ISL_2553109 | Lighthouse Lab in Alderley Park | Wellcome Sanger Institute for the COVID-19 Genomics UK (COG-UK) Consortium | Jacquelyn Wynn, Mairead Hyland, The Lighthouse Lab in Alderley Park and Alex Alderton, Roberto Amato, Jeffrey Barrett, Sonia Goncalves, Ewan Harrison, David K. Jackson, Ian Johnston, Dominic Kwiatkowski, Cordelia Langford, John Sillitoe on behalf of the Wellcome Sanger Institute COVID-19 Surveillance Team |
| EPI_ISL_2553110, EPI_ISL_2553111 | Lighthouse Lab in Milton Keynes | Wellcome Sanger Institute for the COVID-19 Genomics UK (COG-UK) Consortium | The Lighthouse Lab in Milton Keynes and Alex Alderton, Roberto Amato, Jeffrey Barrett, Sonia Goncalves, Ewan Harrison, David K. Jackson, Ian Johnston, Dominic Kwiatkowski, Cordelia Langford, John Sillitoe on behalf of the Wellcome Sanger Institute COVID-19 Surveillance Team |
| EPI_ISL_2553113 | Lighthouse Lab in Alderley Park | Wellcome Sanger Institute for the COVID-19 Genomics UK (COG-UK) Consortium | Jacquelyn Wynn, Mairead Hyland, The Lighthouse Lab in Alderley Park and Alex Alderton, Roberto Amato, Jeffrey Barrett, Sonia Goncalves, Ewan Harrison, David K. Jackson, Ian Johnston, Dominic Kwiatkowski, Cordelia Langford, John Sillitoe on behalf of the Wellcome Sanger Institute COVID-19 Surveillance Team |
| EPI_ISL_2553114 | Lighthouse Lab in Milton Keynes | Wellcome Sanger Institute for the COVID-19 Genomics UK (COG-UK) Consortium | The Lighthouse Lab in Milton Keynes and Alex Alderton, Roberto Amato, Jeffrey Barrett, Sonia Goncalves, Ewan Harrison, David K. Jackson, Ian Johnston, Dominic Kwiatkowski, Cordelia Langford, John Sillitoe on behalf of the Wellcome Sanger Institute COVID-19 Surveillance Team |
| EPI_ISL_2553115 | Lighthouse Lab in Glasgow | Wellcome Sanger Institute for the COVID-19 Genomics UK (COG-UK) Consortium | Harper VanSteenhouse, Yumi Kasai, David Gray, Carol Clugston, Anna Dominiczak and Alex Alderton, Roberto Amato, Jeffrey Barrett, Sonia Goncalves, Ewan Harrison, David K. Jackson, Ian Johnston, Dominic Kwiatkowski, Cordelia Langford, John Sillitoe on behalf of the Wellcome Sanger Institute COVID-19 Surveillance Team |
| EPI_ISL_2553116 | Lighthouse Lab in Alderley Park | Wellcome Sanger Institute for the COVID-19 Genomics UK (COG-UK) Consortium | Jacquelyn Wynn, Mairead Hyland, The Lighthouse Lab in Alderley Park and Alex Alderton, Roberto Amato, Jeffrey Barrett, Sonia Goncalves, Ewan Harrison, David K. Jackson, Ian Johnston, Dominic Kwiatkowski, Cordelia Langford, John Sillitoe on behalf of the Wellcome Sanger Institute COVID-19 Surveillance Team |
| EPI_ISL_2553118 | Lighthouse Lab in Glasgow | Wellcome Sanger Institute for the COVID-19 Genomics UK (COG-UK) Consortium | Harper VanSteenhouse, Yumi Kasai, David Gray, Carol Clugston, Anna Dominiczak and Alex Alderton, Roberto Amato, Jeffrey Barrett, Sonia Goncalves, Ewan Harrison, David K. Jackson, Ian Johnston, Dominic Kwiatkowski, Cordelia Langford, John Sillitoe on behalf of the Wellcome Sanger Institute COVID-19 Surveillance Team |
| EPI_ISL_2553119 | Lighthouse Lab in Alderley Park | Wellcome Sanger Institute for the COVID-19 Genomics UK (COG-UK) Consortium | Jacquelyn Wynn, Mairead Hyland, The Lighthouse Lab in Alderley Park and Alex Alderton, Roberto Amato, Jeffrey Barrett, Sonia Goncalves, Ewan Harrison, David K. Jackson, Ian Johnston, Dominic Kwiatkowski, Cordelia Langford, John Sillitoe on behalf of the Wellcome Sanger Institute COVID-19 Surveillance Team |
| EPI_ISL_2553120, EPI_ISL_2553121, EPI_ISL_2553122 | Lighthouse Lab in Glasgow | Wellcome Sanger Institute for the COVID-19 Genomics UK (COG-UK) Consortium | Harper VanSteenhouse, Yumi Kasai, David Gray, Carol Clugston, Anna Dominiczak and Alex Alderton, Roberto Amato, Jeffrey Barrett, Sonia Goncalves, Ewan Harrison, David K. Jackson, Ian Johnston, Dominic Kwiatkowski, Cordelia Langford, John Sillitoe on behalf of the Wellcome Sanger Institute COVID-19 Surveillance Team |
| EPI_ISL_2553124, EPI_ISL_2553125 | Lighthouse Lab in Alderley Park | Wellcome Sanger Institute for the COVID-19 Genomics UK (COG-UK) Consortium | Jacquelyn Wynn, Mairead Hyland, The Lighthouse Lab in Alderley Park and Alex Alderton, Roberto Amato, Jeffrey Barrett, Sonia Goncalves, Ewan Harrison, David K. Jackson, Ian Johnston, Dominic Kwiatkowski, Cordelia Langford, John Sillitoe on behalf of the Wellcome Sanger Institute COVID-19 Surveillance Team |
| EPI_ISL_2553126, EPI_ISL_2553127, EPI_ISL_2553128 | Lighthouse Lab in Milton Keynes | Wellcome Sanger Institute for the COVID-19 Genomics UK (COG-UK) Consortium | The Lighthouse Lab in Milton Keynes and Alex Alderton, Roberto Amato, Jeffrey Barrett, Sonia Goncalves, Ewan Harrison, David K. Jackson, Ian Johnston, Dominic Kwiatkowski, Cordelia Langford, John Sillitoe on behalf of the Wellcome Sanger Institute COVID-19 Surveillance Team |
| EPI_ISL_2553129, EPI_ISL_2553130 | Lighthouse Lab in Glasgow | Wellcome Sanger Institute for the COVID-19 Genomics UK (COG-UK) Consortium | Harper VanSteenhouse, Yumi Kasai, David Gray, Carol Clugston, Anna Dominiczak and Alex Alderton, Roberto Amato, Jeffrey Barrett, Sonia Goncalves, Ewan Harrison, David K. Jackson, Ian Johnston, Dominic Kwiatkowski, Cordelia Langford, John Sillitoe on behalf of the Wellcome Sanger Institute COVID-19 Surveillance Team |
| EPI_ISL_2553131, EPI_ISL_2553132, EPI_ISL_2553133, EPI_ISL_2553134, EPI_ISL_2553137 | Lighthouse Lab in Alderley Park | Wellcome Sanger Institute for the COVID-19 Genomics UK (COG-UK) Consortium | Jacquelyn Wynn, Mairead Hyland, The Lighthouse Lab in Alderley Park and Alex Alderton, Roberto Amato, Jeffrey Barrett, Sonia Goncalves, Ewan Harrison, David K. Jackson, Ian Johnston, Dominic Kwiatkowski, Cordelia Langford, John Sillitoe on behalf of the Wellcome Sanger Institute COVID-19 Surveillance Team |
| EPI_ISL_2553139, EPI_ISL_2553141 | Lighthouse Lab in Glasgow | Wellcome Sanger Institute for the COVID-19 Genomics UK (COG-UK) Consortium | Harper VanSteenhouse, Yumi Kasai, David Gray, Carol Clugston, Anna Dominiczak and Alex Alderton, Roberto Amato, Jeffrey Barrett, Sonia Goncalves, Ewan Harrison, David K. Jackson, Ian Johnston, Dominic Kwiatkowski, Cordelia Langford, John Sillitoe on behalf of the Wellcome Sanger Institute COVID-19 Surveillance Team |
| EPI_ISL_2553142 | Lighthouse Lab in Milton Keynes | Wellcome Sanger Institute for the COVID-19 Genomics UK (COG-UK) Consortium | The Lighthouse Lab in Milton Keynes and Alex Alderton, Roberto Amato, Jeffrey Barrett, Sonia Goncalves, Ewan Harrison, David K. Jackson, Ian Johnston, Dominic Kwiatkowski, Cordelia Langford, John Sillitoe on behalf of the Wellcome Sanger Institute COVID-19 Surveillance Team |
| EPI_ISL_2553143 | Lighthouse Lab in Alderley Park | Wellcome Sanger Institute for the COVID-19 Genomics UK (COG-UK) Consortium | Jacquelyn Wynn, Mairead Hyland, The Lighthouse Lab in Alderley Park and Alex Alderton, Roberto Amato, Jeffrey Barrett, Sonia Goncalves, Ewan Harrison, David K. Jackson, Ian Johnston, Dominic Kwiatkowski, Cordelia Langford, John Sillitoe on behalf of the Wellcome Sanger Institute COVID-19 Surveillance Team |
| EPI_ISL_2553144 | Lighthouse Lab in Glasgow | Wellcome Sanger Institute for the COVID-19 Genomics UK (COG-UK) Consortium | Harper VanSteenhouse, Yumi Kasai, David Gray, Carol Clugston, Anna Dominiczak and Alex Alderton, Roberto Amato, Jeffrey Barrett, Sonia Goncalves, Ewan Harrison, David K. Jackson, Ian Johnston, Dominic Kwiatkowski, Cordelia Langford, John Sillitoe on behalf of the Wellcome Sanger Institute COVID-19 Surveillance Team |
| EPI_ISL_2553145, EPI_ISL_2553146 | Lighthouse Lab in Alderley Park | Wellcome Sanger Institute for the COVID-19 Genomics UK (COG-UK) Consortium | Jacquelyn Wynn, Mairead Hyland, The Lighthouse Lab in Alderley Park and Alex Alderton, Roberto Amato, Jeffrey Barrett, Sonia Goncalves, Ewan Harrison, David K. Jackson, Ian Johnston, Dominic Kwiatkowski, Cordelia Langford, John Sillitoe on behalf of the Wellcome Sanger Institute COVID-19 Surveillance Team |
| EPI_ISL_2553147 | Lighthouse Lab in Milton Keynes | Wellcome Sanger Institute for the COVID-19 Genomics UK (COG-UK) Consortium | The Lighthouse Lab in Milton Keynes and Alex Alderton, Roberto Amato, Jeffrey Barrett, Sonia Goncalves, Ewan Harrison, David K. Jackson, Ian Johnston, Dominic Kwiatkowski, Cordelia Langford, John Sillitoe on behalf of the Wellcome Sanger Institute COVID-19 Surveillance Team |
| EPI_ISL_2553148, EPI_ISL_2553149, EPI_ISL_2553150, EPI_ISL_2553151 | Lighthouse Lab in Glasgow | Wellcome Sanger Institute for the COVID-19 Genomics UK (COG-UK) Consortium | Harper VanSteenhouse, Yumi Kasai, David Gray, Carol Clugston, Anna Dominiczak and Alex Alderton, Roberto Amato, Jeffrey Barrett, Sonia Goncalves, Ewan Harrison, David K. Jackson, Ian Johnston, Dominic Kwiatkowski, Cordelia Langford, John Sillitoe on behalf of the Wellcome Sanger Institute COVID-19 Surveillance Team |
| EPI_ISL_2553153 | Lighthouse Lab in Alderley Park | Wellcome Sanger Institute for the COVID-19 Genomics UK (COG-UK) Consortium | Jacquelyn Wynn, Mairead Hyland, The Lighthouse Lab in Alderley Park and Alex Alderton, Roberto Amato, Jeffrey Barrett, Sonia Goncalves, Ewan Harrison, David K. Jackson, Ian Johnston, Dominic Kwiatkowski, Cordelia Langford, John Sillitoe on behalf of the Wellcome Sanger Institute COVID-19 Surveillance Team |
| EPI_ISL_2553154 | Lighthouse Lab in Milton Keynes | Wellcome Sanger Institute for the COVID-19 Genomics UK (COG-UK) Consortium | The Lighthouse Lab in Milton Keynes and Alex Alderton, Roberto Amato, Jeffrey Barrett, Sonia Goncalves, Ewan Harrison, David K. Jackson, Ian Johnston, Dominic Kwiatkowski, Cordelia Langford, John Sillitoe on behalf of the Wellcome Sanger Institute COVID-19 Surveillance Team |
| EPI_ISL_2553155 | Lighthouse Lab in Glasgow | Wellcome Sanger Institute for the COVID-19 Genomics UK (COG-UK) Consortium | Harper VanSteenhouse, Yumi Kasai, David Gray, Carol Clugston, Anna Dominiczak and Alex Alderton, Roberto Amato, Jeffrey Barrett, Sonia Goncalves, Ewan Harrison, David K. Jackson, Ian Johnston, Dominic Kwiatkowski, Cordelia Langford, John Sillitoe on behalf of the Wellcome Sanger Institute COVID-19 Surveillance Team |
| EPI_ISL_2553157 | Lighthouse Lab in Milton Keynes | Wellcome Sanger Institute for the COVID-19 Genomics UK (COG-UK) Consortium | The Lighthouse Lab in Milton Keynes and Alex Alderton, Roberto Amato, Jeffrey Barrett, Sonia Goncalves, Ewan Harrison, David K. Jackson, Ian Johnston, Dominic Kwiatkowski, Cordelia Langford, John Sillitoe on behalf of the Wellcome Sanger Institute COVID-19 Surveillance Team |
| EPI_ISL_2553158, EPI_ISL_2553159, EPI_ISL_2553160 | Lighthouse Lab in Glasgow | Wellcome Sanger Institute for the COVID-19 Genomics UK (COG-UK) Consortium | Harper VanSteenhouse, Yumi Kasai, David Gray, Carol Clugston, Anna Dominiczak and Alex Alderton, Roberto Amato, Jeffrey Barrett, Sonia Goncalves, Ewan Harrison, David K. Jackson, Ian Johnston, Dominic Kwiatkowski, Cordelia Langford, John Sillitoe on behalf of the Wellcome Sanger Institute COVID-19 Surveillance Team |
| EPI_ISL_2553161 | Lighthouse Lab in Milton Keynes | Wellcome Sanger Institute for the COVID-19 Genomics UK (COG-UK) Consortium | The Lighthouse Lab in Milton Keynes and Alex Alderton, Roberto Amato, Jeffrey Barrett, Sonia Goncalves, Ewan Harrison, David K. Jackson, Ian Johnston, Dominic Kwiatkowski, Cordelia Langford, John Sillitoe on behalf of the Wellcome Sanger Institute COVID-19 Surveillance Team |
| EPI_ISL_2553162 | Lighthouse Lab in Glasgow | Wellcome Sanger Institute for the COVID-19 Genomics UK (COG-UK) Consortium | Harper VanSteenhouse, Yumi Kasai, David Gray, Carol Clugston, Anna Dominiczak and Alex Alderton, Roberto Amato, Jeffrey Barrett, Sonia Goncalves, Ewan Harrison, David K. Jackson, Ian Johnston, Dominic Kwiatkowski, Cordelia Langford, John Sillitoe on behalf of the Wellcome Sanger Institute COVID-19 Surveillance Team |

| | | | |
|---|---|---|---|
| EPI_ISL_2553163, EPI_ISL_2553164 | Lighthouse Lab in Milton Keynes | Wellcome Sanger Institute for the COVID-19 Genomics UK (COG-UK) Consortium | The Lighthouse Lab in Milton Keynes and Alex Alderton, Roberto Amato, Jeffrey Barrett, Sonia Goncalves, Ewan Harrison, David K. Jackson, Ian Johnston, Dominic Kwiatkowski, Cordelia Langford, John Sillitoe on behalf of the Wellcome Sanger Institute COVID-19 Surveillance Team |
| EPI_ISL_2553165 | Lighthouse Lab in Glasgow | Wellcome Sanger Institute for the COVID-19 Genomics UK (COG-UK) Consortium | Harper VanSteenhouse, Yumi Kasai, David Gray, Carol Clugston, Anna Dominiczak and Alex Alderton, Roberto Amato, Jeffrey Barrett, Sonia Goncalves, Ewan Harrison, David K. Jackson, Ian Johnston, Dominic Kwiatkowski, Cordelia Langford, John Sillitoe on behalf of the Wellcome Sanger Institute COVID-19 Surveillance Team |
| EPI_ISL_2553166 | Lighthouse Lab in Milton Keynes | Wellcome Sanger Institute for the COVID-19 Genomics UK (COG-UK) Consortium | The Lighthouse Lab in Milton Keynes and Alex Alderton, Roberto Amato, Jeffrey Barrett, Sonia Goncalves, Ewan Harrison, David K. Jackson, Ian Johnston, Dominic Kwiatkowski, Cordelia Langford, John Sillitoe on behalf of the Wellcome Sanger Institute COVID-19 Surveillance Team |
| EPI_ISL_2553167 | Lighthouse Lab in Glasgow | Wellcome Sanger Institute for the COVID-19 Genomics UK (COG-UK) Consortium | Harper VanSteenhouse, Yumi Kasai, David Gray, Carol Clugston, Anna Dominiczak and Alex Alderton, Roberto Amato, Jeffrey Barrett, Sonia Goncalves, Ewan Harrison, David K. Jackson, Ian Johnston, Dominic Kwiatkowski, Cordelia Langford, John Sillitoe on behalf of the Wellcome Sanger Institute COVID-19 Surveillance Team |
| EPI_ISL_2553168 | Lighthouse Lab in Milton Keynes | Wellcome Sanger Institute for the COVID-19 Genomics UK (COG-UK) Consortium | The Lighthouse Lab in Milton Keynes and Alex Alderton, Roberto Amato, Jeffrey Barrett, Sonia Goncalves, Ewan Harrison, David K. Jackson, Ian Johnston, Dominic Kwiatkowski, Cordelia Langford, John Sillitoe on behalf of the Wellcome Sanger Institute COVID-19 Surveillance Team |
| EPI_ISL_2553169, EPI_ISL_2553170 | Lighthouse Lab in Alderley Park | Wellcome Sanger Institute for the COVID-19 Genomics UK (COG-UK) Consortium | Jacquelyn Wynn, Mairead Hyland, The Lighthouse Lab in Alderley Park and Alex Alderton, Roberto Amato, Jeffrey Barrett, Sonia Goncalves, Ewan Harrison, David K. Jackson, Ian Johnston, Dominic Kwiatkowski, Cordelia Langford, John Sillitoe on behalf of the Wellcome Sanger Institute COVID-19 Surveillance Team |
| EPI_ISL_2553171 | Lighthouse Lab in Glasgow | Wellcome Sanger Institute for the COVID-19 Genomics UK (COG-UK) Consortium | Harper VanSteenhouse, Yumi Kasai, David Gray, Carol Clugston, Anna Dominiczak and Alex Alderton, Roberto Amato, Jeffrey Barrett, Sonia Goncalves, Ewan Harrison, David K. Jackson, Ian Johnston, Dominic Kwiatkowski, Cordelia Langford, John Sillitoe on behalf of the Wellcome Sanger Institute COVID-19 Surveillance Team |
| EPI_ISL_2553172, EPI_ISL_2553174 | Lighthouse Lab in Alderley Park | Wellcome Sanger Institute for the COVID-19 Genomics UK (COG-UK) Consortium | Jacquelyn Wynn, Mairead Hyland, The Lighthouse Lab in Alderley Park and Alex Alderton, Roberto Amato, Jeffrey Barrett, Sonia Goncalves, Ewan Harrison, David K. Jackson, Ian Johnston, Dominic Kwiatkowski, Cordelia Langford, John Sillitoe on behalf of the Wellcome Sanger Institute COVID-19 Surveillance Team |
| EPI_ISL_2553175, EPI_ISL_2553176 | Lighthouse Lab in Glasgow | Wellcome Sanger Institute for the COVID-19 Genomics UK (COG-UK) Consortium | Harper VanSteenhouse, Yumi Kasai, David Gray, Carol Clugston, Anna Dominiczak and Alex Alderton, Roberto Amato, Jeffrey Barrett, Sonia Goncalves, Ewan Harrison, David K. Jackson, Ian Johnston, Dominic Kwiatkowski, Cordelia Langford, John Sillitoe on behalf of the Wellcome Sanger Institute COVID-19 Surveillance Team |
| EPI_ISL_2553177 | Lighthouse Lab in Alderley Park | Wellcome Sanger Institute for the COVID-19 Genomics UK (COG-UK) Consortium | Jacquelyn Wynn, Mairead Hyland, The Lighthouse Lab in Alderley Park and Alex Alderton, Roberto Amato, Jeffrey Barrett, Sonia Goncalves, Ewan Harrison, David K. Jackson, Ian Johnston, Dominic Kwiatkowski, Cordelia Langford, John Sillitoe on behalf of the Wellcome Sanger Institute COVID-19 Surveillance Team |
| EPI_ISL_2553178 | Lighthouse Lab in Glasgow | Wellcome Sanger Institute for the COVID-19 Genomics UK (COG-UK) Consortium | Harper VanSteenhouse, Yumi Kasai, David Gray, Carol Clugston, Anna Dominiczak and Alex Alderton, Roberto Amato, Jeffrey Barrett, Sonia Goncalves, Ewan Harrison, David K. Jackson, Ian Johnston, Dominic Kwiatkowski, Cordelia Langford, John Sillitoe on behalf of the Wellcome Sanger Institute COVID-19 Surveillance Team |
| EPI_ISL_2553179 | Lighthouse Lab in Milton Keynes | Wellcome Sanger Institute for the COVID-19 Genomics UK (COG-UK) Consortium | The Lighthouse Lab in Milton Keynes and Alex Alderton, Roberto Amato, Jeffrey Barrett, Sonia Goncalves, Ewan Harrison, David K. Jackson, Ian Johnston, Dominic Kwiatkowski, Cordelia Langford, John Sillitoe on behalf of the Wellcome Sanger Institute COVID-19 Surveillance Team |
| EPI_ISL_2553180 | Lighthouse Lab in Glasgow | Wellcome Sanger Institute for the COVID-19 Genomics UK (COG-UK) Consortium | Harper VanSteenhouse, Yumi Kasai, David Gray, Carol Clugston, Anna Dominiczak and Alex Alderton, Roberto Amato, Jeffrey Barrett, Sonia Goncalves, Ewan Harrison, David K. Jackson, Ian Johnston, Dominic Kwiatkowski, Cordelia Langford, John Sillitoe on behalf of the Wellcome Sanger Institute COVID-19 Surveillance Team |
| EPI_ISL_2553181 | Lighthouse Lab in Milton Keynes | Wellcome Sanger Institute for the COVID-19 Genomics UK (COG-UK) Consortium | The Lighthouse Lab in Milton Keynes and Alex Alderton, Roberto Amato, Jeffrey Barrett, Sonia Goncalves, Ewan Harrison, David K. Jackson, Ian Johnston, Dominic Kwiatkowski, Cordelia Langford, John Sillitoe on behalf of the Wellcome Sanger Institute COVID-19 Surveillance Team |
| EPI_ISL_2553182 | Lighthouse Lab in Glasgow | Wellcome Sanger Institute for the COVID-19 Genomics UK (COG-UK) Consortium | Harper VanSteenhouse, Yumi Kasai, David Gray, Carol Clugston, Anna Dominiczak and Alex Alderton, Roberto Amato, Jeffrey Barrett, Sonia Goncalves, Ewan Harrison, David K. Jackson, Ian Johnston, Dominic Kwiatkowski, Cordelia Langford, John Sillitoe on behalf of the Wellcome Sanger Institute COVID-19 Surveillance Team |
| EPI_ISL_2553183 | Lighthouse Lab in Alderley Park | Wellcome Sanger Institute for the COVID-19 Genomics UK (COG-UK) Consortium | Jacquelyn Wynn, Mairead Hyland, The Lighthouse Lab in Alderley Park and Alex Alderton, Roberto Amato, Jeffrey Barrett, Sonia Goncalves, Ewan Harrison, David K. Jackson, Ian Johnston, Dominic Kwiatkowski, Cordelia Langford, John Sillitoe on behalf of the Wellcome Sanger Institute COVID-19 Surveillance Team |
| EPI_ISL_2553184, EPI_ISL_2553185, EPI_ISL_2553186, EPI_ISL_2553187 | Lighthouse Lab in Glasgow | Wellcome Sanger Institute for the COVID-19 Genomics UK (COG-UK) Consortium | Harper VanSteenhouse, Yumi Kasai, David Gray, Carol Clugston, Anna Dominiczak and Alex Alderton, Roberto Amato, Jeffrey Barrett, Sonia Goncalves, Ewan Harrison, David K. Jackson, Ian Johnston, Dominic Kwiatkowski, Cordelia Langford, John Sillitoe on behalf of the Wellcome Sanger Institute COVID-19 Surveillance Team |
| EPI_ISL_2553188, EPI_ISL_2553189, EPI_ISL_2553190 | Lighthouse Lab in Alderley Park | Wellcome Sanger Institute for the COVID-19 Genomics UK (COG-UK) Consortium | Jacquelyn Wynn, Mairead Hyland, The Lighthouse Lab in Alderley Park and Alex Alderton, Roberto Amato, Jeffrey Barrett, Sonia Goncalves, Ewan Harrison, David K. Jackson, Ian Johnston, Dominic Kwiatkowski, Cordelia Langford, John Sillitoe on behalf of the Wellcome Sanger Institute COVID-19 Surveillance Team |
| EPI_ISL_2553192 | Lighthouse Lab in Glasgow | Wellcome Sanger Institute for the COVID-19 Genomics UK (COG-UK) Consortium | Harper VanSteenhouse, Yumi Kasai, David Gray, Carol Clugston, Anna Dominiczak and Alex Alderton, Roberto Amato, Jeffrey Barrett, Sonia Goncalves, Ewan Harrison, David K. Jackson, Ian Johnston, Dominic Kwiatkowski, Cordelia Langford, John Sillitoe on behalf of the Wellcome Sanger Institute COVID-19 Surveillance Team |
| EPI_ISL_2553193, EPI_ISL_2553194, EPI_ISL_2553195, EPI_ISL_2553197 | Lighthouse Lab in Milton Keynes | Wellcome Sanger Institute for the COVID-19 Genomics UK (COG-UK) Consortium | The Lighthouse Lab in Milton Keynes and Alex Alderton, Roberto Amato, Jeffrey Barrett, Sonia Goncalves, Ewan Harrison, David K. Jackson, Ian Johnston, Dominic Kwiatkowski, Cordelia Langford, John Sillitoe on behalf of the Wellcome Sanger Institute COVID-19 Surveillance Team |
| EPI_ISL_2553198 | Lighthouse Lab in Glasgow | Wellcome Sanger Institute for the COVID-19 Genomics UK (COG-UK) Consortium | Harper VanSteenhouse, Yumi Kasai, David Gray, Carol Clugston, Anna Dominiczak and Alex Alderton, Roberto Amato, Jeffrey Barrett, Sonia Goncalves, Ewan Harrison, David K. Jackson, Ian Johnston, Dominic Kwiatkowski, Cordelia Langford, John Sillitoe on behalf of the Wellcome Sanger Institute COVID-19 Surveillance Team |
| EPI_ISL_2553199 | Lighthouse Lab in Milton Keynes | Wellcome Sanger Institute for the COVID-19 Genomics UK (COG-UK) Consortium | The Lighthouse Lab in Milton Keynes and Alex Alderton, Roberto Amato, Jeffrey Barrett, Sonia Goncalves, Ewan Harrison, David K. Jackson, Ian Johnston, Dominic Kwiatkowski, Cordelia Langford, John Sillitoe on behalf of the Wellcome Sanger Institute COVID-19 Surveillance Team |
| EPI_ISL_2553200 | Lighthouse Lab in Glasgow | Wellcome Sanger Institute for the COVID-19 Genomics UK (COG-UK) Consortium | Harper VanSteenhouse, Yumi Kasai, David Gray, Carol Clugston, Anna Dominiczak and Alex Alderton, Roberto Amato, Jeffrey Barrett, Sonia Goncalves, Ewan Harrison, David K. Jackson, Ian Johnston, Dominic Kwiatkowski, Cordelia Langford, John Sillitoe on behalf of the Wellcome Sanger Institute COVID-19 Surveillance Team |
| EPI_ISL_2553202 | Lighthouse Lab in Alderley Park | Wellcome Sanger Institute for the COVID-19 Genomics UK (COG-UK) Consortium | Jacquelyn Wynn, Mairead Hyland, The Lighthouse Lab in Alderley Park and Alex Alderton, Roberto Amato, Jeffrey Barrett, Sonia Goncalves, Ewan Harrison, David K. Jackson, Ian Johnston, Dominic Kwiatkowski, Cordelia Langford, John Sillitoe on behalf of the Wellcome Sanger Institute COVID-19 Surveillance Team |
| EPI_ISL_2553204 | Lighthouse Lab in Milton Keynes | Wellcome Sanger Institute for the COVID-19 Genomics UK (COG-UK) Consortium | The Lighthouse Lab in Milton Keynes and Alex Alderton, Roberto Amato, Jeffrey Barrett, Sonia Goncalves, Ewan Harrison, David K. Jackson, Ian Johnston, Dominic Kwiatkowski, Cordelia Langford, John Sillitoe on behalf of the Wellcome Sanger Institute COVID-19 Surveillance Team |
| EPI_ISL_2553205, EPI_ISL_2553206, EPI_ISL_2553207, EPI_ISL_2553209 | Lighthouse Lab in Glasgow | Wellcome Sanger Institute for the COVID-19 Genomics UK (COG-UK) Consortium | Harper VanSteenhouse, Yumi Kasai, David Gray, Carol Clugston, Anna Dominiczak and Alex Alderton, Roberto Amato, Jeffrey Barrett, Sonia Goncalves, Ewan Harrison, David K. Jackson, Ian Johnston, Dominic Kwiatkowski, Cordelia Langford, John Sillitoe on behalf of the Wellcome Sanger Institute COVID-19 Surveillance Team |
| EPI_ISL_2553210 | Lighthouse Lab in Milton Keynes | Wellcome Sanger Institute for the COVID-19 Genomics UK (COG-UK) Consortium | The Lighthouse Lab in Milton Keynes and Alex Alderton, Roberto Amato, Jeffrey Barrett, Sonia Goncalves, Ewan Harrison, David K. Jackson, Ian Johnston, Dominic Kwiatkowski, Cordelia Langford, John Sillitoe on behalf of the Wellcome Sanger Institute COVID-19 Surveillance Team |
| EPI_ISL_2553211, EPI_ISL_2553214, EPI_ISL_2553215 | Lighthouse Lab in Glasgow | Wellcome Sanger Institute for the COVID-19 Genomics UK (COG-UK) Consortium | Harper VanSteenhouse, Yumi Kasai, David Gray, Carol Clugston, Anna Dominiczak and Alex Alderton, Roberto Amato, Jeffrey Barrett, Sonia Goncalves, Ewan Harrison, David K. Jackson, Ian Johnston, Dominic Kwiatkowski, Cordelia Langford, John Sillitoe on behalf of the Wellcome Sanger Institute COVID-19 Surveillance Team |
| EPI_ISL_2553216 | Lighthouse Lab in Alderley Park | Wellcome Sanger Institute for the COVID-19 Genomics UK (COG-UK) Consortium | Jacquelyn Wynn, Mairead Hyland, The Lighthouse Lab in Alderley Park and Alex Alderton, Roberto Amato, Jeffrey Barrett, Sonia Goncalves, Ewan Harrison, David K. Jackson, Ian Johnston, Dominic Kwiatkowski, Cordelia Langford, John Sillitoe on behalf of the Wellcome Sanger Institute COVID-19 Surveillance Team |
| EPI_ISL_2553217, EPI_ISL_2553218 | Lighthouse Lab in Glasgow | Wellcome Sanger Institute for the COVID-19 Genomics UK (COG-UK) Consortium | Harper VanSteenhouse, Yumi Kasai, David Gray, Carol Clugston, Anna Dominiczak and Alex Alderton, Roberto Amato, Jeffrey Barrett, Sonia Goncalves, Ewan Harrison, David K. Jackson, Ian Johnston, Dominic Kwiatkowski, Cordelia Langford, John Sillitoe on behalf of the Wellcome Sanger Institute COVID-19 Surveillance Team |
| EPI_ISL_2553219 | Lighthouse Lab in Alderley Park | Wellcome Sanger Institute for the COVID-19 Genomics UK (COG-UK) Consortium | Jacquelyn Wynn, Mairead Hyland, The Lighthouse Lab in Alderley Park and Alex Alderton, Roberto Amato, Jeffrey Barrett, Sonia Goncalves, Ewan Harrison, David K. Jackson, Ian Johnston, Dominic Kwiatkowski, Cordelia Langford, John Sillitoe on behalf of the Wellcome Sanger Institute COVID-19 Surveillance Team |
| EPI_ISL_2553220, EPI_ISL_2553221, EPI_ISL_2553222 | Lighthouse Lab in Glasgow | Wellcome Sanger Institute for the COVID-19 Genomics UK (COG-UK) Consortium | Harper VanSteenhouse, Yumi Kasai, David Gray, Carol Clugston, Anna Dominiczak and Alex Alderton, Roberto Amato, Jeffrey Barrett, Sonia Goncalves, Ewan Harrison, David K. Jackson, Ian Johnston, Dominic Kwiatkowski, Cordelia Langford, John Sillitoe on behalf of the Wellcome Sanger Institute COVID-19 Surveillance Team |
| EPI_ISL_2553223 | Lighthouse Lab in Alderley Park | Wellcome Sanger Institute for the COVID-19 Genomics UK (COG-UK) Consortium | Jacquelyn Wynn, Mairead Hyland, The Lighthouse Lab in Alderley Park and Alex Alderton, Roberto Amato, Jeffrey Barrett, Sonia Goncalves, Ewan Harrison, David K. Jackson, Ian Johnston, Dominic Kwiatkowski, Cordelia Langford, John Sillitoe on behalf of the Wellcome Sanger Institute COVID-19 Surveillance Team |
| EPI_ISL_2553224 | Lighthouse Lab in Milton Keynes | Wellcome Sanger Institute for the COVID-19 Genomics UK (COG-UK) Consortium | The Lighthouse Lab in Milton Keynes and Alex Alderton, Roberto Amato, Jeffrey Barrett, Sonia Goncalves, Ewan Harrison, David K. Jackson, Ian Johnston, Dominic Kwiatkowski, Cordelia Langford, John Sillitoe on behalf of the Wellcome Sanger Institute COVID-19 Surveillance Team |
| EPI_ISL_2553225 | Lighthouse Lab in Glasgow | Wellcome Sanger Institute for the COVID-19 Genomics UK (COG-UK) Consortium | Harper VanSteenhouse, Yumi Kasai, David Gray, Carol Clugston, Anna Dominiczak and Alex Alderton, Roberto Amato, Jeffrey Barrett, Sonia Goncalves, Ewan Harrison, David K. Jackson, Ian Johnston, Dominic Kwiatkowski, Cordelia Langford, John Sillitoe on behalf of the Wellcome Sanger Institute COVID-19 Surveillance Team |
| EPI_ISL_2553228 | Lighthouse Lab in Milton Keynes | Wellcome Sanger Institute for the COVID-19 Genomics UK (COG-UK) Consortium | The Lighthouse Lab in Milton Keynes and Alex Alderton, Roberto Amato, Jeffrey Barrett, Sonia Goncalves, Ewan Harrison, David K. Jackson, Ian Johnston, Dominic Kwiatkowski, Cordelia Langford, John Sillitoe on behalf of the Wellcome Sanger Institute COVID-19 Surveillance Team |
| EPI_ISL_2553229 | Lighthouse Lab in Alderley Park | Wellcome Sanger Institute for the COVID-19 Genomics UK (COG-UK) Consortium | Jacquelyn Wynn, Mairead Hyland, The Lighthouse Lab in Alderley Park and Alex Alderton, Roberto Amato, Jeffrey Barrett, Sonia Goncalves, Ewan Harrison, David K. Jackson, Ian Johnston, Dominic Kwiatkowski, Cordelia Langford, John Sillitoe on behalf of the Wellcome Sanger Institute COVID-19 Surveillance Team |
| EPI_ISL_2553230 | Lighthouse Lab in Milton Keynes | Wellcome Sanger Institute for the COVID-19 Genomics UK (COG-UK) Consortium | The Lighthouse Lab in Milton Keynes and Alex Alderton, Roberto Amato, Jeffrey Barrett, Sonia Goncalves, Ewan Harrison, David K. Jackson, Ian Johnston, Dominic Kwiatkowski, Cordelia Langford, John Sillitoe on behalf of the Wellcome Sanger Institute COVID-19 Surveillance Team |
| EPI_ISL_2553231 | Lighthouse Lab in Alderley Park | Wellcome Sanger Institute for the COVID-19 Genomics UK (COG-UK) Consortium | Jacquelyn Wynn, Mairead Hyland, The Lighthouse Lab in Alderley Park and Alex Alderton, Roberto Amato, Jeffrey Barrett, Sonia Goncalves, Ewan Harrison, David K. Jackson, Ian Johnston, Dominic Kwiatkowski, Cordelia Langford, John Sillitoe on behalf of the Wellcome Sanger Institute COVID-19 Surveillance Team |
| EPI_ISL_2553232, EPI_ISL_2553233, EPI_ISL_2553234 | Lighthouse Lab in Glasgow | Wellcome Sanger Institute for the COVID-19 Genomics UK (COG-UK) Consortium | Harper VanSteenhouse, Yumi Kasai, David Gray, Carol Clugston, Anna Dominiczak and Alex Alderton, Roberto Amato, Jeffrey Barrett, Sonia Goncalves, Ewan Harrison, David K. Jackson, Ian Johnston, Dominic Kwiatkowski, Cordelia Langford, John Sillitoe on behalf of the Wellcome Sanger Institute COVID-19 Surveillance Team |
| EPI_ISL_2553235, EPI_ISL_2553237 | Lighthouse Lab in Milton Keynes | Wellcome Sanger Institute for the COVID-19 Genomics UK (COG-UK) Consortium | The Lighthouse Lab in Milton Keynes and Alex Alderton, Roberto Amato, Jeffrey Barrett, Sonia Goncalves, Ewan Harrison, David K. Jackson, Ian Johnston, Dominic Kwiatkowski, Cordelia Langford, John Sillitoe on behalf of the Wellcome Sanger Institute COVID-19 Surveillance Team |

| | | | |
|---|---|---|---|
| EPI_ISL_2553239 | Lighthouse Lab in Alderley Park | Wellcome Sanger Institute for the COVID-19 Genomics UK (COG-UK) Consortium | Jacquelyn Wynn, Mairead Hyland, The Lighthouse Lab in Alderley Park and Alex Alderton, Roberto Amato, Jeffrey Barrett, Sonia Goncalves, Ewan Harrison, David K. Jackson, Ian Johnston, Dominic Kwiatkowski, Cordelia Langford, John Sillitoe on behalf of the Wellcome Sanger Institute COVID-19 Surveillance Team |
| EPI_ISL_2553240, EPI_ISL_2553241 | Lighthouse Lab in Milton Keynes | Wellcome Sanger Institute for the COVID-19 Genomics UK (COG-UK) Consortium | The Lighthouse Lab in Milton Keynes and Alex Alderton, Roberto Amato, Jeffrey Barrett, Sonia Goncalves, Ewan Harrison, David K. Jackson, Ian Johnston, Dominic Kwiatkowski, Cordelia Langford, John Sillitoe on behalf of the Wellcome Sanger Institute COVID-19 Surveillance Team |
| EPI_ISL_2553242 | Lighthouse Lab in Alderley Park | Wellcome Sanger Institute for the COVID-19 Genomics UK (COG-UK) Consortium | Jacquelyn Wynn, Mairead Hyland, The Lighthouse Lab in Alderley Park and Alex Alderton, Roberto Amato, Jeffrey Barrett, Sonia Goncalves, Ewan Harrison, David K. Jackson, Ian Johnston, Dominic Kwiatkowski, Cordelia Langford, John Sillitoe on behalf of the Wellcome Sanger Institute COVID-19 Surveillance Team |
| EPI_ISL_2553243 | Lighthouse Lab in Glasgow | Wellcome Sanger Institute for the COVID-19 Genomics UK (COG-UK) Consortium | Harper VanSteenhouse, Yumi Kasai, David Gray, Carol Clugston, Anna Dominiczak and Alex Alderton, Roberto Amato, Jeffrey Barrett, Sonia Goncalves, Ewan Harrison, David K. Jackson, Ian Johnston, Dominic Kwiatkowski, Cordelia Langford, John Sillitoe on behalf of the Wellcome Sanger Institute COVID-19 Surveillance Team |
| EPI_ISL_2553244, EPI_ISL_2553245, EPI_ISL_2553246 | Lighthouse Lab in Milton Keynes | Wellcome Sanger Institute for the COVID-19 Genomics UK (COG-UK) Consortium | The Lighthouse Lab in Milton Keynes and Alex Alderton, Roberto Amato, Jeffrey Barrett, Sonia Goncalves, Ewan Harrison, David K. Jackson, Ian Johnston, Dominic Kwiatkowski, Cordelia Langford, John Sillitoe on behalf of the Wellcome Sanger Institute COVID-19 Surveillance Team |
| EPI_ISL_2553247 | Lighthouse Lab in Alderley Park | Wellcome Sanger Institute for the COVID-19 Genomics UK (COG-UK) Consortium | Jacquelyn Wynn, Mairead Hyland, The Lighthouse Lab in Alderley Park and Alex Alderton, Roberto Amato, Jeffrey Barrett, Sonia Goncalves, Ewan Harrison, David K. Jackson, Ian Johnston, Dominic Kwiatkowski, Cordelia Langford, John Sillitoe on behalf of the Wellcome Sanger Institute COVID-19 Surveillance Team |
| EPI_ISL_2553248, EPI_ISL_2553249 | Lighthouse Lab in Glasgow | Wellcome Sanger Institute for the COVID-19 Genomics UK (COG-UK) Consortium | Harper VanSteenhouse, Yumi Kasai, David Gray, Carol Clugston, Anna Dominiczak and Alex Alderton, Roberto Amato, Jeffrey Barrett, Sonia Goncalves, Ewan Harrison, David K. Jackson, Ian Johnston, Dominic Kwiatkowski, Cordelia Langford, John Sillitoe on behalf of the Wellcome Sanger Institute COVID-19 Surveillance Team |
| EPI_ISL_2553250 | Lighthouse Lab in Milton Keynes | Wellcome Sanger Institute for the COVID-19 Genomics UK (COG-UK) Consortium | The Lighthouse Lab in Milton Keynes and Alex Alderton, Roberto Amato, Jeffrey Barrett, Sonia Goncalves, Ewan Harrison, David K. Jackson, Ian Johnston, Dominic Kwiatkowski, Cordelia Langford, John Sillitoe on behalf of the Wellcome Sanger Institute COVID-19 Surveillance Team |
| EPI_ISL_2553251, EPI_ISL_2553252, EPI_ISL_2553254 | Lighthouse Lab in Glasgow | Wellcome Sanger Institute for the COVID-19 Genomics UK (COG-UK) Consortium | Harper VanSteenhouse, Yumi Kasai, David Gray, Carol Clugston, Anna Dominiczak and Alex Alderton, Roberto Amato, Jeffrey Barrett, Sonia Goncalves, Ewan Harrison, David K. Jackson, Ian Johnston, Dominic Kwiatkowski, Cordelia Langford, John Sillitoe on behalf of the Wellcome Sanger Institute COVID-19 Surveillance Team |
| EPI_ISL_2553255 | Lighthouse Lab in Alderley Park | Wellcome Sanger Institute for the COVID-19 Genomics UK (COG-UK) Consortium | Jacquelyn Wynn, Mairead Hyland, The Lighthouse Lab in Alderley Park and Alex Alderton, Roberto Amato, Jeffrey Barrett, Sonia Goncalves, Ewan Harrison, David K. Jackson, Ian Johnston, Dominic Kwiatkowski, Cordelia Langford, John Sillitoe on behalf of the Wellcome Sanger Institute COVID-19 Surveillance Team |
| EPI_ISL_2553256 | Lighthouse Lab in Milton Keynes | Wellcome Sanger Institute for the COVID-19 Genomics UK (COG-UK) Consortium | The Lighthouse Lab in Milton Keynes and Alex Alderton, Roberto Amato, Jeffrey Barrett, Sonia Goncalves, Ewan Harrison, David K. Jackson, Ian Johnston, Dominic Kwiatkowski, Cordelia Langford, John Sillitoe on behalf of the Wellcome Sanger Institute COVID-19 Surveillance Team |
| EPI_ISL_2553257, EPI_ISL_2553258, EPI_ISL_2553259, EPI_ISL_2553260, EPI_ISL_2553261 | Lighthouse Lab in Glasgow | Wellcome Sanger Institute for the COVID-19 Genomics UK (COG-UK) Consortium | Harper VanSteenhouse, Yumi Kasai, David Gray, Carol Clugston, Anna Dominiczak and Alex Alderton, Roberto Amato, Jeffrey Barrett, Sonia Goncalves, Ewan Harrison, David K. Jackson, Ian Johnston, Dominic Kwiatkowski, Cordelia Langford, John Sillitoe on behalf of the Wellcome Sanger Institute COVID-19 Surveillance Team |
| EPI_ISL_2553262 | Lighthouse Lab in Milton Keynes | Wellcome Sanger Institute for the COVID-19 Genomics UK (COG-UK) Consortium | The Lighthouse Lab in Milton Keynes and Alex Alderton, Roberto Amato, Jeffrey Barrett, Sonia Goncalves, Ewan Harrison, David K. Jackson, Ian Johnston, Dominic Kwiatkowski, Cordelia Langford, John Sillitoe on behalf of the Wellcome Sanger Institute COVID-19 Surveillance Team |
| EPI_ISL_2553263, EPI_ISL_2553264 | Lighthouse Lab in Alderley Park | Wellcome Sanger Institute for the COVID-19 Genomics UK (COG-UK) Consortium | Jacquelyn Wynn, Mairead Hyland, The Lighthouse Lab in Alderley Park and Alex Alderton, Roberto Amato, Jeffrey Barrett, Sonia Goncalves, Ewan Harrison, David K. Jackson, Ian Johnston, Dominic Kwiatkowski, Cordelia Langford, John Sillitoe on behalf of the Wellcome Sanger Institute COVID-19 Surveillance Team |
| EPI_ISL_2553265 | Lighthouse Lab in Milton Keynes | Wellcome Sanger Institute for the COVID-19 Genomics UK (COG-UK) Consortium | The Lighthouse Lab in Milton Keynes and Alex Alderton, Roberto Amato, Jeffrey Barrett, Sonia Goncalves, Ewan Harrison, David K. Jackson, Ian Johnston, Dominic Kwiatkowski, Cordelia Langford, John Sillitoe on behalf of the Wellcome Sanger Institute COVID-19 Surveillance Team |
| EPI_ISL_2553266, EPI_ISL_2553267 | Lighthouse Lab in Glasgow | Wellcome Sanger Institute for the COVID-19 Genomics UK (COG-UK) Consortium | Harper VanSteenhouse, Yumi Kasai, David Gray, Carol Clugston, Anna Dominiczak and Alex Alderton, Roberto Amato, Jeffrey Barrett, Sonia Goncalves, Ewan Harrison, David K. Jackson, Ian Johnston, Dominic Kwiatkowski, Cordelia Langford, John Sillitoe on behalf of the Wellcome Sanger Institute COVID-19 Surveillance Team |
| EPI_ISL_2553268 | Lighthouse Lab in Alderley Park | Wellcome Sanger Institute for the COVID-19 Genomics UK (COG-UK) Consortium | Jacquelyn Wynn, Mairead Hyland, The Lighthouse Lab in Alderley Park and Alex Alderton, Roberto Amato, Jeffrey Barrett, Sonia Goncalves, Ewan Harrison, David K. Jackson, Ian Johnston, Dominic Kwiatkowski, Cordelia Langford, John Sillitoe on behalf of the Wellcome Sanger Institute COVID-19 Surveillance Team |
| EPI_ISL_2553269 | Lighthouse Lab in Milton Keynes | Wellcome Sanger Institute for the COVID-19 Genomics UK (COG-UK) Consortium | The Lighthouse Lab in Milton Keynes and Alex Alderton, Roberto Amato, Jeffrey Barrett, Sonia Goncalves, Ewan Harrison, David K. Jackson, Ian Johnston, Dominic Kwiatkowski, Cordelia Langford, John Sillitoe on behalf of the Wellcome Sanger Institute COVID-19 Surveillance Team |
| EPI_ISL_2553270 | Lighthouse Lab in Alderley Park | Wellcome Sanger Institute for the COVID-19 Genomics UK (COG-UK) Consortium | Jacquelyn Wynn, Mairead Hyland, The Lighthouse Lab in Alderley Park and Alex Alderton, Roberto Amato, Jeffrey Barrett, Sonia Goncalves, Ewan Harrison, David K. Jackson, Ian Johnston, Dominic Kwiatkowski, Cordelia Langford, John Sillitoe on behalf of the Wellcome Sanger Institute COVID-19 Surveillance Team |
| EPI_ISL_2553271 | Lighthouse Lab in Milton Keynes | Wellcome Sanger Institute for the COVID-19 Genomics UK (COG-UK) Consortium | The Lighthouse Lab in Milton Keynes and Alex Alderton, Roberto Amato, Jeffrey Barrett, Sonia Goncalves, Ewan Harrison, David K. Jackson, Ian Johnston, Dominic Kwiatkowski, Cordelia Langford, John Sillitoe on behalf of the Wellcome Sanger Institute COVID-19 Surveillance Team |
| EPI_ISL_2553272 | Lighthouse Lab in Alderley Park | Wellcome Sanger Institute for the COVID-19 Genomics UK (COG-UK) Consortium | Jacquelyn Wynn, Mairead Hyland, The Lighthouse Lab in Alderley Park and Alex Alderton, Roberto Amato, Jeffrey Barrett, Sonia Goncalves, Ewan Harrison, David K. Jackson, Ian Johnston, Dominic Kwiatkowski, Cordelia Langford, John Sillitoe on behalf of the Wellcome Sanger Institute COVID-19 Surveillance Team |
| EPI_ISL_2553273 | Lighthouse Lab in Milton Keynes | Wellcome Sanger Institute for the COVID-19 Genomics UK (COG-UK) Consortium | The Lighthouse Lab in Milton Keynes and Alex Alderton, Roberto Amato, Jeffrey Barrett, Sonia Goncalves, Ewan Harrison, David K. Jackson, Ian Johnston, Dominic Kwiatkowski, Cordelia Langford, John Sillitoe on behalf of the Wellcome Sanger Institute COVID-19 Surveillance Team |
| EPI_ISL_2553274 | Lighthouse Lab in Alderley Park | Wellcome Sanger Institute for the COVID-19 Genomics UK (COG-UK) Consortium | Jacquelyn Wynn, Mairead Hyland, The Lighthouse Lab in Alderley Park and Alex Alderton, Roberto Amato, Jeffrey Barrett, Sonia Goncalves, Ewan Harrison, David K. Jackson, Ian Johnston, Dominic Kwiatkowski, Cordelia Langford, John Sillitoe on behalf of the Wellcome Sanger Institute COVID-19 Surveillance Team |
| EPI_ISL_2553275 | Lighthouse Lab in Milton Keynes | Wellcome Sanger Institute for the COVID-19 Genomics UK (COG-UK) Consortium | The Lighthouse Lab in Milton Keynes and Alex Alderton, Roberto Amato, Jeffrey Barrett, Sonia Goncalves, Ewan Harrison, David K. Jackson, Ian Johnston, Dominic Kwiatkowski, Cordelia Langford, John Sillitoe on behalf of the Wellcome Sanger Institute COVID-19 Surveillance Team |
| EPI_ISL_2553276, EPI_ISL_2553277 | Lighthouse Lab in Glasgow | Wellcome Sanger Institute for the COVID-19 Genomics UK (COG-UK) Consortium | Harper VanSteenhouse, Yumi Kasai, David Gray, Carol Clugston, Anna Dominiczak and Alex Alderton, Roberto Amato, Jeffrey Barrett, Sonia Goncalves, Ewan Harrison, David K. Jackson, Ian Johnston, Dominic Kwiatkowski, Cordelia Langford, John Sillitoe on behalf of the Wellcome Sanger Institute COVID-19 Surveillance Team |
| EPI_ISL_2553278 | Lighthouse Lab in Alderley Park | Wellcome Sanger Institute for the COVID-19 Genomics UK (COG-UK) Consortium | Jacquelyn Wynn, Mairead Hyland, The Lighthouse Lab in Alderley Park and Alex Alderton, Roberto Amato, Jeffrey Barrett, Sonia Goncalves, Ewan Harrison, David K. Jackson, Ian Johnston, Dominic Kwiatkowski, Cordelia Langford, John Sillitoe on behalf of the Wellcome Sanger Institute COVID-19 Surveillance Team |
| EPI_ISL_2553279, EPI_ISL_2553280, EPI_ISL_2553281 | Lighthouse Lab in Glasgow | Wellcome Sanger Institute for the COVID-19 Genomics UK (COG-UK) Consortium | Harper VanSteenhouse, Yumi Kasai, David Gray, Carol Clugston, Anna Dominiczak and Alex Alderton, Roberto Amato, Jeffrey Barrett, Sonia Goncalves, Ewan Harrison, David K. Jackson, Ian Johnston, Dominic Kwiatkowski, Cordelia Langford, John Sillitoe on behalf of the Wellcome Sanger Institute COVID-19 Surveillance Team |
| EPI_ISL_2553282 | Lighthouse Lab in Alderley Park | Wellcome Sanger Institute for the COVID-19 Genomics UK (COG-UK) Consortium | Jacquelyn Wynn, Mairead Hyland, The Lighthouse Lab in Alderley Park and Alex Alderton, Roberto Amato, Jeffrey Barrett, Sonia Goncalves, Ewan Harrison, David K. Jackson, Ian Johnston, Dominic Kwiatkowski, Cordelia Langford, John Sillitoe on behalf of the Wellcome Sanger Institute COVID-19 Surveillance Team |
| EPI_ISL_2553283 | Lighthouse Lab in Milton Keynes | Wellcome Sanger Institute for the COVID-19 Genomics UK (COG-UK) Consortium | The Lighthouse Lab in Milton Keynes and Alex Alderton, Roberto Amato, Jeffrey Barrett, Sonia Goncalves, Ewan Harrison, David K. Jackson, Ian Johnston, Dominic Kwiatkowski, Cordelia Langford, John Sillitoe on behalf of the Wellcome Sanger Institute COVID-19 Surveillance Team |
| EPI_ISL_2553284, EPI_ISL_2553285 | Lighthouse Lab in Glasgow | Wellcome Sanger Institute for the COVID-19 Genomics UK (COG-UK) Consortium | Harper VanSteenhouse, Yumi Kasai, David Gray, Carol Clugston, Anna Dominiczak and Alex Alderton, Roberto Amato, Jeffrey Barrett, Sonia Goncalves, Ewan Harrison, David K. Jackson, Ian Johnston, Dominic Kwiatkowski, Cordelia Langford, John Sillitoe on behalf of the Wellcome Sanger Institute COVID-19 Surveillance Team |
| EPI_ISL_2553286 | Lighthouse Lab in Milton Keynes | Wellcome Sanger Institute for the COVID-19 Genomics UK (COG-UK) Consortium | The Lighthouse Lab in Milton Keynes and Alex Alderton, Roberto Amato, Jeffrey Barrett, Sonia Goncalves, Ewan Harrison, David K. Jackson, Ian Johnston, Dominic Kwiatkowski, Cordelia Langford, John Sillitoe on behalf of the Wellcome Sanger Institute COVID-19 Surveillance Team |
| EPI_ISL_2553287, EPI_ISL_2553288 | Lighthouse Lab in Glasgow | Wellcome Sanger Institute for the COVID-19 Genomics UK (COG-UK) Consortium | Harper VanSteenhouse, Yumi Kasai, David Gray, Carol Clugston, Anna Dominiczak and Alex Alderton, Roberto Amato, Jeffrey Barrett, Sonia Goncalves, Ewan Harrison, David K. Jackson, Ian Johnston, Dominic Kwiatkowski, Cordelia Langford, John Sillitoe on behalf of the Wellcome Sanger Institute COVID-19 Surveillance Team |
| EPI_ISL_2553289 | Lighthouse Lab in Milton Keynes | Wellcome Sanger Institute for the COVID-19 Genomics UK (COG-UK) Consortium | The Lighthouse Lab in Milton Keynes and Alex Alderton, Roberto Amato, Jeffrey Barrett, Sonia Goncalves, Ewan Harrison, David K. Jackson, Ian Johnston, Dominic Kwiatkowski, Cordelia Langford, John Sillitoe on behalf of the Wellcome Sanger Institute COVID-19 Surveillance Team |
| EPI_ISL_2553290 | Lighthouse Lab in Alderley Park | Wellcome Sanger Institute for the COVID-19 Genomics UK (COG-UK) Consortium | Jacquelyn Wynn, Mairead Hyland, The Lighthouse Lab in Alderley Park and Alex Alderton, Roberto Amato, Jeffrey Barrett, Sonia Goncalves, Ewan Harrison, David K. Jackson, Ian Johnston, Dominic Kwiatkowski, Cordelia Langford, John Sillitoe on behalf of the Wellcome Sanger Institute COVID-19 Surveillance Team |
| EPI_ISL_2553291 | Lighthouse Lab in Glasgow | Wellcome Sanger Institute for the COVID-19 Genomics UK (COG-UK) Consortium | Harper VanSteenhouse, Yumi Kasai, David Gray, Carol Clugston, Anna Dominiczak and Alex Alderton, Roberto Amato, Jeffrey Barrett, Sonia Goncalves, Ewan Harrison, David K. Jackson, Ian Johnston, Dominic Kwiatkowski, Cordelia Langford, John Sillitoe on behalf of the Wellcome Sanger Institute COVID-19 Surveillance Team |
| EPI_ISL_2553292 | Lighthouse Lab in Milton Keynes | Wellcome Sanger Institute for the COVID-19 Genomics UK (COG-UK) Consortium | The Lighthouse Lab in Milton Keynes and Alex Alderton, Roberto Amato, Jeffrey Barrett, Sonia Goncalves, Ewan Harrison, David K. Jackson, Ian Johnston, Dominic Kwiatkowski, Cordelia Langford, John Sillitoe on behalf of the Wellcome Sanger Institute COVID-19 Surveillance Team |
| EPI_ISL_2553293 | Lighthouse Lab in Glasgow | Wellcome Sanger Institute for the COVID-19 Genomics UK (COG-UK) Consortium | Harper VanSteenhouse, Yumi Kasai, David Gray, Carol Clugston, Anna Dominiczak and Alex Alderton, Roberto Amato, Jeffrey Barrett, Sonia Goncalves, Ewan Harrison, David K. Jackson, Ian Johnston, Dominic Kwiatkowski, Cordelia Langford, John Sillitoe on behalf of the Wellcome Sanger Institute COVID-19 Surveillance Team |
| EPI_ISL_2553294 | Lighthouse Lab in Milton Keynes | Wellcome Sanger Institute for the COVID-19 Genomics UK (COG-UK) Consortium | The Lighthouse Lab in Milton Keynes and Alex Alderton, Roberto Amato, Jeffrey Barrett, Sonia Goncalves, Ewan Harrison, David K. Jackson, Ian Johnston, Dominic Kwiatkowski, Cordelia Langford, John Sillitoe on behalf of the Wellcome Sanger Institute COVID-19 Surveillance Team |
| EPI_ISL_2553295 | Lighthouse Lab in Glasgow | Wellcome Sanger Institute for the COVID-19 Genomics UK (COG-UK) Consortium | Harper VanSteenhouse, Yumi Kasai, David Gray, Carol Clugston, Anna Dominiczak and Alex Alderton, Roberto Amato, Jeffrey Barrett, Sonia Goncalves, Ewan Harrison, David K. Jackson, Ian Johnston, Dominic Kwiatkowski, Cordelia Langford, John Sillitoe on behalf of the Wellcome Sanger Institute COVID-19 Surveillance Team |
| EPI_ISL_2553296 | Lighthouse Lab in Milton Keynes | Wellcome Sanger Institute for the COVID-19 Genomics UK (COG-UK) Consortium | The Lighthouse Lab in Milton Keynes and Alex Alderton, Roberto Amato, Jeffrey Barrett, Sonia Goncalves, Ewan Harrison, David K. Jackson, Ian Johnston, Dominic Kwiatkowski, Cordelia Langford, John Sillitoe on behalf of the Wellcome Sanger Institute COVID-19 Surveillance Team |
| EPI_ISL_2553297 | Lighthouse Lab in Glasgow | Wellcome Sanger Institute for the COVID-19 Genomics UK (COG-UK) Consortium | Harper VanSteenhouse, Yumi Kasai, David Gray, Carol Clugston, Anna Dominiczak and Alex Alderton, Roberto Amato, Jeffrey Barrett, Sonia Goncalves, Ewan Harrison, David K. Jackson, Ian Johnston, Dominic Kwiatkowski, Cordelia Langford, John Sillitoe on behalf of the Wellcome Sanger Institute COVID-19 Surveillance Team |
| EPI_ISL_2553298 | Lighthouse Lab in Milton Keynes | Wellcome Sanger Institute for the COVID-19 Genomics UK (COG-UK) Consortium | The Lighthouse Lab in Milton Keynes and Alex Alderton, Roberto Amato, Jeffrey Barrett, Sonia Goncalves, Ewan Harrison, David K. Jackson, Ian Johnston, Dominic Kwiatkowski, Cordelia Langford, John Sillitoe on behalf of the Wellcome Sanger Institute COVID-19 Surveillance Team |
| EPI_ISL_2553299, | Lighthouse Lab in Glasgow | Wellcome Sanger Institute for the COVID-19 Genomics UK (COG-UK) | Harper VanSteenhouse, Yumi Kasai, David Gray, Carol Clugston, Anna Dominiczak and Alex Alderton, Roberto Amato, Jeffrey Barrett, Sonia Goncalves, Ewan Harrison, David K. Jackson, Ian Johnston, Dominic Kwiatkowski, Cordelia Langford, |

| | | | |
|---|---|---|---|
| EPI_ISL_2553300, EPI_ISL_2553301 | | Consortium | John Sillitoe on behalf of the Wellcome Sanger Institute COVID-19 Surveillance Team |
| EPI_ISL_2553302 | Lighthouse Lab in Milton Keynes | Wellcome Sanger Institute for the COVID-19 Genomics UK (COG-UK) Consortium | The Lighthouse Lab in Milton Keynes and Alex Alderton, Roberto Amato, Jeffrey Barrett, Sonia Goncalves, Ewan Harrison, David K. Jackson, Ian Johnston, Dominic Kwiatkowski, Cordelia Langford, John Sillitoe on behalf of the Wellcome Sanger Institute COVID-19 Surveillance Team |
| EPI_ISL_2553303 | Lighthouse Lab in Alderley Park | Wellcome Sanger Institute for the COVID-19 Genomics UK (COG-UK) Consortium | Jacquelyn Wynn, Mairead Hyland, The Lighthouse Lab in Alderley Park and Alex Alderton, Roberto Amato, Jeffrey Barrett, Sonia Goncalves, Ewan Harrison, David K. Jackson, Ian Johnston, Dominic Kwiatkowski, Cordelia Langford, John Sillitoe on behalf of the Wellcome Sanger Institute COVID-19 Surveillance Team |
| EPI_ISL_2553304 | Lighthouse Lab in Milton Keynes | Wellcome Sanger Institute for the COVID-19 Genomics UK (COG-UK) Consortium | The Lighthouse Lab in Milton Keynes and Alex Alderton, Roberto Amato, Jeffrey Barrett, Sonia Goncalves, Ewan Harrison, David K. Jackson, Ian Johnston, Dominic Kwiatkowski, Cordelia Langford, John Sillitoe on behalf of the Wellcome Sanger Institute COVID-19 Surveillance Team |
| EPI_ISL_2553305 | Lighthouse Lab in Glasgow | Wellcome Sanger Institute for the COVID-19 Genomics UK (COG-UK) Consortium | Harper VanSteenhouse, Yumi Kasai, David Gray, Carol Clugston, Anna Dominiczak and Alex Alderton, Roberto Amato, Jeffrey Barrett, Sonia Goncalves, Ewan Harrison, David K. Jackson, Ian Johnston, Dominic Kwiatkowski, Cordelia Langford, John Sillitoe on behalf of the Wellcome Sanger Institute COVID-19 Surveillance Team |
| EPI_ISL_2553306 | Lighthouse Lab in Alderley Park | Wellcome Sanger Institute for the COVID-19 Genomics UK (COG-UK) Consortium | Jacquelyn Wynn, Mairead Hyland, The Lighthouse Lab in Alderley Park and Alex Alderton, Roberto Amato, Jeffrey Barrett, Sonia Goncalves, Ewan Harrison, David K. Jackson, Ian Johnston, Dominic Kwiatkowski, Cordelia Langford, John Sillitoe on behalf of the Wellcome Sanger Institute COVID-19 Surveillance Team |
| EPI_ISL_2553307, EPI_ISL_2553308 | Lighthouse Lab in Glasgow | Wellcome Sanger Institute for the COVID-19 Genomics UK (COG-UK) Consortium | Harper VanSteenhouse, Yumi Kasai, David Gray, Carol Clugston, Anna Dominiczak and Alex Alderton, Roberto Amato, Jeffrey Barrett, Sonia Goncalves, Ewan Harrison, David K. Jackson, Ian Johnston, Dominic Kwiatkowski, Cordelia Langford, John Sillitoe on behalf of the Wellcome Sanger Institute COVID-19 Surveillance Team |
| EPI_ISL_2553309 | Lighthouse Lab in Alderley Park | Wellcome Sanger Institute for the COVID-19 Genomics UK (COG-UK) Consortium | Jacquelyn Wynn, Mairead Hyland, The Lighthouse Lab in Alderley Park and Alex Alderton, Roberto Amato, Jeffrey Barrett, Sonia Goncalves, Ewan Harrison, David K. Jackson, Ian Johnston, Dominic Kwiatkowski, Cordelia Langford, John Sillitoe on behalf of the Wellcome Sanger Institute COVID-19 Surveillance Team |
| EPI_ISL_2553310, EPI_ISL_2553311 | Lighthouse Lab in Glasgow | Wellcome Sanger Institute for the COVID-19 Genomics UK (COG-UK) Consortium | Harper VanSteenhouse, Yumi Kasai, David Gray, Carol Clugston, Anna Dominiczak and Alex Alderton, Roberto Amato, Jeffrey Barrett, Sonia Goncalves, Ewan Harrison, David K. Jackson, Ian Johnston, Dominic Kwiatkowski, Cordelia Langford, John Sillitoe on behalf of the Wellcome Sanger Institute COVID-19 Surveillance Team |
| EPI_ISL_2553312, EPI_ISL_2553313 | Lighthouse Lab in Milton Keynes | Wellcome Sanger Institute for the COVID-19 Genomics UK (COG-UK) Consortium | The Lighthouse Lab in Milton Keynes and Alex Alderton, Roberto Amato, Jeffrey Barrett, Sonia Goncalves, Ewan Harrison, David K. Jackson, Ian Johnston, Dominic Kwiatkowski, Cordelia Langford, John Sillitoe on behalf of the Wellcome Sanger Institute COVID-19 Surveillance Team |
| EPI_ISL_2553314 | Lighthouse Lab in Alderley Park | Wellcome Sanger Institute for the COVID-19 Genomics UK (COG-UK) Consortium | Jacquelyn Wynn, Mairead Hyland, The Lighthouse Lab in Alderley Park and Alex Alderton, Roberto Amato, Jeffrey Barrett, Sonia Goncalves, Ewan Harrison, David K. Jackson, Ian Johnston, Dominic Kwiatkowski, Cordelia Langford, John Sillitoe on behalf of the Wellcome Sanger Institute COVID-19 Surveillance Team |
| EPI_ISL_2553315 | Lighthouse Lab in Milton Keynes | Wellcome Sanger Institute for the COVID-19 Genomics UK (COG-UK) Consortium | The Lighthouse Lab in Milton Keynes and Alex Alderton, Roberto Amato, Jeffrey Barrett, Sonia Goncalves, Ewan Harrison, David K. Jackson, Ian Johnston, Dominic Kwiatkowski, Cordelia Langford, John Sillitoe on behalf of the Wellcome Sanger Institute COVID-19 Surveillance Team |
| EPI_ISL_2553316 | Lighthouse Lab in Glasgow | Wellcome Sanger Institute for the COVID-19 Genomics UK (COG-UK) Consortium | Harper VanSteenhouse, Yumi Kasai, David Gray, Carol Clugston, Anna Dominiczak and Alex Alderton, Roberto Amato, Jeffrey Barrett, Sonia Goncalves, Ewan Harrison, David K. Jackson, Ian Johnston, Dominic Kwiatkowski, Cordelia Langford, John Sillitoe on behalf of the Wellcome Sanger Institute COVID-19 Surveillance Team |
| EPI_ISL_2553318 | Lighthouse Lab in Alderley Park | Wellcome Sanger Institute for the COVID-19 Genomics UK (COG-UK) Consortium | Jacquelyn Wynn, Mairead Hyland, The Lighthouse Lab in Alderley Park and Alex Alderton, Roberto Amato, Jeffrey Barrett, Sonia Goncalves, Ewan Harrison, David K. Jackson, Ian Johnston, Dominic Kwiatkowski, Cordelia Langford, John Sillitoe on behalf of the Wellcome Sanger Institute COVID-19 Surveillance Team |
| EPI_ISL_2553319 | Lighthouse Lab in Glasgow | Wellcome Sanger Institute for the COVID-19 Genomics UK (COG-UK) Consortium | Harper VanSteenhouse, Yumi Kasai, David Gray, Carol Clugston, Anna Dominiczak and Alex Alderton, Roberto Amato, Jeffrey Barrett, Sonia Goncalves, Ewan Harrison, David K. Jackson, Ian Johnston, Dominic Kwiatkowski, Cordelia Langford, John Sillitoe on behalf of the Wellcome Sanger Institute COVID-19 Surveillance Team |
| EPI_ISL_2553320 | Lighthouse Lab in Milton Keynes | Wellcome Sanger Institute for the COVID-19 Genomics UK (COG-UK) Consortium | The Lighthouse Lab in Milton Keynes and Alex Alderton, Roberto Amato, Jeffrey Barrett, Sonia Goncalves, Ewan Harrison, David K. Jackson, Ian Johnston, Dominic Kwiatkowski, Cordelia Langford, John Sillitoe on behalf of the Wellcome Sanger Institute COVID-19 Surveillance Team |
| EPI_ISL_2553321 | Lighthouse Lab in Alderley Park | Wellcome Sanger Institute for the COVID-19 Genomics UK (COG-UK) Consortium | Jacquelyn Wynn, Mairead Hyland, The Lighthouse Lab in Alderley Park and Alex Alderton, Roberto Amato, Jeffrey Barrett, Sonia Goncalves, Ewan Harrison, David K. Jackson, Ian Johnston, Dominic Kwiatkowski, Cordelia Langford, John Sillitoe on behalf of the Wellcome Sanger Institute COVID-19 Surveillance Team |
| EPI_ISL_2553322 | Lighthouse Lab in Milton Keynes | Wellcome Sanger Institute for the COVID-19 Genomics UK (COG-UK) Consortium | The Lighthouse Lab in Milton Keynes and Alex Alderton, Roberto Amato, Jeffrey Barrett, Sonia Goncalves, Ewan Harrison, David K. Jackson, Ian Johnston, Dominic Kwiatkowski, Cordelia Langford, John Sillitoe on behalf of the Wellcome Sanger Institute COVID-19 Surveillance Team |
| EPI_ISL_2553323, EPI_ISL_2553324 | Lighthouse Lab in Glasgow | Wellcome Sanger Institute for the COVID-19 Genomics UK (COG-UK) Consortium | Harper VanSteenhouse, Yumi Kasai, David Gray, Carol Clugston, Anna Dominiczak and Alex Alderton, Roberto Amato, Jeffrey Barrett, Sonia Goncalves, Ewan Harrison, David K. Jackson, Ian Johnston, Dominic Kwiatkowski, Cordelia Langford, John Sillitoe on behalf of the Wellcome Sanger Institute COVID-19 Surveillance Team |
| EPI_ISL_2553325 | Lighthouse Lab in Milton Keynes | Wellcome Sanger Institute for the COVID-19 Genomics UK (COG-UK) Consortium | The Lighthouse Lab in Milton Keynes and Alex Alderton, Roberto Amato, Jeffrey Barrett, Sonia Goncalves, Ewan Harrison, David K. Jackson, Ian Johnston, Dominic Kwiatkowski, Cordelia Langford, John Sillitoe on behalf of the Wellcome Sanger Institute COVID-19 Surveillance Team |
| EPI_ISL_2553326, EPI_ISL_2553328, EPI_ISL_2553330, EPI_ISL_2553331 | Lighthouse Lab in Glasgow | Wellcome Sanger Institute for the COVID-19 Genomics UK (COG-UK) Consortium | Harper VanSteenhouse, Yumi Kasai, David Gray, Carol Clugston, Anna Dominiczak and Alex Alderton, Roberto Amato, Jeffrey Barrett, Sonia Goncalves, Ewan Harrison, David K. Jackson, Ian Johnston, Dominic Kwiatkowski, Cordelia Langford, John Sillitoe on behalf of the Wellcome Sanger Institute COVID-19 Surveillance Team |
| EPI_ISL_2553332, EPI_ISL_2553333 | Lighthouse Lab in Alderley Park | Wellcome Sanger Institute for the COVID-19 Genomics UK (COG-UK) Consortium | Jacquelyn Wynn, Mairead Hyland, The Lighthouse Lab in Alderley Park and Alex Alderton, Roberto Amato, Jeffrey Barrett, Sonia Goncalves, Ewan Harrison, David K. Jackson, Ian Johnston, Dominic Kwiatkowski, Cordelia Langford, John Sillitoe on behalf of the Wellcome Sanger Institute COVID-19 Surveillance Team |
| EPI_ISL_2553335, EPI_ISL_2553336 | Lighthouse Lab in Glasgow | Wellcome Sanger Institute for the COVID-19 Genomics UK (COG-UK) Consortium | Harper VanSteenhouse, Yumi Kasai, David Gray, Carol Clugston, Anna Dominiczak and Alex Alderton, Roberto Amato, Jeffrey Barrett, Sonia Goncalves, Ewan Harrison, David K. Jackson, Ian Johnston, Dominic Kwiatkowski, Cordelia Langford, John Sillitoe on behalf of the Wellcome Sanger Institute COVID-19 Surveillance Team |
| EPI_ISL_2553337 | Lighthouse Lab in Alderley Park | Wellcome Sanger Institute for the COVID-19 Genomics UK (COG-UK) Consortium | Jacquelyn Wynn, Mairead Hyland, The Lighthouse Lab in Alderley Park and Alex Alderton, Roberto Amato, Jeffrey Barrett, Sonia Goncalves, Ewan Harrison, David K. Jackson, Ian Johnston, Dominic Kwiatkowski, Cordelia Langford, John Sillitoe on behalf of the Wellcome Sanger Institute COVID-19 Surveillance Team |
| EPI_ISL_2553338, EPI_ISL_2553339 | Lighthouse Lab in Milton Keynes | Wellcome Sanger Institute for the COVID-19 Genomics UK (COG-UK) Consortium | The Lighthouse Lab in Milton Keynes and Alex Alderton, Roberto Amato, Jeffrey Barrett, Sonia Goncalves, Ewan Harrison, David K. Jackson, Ian Johnston, Dominic Kwiatkowski, Cordelia Langford, John Sillitoe on behalf of the Wellcome Sanger Institute COVID-19 Surveillance Team |
| EPI_ISL_2553340 | Lighthouse Lab in Alderley Park | Wellcome Sanger Institute for the COVID-19 Genomics UK (COG-UK) Consortium | Jacquelyn Wynn, Mairead Hyland, The Lighthouse Lab in Alderley Park and Alex Alderton, Roberto Amato, Jeffrey Barrett, Sonia Goncalves, Ewan Harrison, David K. Jackson, Ian Johnston, Dominic Kwiatkowski, Cordelia Langford, John Sillitoe on behalf of the Wellcome Sanger Institute COVID-19 Surveillance Team |
| EPI_ISL_2553341, EPI_ISL_2553342, EPI_ISL_2553343, EPI_ISL_2553344 | Lighthouse Lab in Glasgow | Wellcome Sanger Institute for the COVID-19 Genomics UK (COG-UK) Consortium | Harper VanSteenhouse, Yumi Kasai, David Gray, Carol Clugston, Anna Dominiczak and Alex Alderton, Roberto Amato, Jeffrey Barrett, Sonia Goncalves, Ewan Harrison, David K. Jackson, Ian Johnston, Dominic Kwiatkowski, Cordelia Langford, John Sillitoe on behalf of the Wellcome Sanger Institute COVID-19 Surveillance Team |
| EPI_ISL_2553345, EPI_ISL_2553346 | Lighthouse Lab in Alderley Park | Wellcome Sanger Institute for the COVID-19 Genomics UK (COG-UK) Consortium | Jacquelyn Wynn, Mairead Hyland, The Lighthouse Lab in Alderley Park and Alex Alderton, Roberto Amato, Jeffrey Barrett, Sonia Goncalves, Ewan Harrison, David K. Jackson, Ian Johnston, Dominic Kwiatkowski, Cordelia Langford, John Sillitoe on behalf of the Wellcome Sanger Institute COVID-19 Surveillance Team |
| EPI_ISL_2553347 | Lighthouse Lab in Milton Keynes | Wellcome Sanger Institute for the COVID-19 Genomics UK (COG-UK) Consortium | The Lighthouse Lab in Milton Keynes and Alex Alderton, Roberto Amato, Jeffrey Barrett, Sonia Goncalves, Ewan Harrison, David K. Jackson, Ian Johnston, Dominic Kwiatkowski, Cordelia Langford, John Sillitoe on behalf of the Wellcome Sanger Institute COVID-19 Surveillance Team |
| EPI_ISL_2553348, EPI_ISL_2553349 | Lighthouse Lab in Glasgow | Wellcome Sanger Institute for the COVID-19 Genomics UK (COG-UK) Consortium | Harper VanSteenhouse, Yumi Kasai, David Gray, Carol Clugston, Anna Dominiczak and Alex Alderton, Roberto Amato, Jeffrey Barrett, Sonia Goncalves, Ewan Harrison, David K. Jackson, Ian Johnston, Dominic Kwiatkowski, Cordelia Langford, John Sillitoe on behalf of the Wellcome Sanger Institute COVID-19 Surveillance Team |
| EPI_ISL_2553350 | Lighthouse Lab in Alderley Park | Wellcome Sanger Institute for the COVID-19 Genomics UK (COG-UK) Consortium | Jacquelyn Wynn, Mairead Hyland, The Lighthouse Lab in Alderley Park and Alex Alderton, Roberto Amato, Jeffrey Barrett, Sonia Goncalves, Ewan Harrison, David K. Jackson, Ian Johnston, Dominic Kwiatkowski, Cordelia Langford, John Sillitoe on behalf of the Wellcome Sanger Institute COVID-19 Surveillance Team |
| EPI_ISL_2553351 | Lighthouse Lab in Milton Keynes | Wellcome Sanger Institute for the COVID-19 Genomics UK (COG-UK) Consortium | The Lighthouse Lab in Milton Keynes and Alex Alderton, Roberto Amato, Jeffrey Barrett, Sonia Goncalves, Ewan Harrison, David K. Jackson, Ian Johnston, Dominic Kwiatkowski, Cordelia Langford, John Sillitoe on behalf of the Wellcome Sanger Institute COVID-19 Surveillance Team |
| EPI_ISL_2553352 | Lighthouse Lab in Alderley Park | Wellcome Sanger Institute for the COVID-19 Genomics UK (COG-UK) Consortium | Jacquelyn Wynn, Mairead Hyland, The Lighthouse Lab in Alderley Park and Alex Alderton, Roberto Amato, Jeffrey Barrett, Sonia Goncalves, Ewan Harrison, David K. Jackson, Ian Johnston, Dominic Kwiatkowski, Cordelia Langford, John Sillitoe on behalf of the Wellcome Sanger Institute COVID-19 Surveillance Team |
| EPI_ISL_2553353, EPI_ISL_2553354 | Lighthouse Lab in Glasgow | Wellcome Sanger Institute for the COVID-19 Genomics UK (COG-UK) Consortium | Harper VanSteenhouse, Yumi Kasai, David Gray, Carol Clugston, Anna Dominiczak and Alex Alderton, Roberto Amato, Jeffrey Barrett, Sonia Goncalves, Ewan Harrison, David K. Jackson, Ian Johnston, Dominic Kwiatkowski, Cordelia Langford, John Sillitoe on behalf of the Wellcome Sanger Institute COVID-19 Surveillance Team |
| EPI_ISL_2553356 | Lighthouse Lab in Milton Keynes | Wellcome Sanger Institute for the COVID-19 Genomics UK (COG-UK) Consortium | The Lighthouse Lab in Milton Keynes and Alex Alderton, Roberto Amato, Jeffrey Barrett, Sonia Goncalves, Ewan Harrison, David K. Jackson, Ian Johnston, Dominic Kwiatkowski, Cordelia Langford, John Sillitoe on behalf of the Wellcome Sanger Institute COVID-19 Surveillance Team |
| EPI_ISL_2553357, EPI_ISL_2553358 | Lighthouse Lab in Glasgow | Wellcome Sanger Institute for the COVID-19 Genomics UK (COG-UK) Consortium | Harper VanSteenhouse, Yumi Kasai, David Gray, Carol Clugston, Anna Dominiczak and Alex Alderton, Roberto Amato, Jeffrey Barrett, Sonia Goncalves, Ewan Harrison, David K. Jackson, Ian Johnston, Dominic Kwiatkowski, Cordelia Langford, John Sillitoe on behalf of the Wellcome Sanger Institute COVID-19 Surveillance Team |
| EPI_ISL_2553359, EPI_ISL_2553360, EPI_ISL_2553361, EPI_ISL_2553362 | Lighthouse Lab in Milton Keynes | Wellcome Sanger Institute for the COVID-19 Genomics UK (COG-UK) Consortium | The Lighthouse Lab in Milton Keynes and Alex Alderton, Roberto Amato, Jeffrey Barrett, Sonia Goncalves, Ewan Harrison, David K. Jackson, Ian Johnston, Dominic Kwiatkowski, Cordelia Langford, John Sillitoe on behalf of the Wellcome Sanger Institute COVID-19 Surveillance Team |
| EPI_ISL_2553363, EPI_ISL_2553364, EPI_ISL_2553365, EPI_ISL_2553366, EPI_ISL_2553367 | Lighthouse Lab in Glasgow | Wellcome Sanger Institute for the COVID-19 Genomics UK (COG-UK) Consortium | Harper VanSteenhouse, Yumi Kasai, David Gray, Carol Clugston, Anna Dominiczak and Alex Alderton, Roberto Amato, Jeffrey Barrett, Sonia Goncalves, Ewan Harrison, David K. Jackson, Ian Johnston, Dominic Kwiatkowski, Cordelia Langford, John Sillitoe on behalf of the Wellcome Sanger Institute COVID-19 Surveillance Team |
| EPI_ISL_2553368 | Lighthouse Lab in Alderley Park | Wellcome Sanger Institute for the COVID-19 Genomics UK (COG-UK) Consortium | Jacquelyn Wynn, Mairead Hyland, The Lighthouse Lab in Alderley Park and Alex Alderton, Roberto Amato, Jeffrey Barrett, Sonia Goncalves, Ewan Harrison, David K. Jackson, Ian Johnston, Dominic Kwiatkowski, Cordelia Langford, John Sillitoe on behalf of the Wellcome Sanger Institute COVID-19 Surveillance Team |
| EPI_ISL_2553369, EPI_ISL_2553370 | Lighthouse Lab in Glasgow | Wellcome Sanger Institute for the COVID-19 Genomics UK (COG-UK) Consortium | Harper VanSteenhouse, Yumi Kasai, David Gray, Carol Clugston, Anna Dominiczak and Alex Alderton, Roberto Amato, Jeffrey Barrett, Sonia Goncalves, Ewan Harrison, David K. Jackson, Ian Johnston, Dominic Kwiatkowski, Cordelia Langford, John Sillitoe on behalf of the Wellcome Sanger Institute COVID-19 Surveillance Team |
| EPI_ISL_2553371 | Lighthouse Lab in Alderley Park | Wellcome Sanger Institute for the COVID-19 Genomics UK (COG-UK) Consortium | Jacquelyn Wynn, Mairead Hyland, The Lighthouse Lab in Alderley Park and Alex Alderton, Roberto Amato, Jeffrey Barrett, Sonia Goncalves, Ewan Harrison, David K. Jackson, Ian Johnston, Dominic Kwiatkowski, Cordelia Langford, John Sillitoe on behalf of the Wellcome Sanger Institute COVID-19 Surveillance Team |
| EPI_ISL_2553372 | Lighthouse Lab in Glasgow | Wellcome Sanger Institute for the COVID-19 Genomics UK (COG-UK) | Harper VanSteenhouse, Yumi Kasai, David Gray, Carol Clugston, Anna Dominiczak and Alex Alderton, Roberto Amato, Jeffrey Barrett, Sonia Goncalves, Ewan Harrison, David K. Jackson, Ian Johnston, Dominic Kwiatkowski, Cordelia Langford, |

| | | | |
|---|---|---|---|
| | | Consortium | John Sillitoe on behalf of the Wellcome Sanger Institute COVID-19 Surveillance Team |
| EPI_ISL_2553373 | Lighthouse Lab in Alderley Park | Wellcome Sanger Institute for the COVID-19 Genomics UK (COG-UK) Consortium | Jacquelyn Wynn, Mairead Hyland, The Lighthouse Lab in Alderley Park and Alex Alderton, Roberto Amato, Jeffrey Barrett, Sonia Goncalves, Ewan Harrison, David K. Jackson, Ian Johnston, Dominic Kwiatkowski, Cordelia Langford, John Sillitoe on behalf of the Wellcome Sanger Institute COVID-19 Surveillance Team |
| EPI_ISL_2553374 | Lighthouse Lab in Glasgow | Wellcome Sanger Institute for the COVID-19 Genomics UK (COG-UK) Consortium | Harper VanSteenhouse, Yumi Kasai, David Gray, Carol Clugston, Anna Dominiczak and Alex Alderton, Roberto Amato, Jeffrey Barrett, Sonia Goncalves, Ewan Harrison, David K. Jackson, Ian Johnston, Dominic Kwiatkowski, Cordelia Langford, John Sillitoe on behalf of the Wellcome Sanger Institute COVID-19 Surveillance Team |
| EPI_ISL_2553375 | Lighthouse Lab in Alderley Park | Wellcome Sanger Institute for the COVID-19 Genomics UK (COG-UK) Consortium | Jacquelyn Wynn, Mairead Hyland, The Lighthouse Lab in Alderley Park and Alex Alderton, Roberto Amato, Jeffrey Barrett, Sonia Goncalves, Ewan Harrison, David K. Jackson, Ian Johnston, Dominic Kwiatkowski, Cordelia Langford, John Sillitoe on behalf of the Wellcome Sanger Institute COVID-19 Surveillance Team |
| EPI_ISL_2553376, EPI_ISL_2553377, EPI_ISL_2553378 | Lighthouse Lab in Glasgow | Wellcome Sanger Institute for the COVID-19 Genomics UK (COG-UK) Consortium | Harper VanSteenhouse, Yumi Kasai, David Gray, Carol Clugston, Anna Dominiczak and Alex Alderton, Roberto Amato, Jeffrey Barrett, Sonia Goncalves, Ewan Harrison, David K. Jackson, Ian Johnston, Dominic Kwiatkowski, Cordelia Langford, John Sillitoe on behalf of the Wellcome Sanger Institute COVID-19 Surveillance Team |
| EPI_ISL_2553379, EPI_ISL_2553380 | Lighthouse Lab in Milton Keynes | Wellcome Sanger Institute for the COVID-19 Genomics UK (COG-UK) Consortium | The Lighthouse Lab in Milton Keynes and Alex Alderton, Roberto Amato, Jeffrey Barrett, Sonia Goncalves, Ewan Harrison, David K. Jackson, Ian Johnston, Dominic Kwiatkowski, Cordelia Langford, John Sillitoe on behalf of the Wellcome Sanger Institute COVID-19 Surveillance Team |
| EPI_ISL_2553381 | Lighthouse Lab in Alderley Park | Wellcome Sanger Institute for the COVID-19 Genomics UK (COG-UK) Consortium | Jacquelyn Wynn, Mairead Hyland, The Lighthouse Lab in Alderley Park and Alex Alderton, Roberto Amato, Jeffrey Barrett, Sonia Goncalves, Ewan Harrison, David K. Jackson, Ian Johnston, Dominic Kwiatkowski, Cordelia Langford, John Sillitoe on behalf of the Wellcome Sanger Institute COVID-19 Surveillance Team |
| EPI_ISL_2553382 | Lighthouse Lab in Glasgow | Wellcome Sanger Institute for the COVID-19 Genomics UK (COG-UK) Consortium | Harper VanSteenhouse, Yumi Kasai, David Gray, Carol Clugston, Anna Dominiczak and Alex Alderton, Roberto Amato, Jeffrey Barrett, Sonia Goncalves, Ewan Harrison, David K. Jackson, Ian Johnston, Dominic Kwiatkowski, Cordelia Langford, John Sillitoe on behalf of the Wellcome Sanger Institute COVID-19 Surveillance Team |
| EPI_ISL_2553384 | Lighthouse Lab in Milton Keynes | Wellcome Sanger Institute for the COVID-19 Genomics UK (COG-UK) Consortium | The Lighthouse Lab in Milton Keynes and Alex Alderton, Roberto Amato, Jeffrey Barrett, Sonia Goncalves, Ewan Harrison, David K. Jackson, Ian Johnston, Dominic Kwiatkowski, Cordelia Langford, John Sillitoe on behalf of the Wellcome Sanger Institute COVID-19 Surveillance Team |
| EPI_ISL_2553386 | Lighthouse Lab in Alderley Park | Wellcome Sanger Institute for the COVID-19 Genomics UK (COG-UK) Consortium | Jacquelyn Wynn, Mairead Hyland, The Lighthouse Lab in Alderley Park and Alex Alderton, Roberto Amato, Jeffrey Barrett, Sonia Goncalves, Ewan Harrison, David K. Jackson, Ian Johnston, Dominic Kwiatkowski, Cordelia Langford, John Sillitoe on behalf of the Wellcome Sanger Institute COVID-19 Surveillance Team |
| EPI_ISL_2553387 | Lighthouse Lab in Glasgow | Wellcome Sanger Institute for the COVID-19 Genomics UK (COG-UK) Consortium | Harper VanSteenhouse, Yumi Kasai, David Gray, Carol Clugston, Anna Dominiczak and Alex Alderton, Roberto Amato, Jeffrey Barrett, Sonia Goncalves, Ewan Harrison, David K. Jackson, Ian Johnston, Dominic Kwiatkowski, Cordelia Langford, John Sillitoe on behalf of the Wellcome Sanger Institute COVID-19 Surveillance Team |
| EPI_ISL_2553388, EPI_ISL_2553389 | Lighthouse Lab in Milton Keynes | Wellcome Sanger Institute for the COVID-19 Genomics UK (COG-UK) Consortium | The Lighthouse Lab in Milton Keynes and Alex Alderton, Roberto Amato, Jeffrey Barrett, Sonia Goncalves, Ewan Harrison, David K. Jackson, Ian Johnston, Dominic Kwiatkowski, Cordelia Langford, John Sillitoe on behalf of the Wellcome Sanger Institute COVID-19 Surveillance Team |
| EPI_ISL_2553391 | Lighthouse Lab in Alderley Park | Wellcome Sanger Institute for the COVID-19 Genomics UK (COG-UK) Consortium | Jacquelyn Wynn, Mairead Hyland, The Lighthouse Lab in Alderley Park and Alex Alderton, Roberto Amato, Jeffrey Barrett, Sonia Goncalves, Ewan Harrison, David K. Jackson, Ian Johnston, Dominic Kwiatkowski, Cordelia Langford, John Sillitoe on behalf of the Wellcome Sanger Institute COVID-19 Surveillance Team |
| EPI_ISL_2553392, EPI_ISL_2553394 | Lighthouse Lab in Glasgow | Wellcome Sanger Institute for the COVID-19 Genomics UK (COG-UK) Consortium | Harper VanSteenhouse, Yumi Kasai, David Gray, Carol Clugston, Anna Dominiczak and Alex Alderton, Roberto Amato, Jeffrey Barrett, Sonia Goncalves, Ewan Harrison, David K. Jackson, Ian Johnston, Dominic Kwiatkowski, Cordelia Langford, John Sillitoe on behalf of the Wellcome Sanger Institute COVID-19 Surveillance Team |
| EPI_ISL_2553396, EPI_ISL_2553397, EPI_ISL_2553398, EPI_ISL_2553399 | Lighthouse Lab in Milton Keynes | Wellcome Sanger Institute for the COVID-19 Genomics UK (COG-UK) Consortium | The Lighthouse Lab in Milton Keynes and Alex Alderton, Roberto Amato, Jeffrey Barrett, Sonia Goncalves, Ewan Harrison, David K. Jackson, Ian Johnston, Dominic Kwiatkowski, Cordelia Langford, John Sillitoe on behalf of the Wellcome Sanger Institute COVID-19 Surveillance Team |
| EPI_ISL_2553400, EPI_ISL_2553401, EPI_ISL_2553402 | Lighthouse Lab in Glasgow | Wellcome Sanger Institute for the COVID-19 Genomics UK (COG-UK) Consortium | Harper VanSteenhouse, Yumi Kasai, David Gray, Carol Clugston, Anna Dominiczak and Alex Alderton, Roberto Amato, Jeffrey Barrett, Sonia Goncalves, Ewan Harrison, David K. Jackson, Ian Johnston, Dominic Kwiatkowski, Cordelia Langford, John Sillitoe on behalf of the Wellcome Sanger Institute COVID-19 Surveillance Team |
| EPI_ISL_2553403 | Lighthouse Lab in Alderley Park | Wellcome Sanger Institute for the COVID-19 Genomics UK (COG-UK) Consortium | Jacquelyn Wynn, Mairead Hyland, The Lighthouse Lab in Alderley Park and Alex Alderton, Roberto Amato, Jeffrey Barrett, Sonia Goncalves, Ewan Harrison, David K. Jackson, Ian Johnston, Dominic Kwiatkowski, Cordelia Langford, John Sillitoe on behalf of the Wellcome Sanger Institute COVID-19 Surveillance Team |
| EPI_ISL_2553404 | Lighthouse Lab in Glasgow | Wellcome Sanger Institute for the COVID-19 Genomics UK (COG-UK) Consortium | Harper VanSteenhouse, Yumi Kasai, David Gray, Carol Clugston, Anna Dominiczak and Alex Alderton, Roberto Amato, Jeffrey Barrett, Sonia Goncalves, Ewan Harrison, David K. Jackson, Ian Johnston, Dominic Kwiatkowski, Cordelia Langford, John Sillitoe on behalf of the Wellcome Sanger Institute COVID-19 Surveillance Team |
| EPI_ISL_2553405 | Lighthouse Lab in Alderley Park | Wellcome Sanger Institute for the COVID-19 Genomics UK (COG-UK) Consortium | Jacquelyn Wynn, Mairead Hyland, The Lighthouse Lab in Alderley Park and Alex Alderton, Roberto Amato, Jeffrey Barrett, Sonia Goncalves, Ewan Harrison, David K. Jackson, Ian Johnston, Dominic Kwiatkowski, Cordelia Langford, John Sillitoe on behalf of the Wellcome Sanger Institute COVID-19 Surveillance Team |
| EPI_ISL_2553406, EPI_ISL_2553407 | Lighthouse Lab in Glasgow | Wellcome Sanger Institute for the COVID-19 Genomics UK (COG-UK) Consortium | Harper VanSteenhouse, Yumi Kasai, David Gray, Carol Clugston, Anna Dominiczak and Alex Alderton, Roberto Amato, Jeffrey Barrett, Sonia Goncalves, Ewan Harrison, David K. Jackson, Ian Johnston, Dominic Kwiatkowski, Cordelia Langford, John Sillitoe on behalf of the Wellcome Sanger Institute COVID-19 Surveillance Team |
| EPI_ISL_2553408 | Lighthouse Lab in Alderley Park | Wellcome Sanger Institute for the COVID-19 Genomics UK (COG-UK) Consortium | Jacquelyn Wynn, Mairead Hyland, The Lighthouse Lab in Alderley Park and Alex Alderton, Roberto Amato, Jeffrey Barrett, Sonia Goncalves, Ewan Harrison, David K. Jackson, Ian Johnston, Dominic Kwiatkowski, Cordelia Langford, John Sillitoe on behalf of the Wellcome Sanger Institute COVID-19 Surveillance Team |
| EPI_ISL_2553409 | Lighthouse Lab in Milton Keynes | Wellcome Sanger Institute for the COVID-19 Genomics UK (COG-UK) Consortium | The Lighthouse Lab in Milton Keynes and Alex Alderton, Roberto Amato, Jeffrey Barrett, Sonia Goncalves, Ewan Harrison, David K. Jackson, Ian Johnston, Dominic Kwiatkowski, Cordelia Langford, John Sillitoe on behalf of the Wellcome Sanger Institute COVID-19 Surveillance Team |
| EPI_ISL_2553410 | Lighthouse Lab in Alderley Park | Wellcome Sanger Institute for the COVID-19 Genomics UK (COG-UK) Consortium | Jacquelyn Wynn, Mairead Hyland, The Lighthouse Lab in Alderley Park and Alex Alderton, Roberto Amato, Jeffrey Barrett, Sonia Goncalves, Ewan Harrison, David K. Jackson, Ian Johnston, Dominic Kwiatkowski, Cordelia Langford, John Sillitoe on behalf of the Wellcome Sanger Institute COVID-19 Surveillance Team |
| EPI_ISL_2553411 | Lighthouse Lab in Glasgow | Wellcome Sanger Institute for the COVID-19 Genomics UK (COG-UK) Consortium | Harper VanSteenhouse, Yumi Kasai, David Gray, Carol Clugston, Anna Dominiczak and Alex Alderton, Roberto Amato, Jeffrey Barrett, Sonia Goncalves, Ewan Harrison, David K. Jackson, Ian Johnston, Dominic Kwiatkowski, Cordelia Langford, John Sillitoe on behalf of the Wellcome Sanger Institute COVID-19 Surveillance Team |
| EPI_ISL_2553412, EPI_ISL_2553413 | Lighthouse Lab in Milton Keynes | Wellcome Sanger Institute for the COVID-19 Genomics UK (COG-UK) Consortium | The Lighthouse Lab in Milton Keynes and Alex Alderton, Roberto Amato, Jeffrey Barrett, Sonia Goncalves, Ewan Harrison, David K. Jackson, Ian Johnston, Dominic Kwiatkowski, Cordelia Langford, John Sillitoe on behalf of the Wellcome Sanger Institute COVID-19 Surveillance Team |
| EPI_ISL_2553414 | Lighthouse Lab in Alderley Park | Wellcome Sanger Institute for the COVID-19 Genomics UK (COG-UK) Consortium | Jacquelyn Wynn, Mairead Hyland, The Lighthouse Lab in Alderley Park and Alex Alderton, Roberto Amato, Jeffrey Barrett, Sonia Goncalves, Ewan Harrison, David K. Jackson, Ian Johnston, Dominic Kwiatkowski, Cordelia Langford, John Sillitoe on behalf of the Wellcome Sanger Institute COVID-19 Surveillance Team |
| EPI_ISL_2553415 | Lighthouse Lab in Glasgow | Wellcome Sanger Institute for the COVID-19 Genomics UK (COG-UK) Consortium | Harper VanSteenhouse, Yumi Kasai, David Gray, Carol Clugston, Anna Dominiczak and Alex Alderton, Roberto Amato, Jeffrey Barrett, Sonia Goncalves, Ewan Harrison, David K. Jackson, Ian Johnston, Dominic Kwiatkowski, Cordelia Langford, John Sillitoe on behalf of the Wellcome Sanger Institute COVID-19 Surveillance Team |
| EPI_ISL_2553416 | Lighthouse Lab in Milton Keynes | Wellcome Sanger Institute for the COVID-19 Genomics UK (COG-UK) Consortium | The Lighthouse Lab in Milton Keynes and Alex Alderton, Roberto Amato, Jeffrey Barrett, Sonia Goncalves, Ewan Harrison, David K. Jackson, Ian Johnston, Dominic Kwiatkowski, Cordelia Langford, John Sillitoe on behalf of the Wellcome Sanger Institute COVID-19 Surveillance Team |
| EPI_ISL_2553417 | Lighthouse Lab in Glasgow | Wellcome Sanger Institute for the COVID-19 Genomics UK (COG-UK) Consortium | Harper VanSteenhouse, Yumi Kasai, David Gray, Carol Clugston, Anna Dominiczak and Alex Alderton, Roberto Amato, Jeffrey Barrett, Sonia Goncalves, Ewan Harrison, David K. Jackson, Ian Johnston, Dominic Kwiatkowski, Cordelia Langford, John Sillitoe on behalf of the Wellcome Sanger Institute COVID-19 Surveillance Team |
| EPI_ISL_2553418, EPI_ISL_2553419, EPI_ISL_2553420 | Lighthouse Lab in Alderley Park | Wellcome Sanger Institute for the COVID-19 Genomics UK (COG-UK) Consortium | Jacquelyn Wynn, Mairead Hyland, The Lighthouse Lab in Alderley Park and Alex Alderton, Roberto Amato, Jeffrey Barrett, Sonia Goncalves, Ewan Harrison, David K. Jackson, Ian Johnston, Dominic Kwiatkowski, Cordelia Langford, John Sillitoe on behalf of the Wellcome Sanger Institute COVID-19 Surveillance Team |
| EPI_ISL_2553421 | Lighthouse Lab in Milton Keynes | Wellcome Sanger Institute for the COVID-19 Genomics UK (COG-UK) Consortium | The Lighthouse Lab in Milton Keynes and Alex Alderton, Roberto Amato, Jeffrey Barrett, Sonia Goncalves, Ewan Harrison, David K. Jackson, Ian Johnston, Dominic Kwiatkowski, Cordelia Langford, John Sillitoe on behalf of the Wellcome Sanger Institute COVID-19 Surveillance Team |
| EPI_ISL_2553422 | Lighthouse Lab in Alderley Park | Wellcome Sanger Institute for the COVID-19 Genomics UK (COG-UK) Consortium | Jacquelyn Wynn, Mairead Hyland, The Lighthouse Lab in Alderley Park and Alex Alderton, Roberto Amato, Jeffrey Barrett, Sonia Goncalves, Ewan Harrison, David K. Jackson, Ian Johnston, Dominic Kwiatkowski, Cordelia Langford, John Sillitoe on behalf of the Wellcome Sanger Institute COVID-19 Surveillance Team |
| EPI_ISL_2553423 | Lighthouse Lab in Glasgow | Wellcome Sanger Institute for the COVID-19 Genomics UK (COG-UK) Consortium | Harper VanSteenhouse, Yumi Kasai, David Gray, Carol Clugston, Anna Dominiczak and Alex Alderton, Roberto Amato, Jeffrey Barrett, Sonia Goncalves, Ewan Harrison, David K. Jackson, Ian Johnston, Dominic Kwiatkowski, Cordelia Langford, John Sillitoe on behalf of the Wellcome Sanger Institute COVID-19 Surveillance Team |
| EPI_ISL_2553425 | Lighthouse Lab in Alderley Park | Wellcome Sanger Institute for the COVID-19 Genomics UK (COG-UK) Consortium | Jacquelyn Wynn, Mairead Hyland, The Lighthouse Lab in Alderley Park and Alex Alderton, Roberto Amato, Jeffrey Barrett, Sonia Goncalves, Ewan Harrison, David K. Jackson, Ian Johnston, Dominic Kwiatkowski, Cordelia Langford, John Sillitoe on behalf of the Wellcome Sanger Institute COVID-19 Surveillance Team |
| EPI_ISL_2553426 | Lighthouse Lab in Milton Keynes | Wellcome Sanger Institute for the COVID-19 Genomics UK (COG-UK) Consortium | The Lighthouse Lab in Milton Keynes and Alex Alderton, Roberto Amato, Jeffrey Barrett, Sonia Goncalves, Ewan Harrison, David K. Jackson, Ian Johnston, Dominic Kwiatkowski, Cordelia Langford, John Sillitoe on behalf of the Wellcome Sanger Institute COVID-19 Surveillance Team |
| EPI_ISL_2553428 | Lighthouse Lab in Glasgow | Wellcome Sanger Institute for the COVID-19 Genomics UK (COG-UK) Consortium | Harper VanSteenhouse, Yumi Kasai, David Gray, Carol Clugston, Anna Dominiczak and Alex Alderton, Roberto Amato, Jeffrey Barrett, Sonia Goncalves, Ewan Harrison, David K. Jackson, Ian Johnston, Dominic Kwiatkowski, Cordelia Langford, John Sillitoe on behalf of the Wellcome Sanger Institute COVID-19 Surveillance Team |
| EPI_ISL_2553429 | Lighthouse Lab in Milton Keynes | Wellcome Sanger Institute for the COVID-19 Genomics UK (COG-UK) Consortium | The Lighthouse Lab in Milton Keynes and Alex Alderton, Roberto Amato, Jeffrey Barrett, Sonia Goncalves, Ewan Harrison, David K. Jackson, Ian Johnston, Dominic Kwiatkowski, Cordelia Langford, John Sillitoe on behalf of the Wellcome Sanger Institute COVID-19 Surveillance Team |
| EPI_ISL_2553430, EPI_ISL_2553432, EPI_ISL_2553433 | Lighthouse Lab in Glasgow | Wellcome Sanger Institute for the COVID-19 Genomics UK (COG-UK) Consortium | Harper VanSteenhouse, Yumi Kasai, David Gray, Carol Clugston, Anna Dominiczak and Alex Alderton, Roberto Amato, Jeffrey Barrett, Sonia Goncalves, Ewan Harrison, David K. Jackson, Ian Johnston, Dominic Kwiatkowski, Cordelia Langford, John Sillitoe on behalf of the Wellcome Sanger Institute COVID-19 Surveillance Team |
| EPI_ISL_2553434 | Lighthouse Lab in Milton Keynes | Wellcome Sanger Institute for the COVID-19 Genomics UK (COG-UK) Consortium | The Lighthouse Lab in Milton Keynes and Alex Alderton, Roberto Amato, Jeffrey Barrett, Sonia Goncalves, Ewan Harrison, David K. Jackson, Ian Johnston, Dominic Kwiatkowski, Cordelia Langford, John Sillitoe on behalf of the Wellcome Sanger Institute COVID-19 Surveillance Team |
| EPI_ISL_2553435 | Lighthouse Lab in Glasgow | Wellcome Sanger Institute for the COVID-19 Genomics UK (COG-UK) Consortium | Harper VanSteenhouse, Yumi Kasai, David Gray, Carol Clugston, Anna Dominiczak and Alex Alderton, Roberto Amato, Jeffrey Barrett, Sonia Goncalves, Ewan Harrison, David K. Jackson, Ian Johnston, Dominic Kwiatkowski, Cordelia Langford, John Sillitoe on behalf of the Wellcome Sanger Institute COVID-19 Surveillance Team |
| EPI_ISL_2553437, EPI_ISL_2553438, EPI_ISL_2553439, EPI_ISL_2553440 | Lighthouse Lab in Alderley Park | Wellcome Sanger Institute for the COVID-19 Genomics UK (COG-UK) Consortium | Jacquelyn Wynn, Mairead Hyland, The Lighthouse Lab in Alderley Park and Alex Alderton, Roberto Amato, Jeffrey Barrett, Sonia Goncalves, Ewan Harrison, David K. Jackson, Ian Johnston, Dominic Kwiatkowski, Cordelia Langford, John Sillitoe on behalf of the Wellcome Sanger Institute COVID-19 Surveillance Team |
| EPI_ISL_2553441 | Lighthouse Lab in Glasgow | Wellcome Sanger Institute for the COVID-19 Genomics UK (COG-UK) Consortium | Harper VanSteenhouse, Yumi Kasai, David Gray, Carol Clugston, Anna Dominiczak and Alex Alderton, Roberto Amato, Jeffrey Barrett, Sonia Goncalves, Ewan Harrison, David K. Jackson, Ian Johnston, Dominic Kwiatkowski, Cordelia Langford, John Sillitoe on behalf of the Wellcome Sanger Institute COVID-19 Surveillance Team |

| | | | |
|---|---|---|---|
| EPI_ISL_2553442, EPI_ISL_2553443, EPI_ISL_2553444, EPI_ISL_2553445, EPI_ISL_2553446, EPI_ISL_2553447, EPI_ISL_2553448, EPI_ISL_2553449 | Lighthouse Lab in Alderley Park | Wellcome Sanger Institute for the COVID-19 Genomics UK (COG-UK) Consortium | Jacquelyn Wynn, Mairead Hyland, The Lighthouse Lab in Alderley Park and Alex Alderton, Roberto Amato, Jeffrey Barrett, Sonia Goncalves, Ewan Harrison, David K. Jackson, Ian Johnston, Dominic Kwiatkowski, Cordelia Langford, John Sillitoe on behalf of the Wellcome Sanger Institute COVID-19 Surveillance Team |
| EPI_ISL_2553450 | Lighthouse Lab in Glasgow | Wellcome Sanger Institute for the COVID-19 Genomics UK (COG-UK) Consortium | Harper VanSteenhouse, Yumi Kasai, David Gray, Carol Clugston, Anna Dominiczak and Alex Alderton, Roberto Amato, Jeffrey Barrett, Sonia Goncalves, Ewan Harrison, David K. Jackson, Ian Johnston, Dominic Kwiatkowski, Cordelia Langford, John Sillitoe on behalf of the Wellcome Sanger Institute COVID-19 Surveillance Team |
| EPI_ISL_2553451, EPI_ISL_2553452, EPI_ISL_2553453, EPI_ISL_2553454, EPI_ISL_2553455, EPI_ISL_2553456 | Lighthouse Lab in Alderley Park | Wellcome Sanger Institute for the COVID-19 Genomics UK (COG-UK) Consortium | Jacquelyn Wynn, Mairead Hyland, The Lighthouse Lab in Alderley Park and Alex Alderton, Roberto Amato, Jeffrey Barrett, Sonia Goncalves, Ewan Harrison, David K. Jackson, Ian Johnston, Dominic Kwiatkowski, Cordelia Langford, John Sillitoe on behalf of the Wellcome Sanger Institute COVID-19 Surveillance Team |
| EPI_ISL_2553458 | Lighthouse Lab in Glasgow | Wellcome Sanger Institute for the COVID-19 Genomics UK (COG-UK) Consortium | Harper VanSteenhouse, Yumi Kasai, David Gray, Carol Clugston, Anna Dominiczak and Alex Alderton, Roberto Amato, Jeffrey Barrett, Sonia Goncalves, Ewan Harrison, David K. Jackson, Ian Johnston, Dominic Kwiatkowski, Cordelia Langford, John Sillitoe on behalf of the Wellcome Sanger Institute COVID-19 Surveillance Team |
| EPI_ISL_2553459, EPI_ISL_2553460, EPI_ISL_2553461 | Lighthouse Lab in Alderley Park | Wellcome Sanger Institute for the COVID-19 Genomics UK (COG-UK) Consortium | Jacquelyn Wynn, Mairead Hyland, The Lighthouse Lab in Alderley Park and Alex Alderton, Roberto Amato, Jeffrey Barrett, Sonia Goncalves, Ewan Harrison, David K. Jackson, Ian Johnston, Dominic Kwiatkowski, Cordelia Langford, John Sillitoe on behalf of the Wellcome Sanger Institute COVID-19 Surveillance Team |
| EPI_ISL_2553462 | Lighthouse Lab in Glasgow | Wellcome Sanger Institute for the COVID-19 Genomics UK (COG-UK) Consortium | Harper VanSteenhouse, Yumi Kasai, David Gray, Carol Clugston, Anna Dominiczak and Alex Alderton, Roberto Amato, Jeffrey Barrett, Sonia Goncalves, Ewan Harrison, David K. Jackson, Ian Johnston, Dominic Kwiatkowski, Cordelia Langford, John Sillitoe on behalf of the Wellcome Sanger Institute COVID-19 Surveillance Team |
| EPI_ISL_2553463, EPI_ISL_2553464 | Lighthouse Lab in Alderley Park | Wellcome Sanger Institute for the COVID-19 Genomics UK (COG-UK) Consortium | Jacquelyn Wynn, Mairead Hyland, The Lighthouse Lab in Alderley Park and Alex Alderton, Roberto Amato, Jeffrey Barrett, Sonia Goncalves, Ewan Harrison, David K. Jackson, Ian Johnston, Dominic Kwiatkowski, Cordelia Langford, John Sillitoe on behalf of the Wellcome Sanger Institute COVID-19 Surveillance Team |
| EPI_ISL_2553465, EPI_ISL_2553466, EPI_ISL_2553467 | Lighthouse Lab in Glasgow | Wellcome Sanger Institute for the COVID-19 Genomics UK (COG-UK) Consortium | Harper VanSteenhouse, Yumi Kasai, David Gray, Carol Clugston, Anna Dominiczak and Alex Alderton, Roberto Amato, Jeffrey Barrett, Sonia Goncalves, Ewan Harrison, David K. Jackson, Ian Johnston, Dominic Kwiatkowski, Cordelia Langford, John Sillitoe on behalf of the Wellcome Sanger Institute COVID-19 Surveillance Team |
| EPI_ISL_2553468, EPI_ISL_2553469, EPI_ISL_2553470, EPI_ISL_2553471, EPI_ISL_2553472, EPI_ISL_2553473, EPI_ISL_2553474, EPI_ISL_2553475, EPI_ISL_2553476, EPI_ISL_2553477, EPI_ISL_2553478, EPI_ISL_2553479, EPI_ISL_2553480, EPI_ISL_2553481, EPI_ISL_2553482, EPI_ISL_2553483, EPI_ISL_2553484, EPI_ISL_2553485, EPI_ISL_2553486, EPI_ISL_2553487, EPI_ISL_2553488, EPI_ISL_2553489, EPI_ISL_2553490 | | | |
| see above | Lighthouse Lab in Alderley Park | Wellcome Sanger Institute for the COVID-19 Genomics UK (COG-UK) Consortium | Jacquelyn Wynn, Mairead Hyland, The Lighthouse Lab in Alderley Park and Alex Alderton, Roberto Amato, Jeffrey Barrett, Sonia Goncalves, Ewan Harrison, David K. Jackson, Ian Johnston, Dominic Kwiatkowski, Cordelia Langford, John Sillitoe on behalf of the Wellcome Sanger Institute COVID-19 Surveillance Team |
| EPI_ISL_2553491 | Lighthouse Lab in Glasgow | Wellcome Sanger Institute for the COVID-19 Genomics UK (COG-UK) Consortium | Harper VanSteenhouse, Yumi Kasai, David Gray, Carol Clugston, Anna Dominiczak and Alex Alderton, Roberto Amato, Jeffrey Barrett, Sonia Goncalves, Ewan Harrison, David K. Jackson, Ian Johnston, Dominic Kwiatkowski, Cordelia Langford, John Sillitoe on behalf of the Wellcome Sanger Institute COVID-19 Surveillance Team |
| EPI_ISL_2553492, EPI_ISL_2553494 | Lighthouse Lab in Alderley Park | Wellcome Sanger Institute for the COVID-19 Genomics UK (COG-UK) Consortium | Jacquelyn Wynn, Mairead Hyland, The Lighthouse Lab in Alderley Park and Alex Alderton, Roberto Amato, Jeffrey Barrett, Sonia Goncalves, Ewan Harrison, David K. Jackson, Ian Johnston, Dominic Kwiatkowski, Cordelia Langford, John Sillitoe on behalf of the Wellcome Sanger Institute COVID-19 Surveillance Team |
| EPI_ISL_2553495 | Lighthouse Lab in Glasgow | Wellcome Sanger Institute for the COVID-19 Genomics UK (COG-UK) Consortium | Harper VanSteenhouse, Yumi Kasai, David Gray, Carol Clugston, Anna Dominiczak and Alex Alderton, Roberto Amato, Jeffrey Barrett, Sonia Goncalves, Ewan Harrison, David K. Jackson, Ian Johnston, Dominic Kwiatkowski, Cordelia Langford, John Sillitoe on behalf of the Wellcome Sanger Institute COVID-19 Surveillance Team |
| EPI_ISL_2553496, EPI_ISL_2553497 | Lighthouse Lab in Alderley Park | Wellcome Sanger Institute for the COVID-19 Genomics UK (COG-UK) Consortium | Jacquelyn Wynn, Mairead Hyland, The Lighthouse Lab in Alderley Park and Alex Alderton, Roberto Amato, Jeffrey Barrett, Sonia Goncalves, Ewan Harrison, David K. Jackson, Ian Johnston, Dominic Kwiatkowski, Cordelia Langford, John Sillitoe on behalf of the Wellcome Sanger Institute COVID-19 Surveillance Team |
| EPI_ISL_2553498 | Lighthouse Lab in Glasgow | Wellcome Sanger Institute for the COVID-19 Genomics UK (COG-UK) Consortium | Harper VanSteenhouse, Yumi Kasai, David Gray, Carol Clugston, Anna Dominiczak and Alex Alderton, Roberto Amato, Jeffrey Barrett, Sonia Goncalves, Ewan Harrison, David K. Jackson, Ian Johnston, Dominic Kwiatkowski, Cordelia Langford, John Sillitoe on behalf of the Wellcome Sanger Institute COVID-19 Surveillance Team |
| EPI_ISL_2553499, EPI_ISL_2553500 | Lighthouse Lab in Alderley Park | Wellcome Sanger Institute for the COVID-19 Genomics UK (COG-UK) Consortium | Jacquelyn Wynn, Mairead Hyland, The Lighthouse Lab in Alderley Park and Alex Alderton, Roberto Amato, Jeffrey Barrett, Sonia Goncalves, Ewan Harrison, David K. Jackson, Ian Johnston, Dominic Kwiatkowski, Cordelia Langford, John Sillitoe on behalf of the Wellcome Sanger Institute COVID-19 Surveillance Team |
| EPI_ISL_2553501 | Lighthouse Lab in Glasgow | Wellcome Sanger Institute for the COVID-19 Genomics UK (COG-UK) Consortium | Harper VanSteenhouse, Yumi Kasai, David Gray, Carol Clugston, Anna Dominiczak and Alex Alderton, Roberto Amato, Jeffrey Barrett, Sonia Goncalves, Ewan Harrison, David K. Jackson, Ian Johnston, Dominic Kwiatkowski, Cordelia Langford, John Sillitoe on behalf of the Wellcome Sanger Institute COVID-19 Surveillance Team |
| EPI_ISL_2553502, EPI_ISL_2553503, EPI_ISL_2553504, EPI_ISL_2553505, EPI_ISL_2553506 | Lighthouse Lab in Alderley Park | Wellcome Sanger Institute for the COVID-19 Genomics UK (COG-UK) Consortium | Jacquelyn Wynn, Mairead Hyland, The Lighthouse Lab in Alderley Park and Alex Alderton, Roberto Amato, Jeffrey Barrett, Sonia Goncalves, Ewan Harrison, David K. Jackson, Ian Johnston, Dominic Kwiatkowski, Cordelia Langford, John Sillitoe on behalf of the Wellcome Sanger Institute COVID-19 Surveillance Team |
| EPI_ISL_2553507, EPI_ISL_2553508 | Lighthouse Lab in Glasgow | Wellcome Sanger Institute for the COVID-19 Genomics UK (COG-UK) Consortium | Harper VanSteenhouse, Yumi Kasai, David Gray, Carol Clugston, Anna Dominiczak and Alex Alderton, Roberto Amato, Jeffrey Barrett, Sonia Goncalves, Ewan Harrison, David K. Jackson, Ian Johnston, Dominic Kwiatkowski, Cordelia Langford, John Sillitoe on behalf of the Wellcome Sanger Institute COVID-19 Surveillance Team |
| EPI_ISL_2553509, EPI_ISL_2553510, EPI_ISL_2553511, EPI_ISL_2553512, EPI_ISL_2553513, EPI_ISL_2553514, EPI_ISL_2553515, EPI_ISL_2553516, EPI_ISL_2553518, EPI_ISL_2553519, EPI_ISL_2553520, EPI_ISL_2553521, EPI_ISL_2553522, EPI_ISL_2553523, EPI_ISL_2553525, EPI_ISL_2553526, EPI_ISL_2553527 | | | |
| see above | Lighthouse Lab in Alderley Park | Wellcome Sanger Institute for the COVID-19 Genomics UK (COG-UK) Consortium | Jacquelyn Wynn, Mairead Hyland, The Lighthouse Lab in Alderley Park and Alex Alderton, Roberto Amato, Jeffrey Barrett, Sonia Goncalves, Ewan Harrison, David K. Jackson, Ian Johnston, Dominic Kwiatkowski, Cordelia Langford, John Sillitoe on behalf of the Wellcome Sanger Institute COVID-19 Surveillance Team |
| EPI_ISL_2553528, EPI_ISL_2553529, EPI_ISL_2553530, EPI_ISL_2553531 | Lighthouse Lab in Glasgow | Wellcome Sanger Institute for the COVID-19 Genomics UK (COG-UK) Consortium | Harper VanSteenhouse, Yumi Kasai, David Gray, Carol Clugston, Anna Dominiczak and Alex Alderton, Roberto Amato, Jeffrey Barrett, Sonia Goncalves, Ewan Harrison, David K. Jackson, Ian Johnston, Dominic Kwiatkowski, Cordelia Langford, John Sillitoe on behalf of the Wellcome Sanger Institute COVID-19 Surveillance Team |
| EPI_ISL_2553533, EPI_ISL_2553534 | Lighthouse Lab in Alderley Park | Wellcome Sanger Institute for the COVID-19 Genomics UK (COG-UK) Consortium | Jacquelyn Wynn, Mairead Hyland, The Lighthouse Lab in Alderley Park and Alex Alderton, Roberto Amato, Jeffrey Barrett, Sonia Goncalves, Ewan Harrison, David K. Jackson, Ian Johnston, Dominic Kwiatkowski, Cordelia Langford, John Sillitoe on behalf of the Wellcome Sanger Institute COVID-19 Surveillance Team |
| EPI_ISL_2553535, EPI_ISL_2553536, EPI_ISL_2553537 | Lighthouse Lab in Glasgow | Wellcome Sanger Institute for the COVID-19 Genomics UK (COG-UK) Consortium | Harper VanSteenhouse, Yumi Kasai, David Gray, Carol Clugston, Anna Dominiczak and Alex Alderton, Roberto Amato, Jeffrey Barrett, Sonia Goncalves, Ewan Harrison, David K. Jackson, Ian Johnston, Dominic Kwiatkowski, Cordelia Langford, John Sillitoe on behalf of the Wellcome Sanger Institute COVID-19 Surveillance Team |
| EPI_ISL_2553538 | Lighthouse Lab in Alderley Park | Wellcome Sanger Institute for the COVID-19 Genomics UK (COG-UK) Consortium | Jacquelyn Wynn, Mairead Hyland, The Lighthouse Lab in Alderley Park and Alex Alderton, Roberto Amato, Jeffrey Barrett, Sonia Goncalves, Ewan Harrison, David K. Jackson, Ian Johnston, Dominic Kwiatkowski, Cordelia Langford, John Sillitoe on behalf of the Wellcome Sanger Institute COVID-19 Surveillance Team |
| EPI_ISL_2553539 | Lighthouse Lab in Glasgow | Wellcome Sanger Institute for the COVID-19 Genomics UK (COG-UK) Consortium | Harper VanSteenhouse, Yumi Kasai, David Gray, Carol Clugston, Anna Dominiczak and Alex Alderton, Roberto Amato, Jeffrey Barrett, Sonia Goncalves, Ewan Harrison, David K. Jackson, Ian Johnston, Dominic Kwiatkowski, Cordelia Langford, John Sillitoe on behalf of the Wellcome Sanger Institute COVID-19 Surveillance Team |
| EPI_ISL_2553540, EPI_ISL_2553541, EPI_ISL_2553542, EPI_ISL_2553543, EPI_ISL_2553544, EPI_ISL_2553545 | Lighthouse Lab in Alderley Park | Wellcome Sanger Institute for the COVID-19 Genomics UK (COG-UK) Consortium | Jacquelyn Wynn, Mairead Hyland, The Lighthouse Lab in Alderley Park and Alex Alderton, Roberto Amato, Jeffrey Barrett, Sonia Goncalves, Ewan Harrison, David K. Jackson, Ian Johnston, Dominic Kwiatkowski, Cordelia Langford, John Sillitoe on behalf of the Wellcome Sanger Institute COVID-19 Surveillance Team |
| EPI_ISL_2553546 | Lighthouse Lab in Glasgow | Wellcome Sanger Institute for the COVID-19 Genomics UK (COG-UK) Consortium | Harper VanSteenhouse, Yumi Kasai, David Gray, Carol Clugston, Anna Dominiczak and Alex Alderton, Roberto Amato, Jeffrey Barrett, Sonia Goncalves, Ewan Harrison, David K. Jackson, Ian Johnston, Dominic Kwiatkowski, Cordelia Langford, John Sillitoe on behalf of the Wellcome Sanger Institute COVID-19 Surveillance Team |
| EPI_ISL_2553547, EPI_ISL_2553548 | Lighthouse Lab in Alderley Park | Wellcome Sanger Institute for the COVID-19 Genomics UK (COG-UK) Consortium | Jacquelyn Wynn, Mairead Hyland, The Lighthouse Lab in Alderley Park and Alex Alderton, Roberto Amato, Jeffrey Barrett, Sonia Goncalves, Ewan Harrison, David K. Jackson, Ian Johnston, Dominic Kwiatkowski, Cordelia Langford, John Sillitoe on behalf of the Wellcome Sanger Institute COVID-19 Surveillance Team |
| EPI_ISL_2553549 | Lighthouse Lab in Glasgow | Wellcome Sanger Institute for the COVID-19 Genomics UK (COG-UK) Consortium | Harper VanSteenhouse, Yumi Kasai, David Gray, Carol Clugston, Anna Dominiczak and Alex Alderton, Roberto Amato, Jeffrey Barrett, Sonia Goncalves, Ewan Harrison, David K. Jackson, Ian Johnston, Dominic Kwiatkowski, Cordelia Langford, John Sillitoe on behalf of the Wellcome Sanger Institute COVID-19 Surveillance Team |
| EPI_ISL_2553550 | Lighthouse Lab in Alderley Park | Wellcome Sanger Institute for the COVID-19 Genomics UK (COG-UK) Consortium | Jacquelyn Wynn, Mairead Hyland, The Lighthouse Lab in Alderley Park and Alex Alderton, Roberto Amato, Jeffrey Barrett, Sonia Goncalves, Ewan Harrison, David K. Jackson, Ian Johnston, Dominic Kwiatkowski, Cordelia Langford, John Sillitoe on behalf of the Wellcome Sanger Institute COVID-19 Surveillance Team |
| EPI_ISL_2553551 | Lighthouse Lab in Glasgow | Wellcome Sanger Institute for the COVID-19 Genomics UK (COG-UK) Consortium | Harper VanSteenhouse, Yumi Kasai, David Gray, Carol Clugston, Anna Dominiczak and Alex Alderton, Roberto Amato, Jeffrey Barrett, Sonia Goncalves, Ewan Harrison, David K. Jackson, Ian Johnston, Dominic Kwiatkowski, Cordelia Langford, John Sillitoe on behalf of the Wellcome Sanger Institute COVID-19 Surveillance Team |
| EPI_ISL_2553552, EPI_ISL_2553553, EPI_ISL_2553554, EPI_ISL_2553555, EPI_ISL_2553556, EPI_ISL_2553557, EPI_ISL_2553558, EPI_ISL_2553559, EPI_ISL_2553560, EPI_ISL_2553561, EPI_ISL_2553562, EPI_ISL_2553564 | | | |
| see above | Lighthouse Lab in Alderley Park | Wellcome Sanger Institute for the COVID-19 Genomics UK (COG-UK) Consortium | Jacquelyn Wynn, Mairead Hyland, The Lighthouse Lab in Alderley Park and Alex Alderton, Roberto Amato, Jeffrey Barrett, Sonia Goncalves, Ewan Harrison, David K. Jackson, Ian Johnston, Dominic Kwiatkowski, Cordelia Langford, John Sillitoe on behalf of the Wellcome Sanger Institute COVID-19 Surveillance Team |
| EPI_ISL_2553565, EPI_ISL_2553566 | Lighthouse Lab in Glasgow | Wellcome Sanger Institute for the COVID-19 Genomics UK (COG-UK) Consortium | Harper VanSteenhouse, Yumi Kasai, David Gray, Carol Clugston, Anna Dominiczak and Alex Alderton, Roberto Amato, Jeffrey Barrett, Sonia Goncalves, Ewan Harrison, David K. Jackson, Ian Johnston, Dominic Kwiatkowski, Cordelia Langford, John Sillitoe on behalf of the Wellcome Sanger Institute COVID-19 Surveillance Team |
| EPI_ISL_2553567, EPI_ISL_2553568 | Lighthouse Lab in Alderley Park | Wellcome Sanger Institute for the COVID-19 Genomics UK (COG-UK) Consortium | Jacquelyn Wynn, Mairead Hyland, The Lighthouse Lab in Alderley Park and Alex Alderton, Roberto Amato, Jeffrey Barrett, Sonia Goncalves, Ewan Harrison, David K. Jackson, Ian Johnston, Dominic Kwiatkowski, Cordelia Langford, John Sillitoe on behalf of the Wellcome Sanger Institute COVID-19 Surveillance Team |
| EPI_ISL_2553569 | Lighthouse Lab in Glasgow | Wellcome Sanger Institute for the COVID-19 Genomics UK (COG-UK) Consortium | Harper VanSteenhouse, Yumi Kasai, David Gray, Carol Clugston, Anna Dominiczak and Alex Alderton, Roberto Amato, Jeffrey Barrett, Sonia Goncalves, Ewan Harrison, David K. Jackson, Ian Johnston, Dominic Kwiatkowski, Cordelia Langford, John Sillitoe on behalf of the Wellcome Sanger Institute COVID-19 Surveillance Team |

| | | | |
|---|---|---|---|
| EPI_ISL_2553570 | Lighthouse Lab in Alderley Park | Wellcome Sanger Institute for the COVID-19 Genomics UK (COG-UK) Consortium | Jacquelyn Wynn, Mairead Hyland, The Lighthouse Lab in Alderley Park and Alex Alderton, Roberto Amato, Jeffrey Barrett, Sonia Goncalves, Ewan Harrison, David K. Jackson, Ian Johnston, Dominic Kwiatkowski, Cordelia Langford, John Sillitoe on behalf of the Wellcome Sanger Institute COVID-19 Surveillance Team |
| EPI_ISL_2553571 | Lighthouse Lab in Glasgow | Wellcome Sanger Institute for the COVID-19 Genomics UK (COG-UK) Consortium | Harper VanSteenhouse, Yumi Kasai, David Gray, Carol Clugston, Anna Dominiczak and Alex Alderton, Roberto Amato, Jeffrey Barrett, Sonia Goncalves, Ewan Harrison, David K. Jackson, Ian Johnston, Dominic Kwiatkowski, Cordelia Langford, John Sillitoe on behalf of the Wellcome Sanger Institute COVID-19 Surveillance Team |
| EPI_ISL_2553572 | Lighthouse Lab in Alderley Park | Wellcome Sanger Institute for the COVID-19 Genomics UK (COG-UK) Consortium | Jacquelyn Wynn, Mairead Hyland, The Lighthouse Lab in Alderley Park and Alex Alderton, Roberto Amato, Jeffrey Barrett, Sonia Goncalves, Ewan Harrison, David K. Jackson, Ian Johnston, Dominic Kwiatkowski, Cordelia Langford, John Sillitoe on behalf of the Wellcome Sanger Institute COVID-19 Surveillance Team |
| EPI_ISL_2553573 | Lighthouse Lab in Glasgow | Wellcome Sanger Institute for the COVID-19 Genomics UK (COG-UK) Consortium | Harper VanSteenhouse, Yumi Kasai, David Gray, Carol Clugston, Anna Dominiczak and Alex Alderton, Roberto Amato, Jeffrey Barrett, Sonia Goncalves, Ewan Harrison, David K. Jackson, Ian Johnston, Dominic Kwiatkowski, Cordelia Langford, John Sillitoe on behalf of the Wellcome Sanger Institute COVID-19 Surveillance Team |
| EPI_ISL_2553574, EPI_ISL_2553575, EPI_ISL_2553576, EPI_ISL_2553577, EPI_ISL_2553578, EPI_ISL_2553579, EPI_ISL_2553580, EPI_ISL_2553581, EPI_ISL_2553582, EPI_ISL_2553584, EPI_ISL_2553585, EPI_ISL_2553586, EPI_ISL_2553587, EPI_ISL_2553588, EPI_ISL_2553589, EPI_ISL_2553590, EPI_ISL_2553591 | | | |
| see above | Lighthouse Lab in Alderley Park | Wellcome Sanger Institute for the COVID-19 Genomics UK (COG-UK) Consortium | Jacquelyn Wynn, Mairead Hyland, The Lighthouse Lab in Alderley Park and Alex Alderton, Roberto Amato, Jeffrey Barrett, Sonia Goncalves, Ewan Harrison, David K. Jackson, Ian Johnston, Dominic Kwiatkowski, Cordelia Langford, John Sillitoe on behalf of the Wellcome Sanger Institute COVID-19 Surveillance Team |
| EPI_ISL_2553592, EPI_ISL_2553593 | Lighthouse Lab in Glasgow | Wellcome Sanger Institute for the COVID-19 Genomics UK (COG-UK) Consortium | Harper VanSteenhouse, Yumi Kasai, David Gray, Carol Clugston, Anna Dominiczak and Alex Alderton, Roberto Amato, Jeffrey Barrett, Sonia Goncalves, Ewan Harrison, David K. Jackson, Ian Johnston, Dominic Kwiatkowski, Cordelia Langford, John Sillitoe on behalf of the Wellcome Sanger Institute COVID-19 Surveillance Team |
| EPI_ISL_2553594 | Lighthouse Lab in Alderley Park | Wellcome Sanger Institute for the COVID-19 Genomics UK (COG-UK) Consortium | Jacquelyn Wynn, Mairead Hyland, The Lighthouse Lab in Alderley Park and Alex Alderton, Roberto Amato, Jeffrey Barrett, Sonia Goncalves, Ewan Harrison, David K. Jackson, Ian Johnston, Dominic Kwiatkowski, Cordelia Langford, John Sillitoe on behalf of the Wellcome Sanger Institute COVID-19 Surveillance Team |
| EPI_ISL_2553595, EPI_ISL_2553596 | Lighthouse Lab in Glasgow | Wellcome Sanger Institute for the COVID-19 Genomics UK (COG-UK) Consortium | Harper VanSteenhouse, Yumi Kasai, David Gray, Carol Clugston, Anna Dominiczak and Alex Alderton, Roberto Amato, Jeffrey Barrett, Sonia Goncalves, Ewan Harrison, David K. Jackson, Ian Johnston, Dominic Kwiatkowski, Cordelia Langford, John Sillitoe on behalf of the Wellcome Sanger Institute COVID-19 Surveillance Team |
| EPI_ISL_2553597, EPI_ISL_2553598, EPI_ISL_2553599 | Lighthouse Lab in Alderley Park | Wellcome Sanger Institute for the COVID-19 Genomics UK (COG-UK) Consortium | Jacquelyn Wynn, Mairead Hyland, The Lighthouse Lab in Alderley Park and Alex Alderton, Roberto Amato, Jeffrey Barrett, Sonia Goncalves, Ewan Harrison, David K. Jackson, Ian Johnston, Dominic Kwiatkowski, Cordelia Langford, John Sillitoe on behalf of the Wellcome Sanger Institute COVID-19 Surveillance Team |
| EPI_ISL_2553600 | Lighthouse Lab in Glasgow | Wellcome Sanger Institute for the COVID-19 Genomics UK (COG-UK) Consortium | Harper VanSteenhouse, Yumi Kasai, David Gray, Carol Clugston, Anna Dominiczak and Alex Alderton, Roberto Amato, Jeffrey Barrett, Sonia Goncalves, Ewan Harrison, David K. Jackson, Ian Johnston, Dominic Kwiatkowski, Cordelia Langford, John Sillitoe on behalf of the Wellcome Sanger Institute COVID-19 Surveillance Team |
| EPI_ISL_2553601, EPI_ISL_2553602, EPI_ISL_2553603, EPI_ISL_2553604, EPI_ISL_2553605, EPI_ISL_2553606, EPI_ISL_2553607, EPI_ISL_2553608, EPI_ISL_2553609, EPI_ISL_2553610, EPI_ISL_2553611, EPI_ISL_2553612, EPI_ISL_2553613 | | | |
| see above | Lighthouse Lab in Alderley Park | Wellcome Sanger Institute for the COVID-19 Genomics UK (COG-UK) Consortium | Jacquelyn Wynn, Mairead Hyland, The Lighthouse Lab in Alderley Park and Alex Alderton, Roberto Amato, Jeffrey Barrett, Sonia Goncalves, Ewan Harrison, David K. Jackson, Ian Johnston, Dominic Kwiatkowski, Cordelia Langford, John Sillitoe on behalf of the Wellcome Sanger Institute COVID-19 Surveillance Team |
| EPI_ISL_2553614 | Lighthouse Lab in Glasgow | Wellcome Sanger Institute for the COVID-19 Genomics UK (COG-UK) Consortium | Harper VanSteenhouse, Yumi Kasai, David Gray, Carol Clugston, Anna Dominiczak and Alex Alderton, Roberto Amato, Jeffrey Barrett, Sonia Goncalves, Ewan Harrison, David K. Jackson, Ian Johnston, Dominic Kwiatkowski, Cordelia Langford, John Sillitoe on behalf of the Wellcome Sanger Institute COVID-19 Surveillance Team |
| EPI_ISL_2553615, EPI_ISL_2553616, EPI_ISL_2553617, EPI_ISL_2553618, EPI_ISL_2553619, EPI_ISL_2553620, EPI_ISL_2553621, EPI_ISL_2553622 | Lighthouse Lab in Alderley Park | Wellcome Sanger Institute for the COVID-19 Genomics UK (COG-UK) Consortium | Jacquelyn Wynn, Mairead Hyland, The Lighthouse Lab in Alderley Park and Alex Alderton, Roberto Amato, Jeffrey Barrett, Sonia Goncalves, Ewan Harrison, David K. Jackson, Ian Johnston, Dominic Kwiatkowski, Cordelia Langford, John Sillitoe on behalf of the Wellcome Sanger Institute COVID-19 Surveillance Team |
| EPI_ISL_2553623 | Lighthouse Lab in Glasgow | Wellcome Sanger Institute for the COVID-19 Genomics UK (COG-UK) Consortium | Harper VanSteenhouse, Yumi Kasai, David Gray, Carol Clugston, Anna Dominiczak and Alex Alderton, Roberto Amato, Jeffrey Barrett, Sonia Goncalves, Ewan Harrison, David K. Jackson, Ian Johnston, Dominic Kwiatkowski, Cordelia Langford, John Sillitoe on behalf of the Wellcome Sanger Institute COVID-19 Surveillance Team |
| EPI_ISL_2553625 | Lighthouse Lab in Alderley Park | Wellcome Sanger Institute for the COVID-19 Genomics UK (COG-UK) Consortium | Jacquelyn Wynn, Mairead Hyland, The Lighthouse Lab in Alderley Park and Alex Alderton, Roberto Amato, Jeffrey Barrett, Sonia Goncalves, Ewan Harrison, David K. Jackson, Ian Johnston, Dominic Kwiatkowski, Cordelia Langford, John Sillitoe on behalf of the Wellcome Sanger Institute COVID-19 Surveillance Team |
| EPI_ISL_2553626, EPI_ISL_2553627 | Lighthouse Lab in Glasgow | Wellcome Sanger Institute for the COVID-19 Genomics UK (COG-UK) Consortium | Harper VanSteenhouse, Yumi Kasai, David Gray, Carol Clugston, Anna Dominiczak and Alex Alderton, Roberto Amato, Jeffrey Barrett, Sonia Goncalves, Ewan Harrison, David K. Jackson, Ian Johnston, Dominic Kwiatkowski, Cordelia Langford, John Sillitoe on behalf of the Wellcome Sanger Institute COVID-19 Surveillance Team |
| EPI_ISL_2553629 | Lighthouse Lab in Alderley Park | Wellcome Sanger Institute for the COVID-19 Genomics UK (COG-UK) Consortium | Jacquelyn Wynn, Mairead Hyland, The Lighthouse Lab in Alderley Park and Alex Alderton, Roberto Amato, Jeffrey Barrett, Sonia Goncalves, Ewan Harrison, David K. Jackson, Ian Johnston, Dominic Kwiatkowski, Cordelia Langford, John Sillitoe on behalf of the Wellcome Sanger Institute COVID-19 Surveillance Team |
| EPI_ISL_2553630 | Lighthouse Lab in Glasgow | Wellcome Sanger Institute for the COVID-19 Genomics UK (COG-UK) Consortium | Harper VanSteenhouse, Yumi Kasai, David Gray, Carol Clugston, Anna Dominiczak and Alex Alderton, Roberto Amato, Jeffrey Barrett, Sonia Goncalves, Ewan Harrison, David K. Jackson, Ian Johnston, Dominic Kwiatkowski, Cordelia Langford, John Sillitoe on behalf of the Wellcome Sanger Institute COVID-19 Surveillance Team |
| EPI_ISL_2553631 | Lighthouse Lab in Alderley Park | Wellcome Sanger Institute for the COVID-19 Genomics UK (COG-UK) Consortium | Jacquelyn Wynn, Mairead Hyland, The Lighthouse Lab in Alderley Park and Alex Alderton, Roberto Amato, Jeffrey Barrett, Sonia Goncalves, Ewan Harrison, David K. Jackson, Ian Johnston, Dominic Kwiatkowski, Cordelia Langford, John Sillitoe on behalf of the Wellcome Sanger Institute COVID-19 Surveillance Team |
| EPI_ISL_2553632 | Lighthouse Lab in Glasgow | Wellcome Sanger Institute for the COVID-19 Genomics UK (COG-UK) Consortium | Harper VanSteenhouse, Yumi Kasai, David Gray, Carol Clugston, Anna Dominiczak and Alex Alderton, Roberto Amato, Jeffrey Barrett, Sonia Goncalves, Ewan Harrison, David K. Jackson, Ian Johnston, Dominic Kwiatkowski, Cordelia Langford, John Sillitoe on behalf of the Wellcome Sanger Institute COVID-19 Surveillance Team |
| EPI_ISL_2553633, EPI_ISL_2553634 | Lighthouse Lab in Alderley Park | Wellcome Sanger Institute for the COVID-19 Genomics UK (COG-UK) Consortium | Jacquelyn Wynn, Mairead Hyland, The Lighthouse Lab in Alderley Park and Alex Alderton, Roberto Amato, Jeffrey Barrett, Sonia Goncalves, Ewan Harrison, David K. Jackson, Ian Johnston, Dominic Kwiatkowski, Cordelia Langford, John Sillitoe on behalf of the Wellcome Sanger Institute COVID-19 Surveillance Team |
| EPI_ISL_2553635 | Lighthouse Lab in Glasgow | Wellcome Sanger Institute for the COVID-19 Genomics UK (COG-UK) Consortium | Harper VanSteenhouse, Yumi Kasai, David Gray, Carol Clugston, Anna Dominiczak and Alex Alderton, Roberto Amato, Jeffrey Barrett, Sonia Goncalves, Ewan Harrison, David K. Jackson, Ian Johnston, Dominic Kwiatkowski, Cordelia Langford, John Sillitoe on behalf of the Wellcome Sanger Institute COVID-19 Surveillance Team |
| EPI_ISL_2553636 | Lighthouse Lab in Alderley Park | Wellcome Sanger Institute for the COVID-19 Genomics UK (COG-UK) Consortium | Jacquelyn Wynn, Mairead Hyland, The Lighthouse Lab in Alderley Park and Alex Alderton, Roberto Amato, Jeffrey Barrett, Sonia Goncalves, Ewan Harrison, David K. Jackson, Ian Johnston, Dominic Kwiatkowski, Cordelia Langford, John Sillitoe on behalf of the Wellcome Sanger Institute COVID-19 Surveillance Team |
| EPI_ISL_2553637 | Lighthouse Lab in Glasgow | Wellcome Sanger Institute for the COVID-19 Genomics UK (COG-UK) Consortium | Harper VanSteenhouse, Yumi Kasai, David Gray, Carol Clugston, Anna Dominiczak and Alex Alderton, Roberto Amato, Jeffrey Barrett, Sonia Goncalves, Ewan Harrison, David K. Jackson, Ian Johnston, Dominic Kwiatkowski, Cordelia Langford, John Sillitoe on behalf of the Wellcome Sanger Institute COVID-19 Surveillance Team |
| EPI_ISL_2553638, EPI_ISL_2553639, EPI_ISL_2553640 | Lighthouse Lab in Alderley Park | Wellcome Sanger Institute for the COVID-19 Genomics UK (COG-UK) Consortium | Jacquelyn Wynn, Mairead Hyland, The Lighthouse Lab in Alderley Park and Alex Alderton, Roberto Amato, Jeffrey Barrett, Sonia Goncalves, Ewan Harrison, David K. Jackson, Ian Johnston, Dominic Kwiatkowski, Cordelia Langford, John Sillitoe on behalf of the Wellcome Sanger Institute COVID-19 Surveillance Team |
| EPI_ISL_2553641 | Lighthouse Lab in Glasgow | Wellcome Sanger Institute for the COVID-19 Genomics UK (COG-UK) Consortium | Harper VanSteenhouse, Yumi Kasai, David Gray, Carol Clugston, Anna Dominiczak and Alex Alderton, Roberto Amato, Jeffrey Barrett, Sonia Goncalves, Ewan Harrison, David K. Jackson, Ian Johnston, Dominic Kwiatkowski, Cordelia Langford, John Sillitoe on behalf of the Wellcome Sanger Institute COVID-19 Surveillance Team |
| EPI_ISL_2553642 | Lighthouse Lab in Alderley Park | Wellcome Sanger Institute for the COVID-19 Genomics UK (COG-UK) Consortium | Jacquelyn Wynn, Mairead Hyland, The Lighthouse Lab in Alderley Park and Alex Alderton, Roberto Amato, Jeffrey Barrett, Sonia Goncalves, Ewan Harrison, David K. Jackson, Ian Johnston, Dominic Kwiatkowski, Cordelia Langford, John Sillitoe on behalf of the Wellcome Sanger Institute COVID-19 Surveillance Team |
| EPI_ISL_2553643 | Lighthouse Lab in Glasgow | Wellcome Sanger Institute for the COVID-19 Genomics UK (COG-UK) Consortium | Harper VanSteenhouse, Yumi Kasai, David Gray, Carol Clugston, Anna Dominiczak and Alex Alderton, Roberto Amato, Jeffrey Barrett, Sonia Goncalves, Ewan Harrison, David K. Jackson, Ian Johnston, Dominic Kwiatkowski, Cordelia Langford, John Sillitoe on behalf of the Wellcome Sanger Institute COVID-19 Surveillance Team |
| EPI_ISL_2553644 | Lighthouse Lab in Alderley Park | Wellcome Sanger Institute for the COVID-19 Genomics UK (COG-UK) Consortium | Jacquelyn Wynn, Mairead Hyland, The Lighthouse Lab in Alderley Park and Alex Alderton, Roberto Amato, Jeffrey Barrett, Sonia Goncalves, Ewan Harrison, David K. Jackson, Ian Johnston, Dominic Kwiatkowski, Cordelia Langford, John Sillitoe on behalf of the Wellcome Sanger Institute COVID-19 Surveillance Team |
| EPI_ISL_2553646 | Lighthouse Lab in Glasgow | Wellcome Sanger Institute for the COVID-19 Genomics UK (COG-UK) Consortium | Harper VanSteenhouse, Yumi Kasai, David Gray, Carol Clugston, Anna Dominiczak and Alex Alderton, Roberto Amato, Jeffrey Barrett, Sonia Goncalves, Ewan Harrison, David K. Jackson, Ian Johnston, Dominic Kwiatkowski, Cordelia Langford, John Sillitoe on behalf of the Wellcome Sanger Institute COVID-19 Surveillance Team |
| EPI_ISL_2553647, EPI_ISL_2553648 | Lighthouse Lab in Alderley Park | Wellcome Sanger Institute for the COVID-19 Genomics UK (COG-UK) Consortium | Jacquelyn Wynn, Mairead Hyland, The Lighthouse Lab in Alderley Park and Alex Alderton, Roberto Amato, Jeffrey Barrett, Sonia Goncalves, Ewan Harrison, David K. Jackson, Ian Johnston, Dominic Kwiatkowski, Cordelia Langford, John Sillitoe on behalf of the Wellcome Sanger Institute COVID-19 Surveillance Team |
| EPI_ISL_2553649 | Lighthouse Lab in Glasgow | Wellcome Sanger Institute for the COVID-19 Genomics UK (COG-UK) Consortium | Harper VanSteenhouse, Yumi Kasai, David Gray, Carol Clugston, Anna Dominiczak and Alex Alderton, Roberto Amato, Jeffrey Barrett, Sonia Goncalves, Ewan Harrison, David K. Jackson, Ian Johnston, Dominic Kwiatkowski, Cordelia Langford, John Sillitoe on behalf of the Wellcome Sanger Institute COVID-19 Surveillance Team |
| EPI_ISL_2553650, EPI_ISL_2553651, EPI_ISL_2553652, EPI_ISL_2553653, EPI_ISL_2553654, EPI_ISL_2553655, EPI_ISL_2553656 | Lighthouse Lab in Alderley Park | Wellcome Sanger Institute for the COVID-19 Genomics UK (COG-UK) Consortium | Jacquelyn Wynn, Mairead Hyland, The Lighthouse Lab in Alderley Park and Alex Alderton, Roberto Amato, Jeffrey Barrett, Sonia Goncalves, Ewan Harrison, David K. Jackson, Ian Johnston, Dominic Kwiatkowski, Cordelia Langford, John Sillitoe on behalf of the Wellcome Sanger Institute COVID-19 Surveillance Team |
| EPI_ISL_2553657 | Lighthouse Lab in Glasgow | Wellcome Sanger Institute for the COVID-19 Genomics UK (COG-UK) Consortium | Harper VanSteenhouse, Yumi Kasai, David Gray, Carol Clugston, Anna Dominiczak and Alex Alderton, Roberto Amato, Jeffrey Barrett, Sonia Goncalves, Ewan Harrison, David K. Jackson, Ian Johnston, Dominic Kwiatkowski, Cordelia Langford, John Sillitoe on behalf of the Wellcome Sanger Institute COVID-19 Surveillance Team |
| EPI_ISL_2553658, EPI_ISL_2553659, EPI_ISL_2553660, EPI_ISL_2553661, EPI_ISL_2553662, EPI_ISL_2553663, EPI_ISL_2553664 | Lighthouse Lab in Alderley Park | Wellcome Sanger Institute for the COVID-19 Genomics UK (COG-UK) Consortium | Jacquelyn Wynn, Mairead Hyland, The Lighthouse Lab in Alderley Park and Alex Alderton, Roberto Amato, Jeffrey Barrett, Sonia Goncalves, Ewan Harrison, David K. Jackson, Ian Johnston, Dominic Kwiatkowski, Cordelia Langford, John Sillitoe on behalf of the Wellcome Sanger Institute COVID-19 Surveillance Team |
| EPI_ISL_2553665 | Lighthouse Lab in Glasgow | Wellcome Sanger Institute for the COVID-19 Genomics UK (COG-UK) Consortium | Harper VanSteenhouse, Yumi Kasai, David Gray, Carol Clugston, Anna Dominiczak and Alex Alderton, Roberto Amato, Jeffrey Barrett, Sonia Goncalves, Ewan Harrison, David K. Jackson, Ian Johnston, Dominic Kwiatkowski, Cordelia Langford, John Sillitoe on behalf of the Wellcome Sanger Institute COVID-19 Surveillance Team |

| | | | |
|---|---|---|---|
| EPI_ISL_2553667, EPI_ISL_2553668, EPI_ISL_2553669, EPI_ISL_2553670 | Lighthouse Lab in Alderley Park | Wellcome Sanger Institute for the COVID-19 Genomics UK (COG-UK) Consortium | Jacquelyn Wynn, Mairead Hyland, The Lighthouse Lab in Alderley Park and Alex Alderton, Roberto Amato, Jeffrey Barrett, Sonia Goncalves, Ewan Harrison, David K. Jackson, Ian Johnston, Dominic Kwiatkowski, Cordelia Langford, John Sillitoe on behalf of the Wellcome Sanger Institute COVID-19 Surveillance Team |
| EPI_ISL_2553671 | Lighthouse Lab in Glasgow | Wellcome Sanger Institute for the COVID-19 Genomics UK (COG-UK) Consortium | Harper VanSteenhouse, Yumi Kasai, David Gray, Carol Clugston, Anna Dominiczak and Alex Alderton, Roberto Amato, Jeffrey Barrett, Sonia Goncalves, Ewan Harrison, David K. Jackson, Ian Johnston, Dominic Kwiatkowski, Cordelia Langford, John Sillitoe on behalf of the Wellcome Sanger Institute COVID-19 Surveillance Team |
| EPI_ISL_2553672 | Lighthouse Lab in Alderley Park | Wellcome Sanger Institute for the COVID-19 Genomics UK (COG-UK) Consortium | Jacquelyn Wynn, Mairead Hyland, The Lighthouse Lab in Alderley Park and Alex Alderton, Roberto Amato, Jeffrey Barrett, Sonia Goncalves, Ewan Harrison, David K. Jackson, Ian Johnston, Dominic Kwiatkowski, Cordelia Langford, John Sillitoe on behalf of the Wellcome Sanger Institute COVID-19 Surveillance Team |
| EPI_ISL_2553673 | Lighthouse Lab in Glasgow | Wellcome Sanger Institute for the COVID-19 Genomics UK (COG-UK) Consortium | Harper VanSteenhouse, Yumi Kasai, David Gray, Carol Clugston, Anna Dominiczak and Alex Alderton, Roberto Amato, Jeffrey Barrett, Sonia Goncalves, Ewan Harrison, David K. Jackson, Ian Johnston, Dominic Kwiatkowski, Cordelia Langford, John Sillitoe on behalf of the Wellcome Sanger Institute COVID-19 Surveillance Team |
| EPI_ISL_2553674 | Lighthouse Lab in Alderley Park | Wellcome Sanger Institute for the COVID-19 Genomics UK (COG-UK) Consortium | Jacquelyn Wynn, Mairead Hyland, The Lighthouse Lab in Alderley Park and Alex Alderton, Roberto Amato, Jeffrey Barrett, Sonia Goncalves, Ewan Harrison, David K. Jackson, Ian Johnston, Dominic Kwiatkowski, Cordelia Langford, John Sillitoe on behalf of the Wellcome Sanger Institute COVID-19 Surveillance Team |
| EPI_ISL_2553675 | Lighthouse Lab in Glasgow | Wellcome Sanger Institute for the COVID-19 Genomics UK (COG-UK) Consortium | Harper VanSteenhouse, Yumi Kasai, David Gray, Carol Clugston, Anna Dominiczak and Alex Alderton, Roberto Amato, Jeffrey Barrett, Sonia Goncalves, Ewan Harrison, David K. Jackson, Ian Johnston, Dominic Kwiatkowski, Cordelia Langford, John Sillitoe on behalf of the Wellcome Sanger Institute COVID-19 Surveillance Team |
| EPI_ISL_2553676, EPI_ISL_2553677, EPI_ISL_2553678, EPI_ISL_2553679 | Lighthouse Lab in Alderley Park | Wellcome Sanger Institute for the COVID-19 Genomics UK (COG-UK) Consortium | Jacquelyn Wynn, Mairead Hyland, The Lighthouse Lab in Alderley Park and Alex Alderton, Roberto Amato, Jeffrey Barrett, Sonia Goncalves, Ewan Harrison, David K. Jackson, Ian Johnston, Dominic Kwiatkowski, Cordelia Langford, John Sillitoe on behalf of the Wellcome Sanger Institute COVID-19 Surveillance Team |
| EPI_ISL_2553680 | Lighthouse Lab in Glasgow | Wellcome Sanger Institute for the COVID-19 Genomics UK (COG-UK) Consortium | Harper VanSteenhouse, Yumi Kasai, David Gray, Carol Clugston, Anna Dominiczak and Alex Alderton, Roberto Amato, Jeffrey Barrett, Sonia Goncalves, Ewan Harrison, David K. Jackson, Ian Johnston, Dominic Kwiatkowski, Cordelia Langford, John Sillitoe on behalf of the Wellcome Sanger Institute COVID-19 Surveillance Team |
| EPI_ISL_2553681 | Lighthouse Lab in Alderley Park | Wellcome Sanger Institute for the COVID-19 Genomics UK (COG-UK) Consortium | Jacquelyn Wynn, Mairead Hyland, The Lighthouse Lab in Alderley Park and Alex Alderton, Roberto Amato, Jeffrey Barrett, Sonia Goncalves, Ewan Harrison, David K. Jackson, Ian Johnston, Dominic Kwiatkowski, Cordelia Langford, John Sillitoe on behalf of the Wellcome Sanger Institute COVID-19 Surveillance Team |
| EPI_ISL_2553682 | Lighthouse Lab in Glasgow | Wellcome Sanger Institute for the COVID-19 Genomics UK (COG-UK) Consortium | Harper VanSteenhouse, Yumi Kasai, David Gray, Carol Clugston, Anna Dominiczak and Alex Alderton, Roberto Amato, Jeffrey Barrett, Sonia Goncalves, Ewan Harrison, David K. Jackson, Ian Johnston, Dominic Kwiatkowski, Cordelia Langford, John Sillitoe on behalf of the Wellcome Sanger Institute COVID-19 Surveillance Team |
| EPI_ISL_2553683, EPI_ISL_2553684 | Lighthouse Lab in Alderley Park | Wellcome Sanger Institute for the COVID-19 Genomics UK (COG-UK) Consortium | Jacquelyn Wynn, Mairead Hyland, The Lighthouse Lab in Alderley Park and Alex Alderton, Roberto Amato, Jeffrey Barrett, Sonia Goncalves, Ewan Harrison, David K. Jackson, Ian Johnston, Dominic Kwiatkowski, Cordelia Langford, John Sillitoe on behalf of the Wellcome Sanger Institute COVID-19 Surveillance Team |
| EPI_ISL_2553685 | Lighthouse Lab in Glasgow | Wellcome Sanger Institute for the COVID-19 Genomics UK (COG-UK) Consortium | Harper VanSteenhouse, Yumi Kasai, David Gray, Carol Clugston, Anna Dominiczak and Alex Alderton, Roberto Amato, Jeffrey Barrett, Sonia Goncalves, Ewan Harrison, David K. Jackson, Ian Johnston, Dominic Kwiatkowski, Cordelia Langford, John Sillitoe on behalf of the Wellcome Sanger Institute COVID-19 Surveillance Team |
| EPI_ISL_2553686 | Lighthouse Lab in Alderley Park | Wellcome Sanger Institute for the COVID-19 Genomics UK (COG-UK) Consortium | Jacquelyn Wynn, Mairead Hyland, The Lighthouse Lab in Alderley Park and Alex Alderton, Roberto Amato, Jeffrey Barrett, Sonia Goncalves, Ewan Harrison, David K. Jackson, Ian Johnston, Dominic Kwiatkowski, Cordelia Langford, John Sillitoe on behalf of the Wellcome Sanger Institute COVID-19 Surveillance Team |
| EPI_ISL_2553687 | Lighthouse Lab in Glasgow | Wellcome Sanger Institute for the COVID-19 Genomics UK (COG-UK) Consortium | Harper VanSteenhouse, Yumi Kasai, David Gray, Carol Clugston, Anna Dominiczak and Alex Alderton, Roberto Amato, Jeffrey Barrett, Sonia Goncalves, Ewan Harrison, David K. Jackson, Ian Johnston, Dominic Kwiatkowski, Cordelia Langford, John Sillitoe on behalf of the Wellcome Sanger Institute COVID-19 Surveillance Team |
| EPI_ISL_2553688, EPI_ISL_2553689 | Lighthouse Lab in Alderley Park | Wellcome Sanger Institute for the COVID-19 Genomics UK (COG-UK) Consortium | Jacquelyn Wynn, Mairead Hyland, The Lighthouse Lab in Alderley Park and Alex Alderton, Roberto Amato, Jeffrey Barrett, Sonia Goncalves, Ewan Harrison, David K. Jackson, Ian Johnston, Dominic Kwiatkowski, Cordelia Langford, John Sillitoe on behalf of the Wellcome Sanger Institute COVID-19 Surveillance Team |
| EPI_ISL_2553690, EPI_ISL_2553691, EPI_ISL_2553692 | Lighthouse Lab in Glasgow | Wellcome Sanger Institute for the COVID-19 Genomics UK (COG-UK) Consortium | Harper VanSteenhouse, Yumi Kasai, David Gray, Carol Clugston, Anna Dominiczak and Alex Alderton, Roberto Amato, Jeffrey Barrett, Sonia Goncalves, Ewan Harrison, David K. Jackson, Ian Johnston, Dominic Kwiatkowski, Cordelia Langford, John Sillitoe on behalf of the Wellcome Sanger Institute COVID-19 Surveillance Team |
| EPI_ISL_2553693, EPI_ISL_2553694, EPI_ISL_2553695, EPI_ISL_2553696, EPI_ISL_2553697, EPI_ISL_2553698 | Lighthouse Lab in Alderley Park | Wellcome Sanger Institute for the COVID-19 Genomics UK (COG-UK) Consortium | Jacquelyn Wynn, Mairead Hyland, The Lighthouse Lab in Alderley Park and Alex Alderton, Roberto Amato, Jeffrey Barrett, Sonia Goncalves, Ewan Harrison, David K. Jackson, Ian Johnston, Dominic Kwiatkowski, Cordelia Langford, John Sillitoe on behalf of the Wellcome Sanger Institute COVID-19 Surveillance Team |
| EPI_ISL_2553699 | Lighthouse Lab in Glasgow | Wellcome Sanger Institute for the COVID-19 Genomics UK (COG-UK) Consortium | Harper VanSteenhouse, Yumi Kasai, David Gray, Carol Clugston, Anna Dominiczak and Alex Alderton, Roberto Amato, Jeffrey Barrett, Sonia Goncalves, Ewan Harrison, David K. Jackson, Ian Johnston, Dominic Kwiatkowski, Cordelia Langford, John Sillitoe on behalf of the Wellcome Sanger Institute COVID-19 Surveillance Team |
| EPI_ISL_2553701, EPI_ISL_2553702, EPI_ISL_2553703, EPI_ISL_2553704, EPI_ISL_2553705 | Lighthouse Lab in Alderley Park | Wellcome Sanger Institute for the COVID-19 Genomics UK (COG-UK) Consortium | Jacquelyn Wynn, Mairead Hyland, The Lighthouse Lab in Alderley Park and Alex Alderton, Roberto Amato, Jeffrey Barrett, Sonia Goncalves, Ewan Harrison, David K. Jackson, Ian Johnston, Dominic Kwiatkowski, Cordelia Langford, John Sillitoe on behalf of the Wellcome Sanger Institute COVID-19 Surveillance Team |
| EPI_ISL_2553706 | Lighthouse Lab in Glasgow | Wellcome Sanger Institute for the COVID-19 Genomics UK (COG-UK) Consortium | Harper VanSteenhouse, Yumi Kasai, David Gray, Carol Clugston, Anna Dominiczak and Alex Alderton, Roberto Amato, Jeffrey Barrett, Sonia Goncalves, Ewan Harrison, David K. Jackson, Ian Johnston, Dominic Kwiatkowski, Cordelia Langford, John Sillitoe on behalf of the Wellcome Sanger Institute COVID-19 Surveillance Team |
| EPI_ISL_2553707, EPI_ISL_2553708, EPI_ISL_2553709 | Lighthouse Lab in Alderley Park | Wellcome Sanger Institute for the COVID-19 Genomics UK (COG-UK) Consortium | Jacquelyn Wynn, Mairead Hyland, The Lighthouse Lab in Alderley Park and Alex Alderton, Roberto Amato, Jeffrey Barrett, Sonia Goncalves, Ewan Harrison, David K. Jackson, Ian Johnston, Dominic Kwiatkowski, Cordelia Langford, John Sillitoe on behalf of the Wellcome Sanger Institute COVID-19 Surveillance Team |
| EPI_ISL_2553710 | Lighthouse Lab in Glasgow | Wellcome Sanger Institute for the COVID-19 Genomics UK (COG-UK) Consortium | Harper VanSteenhouse, Yumi Kasai, David Gray, Carol Clugston, Anna Dominiczak and Alex Alderton, Roberto Amato, Jeffrey Barrett, Sonia Goncalves, Ewan Harrison, David K. Jackson, Ian Johnston, Dominic Kwiatkowski, Cordelia Langford, John Sillitoe on behalf of the Wellcome Sanger Institute COVID-19 Surveillance Team |
| EPI_ISL_2553711, EPI_ISL_2553712, EPI_ISL_2553713, EPI_ISL_2553714 | Lighthouse Lab in Alderley Park | Wellcome Sanger Institute for the COVID-19 Genomics UK (COG-UK) Consortium | Jacquelyn Wynn, Mairead Hyland, The Lighthouse Lab in Alderley Park and Alex Alderton, Roberto Amato, Jeffrey Barrett, Sonia Goncalves, Ewan Harrison, David K. Jackson, Ian Johnston, Dominic Kwiatkowski, Cordelia Langford, John Sillitoe on behalf of the Wellcome Sanger Institute COVID-19 Surveillance Team |
| EPI_ISL_2553716, EPI_ISL_2553717 | Lighthouse Lab in Glasgow | Wellcome Sanger Institute for the COVID-19 Genomics UK (COG-UK) Consortium | Harper VanSteenhouse, Yumi Kasai, David Gray, Carol Clugston, Anna Dominiczak and Alex Alderton, Roberto Amato, Jeffrey Barrett, Sonia Goncalves, Ewan Harrison, David K. Jackson, Ian Johnston, Dominic Kwiatkowski, Cordelia Langford, John Sillitoe on behalf of the Wellcome Sanger Institute COVID-19 Surveillance Team |
| EPI_ISL_2553718, EPI_ISL_2553719, EPI_ISL_2553720, EPI_ISL_2553721, EPI_ISL_2553722, EPI_ISL_2553723, EPI_ISL_2553724, EPI_ISL_2553726, EPI_ISL_2553727 | Lighthouse Lab in Alderley Park | Wellcome Sanger Institute for the COVID-19 Genomics UK (COG-UK) Consortium | Jacquelyn Wynn, Mairead Hyland, The Lighthouse Lab in Alderley Park and Alex Alderton, Roberto Amato, Jeffrey Barrett, Sonia Goncalves, Ewan Harrison, David K. Jackson, Ian Johnston, Dominic Kwiatkowski, Cordelia Langford, John Sillitoe on behalf of the Wellcome Sanger Institute COVID-19 Surveillance Team |
| EPI_ISL_2553728 | Lighthouse Lab in Glasgow | Wellcome Sanger Institute for the COVID-19 Genomics UK (COG-UK) Consortium | Harper VanSteenhouse, Yumi Kasai, David Gray, Carol Clugston, Anna Dominiczak and Alex Alderton, Roberto Amato, Jeffrey Barrett, Sonia Goncalves, Ewan Harrison, David K. Jackson, Ian Johnston, Dominic Kwiatkowski, Cordelia Langford, John Sillitoe on behalf of the Wellcome Sanger Institute COVID-19 Surveillance Team |
| EPI_ISL_2553729, EPI_ISL_2553730, EPI_ISL_2553731, EPI_ISL_2553732, EPI_ISL_2553733, EPI_ISL_2553734, EPI_ISL_2553735, EPI_ISL_2553736, EPI_ISL_2553737, EPI_ISL_2553738, EPI_ISL_2553740 | | | |
| see above | Lighthouse Lab in Alderley Park | Wellcome Sanger Institute for the COVID-19 Genomics UK (COG-UK) Consortium | Jacquelyn Wynn, Mairead Hyland, The Lighthouse Lab in Alderley Park and Alex Alderton, Roberto Amato, Jeffrey Barrett, Sonia Goncalves, Ewan Harrison, David K. Jackson, Ian Johnston, Dominic Kwiatkowski, Cordelia Langford, John Sillitoe on behalf of the Wellcome Sanger Institute COVID-19 Surveillance Team |
| EPI_ISL_2553741, EPI_ISL_2553742, EPI_ISL_2553743 | Lighthouse Lab in Glasgow | Wellcome Sanger Institute for the COVID-19 Genomics UK (COG-UK) Consortium | Harper VanSteenhouse, Yumi Kasai, David Gray, Carol Clugston, Anna Dominiczak and Alex Alderton, Roberto Amato, Jeffrey Barrett, Sonia Goncalves, Ewan Harrison, David K. Jackson, Ian Johnston, Dominic Kwiatkowski, Cordelia Langford, John Sillitoe on behalf of the Wellcome Sanger Institute COVID-19 Surveillance Team |
| EPI_ISL_2553744, EPI_ISL_2553745, EPI_ISL_2553746 | Lighthouse Lab in Alderley Park | Wellcome Sanger Institute for the COVID-19 Genomics UK (COG-UK) Consortium | Jacquelyn Wynn, Mairead Hyland, The Lighthouse Lab in Alderley Park and Alex Alderton, Roberto Amato, Jeffrey Barrett, Sonia Goncalves, Ewan Harrison, David K. Jackson, Ian Johnston, Dominic Kwiatkowski, Cordelia Langford, John Sillitoe on behalf of the Wellcome Sanger Institute COVID-19 Surveillance Team |
| EPI_ISL_2553747 | Lighthouse Lab in Glasgow | Wellcome Sanger Institute for the COVID-19 Genomics UK (COG-UK) Consortium | Harper VanSteenhouse, Yumi Kasai, David Gray, Carol Clugston, Anna Dominiczak and Alex Alderton, Roberto Amato, Jeffrey Barrett, Sonia Goncalves, Ewan Harrison, David K. Jackson, Ian Johnston, Dominic Kwiatkowski, Cordelia Langford, John Sillitoe on behalf of the Wellcome Sanger Institute COVID-19 Surveillance Team |
| EPI_ISL_2553748 | Lighthouse Lab in Alderley Park | Wellcome Sanger Institute for the COVID-19 Genomics UK (COG-UK) Consortium | Jacquelyn Wynn, Mairead Hyland, The Lighthouse Lab in Alderley Park and Alex Alderton, Roberto Amato, Jeffrey Barrett, Sonia Goncalves, Ewan Harrison, David K. Jackson, Ian Johnston, Dominic Kwiatkowski, Cordelia Langford, John Sillitoe on behalf of the Wellcome Sanger Institute COVID-19 Surveillance Team |
| EPI_ISL_2553749 | Lighthouse Lab in Glasgow | Wellcome Sanger Institute for the COVID-19 Genomics UK (COG-UK) Consortium | Harper VanSteenhouse, Yumi Kasai, David Gray, Carol Clugston, Anna Dominiczak and Alex Alderton, Roberto Amato, Jeffrey Barrett, Sonia Goncalves, Ewan Harrison, David K. Jackson, Ian Johnston, Dominic Kwiatkowski, Cordelia Langford, John Sillitoe on behalf of the Wellcome Sanger Institute COVID-19 Surveillance Team |
| EPI_ISL_2553751, EPI_ISL_2553752, EPI_ISL_2553753, EPI_ISL_2553754, EPI_ISL_2553756, EPI_ISL_2553757, EPI_ISL_2553758 | Lighthouse Lab in Alderley Park | Wellcome Sanger Institute for the COVID-19 Genomics UK (COG-UK) Consortium | Jacquelyn Wynn, Mairead Hyland, The Lighthouse Lab in Alderley Park and Alex Alderton, Roberto Amato, Jeffrey Barrett, Sonia Goncalves, Ewan Harrison, David K. Jackson, Ian Johnston, Dominic Kwiatkowski, Cordelia Langford, John Sillitoe on behalf of the Wellcome Sanger Institute COVID-19 Surveillance Team |

| | | | |
|---|---|---|---|
| EPI_ISL_2553759 | Lighthouse Lab in Glasgow | Wellcome Sanger Institute for the COVID-19 Genomics UK (COG-UK) Consortium | Harper VanSteenhouse, Yumi Kasai, David Gray, Carol Clugston, Anna Dominiczak and Alex Alderton, Roberto Amato, Jeffrey Barrett, Sonia Goncalves, Ewan Harrison, David K. Jackson, Ian Johnston, Dominic Kwiatkowski, Cordelia Langford, John Sillitoe on behalf of the Wellcome Sanger Institute COVID-19 Surveillance Team |
| EPI_ISL_2553760, EPI_ISL_2553761 | Lighthouse Lab in Alderley Park | Wellcome Sanger Institute for the COVID-19 Genomics UK (COG-UK) Consortium | Jacquelyn Wynn, Mairead Hyland, The Lighthouse Lab in Alderley Park and Alex Alderton, Roberto Amato, Jeffrey Barrett, Sonia Goncalves, Ewan Harrison, David K. Jackson, Ian Johnston, Dominic Kwiatkowski, Cordelia Langford, John Sillitoe on behalf of the Wellcome Sanger Institute COVID-19 Surveillance Team |
| EPI_ISL_2553762 | Lighthouse Lab in Glasgow | Wellcome Sanger Institute for the COVID-19 Genomics UK (COG-UK) Consortium | Harper VanSteenhouse, Yumi Kasai, David Gray, Carol Clugston, Anna Dominiczak and Alex Alderton, Roberto Amato, Jeffrey Barrett, Sonia Goncalves, Ewan Harrison, David K. Jackson, Ian Johnston, Dominic Kwiatkowski, Cordelia Langford, John Sillitoe on behalf of the Wellcome Sanger Institute COVID-19 Surveillance Team |
| EPI_ISL_2553763 | Lighthouse Lab in Alderley Park | Wellcome Sanger Institute for the COVID-19 Genomics UK (COG-UK) Consortium | Jacquelyn Wynn, Mairead Hyland, The Lighthouse Lab in Alderley Park and Alex Alderton, Roberto Amato, Jeffrey Barrett, Sonia Goncalves, Ewan Harrison, David K. Jackson, Ian Johnston, Dominic Kwiatkowski, Cordelia Langford, John Sillitoe on behalf of the Wellcome Sanger Institute COVID-19 Surveillance Team |
| EPI_ISL_2553764 | Lighthouse Lab in Glasgow | Wellcome Sanger Institute for the COVID-19 Genomics UK (COG-UK) Consortium | Harper VanSteenhouse, Yumi Kasai, David Gray, Carol Clugston, Anna Dominiczak and Alex Alderton, Roberto Amato, Jeffrey Barrett, Sonia Goncalves, Ewan Harrison, David K. Jackson, Ian Johnston, Dominic Kwiatkowski, Cordelia Langford, John Sillitoe on behalf of the Wellcome Sanger Institute COVID-19 Surveillance Team |
| EPI_ISL_2553765, EPI_ISL_2553766, EPI_ISL_2553767, EPI_ISL_2553768, EPI_ISL_2553769, EPI_ISL_2553770, EPI_ISL_2553771 | Lighthouse Lab in Alderley Park | Wellcome Sanger Institute for the COVID-19 Genomics UK (COG-UK) Consortium | Jacquelyn Wynn, Mairead Hyland, The Lighthouse Lab in Alderley Park and Alex Alderton, Roberto Amato, Jeffrey Barrett, Sonia Goncalves, Ewan Harrison, David K. Jackson, Ian Johnston, Dominic Kwiatkowski, Cordelia Langford, John Sillitoe on behalf of the Wellcome Sanger Institute COVID-19 Surveillance Team |
| EPI_ISL_2553772 | Lighthouse Lab in Glasgow | Wellcome Sanger Institute for the COVID-19 Genomics UK (COG-UK) Consortium | Harper VanSteenhouse, Yumi Kasai, David Gray, Carol Clugston, Anna Dominiczak and Alex Alderton, Roberto Amato, Jeffrey Barrett, Sonia Goncalves, Ewan Harrison, David K. Jackson, Ian Johnston, Dominic Kwiatkowski, Cordelia Langford, John Sillitoe on behalf of the Wellcome Sanger Institute COVID-19 Surveillance Team |
| EPI_ISL_2553773 | Lighthouse Lab in Alderley Park | Wellcome Sanger Institute for the COVID-19 Genomics UK (COG-UK) Consortium | Jacquelyn Wynn, Mairead Hyland, The Lighthouse Lab in Alderley Park and Alex Alderton, Roberto Amato, Jeffrey Barrett, Sonia Goncalves, Ewan Harrison, David K. Jackson, Ian Johnston, Dominic Kwiatkowski, Cordelia Langford, John Sillitoe on behalf of the Wellcome Sanger Institute COVID-19 Surveillance Team |
| EPI_ISL_2553774, EPI_ISL_2553775, EPI_ISL_2553776 | Lighthouse Lab in Glasgow | Wellcome Sanger Institute for the COVID-19 Genomics UK (COG-UK) Consortium | Harper VanSteenhouse, Yumi Kasai, David Gray, Carol Clugston, Anna Dominiczak and Alex Alderton, Roberto Amato, Jeffrey Barrett, Sonia Goncalves, Ewan Harrison, David K. Jackson, Ian Johnston, Dominic Kwiatkowski, Cordelia Langford, John Sillitoe on behalf of the Wellcome Sanger Institute COVID-19 Surveillance Team |
| EPI_ISL_2553777, EPI_ISL_2553778 | Lighthouse Lab in Alderley Park | Wellcome Sanger Institute for the COVID-19 Genomics UK (COG-UK) Consortium | Jacquelyn Wynn, Mairead Hyland, The Lighthouse Lab in Alderley Park and Alex Alderton, Roberto Amato, Jeffrey Barrett, Sonia Goncalves, Ewan Harrison, David K. Jackson, Ian Johnston, Dominic Kwiatkowski, Cordelia Langford, John Sillitoe on behalf of the Wellcome Sanger Institute COVID-19 Surveillance Team |
| EPI_ISL_2553779, EPI_ISL_2553780 | Lighthouse Lab in Glasgow | Wellcome Sanger Institute for the COVID-19 Genomics UK (COG-UK) Consortium | Harper VanSteenhouse, Yumi Kasai, David Gray, Carol Clugston, Anna Dominiczak and Alex Alderton, Roberto Amato, Jeffrey Barrett, Sonia Goncalves, Ewan Harrison, David K. Jackson, Ian Johnston, Dominic Kwiatkowski, Cordelia Langford, John Sillitoe on behalf of the Wellcome Sanger Institute COVID-19 Surveillance Team |
| EPI_ISL_2553781, EPI_ISL_2553782, EPI_ISL_2553783, EPI_ISL_2553784, EPI_ISL_2553785, EPI_ISL_2553786, EPI_ISL_2553787, EPI_ISL_2553788, EPI_ISL_2553789 | Lighthouse Lab in Alderley Park | Wellcome Sanger Institute for the COVID-19 Genomics UK (COG-UK) Consortium | Jacquelyn Wynn, Mairead Hyland, The Lighthouse Lab in Alderley Park and Alex Alderton, Roberto Amato, Jeffrey Barrett, Sonia Goncalves, Ewan Harrison, David K. Jackson, Ian Johnston, Dominic Kwiatkowski, Cordelia Langford, John Sillitoe on behalf of the Wellcome Sanger Institute COVID-19 Surveillance Team |
| EPI_ISL_2553790, EPI_ISL_2553791 | Health Services Laboratories | Wellcome Sanger Institute for the COVID-19 Genomics UK (COG-UK) Consortium | Health Services Laboratories and Alex Alderton, Roberto Amato, Jeffrey Barrett, Sonia Goncalves, Ewan Harrison, David K. Jackson, Ian Johnston, Dominic Kwiatkowski, Cordelia Langford, John Sillitoe on behalf of the Wellcome Sanger Institute COVID-19 Surveillance Team |
| EPI_ISL_2553792 | Lighthouse Lab in Alderley Park | Wellcome Sanger Institute for the COVID-19 Genomics UK (COG-UK) Consortium | Jacquelyn Wynn, Mairead Hyland, The Lighthouse Lab in Alderley Park and Alex Alderton, Roberto Amato, Jeffrey Barrett, Sonia Goncalves, Ewan Harrison, David K. Jackson, Ian Johnston, Dominic Kwiatkowski, Cordelia Langford, John Sillitoe on behalf of the Wellcome Sanger Institute COVID-19 Surveillance Team |
| EPI_ISL_2553793 | Lighthouse Lab in Milton Keynes | Wellcome Sanger Institute for the COVID-19 Genomics UK (COG-UK) Consortium | The Lighthouse Lab in Milton Keynes and Alex Alderton, Roberto Amato, Jeffrey Barrett, Sonia Goncalves, Ewan Harrison, David K. Jackson, Ian Johnston, Dominic Kwiatkowski, Cordelia Langford, John Sillitoe on behalf of the Wellcome Sanger Institute COVID-19 Surveillance Team |
| EPI_ISL_2553794 | Lighthouse Lab in Alderley Park | Wellcome Sanger Institute for the COVID-19 Genomics UK (COG-UK) Consortium | Jacquelyn Wynn, Mairead Hyland, The Lighthouse Lab in Alderley Park and Alex Alderton, Roberto Amato, Jeffrey Barrett, Sonia Goncalves, Ewan Harrison, David K. Jackson, Ian Johnston, Dominic Kwiatkowski, Cordelia Langford, John Sillitoe on behalf of the Wellcome Sanger Institute COVID-19 Surveillance Team |
| EPI_ISL_2553795, EPI_ISL_2553796 | Lighthouse Lab in Milton Keynes | Wellcome Sanger Institute for the COVID-19 Genomics UK (COG-UK) Consortium | The Lighthouse Lab in Milton Keynes and Alex Alderton, Roberto Amato, Jeffrey Barrett, Sonia Goncalves, Ewan Harrison, David K. Jackson, Ian Johnston, Dominic Kwiatkowski, Cordelia Langford, John Sillitoe on behalf of the Wellcome Sanger Institute COVID-19 Surveillance Team |
| EPI_ISL_2553797 | Health Services Laboratories | Wellcome Sanger Institute for the COVID-19 Genomics UK (COG-UK) Consortium | Health Services Laboratories and Alex Alderton, Roberto Amato, Jeffrey Barrett, Sonia Goncalves, Ewan Harrison, David K. Jackson, Ian Johnston, Dominic Kwiatkowski, Cordelia Langford, John Sillitoe on behalf of the Wellcome Sanger Institute COVID-19 Surveillance Team |
| EPI_ISL_2553798 | Lighthouse Lab in Milton Keynes | Wellcome Sanger Institute for the COVID-19 Genomics UK (COG-UK) Consortium | The Lighthouse Lab in Milton Keynes and Alex Alderton, Roberto Amato, Jeffrey Barrett, Sonia Goncalves, Ewan Harrison, David K. Jackson, Ian Johnston, Dominic Kwiatkowski, Cordelia Langford, John Sillitoe on behalf of the Wellcome Sanger Institute COVID-19 Surveillance Team |
| EPI_ISL_2553799 | Lighthouse Lab in Alderley Park | Wellcome Sanger Institute for the COVID-19 Genomics UK (COG-UK) Consortium | Jacquelyn Wynn, Mairead Hyland, The Lighthouse Lab in Alderley Park and Alex Alderton, Roberto Amato, Jeffrey Barrett, Sonia Goncalves, Ewan Harrison, David K. Jackson, Ian Johnston, Dominic Kwiatkowski, Cordelia Langford, John Sillitoe on behalf of the Wellcome Sanger Institute COVID-19 Surveillance Team |
| EPI_ISL_2553800 | Health Services Laboratories | Wellcome Sanger Institute for the COVID-19 Genomics UK (COG-UK) Consortium | Health Services Laboratories and Alex Alderton, Roberto Amato, Jeffrey Barrett, Sonia Goncalves, Ewan Harrison, David K. Jackson, Ian Johnston, Dominic Kwiatkowski, Cordelia Langford, John Sillitoe on behalf of the Wellcome Sanger Institute COVID-19 Surveillance Team |
| EPI_ISL_2553802 | Lighthouse Lab in Alderley Park | Wellcome Sanger Institute for the COVID-19 Genomics UK (COG-UK) Consortium | Jacquelyn Wynn, Mairead Hyland, The Lighthouse Lab in Alderley Park and Alex Alderton, Roberto Amato, Jeffrey Barrett, Sonia Goncalves, Ewan Harrison, David K. Jackson, Ian Johnston, Dominic Kwiatkowski, Cordelia Langford, John Sillitoe on behalf of the Wellcome Sanger Institute COVID-19 Surveillance Team |
| EPI_ISL_2553803, EPI_ISL_2553804, EPI_ISL_2553805 | Health Services Laboratories | Wellcome Sanger Institute for the COVID-19 Genomics UK (COG-UK) Consortium | Health Services Laboratories and Alex Alderton, Roberto Amato, Jeffrey Barrett, Sonia Goncalves, Ewan Harrison, David K. Jackson, Ian Johnston, Dominic Kwiatkowski, Cordelia Langford, John Sillitoe on behalf of the Wellcome Sanger Institute COVID-19 Surveillance Team |
| EPI_ISL_2553806 | Lighthouse Lab in Alderley Park | Wellcome Sanger Institute for the COVID-19 Genomics UK (COG-UK) Consortium | Jacquelyn Wynn, Mairead Hyland, The Lighthouse Lab in Alderley Park and Alex Alderton, Roberto Amato, Jeffrey Barrett, Sonia Goncalves, Ewan Harrison, David K. Jackson, Ian Johnston, Dominic Kwiatkowski, Cordelia Langford, John Sillitoe on behalf of the Wellcome Sanger Institute COVID-19 Surveillance Team |
| EPI_ISL_2553807 | Health Services Laboratories | Wellcome Sanger Institute for the COVID-19 Genomics UK (COG-UK) Consortium | Health Services Laboratories and Alex Alderton, Roberto Amato, Jeffrey Barrett, Sonia Goncalves, Ewan Harrison, David K. Jackson, Ian Johnston, Dominic Kwiatkowski, Cordelia Langford, John Sillitoe on behalf of the Wellcome Sanger Institute COVID-19 Surveillance Team |
| EPI_ISL_2553808, EPI_ISL_2553809 | Lighthouse Lab in Milton Keynes | Wellcome Sanger Institute for the COVID-19 Genomics UK (COG-UK) Consortium | The Lighthouse Lab in Milton Keynes and Alex Alderton, Roberto Amato, Jeffrey Barrett, Sonia Goncalves, Ewan Harrison, David K. Jackson, Ian Johnston, Dominic Kwiatkowski, Cordelia Langford, John Sillitoe on behalf of the Wellcome Sanger Institute COVID-19 Surveillance Team |
| EPI_ISL_2553810, EPI_ISL_2553811 | Health Services Laboratories | Wellcome Sanger Institute for the COVID-19 Genomics UK (COG-UK) Consortium | Health Services Laboratories and Alex Alderton, Roberto Amato, Jeffrey Barrett, Sonia Goncalves, Ewan Harrison, David K. Jackson, Ian Johnston, Dominic Kwiatkowski, Cordelia Langford, John Sillitoe on behalf of the Wellcome Sanger Institute COVID-19 Surveillance Team |
| EPI_ISL_2553812 | Lighthouse Lab in Alderley Park | Wellcome Sanger Institute for the COVID-19 Genomics UK (COG-UK) Consortium | Jacquelyn Wynn, Mairead Hyland, The Lighthouse Lab in Alderley Park and Alex Alderton, Roberto Amato, Jeffrey Barrett, Sonia Goncalves, Ewan Harrison, David K. Jackson, Ian Johnston, Dominic Kwiatkowski, Cordelia Langford, John Sillitoe on behalf of the Wellcome Sanger Institute COVID-19 Surveillance Team |
| EPI_ISL_2553813 | Health Services Laboratories | Wellcome Sanger Institute for the COVID-19 Genomics UK (COG-UK) Consortium | Health Services Laboratories and Alex Alderton, Roberto Amato, Jeffrey Barrett, Sonia Goncalves, Ewan Harrison, David K. Jackson, Ian Johnston, Dominic Kwiatkowski, Cordelia Langford, John Sillitoe on behalf of the Wellcome Sanger Institute COVID-19 Surveillance Team |
| EPI_ISL_2553814, EPI_ISL_2553815, EPI_ISL_2553816 | Lighthouse Lab in Milton Keynes | Wellcome Sanger Institute for the COVID-19 Genomics UK (COG-UK) Consortium | The Lighthouse Lab in Milton Keynes and Alex Alderton, Roberto Amato, Jeffrey Barrett, Sonia Goncalves, Ewan Harrison, David K. Jackson, Ian Johnston, Dominic Kwiatkowski, Cordelia Langford, John Sillitoe on behalf of the Wellcome Sanger Institute COVID-19 Surveillance Team |
| EPI_ISL_2553817 | Health Services Laboratories | Wellcome Sanger Institute for the COVID-19 Genomics UK (COG-UK) Consortium | Health Services Laboratories and Alex Alderton, Roberto Amato, Jeffrey Barrett, Sonia Goncalves, Ewan Harrison, David K. Jackson, Ian Johnston, Dominic Kwiatkowski, Cordelia Langford, John Sillitoe on behalf of the Wellcome Sanger Institute COVID-19 Surveillance Team |
| EPI_ISL_2553818 | Lighthouse Lab in Milton Keynes | Wellcome Sanger Institute for the COVID-19 Genomics UK (COG-UK) Consortium | The Lighthouse Lab in Milton Keynes and Alex Alderton, Roberto Amato, Jeffrey Barrett, Sonia Goncalves, Ewan Harrison, David K. Jackson, Ian Johnston, Dominic Kwiatkowski, Cordelia Langford, John Sillitoe on behalf of the Wellcome Sanger Institute COVID-19 Surveillance Team |
| EPI_ISL_2553819 | Lighthouse Lab in Alderley Park | Wellcome Sanger Institute for the COVID-19 Genomics UK (COG-UK) Consortium | Jacquelyn Wynn, Mairead Hyland, The Lighthouse Lab in Alderley Park and Alex Alderton, Roberto Amato, Jeffrey Barrett, Sonia Goncalves, Ewan Harrison, David K. Jackson, Ian Johnston, Dominic Kwiatkowski, Cordelia Langford, John Sillitoe on behalf of the Wellcome Sanger Institute COVID-19 Surveillance Team |
| EPI_ISL_2553821, EPI_ISL_2553822, EPI_ISL_2553824 | Health Services Laboratories | Wellcome Sanger Institute for the COVID-19 Genomics UK (COG-UK) Consortium | Health Services Laboratories and Alex Alderton, Roberto Amato, Jeffrey Barrett, Sonia Goncalves, Ewan Harrison, David K. Jackson, Ian Johnston, Dominic Kwiatkowski, Cordelia Langford, John Sillitoe on behalf of the Wellcome Sanger Institute COVID-19 Surveillance Team |
| EPI_ISL_2553825 | Lighthouse Lab in Alderley Park | Wellcome Sanger Institute for the COVID-19 Genomics UK (COG-UK) Consortium | Jacquelyn Wynn, Mairead Hyland, The Lighthouse Lab in Alderley Park and Alex Alderton, Roberto Amato, Jeffrey Barrett, Sonia Goncalves, Ewan Harrison, David K. Jackson, Ian Johnston, Dominic Kwiatkowski, Cordelia Langford, John Sillitoe on behalf of the Wellcome Sanger Institute COVID-19 Surveillance Team |
| EPI_ISL_2553826 | Health Services Laboratories | Wellcome Sanger Institute for the COVID-19 Genomics UK (COG-UK) Consortium | Health Services Laboratories and Alex Alderton, Roberto Amato, Jeffrey Barrett, Sonia Goncalves, Ewan Harrison, David K. Jackson, Ian Johnston, Dominic Kwiatkowski, Cordelia Langford, John Sillitoe on behalf of the Wellcome Sanger Institute COVID-19 Surveillance Team |
| EPI_ISL_2553827, EPI_ISL_2553828, EPI_ISL_2553829 | Lighthouse Lab in Milton Keynes | Wellcome Sanger Institute for the COVID-19 Genomics UK (COG-UK) Consortium | The Lighthouse Lab in Milton Keynes and Alex Alderton, Roberto Amato, Jeffrey Barrett, Sonia Goncalves, Ewan Harrison, David K. Jackson, Ian Johnston, Dominic Kwiatkowski, Cordelia Langford, John Sillitoe on behalf of the Wellcome Sanger Institute COVID-19 Surveillance Team |
| EPI_ISL_2553830 | Lighthouse Lab in Alderley Park | Wellcome Sanger Institute for the COVID-19 Genomics UK (COG-UK) Consortium | Jacquelyn Wynn, Mairead Hyland, The Lighthouse Lab in Alderley Park and Alex Alderton, Roberto Amato, Jeffrey Barrett, Sonia Goncalves, Ewan Harrison, David K. Jackson, Ian Johnston, Dominic Kwiatkowski, Cordelia Langford, John Sillitoe on behalf of the Wellcome Sanger Institute COVID-19 Surveillance Team |

| | | | |
|---|---|---|---|
| EPI_ISL_2553831, EPI_ISL_2553832 | Lighthouse Lab in Milton Keynes | Wellcome Sanger Institute for the COVID-19 Genomics UK (COG-UK) Consortium | The Lighthouse Lab in Milton Keynes and Alex Alderton, Roberto Amato, Jeffrey Barrett, Sonia Goncalves, Ewan Harrison, David K. Jackson, Ian Johnston, Dominic Kwiatkowski, Cordelia Langford, John Sillitoe on behalf of the Wellcome Sanger Institute COVID-19 Surveillance Team |
| EPI_ISL_2553833, EPI_ISL_2553834 | Health Services Laboratories | Wellcome Sanger Institute for the COVID-19 Genomics UK (COG-UK) Consortium | Health Services Laboratories and Alex Alderton, Roberto Amato, Jeffrey Barrett, Sonia Goncalves, Ewan Harrison, David K. Jackson, Ian Johnston, Dominic Kwiatkowski, Cordelia Langford, John Sillitoe on behalf of the Wellcome Sanger Institute COVID-19 Surveillance Team |
| EPI_ISL_2553835 | Lighthouse Lab in Milton Keynes | Wellcome Sanger Institute for the COVID-19 Genomics UK (COG-UK) Consortium | The Lighthouse Lab in Milton Keynes and Alex Alderton, Roberto Amato, Jeffrey Barrett, Sonia Goncalves, Ewan Harrison, David K. Jackson, Ian Johnston, Dominic Kwiatkowski, Cordelia Langford, John Sillitoe on behalf of the Wellcome Sanger Institute COVID-19 Surveillance Team |
| EPI_ISL_2553836 | Lighthouse Lab in Alderley Park | Wellcome Sanger Institute for the COVID-19 Genomics UK (COG-UK) Consortium | Jacquelyn Wynn, Mairead Hyland, The Lighthouse Lab in Alderley Park and Alex Alderton, Roberto Amato, Jeffrey Barrett, Sonia Goncalves, Ewan Harrison, David K. Jackson, Ian Johnston, Dominic Kwiatkowski, Cordelia Langford, John Sillitoe on behalf of the Wellcome Sanger Institute COVID-19 Surveillance Team |
| EPI_ISL_2553837 | Lighthouse Lab in Milton Keynes | Wellcome Sanger Institute for the COVID-19 Genomics UK (COG-UK) Consortium | The Lighthouse Lab in Milton Keynes and Alex Alderton, Roberto Amato, Jeffrey Barrett, Sonia Goncalves, Ewan Harrison, David K. Jackson, Ian Johnston, Dominic Kwiatkowski, Cordelia Langford, John Sillitoe on behalf of the Wellcome Sanger Institute COVID-19 Surveillance Team |
| EPI_ISL_2553838, EPI_ISL_2553839 | Health Services Laboratories | Wellcome Sanger Institute for the COVID-19 Genomics UK (COG-UK) Consortium | Health Services Laboratories and Alex Alderton, Roberto Amato, Jeffrey Barrett, Sonia Goncalves, Ewan Harrison, David K. Jackson, Ian Johnston, Dominic Kwiatkowski, Cordelia Langford, John Sillitoe on behalf of the Wellcome Sanger Institute COVID-19 Surveillance Team |
| EPI_ISL_2553840 | Lighthouse Lab in Milton Keynes | Wellcome Sanger Institute for the COVID-19 Genomics UK (COG-UK) Consortium | The Lighthouse Lab in Milton Keynes and Alex Alderton, Roberto Amato, Jeffrey Barrett, Sonia Goncalves, Ewan Harrison, David K. Jackson, Ian Johnston, Dominic Kwiatkowski, Cordelia Langford, John Sillitoe on behalf of the Wellcome Sanger Institute COVID-19 Surveillance Team |
| EPI_ISL_2553841 | Health Services Laboratories | Wellcome Sanger Institute for the COVID-19 Genomics UK (COG-UK) Consortium | Health Services Laboratories and Alex Alderton, Roberto Amato, Jeffrey Barrett, Sonia Goncalves, Ewan Harrison, David K. Jackson, Ian Johnston, Dominic Kwiatkowski, Cordelia Langford, John Sillitoe on behalf of the Wellcome Sanger Institute COVID-19 Surveillance Team |
| EPI_ISL_2553842 | Lighthouse Lab in Alderley Park | Wellcome Sanger Institute for the COVID-19 Genomics UK (COG-UK) Consortium | Jacquelyn Wynn, Mairead Hyland, The Lighthouse Lab in Alderley Park and Alex Alderton, Roberto Amato, Jeffrey Barrett, Sonia Goncalves, Ewan Harrison, David K. Jackson, Ian Johnston, Dominic Kwiatkowski, Cordelia Langford, John Sillitoe on behalf of the Wellcome Sanger Institute COVID-19 Surveillance Team |
| EPI_ISL_2553843, EPI_ISL_2553844 | Health Services Laboratories | Wellcome Sanger Institute for the COVID-19 Genomics UK (COG-UK) Consortium | Health Services Laboratories and Alex Alderton, Roberto Amato, Jeffrey Barrett, Sonia Goncalves, Ewan Harrison, David K. Jackson, Ian Johnston, Dominic Kwiatkowski, Cordelia Langford, John Sillitoe on behalf of the Wellcome Sanger Institute COVID-19 Surveillance Team |
| EPI_ISL_2553845 | Lighthouse Lab in Alderley Park | Wellcome Sanger Institute for the COVID-19 Genomics UK (COG-UK) Consortium | Jacquelyn Wynn, Mairead Hyland, The Lighthouse Lab in Alderley Park and Alex Alderton, Roberto Amato, Jeffrey Barrett, Sonia Goncalves, Ewan Harrison, David K. Jackson, Ian Johnston, Dominic Kwiatkowski, Cordelia Langford, John Sillitoe on behalf of the Wellcome Sanger Institute COVID-19 Surveillance Team |
| EPI_ISL_2553846 | Lighthouse Lab in Milton Keynes | Wellcome Sanger Institute for the COVID-19 Genomics UK (COG-UK) Consortium | The Lighthouse Lab in Milton Keynes and Alex Alderton, Roberto Amato, Jeffrey Barrett, Sonia Goncalves, Ewan Harrison, David K. Jackson, Ian Johnston, Dominic Kwiatkowski, Cordelia Langford, John Sillitoe on behalf of the Wellcome Sanger Institute COVID-19 Surveillance Team |
| EPI_ISL_2553847 | Health Services Laboratories | Wellcome Sanger Institute for the COVID-19 Genomics UK (COG-UK) Consortium | Health Services Laboratories and Alex Alderton, Roberto Amato, Jeffrey Barrett, Sonia Goncalves, Ewan Harrison, David K. Jackson, Ian Johnston, Dominic Kwiatkowski, Cordelia Langford, John Sillitoe on behalf of the Wellcome Sanger Institute COVID-19 Surveillance Team |
| EPI_ISL_2553848 | Lighthouse Lab in Milton Keynes | Wellcome Sanger Institute for the COVID-19 Genomics UK (COG-UK) Consortium | The Lighthouse Lab in Milton Keynes and Alex Alderton, Roberto Amato, Jeffrey Barrett, Sonia Goncalves, Ewan Harrison, David K. Jackson, Ian Johnston, Dominic Kwiatkowski, Cordelia Langford, John Sillitoe on behalf of the Wellcome Sanger Institute COVID-19 Surveillance Team |
| EPI_ISL_2553849 | Health Services Laboratories | Wellcome Sanger Institute for the COVID-19 Genomics UK (COG-UK) Consortium | Health Services Laboratories and Alex Alderton, Roberto Amato, Jeffrey Barrett, Sonia Goncalves, Ewan Harrison, David K. Jackson, Ian Johnston, Dominic Kwiatkowski, Cordelia Langford, John Sillitoe on behalf of the Wellcome Sanger Institute COVID-19 Surveillance Team |
| EPI_ISL_2553850 | Lighthouse Lab in Alderley Park | Wellcome Sanger Institute for the COVID-19 Genomics UK (COG-UK) Consortium | Jacquelyn Wynn, Mairead Hyland, The Lighthouse Lab in Alderley Park and Alex Alderton, Roberto Amato, Jeffrey Barrett, Sonia Goncalves, Ewan Harrison, David K. Jackson, Ian Johnston, Dominic Kwiatkowski, Cordelia Langford, John Sillitoe on behalf of the Wellcome Sanger Institute COVID-19 Surveillance Team |
| EPI_ISL_2553851 | Health Services Laboratories | Wellcome Sanger Institute for the COVID-19 Genomics UK (COG-UK) Consortium | Health Services Laboratories and Alex Alderton, Roberto Amato, Jeffrey Barrett, Sonia Goncalves, Ewan Harrison, David K. Jackson, Ian Johnston, Dominic Kwiatkowski, Cordelia Langford, John Sillitoe on behalf of the Wellcome Sanger Institute COVID-19 Surveillance Team |
| EPI_ISL_2553852, EPI_ISL_2553854 | Lighthouse Lab in Milton Keynes | Wellcome Sanger Institute for the COVID-19 Genomics UK (COG-UK) Consortium | The Lighthouse Lab in Milton Keynes and Alex Alderton, Roberto Amato, Jeffrey Barrett, Sonia Goncalves, Ewan Harrison, David K. Jackson, Ian Johnston, Dominic Kwiatkowski, Cordelia Langford, John Sillitoe on behalf of the Wellcome Sanger Institute COVID-19 Surveillance Team |
| EPI_ISL_2553855 | Lighthouse Lab in Alderley Park | Wellcome Sanger Institute for the COVID-19 Genomics UK (COG-UK) Consortium | Jacquelyn Wynn, Mairead Hyland, The Lighthouse Lab in Alderley Park and Alex Alderton, Roberto Amato, Jeffrey Barrett, Sonia Goncalves, Ewan Harrison, David K. Jackson, Ian Johnston, Dominic Kwiatkowski, Cordelia Langford, John Sillitoe on behalf of the Wellcome Sanger Institute COVID-19 Surveillance Team |
| EPI_ISL_2553856, EPI_ISL_2553857, EPI_ISL_2553858 | Health Services Laboratories | Wellcome Sanger Institute for the COVID-19 Genomics UK (COG-UK) Consortium | Health Services Laboratories and Alex Alderton, Roberto Amato, Jeffrey Barrett, Sonia Goncalves, Ewan Harrison, David K. Jackson, Ian Johnston, Dominic Kwiatkowski, Cordelia Langford, John Sillitoe on behalf of the Wellcome Sanger Institute COVID-19 Surveillance Team |
| EPI_ISL_2553859 | Lighthouse Lab in Milton Keynes | Wellcome Sanger Institute for the COVID-19 Genomics UK (COG-UK) Consortium | The Lighthouse Lab in Milton Keynes and Alex Alderton, Roberto Amato, Jeffrey Barrett, Sonia Goncalves, Ewan Harrison, David K. Jackson, Ian Johnston, Dominic Kwiatkowski, Cordelia Langford, John Sillitoe on behalf of the Wellcome Sanger Institute COVID-19 Surveillance Team |
| EPI_ISL_2553860 | Health Services Laboratories | Wellcome Sanger Institute for the COVID-19 Genomics UK (COG-UK) Consortium | Health Services Laboratories and Alex Alderton, Roberto Amato, Jeffrey Barrett, Sonia Goncalves, Ewan Harrison, David K. Jackson, Ian Johnston, Dominic Kwiatkowski, Cordelia Langford, John Sillitoe on behalf of the Wellcome Sanger Institute COVID-19 Surveillance Team |
| EPI_ISL_2553861 | Lighthouse Lab in Milton Keynes | Wellcome Sanger Institute for the COVID-19 Genomics UK (COG-UK) Consortium | The Lighthouse Lab in Milton Keynes and Alex Alderton, Roberto Amato, Jeffrey Barrett, Sonia Goncalves, Ewan Harrison, David K. Jackson, Ian Johnston, Dominic Kwiatkowski, Cordelia Langford, John Sillitoe on behalf of the Wellcome Sanger Institute COVID-19 Surveillance Team |
| EPI_ISL_2553862 | Health Services Laboratories | Wellcome Sanger Institute for the COVID-19 Genomics UK (COG-UK) Consortium | Health Services Laboratories and Alex Alderton, Roberto Amato, Jeffrey Barrett, Sonia Goncalves, Ewan Harrison, David K. Jackson, Ian Johnston, Dominic Kwiatkowski, Cordelia Langford, John Sillitoe on behalf of the Wellcome Sanger Institute COVID-19 Surveillance Team |
| EPI_ISL_2553863, EPI_ISL_2553864 | Lighthouse Lab in Milton Keynes | Wellcome Sanger Institute for the COVID-19 Genomics UK (COG-UK) Consortium | The Lighthouse Lab in Milton Keynes and Alex Alderton, Roberto Amato, Jeffrey Barrett, Sonia Goncalves, Ewan Harrison, David K. Jackson, Ian Johnston, Dominic Kwiatkowski, Cordelia Langford, John Sillitoe on behalf of the Wellcome Sanger Institute COVID-19 Surveillance Team |
| EPI_ISL_2553865, EPI_ISL_2553866 | Health Services Laboratories | Wellcome Sanger Institute for the COVID-19 Genomics UK (COG-UK) Consortium | Health Services Laboratories and Alex Alderton, Roberto Amato, Jeffrey Barrett, Sonia Goncalves, Ewan Harrison, David K. Jackson, Ian Johnston, Dominic Kwiatkowski, Cordelia Langford, John Sillitoe on behalf of the Wellcome Sanger Institute COVID-19 Surveillance Team |
| EPI_ISL_2553867 | Lighthouse Lab in Alderley Park | Wellcome Sanger Institute for the COVID-19 Genomics UK (COG-UK) Consortium | Jacquelyn Wynn, Mairead Hyland, The Lighthouse Lab in Alderley Park and Alex Alderton, Roberto Amato, Jeffrey Barrett, Sonia Goncalves, Ewan Harrison, David K. Jackson, Ian Johnston, Dominic Kwiatkowski, Cordelia Langford, John Sillitoe on behalf of the Wellcome Sanger Institute COVID-19 Surveillance Team |
| EPI_ISL_2553868 | Lighthouse Lab in Milton Keynes | Wellcome Sanger Institute for the COVID-19 Genomics UK (COG-UK) Consortium | The Lighthouse Lab in Milton Keynes and Alex Alderton, Roberto Amato, Jeffrey Barrett, Sonia Goncalves, Ewan Harrison, David K. Jackson, Ian Johnston, Dominic Kwiatkowski, Cordelia Langford, John Sillitoe on behalf of the Wellcome Sanger Institute COVID-19 Surveillance Team |
| EPI_ISL_2553869 | Health Services Laboratories | Wellcome Sanger Institute for the COVID-19 Genomics UK (COG-UK) Consortium | Health Services Laboratories and Alex Alderton, Roberto Amato, Jeffrey Barrett, Sonia Goncalves, Ewan Harrison, David K. Jackson, Ian Johnston, Dominic Kwiatkowski, Cordelia Langford, John Sillitoe on behalf of the Wellcome Sanger Institute COVID-19 Surveillance Team |
| EPI_ISL_2553870, EPI_ISL_2553871 | Lighthouse Lab in Alderley Park | Wellcome Sanger Institute for the COVID-19 Genomics UK (COG-UK) Consortium | Jacquelyn Wynn, Mairead Hyland, The Lighthouse Lab in Alderley Park and Alex Alderton, Roberto Amato, Jeffrey Barrett, Sonia Goncalves, Ewan Harrison, David K. Jackson, Ian Johnston, Dominic Kwiatkowski, Cordelia Langford, John Sillitoe on behalf of the Wellcome Sanger Institute COVID-19 Surveillance Team |
| EPI_ISL_2553872 | Health Services Laboratories | Wellcome Sanger Institute for the COVID-19 Genomics UK (COG-UK) Consortium | Health Services Laboratories and Alex Alderton, Roberto Amato, Jeffrey Barrett, Sonia Goncalves, Ewan Harrison, David K. Jackson, Ian Johnston, Dominic Kwiatkowski, Cordelia Langford, John Sillitoe on behalf of the Wellcome Sanger Institute COVID-19 Surveillance Team |
| EPI_ISL_2553873 | Lighthouse Lab in Milton Keynes | Wellcome Sanger Institute for the COVID-19 Genomics UK (COG-UK) Consortium | The Lighthouse Lab in Milton Keynes and Alex Alderton, Roberto Amato, Jeffrey Barrett, Sonia Goncalves, Ewan Harrison, David K. Jackson, Ian Johnston, Dominic Kwiatkowski, Cordelia Langford, John Sillitoe on behalf of the Wellcome Sanger Institute COVID-19 Surveillance Team |
| EPI_ISL_2553874, EPI_ISL_2553875, EPI_ISL_2553876, EPI_ISL_2553877 | Health Services Laboratories | Wellcome Sanger Institute for the COVID-19 Genomics UK (COG-UK) Consortium | Health Services Laboratories and Alex Alderton, Roberto Amato, Jeffrey Barrett, Sonia Goncalves, Ewan Harrison, David K. Jackson, Ian Johnston, Dominic Kwiatkowski, Cordelia Langford, John Sillitoe on behalf of the Wellcome Sanger Institute COVID-19 Surveillance Team |
| EPI_ISL_2553878 | Lighthouse Lab in Alderley Park | Wellcome Sanger Institute for the COVID-19 Genomics UK (COG-UK) Consortium | Jacquelyn Wynn, Mairead Hyland, The Lighthouse Lab in Alderley Park and Alex Alderton, Roberto Amato, Jeffrey Barrett, Sonia Goncalves, Ewan Harrison, David K. Jackson, Ian Johnston, Dominic Kwiatkowski, Cordelia Langford, John Sillitoe on behalf of the Wellcome Sanger Institute COVID-19 Surveillance Team |
| EPI_ISL_2553879 | Health Services Laboratories | Wellcome Sanger Institute for the COVID-19 Genomics UK (COG-UK) Consortium | Health Services Laboratories and Alex Alderton, Roberto Amato, Jeffrey Barrett, Sonia Goncalves, Ewan Harrison, David K. Jackson, Ian Johnston, Dominic Kwiatkowski, Cordelia Langford, John Sillitoe on behalf of the Wellcome Sanger Institute COVID-19 Surveillance Team |
| EPI_ISL_2553881 | Lighthouse Lab in Alderley Park | Wellcome Sanger Institute for the COVID-19 Genomics UK (COG-UK) Consortium | Jacquelyn Wynn, Mairead Hyland, The Lighthouse Lab in Alderley Park and Alex Alderton, Roberto Amato, Jeffrey Barrett, Sonia Goncalves, Ewan Harrison, David K. Jackson, Ian Johnston, Dominic Kwiatkowski, Cordelia Langford, John Sillitoe on behalf of the Wellcome Sanger Institute COVID-19 Surveillance Team |
| EPI_ISL_2553882, EPI_ISL_2553883, EPI_ISL_2553884 | Health Services Laboratories | Wellcome Sanger Institute for the COVID-19 Genomics UK (COG-UK) Consortium | Health Services Laboratories and Alex Alderton, Roberto Amato, Jeffrey Barrett, Sonia Goncalves, Ewan Harrison, David K. Jackson, Ian Johnston, Dominic Kwiatkowski, Cordelia Langford, John Sillitoe on behalf of the Wellcome Sanger Institute COVID-19 Surveillance Team |
| EPI_ISL_2553885, EPI_ISL_2553886 | Lighthouse Lab in Milton Keynes | Wellcome Sanger Institute for the COVID-19 Genomics UK (COG-UK) Consortium | The Lighthouse Lab in Milton Keynes and Alex Alderton, Roberto Amato, Jeffrey Barrett, Sonia Goncalves, Ewan Harrison, David K. Jackson, Ian Johnston, Dominic Kwiatkowski, Cordelia Langford, John Sillitoe on behalf of the Wellcome Sanger Institute COVID-19 Surveillance Team |
| EPI_ISL_2553887, EPI_ISL_2553889 | Lighthouse Lab in Alderley Park | Wellcome Sanger Institute for the COVID-19 Genomics UK (COG-UK) Consortium | Jacquelyn Wynn, Mairead Hyland, The Lighthouse Lab in Alderley Park and Alex Alderton, Roberto Amato, Jeffrey Barrett, Sonia Goncalves, Ewan Harrison, David K. Jackson, Ian Johnston, Dominic Kwiatkowski, Cordelia Langford, John Sillitoe on behalf of the Wellcome Sanger Institute COVID-19 Surveillance Team |
| EPI_ISL_2553890 | Health Services Laboratories | Wellcome Sanger Institute for the COVID-19 Genomics UK (COG-UK) Consortium | Health Services Laboratories and Alex Alderton, Roberto Amato, Jeffrey Barrett, Sonia Goncalves, Ewan Harrison, David K. Jackson, Ian Johnston, Dominic Kwiatkowski, Cordelia Langford, John Sillitoe on behalf of the Wellcome Sanger Institute COVID-19 Surveillance Team |
| EPI_ISL_2553892, EPI_ISL_2553893 | Lighthouse Lab in Milton Keynes | Wellcome Sanger Institute for the COVID-19 Genomics UK (COG-UK) Consortium | The Lighthouse Lab in Milton Keynes and Alex Alderton, Roberto Amato, Jeffrey Barrett, Sonia Goncalves, Ewan Harrison, David K. Jackson, Ian Johnston, Dominic Kwiatkowski, Cordelia Langford, John Sillitoe on behalf of the Wellcome Sanger Institute COVID-19 Surveillance Team |
| EPI_ISL_2553894, EPI_ISL_2553895 | Health Services Laboratories | Wellcome Sanger Institute for the COVID-19 Genomics UK (COG-UK) Consortium | Health Services Laboratories and Alex Alderton, Roberto Amato, Jeffrey Barrett, Sonia Goncalves, Ewan Harrison, David K. Jackson, Ian Johnston, Dominic Kwiatkowski, Cordelia Langford, John Sillitoe on behalf of the Wellcome Sanger Institute COVID-19 Surveillance Team |
| EPI_ISL_2553896 | Lighthouse Lab in Alderley Park | Wellcome Sanger Institute for the COVID-19 Genomics UK (COG-UK) Consortium | Jacquelyn Wynn, Mairead Hyland, The Lighthouse Lab in Alderley Park and Alex Alderton, Roberto Amato, Jeffrey Barrett, Sonia Goncalves, Ewan Harrison, David K. Jackson, Ian Johnston, Dominic Kwiatkowski, Cordelia Langford, John Sillitoe on behalf of the Wellcome Sanger Institute COVID-19 Surveillance Team |
| EPI_ISL_2553897, | Health Services Laboratories | Wellcome Sanger Institute for the COVID-19 Genomics UK (COG-UK) | Health Services Laboratories and Alex Alderton, Roberto Amato, Jeffrey Barrett, Sonia Goncalves, Ewan Harrison, David K. Jackson, Ian Johnston, Dominic Kwiatkowski, Cordelia Langford, John Sillitoe on behalf of the Wellcome Sanger Institute |

| | | | |
|---|---|---|---|
| EPI_ISL_2553898, EPI_ISL_2553899 | | Consortium | COVID-19 Surveillance Team |
| EPI_ISL_2553900 | Lighthouse Lab in Alderley Park | Wellcome Sanger Institute for the COVID-19 Genomics UK (COG-UK) Consortium | Jacquelyn Wynn, Mairead Hyland, The Lighthouse Lab in Alderley Park and Alex Alderton, Roberto Amato, Jeffrey Barrett, Sonia Goncalves, Ewan Harrison, David K. Jackson, Ian Johnston, Dominic Kwiatkowski, Cordelia Langford, John Sillitoe on behalf of the Wellcome Sanger Institute COVID-19 Surveillance Team |
| EPI_ISL_2553901, EPI_ISL_2553902 | Health Services Laboratories | Wellcome Sanger Institute for the COVID-19 Genomics UK (COG-UK) Consortium | Health Services Laboratories and Alex Alderton, Roberto Amato, Jeffrey Barrett, Sonia Goncalves, Ewan Harrison, David K. Jackson, Ian Johnston, Dominic Kwiatkowski, Cordelia Langford, John Sillitoe on behalf of the Wellcome Sanger Institute COVID-19 Surveillance Team |
| EPI_ISL_2553903, EPI_ISL_2553904 | Lighthouse Lab in Alderley Park | Wellcome Sanger Institute for the COVID-19 Genomics UK (COG-UK) Consortium | Jacquelyn Wynn, Mairead Hyland, The Lighthouse Lab in Alderley Park and Alex Alderton, Roberto Amato, Jeffrey Barrett, Sonia Goncalves, Ewan Harrison, David K. Jackson, Ian Johnston, Dominic Kwiatkowski, Cordelia Langford, John Sillitoe on behalf of the Wellcome Sanger Institute COVID-19 Surveillance Team |
| EPI_ISL_2553905 | Lighthouse Lab in Milton Keynes | Wellcome Sanger Institute for the COVID-19 Genomics UK (COG-UK) Consortium | The Lighthouse Lab in Milton Keynes and Alex Alderton, Roberto Amato, Jeffrey Barrett, Sonia Goncalves, Ewan Harrison, David K. Jackson, Ian Johnston, Dominic Kwiatkowski, Cordelia Langford, John Sillitoe on behalf of the Wellcome Sanger Institute COVID-19 Surveillance Team |
| EPI_ISL_2553906, EPI_ISL_2553907, EPI_ISL_2553908 | Health Services Laboratories | Wellcome Sanger Institute for the COVID-19 Genomics UK (COG-UK) Consortium | Health Services Laboratories and Alex Alderton, Roberto Amato, Jeffrey Barrett, Sonia Goncalves, Ewan Harrison, David K. Jackson, Ian Johnston, Dominic Kwiatkowski, Cordelia Langford, John Sillitoe on behalf of the Wellcome Sanger Institute COVID-19 Surveillance Team |
| EPI_ISL_2553909 | Lighthouse Lab in Milton Keynes | Wellcome Sanger Institute for the COVID-19 Genomics UK (COG-UK) Consortium | The Lighthouse Lab in Milton Keynes and Alex Alderton, Roberto Amato, Jeffrey Barrett, Sonia Goncalves, Ewan Harrison, David K. Jackson, Ian Johnston, Dominic Kwiatkowski, Cordelia Langford, John Sillitoe on behalf of the Wellcome Sanger Institute COVID-19 Surveillance Team |
| EPI_ISL_2553910, EPI_ISL_2553911, EPI_ISL_2553912, EPI_ISL_2553913 | Health Services Laboratories | Wellcome Sanger Institute for the COVID-19 Genomics UK (COG-UK) Consortium | Health Services Laboratories and Alex Alderton, Roberto Amato, Jeffrey Barrett, Sonia Goncalves, Ewan Harrison, David K. Jackson, Ian Johnston, Dominic Kwiatkowski, Cordelia Langford, John Sillitoe on behalf of the Wellcome Sanger Institute COVID-19 Surveillance Team |
| EPI_ISL_2553914 | Lighthouse Lab in Alderley Park | Wellcome Sanger Institute for the COVID-19 Genomics UK (COG-UK) Consortium | Jacquelyn Wynn, Mairead Hyland, The Lighthouse Lab in Alderley Park and Alex Alderton, Roberto Amato, Jeffrey Barrett, Sonia Goncalves, Ewan Harrison, David K. Jackson, Ian Johnston, Dominic Kwiatkowski, Cordelia Langford, John Sillitoe on behalf of the Wellcome Sanger Institute COVID-19 Surveillance Team |
| EPI_ISL_2553915 | Lighthouse Lab in Milton Keynes | Wellcome Sanger Institute for the COVID-19 Genomics UK (COG-UK) Consortium | The Lighthouse Lab in Milton Keynes and Alex Alderton, Roberto Amato, Jeffrey Barrett, Sonia Goncalves, Ewan Harrison, David K. Jackson, Ian Johnston, Dominic Kwiatkowski, Cordelia Langford, John Sillitoe on behalf of the Wellcome Sanger Institute COVID-19 Surveillance Team |
| EPI_ISL_2553916 | Lighthouse Lab in Alderley Park | Wellcome Sanger Institute for the COVID-19 Genomics UK (COG-UK) Consortium | Jacquelyn Wynn, Mairead Hyland, The Lighthouse Lab in Alderley Park and Alex Alderton, Roberto Amato, Jeffrey Barrett, Sonia Goncalves, Ewan Harrison, David K. Jackson, Ian Johnston, Dominic Kwiatkowski, Cordelia Langford, John Sillitoe on behalf of the Wellcome Sanger Institute COVID-19 Surveillance Team |
| EPI_ISL_2553917 | Health Services Laboratories | Wellcome Sanger Institute for the COVID-19 Genomics UK (COG-UK) Consortium | Health Services Laboratories and Alex Alderton, Roberto Amato, Jeffrey Barrett, Sonia Goncalves, Ewan Harrison, David K. Jackson, Ian Johnston, Dominic Kwiatkowski, Cordelia Langford, John Sillitoe on behalf of the Wellcome Sanger Institute COVID-19 Surveillance Team |
| EPI_ISL_2553918 | Lighthouse Lab in Milton Keynes | Wellcome Sanger Institute for the COVID-19 Genomics UK (COG-UK) Consortium | The Lighthouse Lab in Milton Keynes and Alex Alderton, Roberto Amato, Jeffrey Barrett, Sonia Goncalves, Ewan Harrison, David K. Jackson, Ian Johnston, Dominic Kwiatkowski, Cordelia Langford, John Sillitoe on behalf of the Wellcome Sanger Institute COVID-19 Surveillance Team |
| EPI_ISL_2553920 | Health Services Laboratories | Wellcome Sanger Institute for the COVID-19 Genomics UK (COG-UK) Consortium | Health Services Laboratories and Alex Alderton, Roberto Amato, Jeffrey Barrett, Sonia Goncalves, Ewan Harrison, David K. Jackson, Ian Johnston, Dominic Kwiatkowski, Cordelia Langford, John Sillitoe on behalf of the Wellcome Sanger Institute COVID-19 Surveillance Team |
| EPI_ISL_2553921 | Lighthouse Lab in Alderley Park | Wellcome Sanger Institute for the COVID-19 Genomics UK (COG-UK) Consortium | Jacquelyn Wynn, Mairead Hyland, The Lighthouse Lab in Alderley Park and Alex Alderton, Roberto Amato, Jeffrey Barrett, Sonia Goncalves, Ewan Harrison, David K. Jackson, Ian Johnston, Dominic Kwiatkowski, Cordelia Langford, John Sillitoe on behalf of the Wellcome Sanger Institute COVID-19 Surveillance Team |
| EPI_ISL_2553922 | Health Services Laboratories | Wellcome Sanger Institute for the COVID-19 Genomics UK (COG-UK) Consortium | Health Services Laboratories and Alex Alderton, Roberto Amato, Jeffrey Barrett, Sonia Goncalves, Ewan Harrison, David K. Jackson, Ian Johnston, Dominic Kwiatkowski, Cordelia Langford, John Sillitoe on behalf of the Wellcome Sanger Institute COVID-19 Surveillance Team |
| EPI_ISL_2553923 | Lighthouse Lab in Milton Keynes | Wellcome Sanger Institute for the COVID-19 Genomics UK (COG-UK) Consortium | The Lighthouse Lab in Milton Keynes and Alex Alderton, Roberto Amato, Jeffrey Barrett, Sonia Goncalves, Ewan Harrison, David K. Jackson, Ian Johnston, Dominic Kwiatkowski, Cordelia Langford, John Sillitoe on behalf of the Wellcome Sanger Institute COVID-19 Surveillance Team |
| EPI_ISL_2553924, EPI_ISL_2553925, EPI_ISL_2553926 | Health Services Laboratories | Wellcome Sanger Institute for the COVID-19 Genomics UK (COG-UK) Consortium | Health Services Laboratories and Alex Alderton, Roberto Amato, Jeffrey Barrett, Sonia Goncalves, Ewan Harrison, David K. Jackson, Ian Johnston, Dominic Kwiatkowski, Cordelia Langford, John Sillitoe on behalf of the Wellcome Sanger Institute COVID-19 Surveillance Team |
| EPI_ISL_2553927 | Lighthouse Lab in Alderley Park | Wellcome Sanger Institute for the COVID-19 Genomics UK (COG-UK) Consortium | Jacquelyn Wynn, Mairead Hyland, The Lighthouse Lab in Alderley Park and Alex Alderton, Roberto Amato, Jeffrey Barrett, Sonia Goncalves, Ewan Harrison, David K. Jackson, Ian Johnston, Dominic Kwiatkowski, Cordelia Langford, John Sillitoe on behalf of the Wellcome Sanger Institute COVID-19 Surveillance Team |
| EPI_ISL_2553928 | Health Services Laboratories | Wellcome Sanger Institute for the COVID-19 Genomics UK (COG-UK) Consortium | Health Services Laboratories and Alex Alderton, Roberto Amato, Jeffrey Barrett, Sonia Goncalves, Ewan Harrison, David K. Jackson, Ian Johnston, Dominic Kwiatkowski, Cordelia Langford, John Sillitoe on behalf of the Wellcome Sanger Institute COVID-19 Surveillance Team |
| EPI_ISL_2553929 | Lighthouse Lab in Alderley Park | Wellcome Sanger Institute for the COVID-19 Genomics UK (COG-UK) Consortium | Jacquelyn Wynn, Mairead Hyland, The Lighthouse Lab in Alderley Park and Alex Alderton, Roberto Amato, Jeffrey Barrett, Sonia Goncalves, Ewan Harrison, David K. Jackson, Ian Johnston, Dominic Kwiatkowski, Cordelia Langford, John Sillitoe on behalf of the Wellcome Sanger Institute COVID-19 Surveillance Team |
| EPI_ISL_2553930, EPI_ISL_2553931 | Health Services Laboratories | Wellcome Sanger Institute for the COVID-19 Genomics UK (COG-UK) Consortium | Health Services Laboratories and Alex Alderton, Roberto Amato, Jeffrey Barrett, Sonia Goncalves, Ewan Harrison, David K. Jackson, Ian Johnston, Dominic Kwiatkowski, Cordelia Langford, John Sillitoe on behalf of the Wellcome Sanger Institute COVID-19 Surveillance Team |
| EPI_ISL_2553932 | Lighthouse Lab in Alderley Park | Wellcome Sanger Institute for the COVID-19 Genomics UK (COG-UK) Consortium | Jacquelyn Wynn, Mairead Hyland, The Lighthouse Lab in Alderley Park and Alex Alderton, Roberto Amato, Jeffrey Barrett, Sonia Goncalves, Ewan Harrison, David K. Jackson, Ian Johnston, Dominic Kwiatkowski, Cordelia Langford, John Sillitoe on behalf of the Wellcome Sanger Institute COVID-19 Surveillance Team |
| EPI_ISL_2553933 | Lighthouse Lab in Milton Keynes | Wellcome Sanger Institute for the COVID-19 Genomics UK (COG-UK) Consortium | The Lighthouse Lab in Milton Keynes and Alex Alderton, Roberto Amato, Jeffrey Barrett, Sonia Goncalves, Ewan Harrison, David K. Jackson, Ian Johnston, Dominic Kwiatkowski, Cordelia Langford, John Sillitoe on behalf of the Wellcome Sanger Institute COVID-19 Surveillance Team |
| EPI_ISL_2553934, EPI_ISL_2553935, EPI_ISL_2553936, EPI_ISL_2553937 | Health Services Laboratories | Wellcome Sanger Institute for the COVID-19 Genomics UK (COG-UK) Consortium | Health Services Laboratories and Alex Alderton, Roberto Amato, Jeffrey Barrett, Sonia Goncalves, Ewan Harrison, David K. Jackson, Ian Johnston, Dominic Kwiatkowski, Cordelia Langford, John Sillitoe on behalf of the Wellcome Sanger Institute COVID-19 Surveillance Team |
| EPI_ISL_2553938 | Lighthouse Lab in Alderley Park | Wellcome Sanger Institute for the COVID-19 Genomics UK (COG-UK) Consortium | Jacquelyn Wynn, Mairead Hyland, The Lighthouse Lab in Alderley Park and Alex Alderton, Roberto Amato, Jeffrey Barrett, Sonia Goncalves, Ewan Harrison, David K. Jackson, Ian Johnston, Dominic Kwiatkowski, Cordelia Langford, John Sillitoe on behalf of the Wellcome Sanger Institute COVID-19 Surveillance Team |
| EPI_ISL_2553939, EPI_ISL_2553940, EPI_ISL_2553941 | Health Services Laboratories | Wellcome Sanger Institute for the COVID-19 Genomics UK (COG-UK) Consortium | Health Services Laboratories and Alex Alderton, Roberto Amato, Jeffrey Barrett, Sonia Goncalves, Ewan Harrison, David K. Jackson, Ian Johnston, Dominic Kwiatkowski, Cordelia Langford, John Sillitoe on behalf of the Wellcome Sanger Institute COVID-19 Surveillance Team |
| EPI_ISL_2553942 | Lighthouse Lab in Alderley Park | Wellcome Sanger Institute for the COVID-19 Genomics UK (COG-UK) Consortium | Jacquelyn Wynn, Mairead Hyland, The Lighthouse Lab in Alderley Park and Alex Alderton, Roberto Amato, Jeffrey Barrett, Sonia Goncalves, Ewan Harrison, David K. Jackson, Ian Johnston, Dominic Kwiatkowski, Cordelia Langford, John Sillitoe on behalf of the Wellcome Sanger Institute COVID-19 Surveillance Team |
| EPI_ISL_2553943 | Health Services Laboratories | Wellcome Sanger Institute for the COVID-19 Genomics UK (COG-UK) Consortium | Health Services Laboratories and Alex Alderton, Roberto Amato, Jeffrey Barrett, Sonia Goncalves, Ewan Harrison, David K. Jackson, Ian Johnston, Dominic Kwiatkowski, Cordelia Langford, John Sillitoe on behalf of the Wellcome Sanger Institute COVID-19 Surveillance Team |
| EPI_ISL_2553944 | Lighthouse Lab in Milton Keynes | Wellcome Sanger Institute for the COVID-19 Genomics UK (COG-UK) Consortium | The Lighthouse Lab in Milton Keynes and Alex Alderton, Roberto Amato, Jeffrey Barrett, Sonia Goncalves, Ewan Harrison, David K. Jackson, Ian Johnston, Dominic Kwiatkowski, Cordelia Langford, John Sillitoe on behalf of the Wellcome Sanger Institute COVID-19 Surveillance Team |
| EPI_ISL_2553945, EPI_ISL_2553946 | Health Services Laboratories | Wellcome Sanger Institute for the COVID-19 Genomics UK (COG-UK) Consortium | Health Services Laboratories and Alex Alderton, Roberto Amato, Jeffrey Barrett, Sonia Goncalves, Ewan Harrison, David K. Jackson, Ian Johnston, Dominic Kwiatkowski, Cordelia Langford, John Sillitoe on behalf of the Wellcome Sanger Institute COVID-19 Surveillance Team |
| EPI_ISL_2553947 | Lighthouse Lab in Milton Keynes | Wellcome Sanger Institute for the COVID-19 Genomics UK (COG-UK) Consortium | The Lighthouse Lab in Milton Keynes and Alex Alderton, Roberto Amato, Jeffrey Barrett, Sonia Goncalves, Ewan Harrison, David K. Jackson, Ian Johnston, Dominic Kwiatkowski, Cordelia Langford, John Sillitoe on behalf of the Wellcome Sanger Institute COVID-19 Surveillance Team |
| EPI_ISL_2553948 | Health Services Laboratories | Wellcome Sanger Institute for the COVID-19 Genomics UK (COG-UK) Consortium | Health Services Laboratories and Alex Alderton, Roberto Amato, Jeffrey Barrett, Sonia Goncalves, Ewan Harrison, David K. Jackson, Ian Johnston, Dominic Kwiatkowski, Cordelia Langford, John Sillitoe on behalf of the Wellcome Sanger Institute COVID-19 Surveillance Team |
| EPI_ISL_2553949, EPI_ISL_2553950 | Lighthouse Lab in Alderley Park | Wellcome Sanger Institute for the COVID-19 Genomics UK (COG-UK) Consortium | Jacquelyn Wynn, Mairead Hyland, The Lighthouse Lab in Alderley Park and Alex Alderton, Roberto Amato, Jeffrey Barrett, Sonia Goncalves, Ewan Harrison, David K. Jackson, Ian Johnston, Dominic Kwiatkowski, Cordelia Langford, John Sillitoe on behalf of the Wellcome Sanger Institute COVID-19 Surveillance Team |
| EPI_ISL_2553951 | Health Services Laboratories | Wellcome Sanger Institute for the COVID-19 Genomics UK (COG-UK) Consortium | Health Services Laboratories and Alex Alderton, Roberto Amato, Jeffrey Barrett, Sonia Goncalves, Ewan Harrison, David K. Jackson, Ian Johnston, Dominic Kwiatkowski, Cordelia Langford, John Sillitoe on behalf of the Wellcome Sanger Institute COVID-19 Surveillance Team |
| EPI_ISL_2553952 | Lighthouse Lab in Alderley Park | Wellcome Sanger Institute for the COVID-19 Genomics UK (COG-UK) Consortium | Jacquelyn Wynn, Mairead Hyland, The Lighthouse Lab in Alderley Park and Alex Alderton, Roberto Amato, Jeffrey Barrett, Sonia Goncalves, Ewan Harrison, David K. Jackson, Ian Johnston, Dominic Kwiatkowski, Cordelia Langford, John Sillitoe on behalf of the Wellcome Sanger Institute COVID-19 Surveillance Team |
| EPI_ISL_2553953, EPI_ISL_2553954 | Health Services Laboratories | Wellcome Sanger Institute for the COVID-19 Genomics UK (COG-UK) Consortium | Health Services Laboratories and Alex Alderton, Roberto Amato, Jeffrey Barrett, Sonia Goncalves, Ewan Harrison, David K. Jackson, Ian Johnston, Dominic Kwiatkowski, Cordelia Langford, John Sillitoe on behalf of the Wellcome Sanger Institute COVID-19 Surveillance Team |
| EPI_ISL_2553955 | Lighthouse Lab in Milton Keynes | Wellcome Sanger Institute for the COVID-19 Genomics UK (COG-UK) Consortium | The Lighthouse Lab in Milton Keynes and Alex Alderton, Roberto Amato, Jeffrey Barrett, Sonia Goncalves, Ewan Harrison, David K. Jackson, Ian Johnston, Dominic Kwiatkowski, Cordelia Langford, John Sillitoe on behalf of the Wellcome Sanger Institute COVID-19 Surveillance Team |
| EPI_ISL_2553956, EPI_ISL_2553957 | Health Services Laboratories | Wellcome Sanger Institute for the COVID-19 Genomics UK (COG-UK) Consortium | Health Services Laboratories and Alex Alderton, Roberto Amato, Jeffrey Barrett, Sonia Goncalves, Ewan Harrison, David K. Jackson, Ian Johnston, Dominic Kwiatkowski, Cordelia Langford, John Sillitoe on behalf of the Wellcome Sanger Institute COVID-19 Surveillance Team |
| EPI_ISL_2553958 | Lighthouse Lab in Milton Keynes | Wellcome Sanger Institute for the COVID-19 Genomics UK (COG-UK) Consortium | The Lighthouse Lab in Milton Keynes and Alex Alderton, Roberto Amato, Jeffrey Barrett, Sonia Goncalves, Ewan Harrison, David K. Jackson, Ian Johnston, Dominic Kwiatkowski, Cordelia Langford, John Sillitoe on behalf of the Wellcome Sanger Institute COVID-19 Surveillance Team |
| EPI_ISL_2553959 | Health Services Laboratories | Wellcome Sanger Institute for the COVID-19 Genomics UK (COG-UK) Consortium | Health Services Laboratories and Alex Alderton, Roberto Amato, Jeffrey Barrett, Sonia Goncalves, Ewan Harrison, David K. Jackson, Ian Johnston, Dominic Kwiatkowski, Cordelia Langford, John Sillitoe on behalf of the Wellcome Sanger Institute COVID-19 Surveillance Team |
| EPI_ISL_2553960 | Lighthouse Lab in Milton Keynes | Wellcome Sanger Institute for the COVID-19 Genomics UK (COG-UK) Consortium | The Lighthouse Lab in Milton Keynes and Alex Alderton, Roberto Amato, Jeffrey Barrett, Sonia Goncalves, Ewan Harrison, David K. Jackson, Ian Johnston, Dominic Kwiatkowski, Cordelia Langford, John Sillitoe on behalf of the Wellcome Sanger Institute COVID-19 Surveillance Team |
| EPI_ISL_2553961, | Health Services Laboratories | Wellcome Sanger Institute for the COVID-19 Genomics UK (COG-UK) | Health Services Laboratories and Alex Alderton, Roberto Amato, Jeffrey Barrett, Sonia Goncalves, Ewan Harrison, David K. Jackson, Ian Johnston, Dominic Kwiatkowski, Cordelia Langford, John Sillitoe on behalf of the Wellcome Sanger Institute |

| | | | |
|---|---|---|---|
| EPI_ISL_2553962 | | Consortium | COVID-19 Surveillance Team |
| EPI_ISL_2553963 | Lighthouse Lab in Alderley Park | Wellcome Sanger Institute for the COVID-19 Genomics UK (COG-UK) Consortium | Jacquelyn Wynn, Mairead Hyland, The Lighthouse Lab in Alderley Park and Alex Alderton, Roberto Amato, Jeffrey Barrett, Sonia Goncalves, Ewan Harrison, David K. Jackson, Ian Johnston, Dominic Kwiatkowski, Cordelia Langford, John Sillitoe on behalf of the Wellcome Sanger Institute COVID-19 Surveillance Team |
| EPI_ISL_2553964 | Lighthouse Lab in Milton Keynes | Wellcome Sanger Institute for the COVID-19 Genomics UK (COG-UK) Consortium | The Lighthouse Lab in Milton Keynes and Alex Alderton, Roberto Amato, Jeffrey Barrett, Sonia Goncalves, Ewan Harrison, David K. Jackson, Ian Johnston, Dominic Kwiatkowski, Cordelia Langford, John Sillitoe on behalf of the Wellcome Sanger Institute COVID-19 Surveillance Team |
| EPI_ISL_2553965 | Health Services Laboratories | Wellcome Sanger Institute for the COVID-19 Genomics UK (COG-UK) Consortium | Health Services Laboratories and Alex Alderton, Roberto Amato, Jeffrey Barrett, Sonia Goncalves, Ewan Harrison, David K. Jackson, Ian Johnston, Dominic Kwiatkowski, Cordelia Langford, John Sillitoe on behalf of the Wellcome Sanger Institute COVID-19 Surveillance Team |
| EPI_ISL_2553966 | Lighthouse Lab in Alderley Park | Wellcome Sanger Institute for the COVID-19 Genomics UK (COG-UK) Consortium | Jacquelyn Wynn, Mairead Hyland, The Lighthouse Lab in Alderley Park and Alex Alderton, Roberto Amato, Jeffrey Barrett, Sonia Goncalves, Ewan Harrison, David K. Jackson, Ian Johnston, Dominic Kwiatkowski, Cordelia Langford, John Sillitoe on behalf of the Wellcome Sanger Institute COVID-19 Surveillance Team |
| EPI_ISL_2553967 | Health Services Laboratories | Wellcome Sanger Institute for the COVID-19 Genomics UK (COG-UK) Consortium | Health Services Laboratories and Alex Alderton, Roberto Amato, Jeffrey Barrett, Sonia Goncalves, Ewan Harrison, David K. Jackson, Ian Johnston, Dominic Kwiatkowski, Cordelia Langford, John Sillitoe on behalf of the Wellcome Sanger Institute COVID-19 Surveillance Team |
| EPI_ISL_2553968, EPI_ISL_2553969, EPI_ISL_2553970 | Lighthouse Lab in Milton Keynes | Wellcome Sanger Institute for the COVID-19 Genomics UK (COG-UK) Consortium | The Lighthouse Lab in Milton Keynes and Alex Alderton, Roberto Amato, Jeffrey Barrett, Sonia Goncalves, Ewan Harrison, David K. Jackson, Ian Johnston, Dominic Kwiatkowski, Cordelia Langford, John Sillitoe on behalf of the Wellcome Sanger Institute COVID-19 Surveillance Team |
| EPI_ISL_2553971 | Lighthouse Lab in Alderley Park | Wellcome Sanger Institute for the COVID-19 Genomics UK (COG-UK) Consortium | Jacquelyn Wynn, Mairead Hyland, The Lighthouse Lab in Alderley Park and Alex Alderton, Roberto Amato, Jeffrey Barrett, Sonia Goncalves, Ewan Harrison, David K. Jackson, Ian Johnston, Dominic Kwiatkowski, Cordelia Langford, John Sillitoe on behalf of the Wellcome Sanger Institute COVID-19 Surveillance Team |
| EPI_ISL_2553972, EPI_ISL_2553974, EPI_ISL_2553975 | Lighthouse Lab in Milton Keynes | Wellcome Sanger Institute for the COVID-19 Genomics UK (COG-UK) Consortium | The Lighthouse Lab in Milton Keynes and Alex Alderton, Roberto Amato, Jeffrey Barrett, Sonia Goncalves, Ewan Harrison, David K. Jackson, Ian Johnston, Dominic Kwiatkowski, Cordelia Langford, John Sillitoe on behalf of the Wellcome Sanger Institute COVID-19 Surveillance Team |
| EPI_ISL_2553976, EPI_ISL_2553977, EPI_ISL_2553978 | Lighthouse Lab in Alderley Park | Wellcome Sanger Institute for the COVID-19 Genomics UK (COG-UK) Consortium | Jacquelyn Wynn, Mairead Hyland, The Lighthouse Lab in Alderley Park and Alex Alderton, Roberto Amato, Jeffrey Barrett, Sonia Goncalves, Ewan Harrison, David K. Jackson, Ian Johnston, Dominic Kwiatkowski, Cordelia Langford, John Sillitoe on behalf of the Wellcome Sanger Institute COVID-19 Surveillance Team |
| EPI_ISL_2553979 | Health Services Laboratories | Wellcome Sanger Institute for the COVID-19 Genomics UK (COG-UK) Consortium | Health Services Laboratories and Alex Alderton, Roberto Amato, Jeffrey Barrett, Sonia Goncalves, Ewan Harrison, David K. Jackson, Ian Johnston, Dominic Kwiatkowski, Cordelia Langford, John Sillitoe on behalf of the Wellcome Sanger Institute COVID-19 Surveillance Team |
| EPI_ISL_2553980 | Lighthouse Lab in Alderley Park | Wellcome Sanger Institute for the COVID-19 Genomics UK (COG-UK) Consortium | Jacquelyn Wynn, Mairead Hyland, The Lighthouse Lab in Alderley Park and Alex Alderton, Roberto Amato, Jeffrey Barrett, Sonia Goncalves, Ewan Harrison, David K. Jackson, Ian Johnston, Dominic Kwiatkowski, Cordelia Langford, John Sillitoe on behalf of the Wellcome Sanger Institute COVID-19 Surveillance Team |
| EPI_ISL_2553981, EPI_ISL_2553982, EPI_ISL_2553983 | Health Services Laboratories | Wellcome Sanger Institute for the COVID-19 Genomics UK (COG-UK) Consortium | Health Services Laboratories and Alex Alderton, Roberto Amato, Jeffrey Barrett, Sonia Goncalves, Ewan Harrison, David K. Jackson, Ian Johnston, Dominic Kwiatkowski, Cordelia Langford, John Sillitoe on behalf of the Wellcome Sanger Institute COVID-19 Surveillance Team |
| EPI_ISL_2553984 | Lighthouse Lab in Alderley Park | Wellcome Sanger Institute for the COVID-19 Genomics UK (COG-UK) Consortium | Jacquelyn Wynn, Mairead Hyland, The Lighthouse Lab in Alderley Park and Alex Alderton, Roberto Amato, Jeffrey Barrett, Sonia Goncalves, Ewan Harrison, David K. Jackson, Ian Johnston, Dominic Kwiatkowski, Cordelia Langford, John Sillitoe on behalf of the Wellcome Sanger Institute COVID-19 Surveillance Team |
| EPI_ISL_2553985, EPI_ISL_2553986, EPI_ISL_2553987 | Health Services Laboratories | Wellcome Sanger Institute for the COVID-19 Genomics UK (COG-UK) Consortium | Health Services Laboratories and Alex Alderton, Roberto Amato, Jeffrey Barrett, Sonia Goncalves, Ewan Harrison, David K. Jackson, Ian Johnston, Dominic Kwiatkowski, Cordelia Langford, John Sillitoe on behalf of the Wellcome Sanger Institute COVID-19 Surveillance Team |
| EPI_ISL_2553988 | Lighthouse Lab in Milton Keynes | Wellcome Sanger Institute for the COVID-19 Genomics UK (COG-UK) Consortium | The Lighthouse Lab in Milton Keynes and Alex Alderton, Roberto Amato, Jeffrey Barrett, Sonia Goncalves, Ewan Harrison, David K. Jackson, Ian Johnston, Dominic Kwiatkowski, Cordelia Langford, John Sillitoe on behalf of the Wellcome Sanger Institute COVID-19 Surveillance Team |
| EPI_ISL_2553989, EPI_ISL_2553990, EPI_ISL_2553991 | Health Services Laboratories | Wellcome Sanger Institute for the COVID-19 Genomics UK (COG-UK) Consortium | Health Services Laboratories and Alex Alderton, Roberto Amato, Jeffrey Barrett, Sonia Goncalves, Ewan Harrison, David K. Jackson, Ian Johnston, Dominic Kwiatkowski, Cordelia Langford, John Sillitoe on behalf of the Wellcome Sanger Institute COVID-19 Surveillance Team |
| EPI_ISL_2553992 | Lighthouse Lab in Alderley Park | Wellcome Sanger Institute for the COVID-19 Genomics UK (COG-UK) Consortium | Jacquelyn Wynn, Mairead Hyland, The Lighthouse Lab in Alderley Park and Alex Alderton, Roberto Amato, Jeffrey Barrett, Sonia Goncalves, Ewan Harrison, David K. Jackson, Ian Johnston, Dominic Kwiatkowski, Cordelia Langford, John Sillitoe on behalf of the Wellcome Sanger Institute COVID-19 Surveillance Team |
| EPI_ISL_2553993 | Lighthouse Lab in Milton Keynes | Wellcome Sanger Institute for the COVID-19 Genomics UK (COG-UK) Consortium | The Lighthouse Lab in Milton Keynes and Alex Alderton, Roberto Amato, Jeffrey Barrett, Sonia Goncalves, Ewan Harrison, David K. Jackson, Ian Johnston, Dominic Kwiatkowski, Cordelia Langford, John Sillitoe on behalf of the Wellcome Sanger Institute COVID-19 Surveillance Team |
| EPI_ISL_2553994, EPI_ISL_2553995 | Health Services Laboratories | Wellcome Sanger Institute for the COVID-19 Genomics UK (COG-UK) Consortium | Health Services Laboratories and Alex Alderton, Roberto Amato, Jeffrey Barrett, Sonia Goncalves, Ewan Harrison, David K. Jackson, Ian Johnston, Dominic Kwiatkowski, Cordelia Langford, John Sillitoe on behalf of the Wellcome Sanger Institute COVID-19 Surveillance Team |
| EPI_ISL_2553996 | Lighthouse Lab in Milton Keynes | Wellcome Sanger Institute for the COVID-19 Genomics UK (COG-UK) Consortium | The Lighthouse Lab in Milton Keynes and Alex Alderton, Roberto Amato, Jeffrey Barrett, Sonia Goncalves, Ewan Harrison, David K. Jackson, Ian Johnston, Dominic Kwiatkowski, Cordelia Langford, John Sillitoe on behalf of the Wellcome Sanger Institute COVID-19 Surveillance Team |
| EPI_ISL_2553997 | Health Services Laboratories | Wellcome Sanger Institute for the COVID-19 Genomics UK (COG-UK) Consortium | Health Services Laboratories and Alex Alderton, Roberto Amato, Jeffrey Barrett, Sonia Goncalves, Ewan Harrison, David K. Jackson, Ian Johnston, Dominic Kwiatkowski, Cordelia Langford, John Sillitoe on behalf of the Wellcome Sanger Institute COVID-19 Surveillance Team |
| EPI_ISL_2553998, EPI_ISL_2553999 | Lighthouse Lab in Milton Keynes | Wellcome Sanger Institute for the COVID-19 Genomics UK (COG-UK) Consortium | The Lighthouse Lab in Milton Keynes and Alex Alderton, Roberto Amato, Jeffrey Barrett, Sonia Goncalves, Ewan Harrison, David K. Jackson, Ian Johnston, Dominic Kwiatkowski, Cordelia Langford, John Sillitoe on behalf of the Wellcome Sanger Institute COVID-19 Surveillance Team |
| EPI_ISL_2554000 | Health Services Laboratories | Wellcome Sanger Institute for the COVID-19 Genomics UK (COG-UK) Consortium | Health Services Laboratories and Alex Alderton, Roberto Amato, Jeffrey Barrett, Sonia Goncalves, Ewan Harrison, David K. Jackson, Ian Johnston, Dominic Kwiatkowski, Cordelia Langford, John Sillitoe on behalf of the Wellcome Sanger Institute COVID-19 Surveillance Team |
| EPI_ISL_2554001 | Lighthouse Lab in Alderley Park | Wellcome Sanger Institute for the COVID-19 Genomics UK (COG-UK) Consortium | Jacquelyn Wynn, Mairead Hyland, The Lighthouse Lab in Alderley Park and Alex Alderton, Roberto Amato, Jeffrey Barrett, Sonia Goncalves, Ewan Harrison, David K. Jackson, Ian Johnston, Dominic Kwiatkowski, Cordelia Langford, John Sillitoe on behalf of the Wellcome Sanger Institute COVID-19 Surveillance Team |
| EPI_ISL_2554002, EPI_ISL_2554003 | Health Services Laboratories | Wellcome Sanger Institute for the COVID-19 Genomics UK (COG-UK) Consortium | Health Services Laboratories and Alex Alderton, Roberto Amato, Jeffrey Barrett, Sonia Goncalves, Ewan Harrison, David K. Jackson, Ian Johnston, Dominic Kwiatkowski, Cordelia Langford, John Sillitoe on behalf of the Wellcome Sanger Institute COVID-19 Surveillance Team |
| EPI_ISL_2554004 | Lighthouse Lab in Milton Keynes | Wellcome Sanger Institute for the COVID-19 Genomics UK (COG-UK) Consortium | The Lighthouse Lab in Milton Keynes and Alex Alderton, Roberto Amato, Jeffrey Barrett, Sonia Goncalves, Ewan Harrison, David K. Jackson, Ian Johnston, Dominic Kwiatkowski, Cordelia Langford, John Sillitoe on behalf of the Wellcome Sanger Institute COVID-19 Surveillance Team |
| EPI_ISL_2554005 | Health Services Laboratories | Wellcome Sanger Institute for the COVID-19 Genomics UK (COG-UK) Consortium | Health Services Laboratories and Alex Alderton, Roberto Amato, Jeffrey Barrett, Sonia Goncalves, Ewan Harrison, David K. Jackson, Ian Johnston, Dominic Kwiatkowski, Cordelia Langford, John Sillitoe on behalf of the Wellcome Sanger Institute COVID-19 Surveillance Team |
| EPI_ISL_2554006 | Lighthouse Lab in Milton Keynes | Wellcome Sanger Institute for the COVID-19 Genomics UK (COG-UK) Consortium | The Lighthouse Lab in Milton Keynes and Alex Alderton, Roberto Amato, Jeffrey Barrett, Sonia Goncalves, Ewan Harrison, David K. Jackson, Ian Johnston, Dominic Kwiatkowski, Cordelia Langford, John Sillitoe on behalf of the Wellcome Sanger Institute COVID-19 Surveillance Team |
| EPI_ISL_2554007 | Lighthouse Lab in Alderley Park | Wellcome Sanger Institute for the COVID-19 Genomics UK (COG-UK) Consortium | Jacquelyn Wynn, Mairead Hyland, The Lighthouse Lab in Alderley Park and Alex Alderton, Roberto Amato, Jeffrey Barrett, Sonia Goncalves, Ewan Harrison, David K. Jackson, Ian Johnston, Dominic Kwiatkowski, Cordelia Langford, John Sillitoe on behalf of the Wellcome Sanger Institute COVID-19 Surveillance Team |
| EPI_ISL_2554008 | Lighthouse Lab in Milton Keynes | Wellcome Sanger Institute for the COVID-19 Genomics UK (COG-UK) Consortium | The Lighthouse Lab in Milton Keynes and Alex Alderton, Roberto Amato, Jeffrey Barrett, Sonia Goncalves, Ewan Harrison, David K. Jackson, Ian Johnston, Dominic Kwiatkowski, Cordelia Langford, John Sillitoe on behalf of the Wellcome Sanger Institute COVID-19 Surveillance Team |
| EPI_ISL_2554009, EPI_ISL_2554010, EPI_ISL_2554011, EPI_ISL_2554012 | Health Services Laboratories | Wellcome Sanger Institute for the COVID-19 Genomics UK (COG-UK) Consortium | Health Services Laboratories and Alex Alderton, Roberto Amato, Jeffrey Barrett, Sonia Goncalves, Ewan Harrison, David K. Jackson, Ian Johnston, Dominic Kwiatkowski, Cordelia Langford, John Sillitoe on behalf of the Wellcome Sanger Institute COVID-19 Surveillance Team |
| EPI_ISL_2554013 | Lighthouse Lab in Milton Keynes | Wellcome Sanger Institute for the COVID-19 Genomics UK (COG-UK) Consortium | The Lighthouse Lab in Milton Keynes and Alex Alderton, Roberto Amato, Jeffrey Barrett, Sonia Goncalves, Ewan Harrison, David K. Jackson, Ian Johnston, Dominic Kwiatkowski, Cordelia Langford, John Sillitoe on behalf of the Wellcome Sanger Institute COVID-19 Surveillance Team |
| EPI_ISL_2554014 | Health Services Laboratories | Wellcome Sanger Institute for the COVID-19 Genomics UK (COG-UK) Consortium | Health Services Laboratories and Alex Alderton, Roberto Amato, Jeffrey Barrett, Sonia Goncalves, Ewan Harrison, David K. Jackson, Ian Johnston, Dominic Kwiatkowski, Cordelia Langford, John Sillitoe on behalf of the Wellcome Sanger Institute COVID-19 Surveillance Team |
| EPI_ISL_2554015 | Lighthouse Lab in Alderley Park | Wellcome Sanger Institute for the COVID-19 Genomics UK (COG-UK) Consortium | Jacquelyn Wynn, Mairead Hyland, The Lighthouse Lab in Alderley Park and Alex Alderton, Roberto Amato, Jeffrey Barrett, Sonia Goncalves, Ewan Harrison, David K. Jackson, Ian Johnston, Dominic Kwiatkowski, Cordelia Langford, John Sillitoe on behalf of the Wellcome Sanger Institute COVID-19 Surveillance Team |
| EPI_ISL_2554016 | Health Services Laboratories | Wellcome Sanger Institute for the COVID-19 Genomics UK (COG-UK) Consortium | Health Services Laboratories and Alex Alderton, Roberto Amato, Jeffrey Barrett, Sonia Goncalves, Ewan Harrison, David K. Jackson, Ian Johnston, Dominic Kwiatkowski, Cordelia Langford, John Sillitoe on behalf of the Wellcome Sanger Institute COVID-19 Surveillance Team |
| EPI_ISL_2554017 | Lighthouse Lab in Milton Keynes | Wellcome Sanger Institute for the COVID-19 Genomics UK (COG-UK) Consortium | The Lighthouse Lab in Milton Keynes and Alex Alderton, Roberto Amato, Jeffrey Barrett, Sonia Goncalves, Ewan Harrison, David K. Jackson, Ian Johnston, Dominic Kwiatkowski, Cordelia Langford, John Sillitoe on behalf of the Wellcome Sanger Institute COVID-19 Surveillance Team |
| EPI_ISL_2554018 | Health Services Laboratories | Wellcome Sanger Institute for the COVID-19 Genomics UK (COG-UK) Consortium | Health Services Laboratories and Alex Alderton, Roberto Amato, Jeffrey Barrett, Sonia Goncalves, Ewan Harrison, David K. Jackson, Ian Johnston, Dominic Kwiatkowski, Cordelia Langford, John Sillitoe on behalf of the Wellcome Sanger Institute COVID-19 Surveillance Team |
| EPI_ISL_2554019 | Lighthouse Lab in Alderley Park | Wellcome Sanger Institute for the COVID-19 Genomics UK (COG-UK) Consortium | Jacquelyn Wynn, Mairead Hyland, The Lighthouse Lab in Alderley Park and Alex Alderton, Roberto Amato, Jeffrey Barrett, Sonia Goncalves, Ewan Harrison, David K. Jackson, Ian Johnston, Dominic Kwiatkowski, Cordelia Langford, John Sillitoe on behalf of the Wellcome Sanger Institute COVID-19 Surveillance Team |
| EPI_ISL_2554020 | Health Services Laboratories | Wellcome Sanger Institute for the COVID-19 Genomics UK (COG-UK) Consortium | Health Services Laboratories and Alex Alderton, Roberto Amato, Jeffrey Barrett, Sonia Goncalves, Ewan Harrison, David K. Jackson, Ian Johnston, Dominic Kwiatkowski, Cordelia Langford, John Sillitoe on behalf of the Wellcome Sanger Institute COVID-19 Surveillance Team |
| EPI_ISL_2554021 | Lighthouse Lab in Alderley Park | Wellcome Sanger Institute for the COVID-19 Genomics UK (COG-UK) Consortium | Jacquelyn Wynn, Mairead Hyland, The Lighthouse Lab in Alderley Park and Alex Alderton, Roberto Amato, Jeffrey Barrett, Sonia Goncalves, Ewan Harrison, David K. Jackson, Ian Johnston, Dominic Kwiatkowski, Cordelia Langford, John Sillitoe on behalf of the Wellcome Sanger Institute COVID-19 Surveillance Team |
| EPI_ISL_2554022 | Health Services Laboratories | Wellcome Sanger Institute for the COVID-19 Genomics UK (COG-UK) Consortium | Health Services Laboratories and Alex Alderton, Roberto Amato, Jeffrey Barrett, Sonia Goncalves, Ewan Harrison, David K. Jackson, Ian Johnston, Dominic Kwiatkowski, Cordelia Langford, John Sillitoe on behalf of the Wellcome Sanger Institute COVID-19 Surveillance Team |
| EPI_ISL_2554023 | Lighthouse Lab in Alderley Park | Wellcome Sanger Institute for the COVID-19 Genomics UK (COG-UK) | Jacquelyn Wynn, Mairead Hyland, The Lighthouse Lab in Alderley Park and Alex Alderton, Roberto Amato, Jeffrey Barrett, Sonia Goncalves, Ewan Harrison, David K. Jackson, Ian Johnston, Dominic Kwiatkowski, Cordelia Langford, John Sillitoe on |

| | | | |
|---|---|---|---|
| | | Consortium | behalf of the Wellcome Sanger Institute COVID-19 Surveillance Team |
| EPI_ISL_2554024, EPI_ISL_2554025 | Lighthouse Lab in Milton Keynes | Wellcome Sanger Institute for the COVID-19 Genomics UK (COG-UK) Consortium | The Lighthouse Lab in Milton Keynes and Alex Alderton, Roberto Amato, Jeffrey Barrett, Sonia Goncalves, Ewan Harrison, David K. Jackson, Ian Johnston, Dominic Kwiatkowski, Cordelia Langford, John Sillitoe on behalf of the Wellcome Sanger Institute COVID-19 Surveillance Team |
| EPI_ISL_2554026 | Lighthouse Lab in Alderley Park | Wellcome Sanger Institute for the COVID-19 Genomics UK (COG-UK) Consortium | Jacquelyn Wynn, Mairead Hyland, The Lighthouse Lab in Alderley Park and Alex Alderton, Roberto Amato, Jeffrey Barrett, Sonia Goncalves, Ewan Harrison, David K. Jackson, Ian Johnston, Dominic Kwiatkowski, Cordelia Langford, John Sillitoe on behalf of the Wellcome Sanger Institute COVID-19 Surveillance Team |
| EPI_ISL_2554027 | Lighthouse Lab in Milton Keynes | Wellcome Sanger Institute for the COVID-19 Genomics UK (COG-UK) Consortium | The Lighthouse Lab in Milton Keynes and Alex Alderton, Roberto Amato, Jeffrey Barrett, Sonia Goncalves, Ewan Harrison, David K. Jackson, Ian Johnston, Dominic Kwiatkowski, Cordelia Langford, John Sillitoe on behalf of the Wellcome Sanger Institute COVID-19 Surveillance Team |
| EPI_ISL_2554028, EPI_ISL_2554029 | Health Services Laboratories | Wellcome Sanger Institute for the COVID-19 Genomics UK (COG-UK) Consortium | Health Services Laboratories and Alex Alderton, Roberto Amato, Jeffrey Barrett, Sonia Goncalves, Ewan Harrison, David K. Jackson, Ian Johnston, Dominic Kwiatkowski, Cordelia Langford, John Sillitoe on behalf of the Wellcome Sanger Institute COVID-19 Surveillance Team |
| EPI_ISL_2554030 | Lighthouse Lab in Milton Keynes | Wellcome Sanger Institute for the COVID-19 Genomics UK (COG-UK) Consortium | The Lighthouse Lab in Milton Keynes and Alex Alderton, Roberto Amato, Jeffrey Barrett, Sonia Goncalves, Ewan Harrison, David K. Jackson, Ian Johnston, Dominic Kwiatkowski, Cordelia Langford, John Sillitoe on behalf of the Wellcome Sanger Institute COVID-19 Surveillance Team |
| EPI_ISL_2554031 | Health Services Laboratories | Wellcome Sanger Institute for the COVID-19 Genomics UK (COG-UK) Consortium | Health Services Laboratories and Alex Alderton, Roberto Amato, Jeffrey Barrett, Sonia Goncalves, Ewan Harrison, David K. Jackson, Ian Johnston, Dominic Kwiatkowski, Cordelia Langford, John Sillitoe on behalf of the Wellcome Sanger Institute COVID-19 Surveillance Team |
| EPI_ISL_2554032 | Lighthouse Lab in Alderley Park | Wellcome Sanger Institute for the COVID-19 Genomics UK (COG-UK) Consortium | Jacquelyn Wynn, Mairead Hyland, The Lighthouse Lab in Alderley Park and Alex Alderton, Roberto Amato, Jeffrey Barrett, Sonia Goncalves, Ewan Harrison, David K. Jackson, Ian Johnston, Dominic Kwiatkowski, Cordelia Langford, John Sillitoe on behalf of the Wellcome Sanger Institute COVID-19 Surveillance Team |
| EPI_ISL_2554034 | Lighthouse Lab in Milton Keynes | Wellcome Sanger Institute for the COVID-19 Genomics UK (COG-UK) Consortium | The Lighthouse Lab in Milton Keynes and Alex Alderton, Roberto Amato, Jeffrey Barrett, Sonia Goncalves, Ewan Harrison, David K. Jackson, Ian Johnston, Dominic Kwiatkowski, Cordelia Langford, John Sillitoe on behalf of the Wellcome Sanger Institute COVID-19 Surveillance Team |
| EPI_ISL_2554035 | Health Services Laboratories | Wellcome Sanger Institute for the COVID-19 Genomics UK (COG-UK) Consortium | Health Services Laboratories and Alex Alderton, Roberto Amato, Jeffrey Barrett, Sonia Goncalves, Ewan Harrison, David K. Jackson, Ian Johnston, Dominic Kwiatkowski, Cordelia Langford, John Sillitoe on behalf of the Wellcome Sanger Institute COVID-19 Surveillance Team |
| EPI_ISL_2554036 | Lighthouse Lab in Milton Keynes | Wellcome Sanger Institute for the COVID-19 Genomics UK (COG-UK) Consortium | The Lighthouse Lab in Milton Keynes and Alex Alderton, Roberto Amato, Jeffrey Barrett, Sonia Goncalves, Ewan Harrison, David K. Jackson, Ian Johnston, Dominic Kwiatkowski, Cordelia Langford, John Sillitoe on behalf of the Wellcome Sanger Institute COVID-19 Surveillance Team |
| EPI_ISL_2554037 | Health Services Laboratories | Wellcome Sanger Institute for the COVID-19 Genomics UK (COG-UK) Consortium | Health Services Laboratories and Alex Alderton, Roberto Amato, Jeffrey Barrett, Sonia Goncalves, Ewan Harrison, David K. Jackson, Ian Johnston, Dominic Kwiatkowski, Cordelia Langford, John Sillitoe on behalf of the Wellcome Sanger Institute COVID-19 Surveillance Team |
| EPI_ISL_2554038 | Lighthouse Lab in Milton Keynes | Wellcome Sanger Institute for the COVID-19 Genomics UK (COG-UK) Consortium | The Lighthouse Lab in Milton Keynes and Alex Alderton, Roberto Amato, Jeffrey Barrett, Sonia Goncalves, Ewan Harrison, David K. Jackson, Ian Johnston, Dominic Kwiatkowski, Cordelia Langford, John Sillitoe on behalf of the Wellcome Sanger Institute COVID-19 Surveillance Team |
| EPI_ISL_2554039, EPI_ISL_2554040, EPI_ISL_2554041, EPI_ISL_2554042, EPI_ISL_2554043 | Lighthouse Lab in Alderley Park | Wellcome Sanger Institute for the COVID-19 Genomics UK (COG-UK) Consortium | Jacquelyn Wynn, Mairead Hyland, The Lighthouse Lab in Alderley Park and Alex Alderton, Roberto Amato, Jeffrey Barrett, Sonia Goncalves, Ewan Harrison, David K. Jackson, Ian Johnston, Dominic Kwiatkowski, Cordelia Langford, John Sillitoe on behalf of the Wellcome Sanger Institute COVID-19 Surveillance Team |
| EPI_ISL_2554044, EPI_ISL_2554045, EPI_ISL_2554046 | Health Services Laboratories | Wellcome Sanger Institute for the COVID-19 Genomics UK (COG-UK) Consortium | Health Services Laboratories and Alex Alderton, Roberto Amato, Jeffrey Barrett, Sonia Goncalves, Ewan Harrison, David K. Jackson, Ian Johnston, Dominic Kwiatkowski, Cordelia Langford, John Sillitoe on behalf of the Wellcome Sanger Institute COVID-19 Surveillance Team |
| EPI_ISL_2554047 | Lighthouse Lab in Alderley Park | Wellcome Sanger Institute for the COVID-19 Genomics UK (COG-UK) Consortium | Jacquelyn Wynn, Mairead Hyland, The Lighthouse Lab in Alderley Park and Alex Alderton, Roberto Amato, Jeffrey Barrett, Sonia Goncalves, Ewan Harrison, David K. Jackson, Ian Johnston, Dominic Kwiatkowski, Cordelia Langford, John Sillitoe on behalf of the Wellcome Sanger Institute COVID-19 Surveillance Team |
| EPI_ISL_2554048 | Health Services Laboratories | Wellcome Sanger Institute for the COVID-19 Genomics UK (COG-UK) Consortium | Health Services Laboratories and Alex Alderton, Roberto Amato, Jeffrey Barrett, Sonia Goncalves, Ewan Harrison, David K. Jackson, Ian Johnston, Dominic Kwiatkowski, Cordelia Langford, John Sillitoe on behalf of the Wellcome Sanger Institute COVID-19 Surveillance Team |
| EPI_ISL_2554049, EPI_ISL_2554050, EPI_ISL_2554051 | Lighthouse Lab in Milton Keynes | Wellcome Sanger Institute for the COVID-19 Genomics UK (COG-UK) Consortium | The Lighthouse Lab in Milton Keynes and Alex Alderton, Roberto Amato, Jeffrey Barrett, Sonia Goncalves, Ewan Harrison, David K. Jackson, Ian Johnston, Dominic Kwiatkowski, Cordelia Langford, John Sillitoe on behalf of the Wellcome Sanger Institute COVID-19 Surveillance Team |
| EPI_ISL_2554052 | Lighthouse Lab in Alderley Park | Wellcome Sanger Institute for the COVID-19 Genomics UK (COG-UK) Consortium | Jacquelyn Wynn, Mairead Hyland, The Lighthouse Lab in Alderley Park and Alex Alderton, Roberto Amato, Jeffrey Barrett, Sonia Goncalves, Ewan Harrison, David K. Jackson, Ian Johnston, Dominic Kwiatkowski, Cordelia Langford, John Sillitoe on behalf of the Wellcome Sanger Institute COVID-19 Surveillance Team |
| EPI_ISL_2554054 | Health Services Laboratories | Wellcome Sanger Institute for the COVID-19 Genomics UK (COG-UK) Consortium | Health Services Laboratories and Alex Alderton, Roberto Amato, Jeffrey Barrett, Sonia Goncalves, Ewan Harrison, David K. Jackson, Ian Johnston, Dominic Kwiatkowski, Cordelia Langford, John Sillitoe on behalf of the Wellcome Sanger Institute COVID-19 Surveillance Team |
| EPI_ISL_2554055, EPI_ISL_2554056 | Lighthouse Lab in Alderley Park | Wellcome Sanger Institute for the COVID-19 Genomics UK (COG-UK) Consortium | Jacquelyn Wynn, Mairead Hyland, The Lighthouse Lab in Alderley Park and Alex Alderton, Roberto Amato, Jeffrey Barrett, Sonia Goncalves, Ewan Harrison, David K. Jackson, Ian Johnston, Dominic Kwiatkowski, Cordelia Langford, John Sillitoe on behalf of the Wellcome Sanger Institute COVID-19 Surveillance Team |
| EPI_ISL_2554057 | Health Services Laboratories | Wellcome Sanger Institute for the COVID-19 Genomics UK (COG-UK) Consortium | Health Services Laboratories and Alex Alderton, Roberto Amato, Jeffrey Barrett, Sonia Goncalves, Ewan Harrison, David K. Jackson, Ian Johnston, Dominic Kwiatkowski, Cordelia Langford, John Sillitoe on behalf of the Wellcome Sanger Institute COVID-19 Surveillance Team |
| EPI_ISL_2554058, EPI_ISL_2554059, EPI_ISL_2554060 | Lighthouse Lab in Milton Keynes | Wellcome Sanger Institute for the COVID-19 Genomics UK (COG-UK) Consortium | The Lighthouse Lab in Milton Keynes and Alex Alderton, Roberto Amato, Jeffrey Barrett, Sonia Goncalves, Ewan Harrison, David K. Jackson, Ian Johnston, Dominic Kwiatkowski, Cordelia Langford, John Sillitoe on behalf of the Wellcome Sanger Institute COVID-19 Surveillance Team |
| EPI_ISL_2554061, EPI_ISL_2554062 | Health Services Laboratories | Wellcome Sanger Institute for the COVID-19 Genomics UK (COG-UK) Consortium | Health Services Laboratories and Alex Alderton, Roberto Amato, Jeffrey Barrett, Sonia Goncalves, Ewan Harrison, David K. Jackson, Ian Johnston, Dominic Kwiatkowski, Cordelia Langford, John Sillitoe on behalf of the Wellcome Sanger Institute COVID-19 Surveillance Team |
| EPI_ISL_2554063, EPI_ISL_2554064 | Lighthouse Lab in Milton Keynes | Wellcome Sanger Institute for the COVID-19 Genomics UK (COG-UK) Consortium | The Lighthouse Lab in Milton Keynes and Alex Alderton, Roberto Amato, Jeffrey Barrett, Sonia Goncalves, Ewan Harrison, David K. Jackson, Ian Johnston, Dominic Kwiatkowski, Cordelia Langford, John Sillitoe on behalf of the Wellcome Sanger Institute COVID-19 Surveillance Team |
| EPI_ISL_2554065 | Lighthouse Lab in Alderley Park | Wellcome Sanger Institute for the COVID-19 Genomics UK (COG-UK) Consortium | Jacquelyn Wynn, Mairead Hyland, The Lighthouse Lab in Alderley Park and Alex Alderton, Roberto Amato, Jeffrey Barrett, Sonia Goncalves, Ewan Harrison, David K. Jackson, Ian Johnston, Dominic Kwiatkowski, Cordelia Langford, John Sillitoe on behalf of the Wellcome Sanger Institute COVID-19 Surveillance Team |
| EPI_ISL_2554066 | Lighthouse Lab in Milton Keynes | Wellcome Sanger Institute for the COVID-19 Genomics UK (COG-UK) Consortium | The Lighthouse Lab in Milton Keynes and Alex Alderton, Roberto Amato, Jeffrey Barrett, Sonia Goncalves, Ewan Harrison, David K. Jackson, Ian Johnston, Dominic Kwiatkowski, Cordelia Langford, John Sillitoe on behalf of the Wellcome Sanger Institute COVID-19 Surveillance Team |
| EPI_ISL_2554067 | Health Services Laboratories | Wellcome Sanger Institute for the COVID-19 Genomics UK (COG-UK) Consortium | Health Services Laboratories and Alex Alderton, Roberto Amato, Jeffrey Barrett, Sonia Goncalves, Ewan Harrison, David K. Jackson, Ian Johnston, Dominic Kwiatkowski, Cordelia Langford, John Sillitoe on behalf of the Wellcome Sanger Institute COVID-19 Surveillance Team |
| EPI_ISL_2554068 | Lighthouse Lab in Alderley Park | Wellcome Sanger Institute for the COVID-19 Genomics UK (COG-UK) Consortium | Jacquelyn Wynn, Mairead Hyland, The Lighthouse Lab in Alderley Park and Alex Alderton, Roberto Amato, Jeffrey Barrett, Sonia Goncalves, Ewan Harrison, David K. Jackson, Ian Johnston, Dominic Kwiatkowski, Cordelia Langford, John Sillitoe on behalf of the Wellcome Sanger Institute COVID-19 Surveillance Team |
| EPI_ISL_2554069 | Health Services Laboratories | Wellcome Sanger Institute for the COVID-19 Genomics UK (COG-UK) Consortium | Health Services Laboratories and Alex Alderton, Roberto Amato, Jeffrey Barrett, Sonia Goncalves, Ewan Harrison, David K. Jackson, Ian Johnston, Dominic Kwiatkowski, Cordelia Langford, John Sillitoe on behalf of the Wellcome Sanger Institute COVID-19 Surveillance Team |
| EPI_ISL_2554070, EPI_ISL_2554071 | Lighthouse Lab in Milton Keynes | Wellcome Sanger Institute for the COVID-19 Genomics UK (COG-UK) Consortium | The Lighthouse Lab in Milton Keynes and Alex Alderton, Roberto Amato, Jeffrey Barrett, Sonia Goncalves, Ewan Harrison, David K. Jackson, Ian Johnston, Dominic Kwiatkowski, Cordelia Langford, John Sillitoe on behalf of the Wellcome Sanger Institute COVID-19 Surveillance Team |
| EPI_ISL_2554072, EPI_ISL_2554073, EPI_ISL_2554074, EPI_ISL_2554075 | Health Services Laboratories | Wellcome Sanger Institute for the COVID-19 Genomics UK (COG-UK) Consortium | Health Services Laboratories and Alex Alderton, Roberto Amato, Jeffrey Barrett, Sonia Goncalves, Ewan Harrison, David K. Jackson, Ian Johnston, Dominic Kwiatkowski, Cordelia Langford, John Sillitoe on behalf of the Wellcome Sanger Institute COVID-19 Surveillance Team |
| EPI_ISL_2554076, EPI_ISL_2554077, EPI_ISL_2554078 | Lighthouse Lab in Milton Keynes | Wellcome Sanger Institute for the COVID-19 Genomics UK (COG-UK) Consortium | The Lighthouse Lab in Milton Keynes and Alex Alderton, Roberto Amato, Jeffrey Barrett, Sonia Goncalves, Ewan Harrison, David K. Jackson, Ian Johnston, Dominic Kwiatkowski, Cordelia Langford, John Sillitoe on behalf of the Wellcome Sanger Institute COVID-19 Surveillance Team |
| EPI_ISL_2554079, EPI_ISL_2554080, EPI_ISL_2554081, EPI_ISL_2554082 | Health Services Laboratories | Wellcome Sanger Institute for the COVID-19 Genomics UK (COG-UK) Consortium | Health Services Laboratories and Alex Alderton, Roberto Amato, Jeffrey Barrett, Sonia Goncalves, Ewan Harrison, David K. Jackson, Ian Johnston, Dominic Kwiatkowski, Cordelia Langford, John Sillitoe on behalf of the Wellcome Sanger Institute COVID-19 Surveillance Team |
| EPI_ISL_2554111 | Lighthouse Lab in Milton Keynes | Wellcome Sanger Institute for the COVID-19 Genomics UK (COG-UK) Consortium | The Lighthouse Lab in Milton Keynes and Alex Alderton, Roberto Amato, Jeffrey Barrett, Sonia Goncalves, Ewan Harrison, David K. Jackson, Ian Johnston, Dominic Kwiatkowski, Cordelia Langford, John Sillitoe on behalf of the Wellcome Sanger Institute COVID-19 Surveillance Team |
| EPI_ISL_2554112, EPI_ISL_2554113 | Lighthouse Lab in Alderley Park | Wellcome Sanger Institute for the COVID-19 Genomics UK (COG-UK) Consortium | Jacquelyn Wynn, Mairead Hyland, The Lighthouse Lab in Alderley Park and Alex Alderton, Roberto Amato, Jeffrey Barrett, Sonia Goncalves, Ewan Harrison, David K. Jackson, Ian Johnston, Dominic Kwiatkowski, Cordelia Langford, John Sillitoe on behalf of the Wellcome Sanger Institute COVID-19 Surveillance Team |
| EPI_ISL_2554114, EPI_ISL_2554115 | Lighthouse Lab in Milton Keynes | Wellcome Sanger Institute for the COVID-19 Genomics UK (COG-UK) Consortium | The Lighthouse Lab in Milton Keynes and Alex Alderton, Roberto Amato, Jeffrey Barrett, Sonia Goncalves, Ewan Harrison, David K. Jackson, Ian Johnston, Dominic Kwiatkowski, Cordelia Langford, John Sillitoe on behalf of the Wellcome Sanger Institute COVID-19 Surveillance Team |
| EPI_ISL_2554116, EPI_ISL_2554117 | Health Services Laboratories | Wellcome Sanger Institute for the COVID-19 Genomics UK (COG-UK) Consortium | Health Services Laboratories and Alex Alderton, Roberto Amato, Jeffrey Barrett, Sonia Goncalves, Ewan Harrison, David K. Jackson, Ian Johnston, Dominic Kwiatkowski, Cordelia Langford, John Sillitoe on behalf of the Wellcome Sanger Institute COVID-19 Surveillance Team |
| EPI_ISL_2554118 | Lighthouse Lab in Alderley Park | Wellcome Sanger Institute for the COVID-19 Genomics UK (COG-UK) Consortium | Jacquelyn Wynn, Mairead Hyland, The Lighthouse Lab in Alderley Park and Alex Alderton, Roberto Amato, Jeffrey Barrett, Sonia Goncalves, Ewan Harrison, David K. Jackson, Ian Johnston, Dominic Kwiatkowski, Cordelia Langford, John Sillitoe on behalf of the Wellcome Sanger Institute COVID-19 Surveillance Team |
| EPI_ISL_2554119 | Lighthouse Lab in Milton Keynes | Wellcome Sanger Institute for the COVID-19 Genomics UK (COG-UK) Consortium | The Lighthouse Lab in Milton Keynes and Alex Alderton, Roberto Amato, Jeffrey Barrett, Sonia Goncalves, Ewan Harrison, David K. Jackson, Ian Johnston, Dominic Kwiatkowski, Cordelia Langford, John Sillitoe on behalf of the Wellcome Sanger Institute COVID-19 Surveillance Team |
| EPI_ISL_2554120 | Health Services Laboratories | Wellcome Sanger Institute for the COVID-19 Genomics UK (COG-UK) Consortium | Health Services Laboratories and Alex Alderton, Roberto Amato, Jeffrey Barrett, Sonia Goncalves, Ewan Harrison, David K. Jackson, Ian Johnston, Dominic Kwiatkowski, Cordelia Langford, John Sillitoe on behalf of the Wellcome Sanger Institute COVID-19 Surveillance Team |

| | | | |
|---|---|---|---|
| EPI_ISL_2554121 | Lighthouse Lab in Milton Keynes | Wellcome Sanger Institute for the COVID-19 Genomics UK (COG-UK) Consortium | The Lighthouse Lab in Milton Keynes and Alex Alderton, Roberto Amato, Jeffrey Barrett, Sonia Goncalves, Ewan Harrison, David K. Jackson, Ian Johnston, Dominic Kwiatkowski, Cordelia Langford, John Sillitoe on behalf of the Wellcome Sanger Institute COVID-19 Surveillance Team |
| EPI_ISL_2554122, EPI_ISL_2554123, EPI_ISL_2554124 | Health Services Laboratories | Wellcome Sanger Institute for the COVID-19 Genomics UK (COG-UK) Consortium | Health Services Laboratories and Alex Alderton, Roberto Amato, Jeffrey Barrett, Sonia Goncalves, Ewan Harrison, David K. Jackson, Ian Johnston, Dominic Kwiatkowski, Cordelia Langford, John Sillitoe on behalf of the Wellcome Sanger Institute COVID-19 Surveillance Team |
| EPI_ISL_2554125, EPI_ISL_2554126 | Lighthouse Lab in Alderley Park | Wellcome Sanger Institute for the COVID-19 Genomics UK (COG-UK) Consortium | Jacquelyn Wynn, Mairead Hyland, The Lighthouse Lab in Alderley Park and Alex Alderton, Roberto Amato, Jeffrey Barrett, Sonia Goncalves, Ewan Harrison, David K. Jackson, Ian Johnston, Dominic Kwiatkowski, Cordelia Langford, John Sillitoe on behalf of the Wellcome Sanger Institute COVID-19 Surveillance Team |
| EPI_ISL_2554127, EPI_ISL_2554128 | Health Services Laboratories | Wellcome Sanger Institute for the COVID-19 Genomics UK (COG-UK) Consortium | Health Services Laboratories and Alex Alderton, Roberto Amato, Jeffrey Barrett, Sonia Goncalves, Ewan Harrison, David K. Jackson, Ian Johnston, Dominic Kwiatkowski, Cordelia Langford, John Sillitoe on behalf of the Wellcome Sanger Institute COVID-19 Surveillance Team |
| EPI_ISL_2554129 | Lighthouse Lab in Alderley Park | Wellcome Sanger Institute for the COVID-19 Genomics UK (COG-UK) Consortium | Jacquelyn Wynn, Mairead Hyland, The Lighthouse Lab in Alderley Park and Alex Alderton, Roberto Amato, Jeffrey Barrett, Sonia Goncalves, Ewan Harrison, David K. Jackson, Ian Johnston, Dominic Kwiatkowski, Cordelia Langford, John Sillitoe on behalf of the Wellcome Sanger Institute COVID-19 Surveillance Team |
| EPI_ISL_2554130, EPI_ISL_2554131 | Health Services Laboratories | Wellcome Sanger Institute for the COVID-19 Genomics UK (COG-UK) Consortium | Health Services Laboratories and Alex Alderton, Roberto Amato, Jeffrey Barrett, Sonia Goncalves, Ewan Harrison, David K. Jackson, Ian Johnston, Dominic Kwiatkowski, Cordelia Langford, John Sillitoe on behalf of the Wellcome Sanger Institute COVID-19 Surveillance Team |
| EPI_ISL_2554132 | Lighthouse Lab in Alderley Park | Wellcome Sanger Institute for the COVID-19 Genomics UK (COG-UK) Consortium | Jacquelyn Wynn, Mairead Hyland, The Lighthouse Lab in Alderley Park and Alex Alderton, Roberto Amato, Jeffrey Barrett, Sonia Goncalves, Ewan Harrison, David K. Jackson, Ian Johnston, Dominic Kwiatkowski, Cordelia Langford, John Sillitoe on behalf of the Wellcome Sanger Institute COVID-19 Surveillance Team |
| EPI_ISL_2554133 | Lighthouse Lab in Milton Keynes | Wellcome Sanger Institute for the COVID-19 Genomics UK (COG-UK) Consortium | The Lighthouse Lab in Milton Keynes and Alex Alderton, Roberto Amato, Jeffrey Barrett, Sonia Goncalves, Ewan Harrison, David K. Jackson, Ian Johnston, Dominic Kwiatkowski, Cordelia Langford, John Sillitoe on behalf of the Wellcome Sanger Institute COVID-19 Surveillance Team |
| EPI_ISL_2554134, EPI_ISL_2554136 | Health Services Laboratories | Wellcome Sanger Institute for the COVID-19 Genomics UK (COG-UK) Consortium | Health Services Laboratories and Alex Alderton, Roberto Amato, Jeffrey Barrett, Sonia Goncalves, Ewan Harrison, David K. Jackson, Ian Johnston, Dominic Kwiatkowski, Cordelia Langford, John Sillitoe on behalf of the Wellcome Sanger Institute COVID-19 Surveillance Team |
| EPI_ISL_2554137, EPI_ISL_2554138, EPI_ISL_2554139 | Lighthouse Lab in Milton Keynes | Wellcome Sanger Institute for the COVID-19 Genomics UK (COG-UK) Consortium | The Lighthouse Lab in Milton Keynes and Alex Alderton, Roberto Amato, Jeffrey Barrett, Sonia Goncalves, Ewan Harrison, David K. Jackson, Ian Johnston, Dominic Kwiatkowski, Cordelia Langford, John Sillitoe on behalf of the Wellcome Sanger Institute COVID-19 Surveillance Team |
| EPI_ISL_2554140 | Health Services Laboratories | Wellcome Sanger Institute for the COVID-19 Genomics UK (COG-UK) Consortium | Health Services Laboratories and Alex Alderton, Roberto Amato, Jeffrey Barrett, Sonia Goncalves, Ewan Harrison, David K. Jackson, Ian Johnston, Dominic Kwiatkowski, Cordelia Langford, John Sillitoe on behalf of the Wellcome Sanger Institute COVID-19 Surveillance Team |
| EPI_ISL_2554141 | Lighthouse Laboratory Plymouth | Wellcome Sanger Institute for the COVID-19 Genomics UK (COG-UK) Consortium | Lighthouse Laboratory Plymouth and Alex Alderton, Roberto Amato, Jeffrey Barrett, Sonia Goncalves, Ewan Harrison, David K. Jackson, Ian Johnston, Dominic Kwiatkowski, Cordelia Langford, John Sillitoe on behalf of the Wellcome Sanger Institute COVID-19 Surveillance Team |
| EPI_ISL_2554142, EPI_ISL_2554143 | Lighthouse Lab in Alderley Park | Wellcome Sanger Institute for the COVID-19 Genomics UK (COG-UK) Consortium | Jacquelyn Wynn, Mairead Hyland, The Lighthouse Lab in Alderley Park and Alex Alderton, Roberto Amato, Jeffrey Barrett, Sonia Goncalves, Ewan Harrison, David K. Jackson, Ian Johnston, Dominic Kwiatkowski, Cordelia Langford, John Sillitoe on behalf of the Wellcome Sanger Institute COVID-19 Surveillance Team |
| EPI_ISL_2554144 | Lighthouse Lab in Glasgow | Wellcome Sanger Institute for the COVID-19 Genomics UK (COG-UK) Consortium | Harper VanSteenhouse, Yumi Kasai, David Gray, Carol Clugston, Anna Dominiczak and Alex Alderton, Roberto Amato, Jeffrey Barrett, Sonia Goncalves, Ewan Harrison, David K. Jackson, Ian Johnston, Dominic Kwiatkowski, Cordelia Langford, John Sillitoe on behalf of the Wellcome Sanger Institute COVID-19 Surveillance Team |
| EPI_ISL_2554145 | Lighthouse Laboratory Plymouth | Wellcome Sanger Institute for the COVID-19 Genomics UK (COG-UK) Consortium | Lighthouse Laboratory Plymouth and Alex Alderton, Roberto Amato, Jeffrey Barrett, Sonia Goncalves, Ewan Harrison, David K. Jackson, Ian Johnston, Dominic Kwiatkowski, Cordelia Langford, John Sillitoe on behalf of the Wellcome Sanger Institute COVID-19 Surveillance Team |
| EPI_ISL_2554146 | Lighthouse Lab in Glasgow | Wellcome Sanger Institute for the COVID-19 Genomics UK (COG-UK) Consortium | Harper VanSteenhouse, Yumi Kasai, David Gray, Carol Clugston, Anna Dominiczak and Alex Alderton, Roberto Amato, Jeffrey Barrett, Sonia Goncalves, Ewan Harrison, David K. Jackson, Ian Johnston, Dominic Kwiatkowski, Cordelia Langford, John Sillitoe on behalf of the Wellcome Sanger Institute COVID-19 Surveillance Team |
| EPI_ISL_2554147, EPI_ISL_2554148, EPI_ISL_2554149 | Lighthouse Lab in Alderley Park | Wellcome Sanger Institute for the COVID-19 Genomics UK (COG-UK) Consortium | Jacquelyn Wynn, Mairead Hyland, The Lighthouse Lab in Alderley Park and Alex Alderton, Roberto Amato, Jeffrey Barrett, Sonia Goncalves, Ewan Harrison, David K. Jackson, Ian Johnston, Dominic Kwiatkowski, Cordelia Langford, John Sillitoe on behalf of the Wellcome Sanger Institute COVID-19 Surveillance Team |
| EPI_ISL_2554151 | Lighthouse Laboratory Plymouth | Wellcome Sanger Institute for the COVID-19 Genomics UK (COG-UK) Consortium | Lighthouse Laboratory Plymouth and Alex Alderton, Roberto Amato, Jeffrey Barrett, Sonia Goncalves, Ewan Harrison, David K. Jackson, Ian Johnston, Dominic Kwiatkowski, Cordelia Langford, John Sillitoe on behalf of the Wellcome Sanger Institute COVID-19 Surveillance Team |
| EPI_ISL_2554152, EPI_ISL_2554153 | Lighthouse Lab in Glasgow | Wellcome Sanger Institute for the COVID-19 Genomics UK (COG-UK) Consortium | Harper VanSteenhouse, Yumi Kasai, David Gray, Carol Clugston, Anna Dominiczak and Alex Alderton, Roberto Amato, Jeffrey Barrett, Sonia Goncalves, Ewan Harrison, David K. Jackson, Ian Johnston, Dominic Kwiatkowski, Cordelia Langford, John Sillitoe on behalf of the Wellcome Sanger Institute COVID-19 Surveillance Team |
| EPI_ISL_2554154 | Lighthouse Lab in Alderley Park | Wellcome Sanger Institute for the COVID-19 Genomics UK (COG-UK) Consortium | Jacquelyn Wynn, Mairead Hyland, The Lighthouse Lab in Alderley Park and Alex Alderton, Roberto Amato, Jeffrey Barrett, Sonia Goncalves, Ewan Harrison, David K. Jackson, Ian Johnston, Dominic Kwiatkowski, Cordelia Langford, John Sillitoe on behalf of the Wellcome Sanger Institute COVID-19 Surveillance Team |
| EPI_ISL_2554155 | Lighthouse Laboratory Plymouth | Wellcome Sanger Institute for the COVID-19 Genomics UK (COG-UK) Consortium | Lighthouse Laboratory Plymouth and Alex Alderton, Roberto Amato, Jeffrey Barrett, Sonia Goncalves, Ewan Harrison, David K. Jackson, Ian Johnston, Dominic Kwiatkowski, Cordelia Langford, John Sillitoe on behalf of the Wellcome Sanger Institute COVID-19 Surveillance Team |
| EPI_ISL_2554156, EPI_ISL_2554157, EPI_ISL_2554158 | Lighthouse Lab in Alderley Park | Wellcome Sanger Institute for the COVID-19 Genomics UK (COG-UK) Consortium | Jacquelyn Wynn, Mairead Hyland, The Lighthouse Lab in Alderley Park and Alex Alderton, Roberto Amato, Jeffrey Barrett, Sonia Goncalves, Ewan Harrison, David K. Jackson, Ian Johnston, Dominic Kwiatkowski, Cordelia Langford, John Sillitoe on behalf of the Wellcome Sanger Institute COVID-19 Surveillance Team |
| EPI_ISL_2554159 | Lighthouse Laboratory Plymouth | Wellcome Sanger Institute for the COVID-19 Genomics UK (COG-UK) Consortium | Lighthouse Laboratory Plymouth and Alex Alderton, Roberto Amato, Jeffrey Barrett, Sonia Goncalves, Ewan Harrison, David K. Jackson, Ian Johnston, Dominic Kwiatkowski, Cordelia Langford, John Sillitoe on behalf of the Wellcome Sanger Institute COVID-19 Surveillance Team |
| EPI_ISL_2554160 | Lighthouse Lab in Glasgow | Wellcome Sanger Institute for the COVID-19 Genomics UK (COG-UK) Consortium | Harper VanSteenhouse, Yumi Kasai, David Gray, Carol Clugston, Anna Dominiczak and Alex Alderton, Roberto Amato, Jeffrey Barrett, Sonia Goncalves, Ewan Harrison, David K. Jackson, Ian Johnston, Dominic Kwiatkowski, Cordelia Langford, John Sillitoe on behalf of the Wellcome Sanger Institute COVID-19 Surveillance Team |
| EPI_ISL_2554161 | Lighthouse Laboratory Plymouth | Wellcome Sanger Institute for the COVID-19 Genomics UK (COG-UK) Consortium | Lighthouse Laboratory Plymouth and Alex Alderton, Roberto Amato, Jeffrey Barrett, Sonia Goncalves, Ewan Harrison, David K. Jackson, Ian Johnston, Dominic Kwiatkowski, Cordelia Langford, John Sillitoe on behalf of the Wellcome Sanger Institute COVID-19 Surveillance Team |
| EPI_ISL_2554162 | Lighthouse Lab in Alderley Park | Wellcome Sanger Institute for the COVID-19 Genomics UK (COG-UK) Consortium | Jacquelyn Wynn, Mairead Hyland, The Lighthouse Lab in Alderley Park and Alex Alderton, Roberto Amato, Jeffrey Barrett, Sonia Goncalves, Ewan Harrison, David K. Jackson, Ian Johnston, Dominic Kwiatkowski, Cordelia Langford, John Sillitoe on behalf of the Wellcome Sanger Institute COVID-19 Surveillance Team |
| EPI_ISL_2554163 | Lighthouse Lab in Glasgow | Wellcome Sanger Institute for the COVID-19 Genomics UK (COG-UK) Consortium | Harper VanSteenhouse, Yumi Kasai, David Gray, Carol Clugston, Anna Dominiczak and Alex Alderton, Roberto Amato, Jeffrey Barrett, Sonia Goncalves, Ewan Harrison, David K. Jackson, Ian Johnston, Dominic Kwiatkowski, Cordelia Langford, John Sillitoe on behalf of the Wellcome Sanger Institute COVID-19 Surveillance Team |
| EPI_ISL_2554164 | Lighthouse Lab in Alderley Park | Wellcome Sanger Institute for the COVID-19 Genomics UK (COG-UK) Consortium | Jacquelyn Wynn, Mairead Hyland, The Lighthouse Lab in Alderley Park and Alex Alderton, Roberto Amato, Jeffrey Barrett, Sonia Goncalves, Ewan Harrison, David K. Jackson, Ian Johnston, Dominic Kwiatkowski, Cordelia Langford, John Sillitoe on behalf of the Wellcome Sanger Institute COVID-19 Surveillance Team |
| EPI_ISL_2554165 | Lighthouse Lab in Glasgow | Wellcome Sanger Institute for the COVID-19 Genomics UK (COG-UK) Consortium | Harper VanSteenhouse, Yumi Kasai, David Gray, Carol Clugston, Anna Dominiczak and Alex Alderton, Roberto Amato, Jeffrey Barrett, Sonia Goncalves, Ewan Harrison, David K. Jackson, Ian Johnston, Dominic Kwiatkowski, Cordelia Langford, John Sillitoe on behalf of the Wellcome Sanger Institute COVID-19 Surveillance Team |
| EPI_ISL_2554166, EPI_ISL_2554167, EPI_ISL_2554168, EPI_ISL_2554169, EPI_ISL_2554170, EPI_ISL_2554172 | Lighthouse Lab in Alderley Park | Wellcome Sanger Institute for the COVID-19 Genomics UK (COG-UK) Consortium | Jacquelyn Wynn, Mairead Hyland, The Lighthouse Lab in Alderley Park and Alex Alderton, Roberto Amato, Jeffrey Barrett, Sonia Goncalves, Ewan Harrison, David K. Jackson, Ian Johnston, Dominic Kwiatkowski, Cordelia Langford, John Sillitoe on behalf of the Wellcome Sanger Institute COVID-19 Surveillance Team |
| EPI_ISL_2554173 | Lighthouse Laboratory Plymouth | Wellcome Sanger Institute for the COVID-19 Genomics UK (COG-UK) Consortium | Lighthouse Laboratory Plymouth and Alex Alderton, Roberto Amato, Jeffrey Barrett, Sonia Goncalves, Ewan Harrison, David K. Jackson, Ian Johnston, Dominic Kwiatkowski, Cordelia Langford, John Sillitoe on behalf of the Wellcome Sanger Institute COVID-19 Surveillance Team |
| EPI_ISL_2554175, EPI_ISL_2554176 | Lighthouse Lab in Glasgow | Wellcome Sanger Institute for the COVID-19 Genomics UK (COG-UK) Consortium | Harper VanSteenhouse, Yumi Kasai, David Gray, Carol Clugston, Anna Dominiczak and Alex Alderton, Roberto Amato, Jeffrey Barrett, Sonia Goncalves, Ewan Harrison, David K. Jackson, Ian Johnston, Dominic Kwiatkowski, Cordelia Langford, John Sillitoe on behalf of the Wellcome Sanger Institute COVID-19 Surveillance Team |
| EPI_ISL_2554177, EPI_ISL_2554178, EPI_ISL_2554179 | Lighthouse Lab in Alderley Park | Wellcome Sanger Institute for the COVID-19 Genomics UK (COG-UK) Consortium | Jacquelyn Wynn, Mairead Hyland, The Lighthouse Lab in Alderley Park and Alex Alderton, Roberto Amato, Jeffrey Barrett, Sonia Goncalves, Ewan Harrison, David K. Jackson, Ian Johnston, Dominic Kwiatkowski, Cordelia Langford, John Sillitoe on behalf of the Wellcome Sanger Institute COVID-19 Surveillance Team |
| EPI_ISL_2554180 | Lighthouse Lab in Glasgow | Wellcome Sanger Institute for the COVID-19 Genomics UK (COG-UK) Consortium | Harper VanSteenhouse, Yumi Kasai, David Gray, Carol Clugston, Anna Dominiczak and Alex Alderton, Roberto Amato, Jeffrey Barrett, Sonia Goncalves, Ewan Harrison, David K. Jackson, Ian Johnston, Dominic Kwiatkowski, Cordelia Langford, John Sillitoe on behalf of the Wellcome Sanger Institute COVID-19 Surveillance Team |
| EPI_ISL_2554181, EPI_ISL_2554182, EPI_ISL_2554184 | Lighthouse Lab in Alderley Park | Wellcome Sanger Institute for the COVID-19 Genomics UK (COG-UK) Consortium | Jacquelyn Wynn, Mairead Hyland, The Lighthouse Lab in Alderley Park and Alex Alderton, Roberto Amato, Jeffrey Barrett, Sonia Goncalves, Ewan Harrison, David K. Jackson, Ian Johnston, Dominic Kwiatkowski, Cordelia Langford, John Sillitoe on behalf of the Wellcome Sanger Institute COVID-19 Surveillance Team |
| EPI_ISL_2554185 | Lighthouse Laboratory Plymouth | Wellcome Sanger Institute for the COVID-19 Genomics UK (COG-UK) Consortium | Lighthouse Laboratory Plymouth and Alex Alderton, Roberto Amato, Jeffrey Barrett, Sonia Goncalves, Ewan Harrison, David K. Jackson, Ian Johnston, Dominic Kwiatkowski, Cordelia Langford, John Sillitoe on behalf of the Wellcome Sanger Institute COVID-19 Surveillance Team |
| EPI_ISL_2554186, EPI_ISL_2554187 | Lighthouse Lab in Glasgow | Wellcome Sanger Institute for the COVID-19 Genomics UK (COG-UK) Consortium | Harper VanSteenhouse, Yumi Kasai, David Gray, Carol Clugston, Anna Dominiczak and Alex Alderton, Roberto Amato, Jeffrey Barrett, Sonia Goncalves, Ewan Harrison, David K. Jackson, Ian Johnston, Dominic Kwiatkowski, Cordelia Langford, John Sillitoe on behalf of the Wellcome Sanger Institute COVID-19 Surveillance Team |
| EPI_ISL_2554188, EPI_ISL_2554189, EPI_ISL_2554190, EPI_ISL_2554191, EPI_ISL_2554192 | Lighthouse Lab in Alderley Park | Wellcome Sanger Institute for the COVID-19 Genomics UK (COG-UK) Consortium | Jacquelyn Wynn, Mairead Hyland, The Lighthouse Lab in Alderley Park and Alex Alderton, Roberto Amato, Jeffrey Barrett, Sonia Goncalves, Ewan Harrison, David K. Jackson, Ian Johnston, Dominic Kwiatkowski, Cordelia Langford, John Sillitoe on behalf of the Wellcome Sanger Institute COVID-19 Surveillance Team |
| EPI_ISL_2554193 | Lighthouse Lab in Glasgow | Wellcome Sanger Institute for the COVID-19 Genomics UK (COG-UK) Consortium | Harper VanSteenhouse, Yumi Kasai, David Gray, Carol Clugston, Anna Dominiczak and Alex Alderton, Roberto Amato, Jeffrey Barrett, Sonia Goncalves, Ewan Harrison, David K. Jackson, Ian Johnston, Dominic Kwiatkowski, Cordelia Langford, John Sillitoe on behalf of the Wellcome Sanger Institute COVID-19 Surveillance Team |
| EPI_ISL_2554194 | Lighthouse Laboratory Plymouth | Wellcome Sanger Institute for the COVID-19 Genomics UK (COG-UK) Consortium | Lighthouse Laboratory Plymouth and Alex Alderton, Roberto Amato, Jeffrey Barrett, Sonia Goncalves, Ewan Harrison, David K. Jackson, Ian Johnston, Dominic Kwiatkowski, Cordelia Langford, John Sillitoe on behalf of the Wellcome Sanger Institute COVID-19 Surveillance Team |

| | | | |
|---|---|---|---|
| EPI_ISL_2554195 | Lighthouse Lab in Alderley Park | Wellcome Sanger Institute for the COVID-19 Genomics UK (COG-UK) Consortium | Jacquelyn Wynn, Mairead Hyland, The Lighthouse Lab in Alderley Park and Alex Alderton, Roberto Amato, Jeffrey Barrett, Sonia Goncalves, Ewan Harrison, David K. Jackson, Ian Johnston, Dominic Kwiatkowski, Cordelia Langford, John Sillitoe on behalf of the Wellcome Sanger Institute COVID-19 Surveillance Team |
| EPI_ISL_2554196 | Lighthouse Lab in Glasgow | Wellcome Sanger Institute for the COVID-19 Genomics UK (COG-UK) Consortium | Harper VanSteenhouse, Yumi Kasai, David Gray, Carol Clugston, Anna Dominiczak and Alex Alderton, Roberto Amato, Jeffrey Barrett, Sonia Goncalves, Ewan Harrison, David K. Jackson, Ian Johnston, Dominic Kwiatkowski, Cordelia Langford, John Sillitoe on behalf of the Wellcome Sanger Institute COVID-19 Surveillance Team |
| EPI_ISL_2554197, EPI_ISL_2554198, EPI_ISL_2554199, EPI_ISL_2554200, EPI_ISL_2554201, EPI_ISL_2554202, EPI_ISL_2554203 | Lighthouse Lab in Alderley Park | Wellcome Sanger Institute for the COVID-19 Genomics UK (COG-UK) Consortium | Jacquelyn Wynn, Mairead Hyland, The Lighthouse Lab in Alderley Park and Alex Alderton, Roberto Amato, Jeffrey Barrett, Sonia Goncalves, Ewan Harrison, David K. Jackson, Ian Johnston, Dominic Kwiatkowski, Cordelia Langford, John Sillitoe on behalf of the Wellcome Sanger Institute COVID-19 Surveillance Team |
| EPI_ISL_2554204 | Lighthouse Laboratory Plymouth | Wellcome Sanger Institute for the COVID-19 Genomics UK (COG-UK) Consortium | Lighthouse Laboratory Plymouth and Alex Alderton, Roberto Amato, Jeffrey Barrett, Sonia Goncalves, Ewan Harrison, David K. Jackson, Ian Johnston, Dominic Kwiatkowski, Cordelia Langford, John Sillitoe on behalf of the Wellcome Sanger Institute COVID-19 Surveillance Team |
| EPI_ISL_2554205 | Lighthouse Lab in Alderley Park | Wellcome Sanger Institute for the COVID-19 Genomics UK (COG-UK) Consortium | Jacquelyn Wynn, Mairead Hyland, The Lighthouse Lab in Alderley Park and Alex Alderton, Roberto Amato, Jeffrey Barrett, Sonia Goncalves, Ewan Harrison, David K. Jackson, Ian Johnston, Dominic Kwiatkowski, Cordelia Langford, John Sillitoe on behalf of the Wellcome Sanger Institute COVID-19 Surveillance Team |
| EPI_ISL_2554206 | Lighthouse Laboratory Plymouth | Wellcome Sanger Institute for the COVID-19 Genomics UK (COG-UK) Consortium | Lighthouse Laboratory Plymouth and Alex Alderton, Roberto Amato, Jeffrey Barrett, Sonia Goncalves, Ewan Harrison, David K. Jackson, Ian Johnston, Dominic Kwiatkowski, Cordelia Langford, John Sillitoe on behalf of the Wellcome Sanger Institute COVID-19 Surveillance Team |
| EPI_ISL_2554207 | Lighthouse Lab in Alderley Park | Wellcome Sanger Institute for the COVID-19 Genomics UK (COG-UK) Consortium | Jacquelyn Wynn, Mairead Hyland, The Lighthouse Lab in Alderley Park and Alex Alderton, Roberto Amato, Jeffrey Barrett, Sonia Goncalves, Ewan Harrison, David K. Jackson, Ian Johnston, Dominic Kwiatkowski, Cordelia Langford, John Sillitoe on behalf of the Wellcome Sanger Institute COVID-19 Surveillance Team |
| EPI_ISL_2554208, EPI_ISL_2554209 | Lighthouse Laboratory Plymouth | Wellcome Sanger Institute for the COVID-19 Genomics UK (COG-UK) Consortium | Lighthouse Laboratory Plymouth and Alex Alderton, Roberto Amato, Jeffrey Barrett, Sonia Goncalves, Ewan Harrison, David K. Jackson, Ian Johnston, Dominic Kwiatkowski, Cordelia Langford, John Sillitoe on behalf of the Wellcome Sanger Institute COVID-19 Surveillance Team |
| EPI_ISL_2554210, EPI_ISL_2554211 | Lighthouse Lab in Alderley Park | Wellcome Sanger Institute for the COVID-19 Genomics UK (COG-UK) Consortium | Jacquelyn Wynn, Mairead Hyland, The Lighthouse Lab in Alderley Park and Alex Alderton, Roberto Amato, Jeffrey Barrett, Sonia Goncalves, Ewan Harrison, David K. Jackson, Ian Johnston, Dominic Kwiatkowski, Cordelia Langford, John Sillitoe on behalf of the Wellcome Sanger Institute COVID-19 Surveillance Team |
| EPI_ISL_2554212, EPI_ISL_2554213 | Lighthouse Laboratory Plymouth | Wellcome Sanger Institute for the COVID-19 Genomics UK (COG-UK) Consortium | Lighthouse Laboratory Plymouth and Alex Alderton, Roberto Amato, Jeffrey Barrett, Sonia Goncalves, Ewan Harrison, David K. Jackson, Ian Johnston, Dominic Kwiatkowski, Cordelia Langford, John Sillitoe on behalf of the Wellcome Sanger Institute COVID-19 Surveillance Team |
| EPI_ISL_2554214 | Lighthouse Lab in Glasgow | Wellcome Sanger Institute for the COVID-19 Genomics UK (COG-UK) Consortium | Harper VanSteenhouse, Yumi Kasai, David Gray, Carol Clugston, Anna Dominiczak and Alex Alderton, Roberto Amato, Jeffrey Barrett, Sonia Goncalves, Ewan Harrison, David K. Jackson, Ian Johnston, Dominic Kwiatkowski, Cordelia Langford, John Sillitoe on behalf of the Wellcome Sanger Institute COVID-19 Surveillance Team |
| EPI_ISL_2554215, EPI_ISL_2554216 | Lighthouse Lab in Alderley Park | Wellcome Sanger Institute for the COVID-19 Genomics UK (COG-UK) Consortium | Jacquelyn Wynn, Mairead Hyland, The Lighthouse Lab in Alderley Park and Alex Alderton, Roberto Amato, Jeffrey Barrett, Sonia Goncalves, Ewan Harrison, David K. Jackson, Ian Johnston, Dominic Kwiatkowski, Cordelia Langford, John Sillitoe on behalf of the Wellcome Sanger Institute COVID-19 Surveillance Team |
| EPI_ISL_2554217 | Lighthouse Laboratory Plymouth | Wellcome Sanger Institute for the COVID-19 Genomics UK (COG-UK) Consortium | Lighthouse Laboratory Plymouth and Alex Alderton, Roberto Amato, Jeffrey Barrett, Sonia Goncalves, Ewan Harrison, David K. Jackson, Ian Johnston, Dominic Kwiatkowski, Cordelia Langford, John Sillitoe on behalf of the Wellcome Sanger Institute COVID-19 Surveillance Team |
| EPI_ISL_2554218 | Lighthouse Lab in Alderley Park | Wellcome Sanger Institute for the COVID-19 Genomics UK (COG-UK) Consortium | Jacquelyn Wynn, Mairead Hyland, The Lighthouse Lab in Alderley Park and Alex Alderton, Roberto Amato, Jeffrey Barrett, Sonia Goncalves, Ewan Harrison, David K. Jackson, Ian Johnston, Dominic Kwiatkowski, Cordelia Langford, John Sillitoe on behalf of the Wellcome Sanger Institute COVID-19 Surveillance Team |
| EPI_ISL_2554219, EPI_ISL_2554220 | Lighthouse Lab in Glasgow | Wellcome Sanger Institute for the COVID-19 Genomics UK (COG-UK) Consortium | Harper VanSteenhouse, Yumi Kasai, David Gray, Carol Clugston, Anna Dominiczak and Alex Alderton, Roberto Amato, Jeffrey Barrett, Sonia Goncalves, Ewan Harrison, David K. Jackson, Ian Johnston, Dominic Kwiatkowski, Cordelia Langford, John Sillitoe on behalf of the Wellcome Sanger Institute COVID-19 Surveillance Team |
| EPI_ISL_2554221 | Lighthouse Lab in Alderley Park | Wellcome Sanger Institute for the COVID-19 Genomics UK (COG-UK) Consortium | Jacquelyn Wynn, Mairead Hyland, The Lighthouse Lab in Alderley Park and Alex Alderton, Roberto Amato, Jeffrey Barrett, Sonia Goncalves, Ewan Harrison, David K. Jackson, Ian Johnston, Dominic Kwiatkowski, Cordelia Langford, John Sillitoe on behalf of the Wellcome Sanger Institute COVID-19 Surveillance Team |
| EPI_ISL_2554222 | Lighthouse Lab in Glasgow | Wellcome Sanger Institute for the COVID-19 Genomics UK (COG-UK) Consortium | Harper VanSteenhouse, Yumi Kasai, David Gray, Carol Clugston, Anna Dominiczak and Alex Alderton, Roberto Amato, Jeffrey Barrett, Sonia Goncalves, Ewan Harrison, David K. Jackson, Ian Johnston, Dominic Kwiatkowski, Cordelia Langford, John Sillitoe on behalf of the Wellcome Sanger Institute COVID-19 Surveillance Team |
| EPI_ISL_2554223, EPI_ISL_2554224, EPI_ISL_2554225 | Lighthouse Lab in Alderley Park | Wellcome Sanger Institute for the COVID-19 Genomics UK (COG-UK) Consortium | Jacquelyn Wynn, Mairead Hyland, The Lighthouse Lab in Alderley Park and Alex Alderton, Roberto Amato, Jeffrey Barrett, Sonia Goncalves, Ewan Harrison, David K. Jackson, Ian Johnston, Dominic Kwiatkowski, Cordelia Langford, John Sillitoe on behalf of the Wellcome Sanger Institute COVID-19 Surveillance Team |
| EPI_ISL_2554226 | Lighthouse Lab in Glasgow | Wellcome Sanger Institute for the COVID-19 Genomics UK (COG-UK) Consortium | Harper VanSteenhouse, Yumi Kasai, David Gray, Carol Clugston, Anna Dominiczak and Alex Alderton, Roberto Amato, Jeffrey Barrett, Sonia Goncalves, Ewan Harrison, David K. Jackson, Ian Johnston, Dominic Kwiatkowski, Cordelia Langford, John Sillitoe on behalf of the Wellcome Sanger Institute COVID-19 Surveillance Team |
| EPI_ISL_2554227 | Lighthouse Lab in Alderley Park | Wellcome Sanger Institute for the COVID-19 Genomics UK (COG-UK) Consortium | Jacquelyn Wynn, Mairead Hyland, The Lighthouse Lab in Alderley Park and Alex Alderton, Roberto Amato, Jeffrey Barrett, Sonia Goncalves, Ewan Harrison, David K. Jackson, Ian Johnston, Dominic Kwiatkowski, Cordelia Langford, John Sillitoe on behalf of the Wellcome Sanger Institute COVID-19 Surveillance Team |
| EPI_ISL_2554228 | Lighthouse Lab in Glasgow | Wellcome Sanger Institute for the COVID-19 Genomics UK (COG-UK) Consortium | Harper VanSteenhouse, Yumi Kasai, David Gray, Carol Clugston, Anna Dominiczak and Alex Alderton, Roberto Amato, Jeffrey Barrett, Sonia Goncalves, Ewan Harrison, David K. Jackson, Ian Johnston, Dominic Kwiatkowski, Cordelia Langford, John Sillitoe on behalf of the Wellcome Sanger Institute COVID-19 Surveillance Team |
| EPI_ISL_2554229, EPI_ISL_2554230 | Lighthouse Lab in Alderley Park | Wellcome Sanger Institute for the COVID-19 Genomics UK (COG-UK) Consortium | Jacquelyn Wynn, Mairead Hyland, The Lighthouse Lab in Alderley Park and Alex Alderton, Roberto Amato, Jeffrey Barrett, Sonia Goncalves, Ewan Harrison, David K. Jackson, Ian Johnston, Dominic Kwiatkowski, Cordelia Langford, John Sillitoe on behalf of the Wellcome Sanger Institute COVID-19 Surveillance Team |
| EPI_ISL_2554231, EPI_ISL_2554232 | Lighthouse Laboratory Plymouth | Wellcome Sanger Institute for the COVID-19 Genomics UK (COG-UK) Consortium | Lighthouse Laboratory Plymouth and Alex Alderton, Roberto Amato, Jeffrey Barrett, Sonia Goncalves, Ewan Harrison, David K. Jackson, Ian Johnston, Dominic Kwiatkowski, Cordelia Langford, John Sillitoe on behalf of the Wellcome Sanger Institute COVID-19 Surveillance Team |
| EPI_ISL_2554233 | Lighthouse Lab in Alderley Park | Wellcome Sanger Institute for the COVID-19 Genomics UK (COG-UK) Consortium | Jacquelyn Wynn, Mairead Hyland, The Lighthouse Lab in Alderley Park and Alex Alderton, Roberto Amato, Jeffrey Barrett, Sonia Goncalves, Ewan Harrison, David K. Jackson, Ian Johnston, Dominic Kwiatkowski, Cordelia Langford, John Sillitoe on behalf of the Wellcome Sanger Institute COVID-19 Surveillance Team |
| EPI_ISL_2554235 | Lighthouse Laboratory Plymouth | Wellcome Sanger Institute for the COVID-19 Genomics UK (COG-UK) Consortium | Lighthouse Laboratory Plymouth and Alex Alderton, Roberto Amato, Jeffrey Barrett, Sonia Goncalves, Ewan Harrison, David K. Jackson, Ian Johnston, Dominic Kwiatkowski, Cordelia Langford, John Sillitoe on behalf of the Wellcome Sanger Institute COVID-19 Surveillance Team |
| EPI_ISL_2554236 | Lighthouse Lab in Alderley Park | Wellcome Sanger Institute for the COVID-19 Genomics UK (COG-UK) Consortium | Jacquelyn Wynn, Mairead Hyland, The Lighthouse Lab in Alderley Park and Alex Alderton, Roberto Amato, Jeffrey Barrett, Sonia Goncalves, Ewan Harrison, David K. Jackson, Ian Johnston, Dominic Kwiatkowski, Cordelia Langford, John Sillitoe on behalf of the Wellcome Sanger Institute COVID-19 Surveillance Team |
| EPI_ISL_2554237 | Lighthouse Lab in Glasgow | Wellcome Sanger Institute for the COVID-19 Genomics UK (COG-UK) Consortium | Harper VanSteenhouse, Yumi Kasai, David Gray, Carol Clugston, Anna Dominiczak and Alex Alderton, Roberto Amato, Jeffrey Barrett, Sonia Goncalves, Ewan Harrison, David K. Jackson, Ian Johnston, Dominic Kwiatkowski, Cordelia Langford, John Sillitoe on behalf of the Wellcome Sanger Institute COVID-19 Surveillance Team |
| EPI_ISL_2554238, EPI_ISL_2554239, EPI_ISL_2554240, EPI_ISL_2554241, EPI_ISL_2554242 | Lighthouse Lab in Alderley Park | Wellcome Sanger Institute for the COVID-19 Genomics UK (COG-UK) Consortium | Jacquelyn Wynn, Mairead Hyland, The Lighthouse Lab in Alderley Park and Alex Alderton, Roberto Amato, Jeffrey Barrett, Sonia Goncalves, Ewan Harrison, David K. Jackson, Ian Johnston, Dominic Kwiatkowski, Cordelia Langford, John Sillitoe on behalf of the Wellcome Sanger Institute COVID-19 Surveillance Team |
| EPI_ISL_2554243, EPI_ISL_2554244 | Lighthouse Lab in Glasgow | Wellcome Sanger Institute for the COVID-19 Genomics UK (COG-UK) Consortium | Harper VanSteenhouse, Yumi Kasai, David Gray, Carol Clugston, Anna Dominiczak and Alex Alderton, Roberto Amato, Jeffrey Barrett, Sonia Goncalves, Ewan Harrison, David K. Jackson, Ian Johnston, Dominic Kwiatkowski, Cordelia Langford, John Sillitoe on behalf of the Wellcome Sanger Institute COVID-19 Surveillance Team |
| EPI_ISL_2554245, EPI_ISL_2554246, EPI_ISL_2554247 | Lighthouse Lab in Alderley Park | Wellcome Sanger Institute for the COVID-19 Genomics UK (COG-UK) Consortium | Jacquelyn Wynn, Mairead Hyland, The Lighthouse Lab in Alderley Park and Alex Alderton, Roberto Amato, Jeffrey Barrett, Sonia Goncalves, Ewan Harrison, David K. Jackson, Ian Johnston, Dominic Kwiatkowski, Cordelia Langford, John Sillitoe on behalf of the Wellcome Sanger Institute COVID-19 Surveillance Team |
| EPI_ISL_2554248 | Lighthouse Lab in Glasgow | Wellcome Sanger Institute for the COVID-19 Genomics UK (COG-UK) Consortium | Harper VanSteenhouse, Yumi Kasai, David Gray, Carol Clugston, Anna Dominiczak and Alex Alderton, Roberto Amato, Jeffrey Barrett, Sonia Goncalves, Ewan Harrison, David K. Jackson, Ian Johnston, Dominic Kwiatkowski, Cordelia Langford, John Sillitoe on behalf of the Wellcome Sanger Institute COVID-19 Surveillance Team |
| EPI_ISL_2554249, EPI_ISL_2554250, EPI_ISL_2554251, EPI_ISL_2554252, EPI_ISL_2554253, EPI_ISL_2554254, EPI_ISL_2554255, EPI_ISL_2554256 | Lighthouse Lab in Alderley Park | Wellcome Sanger Institute for the COVID-19 Genomics UK (COG-UK) Consortium | Jacquelyn Wynn, Mairead Hyland, The Lighthouse Lab in Alderley Park and Alex Alderton, Roberto Amato, Jeffrey Barrett, Sonia Goncalves, Ewan Harrison, David K. Jackson, Ian Johnston, Dominic Kwiatkowski, Cordelia Langford, John Sillitoe on behalf of the Wellcome Sanger Institute COVID-19 Surveillance Team |
| EPI_ISL_2554257 | Lighthouse Lab in Glasgow | Wellcome Sanger Institute for the COVID-19 Genomics UK (COG-UK) Consortium | Harper VanSteenhouse, Yumi Kasai, David Gray, Carol Clugston, Anna Dominiczak and Alex Alderton, Roberto Amato, Jeffrey Barrett, Sonia Goncalves, Ewan Harrison, David K. Jackson, Ian Johnston, Dominic Kwiatkowski, Cordelia Langford, John Sillitoe on behalf of the Wellcome Sanger Institute COVID-19 Surveillance Team |
| EPI_ISL_2554258, EPI_ISL_2554259 | Lighthouse Lab in Alderley Park | Wellcome Sanger Institute for the COVID-19 Genomics UK (COG-UK) Consortium | Jacquelyn Wynn, Mairead Hyland, The Lighthouse Lab in Alderley Park and Alex Alderton, Roberto Amato, Jeffrey Barrett, Sonia Goncalves, Ewan Harrison, David K. Jackson, Ian Johnston, Dominic Kwiatkowski, Cordelia Langford, John Sillitoe on behalf of the Wellcome Sanger Institute COVID-19 Surveillance Team |
| EPI_ISL_2554260 | Lighthouse Lab in Glasgow | Wellcome Sanger Institute for the COVID-19 Genomics UK (COG-UK) Consortium | Harper VanSteenhouse, Yumi Kasai, David Gray, Carol Clugston, Anna Dominiczak and Alex Alderton, Roberto Amato, Jeffrey Barrett, Sonia Goncalves, Ewan Harrison, David K. Jackson, Ian Johnston, Dominic Kwiatkowski, Cordelia Langford, John Sillitoe on behalf of the Wellcome Sanger Institute COVID-19 Surveillance Team |
| EPI_ISL_2554261 | Lighthouse Lab in Alderley Park | Wellcome Sanger Institute for the COVID-19 Genomics UK (COG-UK) Consortium | Jacquelyn Wynn, Mairead Hyland, The Lighthouse Lab in Alderley Park and Alex Alderton, Roberto Amato, Jeffrey Barrett, Sonia Goncalves, Ewan Harrison, David K. Jackson, Ian Johnston, Dominic Kwiatkowski, Cordelia Langford, John Sillitoe on behalf of the Wellcome Sanger Institute COVID-19 Surveillance Team |
| EPI_ISL_2554262 | Lighthouse Lab in Glasgow | Wellcome Sanger Institute for the COVID-19 Genomics UK (COG-UK) Consortium | Harper VanSteenhouse, Yumi Kasai, David Gray, Carol Clugston, Anna Dominiczak and Alex Alderton, Roberto Amato, Jeffrey Barrett, Sonia Goncalves, Ewan Harrison, David K. Jackson, Ian Johnston, Dominic Kwiatkowski, Cordelia Langford, John Sillitoe on behalf of the Wellcome Sanger Institute COVID-19 Surveillance Team |
| EPI_ISL_2554263 | Lighthouse Laboratory Plymouth | Wellcome Sanger Institute for the COVID-19 Genomics UK (COG-UK) Consortium | Lighthouse Laboratory Plymouth and Alex Alderton, Roberto Amato, Jeffrey Barrett, Sonia Goncalves, Ewan Harrison, David K. Jackson, Ian Johnston, Dominic Kwiatkowski, Cordelia Langford, John Sillitoe on behalf of the Wellcome Sanger Institute COVID-19 Surveillance Team |
| EPI_ISL_2554264, | Lighthouse Lab in Alderley Park | Wellcome Sanger Institute for the COVID-19 Genomics UK (COG-UK) | Jacquelyn Wynn, Mairead Hyland, The Lighthouse Lab in Alderley Park and Alex Alderton, Roberto Amato, Jeffrey Barrett, Sonia Goncalves, Ewan Harrison, David K. Jackson, Ian Johnston, Dominic Kwiatkowski, Cordelia Langford, John Sillitoe on |

| | | | |
|---|---|---|---|
| EPI_ISL_2554265 | | Consortium | behalf of the Wellcome Sanger Institute COVID-19 Surveillance Team |
| EPI_ISL_2554266 | Lighthouse Laboratory Plymouth | Wellcome Sanger Institute for the COVID-19 Genomics UK (COG-UK) Consortium | Lighthouse Laboratory Plymouth and Alex Alderton, Roberto Amato, Jeffrey Barrett, Sonia Goncalves, Ewan Harrison, David K. Jackson, Ian Johnston, Dominic Kwiatkowski, Cordelia Langford, John Sillitoe on behalf of the Wellcome Sanger Institute COVID-19 Surveillance Team |
| EPI_ISL_2554267 | Lighthouse Lab in Alderley Park | Wellcome Sanger Institute for the COVID-19 Genomics UK (COG-UK) Consortium | Jacquelyn Wynn, Mairead Hyland, The Lighthouse Lab in Alderley Park and Alex Alderton, Roberto Amato, Jeffrey Barrett, Sonia Goncalves, Ewan Harrison, David K. Jackson, Ian Johnston, Dominic Kwiatkowski, Cordelia Langford, John Sillitoe on behalf of the Wellcome Sanger Institute COVID-19 Surveillance Team |
| EPI_ISL_2554268 | Lighthouse Laboratory Plymouth | Wellcome Sanger Institute for the COVID-19 Genomics UK (COG-UK) Consortium | Lighthouse Laboratory Plymouth and Alex Alderton, Roberto Amato, Jeffrey Barrett, Sonia Goncalves, Ewan Harrison, David K. Jackson, Ian Johnston, Dominic Kwiatkowski, Cordelia Langford, John Sillitoe on behalf of the Wellcome Sanger Institute COVID-19 Surveillance Team |
| EPI_ISL_2554269, EPI_ISL_2554271 | Lighthouse Lab in Glasgow | Wellcome Sanger Institute for the COVID-19 Genomics UK (COG-UK) Consortium | Harper VanSteenhouse, Yumi Kasai, David Gray, Carol Clugston, Anna Dominiczak and Alex Alderton, Roberto Amato, Jeffrey Barrett, Sonia Goncalves, Ewan Harrison, David K. Jackson, Ian Johnston, Dominic Kwiatkowski, Cordelia Langford, John Sillitoe on behalf of the Wellcome Sanger Institute COVID-19 Surveillance Team |
| EPI_ISL_2554272 | Lighthouse Lab in Alderley Park | Wellcome Sanger Institute for the COVID-19 Genomics UK (COG-UK) Consortium | Jacquelyn Wynn, Mairead Hyland, The Lighthouse Lab in Alderley Park and Alex Alderton, Roberto Amato, Jeffrey Barrett, Sonia Goncalves, Ewan Harrison, David K. Jackson, Ian Johnston, Dominic Kwiatkowski, Cordelia Langford, John Sillitoe on behalf of the Wellcome Sanger Institute COVID-19 Surveillance Team |
| EPI_ISL_2554273 | Lighthouse Lab in Glasgow | Wellcome Sanger Institute for the COVID-19 Genomics UK (COG-UK) Consortium | Harper VanSteenhouse, Yumi Kasai, David Gray, Carol Clugston, Anna Dominiczak and Alex Alderton, Roberto Amato, Jeffrey Barrett, Sonia Goncalves, Ewan Harrison, David K. Jackson, Ian Johnston, Dominic Kwiatkowski, Cordelia Langford, John Sillitoe on behalf of the Wellcome Sanger Institute COVID-19 Surveillance Team |
| EPI_ISL_2554274 | Lighthouse Laboratory Plymouth | Wellcome Sanger Institute for the COVID-19 Genomics UK (COG-UK) Consortium | Lighthouse Laboratory Plymouth and Alex Alderton, Roberto Amato, Jeffrey Barrett, Sonia Goncalves, Ewan Harrison, David K. Jackson, Ian Johnston, Dominic Kwiatkowski, Cordelia Langford, John Sillitoe on behalf of the Wellcome Sanger Institute COVID-19 Surveillance Team |
| EPI_ISL_2554275 | Lighthouse Lab in Alderley Park | Wellcome Sanger Institute for the COVID-19 Genomics UK (COG-UK) Consortium | Jacquelyn Wynn, Mairead Hyland, The Lighthouse Lab in Alderley Park and Alex Alderton, Roberto Amato, Jeffrey Barrett, Sonia Goncalves, Ewan Harrison, David K. Jackson, Ian Johnston, Dominic Kwiatkowski, Cordelia Langford, John Sillitoe on behalf of the Wellcome Sanger Institute COVID-19 Surveillance Team |
| EPI_ISL_2554276, EPI_ISL_2554277 | Lighthouse Laboratory Plymouth | Wellcome Sanger Institute for the COVID-19 Genomics UK (COG-UK) Consortium | Lighthouse Laboratory Plymouth and Alex Alderton, Roberto Amato, Jeffrey Barrett, Sonia Goncalves, Ewan Harrison, David K. Jackson, Ian Johnston, Dominic Kwiatkowski, Cordelia Langford, John Sillitoe on behalf of the Wellcome Sanger Institute COVID-19 Surveillance Team |
| EPI_ISL_2554278 | Lighthouse Lab in Alderley Park | Wellcome Sanger Institute for the COVID-19 Genomics UK (COG-UK) Consortium | Jacquelyn Wynn, Mairead Hyland, The Lighthouse Lab in Alderley Park and Alex Alderton, Roberto Amato, Jeffrey Barrett, Sonia Goncalves, Ewan Harrison, David K. Jackson, Ian Johnston, Dominic Kwiatkowski, Cordelia Langford, John Sillitoe on behalf of the Wellcome Sanger Institute COVID-19 Surveillance Team |
| EPI_ISL_2554280 | Lighthouse Laboratory Plymouth | Wellcome Sanger Institute for the COVID-19 Genomics UK (COG-UK) Consortium | Lighthouse Laboratory Plymouth and Alex Alderton, Roberto Amato, Jeffrey Barrett, Sonia Goncalves, Ewan Harrison, David K. Jackson, Ian Johnston, Dominic Kwiatkowski, Cordelia Langford, John Sillitoe on behalf of the Wellcome Sanger Institute COVID-19 Surveillance Team |
| EPI_ISL_2554281 | Lighthouse Lab in Glasgow | Wellcome Sanger Institute for the COVID-19 Genomics UK (COG-UK) Consortium | Harper VanSteenhouse, Yumi Kasai, David Gray, Carol Clugston, Anna Dominiczak and Alex Alderton, Roberto Amato, Jeffrey Barrett, Sonia Goncalves, Ewan Harrison, David K. Jackson, Ian Johnston, Dominic Kwiatkowski, Cordelia Langford, John Sillitoe on behalf of the Wellcome Sanger Institute COVID-19 Surveillance Team |
| EPI_ISL_2554282 | Lighthouse Lab in Alderley Park | Wellcome Sanger Institute for the COVID-19 Genomics UK (COG-UK) Consortium | Jacquelyn Wynn, Mairead Hyland, The Lighthouse Lab in Alderley Park and Alex Alderton, Roberto Amato, Jeffrey Barrett, Sonia Goncalves, Ewan Harrison, David K. Jackson, Ian Johnston, Dominic Kwiatkowski, Cordelia Langford, John Sillitoe on behalf of the Wellcome Sanger Institute COVID-19 Surveillance Team |
| EPI_ISL_2554283 | Lighthouse Laboratory Plymouth | Wellcome Sanger Institute for the COVID-19 Genomics UK (COG-UK) Consortium | Lighthouse Laboratory Plymouth and Alex Alderton, Roberto Amato, Jeffrey Barrett, Sonia Goncalves, Ewan Harrison, David K. Jackson, Ian Johnston, Dominic Kwiatkowski, Cordelia Langford, John Sillitoe on behalf of the Wellcome Sanger Institute COVID-19 Surveillance Team |
| EPI_ISL_2554284 | Lighthouse Lab in Alderley Park | Wellcome Sanger Institute for the COVID-19 Genomics UK (COG-UK) Consortium | Jacquelyn Wynn, Mairead Hyland, The Lighthouse Lab in Alderley Park and Alex Alderton, Roberto Amato, Jeffrey Barrett, Sonia Goncalves, Ewan Harrison, David K. Jackson, Ian Johnston, Dominic Kwiatkowski, Cordelia Langford, John Sillitoe on behalf of the Wellcome Sanger Institute COVID-19 Surveillance Team |
| EPI_ISL_2554285 | Lighthouse Laboratory Plymouth | Wellcome Sanger Institute for the COVID-19 Genomics UK (COG-UK) Consortium | Lighthouse Laboratory Plymouth and Alex Alderton, Roberto Amato, Jeffrey Barrett, Sonia Goncalves, Ewan Harrison, David K. Jackson, Ian Johnston, Dominic Kwiatkowski, Cordelia Langford, John Sillitoe on behalf of the Wellcome Sanger Institute COVID-19 Surveillance Team |
| EPI_ISL_2554286, EPI_ISL_2554287 | Lighthouse Lab in Alderley Park | Wellcome Sanger Institute for the COVID-19 Genomics UK (COG-UK) Consortium | Jacquelyn Wynn, Mairead Hyland, The Lighthouse Lab in Alderley Park and Alex Alderton, Roberto Amato, Jeffrey Barrett, Sonia Goncalves, Ewan Harrison, David K. Jackson, Ian Johnston, Dominic Kwiatkowski, Cordelia Langford, John Sillitoe on behalf of the Wellcome Sanger Institute COVID-19 Surveillance Team |
| EPI_ISL_2554288 | Lighthouse Lab in Glasgow | Wellcome Sanger Institute for the COVID-19 Genomics UK (COG-UK) Consortium | Harper VanSteenhouse, Yumi Kasai, David Gray, Carol Clugston, Anna Dominiczak and Alex Alderton, Roberto Amato, Jeffrey Barrett, Sonia Goncalves, Ewan Harrison, David K. Jackson, Ian Johnston, Dominic Kwiatkowski, Cordelia Langford, John Sillitoe on behalf of the Wellcome Sanger Institute COVID-19 Surveillance Team |
| EPI_ISL_2554289, EPI_ISL_2554290, EPI_ISL_2554291, EPI_ISL_2554292, EPI_ISL_2554293, EPI_ISL_2554294 | Lighthouse Lab in Alderley Park | Wellcome Sanger Institute for the COVID-19 Genomics UK (COG-UK) Consortium | Jacquelyn Wynn, Mairead Hyland, The Lighthouse Lab in Alderley Park and Alex Alderton, Roberto Amato, Jeffrey Barrett, Sonia Goncalves, Ewan Harrison, David K. Jackson, Ian Johnston, Dominic Kwiatkowski, Cordelia Langford, John Sillitoe on behalf of the Wellcome Sanger Institute COVID-19 Surveillance Team |
| EPI_ISL_2554295, EPI_ISL_2554296, EPI_ISL_2554297, EPI_ISL_2554298 | Lighthouse Laboratory Plymouth | Wellcome Sanger Institute for the COVID-19 Genomics UK (COG-UK) Consortium | Lighthouse Laboratory Plymouth and Alex Alderton, Roberto Amato, Jeffrey Barrett, Sonia Goncalves, Ewan Harrison, David K. Jackson, Ian Johnston, Dominic Kwiatkowski, Cordelia Langford, John Sillitoe on behalf of the Wellcome Sanger Institute COVID-19 Surveillance Team |
| EPI_ISL_2554299 | Lighthouse Lab in Alderley Park | Wellcome Sanger Institute for the COVID-19 Genomics UK (COG-UK) Consortium | Jacquelyn Wynn, Mairead Hyland, The Lighthouse Lab in Alderley Park and Alex Alderton, Roberto Amato, Jeffrey Barrett, Sonia Goncalves, Ewan Harrison, David K. Jackson, Ian Johnston, Dominic Kwiatkowski, Cordelia Langford, John Sillitoe on behalf of the Wellcome Sanger Institute COVID-19 Surveillance Team |
| EPI_ISL_2554300, EPI_ISL_2554301 | Lighthouse Lab in Glasgow | Wellcome Sanger Institute for the COVID-19 Genomics UK (COG-UK) Consortium | Harper VanSteenhouse, Yumi Kasai, David Gray, Carol Clugston, Anna Dominiczak and Alex Alderton, Roberto Amato, Jeffrey Barrett, Sonia Goncalves, Ewan Harrison, David K. Jackson, Ian Johnston, Dominic Kwiatkowski, Cordelia Langford, John Sillitoe on behalf of the Wellcome Sanger Institute COVID-19 Surveillance Team |
| EPI_ISL_2554302, EPI_ISL_2554303 | Lighthouse Laboratory Plymouth | Wellcome Sanger Institute for the COVID-19 Genomics UK (COG-UK) Consortium | Lighthouse Laboratory Plymouth and Alex Alderton, Roberto Amato, Jeffrey Barrett, Sonia Goncalves, Ewan Harrison, David K. Jackson, Ian Johnston, Dominic Kwiatkowski, Cordelia Langford, John Sillitoe on behalf of the Wellcome Sanger Institute COVID-19 Surveillance Team |
| EPI_ISL_2554304 | Lighthouse Lab in Alderley Park | Wellcome Sanger Institute for the COVID-19 Genomics UK (COG-UK) Consortium | Jacquelyn Wynn, Mairead Hyland, The Lighthouse Lab in Alderley Park and Alex Alderton, Roberto Amato, Jeffrey Barrett, Sonia Goncalves, Ewan Harrison, David K. Jackson, Ian Johnston, Dominic Kwiatkowski, Cordelia Langford, John Sillitoe on behalf of the Wellcome Sanger Institute COVID-19 Surveillance Team |
| EPI_ISL_2554305, EPI_ISL_2554307, EPI_ISL_2554308 | Lighthouse Laboratory Plymouth | Wellcome Sanger Institute for the COVID-19 Genomics UK (COG-UK) Consortium | Lighthouse Laboratory Plymouth and Alex Alderton, Roberto Amato, Jeffrey Barrett, Sonia Goncalves, Ewan Harrison, David K. Jackson, Ian Johnston, Dominic Kwiatkowski, Cordelia Langford, John Sillitoe on behalf of the Wellcome Sanger Institute COVID-19 Surveillance Team |
| EPI_ISL_2554309 | Lighthouse Lab in Alderley Park | Wellcome Sanger Institute for the COVID-19 Genomics UK (COG-UK) Consortium | Jacquelyn Wynn, Mairead Hyland, The Lighthouse Lab in Alderley Park and Alex Alderton, Roberto Amato, Jeffrey Barrett, Sonia Goncalves, Ewan Harrison, David K. Jackson, Ian Johnston, Dominic Kwiatkowski, Cordelia Langford, John Sillitoe on behalf of the Wellcome Sanger Institute COVID-19 Surveillance Team |
| EPI_ISL_2554310 | Lighthouse Laboratory Plymouth | Wellcome Sanger Institute for the COVID-19 Genomics UK (COG-UK) Consortium | Lighthouse Laboratory Plymouth and Alex Alderton, Roberto Amato, Jeffrey Barrett, Sonia Goncalves, Ewan Harrison, David K. Jackson, Ian Johnston, Dominic Kwiatkowski, Cordelia Langford, John Sillitoe on behalf of the Wellcome Sanger Institute COVID-19 Surveillance Team |
| EPI_ISL_2554312 | Lighthouse Lab in Glasgow | Wellcome Sanger Institute for the COVID-19 Genomics UK (COG-UK) Consortium | Harper VanSteenhouse, Yumi Kasai, David Gray, Carol Clugston, Anna Dominiczak and Alex Alderton, Roberto Amato, Jeffrey Barrett, Sonia Goncalves, Ewan Harrison, David K. Jackson, Ian Johnston, Dominic Kwiatkowski, Cordelia Langford, John Sillitoe on behalf of the Wellcome Sanger Institute COVID-19 Surveillance Team |
| EPI_ISL_2554313 | Lighthouse Lab in Alderley Park | Wellcome Sanger Institute for the COVID-19 Genomics UK (COG-UK) Consortium | Jacquelyn Wynn, Mairead Hyland, The Lighthouse Lab in Alderley Park and Alex Alderton, Roberto Amato, Jeffrey Barrett, Sonia Goncalves, Ewan Harrison, David K. Jackson, Ian Johnston, Dominic Kwiatkowski, Cordelia Langford, John Sillitoe on behalf of the Wellcome Sanger Institute COVID-19 Surveillance Team |
| EPI_ISL_2554314 | Lighthouse Lab in Glasgow | Wellcome Sanger Institute for the COVID-19 Genomics UK (COG-UK) Consortium | Harper VanSteenhouse, Yumi Kasai, David Gray, Carol Clugston, Anna Dominiczak and Alex Alderton, Roberto Amato, Jeffrey Barrett, Sonia Goncalves, Ewan Harrison, David K. Jackson, Ian Johnston, Dominic Kwiatkowski, Cordelia Langford, John Sillitoe on behalf of the Wellcome Sanger Institute COVID-19 Surveillance Team |
| EPI_ISL_2554315, EPI_ISL_2554316, EPI_ISL_2554317 | Lighthouse Laboratory Plymouth | Wellcome Sanger Institute for the COVID-19 Genomics UK (COG-UK) Consortium | Lighthouse Laboratory Plymouth and Alex Alderton, Roberto Amato, Jeffrey Barrett, Sonia Goncalves, Ewan Harrison, David K. Jackson, Ian Johnston, Dominic Kwiatkowski, Cordelia Langford, John Sillitoe on behalf of the Wellcome Sanger Institute COVID-19 Surveillance Team |
| EPI_ISL_2554318 | Lighthouse Lab in Glasgow | Wellcome Sanger Institute for the COVID-19 Genomics UK (COG-UK) Consortium | Harper VanSteenhouse, Yumi Kasai, David Gray, Carol Clugston, Anna Dominiczak and Alex Alderton, Roberto Amato, Jeffrey Barrett, Sonia Goncalves, Ewan Harrison, David K. Jackson, Ian Johnston, Dominic Kwiatkowski, Cordelia Langford, John Sillitoe on behalf of the Wellcome Sanger Institute COVID-19 Surveillance Team |
| EPI_ISL_2554319 | Lighthouse Laboratory Plymouth | Wellcome Sanger Institute for the COVID-19 Genomics UK (COG-UK) Consortium | Lighthouse Laboratory Plymouth and Alex Alderton, Roberto Amato, Jeffrey Barrett, Sonia Goncalves, Ewan Harrison, David K. Jackson, Ian Johnston, Dominic Kwiatkowski, Cordelia Langford, John Sillitoe on behalf of the Wellcome Sanger Institute COVID-19 Surveillance Team |
| EPI_ISL_2554320, EPI_ISL_2554321, EPI_ISL_2554322, EPI_ISL_2554323 | Lighthouse Lab in Glasgow | Wellcome Sanger Institute for the COVID-19 Genomics UK (COG-UK) Consortium | Harper VanSteenhouse, Yumi Kasai, David Gray, Carol Clugston, Anna Dominiczak and Alex Alderton, Roberto Amato, Jeffrey Barrett, Sonia Goncalves, Ewan Harrison, David K. Jackson, Ian Johnston, Dominic Kwiatkowski, Cordelia Langford, John Sillitoe on behalf of the Wellcome Sanger Institute COVID-19 Surveillance Team |
| EPI_ISL_2554324 | Lighthouse Laboratory Plymouth | Wellcome Sanger Institute for the COVID-19 Genomics UK (COG-UK) Consortium | Lighthouse Laboratory Plymouth and Alex Alderton, Roberto Amato, Jeffrey Barrett, Sonia Goncalves, Ewan Harrison, David K. Jackson, Ian Johnston, Dominic Kwiatkowski, Cordelia Langford, John Sillitoe on behalf of the Wellcome Sanger Institute COVID-19 Surveillance Team |
| EPI_ISL_2554325, EPI_ISL_2554326, EPI_ISL_2554328, EPI_ISL_2554329 | Lighthouse Lab in Alderley Park | Wellcome Sanger Institute for the COVID-19 Genomics UK (COG-UK) Consortium | Jacquelyn Wynn, Mairead Hyland, The Lighthouse Lab in Alderley Park and Alex Alderton, Roberto Amato, Jeffrey Barrett, Sonia Goncalves, Ewan Harrison, David K. Jackson, Ian Johnston, Dominic Kwiatkowski, Cordelia Langford, John Sillitoe on behalf of the Wellcome Sanger Institute COVID-19 Surveillance Team |
| EPI_ISL_2554330 | Lighthouse Lab in Glasgow | Wellcome Sanger Institute for the COVID-19 Genomics UK (COG-UK) Consortium | Harper VanSteenhouse, Yumi Kasai, David Gray, Carol Clugston, Anna Dominiczak and Alex Alderton, Roberto Amato, Jeffrey Barrett, Sonia Goncalves, Ewan Harrison, David K. Jackson, Ian Johnston, Dominic Kwiatkowski, Cordelia Langford, John Sillitoe on behalf of the Wellcome Sanger Institute COVID-19 Surveillance Team |
| EPI_ISL_2554331, EPI_ISL_2554332 | Lighthouse Laboratory Plymouth | Wellcome Sanger Institute for the COVID-19 Genomics UK (COG-UK) Consortium | Lighthouse Laboratory Plymouth and Alex Alderton, Roberto Amato, Jeffrey Barrett, Sonia Goncalves, Ewan Harrison, David K. Jackson, Ian Johnston, Dominic Kwiatkowski, Cordelia Langford, John Sillitoe on behalf of the Wellcome Sanger Institute COVID-19 Surveillance Team |
| EPI_ISL_2554334, EPI_ISL_2554337 | Lighthouse Lab in Alderley Park | Wellcome Sanger Institute for the COVID-19 Genomics UK (COG-UK) Consortium | Jacquelyn Wynn, Mairead Hyland, The Lighthouse Lab in Alderley Park and Alex Alderton, Roberto Amato, Jeffrey Barrett, Sonia Goncalves, Ewan Harrison, David K. Jackson, Ian Johnston, Dominic Kwiatkowski, Cordelia Langford, John Sillitoe on behalf of the Wellcome Sanger Institute COVID-19 Surveillance Team |
| EPI_ISL_2554338 | Lighthouse Laboratory Plymouth | Wellcome Sanger Institute for the COVID-19 Genomics UK (COG-UK) | Lighthouse Laboratory Plymouth and Alex Alderton, Roberto Amato, Jeffrey Barrett, Sonia Goncalves, Ewan Harrison, David K. Jackson, Ian Johnston, Dominic Kwiatkowski, Cordelia Langford, John Sillitoe on behalf of the Wellcome Sanger |

| | | | |
|---|---|---|---|
| | | Consortium | Institute COVID-19 Surveillance Team |
| EPI_ISL_2554339 | Lighthouse Lab in Alderley Park | Wellcome Sanger Institute for the COVID-19 Genomics UK (COG-UK) Consortium | Jacquelyn Wynn, Mairead Hyland, The Lighthouse Lab in Alderley Park and Alex Alderton, Roberto Amato, Jeffrey Barrett, Sonia Goncalves, Ewan Harrison, David K. Jackson, Ian Johnston, Dominic Kwiatkowski, Cordelia Langford, John Sillitoe on behalf of the Wellcome Sanger Institute COVID-19 Surveillance Team |
| EPI_ISL_2554340 | Lighthouse Laboratory Plymouth | Wellcome Sanger Institute for the COVID-19 Genomics UK (COG-UK) Consortium | Lighthouse Laboratory Plymouth and Alex Alderton, Roberto Amato, Jeffrey Barrett, Sonia Goncalves, Ewan Harrison, David K. Jackson, Ian Johnston, Dominic Kwiatkowski, Cordelia Langford, John Sillitoe on behalf of the Wellcome Sanger Institute COVID-19 Surveillance Team |
| EPI_ISL_2554341, EPI_ISL_2554342, EPI_ISL_2554344, EPI_ISL_2554345, EPI_ISL_2554346 | Lighthouse Lab in Alderley Park | Wellcome Sanger Institute for the COVID-19 Genomics UK (COG-UK) Consortium | Jacquelyn Wynn, Mairead Hyland, The Lighthouse Lab in Alderley Park and Alex Alderton, Roberto Amato, Jeffrey Barrett, Sonia Goncalves, Ewan Harrison, David K. Jackson, Ian Johnston, Dominic Kwiatkowski, Cordelia Langford, John Sillitoe on behalf of the Wellcome Sanger Institute COVID-19 Surveillance Team |
| EPI_ISL_2554347, EPI_ISL_2554348 | Lighthouse Lab in Glasgow | Wellcome Sanger Institute for the COVID-19 Genomics UK (COG-UK) Consortium | Harper VanSteenhouse, Yumi Kasai, David Gray, Carol Clugston, Anna Dominiczak and Alex Alderton, Roberto Amato, Jeffrey Barrett, Sonia Goncalves, Ewan Harrison, David K. Jackson, Ian Johnston, Dominic Kwiatkowski, Cordelia Langford, John Sillitoe on behalf of the Wellcome Sanger Institute COVID-19 Surveillance Team |
| EPI_ISL_2554349 | Lighthouse Lab in Alderley Park | Wellcome Sanger Institute for the COVID-19 Genomics UK (COG-UK) Consortium | Jacquelyn Wynn, Mairead Hyland, The Lighthouse Lab in Alderley Park and Alex Alderton, Roberto Amato, Jeffrey Barrett, Sonia Goncalves, Ewan Harrison, David K. Jackson, Ian Johnston, Dominic Kwiatkowski, Cordelia Langford, John Sillitoe on behalf of the Wellcome Sanger Institute COVID-19 Surveillance Team |
| EPI_ISL_2554350 | Lighthouse Laboratory Plymouth | Wellcome Sanger Institute for the COVID-19 Genomics UK (COG-UK) Consortium | Lighthouse Laboratory Plymouth and Alex Alderton, Roberto Amato, Jeffrey Barrett, Sonia Goncalves, Ewan Harrison, David K. Jackson, Ian Johnston, Dominic Kwiatkowski, Cordelia Langford, John Sillitoe on behalf of the Wellcome Sanger Institute COVID-19 Surveillance Team |
| EPI_ISL_2554351 | Lighthouse Lab in Alderley Park | Wellcome Sanger Institute for the COVID-19 Genomics UK (COG-UK) Consortium | Jacquelyn Wynn, Mairead Hyland, The Lighthouse Lab in Alderley Park and Alex Alderton, Roberto Amato, Jeffrey Barrett, Sonia Goncalves, Ewan Harrison, David K. Jackson, Ian Johnston, Dominic Kwiatkowski, Cordelia Langford, John Sillitoe on behalf of the Wellcome Sanger Institute COVID-19 Surveillance Team |
| EPI_ISL_2554352, EPI_ISL_2554353, EPI_ISL_2554354 | Lighthouse Laboratory Plymouth | Wellcome Sanger Institute for the COVID-19 Genomics UK (COG-UK) Consortium | Lighthouse Laboratory Plymouth and Alex Alderton, Roberto Amato, Jeffrey Barrett, Sonia Goncalves, Ewan Harrison, David K. Jackson, Ian Johnston, Dominic Kwiatkowski, Cordelia Langford, John Sillitoe on behalf of the Wellcome Sanger Institute COVID-19 Surveillance Team |
| EPI_ISL_2554355, EPI_ISL_2554356 | Lighthouse Lab in Glasgow | Wellcome Sanger Institute for the COVID-19 Genomics UK (COG-UK) Consortium | Harper VanSteenhouse, Yumi Kasai, David Gray, Carol Clugston, Anna Dominiczak and Alex Alderton, Roberto Amato, Jeffrey Barrett, Sonia Goncalves, Ewan Harrison, David K. Jackson, Ian Johnston, Dominic Kwiatkowski, Cordelia Langford, John Sillitoe on behalf of the Wellcome Sanger Institute COVID-19 Surveillance Team |
| EPI_ISL_2554357, EPI_ISL_2554358 | Lighthouse Lab in Alderley Park | Wellcome Sanger Institute for the COVID-19 Genomics UK (COG-UK) Consortium | Jacquelyn Wynn, Mairead Hyland, The Lighthouse Lab in Alderley Park and Alex Alderton, Roberto Amato, Jeffrey Barrett, Sonia Goncalves, Ewan Harrison, David K. Jackson, Ian Johnston, Dominic Kwiatkowski, Cordelia Langford, John Sillitoe on behalf of the Wellcome Sanger Institute COVID-19 Surveillance Team |
| EPI_ISL_2554359 | Lighthouse Laboratory Plymouth | Wellcome Sanger Institute for the COVID-19 Genomics UK (COG-UK) Consortium | Lighthouse Laboratory Plymouth and Alex Alderton, Roberto Amato, Jeffrey Barrett, Sonia Goncalves, Ewan Harrison, David K. Jackson, Ian Johnston, Dominic Kwiatkowski, Cordelia Langford, John Sillitoe on behalf of the Wellcome Sanger Institute COVID-19 Surveillance Team |
| EPI_ISL_2554360, EPI_ISL_2554361, EPI_ISL_2554362 | Lighthouse Lab in Glasgow | Wellcome Sanger Institute for the COVID-19 Genomics UK (COG-UK) Consortium | Harper VanSteenhouse, Yumi Kasai, David Gray, Carol Clugston, Anna Dominiczak and Alex Alderton, Roberto Amato, Jeffrey Barrett, Sonia Goncalves, Ewan Harrison, David K. Jackson, Ian Johnston, Dominic Kwiatkowski, Cordelia Langford, John Sillitoe on behalf of the Wellcome Sanger Institute COVID-19 Surveillance Team |
| EPI_ISL_2554363, EPI_ISL_2554364 | Lighthouse Lab in Alderley Park | Wellcome Sanger Institute for the COVID-19 Genomics UK (COG-UK) Consortium | Jacquelyn Wynn, Mairead Hyland, The Lighthouse Lab in Alderley Park and Alex Alderton, Roberto Amato, Jeffrey Barrett, Sonia Goncalves, Ewan Harrison, David K. Jackson, Ian Johnston, Dominic Kwiatkowski, Cordelia Langford, John Sillitoe on behalf of the Wellcome Sanger Institute COVID-19 Surveillance Team |
| EPI_ISL_2554365 | Lighthouse Lab in Glasgow | Wellcome Sanger Institute for the COVID-19 Genomics UK (COG-UK) Consortium | Harper VanSteenhouse, Yumi Kasai, David Gray, Carol Clugston, Anna Dominiczak and Alex Alderton, Roberto Amato, Jeffrey Barrett, Sonia Goncalves, Ewan Harrison, David K. Jackson, Ian Johnston, Dominic Kwiatkowski, Cordelia Langford, John Sillitoe on behalf of the Wellcome Sanger Institute COVID-19 Surveillance Team |
| EPI_ISL_2554366 | Lighthouse Laboratory Plymouth | Wellcome Sanger Institute for the COVID-19 Genomics UK (COG-UK) Consortium | Lighthouse Laboratory Plymouth and Alex Alderton, Roberto Amato, Jeffrey Barrett, Sonia Goncalves, Ewan Harrison, David K. Jackson, Ian Johnston, Dominic Kwiatkowski, Cordelia Langford, John Sillitoe on behalf of the Wellcome Sanger Institute COVID-19 Surveillance Team |
| EPI_ISL_2554367 | Lighthouse Lab in Glasgow | Wellcome Sanger Institute for the COVID-19 Genomics UK (COG-UK) Consortium | Harper VanSteenhouse, Yumi Kasai, David Gray, Carol Clugston, Anna Dominiczak and Alex Alderton, Roberto Amato, Jeffrey Barrett, Sonia Goncalves, Ewan Harrison, David K. Jackson, Ian Johnston, Dominic Kwiatkowski, Cordelia Langford, John Sillitoe on behalf of the Wellcome Sanger Institute COVID-19 Surveillance Team |
| EPI_ISL_2554368, EPI_ISL_2554370 | Lighthouse Lab in Alderley Park | Wellcome Sanger Institute for the COVID-19 Genomics UK (COG-UK) Consortium | Jacquelyn Wynn, Mairead Hyland, The Lighthouse Lab in Alderley Park and Alex Alderton, Roberto Amato, Jeffrey Barrett, Sonia Goncalves, Ewan Harrison, David K. Jackson, Ian Johnston, Dominic Kwiatkowski, Cordelia Langford, John Sillitoe on behalf of the Wellcome Sanger Institute COVID-19 Surveillance Team |
| EPI_ISL_2554371 | Lighthouse Laboratory Plymouth | Wellcome Sanger Institute for the COVID-19 Genomics UK (COG-UK) Consortium | Lighthouse Laboratory Plymouth and Alex Alderton, Roberto Amato, Jeffrey Barrett, Sonia Goncalves, Ewan Harrison, David K. Jackson, Ian Johnston, Dominic Kwiatkowski, Cordelia Langford, John Sillitoe on behalf of the Wellcome Sanger Institute COVID-19 Surveillance Team |
| EPI_ISL_2554372 | Lighthouse Lab in Glasgow | Wellcome Sanger Institute for the COVID-19 Genomics UK (COG-UK) Consortium | Harper VanSteenhouse, Yumi Kasai, David Gray, Carol Clugston, Anna Dominiczak and Alex Alderton, Roberto Amato, Jeffrey Barrett, Sonia Goncalves, Ewan Harrison, David K. Jackson, Ian Johnston, Dominic Kwiatkowski, Cordelia Langford, John Sillitoe on behalf of the Wellcome Sanger Institute COVID-19 Surveillance Team |
| EPI_ISL_2554373 | Lighthouse Lab in Alderley Park | Wellcome Sanger Institute for the COVID-19 Genomics UK (COG-UK) Consortium | Jacquelyn Wynn, Mairead Hyland, The Lighthouse Lab in Alderley Park and Alex Alderton, Roberto Amato, Jeffrey Barrett, Sonia Goncalves, Ewan Harrison, David K. Jackson, Ian Johnston, Dominic Kwiatkowski, Cordelia Langford, John Sillitoe on behalf of the Wellcome Sanger Institute COVID-19 Surveillance Team |
| EPI_ISL_2554374 | Lighthouse Lab in Glasgow | Wellcome Sanger Institute for the COVID-19 Genomics UK (COG-UK) Consortium | Harper VanSteenhouse, Yumi Kasai, David Gray, Carol Clugston, Anna Dominiczak and Alex Alderton, Roberto Amato, Jeffrey Barrett, Sonia Goncalves, Ewan Harrison, David K. Jackson, Ian Johnston, Dominic Kwiatkowski, Cordelia Langford, John Sillitoe on behalf of the Wellcome Sanger Institute COVID-19 Surveillance Team |
| EPI_ISL_2554375 | Lighthouse Laboratory Plymouth | Wellcome Sanger Institute for the COVID-19 Genomics UK (COG-UK) Consortium | Lighthouse Laboratory Plymouth and Alex Alderton, Roberto Amato, Jeffrey Barrett, Sonia Goncalves, Ewan Harrison, David K. Jackson, Ian Johnston, Dominic Kwiatkowski, Cordelia Langford, John Sillitoe on behalf of the Wellcome Sanger Institute COVID-19 Surveillance Team |
| EPI_ISL_2554377, EPI_ISL_2554378, EPI_ISL_2554379, EPI_ISL_2554380 | Lighthouse Lab in Alderley Park | Wellcome Sanger Institute for the COVID-19 Genomics UK (COG-UK) Consortium | Jacquelyn Wynn, Mairead Hyland, The Lighthouse Lab in Alderley Park and Alex Alderton, Roberto Amato, Jeffrey Barrett, Sonia Goncalves, Ewan Harrison, David K. Jackson, Ian Johnston, Dominic Kwiatkowski, Cordelia Langford, John Sillitoe on behalf of the Wellcome Sanger Institute COVID-19 Surveillance Team |
| EPI_ISL_2554382 | Lighthouse Lab in Glasgow | Wellcome Sanger Institute for the COVID-19 Genomics UK (COG-UK) Consortium | Harper VanSteenhouse, Yumi Kasai, David Gray, Carol Clugston, Anna Dominiczak and Alex Alderton, Roberto Amato, Jeffrey Barrett, Sonia Goncalves, Ewan Harrison, David K. Jackson, Ian Johnston, Dominic Kwiatkowski, Cordelia Langford, John Sillitoe on behalf of the Wellcome Sanger Institute COVID-19 Surveillance Team |
| EPI_ISL_2554383, EPI_ISL_2554384 | Lighthouse Lab in Alderley Park | Wellcome Sanger Institute for the COVID-19 Genomics UK (COG-UK) Consortium | Jacquelyn Wynn, Mairead Hyland, The Lighthouse Lab in Alderley Park and Alex Alderton, Roberto Amato, Jeffrey Barrett, Sonia Goncalves, Ewan Harrison, David K. Jackson, Ian Johnston, Dominic Kwiatkowski, Cordelia Langford, John Sillitoe on behalf of the Wellcome Sanger Institute COVID-19 Surveillance Team |
| EPI_ISL_2554385, EPI_ISL_2554386 | Lighthouse Laboratory Plymouth | Wellcome Sanger Institute for the COVID-19 Genomics UK (COG-UK) Consortium | Lighthouse Laboratory Plymouth and Alex Alderton, Roberto Amato, Jeffrey Barrett, Sonia Goncalves, Ewan Harrison, David K. Jackson, Ian Johnston, Dominic Kwiatkowski, Cordelia Langford, John Sillitoe on behalf of the Wellcome Sanger Institute COVID-19 Surveillance Team |
| EPI_ISL_2554387 | Lighthouse Lab in Glasgow | Wellcome Sanger Institute for the COVID-19 Genomics UK (COG-UK) Consortium | Harper VanSteenhouse, Yumi Kasai, David Gray, Carol Clugston, Anna Dominiczak and Alex Alderton, Roberto Amato, Jeffrey Barrett, Sonia Goncalves, Ewan Harrison, David K. Jackson, Ian Johnston, Dominic Kwiatkowski, Cordelia Langford, John Sillitoe on behalf of the Wellcome Sanger Institute COVID-19 Surveillance Team |
| EPI_ISL_2554388, EPI_ISL_2554389, EPI_ISL_2554390 | Lighthouse Lab in Alderley Park | Wellcome Sanger Institute for the COVID-19 Genomics UK (COG-UK) Consortium | Jacquelyn Wynn, Mairead Hyland, The Lighthouse Lab in Alderley Park and Alex Alderton, Roberto Amato, Jeffrey Barrett, Sonia Goncalves, Ewan Harrison, David K. Jackson, Ian Johnston, Dominic Kwiatkowski, Cordelia Langford, John Sillitoe on behalf of the Wellcome Sanger Institute COVID-19 Surveillance Team |
| EPI_ISL_2554391 | Lighthouse Laboratory Plymouth | Wellcome Sanger Institute for the COVID-19 Genomics UK (COG-UK) Consortium | Lighthouse Laboratory Plymouth and Alex Alderton, Roberto Amato, Jeffrey Barrett, Sonia Goncalves, Ewan Harrison, David K. Jackson, Ian Johnston, Dominic Kwiatkowski, Cordelia Langford, John Sillitoe on behalf of the Wellcome Sanger Institute COVID-19 Surveillance Team |
| EPI_ISL_2554392 | Lighthouse Lab in Glasgow | Wellcome Sanger Institute for the COVID-19 Genomics UK (COG-UK) Consortium | Harper VanSteenhouse, Yumi Kasai, David Gray, Carol Clugston, Anna Dominiczak and Alex Alderton, Roberto Amato, Jeffrey Barrett, Sonia Goncalves, Ewan Harrison, David K. Jackson, Ian Johnston, Dominic Kwiatkowski, Cordelia Langford, John Sillitoe on behalf of the Wellcome Sanger Institute COVID-19 Surveillance Team |
| EPI_ISL_2554393, EPI_ISL_2554394 | Lighthouse Laboratory Plymouth | Wellcome Sanger Institute for the COVID-19 Genomics UK (COG-UK) Consortium | Lighthouse Laboratory Plymouth and Alex Alderton, Roberto Amato, Jeffrey Barrett, Sonia Goncalves, Ewan Harrison, David K. Jackson, Ian Johnston, Dominic Kwiatkowski, Cordelia Langford, John Sillitoe on behalf of the Wellcome Sanger Institute COVID-19 Surveillance Team |
| EPI_ISL_2554395 | Lighthouse Lab in Glasgow | Wellcome Sanger Institute for the COVID-19 Genomics UK (COG-UK) Consortium | Harper VanSteenhouse, Yumi Kasai, David Gray, Carol Clugston, Anna Dominiczak and Alex Alderton, Roberto Amato, Jeffrey Barrett, Sonia Goncalves, Ewan Harrison, David K. Jackson, Ian Johnston, Dominic Kwiatkowski, Cordelia Langford, John Sillitoe on behalf of the Wellcome Sanger Institute COVID-19 Surveillance Team |
| EPI_ISL_2554397 | Lighthouse Lab in Alderley Park | Wellcome Sanger Institute for the COVID-19 Genomics UK (COG-UK) Consortium | Jacquelyn Wynn, Mairead Hyland, The Lighthouse Lab in Alderley Park and Alex Alderton, Roberto Amato, Jeffrey Barrett, Sonia Goncalves, Ewan Harrison, David K. Jackson, Ian Johnston, Dominic Kwiatkowski, Cordelia Langford, John Sillitoe on behalf of the Wellcome Sanger Institute COVID-19 Surveillance Team |
| EPI_ISL_2554398 | Lighthouse Lab in Glasgow | Wellcome Sanger Institute for the COVID-19 Genomics UK (COG-UK) Consortium | Harper VanSteenhouse, Yumi Kasai, David Gray, Carol Clugston, Anna Dominiczak and Alex Alderton, Roberto Amato, Jeffrey Barrett, Sonia Goncalves, Ewan Harrison, David K. Jackson, Ian Johnston, Dominic Kwiatkowski, Cordelia Langford, John Sillitoe on behalf of the Wellcome Sanger Institute COVID-19 Surveillance Team |
| EPI_ISL_2554399 | Lighthouse Laboratory Plymouth | Wellcome Sanger Institute for the COVID-19 Genomics UK (COG-UK) Consortium | Lighthouse Laboratory Plymouth and Alex Alderton, Roberto Amato, Jeffrey Barrett, Sonia Goncalves, Ewan Harrison, David K. Jackson, Ian Johnston, Dominic Kwiatkowski, Cordelia Langford, John Sillitoe on behalf of the Wellcome Sanger Institute COVID-19 Surveillance Team |
| EPI_ISL_2554401 | Lighthouse Lab in Glasgow | Wellcome Sanger Institute for the COVID-19 Genomics UK (COG-UK) Consortium | Harper VanSteenhouse, Yumi Kasai, David Gray, Carol Clugston, Anna Dominiczak and Alex Alderton, Roberto Amato, Jeffrey Barrett, Sonia Goncalves, Ewan Harrison, David K. Jackson, Ian Johnston, Dominic Kwiatkowski, Cordelia Langford, John Sillitoe on behalf of the Wellcome Sanger Institute COVID-19 Surveillance Team |
| EPI_ISL_2554403 | Lighthouse Lab in Alderley Park | Wellcome Sanger Institute for the COVID-19 Genomics UK (COG-UK) Consortium | Jacquelyn Wynn, Mairead Hyland, The Lighthouse Lab in Alderley Park and Alex Alderton, Roberto Amato, Jeffrey Barrett, Sonia Goncalves, Ewan Harrison, David K. Jackson, Ian Johnston, Dominic Kwiatkowski, Cordelia Langford, John Sillitoe on behalf of the Wellcome Sanger Institute COVID-19 Surveillance Team |
| EPI_ISL_2554404 | Lighthouse Lab in Glasgow | Wellcome Sanger Institute for the COVID-19 Genomics UK (COG-UK) Consortium | Harper VanSteenhouse, Yumi Kasai, David Gray, Carol Clugston, Anna Dominiczak and Alex Alderton, Roberto Amato, Jeffrey Barrett, Sonia Goncalves, Ewan Harrison, David K. Jackson, Ian Johnston, Dominic Kwiatkowski, Cordelia Langford, John Sillitoe on behalf of the Wellcome Sanger Institute COVID-19 Surveillance Team |
| EPI_ISL_2554405 | Lighthouse Lab in Alderley Park | Wellcome Sanger Institute for the COVID-19 Genomics UK (COG-UK) Consortium | Jacquelyn Wynn, Mairead Hyland, The Lighthouse Lab in Alderley Park and Alex Alderton, Roberto Amato, Jeffrey Barrett, Sonia Goncalves, Ewan Harrison, David K. Jackson, Ian Johnston, Dominic Kwiatkowski, Cordelia Langford, John Sillitoe on behalf of the Wellcome Sanger Institute COVID-19 Surveillance Team |
| EPI_ISL_2554406, EPI_ISL_2554407 | Lighthouse Laboratory Plymouth | Wellcome Sanger Institute for the COVID-19 Genomics UK (COG-UK) Consortium | Lighthouse Laboratory Plymouth and Alex Alderton, Roberto Amato, Jeffrey Barrett, Sonia Goncalves, Ewan Harrison, David K. Jackson, Ian Johnston, Dominic Kwiatkowski, Cordelia Langford, John Sillitoe on behalf of the Wellcome Sanger Institute COVID-19 Surveillance Team |
| EPI_ISL_2554408, EPI_ISL_2554410 | Lighthouse Lab in Alderley Park | Wellcome Sanger Institute for the COVID-19 Genomics UK (COG-UK) Consortium | Jacquelyn Wynn, Mairead Hyland, The Lighthouse Lab in Alderley Park and Alex Alderton, Roberto Amato, Jeffrey Barrett, Sonia Goncalves, Ewan Harrison, David K. Jackson, Ian Johnston, Dominic Kwiatkowski, Cordelia Langford, John Sillitoe on behalf of the Wellcome Sanger Institute COVID-19 Surveillance Team |
| EPI_ISL_2554411, | Lighthouse Lab in Glasgow | Wellcome Sanger Institute for the COVID-19 Genomics UK (COG-UK) | Harper VanSteenhouse, Yumi Kasai, David Gray, Carol Clugston, Anna Dominiczak and Alex Alderton, Roberto Amato, Jeffrey Barrett, Sonia Goncalves, Ewan Harrison, David K. Jackson, Ian Johnston, Dominic Kwiatkowski, Cordelia Langford, |

| | | | |
|---|---|---|---|
| EPI_ISL_2554412 | | Consortium | John Sillitoe on behalf of the Wellcome Sanger Institute COVID-19 Surveillance Team |
| EPI_ISL_2554413 | Lighthouse Laboratory Plymouth | Wellcome Sanger Institute for the COVID-19 Genomics UK (COG-UK) Consortium | Lighthouse Laboratory Plymouth and Alex Alderton, Roberto Amato, Jeffrey Barrett, Sonia Goncalves, Ewan Harrison, David K. Jackson, Ian Johnston, Dominic Kwiatkowski, Cordelia Langford, John Sillitoe on behalf of the Wellcome Sanger Institute COVID-19 Surveillance Team |
| EPI_ISL_2554414 | Lighthouse Lab in Alderley Park | Wellcome Sanger Institute for the COVID-19 Genomics UK (COG-UK) Consortium | Jacquelyn Wynn, Mairead Hyland, The Lighthouse Lab in Alderley Park and Alex Alderton, Roberto Amato, Jeffrey Barrett, Sonia Goncalves, Ewan Harrison, David K. Jackson, Ian Johnston, Dominic Kwiatkowski, Cordelia Langford, John Sillitoe on behalf of the Wellcome Sanger Institute COVID-19 Surveillance Team |
| EPI_ISL_2554415 | Lighthouse Laboratory Plymouth | Wellcome Sanger Institute for the COVID-19 Genomics UK (COG-UK) Consortium | Lighthouse Laboratory Plymouth and Alex Alderton, Roberto Amato, Jeffrey Barrett, Sonia Goncalves, Ewan Harrison, David K. Jackson, Ian Johnston, Dominic Kwiatkowski, Cordelia Langford, John Sillitoe on behalf of the Wellcome Sanger Institute COVID-19 Surveillance Team |
| EPI_ISL_2554416 | Lighthouse Lab in Glasgow | Wellcome Sanger Institute for the COVID-19 Genomics UK (COG-UK) Consortium | Harper VanSteenhouse, Yumi Kasai, David Gray, Carol Clugston, Anna Dominiczak and Alex Alderton, Roberto Amato, Jeffrey Barrett, Sonia Goncalves, Ewan Harrison, David K. Jackson, Ian Johnston, Dominic Kwiatkowski, Cordelia Langford, John Sillitoe on behalf of the Wellcome Sanger Institute COVID-19 Surveillance Team |
| EPI_ISL_2554417, EPI_ISL_2554418, EPI_ISL_2554420 | Lighthouse Lab in Alderley Park | Wellcome Sanger Institute for the COVID-19 Genomics UK (COG-UK) Consortium | Jacquelyn Wynn, Mairead Hyland, The Lighthouse Lab in Alderley Park and Alex Alderton, Roberto Amato, Jeffrey Barrett, Sonia Goncalves, Ewan Harrison, David K. Jackson, Ian Johnston, Dominic Kwiatkowski, Cordelia Langford, John Sillitoe on behalf of the Wellcome Sanger Institute COVID-19 Surveillance Team |
| EPI_ISL_2554421, EPI_ISL_2554422 | Lighthouse Lab in Glasgow | Wellcome Sanger Institute for the COVID-19 Genomics UK (COG-UK) Consortium | Harper VanSteenhouse, Yumi Kasai, David Gray, Carol Clugston, Anna Dominiczak and Alex Alderton, Roberto Amato, Jeffrey Barrett, Sonia Goncalves, Ewan Harrison, David K. Jackson, Ian Johnston, Dominic Kwiatkowski, Cordelia Langford, John Sillitoe on behalf of the Wellcome Sanger Institute COVID-19 Surveillance Team |
| EPI_ISL_2554424 | Lighthouse Lab in Alderley Park | Wellcome Sanger Institute for the COVID-19 Genomics UK (COG-UK) Consortium | Jacquelyn Wynn, Mairead Hyland, The Lighthouse Lab in Alderley Park and Alex Alderton, Roberto Amato, Jeffrey Barrett, Sonia Goncalves, Ewan Harrison, David K. Jackson, Ian Johnston, Dominic Kwiatkowski, Cordelia Langford, John Sillitoe on behalf of the Wellcome Sanger Institute COVID-19 Surveillance Team |
| EPI_ISL_2554425 | Lighthouse Lab in Glasgow | Wellcome Sanger Institute for the COVID-19 Genomics UK (COG-UK) Consortium | Harper VanSteenhouse, Yumi Kasai, David Gray, Carol Clugston, Anna Dominiczak and Alex Alderton, Roberto Amato, Jeffrey Barrett, Sonia Goncalves, Ewan Harrison, David K. Jackson, Ian Johnston, Dominic Kwiatkowski, Cordelia Langford, John Sillitoe on behalf of the Wellcome Sanger Institute COVID-19 Surveillance Team |
| EPI_ISL_2554426, EPI_ISL_2554427, EPI_ISL_2554428, EPI_ISL_2554429, EPI_ISL_2554430 | Lighthouse Lab in Alderley Park | Wellcome Sanger Institute for the COVID-19 Genomics UK (COG-UK) Consortium | Jacquelyn Wynn, Mairead Hyland, The Lighthouse Lab in Alderley Park and Alex Alderton, Roberto Amato, Jeffrey Barrett, Sonia Goncalves, Ewan Harrison, David K. Jackson, Ian Johnston, Dominic Kwiatkowski, Cordelia Langford, John Sillitoe on behalf of the Wellcome Sanger Institute COVID-19 Surveillance Team |
| EPI_ISL_2554431 | Lighthouse Lab in Glasgow | Wellcome Sanger Institute for the COVID-19 Genomics UK (COG-UK) Consortium | Harper VanSteenhouse, Yumi Kasai, David Gray, Carol Clugston, Anna Dominiczak and Alex Alderton, Roberto Amato, Jeffrey Barrett, Sonia Goncalves, Ewan Harrison, David K. Jackson, Ian Johnston, Dominic Kwiatkowski, Cordelia Langford, John Sillitoe on behalf of the Wellcome Sanger Institute COVID-19 Surveillance Team |
| EPI_ISL_2554432 | Lighthouse Lab in Alderley Park | Wellcome Sanger Institute for the COVID-19 Genomics UK (COG-UK) Consortium | Jacquelyn Wynn, Mairead Hyland, The Lighthouse Lab in Alderley Park and Alex Alderton, Roberto Amato, Jeffrey Barrett, Sonia Goncalves, Ewan Harrison, David K. Jackson, Ian Johnston, Dominic Kwiatkowski, Cordelia Langford, John Sillitoe on behalf of the Wellcome Sanger Institute COVID-19 Surveillance Team |
| EPI_ISL_2554433 | Lighthouse Lab in Glasgow | Wellcome Sanger Institute for the COVID-19 Genomics UK (COG-UK) Consortium | Harper VanSteenhouse, Yumi Kasai, David Gray, Carol Clugston, Anna Dominiczak and Alex Alderton, Roberto Amato, Jeffrey Barrett, Sonia Goncalves, Ewan Harrison, David K. Jackson, Ian Johnston, Dominic Kwiatkowski, Cordelia Langford, John Sillitoe on behalf of the Wellcome Sanger Institute COVID-19 Surveillance Team |
| EPI_ISL_2554434 | Lighthouse Lab in Alderley Park | Wellcome Sanger Institute for the COVID-19 Genomics UK (COG-UK) Consortium | Jacquelyn Wynn, Mairead Hyland, The Lighthouse Lab in Alderley Park and Alex Alderton, Roberto Amato, Jeffrey Barrett, Sonia Goncalves, Ewan Harrison, David K. Jackson, Ian Johnston, Dominic Kwiatkowski, Cordelia Langford, John Sillitoe on behalf of the Wellcome Sanger Institute COVID-19 Surveillance Team |
| EPI_ISL_2554436, EPI_ISL_2554437 | Lighthouse Laboratory Plymouth | Wellcome Sanger Institute for the COVID-19 Genomics UK (COG-UK) Consortium | Lighthouse Laboratory Plymouth and Alex Alderton, Roberto Amato, Jeffrey Barrett, Sonia Goncalves, Ewan Harrison, David K. Jackson, Ian Johnston, Dominic Kwiatkowski, Cordelia Langford, John Sillitoe on behalf of the Wellcome Sanger Institute COVID-19 Surveillance Team |
| EPI_ISL_2554438 | Lighthouse Lab in Glasgow | Wellcome Sanger Institute for the COVID-19 Genomics UK (COG-UK) Consortium | Harper VanSteenhouse, Yumi Kasai, David Gray, Carol Clugston, Anna Dominiczak and Alex Alderton, Roberto Amato, Jeffrey Barrett, Sonia Goncalves, Ewan Harrison, David K. Jackson, Ian Johnston, Dominic Kwiatkowski, Cordelia Langford, John Sillitoe on behalf of the Wellcome Sanger Institute COVID-19 Surveillance Team |
| EPI_ISL_2554439 | Lighthouse Lab in Alderley Park | Wellcome Sanger Institute for the COVID-19 Genomics UK (COG-UK) Consortium | Jacquelyn Wynn, Mairead Hyland, The Lighthouse Lab in Alderley Park and Alex Alderton, Roberto Amato, Jeffrey Barrett, Sonia Goncalves, Ewan Harrison, David K. Jackson, Ian Johnston, Dominic Kwiatkowski, Cordelia Langford, John Sillitoe on behalf of the Wellcome Sanger Institute COVID-19 Surveillance Team |
| EPI_ISL_2554440, EPI_ISL_2554441 | Lighthouse Lab in Glasgow | Wellcome Sanger Institute for the COVID-19 Genomics UK (COG-UK) Consortium | Harper VanSteenhouse, Yumi Kasai, David Gray, Carol Clugston, Anna Dominiczak and Alex Alderton, Roberto Amato, Jeffrey Barrett, Sonia Goncalves, Ewan Harrison, David K. Jackson, Ian Johnston, Dominic Kwiatkowski, Cordelia Langford, John Sillitoe on behalf of the Wellcome Sanger Institute COVID-19 Surveillance Team |
| EPI_ISL_2554442, EPI_ISL_2554443, EPI_ISL_2554444, EPI_ISL_2554445, EPI_ISL_2554446 | Lighthouse Lab in Alderley Park | Wellcome Sanger Institute for the COVID-19 Genomics UK (COG-UK) Consortium | Jacquelyn Wynn, Mairead Hyland, The Lighthouse Lab in Alderley Park and Alex Alderton, Roberto Amato, Jeffrey Barrett, Sonia Goncalves, Ewan Harrison, David K. Jackson, Ian Johnston, Dominic Kwiatkowski, Cordelia Langford, John Sillitoe on behalf of the Wellcome Sanger Institute COVID-19 Surveillance Team |
| EPI_ISL_2554447 | Lighthouse Lab in Glasgow | Wellcome Sanger Institute for the COVID-19 Genomics UK (COG-UK) Consortium | Harper VanSteenhouse, Yumi Kasai, David Gray, Carol Clugston, Anna Dominiczak and Alex Alderton, Roberto Amato, Jeffrey Barrett, Sonia Goncalves, Ewan Harrison, David K. Jackson, Ian Johnston, Dominic Kwiatkowski, Cordelia Langford, John Sillitoe on behalf of the Wellcome Sanger Institute COVID-19 Surveillance Team |
| EPI_ISL_2554448 | Lighthouse Lab in Alderley Park | Wellcome Sanger Institute for the COVID-19 Genomics UK (COG-UK) Consortium | Jacquelyn Wynn, Mairead Hyland, The Lighthouse Lab in Alderley Park and Alex Alderton, Roberto Amato, Jeffrey Barrett, Sonia Goncalves, Ewan Harrison, David K. Jackson, Ian Johnston, Dominic Kwiatkowski, Cordelia Langford, John Sillitoe on behalf of the Wellcome Sanger Institute COVID-19 Surveillance Team |
| EPI_ISL_2554449 | Lighthouse Lab in Glasgow | Wellcome Sanger Institute for the COVID-19 Genomics UK (COG-UK) Consortium | Harper VanSteenhouse, Yumi Kasai, David Gray, Carol Clugston, Anna Dominiczak and Alex Alderton, Roberto Amato, Jeffrey Barrett, Sonia Goncalves, Ewan Harrison, David K. Jackson, Ian Johnston, Dominic Kwiatkowski, Cordelia Langford, John Sillitoe on behalf of the Wellcome Sanger Institute COVID-19 Surveillance Team |
| EPI_ISL_2554450 | Lighthouse Laboratory Plymouth | Wellcome Sanger Institute for the COVID-19 Genomics UK (COG-UK) Consortium | Lighthouse Laboratory Plymouth and Alex Alderton, Roberto Amato, Jeffrey Barrett, Sonia Goncalves, Ewan Harrison, David K. Jackson, Ian Johnston, Dominic Kwiatkowski, Cordelia Langford, John Sillitoe on behalf of the Wellcome Sanger Institute COVID-19 Surveillance Team |
| EPI_ISL_2554451, EPI_ISL_2554452 | Lighthouse Lab in Alderley Park | Wellcome Sanger Institute for the COVID-19 Genomics UK (COG-UK) Consortium | Jacquelyn Wynn, Mairead Hyland, The Lighthouse Lab in Alderley Park and Alex Alderton, Roberto Amato, Jeffrey Barrett, Sonia Goncalves, Ewan Harrison, David K. Jackson, Ian Johnston, Dominic Kwiatkowski, Cordelia Langford, John Sillitoe on behalf of the Wellcome Sanger Institute COVID-19 Surveillance Team |
| EPI_ISL_2554453 | Lighthouse Laboratory Plymouth | Wellcome Sanger Institute for the COVID-19 Genomics UK (COG-UK) Consortium | Lighthouse Laboratory Plymouth and Alex Alderton, Roberto Amato, Jeffrey Barrett, Sonia Goncalves, Ewan Harrison, David K. Jackson, Ian Johnston, Dominic Kwiatkowski, Cordelia Langford, John Sillitoe on behalf of the Wellcome Sanger Institute COVID-19 Surveillance Team |
| EPI_ISL_2554454 | Lighthouse Lab in Alderley Park | Wellcome Sanger Institute for the COVID-19 Genomics UK (COG-UK) Consortium | Jacquelyn Wynn, Mairead Hyland, The Lighthouse Lab in Alderley Park and Alex Alderton, Roberto Amato, Jeffrey Barrett, Sonia Goncalves, Ewan Harrison, David K. Jackson, Ian Johnston, Dominic Kwiatkowski, Cordelia Langford, John Sillitoe on behalf of the Wellcome Sanger Institute COVID-19 Surveillance Team |
| EPI_ISL_2554455 | Lighthouse Laboratory Plymouth | Wellcome Sanger Institute for the COVID-19 Genomics UK (COG-UK) Consortium | Lighthouse Laboratory Plymouth and Alex Alderton, Roberto Amato, Jeffrey Barrett, Sonia Goncalves, Ewan Harrison, David K. Jackson, Ian Johnston, Dominic Kwiatkowski, Cordelia Langford, John Sillitoe on behalf of the Wellcome Sanger Institute COVID-19 Surveillance Team |
| EPI_ISL_2554456 | Lighthouse Lab in Glasgow | Wellcome Sanger Institute for the COVID-19 Genomics UK (COG-UK) Consortium | Harper VanSteenhouse, Yumi Kasai, David Gray, Carol Clugston, Anna Dominiczak and Alex Alderton, Roberto Amato, Jeffrey Barrett, Sonia Goncalves, Ewan Harrison, David K. Jackson, Ian Johnston, Dominic Kwiatkowski, Cordelia Langford, John Sillitoe on behalf of the Wellcome Sanger Institute COVID-19 Surveillance Team |
| EPI_ISL_2554457 | Lighthouse Lab in Alderley Park | Wellcome Sanger Institute for the COVID-19 Genomics UK (COG-UK) Consortium | Jacquelyn Wynn, Mairead Hyland, The Lighthouse Lab in Alderley Park and Alex Alderton, Roberto Amato, Jeffrey Barrett, Sonia Goncalves, Ewan Harrison, David K. Jackson, Ian Johnston, Dominic Kwiatkowski, Cordelia Langford, John Sillitoe on behalf of the Wellcome Sanger Institute COVID-19 Surveillance Team |
| EPI_ISL_2554458 | Lighthouse Laboratory Plymouth | Wellcome Sanger Institute for the COVID-19 Genomics UK (COG-UK) Consortium | Lighthouse Laboratory Plymouth and Alex Alderton, Roberto Amato, Jeffrey Barrett, Sonia Goncalves, Ewan Harrison, David K. Jackson, Ian Johnston, Dominic Kwiatkowski, Cordelia Langford, John Sillitoe on behalf of the Wellcome Sanger Institute COVID-19 Surveillance Team |
| EPI_ISL_2554459, EPI_ISL_2554460 | Lighthouse Lab in Alderley Park | Wellcome Sanger Institute for the COVID-19 Genomics UK (COG-UK) Consortium | Jacquelyn Wynn, Mairead Hyland, The Lighthouse Lab in Alderley Park and Alex Alderton, Roberto Amato, Jeffrey Barrett, Sonia Goncalves, Ewan Harrison, David K. Jackson, Ian Johnston, Dominic Kwiatkowski, Cordelia Langford, John Sillitoe on behalf of the Wellcome Sanger Institute COVID-19 Surveillance Team |
| EPI_ISL_2554461, EPI_ISL_2554462, EPI_ISL_2554463 | Lighthouse Laboratory Plymouth | Wellcome Sanger Institute for the COVID-19 Genomics UK (COG-UK) Consortium | Lighthouse Laboratory Plymouth and Alex Alderton, Roberto Amato, Jeffrey Barrett, Sonia Goncalves, Ewan Harrison, David K. Jackson, Ian Johnston, Dominic Kwiatkowski, Cordelia Langford, John Sillitoe on behalf of the Wellcome Sanger Institute COVID-19 Surveillance Team |
| EPI_ISL_2554464 | Lighthouse Lab in Glasgow | Wellcome Sanger Institute for the COVID-19 Genomics UK (COG-UK) Consortium | Harper VanSteenhouse, Yumi Kasai, David Gray, Carol Clugston, Anna Dominiczak and Alex Alderton, Roberto Amato, Jeffrey Barrett, Sonia Goncalves, Ewan Harrison, David K. Jackson, Ian Johnston, Dominic Kwiatkowski, Cordelia Langford, John Sillitoe on behalf of the Wellcome Sanger Institute COVID-19 Surveillance Team |
| EPI_ISL_2554465 | Lighthouse Laboratory Plymouth | Wellcome Sanger Institute for the COVID-19 Genomics UK (COG-UK) Consortium | Lighthouse Laboratory Plymouth and Alex Alderton, Roberto Amato, Jeffrey Barrett, Sonia Goncalves, Ewan Harrison, David K. Jackson, Ian Johnston, Dominic Kwiatkowski, Cordelia Langford, John Sillitoe on behalf of the Wellcome Sanger Institute COVID-19 Surveillance Team |
| EPI_ISL_2554466 | Lighthouse Lab in Alderley Park | Wellcome Sanger Institute for the COVID-19 Genomics UK (COG-UK) Consortium | Jacquelyn Wynn, Mairead Hyland, The Lighthouse Lab in Alderley Park and Alex Alderton, Roberto Amato, Jeffrey Barrett, Sonia Goncalves, Ewan Harrison, David K. Jackson, Ian Johnston, Dominic Kwiatkowski, Cordelia Langford, John Sillitoe on behalf of the Wellcome Sanger Institute COVID-19 Surveillance Team |
| EPI_ISL_2554467 | Lighthouse Laboratory Plymouth | Wellcome Sanger Institute for the COVID-19 Genomics UK (COG-UK) Consortium | Lighthouse Laboratory Plymouth and Alex Alderton, Roberto Amato, Jeffrey Barrett, Sonia Goncalves, Ewan Harrison, David K. Jackson, Ian Johnston, Dominic Kwiatkowski, Cordelia Langford, John Sillitoe on behalf of the Wellcome Sanger Institute COVID-19 Surveillance Team |
| EPI_ISL_2554468, EPI_ISL_2554469 | Lighthouse Lab in Alderley Park | Wellcome Sanger Institute for the COVID-19 Genomics UK (COG-UK) Consortium | Jacquelyn Wynn, Mairead Hyland, The Lighthouse Lab in Alderley Park and Alex Alderton, Roberto Amato, Jeffrey Barrett, Sonia Goncalves, Ewan Harrison, David K. Jackson, Ian Johnston, Dominic Kwiatkowski, Cordelia Langford, John Sillitoe on behalf of the Wellcome Sanger Institute COVID-19 Surveillance Team |
| EPI_ISL_2554470 | Lighthouse Lab in Glasgow | Wellcome Sanger Institute for the COVID-19 Genomics UK (COG-UK) Consortium | Harper VanSteenhouse, Yumi Kasai, David Gray, Carol Clugston, Anna Dominiczak and Alex Alderton, Roberto Amato, Jeffrey Barrett, Sonia Goncalves, Ewan Harrison, David K. Jackson, Ian Johnston, Dominic Kwiatkowski, Cordelia Langford, John Sillitoe on behalf of the Wellcome Sanger Institute COVID-19 Surveillance Team |
| EPI_ISL_2554471, EPI_ISL_2554472, EPI_ISL_2554473 | Lighthouse Lab in Alderley Park | Wellcome Sanger Institute for the COVID-19 Genomics UK (COG-UK) Consortium | Jacquelyn Wynn, Mairead Hyland, The Lighthouse Lab in Alderley Park and Alex Alderton, Roberto Amato, Jeffrey Barrett, Sonia Goncalves, Ewan Harrison, David K. Jackson, Ian Johnston, Dominic Kwiatkowski, Cordelia Langford, John Sillitoe on behalf of the Wellcome Sanger Institute COVID-19 Surveillance Team |
| EPI_ISL_2554474, EPI_ISL_2554475, EPI_ISL_2554476, EPI_ISL_2554478 | Lighthouse Lab in Glasgow | Wellcome Sanger Institute for the COVID-19 Genomics UK (COG-UK) Consortium | Harper VanSteenhouse, Yumi Kasai, David Gray, Carol Clugston, Anna Dominiczak and Alex Alderton, Roberto Amato, Jeffrey Barrett, Sonia Goncalves, Ewan Harrison, David K. Jackson, Ian Johnston, Dominic Kwiatkowski, Cordelia Langford, John Sillitoe on behalf of the Wellcome Sanger Institute COVID-19 Surveillance Team |
| EPI_ISL_2554479, EPI_ISL_2554480, | Lighthouse Lab in Alderley Park | Wellcome Sanger Institute for the COVID-19 Genomics UK (COG-UK) Consortium | Jacquelyn Wynn, Mairead Hyland, The Lighthouse Lab in Alderley Park and Alex Alderton, Roberto Amato, Jeffrey Barrett, Sonia Goncalves, Ewan Harrison, David K. Jackson, Ian Johnston, Dominic Kwiatkowski, Cordelia Langford, John Sillitoe on behalf of the Wellcome Sanger Institute COVID-19 Surveillance Team |

| | | | |
|---|---|---|---|
| EPI_ISL_2554482 | | | |
| EPI_ISL_2554483 | Lighthouse Lab in Glasgow | Wellcome Sanger Institute for the COVID-19 Genomics UK (COG-UK) Consortium | Harper VanSteenhouse, Yumi Kasai, David Gray, Carol Clugston, Anna Dominiczak and Alex Alderton, Roberto Amato, Jeffrey Barrett, Sonia Goncalves, Ewan Harrison, David K. Jackson, Ian Johnston, Dominic Kwiatkowski, Cordelia Langford, John Sillitoe on behalf of the Wellcome Sanger Institute COVID-19 Surveillance Team |
| EPI_ISL_2554484 | Lighthouse Lab in Alderley Park | Wellcome Sanger Institute for the COVID-19 Genomics UK (COG-UK) Consortium | Jacquelyn Wynn, Mairead Hyland, The Lighthouse Lab in Alderley Park and Alex Alderton, Roberto Amato, Jeffrey Barrett, Sonia Goncalves, Ewan Harrison, David K. Jackson, Ian Johnston, Dominic Kwiatkowski, Cordelia Langford, John Sillitoe on behalf of the Wellcome Sanger Institute COVID-19 Surveillance Team |
| EPI_ISL_2554485 | Lighthouse Lab in Glasgow | Wellcome Sanger Institute for the COVID-19 Genomics UK (COG-UK) Consortium | Harper VanSteenhouse, Yumi Kasai, David Gray, Carol Clugston, Anna Dominiczak and Alex Alderton, Roberto Amato, Jeffrey Barrett, Sonia Goncalves, Ewan Harrison, David K. Jackson, Ian Johnston, Dominic Kwiatkowski, Cordelia Langford, John Sillitoe on behalf of the Wellcome Sanger Institute COVID-19 Surveillance Team |
| EPI_ISL_2554486 | Lighthouse Lab in Alderley Park | Wellcome Sanger Institute for the COVID-19 Genomics UK (COG-UK) Consortium | Jacquelyn Wynn, Mairead Hyland, The Lighthouse Lab in Alderley Park and Alex Alderton, Roberto Amato, Jeffrey Barrett, Sonia Goncalves, Ewan Harrison, David K. Jackson, Ian Johnston, Dominic Kwiatkowski, Cordelia Langford, John Sillitoe on behalf of the Wellcome Sanger Institute COVID-19 Surveillance Team |
| EPI_ISL_2554487 | Lighthouse Lab in Glasgow | Wellcome Sanger Institute for the COVID-19 Genomics UK (COG-UK) Consortium | Harper VanSteenhouse, Yumi Kasai, David Gray, Carol Clugston, Anna Dominiczak and Alex Alderton, Roberto Amato, Jeffrey Barrett, Sonia Goncalves, Ewan Harrison, David K. Jackson, Ian Johnston, Dominic Kwiatkowski, Cordelia Langford, John Sillitoe on behalf of the Wellcome Sanger Institute COVID-19 Surveillance Team |
| EPI_ISL_2554488, EPI_ISL_2554489, EPI_ISL_2554490 | Lighthouse Lab in Alderley Park | Wellcome Sanger Institute for the COVID-19 Genomics UK (COG-UK) Consortium | Jacquelyn Wynn, Mairead Hyland, The Lighthouse Lab in Alderley Park and Alex Alderton, Roberto Amato, Jeffrey Barrett, Sonia Goncalves, Ewan Harrison, David K. Jackson, Ian Johnston, Dominic Kwiatkowski, Cordelia Langford, John Sillitoe on behalf of the Wellcome Sanger Institute COVID-19 Surveillance Team |
| EPI_ISL_2554491 | Lighthouse Lab in Glasgow | Wellcome Sanger Institute for the COVID-19 Genomics UK (COG-UK) Consortium | Harper VanSteenhouse, Yumi Kasai, David Gray, Carol Clugston, Anna Dominiczak and Alex Alderton, Roberto Amato, Jeffrey Barrett, Sonia Goncalves, Ewan Harrison, David K. Jackson, Ian Johnston, Dominic Kwiatkowski, Cordelia Langford, John Sillitoe on behalf of the Wellcome Sanger Institute COVID-19 Surveillance Team |
| EPI_ISL_2554492 | Lighthouse Laboratory Plymouth | Wellcome Sanger Institute for the COVID-19 Genomics UK (COG-UK) Consortium | Lighthouse Laboratory Plymouth and Alex Alderton, Roberto Amato, Jeffrey Barrett, Sonia Goncalves, Ewan Harrison, David K. Jackson, Ian Johnston, Dominic Kwiatkowski, Cordelia Langford, John Sillitoe on behalf of the Wellcome Sanger Institute COVID-19 Surveillance Team |
| EPI_ISL_2554493 | Lighthouse Lab in Milton Keynes | Wellcome Sanger Institute for the COVID-19 Genomics UK (COG-UK) Consortium | The Lighthouse Lab in Milton Keynes and Alex Alderton, Roberto Amato, Jeffrey Barrett, Sonia Goncalves, Ewan Harrison, David K. Jackson, Ian Johnston, Dominic Kwiatkowski, Cordelia Langford, John Sillitoe on behalf of the Wellcome Sanger Institute COVID-19 Surveillance Team |
| EPI_ISL_2554495, EPI_ISL_2554496 | Lighthouse Lab in Glasgow | Wellcome Sanger Institute for the COVID-19 Genomics UK (COG-UK) Consortium | Harper VanSteenhouse, Yumi Kasai, David Gray, Carol Clugston, Anna Dominiczak and Alex Alderton, Roberto Amato, Jeffrey Barrett, Sonia Goncalves, Ewan Harrison, David K. Jackson, Ian Johnston, Dominic Kwiatkowski, Cordelia Langford, John Sillitoe on behalf of the Wellcome Sanger Institute COVID-19 Surveillance Team |
| EPI_ISL_2554497, EPI_ISL_2554498 | Lighthouse Lab in Milton Keynes | Wellcome Sanger Institute for the COVID-19 Genomics UK (COG-UK) Consortium | The Lighthouse Lab in Milton Keynes and Alex Alderton, Roberto Amato, Jeffrey Barrett, Sonia Goncalves, Ewan Harrison, David K. Jackson, Ian Johnston, Dominic Kwiatkowski, Cordelia Langford, John Sillitoe on behalf of the Wellcome Sanger Institute COVID-19 Surveillance Team |
| EPI_ISL_2554499, EPI_ISL_2554500 | Lighthouse Lab in Glasgow | Wellcome Sanger Institute for the COVID-19 Genomics UK (COG-UK) Consortium | Harper VanSteenhouse, Yumi Kasai, David Gray, Carol Clugston, Anna Dominiczak and Alex Alderton, Roberto Amato, Jeffrey Barrett, Sonia Goncalves, Ewan Harrison, David K. Jackson, Ian Johnston, Dominic Kwiatkowski, Cordelia Langford, John Sillitoe on behalf of the Wellcome Sanger Institute COVID-19 Surveillance Team |
| EPI_ISL_2554501 | Lighthouse Lab in Milton Keynes | Wellcome Sanger Institute for the COVID-19 Genomics UK (COG-UK) Consortium | The Lighthouse Lab in Milton Keynes and Alex Alderton, Roberto Amato, Jeffrey Barrett, Sonia Goncalves, Ewan Harrison, David K. Jackson, Ian Johnston, Dominic Kwiatkowski, Cordelia Langford, John Sillitoe on behalf of the Wellcome Sanger Institute COVID-19 Surveillance Team |
| EPI_ISL_2554503 | Lighthouse Lab in Glasgow | Wellcome Sanger Institute for the COVID-19 Genomics UK (COG-UK) Consortium | Harper VanSteenhouse, Yumi Kasai, David Gray, Carol Clugston, Anna Dominiczak and Alex Alderton, Roberto Amato, Jeffrey Barrett, Sonia Goncalves, Ewan Harrison, David K. Jackson, Ian Johnston, Dominic Kwiatkowski, Cordelia Langford, John Sillitoe on behalf of the Wellcome Sanger Institute COVID-19 Surveillance Team |
| EPI_ISL_2554504, EPI_ISL_2554506 | Lighthouse Lab in Milton Keynes | Wellcome Sanger Institute for the COVID-19 Genomics UK (COG-UK) Consortium | The Lighthouse Lab in Milton Keynes and Alex Alderton, Roberto Amato, Jeffrey Barrett, Sonia Goncalves, Ewan Harrison, David K. Jackson, Ian Johnston, Dominic Kwiatkowski, Cordelia Langford, John Sillitoe on behalf of the Wellcome Sanger Institute COVID-19 Surveillance Team |
| EPI_ISL_2554507, EPI_ISL_2554508 | Lighthouse Lab in Glasgow | Wellcome Sanger Institute for the COVID-19 Genomics UK (COG-UK) Consortium | Harper VanSteenhouse, Yumi Kasai, David Gray, Carol Clugston, Anna Dominiczak and Alex Alderton, Roberto Amato, Jeffrey Barrett, Sonia Goncalves, Ewan Harrison, David K. Jackson, Ian Johnston, Dominic Kwiatkowski, Cordelia Langford, John Sillitoe on behalf of the Wellcome Sanger Institute COVID-19 Surveillance Team |
| EPI_ISL_2554509, EPI_ISL_2554510 | Lighthouse Lab in Milton Keynes | Wellcome Sanger Institute for the COVID-19 Genomics UK (COG-UK) Consortium | The Lighthouse Lab in Milton Keynes and Alex Alderton, Roberto Amato, Jeffrey Barrett, Sonia Goncalves, Ewan Harrison, David K. Jackson, Ian Johnston, Dominic Kwiatkowski, Cordelia Langford, John Sillitoe on behalf of the Wellcome Sanger Institute COVID-19 Surveillance Team |
| EPI_ISL_2554511 | Lighthouse Lab in Glasgow | Wellcome Sanger Institute for the COVID-19 Genomics UK (COG-UK) Consortium | Harper VanSteenhouse, Yumi Kasai, David Gray, Carol Clugston, Anna Dominiczak and Alex Alderton, Roberto Amato, Jeffrey Barrett, Sonia Goncalves, Ewan Harrison, David K. Jackson, Ian Johnston, Dominic Kwiatkowski, Cordelia Langford, John Sillitoe on behalf of the Wellcome Sanger Institute COVID-19 Surveillance Team |
| EPI_ISL_2554513 | Lighthouse Lab in Milton Keynes | Wellcome Sanger Institute for the COVID-19 Genomics UK (COG-UK) Consortium | The Lighthouse Lab in Milton Keynes and Alex Alderton, Roberto Amato, Jeffrey Barrett, Sonia Goncalves, Ewan Harrison, David K. Jackson, Ian Johnston, Dominic Kwiatkowski, Cordelia Langford, John Sillitoe on behalf of the Wellcome Sanger Institute COVID-19 Surveillance Team |
| EPI_ISL_2554515, EPI_ISL_2554516 | Lighthouse Lab in Glasgow | Wellcome Sanger Institute for the COVID-19 Genomics UK (COG-UK) Consortium | Harper VanSteenhouse, Yumi Kasai, David Gray, Carol Clugston, Anna Dominiczak and Alex Alderton, Roberto Amato, Jeffrey Barrett, Sonia Goncalves, Ewan Harrison, David K. Jackson, Ian Johnston, Dominic Kwiatkowski, Cordelia Langford, John Sillitoe on behalf of the Wellcome Sanger Institute COVID-19 Surveillance Team |
| EPI_ISL_2554518, EPI_ISL_2554519, EPI_ISL_2554521 | Lighthouse Lab in Milton Keynes | Wellcome Sanger Institute for the COVID-19 Genomics UK (COG-UK) Consortium | The Lighthouse Lab in Milton Keynes and Alex Alderton, Roberto Amato, Jeffrey Barrett, Sonia Goncalves, Ewan Harrison, David K. Jackson, Ian Johnston, Dominic Kwiatkowski, Cordelia Langford, John Sillitoe on behalf of the Wellcome Sanger Institute COVID-19 Surveillance Team |
| EPI_ISL_2554522 | Lighthouse Lab in Glasgow | Wellcome Sanger Institute for the COVID-19 Genomics UK (COG-UK) Consortium | Harper VanSteenhouse, Yumi Kasai, David Gray, Carol Clugston, Anna Dominiczak and Alex Alderton, Roberto Amato, Jeffrey Barrett, Sonia Goncalves, Ewan Harrison, David K. Jackson, Ian Johnston, Dominic Kwiatkowski, Cordelia Langford, John Sillitoe on behalf of the Wellcome Sanger Institute COVID-19 Surveillance Team |
| EPI_ISL_2554523, EPI_ISL_2554524 | Lighthouse Lab in Milton Keynes | Wellcome Sanger Institute for the COVID-19 Genomics UK (COG-UK) Consortium | The Lighthouse Lab in Milton Keynes and Alex Alderton, Roberto Amato, Jeffrey Barrett, Sonia Goncalves, Ewan Harrison, David K. Jackson, Ian Johnston, Dominic Kwiatkowski, Cordelia Langford, John Sillitoe on behalf of the Wellcome Sanger Institute COVID-19 Surveillance Team |
| EPI_ISL_2554525 | Lighthouse Lab in Glasgow | Wellcome Sanger Institute for the COVID-19 Genomics UK (COG-UK) Consortium | Harper VanSteenhouse, Yumi Kasai, David Gray, Carol Clugston, Anna Dominiczak and Alex Alderton, Roberto Amato, Jeffrey Barrett, Sonia Goncalves, Ewan Harrison, David K. Jackson, Ian Johnston, Dominic Kwiatkowski, Cordelia Langford, John Sillitoe on behalf of the Wellcome Sanger Institute COVID-19 Surveillance Team |
| EPI_ISL_2554526 | Lighthouse Lab in Milton Keynes | Wellcome Sanger Institute for the COVID-19 Genomics UK (COG-UK) Consortium | The Lighthouse Lab in Milton Keynes and Alex Alderton, Roberto Amato, Jeffrey Barrett, Sonia Goncalves, Ewan Harrison, David K. Jackson, Ian Johnston, Dominic Kwiatkowski, Cordelia Langford, John Sillitoe on behalf of the Wellcome Sanger Institute COVID-19 Surveillance Team |
| EPI_ISL_2554528, EPI_ISL_2554529 | Lighthouse Lab in Glasgow | Wellcome Sanger Institute for the COVID-19 Genomics UK (COG-UK) Consortium | Harper VanSteenhouse, Yumi Kasai, David Gray, Carol Clugston, Anna Dominiczak and Alex Alderton, Roberto Amato, Jeffrey Barrett, Sonia Goncalves, Ewan Harrison, David K. Jackson, Ian Johnston, Dominic Kwiatkowski, Cordelia Langford, John Sillitoe on behalf of the Wellcome Sanger Institute COVID-19 Surveillance Team |
| EPI_ISL_2554530, EPI_ISL_2554532, EPI_ISL_2554533, EPI_ISL_2554534, EPI_ISL_2554535, EPI_ISL_2554536 | Lighthouse Lab in Milton Keynes | Wellcome Sanger Institute for the COVID-19 Genomics UK (COG-UK) Consortium | The Lighthouse Lab in Milton Keynes and Alex Alderton, Roberto Amato, Jeffrey Barrett, Sonia Goncalves, Ewan Harrison, David K. Jackson, Ian Johnston, Dominic Kwiatkowski, Cordelia Langford, John Sillitoe on behalf of the Wellcome Sanger Institute COVID-19 Surveillance Team |
| EPI_ISL_2554537, EPI_ISL_2554538 | Lighthouse Lab in Glasgow | Wellcome Sanger Institute for the COVID-19 Genomics UK (COG-UK) Consortium | Harper VanSteenhouse, Yumi Kasai, David Gray, Carol Clugston, Anna Dominiczak and Alex Alderton, Roberto Amato, Jeffrey Barrett, Sonia Goncalves, Ewan Harrison, David K. Jackson, Ian Johnston, Dominic Kwiatkowski, Cordelia Langford, John Sillitoe on behalf of the Wellcome Sanger Institute COVID-19 Surveillance Team |
| EPI_ISL_2554539, EPI_ISL_2554540, EPI_ISL_2554541 | Lighthouse Lab in Milton Keynes | Wellcome Sanger Institute for the COVID-19 Genomics UK (COG-UK) Consortium | The Lighthouse Lab in Milton Keynes and Alex Alderton, Roberto Amato, Jeffrey Barrett, Sonia Goncalves, Ewan Harrison, David K. Jackson, Ian Johnston, Dominic Kwiatkowski, Cordelia Langford, John Sillitoe on behalf of the Wellcome Sanger Institute COVID-19 Surveillance Team |
| EPI_ISL_2554542 | Lighthouse Lab in Glasgow | Wellcome Sanger Institute for the COVID-19 Genomics UK (COG-UK) Consortium | Harper VanSteenhouse, Yumi Kasai, David Gray, Carol Clugston, Anna Dominiczak and Alex Alderton, Roberto Amato, Jeffrey Barrett, Sonia Goncalves, Ewan Harrison, David K. Jackson, Ian Johnston, Dominic Kwiatkowski, Cordelia Langford, John Sillitoe on behalf of the Wellcome Sanger Institute COVID-19 Surveillance Team |
| EPI_ISL_2554543, EPI_ISL_2554544, EPI_ISL_2554545 | Lighthouse Lab in Milton Keynes | Wellcome Sanger Institute for the COVID-19 Genomics UK (COG-UK) Consortium | The Lighthouse Lab in Milton Keynes and Alex Alderton, Roberto Amato, Jeffrey Barrett, Sonia Goncalves, Ewan Harrison, David K. Jackson, Ian Johnston, Dominic Kwiatkowski, Cordelia Langford, John Sillitoe on behalf of the Wellcome Sanger Institute COVID-19 Surveillance Team |
| EPI_ISL_2554546, EPI_ISL_2554547, EPI_ISL_2554549 | Lighthouse Lab in Glasgow | Wellcome Sanger Institute for the COVID-19 Genomics UK (COG-UK) Consortium | Harper VanSteenhouse, Yumi Kasai, David Gray, Carol Clugston, Anna Dominiczak and Alex Alderton, Roberto Amato, Jeffrey Barrett, Sonia Goncalves, Ewan Harrison, David K. Jackson, Ian Johnston, Dominic Kwiatkowski, Cordelia Langford, John Sillitoe on behalf of the Wellcome Sanger Institute COVID-19 Surveillance Team |
| EPI_ISL_2554550, EPI_ISL_2554551, EPI_ISL_2554552, EPI_ISL_2554553 | Lighthouse Lab in Milton Keynes | Wellcome Sanger Institute for the COVID-19 Genomics UK (COG-UK) Consortium | The Lighthouse Lab in Milton Keynes and Alex Alderton, Roberto Amato, Jeffrey Barrett, Sonia Goncalves, Ewan Harrison, David K. Jackson, Ian Johnston, Dominic Kwiatkowski, Cordelia Langford, John Sillitoe on behalf of the Wellcome Sanger Institute COVID-19 Surveillance Team |
| EPI_ISL_2554554 | Lighthouse Lab in Glasgow | Wellcome Sanger Institute for the COVID-19 Genomics UK (COG-UK) Consortium | Harper VanSteenhouse, Yumi Kasai, David Gray, Carol Clugston, Anna Dominiczak and Alex Alderton, Roberto Amato, Jeffrey Barrett, Sonia Goncalves, Ewan Harrison, David K. Jackson, Ian Johnston, Dominic Kwiatkowski, Cordelia Langford, John Sillitoe on behalf of the Wellcome Sanger Institute COVID-19 Surveillance Team |
| EPI_ISL_2554555, EPI_ISL_2554556, EPI_ISL_2554557, EPI_ISL_2554558 | Lighthouse Lab in Milton Keynes | Wellcome Sanger Institute for the COVID-19 Genomics UK (COG-UK) Consortium | The Lighthouse Lab in Milton Keynes and Alex Alderton, Roberto Amato, Jeffrey Barrett, Sonia Goncalves, Ewan Harrison, David K. Jackson, Ian Johnston, Dominic Kwiatkowski, Cordelia Langford, John Sillitoe on behalf of the Wellcome Sanger Institute COVID-19 Surveillance Team |
| EPI_ISL_2554559, EPI_ISL_2554560, EPI_ISL_2554561, EPI_ISL_2554562 | Lighthouse Lab in Glasgow | Wellcome Sanger Institute for the COVID-19 Genomics UK (COG-UK) Consortium | Harper VanSteenhouse, Yumi Kasai, David Gray, Carol Clugston, Anna Dominiczak and Alex Alderton, Roberto Amato, Jeffrey Barrett, Sonia Goncalves, Ewan Harrison, David K. Jackson, Ian Johnston, Dominic Kwiatkowski, Cordelia Langford, John Sillitoe on behalf of the Wellcome Sanger Institute COVID-19 Surveillance Team |
| EPI_ISL_2554563 | Lighthouse Lab in Milton Keynes | Wellcome Sanger Institute for the COVID-19 Genomics UK (COG-UK) Consortium | The Lighthouse Lab in Milton Keynes and Alex Alderton, Roberto Amato, Jeffrey Barrett, Sonia Goncalves, Ewan Harrison, David K. Jackson, Ian Johnston, Dominic Kwiatkowski, Cordelia Langford, John Sillitoe on behalf of the Wellcome Sanger Institute COVID-19 Surveillance Team |
| EPI_ISL_2554564, EPI_ISL_2554565, EPI_ISL_2554566, | Lighthouse Lab in Glasgow | Wellcome Sanger Institute for the COVID-19 Genomics UK (COG-UK) Consortium | Harper VanSteenhouse, Yumi Kasai, David Gray, Carol Clugston, Anna Dominiczak and Alex Alderton, Roberto Amato, Jeffrey Barrett, Sonia Goncalves, Ewan Harrison, David K. Jackson, Ian Johnston, Dominic Kwiatkowski, Cordelia Langford, John Sillitoe on behalf of the Wellcome Sanger Institute COVID-19 Surveillance Team |

| | | | |
|---|---|---|---|
| EPI_ISL_2554567 | | | |
| EPI_ISL_2554568 | Lighthouse Lab in Milton Keynes | Wellcome Sanger Institute for the COVID-19 Genomics UK (COG-UK) Consortium | The Lighthouse Lab in Milton Keynes and Alex Alderton, Roberto Amato, Jeffrey Barrett, Sonia Goncalves, Ewan Harrison, David K. Jackson, Ian Johnston, Dominic Kwiatkowski, Cordelia Langford, John Sillitoe on behalf of the Wellcome Sanger Institute COVID-19 Surveillance Team |
| EPI_ISL_2554569 | Lighthouse Lab in Glasgow | Wellcome Sanger Institute for the COVID-19 Genomics UK (COG-UK) Consortium | Harper VanSteenhouse, Yumi Kasai, David Gray, Carol Clugston, Anna Dominiczak and Alex Alderton, Roberto Amato, Jeffrey Barrett, Sonia Goncalves, Ewan Harrison, David K. Jackson, Ian Johnston, Dominic Kwiatkowski, Cordelia Langford, John Sillitoe on behalf of the Wellcome Sanger Institute COVID-19 Surveillance Team |
| EPI_ISL_2554570 | Lighthouse Lab in Milton Keynes | Wellcome Sanger Institute for the COVID-19 Genomics UK (COG-UK) Consortium | The Lighthouse Lab in Milton Keynes and Alex Alderton, Roberto Amato, Jeffrey Barrett, Sonia Goncalves, Ewan Harrison, David K. Jackson, Ian Johnston, Dominic Kwiatkowski, Cordelia Langford, John Sillitoe on behalf of the Wellcome Sanger Institute COVID-19 Surveillance Team |
| EPI_ISL_2554571, EPI_ISL_2554572, EPI_ISL_2554573, EPI_ISL_2554574 | Lighthouse Lab in Glasgow | Wellcome Sanger Institute for the COVID-19 Genomics UK (COG-UK) Consortium | Harper VanSteenhouse, Yumi Kasai, David Gray, Carol Clugston, Anna Dominiczak and Alex Alderton, Roberto Amato, Jeffrey Barrett, Sonia Goncalves, Ewan Harrison, David K. Jackson, Ian Johnston, Dominic Kwiatkowski, Cordelia Langford, John Sillitoe on behalf of the Wellcome Sanger Institute COVID-19 Surveillance Team |
| EPI_ISL_2554575 | Lighthouse Lab in Milton Keynes | Wellcome Sanger Institute for the COVID-19 Genomics UK (COG-UK) Consortium | The Lighthouse Lab in Milton Keynes and Alex Alderton, Roberto Amato, Jeffrey Barrett, Sonia Goncalves, Ewan Harrison, David K. Jackson, Ian Johnston, Dominic Kwiatkowski, Cordelia Langford, John Sillitoe on behalf of the Wellcome Sanger Institute COVID-19 Surveillance Team |
| EPI_ISL_2554576 | Lighthouse Lab in Glasgow | Wellcome Sanger Institute for the COVID-19 Genomics UK (COG-UK) Consortium | Harper VanSteenhouse, Yumi Kasai, David Gray, Carol Clugston, Anna Dominiczak and Alex Alderton, Roberto Amato, Jeffrey Barrett, Sonia Goncalves, Ewan Harrison, David K. Jackson, Ian Johnston, Dominic Kwiatkowski, Cordelia Langford, John Sillitoe on behalf of the Wellcome Sanger Institute COVID-19 Surveillance Team |
| EPI_ISL_2554577 | Lighthouse Lab in Milton Keynes | Wellcome Sanger Institute for the COVID-19 Genomics UK (COG-UK) Consortium | The Lighthouse Lab in Milton Keynes and Alex Alderton, Roberto Amato, Jeffrey Barrett, Sonia Goncalves, Ewan Harrison, David K. Jackson, Ian Johnston, Dominic Kwiatkowski, Cordelia Langford, John Sillitoe on behalf of the Wellcome Sanger Institute COVID-19 Surveillance Team |
| EPI_ISL_2554578 | Lighthouse Lab in Glasgow | Wellcome Sanger Institute for the COVID-19 Genomics UK (COG-UK) Consortium | Harper VanSteenhouse, Yumi Kasai, David Gray, Carol Clugston, Anna Dominiczak and Alex Alderton, Roberto Amato, Jeffrey Barrett, Sonia Goncalves, Ewan Harrison, David K. Jackson, Ian Johnston, Dominic Kwiatkowski, Cordelia Langford, John Sillitoe on behalf of the Wellcome Sanger Institute COVID-19 Surveillance Team |
| EPI_ISL_2554579, EPI_ISL_2554580 | Lighthouse Lab in Milton Keynes | Wellcome Sanger Institute for the COVID-19 Genomics UK (COG-UK) Consortium | The Lighthouse Lab in Milton Keynes and Alex Alderton, Roberto Amato, Jeffrey Barrett, Sonia Goncalves, Ewan Harrison, David K. Jackson, Ian Johnston, Dominic Kwiatkowski, Cordelia Langford, John Sillitoe on behalf of the Wellcome Sanger Institute COVID-19 Surveillance Team |
| EPI_ISL_2554581 | Lighthouse Lab in Glasgow | Wellcome Sanger Institute for the COVID-19 Genomics UK (COG-UK) Consortium | Harper VanSteenhouse, Yumi Kasai, David Gray, Carol Clugston, Anna Dominiczak and Alex Alderton, Roberto Amato, Jeffrey Barrett, Sonia Goncalves, Ewan Harrison, David K. Jackson, Ian Johnston, Dominic Kwiatkowski, Cordelia Langford, John Sillitoe on behalf of the Wellcome Sanger Institute COVID-19 Surveillance Team |
| EPI_ISL_2554582, EPI_ISL_2554583 | Lighthouse Lab in Milton Keynes | Wellcome Sanger Institute for the COVID-19 Genomics UK (COG-UK) Consortium | The Lighthouse Lab in Milton Keynes and Alex Alderton, Roberto Amato, Jeffrey Barrett, Sonia Goncalves, Ewan Harrison, David K. Jackson, Ian Johnston, Dominic Kwiatkowski, Cordelia Langford, John Sillitoe on behalf of the Wellcome Sanger Institute COVID-19 Surveillance Team |
| EPI_ISL_2554584 | Lighthouse Lab in Glasgow | Wellcome Sanger Institute for the COVID-19 Genomics UK (COG-UK) Consortium | Harper VanSteenhouse, Yumi Kasai, David Gray, Carol Clugston, Anna Dominiczak and Alex Alderton, Roberto Amato, Jeffrey Barrett, Sonia Goncalves, Ewan Harrison, David K. Jackson, Ian Johnston, Dominic Kwiatkowski, Cordelia Langford, John Sillitoe on behalf of the Wellcome Sanger Institute COVID-19 Surveillance Team |
| EPI_ISL_2554585, EPI_ISL_2554586, EPI_ISL_2554587 | Lighthouse Lab in Milton Keynes | Wellcome Sanger Institute for the COVID-19 Genomics UK (COG-UK) Consortium | The Lighthouse Lab in Milton Keynes and Alex Alderton, Roberto Amato, Jeffrey Barrett, Sonia Goncalves, Ewan Harrison, David K. Jackson, Ian Johnston, Dominic Kwiatkowski, Cordelia Langford, John Sillitoe on behalf of the Wellcome Sanger Institute COVID-19 Surveillance Team |
| EPI_ISL_2554588 | Lighthouse Lab in Glasgow | Wellcome Sanger Institute for the COVID-19 Genomics UK (COG-UK) Consortium | Harper VanSteenhouse, Yumi Kasai, David Gray, Carol Clugston, Anna Dominiczak and Alex Alderton, Roberto Amato, Jeffrey Barrett, Sonia Goncalves, Ewan Harrison, David K. Jackson, Ian Johnston, Dominic Kwiatkowski, Cordelia Langford, John Sillitoe on behalf of the Wellcome Sanger Institute COVID-19 Surveillance Team |
| EPI_ISL_2554589 | Lighthouse Lab in Milton Keynes | Wellcome Sanger Institute for the COVID-19 Genomics UK (COG-UK) Consortium | The Lighthouse Lab in Milton Keynes and Alex Alderton, Roberto Amato, Jeffrey Barrett, Sonia Goncalves, Ewan Harrison, David K. Jackson, Ian Johnston, Dominic Kwiatkowski, Cordelia Langford, John Sillitoe on behalf of the Wellcome Sanger Institute COVID-19 Surveillance Team |
| EPI_ISL_2554590 | Lighthouse Lab in Glasgow | Wellcome Sanger Institute for the COVID-19 Genomics UK (COG-UK) Consortium | Harper VanSteenhouse, Yumi Kasai, David Gray, Carol Clugston, Anna Dominiczak and Alex Alderton, Roberto Amato, Jeffrey Barrett, Sonia Goncalves, Ewan Harrison, David K. Jackson, Ian Johnston, Dominic Kwiatkowski, Cordelia Langford, John Sillitoe on behalf of the Wellcome Sanger Institute COVID-19 Surveillance Team |
| EPI_ISL_2554591 | Lighthouse Lab in Milton Keynes | Wellcome Sanger Institute for the COVID-19 Genomics UK (COG-UK) Consortium | The Lighthouse Lab in Milton Keynes and Alex Alderton, Roberto Amato, Jeffrey Barrett, Sonia Goncalves, Ewan Harrison, David K. Jackson, Ian Johnston, Dominic Kwiatkowski, Cordelia Langford, John Sillitoe on behalf of the Wellcome Sanger Institute COVID-19 Surveillance Team |
| EPI_ISL_2554592, EPI_ISL_2554593 | Lighthouse Lab in Glasgow | Wellcome Sanger Institute for the COVID-19 Genomics UK (COG-UK) Consortium | Harper VanSteenhouse, Yumi Kasai, David Gray, Carol Clugston, Anna Dominiczak and Alex Alderton, Roberto Amato, Jeffrey Barrett, Sonia Goncalves, Ewan Harrison, David K. Jackson, Ian Johnston, Dominic Kwiatkowski, Cordelia Langford, John Sillitoe on behalf of the Wellcome Sanger Institute COVID-19 Surveillance Team |
| EPI_ISL_2554594, EPI_ISL_2554595 | Lighthouse Lab in Milton Keynes | Wellcome Sanger Institute for the COVID-19 Genomics UK (COG-UK) Consortium | The Lighthouse Lab in Milton Keynes and Alex Alderton, Roberto Amato, Jeffrey Barrett, Sonia Goncalves, Ewan Harrison, David K. Jackson, Ian Johnston, Dominic Kwiatkowski, Cordelia Langford, John Sillitoe on behalf of the Wellcome Sanger Institute COVID-19 Surveillance Team |
| EPI_ISL_2554597, EPI_ISL_2554598, EPI_ISL_2554600, EPI_ISL_2554601, EPI_ISL_2554602, EPI_ISL_2554603 | Lighthouse Lab in Glasgow | Wellcome Sanger Institute for the COVID-19 Genomics UK (COG-UK) Consortium | Harper VanSteenhouse, Yumi Kasai, David Gray, Carol Clugston, Anna Dominiczak and Alex Alderton, Roberto Amato, Jeffrey Barrett, Sonia Goncalves, Ewan Harrison, David K. Jackson, Ian Johnston, Dominic Kwiatkowski, Cordelia Langford, John Sillitoe on behalf of the Wellcome Sanger Institute COVID-19 Surveillance Team |
| EPI_ISL_2554604, EPI_ISL_2554605, EPI_ISL_2554606 | Lighthouse Lab in Milton Keynes | Wellcome Sanger Institute for the COVID-19 Genomics UK (COG-UK) Consortium | The Lighthouse Lab in Milton Keynes and Alex Alderton, Roberto Amato, Jeffrey Barrett, Sonia Goncalves, Ewan Harrison, David K. Jackson, Ian Johnston, Dominic Kwiatkowski, Cordelia Langford, John Sillitoe on behalf of the Wellcome Sanger Institute COVID-19 Surveillance Team |
| EPI_ISL_2554608 | Lighthouse Lab in Glasgow | Wellcome Sanger Institute for the COVID-19 Genomics UK (COG-UK) Consortium | Harper VanSteenhouse, Yumi Kasai, David Gray, Carol Clugston, Anna Dominiczak and Alex Alderton, Roberto Amato, Jeffrey Barrett, Sonia Goncalves, Ewan Harrison, David K. Jackson, Ian Johnston, Dominic Kwiatkowski, Cordelia Langford, John Sillitoe on behalf of the Wellcome Sanger Institute COVID-19 Surveillance Team |
| EPI_ISL_2554609, EPI_ISL_2554610 | Lighthouse Lab in Milton Keynes | Wellcome Sanger Institute for the COVID-19 Genomics UK (COG-UK) Consortium | The Lighthouse Lab in Milton Keynes and Alex Alderton, Roberto Amato, Jeffrey Barrett, Sonia Goncalves, Ewan Harrison, David K. Jackson, Ian Johnston, Dominic Kwiatkowski, Cordelia Langford, John Sillitoe on behalf of the Wellcome Sanger Institute COVID-19 Surveillance Team |
| EPI_ISL_2554611 | Lighthouse Lab in Glasgow | Wellcome Sanger Institute for the COVID-19 Genomics UK (COG-UK) Consortium | Harper VanSteenhouse, Yumi Kasai, David Gray, Carol Clugston, Anna Dominiczak and Alex Alderton, Roberto Amato, Jeffrey Barrett, Sonia Goncalves, Ewan Harrison, David K. Jackson, Ian Johnston, Dominic Kwiatkowski, Cordelia Langford, John Sillitoe on behalf of the Wellcome Sanger Institute COVID-19 Surveillance Team |
| EPI_ISL_2554612 | Lighthouse Lab in Milton Keynes | Wellcome Sanger Institute for the COVID-19 Genomics UK (COG-UK) Consortium | The Lighthouse Lab in Milton Keynes and Alex Alderton, Roberto Amato, Jeffrey Barrett, Sonia Goncalves, Ewan Harrison, David K. Jackson, Ian Johnston, Dominic Kwiatkowski, Cordelia Langford, John Sillitoe on behalf of the Wellcome Sanger Institute COVID-19 Surveillance Team |
| EPI_ISL_2554613 | Lighthouse Lab in Glasgow | Wellcome Sanger Institute for the COVID-19 Genomics UK (COG-UK) Consortium | Harper VanSteenhouse, Yumi Kasai, David Gray, Carol Clugston, Anna Dominiczak and Alex Alderton, Roberto Amato, Jeffrey Barrett, Sonia Goncalves, Ewan Harrison, David K. Jackson, Ian Johnston, Dominic Kwiatkowski, Cordelia Langford, John Sillitoe on behalf of the Wellcome Sanger Institute COVID-19 Surveillance Team |
| EPI_ISL_2554614 | Lighthouse Lab in Milton Keynes | Wellcome Sanger Institute for the COVID-19 Genomics UK (COG-UK) Consortium | The Lighthouse Lab in Milton Keynes and Alex Alderton, Roberto Amato, Jeffrey Barrett, Sonia Goncalves, Ewan Harrison, David K. Jackson, Ian Johnston, Dominic Kwiatkowski, Cordelia Langford, John Sillitoe on behalf of the Wellcome Sanger Institute COVID-19 Surveillance Team |
| EPI_ISL_2554615, EPI_ISL_2554616, EPI_ISL_2554618, EPI_ISL_2554619 | Lighthouse Lab in Glasgow | Wellcome Sanger Institute for the COVID-19 Genomics UK (COG-UK) Consortium | Harper VanSteenhouse, Yumi Kasai, David Gray, Carol Clugston, Anna Dominiczak and Alex Alderton, Roberto Amato, Jeffrey Barrett, Sonia Goncalves, Ewan Harrison, David K. Jackson, Ian Johnston, Dominic Kwiatkowski, Cordelia Langford, John Sillitoe on behalf of the Wellcome Sanger Institute COVID-19 Surveillance Team |
| EPI_ISL_2554620, EPI_ISL_2554621 | Lighthouse Lab in Milton Keynes | Wellcome Sanger Institute for the COVID-19 Genomics UK (COG-UK) Consortium | The Lighthouse Lab in Milton Keynes and Alex Alderton, Roberto Amato, Jeffrey Barrett, Sonia Goncalves, Ewan Harrison, David K. Jackson, Ian Johnston, Dominic Kwiatkowski, Cordelia Langford, John Sillitoe on behalf of the Wellcome Sanger Institute COVID-19 Surveillance Team |
| EPI_ISL_2554622 | Lighthouse Lab in Glasgow | Wellcome Sanger Institute for the COVID-19 Genomics UK (COG-UK) Consortium | Harper VanSteenhouse, Yumi Kasai, David Gray, Carol Clugston, Anna Dominiczak and Alex Alderton, Roberto Amato, Jeffrey Barrett, Sonia Goncalves, Ewan Harrison, David K. Jackson, Ian Johnston, Dominic Kwiatkowski, Cordelia Langford, John Sillitoe on behalf of the Wellcome Sanger Institute COVID-19 Surveillance Team |
| EPI_ISL_2554623, EPI_ISL_2554624 | Lighthouse Lab in Milton Keynes | Wellcome Sanger Institute for the COVID-19 Genomics UK (COG-UK) Consortium | The Lighthouse Lab in Milton Keynes and Alex Alderton, Roberto Amato, Jeffrey Barrett, Sonia Goncalves, Ewan Harrison, David K. Jackson, Ian Johnston, Dominic Kwiatkowski, Cordelia Langford, John Sillitoe on behalf of the Wellcome Sanger Institute COVID-19 Surveillance Team |
| EPI_ISL_2554625, EPI_ISL_2554626, EPI_ISL_2554627 | Lighthouse Lab in Glasgow | Wellcome Sanger Institute for the COVID-19 Genomics UK (COG-UK) Consortium | Harper VanSteenhouse, Yumi Kasai, David Gray, Carol Clugston, Anna Dominiczak and Alex Alderton, Roberto Amato, Jeffrey Barrett, Sonia Goncalves, Ewan Harrison, David K. Jackson, Ian Johnston, Dominic Kwiatkowski, Cordelia Langford, John Sillitoe on behalf of the Wellcome Sanger Institute COVID-19 Surveillance Team |
| EPI_ISL_2554628, EPI_ISL_2554629 | Lighthouse Lab in Milton Keynes | Wellcome Sanger Institute for the COVID-19 Genomics UK (COG-UK) Consortium | The Lighthouse Lab in Milton Keynes and Alex Alderton, Roberto Amato, Jeffrey Barrett, Sonia Goncalves, Ewan Harrison, David K. Jackson, Ian Johnston, Dominic Kwiatkowski, Cordelia Langford, John Sillitoe on behalf of the Wellcome Sanger Institute COVID-19 Surveillance Team |
| EPI_ISL_2554630 | Lighthouse Lab in Glasgow | Wellcome Sanger Institute for the COVID-19 Genomics UK (COG-UK) Consortium | Harper VanSteenhouse, Yumi Kasai, David Gray, Carol Clugston, Anna Dominiczak and Alex Alderton, Roberto Amato, Jeffrey Barrett, Sonia Goncalves, Ewan Harrison, David K. Jackson, Ian Johnston, Dominic Kwiatkowski, Cordelia Langford, John Sillitoe on behalf of the Wellcome Sanger Institute COVID-19 Surveillance Team |
| EPI_ISL_2554631 | Lighthouse Lab in Milton Keynes | Wellcome Sanger Institute for the COVID-19 Genomics UK (COG-UK) Consortium | The Lighthouse Lab in Milton Keynes and Alex Alderton, Roberto Amato, Jeffrey Barrett, Sonia Goncalves, Ewan Harrison, David K. Jackson, Ian Johnston, Dominic Kwiatkowski, Cordelia Langford, John Sillitoe on behalf of the Wellcome Sanger Institute COVID-19 Surveillance Team |
| EPI_ISL_2554632 | Lighthouse Lab in Glasgow | Wellcome Sanger Institute for the COVID-19 Genomics UK (COG-UK) Consortium | Harper VanSteenhouse, Yumi Kasai, David Gray, Carol Clugston, Anna Dominiczak and Alex Alderton, Roberto Amato, Jeffrey Barrett, Sonia Goncalves, Ewan Harrison, David K. Jackson, Ian Johnston, Dominic Kwiatkowski, Cordelia Langford, John Sillitoe on behalf of the Wellcome Sanger Institute COVID-19 Surveillance Team |
| EPI_ISL_2554633 | Lighthouse Lab in Milton Keynes | Wellcome Sanger Institute for the COVID-19 Genomics UK (COG-UK) Consortium | The Lighthouse Lab in Milton Keynes and Alex Alderton, Roberto Amato, Jeffrey Barrett, Sonia Goncalves, Ewan Harrison, David K. Jackson, Ian Johnston, Dominic Kwiatkowski, Cordelia Langford, John Sillitoe on behalf of the Wellcome Sanger Institute COVID-19 Surveillance Team |
| EPI_ISL_2554634 | Lighthouse Lab in Glasgow | Wellcome Sanger Institute for the COVID-19 Genomics UK (COG-UK) Consortium | Harper VanSteenhouse, Yumi Kasai, David Gray, Carol Clugston, Anna Dominiczak and Alex Alderton, Roberto Amato, Jeffrey Barrett, Sonia Goncalves, Ewan Harrison, David K. Jackson, Ian Johnston, Dominic Kwiatkowski, Cordelia Langford, John Sillitoe on behalf of the Wellcome Sanger Institute COVID-19 Surveillance Team |
| EPI_ISL_2554635, EPI_ISL_2554636, EPI_ISL_2554638, EPI_ISL_2554639, | Lighthouse Lab in Milton Keynes | Wellcome Sanger Institute for the COVID-19 Genomics UK (COG-UK) Consortium | The Lighthouse Lab in Milton Keynes and Alex Alderton, Roberto Amato, Jeffrey Barrett, Sonia Goncalves, Ewan Harrison, David K. Jackson, Ian Johnston, Dominic Kwiatkowski, Cordelia Langford, John Sillitoe on behalf of the Wellcome Sanger Institute COVID-19 Surveillance Team |

| | | | |
|---|---|---|---|
| EPI_ISL_2554640, EPI_ISL_2554641 | | | |
| EPI_ISL_2554642, EPI_ISL_2554643 | Lighthouse Lab in Glasgow | Wellcome Sanger Institute for the COVID-19 Genomics UK (COG-UK) Consortium | Harper VanSteenhouse, Yumi Kasai, David Gray, Carol Clugston, Anna Dominiczak and Alex Alderton, Roberto Amato, Jeffrey Barrett, Sonia Goncalves, Ewan Harrison, David K. Jackson, Ian Johnston, Dominic Kwiatkowski, Cordelia Langford, John Sillitoe on behalf of the Wellcome Sanger Institute COVID-19 Surveillance Team |
| EPI_ISL_2554644 | Lighthouse Lab in Milton Keynes | Wellcome Sanger Institute for the COVID-19 Genomics UK (COG-UK) Consortium | The Lighthouse Lab in Milton Keynes and Alex Alderton, Roberto Amato, Jeffrey Barrett, Sonia Goncalves, Ewan Harrison, David K. Jackson, Ian Johnston, Dominic Kwiatkowski, Cordelia Langford, John Sillitoe on behalf of the Wellcome Sanger Institute COVID-19 Surveillance Team |
| EPI_ISL_2554645, EPI_ISL_2554646 | Lighthouse Lab in Glasgow | Wellcome Sanger Institute for the COVID-19 Genomics UK (COG-UK) Consortium | Harper VanSteenhouse, Yumi Kasai, David Gray, Carol Clugston, Anna Dominiczak and Alex Alderton, Roberto Amato, Jeffrey Barrett, Sonia Goncalves, Ewan Harrison, David K. Jackson, Ian Johnston, Dominic Kwiatkowski, Cordelia Langford, John Sillitoe on behalf of the Wellcome Sanger Institute COVID-19 Surveillance Team |
| EPI_ISL_2554647, EPI_ISL_2554648 | Lighthouse Lab in Milton Keynes | Wellcome Sanger Institute for the COVID-19 Genomics UK (COG-UK) Consortium | The Lighthouse Lab in Milton Keynes and Alex Alderton, Roberto Amato, Jeffrey Barrett, Sonia Goncalves, Ewan Harrison, David K. Jackson, Ian Johnston, Dominic Kwiatkowski, Cordelia Langford, John Sillitoe on behalf of the Wellcome Sanger Institute COVID-19 Surveillance Team |
| EPI_ISL_2554649, EPI_ISL_2554650, EPI_ISL_2554651, EPI_ISL_2554652 | Lighthouse Lab in Glasgow | Wellcome Sanger Institute for the COVID-19 Genomics UK (COG-UK) Consortium | Harper VanSteenhouse, Yumi Kasai, David Gray, Carol Clugston, Anna Dominiczak and Alex Alderton, Roberto Amato, Jeffrey Barrett, Sonia Goncalves, Ewan Harrison, David K. Jackson, Ian Johnston, Dominic Kwiatkowski, Cordelia Langford, John Sillitoe on behalf of the Wellcome Sanger Institute COVID-19 Surveillance Team |
| EPI_ISL_2554653 | Lighthouse Lab in Milton Keynes | Wellcome Sanger Institute for the COVID-19 Genomics UK (COG-UK) Consortium | The Lighthouse Lab in Milton Keynes and Alex Alderton, Roberto Amato, Jeffrey Barrett, Sonia Goncalves, Ewan Harrison, David K. Jackson, Ian Johnston, Dominic Kwiatkowski, Cordelia Langford, John Sillitoe on behalf of the Wellcome Sanger Institute COVID-19 Surveillance Team |
| EPI_ISL_2554654 | Lighthouse Lab in Glasgow | Wellcome Sanger Institute for the COVID-19 Genomics UK (COG-UK) Consortium | Harper VanSteenhouse, Yumi Kasai, David Gray, Carol Clugston, Anna Dominiczak and Alex Alderton, Roberto Amato, Jeffrey Barrett, Sonia Goncalves, Ewan Harrison, David K. Jackson, Ian Johnston, Dominic Kwiatkowski, Cordelia Langford, John Sillitoe on behalf of the Wellcome Sanger Institute COVID-19 Surveillance Team |
| EPI_ISL_2554655, EPI_ISL_2554656 | Lighthouse Lab in Milton Keynes | Wellcome Sanger Institute for the COVID-19 Genomics UK (COG-UK) Consortium | The Lighthouse Lab in Milton Keynes and Alex Alderton, Roberto Amato, Jeffrey Barrett, Sonia Goncalves, Ewan Harrison, David K. Jackson, Ian Johnston, Dominic Kwiatkowski, Cordelia Langford, John Sillitoe on behalf of the Wellcome Sanger Institute COVID-19 Surveillance Team |
| EPI_ISL_2554657, EPI_ISL_2554658 | Lighthouse Lab in Glasgow | Wellcome Sanger Institute for the COVID-19 Genomics UK (COG-UK) Consortium | Harper VanSteenhouse, Yumi Kasai, David Gray, Carol Clugston, Anna Dominiczak and Alex Alderton, Roberto Amato, Jeffrey Barrett, Sonia Goncalves, Ewan Harrison, David K. Jackson, Ian Johnston, Dominic Kwiatkowski, Cordelia Langford, John Sillitoe on behalf of the Wellcome Sanger Institute COVID-19 Surveillance Team |
| EPI_ISL_2554659 | Lighthouse Lab in Milton Keynes | Wellcome Sanger Institute for the COVID-19 Genomics UK (COG-UK) Consortium | The Lighthouse Lab in Milton Keynes and Alex Alderton, Roberto Amato, Jeffrey Barrett, Sonia Goncalves, Ewan Harrison, David K. Jackson, Ian Johnston, Dominic Kwiatkowski, Cordelia Langford, John Sillitoe on behalf of the Wellcome Sanger Institute COVID-19 Surveillance Team |
| EPI_ISL_2554662, EPI_ISL_2554663, EPI_ISL_2554664 | Lighthouse Lab in Glasgow | Wellcome Sanger Institute for the COVID-19 Genomics UK (COG-UK) Consortium | Harper VanSteenhouse, Yumi Kasai, David Gray, Carol Clugston, Anna Dominiczak and Alex Alderton, Roberto Amato, Jeffrey Barrett, Sonia Goncalves, Ewan Harrison, David K. Jackson, Ian Johnston, Dominic Kwiatkowski, Cordelia Langford, John Sillitoe on behalf of the Wellcome Sanger Institute COVID-19 Surveillance Team |
| EPI_ISL_2554665 | Lighthouse Lab in Milton Keynes | Wellcome Sanger Institute for the COVID-19 Genomics UK (COG-UK) Consortium | The Lighthouse Lab in Milton Keynes and Alex Alderton, Roberto Amato, Jeffrey Barrett, Sonia Goncalves, Ewan Harrison, David K. Jackson, Ian Johnston, Dominic Kwiatkowski, Cordelia Langford, John Sillitoe on behalf of the Wellcome Sanger Institute COVID-19 Surveillance Team |
| EPI_ISL_2554666, EPI_ISL_2554667 | Lighthouse Lab in Glasgow | Wellcome Sanger Institute for the COVID-19 Genomics UK (COG-UK) Consortium | Harper VanSteenhouse, Yumi Kasai, David Gray, Carol Clugston, Anna Dominiczak and Alex Alderton, Roberto Amato, Jeffrey Barrett, Sonia Goncalves, Ewan Harrison, David K. Jackson, Ian Johnston, Dominic Kwiatkowski, Cordelia Langford, John Sillitoe on behalf of the Wellcome Sanger Institute COVID-19 Surveillance Team |
| EPI_ISL_2554668, EPI_ISL_2554669, EPI_ISL_2554670 | Lighthouse Lab in Milton Keynes | Wellcome Sanger Institute for the COVID-19 Genomics UK (COG-UK) Consortium | The Lighthouse Lab in Milton Keynes and Alex Alderton, Roberto Amato, Jeffrey Barrett, Sonia Goncalves, Ewan Harrison, David K. Jackson, Ian Johnston, Dominic Kwiatkowski, Cordelia Langford, John Sillitoe on behalf of the Wellcome Sanger Institute COVID-19 Surveillance Team |
| EPI_ISL_2554671, EPI_ISL_2554672, EPI_ISL_2554673, EPI_ISL_2554674 | Lighthouse Lab in Glasgow | Wellcome Sanger Institute for the COVID-19 Genomics UK (COG-UK) Consortium | Harper VanSteenhouse, Yumi Kasai, David Gray, Carol Clugston, Anna Dominiczak and Alex Alderton, Roberto Amato, Jeffrey Barrett, Sonia Goncalves, Ewan Harrison, David K. Jackson, Ian Johnston, Dominic Kwiatkowski, Cordelia Langford, John Sillitoe on behalf of the Wellcome Sanger Institute COVID-19 Surveillance Team |
| EPI_ISL_2554675, EPI_ISL_2554676 | Lighthouse Lab in Milton Keynes | Wellcome Sanger Institute for the COVID-19 Genomics UK (COG-UK) Consortium | The Lighthouse Lab in Milton Keynes and Alex Alderton, Roberto Amato, Jeffrey Barrett, Sonia Goncalves, Ewan Harrison, David K. Jackson, Ian Johnston, Dominic Kwiatkowski, Cordelia Langford, John Sillitoe on behalf of the Wellcome Sanger Institute COVID-19 Surveillance Team |
| EPI_ISL_2554677 | Lighthouse Lab in Glasgow | Wellcome Sanger Institute for the COVID-19 Genomics UK (COG-UK) Consortium | Harper VanSteenhouse, Yumi Kasai, David Gray, Carol Clugston, Anna Dominiczak and Alex Alderton, Roberto Amato, Jeffrey Barrett, Sonia Goncalves, Ewan Harrison, David K. Jackson, Ian Johnston, Dominic Kwiatkowski, Cordelia Langford, John Sillitoe on behalf of the Wellcome Sanger Institute COVID-19 Surveillance Team |
| EPI_ISL_2554678 | Lighthouse Lab in Milton Keynes | Wellcome Sanger Institute for the COVID-19 Genomics UK (COG-UK) Consortium | The Lighthouse Lab in Milton Keynes and Alex Alderton, Roberto Amato, Jeffrey Barrett, Sonia Goncalves, Ewan Harrison, David K. Jackson, Ian Johnston, Dominic Kwiatkowski, Cordelia Langford, John Sillitoe on behalf of the Wellcome Sanger Institute COVID-19 Surveillance Team |
| EPI_ISL_2554679 | Lighthouse Lab in Glasgow | Wellcome Sanger Institute for the COVID-19 Genomics UK (COG-UK) Consortium | Harper VanSteenhouse, Yumi Kasai, David Gray, Carol Clugston, Anna Dominiczak and Alex Alderton, Roberto Amato, Jeffrey Barrett, Sonia Goncalves, Ewan Harrison, David K. Jackson, Ian Johnston, Dominic Kwiatkowski, Cordelia Langford, John Sillitoe on behalf of the Wellcome Sanger Institute COVID-19 Surveillance Team |
| EPI_ISL_2554680 | Lighthouse Lab in Milton Keynes | Wellcome Sanger Institute for the COVID-19 Genomics UK (COG-UK) Consortium | The Lighthouse Lab in Milton Keynes and Alex Alderton, Roberto Amato, Jeffrey Barrett, Sonia Goncalves, Ewan Harrison, David K. Jackson, Ian Johnston, Dominic Kwiatkowski, Cordelia Langford, John Sillitoe on behalf of the Wellcome Sanger Institute COVID-19 Surveillance Team |
| EPI_ISL_2554681, EPI_ISL_2554682, EPI_ISL_2554683 | Lighthouse Lab in Glasgow | Wellcome Sanger Institute for the COVID-19 Genomics UK (COG-UK) Consortium | Harper VanSteenhouse, Yumi Kasai, David Gray, Carol Clugston, Anna Dominiczak and Alex Alderton, Roberto Amato, Jeffrey Barrett, Sonia Goncalves, Ewan Harrison, David K. Jackson, Ian Johnston, Dominic Kwiatkowski, Cordelia Langford, John Sillitoe on behalf of the Wellcome Sanger Institute COVID-19 Surveillance Team |
| EPI_ISL_2554684, EPI_ISL_2554687 | Lighthouse Lab in Milton Keynes | Wellcome Sanger Institute for the COVID-19 Genomics UK (COG-UK) Consortium | The Lighthouse Lab in Milton Keynes and Alex Alderton, Roberto Amato, Jeffrey Barrett, Sonia Goncalves, Ewan Harrison, David K. Jackson, Ian Johnston, Dominic Kwiatkowski, Cordelia Langford, John Sillitoe on behalf of the Wellcome Sanger Institute COVID-19 Surveillance Team |
| EPI_ISL_2554688 | Lighthouse Lab in Glasgow | Wellcome Sanger Institute for the COVID-19 Genomics UK (COG-UK) Consortium | Harper VanSteenhouse, Yumi Kasai, David Gray, Carol Clugston, Anna Dominiczak and Alex Alderton, Roberto Amato, Jeffrey Barrett, Sonia Goncalves, Ewan Harrison, David K. Jackson, Ian Johnston, Dominic Kwiatkowski, Cordelia Langford, John Sillitoe on behalf of the Wellcome Sanger Institute COVID-19 Surveillance Team |
| EPI_ISL_2554689 | Lighthouse Lab in Milton Keynes | Wellcome Sanger Institute for the COVID-19 Genomics UK (COG-UK) Consortium | The Lighthouse Lab in Milton Keynes and Alex Alderton, Roberto Amato, Jeffrey Barrett, Sonia Goncalves, Ewan Harrison, David K. Jackson, Ian Johnston, Dominic Kwiatkowski, Cordelia Langford, John Sillitoe on behalf of the Wellcome Sanger Institute COVID-19 Surveillance Team |
| EPI_ISL_2554691, EPI_ISL_2554692 | Lighthouse Lab in Glasgow | Wellcome Sanger Institute for the COVID-19 Genomics UK (COG-UK) Consortium | Harper VanSteenhouse, Yumi Kasai, David Gray, Carol Clugston, Anna Dominiczak and Alex Alderton, Roberto Amato, Jeffrey Barrett, Sonia Goncalves, Ewan Harrison, David K. Jackson, Ian Johnston, Dominic Kwiatkowski, Cordelia Langford, John Sillitoe on behalf of the Wellcome Sanger Institute COVID-19 Surveillance Team |
| EPI_ISL_2554693, EPI_ISL_2554694 | Lighthouse Lab in Milton Keynes | Wellcome Sanger Institute for the COVID-19 Genomics UK (COG-UK) Consortium | The Lighthouse Lab in Milton Keynes and Alex Alderton, Roberto Amato, Jeffrey Barrett, Sonia Goncalves, Ewan Harrison, David K. Jackson, Ian Johnston, Dominic Kwiatkowski, Cordelia Langford, John Sillitoe on behalf of the Wellcome Sanger Institute COVID-19 Surveillance Team |
| EPI_ISL_2554695 | Lighthouse Lab in Glasgow | Wellcome Sanger Institute for the COVID-19 Genomics UK (COG-UK) Consortium | Harper VanSteenhouse, Yumi Kasai, David Gray, Carol Clugston, Anna Dominiczak and Alex Alderton, Roberto Amato, Jeffrey Barrett, Sonia Goncalves, Ewan Harrison, David K. Jackson, Ian Johnston, Dominic Kwiatkowski, Cordelia Langford, John Sillitoe on behalf of the Wellcome Sanger Institute COVID-19 Surveillance Team |
| EPI_ISL_2554696 | Lighthouse Lab in Milton Keynes | Wellcome Sanger Institute for the COVID-19 Genomics UK (COG-UK) Consortium | The Lighthouse Lab in Milton Keynes and Alex Alderton, Roberto Amato, Jeffrey Barrett, Sonia Goncalves, Ewan Harrison, David K. Jackson, Ian Johnston, Dominic Kwiatkowski, Cordelia Langford, John Sillitoe on behalf of the Wellcome Sanger Institute COVID-19 Surveillance Team |
| EPI_ISL_2554698 | Lighthouse Lab in Glasgow | Wellcome Sanger Institute for the COVID-19 Genomics UK (COG-UK) Consortium | Harper VanSteenhouse, Yumi Kasai, David Gray, Carol Clugston, Anna Dominiczak and Alex Alderton, Roberto Amato, Jeffrey Barrett, Sonia Goncalves, Ewan Harrison, David K. Jackson, Ian Johnston, Dominic Kwiatkowski, Cordelia Langford, John Sillitoe on behalf of the Wellcome Sanger Institute COVID-19 Surveillance Team |
| EPI_ISL_2554700 | Lighthouse Lab in Milton Keynes | Wellcome Sanger Institute for the COVID-19 Genomics UK (COG-UK) Consortium | The Lighthouse Lab in Milton Keynes and Alex Alderton, Roberto Amato, Jeffrey Barrett, Sonia Goncalves, Ewan Harrison, David K. Jackson, Ian Johnston, Dominic Kwiatkowski, Cordelia Langford, John Sillitoe on behalf of the Wellcome Sanger Institute COVID-19 Surveillance Team |
| EPI_ISL_2554703 | Lighthouse Lab in Glasgow | Wellcome Sanger Institute for the COVID-19 Genomics UK (COG-UK) Consortium | Harper VanSteenhouse, Yumi Kasai, David Gray, Carol Clugston, Anna Dominiczak and Alex Alderton, Roberto Amato, Jeffrey Barrett, Sonia Goncalves, Ewan Harrison, David K. Jackson, Ian Johnston, Dominic Kwiatkowski, Cordelia Langford, John Sillitoe on behalf of the Wellcome Sanger Institute COVID-19 Surveillance Team |
| EPI_ISL_2554704, EPI_ISL_2554705 | Lighthouse Lab in Milton Keynes | Wellcome Sanger Institute for the COVID-19 Genomics UK (COG-UK) Consortium | The Lighthouse Lab in Milton Keynes and Alex Alderton, Roberto Amato, Jeffrey Barrett, Sonia Goncalves, Ewan Harrison, David K. Jackson, Ian Johnston, Dominic Kwiatkowski, Cordelia Langford, John Sillitoe on behalf of the Wellcome Sanger Institute COVID-19 Surveillance Team |
| EPI_ISL_2554706, EPI_ISL_2554707 | Lighthouse Lab in Glasgow | Wellcome Sanger Institute for the COVID-19 Genomics UK (COG-UK) Consortium | Harper VanSteenhouse, Yumi Kasai, David Gray, Carol Clugston, Anna Dominiczak and Alex Alderton, Roberto Amato, Jeffrey Barrett, Sonia Goncalves, Ewan Harrison, David K. Jackson, Ian Johnston, Dominic Kwiatkowski, Cordelia Langford, John Sillitoe on behalf of the Wellcome Sanger Institute COVID-19 Surveillance Team |
| EPI_ISL_2554708, EPI_ISL_2554710, EPI_ISL_2554711, EPI_ISL_2554712 | Lighthouse Lab in Milton Keynes | Wellcome Sanger Institute for the COVID-19 Genomics UK (COG-UK) Consortium | The Lighthouse Lab in Milton Keynes and Alex Alderton, Roberto Amato, Jeffrey Barrett, Sonia Goncalves, Ewan Harrison, David K. Jackson, Ian Johnston, Dominic Kwiatkowski, Cordelia Langford, John Sillitoe on behalf of the Wellcome Sanger Institute COVID-19 Surveillance Team |
| EPI_ISL_2554713, EPI_ISL_2554714 | Lighthouse Lab in Glasgow | Wellcome Sanger Institute for the COVID-19 Genomics UK (COG-UK) Consortium | Harper VanSteenhouse, Yumi Kasai, David Gray, Carol Clugston, Anna Dominiczak and Alex Alderton, Roberto Amato, Jeffrey Barrett, Sonia Goncalves, Ewan Harrison, David K. Jackson, Ian Johnston, Dominic Kwiatkowski, Cordelia Langford, John Sillitoe on behalf of the Wellcome Sanger Institute COVID-19 Surveillance Team |
| EPI_ISL_2554715 | Lighthouse Lab in Milton Keynes | Wellcome Sanger Institute for the COVID-19 Genomics UK (COG-UK) Consortium | The Lighthouse Lab in Milton Keynes and Alex Alderton, Roberto Amato, Jeffrey Barrett, Sonia Goncalves, Ewan Harrison, David K. Jackson, Ian Johnston, Dominic Kwiatkowski, Cordelia Langford, John Sillitoe on behalf of the Wellcome Sanger Institute COVID-19 Surveillance Team |
| EPI_ISL_2554716, EPI_ISL_2554717, EPI_ISL_2554719 | Lighthouse Lab in Glasgow | Wellcome Sanger Institute for the COVID-19 Genomics UK (COG-UK) Consortium | Harper VanSteenhouse, Yumi Kasai, David Gray, Carol Clugston, Anna Dominiczak and Alex Alderton, Roberto Amato, Jeffrey Barrett, Sonia Goncalves, Ewan Harrison, David K. Jackson, Ian Johnston, Dominic Kwiatkowski, Cordelia Langford, John Sillitoe on behalf of the Wellcome Sanger Institute COVID-19 Surveillance Team |
| EPI_ISL_2554720 | Lighthouse Lab in Milton Keynes | Wellcome Sanger Institute for the COVID-19 Genomics UK (COG-UK) Consortium | The Lighthouse Lab in Milton Keynes and Alex Alderton, Roberto Amato, Jeffrey Barrett, Sonia Goncalves, Ewan Harrison, David K. Jackson, Ian Johnston, Dominic Kwiatkowski, Cordelia Langford, John Sillitoe on behalf of the Wellcome Sanger Institute COVID-19 Surveillance Team |
| EPI_ISL_2554721 | Lighthouse Lab in Glasgow | Wellcome Sanger Institute for the COVID-19 Genomics UK (COG-UK) Consortium | Harper VanSteenhouse, Yumi Kasai, David Gray, Carol Clugston, Anna Dominiczak and Alex Alderton, Roberto Amato, Jeffrey Barrett, Sonia Goncalves, Ewan Harrison, David K. Jackson, Ian Johnston, Dominic Kwiatkowski, Cordelia Langford, John Sillitoe on behalf of the Wellcome Sanger Institute COVID-19 Surveillance Team |
| EPI_ISL_2554722 | Lighthouse Lab in Milton Keynes | Wellcome Sanger Institute for the COVID-19 Genomics UK (COG-UK) Consortium | The Lighthouse Lab in Milton Keynes and Alex Alderton, Roberto Amato, Jeffrey Barrett, Sonia Goncalves, Ewan Harrison, David K. Jackson, Ian Johnston, Dominic Kwiatkowski, Cordelia Langford, John Sillitoe on behalf of the Wellcome Sanger Institute COVID-19 Surveillance Team |

| | | | |
|---|---|---|---|
| EPI_ISL_2554723 | Lighthouse Lab in Glasgow | Wellcome Sanger Institute for the COVID-19 Genomics UK (COG-UK) Consortium | Harper VanSteenhouse, Yumi Kasai, David Gray, Carol Clugston, Anna Dominiczak and Alex Alderton, Roberto Amato, Jeffrey Barrett, Sonia Goncalves, Ewan Harrison, David K. Jackson, Ian Johnston, Dominic Kwiatkowski, Cordelia Langford, John Sillitoe on behalf of the Wellcome Sanger Institute COVID-19 Surveillance Team |
| EPI_ISL_2554724, EPI_ISL_2554725, EPI_ISL_2554726, EPI_ISL_2554727, EPI_ISL_2554728 | Lighthouse Lab in Milton Keynes | Wellcome Sanger Institute for the COVID-19 Genomics UK (COG-UK) Consortium | The Lighthouse Lab in Milton Keynes and Alex Alderton, Roberto Amato, Jeffrey Barrett, Sonia Goncalves, Ewan Harrison, David K. Jackson, Ian Johnston, Dominic Kwiatkowski, Cordelia Langford, John Sillitoe on behalf of the Wellcome Sanger Institute COVID-19 Surveillance Team |
| EPI_ISL_2554732 | Lighthouse Lab in Glasgow | Wellcome Sanger Institute for the COVID-19 Genomics UK (COG-UK) Consortium | Harper VanSteenhouse, Yumi Kasai, David Gray, Carol Clugston, Anna Dominiczak and Alex Alderton, Roberto Amato, Jeffrey Barrett, Sonia Goncalves, Ewan Harrison, David K. Jackson, Ian Johnston, Dominic Kwiatkowski, Cordelia Langford, John Sillitoe on behalf of the Wellcome Sanger Institute COVID-19 Surveillance Team |
| EPI_ISL_2554733, EPI_ISL_2554734, EPI_ISL_2554735 | Lighthouse Lab in Milton Keynes | Wellcome Sanger Institute for the COVID-19 Genomics UK (COG-UK) Consortium | The Lighthouse Lab in Milton Keynes and Alex Alderton, Roberto Amato, Jeffrey Barrett, Sonia Goncalves, Ewan Harrison, David K. Jackson, Ian Johnston, Dominic Kwiatkowski, Cordelia Langford, John Sillitoe on behalf of the Wellcome Sanger Institute COVID-19 Surveillance Team |
| EPI_ISL_2554736, EPI_ISL_2554737 | Lighthouse Lab in Glasgow | Wellcome Sanger Institute for the COVID-19 Genomics UK (COG-UK) Consortium | Harper VanSteenhouse, Yumi Kasai, David Gray, Carol Clugston, Anna Dominiczak and Alex Alderton, Roberto Amato, Jeffrey Barrett, Sonia Goncalves, Ewan Harrison, David K. Jackson, Ian Johnston, Dominic Kwiatkowski, Cordelia Langford, John Sillitoe on behalf of the Wellcome Sanger Institute COVID-19 Surveillance Team |
| EPI_ISL_2554738, EPI_ISL_2554739, EPI_ISL_2554740 | Lighthouse Lab in Milton Keynes | Wellcome Sanger Institute for the COVID-19 Genomics UK (COG-UK) Consortium | The Lighthouse Lab in Milton Keynes and Alex Alderton, Roberto Amato, Jeffrey Barrett, Sonia Goncalves, Ewan Harrison, David K. Jackson, Ian Johnston, Dominic Kwiatkowski, Cordelia Langford, John Sillitoe on behalf of the Wellcome Sanger Institute COVID-19 Surveillance Team |
| EPI_ISL_2554741 | Lighthouse Lab in Glasgow | Wellcome Sanger Institute for the COVID-19 Genomics UK (COG-UK) Consortium | Harper VanSteenhouse, Yumi Kasai, David Gray, Carol Clugston, Anna Dominiczak and Alex Alderton, Roberto Amato, Jeffrey Barrett, Sonia Goncalves, Ewan Harrison, David K. Jackson, Ian Johnston, Dominic Kwiatkowski, Cordelia Langford, John Sillitoe on behalf of the Wellcome Sanger Institute COVID-19 Surveillance Team |
| EPI_ISL_2554742 | Lighthouse Lab in Milton Keynes | Wellcome Sanger Institute for the COVID-19 Genomics UK (COG-UK) Consortium | The Lighthouse Lab in Milton Keynes and Alex Alderton, Roberto Amato, Jeffrey Barrett, Sonia Goncalves, Ewan Harrison, David K. Jackson, Ian Johnston, Dominic Kwiatkowski, Cordelia Langford, John Sillitoe on behalf of the Wellcome Sanger Institute COVID-19 Surveillance Team |
| EPI_ISL_2554744, EPI_ISL_2554745 | Lighthouse Lab in Glasgow | Wellcome Sanger Institute for the COVID-19 Genomics UK (COG-UK) Consortium | Harper VanSteenhouse, Yumi Kasai, David Gray, Carol Clugston, Anna Dominiczak and Alex Alderton, Roberto Amato, Jeffrey Barrett, Sonia Goncalves, Ewan Harrison, David K. Jackson, Ian Johnston, Dominic Kwiatkowski, Cordelia Langford, John Sillitoe on behalf of the Wellcome Sanger Institute COVID-19 Surveillance Team |
| EPI_ISL_2554746, EPI_ISL_2554747 | Lighthouse Lab in Milton Keynes | Wellcome Sanger Institute for the COVID-19 Genomics UK (COG-UK) Consortium | The Lighthouse Lab in Milton Keynes and Alex Alderton, Roberto Amato, Jeffrey Barrett, Sonia Goncalves, Ewan Harrison, David K. Jackson, Ian Johnston, Dominic Kwiatkowski, Cordelia Langford, John Sillitoe on behalf of the Wellcome Sanger Institute COVID-19 Surveillance Team |
| EPI_ISL_2554748, EPI_ISL_2554749, EPI_ISL_2554750, EPI_ISL_2554751 | Lighthouse Lab in Glasgow | Wellcome Sanger Institute for the COVID-19 Genomics UK (COG-UK) Consortium | Harper VanSteenhouse, Yumi Kasai, David Gray, Carol Clugston, Anna Dominiczak and Alex Alderton, Roberto Amato, Jeffrey Barrett, Sonia Goncalves, Ewan Harrison, David K. Jackson, Ian Johnston, Dominic Kwiatkowski, Cordelia Langford, John Sillitoe on behalf of the Wellcome Sanger Institute COVID-19 Surveillance Team |
| EPI_ISL_2554752, EPI_ISL_2554753 | Lighthouse Lab in Milton Keynes | Wellcome Sanger Institute for the COVID-19 Genomics UK (COG-UK) Consortium | The Lighthouse Lab in Milton Keynes and Alex Alderton, Roberto Amato, Jeffrey Barrett, Sonia Goncalves, Ewan Harrison, David K. Jackson, Ian Johnston, Dominic Kwiatkowski, Cordelia Langford, John Sillitoe on behalf of the Wellcome Sanger Institute COVID-19 Surveillance Team |
| EPI_ISL_2554754 | Lighthouse Lab in Glasgow | Wellcome Sanger Institute for the COVID-19 Genomics UK (COG-UK) Consortium | Harper VanSteenhouse, Yumi Kasai, David Gray, Carol Clugston, Anna Dominiczak and Alex Alderton, Roberto Amato, Jeffrey Barrett, Sonia Goncalves, Ewan Harrison, David K. Jackson, Ian Johnston, Dominic Kwiatkowski, Cordelia Langford, John Sillitoe on behalf of the Wellcome Sanger Institute COVID-19 Surveillance Team |
| EPI_ISL_2554755 | Lighthouse Lab in Milton Keynes | Wellcome Sanger Institute for the COVID-19 Genomics UK (COG-UK) Consortium | The Lighthouse Lab in Milton Keynes and Alex Alderton, Roberto Amato, Jeffrey Barrett, Sonia Goncalves, Ewan Harrison, David K. Jackson, Ian Johnston, Dominic Kwiatkowski, Cordelia Langford, John Sillitoe on behalf of the Wellcome Sanger Institute COVID-19 Surveillance Team |
| EPI_ISL_2554756 | Lighthouse Lab in Glasgow | Wellcome Sanger Institute for the COVID-19 Genomics UK (COG-UK) Consortium | Harper VanSteenhouse, Yumi Kasai, David Gray, Carol Clugston, Anna Dominiczak and Alex Alderton, Roberto Amato, Jeffrey Barrett, Sonia Goncalves, Ewan Harrison, David K. Jackson, Ian Johnston, Dominic Kwiatkowski, Cordelia Langford, John Sillitoe on behalf of the Wellcome Sanger Institute COVID-19 Surveillance Team |
| EPI_ISL_2554757, EPI_ISL_2554759, EPI_ISL_2554760, EPI_ISL_2554761, EPI_ISL_2554762 | Lighthouse Lab in Milton Keynes | Wellcome Sanger Institute for the COVID-19 Genomics UK (COG-UK) Consortium | The Lighthouse Lab in Milton Keynes and Alex Alderton, Roberto Amato, Jeffrey Barrett, Sonia Goncalves, Ewan Harrison, David K. Jackson, Ian Johnston, Dominic Kwiatkowski, Cordelia Langford, John Sillitoe on behalf of the Wellcome Sanger Institute COVID-19 Surveillance Team |
| EPI_ISL_2554763 | Lighthouse Lab in Glasgow | Wellcome Sanger Institute for the COVID-19 Genomics UK (COG-UK) Consortium | Harper VanSteenhouse, Yumi Kasai, David Gray, Carol Clugston, Anna Dominiczak and Alex Alderton, Roberto Amato, Jeffrey Barrett, Sonia Goncalves, Ewan Harrison, David K. Jackson, Ian Johnston, Dominic Kwiatkowski, Cordelia Langford, John Sillitoe on behalf of the Wellcome Sanger Institute COVID-19 Surveillance Team |
| EPI_ISL_2554764, EPI_ISL_2554765, EPI_ISL_2554766, EPI_ISL_2554767, EPI_ISL_2554768, EPI_ISL_2554769 | Lighthouse Lab in Milton Keynes | Wellcome Sanger Institute for the COVID-19 Genomics UK (COG-UK) Consortium | The Lighthouse Lab in Milton Keynes and Alex Alderton, Roberto Amato, Jeffrey Barrett, Sonia Goncalves, Ewan Harrison, David K. Jackson, Ian Johnston, Dominic Kwiatkowski, Cordelia Langford, John Sillitoe on behalf of the Wellcome Sanger Institute COVID-19 Surveillance Team |
| EPI_ISL_2554770 | Lighthouse Lab in Glasgow | Wellcome Sanger Institute for the COVID-19 Genomics UK (COG-UK) Consortium | Harper VanSteenhouse, Yumi Kasai, David Gray, Carol Clugston, Anna Dominiczak and Alex Alderton, Roberto Amato, Jeffrey Barrett, Sonia Goncalves, Ewan Harrison, David K. Jackson, Ian Johnston, Dominic Kwiatkowski, Cordelia Langford, John Sillitoe on behalf of the Wellcome Sanger Institute COVID-19 Surveillance Team |
| EPI_ISL_2554771, EPI_ISL_2554772, EPI_ISL_2554773, EPI_ISL_2554774, EPI_ISL_2554775, EPI_ISL_2554776 | Lighthouse Lab in Milton Keynes | Wellcome Sanger Institute for the COVID-19 Genomics UK (COG-UK) Consortium | The Lighthouse Lab in Milton Keynes and Alex Alderton, Roberto Amato, Jeffrey Barrett, Sonia Goncalves, Ewan Harrison, David K. Jackson, Ian Johnston, Dominic Kwiatkowski, Cordelia Langford, John Sillitoe on behalf of the Wellcome Sanger Institute COVID-19 Surveillance Team |
| EPI_ISL_2554777 | Lighthouse Lab in Glasgow | Wellcome Sanger Institute for the COVID-19 Genomics UK (COG-UK) Consortium | Harper VanSteenhouse, Yumi Kasai, David Gray, Carol Clugston, Anna Dominiczak and Alex Alderton, Roberto Amato, Jeffrey Barrett, Sonia Goncalves, Ewan Harrison, David K. Jackson, Ian Johnston, Dominic Kwiatkowski, Cordelia Langford, John Sillitoe on behalf of the Wellcome Sanger Institute COVID-19 Surveillance Team |
| EPI_ISL_2554778 | Lighthouse Lab in Milton Keynes | Wellcome Sanger Institute for the COVID-19 Genomics UK (COG-UK) Consortium | The Lighthouse Lab in Milton Keynes and Alex Alderton, Roberto Amato, Jeffrey Barrett, Sonia Goncalves, Ewan Harrison, David K. Jackson, Ian Johnston, Dominic Kwiatkowski, Cordelia Langford, John Sillitoe on behalf of the Wellcome Sanger Institute COVID-19 Surveillance Team |
| EPI_ISL_2554779 | Lighthouse Lab in Glasgow | Wellcome Sanger Institute for the COVID-19 Genomics UK (COG-UK) Consortium | Harper VanSteenhouse, Yumi Kasai, David Gray, Carol Clugston, Anna Dominiczak and Alex Alderton, Roberto Amato, Jeffrey Barrett, Sonia Goncalves, Ewan Harrison, David K. Jackson, Ian Johnston, Dominic Kwiatkowski, Cordelia Langford, John Sillitoe on behalf of the Wellcome Sanger Institute COVID-19 Surveillance Team |
| EPI_ISL_2554781 | Lighthouse Lab in Milton Keynes | Wellcome Sanger Institute for the COVID-19 Genomics UK (COG-UK) Consortium | The Lighthouse Lab in Milton Keynes and Alex Alderton, Roberto Amato, Jeffrey Barrett, Sonia Goncalves, Ewan Harrison, David K. Jackson, Ian Johnston, Dominic Kwiatkowski, Cordelia Langford, John Sillitoe on behalf of the Wellcome Sanger Institute COVID-19 Surveillance Team |
| EPI_ISL_2554782 | Lighthouse Lab in Glasgow | Wellcome Sanger Institute for the COVID-19 Genomics UK (COG-UK) Consortium | Harper VanSteenhouse, Yumi Kasai, David Gray, Carol Clugston, Anna Dominiczak and Alex Alderton, Roberto Amato, Jeffrey Barrett, Sonia Goncalves, Ewan Harrison, David K. Jackson, Ian Johnston, Dominic Kwiatkowski, Cordelia Langford, John Sillitoe on behalf of the Wellcome Sanger Institute COVID-19 Surveillance Team |
| EPI_ISL_2554783 | Lighthouse Lab in Milton Keynes | Wellcome Sanger Institute for the COVID-19 Genomics UK (COG-UK) Consortium | The Lighthouse Lab in Milton Keynes and Alex Alderton, Roberto Amato, Jeffrey Barrett, Sonia Goncalves, Ewan Harrison, David K. Jackson, Ian Johnston, Dominic Kwiatkowski, Cordelia Langford, John Sillitoe on behalf of the Wellcome Sanger Institute COVID-19 Surveillance Team |
| EPI_ISL_2554784, EPI_ISL_2554785, EPI_ISL_2554786, EPI_ISL_2554787 | Lighthouse Lab in Glasgow | Wellcome Sanger Institute for the COVID-19 Genomics UK (COG-UK) Consortium | Harper VanSteenhouse, Yumi Kasai, David Gray, Carol Clugston, Anna Dominiczak and Alex Alderton, Roberto Amato, Jeffrey Barrett, Sonia Goncalves, Ewan Harrison, David K. Jackson, Ian Johnston, Dominic Kwiatkowski, Cordelia Langford, John Sillitoe on behalf of the Wellcome Sanger Institute COVID-19 Surveillance Team |
| EPI_ISL_2554788 | Lighthouse Lab in Milton Keynes | Wellcome Sanger Institute for the COVID-19 Genomics UK (COG-UK) Consortium | The Lighthouse Lab in Milton Keynes and Alex Alderton, Roberto Amato, Jeffrey Barrett, Sonia Goncalves, Ewan Harrison, David K. Jackson, Ian Johnston, Dominic Kwiatkowski, Cordelia Langford, John Sillitoe on behalf of the Wellcome Sanger Institute COVID-19 Surveillance Team |
| EPI_ISL_2554789 | Lighthouse Lab in Glasgow | Wellcome Sanger Institute for the COVID-19 Genomics UK (COG-UK) Consortium | Harper VanSteenhouse, Yumi Kasai, David Gray, Carol Clugston, Anna Dominiczak and Alex Alderton, Roberto Amato, Jeffrey Barrett, Sonia Goncalves, Ewan Harrison, David K. Jackson, Ian Johnston, Dominic Kwiatkowski, Cordelia Langford, John Sillitoe on behalf of the Wellcome Sanger Institute COVID-19 Surveillance Team |
| EPI_ISL_2554790, EPI_ISL_2554791 | Lighthouse Lab in Milton Keynes | Wellcome Sanger Institute for the COVID-19 Genomics UK (COG-UK) Consortium | The Lighthouse Lab in Milton Keynes and Alex Alderton, Roberto Amato, Jeffrey Barrett, Sonia Goncalves, Ewan Harrison, David K. Jackson, Ian Johnston, Dominic Kwiatkowski, Cordelia Langford, John Sillitoe on behalf of the Wellcome Sanger Institute COVID-19 Surveillance Team |
| EPI_ISL_2554792 | Lighthouse Lab in Glasgow | Wellcome Sanger Institute for the COVID-19 Genomics UK (COG-UK) Consortium | Harper VanSteenhouse, Yumi Kasai, David Gray, Carol Clugston, Anna Dominiczak and Alex Alderton, Roberto Amato, Jeffrey Barrett, Sonia Goncalves, Ewan Harrison, David K. Jackson, Ian Johnston, Dominic Kwiatkowski, Cordelia Langford, John Sillitoe on behalf of the Wellcome Sanger Institute COVID-19 Surveillance Team |
| EPI_ISL_2554793, EPI_ISL_2554794 | Lighthouse Lab in Milton Keynes | Wellcome Sanger Institute for the COVID-19 Genomics UK (COG-UK) Consortium | The Lighthouse Lab in Milton Keynes and Alex Alderton, Roberto Amato, Jeffrey Barrett, Sonia Goncalves, Ewan Harrison, David K. Jackson, Ian Johnston, Dominic Kwiatkowski, Cordelia Langford, John Sillitoe on behalf of the Wellcome Sanger Institute COVID-19 Surveillance Team |
| EPI_ISL_2554796, EPI_ISL_2554797, EPI_ISL_2554798 | Lighthouse Lab in Glasgow | Wellcome Sanger Institute for the COVID-19 Genomics UK (COG-UK) Consortium | Harper VanSteenhouse, Yumi Kasai, David Gray, Carol Clugston, Anna Dominiczak and Alex Alderton, Roberto Amato, Jeffrey Barrett, Sonia Goncalves, Ewan Harrison, David K. Jackson, Ian Johnston, Dominic Kwiatkowski, Cordelia Langford, John Sillitoe on behalf of the Wellcome Sanger Institute COVID-19 Surveillance Team |
| EPI_ISL_2554799, EPI_ISL_2554800 | Lighthouse Lab in Milton Keynes | Wellcome Sanger Institute for the COVID-19 Genomics UK (COG-UK) Consortium | The Lighthouse Lab in Milton Keynes and Alex Alderton, Roberto Amato, Jeffrey Barrett, Sonia Goncalves, Ewan Harrison, David K. Jackson, Ian Johnston, Dominic Kwiatkowski, Cordelia Langford, John Sillitoe on behalf of the Wellcome Sanger Institute COVID-19 Surveillance Team |
| EPI_ISL_2554801, EPI_ISL_2554802, EPI_ISL_2554803 | Lighthouse Lab in Glasgow | Wellcome Sanger Institute for the COVID-19 Genomics UK (COG-UK) Consortium | Harper VanSteenhouse, Yumi Kasai, David Gray, Carol Clugston, Anna Dominiczak and Alex Alderton, Roberto Amato, Jeffrey Barrett, Sonia Goncalves, Ewan Harrison, David K. Jackson, Ian Johnston, Dominic Kwiatkowski, Cordelia Langford, John Sillitoe on behalf of the Wellcome Sanger Institute COVID-19 Surveillance Team |
| EPI_ISL_2554804, EPI_ISL_2554805, | Lighthouse Lab in Milton Keynes | Wellcome Sanger Institute for the COVID-19 Genomics UK (COG-UK) Consortium | The Lighthouse Lab in Milton Keynes and Alex Alderton, Roberto Amato, Jeffrey Barrett, Sonia Goncalves, Ewan Harrison, David K. Jackson, Ian Johnston, Dominic Kwiatkowski, Cordelia Langford, John Sillitoe on behalf of the Wellcome Sanger Institute COVID-19 Surveillance Team |

| | | | |
|---|---|---|---|
| EPI_ISL_2554806 | | | |
| EPI_ISL_2554807 | Lighthouse Lab in Glasgow | Wellcome Sanger Institute for the COVID-19 Genomics UK (COG-UK) Consortium | Harper VanSteenhouse, Yumi Kasai, David Gray, Carol Clugston, Anna Dominiczak and Alex Alderton, Roberto Amato, Jeffrey Barrett, Sonia Goncalves, Ewan Harrison, David K. Jackson, Ian Johnston, Dominic Kwiatkowski, Cordelia Langford, John Sillitoe on behalf of the Wellcome Sanger Institute COVID-19 Surveillance Team |
| EPI_ISL_2554808 | Lighthouse Lab in Milton Keynes | Wellcome Sanger Institute for the COVID-19 Genomics UK (COG-UK) Consortium | The Lighthouse Lab in Milton Keynes and Alex Alderton, Roberto Amato, Jeffrey Barrett, Sonia Goncalves, Ewan Harrison, David K. Jackson, Ian Johnston, Dominic Kwiatkowski, Cordelia Langford, John Sillitoe on behalf of the Wellcome Sanger Institute COVID-19 Surveillance Team |
| EPI_ISL_2554809, EPI_ISL_2554810 | Lighthouse Lab in Glasgow | Wellcome Sanger Institute for the COVID-19 Genomics UK (COG-UK) Consortium | Harper VanSteenhouse, Yumi Kasai, David Gray, Carol Clugston, Anna Dominiczak and Alex Alderton, Roberto Amato, Jeffrey Barrett, Sonia Goncalves, Ewan Harrison, David K. Jackson, Ian Johnston, Dominic Kwiatkowski, Cordelia Langford, John Sillitoe on behalf of the Wellcome Sanger Institute COVID-19 Surveillance Team |
| EPI_ISL_2554811 | Lighthouse Lab in Milton Keynes | Wellcome Sanger Institute for the COVID-19 Genomics UK (COG-UK) Consortium | The Lighthouse Lab in Milton Keynes and Alex Alderton, Roberto Amato, Jeffrey Barrett, Sonia Goncalves, Ewan Harrison, David K. Jackson, Ian Johnston, Dominic Kwiatkowski, Cordelia Langford, John Sillitoe on behalf of the Wellcome Sanger Institute COVID-19 Surveillance Team |
| EPI_ISL_2554812, EPI_ISL_2554813, EPI_ISL_2554814 | Lighthouse Lab in Glasgow | Wellcome Sanger Institute for the COVID-19 Genomics UK (COG-UK) Consortium | Harper VanSteenhouse, Yumi Kasai, David Gray, Carol Clugston, Anna Dominiczak and Alex Alderton, Roberto Amato, Jeffrey Barrett, Sonia Goncalves, Ewan Harrison, David K. Jackson, Ian Johnston, Dominic Kwiatkowski, Cordelia Langford, John Sillitoe on behalf of the Wellcome Sanger Institute COVID-19 Surveillance Team |
| EPI_ISL_2554815, EPI_ISL_2554817, EPI_ISL_2554818 | Lighthouse Lab in Milton Keynes | Wellcome Sanger Institute for the COVID-19 Genomics UK (COG-UK) Consortium | The Lighthouse Lab in Milton Keynes and Alex Alderton, Roberto Amato, Jeffrey Barrett, Sonia Goncalves, Ewan Harrison, David K. Jackson, Ian Johnston, Dominic Kwiatkowski, Cordelia Langford, John Sillitoe on behalf of the Wellcome Sanger Institute COVID-19 Surveillance Team |
| EPI_ISL_2554819, EPI_ISL_2554820 | Lighthouse Lab in Glasgow | Wellcome Sanger Institute for the COVID-19 Genomics UK (COG-UK) Consortium | Harper VanSteenhouse, Yumi Kasai, David Gray, Carol Clugston, Anna Dominiczak and Alex Alderton, Roberto Amato, Jeffrey Barrett, Sonia Goncalves, Ewan Harrison, David K. Jackson, Ian Johnston, Dominic Kwiatkowski, Cordelia Langford, John Sillitoe on behalf of the Wellcome Sanger Institute COVID-19 Surveillance Team |
| EPI_ISL_2554821 | Lighthouse Lab in Milton Keynes | Wellcome Sanger Institute for the COVID-19 Genomics UK (COG-UK) Consortium | The Lighthouse Lab in Milton Keynes and Alex Alderton, Roberto Amato, Jeffrey Barrett, Sonia Goncalves, Ewan Harrison, David K. Jackson, Ian Johnston, Dominic Kwiatkowski, Cordelia Langford, John Sillitoe on behalf of the Wellcome Sanger Institute COVID-19 Surveillance Team |
| EPI_ISL_2554822, EPI_ISL_2554823, EPI_ISL_2554824, EPI_ISL_2554825, EPI_ISL_2554826, EPI_ISL_2554827 | Lighthouse Lab in Glasgow | Wellcome Sanger Institute for the COVID-19 Genomics UK (COG-UK) Consortium | Harper VanSteenhouse, Yumi Kasai, David Gray, Carol Clugston, Anna Dominiczak and Alex Alderton, Roberto Amato, Jeffrey Barrett, Sonia Goncalves, Ewan Harrison, David K. Jackson, Ian Johnston, Dominic Kwiatkowski, Cordelia Langford, John Sillitoe on behalf of the Wellcome Sanger Institute COVID-19 Surveillance Team |
| EPI_ISL_2554828, EPI_ISL_2554829 | Lighthouse Lab in Milton Keynes | Wellcome Sanger Institute for the COVID-19 Genomics UK (COG-UK) Consortium | The Lighthouse Lab in Milton Keynes and Alex Alderton, Roberto Amato, Jeffrey Barrett, Sonia Goncalves, Ewan Harrison, David K. Jackson, Ian Johnston, Dominic Kwiatkowski, Cordelia Langford, John Sillitoe on behalf of the Wellcome Sanger Institute COVID-19 Surveillance Team |
| EPI_ISL_2554830 | Lighthouse Lab in Glasgow | Wellcome Sanger Institute for the COVID-19 Genomics UK (COG-UK) Consortium | Harper VanSteenhouse, Yumi Kasai, David Gray, Carol Clugston, Anna Dominiczak and Alex Alderton, Roberto Amato, Jeffrey Barrett, Sonia Goncalves, Ewan Harrison, David K. Jackson, Ian Johnston, Dominic Kwiatkowski, Cordelia Langford, John Sillitoe on behalf of the Wellcome Sanger Institute COVID-19 Surveillance Team |
| EPI_ISL_2554831, EPI_ISL_2554832, EPI_ISL_2554833 | Lighthouse Lab in Milton Keynes | Wellcome Sanger Institute for the COVID-19 Genomics UK (COG-UK) Consortium | The Lighthouse Lab in Milton Keynes and Alex Alderton, Roberto Amato, Jeffrey Barrett, Sonia Goncalves, Ewan Harrison, David K. Jackson, Ian Johnston, Dominic Kwiatkowski, Cordelia Langford, John Sillitoe on behalf of the Wellcome Sanger Institute COVID-19 Surveillance Team |
| EPI_ISL_2554834 | Lighthouse Lab in Glasgow | Wellcome Sanger Institute for the COVID-19 Genomics UK (COG-UK) Consortium | Harper VanSteenhouse, Yumi Kasai, David Gray, Carol Clugston, Anna Dominiczak and Alex Alderton, Roberto Amato, Jeffrey Barrett, Sonia Goncalves, Ewan Harrison, David K. Jackson, Ian Johnston, Dominic Kwiatkowski, Cordelia Langford, John Sillitoe on behalf of the Wellcome Sanger Institute COVID-19 Surveillance Team |
| EPI_ISL_2554835 | Lighthouse Lab in Milton Keynes | Wellcome Sanger Institute for the COVID-19 Genomics UK (COG-UK) Consortium | The Lighthouse Lab in Milton Keynes and Alex Alderton, Roberto Amato, Jeffrey Barrett, Sonia Goncalves, Ewan Harrison, David K. Jackson, Ian Johnston, Dominic Kwiatkowski, Cordelia Langford, John Sillitoe on behalf of the Wellcome Sanger Institute COVID-19 Surveillance Team |
| EPI_ISL_2554836 | Lighthouse Lab in Glasgow | Wellcome Sanger Institute for the COVID-19 Genomics UK (COG-UK) Consortium | Harper VanSteenhouse, Yumi Kasai, David Gray, Carol Clugston, Anna Dominiczak and Alex Alderton, Roberto Amato, Jeffrey Barrett, Sonia Goncalves, Ewan Harrison, David K. Jackson, Ian Johnston, Dominic Kwiatkowski, Cordelia Langford, John Sillitoe on behalf of the Wellcome Sanger Institute COVID-19 Surveillance Team |
| EPI_ISL_2554837 | Lighthouse Lab in Milton Keynes | Wellcome Sanger Institute for the COVID-19 Genomics UK (COG-UK) Consortium | The Lighthouse Lab in Milton Keynes and Alex Alderton, Roberto Amato, Jeffrey Barrett, Sonia Goncalves, Ewan Harrison, David K. Jackson, Ian Johnston, Dominic Kwiatkowski, Cordelia Langford, John Sillitoe on behalf of the Wellcome Sanger Institute COVID-19 Surveillance Team |
| EPI_ISL_2554839, EPI_ISL_2554840, EPI_ISL_2554841, EPI_ISL_2554842, EPI_ISL_2554843, EPI_ISL_2554845 | Lighthouse Lab in Glasgow | Wellcome Sanger Institute for the COVID-19 Genomics UK (COG-UK) Consortium | Harper VanSteenhouse, Yumi Kasai, David Gray, Carol Clugston, Anna Dominiczak and Alex Alderton, Roberto Amato, Jeffrey Barrett, Sonia Goncalves, Ewan Harrison, David K. Jackson, Ian Johnston, Dominic Kwiatkowski, Cordelia Langford, John Sillitoe on behalf of the Wellcome Sanger Institute COVID-19 Surveillance Team |
| EPI_ISL_2554846, EPI_ISL_2554847 | Lighthouse Lab in Alderley Park | Wellcome Sanger Institute for the COVID-19 Genomics UK (COG-UK) Consortium | Jacquelyn Wynn, Mairead Hyland, The Lighthouse Lab in Alderley Park and Alex Alderton, Roberto Amato, Jeffrey Barrett, Sonia Goncalves, Ewan Harrison, David K. Jackson, Ian Johnston, Dominic Kwiatkowski, Cordelia Langford, John Sillitoe on behalf of the Wellcome Sanger Institute COVID-19 Surveillance Team |
| EPI_ISL_2554848, EPI_ISL_2554849, EPI_ISL_2554850, EPI_ISL_2554851, EPI_ISL_2554852, EPI_ISL_2554853, EPI_ISL_2554854, EPI_ISL_2554855, EPI_ISL_2554857, EPI_ISL_2554858, EPI_ISL_2554859, EPI_ISL_2554861, EPI_ISL_2554862, EPI_ISL_2554864, EPI_ISL_2554865, EPI_ISL_2554866, EPI_ISL_2554867, EPI_ISL_2554868, EPI_ISL_2554869, EPI_ISL_2554870, EPI_ISL_2554871, EPI_ISL_2554872, EPI_ISL_2554873, EPI_ISL_2554874, EPI_ISL_2554876, EPI_ISL_2554877, EPI_ISL_2554878, EPI_ISL_2554880, EPI_ISL_2554881, EPI_ISL_2554882, EPI_ISL_2554884, EPI_ISL_2554886, EPI_ISL_2554887, EPI_ISL_2554888, EPI_ISL_2554889, EPI_ISL_2554893, EPI_ISL_2554894, EPI_ISL_2554895, EPI_ISL_2554896, EPI_ISL_2554897, EPI_ISL_2554898, EPI_ISL_2554902, EPI_ISL_2554904, EPI_ISL_2554905, EPI_ISL_2554906, EPI_ISL_2554907, EPI_ISL_2554908, EPI_ISL_2554909, EPI_ISL_2554910, EPI_ISL_2554911, EPI_ISL_2554912, EPI_ISL_2554915, EPI_ISL_2554917, EPI_ISL_2554918, EPI_ISL_2554919, EPI_ISL_2554920, EPI_ISL_2554921, EPI_ISL_2554922, EPI_ISL_2554923, EPI_ISL_2554924, EPI_ISL_2554927, EPI_ISL_2554928, EPI_ISL_2554929, EPI_ISL_2554930, EPI_ISL_2554931, EPI_ISL_2554932, EPI_ISL_2554933, EPI_ISL_2554936, EPI_ISL_2554937, EPI_ISL_2554938, EPI_ISL_2554939, EPI_ISL_2554940, EPI_ISL_2554941, EPI_ISL_2554942, EPI_ISL_2554943, EPI_ISL_2554944, EPI_ISL_2554945, EPI_ISL_2554947, EPI_ISL_2554949, EPI_ISL_2554950, EPI_ISL_2554951, EPI_ISL_2554952, EPI_ISL_2554953, EPI_ISL_2554955, EPI_ISL_2554956, EPI_ISL_2554957, EPI_ISL_2554960, EPI_ISL_2554961, EPI_ISL_2554963, EPI_ISL_2554964, EPI_ISL_2554966, EPI_ISL_2554967, EPI_ISL_2554968, EPI_ISL_2554969, EPI_ISL_2554970, EPI_ISL_2554971, EPI_ISL_2554972, EPI_ISL_2554973, EPI_ISL_2554974, EPI_ISL_2554975, EPI_ISL_2554976, EPI_ISL_2554977, EPI_ISL_2554978, EPI_ISL_2554979, EPI_ISL_2554980, EPI_ISL_2554981, EPI_ISL_2554982, EPI_ISL_2554983, EPI_ISL_2554984, EPI_ISL_2554987, EPI_ISL_2554988, EPI_ISL_2554989, EPI_ISL_2554990, EPI_ISL_2554992, EPI_ISL_2554993, EPI_ISL_2554995, EPI_ISL_2554996, EPI_ISL_2554997, EPI_ISL_2554998, EPI_ISL_2554999, EPI_ISL_2555000, EPI_ISL_2555001 | | | |
| see above | Lighthouse Lab in Glasgow | Wellcome Sanger Institute for the COVID-19 Genomics UK (COG-UK) Consortium | Harper VanSteenhouse, Yumi Kasai, David Gray, Carol Clugston, Anna Dominiczak and Alex Alderton, Roberto Amato, Jeffrey Barrett, Sonia Goncalves, Ewan Harrison, David K. Jackson, Ian Johnston, Dominic Kwiatkowski, Cordelia Langford, John Sillitoe on behalf of the Wellcome Sanger Institute COVID-19 Surveillance Team |
| EPI_ISL_2555002 | Lighthouse Lab in Alderley Park | Wellcome Sanger Institute for the COVID-19 Genomics UK (COG-UK) Consortium | Jacquelyn Wynn, Mairead Hyland, The Lighthouse Lab in Alderley Park and Alex Alderton, Roberto Amato, Jeffrey Barrett, Sonia Goncalves, Ewan Harrison, David K. Jackson, Ian Johnston, Dominic Kwiatkowski, Cordelia Langford, John Sillitoe on behalf of the Wellcome Sanger Institute COVID-19 Surveillance Team |
| EPI_ISL_2555004 | Health Services Laboratories | Wellcome Sanger Institute for the COVID-19 Genomics UK (COG-UK) Consortium | Health Services Laboratories and Alex Alderton, Roberto Amato, Jeffrey Barrett, Sonia Goncalves, Ewan Harrison, David K. Jackson, Ian Johnston, Dominic Kwiatkowski, Cordelia Langford, John Sillitoe on behalf of the Wellcome Sanger Institute COVID-19 Surveillance Team |
| EPI_ISL_2555005 | Lighthouse Lab in Alderley Park | Wellcome Sanger Institute for the COVID-19 Genomics UK (COG-UK) Consortium | Jacquelyn Wynn, Mairead Hyland, The Lighthouse Lab in Alderley Park and Alex Alderton, Roberto Amato, Jeffrey Barrett, Sonia Goncalves, Ewan Harrison, David K. Jackson, Ian Johnston, Dominic Kwiatkowski, Cordelia Langford, John Sillitoe on behalf of the Wellcome Sanger Institute COVID-19 Surveillance Team |
| EPI_ISL_2555006 | Health Services Laboratories | Wellcome Sanger Institute for the COVID-19 Genomics UK (COG-UK) Consortium | Health Services Laboratories and Alex Alderton, Roberto Amato, Jeffrey Barrett, Sonia Goncalves, Ewan Harrison, David K. Jackson, Ian Johnston, Dominic Kwiatkowski, Cordelia Langford, John Sillitoe on behalf of the Wellcome Sanger Institute COVID-19 Surveillance Team |
| EPI_ISL_2555007 | Lighthouse Lab in Alderley Park | Wellcome Sanger Institute for the COVID-19 Genomics UK (COG-UK) Consortium | Jacquelyn Wynn, Mairead Hyland, The Lighthouse Lab in Alderley Park and Alex Alderton, Roberto Amato, Jeffrey Barrett, Sonia Goncalves, Ewan Harrison, David K. Jackson, Ian Johnston, Dominic Kwiatkowski, Cordelia Langford, John Sillitoe on behalf of the Wellcome Sanger Institute COVID-19 Surveillance Team |
| EPI_ISL_2555009, EPI_ISL_2555010, EPI_ISL_2555011, EPI_ISL_2555012, EPI_ISL_2555013, EPI_ISL_2555014, EPI_ISL_2555015, EPI_ISL_2555016, EPI_ISL_2555017, EPI_ISL_2555019, EPI_ISL_2555020, EPI_ISL_2555021, EPI_ISL_2555024, EPI_ISL_2555025, EPI_ISL_2555026, EPI_ISL_2555027, EPI_ISL_2555028, EPI_ISL_2555029, EPI_ISL_2555030, EPI_ISL_2555031, EPI_ISL_2555032, EPI_ISL_2555033, EPI_ISL_2555034, EPI_ISL_2555035, EPI_ISL_2555036, EPI_ISL_2555037, EPI_ISL_2555038, EPI_ISL_2555039, EPI_ISL_2555040, EPI_ISL_2555041, EPI_ISL_2555043, EPI_ISL_2555044, EPI_ISL_2555045, EPI_ISL_2555046, EPI_ISL_2555049, EPI_ISL_2555050, EPI_ISL_2555051, EPI_ISL_2555052, EPI_ISL_2555053 | | | |
| see above | Lighthouse Lab in Glasgow | Wellcome Sanger Institute for the COVID-19 Genomics UK (COG-UK) Consortium | Harper VanSteenhouse, Yumi Kasai, David Gray, Carol Clugston, Anna Dominiczak and Alex Alderton, Roberto Amato, Jeffrey Barrett, Sonia Goncalves, Ewan Harrison, David K. Jackson, Ian Johnston, Dominic Kwiatkowski, Cordelia Langford, John Sillitoe on behalf of the Wellcome Sanger Institute COVID-19 Surveillance Team |
| EPI_ISL_2555054, EPI_ISL_2555055 | Lighthouse Lab in Milton Keynes | Wellcome Sanger Institute for the COVID-19 Genomics UK (COG-UK) Consortium | The Lighthouse Lab in Milton Keynes and Alex Alderton, Roberto Amato, Jeffrey Barrett, Sonia Goncalves, Ewan Harrison, David K. Jackson, Ian Johnston, Dominic Kwiatkowski, Cordelia Langford, John Sillitoe on behalf of the Wellcome Sanger Institute COVID-19 Surveillance Team |
| EPI_ISL_2555057 | Health Services Laboratories | Wellcome Sanger Institute for the COVID-19 Genomics UK (COG-UK) Consortium | Health Services Laboratories and Alex Alderton, Roberto Amato, Jeffrey Barrett, Sonia Goncalves, Ewan Harrison, David K. Jackson, Ian Johnston, Dominic Kwiatkowski, Cordelia Langford, John Sillitoe on behalf of the Wellcome Sanger Institute COVID-19 Surveillance Team |
| EPI_ISL_2555058, EPI_ISL_2555060, EPI_ISL_2555061, EPI_ISL_2555062, EPI_ISL_2555063, EPI_ISL_2555065, EPI_ISL_2555066, EPI_ISL_2555067, EPI_ISL_2555068, EPI_ISL_2555069, EPI_ISL_2555070, EPI_ISL_2555072, EPI_ISL_2555073, EPI_ISL_2555074, EPI_ISL_2555075, EPI_ISL_2555076, EPI_ISL_2555077, EPI_ISL_2555078, EPI_ISL_2555080, EPI_ISL_2555081, EPI_ISL_2555082, EPI_ISL_2555083, EPI_ISL_2555085, EPI_ISL_2555086, EPI_ISL_2555087, EPI_ISL_2555088, EPI_ISL_2555090, EPI_ISL_2555091, EPI_ISL_2555093, EPI_ISL_2555094, EPI_ISL_2555095, EPI_ISL_2555096, EPI_ISL_2555097, EPI_ISL_2555098, EPI_ISL_2555099, EPI_ISL_2555100, EPI_ISL_2555101, EPI_ISL_2555102, EPI_ISL_2555103, EPI_ISL_2555104, EPI_ISL_2555105, EPI_ISL_2555106, EPI_ISL_2555107, EPI_ISL_2555109, EPI_ISL_2555110, EPI_ISL_2555111, EPI_ISL_2555113, EPI_ISL_2555114, EPI_ISL_2555115, EPI_ISL_2555116, EPI_ISL_2555117, EPI_ISL_2555118, EPI_ISL_2555119, EPI_ISL_2555121, EPI_ISL_2555122, EPI_ISL_2555123, EPI_ISL_2555125, EPI_ISL_2555127, EPI_ISL_2555128, EPI_ISL_2555129, EPI_ISL_2555131, EPI_ISL_2555132, EPI_ISL_2555133, EPI_ISL_2555134, EPI_ISL_2555135, EPI_ISL_2555136, EPI_ISL_2555137, EPI_ISL_2555138, EPI_ISL_2555140, EPI_ISL_2555141, EPI_ISL_2555143, EPI_ISL_2555144, EPI_ISL_2555145, EPI_ISL_2555147, EPI_ISL_2555148, EPI_ISL_2555150, EPI_ISL_2555152, EPI_ISL_2555153, EPI_ISL_2555154, EPI_ISL_2555156, EPI_ISL_2555157, EPI_ISL_2555158, EPI_ISL_2555161, EPI_ISL_2555162, EPI_ISL_2555163, EPI_ISL_2555165, EPI_ISL_2555166, EPI_ISL_2555167, EPI_ISL_2555168, EPI_ISL_2555170, EPI_ISL_2555171, EPI_ISL_2555174, EPI_ISL_2555175, EPI_ISL_2555176, EPI_ISL_2555177, EPI_ISL_2555178, EPI_ISL_2555179, EPI_ISL_2555180, EPI_ISL_2555181, EPI_ISL_2555182, EPI_ISL_2555183, EPI_ISL_2555186, EPI_ISL_2555187, EPI_ISL_2555188, EPI_ISL_2555190, EPI_ISL_2555194, EPI_ISL_2555195, EPI_ISL_2555196, EPI_ISL_2555197, EPI_ISL_2555200, EPI_ISL_2555201, EPI_ISL_2555202, EPI_ISL_2555203, EPI_ISL_2555204, EPI_ISL_2555205, EPI_ISL_2555208, EPI_ISL_2555209, EPI_ISL_2555210, EPI_ISL_2555212, EPI_ISL_2555213, EPI_ISL_2555214, EPI_ISL_2555215, EPI_ISL_2555216, EPI_ISL_2555217, EPI_ISL_2555219, EPI_ISL_2555220, EPI_ISL_2555222, EPI_ISL_2555223, EPI_ISL_2555224, EPI_ISL_2555225, EPI_ISL_2555226, EPI_ISL_2555227, EPI_ISL_2555228, EPI_ISL_2555229, EPI_ISL_2555231, EPI_ISL_2555232, EPI_ISL_2555233, EPI_ISL_2555234, EPI_ISL_2555236, EPI_ISL_2555238, EPI_ISL_2555239, EPI_ISL_2555240, EPI_ISL_2555241, EPI_ISL_2555242, EPI_ISL_2555243, EPI_ISL_2555244, EPI_ISL_2555245, EPI_ISL_2555246, EPI_ISL_2555247, EPI_ISL_2555248, EPI_ISL_2555249, EPI_ISL_2555250, EPI_ISL_2555251, EPI_ISL_2555254, EPI_ISL_2555255, EPI_ISL_2555256, EPI_ISL_2555257, EPI_ISL_2555258, EPI_ISL_2555259, EPI_ISL_2555260, EPI_ISL_2555262, EPI_ISL_2555265, EPI_ISL_2555266, EPI_ISL_2555267, EPI_ISL_2555269, EPI_ISL_2555271, EPI_ISL_2555272, EPI_ISL_2555273, EPI_ISL_2555275, EPI_ISL_2555276, EPI_ISL_2555277, EPI_ISL_2555281, EPI_ISL_2555282, EPI_ISL_2555283, EPI_ISL_2555284, EPI_ISL_2555287, EPI_ISL_2555288, EPI_ISL_2555289, EPI_ISL_2555290, EPI_ISL_2555291, EPI_ISL_2555292, EPI_ISL_2555294, EPI_ISL_2555295, EPI_ISL_2555296, EPI_ISL_2555297, EPI_ISL_2555299, EPI_ISL_2555300, EPI_ISL_2555301, EPI_ISL_2555302, EPI_ISL_2555303, EPI_ISL_2555304, EPI_ISL_2555305, EPI_ISL_2555307, EPI_ISL_2555308, EPI_ISL_2555309, EPI_ISL_2555311, EPI_ISL_2555312, EPI_ISL_2555313, EPI_ISL_2555314, EPI_ISL_2555315, EPI_ISL_2555316, EPI_ISL_2555317, EPI_ISL_2555318, EPI_ISL_2555319, EPI_ISL_2555320, EPI_ISL_2555321, EPI_ISL_2555322, EPI_ISL_2555323, EPI_ISL_2555324, EPI_ISL_2555325, EPI_ISL_2555326, EPI_ISL_2555327, EPI_ISL_2555328, EPI_ISL_2555329, EPI_ISL_2555330, EPI_ISL_2555331, EPI_ISL_2555332, EPI_ISL_2555333, EPI_ISL_2555334, EPI_ISL_2555335, EPI_ISL_2555337, EPI_ISL_2555338, EPI_ISL_2555339, EPI_ISL_2555340, EPI_ISL_2555341, EPI_ISL_2555342, EPI_ISL_2555343, EPI_ISL_2555344, EPI_ISL_2555345, EPI_ISL_2555346, EPI_ISL_2555348, EPI_ISL_2555349, EPI_ISL_2555350, EPI_ISL_2555351, EPI_ISL_2555352, EPI_ISL_2555353, EPI_ISL_2555354, EPI_ISL_2555355, EPI_ISL_2555356, EPI_ISL_2555358, EPI_ISL_2555359, EPI_ISL_2555360, EPI_ISL_2555361, EPI_ISL_2555362, EPI_ISL_2555364, EPI_ISL_2555366, EPI_ISL_2555367, EPI_ISL_2555368, EPI_ISL_2555369, EPI_ISL_2555370, EPI_ISL_2555371, EPI_ISL_2555372, EPI_ISL_2555373, EPI_ISL_2555374, EPI_ISL_2555376, EPI_ISL_2555377, EPI_ISL_2555378, EPI_ISL_2555379, EPI_ISL_2555380, EPI_ISL_2555382, EPI_ISL_2555383, EPI_ISL_2555384, EPI_ISL_2555385, EPI_ISL_2555386, EPI_ISL_2555387, EPI_ISL_2555389, EPI_ISL_2555390, EPI_ISL_2555391, EPI_ISL_2555392, EPI_ISL_2555393, EPI_ISL_2555394, EPI_ISL_2555395, EPI_ISL_2555396, EPI_ISL_2555397, EPI_ISL_2555399, EPI_ISL_2555400, EPI_ISL_2555401, EPI_ISL_2555402, EPI_ISL_2555403, EPI_ISL_2555405, EPI_ISL_2555407, EPI_ISL_2555408, EPI_ISL_2555409, EPI_ISL_2555410, EPI_ISL_2555411, EPI_ISL_2555412, EPI_ISL_2555413, EPI_ISL_2555414, EPI_ISL_2555415, EPI_ISL_2555416, EPI_ISL_2555417, EPI_ISL_2555418, EPI_ISL_2555419, EPI_ISL_2555420, EPI_ISL_2555421, EPI_ISL_2555422, EPI_ISL_2555423, EPI_ISL_2555424, EPI_ISL_2555425, EPI_ISL_2555426, EPI_ISL_2555427, EPI_ISL_2555428, EPI_ISL_2555429, EPI_ISL_2555430, EPI_ISL_2555431, EPI_ISL_2555432, EPI_ISL_2555434, EPI_ISL_2555435, EPI_ISL_2555436, EPI_ISL_2555437, EPI_ISL_2555438, EPI_ISL_2555440, EPI_ISL_2555443, EPI_ISL_2555444, EPI_ISL_2555445, EPI_ISL_2555446, EPI_ISL_2555449, EPI_ISL_2555450, EPI_ISL_2555451, EPI_ISL_2555452, EPI_ISL_2555454, EPI_ISL_2555455, EPI_ISL_2555456, EPI_ISL_2555457, EPI_ISL_2555458, EPI_ISL_2555459, EPI_ISL_2555460, EPI_ISL_2555461, EPI_ISL_2555462, EPI_ISL_2555463, EPI_ISL_2555465, EPI_ISL_2555466, EPI_ISL_2555467, EPI_ISL_2555468, EPI_ISL_2555469, EPI_ISL_2555470, EPI_ISL_2555471, EPI_ISL_2555472, EPI_ISL_2555473, EPI_ISL_2555474, EPI_ISL_2555475, EPI_ISL_2555476, EPI_ISL_2555477, EPI_ISL_2555478, EPI_ISL_2555479, EPI_ISL_2555480, EPI_ISL_2555481, EPI_ISL_2555482, EPI_ISL_2555483, EPI_ISL_2555485, EPI_ISL_2555486, EPI_ISL_2555487, EPI_ISL_2555489, EPI_ISL_2555490, EPI_ISL_2555491, EPI_ISL_2555492, EPI_ISL_2555493, EPI_ISL_2555495, EPI_ISL_2555497, EPI_ISL_2555498, EPI_ISL_2555501, EPI_ISL_2555503, EPI_ISL_2555504, EPI_ISL_2555505, EPI_ISL_2555507, EPI_ISL_2555508, EPI_ISL_2555509, EPI_ISL_2555511, EPI_ISL_2555512, EPI_ISL_2555513, EPI_ISL_2555514, EPI_ISL_2555515, EPI_ISL_2555516, EPI_ISL_2555517, EPI_ISL_2555518, EPI_ISL_2555519, EPI_ISL_2555521, EPI_ISL_2555522, EPI_ISL_2555525 | | | |
| see above | Lighthouse Lab in Glasgow | Wellcome Sanger Institute for the COVID-19 Genomics UK (COG-UK) Consortium | Harper VanSteenhouse, Yumi Kasai, David Gray, Carol Clugston, Anna Dominiczak and Alex Alderton, Roberto Amato, Jeffrey Barrett, Sonia Goncalves, Ewan Harrison, David K. Jackson, Ian Johnston, Dominic Kwiatkowski, Cordelia Langford, John Sillitoe on behalf of the Wellcome Sanger Institute COVID-19 Surveillance Team |
| EPI_ISL_2555526 | Lighthouse Lab in Alderley Park | Wellcome Sanger Institute for the COVID-19 Genomics UK (COG-UK) | Jacquelyn Wynn, Mairead Hyland, The Lighthouse Lab in Alderley Park and Alex Alderton, Roberto Amato, Jeffrey Barrett, Sonia Goncalves, Ewan Harrison, David K. Jackson, Ian Johnston, Dominic Kwiatkowski, Cordelia Langford, John Sillitoe on |

| | | | |
|---|---|---|---|
| | | Consortium | behalf of the Wellcome Sanger Institute COVID-19 Surveillance Team |
| EPI_ISL_2555551, EPI_ISL_2555552, EPI_ISL_2555553, EPI_ISL_2555554, EPI_ISL_2555555, EPI_ISL_2555558, EPI_ISL_2555562, EPI_ISL_2555563, EPI_ISL_2555564, EPI_ISL_2555565, EPI_ISL_2555566, EPI_ISL_2555567, EPI_ISL_2555568, EPI_ISL_2555569, EPI_ISL_2555571, EPI_ISL_2555575, EPI_ISL_2555576, EPI_ISL_2555577, EPI_ISL_2555580, EPI_ISL_2555581, EPI_ISL_2555583, EPI_ISL_2555584, EPI_ISL_2555585, EPI_ISL_2555586, EPI_ISL_2555587, EPI_ISL_2555588, EPI_ISL_2555589, EPI_ISL_2555590, EPI_ISL_2555592, EPI_ISL_2555594, EPI_ISL_2555596, EPI_ISL_2555600, EPI_ISL_2555606, EPI_ISL_2555607, EPI_ISL_2555608, EPI_ISL_2555609, EPI_ISL_2555611, EPI_ISL_2555612, EPI_ISL_2555613, EPI_ISL_2555617, EPI_ISL_2555618, EPI_ISL_2555619, EPI_ISL_2555620, EPI_ISL_2555622, EPI_ISL_2555623, EPI_ISL_2555624, EPI_ISL_2555625, EPI_ISL_2555626, EPI_ISL_2555627, EPI_ISL_2555628, EPI_ISL_2555629, EPI_ISL_2555630, EPI_ISL_2555631, EPI_ISL_2555632, EPI_ISL_2555634, EPI_ISL_2555635, EPI_ISL_2555636, EPI_ISL_2555637, EPI_ISL_2555640, EPI_ISL_2555642, EPI_ISL_2555643, EPI_ISL_2555645, EPI_ISL_2555646, EPI_ISL_2555648, EPI_ISL_2555650, EPI_ISL_2555651, EPI_ISL_2555652, EPI_ISL_2555653, EPI_ISL_2555655, EPI_ISL_2555656, EPI_ISL_2555657, EPI_ISL_2555658 | | | |
| see above | National Centre for Disease Control (NCDC) Biotechnology Division, Delhi | NCDC Delhi, Biotechnology Division INSACOG | Hemlata Lall, Manoj K Singh, Radhakrishnan V. S, Robin Marwal, Mahesh S Dhar, Kalaiarasan Ponnusamy, Meena Datta, Priyanka Singh, Uma Sharma, Hema Gogia, Preeti Madan, Sandhya Kabra, Sujeet K Singh, Partha Rakshit |
| EPI_ISL_2555672, EPI_ISL_2555696 | ICMR-National Institute of Virology | NCDC Delhi, Biotechnology Division | V.Vipat,Meena Datta,S.Jadhav,Priyanka Singh,U.Saha ,M.Das,Uma Sharma,G.Divekar,T.Sanjeevi,Manoj K Singh,S Jadhav,Radhakrishnan V. S,A Walimbe ,Robin Marwal,V Malik,Mahesh S Dhar,K.Iyengar,Kalaiarasan Ponnusamy,K P.Shinde,Hemlata Lall,K.Korabu,Z. Sayyed,Hema Gogia, S.Shelkande,R.Verma,Preeti Madan,S.Bhorekar,B.Apoorva,Sandhya Kabra,A.Titkare,H.Kengale,Sujeet K Singh,V.Autude,A.Awhale,Partha Rakshit,M L Choudhary |
| EPI_ISL_2555701, EPI_ISL_2555714, EPI_ISL_2555716, EPI_ISL_2555720, EPI_ISL_2555724, EPI_ISL_2555732 | ICMR-National Institute of Virology | NCDC Delhi, Biotechnology Division | Meena Datta,V.Vipat,S.Jadhav,Priyanka Singh,U.Saha ,M.Das,Uma Sharma,G.Divekar,T.Sanjeevi,Manoj K Singh,S Jadhav,Radhakrishnan V. S,A Walimbe ,Robin Marwal,V Malik,Mahesh S Dhar,K.Iyengar,Kalaiarasan Ponnusamy,K P.Shinde,Hemlata Lall,K.Korabu,Z. Sayyed,Hema Gogia, S.Shelkande,R.Verma,Preeti Madan,S.Bhorekar,B.Apoorva,Sandhya Kabra,A.Titkare,H.Kengale,Sujeet K Singh,V.Autude,A.Awhale,M L Choudhary,Partha Rakshit |
| EPI_ISL_2555733, EPI_ISL_2555735, EPI_ISL_2555736, EPI_ISL_2555741, EPI_ISL_2555742, EPI_ISL_2555743, EPI_ISL_2555744, EPI_ISL_2555745, EPI_ISL_2555747, EPI_ISL_2555750, EPI_ISL_2555751, EPI_ISL_2555756, EPI_ISL_2555757, EPI_ISL_2555758, EPI_ISL_2555760, EPI_ISL_2555761 | | | |
| see above | National Centre for Disease Control (NCDC) Biotechnology Division, Delhi | NCDC Delhi, Biotechnology Division INSACOG | Manoj K Singh, Radhakrishnan V. S, Robin Marwal, Mahesh S Dhar, Kalaiarasan Ponnusamy, Meena Datta, Priyanka Singh, Uma Sharma, Hemlata Lall, Hema Gogia, Preeti Madan, Sandhya Kabra, Sujeet K Singh, Partha Rakshit |
| EPI_ISL_2555762, EPI_ISL_2555763, EPI_ISL_2555764, EPI_ISL_2555765, EPI_ISL_2555766, EPI_ISL_2555767, EPI_ISL_2555769, EPI_ISL_2555773, EPI_ISL_2555774, EPI_ISL_2555777, EPI_ISL_2555778, EPI_ISL_2555779, EPI_ISL_2555780, EPI_ISL_2555782, EPI_ISL_2555783, EPI_ISL_2555784, EPI_ISL_2555787, EPI_ISL_2555789, EPI_ISL_2555790, EPI_ISL_2555792, EPI_ISL_2555793, EPI_ISL_2555796, EPI_ISL_2555807, EPI_ISL_2555808, EPI_ISL_2555811, EPI_ISL_2555812, EPI_ISL_2555814, EPI_ISL_2555815, EPI_ISL_2555816, EPI_ISL_2555817, EPI_ISL_2555818, EPI_ISL_2555819, EPI_ISL_2555820 | | | |
| see above | National Centre for Disease Control (NCDC) Biotechnology Division, Delhi | NCDC Delhi, Biotechnology Division INSACOG | Radhakrishnan V. S, Robin Marwal, Mahesh S Dhar, Kalaiarasan Ponnusamy, Meena Datta, Priyanka Singh, Uma Sharma, Hemlata Lall, Manoj K Singh, Hema Gogia, Preeti Madan, Sandhya Kabra, Sujeet K Singh, Partha Rakshit |
| EPI_ISL_2555821, EPI_ISL_2555822, EPI_ISL_2555827, EPI_ISL_2555829, EPI_ISL_2555830, EPI_ISL_2555833, EPI_ISL_2555843, EPI_ISL_2555849, EPI_ISL_2555851, EPI_ISL_2555852, EPI_ISL_2555853, EPI_ISL_2555857, EPI_ISL_2555859, EPI_ISL_2555863, EPI_ISL_2555865, EPI_ISL_2555866, EPI_ISL_2555867, EPI_ISL_2555868, EPI_ISL_2555869, EPI_ISL_2555870, EPI_ISL_2555872 | | | |
| see above | National Centre for Disease Control (NCDC) Biotechnology Division, Delhi | NCDC Delhi, Biotechnology Division INSACOG | Robin Marwal, Mahesh S Dhar, Kalaiarasan Ponnusamy, Meena Datta, Priyanka Singh, Uma Sharma, Hemlata Lall, Manoj K Singh, Radhakrishnan V. S, Hema Gogia, Preeti Madan, Sandhya Kabra, Sujeet K Singh, Partha Rakshit |
| EPI_ISL_2555874, EPI_ISL_2555877, EPI_ISL_2555878, EPI_ISL_2555879, EPI_ISL_2555880, EPI_ISL_2555881, EPI_ISL_2555883, EPI_ISL_2555885, EPI_ISL_2555886, EPI_ISL_2555889, EPI_ISL_2555894, EPI_ISL_2555896, EPI_ISL_2555899, EPI_ISL_2555900, EPI_ISL_2555909, EPI_ISL_2555911, EPI_ISL_2555929, EPI_ISL_2555931, EPI_ISL_2555941, EPI_ISL_2555943, EPI_ISL_2555947, EPI_ISL_2555950, EPI_ISL_2555951 | | | |
| see above | National Centre for Disease Control (NCDC) Biotechnology Division, Delhi | NCDC Delhi, Biotechnology Division INSACOG | Mahesh S Dhar, Kalaiarasan Ponnusamy, Meena Datta, Priyanka Singh, Uma Sharma, Hemlata Lall, Manoj K Singh, Radhakrishnan V. S, Robin Marwal, Hema Gogia, Preeti Madan, Sandhya Kabra, Sujeet K Singh, Partha Rakshit |
| EPI_ISL_2555966, EPI_ISL_2555968, EPI_ISL_2555969, EPI_ISL_2555971, EPI_ISL_2555972, EPI_ISL_2555976, EPI_ISL_2555978, EPI_ISL_2555979, EPI_ISL_2555980, EPI_ISL_2555984 | National Centre for Disease Control (NCDC) Biotechnology Division, Delhi | NCDC Delhi, Biotechnology Division INSACOG | Kalaiarasan Ponnusamy, Meena Datta, Priyanka Singh, Uma Sharma, Hemlata Lall, Manoj K Singh, Radhakrishnan V. S, Robin Marwal, Mahesh S Dhar, Hema Gogia, Preeti Madan, Sandhya Kabra, Sujeet K Singh, Partha Rakshit |
| EPI_ISL_2555989, EPI_ISL_2555995, EPI_ISL_2555997, EPI_ISL_2555998, EPI_ISL_2555999, EPI_ISL_2556006, EPI_ISL_2556012, EPI_ISL_2556014, EPI_ISL_2556022, EPI_ISL_2556027, EPI_ISL_2556030, EPI_ISL_2556034, EPI_ISL_2556039, EPI_ISL_2556040, EPI_ISL_2556043, EPI_ISL_2556044 | | | |
| see above | National Centre for Disease Control (NCDC) Biotechnology Division, Delhi | NCDC Delhi, Biotechnology Division INSACOG | Meena Datta, Priyanka Singh, Uma Sharma, Hemlata Lall, Manoj K Singh, Radhakrishnan V. S, Robin Marwal, Mahesh S Dhar, Kalaiarasan Ponnusamy, Hema Gogia, Preeti Madan, Sandhya Kabra, Sujeet K Singh, Partha Rakshit |
| EPI_ISL_2556047, EPI_ISL_2556050, EPI_ISL_2556052, EPI_ISL_2556053, EPI_ISL_2556055, EPI_ISL_2556060, EPI_ISL_2556061, EPI_ISL_2556062, EPI_ISL_2556066, EPI_ISL_2556067, EPI_ISL_2556071, EPI_ISL_2556073, EPI_ISL_2556074, EPI_ISL_2556075, EPI_ISL_2556076, EPI_ISL_2556077, EPI_ISL_2556078, EPI_ISL_2556079, EPI_ISL_2556080, EPI_ISL_2556081, EPI_ISL_2556085, EPI_ISL_2556086, EPI_ISL_2556088, EPI_ISL_2556090, EPI_ISL_2556091, EPI_ISL_2556092, EPI_ISL_2556093, EPI_ISL_2556094, EPI_ISL_2556095, EPI_ISL_2556099, EPI_ISL_2556100, EPI_ISL_2556101, EPI_ISL_2556102, EPI_ISL_2556104, EPI_ISL_2556105, EPI_ISL_2556106, EPI_ISL_2556108, EPI_ISL_2556109, EPI_ISL_2556110 | | | |
| see above | National Centre for Disease Control (NCDC) Biotechnology Division, Delhi | NCDC Delhi, Biotechnology Division INSACOG | Priyanka Singh, Uma Sharma, Hemlata Lall, Manoj K Singh, Radhakrishnan V. S, Robin Marwal, Mahesh S Dhar, Kalaiarasan Ponnusamy, Meena Datta, Hema Gogia, Preeti Madan, Sandhya Kabra, Sujeet K Singh, Partha Rakshit |
| EPI_ISL_2556187 | National Centre for Disease Control (NCDC) Biotechnology Division, Delhi | NCDC Delhi, Biotechnology Division INSACOG | Robin Marwal, Mahesh S Dhar, Kalaiarasan Ponnusamy, Meena Datta, Priyanka Singh, Uma Sharma, Hemlata Lall, Manoj K Singh, Radhakrishnan V. S, Hema Gogia, Preeti Madan, Sandhya Kabra, Sujeet K Singh, Partha Rakshit |
| EPI_ISL_2556200 | National Centre for Disease Control (NCDC) Biotechnology Division, Delhi | NCDC Delhi, Biotechnology Division INSACOG | Mahesh S Dhar, Kalaiarasan Ponnusamy, Meena Datta, Priyanka Singh, Uma Sharma, Hemlata Lall, Manoj K Singh, Radhakrishnan V. S, Robin Marwal, Hema Gogia, Preeti Madan, Sandhya Kabra, Sujeet K Singh, Partha Rakshit |
| EPI_ISL_2556447, EPI_ISL_2556448, EPI_ISL_2556524, EPI_ISL_2556527, EPI_ISL_2556566, EPI_ISL_2556582, EPI_ISL_2556583, EPI_ISL_2556605, EPI_ISL_2556857 | National Centre for Disease Control (NCDC) Biotechnology Division, Delhi | NCDC Delhi, Biotechnology Division INSACOG | Kalaiarasan Ponnusamy, Meena Datta, Priyanka Singh, Uma Sharma, Hemlata Lall, Manoj K Singh, Radhakrishnan V. S, Robin Marwal, Mahesh S Dhar, Hema Gogia, Preeti Madan, Sandhya Kabra, Sujeet K Singh, Partha Rakshit |
| EPI_ISL_2556971, EPI_ISL_2556976, EPI_ISL_2556979, EPI_ISL_2556999, EPI_ISL_2557000, EPI_ISL_2557002, EPI_ISL_2557021, EPI_ISL_2557032 | National Centre for Disease Control (NCDC) Biotechnology Division, Delhi | NCDC Delhi, Biotechnology Division INSACOG | Meena Datta, Priyanka Singh, Uma Sharma, Hemlata Lall, Manoj K Singh, Radhakrishnan V. S, Robin Marwal, Mahesh S Dhar, Kalaiarasan Ponnusamy, Hema Gogia, Preeti Madan, Sandhya Kabra, Sujeet K Singh, Partha Rakshit |
| EPI_ISL_2557049 | Washington State Department of Health Public Health Laboratories | Washington State Department of Health Public Health Laboratories | Drew MacKellar, Philip Dykema, Denny Russell, Joenice Gonzalez, Hannah Gray, Geoff Melly, Vanessa De Los Santos, Darren Lucas, JohnAric Peterson, Avi Singh, Rebecca Cao |
| EPI_ISL_2557085, EPI_ISL_2557094, EPI_ISL_2557096, EPI_ISL_2557104, EPI_ISL_2557110, EPI_ISL_2557113, EPI_ISL_2557122, EPI_ISL_2557123, EPI_ISL_2557130, EPI_ISL_2557132, EPI_ISL_2557144 | | | |
| see above | NJDOH, Public Health and Environmental Laboratories | NJ_PHEL | Lindsey Bodnar, Shiv K. Verma, Ryan Pachucki, Chelsea San Filippo, Dana Woell, Allison Roder, Byeong Jeong |
| EPI_ISL_2557176, EPI_ISL_2557177 | Fimlab Laboratoriot Oy Tampere | Expert Microbiology, National Institute for Health and Welfare | Soile Blomqvist, Jani Halkilahti, Kirsi Liitsola, Haider al-Hello, Päivi Laurila, Erika Lindh, Teemu Smura, Ravi Kant, Phuoc Truong, Olli Vapalahti, Sari Hannula, Pekka Ellonen, Niina Ikonen, Carita Savolainen-Kopra |
| EPI_ISL_2557178 | SYNLAB Suomi | Expert Microbiology, National Institute for Health and Welfare | Soile Blomqvist, Jani Halkilahti, Kirsi Liitsola, Haider al-Hello, Päivi Laurila, Erika Lindh, Teemu Smura, Ravi Kant, Phuoc Truong, Olli Vapalahti, Sari Hannula, Pekka Ellonen, Niina Ikonen, Carita Savolainen-Kopra |
| EPI_ISL_2557179, EPI_ISL_2557180, EPI_ISL_2557182, EPI_ISL_2557183, EPI_ISL_2557184, EPI_ISL_2557185, EPI_ISL_2557186 | Fimlab Laboratoriot Oy Tampere | Expert Microbiology, National Institute for Health and Welfare | Soile Blomqvist, Jani Halkilahti, Kirsi Liitsola, Haider al-Hello, Päivi Laurila, Erika Lindh, Teemu Smura, Ravi Kant, Phuoc Truong, Olli Vapalahti, Sari Hannula, Pekka Ellonen, Niina Ikonen, Carita Savolainen-Kopra |
| EPI_ISL_2557188, EPI_ISL_2557189, EPI_ISL_2557190, EPI_ISL_2557191, EPI_ISL_2557192 | SYNLAB Suomi | Expert Microbiology, National Institute for Health and Welfare | Soile Blomqvist, Jani Halkilahti, Kirsi Liitsola, Haider al-Hello, Päivi Laurila, Erika Lindh, Teemu Smura, Ravi Kant, Phuoc Truong, Olli Vapalahti, Sari Hannula, Pekka Ellonen, Niina Ikonen, Carita Savolainen-Kopra |
| EPI_ISL_2557193, EPI_ISL_2557194, EPI_ISL_2557195, EPI_ISL_2557196, EPI_ISL_2557197, EPI_ISL_2557198, EPI_ISL_2557199, EPI_ISL_2557200, EPI_ISL_2557201, EPI_ISL_2557202, EPI_ISL_2557203, EPI_ISL_2557204, EPI_ISL_2557205, EPI_ISL_2557206, EPI_ISL_2557207, EPI_ISL_2557208, EPI_ISL_2557209, EPI_ISL_2557210 | | | |
| see above | Fimlab Laboratoriot Oy Tampere | Expert Microbiology, National Institute for Health and Welfare | Soile Blomqvist, Jani Halkilahti, Kirsi Liitsola, Haider al-Hello, Päivi Laurila, Erika Lindh, Teemu Smura, Ravi Kant, Phuoc Truong, Olli Vapalahti, Sari Hannula, Pekka Ellonen, Niina Ikonen, Carita Savolainen-Kopra |
| EPI_ISL_2557213, EPI_ISL_2557216, EPI_ISL_2557220, EPI_ISL_2557234, EPI_ISL_2557240 | IN State Department of Health Laboratory Services | IN State Department of Health Laboratory Services | Cassandra Campion, Jamie Yeadon, Brian Pope, Lixia Liu, Kyle Brownlee, Melissa Hindenlang, Mark Glazier |
| EPI_ISL_2557455, EPI_ISL_2557467, EPI_ISL_2557507, | Infinity Biologix | Centers for Disease Control and Prevention Division of Viral Diseases, Pathogen Discovery | Dakota Howard, Dhwani Batra, Peter W. Cook, Kara Moser, Adrian Paskey, Jason Caravas, Benjamin Rambo-Martin, Shatavia Morrison, Christopher Gulvick, Scott Sammons, Yvette Unoarumhi, Darlene Wagner, Matthew Schmerer, Christian Bixby, Yihe Wang, Jonathan Schultz, Chirayu Goswami, Russ Hager, Robin Grimwood, Clinton R. Paden, Duncan MacCannell |

| | | | |
|---|---|---|---|
| EPI_ISL_2557509, EPI_ISL_2557616 | | | |
| EPI_ISL_2557816, EPI_ISL_2557817, EPI_ISL_2557818, EPI_ISL_2557819, EPI_ISL_2557820, EPI_ISL_2557821, EPI_ISL_2557822, EPI_ISL_2557823, EPI_ISL_2557824 | Michigan Department of Health and Human Services, Bureau of Laboratories | Michigan Department of Health and Human Services, Bureau of Laboratories | Blankenship HM, Riner D, Soehnlen MK |
| EPI_ISL_2558004, EPI_ISL_2558005, EPI_ISL_2558006, EPI_ISL_2558007, EPI_ISL_2558042 | Servicio de Microbiologia Hospital Ramon y Cajal | Servicio de Microbiologia Hospital Ramon y Cajal | Ponce M, Galan JC, Martinez L. Abreu M, y Gonzalez-Alba JM |
| EPI_ISL_2558052 | Philippine Red Cross - Clark Molecular Laboratory | Philippine Genome Center | Francis A. Tablizo, Kenneth M. Kim, Maria Sofia L. Yangzon, Elcid Aaron R. Pangilinan, Renato Jacinto Q. Mantaring, Benedict A. Maralit, Marc Edsel C. Ayes, Eva Maria Cutiongco-de la Paz, Alethea R. de Guzman, Jan Michael C. Yap, Jo-Hannah S. Llames, Sheila Mae M. Araiza, Kris P. Punayan, Irish Coleen A. Asin, Candice Francheska B. Tambaoan, Asia Louisa U. Chong, Karol Sophia Agape R. Padilla, Rianna Patricia S. Cruz, Carlo M. Lapid, El King D. Morado, Joshua Gregor A. Dizon, Razel Nikka M. Hao, Arianne A. Zamora, Devon Ray Pacial, Juan Antonio R. Magalang, Marissa Alejandria, Celia Carlos, Anna Ong-Lim, Edsel Maurice Salvaña, John Q. Wong, Jaime C. Montoya, Maria Rosario Singh-Vergeire and Cynthia P. Saloma |
| EPI_ISL_2558053 | First Aide Diagnostic Center | Philippine Genome Center | Francis A. Tablizo, Kenneth M. Kim, Maria Sofia L. Yangzon, Elcid Aaron R. Pangilinan, Renato Jacinto Q. Mantaring, Benedict A. Maralit, Marc Edsel C. Ayes, Eva Maria Cutiongco-de la Paz, Alethea R. de Guzman, Jan Michael C. Yap, Jo-Hannah S. Llames, Sheila Mae M. Araiza, Kris P. Punayan, Irish Coleen A. Asin, Candice Francheska B. Tambaoan, Asia Louisa U. Chong, Karol Sophia Agape R. Padilla, Rianna Patricia S. Cruz, Carlo M. Lapid, El King D. Morado, Joshua Gregor A. Dizon, Razel Nikka M. Hao, Arianne A. Zamora, Devon Ray Pacial, Juan Antonio R. Magalang, Marissa Alejandria, Celia Carlos, Anna Ong-Lim, Edsel Maurice Salvaña, John Q. Wong, Jaime C. Montoya, Maria Rosario Singh-Vergeire and Cynthia P. Saloma |
| EPI_ISL_2558054 | Bureau Of Quarantine (BOQ) | Philippine Genome Center | Francis A. Tablizo, Kenneth M. Kim, Maria Sofia L. Yangzon, Elcid Aaron R. Pangilinan, Renato Jacinto Q. Mantaring, Benedict A. Maralit, Marc Edsel C. Ayes, Eva Maria Cutiongco-de la Paz, Alethea R. de Guzman, Jan Michael C. Yap, Jo-Hannah S. Llames, Sheila Mae M. Araiza, Kris P. Punayan, Irish Coleen A. Asin, Candice Francheska B. Tambaoan, Asia Louisa U. Chong, Karol Sophia Agape R. Padilla, Rianna Patricia S. Cruz, Carlo M. Lapid, El King D. Morado, Joshua Gregor A. Dizon, Razel Nikka M. Hao, Arianne A. Zamora, Devon Ray Pacial, Juan Antonio R. Magalang, Marissa Alejandria, Celia Carlos, Anna Ong-Lim, Edsel Maurice Salvaña, John Q. Wong, Jaime C. Montoya, Maria Rosario Singh-Vergeire and Cynthia P. Saloma |
| EPI_ISL_2558055 | University of the Philippines National Institutes of Health (UP NIH) | Philippine Genome Center | Francis A. Tablizo, Kenneth M. Kim, Maria Sofia L. Yangzon, Elcid Aaron R. Pangilinan, Renato Jacinto Q. Mantaring, Benedict A. Maralit, Marc Edsel C. Ayes, Eva Maria Cutiongco-de la Paz, Alethea R. de Guzman, Jan Michael C. Yap, Jo-Hannah S. Llames, Sheila Mae M. Araiza, Kris P. Punayan, Irish Coleen A. Asin, Candice Francheska B. Tambaoan, Asia Louisa U. Chong, Karol Sophia Agape R. Padilla, Rianna Patricia S. Cruz, Carlo M. Lapid, El King D. Morado, Joshua Gregor A. Dizon, Razel Nikka M. Hao, Arianne A. Zamora, Devon Ray Pacial, Juan Antonio R. Magalang, Marissa Alejandria, Celia Carlos, Anna Ong-Lim, Edsel Maurice Salvaña, John Q. Wong, Jaime C. Montoya, Maria Rosario Singh-Vergeire and Cynthia P. Saloma |
| EPI_ISL_2558056 | Bureau Of Quarantine (BOQ) | Philippine Genome Center | Francis A. Tablizo, Kenneth M. Kim, Maria Sofia L. Yangzon, Elcid Aaron R. Pangilinan, Renato Jacinto Q. Mantaring, Benedict A. Maralit, Marc Edsel C. Ayes, Eva Maria Cutiongco-de la Paz, Alethea R. de Guzman, Jan Michael C. Yap, Jo-Hannah S. Llames, Sheila Mae M. Araiza, Kris P. Punayan, Irish Coleen A. Asin, Candice Francheska B. Tambaoan, Asia Louisa U. Chong, Karol Sophia Agape R. Padilla, Rianna Patricia S. Cruz, Carlo M. Lapid, El King D. Morado, Joshua Gregor A. Dizon, Razel Nikka M. Hao, Arianne A. Zamora, Devon Ray Pacial, Juan Antonio R. Magalang, Marissa Alejandria, Celia Carlos, Anna Ong-Lim, Edsel Maurice Salvaña, John Q. Wong, Jaime C. Montoya, Maria Rosario Singh-Vergeire and Cynthia P. Saloma |
| EPI_ISL_2558057 | First Aide Diagnostic Center | Philippine Genome Center | Francis A. Tablizo, Kenneth M. Kim, Maria Sofia L. Yangzon, Elcid Aaron R. Pangilinan, Renato Jacinto Q. Mantaring, Benedict A. Maralit, Marc Edsel C. Ayes, Eva Maria Cutiongco-de la Paz, Alethea R. de Guzman, Jan Michael C. Yap, Jo-Hannah S. Llames, Sheila Mae M. Araiza, Kris P. Punayan, Irish Coleen A. Asin, Candice Francheska B. Tambaoan, Asia Louisa U. Chong, Karol Sophia Agape R. Padilla, Rianna Patricia S. Cruz, Carlo M. Lapid, El King D. Morado, Joshua Gregor A. Dizon, Razel Nikka M. Hao, Arianne A. Zamora, Devon Ray Pacial, Juan Antonio R. Magalang, Marissa Alejandria, Celia Carlos, Anna Ong-Lim, Edsel Maurice Salvaña, John Q. Wong, Jaime C. Montoya, Maria Rosario Singh-Vergeire and Cynthia P. Saloma |
| EPI_ISL_2558058, EPI_ISL_2558059 | Bureau Of Quarantine (BOQ) | Philippine Genome Center | Francis A. Tablizo, Kenneth M. Kim, Maria Sofia L. Yangzon, Elcid Aaron R. Pangilinan, Renato Jacinto Q. Mantaring, Benedict A. Maralit, Marc Edsel C. Ayes, Eva Maria Cutiongco-de la Paz, Alethea R. de Guzman, Jan Michael C. Yap, Jo-Hannah S. Llames, Sheila Mae M. Araiza, Kris P. Punayan, Irish Coleen A. Asin, Candice Francheska B. Tambaoan, Asia Louisa U. Chong, Karol Sophia Agape R. Padilla, Rianna Patricia S. Cruz, Carlo M. Lapid, El King D. Morado, Joshua Gregor A. Dizon, Razel Nikka M. Hao, Arianne A. Zamora, Devon Ray Pacial, Juan Antonio R. Magalang, Marissa Alejandria, Celia Carlos, Anna Ong-Lim, Edsel Maurice Salvaña, John Q. Wong, Jaime C. Montoya, Maria Rosario Singh-Vergeire and Cynthia P. Saloma |
| EPI_ISL_2558060 | University of the Philippines National Institutes of Health (UP NIH) | Philippine Genome Center | Francis A. Tablizo, Kenneth M. Kim, Maria Sofia L. Yangzon, Elcid Aaron R. Pangilinan, Renato Jacinto Q. Mantaring, Benedict A. Maralit, Marc Edsel C. Ayes, Eva Maria Cutiongco-de la Paz, Alethea R. de Guzman, Jan Michael C. Yap, Jo-Hannah S. Llames, Sheila Mae M. Araiza, Kris P. Punayan, Irish Coleen A. Asin, Candice Francheska B. Tambaoan, Asia Louisa U. Chong, Karol Sophia Agape R. Padilla, Rianna Patricia S. Cruz, Carlo M. Lapid, El King D. Morado, Joshua Gregor A. Dizon, Razel Nikka M. Hao, Arianne A. Zamora, Devon Ray Pacial, Juan Antonio R. Magalang, Marissa Alejandria, Celia Carlos, Anna Ong-Lim, Edsel Maurice Salvaña, John Q. Wong, Jaime C. Montoya, Maria Rosario Singh-Vergeire and Cynthia P. Saloma |
| EPI_ISL_2558061, EPI_ISL_2558062, EPI_ISL_2558063 | Bureau Of Quarantine (BOQ) | Philippine Genome Center | Francis A. Tablizo, Kenneth M. Kim, Maria Sofia L. Yangzon, Elcid Aaron R. Pangilinan, Renato Jacinto Q. Mantaring, Benedict A. Maralit, Marc Edsel C. Ayes, Eva Maria Cutiongco-de la Paz, Alethea R. de Guzman, Jan Michael C. Yap, Jo-Hannah S. Llames, Sheila Mae M. Araiza, Kris P. Punayan, Irish Coleen A. Asin, Candice Francheska B. Tambaoan, Asia Louisa U. Chong, Karol Sophia Agape R. Padilla, Rianna Patricia S. Cruz, Carlo M. Lapid, El King D. Morado, Joshua Gregor A. Dizon, Razel Nikka M. Hao, Arianne A. Zamora, Devon Ray Pacial, Juan Antonio R. Magalang, Marissa Alejandria, Celia Carlos, Anna Ong-Lim, Edsel Maurice Salvaña, John Q. Wong, Jaime C. Montoya, Maria Rosario Singh-Vergeire and Cynthia P. Saloma |
| EPI_ISL_2558139, EPI_ISL_2558195, EPI_ISL_2558352, EPI_ISL_2558380, EPI_ISL_2558570, EPI_ISL_2558595, EPI_ISL_2558639, EPI_ISL_2558644, EPI_ISL_2558667 | Wisconsin State Laboratory of Hygiene Communicable Disease Division | Wisconsin State Laboratory of Hygiene Communicable Disease Division | Abigail C. Shockey, Alicia J. Mooney, Erika M. Hanson, Tonya Danz, Richard Griesser, Sara Wagner, Kelsey R. Florek |
| EPI_ISL_2558695, EPI_ISL_2558696, EPI_ISL_2558698 | Wichita State University Molecular Diagnotic Lab | Kansas Health and Environmental Lab | Katherine Wiggins, Mike Grose, Jonathan Barnell, Ben Olsen, and Phil Adam |
| EPI_ISL_2558699, EPI_ISL_2558700, EPI_ISL_2558703, EPI_ISL_2558704 | Kansas Health and Environmental Lab | Kansas Health and Environmental Lab | Katherine Wiggins, Mike Grose, Jonathan Barnell, Ben Olsen, and Phil Adam |
| EPI_ISL_2558705, EPI_ISL_2558706, EPI_ISL_2558707, EPI_ISL_2558708, EPI_ISL_2558709 | Balai Besar Penelitian dan Pengembangan dan Reservoir Penyakit, Salatiga | Genomik Solidaritas Indonesia Laboratorium | Vania Gavrila Wikasa, Carissa Sintca Wijaya, Dhahlia Agustina Cahyono, Alfin Mohammad Abdillah, Normastuti Adhini Tantyo, Yudistira Aditya Rahmat, Joko Waluyo, Annisa Muthiah Sukirman, Gracia Felias Enos Korompis, Pratiwi Sudarmono, Budi Wiweko, Louisa Markus, Anuraj Shankar, Meutia Ayuputeri Kumaheri, Anindya Pradipta Susanto, Ariel Pradipta |
| EPI_ISL_2558710 | Balai Besar Penelitian dan Pengembangan dan Reservoir Penyakit, Salatiga | Genomik Solidaritas Indonesia Laboratorium | Carissa Sintca Wijaya, Vania Gavrila Wikasa, Dhahlia Agustina Cahyono, Alfin Mohammad Abdillah, Normastuti Adhini Tantyo, Yudistira Aditya Rahmat, Joko Waluyo, Annisa Muthiah Sukirman, Gracia Felias Enos Korompis, Pratiwi Sudarmono, Budi Wiweko, Louisa Markus, Anuraj Shankar, Meutia Ayuputeri Kumaheri, Anindya Pradipta Susanto, Ariel Pradipta |
| EPI_ISL_2558711 | Balai Besar Penelitian dan Pengembangan dan Reservoir Penyakit, Salatiga | Genomik Solidaritas Indonesia Laboratorium | Vania Gavrila Wikasa, Carissa Sintca Wijaya, Dhahlia Agustina Cahyono, Alfin Mohammad Abdillah, Normastuti Adhini Tantyo, Yudistira Aditya Rahmat, Joko Waluyo, Annisa Muthiah Sukirman, Gracia Felias Enos Korompis, Pratiwi Sudarmono, Budi Wiweko, Louisa Markus, Anuraj Shankar, Meutia Ayuputeri Kumaheri, Anindya Pradipta Susanto, Ariel Pradipta |
| EPI_ISL_2558712 | Balai Besar Penelitian dan Pengembangan dan Reservoir Penyakit, Salatiga | Genomik Solidaritas Indonesia Laboratorium | Carissa Sintca Wijaya, Vania Gavrila Wikasa, Dhahlia Agustina Cahyono, Alfin Mohammad Abdillah, Normastuti Adhini Tantyo, Yudistira Aditya Rahmat, Joko Waluyo, Annisa Muthiah Sukirman, Gracia Felias Enos Korompis, Pratiwi Sudarmono, Budi Wiweko, Louisa Markus, Anuraj Shankar, Meutia Ayuputeri Kumaheri, Anindya Pradipta Susanto, Ariel Pradipta |
| EPI_ISL_2558714, EPI_ISL_2558716, EPI_ISL_2558717 | Balai Besar Penelitian dan Pengembangan dan Reservoir Penyakit, Salatiga | Genomik Solidaritas Indonesia Laboratorium | Vania Gavrila Wikasa, Carissa Sintca Wijaya, Dhahlia Agustina Cahyono, Alfin Mohammad Abdillah, Normastuti Adhini Tantyo, Yudistira Aditya Rahmat, Joko Waluyo, Annisa Muthiah Sukirman, Gracia Felias Enos Korompis, Pratiwi Sudarmono, Budi Wiweko, Louisa Markus, Anuraj Shankar, Meutia Ayuputeri Kumaheri, Anindya Pradipta Susanto, Ariel Pradipta |
| EPI_ISL_2558721, EPI_ISL_2558722, EPI_ISL_2558723, EPI_ISL_2558724, EPI_ISL_2558725 | Balai Besar Penelitian dan Pengembangan dan Reservoir Penyakit, Salatiga | Genomik Solidaritas Indonesia Laboratorium | Carissa Sintca Wijaya, Vania Gavrila Wikasa, Dhahlia Agustina Cahyono, Alfin Mohammad Abdillah, Normastuti Adhini Tantyo, Yudistira Aditya Rahmat, Joko Waluyo, Annisa Muthiah Sukirman, Gracia Felias Enos Korompis, Pratiwi Sudarmono, Budi Wiweko, Louisa Markus, Anuraj Shankar, Meutia Ayuputeri Kumaheri, Anindya Pradipta Susanto, Ariel Pradipta |
| EPI_ISL_2558726, EPI_ISL_2558727, EPI_ISL_2558728, EPI_ISL_2558729, EPI_ISL_2558730, EPI_ISL_2558731 | Balai Besar Penelitian dan Pengembangan dan Reservoir Penyakit, Salatiga | Genomik Solidaritas Indonesia Laboratorium | Vania Gavrila Wikasa, Carissa Sintca Wijaya, Dhahlia Agustina Cahyono, Alfin Mohammad Abdillah, Normastuti Adhini Tantyo, Yudistira Aditya Rahmat, Joko Waluyo, Annisa Muthiah Sukirman, Gracia Felias Enos Korompis, Pratiwi Sudarmono, Budi Wiweko, Louisa Markus, Anuraj Shankar, Meutia Ayuputeri Kumaheri, Anindya Pradipta Susanto, Ariel Pradipta |
| EPI_ISL_2558736 | Balai Besar Penelitian dan Pengembangan dan Reservoir Penyakit, Salatiga | Genomik Solidaritas Indonesia Laboratorium | Carissa Sintca Wijaya, Vania Gavrila Wikasa, Dhahlia Agustina Cahyono, Alfin Mohammad Abdillah, Normastuti Adhini Tantyo, Yudistira Aditya Rahmat, Joko Waluyo, Annisa Muthiah Sukirman, Gracia Felias Enos Korompis, Pratiwi Sudarmono, Budi Wiweko, Louisa Markus, Anuraj Shankar, Meutia Ayuputeri Kumaheri, Anindya Pradipta Susanto, Ariel Pradipta |
| EPI_ISL_2558737 | Balai Besar Penelitian dan Pengembangan dan Reservoir Penyakit, Salatiga | Genomik Solidaritas Indonesia Laboratorium | Vania Gavrila Wikasa, Carissa Sintca Wijaya, Dhahlia Agustina Cahyono, Alfin Mohammad Abdillah, Normastuti Adhini Tantyo, Yudistira Aditya Rahmat, Joko Waluyo, Annisa Muthiah Sukirman, Gracia Felias Enos Korompis, Pratiwi Sudarmono, Budi Wiweko, Louisa Markus, Anuraj Shankar, Meutia Ayuputeri Kumaheri, Anindya Pradipta Susanto, Ariel Pradipta |
| EPI_ISL_2559110, EPI_ISL_2559115, EPI_ISL_2559122 | Wyoming Public Health Laboratory | Wyoming Public Health Laboratory | Jim Mildenberger, Taylor Fearing, Channing Weber, Ashley Norberg, Chayse Rowley, Marley Goetz, Brian Dominguez, Elliot Thomasson, Sam Britz, Cari Sloma, Robert Petit, and Rob Christensen |
| EPI_ISL_2561913 | Willis-Knighton Medical Center Hospital Laboratory | LSUHS Emerging Viral Threat Laboratory | Lorie M. Atkins, April N. Johnson, Gregory L. Ware, Alexander Mijalis, Jeremy P. Kamil, Jennifer L. Carroll, Maarten Van Diest, Rona S. Scott, Andrew D. Yurochko, Christopher G. Kevil, John A. Vanchiere, Joseph A. Bocchini |
| EPI_ISL_2561963 | LSUHS Emerging Viral Threat Laboratory | LSUHS Emerging Viral Threat Laboratory | Maarten Van Diest, Rona S. Scott, Gregory L. Ware, Alexander Mijalis,Jeremy P. Kamil,Jennifer L. Carroll, Andrew D. Yurochko, Christopher G. Kevil, John A. Vanchiere |
| EPI_ISL_2561981, EPI_ISL_2561982, EPI_ISL_2561983, EPI_ISL_2561993, EPI_ISL_2561997, EPI_ISL_2561998, EPI_ISL_2561999, EPI_ISL_2562003, EPI_ISL_2562006, EPI_ISL_2562010, EPI_ISL_2562012 | | | |
| see above | cerballiance-IDF | Cerba lab | Emmanuel Lecorche, Sabine Trombert-Paolantoni, Aude Lessenne, Kader Merah, Thierry Collin, Laura Verdurme, Patrice Herisson, Stéphanie Haïm-Boukobza, Bénédicte Roquebert |
| EPI_ISL_2562068, EPI_ISL_2562069, | National Institute of Public Health | State Veterinary Institute Prague | Nagy,A;Jirincova,H;Suri,T;Trnka,D;Vecerova,J |

| | | | |
|---|---|---|---|
| EPI_ISL_2562070, EPI_ISL_2562071, EPI_ISL_2562072, EPI_ISL_2562073, EPI_ISL_2562075, EPI_ISL_2562079 | | | |
| EPI_ISL_2562085 | GMERS Government Medical College (GMC), Gotri | Gujarat Biotechnology Research Centre | Ramesh Pandit,Janvi Raval,Zarna Patel,Nitin Savaliya,Dinesh Kumar,Twinkle Soni,Sonal Sharma ,Zuber Saiyed,Bithika Duttaroy,Umang Mishra,Nitesh Shah,Chaitanya Joshi,Madhvi Joshi |
| EPI_ISL_2562086 | GMERS Government Medical College (GMC), Gotri | Gujarat Biotechnology Research Centre | Janvi Raval,Zarna Patel,Nitin Savaliya,Dinesh Kumar,Twinkle Soni,Sonal Sharma ,Zuber Saiyed,Ramesh Pandit,Bithika Duttaroy,Umang Mishra,Nitesh Shah,Chaitanya Joshi,Madhvi Joshi |
| EPI_ISL_2562087 | GMERS Government Medical College (GMC), Gotri | Gujarat Biotechnology Research Centre | Zarna Patel,Nitin Savaliya,Dinesh Kumar,Twinkle Soni,Sonal Sharma ,Zuber Saiyed,Ramesh Pandit,Janvi Raval,Bithika Duttaroy,Umang Mishra,Nitesh Shah,Chaitanya Joshi,Madhvi Joshi |
| EPI_ISL_2562088 | GMERS Government Medical College (GMC), Gotri | Gujarat Biotechnology Research Centre | Nitin Savaliya,Dinesh Kumar,Twinkle Soni,Sonal Sharma ,Zuber Saiyed,Ramesh Pandit,Janvi Raval,Zarna Patel,Bithika Duttaroy,Umang Mishra,Nitesh Shah,Chaitanya Joshi,Madhvi Joshi |
| EPI_ISL_2562089 | GMERS Government Medical College (GMC), Gotri | Gujarat Biotechnology Research Centre | Dinesh Kumar,Twinkle Soni,Sonal Sharma ,Zuber Saiyed,Ramesh Pandit,Janvi Raval,Zarna Patel,Nitin Savaliya,Bithika Duttaroy,Umang Mishra,Nitesh Shah,Chaitanya Joshi,Madhvi Joshi |
| EPI_ISL_2562091 | GMERS Government Medical College (GMC), Gotri | Gujarat Biotechnology Research Centre | Sonal Sharma ,Zuber Saiyed,Ramesh Pandit,Janvi Raval,Zarna Patel,Nitin Savaliya,Dinesh Kumar,Twinkle Soni,Bithika Duttaroy,Umang Mishra,Nitesh Shah,Chaitanya Joshi,Madhvi Joshi |
| EPI_ISL_2562092 | GMERS Government Medical College (GMC), Gotri | Gujarat Biotechnology Research Centre | Zuber Saiyed,Ramesh Pandit,Janvi Raval,Zarna Patel,Nitin Savaliya,Dinesh Kumar,Twinkle Soni,Sonal Sharma ,Bithika Duttaroy,Umang Mishra,Nitesh Shah,Chaitanya Joshi,Madhvi Joshi |
| EPI_ISL_2562093 | GMERS Government Medical College (GMC), Gotri | Gujarat Biotechnology Research Centre | Ramesh Pandit,Janvi Raval,Zarna Patel,Nitin Savaliya,Dinesh Kumar,Twinkle Soni,Sonal Sharma ,Zuber Saiyed,Bithika Duttaroy,Umang Mishra,Nitesh Shah,Chaitanya Joshi,Madhvi Joshi |
| EPI_ISL_2562094 | GMERS Government Medical College (GMC), Gotri | Gujarat Biotechnology Research Centre | Janvi Raval,Zarna Patel,Nitin Savaliya,Dinesh Kumar,Twinkle Soni,Sonal Sharma ,Zuber Saiyed,Ramesh Pandit,Bithika Duttaroy,Umang Mishra,Nitesh Shah,Chaitanya Joshi,Madhvi Joshi |
| EPI_ISL_2562095 | GMERS Government Medical College (GMC), Gotri | Gujarat Biotechnology Research Centre | Zarna Patel,Nitin Savaliya,Dinesh Kumar,Twinkle Soni,Sonal Sharma ,Zuber Saiyed,Ramesh Pandit,Janvi Raval,Bithika Duttaroy,Umang Mishra,Nitesh Shah,Chaitanya Joshi,Madhvi Joshi |
| EPI_ISL_2562096 | GMERS Government Medical College (GMC), Gotri | Gujarat Biotechnology Research Centre | Nitin Savaliya,Dinesh Kumar,Twinkle Soni,Sonal Sharma ,Zuber Saiyed,Ramesh Pandit,Janvi Raval,Zarna Patel,Bithika Duttaroy,Umang Mishra,Nitesh Shah,Chaitanya Joshi,Madhvi Joshi |
| EPI_ISL_2562097 | GMERS Government Medical College (GMC), Gotri | Gujarat Biotechnology Research Centre | Dinesh Kumar,Twinkle Soni,Sonal Sharma ,Zuber Saiyed,Ramesh Pandit,Janvi Raval,Zarna Patel,Nitin Savaliya,Bithika Duttaroy,Umang Mishra,Nitesh Shah,Chaitanya Joshi,Madhvi Joshi |
| EPI_ISL_2562098 | GMERS Government Medical College (GMC), Gotri | Gujarat Biotechnology Research Centre | Twinkle Soni,Sonal Sharma ,Zuber Saiyed,Ramesh Pandit,Janvi Raval,Zarna Patel,Nitin Savaliya,Dinesh Kumar,Bithika Duttaroy,Umang Mishra,Nitesh Shah,Chaitanya Joshi,Madhvi Joshi |
| EPI_ISL_2562099 | GMERS Government Medical College (GMC), Gotri | Gujarat Biotechnology Research Centre | Sonal Sharma ,Zuber Saiyed,Ramesh Pandit,Janvi Raval,Zarna Patel,Nitin Savaliya,Dinesh Kumar,Twinkle Soni,Bithika Duttaroy,Umang Mishra,Nitesh Shah,Chaitanya Joshi,Madhvi Joshi |
| EPI_ISL_2562100 | GMERS Government Medical College (GMC), Gotri | Gujarat Biotechnology Research Centre | Zuber Saiyed,Ramesh Pandit,Janvi Raval,Zarna Patel,Nitin Savaliya,Dinesh Kumar,Twinkle Soni,Sonal Sharma ,Bithika Duttaroy,Umang Mishra,Nitesh Shah,Chaitanya Joshi,Madhvi Joshi |
| EPI_ISL_2562101 | GMERS Government Medical College (GMC), Gotri | Gujarat Biotechnology Research Centre | Ramesh Pandit,Janvi Raval,Zarna Patel,Nitin Savaliya,Dinesh Kumar,Twinkle Soni,Sonal Sharma ,Zuber Saiyed,Bithika Duttaroy,Umang Mishra,Nitesh Shah,Chaitanya Joshi,Madhvi Joshi |
| EPI_ISL_2562102 | GMERS Government Medical College (GMC), Gotri | Gujarat Biotechnology Research Centre | Janvi Raval,Zarna Patel,Nitin Savaliya,Dinesh Kumar,Twinkle Soni,Sonal Sharma ,Zuber Saiyed,Ramesh Pandit,Bithika Duttaroy,Umang Mishra,Nitesh Shah,Chaitanya Joshi,Madhvi Joshi |
| EPI_ISL_2562103 | GMERS Government Medical College (GMC), Gotri | Gujarat Biotechnology Research Centre | Zarna Patel,Nitin Savaliya,Dinesh Kumar,Twinkle Soni,Sonal Sharma ,Zuber Saiyed,Ramesh Pandit,Janvi Raval,Bithika Duttaroy,Umang Mishra,Nitesh Shah,Chaitanya Joshi,Madhvi Joshi |
| EPI_ISL_2562104 | GMERS Government Medical College (GMC), Gotri | Gujarat Biotechnology Research Centre | Nitin Savaliya,Dinesh Kumar,Twinkle Soni,Sonal Sharma ,Zuber Saiyed,Ramesh Pandit,Janvi Raval,Zarna Patel,Bithika Duttaroy,Umang Mishra,Nitesh Shah,Chaitanya Joshi,Madhvi Joshi |
| EPI_ISL_2562105 | GMERS Government Medical College (GMC), Gotri | Gujarat Biotechnology Research Centre | Dinesh Kumar,Twinkle Soni,Sonal Sharma ,Zuber Saiyed,Ramesh Pandit,Janvi Raval,Zarna Patel,Nitin Savaliya,Bithika Duttaroy,Umang Mishra,Nitesh Shah,Chaitanya Joshi,Madhvi Joshi |
| EPI_ISL_2562106 | GMERS Government Medical College (GMC), Gotri | Gujarat Biotechnology Research Centre | Twinkle Soni,Sonal Sharma ,Zuber Saiyed,Ramesh Pandit,Janvi Raval,Zarna Patel,Nitin Savaliya,Dinesh Kumar,Bithika Duttaroy,Umang Mishra,Nitesh Shah,Chaitanya Joshi,Madhvi Joshi |
| EPI_ISL_2562107 | GMERS Government Medical College (GMC), Gotri | Gujarat Biotechnology Research Centre | Sonal Sharma ,Zuber Saiyed,Ramesh Pandit,Janvi Raval,Zarna Patel,Nitin Savaliya,Dinesh Kumar,Twinkle Soni,Bithika Duttaroy,Umang Mishra,Nitesh Shah,Chaitanya Joshi,Madhvi Joshi |
| EPI_ISL_2562108 | GMERS Government Medical College (GMC), Gotri | Gujarat Biotechnology Research Centre | Zuber Saiyed,Ramesh Pandit,Janvi Raval,Zarna Patel,Nitin Savaliya,Dinesh Kumar,Twinkle Soni,Sonal Sharma ,Bithika Duttaroy,Umang Mishra,Nitesh Shah,Chaitanya Joshi,Madhvi Joshi |
| EPI_ISL_2562109 | GMERS Government Medical College (GMC), Gotri | Gujarat Biotechnology Research Centre | Ramesh Pandit,Janvi Raval,Zarna Patel,Nitin Savaliya,Dinesh Kumar,Twinkle Soni,Sonal Sharma ,Zuber Saiyed,Bithika Duttaroy,Umang Mishra,Nitesh Shah,Chaitanya Joshi,Madhvi Joshi |
| EPI_ISL_2562110 | GMERS Government Medical College (GMC), Gotri | Gujarat Biotechnology Research Centre | Janvi Raval,Zarna Patel,Nitin Savaliya,Dinesh Kumar,Twinkle Soni,Sonal Sharma ,Zuber Saiyed,Ramesh Pandit,Bithika Duttaroy,Umang Mishra,Nitesh Shah,Chaitanya Joshi,Madhvi Joshi |
| EPI_ISL_2562111 | GMERS Government Medical College (GMC), Gotri | Gujarat Biotechnology Research Centre | Zarna Patel,Nitin Savaliya,Dinesh Kumar,Twinkle Soni,Sonal Sharma ,Zuber Saiyed,Ramesh Pandit,Janvi Raval,Bithika Duttaroy,Umang Mishra,Nitesh Shah,Chaitanya Joshi,Madhvi Joshi |
| EPI_ISL_2562268 | Laboratorio de Referencial Nacional de Virus Respiratorios | Laboratorio de Referencial Nacional de Virus Respiratorios | Carlos Padilla Rojas, Veronica Hurtado Vela, Orson Mestanza Millones, Iris Silva Molina, Luis Barcena Flores, Sandra Morales Ruiz, Steve Acedo Lazo, Omar Caceres Rey, Marco Galarza Perez, Henri Bailon Calderon, Nancy Rojas Serrano, Priscila Lope Pari, Lely Solari Zerpa, Joseph Huayra Niquen |
| EPI_ISL_2562474, EPI_ISL_2562478, EPI_ISL_2562479, EPI_ISL_2562480, EPI_ISL_2562483 | Creighton University Medical Center | Creighton COVID Consortium | Holly Stessman, Michael Belshan, Richard Goering, Cynthia Watson, Jake Siedlik |
| EPI_ISL_2562777, EPI_ISL_2562780, EPI_ISL_2562783, EPI_ISL_2562784, EPI_ISL_2562785, EPI_ISL_2562786, EPI_ISL_2562787, EPI_ISL_2562788, EPI_ISL_2562790, EPI_ISL_2562791 | TriCore Reference Laboratories | Center for Global Health, University of New Mexico Health Sciences Center | Daryl Domman, Kurt Schwalm, Valerie Morley, Cecilia Thompson, Kendra Pesko, Karissa Culbreath, Darrell Dinwiddie |
| EPI_ISL_2565774, EPI_ISL_2565777, EPI_ISL_2565778, EPI_ISL_2565779 | TXDSHS | TXDSHS | Rashmi Tuladhar, Bonnie Oh, Jenny Zhang, Maliha Rahman, Mayela Pedrueza, Anita Pokharel, Karen Bobier, Lorraine Rodriguez, Myong Koag, Chun Wang, Rachel Lee, Grace Kubin |
| EPI_ISL_2565876 | Minnesota Department of Health, Public Health Laboratory | Minnesota Department of Health, Public Health Laboratory | Alexandra Lorentz, Jacob Garfin, Matt Plumb, and Xiong Wang |
| EPI_ISL_2565914 | North Dakota Department of Health, Public Health Laboratory | North Dakota Department of Health, Public Health Laboratory | Lisa Wingerter |
| EPI_ISL_2565983 | Genome Information Research Center, Research Institute for Microbial Diseases, Osaka University | Genome Information Research Center, Research Institute for Microbial Diseases, Osaka University | Daisuke Motooka, Kentaro Tanaka, Hiroya Oki, and Shota Nakamura |
| EPI_ISL_2566121 | Nevada State Public Health Laboratory | Nevada State Public Health Laboratory | Andrew Gorzalski, Mark Pandori |
| EPI_ISL_2566200 | Botswana Harvard HIV Reference Laboratory | Botswana Harvard HIV Reference Laboratory | Sikhulile Moyo, Wonderful T. Choga, Pamela Smith-Lawrence, Kgomotso Moruisi, Dorcas Maruapula, Keoratile Ntshambiwa, Thongbotho Mphoyakgosi, Boitumelo Zuze, Botshelo Radibe, Legodile Kooepile, Ontlametse T. Bareng, Roger Shapiro, Shahin Lockman, Joseph Makhema, Madisa Mine, Mosepele Mosepele, Simani Gaseitsiwe |
| EPI_ISL_2566245 | Wyoming Public Health Laboratory | Wyoming Public Health Laboratory | Jim Mildenberger, Taylor Fearing, Channing Weber, Ashley Norberg, Chayse Rowley, Marley Goetz, Brian Dominguez, Elliot Thomasson, Sam Britz, Cari Sloma, Robert Petit, and Rob Christensen |
| EPI_ISL_2566444, EPI_ISL_2566445 | Singapore General Hospital | Department of Microbiology | Nurdyana Abdul Rahman, Kenneth Xin Long Chan, Chayaporn Suphavilai, Kun Lee Lim, Sui Sin Goh, Kian Sing Chan, Lynette Oon, Kern Rei Chng, James Sim Heng Chiak, Niranjan Nagarajan, Karrie Ko |
| EPI_ISL_2566447, EPI_ISL_2566459, EPI_ISL_2566460 | National Institute of Infectious Diseases-Prof. Dr. Matei Bals Molecular Diagnostics Laboratory | National Institute of Infectious Diseases-Prof. Dr. Matei Bals Molecular Diagnostics Laboratory | Corina Casangiu, Leontina Banica, Marius Surleac, Petre Milu, Robert Hohan, Simona Paraschiv, Dan Otelea |
| EPI_ISL_2566565, EPI_ISL_2566579 | MKK Mühlenkreiskliniken, Johannes Wesling Klinikum Minden | Hannover Medical School, Institute of Virology | Lars Steinbrück |
| EPI_ISL_2566597, EPI_ISL_2566598 | Hospital Universitari Joan XXIII | Hospital Universitari Joan XXIII | Gemma Recio, Cristina Gutiérrez, Clara Benavent, Carla Martín, Margarida Terrón, Ester Picó, Natalia Bastón |
| EPI_ISL_2566615, EPI_ISL_2566631, EPI_ISL_2566634, EPI_ISL_2566635, EPI_ISL_2566659, EPI_ISL_2566660, EPI_ISL_2566661 | Shamir Medical Center (Asaf Harofe) | Shamir Medical Center (Asaf Harofe) | Nir Rainy,Chen Weiner,Reut Sorek Abramovich,Yevgeni Yegorov,Anna Vishnevsky,Patricia Benveniste-Lekovitz,Abu Hamad Ramzia,Adina Bar Chaim |
| EPI_ISL_2566734 | WSSE Gorzów Wlkp. | 1. National Institute of Public Health - National Institute of Hygiene; 2. Eurofins Genomics Europe Sequencing GmbH | Wołkowicz Tomasz, Zacharczuk Katarzyna, Sadkowska-Todys Małgorzata, Gierczyński Rafał, Eurofins Genomics Europe Sequencing Team, ECDC COVID-19 WGS support team |
| EPI_ISL_2566903 | Laboratorium Mikrobiologiczne RSS w Grudziądzu | 1. National Institute of Public Health - National Institute of Hygiene; 2. Eurofins Genomics Europe Sequencing GmbH | Wołkowicz Tomasz, Zacharczuk Katarzyna, Sadkowska-Todys Małgorzata, Gierczyński Rafał, Eurofins Genomics Europe Sequencing Team, ECDC COVID-19 WGS support team |
| EPI_ISL_2566972 | Dept. of Microbiology and Infection Control, Akershus University Hospital HF | Dept. of Microbiology and Infection Control, Akershus University Hospital HF | Hege Vangstein Aamot, Alexander Hesselberg Løvestad |
| EPI_ISL_2567003 | AZ Klina | AZ Klina | Carl Vael - Lynsey Berckmans |
| EPI_ISL_2567007, EPI_ISL_2567021, EPI_ISL_2567032, EPI_ISL_2567040, EPI_ISL_2567042, EPI_ISL_2567050, | Shamir Medical Center (Asaf Harofe) | Shamir Medical Center (Asaf Harofe) | Nir Rainy,Chen Weiner,Reut Sorek Abramovich,Yevgeni Yegorov,Anna Vishnevsky,Patricia Benveniste-Lekovitz,Abu Hamad Ramzia,Adina Bar Chaim |

| | | | |
|---|---|---|---|
| EPI_ISL_2567071 | | | |
| EPI_ISL_2567385, EPI_ISL_2567390 | LBM UNIBIO VALENTIN | CNR Virus des Infections Respiratoires - France SUD | Antonin Bal, Gregory Destras, Gwendolyne Burfin, Hadrien Regue, Quentin Semanas, Martine Valette, Bruno Lina, Laurence Josset |
| EPI_ISL_2567439, EPI_ISL_2567441, EPI_ISL_2567444, EPI_ISL_2567452 | MIRIALIS CLUSES BECHET | CNR Virus des Infections Respiratoires - France SUD | Antonin Bal, Gregory Destras, Gwendolyne Burfin, Hadrien Regue, Quentin Semanas, Martine Valette, Bruno Lina, Laurence Josset |
| EPI_ISL_2567490, EPI_ISL_2567491, EPI_ISL_2567492, EPI_ISL_2567493, EPI_ISL_2567494, EPI_ISL_2567495 | LABORATOIRE FORTE BIO ST PAUL LES DAX | CNR Virus des Infections Respiratoires - France SUD | Antonin Bal, Gregory Destras, Gwendolyne Burfin, Hadrien Regue, Quentin Semanas, Martine Valette, Bruno Lina, Laurence Josset |
| EPI_ISL_2567524, EPI_ISL_2567535 | CNR Virus des Infections Respiratoires - France SUD | CNR Virus des Infections Respiratoires - France SUD | Antonin Bal, Gregory Destras, Gwendolyne Burfin, Hadrien Regue, Quentin Semanas, Martine Valette, Bruno Lina, Laurence Josset |
| EPI_ISL_2567635, EPI_ISL_2567636, EPI_ISL_2567637, EPI_ISL_2567638, EPI_ISL_2567639, EPI_ISL_2567640, EPI_ISL_2567641, EPI_ISL_2567642, EPI_ISL_2567643, EPI_ISL_2567644, EPI_ISL_2567645, EPI_ISL_2567646, EPI_ISL_2567647, EPI_ISL_2567648, EPI_ISL_2567649, EPI_ISL_2567650, EPI_ISL_2567651, EPI_ISL_2567652, EPI_ISL_2567653, EPI_ISL_2567654, EPI_ISL_2567655, EPI_ISL_2567656, EPI_ISL_2567657, EPI_ISL_2567658, EPI_ISL_2567659, EPI_ISL_2567661, EPI_ISL_2567662, EPI_ISL_2567663, EPI_ISL_2567664, EPI_ISL_2567665, EPI_ISL_2567666, EPI_ISL_2567667, EPI_ISL_2567669, EPI_ISL_2567670, EPI_ISL_2567671, EPI_ISL_2567672, EPI_ISL_2567673, EPI_ISL_2567674, EPI_ISL_2567675, EPI_ISL_2567677, EPI_ISL_2567678, EPI_ISL_2567679, EPI_ISL_2567680, EPI_ISL_2567681, EPI_ISL_2567682, EPI_ISL_2567683, EPI_ISL_2567684, EPI_ISL_2567685, EPI_ISL_2567686, EPI_ISL_2567687, EPI_ISL_2567688, EPI_ISL_2567689, EPI_ISL_2567690, EPI_ISL_2567691, EPI_ISL_2567692, EPI_ISL_2567693, EPI_ISL_2567694, EPI_ISL_2567695, EPI_ISL_2567698, EPI_ISL_2567699, EPI_ISL_2567700, EPI_ISL_2567701, EPI_ISL_2567702, EPI_ISL_2567703, EPI_ISL_2567704, EPI_ISL_2567706, EPI_ISL_2567707, EPI_ISL_2567708, EPI_ISL_2567709, EPI_ISL_2567710, EPI_ISL_2567711, EPI_ISL_2567712, EPI_ISL_2567713, EPI_ISL_2567714, EPI_ISL_2567715, EPI_ISL_2567716, EPI_ISL_2567717, EPI_ISL_2567719, EPI_ISL_2567720, EPI_ISL_2567721, EPI_ISL_2567722, EPI_ISL_2567723, EPI_ISL_2567724, EPI_ISL_2567725, EPI_ISL_2567727, EPI_ISL_2567728, EPI_ISL_2567729, EPI_ISL_2567730, EPI_ISL_2567731, EPI_ISL_2567732, EPI_ISL_2567733, EPI_ISL_2567734, EPI_ISL_2567735, EPI_ISL_2567736, EPI_ISL_2567737, EPI_ISL_2567738, EPI_ISL_2567739, EPI_ISL_2567740, EPI_ISL_2567741, EPI_ISL_2567742, EPI_ISL_2567743, EPI_ISL_2567744, EPI_ISL_2567746, EPI_ISL_2567747, EPI_ISL_2567748, EPI_ISL_2567749, EPI_ISL_2567751, EPI_ISL_2567752, EPI_ISL_2567753, EPI_ISL_2567754, EPI_ISL_2567755, EPI_ISL_2567756, EPI_ISL_2567757, EPI_ISL_2567758, EPI_ISL_2567759, EPI_ISL_2567760, EPI_ISL_2567761, EPI_ISL_2567762, EPI_ISL_2567763, EPI_ISL_2567764, EPI_ISL_2567765, EPI_ISL_2567766, EPI_ISL_2567767, EPI_ISL_2567768, EPI_ISL_2567769, EPI_ISL_2567770, EPI_ISL_2567771, EPI_ISL_2567772, EPI_ISL_2567773, EPI_ISL_2567774, EPI_ISL_2567775, EPI_ISL_2567776, EPI_ISL_2567778, EPI_ISL_2567779, EPI_ISL_2567780, EPI_ISL_2567781, EPI_ISL_2567782, EPI_ISL_2567783, EPI_ISL_2567784, EPI_ISL_2567785, EPI_ISL_2567786, EPI_ISL_2567788, EPI_ISL_2567790, EPI_ISL_2567791, EPI_ISL_2567792, EPI_ISL_2567793, EPI_ISL_2567794, EPI_ISL_2567795, EPI_ISL_2567796, EPI_ISL_2567797, EPI_ISL_2567798, EPI_ISL_2567799, EPI_ISL_2567801, EPI_ISL_2567802, EPI_ISL_2567803, EPI_ISL_2567804, EPI_ISL_2567805, EPI_ISL_2567806, EPI_ISL_2567807, EPI_ISL_2567808, EPI_ISL_2567809, EPI_ISL_2567810, EPI_ISL_2567811, EPI_ISL_2567812, EPI_ISL_2567813, EPI_ISL_2567814, EPI_ISL_2567815, EPI_ISL_2567816, EPI_ISL_2567817, EPI_ISL_2567818, EPI_ISL_2567819, EPI_ISL_2567820, EPI_ISL_2567821, EPI_ISL_2567822, EPI_ISL_2567823, EPI_ISL_2567824, EPI_ISL_2567825, EPI_ISL_2567826, EPI_ISL_2567827, EPI_ISL_2567828, EPI_ISL_2567829, EPI_ISL_2567831, EPI_ISL_2567832, EPI_ISL_2567833, EPI_ISL_2567834, EPI_ISL_2567836, EPI_ISL_2567837, EPI_ISL_2567838, EPI_ISL_2567839, EPI_ISL_2567840, EPI_ISL_2567841, EPI_ISL_2567842, EPI_ISL_2567843, EPI_ISL_2567844, EPI_ISL_2567845, EPI_ISL_2567846, EPI_ISL_2567847, EPI_ISL_2567848, EPI_ISL_2567849, EPI_ISL_2567850, EPI_ISL_2567851, EPI_ISL_2567852, EPI_ISL_2567853, EPI_ISL_2567854, EPI_ISL_2567855, EPI_ISL_2567856, EPI_ISL_2567857, EPI_ISL_2567858, EPI_ISL_2567859, EPI_ISL_2567860, EPI_ISL_2567861, EPI_ISL_2567862, EPI_ISL_2567863, EPI_ISL_2567864, EPI_ISL_2567865, EPI_ISL_2567866, EPI_ISL_2567867, EPI_ISL_2567868, EPI_ISL_2567869, EPI_ISL_2567870, EPI_ISL_2567871, EPI_ISL_2567872, EPI_ISL_2567874, EPI_ISL_2567875, EPI_ISL_2567877, EPI_ISL_2567878, EPI_ISL_2567879, EPI_ISL_2567880, EPI_ISL_2567881, EPI_ISL_2567882, EPI_ISL_2567883, EPI_ISL_2567884, EPI_ISL_2567885, EPI_ISL_2567886, EPI_ISL_2567888, EPI_ISL_2567889, EPI_ISL_2567890, EPI_ISL_2567891, EPI_ISL_2567892, EPI_ISL_2567894, EPI_ISL_2567895, EPI_ISL_2567896, EPI_ISL_2567897, EPI_ISL_2567898, EPI_ISL_2567899, EPI_ISL_2567900, EPI_ISL_2567901, EPI_ISL_2567902, EPI_ISL_2567903, EPI_ISL_2567904, EPI_ISL_2567905, EPI_ISL_2567906, EPI_ISL_2567907, EPI_ISL_2567908, EPI_ISL_2567909, EPI_ISL_2567910, EPI_ISL_2567911, EPI_ISL_2567913, EPI_ISL_2567914, EPI_ISL_2567915, EPI_ISL_2567916, EPI_ISL_2567917, EPI_ISL_2567918, EPI_ISL_2567919, EPI_ISL_2567920, EPI_ISL_2567921, EPI_ISL_2567922, EPI_ISL_2567923, EPI_ISL_2567928, EPI_ISL_2567929, EPI_ISL_2567930, EPI_ISL_2567931, EPI_ISL_2567932, EPI_ISL_2567933, EPI_ISL_2567934, EPI_ISL_2567935, EPI_ISL_2567936, EPI_ISL_2567937, EPI_ISL_2567938, EPI_ISL_2567939, EPI_ISL_2567940, EPI_ISL_2567941, EPI_ISL_2567942, EPI_ISL_2567943, EPI_ISL_2567944, EPI_ISL_2567945, EPI_ISL_2567946 | | | |
| see above | Lighthouse Lab in Alderley Park | Wellcome Sanger Institute for the COVID-19 Genomics UK (COG-UK) Consortium | Jacquelyn Wynn, Mairead Hyland, The Lighthouse Lab in Alderley Park and Alex Alderton, Roberto Amato, Jeffrey Barrett, Sonia Goncalves, Ewan Harrison, David K. Jackson, Ian Johnston, Dominic Kwiatkowski, Cordelia Langford, John Sillitoe on behalf of the Wellcome Sanger Institute COVID-19 Surveillance Team |
| EPI_ISL_2567947 | Lighthouse Lab in Glasgow | Wellcome Sanger Institute for the COVID-19 Genomics UK (COG-UK) Consortium | Harper VanSteenhouse, Yumi Kasai, David Gray, Carol Clugston, Anna Dominiczak and Alex Alderton, Roberto Amato, Jeffrey Barrett, Sonia Goncalves, Ewan Harrison, David K. Jackson, Ian Johnston, Dominic Kwiatkowski, Cordelia Langford, John Sillitoe on behalf of the Wellcome Sanger Institute COVID-19 Surveillance Team |
| EPI_ISL_2567948, EPI_ISL_2567949 | Lighthouse Lab in Alderley Park | Wellcome Sanger Institute for the COVID-19 Genomics UK (COG-UK) Consortium | Jacquelyn Wynn, Mairead Hyland, The Lighthouse Lab in Alderley Park and Alex Alderton, Roberto Amato, Jeffrey Barrett, Sonia Goncalves, Ewan Harrison, David K. Jackson, Ian Johnston, Dominic Kwiatkowski, Cordelia Langford, John Sillitoe on behalf of the Wellcome Sanger Institute COVID-19 Surveillance Team |
| EPI_ISL_2567950 | Lighthouse Lab in Glasgow | Wellcome Sanger Institute for the COVID-19 Genomics UK (COG-UK) Consortium | Harper VanSteenhouse, Yumi Kasai, David Gray, Carol Clugston, Anna Dominiczak and Alex Alderton, Roberto Amato, Jeffrey Barrett, Sonia Goncalves, Ewan Harrison, David K. Jackson, Ian Johnston, Dominic Kwiatkowski, Cordelia Langford, John Sillitoe on behalf of the Wellcome Sanger Institute COVID-19 Surveillance Team |
| EPI_ISL_2567951 | Lighthouse Lab in Alderley Park | Wellcome Sanger Institute for the COVID-19 Genomics UK (COG-UK) Consortium | Jacquelyn Wynn, Mairead Hyland, The Lighthouse Lab in Alderley Park and Alex Alderton, Roberto Amato, Jeffrey Barrett, Sonia Goncalves, Ewan Harrison, David K. Jackson, Ian Johnston, Dominic Kwiatkowski, Cordelia Langford, John Sillitoe on behalf of the Wellcome Sanger Institute COVID-19 Surveillance Team |
| EPI_ISL_2567952 | Lighthouse Lab in Glasgow | Wellcome Sanger Institute for the COVID-19 Genomics UK (COG-UK) Consortium | Harper VanSteenhouse, Yumi Kasai, David Gray, Carol Clugston, Anna Dominiczak and Alex Alderton, Roberto Amato, Jeffrey Barrett, Sonia Goncalves, Ewan Harrison, David K. Jackson, Ian Johnston, Dominic Kwiatkowski, Cordelia Langford, John Sillitoe on behalf of the Wellcome Sanger Institute COVID-19 Surveillance Team |
| EPI_ISL_2567953, EPI_ISL_2567954 | Lighthouse Lab in Alderley Park | Wellcome Sanger Institute for the COVID-19 Genomics UK (COG-UK) Consortium | Jacquelyn Wynn, Mairead Hyland, The Lighthouse Lab in Alderley Park and Alex Alderton, Roberto Amato, Jeffrey Barrett, Sonia Goncalves, Ewan Harrison, David K. Jackson, Ian Johnston, Dominic Kwiatkowski, Cordelia Langford, John Sillitoe on behalf of the Wellcome Sanger Institute COVID-19 Surveillance Team |
| EPI_ISL_2567955 | Lighthouse Lab in Glasgow | Wellcome Sanger Institute for the COVID-19 Genomics UK (COG-UK) Consortium | Harper VanSteenhouse, Yumi Kasai, David Gray, Carol Clugston, Anna Dominiczak and Alex Alderton, Roberto Amato, Jeffrey Barrett, Sonia Goncalves, Ewan Harrison, David K. Jackson, Ian Johnston, Dominic Kwiatkowski, Cordelia Langford, John Sillitoe on behalf of the Wellcome Sanger Institute COVID-19 Surveillance Team |
| EPI_ISL_2567956, EPI_ISL_2567957 | Lighthouse Lab in Alderley Park | Wellcome Sanger Institute for the COVID-19 Genomics UK (COG-UK) Consortium | Jacquelyn Wynn, Mairead Hyland, The Lighthouse Lab in Alderley Park and Alex Alderton, Roberto Amato, Jeffrey Barrett, Sonia Goncalves, Ewan Harrison, David K. Jackson, Ian Johnston, Dominic Kwiatkowski, Cordelia Langford, John Sillitoe on behalf of the Wellcome Sanger Institute COVID-19 Surveillance Team |
| EPI_ISL_2567958 | Lighthouse Lab in Glasgow | Wellcome Sanger Institute for the COVID-19 Genomics UK (COG-UK) Consortium | Harper VanSteenhouse, Yumi Kasai, David Gray, Carol Clugston, Anna Dominiczak and Alex Alderton, Roberto Amato, Jeffrey Barrett, Sonia Goncalves, Ewan Harrison, David K. Jackson, Ian Johnston, Dominic Kwiatkowski, Cordelia Langford, John Sillitoe on behalf of the Wellcome Sanger Institute COVID-19 Surveillance Team |
| EPI_ISL_2567959 | Lighthouse Lab in Alderley Park | Wellcome Sanger Institute for the COVID-19 Genomics UK (COG-UK) Consortium | Jacquelyn Wynn, Mairead Hyland, The Lighthouse Lab in Alderley Park and Alex Alderton, Roberto Amato, Jeffrey Barrett, Sonia Goncalves, Ewan Harrison, David K. Jackson, Ian Johnston, Dominic Kwiatkowski, Cordelia Langford, John Sillitoe on behalf of the Wellcome Sanger Institute COVID-19 Surveillance Team |
| EPI_ISL_2567960 | Lighthouse Lab in Glasgow | Wellcome Sanger Institute for the COVID-19 Genomics UK (COG-UK) Consortium | Harper VanSteenhouse, Yumi Kasai, David Gray, Carol Clugston, Anna Dominiczak and Alex Alderton, Roberto Amato, Jeffrey Barrett, Sonia Goncalves, Ewan Harrison, David K. Jackson, Ian Johnston, Dominic Kwiatkowski, Cordelia Langford, John Sillitoe on behalf of the Wellcome Sanger Institute COVID-19 Surveillance Team |
| EPI_ISL_2567961 | Lighthouse Lab in Alderley Park | Wellcome Sanger Institute for the COVID-19 Genomics UK (COG-UK) Consortium | Jacquelyn Wynn, Mairead Hyland, The Lighthouse Lab in Alderley Park and Alex Alderton, Roberto Amato, Jeffrey Barrett, Sonia Goncalves, Ewan Harrison, David K. Jackson, Ian Johnston, Dominic Kwiatkowski, Cordelia Langford, John Sillitoe on behalf of the Wellcome Sanger Institute COVID-19 Surveillance Team |
| EPI_ISL_2567962, EPI_ISL_2567963 | Lighthouse Lab in Glasgow | Wellcome Sanger Institute for the COVID-19 Genomics UK (COG-UK) Consortium | Harper VanSteenhouse, Yumi Kasai, David Gray, Carol Clugston, Anna Dominiczak and Alex Alderton, Roberto Amato, Jeffrey Barrett, Sonia Goncalves, Ewan Harrison, David K. Jackson, Ian Johnston, Dominic Kwiatkowski, Cordelia Langford, John Sillitoe on behalf of the Wellcome Sanger Institute COVID-19 Surveillance Team |
| EPI_ISL_2567964, EPI_ISL_2567965 | Lighthouse Lab in Alderley Park | Wellcome Sanger Institute for the COVID-19 Genomics UK (COG-UK) Consortium | Jacquelyn Wynn, Mairead Hyland, The Lighthouse Lab in Alderley Park and Alex Alderton, Roberto Amato, Jeffrey Barrett, Sonia Goncalves, Ewan Harrison, David K. Jackson, Ian Johnston, Dominic Kwiatkowski, Cordelia Langford, John Sillitoe on behalf of the Wellcome Sanger Institute COVID-19 Surveillance Team |
| EPI_ISL_2567966, EPI_ISL_2567967 | Lighthouse Lab in Glasgow | Wellcome Sanger Institute for the COVID-19 Genomics UK (COG-UK) Consortium | Harper VanSteenhouse, Yumi Kasai, David Gray, Carol Clugston, Anna Dominiczak and Alex Alderton, Roberto Amato, Jeffrey Barrett, Sonia Goncalves, Ewan Harrison, David K. Jackson, Ian Johnston, Dominic Kwiatkowski, Cordelia Langford, John Sillitoe on behalf of the Wellcome Sanger Institute COVID-19 Surveillance Team |
| EPI_ISL_2567968, EPI_ISL_2567969 | Lighthouse Lab in Alderley Park | Wellcome Sanger Institute for the COVID-19 Genomics UK (COG-UK) Consortium | Jacquelyn Wynn, Mairead Hyland, The Lighthouse Lab in Alderley Park and Alex Alderton, Roberto Amato, Jeffrey Barrett, Sonia Goncalves, Ewan Harrison, David K. Jackson, Ian Johnston, Dominic Kwiatkowski, Cordelia Langford, John Sillitoe on behalf of the Wellcome Sanger Institute COVID-19 Surveillance Team |
| EPI_ISL_2567970, EPI_ISL_2567971 | Lighthouse Lab in Glasgow | Wellcome Sanger Institute for the COVID-19 Genomics UK (COG-UK) Consortium | Harper VanSteenhouse, Yumi Kasai, David Gray, Carol Clugston, Anna Dominiczak and Alex Alderton, Roberto Amato, Jeffrey Barrett, Sonia Goncalves, Ewan Harrison, David K. Jackson, Ian Johnston, Dominic Kwiatkowski, Cordelia Langford, John Sillitoe on behalf of the Wellcome Sanger Institute COVID-19 Surveillance Team |
| EPI_ISL_2567972 | Lighthouse Lab in Alderley Park | Wellcome Sanger Institute for the COVID-19 Genomics UK (COG-UK) Consortium | Jacquelyn Wynn, Mairead Hyland, The Lighthouse Lab in Alderley Park and Alex Alderton, Roberto Amato, Jeffrey Barrett, Sonia Goncalves, Ewan Harrison, David K. Jackson, Ian Johnston, Dominic Kwiatkowski, Cordelia Langford, John Sillitoe on behalf of the Wellcome Sanger Institute COVID-19 Surveillance Team |
| EPI_ISL_2567973, EPI_ISL_2567974, EPI_ISL_2567975 | Lighthouse Lab in Glasgow | Wellcome Sanger Institute for the COVID-19 Genomics UK (COG-UK) Consortium | Harper VanSteenhouse, Yumi Kasai, David Gray, Carol Clugston, Anna Dominiczak and Alex Alderton, Roberto Amato, Jeffrey Barrett, Sonia Goncalves, Ewan Harrison, David K. Jackson, Ian Johnston, Dominic Kwiatkowski, Cordelia Langford, John Sillitoe on behalf of the Wellcome Sanger Institute COVID-19 Surveillance Team |
| EPI_ISL_2567976, EPI_ISL_2567978 | Lighthouse Lab in Alderley Park | Wellcome Sanger Institute for the COVID-19 Genomics UK (COG-UK) Consortium | Jacquelyn Wynn, Mairead Hyland, The Lighthouse Lab in Alderley Park and Alex Alderton, Roberto Amato, Jeffrey Barrett, Sonia Goncalves, Ewan Harrison, David K. Jackson, Ian Johnston, Dominic Kwiatkowski, Cordelia Langford, John Sillitoe on behalf of the Wellcome Sanger Institute COVID-19 Surveillance Team |
| EPI_ISL_2567979, EPI_ISL_2567980, EPI_ISL_2567981, EPI_ISL_2567982, EPI_ISL_2567983 | Lighthouse Lab in Glasgow | Wellcome Sanger Institute for the COVID-19 Genomics UK (COG-UK) Consortium | Harper VanSteenhouse, Yumi Kasai, David Gray, Carol Clugston, Anna Dominiczak and Alex Alderton, Roberto Amato, Jeffrey Barrett, Sonia Goncalves, Ewan Harrison, David K. Jackson, Ian Johnston, Dominic Kwiatkowski, Cordelia Langford, John Sillitoe on behalf of the Wellcome Sanger Institute COVID-19 Surveillance Team |
| EPI_ISL_2567984 | Lighthouse Lab in Alderley Park | Wellcome Sanger Institute for the COVID-19 Genomics UK (COG-UK) Consortium | Jacquelyn Wynn, Mairead Hyland, The Lighthouse Lab in Alderley Park and Alex Alderton, Roberto Amato, Jeffrey Barrett, Sonia Goncalves, Ewan Harrison, David K. Jackson, Ian Johnston, Dominic Kwiatkowski, Cordelia Langford, John Sillitoe on behalf of the Wellcome Sanger Institute COVID-19 Surveillance Team |
| EPI_ISL_2567985 | Lighthouse Lab in Glasgow | Wellcome Sanger Institute for the COVID-19 Genomics UK (COG-UK) Consortium | Harper VanSteenhouse, Yumi Kasai, David Gray, Carol Clugston, Anna Dominiczak and Alex Alderton, Roberto Amato, Jeffrey Barrett, Sonia Goncalves, Ewan Harrison, David K. Jackson, Ian Johnston, Dominic Kwiatkowski, Cordelia Langford, John Sillitoe on behalf of the Wellcome Sanger Institute COVID-19 Surveillance Team |
| EPI_ISL_2567986, EPI_ISL_2567987 | Lighthouse Lab in Alderley Park | Wellcome Sanger Institute for the COVID-19 Genomics UK (COG-UK) Consortium | Jacquelyn Wynn, Mairead Hyland, The Lighthouse Lab in Alderley Park and Alex Alderton, Roberto Amato, Jeffrey Barrett, Sonia Goncalves, Ewan Harrison, David K. Jackson, Ian Johnston, Dominic Kwiatkowski, Cordelia Langford, John Sillitoe on behalf of the Wellcome Sanger Institute COVID-19 Surveillance Team |
| EPI_ISL_2567988 | Lighthouse Lab in Glasgow | Wellcome Sanger Institute for the COVID-19 Genomics UK (COG-UK) Consortium | Harper VanSteenhouse, Yumi Kasai, David Gray, Carol Clugston, Anna Dominiczak and Alex Alderton, Roberto Amato, Jeffrey Barrett, Sonia Goncalves, Ewan Harrison, David K. Jackson, Ian Johnston, Dominic Kwiatkowski, Cordelia Langford, John Sillitoe on behalf of the Wellcome Sanger Institute COVID-19 Surveillance Team |
| EPI_ISL_2567989, EPI_ISL_2567990 | Lighthouse Lab in Alderley Park | Wellcome Sanger Institute for the COVID-19 Genomics UK (COG-UK) Consortium | Jacquelyn Wynn, Mairead Hyland, The Lighthouse Lab in Alderley Park and Alex Alderton, Roberto Amato, Jeffrey Barrett, Sonia Goncalves, Ewan Harrison, David K. Jackson, Ian Johnston, Dominic Kwiatkowski, Cordelia Langford, John Sillitoe on behalf of the Wellcome Sanger Institute COVID-19 Surveillance Team |
| EPI_ISL_2567991 | Lighthouse Lab in Glasgow | Wellcome Sanger Institute for the COVID-19 Genomics UK (COG-UK) Consortium | Harper VanSteenhouse, Yumi Kasai, David Gray, Carol Clugston, Anna Dominiczak and Alex Alderton, Roberto Amato, Jeffrey Barrett, Sonia Goncalves, Ewan Harrison, David K. Jackson, Ian Johnston, Dominic Kwiatkowski, Cordelia Langford, John Sillitoe on behalf of the Wellcome Sanger Institute COVID-19 Surveillance Team |
| EPI_ISL_2567992 | Lighthouse Lab in Alderley Park | Wellcome Sanger Institute for the COVID-19 Genomics UK (COG-UK) Consortium | Jacquelyn Wynn, Mairead Hyland, The Lighthouse Lab in Alderley Park and Alex Alderton, Roberto Amato, Jeffrey Barrett, Sonia Goncalves, Ewan Harrison, David K. Jackson, Ian Johnston, Dominic Kwiatkowski, Cordelia Langford, John Sillitoe on behalf of the Wellcome Sanger Institute COVID-19 Surveillance Team |
| EPI_ISL_2567993 | Lighthouse Lab in Glasgow | Wellcome Sanger Institute for the COVID-19 Genomics UK (COG-UK) Consortium | Harper VanSteenhouse, Yumi Kasai, David Gray, Carol Clugston, Anna Dominiczak and Alex Alderton, Roberto Amato, Jeffrey Barrett, Sonia Goncalves, Ewan Harrison, David K. Jackson, Ian Johnston, Dominic Kwiatkowski, Cordelia Langford, John Sillitoe on behalf of the Wellcome Sanger Institute COVID-19 Surveillance Team |
| EPI_ISL_2567994 | Lighthouse Lab in Alderley Park | Wellcome Sanger Institute for the COVID-19 Genomics UK (COG-UK) | Jacquelyn Wynn, Mairead Hyland, The Lighthouse Lab in Alderley Park and Alex Alderton, Roberto Amato, Jeffrey Barrett, Sonia Goncalves, Ewan Harrison, David K. Jackson, Ian Johnston, Dominic Kwiatkowski, Cordelia Langford, John Sillitoe on |

| | | | |
|---|---|---|---|
| | | Consortium | behalf of the Wellcome Sanger Institute COVID-19 Surveillance Team |
| EPI_ISL_2567995, EPI_ISL_2567996, EPI_ISL_2567997 | Lighthouse Lab in Glasgow | Wellcome Sanger Institute for the COVID-19 Genomics UK (COG-UK) Consortium | Harper VanSteenhouse, Yumi Kasai, David Gray, Carol Clugston, Anna Dominiczak and Alex Alderton, Roberto Amato, Jeffrey Barrett, Sonia Goncalves, Ewan Harrison, David K. Jackson, Ian Johnston, Dominic Kwiatkowski, Cordelia Langford, John Sillitoe on behalf of the Wellcome Sanger Institute COVID-19 Surveillance Team |
| EPI_ISL_2567998 | Lighthouse Lab in Alderley Park | Wellcome Sanger Institute for the COVID-19 Genomics UK (COG-UK) Consortium | Jacquelyn Wynn, Mairead Hyland, The Lighthouse Lab in Alderley Park and Alex Alderton, Roberto Amato, Jeffrey Barrett, Sonia Goncalves, Ewan Harrison, David K. Jackson, Ian Johnston, Dominic Kwiatkowski, Cordelia Langford, John Sillitoe on behalf of the Wellcome Sanger Institute COVID-19 Surveillance Team |
| EPI_ISL_2567999 | Lighthouse Lab in Glasgow | Wellcome Sanger Institute for the COVID-19 Genomics UK (COG-UK) Consortium | Harper VanSteenhouse, Yumi Kasai, David Gray, Carol Clugston, Anna Dominiczak and Alex Alderton, Roberto Amato, Jeffrey Barrett, Sonia Goncalves, Ewan Harrison, David K. Jackson, Ian Johnston, Dominic Kwiatkowski, Cordelia Langford, John Sillitoe on behalf of the Wellcome Sanger Institute COVID-19 Surveillance Team |
| EPI_ISL_2568000 | Lighthouse Lab in Alderley Park | Wellcome Sanger Institute for the COVID-19 Genomics UK (COG-UK) Consortium | Jacquelyn Wynn, Mairead Hyland, The Lighthouse Lab in Alderley Park and Alex Alderton, Roberto Amato, Jeffrey Barrett, Sonia Goncalves, Ewan Harrison, David K. Jackson, Ian Johnston, Dominic Kwiatkowski, Cordelia Langford, John Sillitoe on behalf of the Wellcome Sanger Institute COVID-19 Surveillance Team |
| EPI_ISL_2568001, EPI_ISL_2568002 | Lighthouse Lab in Glasgow | Wellcome Sanger Institute for the COVID-19 Genomics UK (COG-UK) Consortium | Harper VanSteenhouse, Yumi Kasai, David Gray, Carol Clugston, Anna Dominiczak and Alex Alderton, Roberto Amato, Jeffrey Barrett, Sonia Goncalves, Ewan Harrison, David K. Jackson, Ian Johnston, Dominic Kwiatkowski, Cordelia Langford, John Sillitoe on behalf of the Wellcome Sanger Institute COVID-19 Surveillance Team |
| EPI_ISL_2568003 | Lighthouse Lab in Alderley Park | Wellcome Sanger Institute for the COVID-19 Genomics UK (COG-UK) Consortium | Jacquelyn Wynn, Mairead Hyland, The Lighthouse Lab in Alderley Park and Alex Alderton, Roberto Amato, Jeffrey Barrett, Sonia Goncalves, Ewan Harrison, David K. Jackson, Ian Johnston, Dominic Kwiatkowski, Cordelia Langford, John Sillitoe on behalf of the Wellcome Sanger Institute COVID-19 Surveillance Team |
| EPI_ISL_2568004 | Lighthouse Lab in Glasgow | Wellcome Sanger Institute for the COVID-19 Genomics UK (COG-UK) Consortium | Harper VanSteenhouse, Yumi Kasai, David Gray, Carol Clugston, Anna Dominiczak and Alex Alderton, Roberto Amato, Jeffrey Barrett, Sonia Goncalves, Ewan Harrison, David K. Jackson, Ian Johnston, Dominic Kwiatkowski, Cordelia Langford, John Sillitoe on behalf of the Wellcome Sanger Institute COVID-19 Surveillance Team |
| EPI_ISL_2568005, EPI_ISL_2568006 | Lighthouse Lab in Alderley Park | Wellcome Sanger Institute for the COVID-19 Genomics UK (COG-UK) Consortium | Jacquelyn Wynn, Mairead Hyland, The Lighthouse Lab in Alderley Park and Alex Alderton, Roberto Amato, Jeffrey Barrett, Sonia Goncalves, Ewan Harrison, David K. Jackson, Ian Johnston, Dominic Kwiatkowski, Cordelia Langford, John Sillitoe on behalf of the Wellcome Sanger Institute COVID-19 Surveillance Team |
| EPI_ISL_2568008, EPI_ISL_2568009, EPI_ISL_2568010 | Lighthouse Lab in Glasgow | Wellcome Sanger Institute for the COVID-19 Genomics UK (COG-UK) Consortium | Harper VanSteenhouse, Yumi Kasai, David Gray, Carol Clugston, Anna Dominiczak and Alex Alderton, Roberto Amato, Jeffrey Barrett, Sonia Goncalves, Ewan Harrison, David K. Jackson, Ian Johnston, Dominic Kwiatkowski, Cordelia Langford, John Sillitoe on behalf of the Wellcome Sanger Institute COVID-19 Surveillance Team |
| EPI_ISL_2568011, EPI_ISL_2568012 | Lighthouse Lab in Alderley Park | Wellcome Sanger Institute for the COVID-19 Genomics UK (COG-UK) Consortium | Jacquelyn Wynn, Mairead Hyland, The Lighthouse Lab in Alderley Park and Alex Alderton, Roberto Amato, Jeffrey Barrett, Sonia Goncalves, Ewan Harrison, David K. Jackson, Ian Johnston, Dominic Kwiatkowski, Cordelia Langford, John Sillitoe on behalf of the Wellcome Sanger Institute COVID-19 Surveillance Team |
| EPI_ISL_2568013 | Lighthouse Lab in Glasgow | Wellcome Sanger Institute for the COVID-19 Genomics UK (COG-UK) Consortium | Harper VanSteenhouse, Yumi Kasai, David Gray, Carol Clugston, Anna Dominiczak and Alex Alderton, Roberto Amato, Jeffrey Barrett, Sonia Goncalves, Ewan Harrison, David K. Jackson, Ian Johnston, Dominic Kwiatkowski, Cordelia Langford, John Sillitoe on behalf of the Wellcome Sanger Institute COVID-19 Surveillance Team |
| EPI_ISL_2568014 | Lighthouse Lab in Alderley Park | Wellcome Sanger Institute for the COVID-19 Genomics UK (COG-UK) Consortium | Jacquelyn Wynn, Mairead Hyland, The Lighthouse Lab in Alderley Park and Alex Alderton, Roberto Amato, Jeffrey Barrett, Sonia Goncalves, Ewan Harrison, David K. Jackson, Ian Johnston, Dominic Kwiatkowski, Cordelia Langford, John Sillitoe on behalf of the Wellcome Sanger Institute COVID-19 Surveillance Team |
| EPI_ISL_2568015 | Lighthouse Lab in Glasgow | Wellcome Sanger Institute for the COVID-19 Genomics UK (COG-UK) Consortium | Harper VanSteenhouse, Yumi Kasai, David Gray, Carol Clugston, Anna Dominiczak and Alex Alderton, Roberto Amato, Jeffrey Barrett, Sonia Goncalves, Ewan Harrison, David K. Jackson, Ian Johnston, Dominic Kwiatkowski, Cordelia Langford, John Sillitoe on behalf of the Wellcome Sanger Institute COVID-19 Surveillance Team |
| EPI_ISL_2568016 | Lighthouse Lab in Alderley Park | Wellcome Sanger Institute for the COVID-19 Genomics UK (COG-UK) Consortium | Jacquelyn Wynn, Mairead Hyland, The Lighthouse Lab in Alderley Park and Alex Alderton, Roberto Amato, Jeffrey Barrett, Sonia Goncalves, Ewan Harrison, David K. Jackson, Ian Johnston, Dominic Kwiatkowski, Cordelia Langford, John Sillitoe on behalf of the Wellcome Sanger Institute COVID-19 Surveillance Team |
| EPI_ISL_2568017 | Lighthouse Lab in Glasgow | Wellcome Sanger Institute for the COVID-19 Genomics UK (COG-UK) Consortium | Harper VanSteenhouse, Yumi Kasai, David Gray, Carol Clugston, Anna Dominiczak and Alex Alderton, Roberto Amato, Jeffrey Barrett, Sonia Goncalves, Ewan Harrison, David K. Jackson, Ian Johnston, Dominic Kwiatkowski, Cordelia Langford, John Sillitoe on behalf of the Wellcome Sanger Institute COVID-19 Surveillance Team |
| EPI_ISL_2568018, EPI_ISL_2568019, EPI_ISL_2568020, EPI_ISL_2568021, EPI_ISL_2568022, EPI_ISL_2568023 | Lighthouse Lab in Alderley Park | Wellcome Sanger Institute for the COVID-19 Genomics UK (COG-UK) Consortium | Jacquelyn Wynn, Mairead Hyland, The Lighthouse Lab in Alderley Park and Alex Alderton, Roberto Amato, Jeffrey Barrett, Sonia Goncalves, Ewan Harrison, David K. Jackson, Ian Johnston, Dominic Kwiatkowski, Cordelia Langford, John Sillitoe on behalf of the Wellcome Sanger Institute COVID-19 Surveillance Team |
| EPI_ISL_2568024, EPI_ISL_2568025 | Lighthouse Lab in Glasgow | Wellcome Sanger Institute for the COVID-19 Genomics UK (COG-UK) Consortium | Harper VanSteenhouse, Yumi Kasai, David Gray, Carol Clugston, Anna Dominiczak and Alex Alderton, Roberto Amato, Jeffrey Barrett, Sonia Goncalves, Ewan Harrison, David K. Jackson, Ian Johnston, Dominic Kwiatkowski, Cordelia Langford, John Sillitoe on behalf of the Wellcome Sanger Institute COVID-19 Surveillance Team |
| EPI_ISL_2568026, EPI_ISL_2568027 | Lighthouse Lab in Alderley Park | Wellcome Sanger Institute for the COVID-19 Genomics UK (COG-UK) Consortium | Jacquelyn Wynn, Mairead Hyland, The Lighthouse Lab in Alderley Park and Alex Alderton, Roberto Amato, Jeffrey Barrett, Sonia Goncalves, Ewan Harrison, David K. Jackson, Ian Johnston, Dominic Kwiatkowski, Cordelia Langford, John Sillitoe on behalf of the Wellcome Sanger Institute COVID-19 Surveillance Team |
| EPI_ISL_2568028, EPI_ISL_2568030 | Lighthouse Lab in Glasgow | Wellcome Sanger Institute for the COVID-19 Genomics UK (COG-UK) Consortium | Harper VanSteenhouse, Yumi Kasai, David Gray, Carol Clugston, Anna Dominiczak and Alex Alderton, Roberto Amato, Jeffrey Barrett, Sonia Goncalves, Ewan Harrison, David K. Jackson, Ian Johnston, Dominic Kwiatkowski, Cordelia Langford, John Sillitoe on behalf of the Wellcome Sanger Institute COVID-19 Surveillance Team |
| EPI_ISL_2568031 | Lighthouse Lab in Alderley Park | Wellcome Sanger Institute for the COVID-19 Genomics UK (COG-UK) Consortium | Jacquelyn Wynn, Mairead Hyland, The Lighthouse Lab in Alderley Park and Alex Alderton, Roberto Amato, Jeffrey Barrett, Sonia Goncalves, Ewan Harrison, David K. Jackson, Ian Johnston, Dominic Kwiatkowski, Cordelia Langford, John Sillitoe on behalf of the Wellcome Sanger Institute COVID-19 Surveillance Team |
| EPI_ISL_2568032, EPI_ISL_2568033 | Lighthouse Lab in Glasgow | Wellcome Sanger Institute for the COVID-19 Genomics UK (COG-UK) Consortium | Harper VanSteenhouse, Yumi Kasai, David Gray, Carol Clugston, Anna Dominiczak and Alex Alderton, Roberto Amato, Jeffrey Barrett, Sonia Goncalves, Ewan Harrison, David K. Jackson, Ian Johnston, Dominic Kwiatkowski, Cordelia Langford, John Sillitoe on behalf of the Wellcome Sanger Institute COVID-19 Surveillance Team |
| EPI_ISL_2568034 | Lighthouse Lab in Alderley Park | Wellcome Sanger Institute for the COVID-19 Genomics UK (COG-UK) Consortium | Jacquelyn Wynn, Mairead Hyland, The Lighthouse Lab in Alderley Park and Alex Alderton, Roberto Amato, Jeffrey Barrett, Sonia Goncalves, Ewan Harrison, David K. Jackson, Ian Johnston, Dominic Kwiatkowski, Cordelia Langford, John Sillitoe on behalf of the Wellcome Sanger Institute COVID-19 Surveillance Team |
| EPI_ISL_2568035 | Lighthouse Lab in Glasgow | Wellcome Sanger Institute for the COVID-19 Genomics UK (COG-UK) Consortium | Harper VanSteenhouse, Yumi Kasai, David Gray, Carol Clugston, Anna Dominiczak and Alex Alderton, Roberto Amato, Jeffrey Barrett, Sonia Goncalves, Ewan Harrison, David K. Jackson, Ian Johnston, Dominic Kwiatkowski, Cordelia Langford, John Sillitoe on behalf of the Wellcome Sanger Institute COVID-19 Surveillance Team |
| EPI_ISL_2568036, EPI_ISL_2568037 | Lighthouse Lab in Alderley Park | Wellcome Sanger Institute for the COVID-19 Genomics UK (COG-UK) Consortium | Jacquelyn Wynn, Mairead Hyland, The Lighthouse Lab in Alderley Park and Alex Alderton, Roberto Amato, Jeffrey Barrett, Sonia Goncalves, Ewan Harrison, David K. Jackson, Ian Johnston, Dominic Kwiatkowski, Cordelia Langford, John Sillitoe on behalf of the Wellcome Sanger Institute COVID-19 Surveillance Team |
| EPI_ISL_2568038, EPI_ISL_2568039 | Lighthouse Lab in Glasgow | Wellcome Sanger Institute for the COVID-19 Genomics UK (COG-UK) Consortium | Harper VanSteenhouse, Yumi Kasai, David Gray, Carol Clugston, Anna Dominiczak and Alex Alderton, Roberto Amato, Jeffrey Barrett, Sonia Goncalves, Ewan Harrison, David K. Jackson, Ian Johnston, Dominic Kwiatkowski, Cordelia Langford, John Sillitoe on behalf of the Wellcome Sanger Institute COVID-19 Surveillance Team |
| EPI_ISL_2568040 | Lighthouse Lab in Alderley Park | Wellcome Sanger Institute for the COVID-19 Genomics UK (COG-UK) Consortium | Jacquelyn Wynn, Mairead Hyland, The Lighthouse Lab in Alderley Park and Alex Alderton, Roberto Amato, Jeffrey Barrett, Sonia Goncalves, Ewan Harrison, David K. Jackson, Ian Johnston, Dominic Kwiatkowski, Cordelia Langford, John Sillitoe on behalf of the Wellcome Sanger Institute COVID-19 Surveillance Team |
| EPI_ISL_2568041, EPI_ISL_2568042, EPI_ISL_2568044, EPI_ISL_2568045 | Lighthouse Lab in Glasgow | Wellcome Sanger Institute for the COVID-19 Genomics UK (COG-UK) Consortium | Harper VanSteenhouse, Yumi Kasai, David Gray, Carol Clugston, Anna Dominiczak and Alex Alderton, Roberto Amato, Jeffrey Barrett, Sonia Goncalves, Ewan Harrison, David K. Jackson, Ian Johnston, Dominic Kwiatkowski, Cordelia Langford, John Sillitoe on behalf of the Wellcome Sanger Institute COVID-19 Surveillance Team |
| EPI_ISL_2568046, EPI_ISL_2568047 | Lighthouse Lab in Alderley Park | Wellcome Sanger Institute for the COVID-19 Genomics UK (COG-UK) Consortium | Jacquelyn Wynn, Mairead Hyland, The Lighthouse Lab in Alderley Park and Alex Alderton, Roberto Amato, Jeffrey Barrett, Sonia Goncalves, Ewan Harrison, David K. Jackson, Ian Johnston, Dominic Kwiatkowski, Cordelia Langford, John Sillitoe on behalf of the Wellcome Sanger Institute COVID-19 Surveillance Team |
| EPI_ISL_2568048, EPI_ISL_2568049 | Lighthouse Lab in Glasgow | Wellcome Sanger Institute for the COVID-19 Genomics UK (COG-UK) Consortium | Harper VanSteenhouse, Yumi Kasai, David Gray, Carol Clugston, Anna Dominiczak and Alex Alderton, Roberto Amato, Jeffrey Barrett, Sonia Goncalves, Ewan Harrison, David K. Jackson, Ian Johnston, Dominic Kwiatkowski, Cordelia Langford, John Sillitoe on behalf of the Wellcome Sanger Institute COVID-19 Surveillance Team |
| EPI_ISL_2568050, EPI_ISL_2568051, EPI_ISL_2568052 | Lighthouse Lab in Alderley Park | Wellcome Sanger Institute for the COVID-19 Genomics UK (COG-UK) Consortium | Jacquelyn Wynn, Mairead Hyland, The Lighthouse Lab in Alderley Park and Alex Alderton, Roberto Amato, Jeffrey Barrett, Sonia Goncalves, Ewan Harrison, David K. Jackson, Ian Johnston, Dominic Kwiatkowski, Cordelia Langford, John Sillitoe on behalf of the Wellcome Sanger Institute COVID-19 Surveillance Team |
| EPI_ISL_2568053, EPI_ISL_2568054 | Lighthouse Lab in Glasgow | Wellcome Sanger Institute for the COVID-19 Genomics UK (COG-UK) Consortium | Harper VanSteenhouse, Yumi Kasai, David Gray, Carol Clugston, Anna Dominiczak and Alex Alderton, Roberto Amato, Jeffrey Barrett, Sonia Goncalves, Ewan Harrison, David K. Jackson, Ian Johnston, Dominic Kwiatkowski, Cordelia Langford, John Sillitoe on behalf of the Wellcome Sanger Institute COVID-19 Surveillance Team |
| EPI_ISL_2568055, EPI_ISL_2568056 | Lighthouse Lab in Alderley Park | Wellcome Sanger Institute for the COVID-19 Genomics UK (COG-UK) Consortium | Jacquelyn Wynn, Mairead Hyland, The Lighthouse Lab in Alderley Park and Alex Alderton, Roberto Amato, Jeffrey Barrett, Sonia Goncalves, Ewan Harrison, David K. Jackson, Ian Johnston, Dominic Kwiatkowski, Cordelia Langford, John Sillitoe on behalf of the Wellcome Sanger Institute COVID-19 Surveillance Team |
| EPI_ISL_2568057 | Lighthouse Lab in Glasgow | Wellcome Sanger Institute for the COVID-19 Genomics UK (COG-UK) Consortium | Harper VanSteenhouse, Yumi Kasai, David Gray, Carol Clugston, Anna Dominiczak and Alex Alderton, Roberto Amato, Jeffrey Barrett, Sonia Goncalves, Ewan Harrison, David K. Jackson, Ian Johnston, Dominic Kwiatkowski, Cordelia Langford, John Sillitoe on behalf of the Wellcome Sanger Institute COVID-19 Surveillance Team |
| EPI_ISL_2568058 | Lighthouse Lab in Alderley Park | Wellcome Sanger Institute for the COVID-19 Genomics UK (COG-UK) Consortium | Jacquelyn Wynn, Mairead Hyland, The Lighthouse Lab in Alderley Park and Alex Alderton, Roberto Amato, Jeffrey Barrett, Sonia Goncalves, Ewan Harrison, David K. Jackson, Ian Johnston, Dominic Kwiatkowski, Cordelia Langford, John Sillitoe on behalf of the Wellcome Sanger Institute COVID-19 Surveillance Team |
| EPI_ISL_2568059, EPI_ISL_2568060 | Lighthouse Lab in Glasgow | Wellcome Sanger Institute for the COVID-19 Genomics UK (COG-UK) Consortium | Harper VanSteenhouse, Yumi Kasai, David Gray, Carol Clugston, Anna Dominiczak and Alex Alderton, Roberto Amato, Jeffrey Barrett, Sonia Goncalves, Ewan Harrison, David K. Jackson, Ian Johnston, Dominic Kwiatkowski, Cordelia Langford, John Sillitoe on behalf of the Wellcome Sanger Institute COVID-19 Surveillance Team |
| EPI_ISL_2568061 | Lighthouse Lab in Alderley Park | Wellcome Sanger Institute for the COVID-19 Genomics UK (COG-UK) Consortium | Jacquelyn Wynn, Mairead Hyland, The Lighthouse Lab in Alderley Park and Alex Alderton, Roberto Amato, Jeffrey Barrett, Sonia Goncalves, Ewan Harrison, David K. Jackson, Ian Johnston, Dominic Kwiatkowski, Cordelia Langford, John Sillitoe on behalf of the Wellcome Sanger Institute COVID-19 Surveillance Team |
| EPI_ISL_2568062 | Lighthouse Lab in Glasgow | Wellcome Sanger Institute for the COVID-19 Genomics UK (COG-UK) Consortium | Harper VanSteenhouse, Yumi Kasai, David Gray, Carol Clugston, Anna Dominiczak and Alex Alderton, Roberto Amato, Jeffrey Barrett, Sonia Goncalves, Ewan Harrison, David K. Jackson, Ian Johnston, Dominic Kwiatkowski, Cordelia Langford, John Sillitoe on behalf of the Wellcome Sanger Institute COVID-19 Surveillance Team |
| EPI_ISL_2568063, EPI_ISL_2568064 | Lighthouse Lab in Alderley Park | Wellcome Sanger Institute for the COVID-19 Genomics UK (COG-UK) Consortium | Jacquelyn Wynn, Mairead Hyland, The Lighthouse Lab in Alderley Park and Alex Alderton, Roberto Amato, Jeffrey Barrett, Sonia Goncalves, Ewan Harrison, David K. Jackson, Ian Johnston, Dominic Kwiatkowski, Cordelia Langford, John Sillitoe on behalf of the Wellcome Sanger Institute COVID-19 Surveillance Team |
| EPI_ISL_2568065, EPI_ISL_2568066, EPI_ISL_2568067, EPI_ISL_2568068 | Lighthouse Lab in Glasgow | Wellcome Sanger Institute for the COVID-19 Genomics UK (COG-UK) Consortium | Harper VanSteenhouse, Yumi Kasai, David Gray, Carol Clugston, Anna Dominiczak and Alex Alderton, Roberto Amato, Jeffrey Barrett, Sonia Goncalves, Ewan Harrison, David K. Jackson, Ian Johnston, Dominic Kwiatkowski, Cordelia Langford, John Sillitoe on behalf of the Wellcome Sanger Institute COVID-19 Surveillance Team |
| EPI_ISL_2568069, EPI_ISL_2568070, EPI_ISL_2568071, | Lighthouse Lab in Alderley Park | Wellcome Sanger Institute for the COVID-19 Genomics UK (COG-UK) Consortium | Jacquelyn Wynn, Mairead Hyland, The Lighthouse Lab in Alderley Park and Alex Alderton, Roberto Amato, Jeffrey Barrett, Sonia Goncalves, Ewan Harrison, David K. Jackson, Ian Johnston, Dominic Kwiatkowski, Cordelia Langford, John Sillitoe on behalf of the Wellcome Sanger Institute COVID-19 Surveillance Team |

| | | | |
|---|---|---|---|
| EPI_ISL_2568072, EPI_ISL_2568073 | | | |
| EPI_ISL_2568074, EPI_ISL_2568075 | Lighthouse Lab in Glasgow | Wellcome Sanger Institute for the COVID-19 Genomics UK (COG-UK) Consortium | Harper VanSteenhouse, Yumi Kasai, David Gray, Carol Clugston, Anna Dominiczak and Alex Alderton, Roberto Amato, Jeffrey Barrett, Sonia Goncalves, Ewan Harrison, David K. Jackson, Ian Johnston, Dominic Kwiatkowski, Cordelia Langford, John Sillitoe on behalf of the Wellcome Sanger Institute COVID-19 Surveillance Team |
| EPI_ISL_2568076 | Lighthouse Lab in Alderley Park | Wellcome Sanger Institute for the COVID-19 Genomics UK (COG-UK) Consortium | Jacquelyn Wynn, Mairead Hyland, The Lighthouse Lab in Alderley Park and Alex Alderton, Roberto Amato, Jeffrey Barrett, Sonia Goncalves, Ewan Harrison, David K. Jackson, Ian Johnston, Dominic Kwiatkowski, Cordelia Langford, John Sillitoe on behalf of the Wellcome Sanger Institute COVID-19 Surveillance Team |
| EPI_ISL_2568077, EPI_ISL_2568078, EPI_ISL_2568079, EPI_ISL_2568080, EPI_ISL_2568082 | Lighthouse Lab in Glasgow | Wellcome Sanger Institute for the COVID-19 Genomics UK (COG-UK) Consortium | Harper VanSteenhouse, Yumi Kasai, David Gray, Carol Clugston, Anna Dominiczak and Alex Alderton, Roberto Amato, Jeffrey Barrett, Sonia Goncalves, Ewan Harrison, David K. Jackson, Ian Johnston, Dominic Kwiatkowski, Cordelia Langford, John Sillitoe on behalf of the Wellcome Sanger Institute COVID-19 Surveillance Team |
| EPI_ISL_2568085 | Lighthouse Lab in Alderley Park | Wellcome Sanger Institute for the COVID-19 Genomics UK (COG-UK) Consortium | Jacquelyn Wynn, Mairead Hyland, The Lighthouse Lab in Alderley Park and Alex Alderton, Roberto Amato, Jeffrey Barrett, Sonia Goncalves, Ewan Harrison, David K. Jackson, Ian Johnston, Dominic Kwiatkowski, Cordelia Langford, John Sillitoe on behalf of the Wellcome Sanger Institute COVID-19 Surveillance Team |
| EPI_ISL_2568086 | Lighthouse Lab in Glasgow | Wellcome Sanger Institute for the COVID-19 Genomics UK (COG-UK) Consortium | Harper VanSteenhouse, Yumi Kasai, David Gray, Carol Clugston, Anna Dominiczak and Alex Alderton, Roberto Amato, Jeffrey Barrett, Sonia Goncalves, Ewan Harrison, David K. Jackson, Ian Johnston, Dominic Kwiatkowski, Cordelia Langford, John Sillitoe on behalf of the Wellcome Sanger Institute COVID-19 Surveillance Team |
| EPI_ISL_2568087 | Lighthouse Lab in Alderley Park | Wellcome Sanger Institute for the COVID-19 Genomics UK (COG-UK) Consortium | Jacquelyn Wynn, Mairead Hyland, The Lighthouse Lab in Alderley Park and Alex Alderton, Roberto Amato, Jeffrey Barrett, Sonia Goncalves, Ewan Harrison, David K. Jackson, Ian Johnston, Dominic Kwiatkowski, Cordelia Langford, John Sillitoe on behalf of the Wellcome Sanger Institute COVID-19 Surveillance Team |
| EPI_ISL_2568088 | Lighthouse Lab in Glasgow | Wellcome Sanger Institute for the COVID-19 Genomics UK (COG-UK) Consortium | Harper VanSteenhouse, Yumi Kasai, David Gray, Carol Clugston, Anna Dominiczak and Alex Alderton, Roberto Amato, Jeffrey Barrett, Sonia Goncalves, Ewan Harrison, David K. Jackson, Ian Johnston, Dominic Kwiatkowski, Cordelia Langford, John Sillitoe on behalf of the Wellcome Sanger Institute COVID-19 Surveillance Team |
| EPI_ISL_2568089, EPI_ISL_2568090 | Lighthouse Lab in Alderley Park | Wellcome Sanger Institute for the COVID-19 Genomics UK (COG-UK) Consortium | Jacquelyn Wynn, Mairead Hyland, The Lighthouse Lab in Alderley Park and Alex Alderton, Roberto Amato, Jeffrey Barrett, Sonia Goncalves, Ewan Harrison, David K. Jackson, Ian Johnston, Dominic Kwiatkowski, Cordelia Langford, John Sillitoe on behalf of the Wellcome Sanger Institute COVID-19 Surveillance Team |
| EPI_ISL_2568091 | Lighthouse Lab in Glasgow | Wellcome Sanger Institute for the COVID-19 Genomics UK (COG-UK) Consortium | Harper VanSteenhouse, Yumi Kasai, David Gray, Carol Clugston, Anna Dominiczak and Alex Alderton, Roberto Amato, Jeffrey Barrett, Sonia Goncalves, Ewan Harrison, David K. Jackson, Ian Johnston, Dominic Kwiatkowski, Cordelia Langford, John Sillitoe on behalf of the Wellcome Sanger Institute COVID-19 Surveillance Team |
| EPI_ISL_2568092 | Lighthouse Lab in Alderley Park | Wellcome Sanger Institute for the COVID-19 Genomics UK (COG-UK) Consortium | Jacquelyn Wynn, Mairead Hyland, The Lighthouse Lab in Alderley Park and Alex Alderton, Roberto Amato, Jeffrey Barrett, Sonia Goncalves, Ewan Harrison, David K. Jackson, Ian Johnston, Dominic Kwiatkowski, Cordelia Langford, John Sillitoe on behalf of the Wellcome Sanger Institute COVID-19 Surveillance Team |
| EPI_ISL_2568093, EPI_ISL_2568094 | Lighthouse Lab in Glasgow | Wellcome Sanger Institute for the COVID-19 Genomics UK (COG-UK) Consortium | Harper VanSteenhouse, Yumi Kasai, David Gray, Carol Clugston, Anna Dominiczak and Alex Alderton, Roberto Amato, Jeffrey Barrett, Sonia Goncalves, Ewan Harrison, David K. Jackson, Ian Johnston, Dominic Kwiatkowski, Cordelia Langford, John Sillitoe on behalf of the Wellcome Sanger Institute COVID-19 Surveillance Team |
| EPI_ISL_2568095 | Lighthouse Lab in Alderley Park | Wellcome Sanger Institute for the COVID-19 Genomics UK (COG-UK) Consortium | Jacquelyn Wynn, Mairead Hyland, The Lighthouse Lab in Alderley Park and Alex Alderton, Roberto Amato, Jeffrey Barrett, Sonia Goncalves, Ewan Harrison, David K. Jackson, Ian Johnston, Dominic Kwiatkowski, Cordelia Langford, John Sillitoe on behalf of the Wellcome Sanger Institute COVID-19 Surveillance Team |
| EPI_ISL_2568096 | Lighthouse Lab in Glasgow | Wellcome Sanger Institute for the COVID-19 Genomics UK (COG-UK) Consortium | Harper VanSteenhouse, Yumi Kasai, David Gray, Carol Clugston, Anna Dominiczak and Alex Alderton, Roberto Amato, Jeffrey Barrett, Sonia Goncalves, Ewan Harrison, David K. Jackson, Ian Johnston, Dominic Kwiatkowski, Cordelia Langford, John Sillitoe on behalf of the Wellcome Sanger Institute COVID-19 Surveillance Team |
| EPI_ISL_2568097 | Lighthouse Lab in Alderley Park | Wellcome Sanger Institute for the COVID-19 Genomics UK (COG-UK) Consortium | Jacquelyn Wynn, Mairead Hyland, The Lighthouse Lab in Alderley Park and Alex Alderton, Roberto Amato, Jeffrey Barrett, Sonia Goncalves, Ewan Harrison, David K. Jackson, Ian Johnston, Dominic Kwiatkowski, Cordelia Langford, John Sillitoe on behalf of the Wellcome Sanger Institute COVID-19 Surveillance Team |
| EPI_ISL_2568098, EPI_ISL_2568099 | Lighthouse Lab in Glasgow | Wellcome Sanger Institute for the COVID-19 Genomics UK (COG-UK) Consortium | Harper VanSteenhouse, Yumi Kasai, David Gray, Carol Clugston, Anna Dominiczak and Alex Alderton, Roberto Amato, Jeffrey Barrett, Sonia Goncalves, Ewan Harrison, David K. Jackson, Ian Johnston, Dominic Kwiatkowski, Cordelia Langford, John Sillitoe on behalf of the Wellcome Sanger Institute COVID-19 Surveillance Team |
| EPI_ISL_2568100, EPI_ISL_2568101 | Lighthouse Lab in Alderley Park | Wellcome Sanger Institute for the COVID-19 Genomics UK (COG-UK) Consortium | Jacquelyn Wynn, Mairead Hyland, The Lighthouse Lab in Alderley Park and Alex Alderton, Roberto Amato, Jeffrey Barrett, Sonia Goncalves, Ewan Harrison, David K. Jackson, Ian Johnston, Dominic Kwiatkowski, Cordelia Langford, John Sillitoe on behalf of the Wellcome Sanger Institute COVID-19 Surveillance Team |
| EPI_ISL_2568102, EPI_ISL_2568103 | Lighthouse Lab in Glasgow | Wellcome Sanger Institute for the COVID-19 Genomics UK (COG-UK) Consortium | Harper VanSteenhouse, Yumi Kasai, David Gray, Carol Clugston, Anna Dominiczak and Alex Alderton, Roberto Amato, Jeffrey Barrett, Sonia Goncalves, Ewan Harrison, David K. Jackson, Ian Johnston, Dominic Kwiatkowski, Cordelia Langford, John Sillitoe on behalf of the Wellcome Sanger Institute COVID-19 Surveillance Team |
| EPI_ISL_2568104 | Lighthouse Lab in Alderley Park | Wellcome Sanger Institute for the COVID-19 Genomics UK (COG-UK) Consortium | Jacquelyn Wynn, Mairead Hyland, The Lighthouse Lab in Alderley Park and Alex Alderton, Roberto Amato, Jeffrey Barrett, Sonia Goncalves, Ewan Harrison, David K. Jackson, Ian Johnston, Dominic Kwiatkowski, Cordelia Langford, John Sillitoe on behalf of the Wellcome Sanger Institute COVID-19 Surveillance Team |
| EPI_ISL_2568106, EPI_ISL_2568107, EPI_ISL_2568108 | Lighthouse Lab in Glasgow | Wellcome Sanger Institute for the COVID-19 Genomics UK (COG-UK) Consortium | Harper VanSteenhouse, Yumi Kasai, David Gray, Carol Clugston, Anna Dominiczak and Alex Alderton, Roberto Amato, Jeffrey Barrett, Sonia Goncalves, Ewan Harrison, David K. Jackson, Ian Johnston, Dominic Kwiatkowski, Cordelia Langford, John Sillitoe on behalf of the Wellcome Sanger Institute COVID-19 Surveillance Team |
| EPI_ISL_2568109 | Lighthouse Lab in Milton Keynes | Wellcome Sanger Institute for the COVID-19 Genomics UK (COG-UK) Consortium | The Lighthouse Lab in Milton Keynes and Alex Alderton, Roberto Amato, Jeffrey Barrett, Sonia Goncalves, Ewan Harrison, David K. Jackson, Ian Johnston, Dominic Kwiatkowski, Cordelia Langford, John Sillitoe on behalf of the Wellcome Sanger Institute COVID-19 Surveillance Team |
| EPI_ISL_2568110 | Lighthouse Lab in Alderley Park | Wellcome Sanger Institute for the COVID-19 Genomics UK (COG-UK) Consortium | Jacquelyn Wynn, Mairead Hyland, The Lighthouse Lab in Alderley Park and Alex Alderton, Roberto Amato, Jeffrey Barrett, Sonia Goncalves, Ewan Harrison, David K. Jackson, Ian Johnston, Dominic Kwiatkowski, Cordelia Langford, John Sillitoe on behalf of the Wellcome Sanger Institute COVID-19 Surveillance Team |
| EPI_ISL_2568111 | Lighthouse Lab in Glasgow | Wellcome Sanger Institute for the COVID-19 Genomics UK (COG-UK) Consortium | Harper VanSteenhouse, Yumi Kasai, David Gray, Carol Clugston, Anna Dominiczak and Alex Alderton, Roberto Amato, Jeffrey Barrett, Sonia Goncalves, Ewan Harrison, David K. Jackson, Ian Johnston, Dominic Kwiatkowski, Cordelia Langford, John Sillitoe on behalf of the Wellcome Sanger Institute COVID-19 Surveillance Team |
| EPI_ISL_2568112 | Lighthouse Lab in Alderley Park | Wellcome Sanger Institute for the COVID-19 Genomics UK (COG-UK) Consortium | Jacquelyn Wynn, Mairead Hyland, The Lighthouse Lab in Alderley Park and Alex Alderton, Roberto Amato, Jeffrey Barrett, Sonia Goncalves, Ewan Harrison, David K. Jackson, Ian Johnston, Dominic Kwiatkowski, Cordelia Langford, John Sillitoe on behalf of the Wellcome Sanger Institute COVID-19 Surveillance Team |
| EPI_ISL_2568113, EPI_ISL_2568114 | Lighthouse Lab in Milton Keynes | Wellcome Sanger Institute for the COVID-19 Genomics UK (COG-UK) Consortium | The Lighthouse Lab in Milton Keynes and Alex Alderton, Roberto Amato, Jeffrey Barrett, Sonia Goncalves, Ewan Harrison, David K. Jackson, Ian Johnston, Dominic Kwiatkowski, Cordelia Langford, John Sillitoe on behalf of the Wellcome Sanger Institute COVID-19 Surveillance Team |
| EPI_ISL_2568115 | Lighthouse Lab in Alderley Park | Wellcome Sanger Institute for the COVID-19 Genomics UK (COG-UK) Consortium | Jacquelyn Wynn, Mairead Hyland, The Lighthouse Lab in Alderley Park and Alex Alderton, Roberto Amato, Jeffrey Barrett, Sonia Goncalves, Ewan Harrison, David K. Jackson, Ian Johnston, Dominic Kwiatkowski, Cordelia Langford, John Sillitoe on behalf of the Wellcome Sanger Institute COVID-19 Surveillance Team |
| EPI_ISL_2568116 | Lighthouse Lab in Milton Keynes | Wellcome Sanger Institute for the COVID-19 Genomics UK (COG-UK) Consortium | The Lighthouse Lab in Milton Keynes and Alex Alderton, Roberto Amato, Jeffrey Barrett, Sonia Goncalves, Ewan Harrison, David K. Jackson, Ian Johnston, Dominic Kwiatkowski, Cordelia Langford, John Sillitoe on behalf of the Wellcome Sanger Institute COVID-19 Surveillance Team |
| EPI_ISL_2568117, EPI_ISL_2568118, EPI_ISL_2568119 | Lighthouse Lab in Alderley Park | Wellcome Sanger Institute for the COVID-19 Genomics UK (COG-UK) Consortium | Jacquelyn Wynn, Mairead Hyland, The Lighthouse Lab in Alderley Park and Alex Alderton, Roberto Amato, Jeffrey Barrett, Sonia Goncalves, Ewan Harrison, David K. Jackson, Ian Johnston, Dominic Kwiatkowski, Cordelia Langford, John Sillitoe on behalf of the Wellcome Sanger Institute COVID-19 Surveillance Team |
| EPI_ISL_2568120, EPI_ISL_2568121 | Lighthouse Lab in Milton Keynes | Wellcome Sanger Institute for the COVID-19 Genomics UK (COG-UK) Consortium | The Lighthouse Lab in Milton Keynes and Alex Alderton, Roberto Amato, Jeffrey Barrett, Sonia Goncalves, Ewan Harrison, David K. Jackson, Ian Johnston, Dominic Kwiatkowski, Cordelia Langford, John Sillitoe on behalf of the Wellcome Sanger Institute COVID-19 Surveillance Team |
| EPI_ISL_2568122 | Lighthouse Lab in Alderley Park | Wellcome Sanger Institute for the COVID-19 Genomics UK (COG-UK) Consortium | Jacquelyn Wynn, Mairead Hyland, The Lighthouse Lab in Alderley Park and Alex Alderton, Roberto Amato, Jeffrey Barrett, Sonia Goncalves, Ewan Harrison, David K. Jackson, Ian Johnston, Dominic Kwiatkowski, Cordelia Langford, John Sillitoe on behalf of the Wellcome Sanger Institute COVID-19 Surveillance Team |
| EPI_ISL_2568123 | Lighthouse Lab in Milton Keynes | Wellcome Sanger Institute for the COVID-19 Genomics UK (COG-UK) Consortium | The Lighthouse Lab in Milton Keynes and Alex Alderton, Roberto Amato, Jeffrey Barrett, Sonia Goncalves, Ewan Harrison, David K. Jackson, Ian Johnston, Dominic Kwiatkowski, Cordelia Langford, John Sillitoe on behalf of the Wellcome Sanger Institute COVID-19 Surveillance Team |
| EPI_ISL_2568124 | Lighthouse Lab in Alderley Park | Wellcome Sanger Institute for the COVID-19 Genomics UK (COG-UK) Consortium | Jacquelyn Wynn, Mairead Hyland, The Lighthouse Lab in Alderley Park and Alex Alderton, Roberto Amato, Jeffrey Barrett, Sonia Goncalves, Ewan Harrison, David K. Jackson, Ian Johnston, Dominic Kwiatkowski, Cordelia Langford, John Sillitoe on behalf of the Wellcome Sanger Institute COVID-19 Surveillance Team |
| EPI_ISL_2568125 | Lighthouse Lab in Glasgow | Wellcome Sanger Institute for the COVID-19 Genomics UK (COG-UK) Consortium | Harper VanSteenhouse, Yumi Kasai, David Gray, Carol Clugston, Anna Dominiczak and Alex Alderton, Roberto Amato, Jeffrey Barrett, Sonia Goncalves, Ewan Harrison, David K. Jackson, Ian Johnston, Dominic Kwiatkowski, Cordelia Langford, John Sillitoe on behalf of the Wellcome Sanger Institute COVID-19 Surveillance Team |
| EPI_ISL_2568126 | Lighthouse Lab in Alderley Park | Wellcome Sanger Institute for the COVID-19 Genomics UK (COG-UK) Consortium | Jacquelyn Wynn, Mairead Hyland, The Lighthouse Lab in Alderley Park and Alex Alderton, Roberto Amato, Jeffrey Barrett, Sonia Goncalves, Ewan Harrison, David K. Jackson, Ian Johnston, Dominic Kwiatkowski, Cordelia Langford, John Sillitoe on behalf of the Wellcome Sanger Institute COVID-19 Surveillance Team |
| EPI_ISL_2568127 | Lighthouse Lab in Milton Keynes | Wellcome Sanger Institute for the COVID-19 Genomics UK (COG-UK) Consortium | The Lighthouse Lab in Milton Keynes and Alex Alderton, Roberto Amato, Jeffrey Barrett, Sonia Goncalves, Ewan Harrison, David K. Jackson, Ian Johnston, Dominic Kwiatkowski, Cordelia Langford, John Sillitoe on behalf of the Wellcome Sanger Institute COVID-19 Surveillance Team |
| EPI_ISL_2568128 | Lighthouse Lab in Alderley Park | Wellcome Sanger Institute for the COVID-19 Genomics UK (COG-UK) Consortium | Jacquelyn Wynn, Mairead Hyland, The Lighthouse Lab in Alderley Park and Alex Alderton, Roberto Amato, Jeffrey Barrett, Sonia Goncalves, Ewan Harrison, David K. Jackson, Ian Johnston, Dominic Kwiatkowski, Cordelia Langford, John Sillitoe on behalf of the Wellcome Sanger Institute COVID-19 Surveillance Team |
| EPI_ISL_2568129 | Lighthouse Lab in Glasgow | Wellcome Sanger Institute for the COVID-19 Genomics UK (COG-UK) Consortium | Harper VanSteenhouse, Yumi Kasai, David Gray, Carol Clugston, Anna Dominiczak and Alex Alderton, Roberto Amato, Jeffrey Barrett, Sonia Goncalves, Ewan Harrison, David K. Jackson, Ian Johnston, Dominic Kwiatkowski, Cordelia Langford, John Sillitoe on behalf of the Wellcome Sanger Institute COVID-19 Surveillance Team |
| EPI_ISL_2568130, EPI_ISL_2568131, EPI_ISL_2568132 | Lighthouse Lab in Milton Keynes | Wellcome Sanger Institute for the COVID-19 Genomics UK (COG-UK) Consortium | The Lighthouse Lab in Milton Keynes and Alex Alderton, Roberto Amato, Jeffrey Barrett, Sonia Goncalves, Ewan Harrison, David K. Jackson, Ian Johnston, Dominic Kwiatkowski, Cordelia Langford, John Sillitoe on behalf of the Wellcome Sanger Institute COVID-19 Surveillance Team |
| EPI_ISL_2568133 | Lighthouse Lab in Alderley Park | Wellcome Sanger Institute for the COVID-19 Genomics UK (COG-UK) Consortium | Jacquelyn Wynn, Mairead Hyland, The Lighthouse Lab in Alderley Park and Alex Alderton, Roberto Amato, Jeffrey Barrett, Sonia Goncalves, Ewan Harrison, David K. Jackson, Ian Johnston, Dominic Kwiatkowski, Cordelia Langford, John Sillitoe on behalf of the Wellcome Sanger Institute COVID-19 Surveillance Team |
| EPI_ISL_2568134 | Lighthouse Lab in Glasgow | Wellcome Sanger Institute for the COVID-19 Genomics UK (COG-UK) Consortium | Harper VanSteenhouse, Yumi Kasai, David Gray, Carol Clugston, Anna Dominiczak and Alex Alderton, Roberto Amato, Jeffrey Barrett, Sonia Goncalves, Ewan Harrison, David K. Jackson, Ian Johnston, Dominic Kwiatkowski, Cordelia Langford, John Sillitoe on behalf of the Wellcome Sanger Institute COVID-19 Surveillance Team |
| EPI_ISL_2568136, EPI_ISL_2568137 | Lighthouse Lab in Alderley Park | Wellcome Sanger Institute for the COVID-19 Genomics UK (COG-UK) Consortium | Jacquelyn Wynn, Mairead Hyland, The Lighthouse Lab in Alderley Park and Alex Alderton, Roberto Amato, Jeffrey Barrett, Sonia Goncalves, Ewan Harrison, David K. Jackson, Ian Johnston, Dominic Kwiatkowski, Cordelia Langford, John Sillitoe on behalf of the Wellcome Sanger Institute COVID-19 Surveillance Team |
| EPI_ISL_2568138 | Lighthouse Lab in Glasgow | Wellcome Sanger Institute for the COVID-19 Genomics UK (COG-UK) Consortium | Harper VanSteenhouse, Yumi Kasai, David Gray, Carol Clugston, Anna Dominiczak and Alex Alderton, Roberto Amato, Jeffrey Barrett, Sonia Goncalves, Ewan Harrison, David K. Jackson, Ian Johnston, Dominic Kwiatkowski, Cordelia Langford, John Sillitoe on behalf of the Wellcome Sanger Institute COVID-19 Surveillance Team |
| EPI_ISL_2568140, | Lighthouse Lab in Alderley Park | Wellcome Sanger Institute for the COVID-19 Genomics UK (COG-UK) | Jacquelyn Wynn, Mairead Hyland, The Lighthouse Lab in Alderley Park and Alex Alderton, Roberto Amato, Jeffrey Barrett, Sonia Goncalves, Ewan Harrison, David K. Jackson, Ian Johnston, Dominic Kwiatkowski, Cordelia Langford, John Sillitoe on |

| | | | |
|---|---|---|---|
| EPI_ISL_2568141, EPI_ISL_2568142, EPI_ISL_2568143, EPI_ISL_2568145 | | Consortium | behalf of the Wellcome Sanger Institute COVID-19 Surveillance Team |
| EPI_ISL_2568146 | Lighthouse Lab in Milton Keynes | Wellcome Sanger Institute for the COVID-19 Genomics UK (COG-UK) Consortium | The Lighthouse Lab in Milton Keynes and Alex Alderton, Roberto Amato, Jeffrey Barrett, Sonia Goncalves, Ewan Harrison, David K. Jackson, Ian Johnston, Dominic Kwiatkowski, Cordelia Langford, John Sillitoe on behalf of the Wellcome Sanger Institute COVID-19 Surveillance Team |
| EPI_ISL_2568147 | Lighthouse Lab in Alderley Park | Wellcome Sanger Institute for the COVID-19 Genomics UK (COG-UK) Consortium | Jacquelyn Wynn, Mairead Hyland, The Lighthouse Lab in Alderley Park and Alex Alderton, Roberto Amato, Jeffrey Barrett, Sonia Goncalves, Ewan Harrison, David K. Jackson, Ian Johnston, Dominic Kwiatkowski, Cordelia Langford, John Sillitoe on behalf of the Wellcome Sanger Institute COVID-19 Surveillance Team |
| EPI_ISL_2568148, EPI_ISL_2568149 | Lighthouse Lab in Milton Keynes | Wellcome Sanger Institute for the COVID-19 Genomics UK (COG-UK) Consortium | The Lighthouse Lab in Milton Keynes and Alex Alderton, Roberto Amato, Jeffrey Barrett, Sonia Goncalves, Ewan Harrison, David K. Jackson, Ian Johnston, Dominic Kwiatkowski, Cordelia Langford, John Sillitoe on behalf of the Wellcome Sanger Institute COVID-19 Surveillance Team |
| EPI_ISL_2568150, EPI_ISL_2568151, EPI_ISL_2568152, EPI_ISL_2568153, EPI_ISL_2568154, EPI_ISL_2568155 | Lighthouse Lab in Alderley Park | Wellcome Sanger Institute for the COVID-19 Genomics UK (COG-UK) Consortium | Jacquelyn Wynn, Mairead Hyland, The Lighthouse Lab in Alderley Park and Alex Alderton, Roberto Amato, Jeffrey Barrett, Sonia Goncalves, Ewan Harrison, David K. Jackson, Ian Johnston, Dominic Kwiatkowski, Cordelia Langford, John Sillitoe on behalf of the Wellcome Sanger Institute COVID-19 Surveillance Team |
| EPI_ISL_2568156, EPI_ISL_2568157 | Lighthouse Lab in Milton Keynes | Wellcome Sanger Institute for the COVID-19 Genomics UK (COG-UK) Consortium | The Lighthouse Lab in Milton Keynes and Alex Alderton, Roberto Amato, Jeffrey Barrett, Sonia Goncalves, Ewan Harrison, David K. Jackson, Ian Johnston, Dominic Kwiatkowski, Cordelia Langford, John Sillitoe on behalf of the Wellcome Sanger Institute COVID-19 Surveillance Team |
| EPI_ISL_2568158, EPI_ISL_2568159, EPI_ISL_2568160, EPI_ISL_2568161 | Lighthouse Lab in Alderley Park | Wellcome Sanger Institute for the COVID-19 Genomics UK (COG-UK) Consortium | Jacquelyn Wynn, Mairead Hyland, The Lighthouse Lab in Alderley Park and Alex Alderton, Roberto Amato, Jeffrey Barrett, Sonia Goncalves, Ewan Harrison, David K. Jackson, Ian Johnston, Dominic Kwiatkowski, Cordelia Langford, John Sillitoe on behalf of the Wellcome Sanger Institute COVID-19 Surveillance Team |
| EPI_ISL_2568162 | Lighthouse Lab in Glasgow | Wellcome Sanger Institute for the COVID-19 Genomics UK (COG-UK) Consortium | Harper VanSteenhouse, Yumi Kasai, David Gray, Carol Clugston, Anna Dominiczak and Alex Alderton, Roberto Amato, Jeffrey Barrett, Sonia Goncalves, Ewan Harrison, David K. Jackson, Ian Johnston, Dominic Kwiatkowski, Cordelia Langford, John Sillitoe on behalf of the Wellcome Sanger Institute COVID-19 Surveillance Team |
| EPI_ISL_2568163, EPI_ISL_2568164, EPI_ISL_2568165, EPI_ISL_2568166, EPI_ISL_2568168, EPI_ISL_2568170 | Lighthouse Lab in Alderley Park | Wellcome Sanger Institute for the COVID-19 Genomics UK (COG-UK) Consortium | Jacquelyn Wynn, Mairead Hyland, The Lighthouse Lab in Alderley Park and Alex Alderton, Roberto Amato, Jeffrey Barrett, Sonia Goncalves, Ewan Harrison, David K. Jackson, Ian Johnston, Dominic Kwiatkowski, Cordelia Langford, John Sillitoe on behalf of the Wellcome Sanger Institute COVID-19 Surveillance Team |
| EPI_ISL_2568171 | Lighthouse Lab in Milton Keynes | Wellcome Sanger Institute for the COVID-19 Genomics UK (COG-UK) Consortium | The Lighthouse Lab in Milton Keynes and Alex Alderton, Roberto Amato, Jeffrey Barrett, Sonia Goncalves, Ewan Harrison, David K. Jackson, Ian Johnston, Dominic Kwiatkowski, Cordelia Langford, John Sillitoe on behalf of the Wellcome Sanger Institute COVID-19 Surveillance Team |
| EPI_ISL_2568172, EPI_ISL_2568173, EPI_ISL_2568174 | Lighthouse Lab in Alderley Park | Wellcome Sanger Institute for the COVID-19 Genomics UK (COG-UK) Consortium | Jacquelyn Wynn, Mairead Hyland, The Lighthouse Lab in Alderley Park and Alex Alderton, Roberto Amato, Jeffrey Barrett, Sonia Goncalves, Ewan Harrison, David K. Jackson, Ian Johnston, Dominic Kwiatkowski, Cordelia Langford, John Sillitoe on behalf of the Wellcome Sanger Institute COVID-19 Surveillance Team |
| EPI_ISL_2568175, EPI_ISL_2568176, EPI_ISL_2568178 | Lighthouse Lab in Milton Keynes | Wellcome Sanger Institute for the COVID-19 Genomics UK (COG-UK) Consortium | The Lighthouse Lab in Milton Keynes and Alex Alderton, Roberto Amato, Jeffrey Barrett, Sonia Goncalves, Ewan Harrison, David K. Jackson, Ian Johnston, Dominic Kwiatkowski, Cordelia Langford, John Sillitoe on behalf of the Wellcome Sanger Institute COVID-19 Surveillance Team |
| EPI_ISL_2568179, EPI_ISL_2568180, EPI_ISL_2568181 | Lighthouse Lab in Alderley Park | Wellcome Sanger Institute for the COVID-19 Genomics UK (COG-UK) Consortium | Jacquelyn Wynn, Mairead Hyland, The Lighthouse Lab in Alderley Park and Alex Alderton, Roberto Amato, Jeffrey Barrett, Sonia Goncalves, Ewan Harrison, David K. Jackson, Ian Johnston, Dominic Kwiatkowski, Cordelia Langford, John Sillitoe on behalf of the Wellcome Sanger Institute COVID-19 Surveillance Team |
| EPI_ISL_2568182 | Lighthouse Lab in Milton Keynes | Wellcome Sanger Institute for the COVID-19 Genomics UK (COG-UK) Consortium | The Lighthouse Lab in Milton Keynes and Alex Alderton, Roberto Amato, Jeffrey Barrett, Sonia Goncalves, Ewan Harrison, David K. Jackson, Ian Johnston, Dominic Kwiatkowski, Cordelia Langford, John Sillitoe on behalf of the Wellcome Sanger Institute COVID-19 Surveillance Team |
| EPI_ISL_2568183, EPI_ISL_2568184, EPI_ISL_2568185, EPI_ISL_2568186, EPI_ISL_2568187 | Lighthouse Lab in Alderley Park | Wellcome Sanger Institute for the COVID-19 Genomics UK (COG-UK) Consortium | Jacquelyn Wynn, Mairead Hyland, The Lighthouse Lab in Alderley Park and Alex Alderton, Roberto Amato, Jeffrey Barrett, Sonia Goncalves, Ewan Harrison, David K. Jackson, Ian Johnston, Dominic Kwiatkowski, Cordelia Langford, John Sillitoe on behalf of the Wellcome Sanger Institute COVID-19 Surveillance Team |
| EPI_ISL_2568188 | Lighthouse Lab in Milton Keynes | Wellcome Sanger Institute for the COVID-19 Genomics UK (COG-UK) Consortium | The Lighthouse Lab in Milton Keynes and Alex Alderton, Roberto Amato, Jeffrey Barrett, Sonia Goncalves, Ewan Harrison, David K. Jackson, Ian Johnston, Dominic Kwiatkowski, Cordelia Langford, John Sillitoe on behalf of the Wellcome Sanger Institute COVID-19 Surveillance Team |
| EPI_ISL_2568189 | Lighthouse Lab in Alderley Park | Wellcome Sanger Institute for the COVID-19 Genomics UK (COG-UK) Consortium | Jacquelyn Wynn, Mairead Hyland, The Lighthouse Lab in Alderley Park and Alex Alderton, Roberto Amato, Jeffrey Barrett, Sonia Goncalves, Ewan Harrison, David K. Jackson, Ian Johnston, Dominic Kwiatkowski, Cordelia Langford, John Sillitoe on behalf of the Wellcome Sanger Institute COVID-19 Surveillance Team |
| EPI_ISL_2568190, EPI_ISL_2568192 | Lighthouse Lab in Milton Keynes | Wellcome Sanger Institute for the COVID-19 Genomics UK (COG-UK) Consortium | The Lighthouse Lab in Milton Keynes and Alex Alderton, Roberto Amato, Jeffrey Barrett, Sonia Goncalves, Ewan Harrison, David K. Jackson, Ian Johnston, Dominic Kwiatkowski, Cordelia Langford, John Sillitoe on behalf of the Wellcome Sanger Institute COVID-19 Surveillance Team |
| EPI_ISL_2568193 | Lighthouse Lab in Alderley Park | Wellcome Sanger Institute for the COVID-19 Genomics UK (COG-UK) Consortium | Jacquelyn Wynn, Mairead Hyland, The Lighthouse Lab in Alderley Park and Alex Alderton, Roberto Amato, Jeffrey Barrett, Sonia Goncalves, Ewan Harrison, David K. Jackson, Ian Johnston, Dominic Kwiatkowski, Cordelia Langford, John Sillitoe on behalf of the Wellcome Sanger Institute COVID-19 Surveillance Team |
| EPI_ISL_2568195 | Lighthouse Lab in Milton Keynes | Wellcome Sanger Institute for the COVID-19 Genomics UK (COG-UK) Consortium | The Lighthouse Lab in Milton Keynes and Alex Alderton, Roberto Amato, Jeffrey Barrett, Sonia Goncalves, Ewan Harrison, David K. Jackson, Ian Johnston, Dominic Kwiatkowski, Cordelia Langford, John Sillitoe on behalf of the Wellcome Sanger Institute COVID-19 Surveillance Team |
| EPI_ISL_2568196, EPI_ISL_2568197, EPI_ISL_2568198, EPI_ISL_2568199, EPI_ISL_2568200 | Lighthouse Lab in Alderley Park | Wellcome Sanger Institute for the COVID-19 Genomics UK (COG-UK) Consortium | Jacquelyn Wynn, Mairead Hyland, The Lighthouse Lab in Alderley Park and Alex Alderton, Roberto Amato, Jeffrey Barrett, Sonia Goncalves, Ewan Harrison, David K. Jackson, Ian Johnston, Dominic Kwiatkowski, Cordelia Langford, John Sillitoe on behalf of the Wellcome Sanger Institute COVID-19 Surveillance Team |
| EPI_ISL_2568202 | Lighthouse Lab in Milton Keynes | Wellcome Sanger Institute for the COVID-19 Genomics UK (COG-UK) Consortium | The Lighthouse Lab in Milton Keynes and Alex Alderton, Roberto Amato, Jeffrey Barrett, Sonia Goncalves, Ewan Harrison, David K. Jackson, Ian Johnston, Dominic Kwiatkowski, Cordelia Langford, John Sillitoe on behalf of the Wellcome Sanger Institute COVID-19 Surveillance Team |
| EPI_ISL_2568203, EPI_ISL_2568204, EPI_ISL_2568205, EPI_ISL_2568206 | Lighthouse Lab in Alderley Park | Wellcome Sanger Institute for the COVID-19 Genomics UK (COG-UK) Consortium | Jacquelyn Wynn, Mairead Hyland, The Lighthouse Lab in Alderley Park and Alex Alderton, Roberto Amato, Jeffrey Barrett, Sonia Goncalves, Ewan Harrison, David K. Jackson, Ian Johnston, Dominic Kwiatkowski, Cordelia Langford, John Sillitoe on behalf of the Wellcome Sanger Institute COVID-19 Surveillance Team |
| EPI_ISL_2568207 | Lighthouse Lab in Milton Keynes | Wellcome Sanger Institute for the COVID-19 Genomics UK (COG-UK) Consortium | The Lighthouse Lab in Milton Keynes and Alex Alderton, Roberto Amato, Jeffrey Barrett, Sonia Goncalves, Ewan Harrison, David K. Jackson, Ian Johnston, Dominic Kwiatkowski, Cordelia Langford, John Sillitoe on behalf of the Wellcome Sanger Institute COVID-19 Surveillance Team |
| EPI_ISL_2568208, EPI_ISL_2568209 | Lighthouse Lab in Alderley Park | Wellcome Sanger Institute for the COVID-19 Genomics UK (COG-UK) Consortium | Jacquelyn Wynn, Mairead Hyland, The Lighthouse Lab in Alderley Park and Alex Alderton, Roberto Amato, Jeffrey Barrett, Sonia Goncalves, Ewan Harrison, David K. Jackson, Ian Johnston, Dominic Kwiatkowski, Cordelia Langford, John Sillitoe on behalf of the Wellcome Sanger Institute COVID-19 Surveillance Team |
| EPI_ISL_2568210 | Lighthouse Lab in Milton Keynes | Wellcome Sanger Institute for the COVID-19 Genomics UK (COG-UK) Consortium | The Lighthouse Lab in Milton Keynes and Alex Alderton, Roberto Amato, Jeffrey Barrett, Sonia Goncalves, Ewan Harrison, David K. Jackson, Ian Johnston, Dominic Kwiatkowski, Cordelia Langford, John Sillitoe on behalf of the Wellcome Sanger Institute COVID-19 Surveillance Team |
| EPI_ISL_2568211, EPI_ISL_2568212, EPI_ISL_2568213, EPI_ISL_2568214, EPI_ISL_2568215, EPI_ISL_2568216 | Lighthouse Lab in Alderley Park | Wellcome Sanger Institute for the COVID-19 Genomics UK (COG-UK) Consortium | Jacquelyn Wynn, Mairead Hyland, The Lighthouse Lab in Alderley Park and Alex Alderton, Roberto Amato, Jeffrey Barrett, Sonia Goncalves, Ewan Harrison, David K. Jackson, Ian Johnston, Dominic Kwiatkowski, Cordelia Langford, John Sillitoe on behalf of the Wellcome Sanger Institute COVID-19 Surveillance Team |
| EPI_ISL_2568217 | Lighthouse Lab in Glasgow | Wellcome Sanger Institute for the COVID-19 Genomics UK (COG-UK) Consortium | Harper VanSteenhouse, Yumi Kasai, David Gray, Carol Clugston, Anna Dominiczak and Alex Alderton, Roberto Amato, Jeffrey Barrett, Sonia Goncalves, Ewan Harrison, David K. Jackson, Ian Johnston, Dominic Kwiatkowski, Cordelia Langford, John Sillitoe on behalf of the Wellcome Sanger Institute COVID-19 Surveillance Team |
| EPI_ISL_2568218 | Lighthouse Lab in Milton Keynes | Wellcome Sanger Institute for the COVID-19 Genomics UK (COG-UK) Consortium | The Lighthouse Lab in Milton Keynes and Alex Alderton, Roberto Amato, Jeffrey Barrett, Sonia Goncalves, Ewan Harrison, David K. Jackson, Ian Johnston, Dominic Kwiatkowski, Cordelia Langford, John Sillitoe on behalf of the Wellcome Sanger Institute COVID-19 Surveillance Team |
| EPI_ISL_2568219 | Lighthouse Lab in Glasgow | Wellcome Sanger Institute for the COVID-19 Genomics UK (COG-UK) Consortium | Harper VanSteenhouse, Yumi Kasai, David Gray, Carol Clugston, Anna Dominiczak and Alex Alderton, Roberto Amato, Jeffrey Barrett, Sonia Goncalves, Ewan Harrison, David K. Jackson, Ian Johnston, Dominic Kwiatkowski, Cordelia Langford, John Sillitoe on behalf of the Wellcome Sanger Institute COVID-19 Surveillance Team |
| EPI_ISL_2568220, EPI_ISL_2568221 | Lighthouse Lab in Alderley Park | Wellcome Sanger Institute for the COVID-19 Genomics UK (COG-UK) Consortium | Jacquelyn Wynn, Mairead Hyland, The Lighthouse Lab in Alderley Park and Alex Alderton, Roberto Amato, Jeffrey Barrett, Sonia Goncalves, Ewan Harrison, David K. Jackson, Ian Johnston, Dominic Kwiatkowski, Cordelia Langford, John Sillitoe on behalf of the Wellcome Sanger Institute COVID-19 Surveillance Team |
| EPI_ISL_2568222 | Lighthouse Lab in Milton Keynes | Wellcome Sanger Institute for the COVID-19 Genomics UK (COG-UK) Consortium | The Lighthouse Lab in Milton Keynes and Alex Alderton, Roberto Amato, Jeffrey Barrett, Sonia Goncalves, Ewan Harrison, David K. Jackson, Ian Johnston, Dominic Kwiatkowski, Cordelia Langford, John Sillitoe on behalf of the Wellcome Sanger Institute COVID-19 Surveillance Team |
| EPI_ISL_2568223, EPI_ISL_2568224, EPI_ISL_2568225, EPI_ISL_2568226 | Lighthouse Lab in Alderley Park | Wellcome Sanger Institute for the COVID-19 Genomics UK (COG-UK) Consortium | Jacquelyn Wynn, Mairead Hyland, The Lighthouse Lab in Alderley Park and Alex Alderton, Roberto Amato, Jeffrey Barrett, Sonia Goncalves, Ewan Harrison, David K. Jackson, Ian Johnston, Dominic Kwiatkowski, Cordelia Langford, John Sillitoe on behalf of the Wellcome Sanger Institute COVID-19 Surveillance Team |

| | | | |
|---|---|---|---|
| EPI_ISL_2568227, EPI_ISL_2568228, EPI_ISL_2568229, EPI_ISL_2568230 | Lighthouse Lab in Milton Keynes | Wellcome Sanger Institute for the COVID-19 Genomics UK (COG-UK) Consortium | The Lighthouse Lab in Milton Keynes and Alex Alderton, Roberto Amato, Jeffrey Barrett, Sonia Goncalves, Ewan Harrison, David K. Jackson, Ian Johnston, Dominic Kwiatkowski, Cordelia Langford, John Sillitoe on behalf of the Wellcome Sanger Institute COVID-19 Surveillance Team |
| EPI_ISL_2568231, EPI_ISL_2568232 | Lighthouse Lab in Alderley Park | Wellcome Sanger Institute for the COVID-19 Genomics UK (COG-UK) Consortium | Jacquelyn Wynn, Mairead Hyland, The Lighthouse Lab in Alderley Park and Alex Alderton, Roberto Amato, Jeffrey Barrett, Sonia Goncalves, Ewan Harrison, David K. Jackson, Ian Johnston, Dominic Kwiatkowski, Cordelia Langford, John Sillitoe on behalf of the Wellcome Sanger Institute COVID-19 Surveillance Team |
| EPI_ISL_2568233 | Lighthouse Lab in Milton Keynes | Wellcome Sanger Institute for the COVID-19 Genomics UK (COG-UK) Consortium | The Lighthouse Lab in Milton Keynes and Alex Alderton, Roberto Amato, Jeffrey Barrett, Sonia Goncalves, Ewan Harrison, David K. Jackson, Ian Johnston, Dominic Kwiatkowski, Cordelia Langford, John Sillitoe on behalf of the Wellcome Sanger Institute COVID-19 Surveillance Team |
| EPI_ISL_2568234 | Lighthouse Lab in Alderley Park | Wellcome Sanger Institute for the COVID-19 Genomics UK (COG-UK) Consortium | Jacquelyn Wynn, Mairead Hyland, The Lighthouse Lab in Alderley Park and Alex Alderton, Roberto Amato, Jeffrey Barrett, Sonia Goncalves, Ewan Harrison, David K. Jackson, Ian Johnston, Dominic Kwiatkowski, Cordelia Langford, John Sillitoe on behalf of the Wellcome Sanger Institute COVID-19 Surveillance Team |
| EPI_ISL_2568236 | Lighthouse Lab in Milton Keynes | Wellcome Sanger Institute for the COVID-19 Genomics UK (COG-UK) Consortium | The Lighthouse Lab in Milton Keynes and Alex Alderton, Roberto Amato, Jeffrey Barrett, Sonia Goncalves, Ewan Harrison, David K. Jackson, Ian Johnston, Dominic Kwiatkowski, Cordelia Langford, John Sillitoe on behalf of the Wellcome Sanger Institute COVID-19 Surveillance Team |
| EPI_ISL_2568237, EPI_ISL_2568238, EPI_ISL_2568239, EPI_ISL_2568240, EPI_ISL_2568241, EPI_ISL_2568242 | Lighthouse Lab in Alderley Park | Wellcome Sanger Institute for the COVID-19 Genomics UK (COG-UK) Consortium | Jacquelyn Wynn, Mairead Hyland, The Lighthouse Lab in Alderley Park and Alex Alderton, Roberto Amato, Jeffrey Barrett, Sonia Goncalves, Ewan Harrison, David K. Jackson, Ian Johnston, Dominic Kwiatkowski, Cordelia Langford, John Sillitoe on behalf of the Wellcome Sanger Institute COVID-19 Surveillance Team |
| EPI_ISL_2568243 | Lighthouse Lab in Milton Keynes | Wellcome Sanger Institute for the COVID-19 Genomics UK (COG-UK) Consortium | The Lighthouse Lab in Milton Keynes and Alex Alderton, Roberto Amato, Jeffrey Barrett, Sonia Goncalves, Ewan Harrison, David K. Jackson, Ian Johnston, Dominic Kwiatkowski, Cordelia Langford, John Sillitoe on behalf of the Wellcome Sanger Institute COVID-19 Surveillance Team |
| EPI_ISL_2568245, EPI_ISL_2568246 | Lighthouse Lab in Alderley Park | Wellcome Sanger Institute for the COVID-19 Genomics UK (COG-UK) Consortium | Jacquelyn Wynn, Mairead Hyland, The Lighthouse Lab in Alderley Park and Alex Alderton, Roberto Amato, Jeffrey Barrett, Sonia Goncalves, Ewan Harrison, David K. Jackson, Ian Johnston, Dominic Kwiatkowski, Cordelia Langford, John Sillitoe on behalf of the Wellcome Sanger Institute COVID-19 Surveillance Team |
| EPI_ISL_2568247 | Lighthouse Lab in Milton Keynes | Wellcome Sanger Institute for the COVID-19 Genomics UK (COG-UK) Consortium | The Lighthouse Lab in Milton Keynes and Alex Alderton, Roberto Amato, Jeffrey Barrett, Sonia Goncalves, Ewan Harrison, David K. Jackson, Ian Johnston, Dominic Kwiatkowski, Cordelia Langford, John Sillitoe on behalf of the Wellcome Sanger Institute COVID-19 Surveillance Team |
| EPI_ISL_2568248 | Lighthouse Lab in Alderley Park | Wellcome Sanger Institute for the COVID-19 Genomics UK (COG-UK) Consortium | Jacquelyn Wynn, Mairead Hyland, The Lighthouse Lab in Alderley Park and Alex Alderton, Roberto Amato, Jeffrey Barrett, Sonia Goncalves, Ewan Harrison, David K. Jackson, Ian Johnston, Dominic Kwiatkowski, Cordelia Langford, John Sillitoe on behalf of the Wellcome Sanger Institute COVID-19 Surveillance Team |
| EPI_ISL_2568249 | Lighthouse Lab in Milton Keynes | Wellcome Sanger Institute for the COVID-19 Genomics UK (COG-UK) Consortium | The Lighthouse Lab in Milton Keynes and Alex Alderton, Roberto Amato, Jeffrey Barrett, Sonia Goncalves, Ewan Harrison, David K. Jackson, Ian Johnston, Dominic Kwiatkowski, Cordelia Langford, John Sillitoe on behalf of the Wellcome Sanger Institute COVID-19 Surveillance Team |
| EPI_ISL_2568250 | Lighthouse Lab in Alderley Park | Wellcome Sanger Institute for the COVID-19 Genomics UK (COG-UK) Consortium | Jacquelyn Wynn, Mairead Hyland, The Lighthouse Lab in Alderley Park and Alex Alderton, Roberto Amato, Jeffrey Barrett, Sonia Goncalves, Ewan Harrison, David K. Jackson, Ian Johnston, Dominic Kwiatkowski, Cordelia Langford, John Sillitoe on behalf of the Wellcome Sanger Institute COVID-19 Surveillance Team |
| EPI_ISL_2568251 | Lighthouse Lab in Milton Keynes | Wellcome Sanger Institute for the COVID-19 Genomics UK (COG-UK) Consortium | The Lighthouse Lab in Milton Keynes and Alex Alderton, Roberto Amato, Jeffrey Barrett, Sonia Goncalves, Ewan Harrison, David K. Jackson, Ian Johnston, Dominic Kwiatkowski, Cordelia Langford, John Sillitoe on behalf of the Wellcome Sanger Institute COVID-19 Surveillance Team |
| EPI_ISL_2568252 | Lighthouse Lab in Alderley Park | Wellcome Sanger Institute for the COVID-19 Genomics UK (COG-UK) Consortium | Jacquelyn Wynn, Mairead Hyland, The Lighthouse Lab in Alderley Park and Alex Alderton, Roberto Amato, Jeffrey Barrett, Sonia Goncalves, Ewan Harrison, David K. Jackson, Ian Johnston, Dominic Kwiatkowski, Cordelia Langford, John Sillitoe on behalf of the Wellcome Sanger Institute COVID-19 Surveillance Team |
| EPI_ISL_2568253 | Lighthouse Lab in Milton Keynes | Wellcome Sanger Institute for the COVID-19 Genomics UK (COG-UK) Consortium | The Lighthouse Lab in Milton Keynes and Alex Alderton, Roberto Amato, Jeffrey Barrett, Sonia Goncalves, Ewan Harrison, David K. Jackson, Ian Johnston, Dominic Kwiatkowski, Cordelia Langford, John Sillitoe on behalf of the Wellcome Sanger Institute COVID-19 Surveillance Team |
| EPI_ISL_2568254 | Lighthouse Lab in Alderley Park | Wellcome Sanger Institute for the COVID-19 Genomics UK (COG-UK) Consortium | Jacquelyn Wynn, Mairead Hyland, The Lighthouse Lab in Alderley Park and Alex Alderton, Roberto Amato, Jeffrey Barrett, Sonia Goncalves, Ewan Harrison, David K. Jackson, Ian Johnston, Dominic Kwiatkowski, Cordelia Langford, John Sillitoe on behalf of the Wellcome Sanger Institute COVID-19 Surveillance Team |
| EPI_ISL_2568255 | Lighthouse Lab in Glasgow | Wellcome Sanger Institute for the COVID-19 Genomics UK (COG-UK) Consortium | Harper VanSteenhouse, Yumi Kasai, David Gray, Carol Clugston, Anna Dominiczak and Alex Alderton, Roberto Amato, Jeffrey Barrett, Sonia Goncalves, Ewan Harrison, David K. Jackson, Ian Johnston, Dominic Kwiatkowski, Cordelia Langford, John Sillitoe on behalf of the Wellcome Sanger Institute COVID-19 Surveillance Team |
| EPI_ISL_2568256, EPI_ISL_2568257 | Lighthouse Lab in Milton Keynes | Wellcome Sanger Institute for the COVID-19 Genomics UK (COG-UK) Consortium | The Lighthouse Lab in Milton Keynes and Alex Alderton, Roberto Amato, Jeffrey Barrett, Sonia Goncalves, Ewan Harrison, David K. Jackson, Ian Johnston, Dominic Kwiatkowski, Cordelia Langford, John Sillitoe on behalf of the Wellcome Sanger Institute COVID-19 Surveillance Team |
| EPI_ISL_2568259, EPI_ISL_2568260, EPI_ISL_2568261, EPI_ISL_2568262 | Lighthouse Lab in Alderley Park | Wellcome Sanger Institute for the COVID-19 Genomics UK (COG-UK) Consortium | Jacquelyn Wynn, Mairead Hyland, The Lighthouse Lab in Alderley Park and Alex Alderton, Roberto Amato, Jeffrey Barrett, Sonia Goncalves, Ewan Harrison, David K. Jackson, Ian Johnston, Dominic Kwiatkowski, Cordelia Langford, John Sillitoe on behalf of the Wellcome Sanger Institute COVID-19 Surveillance Team |
| EPI_ISL_2568263, EPI_ISL_2568265 | Lighthouse Lab in Milton Keynes | Wellcome Sanger Institute for the COVID-19 Genomics UK (COG-UK) Consortium | The Lighthouse Lab in Milton Keynes and Alex Alderton, Roberto Amato, Jeffrey Barrett, Sonia Goncalves, Ewan Harrison, David K. Jackson, Ian Johnston, Dominic Kwiatkowski, Cordelia Langford, John Sillitoe on behalf of the Wellcome Sanger Institute COVID-19 Surveillance Team |
| EPI_ISL_2568266 | Lighthouse Lab in Glasgow | Wellcome Sanger Institute for the COVID-19 Genomics UK (COG-UK) Consortium | Harper VanSteenhouse, Yumi Kasai, David Gray, Carol Clugston, Anna Dominiczak and Alex Alderton, Roberto Amato, Jeffrey Barrett, Sonia Goncalves, Ewan Harrison, David K. Jackson, Ian Johnston, Dominic Kwiatkowski, Cordelia Langford, John Sillitoe on behalf of the Wellcome Sanger Institute COVID-19 Surveillance Team |
| EPI_ISL_2568267, EPI_ISL_2568268, EPI_ISL_2568269, EPI_ISL_2568270 | Lighthouse Lab in Alderley Park | Wellcome Sanger Institute for the COVID-19 Genomics UK (COG-UK) Consortium | Jacquelyn Wynn, Mairead Hyland, The Lighthouse Lab in Alderley Park and Alex Alderton, Roberto Amato, Jeffrey Barrett, Sonia Goncalves, Ewan Harrison, David K. Jackson, Ian Johnston, Dominic Kwiatkowski, Cordelia Langford, John Sillitoe on behalf of the Wellcome Sanger Institute COVID-19 Surveillance Team |
| EPI_ISL_2568271 | Lighthouse Lab in Milton Keynes | Wellcome Sanger Institute for the COVID-19 Genomics UK (COG-UK) Consortium | The Lighthouse Lab in Milton Keynes and Alex Alderton, Roberto Amato, Jeffrey Barrett, Sonia Goncalves, Ewan Harrison, David K. Jackson, Ian Johnston, Dominic Kwiatkowski, Cordelia Langford, John Sillitoe on behalf of the Wellcome Sanger Institute COVID-19 Surveillance Team |
| EPI_ISL_2568272, EPI_ISL_2568273 | Lighthouse Lab in Alderley Park | Wellcome Sanger Institute for the COVID-19 Genomics UK (COG-UK) Consortium | Jacquelyn Wynn, Mairead Hyland, The Lighthouse Lab in Alderley Park and Alex Alderton, Roberto Amato, Jeffrey Barrett, Sonia Goncalves, Ewan Harrison, David K. Jackson, Ian Johnston, Dominic Kwiatkowski, Cordelia Langford, John Sillitoe on behalf of the Wellcome Sanger Institute COVID-19 Surveillance Team |
| EPI_ISL_2568274, EPI_ISL_2568275, EPI_ISL_2568277, EPI_ISL_2568278 | Lighthouse Lab in Milton Keynes | Wellcome Sanger Institute for the COVID-19 Genomics UK (COG-UK) Consortium | The Lighthouse Lab in Milton Keynes and Alex Alderton, Roberto Amato, Jeffrey Barrett, Sonia Goncalves, Ewan Harrison, David K. Jackson, Ian Johnston, Dominic Kwiatkowski, Cordelia Langford, John Sillitoe on behalf of the Wellcome Sanger Institute COVID-19 Surveillance Team |
| EPI_ISL_2568279, EPI_ISL_2568280 | Lighthouse Lab in Glasgow | Wellcome Sanger Institute for the COVID-19 Genomics UK (COG-UK) Consortium | Harper VanSteenhouse, Yumi Kasai, David Gray, Carol Clugston, Anna Dominiczak and Alex Alderton, Roberto Amato, Jeffrey Barrett, Sonia Goncalves, Ewan Harrison, David K. Jackson, Ian Johnston, Dominic Kwiatkowski, Cordelia Langford, John Sillitoe on behalf of the Wellcome Sanger Institute COVID-19 Surveillance Team |
| EPI_ISL_2568281 | Lighthouse Lab in Milton Keynes | Wellcome Sanger Institute for the COVID-19 Genomics UK (COG-UK) Consortium | The Lighthouse Lab in Milton Keynes and Alex Alderton, Roberto Amato, Jeffrey Barrett, Sonia Goncalves, Ewan Harrison, David K. Jackson, Ian Johnston, Dominic Kwiatkowski, Cordelia Langford, John Sillitoe on behalf of the Wellcome Sanger Institute COVID-19 Surveillance Team |
| EPI_ISL_2568282, EPI_ISL_2568283 | Lighthouse Lab in Alderley Park | Wellcome Sanger Institute for the COVID-19 Genomics UK (COG-UK) Consortium | Jacquelyn Wynn, Mairead Hyland, The Lighthouse Lab in Alderley Park and Alex Alderton, Roberto Amato, Jeffrey Barrett, Sonia Goncalves, Ewan Harrison, David K. Jackson, Ian Johnston, Dominic Kwiatkowski, Cordelia Langford, John Sillitoe on behalf of the Wellcome Sanger Institute COVID-19 Surveillance Team |
| EPI_ISL_2568284 | Lighthouse Lab in Milton Keynes | Wellcome Sanger Institute for the COVID-19 Genomics UK (COG-UK) Consortium | The Lighthouse Lab in Milton Keynes and Alex Alderton, Roberto Amato, Jeffrey Barrett, Sonia Goncalves, Ewan Harrison, David K. Jackson, Ian Johnston, Dominic Kwiatkowski, Cordelia Langford, John Sillitoe on behalf of the Wellcome Sanger Institute COVID-19 Surveillance Team |
| EPI_ISL_2568285 | Lighthouse Lab in Alderley Park | Wellcome Sanger Institute for the COVID-19 Genomics UK (COG-UK) Consortium | Jacquelyn Wynn, Mairead Hyland, The Lighthouse Lab in Alderley Park and Alex Alderton, Roberto Amato, Jeffrey Barrett, Sonia Goncalves, Ewan Harrison, David K. Jackson, Ian Johnston, Dominic Kwiatkowski, Cordelia Langford, John Sillitoe on behalf of the Wellcome Sanger Institute COVID-19 Surveillance Team |
| EPI_ISL_2568286, EPI_ISL_2568287 | Lighthouse Lab in Milton Keynes | Wellcome Sanger Institute for the COVID-19 Genomics UK (COG-UK) Consortium | The Lighthouse Lab in Milton Keynes and Alex Alderton, Roberto Amato, Jeffrey Barrett, Sonia Goncalves, Ewan Harrison, David K. Jackson, Ian Johnston, Dominic Kwiatkowski, Cordelia Langford, John Sillitoe on behalf of the Wellcome Sanger Institute COVID-19 Surveillance Team |
| EPI_ISL_2568288, EPI_ISL_2568289 | Lighthouse Lab in Alderley Park | Wellcome Sanger Institute for the COVID-19 Genomics UK (COG-UK) Consortium | Jacquelyn Wynn, Mairead Hyland, The Lighthouse Lab in Alderley Park and Alex Alderton, Roberto Amato, Jeffrey Barrett, Sonia Goncalves, Ewan Harrison, David K. Jackson, Ian Johnston, Dominic Kwiatkowski, Cordelia Langford, John Sillitoe on behalf of the Wellcome Sanger Institute COVID-19 Surveillance Team |
| EPI_ISL_2568290 | Lighthouse Lab in Milton Keynes | Wellcome Sanger Institute for the COVID-19 Genomics UK (COG-UK) Consortium | The Lighthouse Lab in Milton Keynes and Alex Alderton, Roberto Amato, Jeffrey Barrett, Sonia Goncalves, Ewan Harrison, David K. Jackson, Ian Johnston, Dominic Kwiatkowski, Cordelia Langford, John Sillitoe on behalf of the Wellcome Sanger Institute COVID-19 Surveillance Team |
| EPI_ISL_2568292, EPI_ISL_2568293, EPI_ISL_2568294, EPI_ISL_2568295 | Lighthouse Lab in Alderley Park | Wellcome Sanger Institute for the COVID-19 Genomics UK (COG-UK) Consortium | Jacquelyn Wynn, Mairead Hyland, The Lighthouse Lab in Alderley Park and Alex Alderton, Roberto Amato, Jeffrey Barrett, Sonia Goncalves, Ewan Harrison, David K. Jackson, Ian Johnston, Dominic Kwiatkowski, Cordelia Langford, John Sillitoe on behalf of the Wellcome Sanger Institute COVID-19 Surveillance Team |
| EPI_ISL_2568296, EPI_ISL_2568297 | Lighthouse Lab in Milton Keynes | Wellcome Sanger Institute for the COVID-19 Genomics UK (COG-UK) Consortium | The Lighthouse Lab in Milton Keynes and Alex Alderton, Roberto Amato, Jeffrey Barrett, Sonia Goncalves, Ewan Harrison, David K. Jackson, Ian Johnston, Dominic Kwiatkowski, Cordelia Langford, John Sillitoe on behalf of the Wellcome Sanger Institute COVID-19 Surveillance Team |
| EPI_ISL_2568298 | Lighthouse Lab in Alderley Park | Wellcome Sanger Institute for the COVID-19 Genomics UK (COG-UK) Consortium | Jacquelyn Wynn, Mairead Hyland, The Lighthouse Lab in Alderley Park and Alex Alderton, Roberto Amato, Jeffrey Barrett, Sonia Goncalves, Ewan Harrison, David K. Jackson, Ian Johnston, Dominic Kwiatkowski, Cordelia Langford, John Sillitoe on behalf of the Wellcome Sanger Institute COVID-19 Surveillance Team |
| EPI_ISL_2568299 | Lighthouse Lab in Milton Keynes | Wellcome Sanger Institute for the COVID-19 Genomics UK (COG-UK) Consortium | The Lighthouse Lab in Milton Keynes and Alex Alderton, Roberto Amato, Jeffrey Barrett, Sonia Goncalves, Ewan Harrison, David K. Jackson, Ian Johnston, Dominic Kwiatkowski, Cordelia Langford, John Sillitoe on behalf of the Wellcome Sanger Institute COVID-19 Surveillance Team |
| EPI_ISL_2568300 | Lighthouse Lab in Alderley Park | Wellcome Sanger Institute for the COVID-19 Genomics UK (COG-UK) Consortium | Jacquelyn Wynn, Mairead Hyland, The Lighthouse Lab in Alderley Park and Alex Alderton, Roberto Amato, Jeffrey Barrett, Sonia Goncalves, Ewan Harrison, David K. Jackson, Ian Johnston, Dominic Kwiatkowski, Cordelia Langford, John Sillitoe on behalf of the Wellcome Sanger Institute COVID-19 Surveillance Team |
| EPI_ISL_2568301, EPI_ISL_2568302 | Lighthouse Lab in Milton Keynes | Wellcome Sanger Institute for the COVID-19 Genomics UK (COG-UK) Consortium | The Lighthouse Lab in Milton Keynes and Alex Alderton, Roberto Amato, Jeffrey Barrett, Sonia Goncalves, Ewan Harrison, David K. Jackson, Ian Johnston, Dominic Kwiatkowski, Cordelia Langford, John Sillitoe on behalf of the Wellcome Sanger Institute COVID-19 Surveillance Team |

| | | | |
|---|---|---|---|
| EPI_ISL_2568303, EPI_ISL_2568304, EPI_ISL_2568305, EPI_ISL_2568306 | Lighthouse Lab in Alderley Park | Wellcome Sanger Institute for the COVID-19 Genomics UK (COG-UK) Consortium | Jacquelyn Wynn, Mairead Hyland, The Lighthouse Lab in Alderley Park and Alex Alderton, Roberto Amato, Jeffrey Barrett, Sonia Goncalves, Ewan Harrison, David K. Jackson, Ian Johnston, Dominic Kwiatkowski, Cordelia Langford, John Sillitoe on behalf of the Wellcome Sanger Institute COVID-19 Surveillance Team |
| EPI_ISL_2568309, EPI_ISL_2568311 | Lighthouse Lab in Milton Keynes | Wellcome Sanger Institute for the COVID-19 Genomics UK (COG-UK) Consortium | The Lighthouse Lab in Milton Keynes and Alex Alderton, Roberto Amato, Jeffrey Barrett, Sonia Goncalves, Ewan Harrison, David K. Jackson, Ian Johnston, Dominic Kwiatkowski, Cordelia Langford, John Sillitoe on behalf of the Wellcome Sanger Institute COVID-19 Surveillance Team |
| EPI_ISL_2568312, EPI_ISL_2568313, EPI_ISL_2568314, EPI_ISL_2568315, EPI_ISL_2568316, EPI_ISL_2568317 | Lighthouse Lab in Alderley Park | Wellcome Sanger Institute for the COVID-19 Genomics UK (COG-UK) Consortium | Jacquelyn Wynn, Mairead Hyland, The Lighthouse Lab in Alderley Park and Alex Alderton, Roberto Amato, Jeffrey Barrett, Sonia Goncalves, Ewan Harrison, David K. Jackson, Ian Johnston, Dominic Kwiatkowski, Cordelia Langford, John Sillitoe on behalf of the Wellcome Sanger Institute COVID-19 Surveillance Team |
| EPI_ISL_2568318 | Lighthouse Lab in Milton Keynes | Wellcome Sanger Institute for the COVID-19 Genomics UK (COG-UK) Consortium | The Lighthouse Lab in Milton Keynes and Alex Alderton, Roberto Amato, Jeffrey Barrett, Sonia Goncalves, Ewan Harrison, David K. Jackson, Ian Johnston, Dominic Kwiatkowski, Cordelia Langford, John Sillitoe on behalf of the Wellcome Sanger Institute COVID-19 Surveillance Team |
| EPI_ISL_2568319, EPI_ISL_2568321 | Lighthouse Lab in Alderley Park | Wellcome Sanger Institute for the COVID-19 Genomics UK (COG-UK) Consortium | Jacquelyn Wynn, Mairead Hyland, The Lighthouse Lab in Alderley Park and Alex Alderton, Roberto Amato, Jeffrey Barrett, Sonia Goncalves, Ewan Harrison, David K. Jackson, Ian Johnston, Dominic Kwiatkowski, Cordelia Langford, John Sillitoe on behalf of the Wellcome Sanger Institute COVID-19 Surveillance Team |
| EPI_ISL_2568322, EPI_ISL_2568324, EPI_ISL_2568325, EPI_ISL_2568326 | Lighthouse Lab in Milton Keynes | Wellcome Sanger Institute for the COVID-19 Genomics UK (COG-UK) Consortium | The Lighthouse Lab in Milton Keynes and Alex Alderton, Roberto Amato, Jeffrey Barrett, Sonia Goncalves, Ewan Harrison, David K. Jackson, Ian Johnston, Dominic Kwiatkowski, Cordelia Langford, John Sillitoe on behalf of the Wellcome Sanger Institute COVID-19 Surveillance Team |
| EPI_ISL_2568327 | Lighthouse Lab in Alderley Park | Wellcome Sanger Institute for the COVID-19 Genomics UK (COG-UK) Consortium | Jacquelyn Wynn, Mairead Hyland, The Lighthouse Lab in Alderley Park and Alex Alderton, Roberto Amato, Jeffrey Barrett, Sonia Goncalves, Ewan Harrison, David K. Jackson, Ian Johnston, Dominic Kwiatkowski, Cordelia Langford, John Sillitoe on behalf of the Wellcome Sanger Institute COVID-19 Surveillance Team |
| EPI_ISL_2568329 | Lighthouse Lab in Milton Keynes | Wellcome Sanger Institute for the COVID-19 Genomics UK (COG-UK) Consortium | The Lighthouse Lab in Milton Keynes and Alex Alderton, Roberto Amato, Jeffrey Barrett, Sonia Goncalves, Ewan Harrison, David K. Jackson, Ian Johnston, Dominic Kwiatkowski, Cordelia Langford, John Sillitoe on behalf of the Wellcome Sanger Institute COVID-19 Surveillance Team |
| EPI_ISL_2568330, EPI_ISL_2568331, EPI_ISL_2568332, EPI_ISL_2568333, EPI_ISL_2568334, EPI_ISL_2568335, EPI_ISL_2568336 | Lighthouse Lab in Alderley Park | Wellcome Sanger Institute for the COVID-19 Genomics UK (COG-UK) Consortium | Jacquelyn Wynn, Mairead Hyland, The Lighthouse Lab in Alderley Park and Alex Alderton, Roberto Amato, Jeffrey Barrett, Sonia Goncalves, Ewan Harrison, David K. Jackson, Ian Johnston, Dominic Kwiatkowski, Cordelia Langford, John Sillitoe on behalf of the Wellcome Sanger Institute COVID-19 Surveillance Team |
| EPI_ISL_2568337 | Lighthouse Lab in Milton Keynes | Wellcome Sanger Institute for the COVID-19 Genomics UK (COG-UK) Consortium | The Lighthouse Lab in Milton Keynes and Alex Alderton, Roberto Amato, Jeffrey Barrett, Sonia Goncalves, Ewan Harrison, David K. Jackson, Ian Johnston, Dominic Kwiatkowski, Cordelia Langford, John Sillitoe on behalf of the Wellcome Sanger Institute COVID-19 Surveillance Team |
| EPI_ISL_2568338, EPI_ISL_2568339, EPI_ISL_2568340, EPI_ISL_2568341, EPI_ISL_2568342, EPI_ISL_2568343, EPI_ISL_2568344, EPI_ISL_2568345 | Lighthouse Lab in Alderley Park | Wellcome Sanger Institute for the COVID-19 Genomics UK (COG-UK) Consortium | Jacquelyn Wynn, Mairead Hyland, The Lighthouse Lab in Alderley Park and Alex Alderton, Roberto Amato, Jeffrey Barrett, Sonia Goncalves, Ewan Harrison, David K. Jackson, Ian Johnston, Dominic Kwiatkowski, Cordelia Langford, John Sillitoe on behalf of the Wellcome Sanger Institute COVID-19 Surveillance Team |
| EPI_ISL_2568346 | Lighthouse Lab in Glasgow | Wellcome Sanger Institute for the COVID-19 Genomics UK (COG-UK) Consortium | Harper VanSteenhouse, Yumi Kasai, David Gray, Carol Clugston, Anna Dominiczak and Alex Alderton, Roberto Amato, Jeffrey Barrett, Sonia Goncalves, Ewan Harrison, David K. Jackson, Ian Johnston, Dominic Kwiatkowski, Cordelia Langford, John Sillitoe on behalf of the Wellcome Sanger Institute COVID-19 Surveillance Team |
| EPI_ISL_2568347, EPI_ISL_2568348, EPI_ISL_2568349, EPI_ISL_2568350, EPI_ISL_2568351 | Lighthouse Lab in Alderley Park | Wellcome Sanger Institute for the COVID-19 Genomics UK (COG-UK) Consortium | Jacquelyn Wynn, Mairead Hyland, The Lighthouse Lab in Alderley Park and Alex Alderton, Roberto Amato, Jeffrey Barrett, Sonia Goncalves, Ewan Harrison, David K. Jackson, Ian Johnston, Dominic Kwiatkowski, Cordelia Langford, John Sillitoe on behalf of the Wellcome Sanger Institute COVID-19 Surveillance Team |
| EPI_ISL_2568352, EPI_ISL_2568353 | Lighthouse Lab in Milton Keynes | Wellcome Sanger Institute for the COVID-19 Genomics UK (COG-UK) Consortium | The Lighthouse Lab in Milton Keynes and Alex Alderton, Roberto Amato, Jeffrey Barrett, Sonia Goncalves, Ewan Harrison, David K. Jackson, Ian Johnston, Dominic Kwiatkowski, Cordelia Langford, John Sillitoe on behalf of the Wellcome Sanger Institute COVID-19 Surveillance Team |
| EPI_ISL_2568354, EPI_ISL_2568355, EPI_ISL_2568356, EPI_ISL_2568357, EPI_ISL_2568358, EPI_ISL_2568359, EPI_ISL_2568360, EPI_ISL_2568361 | Lighthouse Lab in Alderley Park | Wellcome Sanger Institute for the COVID-19 Genomics UK (COG-UK) Consortium | Jacquelyn Wynn, Mairead Hyland, The Lighthouse Lab in Alderley Park and Alex Alderton, Roberto Amato, Jeffrey Barrett, Sonia Goncalves, Ewan Harrison, David K. Jackson, Ian Johnston, Dominic Kwiatkowski, Cordelia Langford, John Sillitoe on behalf of the Wellcome Sanger Institute COVID-19 Surveillance Team |
| EPI_ISL_2568362 | Lighthouse Lab in Milton Keynes | Wellcome Sanger Institute for the COVID-19 Genomics UK (COG-UK) Consortium | The Lighthouse Lab in Milton Keynes and Alex Alderton, Roberto Amato, Jeffrey Barrett, Sonia Goncalves, Ewan Harrison, David K. Jackson, Ian Johnston, Dominic Kwiatkowski, Cordelia Langford, John Sillitoe on behalf of the Wellcome Sanger Institute COVID-19 Surveillance Team |
| EPI_ISL_2568363, EPI_ISL_2568364, EPI_ISL_2568365 | Lighthouse Lab in Alderley Park | Wellcome Sanger Institute for the COVID-19 Genomics UK (COG-UK) Consortium | Jacquelyn Wynn, Mairead Hyland, The Lighthouse Lab in Alderley Park and Alex Alderton, Roberto Amato, Jeffrey Barrett, Sonia Goncalves, Ewan Harrison, David K. Jackson, Ian Johnston, Dominic Kwiatkowski, Cordelia Langford, John Sillitoe on behalf of the Wellcome Sanger Institute COVID-19 Surveillance Team |
| EPI_ISL_2568366, EPI_ISL_2568367 | Lighthouse Lab in Milton Keynes | Wellcome Sanger Institute for the COVID-19 Genomics UK (COG-UK) Consortium | The Lighthouse Lab in Milton Keynes and Alex Alderton, Roberto Amato, Jeffrey Barrett, Sonia Goncalves, Ewan Harrison, David K. Jackson, Ian Johnston, Dominic Kwiatkowski, Cordelia Langford, John Sillitoe on behalf of the Wellcome Sanger Institute COVID-19 Surveillance Team |
| EPI_ISL_2568368, EPI_ISL_2568369 | Lighthouse Lab in Alderley Park | Wellcome Sanger Institute for the COVID-19 Genomics UK (COG-UK) Consortium | Jacquelyn Wynn, Mairead Hyland, The Lighthouse Lab in Alderley Park and Alex Alderton, Roberto Amato, Jeffrey Barrett, Sonia Goncalves, Ewan Harrison, David K. Jackson, Ian Johnston, Dominic Kwiatkowski, Cordelia Langford, John Sillitoe on behalf of the Wellcome Sanger Institute COVID-19 Surveillance Team |
| EPI_ISL_2568370 | Lighthouse Lab in Milton Keynes | Wellcome Sanger Institute for the COVID-19 Genomics UK (COG-UK) Consortium | The Lighthouse Lab in Milton Keynes and Alex Alderton, Roberto Amato, Jeffrey Barrett, Sonia Goncalves, Ewan Harrison, David K. Jackson, Ian Johnston, Dominic Kwiatkowski, Cordelia Langford, John Sillitoe on behalf of the Wellcome Sanger Institute COVID-19 Surveillance Team |
| EPI_ISL_2568371 | Lighthouse Lab in Alderley Park | Wellcome Sanger Institute for the COVID-19 Genomics UK (COG-UK) Consortium | Jacquelyn Wynn, Mairead Hyland, The Lighthouse Lab in Alderley Park and Alex Alderton, Roberto Amato, Jeffrey Barrett, Sonia Goncalves, Ewan Harrison, David K. Jackson, Ian Johnston, Dominic Kwiatkowski, Cordelia Langford, John Sillitoe on behalf of the Wellcome Sanger Institute COVID-19 Surveillance Team |
| EPI_ISL_2568372 | Lighthouse Lab in Milton Keynes | Wellcome Sanger Institute for the COVID-19 Genomics UK (COG-UK) Consortium | The Lighthouse Lab in Milton Keynes and Alex Alderton, Roberto Amato, Jeffrey Barrett, Sonia Goncalves, Ewan Harrison, David K. Jackson, Ian Johnston, Dominic Kwiatkowski, Cordelia Langford, John Sillitoe on behalf of the Wellcome Sanger Institute COVID-19 Surveillance Team |
| EPI_ISL_2568373, EPI_ISL_2568374 | Lighthouse Lab in Alderley Park | Wellcome Sanger Institute for the COVID-19 Genomics UK (COG-UK) Consortium | Jacquelyn Wynn, Mairead Hyland, The Lighthouse Lab in Alderley Park and Alex Alderton, Roberto Amato, Jeffrey Barrett, Sonia Goncalves, Ewan Harrison, David K. Jackson, Ian Johnston, Dominic Kwiatkowski, Cordelia Langford, John Sillitoe on behalf of the Wellcome Sanger Institute COVID-19 Surveillance Team |
| EPI_ISL_2568375 | Lighthouse Lab in Glasgow | Wellcome Sanger Institute for the COVID-19 Genomics UK (COG-UK) Consortium | Harper VanSteenhouse, Yumi Kasai, David Gray, Carol Clugston, Anna Dominiczak and Alex Alderton, Roberto Amato, Jeffrey Barrett, Sonia Goncalves, Ewan Harrison, David K. Jackson, Ian Johnston, Dominic Kwiatkowski, Cordelia Langford, John Sillitoe on behalf of the Wellcome Sanger Institute COVID-19 Surveillance Team |
| EPI_ISL_2568376, EPI_ISL_2568377, EPI_ISL_2568378, EPI_ISL_2568379, EPI_ISL_2568380, EPI_ISL_2568381, EPI_ISL_2568382, EPI_ISL_2568383, EPI_ISL_2568384, EPI_ISL_2568385 | Lighthouse Lab in Alderley Park | Wellcome Sanger Institute for the COVID-19 Genomics UK (COG-UK) Consortium | Jacquelyn Wynn, Mairead Hyland, The Lighthouse Lab in Alderley Park and Alex Alderton, Roberto Amato, Jeffrey Barrett, Sonia Goncalves, Ewan Harrison, David K. Jackson, Ian Johnston, Dominic Kwiatkowski, Cordelia Langford, John Sillitoe on behalf of the Wellcome Sanger Institute COVID-19 Surveillance Team |
| EPI_ISL_2568386, EPI_ISL_2568387, EPI_ISL_2568388 | Lighthouse Lab in Milton Keynes | Wellcome Sanger Institute for the COVID-19 Genomics UK (COG-UK) Consortium | The Lighthouse Lab in Milton Keynes and Alex Alderton, Roberto Amato, Jeffrey Barrett, Sonia Goncalves, Ewan Harrison, David K. Jackson, Ian Johnston, Dominic Kwiatkowski, Cordelia Langford, John Sillitoe on behalf of the Wellcome Sanger Institute COVID-19 Surveillance Team |
| EPI_ISL_2568389 | Lighthouse Lab in Alderley Park | Wellcome Sanger Institute for the COVID-19 Genomics UK (COG-UK) Consortium | Jacquelyn Wynn, Mairead Hyland, The Lighthouse Lab in Alderley Park and Alex Alderton, Roberto Amato, Jeffrey Barrett, Sonia Goncalves, Ewan Harrison, David K. Jackson, Ian Johnston, Dominic Kwiatkowski, Cordelia Langford, John Sillitoe on behalf of the Wellcome Sanger Institute COVID-19 Surveillance Team |
| EPI_ISL_2568390 | Lighthouse Lab in Milton Keynes | Wellcome Sanger Institute for the COVID-19 Genomics UK (COG-UK) Consortium | The Lighthouse Lab in Milton Keynes and Alex Alderton, Roberto Amato, Jeffrey Barrett, Sonia Goncalves, Ewan Harrison, David K. Jackson, Ian Johnston, Dominic Kwiatkowski, Cordelia Langford, John Sillitoe on behalf of the Wellcome Sanger Institute COVID-19 Surveillance Team |
| EPI_ISL_2568391 | Lighthouse Lab in Glasgow | Wellcome Sanger Institute for the COVID-19 Genomics UK (COG-UK) Consortium | Harper VanSteenhouse, Yumi Kasai, David Gray, Carol Clugston, Anna Dominiczak and Alex Alderton, Roberto Amato, Jeffrey Barrett, Sonia Goncalves, Ewan Harrison, David K. Jackson, Ian Johnston, Dominic Kwiatkowski, Cordelia Langford, John Sillitoe on behalf of the Wellcome Sanger Institute COVID-19 Surveillance Team |
| EPI_ISL_2568392 | Lighthouse Lab in Alderley Park | Wellcome Sanger Institute for the COVID-19 Genomics UK (COG-UK) Consortium | Jacquelyn Wynn, Mairead Hyland, The Lighthouse Lab in Alderley Park and Alex Alderton, Roberto Amato, Jeffrey Barrett, Sonia Goncalves, Ewan Harrison, David K. Jackson, Ian Johnston, Dominic Kwiatkowski, Cordelia Langford, John Sillitoe on |

| | | | |
|---|---|---|---|
| | | Consortium | behalf of the Wellcome Sanger Institute COVID-19 Surveillance Team |
| EPI_ISL_2568393 | Lighthouse Lab in Milton Keynes | Wellcome Sanger Institute for the COVID-19 Genomics UK (COG-UK) Consortium | The Lighthouse Lab in Milton Keynes and Alex Alderton, Roberto Amato, Jeffrey Barrett, Sonia Goncalves, Ewan Harrison, David K. Jackson, Ian Johnston, Dominic Kwiatkowski, Cordelia Langford, John Sillitoe on behalf of the Wellcome Sanger Institute COVID-19 Surveillance Team |
| EPI_ISL_2568394 | Lighthouse Lab in Alderley Park | Wellcome Sanger Institute for the COVID-19 Genomics UK (COG-UK) Consortium | Jacquelyn Wynn, Mairead Hyland, The Lighthouse Lab in Alderley Park and Alex Alderton, Roberto Amato, Jeffrey Barrett, Sonia Goncalves, Ewan Harrison, David K. Jackson, Ian Johnston, Dominic Kwiatkowski, Cordelia Langford, John Sillitoe on behalf of the Wellcome Sanger Institute COVID-19 Surveillance Team |
| EPI_ISL_2568395 | Lighthouse Lab in Glasgow | Wellcome Sanger Institute for the COVID-19 Genomics UK (COG-UK) Consortium | Harper VanSteenhouse, Yumi Kasai, David Gray, Carol Clugston, Anna Dominiczak and Alex Alderton, Roberto Amato, Jeffrey Barrett, Sonia Goncalves, Ewan Harrison, David K. Jackson, Ian Johnston, Dominic Kwiatkowski, Cordelia Langford, John Sillitoe on behalf of the Wellcome Sanger Institute COVID-19 Surveillance Team |
| EPI_ISL_2568397 | Lighthouse Lab in Milton Keynes | Wellcome Sanger Institute for the COVID-19 Genomics UK (COG-UK) Consortium | The Lighthouse Lab in Milton Keynes and Alex Alderton, Roberto Amato, Jeffrey Barrett, Sonia Goncalves, Ewan Harrison, David K. Jackson, Ian Johnston, Dominic Kwiatkowski, Cordelia Langford, John Sillitoe on behalf of the Wellcome Sanger Institute COVID-19 Surveillance Team |
| EPI_ISL_2568398, EPI_ISL_2568399, EPI_ISL_2568400, EPI_ISL_2568401, EPI_ISL_2568402 | Lighthouse Lab in Alderley Park | Wellcome Sanger Institute for the COVID-19 Genomics UK (COG-UK) Consortium | Jacquelyn Wynn, Mairead Hyland, The Lighthouse Lab in Alderley Park and Alex Alderton, Roberto Amato, Jeffrey Barrett, Sonia Goncalves, Ewan Harrison, David K. Jackson, Ian Johnston, Dominic Kwiatkowski, Cordelia Langford, John Sillitoe on behalf of the Wellcome Sanger Institute COVID-19 Surveillance Team |
| EPI_ISL_2568403 | Lighthouse Lab in Glasgow | Wellcome Sanger Institute for the COVID-19 Genomics UK (COG-UK) Consortium | Harper VanSteenhouse, Yumi Kasai, David Gray, Carol Clugston, Anna Dominiczak and Alex Alderton, Roberto Amato, Jeffrey Barrett, Sonia Goncalves, Ewan Harrison, David K. Jackson, Ian Johnston, Dominic Kwiatkowski, Cordelia Langford, John Sillitoe on behalf of the Wellcome Sanger Institute COVID-19 Surveillance Team |
| EPI_ISL_2568404, EPI_ISL_2568406, EPI_ISL_2568407, EPI_ISL_2568408, EPI_ISL_2568409, EPI_ISL_2568410, EPI_ISL_2568411, EPI_ISL_2568412 | Lighthouse Lab in Alderley Park | Wellcome Sanger Institute for the COVID-19 Genomics UK (COG-UK) Consortium | Jacquelyn Wynn, Mairead Hyland, The Lighthouse Lab in Alderley Park and Alex Alderton, Roberto Amato, Jeffrey Barrett, Sonia Goncalves, Ewan Harrison, David K. Jackson, Ian Johnston, Dominic Kwiatkowski, Cordelia Langford, John Sillitoe on behalf of the Wellcome Sanger Institute COVID-19 Surveillance Team |
| EPI_ISL_2568413, EPI_ISL_2568414, EPI_ISL_2568415 | Lighthouse Lab in Glasgow | Wellcome Sanger Institute for the COVID-19 Genomics UK (COG-UK) Consortium | Harper VanSteenhouse, Yumi Kasai, David Gray, Carol Clugston, Anna Dominiczak and Alex Alderton, Roberto Amato, Jeffrey Barrett, Sonia Goncalves, Ewan Harrison, David K. Jackson, Ian Johnston, Dominic Kwiatkowski, Cordelia Langford, John Sillitoe on behalf of the Wellcome Sanger Institute COVID-19 Surveillance Team |
| EPI_ISL_2568416 | Lighthouse Lab in Alderley Park | Wellcome Sanger Institute for the COVID-19 Genomics UK (COG-UK) Consortium | Jacquelyn Wynn, Mairead Hyland, The Lighthouse Lab in Alderley Park and Alex Alderton, Roberto Amato, Jeffrey Barrett, Sonia Goncalves, Ewan Harrison, David K. Jackson, Ian Johnston, Dominic Kwiatkowski, Cordelia Langford, John Sillitoe on behalf of the Wellcome Sanger Institute COVID-19 Surveillance Team |
| EPI_ISL_2568417, EPI_ISL_2568418 | Lighthouse Lab in Glasgow | Wellcome Sanger Institute for the COVID-19 Genomics UK (COG-UK) Consortium | Harper VanSteenhouse, Yumi Kasai, David Gray, Carol Clugston, Anna Dominiczak and Alex Alderton, Roberto Amato, Jeffrey Barrett, Sonia Goncalves, Ewan Harrison, David K. Jackson, Ian Johnston, Dominic Kwiatkowski, Cordelia Langford, John Sillitoe on behalf of the Wellcome Sanger Institute COVID-19 Surveillance Team |
| EPI_ISL_2568419 | Lighthouse Lab in Alderley Park | Wellcome Sanger Institute for the COVID-19 Genomics UK (COG-UK) Consortium | Jacquelyn Wynn, Mairead Hyland, The Lighthouse Lab in Alderley Park and Alex Alderton, Roberto Amato, Jeffrey Barrett, Sonia Goncalves, Ewan Harrison, David K. Jackson, Ian Johnston, Dominic Kwiatkowski, Cordelia Langford, John Sillitoe on behalf of the Wellcome Sanger Institute COVID-19 Surveillance Team |
| EPI_ISL_2568420 | Lighthouse Lab in Glasgow | Wellcome Sanger Institute for the COVID-19 Genomics UK (COG-UK) Consortium | Harper VanSteenhouse, Yumi Kasai, David Gray, Carol Clugston, Anna Dominiczak and Alex Alderton, Roberto Amato, Jeffrey Barrett, Sonia Goncalves, Ewan Harrison, David K. Jackson, Ian Johnston, Dominic Kwiatkowski, Cordelia Langford, John Sillitoe on behalf of the Wellcome Sanger Institute COVID-19 Surveillance Team |
| EPI_ISL_2568421 | Lighthouse Lab in Alderley Park | Wellcome Sanger Institute for the COVID-19 Genomics UK (COG-UK) Consortium | Jacquelyn Wynn, Mairead Hyland, The Lighthouse Lab in Alderley Park and Alex Alderton, Roberto Amato, Jeffrey Barrett, Sonia Goncalves, Ewan Harrison, David K. Jackson, Ian Johnston, Dominic Kwiatkowski, Cordelia Langford, John Sillitoe on behalf of the Wellcome Sanger Institute COVID-19 Surveillance Team |
| EPI_ISL_2568422 | Lighthouse Lab in Glasgow | Wellcome Sanger Institute for the COVID-19 Genomics UK (COG-UK) Consortium | Harper VanSteenhouse, Yumi Kasai, David Gray, Carol Clugston, Anna Dominiczak and Alex Alderton, Roberto Amato, Jeffrey Barrett, Sonia Goncalves, Ewan Harrison, David K. Jackson, Ian Johnston, Dominic Kwiatkowski, Cordelia Langford, John Sillitoe on behalf of the Wellcome Sanger Institute COVID-19 Surveillance Team |
| EPI_ISL_2568423 | Lighthouse Lab in Alderley Park | Wellcome Sanger Institute for the COVID-19 Genomics UK (COG-UK) Consortium | Jacquelyn Wynn, Mairead Hyland, The Lighthouse Lab in Alderley Park and Alex Alderton, Roberto Amato, Jeffrey Barrett, Sonia Goncalves, Ewan Harrison, David K. Jackson, Ian Johnston, Dominic Kwiatkowski, Cordelia Langford, John Sillitoe on behalf of the Wellcome Sanger Institute COVID-19 Surveillance Team |
| EPI_ISL_2568424, EPI_ISL_2568425 | Lighthouse Lab in Glasgow | Wellcome Sanger Institute for the COVID-19 Genomics UK (COG-UK) Consortium | Harper VanSteenhouse, Yumi Kasai, David Gray, Carol Clugston, Anna Dominiczak and Alex Alderton, Roberto Amato, Jeffrey Barrett, Sonia Goncalves, Ewan Harrison, David K. Jackson, Ian Johnston, Dominic Kwiatkowski, Cordelia Langford, John Sillitoe on behalf of the Wellcome Sanger Institute COVID-19 Surveillance Team |
| EPI_ISL_2568426 | Lighthouse Lab in Alderley Park | Wellcome Sanger Institute for the COVID-19 Genomics UK (COG-UK) Consortium | Jacquelyn Wynn, Mairead Hyland, The Lighthouse Lab in Alderley Park and Alex Alderton, Roberto Amato, Jeffrey Barrett, Sonia Goncalves, Ewan Harrison, David K. Jackson, Ian Johnston, Dominic Kwiatkowski, Cordelia Langford, John Sillitoe on behalf of the Wellcome Sanger Institute COVID-19 Surveillance Team |
| EPI_ISL_2568427 | Lighthouse Lab in Glasgow | Wellcome Sanger Institute for the COVID-19 Genomics UK (COG-UK) Consortium | Harper VanSteenhouse, Yumi Kasai, David Gray, Carol Clugston, Anna Dominiczak and Alex Alderton, Roberto Amato, Jeffrey Barrett, Sonia Goncalves, Ewan Harrison, David K. Jackson, Ian Johnston, Dominic Kwiatkowski, Cordelia Langford, John Sillitoe on behalf of the Wellcome Sanger Institute COVID-19 Surveillance Team |
| EPI_ISL_2568428, EPI_ISL_2568429, EPI_ISL_2568430 | Lighthouse Lab in Alderley Park | Wellcome Sanger Institute for the COVID-19 Genomics UK (COG-UK) Consortium | Jacquelyn Wynn, Mairead Hyland, The Lighthouse Lab in Alderley Park and Alex Alderton, Roberto Amato, Jeffrey Barrett, Sonia Goncalves, Ewan Harrison, David K. Jackson, Ian Johnston, Dominic Kwiatkowski, Cordelia Langford, John Sillitoe on behalf of the Wellcome Sanger Institute COVID-19 Surveillance Team |
| EPI_ISL_2568431 | Lighthouse Lab in Glasgow | Wellcome Sanger Institute for the COVID-19 Genomics UK (COG-UK) Consortium | Harper VanSteenhouse, Yumi Kasai, David Gray, Carol Clugston, Anna Dominiczak and Alex Alderton, Roberto Amato, Jeffrey Barrett, Sonia Goncalves, Ewan Harrison, David K. Jackson, Ian Johnston, Dominic Kwiatkowski, Cordelia Langford, John Sillitoe on behalf of the Wellcome Sanger Institute COVID-19 Surveillance Team |
| EPI_ISL_2568432, EPI_ISL_2568433, EPI_ISL_2568434, EPI_ISL_2568435 | Lighthouse Lab in Alderley Park | Wellcome Sanger Institute for the COVID-19 Genomics UK (COG-UK) Consortium | Jacquelyn Wynn, Mairead Hyland, The Lighthouse Lab in Alderley Park and Alex Alderton, Roberto Amato, Jeffrey Barrett, Sonia Goncalves, Ewan Harrison, David K. Jackson, Ian Johnston, Dominic Kwiatkowski, Cordelia Langford, John Sillitoe on behalf of the Wellcome Sanger Institute COVID-19 Surveillance Team |
| EPI_ISL_2568436, EPI_ISL_2568437 | Lighthouse Lab in Glasgow | Wellcome Sanger Institute for the COVID-19 Genomics UK (COG-UK) Consortium | Harper VanSteenhouse, Yumi Kasai, David Gray, Carol Clugston, Anna Dominiczak and Alex Alderton, Roberto Amato, Jeffrey Barrett, Sonia Goncalves, Ewan Harrison, David K. Jackson, Ian Johnston, Dominic Kwiatkowski, Cordelia Langford, John Sillitoe on behalf of the Wellcome Sanger Institute COVID-19 Surveillance Team |
| EPI_ISL_2568438, EPI_ISL_2568439, EPI_ISL_2568440 | Lighthouse Lab in Alderley Park | Wellcome Sanger Institute for the COVID-19 Genomics UK (COG-UK) Consortium | Jacquelyn Wynn, Mairead Hyland, The Lighthouse Lab in Alderley Park and Alex Alderton, Roberto Amato, Jeffrey Barrett, Sonia Goncalves, Ewan Harrison, David K. Jackson, Ian Johnston, Dominic Kwiatkowski, Cordelia Langford, John Sillitoe on behalf of the Wellcome Sanger Institute COVID-19 Surveillance Team |
| EPI_ISL_2568441 | Lighthouse Lab in Glasgow | Wellcome Sanger Institute for the COVID-19 Genomics UK (COG-UK) Consortium | Harper VanSteenhouse, Yumi Kasai, David Gray, Carol Clugston, Anna Dominiczak and Alex Alderton, Roberto Amato, Jeffrey Barrett, Sonia Goncalves, Ewan Harrison, David K. Jackson, Ian Johnston, Dominic Kwiatkowski, Cordelia Langford, John Sillitoe on behalf of the Wellcome Sanger Institute COVID-19 Surveillance Team |
| EPI_ISL_2568442, EPI_ISL_2568443, EPI_ISL_2568444, EPI_ISL_2568445, EPI_ISL_2568446, EPI_ISL_2568447 | Lighthouse Lab in Alderley Park | Wellcome Sanger Institute for the COVID-19 Genomics UK (COG-UK) Consortium | Jacquelyn Wynn, Mairead Hyland, The Lighthouse Lab in Alderley Park and Alex Alderton, Roberto Amato, Jeffrey Barrett, Sonia Goncalves, Ewan Harrison, David K. Jackson, Ian Johnston, Dominic Kwiatkowski, Cordelia Langford, John Sillitoe on behalf of the Wellcome Sanger Institute COVID-19 Surveillance Team |
| EPI_ISL_2568448 | Lighthouse Lab in Glasgow | Wellcome Sanger Institute for the COVID-19 Genomics UK (COG-UK) Consortium | Harper VanSteenhouse, Yumi Kasai, David Gray, Carol Clugston, Anna Dominiczak and Alex Alderton, Roberto Amato, Jeffrey Barrett, Sonia Goncalves, Ewan Harrison, David K. Jackson, Ian Johnston, Dominic Kwiatkowski, Cordelia Langford, John Sillitoe on behalf of the Wellcome Sanger Institute COVID-19 Surveillance Team |
| EPI_ISL_2568449 | Lighthouse Lab in Alderley Park | Wellcome Sanger Institute for the COVID-19 Genomics UK (COG-UK) Consortium | Jacquelyn Wynn, Mairead Hyland, The Lighthouse Lab in Alderley Park and Alex Alderton, Roberto Amato, Jeffrey Barrett, Sonia Goncalves, Ewan Harrison, David K. Jackson, Ian Johnston, Dominic Kwiatkowski, Cordelia Langford, John Sillitoe on behalf of the Wellcome Sanger Institute COVID-19 Surveillance Team |
| EPI_ISL_2568450 | Lighthouse Lab in Glasgow | Wellcome Sanger Institute for the COVID-19 Genomics UK (COG-UK) Consortium | Harper VanSteenhouse, Yumi Kasai, David Gray, Carol Clugston, Anna Dominiczak and Alex Alderton, Roberto Amato, Jeffrey Barrett, Sonia Goncalves, Ewan Harrison, David K. Jackson, Ian Johnston, Dominic Kwiatkowski, Cordelia Langford, John Sillitoe on behalf of the Wellcome Sanger Institute COVID-19 Surveillance Team |
| EPI_ISL_2568451, EPI_ISL_2568452, EPI_ISL_2568453, EPI_ISL_2568454, EPI_ISL_2568455 | Lighthouse Lab in Alderley Park | Wellcome Sanger Institute for the COVID-19 Genomics UK (COG-UK) Consortium | Jacquelyn Wynn, Mairead Hyland, The Lighthouse Lab in Alderley Park and Alex Alderton, Roberto Amato, Jeffrey Barrett, Sonia Goncalves, Ewan Harrison, David K. Jackson, Ian Johnston, Dominic Kwiatkowski, Cordelia Langford, John Sillitoe on behalf of the Wellcome Sanger Institute COVID-19 Surveillance Team |
| EPI_ISL_2568456 | Lighthouse Lab in Glasgow | Wellcome Sanger Institute for the COVID-19 Genomics UK (COG-UK) Consortium | Harper VanSteenhouse, Yumi Kasai, David Gray, Carol Clugston, Anna Dominiczak and Alex Alderton, Roberto Amato, Jeffrey Barrett, Sonia Goncalves, Ewan Harrison, David K. Jackson, Ian Johnston, Dominic Kwiatkowski, Cordelia Langford, John Sillitoe on behalf of the Wellcome Sanger Institute COVID-19 Surveillance Team |
| EPI_ISL_2568457, EPI_ISL_2568458, EPI_ISL_2568459, EPI_ISL_2568460, EPI_ISL_2568461 | Lighthouse Lab in Alderley Park | Wellcome Sanger Institute for the COVID-19 Genomics UK (COG-UK) Consortium | Jacquelyn Wynn, Mairead Hyland, The Lighthouse Lab in Alderley Park and Alex Alderton, Roberto Amato, Jeffrey Barrett, Sonia Goncalves, Ewan Harrison, David K. Jackson, Ian Johnston, Dominic Kwiatkowski, Cordelia Langford, John Sillitoe on behalf of the Wellcome Sanger Institute COVID-19 Surveillance Team |
| EPI_ISL_2568462, EPI_ISL_2568463 | Lighthouse Lab in Glasgow | Wellcome Sanger Institute for the COVID-19 Genomics UK (COG-UK) Consortium | Harper VanSteenhouse, Yumi Kasai, David Gray, Carol Clugston, Anna Dominiczak and Alex Alderton, Roberto Amato, Jeffrey Barrett, Sonia Goncalves, Ewan Harrison, David K. Jackson, Ian Johnston, Dominic Kwiatkowski, Cordelia Langford, John Sillitoe on behalf of the Wellcome Sanger Institute COVID-19 Surveillance Team |
| EPI_ISL_2568464 | Lighthouse Lab in Alderley Park | Wellcome Sanger Institute for the COVID-19 Genomics UK (COG-UK) Consortium | Jacquelyn Wynn, Mairead Hyland, The Lighthouse Lab in Alderley Park and Alex Alderton, Roberto Amato, Jeffrey Barrett, Sonia Goncalves, Ewan Harrison, David K. Jackson, Ian Johnston, Dominic Kwiatkowski, Cordelia Langford, John Sillitoe on behalf of the Wellcome Sanger Institute COVID-19 Surveillance Team |
| EPI_ISL_2568465 | Lighthouse Lab in Glasgow | Wellcome Sanger Institute for the COVID-19 Genomics UK (COG-UK) Consortium | Harper VanSteenhouse, Yumi Kasai, David Gray, Carol Clugston, Anna Dominiczak and Alex Alderton, Roberto Amato, Jeffrey Barrett, Sonia Goncalves, Ewan Harrison, David K. Jackson, Ian Johnston, Dominic Kwiatkowski, Cordelia Langford, John Sillitoe on behalf of the Wellcome Sanger Institute COVID-19 Surveillance Team |
| EPI_ISL_2568466 | Lighthouse Lab in Alderley Park | Wellcome Sanger Institute for the COVID-19 Genomics UK (COG-UK) | Jacquelyn Wynn, Mairead Hyland, The Lighthouse Lab in Alderley Park and Alex Alderton, Roberto Amato, Jeffrey Barrett, Sonia Goncalves, Ewan Harrison, David K. Jackson, Ian Johnston, Dominic Kwiatkowski, Cordelia Langford, John Sillitoe on |

| | | | |
|---|---|---|---|
| | | Consortium | behalf of the Wellcome Sanger Institute COVID-19 Surveillance Team |
| EPI_ISL_2568467 | Lighthouse Lab in Glasgow | Wellcome Sanger Institute for the COVID-19 Genomics UK (COG-UK) Consortium | Harper VanSteenhouse, Yumi Kasai, David Gray, Carol Clugston, Anna Dominiczak and Alex Alderton, Roberto Amato, Jeffrey Barrett, Sonia Goncalves, Ewan Harrison, David K. Jackson, Ian Johnston, Dominic Kwiatkowski, Cordelia Langford, John Sillitoe on behalf of the Wellcome Sanger Institute COVID-19 Surveillance Team |
| EPI_ISL_2568468, EPI_ISL_2568469, EPI_ISL_2568470, EPI_ISL_2568471 | Lighthouse Lab in Alderley Park | Wellcome Sanger Institute for the COVID-19 Genomics UK (COG-UK) Consortium | Jacquelyn Wynn, Mairead Hyland, The Lighthouse Lab in Alderley Park and Alex Alderton, Roberto Amato, Jeffrey Barrett, Sonia Goncalves, Ewan Harrison, David K. Jackson, Ian Johnston, Dominic Kwiatkowski, Cordelia Langford, John Sillitoe on behalf of the Wellcome Sanger Institute COVID-19 Surveillance Team |
| EPI_ISL_2568472 | Lighthouse Lab in Glasgow | Wellcome Sanger Institute for the COVID-19 Genomics UK (COG-UK) Consortium | Harper VanSteenhouse, Yumi Kasai, David Gray, Carol Clugston, Anna Dominiczak and Alex Alderton, Roberto Amato, Jeffrey Barrett, Sonia Goncalves, Ewan Harrison, David K. Jackson, Ian Johnston, Dominic Kwiatkowski, Cordelia Langford, John Sillitoe on behalf of the Wellcome Sanger Institute COVID-19 Surveillance Team |
| EPI_ISL_2568473 | Lighthouse Lab in Alderley Park | Wellcome Sanger Institute for the COVID-19 Genomics UK (COG-UK) Consortium | Jacquelyn Wynn, Mairead Hyland, The Lighthouse Lab in Alderley Park and Alex Alderton, Roberto Amato, Jeffrey Barrett, Sonia Goncalves, Ewan Harrison, David K. Jackson, Ian Johnston, Dominic Kwiatkowski, Cordelia Langford, John Sillitoe on behalf of the Wellcome Sanger Institute COVID-19 Surveillance Team |
| EPI_ISL_2568474 | Lighthouse Lab in Glasgow | Wellcome Sanger Institute for the COVID-19 Genomics UK (COG-UK) Consortium | Harper VanSteenhouse, Yumi Kasai, David Gray, Carol Clugston, Anna Dominiczak and Alex Alderton, Roberto Amato, Jeffrey Barrett, Sonia Goncalves, Ewan Harrison, David K. Jackson, Ian Johnston, Dominic Kwiatkowski, Cordelia Langford, John Sillitoe on behalf of the Wellcome Sanger Institute COVID-19 Surveillance Team |
| EPI_ISL_2568475 | Lighthouse Lab in Alderley Park | Wellcome Sanger Institute for the COVID-19 Genomics UK (COG-UK) Consortium | Jacquelyn Wynn, Mairead Hyland, The Lighthouse Lab in Alderley Park and Alex Alderton, Roberto Amato, Jeffrey Barrett, Sonia Goncalves, Ewan Harrison, David K. Jackson, Ian Johnston, Dominic Kwiatkowski, Cordelia Langford, John Sillitoe on behalf of the Wellcome Sanger Institute COVID-19 Surveillance Team |
| EPI_ISL_2568476 | Lighthouse Lab in Glasgow | Wellcome Sanger Institute for the COVID-19 Genomics UK (COG-UK) Consortium | Harper VanSteenhouse, Yumi Kasai, David Gray, Carol Clugston, Anna Dominiczak and Alex Alderton, Roberto Amato, Jeffrey Barrett, Sonia Goncalves, Ewan Harrison, David K. Jackson, Ian Johnston, Dominic Kwiatkowski, Cordelia Langford, John Sillitoe on behalf of the Wellcome Sanger Institute COVID-19 Surveillance Team |
| EPI_ISL_2568477 | Lighthouse Lab in Alderley Park | Wellcome Sanger Institute for the COVID-19 Genomics UK (COG-UK) Consortium | Jacquelyn Wynn, Mairead Hyland, The Lighthouse Lab in Alderley Park and Alex Alderton, Roberto Amato, Jeffrey Barrett, Sonia Goncalves, Ewan Harrison, David K. Jackson, Ian Johnston, Dominic Kwiatkowski, Cordelia Langford, John Sillitoe on behalf of the Wellcome Sanger Institute COVID-19 Surveillance Team |
| EPI_ISL_2568478, EPI_ISL_2568479 | Lighthouse Lab in Glasgow | Wellcome Sanger Institute for the COVID-19 Genomics UK (COG-UK) Consortium | Harper VanSteenhouse, Yumi Kasai, David Gray, Carol Clugston, Anna Dominiczak and Alex Alderton, Roberto Amato, Jeffrey Barrett, Sonia Goncalves, Ewan Harrison, David K. Jackson, Ian Johnston, Dominic Kwiatkowski, Cordelia Langford, John Sillitoe on behalf of the Wellcome Sanger Institute COVID-19 Surveillance Team |
| EPI_ISL_2568480, EPI_ISL_2568481, EPI_ISL_2568482, EPI_ISL_2568483, EPI_ISL_2568484, EPI_ISL_2568485, EPI_ISL_2568486, EPI_ISL_2568487, EPI_ISL_2568488, EPI_ISL_2568489, EPI_ISL_2568490, EPI_ISL_2568491, EPI_ISL_2568492, EPI_ISL_2568493, EPI_ISL_2568494, EPI_ISL_2568495, EPI_ISL_2568496 | | | |
| see above | Lighthouse Lab in Alderley Park | Wellcome Sanger Institute for the COVID-19 Genomics UK (COG-UK) Consortium | Jacquelyn Wynn, Mairead Hyland, The Lighthouse Lab in Alderley Park and Alex Alderton, Roberto Amato, Jeffrey Barrett, Sonia Goncalves, Ewan Harrison, David K. Jackson, Ian Johnston, Dominic Kwiatkowski, Cordelia Langford, John Sillitoe on behalf of the Wellcome Sanger Institute COVID-19 Surveillance Team |
| EPI_ISL_2568497 | Lighthouse Lab in Glasgow | Wellcome Sanger Institute for the COVID-19 Genomics UK (COG-UK) Consortium | Harper VanSteenhouse, Yumi Kasai, David Gray, Carol Clugston, Anna Dominiczak and Alex Alderton, Roberto Amato, Jeffrey Barrett, Sonia Goncalves, Ewan Harrison, David K. Jackson, Ian Johnston, Dominic Kwiatkowski, Cordelia Langford, John Sillitoe on behalf of the Wellcome Sanger Institute COVID-19 Surveillance Team |
| EPI_ISL_2568498, EPI_ISL_2568499, EPI_ISL_2568500 | Lighthouse Lab in Alderley Park | Wellcome Sanger Institute for the COVID-19 Genomics UK (COG-UK) Consortium | Jacquelyn Wynn, Mairead Hyland, The Lighthouse Lab in Alderley Park and Alex Alderton, Roberto Amato, Jeffrey Barrett, Sonia Goncalves, Ewan Harrison, David K. Jackson, Ian Johnston, Dominic Kwiatkowski, Cordelia Langford, John Sillitoe on behalf of the Wellcome Sanger Institute COVID-19 Surveillance Team |
| EPI_ISL_2568502 | Lighthouse Lab in Glasgow | Wellcome Sanger Institute for the COVID-19 Genomics UK (COG-UK) Consortium | Harper VanSteenhouse, Yumi Kasai, David Gray, Carol Clugston, Anna Dominiczak and Alex Alderton, Roberto Amato, Jeffrey Barrett, Sonia Goncalves, Ewan Harrison, David K. Jackson, Ian Johnston, Dominic Kwiatkowski, Cordelia Langford, John Sillitoe on behalf of the Wellcome Sanger Institute COVID-19 Surveillance Team |
| EPI_ISL_2568503, EPI_ISL_2568504, EPI_ISL_2568505, EPI_ISL_2568506, EPI_ISL_2568507, EPI_ISL_2568508 | Lighthouse Lab in Alderley Park | Wellcome Sanger Institute for the COVID-19 Genomics UK (COG-UK) Consortium | Jacquelyn Wynn, Mairead Hyland, The Lighthouse Lab in Alderley Park and Alex Alderton, Roberto Amato, Jeffrey Barrett, Sonia Goncalves, Ewan Harrison, David K. Jackson, Ian Johnston, Dominic Kwiatkowski, Cordelia Langford, John Sillitoe on behalf of the Wellcome Sanger Institute COVID-19 Surveillance Team |
| EPI_ISL_2568509 | Lighthouse Lab in Glasgow | Wellcome Sanger Institute for the COVID-19 Genomics UK (COG-UK) Consortium | Harper VanSteenhouse, Yumi Kasai, David Gray, Carol Clugston, Anna Dominiczak and Alex Alderton, Roberto Amato, Jeffrey Barrett, Sonia Goncalves, Ewan Harrison, David K. Jackson, Ian Johnston, Dominic Kwiatkowski, Cordelia Langford, John Sillitoe on behalf of the Wellcome Sanger Institute COVID-19 Surveillance Team |
| EPI_ISL_2568510, EPI_ISL_2568511 | Lighthouse Lab in Alderley Park | Wellcome Sanger Institute for the COVID-19 Genomics UK (COG-UK) Consortium | Jacquelyn Wynn, Mairead Hyland, The Lighthouse Lab in Alderley Park and Alex Alderton, Roberto Amato, Jeffrey Barrett, Sonia Goncalves, Ewan Harrison, David K. Jackson, Ian Johnston, Dominic Kwiatkowski, Cordelia Langford, John Sillitoe on behalf of the Wellcome Sanger Institute COVID-19 Surveillance Team |
| EPI_ISL_2568512 | Lighthouse Lab in Glasgow | Wellcome Sanger Institute for the COVID-19 Genomics UK (COG-UK) Consortium | Harper VanSteenhouse, Yumi Kasai, David Gray, Carol Clugston, Anna Dominiczak and Alex Alderton, Roberto Amato, Jeffrey Barrett, Sonia Goncalves, Ewan Harrison, David K. Jackson, Ian Johnston, Dominic Kwiatkowski, Cordelia Langford, John Sillitoe on behalf of the Wellcome Sanger Institute COVID-19 Surveillance Team |
| EPI_ISL_2568513, EPI_ISL_2568514 | Lighthouse Lab in Alderley Park | Wellcome Sanger Institute for the COVID-19 Genomics UK (COG-UK) Consortium | Jacquelyn Wynn, Mairead Hyland, The Lighthouse Lab in Alderley Park and Alex Alderton, Roberto Amato, Jeffrey Barrett, Sonia Goncalves, Ewan Harrison, David K. Jackson, Ian Johnston, Dominic Kwiatkowski, Cordelia Langford, John Sillitoe on behalf of the Wellcome Sanger Institute COVID-19 Surveillance Team |
| EPI_ISL_2568515 | Lighthouse Lab in Glasgow | Wellcome Sanger Institute for the COVID-19 Genomics UK (COG-UK) Consortium | Harper VanSteenhouse, Yumi Kasai, David Gray, Carol Clugston, Anna Dominiczak and Alex Alderton, Roberto Amato, Jeffrey Barrett, Sonia Goncalves, Ewan Harrison, David K. Jackson, Ian Johnston, Dominic Kwiatkowski, Cordelia Langford, John Sillitoe on behalf of the Wellcome Sanger Institute COVID-19 Surveillance Team |
| EPI_ISL_2568516, EPI_ISL_2568517, EPI_ISL_2568518 | Lighthouse Lab in Alderley Park | Wellcome Sanger Institute for the COVID-19 Genomics UK (COG-UK) Consortium | Jacquelyn Wynn, Mairead Hyland, The Lighthouse Lab in Alderley Park and Alex Alderton, Roberto Amato, Jeffrey Barrett, Sonia Goncalves, Ewan Harrison, David K. Jackson, Ian Johnston, Dominic Kwiatkowski, Cordelia Langford, John Sillitoe on behalf of the Wellcome Sanger Institute COVID-19 Surveillance Team |
| EPI_ISL_2568519 | Lighthouse Lab in Glasgow | Wellcome Sanger Institute for the COVID-19 Genomics UK (COG-UK) Consortium | Harper VanSteenhouse, Yumi Kasai, David Gray, Carol Clugston, Anna Dominiczak and Alex Alderton, Roberto Amato, Jeffrey Barrett, Sonia Goncalves, Ewan Harrison, David K. Jackson, Ian Johnston, Dominic Kwiatkowski, Cordelia Langford, John Sillitoe on behalf of the Wellcome Sanger Institute COVID-19 Surveillance Team |
| EPI_ISL_2568520, EPI_ISL_2568521 | Lighthouse Lab in Alderley Park | Wellcome Sanger Institute for the COVID-19 Genomics UK (COG-UK) Consortium | Jacquelyn Wynn, Mairead Hyland, The Lighthouse Lab in Alderley Park and Alex Alderton, Roberto Amato, Jeffrey Barrett, Sonia Goncalves, Ewan Harrison, David K. Jackson, Ian Johnston, Dominic Kwiatkowski, Cordelia Langford, John Sillitoe on behalf of the Wellcome Sanger Institute COVID-19 Surveillance Team |
| EPI_ISL_2568522 | Lighthouse Lab in Glasgow | Wellcome Sanger Institute for the COVID-19 Genomics UK (COG-UK) Consortium | Harper VanSteenhouse, Yumi Kasai, David Gray, Carol Clugston, Anna Dominiczak and Alex Alderton, Roberto Amato, Jeffrey Barrett, Sonia Goncalves, Ewan Harrison, David K. Jackson, Ian Johnston, Dominic Kwiatkowski, Cordelia Langford, John Sillitoe on behalf of the Wellcome Sanger Institute COVID-19 Surveillance Team |
| EPI_ISL_2568523, EPI_ISL_2568524, EPI_ISL_2568525, EPI_ISL_2568526, EPI_ISL_2568527, EPI_ISL_2568528, EPI_ISL_2568530, EPI_ISL_2568531, EPI_ISL_2568532, EPI_ISL_2568533 | Lighthouse Lab in Alderley Park | Wellcome Sanger Institute for the COVID-19 Genomics UK (COG-UK) Consortium | Jacquelyn Wynn, Mairead Hyland, The Lighthouse Lab in Alderley Park and Alex Alderton, Roberto Amato, Jeffrey Barrett, Sonia Goncalves, Ewan Harrison, David K. Jackson, Ian Johnston, Dominic Kwiatkowski, Cordelia Langford, John Sillitoe on behalf of the Wellcome Sanger Institute COVID-19 Surveillance Team |
| EPI_ISL_2568534 | Lighthouse Lab in Glasgow | Wellcome Sanger Institute for the COVID-19 Genomics UK (COG-UK) Consortium | Harper VanSteenhouse, Yumi Kasai, David Gray, Carol Clugston, Anna Dominiczak and Alex Alderton, Roberto Amato, Jeffrey Barrett, Sonia Goncalves, Ewan Harrison, David K. Jackson, Ian Johnston, Dominic Kwiatkowski, Cordelia Langford, John Sillitoe on behalf of the Wellcome Sanger Institute COVID-19 Surveillance Team |
| EPI_ISL_2568535, EPI_ISL_2568536, EPI_ISL_2568537 | Lighthouse Lab in Alderley Park | Wellcome Sanger Institute for the COVID-19 Genomics UK (COG-UK) Consortium | Jacquelyn Wynn, Mairead Hyland, The Lighthouse Lab in Alderley Park and Alex Alderton, Roberto Amato, Jeffrey Barrett, Sonia Goncalves, Ewan Harrison, David K. Jackson, Ian Johnston, Dominic Kwiatkowski, Cordelia Langford, John Sillitoe on behalf of the Wellcome Sanger Institute COVID-19 Surveillance Team |
| EPI_ISL_2568538 | Lighthouse Lab in Glasgow | Wellcome Sanger Institute for the COVID-19 Genomics UK (COG-UK) Consortium | Harper VanSteenhouse, Yumi Kasai, David Gray, Carol Clugston, Anna Dominiczak and Alex Alderton, Roberto Amato, Jeffrey Barrett, Sonia Goncalves, Ewan Harrison, David K. Jackson, Ian Johnston, Dominic Kwiatkowski, Cordelia Langford, John Sillitoe on behalf of the Wellcome Sanger Institute COVID-19 Surveillance Team |
| EPI_ISL_2568540, EPI_ISL_2568541, EPI_ISL_2568542, EPI_ISL_2568543, EPI_ISL_2568545, EPI_ISL_2568547, EPI_ISL_2568548, EPI_ISL_2568549, EPI_ISL_2568550, EPI_ISL_2568551, EPI_ISL_2568552, EPI_ISL_2568553 | | | |
| see above | Lighthouse Lab in Alderley Park | Wellcome Sanger Institute for the COVID-19 Genomics UK (COG-UK) Consortium | Jacquelyn Wynn, Mairead Hyland, The Lighthouse Lab in Alderley Park and Alex Alderton, Roberto Amato, Jeffrey Barrett, Sonia Goncalves, Ewan Harrison, David K. Jackson, Ian Johnston, Dominic Kwiatkowski, Cordelia Langford, John Sillitoe on behalf of the Wellcome Sanger Institute COVID-19 Surveillance Team |
| EPI_ISL_2568554 | Lighthouse Lab in Glasgow | Wellcome Sanger Institute for the COVID-19 Genomics UK (COG-UK) Consortium | Harper VanSteenhouse, Yumi Kasai, David Gray, Carol Clugston, Anna Dominiczak and Alex Alderton, Roberto Amato, Jeffrey Barrett, Sonia Goncalves, Ewan Harrison, David K. Jackson, Ian Johnston, Dominic Kwiatkowski, Cordelia Langford, John Sillitoe on behalf of the Wellcome Sanger Institute COVID-19 Surveillance Team |
| EPI_ISL_2568555, EPI_ISL_2568556, EPI_ISL_2568558 | Lighthouse Lab in Alderley Park | Wellcome Sanger Institute for the COVID-19 Genomics UK (COG-UK) Consortium | Jacquelyn Wynn, Mairead Hyland, The Lighthouse Lab in Alderley Park and Alex Alderton, Roberto Amato, Jeffrey Barrett, Sonia Goncalves, Ewan Harrison, David K. Jackson, Ian Johnston, Dominic Kwiatkowski, Cordelia Langford, John Sillitoe on behalf of the Wellcome Sanger Institute COVID-19 Surveillance Team |
| EPI_ISL_2568559 | Lighthouse Lab in Glasgow | Wellcome Sanger Institute for the COVID-19 Genomics UK (COG-UK) Consortium | Harper VanSteenhouse, Yumi Kasai, David Gray, Carol Clugston, Anna Dominiczak and Alex Alderton, Roberto Amato, Jeffrey Barrett, Sonia Goncalves, Ewan Harrison, David K. Jackson, Ian Johnston, Dominic Kwiatkowski, Cordelia Langford, John Sillitoe on behalf of the Wellcome Sanger Institute COVID-19 Surveillance Team |
| EPI_ISL_2568560, EPI_ISL_2568561, EPI_ISL_2568562 | Lighthouse Lab in Alderley Park | Wellcome Sanger Institute for the COVID-19 Genomics UK (COG-UK) Consortium | Jacquelyn Wynn, Mairead Hyland, The Lighthouse Lab in Alderley Park and Alex Alderton, Roberto Amato, Jeffrey Barrett, Sonia Goncalves, Ewan Harrison, David K. Jackson, Ian Johnston, Dominic Kwiatkowski, Cordelia Langford, John Sillitoe on behalf of the Wellcome Sanger Institute COVID-19 Surveillance Team |
| EPI_ISL_2568563 | Lighthouse Lab in Glasgow | Wellcome Sanger Institute for the COVID-19 Genomics UK (COG-UK) Consortium | Harper VanSteenhouse, Yumi Kasai, David Gray, Carol Clugston, Anna Dominiczak and Alex Alderton, Roberto Amato, Jeffrey Barrett, Sonia Goncalves, Ewan Harrison, David K. Jackson, Ian Johnston, Dominic Kwiatkowski, Cordelia Langford, John Sillitoe on behalf of the Wellcome Sanger Institute COVID-19 Surveillance Team |
| EPI_ISL_2568564, EPI_ISL_2568565 | Lighthouse Lab in Alderley Park | Wellcome Sanger Institute for the COVID-19 Genomics UK (COG-UK) Consortium | Jacquelyn Wynn, Mairead Hyland, The Lighthouse Lab in Alderley Park and Alex Alderton, Roberto Amato, Jeffrey Barrett, Sonia Goncalves, Ewan Harrison, David K. Jackson, Ian Johnston, Dominic Kwiatkowski, Cordelia Langford, John Sillitoe on behalf of the Wellcome Sanger Institute COVID-19 Surveillance Team |
| EPI_ISL_2568566, EPI_ISL_2568567, EPI_ISL_2568568 | Lighthouse Lab in Glasgow | Wellcome Sanger Institute for the COVID-19 Genomics UK (COG-UK) Consortium | Harper VanSteenhouse, Yumi Kasai, David Gray, Carol Clugston, Anna Dominiczak and Alex Alderton, Roberto Amato, Jeffrey Barrett, Sonia Goncalves, Ewan Harrison, David K. Jackson, Ian Johnston, Dominic Kwiatkowski, Cordelia Langford, John Sillitoe on behalf of the Wellcome Sanger Institute COVID-19 Surveillance Team |

| | | | |
|---|---|---|---|
| EPI_ISL_2568569, EPI_ISL_2568570, EPI_ISL_2568571, EPI_ISL_2568572, EPI_ISL_2568573, EPI_ISL_2568574, EPI_ISL_2568575, EPI_ISL_2568576, EPI_ISL_2568577, EPI_ISL_2568578 | Lighthouse Lab in Alderley Park | Wellcome Sanger Institute for the COVID-19 Genomics UK (COG-UK) Consortium | Jacquelyn Wynn, Mairead Hyland, The Lighthouse Lab in Alderley Park and Alex Alderton, Roberto Amato, Jeffrey Barrett, Sonia Goncalves, Ewan Harrison, David K. Jackson, Ian Johnston, Dominic Kwiatkowski, Cordelia Langford, John Sillitoe on behalf of the Wellcome Sanger Institute COVID-19 Surveillance Team |
| EPI_ISL_2568579, EPI_ISL_2568580, EPI_ISL_2568581 | Lighthouse Lab in Glasgow | Wellcome Sanger Institute for the COVID-19 Genomics UK (COG-UK) Consortium | Harper VanSteenhouse, Yumi Kasai, David Gray, Carol Clugston, Anna Dominiczak and Alex Alderton, Roberto Amato, Jeffrey Barrett, Sonia Goncalves, Ewan Harrison, David K. Jackson, Ian Johnston, Dominic Kwiatkowski, Cordelia Langford, John Sillitoe on behalf of the Wellcome Sanger Institute COVID-19 Surveillance Team |
| EPI_ISL_2568582 | Lighthouse Lab in Alderley Park | Wellcome Sanger Institute for the COVID-19 Genomics UK (COG-UK) Consortium | Jacquelyn Wynn, Mairead Hyland, The Lighthouse Lab in Alderley Park and Alex Alderton, Roberto Amato, Jeffrey Barrett, Sonia Goncalves, Ewan Harrison, David K. Jackson, Ian Johnston, Dominic Kwiatkowski, Cordelia Langford, John Sillitoe on behalf of the Wellcome Sanger Institute COVID-19 Surveillance Team |
| EPI_ISL_2568583 | Lighthouse Lab in Glasgow | Wellcome Sanger Institute for the COVID-19 Genomics UK (COG-UK) Consortium | Harper VanSteenhouse, Yumi Kasai, David Gray, Carol Clugston, Anna Dominiczak and Alex Alderton, Roberto Amato, Jeffrey Barrett, Sonia Goncalves, Ewan Harrison, David K. Jackson, Ian Johnston, Dominic Kwiatkowski, Cordelia Langford, John Sillitoe on behalf of the Wellcome Sanger Institute COVID-19 Surveillance Team |
| EPI_ISL_2568584, EPI_ISL_2568585 | Lighthouse Lab in Alderley Park | Wellcome Sanger Institute for the COVID-19 Genomics UK (COG-UK) Consortium | Jacquelyn Wynn, Mairead Hyland, The Lighthouse Lab in Alderley Park and Alex Alderton, Roberto Amato, Jeffrey Barrett, Sonia Goncalves, Ewan Harrison, David K. Jackson, Ian Johnston, Dominic Kwiatkowski, Cordelia Langford, John Sillitoe on behalf of the Wellcome Sanger Institute COVID-19 Surveillance Team |
| EPI_ISL_2568586, EPI_ISL_2568587, EPI_ISL_2568588 | Lighthouse Lab in Glasgow | Wellcome Sanger Institute for the COVID-19 Genomics UK (COG-UK) Consortium | Harper VanSteenhouse, Yumi Kasai, David Gray, Carol Clugston, Anna Dominiczak and Alex Alderton, Roberto Amato, Jeffrey Barrett, Sonia Goncalves, Ewan Harrison, David K. Jackson, Ian Johnston, Dominic Kwiatkowski, Cordelia Langford, John Sillitoe on behalf of the Wellcome Sanger Institute COVID-19 Surveillance Team |
| EPI_ISL_2568589, EPI_ISL_2568590, EPI_ISL_2568591, EPI_ISL_2568592 | Lighthouse Lab in Alderley Park | Wellcome Sanger Institute for the COVID-19 Genomics UK (COG-UK) Consortium | Jacquelyn Wynn, Mairead Hyland, The Lighthouse Lab in Alderley Park and Alex Alderton, Roberto Amato, Jeffrey Barrett, Sonia Goncalves, Ewan Harrison, David K. Jackson, Ian Johnston, Dominic Kwiatkowski, Cordelia Langford, John Sillitoe on behalf of the Wellcome Sanger Institute COVID-19 Surveillance Team |
| EPI_ISL_2568593 | Lighthouse Lab in Glasgow | Wellcome Sanger Institute for the COVID-19 Genomics UK (COG-UK) Consortium | Harper VanSteenhouse, Yumi Kasai, David Gray, Carol Clugston, Anna Dominiczak and Alex Alderton, Roberto Amato, Jeffrey Barrett, Sonia Goncalves, Ewan Harrison, David K. Jackson, Ian Johnston, Dominic Kwiatkowski, Cordelia Langford, John Sillitoe on behalf of the Wellcome Sanger Institute COVID-19 Surveillance Team |
| EPI_ISL_2568594 | Lighthouse Lab in Alderley Park | Wellcome Sanger Institute for the COVID-19 Genomics UK (COG-UK) Consortium | Jacquelyn Wynn, Mairead Hyland, The Lighthouse Lab in Alderley Park and Alex Alderton, Roberto Amato, Jeffrey Barrett, Sonia Goncalves, Ewan Harrison, David K. Jackson, Ian Johnston, Dominic Kwiatkowski, Cordelia Langford, John Sillitoe on behalf of the Wellcome Sanger Institute COVID-19 Surveillance Team |
| EPI_ISL_2568595 | Lighthouse Lab in Glasgow | Wellcome Sanger Institute for the COVID-19 Genomics UK (COG-UK) Consortium | Harper VanSteenhouse, Yumi Kasai, David Gray, Carol Clugston, Anna Dominiczak and Alex Alderton, Roberto Amato, Jeffrey Barrett, Sonia Goncalves, Ewan Harrison, David K. Jackson, Ian Johnston, Dominic Kwiatkowski, Cordelia Langford, John Sillitoe on behalf of the Wellcome Sanger Institute COVID-19 Surveillance Team |
| EPI_ISL_2568596, EPI_ISL_2568598, EPI_ISL_2568599, EPI_ISL_2568600, EPI_ISL_2568601, EPI_ISL_2568602 | Lighthouse Lab in Alderley Park | Wellcome Sanger Institute for the COVID-19 Genomics UK (COG-UK) Consortium | Jacquelyn Wynn, Mairead Hyland, The Lighthouse Lab in Alderley Park and Alex Alderton, Roberto Amato, Jeffrey Barrett, Sonia Goncalves, Ewan Harrison, David K. Jackson, Ian Johnston, Dominic Kwiatkowski, Cordelia Langford, John Sillitoe on behalf of the Wellcome Sanger Institute COVID-19 Surveillance Team |
| EPI_ISL_2568603 | Lighthouse Lab in Glasgow | Wellcome Sanger Institute for the COVID-19 Genomics UK (COG-UK) Consortium | Harper VanSteenhouse, Yumi Kasai, David Gray, Carol Clugston, Anna Dominiczak and Alex Alderton, Roberto Amato, Jeffrey Barrett, Sonia Goncalves, Ewan Harrison, David K. Jackson, Ian Johnston, Dominic Kwiatkowski, Cordelia Langford, John Sillitoe on behalf of the Wellcome Sanger Institute COVID-19 Surveillance Team |
| EPI_ISL_2568604, EPI_ISL_2568605, EPI_ISL_2568607, EPI_ISL_2568608, EPI_ISL_2568609, EPI_ISL_2568610 | Lighthouse Lab in Alderley Park | Wellcome Sanger Institute for the COVID-19 Genomics UK (COG-UK) Consortium | Jacquelyn Wynn, Mairead Hyland, The Lighthouse Lab in Alderley Park and Alex Alderton, Roberto Amato, Jeffrey Barrett, Sonia Goncalves, Ewan Harrison, David K. Jackson, Ian Johnston, Dominic Kwiatkowski, Cordelia Langford, John Sillitoe on behalf of the Wellcome Sanger Institute COVID-19 Surveillance Team |
| EPI_ISL_2568611 | Lighthouse Lab in Glasgow | Wellcome Sanger Institute for the COVID-19 Genomics UK (COG-UK) Consortium | Harper VanSteenhouse, Yumi Kasai, David Gray, Carol Clugston, Anna Dominiczak and Alex Alderton, Roberto Amato, Jeffrey Barrett, Sonia Goncalves, Ewan Harrison, David K. Jackson, Ian Johnston, Dominic Kwiatkowski, Cordelia Langford, John Sillitoe on behalf of the Wellcome Sanger Institute COVID-19 Surveillance Team |
| EPI_ISL_2568612, EPI_ISL_2568613, EPI_ISL_2568614, EPI_ISL_2568615, EPI_ISL_2568616, EPI_ISL_2568617, EPI_ISL_2568618 | Lighthouse Lab in Alderley Park | Wellcome Sanger Institute for the COVID-19 Genomics UK (COG-UK) Consortium | Jacquelyn Wynn, Mairead Hyland, The Lighthouse Lab in Alderley Park and Alex Alderton, Roberto Amato, Jeffrey Barrett, Sonia Goncalves, Ewan Harrison, David K. Jackson, Ian Johnston, Dominic Kwiatkowski, Cordelia Langford, John Sillitoe on behalf of the Wellcome Sanger Institute COVID-19 Surveillance Team |
| EPI_ISL_2568619 | Lighthouse Lab in Glasgow | Wellcome Sanger Institute for the COVID-19 Genomics UK (COG-UK) Consortium | Harper VanSteenhouse, Yumi Kasai, David Gray, Carol Clugston, Anna Dominiczak and Alex Alderton, Roberto Amato, Jeffrey Barrett, Sonia Goncalves, Ewan Harrison, David K. Jackson, Ian Johnston, Dominic Kwiatkowski, Cordelia Langford, John Sillitoe on behalf of the Wellcome Sanger Institute COVID-19 Surveillance Team |
| EPI_ISL_2568620, EPI_ISL_2568621, EPI_ISL_2568622 | Lighthouse Lab in Alderley Park | Wellcome Sanger Institute for the COVID-19 Genomics UK (COG-UK) Consortium | Jacquelyn Wynn, Mairead Hyland, The Lighthouse Lab in Alderley Park and Alex Alderton, Roberto Amato, Jeffrey Barrett, Sonia Goncalves, Ewan Harrison, David K. Jackson, Ian Johnston, Dominic Kwiatkowski, Cordelia Langford, John Sillitoe on behalf of the Wellcome Sanger Institute COVID-19 Surveillance Team |
| EPI_ISL_2568623 | Lighthouse Lab in Glasgow | Wellcome Sanger Institute for the COVID-19 Genomics UK (COG-UK) Consortium | Harper VanSteenhouse, Yumi Kasai, David Gray, Carol Clugston, Anna Dominiczak and Alex Alderton, Roberto Amato, Jeffrey Barrett, Sonia Goncalves, Ewan Harrison, David K. Jackson, Ian Johnston, Dominic Kwiatkowski, Cordelia Langford, John Sillitoe on behalf of the Wellcome Sanger Institute COVID-19 Surveillance Team |
| EPI_ISL_2568624, EPI_ISL_2568625, EPI_ISL_2568626, EPI_ISL_2568627 | Lighthouse Lab in Alderley Park | Wellcome Sanger Institute for the COVID-19 Genomics UK (COG-UK) Consortium | Jacquelyn Wynn, Mairead Hyland, The Lighthouse Lab in Alderley Park and Alex Alderton, Roberto Amato, Jeffrey Barrett, Sonia Goncalves, Ewan Harrison, David K. Jackson, Ian Johnston, Dominic Kwiatkowski, Cordelia Langford, John Sillitoe on behalf of the Wellcome Sanger Institute COVID-19 Surveillance Team |
| EPI_ISL_2568628 | Lighthouse Lab in Glasgow | Wellcome Sanger Institute for the COVID-19 Genomics UK (COG-UK) Consortium | Harper VanSteenhouse, Yumi Kasai, David Gray, Carol Clugston, Anna Dominiczak and Alex Alderton, Roberto Amato, Jeffrey Barrett, Sonia Goncalves, Ewan Harrison, David K. Jackson, Ian Johnston, Dominic Kwiatkowski, Cordelia Langford, John Sillitoe on behalf of the Wellcome Sanger Institute COVID-19 Surveillance Team |
| EPI_ISL_2568629, EPI_ISL_2568630, EPI_ISL_2568631, EPI_ISL_2568632 | Lighthouse Lab in Alderley Park | Wellcome Sanger Institute for the COVID-19 Genomics UK (COG-UK) Consortium | Jacquelyn Wynn, Mairead Hyland, The Lighthouse Lab in Alderley Park and Alex Alderton, Roberto Amato, Jeffrey Barrett, Sonia Goncalves, Ewan Harrison, David K. Jackson, Ian Johnston, Dominic Kwiatkowski, Cordelia Langford, John Sillitoe on behalf of the Wellcome Sanger Institute COVID-19 Surveillance Team |
| EPI_ISL_2568633, EPI_ISL_2568634 | Lighthouse Lab in Glasgow | Wellcome Sanger Institute for the COVID-19 Genomics UK (COG-UK) Consortium | Harper VanSteenhouse, Yumi Kasai, David Gray, Carol Clugston, Anna Dominiczak and Alex Alderton, Roberto Amato, Jeffrey Barrett, Sonia Goncalves, Ewan Harrison, David K. Jackson, Ian Johnston, Dominic Kwiatkowski, Cordelia Langford, John Sillitoe on behalf of the Wellcome Sanger Institute COVID-19 Surveillance Team |
| EPI_ISL_2568635 | Lighthouse Lab in Alderley Park | Wellcome Sanger Institute for the COVID-19 Genomics UK (COG-UK) Consortium | Jacquelyn Wynn, Mairead Hyland, The Lighthouse Lab in Alderley Park and Alex Alderton, Roberto Amato, Jeffrey Barrett, Sonia Goncalves, Ewan Harrison, David K. Jackson, Ian Johnston, Dominic Kwiatkowski, Cordelia Langford, John Sillitoe on behalf of the Wellcome Sanger Institute COVID-19 Surveillance Team |
| EPI_ISL_2568636 | Lighthouse Lab in Glasgow | Wellcome Sanger Institute for the COVID-19 Genomics UK (COG-UK) Consortium | Harper VanSteenhouse, Yumi Kasai, David Gray, Carol Clugston, Anna Dominiczak and Alex Alderton, Roberto Amato, Jeffrey Barrett, Sonia Goncalves, Ewan Harrison, David K. Jackson, Ian Johnston, Dominic Kwiatkowski, Cordelia Langford, John Sillitoe on behalf of the Wellcome Sanger Institute COVID-19 Surveillance Team |
| EPI_ISL_2568637, EPI_ISL_2568638, EPI_ISL_2568639, EPI_ISL_2568640 | Lighthouse Lab in Alderley Park | Wellcome Sanger Institute for the COVID-19 Genomics UK (COG-UK) Consortium | Jacquelyn Wynn, Mairead Hyland, The Lighthouse Lab in Alderley Park and Alex Alderton, Roberto Amato, Jeffrey Barrett, Sonia Goncalves, Ewan Harrison, David K. Jackson, Ian Johnston, Dominic Kwiatkowski, Cordelia Langford, John Sillitoe on behalf of the Wellcome Sanger Institute COVID-19 Surveillance Team |
| EPI_ISL_2568641 | Lighthouse Lab in Glasgow | Wellcome Sanger Institute for the COVID-19 Genomics UK (COG-UK) Consortium | Harper VanSteenhouse, Yumi Kasai, David Gray, Carol Clugston, Anna Dominiczak and Alex Alderton, Roberto Amato, Jeffrey Barrett, Sonia Goncalves, Ewan Harrison, David K. Jackson, Ian Johnston, Dominic Kwiatkowski, Cordelia Langford, John Sillitoe on behalf of the Wellcome Sanger Institute COVID-19 Surveillance Team |
| EPI_ISL_2568642, EPI_ISL_2568643, EPI_ISL_2568644 | Lighthouse Lab in Alderley Park | Wellcome Sanger Institute for the COVID-19 Genomics UK (COG-UK) Consortium | Jacquelyn Wynn, Mairead Hyland, The Lighthouse Lab in Alderley Park and Alex Alderton, Roberto Amato, Jeffrey Barrett, Sonia Goncalves, Ewan Harrison, David K. Jackson, Ian Johnston, Dominic Kwiatkowski, Cordelia Langford, John Sillitoe on behalf of the Wellcome Sanger Institute COVID-19 Surveillance Team |
| EPI_ISL_2568645 | Lighthouse Lab in Glasgow | Wellcome Sanger Institute for the COVID-19 Genomics UK (COG-UK) Consortium | Harper VanSteenhouse, Yumi Kasai, David Gray, Carol Clugston, Anna Dominiczak and Alex Alderton, Roberto Amato, Jeffrey Barrett, Sonia Goncalves, Ewan Harrison, David K. Jackson, Ian Johnston, Dominic Kwiatkowski, Cordelia Langford, John Sillitoe on behalf of the Wellcome Sanger Institute COVID-19 Surveillance Team |
| EPI_ISL_2568646 | Lighthouse Lab in Alderley Park | Wellcome Sanger Institute for the COVID-19 Genomics UK (COG-UK) Consortium | Jacquelyn Wynn, Mairead Hyland, The Lighthouse Lab in Alderley Park and Alex Alderton, Roberto Amato, Jeffrey Barrett, Sonia Goncalves, Ewan Harrison, David K. Jackson, Ian Johnston, Dominic Kwiatkowski, Cordelia Langford, John Sillitoe on behalf of the Wellcome Sanger Institute COVID-19 Surveillance Team |
| EPI_ISL_2568647 | Lighthouse Lab in Glasgow | Wellcome Sanger Institute for the COVID-19 Genomics UK (COG-UK) Consortium | Harper VanSteenhouse, Yumi Kasai, David Gray, Carol Clugston, Anna Dominiczak and Alex Alderton, Roberto Amato, Jeffrey Barrett, Sonia Goncalves, Ewan Harrison, David K. Jackson, Ian Johnston, Dominic Kwiatkowski, Cordelia Langford, John Sillitoe on behalf of the Wellcome Sanger Institute COVID-19 Surveillance Team |
| EPI_ISL_2568648, EPI_ISL_2568649 | Lighthouse Lab in Alderley Park | Wellcome Sanger Institute for the COVID-19 Genomics UK (COG-UK) Consortium | Jacquelyn Wynn, Mairead Hyland, The Lighthouse Lab in Alderley Park and Alex Alderton, Roberto Amato, Jeffrey Barrett, Sonia Goncalves, Ewan Harrison, David K. Jackson, Ian Johnston, Dominic Kwiatkowski, Cordelia Langford, John Sillitoe on behalf of the Wellcome Sanger Institute COVID-19 Surveillance Team |
| EPI_ISL_2568650 | Lighthouse Lab in Glasgow | Wellcome Sanger Institute for the COVID-19 Genomics UK (COG-UK) | Harper VanSteenhouse, Yumi Kasai, David Gray, Carol Clugston, Anna Dominiczak and Alex Alderton, Roberto Amato, Jeffrey Barrett, Sonia Goncalves, Ewan Harrison, David K. Jackson, Ian Johnston, Dominic Kwiatkowski, Cordelia Langford, |

| | | | |
|---|---|---|---|
| | | Consortium | John Sillitoe on behalf of the Wellcome Sanger Institute COVID-19 Surveillance Team |
| EPI_ISL_2568651 | Lighthouse Lab in Milton Keynes | Wellcome Sanger Institute for the COVID-19 Genomics UK (COG-UK) Consortium | The Lighthouse Lab in Milton Keynes and Alex Alderton, Roberto Amato, Jeffrey Barrett, Sonia Goncalves, Ewan Harrison, David K. Jackson, Ian Johnston, Dominic Kwiatkowski, Cordelia Langford, John Sillitoe on behalf of the Wellcome Sanger Institute COVID-19 Surveillance Team |
| EPI_ISL_2568652, EPI_ISL_2568653, EPI_ISL_2568654, EPI_ISL_2568655, EPI_ISL_2568656, EPI_ISL_2568657 | Lighthouse Lab in Alderley Park | Wellcome Sanger Institute for the COVID-19 Genomics UK (COG-UK) Consortium | Jacquelyn Wynn, Mairead Hyland, The Lighthouse Lab in Alderley Park and Alex Alderton, Roberto Amato, Jeffrey Barrett, Sonia Goncalves, Ewan Harrison, David K. Jackson, Ian Johnston, Dominic Kwiatkowski, Cordelia Langford, John Sillitoe on behalf of the Wellcome Sanger Institute COVID-19 Surveillance Team |
| EPI_ISL_2568658, EPI_ISL_2568659 | Lighthouse Lab in Milton Keynes | Wellcome Sanger Institute for the COVID-19 Genomics UK (COG-UK) Consortium | The Lighthouse Lab in Milton Keynes and Alex Alderton, Roberto Amato, Jeffrey Barrett, Sonia Goncalves, Ewan Harrison, David K. Jackson, Ian Johnston, Dominic Kwiatkowski, Cordelia Langford, John Sillitoe on behalf of the Wellcome Sanger Institute COVID-19 Surveillance Team |
| EPI_ISL_2568660, EPI_ISL_2568661 | Lighthouse Lab in Alderley Park | Wellcome Sanger Institute for the COVID-19 Genomics UK (COG-UK) Consortium | Jacquelyn Wynn, Mairead Hyland, The Lighthouse Lab in Alderley Park and Alex Alderton, Roberto Amato, Jeffrey Barrett, Sonia Goncalves, Ewan Harrison, David K. Jackson, Ian Johnston, Dominic Kwiatkowski, Cordelia Langford, John Sillitoe on behalf of the Wellcome Sanger Institute COVID-19 Surveillance Team |
| EPI_ISL_2568662 | Lighthouse Lab in Milton Keynes | Wellcome Sanger Institute for the COVID-19 Genomics UK (COG-UK) Consortium | The Lighthouse Lab in Milton Keynes and Alex Alderton, Roberto Amato, Jeffrey Barrett, Sonia Goncalves, Ewan Harrison, David K. Jackson, Ian Johnston, Dominic Kwiatkowski, Cordelia Langford, John Sillitoe on behalf of the Wellcome Sanger Institute COVID-19 Surveillance Team |
| EPI_ISL_2568663 | Lighthouse Laboratory Plymouth | Wellcome Sanger Institute for the COVID-19 Genomics UK (COG-UK) Consortium | Lighthouse Laboratory Plymouth and Alex Alderton, Roberto Amato, Jeffrey Barrett, Sonia Goncalves, Ewan Harrison, David K. Jackson, Ian Johnston, Dominic Kwiatkowski, Cordelia Langford, John Sillitoe on behalf of the Wellcome Sanger Institute COVID-19 Surveillance Team |
| EPI_ISL_2568664 | Lighthouse Lab in Alderley Park | Wellcome Sanger Institute for the COVID-19 Genomics UK (COG-UK) Consortium | Jacquelyn Wynn, Mairead Hyland, The Lighthouse Lab in Alderley Park and Alex Alderton, Roberto Amato, Jeffrey Barrett, Sonia Goncalves, Ewan Harrison, David K. Jackson, Ian Johnston, Dominic Kwiatkowski, Cordelia Langford, John Sillitoe on behalf of the Wellcome Sanger Institute COVID-19 Surveillance Team |
| EPI_ISL_2568666, EPI_ISL_2568667, EPI_ISL_2568668 | Lighthouse Laboratory Plymouth | Wellcome Sanger Institute for the COVID-19 Genomics UK (COG-UK) Consortium | Lighthouse Laboratory Plymouth and Alex Alderton, Roberto Amato, Jeffrey Barrett, Sonia Goncalves, Ewan Harrison, David K. Jackson, Ian Johnston, Dominic Kwiatkowski, Cordelia Langford, John Sillitoe on behalf of the Wellcome Sanger Institute COVID-19 Surveillance Team |
| EPI_ISL_2568670, EPI_ISL_2568671 | Lighthouse Lab in Milton Keynes | Wellcome Sanger Institute for the COVID-19 Genomics UK (COG-UK) Consortium | The Lighthouse Lab in Milton Keynes and Alex Alderton, Roberto Amato, Jeffrey Barrett, Sonia Goncalves, Ewan Harrison, David K. Jackson, Ian Johnston, Dominic Kwiatkowski, Cordelia Langford, John Sillitoe on behalf of the Wellcome Sanger Institute COVID-19 Surveillance Team |
| EPI_ISL_2568672 | Lighthouse Lab in Alderley Park | Wellcome Sanger Institute for the COVID-19 Genomics UK (COG-UK) Consortium | Jacquelyn Wynn, Mairead Hyland, The Lighthouse Lab in Alderley Park and Alex Alderton, Roberto Amato, Jeffrey Barrett, Sonia Goncalves, Ewan Harrison, David K. Jackson, Ian Johnston, Dominic Kwiatkowski, Cordelia Langford, John Sillitoe on behalf of the Wellcome Sanger Institute COVID-19 Surveillance Team |
| EPI_ISL_2568673, EPI_ISL_2568674 | Lighthouse Laboratory Plymouth | Wellcome Sanger Institute for the COVID-19 Genomics UK (COG-UK) Consortium | Lighthouse Laboratory Plymouth and Alex Alderton, Roberto Amato, Jeffrey Barrett, Sonia Goncalves, Ewan Harrison, David K. Jackson, Ian Johnston, Dominic Kwiatkowski, Cordelia Langford, John Sillitoe on behalf of the Wellcome Sanger Institute COVID-19 Surveillance Team |
| EPI_ISL_2568675 | Lighthouse Lab in Alderley Park | Wellcome Sanger Institute for the COVID-19 Genomics UK (COG-UK) Consortium | Jacquelyn Wynn, Mairead Hyland, The Lighthouse Lab in Alderley Park and Alex Alderton, Roberto Amato, Jeffrey Barrett, Sonia Goncalves, Ewan Harrison, David K. Jackson, Ian Johnston, Dominic Kwiatkowski, Cordelia Langford, John Sillitoe on behalf of the Wellcome Sanger Institute COVID-19 Surveillance Team |
| EPI_ISL_2568676 | Lighthouse Lab in Milton Keynes | Wellcome Sanger Institute for the COVID-19 Genomics UK (COG-UK) Consortium | The Lighthouse Lab in Milton Keynes and Alex Alderton, Roberto Amato, Jeffrey Barrett, Sonia Goncalves, Ewan Harrison, David K. Jackson, Ian Johnston, Dominic Kwiatkowski, Cordelia Langford, John Sillitoe on behalf of the Wellcome Sanger Institute COVID-19 Surveillance Team |
| EPI_ISL_2568677 | Lighthouse Laboratory Plymouth | Wellcome Sanger Institute for the COVID-19 Genomics UK (COG-UK) Consortium | Lighthouse Laboratory Plymouth and Alex Alderton, Roberto Amato, Jeffrey Barrett, Sonia Goncalves, Ewan Harrison, David K. Jackson, Ian Johnston, Dominic Kwiatkowski, Cordelia Langford, John Sillitoe on behalf of the Wellcome Sanger Institute COVID-19 Surveillance Team |
| EPI_ISL_2568678 | Lighthouse Lab in Alderley Park | Wellcome Sanger Institute for the COVID-19 Genomics UK (COG-UK) Consortium | Jacquelyn Wynn, Mairead Hyland, The Lighthouse Lab in Alderley Park and Alex Alderton, Roberto Amato, Jeffrey Barrett, Sonia Goncalves, Ewan Harrison, David K. Jackson, Ian Johnston, Dominic Kwiatkowski, Cordelia Langford, John Sillitoe on behalf of the Wellcome Sanger Institute COVID-19 Surveillance Team |
| EPI_ISL_2568679 | Lighthouse Lab in Milton Keynes | Wellcome Sanger Institute for the COVID-19 Genomics UK (COG-UK) Consortium | The Lighthouse Lab in Milton Keynes and Alex Alderton, Roberto Amato, Jeffrey Barrett, Sonia Goncalves, Ewan Harrison, David K. Jackson, Ian Johnston, Dominic Kwiatkowski, Cordelia Langford, John Sillitoe on behalf of the Wellcome Sanger Institute COVID-19 Surveillance Team |
| EPI_ISL_2568680 | Lighthouse Laboratory Plymouth | Wellcome Sanger Institute for the COVID-19 Genomics UK (COG-UK) Consortium | Lighthouse Laboratory Plymouth and Alex Alderton, Roberto Amato, Jeffrey Barrett, Sonia Goncalves, Ewan Harrison, David K. Jackson, Ian Johnston, Dominic Kwiatkowski, Cordelia Langford, John Sillitoe on behalf of the Wellcome Sanger Institute COVID-19 Surveillance Team |
| EPI_ISL_2568681, EPI_ISL_2568683 | Lighthouse Lab in Milton Keynes | Wellcome Sanger Institute for the COVID-19 Genomics UK (COG-UK) Consortium | The Lighthouse Lab in Milton Keynes and Alex Alderton, Roberto Amato, Jeffrey Barrett, Sonia Goncalves, Ewan Harrison, David K. Jackson, Ian Johnston, Dominic Kwiatkowski, Cordelia Langford, John Sillitoe on behalf of the Wellcome Sanger Institute COVID-19 Surveillance Team |
| EPI_ISL_2568684 | Lighthouse Lab in Alderley Park | Wellcome Sanger Institute for the COVID-19 Genomics UK (COG-UK) Consortium | Jacquelyn Wynn, Mairead Hyland, The Lighthouse Lab in Alderley Park and Alex Alderton, Roberto Amato, Jeffrey Barrett, Sonia Goncalves, Ewan Harrison, David K. Jackson, Ian Johnston, Dominic Kwiatkowski, Cordelia Langford, John Sillitoe on behalf of the Wellcome Sanger Institute COVID-19 Surveillance Team |
| EPI_ISL_2568685, EPI_ISL_2568686, EPI_ISL_2568687 | Lighthouse Lab in Milton Keynes | Wellcome Sanger Institute for the COVID-19 Genomics UK (COG-UK) Consortium | The Lighthouse Lab in Milton Keynes and Alex Alderton, Roberto Amato, Jeffrey Barrett, Sonia Goncalves, Ewan Harrison, David K. Jackson, Ian Johnston, Dominic Kwiatkowski, Cordelia Langford, John Sillitoe on behalf of the Wellcome Sanger Institute COVID-19 Surveillance Team |
| EPI_ISL_2568688 | Lighthouse Lab in Alderley Park | Wellcome Sanger Institute for the COVID-19 Genomics UK (COG-UK) Consortium | Jacquelyn Wynn, Mairead Hyland, The Lighthouse Lab in Alderley Park and Alex Alderton, Roberto Amato, Jeffrey Barrett, Sonia Goncalves, Ewan Harrison, David K. Jackson, Ian Johnston, Dominic Kwiatkowski, Cordelia Langford, John Sillitoe on behalf of the Wellcome Sanger Institute COVID-19 Surveillance Team |
| EPI_ISL_2568689, EPI_ISL_2568690, EPI_ISL_2568691 | Lighthouse Laboratory Plymouth | Wellcome Sanger Institute for the COVID-19 Genomics UK (COG-UK) Consortium | Lighthouse Laboratory Plymouth and Alex Alderton, Roberto Amato, Jeffrey Barrett, Sonia Goncalves, Ewan Harrison, David K. Jackson, Ian Johnston, Dominic Kwiatkowski, Cordelia Langford, John Sillitoe on behalf of the Wellcome Sanger Institute COVID-19 Surveillance Team |
| EPI_ISL_2568692 | Lighthouse Lab in Milton Keynes | Wellcome Sanger Institute for the COVID-19 Genomics UK (COG-UK) Consortium | The Lighthouse Lab in Milton Keynes and Alex Alderton, Roberto Amato, Jeffrey Barrett, Sonia Goncalves, Ewan Harrison, David K. Jackson, Ian Johnston, Dominic Kwiatkowski, Cordelia Langford, John Sillitoe on behalf of the Wellcome Sanger Institute COVID-19 Surveillance Team |
| EPI_ISL_2568693 | Lighthouse Lab in Alderley Park | Wellcome Sanger Institute for the COVID-19 Genomics UK (COG-UK) Consortium | Jacquelyn Wynn, Mairead Hyland, The Lighthouse Lab in Alderley Park and Alex Alderton, Roberto Amato, Jeffrey Barrett, Sonia Goncalves, Ewan Harrison, David K. Jackson, Ian Johnston, Dominic Kwiatkowski, Cordelia Langford, John Sillitoe on behalf of the Wellcome Sanger Institute COVID-19 Surveillance Team |
| EPI_ISL_2568696 | Lighthouse Laboratory Plymouth | Wellcome Sanger Institute for the COVID-19 Genomics UK (COG-UK) Consortium | Lighthouse Laboratory Plymouth and Alex Alderton, Roberto Amato, Jeffrey Barrett, Sonia Goncalves, Ewan Harrison, David K. Jackson, Ian Johnston, Dominic Kwiatkowski, Cordelia Langford, John Sillitoe on behalf of the Wellcome Sanger Institute COVID-19 Surveillance Team |
| EPI_ISL_2568697 | Lighthouse Lab in Milton Keynes | Wellcome Sanger Institute for the COVID-19 Genomics UK (COG-UK) Consortium | The Lighthouse Lab in Milton Keynes and Alex Alderton, Roberto Amato, Jeffrey Barrett, Sonia Goncalves, Ewan Harrison, David K. Jackson, Ian Johnston, Dominic Kwiatkowski, Cordelia Langford, John Sillitoe on behalf of the Wellcome Sanger Institute COVID-19 Surveillance Team |
| EPI_ISL_2568698 | Lighthouse Lab in Alderley Park | Wellcome Sanger Institute for the COVID-19 Genomics UK (COG-UK) Consortium | Jacquelyn Wynn, Mairead Hyland, The Lighthouse Lab in Alderley Park and Alex Alderton, Roberto Amato, Jeffrey Barrett, Sonia Goncalves, Ewan Harrison, David K. Jackson, Ian Johnston, Dominic Kwiatkowski, Cordelia Langford, John Sillitoe on behalf of the Wellcome Sanger Institute COVID-19 Surveillance Team |
| EPI_ISL_2568699, EPI_ISL_2568700, EPI_ISL_2568701, EPI_ISL_2568702, EPI_ISL_2568703 | Lighthouse Lab in Milton Keynes | Wellcome Sanger Institute for the COVID-19 Genomics UK (COG-UK) Consortium | The Lighthouse Lab in Milton Keynes and Alex Alderton, Roberto Amato, Jeffrey Barrett, Sonia Goncalves, Ewan Harrison, David K. Jackson, Ian Johnston, Dominic Kwiatkowski, Cordelia Langford, John Sillitoe on behalf of the Wellcome Sanger Institute COVID-19 Surveillance Team |
| EPI_ISL_2568704 | Lighthouse Laboratory Plymouth | Wellcome Sanger Institute for the COVID-19 Genomics UK (COG-UK) Consortium | Lighthouse Laboratory Plymouth and Alex Alderton, Roberto Amato, Jeffrey Barrett, Sonia Goncalves, Ewan Harrison, David K. Jackson, Ian Johnston, Dominic Kwiatkowski, Cordelia Langford, John Sillitoe on behalf of the Wellcome Sanger Institute COVID-19 Surveillance Team |
| EPI_ISL_2568705, EPI_ISL_2568706, EPI_ISL_2568707 | Lighthouse Lab in Milton Keynes | Wellcome Sanger Institute for the COVID-19 Genomics UK (COG-UK) Consortium | The Lighthouse Lab in Milton Keynes and Alex Alderton, Roberto Amato, Jeffrey Barrett, Sonia Goncalves, Ewan Harrison, David K. Jackson, Ian Johnston, Dominic Kwiatkowski, Cordelia Langford, John Sillitoe on behalf of the Wellcome Sanger Institute COVID-19 Surveillance Team |
| EPI_ISL_2568708 | Lighthouse Laboratory Plymouth | Wellcome Sanger Institute for the COVID-19 Genomics UK (COG-UK) Consortium | Lighthouse Laboratory Plymouth and Alex Alderton, Roberto Amato, Jeffrey Barrett, Sonia Goncalves, Ewan Harrison, David K. Jackson, Ian Johnston, Dominic Kwiatkowski, Cordelia Langford, John Sillitoe on behalf of the Wellcome Sanger Institute COVID-19 Surveillance Team |
| EPI_ISL_2568709, EPI_ISL_2568710 | Lighthouse Lab in Milton Keynes | Wellcome Sanger Institute for the COVID-19 Genomics UK (COG-UK) Consortium | The Lighthouse Lab in Milton Keynes and Alex Alderton, Roberto Amato, Jeffrey Barrett, Sonia Goncalves, Ewan Harrison, David K. Jackson, Ian Johnston, Dominic Kwiatkowski, Cordelia Langford, John Sillitoe on behalf of the Wellcome Sanger Institute COVID-19 Surveillance Team |
| EPI_ISL_2568711 | Lighthouse Lab in Alderley Park | Wellcome Sanger Institute for the COVID-19 Genomics UK (COG-UK) Consortium | Jacquelyn Wynn, Mairead Hyland, The Lighthouse Lab in Alderley Park and Alex Alderton, Roberto Amato, Jeffrey Barrett, Sonia Goncalves, Ewan Harrison, David K. Jackson, Ian Johnston, Dominic Kwiatkowski, Cordelia Langford, John Sillitoe on behalf of the Wellcome Sanger Institute COVID-19 Surveillance Team |
| EPI_ISL_2568712, EPI_ISL_2568713, EPI_ISL_2568714 | Lighthouse Lab in Milton Keynes | Wellcome Sanger Institute for the COVID-19 Genomics UK (COG-UK) Consortium | The Lighthouse Lab in Milton Keynes and Alex Alderton, Roberto Amato, Jeffrey Barrett, Sonia Goncalves, Ewan Harrison, David K. Jackson, Ian Johnston, Dominic Kwiatkowski, Cordelia Langford, John Sillitoe on behalf of the Wellcome Sanger Institute COVID-19 Surveillance Team |
| EPI_ISL_2568715, EPI_ISL_2568716 | Lighthouse Laboratory Plymouth | Wellcome Sanger Institute for the COVID-19 Genomics UK (COG-UK) Consortium | Lighthouse Laboratory Plymouth and Alex Alderton, Roberto Amato, Jeffrey Barrett, Sonia Goncalves, Ewan Harrison, David K. Jackson, Ian Johnston, Dominic Kwiatkowski, Cordelia Langford, John Sillitoe on behalf of the Wellcome Sanger Institute COVID-19 Surveillance Team |
| EPI_ISL_2568717, EPI_ISL_2568718 | Lighthouse Lab in Milton Keynes | Wellcome Sanger Institute for the COVID-19 Genomics UK (COG-UK) Consortium | The Lighthouse Lab in Milton Keynes and Alex Alderton, Roberto Amato, Jeffrey Barrett, Sonia Goncalves, Ewan Harrison, David K. Jackson, Ian Johnston, Dominic Kwiatkowski, Cordelia Langford, John Sillitoe on behalf of the Wellcome Sanger Institute COVID-19 Surveillance Team |
| EPI_ISL_2568719 | Lighthouse Lab in Alderley Park | Wellcome Sanger Institute for the COVID-19 Genomics UK (COG-UK) Consortium | Jacquelyn Wynn, Mairead Hyland, The Lighthouse Lab in Alderley Park and Alex Alderton, Roberto Amato, Jeffrey Barrett, Sonia Goncalves, Ewan Harrison, David K. Jackson, Ian Johnston, Dominic Kwiatkowski, Cordelia Langford, John Sillitoe on behalf of the Wellcome Sanger Institute COVID-19 Surveillance Team |
| EPI_ISL_2568720, EPI_ISL_2568721 | Lighthouse Laboratory Plymouth | Wellcome Sanger Institute for the COVID-19 Genomics UK (COG-UK) Consortium | Lighthouse Laboratory Plymouth and Alex Alderton, Roberto Amato, Jeffrey Barrett, Sonia Goncalves, Ewan Harrison, David K. Jackson, Ian Johnston, Dominic Kwiatkowski, Cordelia Langford, John Sillitoe on behalf of the Wellcome Sanger Institute COVID-19 Surveillance Team |
| EPI_ISL_2568722, EPI_ISL_2568723 | Lighthouse Lab in Milton Keynes | Wellcome Sanger Institute for the COVID-19 Genomics UK (COG-UK) Consortium | The Lighthouse Lab in Milton Keynes and Alex Alderton, Roberto Amato, Jeffrey Barrett, Sonia Goncalves, Ewan Harrison, David K. Jackson, Ian Johnston, Dominic Kwiatkowski, Cordelia Langford, John Sillitoe on behalf of the Wellcome Sanger Institute COVID-19 Surveillance Team |
| EPI_ISL_2568724, | Lighthouse Laboratory Plymouth | Wellcome Sanger Institute for the COVID-19 Genomics UK (COG-UK) | Lighthouse Laboratory Plymouth and Alex Alderton, Roberto Amato, Jeffrey Barrett, Sonia Goncalves, Ewan Harrison, David K. Jackson, Ian Johnston, Dominic Kwiatkowski, Cordelia Langford, John Sillitoe on behalf of the Wellcome Sanger |

| | | | |
|---|---|---|---|
| EPI_ISL_2568725 | | Consortium | Institute COVID-19 Surveillance Team |
| EPI_ISL_2568726 | Lighthouse Lab in Milton Keynes | Wellcome Sanger Institute for the COVID-19 Genomics UK (COG-UK) Consortium | The Lighthouse Lab in Milton Keynes and Alex Alderton, Roberto Amato, Jeffrey Barrett, Sonia Goncalves, Ewan Harrison, David K. Jackson, Ian Johnston, Dominic Kwiatkowski, Cordelia Langford, John Sillitoe on behalf of the Wellcome Sanger Institute COVID-19 Surveillance Team |
| EPI_ISL_2568728, EPI_ISL_2568729 | Lighthouse Laboratory Plymouth | Wellcome Sanger Institute for the COVID-19 Genomics UK (COG-UK) Consortium | Lighthouse Laboratory Plymouth and Alex Alderton, Roberto Amato, Jeffrey Barrett, Sonia Goncalves, Ewan Harrison, David K. Jackson, Ian Johnston, Dominic Kwiatkowski, Cordelia Langford, John Sillitoe on behalf of the Wellcome Sanger Institute COVID-19 Surveillance Team |
| EPI_ISL_2568730 | Lighthouse Lab in Alderley Park | Wellcome Sanger Institute for the COVID-19 Genomics UK (COG-UK) Consortium | Jacquelyn Wynn, Mairead Hyland, The Lighthouse Lab in Alderley Park and Alex Alderton, Roberto Amato, Jeffrey Barrett, Sonia Goncalves, Ewan Harrison, David K. Jackson, Ian Johnston, Dominic Kwiatkowski, Cordelia Langford, John Sillitoe on behalf of the Wellcome Sanger Institute COVID-19 Surveillance Team |
| EPI_ISL_2568731, EPI_ISL_2568732, EPI_ISL_2568733, EPI_ISL_2568734, EPI_ISL_2568735, EPI_ISL_2568736, EPI_ISL_2568737, EPI_ISL_2568738 | Lighthouse Lab in Milton Keynes | Wellcome Sanger Institute for the COVID-19 Genomics UK (COG-UK) Consortium | The Lighthouse Lab in Milton Keynes and Alex Alderton, Roberto Amato, Jeffrey Barrett, Sonia Goncalves, Ewan Harrison, David K. Jackson, Ian Johnston, Dominic Kwiatkowski, Cordelia Langford, John Sillitoe on behalf of the Wellcome Sanger Institute COVID-19 Surveillance Team |
| EPI_ISL_2568739 | Lighthouse Laboratory Plymouth | Wellcome Sanger Institute for the COVID-19 Genomics UK (COG-UK) Consortium | Lighthouse Laboratory Plymouth and Alex Alderton, Roberto Amato, Jeffrey Barrett, Sonia Goncalves, Ewan Harrison, David K. Jackson, Ian Johnston, Dominic Kwiatkowski, Cordelia Langford, John Sillitoe on behalf of the Wellcome Sanger Institute COVID-19 Surveillance Team |
| EPI_ISL_2568740 | Lighthouse Lab in Alderley Park | Wellcome Sanger Institute for the COVID-19 Genomics UK (COG-UK) Consortium | Jacquelyn Wynn, Mairead Hyland, The Lighthouse Lab in Alderley Park and Alex Alderton, Roberto Amato, Jeffrey Barrett, Sonia Goncalves, Ewan Harrison, David K. Jackson, Ian Johnston, Dominic Kwiatkowski, Cordelia Langford, John Sillitoe on behalf of the Wellcome Sanger Institute COVID-19 Surveillance Team |
| EPI_ISL_2568741 | Lighthouse Laboratory Plymouth | Wellcome Sanger Institute for the COVID-19 Genomics UK (COG-UK) Consortium | Lighthouse Laboratory Plymouth and Alex Alderton, Roberto Amato, Jeffrey Barrett, Sonia Goncalves, Ewan Harrison, David K. Jackson, Ian Johnston, Dominic Kwiatkowski, Cordelia Langford, John Sillitoe on behalf of the Wellcome Sanger Institute COVID-19 Surveillance Team |
| EPI_ISL_2568742, EPI_ISL_2568743 | Lighthouse Lab in Milton Keynes | Wellcome Sanger Institute for the COVID-19 Genomics UK (COG-UK) Consortium | The Lighthouse Lab in Milton Keynes and Alex Alderton, Roberto Amato, Jeffrey Barrett, Sonia Goncalves, Ewan Harrison, David K. Jackson, Ian Johnston, Dominic Kwiatkowski, Cordelia Langford, John Sillitoe on behalf of the Wellcome Sanger Institute COVID-19 Surveillance Team |
| EPI_ISL_2568744 | Lighthouse Laboratory Plymouth | Wellcome Sanger Institute for the COVID-19 Genomics UK (COG-UK) Consortium | Lighthouse Laboratory Plymouth and Alex Alderton, Roberto Amato, Jeffrey Barrett, Sonia Goncalves, Ewan Harrison, David K. Jackson, Ian Johnston, Dominic Kwiatkowski, Cordelia Langford, John Sillitoe on behalf of the Wellcome Sanger Institute COVID-19 Surveillance Team |
| EPI_ISL_2568745 | Lighthouse Lab in Milton Keynes | Wellcome Sanger Institute for the COVID-19 Genomics UK (COG-UK) Consortium | The Lighthouse Lab in Milton Keynes and Alex Alderton, Roberto Amato, Jeffrey Barrett, Sonia Goncalves, Ewan Harrison, David K. Jackson, Ian Johnston, Dominic Kwiatkowski, Cordelia Langford, John Sillitoe on behalf of the Wellcome Sanger Institute COVID-19 Surveillance Team |
| EPI_ISL_2568746 | Lighthouse Lab in Alderley Park | Wellcome Sanger Institute for the COVID-19 Genomics UK (COG-UK) Consortium | Jacquelyn Wynn, Mairead Hyland, The Lighthouse Lab in Alderley Park and Alex Alderton, Roberto Amato, Jeffrey Barrett, Sonia Goncalves, Ewan Harrison, David K. Jackson, Ian Johnston, Dominic Kwiatkowski, Cordelia Langford, John Sillitoe on behalf of the Wellcome Sanger Institute COVID-19 Surveillance Team |
| EPI_ISL_2568747, EPI_ISL_2568748, EPI_ISL_2568749, EPI_ISL_2568750 | Lighthouse Lab in Milton Keynes | Wellcome Sanger Institute for the COVID-19 Genomics UK (COG-UK) Consortium | The Lighthouse Lab in Milton Keynes and Alex Alderton, Roberto Amato, Jeffrey Barrett, Sonia Goncalves, Ewan Harrison, David K. Jackson, Ian Johnston, Dominic Kwiatkowski, Cordelia Langford, John Sillitoe on behalf of the Wellcome Sanger Institute COVID-19 Surveillance Team |
| EPI_ISL_2568751 | Lighthouse Laboratory Plymouth | Wellcome Sanger Institute for the COVID-19 Genomics UK (COG-UK) Consortium | Lighthouse Laboratory Plymouth and Alex Alderton, Roberto Amato, Jeffrey Barrett, Sonia Goncalves, Ewan Harrison, David K. Jackson, Ian Johnston, Dominic Kwiatkowski, Cordelia Langford, John Sillitoe on behalf of the Wellcome Sanger Institute COVID-19 Surveillance Team |
| EPI_ISL_2568752, EPI_ISL_2568753 | Lighthouse Lab in Alderley Park | Wellcome Sanger Institute for the COVID-19 Genomics UK (COG-UK) Consortium | Jacquelyn Wynn, Mairead Hyland, The Lighthouse Lab in Alderley Park and Alex Alderton, Roberto Amato, Jeffrey Barrett, Sonia Goncalves, Ewan Harrison, David K. Jackson, Ian Johnston, Dominic Kwiatkowski, Cordelia Langford, John Sillitoe on behalf of the Wellcome Sanger Institute COVID-19 Surveillance Team |
| EPI_ISL_2568754, EPI_ISL_2568755 | Lighthouse Lab in Milton Keynes | Wellcome Sanger Institute for the COVID-19 Genomics UK (COG-UK) Consortium | The Lighthouse Lab in Milton Keynes and Alex Alderton, Roberto Amato, Jeffrey Barrett, Sonia Goncalves, Ewan Harrison, David K. Jackson, Ian Johnston, Dominic Kwiatkowski, Cordelia Langford, John Sillitoe on behalf of the Wellcome Sanger Institute COVID-19 Surveillance Team |
| EPI_ISL_2568756 | Lighthouse Lab in Alderley Park | Wellcome Sanger Institute for the COVID-19 Genomics UK (COG-UK) Consortium | Jacquelyn Wynn, Mairead Hyland, The Lighthouse Lab in Alderley Park and Alex Alderton, Roberto Amato, Jeffrey Barrett, Sonia Goncalves, Ewan Harrison, David K. Jackson, Ian Johnston, Dominic Kwiatkowski, Cordelia Langford, John Sillitoe on behalf of the Wellcome Sanger Institute COVID-19 Surveillance Team |
| EPI_ISL_2568757 | Lighthouse Lab in Milton Keynes | Wellcome Sanger Institute for the COVID-19 Genomics UK (COG-UK) Consortium | The Lighthouse Lab in Milton Keynes and Alex Alderton, Roberto Amato, Jeffrey Barrett, Sonia Goncalves, Ewan Harrison, David K. Jackson, Ian Johnston, Dominic Kwiatkowski, Cordelia Langford, John Sillitoe on behalf of the Wellcome Sanger Institute COVID-19 Surveillance Team |
| EPI_ISL_2568758, EPI_ISL_2568760 | Lighthouse Lab in Alderley Park | Wellcome Sanger Institute for the COVID-19 Genomics UK (COG-UK) Consortium | Jacquelyn Wynn, Mairead Hyland, The Lighthouse Lab in Alderley Park and Alex Alderton, Roberto Amato, Jeffrey Barrett, Sonia Goncalves, Ewan Harrison, David K. Jackson, Ian Johnston, Dominic Kwiatkowski, Cordelia Langford, John Sillitoe on behalf of the Wellcome Sanger Institute COVID-19 Surveillance Team |
| EPI_ISL_2568761 | Lighthouse Laboratory Plymouth | Wellcome Sanger Institute for the COVID-19 Genomics UK (COG-UK) Consortium | Lighthouse Laboratory Plymouth and Alex Alderton, Roberto Amato, Jeffrey Barrett, Sonia Goncalves, Ewan Harrison, David K. Jackson, Ian Johnston, Dominic Kwiatkowski, Cordelia Langford, John Sillitoe on behalf of the Wellcome Sanger Institute COVID-19 Surveillance Team |
| EPI_ISL_2568762, EPI_ISL_2568763 | Lighthouse Lab in Milton Keynes | Wellcome Sanger Institute for the COVID-19 Genomics UK (COG-UK) Consortium | The Lighthouse Lab in Milton Keynes and Alex Alderton, Roberto Amato, Jeffrey Barrett, Sonia Goncalves, Ewan Harrison, David K. Jackson, Ian Johnston, Dominic Kwiatkowski, Cordelia Langford, John Sillitoe on behalf of the Wellcome Sanger Institute COVID-19 Surveillance Team |
| EPI_ISL_2568764 | Lighthouse Lab in Alderley Park | Wellcome Sanger Institute for the COVID-19 Genomics UK (COG-UK) Consortium | Jacquelyn Wynn, Mairead Hyland, The Lighthouse Lab in Alderley Park and Alex Alderton, Roberto Amato, Jeffrey Barrett, Sonia Goncalves, Ewan Harrison, David K. Jackson, Ian Johnston, Dominic Kwiatkowski, Cordelia Langford, John Sillitoe on behalf of the Wellcome Sanger Institute COVID-19 Surveillance Team |
| EPI_ISL_2568765, EPI_ISL_2568766 | Lighthouse Lab in Milton Keynes | Wellcome Sanger Institute for the COVID-19 Genomics UK (COG-UK) Consortium | The Lighthouse Lab in Milton Keynes and Alex Alderton, Roberto Amato, Jeffrey Barrett, Sonia Goncalves, Ewan Harrison, David K. Jackson, Ian Johnston, Dominic Kwiatkowski, Cordelia Langford, John Sillitoe on behalf of the Wellcome Sanger Institute COVID-19 Surveillance Team |
| EPI_ISL_2568767 | Lighthouse Laboratory Plymouth | Wellcome Sanger Institute for the COVID-19 Genomics UK (COG-UK) Consortium | Lighthouse Laboratory Plymouth and Alex Alderton, Roberto Amato, Jeffrey Barrett, Sonia Goncalves, Ewan Harrison, David K. Jackson, Ian Johnston, Dominic Kwiatkowski, Cordelia Langford, John Sillitoe on behalf of the Wellcome Sanger Institute COVID-19 Surveillance Team |
| EPI_ISL_2568768, EPI_ISL_2568769, EPI_ISL_2568770, EPI_ISL_2568771, EPI_ISL_2568772, EPI_ISL_2568773, EPI_ISL_2568774 | Lighthouse Lab in Milton Keynes | Wellcome Sanger Institute for the COVID-19 Genomics UK (COG-UK) Consortium | The Lighthouse Lab in Milton Keynes and Alex Alderton, Roberto Amato, Jeffrey Barrett, Sonia Goncalves, Ewan Harrison, David K. Jackson, Ian Johnston, Dominic Kwiatkowski, Cordelia Langford, John Sillitoe on behalf of the Wellcome Sanger Institute COVID-19 Surveillance Team |
| EPI_ISL_2568775, EPI_ISL_2568776 | Lighthouse Lab in Alderley Park | Wellcome Sanger Institute for the COVID-19 Genomics UK (COG-UK) Consortium | Jacquelyn Wynn, Mairead Hyland, The Lighthouse Lab in Alderley Park and Alex Alderton, Roberto Amato, Jeffrey Barrett, Sonia Goncalves, Ewan Harrison, David K. Jackson, Ian Johnston, Dominic Kwiatkowski, Cordelia Langford, John Sillitoe on behalf of the Wellcome Sanger Institute COVID-19 Surveillance Team |
| EPI_ISL_2568777, EPI_ISL_2568778 | Lighthouse Lab in Milton Keynes | Wellcome Sanger Institute for the COVID-19 Genomics UK (COG-UK) Consortium | The Lighthouse Lab in Milton Keynes and Alex Alderton, Roberto Amato, Jeffrey Barrett, Sonia Goncalves, Ewan Harrison, David K. Jackson, Ian Johnston, Dominic Kwiatkowski, Cordelia Langford, John Sillitoe on behalf of the Wellcome Sanger Institute COVID-19 Surveillance Team |
| EPI_ISL_2568779 | Lighthouse Laboratory Plymouth | Wellcome Sanger Institute for the COVID-19 Genomics UK (COG-UK) Consortium | Lighthouse Laboratory Plymouth and Alex Alderton, Roberto Amato, Jeffrey Barrett, Sonia Goncalves, Ewan Harrison, David K. Jackson, Ian Johnston, Dominic Kwiatkowski, Cordelia Langford, John Sillitoe on behalf of the Wellcome Sanger Institute COVID-19 Surveillance Team |
| EPI_ISL_2568780 | Lighthouse Lab in Alderley Park | Wellcome Sanger Institute for the COVID-19 Genomics UK (COG-UK) Consortium | Jacquelyn Wynn, Mairead Hyland, The Lighthouse Lab in Alderley Park and Alex Alderton, Roberto Amato, Jeffrey Barrett, Sonia Goncalves, Ewan Harrison, David K. Jackson, Ian Johnston, Dominic Kwiatkowski, Cordelia Langford, John Sillitoe on behalf of the Wellcome Sanger Institute COVID-19 Surveillance Team |
| EPI_ISL_2568781 | Lighthouse Lab in Milton Keynes | Wellcome Sanger Institute for the COVID-19 Genomics UK (COG-UK) Consortium | The Lighthouse Lab in Milton Keynes and Alex Alderton, Roberto Amato, Jeffrey Barrett, Sonia Goncalves, Ewan Harrison, David K. Jackson, Ian Johnston, Dominic Kwiatkowski, Cordelia Langford, John Sillitoe on behalf of the Wellcome Sanger Institute COVID-19 Surveillance Team |
| EPI_ISL_2568782 | Lighthouse Lab in Alderley Park | Wellcome Sanger Institute for the COVID-19 Genomics UK (COG-UK) Consortium | Jacquelyn Wynn, Mairead Hyland, The Lighthouse Lab in Alderley Park and Alex Alderton, Roberto Amato, Jeffrey Barrett, Sonia Goncalves, Ewan Harrison, David K. Jackson, Ian Johnston, Dominic Kwiatkowski, Cordelia Langford, John Sillitoe on behalf of the Wellcome Sanger Institute COVID-19 Surveillance Team |
| EPI_ISL_2568783 | Lighthouse Lab in Milton Keynes | Wellcome Sanger Institute for the COVID-19 Genomics UK (COG-UK) Consortium | The Lighthouse Lab in Milton Keynes and Alex Alderton, Roberto Amato, Jeffrey Barrett, Sonia Goncalves, Ewan Harrison, David K. Jackson, Ian Johnston, Dominic Kwiatkowski, Cordelia Langford, John Sillitoe on behalf of the Wellcome Sanger Institute COVID-19 Surveillance Team |
| EPI_ISL_2568784 | Lighthouse Lab in Alderley Park | Wellcome Sanger Institute for the COVID-19 Genomics UK (COG-UK) Consortium | Jacquelyn Wynn, Mairead Hyland, The Lighthouse Lab in Alderley Park and Alex Alderton, Roberto Amato, Jeffrey Barrett, Sonia Goncalves, Ewan Harrison, David K. Jackson, Ian Johnston, Dominic Kwiatkowski, Cordelia Langford, John Sillitoe on behalf of the Wellcome Sanger Institute COVID-19 Surveillance Team |
| EPI_ISL_2568785 | Lighthouse Laboratory Plymouth | Wellcome Sanger Institute for the COVID-19 Genomics UK (COG-UK) Consortium | Lighthouse Laboratory Plymouth and Alex Alderton, Roberto Amato, Jeffrey Barrett, Sonia Goncalves, Ewan Harrison, David K. Jackson, Ian Johnston, Dominic Kwiatkowski, Cordelia Langford, John Sillitoe on behalf of the Wellcome Sanger Institute COVID-19 Surveillance Team |
| EPI_ISL_2568786 | Lighthouse Lab in Milton Keynes | Wellcome Sanger Institute for the COVID-19 Genomics UK (COG-UK) Consortium | The Lighthouse Lab in Milton Keynes and Alex Alderton, Roberto Amato, Jeffrey Barrett, Sonia Goncalves, Ewan Harrison, David K. Jackson, Ian Johnston, Dominic Kwiatkowski, Cordelia Langford, John Sillitoe on behalf of the Wellcome Sanger Institute COVID-19 Surveillance Team |
| EPI_ISL_2568787 | Lighthouse Lab in Alderley Park | Wellcome Sanger Institute for the COVID-19 Genomics UK (COG-UK) Consortium | Jacquelyn Wynn, Mairead Hyland, The Lighthouse Lab in Alderley Park and Alex Alderton, Roberto Amato, Jeffrey Barrett, Sonia Goncalves, Ewan Harrison, David K. Jackson, Ian Johnston, Dominic Kwiatkowski, Cordelia Langford, John Sillitoe on behalf of the Wellcome Sanger Institute COVID-19 Surveillance Team |
| EPI_ISL_2568789 | Lighthouse Laboratory Plymouth | Wellcome Sanger Institute for the COVID-19 Genomics UK (COG-UK) Consortium | Lighthouse Laboratory Plymouth and Alex Alderton, Roberto Amato, Jeffrey Barrett, Sonia Goncalves, Ewan Harrison, David K. Jackson, Ian Johnston, Dominic Kwiatkowski, Cordelia Langford, John Sillitoe on behalf of the Wellcome Sanger Institute COVID-19 Surveillance Team |
| EPI_ISL_2568790, EPI_ISL_2568791 | Lighthouse Lab in Milton Keynes | Wellcome Sanger Institute for the COVID-19 Genomics UK (COG-UK) Consortium | The Lighthouse Lab in Milton Keynes and Alex Alderton, Roberto Amato, Jeffrey Barrett, Sonia Goncalves, Ewan Harrison, David K. Jackson, Ian Johnston, Dominic Kwiatkowski, Cordelia Langford, John Sillitoe on behalf of the Wellcome Sanger Institute COVID-19 Surveillance Team |
| EPI_ISL_2568792 | Lighthouse Laboratory Plymouth | Wellcome Sanger Institute for the COVID-19 Genomics UK (COG-UK) Consortium | Lighthouse Laboratory Plymouth and Alex Alderton, Roberto Amato, Jeffrey Barrett, Sonia Goncalves, Ewan Harrison, David K. Jackson, Ian Johnston, Dominic Kwiatkowski, Cordelia Langford, John Sillitoe on behalf of the Wellcome Sanger Institute COVID-19 Surveillance Team |
| EPI_ISL_2568793, EPI_ISL_2568795 | Lighthouse Lab in Milton Keynes | Wellcome Sanger Institute for the COVID-19 Genomics UK (COG-UK) Consortium | The Lighthouse Lab in Milton Keynes and Alex Alderton, Roberto Amato, Jeffrey Barrett, Sonia Goncalves, Ewan Harrison, David K. Jackson, Ian Johnston, Dominic Kwiatkowski, Cordelia Langford, John Sillitoe on behalf of the Wellcome Sanger Institute COVID-19 Surveillance Team |
| EPI_ISL_2568796 | Lighthouse Laboratory Plymouth | Wellcome Sanger Institute for the COVID-19 Genomics UK (COG-UK) | Lighthouse Laboratory Plymouth and Alex Alderton, Roberto Amato, Jeffrey Barrett, Sonia Goncalves, Ewan Harrison, David K. Jackson, Ian Johnston, Dominic Kwiatkowski, Cordelia Langford, John Sillitoe on behalf of the Wellcome Sanger |

| | | | |
|---|---|---|---|
| | | Consortium | Institute COVID-19 Surveillance Team |
| EPI_ISL_2568798, EPI_ISL_2568799, EPI_ISL_2568800 | Lighthouse Lab in Milton Keynes | Wellcome Sanger Institute for the COVID-19 Genomics UK (COG-UK) Consortium | The Lighthouse Lab in Milton Keynes and Alex Alderton, Roberto Amato, Jeffrey Barrett, Sonia Goncalves, Ewan Harrison, David K. Jackson, Ian Johnston, Dominic Kwiatkowski, Cordelia Langford, John Sillitoe on behalf of the Wellcome Sanger Institute COVID-19 Surveillance Team |
| EPI_ISL_2568801, EPI_ISL_2568802 | Lighthouse Laboratory Plymouth | Wellcome Sanger Institute for the COVID-19 Genomics UK (COG-UK) Consortium | Lighthouse Laboratory Plymouth and Alex Alderton, Roberto Amato, Jeffrey Barrett, Sonia Goncalves, Ewan Harrison, David K. Jackson, Ian Johnston, Dominic Kwiatkowski, Cordelia Langford, John Sillitoe on behalf of the Wellcome Sanger Institute COVID-19 Surveillance Team |
| EPI_ISL_2568803 | Lighthouse Lab in Milton Keynes | Wellcome Sanger Institute for the COVID-19 Genomics UK (COG-UK) Consortium | The Lighthouse Lab in Milton Keynes and Alex Alderton, Roberto Amato, Jeffrey Barrett, Sonia Goncalves, Ewan Harrison, David K. Jackson, Ian Johnston, Dominic Kwiatkowski, Cordelia Langford, John Sillitoe on behalf of the Wellcome Sanger Institute COVID-19 Surveillance Team |
| EPI_ISL_2568804 | Lighthouse Lab in Alderley Park | Wellcome Sanger Institute for the COVID-19 Genomics UK (COG-UK) Consortium | Jacquelyn Wynn, Mairead Hyland, The Lighthouse Lab in Alderley Park and Alex Alderton, Roberto Amato, Jeffrey Barrett, Sonia Goncalves, Ewan Harrison, David K. Jackson, Ian Johnston, Dominic Kwiatkowski, Cordelia Langford, John Sillitoe on behalf of the Wellcome Sanger Institute COVID-19 Surveillance Team |
| EPI_ISL_2568806, EPI_ISL_2568807, EPI_ISL_2568808 | Lighthouse Laboratory Plymouth | Wellcome Sanger Institute for the COVID-19 Genomics UK (COG-UK) Consortium | Lighthouse Laboratory Plymouth and Alex Alderton, Roberto Amato, Jeffrey Barrett, Sonia Goncalves, Ewan Harrison, David K. Jackson, Ian Johnston, Dominic Kwiatkowski, Cordelia Langford, John Sillitoe on behalf of the Wellcome Sanger Institute COVID-19 Surveillance Team |
| EPI_ISL_2568809 | Lighthouse Lab in Milton Keynes | Wellcome Sanger Institute for the COVID-19 Genomics UK (COG-UK) Consortium | The Lighthouse Lab in Milton Keynes and Alex Alderton, Roberto Amato, Jeffrey Barrett, Sonia Goncalves, Ewan Harrison, David K. Jackson, Ian Johnston, Dominic Kwiatkowski, Cordelia Langford, John Sillitoe on behalf of the Wellcome Sanger Institute COVID-19 Surveillance Team |
| EPI_ISL_2568810, EPI_ISL_2568812 | Lighthouse Lab in Alderley Park | Wellcome Sanger Institute for the COVID-19 Genomics UK (COG-UK) Consortium | Jacquelyn Wynn, Mairead Hyland, The Lighthouse Lab in Alderley Park and Alex Alderton, Roberto Amato, Jeffrey Barrett, Sonia Goncalves, Ewan Harrison, David K. Jackson, Ian Johnston, Dominic Kwiatkowski, Cordelia Langford, John Sillitoe on behalf of the Wellcome Sanger Institute COVID-19 Surveillance Team |
| EPI_ISL_2568813, EPI_ISL_2568814 | Lighthouse Lab in Milton Keynes | Wellcome Sanger Institute for the COVID-19 Genomics UK (COG-UK) Consortium | The Lighthouse Lab in Milton Keynes and Alex Alderton, Roberto Amato, Jeffrey Barrett, Sonia Goncalves, Ewan Harrison, David K. Jackson, Ian Johnston, Dominic Kwiatkowski, Cordelia Langford, John Sillitoe on behalf of the Wellcome Sanger Institute COVID-19 Surveillance Team |
| EPI_ISL_2568815 | Lighthouse Laboratory Plymouth | Wellcome Sanger Institute for the COVID-19 Genomics UK (COG-UK) Consortium | Lighthouse Laboratory Plymouth and Alex Alderton, Roberto Amato, Jeffrey Barrett, Sonia Goncalves, Ewan Harrison, David K. Jackson, Ian Johnston, Dominic Kwiatkowski, Cordelia Langford, John Sillitoe on behalf of the Wellcome Sanger Institute COVID-19 Surveillance Team |
| EPI_ISL_2568817, EPI_ISL_2568818, EPI_ISL_2568819 | Lighthouse Lab in Milton Keynes | Wellcome Sanger Institute for the COVID-19 Genomics UK (COG-UK) Consortium | The Lighthouse Lab in Milton Keynes and Alex Alderton, Roberto Amato, Jeffrey Barrett, Sonia Goncalves, Ewan Harrison, David K. Jackson, Ian Johnston, Dominic Kwiatkowski, Cordelia Langford, John Sillitoe on behalf of the Wellcome Sanger Institute COVID-19 Surveillance Team |
| EPI_ISL_2568820 | Lighthouse Laboratory Plymouth | Wellcome Sanger Institute for the COVID-19 Genomics UK (COG-UK) Consortium | Lighthouse Laboratory Plymouth and Alex Alderton, Roberto Amato, Jeffrey Barrett, Sonia Goncalves, Ewan Harrison, David K. Jackson, Ian Johnston, Dominic Kwiatkowski, Cordelia Langford, John Sillitoe on behalf of the Wellcome Sanger Institute COVID-19 Surveillance Team |
| EPI_ISL_2568821, EPI_ISL_2568822 | Lighthouse Lab in Milton Keynes | Wellcome Sanger Institute for the COVID-19 Genomics UK (COG-UK) Consortium | The Lighthouse Lab in Milton Keynes and Alex Alderton, Roberto Amato, Jeffrey Barrett, Sonia Goncalves, Ewan Harrison, David K. Jackson, Ian Johnston, Dominic Kwiatkowski, Cordelia Langford, John Sillitoe on behalf of the Wellcome Sanger Institute COVID-19 Surveillance Team |
| EPI_ISL_2568823 | Lighthouse Lab in Alderley Park | Wellcome Sanger Institute for the COVID-19 Genomics UK (COG-UK) Consortium | Jacquelyn Wynn, Mairead Hyland, The Lighthouse Lab in Alderley Park and Alex Alderton, Roberto Amato, Jeffrey Barrett, Sonia Goncalves, Ewan Harrison, David K. Jackson, Ian Johnston, Dominic Kwiatkowski, Cordelia Langford, John Sillitoe on behalf of the Wellcome Sanger Institute COVID-19 Surveillance Team |
| EPI_ISL_2568824, EPI_ISL_2568825, EPI_ISL_2568826, EPI_ISL_2568827, EPI_ISL_2568828 | Lighthouse Lab in Milton Keynes | Wellcome Sanger Institute for the COVID-19 Genomics UK (COG-UK) Consortium | The Lighthouse Lab in Milton Keynes and Alex Alderton, Roberto Amato, Jeffrey Barrett, Sonia Goncalves, Ewan Harrison, David K. Jackson, Ian Johnston, Dominic Kwiatkowski, Cordelia Langford, John Sillitoe on behalf of the Wellcome Sanger Institute COVID-19 Surveillance Team |
| EPI_ISL_2568829 | Lighthouse Lab in Alderley Park | Wellcome Sanger Institute for the COVID-19 Genomics UK (COG-UK) Consortium | Jacquelyn Wynn, Mairead Hyland, The Lighthouse Lab in Alderley Park and Alex Alderton, Roberto Amato, Jeffrey Barrett, Sonia Goncalves, Ewan Harrison, David K. Jackson, Ian Johnston, Dominic Kwiatkowski, Cordelia Langford, John Sillitoe on behalf of the Wellcome Sanger Institute COVID-19 Surveillance Team |
| EPI_ISL_2568830, EPI_ISL_2568831 | Lighthouse Lab in Milton Keynes | Wellcome Sanger Institute for the COVID-19 Genomics UK (COG-UK) Consortium | The Lighthouse Lab in Milton Keynes and Alex Alderton, Roberto Amato, Jeffrey Barrett, Sonia Goncalves, Ewan Harrison, David K. Jackson, Ian Johnston, Dominic Kwiatkowski, Cordelia Langford, John Sillitoe on behalf of the Wellcome Sanger Institute COVID-19 Surveillance Team |
| EPI_ISL_2568832 | Lighthouse Laboratory Plymouth | Wellcome Sanger Institute for the COVID-19 Genomics UK (COG-UK) Consortium | Lighthouse Laboratory Plymouth and Alex Alderton, Roberto Amato, Jeffrey Barrett, Sonia Goncalves, Ewan Harrison, David K. Jackson, Ian Johnston, Dominic Kwiatkowski, Cordelia Langford, John Sillitoe on behalf of the Wellcome Sanger Institute COVID-19 Surveillance Team |
| EPI_ISL_2568833, EPI_ISL_2568834, EPI_ISL_2568835, EPI_ISL_2568836 | Lighthouse Lab in Milton Keynes | Wellcome Sanger Institute for the COVID-19 Genomics UK (COG-UK) Consortium | The Lighthouse Lab in Milton Keynes and Alex Alderton, Roberto Amato, Jeffrey Barrett, Sonia Goncalves, Ewan Harrison, David K. Jackson, Ian Johnston, Dominic Kwiatkowski, Cordelia Langford, John Sillitoe on behalf of the Wellcome Sanger Institute COVID-19 Surveillance Team |
| EPI_ISL_2568837 | Lighthouse Laboratory Plymouth | Wellcome Sanger Institute for the COVID-19 Genomics UK (COG-UK) Consortium | Lighthouse Laboratory Plymouth and Alex Alderton, Roberto Amato, Jeffrey Barrett, Sonia Goncalves, Ewan Harrison, David K. Jackson, Ian Johnston, Dominic Kwiatkowski, Cordelia Langford, John Sillitoe on behalf of the Wellcome Sanger Institute COVID-19 Surveillance Team |
| EPI_ISL_2568838 | Lighthouse Lab in Milton Keynes | Wellcome Sanger Institute for the COVID-19 Genomics UK (COG-UK) Consortium | The Lighthouse Lab in Milton Keynes and Alex Alderton, Roberto Amato, Jeffrey Barrett, Sonia Goncalves, Ewan Harrison, David K. Jackson, Ian Johnston, Dominic Kwiatkowski, Cordelia Langford, John Sillitoe on behalf of the Wellcome Sanger Institute COVID-19 Surveillance Team |
| EPI_ISL_2568839, EPI_ISL_2568840 | Lighthouse Laboratory Plymouth | Wellcome Sanger Institute for the COVID-19 Genomics UK (COG-UK) Consortium | Lighthouse Laboratory Plymouth and Alex Alderton, Roberto Amato, Jeffrey Barrett, Sonia Goncalves, Ewan Harrison, David K. Jackson, Ian Johnston, Dominic Kwiatkowski, Cordelia Langford, John Sillitoe on behalf of the Wellcome Sanger Institute COVID-19 Surveillance Team |
| EPI_ISL_2568841, EPI_ISL_2568842 | Lighthouse Lab in Alderley Park | Wellcome Sanger Institute for the COVID-19 Genomics UK (COG-UK) Consortium | Jacquelyn Wynn, Mairead Hyland, The Lighthouse Lab in Alderley Park and Alex Alderton, Roberto Amato, Jeffrey Barrett, Sonia Goncalves, Ewan Harrison, David K. Jackson, Ian Johnston, Dominic Kwiatkowski, Cordelia Langford, John Sillitoe on behalf of the Wellcome Sanger Institute COVID-19 Surveillance Team |
| EPI_ISL_2568843 | Lighthouse Lab in Milton Keynes | Wellcome Sanger Institute for the COVID-19 Genomics UK (COG-UK) Consortium | The Lighthouse Lab in Milton Keynes and Alex Alderton, Roberto Amato, Jeffrey Barrett, Sonia Goncalves, Ewan Harrison, David K. Jackson, Ian Johnston, Dominic Kwiatkowski, Cordelia Langford, John Sillitoe on behalf of the Wellcome Sanger Institute COVID-19 Surveillance Team |
| EPI_ISL_2568844 | Lighthouse Laboratory Plymouth | Wellcome Sanger Institute for the COVID-19 Genomics UK (COG-UK) Consortium | Lighthouse Laboratory Plymouth and Alex Alderton, Roberto Amato, Jeffrey Barrett, Sonia Goncalves, Ewan Harrison, David K. Jackson, Ian Johnston, Dominic Kwiatkowski, Cordelia Langford, John Sillitoe on behalf of the Wellcome Sanger Institute COVID-19 Surveillance Team |
| EPI_ISL_2568845, EPI_ISL_2568846 | Lighthouse Lab in Alderley Park | Wellcome Sanger Institute for the COVID-19 Genomics UK (COG-UK) Consortium | Jacquelyn Wynn, Mairead Hyland, The Lighthouse Lab in Alderley Park and Alex Alderton, Roberto Amato, Jeffrey Barrett, Sonia Goncalves, Ewan Harrison, David K. Jackson, Ian Johnston, Dominic Kwiatkowski, Cordelia Langford, John Sillitoe on behalf of the Wellcome Sanger Institute COVID-19 Surveillance Team |
| EPI_ISL_2568847 | Lighthouse Laboratory Plymouth | Wellcome Sanger Institute for the COVID-19 Genomics UK (COG-UK) Consortium | Lighthouse Laboratory Plymouth and Alex Alderton, Roberto Amato, Jeffrey Barrett, Sonia Goncalves, Ewan Harrison, David K. Jackson, Ian Johnston, Dominic Kwiatkowski, Cordelia Langford, John Sillitoe on behalf of the Wellcome Sanger Institute COVID-19 Surveillance Team |
| EPI_ISL_2568848 | Lighthouse Lab in Milton Keynes | Wellcome Sanger Institute for the COVID-19 Genomics UK (COG-UK) Consortium | The Lighthouse Lab in Milton Keynes and Alex Alderton, Roberto Amato, Jeffrey Barrett, Sonia Goncalves, Ewan Harrison, David K. Jackson, Ian Johnston, Dominic Kwiatkowski, Cordelia Langford, John Sillitoe on behalf of the Wellcome Sanger Institute COVID-19 Surveillance Team |
| EPI_ISL_2568849, EPI_ISL_2568850, EPI_ISL_2568851 | Lighthouse Lab in Alderley Park | Wellcome Sanger Institute for the COVID-19 Genomics UK (COG-UK) Consortium | Jacquelyn Wynn, Mairead Hyland, The Lighthouse Lab in Alderley Park and Alex Alderton, Roberto Amato, Jeffrey Barrett, Sonia Goncalves, Ewan Harrison, David K. Jackson, Ian Johnston, Dominic Kwiatkowski, Cordelia Langford, John Sillitoe on behalf of the Wellcome Sanger Institute COVID-19 Surveillance Team |
| EPI_ISL_2568852 | Lighthouse Laboratory Plymouth | Wellcome Sanger Institute for the COVID-19 Genomics UK (COG-UK) Consortium | Lighthouse Laboratory Plymouth and Alex Alderton, Roberto Amato, Jeffrey Barrett, Sonia Goncalves, Ewan Harrison, David K. Jackson, Ian Johnston, Dominic Kwiatkowski, Cordelia Langford, John Sillitoe on behalf of the Wellcome Sanger Institute COVID-19 Surveillance Team |
| EPI_ISL_2568853 | Lighthouse Lab in Milton Keynes | Wellcome Sanger Institute for the COVID-19 Genomics UK (COG-UK) Consortium | The Lighthouse Lab in Milton Keynes and Alex Alderton, Roberto Amato, Jeffrey Barrett, Sonia Goncalves, Ewan Harrison, David K. Jackson, Ian Johnston, Dominic Kwiatkowski, Cordelia Langford, John Sillitoe on behalf of the Wellcome Sanger Institute COVID-19 Surveillance Team |
| EPI_ISL_2568854 | Lighthouse Lab in Alderley Park | Wellcome Sanger Institute for the COVID-19 Genomics UK (COG-UK) Consortium | Jacquelyn Wynn, Mairead Hyland, The Lighthouse Lab in Alderley Park and Alex Alderton, Roberto Amato, Jeffrey Barrett, Sonia Goncalves, Ewan Harrison, David K. Jackson, Ian Johnston, Dominic Kwiatkowski, Cordelia Langford, John Sillitoe on behalf of the Wellcome Sanger Institute COVID-19 Surveillance Team |
| EPI_ISL_2568855 | Lighthouse Laboratory Plymouth | Wellcome Sanger Institute for the COVID-19 Genomics UK (COG-UK) Consortium | Lighthouse Laboratory Plymouth and Alex Alderton, Roberto Amato, Jeffrey Barrett, Sonia Goncalves, Ewan Harrison, David K. Jackson, Ian Johnston, Dominic Kwiatkowski, Cordelia Langford, John Sillitoe on behalf of the Wellcome Sanger Institute COVID-19 Surveillance Team |
| EPI_ISL_2568856 | Lighthouse Lab in Alderley Park | Wellcome Sanger Institute for the COVID-19 Genomics UK (COG-UK) Consortium | Jacquelyn Wynn, Mairead Hyland, The Lighthouse Lab in Alderley Park and Alex Alderton, Roberto Amato, Jeffrey Barrett, Sonia Goncalves, Ewan Harrison, David K. Jackson, Ian Johnston, Dominic Kwiatkowski, Cordelia Langford, John Sillitoe on behalf of the Wellcome Sanger Institute COVID-19 Surveillance Team |
| EPI_ISL_2568857, EPI_ISL_2568859, EPI_ISL_2568860, EPI_ISL_2568861, EPI_ISL_2568862 | Lighthouse Lab in Milton Keynes | Wellcome Sanger Institute for the COVID-19 Genomics UK (COG-UK) Consortium | The Lighthouse Lab in Milton Keynes and Alex Alderton, Roberto Amato, Jeffrey Barrett, Sonia Goncalves, Ewan Harrison, David K. Jackson, Ian Johnston, Dominic Kwiatkowski, Cordelia Langford, John Sillitoe on behalf of the Wellcome Sanger Institute COVID-19 Surveillance Team |
| EPI_ISL_2568863 | Lighthouse Laboratory Plymouth | Wellcome Sanger Institute for the COVID-19 Genomics UK (COG-UK) Consortium | Lighthouse Laboratory Plymouth and Alex Alderton, Roberto Amato, Jeffrey Barrett, Sonia Goncalves, Ewan Harrison, David K. Jackson, Ian Johnston, Dominic Kwiatkowski, Cordelia Langford, John Sillitoe on behalf of the Wellcome Sanger Institute COVID-19 Surveillance Team |
| EPI_ISL_2568864 | Lighthouse Lab in Alderley Park | Wellcome Sanger Institute for the COVID-19 Genomics UK (COG-UK) Consortium | Jacquelyn Wynn, Mairead Hyland, The Lighthouse Lab in Alderley Park and Alex Alderton, Roberto Amato, Jeffrey Barrett, Sonia Goncalves, Ewan Harrison, David K. Jackson, Ian Johnston, Dominic Kwiatkowski, Cordelia Langford, John Sillitoe on behalf of the Wellcome Sanger Institute COVID-19 Surveillance Team |
| EPI_ISL_2568865, EPI_ISL_2568866, EPI_ISL_2568867 | Lighthouse Lab in Milton Keynes | Wellcome Sanger Institute for the COVID-19 Genomics UK (COG-UK) Consortium | The Lighthouse Lab in Milton Keynes and Alex Alderton, Roberto Amato, Jeffrey Barrett, Sonia Goncalves, Ewan Harrison, David K. Jackson, Ian Johnston, Dominic Kwiatkowski, Cordelia Langford, John Sillitoe on behalf of the Wellcome Sanger Institute COVID-19 Surveillance Team |
| EPI_ISL_2568868, EPI_ISL_2568869 | Lighthouse Laboratory Plymouth | Wellcome Sanger Institute for the COVID-19 Genomics UK (COG-UK) Consortium | Lighthouse Laboratory Plymouth and Alex Alderton, Roberto Amato, Jeffrey Barrett, Sonia Goncalves, Ewan Harrison, David K. Jackson, Ian Johnston, Dominic Kwiatkowski, Cordelia Langford, John Sillitoe on behalf of the Wellcome Sanger Institute COVID-19 Surveillance Team |
| EPI_ISL_2568870 | Lighthouse Lab in Alderley Park | Wellcome Sanger Institute for the COVID-19 Genomics UK (COG-UK) Consortium | Jacquelyn Wynn, Mairead Hyland, The Lighthouse Lab in Alderley Park and Alex Alderton, Roberto Amato, Jeffrey Barrett, Sonia Goncalves, Ewan Harrison, David K. Jackson, Ian Johnston, Dominic Kwiatkowski, Cordelia Langford, John Sillitoe on behalf of the Wellcome Sanger Institute COVID-19 Surveillance Team |
| EPI_ISL_2568871 | Lighthouse Lab in Milton Keynes | Wellcome Sanger Institute for the COVID-19 Genomics UK (COG-UK) | The Lighthouse Lab in Milton Keynes and Alex Alderton, Roberto Amato, Jeffrey Barrett, Sonia Goncalves, Ewan Harrison, David K. Jackson, Ian Johnston, Dominic Kwiatkowski, Cordelia Langford, John Sillitoe on behalf of the Wellcome Sanger |

| | | | |
|---|---|---|---|
| | | Consortium | Institute COVID-19 Surveillance Team |
| EPI_ISL_2568872 | Lighthouse Lab in Alderley Park | Wellcome Sanger Institute for the COVID-19 Genomics UK (COG-UK) Consortium | Jacquelyn Wynn, Mairead Hyland, The Lighthouse Lab in Alderley Park and Alex Alderton, Roberto Amato, Jeffrey Barrett, Sonia Goncalves, Ewan Harrison, David K. Jackson, Ian Johnston, Dominic Kwiatkowski, Cordelia Langford, John Sillitoe on behalf of the Wellcome Sanger Institute COVID-19 Surveillance Team |
| EPI_ISL_2568873 | Lighthouse Laboratory Plymouth | Wellcome Sanger Institute for the COVID-19 Genomics UK (COG-UK) Consortium | Lighthouse Laboratory Plymouth and Alex Alderton, Roberto Amato, Jeffrey Barrett, Sonia Goncalves, Ewan Harrison, David K. Jackson, Ian Johnston, Dominic Kwiatkowski, Cordelia Langford, John Sillitoe on behalf of the Wellcome Sanger Institute COVID-19 Surveillance Team |
| EPI_ISL_2568874 | Lighthouse Lab in Alderley Park | Wellcome Sanger Institute for the COVID-19 Genomics UK (COG-UK) Consortium | Jacquelyn Wynn, Mairead Hyland, The Lighthouse Lab in Alderley Park and Alex Alderton, Roberto Amato, Jeffrey Barrett, Sonia Goncalves, Ewan Harrison, David K. Jackson, Ian Johnston, Dominic Kwiatkowski, Cordelia Langford, John Sillitoe on behalf of the Wellcome Sanger Institute COVID-19 Surveillance Team |
| EPI_ISL_2568875 | Lighthouse Lab in Milton Keynes | Wellcome Sanger Institute for the COVID-19 Genomics UK (COG-UK) Consortium | The Lighthouse Lab in Milton Keynes and Alex Alderton, Roberto Amato, Jeffrey Barrett, Sonia Goncalves, Ewan Harrison, David K. Jackson, Ian Johnston, Dominic Kwiatkowski, Cordelia Langford, John Sillitoe on behalf of the Wellcome Sanger Institute COVID-19 Surveillance Team |
| EPI_ISL_2568876 | Lighthouse Laboratory Plymouth | Wellcome Sanger Institute for the COVID-19 Genomics UK (COG-UK) Consortium | Lighthouse Laboratory Plymouth and Alex Alderton, Roberto Amato, Jeffrey Barrett, Sonia Goncalves, Ewan Harrison, David K. Jackson, Ian Johnston, Dominic Kwiatkowski, Cordelia Langford, John Sillitoe on behalf of the Wellcome Sanger Institute COVID-19 Surveillance Team |
| EPI_ISL_2568877, EPI_ISL_2568878, EPI_ISL_2568879 | Lighthouse Lab in Alderley Park | Wellcome Sanger Institute for the COVID-19 Genomics UK (COG-UK) Consortium | Jacquelyn Wynn, Mairead Hyland, The Lighthouse Lab in Alderley Park and Alex Alderton, Roberto Amato, Jeffrey Barrett, Sonia Goncalves, Ewan Harrison, David K. Jackson, Ian Johnston, Dominic Kwiatkowski, Cordelia Langford, John Sillitoe on behalf of the Wellcome Sanger Institute COVID-19 Surveillance Team |
| EPI_ISL_2568880 | Lighthouse Lab in Milton Keynes | Wellcome Sanger Institute for the COVID-19 Genomics UK (COG-UK) Consortium | The Lighthouse Lab in Milton Keynes and Alex Alderton, Roberto Amato, Jeffrey Barrett, Sonia Goncalves, Ewan Harrison, David K. Jackson, Ian Johnston, Dominic Kwiatkowski, Cordelia Langford, John Sillitoe on behalf of the Wellcome Sanger Institute COVID-19 Surveillance Team |
| EPI_ISL_2568881 | Lighthouse Laboratory Plymouth | Wellcome Sanger Institute for the COVID-19 Genomics UK (COG-UK) Consortium | Lighthouse Laboratory Plymouth and Alex Alderton, Roberto Amato, Jeffrey Barrett, Sonia Goncalves, Ewan Harrison, David K. Jackson, Ian Johnston, Dominic Kwiatkowski, Cordelia Langford, John Sillitoe on behalf of the Wellcome Sanger Institute COVID-19 Surveillance Team |
| EPI_ISL_2568882 | Lighthouse Lab in Milton Keynes | Wellcome Sanger Institute for the COVID-19 Genomics UK (COG-UK) Consortium | The Lighthouse Lab in Milton Keynes and Alex Alderton, Roberto Amato, Jeffrey Barrett, Sonia Goncalves, Ewan Harrison, David K. Jackson, Ian Johnston, Dominic Kwiatkowski, Cordelia Langford, John Sillitoe on behalf of the Wellcome Sanger Institute COVID-19 Surveillance Team |
| EPI_ISL_2568883, EPI_ISL_2568884 | Lighthouse Lab in Alderley Park | Wellcome Sanger Institute for the COVID-19 Genomics UK (COG-UK) Consortium | Jacquelyn Wynn, Mairead Hyland, The Lighthouse Lab in Alderley Park and Alex Alderton, Roberto Amato, Jeffrey Barrett, Sonia Goncalves, Ewan Harrison, David K. Jackson, Ian Johnston, Dominic Kwiatkowski, Cordelia Langford, John Sillitoe on behalf of the Wellcome Sanger Institute COVID-19 Surveillance Team |
| EPI_ISL_2568885, EPI_ISL_2568886 | Lighthouse Lab in Milton Keynes | Wellcome Sanger Institute for the COVID-19 Genomics UK (COG-UK) Consortium | The Lighthouse Lab in Milton Keynes and Alex Alderton, Roberto Amato, Jeffrey Barrett, Sonia Goncalves, Ewan Harrison, David K. Jackson, Ian Johnston, Dominic Kwiatkowski, Cordelia Langford, John Sillitoe on behalf of the Wellcome Sanger Institute COVID-19 Surveillance Team |
| EPI_ISL_2568887 | Lighthouse Laboratory Plymouth | Wellcome Sanger Institute for the COVID-19 Genomics UK (COG-UK) Consortium | Lighthouse Laboratory Plymouth and Alex Alderton, Roberto Amato, Jeffrey Barrett, Sonia Goncalves, Ewan Harrison, David K. Jackson, Ian Johnston, Dominic Kwiatkowski, Cordelia Langford, John Sillitoe on behalf of the Wellcome Sanger Institute COVID-19 Surveillance Team |
| EPI_ISL_2568888 | Lighthouse Lab in Milton Keynes | Wellcome Sanger Institute for the COVID-19 Genomics UK (COG-UK) Consortium | The Lighthouse Lab in Milton Keynes and Alex Alderton, Roberto Amato, Jeffrey Barrett, Sonia Goncalves, Ewan Harrison, David K. Jackson, Ian Johnston, Dominic Kwiatkowski, Cordelia Langford, John Sillitoe on behalf of the Wellcome Sanger Institute COVID-19 Surveillance Team |
| EPI_ISL_2568889 | Lighthouse Laboratory Plymouth | Wellcome Sanger Institute for the COVID-19 Genomics UK (COG-UK) Consortium | Lighthouse Laboratory Plymouth and Alex Alderton, Roberto Amato, Jeffrey Barrett, Sonia Goncalves, Ewan Harrison, David K. Jackson, Ian Johnston, Dominic Kwiatkowski, Cordelia Langford, John Sillitoe on behalf of the Wellcome Sanger Institute COVID-19 Surveillance Team |
| EPI_ISL_2568890, EPI_ISL_2568891 | Lighthouse Lab in Alderley Park | Wellcome Sanger Institute for the COVID-19 Genomics UK (COG-UK) Consortium | Jacquelyn Wynn, Mairead Hyland, The Lighthouse Lab in Alderley Park and Alex Alderton, Roberto Amato, Jeffrey Barrett, Sonia Goncalves, Ewan Harrison, David K. Jackson, Ian Johnston, Dominic Kwiatkowski, Cordelia Langford, John Sillitoe on behalf of the Wellcome Sanger Institute COVID-19 Surveillance Team |
| EPI_ISL_2568892 | Lighthouse Laboratory Plymouth | Wellcome Sanger Institute for the COVID-19 Genomics UK (COG-UK) Consortium | Lighthouse Laboratory Plymouth and Alex Alderton, Roberto Amato, Jeffrey Barrett, Sonia Goncalves, Ewan Harrison, David K. Jackson, Ian Johnston, Dominic Kwiatkowski, Cordelia Langford, John Sillitoe on behalf of the Wellcome Sanger Institute COVID-19 Surveillance Team |
| EPI_ISL_2568893 | Lighthouse Lab in Milton Keynes | Wellcome Sanger Institute for the COVID-19 Genomics UK (COG-UK) Consortium | The Lighthouse Lab in Milton Keynes and Alex Alderton, Roberto Amato, Jeffrey Barrett, Sonia Goncalves, Ewan Harrison, David K. Jackson, Ian Johnston, Dominic Kwiatkowski, Cordelia Langford, John Sillitoe on behalf of the Wellcome Sanger Institute COVID-19 Surveillance Team |
| EPI_ISL_2568894, EPI_ISL_2568895, EPI_ISL_2568896 | Lighthouse Lab in Alderley Park | Wellcome Sanger Institute for the COVID-19 Genomics UK (COG-UK) Consortium | Jacquelyn Wynn, Mairead Hyland, The Lighthouse Lab in Alderley Park and Alex Alderton, Roberto Amato, Jeffrey Barrett, Sonia Goncalves, Ewan Harrison, David K. Jackson, Ian Johnston, Dominic Kwiatkowski, Cordelia Langford, John Sillitoe on behalf of the Wellcome Sanger Institute COVID-19 Surveillance Team |
| EPI_ISL_2568897 | Lighthouse Laboratory Plymouth | Wellcome Sanger Institute for the COVID-19 Genomics UK (COG-UK) Consortium | Lighthouse Laboratory Plymouth and Alex Alderton, Roberto Amato, Jeffrey Barrett, Sonia Goncalves, Ewan Harrison, David K. Jackson, Ian Johnston, Dominic Kwiatkowski, Cordelia Langford, John Sillitoe on behalf of the Wellcome Sanger Institute COVID-19 Surveillance Team |
| EPI_ISL_2568899 | Lighthouse Lab in Milton Keynes | Wellcome Sanger Institute for the COVID-19 Genomics UK (COG-UK) Consortium | The Lighthouse Lab in Milton Keynes and Alex Alderton, Roberto Amato, Jeffrey Barrett, Sonia Goncalves, Ewan Harrison, David K. Jackson, Ian Johnston, Dominic Kwiatkowski, Cordelia Langford, John Sillitoe on behalf of the Wellcome Sanger Institute COVID-19 Surveillance Team |
| EPI_ISL_2568900 | Lighthouse Laboratory Plymouth | Wellcome Sanger Institute for the COVID-19 Genomics UK (COG-UK) Consortium | Lighthouse Laboratory Plymouth and Alex Alderton, Roberto Amato, Jeffrey Barrett, Sonia Goncalves, Ewan Harrison, David K. Jackson, Ian Johnston, Dominic Kwiatkowski, Cordelia Langford, John Sillitoe on behalf of the Wellcome Sanger Institute COVID-19 Surveillance Team |
| EPI_ISL_2568901 | Lighthouse Lab in Alderley Park | Wellcome Sanger Institute for the COVID-19 Genomics UK (COG-UK) Consortium | Jacquelyn Wynn, Mairead Hyland, The Lighthouse Lab in Alderley Park and Alex Alderton, Roberto Amato, Jeffrey Barrett, Sonia Goncalves, Ewan Harrison, David K. Jackson, Ian Johnston, Dominic Kwiatkowski, Cordelia Langford, John Sillitoe on behalf of the Wellcome Sanger Institute COVID-19 Surveillance Team |
| EPI_ISL_2568902 | Lighthouse Lab in Milton Keynes | Wellcome Sanger Institute for the COVID-19 Genomics UK (COG-UK) Consortium | The Lighthouse Lab in Milton Keynes and Alex Alderton, Roberto Amato, Jeffrey Barrett, Sonia Goncalves, Ewan Harrison, David K. Jackson, Ian Johnston, Dominic Kwiatkowski, Cordelia Langford, John Sillitoe on behalf of the Wellcome Sanger Institute COVID-19 Surveillance Team |
| EPI_ISL_2568903, EPI_ISL_2568904, EPI_ISL_2568905, EPI_ISL_2568906, EPI_ISL_2568907 | Lighthouse Lab in Alderley Park | Wellcome Sanger Institute for the COVID-19 Genomics UK (COG-UK) Consortium | Jacquelyn Wynn, Mairead Hyland, The Lighthouse Lab in Alderley Park and Alex Alderton, Roberto Amato, Jeffrey Barrett, Sonia Goncalves, Ewan Harrison, David K. Jackson, Ian Johnston, Dominic Kwiatkowski, Cordelia Langford, John Sillitoe on behalf of the Wellcome Sanger Institute COVID-19 Surveillance Team |
| EPI_ISL_2568908, EPI_ISL_2568909, EPI_ISL_2568910 | Lighthouse Lab in Milton Keynes | Wellcome Sanger Institute for the COVID-19 Genomics UK (COG-UK) Consortium | The Lighthouse Lab in Milton Keynes and Alex Alderton, Roberto Amato, Jeffrey Barrett, Sonia Goncalves, Ewan Harrison, David K. Jackson, Ian Johnston, Dominic Kwiatkowski, Cordelia Langford, John Sillitoe on behalf of the Wellcome Sanger Institute COVID-19 Surveillance Team |
| EPI_ISL_2568911 | Lighthouse Lab in Alderley Park | Wellcome Sanger Institute for the COVID-19 Genomics UK (COG-UK) Consortium | Jacquelyn Wynn, Mairead Hyland, The Lighthouse Lab in Alderley Park and Alex Alderton, Roberto Amato, Jeffrey Barrett, Sonia Goncalves, Ewan Harrison, David K. Jackson, Ian Johnston, Dominic Kwiatkowski, Cordelia Langford, John Sillitoe on behalf of the Wellcome Sanger Institute COVID-19 Surveillance Team |
| EPI_ISL_2568912 | Lighthouse Laboratory Plymouth | Wellcome Sanger Institute for the COVID-19 Genomics UK (COG-UK) Consortium | Lighthouse Laboratory Plymouth and Alex Alderton, Roberto Amato, Jeffrey Barrett, Sonia Goncalves, Ewan Harrison, David K. Jackson, Ian Johnston, Dominic Kwiatkowski, Cordelia Langford, John Sillitoe on behalf of the Wellcome Sanger Institute COVID-19 Surveillance Team |
| EPI_ISL_2568913 | Lighthouse Lab in Alderley Park | Wellcome Sanger Institute for the COVID-19 Genomics UK (COG-UK) Consortium | Jacquelyn Wynn, Mairead Hyland, The Lighthouse Lab in Alderley Park and Alex Alderton, Roberto Amato, Jeffrey Barrett, Sonia Goncalves, Ewan Harrison, David K. Jackson, Ian Johnston, Dominic Kwiatkowski, Cordelia Langford, John Sillitoe on behalf of the Wellcome Sanger Institute COVID-19 Surveillance Team |
| EPI_ISL_2568914 | Lighthouse Laboratory Plymouth | Wellcome Sanger Institute for the COVID-19 Genomics UK (COG-UK) Consortium | Lighthouse Laboratory Plymouth and Alex Alderton, Roberto Amato, Jeffrey Barrett, Sonia Goncalves, Ewan Harrison, David K. Jackson, Ian Johnston, Dominic Kwiatkowski, Cordelia Langford, John Sillitoe on behalf of the Wellcome Sanger Institute COVID-19 Surveillance Team |
| EPI_ISL_2568915 | Lighthouse Lab in Alderley Park | Wellcome Sanger Institute for the COVID-19 Genomics UK (COG-UK) Consortium | Jacquelyn Wynn, Mairead Hyland, The Lighthouse Lab in Alderley Park and Alex Alderton, Roberto Amato, Jeffrey Barrett, Sonia Goncalves, Ewan Harrison, David K. Jackson, Ian Johnston, Dominic Kwiatkowski, Cordelia Langford, John Sillitoe on behalf of the Wellcome Sanger Institute COVID-19 Surveillance Team |
| EPI_ISL_2568916 | Lighthouse Laboratory Plymouth | Wellcome Sanger Institute for the COVID-19 Genomics UK (COG-UK) Consortium | Lighthouse Laboratory Plymouth and Alex Alderton, Roberto Amato, Jeffrey Barrett, Sonia Goncalves, Ewan Harrison, David K. Jackson, Ian Johnston, Dominic Kwiatkowski, Cordelia Langford, John Sillitoe on behalf of the Wellcome Sanger Institute COVID-19 Surveillance Team |
| EPI_ISL_2568917 | Lighthouse Lab in Alderley Park | Wellcome Sanger Institute for the COVID-19 Genomics UK (COG-UK) Consortium | Jacquelyn Wynn, Mairead Hyland, The Lighthouse Lab in Alderley Park and Alex Alderton, Roberto Amato, Jeffrey Barrett, Sonia Goncalves, Ewan Harrison, David K. Jackson, Ian Johnston, Dominic Kwiatkowski, Cordelia Langford, John Sillitoe on behalf of the Wellcome Sanger Institute COVID-19 Surveillance Team |
| EPI_ISL_2568918, EPI_ISL_2568920 | Lighthouse Laboratory Plymouth | Wellcome Sanger Institute for the COVID-19 Genomics UK (COG-UK) Consortium | Lighthouse Laboratory Plymouth and Alex Alderton, Roberto Amato, Jeffrey Barrett, Sonia Goncalves, Ewan Harrison, David K. Jackson, Ian Johnston, Dominic Kwiatkowski, Cordelia Langford, John Sillitoe on behalf of the Wellcome Sanger Institute COVID-19 Surveillance Team |
| EPI_ISL_2568922 | Lighthouse Lab in Milton Keynes | Wellcome Sanger Institute for the COVID-19 Genomics UK (COG-UK) Consortium | The Lighthouse Lab in Milton Keynes and Alex Alderton, Roberto Amato, Jeffrey Barrett, Sonia Goncalves, Ewan Harrison, David K. Jackson, Ian Johnston, Dominic Kwiatkowski, Cordelia Langford, John Sillitoe on behalf of the Wellcome Sanger Institute COVID-19 Surveillance Team |
| EPI_ISL_2568924 | Lighthouse Lab in Alderley Park | Wellcome Sanger Institute for the COVID-19 Genomics UK (COG-UK) Consortium | Jacquelyn Wynn, Mairead Hyland, The Lighthouse Lab in Alderley Park and Alex Alderton, Roberto Amato, Jeffrey Barrett, Sonia Goncalves, Ewan Harrison, David K. Jackson, Ian Johnston, Dominic Kwiatkowski, Cordelia Langford, John Sillitoe on behalf of the Wellcome Sanger Institute COVID-19 Surveillance Team |
| EPI_ISL_2568925 | Lighthouse Lab in Milton Keynes | Wellcome Sanger Institute for the COVID-19 Genomics UK (COG-UK) Consortium | The Lighthouse Lab in Milton Keynes and Alex Alderton, Roberto Amato, Jeffrey Barrett, Sonia Goncalves, Ewan Harrison, David K. Jackson, Ian Johnston, Dominic Kwiatkowski, Cordelia Langford, John Sillitoe on behalf of the Wellcome Sanger Institute COVID-19 Surveillance Team |
| EPI_ISL_2568926 | Lighthouse Laboratory Plymouth | Wellcome Sanger Institute for the COVID-19 Genomics UK (COG-UK) Consortium | Lighthouse Laboratory Plymouth and Alex Alderton, Roberto Amato, Jeffrey Barrett, Sonia Goncalves, Ewan Harrison, David K. Jackson, Ian Johnston, Dominic Kwiatkowski, Cordelia Langford, John Sillitoe on behalf of the Wellcome Sanger Institute COVID-19 Surveillance Team |
| EPI_ISL_2568927 | Lighthouse Lab in Milton Keynes | Wellcome Sanger Institute for the COVID-19 Genomics UK (COG-UK) Consortium | The Lighthouse Lab in Milton Keynes and Alex Alderton, Roberto Amato, Jeffrey Barrett, Sonia Goncalves, Ewan Harrison, David K. Jackson, Ian Johnston, Dominic Kwiatkowski, Cordelia Langford, John Sillitoe on behalf of the Wellcome Sanger Institute COVID-19 Surveillance Team |
| EPI_ISL_2568928 | Lighthouse Lab in Alderley Park | Wellcome Sanger Institute for the COVID-19 Genomics UK (COG-UK) Consortium | Jacquelyn Wynn, Mairead Hyland, The Lighthouse Lab in Alderley Park and Alex Alderton, Roberto Amato, Jeffrey Barrett, Sonia Goncalves, Ewan Harrison, David K. Jackson, Ian Johnston, Dominic Kwiatkowski, Cordelia Langford, John Sillitoe on behalf of the Wellcome Sanger Institute COVID-19 Surveillance Team |
| EPI_ISL_2568929, EPI_ISL_2568930 | Lighthouse Laboratory Plymouth | Wellcome Sanger Institute for the COVID-19 Genomics UK (COG-UK) Consortium | Lighthouse Laboratory Plymouth and Alex Alderton, Roberto Amato, Jeffrey Barrett, Sonia Goncalves, Ewan Harrison, David K. Jackson, Ian Johnston, Dominic Kwiatkowski, Cordelia Langford, John Sillitoe on behalf of the Wellcome Sanger Institute COVID-19 Surveillance Team |
| EPI_ISL_2568931, EPI_ISL_2568932 | Lighthouse Lab in Alderley Park | Wellcome Sanger Institute for the COVID-19 Genomics UK (COG-UK) Consortium | Jacquelyn Wynn, Mairead Hyland, The Lighthouse Lab in Alderley Park and Alex Alderton, Roberto Amato, Jeffrey Barrett, Sonia Goncalves, Ewan Harrison, David K. Jackson, Ian Johnston, Dominic Kwiatkowski, Cordelia Langford, John Sillitoe on behalf of the Wellcome Sanger Institute COVID-19 Surveillance Team |
| EPI_ISL_2568933 | Lighthouse Laboratory Plymouth | Wellcome Sanger Institute for the COVID-19 Genomics UK (COG-UK) Consortium | Lighthouse Laboratory Plymouth and Alex Alderton, Roberto Amato, Jeffrey Barrett, Sonia Goncalves, Ewan Harrison, David K. Jackson, Ian Johnston, Dominic Kwiatkowski, Cordelia Langford, John Sillitoe on behalf of the Wellcome Sanger Institute COVID-19 Surveillance Team |

| | | | |
|---|---|---|---|
| EPI_ISL_2568934 | Lighthouse Lab in Alderley Park | Wellcome Sanger Institute for the COVID-19 Genomics UK (COG-UK) Consortium | Jacquelyn Wynn, Mairead Hyland, The Lighthouse Lab in Alderley Park and Alex Alderton, Roberto Amato, Jeffrey Barrett, Sonia Goncalves, Ewan Harrison, David K. Jackson, Ian Johnston, Dominic Kwiatkowski, Cordelia Langford, John Sillitoe on behalf of the Wellcome Sanger Institute COVID-19 Surveillance Team |
| EPI_ISL_2568935 | Lighthouse Lab in Milton Keynes | Wellcome Sanger Institute for the COVID-19 Genomics UK (COG-UK) Consortium | The Lighthouse Lab in Milton Keynes and Alex Alderton, Roberto Amato, Jeffrey Barrett, Sonia Goncalves, Ewan Harrison, David K. Jackson, Ian Johnston, Dominic Kwiatkowski, Cordelia Langford, John Sillitoe on behalf of the Wellcome Sanger Institute COVID-19 Surveillance Team |
| EPI_ISL_2568936 | Lighthouse Laboratory Plymouth | Wellcome Sanger Institute for the COVID-19 Genomics UK (COG-UK) Consortium | Lighthouse Laboratory Plymouth and Alex Alderton, Roberto Amato, Jeffrey Barrett, Sonia Goncalves, Ewan Harrison, David K. Jackson, Ian Johnston, Dominic Kwiatkowski, Cordelia Langford, John Sillitoe on behalf of the Wellcome Sanger Institute COVID-19 Surveillance Team |
| EPI_ISL_2568937 | Lighthouse Lab in Alderley Park | Wellcome Sanger Institute for the COVID-19 Genomics UK (COG-UK) Consortium | Jacquelyn Wynn, Mairead Hyland, The Lighthouse Lab in Alderley Park and Alex Alderton, Roberto Amato, Jeffrey Barrett, Sonia Goncalves, Ewan Harrison, David K. Jackson, Ian Johnston, Dominic Kwiatkowski, Cordelia Langford, John Sillitoe on behalf of the Wellcome Sanger Institute COVID-19 Surveillance Team |
| EPI_ISL_2568938 | Lighthouse Lab in Milton Keynes | Wellcome Sanger Institute for the COVID-19 Genomics UK (COG-UK) Consortium | The Lighthouse Lab in Milton Keynes and Alex Alderton, Roberto Amato, Jeffrey Barrett, Sonia Goncalves, Ewan Harrison, David K. Jackson, Ian Johnston, Dominic Kwiatkowski, Cordelia Langford, John Sillitoe on behalf of the Wellcome Sanger Institute COVID-19 Surveillance Team |
| EPI_ISL_2568939, EPI_ISL_2568940, EPI_ISL_2568941 | Lighthouse Lab in Alderley Park | Wellcome Sanger Institute for the COVID-19 Genomics UK (COG-UK) Consortium | Jacquelyn Wynn, Mairead Hyland, The Lighthouse Lab in Alderley Park and Alex Alderton, Roberto Amato, Jeffrey Barrett, Sonia Goncalves, Ewan Harrison, David K. Jackson, Ian Johnston, Dominic Kwiatkowski, Cordelia Langford, John Sillitoe on behalf of the Wellcome Sanger Institute COVID-19 Surveillance Team |
| EPI_ISL_2568942, EPI_ISL_2568943 | Lighthouse Lab in Milton Keynes | Wellcome Sanger Institute for the COVID-19 Genomics UK (COG-UK) Consortium | The Lighthouse Lab in Milton Keynes and Alex Alderton, Roberto Amato, Jeffrey Barrett, Sonia Goncalves, Ewan Harrison, David K. Jackson, Ian Johnston, Dominic Kwiatkowski, Cordelia Langford, John Sillitoe on behalf of the Wellcome Sanger Institute COVID-19 Surveillance Team |
| EPI_ISL_2568944 | Lighthouse Laboratory Plymouth | Wellcome Sanger Institute for the COVID-19 Genomics UK (COG-UK) Consortium | Lighthouse Laboratory Plymouth and Alex Alderton, Roberto Amato, Jeffrey Barrett, Sonia Goncalves, Ewan Harrison, David K. Jackson, Ian Johnston, Dominic Kwiatkowski, Cordelia Langford, John Sillitoe on behalf of the Wellcome Sanger Institute COVID-19 Surveillance Team |
| EPI_ISL_2568945 | Lighthouse Lab in Alderley Park | Wellcome Sanger Institute for the COVID-19 Genomics UK (COG-UK) Consortium | Jacquelyn Wynn, Mairead Hyland, The Lighthouse Lab in Alderley Park and Alex Alderton, Roberto Amato, Jeffrey Barrett, Sonia Goncalves, Ewan Harrison, David K. Jackson, Ian Johnston, Dominic Kwiatkowski, Cordelia Langford, John Sillitoe on behalf of the Wellcome Sanger Institute COVID-19 Surveillance Team |
| EPI_ISL_2568946, EPI_ISL_2568947 | Lighthouse Lab in Milton Keynes | Wellcome Sanger Institute for the COVID-19 Genomics UK (COG-UK) Consortium | The Lighthouse Lab in Milton Keynes and Alex Alderton, Roberto Amato, Jeffrey Barrett, Sonia Goncalves, Ewan Harrison, David K. Jackson, Ian Johnston, Dominic Kwiatkowski, Cordelia Langford, John Sillitoe on behalf of the Wellcome Sanger Institute COVID-19 Surveillance Team |
| EPI_ISL_2568948, EPI_ISL_2568949 | Lighthouse Lab in Alderley Park | Wellcome Sanger Institute for the COVID-19 Genomics UK (COG-UK) Consortium | Jacquelyn Wynn, Mairead Hyland, The Lighthouse Lab in Alderley Park and Alex Alderton, Roberto Amato, Jeffrey Barrett, Sonia Goncalves, Ewan Harrison, David K. Jackson, Ian Johnston, Dominic Kwiatkowski, Cordelia Langford, John Sillitoe on behalf of the Wellcome Sanger Institute COVID-19 Surveillance Team |
| EPI_ISL_2568950 | Lighthouse Laboratory Plymouth | Wellcome Sanger Institute for the COVID-19 Genomics UK (COG-UK) Consortium | Lighthouse Laboratory Plymouth and Alex Alderton, Roberto Amato, Jeffrey Barrett, Sonia Goncalves, Ewan Harrison, David K. Jackson, Ian Johnston, Dominic Kwiatkowski, Cordelia Langford, John Sillitoe on behalf of the Wellcome Sanger Institute COVID-19 Surveillance Team |
| EPI_ISL_2568951 | Lighthouse Lab in Milton Keynes | Wellcome Sanger Institute for the COVID-19 Genomics UK (COG-UK) Consortium | The Lighthouse Lab in Milton Keynes and Alex Alderton, Roberto Amato, Jeffrey Barrett, Sonia Goncalves, Ewan Harrison, David K. Jackson, Ian Johnston, Dominic Kwiatkowski, Cordelia Langford, John Sillitoe on behalf of the Wellcome Sanger Institute COVID-19 Surveillance Team |
| EPI_ISL_2568952 | Lighthouse Lab in Alderley Park | Wellcome Sanger Institute for the COVID-19 Genomics UK (COG-UK) Consortium | Jacquelyn Wynn, Mairead Hyland, The Lighthouse Lab in Alderley Park and Alex Alderton, Roberto Amato, Jeffrey Barrett, Sonia Goncalves, Ewan Harrison, David K. Jackson, Ian Johnston, Dominic Kwiatkowski, Cordelia Langford, John Sillitoe on behalf of the Wellcome Sanger Institute COVID-19 Surveillance Team |
| EPI_ISL_2568953, EPI_ISL_2568955 | Lighthouse Lab in Milton Keynes | Wellcome Sanger Institute for the COVID-19 Genomics UK (COG-UK) Consortium | The Lighthouse Lab in Milton Keynes and Alex Alderton, Roberto Amato, Jeffrey Barrett, Sonia Goncalves, Ewan Harrison, David K. Jackson, Ian Johnston, Dominic Kwiatkowski, Cordelia Langford, John Sillitoe on behalf of the Wellcome Sanger Institute COVID-19 Surveillance Team |
| EPI_ISL_2568956 | Lighthouse Laboratory Plymouth | Wellcome Sanger Institute for the COVID-19 Genomics UK (COG-UK) Consortium | Lighthouse Laboratory Plymouth and Alex Alderton, Roberto Amato, Jeffrey Barrett, Sonia Goncalves, Ewan Harrison, David K. Jackson, Ian Johnston, Dominic Kwiatkowski, Cordelia Langford, John Sillitoe on behalf of the Wellcome Sanger Institute COVID-19 Surveillance Team |
| EPI_ISL_2568957 | Lighthouse Lab in Milton Keynes | Wellcome Sanger Institute for the COVID-19 Genomics UK (COG-UK) Consortium | The Lighthouse Lab in Milton Keynes and Alex Alderton, Roberto Amato, Jeffrey Barrett, Sonia Goncalves, Ewan Harrison, David K. Jackson, Ian Johnston, Dominic Kwiatkowski, Cordelia Langford, John Sillitoe on behalf of the Wellcome Sanger Institute COVID-19 Surveillance Team |
| EPI_ISL_2568958 | Lighthouse Laboratory Plymouth | Wellcome Sanger Institute for the COVID-19 Genomics UK (COG-UK) Consortium | Lighthouse Laboratory Plymouth and Alex Alderton, Roberto Amato, Jeffrey Barrett, Sonia Goncalves, Ewan Harrison, David K. Jackson, Ian Johnston, Dominic Kwiatkowski, Cordelia Langford, John Sillitoe on behalf of the Wellcome Sanger Institute COVID-19 Surveillance Team |
| EPI_ISL_2568959 | Lighthouse Lab in Alderley Park | Wellcome Sanger Institute for the COVID-19 Genomics UK (COG-UK) Consortium | Jacquelyn Wynn, Mairead Hyland, The Lighthouse Lab in Alderley Park and Alex Alderton, Roberto Amato, Jeffrey Barrett, Sonia Goncalves, Ewan Harrison, David K. Jackson, Ian Johnston, Dominic Kwiatkowski, Cordelia Langford, John Sillitoe on behalf of the Wellcome Sanger Institute COVID-19 Surveillance Team |
| EPI_ISL_2568960, EPI_ISL_2568961 | Lighthouse Laboratory Plymouth | Wellcome Sanger Institute for the COVID-19 Genomics UK (COG-UK) Consortium | Lighthouse Laboratory Plymouth and Alex Alderton, Roberto Amato, Jeffrey Barrett, Sonia Goncalves, Ewan Harrison, David K. Jackson, Ian Johnston, Dominic Kwiatkowski, Cordelia Langford, John Sillitoe on behalf of the Wellcome Sanger Institute COVID-19 Surveillance Team |
| EPI_ISL_2568963 | Lighthouse Lab in Alderley Park | Wellcome Sanger Institute for the COVID-19 Genomics UK (COG-UK) Consortium | Jacquelyn Wynn, Mairead Hyland, The Lighthouse Lab in Alderley Park and Alex Alderton, Roberto Amato, Jeffrey Barrett, Sonia Goncalves, Ewan Harrison, David K. Jackson, Ian Johnston, Dominic Kwiatkowski, Cordelia Langford, John Sillitoe on behalf of the Wellcome Sanger Institute COVID-19 Surveillance Team |
| EPI_ISL_2568964 | Lighthouse Lab in Milton Keynes | Wellcome Sanger Institute for the COVID-19 Genomics UK (COG-UK) Consortium | The Lighthouse Lab in Milton Keynes and Alex Alderton, Roberto Amato, Jeffrey Barrett, Sonia Goncalves, Ewan Harrison, David K. Jackson, Ian Johnston, Dominic Kwiatkowski, Cordelia Langford, John Sillitoe on behalf of the Wellcome Sanger Institute COVID-19 Surveillance Team |
| EPI_ISL_2568965 | Lighthouse Lab in Alderley Park | Wellcome Sanger Institute for the COVID-19 Genomics UK (COG-UK) Consortium | Jacquelyn Wynn, Mairead Hyland, The Lighthouse Lab in Alderley Park and Alex Alderton, Roberto Amato, Jeffrey Barrett, Sonia Goncalves, Ewan Harrison, David K. Jackson, Ian Johnston, Dominic Kwiatkowski, Cordelia Langford, John Sillitoe on behalf of the Wellcome Sanger Institute COVID-19 Surveillance Team |
| EPI_ISL_2568966 | Lighthouse Lab in Milton Keynes | Wellcome Sanger Institute for the COVID-19 Genomics UK (COG-UK) Consortium | The Lighthouse Lab in Milton Keynes and Alex Alderton, Roberto Amato, Jeffrey Barrett, Sonia Goncalves, Ewan Harrison, David K. Jackson, Ian Johnston, Dominic Kwiatkowski, Cordelia Langford, John Sillitoe on behalf of the Wellcome Sanger Institute COVID-19 Surveillance Team |
| EPI_ISL_2568967 | Lighthouse Lab in Alderley Park | Wellcome Sanger Institute for the COVID-19 Genomics UK (COG-UK) Consortium | Jacquelyn Wynn, Mairead Hyland, The Lighthouse Lab in Alderley Park and Alex Alderton, Roberto Amato, Jeffrey Barrett, Sonia Goncalves, Ewan Harrison, David K. Jackson, Ian Johnston, Dominic Kwiatkowski, Cordelia Langford, John Sillitoe on behalf of the Wellcome Sanger Institute COVID-19 Surveillance Team |
| EPI_ISL_2568968, EPI_ISL_2568969 | Lighthouse Lab in Milton Keynes | Wellcome Sanger Institute for the COVID-19 Genomics UK (COG-UK) Consortium | The Lighthouse Lab in Milton Keynes and Alex Alderton, Roberto Amato, Jeffrey Barrett, Sonia Goncalves, Ewan Harrison, David K. Jackson, Ian Johnston, Dominic Kwiatkowski, Cordelia Langford, John Sillitoe on behalf of the Wellcome Sanger Institute COVID-19 Surveillance Team |
| EPI_ISL_2568970 | Lighthouse Lab in Alderley Park | Wellcome Sanger Institute for the COVID-19 Genomics UK (COG-UK) Consortium | Jacquelyn Wynn, Mairead Hyland, The Lighthouse Lab in Alderley Park and Alex Alderton, Roberto Amato, Jeffrey Barrett, Sonia Goncalves, Ewan Harrison, David K. Jackson, Ian Johnston, Dominic Kwiatkowski, Cordelia Langford, John Sillitoe on behalf of the Wellcome Sanger Institute COVID-19 Surveillance Team |
| EPI_ISL_2568971, EPI_ISL_2568972 | Lighthouse Lab in Milton Keynes | Wellcome Sanger Institute for the COVID-19 Genomics UK (COG-UK) Consortium | The Lighthouse Lab in Milton Keynes and Alex Alderton, Roberto Amato, Jeffrey Barrett, Sonia Goncalves, Ewan Harrison, David K. Jackson, Ian Johnston, Dominic Kwiatkowski, Cordelia Langford, John Sillitoe on behalf of the Wellcome Sanger Institute COVID-19 Surveillance Team |
| EPI_ISL_2568973 | Lighthouse Laboratory Plymouth | Wellcome Sanger Institute for the COVID-19 Genomics UK (COG-UK) Consortium | Lighthouse Laboratory Plymouth and Alex Alderton, Roberto Amato, Jeffrey Barrett, Sonia Goncalves, Ewan Harrison, David K. Jackson, Ian Johnston, Dominic Kwiatkowski, Cordelia Langford, John Sillitoe on behalf of the Wellcome Sanger Institute COVID-19 Surveillance Team |
| EPI_ISL_2568974, EPI_ISL_2568975 | Lighthouse Lab in Milton Keynes | Wellcome Sanger Institute for the COVID-19 Genomics UK (COG-UK) Consortium | The Lighthouse Lab in Milton Keynes and Alex Alderton, Roberto Amato, Jeffrey Barrett, Sonia Goncalves, Ewan Harrison, David K. Jackson, Ian Johnston, Dominic Kwiatkowski, Cordelia Langford, John Sillitoe on behalf of the Wellcome Sanger Institute COVID-19 Surveillance Team |
| EPI_ISL_2568976 | Lighthouse Lab in Alderley Park | Wellcome Sanger Institute for the COVID-19 Genomics UK (COG-UK) Consortium | Jacquelyn Wynn, Mairead Hyland, The Lighthouse Lab in Alderley Park and Alex Alderton, Roberto Amato, Jeffrey Barrett, Sonia Goncalves, Ewan Harrison, David K. Jackson, Ian Johnston, Dominic Kwiatkowski, Cordelia Langford, John Sillitoe on behalf of the Wellcome Sanger Institute COVID-19 Surveillance Team |
| EPI_ISL_2568977 | Lighthouse Lab in Milton Keynes | Wellcome Sanger Institute for the COVID-19 Genomics UK (COG-UK) Consortium | The Lighthouse Lab in Milton Keynes and Alex Alderton, Roberto Amato, Jeffrey Barrett, Sonia Goncalves, Ewan Harrison, David K. Jackson, Ian Johnston, Dominic Kwiatkowski, Cordelia Langford, John Sillitoe on behalf of the Wellcome Sanger Institute COVID-19 Surveillance Team |
| EPI_ISL_2568978 | Lighthouse Lab in Alderley Park | Wellcome Sanger Institute for the COVID-19 Genomics UK (COG-UK) Consortium | Jacquelyn Wynn, Mairead Hyland, The Lighthouse Lab in Alderley Park and Alex Alderton, Roberto Amato, Jeffrey Barrett, Sonia Goncalves, Ewan Harrison, David K. Jackson, Ian Johnston, Dominic Kwiatkowski, Cordelia Langford, John Sillitoe on behalf of the Wellcome Sanger Institute COVID-19 Surveillance Team |
| EPI_ISL_2568979 | Lighthouse Laboratory Plymouth | Wellcome Sanger Institute for the COVID-19 Genomics UK (COG-UK) Consortium | Lighthouse Laboratory Plymouth and Alex Alderton, Roberto Amato, Jeffrey Barrett, Sonia Goncalves, Ewan Harrison, David K. Jackson, Ian Johnston, Dominic Kwiatkowski, Cordelia Langford, John Sillitoe on behalf of the Wellcome Sanger Institute COVID-19 Surveillance Team |
| EPI_ISL_2568980 | Lighthouse Lab in Milton Keynes | Wellcome Sanger Institute for the COVID-19 Genomics UK (COG-UK) Consortium | The Lighthouse Lab in Milton Keynes and Alex Alderton, Roberto Amato, Jeffrey Barrett, Sonia Goncalves, Ewan Harrison, David K. Jackson, Ian Johnston, Dominic Kwiatkowski, Cordelia Langford, John Sillitoe on behalf of the Wellcome Sanger Institute COVID-19 Surveillance Team |
| EPI_ISL_2568981, EPI_ISL_2568982 | Lighthouse Lab in Alderley Park | Wellcome Sanger Institute for the COVID-19 Genomics UK (COG-UK) Consortium | Jacquelyn Wynn, Mairead Hyland, The Lighthouse Lab in Alderley Park and Alex Alderton, Roberto Amato, Jeffrey Barrett, Sonia Goncalves, Ewan Harrison, David K. Jackson, Ian Johnston, Dominic Kwiatkowski, Cordelia Langford, John Sillitoe on behalf of the Wellcome Sanger Institute COVID-19 Surveillance Team |
| EPI_ISL_2568984, EPI_ISL_2568985 | Lighthouse Lab in Milton Keynes | Wellcome Sanger Institute for the COVID-19 Genomics UK (COG-UK) Consortium | The Lighthouse Lab in Milton Keynes and Alex Alderton, Roberto Amato, Jeffrey Barrett, Sonia Goncalves, Ewan Harrison, David K. Jackson, Ian Johnston, Dominic Kwiatkowski, Cordelia Langford, John Sillitoe on behalf of the Wellcome Sanger Institute COVID-19 Surveillance Team |
| EPI_ISL_2568986, EPI_ISL_2568987 | Lighthouse Lab in Alderley Park | Wellcome Sanger Institute for the COVID-19 Genomics UK (COG-UK) Consortium | Jacquelyn Wynn, Mairead Hyland, The Lighthouse Lab in Alderley Park and Alex Alderton, Roberto Amato, Jeffrey Barrett, Sonia Goncalves, Ewan Harrison, David K. Jackson, Ian Johnston, Dominic Kwiatkowski, Cordelia Langford, John Sillitoe on behalf of the Wellcome Sanger Institute COVID-19 Surveillance Team |
| EPI_ISL_2568988 | Lighthouse Laboratory Plymouth | Wellcome Sanger Institute for the COVID-19 Genomics UK (COG-UK) Consortium | Lighthouse Laboratory Plymouth and Alex Alderton, Roberto Amato, Jeffrey Barrett, Sonia Goncalves, Ewan Harrison, David K. Jackson, Ian Johnston, Dominic Kwiatkowski, Cordelia Langford, John Sillitoe on behalf of the Wellcome Sanger Institute COVID-19 Surveillance Team |
| EPI_ISL_2568989 | Lighthouse Lab in Alderley Park | Wellcome Sanger Institute for the COVID-19 Genomics UK (COG-UK) Consortium | Jacquelyn Wynn, Mairead Hyland, The Lighthouse Lab in Alderley Park and Alex Alderton, Roberto Amato, Jeffrey Barrett, Sonia Goncalves, Ewan Harrison, David K. Jackson, Ian Johnston, Dominic Kwiatkowski, Cordelia Langford, John Sillitoe on behalf of the Wellcome Sanger Institute COVID-19 Surveillance Team |
| EPI_ISL_2568990, EPI_ISL_2568992, EPI_ISL_2568993 | Lighthouse Lab in Milton Keynes | Wellcome Sanger Institute for the COVID-19 Genomics UK (COG-UK) Consortium | The Lighthouse Lab in Milton Keynes and Alex Alderton, Roberto Amato, Jeffrey Barrett, Sonia Goncalves, Ewan Harrison, David K. Jackson, Ian Johnston, Dominic Kwiatkowski, Cordelia Langford, John Sillitoe on behalf of the Wellcome Sanger Institute COVID-19 Surveillance Team |
| EPI_ISL_2568994 | Lighthouse Laboratory Plymouth | Wellcome Sanger Institute for the COVID-19 Genomics UK (COG-UK) Consortium | Lighthouse Laboratory Plymouth and Alex Alderton, Roberto Amato, Jeffrey Barrett, Sonia Goncalves, Ewan Harrison, David K. Jackson, Ian Johnston, Dominic Kwiatkowski, Cordelia Langford, John Sillitoe on behalf of the Wellcome Sanger Institute COVID-19 Surveillance Team |
| EPI_ISL_2568995, EPI_ISL_2568996, EPI_ISL_2568997, EPI_ISL_2568998 | Lighthouse Lab in Milton Keynes | Wellcome Sanger Institute for the COVID-19 Genomics UK (COG-UK) Consortium | The Lighthouse Lab in Milton Keynes and Alex Alderton, Roberto Amato, Jeffrey Barrett, Sonia Goncalves, Ewan Harrison, David K. Jackson, Ian Johnston, Dominic Kwiatkowski, Cordelia Langford, John Sillitoe on behalf of the Wellcome Sanger Institute COVID-19 Surveillance Team |
| EPI_ISL_2568999, | Lighthouse Lab in Alderley Park | Wellcome Sanger Institute for the COVID-19 Genomics UK (COG-UK) | Jacquelyn Wynn, Mairead Hyland, The Lighthouse Lab in Alderley Park and Alex Alderton, Roberto Amato, Jeffrey Barrett, Sonia Goncalves, Ewan Harrison, David K. Jackson, Ian Johnston, Dominic Kwiatkowski, Cordelia Langford, John Sillitoe on |

| | | | |
|---|---|---|---|
| EPI_ISL_2569000, EPI_ISL_2569001 | | Consortium | behalf of the Wellcome Sanger Institute COVID-19 Surveillance Team |
| EPI_ISL_2569002 | Lighthouse Laboratory Plymouth | Wellcome Sanger Institute for the COVID-19 Genomics UK (COG-UK) Consortium | Lighthouse Laboratory Plymouth and Alex Alderton, Roberto Amato, Jeffrey Barrett, Sonia Goncalves, Ewan Harrison, David K. Jackson, Ian Johnston, Dominic Kwiatkowski, Cordelia Langford, John Sillitoe on behalf of the Wellcome Sanger Institute COVID-19 Surveillance Team |
| EPI_ISL_2569003 | Lighthouse Lab in Alderley Park | Wellcome Sanger Institute for the COVID-19 Genomics UK (COG-UK) Consortium | Jacquelyn Wynn, Mairead Hyland, The Lighthouse Lab in Alderley Park and Alex Alderton, Roberto Amato, Jeffrey Barrett, Sonia Goncalves, Ewan Harrison, David K. Jackson, Ian Johnston, Dominic Kwiatkowski, Cordelia Langford, John Sillitoe on behalf of the Wellcome Sanger Institute COVID-19 Surveillance Team |
| EPI_ISL_2569004 | Lighthouse Lab in Milton Keynes | Wellcome Sanger Institute for the COVID-19 Genomics UK (COG-UK) Consortium | The Lighthouse Lab in Milton Keynes and Alex Alderton, Roberto Amato, Jeffrey Barrett, Sonia Goncalves, Ewan Harrison, David K. Jackson, Ian Johnston, Dominic Kwiatkowski, Cordelia Langford, John Sillitoe on behalf of the Wellcome Sanger Institute COVID-19 Surveillance Team |
| EPI_ISL_2569005 | Lighthouse Laboratory Plymouth | Wellcome Sanger Institute for the COVID-19 Genomics UK (COG-UK) Consortium | Lighthouse Laboratory Plymouth and Alex Alderton, Roberto Amato, Jeffrey Barrett, Sonia Goncalves, Ewan Harrison, David K. Jackson, Ian Johnston, Dominic Kwiatkowski, Cordelia Langford, John Sillitoe on behalf of the Wellcome Sanger Institute COVID-19 Surveillance Team |
| EPI_ISL_2569006 | Lighthouse Lab in Alderley Park | Wellcome Sanger Institute for the COVID-19 Genomics UK (COG-UK) Consortium | Jacquelyn Wynn, Mairead Hyland, The Lighthouse Lab in Alderley Park and Alex Alderton, Roberto Amato, Jeffrey Barrett, Sonia Goncalves, Ewan Harrison, David K. Jackson, Ian Johnston, Dominic Kwiatkowski, Cordelia Langford, John Sillitoe on behalf of the Wellcome Sanger Institute COVID-19 Surveillance Team |
| EPI_ISL_2569007, EPI_ISL_2569008, EPI_ISL_2569009 | Lighthouse Lab in Milton Keynes | Wellcome Sanger Institute for the COVID-19 Genomics UK (COG-UK) Consortium | The Lighthouse Lab in Milton Keynes and Alex Alderton, Roberto Amato, Jeffrey Barrett, Sonia Goncalves, Ewan Harrison, David K. Jackson, Ian Johnston, Dominic Kwiatkowski, Cordelia Langford, John Sillitoe on behalf of the Wellcome Sanger Institute COVID-19 Surveillance Team |
| EPI_ISL_2569010 | Lighthouse Lab in Alderley Park | Wellcome Sanger Institute for the COVID-19 Genomics UK (COG-UK) Consortium | Jacquelyn Wynn, Mairead Hyland, The Lighthouse Lab in Alderley Park and Alex Alderton, Roberto Amato, Jeffrey Barrett, Sonia Goncalves, Ewan Harrison, David K. Jackson, Ian Johnston, Dominic Kwiatkowski, Cordelia Langford, John Sillitoe on behalf of the Wellcome Sanger Institute COVID-19 Surveillance Team |
| EPI_ISL_2569011 | Lighthouse Laboratory Plymouth | Wellcome Sanger Institute for the COVID-19 Genomics UK (COG-UK) Consortium | Lighthouse Laboratory Plymouth and Alex Alderton, Roberto Amato, Jeffrey Barrett, Sonia Goncalves, Ewan Harrison, David K. Jackson, Ian Johnston, Dominic Kwiatkowski, Cordelia Langford, John Sillitoe on behalf of the Wellcome Sanger Institute COVID-19 Surveillance Team |
| EPI_ISL_2569012 | Lighthouse Lab in Alderley Park | Wellcome Sanger Institute for the COVID-19 Genomics UK (COG-UK) Consortium | Jacquelyn Wynn, Mairead Hyland, The Lighthouse Lab in Alderley Park and Alex Alderton, Roberto Amato, Jeffrey Barrett, Sonia Goncalves, Ewan Harrison, David K. Jackson, Ian Johnston, Dominic Kwiatkowski, Cordelia Langford, John Sillitoe on behalf of the Wellcome Sanger Institute COVID-19 Surveillance Team |
| EPI_ISL_2569013, EPI_ISL_2569014 | Lighthouse Laboratory Plymouth | Wellcome Sanger Institute for the COVID-19 Genomics UK (COG-UK) Consortium | Lighthouse Laboratory Plymouth and Alex Alderton, Roberto Amato, Jeffrey Barrett, Sonia Goncalves, Ewan Harrison, David K. Jackson, Ian Johnston, Dominic Kwiatkowski, Cordelia Langford, John Sillitoe on behalf of the Wellcome Sanger Institute COVID-19 Surveillance Team |
| EPI_ISL_2569015 | Lighthouse Lab in Alderley Park | Wellcome Sanger Institute for the COVID-19 Genomics UK (COG-UK) Consortium | Jacquelyn Wynn, Mairead Hyland, The Lighthouse Lab in Alderley Park and Alex Alderton, Roberto Amato, Jeffrey Barrett, Sonia Goncalves, Ewan Harrison, David K. Jackson, Ian Johnston, Dominic Kwiatkowski, Cordelia Langford, John Sillitoe on behalf of the Wellcome Sanger Institute COVID-19 Surveillance Team |
| EPI_ISL_2569016 | Lighthouse Lab in Glasgow | Wellcome Sanger Institute for the COVID-19 Genomics UK (COG-UK) Consortium | Harper VanSteenhouse, Yumi Kasai, David Gray, Carol Clugston, Anna Dominiczak and Alex Alderton, Roberto Amato, Jeffrey Barrett, Sonia Goncalves, Ewan Harrison, David K. Jackson, Ian Johnston, Dominic Kwiatkowski, Cordelia Langford, John Sillitoe on behalf of the Wellcome Sanger Institute COVID-19 Surveillance Team |
| EPI_ISL_2569017 | Lighthouse Lab in Alderley Park | Wellcome Sanger Institute for the COVID-19 Genomics UK (COG-UK) Consortium | Jacquelyn Wynn, Mairead Hyland, The Lighthouse Lab in Alderley Park and Alex Alderton, Roberto Amato, Jeffrey Barrett, Sonia Goncalves, Ewan Harrison, David K. Jackson, Ian Johnston, Dominic Kwiatkowski, Cordelia Langford, John Sillitoe on behalf of the Wellcome Sanger Institute COVID-19 Surveillance Team |
| EPI_ISL_2569018, EPI_ISL_2569019 | Lighthouse Lab in Milton Keynes | Wellcome Sanger Institute for the COVID-19 Genomics UK (COG-UK) Consortium | The Lighthouse Lab in Milton Keynes and Alex Alderton, Roberto Amato, Jeffrey Barrett, Sonia Goncalves, Ewan Harrison, David K. Jackson, Ian Johnston, Dominic Kwiatkowski, Cordelia Langford, John Sillitoe on behalf of the Wellcome Sanger Institute COVID-19 Surveillance Team |
| EPI_ISL_2569020 | Lighthouse Lab in Glasgow | Wellcome Sanger Institute for the COVID-19 Genomics UK (COG-UK) Consortium | Harper VanSteenhouse, Yumi Kasai, David Gray, Carol Clugston, Anna Dominiczak and Alex Alderton, Roberto Amato, Jeffrey Barrett, Sonia Goncalves, Ewan Harrison, David K. Jackson, Ian Johnston, Dominic Kwiatkowski, Cordelia Langford, John Sillitoe on behalf of the Wellcome Sanger Institute COVID-19 Surveillance Team |
| EPI_ISL_2569021, EPI_ISL_2569022 | Lighthouse Laboratory Plymouth | Wellcome Sanger Institute for the COVID-19 Genomics UK (COG-UK) Consortium | Lighthouse Laboratory Plymouth and Alex Alderton, Roberto Amato, Jeffrey Barrett, Sonia Goncalves, Ewan Harrison, David K. Jackson, Ian Johnston, Dominic Kwiatkowski, Cordelia Langford, John Sillitoe on behalf of the Wellcome Sanger Institute COVID-19 Surveillance Team |
| EPI_ISL_2569023 | Lighthouse Lab in Glasgow | Wellcome Sanger Institute for the COVID-19 Genomics UK (COG-UK) Consortium | Harper VanSteenhouse, Yumi Kasai, David Gray, Carol Clugston, Anna Dominiczak and Alex Alderton, Roberto Amato, Jeffrey Barrett, Sonia Goncalves, Ewan Harrison, David K. Jackson, Ian Johnston, Dominic Kwiatkowski, Cordelia Langford, John Sillitoe on behalf of the Wellcome Sanger Institute COVID-19 Surveillance Team |
| EPI_ISL_2569024, EPI_ISL_2569025 | Lighthouse Lab in Milton Keynes | Wellcome Sanger Institute for the COVID-19 Genomics UK (COG-UK) Consortium | The Lighthouse Lab in Milton Keynes and Alex Alderton, Roberto Amato, Jeffrey Barrett, Sonia Goncalves, Ewan Harrison, David K. Jackson, Ian Johnston, Dominic Kwiatkowski, Cordelia Langford, John Sillitoe on behalf of the Wellcome Sanger Institute COVID-19 Surveillance Team |
| EPI_ISL_2569026 | Lighthouse Lab in Alderley Park | Wellcome Sanger Institute for the COVID-19 Genomics UK (COG-UK) Consortium | Jacquelyn Wynn, Mairead Hyland, The Lighthouse Lab in Alderley Park and Alex Alderton, Roberto Amato, Jeffrey Barrett, Sonia Goncalves, Ewan Harrison, David K. Jackson, Ian Johnston, Dominic Kwiatkowski, Cordelia Langford, John Sillitoe on behalf of the Wellcome Sanger Institute COVID-19 Surveillance Team |
| EPI_ISL_2569027 | Lighthouse Lab in Glasgow | Wellcome Sanger Institute for the COVID-19 Genomics UK (COG-UK) Consortium | Harper VanSteenhouse, Yumi Kasai, David Gray, Carol Clugston, Anna Dominiczak and Alex Alderton, Roberto Amato, Jeffrey Barrett, Sonia Goncalves, Ewan Harrison, David K. Jackson, Ian Johnston, Dominic Kwiatkowski, Cordelia Langford, John Sillitoe on behalf of the Wellcome Sanger Institute COVID-19 Surveillance Team |
| EPI_ISL_2569028 | Lighthouse Lab in Milton Keynes | Wellcome Sanger Institute for the COVID-19 Genomics UK (COG-UK) Consortium | The Lighthouse Lab in Milton Keynes and Alex Alderton, Roberto Amato, Jeffrey Barrett, Sonia Goncalves, Ewan Harrison, David K. Jackson, Ian Johnston, Dominic Kwiatkowski, Cordelia Langford, John Sillitoe on behalf of the Wellcome Sanger Institute COVID-19 Surveillance Team |
| EPI_ISL_2569029 | Lighthouse Lab in Alderley Park | Wellcome Sanger Institute for the COVID-19 Genomics UK (COG-UK) Consortium | Jacquelyn Wynn, Mairead Hyland, The Lighthouse Lab in Alderley Park and Alex Alderton, Roberto Amato, Jeffrey Barrett, Sonia Goncalves, Ewan Harrison, David K. Jackson, Ian Johnston, Dominic Kwiatkowski, Cordelia Langford, John Sillitoe on behalf of the Wellcome Sanger Institute COVID-19 Surveillance Team |
| EPI_ISL_2569030 | Lighthouse Laboratory Plymouth | Wellcome Sanger Institute for the COVID-19 Genomics UK (COG-UK) Consortium | Lighthouse Laboratory Plymouth and Alex Alderton, Roberto Amato, Jeffrey Barrett, Sonia Goncalves, Ewan Harrison, David K. Jackson, Ian Johnston, Dominic Kwiatkowski, Cordelia Langford, John Sillitoe on behalf of the Wellcome Sanger Institute COVID-19 Surveillance Team |
| EPI_ISL_2569031 | Lighthouse Lab in Alderley Park | Wellcome Sanger Institute for the COVID-19 Genomics UK (COG-UK) Consortium | Jacquelyn Wynn, Mairead Hyland, The Lighthouse Lab in Alderley Park and Alex Alderton, Roberto Amato, Jeffrey Barrett, Sonia Goncalves, Ewan Harrison, David K. Jackson, Ian Johnston, Dominic Kwiatkowski, Cordelia Langford, John Sillitoe on behalf of the Wellcome Sanger Institute COVID-19 Surveillance Team |
| EPI_ISL_2569032 | Lighthouse Lab in Glasgow | Wellcome Sanger Institute for the COVID-19 Genomics UK (COG-UK) Consortium | Harper VanSteenhouse, Yumi Kasai, David Gray, Carol Clugston, Anna Dominiczak and Alex Alderton, Roberto Amato, Jeffrey Barrett, Sonia Goncalves, Ewan Harrison, David K. Jackson, Ian Johnston, Dominic Kwiatkowski, Cordelia Langford, John Sillitoe on behalf of the Wellcome Sanger Institute COVID-19 Surveillance Team |
| EPI_ISL_2569033 | Lighthouse Lab in Milton Keynes | Wellcome Sanger Institute for the COVID-19 Genomics UK (COG-UK) Consortium | The Lighthouse Lab in Milton Keynes and Alex Alderton, Roberto Amato, Jeffrey Barrett, Sonia Goncalves, Ewan Harrison, David K. Jackson, Ian Johnston, Dominic Kwiatkowski, Cordelia Langford, John Sillitoe on behalf of the Wellcome Sanger Institute COVID-19 Surveillance Team |
| EPI_ISL_2569035 | Lighthouse Lab in Glasgow | Wellcome Sanger Institute for the COVID-19 Genomics UK (COG-UK) Consortium | Harper VanSteenhouse, Yumi Kasai, David Gray, Carol Clugston, Anna Dominiczak and Alex Alderton, Roberto Amato, Jeffrey Barrett, Sonia Goncalves, Ewan Harrison, David K. Jackson, Ian Johnston, Dominic Kwiatkowski, Cordelia Langford, John Sillitoe on behalf of the Wellcome Sanger Institute COVID-19 Surveillance Team |
| EPI_ISL_2569036 | Lighthouse Lab in Milton Keynes | Wellcome Sanger Institute for the COVID-19 Genomics UK (COG-UK) Consortium | The Lighthouse Lab in Milton Keynes and Alex Alderton, Roberto Amato, Jeffrey Barrett, Sonia Goncalves, Ewan Harrison, David K. Jackson, Ian Johnston, Dominic Kwiatkowski, Cordelia Langford, John Sillitoe on behalf of the Wellcome Sanger Institute COVID-19 Surveillance Team |
| EPI_ISL_2569037 | Lighthouse Laboratory Plymouth | Wellcome Sanger Institute for the COVID-19 Genomics UK (COG-UK) Consortium | Lighthouse Laboratory Plymouth and Alex Alderton, Roberto Amato, Jeffrey Barrett, Sonia Goncalves, Ewan Harrison, David K. Jackson, Ian Johnston, Dominic Kwiatkowski, Cordelia Langford, John Sillitoe on behalf of the Wellcome Sanger Institute COVID-19 Surveillance Team |
| EPI_ISL_2569038, EPI_ISL_2569039 | Lighthouse Lab in Alderley Park | Wellcome Sanger Institute for the COVID-19 Genomics UK (COG-UK) Consortium | Jacquelyn Wynn, Mairead Hyland, The Lighthouse Lab in Alderley Park and Alex Alderton, Roberto Amato, Jeffrey Barrett, Sonia Goncalves, Ewan Harrison, David K. Jackson, Ian Johnston, Dominic Kwiatkowski, Cordelia Langford, John Sillitoe on behalf of the Wellcome Sanger Institute COVID-19 Surveillance Team |
| EPI_ISL_2569040 | Lighthouse Laboratory Plymouth | Wellcome Sanger Institute for the COVID-19 Genomics UK (COG-UK) Consortium | Lighthouse Laboratory Plymouth and Alex Alderton, Roberto Amato, Jeffrey Barrett, Sonia Goncalves, Ewan Harrison, David K. Jackson, Ian Johnston, Dominic Kwiatkowski, Cordelia Langford, John Sillitoe on behalf of the Wellcome Sanger Institute COVID-19 Surveillance Team |
| EPI_ISL_2569041 | Lighthouse Lab in Alderley Park | Wellcome Sanger Institute for the COVID-19 Genomics UK (COG-UK) Consortium | Jacquelyn Wynn, Mairead Hyland, The Lighthouse Lab in Alderley Park and Alex Alderton, Roberto Amato, Jeffrey Barrett, Sonia Goncalves, Ewan Harrison, David K. Jackson, Ian Johnston, Dominic Kwiatkowski, Cordelia Langford, John Sillitoe on behalf of the Wellcome Sanger Institute COVID-19 Surveillance Team |
| EPI_ISL_2569042 | Lighthouse Lab in Milton Keynes | Wellcome Sanger Institute for the COVID-19 Genomics UK (COG-UK) Consortium | The Lighthouse Lab in Milton Keynes and Alex Alderton, Roberto Amato, Jeffrey Barrett, Sonia Goncalves, Ewan Harrison, David K. Jackson, Ian Johnston, Dominic Kwiatkowski, Cordelia Langford, John Sillitoe on behalf of the Wellcome Sanger Institute COVID-19 Surveillance Team |
| EPI_ISL_2569043, EPI_ISL_2569044 | Lighthouse Lab in Alderley Park | Wellcome Sanger Institute for the COVID-19 Genomics UK (COG-UK) Consortium | Jacquelyn Wynn, Mairead Hyland, The Lighthouse Lab in Alderley Park and Alex Alderton, Roberto Amato, Jeffrey Barrett, Sonia Goncalves, Ewan Harrison, David K. Jackson, Ian Johnston, Dominic Kwiatkowski, Cordelia Langford, John Sillitoe on behalf of the Wellcome Sanger Institute COVID-19 Surveillance Team |
| EPI_ISL_2569045 | Lighthouse Lab in Milton Keynes | Wellcome Sanger Institute for the COVID-19 Genomics UK (COG-UK) Consortium | The Lighthouse Lab in Milton Keynes and Alex Alderton, Roberto Amato, Jeffrey Barrett, Sonia Goncalves, Ewan Harrison, David K. Jackson, Ian Johnston, Dominic Kwiatkowski, Cordelia Langford, John Sillitoe on behalf of the Wellcome Sanger Institute COVID-19 Surveillance Team |
| EPI_ISL_2569046 | Lighthouse Lab in Glasgow | Wellcome Sanger Institute for the COVID-19 Genomics UK (COG-UK) Consortium | Harper VanSteenhouse, Yumi Kasai, David Gray, Carol Clugston, Anna Dominiczak and Alex Alderton, Roberto Amato, Jeffrey Barrett, Sonia Goncalves, Ewan Harrison, David K. Jackson, Ian Johnston, Dominic Kwiatkowski, Cordelia Langford, John Sillitoe on behalf of the Wellcome Sanger Institute COVID-19 Surveillance Team |
| EPI_ISL_2569047 | Lighthouse Lab in Milton Keynes | Wellcome Sanger Institute for the COVID-19 Genomics UK (COG-UK) Consortium | The Lighthouse Lab in Milton Keynes and Alex Alderton, Roberto Amato, Jeffrey Barrett, Sonia Goncalves, Ewan Harrison, David K. Jackson, Ian Johnston, Dominic Kwiatkowski, Cordelia Langford, John Sillitoe on behalf of the Wellcome Sanger Institute COVID-19 Surveillance Team |
| EPI_ISL_2569048, EPI_ISL_2569049, EPI_ISL_2569050 | Lighthouse Lab in Alderley Park | Wellcome Sanger Institute for the COVID-19 Genomics UK (COG-UK) Consortium | Jacquelyn Wynn, Mairead Hyland, The Lighthouse Lab in Alderley Park and Alex Alderton, Roberto Amato, Jeffrey Barrett, Sonia Goncalves, Ewan Harrison, David K. Jackson, Ian Johnston, Dominic Kwiatkowski, Cordelia Langford, John Sillitoe on behalf of the Wellcome Sanger Institute COVID-19 Surveillance Team |
| EPI_ISL_2569051 | Lighthouse Laboratory Plymouth | Wellcome Sanger Institute for the COVID-19 Genomics UK (COG-UK) Consortium | Lighthouse Laboratory Plymouth and Alex Alderton, Roberto Amato, Jeffrey Barrett, Sonia Goncalves, Ewan Harrison, David K. Jackson, Ian Johnston, Dominic Kwiatkowski, Cordelia Langford, John Sillitoe on behalf of the Wellcome Sanger Institute COVID-19 Surveillance Team |
| EPI_ISL_2569052 | Lighthouse Lab in Alderley Park | Wellcome Sanger Institute for the COVID-19 Genomics UK (COG-UK) Consortium | Jacquelyn Wynn, Mairead Hyland, The Lighthouse Lab in Alderley Park and Alex Alderton, Roberto Amato, Jeffrey Barrett, Sonia Goncalves, Ewan Harrison, David K. Jackson, Ian Johnston, Dominic Kwiatkowski, Cordelia Langford, John Sillitoe on behalf of the Wellcome Sanger Institute COVID-19 Surveillance Team |
| EPI_ISL_2569053 | Lighthouse Lab in Milton Keynes | Wellcome Sanger Institute for the COVID-19 Genomics UK (COG-UK) Consortium | The Lighthouse Lab in Milton Keynes and Alex Alderton, Roberto Amato, Jeffrey Barrett, Sonia Goncalves, Ewan Harrison, David K. Jackson, Ian Johnston, Dominic Kwiatkowski, Cordelia Langford, John Sillitoe on behalf of the Wellcome Sanger Institute COVID-19 Surveillance Team |
| EPI_ISL_2569054 | Lighthouse Lab in Glasgow | Wellcome Sanger Institute for the COVID-19 Genomics UK (COG-UK) Consortium | Harper VanSteenhouse, Yumi Kasai, David Gray, Carol Clugston, Anna Dominiczak and Alex Alderton, Roberto Amato, Jeffrey Barrett, Sonia Goncalves, Ewan Harrison, David K. Jackson, Ian Johnston, Dominic Kwiatkowski, Cordelia Langford, John Sillitoe on behalf of the Wellcome Sanger Institute COVID-19 Surveillance Team |
| EPI_ISL_2569055, EPI_ISL_2569056 | Lighthouse Laboratory Plymouth | Wellcome Sanger Institute for the COVID-19 Genomics UK (COG-UK) Consortium | Lighthouse Laboratory Plymouth and Alex Alderton, Roberto Amato, Jeffrey Barrett, Sonia Goncalves, Ewan Harrison, David K. Jackson, Ian Johnston, Dominic Kwiatkowski, Cordelia Langford, John Sillitoe on behalf of the Wellcome Sanger Institute COVID-19 Surveillance Team |

| | | | |
|---|---|---|---|
| EPI_ISL_2569057 | Lighthouse Lab in Alderley Park | Wellcome Sanger Institute for the COVID-19 Genomics UK (COG-UK) Consortium | Jacquelyn Wynn, Mairead Hyland, The Lighthouse Lab in Alderley Park and Alex Alderton, Roberto Amato, Jeffrey Barrett, Sonia Goncalves, Ewan Harrison, David K. Jackson, Ian Johnston, Dominic Kwiatkowski, Cordelia Langford, John Sillitoe on behalf of the Wellcome Sanger Institute COVID-19 Surveillance Team |
| EPI_ISL_2569058, EPI_ISL_2569059 | Lighthouse Lab in Milton Keynes | Wellcome Sanger Institute for the COVID-19 Genomics UK (COG-UK) Consortium | The Lighthouse Lab in Milton Keynes and Alex Alderton, Roberto Amato, Jeffrey Barrett, Sonia Goncalves, Ewan Harrison, David K. Jackson, Ian Johnston, Dominic Kwiatkowski, Cordelia Langford, John Sillitoe on behalf of the Wellcome Sanger Institute COVID-19 Surveillance Team |
| EPI_ISL_2569060 | Lighthouse Laboratory Plymouth | Wellcome Sanger Institute for the COVID-19 Genomics UK (COG-UK) Consortium | Lighthouse Laboratory Plymouth and Alex Alderton, Roberto Amato, Jeffrey Barrett, Sonia Goncalves, Ewan Harrison, David K. Jackson, Ian Johnston, Dominic Kwiatkowski, Cordelia Langford, John Sillitoe on behalf of the Wellcome Sanger Institute COVID-19 Surveillance Team |
| EPI_ISL_2569061 | Lighthouse Lab in Alderley Park | Wellcome Sanger Institute for the COVID-19 Genomics UK (COG-UK) Consortium | Jacquelyn Wynn, Mairead Hyland, The Lighthouse Lab in Alderley Park and Alex Alderton, Roberto Amato, Jeffrey Barrett, Sonia Goncalves, Ewan Harrison, David K. Jackson, Ian Johnston, Dominic Kwiatkowski, Cordelia Langford, John Sillitoe on behalf of the Wellcome Sanger Institute COVID-19 Surveillance Team |
| EPI_ISL_2569062 | Lighthouse Lab in Milton Keynes | Wellcome Sanger Institute for the COVID-19 Genomics UK (COG-UK) Consortium | The Lighthouse Lab in Milton Keynes and Alex Alderton, Roberto Amato, Jeffrey Barrett, Sonia Goncalves, Ewan Harrison, David K. Jackson, Ian Johnston, Dominic Kwiatkowski, Cordelia Langford, John Sillitoe on behalf of the Wellcome Sanger Institute COVID-19 Surveillance Team |
| EPI_ISL_2569063 | Lighthouse Laboratory Plymouth | Wellcome Sanger Institute for the COVID-19 Genomics UK (COG-UK) Consortium | Lighthouse Laboratory Plymouth and Alex Alderton, Roberto Amato, Jeffrey Barrett, Sonia Goncalves, Ewan Harrison, David K. Jackson, Ian Johnston, Dominic Kwiatkowski, Cordelia Langford, John Sillitoe on behalf of the Wellcome Sanger Institute COVID-19 Surveillance Team |
| EPI_ISL_2569064 | Lighthouse Lab in Milton Keynes | Wellcome Sanger Institute for the COVID-19 Genomics UK (COG-UK) Consortium | The Lighthouse Lab in Milton Keynes and Alex Alderton, Roberto Amato, Jeffrey Barrett, Sonia Goncalves, Ewan Harrison, David K. Jackson, Ian Johnston, Dominic Kwiatkowski, Cordelia Langford, John Sillitoe on behalf of the Wellcome Sanger Institute COVID-19 Surveillance Team |
| EPI_ISL_2569065 | Lighthouse Lab in Alderley Park | Wellcome Sanger Institute for the COVID-19 Genomics UK (COG-UK) Consortium | Jacquelyn Wynn, Mairead Hyland, The Lighthouse Lab in Alderley Park and Alex Alderton, Roberto Amato, Jeffrey Barrett, Sonia Goncalves, Ewan Harrison, David K. Jackson, Ian Johnston, Dominic Kwiatkowski, Cordelia Langford, John Sillitoe on behalf of the Wellcome Sanger Institute COVID-19 Surveillance Team |
| EPI_ISL_2569066 | Lighthouse Lab in Glasgow | Wellcome Sanger Institute for the COVID-19 Genomics UK (COG-UK) Consortium | Harper VanSteenhouse, Yumi Kasai, David Gray, Carol Clugston, Anna Dominiczak and Alex Alderton, Roberto Amato, Jeffrey Barrett, Sonia Goncalves, Ewan Harrison, David K. Jackson, Ian Johnston, Dominic Kwiatkowski, Cordelia Langford, John Sillitoe on behalf of the Wellcome Sanger Institute COVID-19 Surveillance Team |
| EPI_ISL_2569067 | Lighthouse Laboratory Plymouth | Wellcome Sanger Institute for the COVID-19 Genomics UK (COG-UK) Consortium | Lighthouse Laboratory Plymouth and Alex Alderton, Roberto Amato, Jeffrey Barrett, Sonia Goncalves, Ewan Harrison, David K. Jackson, Ian Johnston, Dominic Kwiatkowski, Cordelia Langford, John Sillitoe on behalf of the Wellcome Sanger Institute COVID-19 Surveillance Team |
| EPI_ISL_2569069 | Lighthouse Lab in Milton Keynes | Wellcome Sanger Institute for the COVID-19 Genomics UK (COG-UK) Consortium | The Lighthouse Lab in Milton Keynes and Alex Alderton, Roberto Amato, Jeffrey Barrett, Sonia Goncalves, Ewan Harrison, David K. Jackson, Ian Johnston, Dominic Kwiatkowski, Cordelia Langford, John Sillitoe on behalf of the Wellcome Sanger Institute COVID-19 Surveillance Team |
| EPI_ISL_2569070 | Lighthouse Lab in Glasgow | Wellcome Sanger Institute for the COVID-19 Genomics UK (COG-UK) Consortium | Harper VanSteenhouse, Yumi Kasai, David Gray, Carol Clugston, Anna Dominiczak and Alex Alderton, Roberto Amato, Jeffrey Barrett, Sonia Goncalves, Ewan Harrison, David K. Jackson, Ian Johnston, Dominic Kwiatkowski, Cordelia Langford, John Sillitoe on behalf of the Wellcome Sanger Institute COVID-19 Surveillance Team |
| EPI_ISL_2569071 | Lighthouse Lab in Milton Keynes | Wellcome Sanger Institute for the COVID-19 Genomics UK (COG-UK) Consortium | The Lighthouse Lab in Milton Keynes and Alex Alderton, Roberto Amato, Jeffrey Barrett, Sonia Goncalves, Ewan Harrison, David K. Jackson, Ian Johnston, Dominic Kwiatkowski, Cordelia Langford, John Sillitoe on behalf of the Wellcome Sanger Institute COVID-19 Surveillance Team |
| EPI_ISL_2569072 | Lighthouse Lab in Glasgow | Wellcome Sanger Institute for the COVID-19 Genomics UK (COG-UK) Consortium | Harper VanSteenhouse, Yumi Kasai, David Gray, Carol Clugston, Anna Dominiczak and Alex Alderton, Roberto Amato, Jeffrey Barrett, Sonia Goncalves, Ewan Harrison, David K. Jackson, Ian Johnston, Dominic Kwiatkowski, Cordelia Langford, John Sillitoe on behalf of the Wellcome Sanger Institute COVID-19 Surveillance Team |
| EPI_ISL_2569073 | Lighthouse Laboratory Plymouth | Wellcome Sanger Institute for the COVID-19 Genomics UK (COG-UK) Consortium | Lighthouse Laboratory Plymouth and Alex Alderton, Roberto Amato, Jeffrey Barrett, Sonia Goncalves, Ewan Harrison, David K. Jackson, Ian Johnston, Dominic Kwiatkowski, Cordelia Langford, John Sillitoe on behalf of the Wellcome Sanger Institute COVID-19 Surveillance Team |
| EPI_ISL_2569074 | Lighthouse Lab in Milton Keynes | Wellcome Sanger Institute for the COVID-19 Genomics UK (COG-UK) Consortium | The Lighthouse Lab in Milton Keynes and Alex Alderton, Roberto Amato, Jeffrey Barrett, Sonia Goncalves, Ewan Harrison, David K. Jackson, Ian Johnston, Dominic Kwiatkowski, Cordelia Langford, John Sillitoe on behalf of the Wellcome Sanger Institute COVID-19 Surveillance Team |
| EPI_ISL_2569075 | Lighthouse Lab in Alderley Park | Wellcome Sanger Institute for the COVID-19 Genomics UK (COG-UK) Consortium | Jacquelyn Wynn, Mairead Hyland, The Lighthouse Lab in Alderley Park and Alex Alderton, Roberto Amato, Jeffrey Barrett, Sonia Goncalves, Ewan Harrison, David K. Jackson, Ian Johnston, Dominic Kwiatkowski, Cordelia Langford, John Sillitoe on behalf of the Wellcome Sanger Institute COVID-19 Surveillance Team |
| EPI_ISL_2569076, EPI_ISL_2569077 | Lighthouse Laboratory Plymouth | Wellcome Sanger Institute for the COVID-19 Genomics UK (COG-UK) Consortium | Lighthouse Laboratory Plymouth and Alex Alderton, Roberto Amato, Jeffrey Barrett, Sonia Goncalves, Ewan Harrison, David K. Jackson, Ian Johnston, Dominic Kwiatkowski, Cordelia Langford, John Sillitoe on behalf of the Wellcome Sanger Institute COVID-19 Surveillance Team |
| EPI_ISL_2569078 | Lighthouse Lab in Glasgow | Wellcome Sanger Institute for the COVID-19 Genomics UK (COG-UK) Consortium | Harper VanSteenhouse, Yumi Kasai, David Gray, Carol Clugston, Anna Dominiczak and Alex Alderton, Roberto Amato, Jeffrey Barrett, Sonia Goncalves, Ewan Harrison, David K. Jackson, Ian Johnston, Dominic Kwiatkowski, Cordelia Langford, John Sillitoe on behalf of the Wellcome Sanger Institute COVID-19 Surveillance Team |
| EPI_ISL_2569079 | Lighthouse Lab in Milton Keynes | Wellcome Sanger Institute for the COVID-19 Genomics UK (COG-UK) Consortium | The Lighthouse Lab in Milton Keynes and Alex Alderton, Roberto Amato, Jeffrey Barrett, Sonia Goncalves, Ewan Harrison, David K. Jackson, Ian Johnston, Dominic Kwiatkowski, Cordelia Langford, John Sillitoe on behalf of the Wellcome Sanger Institute COVID-19 Surveillance Team |
| EPI_ISL_2569080 | Lighthouse Lab in Glasgow | Wellcome Sanger Institute for the COVID-19 Genomics UK (COG-UK) Consortium | Harper VanSteenhouse, Yumi Kasai, David Gray, Carol Clugston, Anna Dominiczak and Alex Alderton, Roberto Amato, Jeffrey Barrett, Sonia Goncalves, Ewan Harrison, David K. Jackson, Ian Johnston, Dominic Kwiatkowski, Cordelia Langford, John Sillitoe on behalf of the Wellcome Sanger Institute COVID-19 Surveillance Team |
| EPI_ISL_2569081, EPI_ISL_2569082 | Lighthouse Lab in Milton Keynes | Wellcome Sanger Institute for the COVID-19 Genomics UK (COG-UK) Consortium | The Lighthouse Lab in Milton Keynes and Alex Alderton, Roberto Amato, Jeffrey Barrett, Sonia Goncalves, Ewan Harrison, David K. Jackson, Ian Johnston, Dominic Kwiatkowski, Cordelia Langford, John Sillitoe on behalf of the Wellcome Sanger Institute COVID-19 Surveillance Team |
| EPI_ISL_2569083 | Lighthouse Laboratory Plymouth | Wellcome Sanger Institute for the COVID-19 Genomics UK (COG-UK) Consortium | Lighthouse Laboratory Plymouth and Alex Alderton, Roberto Amato, Jeffrey Barrett, Sonia Goncalves, Ewan Harrison, David K. Jackson, Ian Johnston, Dominic Kwiatkowski, Cordelia Langford, John Sillitoe on behalf of the Wellcome Sanger Institute COVID-19 Surveillance Team |
| EPI_ISL_2569084 | Lighthouse Lab in Milton Keynes | Wellcome Sanger Institute for the COVID-19 Genomics UK (COG-UK) Consortium | The Lighthouse Lab in Milton Keynes and Alex Alderton, Roberto Amato, Jeffrey Barrett, Sonia Goncalves, Ewan Harrison, David K. Jackson, Ian Johnston, Dominic Kwiatkowski, Cordelia Langford, John Sillitoe on behalf of the Wellcome Sanger Institute COVID-19 Surveillance Team |
| EPI_ISL_2569085, EPI_ISL_2569086 | Lighthouse Lab in Alderley Park | Wellcome Sanger Institute for the COVID-19 Genomics UK (COG-UK) Consortium | Jacquelyn Wynn, Mairead Hyland, The Lighthouse Lab in Alderley Park and Alex Alderton, Roberto Amato, Jeffrey Barrett, Sonia Goncalves, Ewan Harrison, David K. Jackson, Ian Johnston, Dominic Kwiatkowski, Cordelia Langford, John Sillitoe on behalf of the Wellcome Sanger Institute COVID-19 Surveillance Team |
| EPI_ISL_2569088 | Lighthouse Lab in Milton Keynes | Wellcome Sanger Institute for the COVID-19 Genomics UK (COG-UK) Consortium | The Lighthouse Lab in Milton Keynes and Alex Alderton, Roberto Amato, Jeffrey Barrett, Sonia Goncalves, Ewan Harrison, David K. Jackson, Ian Johnston, Dominic Kwiatkowski, Cordelia Langford, John Sillitoe on behalf of the Wellcome Sanger Institute COVID-19 Surveillance Team |
| EPI_ISL_2569089 | Lighthouse Lab in Alderley Park | Wellcome Sanger Institute for the COVID-19 Genomics UK (COG-UK) Consortium | Jacquelyn Wynn, Mairead Hyland, The Lighthouse Lab in Alderley Park and Alex Alderton, Roberto Amato, Jeffrey Barrett, Sonia Goncalves, Ewan Harrison, David K. Jackson, Ian Johnston, Dominic Kwiatkowski, Cordelia Langford, John Sillitoe on behalf of the Wellcome Sanger Institute COVID-19 Surveillance Team |
| EPI_ISL_2569090 | Lighthouse Lab in Glasgow | Wellcome Sanger Institute for the COVID-19 Genomics UK (COG-UK) Consortium | Harper VanSteenhouse, Yumi Kasai, David Gray, Carol Clugston, Anna Dominiczak and Alex Alderton, Roberto Amato, Jeffrey Barrett, Sonia Goncalves, Ewan Harrison, David K. Jackson, Ian Johnston, Dominic Kwiatkowski, Cordelia Langford, John Sillitoe on behalf of the Wellcome Sanger Institute COVID-19 Surveillance Team |
| EPI_ISL_2569091 | Lighthouse Lab in Milton Keynes | Wellcome Sanger Institute for the COVID-19 Genomics UK (COG-UK) Consortium | The Lighthouse Lab in Milton Keynes and Alex Alderton, Roberto Amato, Jeffrey Barrett, Sonia Goncalves, Ewan Harrison, David K. Jackson, Ian Johnston, Dominic Kwiatkowski, Cordelia Langford, John Sillitoe on behalf of the Wellcome Sanger Institute COVID-19 Surveillance Team |
| EPI_ISL_2569092 | Lighthouse Lab in Glasgow | Wellcome Sanger Institute for the COVID-19 Genomics UK (COG-UK) Consortium | Harper VanSteenhouse, Yumi Kasai, David Gray, Carol Clugston, Anna Dominiczak and Alex Alderton, Roberto Amato, Jeffrey Barrett, Sonia Goncalves, Ewan Harrison, David K. Jackson, Ian Johnston, Dominic Kwiatkowski, Cordelia Langford, John Sillitoe on behalf of the Wellcome Sanger Institute COVID-19 Surveillance Team |
| EPI_ISL_2569093 | Lighthouse Lab in Milton Keynes | Wellcome Sanger Institute for the COVID-19 Genomics UK (COG-UK) Consortium | The Lighthouse Lab in Milton Keynes and Alex Alderton, Roberto Amato, Jeffrey Barrett, Sonia Goncalves, Ewan Harrison, David K. Jackson, Ian Johnston, Dominic Kwiatkowski, Cordelia Langford, John Sillitoe on behalf of the Wellcome Sanger Institute COVID-19 Surveillance Team |
| EPI_ISL_2569094 | Lighthouse Lab in Glasgow | Wellcome Sanger Institute for the COVID-19 Genomics UK (COG-UK) Consortium | Harper VanSteenhouse, Yumi Kasai, David Gray, Carol Clugston, Anna Dominiczak and Alex Alderton, Roberto Amato, Jeffrey Barrett, Sonia Goncalves, Ewan Harrison, David K. Jackson, Ian Johnston, Dominic Kwiatkowski, Cordelia Langford, John Sillitoe on behalf of the Wellcome Sanger Institute COVID-19 Surveillance Team |
| EPI_ISL_2569095 | Lighthouse Laboratory Plymouth | Wellcome Sanger Institute for the COVID-19 Genomics UK (COG-UK) Consortium | Lighthouse Laboratory Plymouth and Alex Alderton, Roberto Amato, Jeffrey Barrett, Sonia Goncalves, Ewan Harrison, David K. Jackson, Ian Johnston, Dominic Kwiatkowski, Cordelia Langford, John Sillitoe on behalf of the Wellcome Sanger Institute COVID-19 Surveillance Team |
| EPI_ISL_2569096 | Lighthouse Lab in Alderley Park | Wellcome Sanger Institute for the COVID-19 Genomics UK (COG-UK) Consortium | Jacquelyn Wynn, Mairead Hyland, The Lighthouse Lab in Alderley Park and Alex Alderton, Roberto Amato, Jeffrey Barrett, Sonia Goncalves, Ewan Harrison, David K. Jackson, Ian Johnston, Dominic Kwiatkowski, Cordelia Langford, John Sillitoe on behalf of the Wellcome Sanger Institute COVID-19 Surveillance Team |
| EPI_ISL_2569097 | Lighthouse Laboratory Plymouth | Wellcome Sanger Institute for the COVID-19 Genomics UK (COG-UK) Consortium | Lighthouse Laboratory Plymouth and Alex Alderton, Roberto Amato, Jeffrey Barrett, Sonia Goncalves, Ewan Harrison, David K. Jackson, Ian Johnston, Dominic Kwiatkowski, Cordelia Langford, John Sillitoe on behalf of the Wellcome Sanger Institute COVID-19 Surveillance Team |
| EPI_ISL_2569098 | Lighthouse Lab in Glasgow | Wellcome Sanger Institute for the COVID-19 Genomics UK (COG-UK) Consortium | Harper VanSteenhouse, Yumi Kasai, David Gray, Carol Clugston, Anna Dominiczak and Alex Alderton, Roberto Amato, Jeffrey Barrett, Sonia Goncalves, Ewan Harrison, David K. Jackson, Ian Johnston, Dominic Kwiatkowski, Cordelia Langford, John Sillitoe on behalf of the Wellcome Sanger Institute COVID-19 Surveillance Team |
| EPI_ISL_2569099 | Lighthouse Laboratory Plymouth | Wellcome Sanger Institute for the COVID-19 Genomics UK (COG-UK) Consortium | Lighthouse Laboratory Plymouth and Alex Alderton, Roberto Amato, Jeffrey Barrett, Sonia Goncalves, Ewan Harrison, David K. Jackson, Ian Johnston, Dominic Kwiatkowski, Cordelia Langford, John Sillitoe on behalf of the Wellcome Sanger Institute COVID-19 Surveillance Team |
| EPI_ISL_2569100 | Lighthouse Lab in Glasgow | Wellcome Sanger Institute for the COVID-19 Genomics UK (COG-UK) Consortium | Harper VanSteenhouse, Yumi Kasai, David Gray, Carol Clugston, Anna Dominiczak and Alex Alderton, Roberto Amato, Jeffrey Barrett, Sonia Goncalves, Ewan Harrison, David K. Jackson, Ian Johnston, Dominic Kwiatkowski, Cordelia Langford, John Sillitoe on behalf of the Wellcome Sanger Institute COVID-19 Surveillance Team |
| EPI_ISL_2569101, EPI_ISL_2569102 | Lighthouse Lab in Milton Keynes | Wellcome Sanger Institute for the COVID-19 Genomics UK (COG-UK) Consortium | The Lighthouse Lab in Milton Keynes and Alex Alderton, Roberto Amato, Jeffrey Barrett, Sonia Goncalves, Ewan Harrison, David K. Jackson, Ian Johnston, Dominic Kwiatkowski, Cordelia Langford, John Sillitoe on behalf of the Wellcome Sanger Institute COVID-19 Surveillance Team |
| EPI_ISL_2569103 | Lighthouse Lab in Glasgow | Wellcome Sanger Institute for the COVID-19 Genomics UK (COG-UK) Consortium | Harper VanSteenhouse, Yumi Kasai, David Gray, Carol Clugston, Anna Dominiczak and Alex Alderton, Roberto Amato, Jeffrey Barrett, Sonia Goncalves, Ewan Harrison, David K. Jackson, Ian Johnston, Dominic Kwiatkowski, Cordelia Langford, John Sillitoe on behalf of the Wellcome Sanger Institute COVID-19 Surveillance Team |
| EPI_ISL_2569104 | Lighthouse Laboratory Plymouth | Wellcome Sanger Institute for the COVID-19 Genomics UK (COG-UK) Consortium | Lighthouse Laboratory Plymouth and Alex Alderton, Roberto Amato, Jeffrey Barrett, Sonia Goncalves, Ewan Harrison, David K. Jackson, Ian Johnston, Dominic Kwiatkowski, Cordelia Langford, John Sillitoe on behalf of the Wellcome Sanger Institute COVID-19 Surveillance Team |
| EPI_ISL_2569105 | Lighthouse Lab in Milton Keynes | Wellcome Sanger Institute for the COVID-19 Genomics UK (COG-UK) Consortium | The Lighthouse Lab in Milton Keynes and Alex Alderton, Roberto Amato, Jeffrey Barrett, Sonia Goncalves, Ewan Harrison, David K. Jackson, Ian Johnston, Dominic Kwiatkowski, Cordelia Langford, John Sillitoe on behalf of the Wellcome Sanger Institute COVID-19 Surveillance Team |
| EPI_ISL_2569106, EPI_ISL_2569107 | Lighthouse Laboratory Plymouth | Wellcome Sanger Institute for the COVID-19 Genomics UK (COG-UK) Consortium | Lighthouse Laboratory Plymouth and Alex Alderton, Roberto Amato, Jeffrey Barrett, Sonia Goncalves, Ewan Harrison, David K. Jackson, Ian Johnston, Dominic Kwiatkowski, Cordelia Langford, John Sillitoe on behalf of the Wellcome Sanger Institute COVID-19 Surveillance Team |
| EPI_ISL_2569108, EPI_ISL_2569109, EPI_ISL_2569110 | Lighthouse Lab in Milton Keynes | Wellcome Sanger Institute for the COVID-19 Genomics UK (COG-UK) Consortium | The Lighthouse Lab in Milton Keynes and Alex Alderton, Roberto Amato, Jeffrey Barrett, Sonia Goncalves, Ewan Harrison, David K. Jackson, Ian Johnston, Dominic Kwiatkowski, Cordelia Langford, John Sillitoe on behalf of the Wellcome Sanger Institute COVID-19 Surveillance Team |
| EPI_ISL_2569111 | Lighthouse Lab in Glasgow | Wellcome Sanger Institute for the COVID-19 Genomics UK (COG-UK) | Harper VanSteenhouse, Yumi Kasai, David Gray, Carol Clugston, Anna Dominiczak and Alex Alderton, Roberto Amato, Jeffrey Barrett, Sonia Goncalves, Ewan Harrison, David K. Jackson, Ian Johnston, Dominic Kwiatkowski, Cordelia Langford, |

| | | | |
|---|---|---|---|
| | | Consortium | John Sillitoe on behalf of the Wellcome Sanger Institute COVID-19 Surveillance Team |
| EPI_ISL_2569112 | Lighthouse Laboratory Plymouth | Wellcome Sanger Institute for the COVID-19 Genomics UK (COG-UK) Consortium | Lighthouse Laboratory Plymouth and Alex Alderton, Roberto Amato, Jeffrey Barrett, Sonia Goncalves, Ewan Harrison, David K. Jackson, Ian Johnston, Dominic Kwiatkowski, Cordelia Langford, John Sillitoe on behalf of the Wellcome Sanger Institute COVID-19 Surveillance Team |
| EPI_ISL_2569113 | Lighthouse Lab in Alderley Park | Wellcome Sanger Institute for the COVID-19 Genomics UK (COG-UK) Consortium | Jacquelyn Wynn, Mairead Hyland, The Lighthouse Lab in Alderley Park and Alex Alderton, Roberto Amato, Jeffrey Barrett, Sonia Goncalves, Ewan Harrison, David K. Jackson, Ian Johnston, Dominic Kwiatkowski, Cordelia Langford, John Sillitoe on behalf of the Wellcome Sanger Institute COVID-19 Surveillance Team |
| EPI_ISL_2569114 | Lighthouse Laboratory Plymouth | Wellcome Sanger Institute for the COVID-19 Genomics UK (COG-UK) Consortium | Lighthouse Laboratory Plymouth and Alex Alderton, Roberto Amato, Jeffrey Barrett, Sonia Goncalves, Ewan Harrison, David K. Jackson, Ian Johnston, Dominic Kwiatkowski, Cordelia Langford, John Sillitoe on behalf of the Wellcome Sanger Institute COVID-19 Surveillance Team |
| EPI_ISL_2569115 | Lighthouse Lab in Milton Keynes | Wellcome Sanger Institute for the COVID-19 Genomics UK (COG-UK) Consortium | The Lighthouse Lab in Milton Keynes and Alex Alderton, Roberto Amato, Jeffrey Barrett, Sonia Goncalves, Ewan Harrison, David K. Jackson, Ian Johnston, Dominic Kwiatkowski, Cordelia Langford, John Sillitoe on behalf of the Wellcome Sanger Institute COVID-19 Surveillance Team |
| EPI_ISL_2569116 | Lighthouse Laboratory Plymouth | Wellcome Sanger Institute for the COVID-19 Genomics UK (COG-UK) Consortium | Lighthouse Laboratory Plymouth and Alex Alderton, Roberto Amato, Jeffrey Barrett, Sonia Goncalves, Ewan Harrison, David K. Jackson, Ian Johnston, Dominic Kwiatkowski, Cordelia Langford, John Sillitoe on behalf of the Wellcome Sanger Institute COVID-19 Surveillance Team |
| EPI_ISL_2569117 | Lighthouse Lab in Alderley Park | Wellcome Sanger Institute for the COVID-19 Genomics UK (COG-UK) Consortium | Jacquelyn Wynn, Mairead Hyland, The Lighthouse Lab in Alderley Park and Alex Alderton, Roberto Amato, Jeffrey Barrett, Sonia Goncalves, Ewan Harrison, David K. Jackson, Ian Johnston, Dominic Kwiatkowski, Cordelia Langford, John Sillitoe on behalf of the Wellcome Sanger Institute COVID-19 Surveillance Team |
| EPI_ISL_2569118, EPI_ISL_2569119 | Lighthouse Lab in Milton Keynes | Wellcome Sanger Institute for the COVID-19 Genomics UK (COG-UK) Consortium | The Lighthouse Lab in Milton Keynes and Alex Alderton, Roberto Amato, Jeffrey Barrett, Sonia Goncalves, Ewan Harrison, David K. Jackson, Ian Johnston, Dominic Kwiatkowski, Cordelia Langford, John Sillitoe on behalf of the Wellcome Sanger Institute COVID-19 Surveillance Team |
| EPI_ISL_2569121 | Lighthouse Laboratory Plymouth | Wellcome Sanger Institute for the COVID-19 Genomics UK (COG-UK) Consortium | Lighthouse Laboratory Plymouth and Alex Alderton, Roberto Amato, Jeffrey Barrett, Sonia Goncalves, Ewan Harrison, David K. Jackson, Ian Johnston, Dominic Kwiatkowski, Cordelia Langford, John Sillitoe on behalf of the Wellcome Sanger Institute COVID-19 Surveillance Team |
| EPI_ISL_2569122, EPI_ISL_2569124 | Lighthouse Lab in Alderley Park | Wellcome Sanger Institute for the COVID-19 Genomics UK (COG-UK) Consortium | Jacquelyn Wynn, Mairead Hyland, The Lighthouse Lab in Alderley Park and Alex Alderton, Roberto Amato, Jeffrey Barrett, Sonia Goncalves, Ewan Harrison, David K. Jackson, Ian Johnston, Dominic Kwiatkowski, Cordelia Langford, John Sillitoe on behalf of the Wellcome Sanger Institute COVID-19 Surveillance Team |
| EPI_ISL_2569125 | Lighthouse Lab in Glasgow | Wellcome Sanger Institute for the COVID-19 Genomics UK (COG-UK) Consortium | Harper VanSteenhouse, Yumi Kasai, David Gray, Carol Clugston, Anna Dominiczak and Alex Alderton, Roberto Amato, Jeffrey Barrett, Sonia Goncalves, Ewan Harrison, David K. Jackson, Ian Johnston, Dominic Kwiatkowski, Cordelia Langford, John Sillitoe on behalf of the Wellcome Sanger Institute COVID-19 Surveillance Team |
| EPI_ISL_2569126 | Lighthouse Laboratory Plymouth | Wellcome Sanger Institute for the COVID-19 Genomics UK (COG-UK) Consortium | Lighthouse Laboratory Plymouth and Alex Alderton, Roberto Amato, Jeffrey Barrett, Sonia Goncalves, Ewan Harrison, David K. Jackson, Ian Johnston, Dominic Kwiatkowski, Cordelia Langford, John Sillitoe on behalf of the Wellcome Sanger Institute COVID-19 Surveillance Team |
| EPI_ISL_2569127, EPI_ISL_2569128, EPI_ISL_2569129, EPI_ISL_2569130 | Lighthouse Lab in Alderley Park | Wellcome Sanger Institute for the COVID-19 Genomics UK (COG-UK) Consortium | Jacquelyn Wynn, Mairead Hyland, The Lighthouse Lab in Alderley Park and Alex Alderton, Roberto Amato, Jeffrey Barrett, Sonia Goncalves, Ewan Harrison, David K. Jackson, Ian Johnston, Dominic Kwiatkowski, Cordelia Langford, John Sillitoe on behalf of the Wellcome Sanger Institute COVID-19 Surveillance Team |
| EPI_ISL_2569131, EPI_ISL_2569132 | Lighthouse Laboratory Plymouth | Wellcome Sanger Institute for the COVID-19 Genomics UK (COG-UK) Consortium | Lighthouse Laboratory Plymouth and Alex Alderton, Roberto Amato, Jeffrey Barrett, Sonia Goncalves, Ewan Harrison, David K. Jackson, Ian Johnston, Dominic Kwiatkowski, Cordelia Langford, John Sillitoe on behalf of the Wellcome Sanger Institute COVID-19 Surveillance Team |
| EPI_ISL_2569133 | Lighthouse Lab in Alderley Park | Wellcome Sanger Institute for the COVID-19 Genomics UK (COG-UK) Consortium | Jacquelyn Wynn, Mairead Hyland, The Lighthouse Lab in Alderley Park and Alex Alderton, Roberto Amato, Jeffrey Barrett, Sonia Goncalves, Ewan Harrison, David K. Jackson, Ian Johnston, Dominic Kwiatkowski, Cordelia Langford, John Sillitoe on behalf of the Wellcome Sanger Institute COVID-19 Surveillance Team |
| EPI_ISL_2569134, EPI_ISL_2569135 | Lighthouse Lab in Glasgow | Wellcome Sanger Institute for the COVID-19 Genomics UK (COG-UK) Consortium | Harper VanSteenhouse, Yumi Kasai, David Gray, Carol Clugston, Anna Dominiczak and Alex Alderton, Roberto Amato, Jeffrey Barrett, Sonia Goncalves, Ewan Harrison, David K. Jackson, Ian Johnston, Dominic Kwiatkowski, Cordelia Langford, John Sillitoe on behalf of the Wellcome Sanger Institute COVID-19 Surveillance Team |
| EPI_ISL_2569136 | Lighthouse Lab in Milton Keynes | Wellcome Sanger Institute for the COVID-19 Genomics UK (COG-UK) Consortium | The Lighthouse Lab in Milton Keynes and Alex Alderton, Roberto Amato, Jeffrey Barrett, Sonia Goncalves, Ewan Harrison, David K. Jackson, Ian Johnston, Dominic Kwiatkowski, Cordelia Langford, John Sillitoe on behalf of the Wellcome Sanger Institute COVID-19 Surveillance Team |
| EPI_ISL_2569137 | Lighthouse Lab in Glasgow | Wellcome Sanger Institute for the COVID-19 Genomics UK (COG-UK) Consortium | Harper VanSteenhouse, Yumi Kasai, David Gray, Carol Clugston, Anna Dominiczak and Alex Alderton, Roberto Amato, Jeffrey Barrett, Sonia Goncalves, Ewan Harrison, David K. Jackson, Ian Johnston, Dominic Kwiatkowski, Cordelia Langford, John Sillitoe on behalf of the Wellcome Sanger Institute COVID-19 Surveillance Team |
| EPI_ISL_2569139 | Lighthouse Lab in Alderley Park | Wellcome Sanger Institute for the COVID-19 Genomics UK (COG-UK) Consortium | Jacquelyn Wynn, Mairead Hyland, The Lighthouse Lab in Alderley Park and Alex Alderton, Roberto Amato, Jeffrey Barrett, Sonia Goncalves, Ewan Harrison, David K. Jackson, Ian Johnston, Dominic Kwiatkowski, Cordelia Langford, John Sillitoe on behalf of the Wellcome Sanger Institute COVID-19 Surveillance Team |
| EPI_ISL_2569140, EPI_ISL_2569141 | Lighthouse Lab in Milton Keynes | Wellcome Sanger Institute for the COVID-19 Genomics UK (COG-UK) Consortium | The Lighthouse Lab in Milton Keynes and Alex Alderton, Roberto Amato, Jeffrey Barrett, Sonia Goncalves, Ewan Harrison, David K. Jackson, Ian Johnston, Dominic Kwiatkowski, Cordelia Langford, John Sillitoe on behalf of the Wellcome Sanger Institute COVID-19 Surveillance Team |
| EPI_ISL_2569143 | Lighthouse Lab in Glasgow | Wellcome Sanger Institute for the COVID-19 Genomics UK (COG-UK) Consortium | Harper VanSteenhouse, Yumi Kasai, David Gray, Carol Clugston, Anna Dominiczak and Alex Alderton, Roberto Amato, Jeffrey Barrett, Sonia Goncalves, Ewan Harrison, David K. Jackson, Ian Johnston, Dominic Kwiatkowski, Cordelia Langford, John Sillitoe on behalf of the Wellcome Sanger Institute COVID-19 Surveillance Team |
| EPI_ISL_2569144, EPI_ISL_2569145 | Lighthouse Lab in Milton Keynes | Wellcome Sanger Institute for the COVID-19 Genomics UK (COG-UK) Consortium | The Lighthouse Lab in Milton Keynes and Alex Alderton, Roberto Amato, Jeffrey Barrett, Sonia Goncalves, Ewan Harrison, David K. Jackson, Ian Johnston, Dominic Kwiatkowski, Cordelia Langford, John Sillitoe on behalf of the Wellcome Sanger Institute COVID-19 Surveillance Team |
| EPI_ISL_2569146 | Lighthouse Laboratory Plymouth | Wellcome Sanger Institute for the COVID-19 Genomics UK (COG-UK) Consortium | Lighthouse Laboratory Plymouth and Alex Alderton, Roberto Amato, Jeffrey Barrett, Sonia Goncalves, Ewan Harrison, David K. Jackson, Ian Johnston, Dominic Kwiatkowski, Cordelia Langford, John Sillitoe on behalf of the Wellcome Sanger Institute COVID-19 Surveillance Team |
| EPI_ISL_2569147 | Lighthouse Lab in Glasgow | Wellcome Sanger Institute for the COVID-19 Genomics UK (COG-UK) Consortium | Harper VanSteenhouse, Yumi Kasai, David Gray, Carol Clugston, Anna Dominiczak and Alex Alderton, Roberto Amato, Jeffrey Barrett, Sonia Goncalves, Ewan Harrison, David K. Jackson, Ian Johnston, Dominic Kwiatkowski, Cordelia Langford, John Sillitoe on behalf of the Wellcome Sanger Institute COVID-19 Surveillance Team |
| EPI_ISL_2569148 | Lighthouse Lab in Milton Keynes | Wellcome Sanger Institute for the COVID-19 Genomics UK (COG-UK) Consortium | The Lighthouse Lab in Milton Keynes and Alex Alderton, Roberto Amato, Jeffrey Barrett, Sonia Goncalves, Ewan Harrison, David K. Jackson, Ian Johnston, Dominic Kwiatkowski, Cordelia Langford, John Sillitoe on behalf of the Wellcome Sanger Institute COVID-19 Surveillance Team |
| EPI_ISL_2569149 | Lighthouse Laboratory Plymouth | Wellcome Sanger Institute for the COVID-19 Genomics UK (COG-UK) Consortium | Lighthouse Laboratory Plymouth and Alex Alderton, Roberto Amato, Jeffrey Barrett, Sonia Goncalves, Ewan Harrison, David K. Jackson, Ian Johnston, Dominic Kwiatkowski, Cordelia Langford, John Sillitoe on behalf of the Wellcome Sanger Institute COVID-19 Surveillance Team |
| EPI_ISL_2569150 | Lighthouse Lab in Milton Keynes | Wellcome Sanger Institute for the COVID-19 Genomics UK (COG-UK) Consortium | The Lighthouse Lab in Milton Keynes and Alex Alderton, Roberto Amato, Jeffrey Barrett, Sonia Goncalves, Ewan Harrison, David K. Jackson, Ian Johnston, Dominic Kwiatkowski, Cordelia Langford, John Sillitoe on behalf of the Wellcome Sanger Institute COVID-19 Surveillance Team |
| EPI_ISL_2569151, EPI_ISL_2569152 | Lighthouse Lab in Alderley Park | Wellcome Sanger Institute for the COVID-19 Genomics UK (COG-UK) Consortium | Jacquelyn Wynn, Mairead Hyland, The Lighthouse Lab in Alderley Park and Alex Alderton, Roberto Amato, Jeffrey Barrett, Sonia Goncalves, Ewan Harrison, David K. Jackson, Ian Johnston, Dominic Kwiatkowski, Cordelia Langford, John Sillitoe on behalf of the Wellcome Sanger Institute COVID-19 Surveillance Team |
| EPI_ISL_2569153, EPI_ISL_2569154 | Lighthouse Laboratory Plymouth | Wellcome Sanger Institute for the COVID-19 Genomics UK (COG-UK) Consortium | Lighthouse Laboratory Plymouth and Alex Alderton, Roberto Amato, Jeffrey Barrett, Sonia Goncalves, Ewan Harrison, David K. Jackson, Ian Johnston, Dominic Kwiatkowski, Cordelia Langford, John Sillitoe on behalf of the Wellcome Sanger Institute COVID-19 Surveillance Team |
| EPI_ISL_2569155, EPI_ISL_2569156 | Lighthouse Lab in Milton Keynes | Wellcome Sanger Institute for the COVID-19 Genomics UK (COG-UK) Consortium | The Lighthouse Lab in Milton Keynes and Alex Alderton, Roberto Amato, Jeffrey Barrett, Sonia Goncalves, Ewan Harrison, David K. Jackson, Ian Johnston, Dominic Kwiatkowski, Cordelia Langford, John Sillitoe on behalf of the Wellcome Sanger Institute COVID-19 Surveillance Team |
| EPI_ISL_2569157, EPI_ISL_2569158, EPI_ISL_2569159 | Lighthouse Laboratory Plymouth | Wellcome Sanger Institute for the COVID-19 Genomics UK (COG-UK) Consortium | Lighthouse Laboratory Plymouth and Alex Alderton, Roberto Amato, Jeffrey Barrett, Sonia Goncalves, Ewan Harrison, David K. Jackson, Ian Johnston, Dominic Kwiatkowski, Cordelia Langford, John Sillitoe on behalf of the Wellcome Sanger Institute COVID-19 Surveillance Team |
| EPI_ISL_2569161 | Lighthouse Lab in Alderley Park | Wellcome Sanger Institute for the COVID-19 Genomics UK (COG-UK) Consortium | Jacquelyn Wynn, Mairead Hyland, The Lighthouse Lab in Alderley Park and Alex Alderton, Roberto Amato, Jeffrey Barrett, Sonia Goncalves, Ewan Harrison, David K. Jackson, Ian Johnston, Dominic Kwiatkowski, Cordelia Langford, John Sillitoe on behalf of the Wellcome Sanger Institute COVID-19 Surveillance Team |
| EPI_ISL_2569162, EPI_ISL_2569163, EPI_ISL_2569164 | Lighthouse Laboratory Plymouth | Wellcome Sanger Institute for the COVID-19 Genomics UK (COG-UK) Consortium | Lighthouse Laboratory Plymouth and Alex Alderton, Roberto Amato, Jeffrey Barrett, Sonia Goncalves, Ewan Harrison, David K. Jackson, Ian Johnston, Dominic Kwiatkowski, Cordelia Langford, John Sillitoe on behalf of the Wellcome Sanger Institute COVID-19 Surveillance Team |
| EPI_ISL_2569165 | Lighthouse Lab in Milton Keynes | Wellcome Sanger Institute for the COVID-19 Genomics UK (COG-UK) Consortium | The Lighthouse Lab in Milton Keynes and Alex Alderton, Roberto Amato, Jeffrey Barrett, Sonia Goncalves, Ewan Harrison, David K. Jackson, Ian Johnston, Dominic Kwiatkowski, Cordelia Langford, John Sillitoe on behalf of the Wellcome Sanger Institute COVID-19 Surveillance Team |
| EPI_ISL_2569166, EPI_ISL_2569167 | Lighthouse Lab in Glasgow | Wellcome Sanger Institute for the COVID-19 Genomics UK (COG-UK) Consortium | Harper VanSteenhouse, Yumi Kasai, David Gray, Carol Clugston, Anna Dominiczak and Alex Alderton, Roberto Amato, Jeffrey Barrett, Sonia Goncalves, Ewan Harrison, David K. Jackson, Ian Johnston, Dominic Kwiatkowski, Cordelia Langford, John Sillitoe on behalf of the Wellcome Sanger Institute COVID-19 Surveillance Team |
| EPI_ISL_2569168, EPI_ISL_2569169 | Lighthouse Lab in Alderley Park | Wellcome Sanger Institute for the COVID-19 Genomics UK (COG-UK) Consortium | Jacquelyn Wynn, Mairead Hyland, The Lighthouse Lab in Alderley Park and Alex Alderton, Roberto Amato, Jeffrey Barrett, Sonia Goncalves, Ewan Harrison, David K. Jackson, Ian Johnston, Dominic Kwiatkowski, Cordelia Langford, John Sillitoe on behalf of the Wellcome Sanger Institute COVID-19 Surveillance Team |
| EPI_ISL_2569170 | Lighthouse Lab in Glasgow | Wellcome Sanger Institute for the COVID-19 Genomics UK (COG-UK) Consortium | Harper VanSteenhouse, Yumi Kasai, David Gray, Carol Clugston, Anna Dominiczak and Alex Alderton, Roberto Amato, Jeffrey Barrett, Sonia Goncalves, Ewan Harrison, David K. Jackson, Ian Johnston, Dominic Kwiatkowski, Cordelia Langford, John Sillitoe on behalf of the Wellcome Sanger Institute COVID-19 Surveillance Team |
| EPI_ISL_2569171 | Lighthouse Lab in Alderley Park | Wellcome Sanger Institute for the COVID-19 Genomics UK (COG-UK) Consortium | Jacquelyn Wynn, Mairead Hyland, The Lighthouse Lab in Alderley Park and Alex Alderton, Roberto Amato, Jeffrey Barrett, Sonia Goncalves, Ewan Harrison, David K. Jackson, Ian Johnston, Dominic Kwiatkowski, Cordelia Langford, John Sillitoe on behalf of the Wellcome Sanger Institute COVID-19 Surveillance Team |
| EPI_ISL_2569172 | Lighthouse Lab in Milton Keynes | Wellcome Sanger Institute for the COVID-19 Genomics UK (COG-UK) Consortium | The Lighthouse Lab in Milton Keynes and Alex Alderton, Roberto Amato, Jeffrey Barrett, Sonia Goncalves, Ewan Harrison, David K. Jackson, Ian Johnston, Dominic Kwiatkowski, Cordelia Langford, John Sillitoe on behalf of the Wellcome Sanger Institute COVID-19 Surveillance Team |
| EPI_ISL_2569173 | Lighthouse Laboratory Plymouth | Wellcome Sanger Institute for the COVID-19 Genomics UK (COG-UK) Consortium | Lighthouse Laboratory Plymouth and Alex Alderton, Roberto Amato, Jeffrey Barrett, Sonia Goncalves, Ewan Harrison, David K. Jackson, Ian Johnston, Dominic Kwiatkowski, Cordelia Langford, John Sillitoe on behalf of the Wellcome Sanger Institute COVID-19 Surveillance Team |
| EPI_ISL_2569174 | Lighthouse Lab in Glasgow | Wellcome Sanger Institute for the COVID-19 Genomics UK (COG-UK) Consortium | Harper VanSteenhouse, Yumi Kasai, David Gray, Carol Clugston, Anna Dominiczak and Alex Alderton, Roberto Amato, Jeffrey Barrett, Sonia Goncalves, Ewan Harrison, David K. Jackson, Ian Johnston, Dominic Kwiatkowski, Cordelia Langford, John Sillitoe on behalf of the Wellcome Sanger Institute COVID-19 Surveillance Team |
| EPI_ISL_2569175 | Lighthouse Laboratory Plymouth | Wellcome Sanger Institute for the COVID-19 Genomics UK (COG-UK) Consortium | Lighthouse Laboratory Plymouth and Alex Alderton, Roberto Amato, Jeffrey Barrett, Sonia Goncalves, Ewan Harrison, David K. Jackson, Ian Johnston, Dominic Kwiatkowski, Cordelia Langford, John Sillitoe on behalf of the Wellcome Sanger Institute COVID-19 Surveillance Team |
| EPI_ISL_2569176 | Lighthouse Lab in Milton Keynes | Wellcome Sanger Institute for the COVID-19 Genomics UK (COG-UK) Consortium | The Lighthouse Lab in Milton Keynes and Alex Alderton, Roberto Amato, Jeffrey Barrett, Sonia Goncalves, Ewan Harrison, David K. Jackson, Ian Johnston, Dominic Kwiatkowski, Cordelia Langford, John Sillitoe on behalf of the Wellcome Sanger Institute COVID-19 Surveillance Team |
| EPI_ISL_2569177 | Lighthouse Lab in Glasgow | Wellcome Sanger Institute for the COVID-19 Genomics UK (COG-UK) Consortium | Harper VanSteenhouse, Yumi Kasai, David Gray, Carol Clugston, Anna Dominiczak and Alex Alderton, Roberto Amato, Jeffrey Barrett, Sonia Goncalves, Ewan Harrison, David K. Jackson, Ian Johnston, Dominic Kwiatkowski, Cordelia Langford, John Sillitoe on behalf of the Wellcome Sanger Institute COVID-19 Surveillance Team |

| | | | |
|---|---|---|---|
| EPI_ISL_2569178 | Lighthouse Laboratory Plymouth | Wellcome Sanger Institute for the COVID-19 Genomics UK (COG-UK) Consortium | Lighthouse Laboratory Plymouth and Alex Alderton, Roberto Amato, Jeffrey Barrett, Sonia Goncalves, Ewan Harrison, David K. Jackson, Ian Johnston, Dominic Kwiatkowski, Cordelia Langford, John Sillitoe on behalf of the Wellcome Sanger Institute COVID-19 Surveillance Team |
| EPI_ISL_2569179 | Lighthouse Lab in Glasgow | Wellcome Sanger Institute for the COVID-19 Genomics UK (COG-UK) Consortium | Harper VanSteenhouse, Yumi Kasai, David Gray, Carol Clugston, Anna Dominiczak and Alex Alderton, Roberto Amato, Jeffrey Barrett, Sonia Goncalves, Ewan Harrison, David K. Jackson, Ian Johnston, Dominic Kwiatkowski, Cordelia Langford, John Sillitoe on behalf of the Wellcome Sanger Institute COVID-19 Surveillance Team |
| EPI_ISL_2569180, EPI_ISL_2569181 | Lighthouse Lab in Alderley Park | Wellcome Sanger Institute for the COVID-19 Genomics UK (COG-UK) Consortium | Jacquelyn Wynn, Mairead Hyland, The Lighthouse Lab in Alderley Park and Alex Alderton, Roberto Amato, Jeffrey Barrett, Sonia Goncalves, Ewan Harrison, David K. Jackson, Ian Johnston, Dominic Kwiatkowski, Cordelia Langford, John Sillitoe on behalf of the Wellcome Sanger Institute COVID-19 Surveillance Team |
| EPI_ISL_2569182 | Lighthouse Lab in Milton Keynes | Wellcome Sanger Institute for the COVID-19 Genomics UK (COG-UK) Consortium | The Lighthouse Lab in Milton Keynes and Alex Alderton, Roberto Amato, Jeffrey Barrett, Sonia Goncalves, Ewan Harrison, David K. Jackson, Ian Johnston, Dominic Kwiatkowski, Cordelia Langford, John Sillitoe on behalf of the Wellcome Sanger Institute COVID-19 Surveillance Team |
| EPI_ISL_2569183 | Lighthouse Laboratory Plymouth | Wellcome Sanger Institute for the COVID-19 Genomics UK (COG-UK) Consortium | Lighthouse Laboratory Plymouth and Alex Alderton, Roberto Amato, Jeffrey Barrett, Sonia Goncalves, Ewan Harrison, David K. Jackson, Ian Johnston, Dominic Kwiatkowski, Cordelia Langford, John Sillitoe on behalf of the Wellcome Sanger Institute COVID-19 Surveillance Team |
| EPI_ISL_2569184, EPI_ISL_2569185, EPI_ISL_2569186 | Lighthouse Lab in Alderley Park | Wellcome Sanger Institute for the COVID-19 Genomics UK (COG-UK) Consortium | Jacquelyn Wynn, Mairead Hyland, The Lighthouse Lab in Alderley Park and Alex Alderton, Roberto Amato, Jeffrey Barrett, Sonia Goncalves, Ewan Harrison, David K. Jackson, Ian Johnston, Dominic Kwiatkowski, Cordelia Langford, John Sillitoe on behalf of the Wellcome Sanger Institute COVID-19 Surveillance Team |
| EPI_ISL_2569187 | Lighthouse Laboratory Plymouth | Wellcome Sanger Institute for the COVID-19 Genomics UK (COG-UK) Consortium | Lighthouse Laboratory Plymouth and Alex Alderton, Roberto Amato, Jeffrey Barrett, Sonia Goncalves, Ewan Harrison, David K. Jackson, Ian Johnston, Dominic Kwiatkowski, Cordelia Langford, John Sillitoe on behalf of the Wellcome Sanger Institute COVID-19 Surveillance Team |
| EPI_ISL_2569188, EPI_ISL_2569189 | Lighthouse Lab in Alderley Park | Wellcome Sanger Institute for the COVID-19 Genomics UK (COG-UK) Consortium | Jacquelyn Wynn, Mairead Hyland, The Lighthouse Lab in Alderley Park and Alex Alderton, Roberto Amato, Jeffrey Barrett, Sonia Goncalves, Ewan Harrison, David K. Jackson, Ian Johnston, Dominic Kwiatkowski, Cordelia Langford, John Sillitoe on behalf of the Wellcome Sanger Institute COVID-19 Surveillance Team |
| EPI_ISL_2569190 | Lighthouse Lab in Milton Keynes | Wellcome Sanger Institute for the COVID-19 Genomics UK (COG-UK) Consortium | The Lighthouse Lab in Milton Keynes and Alex Alderton, Roberto Amato, Jeffrey Barrett, Sonia Goncalves, Ewan Harrison, David K. Jackson, Ian Johnston, Dominic Kwiatkowski, Cordelia Langford, John Sillitoe on behalf of the Wellcome Sanger Institute COVID-19 Surveillance Team |
| EPI_ISL_2569191, EPI_ISL_2569192 | Lighthouse Lab in Alderley Park | Wellcome Sanger Institute for the COVID-19 Genomics UK (COG-UK) Consortium | Jacquelyn Wynn, Mairead Hyland, The Lighthouse Lab in Alderley Park and Alex Alderton, Roberto Amato, Jeffrey Barrett, Sonia Goncalves, Ewan Harrison, David K. Jackson, Ian Johnston, Dominic Kwiatkowski, Cordelia Langford, John Sillitoe on behalf of the Wellcome Sanger Institute COVID-19 Surveillance Team |
| EPI_ISL_2569193 | Lighthouse Lab in Milton Keynes | Wellcome Sanger Institute for the COVID-19 Genomics UK (COG-UK) Consortium | The Lighthouse Lab in Milton Keynes and Alex Alderton, Roberto Amato, Jeffrey Barrett, Sonia Goncalves, Ewan Harrison, David K. Jackson, Ian Johnston, Dominic Kwiatkowski, Cordelia Langford, John Sillitoe on behalf of the Wellcome Sanger Institute COVID-19 Surveillance Team |
| EPI_ISL_2569194, EPI_ISL_2569195 | Lighthouse Laboratory Plymouth | Wellcome Sanger Institute for the COVID-19 Genomics UK (COG-UK) Consortium | Lighthouse Laboratory Plymouth and Alex Alderton, Roberto Amato, Jeffrey Barrett, Sonia Goncalves, Ewan Harrison, David K. Jackson, Ian Johnston, Dominic Kwiatkowski, Cordelia Langford, John Sillitoe on behalf of the Wellcome Sanger Institute COVID-19 Surveillance Team |
| EPI_ISL_2569196, EPI_ISL_2569197 | Lighthouse Lab in Milton Keynes | Wellcome Sanger Institute for the COVID-19 Genomics UK (COG-UK) Consortium | The Lighthouse Lab in Milton Keynes and Alex Alderton, Roberto Amato, Jeffrey Barrett, Sonia Goncalves, Ewan Harrison, David K. Jackson, Ian Johnston, Dominic Kwiatkowski, Cordelia Langford, John Sillitoe on behalf of the Wellcome Sanger Institute COVID-19 Surveillance Team |
| EPI_ISL_2569198 | Lighthouse Laboratory Plymouth | Wellcome Sanger Institute for the COVID-19 Genomics UK (COG-UK) Consortium | Lighthouse Laboratory Plymouth and Alex Alderton, Roberto Amato, Jeffrey Barrett, Sonia Goncalves, Ewan Harrison, David K. Jackson, Ian Johnston, Dominic Kwiatkowski, Cordelia Langford, John Sillitoe on behalf of the Wellcome Sanger Institute COVID-19 Surveillance Team |
| EPI_ISL_2569199, EPI_ISL_2569200 | Lighthouse Lab in Milton Keynes | Wellcome Sanger Institute for the COVID-19 Genomics UK (COG-UK) Consortium | The Lighthouse Lab in Milton Keynes and Alex Alderton, Roberto Amato, Jeffrey Barrett, Sonia Goncalves, Ewan Harrison, David K. Jackson, Ian Johnston, Dominic Kwiatkowski, Cordelia Langford, John Sillitoe on behalf of the Wellcome Sanger Institute COVID-19 Surveillance Team |
| EPI_ISL_2569201 | Lighthouse Lab in Alderley Park | Wellcome Sanger Institute for the COVID-19 Genomics UK (COG-UK) Consortium | Jacquelyn Wynn, Mairead Hyland, The Lighthouse Lab in Alderley Park and Alex Alderton, Roberto Amato, Jeffrey Barrett, Sonia Goncalves, Ewan Harrison, David K. Jackson, Ian Johnston, Dominic Kwiatkowski, Cordelia Langford, John Sillitoe on behalf of the Wellcome Sanger Institute COVID-19 Surveillance Team |
| EPI_ISL_2569202 | Lighthouse Lab in Milton Keynes | Wellcome Sanger Institute for the COVID-19 Genomics UK (COG-UK) Consortium | The Lighthouse Lab in Milton Keynes and Alex Alderton, Roberto Amato, Jeffrey Barrett, Sonia Goncalves, Ewan Harrison, David K. Jackson, Ian Johnston, Dominic Kwiatkowski, Cordelia Langford, John Sillitoe on behalf of the Wellcome Sanger Institute COVID-19 Surveillance Team |
| EPI_ISL_2569203 | Lighthouse Lab in Alderley Park | Wellcome Sanger Institute for the COVID-19 Genomics UK (COG-UK) Consortium | Jacquelyn Wynn, Mairead Hyland, The Lighthouse Lab in Alderley Park and Alex Alderton, Roberto Amato, Jeffrey Barrett, Sonia Goncalves, Ewan Harrison, David K. Jackson, Ian Johnston, Dominic Kwiatkowski, Cordelia Langford, John Sillitoe on behalf of the Wellcome Sanger Institute COVID-19 Surveillance Team |
| EPI_ISL_2569204 | Lighthouse Lab in Milton Keynes | Wellcome Sanger Institute for the COVID-19 Genomics UK (COG-UK) Consortium | The Lighthouse Lab in Milton Keynes and Alex Alderton, Roberto Amato, Jeffrey Barrett, Sonia Goncalves, Ewan Harrison, David K. Jackson, Ian Johnston, Dominic Kwiatkowski, Cordelia Langford, John Sillitoe on behalf of the Wellcome Sanger Institute COVID-19 Surveillance Team |
| EPI_ISL_2569205 | Lighthouse Lab in Alderley Park | Wellcome Sanger Institute for the COVID-19 Genomics UK (COG-UK) Consortium | Jacquelyn Wynn, Mairead Hyland, The Lighthouse Lab in Alderley Park and Alex Alderton, Roberto Amato, Jeffrey Barrett, Sonia Goncalves, Ewan Harrison, David K. Jackson, Ian Johnston, Dominic Kwiatkowski, Cordelia Langford, John Sillitoe on behalf of the Wellcome Sanger Institute COVID-19 Surveillance Team |
| EPI_ISL_2569206 | Lighthouse Lab in Glasgow | Wellcome Sanger Institute for the COVID-19 Genomics UK (COG-UK) Consortium | Harper VanSteenhouse, Yumi Kasai, David Gray, Carol Clugston, Anna Dominiczak and Alex Alderton, Roberto Amato, Jeffrey Barrett, Sonia Goncalves, Ewan Harrison, David K. Jackson, Ian Johnston, Dominic Kwiatkowski, Cordelia Langford, John Sillitoe on behalf of the Wellcome Sanger Institute COVID-19 Surveillance Team |
| EPI_ISL_2569207 | Lighthouse Laboratory Plymouth | Wellcome Sanger Institute for the COVID-19 Genomics UK (COG-UK) Consortium | Lighthouse Laboratory Plymouth and Alex Alderton, Roberto Amato, Jeffrey Barrett, Sonia Goncalves, Ewan Harrison, David K. Jackson, Ian Johnston, Dominic Kwiatkowski, Cordelia Langford, John Sillitoe on behalf of the Wellcome Sanger Institute COVID-19 Surveillance Team |
| EPI_ISL_2569208, EPI_ISL_2569209 | Lighthouse Lab in Milton Keynes | Wellcome Sanger Institute for the COVID-19 Genomics UK (COG-UK) Consortium | The Lighthouse Lab in Milton Keynes and Alex Alderton, Roberto Amato, Jeffrey Barrett, Sonia Goncalves, Ewan Harrison, David K. Jackson, Ian Johnston, Dominic Kwiatkowski, Cordelia Langford, John Sillitoe on behalf of the Wellcome Sanger Institute COVID-19 Surveillance Team |
| EPI_ISL_2569211 | Lighthouse Lab in Alderley Park | Wellcome Sanger Institute for the COVID-19 Genomics UK (COG-UK) Consortium | Jacquelyn Wynn, Mairead Hyland, The Lighthouse Lab in Alderley Park and Alex Alderton, Roberto Amato, Jeffrey Barrett, Sonia Goncalves, Ewan Harrison, David K. Jackson, Ian Johnston, Dominic Kwiatkowski, Cordelia Langford, John Sillitoe on behalf of the Wellcome Sanger Institute COVID-19 Surveillance Team |
| EPI_ISL_2569212, EPI_ISL_2569213 | Lighthouse Lab in Milton Keynes | Wellcome Sanger Institute for the COVID-19 Genomics UK (COG-UK) Consortium | The Lighthouse Lab in Milton Keynes and Alex Alderton, Roberto Amato, Jeffrey Barrett, Sonia Goncalves, Ewan Harrison, David K. Jackson, Ian Johnston, Dominic Kwiatkowski, Cordelia Langford, John Sillitoe on behalf of the Wellcome Sanger Institute COVID-19 Surveillance Team |
| EPI_ISL_2569214 | Lighthouse Lab in Glasgow | Wellcome Sanger Institute for the COVID-19 Genomics UK (COG-UK) Consortium | Harper VanSteenhouse, Yumi Kasai, David Gray, Carol Clugston, Anna Dominiczak and Alex Alderton, Roberto Amato, Jeffrey Barrett, Sonia Goncalves, Ewan Harrison, David K. Jackson, Ian Johnston, Dominic Kwiatkowski, Cordelia Langford, John Sillitoe on behalf of the Wellcome Sanger Institute COVID-19 Surveillance Team |
| EPI_ISL_2569215 | Lighthouse Laboratory Plymouth | Wellcome Sanger Institute for the COVID-19 Genomics UK (COG-UK) Consortium | Lighthouse Laboratory Plymouth and Alex Alderton, Roberto Amato, Jeffrey Barrett, Sonia Goncalves, Ewan Harrison, David K. Jackson, Ian Johnston, Dominic Kwiatkowski, Cordelia Langford, John Sillitoe on behalf of the Wellcome Sanger Institute COVID-19 Surveillance Team |
| EPI_ISL_2569216 | Lighthouse Lab in Glasgow | Wellcome Sanger Institute for the COVID-19 Genomics UK (COG-UK) Consortium | Harper VanSteenhouse, Yumi Kasai, David Gray, Carol Clugston, Anna Dominiczak and Alex Alderton, Roberto Amato, Jeffrey Barrett, Sonia Goncalves, Ewan Harrison, David K. Jackson, Ian Johnston, Dominic Kwiatkowski, Cordelia Langford, John Sillitoe on behalf of the Wellcome Sanger Institute COVID-19 Surveillance Team |
| EPI_ISL_2569217, EPI_ISL_2569218 | Lighthouse Laboratory Plymouth | Wellcome Sanger Institute for the COVID-19 Genomics UK (COG-UK) Consortium | Lighthouse Laboratory Plymouth and Alex Alderton, Roberto Amato, Jeffrey Barrett, Sonia Goncalves, Ewan Harrison, David K. Jackson, Ian Johnston, Dominic Kwiatkowski, Cordelia Langford, John Sillitoe on behalf of the Wellcome Sanger Institute COVID-19 Surveillance Team |
| EPI_ISL_2569219 | Lighthouse Lab in Glasgow | Wellcome Sanger Institute for the COVID-19 Genomics UK (COG-UK) Consortium | Harper VanSteenhouse, Yumi Kasai, David Gray, Carol Clugston, Anna Dominiczak and Alex Alderton, Roberto Amato, Jeffrey Barrett, Sonia Goncalves, Ewan Harrison, David K. Jackson, Ian Johnston, Dominic Kwiatkowski, Cordelia Langford, John Sillitoe on behalf of the Wellcome Sanger Institute COVID-19 Surveillance Team |
| EPI_ISL_2569220 | Lighthouse Lab in Alderley Park | Wellcome Sanger Institute for the COVID-19 Genomics UK (COG-UK) Consortium | Jacquelyn Wynn, Mairead Hyland, The Lighthouse Lab in Alderley Park and Alex Alderton, Roberto Amato, Jeffrey Barrett, Sonia Goncalves, Ewan Harrison, David K. Jackson, Ian Johnston, Dominic Kwiatkowski, Cordelia Langford, John Sillitoe on behalf of the Wellcome Sanger Institute COVID-19 Surveillance Team |
| EPI_ISL_2569221 | Lighthouse Laboratory Plymouth | Wellcome Sanger Institute for the COVID-19 Genomics UK (COG-UK) Consortium | Lighthouse Laboratory Plymouth and Alex Alderton, Roberto Amato, Jeffrey Barrett, Sonia Goncalves, Ewan Harrison, David K. Jackson, Ian Johnston, Dominic Kwiatkowski, Cordelia Langford, John Sillitoe on behalf of the Wellcome Sanger Institute COVID-19 Surveillance Team |
| EPI_ISL_2569222 | Lighthouse Lab in Alderley Park | Wellcome Sanger Institute for the COVID-19 Genomics UK (COG-UK) Consortium | Jacquelyn Wynn, Mairead Hyland, The Lighthouse Lab in Alderley Park and Alex Alderton, Roberto Amato, Jeffrey Barrett, Sonia Goncalves, Ewan Harrison, David K. Jackson, Ian Johnston, Dominic Kwiatkowski, Cordelia Langford, John Sillitoe on behalf of the Wellcome Sanger Institute COVID-19 Surveillance Team |
| EPI_ISL_2569223 | Lighthouse Laboratory Plymouth | Wellcome Sanger Institute for the COVID-19 Genomics UK (COG-UK) Consortium | Lighthouse Laboratory Plymouth and Alex Alderton, Roberto Amato, Jeffrey Barrett, Sonia Goncalves, Ewan Harrison, David K. Jackson, Ian Johnston, Dominic Kwiatkowski, Cordelia Langford, John Sillitoe on behalf of the Wellcome Sanger Institute COVID-19 Surveillance Team |
| EPI_ISL_2569224 | Lighthouse Lab in Milton Keynes | Wellcome Sanger Institute for the COVID-19 Genomics UK (COG-UK) Consortium | The Lighthouse Lab in Milton Keynes and Alex Alderton, Roberto Amato, Jeffrey Barrett, Sonia Goncalves, Ewan Harrison, David K. Jackson, Ian Johnston, Dominic Kwiatkowski, Cordelia Langford, John Sillitoe on behalf of the Wellcome Sanger Institute COVID-19 Surveillance Team |
| EPI_ISL_2569225 | Lighthouse Lab in Glasgow | Wellcome Sanger Institute for the COVID-19 Genomics UK (COG-UK) Consortium | Harper VanSteenhouse, Yumi Kasai, David Gray, Carol Clugston, Anna Dominiczak and Alex Alderton, Roberto Amato, Jeffrey Barrett, Sonia Goncalves, Ewan Harrison, David K. Jackson, Ian Johnston, Dominic Kwiatkowski, Cordelia Langford, John Sillitoe on behalf of the Wellcome Sanger Institute COVID-19 Surveillance Team |
| EPI_ISL_2569226 | Lighthouse Lab in Alderley Park | Wellcome Sanger Institute for the COVID-19 Genomics UK (COG-UK) Consortium | Jacquelyn Wynn, Mairead Hyland, The Lighthouse Lab in Alderley Park and Alex Alderton, Roberto Amato, Jeffrey Barrett, Sonia Goncalves, Ewan Harrison, David K. Jackson, Ian Johnston, Dominic Kwiatkowski, Cordelia Langford, John Sillitoe on behalf of the Wellcome Sanger Institute COVID-19 Surveillance Team |
| EPI_ISL_2569227 | Lighthouse Laboratory Plymouth | Wellcome Sanger Institute for the COVID-19 Genomics UK (COG-UK) Consortium | Lighthouse Laboratory Plymouth and Alex Alderton, Roberto Amato, Jeffrey Barrett, Sonia Goncalves, Ewan Harrison, David K. Jackson, Ian Johnston, Dominic Kwiatkowski, Cordelia Langford, John Sillitoe on behalf of the Wellcome Sanger Institute COVID-19 Surveillance Team |
| EPI_ISL_2569228 | Lighthouse Lab in Glasgow | Wellcome Sanger Institute for the COVID-19 Genomics UK (COG-UK) Consortium | Harper VanSteenhouse, Yumi Kasai, David Gray, Carol Clugston, Anna Dominiczak and Alex Alderton, Roberto Amato, Jeffrey Barrett, Sonia Goncalves, Ewan Harrison, David K. Jackson, Ian Johnston, Dominic Kwiatkowski, Cordelia Langford, John Sillitoe on behalf of the Wellcome Sanger Institute COVID-19 Surveillance Team |
| EPI_ISL_2569230 | Lighthouse Lab in Alderley Park | Wellcome Sanger Institute for the COVID-19 Genomics UK (COG-UK) Consortium | Jacquelyn Wynn, Mairead Hyland, The Lighthouse Lab in Alderley Park and Alex Alderton, Roberto Amato, Jeffrey Barrett, Sonia Goncalves, Ewan Harrison, David K. Jackson, Ian Johnston, Dominic Kwiatkowski, Cordelia Langford, John Sillitoe on behalf of the Wellcome Sanger Institute COVID-19 Surveillance Team |
| EPI_ISL_2569232, EPI_ISL_2569233 | Lighthouse Laboratory Plymouth | Wellcome Sanger Institute for the COVID-19 Genomics UK (COG-UK) Consortium | Lighthouse Laboratory Plymouth and Alex Alderton, Roberto Amato, Jeffrey Barrett, Sonia Goncalves, Ewan Harrison, David K. Jackson, Ian Johnston, Dominic Kwiatkowski, Cordelia Langford, John Sillitoe on behalf of the Wellcome Sanger Institute COVID-19 Surveillance Team |
| EPI_ISL_2569234 | Lighthouse Lab in Milton Keynes | Wellcome Sanger Institute for the COVID-19 Genomics UK (COG-UK) Consortium | The Lighthouse Lab in Milton Keynes and Alex Alderton, Roberto Amato, Jeffrey Barrett, Sonia Goncalves, Ewan Harrison, David K. Jackson, Ian Johnston, Dominic Kwiatkowski, Cordelia Langford, John Sillitoe on behalf of the Wellcome Sanger Institute COVID-19 Surveillance Team |
| EPI_ISL_2569235 | Lighthouse Lab in Alderley Park | Wellcome Sanger Institute for the COVID-19 Genomics UK (COG-UK) Consortium | Jacquelyn Wynn, Mairead Hyland, The Lighthouse Lab in Alderley Park and Alex Alderton, Roberto Amato, Jeffrey Barrett, Sonia Goncalves, Ewan Harrison, David K. Jackson, Ian Johnston, Dominic Kwiatkowski, Cordelia Langford, John Sillitoe on behalf of the Wellcome Sanger Institute COVID-19 Surveillance Team |
| EPI_ISL_2569236, EPI_ISL_2569237 | Lighthouse Lab in Milton Keynes | Wellcome Sanger Institute for the COVID-19 Genomics UK (COG-UK) Consortium | The Lighthouse Lab in Milton Keynes and Alex Alderton, Roberto Amato, Jeffrey Barrett, Sonia Goncalves, Ewan Harrison, David K. Jackson, Ian Johnston, Dominic Kwiatkowski, Cordelia Langford, John Sillitoe on behalf of the Wellcome Sanger Institute COVID-19 Surveillance Team |
| EPI_ISL_2569238, | Lighthouse Laboratory Plymouth | Wellcome Sanger Institute for the COVID-19 Genomics UK (COG-UK) | Lighthouse Laboratory Plymouth and Alex Alderton, Roberto Amato, Jeffrey Barrett, Sonia Goncalves, Ewan Harrison, David K. Jackson, Ian Johnston, Dominic Kwiatkowski, Cordelia Langford, John Sillitoe on behalf of the Wellcome Sanger |

| | | | |
|---|---|---|---|
| EPI_ISL_2569239, EPI_ISL_2569240 | | Consortium | Institute COVID-19 Surveillance Team |
| EPI_ISL_2569241 | Lighthouse Lab in Alderley Park | Wellcome Sanger Institute for the COVID-19 Genomics UK (COG-UK) Consortium | Jacquelyn Wynn, Mairead Hyland, The Lighthouse Lab in Alderley Park and Alex Alderton, Roberto Amato, Jeffrey Barrett, Sonia Goncalves, Ewan Harrison, David K. Jackson, Ian Johnston, Dominic Kwiatkowski, Cordelia Langford, John Sillitoe on behalf of the Wellcome Sanger Institute COVID-19 Surveillance Team |
| EPI_ISL_2569242 | Lighthouse Laboratory Plymouth | Wellcome Sanger Institute for the COVID-19 Genomics UK (COG-UK) Consortium | Lighthouse Laboratory Plymouth and Alex Alderton, Roberto Amato, Jeffrey Barrett, Sonia Goncalves, Ewan Harrison, David K. Jackson, Ian Johnston, Dominic Kwiatkowski, Cordelia Langford, John Sillitoe on behalf of the Wellcome Sanger Institute COVID-19 Surveillance Team |
| EPI_ISL_2569243 | Lighthouse Lab in Glasgow | Wellcome Sanger Institute for the COVID-19 Genomics UK (COG-UK) Consortium | Harper VanSteenhouse, Yumi Kasai, David Gray, Carol Clugston, Anna Dominiczak and Alex Alderton, Roberto Amato, Jeffrey Barrett, Sonia Goncalves, Ewan Harrison, David K. Jackson, Ian Johnston, Dominic Kwiatkowski, Cordelia Langford, John Sillitoe on behalf of the Wellcome Sanger Institute COVID-19 Surveillance Team |
| EPI_ISL_2569244 | Lighthouse Lab in Alderley Park | Wellcome Sanger Institute for the COVID-19 Genomics UK (COG-UK) Consortium | Jacquelyn Wynn, Mairead Hyland, The Lighthouse Lab in Alderley Park and Alex Alderton, Roberto Amato, Jeffrey Barrett, Sonia Goncalves, Ewan Harrison, David K. Jackson, Ian Johnston, Dominic Kwiatkowski, Cordelia Langford, John Sillitoe on behalf of the Wellcome Sanger Institute COVID-19 Surveillance Team |
| EPI_ISL_2569245, EPI_ISL_2569246 | Lighthouse Laboratory Plymouth | Wellcome Sanger Institute for the COVID-19 Genomics UK (COG-UK) Consortium | Lighthouse Laboratory Plymouth and Alex Alderton, Roberto Amato, Jeffrey Barrett, Sonia Goncalves, Ewan Harrison, David K. Jackson, Ian Johnston, Dominic Kwiatkowski, Cordelia Langford, John Sillitoe on behalf of the Wellcome Sanger Institute COVID-19 Surveillance Team |
| EPI_ISL_2569248 | Lighthouse Lab in Glasgow | Wellcome Sanger Institute for the COVID-19 Genomics UK (COG-UK) Consortium | Harper VanSteenhouse, Yumi Kasai, David Gray, Carol Clugston, Anna Dominiczak and Alex Alderton, Roberto Amato, Jeffrey Barrett, Sonia Goncalves, Ewan Harrison, David K. Jackson, Ian Johnston, Dominic Kwiatkowski, Cordelia Langford, John Sillitoe on behalf of the Wellcome Sanger Institute COVID-19 Surveillance Team |
| EPI_ISL_2569249 | Lighthouse Lab in Alderley Park | Wellcome Sanger Institute for the COVID-19 Genomics UK (COG-UK) Consortium | Jacquelyn Wynn, Mairead Hyland, The Lighthouse Lab in Alderley Park and Alex Alderton, Roberto Amato, Jeffrey Barrett, Sonia Goncalves, Ewan Harrison, David K. Jackson, Ian Johnston, Dominic Kwiatkowski, Cordelia Langford, John Sillitoe on behalf of the Wellcome Sanger Institute COVID-19 Surveillance Team |
| EPI_ISL_2569250 | Lighthouse Laboratory Plymouth | Wellcome Sanger Institute for the COVID-19 Genomics UK (COG-UK) Consortium | Lighthouse Laboratory Plymouth and Alex Alderton, Roberto Amato, Jeffrey Barrett, Sonia Goncalves, Ewan Harrison, David K. Jackson, Ian Johnston, Dominic Kwiatkowski, Cordelia Langford, John Sillitoe on behalf of the Wellcome Sanger Institute COVID-19 Surveillance Team |
| EPI_ISL_2569251, EPI_ISL_2569252 | Lighthouse Lab in Alderley Park | Wellcome Sanger Institute for the COVID-19 Genomics UK (COG-UK) Consortium | Jacquelyn Wynn, Mairead Hyland, The Lighthouse Lab in Alderley Park and Alex Alderton, Roberto Amato, Jeffrey Barrett, Sonia Goncalves, Ewan Harrison, David K. Jackson, Ian Johnston, Dominic Kwiatkowski, Cordelia Langford, John Sillitoe on behalf of the Wellcome Sanger Institute COVID-19 Surveillance Team |
| EPI_ISL_2569253 | Lighthouse Lab in Glasgow | Wellcome Sanger Institute for the COVID-19 Genomics UK (COG-UK) Consortium | Harper VanSteenhouse, Yumi Kasai, David Gray, Carol Clugston, Anna Dominiczak and Alex Alderton, Roberto Amato, Jeffrey Barrett, Sonia Goncalves, Ewan Harrison, David K. Jackson, Ian Johnston, Dominic Kwiatkowski, Cordelia Langford, John Sillitoe on behalf of the Wellcome Sanger Institute COVID-19 Surveillance Team |
| EPI_ISL_2569254 | Lighthouse Laboratory Plymouth | Wellcome Sanger Institute for the COVID-19 Genomics UK (COG-UK) Consortium | Lighthouse Laboratory Plymouth and Alex Alderton, Roberto Amato, Jeffrey Barrett, Sonia Goncalves, Ewan Harrison, David K. Jackson, Ian Johnston, Dominic Kwiatkowski, Cordelia Langford, John Sillitoe on behalf of the Wellcome Sanger Institute COVID-19 Surveillance Team |
| EPI_ISL_2569255 | Lighthouse Lab in Glasgow | Wellcome Sanger Institute for the COVID-19 Genomics UK (COG-UK) Consortium | Harper VanSteenhouse, Yumi Kasai, David Gray, Carol Clugston, Anna Dominiczak and Alex Alderton, Roberto Amato, Jeffrey Barrett, Sonia Goncalves, Ewan Harrison, David K. Jackson, Ian Johnston, Dominic Kwiatkowski, Cordelia Langford, John Sillitoe on behalf of the Wellcome Sanger Institute COVID-19 Surveillance Team |
| EPI_ISL_2569256 | Lighthouse Lab in Milton Keynes | Wellcome Sanger Institute for the COVID-19 Genomics UK (COG-UK) Consortium | The Lighthouse Lab in Milton Keynes and Alex Alderton, Roberto Amato, Jeffrey Barrett, Sonia Goncalves, Ewan Harrison, David K. Jackson, Ian Johnston, Dominic Kwiatkowski, Cordelia Langford, John Sillitoe on behalf of the Wellcome Sanger Institute COVID-19 Surveillance Team |
| EPI_ISL_2569257, EPI_ISL_2569258 | Lighthouse Lab in Alderley Park | Wellcome Sanger Institute for the COVID-19 Genomics UK (COG-UK) Consortium | Jacquelyn Wynn, Mairead Hyland, The Lighthouse Lab in Alderley Park and Alex Alderton, Roberto Amato, Jeffrey Barrett, Sonia Goncalves, Ewan Harrison, David K. Jackson, Ian Johnston, Dominic Kwiatkowski, Cordelia Langford, John Sillitoe on behalf of the Wellcome Sanger Institute COVID-19 Surveillance Team |
| EPI_ISL_2569259 | Lighthouse Laboratory Plymouth | Wellcome Sanger Institute for the COVID-19 Genomics UK (COG-UK) Consortium | Lighthouse Laboratory Plymouth and Alex Alderton, Roberto Amato, Jeffrey Barrett, Sonia Goncalves, Ewan Harrison, David K. Jackson, Ian Johnston, Dominic Kwiatkowski, Cordelia Langford, John Sillitoe on behalf of the Wellcome Sanger Institute COVID-19 Surveillance Team |
| EPI_ISL_2569260 | Lighthouse Lab in Alderley Park | Wellcome Sanger Institute for the COVID-19 Genomics UK (COG-UK) Consortium | Jacquelyn Wynn, Mairead Hyland, The Lighthouse Lab in Alderley Park and Alex Alderton, Roberto Amato, Jeffrey Barrett, Sonia Goncalves, Ewan Harrison, David K. Jackson, Ian Johnston, Dominic Kwiatkowski, Cordelia Langford, John Sillitoe on behalf of the Wellcome Sanger Institute COVID-19 Surveillance Team |
| EPI_ISL_2569262 | Lighthouse Lab in Glasgow | Wellcome Sanger Institute for the COVID-19 Genomics UK (COG-UK) Consortium | Harper VanSteenhouse, Yumi Kasai, David Gray, Carol Clugston, Anna Dominiczak and Alex Alderton, Roberto Amato, Jeffrey Barrett, Sonia Goncalves, Ewan Harrison, David K. Jackson, Ian Johnston, Dominic Kwiatkowski, Cordelia Langford, John Sillitoe on behalf of the Wellcome Sanger Institute COVID-19 Surveillance Team |
| EPI_ISL_2569263, EPI_ISL_2569264, EPI_ISL_2569265, EPI_ISL_2569266 | Lighthouse Lab in Milton Keynes | Wellcome Sanger Institute for the COVID-19 Genomics UK (COG-UK) Consortium | The Lighthouse Lab in Milton Keynes and Alex Alderton, Roberto Amato, Jeffrey Barrett, Sonia Goncalves, Ewan Harrison, David K. Jackson, Ian Johnston, Dominic Kwiatkowski, Cordelia Langford, John Sillitoe on behalf of the Wellcome Sanger Institute COVID-19 Surveillance Team |
| EPI_ISL_2569267 | Lighthouse Lab in Glasgow | Wellcome Sanger Institute for the COVID-19 Genomics UK (COG-UK) Consortium | Harper VanSteenhouse, Yumi Kasai, David Gray, Carol Clugston, Anna Dominiczak and Alex Alderton, Roberto Amato, Jeffrey Barrett, Sonia Goncalves, Ewan Harrison, David K. Jackson, Ian Johnston, Dominic Kwiatkowski, Cordelia Langford, John Sillitoe on behalf of the Wellcome Sanger Institute COVID-19 Surveillance Team |
| EPI_ISL_2569268 | Lighthouse Laboratory Plymouth | Wellcome Sanger Institute for the COVID-19 Genomics UK (COG-UK) Consortium | Lighthouse Laboratory Plymouth and Alex Alderton, Roberto Amato, Jeffrey Barrett, Sonia Goncalves, Ewan Harrison, David K. Jackson, Ian Johnston, Dominic Kwiatkowski, Cordelia Langford, John Sillitoe on behalf of the Wellcome Sanger Institute COVID-19 Surveillance Team |
| EPI_ISL_2569269, EPI_ISL_2569270 | Lighthouse Lab in Milton Keynes | Wellcome Sanger Institute for the COVID-19 Genomics UK (COG-UK) Consortium | The Lighthouse Lab in Milton Keynes and Alex Alderton, Roberto Amato, Jeffrey Barrett, Sonia Goncalves, Ewan Harrison, David K. Jackson, Ian Johnston, Dominic Kwiatkowski, Cordelia Langford, John Sillitoe on behalf of the Wellcome Sanger Institute COVID-19 Surveillance Team |
| EPI_ISL_2569271, EPI_ISL_2569272 | Lighthouse Laboratory Plymouth | Wellcome Sanger Institute for the COVID-19 Genomics UK (COG-UK) Consortium | Lighthouse Laboratory Plymouth and Alex Alderton, Roberto Amato, Jeffrey Barrett, Sonia Goncalves, Ewan Harrison, David K. Jackson, Ian Johnston, Dominic Kwiatkowski, Cordelia Langford, John Sillitoe on behalf of the Wellcome Sanger Institute COVID-19 Surveillance Team |
| EPI_ISL_2569273, EPI_ISL_2569274, EPI_ISL_2569276, EPI_ISL_2569277 | Lighthouse Lab in Alderley Park | Wellcome Sanger Institute for the COVID-19 Genomics UK (COG-UK) Consortium | Jacquelyn Wynn, Mairead Hyland, The Lighthouse Lab in Alderley Park and Alex Alderton, Roberto Amato, Jeffrey Barrett, Sonia Goncalves, Ewan Harrison, David K. Jackson, Ian Johnston, Dominic Kwiatkowski, Cordelia Langford, John Sillitoe on behalf of the Wellcome Sanger Institute COVID-19 Surveillance Team |
| EPI_ISL_2569278 | Lighthouse Lab in Glasgow | Wellcome Sanger Institute for the COVID-19 Genomics UK (COG-UK) Consortium | Harper VanSteenhouse, Yumi Kasai, David Gray, Carol Clugston, Anna Dominiczak and Alex Alderton, Roberto Amato, Jeffrey Barrett, Sonia Goncalves, Ewan Harrison, David K. Jackson, Ian Johnston, Dominic Kwiatkowski, Cordelia Langford, John Sillitoe on behalf of the Wellcome Sanger Institute COVID-19 Surveillance Team |
| EPI_ISL_2569279 | Lighthouse Laboratory Plymouth | Wellcome Sanger Institute for the COVID-19 Genomics UK (COG-UK) Consortium | Lighthouse Laboratory Plymouth and Alex Alderton, Roberto Amato, Jeffrey Barrett, Sonia Goncalves, Ewan Harrison, David K. Jackson, Ian Johnston, Dominic Kwiatkowski, Cordelia Langford, John Sillitoe on behalf of the Wellcome Sanger Institute COVID-19 Surveillance Team |
| EPI_ISL_2569280 | Lighthouse Lab in Glasgow | Wellcome Sanger Institute for the COVID-19 Genomics UK (COG-UK) Consortium | Harper VanSteenhouse, Yumi Kasai, David Gray, Carol Clugston, Anna Dominiczak and Alex Alderton, Roberto Amato, Jeffrey Barrett, Sonia Goncalves, Ewan Harrison, David K. Jackson, Ian Johnston, Dominic Kwiatkowski, Cordelia Langford, John Sillitoe on behalf of the Wellcome Sanger Institute COVID-19 Surveillance Team |
| EPI_ISL_2569281, EPI_ISL_2569282 | Lighthouse Lab in Alderley Park | Wellcome Sanger Institute for the COVID-19 Genomics UK (COG-UK) Consortium | Jacquelyn Wynn, Mairead Hyland, The Lighthouse Lab in Alderley Park and Alex Alderton, Roberto Amato, Jeffrey Barrett, Sonia Goncalves, Ewan Harrison, David K. Jackson, Ian Johnston, Dominic Kwiatkowski, Cordelia Langford, John Sillitoe on behalf of the Wellcome Sanger Institute COVID-19 Surveillance Team |
| EPI_ISL_2569283 | Lighthouse Lab in Glasgow | Wellcome Sanger Institute for the COVID-19 Genomics UK (COG-UK) Consortium | Harper VanSteenhouse, Yumi Kasai, David Gray, Carol Clugston, Anna Dominiczak and Alex Alderton, Roberto Amato, Jeffrey Barrett, Sonia Goncalves, Ewan Harrison, David K. Jackson, Ian Johnston, Dominic Kwiatkowski, Cordelia Langford, John Sillitoe on behalf of the Wellcome Sanger Institute COVID-19 Surveillance Team |
| EPI_ISL_2569284 | Lighthouse Lab in Milton Keynes | Wellcome Sanger Institute for the COVID-19 Genomics UK (COG-UK) Consortium | The Lighthouse Lab in Milton Keynes and Alex Alderton, Roberto Amato, Jeffrey Barrett, Sonia Goncalves, Ewan Harrison, David K. Jackson, Ian Johnston, Dominic Kwiatkowski, Cordelia Langford, John Sillitoe on behalf of the Wellcome Sanger Institute COVID-19 Surveillance Team |
| EPI_ISL_2569285 | Lighthouse Lab in Alderley Park | Wellcome Sanger Institute for the COVID-19 Genomics UK (COG-UK) Consortium | Jacquelyn Wynn, Mairead Hyland, The Lighthouse Lab in Alderley Park and Alex Alderton, Roberto Amato, Jeffrey Barrett, Sonia Goncalves, Ewan Harrison, David K. Jackson, Ian Johnston, Dominic Kwiatkowski, Cordelia Langford, John Sillitoe on behalf of the Wellcome Sanger Institute COVID-19 Surveillance Team |
| EPI_ISL_2569286, EPI_ISL_2569287 | Lighthouse Lab in Milton Keynes | Wellcome Sanger Institute for the COVID-19 Genomics UK (COG-UK) Consortium | The Lighthouse Lab in Milton Keynes and Alex Alderton, Roberto Amato, Jeffrey Barrett, Sonia Goncalves, Ewan Harrison, David K. Jackson, Ian Johnston, Dominic Kwiatkowski, Cordelia Langford, John Sillitoe on behalf of the Wellcome Sanger Institute COVID-19 Surveillance Team |
| EPI_ISL_2569288 | Lighthouse Lab in Glasgow | Wellcome Sanger Institute for the COVID-19 Genomics UK (COG-UK) Consortium | Harper VanSteenhouse, Yumi Kasai, David Gray, Carol Clugston, Anna Dominiczak and Alex Alderton, Roberto Amato, Jeffrey Barrett, Sonia Goncalves, Ewan Harrison, David K. Jackson, Ian Johnston, Dominic Kwiatkowski, Cordelia Langford, John Sillitoe on behalf of the Wellcome Sanger Institute COVID-19 Surveillance Team |
| EPI_ISL_2569290, EPI_ISL_2569291 | Lighthouse Laboratory Plymouth | Wellcome Sanger Institute for the COVID-19 Genomics UK (COG-UK) Consortium | Lighthouse Laboratory Plymouth and Alex Alderton, Roberto Amato, Jeffrey Barrett, Sonia Goncalves, Ewan Harrison, David K. Jackson, Ian Johnston, Dominic Kwiatkowski, Cordelia Langford, John Sillitoe on behalf of the Wellcome Sanger Institute COVID-19 Surveillance Team |
| EPI_ISL_2569292, EPI_ISL_2569293, EPI_ISL_2569294 | Lighthouse Lab in Alderley Park | Wellcome Sanger Institute for the COVID-19 Genomics UK (COG-UK) Consortium | Jacquelyn Wynn, Mairead Hyland, The Lighthouse Lab in Alderley Park and Alex Alderton, Roberto Amato, Jeffrey Barrett, Sonia Goncalves, Ewan Harrison, David K. Jackson, Ian Johnston, Dominic Kwiatkowski, Cordelia Langford, John Sillitoe on behalf of the Wellcome Sanger Institute COVID-19 Surveillance Team |
| EPI_ISL_2569295 | Lighthouse Lab in Glasgow | Wellcome Sanger Institute for the COVID-19 Genomics UK (COG-UK) Consortium | Harper VanSteenhouse, Yumi Kasai, David Gray, Carol Clugston, Anna Dominiczak and Alex Alderton, Roberto Amato, Jeffrey Barrett, Sonia Goncalves, Ewan Harrison, David K. Jackson, Ian Johnston, Dominic Kwiatkowski, Cordelia Langford, John Sillitoe on behalf of the Wellcome Sanger Institute COVID-19 Surveillance Team |
| EPI_ISL_2569296 | Lighthouse Laboratory Plymouth | Wellcome Sanger Institute for the COVID-19 Genomics UK (COG-UK) Consortium | Lighthouse Laboratory Plymouth and Alex Alderton, Roberto Amato, Jeffrey Barrett, Sonia Goncalves, Ewan Harrison, David K. Jackson, Ian Johnston, Dominic Kwiatkowski, Cordelia Langford, John Sillitoe on behalf of the Wellcome Sanger Institute COVID-19 Surveillance Team |
| EPI_ISL_2569297, EPI_ISL_2569298 | Lighthouse Lab in Milton Keynes | Wellcome Sanger Institute for the COVID-19 Genomics UK (COG-UK) Consortium | The Lighthouse Lab in Milton Keynes and Alex Alderton, Roberto Amato, Jeffrey Barrett, Sonia Goncalves, Ewan Harrison, David K. Jackson, Ian Johnston, Dominic Kwiatkowski, Cordelia Langford, John Sillitoe on behalf of the Wellcome Sanger Institute COVID-19 Surveillance Team |
| EPI_ISL_2569299 | Lighthouse Lab in Alderley Park | Wellcome Sanger Institute for the COVID-19 Genomics UK (COG-UK) Consortium | Jacquelyn Wynn, Mairead Hyland, The Lighthouse Lab in Alderley Park and Alex Alderton, Roberto Amato, Jeffrey Barrett, Sonia Goncalves, Ewan Harrison, David K. Jackson, Ian Johnston, Dominic Kwiatkowski, Cordelia Langford, John Sillitoe on behalf of the Wellcome Sanger Institute COVID-19 Surveillance Team |
| EPI_ISL_2569300 | Lighthouse Lab in Milton Keynes | Wellcome Sanger Institute for the COVID-19 Genomics UK (COG-UK) Consortium | The Lighthouse Lab in Milton Keynes and Alex Alderton, Roberto Amato, Jeffrey Barrett, Sonia Goncalves, Ewan Harrison, David K. Jackson, Ian Johnston, Dominic Kwiatkowski, Cordelia Langford, John Sillitoe on behalf of the Wellcome Sanger Institute COVID-19 Surveillance Team |
| EPI_ISL_2569301 | Lighthouse Lab in Glasgow | Wellcome Sanger Institute for the COVID-19 Genomics UK (COG-UK) Consortium | Harper VanSteenhouse, Yumi Kasai, David Gray, Carol Clugston, Anna Dominiczak and Alex Alderton, Roberto Amato, Jeffrey Barrett, Sonia Goncalves, Ewan Harrison, David K. Jackson, Ian Johnston, Dominic Kwiatkowski, Cordelia Langford, John Sillitoe on behalf of the Wellcome Sanger Institute COVID-19 Surveillance Team |
| EPI_ISL_2569302 | Lighthouse Laboratory Plymouth | Wellcome Sanger Institute for the COVID-19 Genomics UK (COG-UK) Consortium | Lighthouse Laboratory Plymouth and Alex Alderton, Roberto Amato, Jeffrey Barrett, Sonia Goncalves, Ewan Harrison, David K. Jackson, Ian Johnston, Dominic Kwiatkowski, Cordelia Langford, John Sillitoe on behalf of the Wellcome Sanger Institute COVID-19 Surveillance Team |
| EPI_ISL_2569303 | Lighthouse Lab in Alderley Park | Wellcome Sanger Institute for the COVID-19 Genomics UK (COG-UK) Consortium | Jacquelyn Wynn, Mairead Hyland, The Lighthouse Lab in Alderley Park and Alex Alderton, Roberto Amato, Jeffrey Barrett, Sonia Goncalves, Ewan Harrison, David K. Jackson, Ian Johnston, Dominic Kwiatkowski, Cordelia Langford, John Sillitoe on behalf of the Wellcome Sanger Institute COVID-19 Surveillance Team |

| | | | |
|---|---|---|---|
| EPI_ISL_2569304 | Lighthouse Lab in Milton Keynes | Wellcome Sanger Institute for the COVID-19 Genomics UK (COG-UK) Consortium | The Lighthouse Lab in Milton Keynes and Alex Alderton, Roberto Amato, Jeffrey Barrett, Sonia Goncalves, Ewan Harrison, David K. Jackson, Ian Johnston, Dominic Kwiatkowski, Cordelia Langford, John Sillitoe on behalf of the Wellcome Sanger Institute COVID-19 Surveillance Team |
| EPI_ISL_2569305, EPI_ISL_2569306 | Lighthouse Laboratory Plymouth | Wellcome Sanger Institute for the COVID-19 Genomics UK (COG-UK) Consortium | Lighthouse Laboratory Plymouth and Alex Alderton, Roberto Amato, Jeffrey Barrett, Sonia Goncalves, Ewan Harrison, David K. Jackson, Ian Johnston, Dominic Kwiatkowski, Cordelia Langford, John Sillitoe on behalf of the Wellcome Sanger Institute COVID-19 Surveillance Team |
| EPI_ISL_2569307 | Lighthouse Lab in Alderley Park | Wellcome Sanger Institute for the COVID-19 Genomics UK (COG-UK) Consortium | Jacquelyn Wynn, Mairead Hyland, The Lighthouse Lab in Alderley Park and Alex Alderton, Roberto Amato, Jeffrey Barrett, Sonia Goncalves, Ewan Harrison, David K. Jackson, Ian Johnston, Dominic Kwiatkowski, Cordelia Langford, John Sillitoe on behalf of the Wellcome Sanger Institute COVID-19 Surveillance Team |
| EPI_ISL_2569308 | Lighthouse Lab in Milton Keynes | Wellcome Sanger Institute for the COVID-19 Genomics UK (COG-UK) Consortium | The Lighthouse Lab in Milton Keynes and Alex Alderton, Roberto Amato, Jeffrey Barrett, Sonia Goncalves, Ewan Harrison, David K. Jackson, Ian Johnston, Dominic Kwiatkowski, Cordelia Langford, John Sillitoe on behalf of the Wellcome Sanger Institute COVID-19 Surveillance Team |
| EPI_ISL_2569309 | Lighthouse Lab in Alderley Park | Wellcome Sanger Institute for the COVID-19 Genomics UK (COG-UK) Consortium | Jacquelyn Wynn, Mairead Hyland, The Lighthouse Lab in Alderley Park and Alex Alderton, Roberto Amato, Jeffrey Barrett, Sonia Goncalves, Ewan Harrison, David K. Jackson, Ian Johnston, Dominic Kwiatkowski, Cordelia Langford, John Sillitoe on behalf of the Wellcome Sanger Institute COVID-19 Surveillance Team |
| EPI_ISL_2569310 | Lighthouse Laboratory Plymouth | Wellcome Sanger Institute for the COVID-19 Genomics UK (COG-UK) Consortium | Lighthouse Laboratory Plymouth and Alex Alderton, Roberto Amato, Jeffrey Barrett, Sonia Goncalves, Ewan Harrison, David K. Jackson, Ian Johnston, Dominic Kwiatkowski, Cordelia Langford, John Sillitoe on behalf of the Wellcome Sanger Institute COVID-19 Surveillance Team |
| EPI_ISL_2569311 | Lighthouse Lab in Alderley Park | Wellcome Sanger Institute for the COVID-19 Genomics UK (COG-UK) Consortium | Jacquelyn Wynn, Mairead Hyland, The Lighthouse Lab in Alderley Park and Alex Alderton, Roberto Amato, Jeffrey Barrett, Sonia Goncalves, Ewan Harrison, David K. Jackson, Ian Johnston, Dominic Kwiatkowski, Cordelia Langford, John Sillitoe on behalf of the Wellcome Sanger Institute COVID-19 Surveillance Team |
| EPI_ISL_2569312 | Lighthouse Lab in Glasgow | Wellcome Sanger Institute for the COVID-19 Genomics UK (COG-UK) Consortium | Harper VanSteenhouse, Yumi Kasai, David Gray, Carol Clugston, Anna Dominiczak and Alex Alderton, Roberto Amato, Jeffrey Barrett, Sonia Goncalves, Ewan Harrison, David K. Jackson, Ian Johnston, Dominic Kwiatkowski, Cordelia Langford, John Sillitoe on behalf of the Wellcome Sanger Institute COVID-19 Surveillance Team |
| EPI_ISL_2569313 | Lighthouse Laboratory Plymouth | Wellcome Sanger Institute for the COVID-19 Genomics UK (COG-UK) Consortium | Lighthouse Laboratory Plymouth and Alex Alderton, Roberto Amato, Jeffrey Barrett, Sonia Goncalves, Ewan Harrison, David K. Jackson, Ian Johnston, Dominic Kwiatkowski, Cordelia Langford, John Sillitoe on behalf of the Wellcome Sanger Institute COVID-19 Surveillance Team |
| EPI_ISL_2569314 | Lighthouse Lab in Glasgow | Wellcome Sanger Institute for the COVID-19 Genomics UK (COG-UK) Consortium | Harper VanSteenhouse, Yumi Kasai, David Gray, Carol Clugston, Anna Dominiczak and Alex Alderton, Roberto Amato, Jeffrey Barrett, Sonia Goncalves, Ewan Harrison, David K. Jackson, Ian Johnston, Dominic Kwiatkowski, Cordelia Langford, John Sillitoe on behalf of the Wellcome Sanger Institute COVID-19 Surveillance Team |
| EPI_ISL_2569315 | Lighthouse Laboratory Plymouth | Wellcome Sanger Institute for the COVID-19 Genomics UK (COG-UK) Consortium | Lighthouse Laboratory Plymouth and Alex Alderton, Roberto Amato, Jeffrey Barrett, Sonia Goncalves, Ewan Harrison, David K. Jackson, Ian Johnston, Dominic Kwiatkowski, Cordelia Langford, John Sillitoe on behalf of the Wellcome Sanger Institute COVID-19 Surveillance Team |
| EPI_ISL_2569316 | Lighthouse Lab in Alderley Park | Wellcome Sanger Institute for the COVID-19 Genomics UK (COG-UK) Consortium | Jacquelyn Wynn, Mairead Hyland, The Lighthouse Lab in Alderley Park and Alex Alderton, Roberto Amato, Jeffrey Barrett, Sonia Goncalves, Ewan Harrison, David K. Jackson, Ian Johnston, Dominic Kwiatkowski, Cordelia Langford, John Sillitoe on behalf of the Wellcome Sanger Institute COVID-19 Surveillance Team |
| EPI_ISL_2569317 | Lighthouse Lab in Milton Keynes | Wellcome Sanger Institute for the COVID-19 Genomics UK (COG-UK) Consortium | The Lighthouse Lab in Milton Keynes and Alex Alderton, Roberto Amato, Jeffrey Barrett, Sonia Goncalves, Ewan Harrison, David K. Jackson, Ian Johnston, Dominic Kwiatkowski, Cordelia Langford, John Sillitoe on behalf of the Wellcome Sanger Institute COVID-19 Surveillance Team |
| EPI_ISL_2569318, EPI_ISL_2569319, EPI_ISL_2569320, EPI_ISL_2569321, EPI_ISL_2569322, EPI_ISL_2569323, EPI_ISL_2569324, EPI_ISL_2569325, EPI_ISL_2569326, EPI_ISL_2569327, EPI_ISL_2569328, EPI_ISL_2569329, EPI_ISL_2569330, EPI_ISL_2569331, EPI_ISL_2569332, EPI_ISL_2569333, EPI_ISL_2569334, EPI_ISL_2569335, EPI_ISL_2569336, EPI_ISL_2569337, EPI_ISL_2569338, EPI_ISL_2569339, EPI_ISL_2569340, EPI_ISL_2569341, EPI_ISL_2569342, EPI_ISL_2569343, EPI_ISL_2569344, EPI_ISL_2569345, EPI_ISL_2569346, EPI_ISL_2569347, EPI_ISL_2569348, EPI_ISL_2569349, EPI_ISL_2569350, EPI_ISL_2569351, EPI_ISL_2569352, EPI_ISL_2569353, EPI_ISL_2569354, EPI_ISL_2569355, EPI_ISL_2569356, EPI_ISL_2569357, EPI_ISL_2569358, EPI_ISL_2569359, EPI_ISL_2569360, EPI_ISL_2569361, EPI_ISL_2569362, EPI_ISL_2569363, EPI_ISL_2569364, EPI_ISL_2569365, EPI_ISL_2569366, EPI_ISL_2569367, EPI_ISL_2569368, EPI_ISL_2569369, EPI_ISL_2569370, EPI_ISL_2569371, EPI_ISL_2569372, EPI_ISL_2569373, EPI_ISL_2569374, EPI_ISL_2569375, EPI_ISL_2569376, EPI_ISL_2569378, EPI_ISL_2569379, EPI_ISL_2569380, EPI_ISL_2569381, EPI_ISL_2569383, EPI_ISL_2569384, EPI_ISL_2569385, EPI_ISL_2569386, EPI_ISL_2569387, EPI_ISL_2569388, EPI_ISL_2569389, EPI_ISL_2569390, EPI_ISL_2569391, EPI_ISL_2569394, EPI_ISL_2569395, EPI_ISL_2569396, EPI_ISL_2569397, EPI_ISL_2569398, EPI_ISL_2569399, EPI_ISL_2569400, EPI_ISL_2569401, EPI_ISL_2569403, EPI_ISL_2569404, EPI_ISL_2569405, EPI_ISL_2569406, EPI_ISL_2569407, EPI_ISL_2569408, EPI_ISL_2569409, EPI_ISL_2569410, EPI_ISL_2569411, EPI_ISL_2569412, EPI_ISL_2569413, EPI_ISL_2569414, EPI_ISL_2569415, EPI_ISL_2569416, EPI_ISL_2569417, EPI_ISL_2569418, EPI_ISL_2569419, EPI_ISL_2569420, EPI_ISL_2569421, EPI_ISL_2569422, EPI_ISL_2569423, EPI_ISL_2569424, EPI_ISL_2569425, EPI_ISL_2569426, EPI_ISL_2569427, EPI_ISL_2569428, EPI_ISL_2569429, EPI_ISL_2569430, EPI_ISL_2569431, EPI_ISL_2569432, EPI_ISL_2569433, EPI_ISL_2569434, EPI_ISL_2569435, EPI_ISL_2569436, EPI_ISL_2569437, EPI_ISL_2569438, EPI_ISL_2569439, EPI_ISL_2569440, EPI_ISL_2569441, EPI_ISL_2569442, EPI_ISL_2569444, EPI_ISL_2569445, EPI_ISL_2569446, EPI_ISL_2569447, EPI_ISL_2569448, EPI_ISL_2569449, EPI_ISL_2569450, EPI_ISL_2569451, EPI_ISL_2569452, EPI_ISL_2569453, EPI_ISL_2569454, EPI_ISL_2569455, EPI_ISL_2569456, EPI_ISL_2569457, EPI_ISL_2569458, EPI_ISL_2569459, EPI_ISL_2569460, EPI_ISL_2569461, EPI_ISL_2569462, EPI_ISL_2569463, EPI_ISL_2569464, EPI_ISL_2569465, EPI_ISL_2569466, EPI_ISL_2569467, EPI_ISL_2569468, EPI_ISL_2569469, EPI_ISL_2569470, EPI_ISL_2569471, EPI_ISL_2569472, EPI_ISL_2569473, EPI_ISL_2569474, EPI_ISL_2569475, EPI_ISL_2569476, EPI_ISL_2569477, EPI_ISL_2569478, EPI_ISL_2569479, EPI_ISL_2569480, EPI_ISL_2569481, EPI_ISL_2569482, EPI_ISL_2569483, EPI_ISL_2569484, EPI_ISL_2569485, EPI_ISL_2569486, EPI_ISL_2569487, EPI_ISL_2569488, EPI_ISL_2569489, EPI_ISL_2569490, EPI_ISL_2569491, EPI_ISL_2569492, EPI_ISL_2569493, EPI_ISL_2569494, EPI_ISL_2569495, EPI_ISL_2569496, EPI_ISL_2569497, EPI_ISL_2569498, EPI_ISL_2569499, EPI_ISL_2569500, EPI_ISL_2569501, EPI_ISL_2569502, EPI_ISL_2569503, EPI_ISL_2569504, EPI_ISL_2569505, EPI_ISL_2569506, EPI_ISL_2569507, EPI_ISL_2569508, EPI_ISL_2569509, EPI_ISL_2569510, EPI_ISL_2569511, EPI_ISL_2569512, EPI_ISL_2569513, EPI_ISL_2569514, EPI_ISL_2569515, EPI_ISL_2569516, EPI_ISL_2569517, EPI_ISL_2569518, EPI_ISL_2569519, EPI_ISL_2569520, EPI_ISL_2569521, EPI_ISL_2569522, EPI_ISL_2569523, EPI_ISL_2569524, EPI_ISL_2569525, EPI_ISL_2569526, EPI_ISL_2569527, EPI_ISL_2569528, EPI_ISL_2569529, EPI_ISL_2569530, EPI_ISL_2569531, EPI_ISL_2569532, EPI_ISL_2569533, EPI_ISL_2569534, EPI_ISL_2569535, EPI_ISL_2569536, EPI_ISL_2569537, EPI_ISL_2569538, EPI_ISL_2569539, EPI_ISL_2569540, EPI_ISL_2569541, EPI_ISL_2569542, EPI_ISL_2569543, EPI_ISL_2569544, EPI_ISL_2569545, EPI_ISL_2569546, EPI_ISL_2569547, EPI_ISL_2569548, EPI_ISL_2569549, EPI_ISL_2569550, EPI_ISL_2569551, EPI_ISL_2569552, EPI_ISL_2569553, EPI_ISL_2569554, EPI_ISL_2569555, EPI_ISL_2569556, EPI_ISL_2569557, EPI_ISL_2569558, EPI_ISL_2569559, EPI_ISL_2569560, EPI_ISL_2569561, EPI_ISL_2569562, EPI_ISL_2569563, EPI_ISL_2569564, EPI_ISL_2569565, EPI_ISL_2569566, EPI_ISL_2569567, EPI_ISL_2569568, EPI_ISL_2569569, EPI_ISL_2569570, EPI_ISL_2569571, EPI_ISL_2569572, EPI_ISL_2569573, EPI_ISL_2569574, EPI_ISL_2569575, EPI_ISL_2569576, EPI_ISL_2569577, EPI_ISL_2569578, EPI_ISL_2569579, EPI_ISL_2569580, EPI_ISL_2569581, EPI_ISL_2569582, EPI_ISL_2569583, EPI_ISL_2569584, EPI_ISL_2569585, EPI_ISL_2569586, EPI_ISL_2569587, EPI_ISL_2569588, EPI_ISL_2569589, EPI_ISL_2569590, EPI_ISL_2569591, EPI_ISL_2569592, EPI_ISL_2569593, EPI_ISL_2569594, EPI_ISL_2569595, EPI_ISL_2569596, EPI_ISL_2569597, EPI_ISL_2569598, EPI_ISL_2569599, EPI_ISL_2569600, EPI_ISL_2569601, EPI_ISL_2569602, EPI_ISL_2569603, EPI_ISL_2569604, EPI_ISL_2569605, EPI_ISL_2569606, EPI_ISL_2569607, EPI_ISL_2569608, EPI_ISL_2569609, EPI_ISL_2569610, EPI_ISL_2569611, EPI_ISL_2569612, EPI_ISL_2569613, EPI_ISL_2569614, EPI_ISL_2569615, EPI_ISL_2569616, EPI_ISL_2569617, EPI_ISL_2569618, EPI_ISL_2569619, EPI_ISL_2569620, EPI_ISL_2569621, EPI_ISL_2569622, EPI_ISL_2569623, EPI_ISL_2569624, EPI_ISL_2569625, EPI_ISL_2569626, EPI_ISL_2569627, EPI_ISL_2569628, EPI_ISL_2569629, EPI_ISL_2569630, EPI_ISL_2569631, EPI_ISL_2569633, EPI_ISL_2569634, EPI_ISL_2569635, EPI_ISL_2569636, EPI_ISL_2569637, EPI_ISL_2569638, EPI_ISL_2569639, EPI_ISL_2569640, EPI_ISL_2569641, EPI_ISL_2569642, EPI_ISL_2569643, EPI_ISL_2569644, EPI_ISL_2569645, EPI_ISL_2569646, EPI_ISL_2569647, EPI_ISL_2569648, EPI_ISL_2569649, EPI_ISL_2569651, EPI_ISL_2569652, EPI_ISL_2569653, EPI_ISL_2569654, EPI_ISL_2569655, EPI_ISL_2569656, EPI_ISL_2569657, EPI_ISL_2569658, EPI_ISL_2569659, EPI_ISL_2569660, EPI_ISL_2569661, EPI_ISL_2569662 | | | |
| see above | Lighthouse Lab in Alderley Park | Wellcome Sanger Institute for the COVID-19 Genomics UK (COG-UK) Consortium | Jacquelyn Wynn, Mairead Hyland, The Lighthouse Lab in Alderley Park and Alex Alderton, Roberto Amato, Jeffrey Barrett, Sonia Goncalves, Ewan Harrison, David K. Jackson, Ian Johnston, Dominic Kwiatkowski, Cordelia Langford, John Sillitoe on behalf of the Wellcome Sanger Institute COVID-19 Surveillance Team |
| EPI_ISL_2569663 | Lighthouse Lab in Milton Keynes | Wellcome Sanger Institute for the COVID-19 Genomics UK (COG-UK) Consortium | The Lighthouse Lab in Milton Keynes and Alex Alderton, Roberto Amato, Jeffrey Barrett, Sonia Goncalves, Ewan Harrison, David K. Jackson, Ian Johnston, Dominic Kwiatkowski, Cordelia Langford, John Sillitoe on behalf of the Wellcome Sanger Institute COVID-19 Surveillance Team |
| EPI_ISL_2569664 | Lighthouse Lab in Alderley Park | Wellcome Sanger Institute for the COVID-19 Genomics UK (COG-UK) Consortium | Jacquelyn Wynn, Mairead Hyland, The Lighthouse Lab in Alderley Park and Alex Alderton, Roberto Amato, Jeffrey Barrett, Sonia Goncalves, Ewan Harrison, David K. Jackson, Ian Johnston, Dominic Kwiatkowski, Cordelia Langford, John Sillitoe on behalf of the Wellcome Sanger Institute COVID-19 Surveillance Team |
| EPI_ISL_2569665 | Lighthouse Laboratory Plymouth | Wellcome Sanger Institute for the COVID-19 Genomics UK (COG-UK) Consortium | Lighthouse Laboratory Plymouth and Alex Alderton, Roberto Amato, Jeffrey Barrett, Sonia Goncalves, Ewan Harrison, David K. Jackson, Ian Johnston, Dominic Kwiatkowski, Cordelia Langford, John Sillitoe on behalf of the Wellcome Sanger Institute COVID-19 Surveillance Team |
| EPI_ISL_2569666, EPI_ISL_2569667, EPI_ISL_2569668 | Lighthouse Lab in Milton Keynes | Wellcome Sanger Institute for the COVID-19 Genomics UK (COG-UK) Consortium | The Lighthouse Lab in Milton Keynes and Alex Alderton, Roberto Amato, Jeffrey Barrett, Sonia Goncalves, Ewan Harrison, David K. Jackson, Ian Johnston, Dominic Kwiatkowski, Cordelia Langford, John Sillitoe on behalf of the Wellcome Sanger Institute COVID-19 Surveillance Team |
| EPI_ISL_2569669 | Lighthouse Laboratory Plymouth | Wellcome Sanger Institute for the COVID-19 Genomics UK (COG-UK) Consortium | Lighthouse Laboratory Plymouth and Alex Alderton, Roberto Amato, Jeffrey Barrett, Sonia Goncalves, Ewan Harrison, David K. Jackson, Ian Johnston, Dominic Kwiatkowski, Cordelia Langford, John Sillitoe on behalf of the Wellcome Sanger Institute COVID-19 Surveillance Team |
| EPI_ISL_2569670, EPI_ISL_2569671, EPI_ISL_2569672 | Lighthouse Lab in Milton Keynes | Wellcome Sanger Institute for the COVID-19 Genomics UK (COG-UK) Consortium | The Lighthouse Lab in Milton Keynes and Alex Alderton, Roberto Amato, Jeffrey Barrett, Sonia Goncalves, Ewan Harrison, David K. Jackson, Ian Johnston, Dominic Kwiatkowski, Cordelia Langford, John Sillitoe on behalf of the Wellcome Sanger Institute COVID-19 Surveillance Team |
| EPI_ISL_2569673 | Lighthouse Lab in Alderley Park | Wellcome Sanger Institute for the COVID-19 Genomics UK (COG-UK) Consortium | Jacquelyn Wynn, Mairead Hyland, The Lighthouse Lab in Alderley Park and Alex Alderton, Roberto Amato, Jeffrey Barrett, Sonia Goncalves, Ewan Harrison, David K. Jackson, Ian Johnston, Dominic Kwiatkowski, Cordelia Langford, John Sillitoe on behalf of the Wellcome Sanger Institute COVID-19 Surveillance Team |
| EPI_ISL_2569674 | Lighthouse Laboratory Plymouth | Wellcome Sanger Institute for the COVID-19 Genomics UK (COG-UK) Consortium | Lighthouse Laboratory Plymouth and Alex Alderton, Roberto Amato, Jeffrey Barrett, Sonia Goncalves, Ewan Harrison, David K. Jackson, Ian Johnston, Dominic Kwiatkowski, Cordelia Langford, John Sillitoe on behalf of the Wellcome Sanger Institute COVID-19 Surveillance Team |
| EPI_ISL_2569675 | Lighthouse Lab in Alderley Park | Wellcome Sanger Institute for the COVID-19 Genomics UK (COG-UK) Consortium | Jacquelyn Wynn, Mairead Hyland, The Lighthouse Lab in Alderley Park and Alex Alderton, Roberto Amato, Jeffrey Barrett, Sonia Goncalves, Ewan Harrison, David K. Jackson, Ian Johnston, Dominic Kwiatkowski, Cordelia Langford, John Sillitoe on behalf of the Wellcome Sanger Institute COVID-19 Surveillance Team |
| EPI_ISL_2569676 | Lighthouse Lab in Milton Keynes | Wellcome Sanger Institute for the COVID-19 Genomics UK (COG-UK) Consortium | The Lighthouse Lab in Milton Keynes and Alex Alderton, Roberto Amato, Jeffrey Barrett, Sonia Goncalves, Ewan Harrison, David K. Jackson, Ian Johnston, Dominic Kwiatkowski, Cordelia Langford, John Sillitoe on behalf of the Wellcome Sanger Institute COVID-19 Surveillance Team |
| EPI_ISL_2569677, EPI_ISL_2569678 | Lighthouse Lab in Alderley Park | Wellcome Sanger Institute for the COVID-19 Genomics UK (COG-UK) Consortium | Jacquelyn Wynn, Mairead Hyland, The Lighthouse Lab in Alderley Park and Alex Alderton, Roberto Amato, Jeffrey Barrett, Sonia Goncalves, Ewan Harrison, David K. Jackson, Ian Johnston, Dominic Kwiatkowski, Cordelia Langford, John Sillitoe on behalf of the Wellcome Sanger Institute COVID-19 Surveillance Team |
| EPI_ISL_2569679, EPI_ISL_2569680, EPI_ISL_2569681 | Lighthouse Lab in Milton Keynes | Wellcome Sanger Institute for the COVID-19 Genomics UK (COG-UK) Consortium | The Lighthouse Lab in Milton Keynes and Alex Alderton, Roberto Amato, Jeffrey Barrett, Sonia Goncalves, Ewan Harrison, David K. Jackson, Ian Johnston, Dominic Kwiatkowski, Cordelia Langford, John Sillitoe on behalf of the Wellcome Sanger Institute COVID-19 Surveillance Team |
| EPI_ISL_2569682 | Lighthouse Laboratory Plymouth | Wellcome Sanger Institute for the COVID-19 Genomics UK (COG-UK) Consortium | Lighthouse Laboratory Plymouth and Alex Alderton, Roberto Amato, Jeffrey Barrett, Sonia Goncalves, Ewan Harrison, David K. Jackson, Ian Johnston, Dominic Kwiatkowski, Cordelia Langford, John Sillitoe on behalf of the Wellcome Sanger Institute COVID-19 Surveillance Team |
| EPI_ISL_2569683 | Lighthouse Lab in Milton Keynes | Wellcome Sanger Institute for the COVID-19 Genomics UK (COG-UK) Consortium | The Lighthouse Lab in Milton Keynes and Alex Alderton, Roberto Amato, Jeffrey Barrett, Sonia Goncalves, Ewan Harrison, David K. Jackson, Ian Johnston, Dominic Kwiatkowski, Cordelia Langford, John Sillitoe on behalf of the Wellcome Sanger Institute COVID-19 Surveillance Team |
| EPI_ISL_2569684 | Lighthouse Laboratory Plymouth | Wellcome Sanger Institute for the COVID-19 Genomics UK (COG-UK) Consortium | Lighthouse Laboratory Plymouth and Alex Alderton, Roberto Amato, Jeffrey Barrett, Sonia Goncalves, Ewan Harrison, David K. Jackson, Ian Johnston, Dominic Kwiatkowski, Cordelia Langford, John Sillitoe on behalf of the Wellcome Sanger Institute COVID-19 Surveillance Team |
| EPI_ISL_2569685, EPI_ISL_2569686, EPI_ISL_2569687 | Lighthouse Lab in Milton Keynes | Wellcome Sanger Institute for the COVID-19 Genomics UK (COG-UK) Consortium | The Lighthouse Lab in Milton Keynes and Alex Alderton, Roberto Amato, Jeffrey Barrett, Sonia Goncalves, Ewan Harrison, David K. Jackson, Ian Johnston, Dominic Kwiatkowski, Cordelia Langford, John Sillitoe on behalf of the Wellcome Sanger Institute COVID-19 Surveillance Team |
| EPI_ISL_2569688 | Lighthouse Lab in Glasgow | Wellcome Sanger Institute for the COVID-19 Genomics UK (COG-UK) Consortium | Harper VanSteenhouse, Yumi Kasai, David Gray, Carol Clugston, Anna Dominiczak and Alex Alderton, Roberto Amato, Jeffrey Barrett, Sonia Goncalves, Ewan Harrison, David K. Jackson, Ian Johnston, Dominic Kwiatkowski, Cordelia Langford, John Sillitoe on behalf of the Wellcome Sanger Institute COVID-19 Surveillance Team |
| EPI_ISL_2569689, EPI_ISL_2569690 | Lighthouse Lab in Alderley Park | Wellcome Sanger Institute for the COVID-19 Genomics UK (COG-UK) Consortium | Jacquelyn Wynn, Mairead Hyland, The Lighthouse Lab in Alderley Park and Alex Alderton, Roberto Amato, Jeffrey Barrett, Sonia Goncalves, Ewan Harrison, David K. Jackson, Ian Johnston, Dominic Kwiatkowski, Cordelia Langford, John Sillitoe on behalf of the Wellcome Sanger Institute COVID-19 Surveillance Team |
| EPI_ISL_2569691 | Lighthouse Lab in Milton Keynes | Wellcome Sanger Institute for the COVID-19 Genomics UK (COG-UK) Consortium | The Lighthouse Lab in Milton Keynes and Alex Alderton, Roberto Amato, Jeffrey Barrett, Sonia Goncalves, Ewan Harrison, David K. Jackson, Ian Johnston, Dominic Kwiatkowski, Cordelia Langford, John Sillitoe on behalf of the Wellcome Sanger Institute COVID-19 Surveillance Team |
| EPI_ISL_2569692 | Lighthouse Lab in Alderley Park | Wellcome Sanger Institute for the COVID-19 Genomics UK (COG-UK) Consortium | Jacquelyn Wynn, Mairead Hyland, The Lighthouse Lab in Alderley Park and Alex Alderton, Roberto Amato, Jeffrey Barrett, Sonia Goncalves, Ewan Harrison, David K. Jackson, Ian Johnston, Dominic Kwiatkowski, Cordelia Langford, John Sillitoe on behalf of the Wellcome Sanger Institute COVID-19 Surveillance Team |
| EPI_ISL_2569693 | Lighthouse Lab in Milton Keynes | Wellcome Sanger Institute for the COVID-19 Genomics UK (COG-UK) Consortium | The Lighthouse Lab in Milton Keynes and Alex Alderton, Roberto Amato, Jeffrey Barrett, Sonia Goncalves, Ewan Harrison, David K. Jackson, Ian Johnston, Dominic Kwiatkowski, Cordelia Langford, John Sillitoe on behalf of the Wellcome Sanger Institute COVID-19 Surveillance Team |
| EPI_ISL_2569694 | Lighthouse Laboratory Plymouth | Wellcome Sanger Institute for the COVID-19 Genomics UK (COG-UK) Consortium | Lighthouse Laboratory Plymouth and Alex Alderton, Roberto Amato, Jeffrey Barrett, Sonia Goncalves, Ewan Harrison, David K. Jackson, Ian Johnston, Dominic Kwiatkowski, Cordelia Langford, John Sillitoe on behalf of the Wellcome Sanger Institute COVID-19 Surveillance Team |
| EPI_ISL_2569695 | Lighthouse Lab in Alderley Park | Wellcome Sanger Institute for the COVID-19 Genomics UK (COG-UK) | Jacquelyn Wynn, Mairead Hyland, The Lighthouse Lab in Alderley Park and Alex Alderton, Roberto Amato, Jeffrey Barrett, Sonia Goncalves, Ewan Harrison, David K. Jackson, Ian Johnston, Dominic Kwiatkowski, Cordelia Langford, John Sillitoe on |

| | | | |
|---|---|---|---|
| | | Consortium | behalf of the Wellcome Sanger Institute COVID-19 Surveillance Team |
| EPI_ISL_2569696 | Lighthouse Lab in Glasgow | Wellcome Sanger Institute for the COVID-19 Genomics UK (COG-UK) Consortium | Harper VanSteenhouse, Yumi Kasai, David Gray, Carol Clugston, Anna Dominiczak and Alex Alderton, Roberto Amato, Jeffrey Barrett, Sonia Goncalves, Ewan Harrison, David K. Jackson, Ian Johnston, Dominic Kwiatkowski, Cordelia Langford, John Sillitoe on behalf of the Wellcome Sanger Institute COVID-19 Surveillance Team |
| EPI_ISL_2569697, EPI_ISL_2569698 | Lighthouse Lab in Alderley Park | Wellcome Sanger Institute for the COVID-19 Genomics UK (COG-UK) Consortium | Jacquelyn Wynn, Mairead Hyland, The Lighthouse Lab in Alderley Park and Alex Alderton, Roberto Amato, Jeffrey Barrett, Sonia Goncalves, Ewan Harrison, David K. Jackson, Ian Johnston, Dominic Kwiatkowski, Cordelia Langford, John Sillitoe on behalf of the Wellcome Sanger Institute COVID-19 Surveillance Team |
| EPI_ISL_2569699 | Lighthouse Laboratory Plymouth | Wellcome Sanger Institute for the COVID-19 Genomics UK (COG-UK) Consortium | Lighthouse Laboratory Plymouth and Alex Alderton, Roberto Amato, Jeffrey Barrett, Sonia Goncalves, Ewan Harrison, David K. Jackson, Ian Johnston, Dominic Kwiatkowski, Cordelia Langford, John Sillitoe on behalf of the Wellcome Sanger Institute COVID-19 Surveillance Team |
| EPI_ISL_2569700, EPI_ISL_2569701, EPI_ISL_2569702 | Lighthouse Lab in Milton Keynes | Wellcome Sanger Institute for the COVID-19 Genomics UK (COG-UK) Consortium | The Lighthouse Lab in Milton Keynes and Alex Alderton, Roberto Amato, Jeffrey Barrett, Sonia Goncalves, Ewan Harrison, David K. Jackson, Ian Johnston, Dominic Kwiatkowski, Cordelia Langford, John Sillitoe on behalf of the Wellcome Sanger Institute COVID-19 Surveillance Team |
| EPI_ISL_2569703 | Lighthouse Laboratory Plymouth | Wellcome Sanger Institute for the COVID-19 Genomics UK (COG-UK) Consortium | Lighthouse Laboratory Plymouth and Alex Alderton, Roberto Amato, Jeffrey Barrett, Sonia Goncalves, Ewan Harrison, David K. Jackson, Ian Johnston, Dominic Kwiatkowski, Cordelia Langford, John Sillitoe on behalf of the Wellcome Sanger Institute COVID-19 Surveillance Team |
| EPI_ISL_2569704 | Lighthouse Lab in Milton Keynes | Wellcome Sanger Institute for the COVID-19 Genomics UK (COG-UK) Consortium | The Lighthouse Lab in Milton Keynes and Alex Alderton, Roberto Amato, Jeffrey Barrett, Sonia Goncalves, Ewan Harrison, David K. Jackson, Ian Johnston, Dominic Kwiatkowski, Cordelia Langford, John Sillitoe on behalf of the Wellcome Sanger Institute COVID-19 Surveillance Team |
| EPI_ISL_2569705, EPI_ISL_2569706, EPI_ISL_2569707, EPI_ISL_2569708 | Lighthouse Lab in Alderley Park | Wellcome Sanger Institute for the COVID-19 Genomics UK (COG-UK) Consortium | Jacquelyn Wynn, Mairead Hyland, The Lighthouse Lab in Alderley Park and Alex Alderton, Roberto Amato, Jeffrey Barrett, Sonia Goncalves, Ewan Harrison, David K. Jackson, Ian Johnston, Dominic Kwiatkowski, Cordelia Langford, John Sillitoe on behalf of the Wellcome Sanger Institute COVID-19 Surveillance Team |
| EPI_ISL_2569709 | Lighthouse Laboratory Plymouth | Wellcome Sanger Institute for the COVID-19 Genomics UK (COG-UK) Consortium | Lighthouse Laboratory Plymouth and Alex Alderton, Roberto Amato, Jeffrey Barrett, Sonia Goncalves, Ewan Harrison, David K. Jackson, Ian Johnston, Dominic Kwiatkowski, Cordelia Langford, John Sillitoe on behalf of the Wellcome Sanger Institute COVID-19 Surveillance Team |
| EPI_ISL_2569710, EPI_ISL_2569711 | Lighthouse Lab in Alderley Park | Wellcome Sanger Institute for the COVID-19 Genomics UK (COG-UK) Consortium | Jacquelyn Wynn, Mairead Hyland, The Lighthouse Lab in Alderley Park and Alex Alderton, Roberto Amato, Jeffrey Barrett, Sonia Goncalves, Ewan Harrison, David K. Jackson, Ian Johnston, Dominic Kwiatkowski, Cordelia Langford, John Sillitoe on behalf of the Wellcome Sanger Institute COVID-19 Surveillance Team |
| EPI_ISL_2569712, EPI_ISL_2569713 | Lighthouse Lab in Milton Keynes | Wellcome Sanger Institute for the COVID-19 Genomics UK (COG-UK) Consortium | The Lighthouse Lab in Milton Keynes and Alex Alderton, Roberto Amato, Jeffrey Barrett, Sonia Goncalves, Ewan Harrison, David K. Jackson, Ian Johnston, Dominic Kwiatkowski, Cordelia Langford, John Sillitoe on behalf of the Wellcome Sanger Institute COVID-19 Surveillance Team |
| EPI_ISL_2569714 | Lighthouse Lab in Alderley Park | Wellcome Sanger Institute for the COVID-19 Genomics UK (COG-UK) Consortium | Jacquelyn Wynn, Mairead Hyland, The Lighthouse Lab in Alderley Park and Alex Alderton, Roberto Amato, Jeffrey Barrett, Sonia Goncalves, Ewan Harrison, David K. Jackson, Ian Johnston, Dominic Kwiatkowski, Cordelia Langford, John Sillitoe on behalf of the Wellcome Sanger Institute COVID-19 Surveillance Team |
| EPI_ISL_2569715 | Lighthouse Lab in Milton Keynes | Wellcome Sanger Institute for the COVID-19 Genomics UK (COG-UK) Consortium | The Lighthouse Lab in Milton Keynes and Alex Alderton, Roberto Amato, Jeffrey Barrett, Sonia Goncalves, Ewan Harrison, David K. Jackson, Ian Johnston, Dominic Kwiatkowski, Cordelia Langford, John Sillitoe on behalf of the Wellcome Sanger Institute COVID-19 Surveillance Team |
| EPI_ISL_2569716 | Lighthouse Lab in Alderley Park | Wellcome Sanger Institute for the COVID-19 Genomics UK (COG-UK) Consortium | Jacquelyn Wynn, Mairead Hyland, The Lighthouse Lab in Alderley Park and Alex Alderton, Roberto Amato, Jeffrey Barrett, Sonia Goncalves, Ewan Harrison, David K. Jackson, Ian Johnston, Dominic Kwiatkowski, Cordelia Langford, John Sillitoe on behalf of the Wellcome Sanger Institute COVID-19 Surveillance Team |
| EPI_ISL_2569717, EPI_ISL_2569718 | Lighthouse Lab in Milton Keynes | Wellcome Sanger Institute for the COVID-19 Genomics UK (COG-UK) Consortium | The Lighthouse Lab in Milton Keynes and Alex Alderton, Roberto Amato, Jeffrey Barrett, Sonia Goncalves, Ewan Harrison, David K. Jackson, Ian Johnston, Dominic Kwiatkowski, Cordelia Langford, John Sillitoe on behalf of the Wellcome Sanger Institute COVID-19 Surveillance Team |
| EPI_ISL_2569719 | Lighthouse Laboratory Plymouth | Wellcome Sanger Institute for the COVID-19 Genomics UK (COG-UK) Consortium | Lighthouse Laboratory Plymouth and Alex Alderton, Roberto Amato, Jeffrey Barrett, Sonia Goncalves, Ewan Harrison, David K. Jackson, Ian Johnston, Dominic Kwiatkowski, Cordelia Langford, John Sillitoe on behalf of the Wellcome Sanger Institute COVID-19 Surveillance Team |
| EPI_ISL_2569720 | Lighthouse Lab in Milton Keynes | Wellcome Sanger Institute for the COVID-19 Genomics UK (COG-UK) Consortium | The Lighthouse Lab in Milton Keynes and Alex Alderton, Roberto Amato, Jeffrey Barrett, Sonia Goncalves, Ewan Harrison, David K. Jackson, Ian Johnston, Dominic Kwiatkowski, Cordelia Langford, John Sillitoe on behalf of the Wellcome Sanger Institute COVID-19 Surveillance Team |
| EPI_ISL_2569721 | Lighthouse Laboratory Plymouth | Wellcome Sanger Institute for the COVID-19 Genomics UK (COG-UK) Consortium | Lighthouse Laboratory Plymouth and Alex Alderton, Roberto Amato, Jeffrey Barrett, Sonia Goncalves, Ewan Harrison, David K. Jackson, Ian Johnston, Dominic Kwiatkowski, Cordelia Langford, John Sillitoe on behalf of the Wellcome Sanger Institute COVID-19 Surveillance Team |
| EPI_ISL_2569722 | Lighthouse Lab in Milton Keynes | Wellcome Sanger Institute for the COVID-19 Genomics UK (COG-UK) Consortium | The Lighthouse Lab in Milton Keynes and Alex Alderton, Roberto Amato, Jeffrey Barrett, Sonia Goncalves, Ewan Harrison, David K. Jackson, Ian Johnston, Dominic Kwiatkowski, Cordelia Langford, John Sillitoe on behalf of the Wellcome Sanger Institute COVID-19 Surveillance Team |
| EPI_ISL_2569724, EPI_ISL_2569725 | Lighthouse Laboratory Plymouth | Wellcome Sanger Institute for the COVID-19 Genomics UK (COG-UK) Consortium | Lighthouse Laboratory Plymouth and Alex Alderton, Roberto Amato, Jeffrey Barrett, Sonia Goncalves, Ewan Harrison, David K. Jackson, Ian Johnston, Dominic Kwiatkowski, Cordelia Langford, John Sillitoe on behalf of the Wellcome Sanger Institute COVID-19 Surveillance Team |
| EPI_ISL_2569726 | Lighthouse Lab in Alderley Park | Wellcome Sanger Institute for the COVID-19 Genomics UK (COG-UK) Consortium | Jacquelyn Wynn, Mairead Hyland, The Lighthouse Lab in Alderley Park and Alex Alderton, Roberto Amato, Jeffrey Barrett, Sonia Goncalves, Ewan Harrison, David K. Jackson, Ian Johnston, Dominic Kwiatkowski, Cordelia Langford, John Sillitoe on behalf of the Wellcome Sanger Institute COVID-19 Surveillance Team |
| EPI_ISL_2569727 | Lighthouse Laboratory Plymouth | Wellcome Sanger Institute for the COVID-19 Genomics UK (COG-UK) Consortium | Lighthouse Laboratory Plymouth and Alex Alderton, Roberto Amato, Jeffrey Barrett, Sonia Goncalves, Ewan Harrison, David K. Jackson, Ian Johnston, Dominic Kwiatkowski, Cordelia Langford, John Sillitoe on behalf of the Wellcome Sanger Institute COVID-19 Surveillance Team |
| EPI_ISL_2569728, EPI_ISL_2569729 | Lighthouse Lab in Alderley Park | Wellcome Sanger Institute for the COVID-19 Genomics UK (COG-UK) Consortium | Jacquelyn Wynn, Mairead Hyland, The Lighthouse Lab in Alderley Park and Alex Alderton, Roberto Amato, Jeffrey Barrett, Sonia Goncalves, Ewan Harrison, David K. Jackson, Ian Johnston, Dominic Kwiatkowski, Cordelia Langford, John Sillitoe on behalf of the Wellcome Sanger Institute COVID-19 Surveillance Team |
| EPI_ISL_2569730 | Lighthouse Lab in Milton Keynes | Wellcome Sanger Institute for the COVID-19 Genomics UK (COG-UK) Consortium | The Lighthouse Lab in Milton Keynes and Alex Alderton, Roberto Amato, Jeffrey Barrett, Sonia Goncalves, Ewan Harrison, David K. Jackson, Ian Johnston, Dominic Kwiatkowski, Cordelia Langford, John Sillitoe on behalf of the Wellcome Sanger Institute COVID-19 Surveillance Team |
| EPI_ISL_2569731, EPI_ISL_2569732, EPI_ISL_2569733 | Lighthouse Lab in Alderley Park | Wellcome Sanger Institute for the COVID-19 Genomics UK (COG-UK) Consortium | Jacquelyn Wynn, Mairead Hyland, The Lighthouse Lab in Alderley Park and Alex Alderton, Roberto Amato, Jeffrey Barrett, Sonia Goncalves, Ewan Harrison, David K. Jackson, Ian Johnston, Dominic Kwiatkowski, Cordelia Langford, John Sillitoe on behalf of the Wellcome Sanger Institute COVID-19 Surveillance Team |
| EPI_ISL_2569734 | Lighthouse Lab in Milton Keynes | Wellcome Sanger Institute for the COVID-19 Genomics UK (COG-UK) Consortium | The Lighthouse Lab in Milton Keynes and Alex Alderton, Roberto Amato, Jeffrey Barrett, Sonia Goncalves, Ewan Harrison, David K. Jackson, Ian Johnston, Dominic Kwiatkowski, Cordelia Langford, John Sillitoe on behalf of the Wellcome Sanger Institute COVID-19 Surveillance Team |
| EPI_ISL_2569735 | Lighthouse Laboratory Plymouth | Wellcome Sanger Institute for the COVID-19 Genomics UK (COG-UK) Consortium | Lighthouse Laboratory Plymouth and Alex Alderton, Roberto Amato, Jeffrey Barrett, Sonia Goncalves, Ewan Harrison, David K. Jackson, Ian Johnston, Dominic Kwiatkowski, Cordelia Langford, John Sillitoe on behalf of the Wellcome Sanger Institute COVID-19 Surveillance Team |
| EPI_ISL_2569736 | Lighthouse Lab in Alderley Park | Wellcome Sanger Institute for the COVID-19 Genomics UK (COG-UK) Consortium | Jacquelyn Wynn, Mairead Hyland, The Lighthouse Lab in Alderley Park and Alex Alderton, Roberto Amato, Jeffrey Barrett, Sonia Goncalves, Ewan Harrison, David K. Jackson, Ian Johnston, Dominic Kwiatkowski, Cordelia Langford, John Sillitoe on behalf of the Wellcome Sanger Institute COVID-19 Surveillance Team |
| EPI_ISL_2569737, EPI_ISL_2569738 | Lighthouse Lab in Milton Keynes | Wellcome Sanger Institute for the COVID-19 Genomics UK (COG-UK) Consortium | The Lighthouse Lab in Milton Keynes and Alex Alderton, Roberto Amato, Jeffrey Barrett, Sonia Goncalves, Ewan Harrison, David K. Jackson, Ian Johnston, Dominic Kwiatkowski, Cordelia Langford, John Sillitoe on behalf of the Wellcome Sanger Institute COVID-19 Surveillance Team |
| EPI_ISL_2569739 | Lighthouse Laboratory Plymouth | Wellcome Sanger Institute for the COVID-19 Genomics UK (COG-UK) Consortium | Lighthouse Laboratory Plymouth and Alex Alderton, Roberto Amato, Jeffrey Barrett, Sonia Goncalves, Ewan Harrison, David K. Jackson, Ian Johnston, Dominic Kwiatkowski, Cordelia Langford, John Sillitoe on behalf of the Wellcome Sanger Institute COVID-19 Surveillance Team |
| EPI_ISL_2569740 | Lighthouse Lab in Alderley Park | Wellcome Sanger Institute for the COVID-19 Genomics UK (COG-UK) Consortium | Jacquelyn Wynn, Mairead Hyland, The Lighthouse Lab in Alderley Park and Alex Alderton, Roberto Amato, Jeffrey Barrett, Sonia Goncalves, Ewan Harrison, David K. Jackson, Ian Johnston, Dominic Kwiatkowski, Cordelia Langford, John Sillitoe on behalf of the Wellcome Sanger Institute COVID-19 Surveillance Team |
| EPI_ISL_2569741, EPI_ISL_2569742, EPI_ISL_2569743 | Lighthouse Lab in Milton Keynes | Wellcome Sanger Institute for the COVID-19 Genomics UK (COG-UK) Consortium | The Lighthouse Lab in Milton Keynes and Alex Alderton, Roberto Amato, Jeffrey Barrett, Sonia Goncalves, Ewan Harrison, David K. Jackson, Ian Johnston, Dominic Kwiatkowski, Cordelia Langford, John Sillitoe on behalf of the Wellcome Sanger Institute COVID-19 Surveillance Team |
| EPI_ISL_2569744 | Lighthouse Lab in Alderley Park | Wellcome Sanger Institute for the COVID-19 Genomics UK (COG-UK) Consortium | Jacquelyn Wynn, Mairead Hyland, The Lighthouse Lab in Alderley Park and Alex Alderton, Roberto Amato, Jeffrey Barrett, Sonia Goncalves, Ewan Harrison, David K. Jackson, Ian Johnston, Dominic Kwiatkowski, Cordelia Langford, John Sillitoe on behalf of the Wellcome Sanger Institute COVID-19 Surveillance Team |
| EPI_ISL_2569745, EPI_ISL_2569746 | Lighthouse Lab in Milton Keynes | Wellcome Sanger Institute for the COVID-19 Genomics UK (COG-UK) Consortium | The Lighthouse Lab in Milton Keynes and Alex Alderton, Roberto Amato, Jeffrey Barrett, Sonia Goncalves, Ewan Harrison, David K. Jackson, Ian Johnston, Dominic Kwiatkowski, Cordelia Langford, John Sillitoe on behalf of the Wellcome Sanger Institute COVID-19 Surveillance Team |
| EPI_ISL_2569747 | Lighthouse Lab in Alderley Park | Wellcome Sanger Institute for the COVID-19 Genomics UK (COG-UK) Consortium | Jacquelyn Wynn, Mairead Hyland, The Lighthouse Lab in Alderley Park and Alex Alderton, Roberto Amato, Jeffrey Barrett, Sonia Goncalves, Ewan Harrison, David K. Jackson, Ian Johnston, Dominic Kwiatkowski, Cordelia Langford, John Sillitoe on behalf of the Wellcome Sanger Institute COVID-19 Surveillance Team |
| EPI_ISL_2569748 | Lighthouse Lab in Milton Keynes | Wellcome Sanger Institute for the COVID-19 Genomics UK (COG-UK) Consortium | The Lighthouse Lab in Milton Keynes and Alex Alderton, Roberto Amato, Jeffrey Barrett, Sonia Goncalves, Ewan Harrison, David K. Jackson, Ian Johnston, Dominic Kwiatkowski, Cordelia Langford, John Sillitoe on behalf of the Wellcome Sanger Institute COVID-19 Surveillance Team |
| EPI_ISL_2569749 | Lighthouse Lab in Glasgow | Wellcome Sanger Institute for the COVID-19 Genomics UK (COG-UK) Consortium | Harper VanSteenhouse, Yumi Kasai, David Gray, Carol Clugston, Anna Dominiczak and Alex Alderton, Roberto Amato, Jeffrey Barrett, Sonia Goncalves, Ewan Harrison, David K. Jackson, Ian Johnston, Dominic Kwiatkowski, Cordelia Langford, John Sillitoe on behalf of the Wellcome Sanger Institute COVID-19 Surveillance Team |
| EPI_ISL_2569750, EPI_ISL_2569751 | Lighthouse Laboratory Plymouth | Wellcome Sanger Institute for the COVID-19 Genomics UK (COG-UK) Consortium | Lighthouse Laboratory Plymouth and Alex Alderton, Roberto Amato, Jeffrey Barrett, Sonia Goncalves, Ewan Harrison, David K. Jackson, Ian Johnston, Dominic Kwiatkowski, Cordelia Langford, John Sillitoe on behalf of the Wellcome Sanger Institute COVID-19 Surveillance Team |
| EPI_ISL_2569753 | Lighthouse Lab in Alderley Park | Wellcome Sanger Institute for the COVID-19 Genomics UK (COG-UK) Consortium | Jacquelyn Wynn, Mairead Hyland, The Lighthouse Lab in Alderley Park and Alex Alderton, Roberto Amato, Jeffrey Barrett, Sonia Goncalves, Ewan Harrison, David K. Jackson, Ian Johnston, Dominic Kwiatkowski, Cordelia Langford, John Sillitoe on behalf of the Wellcome Sanger Institute COVID-19 Surveillance Team |
| EPI_ISL_2569754, EPI_ISL_2569755, EPI_ISL_2569756 | Lighthouse Lab in Milton Keynes | Wellcome Sanger Institute for the COVID-19 Genomics UK (COG-UK) Consortium | The Lighthouse Lab in Milton Keynes and Alex Alderton, Roberto Amato, Jeffrey Barrett, Sonia Goncalves, Ewan Harrison, David K. Jackson, Ian Johnston, Dominic Kwiatkowski, Cordelia Langford, John Sillitoe on behalf of the Wellcome Sanger Institute COVID-19 Surveillance Team |
| EPI_ISL_2569757 | Lighthouse Lab in Alderley Park | Wellcome Sanger Institute for the COVID-19 Genomics UK (COG-UK) Consortium | Jacquelyn Wynn, Mairead Hyland, The Lighthouse Lab in Alderley Park and Alex Alderton, Roberto Amato, Jeffrey Barrett, Sonia Goncalves, Ewan Harrison, David K. Jackson, Ian Johnston, Dominic Kwiatkowski, Cordelia Langford, John Sillitoe on behalf of the Wellcome Sanger Institute COVID-19 Surveillance Team |
| EPI_ISL_2569758 | Lighthouse Laboratory Plymouth | Wellcome Sanger Institute for the COVID-19 Genomics UK (COG-UK) Consortium | Lighthouse Laboratory Plymouth and Alex Alderton, Roberto Amato, Jeffrey Barrett, Sonia Goncalves, Ewan Harrison, David K. Jackson, Ian Johnston, Dominic Kwiatkowski, Cordelia Langford, John Sillitoe on behalf of the Wellcome Sanger Institute COVID-19 Surveillance Team |
| EPI_ISL_2569759 | Lighthouse Lab in Alderley Park | Wellcome Sanger Institute for the COVID-19 Genomics UK (COG-UK) Consortium | Jacquelyn Wynn, Mairead Hyland, The Lighthouse Lab in Alderley Park and Alex Alderton, Roberto Amato, Jeffrey Barrett, Sonia Goncalves, Ewan Harrison, David K. Jackson, Ian Johnston, Dominic Kwiatkowski, Cordelia Langford, John Sillitoe on behalf of the Wellcome Sanger Institute COVID-19 Surveillance Team |

| | | | |
|---|---|---|---|
| EPI_ISL_2569760, EPI_ISL_2569761 | Lighthouse Lab in Milton Keynes | Wellcome Sanger Institute for the COVID-19 Genomics UK (COG-UK) Consortium | The Lighthouse Lab in Milton Keynes and Alex Alderton, Roberto Amato, Jeffrey Barrett, Sonia Goncalves, Ewan Harrison, David K. Jackson, Ian Johnston, Dominic Kwiatkowski, Cordelia Langford, John Sillitoe on behalf of the Wellcome Sanger Institute COVID-19 Surveillance Team |
| EPI_ISL_2569762, EPI_ISL_2569763 | Lighthouse Laboratory Plymouth | Wellcome Sanger Institute for the COVID-19 Genomics UK (COG-UK) Consortium | Lighthouse Laboratory Plymouth and Alex Alderton, Roberto Amato, Jeffrey Barrett, Sonia Goncalves, Ewan Harrison, David K. Jackson, Ian Johnston, Dominic Kwiatkowski, Cordelia Langford, John Sillitoe on behalf of the Wellcome Sanger Institute COVID-19 Surveillance Team |
| EPI_ISL_2569764 | Lighthouse Lab in Alderley Park | Wellcome Sanger Institute for the COVID-19 Genomics UK (COG-UK) Consortium | Jacquelyn Wynn, Mairead Hyland, The Lighthouse Lab in Alderley Park and Alex Alderton, Roberto Amato, Jeffrey Barrett, Sonia Goncalves, Ewan Harrison, David K. Jackson, Ian Johnston, Dominic Kwiatkowski, Cordelia Langford, John Sillitoe on behalf of the Wellcome Sanger Institute COVID-19 Surveillance Team |
| EPI_ISL_2569765 | Lighthouse Lab in Milton Keynes | Wellcome Sanger Institute for the COVID-19 Genomics UK (COG-UK) Consortium | The Lighthouse Lab in Milton Keynes and Alex Alderton, Roberto Amato, Jeffrey Barrett, Sonia Goncalves, Ewan Harrison, David K. Jackson, Ian Johnston, Dominic Kwiatkowski, Cordelia Langford, John Sillitoe on behalf of the Wellcome Sanger Institute COVID-19 Surveillance Team |
| EPI_ISL_2569766 | Lighthouse Lab in Alderley Park | Wellcome Sanger Institute for the COVID-19 Genomics UK (COG-UK) Consortium | Jacquelyn Wynn, Mairead Hyland, The Lighthouse Lab in Alderley Park and Alex Alderton, Roberto Amato, Jeffrey Barrett, Sonia Goncalves, Ewan Harrison, David K. Jackson, Ian Johnston, Dominic Kwiatkowski, Cordelia Langford, John Sillitoe on behalf of the Wellcome Sanger Institute COVID-19 Surveillance Team |
| EPI_ISL_2569767 | Lighthouse Lab in Milton Keynes | Wellcome Sanger Institute for the COVID-19 Genomics UK (COG-UK) Consortium | The Lighthouse Lab in Milton Keynes and Alex Alderton, Roberto Amato, Jeffrey Barrett, Sonia Goncalves, Ewan Harrison, David K. Jackson, Ian Johnston, Dominic Kwiatkowski, Cordelia Langford, John Sillitoe on behalf of the Wellcome Sanger Institute COVID-19 Surveillance Team |
| EPI_ISL_2569768, EPI_ISL_2569769, EPI_ISL_2569770 | Lighthouse Laboratory Plymouth | Wellcome Sanger Institute for the COVID-19 Genomics UK (COG-UK) Consortium | Lighthouse Laboratory Plymouth and Alex Alderton, Roberto Amato, Jeffrey Barrett, Sonia Goncalves, Ewan Harrison, David K. Jackson, Ian Johnston, Dominic Kwiatkowski, Cordelia Langford, John Sillitoe on behalf of the Wellcome Sanger Institute COVID-19 Surveillance Team |
| EPI_ISL_2569771 | Lighthouse Lab in Alderley Park | Wellcome Sanger Institute for the COVID-19 Genomics UK (COG-UK) Consortium | Jacquelyn Wynn, Mairead Hyland, The Lighthouse Lab in Alderley Park and Alex Alderton, Roberto Amato, Jeffrey Barrett, Sonia Goncalves, Ewan Harrison, David K. Jackson, Ian Johnston, Dominic Kwiatkowski, Cordelia Langford, John Sillitoe on behalf of the Wellcome Sanger Institute COVID-19 Surveillance Team |
| EPI_ISL_2569772 | Lighthouse Laboratory Plymouth | Wellcome Sanger Institute for the COVID-19 Genomics UK (COG-UK) Consortium | Lighthouse Laboratory Plymouth and Alex Alderton, Roberto Amato, Jeffrey Barrett, Sonia Goncalves, Ewan Harrison, David K. Jackson, Ian Johnston, Dominic Kwiatkowski, Cordelia Langford, John Sillitoe on behalf of the Wellcome Sanger Institute COVID-19 Surveillance Team |
| EPI_ISL_2569774, EPI_ISL_2569775 | Lighthouse Lab in Milton Keynes | Wellcome Sanger Institute for the COVID-19 Genomics UK (COG-UK) Consortium | The Lighthouse Lab in Milton Keynes and Alex Alderton, Roberto Amato, Jeffrey Barrett, Sonia Goncalves, Ewan Harrison, David K. Jackson, Ian Johnston, Dominic Kwiatkowski, Cordelia Langford, John Sillitoe on behalf of the Wellcome Sanger Institute COVID-19 Surveillance Team |
| EPI_ISL_2569776 | Lighthouse Lab in Alderley Park | Wellcome Sanger Institute for the COVID-19 Genomics UK (COG-UK) Consortium | Jacquelyn Wynn, Mairead Hyland, The Lighthouse Lab in Alderley Park and Alex Alderton, Roberto Amato, Jeffrey Barrett, Sonia Goncalves, Ewan Harrison, David K. Jackson, Ian Johnston, Dominic Kwiatkowski, Cordelia Langford, John Sillitoe on behalf of the Wellcome Sanger Institute COVID-19 Surveillance Team |
| EPI_ISL_2569777 | Lighthouse Laboratory Plymouth | Wellcome Sanger Institute for the COVID-19 Genomics UK (COG-UK) Consortium | Lighthouse Laboratory Plymouth and Alex Alderton, Roberto Amato, Jeffrey Barrett, Sonia Goncalves, Ewan Harrison, David K. Jackson, Ian Johnston, Dominic Kwiatkowski, Cordelia Langford, John Sillitoe on behalf of the Wellcome Sanger Institute COVID-19 Surveillance Team |
| EPI_ISL_2569778 | Lighthouse Lab in Alderley Park | Wellcome Sanger Institute for the COVID-19 Genomics UK (COG-UK) Consortium | Jacquelyn Wynn, Mairead Hyland, The Lighthouse Lab in Alderley Park and Alex Alderton, Roberto Amato, Jeffrey Barrett, Sonia Goncalves, Ewan Harrison, David K. Jackson, Ian Johnston, Dominic Kwiatkowski, Cordelia Langford, John Sillitoe on behalf of the Wellcome Sanger Institute COVID-19 Surveillance Team |
| EPI_ISL_2569779 | Lighthouse Lab in Milton Keynes | Wellcome Sanger Institute for the COVID-19 Genomics UK (COG-UK) Consortium | The Lighthouse Lab in Milton Keynes and Alex Alderton, Roberto Amato, Jeffrey Barrett, Sonia Goncalves, Ewan Harrison, David K. Jackson, Ian Johnston, Dominic Kwiatkowski, Cordelia Langford, John Sillitoe on behalf of the Wellcome Sanger Institute COVID-19 Surveillance Team |
| EPI_ISL_2569780, EPI_ISL_2569781 | Lighthouse Lab in Alderley Park | Wellcome Sanger Institute for the COVID-19 Genomics UK (COG-UK) Consortium | Jacquelyn Wynn, Mairead Hyland, The Lighthouse Lab in Alderley Park and Alex Alderton, Roberto Amato, Jeffrey Barrett, Sonia Goncalves, Ewan Harrison, David K. Jackson, Ian Johnston, Dominic Kwiatkowski, Cordelia Langford, John Sillitoe on behalf of the Wellcome Sanger Institute COVID-19 Surveillance Team |
| EPI_ISL_2569782, EPI_ISL_2569785 | Lighthouse Laboratory Plymouth | Wellcome Sanger Institute for the COVID-19 Genomics UK (COG-UK) Consortium | Lighthouse Laboratory Plymouth and Alex Alderton, Roberto Amato, Jeffrey Barrett, Sonia Goncalves, Ewan Harrison, David K. Jackson, Ian Johnston, Dominic Kwiatkowski, Cordelia Langford, John Sillitoe on behalf of the Wellcome Sanger Institute COVID-19 Surveillance Team |
| EPI_ISL_2569786 | Lighthouse Lab in Alderley Park | Wellcome Sanger Institute for the COVID-19 Genomics UK (COG-UK) Consortium | Jacquelyn Wynn, Mairead Hyland, The Lighthouse Lab in Alderley Park and Alex Alderton, Roberto Amato, Jeffrey Barrett, Sonia Goncalves, Ewan Harrison, David K. Jackson, Ian Johnston, Dominic Kwiatkowski, Cordelia Langford, John Sillitoe on behalf of the Wellcome Sanger Institute COVID-19 Surveillance Team |
| EPI_ISL_2569787 | Lighthouse Lab in Milton Keynes | Wellcome Sanger Institute for the COVID-19 Genomics UK (COG-UK) Consortium | The Lighthouse Lab in Milton Keynes and Alex Alderton, Roberto Amato, Jeffrey Barrett, Sonia Goncalves, Ewan Harrison, David K. Jackson, Ian Johnston, Dominic Kwiatkowski, Cordelia Langford, John Sillitoe on behalf of the Wellcome Sanger Institute COVID-19 Surveillance Team |
| EPI_ISL_2569788, EPI_ISL_2569789 | Lighthouse Laboratory Plymouth | Wellcome Sanger Institute for the COVID-19 Genomics UK (COG-UK) Consortium | Lighthouse Laboratory Plymouth and Alex Alderton, Roberto Amato, Jeffrey Barrett, Sonia Goncalves, Ewan Harrison, David K. Jackson, Ian Johnston, Dominic Kwiatkowski, Cordelia Langford, John Sillitoe on behalf of the Wellcome Sanger Institute COVID-19 Surveillance Team |
| EPI_ISL_2569790, EPI_ISL_2569791 | Lighthouse Lab in Milton Keynes | Wellcome Sanger Institute for the COVID-19 Genomics UK (COG-UK) Consortium | The Lighthouse Lab in Milton Keynes and Alex Alderton, Roberto Amato, Jeffrey Barrett, Sonia Goncalves, Ewan Harrison, David K. Jackson, Ian Johnston, Dominic Kwiatkowski, Cordelia Langford, John Sillitoe on behalf of the Wellcome Sanger Institute COVID-19 Surveillance Team |
| EPI_ISL_2569792 | Lighthouse Lab in Alderley Park | Wellcome Sanger Institute for the COVID-19 Genomics UK (COG-UK) Consortium | Jacquelyn Wynn, Mairead Hyland, The Lighthouse Lab in Alderley Park and Alex Alderton, Roberto Amato, Jeffrey Barrett, Sonia Goncalves, Ewan Harrison, David K. Jackson, Ian Johnston, Dominic Kwiatkowski, Cordelia Langford, John Sillitoe on behalf of the Wellcome Sanger Institute COVID-19 Surveillance Team |
| EPI_ISL_2569793, EPI_ISL_2569794 | Lighthouse Lab in Milton Keynes | Wellcome Sanger Institute for the COVID-19 Genomics UK (COG-UK) Consortium | The Lighthouse Lab in Milton Keynes and Alex Alderton, Roberto Amato, Jeffrey Barrett, Sonia Goncalves, Ewan Harrison, David K. Jackson, Ian Johnston, Dominic Kwiatkowski, Cordelia Langford, John Sillitoe on behalf of the Wellcome Sanger Institute COVID-19 Surveillance Team |
| EPI_ISL_2569795 | Lighthouse Lab in Alderley Park | Wellcome Sanger Institute for the COVID-19 Genomics UK (COG-UK) Consortium | Jacquelyn Wynn, Mairead Hyland, The Lighthouse Lab in Alderley Park and Alex Alderton, Roberto Amato, Jeffrey Barrett, Sonia Goncalves, Ewan Harrison, David K. Jackson, Ian Johnston, Dominic Kwiatkowski, Cordelia Langford, John Sillitoe on behalf of the Wellcome Sanger Institute COVID-19 Surveillance Team |
| EPI_ISL_2569796 | Lighthouse Laboratory Plymouth | Wellcome Sanger Institute for the COVID-19 Genomics UK (COG-UK) Consortium | Lighthouse Laboratory Plymouth and Alex Alderton, Roberto Amato, Jeffrey Barrett, Sonia Goncalves, Ewan Harrison, David K. Jackson, Ian Johnston, Dominic Kwiatkowski, Cordelia Langford, John Sillitoe on behalf of the Wellcome Sanger Institute COVID-19 Surveillance Team |
| EPI_ISL_2569797 | Lighthouse Lab in Alderley Park | Wellcome Sanger Institute for the COVID-19 Genomics UK (COG-UK) Consortium | Jacquelyn Wynn, Mairead Hyland, The Lighthouse Lab in Alderley Park and Alex Alderton, Roberto Amato, Jeffrey Barrett, Sonia Goncalves, Ewan Harrison, David K. Jackson, Ian Johnston, Dominic Kwiatkowski, Cordelia Langford, John Sillitoe on behalf of the Wellcome Sanger Institute COVID-19 Surveillance Team |
| EPI_ISL_2569798 | Lighthouse Lab in Milton Keynes | Wellcome Sanger Institute for the COVID-19 Genomics UK (COG-UK) Consortium | The Lighthouse Lab in Milton Keynes and Alex Alderton, Roberto Amato, Jeffrey Barrett, Sonia Goncalves, Ewan Harrison, David K. Jackson, Ian Johnston, Dominic Kwiatkowski, Cordelia Langford, John Sillitoe on behalf of the Wellcome Sanger Institute COVID-19 Surveillance Team |
| EPI_ISL_2569799 | Lighthouse Laboratory Plymouth | Wellcome Sanger Institute for the COVID-19 Genomics UK (COG-UK) Consortium | Lighthouse Laboratory Plymouth and Alex Alderton, Roberto Amato, Jeffrey Barrett, Sonia Goncalves, Ewan Harrison, David K. Jackson, Ian Johnston, Dominic Kwiatkowski, Cordelia Langford, John Sillitoe on behalf of the Wellcome Sanger Institute COVID-19 Surveillance Team |
| EPI_ISL_2569800 | Lighthouse Lab in Milton Keynes | Wellcome Sanger Institute for the COVID-19 Genomics UK (COG-UK) Consortium | The Lighthouse Lab in Milton Keynes and Alex Alderton, Roberto Amato, Jeffrey Barrett, Sonia Goncalves, Ewan Harrison, David K. Jackson, Ian Johnston, Dominic Kwiatkowski, Cordelia Langford, John Sillitoe on behalf of the Wellcome Sanger Institute COVID-19 Surveillance Team |
| EPI_ISL_2569801, EPI_ISL_2569802, EPI_ISL_2569803 | Lighthouse Lab in Alderley Park | Wellcome Sanger Institute for the COVID-19 Genomics UK (COG-UK) Consortium | Jacquelyn Wynn, Mairead Hyland, The Lighthouse Lab in Alderley Park and Alex Alderton, Roberto Amato, Jeffrey Barrett, Sonia Goncalves, Ewan Harrison, David K. Jackson, Ian Johnston, Dominic Kwiatkowski, Cordelia Langford, John Sillitoe on behalf of the Wellcome Sanger Institute COVID-19 Surveillance Team |
| EPI_ISL_2569804, EPI_ISL_2569805, EPI_ISL_2569806, EPI_ISL_2569807 | Lighthouse Laboratory Plymouth | Wellcome Sanger Institute for the COVID-19 Genomics UK (COG-UK) Consortium | Lighthouse Laboratory Plymouth and Alex Alderton, Roberto Amato, Jeffrey Barrett, Sonia Goncalves, Ewan Harrison, David K. Jackson, Ian Johnston, Dominic Kwiatkowski, Cordelia Langford, John Sillitoe on behalf of the Wellcome Sanger Institute COVID-19 Surveillance Team |
| EPI_ISL_2569808, EPI_ISL_2569809, EPI_ISL_2569810 | Lighthouse Lab in Milton Keynes | Wellcome Sanger Institute for the COVID-19 Genomics UK (COG-UK) Consortium | The Lighthouse Lab in Milton Keynes and Alex Alderton, Roberto Amato, Jeffrey Barrett, Sonia Goncalves, Ewan Harrison, David K. Jackson, Ian Johnston, Dominic Kwiatkowski, Cordelia Langford, John Sillitoe on behalf of the Wellcome Sanger Institute COVID-19 Surveillance Team |
| EPI_ISL_2569811 | Lighthouse Lab in Alderley Park | Wellcome Sanger Institute for the COVID-19 Genomics UK (COG-UK) Consortium | Jacquelyn Wynn, Mairead Hyland, The Lighthouse Lab in Alderley Park and Alex Alderton, Roberto Amato, Jeffrey Barrett, Sonia Goncalves, Ewan Harrison, David K. Jackson, Ian Johnston, Dominic Kwiatkowski, Cordelia Langford, John Sillitoe on behalf of the Wellcome Sanger Institute COVID-19 Surveillance Team |
| EPI_ISL_2569812 | Lighthouse Lab in Milton Keynes | Wellcome Sanger Institute for the COVID-19 Genomics UK (COG-UK) Consortium | The Lighthouse Lab in Milton Keynes and Alex Alderton, Roberto Amato, Jeffrey Barrett, Sonia Goncalves, Ewan Harrison, David K. Jackson, Ian Johnston, Dominic Kwiatkowski, Cordelia Langford, John Sillitoe on behalf of the Wellcome Sanger Institute COVID-19 Surveillance Team |
| EPI_ISL_2569813, EPI_ISL_2569814 | Lighthouse Lab in Alderley Park | Wellcome Sanger Institute for the COVID-19 Genomics UK (COG-UK) Consortium | Jacquelyn Wynn, Mairead Hyland, The Lighthouse Lab in Alderley Park and Alex Alderton, Roberto Amato, Jeffrey Barrett, Sonia Goncalves, Ewan Harrison, David K. Jackson, Ian Johnston, Dominic Kwiatkowski, Cordelia Langford, John Sillitoe on behalf of the Wellcome Sanger Institute COVID-19 Surveillance Team |
| EPI_ISL_2569815 | Lighthouse Lab in Milton Keynes | Wellcome Sanger Institute for the COVID-19 Genomics UK (COG-UK) Consortium | The Lighthouse Lab in Milton Keynes and Alex Alderton, Roberto Amato, Jeffrey Barrett, Sonia Goncalves, Ewan Harrison, David K. Jackson, Ian Johnston, Dominic Kwiatkowski, Cordelia Langford, John Sillitoe on behalf of the Wellcome Sanger Institute COVID-19 Surveillance Team |
| EPI_ISL_2569816, EPI_ISL_2569817 | Lighthouse Laboratory Plymouth | Wellcome Sanger Institute for the COVID-19 Genomics UK (COG-UK) Consortium | Lighthouse Laboratory Plymouth and Alex Alderton, Roberto Amato, Jeffrey Barrett, Sonia Goncalves, Ewan Harrison, David K. Jackson, Ian Johnston, Dominic Kwiatkowski, Cordelia Langford, John Sillitoe on behalf of the Wellcome Sanger Institute COVID-19 Surveillance Team |
| EPI_ISL_2569818 | Lighthouse Lab in Milton Keynes | Wellcome Sanger Institute for the COVID-19 Genomics UK (COG-UK) Consortium | The Lighthouse Lab in Milton Keynes and Alex Alderton, Roberto Amato, Jeffrey Barrett, Sonia Goncalves, Ewan Harrison, David K. Jackson, Ian Johnston, Dominic Kwiatkowski, Cordelia Langford, John Sillitoe on behalf of the Wellcome Sanger Institute COVID-19 Surveillance Team |
| EPI_ISL_2569819 | Lighthouse Laboratory Plymouth | Wellcome Sanger Institute for the COVID-19 Genomics UK (COG-UK) Consortium | Lighthouse Laboratory Plymouth and Alex Alderton, Roberto Amato, Jeffrey Barrett, Sonia Goncalves, Ewan Harrison, David K. Jackson, Ian Johnston, Dominic Kwiatkowski, Cordelia Langford, John Sillitoe on behalf of the Wellcome Sanger Institute COVID-19 Surveillance Team |
| EPI_ISL_2569820 | Lighthouse Lab in Alderley Park | Wellcome Sanger Institute for the COVID-19 Genomics UK (COG-UK) Consortium | Jacquelyn Wynn, Mairead Hyland, The Lighthouse Lab in Alderley Park and Alex Alderton, Roberto Amato, Jeffrey Barrett, Sonia Goncalves, Ewan Harrison, David K. Jackson, Ian Johnston, Dominic Kwiatkowski, Cordelia Langford, John Sillitoe on behalf of the Wellcome Sanger Institute COVID-19 Surveillance Team |
| EPI_ISL_2569821 | Lighthouse Laboratory Plymouth | Wellcome Sanger Institute for the COVID-19 Genomics UK (COG-UK) Consortium | Lighthouse Laboratory Plymouth and Alex Alderton, Roberto Amato, Jeffrey Barrett, Sonia Goncalves, Ewan Harrison, David K. Jackson, Ian Johnston, Dominic Kwiatkowski, Cordelia Langford, John Sillitoe on behalf of the Wellcome Sanger Institute COVID-19 Surveillance Team |
| EPI_ISL_2569823, EPI_ISL_2569824 | Lighthouse Lab in Alderley Park | Wellcome Sanger Institute for the COVID-19 Genomics UK (COG-UK) Consortium | Jacquelyn Wynn, Mairead Hyland, The Lighthouse Lab in Alderley Park and Alex Alderton, Roberto Amato, Jeffrey Barrett, Sonia Goncalves, Ewan Harrison, David K. Jackson, Ian Johnston, Dominic Kwiatkowski, Cordelia Langford, John Sillitoe on behalf of the Wellcome Sanger Institute COVID-19 Surveillance Team |
| EPI_ISL_2569825 | Lighthouse Laboratory Plymouth | Wellcome Sanger Institute for the COVID-19 Genomics UK (COG-UK) Consortium | Lighthouse Laboratory Plymouth and Alex Alderton, Roberto Amato, Jeffrey Barrett, Sonia Goncalves, Ewan Harrison, David K. Jackson, Ian Johnston, Dominic Kwiatkowski, Cordelia Langford, John Sillitoe on behalf of the Wellcome Sanger Institute COVID-19 Surveillance Team |
| EPI_ISL_2569826 | Lighthouse Lab in Milton Keynes | Wellcome Sanger Institute for the COVID-19 Genomics UK (COG-UK) Consortium | The Lighthouse Lab in Milton Keynes and Alex Alderton, Roberto Amato, Jeffrey Barrett, Sonia Goncalves, Ewan Harrison, David K. Jackson, Ian Johnston, Dominic Kwiatkowski, Cordelia Langford, John Sillitoe on behalf of the Wellcome Sanger Institute COVID-19 Surveillance Team |

| | | | |
|---|---|---|---|
| EPI_ISL_2569827 | Lighthouse Lab in Alderley Park | Wellcome Sanger Institute for the COVID-19 Genomics UK (COG-UK) Consortium | Jacquelyn Wynn, Mairead Hyland, The Lighthouse Lab in Alderley Park and Alex Alderton, Roberto Amato, Jeffrey Barrett, Sonia Goncalves, Ewan Harrison, David K. Jackson, Ian Johnston, Dominic Kwiatkowski, Cordelia Langford, John Sillitoe on behalf of the Wellcome Sanger Institute COVID-19 Surveillance Team |
| EPI_ISL_2569828 | Lighthouse Lab in Milton Keynes | Wellcome Sanger Institute for the COVID-19 Genomics UK (COG-UK) Consortium | The Lighthouse Lab in Milton Keynes and Alex Alderton, Roberto Amato, Jeffrey Barrett, Sonia Goncalves, Ewan Harrison, David K. Jackson, Ian Johnston, Dominic Kwiatkowski, Cordelia Langford, John Sillitoe on behalf of the Wellcome Sanger Institute COVID-19 Surveillance Team |
| EPI_ISL_2569829 | Lighthouse Lab in Alderley Park | Wellcome Sanger Institute for the COVID-19 Genomics UK (COG-UK) Consortium | Jacquelyn Wynn, Mairead Hyland, The Lighthouse Lab in Alderley Park and Alex Alderton, Roberto Amato, Jeffrey Barrett, Sonia Goncalves, Ewan Harrison, David K. Jackson, Ian Johnston, Dominic Kwiatkowski, Cordelia Langford, John Sillitoe on behalf of the Wellcome Sanger Institute COVID-19 Surveillance Team |
| EPI_ISL_2569830 | Lighthouse Lab in Milton Keynes | Wellcome Sanger Institute for the COVID-19 Genomics UK (COG-UK) Consortium | The Lighthouse Lab in Milton Keynes and Alex Alderton, Roberto Amato, Jeffrey Barrett, Sonia Goncalves, Ewan Harrison, David K. Jackson, Ian Johnston, Dominic Kwiatkowski, Cordelia Langford, John Sillitoe on behalf of the Wellcome Sanger Institute COVID-19 Surveillance Team |
| EPI_ISL_2569831 | Lighthouse Lab in Alderley Park | Wellcome Sanger Institute for the COVID-19 Genomics UK (COG-UK) Consortium | Jacquelyn Wynn, Mairead Hyland, The Lighthouse Lab in Alderley Park and Alex Alderton, Roberto Amato, Jeffrey Barrett, Sonia Goncalves, Ewan Harrison, David K. Jackson, Ian Johnston, Dominic Kwiatkowski, Cordelia Langford, John Sillitoe on behalf of the Wellcome Sanger Institute COVID-19 Surveillance Team |
| EPI_ISL_2569832 | Lighthouse Lab in Milton Keynes | Wellcome Sanger Institute for the COVID-19 Genomics UK (COG-UK) Consortium | The Lighthouse Lab in Milton Keynes and Alex Alderton, Roberto Amato, Jeffrey Barrett, Sonia Goncalves, Ewan Harrison, David K. Jackson, Ian Johnston, Dominic Kwiatkowski, Cordelia Langford, John Sillitoe on behalf of the Wellcome Sanger Institute COVID-19 Surveillance Team |
| EPI_ISL_2569834 | Lighthouse Lab in Alderley Park | Wellcome Sanger Institute for the COVID-19 Genomics UK (COG-UK) Consortium | Jacquelyn Wynn, Mairead Hyland, The Lighthouse Lab in Alderley Park and Alex Alderton, Roberto Amato, Jeffrey Barrett, Sonia Goncalves, Ewan Harrison, David K. Jackson, Ian Johnston, Dominic Kwiatkowski, Cordelia Langford, John Sillitoe on behalf of the Wellcome Sanger Institute COVID-19 Surveillance Team |
| EPI_ISL_2569835 | Lighthouse Lab in Milton Keynes | Wellcome Sanger Institute for the COVID-19 Genomics UK (COG-UK) Consortium | The Lighthouse Lab in Milton Keynes and Alex Alderton, Roberto Amato, Jeffrey Barrett, Sonia Goncalves, Ewan Harrison, David K. Jackson, Ian Johnston, Dominic Kwiatkowski, Cordelia Langford, John Sillitoe on behalf of the Wellcome Sanger Institute COVID-19 Surveillance Team |
| EPI_ISL_2569836 | Lighthouse Laboratory Plymouth | Wellcome Sanger Institute for the COVID-19 Genomics UK (COG-UK) Consortium | Lighthouse Laboratory Plymouth and Alex Alderton, Roberto Amato, Jeffrey Barrett, Sonia Goncalves, Ewan Harrison, David K. Jackson, Ian Johnston, Dominic Kwiatkowski, Cordelia Langford, John Sillitoe on behalf of the Wellcome Sanger Institute COVID-19 Surveillance Team |
| EPI_ISL_2569837, EPI_ISL_2569838, EPI_ISL_2569839 | Lighthouse Lab in Alderley Park | Wellcome Sanger Institute for the COVID-19 Genomics UK (COG-UK) Consortium | Jacquelyn Wynn, Mairead Hyland, The Lighthouse Lab in Alderley Park and Alex Alderton, Roberto Amato, Jeffrey Barrett, Sonia Goncalves, Ewan Harrison, David K. Jackson, Ian Johnston, Dominic Kwiatkowski, Cordelia Langford, John Sillitoe on behalf of the Wellcome Sanger Institute COVID-19 Surveillance Team |
| EPI_ISL_2569840 | Lighthouse Lab in Milton Keynes | Wellcome Sanger Institute for the COVID-19 Genomics UK (COG-UK) Consortium | The Lighthouse Lab in Milton Keynes and Alex Alderton, Roberto Amato, Jeffrey Barrett, Sonia Goncalves, Ewan Harrison, David K. Jackson, Ian Johnston, Dominic Kwiatkowski, Cordelia Langford, John Sillitoe on behalf of the Wellcome Sanger Institute COVID-19 Surveillance Team |
| EPI_ISL_2569841 | Lighthouse Laboratory Plymouth | Wellcome Sanger Institute for the COVID-19 Genomics UK (COG-UK) Consortium | Lighthouse Laboratory Plymouth and Alex Alderton, Roberto Amato, Jeffrey Barrett, Sonia Goncalves, Ewan Harrison, David K. Jackson, Ian Johnston, Dominic Kwiatkowski, Cordelia Langford, John Sillitoe on behalf of the Wellcome Sanger Institute COVID-19 Surveillance Team |
| EPI_ISL_2569842, EPI_ISL_2569843 | Lighthouse Lab in Milton Keynes | Wellcome Sanger Institute for the COVID-19 Genomics UK (COG-UK) Consortium | The Lighthouse Lab in Milton Keynes and Alex Alderton, Roberto Amato, Jeffrey Barrett, Sonia Goncalves, Ewan Harrison, David K. Jackson, Ian Johnston, Dominic Kwiatkowski, Cordelia Langford, John Sillitoe on behalf of the Wellcome Sanger Institute COVID-19 Surveillance Team |
| EPI_ISL_2569844, EPI_ISL_2569845 | Lighthouse Laboratory Plymouth | Wellcome Sanger Institute for the COVID-19 Genomics UK (COG-UK) Consortium | Lighthouse Laboratory Plymouth and Alex Alderton, Roberto Amato, Jeffrey Barrett, Sonia Goncalves, Ewan Harrison, David K. Jackson, Ian Johnston, Dominic Kwiatkowski, Cordelia Langford, John Sillitoe on behalf of the Wellcome Sanger Institute COVID-19 Surveillance Team |
| EPI_ISL_2569846 | Lighthouse Lab in Milton Keynes | Wellcome Sanger Institute for the COVID-19 Genomics UK (COG-UK) Consortium | The Lighthouse Lab in Milton Keynes and Alex Alderton, Roberto Amato, Jeffrey Barrett, Sonia Goncalves, Ewan Harrison, David K. Jackson, Ian Johnston, Dominic Kwiatkowski, Cordelia Langford, John Sillitoe on behalf of the Wellcome Sanger Institute COVID-19 Surveillance Team |
| EPI_ISL_2569847 | Lighthouse Lab in Alderley Park | Wellcome Sanger Institute for the COVID-19 Genomics UK (COG-UK) Consortium | Jacquelyn Wynn, Mairead Hyland, The Lighthouse Lab in Alderley Park and Alex Alderton, Roberto Amato, Jeffrey Barrett, Sonia Goncalves, Ewan Harrison, David K. Jackson, Ian Johnston, Dominic Kwiatkowski, Cordelia Langford, John Sillitoe on behalf of the Wellcome Sanger Institute COVID-19 Surveillance Team |
| EPI_ISL_2569848 | Lighthouse Lab in Milton Keynes | Wellcome Sanger Institute for the COVID-19 Genomics UK (COG-UK) Consortium | The Lighthouse Lab in Milton Keynes and Alex Alderton, Roberto Amato, Jeffrey Barrett, Sonia Goncalves, Ewan Harrison, David K. Jackson, Ian Johnston, Dominic Kwiatkowski, Cordelia Langford, John Sillitoe on behalf of the Wellcome Sanger Institute COVID-19 Surveillance Team |
| EPI_ISL_2569849 | Lighthouse Laboratory Plymouth | Wellcome Sanger Institute for the COVID-19 Genomics UK (COG-UK) Consortium | Lighthouse Laboratory Plymouth and Alex Alderton, Roberto Amato, Jeffrey Barrett, Sonia Goncalves, Ewan Harrison, David K. Jackson, Ian Johnston, Dominic Kwiatkowski, Cordelia Langford, John Sillitoe on behalf of the Wellcome Sanger Institute COVID-19 Surveillance Team |
| EPI_ISL_2569850 | Lighthouse Lab in Alderley Park | Wellcome Sanger Institute for the COVID-19 Genomics UK (COG-UK) Consortium | Jacquelyn Wynn, Mairead Hyland, The Lighthouse Lab in Alderley Park and Alex Alderton, Roberto Amato, Jeffrey Barrett, Sonia Goncalves, Ewan Harrison, David K. Jackson, Ian Johnston, Dominic Kwiatkowski, Cordelia Langford, John Sillitoe on behalf of the Wellcome Sanger Institute COVID-19 Surveillance Team |
| EPI_ISL_2569851, EPI_ISL_2569852 | Lighthouse Laboratory Plymouth | Wellcome Sanger Institute for the COVID-19 Genomics UK (COG-UK) Consortium | Lighthouse Laboratory Plymouth and Alex Alderton, Roberto Amato, Jeffrey Barrett, Sonia Goncalves, Ewan Harrison, David K. Jackson, Ian Johnston, Dominic Kwiatkowski, Cordelia Langford, John Sillitoe on behalf of the Wellcome Sanger Institute COVID-19 Surveillance Team |
| EPI_ISL_2569853 | Lighthouse Lab in Milton Keynes | Wellcome Sanger Institute for the COVID-19 Genomics UK (COG-UK) Consortium | The Lighthouse Lab in Milton Keynes and Alex Alderton, Roberto Amato, Jeffrey Barrett, Sonia Goncalves, Ewan Harrison, David K. Jackson, Ian Johnston, Dominic Kwiatkowski, Cordelia Langford, John Sillitoe on behalf of the Wellcome Sanger Institute COVID-19 Surveillance Team |
| EPI_ISL_2569854 | Lighthouse Lab in Alderley Park | Wellcome Sanger Institute for the COVID-19 Genomics UK (COG-UK) Consortium | Jacquelyn Wynn, Mairead Hyland, The Lighthouse Lab in Alderley Park and Alex Alderton, Roberto Amato, Jeffrey Barrett, Sonia Goncalves, Ewan Harrison, David K. Jackson, Ian Johnston, Dominic Kwiatkowski, Cordelia Langford, John Sillitoe on behalf of the Wellcome Sanger Institute COVID-19 Surveillance Team |
| EPI_ISL_2569855 | Lighthouse Lab in Milton Keynes | Wellcome Sanger Institute for the COVID-19 Genomics UK (COG-UK) Consortium | The Lighthouse Lab in Milton Keynes and Alex Alderton, Roberto Amato, Jeffrey Barrett, Sonia Goncalves, Ewan Harrison, David K. Jackson, Ian Johnston, Dominic Kwiatkowski, Cordelia Langford, John Sillitoe on behalf of the Wellcome Sanger Institute COVID-19 Surveillance Team |
| EPI_ISL_2569856, EPI_ISL_2569857 | Lighthouse Lab in Alderley Park | Wellcome Sanger Institute for the COVID-19 Genomics UK (COG-UK) Consortium | Jacquelyn Wynn, Mairead Hyland, The Lighthouse Lab in Alderley Park and Alex Alderton, Roberto Amato, Jeffrey Barrett, Sonia Goncalves, Ewan Harrison, David K. Jackson, Ian Johnston, Dominic Kwiatkowski, Cordelia Langford, John Sillitoe on behalf of the Wellcome Sanger Institute COVID-19 Surveillance Team |
| EPI_ISL_2569858, EPI_ISL_2569859, EPI_ISL_2569860 | Lighthouse Laboratory Plymouth | Wellcome Sanger Institute for the COVID-19 Genomics UK (COG-UK) Consortium | Lighthouse Laboratory Plymouth and Alex Alderton, Roberto Amato, Jeffrey Barrett, Sonia Goncalves, Ewan Harrison, David K. Jackson, Ian Johnston, Dominic Kwiatkowski, Cordelia Langford, John Sillitoe on behalf of the Wellcome Sanger Institute COVID-19 Surveillance Team |
| EPI_ISL_2569861, EPI_ISL_2569862, EPI_ISL_2569863 | Lighthouse Lab in Milton Keynes | Wellcome Sanger Institute for the COVID-19 Genomics UK (COG-UK) Consortium | The Lighthouse Lab in Milton Keynes and Alex Alderton, Roberto Amato, Jeffrey Barrett, Sonia Goncalves, Ewan Harrison, David K. Jackson, Ian Johnston, Dominic Kwiatkowski, Cordelia Langford, John Sillitoe on behalf of the Wellcome Sanger Institute COVID-19 Surveillance Team |
| EPI_ISL_2569864, EPI_ISL_2569865 | Lighthouse Lab in Alderley Park | Wellcome Sanger Institute for the COVID-19 Genomics UK (COG-UK) Consortium | Jacquelyn Wynn, Mairead Hyland, The Lighthouse Lab in Alderley Park and Alex Alderton, Roberto Amato, Jeffrey Barrett, Sonia Goncalves, Ewan Harrison, David K. Jackson, Ian Johnston, Dominic Kwiatkowski, Cordelia Langford, John Sillitoe on behalf of the Wellcome Sanger Institute COVID-19 Surveillance Team |
| EPI_ISL_2569866, EPI_ISL_2569867 | Lighthouse Laboratory Plymouth | Wellcome Sanger Institute for the COVID-19 Genomics UK (COG-UK) Consortium | Lighthouse Laboratory Plymouth and Alex Alderton, Roberto Amato, Jeffrey Barrett, Sonia Goncalves, Ewan Harrison, David K. Jackson, Ian Johnston, Dominic Kwiatkowski, Cordelia Langford, John Sillitoe on behalf of the Wellcome Sanger Institute COVID-19 Surveillance Team |
| EPI_ISL_2569868 | Lighthouse Lab in Alderley Park | Wellcome Sanger Institute for the COVID-19 Genomics UK (COG-UK) Consortium | Jacquelyn Wynn, Mairead Hyland, The Lighthouse Lab in Alderley Park and Alex Alderton, Roberto Amato, Jeffrey Barrett, Sonia Goncalves, Ewan Harrison, David K. Jackson, Ian Johnston, Dominic Kwiatkowski, Cordelia Langford, John Sillitoe on behalf of the Wellcome Sanger Institute COVID-19 Surveillance Team |
| EPI_ISL_2569869 | Lighthouse Laboratory Plymouth | Wellcome Sanger Institute for the COVID-19 Genomics UK (COG-UK) Consortium | Lighthouse Laboratory Plymouth and Alex Alderton, Roberto Amato, Jeffrey Barrett, Sonia Goncalves, Ewan Harrison, David K. Jackson, Ian Johnston, Dominic Kwiatkowski, Cordelia Langford, John Sillitoe on behalf of the Wellcome Sanger Institute COVID-19 Surveillance Team |
| EPI_ISL_2569870 | Lighthouse Lab in Alderley Park | Wellcome Sanger Institute for the COVID-19 Genomics UK (COG-UK) Consortium | Jacquelyn Wynn, Mairead Hyland, The Lighthouse Lab in Alderley Park and Alex Alderton, Roberto Amato, Jeffrey Barrett, Sonia Goncalves, Ewan Harrison, David K. Jackson, Ian Johnston, Dominic Kwiatkowski, Cordelia Langford, John Sillitoe on behalf of the Wellcome Sanger Institute COVID-19 Surveillance Team |
| EPI_ISL_2569871, EPI_ISL_2569872 | Lighthouse Lab in Milton Keynes | Wellcome Sanger Institute for the COVID-19 Genomics UK (COG-UK) Consortium | The Lighthouse Lab in Milton Keynes and Alex Alderton, Roberto Amato, Jeffrey Barrett, Sonia Goncalves, Ewan Harrison, David K. Jackson, Ian Johnston, Dominic Kwiatkowski, Cordelia Langford, John Sillitoe on behalf of the Wellcome Sanger Institute COVID-19 Surveillance Team |
| EPI_ISL_2569873 | Lighthouse Lab in Alderley Park | Wellcome Sanger Institute for the COVID-19 Genomics UK (COG-UK) Consortium | Jacquelyn Wynn, Mairead Hyland, The Lighthouse Lab in Alderley Park and Alex Alderton, Roberto Amato, Jeffrey Barrett, Sonia Goncalves, Ewan Harrison, David K. Jackson, Ian Johnston, Dominic Kwiatkowski, Cordelia Langford, John Sillitoe on behalf of the Wellcome Sanger Institute COVID-19 Surveillance Team |
| EPI_ISL_2569874 | Lighthouse Laboratory Plymouth | Wellcome Sanger Institute for the COVID-19 Genomics UK (COG-UK) Consortium | Lighthouse Laboratory Plymouth and Alex Alderton, Roberto Amato, Jeffrey Barrett, Sonia Goncalves, Ewan Harrison, David K. Jackson, Ian Johnston, Dominic Kwiatkowski, Cordelia Langford, John Sillitoe on behalf of the Wellcome Sanger Institute COVID-19 Surveillance Team |
| EPI_ISL_2569875, EPI_ISL_2569876, EPI_ISL_2569877 | Lighthouse Lab in Milton Keynes | Wellcome Sanger Institute for the COVID-19 Genomics UK (COG-UK) Consortium | The Lighthouse Lab in Milton Keynes and Alex Alderton, Roberto Amato, Jeffrey Barrett, Sonia Goncalves, Ewan Harrison, David K. Jackson, Ian Johnston, Dominic Kwiatkowski, Cordelia Langford, John Sillitoe on behalf of the Wellcome Sanger Institute COVID-19 Surveillance Team |
| EPI_ISL_2569878 | Lighthouse Lab in Alderley Park | Wellcome Sanger Institute for the COVID-19 Genomics UK (COG-UK) Consortium | Jacquelyn Wynn, Mairead Hyland, The Lighthouse Lab in Alderley Park and Alex Alderton, Roberto Amato, Jeffrey Barrett, Sonia Goncalves, Ewan Harrison, David K. Jackson, Ian Johnston, Dominic Kwiatkowski, Cordelia Langford, John Sillitoe on behalf of the Wellcome Sanger Institute COVID-19 Surveillance Team |
| EPI_ISL_2569879 | Lighthouse Lab in Milton Keynes | Wellcome Sanger Institute for the COVID-19 Genomics UK (COG-UK) Consortium | The Lighthouse Lab in Milton Keynes and Alex Alderton, Roberto Amato, Jeffrey Barrett, Sonia Goncalves, Ewan Harrison, David K. Jackson, Ian Johnston, Dominic Kwiatkowski, Cordelia Langford, John Sillitoe on behalf of the Wellcome Sanger Institute COVID-19 Surveillance Team |
| EPI_ISL_2569880 | Lighthouse Laboratory Plymouth | Wellcome Sanger Institute for the COVID-19 Genomics UK (COG-UK) Consortium | Lighthouse Laboratory Plymouth and Alex Alderton, Roberto Amato, Jeffrey Barrett, Sonia Goncalves, Ewan Harrison, David K. Jackson, Ian Johnston, Dominic Kwiatkowski, Cordelia Langford, John Sillitoe on behalf of the Wellcome Sanger Institute COVID-19 Surveillance Team |
| EPI_ISL_2569882 | Lighthouse Lab in Milton Keynes | Wellcome Sanger Institute for the COVID-19 Genomics UK (COG-UK) Consortium | The Lighthouse Lab in Milton Keynes and Alex Alderton, Roberto Amato, Jeffrey Barrett, Sonia Goncalves, Ewan Harrison, David K. Jackson, Ian Johnston, Dominic Kwiatkowski, Cordelia Langford, John Sillitoe on behalf of the Wellcome Sanger Institute COVID-19 Surveillance Team |
| EPI_ISL_2569883 | Lighthouse Lab in Alderley Park | Wellcome Sanger Institute for the COVID-19 Genomics UK (COG-UK) Consortium | Jacquelyn Wynn, Mairead Hyland, The Lighthouse Lab in Alderley Park and Alex Alderton, Roberto Amato, Jeffrey Barrett, Sonia Goncalves, Ewan Harrison, David K. Jackson, Ian Johnston, Dominic Kwiatkowski, Cordelia Langford, John Sillitoe on behalf of the Wellcome Sanger Institute COVID-19 Surveillance Team |
| EPI_ISL_2569884, EPI_ISL_2569885 | Lighthouse Lab in Milton Keynes | Wellcome Sanger Institute for the COVID-19 Genomics UK (COG-UK) Consortium | The Lighthouse Lab in Milton Keynes and Alex Alderton, Roberto Amato, Jeffrey Barrett, Sonia Goncalves, Ewan Harrison, David K. Jackson, Ian Johnston, Dominic Kwiatkowski, Cordelia Langford, John Sillitoe on behalf of the Wellcome Sanger Institute COVID-19 Surveillance Team |
| EPI_ISL_2569886, EPI_ISL_2569887 | Lighthouse Laboratory Plymouth | Wellcome Sanger Institute for the COVID-19 Genomics UK (COG-UK) Consortium | Lighthouse Laboratory Plymouth and Alex Alderton, Roberto Amato, Jeffrey Barrett, Sonia Goncalves, Ewan Harrison, David K. Jackson, Ian Johnston, Dominic Kwiatkowski, Cordelia Langford, John Sillitoe on behalf of the Wellcome Sanger Institute COVID-19 Surveillance Team |
| EPI_ISL_2569888, EPI_ISL_2569889 | Lighthouse Lab in Alderley Park | Wellcome Sanger Institute for the COVID-19 Genomics UK (COG-UK) Consortium | Jacquelyn Wynn, Mairead Hyland, The Lighthouse Lab in Alderley Park and Alex Alderton, Roberto Amato, Jeffrey Barrett, Sonia Goncalves, Ewan Harrison, David K. Jackson, Ian Johnston, Dominic Kwiatkowski, Cordelia Langford, John Sillitoe on behalf of the Wellcome Sanger Institute COVID-19 Surveillance Team |
| EPI_ISL_2569890, | Lighthouse Laboratory Plymouth | Wellcome Sanger Institute for the COVID-19 Genomics UK (COG-UK) | Lighthouse Laboratory Plymouth and Alex Alderton, Roberto Amato, Jeffrey Barrett, Sonia Goncalves, Ewan Harrison, David K. Jackson, Ian Johnston, Dominic Kwiatkowski, Cordelia Langford, John Sillitoe on behalf of the Wellcome Sanger |

| | | | |
|---|---|---|---|
| EPI_ISL_2569891 | | Consortium | Institute COVID-19 Surveillance Team |
| EPI_ISL_2569892 | Lighthouse Lab in Milton Keynes | Wellcome Sanger Institute for the COVID-19 Genomics UK (COG-UK) Consortium | The Lighthouse Lab in Milton Keynes and Alex Alderton, Roberto Amato, Jeffrey Barrett, Sonia Goncalves, Ewan Harrison, David K. Jackson, Ian Johnston, Dominic Kwiatkowski, Cordelia Langford, John Sillitoe on behalf of the Wellcome Sanger Institute COVID-19 Surveillance Team |
| EPI_ISL_2569893 | Lighthouse Laboratory Plymouth | Wellcome Sanger Institute for the COVID-19 Genomics UK (COG-UK) Consortium | Lighthouse Laboratory Plymouth and Alex Alderton, Roberto Amato, Jeffrey Barrett, Sonia Goncalves, Ewan Harrison, David K. Jackson, Ian Johnston, Dominic Kwiatkowski, Cordelia Langford, John Sillitoe on behalf of the Wellcome Sanger Institute COVID-19 Surveillance Team |
| EPI_ISL_2569894, EPI_ISL_2569895, EPI_ISL_2569896 | Lighthouse Lab in Alderley Park | Wellcome Sanger Institute for the COVID-19 Genomics UK (COG-UK) Consortium | Jacquelyn Wynn, Mairead Hyland, The Lighthouse Lab in Alderley Park and Alex Alderton, Roberto Amato, Jeffrey Barrett, Sonia Goncalves, Ewan Harrison, David K. Jackson, Ian Johnston, Dominic Kwiatkowski, Cordelia Langford, John Sillitoe on behalf of the Wellcome Sanger Institute COVID-19 Surveillance Team |
| EPI_ISL_2569897, EPI_ISL_2569898, EPI_ISL_2569899 | Lighthouse Lab in Milton Keynes | Wellcome Sanger Institute for the COVID-19 Genomics UK (COG-UK) Consortium | The Lighthouse Lab in Milton Keynes and Alex Alderton, Roberto Amato, Jeffrey Barrett, Sonia Goncalves, Ewan Harrison, David K. Jackson, Ian Johnston, Dominic Kwiatkowski, Cordelia Langford, John Sillitoe on behalf of the Wellcome Sanger Institute COVID-19 Surveillance Team |
| EPI_ISL_2569900 | Lighthouse Lab in Alderley Park | Wellcome Sanger Institute for the COVID-19 Genomics UK (COG-UK) Consortium | Jacquelyn Wynn, Mairead Hyland, The Lighthouse Lab in Alderley Park and Alex Alderton, Roberto Amato, Jeffrey Barrett, Sonia Goncalves, Ewan Harrison, David K. Jackson, Ian Johnston, Dominic Kwiatkowski, Cordelia Langford, John Sillitoe on behalf of the Wellcome Sanger Institute COVID-19 Surveillance Team |
| EPI_ISL_2569901, EPI_ISL_2569902 | Lighthouse Laboratory Plymouth | Wellcome Sanger Institute for the COVID-19 Genomics UK (COG-UK) Consortium | Lighthouse Laboratory Plymouth and Alex Alderton, Roberto Amato, Jeffrey Barrett, Sonia Goncalves, Ewan Harrison, David K. Jackson, Ian Johnston, Dominic Kwiatkowski, Cordelia Langford, John Sillitoe on behalf of the Wellcome Sanger Institute COVID-19 Surveillance Team |
| EPI_ISL_2569903 | Lighthouse Lab in Milton Keynes | Wellcome Sanger Institute for the COVID-19 Genomics UK (COG-UK) Consortium | The Lighthouse Lab in Milton Keynes and Alex Alderton, Roberto Amato, Jeffrey Barrett, Sonia Goncalves, Ewan Harrison, David K. Jackson, Ian Johnston, Dominic Kwiatkowski, Cordelia Langford, John Sillitoe on behalf of the Wellcome Sanger Institute COVID-19 Surveillance Team |
| EPI_ISL_2569904, EPI_ISL_2569905 | Lighthouse Lab in Alderley Park | Wellcome Sanger Institute for the COVID-19 Genomics UK (COG-UK) Consortium | Jacquelyn Wynn, Mairead Hyland, The Lighthouse Lab in Alderley Park and Alex Alderton, Roberto Amato, Jeffrey Barrett, Sonia Goncalves, Ewan Harrison, David K. Jackson, Ian Johnston, Dominic Kwiatkowski, Cordelia Langford, John Sillitoe on behalf of the Wellcome Sanger Institute COVID-19 Surveillance Team |
| EPI_ISL_2569906 | Lighthouse Laboratory Plymouth | Wellcome Sanger Institute for the COVID-19 Genomics UK (COG-UK) Consortium | Lighthouse Laboratory Plymouth and Alex Alderton, Roberto Amato, Jeffrey Barrett, Sonia Goncalves, Ewan Harrison, David K. Jackson, Ian Johnston, Dominic Kwiatkowski, Cordelia Langford, John Sillitoe on behalf of the Wellcome Sanger Institute COVID-19 Surveillance Team |
| EPI_ISL_2569907 | Lighthouse Lab in Alderley Park | Wellcome Sanger Institute for the COVID-19 Genomics UK (COG-UK) Consortium | Jacquelyn Wynn, Mairead Hyland, The Lighthouse Lab in Alderley Park and Alex Alderton, Roberto Amato, Jeffrey Barrett, Sonia Goncalves, Ewan Harrison, David K. Jackson, Ian Johnston, Dominic Kwiatkowski, Cordelia Langford, John Sillitoe on behalf of the Wellcome Sanger Institute COVID-19 Surveillance Team |
| EPI_ISL_2569909 | Lighthouse Lab in Milton Keynes | Wellcome Sanger Institute for the COVID-19 Genomics UK (COG-UK) Consortium | The Lighthouse Lab in Milton Keynes and Alex Alderton, Roberto Amato, Jeffrey Barrett, Sonia Goncalves, Ewan Harrison, David K. Jackson, Ian Johnston, Dominic Kwiatkowski, Cordelia Langford, John Sillitoe on behalf of the Wellcome Sanger Institute COVID-19 Surveillance Team |
| EPI_ISL_2569910, EPI_ISL_2569911 | Lighthouse Laboratory Plymouth | Wellcome Sanger Institute for the COVID-19 Genomics UK (COG-UK) Consortium | Lighthouse Laboratory Plymouth and Alex Alderton, Roberto Amato, Jeffrey Barrett, Sonia Goncalves, Ewan Harrison, David K. Jackson, Ian Johnston, Dominic Kwiatkowski, Cordelia Langford, John Sillitoe on behalf of the Wellcome Sanger Institute COVID-19 Surveillance Team |
| EPI_ISL_2569912 | Lighthouse Lab in Milton Keynes | Wellcome Sanger Institute for the COVID-19 Genomics UK (COG-UK) Consortium | The Lighthouse Lab in Milton Keynes and Alex Alderton, Roberto Amato, Jeffrey Barrett, Sonia Goncalves, Ewan Harrison, David K. Jackson, Ian Johnston, Dominic Kwiatkowski, Cordelia Langford, John Sillitoe on behalf of the Wellcome Sanger Institute COVID-19 Surveillance Team |
| EPI_ISL_2569913 | Lighthouse Laboratory Plymouth | Wellcome Sanger Institute for the COVID-19 Genomics UK (COG-UK) Consortium | Lighthouse Laboratory Plymouth and Alex Alderton, Roberto Amato, Jeffrey Barrett, Sonia Goncalves, Ewan Harrison, David K. Jackson, Ian Johnston, Dominic Kwiatkowski, Cordelia Langford, John Sillitoe on behalf of the Wellcome Sanger Institute COVID-19 Surveillance Team |
| EPI_ISL_2569914 | Lighthouse Lab in Milton Keynes | Wellcome Sanger Institute for the COVID-19 Genomics UK (COG-UK) Consortium | The Lighthouse Lab in Milton Keynes and Alex Alderton, Roberto Amato, Jeffrey Barrett, Sonia Goncalves, Ewan Harrison, David K. Jackson, Ian Johnston, Dominic Kwiatkowski, Cordelia Langford, John Sillitoe on behalf of the Wellcome Sanger Institute COVID-19 Surveillance Team |
| EPI_ISL_2569915, EPI_ISL_2569916 | Lighthouse Lab in Alderley Park | Wellcome Sanger Institute for the COVID-19 Genomics UK (COG-UK) Consortium | Jacquelyn Wynn, Mairead Hyland, The Lighthouse Lab in Alderley Park and Alex Alderton, Roberto Amato, Jeffrey Barrett, Sonia Goncalves, Ewan Harrison, David K. Jackson, Ian Johnston, Dominic Kwiatkowski, Cordelia Langford, John Sillitoe on behalf of the Wellcome Sanger Institute COVID-19 Surveillance Team |
| EPI_ISL_2569917 | Lighthouse Lab in Milton Keynes | Wellcome Sanger Institute for the COVID-19 Genomics UK (COG-UK) Consortium | The Lighthouse Lab in Milton Keynes and Alex Alderton, Roberto Amato, Jeffrey Barrett, Sonia Goncalves, Ewan Harrison, David K. Jackson, Ian Johnston, Dominic Kwiatkowski, Cordelia Langford, John Sillitoe on behalf of the Wellcome Sanger Institute COVID-19 Surveillance Team |
| EPI_ISL_2569918 | Lighthouse Lab in Alderley Park | Wellcome Sanger Institute for the COVID-19 Genomics UK (COG-UK) Consortium | Jacquelyn Wynn, Mairead Hyland, The Lighthouse Lab in Alderley Park and Alex Alderton, Roberto Amato, Jeffrey Barrett, Sonia Goncalves, Ewan Harrison, David K. Jackson, Ian Johnston, Dominic Kwiatkowski, Cordelia Langford, John Sillitoe on behalf of the Wellcome Sanger Institute COVID-19 Surveillance Team |
| EPI_ISL_2569919 | Lighthouse Lab in Milton Keynes | Wellcome Sanger Institute for the COVID-19 Genomics UK (COG-UK) Consortium | The Lighthouse Lab in Milton Keynes and Alex Alderton, Roberto Amato, Jeffrey Barrett, Sonia Goncalves, Ewan Harrison, David K. Jackson, Ian Johnston, Dominic Kwiatkowski, Cordelia Langford, John Sillitoe on behalf of the Wellcome Sanger Institute COVID-19 Surveillance Team |
| EPI_ISL_2569920 | Lighthouse Lab in Alderley Park | Wellcome Sanger Institute for the COVID-19 Genomics UK (COG-UK) Consortium | Jacquelyn Wynn, Mairead Hyland, The Lighthouse Lab in Alderley Park and Alex Alderton, Roberto Amato, Jeffrey Barrett, Sonia Goncalves, Ewan Harrison, David K. Jackson, Ian Johnston, Dominic Kwiatkowski, Cordelia Langford, John Sillitoe on behalf of the Wellcome Sanger Institute COVID-19 Surveillance Team |
| EPI_ISL_2569921 | Lighthouse Lab in Milton Keynes | Wellcome Sanger Institute for the COVID-19 Genomics UK (COG-UK) Consortium | The Lighthouse Lab in Milton Keynes and Alex Alderton, Roberto Amato, Jeffrey Barrett, Sonia Goncalves, Ewan Harrison, David K. Jackson, Ian Johnston, Dominic Kwiatkowski, Cordelia Langford, John Sillitoe on behalf of the Wellcome Sanger Institute COVID-19 Surveillance Team |
| EPI_ISL_2569922 | Lighthouse Laboratory Plymouth | Wellcome Sanger Institute for the COVID-19 Genomics UK (COG-UK) Consortium | Lighthouse Laboratory Plymouth and Alex Alderton, Roberto Amato, Jeffrey Barrett, Sonia Goncalves, Ewan Harrison, David K. Jackson, Ian Johnston, Dominic Kwiatkowski, Cordelia Langford, John Sillitoe on behalf of the Wellcome Sanger Institute COVID-19 Surveillance Team |
| EPI_ISL_2569923, EPI_ISL_2569924, EPI_ISL_2569925, EPI_ISL_2569926 | Lighthouse Lab in Milton Keynes | Wellcome Sanger Institute for the COVID-19 Genomics UK (COG-UK) Consortium | The Lighthouse Lab in Milton Keynes and Alex Alderton, Roberto Amato, Jeffrey Barrett, Sonia Goncalves, Ewan Harrison, David K. Jackson, Ian Johnston, Dominic Kwiatkowski, Cordelia Langford, John Sillitoe on behalf of the Wellcome Sanger Institute COVID-19 Surveillance Team |
| EPI_ISL_2569927, EPI_ISL_2569928, EPI_ISL_2569929, EPI_ISL_2569930, EPI_ISL_2569931, EPI_ISL_2569932, EPI_ISL_2569933, EPI_ISL_2569934 | Lighthouse Lab in Alderley Park | Wellcome Sanger Institute for the COVID-19 Genomics UK (COG-UK) Consortium | Jacquelyn Wynn, Mairead Hyland, The Lighthouse Lab in Alderley Park and Alex Alderton, Roberto Amato, Jeffrey Barrett, Sonia Goncalves, Ewan Harrison, David K. Jackson, Ian Johnston, Dominic Kwiatkowski, Cordelia Langford, John Sillitoe on behalf of the Wellcome Sanger Institute COVID-19 Surveillance Team |
| EPI_ISL_2569935 | Lighthouse Lab in Milton Keynes | Wellcome Sanger Institute for the COVID-19 Genomics UK (COG-UK) Consortium | The Lighthouse Lab in Milton Keynes and Alex Alderton, Roberto Amato, Jeffrey Barrett, Sonia Goncalves, Ewan Harrison, David K. Jackson, Ian Johnston, Dominic Kwiatkowski, Cordelia Langford, John Sillitoe on behalf of the Wellcome Sanger Institute COVID-19 Surveillance Team |
| EPI_ISL_2569936, EPI_ISL_2569937, EPI_ISL_2569938, EPI_ISL_2569939, EPI_ISL_2569940, EPI_ISL_2569941, EPI_ISL_2569942 | Lighthouse Lab in Alderley Park | Wellcome Sanger Institute for the COVID-19 Genomics UK (COG-UK) Consortium | Jacquelyn Wynn, Mairead Hyland, The Lighthouse Lab in Alderley Park and Alex Alderton, Roberto Amato, Jeffrey Barrett, Sonia Goncalves, Ewan Harrison, David K. Jackson, Ian Johnston, Dominic Kwiatkowski, Cordelia Langford, John Sillitoe on behalf of the Wellcome Sanger Institute COVID-19 Surveillance Team |
| EPI_ISL_2569943 | Lighthouse Lab in Milton Keynes | Wellcome Sanger Institute for the COVID-19 Genomics UK (COG-UK) Consortium | The Lighthouse Lab in Milton Keynes and Alex Alderton, Roberto Amato, Jeffrey Barrett, Sonia Goncalves, Ewan Harrison, David K. Jackson, Ian Johnston, Dominic Kwiatkowski, Cordelia Langford, John Sillitoe on behalf of the Wellcome Sanger Institute COVID-19 Surveillance Team |
| EPI_ISL_2569944, EPI_ISL_2569945, EPI_ISL_2569946, EPI_ISL_2569947, EPI_ISL_2569948, EPI_ISL_2569949 | Lighthouse Lab in Alderley Park | Wellcome Sanger Institute for the COVID-19 Genomics UK (COG-UK) Consortium | Jacquelyn Wynn, Mairead Hyland, The Lighthouse Lab in Alderley Park and Alex Alderton, Roberto Amato, Jeffrey Barrett, Sonia Goncalves, Ewan Harrison, David K. Jackson, Ian Johnston, Dominic Kwiatkowski, Cordelia Langford, John Sillitoe on behalf of the Wellcome Sanger Institute COVID-19 Surveillance Team |
| EPI_ISL_2569950 | Lighthouse Lab in Milton Keynes | Wellcome Sanger Institute for the COVID-19 Genomics UK (COG-UK) Consortium | The Lighthouse Lab in Milton Keynes and Alex Alderton, Roberto Amato, Jeffrey Barrett, Sonia Goncalves, Ewan Harrison, David K. Jackson, Ian Johnston, Dominic Kwiatkowski, Cordelia Langford, John Sillitoe on behalf of the Wellcome Sanger Institute COVID-19 Surveillance Team |
| EPI_ISL_2569951, EPI_ISL_2569953, EPI_ISL_2569954, EPI_ISL_2569955 | Lighthouse Lab in Alderley Park | Wellcome Sanger Institute for the COVID-19 Genomics UK (COG-UK) Consortium | Jacquelyn Wynn, Mairead Hyland, The Lighthouse Lab in Alderley Park and Alex Alderton, Roberto Amato, Jeffrey Barrett, Sonia Goncalves, Ewan Harrison, David K. Jackson, Ian Johnston, Dominic Kwiatkowski, Cordelia Langford, John Sillitoe on behalf of the Wellcome Sanger Institute COVID-19 Surveillance Team |
| EPI_ISL_2569956 | Lighthouse Lab in Milton Keynes | Wellcome Sanger Institute for the COVID-19 Genomics UK (COG-UK) Consortium | The Lighthouse Lab in Milton Keynes and Alex Alderton, Roberto Amato, Jeffrey Barrett, Sonia Goncalves, Ewan Harrison, David K. Jackson, Ian Johnston, Dominic Kwiatkowski, Cordelia Langford, John Sillitoe on behalf of the Wellcome Sanger Institute COVID-19 Surveillance Team |
| EPI_ISL_2569957, EPI_ISL_2569958, EPI_ISL_2569959, EPI_ISL_2569960, EPI_ISL_2569961 | Lighthouse Lab in Alderley Park | Wellcome Sanger Institute for the COVID-19 Genomics UK (COG-UK) Consortium | Jacquelyn Wynn, Mairead Hyland, The Lighthouse Lab in Alderley Park and Alex Alderton, Roberto Amato, Jeffrey Barrett, Sonia Goncalves, Ewan Harrison, David K. Jackson, Ian Johnston, Dominic Kwiatkowski, Cordelia Langford, John Sillitoe on behalf of the Wellcome Sanger Institute COVID-19 Surveillance Team |
| EPI_ISL_2569962 | Lighthouse Lab in Milton Keynes | Wellcome Sanger Institute for the COVID-19 Genomics UK (COG-UK) Consortium | The Lighthouse Lab in Milton Keynes and Alex Alderton, Roberto Amato, Jeffrey Barrett, Sonia Goncalves, Ewan Harrison, David K. Jackson, Ian Johnston, Dominic Kwiatkowski, Cordelia Langford, John Sillitoe on behalf of the Wellcome Sanger Institute COVID-19 Surveillance Team |
| EPI_ISL_2569963, EPI_ISL_2569964, EPI_ISL_2569965, EPI_ISL_2569966, | Lighthouse Lab in Alderley Park | Wellcome Sanger Institute for the COVID-19 Genomics UK (COG-UK) Consortium | Jacquelyn Wynn, Mairead Hyland, The Lighthouse Lab in Alderley Park and Alex Alderton, Roberto Amato, Jeffrey Barrett, Sonia Goncalves, Ewan Harrison, David K. Jackson, Ian Johnston, Dominic Kwiatkowski, Cordelia Langford, John Sillitoe on behalf of the Wellcome Sanger Institute COVID-19 Surveillance Team |

| | | | |
|---|---|---|---|
| EPI_ISL_2569967, EPI_ISL_2569968 | | | |
| EPI_ISL_2569969 | Lighthouse Lab in Milton Keynes | Wellcome Sanger Institute for the COVID-19 Genomics UK (COG-UK) Consortium | The Lighthouse Lab in Milton Keynes and Alex Alderton, Roberto Amato, Jeffrey Barrett, Sonia Goncalves, Ewan Harrison, David K. Jackson, Ian Johnston, Dominic Kwiatkowski, Cordelia Langford, John Sillitoe on behalf of the Wellcome Sanger Institute COVID-19 Surveillance Team |
| EPI_ISL_2569970, EPI_ISL_2569971, EPI_ISL_2569972 | Lighthouse Lab in Alderley Park | Wellcome Sanger Institute for the COVID-19 Genomics UK (COG-UK) Consortium | Jacquelyn Wynn, Mairead Hyland, The Lighthouse Lab in Alderley Park and Alex Alderton, Roberto Amato, Jeffrey Barrett, Sonia Goncalves, Ewan Harrison, David K. Jackson, Ian Johnston, Dominic Kwiatkowski, Cordelia Langford, John Sillitoe on behalf of the Wellcome Sanger Institute COVID-19 Surveillance Team |
| EPI_ISL_2569973 | Lighthouse Lab in Milton Keynes | Wellcome Sanger Institute for the COVID-19 Genomics UK (COG-UK) Consortium | The Lighthouse Lab in Milton Keynes and Alex Alderton, Roberto Amato, Jeffrey Barrett, Sonia Goncalves, Ewan Harrison, David K. Jackson, Ian Johnston, Dominic Kwiatkowski, Cordelia Langford, John Sillitoe on behalf of the Wellcome Sanger Institute COVID-19 Surveillance Team |
| EPI_ISL_2569974, EPI_ISL_2569975 | Lighthouse Lab in Alderley Park | Wellcome Sanger Institute for the COVID-19 Genomics UK (COG-UK) Consortium | Jacquelyn Wynn, Mairead Hyland, The Lighthouse Lab in Alderley Park and Alex Alderton, Roberto Amato, Jeffrey Barrett, Sonia Goncalves, Ewan Harrison, David K. Jackson, Ian Johnston, Dominic Kwiatkowski, Cordelia Langford, John Sillitoe on behalf of the Wellcome Sanger Institute COVID-19 Surveillance Team |
| EPI_ISL_2569976 | Lighthouse Lab in Milton Keynes | Wellcome Sanger Institute for the COVID-19 Genomics UK (COG-UK) Consortium | The Lighthouse Lab in Milton Keynes and Alex Alderton, Roberto Amato, Jeffrey Barrett, Sonia Goncalves, Ewan Harrison, David K. Jackson, Ian Johnston, Dominic Kwiatkowski, Cordelia Langford, John Sillitoe on behalf of the Wellcome Sanger Institute COVID-19 Surveillance Team |
| EPI_ISL_2569977, EPI_ISL_2569978 | Lighthouse Lab in Alderley Park | Wellcome Sanger Institute for the COVID-19 Genomics UK (COG-UK) Consortium | Jacquelyn Wynn, Mairead Hyland, The Lighthouse Lab in Alderley Park and Alex Alderton, Roberto Amato, Jeffrey Barrett, Sonia Goncalves, Ewan Harrison, David K. Jackson, Ian Johnston, Dominic Kwiatkowski, Cordelia Langford, John Sillitoe on behalf of the Wellcome Sanger Institute COVID-19 Surveillance Team |
| EPI_ISL_2569979 | Lighthouse Lab in Milton Keynes | Wellcome Sanger Institute for the COVID-19 Genomics UK (COG-UK) Consortium | The Lighthouse Lab in Milton Keynes and Alex Alderton, Roberto Amato, Jeffrey Barrett, Sonia Goncalves, Ewan Harrison, David K. Jackson, Ian Johnston, Dominic Kwiatkowski, Cordelia Langford, John Sillitoe on behalf of the Wellcome Sanger Institute COVID-19 Surveillance Team |
| EPI_ISL_2569980, EPI_ISL_2569981 | Lighthouse Lab in Alderley Park | Wellcome Sanger Institute for the COVID-19 Genomics UK (COG-UK) Consortium | Jacquelyn Wynn, Mairead Hyland, The Lighthouse Lab in Alderley Park and Alex Alderton, Roberto Amato, Jeffrey Barrett, Sonia Goncalves, Ewan Harrison, David K. Jackson, Ian Johnston, Dominic Kwiatkowski, Cordelia Langford, John Sillitoe on behalf of the Wellcome Sanger Institute COVID-19 Surveillance Team |
| EPI_ISL_2569982 | Lighthouse Lab in Milton Keynes | Wellcome Sanger Institute for the COVID-19 Genomics UK (COG-UK) Consortium | The Lighthouse Lab in Milton Keynes and Alex Alderton, Roberto Amato, Jeffrey Barrett, Sonia Goncalves, Ewan Harrison, David K. Jackson, Ian Johnston, Dominic Kwiatkowski, Cordelia Langford, John Sillitoe on behalf of the Wellcome Sanger Institute COVID-19 Surveillance Team |
| EPI_ISL_2569983, EPI_ISL_2569984, EPI_ISL_2569985, EPI_ISL_2569986, EPI_ISL_2569987, EPI_ISL_2569988, EPI_ISL_2569990, EPI_ISL_2569991, EPI_ISL_2569992, EPI_ISL_2569993, EPI_ISL_2569994 | | | |
| see above | Lighthouse Lab in Alderley Park | Wellcome Sanger Institute for the COVID-19 Genomics UK (COG-UK) Consortium | Jacquelyn Wynn, Mairead Hyland, The Lighthouse Lab in Alderley Park and Alex Alderton, Roberto Amato, Jeffrey Barrett, Sonia Goncalves, Ewan Harrison, David K. Jackson, Ian Johnston, Dominic Kwiatkowski, Cordelia Langford, John Sillitoe on behalf of the Wellcome Sanger Institute COVID-19 Surveillance Team |
| EPI_ISL_2569995 | Lighthouse Lab in Milton Keynes | Wellcome Sanger Institute for the COVID-19 Genomics UK (COG-UK) Consortium | The Lighthouse Lab in Milton Keynes and Alex Alderton, Roberto Amato, Jeffrey Barrett, Sonia Goncalves, Ewan Harrison, David K. Jackson, Ian Johnston, Dominic Kwiatkowski, Cordelia Langford, John Sillitoe on behalf of the Wellcome Sanger Institute COVID-19 Surveillance Team |
| EPI_ISL_2569996, EPI_ISL_2569998 | Lighthouse Lab in Alderley Park | Wellcome Sanger Institute for the COVID-19 Genomics UK (COG-UK) Consortium | Jacquelyn Wynn, Mairead Hyland, The Lighthouse Lab in Alderley Park and Alex Alderton, Roberto Amato, Jeffrey Barrett, Sonia Goncalves, Ewan Harrison, David K. Jackson, Ian Johnston, Dominic Kwiatkowski, Cordelia Langford, John Sillitoe on behalf of the Wellcome Sanger Institute COVID-19 Surveillance Team |
| EPI_ISL_2569999, EPI_ISL_2570000 | Lighthouse Lab in Milton Keynes | Wellcome Sanger Institute for the COVID-19 Genomics UK (COG-UK) Consortium | The Lighthouse Lab in Milton Keynes and Alex Alderton, Roberto Amato, Jeffrey Barrett, Sonia Goncalves, Ewan Harrison, David K. Jackson, Ian Johnston, Dominic Kwiatkowski, Cordelia Langford, John Sillitoe on behalf of the Wellcome Sanger Institute COVID-19 Surveillance Team |
| EPI_ISL_2570001, EPI_ISL_2570002, EPI_ISL_2570003 | Lighthouse Lab in Alderley Park | Wellcome Sanger Institute for the COVID-19 Genomics UK (COG-UK) Consortium | Jacquelyn Wynn, Mairead Hyland, The Lighthouse Lab in Alderley Park and Alex Alderton, Roberto Amato, Jeffrey Barrett, Sonia Goncalves, Ewan Harrison, David K. Jackson, Ian Johnston, Dominic Kwiatkowski, Cordelia Langford, John Sillitoe on behalf of the Wellcome Sanger Institute COVID-19 Surveillance Team |
| EPI_ISL_2570004 | Lighthouse Lab in Milton Keynes | Wellcome Sanger Institute for the COVID-19 Genomics UK (COG-UK) Consortium | The Lighthouse Lab in Milton Keynes and Alex Alderton, Roberto Amato, Jeffrey Barrett, Sonia Goncalves, Ewan Harrison, David K. Jackson, Ian Johnston, Dominic Kwiatkowski, Cordelia Langford, John Sillitoe on behalf of the Wellcome Sanger Institute COVID-19 Surveillance Team |
| EPI_ISL_2570005, EPI_ISL_2570006, EPI_ISL_2570007, EPI_ISL_2570008 | Lighthouse Lab in Alderley Park | Wellcome Sanger Institute for the COVID-19 Genomics UK (COG-UK) Consortium | Jacquelyn Wynn, Mairead Hyland, The Lighthouse Lab in Alderley Park and Alex Alderton, Roberto Amato, Jeffrey Barrett, Sonia Goncalves, Ewan Harrison, David K. Jackson, Ian Johnston, Dominic Kwiatkowski, Cordelia Langford, John Sillitoe on behalf of the Wellcome Sanger Institute COVID-19 Surveillance Team |
| EPI_ISL_2570009 | Lighthouse Lab in Milton Keynes | Wellcome Sanger Institute for the COVID-19 Genomics UK (COG-UK) Consortium | The Lighthouse Lab in Milton Keynes and Alex Alderton, Roberto Amato, Jeffrey Barrett, Sonia Goncalves, Ewan Harrison, David K. Jackson, Ian Johnston, Dominic Kwiatkowski, Cordelia Langford, John Sillitoe on behalf of the Wellcome Sanger Institute COVID-19 Surveillance Team |
| EPI_ISL_2570010, EPI_ISL_2570011 | Lighthouse Lab in Alderley Park | Wellcome Sanger Institute for the COVID-19 Genomics UK (COG-UK) Consortium | Jacquelyn Wynn, Mairead Hyland, The Lighthouse Lab in Alderley Park and Alex Alderton, Roberto Amato, Jeffrey Barrett, Sonia Goncalves, Ewan Harrison, David K. Jackson, Ian Johnston, Dominic Kwiatkowski, Cordelia Langford, John Sillitoe on behalf of the Wellcome Sanger Institute COVID-19 Surveillance Team |
| EPI_ISL_2570012 | Lighthouse Lab in Milton Keynes | Wellcome Sanger Institute for the COVID-19 Genomics UK (COG-UK) Consortium | The Lighthouse Lab in Milton Keynes and Alex Alderton, Roberto Amato, Jeffrey Barrett, Sonia Goncalves, Ewan Harrison, David K. Jackson, Ian Johnston, Dominic Kwiatkowski, Cordelia Langford, John Sillitoe on behalf of the Wellcome Sanger Institute COVID-19 Surveillance Team |
| EPI_ISL_2570013, EPI_ISL_2570014, EPI_ISL_2570015 | Lighthouse Lab in Alderley Park | Wellcome Sanger Institute for the COVID-19 Genomics UK (COG-UK) Consortium | Jacquelyn Wynn, Mairead Hyland, The Lighthouse Lab in Alderley Park and Alex Alderton, Roberto Amato, Jeffrey Barrett, Sonia Goncalves, Ewan Harrison, David K. Jackson, Ian Johnston, Dominic Kwiatkowski, Cordelia Langford, John Sillitoe on behalf of the Wellcome Sanger Institute COVID-19 Surveillance Team |
| EPI_ISL_2570016, EPI_ISL_2570017 | Lighthouse Lab in Milton Keynes | Wellcome Sanger Institute for the COVID-19 Genomics UK (COG-UK) Consortium | The Lighthouse Lab in Milton Keynes and Alex Alderton, Roberto Amato, Jeffrey Barrett, Sonia Goncalves, Ewan Harrison, David K. Jackson, Ian Johnston, Dominic Kwiatkowski, Cordelia Langford, John Sillitoe on behalf of the Wellcome Sanger Institute COVID-19 Surveillance Team |
| EPI_ISL_2570018, EPI_ISL_2570019, EPI_ISL_2570020, EPI_ISL_2570021 | Lighthouse Lab in Alderley Park | Wellcome Sanger Institute for the COVID-19 Genomics UK (COG-UK) Consortium | Jacquelyn Wynn, Mairead Hyland, The Lighthouse Lab in Alderley Park and Alex Alderton, Roberto Amato, Jeffrey Barrett, Sonia Goncalves, Ewan Harrison, David K. Jackson, Ian Johnston, Dominic Kwiatkowski, Cordelia Langford, John Sillitoe on behalf of the Wellcome Sanger Institute COVID-19 Surveillance Team |
| EPI_ISL_2570022 | Lighthouse Lab in Milton Keynes | Wellcome Sanger Institute for the COVID-19 Genomics UK (COG-UK) Consortium | The Lighthouse Lab in Milton Keynes and Alex Alderton, Roberto Amato, Jeffrey Barrett, Sonia Goncalves, Ewan Harrison, David K. Jackson, Ian Johnston, Dominic Kwiatkowski, Cordelia Langford, John Sillitoe on behalf of the Wellcome Sanger Institute COVID-19 Surveillance Team |
| EPI_ISL_2570023, EPI_ISL_2570024, EPI_ISL_2570025 | Lighthouse Lab in Alderley Park | Wellcome Sanger Institute for the COVID-19 Genomics UK (COG-UK) Consortium | Jacquelyn Wynn, Mairead Hyland, The Lighthouse Lab in Alderley Park and Alex Alderton, Roberto Amato, Jeffrey Barrett, Sonia Goncalves, Ewan Harrison, David K. Jackson, Ian Johnston, Dominic Kwiatkowski, Cordelia Langford, John Sillitoe on behalf of the Wellcome Sanger Institute COVID-19 Surveillance Team |
| EPI_ISL_2570026 | Lighthouse Lab in Milton Keynes | Wellcome Sanger Institute for the COVID-19 Genomics UK (COG-UK) Consortium | The Lighthouse Lab in Milton Keynes and Alex Alderton, Roberto Amato, Jeffrey Barrett, Sonia Goncalves, Ewan Harrison, David K. Jackson, Ian Johnston, Dominic Kwiatkowski, Cordelia Langford, John Sillitoe on behalf of the Wellcome Sanger Institute COVID-19 Surveillance Team |
| EPI_ISL_2570027 | Lighthouse Lab in Alderley Park | Wellcome Sanger Institute for the COVID-19 Genomics UK (COG-UK) Consortium | Jacquelyn Wynn, Mairead Hyland, The Lighthouse Lab in Alderley Park and Alex Alderton, Roberto Amato, Jeffrey Barrett, Sonia Goncalves, Ewan Harrison, David K. Jackson, Ian Johnston, Dominic Kwiatkowski, Cordelia Langford, John Sillitoe on behalf of the Wellcome Sanger Institute COVID-19 Surveillance Team |
| EPI_ISL_2570028 | Lighthouse Lab in Milton Keynes | Wellcome Sanger Institute for the COVID-19 Genomics UK (COG-UK) Consortium | The Lighthouse Lab in Milton Keynes and Alex Alderton, Roberto Amato, Jeffrey Barrett, Sonia Goncalves, Ewan Harrison, David K. Jackson, Ian Johnston, Dominic Kwiatkowski, Cordelia Langford, John Sillitoe on behalf of the Wellcome Sanger Institute COVID-19 Surveillance Team |
| EPI_ISL_2570029, EPI_ISL_2570030 | Lighthouse Lab in Alderley Park | Wellcome Sanger Institute for the COVID-19 Genomics UK (COG-UK) Consortium | Jacquelyn Wynn, Mairead Hyland, The Lighthouse Lab in Alderley Park and Alex Alderton, Roberto Amato, Jeffrey Barrett, Sonia Goncalves, Ewan Harrison, David K. Jackson, Ian Johnston, Dominic Kwiatkowski, Cordelia Langford, John Sillitoe on behalf of the Wellcome Sanger Institute COVID-19 Surveillance Team |
| EPI_ISL_2570031, EPI_ISL_2570032, EPI_ISL_2570033 | Lighthouse Lab in Milton Keynes | Wellcome Sanger Institute for the COVID-19 Genomics UK (COG-UK) Consortium | The Lighthouse Lab in Milton Keynes and Alex Alderton, Roberto Amato, Jeffrey Barrett, Sonia Goncalves, Ewan Harrison, David K. Jackson, Ian Johnston, Dominic Kwiatkowski, Cordelia Langford, John Sillitoe on behalf of the Wellcome Sanger Institute COVID-19 Surveillance Team |
| EPI_ISL_2570034, EPI_ISL_2570035, EPI_ISL_2570036, EPI_ISL_2570037, EPI_ISL_2570038 | Lighthouse Lab in Alderley Park | Wellcome Sanger Institute for the COVID-19 Genomics UK (COG-UK) Consortium | Jacquelyn Wynn, Mairead Hyland, The Lighthouse Lab in Alderley Park and Alex Alderton, Roberto Amato, Jeffrey Barrett, Sonia Goncalves, Ewan Harrison, David K. Jackson, Ian Johnston, Dominic Kwiatkowski, Cordelia Langford, John Sillitoe on behalf of the Wellcome Sanger Institute COVID-19 Surveillance Team |
| EPI_ISL_2570039 | Lighthouse Lab in Milton Keynes | Wellcome Sanger Institute for the COVID-19 Genomics UK (COG-UK) Consortium | The Lighthouse Lab in Milton Keynes and Alex Alderton, Roberto Amato, Jeffrey Barrett, Sonia Goncalves, Ewan Harrison, David K. Jackson, Ian Johnston, Dominic Kwiatkowski, Cordelia Langford, John Sillitoe on behalf of the Wellcome Sanger Institute COVID-19 Surveillance Team |
| EPI_ISL_2570040 | Lighthouse Lab in Alderley Park | Wellcome Sanger Institute for the COVID-19 Genomics UK (COG-UK) Consortium | Jacquelyn Wynn, Mairead Hyland, The Lighthouse Lab in Alderley Park and Alex Alderton, Roberto Amato, Jeffrey Barrett, Sonia Goncalves, Ewan Harrison, David K. Jackson, Ian Johnston, Dominic Kwiatkowski, Cordelia Langford, John Sillitoe on behalf of the Wellcome Sanger Institute COVID-19 Surveillance Team |
| EPI_ISL_2570041 | Lighthouse Lab in Milton Keynes | Wellcome Sanger Institute for the COVID-19 Genomics UK (COG-UK) Consortium | The Lighthouse Lab in Milton Keynes and Alex Alderton, Roberto Amato, Jeffrey Barrett, Sonia Goncalves, Ewan Harrison, David K. Jackson, Ian Johnston, Dominic Kwiatkowski, Cordelia Langford, John Sillitoe on behalf of the Wellcome Sanger Institute COVID-19 Surveillance Team |
| EPI_ISL_2570042, EPI_ISL_2570043 | Lighthouse Lab in Alderley Park | Wellcome Sanger Institute for the COVID-19 Genomics UK (COG-UK) Consortium | Jacquelyn Wynn, Mairead Hyland, The Lighthouse Lab in Alderley Park and Alex Alderton, Roberto Amato, Jeffrey Barrett, Sonia Goncalves, Ewan Harrison, David K. Jackson, Ian Johnston, Dominic Kwiatkowski, Cordelia Langford, John Sillitoe on behalf of the Wellcome Sanger Institute COVID-19 Surveillance Team |
| EPI_ISL_2570044 | Lighthouse Lab in Milton Keynes | Wellcome Sanger Institute for the COVID-19 Genomics UK (COG-UK) Consortium | The Lighthouse Lab in Milton Keynes and Alex Alderton, Roberto Amato, Jeffrey Barrett, Sonia Goncalves, Ewan Harrison, David K. Jackson, Ian Johnston, Dominic Kwiatkowski, Cordelia Langford, John Sillitoe on behalf of the Wellcome Sanger Institute COVID-19 Surveillance Team |
| EPI_ISL_2570045, EPI_ISL_2570046 | Lighthouse Lab in Alderley Park | Wellcome Sanger Institute for the COVID-19 Genomics UK (COG-UK) Consortium | Jacquelyn Wynn, Mairead Hyland, The Lighthouse Lab in Alderley Park and Alex Alderton, Roberto Amato, Jeffrey Barrett, Sonia Goncalves, Ewan Harrison, David K. Jackson, Ian Johnston, Dominic Kwiatkowski, Cordelia Langford, John Sillitoe on behalf of the Wellcome Sanger Institute COVID-19 Surveillance Team |
| EPI_ISL_2570049 | Lighthouse Lab in Milton Keynes | Wellcome Sanger Institute for the COVID-19 Genomics UK (COG-UK) Consortium | The Lighthouse Lab in Milton Keynes and Alex Alderton, Roberto Amato, Jeffrey Barrett, Sonia Goncalves, Ewan Harrison, David K. Jackson, Ian Johnston, Dominic Kwiatkowski, Cordelia Langford, John Sillitoe on behalf of the Wellcome Sanger Institute COVID-19 Surveillance Team |
| EPI_ISL_2570050 | Lighthouse Lab in Alderley Park | Wellcome Sanger Institute for the COVID-19 Genomics UK (COG-UK) Consortium | Jacquelyn Wynn, Mairead Hyland, The Lighthouse Lab in Alderley Park and Alex Alderton, Roberto Amato, Jeffrey Barrett, Sonia Goncalves, Ewan Harrison, David K. Jackson, Ian Johnston, Dominic Kwiatkowski, Cordelia Langford, John Sillitoe on behalf of the Wellcome Sanger Institute COVID-19 Surveillance Team |
| EPI_ISL_2570051 | Lighthouse Lab in Milton Keynes | Wellcome Sanger Institute for the COVID-19 Genomics UK (COG-UK) | The Lighthouse Lab in Milton Keynes and Alex Alderton, Roberto Amato, Jeffrey Barrett, Sonia Goncalves, Ewan Harrison, David K. Jackson, Ian Johnston, Dominic Kwiatkowski, Cordelia Langford, John Sillitoe on behalf of the Wellcome Sanger |

| | | | |
|---|---|---|---|
| | | Consortium | Institute COVID-19 Surveillance Team |
| EPI_ISL_2570053, EPI_ISL_2570054, EPI_ISL_2570055, EPI_ISL_2570056, EPI_ISL_2570057, EPI_ISL_2570058, EPI_ISL_2570059, EPI_ISL_2570060 | Lighthouse Lab in Alderley Park | Wellcome Sanger Institute for the COVID-19 Genomics UK (COG-UK) Consortium | Jacquelyn Wynn, Mairead Hyland, The Lighthouse Lab in Alderley Park and Alex Alderton, Roberto Amato, Jeffrey Barrett, Sonia Goncalves, Ewan Harrison, David K. Jackson, Ian Johnston, Dominic Kwiatkowski, Cordelia Langford, John Sillitoe on behalf of the Wellcome Sanger Institute COVID-19 Surveillance Team |
| EPI_ISL_2570061 | Lighthouse Lab in Milton Keynes | Wellcome Sanger Institute for the COVID-19 Genomics UK (COG-UK) Consortium | The Lighthouse Lab in Milton Keynes and Alex Alderton, Roberto Amato, Jeffrey Barrett, Sonia Goncalves, Ewan Harrison, David K. Jackson, Ian Johnston, Dominic Kwiatkowski, Cordelia Langford, John Sillitoe on behalf of the Wellcome Sanger Institute COVID-19 Surveillance Team |
| EPI_ISL_2570062, EPI_ISL_2570063 | Lighthouse Lab in Alderley Park | Wellcome Sanger Institute for the COVID-19 Genomics UK (COG-UK) Consortium | Jacquelyn Wynn, Mairead Hyland, The Lighthouse Lab in Alderley Park and Alex Alderton, Roberto Amato, Jeffrey Barrett, Sonia Goncalves, Ewan Harrison, David K. Jackson, Ian Johnston, Dominic Kwiatkowski, Cordelia Langford, John Sillitoe on behalf of the Wellcome Sanger Institute COVID-19 Surveillance Team |
| EPI_ISL_2570064 | Lighthouse Lab in Milton Keynes | Wellcome Sanger Institute for the COVID-19 Genomics UK (COG-UK) Consortium | The Lighthouse Lab in Milton Keynes and Alex Alderton, Roberto Amato, Jeffrey Barrett, Sonia Goncalves, Ewan Harrison, David K. Jackson, Ian Johnston, Dominic Kwiatkowski, Cordelia Langford, John Sillitoe on behalf of the Wellcome Sanger Institute COVID-19 Surveillance Team |
| EPI_ISL_2570065, EPI_ISL_2570067, EPI_ISL_2570068, EPI_ISL_2570069, EPI_ISL_2570070 | Lighthouse Lab in Alderley Park | Wellcome Sanger Institute for the COVID-19 Genomics UK (COG-UK) Consortium | Jacquelyn Wynn, Mairead Hyland, The Lighthouse Lab in Alderley Park and Alex Alderton, Roberto Amato, Jeffrey Barrett, Sonia Goncalves, Ewan Harrison, David K. Jackson, Ian Johnston, Dominic Kwiatkowski, Cordelia Langford, John Sillitoe on behalf of the Wellcome Sanger Institute COVID-19 Surveillance Team |
| EPI_ISL_2570072, EPI_ISL_2570073, EPI_ISL_2570074, EPI_ISL_2570075 | Lighthouse Lab in Milton Keynes | Wellcome Sanger Institute for the COVID-19 Genomics UK (COG-UK) Consortium | The Lighthouse Lab in Milton Keynes and Alex Alderton, Roberto Amato, Jeffrey Barrett, Sonia Goncalves, Ewan Harrison, David K. Jackson, Ian Johnston, Dominic Kwiatkowski, Cordelia Langford, John Sillitoe on behalf of the Wellcome Sanger Institute COVID-19 Surveillance Team |
| EPI_ISL_2570076, EPI_ISL_2570077, EPI_ISL_2570078 | Lighthouse Lab in Alderley Park | Wellcome Sanger Institute for the COVID-19 Genomics UK (COG-UK) Consortium | Jacquelyn Wynn, Mairead Hyland, The Lighthouse Lab in Alderley Park and Alex Alderton, Roberto Amato, Jeffrey Barrett, Sonia Goncalves, Ewan Harrison, David K. Jackson, Ian Johnston, Dominic Kwiatkowski, Cordelia Langford, John Sillitoe on behalf of the Wellcome Sanger Institute COVID-19 Surveillance Team |
| EPI_ISL_2570079 | Lighthouse Lab in Milton Keynes | Wellcome Sanger Institute for the COVID-19 Genomics UK (COG-UK) Consortium | The Lighthouse Lab in Milton Keynes and Alex Alderton, Roberto Amato, Jeffrey Barrett, Sonia Goncalves, Ewan Harrison, David K. Jackson, Ian Johnston, Dominic Kwiatkowski, Cordelia Langford, John Sillitoe on behalf of the Wellcome Sanger Institute COVID-19 Surveillance Team |
| EPI_ISL_2570080, EPI_ISL_2570081, EPI_ISL_2570082 | Lighthouse Lab in Alderley Park | Wellcome Sanger Institute for the COVID-19 Genomics UK (COG-UK) Consortium | Jacquelyn Wynn, Mairead Hyland, The Lighthouse Lab in Alderley Park and Alex Alderton, Roberto Amato, Jeffrey Barrett, Sonia Goncalves, Ewan Harrison, David K. Jackson, Ian Johnston, Dominic Kwiatkowski, Cordelia Langford, John Sillitoe on behalf of the Wellcome Sanger Institute COVID-19 Surveillance Team |
| EPI_ISL_2570083 | Lighthouse Lab in Milton Keynes | Wellcome Sanger Institute for the COVID-19 Genomics UK (COG-UK) Consortium | The Lighthouse Lab in Milton Keynes and Alex Alderton, Roberto Amato, Jeffrey Barrett, Sonia Goncalves, Ewan Harrison, David K. Jackson, Ian Johnston, Dominic Kwiatkowski, Cordelia Langford, John Sillitoe on behalf of the Wellcome Sanger Institute COVID-19 Surveillance Team |
| EPI_ISL_2570084 | Lighthouse Lab in Alderley Park | Wellcome Sanger Institute for the COVID-19 Genomics UK (COG-UK) Consortium | Jacquelyn Wynn, Mairead Hyland, The Lighthouse Lab in Alderley Park and Alex Alderton, Roberto Amato, Jeffrey Barrett, Sonia Goncalves, Ewan Harrison, David K. Jackson, Ian Johnston, Dominic Kwiatkowski, Cordelia Langford, John Sillitoe on behalf of the Wellcome Sanger Institute COVID-19 Surveillance Team |
| EPI_ISL_2570085 | Lighthouse Lab in Milton Keynes | Wellcome Sanger Institute for the COVID-19 Genomics UK (COG-UK) Consortium | The Lighthouse Lab in Milton Keynes and Alex Alderton, Roberto Amato, Jeffrey Barrett, Sonia Goncalves, Ewan Harrison, David K. Jackson, Ian Johnston, Dominic Kwiatkowski, Cordelia Langford, John Sillitoe on behalf of the Wellcome Sanger Institute COVID-19 Surveillance Team |
| EPI_ISL_2570086, EPI_ISL_2570087, EPI_ISL_2570088, EPI_ISL_2570089, EPI_ISL_2570090, EPI_ISL_2570091, EPI_ISL_2570092 | Lighthouse Lab in Alderley Park | Wellcome Sanger Institute for the COVID-19 Genomics UK (COG-UK) Consortium | Jacquelyn Wynn, Mairead Hyland, The Lighthouse Lab in Alderley Park and Alex Alderton, Roberto Amato, Jeffrey Barrett, Sonia Goncalves, Ewan Harrison, David K. Jackson, Ian Johnston, Dominic Kwiatkowski, Cordelia Langford, John Sillitoe on behalf of the Wellcome Sanger Institute COVID-19 Surveillance Team |
| EPI_ISL_2570093 | Lighthouse Lab in Milton Keynes | Wellcome Sanger Institute for the COVID-19 Genomics UK (COG-UK) Consortium | The Lighthouse Lab in Milton Keynes and Alex Alderton, Roberto Amato, Jeffrey Barrett, Sonia Goncalves, Ewan Harrison, David K. Jackson, Ian Johnston, Dominic Kwiatkowski, Cordelia Langford, John Sillitoe on behalf of the Wellcome Sanger Institute COVID-19 Surveillance Team |
| EPI_ISL_2570094, EPI_ISL_2570095 | Lighthouse Lab in Alderley Park | Wellcome Sanger Institute for the COVID-19 Genomics UK (COG-UK) Consortium | Jacquelyn Wynn, Mairead Hyland, The Lighthouse Lab in Alderley Park and Alex Alderton, Roberto Amato, Jeffrey Barrett, Sonia Goncalves, Ewan Harrison, David K. Jackson, Ian Johnston, Dominic Kwiatkowski, Cordelia Langford, John Sillitoe on behalf of the Wellcome Sanger Institute COVID-19 Surveillance Team |
| EPI_ISL_2570096, EPI_ISL_2570097 | Lighthouse Lab in Milton Keynes | Wellcome Sanger Institute for the COVID-19 Genomics UK (COG-UK) Consortium | The Lighthouse Lab in Milton Keynes and Alex Alderton, Roberto Amato, Jeffrey Barrett, Sonia Goncalves, Ewan Harrison, David K. Jackson, Ian Johnston, Dominic Kwiatkowski, Cordelia Langford, John Sillitoe on behalf of the Wellcome Sanger Institute COVID-19 Surveillance Team |
| EPI_ISL_2570098 | Lighthouse Lab in Alderley Park | Wellcome Sanger Institute for the COVID-19 Genomics UK (COG-UK) Consortium | Jacquelyn Wynn, Mairead Hyland, The Lighthouse Lab in Alderley Park and Alex Alderton, Roberto Amato, Jeffrey Barrett, Sonia Goncalves, Ewan Harrison, David K. Jackson, Ian Johnston, Dominic Kwiatkowski, Cordelia Langford, John Sillitoe on behalf of the Wellcome Sanger Institute COVID-19 Surveillance Team |
| EPI_ISL_2570099 | Lighthouse Lab in Milton Keynes | Wellcome Sanger Institute for the COVID-19 Genomics UK (COG-UK) Consortium | The Lighthouse Lab in Milton Keynes and Alex Alderton, Roberto Amato, Jeffrey Barrett, Sonia Goncalves, Ewan Harrison, David K. Jackson, Ian Johnston, Dominic Kwiatkowski, Cordelia Langford, John Sillitoe on behalf of the Wellcome Sanger Institute COVID-19 Surveillance Team |
| EPI_ISL_2570100, EPI_ISL_2570101, EPI_ISL_2570102, EPI_ISL_2570103 | Lighthouse Lab in Alderley Park | Wellcome Sanger Institute for the COVID-19 Genomics UK (COG-UK) Consortium | Jacquelyn Wynn, Mairead Hyland, The Lighthouse Lab in Alderley Park and Alex Alderton, Roberto Amato, Jeffrey Barrett, Sonia Goncalves, Ewan Harrison, David K. Jackson, Ian Johnston, Dominic Kwiatkowski, Cordelia Langford, John Sillitoe on behalf of the Wellcome Sanger Institute COVID-19 Surveillance Team |
| EPI_ISL_2570104 | Lighthouse Lab in Milton Keynes | Wellcome Sanger Institute for the COVID-19 Genomics UK (COG-UK) Consortium | The Lighthouse Lab in Milton Keynes and Alex Alderton, Roberto Amato, Jeffrey Barrett, Sonia Goncalves, Ewan Harrison, David K. Jackson, Ian Johnston, Dominic Kwiatkowski, Cordelia Langford, John Sillitoe on behalf of the Wellcome Sanger Institute COVID-19 Surveillance Team |
| EPI_ISL_2570105 | Lighthouse Lab in Alderley Park | Wellcome Sanger Institute for the COVID-19 Genomics UK (COG-UK) Consortium | Jacquelyn Wynn, Mairead Hyland, The Lighthouse Lab in Alderley Park and Alex Alderton, Roberto Amato, Jeffrey Barrett, Sonia Goncalves, Ewan Harrison, David K. Jackson, Ian Johnston, Dominic Kwiatkowski, Cordelia Langford, John Sillitoe on behalf of the Wellcome Sanger Institute COVID-19 Surveillance Team |
| EPI_ISL_2570106 | Lighthouse Lab in Milton Keynes | Wellcome Sanger Institute for the COVID-19 Genomics UK (COG-UK) Consortium | The Lighthouse Lab in Milton Keynes and Alex Alderton, Roberto Amato, Jeffrey Barrett, Sonia Goncalves, Ewan Harrison, David K. Jackson, Ian Johnston, Dominic Kwiatkowski, Cordelia Langford, John Sillitoe on behalf of the Wellcome Sanger Institute COVID-19 Surveillance Team |
| EPI_ISL_2570107, EPI_ISL_2570108, EPI_ISL_2570110, EPI_ISL_2570111, EPI_ISL_2570112, EPI_ISL_2570113, EPI_ISL_2570114, EPI_ISL_2570115, EPI_ISL_2570116, EPI_ISL_2570117, EPI_ISL_2570118, EPI_ISL_2570119, EPI_ISL_2570120, EPI_ISL_2570121 | | | |
| see above | Lighthouse Lab in Alderley Park | Wellcome Sanger Institute for the COVID-19 Genomics UK (COG-UK) Consortium | Jacquelyn Wynn, Mairead Hyland, The Lighthouse Lab in Alderley Park and Alex Alderton, Roberto Amato, Jeffrey Barrett, Sonia Goncalves, Ewan Harrison, David K. Jackson, Ian Johnston, Dominic Kwiatkowski, Cordelia Langford, John Sillitoe on behalf of the Wellcome Sanger Institute COVID-19 Surveillance Team |
| EPI_ISL_2570122 | Lighthouse Lab in Milton Keynes | Wellcome Sanger Institute for the COVID-19 Genomics UK (COG-UK) Consortium | The Lighthouse Lab in Milton Keynes and Alex Alderton, Roberto Amato, Jeffrey Barrett, Sonia Goncalves, Ewan Harrison, David K. Jackson, Ian Johnston, Dominic Kwiatkowski, Cordelia Langford, John Sillitoe on behalf of the Wellcome Sanger Institute COVID-19 Surveillance Team |
| EPI_ISL_2570123 | Lighthouse Lab in Alderley Park | Wellcome Sanger Institute for the COVID-19 Genomics UK (COG-UK) Consortium | Jacquelyn Wynn, Mairead Hyland, The Lighthouse Lab in Alderley Park and Alex Alderton, Roberto Amato, Jeffrey Barrett, Sonia Goncalves, Ewan Harrison, David K. Jackson, Ian Johnston, Dominic Kwiatkowski, Cordelia Langford, John Sillitoe on behalf of the Wellcome Sanger Institute COVID-19 Surveillance Team |
| EPI_ISL_2570124 | Lighthouse Lab in Milton Keynes | Wellcome Sanger Institute for the COVID-19 Genomics UK (COG-UK) Consortium | The Lighthouse Lab in Milton Keynes and Alex Alderton, Roberto Amato, Jeffrey Barrett, Sonia Goncalves, Ewan Harrison, David K. Jackson, Ian Johnston, Dominic Kwiatkowski, Cordelia Langford, John Sillitoe on behalf of the Wellcome Sanger Institute COVID-19 Surveillance Team |
| EPI_ISL_2570125, EPI_ISL_2570126, EPI_ISL_2570127, EPI_ISL_2570129, EPI_ISL_2570130 | Lighthouse Lab in Alderley Park | Wellcome Sanger Institute for the COVID-19 Genomics UK (COG-UK) Consortium | Jacquelyn Wynn, Mairead Hyland, The Lighthouse Lab in Alderley Park and Alex Alderton, Roberto Amato, Jeffrey Barrett, Sonia Goncalves, Ewan Harrison, David K. Jackson, Ian Johnston, Dominic Kwiatkowski, Cordelia Langford, John Sillitoe on behalf of the Wellcome Sanger Institute COVID-19 Surveillance Team |
| EPI_ISL_2570131 | Lighthouse Lab in Milton Keynes | Wellcome Sanger Institute for the COVID-19 Genomics UK (COG-UK) Consortium | The Lighthouse Lab in Milton Keynes and Alex Alderton, Roberto Amato, Jeffrey Barrett, Sonia Goncalves, Ewan Harrison, David K. Jackson, Ian Johnston, Dominic Kwiatkowski, Cordelia Langford, John Sillitoe on behalf of the Wellcome Sanger Institute COVID-19 Surveillance Team |
| EPI_ISL_2570132, EPI_ISL_2570133, EPI_ISL_2570134, EPI_ISL_2570135, EPI_ISL_2570136, EPI_ISL_2570137, EPI_ISL_2570138, EPI_ISL_2570139 | Lighthouse Lab in Alderley Park | Wellcome Sanger Institute for the COVID-19 Genomics UK (COG-UK) Consortium | Jacquelyn Wynn, Mairead Hyland, The Lighthouse Lab in Alderley Park and Alex Alderton, Roberto Amato, Jeffrey Barrett, Sonia Goncalves, Ewan Harrison, David K. Jackson, Ian Johnston, Dominic Kwiatkowski, Cordelia Langford, John Sillitoe on behalf of the Wellcome Sanger Institute COVID-19 Surveillance Team |
| EPI_ISL_2570140 | Lighthouse Lab in Milton Keynes | Wellcome Sanger Institute for the COVID-19 Genomics UK (COG-UK) Consortium | The Lighthouse Lab in Milton Keynes and Alex Alderton, Roberto Amato, Jeffrey Barrett, Sonia Goncalves, Ewan Harrison, David K. Jackson, Ian Johnston, Dominic Kwiatkowski, Cordelia Langford, John Sillitoe on behalf of the Wellcome Sanger Institute COVID-19 Surveillance Team |
| EPI_ISL_2570142, EPI_ISL_2570143 | Lighthouse Lab in Alderley Park | Wellcome Sanger Institute for the COVID-19 Genomics UK (COG-UK) Consortium | Jacquelyn Wynn, Mairead Hyland, The Lighthouse Lab in Alderley Park and Alex Alderton, Roberto Amato, Jeffrey Barrett, Sonia Goncalves, Ewan Harrison, David K. Jackson, Ian Johnston, Dominic Kwiatkowski, Cordelia Langford, John Sillitoe on behalf of the Wellcome Sanger Institute COVID-19 Surveillance Team |
| EPI_ISL_2570144 | Lighthouse Lab in Milton Keynes | Wellcome Sanger Institute for the COVID-19 Genomics UK (COG-UK) Consortium | The Lighthouse Lab in Milton Keynes and Alex Alderton, Roberto Amato, Jeffrey Barrett, Sonia Goncalves, Ewan Harrison, David K. Jackson, Ian Johnston, Dominic Kwiatkowski, Cordelia Langford, John Sillitoe on behalf of the Wellcome Sanger Institute COVID-19 Surveillance Team |

| | | | |
|---|---|---|---|
| EPI_ISL_2570146, EPI_ISL_2570147, EPI_ISL_2570148, EPI_ISL_2570149, EPI_ISL_2570150, EPI_ISL_2570151 | Lighthouse Lab in Alderley Park | Wellcome Sanger Institute for the COVID-19 Genomics UK (COG-UK) Consortium | Jacquelyn Wynn, Mairead Hyland, The Lighthouse Lab in Alderley Park and Alex Alderton, Roberto Amato, Jeffrey Barrett, Sonia Goncalves, Ewan Harrison, David K. Jackson, Ian Johnston, Dominic Kwiatkowski, Cordelia Langford, John Sillitoe on behalf of the Wellcome Sanger Institute COVID-19 Surveillance Team |
| EPI_ISL_2570152, EPI_ISL_2570153 | Lighthouse Lab in Milton Keynes | Wellcome Sanger Institute for the COVID-19 Genomics UK (COG-UK) Consortium | The Lighthouse Lab in Milton Keynes and Alex Alderton, Roberto Amato, Jeffrey Barrett, Sonia Goncalves, Ewan Harrison, David K. Jackson, Ian Johnston, Dominic Kwiatkowski, Cordelia Langford, John Sillitoe on behalf of the Wellcome Sanger Institute COVID-19 Surveillance Team |
| EPI_ISL_2570154, EPI_ISL_2570155 | Lighthouse Lab in Alderley Park | Wellcome Sanger Institute for the COVID-19 Genomics UK (COG-UK) Consortium | Jacquelyn Wynn, Mairead Hyland, The Lighthouse Lab in Alderley Park and Alex Alderton, Roberto Amato, Jeffrey Barrett, Sonia Goncalves, Ewan Harrison, David K. Jackson, Ian Johnston, Dominic Kwiatkowski, Cordelia Langford, John Sillitoe on behalf of the Wellcome Sanger Institute COVID-19 Surveillance Team |
| EPI_ISL_2570156 | Lighthouse Lab in Milton Keynes | Wellcome Sanger Institute for the COVID-19 Genomics UK (COG-UK) Consortium | The Lighthouse Lab in Milton Keynes and Alex Alderton, Roberto Amato, Jeffrey Barrett, Sonia Goncalves, Ewan Harrison, David K. Jackson, Ian Johnston, Dominic Kwiatkowski, Cordelia Langford, John Sillitoe on behalf of the Wellcome Sanger Institute COVID-19 Surveillance Team |
| EPI_ISL_2570157, EPI_ISL_2570158, EPI_ISL_2570159, EPI_ISL_2570160, EPI_ISL_2570161, EPI_ISL_2570162, EPI_ISL_2570163, EPI_ISL_2570164, EPI_ISL_2570165 | Lighthouse Lab in Alderley Park | Wellcome Sanger Institute for the COVID-19 Genomics UK (COG-UK) Consortium | Jacquelyn Wynn, Mairead Hyland, The Lighthouse Lab in Alderley Park and Alex Alderton, Roberto Amato, Jeffrey Barrett, Sonia Goncalves, Ewan Harrison, David K. Jackson, Ian Johnston, Dominic Kwiatkowski, Cordelia Langford, John Sillitoe on behalf of the Wellcome Sanger Institute COVID-19 Surveillance Team |
| EPI_ISL_2570166, EPI_ISL_2570167 | Lighthouse Lab in Milton Keynes | Wellcome Sanger Institute for the COVID-19 Genomics UK (COG-UK) Consortium | The Lighthouse Lab in Milton Keynes and Alex Alderton, Roberto Amato, Jeffrey Barrett, Sonia Goncalves, Ewan Harrison, David K. Jackson, Ian Johnston, Dominic Kwiatkowski, Cordelia Langford, John Sillitoe on behalf of the Wellcome Sanger Institute COVID-19 Surveillance Team |
| EPI_ISL_2570168 | Lighthouse Lab in Alderley Park | Wellcome Sanger Institute for the COVID-19 Genomics UK (COG-UK) Consortium | Jacquelyn Wynn, Mairead Hyland, The Lighthouse Lab in Alderley Park and Alex Alderton, Roberto Amato, Jeffrey Barrett, Sonia Goncalves, Ewan Harrison, David K. Jackson, Ian Johnston, Dominic Kwiatkowski, Cordelia Langford, John Sillitoe on behalf of the Wellcome Sanger Institute COVID-19 Surveillance Team |
| EPI_ISL_2570169 | Lighthouse Lab in Milton Keynes | Wellcome Sanger Institute for the COVID-19 Genomics UK (COG-UK) Consortium | The Lighthouse Lab in Milton Keynes and Alex Alderton, Roberto Amato, Jeffrey Barrett, Sonia Goncalves, Ewan Harrison, David K. Jackson, Ian Johnston, Dominic Kwiatkowski, Cordelia Langford, John Sillitoe on behalf of the Wellcome Sanger Institute COVID-19 Surveillance Team |
| EPI_ISL_2570170, EPI_ISL_2570171, EPI_ISL_2570172 | Lighthouse Lab in Alderley Park | Wellcome Sanger Institute for the COVID-19 Genomics UK (COG-UK) Consortium | Jacquelyn Wynn, Mairead Hyland, The Lighthouse Lab in Alderley Park and Alex Alderton, Roberto Amato, Jeffrey Barrett, Sonia Goncalves, Ewan Harrison, David K. Jackson, Ian Johnston, Dominic Kwiatkowski, Cordelia Langford, John Sillitoe on behalf of the Wellcome Sanger Institute COVID-19 Surveillance Team |
| EPI_ISL_2570173, EPI_ISL_2570174, EPI_ISL_2570176 | Lighthouse Lab in Milton Keynes | Wellcome Sanger Institute for the COVID-19 Genomics UK (COG-UK) Consortium | The Lighthouse Lab in Milton Keynes and Alex Alderton, Roberto Amato, Jeffrey Barrett, Sonia Goncalves, Ewan Harrison, David K. Jackson, Ian Johnston, Dominic Kwiatkowski, Cordelia Langford, John Sillitoe on behalf of the Wellcome Sanger Institute COVID-19 Surveillance Team |
| EPI_ISL_2570177, EPI_ISL_2570178 | Lighthouse Lab in Alderley Park | Wellcome Sanger Institute for the COVID-19 Genomics UK (COG-UK) Consortium | Jacquelyn Wynn, Mairead Hyland, The Lighthouse Lab in Alderley Park and Alex Alderton, Roberto Amato, Jeffrey Barrett, Sonia Goncalves, Ewan Harrison, David K. Jackson, Ian Johnston, Dominic Kwiatkowski, Cordelia Langford, John Sillitoe on behalf of the Wellcome Sanger Institute COVID-19 Surveillance Team |
| EPI_ISL_2570179, EPI_ISL_2570180 | Lighthouse Lab in Milton Keynes | Wellcome Sanger Institute for the COVID-19 Genomics UK (COG-UK) Consortium | The Lighthouse Lab in Milton Keynes and Alex Alderton, Roberto Amato, Jeffrey Barrett, Sonia Goncalves, Ewan Harrison, David K. Jackson, Ian Johnston, Dominic Kwiatkowski, Cordelia Langford, John Sillitoe on behalf of the Wellcome Sanger Institute COVID-19 Surveillance Team |
| EPI_ISL_2570181 | Lighthouse Lab in Alderley Park | Wellcome Sanger Institute for the COVID-19 Genomics UK (COG-UK) Consortium | Jacquelyn Wynn, Mairead Hyland, The Lighthouse Lab in Alderley Park and Alex Alderton, Roberto Amato, Jeffrey Barrett, Sonia Goncalves, Ewan Harrison, David K. Jackson, Ian Johnston, Dominic Kwiatkowski, Cordelia Langford, John Sillitoe on behalf of the Wellcome Sanger Institute COVID-19 Surveillance Team |
| EPI_ISL_2570183, EPI_ISL_2570184, EPI_ISL_2570185, EPI_ISL_2570186, EPI_ISL_2570187 | Lighthouse Lab in Milton Keynes | Wellcome Sanger Institute for the COVID-19 Genomics UK (COG-UK) Consortium | The Lighthouse Lab in Milton Keynes and Alex Alderton, Roberto Amato, Jeffrey Barrett, Sonia Goncalves, Ewan Harrison, David K. Jackson, Ian Johnston, Dominic Kwiatkowski, Cordelia Langford, John Sillitoe on behalf of the Wellcome Sanger Institute COVID-19 Surveillance Team |
| EPI_ISL_2570188, EPI_ISL_2570189 | Lighthouse Lab in Alderley Park | Wellcome Sanger Institute for the COVID-19 Genomics UK (COG-UK) Consortium | Jacquelyn Wynn, Mairead Hyland, The Lighthouse Lab in Alderley Park and Alex Alderton, Roberto Amato, Jeffrey Barrett, Sonia Goncalves, Ewan Harrison, David K. Jackson, Ian Johnston, Dominic Kwiatkowski, Cordelia Langford, John Sillitoe on behalf of the Wellcome Sanger Institute COVID-19 Surveillance Team |
| EPI_ISL_2570190 | Lighthouse Lab in Milton Keynes | Wellcome Sanger Institute for the COVID-19 Genomics UK (COG-UK) Consortium | The Lighthouse Lab in Milton Keynes and Alex Alderton, Roberto Amato, Jeffrey Barrett, Sonia Goncalves, Ewan Harrison, David K. Jackson, Ian Johnston, Dominic Kwiatkowski, Cordelia Langford, John Sillitoe on behalf of the Wellcome Sanger Institute COVID-19 Surveillance Team |
| EPI_ISL_2570191 | Lighthouse Lab in Alderley Park | Wellcome Sanger Institute for the COVID-19 Genomics UK (COG-UK) Consortium | Jacquelyn Wynn, Mairead Hyland, The Lighthouse Lab in Alderley Park and Alex Alderton, Roberto Amato, Jeffrey Barrett, Sonia Goncalves, Ewan Harrison, David K. Jackson, Ian Johnston, Dominic Kwiatkowski, Cordelia Langford, John Sillitoe on behalf of the Wellcome Sanger Institute COVID-19 Surveillance Team |
| EPI_ISL_2570192 | Lighthouse Lab in Milton Keynes | Wellcome Sanger Institute for the COVID-19 Genomics UK (COG-UK) Consortium | The Lighthouse Lab in Milton Keynes and Alex Alderton, Roberto Amato, Jeffrey Barrett, Sonia Goncalves, Ewan Harrison, David K. Jackson, Ian Johnston, Dominic Kwiatkowski, Cordelia Langford, John Sillitoe on behalf of the Wellcome Sanger Institute COVID-19 Surveillance Team |
| EPI_ISL_2570193, EPI_ISL_2570195, EPI_ISL_2570196 | Lighthouse Lab in Alderley Park | Wellcome Sanger Institute for the COVID-19 Genomics UK (COG-UK) Consortium | Jacquelyn Wynn, Mairead Hyland, The Lighthouse Lab in Alderley Park and Alex Alderton, Roberto Amato, Jeffrey Barrett, Sonia Goncalves, Ewan Harrison, David K. Jackson, Ian Johnston, Dominic Kwiatkowski, Cordelia Langford, John Sillitoe on behalf of the Wellcome Sanger Institute COVID-19 Surveillance Team |
| EPI_ISL_2570197 | Lighthouse Lab in Milton Keynes | Wellcome Sanger Institute for the COVID-19 Genomics UK (COG-UK) Consortium | The Lighthouse Lab in Milton Keynes and Alex Alderton, Roberto Amato, Jeffrey Barrett, Sonia Goncalves, Ewan Harrison, David K. Jackson, Ian Johnston, Dominic Kwiatkowski, Cordelia Langford, John Sillitoe on behalf of the Wellcome Sanger Institute COVID-19 Surveillance Team |
| EPI_ISL_2570198, EPI_ISL_2570199, EPI_ISL_2570200, EPI_ISL_2570201, EPI_ISL_2570202, EPI_ISL_2570203, EPI_ISL_2570204 | Lighthouse Lab in Alderley Park | Wellcome Sanger Institute for the COVID-19 Genomics UK (COG-UK) Consortium | Jacquelyn Wynn, Mairead Hyland, The Lighthouse Lab in Alderley Park and Alex Alderton, Roberto Amato, Jeffrey Barrett, Sonia Goncalves, Ewan Harrison, David K. Jackson, Ian Johnston, Dominic Kwiatkowski, Cordelia Langford, John Sillitoe on behalf of the Wellcome Sanger Institute COVID-19 Surveillance Team |
| EPI_ISL_2570205 | Lighthouse Lab in Milton Keynes | Wellcome Sanger Institute for the COVID-19 Genomics UK (COG-UK) Consortium | The Lighthouse Lab in Milton Keynes and Alex Alderton, Roberto Amato, Jeffrey Barrett, Sonia Goncalves, Ewan Harrison, David K. Jackson, Ian Johnston, Dominic Kwiatkowski, Cordelia Langford, John Sillitoe on behalf of the Wellcome Sanger Institute COVID-19 Surveillance Team |
| EPI_ISL_2570206, EPI_ISL_2570207 | Lighthouse Lab in Alderley Park | Wellcome Sanger Institute for the COVID-19 Genomics UK (COG-UK) Consortium | Jacquelyn Wynn, Mairead Hyland, The Lighthouse Lab in Alderley Park and Alex Alderton, Roberto Amato, Jeffrey Barrett, Sonia Goncalves, Ewan Harrison, David K. Jackson, Ian Johnston, Dominic Kwiatkowski, Cordelia Langford, John Sillitoe on behalf of the Wellcome Sanger Institute COVID-19 Surveillance Team |
| EPI_ISL_2570208 | Lighthouse Lab in Milton Keynes | Wellcome Sanger Institute for the COVID-19 Genomics UK (COG-UK) Consortium | The Lighthouse Lab in Milton Keynes and Alex Alderton, Roberto Amato, Jeffrey Barrett, Sonia Goncalves, Ewan Harrison, David K. Jackson, Ian Johnston, Dominic Kwiatkowski, Cordelia Langford, John Sillitoe on behalf of the Wellcome Sanger Institute COVID-19 Surveillance Team |
| EPI_ISL_2570209, EPI_ISL_2570210, EPI_ISL_2570211 | Lighthouse Lab in Alderley Park | Wellcome Sanger Institute for the COVID-19 Genomics UK (COG-UK) Consortium | Jacquelyn Wynn, Mairead Hyland, The Lighthouse Lab in Alderley Park and Alex Alderton, Roberto Amato, Jeffrey Barrett, Sonia Goncalves, Ewan Harrison, David K. Jackson, Ian Johnston, Dominic Kwiatkowski, Cordelia Langford, John Sillitoe on behalf of the Wellcome Sanger Institute COVID-19 Surveillance Team |
| EPI_ISL_2570212 | Lighthouse Lab in Milton Keynes | Wellcome Sanger Institute for the COVID-19 Genomics UK (COG-UK) Consortium | The Lighthouse Lab in Milton Keynes and Alex Alderton, Roberto Amato, Jeffrey Barrett, Sonia Goncalves, Ewan Harrison, David K. Jackson, Ian Johnston, Dominic Kwiatkowski, Cordelia Langford, John Sillitoe on behalf of the Wellcome Sanger Institute COVID-19 Surveillance Team |
| EPI_ISL_2570213 | Lighthouse Lab in Alderley Park | Wellcome Sanger Institute for the COVID-19 Genomics UK (COG-UK) Consortium | Jacquelyn Wynn, Mairead Hyland, The Lighthouse Lab in Alderley Park and Alex Alderton, Roberto Amato, Jeffrey Barrett, Sonia Goncalves, Ewan Harrison, David K. Jackson, Ian Johnston, Dominic Kwiatkowski, Cordelia Langford, John Sillitoe on behalf of the Wellcome Sanger Institute COVID-19 Surveillance Team |
| EPI_ISL_2570214 | Lighthouse Lab in Milton Keynes | Wellcome Sanger Institute for the COVID-19 Genomics UK (COG-UK) Consortium | The Lighthouse Lab in Milton Keynes and Alex Alderton, Roberto Amato, Jeffrey Barrett, Sonia Goncalves, Ewan Harrison, David K. Jackson, Ian Johnston, Dominic Kwiatkowski, Cordelia Langford, John Sillitoe on behalf of the Wellcome Sanger Institute COVID-19 Surveillance Team |
| EPI_ISL_2570215 | Lighthouse Lab in Alderley Park | Wellcome Sanger Institute for the COVID-19 Genomics UK (COG-UK) Consortium | Jacquelyn Wynn, Mairead Hyland, The Lighthouse Lab in Alderley Park and Alex Alderton, Roberto Amato, Jeffrey Barrett, Sonia Goncalves, Ewan Harrison, David K. Jackson, Ian Johnston, Dominic Kwiatkowski, Cordelia Langford, John Sillitoe on behalf of the Wellcome Sanger Institute COVID-19 Surveillance Team |
| EPI_ISL_2570216 | Lighthouse Lab in Milton Keynes | Wellcome Sanger Institute for the COVID-19 Genomics UK (COG-UK) Consortium | The Lighthouse Lab in Milton Keynes and Alex Alderton, Roberto Amato, Jeffrey Barrett, Sonia Goncalves, Ewan Harrison, David K. Jackson, Ian Johnston, Dominic Kwiatkowski, Cordelia Langford, John Sillitoe on behalf of the Wellcome Sanger Institute COVID-19 Surveillance Team |
| EPI_ISL_2570217, EPI_ISL_2570218, EPI_ISL_2570219, EPI_ISL_2570220, EPI_ISL_2570221, EPI_ISL_2570222, EPI_ISL_2570223, EPI_ISL_2570224 | Lighthouse Lab in Alderley Park | Wellcome Sanger Institute for the COVID-19 Genomics UK (COG-UK) Consortium | Jacquelyn Wynn, Mairead Hyland, The Lighthouse Lab in Alderley Park and Alex Alderton, Roberto Amato, Jeffrey Barrett, Sonia Goncalves, Ewan Harrison, David K. Jackson, Ian Johnston, Dominic Kwiatkowski, Cordelia Langford, John Sillitoe on behalf of the Wellcome Sanger Institute COVID-19 Surveillance Team |
| EPI_ISL_2570225 | Lighthouse Lab in Milton Keynes | Wellcome Sanger Institute for the COVID-19 Genomics UK (COG-UK) Consortium | The Lighthouse Lab in Milton Keynes and Alex Alderton, Roberto Amato, Jeffrey Barrett, Sonia Goncalves, Ewan Harrison, David K. Jackson, Ian Johnston, Dominic Kwiatkowski, Cordelia Langford, John Sillitoe on behalf of the Wellcome Sanger Institute COVID-19 Surveillance Team |
| EPI_ISL_2570226 | Lighthouse Lab in Alderley Park | Wellcome Sanger Institute for the COVID-19 Genomics UK (COG-UK) Consortium | Jacquelyn Wynn, Mairead Hyland, The Lighthouse Lab in Alderley Park and Alex Alderton, Roberto Amato, Jeffrey Barrett, Sonia Goncalves, Ewan Harrison, David K. Jackson, Ian Johnston, Dominic Kwiatkowski, Cordelia Langford, John Sillitoe on behalf of the Wellcome Sanger Institute COVID-19 Surveillance Team |

| | | | |
|---|---|---|---|
| EPI_ISL_2570227 | Lighthouse Lab in Milton Keynes | Wellcome Sanger Institute for the COVID-19 Genomics UK (COG-UK) Consortium | The Lighthouse Lab in Milton Keynes and Alex Alderton, Roberto Amato, Jeffrey Barrett, Sonia Goncalves, Ewan Harrison, David K. Jackson, Ian Johnston, Dominic Kwiatkowski, Cordelia Langford, John Sillitoe on behalf of the Wellcome Sanger Institute COVID-19 Surveillance Team |
| EPI_ISL_2570228, EPI_ISL_2570229, EPI_ISL_2570230, EPI_ISL_2570231, EPI_ISL_2570232, EPI_ISL_2570233, EPI_ISL_2570234, EPI_ISL_2570235 | Lighthouse Lab in Alderley Park | Wellcome Sanger Institute for the COVID-19 Genomics UK (COG-UK) Consortium | Jacquelyn Wynn, Mairead Hyland, The Lighthouse Lab in Alderley Park and Alex Alderton, Roberto Amato, Jeffrey Barrett, Sonia Goncalves, Ewan Harrison, David K. Jackson, Ian Johnston, Dominic Kwiatkowski, Cordelia Langford, John Sillitoe on behalf of the Wellcome Sanger Institute COVID-19 Surveillance Team |
| EPI_ISL_2570236, EPI_ISL_2570237 | Lighthouse Lab in Milton Keynes | Wellcome Sanger Institute for the COVID-19 Genomics UK (COG-UK) Consortium | The Lighthouse Lab in Milton Keynes and Alex Alderton, Roberto Amato, Jeffrey Barrett, Sonia Goncalves, Ewan Harrison, David K. Jackson, Ian Johnston, Dominic Kwiatkowski, Cordelia Langford, John Sillitoe on behalf of the Wellcome Sanger Institute COVID-19 Surveillance Team |
| EPI_ISL_2570238 | Lighthouse Lab in Alderley Park | Wellcome Sanger Institute for the COVID-19 Genomics UK (COG-UK) Consortium | Jacquelyn Wynn, Mairead Hyland, The Lighthouse Lab in Alderley Park and Alex Alderton, Roberto Amato, Jeffrey Barrett, Sonia Goncalves, Ewan Harrison, David K. Jackson, Ian Johnston, Dominic Kwiatkowski, Cordelia Langford, John Sillitoe on behalf of the Wellcome Sanger Institute COVID-19 Surveillance Team |
| EPI_ISL_2570239 | Lighthouse Lab in Milton Keynes | Wellcome Sanger Institute for the COVID-19 Genomics UK (COG-UK) Consortium | The Lighthouse Lab in Milton Keynes and Alex Alderton, Roberto Amato, Jeffrey Barrett, Sonia Goncalves, Ewan Harrison, David K. Jackson, Ian Johnston, Dominic Kwiatkowski, Cordelia Langford, John Sillitoe on behalf of the Wellcome Sanger Institute COVID-19 Surveillance Team |
| EPI_ISL_2570240, EPI_ISL_2570241 | Lighthouse Lab in Alderley Park | Wellcome Sanger Institute for the COVID-19 Genomics UK (COG-UK) Consortium | Jacquelyn Wynn, Mairead Hyland, The Lighthouse Lab in Alderley Park and Alex Alderton, Roberto Amato, Jeffrey Barrett, Sonia Goncalves, Ewan Harrison, David K. Jackson, Ian Johnston, Dominic Kwiatkowski, Cordelia Langford, John Sillitoe on behalf of the Wellcome Sanger Institute COVID-19 Surveillance Team |
| EPI_ISL_2570242 | Lighthouse Lab in Milton Keynes | Wellcome Sanger Institute for the COVID-19 Genomics UK (COG-UK) Consortium | The Lighthouse Lab in Milton Keynes and Alex Alderton, Roberto Amato, Jeffrey Barrett, Sonia Goncalves, Ewan Harrison, David K. Jackson, Ian Johnston, Dominic Kwiatkowski, Cordelia Langford, John Sillitoe on behalf of the Wellcome Sanger Institute COVID-19 Surveillance Team |
| EPI_ISL_2570243, EPI_ISL_2570244, EPI_ISL_2570245 | Lighthouse Lab in Alderley Park | Wellcome Sanger Institute for the COVID-19 Genomics UK (COG-UK) Consortium | Jacquelyn Wynn, Mairead Hyland, The Lighthouse Lab in Alderley Park and Alex Alderton, Roberto Amato, Jeffrey Barrett, Sonia Goncalves, Ewan Harrison, David K. Jackson, Ian Johnston, Dominic Kwiatkowski, Cordelia Langford, John Sillitoe on behalf of the Wellcome Sanger Institute COVID-19 Surveillance Team |
| EPI_ISL_2570246 | Lighthouse Lab in Milton Keynes | Wellcome Sanger Institute for the COVID-19 Genomics UK (COG-UK) Consortium | The Lighthouse Lab in Milton Keynes and Alex Alderton, Roberto Amato, Jeffrey Barrett, Sonia Goncalves, Ewan Harrison, David K. Jackson, Ian Johnston, Dominic Kwiatkowski, Cordelia Langford, John Sillitoe on behalf of the Wellcome Sanger Institute COVID-19 Surveillance Team |
| EPI_ISL_2570247, EPI_ISL_2570248, EPI_ISL_2570249, EPI_ISL_2570250, EPI_ISL_2570251 | Lighthouse Lab in Alderley Park | Wellcome Sanger Institute for the COVID-19 Genomics UK (COG-UK) Consortium | Jacquelyn Wynn, Mairead Hyland, The Lighthouse Lab in Alderley Park and Alex Alderton, Roberto Amato, Jeffrey Barrett, Sonia Goncalves, Ewan Harrison, David K. Jackson, Ian Johnston, Dominic Kwiatkowski, Cordelia Langford, John Sillitoe on behalf of the Wellcome Sanger Institute COVID-19 Surveillance Team |
| EPI_ISL_2570252 | Lighthouse Lab in Milton Keynes | Wellcome Sanger Institute for the COVID-19 Genomics UK (COG-UK) Consortium | The Lighthouse Lab in Milton Keynes and Alex Alderton, Roberto Amato, Jeffrey Barrett, Sonia Goncalves, Ewan Harrison, David K. Jackson, Ian Johnston, Dominic Kwiatkowski, Cordelia Langford, John Sillitoe on behalf of the Wellcome Sanger Institute COVID-19 Surveillance Team |
| EPI_ISL_2570253, EPI_ISL_2570254, EPI_ISL_2570255, EPI_ISL_2570256, EPI_ISL_2570257, EPI_ISL_2570258, EPI_ISL_2570259 | Lighthouse Lab in Alderley Park | Wellcome Sanger Institute for the COVID-19 Genomics UK (COG-UK) Consortium | Jacquelyn Wynn, Mairead Hyland, The Lighthouse Lab in Alderley Park and Alex Alderton, Roberto Amato, Jeffrey Barrett, Sonia Goncalves, Ewan Harrison, David K. Jackson, Ian Johnston, Dominic Kwiatkowski, Cordelia Langford, John Sillitoe on behalf of the Wellcome Sanger Institute COVID-19 Surveillance Team |
| EPI_ISL_2570260 | Lighthouse Lab in Milton Keynes | Wellcome Sanger Institute for the COVID-19 Genomics UK (COG-UK) Consortium | The Lighthouse Lab in Milton Keynes and Alex Alderton, Roberto Amato, Jeffrey Barrett, Sonia Goncalves, Ewan Harrison, David K. Jackson, Ian Johnston, Dominic Kwiatkowski, Cordelia Langford, John Sillitoe on behalf of the Wellcome Sanger Institute COVID-19 Surveillance Team |
| EPI_ISL_2570261, EPI_ISL_2570262, EPI_ISL_2570263, EPI_ISL_2570264, EPI_ISL_2570265, EPI_ISL_2570266, EPI_ISL_2570267, EPI_ISL_2570268, EPI_ISL_2570269, EPI_ISL_2570270 | Lighthouse Lab in Alderley Park | Wellcome Sanger Institute for the COVID-19 Genomics UK (COG-UK) Consortium | Jacquelyn Wynn, Mairead Hyland, The Lighthouse Lab in Alderley Park and Alex Alderton, Roberto Amato, Jeffrey Barrett, Sonia Goncalves, Ewan Harrison, David K. Jackson, Ian Johnston, Dominic Kwiatkowski, Cordelia Langford, John Sillitoe on behalf of the Wellcome Sanger Institute COVID-19 Surveillance Team |
| EPI_ISL_2570271 | Health Services Laboratories | Wellcome Sanger Institute for the COVID-19 Genomics UK (COG-UK) Consortium | Health Services Laboratories and Alex Alderton, Roberto Amato, Jeffrey Barrett, Sonia Goncalves, Ewan Harrison, David K. Jackson, Ian Johnston, Dominic Kwiatkowski, Cordelia Langford, John Sillitoe on behalf of the Wellcome Sanger Institute COVID-19 Surveillance Team |
| EPI_ISL_2570272, EPI_ISL_2570273, EPI_ISL_2570274 | Lighthouse Lab in Alderley Park | Wellcome Sanger Institute for the COVID-19 Genomics UK (COG-UK) Consortium | Jacquelyn Wynn, Mairead Hyland, The Lighthouse Lab in Alderley Park and Alex Alderton, Roberto Amato, Jeffrey Barrett, Sonia Goncalves, Ewan Harrison, David K. Jackson, Ian Johnston, Dominic Kwiatkowski, Cordelia Langford, John Sillitoe on behalf of the Wellcome Sanger Institute COVID-19 Surveillance Team |
| EPI_ISL_2570275 | Lighthouse Laboratory Plymouth | Wellcome Sanger Institute for the COVID-19 Genomics UK (COG-UK) Consortium | Lighthouse Laboratory Plymouth and Alex Alderton, Roberto Amato, Jeffrey Barrett, Sonia Goncalves, Ewan Harrison, David K. Jackson, Ian Johnston, Dominic Kwiatkowski, Cordelia Langford, John Sillitoe on behalf of the Wellcome Sanger Institute COVID-19 Surveillance Team |
| EPI_ISL_2570276 | Lighthouse Lab in Alderley Park | Wellcome Sanger Institute for the COVID-19 Genomics UK (COG-UK) Consortium | Jacquelyn Wynn, Mairead Hyland, The Lighthouse Lab in Alderley Park and Alex Alderton, Roberto Amato, Jeffrey Barrett, Sonia Goncalves, Ewan Harrison, David K. Jackson, Ian Johnston, Dominic Kwiatkowski, Cordelia Langford, John Sillitoe on behalf of the Wellcome Sanger Institute COVID-19 Surveillance Team |
| EPI_ISL_2570277 | Lighthouse Lab in Glasgow | Wellcome Sanger Institute for the COVID-19 Genomics UK (COG-UK) Consortium | Harper VanSteenhouse, Yumi Kasai, David Gray, Carol Clugston, Anna Dominiczak and Alex Alderton, Roberto Amato, Jeffrey Barrett, Sonia Goncalves, Ewan Harrison, David K. Jackson, Ian Johnston, Dominic Kwiatkowski, Cordelia Langford, John Sillitoe on behalf of the Wellcome Sanger Institute COVID-19 Surveillance Team |
| EPI_ISL_2570279 | Lighthouse Laboratory Plymouth | Wellcome Sanger Institute for the COVID-19 Genomics UK (COG-UK) Consortium | Lighthouse Laboratory Plymouth and Alex Alderton, Roberto Amato, Jeffrey Barrett, Sonia Goncalves, Ewan Harrison, David K. Jackson, Ian Johnston, Dominic Kwiatkowski, Cordelia Langford, John Sillitoe on behalf of the Wellcome Sanger Institute COVID-19 Surveillance Team |
| EPI_ISL_2570280, EPI_ISL_2570281, EPI_ISL_2570282, EPI_ISL_2570283 | Lighthouse Lab in Milton Keynes | Wellcome Sanger Institute for the COVID-19 Genomics UK (COG-UK) Consortium | The Lighthouse Lab in Milton Keynes and Alex Alderton, Roberto Amato, Jeffrey Barrett, Sonia Goncalves, Ewan Harrison, David K. Jackson, Ian Johnston, Dominic Kwiatkowski, Cordelia Langford, John Sillitoe on behalf of the Wellcome Sanger Institute COVID-19 Surveillance Team |
| EPI_ISL_2570284 | Lighthouse Laboratory Plymouth | Wellcome Sanger Institute for the COVID-19 Genomics UK (COG-UK) Consortium | Lighthouse Laboratory Plymouth and Alex Alderton, Roberto Amato, Jeffrey Barrett, Sonia Goncalves, Ewan Harrison, David K. Jackson, Ian Johnston, Dominic Kwiatkowski, Cordelia Langford, John Sillitoe on behalf of the Wellcome Sanger Institute COVID-19 Surveillance Team |
| EPI_ISL_2570285, EPI_ISL_2570286 | Lighthouse Lab in Milton Keynes | Wellcome Sanger Institute for the COVID-19 Genomics UK (COG-UK) Consortium | The Lighthouse Lab in Milton Keynes and Alex Alderton, Roberto Amato, Jeffrey Barrett, Sonia Goncalves, Ewan Harrison, David K. Jackson, Ian Johnston, Dominic Kwiatkowski, Cordelia Langford, John Sillitoe on behalf of the Wellcome Sanger Institute COVID-19 Surveillance Team |
| EPI_ISL_2570287 | Lighthouse Lab in Glasgow | Wellcome Sanger Institute for the COVID-19 Genomics UK (COG-UK) Consortium | Harper VanSteenhouse, Yumi Kasai, David Gray, Carol Clugston, Anna Dominiczak and Alex Alderton, Roberto Amato, Jeffrey Barrett, Sonia Goncalves, Ewan Harrison, David K. Jackson, Ian Johnston, Dominic Kwiatkowski, Cordelia Langford, John Sillitoe on behalf of the Wellcome Sanger Institute COVID-19 Surveillance Team |
| EPI_ISL_2570288 | Lighthouse Laboratory Plymouth | Wellcome Sanger Institute for the COVID-19 Genomics UK (COG-UK) Consortium | Lighthouse Laboratory Plymouth and Alex Alderton, Roberto Amato, Jeffrey Barrett, Sonia Goncalves, Ewan Harrison, David K. Jackson, Ian Johnston, Dominic Kwiatkowski, Cordelia Langford, John Sillitoe on behalf of the Wellcome Sanger Institute COVID-19 Surveillance Team |
| EPI_ISL_2570289, EPI_ISL_2570290 | Lighthouse Lab in Milton Keynes | Wellcome Sanger Institute for the COVID-19 Genomics UK (COG-UK) Consortium | The Lighthouse Lab in Milton Keynes and Alex Alderton, Roberto Amato, Jeffrey Barrett, Sonia Goncalves, Ewan Harrison, David K. Jackson, Ian Johnston, Dominic Kwiatkowski, Cordelia Langford, John Sillitoe on behalf of the Wellcome Sanger Institute COVID-19 Surveillance Team |
| EPI_ISL_2570291, EPI_ISL_2570292 | Lighthouse Lab in Glasgow | Wellcome Sanger Institute for the COVID-19 Genomics UK (COG-UK) Consortium | Harper VanSteenhouse, Yumi Kasai, David Gray, Carol Clugston, Anna Dominiczak and Alex Alderton, Roberto Amato, Jeffrey Barrett, Sonia Goncalves, Ewan Harrison, David K. Jackson, Ian Johnston, Dominic Kwiatkowski, Cordelia Langford, John Sillitoe on behalf of the Wellcome Sanger Institute COVID-19 Surveillance Team |
| EPI_ISL_2570293 | Lighthouse Lab in Alderley Park | Wellcome Sanger Institute for the COVID-19 Genomics UK (COG-UK) Consortium | Jacquelyn Wynn, Mairead Hyland, The Lighthouse Lab in Alderley Park and Alex Alderton, Roberto Amato, Jeffrey Barrett, Sonia Goncalves, Ewan Harrison, David K. Jackson, Ian Johnston, Dominic Kwiatkowski, Cordelia Langford, John Sillitoe on behalf of the Wellcome Sanger Institute COVID-19 Surveillance Team |
| EPI_ISL_2570294 | Lighthouse Lab in Milton Keynes | Wellcome Sanger Institute for the COVID-19 Genomics UK (COG-UK) Consortium | The Lighthouse Lab in Milton Keynes and Alex Alderton, Roberto Amato, Jeffrey Barrett, Sonia Goncalves, Ewan Harrison, David K. Jackson, Ian Johnston, Dominic Kwiatkowski, Cordelia Langford, John Sillitoe on behalf of the Wellcome Sanger Institute COVID-19 Surveillance Team |
| EPI_ISL_2570295, EPI_ISL_2570296 | Lighthouse Laboratory Plymouth | Wellcome Sanger Institute for the COVID-19 Genomics UK (COG-UK) Consortium | Lighthouse Laboratory Plymouth and Alex Alderton, Roberto Amato, Jeffrey Barrett, Sonia Goncalves, Ewan Harrison, David K. Jackson, Ian Johnston, Dominic Kwiatkowski, Cordelia Langford, John Sillitoe on behalf of the Wellcome Sanger Institute COVID-19 Surveillance Team |
| EPI_ISL_2570297 | Lighthouse Lab in Glasgow | Wellcome Sanger Institute for the COVID-19 Genomics UK (COG-UK) Consortium | Harper VanSteenhouse, Yumi Kasai, David Gray, Carol Clugston, Anna Dominiczak and Alex Alderton, Roberto Amato, Jeffrey Barrett, Sonia Goncalves, Ewan Harrison, David K. Jackson, Ian Johnston, Dominic Kwiatkowski, Cordelia Langford, John Sillitoe on behalf of the Wellcome Sanger Institute COVID-19 Surveillance Team |
| EPI_ISL_2570298 | Lighthouse Lab in Alderley Park | Wellcome Sanger Institute for the COVID-19 Genomics UK (COG-UK) Consortium | Jacquelyn Wynn, Mairead Hyland, The Lighthouse Lab in Alderley Park and Alex Alderton, Roberto Amato, Jeffrey Barrett, Sonia Goncalves, Ewan Harrison, David K. Jackson, Ian Johnston, Dominic Kwiatkowski, Cordelia Langford, John Sillitoe on behalf of the Wellcome Sanger Institute COVID-19 Surveillance Team |
| EPI_ISL_2570299 | Lighthouse Laboratory Plymouth | Wellcome Sanger Institute for the COVID-19 Genomics UK (COG-UK) Consortium | Lighthouse Laboratory Plymouth and Alex Alderton, Roberto Amato, Jeffrey Barrett, Sonia Goncalves, Ewan Harrison, David K. Jackson, Ian Johnston, Dominic Kwiatkowski, Cordelia Langford, John Sillitoe on behalf of the Wellcome Sanger Institute COVID-19 Surveillance Team |
| EPI_ISL_2570300, EPI_ISL_2570301 | Lighthouse Lab in Alderley Park | Wellcome Sanger Institute for the COVID-19 Genomics UK (COG-UK) Consortium | Jacquelyn Wynn, Mairead Hyland, The Lighthouse Lab in Alderley Park and Alex Alderton, Roberto Amato, Jeffrey Barrett, Sonia Goncalves, Ewan Harrison, David K. Jackson, Ian Johnston, Dominic Kwiatkowski, Cordelia Langford, John Sillitoe on behalf of the Wellcome Sanger Institute COVID-19 Surveillance Team |
| EPI_ISL_2570302 | Lighthouse Laboratory Plymouth | Wellcome Sanger Institute for the COVID-19 Genomics UK (COG-UK) | Lighthouse Laboratory Plymouth and Alex Alderton, Roberto Amato, Jeffrey Barrett, Sonia Goncalves, Ewan Harrison, David K. Jackson, Ian Johnston, Dominic Kwiatkowski, Cordelia Langford, John Sillitoe on behalf of the Wellcome Sanger |

| | | | |
|---|---|---|---|
| | | Consortium | Institute COVID-19 Surveillance Team |
| EPI_ISL_2570303 | Lighthouse Lab in Milton Keynes | Wellcome Sanger Institute for the COVID-19 Genomics UK (COG-UK) Consortium | The Lighthouse Lab in Milton Keynes and Alex Alderton, Roberto Amato, Jeffrey Barrett, Sonia Goncalves, Ewan Harrison, David K. Jackson, Ian Johnston, Dominic Kwiatkowski, Cordelia Langford, John Sillitoe on behalf of the Wellcome Sanger Institute COVID-19 Surveillance Team |
| EPI_ISL_2570304, EPI_ISL_2570306, EPI_ISL_2570307 | Lighthouse Laboratory Plymouth | Wellcome Sanger Institute for the COVID-19 Genomics UK (COG-UK) Consortium | Lighthouse Laboratory Plymouth and Alex Alderton, Roberto Amato, Jeffrey Barrett, Sonia Goncalves, Ewan Harrison, David K. Jackson, Ian Johnston, Dominic Kwiatkowski, Cordelia Langford, John Sillitoe on behalf of the Wellcome Sanger Institute COVID-19 Surveillance Team |
| EPI_ISL_2570308 | Lighthouse Lab in Alderley Park | Wellcome Sanger Institute for the COVID-19 Genomics UK (COG-UK) Consortium | Jacquelyn Wynn, Mairead Hyland, The Lighthouse Lab in Alderley Park and Alex Alderton, Roberto Amato, Jeffrey Barrett, Sonia Goncalves, Ewan Harrison, David K. Jackson, Ian Johnston, Dominic Kwiatkowski, Cordelia Langford, John Sillitoe on behalf of the Wellcome Sanger Institute COVID-19 Surveillance Team |
| EPI_ISL_2570309 | Lighthouse Lab in Milton Keynes | Wellcome Sanger Institute for the COVID-19 Genomics UK (COG-UK) Consortium | The Lighthouse Lab in Milton Keynes and Alex Alderton, Roberto Amato, Jeffrey Barrett, Sonia Goncalves, Ewan Harrison, David K. Jackson, Ian Johnston, Dominic Kwiatkowski, Cordelia Langford, John Sillitoe on behalf of the Wellcome Sanger Institute COVID-19 Surveillance Team |
| EPI_ISL_2570310 | Lighthouse Lab in Alderley Park | Wellcome Sanger Institute for the COVID-19 Genomics UK (COG-UK) Consortium | Jacquelyn Wynn, Mairead Hyland, The Lighthouse Lab in Alderley Park and Alex Alderton, Roberto Amato, Jeffrey Barrett, Sonia Goncalves, Ewan Harrison, David K. Jackson, Ian Johnston, Dominic Kwiatkowski, Cordelia Langford, John Sillitoe on behalf of the Wellcome Sanger Institute COVID-19 Surveillance Team |
| EPI_ISL_2570311 | Lighthouse Laboratory Plymouth | Wellcome Sanger Institute for the COVID-19 Genomics UK (COG-UK) Consortium | Lighthouse Laboratory Plymouth and Alex Alderton, Roberto Amato, Jeffrey Barrett, Sonia Goncalves, Ewan Harrison, David K. Jackson, Ian Johnston, Dominic Kwiatkowski, Cordelia Langford, John Sillitoe on behalf of the Wellcome Sanger Institute COVID-19 Surveillance Team |
| EPI_ISL_2570312, EPI_ISL_2570313 | Lighthouse Lab in Milton Keynes | Wellcome Sanger Institute for the COVID-19 Genomics UK (COG-UK) Consortium | The Lighthouse Lab in Milton Keynes and Alex Alderton, Roberto Amato, Jeffrey Barrett, Sonia Goncalves, Ewan Harrison, David K. Jackson, Ian Johnston, Dominic Kwiatkowski, Cordelia Langford, John Sillitoe on behalf of the Wellcome Sanger Institute COVID-19 Surveillance Team |
| EPI_ISL_2570314 | Lighthouse Lab in Alderley Park | Wellcome Sanger Institute for the COVID-19 Genomics UK (COG-UK) Consortium | Jacquelyn Wynn, Mairead Hyland, The Lighthouse Lab in Alderley Park and Alex Alderton, Roberto Amato, Jeffrey Barrett, Sonia Goncalves, Ewan Harrison, David K. Jackson, Ian Johnston, Dominic Kwiatkowski, Cordelia Langford, John Sillitoe on behalf of the Wellcome Sanger Institute COVID-19 Surveillance Team |
| EPI_ISL_2570315 | Lighthouse Laboratory Plymouth | Wellcome Sanger Institute for the COVID-19 Genomics UK (COG-UK) Consortium | Lighthouse Laboratory Plymouth and Alex Alderton, Roberto Amato, Jeffrey Barrett, Sonia Goncalves, Ewan Harrison, David K. Jackson, Ian Johnston, Dominic Kwiatkowski, Cordelia Langford, John Sillitoe on behalf of the Wellcome Sanger Institute COVID-19 Surveillance Team |
| EPI_ISL_2570316 | Lighthouse Lab in Milton Keynes | Wellcome Sanger Institute for the COVID-19 Genomics UK (COG-UK) Consortium | The Lighthouse Lab in Milton Keynes and Alex Alderton, Roberto Amato, Jeffrey Barrett, Sonia Goncalves, Ewan Harrison, David K. Jackson, Ian Johnston, Dominic Kwiatkowski, Cordelia Langford, John Sillitoe on behalf of the Wellcome Sanger Institute COVID-19 Surveillance Team |
| EPI_ISL_2570317 | Lighthouse Lab in Alderley Park | Wellcome Sanger Institute for the COVID-19 Genomics UK (COG-UK) Consortium | Jacquelyn Wynn, Mairead Hyland, The Lighthouse Lab in Alderley Park and Alex Alderton, Roberto Amato, Jeffrey Barrett, Sonia Goncalves, Ewan Harrison, David K. Jackson, Ian Johnston, Dominic Kwiatkowski, Cordelia Langford, John Sillitoe on behalf of the Wellcome Sanger Institute COVID-19 Surveillance Team |
| EPI_ISL_2570318, EPI_ISL_2570319, EPI_ISL_2570320, EPI_ISL_2570321, EPI_ISL_2570322 | Lighthouse Laboratory Plymouth | Wellcome Sanger Institute for the COVID-19 Genomics UK (COG-UK) Consortium | Lighthouse Laboratory Plymouth and Alex Alderton, Roberto Amato, Jeffrey Barrett, Sonia Goncalves, Ewan Harrison, David K. Jackson, Ian Johnston, Dominic Kwiatkowski, Cordelia Langford, John Sillitoe on behalf of the Wellcome Sanger Institute COVID-19 Surveillance Team |
| EPI_ISL_2570323 | Lighthouse Lab in Milton Keynes | Wellcome Sanger Institute for the COVID-19 Genomics UK (COG-UK) Consortium | The Lighthouse Lab in Milton Keynes and Alex Alderton, Roberto Amato, Jeffrey Barrett, Sonia Goncalves, Ewan Harrison, David K. Jackson, Ian Johnston, Dominic Kwiatkowski, Cordelia Langford, John Sillitoe on behalf of the Wellcome Sanger Institute COVID-19 Surveillance Team |
| EPI_ISL_2570324 | Lighthouse Lab in Alderley Park | Wellcome Sanger Institute for the COVID-19 Genomics UK (COG-UK) Consortium | Jacquelyn Wynn, Mairead Hyland, The Lighthouse Lab in Alderley Park and Alex Alderton, Roberto Amato, Jeffrey Barrett, Sonia Goncalves, Ewan Harrison, David K. Jackson, Ian Johnston, Dominic Kwiatkowski, Cordelia Langford, John Sillitoe on behalf of the Wellcome Sanger Institute COVID-19 Surveillance Team |
| EPI_ISL_2570325, EPI_ISL_2570326, EPI_ISL_2570327 | Lighthouse Lab in Milton Keynes | Wellcome Sanger Institute for the COVID-19 Genomics UK (COG-UK) Consortium | The Lighthouse Lab in Milton Keynes and Alex Alderton, Roberto Amato, Jeffrey Barrett, Sonia Goncalves, Ewan Harrison, David K. Jackson, Ian Johnston, Dominic Kwiatkowski, Cordelia Langford, John Sillitoe on behalf of the Wellcome Sanger Institute COVID-19 Surveillance Team |
| EPI_ISL_2570328 | Lighthouse Laboratory Plymouth | Wellcome Sanger Institute for the COVID-19 Genomics UK (COG-UK) Consortium | Lighthouse Laboratory Plymouth and Alex Alderton, Roberto Amato, Jeffrey Barrett, Sonia Goncalves, Ewan Harrison, David K. Jackson, Ian Johnston, Dominic Kwiatkowski, Cordelia Langford, John Sillitoe on behalf of the Wellcome Sanger Institute COVID-19 Surveillance Team |
| EPI_ISL_2570329, EPI_ISL_2570330 | Lighthouse Lab in Milton Keynes | Wellcome Sanger Institute for the COVID-19 Genomics UK (COG-UK) Consortium | The Lighthouse Lab in Milton Keynes and Alex Alderton, Roberto Amato, Jeffrey Barrett, Sonia Goncalves, Ewan Harrison, David K. Jackson, Ian Johnston, Dominic Kwiatkowski, Cordelia Langford, John Sillitoe on behalf of the Wellcome Sanger Institute COVID-19 Surveillance Team |
| EPI_ISL_2570331 | Lighthouse Lab in Alderley Park | Wellcome Sanger Institute for the COVID-19 Genomics UK (COG-UK) Consortium | Jacquelyn Wynn, Mairead Hyland, The Lighthouse Lab in Alderley Park and Alex Alderton, Roberto Amato, Jeffrey Barrett, Sonia Goncalves, Ewan Harrison, David K. Jackson, Ian Johnston, Dominic Kwiatkowski, Cordelia Langford, John Sillitoe on behalf of the Wellcome Sanger Institute COVID-19 Surveillance Team |
| EPI_ISL_2570332, EPI_ISL_2570333 | Lighthouse Lab in Milton Keynes | Wellcome Sanger Institute for the COVID-19 Genomics UK (COG-UK) Consortium | The Lighthouse Lab in Milton Keynes and Alex Alderton, Roberto Amato, Jeffrey Barrett, Sonia Goncalves, Ewan Harrison, David K. Jackson, Ian Johnston, Dominic Kwiatkowski, Cordelia Langford, John Sillitoe on behalf of the Wellcome Sanger Institute COVID-19 Surveillance Team |
| EPI_ISL_2570334, EPI_ISL_2570335 | Lighthouse Laboratory Plymouth | Wellcome Sanger Institute for the COVID-19 Genomics UK (COG-UK) Consortium | Lighthouse Laboratory Plymouth and Alex Alderton, Roberto Amato, Jeffrey Barrett, Sonia Goncalves, Ewan Harrison, David K. Jackson, Ian Johnston, Dominic Kwiatkowski, Cordelia Langford, John Sillitoe on behalf of the Wellcome Sanger Institute COVID-19 Surveillance Team |
| EPI_ISL_2570336 | Lighthouse Lab in Glasgow | Wellcome Sanger Institute for the COVID-19 Genomics UK (COG-UK) Consortium | Harper VanSteenhouse, Yumi Kasai, David Gray, Carol Clugston, Anna Dominiczak and Alex Alderton, Roberto Amato, Jeffrey Barrett, Sonia Goncalves, Ewan Harrison, David K. Jackson, Ian Johnston, Dominic Kwiatkowski, Cordelia Langford, John Sillitoe on behalf of the Wellcome Sanger Institute COVID-19 Surveillance Team |
| EPI_ISL_2570337 | Lighthouse Lab in Alderley Park | Wellcome Sanger Institute for the COVID-19 Genomics UK (COG-UK) Consortium | Jacquelyn Wynn, Mairead Hyland, The Lighthouse Lab in Alderley Park and Alex Alderton, Roberto Amato, Jeffrey Barrett, Sonia Goncalves, Ewan Harrison, David K. Jackson, Ian Johnston, Dominic Kwiatkowski, Cordelia Langford, John Sillitoe on behalf of the Wellcome Sanger Institute COVID-19 Surveillance Team |
| EPI_ISL_2570338, EPI_ISL_2570339, EPI_ISL_2570340, EPI_ISL_2570341, EPI_ISL_2570342, EPI_ISL_2570343 | Lighthouse Lab in Milton Keynes | Wellcome Sanger Institute for the COVID-19 Genomics UK (COG-UK) Consortium | The Lighthouse Lab in Milton Keynes and Alex Alderton, Roberto Amato, Jeffrey Barrett, Sonia Goncalves, Ewan Harrison, David K. Jackson, Ian Johnston, Dominic Kwiatkowski, Cordelia Langford, John Sillitoe on behalf of the Wellcome Sanger Institute COVID-19 Surveillance Team |
| EPI_ISL_2570344 | Lighthouse Laboratory Plymouth | Wellcome Sanger Institute for the COVID-19 Genomics UK (COG-UK) Consortium | Lighthouse Laboratory Plymouth and Alex Alderton, Roberto Amato, Jeffrey Barrett, Sonia Goncalves, Ewan Harrison, David K. Jackson, Ian Johnston, Dominic Kwiatkowski, Cordelia Langford, John Sillitoe on behalf of the Wellcome Sanger Institute COVID-19 Surveillance Team |
| EPI_ISL_2570347 | Lighthouse Lab in Alderley Park | Wellcome Sanger Institute for the COVID-19 Genomics UK (COG-UK) Consortium | Jacquelyn Wynn, Mairead Hyland, The Lighthouse Lab in Alderley Park and Alex Alderton, Roberto Amato, Jeffrey Barrett, Sonia Goncalves, Ewan Harrison, David K. Jackson, Ian Johnston, Dominic Kwiatkowski, Cordelia Langford, John Sillitoe on behalf of the Wellcome Sanger Institute COVID-19 Surveillance Team |
| EPI_ISL_2570348, EPI_ISL_2570349 | Lighthouse Lab in Milton Keynes | Wellcome Sanger Institute for the COVID-19 Genomics UK (COG-UK) Consortium | The Lighthouse Lab in Milton Keynes and Alex Alderton, Roberto Amato, Jeffrey Barrett, Sonia Goncalves, Ewan Harrison, David K. Jackson, Ian Johnston, Dominic Kwiatkowski, Cordelia Langford, John Sillitoe on behalf of the Wellcome Sanger Institute COVID-19 Surveillance Team |
| EPI_ISL_2570350 | Lighthouse Lab in Glasgow | Wellcome Sanger Institute for the COVID-19 Genomics UK (COG-UK) Consortium | Harper VanSteenhouse, Yumi Kasai, David Gray, Carol Clugston, Anna Dominiczak and Alex Alderton, Roberto Amato, Jeffrey Barrett, Sonia Goncalves, Ewan Harrison, David K. Jackson, Ian Johnston, Dominic Kwiatkowski, Cordelia Langford, John Sillitoe on behalf of the Wellcome Sanger Institute COVID-19 Surveillance Team |
| EPI_ISL_2570351 | Lighthouse Lab in Milton Keynes | Wellcome Sanger Institute for the COVID-19 Genomics UK (COG-UK) Consortium | The Lighthouse Lab in Milton Keynes and Alex Alderton, Roberto Amato, Jeffrey Barrett, Sonia Goncalves, Ewan Harrison, David K. Jackson, Ian Johnston, Dominic Kwiatkowski, Cordelia Langford, John Sillitoe on behalf of the Wellcome Sanger Institute COVID-19 Surveillance Team |
| EPI_ISL_2570352, EPI_ISL_2570353 | Lighthouse Lab in Alderley Park | Wellcome Sanger Institute for the COVID-19 Genomics UK (COG-UK) Consortium | Jacquelyn Wynn, Mairead Hyland, The Lighthouse Lab in Alderley Park and Alex Alderton, Roberto Amato, Jeffrey Barrett, Sonia Goncalves, Ewan Harrison, David K. Jackson, Ian Johnston, Dominic Kwiatkowski, Cordelia Langford, John Sillitoe on behalf of the Wellcome Sanger Institute COVID-19 Surveillance Team |
| EPI_ISL_2570354, EPI_ISL_2570355 | Lighthouse Laboratory Plymouth | Wellcome Sanger Institute for the COVID-19 Genomics UK (COG-UK) Consortium | Lighthouse Laboratory Plymouth and Alex Alderton, Roberto Amato, Jeffrey Barrett, Sonia Goncalves, Ewan Harrison, David K. Jackson, Ian Johnston, Dominic Kwiatkowski, Cordelia Langford, John Sillitoe on behalf of the Wellcome Sanger Institute COVID-19 Surveillance Team |
| EPI_ISL_2570356, EPI_ISL_2570357, EPI_ISL_2570358, EPI_ISL_2570359, EPI_ISL_2570360, EPI_ISL_2570361, EPI_ISL_2570362, EPI_ISL_2570363, EPI_ISL_2570364 | Lighthouse Lab in Milton Keynes | Wellcome Sanger Institute for the COVID-19 Genomics UK (COG-UK) Consortium | The Lighthouse Lab in Milton Keynes and Alex Alderton, Roberto Amato, Jeffrey Barrett, Sonia Goncalves, Ewan Harrison, David K. Jackson, Ian Johnston, Dominic Kwiatkowski, Cordelia Langford, John Sillitoe on behalf of the Wellcome Sanger Institute COVID-19 Surveillance Team |
| EPI_ISL_2570365, EPI_ISL_2570366 | Lighthouse Laboratory Plymouth | Wellcome Sanger Institute for the COVID-19 Genomics UK (COG-UK) Consortium | Lighthouse Laboratory Plymouth and Alex Alderton, Roberto Amato, Jeffrey Barrett, Sonia Goncalves, Ewan Harrison, David K. Jackson, Ian Johnston, Dominic Kwiatkowski, Cordelia Langford, John Sillitoe on behalf of the Wellcome Sanger Institute COVID-19 Surveillance Team |
| EPI_ISL_2570367 | Lighthouse Lab in Alderley Park | Wellcome Sanger Institute for the COVID-19 Genomics UK (COG-UK) Consortium | Jacquelyn Wynn, Mairead Hyland, The Lighthouse Lab in Alderley Park and Alex Alderton, Roberto Amato, Jeffrey Barrett, Sonia Goncalves, Ewan Harrison, David K. Jackson, Ian Johnston, Dominic Kwiatkowski, Cordelia Langford, John Sillitoe on behalf of the Wellcome Sanger Institute COVID-19 Surveillance Team |
| EPI_ISL_2570368, EPI_ISL_2570370 | Lighthouse Lab in Milton Keynes | Wellcome Sanger Institute for the COVID-19 Genomics UK (COG-UK) Consortium | The Lighthouse Lab in Milton Keynes and Alex Alderton, Roberto Amato, Jeffrey Barrett, Sonia Goncalves, Ewan Harrison, David K. Jackson, Ian Johnston, Dominic Kwiatkowski, Cordelia Langford, John Sillitoe on behalf of the Wellcome Sanger Institute COVID-19 Surveillance Team |
| EPI_ISL_2570371 | Lighthouse Laboratory Plymouth | Wellcome Sanger Institute for the COVID-19 Genomics UK (COG-UK) Consortium | Lighthouse Laboratory Plymouth and Alex Alderton, Roberto Amato, Jeffrey Barrett, Sonia Goncalves, Ewan Harrison, David K. Jackson, Ian Johnston, Dominic Kwiatkowski, Cordelia Langford, John Sillitoe on behalf of the Wellcome Sanger Institute COVID-19 Surveillance Team |
| EPI_ISL_2570372 | Lighthouse Lab in Alderley Park | Wellcome Sanger Institute for the COVID-19 Genomics UK (COG-UK) Consortium | Jacquelyn Wynn, Mairead Hyland, The Lighthouse Lab in Alderley Park and Alex Alderton, Roberto Amato, Jeffrey Barrett, Sonia Goncalves, Ewan Harrison, David K. Jackson, Ian Johnston, Dominic Kwiatkowski, Cordelia Langford, John Sillitoe on behalf of the Wellcome Sanger Institute COVID-19 Surveillance Team |
| EPI_ISL_2570373 | Lighthouse Lab in Glasgow | Wellcome Sanger Institute for the COVID-19 Genomics UK (COG-UK) Consortium | Harper VanSteenhouse, Yumi Kasai, David Gray, Carol Clugston, Anna Dominiczak and Alex Alderton, Roberto Amato, Jeffrey Barrett, Sonia Goncalves, Ewan Harrison, David K. Jackson, Ian Johnston, Dominic Kwiatkowski, Cordelia Langford, John Sillitoe on behalf of the Wellcome Sanger Institute COVID-19 Surveillance Team |
| EPI_ISL_2570374 | Lighthouse Laboratory Plymouth | Wellcome Sanger Institute for the COVID-19 Genomics UK (COG-UK) Consortium | Lighthouse Laboratory Plymouth and Alex Alderton, Roberto Amato, Jeffrey Barrett, Sonia Goncalves, Ewan Harrison, David K. Jackson, Ian Johnston, Dominic Kwiatkowski, Cordelia Langford, John Sillitoe on behalf of the Wellcome Sanger Institute COVID-19 Surveillance Team |

| | | | |
|---|---|---|---|
| EPI_ISL_2570375 | Lighthouse Lab in Milton Keynes | Wellcome Sanger Institute for the COVID-19 Genomics UK (COG-UK) Consortium | The Lighthouse Lab in Milton Keynes and Alex Alderton, Roberto Amato, Jeffrey Barrett, Sonia Goncalves, Ewan Harrison, David K. Jackson, Ian Johnston, Dominic Kwiatkowski, Cordelia Langford, John Sillitoe on behalf of the Wellcome Sanger Institute COVID-19 Surveillance Team |
| EPI_ISL_2570378 | Lighthouse Lab in Glasgow | Wellcome Sanger Institute for the COVID-19 Genomics UK (COG-UK) Consortium | Harper VanSteenhouse, Yumi Kasai, David Gray, Carol Clugston, Anna Dominiczak and Alex Alderton, Roberto Amato, Jeffrey Barrett, Sonia Goncalves, Ewan Harrison, David K. Jackson, Ian Johnston, Dominic Kwiatkowski, Cordelia Langford, John Sillitoe on behalf of the Wellcome Sanger Institute COVID-19 Surveillance Team |
| EPI_ISL_2570379, EPI_ISL_2570380, EPI_ISL_2570381 | Lighthouse Lab in Milton Keynes | Wellcome Sanger Institute for the COVID-19 Genomics UK (COG-UK) Consortium | The Lighthouse Lab in Milton Keynes and Alex Alderton, Roberto Amato, Jeffrey Barrett, Sonia Goncalves, Ewan Harrison, David K. Jackson, Ian Johnston, Dominic Kwiatkowski, Cordelia Langford, John Sillitoe on behalf of the Wellcome Sanger Institute COVID-19 Surveillance Team |
| EPI_ISL_2570382, EPI_ISL_2570383 | Lighthouse Laboratory Plymouth | Wellcome Sanger Institute for the COVID-19 Genomics UK (COG-UK) Consortium | Lighthouse Laboratory Plymouth and Alex Alderton, Roberto Amato, Jeffrey Barrett, Sonia Goncalves, Ewan Harrison, David K. Jackson, Ian Johnston, Dominic Kwiatkowski, Cordelia Langford, John Sillitoe on behalf of the Wellcome Sanger Institute COVID-19 Surveillance Team |
| EPI_ISL_2570384, EPI_ISL_2570385 | Lighthouse Lab in Milton Keynes | Wellcome Sanger Institute for the COVID-19 Genomics UK (COG-UK) Consortium | The Lighthouse Lab in Milton Keynes and Alex Alderton, Roberto Amato, Jeffrey Barrett, Sonia Goncalves, Ewan Harrison, David K. Jackson, Ian Johnston, Dominic Kwiatkowski, Cordelia Langford, John Sillitoe on behalf of the Wellcome Sanger Institute COVID-19 Surveillance Team |
| EPI_ISL_2570386 | Lighthouse Lab in Alderley Park | Wellcome Sanger Institute for the COVID-19 Genomics UK (COG-UK) Consortium | Jacquelyn Wynn, Mairead Hyland, The Lighthouse Lab in Alderley Park and Alex Alderton, Roberto Amato, Jeffrey Barrett, Sonia Goncalves, Ewan Harrison, David K. Jackson, Ian Johnston, Dominic Kwiatkowski, Cordelia Langford, John Sillitoe on behalf of the Wellcome Sanger Institute COVID-19 Surveillance Team |
| EPI_ISL_2570387, EPI_ISL_2570388 | Lighthouse Lab in Milton Keynes | Wellcome Sanger Institute for the COVID-19 Genomics UK (COG-UK) Consortium | The Lighthouse Lab in Milton Keynes and Alex Alderton, Roberto Amato, Jeffrey Barrett, Sonia Goncalves, Ewan Harrison, David K. Jackson, Ian Johnston, Dominic Kwiatkowski, Cordelia Langford, John Sillitoe on behalf of the Wellcome Sanger Institute COVID-19 Surveillance Team |
| EPI_ISL_2570389 | Lighthouse Lab in Alderley Park | Wellcome Sanger Institute for the COVID-19 Genomics UK (COG-UK) Consortium | Jacquelyn Wynn, Mairead Hyland, The Lighthouse Lab in Alderley Park and Alex Alderton, Roberto Amato, Jeffrey Barrett, Sonia Goncalves, Ewan Harrison, David K. Jackson, Ian Johnston, Dominic Kwiatkowski, Cordelia Langford, John Sillitoe on behalf of the Wellcome Sanger Institute COVID-19 Surveillance Team |
| EPI_ISL_2570390, EPI_ISL_2570391 | Lighthouse Lab in Milton Keynes | Wellcome Sanger Institute for the COVID-19 Genomics UK (COG-UK) Consortium | The Lighthouse Lab in Milton Keynes and Alex Alderton, Roberto Amato, Jeffrey Barrett, Sonia Goncalves, Ewan Harrison, David K. Jackson, Ian Johnston, Dominic Kwiatkowski, Cordelia Langford, John Sillitoe on behalf of the Wellcome Sanger Institute COVID-19 Surveillance Team |
| EPI_ISL_2570392 | Lighthouse Lab in Glasgow | Wellcome Sanger Institute for the COVID-19 Genomics UK (COG-UK) Consortium | Harper VanSteenhouse, Yumi Kasai, David Gray, Carol Clugston, Anna Dominiczak and Alex Alderton, Roberto Amato, Jeffrey Barrett, Sonia Goncalves, Ewan Harrison, David K. Jackson, Ian Johnston, Dominic Kwiatkowski, Cordelia Langford, John Sillitoe on behalf of the Wellcome Sanger Institute COVID-19 Surveillance Team |
| EPI_ISL_2570393 | Lighthouse Lab in Milton Keynes | Wellcome Sanger Institute for the COVID-19 Genomics UK (COG-UK) Consortium | The Lighthouse Lab in Milton Keynes and Alex Alderton, Roberto Amato, Jeffrey Barrett, Sonia Goncalves, Ewan Harrison, David K. Jackson, Ian Johnston, Dominic Kwiatkowski, Cordelia Langford, John Sillitoe on behalf of the Wellcome Sanger Institute COVID-19 Surveillance Team |
| EPI_ISL_2570394 | Lighthouse Lab in Glasgow | Wellcome Sanger Institute for the COVID-19 Genomics UK (COG-UK) Consortium | Harper VanSteenhouse, Yumi Kasai, David Gray, Carol Clugston, Anna Dominiczak and Alex Alderton, Roberto Amato, Jeffrey Barrett, Sonia Goncalves, Ewan Harrison, David K. Jackson, Ian Johnston, Dominic Kwiatkowski, Cordelia Langford, John Sillitoe on behalf of the Wellcome Sanger Institute COVID-19 Surveillance Team |
| EPI_ISL_2570395 | Lighthouse Lab in Milton Keynes | Wellcome Sanger Institute for the COVID-19 Genomics UK (COG-UK) Consortium | The Lighthouse Lab in Milton Keynes and Alex Alderton, Roberto Amato, Jeffrey Barrett, Sonia Goncalves, Ewan Harrison, David K. Jackson, Ian Johnston, Dominic Kwiatkowski, Cordelia Langford, John Sillitoe on behalf of the Wellcome Sanger Institute COVID-19 Surveillance Team |
| EPI_ISL_2570396, EPI_ISL_2570397 | Lighthouse Lab in Glasgow | Wellcome Sanger Institute for the COVID-19 Genomics UK (COG-UK) Consortium | Harper VanSteenhouse, Yumi Kasai, David Gray, Carol Clugston, Anna Dominiczak and Alex Alderton, Roberto Amato, Jeffrey Barrett, Sonia Goncalves, Ewan Harrison, David K. Jackson, Ian Johnston, Dominic Kwiatkowski, Cordelia Langford, John Sillitoe on behalf of the Wellcome Sanger Institute COVID-19 Surveillance Team |
| EPI_ISL_2570398, EPI_ISL_2570399 | Lighthouse Laboratory Plymouth | Wellcome Sanger Institute for the COVID-19 Genomics UK (COG-UK) Consortium | Lighthouse Laboratory Plymouth and Alex Alderton, Roberto Amato, Jeffrey Barrett, Sonia Goncalves, Ewan Harrison, David K. Jackson, Ian Johnston, Dominic Kwiatkowski, Cordelia Langford, John Sillitoe on behalf of the Wellcome Sanger Institute COVID-19 Surveillance Team |
| EPI_ISL_2570400 | Lighthouse Lab in Alderley Park | Wellcome Sanger Institute for the COVID-19 Genomics UK (COG-UK) Consortium | Jacquelyn Wynn, Mairead Hyland, The Lighthouse Lab in Alderley Park and Alex Alderton, Roberto Amato, Jeffrey Barrett, Sonia Goncalves, Ewan Harrison, David K. Jackson, Ian Johnston, Dominic Kwiatkowski, Cordelia Langford, John Sillitoe on behalf of the Wellcome Sanger Institute COVID-19 Surveillance Team |
| EPI_ISL_2570401 | Lighthouse Lab in Glasgow | Wellcome Sanger Institute for the COVID-19 Genomics UK (COG-UK) Consortium | Harper VanSteenhouse, Yumi Kasai, David Gray, Carol Clugston, Anna Dominiczak and Alex Alderton, Roberto Amato, Jeffrey Barrett, Sonia Goncalves, Ewan Harrison, David K. Jackson, Ian Johnston, Dominic Kwiatkowski, Cordelia Langford, John Sillitoe on behalf of the Wellcome Sanger Institute COVID-19 Surveillance Team |
| EPI_ISL_2570402 | Lighthouse Lab in Alderley Park | Wellcome Sanger Institute for the COVID-19 Genomics UK (COG-UK) Consortium | Jacquelyn Wynn, Mairead Hyland, The Lighthouse Lab in Alderley Park and Alex Alderton, Roberto Amato, Jeffrey Barrett, Sonia Goncalves, Ewan Harrison, David K. Jackson, Ian Johnston, Dominic Kwiatkowski, Cordelia Langford, John Sillitoe on behalf of the Wellcome Sanger Institute COVID-19 Surveillance Team |
| EPI_ISL_2570403 | Lighthouse Lab in Milton Keynes | Wellcome Sanger Institute for the COVID-19 Genomics UK (COG-UK) Consortium | The Lighthouse Lab in Milton Keynes and Alex Alderton, Roberto Amato, Jeffrey Barrett, Sonia Goncalves, Ewan Harrison, David K. Jackson, Ian Johnston, Dominic Kwiatkowski, Cordelia Langford, John Sillitoe on behalf of the Wellcome Sanger Institute COVID-19 Surveillance Team |
| EPI_ISL_2570404 | Lighthouse Laboratory Plymouth | Wellcome Sanger Institute for the COVID-19 Genomics UK (COG-UK) Consortium | Lighthouse Laboratory Plymouth and Alex Alderton, Roberto Amato, Jeffrey Barrett, Sonia Goncalves, Ewan Harrison, David K. Jackson, Ian Johnston, Dominic Kwiatkowski, Cordelia Langford, John Sillitoe on behalf of the Wellcome Sanger Institute COVID-19 Surveillance Team |
| EPI_ISL_2570405 | Lighthouse Lab in Milton Keynes | Wellcome Sanger Institute for the COVID-19 Genomics UK (COG-UK) Consortium | The Lighthouse Lab in Milton Keynes and Alex Alderton, Roberto Amato, Jeffrey Barrett, Sonia Goncalves, Ewan Harrison, David K. Jackson, Ian Johnston, Dominic Kwiatkowski, Cordelia Langford, John Sillitoe on behalf of the Wellcome Sanger Institute COVID-19 Surveillance Team |
| EPI_ISL_2570406 | Lighthouse Lab in Alderley Park | Wellcome Sanger Institute for the COVID-19 Genomics UK (COG-UK) Consortium | Jacquelyn Wynn, Mairead Hyland, The Lighthouse Lab in Alderley Park and Alex Alderton, Roberto Amato, Jeffrey Barrett, Sonia Goncalves, Ewan Harrison, David K. Jackson, Ian Johnston, Dominic Kwiatkowski, Cordelia Langford, John Sillitoe on behalf of the Wellcome Sanger Institute COVID-19 Surveillance Team |
| EPI_ISL_2570408 | Lighthouse Lab in Milton Keynes | Wellcome Sanger Institute for the COVID-19 Genomics UK (COG-UK) Consortium | The Lighthouse Lab in Milton Keynes and Alex Alderton, Roberto Amato, Jeffrey Barrett, Sonia Goncalves, Ewan Harrison, David K. Jackson, Ian Johnston, Dominic Kwiatkowski, Cordelia Langford, John Sillitoe on behalf of the Wellcome Sanger Institute COVID-19 Surveillance Team |
| EPI_ISL_2570409 | Lighthouse Laboratory Plymouth | Wellcome Sanger Institute for the COVID-19 Genomics UK (COG-UK) Consortium | Lighthouse Laboratory Plymouth and Alex Alderton, Roberto Amato, Jeffrey Barrett, Sonia Goncalves, Ewan Harrison, David K. Jackson, Ian Johnston, Dominic Kwiatkowski, Cordelia Langford, John Sillitoe on behalf of the Wellcome Sanger Institute COVID-19 Surveillance Team |
| EPI_ISL_2570410 | Lighthouse Lab in Milton Keynes | Wellcome Sanger Institute for the COVID-19 Genomics UK (COG-UK) Consortium | The Lighthouse Lab in Milton Keynes and Alex Alderton, Roberto Amato, Jeffrey Barrett, Sonia Goncalves, Ewan Harrison, David K. Jackson, Ian Johnston, Dominic Kwiatkowski, Cordelia Langford, John Sillitoe on behalf of the Wellcome Sanger Institute COVID-19 Surveillance Team |
| EPI_ISL_2570411 | Lighthouse Laboratory Plymouth | Wellcome Sanger Institute for the COVID-19 Genomics UK (COG-UK) Consortium | Lighthouse Laboratory Plymouth and Alex Alderton, Roberto Amato, Jeffrey Barrett, Sonia Goncalves, Ewan Harrison, David K. Jackson, Ian Johnston, Dominic Kwiatkowski, Cordelia Langford, John Sillitoe on behalf of the Wellcome Sanger Institute COVID-19 Surveillance Team |
| EPI_ISL_2570412, EPI_ISL_2570413 | Lighthouse Lab in Milton Keynes | Wellcome Sanger Institute for the COVID-19 Genomics UK (COG-UK) Consortium | The Lighthouse Lab in Milton Keynes and Alex Alderton, Roberto Amato, Jeffrey Barrett, Sonia Goncalves, Ewan Harrison, David K. Jackson, Ian Johnston, Dominic Kwiatkowski, Cordelia Langford, John Sillitoe on behalf of the Wellcome Sanger Institute COVID-19 Surveillance Team |
| EPI_ISL_2570414 | Lighthouse Laboratory Plymouth | Wellcome Sanger Institute for the COVID-19 Genomics UK (COG-UK) Consortium | Lighthouse Laboratory Plymouth and Alex Alderton, Roberto Amato, Jeffrey Barrett, Sonia Goncalves, Ewan Harrison, David K. Jackson, Ian Johnston, Dominic Kwiatkowski, Cordelia Langford, John Sillitoe on behalf of the Wellcome Sanger Institute COVID-19 Surveillance Team |
| EPI_ISL_2570415, EPI_ISL_2570416 | Lighthouse Lab in Milton Keynes | Wellcome Sanger Institute for the COVID-19 Genomics UK (COG-UK) Consortium | The Lighthouse Lab in Milton Keynes and Alex Alderton, Roberto Amato, Jeffrey Barrett, Sonia Goncalves, Ewan Harrison, David K. Jackson, Ian Johnston, Dominic Kwiatkowski, Cordelia Langford, John Sillitoe on behalf of the Wellcome Sanger Institute COVID-19 Surveillance Team |
| EPI_ISL_2570417 | Lighthouse Lab in Alderley Park | Wellcome Sanger Institute for the COVID-19 Genomics UK (COG-UK) Consortium | Jacquelyn Wynn, Mairead Hyland, The Lighthouse Lab in Alderley Park and Alex Alderton, Roberto Amato, Jeffrey Barrett, Sonia Goncalves, Ewan Harrison, David K. Jackson, Ian Johnston, Dominic Kwiatkowski, Cordelia Langford, John Sillitoe on behalf of the Wellcome Sanger Institute COVID-19 Surveillance Team |
| EPI_ISL_2570418 | Lighthouse Lab in Milton Keynes | Wellcome Sanger Institute for the COVID-19 Genomics UK (COG-UK) Consortium | The Lighthouse Lab in Milton Keynes and Alex Alderton, Roberto Amato, Jeffrey Barrett, Sonia Goncalves, Ewan Harrison, David K. Jackson, Ian Johnston, Dominic Kwiatkowski, Cordelia Langford, John Sillitoe on behalf of the Wellcome Sanger Institute COVID-19 Surveillance Team |
| EPI_ISL_2570421, EPI_ISL_2570422 | Lighthouse Laboratory Plymouth | Wellcome Sanger Institute for the COVID-19 Genomics UK (COG-UK) Consortium | Lighthouse Laboratory Plymouth and Alex Alderton, Roberto Amato, Jeffrey Barrett, Sonia Goncalves, Ewan Harrison, David K. Jackson, Ian Johnston, Dominic Kwiatkowski, Cordelia Langford, John Sillitoe on behalf of the Wellcome Sanger Institute COVID-19 Surveillance Team |
| EPI_ISL_2570423, EPI_ISL_2570424 | Lighthouse Lab in Milton Keynes | Wellcome Sanger Institute for the COVID-19 Genomics UK (COG-UK) Consortium | The Lighthouse Lab in Milton Keynes and Alex Alderton, Roberto Amato, Jeffrey Barrett, Sonia Goncalves, Ewan Harrison, David K. Jackson, Ian Johnston, Dominic Kwiatkowski, Cordelia Langford, John Sillitoe on behalf of the Wellcome Sanger Institute COVID-19 Surveillance Team |
| EPI_ISL_2570425, EPI_ISL_2570426 | Lighthouse Laboratory Plymouth | Wellcome Sanger Institute for the COVID-19 Genomics UK (COG-UK) Consortium | Lighthouse Laboratory Plymouth and Alex Alderton, Roberto Amato, Jeffrey Barrett, Sonia Goncalves, Ewan Harrison, David K. Jackson, Ian Johnston, Dominic Kwiatkowski, Cordelia Langford, John Sillitoe on behalf of the Wellcome Sanger Institute COVID-19 Surveillance Team |
| EPI_ISL_2570427 | Lighthouse Lab in Glasgow | Wellcome Sanger Institute for the COVID-19 Genomics UK (COG-UK) Consortium | Harper VanSteenhouse, Yumi Kasai, David Gray, Carol Clugston, Anna Dominiczak and Alex Alderton, Roberto Amato, Jeffrey Barrett, Sonia Goncalves, Ewan Harrison, David K. Jackson, Ian Johnston, Dominic Kwiatkowski, Cordelia Langford, John Sillitoe on behalf of the Wellcome Sanger Institute COVID-19 Surveillance Team |
| EPI_ISL_2570428 | Lighthouse Lab in Milton Keynes | Wellcome Sanger Institute for the COVID-19 Genomics UK (COG-UK) Consortium | The Lighthouse Lab in Milton Keynes and Alex Alderton, Roberto Amato, Jeffrey Barrett, Sonia Goncalves, Ewan Harrison, David K. Jackson, Ian Johnston, Dominic Kwiatkowski, Cordelia Langford, John Sillitoe on behalf of the Wellcome Sanger Institute COVID-19 Surveillance Team |
| EPI_ISL_2570429 | Lighthouse Laboratory Plymouth | Wellcome Sanger Institute for the COVID-19 Genomics UK (COG-UK) Consortium | Lighthouse Laboratory Plymouth and Alex Alderton, Roberto Amato, Jeffrey Barrett, Sonia Goncalves, Ewan Harrison, David K. Jackson, Ian Johnston, Dominic Kwiatkowski, Cordelia Langford, John Sillitoe on behalf of the Wellcome Sanger Institute COVID-19 Surveillance Team |
| EPI_ISL_2570430, EPI_ISL_2570431 | Lighthouse Lab in Milton Keynes | Wellcome Sanger Institute for the COVID-19 Genomics UK (COG-UK) Consortium | The Lighthouse Lab in Milton Keynes and Alex Alderton, Roberto Amato, Jeffrey Barrett, Sonia Goncalves, Ewan Harrison, David K. Jackson, Ian Johnston, Dominic Kwiatkowski, Cordelia Langford, John Sillitoe on behalf of the Wellcome Sanger Institute COVID-19 Surveillance Team |
| EPI_ISL_2570432 | Lighthouse Laboratory Plymouth | Wellcome Sanger Institute for the COVID-19 Genomics UK (COG-UK) Consortium | Lighthouse Laboratory Plymouth and Alex Alderton, Roberto Amato, Jeffrey Barrett, Sonia Goncalves, Ewan Harrison, David K. Jackson, Ian Johnston, Dominic Kwiatkowski, Cordelia Langford, John Sillitoe on behalf of the Wellcome Sanger Institute COVID-19 Surveillance Team |
| EPI_ISL_2570433, EPI_ISL_2570434 | Lighthouse Lab in Milton Keynes | Wellcome Sanger Institute for the COVID-19 Genomics UK (COG-UK) Consortium | The Lighthouse Lab in Milton Keynes and Alex Alderton, Roberto Amato, Jeffrey Barrett, Sonia Goncalves, Ewan Harrison, David K. Jackson, Ian Johnston, Dominic Kwiatkowski, Cordelia Langford, John Sillitoe on behalf of the Wellcome Sanger Institute COVID-19 Surveillance Team |
| EPI_ISL_2570435 | Lighthouse Laboratory Plymouth | Wellcome Sanger Institute for the COVID-19 Genomics UK (COG-UK) Consortium | Lighthouse Laboratory Plymouth and Alex Alderton, Roberto Amato, Jeffrey Barrett, Sonia Goncalves, Ewan Harrison, David K. Jackson, Ian Johnston, Dominic Kwiatkowski, Cordelia Langford, John Sillitoe on behalf of the Wellcome Sanger Institute COVID-19 Surveillance Team |
| EPI_ISL_2570436 | Lighthouse Lab in Glasgow | Wellcome Sanger Institute for the COVID-19 Genomics UK (COG-UK) Consortium | Harper VanSteenhouse, Yumi Kasai, David Gray, Carol Clugston, Anna Dominiczak and Alex Alderton, Roberto Amato, Jeffrey Barrett, Sonia Goncalves, Ewan Harrison, David K. Jackson, Ian Johnston, Dominic Kwiatkowski, Cordelia Langford, John Sillitoe on behalf of the Wellcome Sanger Institute COVID-19 Surveillance Team |
| EPI_ISL_2570437 | Lighthouse Lab in Milton Keynes | Wellcome Sanger Institute for the COVID-19 Genomics UK (COG-UK) Consortium | The Lighthouse Lab in Milton Keynes and Alex Alderton, Roberto Amato, Jeffrey Barrett, Sonia Goncalves, Ewan Harrison, David K. Jackson, Ian Johnston, Dominic Kwiatkowski, Cordelia Langford, John Sillitoe on behalf of the Wellcome Sanger Institute COVID-19 Surveillance Team |
| EPI_ISL_2570438 | Lighthouse Laboratory Plymouth | Wellcome Sanger Institute for the COVID-19 Genomics UK (COG-UK) Consortium | Lighthouse Laboratory Plymouth and Alex Alderton, Roberto Amato, Jeffrey Barrett, Sonia Goncalves, Ewan Harrison, David K. Jackson, Ian Johnston, Dominic Kwiatkowski, Cordelia Langford, John Sillitoe on behalf of the Wellcome Sanger Institute COVID-19 Surveillance Team |
| EPI_ISL_2570439 | Lighthouse Lab in Alderley Park | Wellcome Sanger Institute for the COVID-19 Genomics UK (COG-UK) Consortium | Jacquelyn Wynn, Mairead Hyland, The Lighthouse Lab in Alderley Park and Alex Alderton, Roberto Amato, Jeffrey Barrett, Sonia Goncalves, Ewan Harrison, David K. Jackson, Ian Johnston, Dominic Kwiatkowski, Cordelia Langford, John Sillitoe on behalf of the Wellcome Sanger Institute COVID-19 Surveillance Team |

| | | | |
|---|---|---|---|
| EPI_ISL_2570440 | Lighthouse Lab in Milton Keynes | Wellcome Sanger Institute for the COVID-19 Genomics UK (COG-UK) Consortium | The Lighthouse Lab in Milton Keynes and Alex Alderton, Roberto Amato, Jeffrey Barrett, Sonia Goncalves, Ewan Harrison, David K. Jackson, Ian Johnston, Dominic Kwiatkowski, Cordelia Langford, John Sillitoe on behalf of the Wellcome Sanger Institute COVID-19 Surveillance Team |
| EPI_ISL_2570441 | Lighthouse Lab in Glasgow | Wellcome Sanger Institute for the COVID-19 Genomics UK (COG-UK) Consortium | Harper VanSteenhouse, Yumi Kasai, David Gray, Carol Clugston, Anna Dominiczak and Alex Alderton, Roberto Amato, Jeffrey Barrett, Sonia Goncalves, Ewan Harrison, David K. Jackson, Ian Johnston, Dominic Kwiatkowski, Cordelia Langford, John Sillitoe on behalf of the Wellcome Sanger Institute COVID-19 Surveillance Team |
| EPI_ISL_2570442 | Lighthouse Lab in Alderley Park | Wellcome Sanger Institute for the COVID-19 Genomics UK (COG-UK) Consortium | Jacquelyn Wynn, Mairead Hyland, The Lighthouse Lab in Alderley Park and Alex Alderton, Roberto Amato, Jeffrey Barrett, Sonia Goncalves, Ewan Harrison, David K. Jackson, Ian Johnston, Dominic Kwiatkowski, Cordelia Langford, John Sillitoe on behalf of the Wellcome Sanger Institute COVID-19 Surveillance Team |
| EPI_ISL_2570444 | Lighthouse Laboratory Plymouth | Wellcome Sanger Institute for the COVID-19 Genomics UK (COG-UK) Consortium | Lighthouse Laboratory Plymouth and Alex Alderton, Roberto Amato, Jeffrey Barrett, Sonia Goncalves, Ewan Harrison, David K. Jackson, Ian Johnston, Dominic Kwiatkowski, Cordelia Langford, John Sillitoe on behalf of the Wellcome Sanger Institute COVID-19 Surveillance Team |
| EPI_ISL_2570445 | Lighthouse Lab in Glasgow | Wellcome Sanger Institute for the COVID-19 Genomics UK (COG-UK) Consortium | Harper VanSteenhouse, Yumi Kasai, David Gray, Carol Clugston, Anna Dominiczak and Alex Alderton, Roberto Amato, Jeffrey Barrett, Sonia Goncalves, Ewan Harrison, David K. Jackson, Ian Johnston, Dominic Kwiatkowski, Cordelia Langford, John Sillitoe on behalf of the Wellcome Sanger Institute COVID-19 Surveillance Team |
| EPI_ISL_2570446, EPI_ISL_2570447, EPI_ISL_2570448 | Lighthouse Laboratory Plymouth | Wellcome Sanger Institute for the COVID-19 Genomics UK (COG-UK) Consortium | Lighthouse Laboratory Plymouth and Alex Alderton, Roberto Amato, Jeffrey Barrett, Sonia Goncalves, Ewan Harrison, David K. Jackson, Ian Johnston, Dominic Kwiatkowski, Cordelia Langford, John Sillitoe on behalf of the Wellcome Sanger Institute COVID-19 Surveillance Team |
| EPI_ISL_2570449 | Lighthouse Lab in Milton Keynes | Wellcome Sanger Institute for the COVID-19 Genomics UK (COG-UK) Consortium | The Lighthouse Lab in Milton Keynes and Alex Alderton, Roberto Amato, Jeffrey Barrett, Sonia Goncalves, Ewan Harrison, David K. Jackson, Ian Johnston, Dominic Kwiatkowski, Cordelia Langford, John Sillitoe on behalf of the Wellcome Sanger Institute COVID-19 Surveillance Team |
| EPI_ISL_2570450 | Lighthouse Lab in Glasgow | Wellcome Sanger Institute for the COVID-19 Genomics UK (COG-UK) Consortium | Harper VanSteenhouse, Yumi Kasai, David Gray, Carol Clugston, Anna Dominiczak and Alex Alderton, Roberto Amato, Jeffrey Barrett, Sonia Goncalves, Ewan Harrison, David K. Jackson, Ian Johnston, Dominic Kwiatkowski, Cordelia Langford, John Sillitoe on behalf of the Wellcome Sanger Institute COVID-19 Surveillance Team |
| EPI_ISL_2570451 | Lighthouse Lab in Milton Keynes | Wellcome Sanger Institute for the COVID-19 Genomics UK (COG-UK) Consortium | The Lighthouse Lab in Milton Keynes and Alex Alderton, Roberto Amato, Jeffrey Barrett, Sonia Goncalves, Ewan Harrison, David K. Jackson, Ian Johnston, Dominic Kwiatkowski, Cordelia Langford, John Sillitoe on behalf of the Wellcome Sanger Institute COVID-19 Surveillance Team |
| EPI_ISL_2570453, EPI_ISL_2570454 | Lighthouse Laboratory Plymouth | Wellcome Sanger Institute for the COVID-19 Genomics UK (COG-UK) Consortium | Lighthouse Laboratory Plymouth and Alex Alderton, Roberto Amato, Jeffrey Barrett, Sonia Goncalves, Ewan Harrison, David K. Jackson, Ian Johnston, Dominic Kwiatkowski, Cordelia Langford, John Sillitoe on behalf of the Wellcome Sanger Institute COVID-19 Surveillance Team |
| EPI_ISL_2570455 | Lighthouse Lab in Milton Keynes | Wellcome Sanger Institute for the COVID-19 Genomics UK (COG-UK) Consortium | The Lighthouse Lab in Milton Keynes and Alex Alderton, Roberto Amato, Jeffrey Barrett, Sonia Goncalves, Ewan Harrison, David K. Jackson, Ian Johnston, Dominic Kwiatkowski, Cordelia Langford, John Sillitoe on behalf of the Wellcome Sanger Institute COVID-19 Surveillance Team |
| EPI_ISL_2570456 | Lighthouse Laboratory Plymouth | Wellcome Sanger Institute for the COVID-19 Genomics UK (COG-UK) Consortium | Lighthouse Laboratory Plymouth and Alex Alderton, Roberto Amato, Jeffrey Barrett, Sonia Goncalves, Ewan Harrison, David K. Jackson, Ian Johnston, Dominic Kwiatkowski, Cordelia Langford, John Sillitoe on behalf of the Wellcome Sanger Institute COVID-19 Surveillance Team |
| EPI_ISL_2570457, EPI_ISL_2570458 | Lighthouse Lab in Milton Keynes | Wellcome Sanger Institute for the COVID-19 Genomics UK (COG-UK) Consortium | The Lighthouse Lab in Milton Keynes and Alex Alderton, Roberto Amato, Jeffrey Barrett, Sonia Goncalves, Ewan Harrison, David K. Jackson, Ian Johnston, Dominic Kwiatkowski, Cordelia Langford, John Sillitoe on behalf of the Wellcome Sanger Institute COVID-19 Surveillance Team |
| EPI_ISL_2570459 | Lighthouse Lab in Alderley Park | Wellcome Sanger Institute for the COVID-19 Genomics UK (COG-UK) Consortium | Jacquelyn Wynn, Mairead Hyland, The Lighthouse Lab in Alderley Park and Alex Alderton, Roberto Amato, Jeffrey Barrett, Sonia Goncalves, Ewan Harrison, David K. Jackson, Ian Johnston, Dominic Kwiatkowski, Cordelia Langford, John Sillitoe on behalf of the Wellcome Sanger Institute COVID-19 Surveillance Team |
| EPI_ISL_2570461 | Lighthouse Laboratory Plymouth | Wellcome Sanger Institute for the COVID-19 Genomics UK (COG-UK) Consortium | Lighthouse Laboratory Plymouth and Alex Alderton, Roberto Amato, Jeffrey Barrett, Sonia Goncalves, Ewan Harrison, David K. Jackson, Ian Johnston, Dominic Kwiatkowski, Cordelia Langford, John Sillitoe on behalf of the Wellcome Sanger Institute COVID-19 Surveillance Team |
| EPI_ISL_2570462, EPI_ISL_2570463 | Lighthouse Lab in Milton Keynes | Wellcome Sanger Institute for the COVID-19 Genomics UK (COG-UK) Consortium | The Lighthouse Lab in Milton Keynes and Alex Alderton, Roberto Amato, Jeffrey Barrett, Sonia Goncalves, Ewan Harrison, David K. Jackson, Ian Johnston, Dominic Kwiatkowski, Cordelia Langford, John Sillitoe on behalf of the Wellcome Sanger Institute COVID-19 Surveillance Team |
| EPI_ISL_2570465 | Lighthouse Lab in Alderley Park | Wellcome Sanger Institute for the COVID-19 Genomics UK (COG-UK) Consortium | Jacquelyn Wynn, Mairead Hyland, The Lighthouse Lab in Alderley Park and Alex Alderton, Roberto Amato, Jeffrey Barrett, Sonia Goncalves, Ewan Harrison, David K. Jackson, Ian Johnston, Dominic Kwiatkowski, Cordelia Langford, John Sillitoe on behalf of the Wellcome Sanger Institute COVID-19 Surveillance Team |
| EPI_ISL_2570466 | Lighthouse Laboratory Plymouth | Wellcome Sanger Institute for the COVID-19 Genomics UK (COG-UK) Consortium | Lighthouse Laboratory Plymouth and Alex Alderton, Roberto Amato, Jeffrey Barrett, Sonia Goncalves, Ewan Harrison, David K. Jackson, Ian Johnston, Dominic Kwiatkowski, Cordelia Langford, John Sillitoe on behalf of the Wellcome Sanger Institute COVID-19 Surveillance Team |
| EPI_ISL_2570467, EPI_ISL_2570468 | Lighthouse Lab in Milton Keynes | Wellcome Sanger Institute for the COVID-19 Genomics UK (COG-UK) Consortium | The Lighthouse Lab in Milton Keynes and Alex Alderton, Roberto Amato, Jeffrey Barrett, Sonia Goncalves, Ewan Harrison, David K. Jackson, Ian Johnston, Dominic Kwiatkowski, Cordelia Langford, John Sillitoe on behalf of the Wellcome Sanger Institute COVID-19 Surveillance Team |
| EPI_ISL_2570469, EPI_ISL_2570470, EPI_ISL_2570471 | Lighthouse Laboratory Plymouth | Wellcome Sanger Institute for the COVID-19 Genomics UK (COG-UK) Consortium | Lighthouse Laboratory Plymouth and Alex Alderton, Roberto Amato, Jeffrey Barrett, Sonia Goncalves, Ewan Harrison, David K. Jackson, Ian Johnston, Dominic Kwiatkowski, Cordelia Langford, John Sillitoe on behalf of the Wellcome Sanger Institute COVID-19 Surveillance Team |
| EPI_ISL_2570472 | Lighthouse Lab in Milton Keynes | Wellcome Sanger Institute for the COVID-19 Genomics UK (COG-UK) Consortium | The Lighthouse Lab in Milton Keynes and Alex Alderton, Roberto Amato, Jeffrey Barrett, Sonia Goncalves, Ewan Harrison, David K. Jackson, Ian Johnston, Dominic Kwiatkowski, Cordelia Langford, John Sillitoe on behalf of the Wellcome Sanger Institute COVID-19 Surveillance Team |
| EPI_ISL_2570473, EPI_ISL_2570474, EPI_ISL_2570475 | Lighthouse Lab in Alderley Park | Wellcome Sanger Institute for the COVID-19 Genomics UK (COG-UK) Consortium | Jacquelyn Wynn, Mairead Hyland, The Lighthouse Lab in Alderley Park and Alex Alderton, Roberto Amato, Jeffrey Barrett, Sonia Goncalves, Ewan Harrison, David K. Jackson, Ian Johnston, Dominic Kwiatkowski, Cordelia Langford, John Sillitoe on behalf of the Wellcome Sanger Institute COVID-19 Surveillance Team |
| EPI_ISL_2570476 | Lighthouse Laboratory Plymouth | Wellcome Sanger Institute for the COVID-19 Genomics UK (COG-UK) Consortium | Lighthouse Laboratory Plymouth and Alex Alderton, Roberto Amato, Jeffrey Barrett, Sonia Goncalves, Ewan Harrison, David K. Jackson, Ian Johnston, Dominic Kwiatkowski, Cordelia Langford, John Sillitoe on behalf of the Wellcome Sanger Institute COVID-19 Surveillance Team |
| EPI_ISL_2570477, EPI_ISL_2570478 | Lighthouse Lab in Alderley Park | Wellcome Sanger Institute for the COVID-19 Genomics UK (COG-UK) Consortium | Jacquelyn Wynn, Mairead Hyland, The Lighthouse Lab in Alderley Park and Alex Alderton, Roberto Amato, Jeffrey Barrett, Sonia Goncalves, Ewan Harrison, David K. Jackson, Ian Johnston, Dominic Kwiatkowski, Cordelia Langford, John Sillitoe on behalf of the Wellcome Sanger Institute COVID-19 Surveillance Team |
| EPI_ISL_2570479 | Lighthouse Laboratory Plymouth | Wellcome Sanger Institute for the COVID-19 Genomics UK (COG-UK) Consortium | Lighthouse Laboratory Plymouth and Alex Alderton, Roberto Amato, Jeffrey Barrett, Sonia Goncalves, Ewan Harrison, David K. Jackson, Ian Johnston, Dominic Kwiatkowski, Cordelia Langford, John Sillitoe on behalf of the Wellcome Sanger Institute COVID-19 Surveillance Team |
| EPI_ISL_2570480, EPI_ISL_2570481 | Lighthouse Lab in Alderley Park | Wellcome Sanger Institute for the COVID-19 Genomics UK (COG-UK) Consortium | Jacquelyn Wynn, Mairead Hyland, The Lighthouse Lab in Alderley Park and Alex Alderton, Roberto Amato, Jeffrey Barrett, Sonia Goncalves, Ewan Harrison, David K. Jackson, Ian Johnston, Dominic Kwiatkowski, Cordelia Langford, John Sillitoe on behalf of the Wellcome Sanger Institute COVID-19 Surveillance Team |
| EPI_ISL_2570482, EPI_ISL_2570483 | Lighthouse Laboratory Plymouth | Wellcome Sanger Institute for the COVID-19 Genomics UK (COG-UK) Consortium | Lighthouse Laboratory Plymouth and Alex Alderton, Roberto Amato, Jeffrey Barrett, Sonia Goncalves, Ewan Harrison, David K. Jackson, Ian Johnston, Dominic Kwiatkowski, Cordelia Langford, John Sillitoe on behalf of the Wellcome Sanger Institute COVID-19 Surveillance Team |
| EPI_ISL_2570484 | Lighthouse Lab in Alderley Park | Wellcome Sanger Institute for the COVID-19 Genomics UK (COG-UK) Consortium | Jacquelyn Wynn, Mairead Hyland, The Lighthouse Lab in Alderley Park and Alex Alderton, Roberto Amato, Jeffrey Barrett, Sonia Goncalves, Ewan Harrison, David K. Jackson, Ian Johnston, Dominic Kwiatkowski, Cordelia Langford, John Sillitoe on behalf of the Wellcome Sanger Institute COVID-19 Surveillance Team |
| EPI_ISL_2570485 | Lighthouse Laboratory Plymouth | Wellcome Sanger Institute for the COVID-19 Genomics UK (COG-UK) Consortium | Lighthouse Laboratory Plymouth and Alex Alderton, Roberto Amato, Jeffrey Barrett, Sonia Goncalves, Ewan Harrison, David K. Jackson, Ian Johnston, Dominic Kwiatkowski, Cordelia Langford, John Sillitoe on behalf of the Wellcome Sanger Institute COVID-19 Surveillance Team |
| EPI_ISL_2570486, EPI_ISL_2570487 | Lighthouse Lab in Alderley Park | Wellcome Sanger Institute for the COVID-19 Genomics UK (COG-UK) Consortium | Jacquelyn Wynn, Mairead Hyland, The Lighthouse Lab in Alderley Park and Alex Alderton, Roberto Amato, Jeffrey Barrett, Sonia Goncalves, Ewan Harrison, David K. Jackson, Ian Johnston, Dominic Kwiatkowski, Cordelia Langford, John Sillitoe on behalf of the Wellcome Sanger Institute COVID-19 Surveillance Team |
| EPI_ISL_2570488 | Lighthouse Laboratory Plymouth | Wellcome Sanger Institute for the COVID-19 Genomics UK (COG-UK) Consortium | Lighthouse Laboratory Plymouth and Alex Alderton, Roberto Amato, Jeffrey Barrett, Sonia Goncalves, Ewan Harrison, David K. Jackson, Ian Johnston, Dominic Kwiatkowski, Cordelia Langford, John Sillitoe on behalf of the Wellcome Sanger Institute COVID-19 Surveillance Team |
| EPI_ISL_2570489 | Lighthouse Lab in Alderley Park | Wellcome Sanger Institute for the COVID-19 Genomics UK (COG-UK) Consortium | Jacquelyn Wynn, Mairead Hyland, The Lighthouse Lab in Alderley Park and Alex Alderton, Roberto Amato, Jeffrey Barrett, Sonia Goncalves, Ewan Harrison, David K. Jackson, Ian Johnston, Dominic Kwiatkowski, Cordelia Langford, John Sillitoe on behalf of the Wellcome Sanger Institute COVID-19 Surveillance Team |
| EPI_ISL_2570490 | Lighthouse Laboratory Plymouth | Wellcome Sanger Institute for the COVID-19 Genomics UK (COG-UK) Consortium | Lighthouse Laboratory Plymouth and Alex Alderton, Roberto Amato, Jeffrey Barrett, Sonia Goncalves, Ewan Harrison, David K. Jackson, Ian Johnston, Dominic Kwiatkowski, Cordelia Langford, John Sillitoe on behalf of the Wellcome Sanger Institute COVID-19 Surveillance Team |
| EPI_ISL_2570491, EPI_ISL_2570492, EPI_ISL_2570493, EPI_ISL_2570494, EPI_ISL_2570495, EPI_ISL_2570496, EPI_ISL_2570497 | Lighthouse Lab in Alderley Park | Wellcome Sanger Institute for the COVID-19 Genomics UK (COG-UK) Consortium | Jacquelyn Wynn, Mairead Hyland, The Lighthouse Lab in Alderley Park and Alex Alderton, Roberto Amato, Jeffrey Barrett, Sonia Goncalves, Ewan Harrison, David K. Jackson, Ian Johnston, Dominic Kwiatkowski, Cordelia Langford, John Sillitoe on behalf of the Wellcome Sanger Institute COVID-19 Surveillance Team |
| EPI_ISL_2570498 | Lighthouse Laboratory Plymouth | Wellcome Sanger Institute for the COVID-19 Genomics UK (COG-UK) Consortium | Lighthouse Laboratory Plymouth and Alex Alderton, Roberto Amato, Jeffrey Barrett, Sonia Goncalves, Ewan Harrison, David K. Jackson, Ian Johnston, Dominic Kwiatkowski, Cordelia Langford, John Sillitoe on behalf of the Wellcome Sanger Institute COVID-19 Surveillance Team |
| EPI_ISL_2570499 | Lighthouse Lab in Alderley Park | Wellcome Sanger Institute for the COVID-19 Genomics UK (COG-UK) Consortium | Jacquelyn Wynn, Mairead Hyland, The Lighthouse Lab in Alderley Park and Alex Alderton, Roberto Amato, Jeffrey Barrett, Sonia Goncalves, Ewan Harrison, David K. Jackson, Ian Johnston, Dominic Kwiatkowski, Cordelia Langford, John Sillitoe on behalf of the Wellcome Sanger Institute COVID-19 Surveillance Team |
| EPI_ISL_2570500, EPI_ISL_2570501, EPI_ISL_2570502 | Lighthouse Laboratory Plymouth | Wellcome Sanger Institute for the COVID-19 Genomics UK (COG-UK) Consortium | Lighthouse Laboratory Plymouth and Alex Alderton, Roberto Amato, Jeffrey Barrett, Sonia Goncalves, Ewan Harrison, David K. Jackson, Ian Johnston, Dominic Kwiatkowski, Cordelia Langford, John Sillitoe on behalf of the Wellcome Sanger Institute COVID-19 Surveillance Team |
| EPI_ISL_2570503, EPI_ISL_2570504 | Lighthouse Lab in Alderley Park | Wellcome Sanger Institute for the COVID-19 Genomics UK (COG-UK) Consortium | Jacquelyn Wynn, Mairead Hyland, The Lighthouse Lab in Alderley Park and Alex Alderton, Roberto Amato, Jeffrey Barrett, Sonia Goncalves, Ewan Harrison, David K. Jackson, Ian Johnston, Dominic Kwiatkowski, Cordelia Langford, John Sillitoe on behalf of the Wellcome Sanger Institute COVID-19 Surveillance Team |
| EPI_ISL_2570505 | Lighthouse Laboratory Plymouth | Wellcome Sanger Institute for the COVID-19 Genomics UK (COG-UK) Consortium | Lighthouse Laboratory Plymouth and Alex Alderton, Roberto Amato, Jeffrey Barrett, Sonia Goncalves, Ewan Harrison, David K. Jackson, Ian Johnston, Dominic Kwiatkowski, Cordelia Langford, John Sillitoe on behalf of the Wellcome Sanger Institute COVID-19 Surveillance Team |
| EPI_ISL_2570506, EPI_ISL_2570507, EPI_ISL_2570508 | Lighthouse Lab in Alderley Park | Wellcome Sanger Institute for the COVID-19 Genomics UK (COG-UK) Consortium | Jacquelyn Wynn, Mairead Hyland, The Lighthouse Lab in Alderley Park and Alex Alderton, Roberto Amato, Jeffrey Barrett, Sonia Goncalves, Ewan Harrison, David K. Jackson, Ian Johnston, Dominic Kwiatkowski, Cordelia Langford, John Sillitoe on behalf of the Wellcome Sanger Institute COVID-19 Surveillance Team |
| EPI_ISL_2570509 | Lighthouse Laboratory Plymouth | Wellcome Sanger Institute for the COVID-19 Genomics UK (COG-UK) Consortium | Lighthouse Laboratory Plymouth and Alex Alderton, Roberto Amato, Jeffrey Barrett, Sonia Goncalves, Ewan Harrison, David K. Jackson, Ian Johnston, Dominic Kwiatkowski, Cordelia Langford, John Sillitoe on behalf of the Wellcome Sanger Institute COVID-19 Surveillance Team |

| | | | |
|---|---|---|---|
| EPI_ISL_2570510, EPI_ISL_2570511, EPI_ISL_2570512 | Lighthouse Lab in Alderley Park | Wellcome Sanger Institute for the COVID-19 Genomics UK (COG-UK) Consortium | Jacquelyn Wynn, Mairead Hyland, The Lighthouse Lab in Alderley Park and Alex Alderton, Roberto Amato, Jeffrey Barrett, Sonia Goncalves, Ewan Harrison, David K. Jackson, Ian Johnston, Dominic Kwiatkowski, Cordelia Langford, John Sillitoe on behalf of the Wellcome Sanger Institute COVID-19 Surveillance Team |
| EPI_ISL_2570513 | Lighthouse Laboratory Plymouth | Wellcome Sanger Institute for the COVID-19 Genomics UK (COG-UK) Consortium | Lighthouse Laboratory Plymouth and Alex Alderton, Roberto Amato, Jeffrey Barrett, Sonia Goncalves, Ewan Harrison, David K. Jackson, Ian Johnston, Dominic Kwiatkowski, Cordelia Langford, John Sillitoe on behalf of the Wellcome Sanger Institute COVID-19 Surveillance Team |
| EPI_ISL_2570514, EPI_ISL_2570515, EPI_ISL_2570516 | Lighthouse Lab in Alderley Park | Wellcome Sanger Institute for the COVID-19 Genomics UK (COG-UK) Consortium | Jacquelyn Wynn, Mairead Hyland, The Lighthouse Lab in Alderley Park and Alex Alderton, Roberto Amato, Jeffrey Barrett, Sonia Goncalves, Ewan Harrison, David K. Jackson, Ian Johnston, Dominic Kwiatkowski, Cordelia Langford, John Sillitoe on behalf of the Wellcome Sanger Institute COVID-19 Surveillance Team |
| EPI_ISL_2570517 | Lighthouse Laboratory Plymouth | Wellcome Sanger Institute for the COVID-19 Genomics UK (COG-UK) Consortium | Lighthouse Laboratory Plymouth and Alex Alderton, Roberto Amato, Jeffrey Barrett, Sonia Goncalves, Ewan Harrison, David K. Jackson, Ian Johnston, Dominic Kwiatkowski, Cordelia Langford, John Sillitoe on behalf of the Wellcome Sanger Institute COVID-19 Surveillance Team |
| EPI_ISL_2570518, EPI_ISL_2570519, EPI_ISL_2570520 | Lighthouse Lab in Alderley Park | Wellcome Sanger Institute for the COVID-19 Genomics UK (COG-UK) Consortium | Jacquelyn Wynn, Mairead Hyland, The Lighthouse Lab in Alderley Park and Alex Alderton, Roberto Amato, Jeffrey Barrett, Sonia Goncalves, Ewan Harrison, David K. Jackson, Ian Johnston, Dominic Kwiatkowski, Cordelia Langford, John Sillitoe on behalf of the Wellcome Sanger Institute COVID-19 Surveillance Team |
| EPI_ISL_2570521 | Lighthouse Laboratory Plymouth | Wellcome Sanger Institute for the COVID-19 Genomics UK (COG-UK) Consortium | Lighthouse Laboratory Plymouth and Alex Alderton, Roberto Amato, Jeffrey Barrett, Sonia Goncalves, Ewan Harrison, David K. Jackson, Ian Johnston, Dominic Kwiatkowski, Cordelia Langford, John Sillitoe on behalf of the Wellcome Sanger Institute COVID-19 Surveillance Team |
| EPI_ISL_2570522, EPI_ISL_2570523, EPI_ISL_2570524, EPI_ISL_2570525, EPI_ISL_2570526, EPI_ISL_2570527, EPI_ISL_2570528 | Lighthouse Lab in Alderley Park | Wellcome Sanger Institute for the COVID-19 Genomics UK (COG-UK) Consortium | Jacquelyn Wynn, Mairead Hyland, The Lighthouse Lab in Alderley Park and Alex Alderton, Roberto Amato, Jeffrey Barrett, Sonia Goncalves, Ewan Harrison, David K. Jackson, Ian Johnston, Dominic Kwiatkowski, Cordelia Langford, John Sillitoe on behalf of the Wellcome Sanger Institute COVID-19 Surveillance Team |
| EPI_ISL_2570529 | Lighthouse Laboratory Plymouth | Wellcome Sanger Institute for the COVID-19 Genomics UK (COG-UK) Consortium | Lighthouse Laboratory Plymouth and Alex Alderton, Roberto Amato, Jeffrey Barrett, Sonia Goncalves, Ewan Harrison, David K. Jackson, Ian Johnston, Dominic Kwiatkowski, Cordelia Langford, John Sillitoe on behalf of the Wellcome Sanger Institute COVID-19 Surveillance Team |
| EPI_ISL_2570530, EPI_ISL_2570531, EPI_ISL_2570532, EPI_ISL_2570533, EPI_ISL_2570534, EPI_ISL_2570535, EPI_ISL_2570536 | Lighthouse Lab in Alderley Park | Wellcome Sanger Institute for the COVID-19 Genomics UK (COG-UK) Consortium | Jacquelyn Wynn, Mairead Hyland, The Lighthouse Lab in Alderley Park and Alex Alderton, Roberto Amato, Jeffrey Barrett, Sonia Goncalves, Ewan Harrison, David K. Jackson, Ian Johnston, Dominic Kwiatkowski, Cordelia Langford, John Sillitoe on behalf of the Wellcome Sanger Institute COVID-19 Surveillance Team |
| EPI_ISL_2570537 | Lighthouse Laboratory Plymouth | Wellcome Sanger Institute for the COVID-19 Genomics UK (COG-UK) Consortium | Lighthouse Laboratory Plymouth and Alex Alderton, Roberto Amato, Jeffrey Barrett, Sonia Goncalves, Ewan Harrison, David K. Jackson, Ian Johnston, Dominic Kwiatkowski, Cordelia Langford, John Sillitoe on behalf of the Wellcome Sanger Institute COVID-19 Surveillance Team |
| EPI_ISL_2570538, EPI_ISL_2570539, EPI_ISL_2570540, EPI_ISL_2570541 | Lighthouse Lab in Alderley Park | Wellcome Sanger Institute for the COVID-19 Genomics UK (COG-UK) Consortium | Jacquelyn Wynn, Mairead Hyland, The Lighthouse Lab in Alderley Park and Alex Alderton, Roberto Amato, Jeffrey Barrett, Sonia Goncalves, Ewan Harrison, David K. Jackson, Ian Johnston, Dominic Kwiatkowski, Cordelia Langford, John Sillitoe on behalf of the Wellcome Sanger Institute COVID-19 Surveillance Team |
| EPI_ISL_2570542, EPI_ISL_2570543 | Lighthouse Laboratory Plymouth | Wellcome Sanger Institute for the COVID-19 Genomics UK (COG-UK) Consortium | Lighthouse Laboratory Plymouth and Alex Alderton, Roberto Amato, Jeffrey Barrett, Sonia Goncalves, Ewan Harrison, David K. Jackson, Ian Johnston, Dominic Kwiatkowski, Cordelia Langford, John Sillitoe on behalf of the Wellcome Sanger Institute COVID-19 Surveillance Team |
| EPI_ISL_2570544 | Lighthouse Lab in Alderley Park | Wellcome Sanger Institute for the COVID-19 Genomics UK (COG-UK) Consortium | Jacquelyn Wynn, Mairead Hyland, The Lighthouse Lab in Alderley Park and Alex Alderton, Roberto Amato, Jeffrey Barrett, Sonia Goncalves, Ewan Harrison, David K. Jackson, Ian Johnston, Dominic Kwiatkowski, Cordelia Langford, John Sillitoe on behalf of the Wellcome Sanger Institute COVID-19 Surveillance Team |
| EPI_ISL_2570545 | Lighthouse Laboratory Plymouth | Wellcome Sanger Institute for the COVID-19 Genomics UK (COG-UK) Consortium | Lighthouse Laboratory Plymouth and Alex Alderton, Roberto Amato, Jeffrey Barrett, Sonia Goncalves, Ewan Harrison, David K. Jackson, Ian Johnston, Dominic Kwiatkowski, Cordelia Langford, John Sillitoe on behalf of the Wellcome Sanger Institute COVID-19 Surveillance Team |
| EPI_ISL_2570546, EPI_ISL_2570547, EPI_ISL_2570548, EPI_ISL_2570549, EPI_ISL_2570550 | Lighthouse Lab in Alderley Park | Wellcome Sanger Institute for the COVID-19 Genomics UK (COG-UK) Consortium | Jacquelyn Wynn, Mairead Hyland, The Lighthouse Lab in Alderley Park and Alex Alderton, Roberto Amato, Jeffrey Barrett, Sonia Goncalves, Ewan Harrison, David K. Jackson, Ian Johnston, Dominic Kwiatkowski, Cordelia Langford, John Sillitoe on behalf of the Wellcome Sanger Institute COVID-19 Surveillance Team |
| EPI_ISL_2570552 | Lighthouse Laboratory Plymouth | Wellcome Sanger Institute for the COVID-19 Genomics UK (COG-UK) Consortium | Lighthouse Laboratory Plymouth and Alex Alderton, Roberto Amato, Jeffrey Barrett, Sonia Goncalves, Ewan Harrison, David K. Jackson, Ian Johnston, Dominic Kwiatkowski, Cordelia Langford, John Sillitoe on behalf of the Wellcome Sanger Institute COVID-19 Surveillance Team |
| EPI_ISL_2570553, EPI_ISL_2570554 | Lighthouse Lab in Alderley Park | Wellcome Sanger Institute for the COVID-19 Genomics UK (COG-UK) Consortium | Jacquelyn Wynn, Mairead Hyland, The Lighthouse Lab in Alderley Park and Alex Alderton, Roberto Amato, Jeffrey Barrett, Sonia Goncalves, Ewan Harrison, David K. Jackson, Ian Johnston, Dominic Kwiatkowski, Cordelia Langford, John Sillitoe on behalf of the Wellcome Sanger Institute COVID-19 Surveillance Team |
| EPI_ISL_2570555 | Lighthouse Laboratory Plymouth | Wellcome Sanger Institute for the COVID-19 Genomics UK (COG-UK) Consortium | Lighthouse Laboratory Plymouth and Alex Alderton, Roberto Amato, Jeffrey Barrett, Sonia Goncalves, Ewan Harrison, David K. Jackson, Ian Johnston, Dominic Kwiatkowski, Cordelia Langford, John Sillitoe on behalf of the Wellcome Sanger Institute COVID-19 Surveillance Team |
| EPI_ISL_2570556, EPI_ISL_2570557, EPI_ISL_2570558 | Lighthouse Lab in Alderley Park | Wellcome Sanger Institute for the COVID-19 Genomics UK (COG-UK) Consortium | Jacquelyn Wynn, Mairead Hyland, The Lighthouse Lab in Alderley Park and Alex Alderton, Roberto Amato, Jeffrey Barrett, Sonia Goncalves, Ewan Harrison, David K. Jackson, Ian Johnston, Dominic Kwiatkowski, Cordelia Langford, John Sillitoe on behalf of the Wellcome Sanger Institute COVID-19 Surveillance Team |
| EPI_ISL_2570559 | Lighthouse Laboratory Plymouth | Wellcome Sanger Institute for the COVID-19 Genomics UK (COG-UK) Consortium | Lighthouse Laboratory Plymouth and Alex Alderton, Roberto Amato, Jeffrey Barrett, Sonia Goncalves, Ewan Harrison, David K. Jackson, Ian Johnston, Dominic Kwiatkowski, Cordelia Langford, John Sillitoe on behalf of the Wellcome Sanger Institute COVID-19 Surveillance Team |
| EPI_ISL_2570560, EPI_ISL_2570561, EPI_ISL_2570562, EPI_ISL_2570563, EPI_ISL_2570564 | Lighthouse Lab in Alderley Park | Wellcome Sanger Institute for the COVID-19 Genomics UK (COG-UK) Consortium | Jacquelyn Wynn, Mairead Hyland, The Lighthouse Lab in Alderley Park and Alex Alderton, Roberto Amato, Jeffrey Barrett, Sonia Goncalves, Ewan Harrison, David K. Jackson, Ian Johnston, Dominic Kwiatkowski, Cordelia Langford, John Sillitoe on behalf of the Wellcome Sanger Institute COVID-19 Surveillance Team |
| EPI_ISL_2570565 | Lighthouse Laboratory Plymouth | Wellcome Sanger Institute for the COVID-19 Genomics UK (COG-UK) Consortium | Lighthouse Laboratory Plymouth and Alex Alderton, Roberto Amato, Jeffrey Barrett, Sonia Goncalves, Ewan Harrison, David K. Jackson, Ian Johnston, Dominic Kwiatkowski, Cordelia Langford, John Sillitoe on behalf of the Wellcome Sanger Institute COVID-19 Surveillance Team |
| EPI_ISL_2570566, EPI_ISL_2570567, EPI_ISL_2570568, EPI_ISL_2570569, EPI_ISL_2570570, EPI_ISL_2570571, EPI_ISL_2570572, EPI_ISL_2570573 | Lighthouse Lab in Alderley Park | Wellcome Sanger Institute for the COVID-19 Genomics UK (COG-UK) Consortium | Jacquelyn Wynn, Mairead Hyland, The Lighthouse Lab in Alderley Park and Alex Alderton, Roberto Amato, Jeffrey Barrett, Sonia Goncalves, Ewan Harrison, David K. Jackson, Ian Johnston, Dominic Kwiatkowski, Cordelia Langford, John Sillitoe on behalf of the Wellcome Sanger Institute COVID-19 Surveillance Team |
| EPI_ISL_2570575, EPI_ISL_2570576 | Lighthouse Laboratory Plymouth | Wellcome Sanger Institute for the COVID-19 Genomics UK (COG-UK) Consortium | Lighthouse Laboratory Plymouth and Alex Alderton, Roberto Amato, Jeffrey Barrett, Sonia Goncalves, Ewan Harrison, David K. Jackson, Ian Johnston, Dominic Kwiatkowski, Cordelia Langford, John Sillitoe on behalf of the Wellcome Sanger Institute COVID-19 Surveillance Team |
| EPI_ISL_2570577, EPI_ISL_2570578, EPI_ISL_2570579, EPI_ISL_2570580, EPI_ISL_2570581, EPI_ISL_2570582, EPI_ISL_2570583, EPI_ISL_2570584 | Lighthouse Lab in Alderley Park | Wellcome Sanger Institute for the COVID-19 Genomics UK (COG-UK) Consortium | Jacquelyn Wynn, Mairead Hyland, The Lighthouse Lab in Alderley Park and Alex Alderton, Roberto Amato, Jeffrey Barrett, Sonia Goncalves, Ewan Harrison, David K. Jackson, Ian Johnston, Dominic Kwiatkowski, Cordelia Langford, John Sillitoe on behalf of the Wellcome Sanger Institute COVID-19 Surveillance Team |
| EPI_ISL_2570585 | Lighthouse Laboratory Plymouth | Wellcome Sanger Institute for the COVID-19 Genomics UK (COG-UK) Consortium | Lighthouse Laboratory Plymouth and Alex Alderton, Roberto Amato, Jeffrey Barrett, Sonia Goncalves, Ewan Harrison, David K. Jackson, Ian Johnston, Dominic Kwiatkowski, Cordelia Langford, John Sillitoe on behalf of the Wellcome Sanger Institute COVID-19 Surveillance Team |
| EPI_ISL_2570586, EPI_ISL_2570587, EPI_ISL_2570588 | Lighthouse Lab in Alderley Park | Wellcome Sanger Institute for the COVID-19 Genomics UK (COG-UK) Consortium | Jacquelyn Wynn, Mairead Hyland, The Lighthouse Lab in Alderley Park and Alex Alderton, Roberto Amato, Jeffrey Barrett, Sonia Goncalves, Ewan Harrison, David K. Jackson, Ian Johnston, Dominic Kwiatkowski, Cordelia Langford, John Sillitoe on behalf of the Wellcome Sanger Institute COVID-19 Surveillance Team |
| EPI_ISL_2570589, EPI_ISL_2570590 | Lighthouse Laboratory Plymouth | Wellcome Sanger Institute for the COVID-19 Genomics UK (COG-UK) Consortium | Lighthouse Laboratory Plymouth and Alex Alderton, Roberto Amato, Jeffrey Barrett, Sonia Goncalves, Ewan Harrison, David K. Jackson, Ian Johnston, Dominic Kwiatkowski, Cordelia Langford, John Sillitoe on behalf of the Wellcome Sanger Institute COVID-19 Surveillance Team |
| EPI_ISL_2570591, EPI_ISL_2570592 | Lighthouse Lab in Alderley Park | Wellcome Sanger Institute for the COVID-19 Genomics UK (COG-UK) Consortium | Jacquelyn Wynn, Mairead Hyland, The Lighthouse Lab in Alderley Park and Alex Alderton, Roberto Amato, Jeffrey Barrett, Sonia Goncalves, Ewan Harrison, David K. Jackson, Ian Johnston, Dominic Kwiatkowski, Cordelia Langford, John Sillitoe on behalf of the Wellcome Sanger Institute COVID-19 Surveillance Team |
| EPI_ISL_2570593 | Lighthouse Laboratory Plymouth | Wellcome Sanger Institute for the COVID-19 Genomics UK (COG-UK) Consortium | Lighthouse Laboratory Plymouth and Alex Alderton, Roberto Amato, Jeffrey Barrett, Sonia Goncalves, Ewan Harrison, David K. Jackson, Ian Johnston, Dominic Kwiatkowski, Cordelia Langford, John Sillitoe on behalf of the Wellcome Sanger Institute COVID-19 Surveillance Team |
| EPI_ISL_2570594, | Lighthouse Lab in Alderley Park | Wellcome Sanger Institute for the COVID-19 Genomics UK (COG-UK) | Jacquelyn Wynn, Mairead Hyland, The Lighthouse Lab in Alderley Park and Alex Alderton, Roberto Amato, Jeffrey Barrett, Sonia Goncalves, Ewan Harrison, David K. Jackson, Ian Johnston, Dominic Kwiatkowski, Cordelia Langford, John Sillitoe on |

| | | | |
|---|---|---|---|
| EPI_ISL_2570595, EPI_ISL_2570596, EPI_ISL_2570597 | | Consortium | behalf of the Wellcome Sanger Institute COVID-19 Surveillance Team |
| EPI_ISL_2570598, EPI_ISL_2570599, EPI_ISL_2570600, EPI_ISL_2570601, EPI_ISL_2570602 | Lighthouse Laboratory Plymouth | Wellcome Sanger Institute for the COVID-19 Genomics UK (COG-UK) Consortium | Lighthouse Laboratory Plymouth and Alex Alderton, Roberto Amato, Jeffrey Barrett, Sonia Goncalves, Ewan Harrison, David K. Jackson, Ian Johnston, Dominic Kwiatkowski, Cordelia Langford, John Sillitoe on behalf of the Wellcome Sanger Institute COVID-19 Surveillance Team |
| EPI_ISL_2570603, EPI_ISL_2570604, EPI_ISL_2570605, EPI_ISL_2570606, EPI_ISL_2570607 | Lighthouse Lab in Alderley Park | Wellcome Sanger Institute for the COVID-19 Genomics UK (COG-UK) Consortium | Jacquelyn Wynn, Mairead Hyland, The Lighthouse Lab in Alderley Park and Alex Alderton, Roberto Amato, Jeffrey Barrett, Sonia Goncalves, Ewan Harrison, David K. Jackson, Ian Johnston, Dominic Kwiatkowski, Cordelia Langford, John Sillitoe on behalf of the Wellcome Sanger Institute COVID-19 Surveillance Team |
| EPI_ISL_2570608, EPI_ISL_2570609, EPI_ISL_2570610, EPI_ISL_2570611 | Lighthouse Laboratory Plymouth | Wellcome Sanger Institute for the COVID-19 Genomics UK (COG-UK) Consortium | Lighthouse Laboratory Plymouth and Alex Alderton, Roberto Amato, Jeffrey Barrett, Sonia Goncalves, Ewan Harrison, David K. Jackson, Ian Johnston, Dominic Kwiatkowski, Cordelia Langford, John Sillitoe on behalf of the Wellcome Sanger Institute COVID-19 Surveillance Team |
| EPI_ISL_2570612, EPI_ISL_2570613, EPI_ISL_2570614, EPI_ISL_2570615, EPI_ISL_2570616, EPI_ISL_2570619 | Lighthouse Lab in Alderley Park | Wellcome Sanger Institute for the COVID-19 Genomics UK (COG-UK) Consortium | Jacquelyn Wynn, Mairead Hyland, The Lighthouse Lab in Alderley Park and Alex Alderton, Roberto Amato, Jeffrey Barrett, Sonia Goncalves, Ewan Harrison, David K. Jackson, Ian Johnston, Dominic Kwiatkowski, Cordelia Langford, John Sillitoe on behalf of the Wellcome Sanger Institute COVID-19 Surveillance Team |
| EPI_ISL_2570620, EPI_ISL_2570621 | Lighthouse Laboratory Plymouth | Wellcome Sanger Institute for the COVID-19 Genomics UK (COG-UK) Consortium | Lighthouse Laboratory Plymouth and Alex Alderton, Roberto Amato, Jeffrey Barrett, Sonia Goncalves, Ewan Harrison, David K. Jackson, Ian Johnston, Dominic Kwiatkowski, Cordelia Langford, John Sillitoe on behalf of the Wellcome Sanger Institute COVID-19 Surveillance Team |
| EPI_ISL_2570622, EPI_ISL_2570623, EPI_ISL_2570624, EPI_ISL_2570625, EPI_ISL_2570626, EPI_ISL_2570627, EPI_ISL_2570628, EPI_ISL_2570629, EPI_ISL_2570630, EPI_ISL_2570631, EPI_ISL_2570633 | | | |
| see above | Lighthouse Lab in Alderley Park | Wellcome Sanger Institute for the COVID-19 Genomics UK (COG-UK) Consortium | Jacquelyn Wynn, Mairead Hyland, The Lighthouse Lab in Alderley Park and Alex Alderton, Roberto Amato, Jeffrey Barrett, Sonia Goncalves, Ewan Harrison, David K. Jackson, Ian Johnston, Dominic Kwiatkowski, Cordelia Langford, John Sillitoe on behalf of the Wellcome Sanger Institute COVID-19 Surveillance Team |
| EPI_ISL_2570635 | Lighthouse Laboratory Plymouth | Wellcome Sanger Institute for the COVID-19 Genomics UK (COG-UK) Consortium | Lighthouse Laboratory Plymouth and Alex Alderton, Roberto Amato, Jeffrey Barrett, Sonia Goncalves, Ewan Harrison, David K. Jackson, Ian Johnston, Dominic Kwiatkowski, Cordelia Langford, John Sillitoe on behalf of the Wellcome Sanger Institute COVID-19 Surveillance Team |
| EPI_ISL_2570636, EPI_ISL_2570637 | Lighthouse Lab in Alderley Park | Wellcome Sanger Institute for the COVID-19 Genomics UK (COG-UK) Consortium | Jacquelyn Wynn, Mairead Hyland, The Lighthouse Lab in Alderley Park and Alex Alderton, Roberto Amato, Jeffrey Barrett, Sonia Goncalves, Ewan Harrison, David K. Jackson, Ian Johnston, Dominic Kwiatkowski, Cordelia Langford, John Sillitoe on behalf of the Wellcome Sanger Institute COVID-19 Surveillance Team |
| EPI_ISL_2570638 | Lighthouse Laboratory Plymouth | Wellcome Sanger Institute for the COVID-19 Genomics UK (COG-UK) Consortium | Lighthouse Laboratory Plymouth and Alex Alderton, Roberto Amato, Jeffrey Barrett, Sonia Goncalves, Ewan Harrison, David K. Jackson, Ian Johnston, Dominic Kwiatkowski, Cordelia Langford, John Sillitoe on behalf of the Wellcome Sanger Institute COVID-19 Surveillance Team |
| EPI_ISL_2570639, EPI_ISL_2570640 | Lighthouse Lab in Alderley Park | Wellcome Sanger Institute for the COVID-19 Genomics UK (COG-UK) Consortium | Jacquelyn Wynn, Mairead Hyland, The Lighthouse Lab in Alderley Park and Alex Alderton, Roberto Amato, Jeffrey Barrett, Sonia Goncalves, Ewan Harrison, David K. Jackson, Ian Johnston, Dominic Kwiatkowski, Cordelia Langford, John Sillitoe on behalf of the Wellcome Sanger Institute COVID-19 Surveillance Team |
| EPI_ISL_2570641 | Lighthouse Laboratory Plymouth | Wellcome Sanger Institute for the COVID-19 Genomics UK (COG-UK) Consortium | Lighthouse Laboratory Plymouth and Alex Alderton, Roberto Amato, Jeffrey Barrett, Sonia Goncalves, Ewan Harrison, David K. Jackson, Ian Johnston, Dominic Kwiatkowski, Cordelia Langford, John Sillitoe on behalf of the Wellcome Sanger Institute COVID-19 Surveillance Team |
| EPI_ISL_2570642, EPI_ISL_2570643 | Lighthouse Lab in Alderley Park | Wellcome Sanger Institute for the COVID-19 Genomics UK (COG-UK) Consortium | Jacquelyn Wynn, Mairead Hyland, The Lighthouse Lab in Alderley Park and Alex Alderton, Roberto Amato, Jeffrey Barrett, Sonia Goncalves, Ewan Harrison, David K. Jackson, Ian Johnston, Dominic Kwiatkowski, Cordelia Langford, John Sillitoe on behalf of the Wellcome Sanger Institute COVID-19 Surveillance Team |
| EPI_ISL_2570644 | Lighthouse Laboratory Plymouth | Wellcome Sanger Institute for the COVID-19 Genomics UK (COG-UK) Consortium | Lighthouse Laboratory Plymouth and Alex Alderton, Roberto Amato, Jeffrey Barrett, Sonia Goncalves, Ewan Harrison, David K. Jackson, Ian Johnston, Dominic Kwiatkowski, Cordelia Langford, John Sillitoe on behalf of the Wellcome Sanger Institute COVID-19 Surveillance Team |
| EPI_ISL_2570645 | Lighthouse Lab in Alderley Park | Wellcome Sanger Institute for the COVID-19 Genomics UK (COG-UK) Consortium | Jacquelyn Wynn, Mairead Hyland, The Lighthouse Lab in Alderley Park and Alex Alderton, Roberto Amato, Jeffrey Barrett, Sonia Goncalves, Ewan Harrison, David K. Jackson, Ian Johnston, Dominic Kwiatkowski, Cordelia Langford, John Sillitoe on behalf of the Wellcome Sanger Institute COVID-19 Surveillance Team |
| EPI_ISL_2570647, EPI_ISL_2570648 | Lighthouse Laboratory Plymouth | Wellcome Sanger Institute for the COVID-19 Genomics UK (COG-UK) Consortium | Lighthouse Laboratory Plymouth and Alex Alderton, Roberto Amato, Jeffrey Barrett, Sonia Goncalves, Ewan Harrison, David K. Jackson, Ian Johnston, Dominic Kwiatkowski, Cordelia Langford, John Sillitoe on behalf of the Wellcome Sanger Institute COVID-19 Surveillance Team |
| EPI_ISL_2570649 | Lighthouse Lab in Alderley Park | Wellcome Sanger Institute for the COVID-19 Genomics UK (COG-UK) Consortium | Jacquelyn Wynn, Mairead Hyland, The Lighthouse Lab in Alderley Park and Alex Alderton, Roberto Amato, Jeffrey Barrett, Sonia Goncalves, Ewan Harrison, David K. Jackson, Ian Johnston, Dominic Kwiatkowski, Cordelia Langford, John Sillitoe on behalf of the Wellcome Sanger Institute COVID-19 Surveillance Team |
| EPI_ISL_2570650 | Lighthouse Laboratory Plymouth | Wellcome Sanger Institute for the COVID-19 Genomics UK (COG-UK) Consortium | Lighthouse Laboratory Plymouth and Alex Alderton, Roberto Amato, Jeffrey Barrett, Sonia Goncalves, Ewan Harrison, David K. Jackson, Ian Johnston, Dominic Kwiatkowski, Cordelia Langford, John Sillitoe on behalf of the Wellcome Sanger Institute COVID-19 Surveillance Team |
| EPI_ISL_2570651, EPI_ISL_2570652, EPI_ISL_2570653, EPI_ISL_2570654, EPI_ISL_2570655, EPI_ISL_2570656 | Lighthouse Lab in Alderley Park | Wellcome Sanger Institute for the COVID-19 Genomics UK (COG-UK) Consortium | Jacquelyn Wynn, Mairead Hyland, The Lighthouse Lab in Alderley Park and Alex Alderton, Roberto Amato, Jeffrey Barrett, Sonia Goncalves, Ewan Harrison, David K. Jackson, Ian Johnston, Dominic Kwiatkowski, Cordelia Langford, John Sillitoe on behalf of the Wellcome Sanger Institute COVID-19 Surveillance Team |
| EPI_ISL_2570657 | Lighthouse Laboratory Plymouth | Wellcome Sanger Institute for the COVID-19 Genomics UK (COG-UK) Consortium | Lighthouse Laboratory Plymouth and Alex Alderton, Roberto Amato, Jeffrey Barrett, Sonia Goncalves, Ewan Harrison, David K. Jackson, Ian Johnston, Dominic Kwiatkowski, Cordelia Langford, John Sillitoe on behalf of the Wellcome Sanger Institute COVID-19 Surveillance Team |
| EPI_ISL_2570658 | Lighthouse Lab in Alderley Park | Wellcome Sanger Institute for the COVID-19 Genomics UK (COG-UK) Consortium | Jacquelyn Wynn, Mairead Hyland, The Lighthouse Lab in Alderley Park and Alex Alderton, Roberto Amato, Jeffrey Barrett, Sonia Goncalves, Ewan Harrison, David K. Jackson, Ian Johnston, Dominic Kwiatkowski, Cordelia Langford, John Sillitoe on behalf of the Wellcome Sanger Institute COVID-19 Surveillance Team |
| EPI_ISL_2570659 | Lighthouse Laboratory Plymouth | Wellcome Sanger Institute for the COVID-19 Genomics UK (COG-UK) Consortium | Lighthouse Laboratory Plymouth and Alex Alderton, Roberto Amato, Jeffrey Barrett, Sonia Goncalves, Ewan Harrison, David K. Jackson, Ian Johnston, Dominic Kwiatkowski, Cordelia Langford, John Sillitoe on behalf of the Wellcome Sanger Institute COVID-19 Surveillance Team |
| EPI_ISL_2570660, EPI_ISL_2570661, EPI_ISL_2570662 | Lighthouse Lab in Alderley Park | Wellcome Sanger Institute for the COVID-19 Genomics UK (COG-UK) Consortium | Jacquelyn Wynn, Mairead Hyland, The Lighthouse Lab in Alderley Park and Alex Alderton, Roberto Amato, Jeffrey Barrett, Sonia Goncalves, Ewan Harrison, David K. Jackson, Ian Johnston, Dominic Kwiatkowski, Cordelia Langford, John Sillitoe on behalf of the Wellcome Sanger Institute COVID-19 Surveillance Team |
| EPI_ISL_2570663 | Lighthouse Laboratory Plymouth | Wellcome Sanger Institute for the COVID-19 Genomics UK (COG-UK) Consortium | Lighthouse Laboratory Plymouth and Alex Alderton, Roberto Amato, Jeffrey Barrett, Sonia Goncalves, Ewan Harrison, David K. Jackson, Ian Johnston, Dominic Kwiatkowski, Cordelia Langford, John Sillitoe on behalf of the Wellcome Sanger Institute COVID-19 Surveillance Team |
| EPI_ISL_2570664, EPI_ISL_2570665, EPI_ISL_2570666 | Lighthouse Lab in Alderley Park | Wellcome Sanger Institute for the COVID-19 Genomics UK (COG-UK) Consortium | Jacquelyn Wynn, Mairead Hyland, The Lighthouse Lab in Alderley Park and Alex Alderton, Roberto Amato, Jeffrey Barrett, Sonia Goncalves, Ewan Harrison, David K. Jackson, Ian Johnston, Dominic Kwiatkowski, Cordelia Langford, John Sillitoe on behalf of the Wellcome Sanger Institute COVID-19 Surveillance Team |
| EPI_ISL_2570667 | Lighthouse Laboratory Plymouth | Wellcome Sanger Institute for the COVID-19 Genomics UK (COG-UK) Consortium | Lighthouse Laboratory Plymouth and Alex Alderton, Roberto Amato, Jeffrey Barrett, Sonia Goncalves, Ewan Harrison, David K. Jackson, Ian Johnston, Dominic Kwiatkowski, Cordelia Langford, John Sillitoe on behalf of the Wellcome Sanger Institute COVID-19 Surveillance Team |
| EPI_ISL_2570668, EPI_ISL_2570669, EPI_ISL_2570670, EPI_ISL_2570671, EPI_ISL_2570672 | Lighthouse Lab in Alderley Park | Wellcome Sanger Institute for the COVID-19 Genomics UK (COG-UK) Consortium | Jacquelyn Wynn, Mairead Hyland, The Lighthouse Lab in Alderley Park and Alex Alderton, Roberto Amato, Jeffrey Barrett, Sonia Goncalves, Ewan Harrison, David K. Jackson, Ian Johnston, Dominic Kwiatkowski, Cordelia Langford, John Sillitoe on behalf of the Wellcome Sanger Institute COVID-19 Surveillance Team |
| EPI_ISL_2570673 | Lighthouse Laboratory Plymouth | Wellcome Sanger Institute for the COVID-19 Genomics UK (COG-UK) Consortium | Lighthouse Laboratory Plymouth and Alex Alderton, Roberto Amato, Jeffrey Barrett, Sonia Goncalves, Ewan Harrison, David K. Jackson, Ian Johnston, Dominic Kwiatkowski, Cordelia Langford, John Sillitoe on behalf of the Wellcome Sanger Institute COVID-19 Surveillance Team |
| EPI_ISL_2570674 | Lighthouse Lab in Alderley Park | Wellcome Sanger Institute for the COVID-19 Genomics UK (COG-UK) Consortium | Jacquelyn Wynn, Mairead Hyland, The Lighthouse Lab in Alderley Park and Alex Alderton, Roberto Amato, Jeffrey Barrett, Sonia Goncalves, Ewan Harrison, David K. Jackson, Ian Johnston, Dominic Kwiatkowski, Cordelia Langford, John Sillitoe on behalf of the Wellcome Sanger Institute COVID-19 Surveillance Team |
| EPI_ISL_2570675 | Lighthouse Laboratory Plymouth | Wellcome Sanger Institute for the COVID-19 Genomics UK (COG-UK) Consortium | Lighthouse Laboratory Plymouth and Alex Alderton, Roberto Amato, Jeffrey Barrett, Sonia Goncalves, Ewan Harrison, David K. Jackson, Ian Johnston, Dominic Kwiatkowski, Cordelia Langford, John Sillitoe on behalf of the Wellcome Sanger Institute COVID-19 Surveillance Team |
| EPI_ISL_2570676, EPI_ISL_2570678, EPI_ISL_2570679, EPI_ISL_2570680, EPI_ISL_2570681, EPI_ISL_2570682 | Lighthouse Lab in Alderley Park | Wellcome Sanger Institute for the COVID-19 Genomics UK (COG-UK) Consortium | Jacquelyn Wynn, Mairead Hyland, The Lighthouse Lab in Alderley Park and Alex Alderton, Roberto Amato, Jeffrey Barrett, Sonia Goncalves, Ewan Harrison, David K. Jackson, Ian Johnston, Dominic Kwiatkowski, Cordelia Langford, John Sillitoe on behalf of the Wellcome Sanger Institute COVID-19 Surveillance Team |
| EPI_ISL_2570683, EPI_ISL_2570684 | Lighthouse Laboratory Plymouth | Wellcome Sanger Institute for the COVID-19 Genomics UK (COG-UK) Consortium | Lighthouse Laboratory Plymouth and Alex Alderton, Roberto Amato, Jeffrey Barrett, Sonia Goncalves, Ewan Harrison, David K. Jackson, Ian Johnston, Dominic Kwiatkowski, Cordelia Langford, John Sillitoe on behalf of the Wellcome Sanger Institute COVID-19 Surveillance Team |
| EPI_ISL_2570685, EPI_ISL_2570686, EPI_ISL_2570687, EPI_ISL_2570688, EPI_ISL_2570689, | Lighthouse Lab in Alderley Park | Wellcome Sanger Institute for the COVID-19 Genomics UK (COG-UK) Consortium | Jacquelyn Wynn, Mairead Hyland, The Lighthouse Lab in Alderley Park and Alex Alderton, Roberto Amato, Jeffrey Barrett, Sonia Goncalves, Ewan Harrison, David K. Jackson, Ian Johnston, Dominic Kwiatkowski, Cordelia Langford, John Sillitoe on behalf of the Wellcome Sanger Institute COVID-19 Surveillance Team |

| | | | |
|---|---|---|---|
| EPI_ISL_2570690 | | | |
| EPI_ISL_2570691, EPI_ISL_2570692 | Lighthouse Laboratory Plymouth | Wellcome Sanger Institute for the COVID-19 Genomics UK (COG-UK) Consortium | Lighthouse Laboratory Plymouth and Alex Alderton, Roberto Amato, Jeffrey Barrett, Sonia Goncalves, Ewan Harrison, David K. Jackson, Ian Johnston, Dominic Kwiatkowski, Cordelia Langford, John Sillitoe on behalf of the Wellcome Sanger Institute COVID-19 Surveillance Team |
| EPI_ISL_2570693, EPI_ISL_2570694, EPI_ISL_2570695 | Lighthouse Lab in Alderley Park | Wellcome Sanger Institute for the COVID-19 Genomics UK (COG-UK) Consortium | Jacquelyn Wynn, Mairead Hyland, The Lighthouse Lab in Alderley Park and Alex Alderton, Roberto Amato, Jeffrey Barrett, Sonia Goncalves, Ewan Harrison, David K. Jackson, Ian Johnston, Dominic Kwiatkowski, Cordelia Langford, John Sillitoe on behalf of the Wellcome Sanger Institute COVID-19 Surveillance Team |
| EPI_ISL_2570696 | Lighthouse Laboratory Plymouth | Wellcome Sanger Institute for the COVID-19 Genomics UK (COG-UK) Consortium | Lighthouse Laboratory Plymouth and Alex Alderton, Roberto Amato, Jeffrey Barrett, Sonia Goncalves, Ewan Harrison, David K. Jackson, Ian Johnston, Dominic Kwiatkowski, Cordelia Langford, John Sillitoe on behalf of the Wellcome Sanger Institute COVID-19 Surveillance Team |
| EPI_ISL_2570697, EPI_ISL_2570698, EPI_ISL_2570699, EPI_ISL_2570700, EPI_ISL_2570702, EPI_ISL_2570703 | Lighthouse Lab in Alderley Park | Wellcome Sanger Institute for the COVID-19 Genomics UK (COG-UK) Consortium | Jacquelyn Wynn, Mairead Hyland, The Lighthouse Lab in Alderley Park and Alex Alderton, Roberto Amato, Jeffrey Barrett, Sonia Goncalves, Ewan Harrison, David K. Jackson, Ian Johnston, Dominic Kwiatkowski, Cordelia Langford, John Sillitoe on behalf of the Wellcome Sanger Institute COVID-19 Surveillance Team |
| EPI_ISL_2570704 | Lighthouse Laboratory Plymouth | Wellcome Sanger Institute for the COVID-19 Genomics UK (COG-UK) Consortium | Lighthouse Laboratory Plymouth and Alex Alderton, Roberto Amato, Jeffrey Barrett, Sonia Goncalves, Ewan Harrison, David K. Jackson, Ian Johnston, Dominic Kwiatkowski, Cordelia Langford, John Sillitoe on behalf of the Wellcome Sanger Institute COVID-19 Surveillance Team |
| EPI_ISL_2570706, EPI_ISL_2570707, EPI_ISL_2570708 | Lighthouse Lab in Alderley Park | Wellcome Sanger Institute for the COVID-19 Genomics UK (COG-UK) Consortium | Jacquelyn Wynn, Mairead Hyland, The Lighthouse Lab in Alderley Park and Alex Alderton, Roberto Amato, Jeffrey Barrett, Sonia Goncalves, Ewan Harrison, David K. Jackson, Ian Johnston, Dominic Kwiatkowski, Cordelia Langford, John Sillitoe on behalf of the Wellcome Sanger Institute COVID-19 Surveillance Team |
| EPI_ISL_2570709 | Lighthouse Laboratory Plymouth | Wellcome Sanger Institute for the COVID-19 Genomics UK (COG-UK) Consortium | Lighthouse Laboratory Plymouth and Alex Alderton, Roberto Amato, Jeffrey Barrett, Sonia Goncalves, Ewan Harrison, David K. Jackson, Ian Johnston, Dominic Kwiatkowski, Cordelia Langford, John Sillitoe on behalf of the Wellcome Sanger Institute COVID-19 Surveillance Team |
| EPI_ISL_2570710, EPI_ISL_2570711, EPI_ISL_2570712 | Lighthouse Lab in Alderley Park | Wellcome Sanger Institute for the COVID-19 Genomics UK (COG-UK) Consortium | Jacquelyn Wynn, Mairead Hyland, The Lighthouse Lab in Alderley Park and Alex Alderton, Roberto Amato, Jeffrey Barrett, Sonia Goncalves, Ewan Harrison, David K. Jackson, Ian Johnston, Dominic Kwiatkowski, Cordelia Langford, John Sillitoe on behalf of the Wellcome Sanger Institute COVID-19 Surveillance Team |
| EPI_ISL_2570713 | Lighthouse Laboratory Plymouth | Wellcome Sanger Institute for the COVID-19 Genomics UK (COG-UK) Consortium | Lighthouse Laboratory Plymouth and Alex Alderton, Roberto Amato, Jeffrey Barrett, Sonia Goncalves, Ewan Harrison, David K. Jackson, Ian Johnston, Dominic Kwiatkowski, Cordelia Langford, John Sillitoe on behalf of the Wellcome Sanger Institute COVID-19 Surveillance Team |
| EPI_ISL_2570714, EPI_ISL_2570715, EPI_ISL_2570716 | Lighthouse Lab in Alderley Park | Wellcome Sanger Institute for the COVID-19 Genomics UK (COG-UK) Consortium | Jacquelyn Wynn, Mairead Hyland, The Lighthouse Lab in Alderley Park and Alex Alderton, Roberto Amato, Jeffrey Barrett, Sonia Goncalves, Ewan Harrison, David K. Jackson, Ian Johnston, Dominic Kwiatkowski, Cordelia Langford, John Sillitoe on behalf of the Wellcome Sanger Institute COVID-19 Surveillance Team |
| EPI_ISL_2570717, EPI_ISL_2570718 | Lighthouse Laboratory Plymouth | Wellcome Sanger Institute for the COVID-19 Genomics UK (COG-UK) Consortium | Lighthouse Laboratory Plymouth and Alex Alderton, Roberto Amato, Jeffrey Barrett, Sonia Goncalves, Ewan Harrison, David K. Jackson, Ian Johnston, Dominic Kwiatkowski, Cordelia Langford, John Sillitoe on behalf of the Wellcome Sanger Institute COVID-19 Surveillance Team |
| EPI_ISL_2570719 | Lighthouse Lab in Alderley Park | Wellcome Sanger Institute for the COVID-19 Genomics UK (COG-UK) Consortium | Jacquelyn Wynn, Mairead Hyland, The Lighthouse Lab in Alderley Park and Alex Alderton, Roberto Amato, Jeffrey Barrett, Sonia Goncalves, Ewan Harrison, David K. Jackson, Ian Johnston, Dominic Kwiatkowski, Cordelia Langford, John Sillitoe on behalf of the Wellcome Sanger Institute COVID-19 Surveillance Team |
| EPI_ISL_2570720 | Lighthouse Laboratory Plymouth | Wellcome Sanger Institute for the COVID-19 Genomics UK (COG-UK) Consortium | Lighthouse Laboratory Plymouth and Alex Alderton, Roberto Amato, Jeffrey Barrett, Sonia Goncalves, Ewan Harrison, David K. Jackson, Ian Johnston, Dominic Kwiatkowski, Cordelia Langford, John Sillitoe on behalf of the Wellcome Sanger Institute COVID-19 Surveillance Team |
| EPI_ISL_2570721, EPI_ISL_2570722, EPI_ISL_2570723 | Lighthouse Lab in Alderley Park | Wellcome Sanger Institute for the COVID-19 Genomics UK (COG-UK) Consortium | Jacquelyn Wynn, Mairead Hyland, The Lighthouse Lab in Alderley Park and Alex Alderton, Roberto Amato, Jeffrey Barrett, Sonia Goncalves, Ewan Harrison, David K. Jackson, Ian Johnston, Dominic Kwiatkowski, Cordelia Langford, John Sillitoe on behalf of the Wellcome Sanger Institute COVID-19 Surveillance Team |
| EPI_ISL_2570725 | Lighthouse Laboratory Plymouth | Wellcome Sanger Institute for the COVID-19 Genomics UK (COG-UK) Consortium | Lighthouse Laboratory Plymouth and Alex Alderton, Roberto Amato, Jeffrey Barrett, Sonia Goncalves, Ewan Harrison, David K. Jackson, Ian Johnston, Dominic Kwiatkowski, Cordelia Langford, John Sillitoe on behalf of the Wellcome Sanger Institute COVID-19 Surveillance Team |
| EPI_ISL_2570726, EPI_ISL_2570727, EPI_ISL_2570728 | Lighthouse Lab in Alderley Park | Wellcome Sanger Institute for the COVID-19 Genomics UK (COG-UK) Consortium | Jacquelyn Wynn, Mairead Hyland, The Lighthouse Lab in Alderley Park and Alex Alderton, Roberto Amato, Jeffrey Barrett, Sonia Goncalves, Ewan Harrison, David K. Jackson, Ian Johnston, Dominic Kwiatkowski, Cordelia Langford, John Sillitoe on behalf of the Wellcome Sanger Institute COVID-19 Surveillance Team |
| EPI_ISL_2570729 | Lighthouse Laboratory Plymouth | Wellcome Sanger Institute for the COVID-19 Genomics UK (COG-UK) Consortium | Lighthouse Laboratory Plymouth and Alex Alderton, Roberto Amato, Jeffrey Barrett, Sonia Goncalves, Ewan Harrison, David K. Jackson, Ian Johnston, Dominic Kwiatkowski, Cordelia Langford, John Sillitoe on behalf of the Wellcome Sanger Institute COVID-19 Surveillance Team |
| EPI_ISL_2570730, EPI_ISL_2570732, EPI_ISL_2570733, EPI_ISL_2570734, EPI_ISL_2570735, EPI_ISL_2570736, EPI_ISL_2570737 | Lighthouse Lab in Alderley Park | Wellcome Sanger Institute for the COVID-19 Genomics UK (COG-UK) Consortium | Jacquelyn Wynn, Mairead Hyland, The Lighthouse Lab in Alderley Park and Alex Alderton, Roberto Amato, Jeffrey Barrett, Sonia Goncalves, Ewan Harrison, David K. Jackson, Ian Johnston, Dominic Kwiatkowski, Cordelia Langford, John Sillitoe on behalf of the Wellcome Sanger Institute COVID-19 Surveillance Team |
| EPI_ISL_2570738, EPI_ISL_2570739 | Lighthouse Laboratory Plymouth | Wellcome Sanger Institute for the COVID-19 Genomics UK (COG-UK) Consortium | Lighthouse Laboratory Plymouth and Alex Alderton, Roberto Amato, Jeffrey Barrett, Sonia Goncalves, Ewan Harrison, David K. Jackson, Ian Johnston, Dominic Kwiatkowski, Cordelia Langford, John Sillitoe on behalf of the Wellcome Sanger Institute COVID-19 Surveillance Team |
| EPI_ISL_2570740, EPI_ISL_2570741, EPI_ISL_2570742, EPI_ISL_2570743, EPI_ISL_2570746, EPI_ISL_2570747 | Lighthouse Lab in Alderley Park | Wellcome Sanger Institute for the COVID-19 Genomics UK (COG-UK) Consortium | Jacquelyn Wynn, Mairead Hyland, The Lighthouse Lab in Alderley Park and Alex Alderton, Roberto Amato, Jeffrey Barrett, Sonia Goncalves, Ewan Harrison, David K. Jackson, Ian Johnston, Dominic Kwiatkowski, Cordelia Langford, John Sillitoe on behalf of the Wellcome Sanger Institute COVID-19 Surveillance Team |
| EPI_ISL_2570748 | Lighthouse Laboratory Plymouth | Wellcome Sanger Institute for the COVID-19 Genomics UK (COG-UK) Consortium | Lighthouse Laboratory Plymouth and Alex Alderton, Roberto Amato, Jeffrey Barrett, Sonia Goncalves, Ewan Harrison, David K. Jackson, Ian Johnston, Dominic Kwiatkowski, Cordelia Langford, John Sillitoe on behalf of the Wellcome Sanger Institute COVID-19 Surveillance Team |
| EPI_ISL_2570749, EPI_ISL_2570750 | Lighthouse Lab in Alderley Park | Wellcome Sanger Institute for the COVID-19 Genomics UK (COG-UK) Consortium | Jacquelyn Wynn, Mairead Hyland, The Lighthouse Lab in Alderley Park and Alex Alderton, Roberto Amato, Jeffrey Barrett, Sonia Goncalves, Ewan Harrison, David K. Jackson, Ian Johnston, Dominic Kwiatkowski, Cordelia Langford, John Sillitoe on behalf of the Wellcome Sanger Institute COVID-19 Surveillance Team |
| EPI_ISL_2570751 | Lighthouse Laboratory Plymouth | Wellcome Sanger Institute for the COVID-19 Genomics UK (COG-UK) Consortium | Lighthouse Laboratory Plymouth and Alex Alderton, Roberto Amato, Jeffrey Barrett, Sonia Goncalves, Ewan Harrison, David K. Jackson, Ian Johnston, Dominic Kwiatkowski, Cordelia Langford, John Sillitoe on behalf of the Wellcome Sanger Institute COVID-19 Surveillance Team |
| EPI_ISL_2570752, EPI_ISL_2570753 | Lighthouse Lab in Alderley Park | Wellcome Sanger Institute for the COVID-19 Genomics UK (COG-UK) Consortium | Jacquelyn Wynn, Mairead Hyland, The Lighthouse Lab in Alderley Park and Alex Alderton, Roberto Amato, Jeffrey Barrett, Sonia Goncalves, Ewan Harrison, David K. Jackson, Ian Johnston, Dominic Kwiatkowski, Cordelia Langford, John Sillitoe on behalf of the Wellcome Sanger Institute COVID-19 Surveillance Team |
| EPI_ISL_2570754, EPI_ISL_2570755 | Lighthouse Laboratory Plymouth | Wellcome Sanger Institute for the COVID-19 Genomics UK (COG-UK) Consortium | Lighthouse Laboratory Plymouth and Alex Alderton, Roberto Amato, Jeffrey Barrett, Sonia Goncalves, Ewan Harrison, David K. Jackson, Ian Johnston, Dominic Kwiatkowski, Cordelia Langford, John Sillitoe on behalf of the Wellcome Sanger Institute COVID-19 Surveillance Team |
| EPI_ISL_2570756 | Lighthouse Lab in Alderley Park | Wellcome Sanger Institute for the COVID-19 Genomics UK (COG-UK) Consortium | Jacquelyn Wynn, Mairead Hyland, The Lighthouse Lab in Alderley Park and Alex Alderton, Roberto Amato, Jeffrey Barrett, Sonia Goncalves, Ewan Harrison, David K. Jackson, Ian Johnston, Dominic Kwiatkowski, Cordelia Langford, John Sillitoe on behalf of the Wellcome Sanger Institute COVID-19 Surveillance Team |
| EPI_ISL_2570757 | Lighthouse Laboratory Plymouth | Wellcome Sanger Institute for the COVID-19 Genomics UK (COG-UK) Consortium | Lighthouse Laboratory Plymouth and Alex Alderton, Roberto Amato, Jeffrey Barrett, Sonia Goncalves, Ewan Harrison, David K. Jackson, Ian Johnston, Dominic Kwiatkowski, Cordelia Langford, John Sillitoe on behalf of the Wellcome Sanger Institute COVID-19 Surveillance Team |
| EPI_ISL_2570758, EPI_ISL_2570759, EPI_ISL_2570760 | Lighthouse Lab in Alderley Park | Wellcome Sanger Institute for the COVID-19 Genomics UK (COG-UK) Consortium | Jacquelyn Wynn, Mairead Hyland, The Lighthouse Lab in Alderley Park and Alex Alderton, Roberto Amato, Jeffrey Barrett, Sonia Goncalves, Ewan Harrison, David K. Jackson, Ian Johnston, Dominic Kwiatkowski, Cordelia Langford, John Sillitoe on behalf of the Wellcome Sanger Institute COVID-19 Surveillance Team |
| EPI_ISL_2570761 | Lighthouse Laboratory Plymouth | Wellcome Sanger Institute for the COVID-19 Genomics UK (COG-UK) Consortium | Lighthouse Laboratory Plymouth and Alex Alderton, Roberto Amato, Jeffrey Barrett, Sonia Goncalves, Ewan Harrison, David K. Jackson, Ian Johnston, Dominic Kwiatkowski, Cordelia Langford, John Sillitoe on behalf of the Wellcome Sanger Institute COVID-19 Surveillance Team |
| EPI_ISL_2570762, EPI_ISL_2570763 | Lighthouse Lab in Alderley Park | Wellcome Sanger Institute for the COVID-19 Genomics UK (COG-UK) Consortium | Jacquelyn Wynn, Mairead Hyland, The Lighthouse Lab in Alderley Park and Alex Alderton, Roberto Amato, Jeffrey Barrett, Sonia Goncalves, Ewan Harrison, David K. Jackson, Ian Johnston, Dominic Kwiatkowski, Cordelia Langford, John Sillitoe on behalf of the Wellcome Sanger Institute COVID-19 Surveillance Team |
| EPI_ISL_2570764 | Lighthouse Laboratory Plymouth | Wellcome Sanger Institute for the COVID-19 Genomics UK (COG-UK) Consortium | Lighthouse Laboratory Plymouth and Alex Alderton, Roberto Amato, Jeffrey Barrett, Sonia Goncalves, Ewan Harrison, David K. Jackson, Ian Johnston, Dominic Kwiatkowski, Cordelia Langford, John Sillitoe on behalf of the Wellcome Sanger Institute COVID-19 Surveillance Team |
| EPI_ISL_2570765, EPI_ISL_2570766 | Lighthouse Lab in Alderley Park | Wellcome Sanger Institute for the COVID-19 Genomics UK (COG-UK) Consortium | Jacquelyn Wynn, Mairead Hyland, The Lighthouse Lab in Alderley Park and Alex Alderton, Roberto Amato, Jeffrey Barrett, Sonia Goncalves, Ewan Harrison, David K. Jackson, Ian Johnston, Dominic Kwiatkowski, Cordelia Langford, John Sillitoe on behalf of the Wellcome Sanger Institute COVID-19 Surveillance Team |
| EPI_ISL_2570767 | Lighthouse Laboratory Plymouth | Wellcome Sanger Institute for the COVID-19 Genomics UK (COG-UK) Consortium | Lighthouse Laboratory Plymouth and Alex Alderton, Roberto Amato, Jeffrey Barrett, Sonia Goncalves, Ewan Harrison, David K. Jackson, Ian Johnston, Dominic Kwiatkowski, Cordelia Langford, John Sillitoe on behalf of the Wellcome Sanger Institute COVID-19 Surveillance Team |
| EPI_ISL_2570768, EPI_ISL_2570769 | Lighthouse Lab in Alderley Park | Wellcome Sanger Institute for the COVID-19 Genomics UK (COG-UK) Consortium | Jacquelyn Wynn, Mairead Hyland, The Lighthouse Lab in Alderley Park and Alex Alderton, Roberto Amato, Jeffrey Barrett, Sonia Goncalves, Ewan Harrison, David K. Jackson, Ian Johnston, Dominic Kwiatkowski, Cordelia Langford, John Sillitoe on behalf of the Wellcome Sanger Institute COVID-19 Surveillance Team |
| EPI_ISL_2570770, EPI_ISL_2570771 | Lighthouse Laboratory Plymouth | Wellcome Sanger Institute for the COVID-19 Genomics UK (COG-UK) Consortium | Lighthouse Laboratory Plymouth and Alex Alderton, Roberto Amato, Jeffrey Barrett, Sonia Goncalves, Ewan Harrison, David K. Jackson, Ian Johnston, Dominic Kwiatkowski, Cordelia Langford, John Sillitoe on behalf of the Wellcome Sanger Institute COVID-19 Surveillance Team |
| EPI_ISL_2570772, EPI_ISL_2570773, EPI_ISL_2570774, | Lighthouse Lab in Alderley Park | Wellcome Sanger Institute for the COVID-19 Genomics UK (COG-UK) Consortium | Jacquelyn Wynn, Mairead Hyland, The Lighthouse Lab in Alderley Park and Alex Alderton, Roberto Amato, Jeffrey Barrett, Sonia Goncalves, Ewan Harrison, David K. Jackson, Ian Johnston, Dominic Kwiatkowski, Cordelia Langford, John Sillitoe on behalf of the Wellcome Sanger Institute COVID-19 Surveillance Team |

| | | | |
|---|---|---|---|
| EPI_ISL_2570775 | | | |
| EPI_ISL_2570776 | Lighthouse Laboratory Plymouth | Wellcome Sanger Institute for the COVID-19 Genomics UK (COG-UK) Consortium | Lighthouse Laboratory Plymouth and Alex Alderton, Roberto Amato, Jeffrey Barrett, Sonia Goncalves, Ewan Harrison, David K. Jackson, Ian Johnston, Dominic Kwiatkowski, Cordelia Langford, John Sillitoe on behalf of the Wellcome Sanger Institute COVID-19 Surveillance Team |
| EPI_ISL_2570777 | Lighthouse Lab in Alderley Park | Wellcome Sanger Institute for the COVID-19 Genomics UK (COG-UK) Consortium | Jacquelyn Wynn, Mairead Hyland, The Lighthouse Lab in Alderley Park and Alex Alderton, Roberto Amato, Jeffrey Barrett, Sonia Goncalves, Ewan Harrison, David K. Jackson, Ian Johnston, Dominic Kwiatkowski, Cordelia Langford, John Sillitoe on behalf of the Wellcome Sanger Institute COVID-19 Surveillance Team |
| EPI_ISL_2570778 | Lighthouse Laboratory Plymouth | Wellcome Sanger Institute for the COVID-19 Genomics UK (COG-UK) Consortium | Lighthouse Laboratory Plymouth and Alex Alderton, Roberto Amato, Jeffrey Barrett, Sonia Goncalves, Ewan Harrison, David K. Jackson, Ian Johnston, Dominic Kwiatkowski, Cordelia Langford, John Sillitoe on behalf of the Wellcome Sanger Institute COVID-19 Surveillance Team |
| EPI_ISL_2570779, EPI_ISL_2570780, EPI_ISL_2570781, EPI_ISL_2570782, EPI_ISL_2570783, EPI_ISL_2570784, EPI_ISL_2570785, EPI_ISL_2570786, EPI_ISL_2570787 | Lighthouse Lab in Alderley Park | Wellcome Sanger Institute for the COVID-19 Genomics UK (COG-UK) Consortium | Jacquelyn Wynn, Mairead Hyland, The Lighthouse Lab in Alderley Park and Alex Alderton, Roberto Amato, Jeffrey Barrett, Sonia Goncalves, Ewan Harrison, David K. Jackson, Ian Johnston, Dominic Kwiatkowski, Cordelia Langford, John Sillitoe on behalf of the Wellcome Sanger Institute COVID-19 Surveillance Team |
| EPI_ISL_2570789 | Lighthouse Laboratory Plymouth | Wellcome Sanger Institute for the COVID-19 Genomics UK (COG-UK) Consortium | Lighthouse Laboratory Plymouth and Alex Alderton, Roberto Amato, Jeffrey Barrett, Sonia Goncalves, Ewan Harrison, David K. Jackson, Ian Johnston, Dominic Kwiatkowski, Cordelia Langford, John Sillitoe on behalf of the Wellcome Sanger Institute COVID-19 Surveillance Team |
| EPI_ISL_2570790 | Lighthouse Lab in Alderley Park | Wellcome Sanger Institute for the COVID-19 Genomics UK (COG-UK) Consortium | Jacquelyn Wynn, Mairead Hyland, The Lighthouse Lab in Alderley Park and Alex Alderton, Roberto Amato, Jeffrey Barrett, Sonia Goncalves, Ewan Harrison, David K. Jackson, Ian Johnston, Dominic Kwiatkowski, Cordelia Langford, John Sillitoe on behalf of the Wellcome Sanger Institute COVID-19 Surveillance Team |
| EPI_ISL_2570791 | Lighthouse Laboratory Plymouth | Wellcome Sanger Institute for the COVID-19 Genomics UK (COG-UK) Consortium | Lighthouse Laboratory Plymouth and Alex Alderton, Roberto Amato, Jeffrey Barrett, Sonia Goncalves, Ewan Harrison, David K. Jackson, Ian Johnston, Dominic Kwiatkowski, Cordelia Langford, John Sillitoe on behalf of the Wellcome Sanger Institute COVID-19 Surveillance Team |
| EPI_ISL_2570793, EPI_ISL_2570794, EPI_ISL_2570795, EPI_ISL_2570796, EPI_ISL_2570797, EPI_ISL_2570798, EPI_ISL_2570799 | Lighthouse Lab in Alderley Park | Wellcome Sanger Institute for the COVID-19 Genomics UK (COG-UK) Consortium | Jacquelyn Wynn, Mairead Hyland, The Lighthouse Lab in Alderley Park and Alex Alderton, Roberto Amato, Jeffrey Barrett, Sonia Goncalves, Ewan Harrison, David K. Jackson, Ian Johnston, Dominic Kwiatkowski, Cordelia Langford, John Sillitoe on behalf of the Wellcome Sanger Institute COVID-19 Surveillance Team |
| EPI_ISL_2570800 | Lighthouse Laboratory Plymouth | Wellcome Sanger Institute for the COVID-19 Genomics UK (COG-UK) Consortium | Lighthouse Laboratory Plymouth and Alex Alderton, Roberto Amato, Jeffrey Barrett, Sonia Goncalves, Ewan Harrison, David K. Jackson, Ian Johnston, Dominic Kwiatkowski, Cordelia Langford, John Sillitoe on behalf of the Wellcome Sanger Institute COVID-19 Surveillance Team |
| EPI_ISL_2570801, EPI_ISL_2570802, EPI_ISL_2570803 | Lighthouse Lab in Alderley Park | Wellcome Sanger Institute for the COVID-19 Genomics UK (COG-UK) Consortium | Jacquelyn Wynn, Mairead Hyland, The Lighthouse Lab in Alderley Park and Alex Alderton, Roberto Amato, Jeffrey Barrett, Sonia Goncalves, Ewan Harrison, David K. Jackson, Ian Johnston, Dominic Kwiatkowski, Cordelia Langford, John Sillitoe on behalf of the Wellcome Sanger Institute COVID-19 Surveillance Team |
| EPI_ISL_2570805 | Indrapura Field Hospital Surabaya | Institute of Tropical Disease, Universitas Airlangga | Rima R Prasetya, Krisnoadi Rahardjo, Aldise M Nastri, Jezzy R Dewantari, Febria Rachmanita, Herlin Ferliana, Gatot Soegiarto, Laksmi Wulandari, Resti Yudhawati, Yasuko Mori, Soetjipto, Kazufumi Shimizu, Maria I Lusida |
| EPI_ISL_2570806 | Indrapura Field Hospital Surabaya | Institute of Tropical Disease, Universitas Airlangga | Kazufumi Shimizu, Krisnoadi Rahardjo, Aldise M Nastri, Jezzy R Dewantari, Rima R Prasetya, Febria Rachmanita, Herlin Ferliana, Gatot Soegiarto, Laksmi Wulandari, Resti Yudhawati, Yasuko Mori, Soetjipto, Maria I Lusida |
| EPI_ISL_2570808 | Surabaya Health Office | Institute of Tropical Disease, Universitas Airlangga | Krisnoadi Rahardjo, Aldise M Nastri, Jezzy R Dewantari, Rima R Prasetya, Febria Rachmanita, Herlin Ferliana, Gatot Soegiarto, Laksmi Wulandari, Resti Yudhawati, Yasuko Mori, Soetjipto, Kazufumi Shimizu, Maria I Lusida |
| EPI_ISL_2570809 | Surabaya Health Office | Institute of Tropical Disease, Universitas Airlangga | Aldise M Nastri, Jezzy R Dewantari, Rima R Prasetya, Krisnoadi Rahardjo, Febria Rachmanita, Herlin Ferliana, Gatot Soegiarto, Laksmi Wulandari, Resti Yudhawati, Yasuko Mori, Soetjipto, Kazufumi Shimizu, Maria I Lusida |
| EPI_ISL_2570811 | Surabaya Health Office | Institute of Tropical Disease, Universitas Airlangga | Jezzy R Dewantari, Rima R Prasetya, Krisnoadi Rahardjo, Aldise M Nastri, Febria Rachmanita, Herlin Ferliana, Gatot Soegiarto, Laksmi Wulandari, Resti Yudhawati, Yasuko Mori, Soetjipto, Kazufumi Shimizu, Maria I Lusida |
| EPI_ISL_2574058, EPI_ISL_2574080, EPI_ISL_2574090 | Center for Laboratory Medicine | Center for Laboratory Medicine | Yannick Gerth |
| EPI_ISL_2577027 | IRCCS San Gallicano Dermatological Institute | IRCCS Regina Elena National Cancer Institute | Frauke Goeman,Giulia Orlandi,Eleonora Sperandio,Valentina Ricca,Fabrizio Ensoli,Fulvia Pimpinelli,Maurizio Fanciulli,Giovanni Blandino,Aldo Morrone,Gennaro Ciliberto |
| EPI_ISL_2584380, EPI_ISL_2584418, EPI_ISL_2584419, EPI_ISL_2584480 | Swedish national genomic surveillance program of SARS-CoV-2 | The Public Health Agency of Sweden | Maximilian Riess, Maria Lind Karlberg, Alma Brolund, Swedish national genomic surveillance program of SARS-CoV-2 |
| EPI_ISL_2587054, EPI_ISL_2587069, EPI_ISL_2587140, EPI_ISL_2587399, EPI_ISL_2587642, EPI_ISL_2587661, EPI_ISL_2587754, EPI_ISL_2587768, EPI_ISL_2587770 | SK-Roy Romanow Provincial Laboratory | National Microbiology Laboratory (NML) | Anna Majer, Shari Tyson, Grace Seo, Philip Mabon, Elsie Grudeski, Rhiannon Huzarewich, Russell Mandes, Anneliese Landgraff, Jennifer Tanner, Natalie Knox, Morag Graham, Gary Van Domselaar, Ryan McDonald, Amanda Lang, Rachel DePaulo, Jessica Minion, Nathalie Bastien, Yan Li, Timothy Booth, Darian Hole, Madison Chapel, Kirsten Biggar, CanCOGeN's metadata curation team, Public Health Agency of Canada CanCOGeN team |
| EPI_ISL_2594444 | Wyoming Public Health Laboratory | Wyoming Public Health Laboratory | Jim Mildenberger, Taylor Fearing, Channing Weber, Ashley Norberg, Chayse Rowley, Marley Goetz, Brian Dominguez, Elliot Thomasson, Sam Britz, Cari Sloma, Robert Petit, and Rob Christensen |
| EPI_ISL_2595726 | DSMRC | DSMRC | Khine Zaw Oo, Nay Myo Aung, Ko Ko Win, Phyo Kyaw Aung, Pai Khant Kyaw, Thet Wai Oo, Zaw Win Htun, Kyee Myint, Ko Ko Lwin |
| EPI_ISL_2595727 | Nebraska Public Health Laboratory | NPHL COVID-19 Response Team | NPHL COVID-19 Response Team |
| EPI_ISL_2595746 | MEPHI, Aix Marseille University | MEPHI, Aix Marseille University | Anthony LEVASSEUR |
| EPI_ISL_2595860, EPI_ISL_2596341, EPI_ISL_2596803 | DSMRC | DSMRC | Khine Zaw Oo, Nay Myo Aung, Ko Ko Win, Phyo Kyaw Aung, Pai Khant Kyaw, Thet Wai Oo, Zaw Win Htun, Kyee Myint, Ko Ko Lwin |
| EPI_ISL_2597685, EPI_ISL_2597688, EPI_ISL_2597700 | Hospital of the University of Pennsylvania Molecular Pathology Lab | Bushman Lab - University of Pennsylvania | John K. Everett, Kyle Rodino, Shantan Reddy, Aoife M. Roche, Young Hwang, Scott Sherrill-Mix, Samantha A. Whiteside, Jevon Graham-Wooten, Layla A. Khatib, Ayannah S. Fitzgerald, Arupa Ganguly, Mike Feldman, Brendan Kelly, Ronald G. Collman and Frederic Bushan |
| EPI_ISL_2597810, EPI_ISL_2597815 | Diagnostyka | 1. Tricity SARS-CoV-2 sequencing consortium: University of Gdansk, Medical University of Gdansk, Vaxican Ltd., Invicta Ltd. 2. National Institute of Public Health - National Institute of Hygiene, Warsaw, Poland | Maciej Kosinski, Celina Cybulska, Krystyna Bienkowska Szewczyk, Maciej Grzybek, Karolina Gackowska, Marcin Lubocki, Katarzyna Groth, Lukasz Rabalski, Katarzyna Zacharczuk, Magdalena Nowakowska, Małgorzata Sadkowska-Todys, Tomasz Wołkowicz |
| EPI_ISL_2597819 | UCK | 1. Tricity SARS-CoV-2 sequencing consortium: University of Gdansk, Medical University of Gdansk, Vaxican Ltd., Invicta Ltd. 2. National Institute of Public Health - National Institute of Hygiene, Warsaw, Poland | Maciej Kosinski, Celina Cybulska, Krystyna Bienkowska Szewczyk, Maciej Grzybek, Karolina Gackowska, Marcin Lubocki, Katarzyna Groth, Lukasz Rabalski, Katarzyna Zacharczuk, Magdalena Nowakowska, Małgorzata Sadkowska-Todys, Tomasz Wołkowicz |
| EPI_ISL_2597889 | NZOZ Meditest Diagnostyka Medyczna | 1. Tricity SARS-CoV-2 sequencing consortium: University of Gdansk, Medical University of Gdansk, Vaxican Ltd., Invicta Ltd. 2. National Institute of Public Health - National Institute of Hygiene, Warsaw, Poland | Maciej Kosinski, Celina Cybulska, Krystyna Bienkowska Szewczyk, Maciej Grzybek, Karolina Gackowska, Marcin Lubocki, Katarzyna Groth, Lukasz Rabalski, Katarzyna Zacharczuk, Magdalena Nowakowska, Małgorzata Sadkowska-Todys, Tomasz Wołkowicz |
| EPI_ISL_2597901, EPI_ISL_2597902, EPI_ISL_2597911, EPI_ISL_2597913, EPI_ISL_2597929, EPI_ISL_2597932 | Nebraska Public Health Laboratory | NPHL COVID-19 Response Team | NPHL COVID-19 Response Team |
| EPI_ISL_2597956, EPI_ISL_2597959, EPI_ISL_2597962, EPI_ISL_2597965, EPI_ISL_2597969, EPI_ISL_2597972, EPI_ISL_2597973, EPI_ISL_2597974 | Dianalabs SA | Genesupport | Tanguy ARAUD, Geraldine Jost, Katia Jaton, Nadia Liassine |
| EPI_ISL_2598023, EPI_ISL_2598058, EPI_ISL_2598060, EPI_ISL_2598066, EPI_ISL_2598073, | MD PHL | MD PHL | Maryland Department of Health Laboratories Administration |

| | | | |
|---|---|---|---|
| EPI_ISL_2598074 | | | |
| EPI_ISL_2598088, EPI_ISL_2598092 | TriCore Reference Laboratories | Center for Global Health, University of New Mexico Health Sciences Center | Daryl Domman, Kurt Schwalm, Valerie Morley, Cecilia Thompson, Kendra Pesko, Karissa Culbreath, Darrell Dinwiddie |
| EPI_ISL_2598136 | New Mexico Department of Health Scientific Laboratory | Center for Global Health, University of New Mexico Health Sciences Center | Daryl Domman, Kurt Schwalm, Twila Kunde, Joseph Hicks, Anastacia Griego, Michael Edwards, Valerie Morley, Darrell Dinwiddie |
| EPI_ISL_2598211, EPI_ISL_2598214, EPI_ISL_2598215, EPI_ISL_2598216, EPI_ISL_2598221, EPI_ISL_2598242, EPI_ISL_2598243 | Illinois Department of Public Health - Springfield Lab | Illinois Department of Public Health - Springfield Lab | Bryan Sim, Gordon McCall |
| EPI_ISL_2598296 | PathWest Laboratory Medicine WA | PathWest Laboratory Medicine WA Microbial Surveillance Unit | PathWest Laboratory Medicine WA Microbial Surveillance Unit |
| EPI_ISL_2598388, EPI_ISL_2598392, EPI_ISL_2598397 | Microbiological Diagnostic Unit - Public Health Laboratory (MDU-PHL) | MDU-PHL | Seemann T., Sait, M.L., Sherry, N.L. |
| EPI_ISL_2598420, EPI_ISL_2598467 | Maryland Genomics, Institute for Genome Sciences, University of Maryland School of Medicine | Maryland Genomics, Institute for Genome Sciences, University of Maryland School of Medicine | Tallon, Luke J; Sadzewicz, Lisa D; Humphrys, Mike; Ott, Sandra; Roussey, Holly; Santana-Cruz, Ivette; Vavikolanu, Kranthi; Lim, Jonathan;Hazen, Tracy;Fraser, Claire M; Ravel, Jacques |
| EPI_ISL_2598707, EPI_ISL_2598708, EPI_ISL_2598713, EPI_ISL_2598714, EPI_ISL_2598715, EPI_ISL_2598716, EPI_ISL_2598717, EPI_ISL_2598718, EPI_ISL_2598720, EPI_ISL_2598728, EPI_ISL_2598730, EPI_ISL_2598731, EPI_ISL_2598732, EPI_ISL_2598739, EPI_ISL_2598740, EPI_ISL_2598745, EPI_ISL_2598747, EPI_ISL_2598748, EPI_ISL_2598749, EPI_ISL_2598750, EPI_ISL_2598751, EPI_ISL_2598752, EPI_ISL_2598753, EPI_ISL_2598754, EPI_ISL_2598756, EPI_ISL_2598758, EPI_ISL_2598759, EPI_ISL_2598760, EPI_ISL_2598761, EPI_ISL_2598762, EPI_ISL_2598765, EPI_ISL_2598766, EPI_ISL_2598770, EPI_ISL_2598771, EPI_ISL_2598772, EPI_ISL_2598785, EPI_ISL_2598792, EPI_ISL_2598795, EPI_ISL_2598798, EPI_ISL_2598804, EPI_ISL_2598805 | | | |
| see above | Fulgent Genetics | Centers for Disease Control and Prevention Division of Viral Diseases, Pathogen Discovery | Dakota Howard, Dhwani Batra, Peter W. Cook, Kara Moser, Adrian Paskey, Jason Caravas, Benjamin Rambo-Martin, Shatavia Morrison, Christopher Gulvick, Scott Sammons, Yvette Unoarumhi, Darlene Wagner, Matthew Schmerer, Harry Gao, Mickey Li, John Gao, Joseph Fierro, Benafsh Sapra, Becky Tsai, Yan Meng, Doreen Ng, James Xie, Clinton R. Paden, Duncan MacCannell |
| EPI_ISL_2598863, EPI_ISL_2598917, EPI_ISL_2599076, EPI_ISL_2599162, EPI_ISL_2599233, EPI_ISL_2599267, EPI_ISL_2599575 | Helix/Illumina | Centers for Disease Control and Prevention Division of Viral Diseases, Pathogen Discovery | Dakota Howard, Dhwani Batra, Peter W. Cook, Kara Moser, Adrian Paskey, Jason Caravas, Benjamin Rambo-Martin, Shatavia Morrison, Christopher Gulvick, Scott Sammons, Yvette Unoarumhi, Darlene Wagner, Matthew Schmerer, Eileen de Feo, Jan Antico, Christine Tran, Matthew Tolentino, Shannon Wickline, Kim Gietzen, Brad Sickler, Jingtao Liu, Eric Allen, Phil Febbo, Nicole L. Washington, Simon White, Geraint Levan, Kelly Schiabor Barrett, Elizabeth Cirulli, Alexandre Bolze, Ary Ascencio, Charlotte Rivera-Garcia, Ryan Cho, Jason Nguyen, Sherry Wang, Jimmy Ramirez, Tyler Cassens, Efren Sandoval, Magnus Isaksson, William Lee, David Becker, Marc Laurent, James Lu, Clinton R. Paden, Duncan MacCannell |
| EPI_ISL_2599636, EPI_ISL_2599638, EPI_ISL_2599643, EPI_ISL_2599644, EPI_ISL_2599651, EPI_ISL_2599653, EPI_ISL_2599654, EPI_ISL_2599659, EPI_ISL_2599660, EPI_ISL_2599663, EPI_ISL_2599666, EPI_ISL_2599670, EPI_ISL_2599671, EPI_ISL_2599676, EPI_ISL_2599677, EPI_ISL_2599678, EPI_ISL_2599681, EPI_ISL_2599682, EPI_ISL_2599697, EPI_ISL_2599699, EPI_ISL_2599701, EPI_ISL_2599704, EPI_ISL_2599712, EPI_ISL_2599716, EPI_ISL_2599717, EPI_ISL_2599725, EPI_ISL_2599732, EPI_ISL_2599734, EPI_ISL_2599735, EPI_ISL_2599736, EPI_ISL_2599740, EPI_ISL_2599746, EPI_ISL_2599752, EPI_ISL_2599760, EPI_ISL_2599765, EPI_ISL_2599771, EPI_ISL_2599773, EPI_ISL_2599785, EPI_ISL_2599788, EPI_ISL_2599791 | | | |
| see above | Quest Diagnostics Incorporated | Centers for Disease Control and Prevention Division of Viral Diseases, Pathogen Discovery | Dakota Howard, Dhwani Batra, Peter W. Cook, Kara Moser, Adrian Paskey, Jason Caravas, Benjamin Rambo-Martin, Shatavia Morrison, Christopher Gulvick, Scott Sammons, Yvette Unoarumhi, Darlene Wagner, Matthew Schmerer, S. H. Rosenthal, A. Gerasimova, R. M. Kagan, B. Anderson, M. Hua, Y. Liu, L.E. Bernstein, K.E. Livingston, A. Perez, I. A. Shlyakhter, R. V. Rolando, R. Owen, P. Tanpaiboon, F. Lacbawan, Clinton R. Paden, Duncan MacCannell |
| EPI_ISL_2599793, EPI_ISL_2599798, EPI_ISL_2599800 | Dianalabs SA | Genesupport | Tanguy ARAUD, Geraldine Jost, Katia Jaton, Nadia Liassine |
| EPI_ISL_2599801, EPI_ISL_2599810 | Quest Diagnostics Incorporated | Centers for Disease Control and Prevention Division of Viral Diseases, Pathogen Discovery | Dakota Howard, Dhwani Batra, Peter W. Cook, Kara Moser, Adrian Paskey, Jason Caravas, Benjamin Rambo-Martin, Shatavia Morrison, Christopher Gulvick, Scott Sammons, Yvette Unoarumhi, Darlene Wagner, Matthew Schmerer, S. H. Rosenthal, A. Gerasimova, R. M. Kagan, B. Anderson, M. Hua, Y. Liu, L.E. Bernstein, K.E. Livingston, A. Perez, I. A. Shlyakhter, R. V. Rolando, R. Owen, P. Tanpaiboon, F. Lacbawan, Clinton R. Paden, Duncan MacCannell |
| EPI_ISL_2599815 | Dianalabs SA | Genesupport | Tanguy ARAUD, Geraldine Jost, Katia Jaton, Nadia Liassine |
| EPI_ISL_2599816 | Quest Diagnostics Incorporated | Centers for Disease Control and Prevention Division of Viral Diseases, Pathogen Discovery | Dakota Howard, Dhwani Batra, Peter W. Cook, Kara Moser, Adrian Paskey, Jason Caravas, Benjamin Rambo-Martin, Shatavia Morrison, Christopher Gulvick, Scott Sammons, Yvette Unoarumhi, Darlene Wagner, Matthew Schmerer, S. H. Rosenthal, A. Gerasimova, R. M. Kagan, B. Anderson, M. Hua, Y. Liu, L.E. Bernstein, K.E. Livingston, A. Perez, I. A. Shlyakhter, R. V. Rolando, R. Owen, P. Tanpaiboon, F. Lacbawan, Clinton R. Paden, Duncan MacCannell |
| EPI_ISL_2599818 | Dianalabs SA | Genesupport | Tanguy ARAUD, Geraldine Jost, Katia Jaton, Nadia Liassine |
| EPI_ISL_2599819, EPI_ISL_2599830, EPI_ISL_2599832, EPI_ISL_2599833, EPI_ISL_2599837, EPI_ISL_2599841, EPI_ISL_2599844, EPI_ISL_2599853, EPI_ISL_2599856, EPI_ISL_2599859, EPI_ISL_2599862, EPI_ISL_2599864, EPI_ISL_2599869, EPI_ISL_2599877, EPI_ISL_2599880, EPI_ISL_2599882, EPI_ISL_2599885, EPI_ISL_2599889, EPI_ISL_2599891, EPI_ISL_2599892, EPI_ISL_2599896, EPI_ISL_2599897, EPI_ISL_2599898, EPI_ISL_2599901, EPI_ISL_2599902, EPI_ISL_2599906, EPI_ISL_2599908, EPI_ISL_2599909, EPI_ISL_2599911, EPI_ISL_2599913, EPI_ISL_2599915, EPI_ISL_2599920 | | | |
| see above | Quest Diagnostics Incorporated | Centers for Disease Control and Prevention Division of Viral Diseases, Pathogen Discovery | Dakota Howard, Dhwani Batra, Peter W. Cook, Kara Moser, Adrian Paskey, Jason Caravas, Benjamin Rambo-Martin, Shatavia Morrison, Christopher Gulvick, Scott Sammons, Yvette Unoarumhi, Darlene Wagner, Matthew Schmerer, S. H. Rosenthal, A. Gerasimova, R. M. Kagan, B. Anderson, M. Hua, Y. Liu, L.E. Bernstein, K.E. Livingston, A. Perez, I. A. Shlyakhter, R. V. Rolando, R. Owen, P. Tanpaiboon, F. Lacbawan, Clinton R. Paden, Duncan MacCannell |
| EPI_ISL_2600023, EPI_ISL_2600197 | Aegis Sciences Corporation | Centers for Disease Control and Prevention Division of Viral Diseases, Pathogen Discovery | Dakota Howard, Dhwani Batra, Peter W. Cook, Kara Moser, Adrian Paskey, Jason Caravas, Benjamin Rambo-Martin, Shatavia Morrison, Christopher Gulvick, Scott Sammons, Yvette Unoarumhi, Darlene Wagner, Matthew Schmerer, Cyndi Clark, Patrick Campbell, Rob Case, Vikramsinha Ghorpade, Holly Houdeshell, Ola Kvalvaag, Dillon Nall, Ethan Sanders, Alec Vest, Shaun Westlund, Matthew Hardison, Clinton R. Paden, Duncan MacCannell |
| EPI_ISL_2600347, EPI_ISL_2600349, EPI_ISL_2600350 | Centre for Dengue Research and AICBU, Department of Immunology and Molecular Medicine | Centre for Dengue Research and AICBU, Department of Immunology and Molecular Medicine | Chandima Jeewandara, Deshni Jayathilaka, Dinuka Ariyaratne, Deshan Madhusanka, Diyanath Ranasinghe, Laksiri Gomes, Gathsaurie Neelika Malavige |
| EPI_ISL_2600366, EPI_ISL_2600367, EPI_ISL_2600372 | Institute of Epidemiology, Disease Control and Research (IEDCR) | IEDCR-ideSHi-icddr,b | Hassan Afrad, Sadia Rahman, Manjur Hossain Khan, Firdausi Qadri, Tahmina Shirin |
| EPI_ISL_2600533 | TGen North | TGen North | "Jolene Bowers, Heather Centner, Chris French, Hayley Yaglom, Ashlyn Pfeiffer, Darrin Lemmer, Dave Engelthaler, The Arizona COVID Genomics Union (ACGU)" |
| EPI_ISL_2600537, EPI_ISL_2600538, EPI_ISL_2600539, EPI_ISL_2600540, EPI_ISL_2600541 | Università degli Studi di Perugia | Istituto Zooprofilattico Sperimentale dell'Abruzzo e Molise "G. Caporale" | Mencacci A, Camilloni B, Biagetti M, Giammarioli M, Lorusso A, Marcacci M, Di Domenico M, Ancora M, Curini V, Mangone I, Rinaldi A, Delli Compagni E, Scialabba S, Di Pasquale A, Cammà C, Puglia I, Calistri P, Savini G |
| EPI_ISL_2600780, EPI_ISL_2600781, EPI_ISL_2600782, EPI_ISL_2600783 | Laboratorio Analisi Osp. Città di Castello - Azienda USL Umbria1 | Istituto Zooprofilattico Sperimentale dell'Abruzzo e Molise "G. Caporale" | Malagigi V, Taccani P,Lorusso A, Marcacci M, Di Domenico M, Ancora M, Curini V, Mangone I, Rinaldi A, Delli Compagni E, Scialabba S, Di Pasquale A, Cammà C, Puglia I, Calistri P, Savini G |
| EPI_ISL_2600791 | Azienda Ospedaliera Terni | Istituto Zooprofilattico Sperimentale dell'Abruzzo e Molise "G. Caporale" | Palumbo M, Scaccetti A, Lorusso A, Marcacci M, Di Domenico M, Ancora M, Curini V, Mangone I, Rinaldi A, Delli Compagni E, Scialabba S, Di Pasquale A, Cammà C, Puglia I, Calistri P, Savini G |
| EPI_ISL_2600798 | Azienda USL Umbria 2 | Istituto Zooprofilattico Sperimentale dell'Abruzzo e Molise "G. Caporale" | Proietti A, Pistoni E, Lorusso A, Marcacci M, Di Domenico M, Ancora M, Curini V, Mangone I, Rinaldi A, Delli Compagni E, Scialabba S, Di Pasquale A, Cammà C, Puglia I, Calistri P, Savini G |
| EPI_ISL_2600835, EPI_ISL_2600839, EPI_ISL_2600853 | Hospital Universitari i Politècnic La Fe de València | SeqCOVID-SPAIN consortium/IBV(CSIC) | María Dolores Gómez Ruiz, Eva González Barberá, Ana Gil Brusola, Salvador Giner Almaraz, José Luis López Hontangas and SeqCOVID-SPAIN consortium |
| EPI_ISL_2600912, EPI_ISL_2600915, EPI_ISL_2600916 | Servicio de Microbiología. Hospital Clínico Universitario de Valencia | SeqCOVID-SPAIN consortium/IBV(CSIC) | David Navarro Ortega, Eliseo Albert Vicent, Ignacio Torres and SeqCOVID-SPAIN consortium |
| EPI_ISL_2600929, EPI_ISL_2600931, EPI_ISL_2600932, EPI_ISL_2600933, EPI_ISL_2600936, EPI_ISL_2600949, EPI_ISL_2600950 | Plateforme de testing Namuroise | Plateforme de testing Namuroise | Nicolas GILLIARD, Lesly NYINKEU KEMAMEN; Demars Aurore ; Maschietto Céline ; Nobis Chloé ; Otto Gaetan ; Denis Olivier ; Degosserie Jonathan ; Mullier François |
| EPI_ISL_2600958 | Hematopathology Laboratory, ACTREC, TMC | Hematopathology Laboratory, ACTREC, TMC | Hematopathology Laboratory, ACTREC |
| EPI_ISL_2601045, EPI_ISL_2601046, EPI_ISL_2601047, EPI_ISL_2601048, EPI_ISL_2601050, EPI_ISL_2601051, EPI_ISL_2601052, EPI_ISL_2601054, EPI_ISL_2601057, EPI_ISL_2601059, EPI_ISL_2601063, EPI_ISL_2601065, EPI_ISL_2601069, EPI_ISL_2601078, EPI_ISL_2601079, EPI_ISL_2601080, EPI_ISL_2601084 | | | |
| see above | University Hospitals of Geneva, Laboratory of Virology | HUG, Laboratory of Virology and the Health2030 Genome Center | Samuel Cordey, Ana Rita Goncalves, Laurent Kaiser, Lorenzo Cerutti, Henri Pegeot, Melyssa Elies, Deborah Penet, Keith Harshman, Ioannis Xenarios, Emmanouil Dermitzakis |
| EPI_ISL_2601108, EPI_ISL_2601114, EPI_ISL_2601120, EPI_ISL_2601122, EPI_ISL_2601124, EPI_ISL_2601126, EPI_ISL_2601130, EPI_ISL_2601132, EPI_ISL_2601136, EPI_ISL_2601141, EPI_ISL_2601142, EPI_ISL_2601146, EPI_ISL_2601147, EPI_ISL_2601148, EPI_ISL_2601191, EPI_ISL_2601203, EPI_ISL_2601207, EPI_ISL_2601214, EPI_ISL_2601252, EPI_ISL_2601347, EPI_ISL_2601392, EPI_ISL_2601402 | | | |
| see above | National Virus Reference Laboratory | National Virus Reference Laboratory | Zoe Yandle, Charlene Bennett, Gabriel Gonzalez, Michael Carr, Jonathan Dean, Cillian F De Gascun |
| EPI_ISL_2601661 | RSUPN Cipto Mangunkusumo | Genomik Solidaritas Indonesia Laboratorium | Dhahlia Agustina Cahyono, Alfin Mohammad Abdillah, Eric Tenda, Nuri Dyah Indrasari, Eka Ginanjar, Normastuti Adhini Tantyo, Vania Gavrila Wikasa, Carissa Sintca Wijaya, Ahmad Zoebad Foeady, Annisa Muthiah Sukirman, Gracia Felias Enos Korompis, Pratiwi Sudarmono, Budi Wiweko, Louisa Markus, Anuraj Shankar, Meutia Ayuputeri Kumaheri, Anindya Pradipta Susanto, Ariel Pradipta |
| EPI_ISL_2601662, EPI_ISL_2601663 | RSUPN Cipto Mangunkusumo | Genomik Solidaritas Indonesia Laboratorium | Dhahlia Agustina Cahyono, Normastuti Adhini Tantyo, Eric Tenda, Nuri Dyah Indrasari, Eka Ginanjar, Alfin Mohammad Abdillah, Vania Gavrila Wikasa, Carissa Sintca Wijaya, Ahmad Zoebad Foeady, Annisa Muthiah Sukirman, Gracia Felias Enos Korompis, Pratiwi Sudarmono, Budi Wiweko, Louisa Markus, Anuraj Shankar, Meutia Ayuputeri Kumaheri, Anindya Pradipta Susanto, Ariel Pradipta |
| EPI_ISL_2601664 | RSUPN Cipto Mangunkusumo | Genomik Solidaritas Indonesia Laboratorium | Normastuti Adhini Tantyo, Dhahlia Agustina Cahyono, Eric Tenda, Nuri Dyah Indrasari, Eka Ginanjar, Alfin Mohammad Abdillah, Vania Gavrila Wikasa, Carissa Sintca Wijaya, Ahmad Zoebad Foeady, Annisa Muthiah Sukirman, Gracia Felias Enos Korompis, Pratiwi Sudarmono, Budi Wiweko, Louisa Markus, Anuraj Shankar, Meutia Ayuputeri Kumaheri, Anindya Pradipta Susanto, Ariel Pradipta |
| EPI_ISL_2601665, | RSUPN Cipto Mangunkusumo | Genomik Solidaritas Indonesia Laboratorium | Normastuti Adhini Tantyo, Alfin Mohammad Abdillah, Eric Tenda, Nuri Dyah Indrasari, Eka Ginanjar, Dhahlia Agustina Cahyono, Vania Gavrila Wikasa, Carissa Sintca Wijaya, Ahmad Zoebad Foeady, Annisa Muthiah Sukirman, Gracia Felias Enos |

| | | | |
|---|---|---|---|
| EPI_ISL_2601666 | | | Korompis, Pratiwi Sudarmono, Budi Wiweko, Louisa Markus, Anuraj Shankar, Meutia Ayuputeri Kumaheri, Anindya Pradipta Susanto, Ariel Pradipta |
| EPI_ISL_2601667 | RSUPN Cipto Mangunkusumo | Genomik Solidaritas Indonesia Laboratorium | Alfin Mohammad Abdillah, Normastuti Adhini Tantyo, Eric Tenda, Nuri Dyah Indrasari, Eka Ginanjar, Dhahlia Agustina Cahyono, Vania Gavrila Wikasa, Carissa Sintca Wijaya, Ahmad Zoebad Foeady, Annisa Muthiah Sukirman, Gracia Felias Enos Korompis, Pratiwi Sudarmono, Budi Wiweko, Louisa Markus, Anuraj Shankar, Meutia Ayuputeri Kumaheri, Anindya Pradipta Susanto, Ariel Pradipta |
| EPI_ISL_2601668, EPI_ISL_2601669 | RSUPN Cipto Mangunkusumo | Genomik Solidaritas Indonesia Laboratorium | Alfin Mohammad Abdillah, Dhahlia Agustina Cahyono, Eric Tenda, Nuri Dyah Indrasari, Eka Ginanjar, Normastuti Adhini Tantyo, Vania Gavrila Wikasa, Carissa Sintca Wijaya, Ahmad Zoebad Foeady, Annisa Muthiah Sukirman, Gracia Felias Enos Korompis, Pratiwi Sudarmono, Budi Wiweko, Louisa Markus, Anuraj Shankar, Meutia Ayuputeri Kumaheri, Anindya Pradipta Susanto, Ariel Pradipta |
| EPI_ISL_2601670 | RSUPN Cipto Mangunkusumo | Genomik Solidaritas Indonesia Laboratorium | Alfin Mohammad Abdillah, Normastuti Adhini Tantyo, Eric Tenda, Nuri Dyah Indrasari, Eka Ginanjar, Dhahlia Agustina Cahyono, Vania Gavrila Wikasa, Carissa Sintca Wijaya, Ahmad Zoebad Foeady, Annisa Muthiah Sukirman, Gracia Felias Enos Korompis, Pratiwi Sudarmono, Budi Wiweko, Louisa Markus, Anuraj Shankar, Meutia Ayuputeri Kumaheri, Anindya Pradipta Susanto, Ariel Pradipta |
| EPI_ISL_2601677, EPI_ISL_2601683, EPI_ISL_2601697 | Hospital General Universitario Gregorio Marañón | Hospital General Universitario Gregorio Marañón | Sergio Buenestado Serrano, Pedro Sola Campoy, Laura Pérez-Lago, Cristina Rodriguez-Grande, Marta Herranz Martin, Victor Manuel de la Cueva, Julia Suárez, Pilar Catalán, Patricia Muñoz, Darío García de Viedma |
| EPI_ISL_2601960, EPI_ISL_2601961 | ZOL | Jessa | Berden et al. on behalf of the Jessa_cmdLab |
| EPI_ISL_2601980 | Jessa | Jessa | Berden et al. on behalf of the Jessa_cmdLab |
| EPI_ISL_2602216, EPI_ISL_2602384 | Gravity Diagnostics, LLC | Gravity Diagnostics, LLC | Gravity Diagnostics |
| EPI_ISL_2602420 | Ospedale San Giovanni Evangelista | INMI Lazzaro Spallanzani IRCCS | CEM Gruber, F Santini, G Bonfiglio, O Butera, G Orlandi, E Sperandio, D Di Fusco, D Cerini, E Giombini |
| EPI_ISL_2602432 | Laboratorio di Genetica Medica Ospedale Belcolle | INMI Lazzaro Spallanzani IRCCS | F Messina, CEM Gruber, M Rueca, G Bonfiglio, F De Nicola, A Massacci, F Natoni, G Pessina, E Giombini |
| EPI_ISL_2602435 | Laboratorio di Genetica Medica Ospedale Belcolle | INMI Lazzaro Spallanzani IRCCS | M Rueca, G Bonfiglio, F Messina, CEM Gruber, GF Inger, A Massacci, F Natoni, G Pessina, B Bartolini |
| EPI_ISL_2602618, EPI_ISL_2602677, EPI_ISL_2602679, EPI_ISL_2602703, EPI_ISL_2602712, EPI_ISL_2602713, EPI_ISL_2602716, EPI_ISL_2602720, EPI_ISL_2602733, EPI_ISL_2602734, EPI_ISL_2602735, EPI_ISL_2602736, EPI_ISL_2602739, EPI_ISL_2602748, EPI_ISL_2602750, EPI_ISL_2602759, EPI_ISL_2602761, EPI_ISL_2602763 | | | |
| see above | KEMRI-Wellcome Trust Research Programme,Kilifi | KEMRI-Wellcome Trust Research Programme,Kilifi | Githinji G.,Mohamed K.S.,de Laurent Z.,Mburu M.W.,Matoke D.,Onyango C., |
| EPI_ISL_2602847 | Ospedale di Genzano - ASL RM 6 | INMI Lazzaro Spallanzani IRCCS | F Santini, F Messina, G Bonfiglio, O Butera, G Orlandi, E Sperandio, G Tramini, E Conti, B Bartolini |
| EPI_ISL_2602854 | Ospedale di Genzano - ASL RM 6 | INMI Lazzaro Spallanzani IRCCS | CEM Gruber, F Santini, M Rueca, G Bonfiglio, L Ciuffreda, A Massacci, G Tramini, E Conti, B Bartolini |
| EPI_ISL_2603036, EPI_ISL_2603038, EPI_ISL_2603039, EPI_ISL_2603040, EPI_ISL_2603043, EPI_ISL_2603044, EPI_ISL_2603047, EPI_ISL_2603048, EPI_ISL_2603049, EPI_ISL_2603051, EPI_ISL_2603052, EPI_ISL_2603053, EPI_ISL_2603054, EPI_ISL_2603055, EPI_ISL_2603056, EPI_ISL_2603057, EPI_ISL_2603059, EPI_ISL_2603100, EPI_ISL_2603101 | | | |
| see above | KEMRI-Wellcome Trust Research Programme,Kilifi | KEMRI-Wellcome Trust Research Programme,Kilifi | Githinji G.,Mohamed K.S.,de Laurent Z.,Mburu M.W.,Matoke D.,Onyango C., |
| EPI_ISL_2603202, EPI_ISL_2603209, EPI_ISL_2603221, EPI_ISL_2603222, EPI_ISL_2603228 | Virginia Division of Consolidated Laboratory Services | Virginia Division of Consolidated Laboratory Services | Virginia DCLS |
| EPI_ISL_2603280 | Wyoming Public Health Laboratory | Wyoming Public Health Laboratory | Jim Mildenberger, Taylor Fearing, Channing Weber, Ashley Norberg, Chayse Rowley, Marley Goetz, Brian Dominguez, Elliot Thomasson, Sam Britz, Cari Sloma, Robert Petit, and Rob Christensen |
| EPI_ISL_2603349 | Virginia Division of Consolidated Laboratory Services | Virginia Division of Consolidated Laboratory Services | Virginia DCLS |
| EPI_ISL_2603405, EPI_ISL_2603406 | Wyoming Public Health Laboratory | Wyoming Public Health Laboratory | Jim Mildenberger, Taylor Fearing, Channing Weber, Ashley Norberg, Chayse Rowley, Marley Goetz, Brian Dominguez, Elliot Thomasson, Sam Britz, Cari Sloma, Robert Petit, and Rob Christensen |
| EPI_ISL_2603418, EPI_ISL_2603419, EPI_ISL_2603421 | CHUGA-IBP-laboratoire de Virologie | IBP-laboratoire de virologie | Sylvie Larrat; Patrice Morand; Pascal Poignard; Raphaële Germi; Julien Lupo; Anne Signori -Schmuck; Anne-Karen Faure; Benjamin Nemoz; Aurélie Truffot; Léa Ponderand; Julien Andréani; Alban Caporossi; Hugo Jardin |
| EPI_ISL_2603612, EPI_ISL_2603635, EPI_ISL_2603654 | InDRE | Instituto Nacional de Medicina Genomica | Hidalgo-Miranda A, Mendoza-Vargas A, Reyes-Grajeda JP, Cedro-Tanda A, Gisela Barrera-Badillo, Irma Lopez-Martinez, Jose Ernesto Ramirez González, Gonzalez-Barrera D, Rangel-DeLeon D, Munguia-Garza P, Garcia-Cardenas FJ, Gonzalez-Woge MA, Herrera-Montalvo LA. |
| EPI_ISL_2603666, EPI_ISL_2603667, EPI_ISL_2603690, EPI_ISL_2603704, EPI_ISL_2603708, EPI_ISL_2603718, EPI_ISL_2603768, EPI_ISL_2603774, EPI_ISL_2603800, EPI_ISL_2603801 | Salud Digna | Instituto Nacional de Medicina Genomica | Hidalgo-Miranda A, Cedro-Tanda A, Mendoza-Vargas A, Reyes-Grajeda JP, Abraham Campos-Romero, Moreno-Camacho José Luis, Rodriguez-Gallegos Jorge, Luna-Ruiz Marco, Gonzalez-Barrera D, Rangel-DeLeon D, Munguia-Garza P, Ramirez-Vega O, Escobar-Arrazola, M, Herrera-Montalvo LA. |
| EPI_ISL_2603819, EPI_ISL_2603820, EPI_ISL_2603821, EPI_ISL_2603824 | Queensland Health Forensic and Scientific Services | Queensland Health Forensic and Scientific Services | Son Nguyen |
| EPI_ISL_2604191 | AULSS 2 Marca Trevigiana | Istituto Zooprofilattico Sperimentale delle Venezie | Adelaide Milani, Alessia Schivo, Annalisa Salviato, Elisa Palumbo, Erika Giorgia Quaranta, Luca Tassoni, Ambra Pastori, Edoardo Giussani, Alice Fusaro, Isabella Monne, Calogero Terregino, Antonia Ricci |
| EPI_ISL_2604229, EPI_ISL_2604230, EPI_ISL_2604231, EPI_ISL_2604232 | Montana Public Health Laboratory | Montana Public Health Laboratory | Joy Ritter, Michelle Mozer, Carrie Biskupiak, Deborah Gibson, Michael Dills |
| EPI_ISL_2604822 | Laboratorio di Riferimento Regionale della Sicilia Occidentale per l'Emergenza COVID-19 | Laboratorio di Riferimento Regionale della Sicilia Occidentale per l'Emergenza COVID-19 | Fabio Tramuto, Carmelo Massimo Maida, Daniela Di Naro, Giulia Randazzo, Walter Mazzucco, Giorgio Graziano, Vincenzo Restivo, Claudio Costantino, Francesco Vitale |
| EPI_ISL_2604889, EPI_ISL_2604891, EPI_ISL_2604892, EPI_ISL_2604893, EPI_ISL_2604894, EPI_ISL_2604895, EPI_ISL_2604896, EPI_ISL_2604897, EPI_ISL_2604898, EPI_ISL_2604900, EPI_ISL_2604901, EPI_ISL_2604902, EPI_ISL_2604903, EPI_ISL_2604904, EPI_ISL_2604905, EPI_ISL_2604906, EPI_ISL_2604907, EPI_ISL_2604908, EPI_ISL_2604909, EPI_ISL_2604911 | | | |
| see above | AULSS 2 Marca Trevigiana | Istituto Zooprofilattico Sperimentale delle Venezie | Adelaide Milani, Alessia Schivo, Annalisa Salviato, Elisa Palumbo, Erika Giorgia Quaranta, Luca Tassoni, Ambra Pastori, Edoardo Giussani, Alice Fusaro, Isabella Monne, Calogero Terregino, Antonia Ricci |
| EPI_ISL_2604958 | Inovie AS Biopyrenees | Laboratoire GenBio Clermont Ferrand | Gwladys SEVRIN Pauline LAURENT Florian VERCRUYSSEN |
| EPI_ISL_2604978, EPI_ISL_2605014, EPI_ISL_2605046, EPI_ISL_2605052, EPI_ISL_2605092, EPI_ISL_2605104, EPI_ISL_2605118 | Lab voor klinische biologie | Lab voor klinische biologie | Marija Janevska, Hannelore Hamerlinck, Bruno Verhasselt |
| EPI_ISL_2605126, EPI_ISL_2605128, EPI_ISL_2605129, EPI_ISL_2605130, EPI_ISL_2605132, EPI_ISL_2605133 | Lighthouse Lab in Alderley Park | Wellcome Sanger Institute for the COVID-19 Genomics UK (COG-UK) Consortium | Jacquelyn Wynn, Mairead Hyland, The Lighthouse Lab in Alderley Park and Alex Alderton, Roberto Amato, Jeffrey Barrett, Sonia Goncalves, Ewan Harrison, David K. Jackson, Ian Johnston, Dominic Kwiatkowski, Cordelia Langford, John Sillitoe on behalf of the Wellcome Sanger Institute COVID-19 Surveillance Team |
| EPI_ISL_2605135 | Lighthouse Lab in Glasgow | Wellcome Sanger Institute for the COVID-19 Genomics UK (COG-UK) Consortium | Harper VanSteenhouse, Yumi Kasai, David Gray, Carol Clugston, Anna Dominiczak and Alex Alderton, Roberto Amato, Jeffrey Barrett, Sonia Goncalves, Ewan Harrison, David K. Jackson, Ian Johnston, Dominic Kwiatkowski, Cordelia Langford, John Sillitoe on behalf of the Wellcome Sanger Institute COVID-19 Surveillance Team |
| EPI_ISL_2605137, EPI_ISL_2605138, EPI_ISL_2605140, EPI_ISL_2605141 | Lighthouse Lab in Alderley Park | Wellcome Sanger Institute for the COVID-19 Genomics UK (COG-UK) Consortium | Jacquelyn Wynn, Mairead Hyland, The Lighthouse Lab in Alderley Park and Alex Alderton, Roberto Amato, Jeffrey Barrett, Sonia Goncalves, Ewan Harrison, David K. Jackson, Ian Johnston, Dominic Kwiatkowski, Cordelia Langford, John Sillitoe on behalf of the Wellcome Sanger Institute COVID-19 Surveillance Team |
| EPI_ISL_2605142 | Lighthouse Lab in Glasgow | Wellcome Sanger Institute for the COVID-19 Genomics UK (COG-UK) Consortium | Harper VanSteenhouse, Yumi Kasai, David Gray, Carol Clugston, Anna Dominiczak and Alex Alderton, Roberto Amato, Jeffrey Barrett, Sonia Goncalves, Ewan Harrison, David K. Jackson, Ian Johnston, Dominic Kwiatkowski, Cordelia Langford, John Sillitoe on behalf of the Wellcome Sanger Institute COVID-19 Surveillance Team |
| EPI_ISL_2605144 | Lighthouse Lab in Alderley Park | Wellcome Sanger Institute for the COVID-19 Genomics UK (COG-UK) Consortium | Jacquelyn Wynn, Mairead Hyland, The Lighthouse Lab in Alderley Park and Alex Alderton, Roberto Amato, Jeffrey Barrett, Sonia Goncalves, Ewan Harrison, David K. Jackson, Ian Johnston, Dominic Kwiatkowski, Cordelia Langford, John Sillitoe on behalf of the Wellcome Sanger Institute COVID-19 Surveillance Team |
| EPI_ISL_2605146, EPI_ISL_2605147 | Lighthouse Lab in Glasgow | Wellcome Sanger Institute for the COVID-19 Genomics UK (COG-UK) Consortium | Harper VanSteenhouse, Yumi Kasai, David Gray, Carol Clugston, Anna Dominiczak and Alex Alderton, Roberto Amato, Jeffrey Barrett, Sonia Goncalves, Ewan Harrison, David K. Jackson, Ian Johnston, Dominic Kwiatkowski, Cordelia Langford, John Sillitoe on behalf of the Wellcome Sanger Institute COVID-19 Surveillance Team |
| EPI_ISL_2605149, EPI_ISL_2605150, EPI_ISL_2605152 | Lighthouse Lab in Alderley Park | Wellcome Sanger Institute for the COVID-19 Genomics UK (COG-UK) Consortium | Jacquelyn Wynn, Mairead Hyland, The Lighthouse Lab in Alderley Park and Alex Alderton, Roberto Amato, Jeffrey Barrett, Sonia Goncalves, Ewan Harrison, David K. Jackson, Ian Johnston, Dominic Kwiatkowski, Cordelia Langford, John Sillitoe on behalf of the Wellcome Sanger Institute COVID-19 Surveillance Team |
| EPI_ISL_2605155 | Lighthouse Lab in Glasgow | Wellcome Sanger Institute for the COVID-19 Genomics UK (COG-UK) | Harper VanSteenhouse, Yumi Kasai, David Gray, Carol Clugston, Anna Dominiczak and Alex Alderton, Roberto Amato, Jeffrey Barrett, Sonia Goncalves, Ewan Harrison, David K. Jackson, Ian Johnston, Dominic Kwiatkowski, Cordelia Langford, |

| | | | |
|---|---|---|---|
| | | Consortium | John Sillitoe on behalf of the Wellcome Sanger Institute COVID-19 Surveillance Team |
| EPI_ISL_2605156, EPI_ISL_2605158, EPI_ISL_2605161, EPI_ISL_2605162, EPI_ISL_2605164, EPI_ISL_2605165, EPI_ISL_2605167, EPI_ISL_2605168, EPI_ISL_2605169, EPI_ISL_2605171, EPI_ISL_2605173, EPI_ISL_2605174, EPI_ISL_2605176, EPI_ISL_2605177, EPI_ISL_2605180, EPI_ISL_2605182 | | | |
| see above | Lighthouse Lab in Alderley Park | Wellcome Sanger Institute for the COVID-19 Genomics UK (COG-UK) Consortium | Jacquelyn Wynn, Mairead Hyland, The Lighthouse Lab in Alderley Park and Alex Alderton, Roberto Amato, Jeffrey Barrett, Sonia Goncalves, Ewan Harrison, David K. Jackson, Ian Johnston, Dominic Kwiatkowski, Cordelia Langford, John Sillitoe on behalf of the Wellcome Sanger Institute COVID-19 Surveillance Team |
| EPI_ISL_2605185 | Lighthouse Lab in Glasgow | Wellcome Sanger Institute for the COVID-19 Genomics UK (COG-UK) Consortium | Harper VanSteenhouse, Yumi Kasai, David Gray, Carol Clugston, Anna Dominiczak and Alex Alderton, Roberto Amato, Jeffrey Barrett, Sonia Goncalves, Ewan Harrison, David K. Jackson, Ian Johnston, Dominic Kwiatkowski, Cordelia Langford, John Sillitoe on behalf of the Wellcome Sanger Institute COVID-19 Surveillance Team |
| EPI_ISL_2605186, EPI_ISL_2605188, EPI_ISL_2605189, EPI_ISL_2605191, EPI_ISL_2605192, EPI_ISL_2605194 | Lighthouse Lab in Alderley Park | Wellcome Sanger Institute for the COVID-19 Genomics UK (COG-UK) Consortium | Jacquelyn Wynn, Mairead Hyland, The Lighthouse Lab in Alderley Park and Alex Alderton, Roberto Amato, Jeffrey Barrett, Sonia Goncalves, Ewan Harrison, David K. Jackson, Ian Johnston, Dominic Kwiatkowski, Cordelia Langford, John Sillitoe on behalf of the Wellcome Sanger Institute COVID-19 Surveillance Team |
| EPI_ISL_2605195 | Lighthouse Lab in Glasgow | Wellcome Sanger Institute for the COVID-19 Genomics UK (COG-UK) Consortium | Harper VanSteenhouse, Yumi Kasai, David Gray, Carol Clugston, Anna Dominiczak and Alex Alderton, Roberto Amato, Jeffrey Barrett, Sonia Goncalves, Ewan Harrison, David K. Jackson, Ian Johnston, Dominic Kwiatkowski, Cordelia Langford, John Sillitoe on behalf of the Wellcome Sanger Institute COVID-19 Surveillance Team |
| EPI_ISL_2605197 | Lighthouse Lab in Alderley Park | Wellcome Sanger Institute for the COVID-19 Genomics UK (COG-UK) Consortium | Jacquelyn Wynn, Mairead Hyland, The Lighthouse Lab in Alderley Park and Alex Alderton, Roberto Amato, Jeffrey Barrett, Sonia Goncalves, Ewan Harrison, David K. Jackson, Ian Johnston, Dominic Kwiatkowski, Cordelia Langford, John Sillitoe on behalf of the Wellcome Sanger Institute COVID-19 Surveillance Team |
| EPI_ISL_2605199 | Lighthouse Lab in Glasgow | Wellcome Sanger Institute for the COVID-19 Genomics UK (COG-UK) Consortium | Harper VanSteenhouse, Yumi Kasai, David Gray, Carol Clugston, Anna Dominiczak and Alex Alderton, Roberto Amato, Jeffrey Barrett, Sonia Goncalves, Ewan Harrison, David K. Jackson, Ian Johnston, Dominic Kwiatkowski, Cordelia Langford, John Sillitoe on behalf of the Wellcome Sanger Institute COVID-19 Surveillance Team |
| EPI_ISL_2605200 | Lighthouse Lab in Alderley Park | Wellcome Sanger Institute for the COVID-19 Genomics UK (COG-UK) Consortium | Jacquelyn Wynn, Mairead Hyland, The Lighthouse Lab in Alderley Park and Alex Alderton, Roberto Amato, Jeffrey Barrett, Sonia Goncalves, Ewan Harrison, David K. Jackson, Ian Johnston, Dominic Kwiatkowski, Cordelia Langford, John Sillitoe on behalf of the Wellcome Sanger Institute COVID-19 Surveillance Team |
| EPI_ISL_2605202 | Lighthouse Lab in Glasgow | Wellcome Sanger Institute for the COVID-19 Genomics UK (COG-UK) Consortium | Harper VanSteenhouse, Yumi Kasai, David Gray, Carol Clugston, Anna Dominiczak and Alex Alderton, Roberto Amato, Jeffrey Barrett, Sonia Goncalves, Ewan Harrison, David K. Jackson, Ian Johnston, Dominic Kwiatkowski, Cordelia Langford, John Sillitoe on behalf of the Wellcome Sanger Institute COVID-19 Surveillance Team |
| EPI_ISL_2605203, EPI_ISL_2605205 | Lighthouse Lab in Alderley Park | Wellcome Sanger Institute for the COVID-19 Genomics UK (COG-UK) Consortium | Jacquelyn Wynn, Mairead Hyland, The Lighthouse Lab in Alderley Park and Alex Alderton, Roberto Amato, Jeffrey Barrett, Sonia Goncalves, Ewan Harrison, David K. Jackson, Ian Johnston, Dominic Kwiatkowski, Cordelia Langford, John Sillitoe on behalf of the Wellcome Sanger Institute COVID-19 Surveillance Team |
| EPI_ISL_2605206 | Lighthouse Lab in Glasgow | Wellcome Sanger Institute for the COVID-19 Genomics UK (COG-UK) Consortium | Harper VanSteenhouse, Yumi Kasai, David Gray, Carol Clugston, Anna Dominiczak and Alex Alderton, Roberto Amato, Jeffrey Barrett, Sonia Goncalves, Ewan Harrison, David K. Jackson, Ian Johnston, Dominic Kwiatkowski, Cordelia Langford, John Sillitoe on behalf of the Wellcome Sanger Institute COVID-19 Surveillance Team |
| EPI_ISL_2605208 | Lighthouse Lab in Alderley Park | Wellcome Sanger Institute for the COVID-19 Genomics UK (COG-UK) Consortium | Jacquelyn Wynn, Mairead Hyland, The Lighthouse Lab in Alderley Park and Alex Alderton, Roberto Amato, Jeffrey Barrett, Sonia Goncalves, Ewan Harrison, David K. Jackson, Ian Johnston, Dominic Kwiatkowski, Cordelia Langford, John Sillitoe on behalf of the Wellcome Sanger Institute COVID-19 Surveillance Team |
| EPI_ISL_2605210 | Lighthouse Lab in Glasgow | Wellcome Sanger Institute for the COVID-19 Genomics UK (COG-UK) Consortium | Harper VanSteenhouse, Yumi Kasai, David Gray, Carol Clugston, Anna Dominiczak and Alex Alderton, Roberto Amato, Jeffrey Barrett, Sonia Goncalves, Ewan Harrison, David K. Jackson, Ian Johnston, Dominic Kwiatkowski, Cordelia Langford, John Sillitoe on behalf of the Wellcome Sanger Institute COVID-19 Surveillance Team |
| EPI_ISL_2605211, EPI_ISL_2605213, EPI_ISL_2605214, EPI_ISL_2605216, EPI_ISL_2605217, EPI_ISL_2605219 | Lighthouse Lab in Alderley Park | Wellcome Sanger Institute for the COVID-19 Genomics UK (COG-UK) Consortium | Jacquelyn Wynn, Mairead Hyland, The Lighthouse Lab in Alderley Park and Alex Alderton, Roberto Amato, Jeffrey Barrett, Sonia Goncalves, Ewan Harrison, David K. Jackson, Ian Johnston, Dominic Kwiatkowski, Cordelia Langford, John Sillitoe on behalf of the Wellcome Sanger Institute COVID-19 Surveillance Team |
| EPI_ISL_2605220, EPI_ISL_2605222 | Lighthouse Lab in Glasgow | Wellcome Sanger Institute for the COVID-19 Genomics UK (COG-UK) Consortium | Harper VanSteenhouse, Yumi Kasai, David Gray, Carol Clugston, Anna Dominiczak and Alex Alderton, Roberto Amato, Jeffrey Barrett, Sonia Goncalves, Ewan Harrison, David K. Jackson, Ian Johnston, Dominic Kwiatkowski, Cordelia Langford, John Sillitoe on behalf of the Wellcome Sanger Institute COVID-19 Surveillance Team |
| EPI_ISL_2605223, EPI_ISL_2605225, EPI_ISL_2605226, EPI_ISL_2605227, EPI_ISL_2605229, EPI_ISL_2605230, EPI_ISL_2605232, EPI_ISL_2605233, EPI_ISL_2605235, EPI_ISL_2605236, EPI_ISL_2605238, EPI_ISL_2605239, EPI_ISL_2605241, EPI_ISL_2605242, EPI_ISL_2605243, EPI_ISL_2605245, EPI_ISL_2605246, EPI_ISL_2605248 | | | |
| see above | Lighthouse Lab in Alderley Park | Wellcome Sanger Institute for the COVID-19 Genomics UK (COG-UK) Consortium | Jacquelyn Wynn, Mairead Hyland, The Lighthouse Lab in Alderley Park and Alex Alderton, Roberto Amato, Jeffrey Barrett, Sonia Goncalves, Ewan Harrison, David K. Jackson, Ian Johnston, Dominic Kwiatkowski, Cordelia Langford, John Sillitoe on behalf of the Wellcome Sanger Institute COVID-19 Surveillance Team |
| EPI_ISL_2605249 | Lighthouse Lab in Glasgow | Wellcome Sanger Institute for the COVID-19 Genomics UK (COG-UK) Consortium | Harper VanSteenhouse, Yumi Kasai, David Gray, Carol Clugston, Anna Dominiczak and Alex Alderton, Roberto Amato, Jeffrey Barrett, Sonia Goncalves, Ewan Harrison, David K. Jackson, Ian Johnston, Dominic Kwiatkowski, Cordelia Langford, John Sillitoe on behalf of the Wellcome Sanger Institute COVID-19 Surveillance Team |
| EPI_ISL_2605251, EPI_ISL_2605252, EPI_ISL_2605254, EPI_ISL_2605255, EPI_ISL_2605257, EPI_ISL_2605258 | Lighthouse Lab in Alderley Park | Wellcome Sanger Institute for the COVID-19 Genomics UK (COG-UK) Consortium | Jacquelyn Wynn, Mairead Hyland, The Lighthouse Lab in Alderley Park and Alex Alderton, Roberto Amato, Jeffrey Barrett, Sonia Goncalves, Ewan Harrison, David K. Jackson, Ian Johnston, Dominic Kwiatkowski, Cordelia Langford, John Sillitoe on behalf of the Wellcome Sanger Institute COVID-19 Surveillance Team |
| EPI_ISL_2605260 | Lighthouse Lab in Glasgow | Wellcome Sanger Institute for the COVID-19 Genomics UK (COG-UK) Consortium | Harper VanSteenhouse, Yumi Kasai, David Gray, Carol Clugston, Anna Dominiczak and Alex Alderton, Roberto Amato, Jeffrey Barrett, Sonia Goncalves, Ewan Harrison, David K. Jackson, Ian Johnston, Dominic Kwiatkowski, Cordelia Langford, John Sillitoe on behalf of the Wellcome Sanger Institute COVID-19 Surveillance Team |
| EPI_ISL_2605261 | Lighthouse Lab in Alderley Park | Wellcome Sanger Institute for the COVID-19 Genomics UK (COG-UK) Consortium | Jacquelyn Wynn, Mairead Hyland, The Lighthouse Lab in Alderley Park and Alex Alderton, Roberto Amato, Jeffrey Barrett, Sonia Goncalves, Ewan Harrison, David K. Jackson, Ian Johnston, Dominic Kwiatkowski, Cordelia Langford, John Sillitoe on behalf of the Wellcome Sanger Institute COVID-19 Surveillance Team |
| EPI_ISL_2605263 | Lighthouse Lab in Glasgow | Wellcome Sanger Institute for the COVID-19 Genomics UK (COG-UK) Consortium | Harper VanSteenhouse, Yumi Kasai, David Gray, Carol Clugston, Anna Dominiczak and Alex Alderton, Roberto Amato, Jeffrey Barrett, Sonia Goncalves, Ewan Harrison, David K. Jackson, Ian Johnston, Dominic Kwiatkowski, Cordelia Langford, John Sillitoe on behalf of the Wellcome Sanger Institute COVID-19 Surveillance Team |
| EPI_ISL_2605264, EPI_ISL_2605266, EPI_ISL_2605267, EPI_ISL_2605269 | Lighthouse Lab in Alderley Park | Wellcome Sanger Institute for the COVID-19 Genomics UK (COG-UK) Consortium | Jacquelyn Wynn, Mairead Hyland, The Lighthouse Lab in Alderley Park and Alex Alderton, Roberto Amato, Jeffrey Barrett, Sonia Goncalves, Ewan Harrison, David K. Jackson, Ian Johnston, Dominic Kwiatkowski, Cordelia Langford, John Sillitoe on behalf of the Wellcome Sanger Institute COVID-19 Surveillance Team |
| EPI_ISL_2605270, EPI_ISL_2605274 | Lighthouse Lab in Glasgow | Wellcome Sanger Institute for the COVID-19 Genomics UK (COG-UK) Consortium | Harper VanSteenhouse, Yumi Kasai, David Gray, Carol Clugston, Anna Dominiczak and Alex Alderton, Roberto Amato, Jeffrey Barrett, Sonia Goncalves, Ewan Harrison, David K. Jackson, Ian Johnston, Dominic Kwiatkowski, Cordelia Langford, John Sillitoe on behalf of the Wellcome Sanger Institute COVID-19 Surveillance Team |
| EPI_ISL_2605275, EPI_ISL_2605276, EPI_ISL_2605278, EPI_ISL_2605280, EPI_ISL_2605281, EPI_ISL_2605282, EPI_ISL_2605283, EPI_ISL_2605285, EPI_ISL_2605286, EPI_ISL_2605288, EPI_ISL_2605289 | | | |
| see above | Lighthouse Lab in Alderley Park | Wellcome Sanger Institute for the COVID-19 Genomics UK (COG-UK) Consortium | Jacquelyn Wynn, Mairead Hyland, The Lighthouse Lab in Alderley Park and Alex Alderton, Roberto Amato, Jeffrey Barrett, Sonia Goncalves, Ewan Harrison, David K. Jackson, Ian Johnston, Dominic Kwiatkowski, Cordelia Langford, John Sillitoe on behalf of the Wellcome Sanger Institute COVID-19 Surveillance Team |
| EPI_ISL_2605291 | Lighthouse Lab in Glasgow | Wellcome Sanger Institute for the COVID-19 Genomics UK (COG-UK) Consortium | Harper VanSteenhouse, Yumi Kasai, David Gray, Carol Clugston, Anna Dominiczak and Alex Alderton, Roberto Amato, Jeffrey Barrett, Sonia Goncalves, Ewan Harrison, David K. Jackson, Ian Johnston, Dominic Kwiatkowski, Cordelia Langford, John Sillitoe on behalf of the Wellcome Sanger Institute COVID-19 Surveillance Team |
| EPI_ISL_2605292 | Lighthouse Lab in Alderley Park | Wellcome Sanger Institute for the COVID-19 Genomics UK (COG-UK) Consortium | Jacquelyn Wynn, Mairead Hyland, The Lighthouse Lab in Alderley Park and Alex Alderton, Roberto Amato, Jeffrey Barrett, Sonia Goncalves, Ewan Harrison, David K. Jackson, Ian Johnston, Dominic Kwiatkowski, Cordelia Langford, John Sillitoe on behalf of the Wellcome Sanger Institute COVID-19 Surveillance Team |
| EPI_ISL_2605294 | Lighthouse Lab in Glasgow | Wellcome Sanger Institute for the COVID-19 Genomics UK (COG-UK) Consortium | Harper VanSteenhouse, Yumi Kasai, David Gray, Carol Clugston, Anna Dominiczak and Alex Alderton, Roberto Amato, Jeffrey Barrett, Sonia Goncalves, Ewan Harrison, David K. Jackson, Ian Johnston, Dominic Kwiatkowski, Cordelia Langford, John Sillitoe on behalf of the Wellcome Sanger Institute COVID-19 Surveillance Team |
| EPI_ISL_2605296 | Lighthouse Lab in Alderley Park | Wellcome Sanger Institute for the COVID-19 Genomics UK (COG-UK) Consortium | Jacquelyn Wynn, Mairead Hyland, The Lighthouse Lab in Alderley Park and Alex Alderton, Roberto Amato, Jeffrey Barrett, Sonia Goncalves, Ewan Harrison, David K. Jackson, Ian Johnston, Dominic Kwiatkowski, Cordelia Langford, John Sillitoe on behalf of the Wellcome Sanger Institute COVID-19 Surveillance Team |
| EPI_ISL_2605297 | Lab voor klinische biologie | Lab voor klinische biologie | Marija Janevska, Hannelore Hamerlinck, Bruno Verhasselt |
| EPI_ISL_2605298 | Lighthouse Lab in Alderley Park | Wellcome Sanger Institute for the COVID-19 Genomics UK (COG-UK) Consortium | Jacquelyn Wynn, Mairead Hyland, The Lighthouse Lab in Alderley Park and Alex Alderton, Roberto Amato, Jeffrey Barrett, Sonia Goncalves, Ewan Harrison, David K. Jackson, Ian Johnston, Dominic Kwiatkowski, Cordelia Langford, John Sillitoe on behalf of the Wellcome Sanger Institute COVID-19 Surveillance Team |
| EPI_ISL_2605299 | Lighthouse Lab in Glasgow | Wellcome Sanger Institute for the COVID-19 Genomics UK (COG-UK) Consortium | Harper VanSteenhouse, Yumi Kasai, David Gray, Carol Clugston, Anna Dominiczak and Alex Alderton, Roberto Amato, Jeffrey Barrett, Sonia Goncalves, Ewan Harrison, David K. Jackson, Ian Johnston, Dominic Kwiatkowski, Cordelia Langford, John Sillitoe on behalf of the Wellcome Sanger Institute COVID-19 Surveillance Team |
| EPI_ISL_2605301, EPI_ISL_2605302, EPI_ISL_2605304, EPI_ISL_2605305, EPI_ISL_2605307, EPI_ISL_2605308 | Lighthouse Lab in Alderley Park | Wellcome Sanger Institute for the COVID-19 Genomics UK (COG-UK) Consortium | Jacquelyn Wynn, Mairead Hyland, The Lighthouse Lab in Alderley Park and Alex Alderton, Roberto Amato, Jeffrey Barrett, Sonia Goncalves, Ewan Harrison, David K. Jackson, Ian Johnston, Dominic Kwiatkowski, Cordelia Langford, John Sillitoe on behalf of the Wellcome Sanger Institute COVID-19 Surveillance Team |
| EPI_ISL_2605310, EPI_ISL_2605311 | Lighthouse Lab in Glasgow | Wellcome Sanger Institute for the COVID-19 Genomics UK (COG-UK) Consortium | Harper VanSteenhouse, Yumi Kasai, David Gray, Carol Clugston, Anna Dominiczak and Alex Alderton, Roberto Amato, Jeffrey Barrett, Sonia Goncalves, Ewan Harrison, David K. Jackson, Ian Johnston, Dominic Kwiatkowski, Cordelia Langford, John Sillitoe on behalf of the Wellcome Sanger Institute COVID-19 Surveillance Team |
| EPI_ISL_2605313, EPI_ISL_2605314, EPI_ISL_2605316, EPI_ISL_2605317 | Lighthouse Lab in Alderley Park | Wellcome Sanger Institute for the COVID-19 Genomics UK (COG-UK) Consortium | Jacquelyn Wynn, Mairead Hyland, The Lighthouse Lab in Alderley Park and Alex Alderton, Roberto Amato, Jeffrey Barrett, Sonia Goncalves, Ewan Harrison, David K. Jackson, Ian Johnston, Dominic Kwiatkowski, Cordelia Langford, John Sillitoe on behalf of the Wellcome Sanger Institute COVID-19 Surveillance Team |
| EPI_ISL_2605318 | Lighthouse Lab in Glasgow | Wellcome Sanger Institute for the COVID-19 Genomics UK (COG-UK) Consortium | Harper VanSteenhouse, Yumi Kasai, David Gray, Carol Clugston, Anna Dominiczak and Alex Alderton, Roberto Amato, Jeffrey Barrett, Sonia Goncalves, Ewan Harrison, David K. Jackson, Ian Johnston, Dominic Kwiatkowski, Cordelia Langford, John Sillitoe on behalf of the Wellcome Sanger Institute COVID-19 Surveillance Team |
| EPI_ISL_2605319, EPI_ISL_2605320, | Lighthouse Lab in Alderley Park | Wellcome Sanger Institute for the COVID-19 Genomics UK (COG-UK) Consortium | Jacquelyn Wynn, Mairead Hyland, The Lighthouse Lab in Alderley Park and Alex Alderton, Roberto Amato, Jeffrey Barrett, Sonia Goncalves, Ewan Harrison, David K. Jackson, Ian Johnston, Dominic Kwiatkowski, Cordelia Langford, John Sillitoe on behalf of the Wellcome Sanger Institute COVID-19 Surveillance Team |

| | | | |
|---|---|---|---|
| EPI_ISL_2605321, EPI_ISL_2605322, EPI_ISL_2605324, EPI_ISL_2605325 | | | |
| EPI_ISL_2605326, EPI_ISL_2605330, EPI_ISL_2605333 | Lighthouse Lab in Glasgow | Wellcome Sanger Institute for the COVID-19 Genomics UK (COG-UK) Consortium | Harper VanSteenhouse, Yumi Kasai, David Gray, Carol Clugston, Anna Dominiczak and Alex Alderton, Roberto Amato, Jeffrey Barrett, Sonia Goncalves, Ewan Harrison, David K. Jackson, Ian Johnston, Dominic Kwiatkowski, Cordelia Langford, John Sillitoe on behalf of the Wellcome Sanger Institute COVID-19 Surveillance Team |
| EPI_ISL_2605334, EPI_ISL_2605336, EPI_ISL_2605337, EPI_ISL_2605338, EPI_ISL_2605340 | Lighthouse Lab in Alderley Park | Wellcome Sanger Institute for the COVID-19 Genomics UK (COG-UK) Consortium | Jacquelyn Wynn, Mairead Hyland, The Lighthouse Lab in Alderley Park and Alex Alderton, Roberto Amato, Jeffrey Barrett, Sonia Goncalves, Ewan Harrison, David K. Jackson, Ian Johnston, Dominic Kwiatkowski, Cordelia Langford, John Sillitoe on behalf of the Wellcome Sanger Institute COVID-19 Surveillance Team |
| EPI_ISL_2605342 | Lighthouse Lab in Glasgow | Wellcome Sanger Institute for the COVID-19 Genomics UK (COG-UK) Consortium | Harper VanSteenhouse, Yumi Kasai, David Gray, Carol Clugston, Anna Dominiczak and Alex Alderton, Roberto Amato, Jeffrey Barrett, Sonia Goncalves, Ewan Harrison, David K. Jackson, Ian Johnston, Dominic Kwiatkowski, Cordelia Langford, John Sillitoe on behalf of the Wellcome Sanger Institute COVID-19 Surveillance Team |
| EPI_ISL_2605344, EPI_ISL_2605345, EPI_ISL_2605347, EPI_ISL_2605348, EPI_ISL_2605350, EPI_ISL_2605351, EPI_ISL_2605353, EPI_ISL_2605354 | Lighthouse Lab in Alderley Park | Wellcome Sanger Institute for the COVID-19 Genomics UK (COG-UK) Consortium | Jacquelyn Wynn, Mairead Hyland, The Lighthouse Lab in Alderley Park and Alex Alderton, Roberto Amato, Jeffrey Barrett, Sonia Goncalves, Ewan Harrison, David K. Jackson, Ian Johnston, Dominic Kwiatkowski, Cordelia Langford, John Sillitoe on behalf of the Wellcome Sanger Institute COVID-19 Surveillance Team |
| EPI_ISL_2605355 | Lighthouse Lab in Glasgow | Wellcome Sanger Institute for the COVID-19 Genomics UK (COG-UK) Consortium | Harper VanSteenhouse, Yumi Kasai, David Gray, Carol Clugston, Anna Dominiczak and Alex Alderton, Roberto Amato, Jeffrey Barrett, Sonia Goncalves, Ewan Harrison, David K. Jackson, Ian Johnston, Dominic Kwiatkowski, Cordelia Langford, John Sillitoe on behalf of the Wellcome Sanger Institute COVID-19 Surveillance Team |
| EPI_ISL_2605356 | Lab voor klinische biologie | Lab voor klinische biologie | Marija Janevska, Hannelore Hamerlinck, Bruno Verhasselt |
| EPI_ISL_2605357 | Lighthouse Lab in Alderley Park | Wellcome Sanger Institute for the COVID-19 Genomics UK (COG-UK) Consortium | Jacquelyn Wynn, Mairead Hyland, The Lighthouse Lab in Alderley Park and Alex Alderton, Roberto Amato, Jeffrey Barrett, Sonia Goncalves, Ewan Harrison, David K. Jackson, Ian Johnston, Dominic Kwiatkowski, Cordelia Langford, John Sillitoe on behalf of the Wellcome Sanger Institute COVID-19 Surveillance Team |
| EPI_ISL_2605358 | Lighthouse Lab in Glasgow | Wellcome Sanger Institute for the COVID-19 Genomics UK (COG-UK) Consortium | Harper VanSteenhouse, Yumi Kasai, David Gray, Carol Clugston, Anna Dominiczak and Alex Alderton, Roberto Amato, Jeffrey Barrett, Sonia Goncalves, Ewan Harrison, David K. Jackson, Ian Johnston, Dominic Kwiatkowski, Cordelia Langford, John Sillitoe on behalf of the Wellcome Sanger Institute COVID-19 Surveillance Team |
| EPI_ISL_2605360, EPI_ISL_2605361 | Lighthouse Lab in Alderley Park | Wellcome Sanger Institute for the COVID-19 Genomics UK (COG-UK) Consortium | Jacquelyn Wynn, Mairead Hyland, The Lighthouse Lab in Alderley Park and Alex Alderton, Roberto Amato, Jeffrey Barrett, Sonia Goncalves, Ewan Harrison, David K. Jackson, Ian Johnston, Dominic Kwiatkowski, Cordelia Langford, John Sillitoe on behalf of the Wellcome Sanger Institute COVID-19 Surveillance Team |
| EPI_ISL_2605363 | Lighthouse Lab in Glasgow | Wellcome Sanger Institute for the COVID-19 Genomics UK (COG-UK) Consortium | Harper VanSteenhouse, Yumi Kasai, David Gray, Carol Clugston, Anna Dominiczak and Alex Alderton, Roberto Amato, Jeffrey Barrett, Sonia Goncalves, Ewan Harrison, David K. Jackson, Ian Johnston, Dominic Kwiatkowski, Cordelia Langford, John Sillitoe on behalf of the Wellcome Sanger Institute COVID-19 Surveillance Team |
| EPI_ISL_2605364, EPI_ISL_2605366, EPI_ISL_2605367 | Lighthouse Lab in Alderley Park | Wellcome Sanger Institute for the COVID-19 Genomics UK (COG-UK) Consortium | Jacquelyn Wynn, Mairead Hyland, The Lighthouse Lab in Alderley Park and Alex Alderton, Roberto Amato, Jeffrey Barrett, Sonia Goncalves, Ewan Harrison, David K. Jackson, Ian Johnston, Dominic Kwiatkowski, Cordelia Langford, John Sillitoe on behalf of the Wellcome Sanger Institute COVID-19 Surveillance Team |
| EPI_ISL_2605369, EPI_ISL_2605370 | Lighthouse Lab in Glasgow | Wellcome Sanger Institute for the COVID-19 Genomics UK (COG-UK) Consortium | Harper VanSteenhouse, Yumi Kasai, David Gray, Carol Clugston, Anna Dominiczak and Alex Alderton, Roberto Amato, Jeffrey Barrett, Sonia Goncalves, Ewan Harrison, David K. Jackson, Ian Johnston, Dominic Kwiatkowski, Cordelia Langford, John Sillitoe on behalf of the Wellcome Sanger Institute COVID-19 Surveillance Team |
| EPI_ISL_2605372, EPI_ISL_2605373, EPI_ISL_2605375, EPI_ISL_2605376, EPI_ISL_2605378, EPI_ISL_2605379 | Lighthouse Lab in Alderley Park | Wellcome Sanger Institute for the COVID-19 Genomics UK (COG-UK) Consortium | Jacquelyn Wynn, Mairead Hyland, The Lighthouse Lab in Alderley Park and Alex Alderton, Roberto Amato, Jeffrey Barrett, Sonia Goncalves, Ewan Harrison, David K. Jackson, Ian Johnston, Dominic Kwiatkowski, Cordelia Langford, John Sillitoe on behalf of the Wellcome Sanger Institute COVID-19 Surveillance Team |
| EPI_ISL_2605380 | Lighthouse Lab in Glasgow | Wellcome Sanger Institute for the COVID-19 Genomics UK (COG-UK) Consortium | Harper VanSteenhouse, Yumi Kasai, David Gray, Carol Clugston, Anna Dominiczak and Alex Alderton, Roberto Amato, Jeffrey Barrett, Sonia Goncalves, Ewan Harrison, David K. Jackson, Ian Johnston, Dominic Kwiatkowski, Cordelia Langford, John Sillitoe on behalf of the Wellcome Sanger Institute COVID-19 Surveillance Team |
| EPI_ISL_2605382 | Lighthouse Lab in Alderley Park | Wellcome Sanger Institute for the COVID-19 Genomics UK (COG-UK) Consortium | Jacquelyn Wynn, Mairead Hyland, The Lighthouse Lab in Alderley Park and Alex Alderton, Roberto Amato, Jeffrey Barrett, Sonia Goncalves, Ewan Harrison, David K. Jackson, Ian Johnston, Dominic Kwiatkowski, Cordelia Langford, John Sillitoe on behalf of the Wellcome Sanger Institute COVID-19 Surveillance Team |
| EPI_ISL_2605384, EPI_ISL_2605385 | Lighthouse Lab in Glasgow | Wellcome Sanger Institute for the COVID-19 Genomics UK (COG-UK) Consortium | Harper VanSteenhouse, Yumi Kasai, David Gray, Carol Clugston, Anna Dominiczak and Alex Alderton, Roberto Amato, Jeffrey Barrett, Sonia Goncalves, Ewan Harrison, David K. Jackson, Ian Johnston, Dominic Kwiatkowski, Cordelia Langford, John Sillitoe on behalf of the Wellcome Sanger Institute COVID-19 Surveillance Team |
| EPI_ISL_2605387, EPI_ISL_2605388 | Lighthouse Lab in Alderley Park | Wellcome Sanger Institute for the COVID-19 Genomics UK (COG-UK) Consortium | Jacquelyn Wynn, Mairead Hyland, The Lighthouse Lab in Alderley Park and Alex Alderton, Roberto Amato, Jeffrey Barrett, Sonia Goncalves, Ewan Harrison, David K. Jackson, Ian Johnston, Dominic Kwiatkowski, Cordelia Langford, John Sillitoe on behalf of the Wellcome Sanger Institute COVID-19 Surveillance Team |
| EPI_ISL_2605390 | Lighthouse Lab in Glasgow | Wellcome Sanger Institute for the COVID-19 Genomics UK (COG-UK) Consortium | Harper VanSteenhouse, Yumi Kasai, David Gray, Carol Clugston, Anna Dominiczak and Alex Alderton, Roberto Amato, Jeffrey Barrett, Sonia Goncalves, Ewan Harrison, David K. Jackson, Ian Johnston, Dominic Kwiatkowski, Cordelia Langford, John Sillitoe on behalf of the Wellcome Sanger Institute COVID-19 Surveillance Team |
| EPI_ISL_2605391 | Lighthouse Lab in Alderley Park | Wellcome Sanger Institute for the COVID-19 Genomics UK (COG-UK) Consortium | Jacquelyn Wynn, Mairead Hyland, The Lighthouse Lab in Alderley Park and Alex Alderton, Roberto Amato, Jeffrey Barrett, Sonia Goncalves, Ewan Harrison, David K. Jackson, Ian Johnston, Dominic Kwiatkowski, Cordelia Langford, John Sillitoe on behalf of the Wellcome Sanger Institute COVID-19 Surveillance Team |
| EPI_ISL_2605392 | Lighthouse Lab in Glasgow | Wellcome Sanger Institute for the COVID-19 Genomics UK (COG-UK) Consortium | Harper VanSteenhouse, Yumi Kasai, David Gray, Carol Clugston, Anna Dominiczak and Alex Alderton, Roberto Amato, Jeffrey Barrett, Sonia Goncalves, Ewan Harrison, David K. Jackson, Ian Johnston, Dominic Kwiatkowski, Cordelia Langford, John Sillitoe on behalf of the Wellcome Sanger Institute COVID-19 Surveillance Team |
| EPI_ISL_2605394, EPI_ISL_2605395, EPI_ISL_2605397, EPI_ISL_2605398 | Lighthouse Lab in Alderley Park | Wellcome Sanger Institute for the COVID-19 Genomics UK (COG-UK) Consortium | Jacquelyn Wynn, Mairead Hyland, The Lighthouse Lab in Alderley Park and Alex Alderton, Roberto Amato, Jeffrey Barrett, Sonia Goncalves, Ewan Harrison, David K. Jackson, Ian Johnston, Dominic Kwiatkowski, Cordelia Langford, John Sillitoe on behalf of the Wellcome Sanger Institute COVID-19 Surveillance Team |
| EPI_ISL_2605400 | Lighthouse Lab in Glasgow | Wellcome Sanger Institute for the COVID-19 Genomics UK (COG-UK) Consortium | Harper VanSteenhouse, Yumi Kasai, David Gray, Carol Clugston, Anna Dominiczak and Alex Alderton, Roberto Amato, Jeffrey Barrett, Sonia Goncalves, Ewan Harrison, David K. Jackson, Ian Johnston, Dominic Kwiatkowski, Cordelia Langford, John Sillitoe on behalf of the Wellcome Sanger Institute COVID-19 Surveillance Team |
| EPI_ISL_2605401, EPI_ISL_2605403 | Lighthouse Lab in Alderley Park | Wellcome Sanger Institute for the COVID-19 Genomics UK (COG-UK) Consortium | Jacquelyn Wynn, Mairead Hyland, The Lighthouse Lab in Alderley Park and Alex Alderton, Roberto Amato, Jeffrey Barrett, Sonia Goncalves, Ewan Harrison, David K. Jackson, Ian Johnston, Dominic Kwiatkowski, Cordelia Langford, John Sillitoe on behalf of the Wellcome Sanger Institute COVID-19 Surveillance Team |
| EPI_ISL_2605404 | Lighthouse Lab in Glasgow | Wellcome Sanger Institute for the COVID-19 Genomics UK (COG-UK) Consortium | Harper VanSteenhouse, Yumi Kasai, David Gray, Carol Clugston, Anna Dominiczak and Alex Alderton, Roberto Amato, Jeffrey Barrett, Sonia Goncalves, Ewan Harrison, David K. Jackson, Ian Johnston, Dominic Kwiatkowski, Cordelia Langford, John Sillitoe on behalf of the Wellcome Sanger Institute COVID-19 Surveillance Team |
| EPI_ISL_2605406 | Lighthouse Lab in Alderley Park | Wellcome Sanger Institute for the COVID-19 Genomics UK (COG-UK) Consortium | Jacquelyn Wynn, Mairead Hyland, The Lighthouse Lab in Alderley Park and Alex Alderton, Roberto Amato, Jeffrey Barrett, Sonia Goncalves, Ewan Harrison, David K. Jackson, Ian Johnston, Dominic Kwiatkowski, Cordelia Langford, John Sillitoe on behalf of the Wellcome Sanger Institute COVID-19 Surveillance Team |
| EPI_ISL_2605407, EPI_ISL_2605409 | Lighthouse Lab in Glasgow | Wellcome Sanger Institute for the COVID-19 Genomics UK (COG-UK) Consortium | Harper VanSteenhouse, Yumi Kasai, David Gray, Carol Clugston, Anna Dominiczak and Alex Alderton, Roberto Amato, Jeffrey Barrett, Sonia Goncalves, Ewan Harrison, David K. Jackson, Ian Johnston, Dominic Kwiatkowski, Cordelia Langford, John Sillitoe on behalf of the Wellcome Sanger Institute COVID-19 Surveillance Team |
| EPI_ISL_2605410, EPI_ISL_2605412, EPI_ISL_2605414, EPI_ISL_2605415 | Lighthouse Lab in Alderley Park | Wellcome Sanger Institute for the COVID-19 Genomics UK (COG-UK) Consortium | Jacquelyn Wynn, Mairead Hyland, The Lighthouse Lab in Alderley Park and Alex Alderton, Roberto Amato, Jeffrey Barrett, Sonia Goncalves, Ewan Harrison, David K. Jackson, Ian Johnston, Dominic Kwiatkowski, Cordelia Langford, John Sillitoe on behalf of the Wellcome Sanger Institute COVID-19 Surveillance Team |
| EPI_ISL_2605416 | Lighthouse Lab in Glasgow | Wellcome Sanger Institute for the COVID-19 Genomics UK (COG-UK) Consortium | Harper VanSteenhouse, Yumi Kasai, David Gray, Carol Clugston, Anna Dominiczak and Alex Alderton, Roberto Amato, Jeffrey Barrett, Sonia Goncalves, Ewan Harrison, David K. Jackson, Ian Johnston, Dominic Kwiatkowski, Cordelia Langford, John Sillitoe on behalf of the Wellcome Sanger Institute COVID-19 Surveillance Team |
| EPI_ISL_2605418, EPI_ISL_2605420 | Lighthouse Lab in Alderley Park | Wellcome Sanger Institute for the COVID-19 Genomics UK (COG-UK) Consortium | Jacquelyn Wynn, Mairead Hyland, The Lighthouse Lab in Alderley Park and Alex Alderton, Roberto Amato, Jeffrey Barrett, Sonia Goncalves, Ewan Harrison, David K. Jackson, Ian Johnston, Dominic Kwiatkowski, Cordelia Langford, John Sillitoe on behalf of the Wellcome Sanger Institute COVID-19 Surveillance Team |
| EPI_ISL_2605421 | Lighthouse Lab in Glasgow | Wellcome Sanger Institute for the COVID-19 Genomics UK (COG-UK) Consortium | Harper VanSteenhouse, Yumi Kasai, David Gray, Carol Clugston, Anna Dominiczak and Alex Alderton, Roberto Amato, Jeffrey Barrett, Sonia Goncalves, Ewan Harrison, David K. Jackson, Ian Johnston, Dominic Kwiatkowski, Cordelia Langford, John Sillitoe on behalf of the Wellcome Sanger Institute COVID-19 Surveillance Team |
| EPI_ISL_2605422 | Lighthouse Lab in Alderley Park | Wellcome Sanger Institute for the COVID-19 Genomics UK (COG-UK) Consortium | Jacquelyn Wynn, Mairead Hyland, The Lighthouse Lab in Alderley Park and Alex Alderton, Roberto Amato, Jeffrey Barrett, Sonia Goncalves, Ewan Harrison, David K. Jackson, Ian Johnston, Dominic Kwiatkowski, Cordelia Langford, John Sillitoe on behalf of the Wellcome Sanger Institute COVID-19 Surveillance Team |
| EPI_ISL_2605423 | Lighthouse Lab in Glasgow | Wellcome Sanger Institute for the COVID-19 Genomics UK (COG-UK) Consortium | Harper VanSteenhouse, Yumi Kasai, David Gray, Carol Clugston, Anna Dominiczak and Alex Alderton, Roberto Amato, Jeffrey Barrett, Sonia Goncalves, Ewan Harrison, David K. Jackson, Ian Johnston, Dominic Kwiatkowski, Cordelia Langford, John Sillitoe on behalf of the Wellcome Sanger Institute COVID-19 Surveillance Team |
| EPI_ISL_2605424, EPI_ISL_2605425, EPI_ISL_2605426, EPI_ISL_2605427, EPI_ISL_2605428, EPI_ISL_2605429, EPI_ISL_2605430, EPI_ISL_2605432, EPI_ISL_2605433 | Lighthouse Lab in Alderley Park | Wellcome Sanger Institute for the COVID-19 Genomics UK (COG-UK) Consortium | Jacquelyn Wynn, Mairead Hyland, The Lighthouse Lab in Alderley Park and Alex Alderton, Roberto Amato, Jeffrey Barrett, Sonia Goncalves, Ewan Harrison, David K. Jackson, Ian Johnston, Dominic Kwiatkowski, Cordelia Langford, John Sillitoe on behalf of the Wellcome Sanger Institute COVID-19 Surveillance Team |

| | | | |
|---|---|---|---|
| EPI_ISL_2605435 | Lighthouse Lab in Glasgow | Wellcome Sanger Institute for the COVID-19 Genomics UK (COG-UK) Consortium | Harper VanSteenhouse, Yumi Kasai, David Gray, Carol Clugston, Anna Dominiczak and Alex Alderton, Roberto Amato, Jeffrey Barrett, Sonia Goncalves, Ewan Harrison, David K. Jackson, Ian Johnston, Dominic Kwiatkowski, Cordelia Langford, John Sillitoe on behalf of the Wellcome Sanger Institute COVID-19 Surveillance Team |
| EPI_ISL_2605436 | Lighthouse Lab in Alderley Park | Wellcome Sanger Institute for the COVID-19 Genomics UK (COG-UK) Consortium | Jacquelyn Wynn, Mairead Hyland, The Lighthouse Lab in Alderley Park and Alex Alderton, Roberto Amato, Jeffrey Barrett, Sonia Goncalves, Ewan Harrison, David K. Jackson, Ian Johnston, Dominic Kwiatkowski, Cordelia Langford, John Sillitoe on behalf of the Wellcome Sanger Institute COVID-19 Surveillance Team |
| EPI_ISL_2605437 | Lab voor klinische biologie | Lab voor klinische biologie | Marija Janevska, Hannelore Hamerlinck, Bruno Verhasselt |
| EPI_ISL_2605438, EPI_ISL_2605440, EPI_ISL_2605441, EPI_ISL_2605442 | Lighthouse Lab in Alderley Park | Wellcome Sanger Institute for the COVID-19 Genomics UK (COG-UK) Consortium | Jacquelyn Wynn, Mairead Hyland, The Lighthouse Lab in Alderley Park and Alex Alderton, Roberto Amato, Jeffrey Barrett, Sonia Goncalves, Ewan Harrison, David K. Jackson, Ian Johnston, Dominic Kwiatkowski, Cordelia Langford, John Sillitoe on behalf of the Wellcome Sanger Institute COVID-19 Surveillance Team |
| EPI_ISL_2605444 | Lighthouse Lab in Glasgow | Wellcome Sanger Institute for the COVID-19 Genomics UK (COG-UK) Consortium | Harper VanSteenhouse, Yumi Kasai, David Gray, Carol Clugston, Anna Dominiczak and Alex Alderton, Roberto Amato, Jeffrey Barrett, Sonia Goncalves, Ewan Harrison, David K. Jackson, Ian Johnston, Dominic Kwiatkowski, Cordelia Langford, John Sillitoe on behalf of the Wellcome Sanger Institute COVID-19 Surveillance Team |
| EPI_ISL_2605445, EPI_ISL_2605447, EPI_ISL_2605448, EPI_ISL_2605450, EPI_ISL_2605452, EPI_ISL_2605453, EPI_ISL_2605455 | Lighthouse Lab in Alderley Park | Wellcome Sanger Institute for the COVID-19 Genomics UK (COG-UK) Consortium | Jacquelyn Wynn, Mairead Hyland, The Lighthouse Lab in Alderley Park and Alex Alderton, Roberto Amato, Jeffrey Barrett, Sonia Goncalves, Ewan Harrison, David K. Jackson, Ian Johnston, Dominic Kwiatkowski, Cordelia Langford, John Sillitoe on behalf of the Wellcome Sanger Institute COVID-19 Surveillance Team |
| EPI_ISL_2605456 | Lighthouse Lab in Glasgow | Wellcome Sanger Institute for the COVID-19 Genomics UK (COG-UK) Consortium | Harper VanSteenhouse, Yumi Kasai, David Gray, Carol Clugston, Anna Dominiczak and Alex Alderton, Roberto Amato, Jeffrey Barrett, Sonia Goncalves, Ewan Harrison, David K. Jackson, Ian Johnston, Dominic Kwiatkowski, Cordelia Langford, John Sillitoe on behalf of the Wellcome Sanger Institute COVID-19 Surveillance Team |
| EPI_ISL_2605458 | Lighthouse Lab in Alderley Park | Wellcome Sanger Institute for the COVID-19 Genomics UK (COG-UK) Consortium | Jacquelyn Wynn, Mairead Hyland, The Lighthouse Lab in Alderley Park and Alex Alderton, Roberto Amato, Jeffrey Barrett, Sonia Goncalves, Ewan Harrison, David K. Jackson, Ian Johnston, Dominic Kwiatkowski, Cordelia Langford, John Sillitoe on behalf of the Wellcome Sanger Institute COVID-19 Surveillance Team |
| EPI_ISL_2605460 | Lighthouse Lab in Glasgow | Wellcome Sanger Institute for the COVID-19 Genomics UK (COG-UK) Consortium | Harper VanSteenhouse, Yumi Kasai, David Gray, Carol Clugston, Anna Dominiczak and Alex Alderton, Roberto Amato, Jeffrey Barrett, Sonia Goncalves, Ewan Harrison, David K. Jackson, Ian Johnston, Dominic Kwiatkowski, Cordelia Langford, John Sillitoe on behalf of the Wellcome Sanger Institute COVID-19 Surveillance Team |
| EPI_ISL_2605461, EPI_ISL_2605463, EPI_ISL_2605464, EPI_ISL_2605465 | Lighthouse Lab in Alderley Park | Wellcome Sanger Institute for the COVID-19 Genomics UK (COG-UK) Consortium | Jacquelyn Wynn, Mairead Hyland, The Lighthouse Lab in Alderley Park and Alex Alderton, Roberto Amato, Jeffrey Barrett, Sonia Goncalves, Ewan Harrison, David K. Jackson, Ian Johnston, Dominic Kwiatkowski, Cordelia Langford, John Sillitoe on behalf of the Wellcome Sanger Institute COVID-19 Surveillance Team |
| EPI_ISL_2605467, EPI_ISL_2605468 | Lighthouse Lab in Glasgow | Wellcome Sanger Institute for the COVID-19 Genomics UK (COG-UK) Consortium | Harper VanSteenhouse, Yumi Kasai, David Gray, Carol Clugston, Anna Dominiczak and Alex Alderton, Roberto Amato, Jeffrey Barrett, Sonia Goncalves, Ewan Harrison, David K. Jackson, Ian Johnston, Dominic Kwiatkowski, Cordelia Langford, John Sillitoe on behalf of the Wellcome Sanger Institute COVID-19 Surveillance Team |
| EPI_ISL_2605470, EPI_ISL_2605471, EPI_ISL_2605473, EPI_ISL_2605475, EPI_ISL_2605476, EPI_ISL_2605478, EPI_ISL_2605479, EPI_ISL_2605481 | Lighthouse Lab in Alderley Park | Wellcome Sanger Institute for the COVID-19 Genomics UK (COG-UK) Consortium | Jacquelyn Wynn, Mairead Hyland, The Lighthouse Lab in Alderley Park and Alex Alderton, Roberto Amato, Jeffrey Barrett, Sonia Goncalves, Ewan Harrison, David K. Jackson, Ian Johnston, Dominic Kwiatkowski, Cordelia Langford, John Sillitoe on behalf of the Wellcome Sanger Institute COVID-19 Surveillance Team |
| EPI_ISL_2605482 | Lighthouse Lab in Glasgow | Wellcome Sanger Institute for the COVID-19 Genomics UK (COG-UK) Consortium | Harper VanSteenhouse, Yumi Kasai, David Gray, Carol Clugston, Anna Dominiczak and Alex Alderton, Roberto Amato, Jeffrey Barrett, Sonia Goncalves, Ewan Harrison, David K. Jackson, Ian Johnston, Dominic Kwiatkowski, Cordelia Langford, John Sillitoe on behalf of the Wellcome Sanger Institute COVID-19 Surveillance Team |
| EPI_ISL_2605484, EPI_ISL_2605485, EPI_ISL_2605487, EPI_ISL_2605488, EPI_ISL_2605490, EPI_ISL_2605491, EPI_ISL_2605493, EPI_ISL_2605496, EPI_ISL_2605498 | Lighthouse Lab in Alderley Park | Wellcome Sanger Institute for the COVID-19 Genomics UK (COG-UK) Consortium | Jacquelyn Wynn, Mairead Hyland, The Lighthouse Lab in Alderley Park and Alex Alderton, Roberto Amato, Jeffrey Barrett, Sonia Goncalves, Ewan Harrison, David K. Jackson, Ian Johnston, Dominic Kwiatkowski, Cordelia Langford, John Sillitoe on behalf of the Wellcome Sanger Institute COVID-19 Surveillance Team |
| EPI_ISL_2605499 | Lighthouse Lab in Glasgow | Wellcome Sanger Institute for the COVID-19 Genomics UK (COG-UK) Consortium | Harper VanSteenhouse, Yumi Kasai, David Gray, Carol Clugston, Anna Dominiczak and Alex Alderton, Roberto Amato, Jeffrey Barrett, Sonia Goncalves, Ewan Harrison, David K. Jackson, Ian Johnston, Dominic Kwiatkowski, Cordelia Langford, John Sillitoe on behalf of the Wellcome Sanger Institute COVID-19 Surveillance Team |
| EPI_ISL_2605501 | Lighthouse Lab in Alderley Park | Wellcome Sanger Institute for the COVID-19 Genomics UK (COG-UK) Consortium | Jacquelyn Wynn, Mairead Hyland, The Lighthouse Lab in Alderley Park and Alex Alderton, Roberto Amato, Jeffrey Barrett, Sonia Goncalves, Ewan Harrison, David K. Jackson, Ian Johnston, Dominic Kwiatkowski, Cordelia Langford, John Sillitoe on behalf of the Wellcome Sanger Institute COVID-19 Surveillance Team |
| EPI_ISL_2605502 | Lab voor klinische biologie | Lab voor klinische biologie | Marija Janevska, Hannelore Hamerlinck, Bruno Verhasselt |
| EPI_ISL_2605504 | Lighthouse Lab in Alderley Park | Wellcome Sanger Institute for the COVID-19 Genomics UK (COG-UK) Consortium | Jacquelyn Wynn, Mairead Hyland, The Lighthouse Lab in Alderley Park and Alex Alderton, Roberto Amato, Jeffrey Barrett, Sonia Goncalves, Ewan Harrison, David K. Jackson, Ian Johnston, Dominic Kwiatkowski, Cordelia Langford, John Sillitoe on behalf of the Wellcome Sanger Institute COVID-19 Surveillance Team |
| EPI_ISL_2605505 | Lighthouse Lab in Glasgow | Wellcome Sanger Institute for the COVID-19 Genomics UK (COG-UK) Consortium | Harper VanSteenhouse, Yumi Kasai, David Gray, Carol Clugston, Anna Dominiczak and Alex Alderton, Roberto Amato, Jeffrey Barrett, Sonia Goncalves, Ewan Harrison, David K. Jackson, Ian Johnston, Dominic Kwiatkowski, Cordelia Langford, John Sillitoe on behalf of the Wellcome Sanger Institute COVID-19 Surveillance Team |
| EPI_ISL_2605507, EPI_ISL_2605508, EPI_ISL_2605510, EPI_ISL_2605511, EPI_ISL_2605513, EPI_ISL_2605515, EPI_ISL_2605516, EPI_ISL_2605517, EPI_ISL_2605519, EPI_ISL_2605520, EPI_ISL_2605522, EPI_ISL_2605523, EPI_ISL_2605525, EPI_ISL_2605526, EPI_ISL_2605528, EPI_ISL_2605529, EPI_ISL_2605531, EPI_ISL_2605533, EPI_ISL_2605534, EPI_ISL_2605536, EPI_ISL_2605537, EPI_ISL_2605539, EPI_ISL_2605540, EPI_ISL_2605542, EPI_ISL_2605543 | | | |
| see above | Lighthouse Lab in Alderley Park | Wellcome Sanger Institute for the COVID-19 Genomics UK (COG-UK) Consortium | Jacquelyn Wynn, Mairead Hyland, The Lighthouse Lab in Alderley Park and Alex Alderton, Roberto Amato, Jeffrey Barrett, Sonia Goncalves, Ewan Harrison, David K. Jackson, Ian Johnston, Dominic Kwiatkowski, Cordelia Langford, John Sillitoe on behalf of the Wellcome Sanger Institute COVID-19 Surveillance Team |
| EPI_ISL_2605545 | Lighthouse Lab in Glasgow | Wellcome Sanger Institute for the COVID-19 Genomics UK (COG-UK) Consortium | Harper VanSteenhouse, Yumi Kasai, David Gray, Carol Clugston, Anna Dominiczak and Alex Alderton, Roberto Amato, Jeffrey Barrett, Sonia Goncalves, Ewan Harrison, David K. Jackson, Ian Johnston, Dominic Kwiatkowski, Cordelia Langford, John Sillitoe on behalf of the Wellcome Sanger Institute COVID-19 Surveillance Team |
| EPI_ISL_2605548, EPI_ISL_2605549, EPI_ISL_2605551, EPI_ISL_2605552 | Lighthouse Lab in Alderley Park | Wellcome Sanger Institute for the COVID-19 Genomics UK (COG-UK) Consortium | Jacquelyn Wynn, Mairead Hyland, The Lighthouse Lab in Alderley Park and Alex Alderton, Roberto Amato, Jeffrey Barrett, Sonia Goncalves, Ewan Harrison, David K. Jackson, Ian Johnston, Dominic Kwiatkowski, Cordelia Langford, John Sillitoe on behalf of the Wellcome Sanger Institute COVID-19 Surveillance Team |
| EPI_ISL_2605553 | Lab voor klinische biologie | Lab voor klinische biologie | Marija Janevska, Hannelore Hamerlinck, Bruno Verhasselt |
| EPI_ISL_2605554, EPI_ISL_2605555 | Lighthouse Lab in Glasgow | Wellcome Sanger Institute for the COVID-19 Genomics UK (COG-UK) Consortium | Harper VanSteenhouse, Yumi Kasai, David Gray, Carol Clugston, Anna Dominiczak and Alex Alderton, Roberto Amato, Jeffrey Barrett, Sonia Goncalves, Ewan Harrison, David K. Jackson, Ian Johnston, Dominic Kwiatkowski, Cordelia Langford, John Sillitoe on behalf of the Wellcome Sanger Institute COVID-19 Surveillance Team |
| EPI_ISL_2605558, EPI_ISL_2605560, EPI_ISL_2605561 | Lighthouse Lab in Alderley Park | Wellcome Sanger Institute for the COVID-19 Genomics UK (COG-UK) Consortium | Jacquelyn Wynn, Mairead Hyland, The Lighthouse Lab in Alderley Park and Alex Alderton, Roberto Amato, Jeffrey Barrett, Sonia Goncalves, Ewan Harrison, David K. Jackson, Ian Johnston, Dominic Kwiatkowski, Cordelia Langford, John Sillitoe on behalf of the Wellcome Sanger Institute COVID-19 Surveillance Team |
| EPI_ISL_2605563 | Lighthouse Lab in Glasgow | Wellcome Sanger Institute for the COVID-19 Genomics UK (COG-UK) Consortium | Harper VanSteenhouse, Yumi Kasai, David Gray, Carol Clugston, Anna Dominiczak and Alex Alderton, Roberto Amato, Jeffrey Barrett, Sonia Goncalves, Ewan Harrison, David K. Jackson, Ian Johnston, Dominic Kwiatkowski, Cordelia Langford, John Sillitoe on behalf of the Wellcome Sanger Institute COVID-19 Surveillance Team |
| EPI_ISL_2605564, EPI_ISL_2605566, EPI_ISL_2605567, EPI_ISL_2605569, EPI_ISL_2605570, EPI_ISL_2605572, EPI_ISL_2605573 | Lighthouse Lab in Alderley Park | Wellcome Sanger Institute for the COVID-19 Genomics UK (COG-UK) Consortium | Jacquelyn Wynn, Mairead Hyland, The Lighthouse Lab in Alderley Park and Alex Alderton, Roberto Amato, Jeffrey Barrett, Sonia Goncalves, Ewan Harrison, David K. Jackson, Ian Johnston, Dominic Kwiatkowski, Cordelia Langford, John Sillitoe on behalf of the Wellcome Sanger Institute COVID-19 Surveillance Team |
| EPI_ISL_2605574 | Lighthouse Lab in Glasgow | Wellcome Sanger Institute for the COVID-19 Genomics UK (COG-UK) Consortium | Harper VanSteenhouse, Yumi Kasai, David Gray, Carol Clugston, Anna Dominiczak and Alex Alderton, Roberto Amato, Jeffrey Barrett, Sonia Goncalves, Ewan Harrison, David K. Jackson, Ian Johnston, Dominic Kwiatkowski, Cordelia Langford, John Sillitoe on behalf of the Wellcome Sanger Institute COVID-19 Surveillance Team |
| EPI_ISL_2605576, EPI_ISL_2605578 | Lighthouse Lab in Alderley Park | Wellcome Sanger Institute for the COVID-19 Genomics UK (COG-UK) Consortium | Jacquelyn Wynn, Mairead Hyland, The Lighthouse Lab in Alderley Park and Alex Alderton, Roberto Amato, Jeffrey Barrett, Sonia Goncalves, Ewan Harrison, David K. Jackson, Ian Johnston, Dominic Kwiatkowski, Cordelia Langford, John Sillitoe on behalf of the Wellcome Sanger Institute COVID-19 Surveillance Team |
| EPI_ISL_2605579 | Lab voor klinische biologie | Lab voor klinische biologie | Marija Janevska, Hannelore Hamerlinck, Bruno Verhasselt |
| EPI_ISL_2605580, EPI_ISL_2605581, EPI_ISL_2605583, EPI_ISL_2605584, EPI_ISL_2605586, EPI_ISL_2605588, EPI_ISL_2605589, EPI_ISL_2605591, | Lighthouse Lab in Alderley Park | Wellcome Sanger Institute for the COVID-19 Genomics UK (COG-UK) Consortium | Jacquelyn Wynn, Mairead Hyland, The Lighthouse Lab in Alderley Park and Alex Alderton, Roberto Amato, Jeffrey Barrett, Sonia Goncalves, Ewan Harrison, David K. Jackson, Ian Johnston, Dominic Kwiatkowski, Cordelia Langford, John Sillitoe on behalf of the Wellcome Sanger Institute COVID-19 Surveillance Team |

| | | | |
|---|---|---|---|
| EPI_ISL_2605592 | | | |
| EPI_ISL_2605594, EPI_ISL_2605595 | Lighthouse Lab in Glasgow | Wellcome Sanger Institute for the COVID-19 Genomics UK (COG-UK) Consortium | Harper VanSteenhouse, Yumi Kasai, David Gray, Carol Clugston, Anna Dominiczak and Alex Alderton, Roberto Amato, Jeffrey Barrett, Sonia Goncalves, Ewan Harrison, David K. Jackson, Ian Johnston, Dominic Kwiatkowski, Cordelia Langford, John Sillitoe on behalf of the Wellcome Sanger Institute COVID-19 Surveillance Team |
| EPI_ISL_2605596 | Lab voor klinische biologie | Lab voor klinische biologie | Marija Janevska, Hannelore Hamerlinck, Bruno Verhasselt |
| EPI_ISL_2605597, EPI_ISL_2605600, EPI_ISL_2605601, EPI_ISL_2605603, EPI_ISL_2605606, EPI_ISL_2605607, EPI_ISL_2605609, EPI_ISL_2605611, EPI_ISL_2605612, EPI_ISL_2605614, EPI_ISL_2605615, EPI_ISL_2605617, EPI_ISL_2605618, EPI_ISL_2605620, EPI_ISL_2605621 | | | |
| see above | Lighthouse Lab in Alderley Park | Wellcome Sanger Institute for the COVID-19 Genomics UK (COG-UK) Consortium | Jacquelyn Wynn, Mairead Hyland, The Lighthouse Lab in Alderley Park and Alex Alderton, Roberto Amato, Jeffrey Barrett, Sonia Goncalves, Ewan Harrison, David K. Jackson, Ian Johnston, Dominic Kwiatkowski, Cordelia Langford, John Sillitoe on behalf of the Wellcome Sanger Institute COVID-19 Surveillance Team |
| EPI_ISL_2605622 | Lab voor klinische biologie | Lab voor klinische biologie | Marija Janevska, Hannelore Hamerlinck, Bruno Verhasselt |
| EPI_ISL_2605623, EPI_ISL_2605624, EPI_ISL_2605626, EPI_ISL_2605627 | Lighthouse Lab in Alderley Park | Wellcome Sanger Institute for the COVID-19 Genomics UK (COG-UK) Consortium | Jacquelyn Wynn, Mairead Hyland, The Lighthouse Lab in Alderley Park and Alex Alderton, Roberto Amato, Jeffrey Barrett, Sonia Goncalves, Ewan Harrison, David K. Jackson, Ian Johnston, Dominic Kwiatkowski, Cordelia Langford, John Sillitoe on behalf of the Wellcome Sanger Institute COVID-19 Surveillance Team |
| EPI_ISL_2605629 | Lighthouse Lab in Glasgow | Wellcome Sanger Institute for the COVID-19 Genomics UK (COG-UK) Consortium | Harper VanSteenhouse, Yumi Kasai, David Gray, Carol Clugston, Anna Dominiczak and Alex Alderton, Roberto Amato, Jeffrey Barrett, Sonia Goncalves, Ewan Harrison, David K. Jackson, Ian Johnston, Dominic Kwiatkowski, Cordelia Langford, John Sillitoe on behalf of the Wellcome Sanger Institute COVID-19 Surveillance Team |
| EPI_ISL_2605630, EPI_ISL_2605632, EPI_ISL_2605634, EPI_ISL_2605635, EPI_ISL_2605637 | Lighthouse Lab in Alderley Park | Wellcome Sanger Institute for the COVID-19 Genomics UK (COG-UK) Consortium | Jacquelyn Wynn, Mairead Hyland, The Lighthouse Lab in Alderley Park and Alex Alderton, Roberto Amato, Jeffrey Barrett, Sonia Goncalves, Ewan Harrison, David K. Jackson, Ian Johnston, Dominic Kwiatkowski, Cordelia Langford, John Sillitoe on behalf of the Wellcome Sanger Institute COVID-19 Surveillance Team |
| EPI_ISL_2605638 | Lighthouse Lab in Glasgow | Wellcome Sanger Institute for the COVID-19 Genomics UK (COG-UK) Consortium | Harper VanSteenhouse, Yumi Kasai, David Gray, Carol Clugston, Anna Dominiczak and Alex Alderton, Roberto Amato, Jeffrey Barrett, Sonia Goncalves, Ewan Harrison, David K. Jackson, Ian Johnston, Dominic Kwiatkowski, Cordelia Langford, John Sillitoe on behalf of the Wellcome Sanger Institute COVID-19 Surveillance Team |
| EPI_ISL_2605639, EPI_ISL_2605640 | Lighthouse Lab in Alderley Park | Wellcome Sanger Institute for the COVID-19 Genomics UK (COG-UK) Consortium | Jacquelyn Wynn, Mairead Hyland, The Lighthouse Lab in Alderley Park and Alex Alderton, Roberto Amato, Jeffrey Barrett, Sonia Goncalves, Ewan Harrison, David K. Jackson, Ian Johnston, Dominic Kwiatkowski, Cordelia Langford, John Sillitoe on behalf of the Wellcome Sanger Institute COVID-19 Surveillance Team |
| EPI_ISL_2605642 | Lighthouse Lab in Glasgow | Wellcome Sanger Institute for the COVID-19 Genomics UK (COG-UK) Consortium | Harper VanSteenhouse, Yumi Kasai, David Gray, Carol Clugston, Anna Dominiczak and Alex Alderton, Roberto Amato, Jeffrey Barrett, Sonia Goncalves, Ewan Harrison, David K. Jackson, Ian Johnston, Dominic Kwiatkowski, Cordelia Langford, John Sillitoe on behalf of the Wellcome Sanger Institute COVID-19 Surveillance Team |
| EPI_ISL_2605645, EPI_ISL_2605646 | Lighthouse Lab in Alderley Park | Wellcome Sanger Institute for the COVID-19 Genomics UK (COG-UK) Consortium | Jacquelyn Wynn, Mairead Hyland, The Lighthouse Lab in Alderley Park and Alex Alderton, Roberto Amato, Jeffrey Barrett, Sonia Goncalves, Ewan Harrison, David K. Jackson, Ian Johnston, Dominic Kwiatkowski, Cordelia Langford, John Sillitoe on behalf of the Wellcome Sanger Institute COVID-19 Surveillance Team |
| EPI_ISL_2605647 | Lab voor klinische biologie | Lab voor klinische biologie | Marija Janevska, Hannelore Hamerlinck, Bruno Verhasselt |
| EPI_ISL_2605648, EPI_ISL_2605649, EPI_ISL_2605651, EPI_ISL_2605652, EPI_ISL_2605654, EPI_ISL_2605655 | Lighthouse Lab in Alderley Park | Wellcome Sanger Institute for the COVID-19 Genomics UK (COG-UK) Consortium | Jacquelyn Wynn, Mairead Hyland, The Lighthouse Lab in Alderley Park and Alex Alderton, Roberto Amato, Jeffrey Barrett, Sonia Goncalves, Ewan Harrison, David K. Jackson, Ian Johnston, Dominic Kwiatkowski, Cordelia Langford, John Sillitoe on behalf of the Wellcome Sanger Institute COVID-19 Surveillance Team |
| EPI_ISL_2605656 | Lighthouse Lab in Glasgow | Wellcome Sanger Institute for the COVID-19 Genomics UK (COG-UK) Consortium | Harper VanSteenhouse, Yumi Kasai, David Gray, Carol Clugston, Anna Dominiczak and Alex Alderton, Roberto Amato, Jeffrey Barrett, Sonia Goncalves, Ewan Harrison, David K. Jackson, Ian Johnston, Dominic Kwiatkowski, Cordelia Langford, John Sillitoe on behalf of the Wellcome Sanger Institute COVID-19 Surveillance Team |
| EPI_ISL_2605657, EPI_ISL_2605658, EPI_ISL_2605659, EPI_ISL_2605660, EPI_ISL_2605661, EPI_ISL_2605662, EPI_ISL_2605663, EPI_ISL_2605664, EPI_ISL_2605665, EPI_ISL_2605666, EPI_ISL_2605667, EPI_ISL_2605668, EPI_ISL_2605670, EPI_ISL_2605671, EPI_ISL_2605672, EPI_ISL_2605673, EPI_ISL_2605674, EPI_ISL_2605676, EPI_ISL_2605677, EPI_ISL_2605678, EPI_ISL_2605679, EPI_ISL_2605680, EPI_ISL_2605681, EPI_ISL_2605682, EPI_ISL_2605683, EPI_ISL_2605684, EPI_ISL_2605685, EPI_ISL_2605687, EPI_ISL_2605688, EPI_ISL_2605689, EPI_ISL_2605690, EPI_ISL_2605691, EPI_ISL_2605692, EPI_ISL_2605693, EPI_ISL_2605694, EPI_ISL_2605695, EPI_ISL_2605696, EPI_ISL_2605697, EPI_ISL_2605698, EPI_ISL_2605699, EPI_ISL_2605700, EPI_ISL_2605701, EPI_ISL_2605702, EPI_ISL_2605703, EPI_ISL_2605704, EPI_ISL_2605705, EPI_ISL_2605706, EPI_ISL_2605707, EPI_ISL_2605708, EPI_ISL_2605709 | | | |
| see above | Lighthouse Lab in Alderley Park | Wellcome Sanger Institute for the COVID-19 Genomics UK (COG-UK) Consortium | Jacquelyn Wynn, Mairead Hyland, The Lighthouse Lab in Alderley Park and Alex Alderton, Roberto Amato, Jeffrey Barrett, Sonia Goncalves, Ewan Harrison, David K. Jackson, Ian Johnston, Dominic Kwiatkowski, Cordelia Langford, John Sillitoe on behalf of the Wellcome Sanger Institute COVID-19 Surveillance Team |
| EPI_ISL_2605710 | Lighthouse Lab in Glasgow | Wellcome Sanger Institute for the COVID-19 Genomics UK (COG-UK) Consortium | Harper VanSteenhouse, Yumi Kasai, David Gray, Carol Clugston, Anna Dominiczak and Alex Alderton, Roberto Amato, Jeffrey Barrett, Sonia Goncalves, Ewan Harrison, David K. Jackson, Ian Johnston, Dominic Kwiatkowski, Cordelia Langford, John Sillitoe on behalf of the Wellcome Sanger Institute COVID-19 Surveillance Team |
| EPI_ISL_2605711, EPI_ISL_2605712, EPI_ISL_2605713, EPI_ISL_2605714, EPI_ISL_2605715 | Lighthouse Lab in Alderley Park | Wellcome Sanger Institute for the COVID-19 Genomics UK (COG-UK) Consortium | Jacquelyn Wynn, Mairead Hyland, The Lighthouse Lab in Alderley Park and Alex Alderton, Roberto Amato, Jeffrey Barrett, Sonia Goncalves, Ewan Harrison, David K. Jackson, Ian Johnston, Dominic Kwiatkowski, Cordelia Langford, John Sillitoe on behalf of the Wellcome Sanger Institute COVID-19 Surveillance Team |
| EPI_ISL_2605716 | Lighthouse Lab in Glasgow | Wellcome Sanger Institute for the COVID-19 Genomics UK (COG-UK) Consortium | Harper VanSteenhouse, Yumi Kasai, David Gray, Carol Clugston, Anna Dominiczak and Alex Alderton, Roberto Amato, Jeffrey Barrett, Sonia Goncalves, Ewan Harrison, David K. Jackson, Ian Johnston, Dominic Kwiatkowski, Cordelia Langford, John Sillitoe on behalf of the Wellcome Sanger Institute COVID-19 Surveillance Team |
| EPI_ISL_2605717 | Lighthouse Lab in Alderley Park | Wellcome Sanger Institute for the COVID-19 Genomics UK (COG-UK) Consortium | Jacquelyn Wynn, Mairead Hyland, The Lighthouse Lab in Alderley Park and Alex Alderton, Roberto Amato, Jeffrey Barrett, Sonia Goncalves, Ewan Harrison, David K. Jackson, Ian Johnston, Dominic Kwiatkowski, Cordelia Langford, John Sillitoe on behalf of the Wellcome Sanger Institute COVID-19 Surveillance Team |
| EPI_ISL_2605718 | Lighthouse Lab in Glasgow | Wellcome Sanger Institute for the COVID-19 Genomics UK (COG-UK) Consortium | Harper VanSteenhouse, Yumi Kasai, David Gray, Carol Clugston, Anna Dominiczak and Alex Alderton, Roberto Amato, Jeffrey Barrett, Sonia Goncalves, Ewan Harrison, David K. Jackson, Ian Johnston, Dominic Kwiatkowski, Cordelia Langford, John Sillitoe on behalf of the Wellcome Sanger Institute COVID-19 Surveillance Team |
| EPI_ISL_2605719, EPI_ISL_2605720, EPI_ISL_2605721 | Lighthouse Lab in Alderley Park | Wellcome Sanger Institute for the COVID-19 Genomics UK (COG-UK) Consortium | Jacquelyn Wynn, Mairead Hyland, The Lighthouse Lab in Alderley Park and Alex Alderton, Roberto Amato, Jeffrey Barrett, Sonia Goncalves, Ewan Harrison, David K. Jackson, Ian Johnston, Dominic Kwiatkowski, Cordelia Langford, John Sillitoe on behalf of the Wellcome Sanger Institute COVID-19 Surveillance Team |
| EPI_ISL_2605722 | Lighthouse Lab in Glasgow | Wellcome Sanger Institute for the COVID-19 Genomics UK (COG-UK) Consortium | Harper VanSteenhouse, Yumi Kasai, David Gray, Carol Clugston, Anna Dominiczak and Alex Alderton, Roberto Amato, Jeffrey Barrett, Sonia Goncalves, Ewan Harrison, David K. Jackson, Ian Johnston, Dominic Kwiatkowski, Cordelia Langford, John Sillitoe on behalf of the Wellcome Sanger Institute COVID-19 Surveillance Team |
| EPI_ISL_2605723, EPI_ISL_2605725, EPI_ISL_2605726, EPI_ISL_2605727, EPI_ISL_2605728, EPI_ISL_2605729, EPI_ISL_2605730, EPI_ISL_2605731, EPI_ISL_2605732, EPI_ISL_2605733, EPI_ISL_2605735, EPI_ISL_2605736, EPI_ISL_2605737 | | | |
| see above | Lighthouse Lab in Alderley Park | Wellcome Sanger Institute for the COVID-19 Genomics UK (COG-UK) Consortium | Jacquelyn Wynn, Mairead Hyland, The Lighthouse Lab in Alderley Park and Alex Alderton, Roberto Amato, Jeffrey Barrett, Sonia Goncalves, Ewan Harrison, David K. Jackson, Ian Johnston, Dominic Kwiatkowski, Cordelia Langford, John Sillitoe on behalf of the Wellcome Sanger Institute COVID-19 Surveillance Team |
| EPI_ISL_2605738 | Lighthouse Lab in Glasgow | Wellcome Sanger Institute for the COVID-19 Genomics UK (COG-UK) Consortium | Harper VanSteenhouse, Yumi Kasai, David Gray, Carol Clugston, Anna Dominiczak and Alex Alderton, Roberto Amato, Jeffrey Barrett, Sonia Goncalves, Ewan Harrison, David K. Jackson, Ian Johnston, Dominic Kwiatkowski, Cordelia Langford, John Sillitoe on behalf of the Wellcome Sanger Institute COVID-19 Surveillance Team |
| EPI_ISL_2605739, EPI_ISL_2605740, EPI_ISL_2605741, EPI_ISL_2605742, EPI_ISL_2605743, EPI_ISL_2605745, EPI_ISL_2605746, EPI_ISL_2605747, EPI_ISL_2605748, EPI_ISL_2605749, EPI_ISL_2605750, EPI_ISL_2605751, EPI_ISL_2605752, EPI_ISL_2605753, EPI_ISL_2605754, EPI_ISL_2605755, EPI_ISL_2605756, EPI_ISL_2605757, EPI_ISL_2605758, EPI_ISL_2605759, EPI_ISL_2605760, EPI_ISL_2605761, EPI_ISL_2605762, EPI_ISL_2605763, EPI_ISL_2605764, EPI_ISL_2605765 | | | |
| see above | Lighthouse Lab in Alderley Park | Wellcome Sanger Institute for the COVID-19 Genomics UK (COG-UK) Consortium | Jacquelyn Wynn, Mairead Hyland, The Lighthouse Lab in Alderley Park and Alex Alderton, Roberto Amato, Jeffrey Barrett, Sonia Goncalves, Ewan Harrison, David K. Jackson, Ian Johnston, Dominic Kwiatkowski, Cordelia Langford, John Sillitoe on behalf of the Wellcome Sanger Institute COVID-19 Surveillance Team |
| EPI_ISL_2605766 | Lighthouse Lab in Milton Keynes | Wellcome Sanger Institute for the COVID-19 Genomics UK (COG-UK) Consortium | The Lighthouse Lab in Milton Keynes and Alex Alderton, Roberto Amato, Jeffrey Barrett, Sonia Goncalves, Ewan Harrison, David K. Jackson, Ian Johnston, Dominic Kwiatkowski, Cordelia Langford, John Sillitoe on behalf of the Wellcome Sanger Institute COVID-19 Surveillance Team |
| EPI_ISL_2605767, EPI_ISL_2605768, EPI_ISL_2605769, EPI_ISL_2605770, EPI_ISL_2605771, EPI_ISL_2605772, EPI_ISL_2605773, EPI_ISL_2605774, EPI_ISL_2605775, EPI_ISL_2605776, EPI_ISL_2605777, EPI_ISL_2605778, EPI_ISL_2605779, EPI_ISL_2605780, EPI_ISL_2605781, EPI_ISL_2605782, EPI_ISL_2605783, EPI_ISL_2605784, EPI_ISL_2605785, EPI_ISL_2605786, EPI_ISL_2605787, EPI_ISL_2605788, EPI_ISL_2605789, EPI_ISL_2605790, EPI_ISL_2605791, EPI_ISL_2605792, EPI_ISL_2605793, EPI_ISL_2605794, EPI_ISL_2605795, EPI_ISL_2605796, EPI_ISL_2605797, EPI_ISL_2605798, EPI_ISL_2605799, EPI_ISL_2605800, EPI_ISL_2605801, EPI_ISL_2605802, EPI_ISL_2605803, EPI_ISL_2605804, EPI_ISL_2605805, EPI_ISL_2605806, EPI_ISL_2605807, EPI_ISL_2605808, EPI_ISL_2605809, EPI_ISL_2605810, EPI_ISL_2605811, EPI_ISL_2605812, EPI_ISL_2605813, EPI_ISL_2605814, EPI_ISL_2605816, EPI_ISL_2605817, EPI_ISL_2605818, EPI_ISL_2605819, EPI_ISL_2605820, EPI_ISL_2605821, EPI_ISL_2605822, EPI_ISL_2605823, EPI_ISL_2605824, EPI_ISL_2605825, EPI_ISL_2605826, EPI_ISL_2605827, EPI_ISL_2605828, EPI_ISL_2605829, EPI_ISL_2605830, EPI_ISL_2605831, EPI_ISL_2605832, EPI_ISL_2605833, EPI_ISL_2605834, EPI_ISL_2605835, EPI_ISL_2605836, EPI_ISL_2605837, EPI_ISL_2605838, EPI_ISL_2605839, EPI_ISL_2605840, EPI_ISL_2605841, EPI_ISL_2605842, EPI_ISL_2605843, EPI_ISL_2605844, EPI_ISL_2605845, EPI_ISL_2605846, EPI_ISL_2605847, EPI_ISL_2605848, EPI_ISL_2605849, EPI_ISL_2605850, EPI_ISL_2605851, EPI_ISL_2605852, EPI_ISL_2605853, EPI_ISL_2605854, EPI_ISL_2605855, EPI_ISL_2605856, EPI_ISL_2605857, EPI_ISL_2605858, EPI_ISL_2605859, EPI_ISL_2605860, EPI_ISL_2605861, EPI_ISL_2605862, EPI_ISL_2605863, EPI_ISL_2605864, EPI_ISL_2605865, EPI_ISL_2605866, EPI_ISL_2605867, EPI_ISL_2605868, EPI_ISL_2605869, EPI_ISL_2605870, EPI_ISL_2605872, EPI_ISL_2605873, EPI_ISL_2605874, EPI_ISL_2605875, EPI_ISL_2605876, EPI_ISL_2605878, EPI_ISL_2605879, EPI_ISL_2605881, EPI_ISL_2605882, EPI_ISL_2605883, EPI_ISL_2605884, EPI_ISL_2605886, EPI_ISL_2605887, EPI_ISL_2605888, EPI_ISL_2605889, EPI_ISL_2605890, EPI_ISL_2605891, EPI_ISL_2605892, EPI_ISL_2605893, EPI_ISL_2605894, EPI_ISL_2605895, EPI_ISL_2605896, EPI_ISL_2605897, EPI_ISL_2605898, EPI_ISL_2605899, EPI_ISL_2605900, EPI_ISL_2605901, EPI_ISL_2605902, EPI_ISL_2605903, EPI_ISL_2605904, EPI_ISL_2605905, EPI_ISL_2605906, EPI_ISL_2605907, EPI_ISL_2605909, EPI_ISL_2605910, EPI_ISL_2605911, EPI_ISL_2605912, EPI_ISL_2605913, EPI_ISL_2605914, EPI_ISL_2605916, EPI_ISL_2605917, EPI_ISL_2605918, EPI_ISL_2605919, EPI_ISL_2605920, EPI_ISL_2605921, EPI_ISL_2605922, EPI_ISL_2605923, EPI_ISL_2605924, EPI_ISL_2605925, EPI_ISL_2605926, EPI_ISL_2605927, EPI_ISL_2605928, EPI_ISL_2605929, EPI_ISL_2605930, EPI_ISL_2605931, EPI_ISL_2605932, EPI_ISL_2605933, EPI_ISL_2605934, EPI_ISL_2605935, EPI_ISL_2605936, EPI_ISL_2605937, EPI_ISL_2605938, EPI_ISL_2605939, EPI_ISL_2605940, EPI_ISL_2605941, EPI_ISL_2605942, EPI_ISL_2605943, EPI_ISL_2605944, EPI_ISL_2605945, EPI_ISL_2605946, EPI_ISL_2605947, EPI_ISL_2605948, EPI_ISL_2605949, EPI_ISL_2605950, EPI_ISL_2605951, EPI_ISL_2605952, EPI_ISL_2605953, EPI_ISL_2605954, EPI_ISL_2605955, EPI_ISL_2605956, EPI_ISL_2605957, EPI_ISL_2605958, EPI_ISL_2605959, EPI_ISL_2605960, EPI_ISL_2605961, EPI_ISL_2605962, EPI_ISL_2605963, EPI_ISL_2605964, EPI_ISL_2605965, EPI_ISL_2605966, EPI_ISL_2605967, EPI_ISL_2605968, EPI_ISL_2605969, EPI_ISL_2605970, EPI_ISL_2605972, EPI_ISL_2605973, EPI_ISL_2605974, EPI_ISL_2605975, EPI_ISL_2605976, EPI_ISL_2605977, EPI_ISL_2605978, EPI_ISL_2605979, EPI_ISL_2605980, EPI_ISL_2605981, EPI_ISL_2605982, EPI_ISL_2605983, EPI_ISL_2605984, EPI_ISL_2605985, EPI_ISL_2605986, EPI_ISL_2605987, EPI_ISL_2605988, EPI_ISL_2605989, EPI_ISL_2605990, EPI_ISL_2605991, EPI_ISL_2605992, EPI_ISL_2605993, EPI_ISL_2605994, EPI_ISL_2605995, EPI_ISL_2605996, EPI_ISL_2605997, EPI_ISL_2605998, EPI_ISL_2605999, EPI_ISL_2606000, EPI_ISL_2606001, EPI_ISL_2606002, EPI_ISL_2606003, EPI_ISL_2606004, EPI_ISL_2606005, EPI_ISL_2606006, EPI_ISL_2606007, EPI_ISL_2606008, EPI_ISL_2606009, EPI_ISL_2606010, EPI_ISL_2606011 | | | |
| see above | Lighthouse Lab in Alderley Park | Wellcome Sanger Institute for the COVID-19 Genomics UK (COG-UK) Consortium | Jacquelyn Wynn, Mairead Hyland, The Lighthouse Lab in Alderley Park and Alex Alderton, Roberto Amato, Jeffrey Barrett, Sonia Goncalves, Ewan Harrison, David K. Jackson, Ian Johnston, Dominic Kwiatkowski, Cordelia Langford, John Sillitoe on behalf of the Wellcome Sanger Institute COVID-19 Surveillance Team |
| EPI_ISL_2606012 | Lighthouse Lab in Glasgow | Wellcome Sanger Institute for the COVID-19 Genomics UK (COG-UK) Consortium | Harper VanSteenhouse, Yumi Kasai, David Gray, Carol Clugston, Anna Dominiczak and Alex Alderton, Roberto Amato, Jeffrey Barrett, Sonia Goncalves, Ewan Harrison, David K. Jackson, Ian Johnston, Dominic Kwiatkowski, Cordelia Langford, John Sillitoe on behalf of the Wellcome Sanger Institute COVID-19 Surveillance Team |
| EPI_ISL_2606013, EPI_ISL_2606014, EPI_ISL_2606015, EPI_ISL_2606016, EPI_ISL_2606017, EPI_ISL_2606018, EPI_ISL_2606019, EPI_ISL_2606020, EPI_ISL_2606021, EPI_ISL_2606022 | Lighthouse Lab in Alderley Park | Wellcome Sanger Institute for the COVID-19 Genomics UK (COG-UK) Consortium | Jacquelyn Wynn, Mairead Hyland, The Lighthouse Lab in Alderley Park and Alex Alderton, Roberto Amato, Jeffrey Barrett, Sonia Goncalves, Ewan Harrison, David K. Jackson, Ian Johnston, Dominic Kwiatkowski, Cordelia Langford, John Sillitoe on behalf of the Wellcome Sanger Institute COVID-19 Surveillance Team |

| | | | |
|---|---|---|---|
| EPI_ISL_2606023 | Lighthouse Lab in Milton Keynes | Wellcome Sanger Institute for the COVID-19 Genomics UK (COG-UK) Consortium | The Lighthouse Lab in Milton Keynes and Alex Alderton, Roberto Amato, Jeffrey Barrett, Sonia Goncalves, Ewan Harrison, David K. Jackson, Ian Johnston, Dominic Kwiatkowski, Cordelia Langford, John Sillitoe on behalf of the Wellcome Sanger Institute COVID-19 Surveillance Team |
| EPI_ISL_2606024, EPI_ISL_2606025, EPI_ISL_2606026, EPI_ISL_2606027, EPI_ISL_2606028, EPI_ISL_2606029, EPI_ISL_2606030, EPI_ISL_2606031, EPI_ISL_2606032, EPI_ISL_2606033, EPI_ISL_2606034, EPI_ISL_2606035, EPI_ISL_2606036, EPI_ISL_2606037, EPI_ISL_2606038, EPI_ISL_2606039, EPI_ISL_2606040, EPI_ISL_2606041, EPI_ISL_2606042, EPI_ISL_2606043, EPI_ISL_2606044, EPI_ISL_2606045, EPI_ISL_2606046, EPI_ISL_2606047, EPI_ISL_2606048, EPI_ISL_2606049, EPI_ISL_2606050, EPI_ISL_2606051, EPI_ISL_2606052, EPI_ISL_2606053, EPI_ISL_2606054, EPI_ISL_2606056, EPI_ISL_2606057, EPI_ISL_2606058, EPI_ISL_2606059, EPI_ISL_2606060, EPI_ISL_2606061, EPI_ISL_2606062, EPI_ISL_2606063, EPI_ISL_2606064, EPI_ISL_2606065, EPI_ISL_2606066, EPI_ISL_2606067, EPI_ISL_2606068, EPI_ISL_2606069, EPI_ISL_2606070, EPI_ISL_2606071, EPI_ISL_2606072, EPI_ISL_2606073, EPI_ISL_2606074, EPI_ISL_2606075, EPI_ISL_2606077, EPI_ISL_2606078, EPI_ISL_2606079, EPI_ISL_2606080, EPI_ISL_2606081, EPI_ISL_2606082, EPI_ISL_2606083, EPI_ISL_2606084, EPI_ISL_2606085, EPI_ISL_2606086, EPI_ISL_2606087, EPI_ISL_2606089, EPI_ISL_2606090, EPI_ISL_2606091, EPI_ISL_2606092, EPI_ISL_2606093, EPI_ISL_2606094, EPI_ISL_2606095, EPI_ISL_2606096, EPI_ISL_2606097, EPI_ISL_2606098, EPI_ISL_2606099, EPI_ISL_2606100, EPI_ISL_2606101, EPI_ISL_2606102, EPI_ISL_2606103, EPI_ISL_2606104, EPI_ISL_2606105, EPI_ISL_2606106, EPI_ISL_2606107, EPI_ISL_2606108, EPI_ISL_2606109, EPI_ISL_2606110, EPI_ISL_2606111, EPI_ISL_2606112, EPI_ISL_2606113, EPI_ISL_2606114, EPI_ISL_2606115, EPI_ISL_2606116, EPI_ISL_2606117, EPI_ISL_2606118, EPI_ISL_2606119, EPI_ISL_2606120, EPI_ISL_2606121, EPI_ISL_2606122, EPI_ISL_2606123, EPI_ISL_2606124, EPI_ISL_2606125, EPI_ISL_2606126, EPI_ISL_2606127, EPI_ISL_2606128, EPI_ISL_2606129, EPI_ISL_2606130, EPI_ISL_2606131, EPI_ISL_2606132, EPI_ISL_2606133, EPI_ISL_2606134, EPI_ISL_2606135, EPI_ISL_2606136, EPI_ISL_2606137, EPI_ISL_2606138, EPI_ISL_2606139, EPI_ISL_2606140, EPI_ISL_2606141, EPI_ISL_2606142, EPI_ISL_2606143, EPI_ISL_2606144, EPI_ISL_2606146, EPI_ISL_2606147, EPI_ISL_2606148, EPI_ISL_2606149, EPI_ISL_2606150, EPI_ISL_2606151, EPI_ISL_2606152, EPI_ISL_2606153, EPI_ISL_2606154, EPI_ISL_2606155, EPI_ISL_2606156, EPI_ISL_2606157, EPI_ISL_2606158, EPI_ISL_2606159, EPI_ISL_2606160, EPI_ISL_2606161, EPI_ISL_2606162, EPI_ISL_2606163, EPI_ISL_2606164, EPI_ISL_2606165, EPI_ISL_2606166, EPI_ISL_2606167, EPI_ISL_2606168, EPI_ISL_2606169, EPI_ISL_2606170, EPI_ISL_2606171, EPI_ISL_2606172, EPI_ISL_2606173, EPI_ISL_2606174, EPI_ISL_2606175, EPI_ISL_2606176, EPI_ISL_2606177, EPI_ISL_2606178, EPI_ISL_2606179, EPI_ISL_2606180, EPI_ISL_2606181, EPI_ISL_2606182, EPI_ISL_2606184, EPI_ISL_2606185, EPI_ISL_2606186, EPI_ISL_2606187, EPI_ISL_2606188, EPI_ISL_2606189, EPI_ISL_2606190, EPI_ISL_2606191, EPI_ISL_2606192, EPI_ISL_2606193, EPI_ISL_2606194, EPI_ISL_2606195, EPI_ISL_2606196, EPI_ISL_2606197, EPI_ISL_2606198, EPI_ISL_2606199, EPI_ISL_2606200, EPI_ISL_2606201, EPI_ISL_2606202, EPI_ISL_2606203, EPI_ISL_2606204, EPI_ISL_2606205, EPI_ISL_2606206, EPI_ISL_2606207, EPI_ISL_2606208, EPI_ISL_2606209, EPI_ISL_2606210, EPI_ISL_2606211, EPI_ISL_2606213, EPI_ISL_2606214, EPI_ISL_2606215, EPI_ISL_2606216, EPI_ISL_2606217, EPI_ISL_2606218, EPI_ISL_2606219, EPI_ISL_2606220, EPI_ISL_2606221, EPI_ISL_2606222, EPI_ISL_2606223, EPI_ISL_2606225, EPI_ISL_2606226, EPI_ISL_2606227, EPI_ISL_2606228, EPI_ISL_2606229, EPI_ISL_2606230, EPI_ISL_2606231, EPI_ISL_2606232, EPI_ISL_2606233, EPI_ISL_2606234, EPI_ISL_2606235, EPI_ISL_2606236, EPI_ISL_2606237, EPI_ISL_2606238, EPI_ISL_2606239, EPI_ISL_2606241, EPI_ISL_2606242, EPI_ISL_2606243, EPI_ISL_2606245, EPI_ISL_2606246, EPI_ISL_2606247, EPI_ISL_2606248, EPI_ISL_2606249, EPI_ISL_2606250, EPI_ISL_2606251, EPI_ISL_2606252, EPI_ISL_2606253, EPI_ISL_2606254, EPI_ISL_2606255, EPI_ISL_2606256, EPI_ISL_2606257, EPI_ISL_2606258, EPI_ISL_2606259, EPI_ISL_2606260, EPI_ISL_2606261, EPI_ISL_2606262, EPI_ISL_2606263, EPI_ISL_2606264, EPI_ISL_2606265, EPI_ISL_2606266, EPI_ISL_2606267, EPI_ISL_2606268, EPI_ISL_2606269, EPI_ISL_2606270, EPI_ISL_2606271, EPI_ISL_2606272, EPI_ISL_2606273, EPI_ISL_2606274, EPI_ISL_2606275, EPI_ISL_2606276, EPI_ISL_2606277, EPI_ISL_2606278, EPI_ISL_2606279, EPI_ISL_2606280, EPI_ISL_2606281, EPI_ISL_2606282, EPI_ISL_2606283, EPI_ISL_2606284, EPI_ISL_2606286, EPI_ISL_2606287, EPI_ISL_2606289, EPI_ISL_2606290, EPI_ISL_2606291, EPI_ISL_2606292, EPI_ISL_2606293, EPI_ISL_2606294, EPI_ISL_2606295, EPI_ISL_2606296, EPI_ISL_2606297, EPI_ISL_2606298, EPI_ISL_2606299, EPI_ISL_2606300, EPI_ISL_2606301, EPI_ISL_2606302, EPI_ISL_2606303, EPI_ISL_2606304, EPI_ISL_2606305, EPI_ISL_2606306, EPI_ISL_2606307, EPI_ISL_2606308, EPI_ISL_2606309, EPI_ISL_2606310, EPI_ISL_2606311, EPI_ISL_2606312, EPI_ISL_2606313, EPI_ISL_2606314, EPI_ISL_2606315, EPI_ISL_2606316, EPI_ISL_2606318, EPI_ISL_2606319, EPI_ISL_2606320, EPI_ISL_2606321, EPI_ISL_2606322, EPI_ISL_2606323, EPI_ISL_2606324, EPI_ISL_2606325, EPI_ISL_2606326, EPI_ISL_2606327, EPI_ISL_2606328, EPI_ISL_2606329, EPI_ISL_2606330, EPI_ISL_2606331, EPI_ISL_2606332, EPI_ISL_2606333, EPI_ISL_2606334, EPI_ISL_2606335, EPI_ISL_2606336, EPI_ISL_2606337, EPI_ISL_2606338, EPI_ISL_2606339, EPI_ISL_2606340, EPI_ISL_2606341, EPI_ISL_2606342, EPI_ISL_2606343, EPI_ISL_2606344, EPI_ISL_2606345, EPI_ISL_2606346, EPI_ISL_2606347, EPI_ISL_2606348, EPI_ISL_2606349, EPI_ISL_2606350, EPI_ISL_2606351, EPI_ISL_2606352, EPI_ISL_2606353, EPI_ISL_2606354, EPI_ISL_2606355, EPI_ISL_2606356, EPI_ISL_2606357, EPI_ISL_2606358, EPI_ISL_2606359, EPI_ISL_2606360, EPI_ISL_2606361, EPI_ISL_2606362, EPI_ISL_2606363, EPI_ISL_2606364, EPI_ISL_2606365, EPI_ISL_2606366, EPI_ISL_2606367, EPI_ISL_2606368, EPI_ISL_2606369, EPI_ISL_2606370, EPI_ISL_2606371, EPI_ISL_2606372, EPI_ISL_2606373, EPI_ISL_2606374, EPI_ISL_2606375, EPI_ISL_2606376, EPI_ISL_2606377, EPI_ISL_2606378, EPI_ISL_2606379, EPI_ISL_2606380, EPI_ISL_2606381, EPI_ISL_2606383, EPI_ISL_2606384, EPI_ISL_2606385, EPI_ISL_2606386, EPI_ISL_2606387, EPI_ISL_2606388, EPI_ISL_2606389, EPI_ISL_2606390, EPI_ISL_2606391, EPI_ISL_2606392, EPI_ISL_2606393, EPI_ISL_2606394, EPI_ISL_2606396, EPI_ISL_2606397, EPI_ISL_2606398, EPI_ISL_2606399, EPI_ISL_2606400, EPI_ISL_2606401, EPI_ISL_2606402, EPI_ISL_2606403, EPI_ISL_2606404, EPI_ISL_2606405, EPI_ISL_2606406, EPI_ISL_2606407, EPI_ISL_2606408, EPI_ISL_2606409, EPI_ISL_2606410, EPI_ISL_2606411, EPI_ISL_2606412, EPI_ISL_2606413, EPI_ISL_2606414, EPI_ISL_2606415, EPI_ISL_2606416, EPI_ISL_2606417, EPI_ISL_2606418, EPI_ISL_2606419, EPI_ISL_2606420, EPI_ISL_2606421, EPI_ISL_2606422, EPI_ISL_2606423, EPI_ISL_2606424, EPI_ISL_2606425, EPI_ISL_2606426, EPI_ISL_2606427, EPI_ISL_2606428, EPI_ISL_2606429, EPI_ISL_2606430, EPI_ISL_2606431, EPI_ISL_2606432, EPI_ISL_2606433, EPI_ISL_2606434, EPI_ISL_2606435, EPI_ISL_2606436, EPI_ISL_2606437, EPI_ISL_2606438, EPI_ISL_2606439, EPI_ISL_2606440, EPI_ISL_2606441, EPI_ISL_2606442, EPI_ISL_2606443, EPI_ISL_2606445, EPI_ISL_2606446, EPI_ISL_2606447, EPI_ISL_2606448, EPI_ISL_2606449, EPI_ISL_2606450, EPI_ISL_2606451, EPI_ISL_2606452 | | | |
| see above | Lighthouse Lab in Alderley Park | Wellcome Sanger Institute for the COVID-19 Genomics UK (COG-UK) Consortium | Jacquelyn Wynn, Mairead Hyland, The Lighthouse Lab in Alderley Park and Alex Alderton, Roberto Amato, Jeffrey Barrett, Sonia Goncalves, Ewan Harrison, David K. Jackson, Ian Johnston, Dominic Kwiatkowski, Cordelia Langford, John Sillitoe on behalf of the Wellcome Sanger Institute COVID-19 Surveillance Team |
| EPI_ISL_2606453, EPI_ISL_2606454, EPI_ISL_2606455 | Lighthouse Lab in Milton Keynes | Wellcome Sanger Institute for the COVID-19 Genomics UK (COG-UK) Consortium | The Lighthouse Lab in Milton Keynes and Alex Alderton, Roberto Amato, Jeffrey Barrett, Sonia Goncalves, Ewan Harrison, David K. Jackson, Ian Johnston, Dominic Kwiatkowski, Cordelia Langford, John Sillitoe on behalf of the Wellcome Sanger Institute COVID-19 Surveillance Team |
| EPI_ISL_2606456 | Lighthouse Laboratory Plymouth | Wellcome Sanger Institute for the COVID-19 Genomics UK (COG-UK) Consortium | Lighthouse Laboratory Plymouth and Alex Alderton, Roberto Amato, Jeffrey Barrett, Sonia Goncalves, Ewan Harrison, David K. Jackson, Ian Johnston, Dominic Kwiatkowski, Cordelia Langford, John Sillitoe on behalf of the Wellcome Sanger Institute COVID-19 Surveillance Team |
| EPI_ISL_2606457 | Lighthouse Lab in Glasgow | Wellcome Sanger Institute for the COVID-19 Genomics UK (COG-UK) Consortium | Harper VanSteenhouse, Yumi Kasai, David Gray, Carol Clugston, Anna Dominiczak and Alex Alderton, Roberto Amato, Jeffrey Barrett, Sonia Goncalves, Ewan Harrison, David K. Jackson, Ian Johnston, Dominic Kwiatkowski, Cordelia Langford, John Sillitoe on behalf of the Wellcome Sanger Institute COVID-19 Surveillance Team |
| EPI_ISL_2606458, EPI_ISL_2606459, EPI_ISL_2606460 | Lighthouse Lab in Milton Keynes | Wellcome Sanger Institute for the COVID-19 Genomics UK (COG-UK) Consortium | The Lighthouse Lab in Milton Keynes and Alex Alderton, Roberto Amato, Jeffrey Barrett, Sonia Goncalves, Ewan Harrison, David K. Jackson, Ian Johnston, Dominic Kwiatkowski, Cordelia Langford, John Sillitoe on behalf of the Wellcome Sanger Institute COVID-19 Surveillance Team |
| EPI_ISL_2606461 | Lighthouse Laboratory Plymouth | Wellcome Sanger Institute for the COVID-19 Genomics UK (COG-UK) Consortium | Lighthouse Laboratory Plymouth and Alex Alderton, Roberto Amato, Jeffrey Barrett, Sonia Goncalves, Ewan Harrison, David K. Jackson, Ian Johnston, Dominic Kwiatkowski, Cordelia Langford, John Sillitoe on behalf of the Wellcome Sanger Institute COVID-19 Surveillance Team |
| EPI_ISL_2606462 | Lighthouse Lab in Milton Keynes | Wellcome Sanger Institute for the COVID-19 Genomics UK (COG-UK) Consortium | The Lighthouse Lab in Milton Keynes and Alex Alderton, Roberto Amato, Jeffrey Barrett, Sonia Goncalves, Ewan Harrison, David K. Jackson, Ian Johnston, Dominic Kwiatkowski, Cordelia Langford, John Sillitoe on behalf of the Wellcome Sanger Institute COVID-19 Surveillance Team |
| EPI_ISL_2606463 | Lighthouse Laboratory Plymouth | Wellcome Sanger Institute for the COVID-19 Genomics UK (COG-UK) Consortium | Lighthouse Laboratory Plymouth and Alex Alderton, Roberto Amato, Jeffrey Barrett, Sonia Goncalves, Ewan Harrison, David K. Jackson, Ian Johnston, Dominic Kwiatkowski, Cordelia Langford, John Sillitoe on behalf of the Wellcome Sanger Institute COVID-19 Surveillance Team |
| EPI_ISL_2606464 | Lighthouse Lab in Milton Keynes | Wellcome Sanger Institute for the COVID-19 Genomics UK (COG-UK) Consortium | The Lighthouse Lab in Milton Keynes and Alex Alderton, Roberto Amato, Jeffrey Barrett, Sonia Goncalves, Ewan Harrison, David K. Jackson, Ian Johnston, Dominic Kwiatkowski, Cordelia Langford, John Sillitoe on behalf of the Wellcome Sanger Institute COVID-19 Surveillance Team |
| EPI_ISL_2606465 | Lighthouse Laboratory Plymouth | Wellcome Sanger Institute for the COVID-19 Genomics UK (COG-UK) Consortium | Lighthouse Laboratory Plymouth and Alex Alderton, Roberto Amato, Jeffrey Barrett, Sonia Goncalves, Ewan Harrison, David K. Jackson, Ian Johnston, Dominic Kwiatkowski, Cordelia Langford, John Sillitoe on behalf of the Wellcome Sanger Institute COVID-19 Surveillance Team |
| EPI_ISL_2606466, EPI_ISL_2606467, EPI_ISL_2606468 | Lighthouse Lab in Milton Keynes | Wellcome Sanger Institute for the COVID-19 Genomics UK (COG-UK) Consortium | The Lighthouse Lab in Milton Keynes and Alex Alderton, Roberto Amato, Jeffrey Barrett, Sonia Goncalves, Ewan Harrison, David K. Jackson, Ian Johnston, Dominic Kwiatkowski, Cordelia Langford, John Sillitoe on behalf of the Wellcome Sanger Institute COVID-19 Surveillance Team |
| EPI_ISL_2606469 | Lighthouse Laboratory Plymouth | Wellcome Sanger Institute for the COVID-19 Genomics UK (COG-UK) Consortium | Lighthouse Laboratory Plymouth and Alex Alderton, Roberto Amato, Jeffrey Barrett, Sonia Goncalves, Ewan Harrison, David K. Jackson, Ian Johnston, Dominic Kwiatkowski, Cordelia Langford, John Sillitoe on behalf of the Wellcome Sanger Institute COVID-19 Surveillance Team |
| EPI_ISL_2606470 | Lighthouse Lab in Glasgow | Wellcome Sanger Institute for the COVID-19 Genomics UK (COG-UK) Consortium | Harper VanSteenhouse, Yumi Kasai, David Gray, Carol Clugston, Anna Dominiczak and Alex Alderton, Roberto Amato, Jeffrey Barrett, Sonia Goncalves, Ewan Harrison, David K. Jackson, Ian Johnston, Dominic Kwiatkowski, Cordelia Langford, John Sillitoe on behalf of the Wellcome Sanger Institute COVID-19 Surveillance Team |
| EPI_ISL_2606471, EPI_ISL_2606472, EPI_ISL_2606473 | Lighthouse Lab in Milton Keynes | Wellcome Sanger Institute for the COVID-19 Genomics UK (COG-UK) Consortium | The Lighthouse Lab in Milton Keynes and Alex Alderton, Roberto Amato, Jeffrey Barrett, Sonia Goncalves, Ewan Harrison, David K. Jackson, Ian Johnston, Dominic Kwiatkowski, Cordelia Langford, John Sillitoe on behalf of the Wellcome Sanger Institute COVID-19 Surveillance Team |
| EPI_ISL_2606474, EPI_ISL_2606475 | Lighthouse Laboratory Plymouth | Wellcome Sanger Institute for the COVID-19 Genomics UK (COG-UK) Consortium | Lighthouse Laboratory Plymouth and Alex Alderton, Roberto Amato, Jeffrey Barrett, Sonia Goncalves, Ewan Harrison, David K. Jackson, Ian Johnston, Dominic Kwiatkowski, Cordelia Langford, John Sillitoe on behalf of the Wellcome Sanger Institute COVID-19 Surveillance Team |
| EPI_ISL_2606476, EPI_ISL_2606477, EPI_ISL_2606478 | Lighthouse Lab in Milton Keynes | Wellcome Sanger Institute for the COVID-19 Genomics UK (COG-UK) Consortium | The Lighthouse Lab in Milton Keynes and Alex Alderton, Roberto Amato, Jeffrey Barrett, Sonia Goncalves, Ewan Harrison, David K. Jackson, Ian Johnston, Dominic Kwiatkowski, Cordelia Langford, John Sillitoe on behalf of the Wellcome Sanger Institute COVID-19 Surveillance Team |
| EPI_ISL_2606479 | Lighthouse Lab in Glasgow | Wellcome Sanger Institute for the COVID-19 Genomics UK (COG-UK) Consortium | Harper VanSteenhouse, Yumi Kasai, David Gray, Carol Clugston, Anna Dominiczak and Alex Alderton, Roberto Amato, Jeffrey Barrett, Sonia Goncalves, Ewan Harrison, David K. Jackson, Ian Johnston, Dominic Kwiatkowski, Cordelia Langford, John Sillitoe on behalf of the Wellcome Sanger Institute COVID-19 Surveillance Team |
| EPI_ISL_2606480 | Lighthouse Lab in Milton Keynes | Wellcome Sanger Institute for the COVID-19 Genomics UK (COG-UK) Consortium | The Lighthouse Lab in Milton Keynes and Alex Alderton, Roberto Amato, Jeffrey Barrett, Sonia Goncalves, Ewan Harrison, David K. Jackson, Ian Johnston, Dominic Kwiatkowski, Cordelia Langford, John Sillitoe on behalf of the Wellcome Sanger Institute COVID-19 Surveillance Team |
| EPI_ISL_2606481, EPI_ISL_2606482 | Lighthouse Laboratory Plymouth | Wellcome Sanger Institute for the COVID-19 Genomics UK (COG-UK) Consortium | Lighthouse Laboratory Plymouth and Alex Alderton, Roberto Amato, Jeffrey Barrett, Sonia Goncalves, Ewan Harrison, David K. Jackson, Ian Johnston, Dominic Kwiatkowski, Cordelia Langford, John Sillitoe on behalf of the Wellcome Sanger Institute COVID-19 Surveillance Team |
| EPI_ISL_2606483 | Lighthouse Lab in Milton Keynes | Wellcome Sanger Institute for the COVID-19 Genomics UK (COG-UK) Consortium | The Lighthouse Lab in Milton Keynes and Alex Alderton, Roberto Amato, Jeffrey Barrett, Sonia Goncalves, Ewan Harrison, David K. Jackson, Ian Johnston, Dominic Kwiatkowski, Cordelia Langford, John Sillitoe on behalf of the Wellcome Sanger Institute COVID-19 Surveillance Team |
| EPI_ISL_2606484 | Lighthouse Lab in Glasgow | Wellcome Sanger Institute for the COVID-19 Genomics UK (COG-UK) Consortium | Harper VanSteenhouse, Yumi Kasai, David Gray, Carol Clugston, Anna Dominiczak and Alex Alderton, Roberto Amato, Jeffrey Barrett, Sonia Goncalves, Ewan Harrison, David K. Jackson, Ian Johnston, Dominic Kwiatkowski, Cordelia Langford, John Sillitoe on behalf of the Wellcome Sanger Institute COVID-19 Surveillance Team |
| EPI_ISL_2606485 | Lighthouse Laboratory Plymouth | Wellcome Sanger Institute for the COVID-19 Genomics UK (COG-UK) Consortium | Lighthouse Laboratory Plymouth and Alex Alderton, Roberto Amato, Jeffrey Barrett, Sonia Goncalves, Ewan Harrison, David K. Jackson, Ian Johnston, Dominic Kwiatkowski, Cordelia Langford, John Sillitoe on behalf of the Wellcome Sanger Institute COVID-19 Surveillance Team |
| EPI_ISL_2606486 | Lighthouse Lab in Milton Keynes | Wellcome Sanger Institute for the COVID-19 Genomics UK (COG-UK) Consortium | The Lighthouse Lab in Milton Keynes and Alex Alderton, Roberto Amato, Jeffrey Barrett, Sonia Goncalves, Ewan Harrison, David K. Jackson, Ian Johnston, Dominic Kwiatkowski, Cordelia Langford, John Sillitoe on behalf of the Wellcome Sanger Institute COVID-19 Surveillance Team |
| EPI_ISL_2606487 | Lighthouse Laboratory Plymouth | Wellcome Sanger Institute for the COVID-19 Genomics UK (COG-UK) Consortium | Lighthouse Laboratory Plymouth and Alex Alderton, Roberto Amato, Jeffrey Barrett, Sonia Goncalves, Ewan Harrison, David K. Jackson, Ian Johnston, Dominic Kwiatkowski, Cordelia Langford, John Sillitoe on behalf of the Wellcome Sanger Institute COVID-19 Surveillance Team |
| EPI_ISL_2606488, EPI_ISL_2606489, EPI_ISL_2606490, EPI_ISL_2606491, EPI_ISL_2606492, EPI_ISL_2606493 | Lighthouse Lab in Milton Keynes | Wellcome Sanger Institute for the COVID-19 Genomics UK (COG-UK) Consortium | The Lighthouse Lab in Milton Keynes and Alex Alderton, Roberto Amato, Jeffrey Barrett, Sonia Goncalves, Ewan Harrison, David K. Jackson, Ian Johnston, Dominic Kwiatkowski, Cordelia Langford, John Sillitoe on behalf of the Wellcome Sanger Institute COVID-19 Surveillance Team |
| EPI_ISL_2606494 | Lighthouse Laboratory Plymouth | Wellcome Sanger Institute for the COVID-19 Genomics UK (COG-UK) Consortium | Lighthouse Laboratory Plymouth and Alex Alderton, Roberto Amato, Jeffrey Barrett, Sonia Goncalves, Ewan Harrison, David K. Jackson, Ian Johnston, Dominic Kwiatkowski, Cordelia Langford, John Sillitoe on behalf of the Wellcome Sanger Institute COVID-19 Surveillance Team |
| EPI_ISL_2606495 | Lighthouse Lab in Milton Keynes | Wellcome Sanger Institute for the COVID-19 Genomics UK (COG-UK) Consortium | The Lighthouse Lab in Milton Keynes and Alex Alderton, Roberto Amato, Jeffrey Barrett, Sonia Goncalves, Ewan Harrison, David K. Jackson, Ian Johnston, Dominic Kwiatkowski, Cordelia Langford, John Sillitoe on behalf of the Wellcome Sanger Institute COVID-19 Surveillance Team |
| EPI_ISL_2606496, EPI_ISL_2606497, EPI_ISL_2606498, EPI_ISL_2606500 | Lighthouse Laboratory Plymouth | Wellcome Sanger Institute for the COVID-19 Genomics UK (COG-UK) Consortium | Lighthouse Laboratory Plymouth and Alex Alderton, Roberto Amato, Jeffrey Barrett, Sonia Goncalves, Ewan Harrison, David K. Jackson, Ian Johnston, Dominic Kwiatkowski, Cordelia Langford, John Sillitoe on behalf of the Wellcome Sanger Institute COVID-19 Surveillance Team |
| EPI_ISL_2606501, EPI_ISL_2606502, EPI_ISL_2606503 | Lighthouse Lab in Milton Keynes | Wellcome Sanger Institute for the COVID-19 Genomics UK (COG-UK) Consortium | The Lighthouse Lab in Milton Keynes and Alex Alderton, Roberto Amato, Jeffrey Barrett, Sonia Goncalves, Ewan Harrison, David K. Jackson, Ian Johnston, Dominic Kwiatkowski, Cordelia Langford, John Sillitoe on behalf of the Wellcome Sanger Institute COVID-19 Surveillance Team |
| EPI_ISL_2606504, EPI_ISL_2606505, EPI_ISL_2606506 | Lighthouse Laboratory Plymouth | Wellcome Sanger Institute for the COVID-19 Genomics UK (COG-UK) Consortium | Lighthouse Laboratory Plymouth and Alex Alderton, Roberto Amato, Jeffrey Barrett, Sonia Goncalves, Ewan Harrison, David K. Jackson, Ian Johnston, Dominic Kwiatkowski, Cordelia Langford, John Sillitoe on behalf of the Wellcome Sanger Institute COVID-19 Surveillance Team |
| EPI_ISL_2606507 | Lighthouse Lab in Milton Keynes | Wellcome Sanger Institute for the COVID-19 Genomics UK (COG-UK) | The Lighthouse Lab in Milton Keynes and Alex Alderton, Roberto Amato, Jeffrey Barrett, Sonia Goncalves, Ewan Harrison, David K. Jackson, Ian Johnston, Dominic Kwiatkowski, Cordelia Langford, John Sillitoe on behalf of the Wellcome Sanger |

| | | | |
|---|---|---|---|
| | | Consortium | Institute COVID-19 Surveillance Team |
| EPI_ISL_2606509, EPI_ISL_2606510 | Lighthouse Laboratory Plymouth | Wellcome Sanger Institute for the COVID-19 Genomics UK (COG-UK) Consortium | Lighthouse Laboratory Plymouth and Alex Alderton, Roberto Amato, Jeffrey Barrett, Sonia Goncalves, Ewan Harrison, David K. Jackson, Ian Johnston, Dominic Kwiatkowski, Cordelia Langford, John Sillitoe on behalf of the Wellcome Sanger Institute COVID-19 Surveillance Team |
| EPI_ISL_2606511 | Lighthouse Lab in Milton Keynes | Wellcome Sanger Institute for the COVID-19 Genomics UK (COG-UK) Consortium | The Lighthouse Lab in Milton Keynes and Alex Alderton, Roberto Amato, Jeffrey Barrett, Sonia Goncalves, Ewan Harrison, David K. Jackson, Ian Johnston, Dominic Kwiatkowski, Cordelia Langford, John Sillitoe on behalf of the Wellcome Sanger Institute COVID-19 Surveillance Team |
| EPI_ISL_2606512 | Lighthouse Laboratory Plymouth | Wellcome Sanger Institute for the COVID-19 Genomics UK (COG-UK) Consortium | Lighthouse Laboratory Plymouth and Alex Alderton, Roberto Amato, Jeffrey Barrett, Sonia Goncalves, Ewan Harrison, David K. Jackson, Ian Johnston, Dominic Kwiatkowski, Cordelia Langford, John Sillitoe on behalf of the Wellcome Sanger Institute COVID-19 Surveillance Team |
| EPI_ISL_2606513, EPI_ISL_2606514 | Lighthouse Lab in Milton Keynes | Wellcome Sanger Institute for the COVID-19 Genomics UK (COG-UK) Consortium | The Lighthouse Lab in Milton Keynes and Alex Alderton, Roberto Amato, Jeffrey Barrett, Sonia Goncalves, Ewan Harrison, David K. Jackson, Ian Johnston, Dominic Kwiatkowski, Cordelia Langford, John Sillitoe on behalf of the Wellcome Sanger Institute COVID-19 Surveillance Team |
| EPI_ISL_2606515 | Lighthouse Laboratory Plymouth | Wellcome Sanger Institute for the COVID-19 Genomics UK (COG-UK) Consortium | Lighthouse Laboratory Plymouth and Alex Alderton, Roberto Amato, Jeffrey Barrett, Sonia Goncalves, Ewan Harrison, David K. Jackson, Ian Johnston, Dominic Kwiatkowski, Cordelia Langford, John Sillitoe on behalf of the Wellcome Sanger Institute COVID-19 Surveillance Team |
| EPI_ISL_2606516 | Lighthouse Lab in Milton Keynes | Wellcome Sanger Institute for the COVID-19 Genomics UK (COG-UK) Consortium | The Lighthouse Lab in Milton Keynes and Alex Alderton, Roberto Amato, Jeffrey Barrett, Sonia Goncalves, Ewan Harrison, David K. Jackson, Ian Johnston, Dominic Kwiatkowski, Cordelia Langford, John Sillitoe on behalf of the Wellcome Sanger Institute COVID-19 Surveillance Team |
| EPI_ISL_2606517 | Lighthouse Laboratory Plymouth | Wellcome Sanger Institute for the COVID-19 Genomics UK (COG-UK) Consortium | Lighthouse Laboratory Plymouth and Alex Alderton, Roberto Amato, Jeffrey Barrett, Sonia Goncalves, Ewan Harrison, David K. Jackson, Ian Johnston, Dominic Kwiatkowski, Cordelia Langford, John Sillitoe on behalf of the Wellcome Sanger Institute COVID-19 Surveillance Team |
| EPI_ISL_2606518, EPI_ISL_2606519 | Lighthouse Lab in Milton Keynes | Wellcome Sanger Institute for the COVID-19 Genomics UK (COG-UK) Consortium | The Lighthouse Lab in Milton Keynes and Alex Alderton, Roberto Amato, Jeffrey Barrett, Sonia Goncalves, Ewan Harrison, David K. Jackson, Ian Johnston, Dominic Kwiatkowski, Cordelia Langford, John Sillitoe on behalf of the Wellcome Sanger Institute COVID-19 Surveillance Team |
| EPI_ISL_2606520, EPI_ISL_2606521 | Lighthouse Laboratory Plymouth | Wellcome Sanger Institute for the COVID-19 Genomics UK (COG-UK) Consortium | Lighthouse Laboratory Plymouth and Alex Alderton, Roberto Amato, Jeffrey Barrett, Sonia Goncalves, Ewan Harrison, David K. Jackson, Ian Johnston, Dominic Kwiatkowski, Cordelia Langford, John Sillitoe on behalf of the Wellcome Sanger Institute COVID-19 Surveillance Team |
| EPI_ISL_2606522, EPI_ISL_2606523 | Lighthouse Lab in Milton Keynes | Wellcome Sanger Institute for the COVID-19 Genomics UK (COG-UK) Consortium | The Lighthouse Lab in Milton Keynes and Alex Alderton, Roberto Amato, Jeffrey Barrett, Sonia Goncalves, Ewan Harrison, David K. Jackson, Ian Johnston, Dominic Kwiatkowski, Cordelia Langford, John Sillitoe on behalf of the Wellcome Sanger Institute COVID-19 Surveillance Team |
| EPI_ISL_2606524, EPI_ISL_2606525, EPI_ISL_2606526 | Lighthouse Laboratory Plymouth | Wellcome Sanger Institute for the COVID-19 Genomics UK (COG-UK) Consortium | Lighthouse Laboratory Plymouth and Alex Alderton, Roberto Amato, Jeffrey Barrett, Sonia Goncalves, Ewan Harrison, David K. Jackson, Ian Johnston, Dominic Kwiatkowski, Cordelia Langford, John Sillitoe on behalf of the Wellcome Sanger Institute COVID-19 Surveillance Team |
| EPI_ISL_2606527, EPI_ISL_2606528, EPI_ISL_2606529, EPI_ISL_2606530, EPI_ISL_2606532, EPI_ISL_2606533, EPI_ISL_2606534 | Lighthouse Lab in Milton Keynes | Wellcome Sanger Institute for the COVID-19 Genomics UK (COG-UK) Consortium | The Lighthouse Lab in Milton Keynes and Alex Alderton, Roberto Amato, Jeffrey Barrett, Sonia Goncalves, Ewan Harrison, David K. Jackson, Ian Johnston, Dominic Kwiatkowski, Cordelia Langford, John Sillitoe on behalf of the Wellcome Sanger Institute COVID-19 Surveillance Team |
| EPI_ISL_2606535 | Lighthouse Lab in Glasgow | Wellcome Sanger Institute for the COVID-19 Genomics UK (COG-UK) Consortium | Harper VanSteenhouse, Yumi Kasai, David Gray, Carol Clugston, Anna Dominiczak and Alex Alderton, Roberto Amato, Jeffrey Barrett, Sonia Goncalves, Ewan Harrison, David K. Jackson, Ian Johnston, Dominic Kwiatkowski, Cordelia Langford, John Sillitoe on behalf of the Wellcome Sanger Institute COVID-19 Surveillance Team |
| EPI_ISL_2606536, EPI_ISL_2606537 | Lighthouse Lab in Milton Keynes | Wellcome Sanger Institute for the COVID-19 Genomics UK (COG-UK) Consortium | The Lighthouse Lab in Milton Keynes and Alex Alderton, Roberto Amato, Jeffrey Barrett, Sonia Goncalves, Ewan Harrison, David K. Jackson, Ian Johnston, Dominic Kwiatkowski, Cordelia Langford, John Sillitoe on behalf of the Wellcome Sanger Institute COVID-19 Surveillance Team |
| EPI_ISL_2606538 | Lighthouse Laboratory Plymouth | Wellcome Sanger Institute for the COVID-19 Genomics UK (COG-UK) Consortium | Lighthouse Laboratory Plymouth and Alex Alderton, Roberto Amato, Jeffrey Barrett, Sonia Goncalves, Ewan Harrison, David K. Jackson, Ian Johnston, Dominic Kwiatkowski, Cordelia Langford, John Sillitoe on behalf of the Wellcome Sanger Institute COVID-19 Surveillance Team |
| EPI_ISL_2606539 | Lighthouse Lab in Milton Keynes | Wellcome Sanger Institute for the COVID-19 Genomics UK (COG-UK) Consortium | The Lighthouse Lab in Milton Keynes and Alex Alderton, Roberto Amato, Jeffrey Barrett, Sonia Goncalves, Ewan Harrison, David K. Jackson, Ian Johnston, Dominic Kwiatkowski, Cordelia Langford, John Sillitoe on behalf of the Wellcome Sanger Institute COVID-19 Surveillance Team |
| EPI_ISL_2606540 | Lighthouse Laboratory Plymouth | Wellcome Sanger Institute for the COVID-19 Genomics UK (COG-UK) Consortium | Lighthouse Laboratory Plymouth and Alex Alderton, Roberto Amato, Jeffrey Barrett, Sonia Goncalves, Ewan Harrison, David K. Jackson, Ian Johnston, Dominic Kwiatkowski, Cordelia Langford, John Sillitoe on behalf of the Wellcome Sanger Institute COVID-19 Surveillance Team |
| EPI_ISL_2606541, EPI_ISL_2606542, EPI_ISL_2606543, EPI_ISL_2606544, EPI_ISL_2606545, EPI_ISL_2606546 | Lighthouse Lab in Milton Keynes | Wellcome Sanger Institute for the COVID-19 Genomics UK (COG-UK) Consortium | The Lighthouse Lab in Milton Keynes and Alex Alderton, Roberto Amato, Jeffrey Barrett, Sonia Goncalves, Ewan Harrison, David K. Jackson, Ian Johnston, Dominic Kwiatkowski, Cordelia Langford, John Sillitoe on behalf of the Wellcome Sanger Institute COVID-19 Surveillance Team |
| EPI_ISL_2606547 | Lighthouse Laboratory Plymouth | Wellcome Sanger Institute for the COVID-19 Genomics UK (COG-UK) Consortium | Lighthouse Laboratory Plymouth and Alex Alderton, Roberto Amato, Jeffrey Barrett, Sonia Goncalves, Ewan Harrison, David K. Jackson, Ian Johnston, Dominic Kwiatkowski, Cordelia Langford, John Sillitoe on behalf of the Wellcome Sanger Institute COVID-19 Surveillance Team |
| EPI_ISL_2606548 | Lighthouse Lab in Milton Keynes | Wellcome Sanger Institute for the COVID-19 Genomics UK (COG-UK) Consortium | The Lighthouse Lab in Milton Keynes and Alex Alderton, Roberto Amato, Jeffrey Barrett, Sonia Goncalves, Ewan Harrison, David K. Jackson, Ian Johnston, Dominic Kwiatkowski, Cordelia Langford, John Sillitoe on behalf of the Wellcome Sanger Institute COVID-19 Surveillance Team |
| EPI_ISL_2606549, EPI_ISL_2606550 | Lighthouse Laboratory Plymouth | Wellcome Sanger Institute for the COVID-19 Genomics UK (COG-UK) Consortium | Lighthouse Laboratory Plymouth and Alex Alderton, Roberto Amato, Jeffrey Barrett, Sonia Goncalves, Ewan Harrison, David K. Jackson, Ian Johnston, Dominic Kwiatkowski, Cordelia Langford, John Sillitoe on behalf of the Wellcome Sanger Institute COVID-19 Surveillance Team |
| EPI_ISL_2606551 | Lighthouse Lab in Milton Keynes | Wellcome Sanger Institute for the COVID-19 Genomics UK (COG-UK) Consortium | The Lighthouse Lab in Milton Keynes and Alex Alderton, Roberto Amato, Jeffrey Barrett, Sonia Goncalves, Ewan Harrison, David K. Jackson, Ian Johnston, Dominic Kwiatkowski, Cordelia Langford, John Sillitoe on behalf of the Wellcome Sanger Institute COVID-19 Surveillance Team |
| EPI_ISL_2606552, EPI_ISL_2606553 | Lighthouse Laboratory Plymouth | Wellcome Sanger Institute for the COVID-19 Genomics UK (COG-UK) Consortium | Lighthouse Laboratory Plymouth and Alex Alderton, Roberto Amato, Jeffrey Barrett, Sonia Goncalves, Ewan Harrison, David K. Jackson, Ian Johnston, Dominic Kwiatkowski, Cordelia Langford, John Sillitoe on behalf of the Wellcome Sanger Institute COVID-19 Surveillance Team |
| EPI_ISL_2606554, EPI_ISL_2606555, EPI_ISL_2606556 | Lighthouse Lab in Milton Keynes | Wellcome Sanger Institute for the COVID-19 Genomics UK (COG-UK) Consortium | The Lighthouse Lab in Milton Keynes and Alex Alderton, Roberto Amato, Jeffrey Barrett, Sonia Goncalves, Ewan Harrison, David K. Jackson, Ian Johnston, Dominic Kwiatkowski, Cordelia Langford, John Sillitoe on behalf of the Wellcome Sanger Institute COVID-19 Surveillance Team |
| EPI_ISL_2606557 | Lighthouse Lab in Glasgow | Wellcome Sanger Institute for the COVID-19 Genomics UK (COG-UK) Consortium | Harper VanSteenhouse, Yumi Kasai, David Gray, Carol Clugston, Anna Dominiczak and Alex Alderton, Roberto Amato, Jeffrey Barrett, Sonia Goncalves, Ewan Harrison, David K. Jackson, Ian Johnston, Dominic Kwiatkowski, Cordelia Langford, John Sillitoe on behalf of the Wellcome Sanger Institute COVID-19 Surveillance Team |
| EPI_ISL_2606559, EPI_ISL_2606560, EPI_ISL_2606561, EPI_ISL_2606562, EPI_ISL_2606563, EPI_ISL_2606564 | Lighthouse Lab in Milton Keynes | Wellcome Sanger Institute for the COVID-19 Genomics UK (COG-UK) Consortium | The Lighthouse Lab in Milton Keynes and Alex Alderton, Roberto Amato, Jeffrey Barrett, Sonia Goncalves, Ewan Harrison, David K. Jackson, Ian Johnston, Dominic Kwiatkowski, Cordelia Langford, John Sillitoe on behalf of the Wellcome Sanger Institute COVID-19 Surveillance Team |
| EPI_ISL_2606565 | Lighthouse Laboratory Plymouth | Wellcome Sanger Institute for the COVID-19 Genomics UK (COG-UK) Consortium | Lighthouse Laboratory Plymouth and Alex Alderton, Roberto Amato, Jeffrey Barrett, Sonia Goncalves, Ewan Harrison, David K. Jackson, Ian Johnston, Dominic Kwiatkowski, Cordelia Langford, John Sillitoe on behalf of the Wellcome Sanger Institute COVID-19 Surveillance Team |
| EPI_ISL_2606566, EPI_ISL_2606567 | Lighthouse Lab in Milton Keynes | Wellcome Sanger Institute for the COVID-19 Genomics UK (COG-UK) Consortium | The Lighthouse Lab in Milton Keynes and Alex Alderton, Roberto Amato, Jeffrey Barrett, Sonia Goncalves, Ewan Harrison, David K. Jackson, Ian Johnston, Dominic Kwiatkowski, Cordelia Langford, John Sillitoe on behalf of the Wellcome Sanger Institute COVID-19 Surveillance Team |
| EPI_ISL_2606568, EPI_ISL_2606569 | Lighthouse Laboratory Plymouth | Wellcome Sanger Institute for the COVID-19 Genomics UK (COG-UK) Consortium | Lighthouse Laboratory Plymouth and Alex Alderton, Roberto Amato, Jeffrey Barrett, Sonia Goncalves, Ewan Harrison, David K. Jackson, Ian Johnston, Dominic Kwiatkowski, Cordelia Langford, John Sillitoe on behalf of the Wellcome Sanger Institute COVID-19 Surveillance Team |
| EPI_ISL_2606570 | Lighthouse Lab in Milton Keynes | Wellcome Sanger Institute for the COVID-19 Genomics UK (COG-UK) Consortium | The Lighthouse Lab in Milton Keynes and Alex Alderton, Roberto Amato, Jeffrey Barrett, Sonia Goncalves, Ewan Harrison, David K. Jackson, Ian Johnston, Dominic Kwiatkowski, Cordelia Langford, John Sillitoe on behalf of the Wellcome Sanger Institute COVID-19 Surveillance Team |
| EPI_ISL_2606571, EPI_ISL_2606572, EPI_ISL_2606573 | Lighthouse Laboratory Plymouth | Wellcome Sanger Institute for the COVID-19 Genomics UK (COG-UK) Consortium | Lighthouse Laboratory Plymouth and Alex Alderton, Roberto Amato, Jeffrey Barrett, Sonia Goncalves, Ewan Harrison, David K. Jackson, Ian Johnston, Dominic Kwiatkowski, Cordelia Langford, John Sillitoe on behalf of the Wellcome Sanger Institute COVID-19 Surveillance Team |
| EPI_ISL_2606574, EPI_ISL_2606575 | Lighthouse Lab in Milton Keynes | Wellcome Sanger Institute for the COVID-19 Genomics UK (COG-UK) Consortium | The Lighthouse Lab in Milton Keynes and Alex Alderton, Roberto Amato, Jeffrey Barrett, Sonia Goncalves, Ewan Harrison, David K. Jackson, Ian Johnston, Dominic Kwiatkowski, Cordelia Langford, John Sillitoe on behalf of the Wellcome Sanger Institute COVID-19 Surveillance Team |
| EPI_ISL_2606576, EPI_ISL_2606577 | Lighthouse Laboratory Plymouth | Wellcome Sanger Institute for the COVID-19 Genomics UK (COG-UK) Consortium | Lighthouse Laboratory Plymouth and Alex Alderton, Roberto Amato, Jeffrey Barrett, Sonia Goncalves, Ewan Harrison, David K. Jackson, Ian Johnston, Dominic Kwiatkowski, Cordelia Langford, John Sillitoe on behalf of the Wellcome Sanger Institute COVID-19 Surveillance Team |
| EPI_ISL_2606578 | Lighthouse Lab in Milton Keynes | Wellcome Sanger Institute for the COVID-19 Genomics UK (COG-UK) Consortium | The Lighthouse Lab in Milton Keynes and Alex Alderton, Roberto Amato, Jeffrey Barrett, Sonia Goncalves, Ewan Harrison, David K. Jackson, Ian Johnston, Dominic Kwiatkowski, Cordelia Langford, John Sillitoe on behalf of the Wellcome Sanger Institute COVID-19 Surveillance Team |
| EPI_ISL_2606579 | Lighthouse Laboratory Plymouth | Wellcome Sanger Institute for the COVID-19 Genomics UK (COG-UK) Consortium | Lighthouse Laboratory Plymouth and Alex Alderton, Roberto Amato, Jeffrey Barrett, Sonia Goncalves, Ewan Harrison, David K. Jackson, Ian Johnston, Dominic Kwiatkowski, Cordelia Langford, John Sillitoe on behalf of the Wellcome Sanger Institute COVID-19 Surveillance Team |
| EPI_ISL_2606580, EPI_ISL_2606581, EPI_ISL_2606582 | Lighthouse Lab in Milton Keynes | Wellcome Sanger Institute for the COVID-19 Genomics UK (COG-UK) Consortium | The Lighthouse Lab in Milton Keynes and Alex Alderton, Roberto Amato, Jeffrey Barrett, Sonia Goncalves, Ewan Harrison, David K. Jackson, Ian Johnston, Dominic Kwiatkowski, Cordelia Langford, John Sillitoe on behalf of the Wellcome Sanger Institute COVID-19 Surveillance Team |
| EPI_ISL_2606583 | Lighthouse Laboratory Plymouth | Wellcome Sanger Institute for the COVID-19 Genomics UK (COG-UK) Consortium | Lighthouse Laboratory Plymouth and Alex Alderton, Roberto Amato, Jeffrey Barrett, Sonia Goncalves, Ewan Harrison, David K. Jackson, Ian Johnston, Dominic Kwiatkowski, Cordelia Langford, John Sillitoe on behalf of the Wellcome Sanger Institute COVID-19 Surveillance Team |
| EPI_ISL_2606584, | Lighthouse Lab in Milton Keynes | Wellcome Sanger Institute for the COVID-19 Genomics UK (COG-UK) | The Lighthouse Lab in Milton Keynes and Alex Alderton, Roberto Amato, Jeffrey Barrett, Sonia Goncalves, Ewan Harrison, David K. Jackson, Ian Johnston, Dominic Kwiatkowski, Cordelia Langford, John Sillitoe on behalf of the Wellcome Sanger |

| | | | |
|---|---|---|---|
| EPI_ISL_2606585 | | Consortium | Institute COVID-19 Surveillance Team |
| EPI_ISL_2606586 | Lighthouse Laboratory Plymouth | Wellcome Sanger Institute for the COVID-19 Genomics UK (COG-UK) Consortium | Lighthouse Laboratory Plymouth and Alex Alderton, Roberto Amato, Jeffrey Barrett, Sonia Goncalves, Ewan Harrison, David K. Jackson, Ian Johnston, Dominic Kwiatkowski, Cordelia Langford, John Sillitoe on behalf of the Wellcome Sanger Institute COVID-19 Surveillance Team |
| EPI_ISL_2606587 | Lighthouse Lab in Milton Keynes | Wellcome Sanger Institute for the COVID-19 Genomics UK (COG-UK) Consortium | The Lighthouse Lab in Milton Keynes and Alex Alderton, Roberto Amato, Jeffrey Barrett, Sonia Goncalves, Ewan Harrison, David K. Jackson, Ian Johnston, Dominic Kwiatkowski, Cordelia Langford, John Sillitoe on behalf of the Wellcome Sanger Institute COVID-19 Surveillance Team |
| EPI_ISL_2606588, EPI_ISL_2606589, EPI_ISL_2606590 | Lighthouse Laboratory Plymouth | Wellcome Sanger Institute for the COVID-19 Genomics UK (COG-UK) Consortium | Lighthouse Laboratory Plymouth and Alex Alderton, Roberto Amato, Jeffrey Barrett, Sonia Goncalves, Ewan Harrison, David K. Jackson, Ian Johnston, Dominic Kwiatkowski, Cordelia Langford, John Sillitoe on behalf of the Wellcome Sanger Institute COVID-19 Surveillance Team |
| EPI_ISL_2606591 | Lighthouse Lab in Milton Keynes | Wellcome Sanger Institute for the COVID-19 Genomics UK (COG-UK) Consortium | The Lighthouse Lab in Milton Keynes and Alex Alderton, Roberto Amato, Jeffrey Barrett, Sonia Goncalves, Ewan Harrison, David K. Jackson, Ian Johnston, Dominic Kwiatkowski, Cordelia Langford, John Sillitoe on behalf of the Wellcome Sanger Institute COVID-19 Surveillance Team |
| EPI_ISL_2606592 | Lighthouse Laboratory Plymouth | Wellcome Sanger Institute for the COVID-19 Genomics UK (COG-UK) Consortium | Lighthouse Laboratory Plymouth and Alex Alderton, Roberto Amato, Jeffrey Barrett, Sonia Goncalves, Ewan Harrison, David K. Jackson, Ian Johnston, Dominic Kwiatkowski, Cordelia Langford, John Sillitoe on behalf of the Wellcome Sanger Institute COVID-19 Surveillance Team |
| EPI_ISL_2606593 | Lighthouse Lab in Glasgow | Wellcome Sanger Institute for the COVID-19 Genomics UK (COG-UK) Consortium | Harper VanSteenhouse, Yumi Kasai, David Gray, Carol Clugston, Anna Dominiczak and Alex Alderton, Roberto Amato, Jeffrey Barrett, Sonia Goncalves, Ewan Harrison, David K. Jackson, Ian Johnston, Dominic Kwiatkowski, Cordelia Langford, John Sillitoe on behalf of the Wellcome Sanger Institute COVID-19 Surveillance Team |
| EPI_ISL_2606594 | Lighthouse Lab in Milton Keynes | Wellcome Sanger Institute for the COVID-19 Genomics UK (COG-UK) Consortium | The Lighthouse Lab in Milton Keynes and Alex Alderton, Roberto Amato, Jeffrey Barrett, Sonia Goncalves, Ewan Harrison, David K. Jackson, Ian Johnston, Dominic Kwiatkowski, Cordelia Langford, John Sillitoe on behalf of the Wellcome Sanger Institute COVID-19 Surveillance Team |
| EPI_ISL_2606595 | Lighthouse Laboratory Plymouth | Wellcome Sanger Institute for the COVID-19 Genomics UK (COG-UK) Consortium | Lighthouse Laboratory Plymouth and Alex Alderton, Roberto Amato, Jeffrey Barrett, Sonia Goncalves, Ewan Harrison, David K. Jackson, Ian Johnston, Dominic Kwiatkowski, Cordelia Langford, John Sillitoe on behalf of the Wellcome Sanger Institute COVID-19 Surveillance Team |
| EPI_ISL_2606596 | Lighthouse Lab in Milton Keynes | Wellcome Sanger Institute for the COVID-19 Genomics UK (COG-UK) Consortium | The Lighthouse Lab in Milton Keynes and Alex Alderton, Roberto Amato, Jeffrey Barrett, Sonia Goncalves, Ewan Harrison, David K. Jackson, Ian Johnston, Dominic Kwiatkowski, Cordelia Langford, John Sillitoe on behalf of the Wellcome Sanger Institute COVID-19 Surveillance Team |
| EPI_ISL_2606597 | Lighthouse Laboratory Plymouth | Wellcome Sanger Institute for the COVID-19 Genomics UK (COG-UK) Consortium | Lighthouse Laboratory Plymouth and Alex Alderton, Roberto Amato, Jeffrey Barrett, Sonia Goncalves, Ewan Harrison, David K. Jackson, Ian Johnston, Dominic Kwiatkowski, Cordelia Langford, John Sillitoe on behalf of the Wellcome Sanger Institute COVID-19 Surveillance Team |
| EPI_ISL_2606598, EPI_ISL_2606600, EPI_ISL_2606601 | Lighthouse Lab in Milton Keynes | Wellcome Sanger Institute for the COVID-19 Genomics UK (COG-UK) Consortium | The Lighthouse Lab in Milton Keynes and Alex Alderton, Roberto Amato, Jeffrey Barrett, Sonia Goncalves, Ewan Harrison, David K. Jackson, Ian Johnston, Dominic Kwiatkowski, Cordelia Langford, John Sillitoe on behalf of the Wellcome Sanger Institute COVID-19 Surveillance Team |
| EPI_ISL_2606602 | Lighthouse Laboratory Plymouth | Wellcome Sanger Institute for the COVID-19 Genomics UK (COG-UK) Consortium | Lighthouse Laboratory Plymouth and Alex Alderton, Roberto Amato, Jeffrey Barrett, Sonia Goncalves, Ewan Harrison, David K. Jackson, Ian Johnston, Dominic Kwiatkowski, Cordelia Langford, John Sillitoe on behalf of the Wellcome Sanger Institute COVID-19 Surveillance Team |
| EPI_ISL_2606603, EPI_ISL_2606604 | Lighthouse Lab in Milton Keynes | Wellcome Sanger Institute for the COVID-19 Genomics UK (COG-UK) Consortium | The Lighthouse Lab in Milton Keynes and Alex Alderton, Roberto Amato, Jeffrey Barrett, Sonia Goncalves, Ewan Harrison, David K. Jackson, Ian Johnston, Dominic Kwiatkowski, Cordelia Langford, John Sillitoe on behalf of the Wellcome Sanger Institute COVID-19 Surveillance Team |
| EPI_ISL_2606605 | Lighthouse Laboratory Plymouth | Wellcome Sanger Institute for the COVID-19 Genomics UK (COG-UK) Consortium | Lighthouse Laboratory Plymouth and Alex Alderton, Roberto Amato, Jeffrey Barrett, Sonia Goncalves, Ewan Harrison, David K. Jackson, Ian Johnston, Dominic Kwiatkowski, Cordelia Langford, John Sillitoe on behalf of the Wellcome Sanger Institute COVID-19 Surveillance Team |
| EPI_ISL_2606606, EPI_ISL_2606607, EPI_ISL_2606608, EPI_ISL_2606609, EPI_ISL_2606610 | Lighthouse Lab in Milton Keynes | Wellcome Sanger Institute for the COVID-19 Genomics UK (COG-UK) Consortium | The Lighthouse Lab in Milton Keynes and Alex Alderton, Roberto Amato, Jeffrey Barrett, Sonia Goncalves, Ewan Harrison, David K. Jackson, Ian Johnston, Dominic Kwiatkowski, Cordelia Langford, John Sillitoe on behalf of the Wellcome Sanger Institute COVID-19 Surveillance Team |
| EPI_ISL_2606611 | Lighthouse Laboratory Plymouth | Wellcome Sanger Institute for the COVID-19 Genomics UK (COG-UK) Consortium | Lighthouse Laboratory Plymouth and Alex Alderton, Roberto Amato, Jeffrey Barrett, Sonia Goncalves, Ewan Harrison, David K. Jackson, Ian Johnston, Dominic Kwiatkowski, Cordelia Langford, John Sillitoe on behalf of the Wellcome Sanger Institute COVID-19 Surveillance Team |
| EPI_ISL_2606612, EPI_ISL_2606613, EPI_ISL_2606614, EPI_ISL_2606615, EPI_ISL_2606616, EPI_ISL_2606617, EPI_ISL_2606618 | Lighthouse Lab in Milton Keynes | Wellcome Sanger Institute for the COVID-19 Genomics UK (COG-UK) Consortium | The Lighthouse Lab in Milton Keynes and Alex Alderton, Roberto Amato, Jeffrey Barrett, Sonia Goncalves, Ewan Harrison, David K. Jackson, Ian Johnston, Dominic Kwiatkowski, Cordelia Langford, John Sillitoe on behalf of the Wellcome Sanger Institute COVID-19 Surveillance Team |
| EPI_ISL_2606619 | Lighthouse Laboratory Plymouth | Wellcome Sanger Institute for the COVID-19 Genomics UK (COG-UK) Consortium | Lighthouse Laboratory Plymouth and Alex Alderton, Roberto Amato, Jeffrey Barrett, Sonia Goncalves, Ewan Harrison, David K. Jackson, Ian Johnston, Dominic Kwiatkowski, Cordelia Langford, John Sillitoe on behalf of the Wellcome Sanger Institute COVID-19 Surveillance Team |
| EPI_ISL_2606620, EPI_ISL_2606621, EPI_ISL_2606622, EPI_ISL_2606623 | Lighthouse Lab in Milton Keynes | Wellcome Sanger Institute for the COVID-19 Genomics UK (COG-UK) Consortium | The Lighthouse Lab in Milton Keynes and Alex Alderton, Roberto Amato, Jeffrey Barrett, Sonia Goncalves, Ewan Harrison, David K. Jackson, Ian Johnston, Dominic Kwiatkowski, Cordelia Langford, John Sillitoe on behalf of the Wellcome Sanger Institute COVID-19 Surveillance Team |
| EPI_ISL_2606624 | Lighthouse Lab in Glasgow | Wellcome Sanger Institute for the COVID-19 Genomics UK (COG-UK) Consortium | Harper VanSteenhouse, Yumi Kasai, David Gray, Carol Clugston, Anna Dominiczak and Alex Alderton, Roberto Amato, Jeffrey Barrett, Sonia Goncalves, Ewan Harrison, David K. Jackson, Ian Johnston, Dominic Kwiatkowski, Cordelia Langford, John Sillitoe on behalf of the Wellcome Sanger Institute COVID-19 Surveillance Team |
| EPI_ISL_2606625 | Lighthouse Laboratory Plymouth | Wellcome Sanger Institute for the COVID-19 Genomics UK (COG-UK) Consortium | Lighthouse Laboratory Plymouth and Alex Alderton, Roberto Amato, Jeffrey Barrett, Sonia Goncalves, Ewan Harrison, David K. Jackson, Ian Johnston, Dominic Kwiatkowski, Cordelia Langford, John Sillitoe on behalf of the Wellcome Sanger Institute COVID-19 Surveillance Team |
| EPI_ISL_2606626, EPI_ISL_2606627, EPI_ISL_2606628, EPI_ISL_2606629 | Lighthouse Lab in Milton Keynes | Wellcome Sanger Institute for the COVID-19 Genomics UK (COG-UK) Consortium | The Lighthouse Lab in Milton Keynes and Alex Alderton, Roberto Amato, Jeffrey Barrett, Sonia Goncalves, Ewan Harrison, David K. Jackson, Ian Johnston, Dominic Kwiatkowski, Cordelia Langford, John Sillitoe on behalf of the Wellcome Sanger Institute COVID-19 Surveillance Team |
| EPI_ISL_2606630 | Lighthouse Laboratory Plymouth | Wellcome Sanger Institute for the COVID-19 Genomics UK (COG-UK) Consortium | Lighthouse Laboratory Plymouth and Alex Alderton, Roberto Amato, Jeffrey Barrett, Sonia Goncalves, Ewan Harrison, David K. Jackson, Ian Johnston, Dominic Kwiatkowski, Cordelia Langford, John Sillitoe on behalf of the Wellcome Sanger Institute COVID-19 Surveillance Team |
| EPI_ISL_2606631, EPI_ISL_2606632, EPI_ISL_2606633, EPI_ISL_2606634 | Lighthouse Lab in Milton Keynes | Wellcome Sanger Institute for the COVID-19 Genomics UK (COG-UK) Consortium | The Lighthouse Lab in Milton Keynes and Alex Alderton, Roberto Amato, Jeffrey Barrett, Sonia Goncalves, Ewan Harrison, David K. Jackson, Ian Johnston, Dominic Kwiatkowski, Cordelia Langford, John Sillitoe on behalf of the Wellcome Sanger Institute COVID-19 Surveillance Team |
| EPI_ISL_2606635, EPI_ISL_2606636, EPI_ISL_2606637 | Lighthouse Laboratory Plymouth | Wellcome Sanger Institute for the COVID-19 Genomics UK (COG-UK) Consortium | Lighthouse Laboratory Plymouth and Alex Alderton, Roberto Amato, Jeffrey Barrett, Sonia Goncalves, Ewan Harrison, David K. Jackson, Ian Johnston, Dominic Kwiatkowski, Cordelia Langford, John Sillitoe on behalf of the Wellcome Sanger Institute COVID-19 Surveillance Team |
| EPI_ISL_2606638 | Lighthouse Lab in Milton Keynes | Wellcome Sanger Institute for the COVID-19 Genomics UK (COG-UK) Consortium | The Lighthouse Lab in Milton Keynes and Alex Alderton, Roberto Amato, Jeffrey Barrett, Sonia Goncalves, Ewan Harrison, David K. Jackson, Ian Johnston, Dominic Kwiatkowski, Cordelia Langford, John Sillitoe on behalf of the Wellcome Sanger Institute COVID-19 Surveillance Team |
| EPI_ISL_2606640 | Lighthouse Laboratory Plymouth | Wellcome Sanger Institute for the COVID-19 Genomics UK (COG-UK) Consortium | Lighthouse Laboratory Plymouth and Alex Alderton, Roberto Amato, Jeffrey Barrett, Sonia Goncalves, Ewan Harrison, David K. Jackson, Ian Johnston, Dominic Kwiatkowski, Cordelia Langford, John Sillitoe on behalf of the Wellcome Sanger Institute COVID-19 Surveillance Team |
| EPI_ISL_2606641, EPI_ISL_2606642, EPI_ISL_2606643, EPI_ISL_2606644 | Lighthouse Lab in Milton Keynes | Wellcome Sanger Institute for the COVID-19 Genomics UK (COG-UK) Consortium | The Lighthouse Lab in Milton Keynes and Alex Alderton, Roberto Amato, Jeffrey Barrett, Sonia Goncalves, Ewan Harrison, David K. Jackson, Ian Johnston, Dominic Kwiatkowski, Cordelia Langford, John Sillitoe on behalf of the Wellcome Sanger Institute COVID-19 Surveillance Team |
| EPI_ISL_2606645 | Lighthouse Laboratory Plymouth | Wellcome Sanger Institute for the COVID-19 Genomics UK (COG-UK) Consortium | Lighthouse Laboratory Plymouth and Alex Alderton, Roberto Amato, Jeffrey Barrett, Sonia Goncalves, Ewan Harrison, David K. Jackson, Ian Johnston, Dominic Kwiatkowski, Cordelia Langford, John Sillitoe on behalf of the Wellcome Sanger Institute COVID-19 Surveillance Team |
| EPI_ISL_2606646 | Lighthouse Lab in Milton Keynes | Wellcome Sanger Institute for the COVID-19 Genomics UK (COG-UK) Consortium | The Lighthouse Lab in Milton Keynes and Alex Alderton, Roberto Amato, Jeffrey Barrett, Sonia Goncalves, Ewan Harrison, David K. Jackson, Ian Johnston, Dominic Kwiatkowski, Cordelia Langford, John Sillitoe on behalf of the Wellcome Sanger Institute COVID-19 Surveillance Team |
| EPI_ISL_2606647 | Lighthouse Laboratory Plymouth | Wellcome Sanger Institute for the COVID-19 Genomics UK (COG-UK) Consortium | Lighthouse Laboratory Plymouth and Alex Alderton, Roberto Amato, Jeffrey Barrett, Sonia Goncalves, Ewan Harrison, David K. Jackson, Ian Johnston, Dominic Kwiatkowski, Cordelia Langford, John Sillitoe on behalf of the Wellcome Sanger Institute COVID-19 Surveillance Team |
| EPI_ISL_2606649, EPI_ISL_2606650, EPI_ISL_2606651 | Lighthouse Lab in Milton Keynes | Wellcome Sanger Institute for the COVID-19 Genomics UK (COG-UK) Consortium | The Lighthouse Lab in Milton Keynes and Alex Alderton, Roberto Amato, Jeffrey Barrett, Sonia Goncalves, Ewan Harrison, David K. Jackson, Ian Johnston, Dominic Kwiatkowski, Cordelia Langford, John Sillitoe on behalf of the Wellcome Sanger Institute COVID-19 Surveillance Team |
| EPI_ISL_2606652 | Lighthouse Laboratory Plymouth | Wellcome Sanger Institute for the COVID-19 Genomics UK (COG-UK) Consortium | Lighthouse Laboratory Plymouth and Alex Alderton, Roberto Amato, Jeffrey Barrett, Sonia Goncalves, Ewan Harrison, David K. Jackson, Ian Johnston, Dominic Kwiatkowski, Cordelia Langford, John Sillitoe on behalf of the Wellcome Sanger Institute COVID-19 Surveillance Team |
| EPI_ISL_2606653 | Lighthouse Lab in Milton Keynes | Wellcome Sanger Institute for the COVID-19 Genomics UK (COG-UK) Consortium | The Lighthouse Lab in Milton Keynes and Alex Alderton, Roberto Amato, Jeffrey Barrett, Sonia Goncalves, Ewan Harrison, David K. Jackson, Ian Johnston, Dominic Kwiatkowski, Cordelia Langford, John Sillitoe on behalf of the Wellcome Sanger Institute COVID-19 Surveillance Team |
| EPI_ISL_2606654, EPI_ISL_2606655 | Lighthouse Laboratory Plymouth | Wellcome Sanger Institute for the COVID-19 Genomics UK (COG-UK) Consortium | Lighthouse Laboratory Plymouth and Alex Alderton, Roberto Amato, Jeffrey Barrett, Sonia Goncalves, Ewan Harrison, David K. Jackson, Ian Johnston, Dominic Kwiatkowski, Cordelia Langford, John Sillitoe on behalf of the Wellcome Sanger Institute COVID-19 Surveillance Team |
| EPI_ISL_2606656 | Lighthouse Lab in Milton Keynes | Wellcome Sanger Institute for the COVID-19 Genomics UK (COG-UK) Consortium | The Lighthouse Lab in Milton Keynes and Alex Alderton, Roberto Amato, Jeffrey Barrett, Sonia Goncalves, Ewan Harrison, David K. Jackson, Ian Johnston, Dominic Kwiatkowski, Cordelia Langford, John Sillitoe on behalf of the Wellcome Sanger Institute COVID-19 Surveillance Team |
| EPI_ISL_2606657 | Lighthouse Laboratory Plymouth | Wellcome Sanger Institute for the COVID-19 Genomics UK (COG-UK) | Lighthouse Laboratory Plymouth and Alex Alderton, Roberto Amato, Jeffrey Barrett, Sonia Goncalves, Ewan Harrison, David K. Jackson, Ian Johnston, Dominic Kwiatkowski, Cordelia Langford, John Sillitoe on behalf of the Wellcome Sanger |

| | | | |
|---|---|---|---|
| | | Consortium | Institute COVID-19 Surveillance Team |
| EPI_ISL_2606658, EPI_ISL_2606659, EPI_ISL_2606660, EPI_ISL_2606661, EPI_ISL_2606662, EPI_ISL_2606663 | Lighthouse Lab in Milton Keynes | Wellcome Sanger Institute for the COVID-19 Genomics UK (COG-UK) Consortium | The Lighthouse Lab in Milton Keynes and Alex Alderton, Roberto Amato, Jeffrey Barrett, Sonia Goncalves, Ewan Harrison, David K. Jackson, Ian Johnston, Dominic Kwiatkowski, Cordelia Langford, John Sillitoe on behalf of the Wellcome Sanger Institute COVID-19 Surveillance Team |
| EPI_ISL_2606664, EPI_ISL_2606665 | Lighthouse Laboratory Plymouth | Wellcome Sanger Institute for the COVID-19 Genomics UK (COG-UK) Consortium | Lighthouse Laboratory Plymouth and Alex Alderton, Roberto Amato, Jeffrey Barrett, Sonia Goncalves, Ewan Harrison, David K. Jackson, Ian Johnston, Dominic Kwiatkowski, Cordelia Langford, John Sillitoe on behalf of the Wellcome Sanger Institute COVID-19 Surveillance Team |
| EPI_ISL_2606666, EPI_ISL_2606668 | Lighthouse Lab in Milton Keynes | Wellcome Sanger Institute for the COVID-19 Genomics UK (COG-UK) Consortium | The Lighthouse Lab in Milton Keynes and Alex Alderton, Roberto Amato, Jeffrey Barrett, Sonia Goncalves, Ewan Harrison, David K. Jackson, Ian Johnston, Dominic Kwiatkowski, Cordelia Langford, John Sillitoe on behalf of the Wellcome Sanger Institute COVID-19 Surveillance Team |
| EPI_ISL_2606669 | Lighthouse Laboratory Plymouth | Wellcome Sanger Institute for the COVID-19 Genomics UK (COG-UK) Consortium | Lighthouse Laboratory Plymouth and Alex Alderton, Roberto Amato, Jeffrey Barrett, Sonia Goncalves, Ewan Harrison, David K. Jackson, Ian Johnston, Dominic Kwiatkowski, Cordelia Langford, John Sillitoe on behalf of the Wellcome Sanger Institute COVID-19 Surveillance Team |
| EPI_ISL_2606670 | Lighthouse Lab in Milton Keynes | Wellcome Sanger Institute for the COVID-19 Genomics UK (COG-UK) Consortium | The Lighthouse Lab in Milton Keynes and Alex Alderton, Roberto Amato, Jeffrey Barrett, Sonia Goncalves, Ewan Harrison, David K. Jackson, Ian Johnston, Dominic Kwiatkowski, Cordelia Langford, John Sillitoe on behalf of the Wellcome Sanger Institute COVID-19 Surveillance Team |
| EPI_ISL_2606671 | Lighthouse Laboratory Plymouth | Wellcome Sanger Institute for the COVID-19 Genomics UK (COG-UK) Consortium | Lighthouse Laboratory Plymouth and Alex Alderton, Roberto Amato, Jeffrey Barrett, Sonia Goncalves, Ewan Harrison, David K. Jackson, Ian Johnston, Dominic Kwiatkowski, Cordelia Langford, John Sillitoe on behalf of the Wellcome Sanger Institute COVID-19 Surveillance Team |
| EPI_ISL_2606672 | Lighthouse Lab in Milton Keynes | Wellcome Sanger Institute for the COVID-19 Genomics UK (COG-UK) Consortium | The Lighthouse Lab in Milton Keynes and Alex Alderton, Roberto Amato, Jeffrey Barrett, Sonia Goncalves, Ewan Harrison, David K. Jackson, Ian Johnston, Dominic Kwiatkowski, Cordelia Langford, John Sillitoe on behalf of the Wellcome Sanger Institute COVID-19 Surveillance Team |
| EPI_ISL_2606673, EPI_ISL_2606674 | Lighthouse Laboratory Plymouth | Wellcome Sanger Institute for the COVID-19 Genomics UK (COG-UK) Consortium | Lighthouse Laboratory Plymouth and Alex Alderton, Roberto Amato, Jeffrey Barrett, Sonia Goncalves, Ewan Harrison, David K. Jackson, Ian Johnston, Dominic Kwiatkowski, Cordelia Langford, John Sillitoe on behalf of the Wellcome Sanger Institute COVID-19 Surveillance Team |
| EPI_ISL_2606675, EPI_ISL_2606676, EPI_ISL_2606677 | Lighthouse Lab in Milton Keynes | Wellcome Sanger Institute for the COVID-19 Genomics UK (COG-UK) Consortium | The Lighthouse Lab in Milton Keynes and Alex Alderton, Roberto Amato, Jeffrey Barrett, Sonia Goncalves, Ewan Harrison, David K. Jackson, Ian Johnston, Dominic Kwiatkowski, Cordelia Langford, John Sillitoe on behalf of the Wellcome Sanger Institute COVID-19 Surveillance Team |
| EPI_ISL_2606678 | Lighthouse Laboratory Plymouth | Wellcome Sanger Institute for the COVID-19 Genomics UK (COG-UK) Consortium | Lighthouse Laboratory Plymouth and Alex Alderton, Roberto Amato, Jeffrey Barrett, Sonia Goncalves, Ewan Harrison, David K. Jackson, Ian Johnston, Dominic Kwiatkowski, Cordelia Langford, John Sillitoe on behalf of the Wellcome Sanger Institute COVID-19 Surveillance Team |
| EPI_ISL_2606679, EPI_ISL_2606680, EPI_ISL_2606681 | Lighthouse Lab in Milton Keynes | Wellcome Sanger Institute for the COVID-19 Genomics UK (COG-UK) Consortium | The Lighthouse Lab in Milton Keynes and Alex Alderton, Roberto Amato, Jeffrey Barrett, Sonia Goncalves, Ewan Harrison, David K. Jackson, Ian Johnston, Dominic Kwiatkowski, Cordelia Langford, John Sillitoe on behalf of the Wellcome Sanger Institute COVID-19 Surveillance Team |
| EPI_ISL_2606682 | Lighthouse Laboratory Plymouth | Wellcome Sanger Institute for the COVID-19 Genomics UK (COG-UK) Consortium | Lighthouse Laboratory Plymouth and Alex Alderton, Roberto Amato, Jeffrey Barrett, Sonia Goncalves, Ewan Harrison, David K. Jackson, Ian Johnston, Dominic Kwiatkowski, Cordelia Langford, John Sillitoe on behalf of the Wellcome Sanger Institute COVID-19 Surveillance Team |
| EPI_ISL_2606683, EPI_ISL_2606684 | Lighthouse Lab in Milton Keynes | Wellcome Sanger Institute for the COVID-19 Genomics UK (COG-UK) Consortium | The Lighthouse Lab in Milton Keynes and Alex Alderton, Roberto Amato, Jeffrey Barrett, Sonia Goncalves, Ewan Harrison, David K. Jackson, Ian Johnston, Dominic Kwiatkowski, Cordelia Langford, John Sillitoe on behalf of the Wellcome Sanger Institute COVID-19 Surveillance Team |
| EPI_ISL_2606685 | Lighthouse Laboratory Plymouth | Wellcome Sanger Institute for the COVID-19 Genomics UK (COG-UK) Consortium | Lighthouse Laboratory Plymouth and Alex Alderton, Roberto Amato, Jeffrey Barrett, Sonia Goncalves, Ewan Harrison, David K. Jackson, Ian Johnston, Dominic Kwiatkowski, Cordelia Langford, John Sillitoe on behalf of the Wellcome Sanger Institute COVID-19 Surveillance Team |
| EPI_ISL_2606686 | Lighthouse Lab in Glasgow | Wellcome Sanger Institute for the COVID-19 Genomics UK (COG-UK) Consortium | Harper VanSteenhouse, Yumi Kasai, David Gray, Carol Clugston, Anna Dominiczak and Alex Alderton, Roberto Amato, Jeffrey Barrett, Sonia Goncalves, Ewan Harrison, David K. Jackson, Ian Johnston, Dominic Kwiatkowski, Cordelia Langford, John Sillitoe on behalf of the Wellcome Sanger Institute COVID-19 Surveillance Team |
| EPI_ISL_2606687, EPI_ISL_2606688, EPI_ISL_2606689 | Lighthouse Lab in Milton Keynes | Wellcome Sanger Institute for the COVID-19 Genomics UK (COG-UK) Consortium | The Lighthouse Lab in Milton Keynes and Alex Alderton, Roberto Amato, Jeffrey Barrett, Sonia Goncalves, Ewan Harrison, David K. Jackson, Ian Johnston, Dominic Kwiatkowski, Cordelia Langford, John Sillitoe on behalf of the Wellcome Sanger Institute COVID-19 Surveillance Team |
| EPI_ISL_2606690 | Lighthouse Lab in Glasgow | Wellcome Sanger Institute for the COVID-19 Genomics UK (COG-UK) Consortium | Harper VanSteenhouse, Yumi Kasai, David Gray, Carol Clugston, Anna Dominiczak and Alex Alderton, Roberto Amato, Jeffrey Barrett, Sonia Goncalves, Ewan Harrison, David K. Jackson, Ian Johnston, Dominic Kwiatkowski, Cordelia Langford, John Sillitoe on behalf of the Wellcome Sanger Institute COVID-19 Surveillance Team |
| EPI_ISL_2606691 | Lighthouse Laboratory Plymouth | Wellcome Sanger Institute for the COVID-19 Genomics UK (COG-UK) Consortium | Lighthouse Laboratory Plymouth and Alex Alderton, Roberto Amato, Jeffrey Barrett, Sonia Goncalves, Ewan Harrison, David K. Jackson, Ian Johnston, Dominic Kwiatkowski, Cordelia Langford, John Sillitoe on behalf of the Wellcome Sanger Institute COVID-19 Surveillance Team |
| EPI_ISL_2606692, EPI_ISL_2606693, EPI_ISL_2606694 | Lighthouse Lab in Milton Keynes | Wellcome Sanger Institute for the COVID-19 Genomics UK (COG-UK) Consortium | The Lighthouse Lab in Milton Keynes and Alex Alderton, Roberto Amato, Jeffrey Barrett, Sonia Goncalves, Ewan Harrison, David K. Jackson, Ian Johnston, Dominic Kwiatkowski, Cordelia Langford, John Sillitoe on behalf of the Wellcome Sanger Institute COVID-19 Surveillance Team |
| EPI_ISL_2606695 | Lighthouse Laboratory Plymouth | Wellcome Sanger Institute for the COVID-19 Genomics UK (COG-UK) Consortium | Lighthouse Laboratory Plymouth and Alex Alderton, Roberto Amato, Jeffrey Barrett, Sonia Goncalves, Ewan Harrison, David K. Jackson, Ian Johnston, Dominic Kwiatkowski, Cordelia Langford, John Sillitoe on behalf of the Wellcome Sanger Institute COVID-19 Surveillance Team |
| EPI_ISL_2606696 | Lighthouse Lab in Milton Keynes | Wellcome Sanger Institute for the COVID-19 Genomics UK (COG-UK) Consortium | The Lighthouse Lab in Milton Keynes and Alex Alderton, Roberto Amato, Jeffrey Barrett, Sonia Goncalves, Ewan Harrison, David K. Jackson, Ian Johnston, Dominic Kwiatkowski, Cordelia Langford, John Sillitoe on behalf of the Wellcome Sanger Institute COVID-19 Surveillance Team |
| EPI_ISL_2606697 | Lighthouse Laboratory Plymouth | Wellcome Sanger Institute for the COVID-19 Genomics UK (COG-UK) Consortium | Lighthouse Laboratory Plymouth and Alex Alderton, Roberto Amato, Jeffrey Barrett, Sonia Goncalves, Ewan Harrison, David K. Jackson, Ian Johnston, Dominic Kwiatkowski, Cordelia Langford, John Sillitoe on behalf of the Wellcome Sanger Institute COVID-19 Surveillance Team |
| EPI_ISL_2606698, EPI_ISL_2606699 | Lighthouse Lab in Milton Keynes | Wellcome Sanger Institute for the COVID-19 Genomics UK (COG-UK) Consortium | The Lighthouse Lab in Milton Keynes and Alex Alderton, Roberto Amato, Jeffrey Barrett, Sonia Goncalves, Ewan Harrison, David K. Jackson, Ian Johnston, Dominic Kwiatkowski, Cordelia Langford, John Sillitoe on behalf of the Wellcome Sanger Institute COVID-19 Surveillance Team |
| EPI_ISL_2606700 | Lighthouse Laboratory Plymouth | Wellcome Sanger Institute for the COVID-19 Genomics UK (COG-UK) Consortium | Lighthouse Laboratory Plymouth and Alex Alderton, Roberto Amato, Jeffrey Barrett, Sonia Goncalves, Ewan Harrison, David K. Jackson, Ian Johnston, Dominic Kwiatkowski, Cordelia Langford, John Sillitoe on behalf of the Wellcome Sanger Institute COVID-19 Surveillance Team |
| EPI_ISL_2606701, EPI_ISL_2606702, EPI_ISL_2606703, EPI_ISL_2606705 | Lighthouse Lab in Milton Keynes | Wellcome Sanger Institute for the COVID-19 Genomics UK (COG-UK) Consortium | The Lighthouse Lab in Milton Keynes and Alex Alderton, Roberto Amato, Jeffrey Barrett, Sonia Goncalves, Ewan Harrison, David K. Jackson, Ian Johnston, Dominic Kwiatkowski, Cordelia Langford, John Sillitoe on behalf of the Wellcome Sanger Institute COVID-19 Surveillance Team |
| EPI_ISL_2606706 | Lighthouse Lab in Glasgow | Wellcome Sanger Institute for the COVID-19 Genomics UK (COG-UK) Consortium | Harper VanSteenhouse, Yumi Kasai, David Gray, Carol Clugston, Anna Dominiczak and Alex Alderton, Roberto Amato, Jeffrey Barrett, Sonia Goncalves, Ewan Harrison, David K. Jackson, Ian Johnston, Dominic Kwiatkowski, Cordelia Langford, John Sillitoe on behalf of the Wellcome Sanger Institute COVID-19 Surveillance Team |
| EPI_ISL_2606707, EPI_ISL_2606708, EPI_ISL_2606710, EPI_ISL_2606711 | Lighthouse Lab in Milton Keynes | Wellcome Sanger Institute for the COVID-19 Genomics UK (COG-UK) Consortium | The Lighthouse Lab in Milton Keynes and Alex Alderton, Roberto Amato, Jeffrey Barrett, Sonia Goncalves, Ewan Harrison, David K. Jackson, Ian Johnston, Dominic Kwiatkowski, Cordelia Langford, John Sillitoe on behalf of the Wellcome Sanger Institute COVID-19 Surveillance Team |
| EPI_ISL_2606712 | Lighthouse Lab in Glasgow | Wellcome Sanger Institute for the COVID-19 Genomics UK (COG-UK) Consortium | Harper VanSteenhouse, Yumi Kasai, David Gray, Carol Clugston, Anna Dominiczak and Alex Alderton, Roberto Amato, Jeffrey Barrett, Sonia Goncalves, Ewan Harrison, David K. Jackson, Ian Johnston, Dominic Kwiatkowski, Cordelia Langford, John Sillitoe on behalf of the Wellcome Sanger Institute COVID-19 Surveillance Team |
| EPI_ISL_2606713 | Lighthouse Laboratory Plymouth | Wellcome Sanger Institute for the COVID-19 Genomics UK (COG-UK) Consortium | Lighthouse Laboratory Plymouth and Alex Alderton, Roberto Amato, Jeffrey Barrett, Sonia Goncalves, Ewan Harrison, David K. Jackson, Ian Johnston, Dominic Kwiatkowski, Cordelia Langford, John Sillitoe on behalf of the Wellcome Sanger Institute COVID-19 Surveillance Team |
| EPI_ISL_2606714 | Lighthouse Lab in Milton Keynes | Wellcome Sanger Institute for the COVID-19 Genomics UK (COG-UK) Consortium | The Lighthouse Lab in Milton Keynes and Alex Alderton, Roberto Amato, Jeffrey Barrett, Sonia Goncalves, Ewan Harrison, David K. Jackson, Ian Johnston, Dominic Kwiatkowski, Cordelia Langford, John Sillitoe on behalf of the Wellcome Sanger Institute COVID-19 Surveillance Team |
| EPI_ISL_2606715, EPI_ISL_2606716, EPI_ISL_2606717 | Lighthouse Laboratory Plymouth | Wellcome Sanger Institute for the COVID-19 Genomics UK (COG-UK) Consortium | Lighthouse Laboratory Plymouth and Alex Alderton, Roberto Amato, Jeffrey Barrett, Sonia Goncalves, Ewan Harrison, David K. Jackson, Ian Johnston, Dominic Kwiatkowski, Cordelia Langford, John Sillitoe on behalf of the Wellcome Sanger Institute COVID-19 Surveillance Team |
| EPI_ISL_2606718, EPI_ISL_2606719, EPI_ISL_2606720 | Lighthouse Lab in Milton Keynes | Wellcome Sanger Institute for the COVID-19 Genomics UK (COG-UK) Consortium | The Lighthouse Lab in Milton Keynes and Alex Alderton, Roberto Amato, Jeffrey Barrett, Sonia Goncalves, Ewan Harrison, David K. Jackson, Ian Johnston, Dominic Kwiatkowski, Cordelia Langford, John Sillitoe on behalf of the Wellcome Sanger Institute COVID-19 Surveillance Team |
| EPI_ISL_2606721 | Lighthouse Lab in Glasgow | Wellcome Sanger Institute for the COVID-19 Genomics UK (COG-UK) Consortium | Harper VanSteenhouse, Yumi Kasai, David Gray, Carol Clugston, Anna Dominiczak and Alex Alderton, Roberto Amato, Jeffrey Barrett, Sonia Goncalves, Ewan Harrison, David K. Jackson, Ian Johnston, Dominic Kwiatkowski, Cordelia Langford, John Sillitoe on behalf of the Wellcome Sanger Institute COVID-19 Surveillance Team |
| EPI_ISL_2606722 | Lighthouse Lab in Milton Keynes | Wellcome Sanger Institute for the COVID-19 Genomics UK (COG-UK) Consortium | The Lighthouse Lab in Milton Keynes and Alex Alderton, Roberto Amato, Jeffrey Barrett, Sonia Goncalves, Ewan Harrison, David K. Jackson, Ian Johnston, Dominic Kwiatkowski, Cordelia Langford, John Sillitoe on behalf of the Wellcome Sanger Institute COVID-19 Surveillance Team |
| EPI_ISL_2606723 | Lighthouse Laboratory Plymouth | Wellcome Sanger Institute for the COVID-19 Genomics UK (COG-UK) Consortium | Lighthouse Laboratory Plymouth and Alex Alderton, Roberto Amato, Jeffrey Barrett, Sonia Goncalves, Ewan Harrison, David K. Jackson, Ian Johnston, Dominic Kwiatkowski, Cordelia Langford, John Sillitoe on behalf of the Wellcome Sanger Institute COVID-19 Surveillance Team |
| EPI_ISL_2606724, EPI_ISL_2606725, EPI_ISL_2606726 | Lighthouse Lab in Milton Keynes | Wellcome Sanger Institute for the COVID-19 Genomics UK (COG-UK) Consortium | The Lighthouse Lab in Milton Keynes and Alex Alderton, Roberto Amato, Jeffrey Barrett, Sonia Goncalves, Ewan Harrison, David K. Jackson, Ian Johnston, Dominic Kwiatkowski, Cordelia Langford, John Sillitoe on behalf of the Wellcome Sanger Institute COVID-19 Surveillance Team |
| EPI_ISL_2606727 | Lighthouse Lab in Alderley Park | Wellcome Sanger Institute for the COVID-19 Genomics UK (COG-UK) Consortium | Jacquelyn Wynn, Mairead Hyland, The Lighthouse Lab in Alderley Park and Alex Alderton, Roberto Amato, Jeffrey Barrett, Sonia Goncalves, Ewan Harrison, David K. Jackson, Ian Johnston, Dominic Kwiatkowski, Cordelia Langford, John Sillitoe on behalf of the Wellcome Sanger Institute COVID-19 Surveillance Team |
| EPI_ISL_2606728 | Lighthouse Laboratory Plymouth | Wellcome Sanger Institute for the COVID-19 Genomics UK (COG-UK) Consortium | Lighthouse Laboratory Plymouth and Alex Alderton, Roberto Amato, Jeffrey Barrett, Sonia Goncalves, Ewan Harrison, David K. Jackson, Ian Johnston, Dominic Kwiatkowski, Cordelia Langford, John Sillitoe on behalf of the Wellcome Sanger Institute COVID-19 Surveillance Team |
| EPI_ISL_2606729, | Lighthouse Lab in Alderley Park | Wellcome Sanger Institute for the COVID-19 Genomics UK (COG-UK) | Jacquelyn Wynn, Mairead Hyland, The Lighthouse Lab in Alderley Park and Alex Alderton, Roberto Amato, Jeffrey Barrett, Sonia Goncalves, Ewan Harrison, David K. Jackson, Ian Johnston, Dominic Kwiatkowski, Cordelia Langford, John Sillitoe on |

| | | | |
|---|---|---|---|
| EPI_ISL_2606730 | | Consortium | behalf of the Wellcome Sanger Institute COVID-19 Surveillance Team |
| EPI_ISL_2606731, EPI_ISL_2606732, EPI_ISL_2606733 | Lighthouse Lab in Milton Keynes | Wellcome Sanger Institute for the COVID-19 Genomics UK (COG-UK) Consortium | The Lighthouse Lab in Milton Keynes and Alex Alderton, Roberto Amato, Jeffrey Barrett, Sonia Goncalves, Ewan Harrison, David K. Jackson, Ian Johnston, Dominic Kwiatkowski, Cordelia Langford, John Sillitoe on behalf of the Wellcome Sanger Institute COVID-19 Surveillance Team |
| EPI_ISL_2606734, EPI_ISL_2606735 | Lighthouse Lab in Alderley Park | Wellcome Sanger Institute for the COVID-19 Genomics UK (COG-UK) Consortium | Jacquelyn Wynn, Mairead Hyland, The Lighthouse Lab in Alderley Park and Alex Alderton, Roberto Amato, Jeffrey Barrett, Sonia Goncalves, Ewan Harrison, David K. Jackson, Ian Johnston, Dominic Kwiatkowski, Cordelia Langford, John Sillitoe on behalf of the Wellcome Sanger Institute COVID-19 Surveillance Team |
| EPI_ISL_2606736, EPI_ISL_2606737 | Lighthouse Lab in Glasgow | Wellcome Sanger Institute for the COVID-19 Genomics UK (COG-UK) Consortium | Harper VanSteenhouse, Yumi Kasai, David Gray, Carol Clugston, Anna Dominiczak and Alex Alderton, Roberto Amato, Jeffrey Barrett, Sonia Goncalves, Ewan Harrison, David K. Jackson, Ian Johnston, Dominic Kwiatkowski, Cordelia Langford, John Sillitoe on behalf of the Wellcome Sanger Institute COVID-19 Surveillance Team |
| EPI_ISL_2606738 | Lighthouse Lab in Alderley Park | Wellcome Sanger Institute for the COVID-19 Genomics UK (COG-UK) Consortium | Jacquelyn Wynn, Mairead Hyland, The Lighthouse Lab in Alderley Park and Alex Alderton, Roberto Amato, Jeffrey Barrett, Sonia Goncalves, Ewan Harrison, David K. Jackson, Ian Johnston, Dominic Kwiatkowski, Cordelia Langford, John Sillitoe on behalf of the Wellcome Sanger Institute COVID-19 Surveillance Team |
| EPI_ISL_2606740 | Lighthouse Lab in Glasgow | Wellcome Sanger Institute for the COVID-19 Genomics UK (COG-UK) Consortium | Harper VanSteenhouse, Yumi Kasai, David Gray, Carol Clugston, Anna Dominiczak and Alex Alderton, Roberto Amato, Jeffrey Barrett, Sonia Goncalves, Ewan Harrison, David K. Jackson, Ian Johnston, Dominic Kwiatkowski, Cordelia Langford, John Sillitoe on behalf of the Wellcome Sanger Institute COVID-19 Surveillance Team |
| EPI_ISL_2606741, EPI_ISL_2606742 | Lighthouse Lab in Milton Keynes | Wellcome Sanger Institute for the COVID-19 Genomics UK (COG-UK) Consortium | The Lighthouse Lab in Milton Keynes and Alex Alderton, Roberto Amato, Jeffrey Barrett, Sonia Goncalves, Ewan Harrison, David K. Jackson, Ian Johnston, Dominic Kwiatkowski, Cordelia Langford, John Sillitoe on behalf of the Wellcome Sanger Institute COVID-19 Surveillance Team |
| EPI_ISL_2606743 | Lighthouse Laboratory Plymouth | Wellcome Sanger Institute for the COVID-19 Genomics UK (COG-UK) Consortium | Lighthouse Laboratory Plymouth and Alex Alderton, Roberto Amato, Jeffrey Barrett, Sonia Goncalves, Ewan Harrison, David K. Jackson, Ian Johnston, Dominic Kwiatkowski, Cordelia Langford, John Sillitoe on behalf of the Wellcome Sanger Institute COVID-19 Surveillance Team |
| EPI_ISL_2606744, EPI_ISL_2606745 | Lighthouse Lab in Milton Keynes | Wellcome Sanger Institute for the COVID-19 Genomics UK (COG-UK) Consortium | The Lighthouse Lab in Milton Keynes and Alex Alderton, Roberto Amato, Jeffrey Barrett, Sonia Goncalves, Ewan Harrison, David K. Jackson, Ian Johnston, Dominic Kwiatkowski, Cordelia Langford, John Sillitoe on behalf of the Wellcome Sanger Institute COVID-19 Surveillance Team |
| EPI_ISL_2606746 | Lighthouse Lab in Alderley Park | Wellcome Sanger Institute for the COVID-19 Genomics UK (COG-UK) Consortium | Jacquelyn Wynn, Mairead Hyland, The Lighthouse Lab in Alderley Park and Alex Alderton, Roberto Amato, Jeffrey Barrett, Sonia Goncalves, Ewan Harrison, David K. Jackson, Ian Johnston, Dominic Kwiatkowski, Cordelia Langford, John Sillitoe on behalf of the Wellcome Sanger Institute COVID-19 Surveillance Team |
| EPI_ISL_2606747, EPI_ISL_2606749, EPI_ISL_2606750, EPI_ISL_2606751 | Lighthouse Lab in Milton Keynes | Wellcome Sanger Institute for the COVID-19 Genomics UK (COG-UK) Consortium | The Lighthouse Lab in Milton Keynes and Alex Alderton, Roberto Amato, Jeffrey Barrett, Sonia Goncalves, Ewan Harrison, David K. Jackson, Ian Johnston, Dominic Kwiatkowski, Cordelia Langford, John Sillitoe on behalf of the Wellcome Sanger Institute COVID-19 Surveillance Team |
| EPI_ISL_2606752 | Lighthouse Lab in Alderley Park | Wellcome Sanger Institute for the COVID-19 Genomics UK (COG-UK) Consortium | Jacquelyn Wynn, Mairead Hyland, The Lighthouse Lab in Alderley Park and Alex Alderton, Roberto Amato, Jeffrey Barrett, Sonia Goncalves, Ewan Harrison, David K. Jackson, Ian Johnston, Dominic Kwiatkowski, Cordelia Langford, John Sillitoe on behalf of the Wellcome Sanger Institute COVID-19 Surveillance Team |
| EPI_ISL_2606753 | Lighthouse Lab in Milton Keynes | Wellcome Sanger Institute for the COVID-19 Genomics UK (COG-UK) Consortium | The Lighthouse Lab in Milton Keynes and Alex Alderton, Roberto Amato, Jeffrey Barrett, Sonia Goncalves, Ewan Harrison, David K. Jackson, Ian Johnston, Dominic Kwiatkowski, Cordelia Langford, John Sillitoe on behalf of the Wellcome Sanger Institute COVID-19 Surveillance Team |
| EPI_ISL_2606754 | Lighthouse Lab in Glasgow | Wellcome Sanger Institute for the COVID-19 Genomics UK (COG-UK) Consortium | Harper VanSteenhouse, Yumi Kasai, David Gray, Carol Clugston, Anna Dominiczak and Alex Alderton, Roberto Amato, Jeffrey Barrett, Sonia Goncalves, Ewan Harrison, David K. Jackson, Ian Johnston, Dominic Kwiatkowski, Cordelia Langford, John Sillitoe on behalf of the Wellcome Sanger Institute COVID-19 Surveillance Team |
| EPI_ISL_2606755 | Lighthouse Lab in Alderley Park | Wellcome Sanger Institute for the COVID-19 Genomics UK (COG-UK) Consortium | Jacquelyn Wynn, Mairead Hyland, The Lighthouse Lab in Alderley Park and Alex Alderton, Roberto Amato, Jeffrey Barrett, Sonia Goncalves, Ewan Harrison, David K. Jackson, Ian Johnston, Dominic Kwiatkowski, Cordelia Langford, John Sillitoe on behalf of the Wellcome Sanger Institute COVID-19 Surveillance Team |
| EPI_ISL_2606756 | Lighthouse Lab in Milton Keynes | Wellcome Sanger Institute for the COVID-19 Genomics UK (COG-UK) Consortium | The Lighthouse Lab in Milton Keynes and Alex Alderton, Roberto Amato, Jeffrey Barrett, Sonia Goncalves, Ewan Harrison, David K. Jackson, Ian Johnston, Dominic Kwiatkowski, Cordelia Langford, John Sillitoe on behalf of the Wellcome Sanger Institute COVID-19 Surveillance Team |
| EPI_ISL_2606757 | Lighthouse Lab in Alderley Park | Wellcome Sanger Institute for the COVID-19 Genomics UK (COG-UK) Consortium | Jacquelyn Wynn, Mairead Hyland, The Lighthouse Lab in Alderley Park and Alex Alderton, Roberto Amato, Jeffrey Barrett, Sonia Goncalves, Ewan Harrison, David K. Jackson, Ian Johnston, Dominic Kwiatkowski, Cordelia Langford, John Sillitoe on behalf of the Wellcome Sanger Institute COVID-19 Surveillance Team |
| EPI_ISL_2606758 | Lighthouse Lab in Glasgow | Wellcome Sanger Institute for the COVID-19 Genomics UK (COG-UK) Consortium | Harper VanSteenhouse, Yumi Kasai, David Gray, Carol Clugston, Anna Dominiczak and Alex Alderton, Roberto Amato, Jeffrey Barrett, Sonia Goncalves, Ewan Harrison, David K. Jackson, Ian Johnston, Dominic Kwiatkowski, Cordelia Langford, John Sillitoe on behalf of the Wellcome Sanger Institute COVID-19 Surveillance Team |
| EPI_ISL_2606759 | Lighthouse Lab in Alderley Park | Wellcome Sanger Institute for the COVID-19 Genomics UK (COG-UK) Consortium | Jacquelyn Wynn, Mairead Hyland, The Lighthouse Lab in Alderley Park and Alex Alderton, Roberto Amato, Jeffrey Barrett, Sonia Goncalves, Ewan Harrison, David K. Jackson, Ian Johnston, Dominic Kwiatkowski, Cordelia Langford, John Sillitoe on behalf of the Wellcome Sanger Institute COVID-19 Surveillance Team |
| EPI_ISL_2606760 | Lighthouse Lab in Glasgow | Wellcome Sanger Institute for the COVID-19 Genomics UK (COG-UK) Consortium | Harper VanSteenhouse, Yumi Kasai, David Gray, Carol Clugston, Anna Dominiczak and Alex Alderton, Roberto Amato, Jeffrey Barrett, Sonia Goncalves, Ewan Harrison, David K. Jackson, Ian Johnston, Dominic Kwiatkowski, Cordelia Langford, John Sillitoe on behalf of the Wellcome Sanger Institute COVID-19 Surveillance Team |
| EPI_ISL_2606761, EPI_ISL_2606762 | Lighthouse Lab in Alderley Park | Wellcome Sanger Institute for the COVID-19 Genomics UK (COG-UK) Consortium | Jacquelyn Wynn, Mairead Hyland, The Lighthouse Lab in Alderley Park and Alex Alderton, Roberto Amato, Jeffrey Barrett, Sonia Goncalves, Ewan Harrison, David K. Jackson, Ian Johnston, Dominic Kwiatkowski, Cordelia Langford, John Sillitoe on behalf of the Wellcome Sanger Institute COVID-19 Surveillance Team |
| EPI_ISL_2606763 | Lighthouse Laboratory Plymouth | Wellcome Sanger Institute for the COVID-19 Genomics UK (COG-UK) Consortium | Lighthouse Laboratory Plymouth and Alex Alderton, Roberto Amato, Jeffrey Barrett, Sonia Goncalves, Ewan Harrison, David K. Jackson, Ian Johnston, Dominic Kwiatkowski, Cordelia Langford, John Sillitoe on behalf of the Wellcome Sanger Institute COVID-19 Surveillance Team |
| EPI_ISL_2606764 | Lighthouse Lab in Alderley Park | Wellcome Sanger Institute for the COVID-19 Genomics UK (COG-UK) Consortium | Jacquelyn Wynn, Mairead Hyland, The Lighthouse Lab in Alderley Park and Alex Alderton, Roberto Amato, Jeffrey Barrett, Sonia Goncalves, Ewan Harrison, David K. Jackson, Ian Johnston, Dominic Kwiatkowski, Cordelia Langford, John Sillitoe on behalf of the Wellcome Sanger Institute COVID-19 Surveillance Team |
| EPI_ISL_2606765 | Lighthouse Laboratory Plymouth | Wellcome Sanger Institute for the COVID-19 Genomics UK (COG-UK) Consortium | Lighthouse Laboratory Plymouth and Alex Alderton, Roberto Amato, Jeffrey Barrett, Sonia Goncalves, Ewan Harrison, David K. Jackson, Ian Johnston, Dominic Kwiatkowski, Cordelia Langford, John Sillitoe on behalf of the Wellcome Sanger Institute COVID-19 Surveillance Team |
| EPI_ISL_2606766 | Lighthouse Lab in Alderley Park | Wellcome Sanger Institute for the COVID-19 Genomics UK (COG-UK) Consortium | Jacquelyn Wynn, Mairead Hyland, The Lighthouse Lab in Alderley Park and Alex Alderton, Roberto Amato, Jeffrey Barrett, Sonia Goncalves, Ewan Harrison, David K. Jackson, Ian Johnston, Dominic Kwiatkowski, Cordelia Langford, John Sillitoe on behalf of the Wellcome Sanger Institute COVID-19 Surveillance Team |
| EPI_ISL_2606767 | Lighthouse Lab in Milton Keynes | Wellcome Sanger Institute for the COVID-19 Genomics UK (COG-UK) Consortium | The Lighthouse Lab in Milton Keynes and Alex Alderton, Roberto Amato, Jeffrey Barrett, Sonia Goncalves, Ewan Harrison, David K. Jackson, Ian Johnston, Dominic Kwiatkowski, Cordelia Langford, John Sillitoe on behalf of the Wellcome Sanger Institute COVID-19 Surveillance Team |
| EPI_ISL_2606768 | Lighthouse Lab in Glasgow | Wellcome Sanger Institute for the COVID-19 Genomics UK (COG-UK) Consortium | Harper VanSteenhouse, Yumi Kasai, David Gray, Carol Clugston, Anna Dominiczak and Alex Alderton, Roberto Amato, Jeffrey Barrett, Sonia Goncalves, Ewan Harrison, David K. Jackson, Ian Johnston, Dominic Kwiatkowski, Cordelia Langford, John Sillitoe on behalf of the Wellcome Sanger Institute COVID-19 Surveillance Team |
| EPI_ISL_2606769 | Lighthouse Lab in Alderley Park | Wellcome Sanger Institute for the COVID-19 Genomics UK (COG-UK) Consortium | Jacquelyn Wynn, Mairead Hyland, The Lighthouse Lab in Alderley Park and Alex Alderton, Roberto Amato, Jeffrey Barrett, Sonia Goncalves, Ewan Harrison, David K. Jackson, Ian Johnston, Dominic Kwiatkowski, Cordelia Langford, John Sillitoe on behalf of the Wellcome Sanger Institute COVID-19 Surveillance Team |
| EPI_ISL_2606770, EPI_ISL_2606771 | Lighthouse Lab in Glasgow | Wellcome Sanger Institute for the COVID-19 Genomics UK (COG-UK) Consortium | Harper VanSteenhouse, Yumi Kasai, David Gray, Carol Clugston, Anna Dominiczak and Alex Alderton, Roberto Amato, Jeffrey Barrett, Sonia Goncalves, Ewan Harrison, David K. Jackson, Ian Johnston, Dominic Kwiatkowski, Cordelia Langford, John Sillitoe on behalf of the Wellcome Sanger Institute COVID-19 Surveillance Team |
| EPI_ISL_2606772, EPI_ISL_2606773 | Lighthouse Lab in Alderley Park | Wellcome Sanger Institute for the COVID-19 Genomics UK (COG-UK) Consortium | Jacquelyn Wynn, Mairead Hyland, The Lighthouse Lab in Alderley Park and Alex Alderton, Roberto Amato, Jeffrey Barrett, Sonia Goncalves, Ewan Harrison, David K. Jackson, Ian Johnston, Dominic Kwiatkowski, Cordelia Langford, John Sillitoe on behalf of the Wellcome Sanger Institute COVID-19 Surveillance Team |
| EPI_ISL_2606774, EPI_ISL_2606775 | Lighthouse Lab in Glasgow | Wellcome Sanger Institute for the COVID-19 Genomics UK (COG-UK) Consortium | Harper VanSteenhouse, Yumi Kasai, David Gray, Carol Clugston, Anna Dominiczak and Alex Alderton, Roberto Amato, Jeffrey Barrett, Sonia Goncalves, Ewan Harrison, David K. Jackson, Ian Johnston, Dominic Kwiatkowski, Cordelia Langford, John Sillitoe on behalf of the Wellcome Sanger Institute COVID-19 Surveillance Team |
| EPI_ISL_2606776, EPI_ISL_2606777 | Lighthouse Lab in Alderley Park | Wellcome Sanger Institute for the COVID-19 Genomics UK (COG-UK) Consortium | Jacquelyn Wynn, Mairead Hyland, The Lighthouse Lab in Alderley Park and Alex Alderton, Roberto Amato, Jeffrey Barrett, Sonia Goncalves, Ewan Harrison, David K. Jackson, Ian Johnston, Dominic Kwiatkowski, Cordelia Langford, John Sillitoe on behalf of the Wellcome Sanger Institute COVID-19 Surveillance Team |
| EPI_ISL_2606778 | Lighthouse Lab in Glasgow | Wellcome Sanger Institute for the COVID-19 Genomics UK (COG-UK) Consortium | Harper VanSteenhouse, Yumi Kasai, David Gray, Carol Clugston, Anna Dominiczak and Alex Alderton, Roberto Amato, Jeffrey Barrett, Sonia Goncalves, Ewan Harrison, David K. Jackson, Ian Johnston, Dominic Kwiatkowski, Cordelia Langford, John Sillitoe on behalf of the Wellcome Sanger Institute COVID-19 Surveillance Team |
| EPI_ISL_2606780 | Lighthouse Lab in Alderley Park | Wellcome Sanger Institute for the COVID-19 Genomics UK (COG-UK) Consortium | Jacquelyn Wynn, Mairead Hyland, The Lighthouse Lab in Alderley Park and Alex Alderton, Roberto Amato, Jeffrey Barrett, Sonia Goncalves, Ewan Harrison, David K. Jackson, Ian Johnston, Dominic Kwiatkowski, Cordelia Langford, John Sillitoe on behalf of the Wellcome Sanger Institute COVID-19 Surveillance Team |
| EPI_ISL_2606781 | Lighthouse Lab in Milton Keynes | Wellcome Sanger Institute for the COVID-19 Genomics UK (COG-UK) Consortium | The Lighthouse Lab in Milton Keynes and Alex Alderton, Roberto Amato, Jeffrey Barrett, Sonia Goncalves, Ewan Harrison, David K. Jackson, Ian Johnston, Dominic Kwiatkowski, Cordelia Langford, John Sillitoe on behalf of the Wellcome Sanger Institute COVID-19 Surveillance Team |
| EPI_ISL_2606782, EPI_ISL_2606783, EPI_ISL_2606784 | Lighthouse Lab in Alderley Park | Wellcome Sanger Institute for the COVID-19 Genomics UK (COG-UK) Consortium | Jacquelyn Wynn, Mairead Hyland, The Lighthouse Lab in Alderley Park and Alex Alderton, Roberto Amato, Jeffrey Barrett, Sonia Goncalves, Ewan Harrison, David K. Jackson, Ian Johnston, Dominic Kwiatkowski, Cordelia Langford, John Sillitoe on behalf of the Wellcome Sanger Institute COVID-19 Surveillance Team |
| EPI_ISL_2606785 | Lighthouse Lab in Milton Keynes | Wellcome Sanger Institute for the COVID-19 Genomics UK (COG-UK) Consortium | The Lighthouse Lab in Milton Keynes and Alex Alderton, Roberto Amato, Jeffrey Barrett, Sonia Goncalves, Ewan Harrison, David K. Jackson, Ian Johnston, Dominic Kwiatkowski, Cordelia Langford, John Sillitoe on behalf of the Wellcome Sanger Institute COVID-19 Surveillance Team |
| EPI_ISL_2606786, EPI_ISL_2606788 | Lighthouse Lab in Glasgow | Wellcome Sanger Institute for the COVID-19 Genomics UK (COG-UK) Consortium | Harper VanSteenhouse, Yumi Kasai, David Gray, Carol Clugston, Anna Dominiczak and Alex Alderton, Roberto Amato, Jeffrey Barrett, Sonia Goncalves, Ewan Harrison, David K. Jackson, Ian Johnston, Dominic Kwiatkowski, Cordelia Langford, John Sillitoe on behalf of the Wellcome Sanger Institute COVID-19 Surveillance Team |
| EPI_ISL_2606789 | Lighthouse Lab in Milton Keynes | Wellcome Sanger Institute for the COVID-19 Genomics UK (COG-UK) Consortium | The Lighthouse Lab in Milton Keynes and Alex Alderton, Roberto Amato, Jeffrey Barrett, Sonia Goncalves, Ewan Harrison, David K. Jackson, Ian Johnston, Dominic Kwiatkowski, Cordelia Langford, John Sillitoe on behalf of the Wellcome Sanger Institute COVID-19 Surveillance Team |
| EPI_ISL_2606790 | Lighthouse Lab in Glasgow | Wellcome Sanger Institute for the COVID-19 Genomics UK (COG-UK) Consortium | Harper VanSteenhouse, Yumi Kasai, David Gray, Carol Clugston, Anna Dominiczak and Alex Alderton, Roberto Amato, Jeffrey Barrett, Sonia Goncalves, Ewan Harrison, David K. Jackson, Ian Johnston, Dominic Kwiatkowski, Cordelia Langford, John Sillitoe on behalf of the Wellcome Sanger Institute COVID-19 Surveillance Team |
| EPI_ISL_2606791 | Lighthouse Lab in Milton Keynes | Wellcome Sanger Institute for the COVID-19 Genomics UK (COG-UK) Consortium | The Lighthouse Lab in Milton Keynes and Alex Alderton, Roberto Amato, Jeffrey Barrett, Sonia Goncalves, Ewan Harrison, David K. Jackson, Ian Johnston, Dominic Kwiatkowski, Cordelia Langford, John Sillitoe on behalf of the Wellcome Sanger Institute COVID-19 Surveillance Team |
| EPI_ISL_2606792 | Lighthouse Lab in Alderley Park | Wellcome Sanger Institute for the COVID-19 Genomics UK (COG-UK) Consortium | Jacquelyn Wynn, Mairead Hyland, The Lighthouse Lab in Alderley Park and Alex Alderton, Roberto Amato, Jeffrey Barrett, Sonia Goncalves, Ewan Harrison, David K. Jackson, Ian Johnston, Dominic Kwiatkowski, Cordelia Langford, John Sillitoe on behalf of the Wellcome Sanger Institute COVID-19 Surveillance Team |
| EPI_ISL_2606793 | Lighthouse Lab in Glasgow | Wellcome Sanger Institute for the COVID-19 Genomics UK (COG-UK) Consortium | Harper VanSteenhouse, Yumi Kasai, David Gray, Carol Clugston, Anna Dominiczak and Alex Alderton, Roberto Amato, Jeffrey Barrett, Sonia Goncalves, Ewan Harrison, David K. Jackson, Ian Johnston, Dominic Kwiatkowski, Cordelia Langford, John Sillitoe on behalf of the Wellcome Sanger Institute COVID-19 Surveillance Team |
| EPI_ISL_2606794 | Lighthouse Lab in Milton Keynes | Wellcome Sanger Institute for the COVID-19 Genomics UK (COG-UK) Consortium | The Lighthouse Lab in Milton Keynes and Alex Alderton, Roberto Amato, Jeffrey Barrett, Sonia Goncalves, Ewan Harrison, David K. Jackson, Ian Johnston, Dominic Kwiatkowski, Cordelia Langford, John Sillitoe on behalf of the Wellcome Sanger Institute COVID-19 Surveillance Team |

| | | | |
|---|---|---|---|
| EPI_ISL_2606795 | Lighthouse Laboratory Plymouth | Wellcome Sanger Institute for the COVID-19 Genomics UK (COG-UK) Consortium | Lighthouse Laboratory Plymouth and Alex Alderton, Roberto Amato, Jeffrey Barrett, Sonia Goncalves, Ewan Harrison, David K. Jackson, Ian Johnston, Dominic Kwiatkowski, Cordelia Langford, John Sillitoe on behalf of the Wellcome Sanger Institute COVID-19 Surveillance Team |
| EPI_ISL_2606796 | Lighthouse Lab in Alderley Park | Wellcome Sanger Institute for the COVID-19 Genomics UK (COG-UK) Consortium | Jacquelyn Wynn, Mairead Hyland, The Lighthouse Lab in Alderley Park and Alex Alderton, Roberto Amato, Jeffrey Barrett, Sonia Goncalves, Ewan Harrison, David K. Jackson, Ian Johnston, Dominic Kwiatkowski, Cordelia Langford, John Sillitoe on behalf of the Wellcome Sanger Institute COVID-19 Surveillance Team |
| EPI_ISL_2606797 | Lighthouse Lab in Milton Keynes | Wellcome Sanger Institute for the COVID-19 Genomics UK (COG-UK) Consortium | The Lighthouse Lab in Milton Keynes and Alex Alderton, Roberto Amato, Jeffrey Barrett, Sonia Goncalves, Ewan Harrison, David K. Jackson, Ian Johnston, Dominic Kwiatkowski, Cordelia Langford, John Sillitoe on behalf of the Wellcome Sanger Institute COVID-19 Surveillance Team |
| EPI_ISL_2606798 | Lighthouse Lab in Alderley Park | Wellcome Sanger Institute for the COVID-19 Genomics UK (COG-UK) Consortium | Jacquelyn Wynn, Mairead Hyland, The Lighthouse Lab in Alderley Park and Alex Alderton, Roberto Amato, Jeffrey Barrett, Sonia Goncalves, Ewan Harrison, David K. Jackson, Ian Johnston, Dominic Kwiatkowski, Cordelia Langford, John Sillitoe on behalf of the Wellcome Sanger Institute COVID-19 Surveillance Team |
| EPI_ISL_2606799, EPI_ISL_2606800 | Lighthouse Laboratory Plymouth | Wellcome Sanger Institute for the COVID-19 Genomics UK (COG-UK) Consortium | Lighthouse Laboratory Plymouth and Alex Alderton, Roberto Amato, Jeffrey Barrett, Sonia Goncalves, Ewan Harrison, David K. Jackson, Ian Johnston, Dominic Kwiatkowski, Cordelia Langford, John Sillitoe on behalf of the Wellcome Sanger Institute COVID-19 Surveillance Team |
| EPI_ISL_2606801, EPI_ISL_2606802 | Lighthouse Lab in Alderley Park | Wellcome Sanger Institute for the COVID-19 Genomics UK (COG-UK) Consortium | Jacquelyn Wynn, Mairead Hyland, The Lighthouse Lab in Alderley Park and Alex Alderton, Roberto Amato, Jeffrey Barrett, Sonia Goncalves, Ewan Harrison, David K. Jackson, Ian Johnston, Dominic Kwiatkowski, Cordelia Langford, John Sillitoe on behalf of the Wellcome Sanger Institute COVID-19 Surveillance Team |
| EPI_ISL_2606803 | Lighthouse Lab in Glasgow | Wellcome Sanger Institute for the COVID-19 Genomics UK (COG-UK) Consortium | Harper VanSteenhouse, Yumi Kasai, David Gray, Carol Clugston, Anna Dominiczak and Alex Alderton, Roberto Amato, Jeffrey Barrett, Sonia Goncalves, Ewan Harrison, David K. Jackson, Ian Johnston, Dominic Kwiatkowski, Cordelia Langford, John Sillitoe on behalf of the Wellcome Sanger Institute COVID-19 Surveillance Team |
| EPI_ISL_2606804 | Lighthouse Lab in Milton Keynes | Wellcome Sanger Institute for the COVID-19 Genomics UK (COG-UK) Consortium | The Lighthouse Lab in Milton Keynes and Alex Alderton, Roberto Amato, Jeffrey Barrett, Sonia Goncalves, Ewan Harrison, David K. Jackson, Ian Johnston, Dominic Kwiatkowski, Cordelia Langford, John Sillitoe on behalf of the Wellcome Sanger Institute COVID-19 Surveillance Team |
| EPI_ISL_2606806, EPI_ISL_2606807 | Lighthouse Lab in Glasgow | Wellcome Sanger Institute for the COVID-19 Genomics UK (COG-UK) Consortium | Harper VanSteenhouse, Yumi Kasai, David Gray, Carol Clugston, Anna Dominiczak and Alex Alderton, Roberto Amato, Jeffrey Barrett, Sonia Goncalves, Ewan Harrison, David K. Jackson, Ian Johnston, Dominic Kwiatkowski, Cordelia Langford, John Sillitoe on behalf of the Wellcome Sanger Institute COVID-19 Surveillance Team |
| EPI_ISL_2606808, EPI_ISL_2606809, EPI_ISL_2606810, EPI_ISL_2606811, EPI_ISL_2606812 | Lighthouse Lab in Alderley Park | Wellcome Sanger Institute for the COVID-19 Genomics UK (COG-UK) Consortium | Jacquelyn Wynn, Mairead Hyland, The Lighthouse Lab in Alderley Park and Alex Alderton, Roberto Amato, Jeffrey Barrett, Sonia Goncalves, Ewan Harrison, David K. Jackson, Ian Johnston, Dominic Kwiatkowski, Cordelia Langford, John Sillitoe on behalf of the Wellcome Sanger Institute COVID-19 Surveillance Team |
| EPI_ISL_2606813 | Lighthouse Laboratory Plymouth | Wellcome Sanger Institute for the COVID-19 Genomics UK (COG-UK) Consortium | Lighthouse Laboratory Plymouth and Alex Alderton, Roberto Amato, Jeffrey Barrett, Sonia Goncalves, Ewan Harrison, David K. Jackson, Ian Johnston, Dominic Kwiatkowski, Cordelia Langford, John Sillitoe on behalf of the Wellcome Sanger Institute COVID-19 Surveillance Team |
| EPI_ISL_2606814 | Lighthouse Lab in Alderley Park | Wellcome Sanger Institute for the COVID-19 Genomics UK (COG-UK) Consortium | Jacquelyn Wynn, Mairead Hyland, The Lighthouse Lab in Alderley Park and Alex Alderton, Roberto Amato, Jeffrey Barrett, Sonia Goncalves, Ewan Harrison, David K. Jackson, Ian Johnston, Dominic Kwiatkowski, Cordelia Langford, John Sillitoe on behalf of the Wellcome Sanger Institute COVID-19 Surveillance Team |
| EPI_ISL_2606815 | Lighthouse Laboratory Plymouth | Wellcome Sanger Institute for the COVID-19 Genomics UK (COG-UK) Consortium | Lighthouse Laboratory Plymouth and Alex Alderton, Roberto Amato, Jeffrey Barrett, Sonia Goncalves, Ewan Harrison, David K. Jackson, Ian Johnston, Dominic Kwiatkowski, Cordelia Langford, John Sillitoe on behalf of the Wellcome Sanger Institute COVID-19 Surveillance Team |
| EPI_ISL_2606816, EPI_ISL_2606817 | Lighthouse Lab in Milton Keynes | Wellcome Sanger Institute for the COVID-19 Genomics UK (COG-UK) Consortium | The Lighthouse Lab in Milton Keynes and Alex Alderton, Roberto Amato, Jeffrey Barrett, Sonia Goncalves, Ewan Harrison, David K. Jackson, Ian Johnston, Dominic Kwiatkowski, Cordelia Langford, John Sillitoe on behalf of the Wellcome Sanger Institute COVID-19 Surveillance Team |
| EPI_ISL_2606818 | Lighthouse Laboratory Plymouth | Wellcome Sanger Institute for the COVID-19 Genomics UK (COG-UK) Consortium | Lighthouse Laboratory Plymouth and Alex Alderton, Roberto Amato, Jeffrey Barrett, Sonia Goncalves, Ewan Harrison, David K. Jackson, Ian Johnston, Dominic Kwiatkowski, Cordelia Langford, John Sillitoe on behalf of the Wellcome Sanger Institute COVID-19 Surveillance Team |
| EPI_ISL_2606819 | Lighthouse Lab in Glasgow | Wellcome Sanger Institute for the COVID-19 Genomics UK (COG-UK) Consortium | Harper VanSteenhouse, Yumi Kasai, David Gray, Carol Clugston, Anna Dominiczak and Alex Alderton, Roberto Amato, Jeffrey Barrett, Sonia Goncalves, Ewan Harrison, David K. Jackson, Ian Johnston, Dominic Kwiatkowski, Cordelia Langford, John Sillitoe on behalf of the Wellcome Sanger Institute COVID-19 Surveillance Team |
| EPI_ISL_2606820 | Lighthouse Lab in Alderley Park | Wellcome Sanger Institute for the COVID-19 Genomics UK (COG-UK) Consortium | Jacquelyn Wynn, Mairead Hyland, The Lighthouse Lab in Alderley Park and Alex Alderton, Roberto Amato, Jeffrey Barrett, Sonia Goncalves, Ewan Harrison, David K. Jackson, Ian Johnston, Dominic Kwiatkowski, Cordelia Langford, John Sillitoe on behalf of the Wellcome Sanger Institute COVID-19 Surveillance Team |
| EPI_ISL_2606821 | Lighthouse Lab in Milton Keynes | Wellcome Sanger Institute for the COVID-19 Genomics UK (COG-UK) Consortium | The Lighthouse Lab in Milton Keynes and Alex Alderton, Roberto Amato, Jeffrey Barrett, Sonia Goncalves, Ewan Harrison, David K. Jackson, Ian Johnston, Dominic Kwiatkowski, Cordelia Langford, John Sillitoe on behalf of the Wellcome Sanger Institute COVID-19 Surveillance Team |
| EPI_ISL_2606822 | Lighthouse Lab in Alderley Park | Wellcome Sanger Institute for the COVID-19 Genomics UK (COG-UK) Consortium | Jacquelyn Wynn, Mairead Hyland, The Lighthouse Lab in Alderley Park and Alex Alderton, Roberto Amato, Jeffrey Barrett, Sonia Goncalves, Ewan Harrison, David K. Jackson, Ian Johnston, Dominic Kwiatkowski, Cordelia Langford, John Sillitoe on behalf of the Wellcome Sanger Institute COVID-19 Surveillance Team |
| EPI_ISL_2606823, EPI_ISL_2606824 | Lighthouse Laboratory Plymouth | Wellcome Sanger Institute for the COVID-19 Genomics UK (COG-UK) Consortium | Lighthouse Laboratory Plymouth and Alex Alderton, Roberto Amato, Jeffrey Barrett, Sonia Goncalves, Ewan Harrison, David K. Jackson, Ian Johnston, Dominic Kwiatkowski, Cordelia Langford, John Sillitoe on behalf of the Wellcome Sanger Institute COVID-19 Surveillance Team |
| EPI_ISL_2606825, EPI_ISL_2606826 | Lighthouse Lab in Alderley Park | Wellcome Sanger Institute for the COVID-19 Genomics UK (COG-UK) Consortium | Jacquelyn Wynn, Mairead Hyland, The Lighthouse Lab in Alderley Park and Alex Alderton, Roberto Amato, Jeffrey Barrett, Sonia Goncalves, Ewan Harrison, David K. Jackson, Ian Johnston, Dominic Kwiatkowski, Cordelia Langford, John Sillitoe on behalf of the Wellcome Sanger Institute COVID-19 Surveillance Team |
| EPI_ISL_2606827 | Lighthouse Lab in Milton Keynes | Wellcome Sanger Institute for the COVID-19 Genomics UK (COG-UK) Consortium | The Lighthouse Lab in Milton Keynes and Alex Alderton, Roberto Amato, Jeffrey Barrett, Sonia Goncalves, Ewan Harrison, David K. Jackson, Ian Johnston, Dominic Kwiatkowski, Cordelia Langford, John Sillitoe on behalf of the Wellcome Sanger Institute COVID-19 Surveillance Team |
| EPI_ISL_2606828 | Lighthouse Lab in Alderley Park | Wellcome Sanger Institute for the COVID-19 Genomics UK (COG-UK) Consortium | Jacquelyn Wynn, Mairead Hyland, The Lighthouse Lab in Alderley Park and Alex Alderton, Roberto Amato, Jeffrey Barrett, Sonia Goncalves, Ewan Harrison, David K. Jackson, Ian Johnston, Dominic Kwiatkowski, Cordelia Langford, John Sillitoe on behalf of the Wellcome Sanger Institute COVID-19 Surveillance Team |
| EPI_ISL_2606829 | Lighthouse Lab in Glasgow | Wellcome Sanger Institute for the COVID-19 Genomics UK (COG-UK) Consortium | Harper VanSteenhouse, Yumi Kasai, David Gray, Carol Clugston, Anna Dominiczak and Alex Alderton, Roberto Amato, Jeffrey Barrett, Sonia Goncalves, Ewan Harrison, David K. Jackson, Ian Johnston, Dominic Kwiatkowski, Cordelia Langford, John Sillitoe on behalf of the Wellcome Sanger Institute COVID-19 Surveillance Team |
| EPI_ISL_2606830 | Lighthouse Laboratory Plymouth | Wellcome Sanger Institute for the COVID-19 Genomics UK (COG-UK) Consortium | Lighthouse Laboratory Plymouth and Alex Alderton, Roberto Amato, Jeffrey Barrett, Sonia Goncalves, Ewan Harrison, David K. Jackson, Ian Johnston, Dominic Kwiatkowski, Cordelia Langford, John Sillitoe on behalf of the Wellcome Sanger Institute COVID-19 Surveillance Team |
| EPI_ISL_2606831, EPI_ISL_2606832 | Lighthouse Lab in Alderley Park | Wellcome Sanger Institute for the COVID-19 Genomics UK (COG-UK) Consortium | Jacquelyn Wynn, Mairead Hyland, The Lighthouse Lab in Alderley Park and Alex Alderton, Roberto Amato, Jeffrey Barrett, Sonia Goncalves, Ewan Harrison, David K. Jackson, Ian Johnston, Dominic Kwiatkowski, Cordelia Langford, John Sillitoe on behalf of the Wellcome Sanger Institute COVID-19 Surveillance Team |
| EPI_ISL_2606833, EPI_ISL_2606834 | Lighthouse Laboratory Plymouth | Wellcome Sanger Institute for the COVID-19 Genomics UK (COG-UK) Consortium | Lighthouse Laboratory Plymouth and Alex Alderton, Roberto Amato, Jeffrey Barrett, Sonia Goncalves, Ewan Harrison, David K. Jackson, Ian Johnston, Dominic Kwiatkowski, Cordelia Langford, John Sillitoe on behalf of the Wellcome Sanger Institute COVID-19 Surveillance Team |
| EPI_ISL_2606835 | Lighthouse Lab in Milton Keynes | Wellcome Sanger Institute for the COVID-19 Genomics UK (COG-UK) Consortium | The Lighthouse Lab in Milton Keynes and Alex Alderton, Roberto Amato, Jeffrey Barrett, Sonia Goncalves, Ewan Harrison, David K. Jackson, Ian Johnston, Dominic Kwiatkowski, Cordelia Langford, John Sillitoe on behalf of the Wellcome Sanger Institute COVID-19 Surveillance Team |
| EPI_ISL_2606836, EPI_ISL_2606838 | Lighthouse Lab in Alderley Park | Wellcome Sanger Institute for the COVID-19 Genomics UK (COG-UK) Consortium | Jacquelyn Wynn, Mairead Hyland, The Lighthouse Lab in Alderley Park and Alex Alderton, Roberto Amato, Jeffrey Barrett, Sonia Goncalves, Ewan Harrison, David K. Jackson, Ian Johnston, Dominic Kwiatkowski, Cordelia Langford, John Sillitoe on behalf of the Wellcome Sanger Institute COVID-19 Surveillance Team |
| EPI_ISL_2606839, EPI_ISL_2606840 | Lighthouse Lab in Glasgow | Wellcome Sanger Institute for the COVID-19 Genomics UK (COG-UK) Consortium | Harper VanSteenhouse, Yumi Kasai, David Gray, Carol Clugston, Anna Dominiczak and Alex Alderton, Roberto Amato, Jeffrey Barrett, Sonia Goncalves, Ewan Harrison, David K. Jackson, Ian Johnston, Dominic Kwiatkowski, Cordelia Langford, John Sillitoe on behalf of the Wellcome Sanger Institute COVID-19 Surveillance Team |
| EPI_ISL_2606842 | Lighthouse Lab in Milton Keynes | Wellcome Sanger Institute for the COVID-19 Genomics UK (COG-UK) Consortium | The Lighthouse Lab in Milton Keynes and Alex Alderton, Roberto Amato, Jeffrey Barrett, Sonia Goncalves, Ewan Harrison, David K. Jackson, Ian Johnston, Dominic Kwiatkowski, Cordelia Langford, John Sillitoe on behalf of the Wellcome Sanger Institute COVID-19 Surveillance Team |
| EPI_ISL_2606843 | Lighthouse Lab in Alderley Park | Wellcome Sanger Institute for the COVID-19 Genomics UK (COG-UK) Consortium | Jacquelyn Wynn, Mairead Hyland, The Lighthouse Lab in Alderley Park and Alex Alderton, Roberto Amato, Jeffrey Barrett, Sonia Goncalves, Ewan Harrison, David K. Jackson, Ian Johnston, Dominic Kwiatkowski, Cordelia Langford, John Sillitoe on behalf of the Wellcome Sanger Institute COVID-19 Surveillance Team |
| EPI_ISL_2606844 | Lighthouse Lab in Milton Keynes | Wellcome Sanger Institute for the COVID-19 Genomics UK (COG-UK) Consortium | The Lighthouse Lab in Milton Keynes and Alex Alderton, Roberto Amato, Jeffrey Barrett, Sonia Goncalves, Ewan Harrison, David K. Jackson, Ian Johnston, Dominic Kwiatkowski, Cordelia Langford, John Sillitoe on behalf of the Wellcome Sanger Institute COVID-19 Surveillance Team |
| EPI_ISL_2606845, EPI_ISL_2606846 | Lighthouse Lab in Glasgow | Wellcome Sanger Institute for the COVID-19 Genomics UK (COG-UK) Consortium | Harper VanSteenhouse, Yumi Kasai, David Gray, Carol Clugston, Anna Dominiczak and Alex Alderton, Roberto Amato, Jeffrey Barrett, Sonia Goncalves, Ewan Harrison, David K. Jackson, Ian Johnston, Dominic Kwiatkowski, Cordelia Langford, John Sillitoe on behalf of the Wellcome Sanger Institute COVID-19 Surveillance Team |
| EPI_ISL_2606847 | Lighthouse Lab in Milton Keynes | Wellcome Sanger Institute for the COVID-19 Genomics UK (COG-UK) Consortium | The Lighthouse Lab in Milton Keynes and Alex Alderton, Roberto Amato, Jeffrey Barrett, Sonia Goncalves, Ewan Harrison, David K. Jackson, Ian Johnston, Dominic Kwiatkowski, Cordelia Langford, John Sillitoe on behalf of the Wellcome Sanger Institute COVID-19 Surveillance Team |
| EPI_ISL_2606848 | Lighthouse Lab in Glasgow | Wellcome Sanger Institute for the COVID-19 Genomics UK (COG-UK) Consortium | Harper VanSteenhouse, Yumi Kasai, David Gray, Carol Clugston, Anna Dominiczak and Alex Alderton, Roberto Amato, Jeffrey Barrett, Sonia Goncalves, Ewan Harrison, David K. Jackson, Ian Johnston, Dominic Kwiatkowski, Cordelia Langford, John Sillitoe on behalf of the Wellcome Sanger Institute COVID-19 Surveillance Team |
| EPI_ISL_2606849, EPI_ISL_2606850, EPI_ISL_2606851, EPI_ISL_2606852 | Lighthouse Lab in Milton Keynes | Wellcome Sanger Institute for the COVID-19 Genomics UK (COG-UK) Consortium | The Lighthouse Lab in Milton Keynes and Alex Alderton, Roberto Amato, Jeffrey Barrett, Sonia Goncalves, Ewan Harrison, David K. Jackson, Ian Johnston, Dominic Kwiatkowski, Cordelia Langford, John Sillitoe on behalf of the Wellcome Sanger Institute COVID-19 Surveillance Team |
| EPI_ISL_2606853, EPI_ISL_2606854 | Lighthouse Lab in Alderley Park | Wellcome Sanger Institute for the COVID-19 Genomics UK (COG-UK) Consortium | Jacquelyn Wynn, Mairead Hyland, The Lighthouse Lab in Alderley Park and Alex Alderton, Roberto Amato, Jeffrey Barrett, Sonia Goncalves, Ewan Harrison, David K. Jackson, Ian Johnston, Dominic Kwiatkowski, Cordelia Langford, John Sillitoe on behalf of the Wellcome Sanger Institute COVID-19 Surveillance Team |
| EPI_ISL_2606855, EPI_ISL_2606856 | Lighthouse Laboratory Plymouth | Wellcome Sanger Institute for the COVID-19 Genomics UK (COG-UK) Consortium | Lighthouse Laboratory Plymouth and Alex Alderton, Roberto Amato, Jeffrey Barrett, Sonia Goncalves, Ewan Harrison, David K. Jackson, Ian Johnston, Dominic Kwiatkowski, Cordelia Langford, John Sillitoe on behalf of the Wellcome Sanger Institute COVID-19 Surveillance Team |
| EPI_ISL_2606857, EPI_ISL_2606858 | Lighthouse Lab in Alderley Park | Wellcome Sanger Institute for the COVID-19 Genomics UK (COG-UK) Consortium | Jacquelyn Wynn, Mairead Hyland, The Lighthouse Lab in Alderley Park and Alex Alderton, Roberto Amato, Jeffrey Barrett, Sonia Goncalves, Ewan Harrison, David K. Jackson, Ian Johnston, Dominic Kwiatkowski, Cordelia Langford, John Sillitoe on behalf of the Wellcome Sanger Institute COVID-19 Surveillance Team |
| EPI_ISL_2606859, EPI_ISL_2606860 | Lighthouse Lab in Milton Keynes | Wellcome Sanger Institute for the COVID-19 Genomics UK (COG-UK) Consortium | The Lighthouse Lab in Milton Keynes and Alex Alderton, Roberto Amato, Jeffrey Barrett, Sonia Goncalves, Ewan Harrison, David K. Jackson, Ian Johnston, Dominic Kwiatkowski, Cordelia Langford, John Sillitoe on behalf of the Wellcome Sanger Institute COVID-19 Surveillance Team |
| EPI_ISL_2606861, EPI_ISL_2606862 | Lighthouse Lab in Alderley Park | Wellcome Sanger Institute for the COVID-19 Genomics UK (COG-UK) Consortium | Jacquelyn Wynn, Mairead Hyland, The Lighthouse Lab in Alderley Park and Alex Alderton, Roberto Amato, Jeffrey Barrett, Sonia Goncalves, Ewan Harrison, David K. Jackson, Ian Johnston, Dominic Kwiatkowski, Cordelia Langford, John Sillitoe on behalf of the Wellcome Sanger Institute COVID-19 Surveillance Team |
| EPI_ISL_2606863 | Lighthouse Laboratory Plymouth | Wellcome Sanger Institute for the COVID-19 Genomics UK (COG-UK) Consortium | Lighthouse Laboratory Plymouth and Alex Alderton, Roberto Amato, Jeffrey Barrett, Sonia Goncalves, Ewan Harrison, David K. Jackson, Ian Johnston, Dominic Kwiatkowski, Cordelia Langford, John Sillitoe on behalf of the Wellcome Sanger Institute COVID-19 Surveillance Team |

| | | | |
|---|---|---|---|
| EPI_ISL_2606864 | Lighthouse Lab in Glasgow | Wellcome Sanger Institute for the COVID-19 Genomics UK (COG-UK) Consortium | Harper VanSteenhouse, Yumi Kasai, David Gray, Carol Clugston, Anna Dominiczak and Alex Alderton, Roberto Amato, Jeffrey Barrett, Sonia Goncalves, Ewan Harrison, David K. Jackson, Ian Johnston, Dominic Kwiatkowski, Cordelia Langford, John Sillitoe on behalf of the Wellcome Sanger Institute COVID-19 Surveillance Team |
| EPI_ISL_2606865 | Lighthouse Lab in Milton Keynes | Wellcome Sanger Institute for the COVID-19 Genomics UK (COG-UK) Consortium | The Lighthouse Lab in Milton Keynes and Alex Alderton, Roberto Amato, Jeffrey Barrett, Sonia Goncalves, Ewan Harrison, David K. Jackson, Ian Johnston, Dominic Kwiatkowski, Cordelia Langford, John Sillitoe on behalf of the Wellcome Sanger Institute COVID-19 Surveillance Team |
| EPI_ISL_2606866, EPI_ISL_2606867, EPI_ISL_2606868 | Lighthouse Lab in Alderley Park | Wellcome Sanger Institute for the COVID-19 Genomics UK (COG-UK) Consortium | Jacquelyn Wynn, Mairead Hyland, The Lighthouse Lab in Alderley Park and Alex Alderton, Roberto Amato, Jeffrey Barrett, Sonia Goncalves, Ewan Harrison, David K. Jackson, Ian Johnston, Dominic Kwiatkowski, Cordelia Langford, John Sillitoe on behalf of the Wellcome Sanger Institute COVID-19 Surveillance Team |
| EPI_ISL_2606869 | Lighthouse Laboratory Plymouth | Wellcome Sanger Institute for the COVID-19 Genomics UK (COG-UK) Consortium | Lighthouse Laboratory Plymouth and Alex Alderton, Roberto Amato, Jeffrey Barrett, Sonia Goncalves, Ewan Harrison, David K. Jackson, Ian Johnston, Dominic Kwiatkowski, Cordelia Langford, John Sillitoe on behalf of the Wellcome Sanger Institute COVID-19 Surveillance Team |
| EPI_ISL_2606870 | Lighthouse Lab in Milton Keynes | Wellcome Sanger Institute for the COVID-19 Genomics UK (COG-UK) Consortium | The Lighthouse Lab in Milton Keynes and Alex Alderton, Roberto Amato, Jeffrey Barrett, Sonia Goncalves, Ewan Harrison, David K. Jackson, Ian Johnston, Dominic Kwiatkowski, Cordelia Langford, John Sillitoe on behalf of the Wellcome Sanger Institute COVID-19 Surveillance Team |
| EPI_ISL_2606871 | Lighthouse Lab in Glasgow | Wellcome Sanger Institute for the COVID-19 Genomics UK (COG-UK) Consortium | Harper VanSteenhouse, Yumi Kasai, David Gray, Carol Clugston, Anna Dominiczak and Alex Alderton, Roberto Amato, Jeffrey Barrett, Sonia Goncalves, Ewan Harrison, David K. Jackson, Ian Johnston, Dominic Kwiatkowski, Cordelia Langford, John Sillitoe on behalf of the Wellcome Sanger Institute COVID-19 Surveillance Team |
| EPI_ISL_2606872 | Lighthouse Lab in Milton Keynes | Wellcome Sanger Institute for the COVID-19 Genomics UK (COG-UK) Consortium | The Lighthouse Lab in Milton Keynes and Alex Alderton, Roberto Amato, Jeffrey Barrett, Sonia Goncalves, Ewan Harrison, David K. Jackson, Ian Johnston, Dominic Kwiatkowski, Cordelia Langford, John Sillitoe on behalf of the Wellcome Sanger Institute COVID-19 Surveillance Team |
| EPI_ISL_2606873 | Lighthouse Lab in Alderley Park | Wellcome Sanger Institute for the COVID-19 Genomics UK (COG-UK) Consortium | Jacquelyn Wynn, Mairead Hyland, The Lighthouse Lab in Alderley Park and Alex Alderton, Roberto Amato, Jeffrey Barrett, Sonia Goncalves, Ewan Harrison, David K. Jackson, Ian Johnston, Dominic Kwiatkowski, Cordelia Langford, John Sillitoe on behalf of the Wellcome Sanger Institute COVID-19 Surveillance Team |
| EPI_ISL_2606874 | Lighthouse Laboratory Plymouth | Wellcome Sanger Institute for the COVID-19 Genomics UK (COG-UK) Consortium | Lighthouse Laboratory Plymouth and Alex Alderton, Roberto Amato, Jeffrey Barrett, Sonia Goncalves, Ewan Harrison, David K. Jackson, Ian Johnston, Dominic Kwiatkowski, Cordelia Langford, John Sillitoe on behalf of the Wellcome Sanger Institute COVID-19 Surveillance Team |
| EPI_ISL_2606875 | Lighthouse Lab in Glasgow | Wellcome Sanger Institute for the COVID-19 Genomics UK (COG-UK) Consortium | Harper VanSteenhouse, Yumi Kasai, David Gray, Carol Clugston, Anna Dominiczak and Alex Alderton, Roberto Amato, Jeffrey Barrett, Sonia Goncalves, Ewan Harrison, David K. Jackson, Ian Johnston, Dominic Kwiatkowski, Cordelia Langford, John Sillitoe on behalf of the Wellcome Sanger Institute COVID-19 Surveillance Team |
| EPI_ISL_2606876, EPI_ISL_2606877 | Lighthouse Lab in Alderley Park | Wellcome Sanger Institute for the COVID-19 Genomics UK (COG-UK) Consortium | Jacquelyn Wynn, Mairead Hyland, The Lighthouse Lab in Alderley Park and Alex Alderton, Roberto Amato, Jeffrey Barrett, Sonia Goncalves, Ewan Harrison, David K. Jackson, Ian Johnston, Dominic Kwiatkowski, Cordelia Langford, John Sillitoe on behalf of the Wellcome Sanger Institute COVID-19 Surveillance Team |
| EPI_ISL_2606878, EPI_ISL_2606879 | Lighthouse Lab in Milton Keynes | Wellcome Sanger Institute for the COVID-19 Genomics UK (COG-UK) Consortium | The Lighthouse Lab in Milton Keynes and Alex Alderton, Roberto Amato, Jeffrey Barrett, Sonia Goncalves, Ewan Harrison, David K. Jackson, Ian Johnston, Dominic Kwiatkowski, Cordelia Langford, John Sillitoe on behalf of the Wellcome Sanger Institute COVID-19 Surveillance Team |
| EPI_ISL_2606880, EPI_ISL_2606881 | Lighthouse Lab in Alderley Park | Wellcome Sanger Institute for the COVID-19 Genomics UK (COG-UK) Consortium | Jacquelyn Wynn, Mairead Hyland, The Lighthouse Lab in Alderley Park and Alex Alderton, Roberto Amato, Jeffrey Barrett, Sonia Goncalves, Ewan Harrison, David K. Jackson, Ian Johnston, Dominic Kwiatkowski, Cordelia Langford, John Sillitoe on behalf of the Wellcome Sanger Institute COVID-19 Surveillance Team |
| EPI_ISL_2606882 | Lighthouse Lab in Glasgow | Wellcome Sanger Institute for the COVID-19 Genomics UK (COG-UK) Consortium | Harper VanSteenhouse, Yumi Kasai, David Gray, Carol Clugston, Anna Dominiczak and Alex Alderton, Roberto Amato, Jeffrey Barrett, Sonia Goncalves, Ewan Harrison, David K. Jackson, Ian Johnston, Dominic Kwiatkowski, Cordelia Langford, John Sillitoe on behalf of the Wellcome Sanger Institute COVID-19 Surveillance Team |
| EPI_ISL_2606883 | Lighthouse Lab in Alderley Park | Wellcome Sanger Institute for the COVID-19 Genomics UK (COG-UK) Consortium | Jacquelyn Wynn, Mairead Hyland, The Lighthouse Lab in Alderley Park and Alex Alderton, Roberto Amato, Jeffrey Barrett, Sonia Goncalves, Ewan Harrison, David K. Jackson, Ian Johnston, Dominic Kwiatkowski, Cordelia Langford, John Sillitoe on behalf of the Wellcome Sanger Institute COVID-19 Surveillance Team |
| EPI_ISL_2606884 | Lighthouse Lab in Milton Keynes | Wellcome Sanger Institute for the COVID-19 Genomics UK (COG-UK) Consortium | The Lighthouse Lab in Milton Keynes and Alex Alderton, Roberto Amato, Jeffrey Barrett, Sonia Goncalves, Ewan Harrison, David K. Jackson, Ian Johnston, Dominic Kwiatkowski, Cordelia Langford, John Sillitoe on behalf of the Wellcome Sanger Institute COVID-19 Surveillance Team |
| EPI_ISL_2606885 | Lighthouse Lab in Glasgow | Wellcome Sanger Institute for the COVID-19 Genomics UK (COG-UK) Consortium | Harper VanSteenhouse, Yumi Kasai, David Gray, Carol Clugston, Anna Dominiczak and Alex Alderton, Roberto Amato, Jeffrey Barrett, Sonia Goncalves, Ewan Harrison, David K. Jackson, Ian Johnston, Dominic Kwiatkowski, Cordelia Langford, John Sillitoe on behalf of the Wellcome Sanger Institute COVID-19 Surveillance Team |
| EPI_ISL_2606886 | Lighthouse Lab in Alderley Park | Wellcome Sanger Institute for the COVID-19 Genomics UK (COG-UK) Consortium | Jacquelyn Wynn, Mairead Hyland, The Lighthouse Lab in Alderley Park and Alex Alderton, Roberto Amato, Jeffrey Barrett, Sonia Goncalves, Ewan Harrison, David K. Jackson, Ian Johnston, Dominic Kwiatkowski, Cordelia Langford, John Sillitoe on behalf of the Wellcome Sanger Institute COVID-19 Surveillance Team |
| EPI_ISL_2606887 | Lighthouse Lab in Milton Keynes | Wellcome Sanger Institute for the COVID-19 Genomics UK (COG-UK) Consortium | The Lighthouse Lab in Milton Keynes and Alex Alderton, Roberto Amato, Jeffrey Barrett, Sonia Goncalves, Ewan Harrison, David K. Jackson, Ian Johnston, Dominic Kwiatkowski, Cordelia Langford, John Sillitoe on behalf of the Wellcome Sanger Institute COVID-19 Surveillance Team |
| EPI_ISL_2606888, EPI_ISL_2606889 | Lighthouse Lab in Alderley Park | Wellcome Sanger Institute for the COVID-19 Genomics UK (COG-UK) Consortium | Jacquelyn Wynn, Mairead Hyland, The Lighthouse Lab in Alderley Park and Alex Alderton, Roberto Amato, Jeffrey Barrett, Sonia Goncalves, Ewan Harrison, David K. Jackson, Ian Johnston, Dominic Kwiatkowski, Cordelia Langford, John Sillitoe on behalf of the Wellcome Sanger Institute COVID-19 Surveillance Team |
| EPI_ISL_2606890 | Lighthouse Laboratory Plymouth | Wellcome Sanger Institute for the COVID-19 Genomics UK (COG-UK) Consortium | Lighthouse Laboratory Plymouth and Alex Alderton, Roberto Amato, Jeffrey Barrett, Sonia Goncalves, Ewan Harrison, David K. Jackson, Ian Johnston, Dominic Kwiatkowski, Cordelia Langford, John Sillitoe on behalf of the Wellcome Sanger Institute COVID-19 Surveillance Team |
| EPI_ISL_2606891 | Lighthouse Lab in Alderley Park | Wellcome Sanger Institute for the COVID-19 Genomics UK (COG-UK) Consortium | Jacquelyn Wynn, Mairead Hyland, The Lighthouse Lab in Alderley Park and Alex Alderton, Roberto Amato, Jeffrey Barrett, Sonia Goncalves, Ewan Harrison, David K. Jackson, Ian Johnston, Dominic Kwiatkowski, Cordelia Langford, John Sillitoe on behalf of the Wellcome Sanger Institute COVID-19 Surveillance Team |
| EPI_ISL_2606892 | Lighthouse Lab in Milton Keynes | Wellcome Sanger Institute for the COVID-19 Genomics UK (COG-UK) Consortium | The Lighthouse Lab in Milton Keynes and Alex Alderton, Roberto Amato, Jeffrey Barrett, Sonia Goncalves, Ewan Harrison, David K. Jackson, Ian Johnston, Dominic Kwiatkowski, Cordelia Langford, John Sillitoe on behalf of the Wellcome Sanger Institute COVID-19 Surveillance Team |
| EPI_ISL_2606893, EPI_ISL_2606894 | Lighthouse Lab in Glasgow | Wellcome Sanger Institute for the COVID-19 Genomics UK (COG-UK) Consortium | Harper VanSteenhouse, Yumi Kasai, David Gray, Carol Clugston, Anna Dominiczak and Alex Alderton, Roberto Amato, Jeffrey Barrett, Sonia Goncalves, Ewan Harrison, David K. Jackson, Ian Johnston, Dominic Kwiatkowski, Cordelia Langford, John Sillitoe on behalf of the Wellcome Sanger Institute COVID-19 Surveillance Team |
| EPI_ISL_2606895, EPI_ISL_2606896, EPI_ISL_2606897 | Lighthouse Lab in Alderley Park | Wellcome Sanger Institute for the COVID-19 Genomics UK (COG-UK) Consortium | Jacquelyn Wynn, Mairead Hyland, The Lighthouse Lab in Alderley Park and Alex Alderton, Roberto Amato, Jeffrey Barrett, Sonia Goncalves, Ewan Harrison, David K. Jackson, Ian Johnston, Dominic Kwiatkowski, Cordelia Langford, John Sillitoe on behalf of the Wellcome Sanger Institute COVID-19 Surveillance Team |
| EPI_ISL_2606898 | Lighthouse Lab in Milton Keynes | Wellcome Sanger Institute for the COVID-19 Genomics UK (COG-UK) Consortium | The Lighthouse Lab in Milton Keynes and Alex Alderton, Roberto Amato, Jeffrey Barrett, Sonia Goncalves, Ewan Harrison, David K. Jackson, Ian Johnston, Dominic Kwiatkowski, Cordelia Langford, John Sillitoe on behalf of the Wellcome Sanger Institute COVID-19 Surveillance Team |
| EPI_ISL_2606899 | Lighthouse Lab in Glasgow | Wellcome Sanger Institute for the COVID-19 Genomics UK (COG-UK) Consortium | Harper VanSteenhouse, Yumi Kasai, David Gray, Carol Clugston, Anna Dominiczak and Alex Alderton, Roberto Amato, Jeffrey Barrett, Sonia Goncalves, Ewan Harrison, David K. Jackson, Ian Johnston, Dominic Kwiatkowski, Cordelia Langford, John Sillitoe on behalf of the Wellcome Sanger Institute COVID-19 Surveillance Team |
| EPI_ISL_2606900 | Lighthouse Lab in Alderley Park | Wellcome Sanger Institute for the COVID-19 Genomics UK (COG-UK) Consortium | Jacquelyn Wynn, Mairead Hyland, The Lighthouse Lab in Alderley Park and Alex Alderton, Roberto Amato, Jeffrey Barrett, Sonia Goncalves, Ewan Harrison, David K. Jackson, Ian Johnston, Dominic Kwiatkowski, Cordelia Langford, John Sillitoe on behalf of the Wellcome Sanger Institute COVID-19 Surveillance Team |
| EPI_ISL_2606901 | Lighthouse Laboratory Plymouth | Wellcome Sanger Institute for the COVID-19 Genomics UK (COG-UK) Consortium | Lighthouse Laboratory Plymouth and Alex Alderton, Roberto Amato, Jeffrey Barrett, Sonia Goncalves, Ewan Harrison, David K. Jackson, Ian Johnston, Dominic Kwiatkowski, Cordelia Langford, John Sillitoe on behalf of the Wellcome Sanger Institute COVID-19 Surveillance Team |
| EPI_ISL_2606902 | Lighthouse Lab in Milton Keynes | Wellcome Sanger Institute for the COVID-19 Genomics UK (COG-UK) Consortium | The Lighthouse Lab in Milton Keynes and Alex Alderton, Roberto Amato, Jeffrey Barrett, Sonia Goncalves, Ewan Harrison, David K. Jackson, Ian Johnston, Dominic Kwiatkowski, Cordelia Langford, John Sillitoe on behalf of the Wellcome Sanger Institute COVID-19 Surveillance Team |
| EPI_ISL_2606903 | Lighthouse Lab in Alderley Park | Wellcome Sanger Institute for the COVID-19 Genomics UK (COG-UK) Consortium | Jacquelyn Wynn, Mairead Hyland, The Lighthouse Lab in Alderley Park and Alex Alderton, Roberto Amato, Jeffrey Barrett, Sonia Goncalves, Ewan Harrison, David K. Jackson, Ian Johnston, Dominic Kwiatkowski, Cordelia Langford, John Sillitoe on behalf of the Wellcome Sanger Institute COVID-19 Surveillance Team |
| EPI_ISL_2606904 | Lighthouse Laboratory Plymouth | Wellcome Sanger Institute for the COVID-19 Genomics UK (COG-UK) Consortium | Lighthouse Laboratory Plymouth and Alex Alderton, Roberto Amato, Jeffrey Barrett, Sonia Goncalves, Ewan Harrison, David K. Jackson, Ian Johnston, Dominic Kwiatkowski, Cordelia Langford, John Sillitoe on behalf of the Wellcome Sanger Institute COVID-19 Surveillance Team |
| EPI_ISL_2606905 | Lighthouse Lab in Glasgow | Wellcome Sanger Institute for the COVID-19 Genomics UK (COG-UK) Consortium | Harper VanSteenhouse, Yumi Kasai, David Gray, Carol Clugston, Anna Dominiczak and Alex Alderton, Roberto Amato, Jeffrey Barrett, Sonia Goncalves, Ewan Harrison, David K. Jackson, Ian Johnston, Dominic Kwiatkowski, Cordelia Langford, John Sillitoe on behalf of the Wellcome Sanger Institute COVID-19 Surveillance Team |
| EPI_ISL_2606906, EPI_ISL_2606907 | Lighthouse Lab in Alderley Park | Wellcome Sanger Institute for the COVID-19 Genomics UK (COG-UK) Consortium | Jacquelyn Wynn, Mairead Hyland, The Lighthouse Lab in Alderley Park and Alex Alderton, Roberto Amato, Jeffrey Barrett, Sonia Goncalves, Ewan Harrison, David K. Jackson, Ian Johnston, Dominic Kwiatkowski, Cordelia Langford, John Sillitoe on behalf of the Wellcome Sanger Institute COVID-19 Surveillance Team |
| EPI_ISL_2606908 | Lighthouse Lab in Milton Keynes | Wellcome Sanger Institute for the COVID-19 Genomics UK (COG-UK) Consortium | The Lighthouse Lab in Milton Keynes and Alex Alderton, Roberto Amato, Jeffrey Barrett, Sonia Goncalves, Ewan Harrison, David K. Jackson, Ian Johnston, Dominic Kwiatkowski, Cordelia Langford, John Sillitoe on behalf of the Wellcome Sanger Institute COVID-19 Surveillance Team |
| EPI_ISL_2606909 | Lighthouse Lab in Glasgow | Wellcome Sanger Institute for the COVID-19 Genomics UK (COG-UK) Consortium | Harper VanSteenhouse, Yumi Kasai, David Gray, Carol Clugston, Anna Dominiczak and Alex Alderton, Roberto Amato, Jeffrey Barrett, Sonia Goncalves, Ewan Harrison, David K. Jackson, Ian Johnston, Dominic Kwiatkowski, Cordelia Langford, John Sillitoe on behalf of the Wellcome Sanger Institute COVID-19 Surveillance Team |
| EPI_ISL_2606910 | Lighthouse Lab in Milton Keynes | Wellcome Sanger Institute for the COVID-19 Genomics UK (COG-UK) Consortium | The Lighthouse Lab in Milton Keynes and Alex Alderton, Roberto Amato, Jeffrey Barrett, Sonia Goncalves, Ewan Harrison, David K. Jackson, Ian Johnston, Dominic Kwiatkowski, Cordelia Langford, John Sillitoe on behalf of the Wellcome Sanger Institute COVID-19 Surveillance Team |
| EPI_ISL_2606911 | Lighthouse Lab in Alderley Park | Wellcome Sanger Institute for the COVID-19 Genomics UK (COG-UK) Consortium | Jacquelyn Wynn, Mairead Hyland, The Lighthouse Lab in Alderley Park and Alex Alderton, Roberto Amato, Jeffrey Barrett, Sonia Goncalves, Ewan Harrison, David K. Jackson, Ian Johnston, Dominic Kwiatkowski, Cordelia Langford, John Sillitoe on behalf of the Wellcome Sanger Institute COVID-19 Surveillance Team |
| EPI_ISL_2606912 | Lighthouse Laboratory Plymouth | Wellcome Sanger Institute for the COVID-19 Genomics UK (COG-UK) Consortium | Lighthouse Laboratory Plymouth and Alex Alderton, Roberto Amato, Jeffrey Barrett, Sonia Goncalves, Ewan Harrison, David K. Jackson, Ian Johnston, Dominic Kwiatkowski, Cordelia Langford, John Sillitoe on behalf of the Wellcome Sanger Institute COVID-19 Surveillance Team |
| EPI_ISL_2606913 | Lighthouse Lab in Milton Keynes | Wellcome Sanger Institute for the COVID-19 Genomics UK (COG-UK) Consortium | The Lighthouse Lab in Milton Keynes and Alex Alderton, Roberto Amato, Jeffrey Barrett, Sonia Goncalves, Ewan Harrison, David K. Jackson, Ian Johnston, Dominic Kwiatkowski, Cordelia Langford, John Sillitoe on behalf of the Wellcome Sanger Institute COVID-19 Surveillance Team |
| EPI_ISL_2606914, EPI_ISL_2606915 | Lighthouse Lab in Alderley Park | Wellcome Sanger Institute for the COVID-19 Genomics UK (COG-UK) Consortium | Jacquelyn Wynn, Mairead Hyland, The Lighthouse Lab in Alderley Park and Alex Alderton, Roberto Amato, Jeffrey Barrett, Sonia Goncalves, Ewan Harrison, David K. Jackson, Ian Johnston, Dominic Kwiatkowski, Cordelia Langford, John Sillitoe on behalf of the Wellcome Sanger Institute COVID-19 Surveillance Team |
| EPI_ISL_2606916 | Lighthouse Laboratory Plymouth | Wellcome Sanger Institute for the COVID-19 Genomics UK (COG-UK) Consortium | Lighthouse Laboratory Plymouth and Alex Alderton, Roberto Amato, Jeffrey Barrett, Sonia Goncalves, Ewan Harrison, David K. Jackson, Ian Johnston, Dominic Kwiatkowski, Cordelia Langford, John Sillitoe on behalf of the Wellcome Sanger Institute COVID-19 Surveillance Team |
| EPI_ISL_2606917, EPI_ISL_2606918, EPI_ISL_2606919 | Lighthouse Lab in Alderley Park | Wellcome Sanger Institute for the COVID-19 Genomics UK (COG-UK) Consortium | Jacquelyn Wynn, Mairead Hyland, The Lighthouse Lab in Alderley Park and Alex Alderton, Roberto Amato, Jeffrey Barrett, Sonia Goncalves, Ewan Harrison, David K. Jackson, Ian Johnston, Dominic Kwiatkowski, Cordelia Langford, John Sillitoe on behalf of the Wellcome Sanger Institute COVID-19 Surveillance Team |
| EPI_ISL_2606920 | Lighthouse Lab in Glasgow | Wellcome Sanger Institute for the COVID-19 Genomics UK (COG-UK) | Harper VanSteenhouse, Yumi Kasai, David Gray, Carol Clugston, Anna Dominiczak and Alex Alderton, Roberto Amato, Jeffrey Barrett, Sonia Goncalves, Ewan Harrison, David K. Jackson, Ian Johnston, Dominic Kwiatkowski, Cordelia Langford, |

| | | | |
|---|---|---|---|
| | | Consortium | John Sillitoe on behalf of the Wellcome Sanger Institute COVID-19 Surveillance Team |
| EPI_ISL_2606921, EPI_ISL_2606923, EPI_ISL_2606924 | Lighthouse Lab in Alderley Park | Wellcome Sanger Institute for the COVID-19 Genomics UK (COG-UK) Consortium | Jacquelyn Wynn, Mairead Hyland, The Lighthouse Lab in Alderley Park and Alex Alderton, Roberto Amato, Jeffrey Barrett, Sonia Goncalves, Ewan Harrison, David K. Jackson, Ian Johnston, Dominic Kwiatkowski, Cordelia Langford, John Sillitoe on behalf of the Wellcome Sanger Institute COVID-19 Surveillance Team |
| EPI_ISL_2606925 | Lighthouse Lab in Glasgow | Wellcome Sanger Institute for the COVID-19 Genomics UK (COG-UK) Consortium | Harper VanSteenhouse, Yumi Kasai, David Gray, Carol Clugston, Anna Dominiczak and Alex Alderton, Roberto Amato, Jeffrey Barrett, Sonia Goncalves, Ewan Harrison, David K. Jackson, Ian Johnston, Dominic Kwiatkowski, Cordelia Langford, John Sillitoe on behalf of the Wellcome Sanger Institute COVID-19 Surveillance Team |
| EPI_ISL_2606926 | Lighthouse Lab in Milton Keynes | Wellcome Sanger Institute for the COVID-19 Genomics UK (COG-UK) Consortium | The Lighthouse Lab in Milton Keynes and Alex Alderton, Roberto Amato, Jeffrey Barrett, Sonia Goncalves, Ewan Harrison, David K. Jackson, Ian Johnston, Dominic Kwiatkowski, Cordelia Langford, John Sillitoe on behalf of the Wellcome Sanger Institute COVID-19 Surveillance Team |
| EPI_ISL_2606927 | Lighthouse Laboratory Plymouth | Wellcome Sanger Institute for the COVID-19 Genomics UK (COG-UK) Consortium | Lighthouse Laboratory Plymouth and Alex Alderton, Roberto Amato, Jeffrey Barrett, Sonia Goncalves, Ewan Harrison, David K. Jackson, Ian Johnston, Dominic Kwiatkowski, Cordelia Langford, John Sillitoe on behalf of the Wellcome Sanger Institute COVID-19 Surveillance Team |
| EPI_ISL_2606928, EPI_ISL_2606929 | Lighthouse Lab in Milton Keynes | Wellcome Sanger Institute for the COVID-19 Genomics UK (COG-UK) Consortium | The Lighthouse Lab in Milton Keynes and Alex Alderton, Roberto Amato, Jeffrey Barrett, Sonia Goncalves, Ewan Harrison, David K. Jackson, Ian Johnston, Dominic Kwiatkowski, Cordelia Langford, John Sillitoe on behalf of the Wellcome Sanger Institute COVID-19 Surveillance Team |
| EPI_ISL_2606930, EPI_ISL_2606931 | Lighthouse Lab in Glasgow | Wellcome Sanger Institute for the COVID-19 Genomics UK (COG-UK) Consortium | Harper VanSteenhouse, Yumi Kasai, David Gray, Carol Clugston, Anna Dominiczak and Alex Alderton, Roberto Amato, Jeffrey Barrett, Sonia Goncalves, Ewan Harrison, David K. Jackson, Ian Johnston, Dominic Kwiatkowski, Cordelia Langford, John Sillitoe on behalf of the Wellcome Sanger Institute COVID-19 Surveillance Team |
| EPI_ISL_2606932 | Lighthouse Lab in Milton Keynes | Wellcome Sanger Institute for the COVID-19 Genomics UK (COG-UK) Consortium | The Lighthouse Lab in Milton Keynes and Alex Alderton, Roberto Amato, Jeffrey Barrett, Sonia Goncalves, Ewan Harrison, David K. Jackson, Ian Johnston, Dominic Kwiatkowski, Cordelia Langford, John Sillitoe on behalf of the Wellcome Sanger Institute COVID-19 Surveillance Team |
| EPI_ISL_2606934 | Lighthouse Lab in Alderley Park | Wellcome Sanger Institute for the COVID-19 Genomics UK (COG-UK) Consortium | Jacquelyn Wynn, Mairead Hyland, The Lighthouse Lab in Alderley Park and Alex Alderton, Roberto Amato, Jeffrey Barrett, Sonia Goncalves, Ewan Harrison, David K. Jackson, Ian Johnston, Dominic Kwiatkowski, Cordelia Langford, John Sillitoe on behalf of the Wellcome Sanger Institute COVID-19 Surveillance Team |
| EPI_ISL_2606935 | Lighthouse Lab in Milton Keynes | Wellcome Sanger Institute for the COVID-19 Genomics UK (COG-UK) Consortium | The Lighthouse Lab in Milton Keynes and Alex Alderton, Roberto Amato, Jeffrey Barrett, Sonia Goncalves, Ewan Harrison, David K. Jackson, Ian Johnston, Dominic Kwiatkowski, Cordelia Langford, John Sillitoe on behalf of the Wellcome Sanger Institute COVID-19 Surveillance Team |
| EPI_ISL_2606936 | Lighthouse Laboratory Plymouth | Wellcome Sanger Institute for the COVID-19 Genomics UK (COG-UK) Consortium | Lighthouse Laboratory Plymouth and Alex Alderton, Roberto Amato, Jeffrey Barrett, Sonia Goncalves, Ewan Harrison, David K. Jackson, Ian Johnston, Dominic Kwiatkowski, Cordelia Langford, John Sillitoe on behalf of the Wellcome Sanger Institute COVID-19 Surveillance Team |
| EPI_ISL_2606937, EPI_ISL_2606938 | Lighthouse Lab in Glasgow | Wellcome Sanger Institute for the COVID-19 Genomics UK (COG-UK) Consortium | Harper VanSteenhouse, Yumi Kasai, David Gray, Carol Clugston, Anna Dominiczak and Alex Alderton, Roberto Amato, Jeffrey Barrett, Sonia Goncalves, Ewan Harrison, David K. Jackson, Ian Johnston, Dominic Kwiatkowski, Cordelia Langford, John Sillitoe on behalf of the Wellcome Sanger Institute COVID-19 Surveillance Team |
| EPI_ISL_2606939 | Lighthouse Lab in Milton Keynes | Wellcome Sanger Institute for the COVID-19 Genomics UK (COG-UK) Consortium | The Lighthouse Lab in Milton Keynes and Alex Alderton, Roberto Amato, Jeffrey Barrett, Sonia Goncalves, Ewan Harrison, David K. Jackson, Ian Johnston, Dominic Kwiatkowski, Cordelia Langford, John Sillitoe on behalf of the Wellcome Sanger Institute COVID-19 Surveillance Team |
| EPI_ISL_2606940 | Lighthouse Lab in Alderley Park | Wellcome Sanger Institute for the COVID-19 Genomics UK (COG-UK) Consortium | Jacquelyn Wynn, Mairead Hyland, The Lighthouse Lab in Alderley Park and Alex Alderton, Roberto Amato, Jeffrey Barrett, Sonia Goncalves, Ewan Harrison, David K. Jackson, Ian Johnston, Dominic Kwiatkowski, Cordelia Langford, John Sillitoe on behalf of the Wellcome Sanger Institute COVID-19 Surveillance Team |
| EPI_ISL_2606941 | Lighthouse Lab in Milton Keynes | Wellcome Sanger Institute for the COVID-19 Genomics UK (COG-UK) Consortium | The Lighthouse Lab in Milton Keynes and Alex Alderton, Roberto Amato, Jeffrey Barrett, Sonia Goncalves, Ewan Harrison, David K. Jackson, Ian Johnston, Dominic Kwiatkowski, Cordelia Langford, John Sillitoe on behalf of the Wellcome Sanger Institute COVID-19 Surveillance Team |
| EPI_ISL_2606942 | Lighthouse Laboratory Plymouth | Wellcome Sanger Institute for the COVID-19 Genomics UK (COG-UK) Consortium | Lighthouse Laboratory Plymouth and Alex Alderton, Roberto Amato, Jeffrey Barrett, Sonia Goncalves, Ewan Harrison, David K. Jackson, Ian Johnston, Dominic Kwiatkowski, Cordelia Langford, John Sillitoe on behalf of the Wellcome Sanger Institute COVID-19 Surveillance Team |
| EPI_ISL_2606943 | Lighthouse Lab in Milton Keynes | Wellcome Sanger Institute for the COVID-19 Genomics UK (COG-UK) Consortium | The Lighthouse Lab in Milton Keynes and Alex Alderton, Roberto Amato, Jeffrey Barrett, Sonia Goncalves, Ewan Harrison, David K. Jackson, Ian Johnston, Dominic Kwiatkowski, Cordelia Langford, John Sillitoe on behalf of the Wellcome Sanger Institute COVID-19 Surveillance Team |
| EPI_ISL_2606944 | Lighthouse Lab in Glasgow | Wellcome Sanger Institute for the COVID-19 Genomics UK (COG-UK) Consortium | Harper VanSteenhouse, Yumi Kasai, David Gray, Carol Clugston, Anna Dominiczak and Alex Alderton, Roberto Amato, Jeffrey Barrett, Sonia Goncalves, Ewan Harrison, David K. Jackson, Ian Johnston, Dominic Kwiatkowski, Cordelia Langford, John Sillitoe on behalf of the Wellcome Sanger Institute COVID-19 Surveillance Team |
| EPI_ISL_2606945, EPI_ISL_2606946 | Lighthouse Laboratory Plymouth | Wellcome Sanger Institute for the COVID-19 Genomics UK (COG-UK) Consortium | Lighthouse Laboratory Plymouth and Alex Alderton, Roberto Amato, Jeffrey Barrett, Sonia Goncalves, Ewan Harrison, David K. Jackson, Ian Johnston, Dominic Kwiatkowski, Cordelia Langford, John Sillitoe on behalf of the Wellcome Sanger Institute COVID-19 Surveillance Team |
| EPI_ISL_2606947 | Lighthouse Lab in Milton Keynes | Wellcome Sanger Institute for the COVID-19 Genomics UK (COG-UK) Consortium | The Lighthouse Lab in Milton Keynes and Alex Alderton, Roberto Amato, Jeffrey Barrett, Sonia Goncalves, Ewan Harrison, David K. Jackson, Ian Johnston, Dominic Kwiatkowski, Cordelia Langford, John Sillitoe on behalf of the Wellcome Sanger Institute COVID-19 Surveillance Team |
| EPI_ISL_2606948 | Lighthouse Lab in Alderley Park | Wellcome Sanger Institute for the COVID-19 Genomics UK (COG-UK) Consortium | Jacquelyn Wynn, Mairead Hyland, The Lighthouse Lab in Alderley Park and Alex Alderton, Roberto Amato, Jeffrey Barrett, Sonia Goncalves, Ewan Harrison, David K. Jackson, Ian Johnston, Dominic Kwiatkowski, Cordelia Langford, John Sillitoe on behalf of the Wellcome Sanger Institute COVID-19 Surveillance Team |
| EPI_ISL_2606949, EPI_ISL_2606950 | Lighthouse Laboratory Plymouth | Wellcome Sanger Institute for the COVID-19 Genomics UK (COG-UK) Consortium | Lighthouse Laboratory Plymouth and Alex Alderton, Roberto Amato, Jeffrey Barrett, Sonia Goncalves, Ewan Harrison, David K. Jackson, Ian Johnston, Dominic Kwiatkowski, Cordelia Langford, John Sillitoe on behalf of the Wellcome Sanger Institute COVID-19 Surveillance Team |
| EPI_ISL_2606951, EPI_ISL_2606952, EPI_ISL_2606953 | Lighthouse Lab in Milton Keynes | Wellcome Sanger Institute for the COVID-19 Genomics UK (COG-UK) Consortium | The Lighthouse Lab in Milton Keynes and Alex Alderton, Roberto Amato, Jeffrey Barrett, Sonia Goncalves, Ewan Harrison, David K. Jackson, Ian Johnston, Dominic Kwiatkowski, Cordelia Langford, John Sillitoe on behalf of the Wellcome Sanger Institute COVID-19 Surveillance Team |
| EPI_ISL_2606954 | Lighthouse Lab in Glasgow | Wellcome Sanger Institute for the COVID-19 Genomics UK (COG-UK) Consortium | Harper VanSteenhouse, Yumi Kasai, David Gray, Carol Clugston, Anna Dominiczak and Alex Alderton, Roberto Amato, Jeffrey Barrett, Sonia Goncalves, Ewan Harrison, David K. Jackson, Ian Johnston, Dominic Kwiatkowski, Cordelia Langford, John Sillitoe on behalf of the Wellcome Sanger Institute COVID-19 Surveillance Team |
| EPI_ISL_2606955 | Lighthouse Lab in Milton Keynes | Wellcome Sanger Institute for the COVID-19 Genomics UK (COG-UK) Consortium | The Lighthouse Lab in Milton Keynes and Alex Alderton, Roberto Amato, Jeffrey Barrett, Sonia Goncalves, Ewan Harrison, David K. Jackson, Ian Johnston, Dominic Kwiatkowski, Cordelia Langford, John Sillitoe on behalf of the Wellcome Sanger Institute COVID-19 Surveillance Team |
| EPI_ISL_2606956, EPI_ISL_2606957, EPI_ISL_2606958 | Lighthouse Lab in Alderley Park | Wellcome Sanger Institute for the COVID-19 Genomics UK (COG-UK) Consortium | Jacquelyn Wynn, Mairead Hyland, The Lighthouse Lab in Alderley Park and Alex Alderton, Roberto Amato, Jeffrey Barrett, Sonia Goncalves, Ewan Harrison, David K. Jackson, Ian Johnston, Dominic Kwiatkowski, Cordelia Langford, John Sillitoe on behalf of the Wellcome Sanger Institute COVID-19 Surveillance Team |
| EPI_ISL_2606959, EPI_ISL_2606960 | Lighthouse Lab in Glasgow | Wellcome Sanger Institute for the COVID-19 Genomics UK (COG-UK) Consortium | Harper VanSteenhouse, Yumi Kasai, David Gray, Carol Clugston, Anna Dominiczak and Alex Alderton, Roberto Amato, Jeffrey Barrett, Sonia Goncalves, Ewan Harrison, David K. Jackson, Ian Johnston, Dominic Kwiatkowski, Cordelia Langford, John Sillitoe on behalf of the Wellcome Sanger Institute COVID-19 Surveillance Team |
| EPI_ISL_2606961 | Lighthouse Lab in Alderley Park | Wellcome Sanger Institute for the COVID-19 Genomics UK (COG-UK) Consortium | Jacquelyn Wynn, Mairead Hyland, The Lighthouse Lab in Alderley Park and Alex Alderton, Roberto Amato, Jeffrey Barrett, Sonia Goncalves, Ewan Harrison, David K. Jackson, Ian Johnston, Dominic Kwiatkowski, Cordelia Langford, John Sillitoe on behalf of the Wellcome Sanger Institute COVID-19 Surveillance Team |
| EPI_ISL_2606962 | Lighthouse Laboratory Plymouth | Wellcome Sanger Institute for the COVID-19 Genomics UK (COG-UK) Consortium | Lighthouse Laboratory Plymouth and Alex Alderton, Roberto Amato, Jeffrey Barrett, Sonia Goncalves, Ewan Harrison, David K. Jackson, Ian Johnston, Dominic Kwiatkowski, Cordelia Langford, John Sillitoe on behalf of the Wellcome Sanger Institute COVID-19 Surveillance Team |
| EPI_ISL_2606963, EPI_ISL_2606964 | Lighthouse Lab in Milton Keynes | Wellcome Sanger Institute for the COVID-19 Genomics UK (COG-UK) Consortium | The Lighthouse Lab in Milton Keynes and Alex Alderton, Roberto Amato, Jeffrey Barrett, Sonia Goncalves, Ewan Harrison, David K. Jackson, Ian Johnston, Dominic Kwiatkowski, Cordelia Langford, John Sillitoe on behalf of the Wellcome Sanger Institute COVID-19 Surveillance Team |
| EPI_ISL_2606965 | Lighthouse Lab in Alderley Park | Wellcome Sanger Institute for the COVID-19 Genomics UK (COG-UK) Consortium | Jacquelyn Wynn, Mairead Hyland, The Lighthouse Lab in Alderley Park and Alex Alderton, Roberto Amato, Jeffrey Barrett, Sonia Goncalves, Ewan Harrison, David K. Jackson, Ian Johnston, Dominic Kwiatkowski, Cordelia Langford, John Sillitoe on behalf of the Wellcome Sanger Institute COVID-19 Surveillance Team |
| EPI_ISL_2606966 | Lighthouse Lab in Milton Keynes | Wellcome Sanger Institute for the COVID-19 Genomics UK (COG-UK) Consortium | The Lighthouse Lab in Milton Keynes and Alex Alderton, Roberto Amato, Jeffrey Barrett, Sonia Goncalves, Ewan Harrison, David K. Jackson, Ian Johnston, Dominic Kwiatkowski, Cordelia Langford, John Sillitoe on behalf of the Wellcome Sanger Institute COVID-19 Surveillance Team |
| EPI_ISL_2606967, EPI_ISL_2606968 | Lighthouse Lab in Alderley Park | Wellcome Sanger Institute for the COVID-19 Genomics UK (COG-UK) Consortium | Jacquelyn Wynn, Mairead Hyland, The Lighthouse Lab in Alderley Park and Alex Alderton, Roberto Amato, Jeffrey Barrett, Sonia Goncalves, Ewan Harrison, David K. Jackson, Ian Johnston, Dominic Kwiatkowski, Cordelia Langford, John Sillitoe on behalf of the Wellcome Sanger Institute COVID-19 Surveillance Team |
| EPI_ISL_2606969 | Lighthouse Lab in Milton Keynes | Wellcome Sanger Institute for the COVID-19 Genomics UK (COG-UK) Consortium | The Lighthouse Lab in Milton Keynes and Alex Alderton, Roberto Amato, Jeffrey Barrett, Sonia Goncalves, Ewan Harrison, David K. Jackson, Ian Johnston, Dominic Kwiatkowski, Cordelia Langford, John Sillitoe on behalf of the Wellcome Sanger Institute COVID-19 Surveillance Team |
| EPI_ISL_2606970 | Lighthouse Laboratory Plymouth | Wellcome Sanger Institute for the COVID-19 Genomics UK (COG-UK) Consortium | Lighthouse Laboratory Plymouth and Alex Alderton, Roberto Amato, Jeffrey Barrett, Sonia Goncalves, Ewan Harrison, David K. Jackson, Ian Johnston, Dominic Kwiatkowski, Cordelia Langford, John Sillitoe on behalf of the Wellcome Sanger Institute COVID-19 Surveillance Team |
| EPI_ISL_2606971 | Lighthouse Lab in Glasgow | Wellcome Sanger Institute for the COVID-19 Genomics UK (COG-UK) Consortium | Harper VanSteenhouse, Yumi Kasai, David Gray, Carol Clugston, Anna Dominiczak and Alex Alderton, Roberto Amato, Jeffrey Barrett, Sonia Goncalves, Ewan Harrison, David K. Jackson, Ian Johnston, Dominic Kwiatkowski, Cordelia Langford, John Sillitoe on behalf of the Wellcome Sanger Institute COVID-19 Surveillance Team |
| EPI_ISL_2606972, EPI_ISL_2606973 | Lighthouse Lab in Alderley Park | Wellcome Sanger Institute for the COVID-19 Genomics UK (COG-UK) Consortium | Jacquelyn Wynn, Mairead Hyland, The Lighthouse Lab in Alderley Park and Alex Alderton, Roberto Amato, Jeffrey Barrett, Sonia Goncalves, Ewan Harrison, David K. Jackson, Ian Johnston, Dominic Kwiatkowski, Cordelia Langford, John Sillitoe on behalf of the Wellcome Sanger Institute COVID-19 Surveillance Team |
| EPI_ISL_2606974 | Lighthouse Lab in Milton Keynes | Wellcome Sanger Institute for the COVID-19 Genomics UK (COG-UK) Consortium | The Lighthouse Lab in Milton Keynes and Alex Alderton, Roberto Amato, Jeffrey Barrett, Sonia Goncalves, Ewan Harrison, David K. Jackson, Ian Johnston, Dominic Kwiatkowski, Cordelia Langford, John Sillitoe on behalf of the Wellcome Sanger Institute COVID-19 Surveillance Team |
| EPI_ISL_2606975, EPI_ISL_2606976, EPI_ISL_2606977, EPI_ISL_2606979 | Lighthouse Lab in Alderley Park | Wellcome Sanger Institute for the COVID-19 Genomics UK (COG-UK) Consortium | Jacquelyn Wynn, Mairead Hyland, The Lighthouse Lab in Alderley Park and Alex Alderton, Roberto Amato, Jeffrey Barrett, Sonia Goncalves, Ewan Harrison, David K. Jackson, Ian Johnston, Dominic Kwiatkowski, Cordelia Langford, John Sillitoe on behalf of the Wellcome Sanger Institute COVID-19 Surveillance Team |
| EPI_ISL_2606980 | Lighthouse Laboratory Plymouth | Wellcome Sanger Institute for the COVID-19 Genomics UK (COG-UK) Consortium | Lighthouse Laboratory Plymouth and Alex Alderton, Roberto Amato, Jeffrey Barrett, Sonia Goncalves, Ewan Harrison, David K. Jackson, Ian Johnston, Dominic Kwiatkowski, Cordelia Langford, John Sillitoe on behalf of the Wellcome Sanger Institute COVID-19 Surveillance Team |
| EPI_ISL_2606981 | Lighthouse Lab in Alderley Park | Wellcome Sanger Institute for the COVID-19 Genomics UK (COG-UK) Consortium | Jacquelyn Wynn, Mairead Hyland, The Lighthouse Lab in Alderley Park and Alex Alderton, Roberto Amato, Jeffrey Barrett, Sonia Goncalves, Ewan Harrison, David K. Jackson, Ian Johnston, Dominic Kwiatkowski, Cordelia Langford, John Sillitoe on behalf of the Wellcome Sanger Institute COVID-19 Surveillance Team |
| EPI_ISL_2606982, EPI_ISL_2606983 | Lighthouse Lab in Glasgow | Wellcome Sanger Institute for the COVID-19 Genomics UK (COG-UK) Consortium | Harper VanSteenhouse, Yumi Kasai, David Gray, Carol Clugston, Anna Dominiczak and Alex Alderton, Roberto Amato, Jeffrey Barrett, Sonia Goncalves, Ewan Harrison, David K. Jackson, Ian Johnston, Dominic Kwiatkowski, Cordelia Langford, John Sillitoe on behalf of the Wellcome Sanger Institute COVID-19 Surveillance Team |
| EPI_ISL_2606984 | Lighthouse Lab in Alderley Park | Wellcome Sanger Institute for the COVID-19 Genomics UK (COG-UK) Consortium | Jacquelyn Wynn, Mairead Hyland, The Lighthouse Lab in Alderley Park and Alex Alderton, Roberto Amato, Jeffrey Barrett, Sonia Goncalves, Ewan Harrison, David K. Jackson, Ian Johnston, Dominic Kwiatkowski, Cordelia Langford, John Sillitoe on behalf of the Wellcome Sanger Institute COVID-19 Surveillance Team |
| EPI_ISL_2606985 | Lighthouse Laboratory Plymouth | Wellcome Sanger Institute for the COVID-19 Genomics UK (COG-UK) | Lighthouse Laboratory Plymouth and Alex Alderton, Roberto Amato, Jeffrey Barrett, Sonia Goncalves, Ewan Harrison, David K. Jackson, Ian Johnston, Dominic Kwiatkowski, Cordelia Langford, John Sillitoe on behalf of the Wellcome Sanger |

| | | | |
|---|---|---|---|
| | | Consortium | Institute COVID-19 Surveillance Team |
| EPI_ISL_2606986 | Lighthouse Lab in Alderley Park | Wellcome Sanger Institute for the COVID-19 Genomics UK (COG-UK) Consortium | Jacquelyn Wynn, Mairead Hyland, The Lighthouse Lab in Alderley Park and Alex Alderton, Roberto Amato, Jeffrey Barrett, Sonia Goncalves, Ewan Harrison, David K. Jackson, Ian Johnston, Dominic Kwiatkowski, Cordelia Langford, John Sillitoe on behalf of the Wellcome Sanger Institute COVID-19 Surveillance Team |
| EPI_ISL_2606987 | Lighthouse Lab in Milton Keynes | Wellcome Sanger Institute for the COVID-19 Genomics UK (COG-UK) Consortium | The Lighthouse Lab in Milton Keynes and Alex Alderton, Roberto Amato, Jeffrey Barrett, Sonia Goncalves, Ewan Harrison, David K. Jackson, Ian Johnston, Dominic Kwiatkowski, Cordelia Langford, John Sillitoe on behalf of the Wellcome Sanger Institute COVID-19 Surveillance Team |
| EPI_ISL_2606988 | Lighthouse Lab in Alderley Park | Wellcome Sanger Institute for the COVID-19 Genomics UK (COG-UK) Consortium | Jacquelyn Wynn, Mairead Hyland, The Lighthouse Lab in Alderley Park and Alex Alderton, Roberto Amato, Jeffrey Barrett, Sonia Goncalves, Ewan Harrison, David K. Jackson, Ian Johnston, Dominic Kwiatkowski, Cordelia Langford, John Sillitoe on behalf of the Wellcome Sanger Institute COVID-19 Surveillance Team |
| EPI_ISL_2606989, EPI_ISL_2606990, EPI_ISL_2606991 | Lighthouse Laboratory Plymouth | Wellcome Sanger Institute for the COVID-19 Genomics UK (COG-UK) Consortium | Lighthouse Laboratory Plymouth and Alex Alderton, Roberto Amato, Jeffrey Barrett, Sonia Goncalves, Ewan Harrison, David K. Jackson, Ian Johnston, Dominic Kwiatkowski, Cordelia Langford, John Sillitoe on behalf of the Wellcome Sanger Institute COVID-19 Surveillance Team |
| EPI_ISL_2606992, EPI_ISL_2606993 | Lighthouse Lab in Alderley Park | Wellcome Sanger Institute for the COVID-19 Genomics UK (COG-UK) Consortium | Jacquelyn Wynn, Mairead Hyland, The Lighthouse Lab in Alderley Park and Alex Alderton, Roberto Amato, Jeffrey Barrett, Sonia Goncalves, Ewan Harrison, David K. Jackson, Ian Johnston, Dominic Kwiatkowski, Cordelia Langford, John Sillitoe on behalf of the Wellcome Sanger Institute COVID-19 Surveillance Team |
| EPI_ISL_2606994 | Lighthouse Laboratory Plymouth | Wellcome Sanger Institute for the COVID-19 Genomics UK (COG-UK) Consortium | Lighthouse Laboratory Plymouth and Alex Alderton, Roberto Amato, Jeffrey Barrett, Sonia Goncalves, Ewan Harrison, David K. Jackson, Ian Johnston, Dominic Kwiatkowski, Cordelia Langford, John Sillitoe on behalf of the Wellcome Sanger Institute COVID-19 Surveillance Team |
| EPI_ISL_2606995 | Lighthouse Lab in Glasgow | Wellcome Sanger Institute for the COVID-19 Genomics UK (COG-UK) Consortium | Harper VanSteenhouse, Yumi Kasai, David Gray, Carol Clugston, Anna Dominiczak and Alex Alderton, Roberto Amato, Jeffrey Barrett, Sonia Goncalves, Ewan Harrison, David K. Jackson, Ian Johnston, Dominic Kwiatkowski, Cordelia Langford, John Sillitoe on behalf of the Wellcome Sanger Institute COVID-19 Surveillance Team |
| EPI_ISL_2606996 | Lighthouse Laboratory Plymouth | Wellcome Sanger Institute for the COVID-19 Genomics UK (COG-UK) Consortium | Lighthouse Laboratory Plymouth and Alex Alderton, Roberto Amato, Jeffrey Barrett, Sonia Goncalves, Ewan Harrison, David K. Jackson, Ian Johnston, Dominic Kwiatkowski, Cordelia Langford, John Sillitoe on behalf of the Wellcome Sanger Institute COVID-19 Surveillance Team |
| EPI_ISL_2606997 | Lighthouse Lab in Milton Keynes | Wellcome Sanger Institute for the COVID-19 Genomics UK (COG-UK) Consortium | The Lighthouse Lab in Milton Keynes and Alex Alderton, Roberto Amato, Jeffrey Barrett, Sonia Goncalves, Ewan Harrison, David K. Jackson, Ian Johnston, Dominic Kwiatkowski, Cordelia Langford, John Sillitoe on behalf of the Wellcome Sanger Institute COVID-19 Surveillance Team |
| EPI_ISL_2606998 | Lighthouse Laboratory Plymouth | Wellcome Sanger Institute for the COVID-19 Genomics UK (COG-UK) Consortium | Lighthouse Laboratory Plymouth and Alex Alderton, Roberto Amato, Jeffrey Barrett, Sonia Goncalves, Ewan Harrison, David K. Jackson, Ian Johnston, Dominic Kwiatkowski, Cordelia Langford, John Sillitoe on behalf of the Wellcome Sanger Institute COVID-19 Surveillance Team |
| EPI_ISL_2606999 | Lighthouse Lab in Alderley Park | Wellcome Sanger Institute for the COVID-19 Genomics UK (COG-UK) Consortium | Jacquelyn Wynn, Mairead Hyland, The Lighthouse Lab in Alderley Park and Alex Alderton, Roberto Amato, Jeffrey Barrett, Sonia Goncalves, Ewan Harrison, David K. Jackson, Ian Johnston, Dominic Kwiatkowski, Cordelia Langford, John Sillitoe on behalf of the Wellcome Sanger Institute COVID-19 Surveillance Team |
| EPI_ISL_2607000 | Lighthouse Lab in Milton Keynes | Wellcome Sanger Institute for the COVID-19 Genomics UK (COG-UK) Consortium | The Lighthouse Lab in Milton Keynes and Alex Alderton, Roberto Amato, Jeffrey Barrett, Sonia Goncalves, Ewan Harrison, David K. Jackson, Ian Johnston, Dominic Kwiatkowski, Cordelia Langford, John Sillitoe on behalf of the Wellcome Sanger Institute COVID-19 Surveillance Team |
| EPI_ISL_2607001 | Lighthouse Lab in Alderley Park | Wellcome Sanger Institute for the COVID-19 Genomics UK (COG-UK) Consortium | Jacquelyn Wynn, Mairead Hyland, The Lighthouse Lab in Alderley Park and Alex Alderton, Roberto Amato, Jeffrey Barrett, Sonia Goncalves, Ewan Harrison, David K. Jackson, Ian Johnston, Dominic Kwiatkowski, Cordelia Langford, John Sillitoe on behalf of the Wellcome Sanger Institute COVID-19 Surveillance Team |
| EPI_ISL_2607002 | Lighthouse Lab in Milton Keynes | Wellcome Sanger Institute for the COVID-19 Genomics UK (COG-UK) Consortium | The Lighthouse Lab in Milton Keynes and Alex Alderton, Roberto Amato, Jeffrey Barrett, Sonia Goncalves, Ewan Harrison, David K. Jackson, Ian Johnston, Dominic Kwiatkowski, Cordelia Langford, John Sillitoe on behalf of the Wellcome Sanger Institute COVID-19 Surveillance Team |
| EPI_ISL_2607003 | Lighthouse Laboratory Plymouth | Wellcome Sanger Institute for the COVID-19 Genomics UK (COG-UK) Consortium | Lighthouse Laboratory Plymouth and Alex Alderton, Roberto Amato, Jeffrey Barrett, Sonia Goncalves, Ewan Harrison, David K. Jackson, Ian Johnston, Dominic Kwiatkowski, Cordelia Langford, John Sillitoe on behalf of the Wellcome Sanger Institute COVID-19 Surveillance Team |
| EPI_ISL_2607004 | Lighthouse Lab in Milton Keynes | Wellcome Sanger Institute for the COVID-19 Genomics UK (COG-UK) Consortium | The Lighthouse Lab in Milton Keynes and Alex Alderton, Roberto Amato, Jeffrey Barrett, Sonia Goncalves, Ewan Harrison, David K. Jackson, Ian Johnston, Dominic Kwiatkowski, Cordelia Langford, John Sillitoe on behalf of the Wellcome Sanger Institute COVID-19 Surveillance Team |
| EPI_ISL_2607005 | Lighthouse Lab in Glasgow | Wellcome Sanger Institute for the COVID-19 Genomics UK (COG-UK) Consortium | Harper VanSteenhouse, Yumi Kasai, David Gray, Carol Clugston, Anna Dominiczak and Alex Alderton, Roberto Amato, Jeffrey Barrett, Sonia Goncalves, Ewan Harrison, David K. Jackson, Ian Johnston, Dominic Kwiatkowski, Cordelia Langford, John Sillitoe on behalf of the Wellcome Sanger Institute COVID-19 Surveillance Team |
| EPI_ISL_2607006 | Lighthouse Lab in Alderley Park | Wellcome Sanger Institute for the COVID-19 Genomics UK (COG-UK) Consortium | Jacquelyn Wynn, Mairead Hyland, The Lighthouse Lab in Alderley Park and Alex Alderton, Roberto Amato, Jeffrey Barrett, Sonia Goncalves, Ewan Harrison, David K. Jackson, Ian Johnston, Dominic Kwiatkowski, Cordelia Langford, John Sillitoe on behalf of the Wellcome Sanger Institute COVID-19 Surveillance Team |
| EPI_ISL_2607007 | Lighthouse Lab in Glasgow | Wellcome Sanger Institute for the COVID-19 Genomics UK (COG-UK) Consortium | Harper VanSteenhouse, Yumi Kasai, David Gray, Carol Clugston, Anna Dominiczak and Alex Alderton, Roberto Amato, Jeffrey Barrett, Sonia Goncalves, Ewan Harrison, David K. Jackson, Ian Johnston, Dominic Kwiatkowski, Cordelia Langford, John Sillitoe on behalf of the Wellcome Sanger Institute COVID-19 Surveillance Team |
| EPI_ISL_2607008 | Lighthouse Lab in Alderley Park | Wellcome Sanger Institute for the COVID-19 Genomics UK (COG-UK) Consortium | Jacquelyn Wynn, Mairead Hyland, The Lighthouse Lab in Alderley Park and Alex Alderton, Roberto Amato, Jeffrey Barrett, Sonia Goncalves, Ewan Harrison, David K. Jackson, Ian Johnston, Dominic Kwiatkowski, Cordelia Langford, John Sillitoe on behalf of the Wellcome Sanger Institute COVID-19 Surveillance Team |
| EPI_ISL_2607009 | Lighthouse Laboratory Plymouth | Wellcome Sanger Institute for the COVID-19 Genomics UK (COG-UK) Consortium | Lighthouse Laboratory Plymouth and Alex Alderton, Roberto Amato, Jeffrey Barrett, Sonia Goncalves, Ewan Harrison, David K. Jackson, Ian Johnston, Dominic Kwiatkowski, Cordelia Langford, John Sillitoe on behalf of the Wellcome Sanger Institute COVID-19 Surveillance Team |
| EPI_ISL_2607010 | Lighthouse Lab in Milton Keynes | Wellcome Sanger Institute for the COVID-19 Genomics UK (COG-UK) Consortium | The Lighthouse Lab in Milton Keynes and Alex Alderton, Roberto Amato, Jeffrey Barrett, Sonia Goncalves, Ewan Harrison, David K. Jackson, Ian Johnston, Dominic Kwiatkowski, Cordelia Langford, John Sillitoe on behalf of the Wellcome Sanger Institute COVID-19 Surveillance Team |
| EPI_ISL_2607011 | Lighthouse Lab in Alderley Park | Wellcome Sanger Institute for the COVID-19 Genomics UK (COG-UK) Consortium | Jacquelyn Wynn, Mairead Hyland, The Lighthouse Lab in Alderley Park and Alex Alderton, Roberto Amato, Jeffrey Barrett, Sonia Goncalves, Ewan Harrison, David K. Jackson, Ian Johnston, Dominic Kwiatkowski, Cordelia Langford, John Sillitoe on behalf of the Wellcome Sanger Institute COVID-19 Surveillance Team |
| EPI_ISL_2607012 | Lighthouse Lab in Milton Keynes | Wellcome Sanger Institute for the COVID-19 Genomics UK (COG-UK) Consortium | The Lighthouse Lab in Milton Keynes and Alex Alderton, Roberto Amato, Jeffrey Barrett, Sonia Goncalves, Ewan Harrison, David K. Jackson, Ian Johnston, Dominic Kwiatkowski, Cordelia Langford, John Sillitoe on behalf of the Wellcome Sanger Institute COVID-19 Surveillance Team |
| EPI_ISL_2607013, EPI_ISL_2607014 | Lighthouse Lab in Alderley Park | Wellcome Sanger Institute for the COVID-19 Genomics UK (COG-UK) Consortium | Jacquelyn Wynn, Mairead Hyland, The Lighthouse Lab in Alderley Park and Alex Alderton, Roberto Amato, Jeffrey Barrett, Sonia Goncalves, Ewan Harrison, David K. Jackson, Ian Johnston, Dominic Kwiatkowski, Cordelia Langford, John Sillitoe on behalf of the Wellcome Sanger Institute COVID-19 Surveillance Team |
| EPI_ISL_2607015 | Lighthouse Laboratory Plymouth | Wellcome Sanger Institute for the COVID-19 Genomics UK (COG-UK) Consortium | Lighthouse Laboratory Plymouth and Alex Alderton, Roberto Amato, Jeffrey Barrett, Sonia Goncalves, Ewan Harrison, David K. Jackson, Ian Johnston, Dominic Kwiatkowski, Cordelia Langford, John Sillitoe on behalf of the Wellcome Sanger Institute COVID-19 Surveillance Team |
| EPI_ISL_2607016 | Lighthouse Lab in Glasgow | Wellcome Sanger Institute for the COVID-19 Genomics UK (COG-UK) Consortium | Harper VanSteenhouse, Yumi Kasai, David Gray, Carol Clugston, Anna Dominiczak and Alex Alderton, Roberto Amato, Jeffrey Barrett, Sonia Goncalves, Ewan Harrison, David K. Jackson, Ian Johnston, Dominic Kwiatkowski, Cordelia Langford, John Sillitoe on behalf of the Wellcome Sanger Institute COVID-19 Surveillance Team |
| EPI_ISL_2607017 | Lighthouse Lab in Milton Keynes | Wellcome Sanger Institute for the COVID-19 Genomics UK (COG-UK) Consortium | The Lighthouse Lab in Milton Keynes and Alex Alderton, Roberto Amato, Jeffrey Barrett, Sonia Goncalves, Ewan Harrison, David K. Jackson, Ian Johnston, Dominic Kwiatkowski, Cordelia Langford, John Sillitoe on behalf of the Wellcome Sanger Institute COVID-19 Surveillance Team |
| EPI_ISL_2607018 | Lighthouse Laboratory Plymouth | Wellcome Sanger Institute for the COVID-19 Genomics UK (COG-UK) Consortium | Lighthouse Laboratory Plymouth and Alex Alderton, Roberto Amato, Jeffrey Barrett, Sonia Goncalves, Ewan Harrison, David K. Jackson, Ian Johnston, Dominic Kwiatkowski, Cordelia Langford, John Sillitoe on behalf of the Wellcome Sanger Institute COVID-19 Surveillance Team |
| EPI_ISL_2607019, EPI_ISL_2607020 | Lighthouse Lab in Milton Keynes | Wellcome Sanger Institute for the COVID-19 Genomics UK (COG-UK) Consortium | The Lighthouse Lab in Milton Keynes and Alex Alderton, Roberto Amato, Jeffrey Barrett, Sonia Goncalves, Ewan Harrison, David K. Jackson, Ian Johnston, Dominic Kwiatkowski, Cordelia Langford, John Sillitoe on behalf of the Wellcome Sanger Institute COVID-19 Surveillance Team |
| EPI_ISL_2607021 | Lighthouse Lab in Alderley Park | Wellcome Sanger Institute for the COVID-19 Genomics UK (COG-UK) Consortium | Jacquelyn Wynn, Mairead Hyland, The Lighthouse Lab in Alderley Park and Alex Alderton, Roberto Amato, Jeffrey Barrett, Sonia Goncalves, Ewan Harrison, David K. Jackson, Ian Johnston, Dominic Kwiatkowski, Cordelia Langford, John Sillitoe on behalf of the Wellcome Sanger Institute COVID-19 Surveillance Team |
| EPI_ISL_2607022 | Lighthouse Laboratory Plymouth | Wellcome Sanger Institute for the COVID-19 Genomics UK (COG-UK) Consortium | Lighthouse Laboratory Plymouth and Alex Alderton, Roberto Amato, Jeffrey Barrett, Sonia Goncalves, Ewan Harrison, David K. Jackson, Ian Johnston, Dominic Kwiatkowski, Cordelia Langford, John Sillitoe on behalf of the Wellcome Sanger Institute COVID-19 Surveillance Team |
| EPI_ISL_2607023, EPI_ISL_2607024 | Lighthouse Lab in Milton Keynes | Wellcome Sanger Institute for the COVID-19 Genomics UK (COG-UK) Consortium | The Lighthouse Lab in Milton Keynes and Alex Alderton, Roberto Amato, Jeffrey Barrett, Sonia Goncalves, Ewan Harrison, David K. Jackson, Ian Johnston, Dominic Kwiatkowski, Cordelia Langford, John Sillitoe on behalf of the Wellcome Sanger Institute COVID-19 Surveillance Team |
| EPI_ISL_2607025 | Lighthouse Lab in Alderley Park | Wellcome Sanger Institute for the COVID-19 Genomics UK (COG-UK) Consortium | Jacquelyn Wynn, Mairead Hyland, The Lighthouse Lab in Alderley Park and Alex Alderton, Roberto Amato, Jeffrey Barrett, Sonia Goncalves, Ewan Harrison, David K. Jackson, Ian Johnston, Dominic Kwiatkowski, Cordelia Langford, John Sillitoe on behalf of the Wellcome Sanger Institute COVID-19 Surveillance Team |
| EPI_ISL_2607026 | Lighthouse Laboratory Plymouth | Wellcome Sanger Institute for the COVID-19 Genomics UK (COG-UK) Consortium | Lighthouse Laboratory Plymouth and Alex Alderton, Roberto Amato, Jeffrey Barrett, Sonia Goncalves, Ewan Harrison, David K. Jackson, Ian Johnston, Dominic Kwiatkowski, Cordelia Langford, John Sillitoe on behalf of the Wellcome Sanger Institute COVID-19 Surveillance Team |
| EPI_ISL_2607027 | Lighthouse Lab in Milton Keynes | Wellcome Sanger Institute for the COVID-19 Genomics UK (COG-UK) Consortium | The Lighthouse Lab in Milton Keynes and Alex Alderton, Roberto Amato, Jeffrey Barrett, Sonia Goncalves, Ewan Harrison, David K. Jackson, Ian Johnston, Dominic Kwiatkowski, Cordelia Langford, John Sillitoe on behalf of the Wellcome Sanger Institute COVID-19 Surveillance Team |
| EPI_ISL_2607028, EPI_ISL_2607029, EPI_ISL_2607031 | Lighthouse Lab in Alderley Park | Wellcome Sanger Institute for the COVID-19 Genomics UK (COG-UK) Consortium | Jacquelyn Wynn, Mairead Hyland, The Lighthouse Lab in Alderley Park and Alex Alderton, Roberto Amato, Jeffrey Barrett, Sonia Goncalves, Ewan Harrison, David K. Jackson, Ian Johnston, Dominic Kwiatkowski, Cordelia Langford, John Sillitoe on behalf of the Wellcome Sanger Institute COVID-19 Surveillance Team |
| EPI_ISL_2607032 | Lighthouse Lab in Milton Keynes | Wellcome Sanger Institute for the COVID-19 Genomics UK (COG-UK) Consortium | The Lighthouse Lab in Milton Keynes and Alex Alderton, Roberto Amato, Jeffrey Barrett, Sonia Goncalves, Ewan Harrison, David K. Jackson, Ian Johnston, Dominic Kwiatkowski, Cordelia Langford, John Sillitoe on behalf of the Wellcome Sanger Institute COVID-19 Surveillance Team |
| EPI_ISL_2607033, EPI_ISL_2607034, EPI_ISL_2607035, EPI_ISL_2607036 | Lighthouse Lab in Alderley Park | Wellcome Sanger Institute for the COVID-19 Genomics UK (COG-UK) Consortium | Jacquelyn Wynn, Mairead Hyland, The Lighthouse Lab in Alderley Park and Alex Alderton, Roberto Amato, Jeffrey Barrett, Sonia Goncalves, Ewan Harrison, David K. Jackson, Ian Johnston, Dominic Kwiatkowski, Cordelia Langford, John Sillitoe on behalf of the Wellcome Sanger Institute COVID-19 Surveillance Team |
| EPI_ISL_2607037 | Lighthouse Lab in Milton Keynes | Wellcome Sanger Institute for the COVID-19 Genomics UK (COG-UK) Consortium | The Lighthouse Lab in Milton Keynes and Alex Alderton, Roberto Amato, Jeffrey Barrett, Sonia Goncalves, Ewan Harrison, David K. Jackson, Ian Johnston, Dominic Kwiatkowski, Cordelia Langford, John Sillitoe on behalf of the Wellcome Sanger Institute COVID-19 Surveillance Team |
| EPI_ISL_2607038 | Lighthouse Lab in Alderley Park | Wellcome Sanger Institute for the COVID-19 Genomics UK (COG-UK) Consortium | Jacquelyn Wynn, Mairead Hyland, The Lighthouse Lab in Alderley Park and Alex Alderton, Roberto Amato, Jeffrey Barrett, Sonia Goncalves, Ewan Harrison, David K. Jackson, Ian Johnston, Dominic Kwiatkowski, Cordelia Langford, John Sillitoe on behalf of the Wellcome Sanger Institute COVID-19 Surveillance Team |
| EPI_ISL_2607039 | Lighthouse Lab in Milton Keynes | Wellcome Sanger Institute for the COVID-19 Genomics UK (COG-UK) Consortium | The Lighthouse Lab in Milton Keynes and Alex Alderton, Roberto Amato, Jeffrey Barrett, Sonia Goncalves, Ewan Harrison, David K. Jackson, Ian Johnston, Dominic Kwiatkowski, Cordelia Langford, John Sillitoe on behalf of the Wellcome Sanger Institute COVID-19 Surveillance Team |
| EPI_ISL_2607040 | Lighthouse Lab in Alderley Park | Wellcome Sanger Institute for the COVID-19 Genomics UK (COG-UK) Consortium | Jacquelyn Wynn, Mairead Hyland, The Lighthouse Lab in Alderley Park and Alex Alderton, Roberto Amato, Jeffrey Barrett, Sonia Goncalves, Ewan Harrison, David K. Jackson, Ian Johnston, Dominic Kwiatkowski, Cordelia Langford, John Sillitoe on behalf of the Wellcome Sanger Institute COVID-19 Surveillance Team |

| | | | |
|---|---|---|---|
| EPI_ISL_2607041 | Lighthouse Lab in Milton Keynes | Wellcome Sanger Institute for the COVID-19 Genomics UK (COG-UK) Consortium | The Lighthouse Lab in Milton Keynes and Alex Alderton, Roberto Amato, Jeffrey Barrett, Sonia Goncalves, Ewan Harrison, David K. Jackson, Ian Johnston, Dominic Kwiatkowski, Cordelia Langford, John Sillitoe on behalf of the Wellcome Sanger Institute COVID-19 Surveillance Team |
| EPI_ISL_2607042, EPI_ISL_2607043, EPI_ISL_2607044, EPI_ISL_2607045, EPI_ISL_2607046, EPI_ISL_2607047, EPI_ISL_2607048, EPI_ISL_2607049, EPI_ISL_2607050, EPI_ISL_2607051, EPI_ISL_2607052, EPI_ISL_2607053, EPI_ISL_2607054, EPI_ISL_2607055, EPI_ISL_2607056, EPI_ISL_2607057, EPI_ISL_2607058, EPI_ISL_2607059, EPI_ISL_2607060, EPI_ISL_2607061, EPI_ISL_2607062, EPI_ISL_2607063, EPI_ISL_2607064, EPI_ISL_2607065, EPI_ISL_2607067, EPI_ISL_2607068, EPI_ISL_2607069, EPI_ISL_2607070, EPI_ISL_2607072, EPI_ISL_2607073, EPI_ISL_2607074, EPI_ISL_2607075, EPI_ISL_2607076, EPI_ISL_2607077, EPI_ISL_2607079, EPI_ISL_2607080, EPI_ISL_2607081, EPI_ISL_2607082, EPI_ISL_2607083, EPI_ISL_2607084, EPI_ISL_2607085, EPI_ISL_2607086, EPI_ISL_2607087, EPI_ISL_2607088, EPI_ISL_2607091, EPI_ISL_2607092, EPI_ISL_2607094, EPI_ISL_2607095, EPI_ISL_2607097, EPI_ISL_2607098, EPI_ISL_2607099 | | | |
| see above | Lighthouse Lab in Glasgow | Wellcome Sanger Institute for the COVID-19 Genomics UK (COG-UK) Consortium | Harper VanSteenhouse, Yumi Kasai, David Gray, Carol Clugston, Anna Dominiczak and Alex Alderton, Roberto Amato, Jeffrey Barrett, Sonia Goncalves, Ewan Harrison, David K. Jackson, Ian Johnston, Dominic Kwiatkowski, Cordelia Langford, John Sillitoe on behalf of the Wellcome Sanger Institute COVID-19 Surveillance Team |
| EPI_ISL_2607100 | Lighthouse Lab in Milton Keynes | Wellcome Sanger Institute for the COVID-19 Genomics UK (COG-UK) Consortium | The Lighthouse Lab in Milton Keynes and Alex Alderton, Roberto Amato, Jeffrey Barrett, Sonia Goncalves, Ewan Harrison, David K. Jackson, Ian Johnston, Dominic Kwiatkowski, Cordelia Langford, John Sillitoe on behalf of the Wellcome Sanger Institute COVID-19 Surveillance Team |
| EPI_ISL_2607101, EPI_ISL_2607103, EPI_ISL_2607104, EPI_ISL_2607105, EPI_ISL_2607106, EPI_ISL_2607107, EPI_ISL_2607108, EPI_ISL_2607110, EPI_ISL_2607111, EPI_ISL_2607112, EPI_ISL_2607113, EPI_ISL_2607114, EPI_ISL_2607115, EPI_ISL_2607116, EPI_ISL_2607117, EPI_ISL_2607118, EPI_ISL_2607119, EPI_ISL_2607120, EPI_ISL_2607121, EPI_ISL_2607122, EPI_ISL_2607124, EPI_ISL_2607127, EPI_ISL_2607128, EPI_ISL_2607129, EPI_ISL_2607130, EPI_ISL_2607131, EPI_ISL_2607132, EPI_ISL_2607133, EPI_ISL_2607134, EPI_ISL_2607136, EPI_ISL_2607138, EPI_ISL_2607139, EPI_ISL_2607140, EPI_ISL_2607141, EPI_ISL_2607142, EPI_ISL_2607143, EPI_ISL_2607144, EPI_ISL_2607147, EPI_ISL_2607148, EPI_ISL_2607149, EPI_ISL_2607150, EPI_ISL_2607151, EPI_ISL_2607152, EPI_ISL_2607153, EPI_ISL_2607155, EPI_ISL_2607156, EPI_ISL_2607157, EPI_ISL_2607158, EPI_ISL_2607159, EPI_ISL_2607160, EPI_ISL_2607161, EPI_ISL_2607162, EPI_ISL_2607163, EPI_ISL_2607164, EPI_ISL_2607165, EPI_ISL_2607166, EPI_ISL_2607167, EPI_ISL_2607168, EPI_ISL_2607169, EPI_ISL_2607170, EPI_ISL_2607172, EPI_ISL_2607173, EPI_ISL_2607174, EPI_ISL_2607175, EPI_ISL_2607178, EPI_ISL_2607179, EPI_ISL_2607180, EPI_ISL_2607181, EPI_ISL_2607182, EPI_ISL_2607183, EPI_ISL_2607184, EPI_ISL_2607185, EPI_ISL_2607186, EPI_ISL_2607187, EPI_ISL_2607188, EPI_ISL_2607189, EPI_ISL_2607190, EPI_ISL_2607191, EPI_ISL_2607192, EPI_ISL_2607193, EPI_ISL_2607194, EPI_ISL_2607195, EPI_ISL_2607196, EPI_ISL_2607197, EPI_ISL_2607198, EPI_ISL_2607199, EPI_ISL_2607200, EPI_ISL_2607201, EPI_ISL_2607202, EPI_ISL_2607205, EPI_ISL_2607206, EPI_ISL_2607207, EPI_ISL_2607210, EPI_ISL_2607211, EPI_ISL_2607212, EPI_ISL_2607213, EPI_ISL_2607214, EPI_ISL_2607215, EPI_ISL_2607216, EPI_ISL_2607217, EPI_ISL_2607218, EPI_ISL_2607219, EPI_ISL_2607221, EPI_ISL_2607222, EPI_ISL_2607223, EPI_ISL_2607224, EPI_ISL_2607225, EPI_ISL_2607226, EPI_ISL_2607227, EPI_ISL_2607228, EPI_ISL_2607229, EPI_ISL_2607230, EPI_ISL_2607231, EPI_ISL_2607232, EPI_ISL_2607233, EPI_ISL_2607234, EPI_ISL_2607235, EPI_ISL_2607236, EPI_ISL_2607239, EPI_ISL_2607241, EPI_ISL_2607242, EPI_ISL_2607243, EPI_ISL_2607245, EPI_ISL_2607246, EPI_ISL_2607247, EPI_ISL_2607248, EPI_ISL_2607249, EPI_ISL_2607250 | | | |
| see above | Lighthouse Lab in Glasgow | Wellcome Sanger Institute for the COVID-19 Genomics UK (COG-UK) Consortium | Harper VanSteenhouse, Yumi Kasai, David Gray, Carol Clugston, Anna Dominiczak and Alex Alderton, Roberto Amato, Jeffrey Barrett, Sonia Goncalves, Ewan Harrison, David K. Jackson, Ian Johnston, Dominic Kwiatkowski, Cordelia Langford, John Sillitoe on behalf of the Wellcome Sanger Institute COVID-19 Surveillance Team |
| EPI_ISL_2607251, EPI_ISL_2607252, EPI_ISL_2607253 | Lighthouse Lab in Alderley Park | Wellcome Sanger Institute for the COVID-19 Genomics UK (COG-UK) Consortium | Jacquelyn Wynn, Mairead Hyland, The Lighthouse Lab in Alderley Park and Alex Alderton, Roberto Amato, Jeffrey Barrett, Sonia Goncalves, Ewan Harrison, David K. Jackson, Ian Johnston, Dominic Kwiatkowski, Cordelia Langford, John Sillitoe on behalf of the Wellcome Sanger Institute COVID-19 Surveillance Team |
| EPI_ISL_2607256, EPI_ISL_2607257 | Lighthouse Lab in Milton Keynes | Wellcome Sanger Institute for the COVID-19 Genomics UK (COG-UK) Consortium | The Lighthouse Lab in Milton Keynes and Alex Alderton, Roberto Amato, Jeffrey Barrett, Sonia Goncalves, Ewan Harrison, David K. Jackson, Ian Johnston, Dominic Kwiatkowski, Cordelia Langford, John Sillitoe on behalf of the Wellcome Sanger Institute COVID-19 Surveillance Team |
| EPI_ISL_2607258, EPI_ISL_2607259 | Lighthouse Lab in Alderley Park | Wellcome Sanger Institute for the COVID-19 Genomics UK (COG-UK) Consortium | Jacquelyn Wynn, Mairead Hyland, The Lighthouse Lab in Alderley Park and Alex Alderton, Roberto Amato, Jeffrey Barrett, Sonia Goncalves, Ewan Harrison, David K. Jackson, Ian Johnston, Dominic Kwiatkowski, Cordelia Langford, John Sillitoe on behalf of the Wellcome Sanger Institute COVID-19 Surveillance Team |
| EPI_ISL_2607260, EPI_ISL_2607261, EPI_ISL_2607262, EPI_ISL_2607263, EPI_ISL_2607264, EPI_ISL_2607265, EPI_ISL_2607267, EPI_ISL_2607268, EPI_ISL_2607269, EPI_ISL_2607270, EPI_ISL_2607272, EPI_ISL_2607273, EPI_ISL_2607274, EPI_ISL_2607275, EPI_ISL_2607276, EPI_ISL_2607277, EPI_ISL_2607279, EPI_ISL_2607280, EPI_ISL_2607281, EPI_ISL_2607282, EPI_ISL_2607283, EPI_ISL_2607284, EPI_ISL_2607285, EPI_ISL_2607286, EPI_ISL_2607287, EPI_ISL_2607288, EPI_ISL_2607289, EPI_ISL_2607290, EPI_ISL_2607291, EPI_ISL_2607292, EPI_ISL_2607293, EPI_ISL_2607294, EPI_ISL_2607295, EPI_ISL_2607297, EPI_ISL_2607298, EPI_ISL_2607299, EPI_ISL_2607300, EPI_ISL_2607302, EPI_ISL_2607303, EPI_ISL_2607304, EPI_ISL_2607305, EPI_ISL_2607307, EPI_ISL_2607308, EPI_ISL_2607309, EPI_ISL_2607310, EPI_ISL_2607311, EPI_ISL_2607312, EPI_ISL_2607313, EPI_ISL_2607314, EPI_ISL_2607315, EPI_ISL_2607316, EPI_ISL_2607317, EPI_ISL_2607318, EPI_ISL_2607319, EPI_ISL_2607320, EPI_ISL_2607321, EPI_ISL_2607322, EPI_ISL_2607323, EPI_ISL_2607324, EPI_ISL_2607325, EPI_ISL_2607326, EPI_ISL_2607327, EPI_ISL_2607328, EPI_ISL_2607329, EPI_ISL_2607331, EPI_ISL_2607332, EPI_ISL_2607334, EPI_ISL_2607336, EPI_ISL_2607337, EPI_ISL_2607338, EPI_ISL_2607339, EPI_ISL_2607341, EPI_ISL_2607342, EPI_ISL_2607343, EPI_ISL_2607344, EPI_ISL_2607345, EPI_ISL_2607346, EPI_ISL_2607347, EPI_ISL_2607348, EPI_ISL_2607349, EPI_ISL_2607351, EPI_ISL_2607352, EPI_ISL_2607353, EPI_ISL_2607354, EPI_ISL_2607355, EPI_ISL_2607356, EPI_ISL_2607357, EPI_ISL_2607358, EPI_ISL_2607359, EPI_ISL_2607360, EPI_ISL_2607361, EPI_ISL_2607362, EPI_ISL_2607363, EPI_ISL_2607364, EPI_ISL_2607365, EPI_ISL_2607366, EPI_ISL_2607367, EPI_ISL_2607368, EPI_ISL_2607369, EPI_ISL_2607370, EPI_ISL_2607371, EPI_ISL_2607372, EPI_ISL_2607373, EPI_ISL_2607374, EPI_ISL_2607375, EPI_ISL_2607377, EPI_ISL_2607378, EPI_ISL_2607379, EPI_ISL_2607380, EPI_ISL_2607381, EPI_ISL_2607382, EPI_ISL_2607383, EPI_ISL_2607384, EPI_ISL_2607385, EPI_ISL_2607386, EPI_ISL_2607387, EPI_ISL_2607388, EPI_ISL_2607389, EPI_ISL_2607390, EPI_ISL_2607391, EPI_ISL_2607394, EPI_ISL_2607395, EPI_ISL_2607396, EPI_ISL_2607398, EPI_ISL_2607399, EPI_ISL_2607400, EPI_ISL_2607401, EPI_ISL_2607402, EPI_ISL_2607403, EPI_ISL_2607404, EPI_ISL_2607405, EPI_ISL_2607407, EPI_ISL_2607409, EPI_ISL_2607410, EPI_ISL_2607411, EPI_ISL_2607412, EPI_ISL_2607413, EPI_ISL_2607414, EPI_ISL_2607415, EPI_ISL_2607416, EPI_ISL_2607417, EPI_ISL_2607418, EPI_ISL_2607419, EPI_ISL_2607420, EPI_ISL_2607422, EPI_ISL_2607424, EPI_ISL_2607425, EPI_ISL_2607426, EPI_ISL_2607427, EPI_ISL_2607429, EPI_ISL_2607430, EPI_ISL_2607432, EPI_ISL_2607433, EPI_ISL_2607434, EPI_ISL_2607435, EPI_ISL_2607436, EPI_ISL_2607437, EPI_ISL_2607438, EPI_ISL_2607439, EPI_ISL_2607440, EPI_ISL_2607441, EPI_ISL_2607442, EPI_ISL_2607444, EPI_ISL_2607445, EPI_ISL_2607446, EPI_ISL_2607447, EPI_ISL_2607448, EPI_ISL_2607449, EPI_ISL_2607450, EPI_ISL_2607452, EPI_ISL_2607453, EPI_ISL_2607454, EPI_ISL_2607455, EPI_ISL_2607456, EPI_ISL_2607457, EPI_ISL_2607458, EPI_ISL_2607459, EPI_ISL_2607460, EPI_ISL_2607461, EPI_ISL_2607463, EPI_ISL_2607464, EPI_ISL_2607465, EPI_ISL_2607466, EPI_ISL_2607467, EPI_ISL_2607468, EPI_ISL_2607469, EPI_ISL_2607470, EPI_ISL_2607471, EPI_ISL_2607472, EPI_ISL_2607473, EPI_ISL_2607474, EPI_ISL_2607475, EPI_ISL_2607476, EPI_ISL_2607477, EPI_ISL_2607478, EPI_ISL_2607479, EPI_ISL_2607480, EPI_ISL_2607481, EPI_ISL_2607482, EPI_ISL_2607483, EPI_ISL_2607484, EPI_ISL_2607485, EPI_ISL_2607486, EPI_ISL_2607487, EPI_ISL_2607488, EPI_ISL_2607489, EPI_ISL_2607490, EPI_ISL_2607491, EPI_ISL_2607492, EPI_ISL_2607494, EPI_ISL_2607495, EPI_ISL_2607496, EPI_ISL_2607497, EPI_ISL_2607498, EPI_ISL_2607499, EPI_ISL_2607500, EPI_ISL_2607501, EPI_ISL_2607503 | | | |
| see above | Lighthouse Lab in Glasgow | Wellcome Sanger Institute for the COVID-19 Genomics UK (COG-UK) Consortium | Harper VanSteenhouse, Yumi Kasai, David Gray, Carol Clugston, Anna Dominiczak and Alex Alderton, Roberto Amato, Jeffrey Barrett, Sonia Goncalves, Ewan Harrison, David K. Jackson, Ian Johnston, Dominic Kwiatkowski, Cordelia Langford, John Sillitoe on behalf of the Wellcome Sanger Institute COVID-19 Surveillance Team |
| EPI_ISL_2607504 | Lighthouse Lab in Milton Keynes | Wellcome Sanger Institute for the COVID-19 Genomics UK (COG-UK) Consortium | The Lighthouse Lab in Milton Keynes and Alex Alderton, Roberto Amato, Jeffrey Barrett, Sonia Goncalves, Ewan Harrison, David K. Jackson, Ian Johnston, Dominic Kwiatkowski, Cordelia Langford, John Sillitoe on behalf of the Wellcome Sanger Institute COVID-19 Surveillance Team |
| EPI_ISL_2607506, EPI_ISL_2607507, EPI_ISL_2607508, EPI_ISL_2607509, EPI_ISL_2607510, EPI_ISL_2607512, EPI_ISL_2607513, EPI_ISL_2607514, EPI_ISL_2607515 | Lighthouse Lab in Glasgow | Wellcome Sanger Institute for the COVID-19 Genomics UK (COG-UK) Consortium | Harper VanSteenhouse, Yumi Kasai, David Gray, Carol Clugston, Anna Dominiczak and Alex Alderton, Roberto Amato, Jeffrey Barrett, Sonia Goncalves, Ewan Harrison, David K. Jackson, Ian Johnston, Dominic Kwiatkowski, Cordelia Langford, John Sillitoe on behalf of the Wellcome Sanger Institute COVID-19 Surveillance Team |
| EPI_ISL_2607516 | Lighthouse Lab in Alderley Park | Wellcome Sanger Institute for the COVID-19 Genomics UK (COG-UK) Consortium | Jacquelyn Wynn, Mairead Hyland, The Lighthouse Lab in Alderley Park and Alex Alderton, Roberto Amato, Jeffrey Barrett, Sonia Goncalves, Ewan Harrison, David K. Jackson, Ian Johnston, Dominic Kwiatkowski, Cordelia Langford, John Sillitoe on behalf of the Wellcome Sanger Institute COVID-19 Surveillance Team |
| EPI_ISL_2607517, EPI_ISL_2607518, EPI_ISL_2607519, EPI_ISL_2607520, EPI_ISL_2607521, EPI_ISL_2607522, EPI_ISL_2607523, EPI_ISL_2607525, EPI_ISL_2607526, EPI_ISL_2607527, EPI_ISL_2607528, EPI_ISL_2607529, EPI_ISL_2607530, EPI_ISL_2607531, EPI_ISL_2607532, EPI_ISL_2607533, EPI_ISL_2607534, EPI_ISL_2607535, EPI_ISL_2607537, EPI_ISL_2607538, EPI_ISL_2607539, EPI_ISL_2607540, EPI_ISL_2607541, EPI_ISL_2607543, EPI_ISL_2607544 | | | |
| see above | Lighthouse Lab in Glasgow | Wellcome Sanger Institute for the COVID-19 Genomics UK (COG-UK) Consortium | Harper VanSteenhouse, Yumi Kasai, David Gray, Carol Clugston, Anna Dominiczak and Alex Alderton, Roberto Amato, Jeffrey Barrett, Sonia Goncalves, Ewan Harrison, David K. Jackson, Ian Johnston, Dominic Kwiatkowski, Cordelia Langford, John Sillitoe on behalf of the Wellcome Sanger Institute COVID-19 Surveillance Team |
| EPI_ISL_2607546 | Lighthouse Lab in Alderley Park | Wellcome Sanger Institute for the COVID-19 Genomics UK (COG-UK) Consortium | Jacquelyn Wynn, Mairead Hyland, The Lighthouse Lab in Alderley Park and Alex Alderton, Roberto Amato, Jeffrey Barrett, Sonia Goncalves, Ewan Harrison, David K. Jackson, Ian Johnston, Dominic Kwiatkowski, Cordelia Langford, John Sillitoe on behalf of the Wellcome Sanger Institute COVID-19 Surveillance Team |
| EPI_ISL_2607547, EPI_ISL_2607549, EPI_ISL_2607551, EPI_ISL_2607552, EPI_ISL_2607554, EPI_ISL_2607555, EPI_ISL_2607556, EPI_ISL_2607557, EPI_ISL_2607558, EPI_ISL_2607559, EPI_ISL_2607560, EPI_ISL_2607562, EPI_ISL_2607563, EPI_ISL_2607564, EPI_ISL_2607565, EPI_ISL_2607566, EPI_ISL_2607567, EPI_ISL_2607568, EPI_ISL_2607569, EPI_ISL_2607570, EPI_ISL_2607573, EPI_ISL_2607574, EPI_ISL_2607575, EPI_ISL_2607576, EPI_ISL_2607577, EPI_ISL_2607578, EPI_ISL_2607579, EPI_ISL_2607580, EPI_ISL_2607581, EPI_ISL_2607582, EPI_ISL_2607583, EPI_ISL_2607584, EPI_ISL_2607585, EPI_ISL_2607586, EPI_ISL_2607587, EPI_ISL_2607588, EPI_ISL_2607589, EPI_ISL_2607590, EPI_ISL_2607591, EPI_ISL_2607593, EPI_ISL_2607594, EPI_ISL_2607595, EPI_ISL_2607596, EPI_ISL_2607597, EPI_ISL_2607598, EPI_ISL_2607599, EPI_ISL_2607600, EPI_ISL_2607601, EPI_ISL_2607602, EPI_ISL_2607603, EPI_ISL_2607604, EPI_ISL_2607605, EPI_ISL_2607606, EPI_ISL_2607607, EPI_ISL_2607610, EPI_ISL_2607613, EPI_ISL_2607614, EPI_ISL_2607615, EPI_ISL_2607616, EPI_ISL_2607618, EPI_ISL_2607619, EPI_ISL_2607620, EPI_ISL_2607621, EPI_ISL_2607622, EPI_ISL_2607623, EPI_ISL_2607624, EPI_ISL_2607625, EPI_ISL_2607626, EPI_ISL_2607627, EPI_ISL_2607628, EPI_ISL_2607629, EPI_ISL_2607630, EPI_ISL_2607632, EPI_ISL_2607634, EPI_ISL_2607635 | | | |
| see above | Lighthouse Lab in Glasgow | Wellcome Sanger Institute for the COVID-19 Genomics UK (COG-UK) Consortium | Harper VanSteenhouse, Yumi Kasai, David Gray, Carol Clugston, Anna Dominiczak and Alex Alderton, Roberto Amato, Jeffrey Barrett, Sonia Goncalves, Ewan Harrison, David K. Jackson, Ian Johnston, Dominic Kwiatkowski, Cordelia Langford, John Sillitoe on behalf of the Wellcome Sanger Institute COVID-19 Surveillance Team |
| EPI_ISL_2607636 | Lighthouse Lab in Alderley Park | Wellcome Sanger Institute for the COVID-19 Genomics UK (COG-UK) Consortium | Jacquelyn Wynn, Mairead Hyland, The Lighthouse Lab in Alderley Park and Alex Alderton, Roberto Amato, Jeffrey Barrett, Sonia Goncalves, Ewan Harrison, David K. Jackson, Ian Johnston, Dominic Kwiatkowski, Cordelia Langford, John Sillitoe on behalf of the Wellcome Sanger Institute COVID-19 Surveillance Team |
| EPI_ISL_2607637 | Lighthouse Lab in Milton Keynes | Wellcome Sanger Institute for the COVID-19 Genomics UK (COG-UK) Consortium | The Lighthouse Lab in Milton Keynes and Alex Alderton, Roberto Amato, Jeffrey Barrett, Sonia Goncalves, Ewan Harrison, David K. Jackson, Ian Johnston, Dominic Kwiatkowski, Cordelia Langford, John Sillitoe on behalf of the Wellcome Sanger Institute COVID-19 Surveillance Team |
| EPI_ISL_2607639, EPI_ISL_2607641, EPI_ISL_2607642, EPI_ISL_2607643, EPI_ISL_2607644, EPI_ISL_2607645, EPI_ISL_2607646, EPI_ISL_2607647, EPI_ISL_2607648, EPI_ISL_2607649, EPI_ISL_2607651, EPI_ISL_2607652, EPI_ISL_2607653, EPI_ISL_2607654, EPI_ISL_2607655, EPI_ISL_2607656, EPI_ISL_2607657, EPI_ISL_2607658, EPI_ISL_2607659, EPI_ISL_2607660, EPI_ISL_2607661, EPI_ISL_2607662, EPI_ISL_2607663, EPI_ISL_2607664, EPI_ISL_2607665, EPI_ISL_2607666, EPI_ISL_2607667, EPI_ISL_2607668, EPI_ISL_2607669, EPI_ISL_2607670, EPI_ISL_2607671, EPI_ISL_2607672, EPI_ISL_2607673, EPI_ISL_2607674, EPI_ISL_2607675, EPI_ISL_2607676, EPI_ISL_2607677, EPI_ISL_2607678, EPI_ISL_2607679, EPI_ISL_2607680, EPI_ISL_2607681, EPI_ISL_2607683, EPI_ISL_2607684, EPI_ISL_2607685, EPI_ISL_2607686, EPI_ISL_2607687, EPI_ISL_2607688, EPI_ISL_2607691, EPI_ISL_2607692, EPI_ISL_2607693, EPI_ISL_2607694, EPI_ISL_2607695, EPI_ISL_2607696, EPI_ISL_2607697, EPI_ISL_2607698, EPI_ISL_2607699, EPI_ISL_2607701, EPI_ISL_2607702, EPI_ISL_2607703, EPI_ISL_2607704, EPI_ISL_2607706, EPI_ISL_2607707, EPI_ISL_2607709, EPI_ISL_2607710, EPI_ISL_2607711, EPI_ISL_2607712, EPI_ISL_2607713, EPI_ISL_2607714, EPI_ISL_2607715, EPI_ISL_2607716, EPI_ISL_2607717, EPI_ISL_2607718, EPI_ISL_2607719, EPI_ISL_2607720, EPI_ISL_2607722, EPI_ISL_2607723, EPI_ISL_2607724, EPI_ISL_2607725, EPI_ISL_2607726, EPI_ISL_2607727, EPI_ISL_2607728, EPI_ISL_2607729, EPI_ISL_2607730, EPI_ISL_2607731, EPI_ISL_2607734, EPI_ISL_2607736, EPI_ISL_2607737, EPI_ISL_2607738, EPI_ISL_2607739, EPI_ISL_2607740, EPI_ISL_2607741, EPI_ISL_2607742, EPI_ISL_2607743, EPI_ISL_2607744, EPI_ISL_2607745, EPI_ISL_2607746, EPI_ISL_2607747, EPI_ISL_2607748, EPI_ISL_2607749, EPI_ISL_2607750, EPI_ISL_2607751, EPI_ISL_2607752, EPI_ISL_2607753, EPI_ISL_2607754, EPI_ISL_2607755, EPI_ISL_2607756, EPI_ISL_2607757, EPI_ISL_2607758, EPI_ISL_2607759, EPI_ISL_2607760, EPI_ISL_2607761, EPI_ISL_2607762, EPI_ISL_2607763, EPI_ISL_2607764, EPI_ISL_2607766, EPI_ISL_2607767, EPI_ISL_2607768, EPI_ISL_2607769, EPI_ISL_2607770, EPI_ISL_2607771, EPI_ISL_2607772, EPI_ISL_2607773, EPI_ISL_2607774, EPI_ISL_2607775, EPI_ISL_2607776, EPI_ISL_2607777, EPI_ISL_2607778, EPI_ISL_2607779, EPI_ISL_2607780, EPI_ISL_2607781, EPI_ISL_2607782, EPI_ISL_2607783, EPI_ISL_2607784, EPI_ISL_2607785, EPI_ISL_2607786, EPI_ISL_2607787, EPI_ISL_2607788, EPI_ISL_2607790, EPI_ISL_2607791, EPI_ISL_2607792, EPI_ISL_2607793, EPI_ISL_2607794, EPI_ISL_2607795, EPI_ISL_2607796, EPI_ISL_2607797, EPI_ISL_2607799, EPI_ISL_2607800, EPI_ISL_2607801, EPI_ISL_2607802, EPI_ISL_2607803, EPI_ISL_2607804, EPI_ISL_2607805, EPI_ISL_2607806, EPI_ISL_2607807, EPI_ISL_2607808, EPI_ISL_2607809, EPI_ISL_2607810, EPI_ISL_2607811, EPI_ISL_2607812, EPI_ISL_2607813, EPI_ISL_2607814, EPI_ISL_2607815, EPI_ISL_2607816, EPI_ISL_2607817, EPI_ISL_2607818, EPI_ISL_2607819, EPI_ISL_2607820, EPI_ISL_2607821, EPI_ISL_2607822, EPI_ISL_2607823, EPI_ISL_2607824, EPI_ISL_2607825, EPI_ISL_2607826, EPI_ISL_2607827, EPI_ISL_2607829, EPI_ISL_2607830, EPI_ISL_2607831, EPI_ISL_2607832, EPI_ISL_2607833, EPI_ISL_2607834, EPI_ISL_2607835, EPI_ISL_2607836, EPI_ISL_2607837, EPI_ISL_2607838, EPI_ISL_2607839, EPI_ISL_2607840, EPI_ISL_2607841, EPI_ISL_2607842, EPI_ISL_2607843, EPI_ISL_2607844, EPI_ISL_2607845, EPI_ISL_2607846, EPI_ISL_2607847, EPI_ISL_2607848, EPI_ISL_2607849, EPI_ISL_2607851, EPI_ISL_2607852, EPI_ISL_2607854, EPI_ISL_2607855, EPI_ISL_2607856, EPI_ISL_2607857, EPI_ISL_2607858, EPI_ISL_2607859, EPI_ISL_2607860, EPI_ISL_2607861, EPI_ISL_2607862, EPI_ISL_2607863, EPI_ISL_2607864, EPI_ISL_2607865, EPI_ISL_2607866, EPI_ISL_2607867, EPI_ISL_2607868, EPI_ISL_2607869, EPI_ISL_2607870, EPI_ISL_2607871, EPI_ISL_2607872, EPI_ISL_2607874, EPI_ISL_2607875, EPI_ISL_2607876, EPI_ISL_2607878, EPI_ISL_2607879, EPI_ISL_2607881, EPI_ISL_2607882, EPI_ISL_2607883, EPI_ISL_2607884, EPI_ISL_2607885, EPI_ISL_2607886, EPI_ISL_2607887, EPI_ISL_2607888, EPI_ISL_2607889, EPI_ISL_2607890, EPI_ISL_2607892, EPI_ISL_2607893, EPI_ISL_2607894, EPI_ISL_2607895, EPI_ISL_2607896, EPI_ISL_2607897, EPI_ISL_2607898 | | | |
| see above | Lighthouse Lab in Glasgow | Wellcome Sanger Institute for the COVID-19 Genomics UK (COG-UK) Consortium | Harper VanSteenhouse, Yumi Kasai, David Gray, Carol Clugston, Anna Dominiczak and Alex Alderton, Roberto Amato, Jeffrey Barrett, Sonia Goncalves, Ewan Harrison, David K. Jackson, Ian Johnston, Dominic Kwiatkowski, Cordelia Langford, John Sillitoe on behalf of the Wellcome Sanger Institute COVID-19 Surveillance Team |
| EPI_ISL_2607899, EPI_ISL_2607900 | Lighthouse Lab in Milton Keynes | Wellcome Sanger Institute for the COVID-19 Genomics UK (COG-UK) Consortium | The Lighthouse Lab in Milton Keynes and Alex Alderton, Roberto Amato, Jeffrey Barrett, Sonia Goncalves, Ewan Harrison, David K. Jackson, Ian Johnston, Dominic Kwiatkowski, Cordelia Langford, John Sillitoe on behalf of the Wellcome Sanger Institute COVID-19 Surveillance Team |
| EPI_ISL_2607901, EPI_ISL_2607902 | Lighthouse Lab in Alderley Park | Wellcome Sanger Institute for the COVID-19 Genomics UK (COG-UK) Consortium | Jacquelyn Wynn, Mairead Hyland, The Lighthouse Lab in Alderley Park and Alex Alderton, Roberto Amato, Jeffrey Barrett, Sonia Goncalves, Ewan Harrison, David K. Jackson, Ian Johnston, Dominic Kwiatkowski, Cordelia Langford, John Sillitoe on behalf of the Wellcome Sanger Institute COVID-19 Surveillance Team |
| EPI_ISL_2607903, EPI_ISL_2607904, EPI_ISL_2607905, EPI_ISL_2607906, EPI_ISL_2607907, EPI_ISL_2607908, EPI_ISL_2607909 | Lighthouse Lab in Glasgow | Wellcome Sanger Institute for the COVID-19 Genomics UK (COG-UK) Consortium | Harper VanSteenhouse, Yumi Kasai, David Gray, Carol Clugston, Anna Dominiczak and Alex Alderton, Roberto Amato, Jeffrey Barrett, Sonia Goncalves, Ewan Harrison, David K. Jackson, Ian Johnston, Dominic Kwiatkowski, Cordelia Langford, John Sillitoe on behalf of the Wellcome Sanger Institute COVID-19 Surveillance Team |
| EPI_ISL_2607910 | Lighthouse Laboratory Plymouth | Wellcome Sanger Institute for the COVID-19 Genomics UK (COG-UK) Consortium | Lighthouse Laboratory Plymouth and Alex Alderton, Roberto Amato, Jeffrey Barrett, Sonia Goncalves, Ewan Harrison, David K. Jackson, Ian Johnston, Dominic Kwiatkowski, Cordelia Langford, John Sillitoe on behalf of the Wellcome Sanger Institute COVID-19 Surveillance Team |
| EPI_ISL_2607912 | Lighthouse Lab in Alderley Park | Wellcome Sanger Institute for the COVID-19 Genomics UK (COG-UK) Consortium | Jacquelyn Wynn, Mairead Hyland, The Lighthouse Lab in Alderley Park and Alex Alderton, Roberto Amato, Jeffrey Barrett, Sonia Goncalves, Ewan Harrison, David K. Jackson, Ian Johnston, Dominic Kwiatkowski, Cordelia Langford, John Sillitoe on behalf of the Wellcome Sanger Institute COVID-19 Surveillance Team |
| EPI_ISL_2607913 | Lighthouse Lab in Glasgow | Wellcome Sanger Institute for the COVID-19 Genomics UK (COG-UK) Consortium | Harper VanSteenhouse, Yumi Kasai, David Gray, Carol Clugston, Anna Dominiczak and Alex Alderton, Roberto Amato, Jeffrey Barrett, Sonia Goncalves, Ewan Harrison, David K. Jackson, Ian Johnston, Dominic Kwiatkowski, Cordelia Langford, John Sillitoe on behalf of the Wellcome Sanger Institute COVID-19 Surveillance Team |
| EPI_ISL_2607914 | Lighthouse Lab in Alderley Park | Wellcome Sanger Institute for the COVID-19 Genomics UK (COG-UK) Consortium | Jacquelyn Wynn, Mairead Hyland, The Lighthouse Lab in Alderley Park and Alex Alderton, Roberto Amato, Jeffrey Barrett, Sonia Goncalves, Ewan Harrison, David K. Jackson, Ian Johnston, Dominic Kwiatkowski, Cordelia Langford, John Sillitoe on behalf of the Wellcome Sanger Institute COVID-19 Surveillance Team |

| | | | |
|---|---|---|---|
| EPI_ISL_2607915 | Lighthouse Lab in Milton Keynes | Wellcome Sanger Institute for the COVID-19 Genomics UK (COG-UK) Consortium | The Lighthouse Lab in Milton Keynes and Alex Alderton, Roberto Amato, Jeffrey Barrett, Sonia Goncalves, Ewan Harrison, David K. Jackson, Ian Johnston, Dominic Kwiatkowski, Cordelia Langford, John Sillitoe on behalf of the Wellcome Sanger Institute COVID-19 Surveillance Team |
| EPI_ISL_2607916 | Lighthouse Lab in Glasgow | Wellcome Sanger Institute for the COVID-19 Genomics UK (COG-UK) Consortium | Harper VanSteenhouse, Yumi Kasai, David Gray, Carol Clugston, Anna Dominiczak and Alex Alderton, Roberto Amato, Jeffrey Barrett, Sonia Goncalves, Ewan Harrison, David K. Jackson, Ian Johnston, Dominic Kwiatkowski, Cordelia Langford, John Sillitoe on behalf of the Wellcome Sanger Institute COVID-19 Surveillance Team |
| EPI_ISL_2607917 | Lighthouse Lab in Alderley Park | Wellcome Sanger Institute for the COVID-19 Genomics UK (COG-UK) Consortium | Jacquelyn Wynn, Mairead Hyland, The Lighthouse Lab in Alderley Park and Alex Alderton, Roberto Amato, Jeffrey Barrett, Sonia Goncalves, Ewan Harrison, David K. Jackson, Ian Johnston, Dominic Kwiatkowski, Cordelia Langford, John Sillitoe on behalf of the Wellcome Sanger Institute COVID-19 Surveillance Team |
| EPI_ISL_2607918, EPI_ISL_2607919, EPI_ISL_2607920, EPI_ISL_2607921, EPI_ISL_2607922, EPI_ISL_2607923, EPI_ISL_2607924, EPI_ISL_2607925, EPI_ISL_2607926, EPI_ISL_2607928 | Lighthouse Lab in Glasgow | Wellcome Sanger Institute for the COVID-19 Genomics UK (COG-UK) Consortium | Harper VanSteenhouse, Yumi Kasai, David Gray, Carol Clugston, Anna Dominiczak and Alex Alderton, Roberto Amato, Jeffrey Barrett, Sonia Goncalves, Ewan Harrison, David K. Jackson, Ian Johnston, Dominic Kwiatkowski, Cordelia Langford, John Sillitoe on behalf of the Wellcome Sanger Institute COVID-19 Surveillance Team |
| EPI_ISL_2607929 | Lighthouse Lab in Alderley Park | Wellcome Sanger Institute for the COVID-19 Genomics UK (COG-UK) Consortium | Jacquelyn Wynn, Mairead Hyland, The Lighthouse Lab in Alderley Park and Alex Alderton, Roberto Amato, Jeffrey Barrett, Sonia Goncalves, Ewan Harrison, David K. Jackson, Ian Johnston, Dominic Kwiatkowski, Cordelia Langford, John Sillitoe on behalf of the Wellcome Sanger Institute COVID-19 Surveillance Team |
| EPI_ISL_2607930, EPI_ISL_2607931 | Lighthouse Lab in Glasgow | Wellcome Sanger Institute for the COVID-19 Genomics UK (COG-UK) Consortium | Harper VanSteenhouse, Yumi Kasai, David Gray, Carol Clugston, Anna Dominiczak and Alex Alderton, Roberto Amato, Jeffrey Barrett, Sonia Goncalves, Ewan Harrison, David K. Jackson, Ian Johnston, Dominic Kwiatkowski, Cordelia Langford, John Sillitoe on behalf of the Wellcome Sanger Institute COVID-19 Surveillance Team |
| EPI_ISL_2607932 | Lighthouse Lab in Alderley Park | Wellcome Sanger Institute for the COVID-19 Genomics UK (COG-UK) Consortium | Jacquelyn Wynn, Mairead Hyland, The Lighthouse Lab in Alderley Park and Alex Alderton, Roberto Amato, Jeffrey Barrett, Sonia Goncalves, Ewan Harrison, David K. Jackson, Ian Johnston, Dominic Kwiatkowski, Cordelia Langford, John Sillitoe on behalf of the Wellcome Sanger Institute COVID-19 Surveillance Team |
| EPI_ISL_2607933 | Lighthouse Lab in Glasgow | Wellcome Sanger Institute for the COVID-19 Genomics UK (COG-UK) Consortium | Harper VanSteenhouse, Yumi Kasai, David Gray, Carol Clugston, Anna Dominiczak and Alex Alderton, Roberto Amato, Jeffrey Barrett, Sonia Goncalves, Ewan Harrison, David K. Jackson, Ian Johnston, Dominic Kwiatkowski, Cordelia Langford, John Sillitoe on behalf of the Wellcome Sanger Institute COVID-19 Surveillance Team |
| EPI_ISL_2607934 | Lighthouse Lab in Alderley Park | Wellcome Sanger Institute for the COVID-19 Genomics UK (COG-UK) Consortium | Jacquelyn Wynn, Mairead Hyland, The Lighthouse Lab in Alderley Park and Alex Alderton, Roberto Amato, Jeffrey Barrett, Sonia Goncalves, Ewan Harrison, David K. Jackson, Ian Johnston, Dominic Kwiatkowski, Cordelia Langford, John Sillitoe on behalf of the Wellcome Sanger Institute COVID-19 Surveillance Team |
| EPI_ISL_2607935, EPI_ISL_2607936, EPI_ISL_2607937, EPI_ISL_2607938, EPI_ISL_2607939, EPI_ISL_2607940, EPI_ISL_2607942, EPI_ISL_2607943, EPI_ISL_2607944, EPI_ISL_2607945 | Lighthouse Lab in Glasgow | Wellcome Sanger Institute for the COVID-19 Genomics UK (COG-UK) Consortium | Harper VanSteenhouse, Yumi Kasai, David Gray, Carol Clugston, Anna Dominiczak and Alex Alderton, Roberto Amato, Jeffrey Barrett, Sonia Goncalves, Ewan Harrison, David K. Jackson, Ian Johnston, Dominic Kwiatkowski, Cordelia Langford, John Sillitoe on behalf of the Wellcome Sanger Institute COVID-19 Surveillance Team |
| EPI_ISL_2607946, EPI_ISL_2607947 | Lighthouse Lab in Alderley Park | Wellcome Sanger Institute for the COVID-19 Genomics UK (COG-UK) Consortium | Jacquelyn Wynn, Mairead Hyland, The Lighthouse Lab in Alderley Park and Alex Alderton, Roberto Amato, Jeffrey Barrett, Sonia Goncalves, Ewan Harrison, David K. Jackson, Ian Johnston, Dominic Kwiatkowski, Cordelia Langford, John Sillitoe on behalf of the Wellcome Sanger Institute COVID-19 Surveillance Team |
| EPI_ISL_2607948 | Lighthouse Laboratory Plymouth | Wellcome Sanger Institute for the COVID-19 Genomics UK (COG-UK) Consortium | Lighthouse Laboratory Plymouth and Alex Alderton, Roberto Amato, Jeffrey Barrett, Sonia Goncalves, Ewan Harrison, David K. Jackson, Ian Johnston, Dominic Kwiatkowski, Cordelia Langford, John Sillitoe on behalf of the Wellcome Sanger Institute COVID-19 Surveillance Team |
| EPI_ISL_2607949 | Lighthouse Lab in Glasgow | Wellcome Sanger Institute for the COVID-19 Genomics UK (COG-UK) Consortium | Harper VanSteenhouse, Yumi Kasai, David Gray, Carol Clugston, Anna Dominiczak and Alex Alderton, Roberto Amato, Jeffrey Barrett, Sonia Goncalves, Ewan Harrison, David K. Jackson, Ian Johnston, Dominic Kwiatkowski, Cordelia Langford, John Sillitoe on behalf of the Wellcome Sanger Institute COVID-19 Surveillance Team |
| EPI_ISL_2607951 | Lighthouse Lab in Alderley Park | Wellcome Sanger Institute for the COVID-19 Genomics UK (COG-UK) Consortium | Jacquelyn Wynn, Mairead Hyland, The Lighthouse Lab in Alderley Park and Alex Alderton, Roberto Amato, Jeffrey Barrett, Sonia Goncalves, Ewan Harrison, David K. Jackson, Ian Johnston, Dominic Kwiatkowski, Cordelia Langford, John Sillitoe on behalf of the Wellcome Sanger Institute COVID-19 Surveillance Team |
| EPI_ISL_2607952, EPI_ISL_2607953, EPI_ISL_2607954, EPI_ISL_2607955, EPI_ISL_2607956, EPI_ISL_2607957, EPI_ISL_2607959, EPI_ISL_2607960, EPI_ISL_2607961, EPI_ISL_2607962, EPI_ISL_2607964, EPI_ISL_2607965, EPI_ISL_2607967, EPI_ISL_2607968, EPI_ISL_2607970, EPI_ISL_2607971, EPI_ISL_2607972, EPI_ISL_2607973, EPI_ISL_2607974, EPI_ISL_2607975, EPI_ISL_2607976, EPI_ISL_2607977, EPI_ISL_2607978, EPI_ISL_2607979, EPI_ISL_2607980, EPI_ISL_2607981, EPI_ISL_2607982, EPI_ISL_2607983, EPI_ISL_2607984, EPI_ISL_2607985, EPI_ISL_2607986, EPI_ISL_2607987, EPI_ISL_2607988, EPI_ISL_2607989, EPI_ISL_2607990, EPI_ISL_2607991, EPI_ISL_2607992, EPI_ISL_2607993, EPI_ISL_2607994, EPI_ISL_2607995, EPI_ISL_2607996, EPI_ISL_2607997, EPI_ISL_2607998, EPI_ISL_2608000, EPI_ISL_2608001, EPI_ISL_2608002, EPI_ISL_2608003, EPI_ISL_2608005 | | | |
| see above | Lighthouse Lab in Glasgow | Wellcome Sanger Institute for the COVID-19 Genomics UK (COG-UK) Consortium | Harper VanSteenhouse, Yumi Kasai, David Gray, Carol Clugston, Anna Dominiczak and Alex Alderton, Roberto Amato, Jeffrey Barrett, Sonia Goncalves, Ewan Harrison, David K. Jackson, Ian Johnston, Dominic Kwiatkowski, Cordelia Langford, John Sillitoe on behalf of the Wellcome Sanger Institute COVID-19 Surveillance Team |
| EPI_ISL_2608007 | Lighthouse Lab in Milton Keynes | Wellcome Sanger Institute for the COVID-19 Genomics UK (COG-UK) Consortium | The Lighthouse Lab in Milton Keynes and Alex Alderton, Roberto Amato, Jeffrey Barrett, Sonia Goncalves, Ewan Harrison, David K. Jackson, Ian Johnston, Dominic Kwiatkowski, Cordelia Langford, John Sillitoe on behalf of the Wellcome Sanger Institute COVID-19 Surveillance Team |
| EPI_ISL_2608008, EPI_ISL_2608009, EPI_ISL_2608010, EPI_ISL_2608011, EPI_ISL_2608012, EPI_ISL_2608013, EPI_ISL_2608015, EPI_ISL_2608016, EPI_ISL_2608017, EPI_ISL_2608018, EPI_ISL_2608020, EPI_ISL_2608021, EPI_ISL_2608022, EPI_ISL_2608023, EPI_ISL_2608024, EPI_ISL_2608025, EPI_ISL_2608026, EPI_ISL_2608027, EPI_ISL_2608028, EPI_ISL_2608029, EPI_ISL_2608030, EPI_ISL_2608031, EPI_ISL_2608032, EPI_ISL_2608033 | | | |
| see above | Lighthouse Lab in Glasgow | Wellcome Sanger Institute for the COVID-19 Genomics UK (COG-UK) Consortium | Harper VanSteenhouse, Yumi Kasai, David Gray, Carol Clugston, Anna Dominiczak and Alex Alderton, Roberto Amato, Jeffrey Barrett, Sonia Goncalves, Ewan Harrison, David K. Jackson, Ian Johnston, Dominic Kwiatkowski, Cordelia Langford, John Sillitoe on behalf of the Wellcome Sanger Institute COVID-19 Surveillance Team |
| EPI_ISL_2608034 | Lighthouse Lab in Milton Keynes | Wellcome Sanger Institute for the COVID-19 Genomics UK (COG-UK) Consortium | The Lighthouse Lab in Milton Keynes and Alex Alderton, Roberto Amato, Jeffrey Barrett, Sonia Goncalves, Ewan Harrison, David K. Jackson, Ian Johnston, Dominic Kwiatkowski, Cordelia Langford, John Sillitoe on behalf of the Wellcome Sanger Institute COVID-19 Surveillance Team |
| EPI_ISL_2608035, EPI_ISL_2608036, EPI_ISL_2608037, EPI_ISL_2608040, EPI_ISL_2608041, EPI_ISL_2608043, EPI_ISL_2608044, EPI_ISL_2608045, EPI_ISL_2608046, EPI_ISL_2608047, EPI_ISL_2608048, EPI_ISL_2608049, EPI_ISL_2608050, EPI_ISL_2608051, EPI_ISL_2608052, EPI_ISL_2608053, EPI_ISL_2608055, EPI_ISL_2608056, EPI_ISL_2608057, EPI_ISL_2608058, EPI_ISL_2608059, EPI_ISL_2608060, EPI_ISL_2608061, EPI_ISL_2608062, EPI_ISL_2608063, EPI_ISL_2608064, EPI_ISL_2608065, EPI_ISL_2608066, EPI_ISL_2608067, EPI_ISL_2608068, EPI_ISL_2608069, EPI_ISL_2608070, EPI_ISL_2608071, EPI_ISL_2608072, EPI_ISL_2608073, EPI_ISL_2608074, EPI_ISL_2608075, EPI_ISL_2608076, EPI_ISL_2608077, EPI_ISL_2608078, EPI_ISL_2608079, EPI_ISL_2608080, EPI_ISL_2608081, EPI_ISL_2608082, EPI_ISL_2608083, EPI_ISL_2608084, EPI_ISL_2608085, EPI_ISL_2608086, EPI_ISL_2608087, EPI_ISL_2608088, EPI_ISL_2608089, EPI_ISL_2608090, EPI_ISL_2608091, EPI_ISL_2608092, EPI_ISL_2608093, EPI_ISL_2608094, EPI_ISL_2608096, EPI_ISL_2608097, EPI_ISL_2608098, EPI_ISL_2608099, EPI_ISL_2608100, EPI_ISL_2608101, EPI_ISL_2608102, EPI_ISL_2608103, EPI_ISL_2608104, EPI_ISL_2608105, EPI_ISL_2608107, EPI_ISL_2608108, EPI_ISL_2608110, EPI_ISL_2608112, EPI_ISL_2608115, EPI_ISL_2608116, EPI_ISL_2608117, EPI_ISL_2608118, EPI_ISL_2608120, EPI_ISL_2608121, EPI_ISL_2608122, EPI_ISL_2608123 | | | |
| see above | Lighthouse Lab in Glasgow | Wellcome Sanger Institute for the COVID-19 Genomics UK (COG-UK) Consortium | Harper VanSteenhouse, Yumi Kasai, David Gray, Carol Clugston, Anna Dominiczak and Alex Alderton, Roberto Amato, Jeffrey Barrett, Sonia Goncalves, Ewan Harrison, David K. Jackson, Ian Johnston, Dominic Kwiatkowski, Cordelia Langford, John Sillitoe on behalf of the Wellcome Sanger Institute COVID-19 Surveillance Team |
| EPI_ISL_2608124 | Health Services Laboratories | Wellcome Sanger Institute for the COVID-19 Genomics UK (COG-UK) Consortium | Health Services Laboratories and Alex Alderton, Roberto Amato, Jeffrey Barrett, Sonia Goncalves, Ewan Harrison, David K. Jackson, Ian Johnston, Dominic Kwiatkowski, Cordelia Langford, John Sillitoe on behalf of the Wellcome Sanger Institute COVID-19 Surveillance Team |
| EPI_ISL_2608125, EPI_ISL_2608126, EPI_ISL_2608127, EPI_ISL_2608128, EPI_ISL_2608129, EPI_ISL_2608130, EPI_ISL_2608131, EPI_ISL_2608132, EPI_ISL_2608133, EPI_ISL_2608134, EPI_ISL_2608135, EPI_ISL_2608136, EPI_ISL_2608138, EPI_ISL_2608139, EPI_ISL_2608140, EPI_ISL_2608141, EPI_ISL_2608142, EPI_ISL_2608144, EPI_ISL_2608146, EPI_ISL_2608147, EPI_ISL_2608148, EPI_ISL_2608149, EPI_ISL_2608150, EPI_ISL_2608151, EPI_ISL_2608154, EPI_ISL_2608155, EPI_ISL_2608156, EPI_ISL_2608157, EPI_ISL_2608158, EPI_ISL_2608159, EPI_ISL_2608161, EPI_ISL_2608163, EPI_ISL_2608164, EPI_ISL_2608165, EPI_ISL_2608166, EPI_ISL_2608167, EPI_ISL_2608169, EPI_ISL_2608170, EPI_ISL_2608171, EPI_ISL_2608172, EPI_ISL_2608173, EPI_ISL_2608174, EPI_ISL_2608176, EPI_ISL_2608177, EPI_ISL_2608178, EPI_ISL_2608181, EPI_ISL_2608182, EPI_ISL_2608183, EPI_ISL_2608185, EPI_ISL_2608187, EPI_ISL_2608188, EPI_ISL_2608189, EPI_ISL_2608190, EPI_ISL_2608191, EPI_ISL_2608192, EPI_ISL_2608193, EPI_ISL_2608194, EPI_ISL_2608195, EPI_ISL_2608196, EPI_ISL_2608197, EPI_ISL_2608198, EPI_ISL_2608200, EPI_ISL_2608202, EPI_ISL_2608203, EPI_ISL_2608204, EPI_ISL_2608206, EPI_ISL_2608207 | | | |
| see above | Lighthouse Lab in Glasgow | Wellcome Sanger Institute for the COVID-19 Genomics UK (COG-UK) Consortium | Harper VanSteenhouse, Yumi Kasai, David Gray, Carol Clugston, Anna Dominiczak and Alex Alderton, Roberto Amato, Jeffrey Barrett, Sonia Goncalves, Ewan Harrison, David K. Jackson, Ian Johnston, Dominic Kwiatkowski, Cordelia Langford, John Sillitoe on behalf of the Wellcome Sanger Institute COVID-19 Surveillance Team |
| EPI_ISL_2608208 | Inovie AS MEDILAB 66 | Inovie AS Genbio Clermont-Ferrand | Gwladys SEVRIN Florian VERCRUYSSEN Pauline LAURENT |
| EPI_ISL_2608210, EPI_ISL_2608212, EPI_ISL_2608213, EPI_ISL_2608214, EPI_ISL_2608215, EPI_ISL_2608216, EPI_ISL_2608217, EPI_ISL_2608218, EPI_ISL_2608219, EPI_ISL_2608220, EPI_ISL_2608221, EPI_ISL_2608222, EPI_ISL_2608223, EPI_ISL_2608225, EPI_ISL_2608226, EPI_ISL_2608228, EPI_ISL_2608229, EPI_ISL_2608230, EPI_ISL_2608231, EPI_ISL_2608232, EPI_ISL_2608233, EPI_ISL_2608234, EPI_ISL_2608236, EPI_ISL_2608238, EPI_ISL_2608239, EPI_ISL_2608240, EPI_ISL_2608241, EPI_ISL_2608242, EPI_ISL_2608243, EPI_ISL_2608244, EPI_ISL_2608245, EPI_ISL_2608246, EPI_ISL_2608247, EPI_ISL_2608248, EPI_ISL_2608249, EPI_ISL_2608251, EPI_ISL_2608252, EPI_ISL_2608253, EPI_ISL_2608254, EPI_ISL_2608256, EPI_ISL_2608258, EPI_ISL_2608259, EPI_ISL_2608261, EPI_ISL_2608262, EPI_ISL_2608263, EPI_ISL_2608264, EPI_ISL_2608266, EPI_ISL_2608268, EPI_ISL_2608269, EPI_ISL_2608271, EPI_ISL_2608273, EPI_ISL_2608274, EPI_ISL_2608275, EPI_ISL_2608276, EPI_ISL_2608277, EPI_ISL_2608278, EPI_ISL_2608279, EPI_ISL_2608280, EPI_ISL_2608281, EPI_ISL_2608282, EPI_ISL_2608283 | | | |
| see above | Lighthouse Lab in Glasgow | Wellcome Sanger Institute for the COVID-19 Genomics UK (COG-UK) Consortium | Harper VanSteenhouse, Yumi Kasai, David Gray, Carol Clugston, Anna Dominiczak and Alex Alderton, Roberto Amato, Jeffrey Barrett, Sonia Goncalves, Ewan Harrison, David K. Jackson, Ian Johnston, Dominic Kwiatkowski, Cordelia Langford, John Sillitoe on behalf of the Wellcome Sanger Institute COVID-19 Surveillance Team |
| EPI_ISL_2608285, EPI_ISL_2608286 | Lighthouse Lab in Milton Keynes | Wellcome Sanger Institute for the COVID-19 Genomics UK (COG-UK) Consortium | The Lighthouse Lab in Milton Keynes and Alex Alderton, Roberto Amato, Jeffrey Barrett, Sonia Goncalves, Ewan Harrison, David K. Jackson, Ian Johnston, Dominic Kwiatkowski, Cordelia Langford, John Sillitoe on behalf of the Wellcome Sanger Institute COVID-19 Surveillance Team |
| EPI_ISL_2608316, EPI_ISL_2608317, EPI_ISL_2608319, EPI_ISL_2608324, EPI_ISL_2608330, EPI_ISL_2608333, EPI_ISL_2608334, EPI_ISL_2608336, EPI_ISL_2608340 | Institute for Infectious Diseases | Institute for Infectious Diseases | Stefan Neuenschwander, Christian Baumann, Miguel A Terrazos Miani, Cora SŠgesser, Pascal Bittel, Peter Keller, Franziska Suter-Riniker, Stephen L Leib, Alban Ramette |
| EPI_ISL_2608346 | Medica | Institute of Medical Virology | Verena Kufner, Gabriela Ziltener, Maryam Zaheri, Stefan Schmutz, Annette Audigé, Maria Grünberg, Kevin Steiner, Jon Huder, Cyril Shah, Riccarda Capaul, Guido Bloemberg, Jürg Böni, Michael Huber, Alexandra Trkola |
| EPI_ISL_2608421 | Kampong Cham Rapid Response Team | Virology Unit, Institut Pasteur du Cambodge | Cecile Troupin, Leakhena Pum, Jurre Y Siegers, Ly Sovann, Kraing Sidonn, Yi Sengdoeurn, Chin Savuth, Chau Darapheak, Veasna Duong, Erik A Karlsson |
| EPI_ISL_2608422, EPI_ISL_2608423 | Siem Reap Provinical Laboratory | Virology Unit, Institut Pasteur du Cambodge | Cecile Troupin, Leakhena Pum, Jurre Y Siegers, Ly Sovann, Kraing Sidonn, Yi Sengdoeurn, Chin Savuth, Chau Darapheak, Veasna Duong, Erik A Karlsson |
| EPI_ISL_2608428, EPI_ISL_2608429, EPI_ISL_2608440 | Virology Unit, Institut Pasteur du Cambodge | Virology Unit, Institut Pasteur du Cambodge | Cecile Troupin, Leakhena Pum, Jurre Y Siegers, Ly Sovann, Kraing Sidonn, Yi Sengdoeurn, Chin Savuth, Chau Darapheak, Veasna Duong, Erik A Karlsson |

| | | | |
|---|---|---|---|
| EPI_ISL_2608444 | Kampot Rapid Response Team | Virology Unit, Institut Pasteur du Cambodge | Cecile Troupin, Leakhena Pum, Jurre Y Siegers, Ly Sovann, Kraing Sidonn, Yi Sengdoeurn, Chin Savuth, Chau Darapheak, Veasna Duong, Erik A Karlsson |
| EPI_ISL_2608773, EPI_ISL_2608788, EPI_ISL_2608789, EPI_ISL_2608815, EPI_ISL_2608823, EPI_ISL_2608879 | Department of Virology and Immunology, University of Helsinki and Helsinki University Hospital, Huslab Finland | Department of Virology, Faculty of Medicine, University of Helsinki, Helsinki, Finland | Teemu Smura, Ravi Kant, Phuoc Truong, Hussein Alburkat, Hannimari Kallio-Kokko, Jenni Virtanen, Maija Suvanto, Essi Korhonen, Sari Hannula, Harri Kangas, Hanna Liimatainen, Satu Kurkela, Hanna Jarva, Maija Lappalainen, Pekka Ellonen, Olli Vapalahti |
| EPI_ISL_2609791, EPI_ISL_2609827, EPI_ISL_2610008, EPI_ISL_2610041, EPI_ISL_2610086, EPI_ISL_2610142, EPI_ISL_2610143, EPI_ISL_2610198, EPI_ISL_2610216, EPI_ISL_2610257, EPI_ISL_2610305, EPI_ISL_2610314, EPI_ISL_2610329, EPI_ISL_2610332, EPI_ISL_2610342, EPI_ISL_2610347, EPI_ISL_2610362, EPI_ISL_2610423, EPI_ISL_2610433, EPI_ISL_2610435, EPI_ISL_2610447, EPI_ISL_2610450, EPI_ISL_2610460, EPI_ISL_2610461, EPI_ISL_2610481, EPI_ISL_2610483, EPI_ISL_2610494, EPI_ISL_2610505, EPI_ISL_2610516, EPI_ISL_2610564, EPI_ISL_2610575, EPI_ISL_2610596, EPI_ISL_2610678, EPI_ISL_2610685, EPI_ISL_2610686, EPI_ISL_2610855, EPI_ISL_2610865, EPI_ISL_2610895, EPI_ISL_2610914 | | | |
| see above | Dutch COVID-19 response team | National Institute for Public Health and the Environment (RIVM) | Adam Meijer, Harry Vennema, Dirk Eggink, Jeroen Cremer, Sharon van den Brink, Bas van der Veer, AnneMarie van den Brandt, Lisa Wijsman, Kim Freriks, Ryanne Jaarsma, Euníce Then, Lynn Aarts, Sanne Bos, Melissa van Tuil, Linda van de Nes, Florian Zwagemaker, Dennis Schmitz, Annelies Kroneman, Karim Hajji, Chantal Reusken, on behalf of the national COVID-19 response team |
| EPI_ISL_2610961, EPI_ISL_2610972 | Clinical Virology | Clinical Bacteriology | Tim Roloff, Fanny Wegner, Helena MB Seth-Smith, Alfredo Mari, Karoline Leuzinger, Julia Bielicki, Manuel Battegay, Hans Hirsch, Adrian Egli |
| EPI_ISL_2610974 | Felix-Platter Spital | Clinical Bacteriology | Tim Roloff, Fanny Wegner, Helena MB Seth-Smith, Alfredo Mari, Karoline Leuzinger, Julia Bielicki, Manuel Battegay, Hans Hirsch, Adrian Egli |
| EPI_ISL_2610993, EPI_ISL_2610997, EPI_ISL_2611003 | Clinical Virology | Clinical Bacteriology | Tim Roloff, Fanny Wegner, Helena MB Seth-Smith, Alfredo Mari, Karoline Leuzinger, Julia Bielicki, Manuel Battegay, Hans Hirsch, Adrian Egli |
| EPI_ISL_2611006 | Felix-Platter Spital | Clinical Bacteriology | Tim Roloff, Fanny Wegner, Helena MB Seth-Smith, Alfredo Mari, Karoline Leuzinger, Julia Bielicki, Manuel Battegay, Hans Hirsch, Adrian Egli |
| EPI_ISL_2611028 | Synlab Suisse SA | Clinical Bacteriology | Tim Roloff, Fanny Wegner, Helena MB Seth-Smith, Alfredo Mari, Karoline Leuzinger, Julia Bielicki, Manuel Battegay, Hans Hirsch, Adrian Egli |
| EPI_ISL_2611031 | Labormedizinisches Zentrum Dr Risch | Clinical Bacteriology | Tim Roloff, Fanny Wegner, Helena MB Seth-Smith, Alfredo Mari, Karoline Leuzinger, Julia Bielicki, Nadia Wohlwend,Martin Risch, Lorenz Risch, Manuel Battegay, Hans Hirsch, Adrian Egli |
| EPI_ISL_2611051 | Viollier AG | Department of Biosystems Science and Engineering, ETH Zürich | Christian Beisel, Sarah Nadeau, Chaoran Chen, Ivan Topolsky, Kim Philipp Jablonski, Lara Fuhrmann, Rebecca Denes, Mirjam Feldkamp, Ina Nissen, Natascha Santacroce, Elodie Burcklen, Christiane Beckmann, Maurice Redondo, Olivier Kobel, Christoph Noppen, Niko Beerenwinkel, Tanja Stadler |
| EPI_ISL_2611146, EPI_ISL_2611152, EPI_ISL_2611164, EPI_ISL_2611178, EPI_ISL_2611181, EPI_ISL_2611197, EPI_ISL_2611204, EPI_ISL_2611205, EPI_ISL_2611210, EPI_ISL_2611214, EPI_ISL_2611224, EPI_ISL_2611228, EPI_ISL_2611287, EPI_ISL_2611291, EPI_ISL_2611299, EPI_ISL_2611307, EPI_ISL_2611320, EPI_ISL_2611333, EPI_ISL_2611337, EPI_ISL_2611354, EPI_ISL_2611356, EPI_ISL_2611362, EPI_ISL_2611370, EPI_ISL_2611381, EPI_ISL_2611395, EPI_ISL_2611399, EPI_ISL_2611400, EPI_ISL_2611421, EPI_ISL_2611425, EPI_ISL_2611426, EPI_ISL_2611434, EPI_ISL_2611435, EPI_ISL_2611444, EPI_ISL_2611464, EPI_ISL_2611469, EPI_ISL_2611470, EPI_ISL_2611472, EPI_ISL_2611475, EPI_ISL_2611477, EPI_ISL_2611483, EPI_ISL_2611503, EPI_ISL_2611504, EPI_ISL_2611526, EPI_ISL_2611538, EPI_ISL_2611554, EPI_ISL_2611572, EPI_ISL_2611573, EPI_ISL_2611574, EPI_ISL_2611577, EPI_ISL_2611583, EPI_ISL_2611597, EPI_ISL_2611600, EPI_ISL_2611624, EPI_ISL_2611638, EPI_ISL_2611658, EPI_ISL_2611659 | | | |
| see above | Laboratory Corporation of America | Centers for Disease Control and Prevention Division of Viral Diseases, Pathogen Discovery | Dakota Howard, Dhwani Batra, Peter W. Cook, Kara Moser, Adrian Paskey, Jason Caravas, Benjamin Rambo-Martin, Shatavia Morrison, Christopher Gulvick, Scott Sammons, Yvette Unoarumhi, Darlene Wagner, Matthew Schmerer, Minoo Agarwal, Eyad Almasri, Debbie Boles, Ayla Burns, Nuthawin Charoensri, Oren Cohen, Susan Countryman, Mary Ann Cristobal, Bobbi Croy, Suzanne Dale, Hrushikesh Deshmukh, Amanda Douglas, Vincent Drouillon, Marcia Eisenberg, Howard Engler, Rama Ghatti, Prashant Gupta, Susan Hicks, Jake Humphrey, Lax Iyer, Lisa Pfefferle, Manoj Jain, Matthew Robinson, Mohan Kolli, Brian Krueger, Tim Kuphal, Stanley Letovsky, Michael Levandoski, Craig Lukasik, Jonathan Meltzer, Brian Norvell, Mindy Nye, Scott Parker, Christos Petropoulos, John Pruitt, Steven Ragan, Scott Ryan, Mike Sapeta, Jana Schroth, Suresh Babu Selvaraju, Goran Stevovic, Amanda Suchanek, Andrea Throop, Lyndon Tilson, Thomas Urban, Joe Voshell, Kimberly Wagner, Jonathan Williams, Mary Williamson, Qian Zeng, Tricia Zwiefelhofer, Clinton R. Paden, Duncan MacCannell |
| EPI_ISL_2611662 | PRINCESS MARGARET HOSPITAL | Hong Kong Department of Health | Alan K.L. Tsang, Peter C.W. Yip, Ken H.L. Ng, Edman T.K. Lam, Rickjason C.W. Chan, Dominic N.C. Tsang |
| EPI_ISL_2611665, EPI_ISL_2611667, EPI_ISL_2611668, EPI_ISL_2611671 | HOME QUARANTINE TASKFORCE | Hong Kong Department of Health | Alan K.L. Tsang, Peter C.W. Yip, Ken H.L. Ng, Edman T.K. Lam, Rickjason C.W. Chan, Dominic N.C. Tsang |
| EPI_ISL_2611679, EPI_ISL_2611680, EPI_ISL_2611693, EPI_ISL_2611704, EPI_ISL_2611705, EPI_ISL_2611712, EPI_ISL_2611713, EPI_ISL_2611715, EPI_ISL_2611732, EPI_ISL_2611741, EPI_ISL_2611746, EPI_ISL_2611750, EPI_ISL_2611751, EPI_ISL_2611753, EPI_ISL_2611754, EPI_ISL_2611755, EPI_ISL_2611780, EPI_ISL_2611781, EPI_ISL_2611784, EPI_ISL_2611793, EPI_ISL_2611795, EPI_ISL_2611809 | | | |
| see above | Laboratory Corporation of America | Centers for Disease Control and Prevention Division of Viral Diseases, Pathogen Discovery | Dakota Howard, Dhwani Batra, Peter W. Cook, Kara Moser, Adrian Paskey, Jason Caravas, Benjamin Rambo-Martin, Shatavia Morrison, Christopher Gulvick, Scott Sammons, Yvette Unoarumhi, Darlene Wagner, Matthew Schmerer, Minoo Agarwal, Eyad Almasri, Debbie Boles, Ayla Burns, Nuthawin Charoensri, Oren Cohen, Susan Countryman, Mary Ann Cristobal, Bobbi Croy, Suzanne Dale, Hrushikesh Deshmukh, Amanda Douglas, Vincent Drouillon, Marcia Eisenberg, Howard Engler, Rama Ghatti, Prashant Gupta, Susan Hicks, Jake Humphrey, Lax Iyer, Lisa Pfefferle, Manoj Jain, Matthew Robinson, Mohan Kolli, Brian Krueger, Tim Kuphal, Stanley Letovsky, Michael Levandoski, Craig Lukasik, Jonathan Meltzer, Brian Norvell, Mindy Nye, Scott Parker, Christos Petropoulos, John Pruitt, Steven Ragan, Scott Ryan, Mike Sapeta, Jana Schroth, Suresh Babu Selvaraju, Goran Stevovic, Amanda Suchanek, Andrea Throop, Lyndon Tilson, Thomas Urban, Joe Voshell, Kimberly Wagner, Jonathan Williams, Mary Williamson, Qian Zeng, Tricia Zwiefelhofer, Clinton R. Paden, Duncan MacCannell |
| EPI_ISL_2611847, EPI_ISL_2611849, EPI_ISL_2611850, EPI_ISL_2611855 | Quest Diagnostics Incorporated | Centers for Disease Control and Prevention Division of Viral Diseases, Pathogen Discovery | Dakota Howard, Dhwani Batra, Peter W. Cook, Kara Moser, Adrian Paskey, Jason Caravas, Benjamin Rambo-Martin, Shatavia Morrison, Christopher Gulvick, Scott Sammons, Yvette Unoarumhi, Darlene Wagner, Matthew Schmerer, S. H. Rosenthal, A. Gerasimova, R. M. Kagan, B. Anderson, M. Hua, Y. Liu, L.E. Bernstein, K.E. Livingston, A. Perez, I. A. Shlyakhter, R. V. Rolando, R. Owen, P. Tanpaiboon, F. Lacbawan, Clinton R. Paden, Duncan MacCannell |
| EPI_ISL_2611874 | Klinisk mikrobiologi, Region Västerbotten | CBRN Defence and Security, Swedish Defence Research Agency | Andreas Sjödin, Linda Karlsson, Emelie Näslund Salomonsson, Jonas Näslund, Stina Bäckman, Malin Granberg, Ingrid Dacklin, Anna-Lena Johansson, Kerstin Myrtennäs, David Sundell, Caroline Öhrman, Mats Forsman, Annika Allard, Annika Osterman |
| EPI_ISL_2611978 | Akershus University Hospital, Department for Microbiology and Infectious Disease Control | Norwegian Institute of Public Health, Department of Virology | Kathrine Stene-Johansen, Kamilla Heddeland Instefjord, Hilde Elshaug, Garcia Llorente Ignacio, Jon Bråte, Engebretsen Serina Beate,Pedersen Benedikte Nevjen, Line Victoria Moen, Debech Nadia, Atiya R Ali,Marie Paulsen Madsen, Rasmus Riis Kopperud, Hilde Vollan, Karoline Bragstad, Olav Hungnes |
| EPI_ISL_2611979, EPI_ISL_2611980 | Furst Medical Laboratory | Norwegian Institute of Public Health, Department of Virology | Kathrine Stene-Johansen, Kamilla Heddeland Instefjord, Hilde Elshaug, Garcia Llorente Ignacio, Jon Bråte, Engebretsen Serina Beate,Pedersen Benedikte Nevjen, Line Victoria Moen, Debech Nadia, Atiya R Ali,Marie Paulsen Madsen, Rasmus Riis Kopperud, Hilde Vollan, Karoline Bragstad, Olav Hungnes |
| EPI_ISL_2611987, EPI_ISL_2611988, EPI_ISL_2611989, EPI_ISL_2611990, EPI_ISL_2611991, EPI_ISL_2611992, EPI_ISL_2611993, EPI_ISL_2611994, EPI_ISL_2611995, EPI_ISL_2611996 | Presidio di Brindisi Di Summa - Perrino | Istituto Zooprofilattico Sperimentale della Puglia e della Basilicata | Parisi A., Bianco A., Capozzi L., Del Sambro L., Simone D., Difato L., Pennuzzi G., Santoro A. |
| EPI_ISL_2611998 | Istituto Zooprofilattico Sperimentale della Puglia e della Basilicata | Istituto Zooprofilattico Sperimentale della Puglia e della Basilicata | Parisi A., Bianco A., Capozzi L., Del Sambro L., Simone D., Difato L., Pennuzzi G., Pace L., Galante D. |
| EPI_ISL_2611999, EPI_ISL_2612001, EPI_ISL_2612002 | Ospedale Vito Fazzi | Istituto Zooprofilattico Sperimentale della Puglia e della Basilicata | Parisi A., Bianco A., Capozzi L., Del Sambro L., Simone D., Difato L., Lobreglio G. |
| EPI_ISL_2612004 | Ospedale Santa Caterina Novella | Istituto Zooprofilattico Sperimentale della Puglia e della Basilicata | Parisi A., Bianco A., Capozzi L., Del Sambro L., Simone D., Difato L., Bruno A. R. |
| EPI_ISL_2612005, EPI_ISL_2612006, EPI_ISL_2612007 | Ospedale Vito Fazzi | Istituto Zooprofilattico Sperimentale della Puglia e della Basilicata | Parisi A., Bianco A., Capozzi L., Del Sambro L., Simone D., Difato L., Lobreglio G. |
| EPI_ISL_2612008, EPI_ISL_2612009, EPI_ISL_2612010 | Ospedale Santa Caterina Novella | Istituto Zooprofilattico Sperimentale della Puglia e della Basilicata | Parisi A., Bianco A., Capozzi L., Del Sambro L., Simone D., Difato L., Bruno A. R. |
| EPI_ISL_2612011 | Ospedale Cardinale G. Panico | Istituto Zooprofilattico Sperimentale della Puglia e della Basilicata | Parisi A., Bianco A., Capozzi L., Del Sambro L., Simone D., Difato L., Bruno A. R. |
| EPI_ISL_2612015 | Ospedale Santissima Annunziata | Istituto Zooprofilattico Sperimentale della Puglia e della Basilicata | Parisi A., Bianco A., Capozzi L., Del Sambro L., Simone D., Difato L., Ridolfi D., Giannico A, Tampoia M. |
| EPI_ISL_2612043 | Klinisch Laboratorium ZNA | Klinisch Laboratorium ZNA | Verstrepen et al. |
| EPI_ISL_2612182, EPI_ISL_2612183, EPI_ISL_2612185, EPI_ISL_2612186, EPI_ISL_2612187 | Kansas Health and Environmental Lab | Kansas Health and Environmental Lab | Mike Grose, Katherine Wiggins, Jonathan Barnell, Ben Olsen, and Phil Adam |
| EPI_ISL_2612190, EPI_ISL_2612194 | Wichita State University - Molecular Diagnostics Lab | Kansas Health and Environmental Lab | Mike Grose, Katherine Wiggins, Jonathan Barnell, Ben Olsen, and Phil Adam |
| EPI_ISL_2612300 | DSMRC | DSMRC | Khine Zaw Oo, Nay Myo Aung, Ko Ko Win, Phyo Kyaw Aung, Pai Khant Kyaw, Thet Wai Oo, Zaw Win Htun, Kyee Myint, Ko Ko Lwin |
| EPI_ISL_2612460, EPI_ISL_2612475, EPI_ISL_2612476, EPI_ISL_2612477, EPI_ISL_2612478, EPI_ISL_2612479 | Montana Public Health Laboratory | Montana Public Health Laboratory | Joy Ritter, Michelle Mozer, Carrie Biskupiak, Deborah Gibson, Michael Dills |
| EPI_ISL_2612503, EPI_ISL_2612504, EPI_ISL_2612505, EPI_ISL_2612508 | Nevada State Public Health Laboratory | Nevada State Public Health Laboratory | Andrew Gorzalski, Mark Pandori |

| | | | |
|---|---|---|---|
| EPI_ISL_2612511 | Minnesota Department of Health, Public Health Laboratory | Minnesota Department of Health, Public Health Laboratory | Alexandra Lorentz, Jacob Garfin, Matt Plumb, and Xiong Wang |
| EPI_ISL_2612563 | OCME Office Of Chief Medical Examiner | New York City Public Health Laboratory | Jade Wang, et al. |
| EPI_ISL_2612565 | Labo Analyses Med | National Reference Center for Viruses of Respiratory Infections, Institut Pasteur, Paris | Marion Barbet, Sylvie Behillil, Méline Bizard, Angela Brisebarre, Camille Capel, Vincent Enouf, Louise Lefrançois, Frédéric Lemoine, Christophe Malabat, Corinne Maufrais, Adrien Pain, Etienne Simon-Lorière, Maud Vanpeene, Sylvie Van der Werf |
| EPI_ISL_2612588, EPI_ISL_2612589 | Labo Analyses Med | National Reference Center for Viruses of Respiratory Infections, Institut Pasteur, Paris | Marion Barbet, Sylvie Behillil, Méline Bizard, Angela Brisebarre, Camille Capel, Vincent Enouf, Louise Lefrançois, Frédéric Lemoine, Christophe Malabat, Corinne Maufrais, Etienne Simon-Lorière, Maud Vanpeene, Sylvie Van der Werf ,Nicolas Capron |
| EPI_ISL_2612680 | Labo Analyses Med | National Reference Center for Viruses of Respiratory Infections, Institut Pasteur, Paris | Marion Barbet, Sylvie Behillil, Méline Bizard, Angela Brisebarre, Camille Capel, Vincent Enouf, Louise Lefrançois, Frédéric Lemoine, Christophe Malabat, Corinne Maufrais, Etienne Simon-Lorière, Maud Vanpeene, Sylvie Van der Werf ,Arditi Cocco |
| EPI_ISL_2612770, EPI_ISL_2612771, EPI_ISL_2612772, EPI_ISL_2612773, EPI_ISL_2612774, EPI_ISL_2612775, EPI_ISL_2612776, EPI_ISL_2612777, EPI_ISL_2612786, EPI_ISL_2612787, EPI_ISL_2612788, EPI_ISL_2612789 | | | |
| see above | SYNLAB | GIGA Medical Genomics | Keith Durkin, Maria Artesi, Sébastien Bontems, Raphaël Boreux, Bouchra Boujemla, Nathalie Renotte, Cécile Meex, Pierrette Melin, Marie-Pierre Hayette, Vincent Bours |
| EPI_ISL_2612790 | Department of Clinical Microbiology | GIGA Medical Genomics | Keith Durkin, Maria Artesi, Sébastien Bontems, Raphaël Boreux, Bouchra Boujemla, Nathalie Renotte, Cécile Meex, Pierrette Melin, Marie-Pierre Hayette, Vincent Bours |
| EPI_ISL_2612791, EPI_ISL_2612792 | SYNLAB | GIGA Medical Genomics | Keith Durkin, Maria Artesi, Sébastien Bontems, Raphaël Boreux, Bouchra Boujemla, Nathalie Renotte, Cécile Meex, Pierrette Melin, Marie-Pierre Hayette, Vincent Bours |
| EPI_ISL_2612793 | Department of Clinical Microbiology | GIGA Medical Genomics | Keith Durkin, Maria Artesi, Sébastien Bontems, Raphaël Boreux, Bouchra Boujemla, Nathalie Renotte, Cécile Meex, Pierrette Melin, Marie-Pierre Hayette, Vincent Bours |
| EPI_ISL_2612794 | SYNLAB | GIGA Medical Genomics | Keith Durkin, Maria Artesi, Sébastien Bontems, Raphaël Boreux, Bouchra Boujemla, Nathalie Renotte, Cécile Meex, Pierrette Melin, Marie-Pierre Hayette, Vincent Bours |
| EPI_ISL_2612795 | C.H.C MontLégia | GIGA Medical Genomics | Keith Durkin, Maria Artesi, Sébastien Bontems, Raphaël Boreux, Bouchra Boujemla, Nathalie Renotte, Cécile Meex, Pierrette Melin, Marie-Pierre Hayette, Vincent Bours |
| EPI_ISL_2612796, EPI_ISL_2612797, EPI_ISL_2612798, EPI_ISL_2612805, EPI_ISL_2612837, EPI_ISL_2612838 | SYNLAB | GIGA Medical Genomics | Keith Durkin, Maria Artesi, Sébastien Bontems, Raphaël Boreux, Bouchra Boujemla, Nathalie Renotte, Cécile Meex, Pierrette Melin, Marie-Pierre Hayette, Vincent Bours |
| EPI_ISL_2612839 | University of Liège COVID-19 testing center | GIGA Medical Genomics | Keith Durkin, Maria Artesi, Bouchra Boujemla, Nathalie Renotte, Cécile Meex, Sébastien Bontems, Fabrice Bureau, Laurent Gillet, Wouter Coppieters, Marie-Pierre Hayette, Vincent Bours |
| EPI_ISL_2612904, EPI_ISL_2612906, EPI_ISL_2612907, EPI_ISL_2612909, EPI_ISL_2612910, EPI_ISL_2612911, EPI_ISL_2612912, EPI_ISL_2612913, EPI_ISL_2612914, EPI_ISL_2612915, EPI_ISL_2612916, EPI_ISL_2612917, EPI_ISL_2612922, EPI_ISL_2612923, EPI_ISL_2612926, EPI_ISL_2612927 | | | |
| see above | National Virus Reference Laboratory | National Virus Reference Laboratory | Zoe Yandle, Charlene Bennett, Gabriel Gonzalez, Michael Carr, Jonathan Dean, Cillian F De Gascun |
| EPI_ISL_2612939, EPI_ISL_2613548, EPI_ISL_2613549, EPI_ISL_2613550, EPI_ISL_2613551 | Swedish national genomic surveillance program of SARS-CoV-2 | The Public Health Agency of Sweden | Maximilian Riess, Maria Lind Karlberg, Alma Brolund, Swedish national genomic surveillance program of SARS-CoV-2 |
| EPI_ISL_2613853, EPI_ISL_2613937, EPI_ISL_2614010 | Gravity Diagnostics, LLC | Gravity Diagnostics, LLC | Gravity Diagnostics |
| EPI_ISL_2614397 | Landesamt für Verbraucherschutz Sachsen Anhalt, Magdeburg | Institute of Medical Microbiology and Hospital Hygiene | Prof. Dr. Achim Kaasch, Aljoscha Tersteegen |
| EPI_ISL_2614400, EPI_ISL_2614403 | Lafene Health Center | Kansas State Veterinary Diagnostic Laboratory | Tyler Doerksen, Andreea Lu, Lance Noll, Elizabeth Porter, Emily Cox, Jianfa Bai, Jamie Henningson, Rachel Palinski |
| EPI_ISL_2614404, EPI_ISL_2614405, EPI_ISL_2614409, EPI_ISL_2614410, EPI_ISL_2614411 | Lifebrain Covid Labor GmbH | Dept. of Laboratory Medicine | Harald Esterbauer, Anna Gschaider, Petra Jurkowitsch, Fabian Konig, Sabina Plumer, Robert Strassl, Claudia Weber, Oswald Wagner |
| EPI_ISL_2614413, EPI_ISL_2614414, EPI_ISL_2614415, EPI_ISL_2614416, EPI_ISL_2614417, EPI_ISL_2614418 | Dept. of Laboratory Medicine | Dept. of Laboratory Medicine | Harald Esterbauer, Anna Gschaider, Petra Jurkowitsch, Fabian Konig, Sabina Plumer, Robert Strassl, Claudia Weber, Oswald Wagner |
| EPI_ISL_2614637, EPI_ISL_2614641 | Dr. Yuichiro Oba Department of General Medicine, Osaka General Medical Center | Department of Developmental Medicine, Research Institute, Osaka Women's and Children's Hospital | Itaru Yanagihara, Yukiko Nakura |
| EPI_ISL_2614645 | CSIR Institute of Himalayan Bioresource Technology | CSIR Institute of Himalayan Bioresource Technology | Ravi Kumar, Narendra Tirpude, Vishal Acharya |
| EPI_ISL_2614725 | RS Pelni | Eijkman Institute for Molecular Biology, National Research and Innovation Agency | Willy Agustine, Hidayat Trimarsanto, Lydia V. Panggalo, Iskandar Adnan, Sukma Oktavianthi, Lidwina Priliani, Edison Johar, Frilasita A Yudhaputri, Muhammad Rezki Rasyak, Safarina G Malik, Khin Saw Myint, Amin Soebandrio |
| EPI_ISL_2614726 | RS Pelni | Eijkman Institute for Molecular Biology, National Research and Innovation Agency | Hidayat Trimarsanto, Lydia V. Panggalo, Iskandar Adnan, Sukma Oktavianthi, Lidwina Priliani, Edison Johar, Frilasita A Yudhaputri, Muhammad Rezki Rasyak, Willy Agustine, Safarina G Malik, Khin Saw Myint, Amin Soebandrio |
| EPI_ISL_2614727 | RS Pelni | Eijkman Institute for Molecular Biology, National Research and Innovation Agency | Lydia V. Panggalo, Iskandar Adnan, Sukma Oktavianthi, Lidwina Priliani, Edison Johar, Frilasita A Yudhaputri, Muhammad Rezki Rasyak, Willy Agustine, Hidayat Trimarsanto, Safarina G Malik, Khin Saw Myint, Amin Soebandrio |
| EPI_ISL_2614728 | RS Grha Kedoya | Eijkman Institute for Molecular Biology, National Research and Innovation Agency | Iskandar Adnan, Sukma Oktavianthi, Lidwina Priliani, Edison Johar, Frilasita A Yudhaputri, Muhammad Rezki Rasyak, Willy Agustine, Hidayat Trimarsanto, Lydia V. Panggalo, Safarina G Malik, Khin Saw Myint, Amin Soebandrio |
| EPI_ISL_2614729 | RS Grha Kedoya | Eijkman Institute for Molecular Biology, National Research and Innovation Agency | Sukma Oktavianthi, Lidwina Priliani, Edison Johar, Frilasita A Yudhaputri, Muhammad Rezki Rasyak, Willy Agustine, Hidayat Trimarsanto, Lydia V. Panggalo, Iskandar Adnan, Safarina G Malik, Khin Saw Myint, Amin Soebandrio |
| EPI_ISL_2614730 | EKA Hospital BSD | Eijkman Institute for Molecular Biology, National Research and Innovation Agency | Lidwina Priliani, Edison Johar, Frilasita A Yudhaputri, Muhammad Rezki Rasyak, Willy Agustine, Hidayat Trimarsanto, Lydia V. Panggalo, Iskandar Adnan, Sukma Oktavianthi, Safarina G Malik, Khin Saw Myint, Amin Soebandrio |
| EPI_ISL_2614731 | RS Pelni | Eijkman Institute for Molecular Biology, National Research and Innovation Agency | Iskandar Adnan, Sukma Oktavianthi, Lidwina Priliani, Edison Johar, Frilasita A Yudhaputri, Muhammad Rezki Rasyak, Willy Agustine, Hidayat Trimarsanto, Lydia V. Panggalo, Safarina G Malik, Khin Saw Myint, Amin Soebandrio |
| EPI_ISL_2614732 | RS Sumber Waras | Eijkman Institute for Molecular Biology, National Research and Innovation Agency | Sukma Oktavianthi, Lidwina Priliani, Edison Johar, Frilasita A Yudhaputri, Muhammad Rezki Rasyak, Willy Agustine, Hidayat Trimarsanto, Lydia V. Panggalo, Iskandar Adnan, Safarina G Malik, Khin Saw Myint, Amin Soebandrio |
| EPI_ISL_2614735 | RS Pondok Indah Pondok Indah | Eijkman Institute for Molecular Biology, National Research and Innovation Agency | Frilasita A Yudhaputri, Muhammad Rezki Rasyak, Willy Agustine, Hidayat Trimarsanto, Lydia V. Panggalo, Iskandar Adnan, Sukma Oktavianthi, Lidwina Priliani, Edison Johar, Safarina G Malik, Khin Saw Myint, Amin Soebandrio |
| EPI_ISL_2614736 | RS Hermina Depok | Eijkman Institute for Molecular Biology, National Research and Innovation Agency | Muhammad Rezki Rasyak, Willy Agustine, Hidayat Trimarsanto, Lydia V. Panggalo, Iskandar Adnan, Sukma Oktavianthi, Lidwina Priliani, Edison Johar, Frilasita A Yudhaputri, Safarina G Malik, Khin Saw Myint, Amin Soebandrio |
| EPI_ISL_2614737 | RS Pondok Indah Pondok Indah | Eijkman Institute for Molecular Biology, National Research and Innovation Agency | Willy Agustine, Hidayat Trimarsanto, Lydia V. Panggalo, Iskandar Adnan, Sukma Oktavianthi, Lidwina Priliani, Edison Johar, Frilasita A Yudhaputri, Muhammad Rezki Rasyak, Safarina G Malik, Khin Saw Myint, Amin Soebandrio |
| EPI_ISL_2614738 | RS Pondok Indah Pondok Indah | Eijkman Institute for Molecular Biology, National Research and Innovation Agency | Hidayat Trimarsanto, Lydia V. Panggalo, Iskandar Adnan, Sukma Oktavianthi, Lidwina Priliani, Edison Johar, Frilasita A Yudhaputri, Muhammad Rezki Rasyak, Willy Agustine, Safarina G Malik, Khin Saw Myint, Amin Soebandrio |
| EPI_ISL_2614819, EPI_ISL_2615232 | Nacionalinis maisto ir veterinarijos rizikos vertinimo institutas | National Public Health Surveillance Laboratory | Lukas Zemaitis, Migle Gabrielaite, Jelena Razmuk, Svajune Muralyte, Ana Steponkiene, Lukas Vasionis, Danas Baksa |
| EPI_ISL_2615266, EPI_ISL_2615296, EPI_ISL_2615306, EPI_ISL_2615314, EPI_ISL_2615316, EPI_ISL_2615342, EPI_ISL_2615343, EPI_ISL_2615344, EPI_ISL_2615351, EPI_ISL_2615371, EPI_ISL_2615382, EPI_ISL_2615395, EPI_ISL_2615397 | | | |
| see above | Swedish national genomic surveillance program of SARS-CoV-2 | The Public Health Agency of Sweden | Maximilian Riess, Maria Lind Karlberg, Alma Brolund, Swedish national genomic surveillance program of SARS-CoV-2 |
| EPI_ISL_2615399 | MOH - Jaber Al-Ahmad Hospital (Innovation Research Laboratory) | MOH - Jaber Al-Ahmad Hospital (Innovation Research Laboratory) | Salman Al-Sabah , Mohammad Alghounaim |
| EPI_ISL_2616214 | Austrian Agency for Health and Food Safety (AGES) | Bergthaler laboratory, CeMM Research Center for Molecular Medicine of the Austrian Academy of Sciences | Lukas Endler, Anna Schedl, Fabian Amman, Petr Triska, Matthew Thornton, Thomas Penz, Benedikt Agerer, Maelle Le Moing, Michael Schuster, Bekir Erguner, Jan Laine, Martin Senekowitsch, Christoph Bock, Andreas Bergthaler |
| EPI_ISL_2616523, EPI_ISL_2616543, EPI_ISL_2616586, EPI_ISL_2616663 | Wyoming Public Health Laboratory | Wyoming Public Health Laboratory | Jim Mildenberger, Taylor Fearing, Channing Weber, Ashley Norberg, Chayse Rowley, Marley Goetz, Brian Dominguez, Elliot Thomasson, Sam Britz, Cari Sloma, Robert Petit, and Rob Christensen |
| EPI_ISL_2616773, EPI_ISL_2616774, EPI_ISL_2616775, EPI_ISL_2616776, EPI_ISL_2616777, | Houston Health Dept. | Houston Health Dept. | Ryker Penn, Pamela Brown, Adolpho Lara, Yanlai Lai |

| | | | |
|---|---|---|---|
| EPI_ISL_2616778, EPI_ISL_2616779 | | | |
| EPI_ISL_2616787, EPI_ISL_2616791, EPI_ISL_2616795, EPI_ISL_2616817, EPI_ISL_2616829, EPI_ISL_2616833, EPI_ISL_2616834, EPI_ISL_2616839, EPI_ISL_2616840, EPI_ISL_2616842, EPI_ISL_2616844, EPI_ISL_2616847, EPI_ISL_2616855, EPI_ISL_2616860, EPI_ISL_2616863, EPI_ISL_2616870, EPI_ISL_2616871, EPI_ISL_2616875, EPI_ISL_2616876, EPI_ISL_2616884, EPI_ISL_2616885, EPI_ISL_2616889, EPI_ISL_2616891, EPI_ISL_2616911, EPI_ISL_2616915, EPI_ISL_2616919, EPI_ISL_2616923, EPI_ISL_2616924, EPI_ISL_2616925, EPI_ISL_2616926, EPI_ISL_2616928, EPI_ISL_2616934, EPI_ISL_2616947, EPI_ISL_2616951, EPI_ISL_2616974, EPI_ISL_2616980, EPI_ISL_2616995 | | | |
| see above | Salud Digna | Instituto Nacional de Medicina Genomica | Hidalgo-Miranda A, Cedro-Tanda A, Mendoza-Vargas A, Reyes-Grajeda JP, Abraham Campos-Romero, Moreno-Camacho José Luis, Rodriguez-Gallegos Jorge, Luna-Ruiz Marco, Gonzalez-Barrera D, Rangel-DeLeon D, Munguia-Garza P, Ramirez-Vega O, Escobar-Arrazola, M, Herrera-Montalvo LA. |
| EPI_ISL_2617090 | Africa_CDC - Malawi P6 S2 | CERI, Centre for Epidemic Response and Innvoation, Stellenbosch University and KRISP, KZN Research Innovation and Sequencing Platform, UKZN. | Mvula B, Chilima B, Chiwaula M, Mwangomba W, Panja L, Kasambara W, Auld A, Kim L, Kampira E, Kaba M, Wadonda N, Maida A, Giandhari J, Pillay S, Naidoo Y, Lessells R, Emmanuel SJ, Tegally H, Wilkinson E, de Oliveira T |
| EPI_ISL_2617133, EPI_ISL_2617171, EPI_ISL_2617181, EPI_ISL_2617182 | Instituto Nacional de Investigação em Saúde | CERI, Centre for Epidemic Response and Innvoation, Stellenbosch University and KRISP, KZN Research Innovation and Sequencing Platform, UKZN. | Morais J, Neto Z, Afonso P, Miranda J, David K, Inglês L, Pereira A, Paulo A Carralero RR Paixão JP, Freitas RH, Mufinda M, Lutucuta S, Giandhari J, Pillay S, Naidoo Y, Emmanuel SJ, Tegally H, Wilkinson E, de Oliveira T |
| EPI_ISL_2617435 | Surabaya Health Office | Institute of Tropical Disease, Universitas Airlangga | Rima R Prasetya, Krisnoadi Rahardjo, Aldise M Nastri, Jezzy R Dewantari, Febria Rachmanita, Herlin Ferliana, Gatot Soegiarto, Laksmi Wulandari, Resti Yudhawati, Yasuko Mori, Soetjipto, Kazufumi Shimizu, Maria I Lusida |
| EPI_ISL_2617436 | Surabaya Health Office | Institute of Tropical Disease, Universitas Airlangga | Kazufumi Shimizu, Krisnoadi Rahardjo, Aldise M Nastri, Jezzy R Dewantari, Rima R Prasetya, Febria Rachmanita, Herlin Ferliana, Gatot Soegiarto, Laksmi Wulandari, Resti Yudhawati, Yasuko Mori, Soetjipto, Maria I Lusida |
| EPI_ISL_2617437 | Surabaya Health Office | Institute of Tropical Disease, Universitas Airlangga | Krisnoadi Rahardjo, Aldise M Nastri, Jezzy R Dewantari, Rima R Prasetya, Febria Rachmanita, Herlin Ferliana, Gatot Soegiarto, Laksmi Wulandari, Resti Yudhawati, Yasuko Mori, Soetjipto, Kazufumi Shimizu, Maria I Lusida |
| EPI_ISL_2617439 | Al-Irsyad Hospital | Institute of Tropical Disease, Universitas Airlangga | Jezzy R Dewantari, Rima R Prasetya, Krisnoadi Rahardjo, Aldise M Nastri, Febria Rachmanita, Herlin Ferliana, Gatot Soegiarto, Laksmi Wulandari, Resti Yudhawati, Yasuko Mori, Soetjipto, Kazufumi Shimizu, Maria I Lusida |
| EPI_ISL_2617504 | RSUPP Persahabatan Jakarta | National Institute of Health Research and Development | Hartanti Dian Ikawati, Hana Apsari Pawestri, Subangkit, Kartika Dewi Puspa, Arie Ardiansyah Nugraha, Triyani Soekarso, Krisna Pangesti, Nelly Puspandari, Vivi Setiawaty |
| EPI_ISL_2617513, EPI_ISL_2617516, EPI_ISL_2617517 | RS Umum Medistra | National Institute of Health Research and Development | Hana Apsari Pawestri, Hartanti Dian Ikawati, Subangkit, Kartika Dewi Puspa, Arie Ardiansyah Nugraha, Triyani Soekarso, Krisna Pangesti, Nelly Puspandari, Vivi Setiawaty |
| EPI_ISL_2617518 | RS Hermina Kemayoran | National Institute of Health Research and Development | Hana Apsari Pawestri, Hartanti Dian Ikawati, Subangkit, Kartika Dewi Puspa, Arie Ardiansyah Nugraha, Triyani Soekarso, Krisna Pangesti, Nelly Puspandari, Vivi Setiawaty |
| EPI_ISL_2617523 | RS IMC Bintaro | National Institute of Health Research and Development | Vivi Setiawaty, Hana Apsari Pawestri, Hartanti Dian Ikawati, Subangkit, Kartika Dewi Puspa, Arie Ardiansyah Nugraha, Triyani Soekarso, Krisna Pangesti, Nelly Puspandari |
| EPI_ISL_2617534 | Lab Genomik Solidaritas Indonesia | National Institute of Health Research and Development | Vivi Setiawaty, Hana Apsari Pawestri, Hartanti Dian Ikawati, Subangkit, Kartika Dewi Puspa, Arie Ardiansyah Nugraha, Triyani Soekarso, Krisna Pangesti,Meutia Kumaheri, Nelly Puspandari |
| EPI_ISL_2617583 | City of Milwaukee Health Department Laboratory | City of Milwaukee Health Department Laboratory | Manjeet Khubbar, Amy Bauer, Jennifer Lentz, Samantha Scott, Sanjib Bhattacharyya |
| EPI_ISL_2617631 | Omega Diagnostics at Mounes | Omega Diagnostics at Mounes | Cynthia Corley, Ashleigh McGrail, Latira Haynes-Jacob, Vivek Khare,MD |
| EPI_ISL_2617813, EPI_ISL_2617814, EPI_ISL_2617815, EPI_ISL_2617816, EPI_ISL_2617817, EPI_ISL_2617826 | Pathology and Laboratory Medicine Institute, Cleveland Clinic, Ohio, USA | Pathology and Laboratory Medicine Institute, Cleveland Clinic, Ohio, USA | Concetta Peck, Jennifer Starbuck, Joy Nakitandwe, Kristen McDonnell, David Plunkett, Zheng Jin Tu, Jay Brock, Yu-Wei Cheng, Gary Procop, Daniel Rhoads, Daniel H. Farkas, David Bosler |
| EPI_ISL_2617990, EPI_ISL_2617991, EPI_ISL_2617994, EPI_ISL_2618009, EPI_ISL_2618010, EPI_ISL_2618013, EPI_ISL_2618016, EPI_ISL_2618018, EPI_ISL_2618019, EPI_ISL_2618026, EPI_ISL_2618027, EPI_ISL_2618028, EPI_ISL_2618031, EPI_ISL_2618041, EPI_ISL_2618042, EPI_ISL_2618043, EPI_ISL_2618045, EPI_ISL_2618062, EPI_ISL_2618064, EPI_ISL_2618072, EPI_ISL_2618074, EPI_ISL_2618078, EPI_ISL_2618079, EPI_ISL_2618084, EPI_ISL_2618086, EPI_ISL_2618087, EPI_ISL_2618089, EPI_ISL_2618090, EPI_ISL_2618092, EPI_ISL_2618093, EPI_ISL_2618094, EPI_ISL_2618096, EPI_ISL_2618097, EPI_ISL_2618099, EPI_ISL_2618104, EPI_ISL_2618107, EPI_ISL_2618108, EPI_ISL_2618109, EPI_ISL_2618112 | | | |
| see above | UW Virology Lab | UW Virology Lab | Pavitra Roychoudhury, Hong Xie, Lasata Shrestha, Tien V. Nguyen, Shah Mohamed Bakhash, Michelle Lin, Noah R. Baker, Ricardo Perez, Sean Ellis, Nathan Breit, Robert J. Livingston, Meei-Li Huang, Keith R Jerome, Patrick Mathias, Alexander Greninger |
| EPI_ISL_2618118, EPI_ISL_2618216, EPI_ISL_2618221, EPI_ISL_2618264, EPI_ISL_2618292, EPI_ISL_2618317, EPI_ISL_2618387, EPI_ISL_2618402, EPI_ISL_2618579, EPI_ISL_2618674, EPI_ISL_2618698, EPI_ISL_2618699 | | | |
| see above | UW Virology Lab | UW Virology Lab | Pavitra Roychoudhury, Hong Xie, Lasata Shrestha, Shah Mohamed Bakhash, Tien V. Nguyen, Noah R. Baker, Sean Ellis, Meei-Li Huang, Keith R Jerome, Alexander Greninger |
| EPI_ISL_2618839, EPI_ISL_2618841, EPI_ISL_2618844, EPI_ISL_2618847, EPI_ISL_2618848, EPI_ISL_2618872, EPI_ISL_2618883, EPI_ISL_2618891, EPI_ISL_2618893, EPI_ISL_2618954, EPI_ISL_2618961, EPI_ISL_2618971, EPI_ISL_2618972, EPI_ISL_2618973, EPI_ISL_2618980, EPI_ISL_2618994, EPI_ISL_2618995, EPI_ISL_2618996, EPI_ISL_2619174, EPI_ISL_2619175, EPI_ISL_2619176, EPI_ISL_2619177, EPI_ISL_2619178, EPI_ISL_2619192, EPI_ISL_2619208, EPI_ISL_2619211, EPI_ISL_2619226, EPI_ISL_2619234, EPI_ISL_2619236, EPI_ISL_2619240, EPI_ISL_2619262, EPI_ISL_2619266, EPI_ISL_2619267, EPI_ISL_2619323, EPI_ISL_2619545, EPI_ISL_2619547, EPI_ISL_2619550, EPI_ISL_2619555, EPI_ISL_2619557, EPI_ISL_2619559, EPI_ISL_2619571, EPI_ISL_2619572, EPI_ISL_2619576, EPI_ISL_2619577, EPI_ISL_2619578, EPI_ISL_2619579, EPI_ISL_2619581, EPI_ISL_2619583, EPI_ISL_2619585, EPI_ISL_2619586, EPI_ISL_2619587, EPI_ISL_2619588, EPI_ISL_2619589, EPI_ISL_2619590, EPI_ISL_2619591, EPI_ISL_2619592, EPI_ISL_2619593, EPI_ISL_2619594, EPI_ISL_2619595, EPI_ISL_2619596, EPI_ISL_2619598, EPI_ISL_2619599, EPI_ISL_2619600, EPI_ISL_2619601, EPI_ISL_2619602, EPI_ISL_2619603, EPI_ISL_2619604, EPI_ISL_2619606, EPI_ISL_2619607, EPI_ISL_2619608, EPI_ISL_2619609, EPI_ISL_2619610, EPI_ISL_2619611, EPI_ISL_2619612, EPI_ISL_2619613, EPI_ISL_2619614, EPI_ISL_2619615, EPI_ISL_2619616, EPI_ISL_2619617, EPI_ISL_2619618, EPI_ISL_2619619, EPI_ISL_2619622, EPI_ISL_2619623, EPI_ISL_2619624, EPI_ISL_2619625, EPI_ISL_2619626, EPI_ISL_2619628, EPI_ISL_2619629, EPI_ISL_2619630, EPI_ISL_2619631, EPI_ISL_2619632, EPI_ISL_2619634, EPI_ISL_2619635, EPI_ISL_2619638, EPI_ISL_2619641, EPI_ISL_2619644, EPI_ISL_2619645, EPI_ISL_2619646, EPI_ISL_2619647, EPI_ISL_2619649, EPI_ISL_2619703, EPI_ISL_2619994 | | | |
| see above | Swedish national genomic surveillance program of SARS-CoV-2 | The Public Health Agency of Sweden | Maximilian Riess, Maria Lind Karlberg, Alma Brolund, Swedish national genomic surveillance program of SARS-CoV-2 |
| EPI_ISL_2620719, EPI_ISL_2620720, EPI_ISL_2620721, EPI_ISL_2620722, EPI_ISL_2620723, EPI_ISL_2620724, EPI_ISL_2620725, EPI_ISL_2620726, EPI_ISL_2620727 | Genomik Solidaritas Indonesia Laboratorium | Genomik Solidaritas Indonesia Laboratorium | Dhahlia Agustina Cahyono, Alfin Mohammad Abdillah, Normastuti Adhini Tantyo, Carissa Sintca Wijaya, Vania Gavrila Wikasa, Ahmad Zoebad Foeady, Annisa Muthiah Sukirman, Gracia Felias Enos Korompis, Anuraj Shankar, Meutia Ayuputeri Kumaheri, Ariel Pradipta |
| EPI_ISL_2620735, EPI_ISL_2620737 | Functional Genomics and Proteomics Laboratory (FGPL), Department of Genetic Engineering and Biotechnology, University of Chittagong | Virology Laboratory, International Centre for Diarrhoeal Disease Research, Bangladesh (ICDDR,B) | Dr. Mohammad Al-Forkan, Dr. Sujat Paul, Dr. Laila Khaleda, Mojnu Miah, Rashedul Hasan, Md. Jibran Alam, Amit Datta, Rahee Hasan Chowdhury, Shuva Das, Inzamamul Ismail Shawon, Md. Abdur Rahman, Miftah Mushfique, Amlan Bhattacharjee, Dr. Mustafizur Rahman, Dr. Mohammad Enayet Hossain |
| EPI_ISL_2620738, EPI_ISL_2620739, EPI_ISL_2620740, EPI_ISL_2620741, EPI_ISL_2620742, EPI_ISL_2620743 | State Virus Research and Diagnostic Laboratory (VRDL), AIIMS Raipur | State Virus Research and Diagnostic Laboratory (VRDL), AIIMS Raipur | Pushpendra Singh, Kuldeep Sharma, Priyanka Singh, Somya Sharma, MD Rafiiqullah Khan, Jiyan Chandrawanshi, Sanjay Singh Negi, Anudita Bhargava |
| EPI_ISL_2620826 | Martin Luther King, Jr. Outpatient Center | Los Angeles County Public Health Laboratories | P. Hemarajata et al. |
| EPI_ISL_2620827, EPI_ISL_2620828, EPI_ISL_2620829 | Palmdale Regional Medical Center | Los Angeles County Public Health Laboratories | P. Hemarajata et al. |
| EPI_ISL_2620888, EPI_ISL_2620889 | Iressef Genomics lab | IRESSEF | Souleymane MBOUP, Abdou PADANE , Khadim GUEYE, Diabou Diagne , Samba Ndiour, Papa Alassane DIAW, Birahim Piere NDIAYE, Barada CISSE, Aminata MBOUP, Moustapha MBOW, Ndeye Coumba Toure KANE, Nafisatou LEYE, Gora LO, Ambroise AHOUIDI , Astou Gaye GAYE, Aminata DIA, Yacine DIA |
| EPI_ISL_2620893, EPI_ISL_2620894, EPI_ISL_2620895, EPI_ISL_2620896, EPI_ISL_2620897 | National Institute of Public Health | State Veterinary Institute Prague | Nagy,A;Jirincova,H;Suri,T;Trnka,D;Vecerova,J |
| EPI_ISL_2620951 | Fondazione IRCCS Ca' Granda Ospedale Maggiore Policlinico | Fondazione IRCCS Ca' Granda Ospedale Maggiore Policlinico | Ferruccio Ceriotti, Sara Uceda Renteria |
| EPI_ISL_2621135, EPI_ISL_2621138, EPI_ISL_2621139 | Sedgefield Clinic wc SGE | NHLS/UCT | Arash Iranzadeh, Deelan Doolabh, Lynn Tyers, Bruna Galvao, Innocent Mudau, Marvin Hsiao, Gert Marais, Diana Hardie, Stephen Korsman, Carolyn Williamson |
| EPI_ISL_2621573 | Microbiological Diagnostic Unit - Public Health Laboratory (MDU-PHL) | Microbiological Diagnostic Unit - Public Health Laboratory (MDU-PHL) | Seemann T., Sait, M.L., Sherry, N.L. |
| EPI_ISL_2621633, EPI_ISL_2621654, EPI_ISL_2621655, EPI_ISL_2621658 | Clinical Microbiology, Infection Prevention and Control | Section for Molecular Diagnostics | Björn Hallström, Jonas Björkman |
| EPI_ISL_2621679, EPI_ISL_2621680, EPI_ISL_2621681 | The Caribbean Public Health Agency | Carrington Lab, Department of Preclinical Sciences, Faculty of Medical Sciences, The University of the West Indies | Nikita S. D. Sahadeo, Arianne Brown-Jordan, Sarah Hill, Vernie Ramkissoon, Anushka Ramjag, Kenneth George, Avery Hinds, Karla Georges, Risha Singh, SueMin Nathaniel, Nuno Faria, Oliver Pybus, Christopher Oura, Gabriel Escobar, Christine V. F. Carrington |
| EPI_ISL_2621743, EPI_ISL_2621748, EPI_ISL_2621749, EPI_ISL_2621750, EPI_ISL_2621753, EPI_ISL_2621755, EPI_ISL_2621756, EPI_ISL_2621757, EPI_ISL_2621758, EPI_ISL_2621759, EPI_ISL_2621760, EPI_ISL_2621761, EPI_ISL_2621762, EPI_ISL_2621764, EPI_ISL_2621768, EPI_ISL_2621770 | | | |
| see above | SARS-CoV-2 testing team, National Institute of Infectious Diseases | Pathogen Genomics Center, National Institute of Infectious Diseases | Tsuyoshi Sekizuka, Kentaro Itokawa, Rina Tanaka, Masanori Hashino, Nozomu Hanaoka, Masumichi Saito, Naomi Nojiri, Hazuka Y Furihata, Sana Uchikoba, Hiromizu Takahashi, Tsuguto Fujimoto, Makoto Kuroda |
| EPI_ISL_2621885, EPI_ISL_2621886, EPI_ISL_2621887, EPI_ISL_2621888, EPI_ISL_2621889, EPI_ISL_2621890, EPI_ISL_2621891, EPI_ISL_2621892, EPI_ISL_2621893, EPI_ISL_2621894, EPI_ISL_2621895, EPI_ISL_2621896, EPI_ISL_2621897, EPI_ISL_2621898, EPI_ISL_2621899, EPI_ISL_2621900, EPI_ISL_2621901, EPI_ISL_2621902, EPI_ISL_2621903, EPI_ISL_2621904, EPI_ISL_2621905, EPI_ISL_2621906, EPI_ISL_2621907, EPI_ISL_2621908, EPI_ISL_2621909, EPI_ISL_2621910, EPI_ISL_2621911, EPI_ISL_2621912, EPI_ISL_2621913, EPI_ISL_2621914, EPI_ISL_2621915, EPI_ISL_2621916, EPI_ISL_2621917, EPI_ISL_2621918, EPI_ISL_2621922, EPI_ISL_2621923, EPI_ISL_2621924, EPI_ISL_2621925, EPI_ISL_2621926, EPI_ISL_2621927, EPI_ISL_2621928, EPI_ISL_2621929, EPI_ISL_2621930, EPI_ISL_2621931, EPI_ISL_2621932, EPI_ISL_2621933, EPI_ISL_2621934, EPI_ISL_2621935, EPI_ISL_2621936, EPI_ISL_2621937, EPI_ISL_2621938, EPI_ISL_2621939, EPI_ISL_2621940, EPI_ISL_2621941, EPI_ISL_2621942, EPI_ISL_2621943, EPI_ISL_2621944, EPI_ISL_2621945, EPI_ISL_2621946, EPI_ISL_2621947, EPI_ISL_2621948, EPI_ISL_2621949, EPI_ISL_2621950, EPI_ISL_2621951, EPI_ISL_2621952, EPI_ISL_2621953, EPI_ISL_2621954, EPI_ISL_2621955, EPI_ISL_2621956, EPI_ISL_2621957, EPI_ISL_2621958, EPI_ISL_2621959, EPI_ISL_2621960, EPI_ISL_2621961 | | | |

see above — National Public Health Laboratory, National Centre for Infectious Diseases — National Public Health Laboratory, National Centre for Infectious Diseases — Tze Minn Mak, Zhenyang Zhou, Royce Ang, Lin Cui, Raymond Tzer Pin Lin

EPI_ISL_2622049, EPI_ISL_2622050, EPI_ISL_2622055, EPI_ISL_2622063, EPI_ISL_2622067, EPI_ISL_2622069, EPI_ISL_2622076 — Valais Hospital, Central Institute — Valais Hospital, Central Institute — Alexis Dumoulin, Lorenzo Cerutti, Henri Pegeot, Melyssa Elies, Deborah Penet, Keith Harshman, Ioannis Xenarios, Emmanouil Dermitzakis

EPI_ISL_2622082, EPI_ISL_2622084, EPI_ISL_2622085, EPI_ISL_2622086, EPI_ISL_2622087 — Communicable Disease Laboratory, Public Health Directorate — Communicable Disease Laboratory, Public Health Directorate — Alwasti,H., AlHujairi,Z., AlAbbas,Z., Marhoon,A., Almoamen,G.

EPI_ISL_2622101, EPI_ISL_2622103 — CHUV — Laboratory of genomics and metagenomics — Trestan Pillonel, Damien Jacot, Sébastien Aeby, Gilbert Greub, Claire Bertelli

EPI_ISL_2622176, EPI_ISL_2622177, EPI_ISL_2622178, EPI_ISL_2622179, EPI_ISL_2622180, EPI_ISL_2622181, EPI_ISL_2622182, EPI_ISL_2622183, EPI_ISL_2622184, EPI_ISL_2622185, EPI_ISL_2622186, EPI_ISL_2622187, EPI_ISL_2622190, EPI_ISL_2622191, EPI_ISL_2622192, EPI_ISL_2622193, EPI_ISL_2622195, EPI_ISL_2622196, EPI_ISL_2622197, EPI_ISL_2622198, EPI_ISL_2622199, EPI_ISL_2622200, EPI_ISL_2622201, EPI_ISL_2622202, EPI_ISL_2622203, EPI_ISL_2622204, EPI_ISL_2622205, EPI_ISL_2622206, EPI_ISL_2622207, EPI_ISL_2622208, EPI_ISL_2622209, EPI_ISL_2622210, EPI_ISL_2622211, EPI_ISL_2622212, EPI_ISL_2622213, EPI_ISL_2622214, EPI_ISL_2622215, EPI_ISL_2622216, EPI_ISL_2622217, EPI_ISL_2622218, EPI_ISL_2622219, EPI_ISL_2622220, EPI_ISL_2622221, EPI_ISL_2622222, EPI_ISL_2622223, EPI_ISL_2622224, EPI_ISL_2622225, EPI_ISL_2622226, EPI_ISL_2622227, EPI_ISL_2622228, EPI_ISL_2622229, EPI_ISL_2622230, EPI_ISL_2622231, EPI_ISL_2622232, EPI_ISL_2622233, EPI_ISL_2622234, EPI_ISL_2622235, EPI_ISL_2622236, EPI_ISL_2622237, EPI_ISL_2622238, EPI_ISL_2622239, EPI_ISL_2622240, EPI_ISL_2622241, EPI_ISL_2622242, EPI_ISL_2622243, EPI_ISL_2622244, EPI_ISL_2622245, EPI_ISL_2622246, EPI_ISL_2622247, EPI_ISL_2622248, EPI_ISL_2622249, EPI_ISL_2622250, EPI_ISL_2622251, EPI_ISL_2622252, EPI_ISL_2622253, EPI_ISL_2622254, EPI_ISL_2622255, EPI_ISL_2622256, EPI_ISL_2622257, EPI_ISL_2622258, EPI_ISL_2622259, EPI_ISL_2622260, EPI_ISL_2622261, EPI_ISL_2622262, EPI_ISL_2622263, EPI_ISL_2622264, EPI_ISL_2622265, EPI_ISL_2622266, EPI_ISL_2622267, EPI_ISL_2622268, EPI_ISL_2622269

see above — Department of Pathology, University of Cambridge — COVID-19 Genomics UK (COG-UK) Consortium — Aminu S. Jahun, Yasmin Chaudhry, Iliana Georgana, Myra Hosmillo, Rhys Izuagbe, William L. Hamilton, Martin D. Curran, Surendra Parmar, Ian Goodfellow

EPI_ISL_2622340, EPI_ISL_2622341, EPI_ISL_2622342, EPI_ISL_2622346, EPI_ISL_2622349, EPI_ISL_2622351, EPI_ISL_2622352, EPI_ISL_2622353, EPI_ISL_2622355, EPI_ISL_2622356, EPI_ISL_2622357, EPI_ISL_2622358, EPI_ISL_2622359, EPI_ISL_2622360, EPI_ISL_2622361, EPI_ISL_2622362, EPI_ISL_2622363, EPI_ISL_2622364, EPI_ISL_2622365, EPI_ISL_2622366, EPI_ISL_2622367, EPI_ISL_2622371, EPI_ISL_2622372

see above — Virology Department, Royal Infirmary of Edinburgh, NHS Lothian / School of Biological Sciences, University of Edinburgh — COVID-19 Genomics UK (COG-UK) Consortium — McHugh M, Dewar R, Cotton S, Gallagher A, Maloney D, Rooke S, O'Toole Á, Scher E, Hill V, McCrone JT, Colquhoun R, Yu X, Jackson B, Rambaut A, Templeton K

EPI_ISL_2622377, EPI_ISL_2622379, EPI_ISL_2622380, EPI_ISL_2622381, EPI_ISL_2622382, EPI_ISL_2622383, EPI_ISL_2622384, EPI_ISL_2622385, EPI_ISL_2622387, EPI_ISL_2622388, EPI_ISL_2622389, EPI_ISL_2622391, EPI_ISL_2622392, EPI_ISL_2622393, EPI_ISL_2622396, EPI_ISL_2622397, EPI_ISL_2622398, EPI_ISL_2622399, EPI_ISL_2622400, EPI_ISL_2622402, EPI_ISL_2622403, EPI_ISL_2622404, EPI_ISL_2622406, EPI_ISL_2622407, EPI_ISL_2622408, EPI_ISL_2622409, EPI_ISL_2622411, EPI_ISL_2622412, EPI_ISL_2622413, EPI_ISL_2622415, EPI_ISL_2622416, EPI_ISL_2622417, EPI_ISL_2622418, EPI_ISL_2622419, EPI_ISL_2622421, EPI_ISL_2622425, EPI_ISL_2622427, EPI_ISL_2622431, EPI_ISL_2622432, EPI_ISL_2622434

see above — Liverpool Clinical Laboratories — COVID-19 Genomics UK (COG-UK) Consortium — Sam Haldenby, Alistair Darby, Steve Paterson, Anita Lucaci, Julian Hiscox, M Almsaud, A Alrezaihi, Muhannad Alruwaili, Stuart D Armstrong, Jones Benjamin, Eleanor G Bentley, Anu Chawla, Jordan J Clark, Angela Cowell, Richard Eccles, Isabel García-Dorival, Matthew Gemmell, Alessandro Gerada, PKF Gilmore, Richard Gregory, Ximeng Han, Catherine Hartley, Margaret Hughes, Miren Iturriza-Gomara, James Johnson, L Luu, Jenifer Manson, Charlotte Nelson, Elaine O'Toole, Cassie Olateju, Rebekah Penrice-Randal , Lucille Rainbow, N.P Randle, Trevor Ian Robinson, Parul Sharma, Ghada T Shawli, James P Stewart, Neil Swainston, Ecaterina Vamos, Joanne Watts, Mark Whitehead, Hermione Webster

EPI_ISL_2622435, EPI_ISL_2622437, EPI_ISL_2622438, EPI_ISL_2622439, EPI_ISL_2622440, EPI_ISL_2622441, EPI_ISL_2622442, EPI_ISL_2622443, EPI_ISL_2622444, EPI_ISL_2622445, EPI_ISL_2622446, EPI_ISL_2622447, EPI_ISL_2622448, EPI_ISL_2622449, EPI_ISL_2622450, EPI_ISL_2622451, EPI_ISL_2622452, EPI_ISL_2622453, EPI_ISL_2622454, EPI_ISL_2622455, EPI_ISL_2622456, EPI_ISL_2622457, EPI_ISL_2622458, EPI_ISL_2622459, EPI_ISL_2622460, EPI_ISL_2622461, EPI_ISL_2622462, EPI_ISL_2622463, EPI_ISL_2622464, EPI_ISL_2622465, EPI_ISL_2622466, EPI_ISL_2622467, EPI_ISL_2622468, EPI_ISL_2622469, EPI_ISL_2622470, EPI_ISL_2622471, EPI_ISL_2622472, EPI_ISL_2622473, EPI_ISL_2622474, EPI_ISL_2622475

see above — Barts Health NHS Trust — COVID-19 Genomics UK (COG-UK) Consortium — CUTINO-MOGUEL, Maria-Teresa; HARRINGTON, David; OWOYEMI, Dola; KULASEGARAN-SHYLINI, Raghavendran; BROAD, Claire; KELE, Beatrix

EPI_ISL_2622476, EPI_ISL_2622477, EPI_ISL_2622478, EPI_ISL_2622480, EPI_ISL_2622481, EPI_ISL_2622482, EPI_ISL_2622483, EPI_ISL_2622484, EPI_ISL_2622485, EPI_ISL_2622486, EPI_ISL_2622487, EPI_ISL_2622488, EPI_ISL_2622489, EPI_ISL_2622490, EPI_ISL_2622491, EPI_ISL_2622492, EPI_ISL_2622493, EPI_ISL_2622494, EPI_ISL_2622495, EPI_ISL_2622496, EPI_ISL_2622497, EPI_ISL_2622498, EPI_ISL_2622499, EPI_ISL_2622500, EPI_ISL_2622501, EPI_ISL_2622502, EPI_ISL_2622503, EPI_ISL_2622504, EPI_ISL_2622505, EPI_ISL_2622506, EPI_ISL_2622507, EPI_ISL_2622508, EPI_ISL_2622509, EPI_ISL_2622510, EPI_ISL_2622511, EPI_ISL_2622512, EPI_ISL_2622513, EPI_ISL_2622514, EPI_ISL_2622515, EPI_ISL_2622516, EPI_ISL_2622517, EPI_ISL_2622518, EPI_ISL_2622519, EPI_ISL_2622520, EPI_ISL_2622521, EPI_ISL_2622523, EPI_ISL_2622524, EPI_ISL_2622525, EPI_ISL_2622526, EPI_ISL_2622527, EPI_ISL_2622528, EPI_ISL_2622529, EPI_ISL_2622530, EPI_ISL_2622531, EPI_ISL_2622532, EPI_ISL_2622533, EPI_ISL_2622534, EPI_ISL_2622535, EPI_ISL_2622537, EPI_ISL_2622538, EPI_ISL_2622539, EPI_ISL_2622540, EPI_ISL_2622541, EPI_ISL_2622542, EPI_ISL_2622543, EPI_ISL_2622544, EPI_ISL_2622545, EPI_ISL_2622546, EPI_ISL_2622547, EPI_ISL_2622548, EPI_ISL_2622549, EPI_ISL_2622550, EPI_ISL_2622551, EPI_ISL_2622552, EPI_ISL_2622553, EPI_ISL_2622554, EPI_ISL_2622555, EPI_ISL_2622556, EPI_ISL_2622557, EPI_ISL_2622558, EPI_ISL_2622559, EPI_ISL_2622560, EPI_ISL_2622561, EPI_ISL_2622562, EPI_ISL_2622563, EPI_ISL_2622564, EPI_ISL_2622565, EPI_ISL_2622566, EPI_ISL_2622567, EPI_ISL_2622568, EPI_ISL_2622569, EPI_ISL_2622570, EPI_ISL_2622571, EPI_ISL_2622572, EPI_ISL_2622573, EPI_ISL_2622574, EPI_ISL_2622575, EPI_ISL_2622576, EPI_ISL_2622577, EPI_ISL_2622578, EPI_ISL_2622579, EPI_ISL_2622580, EPI_ISL_2622581, EPI_ISL_2622582, EPI_ISL_2622584, EPI_ISL_2622585, EPI_ISL_2622587, EPI_ISL_2622588, EPI_ISL_2622589, EPI_ISL_2622590, EPI_ISL_2622591, EPI_ISL_2622592, EPI_ISL_2622593, EPI_ISL_2622594, EPI_ISL_2622595, EPI_ISL_2622596, EPI_ISL_2622597, EPI_ISL_2622598, EPI_ISL_2622599, EPI_ISL_2622600, EPI_ISL_2622601, EPI_ISL_2622602, EPI_ISL_2622603, EPI_ISL_2622604, EPI_ISL_2622605, EPI_ISL_2622606, EPI_ISL_2622607, EPI_ISL_2622608, EPI_ISL_2622609, EPI_ISL_2622610, EPI_ISL_2622611, EPI_ISL_2622612, EPI_ISL_2622613, EPI_ISL_2622614, EPI_ISL_2622615, EPI_ISL_2622616, EPI_ISL_2622617, EPI_ISL_2622618, EPI_ISL_2622619, EPI_ISL_2622620, EPI_ISL_2622621, EPI_ISL_2622622, EPI_ISL_2622623, EPI_ISL_2622624, EPI_ISL_2622625, EPI_ISL_2622626, EPI_ISL_2622627, EPI_ISL_2622628, EPI_ISL_2622629, EPI_ISL_2622630, EPI_ISL_2622631, EPI_ISL_2622632, EPI_ISL_2622633, EPI_ISL_2622634, EPI_ISL_2622635, EPI_ISL_2622636, EPI_ISL_2622637, EPI_ISL_2622638, EPI_ISL_2622639, EPI_ISL_2622640, EPI_ISL_2622641, EPI_ISL_2622642, EPI_ISL_2622643, EPI_ISL_2622644, EPI_ISL_2622645, EPI_ISL_2622646, EPI_ISL_2622647, EPI_ISL_2622648, EPI_ISL_2622649, EPI_ISL_2622650, EPI_ISL_2622651, EPI_ISL_2622652, EPI_ISL_2622653, EPI_ISL_2622654, EPI_ISL_2622655, EPI_ISL_2622656, EPI_ISL_2622657, EPI_ISL_2622658, EPI_ISL_2622659, EPI_ISL_2622660, EPI_ISL_2622661, EPI_ISL_2622662, EPI_ISL_2622663, EPI_ISL_2622664, EPI_ISL_2622665, EPI_ISL_2622666, EPI_ISL_2622667, EPI_ISL_2622668, EPI_ISL_2622669, EPI_ISL_2622670, EPI_ISL_2622671, EPI_ISL_2622673, EPI_ISL_2622674, EPI_ISL_2622675, EPI_ISL_2622676, EPI_ISL_2622677, EPI_ISL_2622678, EPI_ISL_2622679, EPI_ISL_2622680, EPI_ISL_2622681, EPI_ISL_2622682, EPI_ISL_2622683, EPI_ISL_2622684, EPI_ISL_2622685, EPI_ISL_2622686, EPI_ISL_2622687, EPI_ISL_2622688, EPI_ISL_2622689, EPI_ISL_2622690, EPI_ISL_2622691, EPI_ISL_2622692, EPI_ISL_2622693, EPI_ISL_2622694, EPI_ISL_2622695, EPI_ISL_2622696, EPI_ISL_2622697, EPI_ISL_2622698, EPI_ISL_2622699, EPI_ISL_2622700, EPI_ISL_2622701, EPI_ISL_2622702, EPI_ISL_2622703, EPI_ISL_2622704, EPI_ISL_2622705, EPI_ISL_2622706, EPI_ISL_2622707, EPI_ISL_2622708, EPI_ISL_2622709, EPI_ISL_2622710, EPI_ISL_2622711, EPI_ISL_2622712, EPI_ISL_2622713, EPI_ISL_2622714, EPI_ISL_2622715, EPI_ISL_2622716, EPI_ISL_2622717, EPI_ISL_2622718, EPI_ISL_2622719, EPI_ISL_2622720, EPI_ISL_2622721, EPI_ISL_2622722, EPI_ISL_2622723, EPI_ISL_2622724, EPI_ISL_2622725, EPI_ISL_2622726, EPI_ISL_2622727, EPI_ISL_2622728, EPI_ISL_2622729, EPI_ISL_2622730, EPI_ISL_2622731, EPI_ISL_2622732, EPI_ISL_2622733, EPI_ISL_2622734, EPI_ISL_2622735, EPI_ISL_2622736, EPI_ISL_2622737, EPI_ISL_2622738, EPI_ISL_2622739, EPI_ISL_2622740, EPI_ISL_2622741, EPI_ISL_2622742, EPI_ISL_2622743, EPI_ISL_2622744, EPI_ISL_2622745, EPI_ISL_2622746, EPI_ISL_2622747, EPI_ISL_2622748, EPI_ISL_2622749, EPI_ISL_2622750, EPI_ISL_2622751, EPI_ISL_2622752, EPI_ISL_2622755, EPI_ISL_2622756, EPI_ISL_2622757, EPI_ISL_2622758, EPI_ISL_2622759, EPI_ISL_2622760, EPI_ISL_2622761, EPI_ISL_2622762, EPI_ISL_2622763, EPI_ISL_2622764, EPI_ISL_2622765, EPI_ISL_2622767, EPI_ISL_2622768, EPI_ISL_2622769, EPI_ISL_2622770, EPI_ISL_2622772, EPI_ISL_2622773, EPI_ISL_2622774, EPI_ISL_2622775, EPI_ISL_2622776, EPI_ISL_2622777, EPI_ISL_2622778, EPI_ISL_2622779, EPI_ISL_2622780, EPI_ISL_2622781, EPI_ISL_2622782, EPI_ISL_2622783, EPI_ISL_2622784, EPI_ISL_2622786, EPI_ISL_2622787, EPI_ISL_2622788, EPI_ISL_2622789, EPI_ISL_2622790, EPI_ISL_2622791, EPI_ISL_2622792, EPI_ISL_2622793, EPI_ISL_2622794, EPI_ISL_2622795, EPI_ISL_2622796, EPI_ISL_2622797, EPI_ISL_2622798, EPI_ISL_2622799, EPI_ISL_2622800, EPI_ISL_2622801, EPI_ISL_2622802, EPI_ISL_2622803, EPI_ISL_2622804, EPI_ISL_2622805, EPI_ISL_2622806, EPI_ISL_2622807, EPI_ISL_2622808, EPI_ISL_2622809, EPI_ISL_2622810, EPI_ISL_2622811, EPI_ISL_2622812, EPI_ISL_2622813, EPI_ISL_2622814, EPI_ISL_2622815, EPI_ISL_2622816, EPI_ISL_2622817, EPI_ISL_2622818, EPI_ISL_2622819, EPI_ISL_2622821, EPI_ISL_2622822, EPI_ISL_2622823, EPI_ISL_2622824, EPI_ISL_2622825, EPI_ISL_2622826, EPI_ISL_2622827, EPI_ISL_2622828, EPI_ISL_2622829, EPI_ISL_2622830, EPI_ISL_2622832, EPI_ISL_2622833, EPI_ISL_2622834, EPI_ISL_2622835, EPI_ISL_2622836, EPI_ISL_2622837, EPI_ISL_2622838, EPI_ISL_2622839, EPI_ISL_2622840, EPI_ISL_2622841, EPI_ISL_2622842, EPI_ISL_2622843, EPI_ISL_2622844, EPI_ISL_2622845, EPI_ISL_2622846, EPI_ISL_2622847, EPI_ISL_2622848, EPI_ISL_2622849, EPI_ISL_2622850, EPI_ISL_2622851, EPI_ISL_2622852, EPI_ISL_2622853, EPI_ISL_2622854, EPI_ISL_2622855, EPI_ISL_2622856, EPI_ISL_2622857, EPI_ISL_2622858, EPI_ISL_2622859, EPI_ISL_2622860, EPI_ISL_2622861, EPI_ISL_2622862, EPI_ISL_2622863, EPI_ISL_2622864, EPI_ISL_2622865, EPI_ISL_2622866, EPI_ISL_2622867, EPI_ISL_2622868, EPI_ISL_2622869, EPI_ISL_2622870, EPI_ISL_2622871, EPI_ISL_2622872, EPI_ISL_2622873, EPI_ISL_2622874, EPI_ISL_2622875, EPI_ISL_2622876, EPI_ISL_2622877, EPI_ISL_2622878, EPI_ISL_2622879, EPI_ISL_2622880, EPI_ISL_2622881, EPI_ISL_2622882, EPI_ISL_2622883, EPI_ISL_2622884, EPI_ISL_2622885, EPI_ISL_2622886, EPI_ISL_2622887, EPI_ISL_2622888, EPI_ISL_2622889, EPI_ISL_2622890, EPI_ISL_2622891, EPI_ISL_2622892, EPI_ISL_2622893, EPI_ISL_2622894, EPI_ISL_2622895, EPI_ISL_2622896, EPI_ISL_2622897, EPI_ISL_2622898, EPI_ISL_2622899, EPI_ISL_2622900, EPI_ISL_2622901, EPI_ISL_2622902, EPI_ISL_2622903, EPI_ISL_2622904, EPI_ISL_2622905, EPI_ISL_2622906, EPI_ISL_2622907, EPI_ISL_2622908, EPI_ISL_2622909, EPI_ISL_2622910, EPI_ISL_2622911, EPI_ISL_2622912, EPI_ISL_2622913, EPI_ISL_2622914, EPI_ISL_2622915, EPI_ISL_2622916, EPI_ISL_2622917, EPI_ISL_2622918, EPI_ISL_2622919, EPI_ISL_2622920, EPI_ISL_2622921, EPI_ISL_2622922, EPI_ISL_2622923, EPI_ISL_2622924, EPI_ISL_2622925, EPI_ISL_2622926, EPI_ISL_2622927, EPI_ISL_2622928, EPI_ISL_2622929, EPI_ISL_2622930, EPI_ISL_2622931, EPI_ISL_2622932, EPI_ISL_2622933, EPI_ISL_2622934, EPI_ISL_2622935, EPI_ISL_2622936, EPI_ISL_2622937, EPI_ISL_2622938, EPI_ISL_2622939, EPI_ISL_2622940, EPI_ISL_2622941, EPI_ISL_2622942, EPI_ISL_2622943, EPI_ISL_2622944, EPI_ISL_2622945, EPI_ISL_2622946, EPI_ISL_2622947, EPI_ISL_2622948, EPI_ISL_2622949, EPI_ISL_2622950, EPI_ISL_2622951, EPI_ISL_2622952, EPI_ISL_2622953, EPI_ISL_2622954, EPI_ISL_2622955, EPI_ISL_2622956, EPI_ISL_2622958, EPI_ISL_2622959, EPI_ISL_2622960, EPI_ISL_2622961, EPI_ISL_2622962, EPI_ISL_2622963, EPI_ISL_2622964, EPI_ISL_2622966, EPI_ISL_2622967, EPI_ISL_2622968, EPI_ISL_2622969, EPI_ISL_2622970, EPI_ISL_2622971, EPI_ISL_2622972, EPI_ISL_2622973, EPI_ISL_2622974, EPI_ISL_2622975, EPI_ISL_2622976, EPI_ISL_2622977, EPI_ISL_2622978, EPI_ISL_2622979, EPI_ISL_2622980, EPI_ISL_2622981, EPI_ISL_2622982, EPI_ISL_2622983, EPI_ISL_2622984, EPI_ISL_2622985, EPI_ISL_2622986, EPI_ISL_2622987, EPI_ISL_2622988, EPI_ISL_2622989, EPI_ISL_2622990, EPI_ISL_2622991, EPI_ISL_2622992, EPI_ISL_2622993, EPI_ISL_2622994, EPI_ISL_2622995, EPI_ISL_2622996, EPI_ISL_2622997, EPI_ISL_2622998, EPI_ISL_2622999, EPI_ISL_2623000, EPI_ISL_2623001, EPI_ISL_2623002, EPI_ISL_2623003, EPI_ISL_2623004, EPI_ISL_2623005, EPI_ISL_2623006, EPI_ISL_2623007, EPI_ISL_2623008, EPI_ISL_2623009, EPI_ISL_2623010, EPI_ISL_2623011, EPI_ISL_2623012, EPI_ISL_2623013, EPI_ISL_2623014, EPI_ISL_2623015, EPI_ISL_2623016, EPI_ISL_2623017, EPI_ISL_2623018, EPI_ISL_2623019, EPI_ISL_2623020, EPI_ISL_2623021, EPI_ISL_2623022, EPI_ISL_2623023, EPI_ISL_2623024, EPI_ISL_2623025, EPI_ISL_2623026, EPI_ISL_2623027, EPI_ISL_2623028, EPI_ISL_2623029, EPI_ISL_2623030, EPI_ISL_2623031, EPI_ISL_2623032, EPI_ISL_2623033, EPI_ISL_2623034, EPI_ISL_2623035, EPI_ISL_2623036, EPI_ISL_2623037, EPI_ISL_2623038, EPI_ISL_2623039, EPI_ISL_2623040, EPI_ISL_2623041, EPI_ISL_2623042, EPI_ISL_2623043, EPI_ISL_2623044, EPI_ISL_2623045, EPI_ISL_2623046, EPI_ISL_2623047, EPI_ISL_2623048, EPI_ISL_2623049, EPI_ISL_2623050, EPI_ISL_2623051, EPI_ISL_2623052, EPI_ISL_2623053, EPI_ISL_2623054, EPI_ISL_2623055, EPI_ISL_2623056, EPI_ISL_2623057, EPI_ISL_2623058, EPI_ISL_2623059, EPI_ISL_2623060, EPI_ISL_2623061, EPI_ISL_2623062, EPI_ISL_2623063, EPI_ISL_2623064, EPI_ISL_2623065, EPI_ISL_2623066, EPI_ISL_2623067, EPI_ISL_2623068, EPI_ISL_2623069, EPI_ISL_2623070, EPI_ISL_2623071, EPI_ISL_2623072, EPI_ISL_2623073, EPI_ISL_2623074, EPI_ISL_2623075, EPI_ISL_2623076, EPI_ISL_2623077, EPI_ISL_2623078, EPI_ISL_2623079, EPI_ISL_2623080, EPI_ISL_2623082, EPI_ISL_2623084, EPI_ISL_2623085, EPI_ISL_2623086, EPI_ISL_2623087, EPI_ISL_2623088, EPI_ISL_2623089, EPI_ISL_2623090, EPI_ISL_2623091, EPI_ISL_2623092, EPI_ISL_2623093, EPI_ISL_2623094, EPI_ISL_2623095, EPI_ISL_2623096, EPI_ISL_2623097, EPI_ISL_2623098, EPI_ISL_2623099, EPI_ISL_2623100, EPI_ISL_2623101, EPI_ISL_2623102, EPI_ISL_2623103, EPI_ISL_2623104, EPI_ISL_2623105, EPI_ISL_2623108, EPI_ISL_2623110, EPI_ISL_2623111, EPI_ISL_2623112, EPI_ISL_2623113, EPI_ISL_2623114, EPI_ISL_2623115, EPI_ISL_2623116, EPI_ISL_2623117, EPI_ISL_2623118, EPI_ISL_2623119, EPI_ISL_2623121, EPI_ISL_2623122, EPI_ISL_2623123, EPI_ISL_2623124

see above — University College London, Great Ormond Street Hospital for Children NHS Foundation Trust, Imperial College Healthcare NHS Trust — COVID-19 Genomics UK (COG-UK) Consortium — Sergi Castellano, Rachel Williams, Mark Kristiansen, Paola Resende Silva, Sunando Roy, Tony Brooks, Helena Tutill, Paola Niola, Patricia Dyal, Charlotte Williams, Leysa Forrest, Yasmin Panchbhaya, Jacqueline Findlay, Samuel Weeks, Julianne Brown, Kathryn Harris, Paul Randell, James Price, Alison Holmes, Judith Breuer

EPI_ISL_2623127, EPI_ISL_2623128, EPI_ISL_2623129, EPI_ISL_2623130, EPI_ISL_2623131, EPI_ISL_2623132, EPI_ISL_2623134, EPI_ISL_2623135, EPI_ISL_2623136, EPI_ISL_2623138, EPI_ISL_2623139, EPI_ISL_2623140, EPI_ISL_2623142, EPI_ISL_2623143, EPI_ISL_2623144, EPI_ISL_2623148, EPI_ISL_2623150, EPI_ISL_2623151, EPI_ISL_2623153, EPI_ISL_2623155, EPI_ISL_2623156, EPI_ISL_2623157, EPI_ISL_2623160, EPI_ISL_2623162, EPI_ISL_2623164, EPI_ISL_2623165, EPI_ISL_2623167, EPI_ISL_2623169, EPI_ISL_2623170, EPI_ISL_2623171, EPI_ISL_2623172, EPI_ISL_2623174, EPI_ISL_2623175, EPI_ISL_2623177, EPI_ISL_2623180, EPI_ISL_2623181, EPI_ISL_2623182, EPI_ISL_2623184, EPI_ISL_2623185, EPI_ISL_2623186, EPI_ISL_2623187, EPI_ISL_2623188, EPI_ISL_2623190, EPI_ISL_2623191, EPI_ISL_2623192, EPI_ISL_2623193, EPI_ISL_2623194, EPI_ISL_2623196, EPI_ISL_2623197, EPI_ISL_2623198, EPI_ISL_2623199, EPI_ISL_2623200, EPI_ISL_2623201, EPI_ISL_2623202, EPI_ISL_2623203, EPI_ISL_2623204, EPI_ISL_2623205, EPI_ISL_2623207, EPI_ISL_2623208, EPI_ISL_2623209, EPI_ISL_2623212, EPI_ISL_2623213, EPI_ISL_2623214, EPI_ISL_2623216, EPI_ISL_2623217, EPI_ISL_2623219, EPI_ISL_2623220, EPI_ISL_2623221, EPI_ISL_2623222, EPI_ISL_2623223, EPI_ISL_2623224, EPI_ISL_2623226, EPI_ISL_2623228, EPI_ISL_2623229, EPI_ISL_2623230, EPI_ISL_2623231, EPI_ISL_2623232, EPI_ISL_2623233, EPI_ISL_2623235, EPI_ISL_2623236, EPI_ISL_2623237, EPI_ISL_2623238, EPI_ISL_2623239, EPI_ISL_2623240, EPI_ISL_2623241, EPI_ISL_2623242, EPI_ISL_2623243, EPI_ISL_2623244, EPI_ISL_2623245, EPI_ISL_2623246, EPI_ISL_2623249, EPI_ISL_2623250, EPI_ISL_2623251, EPI_ISL_2623252, EPI_ISL_2623253, EPI_ISL_2623254, EPI_ISL_2623255, EPI_ISL_2623256, EPI_ISL_2623257, EPI_ISL_2623258, EPI_ISL_2623259, EPI_ISL_2623260, EPI_ISL_2623261, EPI_ISL_2623262, EPI_ISL_2623263, EPI_ISL_2623264, EPI_ISL_2623265, EPI_ISL_2623266, EPI_ISL_2623267, EPI_ISL_2623268, EPI_ISL_2623269, EPI_ISL_2623272, EPI_ISL_2623274, EPI_ISL_2623275, EPI_ISL_2623276, EPI_ISL_2623277, EPI_ISL_2623278, EPI_ISL_2623279, EPI_ISL_2623280, EPI_ISL_2623281, EPI_ISL_2623282, EPI_ISL_2623284, EPI_ISL_2623285, EPI_ISL_2623286, EPI_ISL_2623287, EPI_ISL_2623288, EPI_ISL_2623289, EPI_ISL_2623290, EPI_ISL_2623291, EPI_ISL_2623292, EPI_ISL_2623293, EPI_ISL_2623294, EPI_ISL_2623295, EPI_ISL_2623296, EPI_ISL_2623297, EPI_ISL_2623298, EPI_ISL_2623299, EPI_ISL_2623300, EPI_ISL_2623302, EPI_ISL_2623303, EPI_ISL_2623305, EPI_ISL_2623306, EPI_ISL_2623307, EPI_ISL_2623308, EPI_ISL_2623309, EPI_ISL_2623310, EPI_ISL_2623311, EPI_ISL_2623312, EPI_ISL_2623314, EPI_ISL_2623315, EPI_ISL_2623316, EPI_ISL_2623317, EPI_ISL_2623318, EPI_ISL_2623319, EPI_ISL_2623320, EPI_ISL_2623321, EPI_ISL_2623322, EPI_ISL_2623324, EPI_ISL_2623325, EPI_ISL_2623326, EPI_ISL_2623327, EPI_ISL_2623328, EPI_ISL_2623329, EPI_ISL_2623330, EPI_ISL_2623331, EPI_ISL_2623332, EPI_ISL_2623333, EPI_ISL_2623334, EPI_ISL_2623335, EPI_ISL_2623337, EPI_ISL_2623338, EPI_ISL_2623339, EPI_ISL_2623340, EPI_ISL_2623341, EPI_ISL_2623342, EPI_ISL_2623343, EPI_ISL_2623344, EPI_ISL_2623345, EPI_ISL_2623346, EPI_ISL_2623347, EPI_ISL_2623348, EPI_ISL_2623349, EPI_ISL_2623350, EPI_ISL_2623353, EPI_ISL_2623354, EPI_ISL_2623355, EPI_ISL_2623356, EPI_ISL_2623357, EPI_ISL_2623358, EPI_ISL_2623359, EPI_ISL_2623360, EPI_ISL_2623361, EPI_ISL_2623363, EPI_ISL_2623364, EPI_ISL_2623368, EPI_ISL_2623369, EPI_ISL_2623370, EPI_ISL_2623372, EPI_ISL_2623373, EPI_ISL_2623374, EPI_ISL_2623375, EPI_ISL_2623376, EPI_ISL_2623377, EPI_ISL_2623380, EPI_ISL_2623381, EPI_ISL_2623382, EPI_ISL_2623383, EPI_ISL_2623384, EPI_ISL_2623385, EPI_ISL_2623386, EPI_ISL_2623387, EPI_ISL_2623388, EPI_ISL_2623389, EPI_ISL_2623390, EPI_ISL_2623391, EPI_ISL_2623392, EPI_ISL_2623393, EPI_ISL_2623394, EPI_ISL_2623395, EPI_ISL_2623396, EPI_ISL_2623397, EPI_ISL_2623398, EPI_ISL_2623399, EPI_ISL_2623400, EPI_ISL_2623401, EPI_ISL_2623402, EPI_ISL_2623403, EPI_ISL_2623404, EPI_ISL_2623405, EPI_ISL_2623406, EPI_ISL_2623407, EPI_ISL_2623408, EPI_ISL_2623409, EPI_ISL_2623410, EPI_ISL_2623411, EPI_ISL_2623412, EPI_ISL_2623413, EPI_ISL_2623414, EPI_ISL_2623417, EPI_ISL_2623418, EPI_ISL_2623419, EPI_ISL_2623420, EPI_ISL_2623421, EPI_ISL_2623422, EPI_ISL_2623423, EPI_ISL_2623424, EPI_ISL_2623425, EPI_ISL_2623426, EPI_ISL_2623427, EPI_ISL_2623428, EPI_ISL_2623429, EPI_ISL_2623430, EPI_ISL_2623431, EPI_ISL_2623432, EPI_ISL_2623433, EPI_ISL_2623434, EPI_ISL_2623435, EPI_ISL_2623436, EPI_ISL_2623437, EPI_ISL_2623438, EPI_ISL_2623439, EPI_ISL_2623441, EPI_ISL_2623442, EPI_ISL_2623443, EPI_ISL_2623444, EPI_ISL_2623445, EPI_ISL_2623446, EPI_ISL_2623447, EPI_ISL_2623448, EPI_ISL_2623449, EPI_ISL_2623451, EPI_ISL_2623452, EPI_ISL_2623453, EPI_ISL_2623455, EPI_ISL_2623456, EPI_ISL_2623457, EPI_ISL_2623458, EPI_ISL_2623459, EPI_ISL_2623461, EPI_ISL_2623462, EPI_ISL_2623463, EPI_ISL_2623464, EPI_ISL_2623465, EPI_ISL_2623466, EPI_ISL_2623467, EPI_ISL_2623468, EPI_ISL_2623469, EPI_ISL_2623470, EPI_ISL_2623471, EPI_ISL_2623472, EPI_ISL_2623473, EPI_ISL_2623474, EPI_ISL_2623475, EPI_ISL_2623476, EPI_ISL_2623478, EPI_ISL_2623479, EPI_ISL_2623480, EPI_ISL_2623481, EPI_ISL_2623482, EPI_ISL_2623484, EPI_ISL_2623485, EPI_ISL_2623486, EPI_ISL_2623487, EPI_ISL_2623489, EPI_ISL_2623490, EPI_ISL_2623491, EPI_ISL_2623492, EPI_ISL_2623493, EPI_ISL_2623494, EPI_ISL_2623495, EPI_ISL_2623496, EPI_ISL_2623497, EPI_ISL_2623498, EPI_ISL_2623499, EPI_ISL_2623500, EPI_ISL_2623501, EPI_ISL_2623502, EPI_ISL_2623503, EPI_ISL_2623504, EPI_ISL_2623505, EPI_ISL_2623506, EPI_ISL_2623507, EPI_ISL_2623508, EPI_ISL_2623509, EPI_ISL_2623510, EPI_ISL_2623511, EPI_ISL_2623512, EPI_ISL_2623513, EPI_ISL_2623514, EPI_ISL_2623515, EPI_ISL_2623516, EPI_ISL_2623517, EPI_ISL_2623518, EPI_ISL_2623519, EPI_ISL_2623520, EPI_ISL_2623521, EPI_ISL_2623522, EPI_ISL_2623524, EPI_ISL_2623525, EPI_ISL_2623526, EPI_ISL_2623527, EPI_ISL_2623529, EPI_ISL_2623531, EPI_ISL_2623532, EPI_ISL_2623533, EPI_ISL_2623534, EPI_ISL_2623535, EPI_ISL_2623536, EPI_ISL_2623537, EPI_ISL_2623538, EPI_ISL_2623539, EPI_ISL_2623541, EPI_ISL_2623542, EPI_ISL_2623543, EPI_ISL_2623544, EPI_ISL_2623545, EPI_ISL_2623547, EPI_ISL_2623548, EPI_ISL_2623551, EPI_ISL_2623553, EPI_ISL_2623554, EPI_ISL_2623555, EPI_ISL_2623556, EPI_ISL_2623557, EPI_ISL_2623558, EPI_ISL_2623559, EPI_ISL_2623560, EPI_ISL_2623561, EPI_ISL_2623562, EPI_ISL_2623563, EPI_ISL_2623564, EPI_ISL_2623565, EPI_ISL_2623567, EPI_ISL_2623568, EPI_ISL_2623569, EPI_ISL_2623570, EPI_ISL_2623572, EPI_ISL_2623573, EPI_ISL_2623574, EPI_ISL_2623575, EPI_ISL_2623576, EPI_ISL_2623577, EPI_ISL_2623578, EPI_ISL_2623579, EPI_ISL_2623580, EPI_ISL_2623581, EPI_ISL_2623582, EPI_ISL_2623583, EPI_ISL_2623584, EPI_ISL_2623585, EPI_ISL_2623586, EPI_ISL_2623587, EPI_ISL_2623588, EPI_ISL_2623589, EPI_ISL_2623590, EPI_ISL_2623591, EPI_ISL_2623592, EPI_ISL_2623593, EPI_ISL_2623594, EPI_ISL_2623595, EPI_ISL_2623596, EPI_ISL_2623597, EPI_ISL_2623598, EPI_ISL_2623599, EPI_ISL_2623600, EPI_ISL_2623601, EPI_ISL_2623602, EPI_ISL_2623603, EPI_ISL_2623604, EPI_ISL_2623605, EPI_ISL_2623606, EPI_ISL_2623607, EPI_ISL_2623608, EPI_ISL_2623610, EPI_ISL_2623611, EPI_ISL_2623613, EPI_ISL_2623614, EPI_ISL_2623615, EPI_ISL_2623616, EPI_ISL_2623617, EPI_ISL_2623618, EPI_ISL_2623619, EPI_ISL_2623621, EPI_ISL_2623623, EPI_ISL_2623624, EPI_ISL_2623626, EPI_ISL_2623627, EPI_ISL_2623628, EPI_ISL_2623629, EPI_ISL_2623631, EPI_ISL_2623632, EPI_ISL_2623633, EPI_ISL_2623634, EPI_ISL_2623635, EPI_ISL_2623636, EPI_ISL_2623637, EPI_ISL_2623638, EPI_ISL_2623640, EPI_ISL_2623641, EPI_ISL_2623642, EPI_ISL_2623643, EPI_ISL_2623644, EPI_ISL_2623645, EPI_ISL_2623646, EPI_ISL_2623647

see above — Northumbria University / South Tees Hospitals NHS Foundation Trust / North Cumbria Integrated Care NHS Foundation Trust / North Tees and Hartlepool NHS Foundation Trust / Newcastle Hospitals NHS Foundation Trust — COVID-19 Genomics UK (COG-UK) Consortium — Darren L Smith,Andrew Nelson,Matthew Bashton,Greg R Young,Joshua Loh,John Allan,Mohammad A Tariq,Giles S Holt,Gary Black,Wen C Yew,Lynn Dover,Paul Baker,Steve Liggett,Sarah Essex,Jane Greenaway,Debra Padgett,Clive Graham,Garren Scott,Edward Barton,Emma Swindells,Brendan Payne,Jennifer Collins,Yusri Taha,Gary Eltringham

EPI_ISL_2623648, EPI_ISL_2623649, EPI_ISL_2623650, EPI_ISL_2623651, EPI_ISL_2623652, EPI_ISL_2623653, EPI_ISL_2623654, EPI_ISL_2623655, EPI_ISL_2623656, EPI_ISL_2623657, EPI_ISL_2623658, EPI_ISL_2623659, EPI_ISL_2623660, EPI_ISL_2623661, EPI_ISL_2623662, EPI_ISL_2623663, EPI_ISL_2623664, EPI_ISL_2623665, EPI_ISL_2623668, EPI_ISL_2623669, EPI_ISL_2623670, EPI_ISL_2623671, EPI_ISL_2623672, EPI_ISL_2623673, EPI_ISL_2623674, EPI_ISL_2623675, EPI_ISL_2623676, EPI_ISL_2623678, EPI_ISL_2623680, EPI_ISL_2623681, EPI_ISL_2623682, EPI_ISL_2623683, EPI_ISL_2623684, EPI_ISL_2623685, EPI_ISL_2623686, EPI_ISL_2623687, EPI_ISL_2623688, EPI_ISL_2623689, EPI_ISL_2623690, EPI_ISL_2623691, EPI_ISL_2623692, EPI_ISL_2623693, EPI_ISL_2623694, EPI_ISL_2623695, EPI_ISL_2623697, EPI_ISL_2623698, EPI_ISL_2623699, EPI_ISL_2623700, EPI_ISL_2623701, EPI_ISL_2623702, EPI_ISL_2623703, EPI_ISL_2623704, EPI_ISL_2623705, EPI_ISL_2623709, EPI_ISL_2623710, EPI_ISL_2623711, EPI_ISL_2623712, EPI_ISL_2623713, EPI_ISL_2623714, EPI_ISL_2623715, EPI_ISL_2623716, EPI_ISL_2623717, EPI_ISL_2623718, EPI_ISL_2623719, EPI_ISL_2623720, EPI_ISL_2623721, EPI_ISL_2623722, EPI_ISL_2623724, EPI_ISL_2623725, EPI_ISL_2623726, EPI_ISL_2623727, EPI_ISL_2623728, EPI_ISL_2623729, EPI_ISL_2623730, EPI_ISL_2623731, EPI_ISL_2623732, EPI_ISL_2623733, EPI_ISL_2623734, EPI_ISL_2623735, EPI_ISL_2623736, EPI_ISL_2623737, EPI_ISL_2623738, EPI_ISL_2623740, EPI_ISL_2623742, EPI_ISL_2623743, EPI_ISL_2623744, EPI_ISL_2623745, EPI_ISL_2623746, EPI_ISL_2623747, EPI_ISL_2623748, EPI_ISL_2623749, EPI_ISL_2623750, EPI_ISL_2623751, EPI_ISL_2623752, EPI_ISL_2623753, EPI_ISL_2623754, EPI_ISL_2623755, EPI_ISL_2623756, EPI_ISL_2623757, EPI_ISL_2623759, EPI_ISL_2623760, EPI_ISL_2623761, EPI_ISL_2623762, EPI_ISL_2623763, EPI_ISL_2623764, EPI_ISL_2623765, EPI_ISL_2623766, EPI_ISL_2623767, EPI_ISL_2623768, EPI_ISL_2623769, EPI_ISL_2623770, EPI_ISL_2623772, EPI_ISL_2623773, EPI_ISL_2623774, EPI_ISL_2623776, EPI_ISL_2623778, EPI_ISL_2623779, EPI_ISL_2623780, EPI_ISL_2623781, EPI_ISL_2623782, EPI_ISL_2623783, EPI_ISL_2623784, EPI_ISL_2623785, EPI_ISL_2623786, EPI_ISL_2623787, EPI_ISL_2623788, EPI_ISL_2623789, EPI_ISL_2623790, EPI_ISL_2623791, EPI_ISL_2623792, EPI_ISL_2623793, EPI_ISL_2623794, EPI_ISL_2623798, EPI_ISL_2623799, EPI_ISL_2623800, EPI_ISL_2623801, EPI_ISL_2623802, EPI_ISL_2623803, EPI_ISL_2623804, EPI_ISL_2623805, EPI_ISL_2623806, EPI_ISL_2623807, EPI_ISL_2623808, EPI_ISL_2623809, EPI_ISL_2623810, EPI_ISL_2623811, EPI_ISL_2623812, EPI_ISL_2623813, EPI_ISL_2623814, EPI_ISL_2623815, EPI_ISL_2623816, EPI_ISL_2623817, EPI_ISL_2623818, EPI_ISL_2623819, EPI_ISL_2623820, EPI_ISL_2623822, EPI_ISL_2623823, EPI_ISL_2623824, EPI_ISL_2623825, EPI_ISL_2623826, EPI_ISL_2623827, EPI_ISL_2623828, EPI_ISL_2623829, EPI_ISL_2623830, EPI_ISL_2623831, EPI_ISL_2623832, EPI_ISL_2623833, EPI_ISL_2623834, EPI_ISL_2623835, EPI_ISL_2623836, EPI_ISL_2623837, EPI_ISL_2623838, EPI_ISL_2623839, EPI_ISL_2623840, EPI_ISL_2623841, EPI_ISL_2623843, EPI_ISL_2623844, EPI_ISL_2623845, EPI_ISL_2623846, EPI_ISL_2623847, EPI_ISL_2623848, EPI_ISL_2623849, EPI_ISL_2623850, EPI_ISL_2623851, EPI_ISL_2623852, EPI_ISL_2623854, EPI_ISL_2623855, EPI_ISL_2623856, EPI_ISL_2623858, EPI_ISL_2623859, EPI_ISL_2623861, EPI_ISL_2623862, EPI_ISL_2623863, EPI_ISL_2623865, EPI_ISL_2623866, EPI_ISL_2623867, EPI_ISL_2623868, EPI_ISL_2623869, EPI_ISL_2623870, EPI_ISL_2623871, EPI_ISL_2623872, EPI_ISL_2623873, EPI_ISL_2623874, EPI_ISL_2623875, EPI_ISL_2623876, EPI_ISL_2623877, EPI_ISL_2623878, EPI_ISL_2623879, EPI_ISL_2623880, EPI_ISL_2623881, EPI_ISL_2623882, EPI_ISL_2623883, EPI_ISL_2623884, EPI_ISL_2623885, EPI_ISL_2623886, EPI_ISL_2623887, EPI_ISL_2623888, EPI_ISL_2623889, EPI_ISL_2623890, EPI_ISL_2623891, EPI_ISL_2623892, EPI_ISL_2623893, EPI_ISL_2623894, EPI_ISL_2623895, EPI_ISL_2623896, EPI_ISL_2623897, EPI_ISL_2623898, EPI_ISL_2623899, EPI_ISL_2623900, EPI_ISL_2623901, EPI_ISL_2623902, EPI_ISL_2623903, EPI_ISL_2623904, EPI_ISL_2623905, EPI_ISL_2623906, EPI_ISL_2623907, EPI_ISL_2623908, EPI_ISL_2623909, EPI_ISL_2623910, EPI_ISL_2623911, EPI_ISL_2623912, EPI_ISL_2623913, EPI_ISL_2623914, EPI_ISL_2623915, EPI_ISL_2623916, EPI_ISL_2623917, EPI_ISL_2623918, EPI_ISL_2623920, EPI_ISL_2623921, EPI_ISL_2623923, EPI_ISL_2623926, EPI_ISL_2623927, EPI_ISL_2623928, EPI_ISL_2623929, EPI_ISL_2623930, EPI_ISL_2623932, EPI_ISL_2623933, EPI_ISL_2623934, EPI_ISL_2623936, EPI_ISL_2623937, EPI_ISL_2623939, EPI_ISL_2623940, EPI_ISL_2623941, EPI_ISL_2623943,

| | | | |
|---|---|---|---|
| EPI_ISL_2623944, EPI_ISL_2623945, EPI_ISL_2623946, EPI_ISL_2623947, EPI_ISL_2623948, EPI_ISL_2623949, EPI_ISL_2623951, EPI_ISL_2623952, EPI_ISL_2623953, EPI_ISL_2623954, EPI_ISL_2623957, EPI_ISL_2623958, EPI_ISL_2623960, EPI_ISL_2623961, EPI_ISL_2623962, EPI_ISL_2623963, EPI_ISL_2623964, EPI_ISL_2623966, EPI_ISL_2623967, EPI_ISL_2623968, EPI_ISL_2623969, EPI_ISL_2623970, EPI_ISL_2623971, EPI_ISL_2623972, EPI_ISL_2623973, EPI_ISL_2623974, EPI_ISL_2623975, EPI_ISL_2623976, EPI_ISL_2623977, EPI_ISL_2623978, EPI_ISL_2623979, EPI_ISL_2623980, EPI_ISL_2623981, EPI_ISL_2623982, EPI_ISL_2623983, EPI_ISL_2623984, EPI_ISL_2623985, EPI_ISL_2623986, EPI_ISL_2623987, EPI_ISL_2623988, EPI_ISL_2623989, EPI_ISL_2623991, EPI_ISL_2623992, EPI_ISL_2623993, EPI_ISL_2623995, EPI_ISL_2623996, EPI_ISL_2623998, EPI_ISL_2623999, EPI_ISL_2624000, EPI_ISL_2624002, EPI_ISL_2624003, EPI_ISL_2624006, EPI_ISL_2624007, EPI_ISL_2624008, EPI_ISL_2624010, EPI_ISL_2624011, EPI_ISL_2624013, EPI_ISL_2624017, EPI_ISL_2624019, EPI_ISL_2624020, EPI_ISL_2624023, EPI_ISL_2624026, EPI_ISL_2624027, EPI_ISL_2624028, EPI_ISL_2624029, EPI_ISL_2624030, EPI_ISL_2624031, EPI_ISL_2624032, EPI_ISL_2624033, EPI_ISL_2624034, EPI_ISL_2624035, EPI_ISL_2624036, EPI_ISL_2624037, EPI_ISL_2624039, EPI_ISL_2624040, EPI_ISL_2624041, EPI_ISL_2624042, EPI_ISL_2624043, EPI_ISL_2624044, EPI_ISL_2624045, EPI_ISL_2624046, EPI_ISL_2624047, EPI_ISL_2624049, EPI_ISL_2624050, EPI_ISL_2624053, EPI_ISL_2624054, EPI_ISL_2624055, EPI_ISL_2624056, EPI_ISL_2624057, EPI_ISL_2624061, EPI_ISL_2624062, EPI_ISL_2624063, EPI_ISL_2624064, EPI_ISL_2624065, EPI_ISL_2624067, EPI_ISL_2624068, EPI_ISL_2624069, EPI_ISL_2624070, EPI_ISL_2624071, EPI_ISL_2624072, EPI_ISL_2624075, EPI_ISL_2624077, EPI_ISL_2624078, EPI_ISL_2624080, EPI_ISL_2624083, EPI_ISL_2624085, EPI_ISL_2624086, EPI_ISL_2624087, EPI_ISL_2624088, EPI_ISL_2624089, EPI_ISL_2624090, EPI_ISL_2624093, EPI_ISL_2624094, EPI_ISL_2624095, EPI_ISL_2624096, EPI_ISL_2624097, EPI_ISL_2624098, EPI_ISL_2624099, EPI_ISL_2624100, EPI_ISL_2624101, EPI_ISL_2624102, EPI_ISL_2624103, EPI_ISL_2624104, EPI_ISL_2624105, EPI_ISL_2624107, EPI_ISL_2624108, EPI_ISL_2624109, EPI_ISL_2624110, EPI_ISL_2624111, EPI_ISL_2624112, EPI_ISL_2624113, EPI_ISL_2624114, EPI_ISL_2624115, EPI_ISL_2624116, EPI_ISL_2624117, EPI_ISL_2624118, EPI_ISL_2624119, EPI_ISL_2624120, EPI_ISL_2624121, EPI_ISL_2624123, EPI_ISL_2624125, EPI_ISL_2624126, EPI_ISL_2624127, EPI_ISL_2624128, EPI_ISL_2624129, EPI_ISL_2624130, EPI_ISL_2624131, EPI_ISL_2624132, EPI_ISL_2624133, EPI_ISL_2624134, EPI_ISL_2624135, EPI_ISL_2624136, EPI_ISL_2624137, EPI_ISL_2624138, EPI_ISL_2624140, EPI_ISL_2624141, EPI_ISL_2624142, EPI_ISL_2624143, EPI_ISL_2624144, EPI_ISL_2624145, EPI_ISL_2624146, EPI_ISL_2624147, EPI_ISL_2624148, EPI_ISL_2624149, EPI_ISL_2624150, EPI_ISL_2624151, EPI_ISL_2624152, EPI_ISL_2624154, EPI_ISL_2624155, EPI_ISL_2624156, EPI_ISL_2624157, EPI_ISL_2624158, EPI_ISL_2624159, EPI_ISL_2624160, EPI_ISL_2624162, EPI_ISL_2624165, EPI_ISL_2624169, EPI_ISL_2624170, EPI_ISL_2624176, EPI_ISL_2624177, EPI_ISL_2624181, EPI_ISL_2624182, EPI_ISL_2624183, EPI_ISL_2624184, EPI_ISL_2624185, EPI_ISL_2624186, EPI_ISL_2624187, EPI_ISL_2624188, EPI_ISL_2624190, EPI_ISL_2624191, EPI_ISL_2624192, EPI_ISL_2624193, EPI_ISL_2624194, EPI_ISL_2624195, EPI_ISL_2624196, EPI_ISL_2624198, EPI_ISL_2624199, EPI_ISL_2624200, EPI_ISL_2624201, EPI_ISL_2624202, EPI_ISL_2624203, EPI_ISL_2624204, EPI_ISL_2624205, EPI_ISL_2624207, EPI_ISL_2624208, EPI_ISL_2624211, EPI_ISL_2624212, EPI_ISL_2624214, EPI_ISL_2624215, EPI_ISL_2624216, EPI_ISL_2624217, EPI_ISL_2624218, EPI_ISL_2624219, EPI_ISL_2624220, EPI_ISL_2624221, EPI_ISL_2624222, EPI_ISL_2624223, EPI_ISL_2624224, EPI_ISL_2624226, EPI_ISL_2624227, EPI_ISL_2624228, EPI_ISL_2624229, EPI_ISL_2624230, EPI_ISL_2624231, EPI_ISL_2624232, EPI_ISL_2624235, EPI_ISL_2624236, EPI_ISL_2624237, EPI_ISL_2624238, EPI_ISL_2624239, EPI_ISL_2624241, EPI_ISL_2624242, EPI_ISL_2624243, EPI_ISL_2624244, EPI_ISL_2624245, EPI_ISL_2624247, EPI_ISL_2624248, EPI_ISL_2624249, EPI_ISL_2624250, EPI_ISL_2624253, EPI_ISL_2624254, EPI_ISL_2624255, EPI_ISL_2624256, EPI_ISL_2624257, EPI_ISL_2624258, EPI_ISL_2624261, EPI_ISL_2624262, EPI_ISL_2624263, EPI_ISL_2624264, EPI_ISL_2624265, EPI_ISL_2624266, EPI_ISL_2624268, EPI_ISL_2624269, EPI_ISL_2624271, EPI_ISL_2624272, EPI_ISL_2624273, EPI_ISL_2624274, EPI_ISL_2624275, EPI_ISL_2624276, EPI_ISL_2624277, EPI_ISL_2624278, EPI_ISL_2624279, EPI_ISL_2624280, EPI_ISL_2624281, EPI_ISL_2624282, EPI_ISL_2624283, EPI_ISL_2624286, EPI_ISL_2624288, EPI_ISL_2624290, EPI_ISL_2624291, EPI_ISL_2624292, EPI_ISL_2624293, EPI_ISL_2624294, EPI_ISL_2624295, EPI_ISL_2624296, EPI_ISL_2624297, EPI_ISL_2624298, EPI_ISL_2624299, EPI_ISL_2624300, EPI_ISL_2624302, EPI_ISL_2624304, EPI_ISL_2624305, EPI_ISL_2624306, EPI_ISL_2624307, EPI_ISL_2624308, EPI_ISL_2624311, EPI_ISL_2624313, EPI_ISL_2624314, EPI_ISL_2624315, EPI_ISL_2624316, EPI_ISL_2624317, EPI_ISL_2624319, EPI_ISL_2624321, EPI_ISL_2624322, EPI_ISL_2624323, EPI_ISL_2624325, EPI_ISL_2624326, EPI_ISL_2624327, EPI_ISL_2624328, EPI_ISL_2624329, EPI_ISL_2624330, EPI_ISL_2624331, EPI_ISL_2624332, EPI_ISL_2624333, EPI_ISL_2624334, EPI_ISL_2624335, EPI_ISL_2624336, EPI_ISL_2624337, EPI_ISL_2624338, EPI_ISL_2624339, EPI_ISL_2624340, EPI_ISL_2624341, EPI_ISL_2624342, EPI_ISL_2624343, EPI_ISL_2624344, EPI_ISL_2624345, EPI_ISL_2624346, EPI_ISL_2624347, EPI_ISL_2624348, EPI_ISL_2624349, EPI_ISL_2624350, EPI_ISL_2624351, EPI_ISL_2624353, EPI_ISL_2624354, EPI_ISL_2624355, EPI_ISL_2624358, EPI_ISL_2624359, EPI_ISL_2624360, EPI_ISL_2624361, EPI_ISL_2624362, EPI_ISL_2624365, EPI_ISL_2624366, EPI_ISL_2624367, EPI_ISL_2624368, EPI_ISL_2624369, EPI_ISL_2624370, EPI_ISL_2624371, EPI_ISL_2624372, EPI_ISL_2624373, EPI_ISL_2624374, EPI_ISL_2624375, EPI_ISL_2624376, EPI_ISL_2624377, EPI_ISL_2624378, EPI_ISL_2624379, EPI_ISL_2624380, EPI_ISL_2624384, EPI_ISL_2624386, EPI_ISL_2624387, EPI_ISL_2624388, EPI_ISL_2624390, EPI_ISL_2624392, EPI_ISL_2624393, EPI_ISL_2624394, EPI_ISL_2624395, EPI_ISL_2624396, EPI_ISL_2624397, EPI_ISL_2624398, EPI_ISL_2624400, EPI_ISL_2624401, EPI_ISL_2624402, EPI_ISL_2624404, EPI_ISL_2624405, EPI_ISL_2624406 | | | |
| see above | Quadram Institute Bioscience | COVID-19 Genomics UK (COG-UK) Consortium | Dave J. Baker, Gemma L. Kay, Alp Aydin, Thanh Le-Viet, Steven Rudder, Ana P. Tedim, Anastasia Kolyva, Maria Diaz, Leonardo de Oliveira Martins, Nabil-Fareed Alikhan, Lizzie Meadows, Rachael Stanley, Ngozi Elumogo, Muhammed Yasir, Nicholas M. Thomson, Alexander J Trotter, Rachel Gilroy, Samuel Bloomfield, Claire Stuart, Andrew Bell, Reenesh Prakash, Samir Dervisevic, Alison E. Mather, John Wain, Mark Webber, Andrew J. Page, Justin O'Grady |
| EPI_ISL_2624407, EPI_ISL_2624408, EPI_ISL_2624409, EPI_ISL_2624410, EPI_ISL_2624411, EPI_ISL_2624413, EPI_ISL_2624414, EPI_ISL_2624415, EPI_ISL_2624416, EPI_ISL_2624417, EPI_ISL_2624418, EPI_ISL_2624419, EPI_ISL_2624420, EPI_ISL_2624422, EPI_ISL_2624423, EPI_ISL_2624424, EPI_ISL_2624425, EPI_ISL_2624426, EPI_ISL_2624427, EPI_ISL_2624428, EPI_ISL_2624430, EPI_ISL_2624431, EPI_ISL_2624432, EPI_ISL_2624433, EPI_ISL_2624434, EPI_ISL_2624435, EPI_ISL_2624436, EPI_ISL_2624437, EPI_ISL_2624438, EPI_ISL_2624439, EPI_ISL_2624441, EPI_ISL_2624442, EPI_ISL_2624443, EPI_ISL_2624444, EPI_ISL_2624445, EPI_ISL_2624446, EPI_ISL_2624447, EPI_ISL_2624448, EPI_ISL_2624449, EPI_ISL_2624450, EPI_ISL_2624451, EPI_ISL_2624452, EPI_ISL_2624453, EPI_ISL_2624454, EPI_ISL_2624455, EPI_ISL_2624456, EPI_ISL_2624457, EPI_ISL_2624458, EPI_ISL_2624460, EPI_ISL_2624461, EPI_ISL_2624462, EPI_ISL_2624463, EPI_ISL_2624464, EPI_ISL_2624465, EPI_ISL_2624466, EPI_ISL_2624467, EPI_ISL_2624468, EPI_ISL_2624470, EPI_ISL_2624471, EPI_ISL_2624472, EPI_ISL_2624473, EPI_ISL_2624474, EPI_ISL_2624475, EPI_ISL_2624476, EPI_ISL_2624477, EPI_ISL_2624478, EPI_ISL_2624479, EPI_ISL_2624480, EPI_ISL_2624481, EPI_ISL_2624482, EPI_ISL_2624483, EPI_ISL_2624485, EPI_ISL_2624486, EPI_ISL_2624487, EPI_ISL_2624488, EPI_ISL_2624489, EPI_ISL_2624490, EPI_ISL_2624491, EPI_ISL_2624492, EPI_ISL_2624493, EPI_ISL_2624494, EPI_ISL_2624495, EPI_ISL_2624496, EPI_ISL_2624497, EPI_ISL_2624498, EPI_ISL_2624499, EPI_ISL_2624500, EPI_ISL_2624501, EPI_ISL_2624502, EPI_ISL_2624503, EPI_ISL_2624504, EPI_ISL_2624505, EPI_ISL_2624506, EPI_ISL_2624507, EPI_ISL_2624508, EPI_ISL_2624509, EPI_ISL_2624510, EPI_ISL_2624511, EPI_ISL_2624512, EPI_ISL_2624513, EPI_ISL_2624514, EPI_ISL_2624515, EPI_ISL_2624516, EPI_ISL_2624517, EPI_ISL_2624518, EPI_ISL_2624519, EPI_ISL_2624520, EPI_ISL_2624521, EPI_ISL_2624522, EPI_ISL_2624523, EPI_ISL_2624524 | | | |
| see above | Queens Medical Centre, Clinical Microbiology Department / DeepSeq Nottingham | COVID-19 Genomics UK (COG-UK) Consortium | Gemma Clark, Wendy Smith, Manjinder Khakh, Vicki M Fleming, Michelle M Lister, Hannah Howson-Wells, Jonathan Ball, Timothy Byaruhanga, Jayasree Dey, Emily Park, Jack Hill, Patrick McClure, Joseph Chappell, Theocharis Tsoleridis, Nadine Holmes, Matthew Carlisle, Christopher Moore, Fei Sang, Johnny Debebe, Victoria Wright, Matthew Loose |
| EPI_ISL_2624525, EPI_ISL_2624526, EPI_ISL_2624528, EPI_ISL_2624530, EPI_ISL_2624531, EPI_ISL_2624532, EPI_ISL_2624533, EPI_ISL_2624534, EPI_ISL_2624535, EPI_ISL_2624537, EPI_ISL_2624538, EPI_ISL_2624539, EPI_ISL_2624541, EPI_ISL_2624542, EPI_ISL_2624544, EPI_ISL_2624545, EPI_ISL_2624546, EPI_ISL_2624547, EPI_ISL_2624549, EPI_ISL_2624551, EPI_ISL_2624553, EPI_ISL_2624554, EPI_ISL_2624555, EPI_ISL_2624556, EPI_ISL_2624557, EPI_ISL_2624558, EPI_ISL_2624559, EPI_ISL_2624560, EPI_ISL_2624564, EPI_ISL_2624565, EPI_ISL_2624567, EPI_ISL_2624569, EPI_ISL_2624570, EPI_ISL_2624571, EPI_ISL_2624572, EPI_ISL_2624575, EPI_ISL_2624577, EPI_ISL_2624578, EPI_ISL_2624581, EPI_ISL_2624584, EPI_ISL_2624585, EPI_ISL_2624586 | | | |
| see above | Oxford Viromics, NDM, University of Oxford; Oxford University Hospitals; Basingstoke and North Hampshire Hospital | COVID-19 Genomics UK (COG-UK) Consortium | Tanya Golubchik, David Bonsall, George Macintyre, Amy Trebes, Mariateresa de Cesare, Catrin Moore, Alex Mobbs, Anita Justice, Robert Shaw, Monique Andersson, Timothy Peto, Emma Wise, Nathan Moore, Jessica Lynch, Nick Cortes, Matilde Mori, Stephen Kidd, David Buck, John Todd, Christophe Fraser |
| EPI_ISL_2624587, EPI_ISL_2624588, EPI_ISL_2624589, EPI_ISL_2624590, EPI_ISL_2624591, EPI_ISL_2624592, EPI_ISL_2624593, EPI_ISL_2624594, EPI_ISL_2624595, EPI_ISL_2624596, EPI_ISL_2624597, EPI_ISL_2624598, EPI_ISL_2624599, EPI_ISL_2624600, EPI_ISL_2624601, EPI_ISL_2624602, EPI_ISL_2624603, EPI_ISL_2624604, EPI_ISL_2624605, EPI_ISL_2624606, EPI_ISL_2624607, EPI_ISL_2624608, EPI_ISL_2624609, EPI_ISL_2624611, EPI_ISL_2624613, EPI_ISL_2624614, EPI_ISL_2624615, EPI_ISL_2624616, EPI_ISL_2624617, EPI_ISL_2624618, EPI_ISL_2624619, EPI_ISL_2624620, EPI_ISL_2624621, EPI_ISL_2624622, EPI_ISL_2624623, EPI_ISL_2624624, EPI_ISL_2624625, EPI_ISL_2624626, EPI_ISL_2624627, EPI_ISL_2624628, EPI_ISL_2624629, EPI_ISL_2624630, EPI_ISL_2624631, EPI_ISL_2624632, EPI_ISL_2624633, EPI_ISL_2624634, EPI_ISL_2624635, EPI_ISL_2624636, EPI_ISL_2624637, EPI_ISL_2624639, EPI_ISL_2624640, EPI_ISL_2624641, EPI_ISL_2624642, EPI_ISL_2624643, EPI_ISL_2624644, EPI_ISL_2624645, EPI_ISL_2624647, EPI_ISL_2624648, EPI_ISL_2624649, EPI_ISL_2624650, EPI_ISL_2624652, EPI_ISL_2624653, EPI_ISL_2624654, EPI_ISL_2624655, EPI_ISL_2624656, EPI_ISL_2624657, EPI_ISL_2624658, EPI_ISL_2624659, EPI_ISL_2624660, EPI_ISL_2624661, EPI_ISL_2624662, EPI_ISL_2624664, EPI_ISL_2624665, EPI_ISL_2624666, EPI_ISL_2624667, EPI_ISL_2624668, EPI_ISL_2624669, EPI_ISL_2624670, EPI_ISL_2624671, EPI_ISL_2624672, EPI_ISL_2624674, EPI_ISL_2624675, EPI_ISL_2624677, EPI_ISL_2624678, EPI_ISL_2624679, EPI_ISL_2624680, EPI_ISL_2624681, EPI_ISL_2624682, EPI_ISL_2624683, EPI_ISL_2624684, EPI_ISL_2624685, EPI_ISL_2624686, EPI_ISL_2624689, EPI_ISL_2624690, EPI_ISL_2624691, EPI_ISL_2624692, EPI_ISL_2624693, EPI_ISL_2624694, EPI_ISL_2624695, EPI_ISL_2624697, EPI_ISL_2624698, EPI_ISL_2624699, EPI_ISL_2624700, EPI_ISL_2624701, EPI_ISL_2624702, EPI_ISL_2624703, EPI_ISL_2624704, EPI_ISL_2624705, EPI_ISL_2624706, EPI_ISL_2624707, EPI_ISL_2624708, EPI_ISL_2624709, EPI_ISL_2624710, EPI_ISL_2624711, EPI_ISL_2624712, EPI_ISL_2624713, EPI_ISL_2624714, EPI_ISL_2624715, EPI_ISL_2624716, EPI_ISL_2624717, EPI_ISL_2624718, EPI_ISL_2624719, EPI_ISL_2624720, EPI_ISL_2624721, EPI_ISL_2624722, EPI_ISL_2624723, EPI_ISL_2624724, EPI_ISL_2624725, EPI_ISL_2624730, EPI_ISL_2624731, EPI_ISL_2624734, EPI_ISL_2624735, EPI_ISL_2624736, EPI_ISL_2624737, EPI_ISL_2624739, EPI_ISL_2624740, EPI_ISL_2624741, EPI_ISL_2624743, EPI_ISL_2624744, EPI_ISL_2624746, EPI_ISL_2624747, EPI_ISL_2624748, EPI_ISL_2624749, EPI_ISL_2624750, EPI_ISL_2624751, EPI_ISL_2624752, EPI_ISL_2624753, EPI_ISL_2624754, EPI_ISL_2624755, EPI_ISL_2624757, EPI_ISL_2624759, EPI_ISL_2624760, EPI_ISL_2624761, EPI_ISL_2624762, EPI_ISL_2624763, EPI_ISL_2624765, EPI_ISL_2624766, EPI_ISL_2624767, EPI_ISL_2624768, EPI_ISL_2624769, EPI_ISL_2624770, EPI_ISL_2624771, EPI_ISL_2624772, EPI_ISL_2624773, EPI_ISL_2624774, EPI_ISL_2624775, EPI_ISL_2624776, EPI_ISL_2624778, EPI_ISL_2624780, EPI_ISL_2624781, EPI_ISL_2624782, EPI_ISL_2624786, EPI_ISL_2624787, EPI_ISL_2624789, EPI_ISL_2624791, EPI_ISL_2624792, EPI_ISL_2624793, EPI_ISL_2624794, EPI_ISL_2624796, EPI_ISL_2624797, EPI_ISL_2624798, EPI_ISL_2624804, EPI_ISL_2624805, EPI_ISL_2624806, EPI_ISL_2624807, EPI_ISL_2624808, EPI_ISL_2624810, EPI_ISL_2624811, EPI_ISL_2624812, EPI_ISL_2624813, EPI_ISL_2624814, EPI_ISL_2624815, EPI_ISL_2624816, EPI_ISL_2624817, EPI_ISL_2624818, EPI_ISL_2624819, EPI_ISL_2624821, EPI_ISL_2624822, EPI_ISL_2624823, EPI_ISL_2624824, EPI_ISL_2624825, EPI_ISL_2624826, EPI_ISL_2624827, EPI_ISL_2624828, EPI_ISL_2624829, EPI_ISL_2624830, EPI_ISL_2624831, EPI_ISL_2624832, EPI_ISL_2624833, EPI_ISL_2624834, EPI_ISL_2624835, EPI_ISL_2624836, EPI_ISL_2624837, EPI_ISL_2624838, EPI_ISL_2624842, EPI_ISL_2624843, EPI_ISL_2624844, EPI_ISL_2624845, EPI_ISL_2624847, EPI_ISL_2624848, EPI_ISL_2624849, EPI_ISL_2624850, EPI_ISL_2624851, EPI_ISL_2624852, EPI_ISL_2624853, EPI_ISL_2624854, EPI_ISL_2624855, EPI_ISL_2624856, EPI_ISL_2624857, EPI_ISL_2624858, EPI_ISL_2624859, EPI_ISL_2624860, EPI_ISL_2624861, EPI_ISL_2624862, EPI_ISL_2624863, EPI_ISL_2624864, EPI_ISL_2624865, EPI_ISL_2624866, EPI_ISL_2624867, EPI_ISL_2624868, EPI_ISL_2624869, EPI_ISL_2624870, EPI_ISL_2624871, EPI_ISL_2624872, EPI_ISL_2624873, EPI_ISL_2624874, EPI_ISL_2624875, EPI_ISL_2624876, EPI_ISL_2624877, EPI_ISL_2624881, EPI_ISL_2624883, EPI_ISL_2624884, EPI_ISL_2624888, EPI_ISL_2624889, EPI_ISL_2624891, EPI_ISL_2624892, EPI_ISL_2624893, EPI_ISL_2624894, EPI_ISL_2624896, EPI_ISL_2624898, EPI_ISL_2624899, EPI_ISL_2624900, EPI_ISL_2624901, EPI_ISL_2624902, EPI_ISL_2624903, EPI_ISL_2624904, EPI_ISL_2624905, EPI_ISL_2624906, EPI_ISL_2624908, EPI_ISL_2624909, EPI_ISL_2624910, EPI_ISL_2624912, EPI_ISL_2624914, EPI_ISL_2624915, EPI_ISL_2624917, EPI_ISL_2624918, EPI_ISL_2624919, EPI_ISL_2624920, EPI_ISL_2624921, EPI_ISL_2624922, EPI_ISL_2624923, EPI_ISL_2624925, EPI_ISL_2624926, EPI_ISL_2624929 | | | |
| see above | Originating lab: Wales Specialist Virology Centre Sequencing lab: Pathogen Genomics Unit | Public Health Wales Microbiology Cardiff Wales Specialist Virology Centre | Catherine Moore, Johnathan Evans, Laura Gifford, Malorie Perry, Simon Cottrell, Angela Marchbank, Alec Birchley, Alexander Adams, Amy Gaskin, Bree Gatica-Wilcox, Jason Coombes, Joel Southgate, Lauren Gilbert, Lee Graham, Nicole Pacchiarini, Sara Kumziene-Summerhayes, Sarah Taylor, Sophie Jones, Sara Rey, Matthew Bull, Joanne Watkins, Sally Corden, Tom Connor |
| EPI_ISL_2624930, EPI_ISL_2624931, EPI_ISL_2624932, EPI_ISL_2624933, EPI_ISL_2624934, EPI_ISL_2624935, EPI_ISL_2624936, EPI_ISL_2624937, EPI_ISL_2624938, EPI_ISL_2624939, EPI_ISL_2624940, EPI_ISL_2624941, EPI_ISL_2624942, EPI_ISL_2624943, EPI_ISL_2624944, EPI_ISL_2624945, EPI_ISL_2624946, EPI_ISL_2624947, EPI_ISL_2624948, EPI_ISL_2624949, EPI_ISL_2624950, EPI_ISL_2624951, EPI_ISL_2624952, EPI_ISL_2624953, EPI_ISL_2624954, EPI_ISL_2624955, EPI_ISL_2624956, EPI_ISL_2624957, EPI_ISL_2624958, EPI_ISL_2624959, EPI_ISL_2624960, EPI_ISL_2624961, EPI_ISL_2624962, EPI_ISL_2624963, EPI_ISL_2624964, EPI_ISL_2624965, EPI_ISL_2624966, EPI_ISL_2624968, EPI_ISL_2624969, EPI_ISL_2624970, EPI_ISL_2624971, EPI_ISL_2624972, EPI_ISL_2624973, EPI_ISL_2624974, EPI_ISL_2624975, EPI_ISL_2624976, EPI_ISL_2624977, EPI_ISL_2624978, EPI_ISL_2624979, EPI_ISL_2624981, EPI_ISL_2624982, EPI_ISL_2624983, EPI_ISL_2624984, EPI_ISL_2624985, EPI_ISL_2624986, EPI_ISL_2624987, EPI_ISL_2624988, EPI_ISL_2624989, EPI_ISL_2624990, EPI_ISL_2624991, EPI_ISL_2624992, EPI_ISL_2624993, EPI_ISL_2624994, EPI_ISL_2624995, EPI_ISL_2624996, EPI_ISL_2624997, EPI_ISL_2624998, EPI_ISL_2624999, EPI_ISL_2625000, EPI_ISL_2625001, EPI_ISL_2625002, EPI_ISL_2625003, EPI_ISL_2625004, EPI_ISL_2625005, EPI_ISL_2625006, EPI_ISL_2625007, EPI_ISL_2625008, EPI_ISL_2625009, EPI_ISL_2625010, EPI_ISL_2625011, EPI_ISL_2625012, EPI_ISL_2625013, EPI_ISL_2625014, EPI_ISL_2625015, EPI_ISL_2625016, EPI_ISL_2625017, EPI_ISL_2625018, EPI_ISL_2625019, EPI_ISL_2625020, EPI_ISL_2625022, EPI_ISL_2625023, EPI_ISL_2625024, EPI_ISL_2625025, EPI_ISL_2625026, EPI_ISL_2625027, EPI_ISL_2625028, EPI_ISL_2625029, EPI_ISL_2625030, EPI_ISL_2625031, EPI_ISL_2625032, EPI_ISL_2625033, EPI_ISL_2625034, EPI_ISL_2625035, EPI_ISL_2625036, EPI_ISL_2625037, EPI_ISL_2625038, EPI_ISL_2625039, EPI_ISL_2625040, EPI_ISL_2625041, EPI_ISL_2625042, EPI_ISL_2625043, EPI_ISL_2625044, EPI_ISL_2625045, EPI_ISL_2625046, EPI_ISL_2625047, EPI_ISL_2625048, EPI_ISL_2625049, EPI_ISL_2625050, EPI_ISL_2625051, EPI_ISL_2625052, EPI_ISL_2625053, EPI_ISL_2625054, EPI_ISL_2625055, EPI_ISL_2625057, EPI_ISL_2625058, EPI_ISL_2625059, EPI_ISL_2625060, EPI_ISL_2625061, EPI_ISL_2625062, EPI_ISL_2625063, EPI_ISL_2625064, EPI_ISL_2625065, EPI_ISL_2625066, EPI_ISL_2625068, EPI_ISL_2625069, EPI_ISL_2625070, EPI_ISL_2625071, EPI_ISL_2625072, EPI_ISL_2625073, EPI_ISL_2625074, EPI_ISL_2625075, EPI_ISL_2625076, EPI_ISL_2625077, EPI_ISL_2625078, EPI_ISL_2625079, EPI_ISL_2625080, EPI_ISL_2625081, EPI_ISL_2625082, EPI_ISL_2625083, EPI_ISL_2625084, EPI_ISL_2625085, EPI_ISL_2625086, EPI_ISL_2625087, EPI_ISL_2625088, EPI_ISL_2625089, EPI_ISL_2625090, EPI_ISL_2625091, EPI_ISL_2625092, EPI_ISL_2625093, EPI_ISL_2625094, EPI_ISL_2625095, EPI_ISL_2625096, EPI_ISL_2625097, EPI_ISL_2625098, EPI_ISL_2625099, EPI_ISL_2625100, EPI_ISL_2625101, EPI_ISL_2625102, EPI_ISL_2625103, EPI_ISL_2625104, EPI_ISL_2625105, EPI_ISL_2625106, EPI_ISL_2625107, EPI_ISL_2625108, EPI_ISL_2625109, EPI_ISL_2625110, EPI_ISL_2625111, EPI_ISL_2625112, EPI_ISL_2625113, EPI_ISL_2625114, EPI_ISL_2625115, EPI_ISL_2625116, EPI_ISL_2625117, EPI_ISL_2625118, EPI_ISL_2625119, EPI_ISL_2625120, EPI_ISL_2625121, EPI_ISL_2625122, EPI_ISL_2625123, EPI_ISL_2625124, EPI_ISL_2625125, EPI_ISL_2625126, EPI_ISL_2625127, EPI_ISL_2625128, EPI_ISL_2625129, EPI_ISL_2625130, EPI_ISL_2625131, EPI_ISL_2625132, EPI_ISL_2625133, EPI_ISL_2625134, EPI_ISL_2625135, EPI_ISL_2625136, EPI_ISL_2625137, EPI_ISL_2625138, EPI_ISL_2625140, EPI_ISL_2625141, EPI_ISL_2625142, EPI_ISL_2625143, EPI_ISL_2625144, EPI_ISL_2625145, EPI_ISL_2625146, EPI_ISL_2625147, EPI_ISL_2625148, EPI_ISL_2625149, EPI_ISL_2625150, EPI_ISL_2625151, EPI_ISL_2625152, EPI_ISL_2625153, EPI_ISL_2625154, EPI_ISL_2625155, EPI_ISL_2625156, EPI_ISL_2625157, EPI_ISL_2625158, EPI_ISL_2625159, EPI_ISL_2625160, EPI_ISL_2625161, EPI_ISL_2625162, EPI_ISL_2625163, EPI_ISL_2625164, EPI_ISL_2625165, EPI_ISL_2625166, EPI_ISL_2625167, EPI_ISL_2625168, EPI_ISL_2625169, EPI_ISL_2625170, EPI_ISL_2625171, EPI_ISL_2625172, EPI_ISL_2625173, EPI_ISL_2625174, EPI_ISL_2625175, EPI_ISL_2625176, EPI_ISL_2625177, EPI_ISL_2625178, EPI_ISL_2625179, EPI_ISL_2625180, EPI_ISL_2625181, EPI_ISL_2625182, EPI_ISL_2625183, EPI_ISL_2625184, EPI_ISL_2625185, EPI_ISL_2625186, EPI_ISL_2625187, EPI_ISL_2625188, EPI_ISL_2625189, EPI_ISL_2625190, EPI_ISL_2625191, EPI_ISL_2625192, EPI_ISL_2625193, EPI_ISL_2625194, EPI_ISL_2625195, EPI_ISL_2625196, EPI_ISL_2625197, EPI_ISL_2625198, EPI_ISL_2625199, EPI_ISL_2625200, EPI_ISL_2625201, EPI_ISL_2625202, EPI_ISL_2625203, EPI_ISL_2625204, EPI_ISL_2625205, EPI_ISL_2625206, EPI_ISL_2625208 | | | |
| see above | Centre for Enzyme Innovation, University of Portsmouth / Translational Research Laboratory, Portsmouth Hospitals NHS Trust / Hampshire Hospitals NHS Foundation Trust / Bournemouth University / University Hospitals Dorset NHS Foundation Trust / Brighton and Sussex University Hospitals NHS Trust / Dartford and Gravesham NHS Trust / Isle of Wight NHS Trust / Maidstone and Tunbridge Wells NHS Trust / University Hospital Southampton NHS Foundation Trust | COVID-19 Genomics UK (COG-UK) Consortium | Samuel Robson, Angela Beckett, Salman Goudarzi, Christopher Fearn, Kate Cook, Katie Loveson, Sharon Glaysher, Scott Elliott, Kelly Bicknell, Sarah Wyllie, Allyson Lloyd, Robert Impey, Anoop Chauhan, Stephen Kidd, Nathan Moore, Nick Cortes, Claire Thomas, Anna Mantzaouratou, Sarah Buchan, Magdalena Barrow, Andrew Butt, Liz Sheridan, Jonnie Seymour, Dorian Crudgington, Ben Macklin, Mohammed Hassan-Ibrahim, Cassandra Malone, Benjamin Cogger, Kevin Tucker, Samirakhon Raupova, Rachael Jeremiah, Anibolina Castigador, Emily Macnaughton, Karen Withell, Kordo Saeed, Jacqui Prieto, Adhyana Mahanama, Buddhini Samaraweera, Siona Silvieri, Emanuela Pelosi, Eleri Wilson-Davies, Sarah Jeremiah, Helen Wheeler, Matthew Harvey, Thea Sass, Helen Umpleby, Stephen Aplin |
| EPI_ISL_2625209, EPI_ISL_2625211, EPI_ISL_2625213, EPI_ISL_2625214, EPI_ISL_2625215, EPI_ISL_2625216, EPI_ISL_2625217, EPI_ISL_2625218, EPI_ISL_2625219, EPI_ISL_2625220, EPI_ISL_2625221, EPI_ISL_2625223, EPI_ISL_2625224, EPI_ISL_2625226, EPI_ISL_2625227, EPI_ISL_2625228, EPI_ISL_2625229, EPI_ISL_2625230, EPI_ISL_2625232, EPI_ISL_2625234, EPI_ISL_2625235, EPI_ISL_2625236, EPI_ISL_2625237, EPI_ISL_2625238, EPI_ISL_2625239, EPI_ISL_2625240, EPI_ISL_2625241, EPI_ISL_2625242, EPI_ISL_2625243, EPI_ISL_2625244, EPI_ISL_2625245, EPI_ISL_2625246, EPI_ISL_2625247, EPI_ISL_2625248, EPI_ISL_2625249, EPI_ISL_2625250, EPI_ISL_2625251, EPI_ISL_2625252, EPI_ISL_2625253, EPI_ISL_2625254, EPI_ISL_2625255, EPI_ISL_2625256, EPI_ISL_2625257, EPI_ISL_2625258, EPI_ISL_2625259, EPI_ISL_2625260, EPI_ISL_2625261, EPI_ISL_2625262, EPI_ISL_2625263, EPI_ISL_2625264, EPI_ISL_2625265, EPI_ISL_2625266, EPI_ISL_2625267, EPI_ISL_2625268, EPI_ISL_2625269, EPI_ISL_2625270, EPI_ISL_2625271, EPI_ISL_2625272, EPI_ISL_2625273, EPI_ISL_2625274, EPI_ISL_2625275, EPI_ISL_2625276, EPI_ISL_2625277, EPI_ISL_2625278, EPI_ISL_2625279, EPI_ISL_2625281, EPI_ISL_2625282, EPI_ISL_2625283, EPI_ISL_2625284, EPI_ISL_2625285, EPI_ISL_2625286, EPI_ISL_2625287, EPI_ISL_2625288, EPI_ISL_2625289, EPI_ISL_2625290, EPI_ISL_2625291, EPI_ISL_2625292, EPI_ISL_2625293, EPI_ISL_2625294, EPI_ISL_2625295, EPI_ISL_2625297, EPI_ISL_2625298, EPI_ISL_2625299, EPI_ISL_2625300, EPI_ISL_2625301, EPI_ISL_2625302, EPI_ISL_2625303, EPI_ISL_2625304, EPI_ISL_2625305, EPI_ISL_2625307, EPI_ISL_2625308, EPI_ISL_2625310, EPI_ISL_2625311, EPI_ISL_2625312, EPI_ISL_2625313, EPI_ISL_2625314, EPI_ISL_2625316, EPI_ISL_2625317, EPI_ISL_2625318, EPI_ISL_2625319, EPI_ISL_2625320, EPI_ISL_2625321, EPI_ISL_2625322, EPI_ISL_2625323, EPI_ISL_2625324, EPI_ISL_2625326, EPI_ISL_2625327, EPI_ISL_2625329, EPI_ISL_2625330, EPI_ISL_2625331, EPI_ISL_2625332, EPI_ISL_2625334, EPI_ISL_2625335, EPI_ISL_2625338, EPI_ISL_2625339, EPI_ISL_2625340, EPI_ISL_2625342, EPI_ISL_2625343, EPI_ISL_2625344, EPI_ISL_2625345, EPI_ISL_2625346, EPI_ISL_2625347, EPI_ISL_2625348, EPI_ISL_2625349, EPI_ISL_2625350, EPI_ISL_2625351, EPI_ISL_2625352, EPI_ISL_2625353, EPI_ISL_2625354, EPI_ISL_2625356, EPI_ISL_2625358, EPI_ISL_2625359, EPI_ISL_2625360, EPI_ISL_2625361, EPI_ISL_2625364, EPI_ISL_2625365, EPI_ISL_2625366, EPI_ISL_2625367, EPI_ISL_2625368, EPI_ISL_2625369, EPI_ISL_2625370, EPI_ISL_2625371, EPI_ISL_2625372, EPI_ISL_2625373, EPI_ISL_2625374, EPI_ISL_2625375, EPI_ISL_2625377, EPI_ISL_2625378, EPI_ISL_2625379, EPI_ISL_2625380, EPI_ISL_2625381, EPI_ISL_2625382, EPI_ISL_2625384, EPI_ISL_2625385, EPI_ISL_2625386, EPI_ISL_2625387, EPI_ISL_2625388, EPI_ISL_2625389, EPI_ISL_2625390, EPI_ISL_2625391, EPI_ISL_2625392, EPI_ISL_2625393, EPI_ISL_2625394, EPI_ISL_2625395, EPI_ISL_2625396, EPI_ISL_2625397, EPI_ISL_2625399, EPI_ISL_2625400, EPI_ISL_2625402, EPI_ISL_2625403, EPI_ISL_2625404, EPI_ISL_2625405, EPI_ISL_2625406, EPI_ISL_2625407, EPI_ISL_2625408, EPI_ISL_2625409, EPI_ISL_2625410, EPI_ISL_2625411, EPI_ISL_2625412, EPI_ISL_2625413, EPI_ISL_2625414, EPI_ISL_2625415, EPI_ISL_2625416, EPI_ISL_2625417, EPI_ISL_2625419, EPI_ISL_2625420, EPI_ISL_2625421, EPI_ISL_2625422, EPI_ISL_2625423, EPI_ISL_2625424, EPI_ISL_2625425, EPI_ISL_2625426, EPI_ISL_2625427, EPI_ISL_2625429, EPI_ISL_2625430, EPI_ISL_2625431, EPI_ISL_2625432, EPI_ISL_2625433, EPI_ISL_2625434, EPI_ISL_2625435, EPI_ISL_2625436, EPI_ISL_2625437, EPI_ISL_2625438, EPI_ISL_2625439, EPI_ISL_2625440, EPI_ISL_2625441, EPI_ISL_2625443, EPI_ISL_2625444, EPI_ISL_2625445, EPI_ISL_2625448, EPI_ISL_2625449, EPI_ISL_2625450, EPI_ISL_2625451, EPI_ISL_2625453, EPI_ISL_2625454, EPI_ISL_2625455, EPI_ISL_2625456, EPI_ISL_2625457, EPI_ISL_2625459, EPI_ISL_2625460, EPI_ISL_2625461, EPI_ISL_2625463, EPI_ISL_2625464, EPI_ISL_2625466, EPI_ISL_2625467, EPI_ISL_2625468, EPI_ISL_2625470, EPI_ISL_2625472, EPI_ISL_2625473, EPI_ISL_2625474, EPI_ISL_2625475, EPI_ISL_2625476, EPI_ISL_2625477, EPI_ISL_2625478, EPI_ISL_2625479, EPI_ISL_2625480, EPI_ISL_2625481, EPI_ISL_2625482, EPI_ISL_2625483, EPI_ISL_2625485, EPI_ISL_2625486, EPI_ISL_2625487, EPI_ISL_2625488, EPI_ISL_2625489, EPI_ISL_2625490, EPI_ISL_2625492, EPI_ISL_2625493, EPI_ISL_2625495, EPI_ISL_2625496, EPI_ISL_2625497, EPI_ISL_2625498, EPI_ISL_2625499, EPI_ISL_2625500, EPI_ISL_2625501, EPI_ISL_2625502, EPI_ISL_2625503, EPI_ISL_2625504, EPI_ISL_2625505, EPI_ISL_2625506, EPI_ISL_2625507, EPI_ISL_2625508, EPI_ISL_2625509, EPI_ISL_2625510, EPI_ISL_2625511, EPI_ISL_2625513, EPI_ISL_2625514, EPI_ISL_2625515, EPI_ISL_2625516, EPI_ISL_2625517, EPI_ISL_2625518, EPI_ISL_2625519, EPI_ISL_2625520, EPI_ISL_2625521, EPI_ISL_2625522, EPI_ISL_2625523, EPI_ISL_2625525, EPI_ISL_2625526, EPI_ISL_2625530, EPI_ISL_2625531, EPI_ISL_2625532, EPI_ISL_2625534, EPI_ISL_2625535, EPI_ISL_2625536, EPI_ISL_2625537, EPI_ISL_2625538, EPI_ISL_2625539, EPI_ISL_2625540, EPI_ISL_2625541, EPI_ISL_2625542, EPI_ISL_2625543, EPI_ISL_2625544, EPI_ISL_2625545, EPI_ISL_2625546, EPI_ISL_2625547, EPI_ISL_2625548, EPI_ISL_2625549, EPI_ISL_2625550, EPI_ISL_2625553, EPI_ISL_2625554, EPI_ISL_2625555, EPI_ISL_2625557, EPI_ISL_2625558, EPI_ISL_2625559, EPI_ISL_2625560, EPI_ISL_2625561, EPI_ISL_2625562, EPI_ISL_2625563, EPI_ISL_2625564, EPI_ISL_2625565, EPI_ISL_2625566, EPI_ISL_2625567, EPI_ISL_2625569, EPI_ISL_2625570, EPI_ISL_2625571, EPI_ISL_2625572, EPI_ISL_2625573, EPI_ISL_2625574, EPI_ISL_2625576, EPI_ISL_2625577, EPI_ISL_2625578, EPI_ISL_2625579, EPI_ISL_2625580, EPI_ISL_2625581, EPI_ISL_2625582, EPI_ISL_2625583, EPI_ISL_2625584, EPI_ISL_2625585, EPI_ISL_2625586, EPI_ISL_2625588, EPI_ISL_2625589, EPI_ISL_2625590, EPI_ISL_2625591, EPI_ISL_2625592, EPI_ISL_2625593, EPI_ISL_2625594, EPI_ISL_2625596, EPI_ISL_2625599, EPI_ISL_2625600, EPI_ISL_2625601, EPI_ISL_2625602, EPI_ISL_2625604, EPI_ISL_2625605, EPI_ISL_2625608, EPI_ISL_2625609, EPI_ISL_2625610, EPI_ISL_2625611, EPI_ISL_2625612, EPI_ISL_2625613, EPI_ISL_2625614, EPI_ISL_2625615, EPI_ISL_2625616, EPI_ISL_2625618, EPI_ISL_2625619, EPI_ISL_2625620 | | | |
| see above | Virology Department, Sheffield Teaching Hospitals NHS Foundation Trust/Department of Infection, Immunity and Cardiovascular Disease, The Medical School, University of Sheffield | COVID-19 Genomics UK (COG-UK) Consortium | Thushan de Silva, Matthew Parker, Nikki Smith, Adri Angyal, Rebecca Brown, Luke Green, Rachel Tucker, Paul Parsons, Danielle Groves, Katie Johnson, Laura Carrilero, Alex Keeley, Dave Partridge, Matthew Wyles, Benjamin Lindsey, Mehmet Yavuz, Mohammad Raza, Cariad Evans |
| EPI_ISL_2625622 | Usansolo-Galdakao University Hospital | Cruces University Hospital | Ana Gual-de-Torrella, Izaskun Alejo-Cancho, Mikel Gallego, Ana Belén de la Hoz |
| EPI_ISL_2626010, EPI_ISL_2626012, EPI_ISL_2626015, EPI_ISL_2626021, EPI_ISL_2626023, EPI_ISL_2626028, EPI_ISL_2626031, EPI_ISL_2626032, EPI_ISL_2626037 | Cliniques universitaires Saint-Luc | UCLouvain/IREC/MBLG | Jean Ruelle, Bertrand Bearzatto, Benoit Kabamba Mukadi |
| EPI_ISL_2626062, EPI_ISL_2626067, EPI_ISL_2626068, EPI_ISL_2626072 | Clinique Saint-Pierre Ottignies | UCLouvain/IREC/MBLG | Jean Ruelle, Bertrand Bearzatto, Benoit Kabamba Mukadi |
| EPI_ISL_2626083, EPI_ISL_2626084, EPI_ISL_2626085, EPI_ISL_2626086, EPI_ISL_2626087, EPI_ISL_2626088, EPI_ISL_2626089, EPI_ISL_2626090, EPI_ISL_2626091, EPI_ISL_2626092, EPI_ISL_2626093, EPI_ISL_2626094, EPI_ISL_2626095, EPI_ISL_2626096 | | | |

| | | | |
|---|---|---|---|
| see above | Cliniques universitaires Saint-Luc | UCLouvain/IREC/MBLG | Jean Ruelle, Bertrand Bearzatto, Benoit Kabamba Mukadi |
| EPI_ISL_2626102, EPI_ISL_2626103 | HOSPITAL UNIVERSITARIO SON ESPASES | HOSPITAL UNIVERSITARIO SON ESPASES | Servicio de Microbiología, Hospital Universitario Son Espases |
| EPI_ISL_2626115 | HOSPITAL MATEU ORFILA | HOSPITAL UNIVERSITARIO SON ESPASES | Servicio de Microbiología, Hospital Universitario Son Espases |
| EPI_ISL_2626116 | HOSPITAL MANACOR | HOSPITAL UNIVERSITARIO SON ESPASES | Servicio de Microbiología, Hospital Universitario Son Espases |
| EPI_ISL_2626126 | HOSPITAL MATEU ORFILA | HOSPITAL UNIVERSITARIO SON ESPASES | Servicio de Microbiología, Hospital Universitario Son Espases |
| EPI_ISL_2626132, EPI_ISL_2626141, EPI_ISL_2626144, EPI_ISL_2626150, EPI_ISL_2626151, EPI_ISL_2626152 | HOSPITAL UNIVERSITARIO SON ESPASES | HOSPITAL UNIVERSITARIO SON ESPASES | Servicio de Microbiología, Hospital Universitario Son Espases |
| EPI_ISL_2626153 | HOSPITAL CAN MISSES | HOSPITAL UNIVERSITARIO SON ESPASES | Servicio de Microbiología, Hospital Universitario Son Espases |
| EPI_ISL_2626158 | HOSPITAL UNIVERSITARIO SON ESPASES | HOSPITAL UNIVERSITARIO SON ESPASES | Servicio de Microbiología, Hospital Universitario Son Espases |
| EPI_ISL_2626166 | HOSPITAL MATEU ORFILA | HOSPITAL UNIVERSITARIO SON ESPASES | Servicio de Microbiología, Hospital Universitario Son Espases |
| EPI_ISL_2626173 | HOSPITAL UNIVERSITARIO SON ESPASES | HOSPITAL UNIVERSITARIO SON ESPASES | Servicio de Microbiología, Hospital Universitario Son Espases |
| EPI_ISL_2626176 | HOSPITAL CAN MISSES | HOSPITAL UNIVERSITARIO SON ESPASES | Servicio de Microbiología, Hospital Universitario Son Espases |
| EPI_ISL_2626180 | Usansolo-Galdakao University Hospital | Cruces University Hospital | Ana Gual-de-Torrella, Izaskun Alejo-Cancho, Mikel Gallego, Ana Belén de la Hoz |
| EPI_ISL_2626269, EPI_ISL_2626279, EPI_ISL_2626319, EPI_ISL_2626324, EPI_ISL_2626341, EPI_ISL_2626392, EPI_ISL_2626393, EPI_ISL_2626415, EPI_ISL_2626416, EPI_ISL_2626422, EPI_ISL_2626423, EPI_ISL_2626424, EPI_ISL_2626425, EPI_ISL_2626429 | | | |
| see above | Molecular diagnostic laboratory of Federal Budget Institution of Science "Central Research Institute of Epidemiology" of The Federal Service on Customers' Rights Protection and Human Well-being Surveillance | Group of Genomics and Postgenomic Technologies of Central Research Institute of Epidemiology | Samoilov A.E., Kaptelova V.V., Korneenko E.V., Sinitsyn S.O., Saenko S.S., Nadtoka M.I., Speranskaya A.S., Tivanova E.V., Kondrasheva L.Y., Smirnova Y.S., Akimkin V.G. |
| EPI_ISL_2626465 | MIRIALIS CLUSES BECHET | CNR Virus des Infections Respiratoires - France SUD | Antonin Bal, Gregory Destras, Gwendolyne Burfin, Hadrien Regue, Quentin Semanas, Martine Valette, Bruno Lina, Laurence Josset |
| EPI_ISL_2626595, EPI_ISL_2626596, EPI_ISL_2626597, EPI_ISL_2626598, EPI_ISL_2626599, EPI_ISL_2626600, EPI_ISL_2626601, EPI_ISL_2626602, EPI_ISL_2626603, EPI_ISL_2626604, EPI_ISL_2626605, EPI_ISL_2626606, EPI_ISL_2626607, EPI_ISL_2626608, EPI_ISL_2626609, EPI_ISL_2626610, EPI_ISL_2626611, EPI_ISL_2626612, EPI_ISL_2626613, EPI_ISL_2626614, EPI_ISL_2626615, EPI_ISL_2626616, EPI_ISL_2626617, EPI_ISL_2626618, EPI_ISL_2626619, EPI_ISL_2626620 | | | |
| see above | Genomik Solidaritas Indonesia Laboratorium | Genomik Solidaritas Indonesia Laboratorium | Alfin Mohammad Abdillah, Normastuti Adhini Tantyo, Dhahlia Agustina Cahyono, Carissa Sintca Wijaya, Vania Gavrila Wikasa, Ahmad Zoebad Foeady, Annisa Muthiah Sukirman, Gracia Felias Enos Korompis, Anuraj Shankar, Meutia Ayuputeri Kumaheri, Ariel Pradipta |
| EPI_ISL_2626707, EPI_ISL_2626714, EPI_ISL_2626725, EPI_ISL_2626734, EPI_ISL_2626736, EPI_ISL_2626739, EPI_ISL_2626740, EPI_ISL_2626754, EPI_ISL_2626755 | Alaska State Virology Laboratory | Alaska State Virology Laboratory | Stephanie DeRonde, Elva House, Jacob Zidek, Lisa Smith, Ph.D., Jack Chen, Ph.D. |
| EPI_ISL_2626772, EPI_ISL_2626779, EPI_ISL_2626804, EPI_ISL_2626814, EPI_ISL_2626819, EPI_ISL_2626822, EPI_ISL_2626823, EPI_ISL_2626828, EPI_ISL_2626832, EPI_ISL_2626834 | Baylor Scott & White-Temple | Baylor Scott & White-Temple | Ari Rao, Linden Morales, Kimberly Walker, Marcus Volz, Shelby Johnson |
| EPI_ISL_2626839, EPI_ISL_2626840, EPI_ISL_2626841, EPI_ISL_2626842, EPI_ISL_2626843, EPI_ISL_2626844, EPI_ISL_2626845, EPI_ISL_2626846, EPI_ISL_2626847, EPI_ISL_2626848, EPI_ISL_2626849, EPI_ISL_2626850, EPI_ISL_2626851 | | | |
| see above | Genomik Solidaritas Indonesia Laboratorium | Genomik Solidaritas Indonesia Laboratorium | Normastuti Adhini Tantyo, Dhahlia Agustina Cahyono, Alfin Mohammad Abdillah, Carissa Sintca Wijaya, Vania Gavrila Wikasa, Ahmad Zoebad Foeady, Annisa Muthiah Sukirman, Gracia Felias Enos Korompis, Anuraj Shankar, Meutia Ayuputeri Kumaheri, Ariel Pradipta |
| EPI_ISL_2626854, EPI_ISL_2626858, EPI_ISL_2626861, EPI_ISL_2626867, EPI_ISL_2626869, EPI_ISL_2626870, EPI_ISL_2626890, EPI_ISL_2626894, EPI_ISL_2626900, EPI_ISL_2626901, EPI_ISL_2626902 | | | |
| see above | Platform BIS UZA/UAntwerpen | Labo Klinische Biologie, UZA | Marie Le Mercier, Jasmine Coppens, Basil Britto Xavier, Christine Lammens, Veerle Matheeussen, Herman Goossens |
| EPI_ISL_2626920 | Algemeen Medisch Labo | Labo Klinische Biologie, UZA | Marie Le Mercier, Jasmine Coppens, Basil Britto Xavier, Christine Lammens, Veerle Matheeussen, Herman Goossens |
| EPI_ISL_2626924 | WESTCHESTER MEDICAL CENTER | Wadsworth Center, New York State Department of Health | Kirsten St. George, Daryl M. Lamson, Alexis Russell, Matthew Shudt, Melissa A Leisner, Jonathan Plitnick, Catharine Prussing, Navjot Singh, John Kelly, Erasmus Schneider, Erica Lasek-Nesselquist |
| EPI_ISL_2626947, EPI_ISL_2626955, EPI_ISL_2626969 | TEMPUS LABS INC | Wadsworth Center, New York State Department of Health | Kirsten St. George, Daryl M. Lamson, Alexis Russell, Matthew Shudt, Melissa A Leisner, Jonathan Plitnick, Catharine Prussing, Navjot Singh, John Kelly, Erasmus Schneider, Erica Lasek-Nesselquist |
| EPI_ISL_2626983, EPI_ISL_2627087, EPI_ISL_2627112 | URMC LABS | Wadsworth Center, New York State Department of Health | Kirsten St. George, Daryl M. Lamson, Alexis Russell, Matthew Shudt, Melissa A Leisner, Jonathan Plitnick, Catharine Prussing, Navjot Singh, John Kelly, Erasmus Schneider, Erica Lasek-Nesselquist |
| EPI_ISL_2627139 | WESTCHESTER MEDICAL CENTER | Wadsworth Center, New York State Department of Health | Kirsten St. George, Daryl M. Lamson, Alexis Russell, Matthew Shudt, Melissa A Leisner, Jonathan Plitnick, Catharine Prussing, Navjot Singh, John Kelly, Erasmus Schneider, Erica Lasek-Nesselquist |
| EPI_ISL_2627178 | NORTHWELL HEALTH LABORATORIES | Wadsworth Center, New York State Department of Health | Kirsten St. George, Daryl M. Lamson, Alexis Russell, Matthew Shudt, Melissa A Leisner, Jonathan Plitnick, Catharine Prussing, Navjot Singh, John Kelly, Erasmus Schneider, Erica Lasek-Nesselquist |
| EPI_ISL_2627360, EPI_ISL_2627361, EPI_ISL_2627362, EPI_ISL_2627363, EPI_ISL_2627364, EPI_ISL_2627365, EPI_ISL_2627366, EPI_ISL_2627367, EPI_ISL_2627368, EPI_ISL_2627369, EPI_ISL_2627370, EPI_ISL_2627372, EPI_ISL_2627373, EPI_ISL_2627374, EPI_ISL_2627375, EPI_ISL_2627376, EPI_ISL_2627377, EPI_ISL_2627378, EPI_ISL_2627379, EPI_ISL_2627380, EPI_ISL_2627381, EPI_ISL_2627382, EPI_ISL_2627383, EPI_ISL_2627384, EPI_ISL_2627385, EPI_ISL_2627386, EPI_ISL_2627387, EPI_ISL_2627388, EPI_ISL_2627389, EPI_ISL_2627390, EPI_ISL_2627391, EPI_ISL_2627392, EPI_ISL_2627393, EPI_ISL_2627394 | | | |
| see above | Virology Laboratory, International Centre for Diarrhoeal Disease Research, Bangladesh (ICDDR,B) | Virology Laboratory, International Centre for Diarrhoeal Disease Research, Bangladesh (ICDDR,B) | Mohammad Enayet Hossain, Mojnu Miah, Rashedul Hasan, Md. Mahfuzur Rahman, Mohammed Ziaur Rahman, Mustafizur Rahman |
| EPI_ISL_2627509, EPI_ISL_2627511, EPI_ISL_2627512, EPI_ISL_2627514, EPI_ISL_2627515, EPI_ISL_2627516, EPI_ISL_2627519, EPI_ISL_2627521 | Servicio de Microbiología. Hospital Universitario Doctor Peset | SeqCOVID-SPAIN consortium/IBV(CSIC) | Juan Alberola Enguídanos, Juan José Camarena Miñana, Rosa González Pellicer, José Miguel Nogueira Coito and SeqCOVID-SPAIN consortium |
| EPI_ISL_2627536, EPI_ISL_2627537, EPI_ISL_2627538, EPI_ISL_2627539, EPI_ISL_2627540, EPI_ISL_2627541, EPI_ISL_2627542, EPI_ISL_2627543, EPI_ISL_2627544, EPI_ISL_2627545, EPI_ISL_2627546, EPI_ISL_2627547, EPI_ISL_2627548, EPI_ISL_2627549, EPI_ISL_2627550, EPI_ISL_2627551, EPI_ISL_2627552, EPI_ISL_2627553, EPI_ISL_2627554, EPI_ISL_2627555, EPI_ISL_2627556, EPI_ISL_2627557, EPI_ISL_2627558, EPI_ISL_2627559, EPI_ISL_2627560, EPI_ISL_2627561, EPI_ISL_2627562, EPI_ISL_2627563, EPI_ISL_2627564, EPI_ISL_2627565, EPI_ISL_2627566, EPI_ISL_2627567, EPI_ISL_2627568, EPI_ISL_2627569, EPI_ISL_2627570, EPI_ISL_2627571, EPI_ISL_2627572, EPI_ISL_2627573, EPI_ISL_2627574, EPI_ISL_2627575, EPI_ISL_2627576, EPI_ISL_2627577, EPI_ISL_2627578, EPI_ISL_2627579, EPI_ISL_2627580 | | | |
| see above | National Public Health Laboratory, National Centre for Infectious Diseases | National Public Health Laboratory, National Centre for Infectious Diseases | Tze Minn Mak, Katherine Ching, Zhenyang Zhou, Royce Ang, Lin Cui, Raymond Tzer Pin Lin |
| EPI_ISL_2627588, EPI_ISL_2627615, EPI_ISL_2627636, EPI_ISL_2627686, EPI_ISL_2627728 | National Virus Reference Laboratory | National Virus Reference Laboratory | Zoe Yandle, Charlene Bennett, Gabriel Gonzalez, Michael Carr, Jonathan Dean, Cillian F De Gascun |
| EPI_ISL_2627906 | Orebro University Hospital, Dept Laboratory Medicine, Clinical Microbiology | Orebro University Hospital | Sundqvist M et al |
| EPI_ISL_2627920, EPI_ISL_2627921 | Kansas Health and Environmental Lab | Kansas Health and Environmental Lab | Mike Grose, Katherine Wiggins, Jonathan Barnell, Ben Olsen, and Phil Adam |
| EPI_ISL_2627923, EPI_ISL_2627924 | Pharmacare s.r.l. | AMES Centro Polidiagnostico Strumentale S.r.l. | "Giovanni Savarese, Raffaella Ruggiero, Eloisa Evangelista, Antonella Di Carlo, Luisa Circelli, Luigi D'Amore, Roberto Sirica, Antonio Fico" |
| EPI_ISL_2627925, EPI_ISL_2627926 | Kansas Health and Environmental Lab | Kansas Health and Environmental Lab | Mike Grose, Katherine Wiggins, Jonathan Barnell, Ben Olsen, and Phil Adam |
| EPI_ISL_2627929, | Wichita State University - Molecular Diagnostics Lab | Kansas Health and Environmental Lab | Mike Grose, Katherine Wiggins, Jonathan Barnell, Ben Olsen, and Phil Adam |

| | | | |
|---|---|---|---|
| EPI_ISL_2627931, EPI_ISL_2627932, EPI_ISL_2627933, EPI_ISL_2627934, EPI_ISL_2627935, EPI_ISL_2627936, EPI_ISL_2627937 | | | |
| EPI_ISL_2627990 | Armies | National Reference Center for Viruses of Respiratory Infections, Institut Pasteur, Paris | Marion Barbet, Sylvie Behillil, Méline Bizard, Angela Brisebarre, Camille Capel, Vincent Enouf, Louise Lefrançois, Frédéric Lemoine, Christophe Malabat, Corinne Maufrais, Etienne Simon-Lorière, Maud Vanpeene, Sylvie Van der Werf ,Marine Desroches |
| EPI_ISL_2627997 | Labo Analyses Med | National Reference Center for Viruses of Respiratory Infections, Institut Pasteur, Paris | Marion Barbet, Sylvie Behillil, Méline Bizard, Angela Brisebarre, Camille Capel, Vincent Enouf, Louise Lefrançois, Frédéric Lemoine, Christophe Malabat, Corinne Maufrais, Pierre Lechat, Etienne Simon-Lorière, Maud Vanpeene, Sylvie Van der Werf ,Caroline Devaux |
| EPI_ISL_2628016, EPI_ISL_2628025, EPI_ISL_2628026 | Montana Public Health Laboratory | Montana Public Health Laboratory | Joy Ritter, Michelle Mozer, Carrie Biskupiak, Deborah Gibson, Michael Dills |
| EPI_ISL_2628050, EPI_ISL_2628051, EPI_ISL_2628053, EPI_ISL_2628056, EPI_ISL_2628058, EPI_ISL_2628059, EPI_ISL_2628060 | LHUB-ULB | Labo Klinische Biologie, UZA | Marie Le Mercier, Jasmine Coppens, Basil Britto Xavier, Christine Lammens, Veerle Matheeussen, Herman Goossens |
| EPI_ISL_2628074, EPI_ISL_2628086, EPI_ISL_2628100, EPI_ISL_2628103, EPI_ISL_2628111, EPI_ISL_2628121, EPI_ISL_2628123, EPI_ISL_2628125 | Platform BIS UZA/UAntwerpen | Labo Klinische Biologie, UZA | Marie Le Mercier, Jasmine Coppens, Basil Britto Xavier, Christine Lammens, Veerle Matheeussen, Herman Goossens |
| EPI_ISL_2628127, EPI_ISL_2628128, EPI_ISL_2628130, EPI_ISL_2628137, EPI_ISL_2628138, EPI_ISL_2628141, EPI_ISL_2628144, EPI_ISL_2628145, EPI_ISL_2628147, EPI_ISL_2628150, EPI_ISL_2628153, EPI_ISL_2628155, EPI_ISL_2628156, EPI_ISL_2628158, EPI_ISL_2628159, EPI_ISL_2628161, EPI_ISL_2628164 | | | |
| see above | Hospital Universitari Vall d'Hebron - Vall d'Hebron Institut de Recerca | Hospital Universitari Vall d'Hebron - Vall d'Hebron Institut de Recerca | Cristina Andrés, Maria Piñana, Alejandra González-Sánchez, Damir Garcia-Cehic, Ariadna Rando, Juliana Esperalba, Maria Gema Codina, Carla Castillo, Maria Carmen Martin, Tomàs Pumarola, Josep Quer, Andrés Antón |
| EPI_ISL_2628166 | Hospital Universitari Arnau de Vilanova | Hospital Universitari Vall d'Hebron - Vall d'Hebron Institut de Recerca | Cristina Andrés, Maria Piñana, Alejandra González-Sánchez, Damir Garcia-Cehic, Ariadna Rando, Juliana Esperalba, Maria Gema Codina, Carla Castillo, Maria Carmen Martin, Tomàs Pumarola, Josep Quer, Andrés Antón |
| EPI_ISL_2628174, EPI_ISL_2628176, EPI_ISL_2628179, EPI_ISL_2628180, EPI_ISL_2628184, EPI_ISL_2628187, EPI_ISL_2628189, EPI_ISL_2628190, EPI_ISL_2628192 | Hospital Universitari Vall d'Hebron - Vall d'Hebron Institut de Recerca | Hospital Universitari Vall d'Hebron - Vall d'Hebron Institut de Recerca | Cristina Andrés, Maria Piñana, Alejandra González-Sánchez, Damir Garcia-Cehic, Ariadna Rando, Juliana Esperalba, Maria Gema Codina, Carla Castillo, Maria Carmen Martin, Tomàs Pumarola, Josep Quer, Andrés Antón |
| EPI_ISL_2628286 | Hospital | National Reference Center for Viruses of Respiratory Infections, Institut Pasteur, Paris | Marion Barbet, Sylvie Behillil, Méline Bizard, Angela Brisebarre, Camille Capel, Vincent Enouf, Louise Lefrançois, Frédéric Lemoine, Christophe Malabat, Corinne Maufrais, Etienne Simon-Lorière, Maud Vanpeene, Sylvie Van der Werf ,Marianne Coste-Burel |
| EPI_ISL_2628536, EPI_ISL_2628544 | Sharp HealthCare Laboratory | Andersen lab at Scripps Research | SEARCH Alliance San Diego with Aaron Harding, Jacquelyn Berumen, Cathy Woerle, Liam McGinnis, Art Mendoza, Omid Bakhtar |
| EPI_ISL_2628620, EPI_ISL_2628641, EPI_ISL_2628642, EPI_ISL_2628648, EPI_ISL_2628649, EPI_ISL_2628656, EPI_ISL_2628660, EPI_ISL_2628676, EPI_ISL_2628682, EPI_ISL_2628683, EPI_ISL_2628685, EPI_ISL_2628687, EPI_ISL_2628688, EPI_ISL_2628690, EPI_ISL_2628691, EPI_ISL_2628693, EPI_ISL_2628696, EPI_ISL_2628701, EPI_ISL_2628706, EPI_ISL_2628712, EPI_ISL_2628713, EPI_ISL_2628718, EPI_ISL_2628720, EPI_ISL_2628721, EPI_ISL_2628722, EPI_ISL_2628725, EPI_ISL_2628726, EPI_ISL_2628728, EPI_ISL_2628729, EPI_ISL_2628730, EPI_ISL_2628733, EPI_ISL_2628734, EPI_ISL_2628737, EPI_ISL_2628738, EPI_ISL_2628744, EPI_ISL_2628746, EPI_ISL_2628754, EPI_ISL_2628758 | | | |
| see above | UNILABS | Instituto Nacional de Saude (INSA) | Borges et al |
| EPI_ISL_2628760 | San Diego County Public Health Laboratory | Andersen lab at Scripps Research | SEARCH Alliance San Diego with Ashleigh Murphy, Jovan Shephard, Brett Austin |
| EPI_ISL_2628765, EPI_ISL_2628767, EPI_ISL_2628769, EPI_ISL_2628777, EPI_ISL_2628778, EPI_ISL_2628799, EPI_ISL_2628802, EPI_ISL_2628803, EPI_ISL_2628806, EPI_ISL_2628807, EPI_ISL_2628810, EPI_ISL_2628811, EPI_ISL_2628813, EPI_ISL_2628821, EPI_ISL_2628823, EPI_ISL_2628824, EPI_ISL_2628827, EPI_ISL_2628829, EPI_ISL_2628831, EPI_ISL_2628843, EPI_ISL_2628847, EPI_ISL_2628850, EPI_ISL_2628852, EPI_ISL_2628853, EPI_ISL_2628857, EPI_ISL_2628858, EPI_ISL_2628862, EPI_ISL_2628867, EPI_ISL_2628868, EPI_ISL_2628873, EPI_ISL_2628880, EPI_ISL_2628882, EPI_ISL_2628883, EPI_ISL_2628890, EPI_ISL_2628892, EPI_ISL_2628900, EPI_ISL_2628907, EPI_ISL_2628916, EPI_ISL_2628920, EPI_ISL_2628921, EPI_ISL_2628923, EPI_ISL_2628925, EPI_ISL_2628927, EPI_ISL_2628928, EPI_ISL_2628930, EPI_ISL_2628931, EPI_ISL_2628934, EPI_ISL_2628936, EPI_ISL_2628938, EPI_ISL_2628939, EPI_ISL_2628941, EPI_ISL_2628942, EPI_ISL_2628943, EPI_ISL_2628945, EPI_ISL_2628946, EPI_ISL_2628947, EPI_ISL_2628949, EPI_ISL_2628952, EPI_ISL_2628958, EPI_ISL_2628959, EPI_ISL_2628960, EPI_ISL_2628964, EPI_ISL_2628966, EPI_ISL_2628967, EPI_ISL_2628968, EPI_ISL_2628970, EPI_ISL_2628971, EPI_ISL_2628972, EPI_ISL_2628973, EPI_ISL_2628974, EPI_ISL_2628975, EPI_ISL_2628976, EPI_ISL_2628980, EPI_ISL_2628981, EPI_ISL_2628982, EPI_ISL_2628984, EPI_ISL_2628986, EPI_ISL_2628993, EPI_ISL_2628994 | | | |
| see above | UNILABS | Instituto Nacional de Saude (INSA) | Borges et al |
| EPI_ISL_2629265 | CENTRE HOSPITALIER PIERRE OUDOT | CNR Virus des Infections Respiratoires - France SUD | Antonin Bal, Gregory Destras, Gwendolyne Burfin, Hadrien Regue, Quentin Semanas, Martine Valette, Bruno Lina, Laurence Josset |
| EPI_ISL_2629328 | LABORATOIRE UNILIANS DECINES | CNR Virus des Infections Respiratoires - France SUD | Antonin Bal, Gregory Destras, Gwendolyne Burfin, Hadrien Regue, Quentin Semanas, Martine Valette, Bruno Lina, Laurence Josset |
| EPI_ISL_2629329 | MIRIALIS CLUSES BECHET | CNR Virus des Infections Respiratoires - France SUD | Antonin Bal, Gregory Destras, Gwendolyne Burfin, Hadrien Regue, Quentin Semanas, Martine Valette, Bruno Lina, Laurence Josset |
| EPI_ISL_2629330, EPI_ISL_2629425 | LABORATOIRE UNILIANS DECINES | CNR Virus des Infections Respiratoires - France SUD | Antonin Bal, Gregory Destras, Gwendolyne Burfin, Hadrien Regue, Quentin Semanas, Martine Valette, Bruno Lina, Laurence Josset |
| EPI_ISL_2629445 | LABORATOIRE BIOESTEREL | CNR Virus des Infections Respiratoires - France SUD | Antonin Bal, Gregory Destras, Gwendolyne Burfin, Hadrien Regue, Quentin Semanas, Martine Valette, Bruno Lina, Laurence Josset |
| EPI_ISL_2629457 | DYOMEDEA-LABORATOIRE DE LA SAUVEGARDE | CNR Virus des Infections Respiratoires - France SUD | Antonin Bal, Gregory Destras, Gwendolyne Burfin, Hadrien Regue, Quentin Semanas, Martine Valette, Bruno Lina, Laurence Josset |
| EPI_ISL_2629481, EPI_ISL_2629483, EPI_ISL_2629484 | LABORATOIRE FORTE BIO ST PAUL LES DAX | CNR Virus des Infections Respiratoires - France SUD | Antonin Bal, Gregory Destras, Gwendolyne Burfin, Hadrien Regue, Quentin Semanas, Martine Valette, Bruno Lina, Laurence Josset |
| EPI_ISL_2629491, EPI_ISL_2629496, EPI_ISL_2629497, EPI_ISL_2629501, EPI_ISL_2629502, EPI_ISL_2629509 | AX BIO OCEAN | CNR Virus des Infections Respiratoires - France SUD | Antonin Bal, Gregory Destras, Gwendolyne Burfin, Hadrien Regue, Quentin Semanas, Martine Valette, Bruno Lina, Laurence Josset |
| EPI_ISL_2629511 | MIRIALIS CLUSES BECHET | CNR Virus des Infections Respiratoires - France SUD | Antonin Bal, Gregory Destras, Gwendolyne Burfin, Hadrien Regue, Quentin Semanas, Martine Valette, Bruno Lina, Laurence Josset |
| EPI_ISL_2629561 | LABM DU PRE - LABOMAINE | CNR Virus des Infections Respiratoires - France SUD | Antonin Bal, Gregory Destras, Gwendolyne Burfin, Hadrien Regue, Quentin Semanas, Martine Valette, Bruno Lina, Laurence Josset |
| EPI_ISL_2629567 | CERBALLIANCE - LAMAT | CNR Virus des Infections Respiratoires - France SUD | Antonin Bal, Gregory Destras, Gwendolyne Burfin, Hadrien Regue, Quentin Semanas, Martine Valette, Bruno Lina, Laurence Josset |
| EPI_ISL_2629572, EPI_ISL_2629579 | CNR Virus des Infections Respiratoires - France SUD | CNR Virus des Infections Respiratoires - France SUD | Antonin Bal, Gregory Destras, Gwendolyne Burfin, Hadrien Regue, Quentin Semanas, Martine Valette, Bruno Lina, Laurence Josset |
| EPI_ISL_2629587 | GA Department of Public Health | GA Department of Public Health | Stacy Reeves, Jonathan Edwards, Cynthia Dixey, Tonia Parrott, Aliyah Fields, Taylor Smith |
| EPI_ISL_2629889, EPI_ISL_2629891 | Wyoming Public Health Laboratory | Wyoming Public Health Laboratory | Jim Mildenberger, Taylor Fearing, Channing Weber, Ashley Norberg, Chayse Rowley, Marley Goetz, Brian Dominguez, Elliot Thomasson, Sam Britz, Cari Sloma, Robert Petit, and Rob Christensen |
| EPI_ISL_2629946, EPI_ISL_2629947, EPI_ISL_2629955, EPI_ISL_2629965, EPI_ISL_2629977, EPI_ISL_2629992, EPI_ISL_2629994, EPI_ISL_2629995, EPI_ISL_2629996, EPI_ISL_2629998, EPI_ISL_2630015, EPI_ISL_2630017, EPI_ISL_2630020, EPI_ISL_2630022, EPI_ISL_2630034, EPI_ISL_2630040, EPI_ISL_2630065, EPI_ISL_2630066, EPI_ISL_2630070, EPI_ISL_2630106, EPI_ISL_2630108, EPI_ISL_2630109, EPI_ISL_2630110, EPI_ISL_2630113, EPI_ISL_2630115, EPI_ISL_2630119, EPI_ISL_2630127, EPI_ISL_2630128, EPI_ISL_2630139, EPI_ISL_2630140, EPI_ISL_2630141, EPI_ISL_2630173, EPI_ISL_2630174, EPI_ISL_2630177, EPI_ISL_2630179, EPI_ISL_2630184, EPI_ISL_2630192, EPI_ISL_2630209, EPI_ISL_2630214, EPI_ISL_2630218, EPI_ISL_2630222, EPI_ISL_2630227, EPI_ISL_2630232, EPI_ISL_2630236, EPI_ISL_2630237, EPI_ISL_2630242, EPI_ISL_2630247, EPI_ISL_2630249, EPI_ISL_2630261, EPI_ISL_2630262, EPI_ISL_2630263, EPI_ISL_2630264, EPI_ISL_2630265, EPI_ISL_2630278, EPI_ISL_2630283, EPI_ISL_2630284, EPI_ISL_2630297, EPI_ISL_2630308, EPI_ISL_2630324, EPI_ISL_2630327, EPI_ISL_2630329, EPI_ISL_2630330, EPI_ISL_2630368, EPI_ISL_2630413, EPI_ISL_2630442, EPI_ISL_2630444, EPI_ISL_2630447, EPI_ISL_2630451, EPI_ISL_2630455, EPI_ISL_2630460, EPI_ISL_2630461, EPI_ISL_2630470, EPI_ISL_2630474, EPI_ISL_2630481, EPI_ISL_2630482, EPI_ISL_2630483, EPI_ISL_2630485, EPI_ISL_2630494, EPI_ISL_2630496, EPI_ISL_2630498, EPI_ISL_2630500, EPI_ISL_2630501, EPI_ISL_2630504, EPI_ISL_2630509, EPI_ISL_2630519, EPI_ISL_2630523, EPI_ISL_2630525, EPI_ISL_2630526, EPI_ISL_2630528, EPI_ISL_2630533, EPI_ISL_2630536, EPI_ISL_2630538, EPI_ISL_2630543, EPI_ISL_2630554, EPI_ISL_2630556, EPI_ISL_2630566, EPI_ISL_2630568, EPI_ISL_2630573, EPI_ISL_2630575, EPI_ISL_2630576, EPI_ISL_2630578, EPI_ISL_2630580, EPI_ISL_2630581, EPI_ISL_2630582, EPI_ISL_2630591, EPI_ISL_2630594, EPI_ISL_2630599, EPI_ISL_2630602, EPI_ISL_2630603, EPI_ISL_2630606, EPI_ISL_2630608, EPI_ISL_2630612, EPI_ISL_2630613, EPI_ISL_2630615, EPI_ISL_2630619, EPI_ISL_2630627, EPI_ISL_2630629, EPI_ISL_2630639, EPI_ISL_2630640, EPI_ISL_2630647, EPI_ISL_2630651, EPI_ISL_2630658, EPI_ISL_2630659, EPI_ISL_2630660, EPI_ISL_2630661, EPI_ISL_2630669, EPI_ISL_2630670, EPI_ISL_2630671, EPI_ISL_2630672, EPI_ISL_2630684, EPI_ISL_2630685, EPI_ISL_2630686, EPI_ISL_2630696, EPI_ISL_2630721, EPI_ISL_2630743, EPI_ISL_2630750, EPI_ISL_2630753, EPI_ISL_2630754, EPI_ISL_2630790, EPI_ISL_2630812, EPI_ISL_2630814, EPI_ISL_2630816, EPI_ISL_2630826, EPI_ISL_2630827, EPI_ISL_2630828, EPI_ISL_2630829, EPI_ISL_2630830, EPI_ISL_2630834, EPI_ISL_2630838, EPI_ISL_2630839, EPI_ISL_2630841, EPI_ISL_2630842, EPI_ISL_2630843, EPI_ISL_2630845, EPI_ISL_2630847, EPI_ISL_2630848, EPI_ISL_2630849, EPI_ISL_2630860, EPI_ISL_2630861, EPI_ISL_2630864, EPI_ISL_2630865, EPI_ISL_2630871, EPI_ISL_2630872, EPI_ISL_2630873, EPI_ISL_2630874, EPI_ISL_2630875, EPI_ISL_2630877, EPI_ISL_2630880, EPI_ISL_2630888, EPI_ISL_2630892, EPI_ISL_2630893, EPI_ISL_2630895, EPI_ISL_2630897, EPI_ISL_2630901, EPI_ISL_2630904, EPI_ISL_2630906, EPI_ISL_2630921, EPI_ISL_2630927, EPI_ISL_2630928, EPI_ISL_2630930, EPI_ISL_2630933, EPI_ISL_2630934, EPI_ISL_2630939, EPI_ISL_2630941, EPI_ISL_2630943, EPI_ISL_2630959, EPI_ISL_2630980, EPI_ISL_2630981, EPI_ISL_2630982, EPI_ISL_2630983, EPI_ISL_2630985, EPI_ISL_2630992, EPI_ISL_2630997, EPI_ISL_2630998, EPI_ISL_2631001, EPI_ISL_2631002, EPI_ISL_2631007, EPI_ISL_2631012, EPI_ISL_2631014, EPI_ISL_2631019, EPI_ISL_2631020, EPI_ISL_2631021, EPI_ISL_2631022, EPI_ISL_2631023, EPI_ISL_2631026, EPI_ISL_2631028, EPI_ISL_2631030, EPI_ISL_2631031, EPI_ISL_2631033, EPI_ISL_2631042, EPI_ISL_2631055, EPI_ISL_2631057, EPI_ISL_2631058, EPI_ISL_2631062, EPI_ISL_2631064, EPI_ISL_2631065, EPI_ISL_2631067, EPI_ISL_2631071, EPI_ISL_2631074, EPI_ISL_2631079, EPI_ISL_2631080, EPI_ISL_2631082, EPI_ISL_2631086, EPI_ISL_2631087, EPI_ISL_2631094, EPI_ISL_2631097, EPI_ISL_2631102, EPI_ISL_2631103, EPI_ISL_2631106, EPI_ISL_2631122, EPI_ISL_2631123, EPI_ISL_2631126, EPI_ISL_2631128, EPI_ISL_2631130, EPI_ISL_2631131, EPI_ISL_2631138, EPI_ISL_2631143, EPI_ISL_2631145, EPI_ISL_2631148, EPI_ISL_2631150 | | | |
| see above | Public Health Ontario Laboratory | Public Health Ontario Laboratory | Vanessa G Allen, Philip Banh, Yao Chen, Richard de Borja, Alireza Eshaghi, Nahuel Fittipaldi, Christine Frantz, Jonathan B Gubbay, Jennifer L Guthrie, Lawrence Heisler, Esha Joshi, Michael Laszloffy, Aimin Li, Michael CY Li, Dean Maxwell, Sandeep Nagra, Samir N Patel, Jared Simpson, Karthikeyan Sivaraman, Ashleigh Sullivan, Yogi Sundaravadanam, Sarah Teatero, Andre Villegas, Matthew Watson, Sandra Zittermann |
| EPI_ISL_2631214 | Santa Clara Valley Medical Center | Santa Clara County Public Health Laboratory | Santa Clara County Public Health Department |
| EPI_ISL_2631342, | Pandemic Response Lab - NYC | Pandemic Response Lab, R&D | Henry Lee, Michael Hammerling, Melissa Hopkins, Cybill del Castillo, Shinyoung Clair Kang, William Ward, Pradeep Bugga, Sol Rey, Dylan Law, Katharine Nelson, Haiping Hao, Jon Laurent |

| | | | |
|---|---|---|---|
| EPI_ISL_2631346, EPI_ISL_2631357, EPI_ISL_2631375, EPI_ISL_2631389, EPI_ISL_2631390, EPI_ISL_2631392, EPI_ISL_2631396, EPI_ISL_2631401, EPI_ISL_2631404 | | | |
| EPI_ISL_2631443 | UPTD Labkesda Karawang | Indonesian Institute of Sciences (LIPI)/National Agency of Research and Innovation (BRIN) and National Institute of Health Research and Development (NIHRD) | Herjuno A. Nugroho, Idris, Sri Swasthikawati, Isa Nuryana, Indriawati, Yuliawati, Listiana Oktavia, Gita Syahputra, Andri Wardiana, Syam B. Iryanto, Yayuk Sri Rahayu, Subangkit, Vivi Setiawaty, Puspita Lisdiyanti |
| EPI_ISL_2631444 | UPTD Labkesda Karawang | Indonesian Institute of Sciences (LIPI)/National Agency of Research and Innovation (BRIN) and National Institute of Health Research and Development (NIHRD) | Idris, Sri Swasthikawati, Isa Nuryana, Indriawati, Yuliawati, Listiana Oktavia, Herjuno A. Nugroho, Andri Wardiana, Syam B. Iryanto, Yayuk Sri Rahayu, Subangkit, Vivi Setiawaty Puspita, Sugiyono Saputra, Puspita Lisdiyanti |
| EPI_ISL_2631445 | UPTD Labkesda Karawang | Indonesian Institute of Sciences (LIPI)/National Agency of Research and Innovation (BRIN) and National Institute of Health Research and Development (NIHRD) | Idris, Sri Swasthikawati, Isa Nuryana, Indriawati, Yuliawati, Listiana Oktavia, Herjuno A. Nugroho, Andri Wardiana, Syam B. Iryanto, Yayuk Sri Rahayu, Subangkit, Vivi Setiawaty, Sugiyono Saputra, Puspita Lisdiyanti |
| EPI_ISL_2631446, EPI_ISL_2631447 | UPTD Labkesda Karawang | Indonesian Institute of Sciences (LIPI)/National Agency of Research and Innovation (BRIN) and National Institute of Health Research and Development (NIHRD) | Sri Swasthikawati, Isa Nuryana, Indriawati, Yuliawati, Listiana Oktavia, Herjuno A. Nugroho, Idris, Anggia Prasetyoputri, Syam B. Iryanto, Gita Syahputra, Yayuk Sri Rahayu, Subangkit, Vivi Setiawaty, Sugiyono Saputra, Puspita Lisdiyanti |
| EPI_ISL_2631448, EPI_ISL_2631449 | UPTD Labkesda Karawang | Indonesian Institute of Sciences (LIPI)/National Agency of Research and Innovation (BRIN) and National Institute of Health Research and Development (NIHRD) | Isa Nuryana, Indriawati, Yuliawati, Listiana Oktavia, Herjuno A. Nugroho, Idris, Sri Swasthikawati, Anggia Prasetyoputri, Syam B. Iryanto, Yayuk Sri Rahayu, Subangkit, Vivi Setiawaty, Sugiyono Saputra, Puspita Lisdiyanti |
| EPI_ISL_2631450 | UPTD Labkesda Karawang | Indonesian Institute of Sciences (LIPI)/National Agency of Research and Innovation (BRIN) and National Institute of Health Research and Development (NIHRD) | Indriawati, Yuliawati, Listiana Oktavia, Herjuno A. Nugroho, Idris, Sri Swasthikawati, Isa Nuryana, Ratih A. Ningrum, Anik B. Dharmayanthi, Yayuk Sri Rahayu, Subangkit, Vivi Setiawaty, Puspita Lisdiyanti, Anggia Prasetyoputri |
| EPI_ISL_2631451 | UPTD Labkesda Karawang | Indonesian Institute of Sciences (LIPI)/National Agency of Research and Innovation (BRIN) and National Institute of Health Research and Development (NIHRD) | Yuliawati, Listiana Oktavia, Herjuno A. Nugroho, Idris, Sri Swasthikawati, Isa Nuryana, Indriawati, Ratih A. Ningrum, Anik B. Dharmayanthi, Yayuk Sri Rahayu, Subangkit, Vivi Setiawaty, Puspita Lisdiyanti, Anggia Prasetyoputri |
| EPI_ISL_2631452 | UPTD Labkesda Karawang | Indonesian Institute of Sciences (LIPI)/National Agency of Research and Innovation (BRIN) and National Institute of Health Research and Development (NIHRD) | Yuliawati, Listiana Oktavia, Herjuno A. Nugroho, Idris, Sri Swasthikawati, Isa Nuryana, Indriawati, Ratih A. Ningrum, Anik B. Dharmayanthi, Yayuk Sri Rahayu, Subangkit, Vivi Setiawaty, Puspita Lisdiyanti, Andri Wardiana |
| EPI_ISL_2631453 | UPTD Labkesda Karawang | Indonesian Institute of Sciences (LIPI)/National Agency of Research and Innovation (BRIN) and National Institute of Health Research and Development (NIHRD) | Yuliawati, Listiana Oktavia, Herjuno A. Nugroho, Idris, Sri Swasthikawati, Isa Nuryana, Indriawati, Ratih A. Ningrum, Anik B. Dharmayanthi, Yayuk Sri Rahayu, Subangkit, Vivi Setiawaty Puspita Lisdiyanti, Sugiyono Saputra |
| EPI_ISL_2631454, EPI_ISL_2631455 | RSK Paru Karawang | Indonesian Institute of Sciences (LIPI)/National Agency of Research and Innovation (BRIN) and National Institute of Health Research and Development (NIHRD) | Listiana Oktavia, Herjuno A. Nugroho, Idris, Sri Swasthikawati, Isa Nuryana, Indriawati, Yuliawati, Anggia Prasetyoputri, Eko Wahyu Putro, Syam B. Iryanto, Yayuk Sri Rahayu, Subangkit, Vivi Setiawaty, Puspita Lisdiyanti |
| EPI_ISL_2631456 | RSK Paru Karawang | Indonesian Institute of Sciences (LIPI)/National Agency of Research and Innovation (BRIN) and National Institute of Health Research and Development (NIHRD) | Fahrurrozi, Linda Sukmarini, Nova D. Yanthi, Herjuno A. Nugroho, Idris, Sri Swasthikawati, Anggia Prasetyoputri, Eko Wahyu Putro, Syam B. Iryanto, Yayuk Sri Rahayu, Subangkit, Vivi Setiawaty, Puspita Lisdiyanti |
| EPI_ISL_2631457, EPI_ISL_2631458 | RSK Paru Karawang | Indonesian Institute of Sciences (LIPI)/National Agency of Research and Innovation (BRIN) and National Institute of Health Research and Development (NIHRD) | Fahrurrozi, Linda Sukmarini, Nova D. Yanthi, Herjuno A. Nugroho, Idris, Sri Swasthikawati, Anggia Prasetyoputri, Eko Wahyu Putro, Syam B. Iryanto, Yayuk Sri Rahayu, Subangkit, Vivi Setiawaty, Ratih A. Ningrum, Puspita Lisdiyanti |
| EPI_ISL_2631459 | RSK Paru Karawang | Indonesian Institute of Sciences (LIPI)/National Agency of Research and Innovation (BRIN) and National Institute of Health Research and Development (NIHRD) | Linda Sukmarini, Fahrurrozi, Nova D. Yanthi, Herjuno A. Nugroho, Idris, Sri Swasthikawati, Anggia Prasetyoputri, Ruby Setiawan, Syam B. Iryanto, Yayuk Sri Rahayu, Subangkit, Vivi Setiawaty, Ratih A. Ningrum, Puspita Lisdiyanti |
| EPI_ISL_2631460 | RSK Paru Karawang | Indonesian Institute of Sciences (LIPI)/National Agency of Research and Innovation (BRIN) and National Institute of Health Research and Development (NIHRD) | Linda Sukmarini, Fahrurrozi, Nova D. Yanthi, Herjuno A. Nugroho, Idris, Sri Swasthikawati, Anik B. Dharmayanthi, Ruby Setiawan, Syam B. Iryanto, Yayuk Sri Rahayu, Subangkit, Vivi Setiawaty, Ratih A. Ningrum, Puspita Lisdiyanti |
| EPI_ISL_2631461 | RSK Paru Karawang | Indonesian Institute of Sciences (LIPI)/National Agency of Research and Innovation (BRIN) and National Institute of Health Research and Development (NIHRD) | Linda Sukmarini, Fahrurrozi, Nova D. Yanthi, Herjuno A. Nugroho, Idris, Sri Swasthikawati, Anik B. Dharmayanthi, Ruby Setiawan, Syam B. Iryanto, Yayuk Sri Rahayu, Subangkit, Vivi Setiawaty, Puspita Lisdiyanti |
| EPI_ISL_2631462 | RSK Paru Karawang | Indonesian Institute of Sciences (LIPI)/National Agency of Research and Innovation (BRIN) and National Institute of Health Research and Development (NIHRD) | Linda Sukmarini, Fahrurrozi, Nova D. Yanthi, Herjuno A. Nugroho, Idris, Sri Swasthikawati, Anggia Prasetyoputri, Ruby Setiawan, Syam B. Iryanto, Yayuk Sri Rahayu, Subangkit, Vivi Setiawaty, Puspita Lisdiyanti |
| EPI_ISL_2631463 | RSK Paru Karawang | Indonesian Institute of Sciences (LIPI)/National Agency of Research and Innovation (BRIN) and National Institute of Health Research and Development (NIHRD) | Yuliawati, Listiana Oktavia, Herjuno A. Nugroho, Idris, Sri Swasthikawati, Isa Nuryana, Indriawati, Ratih A. Ningrum, Anik B. Dharmayanthi, Yayuk Sri Rahayu, Subangkit, Vivi Setiawaty Puspita Lisdiyanti, |
| EPI_ISL_2631465, EPI_ISL_2631466, EPI_ISL_2631467 | RS Bayukarta | Indonesian Institute of Sciences (LIPI)/National Agency of Research and Innovation (BRIN) and National Institute of Health Research and Development (NIHRD) | Nova D. Yanthi, Linda Sukmarini, Fahrurrozi, Herjuno A. Nugroho, Idris, Sri Swasthikawati, Anggia Prasetyoputri, Eko Wahyu Putro, Syam B. Iryanto, Yayuk Sri Rahayu, Subangkit, Vivi Setiawaty, Sugiyono Saputra, Puspita Lisdiyanti |
| EPI_ISL_2631468 | RS Bayukarta | Indonesian Institute of Sciences (LIPI)/National Agency of Research and Innovation (BRIN) and National Institute of Health Research and Development (NIHRD) | Listiana Oktavia, Herjuno A. Nugroho, Idris, Sri Swasthikawati, Isa Nuryana, Indriawati, Yuliawati, Anggia Prasetyoputri, Eko Wahyu Putro, Syam B. Iryanto, Yayuk Sri Rahayu, Subangkit, Vivi Setiawaty, Puspita Lisdiyanti |
| EPI_ISL_2631469 | RSUD Karawang | Indonesian Institute of Sciences (LIPI)/National Agency of Research and Innovation (BRIN) and National Institute of Health Research and Development (NIHRD) | Listiana Oktavia, Herjuno A. Nugroho, Idris, Sri Swasthikawati, Isa Nuryana, Indriawati, Yuliawati, Anggia Prasetyoputri, Eko Wahyu Putro, Syam B. Iryanto, Yayuk Sri Rahayu, Subangkit, Vivi Setiawaty, Puspita Lisdiyanti |
| EPI_ISL_2631470 | RSUD Karawang | Indonesian Institute of Sciences (LIPI)/National Agency of Research and Innovation (BRIN) and National Institute of Health Research and Development (NIHRD) | Herjuno A. Nugroho, Idris, Sri Swasthikawati, Isa Nuryana, Indriawati, Yuliawati, Listiana Oktavia, Gita Syahputra, Andri Wardiana, Syam B. Iryanto, Yayuk Sri Rahayu, Subangkit, Vivi Setiawaty Puspita Lisdiyanti |
| EPI_ISL_2631471 | RSUD Karawang | Indonesian Institute of Sciences (LIPI)/National Agency of Research and Innovation (BRIN) and National Institute of Health Research and Development (NIHRD) | Herjuno A. Nugroho, Idris, Sri Swasthikawati, Isa Nuryana, Indriawati, Yuliawati, Listiana Oktavia, Gita Syahputra, Andri Wardiana, Syam B. Iryanto, Yayuk Sri Rahayu, Subangkit, Vivi Setiawaty Puspita Lisdiyanti, |
| EPI_ISL_2631474 | RSUD Karawang | Indonesian Institute of Sciences (LIPI)/National Agency of Research and Innovation (BRIN) and National Institute of Health Research and Development (NIHRD) | Fahrurrozi, Linda Sukmarini, Nova D. Yanthi, Herjuno A. Nugroho, Idris, Sri Swasthikawati, Anggia Prasetyoputri, Eko Wahyu Putro, Syam B. Iryanto, Yayuk Sri Rahayu, Subangkit, Vivi Setiawaty, Puspita Lisdiyanti |
| EPI_ISL_2631475 | RSUD Karawang | Indonesian Institute of Sciences (LIPI)/National Agency of Research and Innovation (BRIN) and National Institute of Health Research and Development (NIHRD) | Isa Nuryana, Indriawati, Yuliawati, Listiana Oktavia, Herjuno A. Nugroho, Idris, Sri Swasthikawati, Anggia Prasetyoputri, Anik B. Dharmayanthi, Yayuk Sri Rahayu, Subangkit, Vivi Setiawaty Puspita Lisdiyanti, Sugiyono Saputra |
| EPI_ISL_2631476, EPI_ISL_2631477 | RSUD Karawang | Indonesian Institute of Sciences (LIPI)/National Agency of Research and Innovation (BRIN) and National Institute of Health Research and Development (NIHRD) | Indriawati, Yuliawati, Listiana Oktavia, Herjuno A. Nugroho, Idris, Sri Swasthikawati, Isa Nuryana, Ratih A. Ningrum, Anik B. Dharmayanthi, Yayuk Sri Rahayu, Subangkit, Vivi Setiawaty, Puspita Lisdiyanti |
| EPI_ISL_2631481 | RS Primaya | Indonesian Institute of Sciences (LIPI)/National Agency of Research and Innovation (BRIN) and National Institute of Health Research and Development (NIHRD) | Hariyatun, Alfi T. Fathurahman, Asep M. Ridwanulloh, Andri Wardiana, Erwin F. Hasrianda, Syam B. Iryanto, Ruby Setiawan, Anik B. Dharmayanthi, Subangkit, Yayuk Sri Rahayu, Vivi Setiawaty, Ratih A. Ningrum, Sugiyono Saputra |
| EPI_ISL_2631482, EPI_ISL_2631483, EPI_ISL_2631484, EPI_ISL_2631485 | RS Primaya | Indonesian Institute of Sciences (LIPI)/National Agency of Research and Innovation (BRIN) and National Institute of Health Research and Development (NIHRD) | Alfi T. Fathurahman, Asep M. Ridwanulloh, Erwin F. Hasrianda, Andri Wardiana, Syam B. Iryanto, Ruby Setiawan, Hariyatun, Anggia Prasetyoputri, Subangkit, Yayuk Sri Rahayu, Vivi Setiawaty, Anik B. Dharmayanthi, Sugiyono Saputra |
| EPI_ISL_2631486 | RS Primaya | Indonesian Institute of Sciences (LIPI)/National Agency of Research and Innovation (BRIN) and National Institute of Health Research and Development (NIHRD) | Asep M. Ridwanulloh, Andri Wardiana, Idris, Erwin F. Hasrianda, Syam B. Iryanto, Ruby Setiawan, Hariyatun, Alfi T. Fathurahman, Anik B. Dharmayanthi, Subangkit, Yayuk Sri Rahayu, Vivi Setiawaty, Anggia Prasetyoputri |
| EPI_ISL_2631487, EPI_ISL_2631488 | Intibios Karawang | Indonesian Institute of Sciences (LIPI)/National Agency of Research and Innovation (BRIN) and National Institute of Health Research and Development (NIHRD) | Asep M. Ridwanulloh, Andri Wardiana, Idris, Erwin F. Hasrianda, Syam B. Iryanto, Ruby Setiawan, Hariyatun, Alfi T. Fathurahman, Anik B. Dharmayanthi, Subangkit, Yayuk Sri Rahayu, Vivi Setiawaty, Anggia Prasetyoputri |

| | | | |
|---|---|---|---|
| EPI_ISL_2631489 | Intibios Karawang | Indonesian Institute of Sciences (LIPI)/National Agency of Research and Innovation (BRIN) and National Institute of Health Research and Development (NIHRD) | Asep M. Ridwanulloh, Andri Wardiana, Syam B. Iryanto, Ruby Setiawan, Hariyatun, Alfi T. Fathurahman, Anik B. Dharmayanthi, Subangkit, Yayuk Sri Rahayu, Vivi Setiawaty, Anggia Prasetyoputri, Puspita Lisdiyanti |
| EPI_ISL_2631490 | Intibios Karawang | Indonesian Institute of Sciences (LIPI)/National Agency of Research and Innovation (BRIN) and National Institute of Health Research and Development (NIHRD) | Andri Wardiana, Syam B. Iryanto, Ade Andriani, Hariyatun, Alfi T. Fathurahman, Idris, Erwin F. Hasrianda, Asep M. Ridwanulloh, Anik B. Dharmayanthi, Subangkit, Yayuk Sri Rahayu, Vivi Setiawaty, Puspita Lisdiyanti |
| EPI_ISL_2631493 | Intibios Karawang | Indonesian Institute of Sciences (LIPI)/National Agency of Research and Innovation (BRIN) and National Institute of Health Research and Development (NIHRD) | Andri Wardiana, Syam B. Iryanto, Erwin F. Hasrianda, Ade Andriani, Hariyatun, Alfi T. Fathurahman, Asep M. Ridwanulloh, Anik B. Dharmayanthi, Subangkit, Yayuk Sri Rahayu, Vivi Setiawaty, Puspita Lisdiyanti |
| EPI_ISL_2631494, EPI_ISL_2631495, EPI_ISL_2631496, EPI_ISL_2631497 | Intibios Karawang | Indonesian Institute of Sciences (LIPI)/National Agency of Research and Innovation (BRIN) and National Institute of Health Research and Development (NIHRD) | Erwin F. Hasrianda, Andri Wardiana, Syam B. Iryanto, Ade Andriani, Hariyatun, Alfi T. Fathurahman, Asep M. Ridwanulloh, Anik B. Dharmayanthi, Subangkit, Yayuk Sri Rahayu, Vivi Setiawaty, Puspita Lisdiyanti, Ratih A. Ningrum |
| EPI_ISL_2631621, EPI_ISL_2631629 | SYNLAB MVZ Weiden | Robert Koch Institute | unknown |
| EPI_ISL_2631631 | SYNLAB MVZ Leinfelden-Echterdingen | Robert Koch Institute | unknown |
| EPI_ISL_2631644 | SYNLAB MVZ Weiden | Robert Koch Institute | unknown |
| EPI_ISL_2631647 | Synlab MVZ Augsburg | Robert Koch Institute | unknown |
| EPI_ISL_2631648 | SYNLAB MVZ Leverkusen | Robert Koch Institute | unknown |
| EPI_ISL_2631649, EPI_ISL_2631657 | SYNLAB MVZ Trier | Robert Koch Institute | unknown |
| EPI_ISL_2631685, EPI_ISL_2631701, EPI_ISL_2631706 | SYNLAB MVZ Weiden | Robert Koch Institute | unknown |
| EPI_ISL_2631816 | MVZ für Laboratoriumsmedizin und Mikrobiologie Koblenz-Mittelrhein (Labor Koblenz) | Robert Koch Institute | unknown |
| EPI_ISL_2631831, EPI_ISL_2631833 | Labor Dr. Wisplinghoff - Köln | Robert Koch Institute | unknown |
| EPI_ISL_2631839 | MVZ für Laboratoriumsmedizin und Mikrobiologie Koblenz-Mittelrhein (Labor Koblenz) | Robert Koch Institute | unknown |
| EPI_ISL_2631872, EPI_ISL_2631875, EPI_ISL_2631885 | Clemenshospital - Alexianer Clemenshospital Münster | Robert Koch Institute | unknown |
| EPI_ISL_2631896, EPI_ISL_2631909, EPI_ISL_2631929 | Eurofins LifeCodexx GmbH | Robert Koch Institute | unknown |
| EPI_ISL_2631935, EPI_ISL_2631962, EPI_ISL_2632011, EPI_ISL_2632017, EPI_ISL_2632018 | MVZ Labor Krone GbR | Robert Koch Institute | unknown |
| EPI_ISL_2632025, EPI_ISL_2632031, EPI_ISL_2632035, EPI_ISL_2632057 | MVZ Dr. Eberhard & Partner Dortmund | Robert Koch Institute | unknown |
| EPI_ISL_2632109 | St. Elisabeth-Gruppe GmbH - Katholische Kliniken Rhein-Ruhr | Robert Koch Institute | unknown |
| EPI_ISL_2632119, EPI_ISL_2632121, EPI_ISL_2632123, EPI_ISL_2632124, EPI_ISL_2632127, EPI_ISL_2632133, EPI_ISL_2632134, EPI_ISL_2632135, EPI_ISL_2632137 | Corona-Testzentrum ifp Institut für Produktqualität GmbH | Robert Koch Institute | unknown |
| EPI_ISL_2632158, EPI_ISL_2632162, EPI_ISL_2632165, EPI_ISL_2632171, EPI_ISL_2632172, EPI_ISL_2632175 | Labor Dr. Heidrich & Kollegen MVZ GmbH Hamburg | Robert Koch Institute | unknown |
| EPI_ISL_2632198, EPI_ISL_2632201, EPI_ISL_2632206, EPI_ISL_2632208, EPI_ISL_2632212, EPI_ISL_2632239 | Medizinische Laboratorien Düsseldorf | Robert Koch Institute | unknown |
| EPI_ISL_2632244 | nordlab - Partnerschaftspraxis für Laboratoriumsmedizin | Robert Koch Institute | unknown |
| EPI_ISL_2632261 | amedes MVZ DIAMEDIS Sennestadt | Robert Koch Institute | unknown |
| EPI_ISL_2632284 | amedes MVZ für Laboratoriumsdiagnostik Raubling GmbH | Robert Koch Institute | unknown |
| EPI_ISL_2632299 | amedes MVZ DIAMEDIS Sennestadt | Robert Koch Institute | unknown |
| EPI_ISL_2632325 | Niedersächsisches Landesgesundheitsamt (NLGA) | Robert Koch Institute | unknown |
| EPI_ISL_2632344 | Bioscientia Labor Wermsdorf | Robert Koch Institute | unknown |
| EPI_ISL_2632377 | LabKom - MVZ Labor Bochum MLB GmbH | Robert Koch Institute | unknown |
| EPI_ISL_2632383 | Labor 28 MVZ GmbH | Robert Koch Institute | unknown |
| EPI_ISL_2632415 | Bioscientia Labor Wermsdorf | Robert Koch Institute | unknown |
| EPI_ISL_2632416 | Sonic - Bioscientia - MVZ Labor Saar GmbH | Robert Koch Institute | unknown |
| EPI_ISL_2632446, EPI_ISL_2632467, EPI_ISL_2632499 | Bioscientia Labor Wermsdorf | Robert Koch Institute | unknown |
| EPI_ISL_2632511, EPI_ISL_2632524 | LabKom - MVZ Labor Bochum MLB GmbH | Robert Koch Institute | unknown |
| EPI_ISL_2632525, EPI_ISL_2632530 | Bioscientia Labor Wermsdorf | Robert Koch Institute | unknown |
| EPI_ISL_2632563 | LabKom - Labor Hannover MVZ GmbH | Robert Koch Institute | unknown |
| EPI_ISL_2632567, EPI_ISL_2632568, EPI_ISL_2632586, EPI_ISL_2632638 | Bioscientia Labor Wermsdorf | Robert Koch Institute | unknown |
| EPI_ISL_2632664, EPI_ISL_2632670, EPI_ISL_2632692, EPI_ISL_2632723 | Limbach - MVZ Humangenetik Ulm | Robert Koch Institute | unknown |

| | | | |
|---|---|---|---|
| EPI_ISL_2632799, EPI_ISL_2632800, EPI_ISL_2632813 | MVZ Labor Dr. Limbach & Kollegen GbR | Robert Koch Institute | unknown |
| EPI_ISL_2632820 | Niedersächsisches Landesgesundheitsamt (NLGA) | Robert Koch Institute | unknown |
| EPI_ISL_2632832, EPI_ISL_2632844 | MVZ Labor Dr. Limbach & Kollegen GbR | Robert Koch Institute | unknown |
| EPI_ISL_2632850, EPI_ISL_2632852, EPI_ISL_2632867 | MVZ für Laboratoriumsmedizin und Mikrobiologie Koblenz-Mittelrhein (Labor Koblenz) | Robert Koch Institute | unknown |
| EPI_ISL_2632871 | MDI Limbach Berlin GmbH; MVZ Labor Berlin | Robert Koch Institute | unknown |
| EPI_ISL_2632872 | MVZ für Laboratoriumsmedizin und Mikrobiologie Koblenz-Mittelrhein (Labor Koblenz) | Robert Koch Institute | unknown |
| EPI_ISL_2632874 | MDI Limbach Berlin GmbH; MVZ Labor Berlin | Robert Koch Institute | unknown |
| EPI_ISL_2632878, EPI_ISL_2632882 | MVZ für Laboratoriumsmedizin und Mikrobiologie Koblenz-Mittelrhein (Labor Koblenz) | Robert Koch Institute | unknown |
| EPI_ISL_2632886 | MDI Limbach Berlin GmbH; MVZ Labor Berlin | Robert Koch Institute | unknown |
| EPI_ISL_2632888, EPI_ISL_2632895 | MVZ für Laboratoriumsmedizin und Mikrobiologie Koblenz-Mittelrhein (Labor Koblenz) | Robert Koch Institute | unknown |
| EPI_ISL_2632919, EPI_ISL_2632938 | MVZ Labor Dr. Limbach & Kollegen GbR | Robert Koch Institute | unknown |
| EPI_ISL_2632959 | Limbach - MVZ Gemeinschaftslabor Suhl | Robert Koch Institute | unknown |
| EPI_ISL_2632963, EPI_ISL_2632977 | MDI Limbach Berlin GmbH; MVZ Labor Berlin | Robert Koch Institute | unknown |
| EPI_ISL_2632988 | SYNLAB MVZ Weiden | Robert Koch Institute | unknown |
| EPI_ISL_2632991 | SYNLAB MVZ Leinfelden-Echterdingen | Robert Koch Institute | unknown |
| EPI_ISL_2632997 | SYNLAB MVZ Trier | Robert Koch Institute | unknown |
| EPI_ISL_2633019 | SYNLAB MVZ Leinfelden-Echterdingen | Robert Koch Institute | unknown |
| EPI_ISL_2633032 | Synlab MVZ Augsburg | Robert Koch Institute | unknown |
| EPI_ISL_2633044, EPI_ISL_2633068, EPI_ISL_2633074, EPI_ISL_2633075 | SYNLAB MVZ Leverkusen | Robert Koch Institute | unknown |
| EPI_ISL_2633092 | SYNLAB MVZ Trier | Robert Koch Institute | unknown |
| EPI_ISL_2633103, EPI_ISL_2633141 | SYNLAB MVZ Weiden | Robert Koch Institute | unknown |
| EPI_ISL_2633160 | SYNLAB MVZ Leverkusen | Robert Koch Institute | unknown |
| EPI_ISL_2633170 | SYNLAB MVZ Weiden | Robert Koch Institute | unknown |
| EPI_ISL_2633171 | SYNLAB MVZ Leinfelden-Echterdingen | Robert Koch Institute | unknown |
| EPI_ISL_2633203 | Synlab MVZ Augsburg | Robert Koch Institute | unknown |
| EPI_ISL_2633206 | SYNLAB MVZ Weiden | Robert Koch Institute | unknown |
| EPI_ISL_2633212 | SYNLAB MVZ Dachau | Robert Koch Institute | unknown |
| EPI_ISL_2633216, EPI_ISL_2633229 | SYNLAB MVZ Weiden | Robert Koch Institute | unknown |
| EPI_ISL_2633233 | SYNLAB MVZ Berlin | Robert Koch Institute | unknown |
| EPI_ISL_2633252, EPI_ISL_2633253 | SYNLAB MVZ Weiden | Robert Koch Institute | unknown |
| EPI_ISL_2633283 | Praxisgemeinschaft für Laboratoriumsmedizin Labor Blumenstraße; Praxis Dr. Sel und Dr. Späte | Robert Koch Institute | unknown |
| EPI_ISL_2633318 | SYNLAB MVZ Weiden | Robert Koch Institute | unknown |
| EPI_ISL_2633340 | SYNLAB MVZ Trier | Robert Koch Institute | unknown |
| EPI_ISL_2633360 | SYNLAB MVZ Weiden | Robert Koch Institute | unknown |
| EPI_ISL_2633364, EPI_ISL_2633373 | SYNLAB MVZ Leinfelden-Echterdingen | Robert Koch Institute | unknown |
| EPI_ISL_2633375 | SYNLAB MVZ Weiden | Robert Koch Institute | unknown |
| EPI_ISL_2633419 | MVZ Dr. Eberhard & Partner Dortmund | Robert Koch Institute | unknown |
| EPI_ISL_2633433, EPI_ISL_2633435 | MVZ für Laboratoriumsmedizin und Mikrobiologie Würzburg | Robert Koch Institute | unknown |
| EPI_ISL_2633502 | CENTOGENE Frankfurt Laboratory: Niederlassung Industriepark Höchst | Robert Koch Institute | unknown |
| EPI_ISL_2633506 | Area of Virology, Serology and Virology Division (SAViD), New South Wales Health Pathology Randwick | Virology Research Laboratory; Area of Virology, Serology and Virology Division (SAViD), New South Wales Health Pathology Randwick | Foster, C.; Au, J.; Ruiz Silva, M.; Deveson, I.; Bull, R.; Van Hal, S.; Rawlinson, W. |
| EPI_ISL_2633512 | CENTOGENE Frankfurt Laboratory: Niederlassung Industriepark Höchst | Robert Koch Institute | unknown |
| EPI_ISL_2633515, EPI_ISL_2633520, EPI_ISL_2633525 | Area of Virology, Serology and Virology Division (SAViD), New South Wales Health Pathology Randwick | Virology Research Laboratory; Area of Virology, Serology and Virology Division (SAViD), New South Wales Health Pathology Randwick | Foster, C.; Au, J.; Ruiz Silva, M.; Deveson, I.; Bull, R.; Van Hal, S.; Rawlinson, W. |
| EPI_ISL_2633549, EPI_ISL_2633557, EPI_ISL_2633565, EPI_ISL_2633566, EPI_ISL_2633569, EPI_ISL_2633577, EPI_ISL_2633581, EPI_ISL_2633595, EPI_ISL_2633596, EPI_ISL_2633602, EPI_ISL_2633606, EPI_ISL_2633614, EPI_ISL_2633637, EPI_ISL_2633639, EPI_ISL_2633652, EPI_ISL_2633664, EPI_ISL_2633688, EPI_ISL_2633709, EPI_ISL_2633711, EPI_ISL_2633712, EPI_ISL_2633713 | | | |
| see above | CENTOGENE Frankfurt Laboratory: Niederlassung Industriepark Höchst | Robert Koch Institute | unknown |
| EPI_ISL_2633742, EPI_ISL_2633744, EPI_ISL_2633747 | Labormedizin Darmstadt | Robert Koch Institute | unknown |
| EPI_ISL_2633751 | Universitätsklinikum Bonn - Institut für Virologie | Robert Koch Institute | unknown |
| EPI_ISL_2633756, EPI_ISL_2633760, EPI_ISL_2633762, EPI_ISL_2633765, EPI_ISL_2633767 | Labormedizin Darmstadt | Robert Koch Institute | unknown |
| EPI_ISL_2633771 | IMD Labor Frankfurt | Robert Koch Institute | unknown |
| EPI_ISL_2633773 | Universitätsklinikum Bonn - Institut für Virologie | Robert Koch Institute | unknown |
| EPI_ISL_2633775 | Labormedizin Darmstadt | Robert Koch Institute | unknown |
| EPI_ISL_2633776 | Universitätsklinikum Bonn - Institut für Virologie | Robert Koch Institute | unknown |
| EPI_ISL_2633780 | IMD Labor Frankfurt | Robert Koch Institute | unknown |
| EPI_ISL_2633782, EPI_ISL_2633784, EPI_ISL_2633789, EPI_ISL_2633794, EPI_ISL_2633797, EPI_ISL_2633800, EPI_ISL_2633801, EPI_ISL_2633802, EPI_ISL_2633804, EPI_ISL_2633805, EPI_ISL_2633811, EPI_ISL_2633812, EPI_ISL_2633813, EPI_ISL_2633816, EPI_ISL_2633817, EPI_ISL_2633818 | | | |
| see above | Labormedizin Darmstadt | Robert Koch Institute | unknown |

| | | | |
|---|---|---|---|
| EPI_ISL_2633823 | Universitätsklinikum Bonn - Institut für Virologie | Robert Koch Institute | unknown |
| EPI_ISL_2633824, EPI_ISL_2633828, EPI_ISL_2633831, EPI_ISL_2633832 | Labormedizin Darmstadt | Robert Koch Institute | unknown |
| EPI_ISL_2633839 | Universitätsklinikum Bonn - Institut für Virologie | Robert Koch Institute | unknown |
| EPI_ISL_2633841, EPI_ISL_2633846, EPI_ISL_2633852, EPI_ISL_2633853 | Labormedizin Darmstadt | Robert Koch Institute | unknown |
| EPI_ISL_2633854 | Universitätsklinikum Bonn - Institut für Virologie | Robert Koch Institute | unknown |
| EPI_ISL_2633860, EPI_ISL_2633866, EPI_ISL_2633873, EPI_ISL_2633891 | Labormedizin Darmstadt | Robert Koch Institute | unknown |
| EPI_ISL_2633901 | Bioscientia Labor Wermsdorf | Robert Koch Institute | unknown |
| EPI_ISL_2633908 | Sonic - Labor Dr. von Froreich GmbH | Robert Koch Institute | unknown |
| EPI_ISL_2633920, EPI_ISL_2633927, EPI_ISL_2633934, EPI_ISL_2634012, EPI_ISL_2634030, EPI_ISL_2634041, EPI_ISL_2634045, EPI_ISL_2634082, EPI_ISL_2634087, EPI_ISL_2634146, EPI_ISL_2634174, EPI_ISL_2634190 | | | |
| see above | Bioscientia Labor Wermsdorf | Robert Koch Institute | unknown |
| EPI_ISL_2634197 | Sonic - Dr. Staber & Kollegen GmbH München | Robert Koch Institute | unknown |
| EPI_ISL_2634202 | Bioscientia Labor Wermsdorf | Robert Koch Institute | unknown |
| EPI_ISL_2634222 | LabKom - Labor Augsburg MVZ GmbH | Robert Koch Institute | unknown |
| EPI_ISL_2634227 | Labor Dr. Schumacher MVZ | Robert Koch Institute | unknown |
| EPI_ISL_2634249 | Bioscientia MVZ Labor Karlsruhe GmbH | Robert Koch Institute | unknown |
| EPI_ISL_2634271 | Sonic - Labor Dr. von Froreich GmbH | Robert Koch Institute | unknown |
| EPI_ISL_2634294 | Bioscientia Labor Wermsdorf | Robert Koch Institute | unknown |
| EPI_ISL_2634385 | Sonic - Dr. Staber & Kollegen GmbH München | Robert Koch Institute | unknown |
| EPI_ISL_2634397 | Sonic - Bioscientia - MVZ Labor Saar GmbH | Robert Koch Institute | unknown |
| EPI_ISL_2634414 | Sonic - Labor Lademannbogen MVZ GmbH | Robert Koch Institute | unknown |
| EPI_ISL_2634448, EPI_ISL_2634452 | Bioscientia Labor Wermsdorf | Robert Koch Institute | unknown |
| EPI_ISL_2634455 | Sonic - Labor Dr. von Froreich GmbH | Robert Koch Institute | unknown |
| EPI_ISL_2634477, EPI_ISL_2634501 | labopart - Medizinische Laboratorien Dresden | Robert Koch Institute | unknown |
| EPI_ISL_2634513, EPI_ISL_2634520, EPI_ISL_2634529, EPI_ISL_2634538, EPI_ISL_2634540, EPI_ISL_2634561, EPI_ISL_2634569, EPI_ISL_2634577, EPI_ISL_2634579, EPI_ISL_2634592, EPI_ISL_2634594, EPI_ISL_2634596, EPI_ISL_2634623, EPI_ISL_2634639, EPI_ISL_2634643, EPI_ISL_2634645, EPI_ISL_2634648, EPI_ISL_2634650, EPI_ISL_2634661, EPI_ISL_2634664, EPI_ISL_2634674, EPI_ISL_2634676, EPI_ISL_2634680, EPI_ISL_2634683, EPI_ISL_2634685 | | | |
| see above | Eurofins LifeCodexx GmbH | Robert Koch Institute | unknown |
| EPI_ISL_2634704, EPI_ISL_2634715 | Clemenshospital - Alexianer Clemenshospital Münster | Robert Koch Institute | unknown |
| EPI_ISL_2634738 | Uniklinikum Carl Gustav Carus an der TU Dresden; Institut für Virologie | Robert Koch Institute | unknown |
| EPI_ISL_2634756, EPI_ISL_2634779 | Landesuntersuchungsanstalt für das Gesundheits- und Veterinärwesen (LUA) Sachsen | Robert Koch Institute | unknown |
| EPI_ISL_2634815 | Medizinisches Labor Ostsachsen MVZ GbR Nebenbetriebsstätte Dresden | Robert Koch Institute | unknown |
| EPI_ISL_2634860, EPI_ISL_2634862, EPI_ISL_2634864, EPI_ISL_2634869, EPI_ISL_2634876, EPI_ISL_2634882, EPI_ISL_2634884, EPI_ISL_2634886, EPI_ISL_2634888, EPI_ISL_2634891, EPI_ISL_2634893, EPI_ISL_2634895, EPI_ISL_2634905, EPI_ISL_2634910, EPI_ISL_2634919, EPI_ISL_2634921 | | | |
| see above | Landesgesundheitsamt Baden-Württemberg | Robert Koch Institute | unknown |
| EPI_ISL_2634925 | Pandemic Response Lab - NYC | Pandemic Response Lab, R&D | Henry Lee, Michael Hammerling, Melissa Hopkins, Cybill del Castillo, Shinyoung Clair Kang, William Ward, Pradeep Bugga, Sol Rey, Dylan Law, Katharine Nelson, Haiping Hao, Jon Laurent |
| EPI_ISL_2634926 | Landesgesundheitsamt Baden-Württemberg | Robert Koch Institute | unknown |
| EPI_ISL_2634927 | Pandemic Response Lab - NYC | Pandemic Response Lab, R&D | Henry Lee, Michael Hammerling, Melissa Hopkins, Cybill del Castillo, Shinyoung Clair Kang, William Ward, Pradeep Bugga, Sol Rey, Dylan Law, Katharine Nelson, Haiping Hao, Jon Laurent |
| EPI_ISL_2634928, EPI_ISL_2634932 | Landesgesundheitsamt Baden-Württemberg | Robert Koch Institute | unknown |
| EPI_ISL_2634933 | Pandemic Response Lab - NYC | Pandemic Response Lab, R&D | Henry Lee, Michael Hammerling, Melissa Hopkins, Cybill del Castillo, Shinyoung Clair Kang, William Ward, Pradeep Bugga, Sol Rey, Dylan Law, Katharine Nelson, Haiping Hao, Jon Laurent |
| EPI_ISL_2634935 | Landesgesundheitsamt Baden-Württemberg | Robert Koch Institute | unknown |
| EPI_ISL_2634938, EPI_ISL_2634940, EPI_ISL_2634942, EPI_ISL_2634945, EPI_ISL_2634947, EPI_ISL_2634949, EPI_ISL_2634951, EPI_ISL_2634952, EPI_ISL_2634958, EPI_ISL_2634963, EPI_ISL_2634969 | | | |
| see above | Pandemic Response Lab - NYC | Pandemic Response Lab, R&D | Henry Lee, Michael Hammerling, Melissa Hopkins, Cybill del Castillo, Shinyoung Clair Kang, William Ward, Pradeep Bugga, Sol Rey, Dylan Law, Katharine Nelson, Haiping Hao, Jon Laurent |
| EPI_ISL_2634978, EPI_ISL_2635003 | Bayerisches Landesamt für Gesundheit und Lebensmittelsicherheit (LGL) | Robert Koch Institute | unknown |
| EPI_ISL_2635088, EPI_ISL_2635091, EPI_ISL_2635098, EPI_ISL_2635100, EPI_ISL_2635102, EPI_ISL_2635104, EPI_ISL_2635105, EPI_ISL_2635109, EPI_ISL_2635111, EPI_ISL_2635115, EPI_ISL_2635116, EPI_ISL_2635118, EPI_ISL_2635120, EPI_ISL_2635122, EPI_ISL_2635124, EPI_ISL_2635126, EPI_ISL_2635127, EPI_ISL_2635130, EPI_ISL_2635131, EPI_ISL_2635133, EPI_ISL_2635135, EPI_ISL_2635137, EPI_ISL_2635138 | | | |
| see above | Missouri State Public Health Laboratory | Missouri State Public Health Laboratory | Matthew Sinn, Joshua Barry, Ashley New |
| EPI_ISL_2635139 | MDI Limbach Berlin GmbH; MVZ Labor Berlin | Robert Koch Institute | unknown |
| EPI_ISL_2635140 | Missouri State Public Health Laboratory | Missouri State Public Health Laboratory | Matthew Sinn, Joshua Barry, Ashley New |
| EPI_ISL_2635141 | MDI Limbach Berlin GmbH; MVZ Labor Berlin | Robert Koch Institute | unknown |
| EPI_ISL_2635144, EPI_ISL_2635147, EPI_ISL_2635149, EPI_ISL_2635151 | Missouri State Public Health Laboratory | Missouri State Public Health Laboratory | Matthew Sinn, Joshua Barry, Ashley New |
| EPI_ISL_2635152 | Limbach - MVZ Labor Eveld & Kollegen Essen | Robert Koch Institute | unknown |
| EPI_ISL_2635156, EPI_ISL_2635158 | Missouri State Public Health Laboratory | Missouri State Public Health Laboratory | Matthew Sinn, Joshua Barry, Ashley New |
| EPI_ISL_2635172 | MVZ Labor Dr. Limbach & Kollegen GbR | Robert Koch Institute | unknown |
| EPI_ISL_2635198, EPI_ISL_2635222, EPI_ISL_2635246, EPI_ISL_2635263, EPI_ISL_2635316 | SYNLAB MVZ Leverkusen | Robert Koch Institute | unknown |
| EPI_ISL_2635341 | SYNLAB MVZ Leinfelden-Echterdingen | Robert Koch Institute | unknown |
| EPI_ISL_2635355, EPI_ISL_2635391 | SYNLAB MVZ Leverkusen | Robert Koch Institute | unknown |
| EPI_ISL_2635404 | SYNLAB MVZ Leinfelden-Echterdingen | Robert Koch Institute | unknown |
| EPI_ISL_2635420 | Synlab MVZ Augsburg | Robert Koch Institute | unknown |
| EPI_ISL_2635421, EPI_ISL_2635426 | SYNLAB MVZ Weiden | Robert Koch Institute | unknown |
| EPI_ISL_2635433 | Synlab MVZ Augsburg | Robert Koch Institute | unknown |

| | | | |
|---|---|---|---|
| EPI_ISL_2635435 | SYNLAB MVZ Dachau | Robert Koch Institute | unknown |
| EPI_ISL_2635437 | SYNLAB MVZ Heidelberg | Robert Koch Institute | unknown |
| EPI_ISL_2635438, EPI_ISL_2635439 | SYNLAB MVZ Weiden | Robert Koch Institute | unknown |
| EPI_ISL_2635440, EPI_ISL_2635445, EPI_ISL_2635447 | SYNLAB MVZ Heidelberg | Robert Koch Institute | unknown |
| EPI_ISL_2635449 | SYNLAB MVZ Weiden | Robert Koch Institute | unknown |
| EPI_ISL_2635451 | SYNLAB MVZ Dachau | Robert Koch Institute | unknown |
| EPI_ISL_2635452 | SYNLAB MVZ Trier | Robert Koch Institute | unknown |
| EPI_ISL_2635455 | SYNLAB MVZ Hamburg | Robert Koch Institute | unknown |
| EPI_ISL_2635461 | SYNLAB MVZ Heidelberg | Robert Koch Institute | unknown |
| EPI_ISL_2635462 | Synlab MVZ Augsburg | Robert Koch Institute | unknown |
| EPI_ISL_2635465, EPI_ISL_2635466 | SYNLAB MVZ Weiden | Robert Koch Institute | unknown |
| EPI_ISL_2635467 | SYNLAB MVZ Hamburg | Robert Koch Institute | unknown |
| EPI_ISL_2635468 | SYNLAB MVZ Trier | Robert Koch Institute | unknown |
| EPI_ISL_2635476, EPI_ISL_2635484 | MVZ Dr. Eberhard & Partner Dortmund | Robert Koch Institute | unknown |
| EPI_ISL_2635490 | MVZ für Laboratoriumsmedizin und Mikrobiologie Würzburg | Robert Koch Institute | unknown |
| EPI_ISL_2635491, EPI_ISL_2635494, EPI_ISL_2635495, EPI_ISL_2635496, EPI_ISL_2635497, EPI_ISL_2635498, EPI_ISL_2635499, EPI_ISL_2635500, EPI_ISL_2635502, EPI_ISL_2635504, EPI_ISL_2635506, EPI_ISL_2635509 | | | |
| see above | IMD - Labor Oderland | Robert Koch Institute | unknown |
| EPI_ISL_2635530, EPI_ISL_2635536, EPI_ISL_2635542, EPI_ISL_2635543, EPI_ISL_2635551, EPI_ISL_2635553, EPI_ISL_2635562 | Labor Becker & Kollegen (Standort München) | Robert Koch Institute | unknown |
| EPI_ISL_2635564 | Landesamt für Verbraucherschutz des Landes Sachsen-Anhalt | Robert Koch Institute | unknown |
| EPI_ISL_2635569, EPI_ISL_2635582, EPI_ISL_2635584, EPI_ISL_2635588, EPI_ISL_2635591, EPI_ISL_2635598 | Medizinische Laboratorien Düsseldorf | Robert Koch Institute | unknown |
| EPI_ISL_2635602 | nordlab - Partnerschaftspraxis für Laboratoriumsmedizin | Robert Koch Institute | unknown |
| EPI_ISL_2635604, EPI_ISL_2635608 | amedes MVZ für Labordiagnostik Rhein-Main | Robert Koch Institute | unknown |
| EPI_ISL_2635611, EPI_ISL_2635629 | nordlab - Partnerschaftspraxis für Laboratoriumsmedizin | Robert Koch Institute | unknown |
| EPI_ISL_2635630 | amedes MVZ für Labordiagnostik Rhein-Main | Robert Koch Institute | unknown |
| EPI_ISL_2635635 | amedes MVZ Hannover | Robert Koch Institute | unknown |
| EPI_ISL_2635640 | amedes MVZ für Labordiagnostik Rhein-Main | Robert Koch Institute | unknown |
| EPI_ISL_2635648 | nordlab - Partnerschaftspraxis für Laboratoriumsmedizin | Robert Koch Institute | unknown |
| EPI_ISL_2635650 | amedes MVZ für Laboratoriumsmedizin Hamburg | Robert Koch Institute | unknown |
| EPI_ISL_2635657, EPI_ISL_2635660 | nordlab - Partnerschaftspraxis für Laboratoriumsmedizin | Robert Koch Institute | unknown |
| EPI_ISL_2635662 | St. Franziskus-Hospital Münster | Robert Koch Institute | unknown |
| EPI_ISL_2635666, EPI_ISL_2635708 | Limbach - MVZ Humangenetik Ulm | Robert Koch Institute | unknown |
| EPI_ISL_2635743 | Limbach - MVZ Labor Eveld & Kollegen Essen | Robert Koch Institute | unknown |
| EPI_ISL_2635757, EPI_ISL_2635758, EPI_ISL_2635759, EPI_ISL_2635760 | Labor Dr. Wisplinghoff - Köln | Robert Koch Institute | unknown |
| EPI_ISL_2635769 | Limbach - MVZ Gemeinschaftslabor Suhl | Robert Koch Institute | unknown |
| EPI_ISL_2635770, EPI_ISL_2635771 | MVZ Labor Dr. Limbach & Kollegen GbR | Robert Koch Institute | unknown |
| EPI_ISL_2635776, EPI_ISL_2635777, EPI_ISL_2635781, EPI_ISL_2635784 | MDI Limbach Berlin GmbH; MVZ Labor Berlin | Robert Koch Institute | unknown |
| EPI_ISL_2635785, EPI_ISL_2635788 | Med. Labor Prof. Schenk Dr. Ansorge & Kollegen | Robert Koch Institute | unknown |
| EPI_ISL_2635811 | MDI Limbach Berlin GmbH; MVZ Labor Berlin | Robert Koch Institute | unknown |
| EPI_ISL_2635814, EPI_ISL_2635815, EPI_ISL_2635816, EPI_ISL_2635817, EPI_ISL_2635818, EPI_ISL_2635819, EPI_ISL_2635820 | Niedersächsisches Landesgesundheitsamt (NLGA) | Robert Koch Institute | unknown |
| EPI_ISL_2635831 | MDI Limbach Berlin GmbH; MVZ Labor Berlin | Robert Koch Institute | unknown |
| EPI_ISL_2635832 | Limbach - MVZ Gemeinschaftslabor Suhl | Robert Koch Institute | unknown |
| EPI_ISL_2635834 | MDI Limbach Berlin GmbH; MVZ Labor Berlin | Robert Koch Institute | unknown |
| EPI_ISL_2635835 | Labor Prof. Dr. G. Enders MVZ GbR | Robert Koch Institute | unknown |
| EPI_ISL_2635836, EPI_ISL_2635838 | MVZ für Laboratoriumsmedizin und Mikrobiologie Koblenz-Mittelrhein (Labor Koblenz) | Robert Koch Institute | unknown |
| EPI_ISL_2635842 | MDI Limbach Berlin GmbH; MVZ Labor Berlin | Robert Koch Institute | unknown |
| EPI_ISL_2635845, EPI_ISL_2635846, EPI_ISL_2635847, EPI_ISL_2635849, EPI_ISL_2635850, EPI_ISL_2635852, EPI_ISL_2635853, EPI_ISL_2635855, EPI_ISL_2635856, EPI_ISL_2635857, EPI_ISL_2635858, EPI_ISL_2635859, EPI_ISL_2635860 | | | |
| see above | Labor Dr. Heidrich & Kollegen MVZ GmbH Hamburg | Robert Koch Institute | unknown |
| EPI_ISL_2635861, EPI_ISL_2635862, EPI_ISL_2635864 | MVZ Labor Dr. Limbach & Kollegen GbR | Robert Koch Institute | unknown |
| EPI_ISL_2635868 | Limbach - MVZ Labor Eveld & Kollegen Essen | Robert Koch Institute | unknown |
| EPI_ISL_2635869, EPI_ISL_2635880, | MVZ Labor Dr. Limbach & Kollegen GbR | Robert Koch Institute | unknown |

| | | | |
|---|---|---|---|
| EPI_ISL_2635884, EPI_ISL_2635885 | | | |
| EPI_ISL_2635893, EPI_ISL_2635894 | Limbach - MVZ Gemeinschaftslabor Suhl | Robert Koch Institute | unknown |
| EPI_ISL_2635896 | MVZ Medizinisches Labor Hannover GmbH | Robert Koch Institute | unknown |
| EPI_ISL_2635897, EPI_ISL_2635898 | MVZ Labor Dr. Limbach & Kollegen GbR | Robert Koch Institute | unknown |
| EPI_ISL_2635903, EPI_ISL_2635905 | Limbach - MVZ Labor Ludwigsburg | Robert Koch Institute | unknown |
| EPI_ISL_2635948 | Helios Universitätsklinikum Wuppertal: Universität Witten/Herdecke: Institut für Medizinische Labordiagnostik | Robert Koch Institute | unknown |
| EPI_ISL_2635974, EPI_ISL_2635975 | Gemeinschaftspraxis für Pathologie Dr. Blume Prof. Dr. Dämmrich Dr. Heine Dr. Dämmrich | Robert Koch Institute | unknown |
| EPI_ISL_2636008 | Laborarztpraxis Dres. med. Walther Weindel & Kollegen | Robert Koch Institute | unknown |
| EPI_ISL_2636034, EPI_ISL_2636059, EPI_ISL_2636071 | Labor ZOTZ\|KLIMAS; MVZ Düsseldorf-Centrum | Robert Koch Institute | unknown |
| EPI_ISL_2636081, EPI_ISL_2636093, EPI_ISL_2636097 | Medizinisch-Diagnostisches Labor Kempten allgäulab | Robert Koch Institute | unknown |
| EPI_ISL_2636126, EPI_ISL_2636129, EPI_ISL_2636134, EPI_ISL_2636139, EPI_ISL_2636146 | Corona-Testzentrum ifp Institut für Produktqualität GmbH | Robert Koch Institute | unknown |
| EPI_ISL_2636161, EPI_ISL_2636166, EPI_ISL_2636189 | Israel Central Virology laboratory | Israel National Consortium for SARS-CoV-2 sequencing | Neta Zuckerman, Efrat Dahan Bucris, Michal Mandelboim, Dana Bar-Ilan, Miranda Geva, Oran Erster, Efrat Glick-Saar, Omri Nayshool, Gideon Rechavi, Ella Mendelson, Orna Mor |
| EPI_ISL_2636195 | MVZ Labor Dr. Reising-Ackermann und Kollegen | Robert Koch Institute | unknown |
| EPI_ISL_2636204 | Israel Central Virology laboratory | Israel National Consortium for SARS-CoV-2 sequencing | Neta Zuckerman, Efrat Dahan Bucris, Michal Mandelboim, Dana Bar-Ilan, Miranda Geva, Oran Erster, Efrat Glick-Saar, Omri Nayshool, Gideon Rechavi, Ella Mendelson, Orna Mor |
| EPI_ISL_2636216 | MVZ Labor Dr. Reising-Ackermann und Kollegen | Robert Koch Institute | unknown |
| EPI_ISL_2636218 | Labor Mönchengladbach MVZ Dr. Stein + Kollegen GbR | Robert Koch Institute | unknown |
| EPI_ISL_2636241 | Israel Central Virology laboratory | Israel National Consortium for SARS-CoV-2 sequencing | Neta Zuckerman, Efrat Dahan Bucris, Michal Mandelboim, Dana Bar-Ilan, Miranda Geva, Oran Erster, Efrat Glick-Saar, Omri Nayshool, Gideon Rechavi, Ella Mendelson, Orna Mor |
| EPI_ISL_2636270 | Diagnosticum - Labor Neukirchen | Robert Koch Institute | unknown |
| EPI_ISL_2636298, EPI_ISL_2636309, EPI_ISL_2636326, EPI_ISL_2636333, EPI_ISL_2636334, EPI_ISL_2636336, EPI_ISL_2636343, EPI_ISL_2636345, EPI_ISL_2636347, EPI_ISL_2636353, EPI_ISL_2636354, EPI_ISL_2636356, EPI_ISL_2636358, EPI_ISL_2636361, EPI_ISL_2636363, EPI_ISL_2636365, EPI_ISL_2636367, EPI_ISL_2636368, EPI_ISL_2636371, EPI_ISL_2636373, EPI_ISL_2636377, EPI_ISL_2636379, EPI_ISL_2636383, EPI_ISL_2636385, EPI_ISL_2636387 | | | |
| see above | Israel Central Virology laboratory | Israel National Consortium for SARS-CoV-2 sequencing | Neta Zuckerman, Efrat Dahan Bucris, Michal Mandelboim, Dana Bar-Ilan, Miranda Geva, Oran Erster, Efrat Glick-Saar, Omri Nayshool, Gideon Rechavi, Ella Mendelson, Orna Mor |
| EPI_ISL_2636388 | Diagnosticum - Labor Neukirchen | Robert Koch Institute | unknown |
| EPI_ISL_2636394, EPI_ISL_2636396, EPI_ISL_2636399 | Israel Central Virology laboratory | Israel National Consortium for SARS-CoV-2 sequencing | Neta Zuckerman, Efrat Dahan Bucris, Michal Mandelboim, Dana Bar-Ilan, Miranda Geva, Oran Erster, Efrat Glick-Saar, Omri Nayshool, Gideon Rechavi, Ella Mendelson, Orna Mor |
| EPI_ISL_2636428, EPI_ISL_2636439 | Diagnosticum - Labor Neukirchen | Robert Koch Institute | unknown |
| EPI_ISL_2636473 | MDI Limbach Berlin GmbH; MVZ Labor Berlin | Robert Koch Institute | unknown |
| EPI_ISL_2636474 | Limbach - MVZ Gemeinschaftslabor Suhl | Robert Koch Institute | unknown |
| EPI_ISL_2636481 | Medizinisches Labor Ostsachsen MVZ GbR Nebenbetriebsstätte Dresden | Robert Koch Institute | unknown |
| EPI_ISL_2636485 | MVZ für Laboratoriumsmedizin und Mikrobiologie Koblenz-Mittelrhein (Labor Koblenz) | Robert Koch Institute | unknown |
| EPI_ISL_2636496 | MDI Limbach Berlin GmbH; MVZ Labor Berlin | Robert Koch Institute | unknown |
| EPI_ISL_2636498 | MVZ Labor Dr. Limbach & Kollegen GbR | Robert Koch Institute | unknown |
| EPI_ISL_2636500, EPI_ISL_2636505, EPI_ISL_2636515 | MDI Limbach Berlin GmbH; MVZ Labor Berlin | Robert Koch Institute | unknown |
| EPI_ISL_2636523 | Limbach - Ganzimmun Diagnostics AG Mainz | Robert Koch Institute | unknown |
| EPI_ISL_2636550, EPI_ISL_2636573, EPI_ISL_2636603, EPI_ISL_2636604, EPI_ISL_2636650, EPI_ISL_2636664, EPI_ISL_2636700, EPI_ISL_2636710, EPI_ISL_2636716 | Diagnosticum - Labor Neukirchen | Robert Koch Institute | unknown |
| EPI_ISL_2636741, EPI_ISL_2636742, EPI_ISL_2636743, EPI_ISL_2636744, EPI_ISL_2636746, EPI_ISL_2636751, EPI_ISL_2636752, EPI_ISL_2636758 | MVZ Labor Dr. Fenner und Kollegen (Standort Hamburg) | Robert Koch Institute | unknown |
| EPI_ISL_2636791, EPI_ISL_2636795, EPI_ISL_2636796, EPI_ISL_2636809, EPI_ISL_2636811, EPI_ISL_2636817, EPI_ISL_2636828, EPI_ISL_2636830, EPI_ISL_2636837, EPI_ISL_2636843, EPI_ISL_2636867 | | | |
| see above | Medizinisch-Diagnostisches Labor Kempten allgäulab | Robert Koch Institute | unknown |
| EPI_ISL_2636876 | MVZ Labor Dr. Limbach & Kollegen GbR | Robert Koch Institute | unknown |
| EPI_ISL_2636880 | Laborarztpraxis Osnabrück | Robert Koch Institute | unknown |
| EPI_ISL_2636885 | Procomcure Biotech Germany GmbH | Robert Koch Institute | unknown |
| EPI_ISL_2636887 | Laborarztpraxis Osnabrück | Robert Koch Institute | unknown |
| EPI_ISL_2636896 | Medizinisches Versorgungszentrum für Labormedizin und Mikrobiologie Ruhr GmbH - mvzlm RUHR GmbH | Robert Koch Institute | unknown |
| EPI_ISL_2636901 | Labor Dr. Spranger | Robert Koch Institute | unknown |
| EPI_ISL_2636904, EPI_ISL_2636905 | Labormedizin Darmstadt | Robert Koch Institute | unknown |
| EPI_ISL_2636906 | Universitätsklinikum des Saarlandes - Institut für Virologie | Robert Koch Institute | unknown |
| EPI_ISL_2636914, EPI_ISL_2636921, EPI_ISL_2636935, EPI_ISL_2636938, EPI_ISL_2636949, EPI_ISL_2636970, | Bioscientia Labor Wermsdorf | Robert Koch Institute | unknown |

| | | | |
|---|---|---|---|
| EPI_ISL_2636979, EPI_ISL_2636986, EPI_ISL_2636988, EPI_ISL_2636993 | | | |
| EPI_ISL_2636998 | LabKom - Labor Hannover MVZ GmbH | Robert Koch Institute | unknown |
| EPI_ISL_2637001, EPI_ISL_2637003, EPI_ISL_2637010, EPI_ISL_2637034 | Bioscientia Labor Wermsdorf | Robert Koch Institute | unknown |
| EPI_ISL_2637035 | Labor 28 MVZ GmbH | Robert Koch Institute | unknown |
| EPI_ISL_2637039, EPI_ISL_2637046, EPI_ISL_2637061 | Bioscientia Labor Wermsdorf | Robert Koch Institute | unknown |
| EPI_ISL_2637069 | Labor Dr. Schumacher MVZ | Robert Koch Institute | unknown |
| EPI_ISL_2637070 | Bioscientia Labor Wermsdorf | Robert Koch Institute | unknown |
| EPI_ISL_2637091 | Sonic - Bioscientia - MVZ Labor Saar GmbH | Robert Koch Institute | unknown |
| EPI_ISL_2637098, EPI_ISL_2637099, EPI_ISL_2637103, EPI_ISL_2637122, EPI_ISL_2637124 | Bioscientia Labor Wermsdorf | Robert Koch Institute | unknown |
| EPI_ISL_2637132 | LabKom - Labor Augsburg MVZ GmbH | Robert Koch Institute | unknown |
| EPI_ISL_2637146, EPI_ISL_2637153, EPI_ISL_2637172, EPI_ISL_2637175, EPI_ISL_2637176 | Bioscientia Labor Wermsdorf | Robert Koch Institute | unknown |
| EPI_ISL_2637177 | LabKom - MVZ Labor Bochum MLB GmbH | Robert Koch Institute | unknown |
| EPI_ISL_2637182 | Bioscientia Labor Wermsdorf | Robert Koch Institute | unknown |
| EPI_ISL_2637184 | Labor 28 MVZ GmbH | Robert Koch Institute | unknown |
| EPI_ISL_2637188 | Bioscientia Labor Wermsdorf | Robert Koch Institute | unknown |
| EPI_ISL_2637190 | Labor 28 MVZ GmbH | Robert Koch Institute | unknown |
| EPI_ISL_2637195 | IMD - Labor Oderland | Robert Koch Institute | unknown |
| EPI_ISL_2637204 | IMD - MVZ Labor Greifswald | Robert Koch Institute | unknown |
| EPI_ISL_2637207 | IMD - Labor Oderland | Robert Koch Institute | unknown |
| EPI_ISL_2637224 | Department of Microbiology, Yamanashi Institute for Public Health and Environment | Genome Analysis Center, Yamanashi Central Hospital | Yosuke Hirotsu |
| EPI_ISL_2637244, EPI_ISL_2637245 | Hospital Universitari Bellvitge | Microbiology Department | Sara Marti, Aida Gonzalez-Diaz, Laura Calatayud, Jordi Niubó, Miguel Fernandez-Huerta, Carmen Ardanuy, Jordi Camara, M Angeles Domínguez |
| EPI_ISL_2637247, EPI_ISL_2637250, EPI_ISL_2637251, EPI_ISL_2637252, EPI_ISL_2637253, EPI_ISL_2637255, EPI_ISL_2637256, EPI_ISL_2637257, EPI_ISL_2637258, EPI_ISL_2637259, EPI_ISL_2637261 | | | |
| see above | CLILAB | Microbiology Department | Sara Marti, Aida Gonzalez-Diaz, Laura Calatayud, Jordi Niubó, Miguel Fernandez-Huerta, Carmen Ardanuy, Jordi Camara, M Angeles Domínguez |
| EPI_ISL_2637262 | Hospital Universitari Bellvitge | Microbiology Department | Sara Marti, Aida Gonzalez-Diaz, Laura Calatayud, Jordi Niubó, Miguel Fernandez-Huerta, Carmen Ardanuy, Jordi Camara, M Angeles Domínguez |
| EPI_ISL_2637263, EPI_ISL_2637264, EPI_ISL_2637265, EPI_ISL_2637267, EPI_ISL_2637268 | CLILAB | Microbiology Department | Sara Marti, Aida Gonzalez-Diaz, Laura Calatayud, Jordi Niubó, Miguel Fernandez-Huerta, Carmen Ardanuy, Jordi Camara, M Angeles Domínguez |
| EPI_ISL_2637269, EPI_ISL_2637270, EPI_ISL_2637276, EPI_ISL_2637278, EPI_ISL_2637280, EPI_ISL_2637281, EPI_ISL_2637282, EPI_ISL_2637283, EPI_ISL_2637284, EPI_ISL_2637285, EPI_ISL_2637286, EPI_ISL_2637287, EPI_ISL_2637288, EPI_ISL_2637289, EPI_ISL_2637290, EPI_ISL_2637291, EPI_ISL_2637292, EPI_ISL_2637293, EPI_ISL_2637294, EPI_ISL_2637296, EPI_ISL_2637298, EPI_ISL_2637299, EPI_ISL_2637301, EPI_ISL_2637302, EPI_ISL_2637303, EPI_ISL_2637304, EPI_ISL_2637305, EPI_ISL_2637307, EPI_ISL_2637308, EPI_ISL_2637309, EPI_ISL_2637310, EPI_ISL_2637311, EPI_ISL_2637312, EPI_ISL_2637319, EPI_ISL_2637322, EPI_ISL_2637323, EPI_ISL_2637327, EPI_ISL_2637332, EPI_ISL_2637333, EPI_ISL_2637334, EPI_ISL_2637335, EPI_ISL_2637336, EPI_ISL_2637337, EPI_ISL_2637349, EPI_ISL_2637350, EPI_ISL_2637351, EPI_ISL_2637355 | | | |
| see above | Hospital Universitari Bellvitge | Microbiology Department | Sara Marti, Aida Gonzalez-Diaz, Laura Calatayud, Jordi Niubó, Miguel Fernandez-Huerta, Carmen Ardanuy, Jordi Camara, M Angeles Domínguez |
| EPI_ISL_2637356 | CLILAB | Microbiology Department | Sara Marti, Aida Gonzalez-Diaz, Laura Calatayud, Jordi Niubó, Miguel Fernandez-Huerta, Carmen Ardanuy, Jordi Camara, M Angeles Domínguez |
| EPI_ISL_2637357, EPI_ISL_2637358, EPI_ISL_2637359 | Hospital Universitari Bellvitge | Microbiology Department | Sara Marti, Aida Gonzalez-Diaz, Laura Calatayud, Jordi Niubó, Miguel Fernandez-Huerta, Carmen Ardanuy, Jordi Camara, M Angeles Domínguez |
| EPI_ISL_2637360, EPI_ISL_2637361, EPI_ISL_2637363, EPI_ISL_2637365, EPI_ISL_2637366 | CLILAB | Microbiology Department | Sara Marti, Aida Gonzalez-Diaz, Laura Calatayud, Jordi Niubó, Miguel Fernandez-Huerta, Carmen Ardanuy, Jordi Camara, M Angeles Domínguez |
| EPI_ISL_2637367, EPI_ISL_2637368, EPI_ISL_2637369 | Hospital Universitari Bellvitge | Microbiology Department | Sara Marti, Aida Gonzalez-Diaz, Laura Calatayud, Jordi Niubó, Miguel Fernandez-Huerta, Carmen Ardanuy, Jordi Camara, M Angeles Domínguez |
| EPI_ISL_2637370, EPI_ISL_2637371 | CLILAB | Microbiology Department | Sara Marti, Aida Gonzalez-Diaz, Laura Calatayud, Jordi Niubó, Miguel Fernandez-Huerta, Carmen Ardanuy, Jordi Camara, M Angeles Domínguez |
| EPI_ISL_2637374, EPI_ISL_2637377, EPI_ISL_2637378, EPI_ISL_2637379, EPI_ISL_2637380 | Hospital Universitari Bellvitge | Microbiology Department | Sara Marti, Aida Gonzalez-Diaz, Laura Calatayud, Jordi Niubó, Miguel Fernandez-Huerta, Carmen Ardanuy, Jordi Camara, M Angeles Domínguez |
| EPI_ISL_2637382 | CLILAB | Microbiology Department | Sara Marti, Aida Gonzalez-Diaz, Laura Calatayud, Jordi Niubó, Miguel Fernandez-Huerta, Carmen Ardanuy, Jordi Camara, M Angeles Domínguez |
| EPI_ISL_2637383, EPI_ISL_2637384 | Hospital Universitari Bellvitge | Microbiology Department | Sara Marti, Aida Gonzalez-Diaz, Laura Calatayud, Jordi Niubó, Miguel Fernandez-Huerta, Carmen Ardanuy, Jordi Camara, M Angeles Domínguez |
| EPI_ISL_2637529, EPI_ISL_2637531 | BROUSSAIS | Department of Virology, Henri Mondor University Hospital, Assistance Publique Hôpitaux de Paris, Université Paris-Est Créteil, INSERM U955 | Christophe Rodriguez, Slim Fourati, Vanessa Demontant, Guillaume Gricourt, Melissa N'Debi, Alexandre Soulier, Elisabeth Trawinski, Jean-Michel Pawlotsky |
| EPI_ISL_2637553 | CHRU STRASBOURG | Department of Virology, Henri Mondor University Hospital, Assistance Publique Hôpitaux de Paris, Université Paris-Est Créteil, INSERM U955 | Christophe Rodriguez, Slim Fourati, Vanessa Demontant, Guillaume Gricourt, Melissa N'Debi, Alexandre Soulier, Elisabeth Trawinski, Jean-Michel Pawlotsky |
| EPI_ISL_2637555, EPI_ISL_2637566, EPI_ISL_2637578, EPI_ISL_2637609 | BROUSSAIS | Department of Virology, Henri Mondor University Hospital, Assistance Publique Hôpitaux de Paris, Université Paris-Est Créteil, INSERM U955 | Christophe Rodriguez, Slim Fourati, Vanessa Demontant, Guillaume Gricourt, Melissa N'Debi, Alexandre Soulier, Elisabeth Trawinski, Jean-Michel Pawlotsky |
| EPI_ISL_2637626 | CHI VILLENEUVE ST GEORGES | Department of Virology, Henri Mondor University Hospital, Assistance Publique Hôpitaux de Paris, Université Paris-Est Créteil, INSERM U955 | Christophe Rodriguez, Slim Fourati, Vanessa Demontant, Guillaume Gricourt, Melissa N'Debi, Alexandre Soulier, Elisabeth Trawinski, Jean-Michel Pawlotsky |
| EPI_ISL_2637631 | BROUSSAIS | Department of Virology, Henri Mondor University Hospital, Assistance Publique Hôpitaux de Paris, Université Paris-Est Créteil, INSERM U955 | Christophe Rodriguez, Slim Fourati, Vanessa Demontant, Guillaume Gricourt, Melissa N'Debi, Alexandre Soulier, Elisabeth Trawinski, Jean-Michel Pawlotsky |
| EPI_ISL_2637648 | CHI VILLENEUVE ST GEORGES | Department of Virology, Henri Mondor University Hospital, Assistance Publique Hôpitaux de Paris, Université Paris-Est Créteil, INSERM U955 | Christophe Rodriguez, Slim Fourati, Vanessa Demontant, Guillaume Gricourt, Melissa N'Debi, Alexandre Soulier, Elisabeth Trawinski, Jean-Michel Pawlotsky |

| | | | |
|---|---|---|---|
| EPI_ISL_2637666 | Hôpital Avicenne | Department of Virology, Henri Mondor University Hospital, Assistance Publique Hôpitaux de Paris, Université Paris-Est Créteil, INSERM U955 | Christophe Rodriguez, Slim Fourati, Vanessa Demontant, Guillaume Gricourt, Melissa N'Debi, Alexandre Soulier, Elisabeth Trawinski, Jean-Michel Pawlotsky |
| EPI_ISL_2637667 | Laboratoire CBM 25 TERRE ROUGE | Department of Virology, Henri Mondor University Hospital, Assistance Publique Hôpitaux de Paris, Université Paris-Est Créteil, INSERM U955 | Christophe Rodriguez, Slim Fourati, Vanessa Demontant, Guillaume Gricourt, Melissa N'Debi, Alexandre Soulier, Elisabeth Trawinski, Jean-Michel Pawlotsky |
| EPI_ISL_2637674 | BROUSSAIS | Department of Virology, Henri Mondor University Hospital, Assistance Publique Hôpitaux de Paris, Université Paris-Est Créteil, INSERM U955 | Christophe Rodriguez, Slim Fourati, Vanessa Demontant, Guillaume Gricourt, Melissa N'Debi, Alexandre Soulier, Elisabeth Trawinski, Jean-Michel Pawlotsky |
| EPI_ISL_2637692 | Hôpital Bichat-Claude Bernard | Department of Virology, Henri Mondor University Hospital, Assistance Publique Hôpitaux de Paris, Université Paris-Est Créteil, INSERM U955 | Christophe Rodriguez, Slim Fourati, Vanessa Demontant, Guillaume Gricourt, Melissa N'Debi, Alexandre Soulier, Elisabeth Trawinski, Jean-Michel Pawlotsky |
| EPI_ISL_2637730 | CH.INTERCOMMUNAL DE CRETEIL | Department of Virology, Henri Mondor University Hospital, Assistance Publique Hôpitaux de Paris, Université Paris-Est Créteil, INSERM U955 | Christophe Rodriguez, Slim Fourati, Vanessa Demontant, Guillaume Gricourt, Melissa N'Debi, Alexandre Soulier, Elisabeth Trawinski, Jean-Michel Pawlotsky |
| EPI_ISL_2637733 | BROUSSAIS | Department of Virology, Henri Mondor University Hospital, Assistance Publique Hôpitaux de Paris, Université Paris-Est Créteil, INSERM U955 | Christophe Rodriguez, Slim Fourati, Vanessa Demontant, Guillaume Gricourt, Melissa N'Debi, Alexandre Soulier, Elisabeth Trawinski, Jean-Michel Pawlotsky |
| EPI_ISL_2637735 | GH A.CHENEVIER-H.MONDOR | Department of Virology, Henri Mondor University Hospital, Assistance Publique Hôpitaux de Paris, Université Paris-Est Créteil, INSERM U955 | Christophe Rodriguez, Slim Fourati, Vanessa Demontant, Guillaume Gricourt, Melissa N'Debi, Alexandre Soulier, Elisabeth Trawinski, Jean-Michel Pawlotsky |
| EPI_ISL_2637771 | CHRU STRASBOURG | Department of Virology, Henri Mondor University Hospital, Assistance Publique Hôpitaux de Paris, Université Paris-Est Créteil, INSERM U955 | Christophe Rodriguez, Slim Fourati, Vanessa Demontant, Guillaume Gricourt, Melissa N'Debi, Alexandre Soulier, Elisabeth Trawinski, Jean-Michel Pawlotsky |
| EPI_ISL_2637778 | BROUSSAIS | Department of Virology, Henri Mondor University Hospital, Assistance Publique Hôpitaux de Paris, Université Paris-Est Créteil, INSERM U955 | Christophe Rodriguez, Slim Fourati, Vanessa Demontant, Guillaume Gricourt, Melissa N'Debi, Alexandre Soulier, Elisabeth Trawinski, Jean-Michel Pawlotsky |
| EPI_ISL_2637784 | Laboratoire LENYS | Department of Virology, Henri Mondor University Hospital, Assistance Publique Hôpitaux de Paris, Université Paris-Est Créteil, INSERM U955 | Christophe Rodriguez, Slim Fourati, Vanessa Demontant, Guillaume Gricourt, Melissa N'Debi, Alexandre Soulier, Elisabeth Trawinski, Jean-Michel Pawlotsky |
| EPI_ISL_2637786 | Hôpital Bichat-Claude Bernard | Department of Virology, Henri Mondor University Hospital, Assistance Publique Hôpitaux de Paris, Université Paris-Est Créteil, INSERM U955 | Christophe Rodriguez, Slim Fourati, Vanessa Demontant, Guillaume Gricourt, Melissa N'Debi, Alexandre Soulier, Elisabeth Trawinski, Jean-Michel Pawlotsky |
| EPI_ISL_2637788 | BROUSSAIS | Department of Virology, Henri Mondor University Hospital, Assistance Publique Hôpitaux de Paris, Université Paris-Est Créteil, INSERM U955 | Christophe Rodriguez, Slim Fourati, Vanessa Demontant, Guillaume Gricourt, Melissa N'Debi, Alexandre Soulier, Elisabeth Trawinski, Jean-Michel Pawlotsky |
| EPI_ISL_2637805 | Plateau technique Biomer Metz Verriers | Department of Virology, Henri Mondor University Hospital, Assistance Publique Hôpitaux de Paris, Université Paris-Est Créteil, INSERM U955 | Christophe Rodriguez, Slim Fourati, Vanessa Demontant, Guillaume Gricourt, Melissa N'Debi, Alexandre Soulier, Elisabeth Trawinski, Jean-Michel Pawlotsky |
| EPI_ISL_2637809 | DYNALAB | Department of Virology, Henri Mondor University Hospital, Assistance Publique Hôpitaux de Paris, Université Paris-Est Créteil, INSERM U955 | Christophe Rodriguez, Slim Fourati, Vanessa Demontant, Guillaume Gricourt, Melissa N'Debi, Alexandre Soulier, Elisabeth Trawinski, Jean-Michel Pawlotsky |
| EPI_ISL_2637861 | Plateau technique Biomer Metz Verriers | Department of Virology, Henri Mondor University Hospital, Assistance Publique Hôpitaux de Paris, Université Paris-Est Créteil, INSERM U955 | Christophe Rodriguez, Slim Fourati, Vanessa Demontant, Guillaume Gricourt, Melissa N'Debi, Alexandre Soulier, Elisabeth Trawinski, Jean-Michel Pawlotsky |
| EPI_ISL_2637877, EPI_ISL_2637905 | SELAS MEDILYS | Department of Virology, Henri Mondor University Hospital, Assistance Publique Hôpitaux de Paris, Université Paris-Est Créteil, INSERM U955 | Christophe Rodriguez, Slim Fourati, Vanessa Demontant, Guillaume Gricourt, Melissa N'Debi, Alexandre Soulier, Elisabeth Trawinski, Jean-Michel Pawlotsky |
| EPI_ISL_2637938 | Biogroup Bio Lam-LCD Saint-Denis | Department of Virology, Henri Mondor University Hospital, Assistance Publique Hôpitaux de Paris, Université Paris-Est Créteil, INSERM U955 | Christophe Rodriguez, Slim Fourati, Vanessa Demontant, Guillaume Gricourt, Melissa N'Debi, Alexandre Soulier, Elisabeth Trawinski, Jean-Michel Pawlotsky |
| EPI_ISL_2638024 | BIOMNIS EUROFINS IVRY | Department of Virology, Henri Mondor University Hospital, Assistance Publique Hôpitaux de Paris, Université Paris-Est Créteil, INSERM U955 | Christophe Rodriguez, Slim Fourati, Vanessa Demontant, Guillaume Gricourt, Melissa N'Debi, Alexandre Soulier, Elisabeth Trawinski, Jean-Michel Pawlotsky |
| EPI_ISL_2638046 | Clinique Saint-Pierre Ottignies | UCLouvain/IREC/MBLG | Jean Ruelle, Bertrand Bearzatto, Benoit Kabamba Mukadi |
| EPI_ISL_2638089 | Medical Laboratories Duesseldorf | Center of Medical Microbiology, Virology, and Hospital Hygiene, University of Duesseldorf | Maximilian Damagnez;Alexander Dilthey;Angelika Helmer;Torsten Houwaart;Lisanna Hülse;Christian Lange;Malte Kohns Vasconcelos;Nadine Lübke;Jessica Nicolai;Klaus Pfeffer;Daniel Strelow;Jörg Timm;Andreas Walker;Tobias Wienemann |
| EPI_ISL_2638098, EPI_ISL_2638100, EPI_ISL_2638101, EPI_ISL_2638104, EPI_ISL_2638108, EPI_ISL_2638110, EPI_ISL_2638111, EPI_ISL_2638112, EPI_ISL_2638113, EPI_ISL_2638114, EPI_ISL_2638116, EPI_ISL_2638117, EPI_ISL_2638118, EPI_ISL_2638119, EPI_ISL_2638121, EPI_ISL_2638122, EPI_ISL_2638123, EPI_ISL_2638126, EPI_ISL_2638127, EPI_ISL_2638128, EPI_ISL_2638129, EPI_ISL_2638130, EPI_ISL_2638131, EPI_ISL_2638132, EPI_ISL_2638133, EPI_ISL_2638134, EPI_ISL_2638135, EPI_ISL_2638138 | | | |
| see above | AULSS 2 Marca Trevigiana | Istituto Zooprofilattico Sperimentale delle Venezie | Adelaide Milani, Alessia Schivo, Annalisa Salviato, Elisa Palumbo, Erika Giorgia Quaranta, Luca Tassoni, Ambra Pastori, Edoardo Giussani, Alice Fusaro, Isabella Monne, Calogero Terregino, Antonia Ricci |
| EPI_ISL_2638161, EPI_ISL_2638162 | Lighthouse Lab in Milton Keynes | Wellcome Sanger Institute for the COVID-19 Genomics UK (COG-UK) Consortium | The Lighthouse Lab in Milton Keynes and Alex Alderton, Roberto Amato, Jeffrey Barrett, Sonia Goncalves, Ewan Harrison, David K. Jackson, Ian Johnston, Dominic Kwiatkowski, Cordelia Langford, John Sillitoe on behalf of the Wellcome Sanger Institute COVID-19 Surveillance Team |
| EPI_ISL_2638163 | Lighthouse Lab in Alderley Park | Wellcome Sanger Institute for the COVID-19 Genomics UK (COG-UK) Consortium | Jacquelyn Wynn, Mairead Hyland, The Lighthouse Lab in Alderley Park and Alex Alderton, Roberto Amato, Jeffrey Barrett, Sonia Goncalves, Ewan Harrison, David K. Jackson, Ian Johnston, Dominic Kwiatkowski, Cordelia Langford, John Sillitoe on behalf of the Wellcome Sanger Institute COVID-19 Surveillance Team |
| EPI_ISL_2638164 | Lighthouse Lab in Milton Keynes | Wellcome Sanger Institute for the COVID-19 Genomics UK (COG-UK) Consortium | The Lighthouse Lab in Milton Keynes and Alex Alderton, Roberto Amato, Jeffrey Barrett, Sonia Goncalves, Ewan Harrison, David K. Jackson, Ian Johnston, Dominic Kwiatkowski, Cordelia Langford, John Sillitoe on behalf of the Wellcome Sanger Institute COVID-19 Surveillance Team |
| EPI_ISL_2638165 | Lighthouse Lab in Glasgow | Wellcome Sanger Institute for the COVID-19 Genomics UK (COG-UK) Consortium | Harper VanSteenhouse, Yumi Kasai, David Gray, Carol Clugston, Anna Dominiczak and Alex Alderton, Roberto Amato, Jeffrey Barrett, Sonia Goncalves, Ewan Harrison, David K. Jackson, Ian Johnston, Dominic Kwiatkowski, Cordelia Langford, John Sillitoe on behalf of the Wellcome Sanger Institute COVID-19 Surveillance Team |
| EPI_ISL_2638166, EPI_ISL_2638167 | Lighthouse Lab in Alderley Park | Wellcome Sanger Institute for the COVID-19 Genomics UK (COG-UK) Consortium | Jacquelyn Wynn, Mairead Hyland, The Lighthouse Lab in Alderley Park and Alex Alderton, Roberto Amato, Jeffrey Barrett, Sonia Goncalves, Ewan Harrison, David K. Jackson, Ian Johnston, Dominic Kwiatkowski, Cordelia Langford, John Sillitoe on behalf of the Wellcome Sanger Institute COVID-19 Surveillance Team |
| EPI_ISL_2638168 | Lighthouse Lab in Milton Keynes | Wellcome Sanger Institute for the COVID-19 Genomics UK (COG-UK) Consortium | The Lighthouse Lab in Milton Keynes and Alex Alderton, Roberto Amato, Jeffrey Barrett, Sonia Goncalves, Ewan Harrison, David K. Jackson, Ian Johnston, Dominic Kwiatkowski, Cordelia Langford, John Sillitoe on behalf of the Wellcome Sanger Institute COVID-19 Surveillance Team |
| EPI_ISL_2638169 | Lighthouse Lab in Alderley Park | Wellcome Sanger Institute for the COVID-19 Genomics UK (COG-UK) Consortium | Jacquelyn Wynn, Mairead Hyland, The Lighthouse Lab in Alderley Park and Alex Alderton, Roberto Amato, Jeffrey Barrett, Sonia Goncalves, Ewan Harrison, David K. Jackson, Ian Johnston, Dominic Kwiatkowski, Cordelia Langford, John Sillitoe on behalf of the Wellcome Sanger Institute COVID-19 Surveillance Team |
| EPI_ISL_2638170 | Lighthouse Lab in Milton Keynes | Wellcome Sanger Institute for the COVID-19 Genomics UK (COG-UK) Consortium | The Lighthouse Lab in Milton Keynes and Alex Alderton, Roberto Amato, Jeffrey Barrett, Sonia Goncalves, Ewan Harrison, David K. Jackson, Ian Johnston, Dominic Kwiatkowski, Cordelia Langford, John Sillitoe on behalf of the Wellcome Sanger Institute COVID-19 Surveillance Team |
| EPI_ISL_2638171 | Lighthouse Lab in Alderley Park | Wellcome Sanger Institute for the COVID-19 Genomics UK (COG-UK) Consortium | Jacquelyn Wynn, Mairead Hyland, The Lighthouse Lab in Alderley Park and Alex Alderton, Roberto Amato, Jeffrey Barrett, Sonia Goncalves, Ewan Harrison, David K. Jackson, Ian Johnston, Dominic Kwiatkowski, Cordelia Langford, John Sillitoe on behalf of the Wellcome Sanger Institute COVID-19 Surveillance Team |
| EPI_ISL_2638172 | Health Services Laboratories | Wellcome Sanger Institute for the COVID-19 Genomics UK (COG-UK) Consortium | Health Services Laboratories and Alex Alderton, Roberto Amato, Jeffrey Barrett, Sonia Goncalves, Ewan Harrison, David K. Jackson, Ian Johnston, Dominic Kwiatkowski, Cordelia Langford, John Sillitoe on behalf of the Wellcome Sanger Institute COVID-19 Surveillance Team |
| EPI_ISL_2638173 | Lighthouse Lab in Alderley Park | Wellcome Sanger Institute for the COVID-19 Genomics UK (COG-UK) Consortium | Jacquelyn Wynn, Mairead Hyland, The Lighthouse Lab in Alderley Park and Alex Alderton, Roberto Amato, Jeffrey Barrett, Sonia Goncalves, Ewan Harrison, David K. Jackson, Ian Johnston, Dominic Kwiatkowski, Cordelia Langford, John Sillitoe on behalf of the Wellcome Sanger Institute COVID-19 Surveillance Team |
| EPI_ISL_2638174 | Lighthouse Lab in Milton Keynes | Wellcome Sanger Institute for the COVID-19 Genomics UK (COG-UK) Consortium | The Lighthouse Lab in Milton Keynes and Alex Alderton, Roberto Amato, Jeffrey Barrett, Sonia Goncalves, Ewan Harrison, David K. Jackson, Ian Johnston, Dominic Kwiatkowski, Cordelia Langford, John Sillitoe on behalf of the Wellcome Sanger Institute COVID-19 Surveillance Team |
| EPI_ISL_2638175 | Health Services Laboratories | Wellcome Sanger Institute for the COVID-19 Genomics UK (COG-UK) Consortium | Health Services Laboratories and Alex Alderton, Roberto Amato, Jeffrey Barrett, Sonia Goncalves, Ewan Harrison, David K. Jackson, Ian Johnston, Dominic Kwiatkowski, Cordelia Langford, John Sillitoe on behalf of the Wellcome Sanger Institute COVID-19 Surveillance Team |
| EPI_ISL_2638176 | Lighthouse Lab in Milton Keynes | Wellcome Sanger Institute for the COVID-19 Genomics UK (COG-UK) Consortium | The Lighthouse Lab in Milton Keynes and Alex Alderton, Roberto Amato, Jeffrey Barrett, Sonia Goncalves, Ewan Harrison, David K. Jackson, Ian Johnston, Dominic Kwiatkowski, Cordelia Langford, John Sillitoe on behalf of the Wellcome Sanger Institute COVID-19 Surveillance Team |
| EPI_ISL_2638177 | Health Services Laboratories | Wellcome Sanger Institute for the COVID-19 Genomics UK (COG-UK) Consortium | Health Services Laboratories and Alex Alderton, Roberto Amato, Jeffrey Barrett, Sonia Goncalves, Ewan Harrison, David K. Jackson, Ian Johnston, Dominic Kwiatkowski, Cordelia Langford, John Sillitoe on behalf of the Wellcome Sanger Institute COVID-19 Surveillance Team |
| EPI_ISL_2638178, EPI_ISL_2638179 | Lighthouse Lab in Alderley Park | Wellcome Sanger Institute for the COVID-19 Genomics UK (COG-UK) Consortium | Jacquelyn Wynn, Mairead Hyland, The Lighthouse Lab in Alderley Park and Alex Alderton, Roberto Amato, Jeffrey Barrett, Sonia Goncalves, Ewan Harrison, David K. Jackson, Ian Johnston, Dominic Kwiatkowski, Cordelia Langford, John Sillitoe on behalf of the Wellcome Sanger Institute COVID-19 Surveillance Team |
| EPI_ISL_2638180 | Lighthouse Lab in Milton Keynes | Wellcome Sanger Institute for the COVID-19 Genomics UK (COG-UK) | The Lighthouse Lab in Milton Keynes and Alex Alderton, Roberto Amato, Jeffrey Barrett, Sonia Goncalves, Ewan Harrison, David K. Jackson, Ian Johnston, Dominic Kwiatkowski, Cordelia Langford, John Sillitoe on behalf of the Wellcome Sanger |

| | | | |
|---|---|---|---|
| | | Consortium | Institute COVID-19 Surveillance Team |
| EPI_ISL_2638181 | Health Services Laboratories | Wellcome Sanger Institute for the COVID-19 Genomics UK (COG-UK) Consortium | Health Services Laboratories and Alex Alderton, Roberto Amato, Jeffrey Barrett, Sonia Goncalves, Ewan Harrison, David K. Jackson, Ian Johnston, Dominic Kwiatkowski, Cordelia Langford, John Sillitoe on behalf of the Wellcome Sanger Institute COVID-19 Surveillance Team |
| EPI_ISL_2638182 | Lighthouse Lab in Alderley Park | Wellcome Sanger Institute for the COVID-19 Genomics UK (COG-UK) Consortium | Jacquelyn Wynn, Mairead Hyland, The Lighthouse Lab in Alderley Park and Alex Alderton, Roberto Amato, Jeffrey Barrett, Sonia Goncalves, Ewan Harrison, David K. Jackson, Ian Johnston, Dominic Kwiatkowski, Cordelia Langford, John Sillitoe on behalf of the Wellcome Sanger Institute COVID-19 Surveillance Team |
| EPI_ISL_2638183 | Lighthouse Lab in Milton Keynes | Wellcome Sanger Institute for the COVID-19 Genomics UK (COG-UK) Consortium | The Lighthouse Lab in Milton Keynes and Alex Alderton, Roberto Amato, Jeffrey Barrett, Sonia Goncalves, Ewan Harrison, David K. Jackson, Ian Johnston, Dominic Kwiatkowski, Cordelia Langford, John Sillitoe on behalf of the Wellcome Sanger Institute COVID-19 Surveillance Team |
| EPI_ISL_2638184, EPI_ISL_2638185 | Lighthouse Lab in Alderley Park | Wellcome Sanger Institute for the COVID-19 Genomics UK (COG-UK) Consortium | Jacquelyn Wynn, Mairead Hyland, The Lighthouse Lab in Alderley Park and Alex Alderton, Roberto Amato, Jeffrey Barrett, Sonia Goncalves, Ewan Harrison, David K. Jackson, Ian Johnston, Dominic Kwiatkowski, Cordelia Langford, John Sillitoe on behalf of the Wellcome Sanger Institute COVID-19 Surveillance Team |
| EPI_ISL_2638186 | Health Services Laboratories | Wellcome Sanger Institute for the COVID-19 Genomics UK (COG-UK) Consortium | Health Services Laboratories and Alex Alderton, Roberto Amato, Jeffrey Barrett, Sonia Goncalves, Ewan Harrison, David K. Jackson, Ian Johnston, Dominic Kwiatkowski, Cordelia Langford, John Sillitoe on behalf of the Wellcome Sanger Institute COVID-19 Surveillance Team |
| EPI_ISL_2638187, EPI_ISL_2638188, EPI_ISL_2638189, EPI_ISL_2638190 | Lighthouse Lab in Alderley Park | Wellcome Sanger Institute for the COVID-19 Genomics UK (COG-UK) Consortium | Jacquelyn Wynn, Mairead Hyland, The Lighthouse Lab in Alderley Park and Alex Alderton, Roberto Amato, Jeffrey Barrett, Sonia Goncalves, Ewan Harrison, David K. Jackson, Ian Johnston, Dominic Kwiatkowski, Cordelia Langford, John Sillitoe on behalf of the Wellcome Sanger Institute COVID-19 Surveillance Team |
| EPI_ISL_2638191 | Lighthouse Lab in Milton Keynes | Wellcome Sanger Institute for the COVID-19 Genomics UK (COG-UK) Consortium | The Lighthouse Lab in Milton Keynes and Alex Alderton, Roberto Amato, Jeffrey Barrett, Sonia Goncalves, Ewan Harrison, David K. Jackson, Ian Johnston, Dominic Kwiatkowski, Cordelia Langford, John Sillitoe on behalf of the Wellcome Sanger Institute COVID-19 Surveillance Team |
| EPI_ISL_2638192, EPI_ISL_2638194, EPI_ISL_2638195 | Lighthouse Lab in Alderley Park | Wellcome Sanger Institute for the COVID-19 Genomics UK (COG-UK) Consortium | Jacquelyn Wynn, Mairead Hyland, The Lighthouse Lab in Alderley Park and Alex Alderton, Roberto Amato, Jeffrey Barrett, Sonia Goncalves, Ewan Harrison, David K. Jackson, Ian Johnston, Dominic Kwiatkowski, Cordelia Langford, John Sillitoe on behalf of the Wellcome Sanger Institute COVID-19 Surveillance Team |
| EPI_ISL_2638196, EPI_ISL_2638197 | Lighthouse Lab in Milton Keynes | Wellcome Sanger Institute for the COVID-19 Genomics UK (COG-UK) Consortium | The Lighthouse Lab in Milton Keynes and Alex Alderton, Roberto Amato, Jeffrey Barrett, Sonia Goncalves, Ewan Harrison, David K. Jackson, Ian Johnston, Dominic Kwiatkowski, Cordelia Langford, John Sillitoe on behalf of the Wellcome Sanger Institute COVID-19 Surveillance Team |
| EPI_ISL_2638198 | Health Services Laboratories | Wellcome Sanger Institute for the COVID-19 Genomics UK (COG-UK) Consortium | Health Services Laboratories and Alex Alderton, Roberto Amato, Jeffrey Barrett, Sonia Goncalves, Ewan Harrison, David K. Jackson, Ian Johnston, Dominic Kwiatkowski, Cordelia Langford, John Sillitoe on behalf of the Wellcome Sanger Institute COVID-19 Surveillance Team |
| EPI_ISL_2638199 | Lighthouse Lab in Milton Keynes | Wellcome Sanger Institute for the COVID-19 Genomics UK (COG-UK) Consortium | The Lighthouse Lab in Milton Keynes and Alex Alderton, Roberto Amato, Jeffrey Barrett, Sonia Goncalves, Ewan Harrison, David K. Jackson, Ian Johnston, Dominic Kwiatkowski, Cordelia Langford, John Sillitoe on behalf of the Wellcome Sanger Institute COVID-19 Surveillance Team |
| EPI_ISL_2638200 | Lighthouse Lab in Alderley Park | Wellcome Sanger Institute for the COVID-19 Genomics UK (COG-UK) Consortium | Jacquelyn Wynn, Mairead Hyland, The Lighthouse Lab in Alderley Park and Alex Alderton, Roberto Amato, Jeffrey Barrett, Sonia Goncalves, Ewan Harrison, David K. Jackson, Ian Johnston, Dominic Kwiatkowski, Cordelia Langford, John Sillitoe on behalf of the Wellcome Sanger Institute COVID-19 Surveillance Team |
| EPI_ISL_2638201, EPI_ISL_2638202 | Lighthouse Lab in Milton Keynes | Wellcome Sanger Institute for the COVID-19 Genomics UK (COG-UK) Consortium | The Lighthouse Lab in Milton Keynes and Alex Alderton, Roberto Amato, Jeffrey Barrett, Sonia Goncalves, Ewan Harrison, David K. Jackson, Ian Johnston, Dominic Kwiatkowski, Cordelia Langford, John Sillitoe on behalf of the Wellcome Sanger Institute COVID-19 Surveillance Team |
| EPI_ISL_2638203, EPI_ISL_2638204, EPI_ISL_2638205, EPI_ISL_2638206, EPI_ISL_2638207, EPI_ISL_2638208 | Lighthouse Lab in Alderley Park | Wellcome Sanger Institute for the COVID-19 Genomics UK (COG-UK) Consortium | Jacquelyn Wynn, Mairead Hyland, The Lighthouse Lab in Alderley Park and Alex Alderton, Roberto Amato, Jeffrey Barrett, Sonia Goncalves, Ewan Harrison, David K. Jackson, Ian Johnston, Dominic Kwiatkowski, Cordelia Langford, John Sillitoe on behalf of the Wellcome Sanger Institute COVID-19 Surveillance Team |
| EPI_ISL_2638209 | Lighthouse Lab in Milton Keynes | Wellcome Sanger Institute for the COVID-19 Genomics UK (COG-UK) Consortium | The Lighthouse Lab in Milton Keynes and Alex Alderton, Roberto Amato, Jeffrey Barrett, Sonia Goncalves, Ewan Harrison, David K. Jackson, Ian Johnston, Dominic Kwiatkowski, Cordelia Langford, John Sillitoe on behalf of the Wellcome Sanger Institute COVID-19 Surveillance Team |
| EPI_ISL_2638210, EPI_ISL_2638211, EPI_ISL_2638213 | Lighthouse Lab in Alderley Park | Wellcome Sanger Institute for the COVID-19 Genomics UK (COG-UK) Consortium | Jacquelyn Wynn, Mairead Hyland, The Lighthouse Lab in Alderley Park and Alex Alderton, Roberto Amato, Jeffrey Barrett, Sonia Goncalves, Ewan Harrison, David K. Jackson, Ian Johnston, Dominic Kwiatkowski, Cordelia Langford, John Sillitoe on behalf of the Wellcome Sanger Institute COVID-19 Surveillance Team |
| EPI_ISL_2638214, EPI_ISL_2638215 | Lighthouse Lab in Milton Keynes | Wellcome Sanger Institute for the COVID-19 Genomics UK (COG-UK) Consortium | The Lighthouse Lab in Milton Keynes and Alex Alderton, Roberto Amato, Jeffrey Barrett, Sonia Goncalves, Ewan Harrison, David K. Jackson, Ian Johnston, Dominic Kwiatkowski, Cordelia Langford, John Sillitoe on behalf of the Wellcome Sanger Institute COVID-19 Surveillance Team |
| EPI_ISL_2638216, EPI_ISL_2638217, EPI_ISL_2638218 | Lighthouse Lab in Alderley Park | Wellcome Sanger Institute for the COVID-19 Genomics UK (COG-UK) Consortium | Jacquelyn Wynn, Mairead Hyland, The Lighthouse Lab in Alderley Park and Alex Alderton, Roberto Amato, Jeffrey Barrett, Sonia Goncalves, Ewan Harrison, David K. Jackson, Ian Johnston, Dominic Kwiatkowski, Cordelia Langford, John Sillitoe on behalf of the Wellcome Sanger Institute COVID-19 Surveillance Team |
| EPI_ISL_2638219 | Lighthouse Lab in Milton Keynes | Wellcome Sanger Institute for the COVID-19 Genomics UK (COG-UK) Consortium | The Lighthouse Lab in Milton Keynes and Alex Alderton, Roberto Amato, Jeffrey Barrett, Sonia Goncalves, Ewan Harrison, David K. Jackson, Ian Johnston, Dominic Kwiatkowski, Cordelia Langford, John Sillitoe on behalf of the Wellcome Sanger Institute COVID-19 Surveillance Team |
| EPI_ISL_2638220 | Lighthouse Lab in Alderley Park | Wellcome Sanger Institute for the COVID-19 Genomics UK (COG-UK) Consortium | Jacquelyn Wynn, Mairead Hyland, The Lighthouse Lab in Alderley Park and Alex Alderton, Roberto Amato, Jeffrey Barrett, Sonia Goncalves, Ewan Harrison, David K. Jackson, Ian Johnston, Dominic Kwiatkowski, Cordelia Langford, John Sillitoe on behalf of the Wellcome Sanger Institute COVID-19 Surveillance Team |
| EPI_ISL_2638221 | Health Services Laboratories | Wellcome Sanger Institute for the COVID-19 Genomics UK (COG-UK) Consortium | Health Services Laboratories and Alex Alderton, Roberto Amato, Jeffrey Barrett, Sonia Goncalves, Ewan Harrison, David K. Jackson, Ian Johnston, Dominic Kwiatkowski, Cordelia Langford, John Sillitoe on behalf of the Wellcome Sanger Institute COVID-19 Surveillance Team |
| EPI_ISL_2638222 | Lighthouse Lab in Alderley Park | Wellcome Sanger Institute for the COVID-19 Genomics UK (COG-UK) Consortium | Jacquelyn Wynn, Mairead Hyland, The Lighthouse Lab in Alderley Park and Alex Alderton, Roberto Amato, Jeffrey Barrett, Sonia Goncalves, Ewan Harrison, David K. Jackson, Ian Johnston, Dominic Kwiatkowski, Cordelia Langford, John Sillitoe on behalf of the Wellcome Sanger Institute COVID-19 Surveillance Team |
| EPI_ISL_2638223 | Lighthouse Lab in Glasgow | Wellcome Sanger Institute for the COVID-19 Genomics UK (COG-UK) Consortium | Harper VanSteenhouse, Yumi Kasai, David Gray, Carol Clugston, Anna Dominiczak and Alex Alderton, Roberto Amato, Jeffrey Barrett, Sonia Goncalves, Ewan Harrison, David K. Jackson, Ian Johnston, Dominic Kwiatkowski, Cordelia Langford, John Sillitoe on behalf of the Wellcome Sanger Institute COVID-19 Surveillance Team |
| EPI_ISL_2638224 | Lighthouse Lab in Alderley Park | Wellcome Sanger Institute for the COVID-19 Genomics UK (COG-UK) Consortium | Jacquelyn Wynn, Mairead Hyland, The Lighthouse Lab in Alderley Park and Alex Alderton, Roberto Amato, Jeffrey Barrett, Sonia Goncalves, Ewan Harrison, David K. Jackson, Ian Johnston, Dominic Kwiatkowski, Cordelia Langford, John Sillitoe on behalf of the Wellcome Sanger Institute COVID-19 Surveillance Team |
| EPI_ISL_2638225, EPI_ISL_2638226 | Lighthouse Lab in Milton Keynes | Wellcome Sanger Institute for the COVID-19 Genomics UK (COG-UK) Consortium | The Lighthouse Lab in Milton Keynes and Alex Alderton, Roberto Amato, Jeffrey Barrett, Sonia Goncalves, Ewan Harrison, David K. Jackson, Ian Johnston, Dominic Kwiatkowski, Cordelia Langford, John Sillitoe on behalf of the Wellcome Sanger Institute COVID-19 Surveillance Team |
| EPI_ISL_2638227, EPI_ISL_2638228 | Lighthouse Lab in Alderley Park | Wellcome Sanger Institute for the COVID-19 Genomics UK (COG-UK) Consortium | Jacquelyn Wynn, Mairead Hyland, The Lighthouse Lab in Alderley Park and Alex Alderton, Roberto Amato, Jeffrey Barrett, Sonia Goncalves, Ewan Harrison, David K. Jackson, Ian Johnston, Dominic Kwiatkowski, Cordelia Langford, John Sillitoe on behalf of the Wellcome Sanger Institute COVID-19 Surveillance Team |
| EPI_ISL_2638229, EPI_ISL_2638230 | Lighthouse Lab in Milton Keynes | Wellcome Sanger Institute for the COVID-19 Genomics UK (COG-UK) Consortium | The Lighthouse Lab in Milton Keynes and Alex Alderton, Roberto Amato, Jeffrey Barrett, Sonia Goncalves, Ewan Harrison, David K. Jackson, Ian Johnston, Dominic Kwiatkowski, Cordelia Langford, John Sillitoe on behalf of the Wellcome Sanger Institute COVID-19 Surveillance Team |
| EPI_ISL_2638231 | Lighthouse Lab in Alderley Park | Wellcome Sanger Institute for the COVID-19 Genomics UK (COG-UK) Consortium | Jacquelyn Wynn, Mairead Hyland, The Lighthouse Lab in Alderley Park and Alex Alderton, Roberto Amato, Jeffrey Barrett, Sonia Goncalves, Ewan Harrison, David K. Jackson, Ian Johnston, Dominic Kwiatkowski, Cordelia Langford, John Sillitoe on behalf of the Wellcome Sanger Institute COVID-19 Surveillance Team |
| EPI_ISL_2638232 | Health Services Laboratories | Wellcome Sanger Institute for the COVID-19 Genomics UK (COG-UK) Consortium | Health Services Laboratories and Alex Alderton, Roberto Amato, Jeffrey Barrett, Sonia Goncalves, Ewan Harrison, David K. Jackson, Ian Johnston, Dominic Kwiatkowski, Cordelia Langford, John Sillitoe on behalf of the Wellcome Sanger Institute COVID-19 Surveillance Team |
| EPI_ISL_2638233, EPI_ISL_2638234, EPI_ISL_2638235, EPI_ISL_2638236 | Lighthouse Lab in Alderley Park | Wellcome Sanger Institute for the COVID-19 Genomics UK (COG-UK) Consortium | Jacquelyn Wynn, Mairead Hyland, The Lighthouse Lab in Alderley Park and Alex Alderton, Roberto Amato, Jeffrey Barrett, Sonia Goncalves, Ewan Harrison, David K. Jackson, Ian Johnston, Dominic Kwiatkowski, Cordelia Langford, John Sillitoe on behalf of the Wellcome Sanger Institute COVID-19 Surveillance Team |
| EPI_ISL_2638237 | Lighthouse Lab in Milton Keynes | Wellcome Sanger Institute for the COVID-19 Genomics UK (COG-UK) Consortium | The Lighthouse Lab in Milton Keynes and Alex Alderton, Roberto Amato, Jeffrey Barrett, Sonia Goncalves, Ewan Harrison, David K. Jackson, Ian Johnston, Dominic Kwiatkowski, Cordelia Langford, John Sillitoe on behalf of the Wellcome Sanger Institute COVID-19 Surveillance Team |
| EPI_ISL_2638238 | Lighthouse Lab in Alderley Park | Wellcome Sanger Institute for the COVID-19 Genomics UK (COG-UK) Consortium | Jacquelyn Wynn, Mairead Hyland, The Lighthouse Lab in Alderley Park and Alex Alderton, Roberto Amato, Jeffrey Barrett, Sonia Goncalves, Ewan Harrison, David K. Jackson, Ian Johnston, Dominic Kwiatkowski, Cordelia Langford, John Sillitoe on behalf of the Wellcome Sanger Institute COVID-19 Surveillance Team |
| EPI_ISL_2638239 | Lighthouse Lab in Milton Keynes | Wellcome Sanger Institute for the COVID-19 Genomics UK (COG-UK) Consortium | The Lighthouse Lab in Milton Keynes and Alex Alderton, Roberto Amato, Jeffrey Barrett, Sonia Goncalves, Ewan Harrison, David K. Jackson, Ian Johnston, Dominic Kwiatkowski, Cordelia Langford, John Sillitoe on behalf of the Wellcome Sanger Institute COVID-19 Surveillance Team |
| EPI_ISL_2638240, EPI_ISL_2638241, EPI_ISL_2638242 | Lighthouse Lab in Alderley Park | Wellcome Sanger Institute for the COVID-19 Genomics UK (COG-UK) Consortium | Jacquelyn Wynn, Mairead Hyland, The Lighthouse Lab in Alderley Park and Alex Alderton, Roberto Amato, Jeffrey Barrett, Sonia Goncalves, Ewan Harrison, David K. Jackson, Ian Johnston, Dominic Kwiatkowski, Cordelia Langford, John Sillitoe on behalf of the Wellcome Sanger Institute COVID-19 Surveillance Team |
| EPI_ISL_2638243, EPI_ISL_2638244 | Lighthouse Lab in Milton Keynes | Wellcome Sanger Institute for the COVID-19 Genomics UK (COG-UK) Consortium | The Lighthouse Lab in Milton Keynes and Alex Alderton, Roberto Amato, Jeffrey Barrett, Sonia Goncalves, Ewan Harrison, David K. Jackson, Ian Johnston, Dominic Kwiatkowski, Cordelia Langford, John Sillitoe on behalf of the Wellcome Sanger Institute COVID-19 Surveillance Team |
| EPI_ISL_2638245, EPI_ISL_2638246, EPI_ISL_2638247, EPI_ISL_2638248 | Lighthouse Lab in Alderley Park | Wellcome Sanger Institute for the COVID-19 Genomics UK (COG-UK) Consortium | Jacquelyn Wynn, Mairead Hyland, The Lighthouse Lab in Alderley Park and Alex Alderton, Roberto Amato, Jeffrey Barrett, Sonia Goncalves, Ewan Harrison, David K. Jackson, Ian Johnston, Dominic Kwiatkowski, Cordelia Langford, John Sillitoe on behalf of the Wellcome Sanger Institute COVID-19 Surveillance Team |
| EPI_ISL_2638249 | Lighthouse Lab in Glasgow | Wellcome Sanger Institute for the COVID-19 Genomics UK (COG-UK) Consortium | Harper VanSteenhouse, Yumi Kasai, David Gray, Carol Clugston, Anna Dominiczak and Alex Alderton, Roberto Amato, Jeffrey Barrett, Sonia Goncalves, Ewan Harrison, David K. Jackson, Ian Johnston, Dominic Kwiatkowski, Cordelia Langford, John Sillitoe on behalf of the Wellcome Sanger Institute COVID-19 Surveillance Team |
| EPI_ISL_2638250, EPI_ISL_2638251, EPI_ISL_2638253 | Lighthouse Lab in Alderley Park | Wellcome Sanger Institute for the COVID-19 Genomics UK (COG-UK) Consortium | Jacquelyn Wynn, Mairead Hyland, The Lighthouse Lab in Alderley Park and Alex Alderton, Roberto Amato, Jeffrey Barrett, Sonia Goncalves, Ewan Harrison, David K. Jackson, Ian Johnston, Dominic Kwiatkowski, Cordelia Langford, John Sillitoe on behalf of the Wellcome Sanger Institute COVID-19 Surveillance Team |

| | | | |
|---|---|---|---|
| EPI_ISL_2638254, EPI_ISL_2638255, EPI_ISL_2638256, EPI_ISL_2638257 | Lighthouse Lab in Milton Keynes | Wellcome Sanger Institute for the COVID-19 Genomics UK (COG-UK) Consortium | The Lighthouse Lab in Milton Keynes and Alex Alderton, Roberto Amato, Jeffrey Barrett, Sonia Goncalves, Ewan Harrison, David K. Jackson, Ian Johnston, Dominic Kwiatkowski, Cordelia Langford, John Sillitoe on behalf of the Wellcome Sanger Institute COVID-19 Surveillance Team |
| EPI_ISL_2638258 | Health Services Laboratories | Wellcome Sanger Institute for the COVID-19 Genomics UK (COG-UK) Consortium | Health Services Laboratories and Alex Alderton, Roberto Amato, Jeffrey Barrett, Sonia Goncalves, Ewan Harrison, David K. Jackson, Ian Johnston, Dominic Kwiatkowski, Cordelia Langford, John Sillitoe on behalf of the Wellcome Sanger Institute COVID-19 Surveillance Team |
| EPI_ISL_2638259 | Lighthouse Lab in Alderley Park | Wellcome Sanger Institute for the COVID-19 Genomics UK (COG-UK) Consortium | Jacquelyn Wynn, Mairead Hyland, The Lighthouse Lab in Alderley Park and Alex Alderton, Roberto Amato, Jeffrey Barrett, Sonia Goncalves, Ewan Harrison, David K. Jackson, Ian Johnston, Dominic Kwiatkowski, Cordelia Langford, John Sillitoe on behalf of the Wellcome Sanger Institute COVID-19 Surveillance Team |
| EPI_ISL_2638260 | Health Services Laboratories | Wellcome Sanger Institute for the COVID-19 Genomics UK (COG-UK) Consortium | Health Services Laboratories and Alex Alderton, Roberto Amato, Jeffrey Barrett, Sonia Goncalves, Ewan Harrison, David K. Jackson, Ian Johnston, Dominic Kwiatkowski, Cordelia Langford, John Sillitoe on behalf of the Wellcome Sanger Institute COVID-19 Surveillance Team |
| EPI_ISL_2638261, EPI_ISL_2638262 | Lighthouse Lab in Milton Keynes | Wellcome Sanger Institute for the COVID-19 Genomics UK (COG-UK) Consortium | The Lighthouse Lab in Milton Keynes and Alex Alderton, Roberto Amato, Jeffrey Barrett, Sonia Goncalves, Ewan Harrison, David K. Jackson, Ian Johnston, Dominic Kwiatkowski, Cordelia Langford, John Sillitoe on behalf of the Wellcome Sanger Institute COVID-19 Surveillance Team |
| EPI_ISL_2638263 | Lighthouse Lab in Alderley Park | Wellcome Sanger Institute for the COVID-19 Genomics UK (COG-UK) Consortium | Jacquelyn Wynn, Mairead Hyland, The Lighthouse Lab in Alderley Park and Alex Alderton, Roberto Amato, Jeffrey Barrett, Sonia Goncalves, Ewan Harrison, David K. Jackson, Ian Johnston, Dominic Kwiatkowski, Cordelia Langford, John Sillitoe on behalf of the Wellcome Sanger Institute COVID-19 Surveillance Team |
| EPI_ISL_2638264, EPI_ISL_2638265, EPI_ISL_2638266 | Lighthouse Lab in Milton Keynes | Wellcome Sanger Institute for the COVID-19 Genomics UK (COG-UK) Consortium | The Lighthouse Lab in Milton Keynes and Alex Alderton, Roberto Amato, Jeffrey Barrett, Sonia Goncalves, Ewan Harrison, David K. Jackson, Ian Johnston, Dominic Kwiatkowski, Cordelia Langford, John Sillitoe on behalf of the Wellcome Sanger Institute COVID-19 Surveillance Team |
| EPI_ISL_2638267 | Lighthouse Lab in Alderley Park | Wellcome Sanger Institute for the COVID-19 Genomics UK (COG-UK) Consortium | Jacquelyn Wynn, Mairead Hyland, The Lighthouse Lab in Alderley Park and Alex Alderton, Roberto Amato, Jeffrey Barrett, Sonia Goncalves, Ewan Harrison, David K. Jackson, Ian Johnston, Dominic Kwiatkowski, Cordelia Langford, John Sillitoe on behalf of the Wellcome Sanger Institute COVID-19 Surveillance Team |
| EPI_ISL_2638268 | Lighthouse Lab in Milton Keynes | Wellcome Sanger Institute for the COVID-19 Genomics UK (COG-UK) Consortium | The Lighthouse Lab in Milton Keynes and Alex Alderton, Roberto Amato, Jeffrey Barrett, Sonia Goncalves, Ewan Harrison, David K. Jackson, Ian Johnston, Dominic Kwiatkowski, Cordelia Langford, John Sillitoe on behalf of the Wellcome Sanger Institute COVID-19 Surveillance Team |
| EPI_ISL_2638269 | Lighthouse Lab in Alderley Park | Wellcome Sanger Institute for the COVID-19 Genomics UK (COG-UK) Consortium | Jacquelyn Wynn, Mairead Hyland, The Lighthouse Lab in Alderley Park and Alex Alderton, Roberto Amato, Jeffrey Barrett, Sonia Goncalves, Ewan Harrison, David K. Jackson, Ian Johnston, Dominic Kwiatkowski, Cordelia Langford, John Sillitoe on behalf of the Wellcome Sanger Institute COVID-19 Surveillance Team |
| EPI_ISL_2638270 | Lighthouse Lab in Milton Keynes | Wellcome Sanger Institute for the COVID-19 Genomics UK (COG-UK) Consortium | The Lighthouse Lab in Milton Keynes and Alex Alderton, Roberto Amato, Jeffrey Barrett, Sonia Goncalves, Ewan Harrison, David K. Jackson, Ian Johnston, Dominic Kwiatkowski, Cordelia Langford, John Sillitoe on behalf of the Wellcome Sanger Institute COVID-19 Surveillance Team |
| EPI_ISL_2638271 | Lighthouse Lab in Glasgow | Wellcome Sanger Institute for the COVID-19 Genomics UK (COG-UK) Consortium | Harper VanSteenhouse, Yumi Kasai, David Gray, Carol Clugston, Anna Dominiczak and Alex Alderton, Roberto Amato, Jeffrey Barrett, Sonia Goncalves, Ewan Harrison, David K. Jackson, Ian Johnston, Dominic Kwiatkowski, Cordelia Langford, John Sillitoe on behalf of the Wellcome Sanger Institute COVID-19 Surveillance Team |
| EPI_ISL_2638272, EPI_ISL_2638273 | Lighthouse Lab in Alderley Park | Wellcome Sanger Institute for the COVID-19 Genomics UK (COG-UK) Consortium | Jacquelyn Wynn, Mairead Hyland, The Lighthouse Lab in Alderley Park and Alex Alderton, Roberto Amato, Jeffrey Barrett, Sonia Goncalves, Ewan Harrison, David K. Jackson, Ian Johnston, Dominic Kwiatkowski, Cordelia Langford, John Sillitoe on behalf of the Wellcome Sanger Institute COVID-19 Surveillance Team |
| EPI_ISL_2638274 | Lighthouse Lab in Milton Keynes | Wellcome Sanger Institute for the COVID-19 Genomics UK (COG-UK) Consortium | The Lighthouse Lab in Milton Keynes and Alex Alderton, Roberto Amato, Jeffrey Barrett, Sonia Goncalves, Ewan Harrison, David K. Jackson, Ian Johnston, Dominic Kwiatkowski, Cordelia Langford, John Sillitoe on behalf of the Wellcome Sanger Institute COVID-19 Surveillance Team |
| EPI_ISL_2638275 | Lighthouse Lab in Alderley Park | Wellcome Sanger Institute for the COVID-19 Genomics UK (COG-UK) Consortium | Jacquelyn Wynn, Mairead Hyland, The Lighthouse Lab in Alderley Park and Alex Alderton, Roberto Amato, Jeffrey Barrett, Sonia Goncalves, Ewan Harrison, David K. Jackson, Ian Johnston, Dominic Kwiatkowski, Cordelia Langford, John Sillitoe on behalf of the Wellcome Sanger Institute COVID-19 Surveillance Team |
| EPI_ISL_2638276, EPI_ISL_2638277 | Lighthouse Lab in Milton Keynes | Wellcome Sanger Institute for the COVID-19 Genomics UK (COG-UK) Consortium | The Lighthouse Lab in Milton Keynes and Alex Alderton, Roberto Amato, Jeffrey Barrett, Sonia Goncalves, Ewan Harrison, David K. Jackson, Ian Johnston, Dominic Kwiatkowski, Cordelia Langford, John Sillitoe on behalf of the Wellcome Sanger Institute COVID-19 Surveillance Team |
| EPI_ISL_2638278, EPI_ISL_2638279 | Lighthouse Lab in Alderley Park | Wellcome Sanger Institute for the COVID-19 Genomics UK (COG-UK) Consortium | Jacquelyn Wynn, Mairead Hyland, The Lighthouse Lab in Alderley Park and Alex Alderton, Roberto Amato, Jeffrey Barrett, Sonia Goncalves, Ewan Harrison, David K. Jackson, Ian Johnston, Dominic Kwiatkowski, Cordelia Langford, John Sillitoe on behalf of the Wellcome Sanger Institute COVID-19 Surveillance Team |
| EPI_ISL_2638280 | Lighthouse Lab in Milton Keynes | Wellcome Sanger Institute for the COVID-19 Genomics UK (COG-UK) Consortium | The Lighthouse Lab in Milton Keynes and Alex Alderton, Roberto Amato, Jeffrey Barrett, Sonia Goncalves, Ewan Harrison, David K. Jackson, Ian Johnston, Dominic Kwiatkowski, Cordelia Langford, John Sillitoe on behalf of the Wellcome Sanger Institute COVID-19 Surveillance Team |
| EPI_ISL_2638281 | Lighthouse Lab in Alderley Park | Wellcome Sanger Institute for the COVID-19 Genomics UK (COG-UK) Consortium | Jacquelyn Wynn, Mairead Hyland, The Lighthouse Lab in Alderley Park and Alex Alderton, Roberto Amato, Jeffrey Barrett, Sonia Goncalves, Ewan Harrison, David K. Jackson, Ian Johnston, Dominic Kwiatkowski, Cordelia Langford, John Sillitoe on behalf of the Wellcome Sanger Institute COVID-19 Surveillance Team |
| EPI_ISL_2638282 | Lighthouse Lab in Milton Keynes | Wellcome Sanger Institute for the COVID-19 Genomics UK (COG-UK) Consortium | The Lighthouse Lab in Milton Keynes and Alex Alderton, Roberto Amato, Jeffrey Barrett, Sonia Goncalves, Ewan Harrison, David K. Jackson, Ian Johnston, Dominic Kwiatkowski, Cordelia Langford, John Sillitoe on behalf of the Wellcome Sanger Institute COVID-19 Surveillance Team |
| EPI_ISL_2638283, EPI_ISL_2638284 | Lighthouse Lab in Alderley Park | Wellcome Sanger Institute for the COVID-19 Genomics UK (COG-UK) Consortium | Jacquelyn Wynn, Mairead Hyland, The Lighthouse Lab in Alderley Park and Alex Alderton, Roberto Amato, Jeffrey Barrett, Sonia Goncalves, Ewan Harrison, David K. Jackson, Ian Johnston, Dominic Kwiatkowski, Cordelia Langford, John Sillitoe on behalf of the Wellcome Sanger Institute COVID-19 Surveillance Team |
| EPI_ISL_2638285 | Lighthouse Lab in Milton Keynes | Wellcome Sanger Institute for the COVID-19 Genomics UK (COG-UK) Consortium | The Lighthouse Lab in Milton Keynes and Alex Alderton, Roberto Amato, Jeffrey Barrett, Sonia Goncalves, Ewan Harrison, David K. Jackson, Ian Johnston, Dominic Kwiatkowski, Cordelia Langford, John Sillitoe on behalf of the Wellcome Sanger Institute COVID-19 Surveillance Team |
| EPI_ISL_2638286 | Lighthouse Lab in Alderley Park | Wellcome Sanger Institute for the COVID-19 Genomics UK (COG-UK) Consortium | Jacquelyn Wynn, Mairead Hyland, The Lighthouse Lab in Alderley Park and Alex Alderton, Roberto Amato, Jeffrey Barrett, Sonia Goncalves, Ewan Harrison, David K. Jackson, Ian Johnston, Dominic Kwiatkowski, Cordelia Langford, John Sillitoe on behalf of the Wellcome Sanger Institute COVID-19 Surveillance Team |
| EPI_ISL_2638287 | Lighthouse Lab in Glasgow | Wellcome Sanger Institute for the COVID-19 Genomics UK (COG-UK) Consortium | Harper VanSteenhouse, Yumi Kasai, David Gray, Carol Clugston, Anna Dominiczak and Alex Alderton, Roberto Amato, Jeffrey Barrett, Sonia Goncalves, Ewan Harrison, David K. Jackson, Ian Johnston, Dominic Kwiatkowski, Cordelia Langford, John Sillitoe on behalf of the Wellcome Sanger Institute COVID-19 Surveillance Team |
| EPI_ISL_2638288, EPI_ISL_2638289, EPI_ISL_2638290 | Lighthouse Lab in Alderley Park | Wellcome Sanger Institute for the COVID-19 Genomics UK (COG-UK) Consortium | Jacquelyn Wynn, Mairead Hyland, The Lighthouse Lab in Alderley Park and Alex Alderton, Roberto Amato, Jeffrey Barrett, Sonia Goncalves, Ewan Harrison, David K. Jackson, Ian Johnston, Dominic Kwiatkowski, Cordelia Langford, John Sillitoe on behalf of the Wellcome Sanger Institute COVID-19 Surveillance Team |
| EPI_ISL_2638291, EPI_ISL_2638292 | Lighthouse Lab in Milton Keynes | Wellcome Sanger Institute for the COVID-19 Genomics UK (COG-UK) Consortium | The Lighthouse Lab in Milton Keynes and Alex Alderton, Roberto Amato, Jeffrey Barrett, Sonia Goncalves, Ewan Harrison, David K. Jackson, Ian Johnston, Dominic Kwiatkowski, Cordelia Langford, John Sillitoe on behalf of the Wellcome Sanger Institute COVID-19 Surveillance Team |
| EPI_ISL_2638293 | Lighthouse Lab in Alderley Park | Wellcome Sanger Institute for the COVID-19 Genomics UK (COG-UK) Consortium | Jacquelyn Wynn, Mairead Hyland, The Lighthouse Lab in Alderley Park and Alex Alderton, Roberto Amato, Jeffrey Barrett, Sonia Goncalves, Ewan Harrison, David K. Jackson, Ian Johnston, Dominic Kwiatkowski, Cordelia Langford, John Sillitoe on behalf of the Wellcome Sanger Institute COVID-19 Surveillance Team |
| EPI_ISL_2638294, EPI_ISL_2638295 | Health Services Laboratories | Wellcome Sanger Institute for the COVID-19 Genomics UK (COG-UK) Consortium | Health Services Laboratories and Alex Alderton, Roberto Amato, Jeffrey Barrett, Sonia Goncalves, Ewan Harrison, David K. Jackson, Ian Johnston, Dominic Kwiatkowski, Cordelia Langford, John Sillitoe on behalf of the Wellcome Sanger Institute COVID-19 Surveillance Team |
| EPI_ISL_2638296 | Lighthouse Lab in Milton Keynes | Wellcome Sanger Institute for the COVID-19 Genomics UK (COG-UK) Consortium | The Lighthouse Lab in Milton Keynes and Alex Alderton, Roberto Amato, Jeffrey Barrett, Sonia Goncalves, Ewan Harrison, David K. Jackson, Ian Johnston, Dominic Kwiatkowski, Cordelia Langford, John Sillitoe on behalf of the Wellcome Sanger Institute COVID-19 Surveillance Team |
| EPI_ISL_2638297 | Lighthouse Lab in Alderley Park | Wellcome Sanger Institute for the COVID-19 Genomics UK (COG-UK) Consortium | Jacquelyn Wynn, Mairead Hyland, The Lighthouse Lab in Alderley Park and Alex Alderton, Roberto Amato, Jeffrey Barrett, Sonia Goncalves, Ewan Harrison, David K. Jackson, Ian Johnston, Dominic Kwiatkowski, Cordelia Langford, John Sillitoe on behalf of the Wellcome Sanger Institute COVID-19 Surveillance Team |
| EPI_ISL_2638299 | Lighthouse Lab in Milton Keynes | Wellcome Sanger Institute for the COVID-19 Genomics UK (COG-UK) Consortium | The Lighthouse Lab in Milton Keynes and Alex Alderton, Roberto Amato, Jeffrey Barrett, Sonia Goncalves, Ewan Harrison, David K. Jackson, Ian Johnston, Dominic Kwiatkowski, Cordelia Langford, John Sillitoe on behalf of the Wellcome Sanger Institute COVID-19 Surveillance Team |
| EPI_ISL_2638300, EPI_ISL_2638301 | Lighthouse Lab in Alderley Park | Wellcome Sanger Institute for the COVID-19 Genomics UK (COG-UK) Consortium | Jacquelyn Wynn, Mairead Hyland, The Lighthouse Lab in Alderley Park and Alex Alderton, Roberto Amato, Jeffrey Barrett, Sonia Goncalves, Ewan Harrison, David K. Jackson, Ian Johnston, Dominic Kwiatkowski, Cordelia Langford, John Sillitoe on behalf of the Wellcome Sanger Institute COVID-19 Surveillance Team |
| EPI_ISL_2638302, EPI_ISL_2638303 | Lighthouse Lab in Milton Keynes | Wellcome Sanger Institute for the COVID-19 Genomics UK (COG-UK) Consortium | The Lighthouse Lab in Milton Keynes and Alex Alderton, Roberto Amato, Jeffrey Barrett, Sonia Goncalves, Ewan Harrison, David K. Jackson, Ian Johnston, Dominic Kwiatkowski, Cordelia Langford, John Sillitoe on behalf of the Wellcome Sanger Institute COVID-19 Surveillance Team |
| EPI_ISL_2638304, EPI_ISL_2638305 | Lighthouse Lab in Alderley Park | Wellcome Sanger Institute for the COVID-19 Genomics UK (COG-UK) Consortium | Jacquelyn Wynn, Mairead Hyland, The Lighthouse Lab in Alderley Park and Alex Alderton, Roberto Amato, Jeffrey Barrett, Sonia Goncalves, Ewan Harrison, David K. Jackson, Ian Johnston, Dominic Kwiatkowski, Cordelia Langford, John Sillitoe on behalf of the Wellcome Sanger Institute COVID-19 Surveillance Team |
| EPI_ISL_2638306 | Lighthouse Lab in Milton Keynes | Wellcome Sanger Institute for the COVID-19 Genomics UK (COG-UK) Consortium | The Lighthouse Lab in Milton Keynes and Alex Alderton, Roberto Amato, Jeffrey Barrett, Sonia Goncalves, Ewan Harrison, David K. Jackson, Ian Johnston, Dominic Kwiatkowski, Cordelia Langford, John Sillitoe on behalf of the Wellcome Sanger Institute COVID-19 Surveillance Team |
| EPI_ISL_2638307 | Lighthouse Lab in Glasgow | Wellcome Sanger Institute for the COVID-19 Genomics UK (COG-UK) Consortium | Harper VanSteenhouse, Yumi Kasai, David Gray, Carol Clugston, Anna Dominiczak and Alex Alderton, Roberto Amato, Jeffrey Barrett, Sonia Goncalves, Ewan Harrison, David K. Jackson, Ian Johnston, Dominic Kwiatkowski, Cordelia Langford, John Sillitoe on behalf of the Wellcome Sanger Institute COVID-19 Surveillance Team |
| EPI_ISL_2638308 | Lighthouse Lab in Milton Keynes | Wellcome Sanger Institute for the COVID-19 Genomics UK (COG-UK) Consortium | The Lighthouse Lab in Milton Keynes and Alex Alderton, Roberto Amato, Jeffrey Barrett, Sonia Goncalves, Ewan Harrison, David K. Jackson, Ian Johnston, Dominic Kwiatkowski, Cordelia Langford, John Sillitoe on behalf of the Wellcome Sanger Institute COVID-19 Surveillance Team |
| EPI_ISL_2638309 | Lighthouse Lab in Alderley Park | Wellcome Sanger Institute for the COVID-19 Genomics UK (COG-UK) Consortium | Jacquelyn Wynn, Mairead Hyland, The Lighthouse Lab in Alderley Park and Alex Alderton, Roberto Amato, Jeffrey Barrett, Sonia Goncalves, Ewan Harrison, David K. Jackson, Ian Johnston, Dominic Kwiatkowski, Cordelia Langford, John Sillitoe on behalf of the Wellcome Sanger Institute COVID-19 Surveillance Team |
| EPI_ISL_2638310 | Lighthouse Lab in Milton Keynes | Wellcome Sanger Institute for the COVID-19 Genomics UK (COG-UK) Consortium | The Lighthouse Lab in Milton Keynes and Alex Alderton, Roberto Amato, Jeffrey Barrett, Sonia Goncalves, Ewan Harrison, David K. Jackson, Ian Johnston, Dominic Kwiatkowski, Cordelia Langford, John Sillitoe on behalf of the Wellcome Sanger Institute COVID-19 Surveillance Team |
| EPI_ISL_2638311, EPI_ISL_2638312 | Lighthouse Lab in Alderley Park | Wellcome Sanger Institute for the COVID-19 Genomics UK (COG-UK) Consortium | Jacquelyn Wynn, Mairead Hyland, The Lighthouse Lab in Alderley Park and Alex Alderton, Roberto Amato, Jeffrey Barrett, Sonia Goncalves, Ewan Harrison, David K. Jackson, Ian Johnston, Dominic Kwiatkowski, Cordelia Langford, John Sillitoe on behalf of the Wellcome Sanger Institute COVID-19 Surveillance Team |
| EPI_ISL_2638313 | Health Services Laboratories | Wellcome Sanger Institute for the COVID-19 Genomics UK (COG-UK) Consortium | Health Services Laboratories and Alex Alderton, Roberto Amato, Jeffrey Barrett, Sonia Goncalves, Ewan Harrison, David K. Jackson, Ian Johnston, Dominic Kwiatkowski, Cordelia Langford, John Sillitoe on behalf of the Wellcome Sanger Institute COVID-19 Surveillance Team |
| EPI_ISL_2638314, EPI_ISL_2638315 | Lighthouse Lab in Alderley Park | Wellcome Sanger Institute for the COVID-19 Genomics UK (COG-UK) Consortium | Jacquelyn Wynn, Mairead Hyland, The Lighthouse Lab in Alderley Park and Alex Alderton, Roberto Amato, Jeffrey Barrett, Sonia Goncalves, Ewan Harrison, David K. Jackson, Ian Johnston, Dominic Kwiatkowski, Cordelia Langford, John Sillitoe on behalf of the Wellcome Sanger Institute COVID-19 Surveillance Team |
| EPI_ISL_2638317, EPI_ISL_2638318, EPI_ISL_2638319 | Lighthouse Lab in Milton Keynes | Wellcome Sanger Institute for the COVID-19 Genomics UK (COG-UK) Consortium | The Lighthouse Lab in Milton Keynes and Alex Alderton, Roberto Amato, Jeffrey Barrett, Sonia Goncalves, Ewan Harrison, David K. Jackson, Ian Johnston, Dominic Kwiatkowski, Cordelia Langford, John Sillitoe on behalf of the Wellcome Sanger Institute COVID-19 Surveillance Team |

| | | | |
|---|---|---|---|
| EPI_ISL_2638320 | Health Services Laboratories | Wellcome Sanger Institute for the COVID-19 Genomics UK (COG-UK) Consortium | Health Services Laboratories and Alex Alderton, Roberto Amato, Jeffrey Barrett, Sonia Goncalves, Ewan Harrison, David K. Jackson, Ian Johnston, Dominic Kwiatkowski, Cordelia Langford, John Sillitoe on behalf of the Wellcome Sanger Institute COVID-19 Surveillance Team |
| EPI_ISL_2638321 | Lighthouse Lab in Alderley Park | Wellcome Sanger Institute for the COVID-19 Genomics UK (COG-UK) Consortium | Jacquelyn Wynn, Mairead Hyland, The Lighthouse Lab in Alderley Park and Alex Alderton, Roberto Amato, Jeffrey Barrett, Sonia Goncalves, Ewan Harrison, David K. Jackson, Ian Johnston, Dominic Kwiatkowski, Cordelia Langford, John Sillitoe on behalf of the Wellcome Sanger Institute COVID-19 Surveillance Team |
| EPI_ISL_2638322 | Lighthouse Lab in Milton Keynes | Wellcome Sanger Institute for the COVID-19 Genomics UK (COG-UK) Consortium | The Lighthouse Lab in Milton Keynes and Alex Alderton, Roberto Amato, Jeffrey Barrett, Sonia Goncalves, Ewan Harrison, David K. Jackson, Ian Johnston, Dominic Kwiatkowski, Cordelia Langford, John Sillitoe on behalf of the Wellcome Sanger Institute COVID-19 Surveillance Team |
| EPI_ISL_2638323 | Lighthouse Lab in Alderley Park | Wellcome Sanger Institute for the COVID-19 Genomics UK (COG-UK) Consortium | Jacquelyn Wynn, Mairead Hyland, The Lighthouse Lab in Alderley Park and Alex Alderton, Roberto Amato, Jeffrey Barrett, Sonia Goncalves, Ewan Harrison, David K. Jackson, Ian Johnston, Dominic Kwiatkowski, Cordelia Langford, John Sillitoe on behalf of the Wellcome Sanger Institute COVID-19 Surveillance Team |
| EPI_ISL_2638324 | Lighthouse Lab in Glasgow | Wellcome Sanger Institute for the COVID-19 Genomics UK (COG-UK) Consortium | Harper VanSteenhouse, Yumi Kasai, David Gray, Carol Clugston, Anna Dominiczak and Alex Alderton, Roberto Amato, Jeffrey Barrett, Sonia Goncalves, Ewan Harrison, David K. Jackson, Ian Johnston, Dominic Kwiatkowski, Cordelia Langford, John Sillitoe on behalf of the Wellcome Sanger Institute COVID-19 Surveillance Team |
| EPI_ISL_2638325, EPI_ISL_2638326 | Lighthouse Lab in Alderley Park | Wellcome Sanger Institute for the COVID-19 Genomics UK (COG-UK) Consortium | Jacquelyn Wynn, Mairead Hyland, The Lighthouse Lab in Alderley Park and Alex Alderton, Roberto Amato, Jeffrey Barrett, Sonia Goncalves, Ewan Harrison, David K. Jackson, Ian Johnston, Dominic Kwiatkowski, Cordelia Langford, John Sillitoe on behalf of the Wellcome Sanger Institute COVID-19 Surveillance Team |
| EPI_ISL_2638327, EPI_ISL_2638328, EPI_ISL_2638329 | Lighthouse Lab in Milton Keynes | Wellcome Sanger Institute for the COVID-19 Genomics UK (COG-UK) Consortium | The Lighthouse Lab in Milton Keynes and Alex Alderton, Roberto Amato, Jeffrey Barrett, Sonia Goncalves, Ewan Harrison, David K. Jackson, Ian Johnston, Dominic Kwiatkowski, Cordelia Langford, John Sillitoe on behalf of the Wellcome Sanger Institute COVID-19 Surveillance Team |
| EPI_ISL_2638330 | Health Services Laboratories | Wellcome Sanger Institute for the COVID-19 Genomics UK (COG-UK) Consortium | Health Services Laboratories and Alex Alderton, Roberto Amato, Jeffrey Barrett, Sonia Goncalves, Ewan Harrison, David K. Jackson, Ian Johnston, Dominic Kwiatkowski, Cordelia Langford, John Sillitoe on behalf of the Wellcome Sanger Institute COVID-19 Surveillance Team |
| EPI_ISL_2638331, EPI_ISL_2638332 | Lighthouse Lab in Alderley Park | Wellcome Sanger Institute for the COVID-19 Genomics UK (COG-UK) Consortium | Jacquelyn Wynn, Mairead Hyland, The Lighthouse Lab in Alderley Park and Alex Alderton, Roberto Amato, Jeffrey Barrett, Sonia Goncalves, Ewan Harrison, David K. Jackson, Ian Johnston, Dominic Kwiatkowski, Cordelia Langford, John Sillitoe on behalf of the Wellcome Sanger Institute COVID-19 Surveillance Team |
| EPI_ISL_2638333 | Lighthouse Lab in Milton Keynes | Wellcome Sanger Institute for the COVID-19 Genomics UK (COG-UK) Consortium | The Lighthouse Lab in Milton Keynes and Alex Alderton, Roberto Amato, Jeffrey Barrett, Sonia Goncalves, Ewan Harrison, David K. Jackson, Ian Johnston, Dominic Kwiatkowski, Cordelia Langford, John Sillitoe on behalf of the Wellcome Sanger Institute COVID-19 Surveillance Team |
| EPI_ISL_2638334 | Lighthouse Lab in Alderley Park | Wellcome Sanger Institute for the COVID-19 Genomics UK (COG-UK) Consortium | Jacquelyn Wynn, Mairead Hyland, The Lighthouse Lab in Alderley Park and Alex Alderton, Roberto Amato, Jeffrey Barrett, Sonia Goncalves, Ewan Harrison, David K. Jackson, Ian Johnston, Dominic Kwiatkowski, Cordelia Langford, John Sillitoe on behalf of the Wellcome Sanger Institute COVID-19 Surveillance Team |
| EPI_ISL_2638335, EPI_ISL_2638336 | Lighthouse Lab in Milton Keynes | Wellcome Sanger Institute for the COVID-19 Genomics UK (COG-UK) Consortium | The Lighthouse Lab in Milton Keynes and Alex Alderton, Roberto Amato, Jeffrey Barrett, Sonia Goncalves, Ewan Harrison, David K. Jackson, Ian Johnston, Dominic Kwiatkowski, Cordelia Langford, John Sillitoe on behalf of the Wellcome Sanger Institute COVID-19 Surveillance Team |
| EPI_ISL_2638337 | Lighthouse Lab in Alderley Park | Wellcome Sanger Institute for the COVID-19 Genomics UK (COG-UK) Consortium | Jacquelyn Wynn, Mairead Hyland, The Lighthouse Lab in Alderley Park and Alex Alderton, Roberto Amato, Jeffrey Barrett, Sonia Goncalves, Ewan Harrison, David K. Jackson, Ian Johnston, Dominic Kwiatkowski, Cordelia Langford, John Sillitoe on behalf of the Wellcome Sanger Institute COVID-19 Surveillance Team |
| EPI_ISL_2638338, EPI_ISL_2638339 | Lighthouse Lab in Milton Keynes | Wellcome Sanger Institute for the COVID-19 Genomics UK (COG-UK) Consortium | The Lighthouse Lab in Milton Keynes and Alex Alderton, Roberto Amato, Jeffrey Barrett, Sonia Goncalves, Ewan Harrison, David K. Jackson, Ian Johnston, Dominic Kwiatkowski, Cordelia Langford, John Sillitoe on behalf of the Wellcome Sanger Institute COVID-19 Surveillance Team |
| EPI_ISL_2638340 | Lighthouse Lab in Alderley Park | Wellcome Sanger Institute for the COVID-19 Genomics UK (COG-UK) Consortium | Jacquelyn Wynn, Mairead Hyland, The Lighthouse Lab in Alderley Park and Alex Alderton, Roberto Amato, Jeffrey Barrett, Sonia Goncalves, Ewan Harrison, David K. Jackson, Ian Johnston, Dominic Kwiatkowski, Cordelia Langford, John Sillitoe on behalf of the Wellcome Sanger Institute COVID-19 Surveillance Team |
| EPI_ISL_2638341, EPI_ISL_2638342 | Lighthouse Lab in Milton Keynes | Wellcome Sanger Institute for the COVID-19 Genomics UK (COG-UK) Consortium | The Lighthouse Lab in Milton Keynes and Alex Alderton, Roberto Amato, Jeffrey Barrett, Sonia Goncalves, Ewan Harrison, David K. Jackson, Ian Johnston, Dominic Kwiatkowski, Cordelia Langford, John Sillitoe on behalf of the Wellcome Sanger Institute COVID-19 Surveillance Team |
| EPI_ISL_2638343, EPI_ISL_2638344 | Health Services Laboratories | Wellcome Sanger Institute for the COVID-19 Genomics UK (COG-UK) Consortium | Health Services Laboratories and Alex Alderton, Roberto Amato, Jeffrey Barrett, Sonia Goncalves, Ewan Harrison, David K. Jackson, Ian Johnston, Dominic Kwiatkowski, Cordelia Langford, John Sillitoe on behalf of the Wellcome Sanger Institute COVID-19 Surveillance Team |
| EPI_ISL_2638345, EPI_ISL_2638346 | Lighthouse Lab in Alderley Park | Wellcome Sanger Institute for the COVID-19 Genomics UK (COG-UK) Consortium | Jacquelyn Wynn, Mairead Hyland, The Lighthouse Lab in Alderley Park and Alex Alderton, Roberto Amato, Jeffrey Barrett, Sonia Goncalves, Ewan Harrison, David K. Jackson, Ian Johnston, Dominic Kwiatkowski, Cordelia Langford, John Sillitoe on behalf of the Wellcome Sanger Institute COVID-19 Surveillance Team |
| EPI_ISL_2638347, EPI_ISL_2638348 | Lighthouse Lab in Milton Keynes | Wellcome Sanger Institute for the COVID-19 Genomics UK (COG-UK) Consortium | The Lighthouse Lab in Milton Keynes and Alex Alderton, Roberto Amato, Jeffrey Barrett, Sonia Goncalves, Ewan Harrison, David K. Jackson, Ian Johnston, Dominic Kwiatkowski, Cordelia Langford, John Sillitoe on behalf of the Wellcome Sanger Institute COVID-19 Surveillance Team |
| EPI_ISL_2638349 | Lighthouse Lab in Glasgow | Wellcome Sanger Institute for the COVID-19 Genomics UK (COG-UK) Consortium | Harper VanSteenhouse, Yumi Kasai, David Gray, Carol Clugston, Anna Dominiczak and Alex Alderton, Roberto Amato, Jeffrey Barrett, Sonia Goncalves, Ewan Harrison, David K. Jackson, Ian Johnston, Dominic Kwiatkowski, Cordelia Langford, John Sillitoe on behalf of the Wellcome Sanger Institute COVID-19 Surveillance Team |
| EPI_ISL_2638350 | Lighthouse Lab in Alderley Park | Wellcome Sanger Institute for the COVID-19 Genomics UK (COG-UK) Consortium | Jacquelyn Wynn, Mairead Hyland, The Lighthouse Lab in Alderley Park and Alex Alderton, Roberto Amato, Jeffrey Barrett, Sonia Goncalves, Ewan Harrison, David K. Jackson, Ian Johnston, Dominic Kwiatkowski, Cordelia Langford, John Sillitoe on behalf of the Wellcome Sanger Institute COVID-19 Surveillance Team |
| EPI_ISL_2638351, EPI_ISL_2638352 | Lighthouse Lab in Milton Keynes | Wellcome Sanger Institute for the COVID-19 Genomics UK (COG-UK) Consortium | The Lighthouse Lab in Milton Keynes and Alex Alderton, Roberto Amato, Jeffrey Barrett, Sonia Goncalves, Ewan Harrison, David K. Jackson, Ian Johnston, Dominic Kwiatkowski, Cordelia Langford, John Sillitoe on behalf of the Wellcome Sanger Institute COVID-19 Surveillance Team |
| EPI_ISL_2638353 | Lighthouse Lab in Alderley Park | Wellcome Sanger Institute for the COVID-19 Genomics UK (COG-UK) Consortium | Jacquelyn Wynn, Mairead Hyland, The Lighthouse Lab in Alderley Park and Alex Alderton, Roberto Amato, Jeffrey Barrett, Sonia Goncalves, Ewan Harrison, David K. Jackson, Ian Johnston, Dominic Kwiatkowski, Cordelia Langford, John Sillitoe on behalf of the Wellcome Sanger Institute COVID-19 Surveillance Team |
| EPI_ISL_2638354 | Health Services Laboratories | Wellcome Sanger Institute for the COVID-19 Genomics UK (COG-UK) Consortium | Health Services Laboratories and Alex Alderton, Roberto Amato, Jeffrey Barrett, Sonia Goncalves, Ewan Harrison, David K. Jackson, Ian Johnston, Dominic Kwiatkowski, Cordelia Langford, John Sillitoe on behalf of the Wellcome Sanger Institute COVID-19 Surveillance Team |
| EPI_ISL_2638355 | Lighthouse Lab in Milton Keynes | Wellcome Sanger Institute for the COVID-19 Genomics UK (COG-UK) Consortium | The Lighthouse Lab in Milton Keynes and Alex Alderton, Roberto Amato, Jeffrey Barrett, Sonia Goncalves, Ewan Harrison, David K. Jackson, Ian Johnston, Dominic Kwiatkowski, Cordelia Langford, John Sillitoe on behalf of the Wellcome Sanger Institute COVID-19 Surveillance Team |
| EPI_ISL_2638356 | Health Services Laboratories | Wellcome Sanger Institute for the COVID-19 Genomics UK (COG-UK) Consortium | Health Services Laboratories and Alex Alderton, Roberto Amato, Jeffrey Barrett, Sonia Goncalves, Ewan Harrison, David K. Jackson, Ian Johnston, Dominic Kwiatkowski, Cordelia Langford, John Sillitoe on behalf of the Wellcome Sanger Institute COVID-19 Surveillance Team |
| EPI_ISL_2638357, EPI_ISL_2638358, EPI_ISL_2638359, EPI_ISL_2638360 | Lighthouse Lab in Milton Keynes | Wellcome Sanger Institute for the COVID-19 Genomics UK (COG-UK) Consortium | The Lighthouse Lab in Milton Keynes and Alex Alderton, Roberto Amato, Jeffrey Barrett, Sonia Goncalves, Ewan Harrison, David K. Jackson, Ian Johnston, Dominic Kwiatkowski, Cordelia Langford, John Sillitoe on behalf of the Wellcome Sanger Institute COVID-19 Surveillance Team |
| EPI_ISL_2638361 | Lighthouse Lab in Alderley Park | Wellcome Sanger Institute for the COVID-19 Genomics UK (COG-UK) Consortium | Jacquelyn Wynn, Mairead Hyland, The Lighthouse Lab in Alderley Park and Alex Alderton, Roberto Amato, Jeffrey Barrett, Sonia Goncalves, Ewan Harrison, David K. Jackson, Ian Johnston, Dominic Kwiatkowski, Cordelia Langford, John Sillitoe on behalf of the Wellcome Sanger Institute COVID-19 Surveillance Team |
| EPI_ISL_2638362 | Lighthouse Lab in Milton Keynes | Wellcome Sanger Institute for the COVID-19 Genomics UK (COG-UK) Consortium | The Lighthouse Lab in Milton Keynes and Alex Alderton, Roberto Amato, Jeffrey Barrett, Sonia Goncalves, Ewan Harrison, David K. Jackson, Ian Johnston, Dominic Kwiatkowski, Cordelia Langford, John Sillitoe on behalf of the Wellcome Sanger Institute COVID-19 Surveillance Team |
| EPI_ISL_2638363, EPI_ISL_2638364 | Lighthouse Lab in Alderley Park | Wellcome Sanger Institute for the COVID-19 Genomics UK (COG-UK) Consortium | Jacquelyn Wynn, Mairead Hyland, The Lighthouse Lab in Alderley Park and Alex Alderton, Roberto Amato, Jeffrey Barrett, Sonia Goncalves, Ewan Harrison, David K. Jackson, Ian Johnston, Dominic Kwiatkowski, Cordelia Langford, John Sillitoe on behalf of the Wellcome Sanger Institute COVID-19 Surveillance Team |
| EPI_ISL_2638365, EPI_ISL_2638366, EPI_ISL_2638367 | Lighthouse Lab in Milton Keynes | Wellcome Sanger Institute for the COVID-19 Genomics UK (COG-UK) Consortium | The Lighthouse Lab in Milton Keynes and Alex Alderton, Roberto Amato, Jeffrey Barrett, Sonia Goncalves, Ewan Harrison, David K. Jackson, Ian Johnston, Dominic Kwiatkowski, Cordelia Langford, John Sillitoe on behalf of the Wellcome Sanger Institute COVID-19 Surveillance Team |
| EPI_ISL_2638368, EPI_ISL_2638369 | Health Services Laboratories | Wellcome Sanger Institute for the COVID-19 Genomics UK (COG-UK) Consortium | Health Services Laboratories and Alex Alderton, Roberto Amato, Jeffrey Barrett, Sonia Goncalves, Ewan Harrison, David K. Jackson, Ian Johnston, Dominic Kwiatkowski, Cordelia Langford, John Sillitoe on behalf of the Wellcome Sanger Institute COVID-19 Surveillance Team |
| EPI_ISL_2638370 | Lighthouse Lab in Alderley Park | Wellcome Sanger Institute for the COVID-19 Genomics UK (COG-UK) Consortium | Jacquelyn Wynn, Mairead Hyland, The Lighthouse Lab in Alderley Park and Alex Alderton, Roberto Amato, Jeffrey Barrett, Sonia Goncalves, Ewan Harrison, David K. Jackson, Ian Johnston, Dominic Kwiatkowski, Cordelia Langford, John Sillitoe on behalf of the Wellcome Sanger Institute COVID-19 Surveillance Team |
| EPI_ISL_2638371, EPI_ISL_2638372 | Lighthouse Lab in Milton Keynes | Wellcome Sanger Institute for the COVID-19 Genomics UK (COG-UK) Consortium | The Lighthouse Lab in Milton Keynes and Alex Alderton, Roberto Amato, Jeffrey Barrett, Sonia Goncalves, Ewan Harrison, David K. Jackson, Ian Johnston, Dominic Kwiatkowski, Cordelia Langford, John Sillitoe on behalf of the Wellcome Sanger Institute COVID-19 Surveillance Team |
| EPI_ISL_2638373, EPI_ISL_2638374, EPI_ISL_2638375 | Lighthouse Lab in Alderley Park | Wellcome Sanger Institute for the COVID-19 Genomics UK (COG-UK) Consortium | Jacquelyn Wynn, Mairead Hyland, The Lighthouse Lab in Alderley Park and Alex Alderton, Roberto Amato, Jeffrey Barrett, Sonia Goncalves, Ewan Harrison, David K. Jackson, Ian Johnston, Dominic Kwiatkowski, Cordelia Langford, John Sillitoe on behalf of the Wellcome Sanger Institute COVID-19 Surveillance Team |
| EPI_ISL_2638376 | Health Services Laboratories | Wellcome Sanger Institute for the COVID-19 Genomics UK (COG-UK) Consortium | Health Services Laboratories and Alex Alderton, Roberto Amato, Jeffrey Barrett, Sonia Goncalves, Ewan Harrison, David K. Jackson, Ian Johnston, Dominic Kwiatkowski, Cordelia Langford, John Sillitoe on behalf of the Wellcome Sanger Institute COVID-19 Surveillance Team |
| EPI_ISL_2638377 | Lighthouse Lab in Glasgow | Wellcome Sanger Institute for the COVID-19 Genomics UK (COG-UK) Consortium | Harper VanSteenhouse, Yumi Kasai, David Gray, Carol Clugston, Anna Dominiczak and Alex Alderton, Roberto Amato, Jeffrey Barrett, Sonia Goncalves, Ewan Harrison, David K. Jackson, Ian Johnston, Dominic Kwiatkowski, Cordelia Langford, John Sillitoe on behalf of the Wellcome Sanger Institute COVID-19 Surveillance Team |
| EPI_ISL_2638378 | Lighthouse Lab in Alderley Park | Wellcome Sanger Institute for the COVID-19 Genomics UK (COG-UK) Consortium | Jacquelyn Wynn, Mairead Hyland, The Lighthouse Lab in Alderley Park and Alex Alderton, Roberto Amato, Jeffrey Barrett, Sonia Goncalves, Ewan Harrison, David K. Jackson, Ian Johnston, Dominic Kwiatkowski, Cordelia Langford, John Sillitoe on behalf of the Wellcome Sanger Institute COVID-19 Surveillance Team |
| EPI_ISL_2638379 | Lighthouse Lab in Milton Keynes | Wellcome Sanger Institute for the COVID-19 Genomics UK (COG-UK) Consortium | The Lighthouse Lab in Milton Keynes and Alex Alderton, Roberto Amato, Jeffrey Barrett, Sonia Goncalves, Ewan Harrison, David K. Jackson, Ian Johnston, Dominic Kwiatkowski, Cordelia Langford, John Sillitoe on behalf of the Wellcome Sanger Institute COVID-19 Surveillance Team |
| EPI_ISL_2638380, EPI_ISL_2638381, EPI_ISL_2638382 | Lighthouse Lab in Alderley Park | Wellcome Sanger Institute for the COVID-19 Genomics UK (COG-UK) Consortium | Jacquelyn Wynn, Mairead Hyland, The Lighthouse Lab in Alderley Park and Alex Alderton, Roberto Amato, Jeffrey Barrett, Sonia Goncalves, Ewan Harrison, David K. Jackson, Ian Johnston, Dominic Kwiatkowski, Cordelia Langford, John Sillitoe on behalf of the Wellcome Sanger Institute COVID-19 Surveillance Team |
| EPI_ISL_2638383 | Lighthouse Lab in Milton Keynes | Wellcome Sanger Institute for the COVID-19 Genomics UK (COG-UK) Consortium | The Lighthouse Lab in Milton Keynes and Alex Alderton, Roberto Amato, Jeffrey Barrett, Sonia Goncalves, Ewan Harrison, David K. Jackson, Ian Johnston, Dominic Kwiatkowski, Cordelia Langford, John Sillitoe on behalf of the Wellcome Sanger Institute COVID-19 Surveillance Team |
| EPI_ISL_2638384, EPI_ISL_2638385 | Lighthouse Lab in Alderley Park | Wellcome Sanger Institute for the COVID-19 Genomics UK (COG-UK) Consortium | Jacquelyn Wynn, Mairead Hyland, The Lighthouse Lab in Alderley Park and Alex Alderton, Roberto Amato, Jeffrey Barrett, Sonia Goncalves, Ewan Harrison, David K. Jackson, Ian Johnston, Dominic Kwiatkowski, Cordelia Langford, John Sillitoe on behalf of the Wellcome Sanger Institute COVID-19 Surveillance Team |
| EPI_ISL_2638386 | Lighthouse Lab in Milton Keynes | Wellcome Sanger Institute for the COVID-19 Genomics UK (COG-UK) Consortium | The Lighthouse Lab in Milton Keynes and Alex Alderton, Roberto Amato, Jeffrey Barrett, Sonia Goncalves, Ewan Harrison, David K. Jackson, Ian Johnston, Dominic Kwiatkowski, Cordelia Langford, John Sillitoe on behalf of the Wellcome Sanger Institute COVID-19 Surveillance Team |

| | | | |
|---|---|---|---|
| EPI_ISL_2638387, EPI_ISL_2638388, EPI_ISL_2638389, EPI_ISL_2638390 | Lighthouse Lab in Alderley Park | Wellcome Sanger Institute for the COVID-19 Genomics UK (COG-UK) Consortium | Jacquelyn Wynn, Mairead Hyland, The Lighthouse Lab in Alderley Park and Alex Alderton, Roberto Amato, Jeffrey Barrett, Sonia Goncalves, Ewan Harrison, David K. Jackson, Ian Johnston, Dominic Kwiatkowski, Cordelia Langford, John Sillitoe on behalf of the Wellcome Sanger Institute COVID-19 Surveillance Team |
| EPI_ISL_2638391, EPI_ISL_2638392, EPI_ISL_2638393, EPI_ISL_2638394 | Lighthouse Lab in Milton Keynes | Wellcome Sanger Institute for the COVID-19 Genomics UK (COG-UK) Consortium | The Lighthouse Lab in Milton Keynes and Alex Alderton, Roberto Amato, Jeffrey Barrett, Sonia Goncalves, Ewan Harrison, David K. Jackson, Ian Johnston, Dominic Kwiatkowski, Cordelia Langford, John Sillitoe on behalf of the Wellcome Sanger Institute COVID-19 Surveillance Team |
| EPI_ISL_2638395, EPI_ISL_2638396 | Lighthouse Laboratory Plymouth | Wellcome Sanger Institute for the COVID-19 Genomics UK (COG-UK) Consortium | Lighthouse Laboratory Plymouth and Alex Alderton, Roberto Amato, Jeffrey Barrett, Sonia Goncalves, Ewan Harrison, David K. Jackson, Ian Johnston, Dominic Kwiatkowski, Cordelia Langford, John Sillitoe on behalf of the Wellcome Sanger Institute COVID-19 Surveillance Team |
| EPI_ISL_2638397 | Lighthouse Lab in Glasgow | Wellcome Sanger Institute for the COVID-19 Genomics UK (COG-UK) Consortium | Harper VanSteenhouse, Yumi Kasai, David Gray, Carol Clugston, Anna Dominiczak and Alex Alderton, Roberto Amato, Jeffrey Barrett, Sonia Goncalves, Ewan Harrison, David K. Jackson, Ian Johnston, Dominic Kwiatkowski, Cordelia Langford, John Sillitoe on behalf of the Wellcome Sanger Institute COVID-19 Surveillance Team |
| EPI_ISL_2638398 | Lighthouse Laboratory Plymouth | Wellcome Sanger Institute for the COVID-19 Genomics UK (COG-UK) Consortium | Lighthouse Laboratory Plymouth and Alex Alderton, Roberto Amato, Jeffrey Barrett, Sonia Goncalves, Ewan Harrison, David K. Jackson, Ian Johnston, Dominic Kwiatkowski, Cordelia Langford, John Sillitoe on behalf of the Wellcome Sanger Institute COVID-19 Surveillance Team |
| EPI_ISL_2638399 | Lighthouse Lab in Glasgow | Wellcome Sanger Institute for the COVID-19 Genomics UK (COG-UK) Consortium | Harper VanSteenhouse, Yumi Kasai, David Gray, Carol Clugston, Anna Dominiczak and Alex Alderton, Roberto Amato, Jeffrey Barrett, Sonia Goncalves, Ewan Harrison, David K. Jackson, Ian Johnston, Dominic Kwiatkowski, Cordelia Langford, John Sillitoe on behalf of the Wellcome Sanger Institute COVID-19 Surveillance Team |
| EPI_ISL_2638400 | Lighthouse Lab in Milton Keynes | Wellcome Sanger Institute for the COVID-19 Genomics UK (COG-UK) Consortium | The Lighthouse Lab in Milton Keynes and Alex Alderton, Roberto Amato, Jeffrey Barrett, Sonia Goncalves, Ewan Harrison, David K. Jackson, Ian Johnston, Dominic Kwiatkowski, Cordelia Langford, John Sillitoe on behalf of the Wellcome Sanger Institute COVID-19 Surveillance Team |
| EPI_ISL_2638401, EPI_ISL_2638402 | Lighthouse Laboratory Plymouth | Wellcome Sanger Institute for the COVID-19 Genomics UK (COG-UK) Consortium | Lighthouse Laboratory Plymouth and Alex Alderton, Roberto Amato, Jeffrey Barrett, Sonia Goncalves, Ewan Harrison, David K. Jackson, Ian Johnston, Dominic Kwiatkowski, Cordelia Langford, John Sillitoe on behalf of the Wellcome Sanger Institute COVID-19 Surveillance Team |
| EPI_ISL_2638404, EPI_ISL_2638405, EPI_ISL_2638406 | Lighthouse Lab in Milton Keynes | Wellcome Sanger Institute for the COVID-19 Genomics UK (COG-UK) Consortium | The Lighthouse Lab in Milton Keynes and Alex Alderton, Roberto Amato, Jeffrey Barrett, Sonia Goncalves, Ewan Harrison, David K. Jackson, Ian Johnston, Dominic Kwiatkowski, Cordelia Langford, John Sillitoe on behalf of the Wellcome Sanger Institute COVID-19 Surveillance Team |
| EPI_ISL_2638407 | Lighthouse Laboratory Plymouth | Wellcome Sanger Institute for the COVID-19 Genomics UK (COG-UK) Consortium | Lighthouse Laboratory Plymouth and Alex Alderton, Roberto Amato, Jeffrey Barrett, Sonia Goncalves, Ewan Harrison, David K. Jackson, Ian Johnston, Dominic Kwiatkowski, Cordelia Langford, John Sillitoe on behalf of the Wellcome Sanger Institute COVID-19 Surveillance Team |
| EPI_ISL_2638408, EPI_ISL_2638409 | Lighthouse Lab in Milton Keynes | Wellcome Sanger Institute for the COVID-19 Genomics UK (COG-UK) Consortium | The Lighthouse Lab in Milton Keynes and Alex Alderton, Roberto Amato, Jeffrey Barrett, Sonia Goncalves, Ewan Harrison, David K. Jackson, Ian Johnston, Dominic Kwiatkowski, Cordelia Langford, John Sillitoe on behalf of the Wellcome Sanger Institute COVID-19 Surveillance Team |
| EPI_ISL_2638410 | Lighthouse Lab in Glasgow | Wellcome Sanger Institute for the COVID-19 Genomics UK (COG-UK) Consortium | Harper VanSteenhouse, Yumi Kasai, David Gray, Carol Clugston, Anna Dominiczak and Alex Alderton, Roberto Amato, Jeffrey Barrett, Sonia Goncalves, Ewan Harrison, David K. Jackson, Ian Johnston, Dominic Kwiatkowski, Cordelia Langford, John Sillitoe on behalf of the Wellcome Sanger Institute COVID-19 Surveillance Team |
| EPI_ISL_2638411 | Lighthouse Lab in Milton Keynes | Wellcome Sanger Institute for the COVID-19 Genomics UK (COG-UK) Consortium | The Lighthouse Lab in Milton Keynes and Alex Alderton, Roberto Amato, Jeffrey Barrett, Sonia Goncalves, Ewan Harrison, David K. Jackson, Ian Johnston, Dominic Kwiatkowski, Cordelia Langford, John Sillitoe on behalf of the Wellcome Sanger Institute COVID-19 Surveillance Team |
| EPI_ISL_2638412, EPI_ISL_2638413 | Lighthouse Laboratory Plymouth | Wellcome Sanger Institute for the COVID-19 Genomics UK (COG-UK) Consortium | Lighthouse Laboratory Plymouth and Alex Alderton, Roberto Amato, Jeffrey Barrett, Sonia Goncalves, Ewan Harrison, David K. Jackson, Ian Johnston, Dominic Kwiatkowski, Cordelia Langford, John Sillitoe on behalf of the Wellcome Sanger Institute COVID-19 Surveillance Team |
| EPI_ISL_2638414 | Lighthouse Lab in Glasgow | Wellcome Sanger Institute for the COVID-19 Genomics UK (COG-UK) Consortium | Harper VanSteenhouse, Yumi Kasai, David Gray, Carol Clugston, Anna Dominiczak and Alex Alderton, Roberto Amato, Jeffrey Barrett, Sonia Goncalves, Ewan Harrison, David K. Jackson, Ian Johnston, Dominic Kwiatkowski, Cordelia Langford, John Sillitoe on behalf of the Wellcome Sanger Institute COVID-19 Surveillance Team |
| EPI_ISL_2638415 | Lighthouse Laboratory Plymouth | Wellcome Sanger Institute for the COVID-19 Genomics UK (COG-UK) Consortium | Lighthouse Laboratory Plymouth and Alex Alderton, Roberto Amato, Jeffrey Barrett, Sonia Goncalves, Ewan Harrison, David K. Jackson, Ian Johnston, Dominic Kwiatkowski, Cordelia Langford, John Sillitoe on behalf of the Wellcome Sanger Institute COVID-19 Surveillance Team |
| EPI_ISL_2638416 | Lighthouse Lab in Glasgow | Wellcome Sanger Institute for the COVID-19 Genomics UK (COG-UK) Consortium | Harper VanSteenhouse, Yumi Kasai, David Gray, Carol Clugston, Anna Dominiczak and Alex Alderton, Roberto Amato, Jeffrey Barrett, Sonia Goncalves, Ewan Harrison, David K. Jackson, Ian Johnston, Dominic Kwiatkowski, Cordelia Langford, John Sillitoe on behalf of the Wellcome Sanger Institute COVID-19 Surveillance Team |
| EPI_ISL_2638417 | Lighthouse Laboratory Plymouth | Wellcome Sanger Institute for the COVID-19 Genomics UK (COG-UK) Consortium | Lighthouse Laboratory Plymouth and Alex Alderton, Roberto Amato, Jeffrey Barrett, Sonia Goncalves, Ewan Harrison, David K. Jackson, Ian Johnston, Dominic Kwiatkowski, Cordelia Langford, John Sillitoe on behalf of the Wellcome Sanger Institute COVID-19 Surveillance Team |
| EPI_ISL_2638419, EPI_ISL_2638420 | Lighthouse Lab in Glasgow | Wellcome Sanger Institute for the COVID-19 Genomics UK (COG-UK) Consortium | Harper VanSteenhouse, Yumi Kasai, David Gray, Carol Clugston, Anna Dominiczak and Alex Alderton, Roberto Amato, Jeffrey Barrett, Sonia Goncalves, Ewan Harrison, David K. Jackson, Ian Johnston, Dominic Kwiatkowski, Cordelia Langford, John Sillitoe on behalf of the Wellcome Sanger Institute COVID-19 Surveillance Team |
| EPI_ISL_2638421 | Lighthouse Laboratory Plymouth | Wellcome Sanger Institute for the COVID-19 Genomics UK (COG-UK) Consortium | Lighthouse Laboratory Plymouth and Alex Alderton, Roberto Amato, Jeffrey Barrett, Sonia Goncalves, Ewan Harrison, David K. Jackson, Ian Johnston, Dominic Kwiatkowski, Cordelia Langford, John Sillitoe on behalf of the Wellcome Sanger Institute COVID-19 Surveillance Team |
| EPI_ISL_2638422 | Lighthouse Lab in Milton Keynes | Wellcome Sanger Institute for the COVID-19 Genomics UK (COG-UK) Consortium | The Lighthouse Lab in Milton Keynes and Alex Alderton, Roberto Amato, Jeffrey Barrett, Sonia Goncalves, Ewan Harrison, David K. Jackson, Ian Johnston, Dominic Kwiatkowski, Cordelia Langford, John Sillitoe on behalf of the Wellcome Sanger Institute COVID-19 Surveillance Team |
| EPI_ISL_2638423 | Lighthouse Lab in Glasgow | Wellcome Sanger Institute for the COVID-19 Genomics UK (COG-UK) Consortium | Harper VanSteenhouse, Yumi Kasai, David Gray, Carol Clugston, Anna Dominiczak and Alex Alderton, Roberto Amato, Jeffrey Barrett, Sonia Goncalves, Ewan Harrison, David K. Jackson, Ian Johnston, Dominic Kwiatkowski, Cordelia Langford, John Sillitoe on behalf of the Wellcome Sanger Institute COVID-19 Surveillance Team |
| EPI_ISL_2638424 | Lighthouse Lab in Milton Keynes | Wellcome Sanger Institute for the COVID-19 Genomics UK (COG-UK) Consortium | The Lighthouse Lab in Milton Keynes and Alex Alderton, Roberto Amato, Jeffrey Barrett, Sonia Goncalves, Ewan Harrison, David K. Jackson, Ian Johnston, Dominic Kwiatkowski, Cordelia Langford, John Sillitoe on behalf of the Wellcome Sanger Institute COVID-19 Surveillance Team |
| EPI_ISL_2638425, EPI_ISL_2638426 | Lighthouse Laboratory Plymouth | Wellcome Sanger Institute for the COVID-19 Genomics UK (COG-UK) Consortium | Lighthouse Laboratory Plymouth and Alex Alderton, Roberto Amato, Jeffrey Barrett, Sonia Goncalves, Ewan Harrison, David K. Jackson, Ian Johnston, Dominic Kwiatkowski, Cordelia Langford, John Sillitoe on behalf of the Wellcome Sanger Institute COVID-19 Surveillance Team |
| EPI_ISL_2638427 | Lighthouse Lab in Glasgow | Wellcome Sanger Institute for the COVID-19 Genomics UK (COG-UK) Consortium | Harper VanSteenhouse, Yumi Kasai, David Gray, Carol Clugston, Anna Dominiczak and Alex Alderton, Roberto Amato, Jeffrey Barrett, Sonia Goncalves, Ewan Harrison, David K. Jackson, Ian Johnston, Dominic Kwiatkowski, Cordelia Langford, John Sillitoe on behalf of the Wellcome Sanger Institute COVID-19 Surveillance Team |
| EPI_ISL_2638428, EPI_ISL_2638429, EPI_ISL_2638430 | Lighthouse Lab in Milton Keynes | Wellcome Sanger Institute for the COVID-19 Genomics UK (COG-UK) Consortium | The Lighthouse Lab in Milton Keynes and Alex Alderton, Roberto Amato, Jeffrey Barrett, Sonia Goncalves, Ewan Harrison, David K. Jackson, Ian Johnston, Dominic Kwiatkowski, Cordelia Langford, John Sillitoe on behalf of the Wellcome Sanger Institute COVID-19 Surveillance Team |
| EPI_ISL_2638431, EPI_ISL_2638433 | Lighthouse Laboratory Plymouth | Wellcome Sanger Institute for the COVID-19 Genomics UK (COG-UK) Consortium | Lighthouse Laboratory Plymouth and Alex Alderton, Roberto Amato, Jeffrey Barrett, Sonia Goncalves, Ewan Harrison, David K. Jackson, Ian Johnston, Dominic Kwiatkowski, Cordelia Langford, John Sillitoe on behalf of the Wellcome Sanger Institute COVID-19 Surveillance Team |
| EPI_ISL_2638434, EPI_ISL_2638435 | Lighthouse Lab in Glasgow | Wellcome Sanger Institute for the COVID-19 Genomics UK (COG-UK) Consortium | Harper VanSteenhouse, Yumi Kasai, David Gray, Carol Clugston, Anna Dominiczak and Alex Alderton, Roberto Amato, Jeffrey Barrett, Sonia Goncalves, Ewan Harrison, David K. Jackson, Ian Johnston, Dominic Kwiatkowski, Cordelia Langford, John Sillitoe on behalf of the Wellcome Sanger Institute COVID-19 Surveillance Team |
| EPI_ISL_2638436, EPI_ISL_2638437 | Lighthouse Lab in Milton Keynes | Wellcome Sanger Institute for the COVID-19 Genomics UK (COG-UK) Consortium | The Lighthouse Lab in Milton Keynes and Alex Alderton, Roberto Amato, Jeffrey Barrett, Sonia Goncalves, Ewan Harrison, David K. Jackson, Ian Johnston, Dominic Kwiatkowski, Cordelia Langford, John Sillitoe on behalf of the Wellcome Sanger Institute COVID-19 Surveillance Team |
| EPI_ISL_2638438, EPI_ISL_2638439 | Lighthouse Laboratory Plymouth | Wellcome Sanger Institute for the COVID-19 Genomics UK (COG-UK) Consortium | Lighthouse Laboratory Plymouth and Alex Alderton, Roberto Amato, Jeffrey Barrett, Sonia Goncalves, Ewan Harrison, David K. Jackson, Ian Johnston, Dominic Kwiatkowski, Cordelia Langford, John Sillitoe on behalf of the Wellcome Sanger Institute COVID-19 Surveillance Team |
| EPI_ISL_2638440 | Lighthouse Lab in Glasgow | Wellcome Sanger Institute for the COVID-19 Genomics UK (COG-UK) Consortium | Harper VanSteenhouse, Yumi Kasai, David Gray, Carol Clugston, Anna Dominiczak and Alex Alderton, Roberto Amato, Jeffrey Barrett, Sonia Goncalves, Ewan Harrison, David K. Jackson, Ian Johnston, Dominic Kwiatkowski, Cordelia Langford, John Sillitoe on behalf of the Wellcome Sanger Institute COVID-19 Surveillance Team |
| EPI_ISL_2638441 | Lighthouse Lab in Milton Keynes | Wellcome Sanger Institute for the COVID-19 Genomics UK (COG-UK) Consortium | The Lighthouse Lab in Milton Keynes and Alex Alderton, Roberto Amato, Jeffrey Barrett, Sonia Goncalves, Ewan Harrison, David K. Jackson, Ian Johnston, Dominic Kwiatkowski, Cordelia Langford, John Sillitoe on behalf of the Wellcome Sanger Institute COVID-19 Surveillance Team |
| EPI_ISL_2638442 | Lighthouse Laboratory Plymouth | Wellcome Sanger Institute for the COVID-19 Genomics UK (COG-UK) Consortium | Lighthouse Laboratory Plymouth and Alex Alderton, Roberto Amato, Jeffrey Barrett, Sonia Goncalves, Ewan Harrison, David K. Jackson, Ian Johnston, Dominic Kwiatkowski, Cordelia Langford, John Sillitoe on behalf of the Wellcome Sanger Institute COVID-19 Surveillance Team |
| EPI_ISL_2638443 | Lighthouse Lab in Glasgow | Wellcome Sanger Institute for the COVID-19 Genomics UK (COG-UK) Consortium | Harper VanSteenhouse, Yumi Kasai, David Gray, Carol Clugston, Anna Dominiczak and Alex Alderton, Roberto Amato, Jeffrey Barrett, Sonia Goncalves, Ewan Harrison, David K. Jackson, Ian Johnston, Dominic Kwiatkowski, Cordelia Langford, John Sillitoe on behalf of the Wellcome Sanger Institute COVID-19 Surveillance Team |
| EPI_ISL_2638444 | Lighthouse Lab in Milton Keynes | Wellcome Sanger Institute for the COVID-19 Genomics UK (COG-UK) Consortium | The Lighthouse Lab in Milton Keynes and Alex Alderton, Roberto Amato, Jeffrey Barrett, Sonia Goncalves, Ewan Harrison, David K. Jackson, Ian Johnston, Dominic Kwiatkowski, Cordelia Langford, John Sillitoe on behalf of the Wellcome Sanger Institute COVID-19 Surveillance Team |
| EPI_ISL_2638445 | Lighthouse Laboratory Plymouth | Wellcome Sanger Institute for the COVID-19 Genomics UK (COG-UK) Consortium | Lighthouse Laboratory Plymouth and Alex Alderton, Roberto Amato, Jeffrey Barrett, Sonia Goncalves, Ewan Harrison, David K. Jackson, Ian Johnston, Dominic Kwiatkowski, Cordelia Langford, John Sillitoe on behalf of the Wellcome Sanger Institute COVID-19 Surveillance Team |
| EPI_ISL_2638446, EPI_ISL_2638447 | Lighthouse Lab in Milton Keynes | Wellcome Sanger Institute for the COVID-19 Genomics UK (COG-UK) Consortium | The Lighthouse Lab in Milton Keynes and Alex Alderton, Roberto Amato, Jeffrey Barrett, Sonia Goncalves, Ewan Harrison, David K. Jackson, Ian Johnston, Dominic Kwiatkowski, Cordelia Langford, John Sillitoe on behalf of the Wellcome Sanger Institute COVID-19 Surveillance Team |
| EPI_ISL_2638448 | Lighthouse Laboratory Plymouth | Wellcome Sanger Institute for the COVID-19 Genomics UK (COG-UK) Consortium | Lighthouse Laboratory Plymouth and Alex Alderton, Roberto Amato, Jeffrey Barrett, Sonia Goncalves, Ewan Harrison, David K. Jackson, Ian Johnston, Dominic Kwiatkowski, Cordelia Langford, John Sillitoe on behalf of the Wellcome Sanger Institute COVID-19 Surveillance Team |
| EPI_ISL_2638449 | Lighthouse Lab in Glasgow | Wellcome Sanger Institute for the COVID-19 Genomics UK (COG-UK) Consortium | Harper VanSteenhouse, Yumi Kasai, David Gray, Carol Clugston, Anna Dominiczak and Alex Alderton, Roberto Amato, Jeffrey Barrett, Sonia Goncalves, Ewan Harrison, David K. Jackson, Ian Johnston, Dominic Kwiatkowski, Cordelia Langford, John Sillitoe on behalf of the Wellcome Sanger Institute COVID-19 Surveillance Team |
| EPI_ISL_2638450 | Lighthouse Lab in Milton Keynes | Wellcome Sanger Institute for the COVID-19 Genomics UK (COG-UK) Consortium | The Lighthouse Lab in Milton Keynes and Alex Alderton, Roberto Amato, Jeffrey Barrett, Sonia Goncalves, Ewan Harrison, David K. Jackson, Ian Johnston, Dominic Kwiatkowski, Cordelia Langford, John Sillitoe on behalf of the Wellcome Sanger Institute COVID-19 Surveillance Team |
| EPI_ISL_2638451 | Lighthouse Laboratory Plymouth | Wellcome Sanger Institute for the COVID-19 Genomics UK (COG-UK) Consortium | Lighthouse Laboratory Plymouth and Alex Alderton, Roberto Amato, Jeffrey Barrett, Sonia Goncalves, Ewan Harrison, David K. Jackson, Ian Johnston, Dominic Kwiatkowski, Cordelia Langford, John Sillitoe on behalf of the Wellcome Sanger Institute COVID-19 Surveillance Team |
| EPI_ISL_2638452 | Lighthouse Lab in Milton Keynes | Wellcome Sanger Institute for the COVID-19 Genomics UK (COG-UK) Consortium | The Lighthouse Lab in Milton Keynes and Alex Alderton, Roberto Amato, Jeffrey Barrett, Sonia Goncalves, Ewan Harrison, David K. Jackson, Ian Johnston, Dominic Kwiatkowski, Cordelia Langford, John Sillitoe on behalf of the Wellcome Sanger Institute COVID-19 Surveillance Team |
| EPI_ISL_2638453, | Lighthouse Lab in Glasgow | Wellcome Sanger Institute for the COVID-19 Genomics UK (COG-UK) | Harper VanSteenhouse, Yumi Kasai, David Gray, Carol Clugston, Anna Dominiczak and Alex Alderton, Roberto Amato, Jeffrey Barrett, Sonia Goncalves, Ewan Harrison, David K. Jackson, Ian Johnston, Dominic Kwiatkowski, Cordelia Langford, |

| | | | |
|---|---|---|---|
| EPI_ISL_2638454 | | Consortium | John Sillitoe on behalf of the Wellcome Sanger Institute COVID-19 Surveillance Team |
| EPI_ISL_2638455 | Lighthouse Laboratory Plymouth | Wellcome Sanger Institute for the COVID-19 Genomics UK (COG-UK) Consortium | Lighthouse Laboratory Plymouth and Alex Alderton, Roberto Amato, Jeffrey Barrett, Sonia Goncalves, Ewan Harrison, David K. Jackson, Ian Johnston, Dominic Kwiatkowski, Cordelia Langford, John Sillitoe on behalf of the Wellcome Sanger Institute COVID-19 Surveillance Team |
| EPI_ISL_2638456 | Lighthouse Lab in Glasgow | Wellcome Sanger Institute for the COVID-19 Genomics UK (COG-UK) Consortium | Harper VanSteenhouse, Yumi Kasai, David Gray, Carol Clugston, Anna Dominiczak and Alex Alderton, Roberto Amato, Jeffrey Barrett, Sonia Goncalves, Ewan Harrison, David K. Jackson, Ian Johnston, Dominic Kwiatkowski, Cordelia Langford, John Sillitoe on behalf of the Wellcome Sanger Institute COVID-19 Surveillance Team |
| EPI_ISL_2638457 | Lighthouse Lab in Milton Keynes | Wellcome Sanger Institute for the COVID-19 Genomics UK (COG-UK) Consortium | The Lighthouse Lab in Milton Keynes and Alex Alderton, Roberto Amato, Jeffrey Barrett, Sonia Goncalves, Ewan Harrison, David K. Jackson, Ian Johnston, Dominic Kwiatkowski, Cordelia Langford, John Sillitoe on behalf of the Wellcome Sanger Institute COVID-19 Surveillance Team |
| EPI_ISL_2638458 | Lighthouse Lab in Glasgow | Wellcome Sanger Institute for the COVID-19 Genomics UK (COG-UK) Consortium | Harper VanSteenhouse, Yumi Kasai, David Gray, Carol Clugston, Anna Dominiczak and Alex Alderton, Roberto Amato, Jeffrey Barrett, Sonia Goncalves, Ewan Harrison, David K. Jackson, Ian Johnston, Dominic Kwiatkowski, Cordelia Langford, John Sillitoe on behalf of the Wellcome Sanger Institute COVID-19 Surveillance Team |
| EPI_ISL_2638459 | Lighthouse Laboratory Plymouth | Wellcome Sanger Institute for the COVID-19 Genomics UK (COG-UK) Consortium | Lighthouse Laboratory Plymouth and Alex Alderton, Roberto Amato, Jeffrey Barrett, Sonia Goncalves, Ewan Harrison, David K. Jackson, Ian Johnston, Dominic Kwiatkowski, Cordelia Langford, John Sillitoe on behalf of the Wellcome Sanger Institute COVID-19 Surveillance Team |
| EPI_ISL_2638460 | Lighthouse Lab in Milton Keynes | Wellcome Sanger Institute for the COVID-19 Genomics UK (COG-UK) Consortium | The Lighthouse Lab in Milton Keynes and Alex Alderton, Roberto Amato, Jeffrey Barrett, Sonia Goncalves, Ewan Harrison, David K. Jackson, Ian Johnston, Dominic Kwiatkowski, Cordelia Langford, John Sillitoe on behalf of the Wellcome Sanger Institute COVID-19 Surveillance Team |
| EPI_ISL_2638461 | Lighthouse Laboratory Plymouth | Wellcome Sanger Institute for the COVID-19 Genomics UK (COG-UK) Consortium | Lighthouse Laboratory Plymouth and Alex Alderton, Roberto Amato, Jeffrey Barrett, Sonia Goncalves, Ewan Harrison, David K. Jackson, Ian Johnston, Dominic Kwiatkowski, Cordelia Langford, John Sillitoe on behalf of the Wellcome Sanger Institute COVID-19 Surveillance Team |
| EPI_ISL_2638462 | Lighthouse Lab in Milton Keynes | Wellcome Sanger Institute for the COVID-19 Genomics UK (COG-UK) Consortium | The Lighthouse Lab in Milton Keynes and Alex Alderton, Roberto Amato, Jeffrey Barrett, Sonia Goncalves, Ewan Harrison, David K. Jackson, Ian Johnston, Dominic Kwiatkowski, Cordelia Langford, John Sillitoe on behalf of the Wellcome Sanger Institute COVID-19 Surveillance Team |
| EPI_ISL_2638463, EPI_ISL_2638464 | Lighthouse Laboratory Plymouth | Wellcome Sanger Institute for the COVID-19 Genomics UK (COG-UK) Consortium | Lighthouse Laboratory Plymouth and Alex Alderton, Roberto Amato, Jeffrey Barrett, Sonia Goncalves, Ewan Harrison, David K. Jackson, Ian Johnston, Dominic Kwiatkowski, Cordelia Langford, John Sillitoe on behalf of the Wellcome Sanger Institute COVID-19 Surveillance Team |
| EPI_ISL_2638465 | Lighthouse Lab in Milton Keynes | Wellcome Sanger Institute for the COVID-19 Genomics UK (COG-UK) Consortium | The Lighthouse Lab in Milton Keynes and Alex Alderton, Roberto Amato, Jeffrey Barrett, Sonia Goncalves, Ewan Harrison, David K. Jackson, Ian Johnston, Dominic Kwiatkowski, Cordelia Langford, John Sillitoe on behalf of the Wellcome Sanger Institute COVID-19 Surveillance Team |
| EPI_ISL_2638466, EPI_ISL_2638467, EPI_ISL_2638468, EPI_ISL_2638469, EPI_ISL_2638470, EPI_ISL_2638471, EPI_ISL_2638472 | Lighthouse Laboratory Plymouth | Wellcome Sanger Institute for the COVID-19 Genomics UK (COG-UK) Consortium | Lighthouse Laboratory Plymouth and Alex Alderton, Roberto Amato, Jeffrey Barrett, Sonia Goncalves, Ewan Harrison, David K. Jackson, Ian Johnston, Dominic Kwiatkowski, Cordelia Langford, John Sillitoe on behalf of the Wellcome Sanger Institute COVID-19 Surveillance Team |
| EPI_ISL_2638473 | Lighthouse Lab in Milton Keynes | Wellcome Sanger Institute for the COVID-19 Genomics UK (COG-UK) Consortium | The Lighthouse Lab in Milton Keynes and Alex Alderton, Roberto Amato, Jeffrey Barrett, Sonia Goncalves, Ewan Harrison, David K. Jackson, Ian Johnston, Dominic Kwiatkowski, Cordelia Langford, John Sillitoe on behalf of the Wellcome Sanger Institute COVID-19 Surveillance Team |
| EPI_ISL_2638474, EPI_ISL_2638475 | Lighthouse Laboratory Plymouth | Wellcome Sanger Institute for the COVID-19 Genomics UK (COG-UK) Consortium | Lighthouse Laboratory Plymouth and Alex Alderton, Roberto Amato, Jeffrey Barrett, Sonia Goncalves, Ewan Harrison, David K. Jackson, Ian Johnston, Dominic Kwiatkowski, Cordelia Langford, John Sillitoe on behalf of the Wellcome Sanger Institute COVID-19 Surveillance Team |
| EPI_ISL_2638476 | Lighthouse Lab in Glasgow | Wellcome Sanger Institute for the COVID-19 Genomics UK (COG-UK) Consortium | Harper VanSteenhouse, Yumi Kasai, David Gray, Carol Clugston, Anna Dominiczak and Alex Alderton, Roberto Amato, Jeffrey Barrett, Sonia Goncalves, Ewan Harrison, David K. Jackson, Ian Johnston, Dominic Kwiatkowski, Cordelia Langford, John Sillitoe on behalf of the Wellcome Sanger Institute COVID-19 Surveillance Team |
| EPI_ISL_2638477, EPI_ISL_2638478 | Lighthouse Lab in Milton Keynes | Wellcome Sanger Institute for the COVID-19 Genomics UK (COG-UK) Consortium | The Lighthouse Lab in Milton Keynes and Alex Alderton, Roberto Amato, Jeffrey Barrett, Sonia Goncalves, Ewan Harrison, David K. Jackson, Ian Johnston, Dominic Kwiatkowski, Cordelia Langford, John Sillitoe on behalf of the Wellcome Sanger Institute COVID-19 Surveillance Team |
| EPI_ISL_2638479, EPI_ISL_2638480 | Lighthouse Lab in Glasgow | Wellcome Sanger Institute for the COVID-19 Genomics UK (COG-UK) Consortium | Harper VanSteenhouse, Yumi Kasai, David Gray, Carol Clugston, Anna Dominiczak and Alex Alderton, Roberto Amato, Jeffrey Barrett, Sonia Goncalves, Ewan Harrison, David K. Jackson, Ian Johnston, Dominic Kwiatkowski, Cordelia Langford, John Sillitoe on behalf of the Wellcome Sanger Institute COVID-19 Surveillance Team |
| EPI_ISL_2638481 | Lighthouse Laboratory Plymouth | Wellcome Sanger Institute for the COVID-19 Genomics UK (COG-UK) Consortium | Lighthouse Laboratory Plymouth and Alex Alderton, Roberto Amato, Jeffrey Barrett, Sonia Goncalves, Ewan Harrison, David K. Jackson, Ian Johnston, Dominic Kwiatkowski, Cordelia Langford, John Sillitoe on behalf of the Wellcome Sanger Institute COVID-19 Surveillance Team |
| EPI_ISL_2638482 | Lighthouse Lab in Glasgow | Wellcome Sanger Institute for the COVID-19 Genomics UK (COG-UK) Consortium | Harper VanSteenhouse, Yumi Kasai, David Gray, Carol Clugston, Anna Dominiczak and Alex Alderton, Roberto Amato, Jeffrey Barrett, Sonia Goncalves, Ewan Harrison, David K. Jackson, Ian Johnston, Dominic Kwiatkowski, Cordelia Langford, John Sillitoe on behalf of the Wellcome Sanger Institute COVID-19 Surveillance Team |
| EPI_ISL_2638483, EPI_ISL_2638484 | Lighthouse Laboratory Plymouth | Wellcome Sanger Institute for the COVID-19 Genomics UK (COG-UK) Consortium | Lighthouse Laboratory Plymouth and Alex Alderton, Roberto Amato, Jeffrey Barrett, Sonia Goncalves, Ewan Harrison, David K. Jackson, Ian Johnston, Dominic Kwiatkowski, Cordelia Langford, John Sillitoe on behalf of the Wellcome Sanger Institute COVID-19 Surveillance Team |
| EPI_ISL_2638485, EPI_ISL_2638486 | Lighthouse Lab in Glasgow | Wellcome Sanger Institute for the COVID-19 Genomics UK (COG-UK) Consortium | Harper VanSteenhouse, Yumi Kasai, David Gray, Carol Clugston, Anna Dominiczak and Alex Alderton, Roberto Amato, Jeffrey Barrett, Sonia Goncalves, Ewan Harrison, David K. Jackson, Ian Johnston, Dominic Kwiatkowski, Cordelia Langford, John Sillitoe on behalf of the Wellcome Sanger Institute COVID-19 Surveillance Team |
| EPI_ISL_2638487 | Lighthouse Lab in Milton Keynes | Wellcome Sanger Institute for the COVID-19 Genomics UK (COG-UK) Consortium | The Lighthouse Lab in Milton Keynes and Alex Alderton, Roberto Amato, Jeffrey Barrett, Sonia Goncalves, Ewan Harrison, David K. Jackson, Ian Johnston, Dominic Kwiatkowski, Cordelia Langford, John Sillitoe on behalf of the Wellcome Sanger Institute COVID-19 Surveillance Team |
| EPI_ISL_2638488, EPI_ISL_2638489 | Lighthouse Laboratory Plymouth | Wellcome Sanger Institute for the COVID-19 Genomics UK (COG-UK) Consortium | Lighthouse Laboratory Plymouth and Alex Alderton, Roberto Amato, Jeffrey Barrett, Sonia Goncalves, Ewan Harrison, David K. Jackson, Ian Johnston, Dominic Kwiatkowski, Cordelia Langford, John Sillitoe on behalf of the Wellcome Sanger Institute COVID-19 Surveillance Team |
| EPI_ISL_2638490 | Lighthouse Lab in Milton Keynes | Wellcome Sanger Institute for the COVID-19 Genomics UK (COG-UK) Consortium | The Lighthouse Lab in Milton Keynes and Alex Alderton, Roberto Amato, Jeffrey Barrett, Sonia Goncalves, Ewan Harrison, David K. Jackson, Ian Johnston, Dominic Kwiatkowski, Cordelia Langford, John Sillitoe on behalf of the Wellcome Sanger Institute COVID-19 Surveillance Team |
| EPI_ISL_2638491, EPI_ISL_2638492, EPI_ISL_2638493 | Lighthouse Laboratory Plymouth | Wellcome Sanger Institute for the COVID-19 Genomics UK (COG-UK) Consortium | Lighthouse Laboratory Plymouth and Alex Alderton, Roberto Amato, Jeffrey Barrett, Sonia Goncalves, Ewan Harrison, David K. Jackson, Ian Johnston, Dominic Kwiatkowski, Cordelia Langford, John Sillitoe on behalf of the Wellcome Sanger Institute COVID-19 Surveillance Team |
| EPI_ISL_2638494 | Lighthouse Lab in Glasgow | Wellcome Sanger Institute for the COVID-19 Genomics UK (COG-UK) Consortium | Harper VanSteenhouse, Yumi Kasai, David Gray, Carol Clugston, Anna Dominiczak and Alex Alderton, Roberto Amato, Jeffrey Barrett, Sonia Goncalves, Ewan Harrison, David K. Jackson, Ian Johnston, Dominic Kwiatkowski, Cordelia Langford, John Sillitoe on behalf of the Wellcome Sanger Institute COVID-19 Surveillance Team |
| EPI_ISL_2638495, EPI_ISL_2638496, EPI_ISL_2638497 | Lighthouse Laboratory Plymouth | Wellcome Sanger Institute for the COVID-19 Genomics UK (COG-UK) Consortium | Lighthouse Laboratory Plymouth and Alex Alderton, Roberto Amato, Jeffrey Barrett, Sonia Goncalves, Ewan Harrison, David K. Jackson, Ian Johnston, Dominic Kwiatkowski, Cordelia Langford, John Sillitoe on behalf of the Wellcome Sanger Institute COVID-19 Surveillance Team |
| EPI_ISL_2638498 | Lighthouse Lab in Glasgow | Wellcome Sanger Institute for the COVID-19 Genomics UK (COG-UK) Consortium | Harper VanSteenhouse, Yumi Kasai, David Gray, Carol Clugston, Anna Dominiczak and Alex Alderton, Roberto Amato, Jeffrey Barrett, Sonia Goncalves, Ewan Harrison, David K. Jackson, Ian Johnston, Dominic Kwiatkowski, Cordelia Langford, John Sillitoe on behalf of the Wellcome Sanger Institute COVID-19 Surveillance Team |
| EPI_ISL_2638499, EPI_ISL_2638500 | Lighthouse Laboratory Plymouth | Wellcome Sanger Institute for the COVID-19 Genomics UK (COG-UK) Consortium | Lighthouse Laboratory Plymouth and Alex Alderton, Roberto Amato, Jeffrey Barrett, Sonia Goncalves, Ewan Harrison, David K. Jackson, Ian Johnston, Dominic Kwiatkowski, Cordelia Langford, John Sillitoe on behalf of the Wellcome Sanger Institute COVID-19 Surveillance Team |
| EPI_ISL_2638501 | Lighthouse Lab in Glasgow | Wellcome Sanger Institute for the COVID-19 Genomics UK (COG-UK) Consortium | Harper VanSteenhouse, Yumi Kasai, David Gray, Carol Clugston, Anna Dominiczak and Alex Alderton, Roberto Amato, Jeffrey Barrett, Sonia Goncalves, Ewan Harrison, David K. Jackson, Ian Johnston, Dominic Kwiatkowski, Cordelia Langford, John Sillitoe on behalf of the Wellcome Sanger Institute COVID-19 Surveillance Team |
| EPI_ISL_2638502 | Lighthouse Laboratory Plymouth | Wellcome Sanger Institute for the COVID-19 Genomics UK (COG-UK) Consortium | Lighthouse Laboratory Plymouth and Alex Alderton, Roberto Amato, Jeffrey Barrett, Sonia Goncalves, Ewan Harrison, David K. Jackson, Ian Johnston, Dominic Kwiatkowski, Cordelia Langford, John Sillitoe on behalf of the Wellcome Sanger Institute COVID-19 Surveillance Team |
| EPI_ISL_2638503 | Lighthouse Lab in Glasgow | Wellcome Sanger Institute for the COVID-19 Genomics UK (COG-UK) Consortium | Harper VanSteenhouse, Yumi Kasai, David Gray, Carol Clugston, Anna Dominiczak and Alex Alderton, Roberto Amato, Jeffrey Barrett, Sonia Goncalves, Ewan Harrison, David K. Jackson, Ian Johnston, Dominic Kwiatkowski, Cordelia Langford, John Sillitoe on behalf of the Wellcome Sanger Institute COVID-19 Surveillance Team |
| EPI_ISL_2638504 | Lighthouse Lab in Milton Keynes | Wellcome Sanger Institute for the COVID-19 Genomics UK (COG-UK) Consortium | The Lighthouse Lab in Milton Keynes and Alex Alderton, Roberto Amato, Jeffrey Barrett, Sonia Goncalves, Ewan Harrison, David K. Jackson, Ian Johnston, Dominic Kwiatkowski, Cordelia Langford, John Sillitoe on behalf of the Wellcome Sanger Institute COVID-19 Surveillance Team |
| EPI_ISL_2638505, EPI_ISL_2638506 | Lighthouse Lab in Glasgow | Wellcome Sanger Institute for the COVID-19 Genomics UK (COG-UK) Consortium | Harper VanSteenhouse, Yumi Kasai, David Gray, Carol Clugston, Anna Dominiczak and Alex Alderton, Roberto Amato, Jeffrey Barrett, Sonia Goncalves, Ewan Harrison, David K. Jackson, Ian Johnston, Dominic Kwiatkowski, Cordelia Langford, John Sillitoe on behalf of the Wellcome Sanger Institute COVID-19 Surveillance Team |
| EPI_ISL_2638507 | Lighthouse Laboratory Plymouth | Wellcome Sanger Institute for the COVID-19 Genomics UK (COG-UK) Consortium | Lighthouse Laboratory Plymouth and Alex Alderton, Roberto Amato, Jeffrey Barrett, Sonia Goncalves, Ewan Harrison, David K. Jackson, Ian Johnston, Dominic Kwiatkowski, Cordelia Langford, John Sillitoe on behalf of the Wellcome Sanger Institute COVID-19 Surveillance Team |
| EPI_ISL_2638508, EPI_ISL_2638509, EPI_ISL_2638510 | Lighthouse Lab in Milton Keynes | Wellcome Sanger Institute for the COVID-19 Genomics UK (COG-UK) Consortium | The Lighthouse Lab in Milton Keynes and Alex Alderton, Roberto Amato, Jeffrey Barrett, Sonia Goncalves, Ewan Harrison, David K. Jackson, Ian Johnston, Dominic Kwiatkowski, Cordelia Langford, John Sillitoe on behalf of the Wellcome Sanger Institute COVID-19 Surveillance Team |
| EPI_ISL_2638511 | Lighthouse Laboratory Plymouth | Wellcome Sanger Institute for the COVID-19 Genomics UK (COG-UK) Consortium | Lighthouse Laboratory Plymouth and Alex Alderton, Roberto Amato, Jeffrey Barrett, Sonia Goncalves, Ewan Harrison, David K. Jackson, Ian Johnston, Dominic Kwiatkowski, Cordelia Langford, John Sillitoe on behalf of the Wellcome Sanger Institute COVID-19 Surveillance Team |
| EPI_ISL_2638512 | Lighthouse Lab in Glasgow | Wellcome Sanger Institute for the COVID-19 Genomics UK (COG-UK) Consortium | Harper VanSteenhouse, Yumi Kasai, David Gray, Carol Clugston, Anna Dominiczak and Alex Alderton, Roberto Amato, Jeffrey Barrett, Sonia Goncalves, Ewan Harrison, David K. Jackson, Ian Johnston, Dominic Kwiatkowski, Cordelia Langford, John Sillitoe on behalf of the Wellcome Sanger Institute COVID-19 Surveillance Team |
| EPI_ISL_2638513, EPI_ISL_2638514, EPI_ISL_2638515 | Lighthouse Lab in Milton Keynes | Wellcome Sanger Institute for the COVID-19 Genomics UK (COG-UK) Consortium | The Lighthouse Lab in Milton Keynes and Alex Alderton, Roberto Amato, Jeffrey Barrett, Sonia Goncalves, Ewan Harrison, David K. Jackson, Ian Johnston, Dominic Kwiatkowski, Cordelia Langford, John Sillitoe on behalf of the Wellcome Sanger Institute COVID-19 Surveillance Team |
| EPI_ISL_2638516 | Lighthouse Lab in Glasgow | Wellcome Sanger Institute for the COVID-19 Genomics UK (COG-UK) Consortium | Harper VanSteenhouse, Yumi Kasai, David Gray, Carol Clugston, Anna Dominiczak and Alex Alderton, Roberto Amato, Jeffrey Barrett, Sonia Goncalves, Ewan Harrison, David K. Jackson, Ian Johnston, Dominic Kwiatkowski, Cordelia Langford, John Sillitoe on behalf of the Wellcome Sanger Institute COVID-19 Surveillance Team |
| EPI_ISL_2638517 | Lighthouse Lab in Milton Keynes | Wellcome Sanger Institute for the COVID-19 Genomics UK (COG-UK) Consortium | The Lighthouse Lab in Milton Keynes and Alex Alderton, Roberto Amato, Jeffrey Barrett, Sonia Goncalves, Ewan Harrison, David K. Jackson, Ian Johnston, Dominic Kwiatkowski, Cordelia Langford, John Sillitoe on behalf of the Wellcome Sanger Institute COVID-19 Surveillance Team |
| EPI_ISL_2638519 | Lighthouse Laboratory Plymouth | Wellcome Sanger Institute for the COVID-19 Genomics UK (COG-UK) Consortium | Lighthouse Laboratory Plymouth and Alex Alderton, Roberto Amato, Jeffrey Barrett, Sonia Goncalves, Ewan Harrison, David K. Jackson, Ian Johnston, Dominic Kwiatkowski, Cordelia Langford, John Sillitoe on behalf of the Wellcome Sanger Institute COVID-19 Surveillance Team |

| | | | |
|---|---|---|---|
| EPI_ISL_2638520 | Lighthouse Lab in Glasgow | Wellcome Sanger Institute for the COVID-19 Genomics UK (COG-UK) Consortium | Harper VanSteenhouse, Yumi Kasai, David Gray, Carol Clugston, Anna Dominiczak and Alex Alderton, Roberto Amato, Jeffrey Barrett, Sonia Goncalves, Ewan Harrison, David K. Jackson, Ian Johnston, Dominic Kwiatkowski, Cordelia Langford, John Sillitoe on behalf of the Wellcome Sanger Institute COVID-19 Surveillance Team |
| EPI_ISL_2638521 | Lighthouse Lab in Milton Keynes | Wellcome Sanger Institute for the COVID-19 Genomics UK (COG-UK) Consortium | The Lighthouse Lab in Milton Keynes and Alex Alderton, Roberto Amato, Jeffrey Barrett, Sonia Goncalves, Ewan Harrison, David K. Jackson, Ian Johnston, Dominic Kwiatkowski, Cordelia Langford, John Sillitoe on behalf of the Wellcome Sanger Institute COVID-19 Surveillance Team |
| EPI_ISL_2638522 | Lighthouse Laboratory Plymouth | Wellcome Sanger Institute for the COVID-19 Genomics UK (COG-UK) Consortium | Lighthouse Laboratory Plymouth and Alex Alderton, Roberto Amato, Jeffrey Barrett, Sonia Goncalves, Ewan Harrison, David K. Jackson, Ian Johnston, Dominic Kwiatkowski, Cordelia Langford, John Sillitoe on behalf of the Wellcome Sanger Institute COVID-19 Surveillance Team |
| EPI_ISL_2638523 | Lighthouse Lab in Milton Keynes | Wellcome Sanger Institute for the COVID-19 Genomics UK (COG-UK) Consortium | The Lighthouse Lab in Milton Keynes and Alex Alderton, Roberto Amato, Jeffrey Barrett, Sonia Goncalves, Ewan Harrison, David K. Jackson, Ian Johnston, Dominic Kwiatkowski, Cordelia Langford, John Sillitoe on behalf of the Wellcome Sanger Institute COVID-19 Surveillance Team |
| EPI_ISL_2638524, EPI_ISL_2638525, EPI_ISL_2638526 | Lighthouse Laboratory Plymouth | Wellcome Sanger Institute for the COVID-19 Genomics UK (COG-UK) Consortium | Lighthouse Laboratory Plymouth and Alex Alderton, Roberto Amato, Jeffrey Barrett, Sonia Goncalves, Ewan Harrison, David K. Jackson, Ian Johnston, Dominic Kwiatkowski, Cordelia Langford, John Sillitoe on behalf of the Wellcome Sanger Institute COVID-19 Surveillance Team |
| EPI_ISL_2638527 | Lighthouse Lab in Glasgow | Wellcome Sanger Institute for the COVID-19 Genomics UK (COG-UK) Consortium | Harper VanSteenhouse, Yumi Kasai, David Gray, Carol Clugston, Anna Dominiczak and Alex Alderton, Roberto Amato, Jeffrey Barrett, Sonia Goncalves, Ewan Harrison, David K. Jackson, Ian Johnston, Dominic Kwiatkowski, Cordelia Langford, John Sillitoe on behalf of the Wellcome Sanger Institute COVID-19 Surveillance Team |
| EPI_ISL_2638528 | Lighthouse Lab in Milton Keynes | Wellcome Sanger Institute for the COVID-19 Genomics UK (COG-UK) Consortium | The Lighthouse Lab in Milton Keynes and Alex Alderton, Roberto Amato, Jeffrey Barrett, Sonia Goncalves, Ewan Harrison, David K. Jackson, Ian Johnston, Dominic Kwiatkowski, Cordelia Langford, John Sillitoe on behalf of the Wellcome Sanger Institute COVID-19 Surveillance Team |
| EPI_ISL_2638529 | Lighthouse Laboratory Plymouth | Wellcome Sanger Institute for the COVID-19 Genomics UK (COG-UK) Consortium | Lighthouse Laboratory Plymouth and Alex Alderton, Roberto Amato, Jeffrey Barrett, Sonia Goncalves, Ewan Harrison, David K. Jackson, Ian Johnston, Dominic Kwiatkowski, Cordelia Langford, John Sillitoe on behalf of the Wellcome Sanger Institute COVID-19 Surveillance Team |
| EPI_ISL_2638530 | Lighthouse Lab in Glasgow | Wellcome Sanger Institute for the COVID-19 Genomics UK (COG-UK) Consortium | Harper VanSteenhouse, Yumi Kasai, David Gray, Carol Clugston, Anna Dominiczak and Alex Alderton, Roberto Amato, Jeffrey Barrett, Sonia Goncalves, Ewan Harrison, David K. Jackson, Ian Johnston, Dominic Kwiatkowski, Cordelia Langford, John Sillitoe on behalf of the Wellcome Sanger Institute COVID-19 Surveillance Team |
| EPI_ISL_2638531, EPI_ISL_2638532, EPI_ISL_2638533 | Lighthouse Laboratory Plymouth | Wellcome Sanger Institute for the COVID-19 Genomics UK (COG-UK) Consortium | Lighthouse Laboratory Plymouth and Alex Alderton, Roberto Amato, Jeffrey Barrett, Sonia Goncalves, Ewan Harrison, David K. Jackson, Ian Johnston, Dominic Kwiatkowski, Cordelia Langford, John Sillitoe on behalf of the Wellcome Sanger Institute COVID-19 Surveillance Team |
| EPI_ISL_2638534 | Lighthouse Lab in Glasgow | Wellcome Sanger Institute for the COVID-19 Genomics UK (COG-UK) Consortium | Harper VanSteenhouse, Yumi Kasai, David Gray, Carol Clugston, Anna Dominiczak and Alex Alderton, Roberto Amato, Jeffrey Barrett, Sonia Goncalves, Ewan Harrison, David K. Jackson, Ian Johnston, Dominic Kwiatkowski, Cordelia Langford, John Sillitoe on behalf of the Wellcome Sanger Institute COVID-19 Surveillance Team |
| EPI_ISL_2638535, EPI_ISL_2638536, EPI_ISL_2638537 | Lighthouse Lab in Milton Keynes | Wellcome Sanger Institute for the COVID-19 Genomics UK (COG-UK) Consortium | The Lighthouse Lab in Milton Keynes and Alex Alderton, Roberto Amato, Jeffrey Barrett, Sonia Goncalves, Ewan Harrison, David K. Jackson, Ian Johnston, Dominic Kwiatkowski, Cordelia Langford, John Sillitoe on behalf of the Wellcome Sanger Institute COVID-19 Surveillance Team |
| EPI_ISL_2638538, EPI_ISL_2638539 | Lighthouse Lab in Glasgow | Wellcome Sanger Institute for the COVID-19 Genomics UK (COG-UK) Consortium | Harper VanSteenhouse, Yumi Kasai, David Gray, Carol Clugston, Anna Dominiczak and Alex Alderton, Roberto Amato, Jeffrey Barrett, Sonia Goncalves, Ewan Harrison, David K. Jackson, Ian Johnston, Dominic Kwiatkowski, Cordelia Langford, John Sillitoe on behalf of the Wellcome Sanger Institute COVID-19 Surveillance Team |
| EPI_ISL_2638540 | Lighthouse Lab in Milton Keynes | Wellcome Sanger Institute for the COVID-19 Genomics UK (COG-UK) Consortium | The Lighthouse Lab in Milton Keynes and Alex Alderton, Roberto Amato, Jeffrey Barrett, Sonia Goncalves, Ewan Harrison, David K. Jackson, Ian Johnston, Dominic Kwiatkowski, Cordelia Langford, John Sillitoe on behalf of the Wellcome Sanger Institute COVID-19 Surveillance Team |
| EPI_ISL_2638541 | Lighthouse Laboratory Plymouth | Wellcome Sanger Institute for the COVID-19 Genomics UK (COG-UK) Consortium | Lighthouse Laboratory Plymouth and Alex Alderton, Roberto Amato, Jeffrey Barrett, Sonia Goncalves, Ewan Harrison, David K. Jackson, Ian Johnston, Dominic Kwiatkowski, Cordelia Langford, John Sillitoe on behalf of the Wellcome Sanger Institute COVID-19 Surveillance Team |
| EPI_ISL_2638542 | Lighthouse Lab in Milton Keynes | Wellcome Sanger Institute for the COVID-19 Genomics UK (COG-UK) Consortium | The Lighthouse Lab in Milton Keynes and Alex Alderton, Roberto Amato, Jeffrey Barrett, Sonia Goncalves, Ewan Harrison, David K. Jackson, Ian Johnston, Dominic Kwiatkowski, Cordelia Langford, John Sillitoe on behalf of the Wellcome Sanger Institute COVID-19 Surveillance Team |
| EPI_ISL_2638543 | Lighthouse Laboratory Plymouth | Wellcome Sanger Institute for the COVID-19 Genomics UK (COG-UK) Consortium | Lighthouse Laboratory Plymouth and Alex Alderton, Roberto Amato, Jeffrey Barrett, Sonia Goncalves, Ewan Harrison, David K. Jackson, Ian Johnston, Dominic Kwiatkowski, Cordelia Langford, John Sillitoe on behalf of the Wellcome Sanger Institute COVID-19 Surveillance Team |
| EPI_ISL_2638544 | Lighthouse Lab in Glasgow | Wellcome Sanger Institute for the COVID-19 Genomics UK (COG-UK) Consortium | Harper VanSteenhouse, Yumi Kasai, David Gray, Carol Clugston, Anna Dominiczak and Alex Alderton, Roberto Amato, Jeffrey Barrett, Sonia Goncalves, Ewan Harrison, David K. Jackson, Ian Johnston, Dominic Kwiatkowski, Cordelia Langford, John Sillitoe on behalf of the Wellcome Sanger Institute COVID-19 Surveillance Team |
| EPI_ISL_2638545 | Lighthouse Lab in Milton Keynes | Wellcome Sanger Institute for the COVID-19 Genomics UK (COG-UK) Consortium | The Lighthouse Lab in Milton Keynes and Alex Alderton, Roberto Amato, Jeffrey Barrett, Sonia Goncalves, Ewan Harrison, David K. Jackson, Ian Johnston, Dominic Kwiatkowski, Cordelia Langford, John Sillitoe on behalf of the Wellcome Sanger Institute COVID-19 Surveillance Team |
| EPI_ISL_2638546, EPI_ISL_2638547 | Lighthouse Laboratory Plymouth | Wellcome Sanger Institute for the COVID-19 Genomics UK (COG-UK) Consortium | Lighthouse Laboratory Plymouth and Alex Alderton, Roberto Amato, Jeffrey Barrett, Sonia Goncalves, Ewan Harrison, David K. Jackson, Ian Johnston, Dominic Kwiatkowski, Cordelia Langford, John Sillitoe on behalf of the Wellcome Sanger Institute COVID-19 Surveillance Team |
| EPI_ISL_2638548 | Lighthouse Lab in Glasgow | Wellcome Sanger Institute for the COVID-19 Genomics UK (COG-UK) Consortium | Harper VanSteenhouse, Yumi Kasai, David Gray, Carol Clugston, Anna Dominiczak and Alex Alderton, Roberto Amato, Jeffrey Barrett, Sonia Goncalves, Ewan Harrison, David K. Jackson, Ian Johnston, Dominic Kwiatkowski, Cordelia Langford, John Sillitoe on behalf of the Wellcome Sanger Institute COVID-19 Surveillance Team |
| EPI_ISL_2638549, EPI_ISL_2638550, EPI_ISL_2638551, EPI_ISL_2638552, EPI_ISL_2638553 | Lighthouse Laboratory Plymouth | Wellcome Sanger Institute for the COVID-19 Genomics UK (COG-UK) Consortium | Lighthouse Laboratory Plymouth and Alex Alderton, Roberto Amato, Jeffrey Barrett, Sonia Goncalves, Ewan Harrison, David K. Jackson, Ian Johnston, Dominic Kwiatkowski, Cordelia Langford, John Sillitoe on behalf of the Wellcome Sanger Institute COVID-19 Surveillance Team |
| EPI_ISL_2638554 | Lighthouse Lab in Milton Keynes | Wellcome Sanger Institute for the COVID-19 Genomics UK (COG-UK) Consortium | The Lighthouse Lab in Milton Keynes and Alex Alderton, Roberto Amato, Jeffrey Barrett, Sonia Goncalves, Ewan Harrison, David K. Jackson, Ian Johnston, Dominic Kwiatkowski, Cordelia Langford, John Sillitoe on behalf of the Wellcome Sanger Institute COVID-19 Surveillance Team |
| EPI_ISL_2638555, EPI_ISL_2638556 | Lighthouse Laboratory Plymouth | Wellcome Sanger Institute for the COVID-19 Genomics UK (COG-UK) Consortium | Lighthouse Laboratory Plymouth and Alex Alderton, Roberto Amato, Jeffrey Barrett, Sonia Goncalves, Ewan Harrison, David K. Jackson, Ian Johnston, Dominic Kwiatkowski, Cordelia Langford, John Sillitoe on behalf of the Wellcome Sanger Institute COVID-19 Surveillance Team |
| EPI_ISL_2638557 | Lighthouse Lab in Milton Keynes | Wellcome Sanger Institute for the COVID-19 Genomics UK (COG-UK) Consortium | The Lighthouse Lab in Milton Keynes and Alex Alderton, Roberto Amato, Jeffrey Barrett, Sonia Goncalves, Ewan Harrison, David K. Jackson, Ian Johnston, Dominic Kwiatkowski, Cordelia Langford, John Sillitoe on behalf of the Wellcome Sanger Institute COVID-19 Surveillance Team |
| EPI_ISL_2638558, EPI_ISL_2638559 | Lighthouse Laboratory Plymouth | Wellcome Sanger Institute for the COVID-19 Genomics UK (COG-UK) Consortium | Lighthouse Laboratory Plymouth and Alex Alderton, Roberto Amato, Jeffrey Barrett, Sonia Goncalves, Ewan Harrison, David K. Jackson, Ian Johnston, Dominic Kwiatkowski, Cordelia Langford, John Sillitoe on behalf of the Wellcome Sanger Institute COVID-19 Surveillance Team |
| EPI_ISL_2638560, EPI_ISL_2638561 | Lighthouse Lab in Milton Keynes | Wellcome Sanger Institute for the COVID-19 Genomics UK (COG-UK) Consortium | The Lighthouse Lab in Milton Keynes and Alex Alderton, Roberto Amato, Jeffrey Barrett, Sonia Goncalves, Ewan Harrison, David K. Jackson, Ian Johnston, Dominic Kwiatkowski, Cordelia Langford, John Sillitoe on behalf of the Wellcome Sanger Institute COVID-19 Surveillance Team |
| EPI_ISL_2638562, EPI_ISL_2638563 | Lighthouse Lab in Glasgow | Wellcome Sanger Institute for the COVID-19 Genomics UK (COG-UK) Consortium | Harper VanSteenhouse, Yumi Kasai, David Gray, Carol Clugston, Anna Dominiczak and Alex Alderton, Roberto Amato, Jeffrey Barrett, Sonia Goncalves, Ewan Harrison, David K. Jackson, Ian Johnston, Dominic Kwiatkowski, Cordelia Langford, John Sillitoe on behalf of the Wellcome Sanger Institute COVID-19 Surveillance Team |
| EPI_ISL_2638564, EPI_ISL_2638565, EPI_ISL_2638566, EPI_ISL_2638567 | Lighthouse Lab in Milton Keynes | Wellcome Sanger Institute for the COVID-19 Genomics UK (COG-UK) Consortium | The Lighthouse Lab in Milton Keynes and Alex Alderton, Roberto Amato, Jeffrey Barrett, Sonia Goncalves, Ewan Harrison, David K. Jackson, Ian Johnston, Dominic Kwiatkowski, Cordelia Langford, John Sillitoe on behalf of the Wellcome Sanger Institute COVID-19 Surveillance Team |
| EPI_ISL_2638568 | Lighthouse Lab in Glasgow | Wellcome Sanger Institute for the COVID-19 Genomics UK (COG-UK) Consortium | Harper VanSteenhouse, Yumi Kasai, David Gray, Carol Clugston, Anna Dominiczak and Alex Alderton, Roberto Amato, Jeffrey Barrett, Sonia Goncalves, Ewan Harrison, David K. Jackson, Ian Johnston, Dominic Kwiatkowski, Cordelia Langford, John Sillitoe on behalf of the Wellcome Sanger Institute COVID-19 Surveillance Team |
| EPI_ISL_2638569 | Lighthouse Laboratory Plymouth | Wellcome Sanger Institute for the COVID-19 Genomics UK (COG-UK) Consortium | Lighthouse Laboratory Plymouth and Alex Alderton, Roberto Amato, Jeffrey Barrett, Sonia Goncalves, Ewan Harrison, David K. Jackson, Ian Johnston, Dominic Kwiatkowski, Cordelia Langford, John Sillitoe on behalf of the Wellcome Sanger Institute COVID-19 Surveillance Team |
| EPI_ISL_2638570, EPI_ISL_2638571 | Lighthouse Lab in Milton Keynes | Wellcome Sanger Institute for the COVID-19 Genomics UK (COG-UK) Consortium | The Lighthouse Lab in Milton Keynes and Alex Alderton, Roberto Amato, Jeffrey Barrett, Sonia Goncalves, Ewan Harrison, David K. Jackson, Ian Johnston, Dominic Kwiatkowski, Cordelia Langford, John Sillitoe on behalf of the Wellcome Sanger Institute COVID-19 Surveillance Team |
| EPI_ISL_2638572 | Lighthouse Lab in Glasgow | Wellcome Sanger Institute for the COVID-19 Genomics UK (COG-UK) Consortium | Harper VanSteenhouse, Yumi Kasai, David Gray, Carol Clugston, Anna Dominiczak and Alex Alderton, Roberto Amato, Jeffrey Barrett, Sonia Goncalves, Ewan Harrison, David K. Jackson, Ian Johnston, Dominic Kwiatkowski, Cordelia Langford, John Sillitoe on behalf of the Wellcome Sanger Institute COVID-19 Surveillance Team |
| EPI_ISL_2638573 | Lighthouse Lab in Milton Keynes | Wellcome Sanger Institute for the COVID-19 Genomics UK (COG-UK) Consortium | The Lighthouse Lab in Milton Keynes and Alex Alderton, Roberto Amato, Jeffrey Barrett, Sonia Goncalves, Ewan Harrison, David K. Jackson, Ian Johnston, Dominic Kwiatkowski, Cordelia Langford, John Sillitoe on behalf of the Wellcome Sanger Institute COVID-19 Surveillance Team |
| EPI_ISL_2638574 | Lighthouse Lab in Glasgow | Wellcome Sanger Institute for the COVID-19 Genomics UK (COG-UK) Consortium | Harper VanSteenhouse, Yumi Kasai, David Gray, Carol Clugston, Anna Dominiczak and Alex Alderton, Roberto Amato, Jeffrey Barrett, Sonia Goncalves, Ewan Harrison, David K. Jackson, Ian Johnston, Dominic Kwiatkowski, Cordelia Langford, John Sillitoe on behalf of the Wellcome Sanger Institute COVID-19 Surveillance Team |
| EPI_ISL_2638575 | Lighthouse Laboratory Plymouth | Wellcome Sanger Institute for the COVID-19 Genomics UK (COG-UK) Consortium | Lighthouse Laboratory Plymouth and Alex Alderton, Roberto Amato, Jeffrey Barrett, Sonia Goncalves, Ewan Harrison, David K. Jackson, Ian Johnston, Dominic Kwiatkowski, Cordelia Langford, John Sillitoe on behalf of the Wellcome Sanger Institute COVID-19 Surveillance Team |
| EPI_ISL_2638576 | Lighthouse Lab in Milton Keynes | Wellcome Sanger Institute for the COVID-19 Genomics UK (COG-UK) Consortium | The Lighthouse Lab in Milton Keynes and Alex Alderton, Roberto Amato, Jeffrey Barrett, Sonia Goncalves, Ewan Harrison, David K. Jackson, Ian Johnston, Dominic Kwiatkowski, Cordelia Langford, John Sillitoe on behalf of the Wellcome Sanger Institute COVID-19 Surveillance Team |
| EPI_ISL_2638577 | Lighthouse Lab in Glasgow | Wellcome Sanger Institute for the COVID-19 Genomics UK (COG-UK) Consortium | Harper VanSteenhouse, Yumi Kasai, David Gray, Carol Clugston, Anna Dominiczak and Alex Alderton, Roberto Amato, Jeffrey Barrett, Sonia Goncalves, Ewan Harrison, David K. Jackson, Ian Johnston, Dominic Kwiatkowski, Cordelia Langford, John Sillitoe on behalf of the Wellcome Sanger Institute COVID-19 Surveillance Team |
| EPI_ISL_2638578, EPI_ISL_2638579, EPI_ISL_2638580 | Lighthouse Lab in Milton Keynes | Wellcome Sanger Institute for the COVID-19 Genomics UK (COG-UK) Consortium | The Lighthouse Lab in Milton Keynes and Alex Alderton, Roberto Amato, Jeffrey Barrett, Sonia Goncalves, Ewan Harrison, David K. Jackson, Ian Johnston, Dominic Kwiatkowski, Cordelia Langford, John Sillitoe on behalf of the Wellcome Sanger Institute COVID-19 Surveillance Team |
| EPI_ISL_2638581 | Lighthouse Lab in Glasgow | Wellcome Sanger Institute for the COVID-19 Genomics UK (COG-UK) Consortium | Harper VanSteenhouse, Yumi Kasai, David Gray, Carol Clugston, Anna Dominiczak and Alex Alderton, Roberto Amato, Jeffrey Barrett, Sonia Goncalves, Ewan Harrison, David K. Jackson, Ian Johnston, Dominic Kwiatkowski, Cordelia Langford, John Sillitoe on behalf of the Wellcome Sanger Institute COVID-19 Surveillance Team |
| EPI_ISL_2638582 | Lighthouse Lab in Milton Keynes | Wellcome Sanger Institute for the COVID-19 Genomics UK (COG-UK) Consortium | The Lighthouse Lab in Milton Keynes and Alex Alderton, Roberto Amato, Jeffrey Barrett, Sonia Goncalves, Ewan Harrison, David K. Jackson, Ian Johnston, Dominic Kwiatkowski, Cordelia Langford, John Sillitoe on behalf of the Wellcome Sanger Institute COVID-19 Surveillance Team |
| EPI_ISL_2638583 | Lighthouse Laboratory Plymouth | Wellcome Sanger Institute for the COVID-19 Genomics UK (COG-UK) | Lighthouse Laboratory Plymouth and Alex Alderton, Roberto Amato, Jeffrey Barrett, Sonia Goncalves, Ewan Harrison, David K. Jackson, Ian Johnston, Dominic Kwiatkowski, Cordelia Langford, John Sillitoe on behalf of the Wellcome Sanger |

| | | | |
|---|---|---|---|
| | | Consortium | Institute COVID-19 Surveillance Team |
| EPI_ISL_2638585 | Lighthouse Lab in Milton Keynes | Wellcome Sanger Institute for the COVID-19 Genomics UK (COG-UK) Consortium | The Lighthouse Lab in Milton Keynes and Alex Alderton, Roberto Amato, Jeffrey Barrett, Sonia Goncalves, Ewan Harrison, David K. Jackson, Ian Johnston, Dominic Kwiatkowski, Cordelia Langford, John Sillitoe on behalf of the Wellcome Sanger Institute COVID-19 Surveillance Team |
| EPI_ISL_2638586 | Lighthouse Lab in Glasgow | Wellcome Sanger Institute for the COVID-19 Genomics UK (COG-UK) Consortium | Harper VanSteenhouse, Yumi Kasai, David Gray, Carol Clugston, Anna Dominiczak and Alex Alderton, Roberto Amato, Jeffrey Barrett, Sonia Goncalves, Ewan Harrison, David K. Jackson, Ian Johnston, Dominic Kwiatkowski, Cordelia Langford, John Sillitoe on behalf of the Wellcome Sanger Institute COVID-19 Surveillance Team |
| EPI_ISL_2638587 | Lighthouse Laboratory Plymouth | Wellcome Sanger Institute for the COVID-19 Genomics UK (COG-UK) Consortium | Lighthouse Laboratory Plymouth and Alex Alderton, Roberto Amato, Jeffrey Barrett, Sonia Goncalves, Ewan Harrison, David K. Jackson, Ian Johnston, Dominic Kwiatkowski, Cordelia Langford, John Sillitoe on behalf of the Wellcome Sanger Institute COVID-19 Surveillance Team |
| EPI_ISL_2638588 | Lighthouse Lab in Milton Keynes | Wellcome Sanger Institute for the COVID-19 Genomics UK (COG-UK) Consortium | The Lighthouse Lab in Milton Keynes and Alex Alderton, Roberto Amato, Jeffrey Barrett, Sonia Goncalves, Ewan Harrison, David K. Jackson, Ian Johnston, Dominic Kwiatkowski, Cordelia Langford, John Sillitoe on behalf of the Wellcome Sanger Institute COVID-19 Surveillance Team |
| EPI_ISL_2638589, EPI_ISL_2638590, EPI_ISL_2638591, EPI_ISL_2638592 | Lighthouse Lab in Glasgow | Wellcome Sanger Institute for the COVID-19 Genomics UK (COG-UK) Consortium | Harper VanSteenhouse, Yumi Kasai, David Gray, Carol Clugston, Anna Dominiczak and Alex Alderton, Roberto Amato, Jeffrey Barrett, Sonia Goncalves, Ewan Harrison, David K. Jackson, Ian Johnston, Dominic Kwiatkowski, Cordelia Langford, John Sillitoe on behalf of the Wellcome Sanger Institute COVID-19 Surveillance Team |
| EPI_ISL_2638593 | Lighthouse Lab in Milton Keynes | Wellcome Sanger Institute for the COVID-19 Genomics UK (COG-UK) Consortium | The Lighthouse Lab in Milton Keynes and Alex Alderton, Roberto Amato, Jeffrey Barrett, Sonia Goncalves, Ewan Harrison, David K. Jackson, Ian Johnston, Dominic Kwiatkowski, Cordelia Langford, John Sillitoe on behalf of the Wellcome Sanger Institute COVID-19 Surveillance Team |
| EPI_ISL_2638594 | Lighthouse Laboratory Plymouth | Wellcome Sanger Institute for the COVID-19 Genomics UK (COG-UK) Consortium | Lighthouse Laboratory Plymouth and Alex Alderton, Roberto Amato, Jeffrey Barrett, Sonia Goncalves, Ewan Harrison, David K. Jackson, Ian Johnston, Dominic Kwiatkowski, Cordelia Langford, John Sillitoe on behalf of the Wellcome Sanger Institute COVID-19 Surveillance Team |
| EPI_ISL_2638595 | Lighthouse Lab in Glasgow | Wellcome Sanger Institute for the COVID-19 Genomics UK (COG-UK) Consortium | Harper VanSteenhouse, Yumi Kasai, David Gray, Carol Clugston, Anna Dominiczak and Alex Alderton, Roberto Amato, Jeffrey Barrett, Sonia Goncalves, Ewan Harrison, David K. Jackson, Ian Johnston, Dominic Kwiatkowski, Cordelia Langford, John Sillitoe on behalf of the Wellcome Sanger Institute COVID-19 Surveillance Team |
| EPI_ISL_2638596 | Lighthouse Laboratory Plymouth | Wellcome Sanger Institute for the COVID-19 Genomics UK (COG-UK) Consortium | Lighthouse Laboratory Plymouth and Alex Alderton, Roberto Amato, Jeffrey Barrett, Sonia Goncalves, Ewan Harrison, David K. Jackson, Ian Johnston, Dominic Kwiatkowski, Cordelia Langford, John Sillitoe on behalf of the Wellcome Sanger Institute COVID-19 Surveillance Team |
| EPI_ISL_2638597 | Lighthouse Lab in Milton Keynes | Wellcome Sanger Institute for the COVID-19 Genomics UK (COG-UK) Consortium | The Lighthouse Lab in Milton Keynes and Alex Alderton, Roberto Amato, Jeffrey Barrett, Sonia Goncalves, Ewan Harrison, David K. Jackson, Ian Johnston, Dominic Kwiatkowski, Cordelia Langford, John Sillitoe on behalf of the Wellcome Sanger Institute COVID-19 Surveillance Team |
| EPI_ISL_2638598, EPI_ISL_2638599 | Lighthouse Laboratory Plymouth | Wellcome Sanger Institute for the COVID-19 Genomics UK (COG-UK) Consortium | Lighthouse Laboratory Plymouth and Alex Alderton, Roberto Amato, Jeffrey Barrett, Sonia Goncalves, Ewan Harrison, David K. Jackson, Ian Johnston, Dominic Kwiatkowski, Cordelia Langford, John Sillitoe on behalf of the Wellcome Sanger Institute COVID-19 Surveillance Team |
| EPI_ISL_2638600 | Lighthouse Lab in Glasgow | Wellcome Sanger Institute for the COVID-19 Genomics UK (COG-UK) Consortium | Harper VanSteenhouse, Yumi Kasai, David Gray, Carol Clugston, Anna Dominiczak and Alex Alderton, Roberto Amato, Jeffrey Barrett, Sonia Goncalves, Ewan Harrison, David K. Jackson, Ian Johnston, Dominic Kwiatkowski, Cordelia Langford, John Sillitoe on behalf of the Wellcome Sanger Institute COVID-19 Surveillance Team |
| EPI_ISL_2638601 | Lighthouse Lab in Milton Keynes | Wellcome Sanger Institute for the COVID-19 Genomics UK (COG-UK) Consortium | The Lighthouse Lab in Milton Keynes and Alex Alderton, Roberto Amato, Jeffrey Barrett, Sonia Goncalves, Ewan Harrison, David K. Jackson, Ian Johnston, Dominic Kwiatkowski, Cordelia Langford, John Sillitoe on behalf of the Wellcome Sanger Institute COVID-19 Surveillance Team |
| EPI_ISL_2638602 | Lighthouse Laboratory Plymouth | Wellcome Sanger Institute for the COVID-19 Genomics UK (COG-UK) Consortium | Lighthouse Laboratory Plymouth and Alex Alderton, Roberto Amato, Jeffrey Barrett, Sonia Goncalves, Ewan Harrison, David K. Jackson, Ian Johnston, Dominic Kwiatkowski, Cordelia Langford, John Sillitoe on behalf of the Wellcome Sanger Institute COVID-19 Surveillance Team |
| EPI_ISL_2638603 | Lighthouse Lab in Milton Keynes | Wellcome Sanger Institute for the COVID-19 Genomics UK (COG-UK) Consortium | The Lighthouse Lab in Milton Keynes and Alex Alderton, Roberto Amato, Jeffrey Barrett, Sonia Goncalves, Ewan Harrison, David K. Jackson, Ian Johnston, Dominic Kwiatkowski, Cordelia Langford, John Sillitoe on behalf of the Wellcome Sanger Institute COVID-19 Surveillance Team |
| EPI_ISL_2638604, EPI_ISL_2638605, EPI_ISL_2638606 | Lighthouse Lab in Glasgow | Wellcome Sanger Institute for the COVID-19 Genomics UK (COG-UK) Consortium | Harper VanSteenhouse, Yumi Kasai, David Gray, Carol Clugston, Anna Dominiczak and Alex Alderton, Roberto Amato, Jeffrey Barrett, Sonia Goncalves, Ewan Harrison, David K. Jackson, Ian Johnston, Dominic Kwiatkowski, Cordelia Langford, John Sillitoe on behalf of the Wellcome Sanger Institute COVID-19 Surveillance Team |
| EPI_ISL_2638607, EPI_ISL_2638608 | Lighthouse Lab in Milton Keynes | Wellcome Sanger Institute for the COVID-19 Genomics UK (COG-UK) Consortium | The Lighthouse Lab in Milton Keynes and Alex Alderton, Roberto Amato, Jeffrey Barrett, Sonia Goncalves, Ewan Harrison, David K. Jackson, Ian Johnston, Dominic Kwiatkowski, Cordelia Langford, John Sillitoe on behalf of the Wellcome Sanger Institute COVID-19 Surveillance Team |
| EPI_ISL_2638609 | Lighthouse Lab in Glasgow | Wellcome Sanger Institute for the COVID-19 Genomics UK (COG-UK) Consortium | Harper VanSteenhouse, Yumi Kasai, David Gray, Carol Clugston, Anna Dominiczak and Alex Alderton, Roberto Amato, Jeffrey Barrett, Sonia Goncalves, Ewan Harrison, David K. Jackson, Ian Johnston, Dominic Kwiatkowski, Cordelia Langford, John Sillitoe on behalf of the Wellcome Sanger Institute COVID-19 Surveillance Team |
| EPI_ISL_2638610, EPI_ISL_2638611, EPI_ISL_2638612, EPI_ISL_2638613, EPI_ISL_2638614 | Lighthouse Lab in Milton Keynes | Wellcome Sanger Institute for the COVID-19 Genomics UK (COG-UK) Consortium | The Lighthouse Lab in Milton Keynes and Alex Alderton, Roberto Amato, Jeffrey Barrett, Sonia Goncalves, Ewan Harrison, David K. Jackson, Ian Johnston, Dominic Kwiatkowski, Cordelia Langford, John Sillitoe on behalf of the Wellcome Sanger Institute COVID-19 Surveillance Team |
| EPI_ISL_2638615, EPI_ISL_2638616, EPI_ISL_2638617, EPI_ISL_2638618, EPI_ISL_2638619 | Lighthouse Lab in Alderley Park | Wellcome Sanger Institute for the COVID-19 Genomics UK (COG-UK) Consortium | Jacquelyn Wynn, Mairead Hyland, The Lighthouse Lab in Alderley Park and Alex Alderton, Roberto Amato, Jeffrey Barrett, Sonia Goncalves, Ewan Harrison, David K. Jackson, Ian Johnston, Dominic Kwiatkowski, Cordelia Langford, John Sillitoe on behalf of the Wellcome Sanger Institute COVID-19 Surveillance Team |
| EPI_ISL_2638620 | Lighthouse Laboratory Plymouth | Wellcome Sanger Institute for the COVID-19 Genomics UK (COG-UK) Consortium | Lighthouse Laboratory Plymouth and Alex Alderton, Roberto Amato, Jeffrey Barrett, Sonia Goncalves, Ewan Harrison, David K. Jackson, Ian Johnston, Dominic Kwiatkowski, Cordelia Langford, John Sillitoe on behalf of the Wellcome Sanger Institute COVID-19 Surveillance Team |
| EPI_ISL_2638621 | Lighthouse Lab in Milton Keynes | Wellcome Sanger Institute for the COVID-19 Genomics UK (COG-UK) Consortium | The Lighthouse Lab in Milton Keynes and Alex Alderton, Roberto Amato, Jeffrey Barrett, Sonia Goncalves, Ewan Harrison, David K. Jackson, Ian Johnston, Dominic Kwiatkowski, Cordelia Langford, John Sillitoe on behalf of the Wellcome Sanger Institute COVID-19 Surveillance Team |
| EPI_ISL_2638622, EPI_ISL_2638623 | Lighthouse Lab in Alderley Park | Wellcome Sanger Institute for the COVID-19 Genomics UK (COG-UK) Consortium | Jacquelyn Wynn, Mairead Hyland, The Lighthouse Lab in Alderley Park and Alex Alderton, Roberto Amato, Jeffrey Barrett, Sonia Goncalves, Ewan Harrison, David K. Jackson, Ian Johnston, Dominic Kwiatkowski, Cordelia Langford, John Sillitoe on behalf of the Wellcome Sanger Institute COVID-19 Surveillance Team |
| EPI_ISL_2638624 | Lighthouse Lab in Milton Keynes | Wellcome Sanger Institute for the COVID-19 Genomics UK (COG-UK) Consortium | The Lighthouse Lab in Milton Keynes and Alex Alderton, Roberto Amato, Jeffrey Barrett, Sonia Goncalves, Ewan Harrison, David K. Jackson, Ian Johnston, Dominic Kwiatkowski, Cordelia Langford, John Sillitoe on behalf of the Wellcome Sanger Institute COVID-19 Surveillance Team |
| EPI_ISL_2638625 | Lighthouse Lab in Glasgow | Wellcome Sanger Institute for the COVID-19 Genomics UK (COG-UK) Consortium | Harper VanSteenhouse, Yumi Kasai, David Gray, Carol Clugston, Anna Dominiczak and Alex Alderton, Roberto Amato, Jeffrey Barrett, Sonia Goncalves, Ewan Harrison, David K. Jackson, Ian Johnston, Dominic Kwiatkowski, Cordelia Langford, John Sillitoe on behalf of the Wellcome Sanger Institute COVID-19 Surveillance Team |
| EPI_ISL_2638626, EPI_ISL_2638627, EPI_ISL_2638628, EPI_ISL_2638629 | Lighthouse Lab in Alderley Park | Wellcome Sanger Institute for the COVID-19 Genomics UK (COG-UK) Consortium | Jacquelyn Wynn, Mairead Hyland, The Lighthouse Lab in Alderley Park and Alex Alderton, Roberto Amato, Jeffrey Barrett, Sonia Goncalves, Ewan Harrison, David K. Jackson, Ian Johnston, Dominic Kwiatkowski, Cordelia Langford, John Sillitoe on behalf of the Wellcome Sanger Institute COVID-19 Surveillance Team |
| EPI_ISL_2638630 | Lighthouse Laboratory Plymouth | Wellcome Sanger Institute for the COVID-19 Genomics UK (COG-UK) Consortium | Lighthouse Laboratory Plymouth and Alex Alderton, Roberto Amato, Jeffrey Barrett, Sonia Goncalves, Ewan Harrison, David K. Jackson, Ian Johnston, Dominic Kwiatkowski, Cordelia Langford, John Sillitoe on behalf of the Wellcome Sanger Institute COVID-19 Surveillance Team |
| EPI_ISL_2638631 | Lighthouse Lab in Milton Keynes | Wellcome Sanger Institute for the COVID-19 Genomics UK (COG-UK) Consortium | The Lighthouse Lab in Milton Keynes and Alex Alderton, Roberto Amato, Jeffrey Barrett, Sonia Goncalves, Ewan Harrison, David K. Jackson, Ian Johnston, Dominic Kwiatkowski, Cordelia Langford, John Sillitoe on behalf of the Wellcome Sanger Institute COVID-19 Surveillance Team |
| EPI_ISL_2638632, EPI_ISL_2638633, EPI_ISL_2638634, EPI_ISL_2638635, EPI_ISL_2638636, EPI_ISL_2638637, EPI_ISL_2638638 | Lighthouse Lab in Alderley Park | Wellcome Sanger Institute for the COVID-19 Genomics UK (COG-UK) Consortium | Jacquelyn Wynn, Mairead Hyland, The Lighthouse Lab in Alderley Park and Alex Alderton, Roberto Amato, Jeffrey Barrett, Sonia Goncalves, Ewan Harrison, David K. Jackson, Ian Johnston, Dominic Kwiatkowski, Cordelia Langford, John Sillitoe on behalf of the Wellcome Sanger Institute COVID-19 Surveillance Team |
| EPI_ISL_2638639, EPI_ISL_2638640 | Lighthouse Lab in Milton Keynes | Wellcome Sanger Institute for the COVID-19 Genomics UK (COG-UK) Consortium | The Lighthouse Lab in Milton Keynes and Alex Alderton, Roberto Amato, Jeffrey Barrett, Sonia Goncalves, Ewan Harrison, David K. Jackson, Ian Johnston, Dominic Kwiatkowski, Cordelia Langford, John Sillitoe on behalf of the Wellcome Sanger Institute COVID-19 Surveillance Team |
| EPI_ISL_2638641, EPI_ISL_2638642, EPI_ISL_2638643, EPI_ISL_2638644 | Lighthouse Lab in Alderley Park | Wellcome Sanger Institute for the COVID-19 Genomics UK (COG-UK) Consortium | Jacquelyn Wynn, Mairead Hyland, The Lighthouse Lab in Alderley Park and Alex Alderton, Roberto Amato, Jeffrey Barrett, Sonia Goncalves, Ewan Harrison, David K. Jackson, Ian Johnston, Dominic Kwiatkowski, Cordelia Langford, John Sillitoe on behalf of the Wellcome Sanger Institute COVID-19 Surveillance Team |
| EPI_ISL_2638645 | Lighthouse Lab in Milton Keynes | Wellcome Sanger Institute for the COVID-19 Genomics UK (COG-UK) Consortium | The Lighthouse Lab in Milton Keynes and Alex Alderton, Roberto Amato, Jeffrey Barrett, Sonia Goncalves, Ewan Harrison, David K. Jackson, Ian Johnston, Dominic Kwiatkowski, Cordelia Langford, John Sillitoe on behalf of the Wellcome Sanger Institute COVID-19 Surveillance Team |
| EPI_ISL_2638646, EPI_ISL_2638647, EPI_ISL_2638648, EPI_ISL_2638649, EPI_ISL_2638650 | Lighthouse Lab in Alderley Park | Wellcome Sanger Institute for the COVID-19 Genomics UK (COG-UK) Consortium | Jacquelyn Wynn, Mairead Hyland, The Lighthouse Lab in Alderley Park and Alex Alderton, Roberto Amato, Jeffrey Barrett, Sonia Goncalves, Ewan Harrison, David K. Jackson, Ian Johnston, Dominic Kwiatkowski, Cordelia Langford, John Sillitoe on behalf of the Wellcome Sanger Institute COVID-19 Surveillance Team |
| EPI_ISL_2638651 | Lighthouse Laboratory Plymouth | Wellcome Sanger Institute for the COVID-19 Genomics UK (COG-UK) Consortium | Lighthouse Laboratory Plymouth and Alex Alderton, Roberto Amato, Jeffrey Barrett, Sonia Goncalves, Ewan Harrison, David K. Jackson, Ian Johnston, Dominic Kwiatkowski, Cordelia Langford, John Sillitoe on behalf of the Wellcome Sanger Institute COVID-19 Surveillance Team |
| EPI_ISL_2638652, EPI_ISL_2638653, EPI_ISL_2638654, EPI_ISL_2638655, EPI_ISL_2638656 | Lighthouse Lab in Alderley Park | Wellcome Sanger Institute for the COVID-19 Genomics UK (COG-UK) Consortium | Jacquelyn Wynn, Mairead Hyland, The Lighthouse Lab in Alderley Park and Alex Alderton, Roberto Amato, Jeffrey Barrett, Sonia Goncalves, Ewan Harrison, David K. Jackson, Ian Johnston, Dominic Kwiatkowski, Cordelia Langford, John Sillitoe on behalf of the Wellcome Sanger Institute COVID-19 Surveillance Team |

| | | | |
|---|---|---|---|
| EPI_ISL_2638657 | Lighthouse Lab in Milton Keynes | Wellcome Sanger Institute for the COVID-19 Genomics UK (COG-UK) Consortium | The Lighthouse Lab in Milton Keynes and Alex Alderton, Roberto Amato, Jeffrey Barrett, Sonia Goncalves, Ewan Harrison, David K. Jackson, Ian Johnston, Dominic Kwiatkowski, Cordelia Langford, John Sillitoe on behalf of the Wellcome Sanger Institute COVID-19 Surveillance Team |
| EPI_ISL_2638658, EPI_ISL_2638659, EPI_ISL_2638661 | Lighthouse Lab in Alderley Park | Wellcome Sanger Institute for the COVID-19 Genomics UK (COG-UK) Consortium | Jacquelyn Wynn, Mairead Hyland, The Lighthouse Lab in Alderley Park and Alex Alderton, Roberto Amato, Jeffrey Barrett, Sonia Goncalves, Ewan Harrison, David K. Jackson, Ian Johnston, Dominic Kwiatkowski, Cordelia Langford, John Sillitoe on behalf of the Wellcome Sanger Institute COVID-19 Surveillance Team |
| EPI_ISL_2638662 | Lighthouse Lab in Milton Keynes | Wellcome Sanger Institute for the COVID-19 Genomics UK (COG-UK) Consortium | The Lighthouse Lab in Milton Keynes and Alex Alderton, Roberto Amato, Jeffrey Barrett, Sonia Goncalves, Ewan Harrison, David K. Jackson, Ian Johnston, Dominic Kwiatkowski, Cordelia Langford, John Sillitoe on behalf of the Wellcome Sanger Institute COVID-19 Surveillance Team |
| EPI_ISL_2638663, EPI_ISL_2638664 | Lighthouse Lab in Alderley Park | Wellcome Sanger Institute for the COVID-19 Genomics UK (COG-UK) Consortium | Jacquelyn Wynn, Mairead Hyland, The Lighthouse Lab in Alderley Park and Alex Alderton, Roberto Amato, Jeffrey Barrett, Sonia Goncalves, Ewan Harrison, David K. Jackson, Ian Johnston, Dominic Kwiatkowski, Cordelia Langford, John Sillitoe on behalf of the Wellcome Sanger Institute COVID-19 Surveillance Team |
| EPI_ISL_2638665 | Lighthouse Laboratory Plymouth | Wellcome Sanger Institute for the COVID-19 Genomics UK (COG-UK) Consortium | Lighthouse Laboratory Plymouth and Alex Alderton, Roberto Amato, Jeffrey Barrett, Sonia Goncalves, Ewan Harrison, David K. Jackson, Ian Johnston, Dominic Kwiatkowski, Cordelia Langford, John Sillitoe on behalf of the Wellcome Sanger Institute COVID-19 Surveillance Team |
| EPI_ISL_2638666, EPI_ISL_2638667 | Lighthouse Lab in Milton Keynes | Wellcome Sanger Institute for the COVID-19 Genomics UK (COG-UK) Consortium | The Lighthouse Lab in Milton Keynes and Alex Alderton, Roberto Amato, Jeffrey Barrett, Sonia Goncalves, Ewan Harrison, David K. Jackson, Ian Johnston, Dominic Kwiatkowski, Cordelia Langford, John Sillitoe on behalf of the Wellcome Sanger Institute COVID-19 Surveillance Team |
| EPI_ISL_2638668, EPI_ISL_2638669 | Lighthouse Lab in Alderley Park | Wellcome Sanger Institute for the COVID-19 Genomics UK (COG-UK) Consortium | Jacquelyn Wynn, Mairead Hyland, The Lighthouse Lab in Alderley Park and Alex Alderton, Roberto Amato, Jeffrey Barrett, Sonia Goncalves, Ewan Harrison, David K. Jackson, Ian Johnston, Dominic Kwiatkowski, Cordelia Langford, John Sillitoe on behalf of the Wellcome Sanger Institute COVID-19 Surveillance Team |
| EPI_ISL_2638670 | Lighthouse Lab in Milton Keynes | Wellcome Sanger Institute for the COVID-19 Genomics UK (COG-UK) Consortium | The Lighthouse Lab in Milton Keynes and Alex Alderton, Roberto Amato, Jeffrey Barrett, Sonia Goncalves, Ewan Harrison, David K. Jackson, Ian Johnston, Dominic Kwiatkowski, Cordelia Langford, John Sillitoe on behalf of the Wellcome Sanger Institute COVID-19 Surveillance Team |
| EPI_ISL_2638671, EPI_ISL_2638672, EPI_ISL_2638673 | Lighthouse Lab in Alderley Park | Wellcome Sanger Institute for the COVID-19 Genomics UK (COG-UK) Consortium | Jacquelyn Wynn, Mairead Hyland, The Lighthouse Lab in Alderley Park and Alex Alderton, Roberto Amato, Jeffrey Barrett, Sonia Goncalves, Ewan Harrison, David K. Jackson, Ian Johnston, Dominic Kwiatkowski, Cordelia Langford, John Sillitoe on behalf of the Wellcome Sanger Institute COVID-19 Surveillance Team |
| EPI_ISL_2638674, EPI_ISL_2638675, EPI_ISL_2638676 | Lighthouse Lab in Glasgow | Wellcome Sanger Institute for the COVID-19 Genomics UK (COG-UK) Consortium | Harper VanSteenhouse, Yumi Kasai, David Gray, Carol Clugston, Anna Dominiczak and Alex Alderton, Roberto Amato, Jeffrey Barrett, Sonia Goncalves, Ewan Harrison, David K. Jackson, Ian Johnston, Dominic Kwiatkowski, Cordelia Langford, John Sillitoe on behalf of the Wellcome Sanger Institute COVID-19 Surveillance Team |
| EPI_ISL_2638677 | Lighthouse Lab in Alderley Park | Wellcome Sanger Institute for the COVID-19 Genomics UK (COG-UK) Consortium | Jacquelyn Wynn, Mairead Hyland, The Lighthouse Lab in Alderley Park and Alex Alderton, Roberto Amato, Jeffrey Barrett, Sonia Goncalves, Ewan Harrison, David K. Jackson, Ian Johnston, Dominic Kwiatkowski, Cordelia Langford, John Sillitoe on behalf of the Wellcome Sanger Institute COVID-19 Surveillance Team |
| EPI_ISL_2638678 | Lighthouse Lab in Milton Keynes | Wellcome Sanger Institute for the COVID-19 Genomics UK (COG-UK) Consortium | The Lighthouse Lab in Milton Keynes and Alex Alderton, Roberto Amato, Jeffrey Barrett, Sonia Goncalves, Ewan Harrison, David K. Jackson, Ian Johnston, Dominic Kwiatkowski, Cordelia Langford, John Sillitoe on behalf of the Wellcome Sanger Institute COVID-19 Surveillance Team |
| EPI_ISL_2638679, EPI_ISL_2638680 | Lighthouse Lab in Alderley Park | Wellcome Sanger Institute for the COVID-19 Genomics UK (COG-UK) Consortium | Jacquelyn Wynn, Mairead Hyland, The Lighthouse Lab in Alderley Park and Alex Alderton, Roberto Amato, Jeffrey Barrett, Sonia Goncalves, Ewan Harrison, David K. Jackson, Ian Johnston, Dominic Kwiatkowski, Cordelia Langford, John Sillitoe on behalf of the Wellcome Sanger Institute COVID-19 Surveillance Team |
| EPI_ISL_2638681 | Lighthouse Lab in Glasgow | Wellcome Sanger Institute for the COVID-19 Genomics UK (COG-UK) Consortium | Harper VanSteenhouse, Yumi Kasai, David Gray, Carol Clugston, Anna Dominiczak and Alex Alderton, Roberto Amato, Jeffrey Barrett, Sonia Goncalves, Ewan Harrison, David K. Jackson, Ian Johnston, Dominic Kwiatkowski, Cordelia Langford, John Sillitoe on behalf of the Wellcome Sanger Institute COVID-19 Surveillance Team |
| EPI_ISL_2638682 | Lighthouse Lab in Milton Keynes | Wellcome Sanger Institute for the COVID-19 Genomics UK (COG-UK) Consortium | The Lighthouse Lab in Milton Keynes and Alex Alderton, Roberto Amato, Jeffrey Barrett, Sonia Goncalves, Ewan Harrison, David K. Jackson, Ian Johnston, Dominic Kwiatkowski, Cordelia Langford, John Sillitoe on behalf of the Wellcome Sanger Institute COVID-19 Surveillance Team |
| EPI_ISL_2638683 | Lighthouse Lab in Glasgow | Wellcome Sanger Institute for the COVID-19 Genomics UK (COG-UK) Consortium | Harper VanSteenhouse, Yumi Kasai, David Gray, Carol Clugston, Anna Dominiczak and Alex Alderton, Roberto Amato, Jeffrey Barrett, Sonia Goncalves, Ewan Harrison, David K. Jackson, Ian Johnston, Dominic Kwiatkowski, Cordelia Langford, John Sillitoe on behalf of the Wellcome Sanger Institute COVID-19 Surveillance Team |
| EPI_ISL_2638684, EPI_ISL_2638685, EPI_ISL_2638686 | Lighthouse Lab in Alderley Park | Wellcome Sanger Institute for the COVID-19 Genomics UK (COG-UK) Consortium | Jacquelyn Wynn, Mairead Hyland, The Lighthouse Lab in Alderley Park and Alex Alderton, Roberto Amato, Jeffrey Barrett, Sonia Goncalves, Ewan Harrison, David K. Jackson, Ian Johnston, Dominic Kwiatkowski, Cordelia Langford, John Sillitoe on behalf of the Wellcome Sanger Institute COVID-19 Surveillance Team |
| EPI_ISL_2638687 | Lighthouse Lab in Milton Keynes | Wellcome Sanger Institute for the COVID-19 Genomics UK (COG-UK) Consortium | The Lighthouse Lab in Milton Keynes and Alex Alderton, Roberto Amato, Jeffrey Barrett, Sonia Goncalves, Ewan Harrison, David K. Jackson, Ian Johnston, Dominic Kwiatkowski, Cordelia Langford, John Sillitoe on behalf of the Wellcome Sanger Institute COVID-19 Surveillance Team |
| EPI_ISL_2638688, EPI_ISL_2638689, EPI_ISL_2638690 | Lighthouse Lab in Alderley Park | Wellcome Sanger Institute for the COVID-19 Genomics UK (COG-UK) Consortium | Jacquelyn Wynn, Mairead Hyland, The Lighthouse Lab in Alderley Park and Alex Alderton, Roberto Amato, Jeffrey Barrett, Sonia Goncalves, Ewan Harrison, David K. Jackson, Ian Johnston, Dominic Kwiatkowski, Cordelia Langford, John Sillitoe on behalf of the Wellcome Sanger Institute COVID-19 Surveillance Team |
| EPI_ISL_2638691, EPI_ISL_2638692 | Lighthouse Lab in Glasgow | Wellcome Sanger Institute for the COVID-19 Genomics UK (COG-UK) Consortium | Harper VanSteenhouse, Yumi Kasai, David Gray, Carol Clugston, Anna Dominiczak and Alex Alderton, Roberto Amato, Jeffrey Barrett, Sonia Goncalves, Ewan Harrison, David K. Jackson, Ian Johnston, Dominic Kwiatkowski, Cordelia Langford, John Sillitoe on behalf of the Wellcome Sanger Institute COVID-19 Surveillance Team |
| EPI_ISL_2638693, EPI_ISL_2638694 | Lighthouse Lab in Alderley Park | Wellcome Sanger Institute for the COVID-19 Genomics UK (COG-UK) Consortium | Jacquelyn Wynn, Mairead Hyland, The Lighthouse Lab in Alderley Park and Alex Alderton, Roberto Amato, Jeffrey Barrett, Sonia Goncalves, Ewan Harrison, David K. Jackson, Ian Johnston, Dominic Kwiatkowski, Cordelia Langford, John Sillitoe on behalf of the Wellcome Sanger Institute COVID-19 Surveillance Team |
| EPI_ISL_2638695, EPI_ISL_2638696 | Lighthouse Lab in Glasgow | Wellcome Sanger Institute for the COVID-19 Genomics UK (COG-UK) Consortium | Harper VanSteenhouse, Yumi Kasai, David Gray, Carol Clugston, Anna Dominiczak and Alex Alderton, Roberto Amato, Jeffrey Barrett, Sonia Goncalves, Ewan Harrison, David K. Jackson, Ian Johnston, Dominic Kwiatkowski, Cordelia Langford, John Sillitoe on behalf of the Wellcome Sanger Institute COVID-19 Surveillance Team |
| EPI_ISL_2638697, EPI_ISL_2638698, EPI_ISL_2638699, EPI_ISL_2638700, EPI_ISL_2638701, EPI_ISL_2638702, EPI_ISL_2638703, EPI_ISL_2638704 | Lighthouse Lab in Alderley Park | Wellcome Sanger Institute for the COVID-19 Genomics UK (COG-UK) Consortium | Jacquelyn Wynn, Mairead Hyland, The Lighthouse Lab in Alderley Park and Alex Alderton, Roberto Amato, Jeffrey Barrett, Sonia Goncalves, Ewan Harrison, David K. Jackson, Ian Johnston, Dominic Kwiatkowski, Cordelia Langford, John Sillitoe on behalf of the Wellcome Sanger Institute COVID-19 Surveillance Team |
| EPI_ISL_2638705, EPI_ISL_2638706 | Lighthouse Lab in Milton Keynes | Wellcome Sanger Institute for the COVID-19 Genomics UK (COG-UK) Consortium | The Lighthouse Lab in Milton Keynes and Alex Alderton, Roberto Amato, Jeffrey Barrett, Sonia Goncalves, Ewan Harrison, David K. Jackson, Ian Johnston, Dominic Kwiatkowski, Cordelia Langford, John Sillitoe on behalf of the Wellcome Sanger Institute COVID-19 Surveillance Team |
| EPI_ISL_2638707, EPI_ISL_2638708, EPI_ISL_2638709, EPI_ISL_2638710 | Lighthouse Lab in Alderley Park | Wellcome Sanger Institute for the COVID-19 Genomics UK (COG-UK) Consortium | Jacquelyn Wynn, Mairead Hyland, The Lighthouse Lab in Alderley Park and Alex Alderton, Roberto Amato, Jeffrey Barrett, Sonia Goncalves, Ewan Harrison, David K. Jackson, Ian Johnston, Dominic Kwiatkowski, Cordelia Langford, John Sillitoe on behalf of the Wellcome Sanger Institute COVID-19 Surveillance Team |
| EPI_ISL_2638712, EPI_ISL_2638713, EPI_ISL_2638714 | Lighthouse Lab in Milton Keynes | Wellcome Sanger Institute for the COVID-19 Genomics UK (COG-UK) Consortium | The Lighthouse Lab in Milton Keynes and Alex Alderton, Roberto Amato, Jeffrey Barrett, Sonia Goncalves, Ewan Harrison, David K. Jackson, Ian Johnston, Dominic Kwiatkowski, Cordelia Langford, John Sillitoe on behalf of the Wellcome Sanger Institute COVID-19 Surveillance Team |
| EPI_ISL_2638715, EPI_ISL_2638716, EPI_ISL_2638717, EPI_ISL_2638718, EPI_ISL_2638719, EPI_ISL_2638720, EPI_ISL_2638721 | Lighthouse Lab in Alderley Park | Wellcome Sanger Institute for the COVID-19 Genomics UK (COG-UK) Consortium | Jacquelyn Wynn, Mairead Hyland, The Lighthouse Lab in Alderley Park and Alex Alderton, Roberto Amato, Jeffrey Barrett, Sonia Goncalves, Ewan Harrison, David K. Jackson, Ian Johnston, Dominic Kwiatkowski, Cordelia Langford, John Sillitoe on behalf of the Wellcome Sanger Institute COVID-19 Surveillance Team |
| EPI_ISL_2638722, EPI_ISL_2638723 | Lighthouse Lab in Milton Keynes | Wellcome Sanger Institute for the COVID-19 Genomics UK (COG-UK) Consortium | The Lighthouse Lab in Milton Keynes and Alex Alderton, Roberto Amato, Jeffrey Barrett, Sonia Goncalves, Ewan Harrison, David K. Jackson, Ian Johnston, Dominic Kwiatkowski, Cordelia Langford, John Sillitoe on behalf of the Wellcome Sanger Institute COVID-19 Surveillance Team |
| EPI_ISL_2638724, EPI_ISL_2638725, EPI_ISL_2638726, EPI_ISL_2638727 | Lighthouse Lab in Alderley Park | Wellcome Sanger Institute for the COVID-19 Genomics UK (COG-UK) Consortium | Jacquelyn Wynn, Mairead Hyland, The Lighthouse Lab in Alderley Park and Alex Alderton, Roberto Amato, Jeffrey Barrett, Sonia Goncalves, Ewan Harrison, David K. Jackson, Ian Johnston, Dominic Kwiatkowski, Cordelia Langford, John Sillitoe on behalf of the Wellcome Sanger Institute COVID-19 Surveillance Team |
| EPI_ISL_2638728, EPI_ISL_2638729, EPI_ISL_2638730, EPI_ISL_2638731 | Lighthouse Lab in Glasgow | Wellcome Sanger Institute for the COVID-19 Genomics UK (COG-UK) Consortium | Harper VanSteenhouse, Yumi Kasai, David Gray, Carol Clugston, Anna Dominiczak and Alex Alderton, Roberto Amato, Jeffrey Barrett, Sonia Goncalves, Ewan Harrison, David K. Jackson, Ian Johnston, Dominic Kwiatkowski, Cordelia Langford, John Sillitoe on behalf of the Wellcome Sanger Institute COVID-19 Surveillance Team |
| EPI_ISL_2638732 | Lighthouse Lab in Milton Keynes | Wellcome Sanger Institute for the COVID-19 Genomics UK (COG-UK) Consortium | The Lighthouse Lab in Milton Keynes and Alex Alderton, Roberto Amato, Jeffrey Barrett, Sonia Goncalves, Ewan Harrison, David K. Jackson, Ian Johnston, Dominic Kwiatkowski, Cordelia Langford, John Sillitoe on behalf of the Wellcome Sanger Institute COVID-19 Surveillance Team |
| EPI_ISL_2638733 | Lighthouse Lab in Alderley Park | Wellcome Sanger Institute for the COVID-19 Genomics UK (COG-UK) Consortium | Jacquelyn Wynn, Mairead Hyland, The Lighthouse Lab in Alderley Park and Alex Alderton, Roberto Amato, Jeffrey Barrett, Sonia Goncalves, Ewan Harrison, David K. Jackson, Ian Johnston, Dominic Kwiatkowski, Cordelia Langford, John Sillitoe on behalf of the Wellcome Sanger Institute COVID-19 Surveillance Team |
| EPI_ISL_2638734 | Lighthouse Lab in Milton Keynes | Wellcome Sanger Institute for the COVID-19 Genomics UK (COG-UK) Consortium | The Lighthouse Lab in Milton Keynes and Alex Alderton, Roberto Amato, Jeffrey Barrett, Sonia Goncalves, Ewan Harrison, David K. Jackson, Ian Johnston, Dominic Kwiatkowski, Cordelia Langford, John Sillitoe on behalf of the Wellcome Sanger Institute COVID-19 Surveillance Team |
| EPI_ISL_2638735, EPI_ISL_2638736 | Lighthouse Lab in Alderley Park | Wellcome Sanger Institute for the COVID-19 Genomics UK (COG-UK) Consortium | Jacquelyn Wynn, Mairead Hyland, The Lighthouse Lab in Alderley Park and Alex Alderton, Roberto Amato, Jeffrey Barrett, Sonia Goncalves, Ewan Harrison, David K. Jackson, Ian Johnston, Dominic Kwiatkowski, Cordelia Langford, John Sillitoe on behalf of the Wellcome Sanger Institute COVID-19 Surveillance Team |
| EPI_ISL_2638737 | Lighthouse Lab in Milton Keynes | Wellcome Sanger Institute for the COVID-19 Genomics UK (COG-UK) Consortium | The Lighthouse Lab in Milton Keynes and Alex Alderton, Roberto Amato, Jeffrey Barrett, Sonia Goncalves, Ewan Harrison, David K. Jackson, Ian Johnston, Dominic Kwiatkowski, Cordelia Langford, John Sillitoe on behalf of the Wellcome Sanger Institute COVID-19 Surveillance Team |
| EPI_ISL_2638738, | Lighthouse Lab in Alderley Park | Wellcome Sanger Institute for the COVID-19 Genomics UK (COG-UK) | Jacquelyn Wynn, Mairead Hyland, The Lighthouse Lab in Alderley Park and Alex Alderton, Roberto Amato, Jeffrey Barrett, Sonia Goncalves, Ewan Harrison, David K. Jackson, Ian Johnston, Dominic Kwiatkowski, Cordelia Langford, John Sillitoe on |

| | | | |
|---|---|---|---|
| EPI_ISL_2638739, EPI_ISL_2638740 | | Consortium | behalf of the Wellcome Sanger Institute COVID-19 Surveillance Team |
| EPI_ISL_2638741 | Lighthouse Lab in Glasgow | Wellcome Sanger Institute for the COVID-19 Genomics UK (COG-UK) Consortium | Harper VanSteenhouse, Yumi Kasai, David Gray, Carol Clugston, Anna Dominiczak and Alex Alderton, Roberto Amato, Jeffrey Barrett, Sonia Goncalves, Ewan Harrison, David K. Jackson, Ian Johnston, Dominic Kwiatkowski, Cordelia Langford, John Sillitoe on behalf of the Wellcome Sanger Institute COVID-19 Surveillance Team |
| EPI_ISL_2638742 | Lighthouse Lab in Alderley Park | Wellcome Sanger Institute for the COVID-19 Genomics UK (COG-UK) Consortium | Jacquelyn Wynn, Mairead Hyland, The Lighthouse Lab in Alderley Park and Alex Alderton, Roberto Amato, Jeffrey Barrett, Sonia Goncalves, Ewan Harrison, David K. Jackson, Ian Johnston, Dominic Kwiatkowski, Cordelia Langford, John Sillitoe on behalf of the Wellcome Sanger Institute COVID-19 Surveillance Team |
| EPI_ISL_2638743 | Lighthouse Lab in Milton Keynes | Wellcome Sanger Institute for the COVID-19 Genomics UK (COG-UK) Consortium | The Lighthouse Lab in Milton Keynes and Alex Alderton, Roberto Amato, Jeffrey Barrett, Sonia Goncalves, Ewan Harrison, David K. Jackson, Ian Johnston, Dominic Kwiatkowski, Cordelia Langford, John Sillitoe on behalf of the Wellcome Sanger Institute COVID-19 Surveillance Team |
| EPI_ISL_2638744 | Lighthouse Laboratory Plymouth | Wellcome Sanger Institute for the COVID-19 Genomics UK (COG-UK) Consortium | Lighthouse Laboratory Plymouth and Alex Alderton, Roberto Amato, Jeffrey Barrett, Sonia Goncalves, Ewan Harrison, David K. Jackson, Ian Johnston, Dominic Kwiatkowski, Cordelia Langford, John Sillitoe on behalf of the Wellcome Sanger Institute COVID-19 Surveillance Team |
| EPI_ISL_2638745 | Lighthouse Lab in Glasgow | Wellcome Sanger Institute for the COVID-19 Genomics UK (COG-UK) Consortium | Harper VanSteenhouse, Yumi Kasai, David Gray, Carol Clugston, Anna Dominiczak and Alex Alderton, Roberto Amato, Jeffrey Barrett, Sonia Goncalves, Ewan Harrison, David K. Jackson, Ian Johnston, Dominic Kwiatkowski, Cordelia Langford, John Sillitoe on behalf of the Wellcome Sanger Institute COVID-19 Surveillance Team |
| EPI_ISL_2638746, EPI_ISL_2638747, EPI_ISL_2638748, EPI_ISL_2638749, EPI_ISL_2638750 | Lighthouse Lab in Alderley Park | Wellcome Sanger Institute for the COVID-19 Genomics UK (COG-UK) Consortium | Jacquelyn Wynn, Mairead Hyland, The Lighthouse Lab in Alderley Park and Alex Alderton, Roberto Amato, Jeffrey Barrett, Sonia Goncalves, Ewan Harrison, David K. Jackson, Ian Johnston, Dominic Kwiatkowski, Cordelia Langford, John Sillitoe on behalf of the Wellcome Sanger Institute COVID-19 Surveillance Team |
| EPI_ISL_2638751 | Lighthouse Lab in Glasgow | Wellcome Sanger Institute for the COVID-19 Genomics UK (COG-UK) Consortium | Harper VanSteenhouse, Yumi Kasai, David Gray, Carol Clugston, Anna Dominiczak and Alex Alderton, Roberto Amato, Jeffrey Barrett, Sonia Goncalves, Ewan Harrison, David K. Jackson, Ian Johnston, Dominic Kwiatkowski, Cordelia Langford, John Sillitoe on behalf of the Wellcome Sanger Institute COVID-19 Surveillance Team |
| EPI_ISL_2638752, EPI_ISL_2638753 | Lighthouse Lab in Alderley Park | Wellcome Sanger Institute for the COVID-19 Genomics UK (COG-UK) Consortium | Jacquelyn Wynn, Mairead Hyland, The Lighthouse Lab in Alderley Park and Alex Alderton, Roberto Amato, Jeffrey Barrett, Sonia Goncalves, Ewan Harrison, David K. Jackson, Ian Johnston, Dominic Kwiatkowski, Cordelia Langford, John Sillitoe on behalf of the Wellcome Sanger Institute COVID-19 Surveillance Team |
| EPI_ISL_2638754 | Lighthouse Lab in Glasgow | Wellcome Sanger Institute for the COVID-19 Genomics UK (COG-UK) Consortium | Harper VanSteenhouse, Yumi Kasai, David Gray, Carol Clugston, Anna Dominiczak and Alex Alderton, Roberto Amato, Jeffrey Barrett, Sonia Goncalves, Ewan Harrison, David K. Jackson, Ian Johnston, Dominic Kwiatkowski, Cordelia Langford, John Sillitoe on behalf of the Wellcome Sanger Institute COVID-19 Surveillance Team |
| EPI_ISL_2638755, EPI_ISL_2638756, EPI_ISL_2638757, EPI_ISL_2638758 | Lighthouse Lab in Alderley Park | Wellcome Sanger Institute for the COVID-19 Genomics UK (COG-UK) Consortium | Jacquelyn Wynn, Mairead Hyland, The Lighthouse Lab in Alderley Park and Alex Alderton, Roberto Amato, Jeffrey Barrett, Sonia Goncalves, Ewan Harrison, David K. Jackson, Ian Johnston, Dominic Kwiatkowski, Cordelia Langford, John Sillitoe on behalf of the Wellcome Sanger Institute COVID-19 Surveillance Team |
| EPI_ISL_2638759, EPI_ISL_2638760 | Lighthouse Lab in Glasgow | Wellcome Sanger Institute for the COVID-19 Genomics UK (COG-UK) Consortium | Harper VanSteenhouse, Yumi Kasai, David Gray, Carol Clugston, Anna Dominiczak and Alex Alderton, Roberto Amato, Jeffrey Barrett, Sonia Goncalves, Ewan Harrison, David K. Jackson, Ian Johnston, Dominic Kwiatkowski, Cordelia Langford, John Sillitoe on behalf of the Wellcome Sanger Institute COVID-19 Surveillance Team |
| EPI_ISL_2638761, EPI_ISL_2638762, EPI_ISL_2638763, EPI_ISL_2638764, EPI_ISL_2638765, EPI_ISL_2638766 | Lighthouse Lab in Alderley Park | Wellcome Sanger Institute for the COVID-19 Genomics UK (COG-UK) Consortium | Jacquelyn Wynn, Mairead Hyland, The Lighthouse Lab in Alderley Park and Alex Alderton, Roberto Amato, Jeffrey Barrett, Sonia Goncalves, Ewan Harrison, David K. Jackson, Ian Johnston, Dominic Kwiatkowski, Cordelia Langford, John Sillitoe on behalf of the Wellcome Sanger Institute COVID-19 Surveillance Team |
| EPI_ISL_2638767 | Lighthouse Lab in Glasgow | Wellcome Sanger Institute for the COVID-19 Genomics UK (COG-UK) Consortium | Harper VanSteenhouse, Yumi Kasai, David Gray, Carol Clugston, Anna Dominiczak and Alex Alderton, Roberto Amato, Jeffrey Barrett, Sonia Goncalves, Ewan Harrison, David K. Jackson, Ian Johnston, Dominic Kwiatkowski, Cordelia Langford, John Sillitoe on behalf of the Wellcome Sanger Institute COVID-19 Surveillance Team |
| EPI_ISL_2638768 | Lighthouse Lab in Alderley Park | Wellcome Sanger Institute for the COVID-19 Genomics UK (COG-UK) Consortium | Jacquelyn Wynn, Mairead Hyland, The Lighthouse Lab in Alderley Park and Alex Alderton, Roberto Amato, Jeffrey Barrett, Sonia Goncalves, Ewan Harrison, David K. Jackson, Ian Johnston, Dominic Kwiatkowski, Cordelia Langford, John Sillitoe on behalf of the Wellcome Sanger Institute COVID-19 Surveillance Team |
| EPI_ISL_2638769 | Lighthouse Lab in Glasgow | Wellcome Sanger Institute for the COVID-19 Genomics UK (COG-UK) Consortium | Harper VanSteenhouse, Yumi Kasai, David Gray, Carol Clugston, Anna Dominiczak and Alex Alderton, Roberto Amato, Jeffrey Barrett, Sonia Goncalves, Ewan Harrison, David K. Jackson, Ian Johnston, Dominic Kwiatkowski, Cordelia Langford, John Sillitoe on behalf of the Wellcome Sanger Institute COVID-19 Surveillance Team |
| EPI_ISL_2638770 | Lighthouse Lab in Alderley Park | Wellcome Sanger Institute for the COVID-19 Genomics UK (COG-UK) Consortium | Jacquelyn Wynn, Mairead Hyland, The Lighthouse Lab in Alderley Park and Alex Alderton, Roberto Amato, Jeffrey Barrett, Sonia Goncalves, Ewan Harrison, David K. Jackson, Ian Johnston, Dominic Kwiatkowski, Cordelia Langford, John Sillitoe on behalf of the Wellcome Sanger Institute COVID-19 Surveillance Team |
| EPI_ISL_2638771 | Lighthouse Lab in Glasgow | Wellcome Sanger Institute for the COVID-19 Genomics UK (COG-UK) Consortium | Harper VanSteenhouse, Yumi Kasai, David Gray, Carol Clugston, Anna Dominiczak and Alex Alderton, Roberto Amato, Jeffrey Barrett, Sonia Goncalves, Ewan Harrison, David K. Jackson, Ian Johnston, Dominic Kwiatkowski, Cordelia Langford, John Sillitoe on behalf of the Wellcome Sanger Institute COVID-19 Surveillance Team |
| EPI_ISL_2638772, EPI_ISL_2638773, EPI_ISL_2638774 | Lighthouse Lab in Alderley Park | Wellcome Sanger Institute for the COVID-19 Genomics UK (COG-UK) Consortium | Jacquelyn Wynn, Mairead Hyland, The Lighthouse Lab in Alderley Park and Alex Alderton, Roberto Amato, Jeffrey Barrett, Sonia Goncalves, Ewan Harrison, David K. Jackson, Ian Johnston, Dominic Kwiatkowski, Cordelia Langford, John Sillitoe on behalf of the Wellcome Sanger Institute COVID-19 Surveillance Team |
| EPI_ISL_2638775 | Lighthouse Lab in Glasgow | Wellcome Sanger Institute for the COVID-19 Genomics UK (COG-UK) Consortium | Harper VanSteenhouse, Yumi Kasai, David Gray, Carol Clugston, Anna Dominiczak and Alex Alderton, Roberto Amato, Jeffrey Barrett, Sonia Goncalves, Ewan Harrison, David K. Jackson, Ian Johnston, Dominic Kwiatkowski, Cordelia Langford, John Sillitoe on behalf of the Wellcome Sanger Institute COVID-19 Surveillance Team |
| EPI_ISL_2638776, EPI_ISL_2638777, EPI_ISL_2638778, EPI_ISL_2638779, EPI_ISL_2638780, EPI_ISL_2638781, EPI_ISL_2638782, EPI_ISL_2638783 | Lighthouse Lab in Alderley Park | Wellcome Sanger Institute for the COVID-19 Genomics UK (COG-UK) Consortium | Jacquelyn Wynn, Mairead Hyland, The Lighthouse Lab in Alderley Park and Alex Alderton, Roberto Amato, Jeffrey Barrett, Sonia Goncalves, Ewan Harrison, David K. Jackson, Ian Johnston, Dominic Kwiatkowski, Cordelia Langford, John Sillitoe on behalf of the Wellcome Sanger Institute COVID-19 Surveillance Team |
| EPI_ISL_2638784 | Lighthouse Lab in Milton Keynes | Wellcome Sanger Institute for the COVID-19 Genomics UK (COG-UK) Consortium | The Lighthouse Lab in Milton Keynes and Alex Alderton, Roberto Amato, Jeffrey Barrett, Sonia Goncalves, Ewan Harrison, David K. Jackson, Ian Johnston, Dominic Kwiatkowski, Cordelia Langford, John Sillitoe on behalf of the Wellcome Sanger Institute COVID-19 Surveillance Team |
| EPI_ISL_2638785, EPI_ISL_2638786, EPI_ISL_2638787, EPI_ISL_2638788 | Lighthouse Lab in Alderley Park | Wellcome Sanger Institute for the COVID-19 Genomics UK (COG-UK) Consortium | Jacquelyn Wynn, Mairead Hyland, The Lighthouse Lab in Alderley Park and Alex Alderton, Roberto Amato, Jeffrey Barrett, Sonia Goncalves, Ewan Harrison, David K. Jackson, Ian Johnston, Dominic Kwiatkowski, Cordelia Langford, John Sillitoe on behalf of the Wellcome Sanger Institute COVID-19 Surveillance Team |
| EPI_ISL_2638789 | Lighthouse Lab in Glasgow | Wellcome Sanger Institute for the COVID-19 Genomics UK (COG-UK) Consortium | Harper VanSteenhouse, Yumi Kasai, David Gray, Carol Clugston, Anna Dominiczak and Alex Alderton, Roberto Amato, Jeffrey Barrett, Sonia Goncalves, Ewan Harrison, David K. Jackson, Ian Johnston, Dominic Kwiatkowski, Cordelia Langford, John Sillitoe on behalf of the Wellcome Sanger Institute COVID-19 Surveillance Team |
| EPI_ISL_2638790, EPI_ISL_2638791, EPI_ISL_2638792, EPI_ISL_2638793 | Lighthouse Lab in Alderley Park | Wellcome Sanger Institute for the COVID-19 Genomics UK (COG-UK) Consortium | Jacquelyn Wynn, Mairead Hyland, The Lighthouse Lab in Alderley Park and Alex Alderton, Roberto Amato, Jeffrey Barrett, Sonia Goncalves, Ewan Harrison, David K. Jackson, Ian Johnston, Dominic Kwiatkowski, Cordelia Langford, John Sillitoe on behalf of the Wellcome Sanger Institute COVID-19 Surveillance Team |
| EPI_ISL_2638794, EPI_ISL_2638795 | Lighthouse Lab in Milton Keynes | Wellcome Sanger Institute for the COVID-19 Genomics UK (COG-UK) Consortium | The Lighthouse Lab in Milton Keynes and Alex Alderton, Roberto Amato, Jeffrey Barrett, Sonia Goncalves, Ewan Harrison, David K. Jackson, Ian Johnston, Dominic Kwiatkowski, Cordelia Langford, John Sillitoe on behalf of the Wellcome Sanger Institute COVID-19 Surveillance Team |
| EPI_ISL_2638796, EPI_ISL_2638797 | Lighthouse Lab in Alderley Park | Wellcome Sanger Institute for the COVID-19 Genomics UK (COG-UK) Consortium | Jacquelyn Wynn, Mairead Hyland, The Lighthouse Lab in Alderley Park and Alex Alderton, Roberto Amato, Jeffrey Barrett, Sonia Goncalves, Ewan Harrison, David K. Jackson, Ian Johnston, Dominic Kwiatkowski, Cordelia Langford, John Sillitoe on behalf of the Wellcome Sanger Institute COVID-19 Surveillance Team |
| EPI_ISL_2638798 | Lighthouse Lab in Milton Keynes | Wellcome Sanger Institute for the COVID-19 Genomics UK (COG-UK) Consortium | The Lighthouse Lab in Milton Keynes and Alex Alderton, Roberto Amato, Jeffrey Barrett, Sonia Goncalves, Ewan Harrison, David K. Jackson, Ian Johnston, Dominic Kwiatkowski, Cordelia Langford, John Sillitoe on behalf of the Wellcome Sanger Institute COVID-19 Surveillance Team |
| EPI_ISL_2638799, EPI_ISL_2638800, EPI_ISL_2638801 | Lighthouse Lab in Alderley Park | Wellcome Sanger Institute for the COVID-19 Genomics UK (COG-UK) Consortium | Jacquelyn Wynn, Mairead Hyland, The Lighthouse Lab in Alderley Park and Alex Alderton, Roberto Amato, Jeffrey Barrett, Sonia Goncalves, Ewan Harrison, David K. Jackson, Ian Johnston, Dominic Kwiatkowski, Cordelia Langford, John Sillitoe on behalf of the Wellcome Sanger Institute COVID-19 Surveillance Team |
| EPI_ISL_2638802 | Lighthouse Lab in Milton Keynes | Wellcome Sanger Institute for the COVID-19 Genomics UK (COG-UK) Consortium | The Lighthouse Lab in Milton Keynes and Alex Alderton, Roberto Amato, Jeffrey Barrett, Sonia Goncalves, Ewan Harrison, David K. Jackson, Ian Johnston, Dominic Kwiatkowski, Cordelia Langford, John Sillitoe on behalf of the Wellcome Sanger Institute COVID-19 Surveillance Team |
| EPI_ISL_2638803 | Lighthouse Lab in Alderley Park | Wellcome Sanger Institute for the COVID-19 Genomics UK (COG-UK) Consortium | Jacquelyn Wynn, Mairead Hyland, The Lighthouse Lab in Alderley Park and Alex Alderton, Roberto Amato, Jeffrey Barrett, Sonia Goncalves, Ewan Harrison, David K. Jackson, Ian Johnston, Dominic Kwiatkowski, Cordelia Langford, John Sillitoe on behalf of the Wellcome Sanger Institute COVID-19 Surveillance Team |
| EPI_ISL_2638804 | Lighthouse Lab in Milton Keynes | Wellcome Sanger Institute for the COVID-19 Genomics UK (COG-UK) Consortium | The Lighthouse Lab in Milton Keynes and Alex Alderton, Roberto Amato, Jeffrey Barrett, Sonia Goncalves, Ewan Harrison, David K. Jackson, Ian Johnston, Dominic Kwiatkowski, Cordelia Langford, John Sillitoe on behalf of the Wellcome Sanger Institute COVID-19 Surveillance Team |
| EPI_ISL_2638805 | Lighthouse Lab in Glasgow | Wellcome Sanger Institute for the COVID-19 Genomics UK (COG-UK) Consortium | Harper VanSteenhouse, Yumi Kasai, David Gray, Carol Clugston, Anna Dominiczak and Alex Alderton, Roberto Amato, Jeffrey Barrett, Sonia Goncalves, Ewan Harrison, David K. Jackson, Ian Johnston, Dominic Kwiatkowski, Cordelia Langford, John Sillitoe on behalf of the Wellcome Sanger Institute COVID-19 Surveillance Team |
| EPI_ISL_2638806 | Lighthouse Lab in Milton Keynes | Wellcome Sanger Institute for the COVID-19 Genomics UK (COG-UK) Consortium | The Lighthouse Lab in Milton Keynes and Alex Alderton, Roberto Amato, Jeffrey Barrett, Sonia Goncalves, Ewan Harrison, David K. Jackson, Ian Johnston, Dominic Kwiatkowski, Cordelia Langford, John Sillitoe on behalf of the Wellcome Sanger Institute COVID-19 Surveillance Team |
| EPI_ISL_2638807 | Lighthouse Lab in Glasgow | Wellcome Sanger Institute for the COVID-19 Genomics UK (COG-UK) Consortium | Harper VanSteenhouse, Yumi Kasai, David Gray, Carol Clugston, Anna Dominiczak and Alex Alderton, Roberto Amato, Jeffrey Barrett, Sonia Goncalves, Ewan Harrison, David K. Jackson, Ian Johnston, Dominic Kwiatkowski, Cordelia Langford, John Sillitoe on behalf of the Wellcome Sanger Institute COVID-19 Surveillance Team |
| EPI_ISL_2638808 | Lighthouse Lab in Alderley Park | Wellcome Sanger Institute for the COVID-19 Genomics UK (COG-UK) Consortium | Jacquelyn Wynn, Mairead Hyland, The Lighthouse Lab in Alderley Park and Alex Alderton, Roberto Amato, Jeffrey Barrett, Sonia Goncalves, Ewan Harrison, David K. Jackson, Ian Johnston, Dominic Kwiatkowski, Cordelia Langford, John Sillitoe on behalf of the Wellcome Sanger Institute COVID-19 Surveillance Team |

| | | | |
|---|---|---|---|
| EPI_ISL_2638809, EPI_ISL_2638810 | Lighthouse Lab in Milton Keynes | Wellcome Sanger Institute for the COVID-19 Genomics UK (COG-UK) Consortium | The Lighthouse Lab in Milton Keynes and Alex Alderton, Roberto Amato, Jeffrey Barrett, Sonia Goncalves, Ewan Harrison, David K. Jackson, Ian Johnston, Dominic Kwiatkowski, Cordelia Langford, John Sillitoe on behalf of the Wellcome Sanger Institute COVID-19 Surveillance Team |
| EPI_ISL_2638811, EPI_ISL_2638812 | Lighthouse Lab in Alderley Park | Wellcome Sanger Institute for the COVID-19 Genomics UK (COG-UK) Consortium | Jacquelyn Wynn, Mairead Hyland, The Lighthouse Lab in Alderley Park and Alex Alderton, Roberto Amato, Jeffrey Barrett, Sonia Goncalves, Ewan Harrison, David K. Jackson, Ian Johnston, Dominic Kwiatkowski, Cordelia Langford, John Sillitoe on behalf of the Wellcome Sanger Institute COVID-19 Surveillance Team |
| EPI_ISL_2638813 | Lighthouse Lab in Glasgow | Wellcome Sanger Institute for the COVID-19 Genomics UK (COG-UK) Consortium | Harper VanSteenhouse, Yumi Kasai, David Gray, Carol Clugston, Anna Dominiczak and Alex Alderton, Roberto Amato, Jeffrey Barrett, Sonia Goncalves, Ewan Harrison, David K. Jackson, Ian Johnston, Dominic Kwiatkowski, Cordelia Langford, John Sillitoe on behalf of the Wellcome Sanger Institute COVID-19 Surveillance Team |
| EPI_ISL_2638814, EPI_ISL_2638815, EPI_ISL_2638816, EPI_ISL_2638817 | Lighthouse Lab in Alderley Park | Wellcome Sanger Institute for the COVID-19 Genomics UK (COG-UK) Consortium | Jacquelyn Wynn, Mairead Hyland, The Lighthouse Lab in Alderley Park and Alex Alderton, Roberto Amato, Jeffrey Barrett, Sonia Goncalves, Ewan Harrison, David K. Jackson, Ian Johnston, Dominic Kwiatkowski, Cordelia Langford, John Sillitoe on behalf of the Wellcome Sanger Institute COVID-19 Surveillance Team |
| EPI_ISL_2638818, EPI_ISL_2638819 | Lighthouse Laboratory Plymouth | Wellcome Sanger Institute for the COVID-19 Genomics UK (COG-UK) Consortium | Lighthouse Laboratory Plymouth and Alex Alderton, Roberto Amato, Jeffrey Barrett, Sonia Goncalves, Ewan Harrison, David K. Jackson, Ian Johnston, Dominic Kwiatkowski, Cordelia Langford, John Sillitoe on behalf of the Wellcome Sanger Institute COVID-19 Surveillance Team |
| EPI_ISL_2638820, EPI_ISL_2638821 | Lighthouse Lab in Alderley Park | Wellcome Sanger Institute for the COVID-19 Genomics UK (COG-UK) Consortium | Jacquelyn Wynn, Mairead Hyland, The Lighthouse Lab in Alderley Park and Alex Alderton, Roberto Amato, Jeffrey Barrett, Sonia Goncalves, Ewan Harrison, David K. Jackson, Ian Johnston, Dominic Kwiatkowski, Cordelia Langford, John Sillitoe on behalf of the Wellcome Sanger Institute COVID-19 Surveillance Team |
| EPI_ISL_2638822 | Lighthouse Lab in Glasgow | Wellcome Sanger Institute for the COVID-19 Genomics UK (COG-UK) Consortium | Harper VanSteenhouse, Yumi Kasai, David Gray, Carol Clugston, Anna Dominiczak and Alex Alderton, Roberto Amato, Jeffrey Barrett, Sonia Goncalves, Ewan Harrison, David K. Jackson, Ian Johnston, Dominic Kwiatkowski, Cordelia Langford, John Sillitoe on behalf of the Wellcome Sanger Institute COVID-19 Surveillance Team |
| EPI_ISL_2638823, EPI_ISL_2638824, EPI_ISL_2638825, EPI_ISL_2638826 | Lighthouse Lab in Milton Keynes | Wellcome Sanger Institute for the COVID-19 Genomics UK (COG-UK) Consortium | The Lighthouse Lab in Milton Keynes and Alex Alderton, Roberto Amato, Jeffrey Barrett, Sonia Goncalves, Ewan Harrison, David K. Jackson, Ian Johnston, Dominic Kwiatkowski, Cordelia Langford, John Sillitoe on behalf of the Wellcome Sanger Institute COVID-19 Surveillance Team |
| EPI_ISL_2638827, EPI_ISL_2638828 | Lighthouse Lab in Alderley Park | Wellcome Sanger Institute for the COVID-19 Genomics UK (COG-UK) Consortium | Jacquelyn Wynn, Mairead Hyland, The Lighthouse Lab in Alderley Park and Alex Alderton, Roberto Amato, Jeffrey Barrett, Sonia Goncalves, Ewan Harrison, David K. Jackson, Ian Johnston, Dominic Kwiatkowski, Cordelia Langford, John Sillitoe on behalf of the Wellcome Sanger Institute COVID-19 Surveillance Team |
| EPI_ISL_2638829 | Lighthouse Lab in Glasgow | Wellcome Sanger Institute for the COVID-19 Genomics UK (COG-UK) Consortium | Harper VanSteenhouse, Yumi Kasai, David Gray, Carol Clugston, Anna Dominiczak and Alex Alderton, Roberto Amato, Jeffrey Barrett, Sonia Goncalves, Ewan Harrison, David K. Jackson, Ian Johnston, Dominic Kwiatkowski, Cordelia Langford, John Sillitoe on behalf of the Wellcome Sanger Institute COVID-19 Surveillance Team |
| EPI_ISL_2638830 | Lighthouse Lab in Milton Keynes | Wellcome Sanger Institute for the COVID-19 Genomics UK (COG-UK) Consortium | The Lighthouse Lab in Milton Keynes and Alex Alderton, Roberto Amato, Jeffrey Barrett, Sonia Goncalves, Ewan Harrison, David K. Jackson, Ian Johnston, Dominic Kwiatkowski, Cordelia Langford, John Sillitoe on behalf of the Wellcome Sanger Institute COVID-19 Surveillance Team |
| EPI_ISL_2638831, EPI_ISL_2638832 | Lighthouse Lab in Alderley Park | Wellcome Sanger Institute for the COVID-19 Genomics UK (COG-UK) Consortium | Jacquelyn Wynn, Mairead Hyland, The Lighthouse Lab in Alderley Park and Alex Alderton, Roberto Amato, Jeffrey Barrett, Sonia Goncalves, Ewan Harrison, David K. Jackson, Ian Johnston, Dominic Kwiatkowski, Cordelia Langford, John Sillitoe on behalf of the Wellcome Sanger Institute COVID-19 Surveillance Team |
| EPI_ISL_2638833, EPI_ISL_2638834 | Lighthouse Lab in Glasgow | Wellcome Sanger Institute for the COVID-19 Genomics UK (COG-UK) Consortium | Harper VanSteenhouse, Yumi Kasai, David Gray, Carol Clugston, Anna Dominiczak and Alex Alderton, Roberto Amato, Jeffrey Barrett, Sonia Goncalves, Ewan Harrison, David K. Jackson, Ian Johnston, Dominic Kwiatkowski, Cordelia Langford, John Sillitoe on behalf of the Wellcome Sanger Institute COVID-19 Surveillance Team |
| EPI_ISL_2638835 | Lighthouse Lab in Milton Keynes | Wellcome Sanger Institute for the COVID-19 Genomics UK (COG-UK) Consortium | The Lighthouse Lab in Milton Keynes and Alex Alderton, Roberto Amato, Jeffrey Barrett, Sonia Goncalves, Ewan Harrison, David K. Jackson, Ian Johnston, Dominic Kwiatkowski, Cordelia Langford, John Sillitoe on behalf of the Wellcome Sanger Institute COVID-19 Surveillance Team |
| EPI_ISL_2638836 | Lighthouse Laboratory Plymouth | Wellcome Sanger Institute for the COVID-19 Genomics UK (COG-UK) Consortium | Lighthouse Laboratory Plymouth and Alex Alderton, Roberto Amato, Jeffrey Barrett, Sonia Goncalves, Ewan Harrison, David K. Jackson, Ian Johnston, Dominic Kwiatkowski, Cordelia Langford, John Sillitoe on behalf of the Wellcome Sanger Institute COVID-19 Surveillance Team |
| EPI_ISL_2638837 | Lighthouse Lab in Milton Keynes | Wellcome Sanger Institute for the COVID-19 Genomics UK (COG-UK) Consortium | The Lighthouse Lab in Milton Keynes and Alex Alderton, Roberto Amato, Jeffrey Barrett, Sonia Goncalves, Ewan Harrison, David K. Jackson, Ian Johnston, Dominic Kwiatkowski, Cordelia Langford, John Sillitoe on behalf of the Wellcome Sanger Institute COVID-19 Surveillance Team |
| EPI_ISL_2638838, EPI_ISL_2638839, EPI_ISL_2638840, EPI_ISL_2638841 | Lighthouse Lab in Alderley Park | Wellcome Sanger Institute for the COVID-19 Genomics UK (COG-UK) Consortium | Jacquelyn Wynn, Mairead Hyland, The Lighthouse Lab in Alderley Park and Alex Alderton, Roberto Amato, Jeffrey Barrett, Sonia Goncalves, Ewan Harrison, David K. Jackson, Ian Johnston, Dominic Kwiatkowski, Cordelia Langford, John Sillitoe on behalf of the Wellcome Sanger Institute COVID-19 Surveillance Team |
| EPI_ISL_2638842 | Lighthouse Laboratory Plymouth | Wellcome Sanger Institute for the COVID-19 Genomics UK (COG-UK) Consortium | Lighthouse Laboratory Plymouth and Alex Alderton, Roberto Amato, Jeffrey Barrett, Sonia Goncalves, Ewan Harrison, David K. Jackson, Ian Johnston, Dominic Kwiatkowski, Cordelia Langford, John Sillitoe on behalf of the Wellcome Sanger Institute COVID-19 Surveillance Team |
| EPI_ISL_2638843 | Lighthouse Lab in Milton Keynes | Wellcome Sanger Institute for the COVID-19 Genomics UK (COG-UK) Consortium | The Lighthouse Lab in Milton Keynes and Alex Alderton, Roberto Amato, Jeffrey Barrett, Sonia Goncalves, Ewan Harrison, David K. Jackson, Ian Johnston, Dominic Kwiatkowski, Cordelia Langford, John Sillitoe on behalf of the Wellcome Sanger Institute COVID-19 Surveillance Team |
| EPI_ISL_2638844 | Lighthouse Lab in Glasgow | Wellcome Sanger Institute for the COVID-19 Genomics UK (COG-UK) Consortium | Harper VanSteenhouse, Yumi Kasai, David Gray, Carol Clugston, Anna Dominiczak and Alex Alderton, Roberto Amato, Jeffrey Barrett, Sonia Goncalves, Ewan Harrison, David K. Jackson, Ian Johnston, Dominic Kwiatkowski, Cordelia Langford, John Sillitoe on behalf of the Wellcome Sanger Institute COVID-19 Surveillance Team |
| EPI_ISL_2638845, EPI_ISL_2638846, EPI_ISL_2638847 | Lighthouse Lab in Alderley Park | Wellcome Sanger Institute for the COVID-19 Genomics UK (COG-UK) Consortium | Jacquelyn Wynn, Mairead Hyland, The Lighthouse Lab in Alderley Park and Alex Alderton, Roberto Amato, Jeffrey Barrett, Sonia Goncalves, Ewan Harrison, David K. Jackson, Ian Johnston, Dominic Kwiatkowski, Cordelia Langford, John Sillitoe on behalf of the Wellcome Sanger Institute COVID-19 Surveillance Team |
| EPI_ISL_2638848 | Lighthouse Lab in Glasgow | Wellcome Sanger Institute for the COVID-19 Genomics UK (COG-UK) Consortium | Harper VanSteenhouse, Yumi Kasai, David Gray, Carol Clugston, Anna Dominiczak and Alex Alderton, Roberto Amato, Jeffrey Barrett, Sonia Goncalves, Ewan Harrison, David K. Jackson, Ian Johnston, Dominic Kwiatkowski, Cordelia Langford, John Sillitoe on behalf of the Wellcome Sanger Institute COVID-19 Surveillance Team |
| EPI_ISL_2638849, EPI_ISL_2638850 | Lighthouse Lab in Alderley Park | Wellcome Sanger Institute for the COVID-19 Genomics UK (COG-UK) Consortium | Jacquelyn Wynn, Mairead Hyland, The Lighthouse Lab in Alderley Park and Alex Alderton, Roberto Amato, Jeffrey Barrett, Sonia Goncalves, Ewan Harrison, David K. Jackson, Ian Johnston, Dominic Kwiatkowski, Cordelia Langford, John Sillitoe on behalf of the Wellcome Sanger Institute COVID-19 Surveillance Team |
| EPI_ISL_2638851 | Lighthouse Lab in Milton Keynes | Wellcome Sanger Institute for the COVID-19 Genomics UK (COG-UK) Consortium | The Lighthouse Lab in Milton Keynes and Alex Alderton, Roberto Amato, Jeffrey Barrett, Sonia Goncalves, Ewan Harrison, David K. Jackson, Ian Johnston, Dominic Kwiatkowski, Cordelia Langford, John Sillitoe on behalf of the Wellcome Sanger Institute COVID-19 Surveillance Team |
| EPI_ISL_2638852, EPI_ISL_2638854 | Lighthouse Lab in Glasgow | Wellcome Sanger Institute for the COVID-19 Genomics UK (COG-UK) Consortium | Harper VanSteenhouse, Yumi Kasai, David Gray, Carol Clugston, Anna Dominiczak and Alex Alderton, Roberto Amato, Jeffrey Barrett, Sonia Goncalves, Ewan Harrison, David K. Jackson, Ian Johnston, Dominic Kwiatkowski, Cordelia Langford, John Sillitoe on behalf of the Wellcome Sanger Institute COVID-19 Surveillance Team |
| EPI_ISL_2638855, EPI_ISL_2638856, EPI_ISL_2638857 | Lighthouse Lab in Alderley Park | Wellcome Sanger Institute for the COVID-19 Genomics UK (COG-UK) Consortium | Jacquelyn Wynn, Mairead Hyland, The Lighthouse Lab in Alderley Park and Alex Alderton, Roberto Amato, Jeffrey Barrett, Sonia Goncalves, Ewan Harrison, David K. Jackson, Ian Johnston, Dominic Kwiatkowski, Cordelia Langford, John Sillitoe on behalf of the Wellcome Sanger Institute COVID-19 Surveillance Team |
| EPI_ISL_2638858, EPI_ISL_2638859 | Lighthouse Lab in Glasgow | Wellcome Sanger Institute for the COVID-19 Genomics UK (COG-UK) Consortium | Harper VanSteenhouse, Yumi Kasai, David Gray, Carol Clugston, Anna Dominiczak and Alex Alderton, Roberto Amato, Jeffrey Barrett, Sonia Goncalves, Ewan Harrison, David K. Jackson, Ian Johnston, Dominic Kwiatkowski, Cordelia Langford, John Sillitoe on behalf of the Wellcome Sanger Institute COVID-19 Surveillance Team |
| EPI_ISL_2638860, EPI_ISL_2638861, EPI_ISL_2638862 | Lighthouse Lab in Milton Keynes | Wellcome Sanger Institute for the COVID-19 Genomics UK (COG-UK) Consortium | The Lighthouse Lab in Milton Keynes and Alex Alderton, Roberto Amato, Jeffrey Barrett, Sonia Goncalves, Ewan Harrison, David K. Jackson, Ian Johnston, Dominic Kwiatkowski, Cordelia Langford, John Sillitoe on behalf of the Wellcome Sanger Institute COVID-19 Surveillance Team |
| EPI_ISL_2638863, EPI_ISL_2638864 | Lighthouse Lab in Alderley Park | Wellcome Sanger Institute for the COVID-19 Genomics UK (COG-UK) Consortium | Jacquelyn Wynn, Mairead Hyland, The Lighthouse Lab in Alderley Park and Alex Alderton, Roberto Amato, Jeffrey Barrett, Sonia Goncalves, Ewan Harrison, David K. Jackson, Ian Johnston, Dominic Kwiatkowski, Cordelia Langford, John Sillitoe on behalf of the Wellcome Sanger Institute COVID-19 Surveillance Team |
| EPI_ISL_2638865 | Lighthouse Lab in Glasgow | Wellcome Sanger Institute for the COVID-19 Genomics UK (COG-UK) Consortium | Harper VanSteenhouse, Yumi Kasai, David Gray, Carol Clugston, Anna Dominiczak and Alex Alderton, Roberto Amato, Jeffrey Barrett, Sonia Goncalves, Ewan Harrison, David K. Jackson, Ian Johnston, Dominic Kwiatkowski, Cordelia Langford, John Sillitoe on behalf of the Wellcome Sanger Institute COVID-19 Surveillance Team |
| EPI_ISL_2638866, EPI_ISL_2638867, EPI_ISL_2638868 | Lighthouse Lab in Alderley Park | Wellcome Sanger Institute for the COVID-19 Genomics UK (COG-UK) Consortium | Jacquelyn Wynn, Mairead Hyland, The Lighthouse Lab in Alderley Park and Alex Alderton, Roberto Amato, Jeffrey Barrett, Sonia Goncalves, Ewan Harrison, David K. Jackson, Ian Johnston, Dominic Kwiatkowski, Cordelia Langford, John Sillitoe on behalf of the Wellcome Sanger Institute COVID-19 Surveillance Team |
| EPI_ISL_2638869 | Lighthouse Lab in Glasgow | Wellcome Sanger Institute for the COVID-19 Genomics UK (COG-UK) Consortium | Harper VanSteenhouse, Yumi Kasai, David Gray, Carol Clugston, Anna Dominiczak and Alex Alderton, Roberto Amato, Jeffrey Barrett, Sonia Goncalves, Ewan Harrison, David K. Jackson, Ian Johnston, Dominic Kwiatkowski, Cordelia Langford, John Sillitoe on behalf of the Wellcome Sanger Institute COVID-19 Surveillance Team |
| EPI_ISL_2638870 | Lighthouse Lab in Milton Keynes | Wellcome Sanger Institute for the COVID-19 Genomics UK (COG-UK) Consortium | The Lighthouse Lab in Milton Keynes and Alex Alderton, Roberto Amato, Jeffrey Barrett, Sonia Goncalves, Ewan Harrison, David K. Jackson, Ian Johnston, Dominic Kwiatkowski, Cordelia Langford, John Sillitoe on behalf of the Wellcome Sanger Institute COVID-19 Surveillance Team |
| EPI_ISL_2638872 | Lighthouse Lab in Alderley Park | Wellcome Sanger Institute for the COVID-19 Genomics UK (COG-UK) Consortium | Jacquelyn Wynn, Mairead Hyland, The Lighthouse Lab in Alderley Park and Alex Alderton, Roberto Amato, Jeffrey Barrett, Sonia Goncalves, Ewan Harrison, David K. Jackson, Ian Johnston, Dominic Kwiatkowski, Cordelia Langford, John Sillitoe on behalf of the Wellcome Sanger Institute COVID-19 Surveillance Team |
| EPI_ISL_2638873 | Lighthouse Laboratory Plymouth | Wellcome Sanger Institute for the COVID-19 Genomics UK (COG-UK) Consortium | Lighthouse Laboratory Plymouth and Alex Alderton, Roberto Amato, Jeffrey Barrett, Sonia Goncalves, Ewan Harrison, David K. Jackson, Ian Johnston, Dominic Kwiatkowski, Cordelia Langford, John Sillitoe on behalf of the Wellcome Sanger Institute COVID-19 Surveillance Team |
| EPI_ISL_2638874 | Lighthouse Lab in Alderley Park | Wellcome Sanger Institute for the COVID-19 Genomics UK (COG-UK) Consortium | Jacquelyn Wynn, Mairead Hyland, The Lighthouse Lab in Alderley Park and Alex Alderton, Roberto Amato, Jeffrey Barrett, Sonia Goncalves, Ewan Harrison, David K. Jackson, Ian Johnston, Dominic Kwiatkowski, Cordelia Langford, John Sillitoe on behalf of the Wellcome Sanger Institute COVID-19 Surveillance Team |
| EPI_ISL_2638876 | Lighthouse Lab in Milton Keynes | Wellcome Sanger Institute for the COVID-19 Genomics UK (COG-UK) Consortium | The Lighthouse Lab in Milton Keynes and Alex Alderton, Roberto Amato, Jeffrey Barrett, Sonia Goncalves, Ewan Harrison, David K. Jackson, Ian Johnston, Dominic Kwiatkowski, Cordelia Langford, John Sillitoe on behalf of the Wellcome Sanger Institute COVID-19 Surveillance Team |
| EPI_ISL_2638877 | Lighthouse Lab in Alderley Park | Wellcome Sanger Institute for the COVID-19 Genomics UK (COG-UK) Consortium | Jacquelyn Wynn, Mairead Hyland, The Lighthouse Lab in Alderley Park and Alex Alderton, Roberto Amato, Jeffrey Barrett, Sonia Goncalves, Ewan Harrison, David K. Jackson, Ian Johnston, Dominic Kwiatkowski, Cordelia Langford, John Sillitoe on behalf of the Wellcome Sanger Institute COVID-19 Surveillance Team |
| EPI_ISL_2638878 | Lighthouse Lab in Milton Keynes | Wellcome Sanger Institute for the COVID-19 Genomics UK (COG-UK) Consortium | The Lighthouse Lab in Milton Keynes and Alex Alderton, Roberto Amato, Jeffrey Barrett, Sonia Goncalves, Ewan Harrison, David K. Jackson, Ian Johnston, Dominic Kwiatkowski, Cordelia Langford, John Sillitoe on behalf of the Wellcome Sanger Institute COVID-19 Surveillance Team |
| EPI_ISL_2638879 | Lighthouse Lab in Alderley Park | Wellcome Sanger Institute for the COVID-19 Genomics UK (COG-UK) Consortium | Jacquelyn Wynn, Mairead Hyland, The Lighthouse Lab in Alderley Park and Alex Alderton, Roberto Amato, Jeffrey Barrett, Sonia Goncalves, Ewan Harrison, David K. Jackson, Ian Johnston, Dominic Kwiatkowski, Cordelia Langford, John Sillitoe on behalf of the Wellcome Sanger Institute COVID-19 Surveillance Team |
| EPI_ISL_2638880 | Lighthouse Lab in Milton Keynes | Wellcome Sanger Institute for the COVID-19 Genomics UK (COG-UK) Consortium | The Lighthouse Lab in Milton Keynes and Alex Alderton, Roberto Amato, Jeffrey Barrett, Sonia Goncalves, Ewan Harrison, David K. Jackson, Ian Johnston, Dominic Kwiatkowski, Cordelia Langford, John Sillitoe on behalf of the Wellcome Sanger Institute COVID-19 Surveillance Team |

| | | | |
|---|---|---|---|
| EPI_ISL_2638881 | Lighthouse Lab in Alderley Park | Wellcome Sanger Institute for the COVID-19 Genomics UK (COG-UK) Consortium | Jacquelyn Wynn, Mairead Hyland, The Lighthouse Lab in Alderley Park and Alex Alderton, Roberto Amato, Jeffrey Barrett, Sonia Goncalves, Ewan Harrison, David K. Jackson, Ian Johnston, Dominic Kwiatkowski, Cordelia Langford, John Sillitoe on behalf of the Wellcome Sanger Institute COVID-19 Surveillance Team |
| EPI_ISL_2638882, EPI_ISL_2638883 | Lighthouse Lab in Milton Keynes | Wellcome Sanger Institute for the COVID-19 Genomics UK (COG-UK) Consortium | The Lighthouse Lab in Milton Keynes and Alex Alderton, Roberto Amato, Jeffrey Barrett, Sonia Goncalves, Ewan Harrison, David K. Jackson, Ian Johnston, Dominic Kwiatkowski, Cordelia Langford, John Sillitoe on behalf of the Wellcome Sanger Institute COVID-19 Surveillance Team |
| EPI_ISL_2638884 | Lighthouse Lab in Alderley Park | Wellcome Sanger Institute for the COVID-19 Genomics UK (COG-UK) Consortium | Jacquelyn Wynn, Mairead Hyland, The Lighthouse Lab in Alderley Park and Alex Alderton, Roberto Amato, Jeffrey Barrett, Sonia Goncalves, Ewan Harrison, David K. Jackson, Ian Johnston, Dominic Kwiatkowski, Cordelia Langford, John Sillitoe on behalf of the Wellcome Sanger Institute COVID-19 Surveillance Team |
| EPI_ISL_2638885, EPI_ISL_2638886 | Lighthouse Lab in Milton Keynes | Wellcome Sanger Institute for the COVID-19 Genomics UK (COG-UK) Consortium | The Lighthouse Lab in Milton Keynes and Alex Alderton, Roberto Amato, Jeffrey Barrett, Sonia Goncalves, Ewan Harrison, David K. Jackson, Ian Johnston, Dominic Kwiatkowski, Cordelia Langford, John Sillitoe on behalf of the Wellcome Sanger Institute COVID-19 Surveillance Team |
| EPI_ISL_2638887 | Lighthouse Lab in Glasgow | Wellcome Sanger Institute for the COVID-19 Genomics UK (COG-UK) Consortium | Harper VanSteenhouse, Yumi Kasai, David Gray, Carol Clugston, Anna Dominiczak and Alex Alderton, Roberto Amato, Jeffrey Barrett, Sonia Goncalves, Ewan Harrison, David K. Jackson, Ian Johnston, Dominic Kwiatkowski, Cordelia Langford, John Sillitoe on behalf of the Wellcome Sanger Institute COVID-19 Surveillance Team |
| EPI_ISL_2638888 | Lighthouse Lab in Milton Keynes | Wellcome Sanger Institute for the COVID-19 Genomics UK (COG-UK) Consortium | The Lighthouse Lab in Milton Keynes and Alex Alderton, Roberto Amato, Jeffrey Barrett, Sonia Goncalves, Ewan Harrison, David K. Jackson, Ian Johnston, Dominic Kwiatkowski, Cordelia Langford, John Sillitoe on behalf of the Wellcome Sanger Institute COVID-19 Surveillance Team |
| EPI_ISL_2638889, EPI_ISL_2638890, EPI_ISL_2638891 | Lighthouse Lab in Alderley Park | Wellcome Sanger Institute for the COVID-19 Genomics UK (COG-UK) Consortium | Jacquelyn Wynn, Mairead Hyland, The Lighthouse Lab in Alderley Park and Alex Alderton, Roberto Amato, Jeffrey Barrett, Sonia Goncalves, Ewan Harrison, David K. Jackson, Ian Johnston, Dominic Kwiatkowski, Cordelia Langford, John Sillitoe on behalf of the Wellcome Sanger Institute COVID-19 Surveillance Team |
| EPI_ISL_2638892, EPI_ISL_2638893 | Lighthouse Lab in Milton Keynes | Wellcome Sanger Institute for the COVID-19 Genomics UK (COG-UK) Consortium | The Lighthouse Lab in Milton Keynes and Alex Alderton, Roberto Amato, Jeffrey Barrett, Sonia Goncalves, Ewan Harrison, David K. Jackson, Ian Johnston, Dominic Kwiatkowski, Cordelia Langford, John Sillitoe on behalf of the Wellcome Sanger Institute COVID-19 Surveillance Team |
| EPI_ISL_2638894 | Lighthouse Laboratory Plymouth | Wellcome Sanger Institute for the COVID-19 Genomics UK (COG-UK) Consortium | Lighthouse Laboratory Plymouth and Alex Alderton, Roberto Amato, Jeffrey Barrett, Sonia Goncalves, Ewan Harrison, David K. Jackson, Ian Johnston, Dominic Kwiatkowski, Cordelia Langford, John Sillitoe on behalf of the Wellcome Sanger Institute COVID-19 Surveillance Team |
| EPI_ISL_2638895, EPI_ISL_2638896, EPI_ISL_2638897, EPI_ISL_2638898 | Lighthouse Lab in Alderley Park | Wellcome Sanger Institute for the COVID-19 Genomics UK (COG-UK) Consortium | Jacquelyn Wynn, Mairead Hyland, The Lighthouse Lab in Alderley Park and Alex Alderton, Roberto Amato, Jeffrey Barrett, Sonia Goncalves, Ewan Harrison, David K. Jackson, Ian Johnston, Dominic Kwiatkowski, Cordelia Langford, John Sillitoe on behalf of the Wellcome Sanger Institute COVID-19 Surveillance Team |
| EPI_ISL_2638899 | Lighthouse Lab in Milton Keynes | Wellcome Sanger Institute for the COVID-19 Genomics UK (COG-UK) Consortium | The Lighthouse Lab in Milton Keynes and Alex Alderton, Roberto Amato, Jeffrey Barrett, Sonia Goncalves, Ewan Harrison, David K. Jackson, Ian Johnston, Dominic Kwiatkowski, Cordelia Langford, John Sillitoe on behalf of the Wellcome Sanger Institute COVID-19 Surveillance Team |
| EPI_ISL_2638900 | Lighthouse Lab in Alderley Park | Wellcome Sanger Institute for the COVID-19 Genomics UK (COG-UK) Consortium | Jacquelyn Wynn, Mairead Hyland, The Lighthouse Lab in Alderley Park and Alex Alderton, Roberto Amato, Jeffrey Barrett, Sonia Goncalves, Ewan Harrison, David K. Jackson, Ian Johnston, Dominic Kwiatkowski, Cordelia Langford, John Sillitoe on behalf of the Wellcome Sanger Institute COVID-19 Surveillance Team |
| EPI_ISL_2638901 | Lighthouse Laboratory Plymouth | Wellcome Sanger Institute for the COVID-19 Genomics UK (COG-UK) Consortium | Lighthouse Laboratory Plymouth and Alex Alderton, Roberto Amato, Jeffrey Barrett, Sonia Goncalves, Ewan Harrison, David K. Jackson, Ian Johnston, Dominic Kwiatkowski, Cordelia Langford, John Sillitoe on behalf of the Wellcome Sanger Institute COVID-19 Surveillance Team |
| EPI_ISL_2638902, EPI_ISL_2638904 | Lighthouse Lab in Glasgow | Wellcome Sanger Institute for the COVID-19 Genomics UK (COG-UK) Consortium | Harper VanSteenhouse, Yumi Kasai, David Gray, Carol Clugston, Anna Dominiczak and Alex Alderton, Roberto Amato, Jeffrey Barrett, Sonia Goncalves, Ewan Harrison, David K. Jackson, Ian Johnston, Dominic Kwiatkowski, Cordelia Langford, John Sillitoe on behalf of the Wellcome Sanger Institute COVID-19 Surveillance Team |
| EPI_ISL_2638905, EPI_ISL_2638906 | Lighthouse Lab in Alderley Park | Wellcome Sanger Institute for the COVID-19 Genomics UK (COG-UK) Consortium | Jacquelyn Wynn, Mairead Hyland, The Lighthouse Lab in Alderley Park and Alex Alderton, Roberto Amato, Jeffrey Barrett, Sonia Goncalves, Ewan Harrison, David K. Jackson, Ian Johnston, Dominic Kwiatkowski, Cordelia Langford, John Sillitoe on behalf of the Wellcome Sanger Institute COVID-19 Surveillance Team |
| EPI_ISL_2638907 | Lighthouse Lab in Milton Keynes | Wellcome Sanger Institute for the COVID-19 Genomics UK (COG-UK) Consortium | The Lighthouse Lab in Milton Keynes and Alex Alderton, Roberto Amato, Jeffrey Barrett, Sonia Goncalves, Ewan Harrison, David K. Jackson, Ian Johnston, Dominic Kwiatkowski, Cordelia Langford, John Sillitoe on behalf of the Wellcome Sanger Institute COVID-19 Surveillance Team |
| EPI_ISL_2638908, EPI_ISL_2638909, EPI_ISL_2638910, EPI_ISL_2638911, EPI_ISL_2638912, EPI_ISL_2638913, EPI_ISL_2638914, EPI_ISL_2638915 | Lighthouse Lab in Alderley Park | Wellcome Sanger Institute for the COVID-19 Genomics UK (COG-UK) Consortium | Jacquelyn Wynn, Mairead Hyland, The Lighthouse Lab in Alderley Park and Alex Alderton, Roberto Amato, Jeffrey Barrett, Sonia Goncalves, Ewan Harrison, David K. Jackson, Ian Johnston, Dominic Kwiatkowski, Cordelia Langford, John Sillitoe on behalf of the Wellcome Sanger Institute COVID-19 Surveillance Team |
| EPI_ISL_2638916, EPI_ISL_2638917, EPI_ISL_2638918, EPI_ISL_2638919, EPI_ISL_2638920 | Lighthouse Laboratory Plymouth | Wellcome Sanger Institute for the COVID-19 Genomics UK (COG-UK) Consortium | Lighthouse Laboratory Plymouth and Alex Alderton, Roberto Amato, Jeffrey Barrett, Sonia Goncalves, Ewan Harrison, David K. Jackson, Ian Johnston, Dominic Kwiatkowski, Cordelia Langford, John Sillitoe on behalf of the Wellcome Sanger Institute COVID-19 Surveillance Team |
| EPI_ISL_2638921 | Lighthouse Lab in Milton Keynes | Wellcome Sanger Institute for the COVID-19 Genomics UK (COG-UK) Consortium | The Lighthouse Lab in Milton Keynes and Alex Alderton, Roberto Amato, Jeffrey Barrett, Sonia Goncalves, Ewan Harrison, David K. Jackson, Ian Johnston, Dominic Kwiatkowski, Cordelia Langford, John Sillitoe on behalf of the Wellcome Sanger Institute COVID-19 Surveillance Team |
| EPI_ISL_2638922, EPI_ISL_2638923 | Lighthouse Laboratory Plymouth | Wellcome Sanger Institute for the COVID-19 Genomics UK (COG-UK) Consortium | Lighthouse Laboratory Plymouth and Alex Alderton, Roberto Amato, Jeffrey Barrett, Sonia Goncalves, Ewan Harrison, David K. Jackson, Ian Johnston, Dominic Kwiatkowski, Cordelia Langford, John Sillitoe on behalf of the Wellcome Sanger Institute COVID-19 Surveillance Team |
| EPI_ISL_2638924 | Lighthouse Lab in Milton Keynes | Wellcome Sanger Institute for the COVID-19 Genomics UK (COG-UK) Consortium | The Lighthouse Lab in Milton Keynes and Alex Alderton, Roberto Amato, Jeffrey Barrett, Sonia Goncalves, Ewan Harrison, David K. Jackson, Ian Johnston, Dominic Kwiatkowski, Cordelia Langford, John Sillitoe on behalf of the Wellcome Sanger Institute COVID-19 Surveillance Team |
| EPI_ISL_2638925 | Lighthouse Lab in Alderley Park | Wellcome Sanger Institute for the COVID-19 Genomics UK (COG-UK) Consortium | Jacquelyn Wynn, Mairead Hyland, The Lighthouse Lab in Alderley Park and Alex Alderton, Roberto Amato, Jeffrey Barrett, Sonia Goncalves, Ewan Harrison, David K. Jackson, Ian Johnston, Dominic Kwiatkowski, Cordelia Langford, John Sillitoe on behalf of the Wellcome Sanger Institute COVID-19 Surveillance Team |
| EPI_ISL_2638926, EPI_ISL_2638927, EPI_ISL_2638928, EPI_ISL_2638929 | Lighthouse Laboratory Plymouth | Wellcome Sanger Institute for the COVID-19 Genomics UK (COG-UK) Consortium | Lighthouse Laboratory Plymouth and Alex Alderton, Roberto Amato, Jeffrey Barrett, Sonia Goncalves, Ewan Harrison, David K. Jackson, Ian Johnston, Dominic Kwiatkowski, Cordelia Langford, John Sillitoe on behalf of the Wellcome Sanger Institute COVID-19 Surveillance Team |
| EPI_ISL_2638930 | Lighthouse Lab in Milton Keynes | Wellcome Sanger Institute for the COVID-19 Genomics UK (COG-UK) Consortium | The Lighthouse Lab in Milton Keynes and Alex Alderton, Roberto Amato, Jeffrey Barrett, Sonia Goncalves, Ewan Harrison, David K. Jackson, Ian Johnston, Dominic Kwiatkowski, Cordelia Langford, John Sillitoe on behalf of the Wellcome Sanger Institute COVID-19 Surveillance Team |
| EPI_ISL_2638931, EPI_ISL_2638932, EPI_ISL_2638933 | Lighthouse Laboratory Plymouth | Wellcome Sanger Institute for the COVID-19 Genomics UK (COG-UK) Consortium | Lighthouse Laboratory Plymouth and Alex Alderton, Roberto Amato, Jeffrey Barrett, Sonia Goncalves, Ewan Harrison, David K. Jackson, Ian Johnston, Dominic Kwiatkowski, Cordelia Langford, John Sillitoe on behalf of the Wellcome Sanger Institute COVID-19 Surveillance Team |
| EPI_ISL_2638934 | Health Services Laboratories | Wellcome Sanger Institute for the COVID-19 Genomics UK (COG-UK) Consortium | Health Services Laboratories and Alex Alderton, Roberto Amato, Jeffrey Barrett, Sonia Goncalves, Ewan Harrison, David K. Jackson, Ian Johnston, Dominic Kwiatkowski, Cordelia Langford, John Sillitoe on behalf of the Wellcome Sanger Institute COVID-19 Surveillance Team |
| EPI_ISL_2638935, EPI_ISL_2638936, EPI_ISL_2638937, EPI_ISL_2638938 | Lighthouse Laboratory Plymouth | Wellcome Sanger Institute for the COVID-19 Genomics UK (COG-UK) Consortium | Lighthouse Laboratory Plymouth and Alex Alderton, Roberto Amato, Jeffrey Barrett, Sonia Goncalves, Ewan Harrison, David K. Jackson, Ian Johnston, Dominic Kwiatkowski, Cordelia Langford, John Sillitoe on behalf of the Wellcome Sanger Institute COVID-19 Surveillance Team |
| EPI_ISL_2638939 | Health Services Laboratories | Wellcome Sanger Institute for the COVID-19 Genomics UK (COG-UK) Consortium | Health Services Laboratories and Alex Alderton, Roberto Amato, Jeffrey Barrett, Sonia Goncalves, Ewan Harrison, David K. Jackson, Ian Johnston, Dominic Kwiatkowski, Cordelia Langford, John Sillitoe on behalf of the Wellcome Sanger Institute COVID-19 Surveillance Team |
| EPI_ISL_2638940, EPI_ISL_2638941, EPI_ISL_2638942 | Lighthouse Laboratory Plymouth | Wellcome Sanger Institute for the COVID-19 Genomics UK (COG-UK) Consortium | Lighthouse Laboratory Plymouth and Alex Alderton, Roberto Amato, Jeffrey Barrett, Sonia Goncalves, Ewan Harrison, David K. Jackson, Ian Johnston, Dominic Kwiatkowski, Cordelia Langford, John Sillitoe on behalf of the Wellcome Sanger Institute COVID-19 Surveillance Team |
| EPI_ISL_2638943 | Lighthouse Lab in Milton Keynes | Wellcome Sanger Institute for the COVID-19 Genomics UK (COG-UK) Consortium | The Lighthouse Lab in Milton Keynes and Alex Alderton, Roberto Amato, Jeffrey Barrett, Sonia Goncalves, Ewan Harrison, David K. Jackson, Ian Johnston, Dominic Kwiatkowski, Cordelia Langford, John Sillitoe on behalf of the Wellcome Sanger Institute COVID-19 Surveillance Team |
| EPI_ISL_2638944 | Lighthouse Laboratory Plymouth | Wellcome Sanger Institute for the COVID-19 Genomics UK (COG-UK) Consortium | Lighthouse Laboratory Plymouth and Alex Alderton, Roberto Amato, Jeffrey Barrett, Sonia Goncalves, Ewan Harrison, David K. Jackson, Ian Johnston, Dominic Kwiatkowski, Cordelia Langford, John Sillitoe on behalf of the Wellcome Sanger Institute COVID-19 Surveillance Team |
| EPI_ISL_2638945 | Lighthouse Lab in Milton Keynes | Wellcome Sanger Institute for the COVID-19 Genomics UK (COG-UK) Consortium | The Lighthouse Lab in Milton Keynes and Alex Alderton, Roberto Amato, Jeffrey Barrett, Sonia Goncalves, Ewan Harrison, David K. Jackson, Ian Johnston, Dominic Kwiatkowski, Cordelia Langford, John Sillitoe on behalf of the Wellcome Sanger Institute COVID-19 Surveillance Team |
| EPI_ISL_2638946 | Lighthouse Laboratory Plymouth | Wellcome Sanger Institute for the COVID-19 Genomics UK (COG-UK) Consortium | Lighthouse Laboratory Plymouth and Alex Alderton, Roberto Amato, Jeffrey Barrett, Sonia Goncalves, Ewan Harrison, David K. Jackson, Ian Johnston, Dominic Kwiatkowski, Cordelia Langford, John Sillitoe on behalf of the Wellcome Sanger Institute COVID-19 Surveillance Team |
| EPI_ISL_2638947 | Health Services Laboratories | Wellcome Sanger Institute for the COVID-19 Genomics UK (COG-UK) Consortium | Health Services Laboratories and Alex Alderton, Roberto Amato, Jeffrey Barrett, Sonia Goncalves, Ewan Harrison, David K. Jackson, Ian Johnston, Dominic Kwiatkowski, Cordelia Langford, John Sillitoe on behalf of the Wellcome Sanger Institute COVID-19 Surveillance Team |
| EPI_ISL_2638948, EPI_ISL_2638949 | Lighthouse Laboratory Plymouth | Wellcome Sanger Institute for the COVID-19 Genomics UK (COG-UK) Consortium | Lighthouse Laboratory Plymouth and Alex Alderton, Roberto Amato, Jeffrey Barrett, Sonia Goncalves, Ewan Harrison, David K. Jackson, Ian Johnston, Dominic Kwiatkowski, Cordelia Langford, John Sillitoe on behalf of the Wellcome Sanger Institute COVID-19 Surveillance Team |
| EPI_ISL_2638950 | Lighthouse Lab in Milton Keynes | Wellcome Sanger Institute for the COVID-19 Genomics UK (COG-UK) Consortium | The Lighthouse Lab in Milton Keynes and Alex Alderton, Roberto Amato, Jeffrey Barrett, Sonia Goncalves, Ewan Harrison, David K. Jackson, Ian Johnston, Dominic Kwiatkowski, Cordelia Langford, John Sillitoe on behalf of the Wellcome Sanger Institute COVID-19 Surveillance Team |
| EPI_ISL_2638951, EPI_ISL_2638952 | Lighthouse Laboratory Plymouth | Wellcome Sanger Institute for the COVID-19 Genomics UK (COG-UK) Consortium | Lighthouse Laboratory Plymouth and Alex Alderton, Roberto Amato, Jeffrey Barrett, Sonia Goncalves, Ewan Harrison, David K. Jackson, Ian Johnston, Dominic Kwiatkowski, Cordelia Langford, John Sillitoe on behalf of the Wellcome Sanger Institute COVID-19 Surveillance Team |
| EPI_ISL_2638953 | Lighthouse Lab in Milton Keynes | Wellcome Sanger Institute for the COVID-19 Genomics UK (COG-UK) | The Lighthouse Lab in Milton Keynes and Alex Alderton, Roberto Amato, Jeffrey Barrett, Sonia Goncalves, Ewan Harrison, David K. Jackson, Ian Johnston, Dominic Kwiatkowski, Cordelia Langford, John Sillitoe on behalf of the Wellcome Sanger |

| | | | |
|---|---|---|---|
| | | Consortium | Institute COVID-19 Surveillance Team |
| EPI_ISL_2638954 | Health Services Laboratories | Wellcome Sanger Institute for the COVID-19 Genomics UK (COG-UK) Consortium | Health Services Laboratories and Alex Alderton, Roberto Amato, Jeffrey Barrett, Sonia Goncalves, Ewan Harrison, David K. Jackson, Ian Johnston, Dominic Kwiatkowski, Cordelia Langford, John Sillitoe on behalf of the Wellcome Sanger Institute COVID-19 Surveillance Team |
| EPI_ISL_2638955 | Lighthouse Laboratory Plymouth | Wellcome Sanger Institute for the COVID-19 Genomics UK (COG-UK) Consortium | Lighthouse Laboratory Plymouth and Alex Alderton, Roberto Amato, Jeffrey Barrett, Sonia Goncalves, Ewan Harrison, David K. Jackson, Ian Johnston, Dominic Kwiatkowski, Cordelia Langford, John Sillitoe on behalf of the Wellcome Sanger Institute COVID-19 Surveillance Team |
| EPI_ISL_2638956 | Health Services Laboratories | Wellcome Sanger Institute for the COVID-19 Genomics UK (COG-UK) Consortium | Health Services Laboratories and Alex Alderton, Roberto Amato, Jeffrey Barrett, Sonia Goncalves, Ewan Harrison, David K. Jackson, Ian Johnston, Dominic Kwiatkowski, Cordelia Langford, John Sillitoe on behalf of the Wellcome Sanger Institute COVID-19 Surveillance Team |
| EPI_ISL_2638957, EPI_ISL_2638958 | Lighthouse Laboratory Plymouth | Wellcome Sanger Institute for the COVID-19 Genomics UK (COG-UK) Consortium | Lighthouse Laboratory Plymouth and Alex Alderton, Roberto Amato, Jeffrey Barrett, Sonia Goncalves, Ewan Harrison, David K. Jackson, Ian Johnston, Dominic Kwiatkowski, Cordelia Langford, John Sillitoe on behalf of the Wellcome Sanger Institute COVID-19 Surveillance Team |
| EPI_ISL_2638959 | Lighthouse Lab in Milton Keynes | Wellcome Sanger Institute for the COVID-19 Genomics UK (COG-UK) Consortium | The Lighthouse Lab in Milton Keynes and Alex Alderton, Roberto Amato, Jeffrey Barrett, Sonia Goncalves, Ewan Harrison, David K. Jackson, Ian Johnston, Dominic Kwiatkowski, Cordelia Langford, John Sillitoe on behalf of the Wellcome Sanger Institute COVID-19 Surveillance Team |
| EPI_ISL_2638960, EPI_ISL_2638961 | Health Services Laboratories | Wellcome Sanger Institute for the COVID-19 Genomics UK (COG-UK) Consortium | Health Services Laboratories and Alex Alderton, Roberto Amato, Jeffrey Barrett, Sonia Goncalves, Ewan Harrison, David K. Jackson, Ian Johnston, Dominic Kwiatkowski, Cordelia Langford, John Sillitoe on behalf of the Wellcome Sanger Institute COVID-19 Surveillance Team |
| EPI_ISL_2638962 | Lighthouse Lab in Alderley Park | Wellcome Sanger Institute for the COVID-19 Genomics UK (COG-UK) Consortium | Jacquelyn Wynn, Mairead Hyland, The Lighthouse Lab in Alderley Park and Alex Alderton, Roberto Amato, Jeffrey Barrett, Sonia Goncalves, Ewan Harrison, David K. Jackson, Ian Johnston, Dominic Kwiatkowski, Cordelia Langford, John Sillitoe on behalf of the Wellcome Sanger Institute COVID-19 Surveillance Team |
| EPI_ISL_2638963, EPI_ISL_2638965, EPI_ISL_2638966, EPI_ISL_2638967 | Lighthouse Laboratory Plymouth | Wellcome Sanger Institute for the COVID-19 Genomics UK (COG-UK) Consortium | Lighthouse Laboratory Plymouth and Alex Alderton, Roberto Amato, Jeffrey Barrett, Sonia Goncalves, Ewan Harrison, David K. Jackson, Ian Johnston, Dominic Kwiatkowski, Cordelia Langford, John Sillitoe on behalf of the Wellcome Sanger Institute COVID-19 Surveillance Team |
| EPI_ISL_2638968 | Health Services Laboratories | Wellcome Sanger Institute for the COVID-19 Genomics UK (COG-UK) Consortium | Health Services Laboratories and Alex Alderton, Roberto Amato, Jeffrey Barrett, Sonia Goncalves, Ewan Harrison, David K. Jackson, Ian Johnston, Dominic Kwiatkowski, Cordelia Langford, John Sillitoe on behalf of the Wellcome Sanger Institute COVID-19 Surveillance Team |
| EPI_ISL_2638969, EPI_ISL_2638971 | Lighthouse Laboratory Plymouth | Wellcome Sanger Institute for the COVID-19 Genomics UK (COG-UK) Consortium | Lighthouse Laboratory Plymouth and Alex Alderton, Roberto Amato, Jeffrey Barrett, Sonia Goncalves, Ewan Harrison, David K. Jackson, Ian Johnston, Dominic Kwiatkowski, Cordelia Langford, John Sillitoe on behalf of the Wellcome Sanger Institute COVID-19 Surveillance Team |
| EPI_ISL_2638972 | Health Services Laboratories | Wellcome Sanger Institute for the COVID-19 Genomics UK (COG-UK) Consortium | Health Services Laboratories and Alex Alderton, Roberto Amato, Jeffrey Barrett, Sonia Goncalves, Ewan Harrison, David K. Jackson, Ian Johnston, Dominic Kwiatkowski, Cordelia Langford, John Sillitoe on behalf of the Wellcome Sanger Institute COVID-19 Surveillance Team |
| EPI_ISL_2638973, EPI_ISL_2638974, EPI_ISL_2638975 | Lighthouse Laboratory Plymouth | Wellcome Sanger Institute for the COVID-19 Genomics UK (COG-UK) Consortium | Lighthouse Laboratory Plymouth and Alex Alderton, Roberto Amato, Jeffrey Barrett, Sonia Goncalves, Ewan Harrison, David K. Jackson, Ian Johnston, Dominic Kwiatkowski, Cordelia Langford, John Sillitoe on behalf of the Wellcome Sanger Institute COVID-19 Surveillance Team |
| EPI_ISL_2638976 | Health Services Laboratories | Wellcome Sanger Institute for the COVID-19 Genomics UK (COG-UK) Consortium | Health Services Laboratories and Alex Alderton, Roberto Amato, Jeffrey Barrett, Sonia Goncalves, Ewan Harrison, David K. Jackson, Ian Johnston, Dominic Kwiatkowski, Cordelia Langford, John Sillitoe on behalf of the Wellcome Sanger Institute COVID-19 Surveillance Team |
| EPI_ISL_2638977 | Lighthouse Laboratory Plymouth | Wellcome Sanger Institute for the COVID-19 Genomics UK (COG-UK) Consortium | Lighthouse Laboratory Plymouth and Alex Alderton, Roberto Amato, Jeffrey Barrett, Sonia Goncalves, Ewan Harrison, David K. Jackson, Ian Johnston, Dominic Kwiatkowski, Cordelia Langford, John Sillitoe on behalf of the Wellcome Sanger Institute COVID-19 Surveillance Team |
| EPI_ISL_2638978, EPI_ISL_2638979 | Health Services Laboratories | Wellcome Sanger Institute for the COVID-19 Genomics UK (COG-UK) Consortium | Health Services Laboratories and Alex Alderton, Roberto Amato, Jeffrey Barrett, Sonia Goncalves, Ewan Harrison, David K. Jackson, Ian Johnston, Dominic Kwiatkowski, Cordelia Langford, John Sillitoe on behalf of the Wellcome Sanger Institute COVID-19 Surveillance Team |
| EPI_ISL_2638980, EPI_ISL_2638981 | Lighthouse Lab in Milton Keynes | Wellcome Sanger Institute for the COVID-19 Genomics UK (COG-UK) Consortium | The Lighthouse Lab in Milton Keynes and Alex Alderton, Roberto Amato, Jeffrey Barrett, Sonia Goncalves, Ewan Harrison, David K. Jackson, Ian Johnston, Dominic Kwiatkowski, Cordelia Langford, John Sillitoe on behalf of the Wellcome Sanger Institute COVID-19 Surveillance Team |
| EPI_ISL_2638982 | Health Services Laboratories | Wellcome Sanger Institute for the COVID-19 Genomics UK (COG-UK) Consortium | Health Services Laboratories and Alex Alderton, Roberto Amato, Jeffrey Barrett, Sonia Goncalves, Ewan Harrison, David K. Jackson, Ian Johnston, Dominic Kwiatkowski, Cordelia Langford, John Sillitoe on behalf of the Wellcome Sanger Institute COVID-19 Surveillance Team |
| EPI_ISL_2638983, EPI_ISL_2638984, EPI_ISL_2638985 | Lighthouse Laboratory Plymouth | Wellcome Sanger Institute for the COVID-19 Genomics UK (COG-UK) Consortium | Lighthouse Laboratory Plymouth and Alex Alderton, Roberto Amato, Jeffrey Barrett, Sonia Goncalves, Ewan Harrison, David K. Jackson, Ian Johnston, Dominic Kwiatkowski, Cordelia Langford, John Sillitoe on behalf of the Wellcome Sanger Institute COVID-19 Surveillance Team |
| EPI_ISL_2638986 | Lighthouse Lab in Milton Keynes | Wellcome Sanger Institute for the COVID-19 Genomics UK (COG-UK) Consortium | The Lighthouse Lab in Milton Keynes and Alex Alderton, Roberto Amato, Jeffrey Barrett, Sonia Goncalves, Ewan Harrison, David K. Jackson, Ian Johnston, Dominic Kwiatkowski, Cordelia Langford, John Sillitoe on behalf of the Wellcome Sanger Institute COVID-19 Surveillance Team |
| EPI_ISL_2638987 | Lighthouse Laboratory Plymouth | Wellcome Sanger Institute for the COVID-19 Genomics UK (COG-UK) Consortium | Lighthouse Laboratory Plymouth and Alex Alderton, Roberto Amato, Jeffrey Barrett, Sonia Goncalves, Ewan Harrison, David K. Jackson, Ian Johnston, Dominic Kwiatkowski, Cordelia Langford, John Sillitoe on behalf of the Wellcome Sanger Institute COVID-19 Surveillance Team |
| EPI_ISL_2638988 | Lighthouse Lab in Milton Keynes | Wellcome Sanger Institute for the COVID-19 Genomics UK (COG-UK) Consortium | The Lighthouse Lab in Milton Keynes and Alex Alderton, Roberto Amato, Jeffrey Barrett, Sonia Goncalves, Ewan Harrison, David K. Jackson, Ian Johnston, Dominic Kwiatkowski, Cordelia Langford, John Sillitoe on behalf of the Wellcome Sanger Institute COVID-19 Surveillance Team |
| EPI_ISL_2638989 | Lighthouse Laboratory Plymouth | Wellcome Sanger Institute for the COVID-19 Genomics UK (COG-UK) Consortium | Lighthouse Laboratory Plymouth and Alex Alderton, Roberto Amato, Jeffrey Barrett, Sonia Goncalves, Ewan Harrison, David K. Jackson, Ian Johnston, Dominic Kwiatkowski, Cordelia Langford, John Sillitoe on behalf of the Wellcome Sanger Institute COVID-19 Surveillance Team |
| EPI_ISL_2638990 | Lighthouse Lab in Milton Keynes | Wellcome Sanger Institute for the COVID-19 Genomics UK (COG-UK) Consortium | The Lighthouse Lab in Milton Keynes and Alex Alderton, Roberto Amato, Jeffrey Barrett, Sonia Goncalves, Ewan Harrison, David K. Jackson, Ian Johnston, Dominic Kwiatkowski, Cordelia Langford, John Sillitoe on behalf of the Wellcome Sanger Institute COVID-19 Surveillance Team |
| EPI_ISL_2638991 | Health Services Laboratories | Wellcome Sanger Institute for the COVID-19 Genomics UK (COG-UK) Consortium | Health Services Laboratories and Alex Alderton, Roberto Amato, Jeffrey Barrett, Sonia Goncalves, Ewan Harrison, David K. Jackson, Ian Johnston, Dominic Kwiatkowski, Cordelia Langford, John Sillitoe on behalf of the Wellcome Sanger Institute COVID-19 Surveillance Team |
| EPI_ISL_2638992 | Lighthouse Lab in Milton Keynes | Wellcome Sanger Institute for the COVID-19 Genomics UK (COG-UK) Consortium | The Lighthouse Lab in Milton Keynes and Alex Alderton, Roberto Amato, Jeffrey Barrett, Sonia Goncalves, Ewan Harrison, David K. Jackson, Ian Johnston, Dominic Kwiatkowski, Cordelia Langford, John Sillitoe on behalf of the Wellcome Sanger Institute COVID-19 Surveillance Team |
| EPI_ISL_2638993 | Lighthouse Laboratory Plymouth | Wellcome Sanger Institute for the COVID-19 Genomics UK (COG-UK) Consortium | Lighthouse Laboratory Plymouth and Alex Alderton, Roberto Amato, Jeffrey Barrett, Sonia Goncalves, Ewan Harrison, David K. Jackson, Ian Johnston, Dominic Kwiatkowski, Cordelia Langford, John Sillitoe on behalf of the Wellcome Sanger Institute COVID-19 Surveillance Team |
| EPI_ISL_2638994, EPI_ISL_2638995 | Health Services Laboratories | Wellcome Sanger Institute for the COVID-19 Genomics UK (COG-UK) Consortium | Health Services Laboratories and Alex Alderton, Roberto Amato, Jeffrey Barrett, Sonia Goncalves, Ewan Harrison, David K. Jackson, Ian Johnston, Dominic Kwiatkowski, Cordelia Langford, John Sillitoe on behalf of the Wellcome Sanger Institute COVID-19 Surveillance Team |
| EPI_ISL_2638996 | Lighthouse Laboratory Plymouth | Wellcome Sanger Institute for the COVID-19 Genomics UK (COG-UK) Consortium | Lighthouse Laboratory Plymouth and Alex Alderton, Roberto Amato, Jeffrey Barrett, Sonia Goncalves, Ewan Harrison, David K. Jackson, Ian Johnston, Dominic Kwiatkowski, Cordelia Langford, John Sillitoe on behalf of the Wellcome Sanger Institute COVID-19 Surveillance Team |
| EPI_ISL_2638997 | Lighthouse Lab in Milton Keynes | Wellcome Sanger Institute for the COVID-19 Genomics UK (COG-UK) Consortium | The Lighthouse Lab in Milton Keynes and Alex Alderton, Roberto Amato, Jeffrey Barrett, Sonia Goncalves, Ewan Harrison, David K. Jackson, Ian Johnston, Dominic Kwiatkowski, Cordelia Langford, John Sillitoe on behalf of the Wellcome Sanger Institute COVID-19 Surveillance Team |
| EPI_ISL_2638998, EPI_ISL_2638999, EPI_ISL_2639000 | Lighthouse Laboratory Plymouth | Wellcome Sanger Institute for the COVID-19 Genomics UK (COG-UK) Consortium | Lighthouse Laboratory Plymouth and Alex Alderton, Roberto Amato, Jeffrey Barrett, Sonia Goncalves, Ewan Harrison, David K. Jackson, Ian Johnston, Dominic Kwiatkowski, Cordelia Langford, John Sillitoe on behalf of the Wellcome Sanger Institute COVID-19 Surveillance Team |
| EPI_ISL_2639001 | Health Services Laboratories | Wellcome Sanger Institute for the COVID-19 Genomics UK (COG-UK) Consortium | Health Services Laboratories and Alex Alderton, Roberto Amato, Jeffrey Barrett, Sonia Goncalves, Ewan Harrison, David K. Jackson, Ian Johnston, Dominic Kwiatkowski, Cordelia Langford, John Sillitoe on behalf of the Wellcome Sanger Institute COVID-19 Surveillance Team |
| EPI_ISL_2639002 | Lighthouse Lab in Milton Keynes | Wellcome Sanger Institute for the COVID-19 Genomics UK (COG-UK) Consortium | The Lighthouse Lab in Milton Keynes and Alex Alderton, Roberto Amato, Jeffrey Barrett, Sonia Goncalves, Ewan Harrison, David K. Jackson, Ian Johnston, Dominic Kwiatkowski, Cordelia Langford, John Sillitoe on behalf of the Wellcome Sanger Institute COVID-19 Surveillance Team |
| EPI_ISL_2639003 | Lighthouse Laboratory Plymouth | Wellcome Sanger Institute for the COVID-19 Genomics UK (COG-UK) Consortium | Lighthouse Laboratory Plymouth and Alex Alderton, Roberto Amato, Jeffrey Barrett, Sonia Goncalves, Ewan Harrison, David K. Jackson, Ian Johnston, Dominic Kwiatkowski, Cordelia Langford, John Sillitoe on behalf of the Wellcome Sanger Institute COVID-19 Surveillance Team |
| EPI_ISL_2639004 | Lighthouse Lab in Milton Keynes | Wellcome Sanger Institute for the COVID-19 Genomics UK (COG-UK) Consortium | The Lighthouse Lab in Milton Keynes and Alex Alderton, Roberto Amato, Jeffrey Barrett, Sonia Goncalves, Ewan Harrison, David K. Jackson, Ian Johnston, Dominic Kwiatkowski, Cordelia Langford, John Sillitoe on behalf of the Wellcome Sanger Institute COVID-19 Surveillance Team |
| EPI_ISL_2639005 | Health Services Laboratories | Wellcome Sanger Institute for the COVID-19 Genomics UK (COG-UK) Consortium | Health Services Laboratories and Alex Alderton, Roberto Amato, Jeffrey Barrett, Sonia Goncalves, Ewan Harrison, David K. Jackson, Ian Johnston, Dominic Kwiatkowski, Cordelia Langford, John Sillitoe on behalf of the Wellcome Sanger Institute COVID-19 Surveillance Team |
| EPI_ISL_2639006, EPI_ISL_2639007, EPI_ISL_2639008, EPI_ISL_2639009 | Lighthouse Laboratory Plymouth | Wellcome Sanger Institute for the COVID-19 Genomics UK (COG-UK) Consortium | Lighthouse Laboratory Plymouth and Alex Alderton, Roberto Amato, Jeffrey Barrett, Sonia Goncalves, Ewan Harrison, David K. Jackson, Ian Johnston, Dominic Kwiatkowski, Cordelia Langford, John Sillitoe on behalf of the Wellcome Sanger Institute COVID-19 Surveillance Team |
| EPI_ISL_2639010, EPI_ISL_2639011 | Lighthouse Lab in Milton Keynes | Wellcome Sanger Institute for the COVID-19 Genomics UK (COG-UK) Consortium | The Lighthouse Lab in Milton Keynes and Alex Alderton, Roberto Amato, Jeffrey Barrett, Sonia Goncalves, Ewan Harrison, David K. Jackson, Ian Johnston, Dominic Kwiatkowski, Cordelia Langford, John Sillitoe on behalf of the Wellcome Sanger Institute COVID-19 Surveillance Team |
| EPI_ISL_2639012 | Lighthouse Lab in Alderley Park | Wellcome Sanger Institute for the COVID-19 Genomics UK (COG-UK) Consortium | Jacquelyn Wynn, Mairead Hyland, The Lighthouse Lab in Alderley Park and Alex Alderton, Roberto Amato, Jeffrey Barrett, Sonia Goncalves, Ewan Harrison, David K. Jackson, Ian Johnston, Dominic Kwiatkowski, Cordelia Langford, John Sillitoe on behalf of the Wellcome Sanger Institute COVID-19 Surveillance Team |
| EPI_ISL_2639013 | Lighthouse Laboratory Plymouth | Wellcome Sanger Institute for the COVID-19 Genomics UK (COG-UK) Consortium | Lighthouse Laboratory Plymouth and Alex Alderton, Roberto Amato, Jeffrey Barrett, Sonia Goncalves, Ewan Harrison, David K. Jackson, Ian Johnston, Dominic Kwiatkowski, Cordelia Langford, John Sillitoe on behalf of the Wellcome Sanger Institute COVID-19 Surveillance Team |
| EPI_ISL_2639014 | Lighthouse Lab in Milton Keynes | Wellcome Sanger Institute for the COVID-19 Genomics UK (COG-UK) Consortium | The Lighthouse Lab in Milton Keynes and Alex Alderton, Roberto Amato, Jeffrey Barrett, Sonia Goncalves, Ewan Harrison, David K. Jackson, Ian Johnston, Dominic Kwiatkowski, Cordelia Langford, John Sillitoe on behalf of the Wellcome Sanger Institute COVID-19 Surveillance Team |
| EPI_ISL_2639015 | Lighthouse Laboratory Plymouth | Wellcome Sanger Institute for the COVID-19 Genomics UK (COG-UK) Consortium | Lighthouse Laboratory Plymouth and Alex Alderton, Roberto Amato, Jeffrey Barrett, Sonia Goncalves, Ewan Harrison, David K. Jackson, Ian Johnston, Dominic Kwiatkowski, Cordelia Langford, John Sillitoe on behalf of the Wellcome Sanger Institute COVID-19 Surveillance Team |
| EPI_ISL_2639016 | Lighthouse Lab in Milton Keynes | Wellcome Sanger Institute for the COVID-19 Genomics UK (COG-UK) Consortium | The Lighthouse Lab in Milton Keynes and Alex Alderton, Roberto Amato, Jeffrey Barrett, Sonia Goncalves, Ewan Harrison, David K. Jackson, Ian Johnston, Dominic Kwiatkowski, Cordelia Langford, John Sillitoe on behalf of the Wellcome Sanger Institute COVID-19 Surveillance Team |

| | | | |
|---|---|---|---|
| EPI_ISL_2639017 | Lighthouse Laboratory Plymouth | Wellcome Sanger Institute for the COVID-19 Genomics UK (COG-UK) Consortium | Lighthouse Laboratory Plymouth and Alex Alderton, Roberto Amato, Jeffrey Barrett, Sonia Goncalves, Ewan Harrison, David K. Jackson, Ian Johnston, Dominic Kwiatkowski, Cordelia Langford, John Sillitoe on behalf of the Wellcome Sanger Institute COVID-19 Surveillance Team |
| EPI_ISL_2639018 | Lighthouse Lab in Milton Keynes | Wellcome Sanger Institute for the COVID-19 Genomics UK (COG-UK) Consortium | The Lighthouse Lab in Milton Keynes and Alex Alderton, Roberto Amato, Jeffrey Barrett, Sonia Goncalves, Ewan Harrison, David K. Jackson, Ian Johnston, Dominic Kwiatkowski, Cordelia Langford, John Sillitoe on behalf of the Wellcome Sanger Institute COVID-19 Surveillance Team |
| EPI_ISL_2639019 | Lighthouse Lab in Alderley Park | Wellcome Sanger Institute for the COVID-19 Genomics UK (COG-UK) Consortium | Jacquelyn Wynn, Mairead Hyland, The Lighthouse Lab in Alderley Park and Alex Alderton, Roberto Amato, Jeffrey Barrett, Sonia Goncalves, Ewan Harrison, David K. Jackson, Ian Johnston, Dominic Kwiatkowski, Cordelia Langford, John Sillitoe on behalf of the Wellcome Sanger Institute COVID-19 Surveillance Team |
| EPI_ISL_2639020 | Lighthouse Lab in Milton Keynes | Wellcome Sanger Institute for the COVID-19 Genomics UK (COG-UK) Consortium | The Lighthouse Lab in Milton Keynes and Alex Alderton, Roberto Amato, Jeffrey Barrett, Sonia Goncalves, Ewan Harrison, David K. Jackson, Ian Johnston, Dominic Kwiatkowski, Cordelia Langford, John Sillitoe on behalf of the Wellcome Sanger Institute COVID-19 Surveillance Team |
| EPI_ISL_2639021 | Lighthouse Laboratory Plymouth | Wellcome Sanger Institute for the COVID-19 Genomics UK (COG-UK) Consortium | Lighthouse Laboratory Plymouth and Alex Alderton, Roberto Amato, Jeffrey Barrett, Sonia Goncalves, Ewan Harrison, David K. Jackson, Ian Johnston, Dominic Kwiatkowski, Cordelia Langford, John Sillitoe on behalf of the Wellcome Sanger Institute COVID-19 Surveillance Team |
| EPI_ISL_2639022 | Health Services Laboratories | Wellcome Sanger Institute for the COVID-19 Genomics UK (COG-UK) Consortium | Health Services Laboratories and Alex Alderton, Roberto Amato, Jeffrey Barrett, Sonia Goncalves, Ewan Harrison, David K. Jackson, Ian Johnston, Dominic Kwiatkowski, Cordelia Langford, John Sillitoe on behalf of the Wellcome Sanger Institute COVID-19 Surveillance Team |
| EPI_ISL_2639023, EPI_ISL_2639024, EPI_ISL_2639025, EPI_ISL_2639026, EPI_ISL_2639027 | Lighthouse Laboratory Plymouth | Wellcome Sanger Institute for the COVID-19 Genomics UK (COG-UK) Consortium | Lighthouse Laboratory Plymouth and Alex Alderton, Roberto Amato, Jeffrey Barrett, Sonia Goncalves, Ewan Harrison, David K. Jackson, Ian Johnston, Dominic Kwiatkowski, Cordelia Langford, John Sillitoe on behalf of the Wellcome Sanger Institute COVID-19 Surveillance Team |
| EPI_ISL_2639028 | Lighthouse Lab in Milton Keynes | Wellcome Sanger Institute for the COVID-19 Genomics UK (COG-UK) Consortium | The Lighthouse Lab in Milton Keynes and Alex Alderton, Roberto Amato, Jeffrey Barrett, Sonia Goncalves, Ewan Harrison, David K. Jackson, Ian Johnston, Dominic Kwiatkowski, Cordelia Langford, John Sillitoe on behalf of the Wellcome Sanger Institute COVID-19 Surveillance Team |
| EPI_ISL_2639030 | Health Services Laboratories | Wellcome Sanger Institute for the COVID-19 Genomics UK (COG-UK) Consortium | Health Services Laboratories and Alex Alderton, Roberto Amato, Jeffrey Barrett, Sonia Goncalves, Ewan Harrison, David K. Jackson, Ian Johnston, Dominic Kwiatkowski, Cordelia Langford, John Sillitoe on behalf of the Wellcome Sanger Institute COVID-19 Surveillance Team |
| EPI_ISL_2639031 | Lighthouse Lab in Milton Keynes | Wellcome Sanger Institute for the COVID-19 Genomics UK (COG-UK) Consortium | The Lighthouse Lab in Milton Keynes and Alex Alderton, Roberto Amato, Jeffrey Barrett, Sonia Goncalves, Ewan Harrison, David K. Jackson, Ian Johnston, Dominic Kwiatkowski, Cordelia Langford, John Sillitoe on behalf of the Wellcome Sanger Institute COVID-19 Surveillance Team |
| EPI_ISL_2639032 | Lighthouse Laboratory Plymouth | Wellcome Sanger Institute for the COVID-19 Genomics UK (COG-UK) Consortium | Lighthouse Laboratory Plymouth and Alex Alderton, Roberto Amato, Jeffrey Barrett, Sonia Goncalves, Ewan Harrison, David K. Jackson, Ian Johnston, Dominic Kwiatkowski, Cordelia Langford, John Sillitoe on behalf of the Wellcome Sanger Institute COVID-19 Surveillance Team |
| EPI_ISL_2639033 | Health Services Laboratories | Wellcome Sanger Institute for the COVID-19 Genomics UK (COG-UK) Consortium | Health Services Laboratories and Alex Alderton, Roberto Amato, Jeffrey Barrett, Sonia Goncalves, Ewan Harrison, David K. Jackson, Ian Johnston, Dominic Kwiatkowski, Cordelia Langford, John Sillitoe on behalf of the Wellcome Sanger Institute COVID-19 Surveillance Team |
| EPI_ISL_2639034 | Lighthouse Lab in Milton Keynes | Wellcome Sanger Institute for the COVID-19 Genomics UK (COG-UK) Consortium | The Lighthouse Lab in Milton Keynes and Alex Alderton, Roberto Amato, Jeffrey Barrett, Sonia Goncalves, Ewan Harrison, David K. Jackson, Ian Johnston, Dominic Kwiatkowski, Cordelia Langford, John Sillitoe on behalf of the Wellcome Sanger Institute COVID-19 Surveillance Team |
| EPI_ISL_2639035 | Lighthouse Laboratory Plymouth | Wellcome Sanger Institute for the COVID-19 Genomics UK (COG-UK) Consortium | Lighthouse Laboratory Plymouth and Alex Alderton, Roberto Amato, Jeffrey Barrett, Sonia Goncalves, Ewan Harrison, David K. Jackson, Ian Johnston, Dominic Kwiatkowski, Cordelia Langford, John Sillitoe on behalf of the Wellcome Sanger Institute COVID-19 Surveillance Team |
| EPI_ISL_2639036 | Health Services Laboratories | Wellcome Sanger Institute for the COVID-19 Genomics UK (COG-UK) Consortium | Health Services Laboratories and Alex Alderton, Roberto Amato, Jeffrey Barrett, Sonia Goncalves, Ewan Harrison, David K. Jackson, Ian Johnston, Dominic Kwiatkowski, Cordelia Langford, John Sillitoe on behalf of the Wellcome Sanger Institute COVID-19 Surveillance Team |
| EPI_ISL_2639037 | Lighthouse Lab in Milton Keynes | Wellcome Sanger Institute for the COVID-19 Genomics UK (COG-UK) Consortium | The Lighthouse Lab in Milton Keynes and Alex Alderton, Roberto Amato, Jeffrey Barrett, Sonia Goncalves, Ewan Harrison, David K. Jackson, Ian Johnston, Dominic Kwiatkowski, Cordelia Langford, John Sillitoe on behalf of the Wellcome Sanger Institute COVID-19 Surveillance Team |
| EPI_ISL_2639038 | Lighthouse Lab in Alderley Park | Wellcome Sanger Institute for the COVID-19 Genomics UK (COG-UK) Consortium | Jacquelyn Wynn, Mairead Hyland, The Lighthouse Lab in Alderley Park and Alex Alderton, Roberto Amato, Jeffrey Barrett, Sonia Goncalves, Ewan Harrison, David K. Jackson, Ian Johnston, Dominic Kwiatkowski, Cordelia Langford, John Sillitoe on behalf of the Wellcome Sanger Institute COVID-19 Surveillance Team |
| EPI_ISL_2639039, EPI_ISL_2639040, EPI_ISL_2639041 | Lighthouse Laboratory Plymouth | Wellcome Sanger Institute for the COVID-19 Genomics UK (COG-UK) Consortium | Lighthouse Laboratory Plymouth and Alex Alderton, Roberto Amato, Jeffrey Barrett, Sonia Goncalves, Ewan Harrison, David K. Jackson, Ian Johnston, Dominic Kwiatkowski, Cordelia Langford, John Sillitoe on behalf of the Wellcome Sanger Institute COVID-19 Surveillance Team |
| EPI_ISL_2639042, EPI_ISL_2639043 | Health Services Laboratories | Wellcome Sanger Institute for the COVID-19 Genomics UK (COG-UK) Consortium | Health Services Laboratories and Alex Alderton, Roberto Amato, Jeffrey Barrett, Sonia Goncalves, Ewan Harrison, David K. Jackson, Ian Johnston, Dominic Kwiatkowski, Cordelia Langford, John Sillitoe on behalf of the Wellcome Sanger Institute COVID-19 Surveillance Team |
| EPI_ISL_2639044 | Lighthouse Laboratory Plymouth | Wellcome Sanger Institute for the COVID-19 Genomics UK (COG-UK) Consortium | Lighthouse Laboratory Plymouth and Alex Alderton, Roberto Amato, Jeffrey Barrett, Sonia Goncalves, Ewan Harrison, David K. Jackson, Ian Johnston, Dominic Kwiatkowski, Cordelia Langford, John Sillitoe on behalf of the Wellcome Sanger Institute COVID-19 Surveillance Team |
| EPI_ISL_2639045 | Lighthouse Lab in Alderley Park | Wellcome Sanger Institute for the COVID-19 Genomics UK (COG-UK) Consortium | Jacquelyn Wynn, Mairead Hyland, The Lighthouse Lab in Alderley Park and Alex Alderton, Roberto Amato, Jeffrey Barrett, Sonia Goncalves, Ewan Harrison, David K. Jackson, Ian Johnston, Dominic Kwiatkowski, Cordelia Langford, John Sillitoe on behalf of the Wellcome Sanger Institute COVID-19 Surveillance Team |
| EPI_ISL_2639046, EPI_ISL_2639047, EPI_ISL_2639048, EPI_ISL_2639049, EPI_ISL_2639050 | Lighthouse Laboratory Plymouth | Wellcome Sanger Institute for the COVID-19 Genomics UK (COG-UK) Consortium | Lighthouse Laboratory Plymouth and Alex Alderton, Roberto Amato, Jeffrey Barrett, Sonia Goncalves, Ewan Harrison, David K. Jackson, Ian Johnston, Dominic Kwiatkowski, Cordelia Langford, John Sillitoe on behalf of the Wellcome Sanger Institute COVID-19 Surveillance Team |
| EPI_ISL_2639051 | Health Services Laboratories | Wellcome Sanger Institute for the COVID-19 Genomics UK (COG-UK) Consortium | Health Services Laboratories and Alex Alderton, Roberto Amato, Jeffrey Barrett, Sonia Goncalves, Ewan Harrison, David K. Jackson, Ian Johnston, Dominic Kwiatkowski, Cordelia Langford, John Sillitoe on behalf of the Wellcome Sanger Institute COVID-19 Surveillance Team |
| EPI_ISL_2639052 | Lighthouse Laboratory Plymouth | Wellcome Sanger Institute for the COVID-19 Genomics UK (COG-UK) Consortium | Lighthouse Laboratory Plymouth and Alex Alderton, Roberto Amato, Jeffrey Barrett, Sonia Goncalves, Ewan Harrison, David K. Jackson, Ian Johnston, Dominic Kwiatkowski, Cordelia Langford, John Sillitoe on behalf of the Wellcome Sanger Institute COVID-19 Surveillance Team |
| EPI_ISL_2639053 | Health Services Laboratories | Wellcome Sanger Institute for the COVID-19 Genomics UK (COG-UK) Consortium | Health Services Laboratories and Alex Alderton, Roberto Amato, Jeffrey Barrett, Sonia Goncalves, Ewan Harrison, David K. Jackson, Ian Johnston, Dominic Kwiatkowski, Cordelia Langford, John Sillitoe on behalf of the Wellcome Sanger Institute COVID-19 Surveillance Team |
| EPI_ISL_2639054 | Lighthouse Lab in Milton Keynes | Wellcome Sanger Institute for the COVID-19 Genomics UK (COG-UK) Consortium | The Lighthouse Lab in Milton Keynes and Alex Alderton, Roberto Amato, Jeffrey Barrett, Sonia Goncalves, Ewan Harrison, David K. Jackson, Ian Johnston, Dominic Kwiatkowski, Cordelia Langford, John Sillitoe on behalf of the Wellcome Sanger Institute COVID-19 Surveillance Team |
| EPI_ISL_2639055, EPI_ISL_2639056 | Lighthouse Laboratory Plymouth | Wellcome Sanger Institute for the COVID-19 Genomics UK (COG-UK) Consortium | Lighthouse Laboratory Plymouth and Alex Alderton, Roberto Amato, Jeffrey Barrett, Sonia Goncalves, Ewan Harrison, David K. Jackson, Ian Johnston, Dominic Kwiatkowski, Cordelia Langford, John Sillitoe on behalf of the Wellcome Sanger Institute COVID-19 Surveillance Team |
| EPI_ISL_2639057 | Lighthouse Lab in Milton Keynes | Wellcome Sanger Institute for the COVID-19 Genomics UK (COG-UK) Consortium | The Lighthouse Lab in Milton Keynes and Alex Alderton, Roberto Amato, Jeffrey Barrett, Sonia Goncalves, Ewan Harrison, David K. Jackson, Ian Johnston, Dominic Kwiatkowski, Cordelia Langford, John Sillitoe on behalf of the Wellcome Sanger Institute COVID-19 Surveillance Team |
| EPI_ISL_2639058 | Lighthouse Laboratory Plymouth | Wellcome Sanger Institute for the COVID-19 Genomics UK (COG-UK) Consortium | Lighthouse Laboratory Plymouth and Alex Alderton, Roberto Amato, Jeffrey Barrett, Sonia Goncalves, Ewan Harrison, David K. Jackson, Ian Johnston, Dominic Kwiatkowski, Cordelia Langford, John Sillitoe on behalf of the Wellcome Sanger Institute COVID-19 Surveillance Team |
| EPI_ISL_2639059 | Health Services Laboratories | Wellcome Sanger Institute for the COVID-19 Genomics UK (COG-UK) Consortium | Health Services Laboratories and Alex Alderton, Roberto Amato, Jeffrey Barrett, Sonia Goncalves, Ewan Harrison, David K. Jackson, Ian Johnston, Dominic Kwiatkowski, Cordelia Langford, John Sillitoe on behalf of the Wellcome Sanger Institute COVID-19 Surveillance Team |
| EPI_ISL_2639060 | Lighthouse Lab in Milton Keynes | Wellcome Sanger Institute for the COVID-19 Genomics UK (COG-UK) Consortium | The Lighthouse Lab in Milton Keynes and Alex Alderton, Roberto Amato, Jeffrey Barrett, Sonia Goncalves, Ewan Harrison, David K. Jackson, Ian Johnston, Dominic Kwiatkowski, Cordelia Langford, John Sillitoe on behalf of the Wellcome Sanger Institute COVID-19 Surveillance Team |
| EPI_ISL_2639061 | Lighthouse Lab in Alderley Park | Wellcome Sanger Institute for the COVID-19 Genomics UK (COG-UK) Consortium | Jacquelyn Wynn, Mairead Hyland, The Lighthouse Lab in Alderley Park and Alex Alderton, Roberto Amato, Jeffrey Barrett, Sonia Goncalves, Ewan Harrison, David K. Jackson, Ian Johnston, Dominic Kwiatkowski, Cordelia Langford, John Sillitoe on behalf of the Wellcome Sanger Institute COVID-19 Surveillance Team |
| EPI_ISL_2639062 | Lighthouse Laboratory Plymouth | Wellcome Sanger Institute for the COVID-19 Genomics UK (COG-UK) Consortium | Lighthouse Laboratory Plymouth and Alex Alderton, Roberto Amato, Jeffrey Barrett, Sonia Goncalves, Ewan Harrison, David K. Jackson, Ian Johnston, Dominic Kwiatkowski, Cordelia Langford, John Sillitoe on behalf of the Wellcome Sanger Institute COVID-19 Surveillance Team |
| EPI_ISL_2639063 | Lighthouse Lab in Milton Keynes | Wellcome Sanger Institute for the COVID-19 Genomics UK (COG-UK) Consortium | The Lighthouse Lab in Milton Keynes and Alex Alderton, Roberto Amato, Jeffrey Barrett, Sonia Goncalves, Ewan Harrison, David K. Jackson, Ian Johnston, Dominic Kwiatkowski, Cordelia Langford, John Sillitoe on behalf of the Wellcome Sanger Institute COVID-19 Surveillance Team |
| EPI_ISL_2639064 | Lighthouse Lab in Alderley Park | Wellcome Sanger Institute for the COVID-19 Genomics UK (COG-UK) Consortium | Jacquelyn Wynn, Mairead Hyland, The Lighthouse Lab in Alderley Park and Alex Alderton, Roberto Amato, Jeffrey Barrett, Sonia Goncalves, Ewan Harrison, David K. Jackson, Ian Johnston, Dominic Kwiatkowski, Cordelia Langford, John Sillitoe on behalf of the Wellcome Sanger Institute COVID-19 Surveillance Team |
| EPI_ISL_2639065 | Lighthouse Laboratory Plymouth | Wellcome Sanger Institute for the COVID-19 Genomics UK (COG-UK) Consortium | Lighthouse Laboratory Plymouth and Alex Alderton, Roberto Amato, Jeffrey Barrett, Sonia Goncalves, Ewan Harrison, David K. Jackson, Ian Johnston, Dominic Kwiatkowski, Cordelia Langford, John Sillitoe on behalf of the Wellcome Sanger Institute COVID-19 Surveillance Team |
| EPI_ISL_2639066 | Health Services Laboratories | Wellcome Sanger Institute for the COVID-19 Genomics UK (COG-UK) Consortium | Health Services Laboratories and Alex Alderton, Roberto Amato, Jeffrey Barrett, Sonia Goncalves, Ewan Harrison, David K. Jackson, Ian Johnston, Dominic Kwiatkowski, Cordelia Langford, John Sillitoe on behalf of the Wellcome Sanger Institute COVID-19 Surveillance Team |
| EPI_ISL_2639067, EPI_ISL_2639068 | Lighthouse Laboratory Plymouth | Wellcome Sanger Institute for the COVID-19 Genomics UK (COG-UK) Consortium | Lighthouse Laboratory Plymouth and Alex Alderton, Roberto Amato, Jeffrey Barrett, Sonia Goncalves, Ewan Harrison, David K. Jackson, Ian Johnston, Dominic Kwiatkowski, Cordelia Langford, John Sillitoe on behalf of the Wellcome Sanger Institute COVID-19 Surveillance Team |
| EPI_ISL_2639069 | Lighthouse Lab in Milton Keynes | Wellcome Sanger Institute for the COVID-19 Genomics UK (COG-UK) Consortium | The Lighthouse Lab in Milton Keynes and Alex Alderton, Roberto Amato, Jeffrey Barrett, Sonia Goncalves, Ewan Harrison, David K. Jackson, Ian Johnston, Dominic Kwiatkowski, Cordelia Langford, John Sillitoe on behalf of the Wellcome Sanger Institute COVID-19 Surveillance Team |
| EPI_ISL_2639070, EPI_ISL_2639071 | Lighthouse Laboratory Plymouth | Wellcome Sanger Institute for the COVID-19 Genomics UK (COG-UK) Consortium | Lighthouse Laboratory Plymouth and Alex Alderton, Roberto Amato, Jeffrey Barrett, Sonia Goncalves, Ewan Harrison, David K. Jackson, Ian Johnston, Dominic Kwiatkowski, Cordelia Langford, John Sillitoe on behalf of the Wellcome Sanger Institute COVID-19 Surveillance Team |
| EPI_ISL_2639072, EPI_ISL_2639073 | Health Services Laboratories | Wellcome Sanger Institute for the COVID-19 Genomics UK (COG-UK) Consortium | Health Services Laboratories and Alex Alderton, Roberto Amato, Jeffrey Barrett, Sonia Goncalves, Ewan Harrison, David K. Jackson, Ian Johnston, Dominic Kwiatkowski, Cordelia Langford, John Sillitoe on behalf of the Wellcome Sanger Institute COVID-19 Surveillance Team |
| EPI_ISL_2639074 | Lighthouse Lab in Milton Keynes | Wellcome Sanger Institute for the COVID-19 Genomics UK (COG-UK) Consortium | The Lighthouse Lab in Milton Keynes and Alex Alderton, Roberto Amato, Jeffrey Barrett, Sonia Goncalves, Ewan Harrison, David K. Jackson, Ian Johnston, Dominic Kwiatkowski, Cordelia Langford, John Sillitoe on behalf of the Wellcome Sanger Institute COVID-19 Surveillance Team |

| | | | |
|---|---|---|---|
| EPI_ISL_2639075, EPI_ISL_2639076, EPI_ISL_2639077, EPI_ISL_2639078, EPI_ISL_2639079, EPI_ISL_2639080, EPI_ISL_2639081, EPI_ISL_2639082, EPI_ISL_2639083 | Lighthouse Laboratory Plymouth | Wellcome Sanger Institute for the COVID-19 Genomics UK (COG-UK) Consortium | Lighthouse Laboratory Plymouth and Alex Alderton, Roberto Amato, Jeffrey Barrett, Sonia Goncalves, Ewan Harrison, David K. Jackson, Ian Johnston, Dominic Kwiatkowski, Cordelia Langford, John Sillitoe on behalf of the Wellcome Sanger Institute COVID-19 Surveillance Team |
| EPI_ISL_2639084 | Health Services Laboratories | Wellcome Sanger Institute for the COVID-19 Genomics UK (COG-UK) Consortium | Health Services Laboratories and Alex Alderton, Roberto Amato, Jeffrey Barrett, Sonia Goncalves, Ewan Harrison, David K. Jackson, Ian Johnston, Dominic Kwiatkowski, Cordelia Langford, John Sillitoe on behalf of the Wellcome Sanger Institute COVID-19 Surveillance Team |
| EPI_ISL_2639085 | Lighthouse Laboratory Plymouth | Wellcome Sanger Institute for the COVID-19 Genomics UK (COG-UK) Consortium | Lighthouse Laboratory Plymouth and Alex Alderton, Roberto Amato, Jeffrey Barrett, Sonia Goncalves, Ewan Harrison, David K. Jackson, Ian Johnston, Dominic Kwiatkowski, Cordelia Langford, John Sillitoe on behalf of the Wellcome Sanger Institute COVID-19 Surveillance Team |
| EPI_ISL_2639086 | Lighthouse Lab in Alderley Park | Wellcome Sanger Institute for the COVID-19 Genomics UK (COG-UK) Consortium | Jacquelyn Wynn, Mairead Hyland, The Lighthouse Lab in Alderley Park and Alex Alderton, Roberto Amato, Jeffrey Barrett, Sonia Goncalves, Ewan Harrison, David K. Jackson, Ian Johnston, Dominic Kwiatkowski, Cordelia Langford, John Sillitoe on behalf of the Wellcome Sanger Institute COVID-19 Surveillance Team |
| EPI_ISL_2639087 | Lighthouse Laboratory Plymouth | Wellcome Sanger Institute for the COVID-19 Genomics UK (COG-UK) Consortium | Lighthouse Laboratory Plymouth and Alex Alderton, Roberto Amato, Jeffrey Barrett, Sonia Goncalves, Ewan Harrison, David K. Jackson, Ian Johnston, Dominic Kwiatkowski, Cordelia Langford, John Sillitoe on behalf of the Wellcome Sanger Institute COVID-19 Surveillance Team |
| EPI_ISL_2639088 | Lighthouse Lab in Milton Keynes | Wellcome Sanger Institute for the COVID-19 Genomics UK (COG-UK) Consortium | The Lighthouse Lab in Milton Keynes and Alex Alderton, Roberto Amato, Jeffrey Barrett, Sonia Goncalves, Ewan Harrison, David K. Jackson, Ian Johnston, Dominic Kwiatkowski, Cordelia Langford, John Sillitoe on behalf of the Wellcome Sanger Institute COVID-19 Surveillance Team |
| EPI_ISL_2639089, EPI_ISL_2639090, EPI_ISL_2639091, EPI_ISL_2639093, EPI_ISL_2639094, EPI_ISL_2639095, EPI_ISL_2639096, EPI_ISL_2639097 | Lighthouse Laboratory Plymouth | Wellcome Sanger Institute for the COVID-19 Genomics UK (COG-UK) Consortium | Lighthouse Laboratory Plymouth and Alex Alderton, Roberto Amato, Jeffrey Barrett, Sonia Goncalves, Ewan Harrison, David K. Jackson, Ian Johnston, Dominic Kwiatkowski, Cordelia Langford, John Sillitoe on behalf of the Wellcome Sanger Institute COVID-19 Surveillance Team |
| EPI_ISL_2639098 | Health Services Laboratories | Wellcome Sanger Institute for the COVID-19 Genomics UK (COG-UK) Consortium | Health Services Laboratories and Alex Alderton, Roberto Amato, Jeffrey Barrett, Sonia Goncalves, Ewan Harrison, David K. Jackson, Ian Johnston, Dominic Kwiatkowski, Cordelia Langford, John Sillitoe on behalf of the Wellcome Sanger Institute COVID-19 Surveillance Team |
| EPI_ISL_2639099 | Lighthouse Lab in Milton Keynes | Wellcome Sanger Institute for the COVID-19 Genomics UK (COG-UK) Consortium | The Lighthouse Lab in Milton Keynes and Alex Alderton, Roberto Amato, Jeffrey Barrett, Sonia Goncalves, Ewan Harrison, David K. Jackson, Ian Johnston, Dominic Kwiatkowski, Cordelia Langford, John Sillitoe on behalf of the Wellcome Sanger Institute COVID-19 Surveillance Team |
| EPI_ISL_2639100 | Lighthouse Laboratory Plymouth | Wellcome Sanger Institute for the COVID-19 Genomics UK (COG-UK) Consortium | Lighthouse Laboratory Plymouth and Alex Alderton, Roberto Amato, Jeffrey Barrett, Sonia Goncalves, Ewan Harrison, David K. Jackson, Ian Johnston, Dominic Kwiatkowski, Cordelia Langford, John Sillitoe on behalf of the Wellcome Sanger Institute COVID-19 Surveillance Team |
| EPI_ISL_2639101 | Health Services Laboratories | Wellcome Sanger Institute for the COVID-19 Genomics UK (COG-UK) Consortium | Health Services Laboratories and Alex Alderton, Roberto Amato, Jeffrey Barrett, Sonia Goncalves, Ewan Harrison, David K. Jackson, Ian Johnston, Dominic Kwiatkowski, Cordelia Langford, John Sillitoe on behalf of the Wellcome Sanger Institute COVID-19 Surveillance Team |
| EPI_ISL_2639102, EPI_ISL_2639103, EPI_ISL_2639104, EPI_ISL_2639105, EPI_ISL_2639106, EPI_ISL_2639107 | Lighthouse Laboratory Plymouth | Wellcome Sanger Institute for the COVID-19 Genomics UK (COG-UK) Consortium | Lighthouse Laboratory Plymouth and Alex Alderton, Roberto Amato, Jeffrey Barrett, Sonia Goncalves, Ewan Harrison, David K. Jackson, Ian Johnston, Dominic Kwiatkowski, Cordelia Langford, John Sillitoe on behalf of the Wellcome Sanger Institute COVID-19 Surveillance Team |
| EPI_ISL_2639108 | Health Services Laboratories | Wellcome Sanger Institute for the COVID-19 Genomics UK (COG-UK) Consortium | Health Services Laboratories and Alex Alderton, Roberto Amato, Jeffrey Barrett, Sonia Goncalves, Ewan Harrison, David K. Jackson, Ian Johnston, Dominic Kwiatkowski, Cordelia Langford, John Sillitoe on behalf of the Wellcome Sanger Institute COVID-19 Surveillance Team |
| EPI_ISL_2639109 | Lighthouse Laboratory Plymouth | Wellcome Sanger Institute for the COVID-19 Genomics UK (COG-UK) Consortium | Lighthouse Laboratory Plymouth and Alex Alderton, Roberto Amato, Jeffrey Barrett, Sonia Goncalves, Ewan Harrison, David K. Jackson, Ian Johnston, Dominic Kwiatkowski, Cordelia Langford, John Sillitoe on behalf of the Wellcome Sanger Institute COVID-19 Surveillance Team |
| EPI_ISL_2639110 | Lighthouse Lab in Milton Keynes | Wellcome Sanger Institute for the COVID-19 Genomics UK (COG-UK) Consortium | The Lighthouse Lab in Milton Keynes and Alex Alderton, Roberto Amato, Jeffrey Barrett, Sonia Goncalves, Ewan Harrison, David K. Jackson, Ian Johnston, Dominic Kwiatkowski, Cordelia Langford, John Sillitoe on behalf of the Wellcome Sanger Institute COVID-19 Surveillance Team |
| EPI_ISL_2639111, EPI_ISL_2639112 | Lighthouse Laboratory Plymouth | Wellcome Sanger Institute for the COVID-19 Genomics UK (COG-UK) Consortium | Lighthouse Laboratory Plymouth and Alex Alderton, Roberto Amato, Jeffrey Barrett, Sonia Goncalves, Ewan Harrison, David K. Jackson, Ian Johnston, Dominic Kwiatkowski, Cordelia Langford, John Sillitoe on behalf of the Wellcome Sanger Institute COVID-19 Surveillance Team |
| EPI_ISL_2639113 | Health Services Laboratories | Wellcome Sanger Institute for the COVID-19 Genomics UK (COG-UK) Consortium | Health Services Laboratories and Alex Alderton, Roberto Amato, Jeffrey Barrett, Sonia Goncalves, Ewan Harrison, David K. Jackson, Ian Johnston, Dominic Kwiatkowski, Cordelia Langford, John Sillitoe on behalf of the Wellcome Sanger Institute COVID-19 Surveillance Team |
| EPI_ISL_2639114, EPI_ISL_2639115 | Lighthouse Laboratory Plymouth | Wellcome Sanger Institute for the COVID-19 Genomics UK (COG-UK) Consortium | Lighthouse Laboratory Plymouth and Alex Alderton, Roberto Amato, Jeffrey Barrett, Sonia Goncalves, Ewan Harrison, David K. Jackson, Ian Johnston, Dominic Kwiatkowski, Cordelia Langford, John Sillitoe on behalf of the Wellcome Sanger Institute COVID-19 Surveillance Team |
| EPI_ISL_2639116 | Health Services Laboratories | Wellcome Sanger Institute for the COVID-19 Genomics UK (COG-UK) Consortium | Health Services Laboratories and Alex Alderton, Roberto Amato, Jeffrey Barrett, Sonia Goncalves, Ewan Harrison, David K. Jackson, Ian Johnston, Dominic Kwiatkowski, Cordelia Langford, John Sillitoe on behalf of the Wellcome Sanger Institute COVID-19 Surveillance Team |
| EPI_ISL_2639117 | Lighthouse Laboratory Plymouth | Wellcome Sanger Institute for the COVID-19 Genomics UK (COG-UK) Consortium | Lighthouse Laboratory Plymouth and Alex Alderton, Roberto Amato, Jeffrey Barrett, Sonia Goncalves, Ewan Harrison, David K. Jackson, Ian Johnston, Dominic Kwiatkowski, Cordelia Langford, John Sillitoe on behalf of the Wellcome Sanger Institute COVID-19 Surveillance Team |
| EPI_ISL_2639118 | Lighthouse Lab in Milton Keynes | Wellcome Sanger Institute for the COVID-19 Genomics UK (COG-UK) Consortium | The Lighthouse Lab in Milton Keynes and Alex Alderton, Roberto Amato, Jeffrey Barrett, Sonia Goncalves, Ewan Harrison, David K. Jackson, Ian Johnston, Dominic Kwiatkowski, Cordelia Langford, John Sillitoe on behalf of the Wellcome Sanger Institute COVID-19 Surveillance Team |
| EPI_ISL_2639119 | Lighthouse Laboratory Plymouth | Wellcome Sanger Institute for the COVID-19 Genomics UK (COG-UK) Consortium | Lighthouse Laboratory Plymouth and Alex Alderton, Roberto Amato, Jeffrey Barrett, Sonia Goncalves, Ewan Harrison, David K. Jackson, Ian Johnston, Dominic Kwiatkowski, Cordelia Langford, John Sillitoe on behalf of the Wellcome Sanger Institute COVID-19 Surveillance Team |
| EPI_ISL_2639120 | Lighthouse Lab in Milton Keynes | Wellcome Sanger Institute for the COVID-19 Genomics UK (COG-UK) Consortium | The Lighthouse Lab in Milton Keynes and Alex Alderton, Roberto Amato, Jeffrey Barrett, Sonia Goncalves, Ewan Harrison, David K. Jackson, Ian Johnston, Dominic Kwiatkowski, Cordelia Langford, John Sillitoe on behalf of the Wellcome Sanger Institute COVID-19 Surveillance Team |
| EPI_ISL_2639121 | Health Services Laboratories | Wellcome Sanger Institute for the COVID-19 Genomics UK (COG-UK) Consortium | Health Services Laboratories and Alex Alderton, Roberto Amato, Jeffrey Barrett, Sonia Goncalves, Ewan Harrison, David K. Jackson, Ian Johnston, Dominic Kwiatkowski, Cordelia Langford, John Sillitoe on behalf of the Wellcome Sanger Institute COVID-19 Surveillance Team |
| EPI_ISL_2639122, EPI_ISL_2639123, EPI_ISL_2639124, EPI_ISL_2639125, EPI_ISL_2639126, EPI_ISL_2639127, EPI_ISL_2639128 | Lighthouse Laboratory Plymouth | Wellcome Sanger Institute for the COVID-19 Genomics UK (COG-UK) Consortium | Lighthouse Laboratory Plymouth and Alex Alderton, Roberto Amato, Jeffrey Barrett, Sonia Goncalves, Ewan Harrison, David K. Jackson, Ian Johnston, Dominic Kwiatkowski, Cordelia Langford, John Sillitoe on behalf of the Wellcome Sanger Institute COVID-19 Surveillance Team |
| EPI_ISL_2639129 | Lighthouse Lab in Alderley Park | Wellcome Sanger Institute for the COVID-19 Genomics UK (COG-UK) Consortium | Jacquelyn Wynn, Mairead Hyland, The Lighthouse Lab in Alderley Park and Alex Alderton, Roberto Amato, Jeffrey Barrett, Sonia Goncalves, Ewan Harrison, David K. Jackson, Ian Johnston, Dominic Kwiatkowski, Cordelia Langford, John Sillitoe on behalf of the Wellcome Sanger Institute COVID-19 Surveillance Team |
| EPI_ISL_2639130 | Health Services Laboratories | Wellcome Sanger Institute for the COVID-19 Genomics UK (COG-UK) Consortium | Health Services Laboratories and Alex Alderton, Roberto Amato, Jeffrey Barrett, Sonia Goncalves, Ewan Harrison, David K. Jackson, Ian Johnston, Dominic Kwiatkowski, Cordelia Langford, John Sillitoe on behalf of the Wellcome Sanger Institute COVID-19 Surveillance Team |
| EPI_ISL_2639131, EPI_ISL_2639132 | Lighthouse Laboratory Plymouth | Wellcome Sanger Institute for the COVID-19 Genomics UK (COG-UK) Consortium | Lighthouse Laboratory Plymouth and Alex Alderton, Roberto Amato, Jeffrey Barrett, Sonia Goncalves, Ewan Harrison, David K. Jackson, Ian Johnston, Dominic Kwiatkowski, Cordelia Langford, John Sillitoe on behalf of the Wellcome Sanger Institute COVID-19 Surveillance Team |
| EPI_ISL_2639133 | Lighthouse Lab in Alderley Park | Wellcome Sanger Institute for the COVID-19 Genomics UK (COG-UK) Consortium | Jacquelyn Wynn, Mairead Hyland, The Lighthouse Lab in Alderley Park and Alex Alderton, Roberto Amato, Jeffrey Barrett, Sonia Goncalves, Ewan Harrison, David K. Jackson, Ian Johnston, Dominic Kwiatkowski, Cordelia Langford, John Sillitoe on behalf of the Wellcome Sanger Institute COVID-19 Surveillance Team |
| EPI_ISL_2639134, EPI_ISL_2639135, EPI_ISL_2639136 | Lighthouse Laboratory Plymouth | Wellcome Sanger Institute for the COVID-19 Genomics UK (COG-UK) Consortium | Lighthouse Laboratory Plymouth and Alex Alderton, Roberto Amato, Jeffrey Barrett, Sonia Goncalves, Ewan Harrison, David K. Jackson, Ian Johnston, Dominic Kwiatkowski, Cordelia Langford, John Sillitoe on behalf of the Wellcome Sanger Institute COVID-19 Surveillance Team |
| EPI_ISL_2639137 | Health Services Laboratories | Wellcome Sanger Institute for the COVID-19 Genomics UK (COG-UK) Consortium | Health Services Laboratories and Alex Alderton, Roberto Amato, Jeffrey Barrett, Sonia Goncalves, Ewan Harrison, David K. Jackson, Ian Johnston, Dominic Kwiatkowski, Cordelia Langford, John Sillitoe on behalf of the Wellcome Sanger Institute COVID-19 Surveillance Team |
| EPI_ISL_2639138, EPI_ISL_2639139 | Lighthouse Laboratory Plymouth | Wellcome Sanger Institute for the COVID-19 Genomics UK (COG-UK) Consortium | Lighthouse Laboratory Plymouth and Alex Alderton, Roberto Amato, Jeffrey Barrett, Sonia Goncalves, Ewan Harrison, David K. Jackson, Ian Johnston, Dominic Kwiatkowski, Cordelia Langford, John Sillitoe on behalf of the Wellcome Sanger Institute COVID-19 Surveillance Team |
| EPI_ISL_2639140 | Lighthouse Lab in Milton Keynes | Wellcome Sanger Institute for the COVID-19 Genomics UK (COG-UK) Consortium | The Lighthouse Lab in Milton Keynes and Alex Alderton, Roberto Amato, Jeffrey Barrett, Sonia Goncalves, Ewan Harrison, David K. Jackson, Ian Johnston, Dominic Kwiatkowski, Cordelia Langford, John Sillitoe on behalf of the Wellcome Sanger Institute COVID-19 Surveillance Team |
| EPI_ISL_2639141 | Health Services Laboratories | Wellcome Sanger Institute for the COVID-19 Genomics UK (COG-UK) Consortium | Health Services Laboratories and Alex Alderton, Roberto Amato, Jeffrey Barrett, Sonia Goncalves, Ewan Harrison, David K. Jackson, Ian Johnston, Dominic Kwiatkowski, Cordelia Langford, John Sillitoe on behalf of the Wellcome Sanger Institute COVID-19 Surveillance Team |
| EPI_ISL_2639142 | Lighthouse Laboratory Plymouth | Wellcome Sanger Institute for the COVID-19 Genomics UK (COG-UK) Consortium | Lighthouse Laboratory Plymouth and Alex Alderton, Roberto Amato, Jeffrey Barrett, Sonia Goncalves, Ewan Harrison, David K. Jackson, Ian Johnston, Dominic Kwiatkowski, Cordelia Langford, John Sillitoe on behalf of the Wellcome Sanger Institute COVID-19 Surveillance Team |
| EPI_ISL_2639143 | Lighthouse Lab in Milton Keynes | Wellcome Sanger Institute for the COVID-19 Genomics UK (COG-UK) | The Lighthouse Lab in Milton Keynes and Alex Alderton, Roberto Amato, Jeffrey Barrett, Sonia Goncalves, Ewan Harrison, David K. Jackson, Ian Johnston, Dominic Kwiatkowski, Cordelia Langford, John Sillitoe on behalf of the Wellcome Sanger |

| | | | |
|---|---|---|---|
| | | Consortium | Institute COVID-19 Surveillance Team |
| EPI_ISL_2639144 | Health Services Laboratories | Wellcome Sanger Institute for the COVID-19 Genomics UK (COG-UK) Consortium | Health Services Laboratories and Alex Alderton, Roberto Amato, Jeffrey Barrett, Sonia Goncalves, Ewan Harrison, David K. Jackson, Ian Johnston, Dominic Kwiatkowski, Cordelia Langford, John Sillitoe on behalf of the Wellcome Sanger Institute COVID-19 Surveillance Team |
| EPI_ISL_2639145 | Lighthouse Lab in Milton Keynes | Wellcome Sanger Institute for the COVID-19 Genomics UK (COG-UK) Consortium | The Lighthouse Lab in Milton Keynes and Alex Alderton, Roberto Amato, Jeffrey Barrett, Sonia Goncalves, Ewan Harrison, David K. Jackson, Ian Johnston, Dominic Kwiatkowski, Cordelia Langford, John Sillitoe on behalf of the Wellcome Sanger Institute COVID-19 Surveillance Team |
| EPI_ISL_2639146 | Lighthouse Lab in Alderley Park | Wellcome Sanger Institute for the COVID-19 Genomics UK (COG-UK) Consortium | Jacquelyn Wynn, Mairead Hyland, The Lighthouse Lab in Alderley Park and Alex Alderton, Roberto Amato, Jeffrey Barrett, Sonia Goncalves, Ewan Harrison, David K. Jackson, Ian Johnston, Dominic Kwiatkowski, Cordelia Langford, John Sillitoe on behalf of the Wellcome Sanger Institute COVID-19 Surveillance Team |
| EPI_ISL_2639147, EPI_ISL_2639148, EPI_ISL_2639149 | Lighthouse Laboratory Plymouth | Wellcome Sanger Institute for the COVID-19 Genomics UK (COG-UK) Consortium | Lighthouse Laboratory Plymouth and Alex Alderton, Roberto Amato, Jeffrey Barrett, Sonia Goncalves, Ewan Harrison, David K. Jackson, Ian Johnston, Dominic Kwiatkowski, Cordelia Langford, John Sillitoe on behalf of the Wellcome Sanger Institute COVID-19 Surveillance Team |
| EPI_ISL_2639150 | Lighthouse Lab in Alderley Park | Wellcome Sanger Institute for the COVID-19 Genomics UK (COG-UK) Consortium | Jacquelyn Wynn, Mairead Hyland, The Lighthouse Lab in Alderley Park and Alex Alderton, Roberto Amato, Jeffrey Barrett, Sonia Goncalves, Ewan Harrison, David K. Jackson, Ian Johnston, Dominic Kwiatkowski, Cordelia Langford, John Sillitoe on behalf of the Wellcome Sanger Institute COVID-19 Surveillance Team |
| EPI_ISL_2639151, EPI_ISL_2639152, EPI_ISL_2639153, EPI_ISL_2639154, EPI_ISL_2639155 | Lighthouse Laboratory Plymouth | Wellcome Sanger Institute for the COVID-19 Genomics UK (COG-UK) Consortium | Lighthouse Laboratory Plymouth and Alex Alderton, Roberto Amato, Jeffrey Barrett, Sonia Goncalves, Ewan Harrison, David K. Jackson, Ian Johnston, Dominic Kwiatkowski, Cordelia Langford, John Sillitoe on behalf of the Wellcome Sanger Institute COVID-19 Surveillance Team |
| EPI_ISL_2639156 | Lighthouse Lab in Milton Keynes | Wellcome Sanger Institute for the COVID-19 Genomics UK (COG-UK) Consortium | The Lighthouse Lab in Milton Keynes and Alex Alderton, Roberto Amato, Jeffrey Barrett, Sonia Goncalves, Ewan Harrison, David K. Jackson, Ian Johnston, Dominic Kwiatkowski, Cordelia Langford, John Sillitoe on behalf of the Wellcome Sanger Institute COVID-19 Surveillance Team |
| EPI_ISL_2639157 | Health Services Laboratories | Wellcome Sanger Institute for the COVID-19 Genomics UK (COG-UK) Consortium | Health Services Laboratories and Alex Alderton, Roberto Amato, Jeffrey Barrett, Sonia Goncalves, Ewan Harrison, David K. Jackson, Ian Johnston, Dominic Kwiatkowski, Cordelia Langford, John Sillitoe on behalf of the Wellcome Sanger Institute COVID-19 Surveillance Team |
| EPI_ISL_2639158, EPI_ISL_2639159 | Lighthouse Laboratory Plymouth | Wellcome Sanger Institute for the COVID-19 Genomics UK (COG-UK) Consortium | Lighthouse Laboratory Plymouth and Alex Alderton, Roberto Amato, Jeffrey Barrett, Sonia Goncalves, Ewan Harrison, David K. Jackson, Ian Johnston, Dominic Kwiatkowski, Cordelia Langford, John Sillitoe on behalf of the Wellcome Sanger Institute COVID-19 Surveillance Team |
| EPI_ISL_2639160 | Health Services Laboratories | Wellcome Sanger Institute for the COVID-19 Genomics UK (COG-UK) Consortium | Health Services Laboratories and Alex Alderton, Roberto Amato, Jeffrey Barrett, Sonia Goncalves, Ewan Harrison, David K. Jackson, Ian Johnston, Dominic Kwiatkowski, Cordelia Langford, John Sillitoe on behalf of the Wellcome Sanger Institute COVID-19 Surveillance Team |
| EPI_ISL_2639161 | Lighthouse Laboratory Plymouth | Wellcome Sanger Institute for the COVID-19 Genomics UK (COG-UK) Consortium | Lighthouse Laboratory Plymouth and Alex Alderton, Roberto Amato, Jeffrey Barrett, Sonia Goncalves, Ewan Harrison, David K. Jackson, Ian Johnston, Dominic Kwiatkowski, Cordelia Langford, John Sillitoe on behalf of the Wellcome Sanger Institute COVID-19 Surveillance Team |
| EPI_ISL_2639162 | Lighthouse Lab in Milton Keynes | Wellcome Sanger Institute for the COVID-19 Genomics UK (COG-UK) Consortium | The Lighthouse Lab in Milton Keynes and Alex Alderton, Roberto Amato, Jeffrey Barrett, Sonia Goncalves, Ewan Harrison, David K. Jackson, Ian Johnston, Dominic Kwiatkowski, Cordelia Langford, John Sillitoe on behalf of the Wellcome Sanger Institute COVID-19 Surveillance Team |
| EPI_ISL_2639163 | Lighthouse Laboratory Plymouth | Wellcome Sanger Institute for the COVID-19 Genomics UK (COG-UK) Consortium | Lighthouse Laboratory Plymouth and Alex Alderton, Roberto Amato, Jeffrey Barrett, Sonia Goncalves, Ewan Harrison, David K. Jackson, Ian Johnston, Dominic Kwiatkowski, Cordelia Langford, John Sillitoe on behalf of the Wellcome Sanger Institute COVID-19 Surveillance Team |
| EPI_ISL_2639164 | Health Services Laboratories | Wellcome Sanger Institute for the COVID-19 Genomics UK (COG-UK) Consortium | Health Services Laboratories and Alex Alderton, Roberto Amato, Jeffrey Barrett, Sonia Goncalves, Ewan Harrison, David K. Jackson, Ian Johnston, Dominic Kwiatkowski, Cordelia Langford, John Sillitoe on behalf of the Wellcome Sanger Institute COVID-19 Surveillance Team |
| EPI_ISL_2639165 | Lighthouse Laboratory Plymouth | Wellcome Sanger Institute for the COVID-19 Genomics UK (COG-UK) Consortium | Lighthouse Laboratory Plymouth and Alex Alderton, Roberto Amato, Jeffrey Barrett, Sonia Goncalves, Ewan Harrison, David K. Jackson, Ian Johnston, Dominic Kwiatkowski, Cordelia Langford, John Sillitoe on behalf of the Wellcome Sanger Institute COVID-19 Surveillance Team |
| EPI_ISL_2639166 | Lighthouse Lab in Alderley Park | Wellcome Sanger Institute for the COVID-19 Genomics UK (COG-UK) Consortium | Jacquelyn Wynn, Mairead Hyland, The Lighthouse Lab in Alderley Park and Alex Alderton, Roberto Amato, Jeffrey Barrett, Sonia Goncalves, Ewan Harrison, David K. Jackson, Ian Johnston, Dominic Kwiatkowski, Cordelia Langford, John Sillitoe on behalf of the Wellcome Sanger Institute COVID-19 Surveillance Team |
| EPI_ISL_2639167 | Lighthouse Laboratory Plymouth | Wellcome Sanger Institute for the COVID-19 Genomics UK (COG-UK) Consortium | Lighthouse Laboratory Plymouth and Alex Alderton, Roberto Amato, Jeffrey Barrett, Sonia Goncalves, Ewan Harrison, David K. Jackson, Ian Johnston, Dominic Kwiatkowski, Cordelia Langford, John Sillitoe on behalf of the Wellcome Sanger Institute COVID-19 Surveillance Team |
| EPI_ISL_2639168 | Health Services Laboratories | Wellcome Sanger Institute for the COVID-19 Genomics UK (COG-UK) Consortium | Health Services Laboratories and Alex Alderton, Roberto Amato, Jeffrey Barrett, Sonia Goncalves, Ewan Harrison, David K. Jackson, Ian Johnston, Dominic Kwiatkowski, Cordelia Langford, John Sillitoe on behalf of the Wellcome Sanger Institute COVID-19 Surveillance Team |
| EPI_ISL_2639169 | Lighthouse Laboratory Plymouth | Wellcome Sanger Institute for the COVID-19 Genomics UK (COG-UK) Consortium | Lighthouse Laboratory Plymouth and Alex Alderton, Roberto Amato, Jeffrey Barrett, Sonia Goncalves, Ewan Harrison, David K. Jackson, Ian Johnston, Dominic Kwiatkowski, Cordelia Langford, John Sillitoe on behalf of the Wellcome Sanger Institute COVID-19 Surveillance Team |
| EPI_ISL_2639170 | Lighthouse Lab in Milton Keynes | Wellcome Sanger Institute for the COVID-19 Genomics UK (COG-UK) Consortium | The Lighthouse Lab in Milton Keynes and Alex Alderton, Roberto Amato, Jeffrey Barrett, Sonia Goncalves, Ewan Harrison, David K. Jackson, Ian Johnston, Dominic Kwiatkowski, Cordelia Langford, John Sillitoe on behalf of the Wellcome Sanger Institute COVID-19 Surveillance Team |
| EPI_ISL_2639171, EPI_ISL_2639172, EPI_ISL_2639173, EPI_ISL_2639174 | Lighthouse Laboratory Plymouth | Wellcome Sanger Institute for the COVID-19 Genomics UK (COG-UK) Consortium | Lighthouse Laboratory Plymouth and Alex Alderton, Roberto Amato, Jeffrey Barrett, Sonia Goncalves, Ewan Harrison, David K. Jackson, Ian Johnston, Dominic Kwiatkowski, Cordelia Langford, John Sillitoe on behalf of the Wellcome Sanger Institute COVID-19 Surveillance Team |
| EPI_ISL_2639175 | Lighthouse Lab in Alderley Park | Wellcome Sanger Institute for the COVID-19 Genomics UK (COG-UK) Consortium | Jacquelyn Wynn, Mairead Hyland, The Lighthouse Lab in Alderley Park and Alex Alderton, Roberto Amato, Jeffrey Barrett, Sonia Goncalves, Ewan Harrison, David K. Jackson, Ian Johnston, Dominic Kwiatkowski, Cordelia Langford, John Sillitoe on behalf of the Wellcome Sanger Institute COVID-19 Surveillance Team |
| EPI_ISL_2639176 | Lighthouse Lab in Milton Keynes | Wellcome Sanger Institute for the COVID-19 Genomics UK (COG-UK) Consortium | The Lighthouse Lab in Milton Keynes and Alex Alderton, Roberto Amato, Jeffrey Barrett, Sonia Goncalves, Ewan Harrison, David K. Jackson, Ian Johnston, Dominic Kwiatkowski, Cordelia Langford, John Sillitoe on behalf of the Wellcome Sanger Institute COVID-19 Surveillance Team |
| EPI_ISL_2639177, EPI_ISL_2639178, EPI_ISL_2639179, EPI_ISL_2639180, EPI_ISL_2639181 | Lighthouse Laboratory Plymouth | Wellcome Sanger Institute for the COVID-19 Genomics UK (COG-UK) Consortium | Lighthouse Laboratory Plymouth and Alex Alderton, Roberto Amato, Jeffrey Barrett, Sonia Goncalves, Ewan Harrison, David K. Jackson, Ian Johnston, Dominic Kwiatkowski, Cordelia Langford, John Sillitoe on behalf of the Wellcome Sanger Institute COVID-19 Surveillance Team |
| EPI_ISL_2639182, EPI_ISL_2639183 | Health Services Laboratories | Wellcome Sanger Institute for the COVID-19 Genomics UK (COG-UK) Consortium | Health Services Laboratories and Alex Alderton, Roberto Amato, Jeffrey Barrett, Sonia Goncalves, Ewan Harrison, David K. Jackson, Ian Johnston, Dominic Kwiatkowski, Cordelia Langford, John Sillitoe on behalf of the Wellcome Sanger Institute COVID-19 Surveillance Team |
| EPI_ISL_2639184, EPI_ISL_2639185, EPI_ISL_2639186 | Lighthouse Laboratory Plymouth | Wellcome Sanger Institute for the COVID-19 Genomics UK (COG-UK) Consortium | Lighthouse Laboratory Plymouth and Alex Alderton, Roberto Amato, Jeffrey Barrett, Sonia Goncalves, Ewan Harrison, David K. Jackson, Ian Johnston, Dominic Kwiatkowski, Cordelia Langford, John Sillitoe on behalf of the Wellcome Sanger Institute COVID-19 Surveillance Team |
| EPI_ISL_2639187 | Health Services Laboratories | Wellcome Sanger Institute for the COVID-19 Genomics UK (COG-UK) Consortium | Health Services Laboratories and Alex Alderton, Roberto Amato, Jeffrey Barrett, Sonia Goncalves, Ewan Harrison, David K. Jackson, Ian Johnston, Dominic Kwiatkowski, Cordelia Langford, John Sillitoe on behalf of the Wellcome Sanger Institute COVID-19 Surveillance Team |
| EPI_ISL_2639188, EPI_ISL_2639189, EPI_ISL_2639190, EPI_ISL_2639191, EPI_ISL_2639192 | Lighthouse Laboratory Plymouth | Wellcome Sanger Institute for the COVID-19 Genomics UK (COG-UK) Consortium | Lighthouse Laboratory Plymouth and Alex Alderton, Roberto Amato, Jeffrey Barrett, Sonia Goncalves, Ewan Harrison, David K. Jackson, Ian Johnston, Dominic Kwiatkowski, Cordelia Langford, John Sillitoe on behalf of the Wellcome Sanger Institute COVID-19 Surveillance Team |
| EPI_ISL_2639193 | Health Services Laboratories | Wellcome Sanger Institute for the COVID-19 Genomics UK (COG-UK) Consortium | Health Services Laboratories and Alex Alderton, Roberto Amato, Jeffrey Barrett, Sonia Goncalves, Ewan Harrison, David K. Jackson, Ian Johnston, Dominic Kwiatkowski, Cordelia Langford, John Sillitoe on behalf of the Wellcome Sanger Institute COVID-19 Surveillance Team |
| EPI_ISL_2639194 | Lighthouse Laboratory Plymouth | Wellcome Sanger Institute for the COVID-19 Genomics UK (COG-UK) Consortium | Lighthouse Laboratory Plymouth and Alex Alderton, Roberto Amato, Jeffrey Barrett, Sonia Goncalves, Ewan Harrison, David K. Jackson, Ian Johnston, Dominic Kwiatkowski, Cordelia Langford, John Sillitoe on behalf of the Wellcome Sanger Institute COVID-19 Surveillance Team |
| EPI_ISL_2639195 | Lighthouse Lab in Milton Keynes | Wellcome Sanger Institute for the COVID-19 Genomics UK (COG-UK) Consortium | The Lighthouse Lab in Milton Keynes and Alex Alderton, Roberto Amato, Jeffrey Barrett, Sonia Goncalves, Ewan Harrison, David K. Jackson, Ian Johnston, Dominic Kwiatkowski, Cordelia Langford, John Sillitoe on behalf of the Wellcome Sanger Institute COVID-19 Surveillance Team |
| EPI_ISL_2639196 | Lighthouse Laboratory Plymouth | Wellcome Sanger Institute for the COVID-19 Genomics UK (COG-UK) Consortium | Lighthouse Laboratory Plymouth and Alex Alderton, Roberto Amato, Jeffrey Barrett, Sonia Goncalves, Ewan Harrison, David K. Jackson, Ian Johnston, Dominic Kwiatkowski, Cordelia Langford, John Sillitoe on behalf of the Wellcome Sanger Institute COVID-19 Surveillance Team |
| EPI_ISL_2639197, EPI_ISL_2639198 | Lighthouse Lab in Milton Keynes | Wellcome Sanger Institute for the COVID-19 Genomics UK (COG-UK) Consortium | The Lighthouse Lab in Milton Keynes and Alex Alderton, Roberto Amato, Jeffrey Barrett, Sonia Goncalves, Ewan Harrison, David K. Jackson, Ian Johnston, Dominic Kwiatkowski, Cordelia Langford, John Sillitoe on behalf of the Wellcome Sanger Institute COVID-19 Surveillance Team |
| EPI_ISL_2639199, EPI_ISL_2639200, EPI_ISL_2639201 | Lighthouse Laboratory Plymouth | Wellcome Sanger Institute for the COVID-19 Genomics UK (COG-UK) Consortium | Lighthouse Laboratory Plymouth and Alex Alderton, Roberto Amato, Jeffrey Barrett, Sonia Goncalves, Ewan Harrison, David K. Jackson, Ian Johnston, Dominic Kwiatkowski, Cordelia Langford, John Sillitoe on behalf of the Wellcome Sanger Institute COVID-19 Surveillance Team |
| EPI_ISL_2639202 | Lighthouse Lab in Milton Keynes | Wellcome Sanger Institute for the COVID-19 Genomics UK (COG-UK) Consortium | The Lighthouse Lab in Milton Keynes and Alex Alderton, Roberto Amato, Jeffrey Barrett, Sonia Goncalves, Ewan Harrison, David K. Jackson, Ian Johnston, Dominic Kwiatkowski, Cordelia Langford, John Sillitoe on behalf of the Wellcome Sanger Institute COVID-19 Surveillance Team |
| EPI_ISL_2639203, EPI_ISL_2639204 | Health Services Laboratories | Wellcome Sanger Institute for the COVID-19 Genomics UK (COG-UK) Consortium | Health Services Laboratories and Alex Alderton, Roberto Amato, Jeffrey Barrett, Sonia Goncalves, Ewan Harrison, David K. Jackson, Ian Johnston, Dominic Kwiatkowski, Cordelia Langford, John Sillitoe on behalf of the Wellcome Sanger Institute COVID-19 Surveillance Team |
| EPI_ISL_2639205, EPI_ISL_2639206 | Lighthouse Laboratory Plymouth | Wellcome Sanger Institute for the COVID-19 Genomics UK (COG-UK) Consortium | Lighthouse Laboratory Plymouth and Alex Alderton, Roberto Amato, Jeffrey Barrett, Sonia Goncalves, Ewan Harrison, David K. Jackson, Ian Johnston, Dominic Kwiatkowski, Cordelia Langford, John Sillitoe on behalf of the Wellcome Sanger Institute COVID-19 Surveillance Team |
| EPI_ISL_2639207, EPI_ISL_2639208 | Health Services Laboratories | Wellcome Sanger Institute for the COVID-19 Genomics UK (COG-UK) Consortium | Health Services Laboratories and Alex Alderton, Roberto Amato, Jeffrey Barrett, Sonia Goncalves, Ewan Harrison, David K. Jackson, Ian Johnston, Dominic Kwiatkowski, Cordelia Langford, John Sillitoe on behalf of the Wellcome Sanger Institute COVID-19 Surveillance Team |
| EPI_ISL_2639209, EPI_ISL_2639210, | Lighthouse Laboratory Plymouth | Wellcome Sanger Institute for the COVID-19 Genomics UK (COG-UK) Consortium | Lighthouse Laboratory Plymouth and Alex Alderton, Roberto Amato, Jeffrey Barrett, Sonia Goncalves, Ewan Harrison, David K. Jackson, Ian Johnston, Dominic Kwiatkowski, Cordelia Langford, John Sillitoe on behalf of the Wellcome Sanger Institute COVID-19 Surveillance Team |

| | | | |
|---|---|---|---|
| EPI_ISL_2639211, EPI_ISL_2639212 | | | |
| EPI_ISL_2639213 | Health Services Laboratories | Wellcome Sanger Institute for the COVID-19 Genomics UK (COG-UK) Consortium | Health Services Laboratories and Alex Alderton, Roberto Amato, Jeffrey Barrett, Sonia Goncalves, Ewan Harrison, David K. Jackson, Ian Johnston, Dominic Kwiatkowski, Cordelia Langford, John Sillitoe on behalf of the Wellcome Sanger Institute COVID-19 Surveillance Team |
| EPI_ISL_2639214 | Lighthouse Laboratory Plymouth | Wellcome Sanger Institute for the COVID-19 Genomics UK (COG-UK) Consortium | Lighthouse Laboratory Plymouth and Alex Alderton, Roberto Amato, Jeffrey Barrett, Sonia Goncalves, Ewan Harrison, David K. Jackson, Ian Johnston, Dominic Kwiatkowski, Cordelia Langford, John Sillitoe on behalf of the Wellcome Sanger Institute COVID-19 Surveillance Team |
| EPI_ISL_2639215 | Lighthouse Lab in Milton Keynes | Wellcome Sanger Institute for the COVID-19 Genomics UK (COG-UK) Consortium | The Lighthouse Lab in Milton Keynes and Alex Alderton, Roberto Amato, Jeffrey Barrett, Sonia Goncalves, Ewan Harrison, David K. Jackson, Ian Johnston, Dominic Kwiatkowski, Cordelia Langford, John Sillitoe on behalf of the Wellcome Sanger Institute COVID-19 Surveillance Team |
| EPI_ISL_2639216, EPI_ISL_2639217, EPI_ISL_2639218, EPI_ISL_2639219, EPI_ISL_2639220, EPI_ISL_2639221, EPI_ISL_2639222 | Lighthouse Laboratory Plymouth | Wellcome Sanger Institute for the COVID-19 Genomics UK (COG-UK) Consortium | Lighthouse Laboratory Plymouth and Alex Alderton, Roberto Amato, Jeffrey Barrett, Sonia Goncalves, Ewan Harrison, David K. Jackson, Ian Johnston, Dominic Kwiatkowski, Cordelia Langford, John Sillitoe on behalf of the Wellcome Sanger Institute COVID-19 Surveillance Team |
| EPI_ISL_2639223 | Lighthouse Lab in Milton Keynes | Wellcome Sanger Institute for the COVID-19 Genomics UK (COG-UK) Consortium | The Lighthouse Lab in Milton Keynes and Alex Alderton, Roberto Amato, Jeffrey Barrett, Sonia Goncalves, Ewan Harrison, David K. Jackson, Ian Johnston, Dominic Kwiatkowski, Cordelia Langford, John Sillitoe on behalf of the Wellcome Sanger Institute COVID-19 Surveillance Team |
| EPI_ISL_2639224 | Lighthouse Laboratory Plymouth | Wellcome Sanger Institute for the COVID-19 Genomics UK (COG-UK) Consortium | Lighthouse Laboratory Plymouth and Alex Alderton, Roberto Amato, Jeffrey Barrett, Sonia Goncalves, Ewan Harrison, David K. Jackson, Ian Johnston, Dominic Kwiatkowski, Cordelia Langford, John Sillitoe on behalf of the Wellcome Sanger Institute COVID-19 Surveillance Team |
| EPI_ISL_2639225 | Lighthouse Lab in Milton Keynes | Wellcome Sanger Institute for the COVID-19 Genomics UK (COG-UK) Consortium | The Lighthouse Lab in Milton Keynes and Alex Alderton, Roberto Amato, Jeffrey Barrett, Sonia Goncalves, Ewan Harrison, David K. Jackson, Ian Johnston, Dominic Kwiatkowski, Cordelia Langford, John Sillitoe on behalf of the Wellcome Sanger Institute COVID-19 Surveillance Team |
| EPI_ISL_2639226 | Lighthouse Laboratory Plymouth | Wellcome Sanger Institute for the COVID-19 Genomics UK (COG-UK) Consortium | Lighthouse Laboratory Plymouth and Alex Alderton, Roberto Amato, Jeffrey Barrett, Sonia Goncalves, Ewan Harrison, David K. Jackson, Ian Johnston, Dominic Kwiatkowski, Cordelia Langford, John Sillitoe on behalf of the Wellcome Sanger Institute COVID-19 Surveillance Team |
| EPI_ISL_2639227 | Lighthouse Lab in Milton Keynes | Wellcome Sanger Institute for the COVID-19 Genomics UK (COG-UK) Consortium | The Lighthouse Lab in Milton Keynes and Alex Alderton, Roberto Amato, Jeffrey Barrett, Sonia Goncalves, Ewan Harrison, David K. Jackson, Ian Johnston, Dominic Kwiatkowski, Cordelia Langford, John Sillitoe on behalf of the Wellcome Sanger Institute COVID-19 Surveillance Team |
| EPI_ISL_2639228 | Health Services Laboratories | Wellcome Sanger Institute for the COVID-19 Genomics UK (COG-UK) Consortium | Health Services Laboratories and Alex Alderton, Roberto Amato, Jeffrey Barrett, Sonia Goncalves, Ewan Harrison, David K. Jackson, Ian Johnston, Dominic Kwiatkowski, Cordelia Langford, John Sillitoe on behalf of the Wellcome Sanger Institute COVID-19 Surveillance Team |
| EPI_ISL_2639229 | Lighthouse Laboratory Plymouth | Wellcome Sanger Institute for the COVID-19 Genomics UK (COG-UK) Consortium | Lighthouse Laboratory Plymouth and Alex Alderton, Roberto Amato, Jeffrey Barrett, Sonia Goncalves, Ewan Harrison, David K. Jackson, Ian Johnston, Dominic Kwiatkowski, Cordelia Langford, John Sillitoe on behalf of the Wellcome Sanger Institute COVID-19 Surveillance Team |
| EPI_ISL_2639230 | Health Services Laboratories | Wellcome Sanger Institute for the COVID-19 Genomics UK (COG-UK) Consortium | Health Services Laboratories and Alex Alderton, Roberto Amato, Jeffrey Barrett, Sonia Goncalves, Ewan Harrison, David K. Jackson, Ian Johnston, Dominic Kwiatkowski, Cordelia Langford, John Sillitoe on behalf of the Wellcome Sanger Institute COVID-19 Surveillance Team |
| EPI_ISL_2639231, EPI_ISL_2639232, EPI_ISL_2639234, EPI_ISL_2639235, EPI_ISL_2639236, EPI_ISL_2639237 | Lighthouse Laboratory Plymouth | Wellcome Sanger Institute for the COVID-19 Genomics UK (COG-UK) Consortium | Lighthouse Laboratory Plymouth and Alex Alderton, Roberto Amato, Jeffrey Barrett, Sonia Goncalves, Ewan Harrison, David K. Jackson, Ian Johnston, Dominic Kwiatkowski, Cordelia Langford, John Sillitoe on behalf of the Wellcome Sanger Institute COVID-19 Surveillance Team |
| EPI_ISL_2639238 | Lighthouse Lab in Alderley Park | Wellcome Sanger Institute for the COVID-19 Genomics UK (COG-UK) Consortium | Jacquelyn Wynn, Mairead Hyland, The Lighthouse Lab in Alderley Park and Alex Alderton, Roberto Amato, Jeffrey Barrett, Sonia Goncalves, Ewan Harrison, David K. Jackson, Ian Johnston, Dominic Kwiatkowski, Cordelia Langford, John Sillitoe on behalf of the Wellcome Sanger Institute COVID-19 Surveillance Team |
| EPI_ISL_2639239, EPI_ISL_2639240 | Lighthouse Laboratory Plymouth | Wellcome Sanger Institute for the COVID-19 Genomics UK (COG-UK) Consortium | Lighthouse Laboratory Plymouth and Alex Alderton, Roberto Amato, Jeffrey Barrett, Sonia Goncalves, Ewan Harrison, David K. Jackson, Ian Johnston, Dominic Kwiatkowski, Cordelia Langford, John Sillitoe on behalf of the Wellcome Sanger Institute COVID-19 Surveillance Team |
| EPI_ISL_2639241 | Health Services Laboratories | Wellcome Sanger Institute for the COVID-19 Genomics UK (COG-UK) Consortium | Health Services Laboratories and Alex Alderton, Roberto Amato, Jeffrey Barrett, Sonia Goncalves, Ewan Harrison, David K. Jackson, Ian Johnston, Dominic Kwiatkowski, Cordelia Langford, John Sillitoe on behalf of the Wellcome Sanger Institute COVID-19 Surveillance Team |
| EPI_ISL_2639242 | Lighthouse Lab in Milton Keynes | Wellcome Sanger Institute for the COVID-19 Genomics UK (COG-UK) Consortium | The Lighthouse Lab in Milton Keynes and Alex Alderton, Roberto Amato, Jeffrey Barrett, Sonia Goncalves, Ewan Harrison, David K. Jackson, Ian Johnston, Dominic Kwiatkowski, Cordelia Langford, John Sillitoe on behalf of the Wellcome Sanger Institute COVID-19 Surveillance Team |
| EPI_ISL_2639243, EPI_ISL_2639244 | Lighthouse Laboratory Plymouth | Wellcome Sanger Institute for the COVID-19 Genomics UK (COG-UK) Consortium | Lighthouse Laboratory Plymouth and Alex Alderton, Roberto Amato, Jeffrey Barrett, Sonia Goncalves, Ewan Harrison, David K. Jackson, Ian Johnston, Dominic Kwiatkowski, Cordelia Langford, John Sillitoe on behalf of the Wellcome Sanger Institute COVID-19 Surveillance Team |
| EPI_ISL_2639245 | Health Services Laboratories | Wellcome Sanger Institute for the COVID-19 Genomics UK (COG-UK) Consortium | Health Services Laboratories and Alex Alderton, Roberto Amato, Jeffrey Barrett, Sonia Goncalves, Ewan Harrison, David K. Jackson, Ian Johnston, Dominic Kwiatkowski, Cordelia Langford, John Sillitoe on behalf of the Wellcome Sanger Institute COVID-19 Surveillance Team |
| EPI_ISL_2639246 | Lighthouse Laboratory Plymouth | Wellcome Sanger Institute for the COVID-19 Genomics UK (COG-UK) Consortium | Lighthouse Laboratory Plymouth and Alex Alderton, Roberto Amato, Jeffrey Barrett, Sonia Goncalves, Ewan Harrison, David K. Jackson, Ian Johnston, Dominic Kwiatkowski, Cordelia Langford, John Sillitoe on behalf of the Wellcome Sanger Institute COVID-19 Surveillance Team |
| EPI_ISL_2639247 | Lighthouse Lab in Milton Keynes | Wellcome Sanger Institute for the COVID-19 Genomics UK (COG-UK) Consortium | The Lighthouse Lab in Milton Keynes and Alex Alderton, Roberto Amato, Jeffrey Barrett, Sonia Goncalves, Ewan Harrison, David K. Jackson, Ian Johnston, Dominic Kwiatkowski, Cordelia Langford, John Sillitoe on behalf of the Wellcome Sanger Institute COVID-19 Surveillance Team |
| EPI_ISL_2639248 | Lighthouse Laboratory Plymouth | Wellcome Sanger Institute for the COVID-19 Genomics UK (COG-UK) Consortium | Lighthouse Laboratory Plymouth and Alex Alderton, Roberto Amato, Jeffrey Barrett, Sonia Goncalves, Ewan Harrison, David K. Jackson, Ian Johnston, Dominic Kwiatkowski, Cordelia Langford, John Sillitoe on behalf of the Wellcome Sanger Institute COVID-19 Surveillance Team |
| EPI_ISL_2639249 | Lighthouse Lab in Alderley Park | Wellcome Sanger Institute for the COVID-19 Genomics UK (COG-UK) Consortium | Jacquelyn Wynn, Mairead Hyland, The Lighthouse Lab in Alderley Park and Alex Alderton, Roberto Amato, Jeffrey Barrett, Sonia Goncalves, Ewan Harrison, David K. Jackson, Ian Johnston, Dominic Kwiatkowski, Cordelia Langford, John Sillitoe on behalf of the Wellcome Sanger Institute COVID-19 Surveillance Team |
| EPI_ISL_2639250, EPI_ISL_2639251 | Lighthouse Laboratory Plymouth | Wellcome Sanger Institute for the COVID-19 Genomics UK (COG-UK) Consortium | Lighthouse Laboratory Plymouth and Alex Alderton, Roberto Amato, Jeffrey Barrett, Sonia Goncalves, Ewan Harrison, David K. Jackson, Ian Johnston, Dominic Kwiatkowski, Cordelia Langford, John Sillitoe on behalf of the Wellcome Sanger Institute COVID-19 Surveillance Team |
| EPI_ISL_2639252 | Lighthouse Lab in Milton Keynes | Wellcome Sanger Institute for the COVID-19 Genomics UK (COG-UK) Consortium | The Lighthouse Lab in Milton Keynes and Alex Alderton, Roberto Amato, Jeffrey Barrett, Sonia Goncalves, Ewan Harrison, David K. Jackson, Ian Johnston, Dominic Kwiatkowski, Cordelia Langford, John Sillitoe on behalf of the Wellcome Sanger Institute COVID-19 Surveillance Team |
| EPI_ISL_2639253 | Lighthouse Laboratory Plymouth | Wellcome Sanger Institute for the COVID-19 Genomics UK (COG-UK) Consortium | Lighthouse Laboratory Plymouth and Alex Alderton, Roberto Amato, Jeffrey Barrett, Sonia Goncalves, Ewan Harrison, David K. Jackson, Ian Johnston, Dominic Kwiatkowski, Cordelia Langford, John Sillitoe on behalf of the Wellcome Sanger Institute COVID-19 Surveillance Team |
| EPI_ISL_2639254 | Lighthouse Lab in Milton Keynes | Wellcome Sanger Institute for the COVID-19 Genomics UK (COG-UK) Consortium | The Lighthouse Lab in Milton Keynes and Alex Alderton, Roberto Amato, Jeffrey Barrett, Sonia Goncalves, Ewan Harrison, David K. Jackson, Ian Johnston, Dominic Kwiatkowski, Cordelia Langford, John Sillitoe on behalf of the Wellcome Sanger Institute COVID-19 Surveillance Team |
| EPI_ISL_2639255 | Lighthouse Laboratory Plymouth | Wellcome Sanger Institute for the COVID-19 Genomics UK (COG-UK) Consortium | Lighthouse Laboratory Plymouth and Alex Alderton, Roberto Amato, Jeffrey Barrett, Sonia Goncalves, Ewan Harrison, David K. Jackson, Ian Johnston, Dominic Kwiatkowski, Cordelia Langford, John Sillitoe on behalf of the Wellcome Sanger Institute COVID-19 Surveillance Team |

EPI_ISL_2639256, EPI_ISL_2639257, EPI_ISL_2639258, EPI_ISL_2639259, EPI_ISL_2639260, EPI_ISL_2639261, EPI_ISL_2639262, EPI_ISL_2639263, EPI_ISL_2639264, EPI_ISL_2639265, EPI_ISL_2639266, EPI_ISL_2639267, EPI_ISL_2639268, EPI_ISL_2639269, EPI_ISL_2639270, EPI_ISL_2639271, EPI_ISL_2639272, EPI_ISL_2639273, EPI_ISL_2639274, EPI_ISL_2639275, EPI_ISL_2639276, EPI_ISL_2639277,
EPI_ISL_2639278, EPI_ISL_2639279, EPI_ISL_2639280, EPI_ISL_2639281, EPI_ISL_2639282, EPI_ISL_2639283, EPI_ISL_2639284, EPI_ISL_2639285, EPI_ISL_2639286, EPI_ISL_2639287, EPI_ISL_2639288, EPI_ISL_2639289, EPI_ISL_2639290, EPI_ISL_2639291, EPI_ISL_2639292, EPI_ISL_2639293, EPI_ISL_2639294, EPI_ISL_2639295, EPI_ISL_2639296, EPI_ISL_2639297, EPI_ISL_2639298, EPI_ISL_2639299,
EPI_ISL_2639300, EPI_ISL_2639301, EPI_ISL_2639302, EPI_ISL_2639303, EPI_ISL_2639304, EPI_ISL_2639305, EPI_ISL_2639306, EPI_ISL_2639307, EPI_ISL_2639308, EPI_ISL_2639309, EPI_ISL_2639310, EPI_ISL_2639311, EPI_ISL_2639312, EPI_ISL_2639313, EPI_ISL_2639314, EPI_ISL_2639315, EPI_ISL_2639316, EPI_ISL_2639317, EPI_ISL_2639318, EPI_ISL_2639319, EPI_ISL_2639320, EPI_ISL_2639321,
EPI_ISL_2639322, EPI_ISL_2639323, EPI_ISL_2639324, EPI_ISL_2639325, EPI_ISL_2639326, EPI_ISL_2639327, EPI_ISL_2639328, EPI_ISL_2639329, EPI_ISL_2639331, EPI_ISL_2639332, EPI_ISL_2639333, EPI_ISL_2639335, EPI_ISL_2639336, EPI_ISL_2639337, EPI_ISL_2639338, EPI_ISL_2639339, EPI_ISL_2639340, EPI_ISL_2639341, EPI_ISL_2639342, EPI_ISL_2639343, EPI_ISL_2639344, EPI_ISL_2639345,
EPI_ISL_2639346, EPI_ISL_2639347, EPI_ISL_2639349, EPI_ISL_2639350, EPI_ISL_2639351, EPI_ISL_2639352, EPI_ISL_2639353, EPI_ISL_2639354, EPI_ISL_2639355, EPI_ISL_2639356, EPI_ISL_2639357, EPI_ISL_2639358, EPI_ISL_2639359, EPI_ISL_2639360, EPI_ISL_2639361, EPI_ISL_2639362, EPI_ISL_2639363, EPI_ISL_2639364, EPI_ISL_2639365, EPI_ISL_2639366, EPI_ISL_2639368, EPI_ISL_2639369,
EPI_ISL_2639370, EPI_ISL_2639371, EPI_ISL_2639372, EPI_ISL_2639373, EPI_ISL_2639374, EPI_ISL_2639375, EPI_ISL_2639376, EPI_ISL_2639377, EPI_ISL_2639378, EPI_ISL_2639379, EPI_ISL_2639380, EPI_ISL_2639381, EPI_ISL_2639382, EPI_ISL_2639383, EPI_ISL_2639384, EPI_ISL_2639385, EPI_ISL_2639386, EPI_ISL_2639387, EPI_ISL_2639388, EPI_ISL_2639389, EPI_ISL_2639390, EPI_ISL_2639391,
EPI_ISL_2639392, EPI_ISL_2639393, EPI_ISL_2639394, EPI_ISL_2639395, EPI_ISL_2639396, EPI_ISL_2639397, EPI_ISL_2639398, EPI_ISL_2639399, EPI_ISL_2639400, EPI_ISL_2639401, EPI_ISL_2639402, EPI_ISL_2639403, EPI_ISL_2639404, EPI_ISL_2639405, EPI_ISL_2639406, EPI_ISL_2639407, EPI_ISL_2639408, EPI_ISL_2639410, EPI_ISL_2639411, EPI_ISL_2639412, EPI_ISL_2639413, EPI_ISL_2639414,
EPI_ISL_2639415, EPI_ISL_2639416, EPI_ISL_2639417, EPI_ISL_2639418, EPI_ISL_2639419, EPI_ISL_2639420, EPI_ISL_2639421, EPI_ISL_2639422, EPI_ISL_2639423, EPI_ISL_2639424, EPI_ISL_2639425, EPI_ISL_2639426, EPI_ISL_2639427, EPI_ISL_2639428, EPI_ISL_2639429, EPI_ISL_2639430, EPI_ISL_2639431, EPI_ISL_2639432, EPI_ISL_2639433, EPI_ISL_2639434, EPI_ISL_2639435, EPI_ISL_2639436,
EPI_ISL_2639437, EPI_ISL_2639438, EPI_ISL_2639439, EPI_ISL_2639440, EPI_ISL_2639441, EPI_ISL_2639442, EPI_ISL_2639443, EPI_ISL_2639444, EPI_ISL_2639445, EPI_ISL_2639446, EPI_ISL_2639447, EPI_ISL_2639448, EPI_ISL_2639449, EPI_ISL_2639450, EPI_ISL_2639451, EPI_ISL_2639452, EPI_ISL_2639453, EPI_ISL_2639454, EPI_ISL_2639455, EPI_ISL_2639456, EPI_ISL_2639457, EPI_ISL_2639458,
EPI_ISL_2639459, EPI_ISL_2639460, EPI_ISL_2639461, EPI_ISL_2639462, EPI_ISL_2639463, EPI_ISL_2639464, EPI_ISL_2639465, EPI_ISL_2639466, EPI_ISL_2639467, EPI_ISL_2639468, EPI_ISL_2639469, EPI_ISL_2639470, EPI_ISL_2639471, EPI_ISL_2639472, EPI_ISL_2639473, EPI_ISL_2639474, EPI_ISL_2639475, EPI_ISL_2639476, EPI_ISL_2639477, EPI_ISL_2639478, EPI_ISL_2639479, EPI_ISL_2639480,
EPI_ISL_2639481, EPI_ISL_2639482, EPI_ISL_2639483, EPI_ISL_2639484, EPI_ISL_2639485, EPI_ISL_2639486, EPI_ISL_2639487, EPI_ISL_2639488, EPI_ISL_2639489, EPI_ISL_2639490, EPI_ISL_2639491, EPI_ISL_2639492, EPI_ISL_2639493, EPI_ISL_2639494, EPI_ISL_2639495, EPI_ISL_2639496, EPI_ISL_2639497, EPI_ISL_2639498, EPI_ISL_2639499, EPI_ISL_2639500, EPI_ISL_2639501, EPI_ISL_2639502,
EPI_ISL_2639503, EPI_ISL_2639504, EPI_ISL_2639505, EPI_ISL_2639506, EPI_ISL_2639507, EPI_ISL_2639508, EPI_ISL_2639509, EPI_ISL_2639510, EPI_ISL_2639511, EPI_ISL_2639512, EPI_ISL_2639513, EPI_ISL_2639514, EPI_ISL_2639515, EPI_ISL_2639516, EPI_ISL_2639517, EPI_ISL_2639518, EPI_ISL_2639519, EPI_ISL_2639520, EPI_ISL_2639521, EPI_ISL_2639522, EPI_ISL_2639524, EPI_ISL_2639525,
EPI_ISL_2639526, EPI_ISL_2639527, EPI_ISL_2639528, EPI_ISL_2639529, EPI_ISL_2639530, EPI_ISL_2639531, EPI_ISL_2639532, EPI_ISL_2639533, EPI_ISL_2639534, EPI_ISL_2639535, EPI_ISL_2639536, EPI_ISL_2639537, EPI_ISL_2639538, EPI_ISL_2639539, EPI_ISL_2639540, EPI_ISL_2639541, EPI_ISL_2639542, EPI_ISL_2639543, EPI_ISL_2639544, EPI_ISL_2639545, EPI_ISL_2639546, EPI_ISL_2639547,
EPI_ISL_2639548, EPI_ISL_2639549, EPI_ISL_2639550, EPI_ISL_2639551, EPI_ISL_2639552, EPI_ISL_2639553, EPI_ISL_2639554, EPI_ISL_2639555, EPI_ISL_2639556, EPI_ISL_2639557, EPI_ISL_2639558, EPI_ISL_2639559, EPI_ISL_2639560, EPI_ISL_2639561, EPI_ISL_2639562, EPI_ISL_2639563, EPI_ISL_2639564, EPI_ISL_2639565, EPI_ISL_2639566, EPI_ISL_2639567, EPI_ISL_2639568, EPI_ISL_2639569,
EPI_ISL_2639570, EPI_ISL_2639571, EPI_ISL_2639572, EPI_ISL_2639573, EPI_ISL_2639574, EPI_ISL_2639575, EPI_ISL_2639576, EPI_ISL_2639577, EPI_ISL_2639578, EPI_ISL_2639579, EPI_ISL_2639580, EPI_ISL_2639581, EPI_ISL_2639582, EPI_ISL_2639583, EPI_ISL_2639585, EPI_ISL_2639586, EPI_ISL_2639587, EPI_ISL_2639588, EPI_ISL_2639589, EPI_ISL_2639590, EPI_ISL_2639591, EPI_ISL_2639592,
EPI_ISL_2639593, EPI_ISL_2639594, EPI_ISL_2639595, EPI_ISL_2639596, EPI_ISL_2639597, EPI_ISL_2639598, EPI_ISL_2639599, EPI_ISL_2639600, EPI_ISL_2639601, EPI_ISL_2639602, EPI_ISL_2639603, EPI_ISL_2639604, EPI_ISL_2639605, EPI_ISL_2639606, EPI_ISL_2639607, EPI_ISL_2639608, EPI_ISL_2639609, EPI_ISL_2639610, EPI_ISL_2639611, EPI_ISL_2639612, EPI_ISL_2639613, EPI_ISL_2639614,
EPI_ISL_2639615, EPI_ISL_2639616, EPI_ISL_2639618, EPI_ISL_2639619

| | | | |
|---|---|---|---|
| see above | Lighthouse Lab in Alderley Park | Wellcome Sanger Institute for the COVID-19 Genomics UK (COG-UK) Consortium | Jacquelyn Wynn, Mairead Hyland, The Lighthouse Lab in Alderley Park and Alex Alderton, Roberto Amato, Jeffrey Barrett, Sonia Goncalves, Ewan Harrison, David K. Jackson, Ian Johnston, Dominic Kwiatkowski, Cordelia Langford, John Sillitoe on behalf of the Wellcome Sanger Institute COVID-19 Surveillance Team |
| EPI_ISL_2639620 | Lighthouse Laboratory Plymouth | Wellcome Sanger Institute for the COVID-19 Genomics UK (COG-UK) Consortium | Lighthouse Laboratory Plymouth and Alex Alderton, Roberto Amato, Jeffrey Barrett, Sonia Goncalves, Ewan Harrison, David K. Jackson, Ian Johnston, Dominic Kwiatkowski, Cordelia Langford, John Sillitoe on behalf of the Wellcome Sanger Institute COVID-19 Surveillance Team |
| EPI_ISL_2639621 | Lighthouse Lab in Glasgow | Wellcome Sanger Institute for the COVID-19 Genomics UK (COG-UK) Consortium | Harper VanSteenhouse, Yumi Kasai, David Gray, Carol Clugston, Anna Dominiczak and Alex Alderton, Roberto Amato, Jeffrey Barrett, Sonia Goncalves, Ewan Harrison, David K. Jackson, Ian Johnston, Dominic Kwiatkowski, Cordelia Langford, John Sillitoe on behalf of the Wellcome Sanger Institute COVID-19 Surveillance Team |
| EPI_ISL_2639622, EPI_ISL_2639623, EPI_ISL_2639624 | Lighthouse Lab in Alderley Park | Wellcome Sanger Institute for the COVID-19 Genomics UK (COG-UK) Consortium | Jacquelyn Wynn, Mairead Hyland, The Lighthouse Lab in Alderley Park and Alex Alderton, Roberto Amato, Jeffrey Barrett, Sonia Goncalves, Ewan Harrison, David K. Jackson, Ian Johnston, Dominic Kwiatkowski, Cordelia Langford, John Sillitoe on behalf of the Wellcome Sanger Institute COVID-19 Surveillance Team |
| EPI_ISL_2639625 | Lighthouse Lab in Glasgow | Wellcome Sanger Institute for the COVID-19 Genomics UK (COG-UK) Consortium | Harper VanSteenhouse, Yumi Kasai, David Gray, Carol Clugston, Anna Dominiczak and Alex Alderton, Roberto Amato, Jeffrey Barrett, Sonia Goncalves, Ewan Harrison, David K. Jackson, Ian Johnston, Dominic Kwiatkowski, Cordelia Langford, John Sillitoe on behalf of the Wellcome Sanger Institute COVID-19 Surveillance Team |

| | | | |
|---|---|---|---|
| EPI_ISL_2639626 | Lighthouse Laboratory Plymouth | Wellcome Sanger Institute for the COVID-19 Genomics UK (COG-UK) Consortium | Lighthouse Laboratory Plymouth and Alex Alderton, Roberto Amato, Jeffrey Barrett, Sonia Goncalves, Ewan Harrison, David K. Jackson, Ian Johnston, Dominic Kwiatkowski, Cordelia Langford, John Sillitoe on behalf of the Wellcome Sanger Institute COVID-19 Surveillance Team |
| EPI_ISL_2639627 | Lighthouse Lab in Alderley Park | Wellcome Sanger Institute for the COVID-19 Genomics UK (COG-UK) Consortium | Jacquelyn Wynn, Mairead Hyland, The Lighthouse Lab in Alderley Park and Alex Alderton, Roberto Amato, Jeffrey Barrett, Sonia Goncalves, Ewan Harrison, David K. Jackson, Ian Johnston, Dominic Kwiatkowski, Cordelia Langford, John Sillitoe on behalf of the Wellcome Sanger Institute COVID-19 Surveillance Team |
| EPI_ISL_2639628, EPI_ISL_2639629 | Lighthouse Laboratory Plymouth | Wellcome Sanger Institute for the COVID-19 Genomics UK (COG-UK) Consortium | Lighthouse Laboratory Plymouth and Alex Alderton, Roberto Amato, Jeffrey Barrett, Sonia Goncalves, Ewan Harrison, David K. Jackson, Ian Johnston, Dominic Kwiatkowski, Cordelia Langford, John Sillitoe on behalf of the Wellcome Sanger Institute COVID-19 Surveillance Team |
| EPI_ISL_2639630 | Lighthouse Lab in Alderley Park | Wellcome Sanger Institute for the COVID-19 Genomics UK (COG-UK) Consortium | Jacquelyn Wynn, Mairead Hyland, The Lighthouse Lab in Alderley Park and Alex Alderton, Roberto Amato, Jeffrey Barrett, Sonia Goncalves, Ewan Harrison, David K. Jackson, Ian Johnston, Dominic Kwiatkowski, Cordelia Langford, John Sillitoe on behalf of the Wellcome Sanger Institute COVID-19 Surveillance Team |
| EPI_ISL_2639631 | Lighthouse Laboratory Plymouth | Wellcome Sanger Institute for the COVID-19 Genomics UK (COG-UK) Consortium | Lighthouse Laboratory Plymouth and Alex Alderton, Roberto Amato, Jeffrey Barrett, Sonia Goncalves, Ewan Harrison, David K. Jackson, Ian Johnston, Dominic Kwiatkowski, Cordelia Langford, John Sillitoe on behalf of the Wellcome Sanger Institute COVID-19 Surveillance Team |
| EPI_ISL_2639632 | Lighthouse Lab in Alderley Park | Wellcome Sanger Institute for the COVID-19 Genomics UK (COG-UK) Consortium | Jacquelyn Wynn, Mairead Hyland, The Lighthouse Lab in Alderley Park and Alex Alderton, Roberto Amato, Jeffrey Barrett, Sonia Goncalves, Ewan Harrison, David K. Jackson, Ian Johnston, Dominic Kwiatkowski, Cordelia Langford, John Sillitoe on behalf of the Wellcome Sanger Institute COVID-19 Surveillance Team |
| EPI_ISL_2639633 | Lighthouse Laboratory Plymouth | Wellcome Sanger Institute for the COVID-19 Genomics UK (COG-UK) Consortium | Lighthouse Laboratory Plymouth and Alex Alderton, Roberto Amato, Jeffrey Barrett, Sonia Goncalves, Ewan Harrison, David K. Jackson, Ian Johnston, Dominic Kwiatkowski, Cordelia Langford, John Sillitoe on behalf of the Wellcome Sanger Institute COVID-19 Surveillance Team |
| EPI_ISL_2639635 | Lighthouse Lab in Alderley Park | Wellcome Sanger Institute for the COVID-19 Genomics UK (COG-UK) Consortium | Jacquelyn Wynn, Mairead Hyland, The Lighthouse Lab in Alderley Park and Alex Alderton, Roberto Amato, Jeffrey Barrett, Sonia Goncalves, Ewan Harrison, David K. Jackson, Ian Johnston, Dominic Kwiatkowski, Cordelia Langford, John Sillitoe on behalf of the Wellcome Sanger Institute COVID-19 Surveillance Team |
| EPI_ISL_2639636 | Lighthouse Laboratory Plymouth | Wellcome Sanger Institute for the COVID-19 Genomics UK (COG-UK) Consortium | Lighthouse Laboratory Plymouth and Alex Alderton, Roberto Amato, Jeffrey Barrett, Sonia Goncalves, Ewan Harrison, David K. Jackson, Ian Johnston, Dominic Kwiatkowski, Cordelia Langford, John Sillitoe on behalf of the Wellcome Sanger Institute COVID-19 Surveillance Team |
| EPI_ISL_2639637, EPI_ISL_2639638, EPI_ISL_2639639 | Lighthouse Lab in Alderley Park | Wellcome Sanger Institute for the COVID-19 Genomics UK (COG-UK) Consortium | Jacquelyn Wynn, Mairead Hyland, The Lighthouse Lab in Alderley Park and Alex Alderton, Roberto Amato, Jeffrey Barrett, Sonia Goncalves, Ewan Harrison, David K. Jackson, Ian Johnston, Dominic Kwiatkowski, Cordelia Langford, John Sillitoe on behalf of the Wellcome Sanger Institute COVID-19 Surveillance Team |
| EPI_ISL_2639640 | Lighthouse Lab in Glasgow | Wellcome Sanger Institute for the COVID-19 Genomics UK (COG-UK) Consortium | Harper VanSteenhouse, Yumi Kasai, David Gray, Carol Clugston, Anna Dominiczak and Alex Alderton, Roberto Amato, Jeffrey Barrett, Sonia Goncalves, Ewan Harrison, David K. Jackson, Ian Johnston, Dominic Kwiatkowski, Cordelia Langford, John Sillitoe on behalf of the Wellcome Sanger Institute COVID-19 Surveillance Team |
| EPI_ISL_2639641, EPI_ISL_2639642, EPI_ISL_2639643, EPI_ISL_2639644, EPI_ISL_2639645, EPI_ISL_2639646, EPI_ISL_2639647 | Lighthouse Lab in Alderley Park | Wellcome Sanger Institute for the COVID-19 Genomics UK (COG-UK) Consortium | Jacquelyn Wynn, Mairead Hyland, The Lighthouse Lab in Alderley Park and Alex Alderton, Roberto Amato, Jeffrey Barrett, Sonia Goncalves, Ewan Harrison, David K. Jackson, Ian Johnston, Dominic Kwiatkowski, Cordelia Langford, John Sillitoe on behalf of the Wellcome Sanger Institute COVID-19 Surveillance Team |
| EPI_ISL_2639648 | Lighthouse Laboratory Plymouth | Wellcome Sanger Institute for the COVID-19 Genomics UK (COG-UK) Consortium | Lighthouse Laboratory Plymouth and Alex Alderton, Roberto Amato, Jeffrey Barrett, Sonia Goncalves, Ewan Harrison, David K. Jackson, Ian Johnston, Dominic Kwiatkowski, Cordelia Langford, John Sillitoe on behalf of the Wellcome Sanger Institute COVID-19 Surveillance Team |
| EPI_ISL_2639649, EPI_ISL_2639651 | Lighthouse Lab in Alderley Park | Wellcome Sanger Institute for the COVID-19 Genomics UK (COG-UK) Consortium | Jacquelyn Wynn, Mairead Hyland, The Lighthouse Lab in Alderley Park and Alex Alderton, Roberto Amato, Jeffrey Barrett, Sonia Goncalves, Ewan Harrison, David K. Jackson, Ian Johnston, Dominic Kwiatkowski, Cordelia Langford, John Sillitoe on behalf of the Wellcome Sanger Institute COVID-19 Surveillance Team |
| EPI_ISL_2639652 | Lighthouse Laboratory Plymouth | Wellcome Sanger Institute for the COVID-19 Genomics UK (COG-UK) Consortium | Lighthouse Laboratory Plymouth and Alex Alderton, Roberto Amato, Jeffrey Barrett, Sonia Goncalves, Ewan Harrison, David K. Jackson, Ian Johnston, Dominic Kwiatkowski, Cordelia Langford, John Sillitoe on behalf of the Wellcome Sanger Institute COVID-19 Surveillance Team |
| EPI_ISL_2639653, EPI_ISL_2639654 | Lighthouse Lab in Alderley Park | Wellcome Sanger Institute for the COVID-19 Genomics UK (COG-UK) Consortium | Jacquelyn Wynn, Mairead Hyland, The Lighthouse Lab in Alderley Park and Alex Alderton, Roberto Amato, Jeffrey Barrett, Sonia Goncalves, Ewan Harrison, David K. Jackson, Ian Johnston, Dominic Kwiatkowski, Cordelia Langford, John Sillitoe on behalf of the Wellcome Sanger Institute COVID-19 Surveillance Team |
| EPI_ISL_2639655, EPI_ISL_2639656 | Lighthouse Laboratory Plymouth | Wellcome Sanger Institute for the COVID-19 Genomics UK (COG-UK) Consortium | Lighthouse Laboratory Plymouth and Alex Alderton, Roberto Amato, Jeffrey Barrett, Sonia Goncalves, Ewan Harrison, David K. Jackson, Ian Johnston, Dominic Kwiatkowski, Cordelia Langford, John Sillitoe on behalf of the Wellcome Sanger Institute COVID-19 Surveillance Team |
| EPI_ISL_2639657 | Lighthouse Lab in Glasgow | Wellcome Sanger Institute for the COVID-19 Genomics UK (COG-UK) Consortium | Harper VanSteenhouse, Yumi Kasai, David Gray, Carol Clugston, Anna Dominiczak and Alex Alderton, Roberto Amato, Jeffrey Barrett, Sonia Goncalves, Ewan Harrison, David K. Jackson, Ian Johnston, Dominic Kwiatkowski, Cordelia Langford, John Sillitoe on behalf of the Wellcome Sanger Institute COVID-19 Surveillance Team |
| EPI_ISL_2639658 | Lighthouse Laboratory Plymouth | Wellcome Sanger Institute for the COVID-19 Genomics UK (COG-UK) Consortium | Lighthouse Laboratory Plymouth and Alex Alderton, Roberto Amato, Jeffrey Barrett, Sonia Goncalves, Ewan Harrison, David K. Jackson, Ian Johnston, Dominic Kwiatkowski, Cordelia Langford, John Sillitoe on behalf of the Wellcome Sanger Institute COVID-19 Surveillance Team |
| EPI_ISL_2639659 | Lighthouse Lab in Glasgow | Wellcome Sanger Institute for the COVID-19 Genomics UK (COG-UK) Consortium | Harper VanSteenhouse, Yumi Kasai, David Gray, Carol Clugston, Anna Dominiczak and Alex Alderton, Roberto Amato, Jeffrey Barrett, Sonia Goncalves, Ewan Harrison, David K. Jackson, Ian Johnston, Dominic Kwiatkowski, Cordelia Langford, John Sillitoe on behalf of the Wellcome Sanger Institute COVID-19 Surveillance Team |
| EPI_ISL_2639660 | Lighthouse Lab in Alderley Park | Wellcome Sanger Institute for the COVID-19 Genomics UK (COG-UK) Consortium | Jacquelyn Wynn, Mairead Hyland, The Lighthouse Lab in Alderley Park and Alex Alderton, Roberto Amato, Jeffrey Barrett, Sonia Goncalves, Ewan Harrison, David K. Jackson, Ian Johnston, Dominic Kwiatkowski, Cordelia Langford, John Sillitoe on behalf of the Wellcome Sanger Institute COVID-19 Surveillance Team |
| EPI_ISL_2639661 | Lighthouse Laboratory Plymouth | Wellcome Sanger Institute for the COVID-19 Genomics UK (COG-UK) Consortium | Lighthouse Laboratory Plymouth and Alex Alderton, Roberto Amato, Jeffrey Barrett, Sonia Goncalves, Ewan Harrison, David K. Jackson, Ian Johnston, Dominic Kwiatkowski, Cordelia Langford, John Sillitoe on behalf of the Wellcome Sanger Institute COVID-19 Surveillance Team |
| EPI_ISL_2639662, EPI_ISL_2639663, EPI_ISL_2639664, EPI_ISL_2639665, EPI_ISL_2639666 | Lighthouse Lab in Alderley Park | Wellcome Sanger Institute for the COVID-19 Genomics UK (COG-UK) Consortium | Jacquelyn Wynn, Mairead Hyland, The Lighthouse Lab in Alderley Park and Alex Alderton, Roberto Amato, Jeffrey Barrett, Sonia Goncalves, Ewan Harrison, David K. Jackson, Ian Johnston, Dominic Kwiatkowski, Cordelia Langford, John Sillitoe on behalf of the Wellcome Sanger Institute COVID-19 Surveillance Team |
| EPI_ISL_2639667 | Lighthouse Lab in Glasgow | Wellcome Sanger Institute for the COVID-19 Genomics UK (COG-UK) Consortium | Harper VanSteenhouse, Yumi Kasai, David Gray, Carol Clugston, Anna Dominiczak and Alex Alderton, Roberto Amato, Jeffrey Barrett, Sonia Goncalves, Ewan Harrison, David K. Jackson, Ian Johnston, Dominic Kwiatkowski, Cordelia Langford, John Sillitoe on behalf of the Wellcome Sanger Institute COVID-19 Surveillance Team |
| EPI_ISL_2639668, EPI_ISL_2639669 | Lighthouse Laboratory Plymouth | Wellcome Sanger Institute for the COVID-19 Genomics UK (COG-UK) Consortium | Lighthouse Laboratory Plymouth and Alex Alderton, Roberto Amato, Jeffrey Barrett, Sonia Goncalves, Ewan Harrison, David K. Jackson, Ian Johnston, Dominic Kwiatkowski, Cordelia Langford, John Sillitoe on behalf of the Wellcome Sanger Institute COVID-19 Surveillance Team |
| EPI_ISL_2639670 | Lighthouse Lab in Alderley Park | Wellcome Sanger Institute for the COVID-19 Genomics UK (COG-UK) Consortium | Jacquelyn Wynn, Mairead Hyland, The Lighthouse Lab in Alderley Park and Alex Alderton, Roberto Amato, Jeffrey Barrett, Sonia Goncalves, Ewan Harrison, David K. Jackson, Ian Johnston, Dominic Kwiatkowski, Cordelia Langford, John Sillitoe on behalf of the Wellcome Sanger Institute COVID-19 Surveillance Team |
| EPI_ISL_2639671 | Lighthouse Laboratory Plymouth | Wellcome Sanger Institute for the COVID-19 Genomics UK (COG-UK) Consortium | Lighthouse Laboratory Plymouth and Alex Alderton, Roberto Amato, Jeffrey Barrett, Sonia Goncalves, Ewan Harrison, David K. Jackson, Ian Johnston, Dominic Kwiatkowski, Cordelia Langford, John Sillitoe on behalf of the Wellcome Sanger Institute COVID-19 Surveillance Team |
| EPI_ISL_2639672, EPI_ISL_2639673 | Lighthouse Lab in Alderley Park | Wellcome Sanger Institute for the COVID-19 Genomics UK (COG-UK) Consortium | Jacquelyn Wynn, Mairead Hyland, The Lighthouse Lab in Alderley Park and Alex Alderton, Roberto Amato, Jeffrey Barrett, Sonia Goncalves, Ewan Harrison, David K. Jackson, Ian Johnston, Dominic Kwiatkowski, Cordelia Langford, John Sillitoe on behalf of the Wellcome Sanger Institute COVID-19 Surveillance Team |
| EPI_ISL_2639674 | Lighthouse Lab in Glasgow | Wellcome Sanger Institute for the COVID-19 Genomics UK (COG-UK) Consortium | Harper VanSteenhouse, Yumi Kasai, David Gray, Carol Clugston, Anna Dominiczak and Alex Alderton, Roberto Amato, Jeffrey Barrett, Sonia Goncalves, Ewan Harrison, David K. Jackson, Ian Johnston, Dominic Kwiatkowski, Cordelia Langford, John Sillitoe on behalf of the Wellcome Sanger Institute COVID-19 Surveillance Team |
| EPI_ISL_2639675, EPI_ISL_2639676 | Lighthouse Lab in Alderley Park | Wellcome Sanger Institute for the COVID-19 Genomics UK (COG-UK) Consortium | Jacquelyn Wynn, Mairead Hyland, The Lighthouse Lab in Alderley Park and Alex Alderton, Roberto Amato, Jeffrey Barrett, Sonia Goncalves, Ewan Harrison, David K. Jackson, Ian Johnston, Dominic Kwiatkowski, Cordelia Langford, John Sillitoe on behalf of the Wellcome Sanger Institute COVID-19 Surveillance Team |
| EPI_ISL_2639677 | Lighthouse Lab in Glasgow | Wellcome Sanger Institute for the COVID-19 Genomics UK (COG-UK) Consortium | Harper VanSteenhouse, Yumi Kasai, David Gray, Carol Clugston, Anna Dominiczak and Alex Alderton, Roberto Amato, Jeffrey Barrett, Sonia Goncalves, Ewan Harrison, David K. Jackson, Ian Johnston, Dominic Kwiatkowski, Cordelia Langford, John Sillitoe on behalf of the Wellcome Sanger Institute COVID-19 Surveillance Team |
| EPI_ISL_2639678, EPI_ISL_2639679, EPI_ISL_2639680, EPI_ISL_2639681 | Lighthouse Lab in Alderley Park | Wellcome Sanger Institute for the COVID-19 Genomics UK (COG-UK) Consortium | Jacquelyn Wynn, Mairead Hyland, The Lighthouse Lab in Alderley Park and Alex Alderton, Roberto Amato, Jeffrey Barrett, Sonia Goncalves, Ewan Harrison, David K. Jackson, Ian Johnston, Dominic Kwiatkowski, Cordelia Langford, John Sillitoe on behalf of the Wellcome Sanger Institute COVID-19 Surveillance Team |
| EPI_ISL_2639682 | Lighthouse Laboratory Plymouth | Wellcome Sanger Institute for the COVID-19 Genomics UK (COG-UK) Consortium | Lighthouse Laboratory Plymouth and Alex Alderton, Roberto Amato, Jeffrey Barrett, Sonia Goncalves, Ewan Harrison, David K. Jackson, Ian Johnston, Dominic Kwiatkowski, Cordelia Langford, John Sillitoe on behalf of the Wellcome Sanger Institute COVID-19 Surveillance Team |
| EPI_ISL_2639683 | Lighthouse Lab in Alderley Park | Wellcome Sanger Institute for the COVID-19 Genomics UK (COG-UK) Consortium | Jacquelyn Wynn, Mairead Hyland, The Lighthouse Lab in Alderley Park and Alex Alderton, Roberto Amato, Jeffrey Barrett, Sonia Goncalves, Ewan Harrison, David K. Jackson, Ian Johnston, Dominic Kwiatkowski, Cordelia Langford, John Sillitoe on behalf of the Wellcome Sanger Institute COVID-19 Surveillance Team |
| EPI_ISL_2639684 | Lighthouse Laboratory Plymouth | Wellcome Sanger Institute for the COVID-19 Genomics UK (COG-UK) Consortium | Lighthouse Laboratory Plymouth and Alex Alderton, Roberto Amato, Jeffrey Barrett, Sonia Goncalves, Ewan Harrison, David K. Jackson, Ian Johnston, Dominic Kwiatkowski, Cordelia Langford, John Sillitoe on behalf of the Wellcome Sanger Institute COVID-19 Surveillance Team |
| EPI_ISL_2639685 | Lighthouse Lab in Alderley Park | Wellcome Sanger Institute for the COVID-19 Genomics UK (COG-UK) Consortium | Jacquelyn Wynn, Mairead Hyland, The Lighthouse Lab in Alderley Park and Alex Alderton, Roberto Amato, Jeffrey Barrett, Sonia Goncalves, Ewan Harrison, David K. Jackson, Ian Johnston, Dominic Kwiatkowski, Cordelia Langford, John Sillitoe on behalf of the Wellcome Sanger Institute COVID-19 Surveillance Team |
| EPI_ISL_2639686 | Lighthouse Laboratory Plymouth | Wellcome Sanger Institute for the COVID-19 Genomics UK (COG-UK) Consortium | Lighthouse Laboratory Plymouth and Alex Alderton, Roberto Amato, Jeffrey Barrett, Sonia Goncalves, Ewan Harrison, David K. Jackson, Ian Johnston, Dominic Kwiatkowski, Cordelia Langford, John Sillitoe on behalf of the Wellcome Sanger Institute COVID-19 Surveillance Team |
| EPI_ISL_2639687, EPI_ISL_2639688 | Lighthouse Lab in Alderley Park | Wellcome Sanger Institute for the COVID-19 Genomics UK (COG-UK) Consortium | Jacquelyn Wynn, Mairead Hyland, The Lighthouse Lab in Alderley Park and Alex Alderton, Roberto Amato, Jeffrey Barrett, Sonia Goncalves, Ewan Harrison, David K. Jackson, Ian Johnston, Dominic Kwiatkowski, Cordelia Langford, John Sillitoe on behalf of the Wellcome Sanger Institute COVID-19 Surveillance Team |
| EPI_ISL_2639689 | Lighthouse Lab in Glasgow | Wellcome Sanger Institute for the COVID-19 Genomics UK (COG-UK) Consortium | Harper VanSteenhouse, Yumi Kasai, David Gray, Carol Clugston, Anna Dominiczak and Alex Alderton, Roberto Amato, Jeffrey Barrett, Sonia Goncalves, Ewan Harrison, David K. Jackson, Ian Johnston, Dominic Kwiatkowski, Cordelia Langford, John Sillitoe on behalf of the Wellcome Sanger Institute COVID-19 Surveillance Team |
| EPI_ISL_2639690 | Lighthouse Laboratory Plymouth | Wellcome Sanger Institute for the COVID-19 Genomics UK (COG-UK) Consortium | Lighthouse Laboratory Plymouth and Alex Alderton, Roberto Amato, Jeffrey Barrett, Sonia Goncalves, Ewan Harrison, David K. Jackson, Ian Johnston, Dominic Kwiatkowski, Cordelia Langford, John Sillitoe on behalf of the Wellcome Sanger Institute COVID-19 Surveillance Team |
| EPI_ISL_2639691, | Lighthouse Lab in Alderley Park | Wellcome Sanger Institute for the COVID-19 Genomics UK (COG-UK) | Jacquelyn Wynn, Mairead Hyland, The Lighthouse Lab in Alderley Park and Alex Alderton, Roberto Amato, Jeffrey Barrett, Sonia Goncalves, Ewan Harrison, David K. Jackson, Ian Johnston, Dominic Kwiatkowski, Cordelia Langford, John Sillitoe on |

| | | | |
|---|---|---|---|
| EPI_ISL_2639692 | | Consortium | behalf of the Wellcome Sanger Institute COVID-19 Surveillance Team |
| EPI_ISL_2639693, EPI_ISL_2639694, EPI_ISL_2639695, EPI_ISL_2639696 | Lighthouse Laboratory Plymouth | Wellcome Sanger Institute for the COVID-19 Genomics UK (COG-UK) Consortium | Lighthouse Laboratory Plymouth and Alex Alderton, Roberto Amato, Jeffrey Barrett, Sonia Goncalves, Ewan Harrison, David K. Jackson, Ian Johnston, Dominic Kwiatkowski, Cordelia Langford, John Sillitoe on behalf of the Wellcome Sanger Institute COVID-19 Surveillance Team |
| EPI_ISL_2639697 | Lighthouse Lab in Glasgow | Wellcome Sanger Institute for the COVID-19 Genomics UK (COG-UK) Consortium | Harper VanSteenhouse, Yumi Kasai, David Gray, Carol Clugston, Anna Dominiczak and Alex Alderton, Roberto Amato, Jeffrey Barrett, Sonia Goncalves, Ewan Harrison, David K. Jackson, Ian Johnston, Dominic Kwiatkowski, Cordelia Langford, John Sillitoe on behalf of the Wellcome Sanger Institute COVID-19 Surveillance Team |
| EPI_ISL_2639698, EPI_ISL_2639699 | Lighthouse Laboratory Plymouth | Wellcome Sanger Institute for the COVID-19 Genomics UK (COG-UK) Consortium | Lighthouse Laboratory Plymouth and Alex Alderton, Roberto Amato, Jeffrey Barrett, Sonia Goncalves, Ewan Harrison, David K. Jackson, Ian Johnston, Dominic Kwiatkowski, Cordelia Langford, John Sillitoe on behalf of the Wellcome Sanger Institute COVID-19 Surveillance Team |
| EPI_ISL_2639700, EPI_ISL_2639701 | Lighthouse Lab in Alderley Park | Wellcome Sanger Institute for the COVID-19 Genomics UK (COG-UK) Consortium | Jacquelyn Wynn, Mairead Hyland, The Lighthouse Lab in Alderley Park and Alex Alderton, Roberto Amato, Jeffrey Barrett, Sonia Goncalves, Ewan Harrison, David K. Jackson, Ian Johnston, Dominic Kwiatkowski, Cordelia Langford, John Sillitoe on behalf of the Wellcome Sanger Institute COVID-19 Surveillance Team |
| EPI_ISL_2639702 | Lighthouse Lab in Glasgow | Wellcome Sanger Institute for the COVID-19 Genomics UK (COG-UK) Consortium | Harper VanSteenhouse, Yumi Kasai, David Gray, Carol Clugston, Anna Dominiczak and Alex Alderton, Roberto Amato, Jeffrey Barrett, Sonia Goncalves, Ewan Harrison, David K. Jackson, Ian Johnston, Dominic Kwiatkowski, Cordelia Langford, John Sillitoe on behalf of the Wellcome Sanger Institute COVID-19 Surveillance Team |
| EPI_ISL_2639703, EPI_ISL_2639704 | Lighthouse Laboratory Plymouth | Wellcome Sanger Institute for the COVID-19 Genomics UK (COG-UK) Consortium | Lighthouse Laboratory Plymouth and Alex Alderton, Roberto Amato, Jeffrey Barrett, Sonia Goncalves, Ewan Harrison, David K. Jackson, Ian Johnston, Dominic Kwiatkowski, Cordelia Langford, John Sillitoe on behalf of the Wellcome Sanger Institute COVID-19 Surveillance Team |
| EPI_ISL_2639705 | Lighthouse Lab in Alderley Park | Wellcome Sanger Institute for the COVID-19 Genomics UK (COG-UK) Consortium | Jacquelyn Wynn, Mairead Hyland, The Lighthouse Lab in Alderley Park and Alex Alderton, Roberto Amato, Jeffrey Barrett, Sonia Goncalves, Ewan Harrison, David K. Jackson, Ian Johnston, Dominic Kwiatkowski, Cordelia Langford, John Sillitoe on behalf of the Wellcome Sanger Institute COVID-19 Surveillance Team |
| EPI_ISL_2639706 | Lighthouse Laboratory Plymouth | Wellcome Sanger Institute for the COVID-19 Genomics UK (COG-UK) Consortium | Lighthouse Laboratory Plymouth and Alex Alderton, Roberto Amato, Jeffrey Barrett, Sonia Goncalves, Ewan Harrison, David K. Jackson, Ian Johnston, Dominic Kwiatkowski, Cordelia Langford, John Sillitoe on behalf of the Wellcome Sanger Institute COVID-19 Surveillance Team |
| EPI_ISL_2639707, EPI_ISL_2639708 | Lighthouse Lab in Alderley Park | Wellcome Sanger Institute for the COVID-19 Genomics UK (COG-UK) Consortium | Jacquelyn Wynn, Mairead Hyland, The Lighthouse Lab in Alderley Park and Alex Alderton, Roberto Amato, Jeffrey Barrett, Sonia Goncalves, Ewan Harrison, David K. Jackson, Ian Johnston, Dominic Kwiatkowski, Cordelia Langford, John Sillitoe on behalf of the Wellcome Sanger Institute COVID-19 Surveillance Team |
| EPI_ISL_2639710 | Lighthouse Lab in Glasgow | Wellcome Sanger Institute for the COVID-19 Genomics UK (COG-UK) Consortium | Harper VanSteenhouse, Yumi Kasai, David Gray, Carol Clugston, Anna Dominiczak and Alex Alderton, Roberto Amato, Jeffrey Barrett, Sonia Goncalves, Ewan Harrison, David K. Jackson, Ian Johnston, Dominic Kwiatkowski, Cordelia Langford, John Sillitoe on behalf of the Wellcome Sanger Institute COVID-19 Surveillance Team |
| EPI_ISL_2639711, EPI_ISL_2639712 | Lighthouse Lab in Alderley Park | Wellcome Sanger Institute for the COVID-19 Genomics UK (COG-UK) Consortium | Jacquelyn Wynn, Mairead Hyland, The Lighthouse Lab in Alderley Park and Alex Alderton, Roberto Amato, Jeffrey Barrett, Sonia Goncalves, Ewan Harrison, David K. Jackson, Ian Johnston, Dominic Kwiatkowski, Cordelia Langford, John Sillitoe on behalf of the Wellcome Sanger Institute COVID-19 Surveillance Team |
| EPI_ISL_2639713 | Lighthouse Lab in Glasgow | Wellcome Sanger Institute for the COVID-19 Genomics UK (COG-UK) Consortium | Harper VanSteenhouse, Yumi Kasai, David Gray, Carol Clugston, Anna Dominiczak and Alex Alderton, Roberto Amato, Jeffrey Barrett, Sonia Goncalves, Ewan Harrison, David K. Jackson, Ian Johnston, Dominic Kwiatkowski, Cordelia Langford, John Sillitoe on behalf of the Wellcome Sanger Institute COVID-19 Surveillance Team |
| EPI_ISL_2639714, EPI_ISL_2639715 | Lighthouse Lab in Alderley Park | Wellcome Sanger Institute for the COVID-19 Genomics UK (COG-UK) Consortium | Jacquelyn Wynn, Mairead Hyland, The Lighthouse Lab in Alderley Park and Alex Alderton, Roberto Amato, Jeffrey Barrett, Sonia Goncalves, Ewan Harrison, David K. Jackson, Ian Johnston, Dominic Kwiatkowski, Cordelia Langford, John Sillitoe on behalf of the Wellcome Sanger Institute COVID-19 Surveillance Team |
| EPI_ISL_2639716 | Lighthouse Lab in Glasgow | Wellcome Sanger Institute for the COVID-19 Genomics UK (COG-UK) Consortium | Harper VanSteenhouse, Yumi Kasai, David Gray, Carol Clugston, Anna Dominiczak and Alex Alderton, Roberto Amato, Jeffrey Barrett, Sonia Goncalves, Ewan Harrison, David K. Jackson, Ian Johnston, Dominic Kwiatkowski, Cordelia Langford, John Sillitoe on behalf of the Wellcome Sanger Institute COVID-19 Surveillance Team |
| EPI_ISL_2639717 | Lighthouse Laboratory Plymouth | Wellcome Sanger Institute for the COVID-19 Genomics UK (COG-UK) Consortium | Lighthouse Laboratory Plymouth and Alex Alderton, Roberto Amato, Jeffrey Barrett, Sonia Goncalves, Ewan Harrison, David K. Jackson, Ian Johnston, Dominic Kwiatkowski, Cordelia Langford, John Sillitoe on behalf of the Wellcome Sanger Institute COVID-19 Surveillance Team |
| EPI_ISL_2639718, EPI_ISL_2639719 | Lighthouse Lab in Alderley Park | Wellcome Sanger Institute for the COVID-19 Genomics UK (COG-UK) Consortium | Jacquelyn Wynn, Mairead Hyland, The Lighthouse Lab in Alderley Park and Alex Alderton, Roberto Amato, Jeffrey Barrett, Sonia Goncalves, Ewan Harrison, David K. Jackson, Ian Johnston, Dominic Kwiatkowski, Cordelia Langford, John Sillitoe on behalf of the Wellcome Sanger Institute COVID-19 Surveillance Team |
| EPI_ISL_2639720, EPI_ISL_2639721 | Lighthouse Laboratory Plymouth | Wellcome Sanger Institute for the COVID-19 Genomics UK (COG-UK) Consortium | Lighthouse Laboratory Plymouth and Alex Alderton, Roberto Amato, Jeffrey Barrett, Sonia Goncalves, Ewan Harrison, David K. Jackson, Ian Johnston, Dominic Kwiatkowski, Cordelia Langford, John Sillitoe on behalf of the Wellcome Sanger Institute COVID-19 Surveillance Team |
| EPI_ISL_2639722 | Lighthouse Lab in Alderley Park | Wellcome Sanger Institute for the COVID-19 Genomics UK (COG-UK) Consortium | Jacquelyn Wynn, Mairead Hyland, The Lighthouse Lab in Alderley Park and Alex Alderton, Roberto Amato, Jeffrey Barrett, Sonia Goncalves, Ewan Harrison, David K. Jackson, Ian Johnston, Dominic Kwiatkowski, Cordelia Langford, John Sillitoe on behalf of the Wellcome Sanger Institute COVID-19 Surveillance Team |
| EPI_ISL_2639723 | Lighthouse Lab in Glasgow | Wellcome Sanger Institute for the COVID-19 Genomics UK (COG-UK) Consortium | Harper VanSteenhouse, Yumi Kasai, David Gray, Carol Clugston, Anna Dominiczak and Alex Alderton, Roberto Amato, Jeffrey Barrett, Sonia Goncalves, Ewan Harrison, David K. Jackson, Ian Johnston, Dominic Kwiatkowski, Cordelia Langford, John Sillitoe on behalf of the Wellcome Sanger Institute COVID-19 Surveillance Team |
| EPI_ISL_2639724, EPI_ISL_2639725, EPI_ISL_2639726 | Lighthouse Lab in Alderley Park | Wellcome Sanger Institute for the COVID-19 Genomics UK (COG-UK) Consortium | Jacquelyn Wynn, Mairead Hyland, The Lighthouse Lab in Alderley Park and Alex Alderton, Roberto Amato, Jeffrey Barrett, Sonia Goncalves, Ewan Harrison, David K. Jackson, Ian Johnston, Dominic Kwiatkowski, Cordelia Langford, John Sillitoe on behalf of the Wellcome Sanger Institute COVID-19 Surveillance Team |
| EPI_ISL_2639727 | Lighthouse Laboratory Plymouth | Wellcome Sanger Institute for the COVID-19 Genomics UK (COG-UK) Consortium | Lighthouse Laboratory Plymouth and Alex Alderton, Roberto Amato, Jeffrey Barrett, Sonia Goncalves, Ewan Harrison, David K. Jackson, Ian Johnston, Dominic Kwiatkowski, Cordelia Langford, John Sillitoe on behalf of the Wellcome Sanger Institute COVID-19 Surveillance Team |
| EPI_ISL_2639728 | Lighthouse Lab in Glasgow | Wellcome Sanger Institute for the COVID-19 Genomics UK (COG-UK) Consortium | Harper VanSteenhouse, Yumi Kasai, David Gray, Carol Clugston, Anna Dominiczak and Alex Alderton, Roberto Amato, Jeffrey Barrett, Sonia Goncalves, Ewan Harrison, David K. Jackson, Ian Johnston, Dominic Kwiatkowski, Cordelia Langford, John Sillitoe on behalf of the Wellcome Sanger Institute COVID-19 Surveillance Team |
| EPI_ISL_2639729, EPI_ISL_2639730 | Lighthouse Lab in Alderley Park | Wellcome Sanger Institute for the COVID-19 Genomics UK (COG-UK) Consortium | Jacquelyn Wynn, Mairead Hyland, The Lighthouse Lab in Alderley Park and Alex Alderton, Roberto Amato, Jeffrey Barrett, Sonia Goncalves, Ewan Harrison, David K. Jackson, Ian Johnston, Dominic Kwiatkowski, Cordelia Langford, John Sillitoe on behalf of the Wellcome Sanger Institute COVID-19 Surveillance Team |
| EPI_ISL_2639732 | Lighthouse Laboratory Plymouth | Wellcome Sanger Institute for the COVID-19 Genomics UK (COG-UK) Consortium | Lighthouse Laboratory Plymouth and Alex Alderton, Roberto Amato, Jeffrey Barrett, Sonia Goncalves, Ewan Harrison, David K. Jackson, Ian Johnston, Dominic Kwiatkowski, Cordelia Langford, John Sillitoe on behalf of the Wellcome Sanger Institute COVID-19 Surveillance Team |
| EPI_ISL_2639733 | Lighthouse Lab in Glasgow | Wellcome Sanger Institute for the COVID-19 Genomics UK (COG-UK) Consortium | Harper VanSteenhouse, Yumi Kasai, David Gray, Carol Clugston, Anna Dominiczak and Alex Alderton, Roberto Amato, Jeffrey Barrett, Sonia Goncalves, Ewan Harrison, David K. Jackson, Ian Johnston, Dominic Kwiatkowski, Cordelia Langford, John Sillitoe on behalf of the Wellcome Sanger Institute COVID-19 Surveillance Team |
| EPI_ISL_2639734, EPI_ISL_2639735, EPI_ISL_2639736, EPI_ISL_2639737 | Lighthouse Lab in Alderley Park | Wellcome Sanger Institute for the COVID-19 Genomics UK (COG-UK) Consortium | Jacquelyn Wynn, Mairead Hyland, The Lighthouse Lab in Alderley Park and Alex Alderton, Roberto Amato, Jeffrey Barrett, Sonia Goncalves, Ewan Harrison, David K. Jackson, Ian Johnston, Dominic Kwiatkowski, Cordelia Langford, John Sillitoe on behalf of the Wellcome Sanger Institute COVID-19 Surveillance Team |
| EPI_ISL_2639738 | Lighthouse Laboratory Plymouth | Wellcome Sanger Institute for the COVID-19 Genomics UK (COG-UK) Consortium | Lighthouse Laboratory Plymouth and Alex Alderton, Roberto Amato, Jeffrey Barrett, Sonia Goncalves, Ewan Harrison, David K. Jackson, Ian Johnston, Dominic Kwiatkowski, Cordelia Langford, John Sillitoe on behalf of the Wellcome Sanger Institute COVID-19 Surveillance Team |
| EPI_ISL_2639739 | Lighthouse Lab in Alderley Park | Wellcome Sanger Institute for the COVID-19 Genomics UK (COG-UK) Consortium | Jacquelyn Wynn, Mairead Hyland, The Lighthouse Lab in Alderley Park and Alex Alderton, Roberto Amato, Jeffrey Barrett, Sonia Goncalves, Ewan Harrison, David K. Jackson, Ian Johnston, Dominic Kwiatkowski, Cordelia Langford, John Sillitoe on behalf of the Wellcome Sanger Institute COVID-19 Surveillance Team |
| EPI_ISL_2639740 | Lighthouse Laboratory Plymouth | Wellcome Sanger Institute for the COVID-19 Genomics UK (COG-UK) Consortium | Lighthouse Laboratory Plymouth and Alex Alderton, Roberto Amato, Jeffrey Barrett, Sonia Goncalves, Ewan Harrison, David K. Jackson, Ian Johnston, Dominic Kwiatkowski, Cordelia Langford, John Sillitoe on behalf of the Wellcome Sanger Institute COVID-19 Surveillance Team |
| EPI_ISL_2639741 | Lighthouse Lab in Glasgow | Wellcome Sanger Institute for the COVID-19 Genomics UK (COG-UK) Consortium | Harper VanSteenhouse, Yumi Kasai, David Gray, Carol Clugston, Anna Dominiczak and Alex Alderton, Roberto Amato, Jeffrey Barrett, Sonia Goncalves, Ewan Harrison, David K. Jackson, Ian Johnston, Dominic Kwiatkowski, Cordelia Langford, John Sillitoe on behalf of the Wellcome Sanger Institute COVID-19 Surveillance Team |
| EPI_ISL_2639742 | Lighthouse Lab in Alderley Park | Wellcome Sanger Institute for the COVID-19 Genomics UK (COG-UK) Consortium | Jacquelyn Wynn, Mairead Hyland, The Lighthouse Lab in Alderley Park and Alex Alderton, Roberto Amato, Jeffrey Barrett, Sonia Goncalves, Ewan Harrison, David K. Jackson, Ian Johnston, Dominic Kwiatkowski, Cordelia Langford, John Sillitoe on behalf of the Wellcome Sanger Institute COVID-19 Surveillance Team |
| EPI_ISL_2639743 | Lighthouse Lab in Glasgow | Wellcome Sanger Institute for the COVID-19 Genomics UK (COG-UK) Consortium | Harper VanSteenhouse, Yumi Kasai, David Gray, Carol Clugston, Anna Dominiczak and Alex Alderton, Roberto Amato, Jeffrey Barrett, Sonia Goncalves, Ewan Harrison, David K. Jackson, Ian Johnston, Dominic Kwiatkowski, Cordelia Langford, John Sillitoe on behalf of the Wellcome Sanger Institute COVID-19 Surveillance Team |
| EPI_ISL_2639744, EPI_ISL_2639745 | Lighthouse Lab in Alderley Park | Wellcome Sanger Institute for the COVID-19 Genomics UK (COG-UK) Consortium | Jacquelyn Wynn, Mairead Hyland, The Lighthouse Lab in Alderley Park and Alex Alderton, Roberto Amato, Jeffrey Barrett, Sonia Goncalves, Ewan Harrison, David K. Jackson, Ian Johnston, Dominic Kwiatkowski, Cordelia Langford, John Sillitoe on behalf of the Wellcome Sanger Institute COVID-19 Surveillance Team |
| EPI_ISL_2639746 | Lighthouse Lab in Glasgow | Wellcome Sanger Institute for the COVID-19 Genomics UK (COG-UK) Consortium | Harper VanSteenhouse, Yumi Kasai, David Gray, Carol Clugston, Anna Dominiczak and Alex Alderton, Roberto Amato, Jeffrey Barrett, Sonia Goncalves, Ewan Harrison, David K. Jackson, Ian Johnston, Dominic Kwiatkowski, Cordelia Langford, John Sillitoe on behalf of the Wellcome Sanger Institute COVID-19 Surveillance Team |
| EPI_ISL_2639747 | Lighthouse Lab in Alderley Park | Wellcome Sanger Institute for the COVID-19 Genomics UK (COG-UK) Consortium | Jacquelyn Wynn, Mairead Hyland, The Lighthouse Lab in Alderley Park and Alex Alderton, Roberto Amato, Jeffrey Barrett, Sonia Goncalves, Ewan Harrison, David K. Jackson, Ian Johnston, Dominic Kwiatkowski, Cordelia Langford, John Sillitoe on behalf of the Wellcome Sanger Institute COVID-19 Surveillance Team |
| EPI_ISL_2639748 | Lighthouse Laboratory Plymouth | Wellcome Sanger Institute for the COVID-19 Genomics UK (COG-UK) Consortium | Lighthouse Laboratory Plymouth and Alex Alderton, Roberto Amato, Jeffrey Barrett, Sonia Goncalves, Ewan Harrison, David K. Jackson, Ian Johnston, Dominic Kwiatkowski, Cordelia Langford, John Sillitoe on behalf of the Wellcome Sanger Institute COVID-19 Surveillance Team |
| EPI_ISL_2639749 | Lighthouse Lab in Alderley Park | Wellcome Sanger Institute for the COVID-19 Genomics UK (COG-UK) Consortium | Jacquelyn Wynn, Mairead Hyland, The Lighthouse Lab in Alderley Park and Alex Alderton, Roberto Amato, Jeffrey Barrett, Sonia Goncalves, Ewan Harrison, David K. Jackson, Ian Johnston, Dominic Kwiatkowski, Cordelia Langford, John Sillitoe on behalf of the Wellcome Sanger Institute COVID-19 Surveillance Team |
| EPI_ISL_2639750 | Lighthouse Lab in Glasgow | Wellcome Sanger Institute for the COVID-19 Genomics UK (COG-UK) Consortium | Harper VanSteenhouse, Yumi Kasai, David Gray, Carol Clugston, Anna Dominiczak and Alex Alderton, Roberto Amato, Jeffrey Barrett, Sonia Goncalves, Ewan Harrison, David K. Jackson, Ian Johnston, Dominic Kwiatkowski, Cordelia Langford, John Sillitoe on behalf of the Wellcome Sanger Institute COVID-19 Surveillance Team |
| EPI_ISL_2639751, EPI_ISL_2639752, EPI_ISL_2639753, EPI_ISL_2639754 | Lighthouse Lab in Alderley Park | Wellcome Sanger Institute for the COVID-19 Genomics UK (COG-UK) Consortium | Jacquelyn Wynn, Mairead Hyland, The Lighthouse Lab in Alderley Park and Alex Alderton, Roberto Amato, Jeffrey Barrett, Sonia Goncalves, Ewan Harrison, David K. Jackson, Ian Johnston, Dominic Kwiatkowski, Cordelia Langford, John Sillitoe on behalf of the Wellcome Sanger Institute COVID-19 Surveillance Team |
| EPI_ISL_2639755 | Lighthouse Laboratory Plymouth | Wellcome Sanger Institute for the COVID-19 Genomics UK (COG-UK) Consortium | Lighthouse Laboratory Plymouth and Alex Alderton, Roberto Amato, Jeffrey Barrett, Sonia Goncalves, Ewan Harrison, David K. Jackson, Ian Johnston, Dominic Kwiatkowski, Cordelia Langford, John Sillitoe on behalf of the Wellcome Sanger Institute COVID-19 Surveillance Team |
| EPI_ISL_2639756 | Lighthouse Lab in Alderley Park | Wellcome Sanger Institute for the COVID-19 Genomics UK (COG-UK) Consortium | Jacquelyn Wynn, Mairead Hyland, The Lighthouse Lab in Alderley Park and Alex Alderton, Roberto Amato, Jeffrey Barrett, Sonia Goncalves, Ewan Harrison, David K. Jackson, Ian Johnston, Dominic Kwiatkowski, Cordelia Langford, John Sillitoe on behalf of the Wellcome Sanger Institute COVID-19 Surveillance Team |

| | | | |
|---|---|---|---|
| EPI_ISL_2639757 | Lighthouse Laboratory Plymouth | Wellcome Sanger Institute for the COVID-19 Genomics UK (COG-UK) Consortium | Lighthouse Laboratory Plymouth and Alex Alderton, Roberto Amato, Jeffrey Barrett, Sonia Goncalves, Ewan Harrison, David K. Jackson, Ian Johnston, Dominic Kwiatkowski, Cordelia Langford, John Sillitoe on behalf of the Wellcome Sanger Institute COVID-19 Surveillance Team |
| EPI_ISL_2639758 | Lighthouse Lab in Alderley Park | Wellcome Sanger Institute for the COVID-19 Genomics UK (COG-UK) Consortium | Jacquelyn Wynn, Mairead Hyland, The Lighthouse Lab in Alderley Park and Alex Alderton, Roberto Amato, Jeffrey Barrett, Sonia Goncalves, Ewan Harrison, David K. Jackson, Ian Johnston, Dominic Kwiatkowski, Cordelia Langford, John Sillitoe on behalf of the Wellcome Sanger Institute COVID-19 Surveillance Team |
| EPI_ISL_2639759 | Lighthouse Laboratory Plymouth | Wellcome Sanger Institute for the COVID-19 Genomics UK (COG-UK) Consortium | Lighthouse Laboratory Plymouth and Alex Alderton, Roberto Amato, Jeffrey Barrett, Sonia Goncalves, Ewan Harrison, David K. Jackson, Ian Johnston, Dominic Kwiatkowski, Cordelia Langford, John Sillitoe on behalf of the Wellcome Sanger Institute COVID-19 Surveillance Team |
| EPI_ISL_2639760 | Lighthouse Lab in Glasgow | Wellcome Sanger Institute for the COVID-19 Genomics UK (COG-UK) Consortium | Harper VanSteenhouse, Yumi Kasai, David Gray, Carol Clugston, Anna Dominiczak and Alex Alderton, Roberto Amato, Jeffrey Barrett, Sonia Goncalves, Ewan Harrison, David K. Jackson, Ian Johnston, Dominic Kwiatkowski, Cordelia Langford, John Sillitoe on behalf of the Wellcome Sanger Institute COVID-19 Surveillance Team |
| EPI_ISL_2639761 | Lighthouse Lab in Alderley Park | Wellcome Sanger Institute for the COVID-19 Genomics UK (COG-UK) Consortium | Jacquelyn Wynn, Mairead Hyland, The Lighthouse Lab in Alderley Park and Alex Alderton, Roberto Amato, Jeffrey Barrett, Sonia Goncalves, Ewan Harrison, David K. Jackson, Ian Johnston, Dominic Kwiatkowski, Cordelia Langford, John Sillitoe on behalf of the Wellcome Sanger Institute COVID-19 Surveillance Team |
| EPI_ISL_2639762 | Lighthouse Laboratory Plymouth | Wellcome Sanger Institute for the COVID-19 Genomics UK (COG-UK) Consortium | Lighthouse Laboratory Plymouth and Alex Alderton, Roberto Amato, Jeffrey Barrett, Sonia Goncalves, Ewan Harrison, David K. Jackson, Ian Johnston, Dominic Kwiatkowski, Cordelia Langford, John Sillitoe on behalf of the Wellcome Sanger Institute COVID-19 Surveillance Team |
| EPI_ISL_2639763 | Lighthouse Lab in Alderley Park | Wellcome Sanger Institute for the COVID-19 Genomics UK (COG-UK) Consortium | Jacquelyn Wynn, Mairead Hyland, The Lighthouse Lab in Alderley Park and Alex Alderton, Roberto Amato, Jeffrey Barrett, Sonia Goncalves, Ewan Harrison, David K. Jackson, Ian Johnston, Dominic Kwiatkowski, Cordelia Langford, John Sillitoe on behalf of the Wellcome Sanger Institute COVID-19 Surveillance Team |
| EPI_ISL_2639764 | Lighthouse Laboratory Plymouth | Wellcome Sanger Institute for the COVID-19 Genomics UK (COG-UK) Consortium | Lighthouse Laboratory Plymouth and Alex Alderton, Roberto Amato, Jeffrey Barrett, Sonia Goncalves, Ewan Harrison, David K. Jackson, Ian Johnston, Dominic Kwiatkowski, Cordelia Langford, John Sillitoe on behalf of the Wellcome Sanger Institute COVID-19 Surveillance Team |
| EPI_ISL_2639765 | Lighthouse Lab in Glasgow | Wellcome Sanger Institute for the COVID-19 Genomics UK (COG-UK) Consortium | Harper VanSteenhouse, Yumi Kasai, David Gray, Carol Clugston, Anna Dominiczak and Alex Alderton, Roberto Amato, Jeffrey Barrett, Sonia Goncalves, Ewan Harrison, David K. Jackson, Ian Johnston, Dominic Kwiatkowski, Cordelia Langford, John Sillitoe on behalf of the Wellcome Sanger Institute COVID-19 Surveillance Team |
| EPI_ISL_2639766, EPI_ISL_2639767, EPI_ISL_2639768 | Lighthouse Lab in Alderley Park | Wellcome Sanger Institute for the COVID-19 Genomics UK (COG-UK) Consortium | Jacquelyn Wynn, Mairead Hyland, The Lighthouse Lab in Alderley Park and Alex Alderton, Roberto Amato, Jeffrey Barrett, Sonia Goncalves, Ewan Harrison, David K. Jackson, Ian Johnston, Dominic Kwiatkowski, Cordelia Langford, John Sillitoe on behalf of the Wellcome Sanger Institute COVID-19 Surveillance Team |
| EPI_ISL_2639769 | Lighthouse Lab in Glasgow | Wellcome Sanger Institute for the COVID-19 Genomics UK (COG-UK) Consortium | Harper VanSteenhouse, Yumi Kasai, David Gray, Carol Clugston, Anna Dominiczak and Alex Alderton, Roberto Amato, Jeffrey Barrett, Sonia Goncalves, Ewan Harrison, David K. Jackson, Ian Johnston, Dominic Kwiatkowski, Cordelia Langford, John Sillitoe on behalf of the Wellcome Sanger Institute COVID-19 Surveillance Team |
| EPI_ISL_2639770, EPI_ISL_2639771, EPI_ISL_2639772 | Lighthouse Lab in Alderley Park | Wellcome Sanger Institute for the COVID-19 Genomics UK (COG-UK) Consortium | Jacquelyn Wynn, Mairead Hyland, The Lighthouse Lab in Alderley Park and Alex Alderton, Roberto Amato, Jeffrey Barrett, Sonia Goncalves, Ewan Harrison, David K. Jackson, Ian Johnston, Dominic Kwiatkowski, Cordelia Langford, John Sillitoe on behalf of the Wellcome Sanger Institute COVID-19 Surveillance Team |
| EPI_ISL_2639773, EPI_ISL_2639774 | Lighthouse Laboratory Plymouth | Wellcome Sanger Institute for the COVID-19 Genomics UK (COG-UK) Consortium | Lighthouse Laboratory Plymouth and Alex Alderton, Roberto Amato, Jeffrey Barrett, Sonia Goncalves, Ewan Harrison, David K. Jackson, Ian Johnston, Dominic Kwiatkowski, Cordelia Langford, John Sillitoe on behalf of the Wellcome Sanger Institute COVID-19 Surveillance Team |
| EPI_ISL_2639775 | Lighthouse Lab in Glasgow | Wellcome Sanger Institute for the COVID-19 Genomics UK (COG-UK) Consortium | Harper VanSteenhouse, Yumi Kasai, David Gray, Carol Clugston, Anna Dominiczak and Alex Alderton, Roberto Amato, Jeffrey Barrett, Sonia Goncalves, Ewan Harrison, David K. Jackson, Ian Johnston, Dominic Kwiatkowski, Cordelia Langford, John Sillitoe on behalf of the Wellcome Sanger Institute COVID-19 Surveillance Team |
| EPI_ISL_2639776 | Lighthouse Lab in Alderley Park | Wellcome Sanger Institute for the COVID-19 Genomics UK (COG-UK) Consortium | Jacquelyn Wynn, Mairead Hyland, The Lighthouse Lab in Alderley Park and Alex Alderton, Roberto Amato, Jeffrey Barrett, Sonia Goncalves, Ewan Harrison, David K. Jackson, Ian Johnston, Dominic Kwiatkowski, Cordelia Langford, John Sillitoe on behalf of the Wellcome Sanger Institute COVID-19 Surveillance Team |
| EPI_ISL_2639777 | Lighthouse Lab in Glasgow | Wellcome Sanger Institute for the COVID-19 Genomics UK (COG-UK) Consortium | Harper VanSteenhouse, Yumi Kasai, David Gray, Carol Clugston, Anna Dominiczak and Alex Alderton, Roberto Amato, Jeffrey Barrett, Sonia Goncalves, Ewan Harrison, David K. Jackson, Ian Johnston, Dominic Kwiatkowski, Cordelia Langford, John Sillitoe on behalf of the Wellcome Sanger Institute COVID-19 Surveillance Team |
| EPI_ISL_2639778, EPI_ISL_2639779, EPI_ISL_2639780 | Lighthouse Lab in Alderley Park | Wellcome Sanger Institute for the COVID-19 Genomics UK (COG-UK) Consortium | Jacquelyn Wynn, Mairead Hyland, The Lighthouse Lab in Alderley Park and Alex Alderton, Roberto Amato, Jeffrey Barrett, Sonia Goncalves, Ewan Harrison, David K. Jackson, Ian Johnston, Dominic Kwiatkowski, Cordelia Langford, John Sillitoe on behalf of the Wellcome Sanger Institute COVID-19 Surveillance Team |
| EPI_ISL_2639781, EPI_ISL_2639782 | Lighthouse Laboratory Plymouth | Wellcome Sanger Institute for the COVID-19 Genomics UK (COG-UK) Consortium | Lighthouse Laboratory Plymouth and Alex Alderton, Roberto Amato, Jeffrey Barrett, Sonia Goncalves, Ewan Harrison, David K. Jackson, Ian Johnston, Dominic Kwiatkowski, Cordelia Langford, John Sillitoe on behalf of the Wellcome Sanger Institute COVID-19 Surveillance Team |
| EPI_ISL_2639783 | Lighthouse Lab in Alderley Park | Wellcome Sanger Institute for the COVID-19 Genomics UK (COG-UK) Consortium | Jacquelyn Wynn, Mairead Hyland, The Lighthouse Lab in Alderley Park and Alex Alderton, Roberto Amato, Jeffrey Barrett, Sonia Goncalves, Ewan Harrison, David K. Jackson, Ian Johnston, Dominic Kwiatkowski, Cordelia Langford, John Sillitoe on behalf of the Wellcome Sanger Institute COVID-19 Surveillance Team |
| EPI_ISL_2639784 | Lighthouse Laboratory Plymouth | Wellcome Sanger Institute for the COVID-19 Genomics UK (COG-UK) Consortium | Lighthouse Laboratory Plymouth and Alex Alderton, Roberto Amato, Jeffrey Barrett, Sonia Goncalves, Ewan Harrison, David K. Jackson, Ian Johnston, Dominic Kwiatkowski, Cordelia Langford, John Sillitoe on behalf of the Wellcome Sanger Institute COVID-19 Surveillance Team |
| EPI_ISL_2639785, EPI_ISL_2639786, EPI_ISL_2639787 | Lighthouse Lab in Alderley Park | Wellcome Sanger Institute for the COVID-19 Genomics UK (COG-UK) Consortium | Jacquelyn Wynn, Mairead Hyland, The Lighthouse Lab in Alderley Park and Alex Alderton, Roberto Amato, Jeffrey Barrett, Sonia Goncalves, Ewan Harrison, David K. Jackson, Ian Johnston, Dominic Kwiatkowski, Cordelia Langford, John Sillitoe on behalf of the Wellcome Sanger Institute COVID-19 Surveillance Team |
| EPI_ISL_2639788 | Lighthouse Laboratory Plymouth | Wellcome Sanger Institute for the COVID-19 Genomics UK (COG-UK) Consortium | Lighthouse Laboratory Plymouth and Alex Alderton, Roberto Amato, Jeffrey Barrett, Sonia Goncalves, Ewan Harrison, David K. Jackson, Ian Johnston, Dominic Kwiatkowski, Cordelia Langford, John Sillitoe on behalf of the Wellcome Sanger Institute COVID-19 Surveillance Team |
| EPI_ISL_2639789 | Lighthouse Lab in Alderley Park | Wellcome Sanger Institute for the COVID-19 Genomics UK (COG-UK) Consortium | Jacquelyn Wynn, Mairead Hyland, The Lighthouse Lab in Alderley Park and Alex Alderton, Roberto Amato, Jeffrey Barrett, Sonia Goncalves, Ewan Harrison, David K. Jackson, Ian Johnston, Dominic Kwiatkowski, Cordelia Langford, John Sillitoe on behalf of the Wellcome Sanger Institute COVID-19 Surveillance Team |
| EPI_ISL_2639790, EPI_ISL_2639791 | Lighthouse Laboratory Plymouth | Wellcome Sanger Institute for the COVID-19 Genomics UK (COG-UK) Consortium | Lighthouse Laboratory Plymouth and Alex Alderton, Roberto Amato, Jeffrey Barrett, Sonia Goncalves, Ewan Harrison, David K. Jackson, Ian Johnston, Dominic Kwiatkowski, Cordelia Langford, John Sillitoe on behalf of the Wellcome Sanger Institute COVID-19 Surveillance Team |
| EPI_ISL_2639792, EPI_ISL_2639793, EPI_ISL_2639794, EPI_ISL_2639795 | Lighthouse Lab in Alderley Park | Wellcome Sanger Institute for the COVID-19 Genomics UK (COG-UK) Consortium | Jacquelyn Wynn, Mairead Hyland, The Lighthouse Lab in Alderley Park and Alex Alderton, Roberto Amato, Jeffrey Barrett, Sonia Goncalves, Ewan Harrison, David K. Jackson, Ian Johnston, Dominic Kwiatkowski, Cordelia Langford, John Sillitoe on behalf of the Wellcome Sanger Institute COVID-19 Surveillance Team |
| EPI_ISL_2639796, EPI_ISL_2639797 | Lighthouse Lab in Glasgow | Wellcome Sanger Institute for the COVID-19 Genomics UK (COG-UK) Consortium | Harper VanSteenhouse, Yumi Kasai, David Gray, Carol Clugston, Anna Dominiczak and Alex Alderton, Roberto Amato, Jeffrey Barrett, Sonia Goncalves, Ewan Harrison, David K. Jackson, Ian Johnston, Dominic Kwiatkowski, Cordelia Langford, John Sillitoe on behalf of the Wellcome Sanger Institute COVID-19 Surveillance Team |
| EPI_ISL_2639798, EPI_ISL_2639799 | Lighthouse Laboratory Plymouth | Wellcome Sanger Institute for the COVID-19 Genomics UK (COG-UK) Consortium | Lighthouse Laboratory Plymouth and Alex Alderton, Roberto Amato, Jeffrey Barrett, Sonia Goncalves, Ewan Harrison, David K. Jackson, Ian Johnston, Dominic Kwiatkowski, Cordelia Langford, John Sillitoe on behalf of the Wellcome Sanger Institute COVID-19 Surveillance Team |
| EPI_ISL_2639800, EPI_ISL_2639801, EPI_ISL_2639802, EPI_ISL_2639803 | Lighthouse Lab in Alderley Park | Wellcome Sanger Institute for the COVID-19 Genomics UK (COG-UK) Consortium | Jacquelyn Wynn, Mairead Hyland, The Lighthouse Lab in Alderley Park and Alex Alderton, Roberto Amato, Jeffrey Barrett, Sonia Goncalves, Ewan Harrison, David K. Jackson, Ian Johnston, Dominic Kwiatkowski, Cordelia Langford, John Sillitoe on behalf of the Wellcome Sanger Institute COVID-19 Surveillance Team |
| EPI_ISL_2639804 | Lighthouse Lab in Glasgow | Wellcome Sanger Institute for the COVID-19 Genomics UK (COG-UK) Consortium | Harper VanSteenhouse, Yumi Kasai, David Gray, Carol Clugston, Anna Dominiczak and Alex Alderton, Roberto Amato, Jeffrey Barrett, Sonia Goncalves, Ewan Harrison, David K. Jackson, Ian Johnston, Dominic Kwiatkowski, Cordelia Langford, John Sillitoe on behalf of the Wellcome Sanger Institute COVID-19 Surveillance Team |
| EPI_ISL_2639805 | Lighthouse Laboratory Plymouth | Wellcome Sanger Institute for the COVID-19 Genomics UK (COG-UK) Consortium | Lighthouse Laboratory Plymouth and Alex Alderton, Roberto Amato, Jeffrey Barrett, Sonia Goncalves, Ewan Harrison, David K. Jackson, Ian Johnston, Dominic Kwiatkowski, Cordelia Langford, John Sillitoe on behalf of the Wellcome Sanger Institute COVID-19 Surveillance Team |
| EPI_ISL_2639806 | Lighthouse Lab in Glasgow | Wellcome Sanger Institute for the COVID-19 Genomics UK (COG-UK) Consortium | Harper VanSteenhouse, Yumi Kasai, David Gray, Carol Clugston, Anna Dominiczak and Alex Alderton, Roberto Amato, Jeffrey Barrett, Sonia Goncalves, Ewan Harrison, David K. Jackson, Ian Johnston, Dominic Kwiatkowski, Cordelia Langford, John Sillitoe on behalf of the Wellcome Sanger Institute COVID-19 Surveillance Team |
| EPI_ISL_2639807, EPI_ISL_2639808, EPI_ISL_2639809, EPI_ISL_2639810 | Lighthouse Lab in Alderley Park | Wellcome Sanger Institute for the COVID-19 Genomics UK (COG-UK) Consortium | Jacquelyn Wynn, Mairead Hyland, The Lighthouse Lab in Alderley Park and Alex Alderton, Roberto Amato, Jeffrey Barrett, Sonia Goncalves, Ewan Harrison, David K. Jackson, Ian Johnston, Dominic Kwiatkowski, Cordelia Langford, John Sillitoe on behalf of the Wellcome Sanger Institute COVID-19 Surveillance Team |
| EPI_ISL_2639811 | Lighthouse Laboratory Plymouth | Wellcome Sanger Institute for the COVID-19 Genomics UK (COG-UK) Consortium | Lighthouse Laboratory Plymouth and Alex Alderton, Roberto Amato, Jeffrey Barrett, Sonia Goncalves, Ewan Harrison, David K. Jackson, Ian Johnston, Dominic Kwiatkowski, Cordelia Langford, John Sillitoe on behalf of the Wellcome Sanger Institute COVID-19 Surveillance Team |
| EPI_ISL_2639812 | Lighthouse Lab in Glasgow | Wellcome Sanger Institute for the COVID-19 Genomics UK (COG-UK) Consortium | Harper VanSteenhouse, Yumi Kasai, David Gray, Carol Clugston, Anna Dominiczak and Alex Alderton, Roberto Amato, Jeffrey Barrett, Sonia Goncalves, Ewan Harrison, David K. Jackson, Ian Johnston, Dominic Kwiatkowski, Cordelia Langford, John Sillitoe on behalf of the Wellcome Sanger Institute COVID-19 Surveillance Team |
| EPI_ISL_2639813 | Lighthouse Laboratory Plymouth | Wellcome Sanger Institute for the COVID-19 Genomics UK (COG-UK) Consortium | Lighthouse Laboratory Plymouth and Alex Alderton, Roberto Amato, Jeffrey Barrett, Sonia Goncalves, Ewan Harrison, David K. Jackson, Ian Johnston, Dominic Kwiatkowski, Cordelia Langford, John Sillitoe on behalf of the Wellcome Sanger Institute COVID-19 Surveillance Team |
| EPI_ISL_2639814, EPI_ISL_2639815, EPI_ISL_2639816, EPI_ISL_2639817 | Lighthouse Lab in Alderley Park | Wellcome Sanger Institute for the COVID-19 Genomics UK (COG-UK) Consortium | Jacquelyn Wynn, Mairead Hyland, The Lighthouse Lab in Alderley Park and Alex Alderton, Roberto Amato, Jeffrey Barrett, Sonia Goncalves, Ewan Harrison, David K. Jackson, Ian Johnston, Dominic Kwiatkowski, Cordelia Langford, John Sillitoe on behalf of the Wellcome Sanger Institute COVID-19 Surveillance Team |
| EPI_ISL_2639818 | Lighthouse Lab in Glasgow | Wellcome Sanger Institute for the COVID-19 Genomics UK (COG-UK) Consortium | Harper VanSteenhouse, Yumi Kasai, David Gray, Carol Clugston, Anna Dominiczak and Alex Alderton, Roberto Amato, Jeffrey Barrett, Sonia Goncalves, Ewan Harrison, David K. Jackson, Ian Johnston, Dominic Kwiatkowski, Cordelia Langford, John Sillitoe on behalf of the Wellcome Sanger Institute COVID-19 Surveillance Team |
| EPI_ISL_2639819 | Lighthouse Lab in Alderley Park | Wellcome Sanger Institute for the COVID-19 Genomics UK (COG-UK) Consortium | Jacquelyn Wynn, Mairead Hyland, The Lighthouse Lab in Alderley Park and Alex Alderton, Roberto Amato, Jeffrey Barrett, Sonia Goncalves, Ewan Harrison, David K. Jackson, Ian Johnston, Dominic Kwiatkowski, Cordelia Langford, John Sillitoe on behalf of the Wellcome Sanger Institute COVID-19 Surveillance Team |
| EPI_ISL_2639820 | Lighthouse Laboratory Plymouth | Wellcome Sanger Institute for the COVID-19 Genomics UK (COG-UK) Consortium | Lighthouse Laboratory Plymouth and Alex Alderton, Roberto Amato, Jeffrey Barrett, Sonia Goncalves, Ewan Harrison, David K. Jackson, Ian Johnston, Dominic Kwiatkowski, Cordelia Langford, John Sillitoe on behalf of the Wellcome Sanger Institute COVID-19 Surveillance Team |
| EPI_ISL_2639821 | Lighthouse Lab in Alderley Park | Wellcome Sanger Institute for the COVID-19 Genomics UK (COG-UK) Consortium | Jacquelyn Wynn, Mairead Hyland, The Lighthouse Lab in Alderley Park and Alex Alderton, Roberto Amato, Jeffrey Barrett, Sonia Goncalves, Ewan Harrison, David K. Jackson, Ian Johnston, Dominic Kwiatkowski, Cordelia Langford, John Sillitoe on behalf of the Wellcome Sanger Institute COVID-19 Surveillance Team |

| | | | |
|---|---|---|---|
| EPI_ISL_2639822 | Lighthouse Laboratory Plymouth | Wellcome Sanger Institute for the COVID-19 Genomics UK (COG-UK) Consortium | Lighthouse Laboratory Plymouth and Alex Alderton, Roberto Amato, Jeffrey Barrett, Sonia Goncalves, Ewan Harrison, David K. Jackson, Ian Johnston, Dominic Kwiatkowski, Cordelia Langford, John Sillitoe on behalf of the Wellcome Sanger Institute COVID-19 Surveillance Team |
| EPI_ISL_2639823, EPI_ISL_2639824, EPI_ISL_2639825, EPI_ISL_2639826, EPI_ISL_2639827 | Lighthouse Lab in Alderley Park | Wellcome Sanger Institute for the COVID-19 Genomics UK (COG-UK) Consortium | Jacquelyn Wynn, Mairead Hyland, The Lighthouse Lab in Alderley Park and Alex Alderton, Roberto Amato, Jeffrey Barrett, Sonia Goncalves, Ewan Harrison, David K. Jackson, Ian Johnston, Dominic Kwiatkowski, Cordelia Langford, John Sillitoe on behalf of the Wellcome Sanger Institute COVID-19 Surveillance Team |
| EPI_ISL_2639828 | Lighthouse Laboratory Plymouth | Wellcome Sanger Institute for the COVID-19 Genomics UK (COG-UK) Consortium | Lighthouse Laboratory Plymouth and Alex Alderton, Roberto Amato, Jeffrey Barrett, Sonia Goncalves, Ewan Harrison, David K. Jackson, Ian Johnston, Dominic Kwiatkowski, Cordelia Langford, John Sillitoe on behalf of the Wellcome Sanger Institute COVID-19 Surveillance Team |
| EPI_ISL_2639829 | Lighthouse Lab in Glasgow | Wellcome Sanger Institute for the COVID-19 Genomics UK (COG-UK) Consortium | Harper VanSteenhouse, Yumi Kasai, David Gray, Carol Clugston, Anna Dominiczak and Alex Alderton, Roberto Amato, Jeffrey Barrett, Sonia Goncalves, Ewan Harrison, David K. Jackson, Ian Johnston, Dominic Kwiatkowski, Cordelia Langford, John Sillitoe on behalf of the Wellcome Sanger Institute COVID-19 Surveillance Team |
| EPI_ISL_2639830, EPI_ISL_2639831 | Lighthouse Lab in Alderley Park | Wellcome Sanger Institute for the COVID-19 Genomics UK (COG-UK) Consortium | Jacquelyn Wynn, Mairead Hyland, The Lighthouse Lab in Alderley Park and Alex Alderton, Roberto Amato, Jeffrey Barrett, Sonia Goncalves, Ewan Harrison, David K. Jackson, Ian Johnston, Dominic Kwiatkowski, Cordelia Langford, John Sillitoe on behalf of the Wellcome Sanger Institute COVID-19 Surveillance Team |
| EPI_ISL_2639832 | Lighthouse Laboratory Plymouth | Wellcome Sanger Institute for the COVID-19 Genomics UK (COG-UK) Consortium | Lighthouse Laboratory Plymouth and Alex Alderton, Roberto Amato, Jeffrey Barrett, Sonia Goncalves, Ewan Harrison, David K. Jackson, Ian Johnston, Dominic Kwiatkowski, Cordelia Langford, John Sillitoe on behalf of the Wellcome Sanger Institute COVID-19 Surveillance Team |
| EPI_ISL_2639833 | Lighthouse Lab in Glasgow | Wellcome Sanger Institute for the COVID-19 Genomics UK (COG-UK) Consortium | Harper VanSteenhouse, Yumi Kasai, David Gray, Carol Clugston, Anna Dominiczak and Alex Alderton, Roberto Amato, Jeffrey Barrett, Sonia Goncalves, Ewan Harrison, David K. Jackson, Ian Johnston, Dominic Kwiatkowski, Cordelia Langford, John Sillitoe on behalf of the Wellcome Sanger Institute COVID-19 Surveillance Team |
| EPI_ISL_2639834 | Lighthouse Lab in Alderley Park | Wellcome Sanger Institute for the COVID-19 Genomics UK (COG-UK) Consortium | Jacquelyn Wynn, Mairead Hyland, The Lighthouse Lab in Alderley Park and Alex Alderton, Roberto Amato, Jeffrey Barrett, Sonia Goncalves, Ewan Harrison, David K. Jackson, Ian Johnston, Dominic Kwiatkowski, Cordelia Langford, John Sillitoe on behalf of the Wellcome Sanger Institute COVID-19 Surveillance Team |
| EPI_ISL_2639835, EPI_ISL_2639836 | Lighthouse Laboratory Plymouth | Wellcome Sanger Institute for the COVID-19 Genomics UK (COG-UK) Consortium | Lighthouse Laboratory Plymouth and Alex Alderton, Roberto Amato, Jeffrey Barrett, Sonia Goncalves, Ewan Harrison, David K. Jackson, Ian Johnston, Dominic Kwiatkowski, Cordelia Langford, John Sillitoe on behalf of the Wellcome Sanger Institute COVID-19 Surveillance Team |
| EPI_ISL_2639837 | Lighthouse Lab in Alderley Park | Wellcome Sanger Institute for the COVID-19 Genomics UK (COG-UK) Consortium | Jacquelyn Wynn, Mairead Hyland, The Lighthouse Lab in Alderley Park and Alex Alderton, Roberto Amato, Jeffrey Barrett, Sonia Goncalves, Ewan Harrison, David K. Jackson, Ian Johnston, Dominic Kwiatkowski, Cordelia Langford, John Sillitoe on behalf of the Wellcome Sanger Institute COVID-19 Surveillance Team |
| EPI_ISL_2639838, EPI_ISL_2639839, EPI_ISL_2639840, EPI_ISL_2639841 | Lighthouse Laboratory Plymouth | Wellcome Sanger Institute for the COVID-19 Genomics UK (COG-UK) Consortium | Lighthouse Laboratory Plymouth and Alex Alderton, Roberto Amato, Jeffrey Barrett, Sonia Goncalves, Ewan Harrison, David K. Jackson, Ian Johnston, Dominic Kwiatkowski, Cordelia Langford, John Sillitoe on behalf of the Wellcome Sanger Institute COVID-19 Surveillance Team |
| EPI_ISL_2639842, EPI_ISL_2639843 | Lighthouse Lab in Alderley Park | Wellcome Sanger Institute for the COVID-19 Genomics UK (COG-UK) Consortium | Jacquelyn Wynn, Mairead Hyland, The Lighthouse Lab in Alderley Park and Alex Alderton, Roberto Amato, Jeffrey Barrett, Sonia Goncalves, Ewan Harrison, David K. Jackson, Ian Johnston, Dominic Kwiatkowski, Cordelia Langford, John Sillitoe on behalf of the Wellcome Sanger Institute COVID-19 Surveillance Team |
| EPI_ISL_2639844 | Lighthouse Laboratory Plymouth | Wellcome Sanger Institute for the COVID-19 Genomics UK (COG-UK) Consortium | Lighthouse Laboratory Plymouth and Alex Alderton, Roberto Amato, Jeffrey Barrett, Sonia Goncalves, Ewan Harrison, David K. Jackson, Ian Johnston, Dominic Kwiatkowski, Cordelia Langford, John Sillitoe on behalf of the Wellcome Sanger Institute COVID-19 Surveillance Team |
| EPI_ISL_2639845, EPI_ISL_2639846 | Lighthouse Lab in Alderley Park | Wellcome Sanger Institute for the COVID-19 Genomics UK (COG-UK) Consortium | Jacquelyn Wynn, Mairead Hyland, The Lighthouse Lab in Alderley Park and Alex Alderton, Roberto Amato, Jeffrey Barrett, Sonia Goncalves, Ewan Harrison, David K. Jackson, Ian Johnston, Dominic Kwiatkowski, Cordelia Langford, John Sillitoe on behalf of the Wellcome Sanger Institute COVID-19 Surveillance Team |
| EPI_ISL_2639847 | Lighthouse Laboratory Plymouth | Wellcome Sanger Institute for the COVID-19 Genomics UK (COG-UK) Consortium | Lighthouse Laboratory Plymouth and Alex Alderton, Roberto Amato, Jeffrey Barrett, Sonia Goncalves, Ewan Harrison, David K. Jackson, Ian Johnston, Dominic Kwiatkowski, Cordelia Langford, John Sillitoe on behalf of the Wellcome Sanger Institute COVID-19 Surveillance Team |
| EPI_ISL_2639848, EPI_ISL_2639849, EPI_ISL_2639850, EPI_ISL_2639851 | Lighthouse Lab in Alderley Park | Wellcome Sanger Institute for the COVID-19 Genomics UK (COG-UK) Consortium | Jacquelyn Wynn, Mairead Hyland, The Lighthouse Lab in Alderley Park and Alex Alderton, Roberto Amato, Jeffrey Barrett, Sonia Goncalves, Ewan Harrison, David K. Jackson, Ian Johnston, Dominic Kwiatkowski, Cordelia Langford, John Sillitoe on behalf of the Wellcome Sanger Institute COVID-19 Surveillance Team |
| EPI_ISL_2639852 | Lighthouse Lab in Glasgow | Wellcome Sanger Institute for the COVID-19 Genomics UK (COG-UK) Consortium | Harper VanSteenhouse, Yumi Kasai, David Gray, Carol Clugston, Anna Dominiczak and Alex Alderton, Roberto Amato, Jeffrey Barrett, Sonia Goncalves, Ewan Harrison, David K. Jackson, Ian Johnston, Dominic Kwiatkowski, Cordelia Langford, John Sillitoe on behalf of the Wellcome Sanger Institute COVID-19 Surveillance Team |
| EPI_ISL_2639853, EPI_ISL_2639854, EPI_ISL_2639855, EPI_ISL_2639856, EPI_ISL_2639857 | Lighthouse Lab in Alderley Park | Wellcome Sanger Institute for the COVID-19 Genomics UK (COG-UK) Consortium | Jacquelyn Wynn, Mairead Hyland, The Lighthouse Lab in Alderley Park and Alex Alderton, Roberto Amato, Jeffrey Barrett, Sonia Goncalves, Ewan Harrison, David K. Jackson, Ian Johnston, Dominic Kwiatkowski, Cordelia Langford, John Sillitoe on behalf of the Wellcome Sanger Institute COVID-19 Surveillance Team |
| EPI_ISL_2639858, EPI_ISL_2639860 | Lighthouse Laboratory Plymouth | Wellcome Sanger Institute for the COVID-19 Genomics UK (COG-UK) Consortium | Lighthouse Laboratory Plymouth and Alex Alderton, Roberto Amato, Jeffrey Barrett, Sonia Goncalves, Ewan Harrison, David K. Jackson, Ian Johnston, Dominic Kwiatkowski, Cordelia Langford, John Sillitoe on behalf of the Wellcome Sanger Institute COVID-19 Surveillance Team |
| EPI_ISL_2639861 | Lighthouse Lab in Alderley Park | Wellcome Sanger Institute for the COVID-19 Genomics UK (COG-UK) Consortium | Jacquelyn Wynn, Mairead Hyland, The Lighthouse Lab in Alderley Park and Alex Alderton, Roberto Amato, Jeffrey Barrett, Sonia Goncalves, Ewan Harrison, David K. Jackson, Ian Johnston, Dominic Kwiatkowski, Cordelia Langford, John Sillitoe on behalf of the Wellcome Sanger Institute COVID-19 Surveillance Team |
| EPI_ISL_2639862, EPI_ISL_2639863, EPI_ISL_2639864 | Lighthouse Lab in Glasgow | Wellcome Sanger Institute for the COVID-19 Genomics UK (COG-UK) Consortium | Harper VanSteenhouse, Yumi Kasai, David Gray, Carol Clugston, Anna Dominiczak and Alex Alderton, Roberto Amato, Jeffrey Barrett, Sonia Goncalves, Ewan Harrison, David K. Jackson, Ian Johnston, Dominic Kwiatkowski, Cordelia Langford, John Sillitoe on behalf of the Wellcome Sanger Institute COVID-19 Surveillance Team |
| EPI_ISL_2639866 | Lighthouse Lab in Alderley Park | Wellcome Sanger Institute for the COVID-19 Genomics UK (COG-UK) Consortium | Jacquelyn Wynn, Mairead Hyland, The Lighthouse Lab in Alderley Park and Alex Alderton, Roberto Amato, Jeffrey Barrett, Sonia Goncalves, Ewan Harrison, David K. Jackson, Ian Johnston, Dominic Kwiatkowski, Cordelia Langford, John Sillitoe on behalf of the Wellcome Sanger Institute COVID-19 Surveillance Team |
| EPI_ISL_2639867 | Lighthouse Laboratory Plymouth | Wellcome Sanger Institute for the COVID-19 Genomics UK (COG-UK) Consortium | Lighthouse Laboratory Plymouth and Alex Alderton, Roberto Amato, Jeffrey Barrett, Sonia Goncalves, Ewan Harrison, David K. Jackson, Ian Johnston, Dominic Kwiatkowski, Cordelia Langford, John Sillitoe on behalf of the Wellcome Sanger Institute COVID-19 Surveillance Team |
| EPI_ISL_2639868, EPI_ISL_2639869, EPI_ISL_2639870, EPI_ISL_2639871 | Lighthouse Lab in Alderley Park | Wellcome Sanger Institute for the COVID-19 Genomics UK (COG-UK) Consortium | Jacquelyn Wynn, Mairead Hyland, The Lighthouse Lab in Alderley Park and Alex Alderton, Roberto Amato, Jeffrey Barrett, Sonia Goncalves, Ewan Harrison, David K. Jackson, Ian Johnston, Dominic Kwiatkowski, Cordelia Langford, John Sillitoe on behalf of the Wellcome Sanger Institute COVID-19 Surveillance Team |
| EPI_ISL_2639872 | Lighthouse Laboratory Plymouth | Wellcome Sanger Institute for the COVID-19 Genomics UK (COG-UK) Consortium | Lighthouse Laboratory Plymouth and Alex Alderton, Roberto Amato, Jeffrey Barrett, Sonia Goncalves, Ewan Harrison, David K. Jackson, Ian Johnston, Dominic Kwiatkowski, Cordelia Langford, John Sillitoe on behalf of the Wellcome Sanger Institute COVID-19 Surveillance Team |
| EPI_ISL_2639873, EPI_ISL_2639874, EPI_ISL_2639875, EPI_ISL_2639876, EPI_ISL_2639877 | Lighthouse Lab in Alderley Park | Wellcome Sanger Institute for the COVID-19 Genomics UK (COG-UK) Consortium | Jacquelyn Wynn, Mairead Hyland, The Lighthouse Lab in Alderley Park and Alex Alderton, Roberto Amato, Jeffrey Barrett, Sonia Goncalves, Ewan Harrison, David K. Jackson, Ian Johnston, Dominic Kwiatkowski, Cordelia Langford, John Sillitoe on behalf of the Wellcome Sanger Institute COVID-19 Surveillance Team |
| EPI_ISL_2639878 | Lighthouse Lab in Glasgow | Wellcome Sanger Institute for the COVID-19 Genomics UK (COG-UK) Consortium | Harper VanSteenhouse, Yumi Kasai, David Gray, Carol Clugston, Anna Dominiczak and Alex Alderton, Roberto Amato, Jeffrey Barrett, Sonia Goncalves, Ewan Harrison, David K. Jackson, Ian Johnston, Dominic Kwiatkowski, Cordelia Langford, John Sillitoe on behalf of the Wellcome Sanger Institute COVID-19 Surveillance Team |
| EPI_ISL_2639879 | Lighthouse Lab in Alderley Park | Wellcome Sanger Institute for the COVID-19 Genomics UK (COG-UK) Consortium | Jacquelyn Wynn, Mairead Hyland, The Lighthouse Lab in Alderley Park and Alex Alderton, Roberto Amato, Jeffrey Barrett, Sonia Goncalves, Ewan Harrison, David K. Jackson, Ian Johnston, Dominic Kwiatkowski, Cordelia Langford, John Sillitoe on behalf of the Wellcome Sanger Institute COVID-19 Surveillance Team |
| EPI_ISL_2639880 | Lighthouse Laboratory Plymouth | Wellcome Sanger Institute for the COVID-19 Genomics UK (COG-UK) Consortium | Lighthouse Laboratory Plymouth and Alex Alderton, Roberto Amato, Jeffrey Barrett, Sonia Goncalves, Ewan Harrison, David K. Jackson, Ian Johnston, Dominic Kwiatkowski, Cordelia Langford, John Sillitoe on behalf of the Wellcome Sanger Institute COVID-19 Surveillance Team |
| EPI_ISL_2639881, EPI_ISL_2639882, EPI_ISL_2639883, EPI_ISL_2639884 | Lighthouse Lab in Alderley Park | Wellcome Sanger Institute for the COVID-19 Genomics UK (COG-UK) Consortium | Jacquelyn Wynn, Mairead Hyland, The Lighthouse Lab in Alderley Park and Alex Alderton, Roberto Amato, Jeffrey Barrett, Sonia Goncalves, Ewan Harrison, David K. Jackson, Ian Johnston, Dominic Kwiatkowski, Cordelia Langford, John Sillitoe on behalf of the Wellcome Sanger Institute COVID-19 Surveillance Team |
| EPI_ISL_2639885 | Lighthouse Laboratory Plymouth | Wellcome Sanger Institute for the COVID-19 Genomics UK (COG-UK) Consortium | Lighthouse Laboratory Plymouth and Alex Alderton, Roberto Amato, Jeffrey Barrett, Sonia Goncalves, Ewan Harrison, David K. Jackson, Ian Johnston, Dominic Kwiatkowski, Cordelia Langford, John Sillitoe on behalf of the Wellcome Sanger Institute COVID-19 Surveillance Team |
| EPI_ISL_2639886 | Lighthouse Lab in Alderley Park | Wellcome Sanger Institute for the COVID-19 Genomics UK (COG-UK) Consortium | Jacquelyn Wynn, Mairead Hyland, The Lighthouse Lab in Alderley Park and Alex Alderton, Roberto Amato, Jeffrey Barrett, Sonia Goncalves, Ewan Harrison, David K. Jackson, Ian Johnston, Dominic Kwiatkowski, Cordelia Langford, John Sillitoe on behalf of the Wellcome Sanger Institute COVID-19 Surveillance Team |
| EPI_ISL_2639887 | Lighthouse Laboratory Plymouth | Wellcome Sanger Institute for the COVID-19 Genomics UK (COG-UK) Consortium | Lighthouse Laboratory Plymouth and Alex Alderton, Roberto Amato, Jeffrey Barrett, Sonia Goncalves, Ewan Harrison, David K. Jackson, Ian Johnston, Dominic Kwiatkowski, Cordelia Langford, John Sillitoe on behalf of the Wellcome Sanger Institute COVID-19 Surveillance Team |
| EPI_ISL_2639888 | Lighthouse Lab in Glasgow | Wellcome Sanger Institute for the COVID-19 Genomics UK (COG-UK) Consortium | Harper VanSteenhouse, Yumi Kasai, David Gray, Carol Clugston, Anna Dominiczak and Alex Alderton, Roberto Amato, Jeffrey Barrett, Sonia Goncalves, Ewan Harrison, David K. Jackson, Ian Johnston, Dominic Kwiatkowski, Cordelia Langford, John Sillitoe on behalf of the Wellcome Sanger Institute COVID-19 Surveillance Team |
| EPI_ISL_2639889, EPI_ISL_2639890 | Lighthouse Laboratory Plymouth | Wellcome Sanger Institute for the COVID-19 Genomics UK (COG-UK) Consortium | Lighthouse Laboratory Plymouth and Alex Alderton, Roberto Amato, Jeffrey Barrett, Sonia Goncalves, Ewan Harrison, David K. Jackson, Ian Johnston, Dominic Kwiatkowski, Cordelia Langford, John Sillitoe on behalf of the Wellcome Sanger Institute COVID-19 Surveillance Team |
| EPI_ISL_2639891, EPI_ISL_2639892, EPI_ISL_2639893 | Lighthouse Lab in Alderley Park | Wellcome Sanger Institute for the COVID-19 Genomics UK (COG-UK) Consortium | Jacquelyn Wynn, Mairead Hyland, The Lighthouse Lab in Alderley Park and Alex Alderton, Roberto Amato, Jeffrey Barrett, Sonia Goncalves, Ewan Harrison, David K. Jackson, Ian Johnston, Dominic Kwiatkowski, Cordelia Langford, John Sillitoe on behalf of the Wellcome Sanger Institute COVID-19 Surveillance Team |
| EPI_ISL_2639894 | Lighthouse Laboratory Plymouth | Wellcome Sanger Institute for the COVID-19 Genomics UK (COG-UK) Consortium | Lighthouse Laboratory Plymouth and Alex Alderton, Roberto Amato, Jeffrey Barrett, Sonia Goncalves, Ewan Harrison, David K. Jackson, Ian Johnston, Dominic Kwiatkowski, Cordelia Langford, John Sillitoe on behalf of the Wellcome Sanger Institute COVID-19 Surveillance Team |
| EPI_ISL_2639896 | Lighthouse Lab in Glasgow | Wellcome Sanger Institute for the COVID-19 Genomics UK (COG-UK) | Harper VanSteenhouse, Yumi Kasai, David Gray, Carol Clugston, Anna Dominiczak and Alex Alderton, Roberto Amato, Jeffrey Barrett, Sonia Goncalves, Ewan Harrison, David K. Jackson, Ian Johnston, Dominic Kwiatkowski, Cordelia Langford, |

| | | | |
|---|---|---|---|
| | | Consortium | John Sillitoe on behalf of the Wellcome Sanger Institute COVID-19 Surveillance Team |
| EPI_ISL_2639897, EPI_ISL_2639898 | Lighthouse Laboratory Plymouth | Wellcome Sanger Institute for the COVID-19 Genomics UK (COG-UK) Consortium | Lighthouse Laboratory Plymouth and Alex Alderton, Roberto Amato, Jeffrey Barrett, Sonia Goncalves, Ewan Harrison, David K. Jackson, Ian Johnston, Dominic Kwiatkowski, Cordelia Langford, John Sillitoe on behalf of the Wellcome Sanger Institute COVID-19 Surveillance Team |
| EPI_ISL_2639899, EPI_ISL_2639900 | Lighthouse Lab in Alderley Park | Wellcome Sanger Institute for the COVID-19 Genomics UK (COG-UK) Consortium | Jacquelyn Wynn, Mairead Hyland, The Lighthouse Lab in Alderley Park and Alex Alderton, Roberto Amato, Jeffrey Barrett, Sonia Goncalves, Ewan Harrison, David K. Jackson, Ian Johnston, Dominic Kwiatkowski, Cordelia Langford, John Sillitoe on behalf of the Wellcome Sanger Institute COVID-19 Surveillance Team |
| EPI_ISL_2639901 | Lighthouse Laboratory Plymouth | Wellcome Sanger Institute for the COVID-19 Genomics UK (COG-UK) Consortium | Lighthouse Laboratory Plymouth and Alex Alderton, Roberto Amato, Jeffrey Barrett, Sonia Goncalves, Ewan Harrison, David K. Jackson, Ian Johnston, Dominic Kwiatkowski, Cordelia Langford, John Sillitoe on behalf of the Wellcome Sanger Institute COVID-19 Surveillance Team |
| EPI_ISL_2639902 | Lighthouse Lab in Glasgow | Wellcome Sanger Institute for the COVID-19 Genomics UK (COG-UK) Consortium | Harper VanSteenhouse, Yumi Kasai, David Gray, Carol Clugston, Anna Dominiczak and Alex Alderton, Roberto Amato, Jeffrey Barrett, Sonia Goncalves, Ewan Harrison, David K. Jackson, Ian Johnston, Dominic Kwiatkowski, Cordelia Langford, John Sillitoe on behalf of the Wellcome Sanger Institute COVID-19 Surveillance Team |
| EPI_ISL_2639903, EPI_ISL_2639904, EPI_ISL_2639905 | Lighthouse Lab in Alderley Park | Wellcome Sanger Institute for the COVID-19 Genomics UK (COG-UK) Consortium | Jacquelyn Wynn, Mairead Hyland, The Lighthouse Lab in Alderley Park and Alex Alderton, Roberto Amato, Jeffrey Barrett, Sonia Goncalves, Ewan Harrison, David K. Jackson, Ian Johnston, Dominic Kwiatkowski, Cordelia Langford, John Sillitoe on behalf of the Wellcome Sanger Institute COVID-19 Surveillance Team |
| EPI_ISL_2639906, EPI_ISL_2639907 | Lighthouse Laboratory Plymouth | Wellcome Sanger Institute for the COVID-19 Genomics UK (COG-UK) Consortium | Lighthouse Laboratory Plymouth and Alex Alderton, Roberto Amato, Jeffrey Barrett, Sonia Goncalves, Ewan Harrison, David K. Jackson, Ian Johnston, Dominic Kwiatkowski, Cordelia Langford, John Sillitoe on behalf of the Wellcome Sanger Institute COVID-19 Surveillance Team |
| EPI_ISL_2639908, EPI_ISL_2639909, EPI_ISL_2639910, EPI_ISL_2639911, EPI_ISL_2639912, EPI_ISL_2639913, EPI_ISL_2639914, EPI_ISL_2639915, EPI_ISL_2639916 | Lighthouse Lab in Alderley Park | Wellcome Sanger Institute for the COVID-19 Genomics UK (COG-UK) Consortium | Jacquelyn Wynn, Mairead Hyland, The Lighthouse Lab in Alderley Park and Alex Alderton, Roberto Amato, Jeffrey Barrett, Sonia Goncalves, Ewan Harrison, David K. Jackson, Ian Johnston, Dominic Kwiatkowski, Cordelia Langford, John Sillitoe on behalf of the Wellcome Sanger Institute COVID-19 Surveillance Team |
| EPI_ISL_2639917 | Lighthouse Laboratory Plymouth | Wellcome Sanger Institute for the COVID-19 Genomics UK (COG-UK) Consortium | Lighthouse Laboratory Plymouth and Alex Alderton, Roberto Amato, Jeffrey Barrett, Sonia Goncalves, Ewan Harrison, David K. Jackson, Ian Johnston, Dominic Kwiatkowski, Cordelia Langford, John Sillitoe on behalf of the Wellcome Sanger Institute COVID-19 Surveillance Team |
| EPI_ISL_2639918, EPI_ISL_2639919 | Lighthouse Lab in Alderley Park | Wellcome Sanger Institute for the COVID-19 Genomics UK (COG-UK) Consortium | Jacquelyn Wynn, Mairead Hyland, The Lighthouse Lab in Alderley Park and Alex Alderton, Roberto Amato, Jeffrey Barrett, Sonia Goncalves, Ewan Harrison, David K. Jackson, Ian Johnston, Dominic Kwiatkowski, Cordelia Langford, John Sillitoe on behalf of the Wellcome Sanger Institute COVID-19 Surveillance Team |
| EPI_ISL_2639920 | Lighthouse Laboratory Plymouth | Wellcome Sanger Institute for the COVID-19 Genomics UK (COG-UK) Consortium | Lighthouse Laboratory Plymouth and Alex Alderton, Roberto Amato, Jeffrey Barrett, Sonia Goncalves, Ewan Harrison, David K. Jackson, Ian Johnston, Dominic Kwiatkowski, Cordelia Langford, John Sillitoe on behalf of the Wellcome Sanger Institute COVID-19 Surveillance Team |
| EPI_ISL_2639921 | Lighthouse Lab in Glasgow | Wellcome Sanger Institute for the COVID-19 Genomics UK (COG-UK) Consortium | Harper VanSteenhouse, Yumi Kasai, David Gray, Carol Clugston, Anna Dominiczak and Alex Alderton, Roberto Amato, Jeffrey Barrett, Sonia Goncalves, Ewan Harrison, David K. Jackson, Ian Johnston, Dominic Kwiatkowski, Cordelia Langford, John Sillitoe on behalf of the Wellcome Sanger Institute COVID-19 Surveillance Team |
| EPI_ISL_2639922 | Lighthouse Laboratory Plymouth | Wellcome Sanger Institute for the COVID-19 Genomics UK (COG-UK) Consortium | Lighthouse Laboratory Plymouth and Alex Alderton, Roberto Amato, Jeffrey Barrett, Sonia Goncalves, Ewan Harrison, David K. Jackson, Ian Johnston, Dominic Kwiatkowski, Cordelia Langford, John Sillitoe on behalf of the Wellcome Sanger Institute COVID-19 Surveillance Team |
| EPI_ISL_2639923 | Lighthouse Lab in Alderley Park | Wellcome Sanger Institute for the COVID-19 Genomics UK (COG-UK) Consortium | Jacquelyn Wynn, Mairead Hyland, The Lighthouse Lab in Alderley Park and Alex Alderton, Roberto Amato, Jeffrey Barrett, Sonia Goncalves, Ewan Harrison, David K. Jackson, Ian Johnston, Dominic Kwiatkowski, Cordelia Langford, John Sillitoe on behalf of the Wellcome Sanger Institute COVID-19 Surveillance Team |
| EPI_ISL_2639924 | Lighthouse Laboratory Plymouth | Wellcome Sanger Institute for the COVID-19 Genomics UK (COG-UK) Consortium | Lighthouse Laboratory Plymouth and Alex Alderton, Roberto Amato, Jeffrey Barrett, Sonia Goncalves, Ewan Harrison, David K. Jackson, Ian Johnston, Dominic Kwiatkowski, Cordelia Langford, John Sillitoe on behalf of the Wellcome Sanger Institute COVID-19 Surveillance Team |
| EPI_ISL_2639925, EPI_ISL_2639926 | Lighthouse Lab in Alderley Park | Wellcome Sanger Institute for the COVID-19 Genomics UK (COG-UK) Consortium | Jacquelyn Wynn, Mairead Hyland, The Lighthouse Lab in Alderley Park and Alex Alderton, Roberto Amato, Jeffrey Barrett, Sonia Goncalves, Ewan Harrison, David K. Jackson, Ian Johnston, Dominic Kwiatkowski, Cordelia Langford, John Sillitoe on behalf of the Wellcome Sanger Institute COVID-19 Surveillance Team |
| EPI_ISL_2639927, EPI_ISL_2639928 | Lighthouse Laboratory Plymouth | Wellcome Sanger Institute for the COVID-19 Genomics UK (COG-UK) Consortium | Lighthouse Laboratory Plymouth and Alex Alderton, Roberto Amato, Jeffrey Barrett, Sonia Goncalves, Ewan Harrison, David K. Jackson, Ian Johnston, Dominic Kwiatkowski, Cordelia Langford, John Sillitoe on behalf of the Wellcome Sanger Institute COVID-19 Surveillance Team |
| EPI_ISL_2639929, EPI_ISL_2639930 | Lighthouse Lab in Alderley Park | Wellcome Sanger Institute for the COVID-19 Genomics UK (COG-UK) Consortium | Jacquelyn Wynn, Mairead Hyland, The Lighthouse Lab in Alderley Park and Alex Alderton, Roberto Amato, Jeffrey Barrett, Sonia Goncalves, Ewan Harrison, David K. Jackson, Ian Johnston, Dominic Kwiatkowski, Cordelia Langford, John Sillitoe on behalf of the Wellcome Sanger Institute COVID-19 Surveillance Team |
| EPI_ISL_2639931, EPI_ISL_2639932, EPI_ISL_2639933, EPI_ISL_2639934, EPI_ISL_2639935, EPI_ISL_2639936, EPI_ISL_2639938, EPI_ISL_2639940, EPI_ISL_2639941, EPI_ISL_2639942, EPI_ISL_2639943, EPI_ISL_2639944, EPI_ISL_2639945, EPI_ISL_2639946, EPI_ISL_2639947, EPI_ISL_2639948, EPI_ISL_2639949, EPI_ISL_2639950, EPI_ISL_2639951, EPI_ISL_2639953, EPI_ISL_2639954, EPI_ISL_2639955, EPI_ISL_2639956, EPI_ISL_2639957, EPI_ISL_2639958, EPI_ISL_2639960, EPI_ISL_2639961, EPI_ISL_2639962, EPI_ISL_2639963, EPI_ISL_2639964, EPI_ISL_2639965, EPI_ISL_2639966, EPI_ISL_2639967, EPI_ISL_2639968, EPI_ISL_2639969, EPI_ISL_2639970, EPI_ISL_2639971, EPI_ISL_2639972, EPI_ISL_2639974, EPI_ISL_2639975, EPI_ISL_2639976, EPI_ISL_2639977, EPI_ISL_2639978, EPI_ISL_2639979, EPI_ISL_2639980, EPI_ISL_2639981, EPI_ISL_2639982, EPI_ISL_2639983, EPI_ISL_2639984, EPI_ISL_2639985, EPI_ISL_2639986, EPI_ISL_2639987, EPI_ISL_2639988, EPI_ISL_2639989, EPI_ISL_2639991, EPI_ISL_2639992, EPI_ISL_2639993, EPI_ISL_2639994, EPI_ISL_2639996, EPI_ISL_2639997, EPI_ISL_2639998, EPI_ISL_2640001, EPI_ISL_2640002, EPI_ISL_2640003, EPI_ISL_2640004, EPI_ISL_2640005, EPI_ISL_2640006, EPI_ISL_2640007, EPI_ISL_2640008, EPI_ISL_2640009, EPI_ISL_2640010, EPI_ISL_2640011, EPI_ISL_2640012, EPI_ISL_2640013, EPI_ISL_2640014, EPI_ISL_2640015, EPI_ISL_2640016, EPI_ISL_2640017, EPI_ISL_2640018, EPI_ISL_2640019, EPI_ISL_2640020, EPI_ISL_2640021, EPI_ISL_2640022, EPI_ISL_2640023, EPI_ISL_2640024, EPI_ISL_2640025, EPI_ISL_2640026, EPI_ISL_2640027, EPI_ISL_2640028, EPI_ISL_2640029, EPI_ISL_2640030, EPI_ISL_2640031, EPI_ISL_2640032, EPI_ISL_2640033, EPI_ISL_2640034, EPI_ISL_2640035, EPI_ISL_2640036, EPI_ISL_2640037, EPI_ISL_2640038, EPI_ISL_2640039, EPI_ISL_2640040, EPI_ISL_2640041, EPI_ISL_2640042, EPI_ISL_2640043, EPI_ISL_2640044, EPI_ISL_2640045, EPI_ISL_2640046, EPI_ISL_2640047, EPI_ISL_2640048, EPI_ISL_2640049, EPI_ISL_2640050, EPI_ISL_2640051, EPI_ISL_2640052, EPI_ISL_2640053, EPI_ISL_2640054, EPI_ISL_2640055, EPI_ISL_2640056, EPI_ISL_2640057, EPI_ISL_2640058, EPI_ISL_2640059, EPI_ISL_2640060, EPI_ISL_2640061, EPI_ISL_2640062, EPI_ISL_2640063, EPI_ISL_2640064, EPI_ISL_2640065, EPI_ISL_2640066, EPI_ISL_2640067, EPI_ISL_2640068 | | | |
| see above | Lighthouse Lab in Glasgow | Wellcome Sanger Institute for the COVID-19 Genomics UK (COG-UK) Consortium | Harper VanSteenhouse, Yumi Kasai, David Gray, Carol Clugston, Anna Dominiczak and Alex Alderton, Roberto Amato, Jeffrey Barrett, Sonia Goncalves, Ewan Harrison, David K. Jackson, Ian Johnston, Dominic Kwiatkowski, Cordelia Langford, John Sillitoe on behalf of the Wellcome Sanger Institute COVID-19 Surveillance Team |
| EPI_ISL_2640069 | Lighthouse Lab in Alderley Park | Wellcome Sanger Institute for the COVID-19 Genomics UK (COG-UK) Consortium | Jacquelyn Wynn, Mairead Hyland, The Lighthouse Lab in Alderley Park and Alex Alderton, Roberto Amato, Jeffrey Barrett, Sonia Goncalves, Ewan Harrison, David K. Jackson, Ian Johnston, Dominic Kwiatkowski, Cordelia Langford, John Sillitoe on behalf of the Wellcome Sanger Institute COVID-19 Surveillance Team |
| EPI_ISL_2640070, EPI_ISL_2640071, EPI_ISL_2640072, EPI_ISL_2640073, EPI_ISL_2640074, EPI_ISL_2640075, EPI_ISL_2640076 | Lighthouse Lab in Glasgow | Wellcome Sanger Institute for the COVID-19 Genomics UK (COG-UK) Consortium | Harper VanSteenhouse, Yumi Kasai, David Gray, Carol Clugston, Anna Dominiczak and Alex Alderton, Roberto Amato, Jeffrey Barrett, Sonia Goncalves, Ewan Harrison, David K. Jackson, Ian Johnston, Dominic Kwiatkowski, Cordelia Langford, John Sillitoe on behalf of the Wellcome Sanger Institute COVID-19 Surveillance Team |
| EPI_ISL_2640078, EPI_ISL_2640079, EPI_ISL_2640080, EPI_ISL_2640081 | Lighthouse Lab in Alderley Park | Wellcome Sanger Institute for the COVID-19 Genomics UK (COG-UK) Consortium | Jacquelyn Wynn, Mairead Hyland, The Lighthouse Lab in Alderley Park and Alex Alderton, Roberto Amato, Jeffrey Barrett, Sonia Goncalves, Ewan Harrison, David K. Jackson, Ian Johnston, Dominic Kwiatkowski, Cordelia Langford, John Sillitoe on behalf of the Wellcome Sanger Institute COVID-19 Surveillance Team |
| EPI_ISL_2640084, EPI_ISL_2640085, EPI_ISL_2640086, EPI_ISL_2640087, EPI_ISL_2640088, EPI_ISL_2640089, EPI_ISL_2640091, EPI_ISL_2640092, EPI_ISL_2640096, EPI_ISL_2640097, EPI_ISL_2640098, EPI_ISL_2640099, EPI_ISL_2640101, EPI_ISL_2640102, EPI_ISL_2640103, EPI_ISL_2640104, EPI_ISL_2640105, EPI_ISL_2640106, EPI_ISL_2640107 | | | |
| see above | Lighthouse Lab in Glasgow | Wellcome Sanger Institute for the COVID-19 Genomics UK (COG-UK) Consortium | Harper VanSteenhouse, Yumi Kasai, David Gray, Carol Clugston, Anna Dominiczak and Alex Alderton, Roberto Amato, Jeffrey Barrett, Sonia Goncalves, Ewan Harrison, David K. Jackson, Ian Johnston, Dominic Kwiatkowski, Cordelia Langford, John Sillitoe on behalf of the Wellcome Sanger Institute COVID-19 Surveillance Team |
| EPI_ISL_2640108 | Lighthouse Laboratory Plymouth | Wellcome Sanger Institute for the COVID-19 Genomics UK (COG-UK) Consortium | Lighthouse Laboratory Plymouth and Alex Alderton, Roberto Amato, Jeffrey Barrett, Sonia Goncalves, Ewan Harrison, David K. Jackson, Ian Johnston, Dominic Kwiatkowski, Cordelia Langford, John Sillitoe on behalf of the Wellcome Sanger Institute COVID-19 Surveillance Team |
| EPI_ISL_2640109, EPI_ISL_2640110, EPI_ISL_2640111, EPI_ISL_2640113, EPI_ISL_2640114, EPI_ISL_2640115, EPI_ISL_2640116, EPI_ISL_2640117, EPI_ISL_2640119, EPI_ISL_2640120, EPI_ISL_2640124, EPI_ISL_2640125, EPI_ISL_2640126, EPI_ISL_2640128, EPI_ISL_2640129, EPI_ISL_2640131, EPI_ISL_2640132, EPI_ISL_2640133, EPI_ISL_2640134, EPI_ISL_2640135, EPI_ISL_2640136, EPI_ISL_2640137, EPI_ISL_2640138, EPI_ISL_2640139, EPI_ISL_2640140, EPI_ISL_2640141, EPI_ISL_2640142, EPI_ISL_2640143, EPI_ISL_2640144, EPI_ISL_2640146, EPI_ISL_2640147, EPI_ISL_2640148, EPI_ISL_2640149, EPI_ISL_2640150, EPI_ISL_2640151, EPI_ISL_2640153, EPI_ISL_2640154, EPI_ISL_2640155, EPI_ISL_2640156, EPI_ISL_2640157, EPI_ISL_2640158, EPI_ISL_2640159, EPI_ISL_2640160 | | | |
| see above | Lighthouse Lab in Glasgow | Wellcome Sanger Institute for the COVID-19 Genomics UK (COG-UK) Consortium | Harper VanSteenhouse, Yumi Kasai, David Gray, Carol Clugston, Anna Dominiczak and Alex Alderton, Roberto Amato, Jeffrey Barrett, Sonia Goncalves, Ewan Harrison, David K. Jackson, Ian Johnston, Dominic Kwiatkowski, Cordelia Langford, John Sillitoe on behalf of the Wellcome Sanger Institute COVID-19 Surveillance Team |
| EPI_ISL_2640161 | Lighthouse Lab in Alderley Park | Wellcome Sanger Institute for the COVID-19 Genomics UK (COG-UK) Consortium | Jacquelyn Wynn, Mairead Hyland, The Lighthouse Lab in Alderley Park and Alex Alderton, Roberto Amato, Jeffrey Barrett, Sonia Goncalves, Ewan Harrison, David K. Jackson, Ian Johnston, Dominic Kwiatkowski, Cordelia Langford, John Sillitoe on behalf of the Wellcome Sanger Institute COVID-19 Surveillance Team |
| EPI_ISL_2640162, EPI_ISL_2640163, EPI_ISL_2640164, EPI_ISL_2640165, EPI_ISL_2640166, EPI_ISL_2640167, EPI_ISL_2640168, EPI_ISL_2640169, EPI_ISL_2640170, EPI_ISL_2640171, EPI_ISL_2640172, EPI_ISL_2640173, EPI_ISL_2640175, EPI_ISL_2640176, EPI_ISL_2640177, EPI_ISL_2640178, EPI_ISL_2640179, EPI_ISL_2640180, EPI_ISL_2640181, EPI_ISL_2640183, EPI_ISL_2640184, EPI_ISL_2640185, EPI_ISL_2640186, EPI_ISL_2640187, EPI_ISL_2640188, EPI_ISL_2640189, EPI_ISL_2640190, EPI_ISL_2640191, EPI_ISL_2640192, EPI_ISL_2640193, EPI_ISL_2640194, EPI_ISL_2640195, EPI_ISL_2640196, EPI_ISL_2640197, EPI_ISL_2640198, EPI_ISL_2640199, EPI_ISL_2640200, EPI_ISL_2640202, EPI_ISL_2640203, EPI_ISL_2640204, EPI_ISL_2640205, EPI_ISL_2640206, EPI_ISL_2640207, EPI_ISL_2640208, EPI_ISL_2640209, EPI_ISL_2640210, EPI_ISL_2640211, EPI_ISL_2640213, EPI_ISL_2640214, EPI_ISL_2640215, EPI_ISL_2640216, EPI_ISL_2640217, EPI_ISL_2640218, EPI_ISL_2640219, EPI_ISL_2640220, EPI_ISL_2640221, EPI_ISL_2640222, EPI_ISL_2640224, EPI_ISL_2640225, EPI_ISL_2640226, EPI_ISL_2640227, EPI_ISL_2640228, EPI_ISL_2640229, EPI_ISL_2640230, EPI_ISL_2640231, EPI_ISL_2640232, EPI_ISL_2640233, EPI_ISL_2640234, EPI_ISL_2640235, EPI_ISL_2640236, EPI_ISL_2640237, EPI_ISL_2640238, EPI_ISL_2640239, EPI_ISL_2640240, EPI_ISL_2640241, EPI_ISL_2640242, EPI_ISL_2640243, EPI_ISL_2640244, EPI_ISL_2640245, EPI_ISL_2640246, EPI_ISL_2640247, EPI_ISL_2640248, EPI_ISL_2640249, EPI_ISL_2640250, EPI_ISL_2640251, EPI_ISL_2640252, EPI_ISL_2640254, EPI_ISL_2640255, EPI_ISL_2640256, EPI_ISL_2640257, EPI_ISL_2640258, EPI_ISL_2640259, EPI_ISL_2640260, EPI_ISL_2640261, EPI_ISL_2640262, EPI_ISL_2640263, EPI_ISL_2640264, EPI_ISL_2640265, EPI_ISL_2640266, EPI_ISL_2640267, EPI_ISL_2640268, EPI_ISL_2640269, EPI_ISL_2640270, EPI_ISL_2640271, EPI_ISL_2640272, EPI_ISL_2640273, EPI_ISL_2640274, EPI_ISL_2640276, EPI_ISL_2640277, EPI_ISL_2640279, EPI_ISL_2640280, EPI_ISL_2640282, EPI_ISL_2640283, EPI_ISL_2640284, EPI_ISL_2640285, EPI_ISL_2640287, EPI_ISL_2640288, EPI_ISL_2640290, EPI_ISL_2640291, EPI_ISL_2640292, EPI_ISL_2640293, EPI_ISL_2640294, EPI_ISL_2640295, EPI_ISL_2640296, EPI_ISL_2640297, EPI_ISL_2640298, EPI_ISL_2640299, EPI_ISL_2640300, EPI_ISL_2640301, EPI_ISL_2640302, EPI_ISL_2640303, EPI_ISL_2640304, EPI_ISL_2640305, EPI_ISL_2640306, EPI_ISL_2640307, EPI_ISL_2640308, EPI_ISL_2640309, EPI_ISL_2640310, EPI_ISL_2640311, EPI_ISL_2640313, EPI_ISL_2640314, EPI_ISL_2640316, EPI_ISL_2640317, EPI_ISL_2640318, EPI_ISL_2640319, EPI_ISL_2640321, EPI_ISL_2640322, EPI_ISL_2640323, EPI_ISL_2640324, EPI_ISL_2640325 | | | |
| see above | Lighthouse Lab in Glasgow | Wellcome Sanger Institute for the COVID-19 Genomics UK (COG-UK) Consortium | Harper VanSteenhouse, Yumi Kasai, David Gray, Carol Clugston, Anna Dominiczak and Alex Alderton, Roberto Amato, Jeffrey Barrett, Sonia Goncalves, Ewan Harrison, David K. Jackson, Ian Johnston, Dominic Kwiatkowski, Cordelia Langford, John Sillitoe on behalf of the Wellcome Sanger Institute COVID-19 Surveillance Team |
| EPI_ISL_2640326 | Lighthouse Lab in Milton Keynes | Wellcome Sanger Institute for the COVID-19 Genomics UK (COG-UK) Consortium | The Lighthouse Lab in Milton Keynes and Alex Alderton, Roberto Amato, Jeffrey Barrett, Sonia Goncalves, Ewan Harrison, David K. Jackson, Ian Johnston, Dominic Kwiatkowski, Cordelia Langford, John Sillitoe on behalf of the Wellcome Sanger Institute COVID-19 Surveillance Team |
| EPI_ISL_2640327, EPI_ISL_2640328 | Lighthouse Lab in Alderley Park | Wellcome Sanger Institute for the COVID-19 Genomics UK (COG-UK) Consortium | Jacquelyn Wynn, Mairead Hyland, The Lighthouse Lab in Alderley Park and Alex Alderton, Roberto Amato, Jeffrey Barrett, Sonia Goncalves, Ewan Harrison, David K. Jackson, Ian Johnston, Dominic Kwiatkowski, Cordelia Langford, John Sillitoe on behalf of the Wellcome Sanger Institute COVID-19 Surveillance Team |
| EPI_ISL_2640329 | Lighthouse Lab in Milton Keynes | Wellcome Sanger Institute for the COVID-19 Genomics UK (COG-UK) Consortium | The Lighthouse Lab in Milton Keynes and Alex Alderton, Roberto Amato, Jeffrey Barrett, Sonia Goncalves, Ewan Harrison, David K. Jackson, Ian Johnston, Dominic Kwiatkowski, Cordelia Langford, John Sillitoe on behalf of the Wellcome Sanger Institute COVID-19 Surveillance Team |
| EPI_ISL_2640330, EPI_ISL_2640331 | Lighthouse Laboratory Plymouth | Wellcome Sanger Institute for the COVID-19 Genomics UK (COG-UK) Consortium | Lighthouse Laboratory Plymouth and Alex Alderton, Roberto Amato, Jeffrey Barrett, Sonia Goncalves, Ewan Harrison, David K. Jackson, Ian Johnston, Dominic Kwiatkowski, Cordelia Langford, John Sillitoe on behalf of the Wellcome Sanger Institute COVID-19 Surveillance Team |
| EPI_ISL_2640332, EPI_ISL_2640333, EPI_ISL_2640334 | Lighthouse Lab in Alderley Park | Wellcome Sanger Institute for the COVID-19 Genomics UK (COG-UK) Consortium | Jacquelyn Wynn, Mairead Hyland, The Lighthouse Lab in Alderley Park and Alex Alderton, Roberto Amato, Jeffrey Barrett, Sonia Goncalves, Ewan Harrison, David K. Jackson, Ian Johnston, Dominic Kwiatkowski, Cordelia Langford, John Sillitoe on behalf of the Wellcome Sanger Institute COVID-19 Surveillance Team |

| | | | |
|---|---|---|---|
| EPI_ISL_2640335 | Lighthouse Laboratory Plymouth | Wellcome Sanger Institute for the COVID-19 Genomics UK (COG-UK) Consortium | Lighthouse Laboratory Plymouth and Alex Alderton, Roberto Amato, Jeffrey Barrett, Sonia Goncalves, Ewan Harrison, David K. Jackson, Ian Johnston, Dominic Kwiatkowski, Cordelia Langford, John Sillitoe on behalf of the Wellcome Sanger Institute COVID-19 Surveillance Team |
| EPI_ISL_2640336 | Lighthouse Lab in Milton Keynes | Wellcome Sanger Institute for the COVID-19 Genomics UK (COG-UK) Consortium | The Lighthouse Lab in Milton Keynes and Alex Alderton, Roberto Amato, Jeffrey Barrett, Sonia Goncalves, Ewan Harrison, David K. Jackson, Ian Johnston, Dominic Kwiatkowski, Cordelia Langford, John Sillitoe on behalf of the Wellcome Sanger Institute COVID-19 Surveillance Team |
| EPI_ISL_2640337, EPI_ISL_2640338, EPI_ISL_2640339, EPI_ISL_2640340 | Lighthouse Lab in Alderley Park | Wellcome Sanger Institute for the COVID-19 Genomics UK (COG-UK) Consortium | Jacquelyn Wynn, Mairead Hyland, The Lighthouse Lab in Alderley Park and Alex Alderton, Roberto Amato, Jeffrey Barrett, Sonia Goncalves, Ewan Harrison, David K. Jackson, Ian Johnston, Dominic Kwiatkowski, Cordelia Langford, John Sillitoe on behalf of the Wellcome Sanger Institute COVID-19 Surveillance Team |
| EPI_ISL_2640341 | Lighthouse Lab in Milton Keynes | Wellcome Sanger Institute for the COVID-19 Genomics UK (COG-UK) Consortium | The Lighthouse Lab in Milton Keynes and Alex Alderton, Roberto Amato, Jeffrey Barrett, Sonia Goncalves, Ewan Harrison, David K. Jackson, Ian Johnston, Dominic Kwiatkowski, Cordelia Langford, John Sillitoe on behalf of the Wellcome Sanger Institute COVID-19 Surveillance Team |
| EPI_ISL_2640342, EPI_ISL_2640343 | Lighthouse Lab in Alderley Park | Wellcome Sanger Institute for the COVID-19 Genomics UK (COG-UK) Consortium | Jacquelyn Wynn, Mairead Hyland, The Lighthouse Lab in Alderley Park and Alex Alderton, Roberto Amato, Jeffrey Barrett, Sonia Goncalves, Ewan Harrison, David K. Jackson, Ian Johnston, Dominic Kwiatkowski, Cordelia Langford, John Sillitoe on behalf of the Wellcome Sanger Institute COVID-19 Surveillance Team |
| EPI_ISL_2640344, EPI_ISL_2640345 | Lighthouse Lab in Milton Keynes | Wellcome Sanger Institute for the COVID-19 Genomics UK (COG-UK) Consortium | The Lighthouse Lab in Milton Keynes and Alex Alderton, Roberto Amato, Jeffrey Barrett, Sonia Goncalves, Ewan Harrison, David K. Jackson, Ian Johnston, Dominic Kwiatkowski, Cordelia Langford, John Sillitoe on behalf of the Wellcome Sanger Institute COVID-19 Surveillance Team |
| EPI_ISL_2640346 | Lighthouse Lab in Alderley Park | Wellcome Sanger Institute for the COVID-19 Genomics UK (COG-UK) Consortium | Jacquelyn Wynn, Mairead Hyland, The Lighthouse Lab in Alderley Park and Alex Alderton, Roberto Amato, Jeffrey Barrett, Sonia Goncalves, Ewan Harrison, David K. Jackson, Ian Johnston, Dominic Kwiatkowski, Cordelia Langford, John Sillitoe on behalf of the Wellcome Sanger Institute COVID-19 Surveillance Team |
| EPI_ISL_2640347, EPI_ISL_2640348 | Lighthouse Lab in Milton Keynes | Wellcome Sanger Institute for the COVID-19 Genomics UK (COG-UK) Consortium | The Lighthouse Lab in Milton Keynes and Alex Alderton, Roberto Amato, Jeffrey Barrett, Sonia Goncalves, Ewan Harrison, David K. Jackson, Ian Johnston, Dominic Kwiatkowski, Cordelia Langford, John Sillitoe on behalf of the Wellcome Sanger Institute COVID-19 Surveillance Team |
| EPI_ISL_2640349 | Lighthouse Lab in Alderley Park | Wellcome Sanger Institute for the COVID-19 Genomics UK (COG-UK) Consortium | Jacquelyn Wynn, Mairead Hyland, The Lighthouse Lab in Alderley Park and Alex Alderton, Roberto Amato, Jeffrey Barrett, Sonia Goncalves, Ewan Harrison, David K. Jackson, Ian Johnston, Dominic Kwiatkowski, Cordelia Langford, John Sillitoe on behalf of the Wellcome Sanger Institute COVID-19 Surveillance Team |
| EPI_ISL_2640350 | Lighthouse Lab in Milton Keynes | Wellcome Sanger Institute for the COVID-19 Genomics UK (COG-UK) Consortium | The Lighthouse Lab in Milton Keynes and Alex Alderton, Roberto Amato, Jeffrey Barrett, Sonia Goncalves, Ewan Harrison, David K. Jackson, Ian Johnston, Dominic Kwiatkowski, Cordelia Langford, John Sillitoe on behalf of the Wellcome Sanger Institute COVID-19 Surveillance Team |
| EPI_ISL_2640351, EPI_ISL_2640352, EPI_ISL_2640353 | Lighthouse Lab in Alderley Park | Wellcome Sanger Institute for the COVID-19 Genomics UK (COG-UK) Consortium | Jacquelyn Wynn, Mairead Hyland, The Lighthouse Lab in Alderley Park and Alex Alderton, Roberto Amato, Jeffrey Barrett, Sonia Goncalves, Ewan Harrison, David K. Jackson, Ian Johnston, Dominic Kwiatkowski, Cordelia Langford, John Sillitoe on behalf of the Wellcome Sanger Institute COVID-19 Surveillance Team |
| EPI_ISL_2640354 | Lighthouse Lab in Milton Keynes | Wellcome Sanger Institute for the COVID-19 Genomics UK (COG-UK) Consortium | The Lighthouse Lab in Milton Keynes and Alex Alderton, Roberto Amato, Jeffrey Barrett, Sonia Goncalves, Ewan Harrison, David K. Jackson, Ian Johnston, Dominic Kwiatkowski, Cordelia Langford, John Sillitoe on behalf of the Wellcome Sanger Institute COVID-19 Surveillance Team |
| EPI_ISL_2640355 | Lighthouse Lab in Alderley Park | Wellcome Sanger Institute for the COVID-19 Genomics UK (COG-UK) Consortium | Jacquelyn Wynn, Mairead Hyland, The Lighthouse Lab in Alderley Park and Alex Alderton, Roberto Amato, Jeffrey Barrett, Sonia Goncalves, Ewan Harrison, David K. Jackson, Ian Johnston, Dominic Kwiatkowski, Cordelia Langford, John Sillitoe on behalf of the Wellcome Sanger Institute COVID-19 Surveillance Team |
| EPI_ISL_2640356 | Lighthouse Lab in Milton Keynes | Wellcome Sanger Institute for the COVID-19 Genomics UK (COG-UK) Consortium | The Lighthouse Lab in Milton Keynes and Alex Alderton, Roberto Amato, Jeffrey Barrett, Sonia Goncalves, Ewan Harrison, David K. Jackson, Ian Johnston, Dominic Kwiatkowski, Cordelia Langford, John Sillitoe on behalf of the Wellcome Sanger Institute COVID-19 Surveillance Team |
| EPI_ISL_2640357, EPI_ISL_2640358, EPI_ISL_2640359 | Lighthouse Lab in Alderley Park | Wellcome Sanger Institute for the COVID-19 Genomics UK (COG-UK) Consortium | Jacquelyn Wynn, Mairead Hyland, The Lighthouse Lab in Alderley Park and Alex Alderton, Roberto Amato, Jeffrey Barrett, Sonia Goncalves, Ewan Harrison, David K. Jackson, Ian Johnston, Dominic Kwiatkowski, Cordelia Langford, John Sillitoe on behalf of the Wellcome Sanger Institute COVID-19 Surveillance Team |
| EPI_ISL_2640360, EPI_ISL_2640361, EPI_ISL_2640362 | Lighthouse Lab in Milton Keynes | Wellcome Sanger Institute for the COVID-19 Genomics UK (COG-UK) Consortium | The Lighthouse Lab in Milton Keynes and Alex Alderton, Roberto Amato, Jeffrey Barrett, Sonia Goncalves, Ewan Harrison, David K. Jackson, Ian Johnston, Dominic Kwiatkowski, Cordelia Langford, John Sillitoe on behalf of the Wellcome Sanger Institute COVID-19 Surveillance Team |
| EPI_ISL_2640364 | Lighthouse Lab in Alderley Park | Wellcome Sanger Institute for the COVID-19 Genomics UK (COG-UK) Consortium | Jacquelyn Wynn, Mairead Hyland, The Lighthouse Lab in Alderley Park and Alex Alderton, Roberto Amato, Jeffrey Barrett, Sonia Goncalves, Ewan Harrison, David K. Jackson, Ian Johnston, Dominic Kwiatkowski, Cordelia Langford, John Sillitoe on behalf of the Wellcome Sanger Institute COVID-19 Surveillance Team |
| EPI_ISL_2640365 | Lighthouse Lab in Milton Keynes | Wellcome Sanger Institute for the COVID-19 Genomics UK (COG-UK) Consortium | The Lighthouse Lab in Milton Keynes and Alex Alderton, Roberto Amato, Jeffrey Barrett, Sonia Goncalves, Ewan Harrison, David K. Jackson, Ian Johnston, Dominic Kwiatkowski, Cordelia Langford, John Sillitoe on behalf of the Wellcome Sanger Institute COVID-19 Surveillance Team |
| EPI_ISL_2640366 | Lighthouse Lab in Alderley Park | Wellcome Sanger Institute for the COVID-19 Genomics UK (COG-UK) Consortium | Jacquelyn Wynn, Mairead Hyland, The Lighthouse Lab in Alderley Park and Alex Alderton, Roberto Amato, Jeffrey Barrett, Sonia Goncalves, Ewan Harrison, David K. Jackson, Ian Johnston, Dominic Kwiatkowski, Cordelia Langford, John Sillitoe on behalf of the Wellcome Sanger Institute COVID-19 Surveillance Team |
| EPI_ISL_2640367, EPI_ISL_2640368, EPI_ISL_2640369, EPI_ISL_2640370, EPI_ISL_2640372, EPI_ISL_2640373 | Lighthouse Lab in Milton Keynes | Wellcome Sanger Institute for the COVID-19 Genomics UK (COG-UK) Consortium | The Lighthouse Lab in Milton Keynes and Alex Alderton, Roberto Amato, Jeffrey Barrett, Sonia Goncalves, Ewan Harrison, David K. Jackson, Ian Johnston, Dominic Kwiatkowski, Cordelia Langford, John Sillitoe on behalf of the Wellcome Sanger Institute COVID-19 Surveillance Team |
| EPI_ISL_2640374, EPI_ISL_2640375, EPI_ISL_2640377, EPI_ISL_2640378 | Lighthouse Lab in Alderley Park | Wellcome Sanger Institute for the COVID-19 Genomics UK (COG-UK) Consortium | Jacquelyn Wynn, Mairead Hyland, The Lighthouse Lab in Alderley Park and Alex Alderton, Roberto Amato, Jeffrey Barrett, Sonia Goncalves, Ewan Harrison, David K. Jackson, Ian Johnston, Dominic Kwiatkowski, Cordelia Langford, John Sillitoe on behalf of the Wellcome Sanger Institute COVID-19 Surveillance Team |
| EPI_ISL_2640379, EPI_ISL_2640380 | Lighthouse Lab in Milton Keynes | Wellcome Sanger Institute for the COVID-19 Genomics UK (COG-UK) Consortium | The Lighthouse Lab in Milton Keynes and Alex Alderton, Roberto Amato, Jeffrey Barrett, Sonia Goncalves, Ewan Harrison, David K. Jackson, Ian Johnston, Dominic Kwiatkowski, Cordelia Langford, John Sillitoe on behalf of the Wellcome Sanger Institute COVID-19 Surveillance Team |
| EPI_ISL_2640381 | Lighthouse Lab in Alderley Park | Wellcome Sanger Institute for the COVID-19 Genomics UK (COG-UK) Consortium | Jacquelyn Wynn, Mairead Hyland, The Lighthouse Lab in Alderley Park and Alex Alderton, Roberto Amato, Jeffrey Barrett, Sonia Goncalves, Ewan Harrison, David K. Jackson, Ian Johnston, Dominic Kwiatkowski, Cordelia Langford, John Sillitoe on behalf of the Wellcome Sanger Institute COVID-19 Surveillance Team |
| EPI_ISL_2640382, EPI_ISL_2640383, EPI_ISL_2640384 | Lighthouse Lab in Milton Keynes | Wellcome Sanger Institute for the COVID-19 Genomics UK (COG-UK) Consortium | The Lighthouse Lab in Milton Keynes and Alex Alderton, Roberto Amato, Jeffrey Barrett, Sonia Goncalves, Ewan Harrison, David K. Jackson, Ian Johnston, Dominic Kwiatkowski, Cordelia Langford, John Sillitoe on behalf of the Wellcome Sanger Institute COVID-19 Surveillance Team |
| EPI_ISL_2640386 | Lighthouse Lab in Alderley Park | Wellcome Sanger Institute for the COVID-19 Genomics UK (COG-UK) Consortium | Jacquelyn Wynn, Mairead Hyland, The Lighthouse Lab in Alderley Park and Alex Alderton, Roberto Amato, Jeffrey Barrett, Sonia Goncalves, Ewan Harrison, David K. Jackson, Ian Johnston, Dominic Kwiatkowski, Cordelia Langford, John Sillitoe on behalf of the Wellcome Sanger Institute COVID-19 Surveillance Team |
| EPI_ISL_2640387, EPI_ISL_2640388 | Lighthouse Lab in Milton Keynes | Wellcome Sanger Institute for the COVID-19 Genomics UK (COG-UK) Consortium | The Lighthouse Lab in Milton Keynes and Alex Alderton, Roberto Amato, Jeffrey Barrett, Sonia Goncalves, Ewan Harrison, David K. Jackson, Ian Johnston, Dominic Kwiatkowski, Cordelia Langford, John Sillitoe on behalf of the Wellcome Sanger Institute COVID-19 Surveillance Team |
| EPI_ISL_2640389 | Lighthouse Lab in Alderley Park | Wellcome Sanger Institute for the COVID-19 Genomics UK (COG-UK) Consortium | Jacquelyn Wynn, Mairead Hyland, The Lighthouse Lab in Alderley Park and Alex Alderton, Roberto Amato, Jeffrey Barrett, Sonia Goncalves, Ewan Harrison, David K. Jackson, Ian Johnston, Dominic Kwiatkowski, Cordelia Langford, John Sillitoe on behalf of the Wellcome Sanger Institute COVID-19 Surveillance Team |
| EPI_ISL_2640390 | Lighthouse Lab in Milton Keynes | Wellcome Sanger Institute for the COVID-19 Genomics UK (COG-UK) Consortium | The Lighthouse Lab in Milton Keynes and Alex Alderton, Roberto Amato, Jeffrey Barrett, Sonia Goncalves, Ewan Harrison, David K. Jackson, Ian Johnston, Dominic Kwiatkowski, Cordelia Langford, John Sillitoe on behalf of the Wellcome Sanger Institute COVID-19 Surveillance Team |
| EPI_ISL_2640391, EPI_ISL_2640392, EPI_ISL_2640393 | Lighthouse Lab in Alderley Park | Wellcome Sanger Institute for the COVID-19 Genomics UK (COG-UK) Consortium | Jacquelyn Wynn, Mairead Hyland, The Lighthouse Lab in Alderley Park and Alex Alderton, Roberto Amato, Jeffrey Barrett, Sonia Goncalves, Ewan Harrison, David K. Jackson, Ian Johnston, Dominic Kwiatkowski, Cordelia Langford, John Sillitoe on behalf of the Wellcome Sanger Institute COVID-19 Surveillance Team |
| EPI_ISL_2640394 | Lighthouse Lab in Milton Keynes | Wellcome Sanger Institute for the COVID-19 Genomics UK (COG-UK) Consortium | The Lighthouse Lab in Milton Keynes and Alex Alderton, Roberto Amato, Jeffrey Barrett, Sonia Goncalves, Ewan Harrison, David K. Jackson, Ian Johnston, Dominic Kwiatkowski, Cordelia Langford, John Sillitoe on behalf of the Wellcome Sanger Institute COVID-19 Surveillance Team |
| EPI_ISL_2640395, EPI_ISL_2640396, EPI_ISL_2640397 | Lighthouse Lab in Alderley Park | Wellcome Sanger Institute for the COVID-19 Genomics UK (COG-UK) Consortium | Jacquelyn Wynn, Mairead Hyland, The Lighthouse Lab in Alderley Park and Alex Alderton, Roberto Amato, Jeffrey Barrett, Sonia Goncalves, Ewan Harrison, David K. Jackson, Ian Johnston, Dominic Kwiatkowski, Cordelia Langford, John Sillitoe on behalf of the Wellcome Sanger Institute COVID-19 Surveillance Team |
| EPI_ISL_2640398, EPI_ISL_2640399, EPI_ISL_2640400, EPI_ISL_2640402, EPI_ISL_2640403, EPI_ISL_2640404, EPI_ISL_2640405 | Lighthouse Lab in Milton Keynes | Wellcome Sanger Institute for the COVID-19 Genomics UK (COG-UK) Consortium | The Lighthouse Lab in Milton Keynes and Alex Alderton, Roberto Amato, Jeffrey Barrett, Sonia Goncalves, Ewan Harrison, David K. Jackson, Ian Johnston, Dominic Kwiatkowski, Cordelia Langford, John Sillitoe on behalf of the Wellcome Sanger Institute COVID-19 Surveillance Team |
| EPI_ISL_2640406 | Lighthouse Lab in Alderley Park | Wellcome Sanger Institute for the COVID-19 Genomics UK (COG-UK) Consortium | Jacquelyn Wynn, Mairead Hyland, The Lighthouse Lab in Alderley Park and Alex Alderton, Roberto Amato, Jeffrey Barrett, Sonia Goncalves, Ewan Harrison, David K. Jackson, Ian Johnston, Dominic Kwiatkowski, Cordelia Langford, John Sillitoe on behalf of the Wellcome Sanger Institute COVID-19 Surveillance Team |
| EPI_ISL_2640407, EPI_ISL_2640408 | Lighthouse Lab in Milton Keynes | Wellcome Sanger Institute for the COVID-19 Genomics UK (COG-UK) Consortium | The Lighthouse Lab in Milton Keynes and Alex Alderton, Roberto Amato, Jeffrey Barrett, Sonia Goncalves, Ewan Harrison, David K. Jackson, Ian Johnston, Dominic Kwiatkowski, Cordelia Langford, John Sillitoe on behalf of the Wellcome Sanger Institute COVID-19 Surveillance Team |
| EPI_ISL_2640409, EPI_ISL_2640410 | Lighthouse Lab in Alderley Park | Wellcome Sanger Institute for the COVID-19 Genomics UK (COG-UK) Consortium | Jacquelyn Wynn, Mairead Hyland, The Lighthouse Lab in Alderley Park and Alex Alderton, Roberto Amato, Jeffrey Barrett, Sonia Goncalves, Ewan Harrison, David K. Jackson, Ian Johnston, Dominic Kwiatkowski, Cordelia Langford, John Sillitoe on behalf of the Wellcome Sanger Institute COVID-19 Surveillance Team |
| EPI_ISL_2640411, EPI_ISL_2640412, EPI_ISL_2640413 | Lighthouse Lab in Milton Keynes | Wellcome Sanger Institute for the COVID-19 Genomics UK (COG-UK) Consortium | The Lighthouse Lab in Milton Keynes and Alex Alderton, Roberto Amato, Jeffrey Barrett, Sonia Goncalves, Ewan Harrison, David K. Jackson, Ian Johnston, Dominic Kwiatkowski, Cordelia Langford, John Sillitoe on behalf of the Wellcome Sanger Institute COVID-19 Surveillance Team |
| EPI_ISL_2640414 | Lighthouse Lab in Alderley Park | Wellcome Sanger Institute for the COVID-19 Genomics UK (COG-UK) Consortium | Jacquelyn Wynn, Mairead Hyland, The Lighthouse Lab in Alderley Park and Alex Alderton, Roberto Amato, Jeffrey Barrett, Sonia Goncalves, Ewan Harrison, David K. Jackson, Ian Johnston, Dominic Kwiatkowski, Cordelia Langford, John Sillitoe on behalf of the Wellcome Sanger Institute COVID-19 Surveillance Team |

| | | | |
|---|---|---|---|
| EPI_ISL_2640416, EPI_ISL_2640417 | Lighthouse Lab in Milton Keynes | Wellcome Sanger Institute for the COVID-19 Genomics UK (COG-UK) Consortium | The Lighthouse Lab in Milton Keynes and Alex Alderton, Roberto Amato, Jeffrey Barrett, Sonia Goncalves, Ewan Harrison, David K. Jackson, Ian Johnston, Dominic Kwiatkowski, Cordelia Langford, John Sillitoe on behalf of the Wellcome Sanger Institute COVID-19 Surveillance Team |
| EPI_ISL_2640418, EPI_ISL_2640419, EPI_ISL_2640420, EPI_ISL_2640423, EPI_ISL_2640424 | Lighthouse Lab in Alderley Park | Wellcome Sanger Institute for the COVID-19 Genomics UK (COG-UK) Consortium | Jacquelyn Wynn, Mairead Hyland, The Lighthouse Lab in Alderley Park and Alex Alderton, Roberto Amato, Jeffrey Barrett, Sonia Goncalves, Ewan Harrison, David K. Jackson, Ian Johnston, Dominic Kwiatkowski, Cordelia Langford, John Sillitoe on behalf of the Wellcome Sanger Institute COVID-19 Surveillance Team |
| EPI_ISL_2640425 | Lighthouse Lab in Milton Keynes | Wellcome Sanger Institute for the COVID-19 Genomics UK (COG-UK) Consortium | The Lighthouse Lab in Milton Keynes and Alex Alderton, Roberto Amato, Jeffrey Barrett, Sonia Goncalves, Ewan Harrison, David K. Jackson, Ian Johnston, Dominic Kwiatkowski, Cordelia Langford, John Sillitoe on behalf of the Wellcome Sanger Institute COVID-19 Surveillance Team |
| EPI_ISL_2640426, EPI_ISL_2640427 | Lighthouse Lab in Alderley Park | Wellcome Sanger Institute for the COVID-19 Genomics UK (COG-UK) Consortium | Jacquelyn Wynn, Mairead Hyland, The Lighthouse Lab in Alderley Park and Alex Alderton, Roberto Amato, Jeffrey Barrett, Sonia Goncalves, Ewan Harrison, David K. Jackson, Ian Johnston, Dominic Kwiatkowski, Cordelia Langford, John Sillitoe on behalf of the Wellcome Sanger Institute COVID-19 Surveillance Team |
| EPI_ISL_2640429, EPI_ISL_2640430 | Lighthouse Lab in Milton Keynes | Wellcome Sanger Institute for the COVID-19 Genomics UK (COG-UK) Consortium | The Lighthouse Lab in Milton Keynes and Alex Alderton, Roberto Amato, Jeffrey Barrett, Sonia Goncalves, Ewan Harrison, David K. Jackson, Ian Johnston, Dominic Kwiatkowski, Cordelia Langford, John Sillitoe on behalf of the Wellcome Sanger Institute COVID-19 Surveillance Team |
| EPI_ISL_2640431 | Lighthouse Lab in Alderley Park | Wellcome Sanger Institute for the COVID-19 Genomics UK (COG-UK) Consortium | Jacquelyn Wynn, Mairead Hyland, The Lighthouse Lab in Alderley Park and Alex Alderton, Roberto Amato, Jeffrey Barrett, Sonia Goncalves, Ewan Harrison, David K. Jackson, Ian Johnston, Dominic Kwiatkowski, Cordelia Langford, John Sillitoe on behalf of the Wellcome Sanger Institute COVID-19 Surveillance Team |
| EPI_ISL_2640432, EPI_ISL_2640433, EPI_ISL_2640435 | Lighthouse Lab in Milton Keynes | Wellcome Sanger Institute for the COVID-19 Genomics UK (COG-UK) Consortium | The Lighthouse Lab in Milton Keynes and Alex Alderton, Roberto Amato, Jeffrey Barrett, Sonia Goncalves, Ewan Harrison, David K. Jackson, Ian Johnston, Dominic Kwiatkowski, Cordelia Langford, John Sillitoe on behalf of the Wellcome Sanger Institute COVID-19 Surveillance Team |
| EPI_ISL_2640436, EPI_ISL_2640437, EPI_ISL_2640438, EPI_ISL_2640439 | Lighthouse Lab in Alderley Park | Wellcome Sanger Institute for the COVID-19 Genomics UK (COG-UK) Consortium | Jacquelyn Wynn, Mairead Hyland, The Lighthouse Lab in Alderley Park and Alex Alderton, Roberto Amato, Jeffrey Barrett, Sonia Goncalves, Ewan Harrison, David K. Jackson, Ian Johnston, Dominic Kwiatkowski, Cordelia Langford, John Sillitoe on behalf of the Wellcome Sanger Institute COVID-19 Surveillance Team |
| EPI_ISL_2640440 | Lighthouse Lab in Milton Keynes | Wellcome Sanger Institute for the COVID-19 Genomics UK (COG-UK) Consortium | The Lighthouse Lab in Milton Keynes and Alex Alderton, Roberto Amato, Jeffrey Barrett, Sonia Goncalves, Ewan Harrison, David K. Jackson, Ian Johnston, Dominic Kwiatkowski, Cordelia Langford, John Sillitoe on behalf of the Wellcome Sanger Institute COVID-19 Surveillance Team |
| EPI_ISL_2640441, EPI_ISL_2640442, EPI_ISL_2640443, EPI_ISL_2640444, EPI_ISL_2640445 | Lighthouse Lab in Alderley Park | Wellcome Sanger Institute for the COVID-19 Genomics UK (COG-UK) Consortium | Jacquelyn Wynn, Mairead Hyland, The Lighthouse Lab in Alderley Park and Alex Alderton, Roberto Amato, Jeffrey Barrett, Sonia Goncalves, Ewan Harrison, David K. Jackson, Ian Johnston, Dominic Kwiatkowski, Cordelia Langford, John Sillitoe on behalf of the Wellcome Sanger Institute COVID-19 Surveillance Team |
| EPI_ISL_2640446 | Lighthouse Lab in Milton Keynes | Wellcome Sanger Institute for the COVID-19 Genomics UK (COG-UK) Consortium | The Lighthouse Lab in Milton Keynes and Alex Alderton, Roberto Amato, Jeffrey Barrett, Sonia Goncalves, Ewan Harrison, David K. Jackson, Ian Johnston, Dominic Kwiatkowski, Cordelia Langford, John Sillitoe on behalf of the Wellcome Sanger Institute COVID-19 Surveillance Team |
| EPI_ISL_2640447 | Lighthouse Lab in Alderley Park | Wellcome Sanger Institute for the COVID-19 Genomics UK (COG-UK) Consortium | Jacquelyn Wynn, Mairead Hyland, The Lighthouse Lab in Alderley Park and Alex Alderton, Roberto Amato, Jeffrey Barrett, Sonia Goncalves, Ewan Harrison, David K. Jackson, Ian Johnston, Dominic Kwiatkowski, Cordelia Langford, John Sillitoe on behalf of the Wellcome Sanger Institute COVID-19 Surveillance Team |
| EPI_ISL_2640448, EPI_ISL_2640449 | Lighthouse Lab in Milton Keynes | Wellcome Sanger Institute for the COVID-19 Genomics UK (COG-UK) Consortium | The Lighthouse Lab in Milton Keynes and Alex Alderton, Roberto Amato, Jeffrey Barrett, Sonia Goncalves, Ewan Harrison, David K. Jackson, Ian Johnston, Dominic Kwiatkowski, Cordelia Langford, John Sillitoe on behalf of the Wellcome Sanger Institute COVID-19 Surveillance Team |
| EPI_ISL_2640450 | Lighthouse Lab in Alderley Park | Wellcome Sanger Institute for the COVID-19 Genomics UK (COG-UK) Consortium | Jacquelyn Wynn, Mairead Hyland, The Lighthouse Lab in Alderley Park and Alex Alderton, Roberto Amato, Jeffrey Barrett, Sonia Goncalves, Ewan Harrison, David K. Jackson, Ian Johnston, Dominic Kwiatkowski, Cordelia Langford, John Sillitoe on behalf of the Wellcome Sanger Institute COVID-19 Surveillance Team |
| EPI_ISL_2640451 | Lighthouse Lab in Milton Keynes | Wellcome Sanger Institute for the COVID-19 Genomics UK (COG-UK) Consortium | The Lighthouse Lab in Milton Keynes and Alex Alderton, Roberto Amato, Jeffrey Barrett, Sonia Goncalves, Ewan Harrison, David K. Jackson, Ian Johnston, Dominic Kwiatkowski, Cordelia Langford, John Sillitoe on behalf of the Wellcome Sanger Institute COVID-19 Surveillance Team |
| EPI_ISL_2640452, EPI_ISL_2640453 | Lighthouse Lab in Alderley Park | Wellcome Sanger Institute for the COVID-19 Genomics UK (COG-UK) Consortium | Jacquelyn Wynn, Mairead Hyland, The Lighthouse Lab in Alderley Park and Alex Alderton, Roberto Amato, Jeffrey Barrett, Sonia Goncalves, Ewan Harrison, David K. Jackson, Ian Johnston, Dominic Kwiatkowski, Cordelia Langford, John Sillitoe on behalf of the Wellcome Sanger Institute COVID-19 Surveillance Team |
| EPI_ISL_2640454 | Lighthouse Lab in Milton Keynes | Wellcome Sanger Institute for the COVID-19 Genomics UK (COG-UK) Consortium | The Lighthouse Lab in Milton Keynes and Alex Alderton, Roberto Amato, Jeffrey Barrett, Sonia Goncalves, Ewan Harrison, David K. Jackson, Ian Johnston, Dominic Kwiatkowski, Cordelia Langford, John Sillitoe on behalf of the Wellcome Sanger Institute COVID-19 Surveillance Team |
| EPI_ISL_2640455 | Lighthouse Lab in Alderley Park | Wellcome Sanger Institute for the COVID-19 Genomics UK (COG-UK) Consortium | Jacquelyn Wynn, Mairead Hyland, The Lighthouse Lab in Alderley Park and Alex Alderton, Roberto Amato, Jeffrey Barrett, Sonia Goncalves, Ewan Harrison, David K. Jackson, Ian Johnston, Dominic Kwiatkowski, Cordelia Langford, John Sillitoe on behalf of the Wellcome Sanger Institute COVID-19 Surveillance Team |
| EPI_ISL_2640456, EPI_ISL_2640457 | Lighthouse Lab in Milton Keynes | Wellcome Sanger Institute for the COVID-19 Genomics UK (COG-UK) Consortium | The Lighthouse Lab in Milton Keynes and Alex Alderton, Roberto Amato, Jeffrey Barrett, Sonia Goncalves, Ewan Harrison, David K. Jackson, Ian Johnston, Dominic Kwiatkowski, Cordelia Langford, John Sillitoe on behalf of the Wellcome Sanger Institute COVID-19 Surveillance Team |
| EPI_ISL_2640458 | Lighthouse Lab in Alderley Park | Wellcome Sanger Institute for the COVID-19 Genomics UK (COG-UK) Consortium | Jacquelyn Wynn, Mairead Hyland, The Lighthouse Lab in Alderley Park and Alex Alderton, Roberto Amato, Jeffrey Barrett, Sonia Goncalves, Ewan Harrison, David K. Jackson, Ian Johnston, Dominic Kwiatkowski, Cordelia Langford, John Sillitoe on behalf of the Wellcome Sanger Institute COVID-19 Surveillance Team |
| EPI_ISL_2640459 | Lighthouse Lab in Milton Keynes | Wellcome Sanger Institute for the COVID-19 Genomics UK (COG-UK) Consortium | The Lighthouse Lab in Milton Keynes and Alex Alderton, Roberto Amato, Jeffrey Barrett, Sonia Goncalves, Ewan Harrison, David K. Jackson, Ian Johnston, Dominic Kwiatkowski, Cordelia Langford, John Sillitoe on behalf of the Wellcome Sanger Institute COVID-19 Surveillance Team |
| EPI_ISL_2640460, EPI_ISL_2640461, EPI_ISL_2640462, EPI_ISL_2640463 | Lighthouse Lab in Alderley Park | Wellcome Sanger Institute for the COVID-19 Genomics UK (COG-UK) Consortium | Jacquelyn Wynn, Mairead Hyland, The Lighthouse Lab in Alderley Park and Alex Alderton, Roberto Amato, Jeffrey Barrett, Sonia Goncalves, Ewan Harrison, David K. Jackson, Ian Johnston, Dominic Kwiatkowski, Cordelia Langford, John Sillitoe on behalf of the Wellcome Sanger Institute COVID-19 Surveillance Team |
| EPI_ISL_2640464, EPI_ISL_2640465 | Lighthouse Lab in Milton Keynes | Wellcome Sanger Institute for the COVID-19 Genomics UK (COG-UK) Consortium | The Lighthouse Lab in Milton Keynes and Alex Alderton, Roberto Amato, Jeffrey Barrett, Sonia Goncalves, Ewan Harrison, David K. Jackson, Ian Johnston, Dominic Kwiatkowski, Cordelia Langford, John Sillitoe on behalf of the Wellcome Sanger Institute COVID-19 Surveillance Team |
| EPI_ISL_2640466 | Lighthouse Lab in Alderley Park | Wellcome Sanger Institute for the COVID-19 Genomics UK (COG-UK) Consortium | Jacquelyn Wynn, Mairead Hyland, The Lighthouse Lab in Alderley Park and Alex Alderton, Roberto Amato, Jeffrey Barrett, Sonia Goncalves, Ewan Harrison, David K. Jackson, Ian Johnston, Dominic Kwiatkowski, Cordelia Langford, John Sillitoe on behalf of the Wellcome Sanger Institute COVID-19 Surveillance Team |
| EPI_ISL_2640467 | Lighthouse Lab in Milton Keynes | Wellcome Sanger Institute for the COVID-19 Genomics UK (COG-UK) Consortium | The Lighthouse Lab in Milton Keynes and Alex Alderton, Roberto Amato, Jeffrey Barrett, Sonia Goncalves, Ewan Harrison, David K. Jackson, Ian Johnston, Dominic Kwiatkowski, Cordelia Langford, John Sillitoe on behalf of the Wellcome Sanger Institute COVID-19 Surveillance Team |
| EPI_ISL_2640468 | Lighthouse Lab in Alderley Park | Wellcome Sanger Institute for the COVID-19 Genomics UK (COG-UK) Consortium | Jacquelyn Wynn, Mairead Hyland, The Lighthouse Lab in Alderley Park and Alex Alderton, Roberto Amato, Jeffrey Barrett, Sonia Goncalves, Ewan Harrison, David K. Jackson, Ian Johnston, Dominic Kwiatkowski, Cordelia Langford, John Sillitoe on behalf of the Wellcome Sanger Institute COVID-19 Surveillance Team |
| EPI_ISL_2640469, EPI_ISL_2640470 | Lighthouse Lab in Milton Keynes | Wellcome Sanger Institute for the COVID-19 Genomics UK (COG-UK) Consortium | The Lighthouse Lab in Milton Keynes and Alex Alderton, Roberto Amato, Jeffrey Barrett, Sonia Goncalves, Ewan Harrison, David K. Jackson, Ian Johnston, Dominic Kwiatkowski, Cordelia Langford, John Sillitoe on behalf of the Wellcome Sanger Institute COVID-19 Surveillance Team |
| EPI_ISL_2640471 | Lighthouse Lab in Alderley Park | Wellcome Sanger Institute for the COVID-19 Genomics UK (COG-UK) Consortium | Jacquelyn Wynn, Mairead Hyland, The Lighthouse Lab in Alderley Park and Alex Alderton, Roberto Amato, Jeffrey Barrett, Sonia Goncalves, Ewan Harrison, David K. Jackson, Ian Johnston, Dominic Kwiatkowski, Cordelia Langford, John Sillitoe on behalf of the Wellcome Sanger Institute COVID-19 Surveillance Team |
| EPI_ISL_2640472 | Lighthouse Lab in Milton Keynes | Wellcome Sanger Institute for the COVID-19 Genomics UK (COG-UK) Consortium | The Lighthouse Lab in Milton Keynes and Alex Alderton, Roberto Amato, Jeffrey Barrett, Sonia Goncalves, Ewan Harrison, David K. Jackson, Ian Johnston, Dominic Kwiatkowski, Cordelia Langford, John Sillitoe on behalf of the Wellcome Sanger Institute COVID-19 Surveillance Team |
| EPI_ISL_2640473, EPI_ISL_2640474 | Lighthouse Lab in Alderley Park | Wellcome Sanger Institute for the COVID-19 Genomics UK (COG-UK) Consortium | Jacquelyn Wynn, Mairead Hyland, The Lighthouse Lab in Alderley Park and Alex Alderton, Roberto Amato, Jeffrey Barrett, Sonia Goncalves, Ewan Harrison, David K. Jackson, Ian Johnston, Dominic Kwiatkowski, Cordelia Langford, John Sillitoe on behalf of the Wellcome Sanger Institute COVID-19 Surveillance Team |
| EPI_ISL_2640475, EPI_ISL_2640476, EPI_ISL_2640477, EPI_ISL_2640478, EPI_ISL_2640479, EPI_ISL_2640480 | Lighthouse Lab in Milton Keynes | Wellcome Sanger Institute for the COVID-19 Genomics UK (COG-UK) Consortium | The Lighthouse Lab in Milton Keynes and Alex Alderton, Roberto Amato, Jeffrey Barrett, Sonia Goncalves, Ewan Harrison, David K. Jackson, Ian Johnston, Dominic Kwiatkowski, Cordelia Langford, John Sillitoe on behalf of the Wellcome Sanger Institute COVID-19 Surveillance Team |
| EPI_ISL_2640481 | Lighthouse Lab in Alderley Park | Wellcome Sanger Institute for the COVID-19 Genomics UK (COG-UK) Consortium | Jacquelyn Wynn, Mairead Hyland, The Lighthouse Lab in Alderley Park and Alex Alderton, Roberto Amato, Jeffrey Barrett, Sonia Goncalves, Ewan Harrison, David K. Jackson, Ian Johnston, Dominic Kwiatkowski, Cordelia Langford, John Sillitoe on behalf of the Wellcome Sanger Institute COVID-19 Surveillance Team |
| EPI_ISL_2640482, EPI_ISL_2640483 | Lighthouse Lab in Milton Keynes | Wellcome Sanger Institute for the COVID-19 Genomics UK (COG-UK) Consortium | The Lighthouse Lab in Milton Keynes and Alex Alderton, Roberto Amato, Jeffrey Barrett, Sonia Goncalves, Ewan Harrison, David K. Jackson, Ian Johnston, Dominic Kwiatkowski, Cordelia Langford, John Sillitoe on behalf of the Wellcome Sanger Institute COVID-19 Surveillance Team |
| EPI_ISL_2640484, EPI_ISL_2640485 | Lighthouse Lab in Alderley Park | Wellcome Sanger Institute for the COVID-19 Genomics UK (COG-UK) Consortium | Jacquelyn Wynn, Mairead Hyland, The Lighthouse Lab in Alderley Park and Alex Alderton, Roberto Amato, Jeffrey Barrett, Sonia Goncalves, Ewan Harrison, David K. Jackson, Ian Johnston, Dominic Kwiatkowski, Cordelia Langford, John Sillitoe on behalf of the Wellcome Sanger Institute COVID-19 Surveillance Team |
| EPI_ISL_2640486, EPI_ISL_2640487, EPI_ISL_2640488 | Lighthouse Lab in Milton Keynes | Wellcome Sanger Institute for the COVID-19 Genomics UK (COG-UK) Consortium | The Lighthouse Lab in Milton Keynes and Alex Alderton, Roberto Amato, Jeffrey Barrett, Sonia Goncalves, Ewan Harrison, David K. Jackson, Ian Johnston, Dominic Kwiatkowski, Cordelia Langford, John Sillitoe on behalf of the Wellcome Sanger Institute COVID-19 Surveillance Team |
| EPI_ISL_2640490, EPI_ISL_2640491 | Lighthouse Lab in Alderley Park | Wellcome Sanger Institute for the COVID-19 Genomics UK (COG-UK) Consortium | Jacquelyn Wynn, Mairead Hyland, The Lighthouse Lab in Alderley Park and Alex Alderton, Roberto Amato, Jeffrey Barrett, Sonia Goncalves, Ewan Harrison, David K. Jackson, Ian Johnston, Dominic Kwiatkowski, Cordelia Langford, John Sillitoe on behalf of the Wellcome Sanger Institute COVID-19 Surveillance Team |
| EPI_ISL_2640492 | Lighthouse Lab in Milton Keynes | Wellcome Sanger Institute for the COVID-19 Genomics UK (COG-UK) Consortium | The Lighthouse Lab in Milton Keynes and Alex Alderton, Roberto Amato, Jeffrey Barrett, Sonia Goncalves, Ewan Harrison, David K. Jackson, Ian Johnston, Dominic Kwiatkowski, Cordelia Langford, John Sillitoe on behalf of the Wellcome Sanger Institute COVID-19 Surveillance Team |
| EPI_ISL_2640493, EPI_ISL_2640494 | Lighthouse Lab in Alderley Park | Wellcome Sanger Institute for the COVID-19 Genomics UK (COG-UK) Consortium | Jacquelyn Wynn, Mairead Hyland, The Lighthouse Lab in Alderley Park and Alex Alderton, Roberto Amato, Jeffrey Barrett, Sonia Goncalves, Ewan Harrison, David K. Jackson, Ian Johnston, Dominic Kwiatkowski, Cordelia Langford, John Sillitoe on behalf of the Wellcome Sanger Institute COVID-19 Surveillance Team |

| | | | |
|---|---|---|---|
| EPI_ISL_2640495, EPI_ISL_2640496, EPI_ISL_2640497, EPI_ISL_2640498 | Lighthouse Lab in Milton Keynes | Wellcome Sanger Institute for the COVID-19 Genomics UK (COG-UK) Consortium | The Lighthouse Lab in Milton Keynes and Alex Alderton, Roberto Amato, Jeffrey Barrett, Sonia Goncalves, Ewan Harrison, David K. Jackson, Ian Johnston, Dominic Kwiatkowski, Cordelia Langford, John Sillitoe on behalf of the Wellcome Sanger Institute COVID-19 Surveillance Team |
| EPI_ISL_2640499, EPI_ISL_2640500 | Lighthouse Lab in Alderley Park | Wellcome Sanger Institute for the COVID-19 Genomics UK (COG-UK) Consortium | Jacquelyn Wynn, Mairead Hyland, The Lighthouse Lab in Alderley Park and Alex Alderton, Roberto Amato, Jeffrey Barrett, Sonia Goncalves, Ewan Harrison, David K. Jackson, Ian Johnston, Dominic Kwiatkowski, Cordelia Langford, John Sillitoe on behalf of the Wellcome Sanger Institute COVID-19 Surveillance Team |
| EPI_ISL_2640501 | Lighthouse Lab in Milton Keynes | Wellcome Sanger Institute for the COVID-19 Genomics UK (COG-UK) Consortium | The Lighthouse Lab in Milton Keynes and Alex Alderton, Roberto Amato, Jeffrey Barrett, Sonia Goncalves, Ewan Harrison, David K. Jackson, Ian Johnston, Dominic Kwiatkowski, Cordelia Langford, John Sillitoe on behalf of the Wellcome Sanger Institute COVID-19 Surveillance Team |
| EPI_ISL_2640502 | Lighthouse Lab in Alderley Park | Wellcome Sanger Institute for the COVID-19 Genomics UK (COG-UK) Consortium | Jacquelyn Wynn, Mairead Hyland, The Lighthouse Lab in Alderley Park and Alex Alderton, Roberto Amato, Jeffrey Barrett, Sonia Goncalves, Ewan Harrison, David K. Jackson, Ian Johnston, Dominic Kwiatkowski, Cordelia Langford, John Sillitoe on behalf of the Wellcome Sanger Institute COVID-19 Surveillance Team |
| EPI_ISL_2640503, EPI_ISL_2640504, EPI_ISL_2640505, EPI_ISL_2640506, EPI_ISL_2640507, EPI_ISL_2640508, EPI_ISL_2640509 | Lighthouse Lab in Milton Keynes | Wellcome Sanger Institute for the COVID-19 Genomics UK (COG-UK) Consortium | The Lighthouse Lab in Milton Keynes and Alex Alderton, Roberto Amato, Jeffrey Barrett, Sonia Goncalves, Ewan Harrison, David K. Jackson, Ian Johnston, Dominic Kwiatkowski, Cordelia Langford, John Sillitoe on behalf of the Wellcome Sanger Institute COVID-19 Surveillance Team |
| EPI_ISL_2640510 | Lighthouse Lab in Alderley Park | Wellcome Sanger Institute for the COVID-19 Genomics UK (COG-UK) Consortium | Jacquelyn Wynn, Mairead Hyland, The Lighthouse Lab in Alderley Park and Alex Alderton, Roberto Amato, Jeffrey Barrett, Sonia Goncalves, Ewan Harrison, David K. Jackson, Ian Johnston, Dominic Kwiatkowski, Cordelia Langford, John Sillitoe on behalf of the Wellcome Sanger Institute COVID-19 Surveillance Team |
| EPI_ISL_2640511 | Lighthouse Lab in Milton Keynes | Wellcome Sanger Institute for the COVID-19 Genomics UK (COG-UK) Consortium | The Lighthouse Lab in Milton Keynes and Alex Alderton, Roberto Amato, Jeffrey Barrett, Sonia Goncalves, Ewan Harrison, David K. Jackson, Ian Johnston, Dominic Kwiatkowski, Cordelia Langford, John Sillitoe on behalf of the Wellcome Sanger Institute COVID-19 Surveillance Team |
| EPI_ISL_2640512, EPI_ISL_2640513 | Lighthouse Lab in Alderley Park | Wellcome Sanger Institute for the COVID-19 Genomics UK (COG-UK) Consortium | Jacquelyn Wynn, Mairead Hyland, The Lighthouse Lab in Alderley Park and Alex Alderton, Roberto Amato, Jeffrey Barrett, Sonia Goncalves, Ewan Harrison, David K. Jackson, Ian Johnston, Dominic Kwiatkowski, Cordelia Langford, John Sillitoe on behalf of the Wellcome Sanger Institute COVID-19 Surveillance Team |
| EPI_ISL_2640514 | Lighthouse Lab in Milton Keynes | Wellcome Sanger Institute for the COVID-19 Genomics UK (COG-UK) Consortium | The Lighthouse Lab in Milton Keynes and Alex Alderton, Roberto Amato, Jeffrey Barrett, Sonia Goncalves, Ewan Harrison, David K. Jackson, Ian Johnston, Dominic Kwiatkowski, Cordelia Langford, John Sillitoe on behalf of the Wellcome Sanger Institute COVID-19 Surveillance Team |
| EPI_ISL_2640515 | Lighthouse Lab in Alderley Park | Wellcome Sanger Institute for the COVID-19 Genomics UK (COG-UK) Consortium | Jacquelyn Wynn, Mairead Hyland, The Lighthouse Lab in Alderley Park and Alex Alderton, Roberto Amato, Jeffrey Barrett, Sonia Goncalves, Ewan Harrison, David K. Jackson, Ian Johnston, Dominic Kwiatkowski, Cordelia Langford, John Sillitoe on behalf of the Wellcome Sanger Institute COVID-19 Surveillance Team |
| EPI_ISL_2640516 | Lighthouse Lab in Milton Keynes | Wellcome Sanger Institute for the COVID-19 Genomics UK (COG-UK) Consortium | The Lighthouse Lab in Milton Keynes and Alex Alderton, Roberto Amato, Jeffrey Barrett, Sonia Goncalves, Ewan Harrison, David K. Jackson, Ian Johnston, Dominic Kwiatkowski, Cordelia Langford, John Sillitoe on behalf of the Wellcome Sanger Institute COVID-19 Surveillance Team |
| EPI_ISL_2640517 | Lighthouse Lab in Alderley Park | Wellcome Sanger Institute for the COVID-19 Genomics UK (COG-UK) Consortium | Jacquelyn Wynn, Mairead Hyland, The Lighthouse Lab in Alderley Park and Alex Alderton, Roberto Amato, Jeffrey Barrett, Sonia Goncalves, Ewan Harrison, David K. Jackson, Ian Johnston, Dominic Kwiatkowski, Cordelia Langford, John Sillitoe on behalf of the Wellcome Sanger Institute COVID-19 Surveillance Team |
| EPI_ISL_2640518 | Lighthouse Lab in Milton Keynes | Wellcome Sanger Institute for the COVID-19 Genomics UK (COG-UK) Consortium | The Lighthouse Lab in Milton Keynes and Alex Alderton, Roberto Amato, Jeffrey Barrett, Sonia Goncalves, Ewan Harrison, David K. Jackson, Ian Johnston, Dominic Kwiatkowski, Cordelia Langford, John Sillitoe on behalf of the Wellcome Sanger Institute COVID-19 Surveillance Team |
| EPI_ISL_2640520, EPI_ISL_2640521 | Lighthouse Lab in Alderley Park | Wellcome Sanger Institute for the COVID-19 Genomics UK (COG-UK) Consortium | Jacquelyn Wynn, Mairead Hyland, The Lighthouse Lab in Alderley Park and Alex Alderton, Roberto Amato, Jeffrey Barrett, Sonia Goncalves, Ewan Harrison, David K. Jackson, Ian Johnston, Dominic Kwiatkowski, Cordelia Langford, John Sillitoe on behalf of the Wellcome Sanger Institute COVID-19 Surveillance Team |
| EPI_ISL_2640522 | Lighthouse Lab in Milton Keynes | Wellcome Sanger Institute for the COVID-19 Genomics UK (COG-UK) Consortium | The Lighthouse Lab in Milton Keynes and Alex Alderton, Roberto Amato, Jeffrey Barrett, Sonia Goncalves, Ewan Harrison, David K. Jackson, Ian Johnston, Dominic Kwiatkowski, Cordelia Langford, John Sillitoe on behalf of the Wellcome Sanger Institute COVID-19 Surveillance Team |
| EPI_ISL_2640523 | Lighthouse Lab in Alderley Park | Wellcome Sanger Institute for the COVID-19 Genomics UK (COG-UK) Consortium | Jacquelyn Wynn, Mairead Hyland, The Lighthouse Lab in Alderley Park and Alex Alderton, Roberto Amato, Jeffrey Barrett, Sonia Goncalves, Ewan Harrison, David K. Jackson, Ian Johnston, Dominic Kwiatkowski, Cordelia Langford, John Sillitoe on behalf of the Wellcome Sanger Institute COVID-19 Surveillance Team |
| EPI_ISL_2640524 | Lighthouse Lab in Milton Keynes | Wellcome Sanger Institute for the COVID-19 Genomics UK (COG-UK) Consortium | The Lighthouse Lab in Milton Keynes and Alex Alderton, Roberto Amato, Jeffrey Barrett, Sonia Goncalves, Ewan Harrison, David K. Jackson, Ian Johnston, Dominic Kwiatkowski, Cordelia Langford, John Sillitoe on behalf of the Wellcome Sanger Institute COVID-19 Surveillance Team |
| EPI_ISL_2640526, EPI_ISL_2640528 | Lighthouse Lab in Alderley Park | Wellcome Sanger Institute for the COVID-19 Genomics UK (COG-UK) Consortium | Jacquelyn Wynn, Mairead Hyland, The Lighthouse Lab in Alderley Park and Alex Alderton, Roberto Amato, Jeffrey Barrett, Sonia Goncalves, Ewan Harrison, David K. Jackson, Ian Johnston, Dominic Kwiatkowski, Cordelia Langford, John Sillitoe on behalf of the Wellcome Sanger Institute COVID-19 Surveillance Team |
| EPI_ISL_2640529 | Lighthouse Lab in Milton Keynes | Wellcome Sanger Institute for the COVID-19 Genomics UK (COG-UK) Consortium | The Lighthouse Lab in Milton Keynes and Alex Alderton, Roberto Amato, Jeffrey Barrett, Sonia Goncalves, Ewan Harrison, David K. Jackson, Ian Johnston, Dominic Kwiatkowski, Cordelia Langford, John Sillitoe on behalf of the Wellcome Sanger Institute COVID-19 Surveillance Team |
| EPI_ISL_2640530, EPI_ISL_2640531 | Lighthouse Lab in Alderley Park | Wellcome Sanger Institute for the COVID-19 Genomics UK (COG-UK) Consortium | Jacquelyn Wynn, Mairead Hyland, The Lighthouse Lab in Alderley Park and Alex Alderton, Roberto Amato, Jeffrey Barrett, Sonia Goncalves, Ewan Harrison, David K. Jackson, Ian Johnston, Dominic Kwiatkowski, Cordelia Langford, John Sillitoe on behalf of the Wellcome Sanger Institute COVID-19 Surveillance Team |
| EPI_ISL_2640532 | Lighthouse Lab in Milton Keynes | Wellcome Sanger Institute for the COVID-19 Genomics UK (COG-UK) Consortium | The Lighthouse Lab in Milton Keynes and Alex Alderton, Roberto Amato, Jeffrey Barrett, Sonia Goncalves, Ewan Harrison, David K. Jackson, Ian Johnston, Dominic Kwiatkowski, Cordelia Langford, John Sillitoe on behalf of the Wellcome Sanger Institute COVID-19 Surveillance Team |
| EPI_ISL_2640533, EPI_ISL_2640534, EPI_ISL_2640535, EPI_ISL_2640536 | Lighthouse Lab in Alderley Park | Wellcome Sanger Institute for the COVID-19 Genomics UK (COG-UK) Consortium | Jacquelyn Wynn, Mairead Hyland, The Lighthouse Lab in Alderley Park and Alex Alderton, Roberto Amato, Jeffrey Barrett, Sonia Goncalves, Ewan Harrison, David K. Jackson, Ian Johnston, Dominic Kwiatkowski, Cordelia Langford, John Sillitoe on behalf of the Wellcome Sanger Institute COVID-19 Surveillance Team |
| EPI_ISL_2640537, EPI_ISL_2640538, EPI_ISL_2640539, EPI_ISL_2640540 | Lighthouse Lab in Milton Keynes | Wellcome Sanger Institute for the COVID-19 Genomics UK (COG-UK) Consortium | The Lighthouse Lab in Milton Keynes and Alex Alderton, Roberto Amato, Jeffrey Barrett, Sonia Goncalves, Ewan Harrison, David K. Jackson, Ian Johnston, Dominic Kwiatkowski, Cordelia Langford, John Sillitoe on behalf of the Wellcome Sanger Institute COVID-19 Surveillance Team |
| EPI_ISL_2640541, EPI_ISL_2640542 | Lighthouse Lab in Alderley Park | Wellcome Sanger Institute for the COVID-19 Genomics UK (COG-UK) Consortium | Jacquelyn Wynn, Mairead Hyland, The Lighthouse Lab in Alderley Park and Alex Alderton, Roberto Amato, Jeffrey Barrett, Sonia Goncalves, Ewan Harrison, David K. Jackson, Ian Johnston, Dominic Kwiatkowski, Cordelia Langford, John Sillitoe on behalf of the Wellcome Sanger Institute COVID-19 Surveillance Team |
| EPI_ISL_2640543 | Lighthouse Lab in Milton Keynes | Wellcome Sanger Institute for the COVID-19 Genomics UK (COG-UK) Consortium | The Lighthouse Lab in Milton Keynes and Alex Alderton, Roberto Amato, Jeffrey Barrett, Sonia Goncalves, Ewan Harrison, David K. Jackson, Ian Johnston, Dominic Kwiatkowski, Cordelia Langford, John Sillitoe on behalf of the Wellcome Sanger Institute COVID-19 Surveillance Team |
| EPI_ISL_2640544 | Lighthouse Lab in Alderley Park | Wellcome Sanger Institute for the COVID-19 Genomics UK (COG-UK) Consortium | Jacquelyn Wynn, Mairead Hyland, The Lighthouse Lab in Alderley Park and Alex Alderton, Roberto Amato, Jeffrey Barrett, Sonia Goncalves, Ewan Harrison, David K. Jackson, Ian Johnston, Dominic Kwiatkowski, Cordelia Langford, John Sillitoe on behalf of the Wellcome Sanger Institute COVID-19 Surveillance Team |
| EPI_ISL_2640545 | Lighthouse Lab in Milton Keynes | Wellcome Sanger Institute for the COVID-19 Genomics UK (COG-UK) Consortium | The Lighthouse Lab in Milton Keynes and Alex Alderton, Roberto Amato, Jeffrey Barrett, Sonia Goncalves, Ewan Harrison, David K. Jackson, Ian Johnston, Dominic Kwiatkowski, Cordelia Langford, John Sillitoe on behalf of the Wellcome Sanger Institute COVID-19 Surveillance Team |
| EPI_ISL_2640546 | Lighthouse Lab in Alderley Park | Wellcome Sanger Institute for the COVID-19 Genomics UK (COG-UK) Consortium | Jacquelyn Wynn, Mairead Hyland, The Lighthouse Lab in Alderley Park and Alex Alderton, Roberto Amato, Jeffrey Barrett, Sonia Goncalves, Ewan Harrison, David K. Jackson, Ian Johnston, Dominic Kwiatkowski, Cordelia Langford, John Sillitoe on behalf of the Wellcome Sanger Institute COVID-19 Surveillance Team |
| EPI_ISL_2640547, EPI_ISL_2640548 | Lighthouse Lab in Milton Keynes | Wellcome Sanger Institute for the COVID-19 Genomics UK (COG-UK) Consortium | The Lighthouse Lab in Milton Keynes and Alex Alderton, Roberto Amato, Jeffrey Barrett, Sonia Goncalves, Ewan Harrison, David K. Jackson, Ian Johnston, Dominic Kwiatkowski, Cordelia Langford, John Sillitoe on behalf of the Wellcome Sanger Institute COVID-19 Surveillance Team |
| EPI_ISL_2640549 | Lighthouse Lab in Alderley Park | Wellcome Sanger Institute for the COVID-19 Genomics UK (COG-UK) Consortium | Jacquelyn Wynn, Mairead Hyland, The Lighthouse Lab in Alderley Park and Alex Alderton, Roberto Amato, Jeffrey Barrett, Sonia Goncalves, Ewan Harrison, David K. Jackson, Ian Johnston, Dominic Kwiatkowski, Cordelia Langford, John Sillitoe on behalf of the Wellcome Sanger Institute COVID-19 Surveillance Team |
| EPI_ISL_2640550, EPI_ISL_2640551 | Lighthouse Lab in Milton Keynes | Wellcome Sanger Institute for the COVID-19 Genomics UK (COG-UK) Consortium | The Lighthouse Lab in Milton Keynes and Alex Alderton, Roberto Amato, Jeffrey Barrett, Sonia Goncalves, Ewan Harrison, David K. Jackson, Ian Johnston, Dominic Kwiatkowski, Cordelia Langford, John Sillitoe on behalf of the Wellcome Sanger Institute COVID-19 Surveillance Team |
| EPI_ISL_2640552 | Lighthouse Lab in Alderley Park | Wellcome Sanger Institute for the COVID-19 Genomics UK (COG-UK) Consortium | Jacquelyn Wynn, Mairead Hyland, The Lighthouse Lab in Alderley Park and Alex Alderton, Roberto Amato, Jeffrey Barrett, Sonia Goncalves, Ewan Harrison, David K. Jackson, Ian Johnston, Dominic Kwiatkowski, Cordelia Langford, John Sillitoe on behalf of the Wellcome Sanger Institute COVID-19 Surveillance Team |
| EPI_ISL_2640553 | Lighthouse Lab in Milton Keynes | Wellcome Sanger Institute for the COVID-19 Genomics UK (COG-UK) Consortium | The Lighthouse Lab in Milton Keynes and Alex Alderton, Roberto Amato, Jeffrey Barrett, Sonia Goncalves, Ewan Harrison, David K. Jackson, Ian Johnston, Dominic Kwiatkowski, Cordelia Langford, John Sillitoe on behalf of the Wellcome Sanger Institute COVID-19 Surveillance Team |
| EPI_ISL_2640554, EPI_ISL_2640556, EPI_ISL_2640557 | Lighthouse Lab in Alderley Park | Wellcome Sanger Institute for the COVID-19 Genomics UK (COG-UK) Consortium | Jacquelyn Wynn, Mairead Hyland, The Lighthouse Lab in Alderley Park and Alex Alderton, Roberto Amato, Jeffrey Barrett, Sonia Goncalves, Ewan Harrison, David K. Jackson, Ian Johnston, Dominic Kwiatkowski, Cordelia Langford, John Sillitoe on behalf of the Wellcome Sanger Institute COVID-19 Surveillance Team |
| EPI_ISL_2640558 | Lighthouse Lab in Milton Keynes | Wellcome Sanger Institute for the COVID-19 Genomics UK (COG-UK) Consortium | The Lighthouse Lab in Milton Keynes and Alex Alderton, Roberto Amato, Jeffrey Barrett, Sonia Goncalves, Ewan Harrison, David K. Jackson, Ian Johnston, Dominic Kwiatkowski, Cordelia Langford, John Sillitoe on behalf of the Wellcome Sanger Institute COVID-19 Surveillance Team |
| EPI_ISL_2640559, EPI_ISL_2640560 | Lighthouse Lab in Alderley Park | Wellcome Sanger Institute for the COVID-19 Genomics UK (COG-UK) Consortium | Jacquelyn Wynn, Mairead Hyland, The Lighthouse Lab in Alderley Park and Alex Alderton, Roberto Amato, Jeffrey Barrett, Sonia Goncalves, Ewan Harrison, David K. Jackson, Ian Johnston, Dominic Kwiatkowski, Cordelia Langford, John Sillitoe on behalf of the Wellcome Sanger Institute COVID-19 Surveillance Team |
| EPI_ISL_2640561 | Lighthouse Lab in Milton Keynes | Wellcome Sanger Institute for the COVID-19 Genomics UK (COG-UK) Consortium | The Lighthouse Lab in Milton Keynes and Alex Alderton, Roberto Amato, Jeffrey Barrett, Sonia Goncalves, Ewan Harrison, David K. Jackson, Ian Johnston, Dominic Kwiatkowski, Cordelia Langford, John Sillitoe on behalf of the Wellcome Sanger Institute COVID-19 Surveillance Team |
| EPI_ISL_2640562 | Lighthouse Lab in Alderley Park | Wellcome Sanger Institute for the COVID-19 Genomics UK (COG-UK) Consortium | Jacquelyn Wynn, Mairead Hyland, The Lighthouse Lab in Alderley Park and Alex Alderton, Roberto Amato, Jeffrey Barrett, Sonia Goncalves, Ewan Harrison, David K. Jackson, Ian Johnston, Dominic Kwiatkowski, Cordelia Langford, John Sillitoe on behalf of the Wellcome Sanger Institute COVID-19 Surveillance Team |
| EPI_ISL_2640564, EPI_ISL_2640565 | Lighthouse Lab in Milton Keynes | Wellcome Sanger Institute for the COVID-19 Genomics UK (COG-UK) Consortium | The Lighthouse Lab in Milton Keynes and Alex Alderton, Roberto Amato, Jeffrey Barrett, Sonia Goncalves, Ewan Harrison, David K. Jackson, Ian Johnston, Dominic Kwiatkowski, Cordelia Langford, John Sillitoe on behalf of the Wellcome Sanger Institute COVID-19 Surveillance Team |
| EPI_ISL_2640566 | Lighthouse Lab in Alderley Park | Wellcome Sanger Institute for the COVID-19 Genomics UK (COG-UK) Consortium | Jacquelyn Wynn, Mairead Hyland, The Lighthouse Lab in Alderley Park and Alex Alderton, Roberto Amato, Jeffrey Barrett, Sonia Goncalves, Ewan Harrison, David K. Jackson, Ian Johnston, Dominic Kwiatkowski, Cordelia Langford, John Sillitoe on behalf of the Wellcome Sanger Institute COVID-19 Surveillance Team |

| | | | |
|---|---|---|---|
| EPI_ISL_2640567, EPI_ISL_2640568 | Lighthouse Lab in Milton Keynes | Wellcome Sanger Institute for the COVID-19 Genomics UK (COG-UK) Consortium | The Lighthouse Lab in Milton Keynes and Alex Alderton, Roberto Amato, Jeffrey Barrett, Sonia Goncalves, Ewan Harrison, David K. Jackson, Ian Johnston, Dominic Kwiatkowski, Cordelia Langford, John Sillitoe on behalf of the Wellcome Sanger Institute COVID-19 Surveillance Team |
| EPI_ISL_2640569, EPI_ISL_2640570 | Lighthouse Lab in Alderley Park | Wellcome Sanger Institute for the COVID-19 Genomics UK (COG-UK) Consortium | Jacquelyn Wynn, Mairead Hyland, The Lighthouse Lab in Alderley Park and Alex Alderton, Roberto Amato, Jeffrey Barrett, Sonia Goncalves, Ewan Harrison, David K. Jackson, Ian Johnston, Dominic Kwiatkowski, Cordelia Langford, John Sillitoe on behalf of the Wellcome Sanger Institute COVID-19 Surveillance Team |
| EPI_ISL_2640571 | Lighthouse Lab in Milton Keynes | Wellcome Sanger Institute for the COVID-19 Genomics UK (COG-UK) Consortium | The Lighthouse Lab in Milton Keynes and Alex Alderton, Roberto Amato, Jeffrey Barrett, Sonia Goncalves, Ewan Harrison, David K. Jackson, Ian Johnston, Dominic Kwiatkowski, Cordelia Langford, John Sillitoe on behalf of the Wellcome Sanger Institute COVID-19 Surveillance Team |
| EPI_ISL_2640572, EPI_ISL_2640573, EPI_ISL_2640574, EPI_ISL_2640575 | Lighthouse Lab in Alderley Park | Wellcome Sanger Institute for the COVID-19 Genomics UK (COG-UK) Consortium | Jacquelyn Wynn, Mairead Hyland, The Lighthouse Lab in Alderley Park and Alex Alderton, Roberto Amato, Jeffrey Barrett, Sonia Goncalves, Ewan Harrison, David K. Jackson, Ian Johnston, Dominic Kwiatkowski, Cordelia Langford, John Sillitoe on behalf of the Wellcome Sanger Institute COVID-19 Surveillance Team |
| EPI_ISL_2640576 | Lighthouse Lab in Milton Keynes | Wellcome Sanger Institute for the COVID-19 Genomics UK (COG-UK) Consortium | The Lighthouse Lab in Milton Keynes and Alex Alderton, Roberto Amato, Jeffrey Barrett, Sonia Goncalves, Ewan Harrison, David K. Jackson, Ian Johnston, Dominic Kwiatkowski, Cordelia Langford, John Sillitoe on behalf of the Wellcome Sanger Institute COVID-19 Surveillance Team |
| EPI_ISL_2640577, EPI_ISL_2640578 | Lighthouse Lab in Alderley Park | Wellcome Sanger Institute for the COVID-19 Genomics UK (COG-UK) Consortium | Jacquelyn Wynn, Mairead Hyland, The Lighthouse Lab in Alderley Park and Alex Alderton, Roberto Amato, Jeffrey Barrett, Sonia Goncalves, Ewan Harrison, David K. Jackson, Ian Johnston, Dominic Kwiatkowski, Cordelia Langford, John Sillitoe on behalf of the Wellcome Sanger Institute COVID-19 Surveillance Team |
| EPI_ISL_2640579 | Lighthouse Lab in Milton Keynes | Wellcome Sanger Institute for the COVID-19 Genomics UK (COG-UK) Consortium | The Lighthouse Lab in Milton Keynes and Alex Alderton, Roberto Amato, Jeffrey Barrett, Sonia Goncalves, Ewan Harrison, David K. Jackson, Ian Johnston, Dominic Kwiatkowski, Cordelia Langford, John Sillitoe on behalf of the Wellcome Sanger Institute COVID-19 Surveillance Team |
| EPI_ISL_2640580, EPI_ISL_2640581 | Lighthouse Lab in Alderley Park | Wellcome Sanger Institute for the COVID-19 Genomics UK (COG-UK) Consortium | Jacquelyn Wynn, Mairead Hyland, The Lighthouse Lab in Alderley Park and Alex Alderton, Roberto Amato, Jeffrey Barrett, Sonia Goncalves, Ewan Harrison, David K. Jackson, Ian Johnston, Dominic Kwiatkowski, Cordelia Langford, John Sillitoe on behalf of the Wellcome Sanger Institute COVID-19 Surveillance Team |
| EPI_ISL_2640582 | Lighthouse Lab in Milton Keynes | Wellcome Sanger Institute for the COVID-19 Genomics UK (COG-UK) Consortium | The Lighthouse Lab in Milton Keynes and Alex Alderton, Roberto Amato, Jeffrey Barrett, Sonia Goncalves, Ewan Harrison, David K. Jackson, Ian Johnston, Dominic Kwiatkowski, Cordelia Langford, John Sillitoe on behalf of the Wellcome Sanger Institute COVID-19 Surveillance Team |
| EPI_ISL_2640583, EPI_ISL_2640584, EPI_ISL_2640585, EPI_ISL_2640586, EPI_ISL_2640587 | Lighthouse Lab in Alderley Park | Wellcome Sanger Institute for the COVID-19 Genomics UK (COG-UK) Consortium | Jacquelyn Wynn, Mairead Hyland, The Lighthouse Lab in Alderley Park and Alex Alderton, Roberto Amato, Jeffrey Barrett, Sonia Goncalves, Ewan Harrison, David K. Jackson, Ian Johnston, Dominic Kwiatkowski, Cordelia Langford, John Sillitoe on behalf of the Wellcome Sanger Institute COVID-19 Surveillance Team |
| EPI_ISL_2640588, EPI_ISL_2640589 | Lighthouse Lab in Milton Keynes | Wellcome Sanger Institute for the COVID-19 Genomics UK (COG-UK) Consortium | The Lighthouse Lab in Milton Keynes and Alex Alderton, Roberto Amato, Jeffrey Barrett, Sonia Goncalves, Ewan Harrison, David K. Jackson, Ian Johnston, Dominic Kwiatkowski, Cordelia Langford, John Sillitoe on behalf of the Wellcome Sanger Institute COVID-19 Surveillance Team |
| EPI_ISL_2640590, EPI_ISL_2640591, EPI_ISL_2640592, EPI_ISL_2640593, EPI_ISL_2640594, EPI_ISL_2640595, EPI_ISL_2640596, EPI_ISL_2640597, EPI_ISL_2640598, EPI_ISL_2640599, EPI_ISL_2640600 | | | |
| see above | Lighthouse Lab in Alderley Park | Wellcome Sanger Institute for the COVID-19 Genomics UK (COG-UK) Consortium | Jacquelyn Wynn, Mairead Hyland, The Lighthouse Lab in Alderley Park and Alex Alderton, Roberto Amato, Jeffrey Barrett, Sonia Goncalves, Ewan Harrison, David K. Jackson, Ian Johnston, Dominic Kwiatkowski, Cordelia Langford, John Sillitoe on behalf of the Wellcome Sanger Institute COVID-19 Surveillance Team |
| EPI_ISL_2640601 | Lighthouse Lab in Milton Keynes | Wellcome Sanger Institute for the COVID-19 Genomics UK (COG-UK) Consortium | The Lighthouse Lab in Milton Keynes and Alex Alderton, Roberto Amato, Jeffrey Barrett, Sonia Goncalves, Ewan Harrison, David K. Jackson, Ian Johnston, Dominic Kwiatkowski, Cordelia Langford, John Sillitoe on behalf of the Wellcome Sanger Institute COVID-19 Surveillance Team |
| EPI_ISL_2640602, EPI_ISL_2640603 | Lighthouse Lab in Alderley Park | Wellcome Sanger Institute for the COVID-19 Genomics UK (COG-UK) Consortium | Jacquelyn Wynn, Mairead Hyland, The Lighthouse Lab in Alderley Park and Alex Alderton, Roberto Amato, Jeffrey Barrett, Sonia Goncalves, Ewan Harrison, David K. Jackson, Ian Johnston, Dominic Kwiatkowski, Cordelia Langford, John Sillitoe on behalf of the Wellcome Sanger Institute COVID-19 Surveillance Team |
| EPI_ISL_2640604 | Lighthouse Lab in Milton Keynes | Wellcome Sanger Institute for the COVID-19 Genomics UK (COG-UK) Consortium | The Lighthouse Lab in Milton Keynes and Alex Alderton, Roberto Amato, Jeffrey Barrett, Sonia Goncalves, Ewan Harrison, David K. Jackson, Ian Johnston, Dominic Kwiatkowski, Cordelia Langford, John Sillitoe on behalf of the Wellcome Sanger Institute COVID-19 Surveillance Team |
| EPI_ISL_2640605 | Lighthouse Lab in Alderley Park | Wellcome Sanger Institute for the COVID-19 Genomics UK (COG-UK) Consortium | Jacquelyn Wynn, Mairead Hyland, The Lighthouse Lab in Alderley Park and Alex Alderton, Roberto Amato, Jeffrey Barrett, Sonia Goncalves, Ewan Harrison, David K. Jackson, Ian Johnston, Dominic Kwiatkowski, Cordelia Langford, John Sillitoe on behalf of the Wellcome Sanger Institute COVID-19 Surveillance Team |
| EPI_ISL_2640606, EPI_ISL_2640607, EPI_ISL_2640608 | Lighthouse Lab in Milton Keynes | Wellcome Sanger Institute for the COVID-19 Genomics UK (COG-UK) Consortium | The Lighthouse Lab in Milton Keynes and Alex Alderton, Roberto Amato, Jeffrey Barrett, Sonia Goncalves, Ewan Harrison, David K. Jackson, Ian Johnston, Dominic Kwiatkowski, Cordelia Langford, John Sillitoe on behalf of the Wellcome Sanger Institute COVID-19 Surveillance Team |
| EPI_ISL_2640609, EPI_ISL_2640610, EPI_ISL_2640611, EPI_ISL_2640612, EPI_ISL_2640613, EPI_ISL_2640614, EPI_ISL_2640615 | Lighthouse Lab in Alderley Park | Wellcome Sanger Institute for the COVID-19 Genomics UK (COG-UK) Consortium | Jacquelyn Wynn, Mairead Hyland, The Lighthouse Lab in Alderley Park and Alex Alderton, Roberto Amato, Jeffrey Barrett, Sonia Goncalves, Ewan Harrison, David K. Jackson, Ian Johnston, Dominic Kwiatkowski, Cordelia Langford, John Sillitoe on behalf of the Wellcome Sanger Institute COVID-19 Surveillance Team |
| EPI_ISL_2640616, EPI_ISL_2640617 | Lighthouse Lab in Milton Keynes | Wellcome Sanger Institute for the COVID-19 Genomics UK (COG-UK) Consortium | The Lighthouse Lab in Milton Keynes and Alex Alderton, Roberto Amato, Jeffrey Barrett, Sonia Goncalves, Ewan Harrison, David K. Jackson, Ian Johnston, Dominic Kwiatkowski, Cordelia Langford, John Sillitoe on behalf of the Wellcome Sanger Institute COVID-19 Surveillance Team |
| EPI_ISL_2640618 | Lighthouse Lab in Alderley Park | Wellcome Sanger Institute for the COVID-19 Genomics UK (COG-UK) Consortium | Jacquelyn Wynn, Mairead Hyland, The Lighthouse Lab in Alderley Park and Alex Alderton, Roberto Amato, Jeffrey Barrett, Sonia Goncalves, Ewan Harrison, David K. Jackson, Ian Johnston, Dominic Kwiatkowski, Cordelia Langford, John Sillitoe on behalf of the Wellcome Sanger Institute COVID-19 Surveillance Team |
| EPI_ISL_2640619 | Lighthouse Lab in Milton Keynes | Wellcome Sanger Institute for the COVID-19 Genomics UK (COG-UK) Consortium | The Lighthouse Lab in Milton Keynes and Alex Alderton, Roberto Amato, Jeffrey Barrett, Sonia Goncalves, Ewan Harrison, David K. Jackson, Ian Johnston, Dominic Kwiatkowski, Cordelia Langford, John Sillitoe on behalf of the Wellcome Sanger Institute COVID-19 Surveillance Team |
| EPI_ISL_2640620, EPI_ISL_2640621, EPI_ISL_2640622 | Lighthouse Lab in Alderley Park | Wellcome Sanger Institute for the COVID-19 Genomics UK (COG-UK) Consortium | Jacquelyn Wynn, Mairead Hyland, The Lighthouse Lab in Alderley Park and Alex Alderton, Roberto Amato, Jeffrey Barrett, Sonia Goncalves, Ewan Harrison, David K. Jackson, Ian Johnston, Dominic Kwiatkowski, Cordelia Langford, John Sillitoe on behalf of the Wellcome Sanger Institute COVID-19 Surveillance Team |
| EPI_ISL_2640623, EPI_ISL_2640624, EPI_ISL_2640625, EPI_ISL_2640626 | Lighthouse Lab in Milton Keynes | Wellcome Sanger Institute for the COVID-19 Genomics UK (COG-UK) Consortium | The Lighthouse Lab in Milton Keynes and Alex Alderton, Roberto Amato, Jeffrey Barrett, Sonia Goncalves, Ewan Harrison, David K. Jackson, Ian Johnston, Dominic Kwiatkowski, Cordelia Langford, John Sillitoe on behalf of the Wellcome Sanger Institute COVID-19 Surveillance Team |
| EPI_ISL_2640627 | Lighthouse Lab in Alderley Park | Wellcome Sanger Institute for the COVID-19 Genomics UK (COG-UK) Consortium | Jacquelyn Wynn, Mairead Hyland, The Lighthouse Lab in Alderley Park and Alex Alderton, Roberto Amato, Jeffrey Barrett, Sonia Goncalves, Ewan Harrison, David K. Jackson, Ian Johnston, Dominic Kwiatkowski, Cordelia Langford, John Sillitoe on behalf of the Wellcome Sanger Institute COVID-19 Surveillance Team |
| EPI_ISL_2640628, EPI_ISL_2640629 | Lighthouse Lab in Milton Keynes | Wellcome Sanger Institute for the COVID-19 Genomics UK (COG-UK) Consortium | The Lighthouse Lab in Milton Keynes and Alex Alderton, Roberto Amato, Jeffrey Barrett, Sonia Goncalves, Ewan Harrison, David K. Jackson, Ian Johnston, Dominic Kwiatkowski, Cordelia Langford, John Sillitoe on behalf of the Wellcome Sanger Institute COVID-19 Surveillance Team |
| EPI_ISL_2640630, EPI_ISL_2640631, EPI_ISL_2640632, EPI_ISL_2640633 | Lighthouse Lab in Alderley Park | Wellcome Sanger Institute for the COVID-19 Genomics UK (COG-UK) Consortium | Jacquelyn Wynn, Mairead Hyland, The Lighthouse Lab in Alderley Park and Alex Alderton, Roberto Amato, Jeffrey Barrett, Sonia Goncalves, Ewan Harrison, David K. Jackson, Ian Johnston, Dominic Kwiatkowski, Cordelia Langford, John Sillitoe on behalf of the Wellcome Sanger Institute COVID-19 Surveillance Team |
| EPI_ISL_2640634, EPI_ISL_2640635 | Lighthouse Lab in Milton Keynes | Wellcome Sanger Institute for the COVID-19 Genomics UK (COG-UK) Consortium | The Lighthouse Lab in Milton Keynes and Alex Alderton, Roberto Amato, Jeffrey Barrett, Sonia Goncalves, Ewan Harrison, David K. Jackson, Ian Johnston, Dominic Kwiatkowski, Cordelia Langford, John Sillitoe on behalf of the Wellcome Sanger Institute COVID-19 Surveillance Team |
| EPI_ISL_2640636, EPI_ISL_2640637 | Lighthouse Lab in Alderley Park | Wellcome Sanger Institute for the COVID-19 Genomics UK (COG-UK) Consortium | Jacquelyn Wynn, Mairead Hyland, The Lighthouse Lab in Alderley Park and Alex Alderton, Roberto Amato, Jeffrey Barrett, Sonia Goncalves, Ewan Harrison, David K. Jackson, Ian Johnston, Dominic Kwiatkowski, Cordelia Langford, John Sillitoe on behalf of the Wellcome Sanger Institute COVID-19 Surveillance Team |
| EPI_ISL_2640638, EPI_ISL_2640639 | Lighthouse Lab in Milton Keynes | Wellcome Sanger Institute for the COVID-19 Genomics UK (COG-UK) Consortium | The Lighthouse Lab in Milton Keynes and Alex Alderton, Roberto Amato, Jeffrey Barrett, Sonia Goncalves, Ewan Harrison, David K. Jackson, Ian Johnston, Dominic Kwiatkowski, Cordelia Langford, John Sillitoe on behalf of the Wellcome Sanger Institute COVID-19 Surveillance Team |
| EPI_ISL_2640640 | Lighthouse Lab in Alderley Park | Wellcome Sanger Institute for the COVID-19 Genomics UK (COG-UK) Consortium | Jacquelyn Wynn, Mairead Hyland, The Lighthouse Lab in Alderley Park and Alex Alderton, Roberto Amato, Jeffrey Barrett, Sonia Goncalves, Ewan Harrison, David K. Jackson, Ian Johnston, Dominic Kwiatkowski, Cordelia Langford, John Sillitoe on behalf of the Wellcome Sanger Institute COVID-19 Surveillance Team |
| EPI_ISL_2640641 | Lighthouse Lab in Milton Keynes | Wellcome Sanger Institute for the COVID-19 Genomics UK (COG-UK) Consortium | The Lighthouse Lab in Milton Keynes and Alex Alderton, Roberto Amato, Jeffrey Barrett, Sonia Goncalves, Ewan Harrison, David K. Jackson, Ian Johnston, Dominic Kwiatkowski, Cordelia Langford, John Sillitoe on behalf of the Wellcome Sanger Institute COVID-19 Surveillance Team |
| EPI_ISL_2640642 | Lighthouse Lab in Alderley Park | Wellcome Sanger Institute for the COVID-19 Genomics UK (COG-UK) Consortium | Jacquelyn Wynn, Mairead Hyland, The Lighthouse Lab in Alderley Park and Alex Alderton, Roberto Amato, Jeffrey Barrett, Sonia Goncalves, Ewan Harrison, David K. Jackson, Ian Johnston, Dominic Kwiatkowski, Cordelia Langford, John Sillitoe on behalf of the Wellcome Sanger Institute COVID-19 Surveillance Team |
| EPI_ISL_2640643, EPI_ISL_2640644 | Lighthouse Lab in Milton Keynes | Wellcome Sanger Institute for the COVID-19 Genomics UK (COG-UK) Consortium | The Lighthouse Lab in Milton Keynes and Alex Alderton, Roberto Amato, Jeffrey Barrett, Sonia Goncalves, Ewan Harrison, David K. Jackson, Ian Johnston, Dominic Kwiatkowski, Cordelia Langford, John Sillitoe on behalf of the Wellcome Sanger Institute COVID-19 Surveillance Team |
| EPI_ISL_2640645, EPI_ISL_2640646 | Lighthouse Lab in Alderley Park | Wellcome Sanger Institute for the COVID-19 Genomics UK (COG-UK) Consortium | Jacquelyn Wynn, Mairead Hyland, The Lighthouse Lab in Alderley Park and Alex Alderton, Roberto Amato, Jeffrey Barrett, Sonia Goncalves, Ewan Harrison, David K. Jackson, Ian Johnston, Dominic Kwiatkowski, Cordelia Langford, John Sillitoe on behalf of the Wellcome Sanger Institute COVID-19 Surveillance Team |
| EPI_ISL_2640647 | Lighthouse Lab in Milton Keynes | Wellcome Sanger Institute for the COVID-19 Genomics UK (COG-UK) Consortium | The Lighthouse Lab in Milton Keynes and Alex Alderton, Roberto Amato, Jeffrey Barrett, Sonia Goncalves, Ewan Harrison, David K. Jackson, Ian Johnston, Dominic Kwiatkowski, Cordelia Langford, John Sillitoe on behalf of the Wellcome Sanger Institute COVID-19 Surveillance Team |
| EPI_ISL_2640648, EPI_ISL_2640649 | Lighthouse Lab in Alderley Park | Wellcome Sanger Institute for the COVID-19 Genomics UK (COG-UK) Consortium | Jacquelyn Wynn, Mairead Hyland, The Lighthouse Lab in Alderley Park and Alex Alderton, Roberto Amato, Jeffrey Barrett, Sonia Goncalves, Ewan Harrison, David K. Jackson, Ian Johnston, Dominic Kwiatkowski, Cordelia Langford, John Sillitoe on behalf of the Wellcome Sanger Institute COVID-19 Surveillance Team |
| EPI_ISL_2640650 | Lighthouse Lab in Milton Keynes | Wellcome Sanger Institute for the COVID-19 Genomics UK (COG-UK) Consortium | The Lighthouse Lab in Milton Keynes and Alex Alderton, Roberto Amato, Jeffrey Barrett, Sonia Goncalves, Ewan Harrison, David K. Jackson, Ian Johnston, Dominic Kwiatkowski, Cordelia Langford, John Sillitoe on behalf of the Wellcome Sanger Institute COVID-19 Surveillance Team |
| EPI_ISL_2640651, EPI_ISL_2640652 | Lighthouse Lab in Alderley Park | Wellcome Sanger Institute for the COVID-19 Genomics UK (COG-UK) Consortium | Jacquelyn Wynn, Mairead Hyland, The Lighthouse Lab in Alderley Park and Alex Alderton, Roberto Amato, Jeffrey Barrett, Sonia Goncalves, Ewan Harrison, David K. Jackson, Ian Johnston, Dominic Kwiatkowski, Cordelia Langford, John Sillitoe on behalf of the Wellcome Sanger Institute COVID-19 Surveillance Team |

| | | | |
|---|---|---|---|
| EPI_ISL_2640653, EPI_ISL_2640655, EPI_ISL_2640656, EPI_ISL_2640657, EPI_ISL_2640658, EPI_ISL_2640659, EPI_ISL_2640660, EPI_ISL_2640661 | Lighthouse Lab in Milton Keynes | Wellcome Sanger Institute for the COVID-19 Genomics UK (COG-UK) Consortium | The Lighthouse Lab in Milton Keynes and Alex Alderton, Roberto Amato, Jeffrey Barrett, Sonia Goncalves, Ewan Harrison, David K. Jackson, Ian Johnston, Dominic Kwiatkowski, Cordelia Langford, John Sillitoe on behalf of the Wellcome Sanger Institute COVID-19 Surveillance Team |
| EPI_ISL_2640662, EPI_ISL_2640663 | Lighthouse Lab in Alderley Park | Wellcome Sanger Institute for the COVID-19 Genomics UK (COG-UK) Consortium | Jacquelyn Wynn, Mairead Hyland, The Lighthouse Lab in Alderley Park and Alex Alderton, Roberto Amato, Jeffrey Barrett, Sonia Goncalves, Ewan Harrison, David K. Jackson, Ian Johnston, Dominic Kwiatkowski, Cordelia Langford, John Sillitoe on behalf of the Wellcome Sanger Institute COVID-19 Surveillance Team |
| EPI_ISL_2640664, EPI_ISL_2640665 | Lighthouse Lab in Milton Keynes | Wellcome Sanger Institute for the COVID-19 Genomics UK (COG-UK) Consortium | The Lighthouse Lab in Milton Keynes and Alex Alderton, Roberto Amato, Jeffrey Barrett, Sonia Goncalves, Ewan Harrison, David K. Jackson, Ian Johnston, Dominic Kwiatkowski, Cordelia Langford, John Sillitoe on behalf of the Wellcome Sanger Institute COVID-19 Surveillance Team |
| EPI_ISL_2640667, EPI_ISL_2640668 | Lighthouse Lab in Alderley Park | Wellcome Sanger Institute for the COVID-19 Genomics UK (COG-UK) Consortium | Jacquelyn Wynn, Mairead Hyland, The Lighthouse Lab in Alderley Park and Alex Alderton, Roberto Amato, Jeffrey Barrett, Sonia Goncalves, Ewan Harrison, David K. Jackson, Ian Johnston, Dominic Kwiatkowski, Cordelia Langford, John Sillitoe on behalf of the Wellcome Sanger Institute COVID-19 Surveillance Team |
| EPI_ISL_2640669, EPI_ISL_2640670 | Lighthouse Lab in Milton Keynes | Wellcome Sanger Institute for the COVID-19 Genomics UK (COG-UK) Consortium | The Lighthouse Lab in Milton Keynes and Alex Alderton, Roberto Amato, Jeffrey Barrett, Sonia Goncalves, Ewan Harrison, David K. Jackson, Ian Johnston, Dominic Kwiatkowski, Cordelia Langford, John Sillitoe on behalf of the Wellcome Sanger Institute COVID-19 Surveillance Team |
| EPI_ISL_2640671 | Lighthouse Lab in Alderley Park | Wellcome Sanger Institute for the COVID-19 Genomics UK (COG-UK) Consortium | Jacquelyn Wynn, Mairead Hyland, The Lighthouse Lab in Alderley Park and Alex Alderton, Roberto Amato, Jeffrey Barrett, Sonia Goncalves, Ewan Harrison, David K. Jackson, Ian Johnston, Dominic Kwiatkowski, Cordelia Langford, John Sillitoe on behalf of the Wellcome Sanger Institute COVID-19 Surveillance Team |
| EPI_ISL_2640672, EPI_ISL_2640673, EPI_ISL_2640674, EPI_ISL_2640675 | Lighthouse Lab in Milton Keynes | Wellcome Sanger Institute for the COVID-19 Genomics UK (COG-UK) Consortium | The Lighthouse Lab in Milton Keynes and Alex Alderton, Roberto Amato, Jeffrey Barrett, Sonia Goncalves, Ewan Harrison, David K. Jackson, Ian Johnston, Dominic Kwiatkowski, Cordelia Langford, John Sillitoe on behalf of the Wellcome Sanger Institute COVID-19 Surveillance Team |
| EPI_ISL_2640676 | Lighthouse Lab in Alderley Park | Wellcome Sanger Institute for the COVID-19 Genomics UK (COG-UK) Consortium | Jacquelyn Wynn, Mairead Hyland, The Lighthouse Lab in Alderley Park and Alex Alderton, Roberto Amato, Jeffrey Barrett, Sonia Goncalves, Ewan Harrison, David K. Jackson, Ian Johnston, Dominic Kwiatkowski, Cordelia Langford, John Sillitoe on behalf of the Wellcome Sanger Institute COVID-19 Surveillance Team |
| EPI_ISL_2640677, EPI_ISL_2640678 | Lighthouse Lab in Glasgow | Wellcome Sanger Institute for the COVID-19 Genomics UK (COG-UK) Consortium | Harper VanSteenhouse, Yumi Kasai, David Gray, Carol Clugston, Anna Dominiczak and Alex Alderton, Roberto Amato, Jeffrey Barrett, Sonia Goncalves, Ewan Harrison, David K. Jackson, Ian Johnston, Dominic Kwiatkowski, Cordelia Langford, John Sillitoe on behalf of the Wellcome Sanger Institute COVID-19 Surveillance Team |
| EPI_ISL_2640679 | Lighthouse Lab in Milton Keynes | Wellcome Sanger Institute for the COVID-19 Genomics UK (COG-UK) Consortium | The Lighthouse Lab in Milton Keynes and Alex Alderton, Roberto Amato, Jeffrey Barrett, Sonia Goncalves, Ewan Harrison, David K. Jackson, Ian Johnston, Dominic Kwiatkowski, Cordelia Langford, John Sillitoe on behalf of the Wellcome Sanger Institute COVID-19 Surveillance Team |
| EPI_ISL_2640680 | Lighthouse Lab in Glasgow | Wellcome Sanger Institute for the COVID-19 Genomics UK (COG-UK) Consortium | Harper VanSteenhouse, Yumi Kasai, David Gray, Carol Clugston, Anna Dominiczak and Alex Alderton, Roberto Amato, Jeffrey Barrett, Sonia Goncalves, Ewan Harrison, David K. Jackson, Ian Johnston, Dominic Kwiatkowski, Cordelia Langford, John Sillitoe on behalf of the Wellcome Sanger Institute COVID-19 Surveillance Team |
| EPI_ISL_2640681 | Lighthouse Lab in Milton Keynes | Wellcome Sanger Institute for the COVID-19 Genomics UK (COG-UK) Consortium | The Lighthouse Lab in Milton Keynes and Alex Alderton, Roberto Amato, Jeffrey Barrett, Sonia Goncalves, Ewan Harrison, David K. Jackson, Ian Johnston, Dominic Kwiatkowski, Cordelia Langford, John Sillitoe on behalf of the Wellcome Sanger Institute COVID-19 Surveillance Team |
| EPI_ISL_2640682 | Lighthouse Lab in Glasgow | Wellcome Sanger Institute for the COVID-19 Genomics UK (COG-UK) Consortium | Harper VanSteenhouse, Yumi Kasai, David Gray, Carol Clugston, Anna Dominiczak and Alex Alderton, Roberto Amato, Jeffrey Barrett, Sonia Goncalves, Ewan Harrison, David K. Jackson, Ian Johnston, Dominic Kwiatkowski, Cordelia Langford, John Sillitoe on behalf of the Wellcome Sanger Institute COVID-19 Surveillance Team |
| EPI_ISL_2640683 | Lighthouse Lab in Milton Keynes | Wellcome Sanger Institute for the COVID-19 Genomics UK (COG-UK) Consortium | The Lighthouse Lab in Milton Keynes and Alex Alderton, Roberto Amato, Jeffrey Barrett, Sonia Goncalves, Ewan Harrison, David K. Jackson, Ian Johnston, Dominic Kwiatkowski, Cordelia Langford, John Sillitoe on behalf of the Wellcome Sanger Institute COVID-19 Surveillance Team |
| EPI_ISL_2640684 | Lighthouse Lab in Alderley Park | Wellcome Sanger Institute for the COVID-19 Genomics UK (COG-UK) Consortium | Jacquelyn Wynn, Mairead Hyland, The Lighthouse Lab in Alderley Park and Alex Alderton, Roberto Amato, Jeffrey Barrett, Sonia Goncalves, Ewan Harrison, David K. Jackson, Ian Johnston, Dominic Kwiatkowski, Cordelia Langford, John Sillitoe on behalf of the Wellcome Sanger Institute COVID-19 Surveillance Team |
| EPI_ISL_2640685 | Lighthouse Lab in Glasgow | Wellcome Sanger Institute for the COVID-19 Genomics UK (COG-UK) Consortium | Harper VanSteenhouse, Yumi Kasai, David Gray, Carol Clugston, Anna Dominiczak and Alex Alderton, Roberto Amato, Jeffrey Barrett, Sonia Goncalves, Ewan Harrison, David K. Jackson, Ian Johnston, Dominic Kwiatkowski, Cordelia Langford, John Sillitoe on behalf of the Wellcome Sanger Institute COVID-19 Surveillance Team |
| EPI_ISL_2640686, EPI_ISL_2640687, EPI_ISL_2640688 | Lighthouse Lab in Alderley Park | Wellcome Sanger Institute for the COVID-19 Genomics UK (COG-UK) Consortium | Jacquelyn Wynn, Mairead Hyland, The Lighthouse Lab in Alderley Park and Alex Alderton, Roberto Amato, Jeffrey Barrett, Sonia Goncalves, Ewan Harrison, David K. Jackson, Ian Johnston, Dominic Kwiatkowski, Cordelia Langford, John Sillitoe on behalf of the Wellcome Sanger Institute COVID-19 Surveillance Team |
| EPI_ISL_2640689 | Lighthouse Lab in Milton Keynes | Wellcome Sanger Institute for the COVID-19 Genomics UK (COG-UK) Consortium | The Lighthouse Lab in Milton Keynes and Alex Alderton, Roberto Amato, Jeffrey Barrett, Sonia Goncalves, Ewan Harrison, David K. Jackson, Ian Johnston, Dominic Kwiatkowski, Cordelia Langford, John Sillitoe on behalf of the Wellcome Sanger Institute COVID-19 Surveillance Team |
| EPI_ISL_2640691 | Lighthouse Lab in Alderley Park | Wellcome Sanger Institute for the COVID-19 Genomics UK (COG-UK) Consortium | Jacquelyn Wynn, Mairead Hyland, The Lighthouse Lab in Alderley Park and Alex Alderton, Roberto Amato, Jeffrey Barrett, Sonia Goncalves, Ewan Harrison, David K. Jackson, Ian Johnston, Dominic Kwiatkowski, Cordelia Langford, John Sillitoe on behalf of the Wellcome Sanger Institute COVID-19 Surveillance Team |
| EPI_ISL_2640692 | Lighthouse Lab in Milton Keynes | Wellcome Sanger Institute for the COVID-19 Genomics UK (COG-UK) Consortium | The Lighthouse Lab in Milton Keynes and Alex Alderton, Roberto Amato, Jeffrey Barrett, Sonia Goncalves, Ewan Harrison, David K. Jackson, Ian Johnston, Dominic Kwiatkowski, Cordelia Langford, John Sillitoe on behalf of the Wellcome Sanger Institute COVID-19 Surveillance Team |
| EPI_ISL_2640693, EPI_ISL_2640694 | Lighthouse Lab in Alderley Park | Wellcome Sanger Institute for the COVID-19 Genomics UK (COG-UK) Consortium | Jacquelyn Wynn, Mairead Hyland, The Lighthouse Lab in Alderley Park and Alex Alderton, Roberto Amato, Jeffrey Barrett, Sonia Goncalves, Ewan Harrison, David K. Jackson, Ian Johnston, Dominic Kwiatkowski, Cordelia Langford, John Sillitoe on behalf of the Wellcome Sanger Institute COVID-19 Surveillance Team |
| EPI_ISL_2640695, EPI_ISL_2640696 | Lighthouse Lab in Glasgow | Wellcome Sanger Institute for the COVID-19 Genomics UK (COG-UK) Consortium | Harper VanSteenhouse, Yumi Kasai, David Gray, Carol Clugston, Anna Dominiczak and Alex Alderton, Roberto Amato, Jeffrey Barrett, Sonia Goncalves, Ewan Harrison, David K. Jackson, Ian Johnston, Dominic Kwiatkowski, Cordelia Langford, John Sillitoe on behalf of the Wellcome Sanger Institute COVID-19 Surveillance Team |
| EPI_ISL_2640697 | Lighthouse Lab in Milton Keynes | Wellcome Sanger Institute for the COVID-19 Genomics UK (COG-UK) Consortium | The Lighthouse Lab in Milton Keynes and Alex Alderton, Roberto Amato, Jeffrey Barrett, Sonia Goncalves, Ewan Harrison, David K. Jackson, Ian Johnston, Dominic Kwiatkowski, Cordelia Langford, John Sillitoe on behalf of the Wellcome Sanger Institute COVID-19 Surveillance Team |
| EPI_ISL_2640698, EPI_ISL_2640699 | Lighthouse Lab in Alderley Park | Wellcome Sanger Institute for the COVID-19 Genomics UK (COG-UK) Consortium | Jacquelyn Wynn, Mairead Hyland, The Lighthouse Lab in Alderley Park and Alex Alderton, Roberto Amato, Jeffrey Barrett, Sonia Goncalves, Ewan Harrison, David K. Jackson, Ian Johnston, Dominic Kwiatkowski, Cordelia Langford, John Sillitoe on behalf of the Wellcome Sanger Institute COVID-19 Surveillance Team |
| EPI_ISL_2640700 | Lighthouse Lab in Glasgow | Wellcome Sanger Institute for the COVID-19 Genomics UK (COG-UK) Consortium | Harper VanSteenhouse, Yumi Kasai, David Gray, Carol Clugston, Anna Dominiczak and Alex Alderton, Roberto Amato, Jeffrey Barrett, Sonia Goncalves, Ewan Harrison, David K. Jackson, Ian Johnston, Dominic Kwiatkowski, Cordelia Langford, John Sillitoe on behalf of the Wellcome Sanger Institute COVID-19 Surveillance Team |
| EPI_ISL_2640701, EPI_ISL_2640702, EPI_ISL_2640703, EPI_ISL_2640704, EPI_ISL_2640705 | Lighthouse Lab in Alderley Park | Wellcome Sanger Institute for the COVID-19 Genomics UK (COG-UK) Consortium | Jacquelyn Wynn, Mairead Hyland, The Lighthouse Lab in Alderley Park and Alex Alderton, Roberto Amato, Jeffrey Barrett, Sonia Goncalves, Ewan Harrison, David K. Jackson, Ian Johnston, Dominic Kwiatkowski, Cordelia Langford, John Sillitoe on behalf of the Wellcome Sanger Institute COVID-19 Surveillance Team |
| EPI_ISL_2640706 | Lighthouse Lab in Glasgow | Wellcome Sanger Institute for the COVID-19 Genomics UK (COG-UK) Consortium | Harper VanSteenhouse, Yumi Kasai, David Gray, Carol Clugston, Anna Dominiczak and Alex Alderton, Roberto Amato, Jeffrey Barrett, Sonia Goncalves, Ewan Harrison, David K. Jackson, Ian Johnston, Dominic Kwiatkowski, Cordelia Langford, John Sillitoe on behalf of the Wellcome Sanger Institute COVID-19 Surveillance Team |
| EPI_ISL_2640707, EPI_ISL_2640708 | Lighthouse Lab in Alderley Park | Wellcome Sanger Institute for the COVID-19 Genomics UK (COG-UK) Consortium | Jacquelyn Wynn, Mairead Hyland, The Lighthouse Lab in Alderley Park and Alex Alderton, Roberto Amato, Jeffrey Barrett, Sonia Goncalves, Ewan Harrison, David K. Jackson, Ian Johnston, Dominic Kwiatkowski, Cordelia Langford, John Sillitoe on behalf of the Wellcome Sanger Institute COVID-19 Surveillance Team |
| EPI_ISL_2640709 | Lighthouse Lab in Milton Keynes | Wellcome Sanger Institute for the COVID-19 Genomics UK (COG-UK) Consortium | The Lighthouse Lab in Milton Keynes and Alex Alderton, Roberto Amato, Jeffrey Barrett, Sonia Goncalves, Ewan Harrison, David K. Jackson, Ian Johnston, Dominic Kwiatkowski, Cordelia Langford, John Sillitoe on behalf of the Wellcome Sanger Institute COVID-19 Surveillance Team |
| EPI_ISL_2640710 | Lighthouse Lab in Alderley Park | Wellcome Sanger Institute for the COVID-19 Genomics UK (COG-UK) Consortium | Jacquelyn Wynn, Mairead Hyland, The Lighthouse Lab in Alderley Park and Alex Alderton, Roberto Amato, Jeffrey Barrett, Sonia Goncalves, Ewan Harrison, David K. Jackson, Ian Johnston, Dominic Kwiatkowski, Cordelia Langford, John Sillitoe on behalf of the Wellcome Sanger Institute COVID-19 Surveillance Team |
| EPI_ISL_2640711, EPI_ISL_2640712 | Lighthouse Lab in Milton Keynes | Wellcome Sanger Institute for the COVID-19 Genomics UK (COG-UK) Consortium | The Lighthouse Lab in Milton Keynes and Alex Alderton, Roberto Amato, Jeffrey Barrett, Sonia Goncalves, Ewan Harrison, David K. Jackson, Ian Johnston, Dominic Kwiatkowski, Cordelia Langford, John Sillitoe on behalf of the Wellcome Sanger Institute COVID-19 Surveillance Team |
| EPI_ISL_2640713 | Lighthouse Lab in Glasgow | Wellcome Sanger Institute for the COVID-19 Genomics UK (COG-UK) Consortium | Harper VanSteenhouse, Yumi Kasai, David Gray, Carol Clugston, Anna Dominiczak and Alex Alderton, Roberto Amato, Jeffrey Barrett, Sonia Goncalves, Ewan Harrison, David K. Jackson, Ian Johnston, Dominic Kwiatkowski, Cordelia Langford, John Sillitoe on behalf of the Wellcome Sanger Institute COVID-19 Surveillance Team |
| EPI_ISL_2640714, EPI_ISL_2640715 | Lighthouse Lab in Alderley Park | Wellcome Sanger Institute for the COVID-19 Genomics UK (COG-UK) Consortium | Jacquelyn Wynn, Mairead Hyland, The Lighthouse Lab in Alderley Park and Alex Alderton, Roberto Amato, Jeffrey Barrett, Sonia Goncalves, Ewan Harrison, David K. Jackson, Ian Johnston, Dominic Kwiatkowski, Cordelia Langford, John Sillitoe on behalf of the Wellcome Sanger Institute COVID-19 Surveillance Team |
| EPI_ISL_2640716, EPI_ISL_2640718 | Lighthouse Lab in Glasgow | Wellcome Sanger Institute for the COVID-19 Genomics UK (COG-UK) Consortium | Harper VanSteenhouse, Yumi Kasai, David Gray, Carol Clugston, Anna Dominiczak and Alex Alderton, Roberto Amato, Jeffrey Barrett, Sonia Goncalves, Ewan Harrison, David K. Jackson, Ian Johnston, Dominic Kwiatkowski, Cordelia Langford, John Sillitoe on behalf of the Wellcome Sanger Institute COVID-19 Surveillance Team |
| EPI_ISL_2640719, EPI_ISL_2640720, EPI_ISL_2640721, EPI_ISL_2640722 | Lighthouse Lab in Alderley Park | Wellcome Sanger Institute for the COVID-19 Genomics UK (COG-UK) Consortium | Jacquelyn Wynn, Mairead Hyland, The Lighthouse Lab in Alderley Park and Alex Alderton, Roberto Amato, Jeffrey Barrett, Sonia Goncalves, Ewan Harrison, David K. Jackson, Ian Johnston, Dominic Kwiatkowski, Cordelia Langford, John Sillitoe on behalf of the Wellcome Sanger Institute COVID-19 Surveillance Team |
| EPI_ISL_2640723 | Lighthouse Lab in Milton Keynes | Wellcome Sanger Institute for the COVID-19 Genomics UK (COG-UK) Consortium | The Lighthouse Lab in Milton Keynes and Alex Alderton, Roberto Amato, Jeffrey Barrett, Sonia Goncalves, Ewan Harrison, David K. Jackson, Ian Johnston, Dominic Kwiatkowski, Cordelia Langford, John Sillitoe on behalf of the Wellcome Sanger Institute COVID-19 Surveillance Team |
| EPI_ISL_2640724 | Lighthouse Lab in Alderley Park | Wellcome Sanger Institute for the COVID-19 Genomics UK (COG-UK) Consortium | Jacquelyn Wynn, Mairead Hyland, The Lighthouse Lab in Alderley Park and Alex Alderton, Roberto Amato, Jeffrey Barrett, Sonia Goncalves, Ewan Harrison, David K. Jackson, Ian Johnston, Dominic Kwiatkowski, Cordelia Langford, John Sillitoe on behalf of the Wellcome Sanger Institute COVID-19 Surveillance Team |
| EPI_ISL_2640726, EPI_ISL_2640727 | Lighthouse Lab in Glasgow | Wellcome Sanger Institute for the COVID-19 Genomics UK (COG-UK) Consortium | Harper VanSteenhouse, Yumi Kasai, David Gray, Carol Clugston, Anna Dominiczak and Alex Alderton, Roberto Amato, Jeffrey Barrett, Sonia Goncalves, Ewan Harrison, David K. Jackson, Ian Johnston, Dominic Kwiatkowski, Cordelia Langford, John Sillitoe on behalf of the Wellcome Sanger Institute COVID-19 Surveillance Team |
| EPI_ISL_2640728, EPI_ISL_2640729 | Lighthouse Lab in Alderley Park | Wellcome Sanger Institute for the COVID-19 Genomics UK (COG-UK) Consortium | Jacquelyn Wynn, Mairead Hyland, The Lighthouse Lab in Alderley Park and Alex Alderton, Roberto Amato, Jeffrey Barrett, Sonia Goncalves, Ewan Harrison, David K. Jackson, Ian Johnston, Dominic Kwiatkowski, Cordelia Langford, John Sillitoe on behalf of the Wellcome Sanger Institute COVID-19 Surveillance Team |
| EPI_ISL_2640730, | Lighthouse Lab in Milton Keynes | Wellcome Sanger Institute for the COVID-19 Genomics UK (COG-UK) | The Lighthouse Lab in Milton Keynes and Alex Alderton, Roberto Amato, Jeffrey Barrett, Sonia Goncalves, Ewan Harrison, David K. Jackson, Ian Johnston, Dominic Kwiatkowski, Cordelia Langford, John Sillitoe on behalf of the Wellcome Sanger |

| | | | |
|---|---|---|---|
| EPI_ISL_2640731 | | Consortium | Institute COVID-19 Surveillance Team |
| EPI_ISL_2640732 | Lighthouse Lab in Alderley Park | Wellcome Sanger Institute for the COVID-19 Genomics UK (COG-UK) Consortium | Jacquelyn Wynn, Mairead Hyland, The Lighthouse Lab in Alderley Park and Alex Alderton, Roberto Amato, Jeffrey Barrett, Sonia Goncalves, Ewan Harrison, David K. Jackson, Ian Johnston, Dominic Kwiatkowski, Cordelia Langford, John Sillitoe on behalf of the Wellcome Sanger Institute COVID-19 Surveillance Team |
| EPI_ISL_2640733 | Lighthouse Lab in Milton Keynes | Wellcome Sanger Institute for the COVID-19 Genomics UK (COG-UK) Consortium | The Lighthouse Lab in Milton Keynes and Alex Alderton, Roberto Amato, Jeffrey Barrett, Sonia Goncalves, Ewan Harrison, David K. Jackson, Ian Johnston, Dominic Kwiatkowski, Cordelia Langford, John Sillitoe on behalf of the Wellcome Sanger Institute COVID-19 Surveillance Team |
| EPI_ISL_2640734, EPI_ISL_2640735, EPI_ISL_2640737, EPI_ISL_2640738 | Lighthouse Lab in Alderley Park | Wellcome Sanger Institute for the COVID-19 Genomics UK (COG-UK) Consortium | Jacquelyn Wynn, Mairead Hyland, The Lighthouse Lab in Alderley Park and Alex Alderton, Roberto Amato, Jeffrey Barrett, Sonia Goncalves, Ewan Harrison, David K. Jackson, Ian Johnston, Dominic Kwiatkowski, Cordelia Langford, John Sillitoe on behalf of the Wellcome Sanger Institute COVID-19 Surveillance Team |
| EPI_ISL_2640739, EPI_ISL_2640740, EPI_ISL_2640741 | Lighthouse Lab in Glasgow | Wellcome Sanger Institute for the COVID-19 Genomics UK (COG-UK) Consortium | Harper VanSteenhouse, Yumi Kasai, David Gray, Carol Clugston, Anna Dominiczak and Alex Alderton, Roberto Amato, Jeffrey Barrett, Sonia Goncalves, Ewan Harrison, David K. Jackson, Ian Johnston, Dominic Kwiatkowski, Cordelia Langford, John Sillitoe on behalf of the Wellcome Sanger Institute COVID-19 Surveillance Team |
| EPI_ISL_2640742, EPI_ISL_2640743 | Lighthouse Lab in Alderley Park | Wellcome Sanger Institute for the COVID-19 Genomics UK (COG-UK) Consortium | Jacquelyn Wynn, Mairead Hyland, The Lighthouse Lab in Alderley Park and Alex Alderton, Roberto Amato, Jeffrey Barrett, Sonia Goncalves, Ewan Harrison, David K. Jackson, Ian Johnston, Dominic Kwiatkowski, Cordelia Langford, John Sillitoe on behalf of the Wellcome Sanger Institute COVID-19 Surveillance Team |
| EPI_ISL_2640744 | Lighthouse Lab in Glasgow | Wellcome Sanger Institute for the COVID-19 Genomics UK (COG-UK) Consortium | Harper VanSteenhouse, Yumi Kasai, David Gray, Carol Clugston, Anna Dominiczak and Alex Alderton, Roberto Amato, Jeffrey Barrett, Sonia Goncalves, Ewan Harrison, David K. Jackson, Ian Johnston, Dominic Kwiatkowski, Cordelia Langford, John Sillitoe on behalf of the Wellcome Sanger Institute COVID-19 Surveillance Team |
| EPI_ISL_2640745, EPI_ISL_2640746, EPI_ISL_2640747, EPI_ISL_2640748, EPI_ISL_2640749, EPI_ISL_2640750 | Lighthouse Lab in Alderley Park | Wellcome Sanger Institute for the COVID-19 Genomics UK (COG-UK) Consortium | Jacquelyn Wynn, Mairead Hyland, The Lighthouse Lab in Alderley Park and Alex Alderton, Roberto Amato, Jeffrey Barrett, Sonia Goncalves, Ewan Harrison, David K. Jackson, Ian Johnston, Dominic Kwiatkowski, Cordelia Langford, John Sillitoe on behalf of the Wellcome Sanger Institute COVID-19 Surveillance Team |
| EPI_ISL_2640751 | Lighthouse Lab in Glasgow | Wellcome Sanger Institute for the COVID-19 Genomics UK (COG-UK) Consortium | Harper VanSteenhouse, Yumi Kasai, David Gray, Carol Clugston, Anna Dominiczak and Alex Alderton, Roberto Amato, Jeffrey Barrett, Sonia Goncalves, Ewan Harrison, David K. Jackson, Ian Johnston, Dominic Kwiatkowski, Cordelia Langford, John Sillitoe on behalf of the Wellcome Sanger Institute COVID-19 Surveillance Team |
| EPI_ISL_2640752, EPI_ISL_2640753, EPI_ISL_2640754 | Lighthouse Lab in Alderley Park | Wellcome Sanger Institute for the COVID-19 Genomics UK (COG-UK) Consortium | Jacquelyn Wynn, Mairead Hyland, The Lighthouse Lab in Alderley Park and Alex Alderton, Roberto Amato, Jeffrey Barrett, Sonia Goncalves, Ewan Harrison, David K. Jackson, Ian Johnston, Dominic Kwiatkowski, Cordelia Langford, John Sillitoe on behalf of the Wellcome Sanger Institute COVID-19 Surveillance Team |
| EPI_ISL_2640755 | Lighthouse Lab in Milton Keynes | Wellcome Sanger Institute for the COVID-19 Genomics UK (COG-UK) Consortium | The Lighthouse Lab in Milton Keynes and Alex Alderton, Roberto Amato, Jeffrey Barrett, Sonia Goncalves, Ewan Harrison, David K. Jackson, Ian Johnston, Dominic Kwiatkowski, Cordelia Langford, John Sillitoe on behalf of the Wellcome Sanger Institute COVID-19 Surveillance Team |
| EPI_ISL_2640756 | Lighthouse Lab in Alderley Park | Wellcome Sanger Institute for the COVID-19 Genomics UK (COG-UK) Consortium | Jacquelyn Wynn, Mairead Hyland, The Lighthouse Lab in Alderley Park and Alex Alderton, Roberto Amato, Jeffrey Barrett, Sonia Goncalves, Ewan Harrison, David K. Jackson, Ian Johnston, Dominic Kwiatkowski, Cordelia Langford, John Sillitoe on behalf of the Wellcome Sanger Institute COVID-19 Surveillance Team |
| EPI_ISL_2640757 | Lighthouse Lab in Milton Keynes | Wellcome Sanger Institute for the COVID-19 Genomics UK (COG-UK) Consortium | The Lighthouse Lab in Milton Keynes and Alex Alderton, Roberto Amato, Jeffrey Barrett, Sonia Goncalves, Ewan Harrison, David K. Jackson, Ian Johnston, Dominic Kwiatkowski, Cordelia Langford, John Sillitoe on behalf of the Wellcome Sanger Institute COVID-19 Surveillance Team |
| EPI_ISL_2640758, EPI_ISL_2640759 | Lighthouse Lab in Alderley Park | Wellcome Sanger Institute for the COVID-19 Genomics UK (COG-UK) Consortium | Jacquelyn Wynn, Mairead Hyland, The Lighthouse Lab in Alderley Park and Alex Alderton, Roberto Amato, Jeffrey Barrett, Sonia Goncalves, Ewan Harrison, David K. Jackson, Ian Johnston, Dominic Kwiatkowski, Cordelia Langford, John Sillitoe on behalf of the Wellcome Sanger Institute COVID-19 Surveillance Team |
| EPI_ISL_2640760 | Lighthouse Lab in Glasgow | Wellcome Sanger Institute for the COVID-19 Genomics UK (COG-UK) Consortium | Harper VanSteenhouse, Yumi Kasai, David Gray, Carol Clugston, Anna Dominiczak and Alex Alderton, Roberto Amato, Jeffrey Barrett, Sonia Goncalves, Ewan Harrison, David K. Jackson, Ian Johnston, Dominic Kwiatkowski, Cordelia Langford, John Sillitoe on behalf of the Wellcome Sanger Institute COVID-19 Surveillance Team |
| EPI_ISL_2640761 | Lighthouse Lab in Alderley Park | Wellcome Sanger Institute for the COVID-19 Genomics UK (COG-UK) Consortium | Jacquelyn Wynn, Mairead Hyland, The Lighthouse Lab in Alderley Park and Alex Alderton, Roberto Amato, Jeffrey Barrett, Sonia Goncalves, Ewan Harrison, David K. Jackson, Ian Johnston, Dominic Kwiatkowski, Cordelia Langford, John Sillitoe on behalf of the Wellcome Sanger Institute COVID-19 Surveillance Team |
| EPI_ISL_2640762, EPI_ISL_2640763 | Lighthouse Lab in Glasgow | Wellcome Sanger Institute for the COVID-19 Genomics UK (COG-UK) Consortium | Harper VanSteenhouse, Yumi Kasai, David Gray, Carol Clugston, Anna Dominiczak and Alex Alderton, Roberto Amato, Jeffrey Barrett, Sonia Goncalves, Ewan Harrison, David K. Jackson, Ian Johnston, Dominic Kwiatkowski, Cordelia Langford, John Sillitoe on behalf of the Wellcome Sanger Institute COVID-19 Surveillance Team |
| EPI_ISL_2640764 | Lighthouse Lab in Alderley Park | Wellcome Sanger Institute for the COVID-19 Genomics UK (COG-UK) Consortium | Jacquelyn Wynn, Mairead Hyland, The Lighthouse Lab in Alderley Park and Alex Alderton, Roberto Amato, Jeffrey Barrett, Sonia Goncalves, Ewan Harrison, David K. Jackson, Ian Johnston, Dominic Kwiatkowski, Cordelia Langford, John Sillitoe on behalf of the Wellcome Sanger Institute COVID-19 Surveillance Team |
| EPI_ISL_2640765 | Lighthouse Lab in Glasgow | Wellcome Sanger Institute for the COVID-19 Genomics UK (COG-UK) Consortium | Harper VanSteenhouse, Yumi Kasai, David Gray, Carol Clugston, Anna Dominiczak and Alex Alderton, Roberto Amato, Jeffrey Barrett, Sonia Goncalves, Ewan Harrison, David K. Jackson, Ian Johnston, Dominic Kwiatkowski, Cordelia Langford, John Sillitoe on behalf of the Wellcome Sanger Institute COVID-19 Surveillance Team |
| EPI_ISL_2640766, EPI_ISL_2640767, EPI_ISL_2640768 | Lighthouse Lab in Alderley Park | Wellcome Sanger Institute for the COVID-19 Genomics UK (COG-UK) Consortium | Jacquelyn Wynn, Mairead Hyland, The Lighthouse Lab in Alderley Park and Alex Alderton, Roberto Amato, Jeffrey Barrett, Sonia Goncalves, Ewan Harrison, David K. Jackson, Ian Johnston, Dominic Kwiatkowski, Cordelia Langford, John Sillitoe on behalf of the Wellcome Sanger Institute COVID-19 Surveillance Team |
| EPI_ISL_2640769 | Lighthouse Lab in Milton Keynes | Wellcome Sanger Institute for the COVID-19 Genomics UK (COG-UK) Consortium | The Lighthouse Lab in Milton Keynes and Alex Alderton, Roberto Amato, Jeffrey Barrett, Sonia Goncalves, Ewan Harrison, David K. Jackson, Ian Johnston, Dominic Kwiatkowski, Cordelia Langford, John Sillitoe on behalf of the Wellcome Sanger Institute COVID-19 Surveillance Team |
| EPI_ISL_2640770 | Lighthouse Lab in Glasgow | Wellcome Sanger Institute for the COVID-19 Genomics UK (COG-UK) Consortium | Harper VanSteenhouse, Yumi Kasai, David Gray, Carol Clugston, Anna Dominiczak and Alex Alderton, Roberto Amato, Jeffrey Barrett, Sonia Goncalves, Ewan Harrison, David K. Jackson, Ian Johnston, Dominic Kwiatkowski, Cordelia Langford, John Sillitoe on behalf of the Wellcome Sanger Institute COVID-19 Surveillance Team |
| EPI_ISL_2640771 | Lighthouse Lab in Milton Keynes | Wellcome Sanger Institute for the COVID-19 Genomics UK (COG-UK) Consortium | The Lighthouse Lab in Milton Keynes and Alex Alderton, Roberto Amato, Jeffrey Barrett, Sonia Goncalves, Ewan Harrison, David K. Jackson, Ian Johnston, Dominic Kwiatkowski, Cordelia Langford, John Sillitoe on behalf of the Wellcome Sanger Institute COVID-19 Surveillance Team |
| EPI_ISL_2640772, EPI_ISL_2640774, EPI_ISL_2640775, EPI_ISL_2640776 | Lighthouse Lab in Alderley Park | Wellcome Sanger Institute for the COVID-19 Genomics UK (COG-UK) Consortium | Jacquelyn Wynn, Mairead Hyland, The Lighthouse Lab in Alderley Park and Alex Alderton, Roberto Amato, Jeffrey Barrett, Sonia Goncalves, Ewan Harrison, David K. Jackson, Ian Johnston, Dominic Kwiatkowski, Cordelia Langford, John Sillitoe on behalf of the Wellcome Sanger Institute COVID-19 Surveillance Team |
| EPI_ISL_2640777 | Lighthouse Lab in Glasgow | Wellcome Sanger Institute for the COVID-19 Genomics UK (COG-UK) Consortium | Harper VanSteenhouse, Yumi Kasai, David Gray, Carol Clugston, Anna Dominiczak and Alex Alderton, Roberto Amato, Jeffrey Barrett, Sonia Goncalves, Ewan Harrison, David K. Jackson, Ian Johnston, Dominic Kwiatkowski, Cordelia Langford, John Sillitoe on behalf of the Wellcome Sanger Institute COVID-19 Surveillance Team |
| EPI_ISL_2640778 | Lighthouse Lab in Alderley Park | Wellcome Sanger Institute for the COVID-19 Genomics UK (COG-UK) Consortium | Jacquelyn Wynn, Mairead Hyland, The Lighthouse Lab in Alderley Park and Alex Alderton, Roberto Amato, Jeffrey Barrett, Sonia Goncalves, Ewan Harrison, David K. Jackson, Ian Johnston, Dominic Kwiatkowski, Cordelia Langford, John Sillitoe on behalf of the Wellcome Sanger Institute COVID-19 Surveillance Team |
| EPI_ISL_2640779, EPI_ISL_2640780 | Lighthouse Lab in Milton Keynes | Wellcome Sanger Institute for the COVID-19 Genomics UK (COG-UK) Consortium | The Lighthouse Lab in Milton Keynes and Alex Alderton, Roberto Amato, Jeffrey Barrett, Sonia Goncalves, Ewan Harrison, David K. Jackson, Ian Johnston, Dominic Kwiatkowski, Cordelia Langford, John Sillitoe on behalf of the Wellcome Sanger Institute COVID-19 Surveillance Team |
| EPI_ISL_2640781 | Lighthouse Lab in Alderley Park | Wellcome Sanger Institute for the COVID-19 Genomics UK (COG-UK) Consortium | Jacquelyn Wynn, Mairead Hyland, The Lighthouse Lab in Alderley Park and Alex Alderton, Roberto Amato, Jeffrey Barrett, Sonia Goncalves, Ewan Harrison, David K. Jackson, Ian Johnston, Dominic Kwiatkowski, Cordelia Langford, John Sillitoe on behalf of the Wellcome Sanger Institute COVID-19 Surveillance Team |
| EPI_ISL_2640782 | Lighthouse Lab in Milton Keynes | Wellcome Sanger Institute for the COVID-19 Genomics UK (COG-UK) Consortium | The Lighthouse Lab in Milton Keynes and Alex Alderton, Roberto Amato, Jeffrey Barrett, Sonia Goncalves, Ewan Harrison, David K. Jackson, Ian Johnston, Dominic Kwiatkowski, Cordelia Langford, John Sillitoe on behalf of the Wellcome Sanger Institute COVID-19 Surveillance Team |
| EPI_ISL_2640783 | Lighthouse Lab in Glasgow | Wellcome Sanger Institute for the COVID-19 Genomics UK (COG-UK) Consortium | Harper VanSteenhouse, Yumi Kasai, David Gray, Carol Clugston, Anna Dominiczak and Alex Alderton, Roberto Amato, Jeffrey Barrett, Sonia Goncalves, Ewan Harrison, David K. Jackson, Ian Johnston, Dominic Kwiatkowski, Cordelia Langford, John Sillitoe on behalf of the Wellcome Sanger Institute COVID-19 Surveillance Team |
| EPI_ISL_2640784 | Lighthouse Lab in Alderley Park | Wellcome Sanger Institute for the COVID-19 Genomics UK (COG-UK) Consortium | Jacquelyn Wynn, Mairead Hyland, The Lighthouse Lab in Alderley Park and Alex Alderton, Roberto Amato, Jeffrey Barrett, Sonia Goncalves, Ewan Harrison, David K. Jackson, Ian Johnston, Dominic Kwiatkowski, Cordelia Langford, John Sillitoe on behalf of the Wellcome Sanger Institute COVID-19 Surveillance Team |
| EPI_ISL_2640785, EPI_ISL_2640786 | Lighthouse Lab in Milton Keynes | Wellcome Sanger Institute for the COVID-19 Genomics UK (COG-UK) Consortium | The Lighthouse Lab in Milton Keynes and Alex Alderton, Roberto Amato, Jeffrey Barrett, Sonia Goncalves, Ewan Harrison, David K. Jackson, Ian Johnston, Dominic Kwiatkowski, Cordelia Langford, John Sillitoe on behalf of the Wellcome Sanger Institute COVID-19 Surveillance Team |
| EPI_ISL_2640787, EPI_ISL_2640788, EPI_ISL_2640789 | Lighthouse Lab in Alderley Park | Wellcome Sanger Institute for the COVID-19 Genomics UK (COG-UK) Consortium | Jacquelyn Wynn, Mairead Hyland, The Lighthouse Lab in Alderley Park and Alex Alderton, Roberto Amato, Jeffrey Barrett, Sonia Goncalves, Ewan Harrison, David K. Jackson, Ian Johnston, Dominic Kwiatkowski, Cordelia Langford, John Sillitoe on behalf of the Wellcome Sanger Institute COVID-19 Surveillance Team |
| EPI_ISL_2640790, EPI_ISL_2640791 | Lighthouse Lab in Milton Keynes | Wellcome Sanger Institute for the COVID-19 Genomics UK (COG-UK) Consortium | The Lighthouse Lab in Milton Keynes and Alex Alderton, Roberto Amato, Jeffrey Barrett, Sonia Goncalves, Ewan Harrison, David K. Jackson, Ian Johnston, Dominic Kwiatkowski, Cordelia Langford, John Sillitoe on behalf of the Wellcome Sanger Institute COVID-19 Surveillance Team |
| EPI_ISL_2640792 | Lighthouse Lab in Glasgow | Wellcome Sanger Institute for the COVID-19 Genomics UK (COG-UK) Consortium | Harper VanSteenhouse, Yumi Kasai, David Gray, Carol Clugston, Anna Dominiczak and Alex Alderton, Roberto Amato, Jeffrey Barrett, Sonia Goncalves, Ewan Harrison, David K. Jackson, Ian Johnston, Dominic Kwiatkowski, Cordelia Langford, John Sillitoe on behalf of the Wellcome Sanger Institute COVID-19 Surveillance Team |
| EPI_ISL_2640793 | Lighthouse Lab in Milton Keynes | Wellcome Sanger Institute for the COVID-19 Genomics UK (COG-UK) Consortium | The Lighthouse Lab in Milton Keynes and Alex Alderton, Roberto Amato, Jeffrey Barrett, Sonia Goncalves, Ewan Harrison, David K. Jackson, Ian Johnston, Dominic Kwiatkowski, Cordelia Langford, John Sillitoe on behalf of the Wellcome Sanger Institute COVID-19 Surveillance Team |
| EPI_ISL_2640794, EPI_ISL_2640795 | Lighthouse Lab in Alderley Park | Wellcome Sanger Institute for the COVID-19 Genomics UK (COG-UK) Consortium | Jacquelyn Wynn, Mairead Hyland, The Lighthouse Lab in Alderley Park and Alex Alderton, Roberto Amato, Jeffrey Barrett, Sonia Goncalves, Ewan Harrison, David K. Jackson, Ian Johnston, Dominic Kwiatkowski, Cordelia Langford, John Sillitoe on behalf of the Wellcome Sanger Institute COVID-19 Surveillance Team |
| EPI_ISL_2640796 | Lighthouse Lab in Milton Keynes | Wellcome Sanger Institute for the COVID-19 Genomics UK (COG-UK) Consortium | The Lighthouse Lab in Milton Keynes and Alex Alderton, Roberto Amato, Jeffrey Barrett, Sonia Goncalves, Ewan Harrison, David K. Jackson, Ian Johnston, Dominic Kwiatkowski, Cordelia Langford, John Sillitoe on behalf of the Wellcome Sanger Institute COVID-19 Surveillance Team |
| EPI_ISL_2640797, EPI_ISL_2640798 | Lighthouse Lab in Glasgow | Wellcome Sanger Institute for the COVID-19 Genomics UK (COG-UK) Consortium | Harper VanSteenhouse, Yumi Kasai, David Gray, Carol Clugston, Anna Dominiczak and Alex Alderton, Roberto Amato, Jeffrey Barrett, Sonia Goncalves, Ewan Harrison, David K. Jackson, Ian Johnston, Dominic Kwiatkowski, Cordelia Langford, John Sillitoe on behalf of the Wellcome Sanger Institute COVID-19 Surveillance Team |
| EPI_ISL_2640799 | Lighthouse Lab in Alderley Park | Wellcome Sanger Institute for the COVID-19 Genomics UK (COG-UK) Consortium | Jacquelyn Wynn, Mairead Hyland, The Lighthouse Lab in Alderley Park and Alex Alderton, Roberto Amato, Jeffrey Barrett, Sonia Goncalves, Ewan Harrison, David K. Jackson, Ian Johnston, Dominic Kwiatkowski, Cordelia Langford, John Sillitoe on behalf of the Wellcome Sanger Institute COVID-19 Surveillance Team |

| | | | |
|---|---|---|---|
| EPI_ISL_2640800 | Lighthouse Lab in Milton Keynes | Wellcome Sanger Institute for the COVID-19 Genomics UK (COG-UK) Consortium | The Lighthouse Lab in Milton Keynes and Alex Alderton, Roberto Amato, Jeffrey Barrett, Sonia Goncalves, Ewan Harrison, David K. Jackson, Ian Johnston, Dominic Kwiatkowski, Cordelia Langford, John Sillitoe on behalf of the Wellcome Sanger Institute COVID-19 Surveillance Team |
| EPI_ISL_2640801 | Lighthouse Lab in Alderley Park | Wellcome Sanger Institute for the COVID-19 Genomics UK (COG-UK) Consortium | Jacquelyn Wynn, Mairead Hyland, The Lighthouse Lab in Alderley Park and Alex Alderton, Roberto Amato, Jeffrey Barrett, Sonia Goncalves, Ewan Harrison, David K. Jackson, Ian Johnston, Dominic Kwiatkowski, Cordelia Langford, John Sillitoe on behalf of the Wellcome Sanger Institute COVID-19 Surveillance Team |
| EPI_ISL_2640802, EPI_ISL_2640803 | Lighthouse Lab in Glasgow | Wellcome Sanger Institute for the COVID-19 Genomics UK (COG-UK) Consortium | Harper VanSteenhouse, Yumi Kasai, David Gray, Carol Clugston, Anna Dominiczak and Alex Alderton, Roberto Amato, Jeffrey Barrett, Sonia Goncalves, Ewan Harrison, David K. Jackson, Ian Johnston, Dominic Kwiatkowski, Cordelia Langford, John Sillitoe on behalf of the Wellcome Sanger Institute COVID-19 Surveillance Team |
| EPI_ISL_2640804 | Lighthouse Lab in Alderley Park | Wellcome Sanger Institute for the COVID-19 Genomics UK (COG-UK) Consortium | Jacquelyn Wynn, Mairead Hyland, The Lighthouse Lab in Alderley Park and Alex Alderton, Roberto Amato, Jeffrey Barrett, Sonia Goncalves, Ewan Harrison, David K. Jackson, Ian Johnston, Dominic Kwiatkowski, Cordelia Langford, John Sillitoe on behalf of the Wellcome Sanger Institute COVID-19 Surveillance Team |
| EPI_ISL_2640806 | Lighthouse Lab in Milton Keynes | Wellcome Sanger Institute for the COVID-19 Genomics UK (COG-UK) Consortium | The Lighthouse Lab in Milton Keynes and Alex Alderton, Roberto Amato, Jeffrey Barrett, Sonia Goncalves, Ewan Harrison, David K. Jackson, Ian Johnston, Dominic Kwiatkowski, Cordelia Langford, John Sillitoe on behalf of the Wellcome Sanger Institute COVID-19 Surveillance Team |
| EPI_ISL_2640807 | Lighthouse Lab in Glasgow | Wellcome Sanger Institute for the COVID-19 Genomics UK (COG-UK) Consortium | Harper VanSteenhouse, Yumi Kasai, David Gray, Carol Clugston, Anna Dominiczak and Alex Alderton, Roberto Amato, Jeffrey Barrett, Sonia Goncalves, Ewan Harrison, David K. Jackson, Ian Johnston, Dominic Kwiatkowski, Cordelia Langford, John Sillitoe on behalf of the Wellcome Sanger Institute COVID-19 Surveillance Team |
| EPI_ISL_2640808 | Lighthouse Lab in Alderley Park | Wellcome Sanger Institute for the COVID-19 Genomics UK (COG-UK) Consortium | Jacquelyn Wynn, Mairead Hyland, The Lighthouse Lab in Alderley Park and Alex Alderton, Roberto Amato, Jeffrey Barrett, Sonia Goncalves, Ewan Harrison, David K. Jackson, Ian Johnston, Dominic Kwiatkowski, Cordelia Langford, John Sillitoe on behalf of the Wellcome Sanger Institute COVID-19 Surveillance Team |
| EPI_ISL_2640809, EPI_ISL_2640810, EPI_ISL_2640811, EPI_ISL_2640812 | Lighthouse Lab in Milton Keynes | Wellcome Sanger Institute for the COVID-19 Genomics UK (COG-UK) Consortium | The Lighthouse Lab in Milton Keynes and Alex Alderton, Roberto Amato, Jeffrey Barrett, Sonia Goncalves, Ewan Harrison, David K. Jackson, Ian Johnston, Dominic Kwiatkowski, Cordelia Langford, John Sillitoe on behalf of the Wellcome Sanger Institute COVID-19 Surveillance Team |
| EPI_ISL_2640813 | Lighthouse Lab in Glasgow | Wellcome Sanger Institute for the COVID-19 Genomics UK (COG-UK) Consortium | Harper VanSteenhouse, Yumi Kasai, David Gray, Carol Clugston, Anna Dominiczak and Alex Alderton, Roberto Amato, Jeffrey Barrett, Sonia Goncalves, Ewan Harrison, David K. Jackson, Ian Johnston, Dominic Kwiatkowski, Cordelia Langford, John Sillitoe on behalf of the Wellcome Sanger Institute COVID-19 Surveillance Team |
| EPI_ISL_2640815, EPI_ISL_2640816, EPI_ISL_2640817, EPI_ISL_2640818, EPI_ISL_2640819 | Lighthouse Lab in Alderley Park | Wellcome Sanger Institute for the COVID-19 Genomics UK (COG-UK) Consortium | Jacquelyn Wynn, Mairead Hyland, The Lighthouse Lab in Alderley Park and Alex Alderton, Roberto Amato, Jeffrey Barrett, Sonia Goncalves, Ewan Harrison, David K. Jackson, Ian Johnston, Dominic Kwiatkowski, Cordelia Langford, John Sillitoe on behalf of the Wellcome Sanger Institute COVID-19 Surveillance Team |
| EPI_ISL_2640820 | Lighthouse Lab in Milton Keynes | Wellcome Sanger Institute for the COVID-19 Genomics UK (COG-UK) Consortium | The Lighthouse Lab in Milton Keynes and Alex Alderton, Roberto Amato, Jeffrey Barrett, Sonia Goncalves, Ewan Harrison, David K. Jackson, Ian Johnston, Dominic Kwiatkowski, Cordelia Langford, John Sillitoe on behalf of the Wellcome Sanger Institute COVID-19 Surveillance Team |
| EPI_ISL_2640821, EPI_ISL_2640822 | Lighthouse Lab in Alderley Park | Wellcome Sanger Institute for the COVID-19 Genomics UK (COG-UK) Consortium | Jacquelyn Wynn, Mairead Hyland, The Lighthouse Lab in Alderley Park and Alex Alderton, Roberto Amato, Jeffrey Barrett, Sonia Goncalves, Ewan Harrison, David K. Jackson, Ian Johnston, Dominic Kwiatkowski, Cordelia Langford, John Sillitoe on behalf of the Wellcome Sanger Institute COVID-19 Surveillance Team |
| EPI_ISL_2640823 | Lighthouse Lab in Glasgow | Wellcome Sanger Institute for the COVID-19 Genomics UK (COG-UK) Consortium | Harper VanSteenhouse, Yumi Kasai, David Gray, Carol Clugston, Anna Dominiczak and Alex Alderton, Roberto Amato, Jeffrey Barrett, Sonia Goncalves, Ewan Harrison, David K. Jackson, Ian Johnston, Dominic Kwiatkowski, Cordelia Langford, John Sillitoe on behalf of the Wellcome Sanger Institute COVID-19 Surveillance Team |
| EPI_ISL_2640824 | Lighthouse Lab in Milton Keynes | Wellcome Sanger Institute for the COVID-19 Genomics UK (COG-UK) Consortium | The Lighthouse Lab in Milton Keynes and Alex Alderton, Roberto Amato, Jeffrey Barrett, Sonia Goncalves, Ewan Harrison, David K. Jackson, Ian Johnston, Dominic Kwiatkowski, Cordelia Langford, John Sillitoe on behalf of the Wellcome Sanger Institute COVID-19 Surveillance Team |
| EPI_ISL_2640825 | Lighthouse Lab in Glasgow | Wellcome Sanger Institute for the COVID-19 Genomics UK (COG-UK) Consortium | Harper VanSteenhouse, Yumi Kasai, David Gray, Carol Clugston, Anna Dominiczak and Alex Alderton, Roberto Amato, Jeffrey Barrett, Sonia Goncalves, Ewan Harrison, David K. Jackson, Ian Johnston, Dominic Kwiatkowski, Cordelia Langford, John Sillitoe on behalf of the Wellcome Sanger Institute COVID-19 Surveillance Team |
| EPI_ISL_2640826, EPI_ISL_2640827 | Lighthouse Lab in Alderley Park | Wellcome Sanger Institute for the COVID-19 Genomics UK (COG-UK) Consortium | Jacquelyn Wynn, Mairead Hyland, The Lighthouse Lab in Alderley Park and Alex Alderton, Roberto Amato, Jeffrey Barrett, Sonia Goncalves, Ewan Harrison, David K. Jackson, Ian Johnston, Dominic Kwiatkowski, Cordelia Langford, John Sillitoe on behalf of the Wellcome Sanger Institute COVID-19 Surveillance Team |
| EPI_ISL_2640828 | Lighthouse Lab in Milton Keynes | Wellcome Sanger Institute for the COVID-19 Genomics UK (COG-UK) Consortium | The Lighthouse Lab in Milton Keynes and Alex Alderton, Roberto Amato, Jeffrey Barrett, Sonia Goncalves, Ewan Harrison, David K. Jackson, Ian Johnston, Dominic Kwiatkowski, Cordelia Langford, John Sillitoe on behalf of the Wellcome Sanger Institute COVID-19 Surveillance Team |
| EPI_ISL_2640829, EPI_ISL_2640830 | Lighthouse Lab in Glasgow | Wellcome Sanger Institute for the COVID-19 Genomics UK (COG-UK) Consortium | Harper VanSteenhouse, Yumi Kasai, David Gray, Carol Clugston, Anna Dominiczak and Alex Alderton, Roberto Amato, Jeffrey Barrett, Sonia Goncalves, Ewan Harrison, David K. Jackson, Ian Johnston, Dominic Kwiatkowski, Cordelia Langford, John Sillitoe on behalf of the Wellcome Sanger Institute COVID-19 Surveillance Team |
| EPI_ISL_2640831, EPI_ISL_2640832, EPI_ISL_2640833 | Lighthouse Lab in Alderley Park | Wellcome Sanger Institute for the COVID-19 Genomics UK (COG-UK) Consortium | Jacquelyn Wynn, Mairead Hyland, The Lighthouse Lab in Alderley Park and Alex Alderton, Roberto Amato, Jeffrey Barrett, Sonia Goncalves, Ewan Harrison, David K. Jackson, Ian Johnston, Dominic Kwiatkowski, Cordelia Langford, John Sillitoe on behalf of the Wellcome Sanger Institute COVID-19 Surveillance Team |
| EPI_ISL_2640834 | Lighthouse Lab in Milton Keynes | Wellcome Sanger Institute for the COVID-19 Genomics UK (COG-UK) Consortium | The Lighthouse Lab in Milton Keynes and Alex Alderton, Roberto Amato, Jeffrey Barrett, Sonia Goncalves, Ewan Harrison, David K. Jackson, Ian Johnston, Dominic Kwiatkowski, Cordelia Langford, John Sillitoe on behalf of the Wellcome Sanger Institute COVID-19 Surveillance Team |
| EPI_ISL_2640835 | Lighthouse Lab in Alderley Park | Wellcome Sanger Institute for the COVID-19 Genomics UK (COG-UK) Consortium | Jacquelyn Wynn, Mairead Hyland, The Lighthouse Lab in Alderley Park and Alex Alderton, Roberto Amato, Jeffrey Barrett, Sonia Goncalves, Ewan Harrison, David K. Jackson, Ian Johnston, Dominic Kwiatkowski, Cordelia Langford, John Sillitoe on behalf of the Wellcome Sanger Institute COVID-19 Surveillance Team |
| EPI_ISL_2640836 | Lighthouse Lab in Glasgow | Wellcome Sanger Institute for the COVID-19 Genomics UK (COG-UK) Consortium | Harper VanSteenhouse, Yumi Kasai, David Gray, Carol Clugston, Anna Dominiczak and Alex Alderton, Roberto Amato, Jeffrey Barrett, Sonia Goncalves, Ewan Harrison, David K. Jackson, Ian Johnston, Dominic Kwiatkowski, Cordelia Langford, John Sillitoe on behalf of the Wellcome Sanger Institute COVID-19 Surveillance Team |
| EPI_ISL_2640837, EPI_ISL_2640838 | Lighthouse Lab in Alderley Park | Wellcome Sanger Institute for the COVID-19 Genomics UK (COG-UK) Consortium | Jacquelyn Wynn, Mairead Hyland, The Lighthouse Lab in Alderley Park and Alex Alderton, Roberto Amato, Jeffrey Barrett, Sonia Goncalves, Ewan Harrison, David K. Jackson, Ian Johnston, Dominic Kwiatkowski, Cordelia Langford, John Sillitoe on behalf of the Wellcome Sanger Institute COVID-19 Surveillance Team |
| EPI_ISL_2640839 | Lighthouse Lab in Milton Keynes | Wellcome Sanger Institute for the COVID-19 Genomics UK (COG-UK) Consortium | The Lighthouse Lab in Milton Keynes and Alex Alderton, Roberto Amato, Jeffrey Barrett, Sonia Goncalves, Ewan Harrison, David K. Jackson, Ian Johnston, Dominic Kwiatkowski, Cordelia Langford, John Sillitoe on behalf of the Wellcome Sanger Institute COVID-19 Surveillance Team |
| EPI_ISL_2640840 | Lighthouse Lab in Glasgow | Wellcome Sanger Institute for the COVID-19 Genomics UK (COG-UK) Consortium | Harper VanSteenhouse, Yumi Kasai, David Gray, Carol Clugston, Anna Dominiczak and Alex Alderton, Roberto Amato, Jeffrey Barrett, Sonia Goncalves, Ewan Harrison, David K. Jackson, Ian Johnston, Dominic Kwiatkowski, Cordelia Langford, John Sillitoe on behalf of the Wellcome Sanger Institute COVID-19 Surveillance Team |
| EPI_ISL_2640841 | Lighthouse Lab in Milton Keynes | Wellcome Sanger Institute for the COVID-19 Genomics UK (COG-UK) Consortium | The Lighthouse Lab in Milton Keynes and Alex Alderton, Roberto Amato, Jeffrey Barrett, Sonia Goncalves, Ewan Harrison, David K. Jackson, Ian Johnston, Dominic Kwiatkowski, Cordelia Langford, John Sillitoe on behalf of the Wellcome Sanger Institute COVID-19 Surveillance Team |
| EPI_ISL_2640842, EPI_ISL_2640843 | Lighthouse Lab in Alderley Park | Wellcome Sanger Institute for the COVID-19 Genomics UK (COG-UK) Consortium | Jacquelyn Wynn, Mairead Hyland, The Lighthouse Lab in Alderley Park and Alex Alderton, Roberto Amato, Jeffrey Barrett, Sonia Goncalves, Ewan Harrison, David K. Jackson, Ian Johnston, Dominic Kwiatkowski, Cordelia Langford, John Sillitoe on behalf of the Wellcome Sanger Institute COVID-19 Surveillance Team |
| EPI_ISL_2640844 | Lighthouse Lab in Glasgow | Wellcome Sanger Institute for the COVID-19 Genomics UK (COG-UK) Consortium | Harper VanSteenhouse, Yumi Kasai, David Gray, Carol Clugston, Anna Dominiczak and Alex Alderton, Roberto Amato, Jeffrey Barrett, Sonia Goncalves, Ewan Harrison, David K. Jackson, Ian Johnston, Dominic Kwiatkowski, Cordelia Langford, John Sillitoe on behalf of the Wellcome Sanger Institute COVID-19 Surveillance Team |
| EPI_ISL_2640845 | Lighthouse Lab in Milton Keynes | Wellcome Sanger Institute for the COVID-19 Genomics UK (COG-UK) Consortium | The Lighthouse Lab in Milton Keynes and Alex Alderton, Roberto Amato, Jeffrey Barrett, Sonia Goncalves, Ewan Harrison, David K. Jackson, Ian Johnston, Dominic Kwiatkowski, Cordelia Langford, John Sillitoe on behalf of the Wellcome Sanger Institute COVID-19 Surveillance Team |
| EPI_ISL_2640846, EPI_ISL_2640847 | Lighthouse Lab in Glasgow | Wellcome Sanger Institute for the COVID-19 Genomics UK (COG-UK) Consortium | Harper VanSteenhouse, Yumi Kasai, David Gray, Carol Clugston, Anna Dominiczak and Alex Alderton, Roberto Amato, Jeffrey Barrett, Sonia Goncalves, Ewan Harrison, David K. Jackson, Ian Johnston, Dominic Kwiatkowski, Cordelia Langford, John Sillitoe on behalf of the Wellcome Sanger Institute COVID-19 Surveillance Team |
| EPI_ISL_2640848 | Lighthouse Lab in Alderley Park | Wellcome Sanger Institute for the COVID-19 Genomics UK (COG-UK) Consortium | Jacquelyn Wynn, Mairead Hyland, The Lighthouse Lab in Alderley Park and Alex Alderton, Roberto Amato, Jeffrey Barrett, Sonia Goncalves, Ewan Harrison, David K. Jackson, Ian Johnston, Dominic Kwiatkowski, Cordelia Langford, John Sillitoe on behalf of the Wellcome Sanger Institute COVID-19 Surveillance Team |
| EPI_ISL_2640849, EPI_ISL_2640850 | Lighthouse Lab in Glasgow | Wellcome Sanger Institute for the COVID-19 Genomics UK (COG-UK) Consortium | Harper VanSteenhouse, Yumi Kasai, David Gray, Carol Clugston, Anna Dominiczak and Alex Alderton, Roberto Amato, Jeffrey Barrett, Sonia Goncalves, Ewan Harrison, David K. Jackson, Ian Johnston, Dominic Kwiatkowski, Cordelia Langford, John Sillitoe on behalf of the Wellcome Sanger Institute COVID-19 Surveillance Team |
| EPI_ISL_2640851 | Lighthouse Lab in Milton Keynes | Wellcome Sanger Institute for the COVID-19 Genomics UK (COG-UK) Consortium | The Lighthouse Lab in Milton Keynes and Alex Alderton, Roberto Amato, Jeffrey Barrett, Sonia Goncalves, Ewan Harrison, David K. Jackson, Ian Johnston, Dominic Kwiatkowski, Cordelia Langford, John Sillitoe on behalf of the Wellcome Sanger Institute COVID-19 Surveillance Team |
| EPI_ISL_2640852 | Lighthouse Lab in Glasgow | Wellcome Sanger Institute for the COVID-19 Genomics UK (COG-UK) Consortium | Harper VanSteenhouse, Yumi Kasai, David Gray, Carol Clugston, Anna Dominiczak and Alex Alderton, Roberto Amato, Jeffrey Barrett, Sonia Goncalves, Ewan Harrison, David K. Jackson, Ian Johnston, Dominic Kwiatkowski, Cordelia Langford, John Sillitoe on behalf of the Wellcome Sanger Institute COVID-19 Surveillance Team |
| EPI_ISL_2640853, EPI_ISL_2640854 | Lighthouse Lab in Alderley Park | Wellcome Sanger Institute for the COVID-19 Genomics UK (COG-UK) Consortium | Jacquelyn Wynn, Mairead Hyland, The Lighthouse Lab in Alderley Park and Alex Alderton, Roberto Amato, Jeffrey Barrett, Sonia Goncalves, Ewan Harrison, David K. Jackson, Ian Johnston, Dominic Kwiatkowski, Cordelia Langford, John Sillitoe on behalf of the Wellcome Sanger Institute COVID-19 Surveillance Team |
| EPI_ISL_2640855 | Lighthouse Lab in Milton Keynes | Wellcome Sanger Institute for the COVID-19 Genomics UK (COG-UK) Consortium | The Lighthouse Lab in Milton Keynes and Alex Alderton, Roberto Amato, Jeffrey Barrett, Sonia Goncalves, Ewan Harrison, David K. Jackson, Ian Johnston, Dominic Kwiatkowski, Cordelia Langford, John Sillitoe on behalf of the Wellcome Sanger Institute COVID-19 Surveillance Team |
| EPI_ISL_2640856, EPI_ISL_2640857 | Lighthouse Lab in Glasgow | Wellcome Sanger Institute for the COVID-19 Genomics UK (COG-UK) Consortium | Harper VanSteenhouse, Yumi Kasai, David Gray, Carol Clugston, Anna Dominiczak and Alex Alderton, Roberto Amato, Jeffrey Barrett, Sonia Goncalves, Ewan Harrison, David K. Jackson, Ian Johnston, Dominic Kwiatkowski, Cordelia Langford, John Sillitoe on behalf of the Wellcome Sanger Institute COVID-19 Surveillance Team |
| EPI_ISL_2640858 | Lighthouse Lab in Milton Keynes | Wellcome Sanger Institute for the COVID-19 Genomics UK (COG-UK) Consortium | The Lighthouse Lab in Milton Keynes and Alex Alderton, Roberto Amato, Jeffrey Barrett, Sonia Goncalves, Ewan Harrison, David K. Jackson, Ian Johnston, Dominic Kwiatkowski, Cordelia Langford, John Sillitoe on behalf of the Wellcome Sanger Institute COVID-19 Surveillance Team |
| EPI_ISL_2640859, EPI_ISL_2640860, EPI_ISL_2640861, EPI_ISL_2640862, EPI_ISL_2640863, EPI_ISL_2640864 | Lighthouse Lab in Alderley Park | Wellcome Sanger Institute for the COVID-19 Genomics UK (COG-UK) Consortium | Jacquelyn Wynn, Mairead Hyland, The Lighthouse Lab in Alderley Park and Alex Alderton, Roberto Amato, Jeffrey Barrett, Sonia Goncalves, Ewan Harrison, David K. Jackson, Ian Johnston, Dominic Kwiatkowski, Cordelia Langford, John Sillitoe on behalf of the Wellcome Sanger Institute COVID-19 Surveillance Team |
| EPI_ISL_2640865 | Lighthouse Lab in Milton Keynes | Wellcome Sanger Institute for the COVID-19 Genomics UK (COG-UK) Consortium | The Lighthouse Lab in Milton Keynes and Alex Alderton, Roberto Amato, Jeffrey Barrett, Sonia Goncalves, Ewan Harrison, David K. Jackson, Ian Johnston, Dominic Kwiatkowski, Cordelia Langford, John Sillitoe on behalf of the Wellcome Sanger Institute COVID-19 Surveillance Team |
| EPI_ISL_2640866 | Lighthouse Lab in Alderley Park | Wellcome Sanger Institute for the COVID-19 Genomics UK (COG-UK) Consortium | Jacquelyn Wynn, Mairead Hyland, The Lighthouse Lab in Alderley Park and Alex Alderton, Roberto Amato, Jeffrey Barrett, Sonia Goncalves, Ewan Harrison, David K. Jackson, Ian Johnston, Dominic Kwiatkowski, Cordelia Langford, John Sillitoe on behalf of the Wellcome Sanger Institute COVID-19 Surveillance Team |

| | | | |
|---|---|---|---|
| EPI_ISL_2640867, EPI_ISL_2640868 | Lighthouse Lab in Milton Keynes | Wellcome Sanger Institute for the COVID-19 Genomics UK (COG-UK) Consortium | The Lighthouse Lab in Milton Keynes and Alex Alderton, Roberto Amato, Jeffrey Barrett, Sonia Goncalves, Ewan Harrison, David K. Jackson, Ian Johnston, Dominic Kwiatkowski, Cordelia Langford, John Sillitoe on behalf of the Wellcome Sanger Institute COVID-19 Surveillance Team |
| EPI_ISL_2640869, EPI_ISL_2640870, EPI_ISL_2640871, EPI_ISL_2640872 | Lighthouse Lab in Alderley Park | Wellcome Sanger Institute for the COVID-19 Genomics UK (COG-UK) Consortium | Jacquelyn Wynn, Mairead Hyland, The Lighthouse Lab in Alderley Park and Alex Alderton, Roberto Amato, Jeffrey Barrett, Sonia Goncalves, Ewan Harrison, David K. Jackson, Ian Johnston, Dominic Kwiatkowski, Cordelia Langford, John Sillitoe on behalf of the Wellcome Sanger Institute COVID-19 Surveillance Team |
| EPI_ISL_2640873, EPI_ISL_2640874 | Lighthouse Lab in Milton Keynes | Wellcome Sanger Institute for the COVID-19 Genomics UK (COG-UK) Consortium | The Lighthouse Lab in Milton Keynes and Alex Alderton, Roberto Amato, Jeffrey Barrett, Sonia Goncalves, Ewan Harrison, David K. Jackson, Ian Johnston, Dominic Kwiatkowski, Cordelia Langford, John Sillitoe on behalf of the Wellcome Sanger Institute COVID-19 Surveillance Team |
| EPI_ISL_2640875 | Lighthouse Lab in Glasgow | Wellcome Sanger Institute for the COVID-19 Genomics UK (COG-UK) Consortium | Harper VanSteenhouse, Yumi Kasai, David Gray, Carol Clugston, Anna Dominiczak and Alex Alderton, Roberto Amato, Jeffrey Barrett, Sonia Goncalves, Ewan Harrison, David K. Jackson, Ian Johnston, Dominic Kwiatkowski, Cordelia Langford, John Sillitoe on behalf of the Wellcome Sanger Institute COVID-19 Surveillance Team |
| EPI_ISL_2640876, EPI_ISL_2640877 | Lighthouse Lab in Alderley Park | Wellcome Sanger Institute for the COVID-19 Genomics UK (COG-UK) Consortium | Jacquelyn Wynn, Mairead Hyland, The Lighthouse Lab in Alderley Park and Alex Alderton, Roberto Amato, Jeffrey Barrett, Sonia Goncalves, Ewan Harrison, David K. Jackson, Ian Johnston, Dominic Kwiatkowski, Cordelia Langford, John Sillitoe on behalf of the Wellcome Sanger Institute COVID-19 Surveillance Team |
| EPI_ISL_2640878 | Lighthouse Lab in Glasgow | Wellcome Sanger Institute for the COVID-19 Genomics UK (COG-UK) Consortium | Harper VanSteenhouse, Yumi Kasai, David Gray, Carol Clugston, Anna Dominiczak and Alex Alderton, Roberto Amato, Jeffrey Barrett, Sonia Goncalves, Ewan Harrison, David K. Jackson, Ian Johnston, Dominic Kwiatkowski, Cordelia Langford, John Sillitoe on behalf of the Wellcome Sanger Institute COVID-19 Surveillance Team |
| EPI_ISL_2640879 | Lighthouse Lab in Alderley Park | Wellcome Sanger Institute for the COVID-19 Genomics UK (COG-UK) Consortium | Jacquelyn Wynn, Mairead Hyland, The Lighthouse Lab in Alderley Park and Alex Alderton, Roberto Amato, Jeffrey Barrett, Sonia Goncalves, Ewan Harrison, David K. Jackson, Ian Johnston, Dominic Kwiatkowski, Cordelia Langford, John Sillitoe on behalf of the Wellcome Sanger Institute COVID-19 Surveillance Team |
| EPI_ISL_2640880 | Lighthouse Lab in Glasgow | Wellcome Sanger Institute for the COVID-19 Genomics UK (COG-UK) Consortium | Harper VanSteenhouse, Yumi Kasai, David Gray, Carol Clugston, Anna Dominiczak and Alex Alderton, Roberto Amato, Jeffrey Barrett, Sonia Goncalves, Ewan Harrison, David K. Jackson, Ian Johnston, Dominic Kwiatkowski, Cordelia Langford, John Sillitoe on behalf of the Wellcome Sanger Institute COVID-19 Surveillance Team |
| EPI_ISL_2640881, EPI_ISL_2640882, EPI_ISL_2640883, EPI_ISL_2640884 | Lighthouse Lab in Alderley Park | Wellcome Sanger Institute for the COVID-19 Genomics UK (COG-UK) Consortium | Jacquelyn Wynn, Mairead Hyland, The Lighthouse Lab in Alderley Park and Alex Alderton, Roberto Amato, Jeffrey Barrett, Sonia Goncalves, Ewan Harrison, David K. Jackson, Ian Johnston, Dominic Kwiatkowski, Cordelia Langford, John Sillitoe on behalf of the Wellcome Sanger Institute COVID-19 Surveillance Team |
| EPI_ISL_2640885 | Lighthouse Lab in Milton Keynes | Wellcome Sanger Institute for the COVID-19 Genomics UK (COG-UK) Consortium | The Lighthouse Lab in Milton Keynes and Alex Alderton, Roberto Amato, Jeffrey Barrett, Sonia Goncalves, Ewan Harrison, David K. Jackson, Ian Johnston, Dominic Kwiatkowski, Cordelia Langford, John Sillitoe on behalf of the Wellcome Sanger Institute COVID-19 Surveillance Team |
| EPI_ISL_2640886 | Lighthouse Lab in Alderley Park | Wellcome Sanger Institute for the COVID-19 Genomics UK (COG-UK) Consortium | Jacquelyn Wynn, Mairead Hyland, The Lighthouse Lab in Alderley Park and Alex Alderton, Roberto Amato, Jeffrey Barrett, Sonia Goncalves, Ewan Harrison, David K. Jackson, Ian Johnston, Dominic Kwiatkowski, Cordelia Langford, John Sillitoe on behalf of the Wellcome Sanger Institute COVID-19 Surveillance Team |
| EPI_ISL_2640887 | Lighthouse Lab in Milton Keynes | Wellcome Sanger Institute for the COVID-19 Genomics UK (COG-UK) Consortium | The Lighthouse Lab in Milton Keynes and Alex Alderton, Roberto Amato, Jeffrey Barrett, Sonia Goncalves, Ewan Harrison, David K. Jackson, Ian Johnston, Dominic Kwiatkowski, Cordelia Langford, John Sillitoe on behalf of the Wellcome Sanger Institute COVID-19 Surveillance Team |
| EPI_ISL_2640889 | Lighthouse Lab in Alderley Park | Wellcome Sanger Institute for the COVID-19 Genomics UK (COG-UK) Consortium | Jacquelyn Wynn, Mairead Hyland, The Lighthouse Lab in Alderley Park and Alex Alderton, Roberto Amato, Jeffrey Barrett, Sonia Goncalves, Ewan Harrison, David K. Jackson, Ian Johnston, Dominic Kwiatkowski, Cordelia Langford, John Sillitoe on behalf of the Wellcome Sanger Institute COVID-19 Surveillance Team |
| EPI_ISL_2640890 | Lighthouse Lab in Milton Keynes | Wellcome Sanger Institute for the COVID-19 Genomics UK (COG-UK) Consortium | The Lighthouse Lab in Milton Keynes and Alex Alderton, Roberto Amato, Jeffrey Barrett, Sonia Goncalves, Ewan Harrison, David K. Jackson, Ian Johnston, Dominic Kwiatkowski, Cordelia Langford, John Sillitoe on behalf of the Wellcome Sanger Institute COVID-19 Surveillance Team |
| EPI_ISL_2640891, EPI_ISL_2640892, EPI_ISL_2640894 | Lighthouse Lab in Alderley Park | Wellcome Sanger Institute for the COVID-19 Genomics UK (COG-UK) Consortium | Jacquelyn Wynn, Mairead Hyland, The Lighthouse Lab in Alderley Park and Alex Alderton, Roberto Amato, Jeffrey Barrett, Sonia Goncalves, Ewan Harrison, David K. Jackson, Ian Johnston, Dominic Kwiatkowski, Cordelia Langford, John Sillitoe on behalf of the Wellcome Sanger Institute COVID-19 Surveillance Team |
| EPI_ISL_2640895 | Lighthouse Lab in Glasgow | Wellcome Sanger Institute for the COVID-19 Genomics UK (COG-UK) Consortium | Harper VanSteenhouse, Yumi Kasai, David Gray, Carol Clugston, Anna Dominiczak and Alex Alderton, Roberto Amato, Jeffrey Barrett, Sonia Goncalves, Ewan Harrison, David K. Jackson, Ian Johnston, Dominic Kwiatkowski, Cordelia Langford, John Sillitoe on behalf of the Wellcome Sanger Institute COVID-19 Surveillance Team |
| EPI_ISL_2640896, EPI_ISL_2640897, EPI_ISL_2640898 | Lighthouse Lab in Alderley Park | Wellcome Sanger Institute for the COVID-19 Genomics UK (COG-UK) Consortium | Jacquelyn Wynn, Mairead Hyland, The Lighthouse Lab in Alderley Park and Alex Alderton, Roberto Amato, Jeffrey Barrett, Sonia Goncalves, Ewan Harrison, David K. Jackson, Ian Johnston, Dominic Kwiatkowski, Cordelia Langford, John Sillitoe on behalf of the Wellcome Sanger Institute COVID-19 Surveillance Team |
| EPI_ISL_2640899 | Lighthouse Lab in Milton Keynes | Wellcome Sanger Institute for the COVID-19 Genomics UK (COG-UK) Consortium | The Lighthouse Lab in Milton Keynes and Alex Alderton, Roberto Amato, Jeffrey Barrett, Sonia Goncalves, Ewan Harrison, David K. Jackson, Ian Johnston, Dominic Kwiatkowski, Cordelia Langford, John Sillitoe on behalf of the Wellcome Sanger Institute COVID-19 Surveillance Team |
| EPI_ISL_2640900, EPI_ISL_2640901 | Lighthouse Lab in Glasgow | Wellcome Sanger Institute for the COVID-19 Genomics UK (COG-UK) Consortium | Harper VanSteenhouse, Yumi Kasai, David Gray, Carol Clugston, Anna Dominiczak and Alex Alderton, Roberto Amato, Jeffrey Barrett, Sonia Goncalves, Ewan Harrison, David K. Jackson, Ian Johnston, Dominic Kwiatkowski, Cordelia Langford, John Sillitoe on behalf of the Wellcome Sanger Institute COVID-19 Surveillance Team |
| EPI_ISL_2640902 | Lighthouse Lab in Alderley Park | Wellcome Sanger Institute for the COVID-19 Genomics UK (COG-UK) Consortium | Jacquelyn Wynn, Mairead Hyland, The Lighthouse Lab in Alderley Park and Alex Alderton, Roberto Amato, Jeffrey Barrett, Sonia Goncalves, Ewan Harrison, David K. Jackson, Ian Johnston, Dominic Kwiatkowski, Cordelia Langford, John Sillitoe on behalf of the Wellcome Sanger Institute COVID-19 Surveillance Team |
| EPI_ISL_2640903 | Lighthouse Lab in Milton Keynes | Wellcome Sanger Institute for the COVID-19 Genomics UK (COG-UK) Consortium | The Lighthouse Lab in Milton Keynes and Alex Alderton, Roberto Amato, Jeffrey Barrett, Sonia Goncalves, Ewan Harrison, David K. Jackson, Ian Johnston, Dominic Kwiatkowski, Cordelia Langford, John Sillitoe on behalf of the Wellcome Sanger Institute COVID-19 Surveillance Team |
| EPI_ISL_2640904 | Lighthouse Lab in Alderley Park | Wellcome Sanger Institute for the COVID-19 Genomics UK (COG-UK) Consortium | Jacquelyn Wynn, Mairead Hyland, The Lighthouse Lab in Alderley Park and Alex Alderton, Roberto Amato, Jeffrey Barrett, Sonia Goncalves, Ewan Harrison, David K. Jackson, Ian Johnston, Dominic Kwiatkowski, Cordelia Langford, John Sillitoe on behalf of the Wellcome Sanger Institute COVID-19 Surveillance Team |
| EPI_ISL_2640905 | Lighthouse Lab in Glasgow | Wellcome Sanger Institute for the COVID-19 Genomics UK (COG-UK) Consortium | Harper VanSteenhouse, Yumi Kasai, David Gray, Carol Clugston, Anna Dominiczak and Alex Alderton, Roberto Amato, Jeffrey Barrett, Sonia Goncalves, Ewan Harrison, David K. Jackson, Ian Johnston, Dominic Kwiatkowski, Cordelia Langford, John Sillitoe on behalf of the Wellcome Sanger Institute COVID-19 Surveillance Team |
| EPI_ISL_2640906 | Lighthouse Lab in Alderley Park | Wellcome Sanger Institute for the COVID-19 Genomics UK (COG-UK) Consortium | Jacquelyn Wynn, Mairead Hyland, The Lighthouse Lab in Alderley Park and Alex Alderton, Roberto Amato, Jeffrey Barrett, Sonia Goncalves, Ewan Harrison, David K. Jackson, Ian Johnston, Dominic Kwiatkowski, Cordelia Langford, John Sillitoe on behalf of the Wellcome Sanger Institute COVID-19 Surveillance Team |
| EPI_ISL_2640907 | Lighthouse Lab in Milton Keynes | Wellcome Sanger Institute for the COVID-19 Genomics UK (COG-UK) Consortium | The Lighthouse Lab in Milton Keynes and Alex Alderton, Roberto Amato, Jeffrey Barrett, Sonia Goncalves, Ewan Harrison, David K. Jackson, Ian Johnston, Dominic Kwiatkowski, Cordelia Langford, John Sillitoe on behalf of the Wellcome Sanger Institute COVID-19 Surveillance Team |
| EPI_ISL_2640908, EPI_ISL_2640909, EPI_ISL_2640910 | Lighthouse Lab in Glasgow | Wellcome Sanger Institute for the COVID-19 Genomics UK (COG-UK) Consortium | Harper VanSteenhouse, Yumi Kasai, David Gray, Carol Clugston, Anna Dominiczak and Alex Alderton, Roberto Amato, Jeffrey Barrett, Sonia Goncalves, Ewan Harrison, David K. Jackson, Ian Johnston, Dominic Kwiatkowski, Cordelia Langford, John Sillitoe on behalf of the Wellcome Sanger Institute COVID-19 Surveillance Team |
| EPI_ISL_2640911, EPI_ISL_2640912 | Lighthouse Lab in Milton Keynes | Wellcome Sanger Institute for the COVID-19 Genomics UK (COG-UK) Consortium | The Lighthouse Lab in Milton Keynes and Alex Alderton, Roberto Amato, Jeffrey Barrett, Sonia Goncalves, Ewan Harrison, David K. Jackson, Ian Johnston, Dominic Kwiatkowski, Cordelia Langford, John Sillitoe on behalf of the Wellcome Sanger Institute COVID-19 Surveillance Team |
| EPI_ISL_2640913 | Lighthouse Lab in Glasgow | Wellcome Sanger Institute for the COVID-19 Genomics UK (COG-UK) Consortium | Harper VanSteenhouse, Yumi Kasai, David Gray, Carol Clugston, Anna Dominiczak and Alex Alderton, Roberto Amato, Jeffrey Barrett, Sonia Goncalves, Ewan Harrison, David K. Jackson, Ian Johnston, Dominic Kwiatkowski, Cordelia Langford, John Sillitoe on behalf of the Wellcome Sanger Institute COVID-19 Surveillance Team |
| EPI_ISL_2640914 | Lighthouse Lab in Alderley Park | Wellcome Sanger Institute for the COVID-19 Genomics UK (COG-UK) Consortium | Jacquelyn Wynn, Mairead Hyland, The Lighthouse Lab in Alderley Park and Alex Alderton, Roberto Amato, Jeffrey Barrett, Sonia Goncalves, Ewan Harrison, David K. Jackson, Ian Johnston, Dominic Kwiatkowski, Cordelia Langford, John Sillitoe on behalf of the Wellcome Sanger Institute COVID-19 Surveillance Team |
| EPI_ISL_2640915 | Lighthouse Lab in Glasgow | Wellcome Sanger Institute for the COVID-19 Genomics UK (COG-UK) Consortium | Harper VanSteenhouse, Yumi Kasai, David Gray, Carol Clugston, Anna Dominiczak and Alex Alderton, Roberto Amato, Jeffrey Barrett, Sonia Goncalves, Ewan Harrison, David K. Jackson, Ian Johnston, Dominic Kwiatkowski, Cordelia Langford, John Sillitoe on behalf of the Wellcome Sanger Institute COVID-19 Surveillance Team |
| EPI_ISL_2640916 | Lighthouse Lab in Milton Keynes | Wellcome Sanger Institute for the COVID-19 Genomics UK (COG-UK) Consortium | The Lighthouse Lab in Milton Keynes and Alex Alderton, Roberto Amato, Jeffrey Barrett, Sonia Goncalves, Ewan Harrison, David K. Jackson, Ian Johnston, Dominic Kwiatkowski, Cordelia Langford, John Sillitoe on behalf of the Wellcome Sanger Institute COVID-19 Surveillance Team |
| EPI_ISL_2640918 | Lighthouse Lab in Alderley Park | Wellcome Sanger Institute for the COVID-19 Genomics UK (COG-UK) Consortium | Jacquelyn Wynn, Mairead Hyland, The Lighthouse Lab in Alderley Park and Alex Alderton, Roberto Amato, Jeffrey Barrett, Sonia Goncalves, Ewan Harrison, David K. Jackson, Ian Johnston, Dominic Kwiatkowski, Cordelia Langford, John Sillitoe on behalf of the Wellcome Sanger Institute COVID-19 Surveillance Team |
| EPI_ISL_2640919 | Lighthouse Lab in Glasgow | Wellcome Sanger Institute for the COVID-19 Genomics UK (COG-UK) Consortium | Harper VanSteenhouse, Yumi Kasai, David Gray, Carol Clugston, Anna Dominiczak and Alex Alderton, Roberto Amato, Jeffrey Barrett, Sonia Goncalves, Ewan Harrison, David K. Jackson, Ian Johnston, Dominic Kwiatkowski, Cordelia Langford, John Sillitoe on behalf of the Wellcome Sanger Institute COVID-19 Surveillance Team |
| EPI_ISL_2640920 | Lighthouse Lab in Alderley Park | Wellcome Sanger Institute for the COVID-19 Genomics UK (COG-UK) Consortium | Jacquelyn Wynn, Mairead Hyland, The Lighthouse Lab in Alderley Park and Alex Alderton, Roberto Amato, Jeffrey Barrett, Sonia Goncalves, Ewan Harrison, David K. Jackson, Ian Johnston, Dominic Kwiatkowski, Cordelia Langford, John Sillitoe on behalf of the Wellcome Sanger Institute COVID-19 Surveillance Team |
| EPI_ISL_2640921, EPI_ISL_2640922, EPI_ISL_2640923 | Lighthouse Lab in Glasgow | Wellcome Sanger Institute for the COVID-19 Genomics UK (COG-UK) Consortium | Harper VanSteenhouse, Yumi Kasai, David Gray, Carol Clugston, Anna Dominiczak and Alex Alderton, Roberto Amato, Jeffrey Barrett, Sonia Goncalves, Ewan Harrison, David K. Jackson, Ian Johnston, Dominic Kwiatkowski, Cordelia Langford, John Sillitoe on behalf of the Wellcome Sanger Institute COVID-19 Surveillance Team |
| EPI_ISL_2640924 | Lighthouse Lab in Milton Keynes | Wellcome Sanger Institute for the COVID-19 Genomics UK (COG-UK) Consortium | The Lighthouse Lab in Milton Keynes and Alex Alderton, Roberto Amato, Jeffrey Barrett, Sonia Goncalves, Ewan Harrison, David K. Jackson, Ian Johnston, Dominic Kwiatkowski, Cordelia Langford, John Sillitoe on behalf of the Wellcome Sanger Institute COVID-19 Surveillance Team |
| EPI_ISL_2640925, EPI_ISL_2640926, EPI_ISL_2640927, EPI_ISL_2640928 | Lighthouse Lab in Alderley Park | Wellcome Sanger Institute for the COVID-19 Genomics UK (COG-UK) Consortium | Jacquelyn Wynn, Mairead Hyland, The Lighthouse Lab in Alderley Park and Alex Alderton, Roberto Amato, Jeffrey Barrett, Sonia Goncalves, Ewan Harrison, David K. Jackson, Ian Johnston, Dominic Kwiatkowski, Cordelia Langford, John Sillitoe on behalf of the Wellcome Sanger Institute COVID-19 Surveillance Team |
| EPI_ISL_2640929 | Lighthouse Lab in Milton Keynes | Wellcome Sanger Institute for the COVID-19 Genomics UK (COG-UK) Consortium | The Lighthouse Lab in Milton Keynes and Alex Alderton, Roberto Amato, Jeffrey Barrett, Sonia Goncalves, Ewan Harrison, David K. Jackson, Ian Johnston, Dominic Kwiatkowski, Cordelia Langford, John Sillitoe on behalf of the Wellcome Sanger Institute COVID-19 Surveillance Team |
| EPI_ISL_2640930 | Lighthouse Lab in Alderley Park | Wellcome Sanger Institute for the COVID-19 Genomics UK (COG-UK) Consortium | Jacquelyn Wynn, Mairead Hyland, The Lighthouse Lab in Alderley Park and Alex Alderton, Roberto Amato, Jeffrey Barrett, Sonia Goncalves, Ewan Harrison, David K. Jackson, Ian Johnston, Dominic Kwiatkowski, Cordelia Langford, John Sillitoe on behalf of the Wellcome Sanger Institute COVID-19 Surveillance Team |
| EPI_ISL_2640931 | Lighthouse Lab in Glasgow | Wellcome Sanger Institute for the COVID-19 Genomics UK (COG-UK) Consortium | Harper VanSteenhouse, Yumi Kasai, David Gray, Carol Clugston, Anna Dominiczak and Alex Alderton, Roberto Amato, Jeffrey Barrett, Sonia Goncalves, Ewan Harrison, David K. Jackson, Ian Johnston, Dominic Kwiatkowski, Cordelia Langford, John Sillitoe on behalf of the Wellcome Sanger Institute COVID-19 Surveillance Team |
| EPI_ISL_2640932 | Lighthouse Lab in Milton Keynes | Wellcome Sanger Institute for the COVID-19 Genomics UK (COG-UK) | The Lighthouse Lab in Milton Keynes and Alex Alderton, Roberto Amato, Jeffrey Barrett, Sonia Goncalves, Ewan Harrison, David K. Jackson, Ian Johnston, Dominic Kwiatkowski, Cordelia Langford, John Sillitoe on behalf of the Wellcome Sanger |

| | | | |
|---|---|---|---|
| | | Consortium | Institute COVID-19 Surveillance Team |
| EPI_ISL_2640934 | Lighthouse Lab in Alderley Park | Wellcome Sanger Institute for the COVID-19 Genomics UK (COG-UK) Consortium | Jacquelyn Wynn, Mairead Hyland, The Lighthouse Lab in Alderley Park and Alex Alderton, Roberto Amato, Jeffrey Barrett, Sonia Goncalves, Ewan Harrison, David K. Jackson, Ian Johnston, Dominic Kwiatkowski, Cordelia Langford, John Sillitoe on behalf of the Wellcome Sanger Institute COVID-19 Surveillance Team |
| EPI_ISL_2640935 | Lighthouse Lab in Glasgow | Wellcome Sanger Institute for the COVID-19 Genomics UK (COG-UK) Consortium | Harper VanSteenhouse, Yumi Kasai, David Gray, Carol Clugston, Anna Dominiczak and Alex Alderton, Roberto Amato, Jeffrey Barrett, Sonia Goncalves, Ewan Harrison, David K. Jackson, Ian Johnston, Dominic Kwiatkowski, Cordelia Langford, John Sillitoe on behalf of the Wellcome Sanger Institute COVID-19 Surveillance Team |
| EPI_ISL_2640936 | Lighthouse Lab in Milton Keynes | Wellcome Sanger Institute for the COVID-19 Genomics UK (COG-UK) Consortium | The Lighthouse Lab in Milton Keynes and Alex Alderton, Roberto Amato, Jeffrey Barrett, Sonia Goncalves, Ewan Harrison, David K. Jackson, Ian Johnston, Dominic Kwiatkowski, Cordelia Langford, John Sillitoe on behalf of the Wellcome Sanger Institute COVID-19 Surveillance Team |
| EPI_ISL_2640937, EPI_ISL_2640938 | Lighthouse Lab in Alderley Park | Wellcome Sanger Institute for the COVID-19 Genomics UK (COG-UK) Consortium | Jacquelyn Wynn, Mairead Hyland, The Lighthouse Lab in Alderley Park and Alex Alderton, Roberto Amato, Jeffrey Barrett, Sonia Goncalves, Ewan Harrison, David K. Jackson, Ian Johnston, Dominic Kwiatkowski, Cordelia Langford, John Sillitoe on behalf of the Wellcome Sanger Institute COVID-19 Surveillance Team |
| EPI_ISL_2640939 | Lighthouse Lab in Milton Keynes | Wellcome Sanger Institute for the COVID-19 Genomics UK (COG-UK) Consortium | The Lighthouse Lab in Milton Keynes and Alex Alderton, Roberto Amato, Jeffrey Barrett, Sonia Goncalves, Ewan Harrison, David K. Jackson, Ian Johnston, Dominic Kwiatkowski, Cordelia Langford, John Sillitoe on behalf of the Wellcome Sanger Institute COVID-19 Surveillance Team |
| EPI_ISL_2640941, EPI_ISL_2640942, EPI_ISL_2640943 | Lighthouse Lab in Alderley Park | Wellcome Sanger Institute for the COVID-19 Genomics UK (COG-UK) Consortium | Jacquelyn Wynn, Mairead Hyland, The Lighthouse Lab in Alderley Park and Alex Alderton, Roberto Amato, Jeffrey Barrett, Sonia Goncalves, Ewan Harrison, David K. Jackson, Ian Johnston, Dominic Kwiatkowski, Cordelia Langford, John Sillitoe on behalf of the Wellcome Sanger Institute COVID-19 Surveillance Team |
| EPI_ISL_2640944 | Lighthouse Lab in Milton Keynes | Wellcome Sanger Institute for the COVID-19 Genomics UK (COG-UK) Consortium | The Lighthouse Lab in Milton Keynes and Alex Alderton, Roberto Amato, Jeffrey Barrett, Sonia Goncalves, Ewan Harrison, David K. Jackson, Ian Johnston, Dominic Kwiatkowski, Cordelia Langford, John Sillitoe on behalf of the Wellcome Sanger Institute COVID-19 Surveillance Team |
| EPI_ISL_2640945 | Lighthouse Lab in Alderley Park | Wellcome Sanger Institute for the COVID-19 Genomics UK (COG-UK) Consortium | Jacquelyn Wynn, Mairead Hyland, The Lighthouse Lab in Alderley Park and Alex Alderton, Roberto Amato, Jeffrey Barrett, Sonia Goncalves, Ewan Harrison, David K. Jackson, Ian Johnston, Dominic Kwiatkowski, Cordelia Langford, John Sillitoe on behalf of the Wellcome Sanger Institute COVID-19 Surveillance Team |
| EPI_ISL_2640946, EPI_ISL_2640947, EPI_ISL_2640949 | Lighthouse Lab in Milton Keynes | Wellcome Sanger Institute for the COVID-19 Genomics UK (COG-UK) Consortium | The Lighthouse Lab in Milton Keynes and Alex Alderton, Roberto Amato, Jeffrey Barrett, Sonia Goncalves, Ewan Harrison, David K. Jackson, Ian Johnston, Dominic Kwiatkowski, Cordelia Langford, John Sillitoe on behalf of the Wellcome Sanger Institute COVID-19 Surveillance Team |
| EPI_ISL_2640950 | Lighthouse Lab in Alderley Park | Wellcome Sanger Institute for the COVID-19 Genomics UK (COG-UK) Consortium | Jacquelyn Wynn, Mairead Hyland, The Lighthouse Lab in Alderley Park and Alex Alderton, Roberto Amato, Jeffrey Barrett, Sonia Goncalves, Ewan Harrison, David K. Jackson, Ian Johnston, Dominic Kwiatkowski, Cordelia Langford, John Sillitoe on behalf of the Wellcome Sanger Institute COVID-19 Surveillance Team |
| EPI_ISL_2640951 | Lighthouse Lab in Glasgow | Wellcome Sanger Institute for the COVID-19 Genomics UK (COG-UK) Consortium | Harper VanSteenhouse, Yumi Kasai, David Gray, Carol Clugston, Anna Dominiczak and Alex Alderton, Roberto Amato, Jeffrey Barrett, Sonia Goncalves, Ewan Harrison, David K. Jackson, Ian Johnston, Dominic Kwiatkowski, Cordelia Langford, John Sillitoe on behalf of the Wellcome Sanger Institute COVID-19 Surveillance Team |
| EPI_ISL_2640952 | Lighthouse Lab in Milton Keynes | Wellcome Sanger Institute for the COVID-19 Genomics UK (COG-UK) Consortium | The Lighthouse Lab in Milton Keynes and Alex Alderton, Roberto Amato, Jeffrey Barrett, Sonia Goncalves, Ewan Harrison, David K. Jackson, Ian Johnston, Dominic Kwiatkowski, Cordelia Langford, John Sillitoe on behalf of the Wellcome Sanger Institute COVID-19 Surveillance Team |
| EPI_ISL_2640953, EPI_ISL_2640954 | Lighthouse Lab in Alderley Park | Wellcome Sanger Institute for the COVID-19 Genomics UK (COG-UK) Consortium | Jacquelyn Wynn, Mairead Hyland, The Lighthouse Lab in Alderley Park and Alex Alderton, Roberto Amato, Jeffrey Barrett, Sonia Goncalves, Ewan Harrison, David K. Jackson, Ian Johnston, Dominic Kwiatkowski, Cordelia Langford, John Sillitoe on behalf of the Wellcome Sanger Institute COVID-19 Surveillance Team |
| EPI_ISL_2640955 | Lighthouse Lab in Glasgow | Wellcome Sanger Institute for the COVID-19 Genomics UK (COG-UK) Consortium | Harper VanSteenhouse, Yumi Kasai, David Gray, Carol Clugston, Anna Dominiczak and Alex Alderton, Roberto Amato, Jeffrey Barrett, Sonia Goncalves, Ewan Harrison, David K. Jackson, Ian Johnston, Dominic Kwiatkowski, Cordelia Langford, John Sillitoe on behalf of the Wellcome Sanger Institute COVID-19 Surveillance Team |
| EPI_ISL_2640956, EPI_ISL_2640957 | Lighthouse Lab in Milton Keynes | Wellcome Sanger Institute for the COVID-19 Genomics UK (COG-UK) Consortium | The Lighthouse Lab in Milton Keynes and Alex Alderton, Roberto Amato, Jeffrey Barrett, Sonia Goncalves, Ewan Harrison, David K. Jackson, Ian Johnston, Dominic Kwiatkowski, Cordelia Langford, John Sillitoe on behalf of the Wellcome Sanger Institute COVID-19 Surveillance Team |
| EPI_ISL_2640958 | Lighthouse Lab in Alderley Park | Wellcome Sanger Institute for the COVID-19 Genomics UK (COG-UK) Consortium | Jacquelyn Wynn, Mairead Hyland, The Lighthouse Lab in Alderley Park and Alex Alderton, Roberto Amato, Jeffrey Barrett, Sonia Goncalves, Ewan Harrison, David K. Jackson, Ian Johnston, Dominic Kwiatkowski, Cordelia Langford, John Sillitoe on behalf of the Wellcome Sanger Institute COVID-19 Surveillance Team |
| EPI_ISL_2640959 | Lighthouse Lab in Milton Keynes | Wellcome Sanger Institute for the COVID-19 Genomics UK (COG-UK) Consortium | The Lighthouse Lab in Milton Keynes and Alex Alderton, Roberto Amato, Jeffrey Barrett, Sonia Goncalves, Ewan Harrison, David K. Jackson, Ian Johnston, Dominic Kwiatkowski, Cordelia Langford, John Sillitoe on behalf of the Wellcome Sanger Institute COVID-19 Surveillance Team |
| EPI_ISL_2640960 | Lighthouse Lab in Glasgow | Wellcome Sanger Institute for the COVID-19 Genomics UK (COG-UK) Consortium | Harper VanSteenhouse, Yumi Kasai, David Gray, Carol Clugston, Anna Dominiczak and Alex Alderton, Roberto Amato, Jeffrey Barrett, Sonia Goncalves, Ewan Harrison, David K. Jackson, Ian Johnston, Dominic Kwiatkowski, Cordelia Langford, John Sillitoe on behalf of the Wellcome Sanger Institute COVID-19 Surveillance Team |
| EPI_ISL_2640961, EPI_ISL_2640962 | Lighthouse Lab in Alderley Park | Wellcome Sanger Institute for the COVID-19 Genomics UK (COG-UK) Consortium | Jacquelyn Wynn, Mairead Hyland, The Lighthouse Lab in Alderley Park and Alex Alderton, Roberto Amato, Jeffrey Barrett, Sonia Goncalves, Ewan Harrison, David K. Jackson, Ian Johnston, Dominic Kwiatkowski, Cordelia Langford, John Sillitoe on behalf of the Wellcome Sanger Institute COVID-19 Surveillance Team |
| EPI_ISL_2640963 | Lighthouse Lab in Glasgow | Wellcome Sanger Institute for the COVID-19 Genomics UK (COG-UK) Consortium | Harper VanSteenhouse, Yumi Kasai, David Gray, Carol Clugston, Anna Dominiczak and Alex Alderton, Roberto Amato, Jeffrey Barrett, Sonia Goncalves, Ewan Harrison, David K. Jackson, Ian Johnston, Dominic Kwiatkowski, Cordelia Langford, John Sillitoe on behalf of the Wellcome Sanger Institute COVID-19 Surveillance Team |
| EPI_ISL_2640964 | Lighthouse Lab in Milton Keynes | Wellcome Sanger Institute for the COVID-19 Genomics UK (COG-UK) Consortium | The Lighthouse Lab in Milton Keynes and Alex Alderton, Roberto Amato, Jeffrey Barrett, Sonia Goncalves, Ewan Harrison, David K. Jackson, Ian Johnston, Dominic Kwiatkowski, Cordelia Langford, John Sillitoe on behalf of the Wellcome Sanger Institute COVID-19 Surveillance Team |
| EPI_ISL_2640965, EPI_ISL_2640966, EPI_ISL_2640968 | Lighthouse Lab in Glasgow | Wellcome Sanger Institute for the COVID-19 Genomics UK (COG-UK) Consortium | Harper VanSteenhouse, Yumi Kasai, David Gray, Carol Clugston, Anna Dominiczak and Alex Alderton, Roberto Amato, Jeffrey Barrett, Sonia Goncalves, Ewan Harrison, David K. Jackson, Ian Johnston, Dominic Kwiatkowski, Cordelia Langford, John Sillitoe on behalf of the Wellcome Sanger Institute COVID-19 Surveillance Team |
| EPI_ISL_2640969, EPI_ISL_2640970 | Lighthouse Lab in Alderley Park | Wellcome Sanger Institute for the COVID-19 Genomics UK (COG-UK) Consortium | Jacquelyn Wynn, Mairead Hyland, The Lighthouse Lab in Alderley Park and Alex Alderton, Roberto Amato, Jeffrey Barrett, Sonia Goncalves, Ewan Harrison, David K. Jackson, Ian Johnston, Dominic Kwiatkowski, Cordelia Langford, John Sillitoe on behalf of the Wellcome Sanger Institute COVID-19 Surveillance Team |
| EPI_ISL_2640971 | Lighthouse Lab in Milton Keynes | Wellcome Sanger Institute for the COVID-19 Genomics UK (COG-UK) Consortium | The Lighthouse Lab in Milton Keynes and Alex Alderton, Roberto Amato, Jeffrey Barrett, Sonia Goncalves, Ewan Harrison, David K. Jackson, Ian Johnston, Dominic Kwiatkowski, Cordelia Langford, John Sillitoe on behalf of the Wellcome Sanger Institute COVID-19 Surveillance Team |
| EPI_ISL_2640972, EPI_ISL_2640973 | Lighthouse Lab in Alderley Park | Wellcome Sanger Institute for the COVID-19 Genomics UK (COG-UK) Consortium | Jacquelyn Wynn, Mairead Hyland, The Lighthouse Lab in Alderley Park and Alex Alderton, Roberto Amato, Jeffrey Barrett, Sonia Goncalves, Ewan Harrison, David K. Jackson, Ian Johnston, Dominic Kwiatkowski, Cordelia Langford, John Sillitoe on behalf of the Wellcome Sanger Institute COVID-19 Surveillance Team |
| EPI_ISL_2640974 | Lighthouse Lab in Glasgow | Wellcome Sanger Institute for the COVID-19 Genomics UK (COG-UK) Consortium | Harper VanSteenhouse, Yumi Kasai, David Gray, Carol Clugston, Anna Dominiczak and Alex Alderton, Roberto Amato, Jeffrey Barrett, Sonia Goncalves, Ewan Harrison, David K. Jackson, Ian Johnston, Dominic Kwiatkowski, Cordelia Langford, John Sillitoe on behalf of the Wellcome Sanger Institute COVID-19 Surveillance Team |
| EPI_ISL_2640975, EPI_ISL_2640976 | Lighthouse Lab in Alderley Park | Wellcome Sanger Institute for the COVID-19 Genomics UK (COG-UK) Consortium | Jacquelyn Wynn, Mairead Hyland, The Lighthouse Lab in Alderley Park and Alex Alderton, Roberto Amato, Jeffrey Barrett, Sonia Goncalves, Ewan Harrison, David K. Jackson, Ian Johnston, Dominic Kwiatkowski, Cordelia Langford, John Sillitoe on behalf of the Wellcome Sanger Institute COVID-19 Surveillance Team |
| EPI_ISL_2640977 | Lighthouse Lab in Glasgow | Wellcome Sanger Institute for the COVID-19 Genomics UK (COG-UK) Consortium | Harper VanSteenhouse, Yumi Kasai, David Gray, Carol Clugston, Anna Dominiczak and Alex Alderton, Roberto Amato, Jeffrey Barrett, Sonia Goncalves, Ewan Harrison, David K. Jackson, Ian Johnston, Dominic Kwiatkowski, Cordelia Langford, John Sillitoe on behalf of the Wellcome Sanger Institute COVID-19 Surveillance Team |
| EPI_ISL_2640978 | Lighthouse Lab in Alderley Park | Wellcome Sanger Institute for the COVID-19 Genomics UK (COG-UK) Consortium | Jacquelyn Wynn, Mairead Hyland, The Lighthouse Lab in Alderley Park and Alex Alderton, Roberto Amato, Jeffrey Barrett, Sonia Goncalves, Ewan Harrison, David K. Jackson, Ian Johnston, Dominic Kwiatkowski, Cordelia Langford, John Sillitoe on behalf of the Wellcome Sanger Institute COVID-19 Surveillance Team |
| EPI_ISL_2640979 | Lighthouse Lab in Milton Keynes | Wellcome Sanger Institute for the COVID-19 Genomics UK (COG-UK) Consortium | The Lighthouse Lab in Milton Keynes and Alex Alderton, Roberto Amato, Jeffrey Barrett, Sonia Goncalves, Ewan Harrison, David K. Jackson, Ian Johnston, Dominic Kwiatkowski, Cordelia Langford, John Sillitoe on behalf of the Wellcome Sanger Institute COVID-19 Surveillance Team |
| EPI_ISL_2640980 | Lighthouse Lab in Alderley Park | Wellcome Sanger Institute for the COVID-19 Genomics UK (COG-UK) Consortium | Jacquelyn Wynn, Mairead Hyland, The Lighthouse Lab in Alderley Park and Alex Alderton, Roberto Amato, Jeffrey Barrett, Sonia Goncalves, Ewan Harrison, David K. Jackson, Ian Johnston, Dominic Kwiatkowski, Cordelia Langford, John Sillitoe on behalf of the Wellcome Sanger Institute COVID-19 Surveillance Team |
| EPI_ISL_2640981 | Lighthouse Lab in Milton Keynes | Wellcome Sanger Institute for the COVID-19 Genomics UK (COG-UK) Consortium | The Lighthouse Lab in Milton Keynes and Alex Alderton, Roberto Amato, Jeffrey Barrett, Sonia Goncalves, Ewan Harrison, David K. Jackson, Ian Johnston, Dominic Kwiatkowski, Cordelia Langford, John Sillitoe on behalf of the Wellcome Sanger Institute COVID-19 Surveillance Team |
| EPI_ISL_2640982 | Lighthouse Lab in Alderley Park | Wellcome Sanger Institute for the COVID-19 Genomics UK (COG-UK) Consortium | Jacquelyn Wynn, Mairead Hyland, The Lighthouse Lab in Alderley Park and Alex Alderton, Roberto Amato, Jeffrey Barrett, Sonia Goncalves, Ewan Harrison, David K. Jackson, Ian Johnston, Dominic Kwiatkowski, Cordelia Langford, John Sillitoe on behalf of the Wellcome Sanger Institute COVID-19 Surveillance Team |
| EPI_ISL_2640983 | Lighthouse Lab in Milton Keynes | Wellcome Sanger Institute for the COVID-19 Genomics UK (COG-UK) Consortium | The Lighthouse Lab in Milton Keynes and Alex Alderton, Roberto Amato, Jeffrey Barrett, Sonia Goncalves, Ewan Harrison, David K. Jackson, Ian Johnston, Dominic Kwiatkowski, Cordelia Langford, John Sillitoe on behalf of the Wellcome Sanger Institute COVID-19 Surveillance Team |
| EPI_ISL_2640984, EPI_ISL_2640985, EPI_ISL_2640986, EPI_ISL_2640987, EPI_ISL_2640988 | Lighthouse Lab in Alderley Park | Wellcome Sanger Institute for the COVID-19 Genomics UK (COG-UK) Consortium | Jacquelyn Wynn, Mairead Hyland, The Lighthouse Lab in Alderley Park and Alex Alderton, Roberto Amato, Jeffrey Barrett, Sonia Goncalves, Ewan Harrison, David K. Jackson, Ian Johnston, Dominic Kwiatkowski, Cordelia Langford, John Sillitoe on behalf of the Wellcome Sanger Institute COVID-19 Surveillance Team |
| EPI_ISL_2640989 | Lighthouse Lab in Milton Keynes | Wellcome Sanger Institute for the COVID-19 Genomics UK (COG-UK) Consortium | The Lighthouse Lab in Milton Keynes and Alex Alderton, Roberto Amato, Jeffrey Barrett, Sonia Goncalves, Ewan Harrison, David K. Jackson, Ian Johnston, Dominic Kwiatkowski, Cordelia Langford, John Sillitoe on behalf of the Wellcome Sanger Institute COVID-19 Surveillance Team |
| EPI_ISL_2640990 | Lighthouse Lab in Alderley Park | Wellcome Sanger Institute for the COVID-19 Genomics UK (COG-UK) Consortium | Jacquelyn Wynn, Mairead Hyland, The Lighthouse Lab in Alderley Park and Alex Alderton, Roberto Amato, Jeffrey Barrett, Sonia Goncalves, Ewan Harrison, David K. Jackson, Ian Johnston, Dominic Kwiatkowski, Cordelia Langford, John Sillitoe on behalf of the Wellcome Sanger Institute COVID-19 Surveillance Team |
| EPI_ISL_2640991 | Lighthouse Lab in Milton Keynes | Wellcome Sanger Institute for the COVID-19 Genomics UK (COG-UK) Consortium | The Lighthouse Lab in Milton Keynes and Alex Alderton, Roberto Amato, Jeffrey Barrett, Sonia Goncalves, Ewan Harrison, David K. Jackson, Ian Johnston, Dominic Kwiatkowski, Cordelia Langford, John Sillitoe on behalf of the Wellcome Sanger Institute COVID-19 Surveillance Team |
| EPI_ISL_2640992 | Lighthouse Lab in Glasgow | Wellcome Sanger Institute for the COVID-19 Genomics UK (COG-UK) Consortium | Harper VanSteenhouse, Yumi Kasai, David Gray, Carol Clugston, Anna Dominiczak and Alex Alderton, Roberto Amato, Jeffrey Barrett, Sonia Goncalves, Ewan Harrison, David K. Jackson, Ian Johnston, Dominic Kwiatkowski, Cordelia Langford, John Sillitoe on behalf of the Wellcome Sanger Institute COVID-19 Surveillance Team |
| EPI_ISL_2640993 | Lighthouse Lab in Milton Keynes | Wellcome Sanger Institute for the COVID-19 Genomics UK (COG-UK) Consortium | The Lighthouse Lab in Milton Keynes and Alex Alderton, Roberto Amato, Jeffrey Barrett, Sonia Goncalves, Ewan Harrison, David K. Jackson, Ian Johnston, Dominic Kwiatkowski, Cordelia Langford, John Sillitoe on behalf of the Wellcome Sanger Institute COVID-19 Surveillance Team |
| EPI_ISL_2640995 | Lighthouse Lab in Alderley Park | Wellcome Sanger Institute for the COVID-19 Genomics UK (COG-UK) Consortium | Jacquelyn Wynn, Mairead Hyland, The Lighthouse Lab in Alderley Park and Alex Alderton, Roberto Amato, Jeffrey Barrett, Sonia Goncalves, Ewan Harrison, David K. Jackson, Ian Johnston, Dominic Kwiatkowski, Cordelia Langford, John Sillitoe on behalf of the Wellcome Sanger Institute COVID-19 Surveillance Team |
| EPI_ISL_2640996, EPI_ISL_2640998 | Lighthouse Lab in Glasgow | Wellcome Sanger Institute for the COVID-19 Genomics UK (COG-UK) Consortium | Harper VanSteenhouse, Yumi Kasai, David Gray, Carol Clugston, Anna Dominiczak and Alex Alderton, Roberto Amato, Jeffrey Barrett, Sonia Goncalves, Ewan Harrison, David K. Jackson, Ian Johnston, Dominic Kwiatkowski, Cordelia Langford, John Sillitoe on behalf of the Wellcome Sanger Institute COVID-19 Surveillance Team |

| | | | |
|---|---|---|---|
| EPI_ISL_2640999 | Lighthouse Lab in Alderley Park | Wellcome Sanger Institute for the COVID-19 Genomics UK (COG-UK) Consortium | Jacquelyn Wynn, Mairead Hyland, The Lighthouse Lab in Alderley Park and Alex Alderton, Roberto Amato, Jeffrey Barrett, Sonia Goncalves, Ewan Harrison, David K. Jackson, Ian Johnston, Dominic Kwiatkowski, Cordelia Langford, John Sillitoe on behalf of the Wellcome Sanger Institute COVID-19 Surveillance Team |
| EPI_ISL_2641000 | Lighthouse Lab in Glasgow | Wellcome Sanger Institute for the COVID-19 Genomics UK (COG-UK) Consortium | Harper VanSteenhouse, Yumi Kasai, David Gray, Carol Clugston, Anna Dominiczak and Alex Alderton, Roberto Amato, Jeffrey Barrett, Sonia Goncalves, Ewan Harrison, David K. Jackson, Ian Johnston, Dominic Kwiatkowski, Cordelia Langford, John Sillitoe on behalf of the Wellcome Sanger Institute COVID-19 Surveillance Team |
| EPI_ISL_2641002 | Lighthouse Lab in Alderley Park | Wellcome Sanger Institute for the COVID-19 Genomics UK (COG-UK) Consortium | Jacquelyn Wynn, Mairead Hyland, The Lighthouse Lab in Alderley Park and Alex Alderton, Roberto Amato, Jeffrey Barrett, Sonia Goncalves, Ewan Harrison, David K. Jackson, Ian Johnston, Dominic Kwiatkowski, Cordelia Langford, John Sillitoe on behalf of the Wellcome Sanger Institute COVID-19 Surveillance Team |
| EPI_ISL_2641003 | Lighthouse Lab in Milton Keynes | Wellcome Sanger Institute for the COVID-19 Genomics UK (COG-UK) Consortium | The Lighthouse Lab in Milton Keynes and Alex Alderton, Roberto Amato, Jeffrey Barrett, Sonia Goncalves, Ewan Harrison, David K. Jackson, Ian Johnston, Dominic Kwiatkowski, Cordelia Langford, John Sillitoe on behalf of the Wellcome Sanger Institute COVID-19 Surveillance Team |
| EPI_ISL_2641004, EPI_ISL_2641005, EPI_ISL_2641007, EPI_ISL_2641008, EPI_ISL_2641009, EPI_ISL_2641010 | Lighthouse Lab in Alderley Park | Wellcome Sanger Institute for the COVID-19 Genomics UK (COG-UK) Consortium | Jacquelyn Wynn, Mairead Hyland, The Lighthouse Lab in Alderley Park and Alex Alderton, Roberto Amato, Jeffrey Barrett, Sonia Goncalves, Ewan Harrison, David K. Jackson, Ian Johnston, Dominic Kwiatkowski, Cordelia Langford, John Sillitoe on behalf of the Wellcome Sanger Institute COVID-19 Surveillance Team |
| EPI_ISL_2641011 | Lighthouse Lab in Milton Keynes | Wellcome Sanger Institute for the COVID-19 Genomics UK (COG-UK) Consortium | The Lighthouse Lab in Milton Keynes and Alex Alderton, Roberto Amato, Jeffrey Barrett, Sonia Goncalves, Ewan Harrison, David K. Jackson, Ian Johnston, Dominic Kwiatkowski, Cordelia Langford, John Sillitoe on behalf of the Wellcome Sanger Institute COVID-19 Surveillance Team |
| EPI_ISL_2641012 | Lighthouse Lab in Glasgow | Wellcome Sanger Institute for the COVID-19 Genomics UK (COG-UK) Consortium | Harper VanSteenhouse, Yumi Kasai, David Gray, Carol Clugston, Anna Dominiczak and Alex Alderton, Roberto Amato, Jeffrey Barrett, Sonia Goncalves, Ewan Harrison, David K. Jackson, Ian Johnston, Dominic Kwiatkowski, Cordelia Langford, John Sillitoe on behalf of the Wellcome Sanger Institute COVID-19 Surveillance Team |
| EPI_ISL_2641013, EPI_ISL_2641014 | Lighthouse Lab in Alderley Park | Wellcome Sanger Institute for the COVID-19 Genomics UK (COG-UK) Consortium | Jacquelyn Wynn, Mairead Hyland, The Lighthouse Lab in Alderley Park and Alex Alderton, Roberto Amato, Jeffrey Barrett, Sonia Goncalves, Ewan Harrison, David K. Jackson, Ian Johnston, Dominic Kwiatkowski, Cordelia Langford, John Sillitoe on behalf of the Wellcome Sanger Institute COVID-19 Surveillance Team |
| EPI_ISL_2641015 | Lighthouse Lab in Milton Keynes | Wellcome Sanger Institute for the COVID-19 Genomics UK (COG-UK) Consortium | The Lighthouse Lab in Milton Keynes and Alex Alderton, Roberto Amato, Jeffrey Barrett, Sonia Goncalves, Ewan Harrison, David K. Jackson, Ian Johnston, Dominic Kwiatkowski, Cordelia Langford, John Sillitoe on behalf of the Wellcome Sanger Institute COVID-19 Surveillance Team |
| EPI_ISL_2641016, EPI_ISL_2641017, EPI_ISL_2641018 | Lighthouse Lab in Alderley Park | Wellcome Sanger Institute for the COVID-19 Genomics UK (COG-UK) Consortium | Jacquelyn Wynn, Mairead Hyland, The Lighthouse Lab in Alderley Park and Alex Alderton, Roberto Amato, Jeffrey Barrett, Sonia Goncalves, Ewan Harrison, David K. Jackson, Ian Johnston, Dominic Kwiatkowski, Cordelia Langford, John Sillitoe on behalf of the Wellcome Sanger Institute COVID-19 Surveillance Team |
| EPI_ISL_2641019 | Lighthouse Lab in Glasgow | Wellcome Sanger Institute for the COVID-19 Genomics UK (COG-UK) Consortium | Harper VanSteenhouse, Yumi Kasai, David Gray, Carol Clugston, Anna Dominiczak and Alex Alderton, Roberto Amato, Jeffrey Barrett, Sonia Goncalves, Ewan Harrison, David K. Jackson, Ian Johnston, Dominic Kwiatkowski, Cordelia Langford, John Sillitoe on behalf of the Wellcome Sanger Institute COVID-19 Surveillance Team |
| EPI_ISL_2641020, EPI_ISL_2641021, EPI_ISL_2641022, EPI_ISL_2641023, EPI_ISL_2641024 | Lighthouse Lab in Alderley Park | Wellcome Sanger Institute for the COVID-19 Genomics UK (COG-UK) Consortium | Jacquelyn Wynn, Mairead Hyland, The Lighthouse Lab in Alderley Park and Alex Alderton, Roberto Amato, Jeffrey Barrett, Sonia Goncalves, Ewan Harrison, David K. Jackson, Ian Johnston, Dominic Kwiatkowski, Cordelia Langford, John Sillitoe on behalf of the Wellcome Sanger Institute COVID-19 Surveillance Team |
| EPI_ISL_2641025 | Lighthouse Lab in Milton Keynes | Wellcome Sanger Institute for the COVID-19 Genomics UK (COG-UK) Consortium | The Lighthouse Lab in Milton Keynes and Alex Alderton, Roberto Amato, Jeffrey Barrett, Sonia Goncalves, Ewan Harrison, David K. Jackson, Ian Johnston, Dominic Kwiatkowski, Cordelia Langford, John Sillitoe on behalf of the Wellcome Sanger Institute COVID-19 Surveillance Team |
| EPI_ISL_2641026 | Lighthouse Lab in Alderley Park | Wellcome Sanger Institute for the COVID-19 Genomics UK (COG-UK) Consortium | Jacquelyn Wynn, Mairead Hyland, The Lighthouse Lab in Alderley Park and Alex Alderton, Roberto Amato, Jeffrey Barrett, Sonia Goncalves, Ewan Harrison, David K. Jackson, Ian Johnston, Dominic Kwiatkowski, Cordelia Langford, John Sillitoe on behalf of the Wellcome Sanger Institute COVID-19 Surveillance Team |
| EPI_ISL_2641027 | Lighthouse Lab in Glasgow | Wellcome Sanger Institute for the COVID-19 Genomics UK (COG-UK) Consortium | Harper VanSteenhouse, Yumi Kasai, David Gray, Carol Clugston, Anna Dominiczak and Alex Alderton, Roberto Amato, Jeffrey Barrett, Sonia Goncalves, Ewan Harrison, David K. Jackson, Ian Johnston, Dominic Kwiatkowski, Cordelia Langford, John Sillitoe on behalf of the Wellcome Sanger Institute COVID-19 Surveillance Team |
| EPI_ISL_2641028, EPI_ISL_2641029 | Lighthouse Lab in Alderley Park | Wellcome Sanger Institute for the COVID-19 Genomics UK (COG-UK) Consortium | Jacquelyn Wynn, Mairead Hyland, The Lighthouse Lab in Alderley Park and Alex Alderton, Roberto Amato, Jeffrey Barrett, Sonia Goncalves, Ewan Harrison, David K. Jackson, Ian Johnston, Dominic Kwiatkowski, Cordelia Langford, John Sillitoe on behalf of the Wellcome Sanger Institute COVID-19 Surveillance Team |
| EPI_ISL_2641030, EPI_ISL_2641031 | Lighthouse Lab in Milton Keynes | Wellcome Sanger Institute for the COVID-19 Genomics UK (COG-UK) Consortium | The Lighthouse Lab in Milton Keynes and Alex Alderton, Roberto Amato, Jeffrey Barrett, Sonia Goncalves, Ewan Harrison, David K. Jackson, Ian Johnston, Dominic Kwiatkowski, Cordelia Langford, John Sillitoe on behalf of the Wellcome Sanger Institute COVID-19 Surveillance Team |
| EPI_ISL_2641032 | Lighthouse Lab in Alderley Park | Wellcome Sanger Institute for the COVID-19 Genomics UK (COG-UK) Consortium | Jacquelyn Wynn, Mairead Hyland, The Lighthouse Lab in Alderley Park and Alex Alderton, Roberto Amato, Jeffrey Barrett, Sonia Goncalves, Ewan Harrison, David K. Jackson, Ian Johnston, Dominic Kwiatkowski, Cordelia Langford, John Sillitoe on behalf of the Wellcome Sanger Institute COVID-19 Surveillance Team |
| EPI_ISL_2641033 | Lighthouse Lab in Milton Keynes | Wellcome Sanger Institute for the COVID-19 Genomics UK (COG-UK) Consortium | The Lighthouse Lab in Milton Keynes and Alex Alderton, Roberto Amato, Jeffrey Barrett, Sonia Goncalves, Ewan Harrison, David K. Jackson, Ian Johnston, Dominic Kwiatkowski, Cordelia Langford, John Sillitoe on behalf of the Wellcome Sanger Institute COVID-19 Surveillance Team |
| EPI_ISL_2641034 | Lighthouse Lab in Alderley Park | Wellcome Sanger Institute for the COVID-19 Genomics UK (COG-UK) Consortium | Jacquelyn Wynn, Mairead Hyland, The Lighthouse Lab in Alderley Park and Alex Alderton, Roberto Amato, Jeffrey Barrett, Sonia Goncalves, Ewan Harrison, David K. Jackson, Ian Johnston, Dominic Kwiatkowski, Cordelia Langford, John Sillitoe on behalf of the Wellcome Sanger Institute COVID-19 Surveillance Team |
| EPI_ISL_2641035 | Lighthouse Lab in Milton Keynes | Wellcome Sanger Institute for the COVID-19 Genomics UK (COG-UK) Consortium | The Lighthouse Lab in Milton Keynes and Alex Alderton, Roberto Amato, Jeffrey Barrett, Sonia Goncalves, Ewan Harrison, David K. Jackson, Ian Johnston, Dominic Kwiatkowski, Cordelia Langford, John Sillitoe on behalf of the Wellcome Sanger Institute COVID-19 Surveillance Team |
| EPI_ISL_2641036, EPI_ISL_2641038 | Lighthouse Lab in Alderley Park | Wellcome Sanger Institute for the COVID-19 Genomics UK (COG-UK) Consortium | Jacquelyn Wynn, Mairead Hyland, The Lighthouse Lab in Alderley Park and Alex Alderton, Roberto Amato, Jeffrey Barrett, Sonia Goncalves, Ewan Harrison, David K. Jackson, Ian Johnston, Dominic Kwiatkowski, Cordelia Langford, John Sillitoe on behalf of the Wellcome Sanger Institute COVID-19 Surveillance Team |
| EPI_ISL_2641039, EPI_ISL_2641041 | Lighthouse Lab in Glasgow | Wellcome Sanger Institute for the COVID-19 Genomics UK (COG-UK) Consortium | Harper VanSteenhouse, Yumi Kasai, David Gray, Carol Clugston, Anna Dominiczak and Alex Alderton, Roberto Amato, Jeffrey Barrett, Sonia Goncalves, Ewan Harrison, David K. Jackson, Ian Johnston, Dominic Kwiatkowski, Cordelia Langford, John Sillitoe on behalf of the Wellcome Sanger Institute COVID-19 Surveillance Team |
| EPI_ISL_2641042 | Lighthouse Lab in Alderley Park | Wellcome Sanger Institute for the COVID-19 Genomics UK (COG-UK) Consortium | Jacquelyn Wynn, Mairead Hyland, The Lighthouse Lab in Alderley Park and Alex Alderton, Roberto Amato, Jeffrey Barrett, Sonia Goncalves, Ewan Harrison, David K. Jackson, Ian Johnston, Dominic Kwiatkowski, Cordelia Langford, John Sillitoe on behalf of the Wellcome Sanger Institute COVID-19 Surveillance Team |
| EPI_ISL_2641043 | Lighthouse Lab in Milton Keynes | Wellcome Sanger Institute for the COVID-19 Genomics UK (COG-UK) Consortium | The Lighthouse Lab in Milton Keynes and Alex Alderton, Roberto Amato, Jeffrey Barrett, Sonia Goncalves, Ewan Harrison, David K. Jackson, Ian Johnston, Dominic Kwiatkowski, Cordelia Langford, John Sillitoe on behalf of the Wellcome Sanger Institute COVID-19 Surveillance Team |
| EPI_ISL_2641044, EPI_ISL_2641045 | Lighthouse Lab in Alderley Park | Wellcome Sanger Institute for the COVID-19 Genomics UK (COG-UK) Consortium | Jacquelyn Wynn, Mairead Hyland, The Lighthouse Lab in Alderley Park and Alex Alderton, Roberto Amato, Jeffrey Barrett, Sonia Goncalves, Ewan Harrison, David K. Jackson, Ian Johnston, Dominic Kwiatkowski, Cordelia Langford, John Sillitoe on behalf of the Wellcome Sanger Institute COVID-19 Surveillance Team |
| EPI_ISL_2641046 | Lighthouse Lab in Glasgow | Wellcome Sanger Institute for the COVID-19 Genomics UK (COG-UK) Consortium | Harper VanSteenhouse, Yumi Kasai, David Gray, Carol Clugston, Anna Dominiczak and Alex Alderton, Roberto Amato, Jeffrey Barrett, Sonia Goncalves, Ewan Harrison, David K. Jackson, Ian Johnston, Dominic Kwiatkowski, Cordelia Langford, John Sillitoe on behalf of the Wellcome Sanger Institute COVID-19 Surveillance Team |
| EPI_ISL_2641047 | Lighthouse Lab in Milton Keynes | Wellcome Sanger Institute for the COVID-19 Genomics UK (COG-UK) Consortium | The Lighthouse Lab in Milton Keynes and Alex Alderton, Roberto Amato, Jeffrey Barrett, Sonia Goncalves, Ewan Harrison, David K. Jackson, Ian Johnston, Dominic Kwiatkowski, Cordelia Langford, John Sillitoe on behalf of the Wellcome Sanger Institute COVID-19 Surveillance Team |
| EPI_ISL_2641048, EPI_ISL_2641049, EPI_ISL_2641050 | Lighthouse Lab in Alderley Park | Wellcome Sanger Institute for the COVID-19 Genomics UK (COG-UK) Consortium | Jacquelyn Wynn, Mairead Hyland, The Lighthouse Lab in Alderley Park and Alex Alderton, Roberto Amato, Jeffrey Barrett, Sonia Goncalves, Ewan Harrison, David K. Jackson, Ian Johnston, Dominic Kwiatkowski, Cordelia Langford, John Sillitoe on behalf of the Wellcome Sanger Institute COVID-19 Surveillance Team |
| EPI_ISL_2641051, EPI_ISL_2641052, EPI_ISL_2641053 | Lighthouse Lab in Milton Keynes | Wellcome Sanger Institute for the COVID-19 Genomics UK (COG-UK) Consortium | The Lighthouse Lab in Milton Keynes and Alex Alderton, Roberto Amato, Jeffrey Barrett, Sonia Goncalves, Ewan Harrison, David K. Jackson, Ian Johnston, Dominic Kwiatkowski, Cordelia Langford, John Sillitoe on behalf of the Wellcome Sanger Institute COVID-19 Surveillance Team |
| EPI_ISL_2641054, EPI_ISL_2641055, EPI_ISL_2641056, EPI_ISL_2641057, EPI_ISL_2641058 | Lighthouse Lab in Alderley Park | Wellcome Sanger Institute for the COVID-19 Genomics UK (COG-UK) Consortium | Jacquelyn Wynn, Mairead Hyland, The Lighthouse Lab in Alderley Park and Alex Alderton, Roberto Amato, Jeffrey Barrett, Sonia Goncalves, Ewan Harrison, David K. Jackson, Ian Johnston, Dominic Kwiatkowski, Cordelia Langford, John Sillitoe on behalf of the Wellcome Sanger Institute COVID-19 Surveillance Team |
| EPI_ISL_2641060 | Lighthouse Lab in Glasgow | Wellcome Sanger Institute for the COVID-19 Genomics UK (COG-UK) Consortium | Harper VanSteenhouse, Yumi Kasai, David Gray, Carol Clugston, Anna Dominiczak and Alex Alderton, Roberto Amato, Jeffrey Barrett, Sonia Goncalves, Ewan Harrison, David K. Jackson, Ian Johnston, Dominic Kwiatkowski, Cordelia Langford, John Sillitoe on behalf of the Wellcome Sanger Institute COVID-19 Surveillance Team |
| EPI_ISL_2641061, EPI_ISL_2641062, EPI_ISL_2641064 | Lighthouse Lab in Milton Keynes | Wellcome Sanger Institute for the COVID-19 Genomics UK (COG-UK) Consortium | The Lighthouse Lab in Milton Keynes and Alex Alderton, Roberto Amato, Jeffrey Barrett, Sonia Goncalves, Ewan Harrison, David K. Jackson, Ian Johnston, Dominic Kwiatkowski, Cordelia Langford, John Sillitoe on behalf of the Wellcome Sanger Institute COVID-19 Surveillance Team |
| EPI_ISL_2641065 | Lighthouse Lab in Glasgow | Wellcome Sanger Institute for the COVID-19 Genomics UK (COG-UK) Consortium | Harper VanSteenhouse, Yumi Kasai, David Gray, Carol Clugston, Anna Dominiczak and Alex Alderton, Roberto Amato, Jeffrey Barrett, Sonia Goncalves, Ewan Harrison, David K. Jackson, Ian Johnston, Dominic Kwiatkowski, Cordelia Langford, John Sillitoe on behalf of the Wellcome Sanger Institute COVID-19 Surveillance Team |
| EPI_ISL_2641066 | Lighthouse Lab in Alderley Park | Wellcome Sanger Institute for the COVID-19 Genomics UK (COG-UK) Consortium | Jacquelyn Wynn, Mairead Hyland, The Lighthouse Lab in Alderley Park and Alex Alderton, Roberto Amato, Jeffrey Barrett, Sonia Goncalves, Ewan Harrison, David K. Jackson, Ian Johnston, Dominic Kwiatkowski, Cordelia Langford, John Sillitoe on behalf of the Wellcome Sanger Institute COVID-19 Surveillance Team |
| EPI_ISL_2641067, EPI_ISL_2641068 | Lighthouse Lab in Glasgow | Wellcome Sanger Institute for the COVID-19 Genomics UK (COG-UK) Consortium | Harper VanSteenhouse, Yumi Kasai, David Gray, Carol Clugston, Anna Dominiczak and Alex Alderton, Roberto Amato, Jeffrey Barrett, Sonia Goncalves, Ewan Harrison, David K. Jackson, Ian Johnston, Dominic Kwiatkowski, Cordelia Langford, John Sillitoe on behalf of the Wellcome Sanger Institute COVID-19 Surveillance Team |
| EPI_ISL_2641069, EPI_ISL_2641070, EPI_ISL_2641071 | Lighthouse Lab in Alderley Park | Wellcome Sanger Institute for the COVID-19 Genomics UK (COG-UK) Consortium | Jacquelyn Wynn, Mairead Hyland, The Lighthouse Lab in Alderley Park and Alex Alderton, Roberto Amato, Jeffrey Barrett, Sonia Goncalves, Ewan Harrison, David K. Jackson, Ian Johnston, Dominic Kwiatkowski, Cordelia Langford, John Sillitoe on behalf of the Wellcome Sanger Institute COVID-19 Surveillance Team |
| EPI_ISL_2641072 | Lighthouse Lab in Milton Keynes | Wellcome Sanger Institute for the COVID-19 Genomics UK (COG-UK) Consortium | The Lighthouse Lab in Milton Keynes and Alex Alderton, Roberto Amato, Jeffrey Barrett, Sonia Goncalves, Ewan Harrison, David K. Jackson, Ian Johnston, Dominic Kwiatkowski, Cordelia Langford, John Sillitoe on behalf of the Wellcome Sanger Institute COVID-19 Surveillance Team |
| EPI_ISL_2641073 | Lighthouse Lab in Alderley Park | Wellcome Sanger Institute for the COVID-19 Genomics UK (COG-UK) Consortium | Jacquelyn Wynn, Mairead Hyland, The Lighthouse Lab in Alderley Park and Alex Alderton, Roberto Amato, Jeffrey Barrett, Sonia Goncalves, Ewan Harrison, David K. Jackson, Ian Johnston, Dominic Kwiatkowski, Cordelia Langford, John Sillitoe on behalf of the Wellcome Sanger Institute COVID-19 Surveillance Team |

| | | | |
|---|---|---|---|
| EPI_ISL_2641074, EPI_ISL_2641075 | Lighthouse Lab in Milton Keynes | Wellcome Sanger Institute for the COVID-19 Genomics UK (COG-UK) Consortium | The Lighthouse Lab in Milton Keynes and Alex Alderton, Roberto Amato, Jeffrey Barrett, Sonia Goncalves, Ewan Harrison, David K. Jackson, Ian Johnston, Dominic Kwiatkowski, Cordelia Langford, John Sillitoe on behalf of the Wellcome Sanger Institute COVID-19 Surveillance Team |
| EPI_ISL_2641076, EPI_ISL_2641077 | Lighthouse Lab in Alderley Park | Wellcome Sanger Institute for the COVID-19 Genomics UK (COG-UK) Consortium | Jacquelyn Wynn, Mairead Hyland, The Lighthouse Lab in Alderley Park and Alex Alderton, Roberto Amato, Jeffrey Barrett, Sonia Goncalves, Ewan Harrison, David K. Jackson, Ian Johnston, Dominic Kwiatkowski, Cordelia Langford, John Sillitoe on behalf of the Wellcome Sanger Institute COVID-19 Surveillance Team |
| EPI_ISL_2641078 | Lighthouse Lab in Milton Keynes | Wellcome Sanger Institute for the COVID-19 Genomics UK (COG-UK) Consortium | The Lighthouse Lab in Milton Keynes and Alex Alderton, Roberto Amato, Jeffrey Barrett, Sonia Goncalves, Ewan Harrison, David K. Jackson, Ian Johnston, Dominic Kwiatkowski, Cordelia Langford, John Sillitoe on behalf of the Wellcome Sanger Institute COVID-19 Surveillance Team |
| EPI_ISL_2641079 | Lighthouse Lab in Alderley Park | Wellcome Sanger Institute for the COVID-19 Genomics UK (COG-UK) Consortium | Jacquelyn Wynn, Mairead Hyland, The Lighthouse Lab in Alderley Park and Alex Alderton, Roberto Amato, Jeffrey Barrett, Sonia Goncalves, Ewan Harrison, David K. Jackson, Ian Johnston, Dominic Kwiatkowski, Cordelia Langford, John Sillitoe on behalf of the Wellcome Sanger Institute COVID-19 Surveillance Team |
| EPI_ISL_2641080 | Lighthouse Lab in Milton Keynes | Wellcome Sanger Institute for the COVID-19 Genomics UK (COG-UK) Consortium | The Lighthouse Lab in Milton Keynes and Alex Alderton, Roberto Amato, Jeffrey Barrett, Sonia Goncalves, Ewan Harrison, David K. Jackson, Ian Johnston, Dominic Kwiatkowski, Cordelia Langford, John Sillitoe on behalf of the Wellcome Sanger Institute COVID-19 Surveillance Team |
| EPI_ISL_2641081 | Lighthouse Lab in Glasgow | Wellcome Sanger Institute for the COVID-19 Genomics UK (COG-UK) Consortium | Harper VanSteenhouse, Yumi Kasai, David Gray, Carol Clugston, Anna Dominiczak and Alex Alderton, Roberto Amato, Jeffrey Barrett, Sonia Goncalves, Ewan Harrison, David K. Jackson, Ian Johnston, Dominic Kwiatkowski, Cordelia Langford, John Sillitoe on behalf of the Wellcome Sanger Institute COVID-19 Surveillance Team |
| EPI_ISL_2641082 | Lighthouse Lab in Alderley Park | Wellcome Sanger Institute for the COVID-19 Genomics UK (COG-UK) Consortium | Jacquelyn Wynn, Mairead Hyland, The Lighthouse Lab in Alderley Park and Alex Alderton, Roberto Amato, Jeffrey Barrett, Sonia Goncalves, Ewan Harrison, David K. Jackson, Ian Johnston, Dominic Kwiatkowski, Cordelia Langford, John Sillitoe on behalf of the Wellcome Sanger Institute COVID-19 Surveillance Team |
| EPI_ISL_2641083, EPI_ISL_2641084 | Lighthouse Lab in Milton Keynes | Wellcome Sanger Institute for the COVID-19 Genomics UK (COG-UK) Consortium | The Lighthouse Lab in Milton Keynes and Alex Alderton, Roberto Amato, Jeffrey Barrett, Sonia Goncalves, Ewan Harrison, David K. Jackson, Ian Johnston, Dominic Kwiatkowski, Cordelia Langford, John Sillitoe on behalf of the Wellcome Sanger Institute COVID-19 Surveillance Team |
| EPI_ISL_2641085 | Lighthouse Lab in Glasgow | Wellcome Sanger Institute for the COVID-19 Genomics UK (COG-UK) Consortium | Harper VanSteenhouse, Yumi Kasai, David Gray, Carol Clugston, Anna Dominiczak and Alex Alderton, Roberto Amato, Jeffrey Barrett, Sonia Goncalves, Ewan Harrison, David K. Jackson, Ian Johnston, Dominic Kwiatkowski, Cordelia Langford, John Sillitoe on behalf of the Wellcome Sanger Institute COVID-19 Surveillance Team |
| EPI_ISL_2641086, EPI_ISL_2641087, EPI_ISL_2641088 | Lighthouse Lab in Alderley Park | Wellcome Sanger Institute for the COVID-19 Genomics UK (COG-UK) Consortium | Jacquelyn Wynn, Mairead Hyland, The Lighthouse Lab in Alderley Park and Alex Alderton, Roberto Amato, Jeffrey Barrett, Sonia Goncalves, Ewan Harrison, David K. Jackson, Ian Johnston, Dominic Kwiatkowski, Cordelia Langford, John Sillitoe on behalf of the Wellcome Sanger Institute COVID-19 Surveillance Team |
| EPI_ISL_2641089 | Lighthouse Lab in Glasgow | Wellcome Sanger Institute for the COVID-19 Genomics UK (COG-UK) Consortium | Harper VanSteenhouse, Yumi Kasai, David Gray, Carol Clugston, Anna Dominiczak and Alex Alderton, Roberto Amato, Jeffrey Barrett, Sonia Goncalves, Ewan Harrison, David K. Jackson, Ian Johnston, Dominic Kwiatkowski, Cordelia Langford, John Sillitoe on behalf of the Wellcome Sanger Institute COVID-19 Surveillance Team |
| EPI_ISL_2641090 | Lighthouse Lab in Alderley Park | Wellcome Sanger Institute for the COVID-19 Genomics UK (COG-UK) Consortium | Jacquelyn Wynn, Mairead Hyland, The Lighthouse Lab in Alderley Park and Alex Alderton, Roberto Amato, Jeffrey Barrett, Sonia Goncalves, Ewan Harrison, David K. Jackson, Ian Johnston, Dominic Kwiatkowski, Cordelia Langford, John Sillitoe on behalf of the Wellcome Sanger Institute COVID-19 Surveillance Team |
| EPI_ISL_2641091, EPI_ISL_2641092, EPI_ISL_2641093 | Lighthouse Lab in Glasgow | Wellcome Sanger Institute for the COVID-19 Genomics UK (COG-UK) Consortium | Harper VanSteenhouse, Yumi Kasai, David Gray, Carol Clugston, Anna Dominiczak and Alex Alderton, Roberto Amato, Jeffrey Barrett, Sonia Goncalves, Ewan Harrison, David K. Jackson, Ian Johnston, Dominic Kwiatkowski, Cordelia Langford, John Sillitoe on behalf of the Wellcome Sanger Institute COVID-19 Surveillance Team |
| EPI_ISL_2641094 | Lighthouse Lab in Milton Keynes | Wellcome Sanger Institute for the COVID-19 Genomics UK (COG-UK) Consortium | The Lighthouse Lab in Milton Keynes and Alex Alderton, Roberto Amato, Jeffrey Barrett, Sonia Goncalves, Ewan Harrison, David K. Jackson, Ian Johnston, Dominic Kwiatkowski, Cordelia Langford, John Sillitoe on behalf of the Wellcome Sanger Institute COVID-19 Surveillance Team |
| EPI_ISL_2641096 | Lighthouse Lab in Glasgow | Wellcome Sanger Institute for the COVID-19 Genomics UK (COG-UK) Consortium | Harper VanSteenhouse, Yumi Kasai, David Gray, Carol Clugston, Anna Dominiczak and Alex Alderton, Roberto Amato, Jeffrey Barrett, Sonia Goncalves, Ewan Harrison, David K. Jackson, Ian Johnston, Dominic Kwiatkowski, Cordelia Langford, John Sillitoe on behalf of the Wellcome Sanger Institute COVID-19 Surveillance Team |
| EPI_ISL_2641097 | Lighthouse Lab in Milton Keynes | Wellcome Sanger Institute for the COVID-19 Genomics UK (COG-UK) Consortium | The Lighthouse Lab in Milton Keynes and Alex Alderton, Roberto Amato, Jeffrey Barrett, Sonia Goncalves, Ewan Harrison, David K. Jackson, Ian Johnston, Dominic Kwiatkowski, Cordelia Langford, John Sillitoe on behalf of the Wellcome Sanger Institute COVID-19 Surveillance Team |
| EPI_ISL_2641098 | Lighthouse Lab in Alderley Park | Wellcome Sanger Institute for the COVID-19 Genomics UK (COG-UK) Consortium | Jacquelyn Wynn, Mairead Hyland, The Lighthouse Lab in Alderley Park and Alex Alderton, Roberto Amato, Jeffrey Barrett, Sonia Goncalves, Ewan Harrison, David K. Jackson, Ian Johnston, Dominic Kwiatkowski, Cordelia Langford, John Sillitoe on behalf of the Wellcome Sanger Institute COVID-19 Surveillance Team |
| EPI_ISL_2641099, EPI_ISL_2641100 | Lighthouse Lab in Milton Keynes | Wellcome Sanger Institute for the COVID-19 Genomics UK (COG-UK) Consortium | The Lighthouse Lab in Milton Keynes and Alex Alderton, Roberto Amato, Jeffrey Barrett, Sonia Goncalves, Ewan Harrison, David K. Jackson, Ian Johnston, Dominic Kwiatkowski, Cordelia Langford, John Sillitoe on behalf of the Wellcome Sanger Institute COVID-19 Surveillance Team |
| EPI_ISL_2641101, EPI_ISL_2641102, EPI_ISL_2641103 | Lighthouse Lab in Alderley Park | Wellcome Sanger Institute for the COVID-19 Genomics UK (COG-UK) Consortium | Jacquelyn Wynn, Mairead Hyland, The Lighthouse Lab in Alderley Park and Alex Alderton, Roberto Amato, Jeffrey Barrett, Sonia Goncalves, Ewan Harrison, David K. Jackson, Ian Johnston, Dominic Kwiatkowski, Cordelia Langford, John Sillitoe on behalf of the Wellcome Sanger Institute COVID-19 Surveillance Team |
| EPI_ISL_2641104 | Lighthouse Lab in Glasgow | Wellcome Sanger Institute for the COVID-19 Genomics UK (COG-UK) Consortium | Harper VanSteenhouse, Yumi Kasai, David Gray, Carol Clugston, Anna Dominiczak and Alex Alderton, Roberto Amato, Jeffrey Barrett, Sonia Goncalves, Ewan Harrison, David K. Jackson, Ian Johnston, Dominic Kwiatkowski, Cordelia Langford, John Sillitoe on behalf of the Wellcome Sanger Institute COVID-19 Surveillance Team |
| EPI_ISL_2641105, EPI_ISL_2641106 | Lighthouse Lab in Alderley Park | Wellcome Sanger Institute for the COVID-19 Genomics UK (COG-UK) Consortium | Jacquelyn Wynn, Mairead Hyland, The Lighthouse Lab in Alderley Park and Alex Alderton, Roberto Amato, Jeffrey Barrett, Sonia Goncalves, Ewan Harrison, David K. Jackson, Ian Johnston, Dominic Kwiatkowski, Cordelia Langford, John Sillitoe on behalf of the Wellcome Sanger Institute COVID-19 Surveillance Team |
| EPI_ISL_2641107 | Lighthouse Lab in Glasgow | Wellcome Sanger Institute for the COVID-19 Genomics UK (COG-UK) Consortium | Harper VanSteenhouse, Yumi Kasai, David Gray, Carol Clugston, Anna Dominiczak and Alex Alderton, Roberto Amato, Jeffrey Barrett, Sonia Goncalves, Ewan Harrison, David K. Jackson, Ian Johnston, Dominic Kwiatkowski, Cordelia Langford, John Sillitoe on behalf of the Wellcome Sanger Institute COVID-19 Surveillance Team |
| EPI_ISL_2641108, EPI_ISL_2641109 | Lighthouse Lab in Alderley Park | Wellcome Sanger Institute for the COVID-19 Genomics UK (COG-UK) Consortium | Jacquelyn Wynn, Mairead Hyland, The Lighthouse Lab in Alderley Park and Alex Alderton, Roberto Amato, Jeffrey Barrett, Sonia Goncalves, Ewan Harrison, David K. Jackson, Ian Johnston, Dominic Kwiatkowski, Cordelia Langford, John Sillitoe on behalf of the Wellcome Sanger Institute COVID-19 Surveillance Team |
| EPI_ISL_2641110, EPI_ISL_2641111, EPI_ISL_2641112, EPI_ISL_2641113 | Lighthouse Lab in Milton Keynes | Wellcome Sanger Institute for the COVID-19 Genomics UK (COG-UK) Consortium | The Lighthouse Lab in Milton Keynes and Alex Alderton, Roberto Amato, Jeffrey Barrett, Sonia Goncalves, Ewan Harrison, David K. Jackson, Ian Johnston, Dominic Kwiatkowski, Cordelia Langford, John Sillitoe on behalf of the Wellcome Sanger Institute COVID-19 Surveillance Team |
| EPI_ISL_2641114, EPI_ISL_2641115, EPI_ISL_2641116 | Lighthouse Lab in Alderley Park | Wellcome Sanger Institute for the COVID-19 Genomics UK (COG-UK) Consortium | Jacquelyn Wynn, Mairead Hyland, The Lighthouse Lab in Alderley Park and Alex Alderton, Roberto Amato, Jeffrey Barrett, Sonia Goncalves, Ewan Harrison, David K. Jackson, Ian Johnston, Dominic Kwiatkowski, Cordelia Langford, John Sillitoe on behalf of the Wellcome Sanger Institute COVID-19 Surveillance Team |
| EPI_ISL_2641118, EPI_ISL_2641119 | Lighthouse Lab in Milton Keynes | Wellcome Sanger Institute for the COVID-19 Genomics UK (COG-UK) Consortium | The Lighthouse Lab in Milton Keynes and Alex Alderton, Roberto Amato, Jeffrey Barrett, Sonia Goncalves, Ewan Harrison, David K. Jackson, Ian Johnston, Dominic Kwiatkowski, Cordelia Langford, John Sillitoe on behalf of the Wellcome Sanger Institute COVID-19 Surveillance Team |
| EPI_ISL_2641120, EPI_ISL_2641121 | Lighthouse Lab in Alderley Park | Wellcome Sanger Institute for the COVID-19 Genomics UK (COG-UK) Consortium | Jacquelyn Wynn, Mairead Hyland, The Lighthouse Lab in Alderley Park and Alex Alderton, Roberto Amato, Jeffrey Barrett, Sonia Goncalves, Ewan Harrison, David K. Jackson, Ian Johnston, Dominic Kwiatkowski, Cordelia Langford, John Sillitoe on behalf of the Wellcome Sanger Institute COVID-19 Surveillance Team |
| EPI_ISL_2641122 | Lighthouse Lab in Glasgow | Wellcome Sanger Institute for the COVID-19 Genomics UK (COG-UK) Consortium | Harper VanSteenhouse, Yumi Kasai, David Gray, Carol Clugston, Anna Dominiczak and Alex Alderton, Roberto Amato, Jeffrey Barrett, Sonia Goncalves, Ewan Harrison, David K. Jackson, Ian Johnston, Dominic Kwiatkowski, Cordelia Langford, John Sillitoe on behalf of the Wellcome Sanger Institute COVID-19 Surveillance Team |
| EPI_ISL_2641123 | Lighthouse Lab in Alderley Park | Wellcome Sanger Institute for the COVID-19 Genomics UK (COG-UK) Consortium | Jacquelyn Wynn, Mairead Hyland, The Lighthouse Lab in Alderley Park and Alex Alderton, Roberto Amato, Jeffrey Barrett, Sonia Goncalves, Ewan Harrison, David K. Jackson, Ian Johnston, Dominic Kwiatkowski, Cordelia Langford, John Sillitoe on behalf of the Wellcome Sanger Institute COVID-19 Surveillance Team |
| EPI_ISL_2641124 | Lighthouse Lab in Glasgow | Wellcome Sanger Institute for the COVID-19 Genomics UK (COG-UK) Consortium | Harper VanSteenhouse, Yumi Kasai, David Gray, Carol Clugston, Anna Dominiczak and Alex Alderton, Roberto Amato, Jeffrey Barrett, Sonia Goncalves, Ewan Harrison, David K. Jackson, Ian Johnston, Dominic Kwiatkowski, Cordelia Langford, John Sillitoe on behalf of the Wellcome Sanger Institute COVID-19 Surveillance Team |
| EPI_ISL_2641126, EPI_ISL_2641127, EPI_ISL_2641128 | Lighthouse Lab in Milton Keynes | Wellcome Sanger Institute for the COVID-19 Genomics UK (COG-UK) Consortium | The Lighthouse Lab in Milton Keynes and Alex Alderton, Roberto Amato, Jeffrey Barrett, Sonia Goncalves, Ewan Harrison, David K. Jackson, Ian Johnston, Dominic Kwiatkowski, Cordelia Langford, John Sillitoe on behalf of the Wellcome Sanger Institute COVID-19 Surveillance Team |
| EPI_ISL_2641129, EPI_ISL_2641131 | Lighthouse Lab in Alderley Park | Wellcome Sanger Institute for the COVID-19 Genomics UK (COG-UK) Consortium | Jacquelyn Wynn, Mairead Hyland, The Lighthouse Lab in Alderley Park and Alex Alderton, Roberto Amato, Jeffrey Barrett, Sonia Goncalves, Ewan Harrison, David K. Jackson, Ian Johnston, Dominic Kwiatkowski, Cordelia Langford, John Sillitoe on behalf of the Wellcome Sanger Institute COVID-19 Surveillance Team |
| EPI_ISL_2641132 | Lighthouse Lab in Glasgow | Wellcome Sanger Institute for the COVID-19 Genomics UK (COG-UK) Consortium | Harper VanSteenhouse, Yumi Kasai, David Gray, Carol Clugston, Anna Dominiczak and Alex Alderton, Roberto Amato, Jeffrey Barrett, Sonia Goncalves, Ewan Harrison, David K. Jackson, Ian Johnston, Dominic Kwiatkowski, Cordelia Langford, John Sillitoe on behalf of the Wellcome Sanger Institute COVID-19 Surveillance Team |
| EPI_ISL_2641133, EPI_ISL_2641134, EPI_ISL_2641135 | Lighthouse Lab in Alderley Park | Wellcome Sanger Institute for the COVID-19 Genomics UK (COG-UK) Consortium | Jacquelyn Wynn, Mairead Hyland, The Lighthouse Lab in Alderley Park and Alex Alderton, Roberto Amato, Jeffrey Barrett, Sonia Goncalves, Ewan Harrison, David K. Jackson, Ian Johnston, Dominic Kwiatkowski, Cordelia Langford, John Sillitoe on behalf of the Wellcome Sanger Institute COVID-19 Surveillance Team |
| EPI_ISL_2641137, EPI_ISL_2641138 | Lighthouse Lab in Glasgow | Wellcome Sanger Institute for the COVID-19 Genomics UK (COG-UK) Consortium | Harper VanSteenhouse, Yumi Kasai, David Gray, Carol Clugston, Anna Dominiczak and Alex Alderton, Roberto Amato, Jeffrey Barrett, Sonia Goncalves, Ewan Harrison, David K. Jackson, Ian Johnston, Dominic Kwiatkowski, Cordelia Langford, John Sillitoe on behalf of the Wellcome Sanger Institute COVID-19 Surveillance Team |
| EPI_ISL_2641139 | Lighthouse Lab in Alderley Park | Wellcome Sanger Institute for the COVID-19 Genomics UK (COG-UK) Consortium | Jacquelyn Wynn, Mairead Hyland, The Lighthouse Lab in Alderley Park and Alex Alderton, Roberto Amato, Jeffrey Barrett, Sonia Goncalves, Ewan Harrison, David K. Jackson, Ian Johnston, Dominic Kwiatkowski, Cordelia Langford, John Sillitoe on behalf of the Wellcome Sanger Institute COVID-19 Surveillance Team |
| EPI_ISL_2641141, EPI_ISL_2641142 | Lighthouse Lab in Glasgow | Wellcome Sanger Institute for the COVID-19 Genomics UK (COG-UK) Consortium | Harper VanSteenhouse, Yumi Kasai, David Gray, Carol Clugston, Anna Dominiczak and Alex Alderton, Roberto Amato, Jeffrey Barrett, Sonia Goncalves, Ewan Harrison, David K. Jackson, Ian Johnston, Dominic Kwiatkowski, Cordelia Langford, John Sillitoe on behalf of the Wellcome Sanger Institute COVID-19 Surveillance Team |
| EPI_ISL_2641143 | Lighthouse Lab in Alderley Park | Wellcome Sanger Institute for the COVID-19 Genomics UK (COG-UK) Consortium | Jacquelyn Wynn, Mairead Hyland, The Lighthouse Lab in Alderley Park and Alex Alderton, Roberto Amato, Jeffrey Barrett, Sonia Goncalves, Ewan Harrison, David K. Jackson, Ian Johnston, Dominic Kwiatkowski, Cordelia Langford, John Sillitoe on behalf of the Wellcome Sanger Institute COVID-19 Surveillance Team |
| EPI_ISL_2641144, EPI_ISL_2641145 | Lighthouse Lab in Glasgow | Wellcome Sanger Institute for the COVID-19 Genomics UK (COG-UK) Consortium | Harper VanSteenhouse, Yumi Kasai, David Gray, Carol Clugston, Anna Dominiczak and Alex Alderton, Roberto Amato, Jeffrey Barrett, Sonia Goncalves, Ewan Harrison, David K. Jackson, Ian Johnston, Dominic Kwiatkowski, Cordelia Langford, John Sillitoe on behalf of the Wellcome Sanger Institute COVID-19 Surveillance Team |
| EPI_ISL_2641146, EPI_ISL_2641147 | Lighthouse Lab in Alderley Park | Wellcome Sanger Institute for the COVID-19 Genomics UK (COG-UK) Consortium | Jacquelyn Wynn, Mairead Hyland, The Lighthouse Lab in Alderley Park and Alex Alderton, Roberto Amato, Jeffrey Barrett, Sonia Goncalves, Ewan Harrison, David K. Jackson, Ian Johnston, Dominic Kwiatkowski, Cordelia Langford, John Sillitoe on behalf of the Wellcome Sanger Institute COVID-19 Surveillance Team |
| EPI_ISL_2641148, EPI_ISL_2641149, EPI_ISL_2641150 | Lighthouse Lab in Milton Keynes | Wellcome Sanger Institute for the COVID-19 Genomics UK (COG-UK) Consortium | The Lighthouse Lab in Milton Keynes and Alex Alderton, Roberto Amato, Jeffrey Barrett, Sonia Goncalves, Ewan Harrison, David K. Jackson, Ian Johnston, Dominic Kwiatkowski, Cordelia Langford, John Sillitoe on behalf of the Wellcome Sanger Institute COVID-19 Surveillance Team |
| EPI_ISL_2641151 | Lighthouse Lab in Alderley Park | Wellcome Sanger Institute for the COVID-19 Genomics UK (COG-UK) | Jacquelyn Wynn, Mairead Hyland, The Lighthouse Lab in Alderley Park and Alex Alderton, Roberto Amato, Jeffrey Barrett, Sonia Goncalves, Ewan Harrison, David K. Jackson, Ian Johnston, Dominic Kwiatkowski, Cordelia Langford, John Sillitoe on |

| | | | |
|---|---|---|---|
| | | Consortium | behalf of the Wellcome Sanger Institute COVID-19 Surveillance Team |
| EPI_ISL_2641152 | Lighthouse Lab in Milton Keynes | Wellcome Sanger Institute for the COVID-19 Genomics UK (COG-UK) Consortium | The Lighthouse Lab in Milton Keynes and Alex Alderton, Roberto Amato, Jeffrey Barrett, Sonia Goncalves, Ewan Harrison, David K. Jackson, Ian Johnston, Dominic Kwiatkowski, Cordelia Langford, John Sillitoe on behalf of the Wellcome Sanger Institute COVID-19 Surveillance Team |
| EPI_ISL_2641153, EPI_ISL_2641154 | Lighthouse Lab in Glasgow | Wellcome Sanger Institute for the COVID-19 Genomics UK (COG-UK) Consortium | Harper VanSteenhouse, Yumi Kasai, David Gray, Carol Clugston, Anna Dominiczak and Alex Alderton, Roberto Amato, Jeffrey Barrett, Sonia Goncalves, Ewan Harrison, David K. Jackson, Ian Johnston, Dominic Kwiatkowski, Cordelia Langford, John Sillitoe on behalf of the Wellcome Sanger Institute COVID-19 Surveillance Team |
| EPI_ISL_2641155 | Lighthouse Lab in Alderley Park | Wellcome Sanger Institute for the COVID-19 Genomics UK (COG-UK) Consortium | Jacquelyn Wynn, Mairead Hyland, The Lighthouse Lab in Alderley Park and Alex Alderton, Roberto Amato, Jeffrey Barrett, Sonia Goncalves, Ewan Harrison, David K. Jackson, Ian Johnston, Dominic Kwiatkowski, Cordelia Langford, John Sillitoe on behalf of the Wellcome Sanger Institute COVID-19 Surveillance Team |
| EPI_ISL_2641156 | Lighthouse Lab in Glasgow | Wellcome Sanger Institute for the COVID-19 Genomics UK (COG-UK) Consortium | Harper VanSteenhouse, Yumi Kasai, David Gray, Carol Clugston, Anna Dominiczak and Alex Alderton, Roberto Amato, Jeffrey Barrett, Sonia Goncalves, Ewan Harrison, David K. Jackson, Ian Johnston, Dominic Kwiatkowski, Cordelia Langford, John Sillitoe on behalf of the Wellcome Sanger Institute COVID-19 Surveillance Team |
| EPI_ISL_2641157, EPI_ISL_2641158 | Lighthouse Lab in Alderley Park | Wellcome Sanger Institute for the COVID-19 Genomics UK (COG-UK) Consortium | Jacquelyn Wynn, Mairead Hyland, The Lighthouse Lab in Alderley Park and Alex Alderton, Roberto Amato, Jeffrey Barrett, Sonia Goncalves, Ewan Harrison, David K. Jackson, Ian Johnston, Dominic Kwiatkowski, Cordelia Langford, John Sillitoe on behalf of the Wellcome Sanger Institute COVID-19 Surveillance Team |
| EPI_ISL_2641159, EPI_ISL_2641160 | Lighthouse Lab in Milton Keynes | Wellcome Sanger Institute for the COVID-19 Genomics UK (COG-UK) Consortium | The Lighthouse Lab in Milton Keynes and Alex Alderton, Roberto Amato, Jeffrey Barrett, Sonia Goncalves, Ewan Harrison, David K. Jackson, Ian Johnston, Dominic Kwiatkowski, Cordelia Langford, John Sillitoe on behalf of the Wellcome Sanger Institute COVID-19 Surveillance Team |
| EPI_ISL_2641161 | Lighthouse Lab in Alderley Park | Wellcome Sanger Institute for the COVID-19 Genomics UK (COG-UK) Consortium | Jacquelyn Wynn, Mairead Hyland, The Lighthouse Lab in Alderley Park and Alex Alderton, Roberto Amato, Jeffrey Barrett, Sonia Goncalves, Ewan Harrison, David K. Jackson, Ian Johnston, Dominic Kwiatkowski, Cordelia Langford, John Sillitoe on behalf of the Wellcome Sanger Institute COVID-19 Surveillance Team |
| EPI_ISL_2641162, EPI_ISL_2641163 | Lighthouse Lab in Milton Keynes | Wellcome Sanger Institute for the COVID-19 Genomics UK (COG-UK) Consortium | The Lighthouse Lab in Milton Keynes and Alex Alderton, Roberto Amato, Jeffrey Barrett, Sonia Goncalves, Ewan Harrison, David K. Jackson, Ian Johnston, Dominic Kwiatkowski, Cordelia Langford, John Sillitoe on behalf of the Wellcome Sanger Institute COVID-19 Surveillance Team |
| EPI_ISL_2641164, EPI_ISL_2641165 | Lighthouse Lab in Glasgow | Wellcome Sanger Institute for the COVID-19 Genomics UK (COG-UK) Consortium | Harper VanSteenhouse, Yumi Kasai, David Gray, Carol Clugston, Anna Dominiczak and Alex Alderton, Roberto Amato, Jeffrey Barrett, Sonia Goncalves, Ewan Harrison, David K. Jackson, Ian Johnston, Dominic Kwiatkowski, Cordelia Langford, John Sillitoe on behalf of the Wellcome Sanger Institute COVID-19 Surveillance Team |
| EPI_ISL_2641166 | Lighthouse Lab in Alderley Park | Wellcome Sanger Institute for the COVID-19 Genomics UK (COG-UK) Consortium | Jacquelyn Wynn, Mairead Hyland, The Lighthouse Lab in Alderley Park and Alex Alderton, Roberto Amato, Jeffrey Barrett, Sonia Goncalves, Ewan Harrison, David K. Jackson, Ian Johnston, Dominic Kwiatkowski, Cordelia Langford, John Sillitoe on behalf of the Wellcome Sanger Institute COVID-19 Surveillance Team |
| EPI_ISL_2641167, EPI_ISL_2641168 | Lighthouse Lab in Milton Keynes | Wellcome Sanger Institute for the COVID-19 Genomics UK (COG-UK) Consortium | The Lighthouse Lab in Milton Keynes and Alex Alderton, Roberto Amato, Jeffrey Barrett, Sonia Goncalves, Ewan Harrison, David K. Jackson, Ian Johnston, Dominic Kwiatkowski, Cordelia Langford, John Sillitoe on behalf of the Wellcome Sanger Institute COVID-19 Surveillance Team |
| EPI_ISL_2641169 | Lighthouse Lab in Glasgow | Wellcome Sanger Institute for the COVID-19 Genomics UK (COG-UK) Consortium | Harper VanSteenhouse, Yumi Kasai, David Gray, Carol Clugston, Anna Dominiczak and Alex Alderton, Roberto Amato, Jeffrey Barrett, Sonia Goncalves, Ewan Harrison, David K. Jackson, Ian Johnston, Dominic Kwiatkowski, Cordelia Langford, John Sillitoe on behalf of the Wellcome Sanger Institute COVID-19 Surveillance Team |
| EPI_ISL_2641170 | Lighthouse Lab in Alderley Park | Wellcome Sanger Institute for the COVID-19 Genomics UK (COG-UK) Consortium | Jacquelyn Wynn, Mairead Hyland, The Lighthouse Lab in Alderley Park and Alex Alderton, Roberto Amato, Jeffrey Barrett, Sonia Goncalves, Ewan Harrison, David K. Jackson, Ian Johnston, Dominic Kwiatkowski, Cordelia Langford, John Sillitoe on behalf of the Wellcome Sanger Institute COVID-19 Surveillance Team |
| EPI_ISL_2641171, EPI_ISL_2641172, EPI_ISL_2641173, EPI_ISL_2641174 | Lighthouse Lab in Milton Keynes | Wellcome Sanger Institute for the COVID-19 Genomics UK (COG-UK) Consortium | The Lighthouse Lab in Milton Keynes and Alex Alderton, Roberto Amato, Jeffrey Barrett, Sonia Goncalves, Ewan Harrison, David K. Jackson, Ian Johnston, Dominic Kwiatkowski, Cordelia Langford, John Sillitoe on behalf of the Wellcome Sanger Institute COVID-19 Surveillance Team |
| EPI_ISL_2641175 | Lighthouse Lab in Alderley Park | Wellcome Sanger Institute for the COVID-19 Genomics UK (COG-UK) Consortium | Jacquelyn Wynn, Mairead Hyland, The Lighthouse Lab in Alderley Park and Alex Alderton, Roberto Amato, Jeffrey Barrett, Sonia Goncalves, Ewan Harrison, David K. Jackson, Ian Johnston, Dominic Kwiatkowski, Cordelia Langford, John Sillitoe on behalf of the Wellcome Sanger Institute COVID-19 Surveillance Team |
| EPI_ISL_2641176, EPI_ISL_2641177, EPI_ISL_2641178, EPI_ISL_2641179 | Lighthouse Lab in Glasgow | Wellcome Sanger Institute for the COVID-19 Genomics UK (COG-UK) Consortium | Harper VanSteenhouse, Yumi Kasai, David Gray, Carol Clugston, Anna Dominiczak and Alex Alderton, Roberto Amato, Jeffrey Barrett, Sonia Goncalves, Ewan Harrison, David K. Jackson, Ian Johnston, Dominic Kwiatkowski, Cordelia Langford, John Sillitoe on behalf of the Wellcome Sanger Institute COVID-19 Surveillance Team |
| EPI_ISL_2641180, EPI_ISL_2641182 | Lighthouse Lab in Alderley Park | Wellcome Sanger Institute for the COVID-19 Genomics UK (COG-UK) Consortium | Jacquelyn Wynn, Mairead Hyland, The Lighthouse Lab in Alderley Park and Alex Alderton, Roberto Amato, Jeffrey Barrett, Sonia Goncalves, Ewan Harrison, David K. Jackson, Ian Johnston, Dominic Kwiatkowski, Cordelia Langford, John Sillitoe on behalf of the Wellcome Sanger Institute COVID-19 Surveillance Team |
| EPI_ISL_2641183 | Lighthouse Lab in Glasgow | Wellcome Sanger Institute for the COVID-19 Genomics UK (COG-UK) Consortium | Harper VanSteenhouse, Yumi Kasai, David Gray, Carol Clugston, Anna Dominiczak and Alex Alderton, Roberto Amato, Jeffrey Barrett, Sonia Goncalves, Ewan Harrison, David K. Jackson, Ian Johnston, Dominic Kwiatkowski, Cordelia Langford, John Sillitoe on behalf of the Wellcome Sanger Institute COVID-19 Surveillance Team |
| EPI_ISL_2641184 | Lighthouse Lab in Alderley Park | Wellcome Sanger Institute for the COVID-19 Genomics UK (COG-UK) Consortium | Jacquelyn Wynn, Mairead Hyland, The Lighthouse Lab in Alderley Park and Alex Alderton, Roberto Amato, Jeffrey Barrett, Sonia Goncalves, Ewan Harrison, David K. Jackson, Ian Johnston, Dominic Kwiatkowski, Cordelia Langford, John Sillitoe on behalf of the Wellcome Sanger Institute COVID-19 Surveillance Team |
| EPI_ISL_2641185 | Lighthouse Lab in Glasgow | Wellcome Sanger Institute for the COVID-19 Genomics UK (COG-UK) Consortium | Harper VanSteenhouse, Yumi Kasai, David Gray, Carol Clugston, Anna Dominiczak and Alex Alderton, Roberto Amato, Jeffrey Barrett, Sonia Goncalves, Ewan Harrison, David K. Jackson, Ian Johnston, Dominic Kwiatkowski, Cordelia Langford, John Sillitoe on behalf of the Wellcome Sanger Institute COVID-19 Surveillance Team |
| EPI_ISL_2641186 | Lighthouse Lab in Alderley Park | Wellcome Sanger Institute for the COVID-19 Genomics UK (COG-UK) Consortium | Jacquelyn Wynn, Mairead Hyland, The Lighthouse Lab in Alderley Park and Alex Alderton, Roberto Amato, Jeffrey Barrett, Sonia Goncalves, Ewan Harrison, David K. Jackson, Ian Johnston, Dominic Kwiatkowski, Cordelia Langford, John Sillitoe on behalf of the Wellcome Sanger Institute COVID-19 Surveillance Team |
| EPI_ISL_2641187 | Lighthouse Lab in Milton Keynes | Wellcome Sanger Institute for the COVID-19 Genomics UK (COG-UK) Consortium | The Lighthouse Lab in Milton Keynes and Alex Alderton, Roberto Amato, Jeffrey Barrett, Sonia Goncalves, Ewan Harrison, David K. Jackson, Ian Johnston, Dominic Kwiatkowski, Cordelia Langford, John Sillitoe on behalf of the Wellcome Sanger Institute COVID-19 Surveillance Team |
| EPI_ISL_2641188 | Lighthouse Lab in Glasgow | Wellcome Sanger Institute for the COVID-19 Genomics UK (COG-UK) Consortium | Harper VanSteenhouse, Yumi Kasai, David Gray, Carol Clugston, Anna Dominiczak and Alex Alderton, Roberto Amato, Jeffrey Barrett, Sonia Goncalves, Ewan Harrison, David K. Jackson, Ian Johnston, Dominic Kwiatkowski, Cordelia Langford, John Sillitoe on behalf of the Wellcome Sanger Institute COVID-19 Surveillance Team |
| EPI_ISL_2641189 | Lighthouse Lab in Alderley Park | Wellcome Sanger Institute for the COVID-19 Genomics UK (COG-UK) Consortium | Jacquelyn Wynn, Mairead Hyland, The Lighthouse Lab in Alderley Park and Alex Alderton, Roberto Amato, Jeffrey Barrett, Sonia Goncalves, Ewan Harrison, David K. Jackson, Ian Johnston, Dominic Kwiatkowski, Cordelia Langford, John Sillitoe on behalf of the Wellcome Sanger Institute COVID-19 Surveillance Team |
| EPI_ISL_2641190 | Lighthouse Lab in Milton Keynes | Wellcome Sanger Institute for the COVID-19 Genomics UK (COG-UK) Consortium | The Lighthouse Lab in Milton Keynes and Alex Alderton, Roberto Amato, Jeffrey Barrett, Sonia Goncalves, Ewan Harrison, David K. Jackson, Ian Johnston, Dominic Kwiatkowski, Cordelia Langford, John Sillitoe on behalf of the Wellcome Sanger Institute COVID-19 Surveillance Team |
| EPI_ISL_2641191 | Lighthouse Lab in Alderley Park | Wellcome Sanger Institute for the COVID-19 Genomics UK (COG-UK) Consortium | Jacquelyn Wynn, Mairead Hyland, The Lighthouse Lab in Alderley Park and Alex Alderton, Roberto Amato, Jeffrey Barrett, Sonia Goncalves, Ewan Harrison, David K. Jackson, Ian Johnston, Dominic Kwiatkowski, Cordelia Langford, John Sillitoe on behalf of the Wellcome Sanger Institute COVID-19 Surveillance Team |
| EPI_ISL_2641192 | Lighthouse Lab in Milton Keynes | Wellcome Sanger Institute for the COVID-19 Genomics UK (COG-UK) Consortium | The Lighthouse Lab in Milton Keynes and Alex Alderton, Roberto Amato, Jeffrey Barrett, Sonia Goncalves, Ewan Harrison, David K. Jackson, Ian Johnston, Dominic Kwiatkowski, Cordelia Langford, John Sillitoe on behalf of the Wellcome Sanger Institute COVID-19 Surveillance Team |
| EPI_ISL_2641193 | Lighthouse Lab in Alderley Park | Wellcome Sanger Institute for the COVID-19 Genomics UK (COG-UK) Consortium | Jacquelyn Wynn, Mairead Hyland, The Lighthouse Lab in Alderley Park and Alex Alderton, Roberto Amato, Jeffrey Barrett, Sonia Goncalves, Ewan Harrison, David K. Jackson, Ian Johnston, Dominic Kwiatkowski, Cordelia Langford, John Sillitoe on behalf of the Wellcome Sanger Institute COVID-19 Surveillance Team |
| EPI_ISL_2641194 | Lighthouse Lab in Milton Keynes | Wellcome Sanger Institute for the COVID-19 Genomics UK (COG-UK) Consortium | The Lighthouse Lab in Milton Keynes and Alex Alderton, Roberto Amato, Jeffrey Barrett, Sonia Goncalves, Ewan Harrison, David K. Jackson, Ian Johnston, Dominic Kwiatkowski, Cordelia Langford, John Sillitoe on behalf of the Wellcome Sanger Institute COVID-19 Surveillance Team |
| EPI_ISL_2641195, EPI_ISL_2641196 | Lighthouse Lab in Glasgow | Wellcome Sanger Institute for the COVID-19 Genomics UK (COG-UK) Consortium | Harper VanSteenhouse, Yumi Kasai, David Gray, Carol Clugston, Anna Dominiczak and Alex Alderton, Roberto Amato, Jeffrey Barrett, Sonia Goncalves, Ewan Harrison, David K. Jackson, Ian Johnston, Dominic Kwiatkowski, Cordelia Langford, John Sillitoe on behalf of the Wellcome Sanger Institute COVID-19 Surveillance Team |
| EPI_ISL_2641197 | Lighthouse Lab in Alderley Park | Wellcome Sanger Institute for the COVID-19 Genomics UK (COG-UK) Consortium | Jacquelyn Wynn, Mairead Hyland, The Lighthouse Lab in Alderley Park and Alex Alderton, Roberto Amato, Jeffrey Barrett, Sonia Goncalves, Ewan Harrison, David K. Jackson, Ian Johnston, Dominic Kwiatkowski, Cordelia Langford, John Sillitoe on behalf of the Wellcome Sanger Institute COVID-19 Surveillance Team |
| EPI_ISL_2641198 | Lighthouse Lab in Glasgow | Wellcome Sanger Institute for the COVID-19 Genomics UK (COG-UK) Consortium | Harper VanSteenhouse, Yumi Kasai, David Gray, Carol Clugston, Anna Dominiczak and Alex Alderton, Roberto Amato, Jeffrey Barrett, Sonia Goncalves, Ewan Harrison, David K. Jackson, Ian Johnston, Dominic Kwiatkowski, Cordelia Langford, John Sillitoe on behalf of the Wellcome Sanger Institute COVID-19 Surveillance Team |
| EPI_ISL_2641199 | Lighthouse Lab in Milton Keynes | Wellcome Sanger Institute for the COVID-19 Genomics UK (COG-UK) Consortium | The Lighthouse Lab in Milton Keynes and Alex Alderton, Roberto Amato, Jeffrey Barrett, Sonia Goncalves, Ewan Harrison, David K. Jackson, Ian Johnston, Dominic Kwiatkowski, Cordelia Langford, John Sillitoe on behalf of the Wellcome Sanger Institute COVID-19 Surveillance Team |
| EPI_ISL_2641200 | Lighthouse Lab in Glasgow | Wellcome Sanger Institute for the COVID-19 Genomics UK (COG-UK) Consortium | Harper VanSteenhouse, Yumi Kasai, David Gray, Carol Clugston, Anna Dominiczak and Alex Alderton, Roberto Amato, Jeffrey Barrett, Sonia Goncalves, Ewan Harrison, David K. Jackson, Ian Johnston, Dominic Kwiatkowski, Cordelia Langford, John Sillitoe on behalf of the Wellcome Sanger Institute COVID-19 Surveillance Team |
| EPI_ISL_2641201, EPI_ISL_2641202, EPI_ISL_2641204, EPI_ISL_2641205, EPI_ISL_2641206, EPI_ISL_2641207 | Lighthouse Lab in Alderley Park | Wellcome Sanger Institute for the COVID-19 Genomics UK (COG-UK) Consortium | Jacquelyn Wynn, Mairead Hyland, The Lighthouse Lab in Alderley Park and Alex Alderton, Roberto Amato, Jeffrey Barrett, Sonia Goncalves, Ewan Harrison, David K. Jackson, Ian Johnston, Dominic Kwiatkowski, Cordelia Langford, John Sillitoe on behalf of the Wellcome Sanger Institute COVID-19 Surveillance Team |
| EPI_ISL_2641208 | Lighthouse Lab in Glasgow | Wellcome Sanger Institute for the COVID-19 Genomics UK (COG-UK) Consortium | Harper VanSteenhouse, Yumi Kasai, David Gray, Carol Clugston, Anna Dominiczak and Alex Alderton, Roberto Amato, Jeffrey Barrett, Sonia Goncalves, Ewan Harrison, David K. Jackson, Ian Johnston, Dominic Kwiatkowski, Cordelia Langford, John Sillitoe on behalf of the Wellcome Sanger Institute COVID-19 Surveillance Team |
| EPI_ISL_2641209, EPI_ISL_2641210, EPI_ISL_2641211, EPI_ISL_2641212 | Lighthouse Lab in Alderley Park | Wellcome Sanger Institute for the COVID-19 Genomics UK (COG-UK) Consortium | Jacquelyn Wynn, Mairead Hyland, The Lighthouse Lab in Alderley Park and Alex Alderton, Roberto Amato, Jeffrey Barrett, Sonia Goncalves, Ewan Harrison, David K. Jackson, Ian Johnston, Dominic Kwiatkowski, Cordelia Langford, John Sillitoe on behalf of the Wellcome Sanger Institute COVID-19 Surveillance Team |
| EPI_ISL_2641213, EPI_ISL_2641214 | Lighthouse Lab in Milton Keynes | Wellcome Sanger Institute for the COVID-19 Genomics UK (COG-UK) Consortium | The Lighthouse Lab in Milton Keynes and Alex Alderton, Roberto Amato, Jeffrey Barrett, Sonia Goncalves, Ewan Harrison, David K. Jackson, Ian Johnston, Dominic Kwiatkowski, Cordelia Langford, John Sillitoe on behalf of the Wellcome Sanger Institute COVID-19 Surveillance Team |
| EPI_ISL_2641215 | Lighthouse Lab in Glasgow | Wellcome Sanger Institute for the COVID-19 Genomics UK (COG-UK) Consortium | Harper VanSteenhouse, Yumi Kasai, David Gray, Carol Clugston, Anna Dominiczak and Alex Alderton, Roberto Amato, Jeffrey Barrett, Sonia Goncalves, Ewan Harrison, David K. Jackson, Ian Johnston, Dominic Kwiatkowski, Cordelia Langford, John Sillitoe on behalf of the Wellcome Sanger Institute COVID-19 Surveillance Team |
| EPI_ISL_2641216 | Lighthouse Lab in Alderley Park | Wellcome Sanger Institute for the COVID-19 Genomics UK (COG-UK) Consortium | Jacquelyn Wynn, Mairead Hyland, The Lighthouse Lab in Alderley Park and Alex Alderton, Roberto Amato, Jeffrey Barrett, Sonia Goncalves, Ewan Harrison, David K. Jackson, Ian Johnston, Dominic Kwiatkowski, Cordelia Langford, John Sillitoe on behalf of the Wellcome Sanger Institute COVID-19 Surveillance Team |

| | | | |
|---|---|---|---|
| EPI_ISL_2641217 | Lighthouse Lab in Glasgow | Wellcome Sanger Institute for the COVID-19 Genomics UK (COG-UK) Consortium | Harper VanSteenhouse, Yumi Kasai, David Gray, Carol Clugston, Anna Dominiczak and Alex Alderton, Roberto Amato, Jeffrey Barrett, Sonia Goncalves, Ewan Harrison, David K. Jackson, Ian Johnston, Dominic Kwiatkowski, Cordelia Langford, John Sillitoe on behalf of the Wellcome Sanger Institute COVID-19 Surveillance Team |
| EPI_ISL_2641218, EPI_ISL_2641219, EPI_ISL_2641220 | Lighthouse Lab in Alderley Park | Wellcome Sanger Institute for the COVID-19 Genomics UK (COG-UK) Consortium | Jacquelyn Wynn, Mairead Hyland, The Lighthouse Lab in Alderley Park and Alex Alderton, Roberto Amato, Jeffrey Barrett, Sonia Goncalves, Ewan Harrison, David K. Jackson, Ian Johnston, Dominic Kwiatkowski, Cordelia Langford, John Sillitoe on behalf of the Wellcome Sanger Institute COVID-19 Surveillance Team |
| EPI_ISL_2641222 | Lighthouse Lab in Milton Keynes | Wellcome Sanger Institute for the COVID-19 Genomics UK (COG-UK) Consortium | The Lighthouse Lab in Milton Keynes and Alex Alderton, Roberto Amato, Jeffrey Barrett, Sonia Goncalves, Ewan Harrison, David K. Jackson, Ian Johnston, Dominic Kwiatkowski, Cordelia Langford, John Sillitoe on behalf of the Wellcome Sanger Institute COVID-19 Surveillance Team |
| EPI_ISL_2641223 | Lighthouse Lab in Alderley Park | Wellcome Sanger Institute for the COVID-19 Genomics UK (COG-UK) Consortium | Jacquelyn Wynn, Mairead Hyland, The Lighthouse Lab in Alderley Park and Alex Alderton, Roberto Amato, Jeffrey Barrett, Sonia Goncalves, Ewan Harrison, David K. Jackson, Ian Johnston, Dominic Kwiatkowski, Cordelia Langford, John Sillitoe on behalf of the Wellcome Sanger Institute COVID-19 Surveillance Team |
| EPI_ISL_2641224 | Lighthouse Lab in Glasgow | Wellcome Sanger Institute for the COVID-19 Genomics UK (COG-UK) Consortium | Harper VanSteenhouse, Yumi Kasai, David Gray, Carol Clugston, Anna Dominiczak and Alex Alderton, Roberto Amato, Jeffrey Barrett, Sonia Goncalves, Ewan Harrison, David K. Jackson, Ian Johnston, Dominic Kwiatkowski, Cordelia Langford, John Sillitoe on behalf of the Wellcome Sanger Institute COVID-19 Surveillance Team |
| EPI_ISL_2641225 | Lighthouse Lab in Alderley Park | Wellcome Sanger Institute for the COVID-19 Genomics UK (COG-UK) Consortium | Jacquelyn Wynn, Mairead Hyland, The Lighthouse Lab in Alderley Park and Alex Alderton, Roberto Amato, Jeffrey Barrett, Sonia Goncalves, Ewan Harrison, David K. Jackson, Ian Johnston, Dominic Kwiatkowski, Cordelia Langford, John Sillitoe on behalf of the Wellcome Sanger Institute COVID-19 Surveillance Team |
| EPI_ISL_2641226 | Lighthouse Lab in Glasgow | Wellcome Sanger Institute for the COVID-19 Genomics UK (COG-UK) Consortium | Harper VanSteenhouse, Yumi Kasai, David Gray, Carol Clugston, Anna Dominiczak and Alex Alderton, Roberto Amato, Jeffrey Barrett, Sonia Goncalves, Ewan Harrison, David K. Jackson, Ian Johnston, Dominic Kwiatkowski, Cordelia Langford, John Sillitoe on behalf of the Wellcome Sanger Institute COVID-19 Surveillance Team |
| EPI_ISL_2641227 | Lighthouse Lab in Milton Keynes | Wellcome Sanger Institute for the COVID-19 Genomics UK (COG-UK) Consortium | The Lighthouse Lab in Milton Keynes and Alex Alderton, Roberto Amato, Jeffrey Barrett, Sonia Goncalves, Ewan Harrison, David K. Jackson, Ian Johnston, Dominic Kwiatkowski, Cordelia Langford, John Sillitoe on behalf of the Wellcome Sanger Institute COVID-19 Surveillance Team |
| EPI_ISL_2641228, EPI_ISL_2641229, EPI_ISL_2641230 | Lighthouse Lab in Glasgow | Wellcome Sanger Institute for the COVID-19 Genomics UK (COG-UK) Consortium | Harper VanSteenhouse, Yumi Kasai, David Gray, Carol Clugston, Anna Dominiczak and Alex Alderton, Roberto Amato, Jeffrey Barrett, Sonia Goncalves, Ewan Harrison, David K. Jackson, Ian Johnston, Dominic Kwiatkowski, Cordelia Langford, John Sillitoe on behalf of the Wellcome Sanger Institute COVID-19 Surveillance Team |
| EPI_ISL_2641232, EPI_ISL_2641233, EPI_ISL_2641234 | Lighthouse Lab in Alderley Park | Wellcome Sanger Institute for the COVID-19 Genomics UK (COG-UK) Consortium | Jacquelyn Wynn, Mairead Hyland, The Lighthouse Lab in Alderley Park and Alex Alderton, Roberto Amato, Jeffrey Barrett, Sonia Goncalves, Ewan Harrison, David K. Jackson, Ian Johnston, Dominic Kwiatkowski, Cordelia Langford, John Sillitoe on behalf of the Wellcome Sanger Institute COVID-19 Surveillance Team |
| EPI_ISL_2641235, EPI_ISL_2641236, EPI_ISL_2641237 | Lighthouse Lab in Milton Keynes | Wellcome Sanger Institute for the COVID-19 Genomics UK (COG-UK) Consortium | The Lighthouse Lab in Milton Keynes and Alex Alderton, Roberto Amato, Jeffrey Barrett, Sonia Goncalves, Ewan Harrison, David K. Jackson, Ian Johnston, Dominic Kwiatkowski, Cordelia Langford, John Sillitoe on behalf of the Wellcome Sanger Institute COVID-19 Surveillance Team |
| EPI_ISL_2641238 | Lighthouse Lab in Glasgow | Wellcome Sanger Institute for the COVID-19 Genomics UK (COG-UK) Consortium | Harper VanSteenhouse, Yumi Kasai, David Gray, Carol Clugston, Anna Dominiczak and Alex Alderton, Roberto Amato, Jeffrey Barrett, Sonia Goncalves, Ewan Harrison, David K. Jackson, Ian Johnston, Dominic Kwiatkowski, Cordelia Langford, John Sillitoe on behalf of the Wellcome Sanger Institute COVID-19 Surveillance Team |
| EPI_ISL_2641239, EPI_ISL_2641240, EPI_ISL_2641241 | Lighthouse Lab in Alderley Park | Wellcome Sanger Institute for the COVID-19 Genomics UK (COG-UK) Consortium | Jacquelyn Wynn, Mairead Hyland, The Lighthouse Lab in Alderley Park and Alex Alderton, Roberto Amato, Jeffrey Barrett, Sonia Goncalves, Ewan Harrison, David K. Jackson, Ian Johnston, Dominic Kwiatkowski, Cordelia Langford, John Sillitoe on behalf of the Wellcome Sanger Institute COVID-19 Surveillance Team |
| EPI_ISL_2641242 | Lighthouse Lab in Glasgow | Wellcome Sanger Institute for the COVID-19 Genomics UK (COG-UK) Consortium | Harper VanSteenhouse, Yumi Kasai, David Gray, Carol Clugston, Anna Dominiczak and Alex Alderton, Roberto Amato, Jeffrey Barrett, Sonia Goncalves, Ewan Harrison, David K. Jackson, Ian Johnston, Dominic Kwiatkowski, Cordelia Langford, John Sillitoe on behalf of the Wellcome Sanger Institute COVID-19 Surveillance Team |
| EPI_ISL_2641243, EPI_ISL_2641244, EPI_ISL_2641245 | Lighthouse Lab in Alderley Park | Wellcome Sanger Institute for the COVID-19 Genomics UK (COG-UK) Consortium | Jacquelyn Wynn, Mairead Hyland, The Lighthouse Lab in Alderley Park and Alex Alderton, Roberto Amato, Jeffrey Barrett, Sonia Goncalves, Ewan Harrison, David K. Jackson, Ian Johnston, Dominic Kwiatkowski, Cordelia Langford, John Sillitoe on behalf of the Wellcome Sanger Institute COVID-19 Surveillance Team |
| EPI_ISL_2641246 | Lighthouse Lab in Glasgow | Wellcome Sanger Institute for the COVID-19 Genomics UK (COG-UK) Consortium | Harper VanSteenhouse, Yumi Kasai, David Gray, Carol Clugston, Anna Dominiczak and Alex Alderton, Roberto Amato, Jeffrey Barrett, Sonia Goncalves, Ewan Harrison, David K. Jackson, Ian Johnston, Dominic Kwiatkowski, Cordelia Langford, John Sillitoe on behalf of the Wellcome Sanger Institute COVID-19 Surveillance Team |
| EPI_ISL_2641247, EPI_ISL_2641249, EPI_ISL_2641250, EPI_ISL_2641251, EPI_ISL_2641252, EPI_ISL_2641253 | Lighthouse Lab in Alderley Park | Wellcome Sanger Institute for the COVID-19 Genomics UK (COG-UK) Consortium | Jacquelyn Wynn, Mairead Hyland, The Lighthouse Lab in Alderley Park and Alex Alderton, Roberto Amato, Jeffrey Barrett, Sonia Goncalves, Ewan Harrison, David K. Jackson, Ian Johnston, Dominic Kwiatkowski, Cordelia Langford, John Sillitoe on behalf of the Wellcome Sanger Institute COVID-19 Surveillance Team |
| EPI_ISL_2641254 | Lighthouse Lab in Milton Keynes | Wellcome Sanger Institute for the COVID-19 Genomics UK (COG-UK) Consortium | The Lighthouse Lab in Milton Keynes and Alex Alderton, Roberto Amato, Jeffrey Barrett, Sonia Goncalves, Ewan Harrison, David K. Jackson, Ian Johnston, Dominic Kwiatkowski, Cordelia Langford, John Sillitoe on behalf of the Wellcome Sanger Institute COVID-19 Surveillance Team |
| EPI_ISL_2641255 | Lighthouse Lab in Alderley Park | Wellcome Sanger Institute for the COVID-19 Genomics UK (COG-UK) Consortium | Jacquelyn Wynn, Mairead Hyland, The Lighthouse Lab in Alderley Park and Alex Alderton, Roberto Amato, Jeffrey Barrett, Sonia Goncalves, Ewan Harrison, David K. Jackson, Ian Johnston, Dominic Kwiatkowski, Cordelia Langford, John Sillitoe on behalf of the Wellcome Sanger Institute COVID-19 Surveillance Team |
| EPI_ISL_2641256 | Lighthouse Lab in Milton Keynes | Wellcome Sanger Institute for the COVID-19 Genomics UK (COG-UK) Consortium | The Lighthouse Lab in Milton Keynes and Alex Alderton, Roberto Amato, Jeffrey Barrett, Sonia Goncalves, Ewan Harrison, David K. Jackson, Ian Johnston, Dominic Kwiatkowski, Cordelia Langford, John Sillitoe on behalf of the Wellcome Sanger Institute COVID-19 Surveillance Team |
| EPI_ISL_2641257 | Lighthouse Lab in Glasgow | Wellcome Sanger Institute for the COVID-19 Genomics UK (COG-UK) Consortium | Harper VanSteenhouse, Yumi Kasai, David Gray, Carol Clugston, Anna Dominiczak and Alex Alderton, Roberto Amato, Jeffrey Barrett, Sonia Goncalves, Ewan Harrison, David K. Jackson, Ian Johnston, Dominic Kwiatkowski, Cordelia Langford, John Sillitoe on behalf of the Wellcome Sanger Institute COVID-19 Surveillance Team |
| EPI_ISL_2641258, EPI_ISL_2641259, EPI_ISL_2641260, EPI_ISL_2641261 | Lighthouse Lab in Alderley Park | Wellcome Sanger Institute for the COVID-19 Genomics UK (COG-UK) Consortium | Jacquelyn Wynn, Mairead Hyland, The Lighthouse Lab in Alderley Park and Alex Alderton, Roberto Amato, Jeffrey Barrett, Sonia Goncalves, Ewan Harrison, David K. Jackson, Ian Johnston, Dominic Kwiatkowski, Cordelia Langford, John Sillitoe on behalf of the Wellcome Sanger Institute COVID-19 Surveillance Team |
| EPI_ISL_2641262 | Lighthouse Lab in Glasgow | Wellcome Sanger Institute for the COVID-19 Genomics UK (COG-UK) Consortium | Harper VanSteenhouse, Yumi Kasai, David Gray, Carol Clugston, Anna Dominiczak and Alex Alderton, Roberto Amato, Jeffrey Barrett, Sonia Goncalves, Ewan Harrison, David K. Jackson, Ian Johnston, Dominic Kwiatkowski, Cordelia Langford, John Sillitoe on behalf of the Wellcome Sanger Institute COVID-19 Surveillance Team |
| EPI_ISL_2641263, EPI_ISL_2641264 | Lighthouse Lab in Milton Keynes | Wellcome Sanger Institute for the COVID-19 Genomics UK (COG-UK) Consortium | The Lighthouse Lab in Milton Keynes and Alex Alderton, Roberto Amato, Jeffrey Barrett, Sonia Goncalves, Ewan Harrison, David K. Jackson, Ian Johnston, Dominic Kwiatkowski, Cordelia Langford, John Sillitoe on behalf of the Wellcome Sanger Institute COVID-19 Surveillance Team |
| EPI_ISL_2641265 | Lighthouse Lab in Glasgow | Wellcome Sanger Institute for the COVID-19 Genomics UK (COG-UK) Consortium | Harper VanSteenhouse, Yumi Kasai, David Gray, Carol Clugston, Anna Dominiczak and Alex Alderton, Roberto Amato, Jeffrey Barrett, Sonia Goncalves, Ewan Harrison, David K. Jackson, Ian Johnston, Dominic Kwiatkowski, Cordelia Langford, John Sillitoe on behalf of the Wellcome Sanger Institute COVID-19 Surveillance Team |
| EPI_ISL_2641266 | Lighthouse Lab in Milton Keynes | Wellcome Sanger Institute for the COVID-19 Genomics UK (COG-UK) Consortium | The Lighthouse Lab in Milton Keynes and Alex Alderton, Roberto Amato, Jeffrey Barrett, Sonia Goncalves, Ewan Harrison, David K. Jackson, Ian Johnston, Dominic Kwiatkowski, Cordelia Langford, John Sillitoe on behalf of the Wellcome Sanger Institute COVID-19 Surveillance Team |
| EPI_ISL_2641267, EPI_ISL_2641268 | Lighthouse Lab in Alderley Park | Wellcome Sanger Institute for the COVID-19 Genomics UK (COG-UK) Consortium | Jacquelyn Wynn, Mairead Hyland, The Lighthouse Lab in Alderley Park and Alex Alderton, Roberto Amato, Jeffrey Barrett, Sonia Goncalves, Ewan Harrison, David K. Jackson, Ian Johnston, Dominic Kwiatkowski, Cordelia Langford, John Sillitoe on behalf of the Wellcome Sanger Institute COVID-19 Surveillance Team |
| EPI_ISL_2641269 | Lighthouse Lab in Glasgow | Wellcome Sanger Institute for the COVID-19 Genomics UK (COG-UK) Consortium | Harper VanSteenhouse, Yumi Kasai, David Gray, Carol Clugston, Anna Dominiczak and Alex Alderton, Roberto Amato, Jeffrey Barrett, Sonia Goncalves, Ewan Harrison, David K. Jackson, Ian Johnston, Dominic Kwiatkowski, Cordelia Langford, John Sillitoe on behalf of the Wellcome Sanger Institute COVID-19 Surveillance Team |
| EPI_ISL_2641270, EPI_ISL_2641271 | Lighthouse Lab in Alderley Park | Wellcome Sanger Institute for the COVID-19 Genomics UK (COG-UK) Consortium | Jacquelyn Wynn, Mairead Hyland, The Lighthouse Lab in Alderley Park and Alex Alderton, Roberto Amato, Jeffrey Barrett, Sonia Goncalves, Ewan Harrison, David K. Jackson, Ian Johnston, Dominic Kwiatkowski, Cordelia Langford, John Sillitoe on behalf of the Wellcome Sanger Institute COVID-19 Surveillance Team |
| EPI_ISL_2641272 | Lighthouse Lab in Milton Keynes | Wellcome Sanger Institute for the COVID-19 Genomics UK (COG-UK) Consortium | The Lighthouse Lab in Milton Keynes and Alex Alderton, Roberto Amato, Jeffrey Barrett, Sonia Goncalves, Ewan Harrison, David K. Jackson, Ian Johnston, Dominic Kwiatkowski, Cordelia Langford, John Sillitoe on behalf of the Wellcome Sanger Institute COVID-19 Surveillance Team |
| EPI_ISL_2641273 | Lighthouse Lab in Alderley Park | Wellcome Sanger Institute for the COVID-19 Genomics UK (COG-UK) Consortium | Jacquelyn Wynn, Mairead Hyland, The Lighthouse Lab in Alderley Park and Alex Alderton, Roberto Amato, Jeffrey Barrett, Sonia Goncalves, Ewan Harrison, David K. Jackson, Ian Johnston, Dominic Kwiatkowski, Cordelia Langford, John Sillitoe on behalf of the Wellcome Sanger Institute COVID-19 Surveillance Team |
| EPI_ISL_2641274 | Lighthouse Lab in Milton Keynes | Wellcome Sanger Institute for the COVID-19 Genomics UK (COG-UK) Consortium | The Lighthouse Lab in Milton Keynes and Alex Alderton, Roberto Amato, Jeffrey Barrett, Sonia Goncalves, Ewan Harrison, David K. Jackson, Ian Johnston, Dominic Kwiatkowski, Cordelia Langford, John Sillitoe on behalf of the Wellcome Sanger Institute COVID-19 Surveillance Team |
| EPI_ISL_2641276 | Lighthouse Lab in Glasgow | Wellcome Sanger Institute for the COVID-19 Genomics UK (COG-UK) Consortium | Harper VanSteenhouse, Yumi Kasai, David Gray, Carol Clugston, Anna Dominiczak and Alex Alderton, Roberto Amato, Jeffrey Barrett, Sonia Goncalves, Ewan Harrison, David K. Jackson, Ian Johnston, Dominic Kwiatkowski, Cordelia Langford, John Sillitoe on behalf of the Wellcome Sanger Institute COVID-19 Surveillance Team |
| EPI_ISL_2641277, EPI_ISL_2641278, EPI_ISL_2641279 | Lighthouse Lab in Alderley Park | Wellcome Sanger Institute for the COVID-19 Genomics UK (COG-UK) Consortium | Jacquelyn Wynn, Mairead Hyland, The Lighthouse Lab in Alderley Park and Alex Alderton, Roberto Amato, Jeffrey Barrett, Sonia Goncalves, Ewan Harrison, David K. Jackson, Ian Johnston, Dominic Kwiatkowski, Cordelia Langford, John Sillitoe on behalf of the Wellcome Sanger Institute COVID-19 Surveillance Team |
| EPI_ISL_2641280, EPI_ISL_2641282, EPI_ISL_2641283 | Lighthouse Lab in Milton Keynes | Wellcome Sanger Institute for the COVID-19 Genomics UK (COG-UK) Consortium | The Lighthouse Lab in Milton Keynes and Alex Alderton, Roberto Amato, Jeffrey Barrett, Sonia Goncalves, Ewan Harrison, David K. Jackson, Ian Johnston, Dominic Kwiatkowski, Cordelia Langford, John Sillitoe on behalf of the Wellcome Sanger Institute COVID-19 Surveillance Team |
| EPI_ISL_2641284, EPI_ISL_2641285 | Lighthouse Lab in Alderley Park | Wellcome Sanger Institute for the COVID-19 Genomics UK (COG-UK) Consortium | Jacquelyn Wynn, Mairead Hyland, The Lighthouse Lab in Alderley Park and Alex Alderton, Roberto Amato, Jeffrey Barrett, Sonia Goncalves, Ewan Harrison, David K. Jackson, Ian Johnston, Dominic Kwiatkowski, Cordelia Langford, John Sillitoe on behalf of the Wellcome Sanger Institute COVID-19 Surveillance Team |
| EPI_ISL_2641286 | Lighthouse Lab in Milton Keynes | Wellcome Sanger Institute for the COVID-19 Genomics UK (COG-UK) Consortium | The Lighthouse Lab in Milton Keynes and Alex Alderton, Roberto Amato, Jeffrey Barrett, Sonia Goncalves, Ewan Harrison, David K. Jackson, Ian Johnston, Dominic Kwiatkowski, Cordelia Langford, John Sillitoe on behalf of the Wellcome Sanger Institute COVID-19 Surveillance Team |
| EPI_ISL_2641287 | Lighthouse Lab in Alderley Park | Wellcome Sanger Institute for the COVID-19 Genomics UK (COG-UK) Consortium | Jacquelyn Wynn, Mairead Hyland, The Lighthouse Lab in Alderley Park and Alex Alderton, Roberto Amato, Jeffrey Barrett, Sonia Goncalves, Ewan Harrison, David K. Jackson, Ian Johnston, Dominic Kwiatkowski, Cordelia Langford, John Sillitoe on behalf of the Wellcome Sanger Institute COVID-19 Surveillance Team |
| EPI_ISL_2641288, EPI_ISL_2641289 | Lighthouse Lab in Milton Keynes | Wellcome Sanger Institute for the COVID-19 Genomics UK (COG-UK) Consortium | The Lighthouse Lab in Milton Keynes and Alex Alderton, Roberto Amato, Jeffrey Barrett, Sonia Goncalves, Ewan Harrison, David K. Jackson, Ian Johnston, Dominic Kwiatkowski, Cordelia Langford, John Sillitoe on behalf of the Wellcome Sanger Institute COVID-19 Surveillance Team |
| EPI_ISL_2641290, | Lighthouse Lab in Alderley Park | Wellcome Sanger Institute for the COVID-19 Genomics UK (COG-UK) | Jacquelyn Wynn, Mairead Hyland, The Lighthouse Lab in Alderley Park and Alex Alderton, Roberto Amato, Jeffrey Barrett, Sonia Goncalves, Ewan Harrison, David K. Jackson, Ian Johnston, Dominic Kwiatkowski, Cordelia Langford, John Sillitoe on |

| | | | |
|---|---|---|---|
| EPI_ISL_2641291, EPI_ISL_2641292 | | Consortium | behalf of the Wellcome Sanger Institute COVID-19 Surveillance Team |
| EPI_ISL_2641293, EPI_ISL_2641294 | Lighthouse Lab in Glasgow | Wellcome Sanger Institute for the COVID-19 Genomics UK (COG-UK) Consortium | Harper VanSteenhouse, Yumi Kasai, David Gray, Carol Clugston, Anna Dominiczak and Alex Alderton, Roberto Amato, Jeffrey Barrett, Sonia Goncalves, Ewan Harrison, David K. Jackson, Ian Johnston, Dominic Kwiatkowski, Cordelia Langford, John Sillitoe on behalf of the Wellcome Sanger Institute COVID-19 Surveillance Team |
| EPI_ISL_2641295, EPI_ISL_2641296 | Lighthouse Lab in Alderley Park | Wellcome Sanger Institute for the COVID-19 Genomics UK (COG-UK) Consortium | Jacquelyn Wynn, Mairead Hyland, The Lighthouse Lab in Alderley Park and Alex Alderton, Roberto Amato, Jeffrey Barrett, Sonia Goncalves, Ewan Harrison, David K. Jackson, Ian Johnston, Dominic Kwiatkowski, Cordelia Langford, John Sillitoe on behalf of the Wellcome Sanger Institute COVID-19 Surveillance Team |
| EPI_ISL_2641298, EPI_ISL_2641299 | Lighthouse Lab in Milton Keynes | Wellcome Sanger Institute for the COVID-19 Genomics UK (COG-UK) Consortium | The Lighthouse Lab in Milton Keynes and Alex Alderton, Roberto Amato, Jeffrey Barrett, Sonia Goncalves, Ewan Harrison, David K. Jackson, Ian Johnston, Dominic Kwiatkowski, Cordelia Langford, John Sillitoe on behalf of the Wellcome Sanger Institute COVID-19 Surveillance Team |
| EPI_ISL_2641300, EPI_ISL_2641301, EPI_ISL_2641302 | Lighthouse Lab in Glasgow | Wellcome Sanger Institute for the COVID-19 Genomics UK (COG-UK) Consortium | Harper VanSteenhouse, Yumi Kasai, David Gray, Carol Clugston, Anna Dominiczak and Alex Alderton, Roberto Amato, Jeffrey Barrett, Sonia Goncalves, Ewan Harrison, David K. Jackson, Ian Johnston, Dominic Kwiatkowski, Cordelia Langford, John Sillitoe on behalf of the Wellcome Sanger Institute COVID-19 Surveillance Team |
| EPI_ISL_2641303, EPI_ISL_2641304 | Lighthouse Lab in Alderley Park | Wellcome Sanger Institute for the COVID-19 Genomics UK (COG-UK) Consortium | Jacquelyn Wynn, Mairead Hyland, The Lighthouse Lab in Alderley Park and Alex Alderton, Roberto Amato, Jeffrey Barrett, Sonia Goncalves, Ewan Harrison, David K. Jackson, Ian Johnston, Dominic Kwiatkowski, Cordelia Langford, John Sillitoe on behalf of the Wellcome Sanger Institute COVID-19 Surveillance Team |
| EPI_ISL_2641305 | Lighthouse Lab in Glasgow | Wellcome Sanger Institute for the COVID-19 Genomics UK (COG-UK) Consortium | Harper VanSteenhouse, Yumi Kasai, David Gray, Carol Clugston, Anna Dominiczak and Alex Alderton, Roberto Amato, Jeffrey Barrett, Sonia Goncalves, Ewan Harrison, David K. Jackson, Ian Johnston, Dominic Kwiatkowski, Cordelia Langford, John Sillitoe on behalf of the Wellcome Sanger Institute COVID-19 Surveillance Team |
| EPI_ISL_2641306 | Lighthouse Lab in Alderley Park | Wellcome Sanger Institute for the COVID-19 Genomics UK (COG-UK) Consortium | Jacquelyn Wynn, Mairead Hyland, The Lighthouse Lab in Alderley Park and Alex Alderton, Roberto Amato, Jeffrey Barrett, Sonia Goncalves, Ewan Harrison, David K. Jackson, Ian Johnston, Dominic Kwiatkowski, Cordelia Langford, John Sillitoe on behalf of the Wellcome Sanger Institute COVID-19 Surveillance Team |
| EPI_ISL_2641308 | Lighthouse Lab in Milton Keynes | Wellcome Sanger Institute for the COVID-19 Genomics UK (COG-UK) Consortium | The Lighthouse Lab in Milton Keynes and Alex Alderton, Roberto Amato, Jeffrey Barrett, Sonia Goncalves, Ewan Harrison, David K. Jackson, Ian Johnston, Dominic Kwiatkowski, Cordelia Langford, John Sillitoe on behalf of the Wellcome Sanger Institute COVID-19 Surveillance Team |
| EPI_ISL_2641309 | Lighthouse Lab in Glasgow | Wellcome Sanger Institute for the COVID-19 Genomics UK (COG-UK) Consortium | Harper VanSteenhouse, Yumi Kasai, David Gray, Carol Clugston, Anna Dominiczak and Alex Alderton, Roberto Amato, Jeffrey Barrett, Sonia Goncalves, Ewan Harrison, David K. Jackson, Ian Johnston, Dominic Kwiatkowski, Cordelia Langford, John Sillitoe on behalf of the Wellcome Sanger Institute COVID-19 Surveillance Team |
| EPI_ISL_2641310, EPI_ISL_2641311, EPI_ISL_2641312 | Lighthouse Lab in Alderley Park | Wellcome Sanger Institute for the COVID-19 Genomics UK (COG-UK) Consortium | Jacquelyn Wynn, Mairead Hyland, The Lighthouse Lab in Alderley Park and Alex Alderton, Roberto Amato, Jeffrey Barrett, Sonia Goncalves, Ewan Harrison, David K. Jackson, Ian Johnston, Dominic Kwiatkowski, Cordelia Langford, John Sillitoe on behalf of the Wellcome Sanger Institute COVID-19 Surveillance Team |
| EPI_ISL_2641313 | Lighthouse Lab in Milton Keynes | Wellcome Sanger Institute for the COVID-19 Genomics UK (COG-UK) Consortium | The Lighthouse Lab in Milton Keynes and Alex Alderton, Roberto Amato, Jeffrey Barrett, Sonia Goncalves, Ewan Harrison, David K. Jackson, Ian Johnston, Dominic Kwiatkowski, Cordelia Langford, John Sillitoe on behalf of the Wellcome Sanger Institute COVID-19 Surveillance Team |
| EPI_ISL_2641314, EPI_ISL_2641317 | Lighthouse Lab in Glasgow | Wellcome Sanger Institute for the COVID-19 Genomics UK (COG-UK) Consortium | Harper VanSteenhouse, Yumi Kasai, David Gray, Carol Clugston, Anna Dominiczak and Alex Alderton, Roberto Amato, Jeffrey Barrett, Sonia Goncalves, Ewan Harrison, David K. Jackson, Ian Johnston, Dominic Kwiatkowski, Cordelia Langford, John Sillitoe on behalf of the Wellcome Sanger Institute COVID-19 Surveillance Team |
| EPI_ISL_2641318 | Lighthouse Lab in Milton Keynes | Wellcome Sanger Institute for the COVID-19 Genomics UK (COG-UK) Consortium | The Lighthouse Lab in Milton Keynes and Alex Alderton, Roberto Amato, Jeffrey Barrett, Sonia Goncalves, Ewan Harrison, David K. Jackson, Ian Johnston, Dominic Kwiatkowski, Cordelia Langford, John Sillitoe on behalf of the Wellcome Sanger Institute COVID-19 Surveillance Team |
| EPI_ISL_2641320 | Lighthouse Lab in Glasgow | Wellcome Sanger Institute for the COVID-19 Genomics UK (COG-UK) Consortium | Harper VanSteenhouse, Yumi Kasai, David Gray, Carol Clugston, Anna Dominiczak and Alex Alderton, Roberto Amato, Jeffrey Barrett, Sonia Goncalves, Ewan Harrison, David K. Jackson, Ian Johnston, Dominic Kwiatkowski, Cordelia Langford, John Sillitoe on behalf of the Wellcome Sanger Institute COVID-19 Surveillance Team |
| EPI_ISL_2641321 | Lighthouse Lab in Alderley Park | Wellcome Sanger Institute for the COVID-19 Genomics UK (COG-UK) Consortium | Jacquelyn Wynn, Mairead Hyland, The Lighthouse Lab in Alderley Park and Alex Alderton, Roberto Amato, Jeffrey Barrett, Sonia Goncalves, Ewan Harrison, David K. Jackson, Ian Johnston, Dominic Kwiatkowski, Cordelia Langford, John Sillitoe on behalf of the Wellcome Sanger Institute COVID-19 Surveillance Team |
| EPI_ISL_2641322 | Lighthouse Lab in Glasgow | Wellcome Sanger Institute for the COVID-19 Genomics UK (COG-UK) Consortium | Harper VanSteenhouse, Yumi Kasai, David Gray, Carol Clugston, Anna Dominiczak and Alex Alderton, Roberto Amato, Jeffrey Barrett, Sonia Goncalves, Ewan Harrison, David K. Jackson, Ian Johnston, Dominic Kwiatkowski, Cordelia Langford, John Sillitoe on behalf of the Wellcome Sanger Institute COVID-19 Surveillance Team |
| EPI_ISL_2641323 | Lighthouse Lab in Alderley Park | Wellcome Sanger Institute for the COVID-19 Genomics UK (COG-UK) Consortium | Jacquelyn Wynn, Mairead Hyland, The Lighthouse Lab in Alderley Park and Alex Alderton, Roberto Amato, Jeffrey Barrett, Sonia Goncalves, Ewan Harrison, David K. Jackson, Ian Johnston, Dominic Kwiatkowski, Cordelia Langford, John Sillitoe on behalf of the Wellcome Sanger Institute COVID-19 Surveillance Team |
| EPI_ISL_2641324, EPI_ISL_2641327 | Lighthouse Lab in Glasgow | Wellcome Sanger Institute for the COVID-19 Genomics UK (COG-UK) Consortium | Harper VanSteenhouse, Yumi Kasai, David Gray, Carol Clugston, Anna Dominiczak and Alex Alderton, Roberto Amato, Jeffrey Barrett, Sonia Goncalves, Ewan Harrison, David K. Jackson, Ian Johnston, Dominic Kwiatkowski, Cordelia Langford, John Sillitoe on behalf of the Wellcome Sanger Institute COVID-19 Surveillance Team |
| EPI_ISL_2641328 | Lighthouse Lab in Alderley Park | Wellcome Sanger Institute for the COVID-19 Genomics UK (COG-UK) Consortium | Jacquelyn Wynn, Mairead Hyland, The Lighthouse Lab in Alderley Park and Alex Alderton, Roberto Amato, Jeffrey Barrett, Sonia Goncalves, Ewan Harrison, David K. Jackson, Ian Johnston, Dominic Kwiatkowski, Cordelia Langford, John Sillitoe on behalf of the Wellcome Sanger Institute COVID-19 Surveillance Team |
| EPI_ISL_2641329 | Lighthouse Lab in Milton Keynes | Wellcome Sanger Institute for the COVID-19 Genomics UK (COG-UK) Consortium | The Lighthouse Lab in Milton Keynes and Alex Alderton, Roberto Amato, Jeffrey Barrett, Sonia Goncalves, Ewan Harrison, David K. Jackson, Ian Johnston, Dominic Kwiatkowski, Cordelia Langford, John Sillitoe on behalf of the Wellcome Sanger Institute COVID-19 Surveillance Team |
| EPI_ISL_2641330 | Lighthouse Lab in Alderley Park | Wellcome Sanger Institute for the COVID-19 Genomics UK (COG-UK) Consortium | Jacquelyn Wynn, Mairead Hyland, The Lighthouse Lab in Alderley Park and Alex Alderton, Roberto Amato, Jeffrey Barrett, Sonia Goncalves, Ewan Harrison, David K. Jackson, Ian Johnston, Dominic Kwiatkowski, Cordelia Langford, John Sillitoe on behalf of the Wellcome Sanger Institute COVID-19 Surveillance Team |
| EPI_ISL_2641331 | Lighthouse Lab in Milton Keynes | Wellcome Sanger Institute for the COVID-19 Genomics UK (COG-UK) Consortium | The Lighthouse Lab in Milton Keynes and Alex Alderton, Roberto Amato, Jeffrey Barrett, Sonia Goncalves, Ewan Harrison, David K. Jackson, Ian Johnston, Dominic Kwiatkowski, Cordelia Langford, John Sillitoe on behalf of the Wellcome Sanger Institute COVID-19 Surveillance Team |
| EPI_ISL_2641332 | Lighthouse Lab in Glasgow | Wellcome Sanger Institute for the COVID-19 Genomics UK (COG-UK) Consortium | Harper VanSteenhouse, Yumi Kasai, David Gray, Carol Clugston, Anna Dominiczak and Alex Alderton, Roberto Amato, Jeffrey Barrett, Sonia Goncalves, Ewan Harrison, David K. Jackson, Ian Johnston, Dominic Kwiatkowski, Cordelia Langford, John Sillitoe on behalf of the Wellcome Sanger Institute COVID-19 Surveillance Team |
| EPI_ISL_2641333, EPI_ISL_2641334 | Lighthouse Lab in Alderley Park | Wellcome Sanger Institute for the COVID-19 Genomics UK (COG-UK) Consortium | Jacquelyn Wynn, Mairead Hyland, The Lighthouse Lab in Alderley Park and Alex Alderton, Roberto Amato, Jeffrey Barrett, Sonia Goncalves, Ewan Harrison, David K. Jackson, Ian Johnston, Dominic Kwiatkowski, Cordelia Langford, John Sillitoe on behalf of the Wellcome Sanger Institute COVID-19 Surveillance Team |
| EPI_ISL_2641335, EPI_ISL_2641336 | Lighthouse Lab in Milton Keynes | Wellcome Sanger Institute for the COVID-19 Genomics UK (COG-UK) Consortium | The Lighthouse Lab in Milton Keynes and Alex Alderton, Roberto Amato, Jeffrey Barrett, Sonia Goncalves, Ewan Harrison, David K. Jackson, Ian Johnston, Dominic Kwiatkowski, Cordelia Langford, John Sillitoe on behalf of the Wellcome Sanger Institute COVID-19 Surveillance Team |
| EPI_ISL_2641337, EPI_ISL_2641338 | Lighthouse Lab in Alderley Park | Wellcome Sanger Institute for the COVID-19 Genomics UK (COG-UK) Consortium | Jacquelyn Wynn, Mairead Hyland, The Lighthouse Lab in Alderley Park and Alex Alderton, Roberto Amato, Jeffrey Barrett, Sonia Goncalves, Ewan Harrison, David K. Jackson, Ian Johnston, Dominic Kwiatkowski, Cordelia Langford, John Sillitoe on behalf of the Wellcome Sanger Institute COVID-19 Surveillance Team |
| EPI_ISL_2641339 | Lighthouse Lab in Glasgow | Wellcome Sanger Institute for the COVID-19 Genomics UK (COG-UK) Consortium | Harper VanSteenhouse, Yumi Kasai, David Gray, Carol Clugston, Anna Dominiczak and Alex Alderton, Roberto Amato, Jeffrey Barrett, Sonia Goncalves, Ewan Harrison, David K. Jackson, Ian Johnston, Dominic Kwiatkowski, Cordelia Langford, John Sillitoe on behalf of the Wellcome Sanger Institute COVID-19 Surveillance Team |
| EPI_ISL_2641340 | Lighthouse Lab in Alderley Park | Wellcome Sanger Institute for the COVID-19 Genomics UK (COG-UK) Consortium | Jacquelyn Wynn, Mairead Hyland, The Lighthouse Lab in Alderley Park and Alex Alderton, Roberto Amato, Jeffrey Barrett, Sonia Goncalves, Ewan Harrison, David K. Jackson, Ian Johnston, Dominic Kwiatkowski, Cordelia Langford, John Sillitoe on behalf of the Wellcome Sanger Institute COVID-19 Surveillance Team |
| EPI_ISL_2641341 | Lighthouse Lab in Glasgow | Wellcome Sanger Institute for the COVID-19 Genomics UK (COG-UK) Consortium | Harper VanSteenhouse, Yumi Kasai, David Gray, Carol Clugston, Anna Dominiczak and Alex Alderton, Roberto Amato, Jeffrey Barrett, Sonia Goncalves, Ewan Harrison, David K. Jackson, Ian Johnston, Dominic Kwiatkowski, Cordelia Langford, John Sillitoe on behalf of the Wellcome Sanger Institute COVID-19 Surveillance Team |
| EPI_ISL_2641342 | Lighthouse Lab in Milton Keynes | Wellcome Sanger Institute for the COVID-19 Genomics UK (COG-UK) Consortium | The Lighthouse Lab in Milton Keynes and Alex Alderton, Roberto Amato, Jeffrey Barrett, Sonia Goncalves, Ewan Harrison, David K. Jackson, Ian Johnston, Dominic Kwiatkowski, Cordelia Langford, John Sillitoe on behalf of the Wellcome Sanger Institute COVID-19 Surveillance Team |
| EPI_ISL_2641343, EPI_ISL_2641344 | Lighthouse Lab in Alderley Park | Wellcome Sanger Institute for the COVID-19 Genomics UK (COG-UK) Consortium | Jacquelyn Wynn, Mairead Hyland, The Lighthouse Lab in Alderley Park and Alex Alderton, Roberto Amato, Jeffrey Barrett, Sonia Goncalves, Ewan Harrison, David K. Jackson, Ian Johnston, Dominic Kwiatkowski, Cordelia Langford, John Sillitoe on behalf of the Wellcome Sanger Institute COVID-19 Surveillance Team |
| EPI_ISL_2641345 | Lighthouse Lab in Glasgow | Wellcome Sanger Institute for the COVID-19 Genomics UK (COG-UK) Consortium | Harper VanSteenhouse, Yumi Kasai, David Gray, Carol Clugston, Anna Dominiczak and Alex Alderton, Roberto Amato, Jeffrey Barrett, Sonia Goncalves, Ewan Harrison, David K. Jackson, Ian Johnston, Dominic Kwiatkowski, Cordelia Langford, John Sillitoe on behalf of the Wellcome Sanger Institute COVID-19 Surveillance Team |
| EPI_ISL_2641346, EPI_ISL_2641347, EPI_ISL_2641348, EPI_ISL_2641349, EPI_ISL_2641350, EPI_ISL_2641352 | Lighthouse Lab in Alderley Park | Wellcome Sanger Institute for the COVID-19 Genomics UK (COG-UK) Consortium | Jacquelyn Wynn, Mairead Hyland, The Lighthouse Lab in Alderley Park and Alex Alderton, Roberto Amato, Jeffrey Barrett, Sonia Goncalves, Ewan Harrison, David K. Jackson, Ian Johnston, Dominic Kwiatkowski, Cordelia Langford, John Sillitoe on behalf of the Wellcome Sanger Institute COVID-19 Surveillance Team |
| EPI_ISL_2641353 | Health Services Laboratories | Wellcome Sanger Institute for the COVID-19 Genomics UK (COG-UK) Consortium | Health Services Laboratories and Alex Alderton, Roberto Amato, Jeffrey Barrett, Sonia Goncalves, Ewan Harrison, David K. Jackson, Ian Johnston, Dominic Kwiatkowski, Cordelia Langford, John Sillitoe on behalf of the Wellcome Sanger Institute COVID-19 Surveillance Team |
| EPI_ISL_2641354, EPI_ISL_2641355 | Lighthouse Lab in Alderley Park | Wellcome Sanger Institute for the COVID-19 Genomics UK (COG-UK) Consortium | Jacquelyn Wynn, Mairead Hyland, The Lighthouse Lab in Alderley Park and Alex Alderton, Roberto Amato, Jeffrey Barrett, Sonia Goncalves, Ewan Harrison, David K. Jackson, Ian Johnston, Dominic Kwiatkowski, Cordelia Langford, John Sillitoe on behalf of the Wellcome Sanger Institute COVID-19 Surveillance Team |
| EPI_ISL_2641356 | Health Services Laboratories | Wellcome Sanger Institute for the COVID-19 Genomics UK (COG-UK) Consortium | Health Services Laboratories and Alex Alderton, Roberto Amato, Jeffrey Barrett, Sonia Goncalves, Ewan Harrison, David K. Jackson, Ian Johnston, Dominic Kwiatkowski, Cordelia Langford, John Sillitoe on behalf of the Wellcome Sanger Institute COVID-19 Surveillance Team |
| EPI_ISL_2641357 | Lighthouse Lab in Alderley Park | Wellcome Sanger Institute for the COVID-19 Genomics UK (COG-UK) Consortium | Jacquelyn Wynn, Mairead Hyland, The Lighthouse Lab in Alderley Park and Alex Alderton, Roberto Amato, Jeffrey Barrett, Sonia Goncalves, Ewan Harrison, David K. Jackson, Ian Johnston, Dominic Kwiatkowski, Cordelia Langford, John Sillitoe on behalf of the Wellcome Sanger Institute COVID-19 Surveillance Team |
| EPI_ISL_2641358 | Health Services Laboratories | Wellcome Sanger Institute for the COVID-19 Genomics UK (COG-UK) Consortium | Health Services Laboratories and Alex Alderton, Roberto Amato, Jeffrey Barrett, Sonia Goncalves, Ewan Harrison, David K. Jackson, Ian Johnston, Dominic Kwiatkowski, Cordelia Langford, John Sillitoe on behalf of the Wellcome Sanger Institute COVID-19 Surveillance Team |
| EPI_ISL_2641359 | Lighthouse Lab in Glasgow | Wellcome Sanger Institute for the COVID-19 Genomics UK (COG-UK) Consortium | Harper VanSteenhouse, Yumi Kasai, David Gray, Carol Clugston, Anna Dominiczak and Alex Alderton, Roberto Amato, Jeffrey Barrett, Sonia Goncalves, Ewan Harrison, David K. Jackson, Ian Johnston, Dominic Kwiatkowski, Cordelia Langford, John Sillitoe on behalf of the Wellcome Sanger Institute COVID-19 Surveillance Team |
| EPI_ISL_2641360 | Health Services Laboratories | Wellcome Sanger Institute for the COVID-19 Genomics UK (COG-UK) Consortium | Health Services Laboratories and Alex Alderton, Roberto Amato, Jeffrey Barrett, Sonia Goncalves, Ewan Harrison, David K. Jackson, Ian Johnston, Dominic Kwiatkowski, Cordelia Langford, John Sillitoe on behalf of the Wellcome Sanger Institute COVID-19 Surveillance Team |
| EPI_ISL_2641361 | Lighthouse Lab in Glasgow | Wellcome Sanger Institute for the COVID-19 Genomics UK (COG-UK) Consortium | Harper VanSteenhouse, Yumi Kasai, David Gray, Carol Clugston, Anna Dominiczak and Alex Alderton, Roberto Amato, Jeffrey Barrett, Sonia Goncalves, Ewan Harrison, David K. Jackson, Ian Johnston, Dominic Kwiatkowski, Cordelia Langford, John Sillitoe on behalf of the Wellcome Sanger Institute COVID-19 Surveillance Team |
| EPI_ISL_2641362, EPI_ISL_2641363 | Lighthouse Lab in Alderley Park | Wellcome Sanger Institute for the COVID-19 Genomics UK (COG-UK) Consortium | Jacquelyn Wynn, Mairead Hyland, The Lighthouse Lab in Alderley Park and Alex Alderton, Roberto Amato, Jeffrey Barrett, Sonia Goncalves, Ewan Harrison, David K. Jackson, Ian Johnston, Dominic Kwiatkowski, Cordelia Langford, John Sillitoe on behalf of the Wellcome Sanger Institute COVID-19 Surveillance Team |

| | | | |
|---|---|---|---|
| EPI_ISL_2641364, EPI_ISL_2641365, EPI_ISL_2641366 | Lighthouse Lab in Glasgow | Wellcome Sanger Institute for the COVID-19 Genomics UK (COG-UK) Consortium | Harper VanSteenhouse, Yumi Kasai, David Gray, Carol Clugston, Anna Dominiczak and Alex Alderton, Roberto Amato, Jeffrey Barrett, Sonia Goncalves, Ewan Harrison, David K. Jackson, Ian Johnston, Dominic Kwiatkowski, Cordelia Langford, John Sillitoe on behalf of the Wellcome Sanger Institute COVID-19 Surveillance Team |
| EPI_ISL_2641367, EPI_ISL_2641368 | Lighthouse Lab in Alderley Park | Wellcome Sanger Institute for the COVID-19 Genomics UK (COG-UK) Consortium | Jacquelyn Wynn, Mairead Hyland, The Lighthouse Lab in Alderley Park and Alex Alderton, Roberto Amato, Jeffrey Barrett, Sonia Goncalves, Ewan Harrison, David K. Jackson, Ian Johnston, Dominic Kwiatkowski, Cordelia Langford, John Sillitoe on behalf of the Wellcome Sanger Institute COVID-19 Surveillance Team |
| EPI_ISL_2641369, EPI_ISL_2641370 | Health Services Laboratories | Wellcome Sanger Institute for the COVID-19 Genomics UK (COG-UK) Consortium | Health Services Laboratories and Alex Alderton, Roberto Amato, Jeffrey Barrett, Sonia Goncalves, Ewan Harrison, David K. Jackson, Ian Johnston, Dominic Kwiatkowski, Cordelia Langford, John Sillitoe on behalf of the Wellcome Sanger Institute COVID-19 Surveillance Team |
| EPI_ISL_2641371, EPI_ISL_2641374, EPI_ISL_2641375 | Lighthouse Lab in Glasgow | Wellcome Sanger Institute for the COVID-19 Genomics UK (COG-UK) Consortium | Harper VanSteenhouse, Yumi Kasai, David Gray, Carol Clugston, Anna Dominiczak and Alex Alderton, Roberto Amato, Jeffrey Barrett, Sonia Goncalves, Ewan Harrison, David K. Jackson, Ian Johnston, Dominic Kwiatkowski, Cordelia Langford, John Sillitoe on behalf of the Wellcome Sanger Institute COVID-19 Surveillance Team |
| EPI_ISL_2641377, EPI_ISL_2641378, EPI_ISL_2641379 | Lighthouse Lab in Alderley Park | Wellcome Sanger Institute for the COVID-19 Genomics UK (COG-UK) Consortium | Jacquelyn Wynn, Mairead Hyland, The Lighthouse Lab in Alderley Park and Alex Alderton, Roberto Amato, Jeffrey Barrett, Sonia Goncalves, Ewan Harrison, David K. Jackson, Ian Johnston, Dominic Kwiatkowski, Cordelia Langford, John Sillitoe on behalf of the Wellcome Sanger Institute COVID-19 Surveillance Team |
| EPI_ISL_2641380 | Lighthouse Lab in Glasgow | Wellcome Sanger Institute for the COVID-19 Genomics UK (COG-UK) Consortium | Harper VanSteenhouse, Yumi Kasai, David Gray, Carol Clugston, Anna Dominiczak and Alex Alderton, Roberto Amato, Jeffrey Barrett, Sonia Goncalves, Ewan Harrison, David K. Jackson, Ian Johnston, Dominic Kwiatkowski, Cordelia Langford, John Sillitoe on behalf of the Wellcome Sanger Institute COVID-19 Surveillance Team |
| EPI_ISL_2641381, EPI_ISL_2641382, EPI_ISL_2641383 | Lighthouse Lab in Alderley Park | Wellcome Sanger Institute for the COVID-19 Genomics UK (COG-UK) Consortium | Jacquelyn Wynn, Mairead Hyland, The Lighthouse Lab in Alderley Park and Alex Alderton, Roberto Amato, Jeffrey Barrett, Sonia Goncalves, Ewan Harrison, David K. Jackson, Ian Johnston, Dominic Kwiatkowski, Cordelia Langford, John Sillitoe on behalf of the Wellcome Sanger Institute COVID-19 Surveillance Team |
| EPI_ISL_2641384 | Health Services Laboratories | Wellcome Sanger Institute for the COVID-19 Genomics UK (COG-UK) Consortium | Health Services Laboratories and Alex Alderton, Roberto Amato, Jeffrey Barrett, Sonia Goncalves, Ewan Harrison, David K. Jackson, Ian Johnston, Dominic Kwiatkowski, Cordelia Langford, John Sillitoe on behalf of the Wellcome Sanger Institute COVID-19 Surveillance Team |
| EPI_ISL_2641385 | Lighthouse Lab in Glasgow | Wellcome Sanger Institute for the COVID-19 Genomics UK (COG-UK) Consortium | Harper VanSteenhouse, Yumi Kasai, David Gray, Carol Clugston, Anna Dominiczak and Alex Alderton, Roberto Amato, Jeffrey Barrett, Sonia Goncalves, Ewan Harrison, David K. Jackson, Ian Johnston, Dominic Kwiatkowski, Cordelia Langford, John Sillitoe on behalf of the Wellcome Sanger Institute COVID-19 Surveillance Team |
| EPI_ISL_2641386, EPI_ISL_2641387 | Lighthouse Lab in Alderley Park | Wellcome Sanger Institute for the COVID-19 Genomics UK (COG-UK) Consortium | Jacquelyn Wynn, Mairead Hyland, The Lighthouse Lab in Alderley Park and Alex Alderton, Roberto Amato, Jeffrey Barrett, Sonia Goncalves, Ewan Harrison, David K. Jackson, Ian Johnston, Dominic Kwiatkowski, Cordelia Langford, John Sillitoe on behalf of the Wellcome Sanger Institute COVID-19 Surveillance Team |
| EPI_ISL_2641388, EPI_ISL_2641389 | Lighthouse Lab in Glasgow | Wellcome Sanger Institute for the COVID-19 Genomics UK (COG-UK) Consortium | Harper VanSteenhouse, Yumi Kasai, David Gray, Carol Clugston, Anna Dominiczak and Alex Alderton, Roberto Amato, Jeffrey Barrett, Sonia Goncalves, Ewan Harrison, David K. Jackson, Ian Johnston, Dominic Kwiatkowski, Cordelia Langford, John Sillitoe on behalf of the Wellcome Sanger Institute COVID-19 Surveillance Team |
| EPI_ISL_2641390 | Health Services Laboratories | Wellcome Sanger Institute for the COVID-19 Genomics UK (COG-UK) Consortium | Health Services Laboratories and Alex Alderton, Roberto Amato, Jeffrey Barrett, Sonia Goncalves, Ewan Harrison, David K. Jackson, Ian Johnston, Dominic Kwiatkowski, Cordelia Langford, John Sillitoe on behalf of the Wellcome Sanger Institute COVID-19 Surveillance Team |
| EPI_ISL_2641391 | Lighthouse Lab in Alderley Park | Wellcome Sanger Institute for the COVID-19 Genomics UK (COG-UK) Consortium | Jacquelyn Wynn, Mairead Hyland, The Lighthouse Lab in Alderley Park and Alex Alderton, Roberto Amato, Jeffrey Barrett, Sonia Goncalves, Ewan Harrison, David K. Jackson, Ian Johnston, Dominic Kwiatkowski, Cordelia Langford, John Sillitoe on behalf of the Wellcome Sanger Institute COVID-19 Surveillance Team |
| EPI_ISL_2641392, EPI_ISL_2641393 | Lighthouse Lab in Glasgow | Wellcome Sanger Institute for the COVID-19 Genomics UK (COG-UK) Consortium | Harper VanSteenhouse, Yumi Kasai, David Gray, Carol Clugston, Anna Dominiczak and Alex Alderton, Roberto Amato, Jeffrey Barrett, Sonia Goncalves, Ewan Harrison, David K. Jackson, Ian Johnston, Dominic Kwiatkowski, Cordelia Langford, John Sillitoe on behalf of the Wellcome Sanger Institute COVID-19 Surveillance Team |
| EPI_ISL_2641394 | Health Services Laboratories | Wellcome Sanger Institute for the COVID-19 Genomics UK (COG-UK) Consortium | Health Services Laboratories and Alex Alderton, Roberto Amato, Jeffrey Barrett, Sonia Goncalves, Ewan Harrison, David K. Jackson, Ian Johnston, Dominic Kwiatkowski, Cordelia Langford, John Sillitoe on behalf of the Wellcome Sanger Institute COVID-19 Surveillance Team |
| EPI_ISL_2641395 | Lighthouse Lab in Alderley Park | Wellcome Sanger Institute for the COVID-19 Genomics UK (COG-UK) Consortium | Jacquelyn Wynn, Mairead Hyland, The Lighthouse Lab in Alderley Park and Alex Alderton, Roberto Amato, Jeffrey Barrett, Sonia Goncalves, Ewan Harrison, David K. Jackson, Ian Johnston, Dominic Kwiatkowski, Cordelia Langford, John Sillitoe on behalf of the Wellcome Sanger Institute COVID-19 Surveillance Team |
| EPI_ISL_2641396, EPI_ISL_2641397 | Lighthouse Lab in Glasgow | Wellcome Sanger Institute for the COVID-19 Genomics UK (COG-UK) Consortium | Harper VanSteenhouse, Yumi Kasai, David Gray, Carol Clugston, Anna Dominiczak and Alex Alderton, Roberto Amato, Jeffrey Barrett, Sonia Goncalves, Ewan Harrison, David K. Jackson, Ian Johnston, Dominic Kwiatkowski, Cordelia Langford, John Sillitoe on behalf of the Wellcome Sanger Institute COVID-19 Surveillance Team |
| EPI_ISL_2641398 | Health Services Laboratories | Wellcome Sanger Institute for the COVID-19 Genomics UK (COG-UK) Consortium | Health Services Laboratories and Alex Alderton, Roberto Amato, Jeffrey Barrett, Sonia Goncalves, Ewan Harrison, David K. Jackson, Ian Johnston, Dominic Kwiatkowski, Cordelia Langford, John Sillitoe on behalf of the Wellcome Sanger Institute COVID-19 Surveillance Team |
| EPI_ISL_2641399, EPI_ISL_2641400 | Lighthouse Lab in Glasgow | Wellcome Sanger Institute for the COVID-19 Genomics UK (COG-UK) Consortium | Harper VanSteenhouse, Yumi Kasai, David Gray, Carol Clugston, Anna Dominiczak and Alex Alderton, Roberto Amato, Jeffrey Barrett, Sonia Goncalves, Ewan Harrison, David K. Jackson, Ian Johnston, Dominic Kwiatkowski, Cordelia Langford, John Sillitoe on behalf of the Wellcome Sanger Institute COVID-19 Surveillance Team |
| EPI_ISL_2641401 | Health Services Laboratories | Wellcome Sanger Institute for the COVID-19 Genomics UK (COG-UK) Consortium | Health Services Laboratories and Alex Alderton, Roberto Amato, Jeffrey Barrett, Sonia Goncalves, Ewan Harrison, David K. Jackson, Ian Johnston, Dominic Kwiatkowski, Cordelia Langford, John Sillitoe on behalf of the Wellcome Sanger Institute COVID-19 Surveillance Team |
| EPI_ISL_2641402 | Lighthouse Lab in Alderley Park | Wellcome Sanger Institute for the COVID-19 Genomics UK (COG-UK) Consortium | Jacquelyn Wynn, Mairead Hyland, The Lighthouse Lab in Alderley Park and Alex Alderton, Roberto Amato, Jeffrey Barrett, Sonia Goncalves, Ewan Harrison, David K. Jackson, Ian Johnston, Dominic Kwiatkowski, Cordelia Langford, John Sillitoe on behalf of the Wellcome Sanger Institute COVID-19 Surveillance Team |
| EPI_ISL_2641403, EPI_ISL_2641404, EPI_ISL_2641405 | Lighthouse Lab in Glasgow | Wellcome Sanger Institute for the COVID-19 Genomics UK (COG-UK) Consortium | Harper VanSteenhouse, Yumi Kasai, David Gray, Carol Clugston, Anna Dominiczak and Alex Alderton, Roberto Amato, Jeffrey Barrett, Sonia Goncalves, Ewan Harrison, David K. Jackson, Ian Johnston, Dominic Kwiatkowski, Cordelia Langford, John Sillitoe on behalf of the Wellcome Sanger Institute COVID-19 Surveillance Team |
| EPI_ISL_2641406 | Lighthouse Lab in Alderley Park | Wellcome Sanger Institute for the COVID-19 Genomics UK (COG-UK) Consortium | Jacquelyn Wynn, Mairead Hyland, The Lighthouse Lab in Alderley Park and Alex Alderton, Roberto Amato, Jeffrey Barrett, Sonia Goncalves, Ewan Harrison, David K. Jackson, Ian Johnston, Dominic Kwiatkowski, Cordelia Langford, John Sillitoe on behalf of the Wellcome Sanger Institute COVID-19 Surveillance Team |
| EPI_ISL_2641407, EPI_ISL_2641409 | Health Services Laboratories | Wellcome Sanger Institute for the COVID-19 Genomics UK (COG-UK) Consortium | Health Services Laboratories and Alex Alderton, Roberto Amato, Jeffrey Barrett, Sonia Goncalves, Ewan Harrison, David K. Jackson, Ian Johnston, Dominic Kwiatkowski, Cordelia Langford, John Sillitoe on behalf of the Wellcome Sanger Institute COVID-19 Surveillance Team |
| EPI_ISL_2641410, EPI_ISL_2641411 | Lighthouse Lab in Glasgow | Wellcome Sanger Institute for the COVID-19 Genomics UK (COG-UK) Consortium | Harper VanSteenhouse, Yumi Kasai, David Gray, Carol Clugston, Anna Dominiczak and Alex Alderton, Roberto Amato, Jeffrey Barrett, Sonia Goncalves, Ewan Harrison, David K. Jackson, Ian Johnston, Dominic Kwiatkowski, Cordelia Langford, John Sillitoe on behalf of the Wellcome Sanger Institute COVID-19 Surveillance Team |
| EPI_ISL_2641412 | Health Services Laboratories | Wellcome Sanger Institute for the COVID-19 Genomics UK (COG-UK) Consortium | Health Services Laboratories and Alex Alderton, Roberto Amato, Jeffrey Barrett, Sonia Goncalves, Ewan Harrison, David K. Jackson, Ian Johnston, Dominic Kwiatkowski, Cordelia Langford, John Sillitoe on behalf of the Wellcome Sanger Institute COVID-19 Surveillance Team |
| EPI_ISL_2641413, EPI_ISL_2641414, EPI_ISL_2641415 | Lighthouse Lab in Alderley Park | Wellcome Sanger Institute for the COVID-19 Genomics UK (COG-UK) Consortium | Jacquelyn Wynn, Mairead Hyland, The Lighthouse Lab in Alderley Park and Alex Alderton, Roberto Amato, Jeffrey Barrett, Sonia Goncalves, Ewan Harrison, David K. Jackson, Ian Johnston, Dominic Kwiatkowski, Cordelia Langford, John Sillitoe on behalf of the Wellcome Sanger Institute COVID-19 Surveillance Team |
| EPI_ISL_2641416, EPI_ISL_2641417, EPI_ISL_2641418 | Lighthouse Lab in Glasgow | Wellcome Sanger Institute for the COVID-19 Genomics UK (COG-UK) Consortium | Harper VanSteenhouse, Yumi Kasai, David Gray, Carol Clugston, Anna Dominiczak and Alex Alderton, Roberto Amato, Jeffrey Barrett, Sonia Goncalves, Ewan Harrison, David K. Jackson, Ian Johnston, Dominic Kwiatkowski, Cordelia Langford, John Sillitoe on behalf of the Wellcome Sanger Institute COVID-19 Surveillance Team |
| EPI_ISL_2641419, EPI_ISL_2641420, EPI_ISL_2641421 | Health Services Laboratories | Wellcome Sanger Institute for the COVID-19 Genomics UK (COG-UK) Consortium | Health Services Laboratories and Alex Alderton, Roberto Amato, Jeffrey Barrett, Sonia Goncalves, Ewan Harrison, David K. Jackson, Ian Johnston, Dominic Kwiatkowski, Cordelia Langford, John Sillitoe on behalf of the Wellcome Sanger Institute COVID-19 Surveillance Team |
| EPI_ISL_2641423 | Lighthouse Lab in Alderley Park | Wellcome Sanger Institute for the COVID-19 Genomics UK (COG-UK) Consortium | Jacquelyn Wynn, Mairead Hyland, The Lighthouse Lab in Alderley Park and Alex Alderton, Roberto Amato, Jeffrey Barrett, Sonia Goncalves, Ewan Harrison, David K. Jackson, Ian Johnston, Dominic Kwiatkowski, Cordelia Langford, John Sillitoe on behalf of the Wellcome Sanger Institute COVID-19 Surveillance Team |
| EPI_ISL_2641424, EPI_ISL_2641425 | Lighthouse Lab in Glasgow | Wellcome Sanger Institute for the COVID-19 Genomics UK (COG-UK) Consortium | Harper VanSteenhouse, Yumi Kasai, David Gray, Carol Clugston, Anna Dominiczak and Alex Alderton, Roberto Amato, Jeffrey Barrett, Sonia Goncalves, Ewan Harrison, David K. Jackson, Ian Johnston, Dominic Kwiatkowski, Cordelia Langford, John Sillitoe on behalf of the Wellcome Sanger Institute COVID-19 Surveillance Team |
| EPI_ISL_2641426 | Health Services Laboratories | Wellcome Sanger Institute for the COVID-19 Genomics UK (COG-UK) Consortium | Health Services Laboratories and Alex Alderton, Roberto Amato, Jeffrey Barrett, Sonia Goncalves, Ewan Harrison, David K. Jackson, Ian Johnston, Dominic Kwiatkowski, Cordelia Langford, John Sillitoe on behalf of the Wellcome Sanger Institute COVID-19 Surveillance Team |
| EPI_ISL_2641427, EPI_ISL_2641428 | Lighthouse Lab in Glasgow | Wellcome Sanger Institute for the COVID-19 Genomics UK (COG-UK) Consortium | Harper VanSteenhouse, Yumi Kasai, David Gray, Carol Clugston, Anna Dominiczak and Alex Alderton, Roberto Amato, Jeffrey Barrett, Sonia Goncalves, Ewan Harrison, David K. Jackson, Ian Johnston, Dominic Kwiatkowski, Cordelia Langford, John Sillitoe on behalf of the Wellcome Sanger Institute COVID-19 Surveillance Team |
| EPI_ISL_2641429 | Health Services Laboratories | Wellcome Sanger Institute for the COVID-19 Genomics UK (COG-UK) Consortium | Health Services Laboratories and Alex Alderton, Roberto Amato, Jeffrey Barrett, Sonia Goncalves, Ewan Harrison, David K. Jackson, Ian Johnston, Dominic Kwiatkowski, Cordelia Langford, John Sillitoe on behalf of the Wellcome Sanger Institute COVID-19 Surveillance Team |
| EPI_ISL_2641430 | Lighthouse Lab in Glasgow | Wellcome Sanger Institute for the COVID-19 Genomics UK (COG-UK) Consortium | Harper VanSteenhouse, Yumi Kasai, David Gray, Carol Clugston, Anna Dominiczak and Alex Alderton, Roberto Amato, Jeffrey Barrett, Sonia Goncalves, Ewan Harrison, David K. Jackson, Ian Johnston, Dominic Kwiatkowski, Cordelia Langford, John Sillitoe on behalf of the Wellcome Sanger Institute COVID-19 Surveillance Team |
| EPI_ISL_2641431, EPI_ISL_2641432 | Health Services Laboratories | Wellcome Sanger Institute for the COVID-19 Genomics UK (COG-UK) Consortium | Health Services Laboratories and Alex Alderton, Roberto Amato, Jeffrey Barrett, Sonia Goncalves, Ewan Harrison, David K. Jackson, Ian Johnston, Dominic Kwiatkowski, Cordelia Langford, John Sillitoe on behalf of the Wellcome Sanger Institute COVID-19 Surveillance Team |
| EPI_ISL_2641433, EPI_ISL_2641434, EPI_ISL_2641435, EPI_ISL_2641438, EPI_ISL_2641439, EPI_ISL_2641441 | Lighthouse Lab in Glasgow | Wellcome Sanger Institute for the COVID-19 Genomics UK (COG-UK) Consortium | Harper VanSteenhouse, Yumi Kasai, David Gray, Carol Clugston, Anna Dominiczak and Alex Alderton, Roberto Amato, Jeffrey Barrett, Sonia Goncalves, Ewan Harrison, David K. Jackson, Ian Johnston, Dominic Kwiatkowski, Cordelia Langford, John Sillitoe on behalf of the Wellcome Sanger Institute COVID-19 Surveillance Team |
| EPI_ISL_2641442 | Lighthouse Lab in Alderley Park | Wellcome Sanger Institute for the COVID-19 Genomics UK (COG-UK) Consortium | Jacquelyn Wynn, Mairead Hyland, The Lighthouse Lab in Alderley Park and Alex Alderton, Roberto Amato, Jeffrey Barrett, Sonia Goncalves, Ewan Harrison, David K. Jackson, Ian Johnston, Dominic Kwiatkowski, Cordelia Langford, John Sillitoe on behalf of the Wellcome Sanger Institute COVID-19 Surveillance Team |
| EPI_ISL_2641443 | Health Services Laboratories | Wellcome Sanger Institute for the COVID-19 Genomics UK (COG-UK) Consortium | Health Services Laboratories and Alex Alderton, Roberto Amato, Jeffrey Barrett, Sonia Goncalves, Ewan Harrison, David K. Jackson, Ian Johnston, Dominic Kwiatkowski, Cordelia Langford, John Sillitoe on behalf of the Wellcome Sanger Institute COVID-19 Surveillance Team |
| EPI_ISL_2641444, EPI_ISL_2641445, EPI_ISL_2641447, | Lighthouse Lab in Glasgow | Wellcome Sanger Institute for the COVID-19 Genomics UK (COG-UK) Consortium | Harper VanSteenhouse, Yumi Kasai, David Gray, Carol Clugston, Anna Dominiczak and Alex Alderton, Roberto Amato, Jeffrey Barrett, Sonia Goncalves, Ewan Harrison, David K. Jackson, Ian Johnston, Dominic Kwiatkowski, Cordelia Langford, John Sillitoe on behalf of the Wellcome Sanger Institute COVID-19 Surveillance Team |

| | | | |
|---|---|---|---|
| EPI_ISL_2641448 | | | |
| EPI_ISL_2641449, EPI_ISL_2641450 | Health Services Laboratories | Wellcome Sanger Institute for the COVID-19 Genomics UK (COG-UK) Consortium | Health Services Laboratories and Alex Alderton, Roberto Amato, Jeffrey Barrett, Sonia Goncalves, Ewan Harrison, David K. Jackson, Ian Johnston, Dominic Kwiatkowski, Cordelia Langford, John Sillitoe on behalf of the Wellcome Sanger Institute COVID-19 Surveillance Team |
| EPI_ISL_2641451, EPI_ISL_2641452, EPI_ISL_2641453, EPI_ISL_2641454, EPI_ISL_2641455, EPI_ISL_2641456, EPI_ISL_2641457 | Lighthouse Lab in Glasgow | Wellcome Sanger Institute for the COVID-19 Genomics UK (COG-UK) Consortium | Harper VanSteenhouse, Yumi Kasai, David Gray, Carol Clugston, Anna Dominiczak and Alex Alderton, Roberto Amato, Jeffrey Barrett, Sonia Goncalves, Ewan Harrison, David K. Jackson, Ian Johnston, Dominic Kwiatkowski, Cordelia Langford, John Sillitoe on behalf of the Wellcome Sanger Institute COVID-19 Surveillance Team |
| EPI_ISL_2641458 | Health Services Laboratories | Wellcome Sanger Institute for the COVID-19 Genomics UK (COG-UK) Consortium | Health Services Laboratories and Alex Alderton, Roberto Amato, Jeffrey Barrett, Sonia Goncalves, Ewan Harrison, David K. Jackson, Ian Johnston, Dominic Kwiatkowski, Cordelia Langford, John Sillitoe on behalf of the Wellcome Sanger Institute COVID-19 Surveillance Team |
| EPI_ISL_2641459 | Lighthouse Lab in Glasgow | Wellcome Sanger Institute for the COVID-19 Genomics UK (COG-UK) Consortium | Harper VanSteenhouse, Yumi Kasai, David Gray, Carol Clugston, Anna Dominiczak and Alex Alderton, Roberto Amato, Jeffrey Barrett, Sonia Goncalves, Ewan Harrison, David K. Jackson, Ian Johnston, Dominic Kwiatkowski, Cordelia Langford, John Sillitoe on behalf of the Wellcome Sanger Institute COVID-19 Surveillance Team |
| EPI_ISL_2641460 | Lighthouse Lab in Alderley Park | Wellcome Sanger Institute for the COVID-19 Genomics UK (COG-UK) Consortium | Jacquelyn Wynn, Mairead Hyland, The Lighthouse Lab in Alderley Park and Alex Alderton, Roberto Amato, Jeffrey Barrett, Sonia Goncalves, Ewan Harrison, David K. Jackson, Ian Johnston, Dominic Kwiatkowski, Cordelia Langford, John Sillitoe on behalf of the Wellcome Sanger Institute COVID-19 Surveillance Team |
| EPI_ISL_2641461 | Lighthouse Lab in Glasgow | Wellcome Sanger Institute for the COVID-19 Genomics UK (COG-UK) Consortium | Harper VanSteenhouse, Yumi Kasai, David Gray, Carol Clugston, Anna Dominiczak and Alex Alderton, Roberto Amato, Jeffrey Barrett, Sonia Goncalves, Ewan Harrison, David K. Jackson, Ian Johnston, Dominic Kwiatkowski, Cordelia Langford, John Sillitoe on behalf of the Wellcome Sanger Institute COVID-19 Surveillance Team |
| EPI_ISL_2641462, EPI_ISL_2641463 | Health Services Laboratories | Wellcome Sanger Institute for the COVID-19 Genomics UK (COG-UK) Consortium | Health Services Laboratories and Alex Alderton, Roberto Amato, Jeffrey Barrett, Sonia Goncalves, Ewan Harrison, David K. Jackson, Ian Johnston, Dominic Kwiatkowski, Cordelia Langford, John Sillitoe on behalf of the Wellcome Sanger Institute COVID-19 Surveillance Team |
| EPI_ISL_2641464, EPI_ISL_2641465 | Lighthouse Lab in Glasgow | Wellcome Sanger Institute for the COVID-19 Genomics UK (COG-UK) Consortium | Harper VanSteenhouse, Yumi Kasai, David Gray, Carol Clugston, Anna Dominiczak and Alex Alderton, Roberto Amato, Jeffrey Barrett, Sonia Goncalves, Ewan Harrison, David K. Jackson, Ian Johnston, Dominic Kwiatkowski, Cordelia Langford, John Sillitoe on behalf of the Wellcome Sanger Institute COVID-19 Surveillance Team |
| EPI_ISL_2641466 | Health Services Laboratories | Wellcome Sanger Institute for the COVID-19 Genomics UK (COG-UK) Consortium | Health Services Laboratories and Alex Alderton, Roberto Amato, Jeffrey Barrett, Sonia Goncalves, Ewan Harrison, David K. Jackson, Ian Johnston, Dominic Kwiatkowski, Cordelia Langford, John Sillitoe on behalf of the Wellcome Sanger Institute COVID-19 Surveillance Team |
| EPI_ISL_2641467 | Lighthouse Lab in Glasgow | Wellcome Sanger Institute for the COVID-19 Genomics UK (COG-UK) Consortium | Harper VanSteenhouse, Yumi Kasai, David Gray, Carol Clugston, Anna Dominiczak and Alex Alderton, Roberto Amato, Jeffrey Barrett, Sonia Goncalves, Ewan Harrison, David K. Jackson, Ian Johnston, Dominic Kwiatkowski, Cordelia Langford, John Sillitoe on behalf of the Wellcome Sanger Institute COVID-19 Surveillance Team |
| EPI_ISL_2641468 | Health Services Laboratories | Wellcome Sanger Institute for the COVID-19 Genomics UK (COG-UK) Consortium | Health Services Laboratories and Alex Alderton, Roberto Amato, Jeffrey Barrett, Sonia Goncalves, Ewan Harrison, David K. Jackson, Ian Johnston, Dominic Kwiatkowski, Cordelia Langford, John Sillitoe on behalf of the Wellcome Sanger Institute COVID-19 Surveillance Team |
| EPI_ISL_2641469 | Lighthouse Lab in Alderley Park | Wellcome Sanger Institute for the COVID-19 Genomics UK (COG-UK) Consortium | Jacquelyn Wynn, Mairead Hyland, The Lighthouse Lab in Alderley Park and Alex Alderton, Roberto Amato, Jeffrey Barrett, Sonia Goncalves, Ewan Harrison, David K. Jackson, Ian Johnston, Dominic Kwiatkowski, Cordelia Langford, John Sillitoe on behalf of the Wellcome Sanger Institute COVID-19 Surveillance Team |
| EPI_ISL_2641470 | Health Services Laboratories | Wellcome Sanger Institute for the COVID-19 Genomics UK (COG-UK) Consortium | Health Services Laboratories and Alex Alderton, Roberto Amato, Jeffrey Barrett, Sonia Goncalves, Ewan Harrison, David K. Jackson, Ian Johnston, Dominic Kwiatkowski, Cordelia Langford, John Sillitoe on behalf of the Wellcome Sanger Institute COVID-19 Surveillance Team |
| EPI_ISL_2641471 | Lighthouse Lab in Glasgow | Wellcome Sanger Institute for the COVID-19 Genomics UK (COG-UK) Consortium | Harper VanSteenhouse, Yumi Kasai, David Gray, Carol Clugston, Anna Dominiczak and Alex Alderton, Roberto Amato, Jeffrey Barrett, Sonia Goncalves, Ewan Harrison, David K. Jackson, Ian Johnston, Dominic Kwiatkowski, Cordelia Langford, John Sillitoe on behalf of the Wellcome Sanger Institute COVID-19 Surveillance Team |
| EPI_ISL_2641472 | Lighthouse Lab in Alderley Park | Wellcome Sanger Institute for the COVID-19 Genomics UK (COG-UK) Consortium | Jacquelyn Wynn, Mairead Hyland, The Lighthouse Lab in Alderley Park and Alex Alderton, Roberto Amato, Jeffrey Barrett, Sonia Goncalves, Ewan Harrison, David K. Jackson, Ian Johnston, Dominic Kwiatkowski, Cordelia Langford, John Sillitoe on behalf of the Wellcome Sanger Institute COVID-19 Surveillance Team |
| EPI_ISL_2641473, EPI_ISL_2641474, EPI_ISL_2641475, EPI_ISL_2641476, EPI_ISL_2641477 | Lighthouse Lab in Glasgow | Wellcome Sanger Institute for the COVID-19 Genomics UK (COG-UK) Consortium | Harper VanSteenhouse, Yumi Kasai, David Gray, Carol Clugston, Anna Dominiczak and Alex Alderton, Roberto Amato, Jeffrey Barrett, Sonia Goncalves, Ewan Harrison, David K. Jackson, Ian Johnston, Dominic Kwiatkowski, Cordelia Langford, John Sillitoe on behalf of the Wellcome Sanger Institute COVID-19 Surveillance Team |
| EPI_ISL_2641480 | Lighthouse Lab in Alderley Park | Wellcome Sanger Institute for the COVID-19 Genomics UK (COG-UK) Consortium | Jacquelyn Wynn, Mairead Hyland, The Lighthouse Lab in Alderley Park and Alex Alderton, Roberto Amato, Jeffrey Barrett, Sonia Goncalves, Ewan Harrison, David K. Jackson, Ian Johnston, Dominic Kwiatkowski, Cordelia Langford, John Sillitoe on behalf of the Wellcome Sanger Institute COVID-19 Surveillance Team |
| EPI_ISL_2641481, EPI_ISL_2641482, EPI_ISL_2641483 | Lighthouse Lab in Glasgow | Wellcome Sanger Institute for the COVID-19 Genomics UK (COG-UK) Consortium | Harper VanSteenhouse, Yumi Kasai, David Gray, Carol Clugston, Anna Dominiczak and Alex Alderton, Roberto Amato, Jeffrey Barrett, Sonia Goncalves, Ewan Harrison, David K. Jackson, Ian Johnston, Dominic Kwiatkowski, Cordelia Langford, John Sillitoe on behalf of the Wellcome Sanger Institute COVID-19 Surveillance Team |
| EPI_ISL_2641484 | Lighthouse Lab in Alderley Park | Wellcome Sanger Institute for the COVID-19 Genomics UK (COG-UK) Consortium | Jacquelyn Wynn, Mairead Hyland, The Lighthouse Lab in Alderley Park and Alex Alderton, Roberto Amato, Jeffrey Barrett, Sonia Goncalves, Ewan Harrison, David K. Jackson, Ian Johnston, Dominic Kwiatkowski, Cordelia Langford, John Sillitoe on behalf of the Wellcome Sanger Institute COVID-19 Surveillance Team |
| EPI_ISL_2641485, EPI_ISL_2641486, EPI_ISL_2641487 | Lighthouse Lab in Glasgow | Wellcome Sanger Institute for the COVID-19 Genomics UK (COG-UK) Consortium | Harper VanSteenhouse, Yumi Kasai, David Gray, Carol Clugston, Anna Dominiczak and Alex Alderton, Roberto Amato, Jeffrey Barrett, Sonia Goncalves, Ewan Harrison, David K. Jackson, Ian Johnston, Dominic Kwiatkowski, Cordelia Langford, John Sillitoe on behalf of the Wellcome Sanger Institute COVID-19 Surveillance Team |
| EPI_ISL_2641488, EPI_ISL_2641489, EPI_ISL_2641490, EPI_ISL_2641491 | Health Services Laboratories | Wellcome Sanger Institute for the COVID-19 Genomics UK (COG-UK) Consortium | Health Services Laboratories and Alex Alderton, Roberto Amato, Jeffrey Barrett, Sonia Goncalves, Ewan Harrison, David K. Jackson, Ian Johnston, Dominic Kwiatkowski, Cordelia Langford, John Sillitoe on behalf of the Wellcome Sanger Institute COVID-19 Surveillance Team |
| EPI_ISL_2641492, EPI_ISL_2641493, EPI_ISL_2641494 | Lighthouse Lab in Alderley Park | Wellcome Sanger Institute for the COVID-19 Genomics UK (COG-UK) Consortium | Jacquelyn Wynn, Mairead Hyland, The Lighthouse Lab in Alderley Park and Alex Alderton, Roberto Amato, Jeffrey Barrett, Sonia Goncalves, Ewan Harrison, David K. Jackson, Ian Johnston, Dominic Kwiatkowski, Cordelia Langford, John Sillitoe on behalf of the Wellcome Sanger Institute COVID-19 Surveillance Team |
| EPI_ISL_2641495 | Health Services Laboratories | Wellcome Sanger Institute for the COVID-19 Genomics UK (COG-UK) Consortium | Health Services Laboratories and Alex Alderton, Roberto Amato, Jeffrey Barrett, Sonia Goncalves, Ewan Harrison, David K. Jackson, Ian Johnston, Dominic Kwiatkowski, Cordelia Langford, John Sillitoe on behalf of the Wellcome Sanger Institute COVID-19 Surveillance Team |
| EPI_ISL_2641496, EPI_ISL_2641497 | Lighthouse Lab in Glasgow | Wellcome Sanger Institute for the COVID-19 Genomics UK (COG-UK) Consortium | Harper VanSteenhouse, Yumi Kasai, David Gray, Carol Clugston, Anna Dominiczak and Alex Alderton, Roberto Amato, Jeffrey Barrett, Sonia Goncalves, Ewan Harrison, David K. Jackson, Ian Johnston, Dominic Kwiatkowski, Cordelia Langford, John Sillitoe on behalf of the Wellcome Sanger Institute COVID-19 Surveillance Team |
| EPI_ISL_2641498, EPI_ISL_2641500 | Lighthouse Lab in Alderley Park | Wellcome Sanger Institute for the COVID-19 Genomics UK (COG-UK) Consortium | Jacquelyn Wynn, Mairead Hyland, The Lighthouse Lab in Alderley Park and Alex Alderton, Roberto Amato, Jeffrey Barrett, Sonia Goncalves, Ewan Harrison, David K. Jackson, Ian Johnston, Dominic Kwiatkowski, Cordelia Langford, John Sillitoe on behalf of the Wellcome Sanger Institute COVID-19 Surveillance Team |
| EPI_ISL_2641501, EPI_ISL_2641502, EPI_ISL_2641503 | Lighthouse Lab in Glasgow | Wellcome Sanger Institute for the COVID-19 Genomics UK (COG-UK) Consortium | Harper VanSteenhouse, Yumi Kasai, David Gray, Carol Clugston, Anna Dominiczak and Alex Alderton, Roberto Amato, Jeffrey Barrett, Sonia Goncalves, Ewan Harrison, David K. Jackson, Ian Johnston, Dominic Kwiatkowski, Cordelia Langford, John Sillitoe on behalf of the Wellcome Sanger Institute COVID-19 Surveillance Team |
| EPI_ISL_2641504 | Health Services Laboratories | Wellcome Sanger Institute for the COVID-19 Genomics UK (COG-UK) Consortium | Health Services Laboratories and Alex Alderton, Roberto Amato, Jeffrey Barrett, Sonia Goncalves, Ewan Harrison, David K. Jackson, Ian Johnston, Dominic Kwiatkowski, Cordelia Langford, John Sillitoe on behalf of the Wellcome Sanger Institute COVID-19 Surveillance Team |
| EPI_ISL_2641505 | Lighthouse Lab in Alderley Park | Wellcome Sanger Institute for the COVID-19 Genomics UK (COG-UK) Consortium | Jacquelyn Wynn, Mairead Hyland, The Lighthouse Lab in Alderley Park and Alex Alderton, Roberto Amato, Jeffrey Barrett, Sonia Goncalves, Ewan Harrison, David K. Jackson, Ian Johnston, Dominic Kwiatkowski, Cordelia Langford, John Sillitoe on behalf of the Wellcome Sanger Institute COVID-19 Surveillance Team |
| EPI_ISL_2641506, EPI_ISL_2641507 | Health Services Laboratories | Wellcome Sanger Institute for the COVID-19 Genomics UK (COG-UK) Consortium | Health Services Laboratories and Alex Alderton, Roberto Amato, Jeffrey Barrett, Sonia Goncalves, Ewan Harrison, David K. Jackson, Ian Johnston, Dominic Kwiatkowski, Cordelia Langford, John Sillitoe on behalf of the Wellcome Sanger Institute COVID-19 Surveillance Team |
| EPI_ISL_2641508, EPI_ISL_2641509 | Lighthouse Lab in Glasgow | Wellcome Sanger Institute for the COVID-19 Genomics UK (COG-UK) Consortium | Harper VanSteenhouse, Yumi Kasai, David Gray, Carol Clugston, Anna Dominiczak and Alex Alderton, Roberto Amato, Jeffrey Barrett, Sonia Goncalves, Ewan Harrison, David K. Jackson, Ian Johnston, Dominic Kwiatkowski, Cordelia Langford, John Sillitoe on behalf of the Wellcome Sanger Institute COVID-19 Surveillance Team |
| EPI_ISL_2641510, EPI_ISL_2641511 | Lighthouse Lab in Alderley Park | Wellcome Sanger Institute for the COVID-19 Genomics UK (COG-UK) Consortium | Jacquelyn Wynn, Mairead Hyland, The Lighthouse Lab in Alderley Park and Alex Alderton, Roberto Amato, Jeffrey Barrett, Sonia Goncalves, Ewan Harrison, David K. Jackson, Ian Johnston, Dominic Kwiatkowski, Cordelia Langford, John Sillitoe on behalf of the Wellcome Sanger Institute COVID-19 Surveillance Team |
| EPI_ISL_2641512 | Health Services Laboratories | Wellcome Sanger Institute for the COVID-19 Genomics UK (COG-UK) Consortium | Health Services Laboratories and Alex Alderton, Roberto Amato, Jeffrey Barrett, Sonia Goncalves, Ewan Harrison, David K. Jackson, Ian Johnston, Dominic Kwiatkowski, Cordelia Langford, John Sillitoe on behalf of the Wellcome Sanger Institute COVID-19 Surveillance Team |
| EPI_ISL_2641513, EPI_ISL_2641514, EPI_ISL_2641515 | Lighthouse Lab in Alderley Park | Wellcome Sanger Institute for the COVID-19 Genomics UK (COG-UK) Consortium | Jacquelyn Wynn, Mairead Hyland, The Lighthouse Lab in Alderley Park and Alex Alderton, Roberto Amato, Jeffrey Barrett, Sonia Goncalves, Ewan Harrison, David K. Jackson, Ian Johnston, Dominic Kwiatkowski, Cordelia Langford, John Sillitoe on behalf of the Wellcome Sanger Institute COVID-19 Surveillance Team |
| EPI_ISL_2641517 | Health Services Laboratories | Wellcome Sanger Institute for the COVID-19 Genomics UK (COG-UK) Consortium | Health Services Laboratories and Alex Alderton, Roberto Amato, Jeffrey Barrett, Sonia Goncalves, Ewan Harrison, David K. Jackson, Ian Johnston, Dominic Kwiatkowski, Cordelia Langford, John Sillitoe on behalf of the Wellcome Sanger Institute COVID-19 Surveillance Team |
| EPI_ISL_2641518 | Lighthouse Lab in Alderley Park | Wellcome Sanger Institute for the COVID-19 Genomics UK (COG-UK) Consortium | Jacquelyn Wynn, Mairead Hyland, The Lighthouse Lab in Alderley Park and Alex Alderton, Roberto Amato, Jeffrey Barrett, Sonia Goncalves, Ewan Harrison, David K. Jackson, Ian Johnston, Dominic Kwiatkowski, Cordelia Langford, John Sillitoe on behalf of the Wellcome Sanger Institute COVID-19 Surveillance Team |
| EPI_ISL_2641519 | Lighthouse Lab in Glasgow | Wellcome Sanger Institute for the COVID-19 Genomics UK (COG-UK) Consortium | Harper VanSteenhouse, Yumi Kasai, David Gray, Carol Clugston, Anna Dominiczak and Alex Alderton, Roberto Amato, Jeffrey Barrett, Sonia Goncalves, Ewan Harrison, David K. Jackson, Ian Johnston, Dominic Kwiatkowski, Cordelia Langford, John Sillitoe on behalf of the Wellcome Sanger Institute COVID-19 Surveillance Team |
| EPI_ISL_2641520 | Lighthouse Lab in Alderley Park | Wellcome Sanger Institute for the COVID-19 Genomics UK (COG-UK) Consortium | Jacquelyn Wynn, Mairead Hyland, The Lighthouse Lab in Alderley Park and Alex Alderton, Roberto Amato, Jeffrey Barrett, Sonia Goncalves, Ewan Harrison, David K. Jackson, Ian Johnston, Dominic Kwiatkowski, Cordelia Langford, John Sillitoe on behalf of the Wellcome Sanger Institute COVID-19 Surveillance Team |
| EPI_ISL_2641521, EPI_ISL_2641522 | Lighthouse Lab in Glasgow | Wellcome Sanger Institute for the COVID-19 Genomics UK (COG-UK) Consortium | Harper VanSteenhouse, Yumi Kasai, David Gray, Carol Clugston, Anna Dominiczak and Alex Alderton, Roberto Amato, Jeffrey Barrett, Sonia Goncalves, Ewan Harrison, David K. Jackson, Ian Johnston, Dominic Kwiatkowski, Cordelia Langford, John Sillitoe on behalf of the Wellcome Sanger Institute COVID-19 Surveillance Team |
| EPI_ISL_2641523, | Health Services Laboratories | Wellcome Sanger Institute for the COVID-19 Genomics UK (COG-UK) | Health Services Laboratories and Alex Alderton, Roberto Amato, Jeffrey Barrett, Sonia Goncalves, Ewan Harrison, David K. Jackson, Ian Johnston, Dominic Kwiatkowski, Cordelia Langford, John Sillitoe on behalf of the Wellcome Sanger Institute |

| | | | |
|---|---|---|---|
| EPI_ISL_2641524 | | Consortium | COVID-19 Surveillance Team |
| EPI_ISL_2641525 | Lighthouse Lab in Alderley Park | Wellcome Sanger Institute for the COVID-19 Genomics UK (COG-UK) Consortium | Jacquelyn Wynn, Mairead Hyland, The Lighthouse Lab in Alderley Park and Alex Alderton, Roberto Amato, Jeffrey Barrett, Sonia Goncalves, Ewan Harrison, David K. Jackson, Ian Johnston, Dominic Kwiatkowski, Cordelia Langford, John Sillitoe on behalf of the Wellcome Sanger Institute COVID-19 Surveillance Team |
| EPI_ISL_2641526, EPI_ISL_2641527 | Lighthouse Lab in Glasgow | Wellcome Sanger Institute for the COVID-19 Genomics UK (COG-UK) Consortium | Harper VanSteenhouse, Yumi Kasai, David Gray, Carol Clugston, Anna Dominiczak and Alex Alderton, Roberto Amato, Jeffrey Barrett, Sonia Goncalves, Ewan Harrison, David K. Jackson, Ian Johnston, Dominic Kwiatkowski, Cordelia Langford, John Sillitoe on behalf of the Wellcome Sanger Institute COVID-19 Surveillance Team |
| EPI_ISL_2641528 | Health Services Laboratories | Wellcome Sanger Institute for the COVID-19 Genomics UK (COG-UK) Consortium | Health Services Laboratories and Alex Alderton, Roberto Amato, Jeffrey Barrett, Sonia Goncalves, Ewan Harrison, David K. Jackson, Ian Johnston, Dominic Kwiatkowski, Cordelia Langford, John Sillitoe on behalf of the Wellcome Sanger Institute COVID-19 Surveillance Team |
| EPI_ISL_2641529 | Lighthouse Lab in Glasgow | Wellcome Sanger Institute for the COVID-19 Genomics UK (COG-UK) Consortium | Harper VanSteenhouse, Yumi Kasai, David Gray, Carol Clugston, Anna Dominiczak and Alex Alderton, Roberto Amato, Jeffrey Barrett, Sonia Goncalves, Ewan Harrison, David K. Jackson, Ian Johnston, Dominic Kwiatkowski, Cordelia Langford, John Sillitoe on behalf of the Wellcome Sanger Institute COVID-19 Surveillance Team |
| EPI_ISL_2641530, EPI_ISL_2641531 | Health Services Laboratories | Wellcome Sanger Institute for the COVID-19 Genomics UK (COG-UK) Consortium | Health Services Laboratories and Alex Alderton, Roberto Amato, Jeffrey Barrett, Sonia Goncalves, Ewan Harrison, David K. Jackson, Ian Johnston, Dominic Kwiatkowski, Cordelia Langford, John Sillitoe on behalf of the Wellcome Sanger Institute COVID-19 Surveillance Team |
| EPI_ISL_2641532 | Lighthouse Lab in Alderley Park | Wellcome Sanger Institute for the COVID-19 Genomics UK (COG-UK) Consortium | Jacquelyn Wynn, Mairead Hyland, The Lighthouse Lab in Alderley Park and Alex Alderton, Roberto Amato, Jeffrey Barrett, Sonia Goncalves, Ewan Harrison, David K. Jackson, Ian Johnston, Dominic Kwiatkowski, Cordelia Langford, John Sillitoe on behalf of the Wellcome Sanger Institute COVID-19 Surveillance Team |
| EPI_ISL_2641533 | Health Services Laboratories | Wellcome Sanger Institute for the COVID-19 Genomics UK (COG-UK) Consortium | Health Services Laboratories and Alex Alderton, Roberto Amato, Jeffrey Barrett, Sonia Goncalves, Ewan Harrison, David K. Jackson, Ian Johnston, Dominic Kwiatkowski, Cordelia Langford, John Sillitoe on behalf of the Wellcome Sanger Institute COVID-19 Surveillance Team |
| EPI_ISL_2641534 | Lighthouse Lab in Alderley Park | Wellcome Sanger Institute for the COVID-19 Genomics UK (COG-UK) Consortium | Jacquelyn Wynn, Mairead Hyland, The Lighthouse Lab in Alderley Park and Alex Alderton, Roberto Amato, Jeffrey Barrett, Sonia Goncalves, Ewan Harrison, David K. Jackson, Ian Johnston, Dominic Kwiatkowski, Cordelia Langford, John Sillitoe on behalf of the Wellcome Sanger Institute COVID-19 Surveillance Team |
| EPI_ISL_2641535 | Health Services Laboratories | Wellcome Sanger Institute for the COVID-19 Genomics UK (COG-UK) Consortium | Health Services Laboratories and Alex Alderton, Roberto Amato, Jeffrey Barrett, Sonia Goncalves, Ewan Harrison, David K. Jackson, Ian Johnston, Dominic Kwiatkowski, Cordelia Langford, John Sillitoe on behalf of the Wellcome Sanger Institute COVID-19 Surveillance Team |
| EPI_ISL_2641536, EPI_ISL_2641537 | Lighthouse Lab in Glasgow | Wellcome Sanger Institute for the COVID-19 Genomics UK (COG-UK) Consortium | Harper VanSteenhouse, Yumi Kasai, David Gray, Carol Clugston, Anna Dominiczak and Alex Alderton, Roberto Amato, Jeffrey Barrett, Sonia Goncalves, Ewan Harrison, David K. Jackson, Ian Johnston, Dominic Kwiatkowski, Cordelia Langford, John Sillitoe on behalf of the Wellcome Sanger Institute COVID-19 Surveillance Team |
| EPI_ISL_2641538 | Health Services Laboratories | Wellcome Sanger Institute for the COVID-19 Genomics UK (COG-UK) Consortium | Health Services Laboratories and Alex Alderton, Roberto Amato, Jeffrey Barrett, Sonia Goncalves, Ewan Harrison, David K. Jackson, Ian Johnston, Dominic Kwiatkowski, Cordelia Langford, John Sillitoe on behalf of the Wellcome Sanger Institute COVID-19 Surveillance Team |
| EPI_ISL_2641539 | Lighthouse Lab in Alderley Park | Wellcome Sanger Institute for the COVID-19 Genomics UK (COG-UK) Consortium | Jacquelyn Wynn, Mairead Hyland, The Lighthouse Lab in Alderley Park and Alex Alderton, Roberto Amato, Jeffrey Barrett, Sonia Goncalves, Ewan Harrison, David K. Jackson, Ian Johnston, Dominic Kwiatkowski, Cordelia Langford, John Sillitoe on behalf of the Wellcome Sanger Institute COVID-19 Surveillance Team |
| EPI_ISL_2641540 | Health Services Laboratories | Wellcome Sanger Institute for the COVID-19 Genomics UK (COG-UK) Consortium | Health Services Laboratories and Alex Alderton, Roberto Amato, Jeffrey Barrett, Sonia Goncalves, Ewan Harrison, David K. Jackson, Ian Johnston, Dominic Kwiatkowski, Cordelia Langford, John Sillitoe on behalf of the Wellcome Sanger Institute COVID-19 Surveillance Team |
| EPI_ISL_2641541, EPI_ISL_2641542, EPI_ISL_2641543 | Lighthouse Lab in Glasgow | Wellcome Sanger Institute for the COVID-19 Genomics UK (COG-UK) Consortium | Harper VanSteenhouse, Yumi Kasai, David Gray, Carol Clugston, Anna Dominiczak and Alex Alderton, Roberto Amato, Jeffrey Barrett, Sonia Goncalves, Ewan Harrison, David K. Jackson, Ian Johnston, Dominic Kwiatkowski, Cordelia Langford, John Sillitoe on behalf of the Wellcome Sanger Institute COVID-19 Surveillance Team |
| EPI_ISL_2641544, EPI_ISL_2641545 | Lighthouse Lab in Alderley Park | Wellcome Sanger Institute for the COVID-19 Genomics UK (COG-UK) Consortium | Jacquelyn Wynn, Mairead Hyland, The Lighthouse Lab in Alderley Park and Alex Alderton, Roberto Amato, Jeffrey Barrett, Sonia Goncalves, Ewan Harrison, David K. Jackson, Ian Johnston, Dominic Kwiatkowski, Cordelia Langford, John Sillitoe on behalf of the Wellcome Sanger Institute COVID-19 Surveillance Team |
| EPI_ISL_2641546 | Lighthouse Lab in Glasgow | Wellcome Sanger Institute for the COVID-19 Genomics UK (COG-UK) Consortium | Harper VanSteenhouse, Yumi Kasai, David Gray, Carol Clugston, Anna Dominiczak and Alex Alderton, Roberto Amato, Jeffrey Barrett, Sonia Goncalves, Ewan Harrison, David K. Jackson, Ian Johnston, Dominic Kwiatkowski, Cordelia Langford, John Sillitoe on behalf of the Wellcome Sanger Institute COVID-19 Surveillance Team |
| EPI_ISL_2641547 | Lighthouse Lab in Alderley Park | Wellcome Sanger Institute for the COVID-19 Genomics UK (COG-UK) Consortium | Jacquelyn Wynn, Mairead Hyland, The Lighthouse Lab in Alderley Park and Alex Alderton, Roberto Amato, Jeffrey Barrett, Sonia Goncalves, Ewan Harrison, David K. Jackson, Ian Johnston, Dominic Kwiatkowski, Cordelia Langford, John Sillitoe on behalf of the Wellcome Sanger Institute COVID-19 Surveillance Team |
| EPI_ISL_2641548 | Lighthouse Lab in Glasgow | Wellcome Sanger Institute for the COVID-19 Genomics UK (COG-UK) Consortium | Harper VanSteenhouse, Yumi Kasai, David Gray, Carol Clugston, Anna Dominiczak and Alex Alderton, Roberto Amato, Jeffrey Barrett, Sonia Goncalves, Ewan Harrison, David K. Jackson, Ian Johnston, Dominic Kwiatkowski, Cordelia Langford, John Sillitoe on behalf of the Wellcome Sanger Institute COVID-19 Surveillance Team |
| EPI_ISL_2641549 | Lighthouse Lab in Alderley Park | Wellcome Sanger Institute for the COVID-19 Genomics UK (COG-UK) Consortium | Jacquelyn Wynn, Mairead Hyland, The Lighthouse Lab in Alderley Park and Alex Alderton, Roberto Amato, Jeffrey Barrett, Sonia Goncalves, Ewan Harrison, David K. Jackson, Ian Johnston, Dominic Kwiatkowski, Cordelia Langford, John Sillitoe on behalf of the Wellcome Sanger Institute COVID-19 Surveillance Team |
| EPI_ISL_2641550, EPI_ISL_2641551 | Lighthouse Lab in Glasgow | Wellcome Sanger Institute for the COVID-19 Genomics UK (COG-UK) Consortium | Harper VanSteenhouse, Yumi Kasai, David Gray, Carol Clugston, Anna Dominiczak and Alex Alderton, Roberto Amato, Jeffrey Barrett, Sonia Goncalves, Ewan Harrison, David K. Jackson, Ian Johnston, Dominic Kwiatkowski, Cordelia Langford, John Sillitoe on behalf of the Wellcome Sanger Institute COVID-19 Surveillance Team |
| EPI_ISL_2641552 | Health Services Laboratories | Wellcome Sanger Institute for the COVID-19 Genomics UK (COG-UK) Consortium | Health Services Laboratories and Alex Alderton, Roberto Amato, Jeffrey Barrett, Sonia Goncalves, Ewan Harrison, David K. Jackson, Ian Johnston, Dominic Kwiatkowski, Cordelia Langford, John Sillitoe on behalf of the Wellcome Sanger Institute COVID-19 Surveillance Team |
| EPI_ISL_2641553 | Lighthouse Lab in Glasgow | Wellcome Sanger Institute for the COVID-19 Genomics UK (COG-UK) Consortium | Harper VanSteenhouse, Yumi Kasai, David Gray, Carol Clugston, Anna Dominiczak and Alex Alderton, Roberto Amato, Jeffrey Barrett, Sonia Goncalves, Ewan Harrison, David K. Jackson, Ian Johnston, Dominic Kwiatkowski, Cordelia Langford, John Sillitoe on behalf of the Wellcome Sanger Institute COVID-19 Surveillance Team |
| EPI_ISL_2641554 | Health Services Laboratories | Wellcome Sanger Institute for the COVID-19 Genomics UK (COG-UK) Consortium | Health Services Laboratories and Alex Alderton, Roberto Amato, Jeffrey Barrett, Sonia Goncalves, Ewan Harrison, David K. Jackson, Ian Johnston, Dominic Kwiatkowski, Cordelia Langford, John Sillitoe on behalf of the Wellcome Sanger Institute COVID-19 Surveillance Team |
| EPI_ISL_2641555 | Lighthouse Lab in Glasgow | Wellcome Sanger Institute for the COVID-19 Genomics UK (COG-UK) Consortium | Harper VanSteenhouse, Yumi Kasai, David Gray, Carol Clugston, Anna Dominiczak and Alex Alderton, Roberto Amato, Jeffrey Barrett, Sonia Goncalves, Ewan Harrison, David K. Jackson, Ian Johnston, Dominic Kwiatkowski, Cordelia Langford, John Sillitoe on behalf of the Wellcome Sanger Institute COVID-19 Surveillance Team |
| EPI_ISL_2641556 | Health Services Laboratories | Wellcome Sanger Institute for the COVID-19 Genomics UK (COG-UK) Consortium | Health Services Laboratories and Alex Alderton, Roberto Amato, Jeffrey Barrett, Sonia Goncalves, Ewan Harrison, David K. Jackson, Ian Johnston, Dominic Kwiatkowski, Cordelia Langford, John Sillitoe on behalf of the Wellcome Sanger Institute COVID-19 Surveillance Team |
| EPI_ISL_2641557 | Lighthouse Lab in Glasgow | Wellcome Sanger Institute for the COVID-19 Genomics UK (COG-UK) Consortium | Harper VanSteenhouse, Yumi Kasai, David Gray, Carol Clugston, Anna Dominiczak and Alex Alderton, Roberto Amato, Jeffrey Barrett, Sonia Goncalves, Ewan Harrison, David K. Jackson, Ian Johnston, Dominic Kwiatkowski, Cordelia Langford, John Sillitoe on behalf of the Wellcome Sanger Institute COVID-19 Surveillance Team |
| EPI_ISL_2641558 | Health Services Laboratories | Wellcome Sanger Institute for the COVID-19 Genomics UK (COG-UK) Consortium | Health Services Laboratories and Alex Alderton, Roberto Amato, Jeffrey Barrett, Sonia Goncalves, Ewan Harrison, David K. Jackson, Ian Johnston, Dominic Kwiatkowski, Cordelia Langford, John Sillitoe on behalf of the Wellcome Sanger Institute COVID-19 Surveillance Team |
| EPI_ISL_2641559 | Lighthouse Lab in Glasgow | Wellcome Sanger Institute for the COVID-19 Genomics UK (COG-UK) Consortium | Harper VanSteenhouse, Yumi Kasai, David Gray, Carol Clugston, Anna Dominiczak and Alex Alderton, Roberto Amato, Jeffrey Barrett, Sonia Goncalves, Ewan Harrison, David K. Jackson, Ian Johnston, Dominic Kwiatkowski, Cordelia Langford, John Sillitoe on behalf of the Wellcome Sanger Institute COVID-19 Surveillance Team |
| EPI_ISL_2641562, EPI_ISL_2641563 | Health Services Laboratories | Wellcome Sanger Institute for the COVID-19 Genomics UK (COG-UK) Consortium | Health Services Laboratories and Alex Alderton, Roberto Amato, Jeffrey Barrett, Sonia Goncalves, Ewan Harrison, David K. Jackson, Ian Johnston, Dominic Kwiatkowski, Cordelia Langford, John Sillitoe on behalf of the Wellcome Sanger Institute COVID-19 Surveillance Team |
| EPI_ISL_2641564, EPI_ISL_2641565 | Lighthouse Lab in Glasgow | Wellcome Sanger Institute for the COVID-19 Genomics UK (COG-UK) Consortium | Harper VanSteenhouse, Yumi Kasai, David Gray, Carol Clugston, Anna Dominiczak and Alex Alderton, Roberto Amato, Jeffrey Barrett, Sonia Goncalves, Ewan Harrison, David K. Jackson, Ian Johnston, Dominic Kwiatkowski, Cordelia Langford, John Sillitoe on behalf of the Wellcome Sanger Institute COVID-19 Surveillance Team |
| EPI_ISL_2641566 | Lighthouse Lab in Alderley Park | Wellcome Sanger Institute for the COVID-19 Genomics UK (COG-UK) Consortium | Jacquelyn Wynn, Mairead Hyland, The Lighthouse Lab in Alderley Park and Alex Alderton, Roberto Amato, Jeffrey Barrett, Sonia Goncalves, Ewan Harrison, David K. Jackson, Ian Johnston, Dominic Kwiatkowski, Cordelia Langford, John Sillitoe on behalf of the Wellcome Sanger Institute COVID-19 Surveillance Team |
| EPI_ISL_2641567 | Health Services Laboratories | Wellcome Sanger Institute for the COVID-19 Genomics UK (COG-UK) Consortium | Health Services Laboratories and Alex Alderton, Roberto Amato, Jeffrey Barrett, Sonia Goncalves, Ewan Harrison, David K. Jackson, Ian Johnston, Dominic Kwiatkowski, Cordelia Langford, John Sillitoe on behalf of the Wellcome Sanger Institute COVID-19 Surveillance Team |
| EPI_ISL_2641568, EPI_ISL_2641569, EPI_ISL_2641571 | Lighthouse Lab in Alderley Park | Wellcome Sanger Institute for the COVID-19 Genomics UK (COG-UK) Consortium | Jacquelyn Wynn, Mairead Hyland, The Lighthouse Lab in Alderley Park and Alex Alderton, Roberto Amato, Jeffrey Barrett, Sonia Goncalves, Ewan Harrison, David K. Jackson, Ian Johnston, Dominic Kwiatkowski, Cordelia Langford, John Sillitoe on behalf of the Wellcome Sanger Institute COVID-19 Surveillance Team |
| EPI_ISL_2641572, EPI_ISL_2641573, EPI_ISL_2641574, EPI_ISL_2641575 | Health Services Laboratories | Wellcome Sanger Institute for the COVID-19 Genomics UK (COG-UK) Consortium | Health Services Laboratories and Alex Alderton, Roberto Amato, Jeffrey Barrett, Sonia Goncalves, Ewan Harrison, David K. Jackson, Ian Johnston, Dominic Kwiatkowski, Cordelia Langford, John Sillitoe on behalf of the Wellcome Sanger Institute COVID-19 Surveillance Team |
| EPI_ISL_2641576 | Lighthouse Lab in Glasgow | Wellcome Sanger Institute for the COVID-19 Genomics UK (COG-UK) Consortium | Harper VanSteenhouse, Yumi Kasai, David Gray, Carol Clugston, Anna Dominiczak and Alex Alderton, Roberto Amato, Jeffrey Barrett, Sonia Goncalves, Ewan Harrison, David K. Jackson, Ian Johnston, Dominic Kwiatkowski, Cordelia Langford, John Sillitoe on behalf of the Wellcome Sanger Institute COVID-19 Surveillance Team |
| EPI_ISL_2641577 | Lighthouse Lab in Alderley Park | Wellcome Sanger Institute for the COVID-19 Genomics UK (COG-UK) Consortium | Jacquelyn Wynn, Mairead Hyland, The Lighthouse Lab in Alderley Park and Alex Alderton, Roberto Amato, Jeffrey Barrett, Sonia Goncalves, Ewan Harrison, David K. Jackson, Ian Johnston, Dominic Kwiatkowski, Cordelia Langford, John Sillitoe on behalf of the Wellcome Sanger Institute COVID-19 Surveillance Team |
| EPI_ISL_2641578, EPI_ISL_2641579, EPI_ISL_2641580, EPI_ISL_2641581 | Lighthouse Lab in Glasgow | Wellcome Sanger Institute for the COVID-19 Genomics UK (COG-UK) Consortium | Harper VanSteenhouse, Yumi Kasai, David Gray, Carol Clugston, Anna Dominiczak and Alex Alderton, Roberto Amato, Jeffrey Barrett, Sonia Goncalves, Ewan Harrison, David K. Jackson, Ian Johnston, Dominic Kwiatkowski, Cordelia Langford, John Sillitoe on behalf of the Wellcome Sanger Institute COVID-19 Surveillance Team |
| EPI_ISL_2641582, EPI_ISL_2641583 | Health Services Laboratories | Wellcome Sanger Institute for the COVID-19 Genomics UK (COG-UK) Consortium | Health Services Laboratories and Alex Alderton, Roberto Amato, Jeffrey Barrett, Sonia Goncalves, Ewan Harrison, David K. Jackson, Ian Johnston, Dominic Kwiatkowski, Cordelia Langford, John Sillitoe on behalf of the Wellcome Sanger Institute COVID-19 Surveillance Team |
| EPI_ISL_2641584 | Lighthouse Lab in Glasgow | Wellcome Sanger Institute for the COVID-19 Genomics UK (COG-UK) Consortium | Harper VanSteenhouse, Yumi Kasai, David Gray, Carol Clugston, Anna Dominiczak and Alex Alderton, Roberto Amato, Jeffrey Barrett, Sonia Goncalves, Ewan Harrison, David K. Jackson, Ian Johnston, Dominic Kwiatkowski, Cordelia Langford, John Sillitoe on behalf of the Wellcome Sanger Institute COVID-19 Surveillance Team |
| EPI_ISL_2641585 | Lighthouse Lab in Alderley Park | Wellcome Sanger Institute for the COVID-19 Genomics UK (COG-UK) Consortium | Jacquelyn Wynn, Mairead Hyland, The Lighthouse Lab in Alderley Park and Alex Alderton, Roberto Amato, Jeffrey Barrett, Sonia Goncalves, Ewan Harrison, David K. Jackson, Ian Johnston, Dominic Kwiatkowski, Cordelia Langford, John Sillitoe on behalf of the Wellcome Sanger Institute COVID-19 Surveillance Team |
| EPI_ISL_2641586, EPI_ISL_2641587 | Health Services Laboratories | Wellcome Sanger Institute for the COVID-19 Genomics UK (COG-UK) Consortium | Health Services Laboratories and Alex Alderton, Roberto Amato, Jeffrey Barrett, Sonia Goncalves, Ewan Harrison, David K. Jackson, Ian Johnston, Dominic Kwiatkowski, Cordelia Langford, John Sillitoe on behalf of the Wellcome Sanger Institute COVID-19 Surveillance Team |
| EPI_ISL_2641588 | Lighthouse Lab in Alderley Park | Wellcome Sanger Institute for the COVID-19 Genomics UK (COG-UK) Consortium | Jacquelyn Wynn, Mairead Hyland, The Lighthouse Lab in Alderley Park and Alex Alderton, Roberto Amato, Jeffrey Barrett, Sonia Goncalves, Ewan Harrison, David K. Jackson, Ian Johnston, Dominic Kwiatkowski, Cordelia Langford, John Sillitoe on behalf of the Wellcome Sanger Institute COVID-19 Surveillance Team |

| | | | |
|---|---|---|---|
| EPI_ISL_2641589 | Lighthouse Lab in Glasgow | Wellcome Sanger Institute for the COVID-19 Genomics UK (COG-UK) Consortium | Harper VanSteenhouse, Yumi Kasai, David Gray, Carol Clugston, Anna Dominiczak and Alex Alderton, Roberto Amato, Jeffrey Barrett, Sonia Goncalves, Ewan Harrison, David K. Jackson, Ian Johnston, Dominic Kwiatkowski, Cordelia Langford, John Sillitoe on behalf of the Wellcome Sanger Institute COVID-19 Surveillance Team |
| EPI_ISL_2641590 | Health Services Laboratories | Wellcome Sanger Institute for the COVID-19 Genomics UK (COG-UK) Consortium | Health Services Laboratories and Alex Alderton, Roberto Amato, Jeffrey Barrett, Sonia Goncalves, Ewan Harrison, David K. Jackson, Ian Johnston, Dominic Kwiatkowski, Cordelia Langford, John Sillitoe on behalf of the Wellcome Sanger Institute COVID-19 Surveillance Team |
| EPI_ISL_2641591 | Lighthouse Lab in Glasgow | Wellcome Sanger Institute for the COVID-19 Genomics UK (COG-UK) Consortium | Harper VanSteenhouse, Yumi Kasai, David Gray, Carol Clugston, Anna Dominiczak and Alex Alderton, Roberto Amato, Jeffrey Barrett, Sonia Goncalves, Ewan Harrison, David K. Jackson, Ian Johnston, Dominic Kwiatkowski, Cordelia Langford, John Sillitoe on behalf of the Wellcome Sanger Institute COVID-19 Surveillance Team |
| EPI_ISL_2641592 | Lighthouse Lab in Alderley Park | Wellcome Sanger Institute for the COVID-19 Genomics UK (COG-UK) Consortium | Jacquelyn Wynn, Mairead Hyland, The Lighthouse Lab in Alderley Park and Alex Alderton, Roberto Amato, Jeffrey Barrett, Sonia Goncalves, Ewan Harrison, David K. Jackson, Ian Johnston, Dominic Kwiatkowski, Cordelia Langford, John Sillitoe on behalf of the Wellcome Sanger Institute COVID-19 Surveillance Team |
| EPI_ISL_2641593 | Health Services Laboratories | Wellcome Sanger Institute for the COVID-19 Genomics UK (COG-UK) Consortium | Health Services Laboratories and Alex Alderton, Roberto Amato, Jeffrey Barrett, Sonia Goncalves, Ewan Harrison, David K. Jackson, Ian Johnston, Dominic Kwiatkowski, Cordelia Langford, John Sillitoe on behalf of the Wellcome Sanger Institute COVID-19 Surveillance Team |
| EPI_ISL_2641594 | Lighthouse Lab in Alderley Park | Wellcome Sanger Institute for the COVID-19 Genomics UK (COG-UK) Consortium | Jacquelyn Wynn, Mairead Hyland, The Lighthouse Lab in Alderley Park and Alex Alderton, Roberto Amato, Jeffrey Barrett, Sonia Goncalves, Ewan Harrison, David K. Jackson, Ian Johnston, Dominic Kwiatkowski, Cordelia Langford, John Sillitoe on behalf of the Wellcome Sanger Institute COVID-19 Surveillance Team |
| EPI_ISL_2641595, EPI_ISL_2641596, EPI_ISL_2641597, EPI_ISL_2641598 | Lighthouse Lab in Glasgow | Wellcome Sanger Institute for the COVID-19 Genomics UK (COG-UK) Consortium | Harper VanSteenhouse, Yumi Kasai, David Gray, Carol Clugston, Anna Dominiczak and Alex Alderton, Roberto Amato, Jeffrey Barrett, Sonia Goncalves, Ewan Harrison, David K. Jackson, Ian Johnston, Dominic Kwiatkowski, Cordelia Langford, John Sillitoe on behalf of the Wellcome Sanger Institute COVID-19 Surveillance Team |
| EPI_ISL_2641599 | Health Services Laboratories | Wellcome Sanger Institute for the COVID-19 Genomics UK (COG-UK) Consortium | Health Services Laboratories and Alex Alderton, Roberto Amato, Jeffrey Barrett, Sonia Goncalves, Ewan Harrison, David K. Jackson, Ian Johnston, Dominic Kwiatkowski, Cordelia Langford, John Sillitoe on behalf of the Wellcome Sanger Institute COVID-19 Surveillance Team |
| EPI_ISL_2641600, EPI_ISL_2641601, EPI_ISL_2641602 | Lighthouse Lab in Glasgow | Wellcome Sanger Institute for the COVID-19 Genomics UK (COG-UK) Consortium | Harper VanSteenhouse, Yumi Kasai, David Gray, Carol Clugston, Anna Dominiczak and Alex Alderton, Roberto Amato, Jeffrey Barrett, Sonia Goncalves, Ewan Harrison, David K. Jackson, Ian Johnston, Dominic Kwiatkowski, Cordelia Langford, John Sillitoe on behalf of the Wellcome Sanger Institute COVID-19 Surveillance Team |
| EPI_ISL_2641603 | Lighthouse Lab in Alderley Park | Wellcome Sanger Institute for the COVID-19 Genomics UK (COG-UK) Consortium | Jacquelyn Wynn, Mairead Hyland, The Lighthouse Lab in Alderley Park and Alex Alderton, Roberto Amato, Jeffrey Barrett, Sonia Goncalves, Ewan Harrison, David K. Jackson, Ian Johnston, Dominic Kwiatkowski, Cordelia Langford, John Sillitoe on behalf of the Wellcome Sanger Institute COVID-19 Surveillance Team |
| EPI_ISL_2641604, EPI_ISL_2641605 | Lighthouse Lab in Glasgow | Wellcome Sanger Institute for the COVID-19 Genomics UK (COG-UK) Consortium | Harper VanSteenhouse, Yumi Kasai, David Gray, Carol Clugston, Anna Dominiczak and Alex Alderton, Roberto Amato, Jeffrey Barrett, Sonia Goncalves, Ewan Harrison, David K. Jackson, Ian Johnston, Dominic Kwiatkowski, Cordelia Langford, John Sillitoe on behalf of the Wellcome Sanger Institute COVID-19 Surveillance Team |
| EPI_ISL_2641606 | Health Services Laboratories | Wellcome Sanger Institute for the COVID-19 Genomics UK (COG-UK) Consortium | Health Services Laboratories and Alex Alderton, Roberto Amato, Jeffrey Barrett, Sonia Goncalves, Ewan Harrison, David K. Jackson, Ian Johnston, Dominic Kwiatkowski, Cordelia Langford, John Sillitoe on behalf of the Wellcome Sanger Institute COVID-19 Surveillance Team |
| EPI_ISL_2641607, EPI_ISL_2641608, EPI_ISL_2641609 | Lighthouse Lab in Glasgow | Wellcome Sanger Institute for the COVID-19 Genomics UK (COG-UK) Consortium | Harper VanSteenhouse, Yumi Kasai, David Gray, Carol Clugston, Anna Dominiczak and Alex Alderton, Roberto Amato, Jeffrey Barrett, Sonia Goncalves, Ewan Harrison, David K. Jackson, Ian Johnston, Dominic Kwiatkowski, Cordelia Langford, John Sillitoe on behalf of the Wellcome Sanger Institute COVID-19 Surveillance Team |
| EPI_ISL_2641610 | Lighthouse Lab in Alderley Park | Wellcome Sanger Institute for the COVID-19 Genomics UK (COG-UK) Consortium | Jacquelyn Wynn, Mairead Hyland, The Lighthouse Lab in Alderley Park and Alex Alderton, Roberto Amato, Jeffrey Barrett, Sonia Goncalves, Ewan Harrison, David K. Jackson, Ian Johnston, Dominic Kwiatkowski, Cordelia Langford, John Sillitoe on behalf of the Wellcome Sanger Institute COVID-19 Surveillance Team |
| EPI_ISL_2641611 | Lighthouse Lab in Glasgow | Wellcome Sanger Institute for the COVID-19 Genomics UK (COG-UK) Consortium | Harper VanSteenhouse, Yumi Kasai, David Gray, Carol Clugston, Anna Dominiczak and Alex Alderton, Roberto Amato, Jeffrey Barrett, Sonia Goncalves, Ewan Harrison, David K. Jackson, Ian Johnston, Dominic Kwiatkowski, Cordelia Langford, John Sillitoe on behalf of the Wellcome Sanger Institute COVID-19 Surveillance Team |
| EPI_ISL_2641612 | Lighthouse Lab in Alderley Park | Wellcome Sanger Institute for the COVID-19 Genomics UK (COG-UK) Consortium | Jacquelyn Wynn, Mairead Hyland, The Lighthouse Lab in Alderley Park and Alex Alderton, Roberto Amato, Jeffrey Barrett, Sonia Goncalves, Ewan Harrison, David K. Jackson, Ian Johnston, Dominic Kwiatkowski, Cordelia Langford, John Sillitoe on behalf of the Wellcome Sanger Institute COVID-19 Surveillance Team |
| EPI_ISL_2641613, EPI_ISL_2641614 | Lighthouse Lab in Glasgow | Wellcome Sanger Institute for the COVID-19 Genomics UK (COG-UK) Consortium | Harper VanSteenhouse, Yumi Kasai, David Gray, Carol Clugston, Anna Dominiczak and Alex Alderton, Roberto Amato, Jeffrey Barrett, Sonia Goncalves, Ewan Harrison, David K. Jackson, Ian Johnston, Dominic Kwiatkowski, Cordelia Langford, John Sillitoe on behalf of the Wellcome Sanger Institute COVID-19 Surveillance Team |
| EPI_ISL_2641615 | Lighthouse Lab in Alderley Park | Wellcome Sanger Institute for the COVID-19 Genomics UK (COG-UK) Consortium | Jacquelyn Wynn, Mairead Hyland, The Lighthouse Lab in Alderley Park and Alex Alderton, Roberto Amato, Jeffrey Barrett, Sonia Goncalves, Ewan Harrison, David K. Jackson, Ian Johnston, Dominic Kwiatkowski, Cordelia Langford, John Sillitoe on behalf of the Wellcome Sanger Institute COVID-19 Surveillance Team |
| EPI_ISL_2641616 | Health Services Laboratories | Wellcome Sanger Institute for the COVID-19 Genomics UK (COG-UK) Consortium | Health Services Laboratories and Alex Alderton, Roberto Amato, Jeffrey Barrett, Sonia Goncalves, Ewan Harrison, David K. Jackson, Ian Johnston, Dominic Kwiatkowski, Cordelia Langford, John Sillitoe on behalf of the Wellcome Sanger Institute COVID-19 Surveillance Team |
| EPI_ISL_2641617 | Lighthouse Lab in Alderley Park | Wellcome Sanger Institute for the COVID-19 Genomics UK (COG-UK) Consortium | Jacquelyn Wynn, Mairead Hyland, The Lighthouse Lab in Alderley Park and Alex Alderton, Roberto Amato, Jeffrey Barrett, Sonia Goncalves, Ewan Harrison, David K. Jackson, Ian Johnston, Dominic Kwiatkowski, Cordelia Langford, John Sillitoe on behalf of the Wellcome Sanger Institute COVID-19 Surveillance Team |
| EPI_ISL_2641618 | Health Services Laboratories | Wellcome Sanger Institute for the COVID-19 Genomics UK (COG-UK) Consortium | Health Services Laboratories and Alex Alderton, Roberto Amato, Jeffrey Barrett, Sonia Goncalves, Ewan Harrison, David K. Jackson, Ian Johnston, Dominic Kwiatkowski, Cordelia Langford, John Sillitoe on behalf of the Wellcome Sanger Institute COVID-19 Surveillance Team |
| EPI_ISL_2641619 | Lighthouse Lab in Glasgow | Wellcome Sanger Institute for the COVID-19 Genomics UK (COG-UK) Consortium | Harper VanSteenhouse, Yumi Kasai, David Gray, Carol Clugston, Anna Dominiczak and Alex Alderton, Roberto Amato, Jeffrey Barrett, Sonia Goncalves, Ewan Harrison, David K. Jackson, Ian Johnston, Dominic Kwiatkowski, Cordelia Langford, John Sillitoe on behalf of the Wellcome Sanger Institute COVID-19 Surveillance Team |
| EPI_ISL_2641620, EPI_ISL_2641621 | Lighthouse Lab in Alderley Park | Wellcome Sanger Institute for the COVID-19 Genomics UK (COG-UK) Consortium | Jacquelyn Wynn, Mairead Hyland, The Lighthouse Lab in Alderley Park and Alex Alderton, Roberto Amato, Jeffrey Barrett, Sonia Goncalves, Ewan Harrison, David K. Jackson, Ian Johnston, Dominic Kwiatkowski, Cordelia Langford, John Sillitoe on behalf of the Wellcome Sanger Institute COVID-19 Surveillance Team |
| EPI_ISL_2641622 | Health Services Laboratories | Wellcome Sanger Institute for the COVID-19 Genomics UK (COG-UK) Consortium | Health Services Laboratories and Alex Alderton, Roberto Amato, Jeffrey Barrett, Sonia Goncalves, Ewan Harrison, David K. Jackson, Ian Johnston, Dominic Kwiatkowski, Cordelia Langford, John Sillitoe on behalf of the Wellcome Sanger Institute COVID-19 Surveillance Team |
| EPI_ISL_2641623 | Lighthouse Lab in Glasgow | Wellcome Sanger Institute for the COVID-19 Genomics UK (COG-UK) Consortium | Harper VanSteenhouse, Yumi Kasai, David Gray, Carol Clugston, Anna Dominiczak and Alex Alderton, Roberto Amato, Jeffrey Barrett, Sonia Goncalves, Ewan Harrison, David K. Jackson, Ian Johnston, Dominic Kwiatkowski, Cordelia Langford, John Sillitoe on behalf of the Wellcome Sanger Institute COVID-19 Surveillance Team |
| EPI_ISL_2641624, EPI_ISL_2641625, EPI_ISL_2641626, EPI_ISL_2641627 | Lighthouse Lab in Alderley Park | Wellcome Sanger Institute for the COVID-19 Genomics UK (COG-UK) Consortium | Jacquelyn Wynn, Mairead Hyland, The Lighthouse Lab in Alderley Park and Alex Alderton, Roberto Amato, Jeffrey Barrett, Sonia Goncalves, Ewan Harrison, David K. Jackson, Ian Johnston, Dominic Kwiatkowski, Cordelia Langford, John Sillitoe on behalf of the Wellcome Sanger Institute COVID-19 Surveillance Team |
| EPI_ISL_2641628, EPI_ISL_2641629 | Health Services Laboratories | Wellcome Sanger Institute for the COVID-19 Genomics UK (COG-UK) Consortium | Health Services Laboratories and Alex Alderton, Roberto Amato, Jeffrey Barrett, Sonia Goncalves, Ewan Harrison, David K. Jackson, Ian Johnston, Dominic Kwiatkowski, Cordelia Langford, John Sillitoe on behalf of the Wellcome Sanger Institute COVID-19 Surveillance Team |
| EPI_ISL_2641630 | Lighthouse Lab in Alderley Park | Wellcome Sanger Institute for the COVID-19 Genomics UK (COG-UK) Consortium | Jacquelyn Wynn, Mairead Hyland, The Lighthouse Lab in Alderley Park and Alex Alderton, Roberto Amato, Jeffrey Barrett, Sonia Goncalves, Ewan Harrison, David K. Jackson, Ian Johnston, Dominic Kwiatkowski, Cordelia Langford, John Sillitoe on behalf of the Wellcome Sanger Institute COVID-19 Surveillance Team |
| EPI_ISL_2641631, EPI_ISL_2641632 | Health Services Laboratories | Wellcome Sanger Institute for the COVID-19 Genomics UK (COG-UK) Consortium | Health Services Laboratories and Alex Alderton, Roberto Amato, Jeffrey Barrett, Sonia Goncalves, Ewan Harrison, David K. Jackson, Ian Johnston, Dominic Kwiatkowski, Cordelia Langford, John Sillitoe on behalf of the Wellcome Sanger Institute COVID-19 Surveillance Team |
| EPI_ISL_2641633, EPI_ISL_2641634 | Lighthouse Lab in Alderley Park | Wellcome Sanger Institute for the COVID-19 Genomics UK (COG-UK) Consortium | Jacquelyn Wynn, Mairead Hyland, The Lighthouse Lab in Alderley Park and Alex Alderton, Roberto Amato, Jeffrey Barrett, Sonia Goncalves, Ewan Harrison, David K. Jackson, Ian Johnston, Dominic Kwiatkowski, Cordelia Langford, John Sillitoe on behalf of the Wellcome Sanger Institute COVID-19 Surveillance Team |
| EPI_ISL_2641635, EPI_ISL_2641636, EPI_ISL_2641637 | Lighthouse Lab in Glasgow | Wellcome Sanger Institute for the COVID-19 Genomics UK (COG-UK) Consortium | Harper VanSteenhouse, Yumi Kasai, David Gray, Carol Clugston, Anna Dominiczak and Alex Alderton, Roberto Amato, Jeffrey Barrett, Sonia Goncalves, Ewan Harrison, David K. Jackson, Ian Johnston, Dominic Kwiatkowski, Cordelia Langford, John Sillitoe on behalf of the Wellcome Sanger Institute COVID-19 Surveillance Team |
| EPI_ISL_2641638, EPI_ISL_2641640 | Lighthouse Lab in Alderley Park | Wellcome Sanger Institute for the COVID-19 Genomics UK (COG-UK) Consortium | Jacquelyn Wynn, Mairead Hyland, The Lighthouse Lab in Alderley Park and Alex Alderton, Roberto Amato, Jeffrey Barrett, Sonia Goncalves, Ewan Harrison, David K. Jackson, Ian Johnston, Dominic Kwiatkowski, Cordelia Langford, John Sillitoe on behalf of the Wellcome Sanger Institute COVID-19 Surveillance Team |
| EPI_ISL_2641641, EPI_ISL_2641642, EPI_ISL_2641643 | Lighthouse Lab in Glasgow | Wellcome Sanger Institute for the COVID-19 Genomics UK (COG-UK) Consortium | Harper VanSteenhouse, Yumi Kasai, David Gray, Carol Clugston, Anna Dominiczak and Alex Alderton, Roberto Amato, Jeffrey Barrett, Sonia Goncalves, Ewan Harrison, David K. Jackson, Ian Johnston, Dominic Kwiatkowski, Cordelia Langford, John Sillitoe on behalf of the Wellcome Sanger Institute COVID-19 Surveillance Team |
| EPI_ISL_2641644 | Lighthouse Lab in Alderley Park | Wellcome Sanger Institute for the COVID-19 Genomics UK (COG-UK) Consortium | Jacquelyn Wynn, Mairead Hyland, The Lighthouse Lab in Alderley Park and Alex Alderton, Roberto Amato, Jeffrey Barrett, Sonia Goncalves, Ewan Harrison, David K. Jackson, Ian Johnston, Dominic Kwiatkowski, Cordelia Langford, John Sillitoe on behalf of the Wellcome Sanger Institute COVID-19 Surveillance Team |
| EPI_ISL_2641645, EPI_ISL_2641646, EPI_ISL_2641647, EPI_ISL_2641649 | Lighthouse Lab in Glasgow | Wellcome Sanger Institute for the COVID-19 Genomics UK (COG-UK) Consortium | Harper VanSteenhouse, Yumi Kasai, David Gray, Carol Clugston, Anna Dominiczak and Alex Alderton, Roberto Amato, Jeffrey Barrett, Sonia Goncalves, Ewan Harrison, David K. Jackson, Ian Johnston, Dominic Kwiatkowski, Cordelia Langford, John Sillitoe on behalf of the Wellcome Sanger Institute COVID-19 Surveillance Team |
| EPI_ISL_2641650 | Health Services Laboratories | Wellcome Sanger Institute for the COVID-19 Genomics UK (COG-UK) Consortium | Health Services Laboratories and Alex Alderton, Roberto Amato, Jeffrey Barrett, Sonia Goncalves, Ewan Harrison, David K. Jackson, Ian Johnston, Dominic Kwiatkowski, Cordelia Langford, John Sillitoe on behalf of the Wellcome Sanger Institute COVID-19 Surveillance Team |
| EPI_ISL_2641651 | Lighthouse Lab in Glasgow | Wellcome Sanger Institute for the COVID-19 Genomics UK (COG-UK) Consortium | Harper VanSteenhouse, Yumi Kasai, David Gray, Carol Clugston, Anna Dominiczak and Alex Alderton, Roberto Amato, Jeffrey Barrett, Sonia Goncalves, Ewan Harrison, David K. Jackson, Ian Johnston, Dominic Kwiatkowski, Cordelia Langford, John Sillitoe on behalf of the Wellcome Sanger Institute COVID-19 Surveillance Team |
| EPI_ISL_2641652, EPI_ISL_2641653 | Lighthouse Lab in Alderley Park | Wellcome Sanger Institute for the COVID-19 Genomics UK (COG-UK) Consortium | Jacquelyn Wynn, Mairead Hyland, The Lighthouse Lab in Alderley Park and Alex Alderton, Roberto Amato, Jeffrey Barrett, Sonia Goncalves, Ewan Harrison, David K. Jackson, Ian Johnston, Dominic Kwiatkowski, Cordelia Langford, John Sillitoe on behalf of the Wellcome Sanger Institute COVID-19 Surveillance Team |
| EPI_ISL_2641654 | Lighthouse Lab in Glasgow | Wellcome Sanger Institute for the COVID-19 Genomics UK (COG-UK) Consortium | Harper VanSteenhouse, Yumi Kasai, David Gray, Carol Clugston, Anna Dominiczak and Alex Alderton, Roberto Amato, Jeffrey Barrett, Sonia Goncalves, Ewan Harrison, David K. Jackson, Ian Johnston, Dominic Kwiatkowski, Cordelia Langford, John Sillitoe on behalf of the Wellcome Sanger Institute COVID-19 Surveillance Team |
| EPI_ISL_2641655 | Lighthouse Lab in Alderley Park | Wellcome Sanger Institute for the COVID-19 Genomics UK (COG-UK) Consortium | Jacquelyn Wynn, Mairead Hyland, The Lighthouse Lab in Alderley Park and Alex Alderton, Roberto Amato, Jeffrey Barrett, Sonia Goncalves, Ewan Harrison, David K. Jackson, Ian Johnston, Dominic Kwiatkowski, Cordelia Langford, John Sillitoe on behalf of the Wellcome Sanger Institute COVID-19 Surveillance Team |
| EPI_ISL_2641656 | Lighthouse Lab in Glasgow | Wellcome Sanger Institute for the COVID-19 Genomics UK (COG-UK) Consortium | Harper VanSteenhouse, Yumi Kasai, David Gray, Carol Clugston, Anna Dominiczak and Alex Alderton, Roberto Amato, Jeffrey Barrett, Sonia Goncalves, Ewan Harrison, David K. Jackson, Ian Johnston, Dominic Kwiatkowski, Cordelia Langford, John Sillitoe on behalf of the Wellcome Sanger Institute COVID-19 Surveillance Team |

| | | | |
|---|---|---|---|
| EPI_ISL_2641657 | Lighthouse Lab in Alderley Park | Wellcome Sanger Institute for the COVID-19 Genomics UK (COG-UK) Consortium | Jacquelyn Wynn, Mairead Hyland, The Lighthouse Lab in Alderley Park and Alex Alderton, Roberto Amato, Jeffrey Barrett, Sonia Goncalves, Ewan Harrison, David K. Jackson, Ian Johnston, Dominic Kwiatkowski, Cordelia Langford, John Sillitoe on behalf of the Wellcome Sanger Institute COVID-19 Surveillance Team |
| EPI_ISL_2641658 | Health Services Laboratories | Wellcome Sanger Institute for the COVID-19 Genomics UK (COG-UK) Consortium | Health Services Laboratories and Alex Alderton, Roberto Amato, Jeffrey Barrett, Sonia Goncalves, Ewan Harrison, David K. Jackson, Ian Johnston, Dominic Kwiatkowski, Cordelia Langford, John Sillitoe on behalf of the Wellcome Sanger Institute COVID-19 Surveillance Team |
| EPI_ISL_2641659 | Lighthouse Lab in Glasgow | Wellcome Sanger Institute for the COVID-19 Genomics UK (COG-UK) Consortium | Harper VanSteenhouse, Yumi Kasai, David Gray, Carol Clugston, Anna Dominiczak and Alex Alderton, Roberto Amato, Jeffrey Barrett, Sonia Goncalves, Ewan Harrison, David K. Jackson, Ian Johnston, Dominic Kwiatkowski, Cordelia Langford, John Sillitoe on behalf of the Wellcome Sanger Institute COVID-19 Surveillance Team |
| EPI_ISL_2641660 | Health Services Laboratories | Wellcome Sanger Institute for the COVID-19 Genomics UK (COG-UK) Consortium | Health Services Laboratories and Alex Alderton, Roberto Amato, Jeffrey Barrett, Sonia Goncalves, Ewan Harrison, David K. Jackson, Ian Johnston, Dominic Kwiatkowski, Cordelia Langford, John Sillitoe on behalf of the Wellcome Sanger Institute COVID-19 Surveillance Team |
| EPI_ISL_2641663 | Lighthouse Lab in Glasgow | Wellcome Sanger Institute for the COVID-19 Genomics UK (COG-UK) Consortium | Harper VanSteenhouse, Yumi Kasai, David Gray, Carol Clugston, Anna Dominiczak and Alex Alderton, Roberto Amato, Jeffrey Barrett, Sonia Goncalves, Ewan Harrison, David K. Jackson, Ian Johnston, Dominic Kwiatkowski, Cordelia Langford, John Sillitoe on behalf of the Wellcome Sanger Institute COVID-19 Surveillance Team |
| EPI_ISL_2641664 | Health Services Laboratories | Wellcome Sanger Institute for the COVID-19 Genomics UK (COG-UK) Consortium | Health Services Laboratories and Alex Alderton, Roberto Amato, Jeffrey Barrett, Sonia Goncalves, Ewan Harrison, David K. Jackson, Ian Johnston, Dominic Kwiatkowski, Cordelia Langford, John Sillitoe on behalf of the Wellcome Sanger Institute COVID-19 Surveillance Team |
| EPI_ISL_2641665 | Lighthouse Lab in Alderley Park | Wellcome Sanger Institute for the COVID-19 Genomics UK (COG-UK) Consortium | Jacquelyn Wynn, Mairead Hyland, The Lighthouse Lab in Alderley Park and Alex Alderton, Roberto Amato, Jeffrey Barrett, Sonia Goncalves, Ewan Harrison, David K. Jackson, Ian Johnston, Dominic Kwiatkowski, Cordelia Langford, John Sillitoe on behalf of the Wellcome Sanger Institute COVID-19 Surveillance Team |
| EPI_ISL_2641666, EPI_ISL_2641667, EPI_ISL_2641668 | Lighthouse Lab in Glasgow | Wellcome Sanger Institute for the COVID-19 Genomics UK (COG-UK) Consortium | Harper VanSteenhouse, Yumi Kasai, David Gray, Carol Clugston, Anna Dominiczak and Alex Alderton, Roberto Amato, Jeffrey Barrett, Sonia Goncalves, Ewan Harrison, David K. Jackson, Ian Johnston, Dominic Kwiatkowski, Cordelia Langford, John Sillitoe on behalf of the Wellcome Sanger Institute COVID-19 Surveillance Team |
| EPI_ISL_2641669, EPI_ISL_2641670 | Lighthouse Lab in Alderley Park | Wellcome Sanger Institute for the COVID-19 Genomics UK (COG-UK) Consortium | Jacquelyn Wynn, Mairead Hyland, The Lighthouse Lab in Alderley Park and Alex Alderton, Roberto Amato, Jeffrey Barrett, Sonia Goncalves, Ewan Harrison, David K. Jackson, Ian Johnston, Dominic Kwiatkowski, Cordelia Langford, John Sillitoe on behalf of the Wellcome Sanger Institute COVID-19 Surveillance Team |
| EPI_ISL_2641671 | Health Services Laboratories | Wellcome Sanger Institute for the COVID-19 Genomics UK (COG-UK) Consortium | Health Services Laboratories and Alex Alderton, Roberto Amato, Jeffrey Barrett, Sonia Goncalves, Ewan Harrison, David K. Jackson, Ian Johnston, Dominic Kwiatkowski, Cordelia Langford, John Sillitoe on behalf of the Wellcome Sanger Institute COVID-19 Surveillance Team |
| EPI_ISL_2641672 | Lighthouse Lab in Glasgow | Wellcome Sanger Institute for the COVID-19 Genomics UK (COG-UK) Consortium | Harper VanSteenhouse, Yumi Kasai, David Gray, Carol Clugston, Anna Dominiczak and Alex Alderton, Roberto Amato, Jeffrey Barrett, Sonia Goncalves, Ewan Harrison, David K. Jackson, Ian Johnston, Dominic Kwiatkowski, Cordelia Langford, John Sillitoe on behalf of the Wellcome Sanger Institute COVID-19 Surveillance Team |
| EPI_ISL_2641673 | Health Services Laboratories | Wellcome Sanger Institute for the COVID-19 Genomics UK (COG-UK) Consortium | Health Services Laboratories and Alex Alderton, Roberto Amato, Jeffrey Barrett, Sonia Goncalves, Ewan Harrison, David K. Jackson, Ian Johnston, Dominic Kwiatkowski, Cordelia Langford, John Sillitoe on behalf of the Wellcome Sanger Institute COVID-19 Surveillance Team |
| EPI_ISL_2641674, EPI_ISL_2641675, EPI_ISL_2641676 | Lighthouse Lab in Glasgow | Wellcome Sanger Institute for the COVID-19 Genomics UK (COG-UK) Consortium | Harper VanSteenhouse, Yumi Kasai, David Gray, Carol Clugston, Anna Dominiczak and Alex Alderton, Roberto Amato, Jeffrey Barrett, Sonia Goncalves, Ewan Harrison, David K. Jackson, Ian Johnston, Dominic Kwiatkowski, Cordelia Langford, John Sillitoe on behalf of the Wellcome Sanger Institute COVID-19 Surveillance Team |
| EPI_ISL_2641677 | Health Services Laboratories | Wellcome Sanger Institute for the COVID-19 Genomics UK (COG-UK) Consortium | Health Services Laboratories and Alex Alderton, Roberto Amato, Jeffrey Barrett, Sonia Goncalves, Ewan Harrison, David K. Jackson, Ian Johnston, Dominic Kwiatkowski, Cordelia Langford, John Sillitoe on behalf of the Wellcome Sanger Institute COVID-19 Surveillance Team |
| EPI_ISL_2641678 | Lighthouse Lab in Glasgow | Wellcome Sanger Institute for the COVID-19 Genomics UK (COG-UK) Consortium | Harper VanSteenhouse, Yumi Kasai, David Gray, Carol Clugston, Anna Dominiczak and Alex Alderton, Roberto Amato, Jeffrey Barrett, Sonia Goncalves, Ewan Harrison, David K. Jackson, Ian Johnston, Dominic Kwiatkowski, Cordelia Langford, John Sillitoe on behalf of the Wellcome Sanger Institute COVID-19 Surveillance Team |
| EPI_ISL_2641679 | Lighthouse Lab in Alderley Park | Wellcome Sanger Institute for the COVID-19 Genomics UK (COG-UK) Consortium | Jacquelyn Wynn, Mairead Hyland, The Lighthouse Lab in Alderley Park and Alex Alderton, Roberto Amato, Jeffrey Barrett, Sonia Goncalves, Ewan Harrison, David K. Jackson, Ian Johnston, Dominic Kwiatkowski, Cordelia Langford, John Sillitoe on behalf of the Wellcome Sanger Institute COVID-19 Surveillance Team |
| EPI_ISL_2641680, EPI_ISL_2641681 | Lighthouse Lab in Glasgow | Wellcome Sanger Institute for the COVID-19 Genomics UK (COG-UK) Consortium | Harper VanSteenhouse, Yumi Kasai, David Gray, Carol Clugston, Anna Dominiczak and Alex Alderton, Roberto Amato, Jeffrey Barrett, Sonia Goncalves, Ewan Harrison, David K. Jackson, Ian Johnston, Dominic Kwiatkowski, Cordelia Langford, John Sillitoe on behalf of the Wellcome Sanger Institute COVID-19 Surveillance Team |
| EPI_ISL_2641682 | Lighthouse Lab in Alderley Park | Wellcome Sanger Institute for the COVID-19 Genomics UK (COG-UK) Consortium | Jacquelyn Wynn, Mairead Hyland, The Lighthouse Lab in Alderley Park and Alex Alderton, Roberto Amato, Jeffrey Barrett, Sonia Goncalves, Ewan Harrison, David K. Jackson, Ian Johnston, Dominic Kwiatkowski, Cordelia Langford, John Sillitoe on behalf of the Wellcome Sanger Institute COVID-19 Surveillance Team |
| EPI_ISL_2641683 | Lighthouse Lab in Glasgow | Wellcome Sanger Institute for the COVID-19 Genomics UK (COG-UK) Consortium | Harper VanSteenhouse, Yumi Kasai, David Gray, Carol Clugston, Anna Dominiczak and Alex Alderton, Roberto Amato, Jeffrey Barrett, Sonia Goncalves, Ewan Harrison, David K. Jackson, Ian Johnston, Dominic Kwiatkowski, Cordelia Langford, John Sillitoe on behalf of the Wellcome Sanger Institute COVID-19 Surveillance Team |
| EPI_ISL_2641684 | Health Services Laboratories | Wellcome Sanger Institute for the COVID-19 Genomics UK (COG-UK) Consortium | Health Services Laboratories and Alex Alderton, Roberto Amato, Jeffrey Barrett, Sonia Goncalves, Ewan Harrison, David K. Jackson, Ian Johnston, Dominic Kwiatkowski, Cordelia Langford, John Sillitoe on behalf of the Wellcome Sanger Institute COVID-19 Surveillance Team |
| EPI_ISL_2641685, EPI_ISL_2641686 | Lighthouse Lab in Glasgow | Wellcome Sanger Institute for the COVID-19 Genomics UK (COG-UK) Consortium | Harper VanSteenhouse, Yumi Kasai, David Gray, Carol Clugston, Anna Dominiczak and Alex Alderton, Roberto Amato, Jeffrey Barrett, Sonia Goncalves, Ewan Harrison, David K. Jackson, Ian Johnston, Dominic Kwiatkowski, Cordelia Langford, John Sillitoe on behalf of the Wellcome Sanger Institute COVID-19 Surveillance Team |
| EPI_ISL_2641687 | Lighthouse Lab in Alderley Park | Wellcome Sanger Institute for the COVID-19 Genomics UK (COG-UK) Consortium | Jacquelyn Wynn, Mairead Hyland, The Lighthouse Lab in Alderley Park and Alex Alderton, Roberto Amato, Jeffrey Barrett, Sonia Goncalves, Ewan Harrison, David K. Jackson, Ian Johnston, Dominic Kwiatkowski, Cordelia Langford, John Sillitoe on behalf of the Wellcome Sanger Institute COVID-19 Surveillance Team |
| EPI_ISL_2641688 | Health Services Laboratories | Wellcome Sanger Institute for the COVID-19 Genomics UK (COG-UK) Consortium | Health Services Laboratories and Alex Alderton, Roberto Amato, Jeffrey Barrett, Sonia Goncalves, Ewan Harrison, David K. Jackson, Ian Johnston, Dominic Kwiatkowski, Cordelia Langford, John Sillitoe on behalf of the Wellcome Sanger Institute COVID-19 Surveillance Team |
| EPI_ISL_2641690, EPI_ISL_2641692, EPI_ISL_2641693, EPI_ISL_2641694, EPI_ISL_2641695, EPI_ISL_2641696 | Lighthouse Lab in Glasgow | Wellcome Sanger Institute for the COVID-19 Genomics UK (COG-UK) Consortium | Harper VanSteenhouse, Yumi Kasai, David Gray, Carol Clugston, Anna Dominiczak and Alex Alderton, Roberto Amato, Jeffrey Barrett, Sonia Goncalves, Ewan Harrison, David K. Jackson, Ian Johnston, Dominic Kwiatkowski, Cordelia Langford, John Sillitoe on behalf of the Wellcome Sanger Institute COVID-19 Surveillance Team |
| EPI_ISL_2641697 | Health Services Laboratories | Wellcome Sanger Institute for the COVID-19 Genomics UK (COG-UK) Consortium | Health Services Laboratories and Alex Alderton, Roberto Amato, Jeffrey Barrett, Sonia Goncalves, Ewan Harrison, David K. Jackson, Ian Johnston, Dominic Kwiatkowski, Cordelia Langford, John Sillitoe on behalf of the Wellcome Sanger Institute COVID-19 Surveillance Team |
| EPI_ISL_2641698, EPI_ISL_2641699, EPI_ISL_2641700, EPI_ISL_2641701, EPI_ISL_2641702 | Lighthouse Lab in Glasgow | Wellcome Sanger Institute for the COVID-19 Genomics UK (COG-UK) Consortium | Harper VanSteenhouse, Yumi Kasai, David Gray, Carol Clugston, Anna Dominiczak and Alex Alderton, Roberto Amato, Jeffrey Barrett, Sonia Goncalves, Ewan Harrison, David K. Jackson, Ian Johnston, Dominic Kwiatkowski, Cordelia Langford, John Sillitoe on behalf of the Wellcome Sanger Institute COVID-19 Surveillance Team |
| EPI_ISL_2641703 | Health Services Laboratories | Wellcome Sanger Institute for the COVID-19 Genomics UK (COG-UK) Consortium | Health Services Laboratories and Alex Alderton, Roberto Amato, Jeffrey Barrett, Sonia Goncalves, Ewan Harrison, David K. Jackson, Ian Johnston, Dominic Kwiatkowski, Cordelia Langford, John Sillitoe on behalf of the Wellcome Sanger Institute COVID-19 Surveillance Team |
| EPI_ISL_2641704, EPI_ISL_2641705 | Lighthouse Lab in Glasgow | Wellcome Sanger Institute for the COVID-19 Genomics UK (COG-UK) Consortium | Harper VanSteenhouse, Yumi Kasai, David Gray, Carol Clugston, Anna Dominiczak and Alex Alderton, Roberto Amato, Jeffrey Barrett, Sonia Goncalves, Ewan Harrison, David K. Jackson, Ian Johnston, Dominic Kwiatkowski, Cordelia Langford, John Sillitoe on behalf of the Wellcome Sanger Institute COVID-19 Surveillance Team |
| EPI_ISL_2641706 | Health Services Laboratories | Wellcome Sanger Institute for the COVID-19 Genomics UK (COG-UK) Consortium | Health Services Laboratories and Alex Alderton, Roberto Amato, Jeffrey Barrett, Sonia Goncalves, Ewan Harrison, David K. Jackson, Ian Johnston, Dominic Kwiatkowski, Cordelia Langford, John Sillitoe on behalf of the Wellcome Sanger Institute COVID-19 Surveillance Team |
| EPI_ISL_2641707, EPI_ISL_2641708 | Lighthouse Lab in Glasgow | Wellcome Sanger Institute for the COVID-19 Genomics UK (COG-UK) Consortium | Harper VanSteenhouse, Yumi Kasai, David Gray, Carol Clugston, Anna Dominiczak and Alex Alderton, Roberto Amato, Jeffrey Barrett, Sonia Goncalves, Ewan Harrison, David K. Jackson, Ian Johnston, Dominic Kwiatkowski, Cordelia Langford, John Sillitoe on behalf of the Wellcome Sanger Institute COVID-19 Surveillance Team |
| EPI_ISL_2641709 | Health Services Laboratories | Wellcome Sanger Institute for the COVID-19 Genomics UK (COG-UK) Consortium | Health Services Laboratories and Alex Alderton, Roberto Amato, Jeffrey Barrett, Sonia Goncalves, Ewan Harrison, David K. Jackson, Ian Johnston, Dominic Kwiatkowski, Cordelia Langford, John Sillitoe on behalf of the Wellcome Sanger Institute COVID-19 Surveillance Team |
| EPI_ISL_2641710, EPI_ISL_2641711 | Lighthouse Lab in Glasgow | Wellcome Sanger Institute for the COVID-19 Genomics UK (COG-UK) Consortium | Harper VanSteenhouse, Yumi Kasai, David Gray, Carol Clugston, Anna Dominiczak and Alex Alderton, Roberto Amato, Jeffrey Barrett, Sonia Goncalves, Ewan Harrison, David K. Jackson, Ian Johnston, Dominic Kwiatkowski, Cordelia Langford, John Sillitoe on behalf of the Wellcome Sanger Institute COVID-19 Surveillance Team |
| EPI_ISL_2641712 | Lighthouse Laboratory Plymouth | Wellcome Sanger Institute for the COVID-19 Genomics UK (COG-UK) Consortium | Lighthouse Laboratory Plymouth and Alex Alderton, Roberto Amato, Jeffrey Barrett, Sonia Goncalves, Ewan Harrison, David K. Jackson, Ian Johnston, Dominic Kwiatkowski, Cordelia Langford, John Sillitoe on behalf of the Wellcome Sanger Institute COVID-19 Surveillance Team |
| EPI_ISL_2641713, EPI_ISL_2641714 | Lighthouse Lab in Alderley Park | Wellcome Sanger Institute for the COVID-19 Genomics UK (COG-UK) Consortium | Jacquelyn Wynn, Mairead Hyland, The Lighthouse Lab in Alderley Park and Alex Alderton, Roberto Amato, Jeffrey Barrett, Sonia Goncalves, Ewan Harrison, David K. Jackson, Ian Johnston, Dominic Kwiatkowski, Cordelia Langford, John Sillitoe on behalf of the Wellcome Sanger Institute COVID-19 Surveillance Team |
| EPI_ISL_2641715, EPI_ISL_2641716, EPI_ISL_2641717 | Lighthouse Lab in Milton Keynes | Wellcome Sanger Institute for the COVID-19 Genomics UK (COG-UK) Consortium | The Lighthouse Lab in Milton Keynes and Alex Alderton, Roberto Amato, Jeffrey Barrett, Sonia Goncalves, Ewan Harrison, David K. Jackson, Ian Johnston, Dominic Kwiatkowski, Cordelia Langford, John Sillitoe on behalf of the Wellcome Sanger Institute COVID-19 Surveillance Team |
| EPI_ISL_2641718 | Lighthouse Lab in Cambridge | Wellcome Sanger Institute for the COVID-19 Genomics UK (COG-UK) Consortium | Rob Howes, The Lighthouse Lab in Cambridge and Alex Alderton, Roberto Amato, Jeffrey Barrett, Sonia Goncalves, Ewan Harrison, David K. Jackson, Ian Johnston, Dominic Kwiatkowski, Cordelia Langford, John Sillitoe on behalf of the Wellcome Sanger Institute COVID-19 Surveillance Team |
| EPI_ISL_2641719 | Lighthouse Lab in Milton Keynes | Wellcome Sanger Institute for the COVID-19 Genomics UK (COG-UK) Consortium | The Lighthouse Lab in Milton Keynes and Alex Alderton, Roberto Amato, Jeffrey Barrett, Sonia Goncalves, Ewan Harrison, David K. Jackson, Ian Johnston, Dominic Kwiatkowski, Cordelia Langford, John Sillitoe on behalf of the Wellcome Sanger Institute COVID-19 Surveillance Team |
| EPI_ISL_2641756, EPI_ISL_2641758, EPI_ISL_2641763, EPI_ISL_2641779, EPI_ISL_2641780, EPI_ISL_2641798, EPI_ISL_2641801, EPI_ISL_2641821, EPI_ISL_2641825, EPI_ISL_2641827, EPI_ISL_2641832, EPI_ISL_2641838, EPI_ISL_2641840, EPI_ISL_2641841, EPI_ISL_2641850, EPI_ISL_2641851, EPI_ISL_2641852, EPI_ISL_2641853, EPI_ISL_2641869, EPI_ISL_2641870, EPI_ISL_2641871, EPI_ISL_2641882, EPI_ISL_2641883, EPI_ISL_2641887, EPI_ISL_2641888, EPI_ISL_2641890, EPI_ISL_2641894, EPI_ISL_2641895, EPI_ISL_2641896, EPI_ISL_2641897, EPI_ISL_2641898, EPI_ISL_2641899, EPI_ISL_2641900, EPI_ISL_2641901, EPI_ISL_2641902, EPI_ISL_2641903, EPI_ISL_2641904, EPI_ISL_2641905, EPI_ISL_2641908, EPI_ISL_2641921, EPI_ISL_2641922, EPI_ISL_2641923, EPI_ISL_2641925, EPI_ISL_2641928, EPI_ISL_2641929, EPI_ISL_2641930, EPI_ISL_2641931, EPI_ISL_2641932, EPI_ISL_2641933, EPI_ISL_2641934, EPI_ISL_2641940, EPI_ISL_2641943, EPI_ISL_2641944, EPI_ISL_2641947, EPI_ISL_2641948, EPI_ISL_2641949, EPI_ISL_2641950, EPI_ISL_2641951, EPI_ISL_2641952, EPI_ISL_2641953, EPI_ISL_2641954, EPI_ISL_2641955, EPI_ISL_2641969, EPI_ISL_2641971, EPI_ISL_2641972, EPI_ISL_2641974, EPI_ISL_2641975, EPI_ISL_2641977, EPI_ISL_2641979, EPI_ISL_2641981, EPI_ISL_2641983, EPI_ISL_2641990, EPI_ISL_2641991, EPI_ISL_2641992, EPI_ISL_2641993, EPI_ISL_2641994, EPI_ISL_2641995, EPI_ISL_2641996, EPI_ISL_2641997, EPI_ISL_2641998, EPI_ISL_2641999, EPI_ISL_2642001 | | | |
| see above | Istituto Zooprofilattico Sperimentale del Mezzogiorno | Telethon Institute of Genetics and Medicine (TIGEM) | Antonio Grimaldi Patrizia Annunziata Francesco Panariello Biancamaria Pierri Claudia Tiberio Teresa Giuliano Valentina Bouche Chiara Colantuono Maria Concetta Cuomo Denise Di Concilio Lucio Di Filippo Anna Manfredi Marcello Salvi Antonio Limone Luigi Atripaldi Pellegrino Cerino Andrea Ballabio Davide Cacchiarelli |
| EPI_ISL_2643672 | Furst Medical Laboratory | Norwegian Institute of Public Health, Department of Virology | Kathrine Stene-Johansen, Kamilla Heddeland Instefjord, Hilde Elshaug, Garcia Llorente Ignacio, Jon Bråte, Engebretsen Serina Beate,Pedersen Benedikte Nevjen, Line Victoria Moen, Debech Nadia, Atiya R Ali,Marie Paulsen Madsen, Rasmus Riis |

| | | | |
|---|---|---|---|
| | | | Kopperud, Hilde Vollan, Karoline Bragstad, Olav Hungnes |
| EPI_ISL_2644451 | Institute of Microbiology and Immunology, Faculty of Medicine, University of Ljubljana | Institute of Microbiology and Immunology, Faculty of Medicine, University of Ljubljana | Alen Suljič, Samo Zakotnik, Tomaž Mark Zorec, Matic Brvar, Doroteja Vlaj, Andraž Celar, Dominika Šturm, Patricija Pozvek, Špela Pleh, Miša Korva, Mario Poljak, Tatjana Avšič – Županc |
| EPI_ISL_2644691 | Medical Microbiology Unit, Department for Laboratory Medicine, Drammen Hospital, Vestre Viken Health Trust, | Norwegian Institute of Public Health, Department of Virology | Kathrine Stene-Johansen, Kamilla Heddeland Instefjord, Hilde Elshaug, Garcia Llorente Ignacio, Jon Bråte, Engebretsen Serina Beate,Pedersen Benedikte Nevjen, Line Victoria Moen, Debech Nadia, Atiya R Ali,Marie Paulsen Madsen, Rasmus Riis Kopperud, Hilde Vollan, Karoline Bragstad, Olav Hungnes |
| EPI_ISL_2644695, EPI_ISL_2644696 | Furst Medical Laboratory | Norwegian Institute of Public Health, Department of Virology | Kathrine Stene-Johansen, Kamilla Heddeland Instefjord, Hilde Elshaug, Garcia Llorente Ignacio, Jon Bråte, Engebretsen Serina Beate,Pedersen Benedikte Nevjen, Line Victoria Moen, Debech Nadia, Atiya R Ali,Marie Paulsen Madsen, Rasmus Riis Kopperud, Hilde Vollan, Karoline Bragstad, Olav Hungnes |
| EPI_ISL_2644701 | Akershus University Hospital, Department for Microbiology and Infectious Disease Control | Norwegian Institute of Public Health, Department of Virology | Kathrine Stene-Johansen, Kamilla Heddeland Instefjord, Hilde Elshaug, Garcia Llorente Ignacio, Jon Bråte, Engebretsen Serina Beate,Pedersen Benedikte Nevjen, Line Victoria Moen, Debech Nadia, Atiya R Ali,Marie Paulsen Madsen, Rasmus Riis Kopperud, Hilde Vollan, Karoline Bragstad, Olav Hungnes |
| EPI_ISL_2644706 | Medical Microbiology Unit, Department for Laboratory Medicine, Drammen Hospital, Vestre Viken Health Trust, | Norwegian Institute of Public Health, Department of Virology | Kathrine Stene-Johansen, Kamilla Heddeland Instefjord, Hilde Elshaug, Garcia Llorente Ignacio, Jon Bråte, Engebretsen Serina Beate,Pedersen Benedikte Nevjen, Line Victoria Moen, Debech Nadia, Atiya R Ali,Marie Paulsen Madsen, Rasmus Riis Kopperud, Hilde Vollan, Karoline Bragstad, Olav Hungnes |
| EPI_ISL_2644715 | Furst Medical Laboratory | Norwegian Institute of Public Health, Department of Virology | Kathrine Stene-Johansen, Kamilla Heddeland Instefjord, Hilde Elshaug, Garcia Llorente Ignacio, Jon Bråte, Engebretsen Serina Beate,Pedersen Benedikte Nevjen, Line Victoria Moen, Debech Nadia, Atiya R Ali,Marie Paulsen Madsen, Rasmus Riis Kopperud, Hilde Vollan, Karoline Bragstad, Olav Hungnes |
| EPI_ISL_2644716 | Dutch COVID-19 response team | Medical Microbiology, Maastricht University Medical Centre | Jozef Dingemans*, Brian van der Veer*, Erik Beuken, Carmen Reumkens, Lieke van Alphen, Christian Hoebe, Paul Savelkoul |
| EPI_ISL_2644777 | Nacionalinis maisto ir veterinarijos rizikos vertinimo institutas | Institute of Biotechnology, Life Sciences Center, Vilnius University | Emilija Vasiliunaite; Milda Norkiene; Danguole Ziogiene; Albertas Timinskas; Alma Gedvilaite |
| EPI_ISL_2644785 | UAB Medicina practica laboratorija | Institute of Biotechnology, Life Sciences Center, Vilnius University | Emilija Vasiliunaite; Milda Norkiene; Danguole Ziogiene; Albertas Timinskas; Alma Gedvilaite |
| EPI_ISL_2644845, EPI_ISL_2644846, EPI_ISL_2644847, EPI_ISL_2644848 | NordLab Kokkola | Expert Microbiology, National Institute for Health and Welfare | Soile Blomqvist, Jani Halkilahti, Kirsi Liitsola, Haider al-Hello, Päivi Laurila, Erika Lindh, Teemu Smura, Ravi Kant, Phuoc Truong, Olli Vapalahti, Sari Hannula, Pekka Ellonen, Niina Ikonen, Carita Savolainen-Kopra |
| EPI_ISL_2644866, EPI_ISL_2644898 | Fimlab Laboratoriot Oy Tampere | Expert Microbiology, National Institute for Health and Welfare | Soile Blomqvist, Jani Halkilahti, Kirsi Liitsola, Haider al-Hello, Päivi Laurila, Erika Lindh, Teemu Smura, Ravi Kant, Phuoc Truong, Olli Vapalahti, Sari Hannula, Pekka Ellonen, Niina Ikonen, Carita Savolainen-Kopra |
| EPI_ISL_2644935, EPI_ISL_2644943 | NordLab Oulu | Expert Microbiology, National Institute for Health and Welfare | Soile Blomqvist, Jani Halkilahti, Kirsi Liitsola, Haider al-Hello, Päivi Laurila, Erika Lindh, Teemu Smura, Ravi Kant, Phuoc Truong, Olli Vapalahti, Sari Hannula, Pekka Ellonen, Niina Ikonen, Carita Savolainen-Kopra |
| EPI_ISL_2644978, EPI_ISL_2644988 | TYKS, Kliininen mikrobiologia | Expert Microbiology, National Institute for Health and Welfare | Soile Blomqvist, Jani Halkilahti, Kirsi Liitsola, Haider al-Hello, Päivi Laurila, Erika Lindh, Teemu Smura, Ravi Kant, Phuoc Truong, Olli Vapalahti, Sari Hannula, Pekka Ellonen, Niina Ikonen, Carita Savolainen-Kopra |
| EPI_ISL_2644992 | SYNLAB Suomi | Expert Microbiology, National Institute for Health and Welfare | Soile Blomqvist, Jani Halkilahti, Kirsi Liitsola, Haider al-Hello, Päivi Laurila, Erika Lindh, Teemu Smura, Ravi Kant, Phuoc Truong, Olli Vapalahti, Sari Hannula, Pekka Ellonen, Niina Ikonen, Carita Savolainen-Kopra |
| EPI_ISL_2645004, EPI_ISL_2645005, EPI_ISL_2645006, EPI_ISL_2645007, EPI_ISL_2645008 | NordLab Kokkola | Expert Microbiology, National Institute for Health and Welfare | Soile Blomqvist, Jani Halkilahti, Kirsi Liitsola, Haider al-Hello, Päivi Laurila, Erika Lindh, Teemu Smura, Ravi Kant, Phuoc Truong, Olli Vapalahti, Sari Hannula, Pekka Ellonen, Niina Ikonen, Carita Savolainen-Kopra |
| EPI_ISL_2645040, EPI_ISL_2645042, EPI_ISL_2645043, EPI_ISL_2645047, EPI_ISL_2645051, EPI_ISL_2645053, EPI_ISL_2645055, EPI_ISL_2645057, EPI_ISL_2645060, EPI_ISL_2645061, EPI_ISL_2645065, EPI_ISL_2645066 | | | |
| see above | NordLab Oulu | Expert Microbiology, National Institute for Health and Welfare | Soile Blomqvist, Jani Halkilahti, Kirsi Liitsola, Haider al-Hello, Päivi Laurila, Erika Lindh, Teemu Smura, Ravi Kant, Phuoc Truong, Olli Vapalahti, Sari Hannula, Pekka Ellonen, Niina Ikonen, Carita Savolainen-Kopra |
| EPI_ISL_2645068, EPI_ISL_2645071, EPI_ISL_2645084, EPI_ISL_2645086, EPI_ISL_2645108, EPI_ISL_2645111, EPI_ISL_2645112, EPI_ISL_2645113 | Fimlab Laboratoriot Oy Tampere | Expert Microbiology, National Institute for Health and Welfare | Soile Blomqvist, Jani Halkilahti, Kirsi Liitsola, Haider al-Hello, Päivi Laurila, Erika Lindh, Teemu Smura, Ravi Kant, Phuoc Truong, Olli Vapalahti, Sari Hannula, Pekka Ellonen, Niina Ikonen, Carita Savolainen-Kopra |
| EPI_ISL_2645119 | Islab, Pohjois-Savon aluelaboratorio | Expert Microbiology, National Institute for Health and Welfare | Soile Blomqvist, Jani Halkilahti, Kirsi Liitsola, Haider al-Hello, Päivi Laurila, Erika Lindh, Teemu Smura, Ravi Kant, Phuoc Truong, Olli Vapalahti, Sari Hannula, Pekka Ellonen, Niina Ikonen, Carita Savolainen-Kopra |
| EPI_ISL_2645129 | Vita Laboratoriot Oy | Expert Microbiology, National Institute for Health and Welfare | Soile Blomqvist, Jani Halkilahti, Kirsi Liitsola, Haider al-Hello, Päivi Laurila, Erika Lindh, Teemu Smura, Ravi Kant, Phuoc Truong, Olli Vapalahti, Sari Hannula, Pekka Ellonen, Niina Ikonen, Carita Savolainen-Kopra |
| EPI_ISL_2645139, EPI_ISL_2645140, EPI_ISL_2645141, EPI_ISL_2645142, EPI_ISL_2645143 | NordLab Kokkola | Expert Microbiology, National Institute for Health and Welfare | Soile Blomqvist, Jani Halkilahti, Kirsi Liitsola, Haider al-Hello, Päivi Laurila, Erika Lindh, Teemu Smura, Ravi Kant, Phuoc Truong, Olli Vapalahti, Sari Hannula, Pekka Ellonen, Niina Ikonen, Carita Savolainen-Kopra |
| EPI_ISL_2645146, EPI_ISL_2645149, EPI_ISL_2645150, EPI_ISL_2645151, EPI_ISL_2645153, EPI_ISL_2645155, EPI_ISL_2645156, EPI_ISL_2645157, EPI_ISL_2645158, EPI_ISL_2645163, EPI_ISL_2645165, EPI_ISL_2645166, EPI_ISL_2645167, EPI_ISL_2645168, EPI_ISL_2645170 | | | |
| see above | NordLab Oulu | Expert Microbiology, National Institute for Health and Welfare | Soile Blomqvist, Jani Halkilahti, Kirsi Liitsola, Haider al-Hello, Päivi Laurila, Erika Lindh, Teemu Smura, Ravi Kant, Phuoc Truong, Olli Vapalahti, Sari Hannula, Pekka Ellonen, Niina Ikonen, Carita Savolainen-Kopra |
| EPI_ISL_2645205, EPI_ISL_2645212, EPI_ISL_2645216, EPI_ISL_2645217, EPI_ISL_2645218 | Fimlab Laboratoriot Oy Tampere | Expert Microbiology, National Institute for Health and Welfare | Soile Blomqvist, Jani Halkilahti, Kirsi Liitsola, Haider al-Hello, Päivi Laurila, Erika Lindh, Teemu Smura, Ravi Kant, Phuoc Truong, Olli Vapalahti, Sari Hannula, Pekka Ellonen, Niina Ikonen, Carita Savolainen-Kopra |
| EPI_ISL_2645273 | Royal GD | Royal GD | Remco Dijkman |
| EPI_ISL_2645281 | SD Public Health Laboratory | Centers for Disease Control and Prevention Division of Viral Diseases, Pathogen Discovery | Mili Sheth, Sarah Nobles, Jasmine Padilla, Mark Burroughs, Shoshona Le, Katie Dillon, Peter Cook, Clinton R. Paden, Dhwani Batra, Krista Queen, Kristen Knipe, Dakota Howard, Yvette Unoarumhi, Darlene Wagner, Matthew Schmerer, Ben L. Rambo-Martin, Kristine Lacek, Sam Shepard, Alison Laufer Halpin, Dave Wentworth, Vivien Dugan, Suxiang Tong, Justin Lee |
| EPI_ISL_2645337 | MI - Michigan Department of Health and Human Services - Bureau of Laboratories | Centers for Disease Control and Prevention Division of Viral Diseases, Pathogen Discovery | Mili Sheth, Sarah Nobles, Jasmine Padilla, Mark Burroughs, Shoshona Le, Katie Dillon, Peter Cook, Clinton R. Paden, Dhwani Batra, Krista Queen, Kristen Knipe, Dakota Howard, Yvette Unoarumhi, Darlene Wagner, Matthew Schmerer, Ben L. Rambo-Martin, Kristine Lacek, Sam Shepard, Alison Laufer Halpin, Dave Wentworth, Vivien Dugan, Suxiang Tong, Justin Lee |
| EPI_ISL_2645411, EPI_ISL_2645412, EPI_ISL_2645413, EPI_ISL_2645414, EPI_ISL_2645415 | Laboratorio Central de Saude Publica do Estado Maranhao (LACEN-MA) | Laboratory of Respiratory Viruses and Measles, Oswaldo Cruz Institute, FIOCRUZ | Paola Resende, Luciana Appolinario, Fernando Motta, Anna Carolina Paixao, Ana Carolina Mendonca, Alice Sampaio Rocha, Taina Venas, Elisa Cavalcante Pereira, Renata Serrano Lopes, Lidio Gonçalves Lima Neto, Marilda Siqueira on behalf of the Fiocruz COVID-19 Genomic Surveillance Network |
| EPI_ISL_2645416 | Laboratorio Central de Saude Publica do Estado Maranhao (LACEN-MA) | Laboratory of Respiratory Viruses and Measles, Oswaldo Cruz Institute, FIOCRUZ | Paola Resende, Alex Pauvolid-Corrêa, Mia Ferreira de Araujo, Ana Beatriz Machado Lima, Luciana Appolinario, Fernando Motta, Anna Carolina Paixao, Ana Carolina Mendonca, Alice Sampaio Rocha, Taina Venas, Elisa Cavalcante Pereira, Renata Serrano Lopes, Lidio Gonçalves Lima Neto, Marilda Siqueira on behalf of the Fiocruz COVID-19 Genomic Surveillance Network |
| EPI_ISL_2645417 | Laboratorio Central de Saude Publica do Estado do Maranhao (LACEN-MA) | Laboratory of Respiratory Viruses and Measles, Oswaldo Cruz Institute, FIOCRUZ | Paola Resende, Luciana Appolinario, Fernando Motta, Anna Carolina Paixao, Ana Carolina Mendonca, Alice Sampaio Rocha, Taina Venas, Elisa Cavalcante Pereira, Renata Serrano Lopes, Lidio Gonçalves Lima Neto, Marilda Siqueira on behalf of the Fiocruz COVID-19 Genomic Surveillance Network |
| EPI_ISL_2645418 | Laboratorio Central de Saude Publica do Estado do Maranhao (LACEN-MA) | Laboratory of Respiratory Viruses and Measles, Oswaldo Cruz Institute, FIOCRUZ | Paola Resende, Alex Pauvolid-Corrêa, Mia Ferreira de Araujo, Ana Beatriz Machado Lima, Luciana Appolinario, Fernando Motta, Anna Carolina Paixao, Ana Carolina Mendonca, Alice Sampaio Rocha, Taina Venas, Elisa Cavalcante Pereira, Renata Serrano Lopes, Lidio Gonçalves Lima Neto, Marilda Siqueira on behalf of the Fiocruz COVID-19 Genomic Surveillance Network |
| EPI_ISL_2645747, EPI_ISL_2645748, EPI_ISL_2645749, EPI_ISL_2645752, EPI_ISL_2645766, EPI_ISL_2645767, EPI_ISL_2645777, EPI_ISL_2645783, EPI_ISL_2645784 | Nebraska Public Health Laboratory | NPHL COVID-19 Response Team | NPHL COVID-19 Response Team |
| EPI_ISL_2645798, EPI_ISL_2645817 | Hospital Universitari Vall d'Hebron - Vall d'Hebron Institut de Recerca | Hospital Universitari Vall d'Hebron - Vall d'Hebron Institut de Recerca | Cristina Andrés, Maria Piñana, Alejandra González-Sánchez, Damir Garcia-Cehic, Ariadna Rando, Juliana Esperalba, Maria Gema Codina, Carla Castillo, Maria Carmen Martin, Tomàs Pumarola, Josep Quer, Andrés Antón |
| EPI_ISL_2645940, EPI_ISL_2645942, EPI_ISL_2645943, EPI_ISL_2645945, EPI_ISL_2645946, EPI_ISL_2645947, EPI_ISL_2645948, EPI_ISL_2645957, EPI_ISL_2645958, EPI_ISL_2645961, EPI_ISL_2645964, EPI_ISL_2645965, EPI_ISL_2645971, EPI_ISL_2645979, EPI_ISL_2645980, EPI_ISL_2645984, EPI_ISL_2645985, EPI_ISL_2645987, EPI_ISL_2645989, EPI_ISL_2645998, EPI_ISL_2645999, EPI_ISL_2646000, EPI_ISL_2646001, EPI_ISL_2646002, EPI_ISL_2646003, EPI_ISL_2646004, EPI_ISL_2646008, EPI_ISL_2646010, EPI_ISL_2646011, EPI_ISL_2646012, EPI_ISL_2646013, EPI_ISL_2646014, EPI_ISL_2646015, EPI_ISL_2646016, EPI_ISL_2646022, EPI_ISL_2646023 | | | |
| see above | UNILABS | Instituto Nacional de Saude (INSA) | Borges et al |
| EPI_ISL_2646026 | INSA | Instituto Nacional de Saude (INSA) | Borges et al |
| EPI_ISL_2646029, EPI_ISL_2646030, EPI_ISL_2646031, EPI_ISL_2646033, EPI_ISL_2646034, EPI_ISL_2646036, EPI_ISL_2646037, EPI_ISL_2646038, EPI_ISL_2646039, EPI_ISL_2646040, EPI_ISL_2646041, EPI_ISL_2646042, EPI_ISL_2646043, EPI_ISL_2646044, EPI_ISL_2646047, EPI_ISL_2646048, EPI_ISL_2646049, EPI_ISL_2646050, EPI_ISL_2646051, EPI_ISL_2646052, EPI_ISL_2646053, EPI_ISL_2646054, EPI_ISL_2646055, EPI_ISL_2646057, EPI_ISL_2646058, EPI_ISL_2646059, EPI_ISL_2646061, EPI_ISL_2646062, EPI_ISL_2646063, EPI_ISL_2646064, EPI_ISL_2646065, EPI_ISL_2646066, EPI_ISL_2646067, EPI_ISL_2646071, EPI_ISL_2646072, EPI_ISL_2646074, EPI_ISL_2646075, EPI_ISL_2646080, EPI_ISL_2646081, EPI_ISL_2646085, EPI_ISL_2646086, EPI_ISL_2646090, EPI_ISL_2646092, EPI_ISL_2646093, EPI_ISL_2646094, EPI_ISL_2646095, EPI_ISL_2646096, EPI_ISL_2646097, EPI_ISL_2646098, EPI_ISL_2646103, EPI_ISL_2646104, EPI_ISL_2646105, EPI_ISL_2646106, EPI_ISL_2646108, EPI_ISL_2646109, EPI_ISL_2646111, EPI_ISL_2646116, EPI_ISL_2646117, EPI_ISL_2646118, EPI_ISL_2646119 | | | |
| see above | UNILABS | Instituto Nacional de Saude (INSA) | Borges et al |
| EPI_ISL_2646219 | cerballiance-IDF | Cerba lab | Emmanuel Lecorche, Sabine Trombert-Paolantoni, Aude Lessenne, Kader Merah, Thierry Collin, Laura Verdurme, Patrice Herisson, Stéphanie Haïm-Boukobza, Bénédicte Roquebert |
| EPI_ISL_2646220, | Exalab | Cerba lab | Emmanuel Lecorche, Sabine Trombert-Paolantoni, Aude Lessenne, Kader Merah, Thierry Collin, Laura Verdurme, Patrice Herisson, Stéphanie Haïm-Boukobza, Bénédicte Roquebert |

| | | | |
|---|---|---|---|
| EPI_ISL_2646221, EPI_ISL_2646222, EPI_ISL_2646223 | | | |
| EPI_ISL_2646224 | cerballiance-IDF | Cerba lab | Emmanuel Lecorche, Sabine Trombert-Paolantoni, Aude Lessenne, Kader Merah, Thierry Collin, Laura Verdurme, Patrice Herisson, Stéphanie Haïm-Boukobza, Bénédicte Roquebert |
| EPI_ISL_2646225, EPI_ISL_2646226, EPI_ISL_2646227, EPI_ISL_2646228, EPI_ISL_2646229, EPI_ISL_2646230, EPI_ISL_2646231 | Exalab | Cerba lab | Emmanuel Lecorche, Sabine Trombert-Paolantoni, Aude Lessenne, Kader Merah, Thierry Collin, Laura Verdurme, Patrice Herisson, Stéphanie Haïm-Boukobza, Bénédicte Roquebert |
| EPI_ISL_2646233, EPI_ISL_2646234, EPI_ISL_2646235, EPI_ISL_2646237, EPI_ISL_2646238 | Cerballiance-IDF | Cerba lab | Emmanuel Lecorche, Sabine Trombert-Paolantoni, Aude Lessenne, Kader Merah, Thierry Collin, Laura Verdurme, Patrice Herisson, Stéphanie Haïm-Boukobza, Bénédicte Roquebert |
| EPI_ISL_2646240 | Cerballiance-Provence | Cerba lab | Emmanuel Lecorche, Sabine Trombert-Paolantoni, Aude Lessenne, Kader Merah, Thierry Collin, Laura Verdurme, Patrice Herisson, Stéphanie Haïm-Boukobza, Bénédicte Roquebert |
| EPI_ISL_2646241 | Cerballiance-Normandie | Cerba lab | Emmanuel Lecorche, Sabine Trombert-Paolantoni, Aude Lessenne, Kader Merah, Thierry Collin, Laura Verdurme, Patrice Herisson, Stéphanie Haïm-Boukobza, Bénédicte Roquebert |
| EPI_ISL_2646242 | Cerballiance-IDF | Cerba lab | Emmanuel Lecorche, Sabine Trombert-Paolantoni, Aude Lessenne, Kader Merah, Thierry Collin, Laura Verdurme, Patrice Herisson, Stéphanie Haïm-Boukobza, Bénédicte Roquebert |
| EPI_ISL_2646243 | Cerballiance-Provence | Cerba lab | Emmanuel Lecorche, Sabine Trombert-Paolantoni, Aude Lessenne, Kader Merah, Thierry Collin, Laura Verdurme, Patrice Herisson, Stéphanie Haïm-Boukobza, Bénédicte Roquebert |
| EPI_ISL_2646245, EPI_ISL_2646246 | Cerballiance-Paris | Cerba lab | Emmanuel Lecorche, Sabine Trombert-Paolantoni, Aude Lessenne, Kader Merah, Thierry Collin, Laura Verdurme, Patrice Herisson, Stéphanie Haïm-Boukobza, Bénédicte Roquebert |
| EPI_ISL_2646247 | Cerballiance-Normandie | Cerba lab | Emmanuel Lecorche, Sabine Trombert-Paolantoni, Aude Lessenne, Kader Merah, Thierry Collin, Laura Verdurme, Patrice Herisson, Stéphanie Haïm-Boukobza, Bénédicte Roquebert |
| EPI_ISL_2646251 | Synlab | Cerba lab | Emmanuel Lecorche, Sabine Trombert-Paolantoni, Aude Lessenne, Kader Merah, Thierry Collin, Laura Verdurme, Patrice Herisson, Stéphanie Haïm-Boukobza, Bénédicte Roquebert |
| EPI_ISL_2646252 | Cerballiance-Paris | Cerba lab | Emmanuel Lecorche, Sabine Trombert-Paolantoni, Aude Lessenne, Kader Merah, Thierry Collin, Laura Verdurme, Patrice Herisson, Stéphanie Haïm-Boukobza, Bénédicte Roquebert |
| EPI_ISL_2646253, EPI_ISL_2646254 | Cerballiance-Pyrénées | Cerba lab | Emmanuel Lecorche, Sabine Trombert-Paolantoni, Aude Lessenne, Kader Merah, Thierry Collin, Laura Verdurme, Patrice Herisson, Stéphanie Haïm-Boukobza, Bénédicte Roquebert |
| EPI_ISL_2646255 | Cerballiance-Paris | Cerba lab | Emmanuel Lecorche, Sabine Trombert-Paolantoni, Aude Lessenne, Kader Merah, Thierry Collin, Laura Verdurme, Patrice Herisson, Stéphanie Haïm-Boukobza, Bénédicte Roquebert |
| EPI_ISL_2646257 | Cerballiance-Pyrénées | Cerba lab | Emmanuel Lecorche, Sabine Trombert-Paolantoni, Aude Lessenne, Kader Merah, Thierry Collin, Laura Verdurme, Patrice Herisson, Stéphanie Haïm-Boukobza, Bénédicte Roquebert |
| EPI_ISL_2646258 | Cerballiance-Paris | Cerba lab | Emmanuel Lecorche, Sabine Trombert-Paolantoni, Aude Lessenne, Kader Merah, Thierry Collin, Laura Verdurme, Patrice Herisson, Stéphanie Haïm-Boukobza, Bénédicte Roquebert |
| EPI_ISL_2646259 | COSEM | Cerba lab | Emmanuel Lecorche, Sabine Trombert-Paolantoni, Aude Lessenne, Kader Merah, Thierry Collin, Laura Verdurme, Patrice Herisson, Stéphanie Haïm-Boukobza, Bénédicte Roquebert |
| EPI_ISL_2646260 | Cerballiance-IDF | Cerba lab | Emmanuel Lecorche, Sabine Trombert-Paolantoni, Aude Lessenne, Kader Merah, Thierry Collin, Laura Verdurme, Patrice Herisson, Stéphanie Haïm-Boukobza, Bénédicte Roquebert |
| EPI_ISL_2646261 | Cerballiance-Paris | Cerba lab | Emmanuel Lecorche, Sabine Trombert-Paolantoni, Aude Lessenne, Kader Merah, Thierry Collin, Laura Verdurme, Patrice Herisson, Stéphanie Haïm-Boukobza, Bénédicte Roquebert |
| EPI_ISL_2646262, EPI_ISL_2646264 | Cerballiance-IDF | Cerba lab | Emmanuel Lecorche, Sabine Trombert-Paolantoni, Aude Lessenne, Kader Merah, Thierry Collin, Laura Verdurme, Patrice Herisson, Stéphanie Haïm-Boukobza, Bénédicte Roquebert |
| EPI_ISL_2646265 | Cerballiance-Paris | Cerba lab | Emmanuel Lecorche, Sabine Trombert-Paolantoni, Aude Lessenne, Kader Merah, Thierry Collin, Laura Verdurme, Patrice Herisson, Stéphanie Haïm-Boukobza, Bénédicte Roquebert |
| EPI_ISL_2646266 | Cerballiance-IDF | Cerba lab | Emmanuel Lecorche, Sabine Trombert-Paolantoni, Aude Lessenne, Kader Merah, Thierry Collin, Laura Verdurme, Patrice Herisson, Stéphanie Haïm-Boukobza, Bénédicte Roquebert |
| EPI_ISL_2646267 | Cerballiance-Paris | Cerba lab | Emmanuel Lecorche, Sabine Trombert-Paolantoni, Aude Lessenne, Kader Merah, Thierry Collin, Laura Verdurme, Patrice Herisson, Stéphanie Haïm-Boukobza, Bénédicte Roquebert |
| EPI_ISL_2646268 | Unilabs | Cerba lab | Emmanuel Lecorche, Sabine Trombert-Paolantoni, Aude Lessenne, Kader Merah, Thierry Collin, Laura Verdurme, Patrice Herisson, Stéphanie Haïm-Boukobza, Bénédicte Roquebert |
| EPI_ISL_2646269 | Cerballiance-IDF | Cerba lab | Emmanuel Lecorche, Sabine Trombert-Paolantoni, Aude Lessenne, Kader Merah, Thierry Collin, Laura Verdurme, Patrice Herisson, Stéphanie Haïm-Boukobza, Bénédicte Roquebert |
| EPI_ISL_2646270 | Cerballiance-HDF | Cerba lab | Emmanuel Lecorche, Sabine Trombert-Paolantoni, Aude Lessenne, Kader Merah, Thierry Collin, Laura Verdurme, Patrice Herisson, Stéphanie Haïm-Boukobza, Bénédicte Roquebert |
| EPI_ISL_2646271 | Kuate | Cerba lab | Emmanuel Lecorche, Sabine Trombert-Paolantoni, Aude Lessenne, Kader Merah, Thierry Collin, Laura Verdurme, Patrice Herisson, Stéphanie Haïm-Boukobza, Bénédicte Roquebert |
| EPI_ISL_2646272 | Hopital Max Fourestier | Cerba lab | Emmanuel Lecorche, Sabine Trombert-Paolantoni, Aude Lessenne, Kader Merah, Thierry Collin, Laura Verdurme, Patrice Herisson, Stéphanie Haïm-Boukobza, Bénédicte Roquebert |
| EPI_ISL_2646273 | Laboratoire Frechin | Cerba lab | Emmanuel Lecorche, Sabine Trombert-Paolantoni, Aude Lessenne, Kader Merah, Thierry Collin, Laura Verdurme, Patrice Herisson, Stéphanie Haïm-Boukobza, Bénédicte Roquebert |
| EPI_ISL_2646274 | Cerballiance-Provence | Cerba lab | Emmanuel Lecorche, Sabine Trombert-Paolantoni, Aude Lessenne, Kader Merah, Thierry Collin, Laura Verdurme, Patrice Herisson, Stéphanie Haïm-Boukobza, Bénédicte Roquebert |
| EPI_ISL_2646275, EPI_ISL_2646276 | Cerballiance-Paris | Cerba lab | Emmanuel Lecorche, Sabine Trombert-Paolantoni, Aude Lessenne, Kader Merah, Thierry Collin, Laura Verdurme, Patrice Herisson, Stéphanie Haïm-Boukobza, Bénédicte Roquebert |
| EPI_ISL_2646277 | Cerballiance-Provence | Cerba lab | Emmanuel Lecorche, Sabine Trombert-Paolantoni, Aude Lessenne, Kader Merah, Thierry Collin, Laura Verdurme, Patrice Herisson, Stéphanie Haïm-Boukobza, Bénédicte Roquebert |
| EPI_ISL_2646278 | Cerballiance-Paris | Cerba lab | Emmanuel Lecorche, Sabine Trombert-Paolantoni, Aude Lessenne, Kader Merah, Thierry Collin, Laura Verdurme, Patrice Herisson, Stéphanie Haïm-Boukobza, Bénédicte Roquebert |
| EPI_ISL_2646279 | Cerballiance-IDF | Cerba lab | Emmanuel Lecorche, Sabine Trombert-Paolantoni, Aude Lessenne, Kader Merah, Thierry Collin, Laura Verdurme, Patrice Herisson, Stéphanie Haïm-Boukobza, Bénédicte Roquebert |
| EPI_ISL_2646302 | ZOL | Jessa | Berden et al. on behalf of Jessa_cmdLab |
| EPI_ISL_2646318 | Jessa | Jessa | Berden et al. on behalf of Jessa_cmdLab |
| EPI_ISL_2646321, EPI_ISL_2646324, EPI_ISL_2646325, EPI_ISL_2646327, EPI_ISL_2646328 | University of Mississippi Medical Center, Department of Pathology | University of Mississippi Medical Center, Molecular and Genomics Core Facility | Ashley C. Johnson, Wenjie Wu, Ithiel J. Frame, Krishna K. Ayyalasomayajula, Michael R. Garrett, D. Ashley Robinson |
| EPI_ISL_2646330, EPI_ISL_2646333, EPI_ISL_2646334, EPI_ISL_2646336, EPI_ISL_2646339, EPI_ISL_2646342, EPI_ISL_2646347, EPI_ISL_2646350, EPI_ISL_2646351 | MS Public Health Laboratory | University of Mississippi Medical Center, Molecular and Genomics Core Facility | Ashley C. Johnson, Wenjie Wu, Ithiel J. Frame, Krishna K. Ayyalasomayajula, Michael R. Garrett, D. Ashley Robinson |
| EPI_ISL_2646362, EPI_ISL_2646364, EPI_ISL_2646408, EPI_ISL_2646411, EPI_ISL_2646437, EPI_ISL_2646445, EPI_ISL_2646499, EPI_ISL_2646503, EPI_ISL_2646524, EPI_ISL_2646542, EPI_ISL_2646550, EPI_ISL_2646554, EPI_ISL_2646557, EPI_ISL_2646579, EPI_ISL_2646602, EPI_ISL_2646608, EPI_ISL_2646611, EPI_ISL_2646617, EPI_ISL_2646627, EPI_ISL_2646631, EPI_ISL_2646639, EPI_ISL_2646650, EPI_ISL_2646651, EPI_ISL_2646660, EPI_ISL_2646676, EPI_ISL_2646687, EPI_ISL_2646688, EPI_ISL_2646695, EPI_ISL_2646705, EPI_ISL_2646709, EPI_ISL_2646716, EPI_ISL_2646723, EPI_ISL_2646762, EPI_ISL_2646813, EPI_ISL_2646829, EPI_ISL_2646856, EPI_ISL_2646923 | | | |
| see above | Mako Medical | Centers for Disease Control and Prevention Division of Viral Diseases, Pathogen Discovery | Dakota Howard, Dhwani Batra, Peter W. Cook, Kara Moser, Adrian Paskey, Jason Caravas, Benjamin Rambo-Martin, Shatavia Morrison, Christopher Gulvick, Scott Sammons, Yvette Unoarumhi, Darlene Wagner, Matthew Schmerer,Matthew Tugwell, Lauren Moon, Clinton R. Paden, Duncan MacCannell |
| EPI_ISL_2646977 | Montana Public Health Laboratory | Montana Public Health Laboratory | Joy Ritter, Michelle Mozer, Carrie Biskupiak, Deborah Gibson, Michael Dills |
| EPI_ISL_2646983 | Mako Medical | Centers for Disease Control and Prevention Division of Viral Diseases, Pathogen Discovery | Dakota Howard, Dhwani Batra, Peter W. Cook, Kara Moser, Adrian Paskey, Jason Caravas, Benjamin Rambo-Martin, Shatavia Morrison, Christopher Gulvick, Scott Sammons, Yvette Unoarumhi, Darlene Wagner, Matthew Schmerer,Matthew Tugwell, Lauren Moon, Clinton R. Paden, Duncan MacCannell |
| EPI_ISL_2647001 | Montana Public Health Laboratory | Montana Public Health Laboratory | Joy Ritter, Michelle Mozer, Carrie Biskupiak, Deborah Gibson, Michael Dills |
| EPI_ISL_2647018, EPI_ISL_2647023, EPI_ISL_2647028, EPI_ISL_2647029, EPI_ISL_2647030, EPI_ISL_2647031, EPI_ISL_2647032, EPI_ISL_2647047, EPI_ISL_2647056, EPI_ISL_2647083, EPI_ISL_2647086, EPI_ISL_2647094, EPI_ISL_2647116, EPI_ISL_2647118, EPI_ISL_2647123, EPI_ISL_2647125, EPI_ISL_2647132, EPI_ISL_2647143, EPI_ISL_2647146, EPI_ISL_2647154, EPI_ISL_2647223, EPI_ISL_2647234, EPI_ISL_2647316, EPI_ISL_2647319, EPI_ISL_2647328, EPI_ISL_2647334, EPI_ISL_2647339, EPI_ISL_2647341, EPI_ISL_2647347, EPI_ISL_2647352, EPI_ISL_2647357, EPI_ISL_2647358, EPI_ISL_2647369, EPI_ISL_2647378, EPI_ISL_2647385, EPI_ISL_2647388, EPI_ISL_2647390, EPI_ISL_2647392, EPI_ISL_2647398, EPI_ISL_2647399, EPI_ISL_2647404, EPI_ISL_2647409, EPI_ISL_2647413, EPI_ISL_2647421, EPI_ISL_2647423, EPI_ISL_2647429, EPI_ISL_2647430, EPI_ISL_2647440, EPI_ISL_2647442, EPI_ISL_2647445, EPI_ISL_2647459, EPI_ISL_2647472, EPI_ISL_2647481, EPI_ISL_2647507, EPI_ISL_2647545, EPI_ISL_2647592, EPI_ISL_2647601, EPI_ISL_2647619, EPI_ISL_2647661, EPI_ISL_2647671, EPI_ISL_2647673, EPI_ISL_2647679, EPI_ISL_2647680, EPI_ISL_2647681, EPI_ISL_2647686, EPI_ISL_2647687, EPI_ISL_2647688, EPI_ISL_2647692, EPI_ISL_2647695, EPI_ISL_2647696, EPI_ISL_2647701, EPI_ISL_2647710, EPI_ISL_2647715, EPI_ISL_2647717, EPI_ISL_2647720, EPI_ISL_2647722, EPI_ISL_2647726, EPI_ISL_2647727, EPI_ISL_2647728, EPI_ISL_2647731, EPI_ISL_2647734, EPI_ISL_2647756, EPI_ISL_2647764, EPI_ISL_2647770, EPI_ISL_2647777, EPI_ISL_2647778, EPI_ISL_2647781, EPI_ISL_2647783, EPI_ISL_2647788, EPI_ISL_2647789, EPI_ISL_2647792, EPI_ISL_2647799, EPI_ISL_2647804, EPI_ISL_2647805, EPI_ISL_2647809, EPI_ISL_2647811, EPI_ISL_2647812, EPI_ISL_2647819, EPI_ISL_2647826, EPI_ISL_2647827, EPI_ISL_2647841, EPI_ISL_2647852, EPI_ISL_2647853, EPI_ISL_2647858, EPI_ISL_2647859, EPI_ISL_2647867, EPI_ISL_2647870, EPI_ISL_2647876, EPI_ISL_2647881, EPI_ISL_2647882, EPI_ISL_2647883, EPI_ISL_2647884, EPI_ISL_2647891, EPI_ISL_2647897, EPI_ISL_2647901, EPI_ISL_2647902, EPI_ISL_2647907, EPI_ISL_2647914, EPI_ISL_2647930, EPI_ISL_2647934, EPI_ISL_2647974, EPI_ISL_2647977, EPI_ISL_2647982, EPI_ISL_2647988 | | | |
| see above | Mako Medical | Centers for Disease Control and Prevention Division of Viral Diseases, Pathogen Discovery | Dakota Howard, Dhwani Batra, Peter W. Cook, Kara Moser, Adrian Paskey, Jason Caravas, Benjamin Rambo-Martin, Shatavia Morrison, Christopher Gulvick, Scott Sammons, Yvette Unoarumhi, Darlene Wagner, Matthew Schmerer,Matthew Tugwell, Lauren Moon, Clinton R. Paden, Duncan MacCannell |
| EPI_ISL_2648221, EPI_ISL_2648222, EPI_ISL_2648223, EPI_ISL_2648224, EPI_ISL_2648226, EPI_ISL_2648228, EPI_ISL_2648232, EPI_ISL_2648235, | Kasane Primary Hospital Laboratory | Botswana Harvard HIV Reference Laboratory | Sikhulile Moyo, Wonderful T. Choga, Dorcas Maruapula, Agnes Karutwaeng, Rose Munyere, Keoratile Ntshambiwa, Thongbotho Mphoyakgosi, Boitumelo Zuze, Botshelo Radibe, Legodile Kooepile, Ontlametse T. Bareng, Roger Shapiro, Shahin Lockman, Joseph Makhema, Madisa Mine, Mosepele Mosepele, Simani Gaseitswe |

| | | | |
|---|---|---|---|
| EPI_ISL_2648236, EPI_ISL_2648237 | | | |
| EPI_ISL_2649677 | National Virus Reference Laboratory | National Virus Reference Laboratory | Zoe Yandle, Charlene Bennett, Gabriel Gonzalez, Michael Carr, Jonathan Dean, Cillian F De Gascun |
| EPI_ISL_2649833, EPI_ISL_2649848, EPI_ISL_2649851, EPI_ISL_2649853, EPI_ISL_2649856, EPI_ISL_2649857, EPI_ISL_2649862, EPI_ISL_2649864, EPI_ISL_2649865, EPI_ISL_2649866, EPI_ISL_2649882 | | | |
| see above | NJDOH, Public Health and Environmental Laboratories | NJ_PHEL | Lindsey Bodnar, Shiv K. Verma, Ryan Pachucki, Chelsea San Filippo, Jacquelyn Deverell, Dana Woell, Allison Roder, Byeong Jeong |
| EPI_ISL_2649992 | Tuanku Ampuan Najihah Hospital, Kuala Pilah | Institute for Medical Research, Infectious Disease Research Centre, National Institutes of Health, Ministry of Health Malaysia | Suppiah J, Kamel K, Mohd Zawawi Z, Azizan MA, Ramly N, Robert F, Thayan R |
| EPI_ISL_2650009 | Laboratorio di Riferimento Regionale della Sicilia Occidentale per l'Emergenza COVID-19 | Laboratorio di Riferimento Regionale della Sicilia Occidentale per l'Emergenza COVID-19 | Fabio Tramuto |
| EPI_ISL_2650011, EPI_ISL_2650012 | Area of Virology, Serology and Virology Division (SAViD), New South Wales Health Pathology Randwick | Virology Research Laboratory; Area of Virology, Serology and Virology Division (SAViD), New South Wales Health Pathology Randwick | Foster, C.; Au, J.; Ruiz Silva, M.; Deveson, I.; Bull, R.; Van Hal, S.; Rawlinson, W. |
| EPI_ISL_2650017, EPI_ISL_2650018 | LabPLUS | Institute of Environmental Science and Research (ESR) | Rachel Boyle, SallyAnn Harbison, Olivia Stroeven, Xiaoyun Ren, Matt Storey, Nikki Freed, Muhammad Faisal, Jing Wang, Hermes Perez, Anja Werno, Antje van der Linden, Arlo Upton, Chris Mansell, David Hammer, Dragana Drinkovic, Gary McAuliffe, Hana Sofia Andersson, James Ussher, Jill Sherwood, Josh Freeman, Julia Howard, Juliet Elvy, Mary DeAlmeida, Matt Blakiston, Matthew Rogers, Max Bloomfield, Michael Addidle, Michelle Balm, Sally Roberts, Sarah Jefferies, Sharmini Muttaiyah, Susan Morpeth, Susan Taylor, Timothy Blackmore, Vani Sathyendran, Veronica Playle, Virginia Hope, Erasmus Smit, Lauren Jelly, Olin Silander, Joep de Ligt |
| EPI_ISL_2650019 | Middlemore Hospital | Institute of Environmental Science and Research (ESR) | Rachel Boyle, SallyAnn Harbison, Olivia Stroeven, Xiaoyun Ren, Matt Storey, Nikki Freed, Muhammad Faisal, Jing Wang, Hermes Perez, Anja Werno, Antje van der Linden, Arlo Upton, Chris Mansell, David Hammer, Dragana Drinkovic, Gary McAuliffe, Hana Sofia Andersson, James Ussher, Jill Sherwood, Josh Freeman, Julia Howard, Juliet Elvy, Mary DeAlmeida, Matt Blakiston, Matthew Rogers, Max Bloomfield, Michael Addidle, Michelle Balm, Sally Roberts, Sarah Jefferies, Sharmini Muttaiyah, Susan Morpeth, Susan Taylor, Timothy Blackmore, Vani Sathyendran, Veronica Playle, Virginia Hope, Erasmus Smit, Lauren Jelly, Olin Silander, Joep de Ligt |
| EPI_ISL_2650020, EPI_ISL_2650021, EPI_ISL_2650022, EPI_ISL_2650023 | LabPLUS | Institute of Environmental Science and Research (ESR) | Rachel Boyle, SallyAnn Harbison, Olivia Stroeven, Xiaoyun Ren, Matt Storey, Nikki Freed, Muhammad Faisal, Jing Wang, Hermes Perez, Anja Werno, Antje van der Linden, Arlo Upton, Chris Mansell, David Hammer, Dragana Drinkovic, Gary McAuliffe, Hana Sofia Andersson, James Ussher, Jill Sherwood, Josh Freeman, Julia Howard, Juliet Elvy, Mary DeAlmeida, Matt Blakiston, Matthew Rogers, Max Bloomfield, Michael Addidle, Michelle Balm, Sally Roberts, Sarah Jefferies, Sharmini Muttaiyah, Susan Morpeth, Susan Taylor, Timothy Blackmore, Vani Sathyendran, Veronica Playle, Virginia Hope, Erasmus Smit, Lauren Jelly, Olin Silander, Joep de Ligt |
| EPI_ISL_2650024 | Middlemore Hospital | Institute of Environmental Science and Research (ESR) | Rachel Boyle, SallyAnn Harbison, Olivia Stroeven, Xiaoyun Ren, Matt Storey, Nikki Freed, Muhammad Faisal, Jing Wang, Hermes Perez, Anja Werno, Antje van der Linden, Arlo Upton, Chris Mansell, David Hammer, Dragana Drinkovic, Gary McAuliffe, Hana Sofia Andersson, James Ussher, Jill Sherwood, Josh Freeman, Julia Howard, Juliet Elvy, Mary DeAlmeida, Matt Blakiston, Matthew Rogers, Max Bloomfield, Michael Addidle, Michelle Balm, Sally Roberts, Sarah Jefferies, Sharmini Muttaiyah, Susan Morpeth, Susan Taylor, Timothy Blackmore, Vani Sathyendran, Veronica Playle, Virginia Hope, Erasmus Smit, Lauren Jelly, Olin Silander, Joep de Ligt |
| EPI_ISL_2650025, EPI_ISL_2650026 | LabPLUS | Institute of Environmental Science and Research (ESR) | Rachel Boyle, SallyAnn Harbison, Olivia Stroeven, Xiaoyun Ren, Matt Storey, Nikki Freed, Muhammad Faisal, Jing Wang, Hermes Perez, Anja Werno, Antje van der Linden, Arlo Upton, Chris Mansell, David Hammer, Dragana Drinkovic, Gary McAuliffe, Hana Sofia Andersson, James Ussher, Jill Sherwood, Josh Freeman, Julia Howard, Juliet Elvy, Mary DeAlmeida, Matt Blakiston, Matthew Rogers, Max Bloomfield, Michael Addidle, Michelle Balm, Sally Roberts, Sarah Jefferies, Sharmini Muttaiyah, Susan Morpeth, Susan Taylor, Timothy Blackmore, Vani Sathyendran, Veronica Playle, Virginia Hope, Erasmus Smit, Lauren Jelly, Olin Silander, Joep de Ligt |
| EPI_ISL_2650134, EPI_ISL_2650143, EPI_ISL_2650146 | Arizona State Public Health Laboratory | Arizona State Public Health Laboratory | Trung Huynh, Jessica Escobar, Katherine Fullerton, Nobuko Fukushima, Matthew Contursi, Stacy White, Linda Getsinger, Victor Waddell |
| EPI_ISL_2650181 | Microbiology Department, Laboratori Clínic Metropolitana Nord. Hospital Universitari Germans Trias i Pujol | Can Ruti SARS-CoV-2 Sequencing Hub (HUGTiP/IrsiCaixa/IGTP) | Marc Noguera-Julian, Pilar Armengol, Ignacio Blanco, Antoni E Bordoy, Francesc Catala-Moll, Pere-Joan Cardona, Maria Casadellà, Cristina Casañ, Gemma Clara, Bonaventura Clotet, Cristina Esteban, Montserrat Giménez, Mercedes Guerrero, Anna Not, Roger Paredes, Mariona Parera, Verónica Saludes, Alba Sánchez, and Elisa Martró on behalf of the Can Ruti SARS-CoV-2 Sequencing Hub |
| EPI_ISL_2650182 | Microbiologia CATLAB | Can Ruti SARS-CoV-2 Sequencing Hub (HUGTiP/IrsiCaixa/IGTP) | Marc Noguera-Julian, Pilar Armengol, Ignacio Blanco, Antoni E Bordoy, Francesc Catala-Moll, Pere-Joan Cardona, Maria Casadellà, Cristina Casañ, Gemma Clara, Bonaventura Clotet, Cristina Esteban, Montserrat Giménez, Mercedes Guerrero, Anna Not, Roger Paredes, Mariona Parera, Verónica Saludes, Alba Sánchez, and Elisa Martró on behalf of the Can Ruti SARS-CoV-2 Sequencing Hub |
| EPI_ISL_2650197, EPI_ISL_2650201 | Microbiology Department, Laboratori Clínic Metropolitana Nord. Hospital Universitari Germans Trias i Pujol | Can Ruti SARS-CoV-2 Sequencing Hub (HUGTiP/IrsiCaixa/IGTP) | Marc Noguera-Julian, Pilar Armengol, Ignacio Blanco, Antoni E Bordoy, Francesc Catala-Moll, Pere-Joan Cardona, Maria Casadellà, Cristina Casañ, Gemma Clara, Bonaventura Clotet, Cristina Esteban, Montserrat Giménez, Mercedes Guerrero, Anna Not, Roger Paredes, Mariona Parera, Verónica Saludes, Alba Sánchez, and Elisa Martró on behalf of the Can Ruti SARS-CoV-2 Sequencing Hub |
| EPI_ISL_2650202 | Microbiologia CATLAB | Can Ruti SARS-CoV-2 Sequencing Hub (HUGTiP/IrsiCaixa/IGTP) | Marc Noguera-Julian, Pilar Armengol, Ignacio Blanco, Antoni E Bordoy, Francesc Catala-Moll, Pere-Joan Cardona, Maria Casadellà, Cristina Casañ, Gemma Clara, Bonaventura Clotet, Cristina Esteban, Montserrat Giménez, Mercedes Guerrero, Anna Not, Roger Paredes, Mariona Parera, Verónica Saludes, Alba Sánchez, and Elisa Martró on behalf of the Can Ruti SARS-CoV-2 Sequencing Hub |
| EPI_ISL_2650208 | Microbiology Department, Laboratori Clínic Metropolitana Nord. Hospital Universitari Germans Trias i Pujol | Can Ruti SARS-CoV-2 Sequencing Hub (HUGTiP/IrsiCaixa/IGTP) | Marc Noguera-Julian, Pilar Armengol, Ignacio Blanco, Antoni E Bordoy, Francesc Catala-Moll, Pere-Joan Cardona, Maria Casadellà, Cristina Casañ, Gemma Clara, Bonaventura Clotet, Cristina Esteban, Montserrat Giménez, Mercedes Guerrero, Anna Not, Roger Paredes, Mariona Parera, Verónica Saludes, Alba Sánchez, and Elisa Martró on behalf of the Can Ruti SARS-CoV-2 Sequencing Hub |
| EPI_ISL_2650211, EPI_ISL_2650219, EPI_ISL_2650220 | Microbiologia CATLAB | Can Ruti SARS-CoV-2 Sequencing Hub (HUGTiP/IrsiCaixa/IGTP) | Marc Noguera-Julian, Pilar Armengol, Ignacio Blanco, Antoni E Bordoy, Francesc Catala-Moll, Pere-Joan Cardona, Maria Casadellà, Cristina Casañ, Gemma Clara, Bonaventura Clotet, Cristina Esteban, Montserrat Giménez, Mercedes Guerrero, Anna Not, Roger Paredes, Mariona Parera, Verónica Saludes, Alba Sánchez, and Elisa Martró on behalf of the Can Ruti SARS-CoV-2 Sequencing Hub |
| EPI_ISL_2650223, EPI_ISL_2650227, EPI_ISL_2650229 | Microbiology Department, Laboratori Clínic Metropolitana Nord. Hospital Universitari Germans Trias i Pujol | Can Ruti SARS-CoV-2 Sequencing Hub (HUGTiP/IrsiCaixa/IGTP) | Marc Noguera-Julian, Pilar Armengol, Ignacio Blanco, Antoni E Bordoy, Francesc Catala-Moll, Pere-Joan Cardona, Maria Casadellà, Cristina Casañ, Gemma Clara, Bonaventura Clotet, Cristina Esteban, Montserrat Giménez, Mercedes Guerrero, Anna Not, Roger Paredes, Mariona Parera, Verónica Saludes, Alba Sánchez, and Elisa Martró on behalf of the Can Ruti SARS-CoV-2 Sequencing Hub |
| EPI_ISL_2650230 | Microbiologia CATLAB | Can Ruti SARS-CoV-2 Sequencing Hub (HUGTiP/IrsiCaixa/IGTP) | Marc Noguera-Julian, Pilar Armengol, Ignacio Blanco, Antoni E Bordoy, Francesc Catala-Moll, Pere-Joan Cardona, Maria Casadellà, Cristina Casañ, Gemma Clara, Bonaventura Clotet, Cristina Esteban, Montserrat Giménez, Mercedes Guerrero, Anna Not, Roger Paredes, Mariona Parera, Verónica Saludes, Alba Sánchez, and Elisa Martró on behalf of the Can Ruti SARS-CoV-2 Sequencing Hub |
| EPI_ISL_2650231, EPI_ISL_2650232, EPI_ISL_2650237 | Microbiology Department, Laboratori Clínic Metropolitana Nord. Hospital Universitari Germans Trias i Pujol | Can Ruti SARS-CoV-2 Sequencing Hub (HUGTiP/IrsiCaixa/IGTP) | Marc Noguera-Julian, Pilar Armengol, Ignacio Blanco, Antoni E Bordoy, Francesc Catala-Moll, Pere-Joan Cardona, Maria Casadellà, Cristina Casañ, Gemma Clara, Bonaventura Clotet, Cristina Esteban, Montserrat Giménez, Mercedes Guerrero, Anna Not, Roger Paredes, Mariona Parera, Verónica Saludes, Alba Sánchez, and Elisa Martró on behalf of the Can Ruti SARS-CoV-2 Sequencing Hub |
| EPI_ISL_2650239, EPI_ISL_2650240 | Microbiologia CATLAB | Can Ruti SARS-CoV-2 Sequencing Hub (HUGTiP/IrsiCaixa/IGTP) | Marc Noguera-Julian, Pilar Armengol, Ignacio Blanco, Antoni E Bordoy, Francesc Catala-Moll, Pere-Joan Cardona, Maria Casadellà, Cristina Casañ, Gemma Clara, Bonaventura Clotet, Cristina Esteban, Montserrat Giménez, Mercedes Guerrero, Anna Not, Roger Paredes, Mariona Parera, Verónica Saludes, Alba Sánchez, and Elisa Martró on behalf of the Can Ruti SARS-CoV-2 Sequencing Hub |
| EPI_ISL_2650241 | Microbiology Department, Laboratori Clínic Metropolitana Nord. Hospital Universitari Germans Trias i Pujol | Can Ruti SARS-CoV-2 Sequencing Hub (HUGTiP/IrsiCaixa/IGTP) | Marc Noguera-Julian, Pilar Armengol, Ignacio Blanco, Antoni E Bordoy, Francesc Catala-Moll, Pere-Joan Cardona, Maria Casadellà, Cristina Casañ, Gemma Clara, Bonaventura Clotet, Cristina Esteban, Montserrat Giménez, Mercedes Guerrero, Anna Not, Roger Paredes, Mariona Parera, Verónica Saludes, Alba Sánchez, and Elisa Martró on behalf of the Can Ruti SARS-CoV-2 Sequencing Hub |
| EPI_ISL_2650247 | Microbiologia CATLAB | Can Ruti SARS-CoV-2 Sequencing Hub (HUGTiP/IrsiCaixa/IGTP) | Marc Noguera-Julian, Pilar Armengol, Ignacio Blanco, Antoni E Bordoy, Francesc Catala-Moll, Pere-Joan Cardona, Maria Casadellà, Cristina Casañ, Gemma Clara, Bonaventura Clotet, Cristina Esteban, Montserrat Giménez, Mercedes Guerrero, Anna Not, Roger Paredes, Mariona Parera, Verónica Saludes, Alba Sánchez, and Elisa Martró on behalf of the Can Ruti SARS-CoV-2 Sequencing Hub |
| EPI_ISL_2650255, EPI_ISL_2650263, EPI_ISL_2650264, EPI_ISL_2650267, EPI_ISL_2650275, EPI_ISL_2650278, EPI_ISL_2650282, EPI_ISL_2650286 | Microbiology Department, Laboratori Clínic Metropolitana Nord. Hospital Universitari Germans Trias i Pujol | Can Ruti SARS-CoV-2 Sequencing Hub (HUGTiP/IrsiCaixa/IGTP) | Marc Noguera-Julian, Pilar Armengol, Ignacio Blanco, Antoni E Bordoy, Francesc Catala-Moll, Pere-Joan Cardona, Maria Casadellà, Cristina Casañ, Gemma Clara, Bonaventura Clotet, Cristina Esteban, Montserrat Giménez, Mercedes Guerrero, Anna Not, Roger Paredes, Mariona Parera, Verónica Saludes, Alba Sánchez, and Elisa Martró on behalf of the Can Ruti SARS-CoV-2 Sequencing Hub |
| EPI_ISL_2650294 | Microbiologia CATLAB | Can Ruti SARS-CoV-2 Sequencing Hub (HUGTiP/IrsiCaixa/IGTP) | Marc Noguera-Julian, Pilar Armengol, Ignacio Blanco, Antoni E Bordoy, Francesc Catala-Moll, Pere-Joan Cardona, Maria Casadellà, Cristina Casañ, Gemma Clara, Bonaventura Clotet, Cristina Esteban, Montserrat Giménez, Mercedes Guerrero, Anna Not, Roger Paredes, Mariona Parera, Verónica Saludes, Alba Sánchez, and Elisa Martró on behalf of the Can Ruti SARS-CoV-2 Sequencing Hub |
| EPI_ISL_2650296, EPI_ISL_2650303, EPI_ISL_2650304 | Microbiology Department, Laboratori Clínic Metropolitana Nord. Hospital Universitari Germans Trias i Pujol | Can Ruti SARS-CoV-2 Sequencing Hub (HUGTiP/IrsiCaixa/IGTP) | Marc Noguera-Julian, Pilar Armengol, Ignacio Blanco, Antoni E Bordoy, Francesc Catala-Moll, Pere-Joan Cardona, Maria Casadellà, Cristina Casañ, Gemma Clara, Bonaventura Clotet, Cristina Esteban, Montserrat Giménez, Mercedes Guerrero, Anna Not, Roger Paredes, Mariona Parera, Verónica Saludes, Alba Sánchez, and Elisa Martró on behalf of the Can Ruti SARS-CoV-2 Sequencing Hub |
| EPI_ISL_2650305 | Microbiologia CATLAB | Can Ruti SARS-CoV-2 Sequencing Hub (HUGTiP/IrsiCaixa/IGTP) | Marc Noguera-Julian, Pilar Armengol, Ignacio Blanco, Antoni E Bordoy, Francesc Catala-Moll, Pere-Joan Cardona, Maria Casadellà, Cristina Casañ, Gemma Clara, Bonaventura Clotet, Cristina Esteban, Montserrat Giménez, Mercedes Guerrero, Anna Not, Roger Paredes, Mariona Parera, Verónica Saludes, Alba Sánchez, and Elisa Martró on behalf of the Can Ruti SARS-CoV-2 Sequencing Hub |
| EPI_ISL_2650306, EPI_ISL_2650308, EPI_ISL_2650310, EPI_ISL_2650315 | Microbiology Department, Laboratori Clínic Metropolitana Nord. Hospital Universitari Germans Trias i Pujol | Can Ruti SARS-CoV-2 Sequencing Hub (HUGTiP/IrsiCaixa/IGTP) | Marc Noguera-Julian, Pilar Armengol, Ignacio Blanco, Antoni E Bordoy, Francesc Catala-Moll, Pere-Joan Cardona, Maria Casadellà, Cristina Casañ, Gemma Clara, Bonaventura Clotet, Cristina Esteban, Montserrat Giménez, Mercedes Guerrero, Anna Not, Roger Paredes, Mariona Parera, Verónica Saludes, Alba Sánchez, and Elisa Martró on behalf of the Can Ruti SARS-CoV-2 Sequencing Hub |
| EPI_ISL_2650317 | Microbiologia CATLAB | Can Ruti SARS-CoV-2 Sequencing Hub (HUGTiP/IrsiCaixa/IGTP) | Marc Noguera-Julian, Pilar Armengol, Ignacio Blanco, Antoni E Bordoy, Francesc Catala-Moll, Pere-Joan Cardona, Maria Casadellà, Cristina Casañ, Gemma Clara, Bonaventura Clotet, Cristina Esteban, Montserrat Giménez, Mercedes Guerrero, Anna Not, Roger Paredes, Mariona Parera, Verónica Saludes, Alba Sánchez, and Elisa Martró on behalf of the Can Ruti SARS-CoV-2 Sequencing Hub |
| EPI_ISL_2650320, EPI_ISL_2650331, EPI_ISL_2650332 | Microbiology Department, Laboratori Clínic Metropolitana Nord. Hospital Universitari Germans Trias i Pujol | Can Ruti SARS-CoV-2 Sequencing Hub (HUGTiP/IrsiCaixa/IGTP) | Marc Noguera-Julian, Pilar Armengol, Ignacio Blanco, Antoni E Bordoy, Francesc Catala-Moll, Pere-Joan Cardona, Maria Casadellà, Cristina Casañ, Gemma Clara, Bonaventura Clotet, Cristina Esteban, Montserrat Giménez, Mercedes Guerrero, Anna Not, Roger Paredes, Mariona Parera, Verónica Saludes, Alba Sánchez, and Elisa Martró on behalf of the Can Ruti SARS-CoV-2 Sequencing Hub |
| EPI_ISL_2650340 | Microbiologia CATLAB | Can Ruti SARS-CoV-2 Sequencing Hub (HUGTiP/IrsiCaixa/IGTP) | Marc Noguera-Julian, Pilar Armengol, Ignacio Blanco, Antoni E Bordoy, Francesc Catala-Moll, Pere-Joan Cardona, Maria Casadellà, Cristina Casañ, Gemma Clara, Bonaventura Clotet, Cristina Esteban, Montserrat Giménez, Mercedes Guerrero, Anna Not, Roger Paredes, Mariona Parera, Verónica Saludes, Alba Sánchez, and Elisa Martró on behalf of the Can Ruti SARS-CoV-2 Sequencing Hub |
| EPI_ISL_2650343 | Microbiology Department, Laboratori Clínic Metropolitana Nord. Hospital Universitari Germans Trias i Pujol | Can Ruti SARS-CoV-2 Sequencing Hub (HUGTiP/IrsiCaixa/IGTP) | Marc Noguera-Julian, Pilar Armengol, Ignacio Blanco, Antoni E Bordoy, Francesc Catala-Moll, Pere-Joan Cardona, Maria Casadellà, Cristina Casañ, Gemma Clara, Bonaventura Clotet, Cristina Esteban, Montserrat Giménez, Mercedes Guerrero, Anna Not, Roger Paredes, Mariona Parera, Verónica Saludes, Alba Sánchez, and Elisa Martró on behalf of the Can Ruti SARS-CoV-2 Sequencing Hub |
| EPI_ISL_2650431 | ASL TO5 | Fondazione del Piemonte per l'Oncologia IRCCS | Antonino Sottile, Silvia Brossa, Paola Marino, Giorgio Giardina |
| EPI_ISL_2650433, EPI_ISL_2650435, EPI_ISL_2650437, EPI_ISL_2650439, EPI_ISL_2650440, EPI_ISL_2650441, EPI_ISL_2650443, EPI_ISL_2650444, EPI_ISL_2650445, EPI_ISL_2650446, EPI_ISL_2650447, EPI_ISL_2650448, EPI_ISL_2650449, EPI_ISL_2650450 | | | |
| see above | Laboratorio di Riferimento Regionale della Sicilia Occidentale per l'Emergenza COVID-19 | Laboratorio di Riferimento Regionale della Sicilia Occidentale per l'Emergenza COVID-19 | Fabio Tramuto, Carmelo Massimo Maida, Daniela Di Naro, Giulia Randazzo, Walter Mazzucco, Giorgio Graziano, Vincenzo Restivo, Claudio Costantino, Francesco Vitale |
| EPI_ISL_2650454, EPI_ISL_2650455, EPI_ISL_2650456, EPI_ISL_2650457, | South Eastern Area Laboratory Services (SEALS) | NSW Health Pathology - Institute of Clinical Pathology and Medical Research; Westmead Hospital; University of Sydney | CIDM-PH et al. |

| | | | |
|---|---|---|---|
| EPI_ISL_2650458, EPI_ISL_2650460, EPI_ISL_2650464, EPI_ISL_2650465, EPI_ISL_2650466 | | | |
| EPI_ISL_2650467 | Laverty Pathology | NSW Health Pathology - Institute of Clinical Pathology and Medical Research; Westmead Hospital; University of Sydney | CIDM-PH et al. |
| EPI_ISL_2650469, EPI_ISL_2650470 | Sydney South West Pathology Service (SSWPS) - Royal Prince Alfred Hospital - NSW Health Pathology | NSW Health Pathology - Institute of Clinical Pathology and Medical Research; Westmead Hospital; University of Sydney | CIDM-PH et al. |
| EPI_ISL_2651048, EPI_ISL_2651049, EPI_ISL_2651076, EPI_ISL_2651095, EPI_ISL_2651096, EPI_ISL_2651109, EPI_ISL_2651112, EPI_ISL_2651114 | Washington State Department of Health Public Health Laboratories | Washington State Department of Health Public Health Laboratories | Drew MacKellar, Philip Dykema, Denny Russell, Joenice Gonzalez, Hannah Gray, Geoff Melly, Vanessa De Los Santos, Darren Lucas, JohnAric Peterson, Avi Singh, Rebecca Cao |
| EPI_ISL_2651115, EPI_ISL_2651116 | Temporary Specimen Collection Centre at the AsiaWorld-Expo | Hong Kong Department of Health | Alan K.L. Tsang, Peter C.W. Yip, Ken H.L. Ng, Edman T.K. Lam, Rickjason C.W. Chan, Dominic N.C. Tsang |
| EPI_ISL_2651117 | North Lantau Hospital | Hong Kong Department of Health | Alan K.L. Tsang, Peter C.W. Yip, Ken H.L. Ng, Edman T.K. Lam, Rickjason C.W. Chan, Dominic N.C. Tsang |
| EPI_ISL_2651118 | HOME QUARANTINE TASKFORCE | Hong Kong Department of Health | Alan K.L. Tsang, Peter C.W. Yip, Ken H.L. Ng, Edman T.K. Lam, Rickjason C.W. Chan, Dominic N.C. Tsang |
| EPI_ISL_2651121 | Temporary Specimen Collection Centre at the AsiaWorld-Expo | Hong Kong Department of Health | Alan K.L. Tsang, Peter C.W. Yip, Ken H.L. Ng, Edman T.K. Lam, Rickjason C.W. Chan, Dominic N.C. Tsang |
| EPI_ISL_2651125 | YAN CHAI HOSPITAL | Hong Kong Department of Health | Alan K.L. Tsang, Peter C.W. Yip, Ken H.L. Ng, Edman T.K. Lam, Rickjason C.W. Chan, Dominic N.C. Tsang |
| EPI_ISL_2651129, EPI_ISL_2651130 | North Lantau Hospital | Hong Kong Department of Health | Alan K.L. Tsang, Peter C.W. Yip, Ken H.L. Ng, Edman T.K. Lam, Rickjason C.W. Chan, Dominic N.C. Tsang |
| EPI_ISL_2651131 | HOME QUARANTINE TASKFORCE | Hong Kong Department of Health | Alan K.L. Tsang, Peter C.W. Yip, Ken H.L. Ng, Edman T.K. Lam, Rickjason C.W. Chan, Dominic N.C. Tsang |
| EPI_ISL_2651134, EPI_ISL_2651135 | Nebraska Public Health Laboratory | NPHL COVID-19 Response Team | NPHL COVID-19 Response Team |
| EPI_ISL_2651240, EPI_ISL_2651241, EPI_ISL_2651242, EPI_ISL_2651243, EPI_ISL_2651244, EPI_ISL_2651245, EPI_ISL_2651246, EPI_ISL_2651247, EPI_ISL_2651248, EPI_ISL_2651249 | Tokyo Metropolitan Institute of Public Health | Tokyo Metropolitan Institute of Public Health | Masaki Hayashi, Takako Yamazaki, Akane Negishi, Hirofumi Miyake, Kenshirou Kuroki, Maya Isogai, Takushi Fujiwara, Mami Nagashima, Jun Suzuki, Kenji Sadamasu |
| EPI_ISL_2651296, EPI_ISL_2651297 | BBMP Urban | INSACOG-KA, NIMHANS | Chitra Pattabiraman, Anson Kunjumon George,Darshan Sreenivas,Pramada Prasad, Ellango Ramasamy, Sony Sharma, Mahesh Kumar.C.S, Ananthapadmanabha Kotambail, Chetan G K, Gautham Arunachal Udupi, Anita S Desai, V Ravi |
| EPI_ISL_2651298 | NIMHANS | INSACOG-KA, NIMHANS | Chitra Pattabiraman, Anson Kunjumon George,Darshan Sreenivas,Pramada Prasad, Ellango Ramasamy, Sony Sharma, Mahesh Kumar.C.S, Ananthapadmanabha Kotambail, Chetan G K, Gautham Arunachal Udupi, Anita S Desai, V Ravi |
| EPI_ISL_2651299, EPI_ISL_2651300, EPI_ISL_2651301, EPI_ISL_2651302 | BBMP Urban | INSACOG-KA, NIMHANS | Chitra Pattabiraman, Anson Kunjumon George,Darshan Sreenivas,Pramada Prasad, Ellango Ramasamy, Sony Sharma, Mahesh Kumar.C.S, Ananthapadmanabha Kotambail, Chetan G K, Gautham Arunachal Udupi, Anita S Desai, V Ravi |
| EPI_ISL_2651303 | Sanjay Gandhi Institute Of Trauma & Orthopaedics Centre | INSACOG-KA, NIMHANS | Chitra Pattabiraman, Anson Kunjumon George,Darshan Sreenivas,Pramada Prasad, Ellango Ramasamy, Sony Sharma, Mahesh Kumar.C.S, Ananthapadmanabha Kotambail, Chetan G K, Gautham Arunachal Udupi, Anita S Desai, V Ravi |
| EPI_ISL_2651304, EPI_ISL_2651305, EPI_ISL_2651306 | BBMP Urban | INSACOG-KA, NIMHANS | Chitra Pattabiraman, Anson Kunjumon George,Darshan Sreenivas,Pramada Prasad, Ellango Ramasamy, Sony Sharma, Mahesh Kumar.C.S, Ananthapadmanabha Kotambail, Chetan G K, Gautham Arunachal Udupi, Anita S Desai, V Ravi |
| EPI_ISL_2651307, EPI_ISL_2651308, EPI_ISL_2651309 | RAILWAY H | INSACOG-KA, NIMHANS | Chitra Pattabiraman, Anson Kunjumon George,Darshan Sreenivas,Pramada Prasad, Ellango Ramasamy, Sony Sharma, Mahesh Kumar.C.S, Ananthapadmanabha Kotambail, Chetan G K, Gautham Arunachal Udupi, Anita S Desai, V Ravi |
| EPI_ISL_2651310, EPI_ISL_2651311 | BBMP Urban | INSACOG-KA, NIMHANS | Chitra Pattabiraman, Anson Kunjumon George,Darshan Sreenivas,Pramada Prasad, Ellango Ramasamy, Sony Sharma, Mahesh Kumar.C.S, Ananthapadmanabha Kotambail, Chetan G K, Gautham Arunachal Udupi, Anita S Desai, V Ravi |
| EPI_ISL_2651312, EPI_ISL_2651313, EPI_ISL_2651314 | NIMHANS | INSACOG-KA, NIMHANS | Chitra Pattabiraman, Anson Kunjumon George,Darshan Sreenivas,Pramada Prasad, Ellango Ramasamy, Sony Sharma, Mahesh Kumar.C.S, Ananthapadmanabha Kotambail, Chetan G K, Gautham Arunachal Udupi, Anita S Desai, V Ravi |
| EPI_ISL_2651315 | BBMP Urban | INSACOG-KA, NIMHANS | Chitra Pattabiraman, Anson Kunjumon George,Darshan Sreenivas,Pramada Prasad, Ellango Ramasamy, Sony Sharma, Mahesh Kumar.C.S, Ananthapadmanabha Kotambail, Chetan G K, Gautham Arunachal Udupi, Anita S Desai, V Ravi |
| EPI_ISL_2651316, EPI_ISL_2651317, EPI_ISL_2651318, EPI_ISL_2651319 | RAILWAY H | INSACOG-KA, NIMHANS | Chitra Pattabiraman, Anson Kunjumon George,Darshan Sreenivas,Pramada Prasad, Ellango Ramasamy, Sony Sharma, Mahesh Kumar.C.S, Ananthapadmanabha Kotambail, Chetan G K, Gautham Arunachal Udupi, Anita S Desai, V Ravi |
| EPI_ISL_2651320 | BBMP Urban | INSACOG-KA, NIMHANS | Chitra Pattabiraman, Anson Kunjumon George,Darshan Sreenivas,Pramada Prasad, Ellango Ramasamy, Sony Sharma, Mahesh Kumar.C.S, Ananthapadmanabha Kotambail, Chetan G K, Gautham Arunachal Udupi, Anita S Desai, V Ravi |
| EPI_ISL_2651321 | NIMHANS | INSACOG-KA, NIMHANS | Chitra Pattabiraman, Anson Kunjumon George,Darshan Sreenivas,Pramada Prasad, Ellango Ramasamy, Sony Sharma, Mahesh Kumar.C.S, Ananthapadmanabha Kotambail, Chetan G K, Gautham Arunachal Udupi, Anita S Desai, V Ravi |
| EPI_ISL_2651322, EPI_ISL_2651323, EPI_ISL_2651324, EPI_ISL_2651325 | RAILWAY H | INSACOG-KA, NIMHANS | Chitra Pattabiraman, Anson Kunjumon George,Darshan Sreenivas,Pramada Prasad, Ellango Ramasamy, Sony Sharma, Mahesh Kumar.C.S, Ananthapadmanabha Kotambail, Chetan G K, Gautham Arunachal Udupi, Anita S Desai, V Ravi |
| EPI_ISL_2651326, EPI_ISL_2651327, EPI_ISL_2651328, EPI_ISL_2651329 | BBMP Urban | INSACOG-KA, NIMHANS | Chitra Pattabiraman, Anson Kunjumon George,Darshan Sreenivas,Pramada Prasad, Ellango Ramasamy, Sony Sharma, Mahesh Kumar.C.S, Ananthapadmanabha Kotambail, Chetan G K, Gautham Arunachal Udupi, Anita S Desai, V Ravi |
| EPI_ISL_2651331 | NIMHANS | INSACOG-KA, NIMHANS | Chitra Pattabiraman, Anson Kunjumon George,Darshan Sreenivas,Pramada Prasad, Ellango Ramasamy, Sony Sharma, Mahesh Kumar.C.S, Ananthapadmanabha Kotambail, Chetan G K, Gautham Arunachal Udupi, Anita S Desai, V Ravi |
| EPI_ISL_2651332, EPI_ISL_2651333, EPI_ISL_2651334, EPI_ISL_2651335, EPI_ISL_2651336, EPI_ISL_2651337, EPI_ISL_2651338, EPI_ISL_2651339, EPI_ISL_2651340 | BBMP Urban | INSACOG-KA, NIMHANS | Chitra Pattabiraman, Anson Kunjumon George,Darshan Sreenivas,Pramada Prasad, Ellango Ramasamy, Sony Sharma, Mahesh Kumar.C.S, Ananthapadmanabha Kotambail, Chetan G K, Gautham Arunachal Udupi, Anita S Desai, V Ravi |
| EPI_ISL_2651341, EPI_ISL_2651342, EPI_ISL_2651343 | RAILWAY H | INSACOG-KA, NIMHANS | Chitra Pattabiraman, Anson Kunjumon George,Darshan Sreenivas,Pramada Prasad, Ellango Ramasamy, Sony Sharma, Mahesh Kumar.C.S, Ananthapadmanabha Kotambail, Chetan G K, Gautham Arunachal Udupi, Anita S Desai, V Ravi |
| EPI_ISL_2651344, EPI_ISL_2651345, EPI_ISL_2651346, EPI_ISL_2651347 | BBMP Urban | INSACOG-KA, NIMHANS | Chitra Pattabiraman, Anson Kunjumon George,Darshan Sreenivas,Pramada Prasad, Ellango Ramasamy, Sony Sharma, Mahesh Kumar.C.S, Ananthapadmanabha Kotambail, Chetan G K, Gautham Arunachal Udupi, Anita S Desai, V Ravi |
| EPI_ISL_2651348 | RAILWAY H | INSACOG-KA, NIMHANS | Chitra Pattabiraman, Anson Kunjumon George,Darshan Sreenivas,Pramada Prasad, Ellango Ramasamy, Sony Sharma, Mahesh Kumar.C.S, Ananthapadmanabha Kotambail, Chetan G K, Gautham Arunachal Udupi, Anita S Desai, V Ravi |
| EPI_ISL_2651349 | BBMP Urban | INSACOG-KA, NIMHANS | Chitra Pattabiraman, Anson Kunjumon George,Darshan Sreenivas,Pramada Prasad, Ellango Ramasamy, Sony Sharma, Mahesh Kumar.C.S, Ananthapadmanabha Kotambail, Chetan G K, Gautham Arunachal Udupi, Anita S Desai, V Ravi |
| EPI_ISL_2651350 | NIMHANS | INSACOG-KA, NIMHANS | Chitra Pattabiraman, Anson Kunjumon George,Darshan Sreenivas,Pramada Prasad, Ellango Ramasamy, Sony Sharma, Mahesh Kumar.C.S, Ananthapadmanabha Kotambail, Chetan G K, Gautham Arunachal Udupi, Anita S Desai, V Ravi |
| EPI_ISL_2651351, EPI_ISL_2651352, EPI_ISL_2651353 | BBMP Urban | INSACOG-KA, NIMHANS | Chitra Pattabiraman, Anson Kunjumon George,Darshan Sreenivas,Pramada Prasad, Ellango Ramasamy, Sony Sharma, Mahesh Kumar.C.S, Ananthapadmanabha Kotambail, Chetan G K, Gautham Arunachal Udupi, Anita S Desai, V Ravi |

| | | | |
|---|---|---|---|
| EPI_ISL_2651354 | Sanjay Gandhi Institute Of Trauma & Orthopaedics Centre | INSACOG-KA, NIMHANS | Chitra Pattabiraman, Anson Kunjumon George,Darshan Sreenivas,Pramada Prasad, Ellango Ramasamy, Sony Sharma, Mahesh Kumar.C.S, Ananthapadmanabha Kotambail, Chetan G K, Gautham Arunachal Udupi, Anita S Desai, V Ravi |
| EPI_ISL_2651355 | NIMHANS | INSACOG-KA, NIMHANS | Chitra Pattabiraman, Anson Kunjumon George,Darshan Sreenivas,Pramada Prasad, Ellango Ramasamy, Sony Sharma, Mahesh Kumar.C.S, Ananthapadmanabha Kotambail, Chetan G K, Gautham Arunachal Udupi, Anita S Desai, V Ravi |
| EPI_ISL_2651356, EPI_ISL_2651357, EPI_ISL_2651358 | BBMP Urban | INSACOG-KA, NIMHANS | Chitra Pattabiraman, Anson Kunjumon George,Darshan Sreenivas,Pramada Prasad, Ellango Ramasamy, Sony Sharma, Mahesh Kumar.C.S, Ananthapadmanabha Kotambail, Chetan G K, Gautham Arunachal Udupi, Anita S Desai, V Ravi |
| EPI_ISL_2651359 | NIMHANS | INSACOG-KA, NIMHANS | Chitra Pattabiraman, Anson Kunjumon George,Darshan Sreenivas,Pramada Prasad, Ellango Ramasamy, Sony Sharma, Mahesh Kumar.C.S, Ananthapadmanabha Kotambail, Chetan G K, Gautham Arunachal Udupi, Anita S Desai, V Ravi |
| EPI_ISL_2651360, EPI_ISL_2651361 | BBMP Urban | INSACOG-KA, NIMHANS | Chitra Pattabiraman, Anson Kunjumon George,Darshan Sreenivas,Pramada Prasad, Ellango Ramasamy, Sony Sharma, Mahesh Kumar.C.S, Ananthapadmanabha Kotambail, Chetan G K, Gautham Arunachal Udupi, Anita S Desai, V Ravi |
| EPI_ISL_2651409 | Lighthouse Lab in Milton Keynes | Wellcome Sanger Institute for the COVID-19 Genomics UK (COG-UK) Consortium | The Lighthouse Lab in Milton Keynes and Alex Alderton, Roberto Amato, Jeffrey Barrett, Sonia Goncalves, Ewan Harrison, David K. Jackson, Ian Johnston, Dominic Kwiatkowski, Cordelia Langford, John Sillitoe on behalf of the Wellcome Sanger Institute COVID-19 Surveillance Team |
| EPI_ISL_2651410 | Lighthouse Lab in Glasgow | Wellcome Sanger Institute for the COVID-19 Genomics UK (COG-UK) Consortium | Harper VanSteenhouse, Yumi Kasai, David Gray, Carol Clugston, Anna Dominiczak and Alex Alderton, Roberto Amato, Jeffrey Barrett, Sonia Goncalves, Ewan Harrison, David K. Jackson, Ian Johnston, Dominic Kwiatkowski, Cordelia Langford, John Sillitoe on behalf of the Wellcome Sanger Institute COVID-19 Surveillance Team |
| EPI_ISL_2651411 | Lighthouse Laboratory Plymouth | Wellcome Sanger Institute for the COVID-19 Genomics UK (COG-UK) Consortium | Lighthouse Laboratory Plymouth and Alex Alderton, Roberto Amato, Jeffrey Barrett, Sonia Goncalves, Ewan Harrison, David K. Jackson, Ian Johnston, Dominic Kwiatkowski, Cordelia Langford, John Sillitoe on behalf of the Wellcome Sanger Institute COVID-19 Surveillance Team |
| EPI_ISL_2651412, EPI_ISL_2651413 | Lighthouse Lab in Glasgow | Wellcome Sanger Institute for the COVID-19 Genomics UK (COG-UK) Consortium | Harper VanSteenhouse, Yumi Kasai, David Gray, Carol Clugston, Anna Dominiczak and Alex Alderton, Roberto Amato, Jeffrey Barrett, Sonia Goncalves, Ewan Harrison, David K. Jackson, Ian Johnston, Dominic Kwiatkowski, Cordelia Langford, John Sillitoe on behalf of the Wellcome Sanger Institute COVID-19 Surveillance Team |
| EPI_ISL_2651414 | Lighthouse Lab in Milton Keynes | Wellcome Sanger Institute for the COVID-19 Genomics UK (COG-UK) Consortium | The Lighthouse Lab in Milton Keynes and Alex Alderton, Roberto Amato, Jeffrey Barrett, Sonia Goncalves, Ewan Harrison, David K. Jackson, Ian Johnston, Dominic Kwiatkowski, Cordelia Langford, John Sillitoe on behalf of the Wellcome Sanger Institute COVID-19 Surveillance Team |
| EPI_ISL_2651415 | Lighthouse Laboratory Plymouth | Wellcome Sanger Institute for the COVID-19 Genomics UK (COG-UK) Consortium | Lighthouse Laboratory Plymouth and Alex Alderton, Roberto Amato, Jeffrey Barrett, Sonia Goncalves, Ewan Harrison, David K. Jackson, Ian Johnston, Dominic Kwiatkowski, Cordelia Langford, John Sillitoe on behalf of the Wellcome Sanger Institute COVID-19 Surveillance Team |
| EPI_ISL_2651416 | Lighthouse Lab in Milton Keynes | Wellcome Sanger Institute for the COVID-19 Genomics UK (COG-UK) Consortium | The Lighthouse Lab in Milton Keynes and Alex Alderton, Roberto Amato, Jeffrey Barrett, Sonia Goncalves, Ewan Harrison, David K. Jackson, Ian Johnston, Dominic Kwiatkowski, Cordelia Langford, John Sillitoe on behalf of the Wellcome Sanger Institute COVID-19 Surveillance Team |
| EPI_ISL_2651417 | Lighthouse Laboratory Plymouth | Wellcome Sanger Institute for the COVID-19 Genomics UK (COG-UK) Consortium | Lighthouse Laboratory Plymouth and Alex Alderton, Roberto Amato, Jeffrey Barrett, Sonia Goncalves, Ewan Harrison, David K. Jackson, Ian Johnston, Dominic Kwiatkowski, Cordelia Langford, John Sillitoe on behalf of the Wellcome Sanger Institute COVID-19 Surveillance Team |
| EPI_ISL_2651418 | Lighthouse Lab in Glasgow | Wellcome Sanger Institute for the COVID-19 Genomics UK (COG-UK) Consortium | Harper VanSteenhouse, Yumi Kasai, David Gray, Carol Clugston, Anna Dominiczak and Alex Alderton, Roberto Amato, Jeffrey Barrett, Sonia Goncalves, Ewan Harrison, David K. Jackson, Ian Johnston, Dominic Kwiatkowski, Cordelia Langford, John Sillitoe on behalf of the Wellcome Sanger Institute COVID-19 Surveillance Team |
| EPI_ISL_2651419 | Lighthouse Lab in Milton Keynes | Wellcome Sanger Institute for the COVID-19 Genomics UK (COG-UK) Consortium | The Lighthouse Lab in Milton Keynes and Alex Alderton, Roberto Amato, Jeffrey Barrett, Sonia Goncalves, Ewan Harrison, David K. Jackson, Ian Johnston, Dominic Kwiatkowski, Cordelia Langford, John Sillitoe on behalf of the Wellcome Sanger Institute COVID-19 Surveillance Team |
| EPI_ISL_2651420 | Lighthouse Laboratory Plymouth | Wellcome Sanger Institute for the COVID-19 Genomics UK (COG-UK) Consortium | Lighthouse Laboratory Plymouth and Alex Alderton, Roberto Amato, Jeffrey Barrett, Sonia Goncalves, Ewan Harrison, David K. Jackson, Ian Johnston, Dominic Kwiatkowski, Cordelia Langford, John Sillitoe on behalf of the Wellcome Sanger Institute COVID-19 Surveillance Team |
| EPI_ISL_2651421 | Lighthouse Lab in Milton Keynes | Wellcome Sanger Institute for the COVID-19 Genomics UK (COG-UK) Consortium | The Lighthouse Lab in Milton Keynes and Alex Alderton, Roberto Amato, Jeffrey Barrett, Sonia Goncalves, Ewan Harrison, David K. Jackson, Ian Johnston, Dominic Kwiatkowski, Cordelia Langford, John Sillitoe on behalf of the Wellcome Sanger Institute COVID-19 Surveillance Team |
| EPI_ISL_2651422 | Lighthouse Laboratory Plymouth | Wellcome Sanger Institute for the COVID-19 Genomics UK (COG-UK) Consortium | Lighthouse Laboratory Plymouth and Alex Alderton, Roberto Amato, Jeffrey Barrett, Sonia Goncalves, Ewan Harrison, David K. Jackson, Ian Johnston, Dominic Kwiatkowski, Cordelia Langford, John Sillitoe on behalf of the Wellcome Sanger Institute COVID-19 Surveillance Team |
| EPI_ISL_2651423 | Lighthouse Lab in Milton Keynes | Wellcome Sanger Institute for the COVID-19 Genomics UK (COG-UK) Consortium | The Lighthouse Lab in Milton Keynes and Alex Alderton, Roberto Amato, Jeffrey Barrett, Sonia Goncalves, Ewan Harrison, David K. Jackson, Ian Johnston, Dominic Kwiatkowski, Cordelia Langford, John Sillitoe on behalf of the Wellcome Sanger Institute COVID-19 Surveillance Team |
| EPI_ISL_2651424 | Lighthouse Laboratory Plymouth | Wellcome Sanger Institute for the COVID-19 Genomics UK (COG-UK) Consortium | Lighthouse Laboratory Plymouth and Alex Alderton, Roberto Amato, Jeffrey Barrett, Sonia Goncalves, Ewan Harrison, David K. Jackson, Ian Johnston, Dominic Kwiatkowski, Cordelia Langford, John Sillitoe on behalf of the Wellcome Sanger Institute COVID-19 Surveillance Team |
| EPI_ISL_2651425, EPI_ISL_2651426 | Lighthouse Lab in Milton Keynes | Wellcome Sanger Institute for the COVID-19 Genomics UK (COG-UK) Consortium | The Lighthouse Lab in Milton Keynes and Alex Alderton, Roberto Amato, Jeffrey Barrett, Sonia Goncalves, Ewan Harrison, David K. Jackson, Ian Johnston, Dominic Kwiatkowski, Cordelia Langford, John Sillitoe on behalf of the Wellcome Sanger Institute COVID-19 Surveillance Team |
| EPI_ISL_2651427 | Lighthouse Lab in Glasgow | Wellcome Sanger Institute for the COVID-19 Genomics UK (COG-UK) Consortium | Harper VanSteenhouse, Yumi Kasai, David Gray, Carol Clugston, Anna Dominiczak and Alex Alderton, Roberto Amato, Jeffrey Barrett, Sonia Goncalves, Ewan Harrison, David K. Jackson, Ian Johnston, Dominic Kwiatkowski, Cordelia Langford, John Sillitoe on behalf of the Wellcome Sanger Institute COVID-19 Surveillance Team |
| EPI_ISL_2651428, EPI_ISL_2651429 | Lighthouse Lab in Milton Keynes | Wellcome Sanger Institute for the COVID-19 Genomics UK (COG-UK) Consortium | The Lighthouse Lab in Milton Keynes and Alex Alderton, Roberto Amato, Jeffrey Barrett, Sonia Goncalves, Ewan Harrison, David K. Jackson, Ian Johnston, Dominic Kwiatkowski, Cordelia Langford, John Sillitoe on behalf of the Wellcome Sanger Institute COVID-19 Surveillance Team |
| EPI_ISL_2651430, EPI_ISL_2651431, EPI_ISL_2651432, EPI_ISL_2651433, EPI_ISL_2651434 | Lighthouse Lab in Glasgow | Wellcome Sanger Institute for the COVID-19 Genomics UK (COG-UK) Consortium | Harper VanSteenhouse, Yumi Kasai, David Gray, Carol Clugston, Anna Dominiczak and Alex Alderton, Roberto Amato, Jeffrey Barrett, Sonia Goncalves, Ewan Harrison, David K. Jackson, Ian Johnston, Dominic Kwiatkowski, Cordelia Langford, John Sillitoe on behalf of the Wellcome Sanger Institute COVID-19 Surveillance Team |
| EPI_ISL_2651435 | Lighthouse Laboratory Plymouth | Wellcome Sanger Institute for the COVID-19 Genomics UK (COG-UK) Consortium | Lighthouse Laboratory Plymouth and Alex Alderton, Roberto Amato, Jeffrey Barrett, Sonia Goncalves, Ewan Harrison, David K. Jackson, Ian Johnston, Dominic Kwiatkowski, Cordelia Langford, John Sillitoe on behalf of the Wellcome Sanger Institute COVID-19 Surveillance Team |
| EPI_ISL_2651436 | Lighthouse Lab in Milton Keynes | Wellcome Sanger Institute for the COVID-19 Genomics UK (COG-UK) Consortium | The Lighthouse Lab in Milton Keynes and Alex Alderton, Roberto Amato, Jeffrey Barrett, Sonia Goncalves, Ewan Harrison, David K. Jackson, Ian Johnston, Dominic Kwiatkowski, Cordelia Langford, John Sillitoe on behalf of the Wellcome Sanger Institute COVID-19 Surveillance Team |
| EPI_ISL_2651437, EPI_ISL_2651438 | Lighthouse Lab in Glasgow | Wellcome Sanger Institute for the COVID-19 Genomics UK (COG-UK) Consortium | Harper VanSteenhouse, Yumi Kasai, David Gray, Carol Clugston, Anna Dominiczak and Alex Alderton, Roberto Amato, Jeffrey Barrett, Sonia Goncalves, Ewan Harrison, David K. Jackson, Ian Johnston, Dominic Kwiatkowski, Cordelia Langford, John Sillitoe on behalf of the Wellcome Sanger Institute COVID-19 Surveillance Team |
| EPI_ISL_2651439 | Lighthouse Laboratory Plymouth | Wellcome Sanger Institute for the COVID-19 Genomics UK (COG-UK) Consortium | Lighthouse Laboratory Plymouth and Alex Alderton, Roberto Amato, Jeffrey Barrett, Sonia Goncalves, Ewan Harrison, David K. Jackson, Ian Johnston, Dominic Kwiatkowski, Cordelia Langford, John Sillitoe on behalf of the Wellcome Sanger Institute COVID-19 Surveillance Team |
| EPI_ISL_2651440, EPI_ISL_2651441, EPI_ISL_2651442 | Lighthouse Lab in Milton Keynes | Wellcome Sanger Institute for the COVID-19 Genomics UK (COG-UK) Consortium | The Lighthouse Lab in Milton Keynes and Alex Alderton, Roberto Amato, Jeffrey Barrett, Sonia Goncalves, Ewan Harrison, David K. Jackson, Ian Johnston, Dominic Kwiatkowski, Cordelia Langford, John Sillitoe on behalf of the Wellcome Sanger Institute COVID-19 Surveillance Team |
| EPI_ISL_2651443, EPI_ISL_2651444 | Lighthouse Laboratory Plymouth | Wellcome Sanger Institute for the COVID-19 Genomics UK (COG-UK) Consortium | Lighthouse Laboratory Plymouth and Alex Alderton, Roberto Amato, Jeffrey Barrett, Sonia Goncalves, Ewan Harrison, David K. Jackson, Ian Johnston, Dominic Kwiatkowski, Cordelia Langford, John Sillitoe on behalf of the Wellcome Sanger Institute COVID-19 Surveillance Team |
| EPI_ISL_2651445, EPI_ISL_2651446, EPI_ISL_2651447 | Lighthouse Lab in Milton Keynes | Wellcome Sanger Institute for the COVID-19 Genomics UK (COG-UK) Consortium | The Lighthouse Lab in Milton Keynes and Alex Alderton, Roberto Amato, Jeffrey Barrett, Sonia Goncalves, Ewan Harrison, David K. Jackson, Ian Johnston, Dominic Kwiatkowski, Cordelia Langford, John Sillitoe on behalf of the Wellcome Sanger Institute COVID-19 Surveillance Team |
| EPI_ISL_2651448 | Lighthouse Laboratory Plymouth | Wellcome Sanger Institute for the COVID-19 Genomics UK (COG-UK) Consortium | Lighthouse Laboratory Plymouth and Alex Alderton, Roberto Amato, Jeffrey Barrett, Sonia Goncalves, Ewan Harrison, David K. Jackson, Ian Johnston, Dominic Kwiatkowski, Cordelia Langford, John Sillitoe on behalf of the Wellcome Sanger Institute COVID-19 Surveillance Team |
| EPI_ISL_2651449 | Lighthouse Lab in Milton Keynes | Wellcome Sanger Institute for the COVID-19 Genomics UK (COG-UK) Consortium | The Lighthouse Lab in Milton Keynes and Alex Alderton, Roberto Amato, Jeffrey Barrett, Sonia Goncalves, Ewan Harrison, David K. Jackson, Ian Johnston, Dominic Kwiatkowski, Cordelia Langford, John Sillitoe on behalf of the Wellcome Sanger Institute COVID-19 Surveillance Team |
| EPI_ISL_2651451 | Lighthouse Lab in Glasgow | Wellcome Sanger Institute for the COVID-19 Genomics UK (COG-UK) Consortium | Harper VanSteenhouse, Yumi Kasai, David Gray, Carol Clugston, Anna Dominiczak and Alex Alderton, Roberto Amato, Jeffrey Barrett, Sonia Goncalves, Ewan Harrison, David K. Jackson, Ian Johnston, Dominic Kwiatkowski, Cordelia Langford, John Sillitoe on behalf of the Wellcome Sanger Institute COVID-19 Surveillance Team |
| EPI_ISL_2651452 | Lighthouse Lab in Milton Keynes | Wellcome Sanger Institute for the COVID-19 Genomics UK (COG-UK) Consortium | The Lighthouse Lab in Milton Keynes and Alex Alderton, Roberto Amato, Jeffrey Barrett, Sonia Goncalves, Ewan Harrison, David K. Jackson, Ian Johnston, Dominic Kwiatkowski, Cordelia Langford, John Sillitoe on behalf of the Wellcome Sanger Institute COVID-19 Surveillance Team |
| EPI_ISL_2651453 | Lighthouse Lab in Glasgow | Wellcome Sanger Institute for the COVID-19 Genomics UK (COG-UK) Consortium | Harper VanSteenhouse, Yumi Kasai, David Gray, Carol Clugston, Anna Dominiczak and Alex Alderton, Roberto Amato, Jeffrey Barrett, Sonia Goncalves, Ewan Harrison, David K. Jackson, Ian Johnston, Dominic Kwiatkowski, Cordelia Langford, John Sillitoe on behalf of the Wellcome Sanger Institute COVID-19 Surveillance Team |
| EPI_ISL_2651454 | Lighthouse Lab in Milton Keynes | Wellcome Sanger Institute for the COVID-19 Genomics UK (COG-UK) Consortium | The Lighthouse Lab in Milton Keynes and Alex Alderton, Roberto Amato, Jeffrey Barrett, Sonia Goncalves, Ewan Harrison, David K. Jackson, Ian Johnston, Dominic Kwiatkowski, Cordelia Langford, John Sillitoe on behalf of the Wellcome Sanger Institute COVID-19 Surveillance Team |
| EPI_ISL_2651455 | Lighthouse Laboratory Plymouth | Wellcome Sanger Institute for the COVID-19 Genomics UK (COG-UK) Consortium | Lighthouse Laboratory Plymouth and Alex Alderton, Roberto Amato, Jeffrey Barrett, Sonia Goncalves, Ewan Harrison, David K. Jackson, Ian Johnston, Dominic Kwiatkowski, Cordelia Langford, John Sillitoe on behalf of the Wellcome Sanger Institute COVID-19 Surveillance Team |
| EPI_ISL_2651456, EPI_ISL_2651457, EPI_ISL_2651458 | Lighthouse Lab in Glasgow | Wellcome Sanger Institute for the COVID-19 Genomics UK (COG-UK) Consortium | Harper VanSteenhouse, Yumi Kasai, David Gray, Carol Clugston, Anna Dominiczak and Alex Alderton, Roberto Amato, Jeffrey Barrett, Sonia Goncalves, Ewan Harrison, David K. Jackson, Ian Johnston, Dominic Kwiatkowski, Cordelia Langford, John Sillitoe on behalf of the Wellcome Sanger Institute COVID-19 Surveillance Team |
| EPI_ISL_2651459, EPI_ISL_2651460 | Lighthouse Laboratory Plymouth | Wellcome Sanger Institute for the COVID-19 Genomics UK (COG-UK) Consortium | Lighthouse Laboratory Plymouth and Alex Alderton, Roberto Amato, Jeffrey Barrett, Sonia Goncalves, Ewan Harrison, David K. Jackson, Ian Johnston, Dominic Kwiatkowski, Cordelia Langford, John Sillitoe on behalf of the Wellcome Sanger Institute COVID-19 Surveillance Team |
| EPI_ISL_2651461, EPI_ISL_2651462, EPI_ISL_2651463 | Lighthouse Lab in Milton Keynes | Wellcome Sanger Institute for the COVID-19 Genomics UK (COG-UK) Consortium | The Lighthouse Lab in Milton Keynes and Alex Alderton, Roberto Amato, Jeffrey Barrett, Sonia Goncalves, Ewan Harrison, David K. Jackson, Ian Johnston, Dominic Kwiatkowski, Cordelia Langford, John Sillitoe on behalf of the Wellcome Sanger Institute COVID-19 Surveillance Team |
| EPI_ISL_2651464 | Lighthouse Laboratory Plymouth | Wellcome Sanger Institute for the COVID-19 Genomics UK (COG-UK) Consortium | Lighthouse Laboratory Plymouth and Alex Alderton, Roberto Amato, Jeffrey Barrett, Sonia Goncalves, Ewan Harrison, David K. Jackson, Ian Johnston, Dominic Kwiatkowski, Cordelia Langford, John Sillitoe on behalf of the Wellcome Sanger Institute COVID-19 Surveillance Team |
| EPI_ISL_2651465, EPI_ISL_2651466 | Lighthouse Lab in Glasgow | Wellcome Sanger Institute for the COVID-19 Genomics UK (COG-UK) Consortium | Harper VanSteenhouse, Yumi Kasai, David Gray, Carol Clugston, Anna Dominiczak and Alex Alderton, Roberto Amato, Jeffrey Barrett, Sonia Goncalves, Ewan Harrison, David K. Jackson, Ian Johnston, Dominic Kwiatkowski, Cordelia Langford, John Sillitoe on behalf of the Wellcome Sanger Institute COVID-19 Surveillance Team |

| | | | |
|---|---|---|---|
| EPI_ISL_2651468 | Lighthouse Laboratory Plymouth | Wellcome Sanger Institute for the COVID-19 Genomics UK (COG-UK) Consortium | Lighthouse Laboratory Plymouth and Alex Alderton, Roberto Amato, Jeffrey Barrett, Sonia Goncalves, Ewan Harrison, David K. Jackson, Ian Johnston, Dominic Kwiatkowski, Cordelia Langford, John Sillitoe on behalf of the Wellcome Sanger Institute COVID-19 Surveillance Team |
| EPI_ISL_2651469 | Lighthouse Lab in Glasgow | Wellcome Sanger Institute for the COVID-19 Genomics UK (COG-UK) Consortium | Harper VanSteenhouse, Yumi Kasai, David Gray, Carol Clugston, Anna Dominiczak and Alex Alderton, Roberto Amato, Jeffrey Barrett, Sonia Goncalves, Ewan Harrison, David K. Jackson, Ian Johnston, Dominic Kwiatkowski, Cordelia Langford, John Sillitoe on behalf of the Wellcome Sanger Institute COVID-19 Surveillance Team |
| EPI_ISL_2651470 | Lighthouse Laboratory Plymouth | Wellcome Sanger Institute for the COVID-19 Genomics UK (COG-UK) Consortium | Lighthouse Laboratory Plymouth and Alex Alderton, Roberto Amato, Jeffrey Barrett, Sonia Goncalves, Ewan Harrison, David K. Jackson, Ian Johnston, Dominic Kwiatkowski, Cordelia Langford, John Sillitoe on behalf of the Wellcome Sanger Institute COVID-19 Surveillance Team |
| EPI_ISL_2651471 | Lighthouse Lab in Milton Keynes | Wellcome Sanger Institute for the COVID-19 Genomics UK (COG-UK) Consortium | The Lighthouse Lab in Milton Keynes and Alex Alderton, Roberto Amato, Jeffrey Barrett, Sonia Goncalves, Ewan Harrison, David K. Jackson, Ian Johnston, Dominic Kwiatkowski, Cordelia Langford, John Sillitoe on behalf of the Wellcome Sanger Institute COVID-19 Surveillance Team |
| EPI_ISL_2651472 | Lighthouse Lab in Glasgow | Wellcome Sanger Institute for the COVID-19 Genomics UK (COG-UK) Consortium | Harper VanSteenhouse, Yumi Kasai, David Gray, Carol Clugston, Anna Dominiczak and Alex Alderton, Roberto Amato, Jeffrey Barrett, Sonia Goncalves, Ewan Harrison, David K. Jackson, Ian Johnston, Dominic Kwiatkowski, Cordelia Langford, John Sillitoe on behalf of the Wellcome Sanger Institute COVID-19 Surveillance Team |
| EPI_ISL_2651473 | Lighthouse Laboratory Plymouth | Wellcome Sanger Institute for the COVID-19 Genomics UK (COG-UK) Consortium | Lighthouse Laboratory Plymouth and Alex Alderton, Roberto Amato, Jeffrey Barrett, Sonia Goncalves, Ewan Harrison, David K. Jackson, Ian Johnston, Dominic Kwiatkowski, Cordelia Langford, John Sillitoe on behalf of the Wellcome Sanger Institute COVID-19 Surveillance Team |
| EPI_ISL_2651474 | Lighthouse Lab in Milton Keynes | Wellcome Sanger Institute for the COVID-19 Genomics UK (COG-UK) Consortium | The Lighthouse Lab in Milton Keynes and Alex Alderton, Roberto Amato, Jeffrey Barrett, Sonia Goncalves, Ewan Harrison, David K. Jackson, Ian Johnston, Dominic Kwiatkowski, Cordelia Langford, John Sillitoe on behalf of the Wellcome Sanger Institute COVID-19 Surveillance Team |
| EPI_ISL_2651475 | Lighthouse Laboratory Plymouth | Wellcome Sanger Institute for the COVID-19 Genomics UK (COG-UK) Consortium | Lighthouse Laboratory Plymouth and Alex Alderton, Roberto Amato, Jeffrey Barrett, Sonia Goncalves, Ewan Harrison, David K. Jackson, Ian Johnston, Dominic Kwiatkowski, Cordelia Langford, John Sillitoe on behalf of the Wellcome Sanger Institute COVID-19 Surveillance Team |
| EPI_ISL_2651476, EPI_ISL_2651477 | Lighthouse Lab in Glasgow | Wellcome Sanger Institute for the COVID-19 Genomics UK (COG-UK) Consortium | Harper VanSteenhouse, Yumi Kasai, David Gray, Carol Clugston, Anna Dominiczak and Alex Alderton, Roberto Amato, Jeffrey Barrett, Sonia Goncalves, Ewan Harrison, David K. Jackson, Ian Johnston, Dominic Kwiatkowski, Cordelia Langford, John Sillitoe on behalf of the Wellcome Sanger Institute COVID-19 Surveillance Team |
| EPI_ISL_2651478 | Lighthouse Lab in Milton Keynes | Wellcome Sanger Institute for the COVID-19 Genomics UK (COG-UK) Consortium | The Lighthouse Lab in Milton Keynes and Alex Alderton, Roberto Amato, Jeffrey Barrett, Sonia Goncalves, Ewan Harrison, David K. Jackson, Ian Johnston, Dominic Kwiatkowski, Cordelia Langford, John Sillitoe on behalf of the Wellcome Sanger Institute COVID-19 Surveillance Team |
| EPI_ISL_2651479 | Lighthouse Laboratory Plymouth | Wellcome Sanger Institute for the COVID-19 Genomics UK (COG-UK) Consortium | Lighthouse Laboratory Plymouth and Alex Alderton, Roberto Amato, Jeffrey Barrett, Sonia Goncalves, Ewan Harrison, David K. Jackson, Ian Johnston, Dominic Kwiatkowski, Cordelia Langford, John Sillitoe on behalf of the Wellcome Sanger Institute COVID-19 Surveillance Team |
| EPI_ISL_2651480, EPI_ISL_2651481, EPI_ISL_2651482 | Lighthouse Lab in Glasgow | Wellcome Sanger Institute for the COVID-19 Genomics UK (COG-UK) Consortium | Harper VanSteenhouse, Yumi Kasai, David Gray, Carol Clugston, Anna Dominiczak and Alex Alderton, Roberto Amato, Jeffrey Barrett, Sonia Goncalves, Ewan Harrison, David K. Jackson, Ian Johnston, Dominic Kwiatkowski, Cordelia Langford, John Sillitoe on behalf of the Wellcome Sanger Institute COVID-19 Surveillance Team |
| EPI_ISL_2651483 | Lighthouse Lab in Milton Keynes | Wellcome Sanger Institute for the COVID-19 Genomics UK (COG-UK) Consortium | The Lighthouse Lab in Milton Keynes and Alex Alderton, Roberto Amato, Jeffrey Barrett, Sonia Goncalves, Ewan Harrison, David K. Jackson, Ian Johnston, Dominic Kwiatkowski, Cordelia Langford, John Sillitoe on behalf of the Wellcome Sanger Institute COVID-19 Surveillance Team |
| EPI_ISL_2651484, EPI_ISL_2651485 | Lighthouse Laboratory Plymouth | Wellcome Sanger Institute for the COVID-19 Genomics UK (COG-UK) Consortium | Lighthouse Laboratory Plymouth and Alex Alderton, Roberto Amato, Jeffrey Barrett, Sonia Goncalves, Ewan Harrison, David K. Jackson, Ian Johnston, Dominic Kwiatkowski, Cordelia Langford, John Sillitoe on behalf of the Wellcome Sanger Institute COVID-19 Surveillance Team |
| EPI_ISL_2651486 | Lighthouse Lab in Glasgow | Wellcome Sanger Institute for the COVID-19 Genomics UK (COG-UK) Consortium | Harper VanSteenhouse, Yumi Kasai, David Gray, Carol Clugston, Anna Dominiczak and Alex Alderton, Roberto Amato, Jeffrey Barrett, Sonia Goncalves, Ewan Harrison, David K. Jackson, Ian Johnston, Dominic Kwiatkowski, Cordelia Langford, John Sillitoe on behalf of the Wellcome Sanger Institute COVID-19 Surveillance Team |
| EPI_ISL_2651487 | Lighthouse Laboratory Plymouth | Wellcome Sanger Institute for the COVID-19 Genomics UK (COG-UK) Consortium | Lighthouse Laboratory Plymouth and Alex Alderton, Roberto Amato, Jeffrey Barrett, Sonia Goncalves, Ewan Harrison, David K. Jackson, Ian Johnston, Dominic Kwiatkowski, Cordelia Langford, John Sillitoe on behalf of the Wellcome Sanger Institute COVID-19 Surveillance Team |
| EPI_ISL_2651488 | Lighthouse Lab in Milton Keynes | Wellcome Sanger Institute for the COVID-19 Genomics UK (COG-UK) Consortium | The Lighthouse Lab in Milton Keynes and Alex Alderton, Roberto Amato, Jeffrey Barrett, Sonia Goncalves, Ewan Harrison, David K. Jackson, Ian Johnston, Dominic Kwiatkowski, Cordelia Langford, John Sillitoe on behalf of the Wellcome Sanger Institute COVID-19 Surveillance Team |
| EPI_ISL_2651489 | Lighthouse Laboratory Plymouth | Wellcome Sanger Institute for the COVID-19 Genomics UK (COG-UK) Consortium | Lighthouse Laboratory Plymouth and Alex Alderton, Roberto Amato, Jeffrey Barrett, Sonia Goncalves, Ewan Harrison, David K. Jackson, Ian Johnston, Dominic Kwiatkowski, Cordelia Langford, John Sillitoe on behalf of the Wellcome Sanger Institute COVID-19 Surveillance Team |
| EPI_ISL_2651490 | Lighthouse Lab in Milton Keynes | Wellcome Sanger Institute for the COVID-19 Genomics UK (COG-UK) Consortium | The Lighthouse Lab in Milton Keynes and Alex Alderton, Roberto Amato, Jeffrey Barrett, Sonia Goncalves, Ewan Harrison, David K. Jackson, Ian Johnston, Dominic Kwiatkowski, Cordelia Langford, John Sillitoe on behalf of the Wellcome Sanger Institute COVID-19 Surveillance Team |
| EPI_ISL_2651491 | Lighthouse Laboratory Plymouth | Wellcome Sanger Institute for the COVID-19 Genomics UK (COG-UK) Consortium | Lighthouse Laboratory Plymouth and Alex Alderton, Roberto Amato, Jeffrey Barrett, Sonia Goncalves, Ewan Harrison, David K. Jackson, Ian Johnston, Dominic Kwiatkowski, Cordelia Langford, John Sillitoe on behalf of the Wellcome Sanger Institute COVID-19 Surveillance Team |
| EPI_ISL_2651492 | Lighthouse Lab in Glasgow | Wellcome Sanger Institute for the COVID-19 Genomics UK (COG-UK) Consortium | Harper VanSteenhouse, Yumi Kasai, David Gray, Carol Clugston, Anna Dominiczak and Alex Alderton, Roberto Amato, Jeffrey Barrett, Sonia Goncalves, Ewan Harrison, David K. Jackson, Ian Johnston, Dominic Kwiatkowski, Cordelia Langford, John Sillitoe on behalf of the Wellcome Sanger Institute COVID-19 Surveillance Team |
| EPI_ISL_2651493, EPI_ISL_2651494 | Lighthouse Lab in Milton Keynes | Wellcome Sanger Institute for the COVID-19 Genomics UK (COG-UK) Consortium | The Lighthouse Lab in Milton Keynes and Alex Alderton, Roberto Amato, Jeffrey Barrett, Sonia Goncalves, Ewan Harrison, David K. Jackson, Ian Johnston, Dominic Kwiatkowski, Cordelia Langford, John Sillitoe on behalf of the Wellcome Sanger Institute COVID-19 Surveillance Team |
| EPI_ISL_2651495, EPI_ISL_2651496, EPI_ISL_2651497 | Lighthouse Laboratory Plymouth | Wellcome Sanger Institute for the COVID-19 Genomics UK (COG-UK) Consortium | Lighthouse Laboratory Plymouth and Alex Alderton, Roberto Amato, Jeffrey Barrett, Sonia Goncalves, Ewan Harrison, David K. Jackson, Ian Johnston, Dominic Kwiatkowski, Cordelia Langford, John Sillitoe on behalf of the Wellcome Sanger Institute COVID-19 Surveillance Team |
| EPI_ISL_2651498 | Lighthouse Lab in Milton Keynes | Wellcome Sanger Institute for the COVID-19 Genomics UK (COG-UK) Consortium | The Lighthouse Lab in Milton Keynes and Alex Alderton, Roberto Amato, Jeffrey Barrett, Sonia Goncalves, Ewan Harrison, David K. Jackson, Ian Johnston, Dominic Kwiatkowski, Cordelia Langford, John Sillitoe on behalf of the Wellcome Sanger Institute COVID-19 Surveillance Team |
| EPI_ISL_2651499, EPI_ISL_2651500 | Lighthouse Lab in Glasgow | Wellcome Sanger Institute for the COVID-19 Genomics UK (COG-UK) Consortium | Harper VanSteenhouse, Yumi Kasai, David Gray, Carol Clugston, Anna Dominiczak and Alex Alderton, Roberto Amato, Jeffrey Barrett, Sonia Goncalves, Ewan Harrison, David K. Jackson, Ian Johnston, Dominic Kwiatkowski, Cordelia Langford, John Sillitoe on behalf of the Wellcome Sanger Institute COVID-19 Surveillance Team |
| EPI_ISL_2651501 | Lighthouse Lab in Milton Keynes | Wellcome Sanger Institute for the COVID-19 Genomics UK (COG-UK) Consortium | The Lighthouse Lab in Milton Keynes and Alex Alderton, Roberto Amato, Jeffrey Barrett, Sonia Goncalves, Ewan Harrison, David K. Jackson, Ian Johnston, Dominic Kwiatkowski, Cordelia Langford, John Sillitoe on behalf of the Wellcome Sanger Institute COVID-19 Surveillance Team |
| EPI_ISL_2651502, EPI_ISL_2651503 | Lighthouse Lab in Glasgow | Wellcome Sanger Institute for the COVID-19 Genomics UK (COG-UK) Consortium | Harper VanSteenhouse, Yumi Kasai, David Gray, Carol Clugston, Anna Dominiczak and Alex Alderton, Roberto Amato, Jeffrey Barrett, Sonia Goncalves, Ewan Harrison, David K. Jackson, Ian Johnston, Dominic Kwiatkowski, Cordelia Langford, John Sillitoe on behalf of the Wellcome Sanger Institute COVID-19 Surveillance Team |
| EPI_ISL_2651504 | Lighthouse Lab in Milton Keynes | Wellcome Sanger Institute for the COVID-19 Genomics UK (COG-UK) Consortium | The Lighthouse Lab in Milton Keynes and Alex Alderton, Roberto Amato, Jeffrey Barrett, Sonia Goncalves, Ewan Harrison, David K. Jackson, Ian Johnston, Dominic Kwiatkowski, Cordelia Langford, John Sillitoe on behalf of the Wellcome Sanger Institute COVID-19 Surveillance Team |
| EPI_ISL_2651505 | Lighthouse Laboratory Plymouth | Wellcome Sanger Institute for the COVID-19 Genomics UK (COG-UK) Consortium | Lighthouse Laboratory Plymouth and Alex Alderton, Roberto Amato, Jeffrey Barrett, Sonia Goncalves, Ewan Harrison, David K. Jackson, Ian Johnston, Dominic Kwiatkowski, Cordelia Langford, John Sillitoe on behalf of the Wellcome Sanger Institute COVID-19 Surveillance Team |
| EPI_ISL_2651506, EPI_ISL_2651507 | Lighthouse Lab in Milton Keynes | Wellcome Sanger Institute for the COVID-19 Genomics UK (COG-UK) Consortium | The Lighthouse Lab in Milton Keynes and Alex Alderton, Roberto Amato, Jeffrey Barrett, Sonia Goncalves, Ewan Harrison, David K. Jackson, Ian Johnston, Dominic Kwiatkowski, Cordelia Langford, John Sillitoe on behalf of the Wellcome Sanger Institute COVID-19 Surveillance Team |
| EPI_ISL_2651508, EPI_ISL_2651509 | Lighthouse Laboratory Plymouth | Wellcome Sanger Institute for the COVID-19 Genomics UK (COG-UK) Consortium | Lighthouse Laboratory Plymouth and Alex Alderton, Roberto Amato, Jeffrey Barrett, Sonia Goncalves, Ewan Harrison, David K. Jackson, Ian Johnston, Dominic Kwiatkowski, Cordelia Langford, John Sillitoe on behalf of the Wellcome Sanger Institute COVID-19 Surveillance Team |
| EPI_ISL_2651510 | Lighthouse Lab in Glasgow | Wellcome Sanger Institute for the COVID-19 Genomics UK (COG-UK) Consortium | Harper VanSteenhouse, Yumi Kasai, David Gray, Carol Clugston, Anna Dominiczak and Alex Alderton, Roberto Amato, Jeffrey Barrett, Sonia Goncalves, Ewan Harrison, David K. Jackson, Ian Johnston, Dominic Kwiatkowski, Cordelia Langford, John Sillitoe on behalf of the Wellcome Sanger Institute COVID-19 Surveillance Team |
| EPI_ISL_2651511 | Lighthouse Lab in Milton Keynes | Wellcome Sanger Institute for the COVID-19 Genomics UK (COG-UK) Consortium | The Lighthouse Lab in Milton Keynes and Alex Alderton, Roberto Amato, Jeffrey Barrett, Sonia Goncalves, Ewan Harrison, David K. Jackson, Ian Johnston, Dominic Kwiatkowski, Cordelia Langford, John Sillitoe on behalf of the Wellcome Sanger Institute COVID-19 Surveillance Team |
| EPI_ISL_2651512, EPI_ISL_2651513, EPI_ISL_2651514 | Lighthouse Laboratory Plymouth | Wellcome Sanger Institute for the COVID-19 Genomics UK (COG-UK) Consortium | Lighthouse Laboratory Plymouth and Alex Alderton, Roberto Amato, Jeffrey Barrett, Sonia Goncalves, Ewan Harrison, David K. Jackson, Ian Johnston, Dominic Kwiatkowski, Cordelia Langford, John Sillitoe on behalf of the Wellcome Sanger Institute COVID-19 Surveillance Team |
| EPI_ISL_2651515, EPI_ISL_2651516 | Lighthouse Lab in Glasgow | Wellcome Sanger Institute for the COVID-19 Genomics UK (COG-UK) Consortium | Harper VanSteenhouse, Yumi Kasai, David Gray, Carol Clugston, Anna Dominiczak and Alex Alderton, Roberto Amato, Jeffrey Barrett, Sonia Goncalves, Ewan Harrison, David K. Jackson, Ian Johnston, Dominic Kwiatkowski, Cordelia Langford, John Sillitoe on behalf of the Wellcome Sanger Institute COVID-19 Surveillance Team |
| EPI_ISL_2651517, EPI_ISL_2651518, EPI_ISL_2651519 | Lighthouse Laboratory Plymouth | Wellcome Sanger Institute for the COVID-19 Genomics UK (COG-UK) Consortium | Lighthouse Laboratory Plymouth and Alex Alderton, Roberto Amato, Jeffrey Barrett, Sonia Goncalves, Ewan Harrison, David K. Jackson, Ian Johnston, Dominic Kwiatkowski, Cordelia Langford, John Sillitoe on behalf of the Wellcome Sanger Institute COVID-19 Surveillance Team |
| EPI_ISL_2651520 | Lighthouse Lab in Milton Keynes | Wellcome Sanger Institute for the COVID-19 Genomics UK (COG-UK) Consortium | The Lighthouse Lab in Milton Keynes and Alex Alderton, Roberto Amato, Jeffrey Barrett, Sonia Goncalves, Ewan Harrison, David K. Jackson, Ian Johnston, Dominic Kwiatkowski, Cordelia Langford, John Sillitoe on behalf of the Wellcome Sanger Institute COVID-19 Surveillance Team |
| EPI_ISL_2651521, EPI_ISL_2651522 | Lighthouse Lab in Glasgow | Wellcome Sanger Institute for the COVID-19 Genomics UK (COG-UK) Consortium | Harper VanSteenhouse, Yumi Kasai, David Gray, Carol Clugston, Anna Dominiczak and Alex Alderton, Roberto Amato, Jeffrey Barrett, Sonia Goncalves, Ewan Harrison, David K. Jackson, Ian Johnston, Dominic Kwiatkowski, Cordelia Langford, John Sillitoe on behalf of the Wellcome Sanger Institute COVID-19 Surveillance Team |
| EPI_ISL_2651523 | Lighthouse Lab in Milton Keynes | Wellcome Sanger Institute for the COVID-19 Genomics UK (COG-UK) Consortium | The Lighthouse Lab in Milton Keynes and Alex Alderton, Roberto Amato, Jeffrey Barrett, Sonia Goncalves, Ewan Harrison, David K. Jackson, Ian Johnston, Dominic Kwiatkowski, Cordelia Langford, John Sillitoe on behalf of the Wellcome Sanger Institute COVID-19 Surveillance Team |
| EPI_ISL_2651524 | Lighthouse Laboratory Plymouth | Wellcome Sanger Institute for the COVID-19 Genomics UK (COG-UK) Consortium | Lighthouse Laboratory Plymouth and Alex Alderton, Roberto Amato, Jeffrey Barrett, Sonia Goncalves, Ewan Harrison, David K. Jackson, Ian Johnston, Dominic Kwiatkowski, Cordelia Langford, John Sillitoe on behalf of the Wellcome Sanger Institute COVID-19 Surveillance Team |
| EPI_ISL_2651525 | Lighthouse Lab in Milton Keynes | Wellcome Sanger Institute for the COVID-19 Genomics UK (COG-UK) Consortium | The Lighthouse Lab in Milton Keynes and Alex Alderton, Roberto Amato, Jeffrey Barrett, Sonia Goncalves, Ewan Harrison, David K. Jackson, Ian Johnston, Dominic Kwiatkowski, Cordelia Langford, John Sillitoe on behalf of the Wellcome Sanger Institute COVID-19 Surveillance Team |
| EPI_ISL_2651526, EPI_ISL_2651527 | Lighthouse Laboratory Plymouth | Wellcome Sanger Institute for the COVID-19 Genomics UK (COG-UK) Consortium | Lighthouse Laboratory Plymouth and Alex Alderton, Roberto Amato, Jeffrey Barrett, Sonia Goncalves, Ewan Harrison, David K. Jackson, Ian Johnston, Dominic Kwiatkowski, Cordelia Langford, John Sillitoe on behalf of the Wellcome Sanger Institute COVID-19 Surveillance Team |
| EPI_ISL_2651528 | Lighthouse Lab in Glasgow | Wellcome Sanger Institute for the COVID-19 Genomics UK (COG-UK) Consortium | Harper VanSteenhouse, Yumi Kasai, David Gray, Carol Clugston, Anna Dominiczak and Alex Alderton, Roberto Amato, Jeffrey Barrett, Sonia Goncalves, Ewan Harrison, David K. Jackson, Ian Johnston, Dominic Kwiatkowski, Cordelia Langford, John Sillitoe on behalf of the Wellcome Sanger Institute COVID-19 Surveillance Team |

| | | | |
|---|---|---|---|
| EPI_ISL_2651529 | Lighthouse Laboratory Plymouth | Wellcome Sanger Institute for the COVID-19 Genomics UK (COG-UK) Consortium | Lighthouse Laboratory Plymouth and Alex Alderton, Roberto Amato, Jeffrey Barrett, Sonia Goncalves, Ewan Harrison, David K. Jackson, Ian Johnston, Dominic Kwiatkowski, Cordelia Langford, John Sillitoe on behalf of the Wellcome Sanger Institute COVID-19 Surveillance Team |
| EPI_ISL_2651530 | Lighthouse Lab in Milton Keynes | Wellcome Sanger Institute for the COVID-19 Genomics UK (COG-UK) Consortium | The Lighthouse Lab in Milton Keynes and Alex Alderton, Roberto Amato, Jeffrey Barrett, Sonia Goncalves, Ewan Harrison, David K. Jackson, Ian Johnston, Dominic Kwiatkowski, Cordelia Langford, John Sillitoe on behalf of the Wellcome Sanger Institute COVID-19 Surveillance Team |
| EPI_ISL_2651531, EPI_ISL_2651532, EPI_ISL_2651533 | Lighthouse Lab in Glasgow | Wellcome Sanger Institute for the COVID-19 Genomics UK (COG-UK) Consortium | Harper VanSteenhouse, Yumi Kasai, David Gray, Carol Clugston, Anna Dominiczak and Alex Alderton, Roberto Amato, Jeffrey Barrett, Sonia Goncalves, Ewan Harrison, David K. Jackson, Ian Johnston, Dominic Kwiatkowski, Cordelia Langford, John Sillitoe on behalf of the Wellcome Sanger Institute COVID-19 Surveillance Team |
| EPI_ISL_2651534, EPI_ISL_2651535 | Lighthouse Laboratory Plymouth | Wellcome Sanger Institute for the COVID-19 Genomics UK (COG-UK) Consortium | Lighthouse Laboratory Plymouth and Alex Alderton, Roberto Amato, Jeffrey Barrett, Sonia Goncalves, Ewan Harrison, David K. Jackson, Ian Johnston, Dominic Kwiatkowski, Cordelia Langford, John Sillitoe on behalf of the Wellcome Sanger Institute COVID-19 Surveillance Team |
| EPI_ISL_2651536, EPI_ISL_2651537 | Lighthouse Lab in Milton Keynes | Wellcome Sanger Institute for the COVID-19 Genomics UK (COG-UK) Consortium | The Lighthouse Lab in Milton Keynes and Alex Alderton, Roberto Amato, Jeffrey Barrett, Sonia Goncalves, Ewan Harrison, David K. Jackson, Ian Johnston, Dominic Kwiatkowski, Cordelia Langford, John Sillitoe on behalf of the Wellcome Sanger Institute COVID-19 Surveillance Team |
| EPI_ISL_2651538, EPI_ISL_2651539 | Lighthouse Lab in Glasgow | Wellcome Sanger Institute for the COVID-19 Genomics UK (COG-UK) Consortium | Harper VanSteenhouse, Yumi Kasai, David Gray, Carol Clugston, Anna Dominiczak and Alex Alderton, Roberto Amato, Jeffrey Barrett, Sonia Goncalves, Ewan Harrison, David K. Jackson, Ian Johnston, Dominic Kwiatkowski, Cordelia Langford, John Sillitoe on behalf of the Wellcome Sanger Institute COVID-19 Surveillance Team |
| EPI_ISL_2651541 | Lighthouse Lab in Milton Keynes | Wellcome Sanger Institute for the COVID-19 Genomics UK (COG-UK) Consortium | The Lighthouse Lab in Milton Keynes and Alex Alderton, Roberto Amato, Jeffrey Barrett, Sonia Goncalves, Ewan Harrison, David K. Jackson, Ian Johnston, Dominic Kwiatkowski, Cordelia Langford, John Sillitoe on behalf of the Wellcome Sanger Institute COVID-19 Surveillance Team |
| EPI_ISL_2651542 | Lighthouse Laboratory Plymouth | Wellcome Sanger Institute for the COVID-19 Genomics UK (COG-UK) Consortium | Lighthouse Laboratory Plymouth and Alex Alderton, Roberto Amato, Jeffrey Barrett, Sonia Goncalves, Ewan Harrison, David K. Jackson, Ian Johnston, Dominic Kwiatkowski, Cordelia Langford, John Sillitoe on behalf of the Wellcome Sanger Institute COVID-19 Surveillance Team |
| EPI_ISL_2651543, EPI_ISL_2651544, EPI_ISL_2651545 | Lighthouse Lab in Milton Keynes | Wellcome Sanger Institute for the COVID-19 Genomics UK (COG-UK) Consortium | The Lighthouse Lab in Milton Keynes and Alex Alderton, Roberto Amato, Jeffrey Barrett, Sonia Goncalves, Ewan Harrison, David K. Jackson, Ian Johnston, Dominic Kwiatkowski, Cordelia Langford, John Sillitoe on behalf of the Wellcome Sanger Institute COVID-19 Surveillance Team |
| EPI_ISL_2651546 | Lighthouse Laboratory Plymouth | Wellcome Sanger Institute for the COVID-19 Genomics UK (COG-UK) Consortium | Lighthouse Laboratory Plymouth and Alex Alderton, Roberto Amato, Jeffrey Barrett, Sonia Goncalves, Ewan Harrison, David K. Jackson, Ian Johnston, Dominic Kwiatkowski, Cordelia Langford, John Sillitoe on behalf of the Wellcome Sanger Institute COVID-19 Surveillance Team |
| EPI_ISL_2651547 | Lighthouse Lab in Glasgow | Wellcome Sanger Institute for the COVID-19 Genomics UK (COG-UK) Consortium | Harper VanSteenhouse, Yumi Kasai, David Gray, Carol Clugston, Anna Dominiczak and Alex Alderton, Roberto Amato, Jeffrey Barrett, Sonia Goncalves, Ewan Harrison, David K. Jackson, Ian Johnston, Dominic Kwiatkowski, Cordelia Langford, John Sillitoe on behalf of the Wellcome Sanger Institute COVID-19 Surveillance Team |
| EPI_ISL_2651548, EPI_ISL_2651549 | Lighthouse Laboratory Plymouth | Wellcome Sanger Institute for the COVID-19 Genomics UK (COG-UK) Consortium | Lighthouse Laboratory Plymouth and Alex Alderton, Roberto Amato, Jeffrey Barrett, Sonia Goncalves, Ewan Harrison, David K. Jackson, Ian Johnston, Dominic Kwiatkowski, Cordelia Langford, John Sillitoe on behalf of the Wellcome Sanger Institute COVID-19 Surveillance Team |
| EPI_ISL_2651550 | Lighthouse Lab in Milton Keynes | Wellcome Sanger Institute for the COVID-19 Genomics UK (COG-UK) Consortium | The Lighthouse Lab in Milton Keynes and Alex Alderton, Roberto Amato, Jeffrey Barrett, Sonia Goncalves, Ewan Harrison, David K. Jackson, Ian Johnston, Dominic Kwiatkowski, Cordelia Langford, John Sillitoe on behalf of the Wellcome Sanger Institute COVID-19 Surveillance Team |
| EPI_ISL_2651551, EPI_ISL_2651552 | Lighthouse Lab in Glasgow | Wellcome Sanger Institute for the COVID-19 Genomics UK (COG-UK) Consortium | Harper VanSteenhouse, Yumi Kasai, David Gray, Carol Clugston, Anna Dominiczak and Alex Alderton, Roberto Amato, Jeffrey Barrett, Sonia Goncalves, Ewan Harrison, David K. Jackson, Ian Johnston, Dominic Kwiatkowski, Cordelia Langford, John Sillitoe on behalf of the Wellcome Sanger Institute COVID-19 Surveillance Team |
| EPI_ISL_2651553, EPI_ISL_2651554 | Lighthouse Lab in Milton Keynes | Wellcome Sanger Institute for the COVID-19 Genomics UK (COG-UK) Consortium | The Lighthouse Lab in Milton Keynes and Alex Alderton, Roberto Amato, Jeffrey Barrett, Sonia Goncalves, Ewan Harrison, David K. Jackson, Ian Johnston, Dominic Kwiatkowski, Cordelia Langford, John Sillitoe on behalf of the Wellcome Sanger Institute COVID-19 Surveillance Team |
| EPI_ISL_2651555 | Lighthouse Laboratory Plymouth | Wellcome Sanger Institute for the COVID-19 Genomics UK (COG-UK) Consortium | Lighthouse Laboratory Plymouth and Alex Alderton, Roberto Amato, Jeffrey Barrett, Sonia Goncalves, Ewan Harrison, David K. Jackson, Ian Johnston, Dominic Kwiatkowski, Cordelia Langford, John Sillitoe on behalf of the Wellcome Sanger Institute COVID-19 Surveillance Team |
| EPI_ISL_2651556, EPI_ISL_2651557 | Lighthouse Lab in Glasgow | Wellcome Sanger Institute for the COVID-19 Genomics UK (COG-UK) Consortium | Harper VanSteenhouse, Yumi Kasai, David Gray, Carol Clugston, Anna Dominiczak and Alex Alderton, Roberto Amato, Jeffrey Barrett, Sonia Goncalves, Ewan Harrison, David K. Jackson, Ian Johnston, Dominic Kwiatkowski, Cordelia Langford, John Sillitoe on behalf of the Wellcome Sanger Institute COVID-19 Surveillance Team |
| EPI_ISL_2651558, EPI_ISL_2651560 | Lighthouse Laboratory Plymouth | Wellcome Sanger Institute for the COVID-19 Genomics UK (COG-UK) Consortium | Lighthouse Laboratory Plymouth and Alex Alderton, Roberto Amato, Jeffrey Barrett, Sonia Goncalves, Ewan Harrison, David K. Jackson, Ian Johnston, Dominic Kwiatkowski, Cordelia Langford, John Sillitoe on behalf of the Wellcome Sanger Institute COVID-19 Surveillance Team |
| EPI_ISL_2651561 | Lighthouse Lab in Glasgow | Wellcome Sanger Institute for the COVID-19 Genomics UK (COG-UK) Consortium | Harper VanSteenhouse, Yumi Kasai, David Gray, Carol Clugston, Anna Dominiczak and Alex Alderton, Roberto Amato, Jeffrey Barrett, Sonia Goncalves, Ewan Harrison, David K. Jackson, Ian Johnston, Dominic Kwiatkowski, Cordelia Langford, John Sillitoe on behalf of the Wellcome Sanger Institute COVID-19 Surveillance Team |
| EPI_ISL_2651562, EPI_ISL_2651563, EPI_ISL_2651564 | Lighthouse Laboratory Plymouth | Wellcome Sanger Institute for the COVID-19 Genomics UK (COG-UK) Consortium | Lighthouse Laboratory Plymouth and Alex Alderton, Roberto Amato, Jeffrey Barrett, Sonia Goncalves, Ewan Harrison, David K. Jackson, Ian Johnston, Dominic Kwiatkowski, Cordelia Langford, John Sillitoe on behalf of the Wellcome Sanger Institute COVID-19 Surveillance Team |
| EPI_ISL_2651565, EPI_ISL_2651566 | Lighthouse Lab in Glasgow | Wellcome Sanger Institute for the COVID-19 Genomics UK (COG-UK) Consortium | Harper VanSteenhouse, Yumi Kasai, David Gray, Carol Clugston, Anna Dominiczak and Alex Alderton, Roberto Amato, Jeffrey Barrett, Sonia Goncalves, Ewan Harrison, David K. Jackson, Ian Johnston, Dominic Kwiatkowski, Cordelia Langford, John Sillitoe on behalf of the Wellcome Sanger Institute COVID-19 Surveillance Team |
| EPI_ISL_2651567 | Lighthouse Lab in Milton Keynes | Wellcome Sanger Institute for the COVID-19 Genomics UK (COG-UK) Consortium | The Lighthouse Lab in Milton Keynes and Alex Alderton, Roberto Amato, Jeffrey Barrett, Sonia Goncalves, Ewan Harrison, David K. Jackson, Ian Johnston, Dominic Kwiatkowski, Cordelia Langford, John Sillitoe on behalf of the Wellcome Sanger Institute COVID-19 Surveillance Team |
| EPI_ISL_2651568 | Lighthouse Laboratory Plymouth | Wellcome Sanger Institute for the COVID-19 Genomics UK (COG-UK) Consortium | Lighthouse Laboratory Plymouth and Alex Alderton, Roberto Amato, Jeffrey Barrett, Sonia Goncalves, Ewan Harrison, David K. Jackson, Ian Johnston, Dominic Kwiatkowski, Cordelia Langford, John Sillitoe on behalf of the Wellcome Sanger Institute COVID-19 Surveillance Team |
| EPI_ISL_2651569, EPI_ISL_2651570 | Lighthouse Lab in Milton Keynes | Wellcome Sanger Institute for the COVID-19 Genomics UK (COG-UK) Consortium | The Lighthouse Lab in Milton Keynes and Alex Alderton, Roberto Amato, Jeffrey Barrett, Sonia Goncalves, Ewan Harrison, David K. Jackson, Ian Johnston, Dominic Kwiatkowski, Cordelia Langford, John Sillitoe on behalf of the Wellcome Sanger Institute COVID-19 Surveillance Team |
| EPI_ISL_2651571, EPI_ISL_2651572 | Lighthouse Lab in Glasgow | Wellcome Sanger Institute for the COVID-19 Genomics UK (COG-UK) Consortium | Harper VanSteenhouse, Yumi Kasai, David Gray, Carol Clugston, Anna Dominiczak and Alex Alderton, Roberto Amato, Jeffrey Barrett, Sonia Goncalves, Ewan Harrison, David K. Jackson, Ian Johnston, Dominic Kwiatkowski, Cordelia Langford, John Sillitoe on behalf of the Wellcome Sanger Institute COVID-19 Surveillance Team |
| EPI_ISL_2651573 | Lighthouse Lab in Milton Keynes | Wellcome Sanger Institute for the COVID-19 Genomics UK (COG-UK) Consortium | The Lighthouse Lab in Milton Keynes and Alex Alderton, Roberto Amato, Jeffrey Barrett, Sonia Goncalves, Ewan Harrison, David K. Jackson, Ian Johnston, Dominic Kwiatkowski, Cordelia Langford, John Sillitoe on behalf of the Wellcome Sanger Institute COVID-19 Surveillance Team |
| EPI_ISL_2651574 | Lighthouse Lab in Glasgow | Wellcome Sanger Institute for the COVID-19 Genomics UK (COG-UK) Consortium | Harper VanSteenhouse, Yumi Kasai, David Gray, Carol Clugston, Anna Dominiczak and Alex Alderton, Roberto Amato, Jeffrey Barrett, Sonia Goncalves, Ewan Harrison, David K. Jackson, Ian Johnston, Dominic Kwiatkowski, Cordelia Langford, John Sillitoe on behalf of the Wellcome Sanger Institute COVID-19 Surveillance Team |
| EPI_ISL_2651575 | Lighthouse Lab in Milton Keynes | Wellcome Sanger Institute for the COVID-19 Genomics UK (COG-UK) Consortium | The Lighthouse Lab in Milton Keynes and Alex Alderton, Roberto Amato, Jeffrey Barrett, Sonia Goncalves, Ewan Harrison, David K. Jackson, Ian Johnston, Dominic Kwiatkowski, Cordelia Langford, John Sillitoe on behalf of the Wellcome Sanger Institute COVID-19 Surveillance Team |
| EPI_ISL_2651576, EPI_ISL_2651577 | Lighthouse Lab in Glasgow | Wellcome Sanger Institute for the COVID-19 Genomics UK (COG-UK) Consortium | Harper VanSteenhouse, Yumi Kasai, David Gray, Carol Clugston, Anna Dominiczak and Alex Alderton, Roberto Amato, Jeffrey Barrett, Sonia Goncalves, Ewan Harrison, David K. Jackson, Ian Johnston, Dominic Kwiatkowski, Cordelia Langford, John Sillitoe on behalf of the Wellcome Sanger Institute COVID-19 Surveillance Team |
| EPI_ISL_2651578, EPI_ISL_2651579 | Lighthouse Laboratory Plymouth | Wellcome Sanger Institute for the COVID-19 Genomics UK (COG-UK) Consortium | Lighthouse Laboratory Plymouth and Alex Alderton, Roberto Amato, Jeffrey Barrett, Sonia Goncalves, Ewan Harrison, David K. Jackson, Ian Johnston, Dominic Kwiatkowski, Cordelia Langford, John Sillitoe on behalf of the Wellcome Sanger Institute COVID-19 Surveillance Team |
| EPI_ISL_2651580 | Lighthouse Lab in Milton Keynes | Wellcome Sanger Institute for the COVID-19 Genomics UK (COG-UK) Consortium | The Lighthouse Lab in Milton Keynes and Alex Alderton, Roberto Amato, Jeffrey Barrett, Sonia Goncalves, Ewan Harrison, David K. Jackson, Ian Johnston, Dominic Kwiatkowski, Cordelia Langford, John Sillitoe on behalf of the Wellcome Sanger Institute COVID-19 Surveillance Team |
| EPI_ISL_2651581 | Lighthouse Laboratory Plymouth | Wellcome Sanger Institute for the COVID-19 Genomics UK (COG-UK) Consortium | Lighthouse Laboratory Plymouth and Alex Alderton, Roberto Amato, Jeffrey Barrett, Sonia Goncalves, Ewan Harrison, David K. Jackson, Ian Johnston, Dominic Kwiatkowski, Cordelia Langford, John Sillitoe on behalf of the Wellcome Sanger Institute COVID-19 Surveillance Team |
| EPI_ISL_2651582 | Lighthouse Lab in Glasgow | Wellcome Sanger Institute for the COVID-19 Genomics UK (COG-UK) Consortium | Harper VanSteenhouse, Yumi Kasai, David Gray, Carol Clugston, Anna Dominiczak and Alex Alderton, Roberto Amato, Jeffrey Barrett, Sonia Goncalves, Ewan Harrison, David K. Jackson, Ian Johnston, Dominic Kwiatkowski, Cordelia Langford, John Sillitoe on behalf of the Wellcome Sanger Institute COVID-19 Surveillance Team |
| EPI_ISL_2651583, EPI_ISL_2651584, EPI_ISL_2651585, EPI_ISL_2651586, EPI_ISL_2651587 | Lighthouse Laboratory Plymouth | Wellcome Sanger Institute for the COVID-19 Genomics UK (COG-UK) Consortium | Lighthouse Laboratory Plymouth and Alex Alderton, Roberto Amato, Jeffrey Barrett, Sonia Goncalves, Ewan Harrison, David K. Jackson, Ian Johnston, Dominic Kwiatkowski, Cordelia Langford, John Sillitoe on behalf of the Wellcome Sanger Institute COVID-19 Surveillance Team |
| EPI_ISL_2651588 | Lighthouse Lab in Glasgow | Wellcome Sanger Institute for the COVID-19 Genomics UK (COG-UK) Consortium | Harper VanSteenhouse, Yumi Kasai, David Gray, Carol Clugston, Anna Dominiczak and Alex Alderton, Roberto Amato, Jeffrey Barrett, Sonia Goncalves, Ewan Harrison, David K. Jackson, Ian Johnston, Dominic Kwiatkowski, Cordelia Langford, John Sillitoe on behalf of the Wellcome Sanger Institute COVID-19 Surveillance Team |
| EPI_ISL_2651589, EPI_ISL_2651590, EPI_ISL_2651591 | Lighthouse Lab in Milton Keynes | Wellcome Sanger Institute for the COVID-19 Genomics UK (COG-UK) Consortium | The Lighthouse Lab in Milton Keynes and Alex Alderton, Roberto Amato, Jeffrey Barrett, Sonia Goncalves, Ewan Harrison, David K. Jackson, Ian Johnston, Dominic Kwiatkowski, Cordelia Langford, John Sillitoe on behalf of the Wellcome Sanger Institute COVID-19 Surveillance Team |
| EPI_ISL_2651592 | Lighthouse Lab in Glasgow | Wellcome Sanger Institute for the COVID-19 Genomics UK (COG-UK) Consortium | Harper VanSteenhouse, Yumi Kasai, David Gray, Carol Clugston, Anna Dominiczak and Alex Alderton, Roberto Amato, Jeffrey Barrett, Sonia Goncalves, Ewan Harrison, David K. Jackson, Ian Johnston, Dominic Kwiatkowski, Cordelia Langford, John Sillitoe on behalf of the Wellcome Sanger Institute COVID-19 Surveillance Team |
| EPI_ISL_2651593 | Lighthouse Lab in Milton Keynes | Wellcome Sanger Institute for the COVID-19 Genomics UK (COG-UK) Consortium | The Lighthouse Lab in Milton Keynes and Alex Alderton, Roberto Amato, Jeffrey Barrett, Sonia Goncalves, Ewan Harrison, David K. Jackson, Ian Johnston, Dominic Kwiatkowski, Cordelia Langford, John Sillitoe on behalf of the Wellcome Sanger Institute COVID-19 Surveillance Team |
| EPI_ISL_2651594, EPI_ISL_2651595 | Lighthouse Lab in Glasgow | Wellcome Sanger Institute for the COVID-19 Genomics UK (COG-UK) Consortium | Harper VanSteenhouse, Yumi Kasai, David Gray, Carol Clugston, Anna Dominiczak and Alex Alderton, Roberto Amato, Jeffrey Barrett, Sonia Goncalves, Ewan Harrison, David K. Jackson, Ian Johnston, Dominic Kwiatkowski, Cordelia Langford, John Sillitoe on behalf of the Wellcome Sanger Institute COVID-19 Surveillance Team |
| EPI_ISL_2651596 | Lighthouse Lab in Milton Keynes | Wellcome Sanger Institute for the COVID-19 Genomics UK (COG-UK) Consortium | The Lighthouse Lab in Milton Keynes and Alex Alderton, Roberto Amato, Jeffrey Barrett, Sonia Goncalves, Ewan Harrison, David K. Jackson, Ian Johnston, Dominic Kwiatkowski, Cordelia Langford, John Sillitoe on behalf of the Wellcome Sanger Institute COVID-19 Surveillance Team |
| EPI_ISL_2651597 | Lighthouse Laboratory Plymouth | Wellcome Sanger Institute for the COVID-19 Genomics UK (COG-UK) Consortium | Lighthouse Laboratory Plymouth and Alex Alderton, Roberto Amato, Jeffrey Barrett, Sonia Goncalves, Ewan Harrison, David K. Jackson, Ian Johnston, Dominic Kwiatkowski, Cordelia Langford, John Sillitoe on behalf of the Wellcome Sanger Institute COVID-19 Surveillance Team |
| EPI_ISL_2651598 | Lighthouse Lab in Glasgow | Wellcome Sanger Institute for the COVID-19 Genomics UK (COG-UK) Consortium | Harper VanSteenhouse, Yumi Kasai, David Gray, Carol Clugston, Anna Dominiczak and Alex Alderton, Roberto Amato, Jeffrey Barrett, Sonia Goncalves, Ewan Harrison, David K. Jackson, Ian Johnston, Dominic Kwiatkowski, Cordelia Langford, John Sillitoe on behalf of the Wellcome Sanger Institute COVID-19 Surveillance Team |
| EPI_ISL_2651599 | Lighthouse Laboratory Plymouth | Wellcome Sanger Institute for the COVID-19 Genomics UK (COG-UK) | Lighthouse Laboratory Plymouth and Alex Alderton, Roberto Amato, Jeffrey Barrett, Sonia Goncalves, Ewan Harrison, David K. Jackson, Ian Johnston, Dominic Kwiatkowski, Cordelia Langford, John Sillitoe on behalf of the Wellcome Sanger |

| | | | |
|---|---|---|---|
| | | Consortium | Institute COVID-19 Surveillance Team |
| EPI_ISL_2651600 | Lighthouse Lab in Milton Keynes | Wellcome Sanger Institute for the COVID-19 Genomics UK (COG-UK) Consortium | The Lighthouse Lab in Milton Keynes and Alex Alderton, Roberto Amato, Jeffrey Barrett, Sonia Goncalves, Ewan Harrison, David K. Jackson, Ian Johnston, Dominic Kwiatkowski, Cordelia Langford, John Sillitoe on behalf of the Wellcome Sanger Institute COVID-19 Surveillance Team |
| EPI_ISL_2651601, EPI_ISL_2651602, EPI_ISL_2651603, EPI_ISL_2651604 | Lighthouse Laboratory Plymouth | Wellcome Sanger Institute for the COVID-19 Genomics UK (COG-UK) Consortium | Lighthouse Laboratory Plymouth and Alex Alderton, Roberto Amato, Jeffrey Barrett, Sonia Goncalves, Ewan Harrison, David K. Jackson, Ian Johnston, Dominic Kwiatkowski, Cordelia Langford, John Sillitoe on behalf of the Wellcome Sanger Institute COVID-19 Surveillance Team |
| EPI_ISL_2651605 | Lighthouse Lab in Glasgow | Wellcome Sanger Institute for the COVID-19 Genomics UK (COG-UK) Consortium | Harper VanSteenhouse, Yumi Kasai, David Gray, Carol Clugston, Anna Dominiczak and Alex Alderton, Roberto Amato, Jeffrey Barrett, Sonia Goncalves, Ewan Harrison, David K. Jackson, Ian Johnston, Dominic Kwiatkowski, Cordelia Langford, John Sillitoe on behalf of the Wellcome Sanger Institute COVID-19 Surveillance Team |
| EPI_ISL_2651608 | Lighthouse Lab in Milton Keynes | Wellcome Sanger Institute for the COVID-19 Genomics UK (COG-UK) Consortium | The Lighthouse Lab in Milton Keynes and Alex Alderton, Roberto Amato, Jeffrey Barrett, Sonia Goncalves, Ewan Harrison, David K. Jackson, Ian Johnston, Dominic Kwiatkowski, Cordelia Langford, John Sillitoe on behalf of the Wellcome Sanger Institute COVID-19 Surveillance Team |
| EPI_ISL_2651609 | Lighthouse Lab in Glasgow | Wellcome Sanger Institute for the COVID-19 Genomics UK (COG-UK) Consortium | Harper VanSteenhouse, Yumi Kasai, David Gray, Carol Clugston, Anna Dominiczak and Alex Alderton, Roberto Amato, Jeffrey Barrett, Sonia Goncalves, Ewan Harrison, David K. Jackson, Ian Johnston, Dominic Kwiatkowski, Cordelia Langford, John Sillitoe on behalf of the Wellcome Sanger Institute COVID-19 Surveillance Team |
| EPI_ISL_2651610 | Lighthouse Laboratory Plymouth | Wellcome Sanger Institute for the COVID-19 Genomics UK (COG-UK) Consortium | Lighthouse Laboratory Plymouth and Alex Alderton, Roberto Amato, Jeffrey Barrett, Sonia Goncalves, Ewan Harrison, David K. Jackson, Ian Johnston, Dominic Kwiatkowski, Cordelia Langford, John Sillitoe on behalf of the Wellcome Sanger Institute COVID-19 Surveillance Team |
| EPI_ISL_2651611 | Lighthouse Lab in Glasgow | Wellcome Sanger Institute for the COVID-19 Genomics UK (COG-UK) Consortium | Harper VanSteenhouse, Yumi Kasai, David Gray, Carol Clugston, Anna Dominiczak and Alex Alderton, Roberto Amato, Jeffrey Barrett, Sonia Goncalves, Ewan Harrison, David K. Jackson, Ian Johnston, Dominic Kwiatkowski, Cordelia Langford, John Sillitoe on behalf of the Wellcome Sanger Institute COVID-19 Surveillance Team |
| EPI_ISL_2651612 | Lighthouse Laboratory Plymouth | Wellcome Sanger Institute for the COVID-19 Genomics UK (COG-UK) Consortium | Lighthouse Laboratory Plymouth and Alex Alderton, Roberto Amato, Jeffrey Barrett, Sonia Goncalves, Ewan Harrison, David K. Jackson, Ian Johnston, Dominic Kwiatkowski, Cordelia Langford, John Sillitoe on behalf of the Wellcome Sanger Institute COVID-19 Surveillance Team |
| EPI_ISL_2651613 | Lighthouse Lab in Glasgow | Wellcome Sanger Institute for the COVID-19 Genomics UK (COG-UK) Consortium | Harper VanSteenhouse, Yumi Kasai, David Gray, Carol Clugston, Anna Dominiczak and Alex Alderton, Roberto Amato, Jeffrey Barrett, Sonia Goncalves, Ewan Harrison, David K. Jackson, Ian Johnston, Dominic Kwiatkowski, Cordelia Langford, John Sillitoe on behalf of the Wellcome Sanger Institute COVID-19 Surveillance Team |
| EPI_ISL_2651614 | Lighthouse Lab in Milton Keynes | Wellcome Sanger Institute for the COVID-19 Genomics UK (COG-UK) Consortium | The Lighthouse Lab in Milton Keynes and Alex Alderton, Roberto Amato, Jeffrey Barrett, Sonia Goncalves, Ewan Harrison, David K. Jackson, Ian Johnston, Dominic Kwiatkowski, Cordelia Langford, John Sillitoe on behalf of the Wellcome Sanger Institute COVID-19 Surveillance Team |
| EPI_ISL_2651615 | Lighthouse Lab in Glasgow | Wellcome Sanger Institute for the COVID-19 Genomics UK (COG-UK) Consortium | Harper VanSteenhouse, Yumi Kasai, David Gray, Carol Clugston, Anna Dominiczak and Alex Alderton, Roberto Amato, Jeffrey Barrett, Sonia Goncalves, Ewan Harrison, David K. Jackson, Ian Johnston, Dominic Kwiatkowski, Cordelia Langford, John Sillitoe on behalf of the Wellcome Sanger Institute COVID-19 Surveillance Team |
| EPI_ISL_2651616 | Lighthouse Lab in Milton Keynes | Wellcome Sanger Institute for the COVID-19 Genomics UK (COG-UK) Consortium | The Lighthouse Lab in Milton Keynes and Alex Alderton, Roberto Amato, Jeffrey Barrett, Sonia Goncalves, Ewan Harrison, David K. Jackson, Ian Johnston, Dominic Kwiatkowski, Cordelia Langford, John Sillitoe on behalf of the Wellcome Sanger Institute COVID-19 Surveillance Team |
| EPI_ISL_2651617, EPI_ISL_2651618 | Lighthouse Laboratory Plymouth | Wellcome Sanger Institute for the COVID-19 Genomics UK (COG-UK) Consortium | Lighthouse Laboratory Plymouth and Alex Alderton, Roberto Amato, Jeffrey Barrett, Sonia Goncalves, Ewan Harrison, David K. Jackson, Ian Johnston, Dominic Kwiatkowski, Cordelia Langford, John Sillitoe on behalf of the Wellcome Sanger Institute COVID-19 Surveillance Team |
| EPI_ISL_2651619, EPI_ISL_2651620 | Lighthouse Lab in Milton Keynes | Wellcome Sanger Institute for the COVID-19 Genomics UK (COG-UK) Consortium | The Lighthouse Lab in Milton Keynes and Alex Alderton, Roberto Amato, Jeffrey Barrett, Sonia Goncalves, Ewan Harrison, David K. Jackson, Ian Johnston, Dominic Kwiatkowski, Cordelia Langford, John Sillitoe on behalf of the Wellcome Sanger Institute COVID-19 Surveillance Team |
| EPI_ISL_2651621 | Lighthouse Laboratory Plymouth | Wellcome Sanger Institute for the COVID-19 Genomics UK (COG-UK) Consortium | Lighthouse Laboratory Plymouth and Alex Alderton, Roberto Amato, Jeffrey Barrett, Sonia Goncalves, Ewan Harrison, David K. Jackson, Ian Johnston, Dominic Kwiatkowski, Cordelia Langford, John Sillitoe on behalf of the Wellcome Sanger Institute COVID-19 Surveillance Team |
| EPI_ISL_2651622 | Lighthouse Lab in Milton Keynes | Wellcome Sanger Institute for the COVID-19 Genomics UK (COG-UK) Consortium | The Lighthouse Lab in Milton Keynes and Alex Alderton, Roberto Amato, Jeffrey Barrett, Sonia Goncalves, Ewan Harrison, David K. Jackson, Ian Johnston, Dominic Kwiatkowski, Cordelia Langford, John Sillitoe on behalf of the Wellcome Sanger Institute COVID-19 Surveillance Team |
| EPI_ISL_2651623 | Lighthouse Lab in Glasgow | Wellcome Sanger Institute for the COVID-19 Genomics UK (COG-UK) Consortium | Harper VanSteenhouse, Yumi Kasai, David Gray, Carol Clugston, Anna Dominiczak and Alex Alderton, Roberto Amato, Jeffrey Barrett, Sonia Goncalves, Ewan Harrison, David K. Jackson, Ian Johnston, Dominic Kwiatkowski, Cordelia Langford, John Sillitoe on behalf of the Wellcome Sanger Institute COVID-19 Surveillance Team |
| EPI_ISL_2651624, EPI_ISL_2651625 | Lighthouse Laboratory Plymouth | Wellcome Sanger Institute for the COVID-19 Genomics UK (COG-UK) Consortium | Lighthouse Laboratory Plymouth and Alex Alderton, Roberto Amato, Jeffrey Barrett, Sonia Goncalves, Ewan Harrison, David K. Jackson, Ian Johnston, Dominic Kwiatkowski, Cordelia Langford, John Sillitoe on behalf of the Wellcome Sanger Institute COVID-19 Surveillance Team |
| EPI_ISL_2651626 | Lighthouse Lab in Milton Keynes | Wellcome Sanger Institute for the COVID-19 Genomics UK (COG-UK) Consortium | The Lighthouse Lab in Milton Keynes and Alex Alderton, Roberto Amato, Jeffrey Barrett, Sonia Goncalves, Ewan Harrison, David K. Jackson, Ian Johnston, Dominic Kwiatkowski, Cordelia Langford, John Sillitoe on behalf of the Wellcome Sanger Institute COVID-19 Surveillance Team |
| EPI_ISL_2651627 | Lighthouse Lab in Glasgow | Wellcome Sanger Institute for the COVID-19 Genomics UK (COG-UK) Consortium | Harper VanSteenhouse, Yumi Kasai, David Gray, Carol Clugston, Anna Dominiczak and Alex Alderton, Roberto Amato, Jeffrey Barrett, Sonia Goncalves, Ewan Harrison, David K. Jackson, Ian Johnston, Dominic Kwiatkowski, Cordelia Langford, John Sillitoe on behalf of the Wellcome Sanger Institute COVID-19 Surveillance Team |
| EPI_ISL_2651628 | Lighthouse Lab in Milton Keynes | Wellcome Sanger Institute for the COVID-19 Genomics UK (COG-UK) Consortium | The Lighthouse Lab in Milton Keynes and Alex Alderton, Roberto Amato, Jeffrey Barrett, Sonia Goncalves, Ewan Harrison, David K. Jackson, Ian Johnston, Dominic Kwiatkowski, Cordelia Langford, John Sillitoe on behalf of the Wellcome Sanger Institute COVID-19 Surveillance Team |
| EPI_ISL_2651629, EPI_ISL_2651630 | Lighthouse Laboratory Plymouth | Wellcome Sanger Institute for the COVID-19 Genomics UK (COG-UK) Consortium | Lighthouse Laboratory Plymouth and Alex Alderton, Roberto Amato, Jeffrey Barrett, Sonia Goncalves, Ewan Harrison, David K. Jackson, Ian Johnston, Dominic Kwiatkowski, Cordelia Langford, John Sillitoe on behalf of the Wellcome Sanger Institute COVID-19 Surveillance Team |
| EPI_ISL_2651631 | Lighthouse Lab in Milton Keynes | Wellcome Sanger Institute for the COVID-19 Genomics UK (COG-UK) Consortium | The Lighthouse Lab in Milton Keynes and Alex Alderton, Roberto Amato, Jeffrey Barrett, Sonia Goncalves, Ewan Harrison, David K. Jackson, Ian Johnston, Dominic Kwiatkowski, Cordelia Langford, John Sillitoe on behalf of the Wellcome Sanger Institute COVID-19 Surveillance Team |
| EPI_ISL_2651632 | Lighthouse Lab in Glasgow | Wellcome Sanger Institute for the COVID-19 Genomics UK (COG-UK) Consortium | Harper VanSteenhouse, Yumi Kasai, David Gray, Carol Clugston, Anna Dominiczak and Alex Alderton, Roberto Amato, Jeffrey Barrett, Sonia Goncalves, Ewan Harrison, David K. Jackson, Ian Johnston, Dominic Kwiatkowski, Cordelia Langford, John Sillitoe on behalf of the Wellcome Sanger Institute COVID-19 Surveillance Team |
| EPI_ISL_2651633 | Lighthouse Laboratory Plymouth | Wellcome Sanger Institute for the COVID-19 Genomics UK (COG-UK) Consortium | Lighthouse Laboratory Plymouth and Alex Alderton, Roberto Amato, Jeffrey Barrett, Sonia Goncalves, Ewan Harrison, David K. Jackson, Ian Johnston, Dominic Kwiatkowski, Cordelia Langford, John Sillitoe on behalf of the Wellcome Sanger Institute COVID-19 Surveillance Team |
| EPI_ISL_2651634 | Lighthouse Lab in Glasgow | Wellcome Sanger Institute for the COVID-19 Genomics UK (COG-UK) Consortium | Harper VanSteenhouse, Yumi Kasai, David Gray, Carol Clugston, Anna Dominiczak and Alex Alderton, Roberto Amato, Jeffrey Barrett, Sonia Goncalves, Ewan Harrison, David K. Jackson, Ian Johnston, Dominic Kwiatkowski, Cordelia Langford, John Sillitoe on behalf of the Wellcome Sanger Institute COVID-19 Surveillance Team |
| EPI_ISL_2651635 | Lighthouse Laboratory Plymouth | Wellcome Sanger Institute for the COVID-19 Genomics UK (COG-UK) Consortium | Lighthouse Laboratory Plymouth and Alex Alderton, Roberto Amato, Jeffrey Barrett, Sonia Goncalves, Ewan Harrison, David K. Jackson, Ian Johnston, Dominic Kwiatkowski, Cordelia Langford, John Sillitoe on behalf of the Wellcome Sanger Institute COVID-19 Surveillance Team |
| EPI_ISL_2651636 | Lighthouse Lab in Milton Keynes | Wellcome Sanger Institute for the COVID-19 Genomics UK (COG-UK) Consortium | The Lighthouse Lab in Milton Keynes and Alex Alderton, Roberto Amato, Jeffrey Barrett, Sonia Goncalves, Ewan Harrison, David K. Jackson, Ian Johnston, Dominic Kwiatkowski, Cordelia Langford, John Sillitoe on behalf of the Wellcome Sanger Institute COVID-19 Surveillance Team |
| EPI_ISL_2651637 | Lighthouse Laboratory Plymouth | Wellcome Sanger Institute for the COVID-19 Genomics UK (COG-UK) Consortium | Lighthouse Laboratory Plymouth and Alex Alderton, Roberto Amato, Jeffrey Barrett, Sonia Goncalves, Ewan Harrison, David K. Jackson, Ian Johnston, Dominic Kwiatkowski, Cordelia Langford, John Sillitoe on behalf of the Wellcome Sanger Institute COVID-19 Surveillance Team |
| EPI_ISL_2651638 | Lighthouse Lab in Glasgow | Wellcome Sanger Institute for the COVID-19 Genomics UK (COG-UK) Consortium | Harper VanSteenhouse, Yumi Kasai, David Gray, Carol Clugston, Anna Dominiczak and Alex Alderton, Roberto Amato, Jeffrey Barrett, Sonia Goncalves, Ewan Harrison, David K. Jackson, Ian Johnston, Dominic Kwiatkowski, Cordelia Langford, John Sillitoe on behalf of the Wellcome Sanger Institute COVID-19 Surveillance Team |
| EPI_ISL_2651639 | Lighthouse Laboratory Plymouth | Wellcome Sanger Institute for the COVID-19 Genomics UK (COG-UK) Consortium | Lighthouse Laboratory Plymouth and Alex Alderton, Roberto Amato, Jeffrey Barrett, Sonia Goncalves, Ewan Harrison, David K. Jackson, Ian Johnston, Dominic Kwiatkowski, Cordelia Langford, John Sillitoe on behalf of the Wellcome Sanger Institute COVID-19 Surveillance Team |
| EPI_ISL_2651640 | Lighthouse Lab in Glasgow | Wellcome Sanger Institute for the COVID-19 Genomics UK (COG-UK) Consortium | Harper VanSteenhouse, Yumi Kasai, David Gray, Carol Clugston, Anna Dominiczak and Alex Alderton, Roberto Amato, Jeffrey Barrett, Sonia Goncalves, Ewan Harrison, David K. Jackson, Ian Johnston, Dominic Kwiatkowski, Cordelia Langford, John Sillitoe on behalf of the Wellcome Sanger Institute COVID-19 Surveillance Team |
| EPI_ISL_2651641 | Lighthouse Lab in Alderley Park | Wellcome Sanger Institute for the COVID-19 Genomics UK (COG-UK) Consortium | Jacquelyn Wynn, Mairead Hyland, The Lighthouse Lab in Alderley Park and Alex Alderton, Roberto Amato, Jeffrey Barrett, Sonia Goncalves, Ewan Harrison, David K. Jackson, Ian Johnston, Dominic Kwiatkowski, Cordelia Langford, John Sillitoe on behalf of the Wellcome Sanger Institute COVID-19 Surveillance Team |
| EPI_ISL_2651642, EPI_ISL_2651643 | Lighthouse Lab in Glasgow | Wellcome Sanger Institute for the COVID-19 Genomics UK (COG-UK) Consortium | Harper VanSteenhouse, Yumi Kasai, David Gray, Carol Clugston, Anna Dominiczak and Alex Alderton, Roberto Amato, Jeffrey Barrett, Sonia Goncalves, Ewan Harrison, David K. Jackson, Ian Johnston, Dominic Kwiatkowski, Cordelia Langford, John Sillitoe on behalf of the Wellcome Sanger Institute COVID-19 Surveillance Team |
| EPI_ISL_2651644, EPI_ISL_2651645, EPI_ISL_2651646, EPI_ISL_2651647, EPI_ISL_2651648, EPI_ISL_2651649 | Lighthouse Lab in Alderley Park | Wellcome Sanger Institute for the COVID-19 Genomics UK (COG-UK) Consortium | Jacquelyn Wynn, Mairead Hyland, The Lighthouse Lab in Alderley Park and Alex Alderton, Roberto Amato, Jeffrey Barrett, Sonia Goncalves, Ewan Harrison, David K. Jackson, Ian Johnston, Dominic Kwiatkowski, Cordelia Langford, John Sillitoe on behalf of the Wellcome Sanger Institute COVID-19 Surveillance Team |
| EPI_ISL_2651650, EPI_ISL_2651651, EPI_ISL_2651652, EPI_ISL_2651653 | Lighthouse Lab in Glasgow | Wellcome Sanger Institute for the COVID-19 Genomics UK (COG-UK) Consortium | Harper VanSteenhouse, Yumi Kasai, David Gray, Carol Clugston, Anna Dominiczak and Alex Alderton, Roberto Amato, Jeffrey Barrett, Sonia Goncalves, Ewan Harrison, David K. Jackson, Ian Johnston, Dominic Kwiatkowski, Cordelia Langford, John Sillitoe on behalf of the Wellcome Sanger Institute COVID-19 Surveillance Team |
| EPI_ISL_2651654 | Lighthouse Lab in Alderley Park | Wellcome Sanger Institute for the COVID-19 Genomics UK (COG-UK) Consortium | Jacquelyn Wynn, Mairead Hyland, The Lighthouse Lab in Alderley Park and Alex Alderton, Roberto Amato, Jeffrey Barrett, Sonia Goncalves, Ewan Harrison, David K. Jackson, Ian Johnston, Dominic Kwiatkowski, Cordelia Langford, John Sillitoe on behalf of the Wellcome Sanger Institute COVID-19 Surveillance Team |
| EPI_ISL_2651656, EPI_ISL_2651657, EPI_ISL_2651658 | Lighthouse Lab in Glasgow | Wellcome Sanger Institute for the COVID-19 Genomics UK (COG-UK) Consortium | Harper VanSteenhouse, Yumi Kasai, David Gray, Carol Clugston, Anna Dominiczak and Alex Alderton, Roberto Amato, Jeffrey Barrett, Sonia Goncalves, Ewan Harrison, David K. Jackson, Ian Johnston, Dominic Kwiatkowski, Cordelia Langford, John Sillitoe on behalf of the Wellcome Sanger Institute COVID-19 Surveillance Team |
| EPI_ISL_2651659 | Lighthouse Lab in Alderley Park | Wellcome Sanger Institute for the COVID-19 Genomics UK (COG-UK) Consortium | Jacquelyn Wynn, Mairead Hyland, The Lighthouse Lab in Alderley Park and Alex Alderton, Roberto Amato, Jeffrey Barrett, Sonia Goncalves, Ewan Harrison, David K. Jackson, Ian Johnston, Dominic Kwiatkowski, Cordelia Langford, John Sillitoe on behalf of the Wellcome Sanger Institute COVID-19 Surveillance Team |
| EPI_ISL_2651660 | Lighthouse Lab in Milton Keynes | Wellcome Sanger Institute for the COVID-19 Genomics UK (COG-UK) Consortium | The Lighthouse Lab in Milton Keynes and Alex Alderton, Roberto Amato, Jeffrey Barrett, Sonia Goncalves, Ewan Harrison, David K. Jackson, Ian Johnston, Dominic Kwiatkowski, Cordelia Langford, John Sillitoe on behalf of the Wellcome Sanger Institute COVID-19 Surveillance Team |
| EPI_ISL_2651661 | Lighthouse Lab in Glasgow | Wellcome Sanger Institute for the COVID-19 Genomics UK (COG-UK) Consortium | Harper VanSteenhouse, Yumi Kasai, David Gray, Carol Clugston, Anna Dominiczak and Alex Alderton, Roberto Amato, Jeffrey Barrett, Sonia Goncalves, Ewan Harrison, David K. Jackson, Ian Johnston, Dominic Kwiatkowski, Cordelia Langford, John Sillitoe on behalf of the Wellcome Sanger Institute COVID-19 Surveillance Team |

| | | | |
|---|---|---|---|
| EPI_ISL_2651662 | Lighthouse Lab in Alderley Park | Wellcome Sanger Institute for the COVID-19 Genomics UK (COG-UK) Consortium | Jacquelyn Wynn, Mairead Hyland, The Lighthouse Lab in Alderley Park and Alex Alderton, Roberto Amato, Jeffrey Barrett, Sonia Goncalves, Ewan Harrison, David K. Jackson, Ian Johnston, Dominic Kwiatkowski, Cordelia Langford, John Sillitoe on behalf of the Wellcome Sanger Institute COVID-19 Surveillance Team |
| EPI_ISL_2651663, EPI_ISL_2651664 | Lighthouse Lab in Glasgow | Wellcome Sanger Institute for the COVID-19 Genomics UK (COG-UK) Consortium | Harper VanSteenhouse, Yumi Kasai, David Gray, Carol Clugston, Anna Dominiczak and Alex Alderton, Roberto Amato, Jeffrey Barrett, Sonia Goncalves, Ewan Harrison, David K. Jackson, Ian Johnston, Dominic Kwiatkowski, Cordelia Langford, John Sillitoe on behalf of the Wellcome Sanger Institute COVID-19 Surveillance Team |
| EPI_ISL_2651665, EPI_ISL_2651666 | Lighthouse Lab in Alderley Park | Wellcome Sanger Institute for the COVID-19 Genomics UK (COG-UK) Consortium | Jacquelyn Wynn, Mairead Hyland, The Lighthouse Lab in Alderley Park and Alex Alderton, Roberto Amato, Jeffrey Barrett, Sonia Goncalves, Ewan Harrison, David K. Jackson, Ian Johnston, Dominic Kwiatkowski, Cordelia Langford, John Sillitoe on behalf of the Wellcome Sanger Institute COVID-19 Surveillance Team |
| EPI_ISL_2651667, EPI_ISL_2651668 | Lighthouse Lab in Glasgow | Wellcome Sanger Institute for the COVID-19 Genomics UK (COG-UK) Consortium | Harper VanSteenhouse, Yumi Kasai, David Gray, Carol Clugston, Anna Dominiczak and Alex Alderton, Roberto Amato, Jeffrey Barrett, Sonia Goncalves, Ewan Harrison, David K. Jackson, Ian Johnston, Dominic Kwiatkowski, Cordelia Langford, John Sillitoe on behalf of the Wellcome Sanger Institute COVID-19 Surveillance Team |
| EPI_ISL_2651669, EPI_ISL_2651671 | Lighthouse Lab in Alderley Park | Wellcome Sanger Institute for the COVID-19 Genomics UK (COG-UK) Consortium | Jacquelyn Wynn, Mairead Hyland, The Lighthouse Lab in Alderley Park and Alex Alderton, Roberto Amato, Jeffrey Barrett, Sonia Goncalves, Ewan Harrison, David K. Jackson, Ian Johnston, Dominic Kwiatkowski, Cordelia Langford, John Sillitoe on behalf of the Wellcome Sanger Institute COVID-19 Surveillance Team |
| EPI_ISL_2651672, EPI_ISL_2651673, EPI_ISL_2651674 | Lighthouse Lab in Glasgow | Wellcome Sanger Institute for the COVID-19 Genomics UK (COG-UK) Consortium | Harper VanSteenhouse, Yumi Kasai, David Gray, Carol Clugston, Anna Dominiczak and Alex Alderton, Roberto Amato, Jeffrey Barrett, Sonia Goncalves, Ewan Harrison, David K. Jackson, Ian Johnston, Dominic Kwiatkowski, Cordelia Langford, John Sillitoe on behalf of the Wellcome Sanger Institute COVID-19 Surveillance Team |
| EPI_ISL_2651675, EPI_ISL_2651676 | Lighthouse Lab in Alderley Park | Wellcome Sanger Institute for the COVID-19 Genomics UK (COG-UK) Consortium | Jacquelyn Wynn, Mairead Hyland, The Lighthouse Lab in Alderley Park and Alex Alderton, Roberto Amato, Jeffrey Barrett, Sonia Goncalves, Ewan Harrison, David K. Jackson, Ian Johnston, Dominic Kwiatkowski, Cordelia Langford, John Sillitoe on behalf of the Wellcome Sanger Institute COVID-19 Surveillance Team |
| EPI_ISL_2651677 | Lighthouse Lab in Glasgow | Wellcome Sanger Institute for the COVID-19 Genomics UK (COG-UK) Consortium | Harper VanSteenhouse, Yumi Kasai, David Gray, Carol Clugston, Anna Dominiczak and Alex Alderton, Roberto Amato, Jeffrey Barrett, Sonia Goncalves, Ewan Harrison, David K. Jackson, Ian Johnston, Dominic Kwiatkowski, Cordelia Langford, John Sillitoe on behalf of the Wellcome Sanger Institute COVID-19 Surveillance Team |
| EPI_ISL_2651678, EPI_ISL_2651680, EPI_ISL_2651681 | Lighthouse Lab in Alderley Park | Wellcome Sanger Institute for the COVID-19 Genomics UK (COG-UK) Consortium | Jacquelyn Wynn, Mairead Hyland, The Lighthouse Lab in Alderley Park and Alex Alderton, Roberto Amato, Jeffrey Barrett, Sonia Goncalves, Ewan Harrison, David K. Jackson, Ian Johnston, Dominic Kwiatkowski, Cordelia Langford, John Sillitoe on behalf of the Wellcome Sanger Institute COVID-19 Surveillance Team |
| EPI_ISL_2651682 | Lighthouse Lab in Glasgow | Wellcome Sanger Institute for the COVID-19 Genomics UK (COG-UK) Consortium | Harper VanSteenhouse, Yumi Kasai, David Gray, Carol Clugston, Anna Dominiczak and Alex Alderton, Roberto Amato, Jeffrey Barrett, Sonia Goncalves, Ewan Harrison, David K. Jackson, Ian Johnston, Dominic Kwiatkowski, Cordelia Langford, John Sillitoe on behalf of the Wellcome Sanger Institute COVID-19 Surveillance Team |
| EPI_ISL_2651683 | Lighthouse Lab in Alderley Park | Wellcome Sanger Institute for the COVID-19 Genomics UK (COG-UK) Consortium | Jacquelyn Wynn, Mairead Hyland, The Lighthouse Lab in Alderley Park and Alex Alderton, Roberto Amato, Jeffrey Barrett, Sonia Goncalves, Ewan Harrison, David K. Jackson, Ian Johnston, Dominic Kwiatkowski, Cordelia Langford, John Sillitoe on behalf of the Wellcome Sanger Institute COVID-19 Surveillance Team |
| EPI_ISL_2651684, EPI_ISL_2651685, EPI_ISL_2651686, EPI_ISL_2651687 | Lighthouse Lab in Glasgow | Wellcome Sanger Institute for the COVID-19 Genomics UK (COG-UK) Consortium | Harper VanSteenhouse, Yumi Kasai, David Gray, Carol Clugston, Anna Dominiczak and Alex Alderton, Roberto Amato, Jeffrey Barrett, Sonia Goncalves, Ewan Harrison, David K. Jackson, Ian Johnston, Dominic Kwiatkowski, Cordelia Langford, John Sillitoe on behalf of the Wellcome Sanger Institute COVID-19 Surveillance Team |
| EPI_ISL_2651688 | Lighthouse Lab in Alderley Park | Wellcome Sanger Institute for the COVID-19 Genomics UK (COG-UK) Consortium | Jacquelyn Wynn, Mairead Hyland, The Lighthouse Lab in Alderley Park and Alex Alderton, Roberto Amato, Jeffrey Barrett, Sonia Goncalves, Ewan Harrison, David K. Jackson, Ian Johnston, Dominic Kwiatkowski, Cordelia Langford, John Sillitoe on behalf of the Wellcome Sanger Institute COVID-19 Surveillance Team |
| EPI_ISL_2651689 | Lighthouse Lab in Glasgow | Wellcome Sanger Institute for the COVID-19 Genomics UK (COG-UK) Consortium | Harper VanSteenhouse, Yumi Kasai, David Gray, Carol Clugston, Anna Dominiczak and Alex Alderton, Roberto Amato, Jeffrey Barrett, Sonia Goncalves, Ewan Harrison, David K. Jackson, Ian Johnston, Dominic Kwiatkowski, Cordelia Langford, John Sillitoe on behalf of the Wellcome Sanger Institute COVID-19 Surveillance Team |
| EPI_ISL_2651690, EPI_ISL_2651691 | Lighthouse Lab in Alderley Park | Wellcome Sanger Institute for the COVID-19 Genomics UK (COG-UK) Consortium | Jacquelyn Wynn, Mairead Hyland, The Lighthouse Lab in Alderley Park and Alex Alderton, Roberto Amato, Jeffrey Barrett, Sonia Goncalves, Ewan Harrison, David K. Jackson, Ian Johnston, Dominic Kwiatkowski, Cordelia Langford, John Sillitoe on behalf of the Wellcome Sanger Institute COVID-19 Surveillance Team |
| EPI_ISL_2651692, EPI_ISL_2651693, EPI_ISL_2651694, EPI_ISL_2651695 | Lighthouse Lab in Glasgow | Wellcome Sanger Institute for the COVID-19 Genomics UK (COG-UK) Consortium | Harper VanSteenhouse, Yumi Kasai, David Gray, Carol Clugston, Anna Dominiczak and Alex Alderton, Roberto Amato, Jeffrey Barrett, Sonia Goncalves, Ewan Harrison, David K. Jackson, Ian Johnston, Dominic Kwiatkowski, Cordelia Langford, John Sillitoe on behalf of the Wellcome Sanger Institute COVID-19 Surveillance Team |
| EPI_ISL_2651696 | Lighthouse Lab in Alderley Park | Wellcome Sanger Institute for the COVID-19 Genomics UK (COG-UK) Consortium | Jacquelyn Wynn, Mairead Hyland, The Lighthouse Lab in Alderley Park and Alex Alderton, Roberto Amato, Jeffrey Barrett, Sonia Goncalves, Ewan Harrison, David K. Jackson, Ian Johnston, Dominic Kwiatkowski, Cordelia Langford, John Sillitoe on behalf of the Wellcome Sanger Institute COVID-19 Surveillance Team |
| EPI_ISL_2651697 | Lighthouse Lab in Glasgow | Wellcome Sanger Institute for the COVID-19 Genomics UK (COG-UK) Consortium | Harper VanSteenhouse, Yumi Kasai, David Gray, Carol Clugston, Anna Dominiczak and Alex Alderton, Roberto Amato, Jeffrey Barrett, Sonia Goncalves, Ewan Harrison, David K. Jackson, Ian Johnston, Dominic Kwiatkowski, Cordelia Langford, John Sillitoe on behalf of the Wellcome Sanger Institute COVID-19 Surveillance Team |
| EPI_ISL_2651698, EPI_ISL_2651699 | Lighthouse Lab in Alderley Park | Wellcome Sanger Institute for the COVID-19 Genomics UK (COG-UK) Consortium | Jacquelyn Wynn, Mairead Hyland, The Lighthouse Lab in Alderley Park and Alex Alderton, Roberto Amato, Jeffrey Barrett, Sonia Goncalves, Ewan Harrison, David K. Jackson, Ian Johnston, Dominic Kwiatkowski, Cordelia Langford, John Sillitoe on behalf of the Wellcome Sanger Institute COVID-19 Surveillance Team |
| EPI_ISL_2651700, EPI_ISL_2651701 | Lighthouse Lab in Glasgow | Wellcome Sanger Institute for the COVID-19 Genomics UK (COG-UK) Consortium | Harper VanSteenhouse, Yumi Kasai, David Gray, Carol Clugston, Anna Dominiczak and Alex Alderton, Roberto Amato, Jeffrey Barrett, Sonia Goncalves, Ewan Harrison, David K. Jackson, Ian Johnston, Dominic Kwiatkowski, Cordelia Langford, John Sillitoe on behalf of the Wellcome Sanger Institute COVID-19 Surveillance Team |
| EPI_ISL_2651702 | Lighthouse Lab in Alderley Park | Wellcome Sanger Institute for the COVID-19 Genomics UK (COG-UK) Consortium | Jacquelyn Wynn, Mairead Hyland, The Lighthouse Lab in Alderley Park and Alex Alderton, Roberto Amato, Jeffrey Barrett, Sonia Goncalves, Ewan Harrison, David K. Jackson, Ian Johnston, Dominic Kwiatkowski, Cordelia Langford, John Sillitoe on behalf of the Wellcome Sanger Institute COVID-19 Surveillance Team |
| EPI_ISL_2651703 | Lighthouse Lab in Glasgow | Wellcome Sanger Institute for the COVID-19 Genomics UK (COG-UK) Consortium | Harper VanSteenhouse, Yumi Kasai, David Gray, Carol Clugston, Anna Dominiczak and Alex Alderton, Roberto Amato, Jeffrey Barrett, Sonia Goncalves, Ewan Harrison, David K. Jackson, Ian Johnston, Dominic Kwiatkowski, Cordelia Langford, John Sillitoe on behalf of the Wellcome Sanger Institute COVID-19 Surveillance Team |
| EPI_ISL_2651704 | Lighthouse Lab in Alderley Park | Wellcome Sanger Institute for the COVID-19 Genomics UK (COG-UK) Consortium | Jacquelyn Wynn, Mairead Hyland, The Lighthouse Lab in Alderley Park and Alex Alderton, Roberto Amato, Jeffrey Barrett, Sonia Goncalves, Ewan Harrison, David K. Jackson, Ian Johnston, Dominic Kwiatkowski, Cordelia Langford, John Sillitoe on behalf of the Wellcome Sanger Institute COVID-19 Surveillance Team |
| EPI_ISL_2651705 | Lighthouse Lab in Milton Keynes | Wellcome Sanger Institute for the COVID-19 Genomics UK (COG-UK) Consortium | The Lighthouse Lab in Milton Keynes and Alex Alderton, Roberto Amato, Jeffrey Barrett, Sonia Goncalves, Ewan Harrison, David K. Jackson, Ian Johnston, Dominic Kwiatkowski, Cordelia Langford, John Sillitoe on behalf of the Wellcome Sanger Institute COVID-19 Surveillance Team |
| EPI_ISL_2651706 | Lighthouse Lab in Alderley Park | Wellcome Sanger Institute for the COVID-19 Genomics UK (COG-UK) Consortium | Jacquelyn Wynn, Mairead Hyland, The Lighthouse Lab in Alderley Park and Alex Alderton, Roberto Amato, Jeffrey Barrett, Sonia Goncalves, Ewan Harrison, David K. Jackson, Ian Johnston, Dominic Kwiatkowski, Cordelia Langford, John Sillitoe on behalf of the Wellcome Sanger Institute COVID-19 Surveillance Team |
| EPI_ISL_2651707, EPI_ISL_2651709, EPI_ISL_2651710, EPI_ISL_2651711, EPI_ISL_2651712, EPI_ISL_2651713, EPI_ISL_2651714 | Lighthouse Lab in Glasgow | Wellcome Sanger Institute for the COVID-19 Genomics UK (COG-UK) Consortium | Harper VanSteenhouse, Yumi Kasai, David Gray, Carol Clugston, Anna Dominiczak and Alex Alderton, Roberto Amato, Jeffrey Barrett, Sonia Goncalves, Ewan Harrison, David K. Jackson, Ian Johnston, Dominic Kwiatkowski, Cordelia Langford, John Sillitoe on behalf of the Wellcome Sanger Institute COVID-19 Surveillance Team |
| EPI_ISL_2651715, EPI_ISL_2651716 | Lighthouse Lab in Alderley Park | Wellcome Sanger Institute for the COVID-19 Genomics UK (COG-UK) Consortium | Jacquelyn Wynn, Mairead Hyland, The Lighthouse Lab in Alderley Park and Alex Alderton, Roberto Amato, Jeffrey Barrett, Sonia Goncalves, Ewan Harrison, David K. Jackson, Ian Johnston, Dominic Kwiatkowski, Cordelia Langford, John Sillitoe on behalf of the Wellcome Sanger Institute COVID-19 Surveillance Team |
| EPI_ISL_2651717, EPI_ISL_2651718 | Lighthouse Lab in Glasgow | Wellcome Sanger Institute for the COVID-19 Genomics UK (COG-UK) Consortium | Harper VanSteenhouse, Yumi Kasai, David Gray, Carol Clugston, Anna Dominiczak and Alex Alderton, Roberto Amato, Jeffrey Barrett, Sonia Goncalves, Ewan Harrison, David K. Jackson, Ian Johnston, Dominic Kwiatkowski, Cordelia Langford, John Sillitoe on behalf of the Wellcome Sanger Institute COVID-19 Surveillance Team |
| EPI_ISL_2651719, EPI_ISL_2651720 | Lighthouse Lab in Alderley Park | Wellcome Sanger Institute for the COVID-19 Genomics UK (COG-UK) Consortium | Jacquelyn Wynn, Mairead Hyland, The Lighthouse Lab in Alderley Park and Alex Alderton, Roberto Amato, Jeffrey Barrett, Sonia Goncalves, Ewan Harrison, David K. Jackson, Ian Johnston, Dominic Kwiatkowski, Cordelia Langford, John Sillitoe on behalf of the Wellcome Sanger Institute COVID-19 Surveillance Team |
| EPI_ISL_2651721, EPI_ISL_2651722, EPI_ISL_2651723, EPI_ISL_2651724, EPI_ISL_2651725 | Lighthouse Lab in Glasgow | Wellcome Sanger Institute for the COVID-19 Genomics UK (COG-UK) Consortium | Harper VanSteenhouse, Yumi Kasai, David Gray, Carol Clugston, Anna Dominiczak and Alex Alderton, Roberto Amato, Jeffrey Barrett, Sonia Goncalves, Ewan Harrison, David K. Jackson, Ian Johnston, Dominic Kwiatkowski, Cordelia Langford, John Sillitoe on behalf of the Wellcome Sanger Institute COVID-19 Surveillance Team |
| EPI_ISL_2651726, EPI_ISL_2651727 | Lighthouse Lab in Alderley Park | Wellcome Sanger Institute for the COVID-19 Genomics UK (COG-UK) Consortium | Jacquelyn Wynn, Mairead Hyland, The Lighthouse Lab in Alderley Park and Alex Alderton, Roberto Amato, Jeffrey Barrett, Sonia Goncalves, Ewan Harrison, David K. Jackson, Ian Johnston, Dominic Kwiatkowski, Cordelia Langford, John Sillitoe on behalf of the Wellcome Sanger Institute COVID-19 Surveillance Team |
| EPI_ISL_2651728, EPI_ISL_2651729 | Lighthouse Lab in Glasgow | Wellcome Sanger Institute for the COVID-19 Genomics UK (COG-UK) Consortium | Harper VanSteenhouse, Yumi Kasai, David Gray, Carol Clugston, Anna Dominiczak and Alex Alderton, Roberto Amato, Jeffrey Barrett, Sonia Goncalves, Ewan Harrison, David K. Jackson, Ian Johnston, Dominic Kwiatkowski, Cordelia Langford, John Sillitoe on behalf of the Wellcome Sanger Institute COVID-19 Surveillance Team |
| EPI_ISL_2651730 | Lighthouse Lab in Alderley Park | Wellcome Sanger Institute for the COVID-19 Genomics UK (COG-UK) Consortium | Jacquelyn Wynn, Mairead Hyland, The Lighthouse Lab in Alderley Park and Alex Alderton, Roberto Amato, Jeffrey Barrett, Sonia Goncalves, Ewan Harrison, David K. Jackson, Ian Johnston, Dominic Kwiatkowski, Cordelia Langford, John Sillitoe on behalf of the Wellcome Sanger Institute COVID-19 Surveillance Team |
| EPI_ISL_2651731 | Lighthouse Lab in Glasgow | Wellcome Sanger Institute for the COVID-19 Genomics UK (COG-UK) Consortium | Harper VanSteenhouse, Yumi Kasai, David Gray, Carol Clugston, Anna Dominiczak and Alex Alderton, Roberto Amato, Jeffrey Barrett, Sonia Goncalves, Ewan Harrison, David K. Jackson, Ian Johnston, Dominic Kwiatkowski, Cordelia Langford, John Sillitoe on behalf of the Wellcome Sanger Institute COVID-19 Surveillance Team |
| EPI_ISL_2651732 | Lighthouse Lab in Alderley Park | Wellcome Sanger Institute for the COVID-19 Genomics UK (COG-UK) Consortium | Jacquelyn Wynn, Mairead Hyland, The Lighthouse Lab in Alderley Park and Alex Alderton, Roberto Amato, Jeffrey Barrett, Sonia Goncalves, Ewan Harrison, David K. Jackson, Ian Johnston, Dominic Kwiatkowski, Cordelia Langford, John Sillitoe on behalf of the Wellcome Sanger Institute COVID-19 Surveillance Team |
| EPI_ISL_2651733, EPI_ISL_2651734, EPI_ISL_2651735, EPI_ISL_2651736, EPI_ISL_2651737, EPI_ISL_2651738, EPI_ISL_2651739, EPI_ISL_2651741 | Lighthouse Lab in Glasgow | Wellcome Sanger Institute for the COVID-19 Genomics UK (COG-UK) Consortium | Harper VanSteenhouse, Yumi Kasai, David Gray, Carol Clugston, Anna Dominiczak and Alex Alderton, Roberto Amato, Jeffrey Barrett, Sonia Goncalves, Ewan Harrison, David K. Jackson, Ian Johnston, Dominic Kwiatkowski, Cordelia Langford, John Sillitoe on behalf of the Wellcome Sanger Institute COVID-19 Surveillance Team |
| EPI_ISL_2651742 | Lighthouse Lab in Alderley Park | Wellcome Sanger Institute for the COVID-19 Genomics UK (COG-UK) Consortium | Jacquelyn Wynn, Mairead Hyland, The Lighthouse Lab in Alderley Park and Alex Alderton, Roberto Amato, Jeffrey Barrett, Sonia Goncalves, Ewan Harrison, David K. Jackson, Ian Johnston, Dominic Kwiatkowski, Cordelia Langford, John Sillitoe on behalf of the Wellcome Sanger Institute COVID-19 Surveillance Team |

| | | | |
|---|---|---|---|
| EPI_ISL_2651743, EPI_ISL_2651744, EPI_ISL_2651745 | Lighthouse Lab in Glasgow | Wellcome Sanger Institute for the COVID-19 Genomics UK (COG-UK) Consortium | Harper VanSteenhouse, Yumi Kasai, David Gray, Carol Clugston, Anna Dominiczak and Alex Alderton, Roberto Amato, Jeffrey Barrett, Sonia Goncalves, Ewan Harrison, David K. Jackson, Ian Johnston, Dominic Kwiatkowski, Cordelia Langford, John Sillitoe on behalf of the Wellcome Sanger Institute COVID-19 Surveillance Team |
| EPI_ISL_2651746 | Lighthouse Lab in Alderley Park | Wellcome Sanger Institute for the COVID-19 Genomics UK (COG-UK) Consortium | Jacquelyn Wynn, Mairead Hyland, The Lighthouse Lab in Alderley Park and Alex Alderton, Roberto Amato, Jeffrey Barrett, Sonia Goncalves, Ewan Harrison, David K. Jackson, Ian Johnston, Dominic Kwiatkowski, Cordelia Langford, John Sillitoe on behalf of the Wellcome Sanger Institute COVID-19 Surveillance Team |
| EPI_ISL_2651747, EPI_ISL_2651748, EPI_ISL_2651749, EPI_ISL_2651750, EPI_ISL_2651751, EPI_ISL_2651752 | Lighthouse Lab in Glasgow | Wellcome Sanger Institute for the COVID-19 Genomics UK (COG-UK) Consortium | Harper VanSteenhouse, Yumi Kasai, David Gray, Carol Clugston, Anna Dominiczak and Alex Alderton, Roberto Amato, Jeffrey Barrett, Sonia Goncalves, Ewan Harrison, David K. Jackson, Ian Johnston, Dominic Kwiatkowski, Cordelia Langford, John Sillitoe on behalf of the Wellcome Sanger Institute COVID-19 Surveillance Team |
| EPI_ISL_2651753, EPI_ISL_2651754, EPI_ISL_2651755, EPI_ISL_2651756 | Lighthouse Lab in Alderley Park | Wellcome Sanger Institute for the COVID-19 Genomics UK (COG-UK) Consortium | Jacquelyn Wynn, Mairead Hyland, The Lighthouse Lab in Alderley Park and Alex Alderton, Roberto Amato, Jeffrey Barrett, Sonia Goncalves, Ewan Harrison, David K. Jackson, Ian Johnston, Dominic Kwiatkowski, Cordelia Langford, John Sillitoe on behalf of the Wellcome Sanger Institute COVID-19 Surveillance Team |
| EPI_ISL_2651757, EPI_ISL_2651758 | Lighthouse Lab in Glasgow | Wellcome Sanger Institute for the COVID-19 Genomics UK (COG-UK) Consortium | Harper VanSteenhouse, Yumi Kasai, David Gray, Carol Clugston, Anna Dominiczak and Alex Alderton, Roberto Amato, Jeffrey Barrett, Sonia Goncalves, Ewan Harrison, David K. Jackson, Ian Johnston, Dominic Kwiatkowski, Cordelia Langford, John Sillitoe on behalf of the Wellcome Sanger Institute COVID-19 Surveillance Team |
| EPI_ISL_2651759, EPI_ISL_2651760 | Lighthouse Lab in Alderley Park | Wellcome Sanger Institute for the COVID-19 Genomics UK (COG-UK) Consortium | Jacquelyn Wynn, Mairead Hyland, The Lighthouse Lab in Alderley Park and Alex Alderton, Roberto Amato, Jeffrey Barrett, Sonia Goncalves, Ewan Harrison, David K. Jackson, Ian Johnston, Dominic Kwiatkowski, Cordelia Langford, John Sillitoe on behalf of the Wellcome Sanger Institute COVID-19 Surveillance Team |
| EPI_ISL_2651761 | Lighthouse Lab in Glasgow | Wellcome Sanger Institute for the COVID-19 Genomics UK (COG-UK) Consortium | Harper VanSteenhouse, Yumi Kasai, David Gray, Carol Clugston, Anna Dominiczak and Alex Alderton, Roberto Amato, Jeffrey Barrett, Sonia Goncalves, Ewan Harrison, David K. Jackson, Ian Johnston, Dominic Kwiatkowski, Cordelia Langford, John Sillitoe on behalf of the Wellcome Sanger Institute COVID-19 Surveillance Team |
| EPI_ISL_2651762, EPI_ISL_2651763 | Lighthouse Lab in Alderley Park | Wellcome Sanger Institute for the COVID-19 Genomics UK (COG-UK) Consortium | Jacquelyn Wynn, Mairead Hyland, The Lighthouse Lab in Alderley Park and Alex Alderton, Roberto Amato, Jeffrey Barrett, Sonia Goncalves, Ewan Harrison, David K. Jackson, Ian Johnston, Dominic Kwiatkowski, Cordelia Langford, John Sillitoe on behalf of the Wellcome Sanger Institute COVID-19 Surveillance Team |
| EPI_ISL_2651764, EPI_ISL_2651765, EPI_ISL_2651766, EPI_ISL_2651767, EPI_ISL_2651768, EPI_ISL_2651769, EPI_ISL_2651770, EPI_ISL_2651771, EPI_ISL_2651772, EPI_ISL_2651773, EPI_ISL_2651774, EPI_ISL_2651775, EPI_ISL_2651776, EPI_ISL_2651777, EPI_ISL_2651778, EPI_ISL_2651779, EPI_ISL_2651780, EPI_ISL_2651781, EPI_ISL_2651782, EPI_ISL_2651783, EPI_ISL_2651784, EPI_ISL_2651785, EPI_ISL_2651787, EPI_ISL_2651788, EPI_ISL_2651789, EPI_ISL_2651791, EPI_ISL_2651792, EPI_ISL_2651793, EPI_ISL_2651795, EPI_ISL_2651796, EPI_ISL_2651797, EPI_ISL_2651798, EPI_ISL_2651799, EPI_ISL_2651800, EPI_ISL_2651801, EPI_ISL_2651802, EPI_ISL_2651803 | | | |
| see above | Lighthouse Lab in Glasgow | Wellcome Sanger Institute for the COVID-19 Genomics UK (COG-UK) Consortium | Harper VanSteenhouse, Yumi Kasai, David Gray, Carol Clugston, Anna Dominiczak and Alex Alderton, Roberto Amato, Jeffrey Barrett, Sonia Goncalves, Ewan Harrison, David K. Jackson, Ian Johnston, Dominic Kwiatkowski, Cordelia Langford, John Sillitoe on behalf of the Wellcome Sanger Institute COVID-19 Surveillance Team |
| EPI_ISL_2651804 | Lighthouse Laboratory Plymouth | Wellcome Sanger Institute for the COVID-19 Genomics UK (COG-UK) Consortium | Lighthouse Laboratory Plymouth and Alex Alderton, Roberto Amato, Jeffrey Barrett, Sonia Goncalves, Ewan Harrison, David K. Jackson, Ian Johnston, Dominic Kwiatkowski, Cordelia Langford, John Sillitoe on behalf of the Wellcome Sanger Institute COVID-19 Surveillance Team |
| EPI_ISL_2651805, EPI_ISL_2651806, EPI_ISL_2651807, EPI_ISL_2651808, EPI_ISL_2651809, EPI_ISL_2651810, EPI_ISL_2651811, EPI_ISL_2651812, EPI_ISL_2651813, EPI_ISL_2651814, EPI_ISL_2651815, EPI_ISL_2651816, EPI_ISL_2651817, EPI_ISL_2651818, EPI_ISL_2651819, EPI_ISL_2651820, EPI_ISL_2651821, EPI_ISL_2651822, EPI_ISL_2651823, EPI_ISL_2651824, EPI_ISL_2651825, EPI_ISL_2651826, EPI_ISL_2651827, EPI_ISL_2651828, EPI_ISL_2651829, EPI_ISL_2651830, EPI_ISL_2651831, EPI_ISL_2651832, EPI_ISL_2651833, EPI_ISL_2651834, EPI_ISL_2651835, EPI_ISL_2651836, EPI_ISL_2651837, EPI_ISL_2651838, EPI_ISL_2651839, EPI_ISL_2651840, EPI_ISL_2651841, EPI_ISL_2651842, EPI_ISL_2651843, EPI_ISL_2651844, EPI_ISL_2651845, EPI_ISL_2651846, EPI_ISL_2651847, EPI_ISL_2651848, EPI_ISL_2651849, EPI_ISL_2651850, EPI_ISL_2651851, EPI_ISL_2651852, EPI_ISL_2651853, EPI_ISL_2651854, EPI_ISL_2651855, EPI_ISL_2651856, EPI_ISL_2651857, EPI_ISL_2651858, EPI_ISL_2651859, EPI_ISL_2651860, EPI_ISL_2651861, EPI_ISL_2651862, EPI_ISL_2651863, EPI_ISL_2651864, EPI_ISL_2651865, EPI_ISL_2651866, EPI_ISL_2651867, EPI_ISL_2651868, EPI_ISL_2651870, EPI_ISL_2651871, EPI_ISL_2651872, EPI_ISL_2651873, EPI_ISL_2651874, EPI_ISL_2651875, EPI_ISL_2651876, EPI_ISL_2651877, EPI_ISL_2651878, EPI_ISL_2651879, EPI_ISL_2651880, EPI_ISL_2651881, EPI_ISL_2651882, EPI_ISL_2651883, EPI_ISL_2651884, EPI_ISL_2651885, EPI_ISL_2651886, EPI_ISL_2651887, EPI_ISL_2651888, EPI_ISL_2651889, EPI_ISL_2651890, EPI_ISL_2651891, EPI_ISL_2651892, EPI_ISL_2651893, EPI_ISL_2651894, EPI_ISL_2651895, EPI_ISL_2651897, EPI_ISL_2651898, EPI_ISL_2651899, EPI_ISL_2651900, EPI_ISL_2651901, EPI_ISL_2651902, EPI_ISL_2651903, EPI_ISL_2651904, EPI_ISL_2651905, EPI_ISL_2651906, EPI_ISL_2651907, EPI_ISL_2651908, EPI_ISL_2651909, EPI_ISL_2651910, EPI_ISL_2651911, EPI_ISL_2651912, EPI_ISL_2651913, EPI_ISL_2651914, EPI_ISL_2651915, EPI_ISL_2651916, EPI_ISL_2651917, EPI_ISL_2651918, EPI_ISL_2651919, EPI_ISL_2651920, EPI_ISL_2651921, EPI_ISL_2651922, EPI_ISL_2651923, EPI_ISL_2651924, EPI_ISL_2651925, EPI_ISL_2651926, EPI_ISL_2651927, EPI_ISL_2651928, EPI_ISL_2651929, EPI_ISL_2651930, EPI_ISL_2651931, EPI_ISL_2651932, EPI_ISL_2651933, EPI_ISL_2651934, EPI_ISL_2651935, EPI_ISL_2651936, EPI_ISL_2651937, EPI_ISL_2651938, EPI_ISL_2651939, EPI_ISL_2651940, EPI_ISL_2651941 | | | |
| see above | Lighthouse Lab in Glasgow | Wellcome Sanger Institute for the COVID-19 Genomics UK (COG-UK) Consortium | Harper VanSteenhouse, Yumi Kasai, David Gray, Carol Clugston, Anna Dominiczak and Alex Alderton, Roberto Amato, Jeffrey Barrett, Sonia Goncalves, Ewan Harrison, David K. Jackson, Ian Johnston, Dominic Kwiatkowski, Cordelia Langford, John Sillitoe on behalf of the Wellcome Sanger Institute COVID-19 Surveillance Team |
| EPI_ISL_2651942 | Lighthouse Lab in Milton Keynes | Wellcome Sanger Institute for the COVID-19 Genomics UK (COG-UK) Consortium | The Lighthouse Lab in Milton Keynes and Alex Alderton, Roberto Amato, Jeffrey Barrett, Sonia Goncalves, Ewan Harrison, David K. Jackson, Ian Johnston, Dominic Kwiatkowski, Cordelia Langford, John Sillitoe on behalf of the Wellcome Sanger Institute COVID-19 Surveillance Team |
| EPI_ISL_2651943 | Lighthouse Laboratory Plymouth | Wellcome Sanger Institute for the COVID-19 Genomics UK (COG-UK) Consortium | Lighthouse Laboratory Plymouth and Alex Alderton, Roberto Amato, Jeffrey Barrett, Sonia Goncalves, Ewan Harrison, David K. Jackson, Ian Johnston, Dominic Kwiatkowski, Cordelia Langford, John Sillitoe on behalf of the Wellcome Sanger Institute COVID-19 Surveillance Team |
| EPI_ISL_2651944 | Lighthouse Lab in Milton Keynes | Wellcome Sanger Institute for the COVID-19 Genomics UK (COG-UK) Consortium | The Lighthouse Lab in Milton Keynes and Alex Alderton, Roberto Amato, Jeffrey Barrett, Sonia Goncalves, Ewan Harrison, David K. Jackson, Ian Johnston, Dominic Kwiatkowski, Cordelia Langford, John Sillitoe on behalf of the Wellcome Sanger Institute COVID-19 Surveillance Team |
| EPI_ISL_2651945 | Lighthouse Laboratory Plymouth | Wellcome Sanger Institute for the COVID-19 Genomics UK (COG-UK) Consortium | Lighthouse Laboratory Plymouth and Alex Alderton, Roberto Amato, Jeffrey Barrett, Sonia Goncalves, Ewan Harrison, David K. Jackson, Ian Johnston, Dominic Kwiatkowski, Cordelia Langford, John Sillitoe on behalf of the Wellcome Sanger Institute COVID-19 Surveillance Team |
| EPI_ISL_2651946, EPI_ISL_2651947, EPI_ISL_2651948 | Lighthouse Lab in Alderley Park | Wellcome Sanger Institute for the COVID-19 Genomics UK (COG-UK) Consortium | Jacquelyn Wynn, Mairead Hyland, The Lighthouse Lab in Alderley Park and Alex Alderton, Roberto Amato, Jeffrey Barrett, Sonia Goncalves, Ewan Harrison, David K. Jackson, Ian Johnston, Dominic Kwiatkowski, Cordelia Langford, John Sillitoe on behalf of the Wellcome Sanger Institute COVID-19 Surveillance Team |
| EPI_ISL_2651949, EPI_ISL_2651950, EPI_ISL_2651951, EPI_ISL_2651952, EPI_ISL_2651953, EPI_ISL_2651956, EPI_ISL_2651957, EPI_ISL_2651958, EPI_ISL_2651959, EPI_ISL_2651960, EPI_ISL_2651961, EPI_ISL_2651963, EPI_ISL_2651964, EPI_ISL_2651965, EPI_ISL_2651966, EPI_ISL_2651967, EPI_ISL_2651968, EPI_ISL_2651969, EPI_ISL_2651971, EPI_ISL_2651972, EPI_ISL_2651973, EPI_ISL_2651974, EPI_ISL_2651975, EPI_ISL_2651976, EPI_ISL_2651977, EPI_ISL_2651978, EPI_ISL_2651979, EPI_ISL_2651980, EPI_ISL_2651981, EPI_ISL_2651982, EPI_ISL_2651983, EPI_ISL_2651984, EPI_ISL_2651986, EPI_ISL_2651987, EPI_ISL_2651988, EPI_ISL_2651989, EPI_ISL_2651990, EPI_ISL_2651991, EPI_ISL_2651992, EPI_ISL_2651993, EPI_ISL_2651994, EPI_ISL_2651995, EPI_ISL_2651996, EPI_ISL_2651997 | | | |
| see above | Lighthouse Lab in Glasgow | Wellcome Sanger Institute for the COVID-19 Genomics UK (COG-UK) Consortium | Harper VanSteenhouse, Yumi Kasai, David Gray, Carol Clugston, Anna Dominiczak and Alex Alderton, Roberto Amato, Jeffrey Barrett, Sonia Goncalves, Ewan Harrison, David K. Jackson, Ian Johnston, Dominic Kwiatkowski, Cordelia Langford, John Sillitoe on behalf of the Wellcome Sanger Institute COVID-19 Surveillance Team |
| EPI_ISL_2651998, EPI_ISL_2651999 | Lighthouse Lab in Alderley Park | Wellcome Sanger Institute for the COVID-19 Genomics UK (COG-UK) Consortium | Jacquelyn Wynn, Mairead Hyland, The Lighthouse Lab in Alderley Park and Alex Alderton, Roberto Amato, Jeffrey Barrett, Sonia Goncalves, Ewan Harrison, David K. Jackson, Ian Johnston, Dominic Kwiatkowski, Cordelia Langford, John Sillitoe on behalf of the Wellcome Sanger Institute COVID-19 Surveillance Team |
| EPI_ISL_2652000 | Lighthouse Lab in Milton Keynes | Wellcome Sanger Institute for the COVID-19 Genomics UK (COG-UK) Consortium | The Lighthouse Lab in Milton Keynes and Alex Alderton, Roberto Amato, Jeffrey Barrett, Sonia Goncalves, Ewan Harrison, David K. Jackson, Ian Johnston, Dominic Kwiatkowski, Cordelia Langford, John Sillitoe on behalf of the Wellcome Sanger Institute COVID-19 Surveillance Team |
| EPI_ISL_2652001 | Lighthouse Lab in Glasgow | Wellcome Sanger Institute for the COVID-19 Genomics UK (COG-UK) Consortium | Harper VanSteenhouse, Yumi Kasai, David Gray, Carol Clugston, Anna Dominiczak and Alex Alderton, Roberto Amato, Jeffrey Barrett, Sonia Goncalves, Ewan Harrison, David K. Jackson, Ian Johnston, Dominic Kwiatkowski, Cordelia Langford, John Sillitoe on behalf of the Wellcome Sanger Institute COVID-19 Surveillance Team |
| EPI_ISL_2652002 | Lighthouse Lab in Alderley Park | Wellcome Sanger Institute for the COVID-19 Genomics UK (COG-UK) Consortium | Jacquelyn Wynn, Mairead Hyland, The Lighthouse Lab in Alderley Park and Alex Alderton, Roberto Amato, Jeffrey Barrett, Sonia Goncalves, Ewan Harrison, David K. Jackson, Ian Johnston, Dominic Kwiatkowski, Cordelia Langford, John Sillitoe on behalf of the Wellcome Sanger Institute COVID-19 Surveillance Team |
| EPI_ISL_2652017, EPI_ISL_2652024, EPI_ISL_2652040 | New Mexico Department of Health Scientific Laboratory | New Mexico Department of Health Scientific Laboratory | Ellie Johnson, D'eldra Malone, Jennifer Benoit, Ratheesh Rajan, Linda Salazar, Keila Gutierrez, Anastacia Griego-Fisher |
| EPI_ISL_2652061, EPI_ISL_2652066 | Hospital Universitari Joan XXIII | Hospital Universitari Joan XXIII | Gemma Recio, Cristina Gutiérrez, Clara Benavent, Carla Martín, Margarida Terrón, Ester Picó, Natalia Bastón |
| EPI_ISL_2652088, EPI_ISL_2652089, EPI_ISL_2652091 | New Mexico Department of Health Scientific Laboratory | New Mexico Department of Health Scientific Laboratory | Ellie Johnson, D'eldra Malone, Jennifer Benoit, Ratheesh Rajan, Linda Salazar, Keila Gutierrez, Anastacia Griego-Fisher |
| EPI_ISL_2652114, EPI_ISL_2652115, EPI_ISL_2652116 | Cerballiance-IDF | Cerba lab | Emmanuel Lecorche, Sabine Trombert-Paolantoni, Aude Lessenne, Kader Merah, Thierry Collin, Laura Verdurme, Patrice Herisson, Stéphanie Haïm-Boukobza, Bénédicte Roquebert |
| EPI_ISL_2652119, EPI_ISL_2652121 | A.S.L. BIELLA | Fondazione del Piemonte per l'Oncologia IRCCS | Antonino Sottile, Silvia Brossa, Paola Marino, Giorgio Giardina |
| EPI_ISL_2652162 | AZ Klina | AZ Klina | Carl Vael - Lynsey Berckmans |
| EPI_ISL_2652214 | Cantacuzino National Military-Medical Institute, Viral Respiratory Infections Laboratory | Cantacuzino Institute Virology | Carmen Cherciu, Luiza Ustea, Nicoleta Paraschiv, Sorin Dinu, Mihaela Oprea, Mihaela Lazar |
| EPI_ISL_2652215 | Clinical Emergency County Hospital, Craiova | Cantacuzino Institute Virology | Carmen Cherciu, Luiza Ustea, Nicoleta Paraschiv, Sorin Dinu, Mihaela Oprea, Mihaela Lazar |
| EPI_ISL_2652216, EPI_ISL_2652217 | Cantacuzino National Military-Medical Institute, Viral Respiratory Infections Laboratory | Cantacuzino Institute Virology | Carmen Cherciu, Luiza Ustea, Nicoleta Paraschiv, Sorin Dinu, Mihaela Oprea, Mihaela Lazar |
| EPI_ISL_2652228, EPI_ISL_2652231, EPI_ISL_2652238 | Institute for Infectious Diseases | Institute for Infectious Diseases | Stefan Neuenschwander, Christian Baumann, Miguel A Terrazos Miani, Cora SÃ¤gesser, Pascal Bittel, Peter Keller, Franziska Suter-Riniker, Stephen L Leib, Alban Ramette |
| EPI_ISL_2652246, EPI_ISL_2652247 | AZ Klina | AZ Klina | Carl Vael - Lynsey Berckmans |
| EPI_ISL_2652249 | Laboratorio de Referencial Nacional de Virus Respiratorios | Laboratorio de Referencial Nacional de Virus Respiratorios | Carlos Padilla Rojas, Veronica Hurtado Vela, Orson Mestanza Millones, Iris Silva Molina, Luis Barcena Flores, Sandra Morales Ruiz, Steve Acedo Lazo, Omar Caceres Rey, Marco Galarza Perez, Henri Bailon Calderon, Nancy Rojas Serrano, Priscila Lope Pari, Lely Solari Zerpa, Joseph Huayra Niquen |
| EPI_ISL_2652262 | Clinical Virology | Clinical Bacteriology | Tim Roloff, Fanny Wegner, Helena MB Seth-Smith, Alfredo Mari, Karoline Leuzinger, Julia Bielicki, Manuel Battegay, Hans Hirsch, Adrian Egli |
| EPI_ISL_2652265 | Kantonsarztamt Solothurn | Clinical Bacteriology | Tim Roloff, Fanny Wegner, Helena MB Seth-Smith, Alfredo Mari, Karoline Leuzinger, Julia Bielicki, Manuel Battegay, Lukas, Fenner, Hans Hirsch, Adrian Egli |
| EPI_ISL_2652269, EPI_ISL_2652273, EPI_ISL_2652276, EPI_ISL_2652277, EPI_ISL_2652281, EPI_ISL_2652282, EPI_ISL_2652283, EPI_ISL_2652289, EPI_ISL_2652293, EPI_ISL_2652299, EPI_ISL_2652300, EPI_ISL_2652304, EPI_ISL_2652312, EPI_ISL_2652314, EPI_ISL_2652316, EPI_ISL_2652317, EPI_ISL_2652322, EPI_ISL_2652325, EPI_ISL_2652326, EPI_ISL_2652328, EPI_ISL_2652331, EPI_ISL_2652332, EPI_ISL_2652334, EPI_ISL_2652335, EPI_ISL_2652337, EPI_ISL_2652339, EPI_ISL_2652341, EPI_ISL_2652342, EPI_ISL_2652343, EPI_ISL_2652345, EPI_ISL_2652347, EPI_ISL_2652349, EPI_ISL_2652356, EPI_ISL_2652357, EPI_ISL_2652359, EPI_ISL_2652360, EPI_ISL_2652362, EPI_ISL_2652363, EPI_ISL_2652365, EPI_ISL_2652366, EPI_ISL_2652367, EPI_ISL_2652369, EPI_ISL_2652371, EPI_ISL_2652374, EPI_ISL_2652375, EPI_ISL_2652376, EPI_ISL_2652378, EPI_ISL_2652379, EPI_ISL_2652381, EPI_ISL_2652387, EPI_ISL_2652388, EPI_ISL_2652390, EPI_ISL_2652395, EPI_ISL_2652397, EPI_ISL_2652398, EPI_ISL_2652399, EPI_ISL_2652401, EPI_ISL_2652402, EPI_ISL_2652403, EPI_ISL_2652404, EPI_ISL_2652405, EPI_ISL_2652406, EPI_ISL_2652407, EPI_ISL_2652408, EPI_ISL_2652409, EPI_ISL_2652411, | | | |

| | | | |
|---|---|---|---|
| EPI_ISL_2652417, EPI_ISL_2652422, EPI_ISL_2652423, EPI_ISL_2652424, EPI_ISL_2652427, EPI_ISL_2652432, EPI_ISL_2652433, EPI_ISL_2652434, EPI_ISL_2652438, EPI_ISL_2652439, EPI_ISL_2652442, EPI_ISL_2652447, EPI_ISL_2652448, EPI_ISL_2652449, EPI_ISL_2652450, EPI_ISL_2652452, EPI_ISL_2652455, EPI_ISL_2652458, EPI_ISL_2652459, EPI_ISL_2652461, EPI_ISL_2652462, EPI_ISL_2652463, EPI_ISL_2652464, EPI_ISL_2652465, EPI_ISL_2652467, EPI_ISL_2652468, EPI_ISL_2652469, EPI_ISL_2652473, EPI_ISL_2652474 | | | |
| see above | Quest Diagnostics Incorporated | Centers for Disease Control and Prevention Division of Viral Diseases, Pathogen Discovery | Dakota Howard, Dhwani Batra, Peter W. Cook, Kara Moser, Adrian Paskey, Jason Caravas, Benjamin Rambo-Martin, Shatavia Morrison, Christopher Gulvick, Scott Sammons, Yvette Unoarumhi, Darlene Wagner, Matthew Schmerer, S. H. Rosenthal, A. Gerasimova, R. M. Kagan, B. Anderson, M. Hua, Y. Liu, L.E. Bernstein, K.E. Livingston, A. Perez, I. A. Shlyakhter, R. V. Rolando, R. Owen, P. Tanpaiboon, F. Lacbawan, Clinton R. Paden, Duncan MacCannell |
| EPI_ISL_2652484, EPI_ISL_2652485, EPI_ISL_2652507 | Minnesota Department of Health, Public Health Laboratory | Minnesota Department of Health, Public Health Laboratory | Alexandra Lorentz, Jacob Garfin, Matt Plumb, and Xiong Wang |
| EPI_ISL_2652571, EPI_ISL_2652644, EPI_ISL_2652646, EPI_ISL_2652668, EPI_ISL_2652849 | Infinity Biologix | Centers for Disease Control and Prevention Division of Viral Diseases, Pathogen Discovery | Dakota Howard, Dhwani Batra, Peter W. Cook, Kara Moser, Adrian Paskey, Jason Caravas, Benjamin Rambo-Martin, Shatavia Morrison, Christopher Gulvick, Scott Sammons, Yvette Unoarumhi, Darlene Wagner, Matthew Schmerer, Christian Bixby, Yihe Wang, Jonathan Schultz, Chirayu Goswami, Russ Hager, Robin Grimwood, Clinton R. Paden, Duncan MacCannell |
| EPI_ISL_2652974 | County of San Luis Obispo Public Health Laboratory | County of San Luis Obispo Public Health Laboratory | Frances Sidhu, Katrina Erwin |
| EPI_ISL_2653001, EPI_ISL_2653060 | Infinity Biologix | Centers for Disease Control and Prevention Division of Viral Diseases, Pathogen Discovery | Dakota Howard, Dhwani Batra, Peter W. Cook, Kara Moser, Adrian Paskey, Jason Caravas, Benjamin Rambo-Martin, Shatavia Morrison, Christopher Gulvick, Scott Sammons, Yvette Unoarumhi, Darlene Wagner, Matthew Schmerer, Christian Bixby, Yihe Wang, Jonathan Schultz, Chirayu Goswami, Russ Hager, Robin Grimwood, Clinton R. Paden, Duncan MacCannell |
| EPI_ISL_2655690, EPI_ISL_2655803, EPI_ISL_2655902, EPI_ISL_2655964, EPI_ISL_2655972, EPI_ISL_2655997, EPI_ISL_2655998, EPI_ISL_2656001, EPI_ISL_2656003, EPI_ISL_2656025, EPI_ISL_2656028, EPI_ISL_2656077, EPI_ISL_2656181, EPI_ISL_2656303, EPI_ISL_2656337, EPI_ISL_2656349, EPI_ISL_2656460, EPI_ISL_2656462, EPI_ISL_2656487, EPI_ISL_2656548, EPI_ISL_2656571, EPI_ISL_2656601, EPI_ISL_2656603, EPI_ISL_2656634, EPI_ISL_2656637, EPI_ISL_2656666, EPI_ISL_2656693, EPI_ISL_2656698, EPI_ISL_2656727, EPI_ISL_2656736, EPI_ISL_2656742, EPI_ISL_2656763, EPI_ISL_2656767, EPI_ISL_2656855, EPI_ISL_2656861, EPI_ISL_2656870, EPI_ISL_2657111, EPI_ISL_2657112, EPI_ISL_2657119 | | | |
| see above | BCCDC Public Health Laboratory | BCCDC Public Health Laboratory | Prystajecky Natalie, Linda Hoang, Dan Fornika, John Tyson, Shannon Russell, Kim Macdonald, Kimia Kamelian, Ana Pacagnella, Corrinne Ng, Loretta Janz, Robert Azana, Mel Krajden |
| EPI_ISL_2657216 | Pracownia Biologii Molekularnej Laboratorium Centralne SPZOZ | 1. Tricity SARS-CoV-2 sequencing consortium: University of Gdansk, Medical University of Gdansk, Vaxican Ltd., Invicta Ltd. 2. National Institute of Public Health - National Institute of Hygiene, Warsaw, Poland | Maciej Kosinski, Celina Cybulska, Krystyna Bienkowska Szewczyk, Maciej Grzybek, Karolina Gackowska, Marcin Lubocki, Katarzyna Groth, Lukasz Rabalski, Katarzyna Zacharczuk, Magdalena Nowakowska, Małgorzata Sadkowska-Todys, Tomasz Wołkowicz |
| EPI_ISL_2657324 | Institute of Virology, Biomedical Research Center of the Slovak Academy of Sciences, Bratislava | Faculty of Natural Sciences, Comenius University, Bratislava | Kristina Borsova, Viktoria Cabanova, Brona Brejova, Viktoria Hodorova, Sabina Fumacova Havlikova, Juraj Kopacek, Martina Lickova, Lubomira Lukacikova, Martina Nebohacova, Monika Slavikova, Tomas Vinar, Boris Klempa, Jozef Nosek |
| EPI_ISL_2657357, EPI_ISL_2657364 | Hospital Universitari Joan XXIII | Hospital Universitari Joan XXIII | Gemma Recio, Cristina Gutiérrez, Clara Benavent, Carla Martín, Margarida Terrón, Ester Picó, Natalia Bastón |
| EPI_ISL_2657378, EPI_ISL_2657380, EPI_ISL_2657383, EPI_ISL_2657384, EPI_ISL_2657385, EPI_ISL_2657386, EPI_ISL_2657388, EPI_ISL_2657389, EPI_ISL_2657390, EPI_ISL_2657393, EPI_ISL_2657394, EPI_ISL_2657397, EPI_ISL_2657404, EPI_ISL_2657457, EPI_ISL_2657458, EPI_ISL_2657473, EPI_ISL_2657474, EPI_ISL_2657475, EPI_ISL_2657476, EPI_ISL_2657477, EPI_ISL_2657478, EPI_ISL_2657479, EPI_ISL_2657480, EPI_ISL_2657481, EPI_ISL_2657482, EPI_ISL_2657483, EPI_ISL_2657484, EPI_ISL_2657485, EPI_ISL_2657486, EPI_ISL_2657487, EPI_ISL_2657488, EPI_ISL_2657490, EPI_ISL_2657491, EPI_ISL_2657492, EPI_ISL_2657494, EPI_ISL_2657495, EPI_ISL_2657497, EPI_ISL_2657498, EPI_ISL_2657499, EPI_ISL_2657500, EPI_ISL_2657505 | | | |
| see above | Austrian Agency for Health and Food Safety (AGES) | Bergthaler laboratory, CeMM Research Center for Molecular Medicine of the Austrian Academy of Sciences | Lukas Endler, Anna Schedl, Fabian Amman, Petr Triska, Matthew Thornton, Thomas Penz, Benedikt Agerer, Maelle Le Moing, Michael Schuster, Bekir Erguner, Jan Laine, Martin Senekowitsch, Christoph Bock, Andreas Bergthaler |
| EPI_ISL_2657533 | Elling group, Institute of Molecular Biotechnology (IMBA) | Bergthaler laboratory, CeMM Research Center for Molecular Medicine of the Austrian Academy of Sciences | Lukas Endler, Anna Schedl, Fabian Amman, Petr Triska, Matthew Thornton, Thomas Penz, Benedikt Agerer, Maelle Le Moing, Michael Schuster, Bekir Erguner, Jan Laine, Martin Senekowitsch, Christoph Bock, Andreas Bergthaler |
| EPI_ISL_2657689, EPI_ISL_2657691 | Wexner Medical Center | The Ohio State University College of Medicine | Koenig,S., Seminetta,J. |
| EPI_ISL_2658150, EPI_ISL_2658151, EPI_ISL_2658152, EPI_ISL_2658153, EPI_ISL_2658154, EPI_ISL_2658156, EPI_ISL_2658157, EPI_ISL_2658158, EPI_ISL_2658159, EPI_ISL_2658160, EPI_ISL_2658161, EPI_ISL_2658162 | | | |
| see above | Kansas Health and Environmental Lab | Kansas Health and Environmental Lab | Mike Grose, Katherine Wiggins, Jonathan Barnell, Ben Olsen, and Phil Adam |
| EPI_ISL_2658186, EPI_ISL_2658187, EPI_ISL_2658190, EPI_ISL_2658191, EPI_ISL_2658219, EPI_ISL_2658223, EPI_ISL_2658225, EPI_ISL_2658228, EPI_ISL_2658239, EPI_ISL_2658247, EPI_ISL_2658249, EPI_ISL_2658251, EPI_ISL_2658254, EPI_ISL_2658256, EPI_ISL_2658259, EPI_ISL_2658260, EPI_ISL_2658262 | | | |
| see above | Servicio de Microbiología. Consorcio Hospital General Universitario de Valencia | SeqCOVID-SPAIN consortium/IBV(CSIC) | María Dolores Ocete, Begoña Fuster Escrivá, Carme Salvador García, Rafael Medina González, Concepción Gimeno Cardona and SeqCOVID-SPAIN consortium |
| EPI_ISL_2658314, EPI_ISL_2658315, EPI_ISL_2658316 | Indian Council of Medical Research-National Institute of Virology, Microbial Containment Complex | Indian Council of Medical Research-National Institute of Virology, Microbial Containment Complex | Pragya D Yadav |
| EPI_ISL_2658317 | Orion Laboratories | Orion Laboratories | Keisha Simoneaux, David Slaughter |
| EPI_ISL_2658330, EPI_ISL_2658339, EPI_ISL_2658344, EPI_ISL_2658345, EPI_ISL_2658346, EPI_ISL_2658347, EPI_ISL_2658352 | UMC Utrecht | UMC Utrecht | Rob Schuurman, Anniek Tanja, Annemarie Wensing, Pascal Mantiri |
| EPI_ISL_2658433 | Furst Medical Laboratory | Norwegian Institute of Public Health, Department of Virology | Kathrine Stene-Johansen, Kamilla Heddeland Instefjord, Hilde Elshaug, Garcia Llorente Ignacio, Jon Bråte, Engebretsen Serina Beate,Pedersen Benedikte Nevjen, Line Victoria Moen, Debech Nadia, Atiya R Ali,Marie Paulsen Madsen, Rasmus Riis Kopperud, Hilde Vollan, Karoline Bragstad, Olav Hungnes |
| EPI_ISL_2658435 | Akershus University Hospital, Department for Microbiology and Infectious Disease Control | Norwegian Institute of Public Health, Department of Virology | Kathrine Stene-Johansen, Kamilla Heddeland Instefjord, Hilde Elshaug, Garcia Llorente Ignacio, Jon Bråte, Engebretsen Serina Beate,Pedersen Benedikte Nevjen, Line Victoria Moen, Debech Nadia, Atiya R Ali,Marie Paulsen Madsen, Rasmus Riis Kopperud, Hilde Vollan, Karoline Bragstad, Olav Hungnes |
| EPI_ISL_2658513 | Basurto University Hospital: Clinical Microbiology Laboratory | Biocruces Bizkaia | Mikel Urrutikoetxea-Gutiérrez, Mª Carmen Nieto Toboso, Estibaliz Ugalde Zarraga, Mikel Gallego Rodrigo, Ana de la Hoz, José Luis Díaz de Tuesta del Arco |
| EPI_ISL_2658598 | Santa Clara County Public Health Laboratory | Chan-Zuckerberg Biohub | CZB Cliahub Consortium |
| EPI_ISL_2658948 | OK Public Health Laboratory, Oklahoma State DOH | Genomics and Discovery, Respiratory Viruses Branch, Division of Viral Diseases, Centers for Disease Control and Prevention | Jing Zhang, Anna Kelleher, Ying Tao, Yan Li, Brian Lynch, Krista Queen, Anna Uehara, Peter Cook, Han Jia Justin Ng, Rachel Marine, Clinton R. Paden, Dhwani Batra, Haibin Wang, Tara Coalter, Jasmine Padilla, Morgan Davis, Mili Sheth, Sarah Nobles, Mark Burroughs, Justin Lee, Adam Retchless, Suxiang Tong |
| EPI_ISL_2658955, EPI_ISL_2658968, EPI_ISL_2658974, EPI_ISL_2658976, EPI_ISL_2658978 | TXDSHS | TXDSHS | Rashmi Tuladhar, Bonnie Oh, Jenny Zhang, Maliha Rahman, Mayela Pedrueza, Anita Pokharel, Karen Bobier, Lorraine Rodriguez, Myong Koag, Chun Wang, Rachel Lee, Grace Kubin |
| EPI_ISL_2658984 | Kaiser Santa Clara | Santa Clara County Public Health Laboratory | Santa Clara County Public Health Department |
| EPI_ISL_2658987, EPI_ISL_2658993, EPI_ISL_2658999, EPI_ISL_2659011, EPI_ISL_2659023, EPI_ISL_2659027, EPI_ISL_2659045, EPI_ISL_2659046, EPI_ISL_2659048 | Santa Clara Valley Medical Center | Santa Clara County Public Health Laboratory | Santa Clara County Public Health Department |
| EPI_ISL_2659051 | Microbiological Diagnostic Unit - Public Health Laboratory (MDU-PHL) | Microbiological Diagnostics Unit Public Health Laboratory | Seemann T., Sait, M.L., Sherry, N.L. |
| EPI_ISL_2659068, EPI_ISL_2659069, EPI_ISL_2659070 | El Camino Hospital | Santa Clara County Public Health Laboratory | Santa Clara County Public Health Department |
| EPI_ISL_2659073 | Good Samaritan Hospital | Santa Clara County Public Health Laboratory | Santa Clara County Public Health Department |
| EPI_ISL_2659079, EPI_ISL_2659090, EPI_ISL_2659099, EPI_ISL_2659101 | Santa Clara Valley Medical Center | Santa Clara County Public Health Laboratory | Santa Clara County Public Health Department |
| EPI_ISL_2659198 | U.O. Microbiologia Laboratorio Unico Centro Servizi AUSL della Romagna | U.O. Microbiologia Laboratorio Unico Centro Servizi AUSL della Romagna | Giorgio Dirani, Silvia Zannoli, Francesca Taddei, Giulia Gatti, Ilaria Poggianti, Vittorio Sambri |
| EPI_ISL_2659676, EPI_ISL_2659700, EPI_ISL_2659710, EPI_ISL_2659714, EPI_ISL_2659731, EPI_ISL_2659752 | MEPHI, Aix Marseille University | MEPHI, Aix Marseille University | Anthony LEVASSEUR |
| EPI_ISL_2660332, EPI_ISL_2660333, EPI_ISL_2660359, | Infectious Disease Program, Broad Institute of Harvard and MIT | Infectious Disease Program, Broad Institute of Harvard and MIT | Lemieux,J.E., Siddle,K.J., Shaw,B., Adams,G., Pierce,V., Turbett,S., Anahtar,M., Branda,J., Slater,D., Harris,J., Lin,A.E., Gladden-Young,A., Lagerborg,K., Rudy,M., DeRuff,K., Carter,A., Normandin,E., Bauer,M., Reilly,S., Tomkins-Tinch,C., Loreth,C., Chaluvadi,S., Neumann,A., Cusick,C., Chapman,S.B., Gnirke,A., Flowers,K., Cerrato,F., Birren,B.W., Gallagher,G., Smole,S., Park,D.J., MacInnis,B.L., Ryan,E., LaRocque,R., Rosenberg,E., Sabeti,P.C. |

| | | | |
|---|---|---|---|
| EPI_ISL_2660391, EPI_ISL_2660396, EPI_ISL_2660425, EPI_ISL_2660444 | | | |
| EPI_ISL_2660698, EPI_ISL_2660699, EPI_ISL_2660700, EPI_ISL_2660701, EPI_ISL_2660702, EPI_ISL_2660703, EPI_ISL_2660704 | AP SSO | CSIR-Centre for Cellular and Molecular Biology-INSACOG | Payel Mukherjee, Lamuk Zaveri, Onkar Kulkarni,Tulasi Nagabandi, Sofia Banu, Ara Sreenivas,Shreekant Verma, Amareshwar Vodapalli ,B Himasri,Valli Nagalakshmi Undamatla, Sumedha Avadhanula, Vidhyadhari Methuku, Priya Nurkurthy, Archana Bharadwaj Siva, Karthik Bharadwaj Tallapaka,Rakesh K Mishra,Divya Tej Sowpati |
| EPI_ISL_2660705, EPI_ISL_2660706, EPI_ISL_2660707, EPI_ISL_2660708, EPI_ISL_2660709, EPI_ISL_2660710, EPI_ISL_2660711, EPI_ISL_2660712, EPI_ISL_2660713, EPI_ISL_2660714 | AP SSO | CSIR-Centre for Cellular and Molecular Biology-INSACOG | Sofia Banu,Payel Mukherjee, Lamuk Zaveri, Onkar Kulkarni,Tulasi Nagabandi, Ara Sreenivas,Shreekant Verma, Amareshwar Vodapalli ,B Himasri,Valli Nagalakshmi Undamatla, Sumedha Avadhanula, Vidhyadhari Methuku, Priya Nurkurthy, Archana Bharadwaj Siva, Karthik Bharadwaj Tallapaka,Rakesh K Mishra,Divya Tej Sowpati |
| EPI_ISL_2660715, EPI_ISL_2660716, EPI_ISL_2660717, EPI_ISL_2660718, EPI_ISL_2660719, EPI_ISL_2660720, EPI_ISL_2660721, EPI_ISL_2660722, EPI_ISL_2660723, EPI_ISL_2660724, EPI_ISL_2660725 | | | |
| see above | AP SSO | CSIR-Centre for Cellular and Molecular Biology-INSACOG | Sumedha Avadhanula, Lamuk Zaveri, Tulasi Nagabandi, Ara Sreenivas,Shreekant Verma, Amareshwar Vodapalli ,B Himasri,Valli Nagalakshmi Undamatla, Vidhyadhari Methuku, Priya Nurkurthy, Payel Mukherjee,Onkar Kulkarni, Sofia Banu, Archana Bharadwaj Siva, Karthik Bharadwaj Tallapaka,Rakesh K Mishra,Divya Tej Sowpati |
| EPI_ISL_2660726, EPI_ISL_2660727, EPI_ISL_2660728, EPI_ISL_2660729, EPI_ISL_2660730, EPI_ISL_2660731, EPI_ISL_2660732, EPI_ISL_2660733, EPI_ISL_2660734 | AP SSO | CSIR-Centre for Cellular and Molecular Biology-INSACOG | Lamuk Zaveri,Tulasi Nagabandi,Payel Mukherjee,Onkar Kulkarni, Sofia Banu, Ara Sreenivas,Shreekant Verma, Amareshwar Vodapalli ,B Himasri,Valli Nagalakshmi Undamatla, Sumedha Avadhanula, Priya Nurkurthy, Vidhyadhari Methuku, Archana Bharadwaj Siva, Karthik Bharadwaj Tallapaka,Rakesh K Mishra,Divya Tej Sowpati |
| EPI_ISL_2660736, EPI_ISL_2660737, EPI_ISL_2660738, EPI_ISL_2660739, EPI_ISL_2660740, EPI_ISL_2660741, EPI_ISL_2660742, EPI_ISL_2660743, EPI_ISL_2660744, EPI_ISL_2660745, EPI_ISL_2660746, EPI_ISL_2660748 | | | |
| see above | AP SSO | CSIR-Centre for Cellular and Molecular Biology-INSACOG | Tulasi Nagabandi,Lamuk Zaveri,Payel Mukherjee,Onkar Kulkarni, Sofia Banu, Ara Sreenivas,Shreekant Verma, Amareshwar Vodapalli ,B Himasri,Valli Nagalakshmi Undamatla, Priya Nurkurthy,Sumedha Avadhanula, Vidhyadhari Methuku, Archana Bharadwaj Siva, Karthik Bharadwaj Tallapaka,Rakesh K Mishra,Divya Tej Sowpati |
| EPI_ISL_2660749, EPI_ISL_2660750, EPI_ISL_2660751, EPI_ISL_2660752, EPI_ISL_2660757, EPI_ISL_2660758, EPI_ISL_2660759 | AP SSO | CSIR-Centre for Cellular and Molecular Biology-INSACOG | Priya Nurkurthy,Ara Sreenivas,Shreekant Verma, Amareshwar Vodapalli ,Lamuk Zaveri,Tulasi Nagabandi,B Himasri,Valli Nagalakshmi Undamatla, Sumedha Avadhanula, Vidhyadhari Methuku,Payel Mukherjee,Onkar Kulkarni, Sofia Banu, Archana Bharadwaj Siva, Karthik Bharadwaj Tallapaka,Rakesh K Mishra,Divya Tej Sowpati |
| EPI_ISL_2660761, EPI_ISL_2660762, EPI_ISL_2660763, EPI_ISL_2660764, EPI_ISL_2660765, EPI_ISL_2660766, EPI_ISL_2660767, EPI_ISL_2660769 | AP SSO | CSIR-Centre for Cellular and Molecular Biology-INSACOG | Ara Sreenivas,Shreekant Verma, Amareshwar Vodapalli ,B Himasri,Valli Nagalakshmi Undamatla,Payel Mukherjee, Lamuk Zaveri, Onkar Kulkarni,Tulasi Nagabandi, Sofia Banu, ,Priya Nurkurthy, Vidhyadhari Methuku, Sumedha Avadhanula, Archana Bharadwaj Siva, Karthik Bharadwaj Tallapaka,Rakesh K Mishra,Divya Tej Sowpati |
| EPI_ISL_2660771, EPI_ISL_2660772, EPI_ISL_2660773, EPI_ISL_2660774, EPI_ISL_2660775, EPI_ISL_2660776, EPI_ISL_2660777, EPI_ISL_2660778, EPI_ISL_2660779, EPI_ISL_2660780, EPI_ISL_2660781, EPI_ISL_2660782, EPI_ISL_2660783, EPI_ISL_2660784, EPI_ISL_2660785, EPI_ISL_2660786 | | | |
| see above | AP SSO | CSIR-Centre for Cellular and Molecular Biology-INSACOG | Shreekant Verma, Amareshwar Vodapalli ,Ara Sreenivas,B Himasri,Valli Nagalakshmi Undamatla,Payel Mukherjee, Lamuk Zaveri, Onkar Kulkarni,Tulasi Nagabandi, Sofia Banu, Sumedha Avadhanula, Priya Nurkurthy, Vidhyadhari Methuku, Archana Bharadwaj Siva, Karthik Bharadwaj Tallapaka,Rakesh K Mishra,Divya Tej Sowpati |
| EPI_ISL_2660787, EPI_ISL_2660788, EPI_ISL_2660789, EPI_ISL_2660790, EPI_ISL_2660791, EPI_ISL_2660792, EPI_ISL_2660793, EPI_ISL_2660794, EPI_ISL_2660795, EPI_ISL_2660796, EPI_ISL_2660797, EPI_ISL_2660798, EPI_ISL_2660799, EPI_ISL_2660800, EPI_ISL_2660801, EPI_ISL_2660802, EPI_ISL_2660803, EPI_ISL_2660804, EPI_ISL_2660805, EPI_ISL_2660806, EPI_ISL_2660807 | | | |
| see above | AP SSO | CSIR-Centre for Cellular and Molecular Biology-INSACOG | Amareshwar Vodapalli,Ara Sreenivas,Shreekant Verma, B Himasri,Valli Nagalakshmi Undamatla,Payel Mukherjee, Lamuk Zaveri, Onkar Kulkarni,Tulasi Nagabandi, Sofia Banu, ,Priya Nurkurthy, Vidhyadhari Methuku, Sumedha Avadhanula, Archana Bharadwaj Siva, Karthik Bharadwaj Tallapaka,Rakesh K Mishra,Divya Tej Sowpati |
| EPI_ISL_2660808, EPI_ISL_2660809, EPI_ISL_2660810, EPI_ISL_2660811, EPI_ISL_2660812, EPI_ISL_2660813, EPI_ISL_2660814, EPI_ISL_2660815 | AP SSO | CSIR-Centre for Cellular and Molecular Biology-INSACOG | B Himasri,Amareshwar Vodapalli,Ara Sreenivas,Shreekant Verma,Valli Nagalakshmi Undamatla,Payel Mukherjee, Lamuk Zaveri, Onkar Kulkarni,Tulasi Nagabandi, Sofia Banu, Vidhyadhari Methuku,Priya Nurkurthy, Sumedha Avadhanula, Archana Bharadwaj Siva, Karthik Bharadwaj Tallapaka,Rakesh K Mishra,Divya Tej Sowpati |
| EPI_ISL_2660816, EPI_ISL_2660818, EPI_ISL_2660819, EPI_ISL_2660820, EPI_ISL_2660821, EPI_ISL_2660822, EPI_ISL_2660823, EPI_ISL_2660824, EPI_ISL_2660825, EPI_ISL_2660826, EPI_ISL_2660827, EPI_ISL_2660828 | | | |
| see above | PASTEUR MOLECULAR LABORATORY - INSACOG MEGHALAYA | CSIR-Centre for Cellular and Molecular Biology-INSACOG | R. Lyngdoh, E Shadap , E Marak,Payel Mukherjee, Lamuk Zaveri, Onkar Kulkarni,Tulasi Nagabandi, Sofia Banu, Ara Sreenivas,Shreekant Verma, Amareshwar Vodapalli ,B Himasri,Valli Nagalakshmi Undamatla, Sumedha Avadhanula, Vidhyadhari Methuku, Priya Nurkurthy, Archana Bharadwaj Siva, Karthik Bharadwaj Tallapaka,Rakesh K Mishra,Divya Tej Sowpati |
| EPI_ISL_2660830, EPI_ISL_2660831, EPI_ISL_2660832, EPI_ISL_2660833, EPI_ISL_2660834, EPI_ISL_2660835, EPI_ISL_2660836, EPI_ISL_2660837 | PASTEUR MOLECULAR LABORATORY - INSACOG MEGHALAYA | CSIR-Centre for Cellular and Molecular Biology-INSACOG | R. Lyngdoh, E Shadap , E Marak,Tulasi Nagabandi,Lamuk Zaveri,Payel Mukherjee,Onkar Kulkarni, Sofia Banu, Ara Sreenivas,Shreekant Verma, Amareshwar Vodapalli ,B Himasri,Valli Nagalakshmi Undamatla, Priya Nurkurthy,Sumedha Avadhanula, Vidhyadhari Methuku, Archana Bharadwaj Siva, Karthik Bharadwaj Tallapaka,Rakesh K Mishra,Divya Tej Sowpati |
| EPI_ISL_2660838, EPI_ISL_2660840, EPI_ISL_2660841, EPI_ISL_2660842, EPI_ISL_2660843, EPI_ISL_2660844 | PASTEUR MOLECULAR LABORATORY - INSACOG MEGHALAYA | CSIR-Centre for Cellular and Molecular Biology-INSACOG | R. Lyngdoh, E Shadap , E Marak,Lamuk Zaveri,Tulasi Nagabandi,Payel Mukherjee,Onkar Kulkarni, Sofia Banu, Ara Sreenivas,Shreekant Verma, Amareshwar Vodapalli ,B Himasri,Valli Nagalakshmi Undamatla, Sumedha Avadhanula, Priya Nurkurthy, Vidhyadhari Methuku, Archana Bharadwaj Siva, Karthik Bharadwaj Tallapaka,Rakesh K Mishra,Divya Tej Sowpati |
| EPI_ISL_2660845, EPI_ISL_2660846, EPI_ISL_2660847, EPI_ISL_2660848, EPI_ISL_2660849, EPI_ISL_2660850, EPI_ISL_2660851, EPI_ISL_2660852, EPI_ISL_2660854, EPI_ISL_2660855 | PASTEUR MOLECULAR LABORATORY - INSACOG MEGHALAYA | CSIR-Centre for Cellular and Molecular Biology-INSACOG | R. Lyngdoh, E Shadap , E Marak,Sofia Banu,Payel Mukherjee, Lamuk Zaveri, Onkar Kulkarni,Tulasi Nagabandi, Ara Sreenivas,Shreekant Verma, Amareshwar Vodapalli ,B Himasri,Valli Nagalakshmi Undamatla, Sumedha Avadhanula, Vidhyadhari Methuku, Priya Nurkurthy, Archana Bharadwaj Siva, Karthik Bharadwaj Tallapaka,Rakesh K Mishra,Divya Tej Sowpati |
| EPI_ISL_2660856, EPI_ISL_2660857, EPI_ISL_2660859, EPI_ISL_2660860, EPI_ISL_2660861, EPI_ISL_2660862, EPI_ISL_2660863, EPI_ISL_2660864 | PASTEUR MOLECULAR LABORATORY - INSACOG MEGHALAYA | CSIR-Centre for Cellular and Molecular Biology-INSACOG | R. Lyngdoh, E Shadap , E Marak,Payel Mukherjee, Lamuk Zaveri, Onkar Kulkarni,Tulasi Nagabandi, Sofia Banu, Ara Sreenivas,Shreekant Verma, Amareshwar Vodapalli ,B Himasri,Valli Nagalakshmi Undamatla, Sumedha Avadhanula, Vidhyadhari Methuku, Priya Nurkurthy, Archana Bharadwaj Siva, Karthik Bharadwaj Tallapaka,Rakesh K Mishra,Divya Tej Sowpati |
| EPI_ISL_2660865, | PASTEUR MOLECULAR LABORATORY - INSACOG MEGHALAYA | CSIR-Centre for Cellular and Molecular Biology-INSACOG | R. Lyngdoh, E Shadap , E Marak,Tulasi Nagabandi,Lamuk Zaveri,Payel Mukherjee,Onkar Kulkarni, Sofia Banu, Ara Sreenivas,Shreekant Verma, Amareshwar Vodapalli ,B Himasri,Valli Nagalakshmi Undamatla, Priya Nurkurthy,Sumedha |

| | | | |
|---|---|---|---|
| EPI_ISL_2660866, EPI_ISL_2660867, EPI_ISL_2660868, EPI_ISL_2660870, EPI_ISL_2660872, EPI_ISL_2660873, EPI_ISL_2660874, EPI_ISL_2660876 | | | Avadhanula, Vidhyadhari Methuku, Archana Bharadwaj Siva, Karthik Bharadwaj Tallapaka,Rakesh K Mishra,Divya Tej Sowpati |
| EPI_ISL_2660877, EPI_ISL_2660878, EPI_ISL_2660879, EPI_ISL_2660880, EPI_ISL_2660881, EPI_ISL_2660882, EPI_ISL_2660883, EPI_ISL_2660884 | PASTEUR MOLECULAR LABORATORY - INSACOG MEGHALAYA | CSIR-Centre for Cellular and Molecular Biology-INSACOG | R. Lyngdoh, E Shadap , E Marak,Lamuk Zaveri,Tulasi Nagabandi,Payel Mukherjee,Onkar Kulkarni, Sofia Banu, Ara Sreenivas,Shreekant Verma, Amareshwar Vodapalli ,B Himasri,Valli Nagalakshmi Undamatla, Sumedha Avadhanula, Priya Nurkurthy, Vidhyadhari Methuku, Archana Bharadwaj Siva, Karthik Bharadwaj Tallapaka,Rakesh K Mishra,Divya Tej Sowpati |
| EPI_ISL_2660885, EPI_ISL_2660886, EPI_ISL_2660887, EPI_ISL_2660888, EPI_ISL_2660889, EPI_ISL_2660890, EPI_ISL_2660891, EPI_ISL_2660892, EPI_ISL_2660894 | PASTEUR MOLECULAR LABORATORY - INSACOG MEGHALAYA | CSIR-Centre for Cellular and Molecular Biology-INSACOG | R. Lyngdoh, E Shadap , E Marak,Sofia Banu,Payel Mukherjee, Lamuk Zaveri, Onkar Kulkarni,Tulasi Nagabandi, Ara Sreenivas,Shreekant Verma, Amareshwar Vodapalli ,B Himasri,Valli Nagalakshmi Undamatla, Sumedha Avadhanula, Vidhyadhari Methuku, Priya Nurkurthy, Archana Bharadwaj Siva, Karthik Bharadwaj Tallapaka,Rakesh K Mishra,Divya Tej Sowpati |
| EPI_ISL_2660896, EPI_ISL_2660897, EPI_ISL_2660898, EPI_ISL_2660899, EPI_ISL_2660900, EPI_ISL_2660901, EPI_ISL_2660902, EPI_ISL_2660903, EPI_ISL_2660904, EPI_ISL_2660905 | PASTEUR MOLECULAR LABORATORY - INSACOG MEGHALAYA | CSIR-Centre for Cellular and Molecular Biology-INSACOG | R. Lyngdoh, E Shadap , E Marak,Payel Mukherjee, Lamuk Zaveri, Onkar Kulkarni,Tulasi Nagabandi, Sofia Banu, Ara Sreenivas,Shreekant Verma, Amareshwar Vodapalli ,B Himasri,Valli Nagalakshmi Undamatla, Sumedha Avadhanula, Vidhyadhari Methuku, Priya Nurkurthy, Archana Bharadwaj Siva, Karthik Bharadwaj Tallapaka,Rakesh K Mishra,Divya Tej Sowpati |
| EPI_ISL_2660906, EPI_ISL_2660907, EPI_ISL_2660908, EPI_ISL_2660909 | PASTEUR MOLECULAR LABORATORY - INSACOG MEGHALAYA | CSIR-Centre for Cellular and Molecular Biology-INSACOG | R. Lyngdoh, E Shadap , E Marak,Tulasi Nagabandi,Lamuk Zaveri,Payel Mukherjee,Onkar Kulkarni, Sofia Banu, Ara Sreenivas,Shreekant Verma, Amareshwar Vodapalli ,B Himasri,Valli Nagalakshmi Undamatla, Priya Nurkurthy,Sumedha Avadhanula, Vidhyadhari Methuku, Archana Bharadwaj Siva, Karthik Bharadwaj Tallapaka,Rakesh K Mishra,Divya Tej Sowpati |
| EPI_ISL_2660993 | CSIR-Centre for Cellular and Molecular Biology | CSIR-Centre for Cellular and Molecular Biology-INSACOG | Shreekant Verma, Amareshwar Vodapalli ,Ara Sreenivas,B Himasri,Valli Nagalakshmi Undamatla,Payel Mukherjee, Lamuk Zaveri, Onkar Kulkarni,Tulasi Nagabandi, Sofia Banu, Sumedha Avadhanula, Priya Nurkurthy, Vidhyadhari Methuku, Archana Bharadwaj Siva, Karthik Bharadwaj Tallapaka,Rakesh K Mishra,Divya Tej Sowpati |
| EPI_ISL_2661022 | CSIR-Centre for Cellular and Molecular Biology | CSIR-Centre for Cellular and Molecular Biology-INSACOG | Amareshwar Vodapalli,Ara Sreenivas,Shreekant Verma, B Himasri,Valli Nagalakshmi Undamatla,Payel Mukherjee, Lamuk Zaveri, Onkar Kulkarni,Tulasi Nagabandi, Sofia Banu, ,Priya Nurkurthy, Vidhyadhari Methuku, Sumedha Avadhanula, Archana Bharadwaj Siva, Karthik Bharadwaj Tallapaka,Rakesh K Mishra,Divya Tej Sowpati |
| EPI_ISL_2661097, EPI_ISL_2661098, EPI_ISL_2661099 | CSIR-Centre for Cellular and Molecular Biology | CSIR-Centre for Cellular and Molecular Biology-INSACOG | Lamuk Zaveri,Tulasi Nagabandi,Payel Mukherjee,Onkar Kulkarni, Sofia Banu, Ara Sreenivas,Shreekant Verma, Amareshwar Vodapalli ,B Himasri,Valli Nagalakshmi Undamatla, Sumedha Avadhanula, Priya Nurkurthy, Vidhyadhari Methuku, Archana Bharadwaj Siva, Karthik Bharadwaj Tallapaka,Rakesh K Mishra,Divya Tej Sowpati |
| EPI_ISL_2661102, EPI_ISL_2661105 | Mapmygenome | CSIR-Centre for Cellular and Molecular Biology-INSACOG | Lamuk Zaveri,Tulasi Nagabandi,Payel Mukherjee,Onkar Kulkarni, Sofia Banu, Ara Sreenivas,Shreekant Verma, Amareshwar Vodapalli ,B Himasri,Valli Nagalakshmi Undamatla, Sumedha Avadhanula, Priya Nurkurthy, Vidhyadhari Methuku, Archana Bharadwaj Siva, Karthik Bharadwaj Tallapaka,Rakesh K Mishra,Divya Tej Sowpati |
| EPI_ISL_2661107, EPI_ISL_2661108, EPI_ISL_2661110, EPI_ISL_2661112, EPI_ISL_2661117, EPI_ISL_2661120, EPI_ISL_2661121 | Mapmygenome | CSIR-Centre for Cellular and Molecular Biology-INSACOG | Tulasi Nagabandi,Lamuk Zaveri,Payel Mukherjee,Onkar Kulkarni, Sofia Banu, Ara Sreenivas,Shreekant Verma, Amareshwar Vodapalli ,B Himasri,Valli Nagalakshmi Undamatla, Priya Nurkurthy,Sumedha Avadhanula, Vidhyadhari Methuku, Archana Bharadwaj Siva, Karthik Bharadwaj Tallapaka,Rakesh K Mishra,Divya Tej Sowpati |
| EPI_ISL_2661126, EPI_ISL_2661127, EPI_ISL_2661128, EPI_ISL_2661130, EPI_ISL_2661131 | Mapmygenome | CSIR-Centre for Cellular and Molecular Biology-INSACOG | Ara Sreenivas,Shreekant Verma, Amareshwar Vodapalli ,B Himasri,Valli Nagalakshmi Undamatla,Payel Mukherjee, Lamuk Zaveri, Onkar Kulkarni,Tulasi Nagabandi, Sofia Banu, ,Priya Nurkurthy, Vidhyadhari Methuku, Sumedha Avadhanula, Archana Bharadwaj Siva, Karthik Bharadwaj Tallapaka,Rakesh K Mishra,Divya Tej Sowpati |
| EPI_ISL_2661134 | CSIR-Centre for Cellular and Molecular Biology | CSIR-Centre for Cellular and Molecular Biology-INSACOG | Ara Sreenivas,Shreekant Verma, Amareshwar Vodapalli ,B Himasri,Valli Nagalakshmi Undamatla,Payel Mukherjee, Lamuk Zaveri, Onkar Kulkarni,Tulasi Nagabandi, Sofia Banu, ,Priya Nurkurthy, Vidhyadhari Methuku, Sumedha Avadhanula, Archana Bharadwaj Siva, Karthik Bharadwaj Tallapaka,Rakesh K Mishra,Divya Tej Sowpati |
| EPI_ISL_2661137, EPI_ISL_2661138, EPI_ISL_2661140, EPI_ISL_2661142 | Mapmygenome | CSIR-Centre for Cellular and Molecular Biology-INSACOG | Ara Sreenivas,Shreekant Verma, Amareshwar Vodapalli ,B Himasri,Valli Nagalakshmi Undamatla,Payel Mukherjee, Lamuk Zaveri, Onkar Kulkarni,Tulasi Nagabandi, Sofia Banu, ,Priya Nurkurthy, Vidhyadhari Methuku, Sumedha Avadhanula, Archana Bharadwaj Siva, Karthik Bharadwaj Tallapaka,Rakesh K Mishra,Divya Tej Sowpati |
| EPI_ISL_2661143, EPI_ISL_2661144, EPI_ISL_2661145 | Mapmygenome | CSIR-Centre for Cellular and Molecular Biology-INSACOG | Shreekant Verma, Amareshwar Vodapalli ,Ara Sreenivas,B Himasri,Valli Nagalakshmi Undamatla,Payel Mukherjee, Lamuk Zaveri, Onkar Kulkarni,Tulasi Nagabandi, Sofia Banu, Sumedha Avadhanula, Priya Nurkurthy, Vidhyadhari Methuku, Archana Bharadwaj Siva, Karthik Bharadwaj Tallapaka,Rakesh K Mishra,Divya Tej Sowpati |
| EPI_ISL_2661155, EPI_ISL_2661156, EPI_ISL_2661157, EPI_ISL_2661158 | CSIR-Centre for Cellular and Molecular Biology | CSIR-Centre for Cellular and Molecular Biology-INSACOG | Shreekant Verma, Amareshwar Vodapalli ,Ara Sreenivas,B Himasri,Valli Nagalakshmi Undamatla,Payel Mukherjee, Lamuk Zaveri, Onkar Kulkarni,Tulasi Nagabandi, Sofia Banu, Sumedha Avadhanula, Priya Nurkurthy, Vidhyadhari Methuku, Archana Bharadwaj Siva, Karthik Bharadwaj Tallapaka,Rakesh K Mishra,Divya Tej Sowpati |
| EPI_ISL_2661159, EPI_ISL_2661160, EPI_ISL_2661161, EPI_ISL_2661162 | CSIR-Centre for Cellular and Molecular Biology | CSIR-Centre for Cellular and Molecular Biology-INSACOG | Sumedha Avadhanula, Lamuk Zaveri, Tulasi Nagabandi, Ara Sreenivas,Shreekant Verma, Amareshwar Vodapalli ,B Himasri,Valli Nagalakshmi Undamatla, Vidhyadhari Methuku, Priya Nurkurthy, Payel Mukherjee,Onkar Kulkarni, Sofia Banu, Archana Bharadwaj Siva, Karthik Bharadwaj Tallapaka,Rakesh K Mishra,Divya Tej Sowpati |
| EPI_ISL_2661163, EPI_ISL_2661164, EPI_ISL_2661165, EPI_ISL_2661166, EPI_ISL_2661167, EPI_ISL_2661168, EPI_ISL_2661169, EPI_ISL_2661170, EPI_ISL_2661171, EPI_ISL_2661172, EPI_ISL_2661173, EPI_ISL_2661174, EPI_ISL_2661175, EPI_ISL_2661176, EPI_ISL_2661177, EPI_ISL_2661178, EPI_ISL_2661179, EPI_ISL_2661180, EPI_ISL_2661181, EPI_ISL_2661182, EPI_ISL_2661183 | | | |
| see above | CSIR-Centre for Cellular and Molecular Biology | CSIR-Centre for Cellular and Molecular Biology-INSACOG | Amareshwar Vodapalli,Ara Sreenivas,Shreekant Verma, B Himasri,Valli Nagalakshmi Undamatla,Payel Mukherjee, Lamuk Zaveri, Onkar Kulkarni,Tulasi Nagabandi, Sofia Banu, ,Priya Nurkurthy, Vidhyadhari Methuku, Sumedha Avadhanula, Archana Bharadwaj Siva, Karthik Bharadwaj Tallapaka,Rakesh K Mishra,Divya Tej Sowpati |
| EPI_ISL_2661184, EPI_ISL_2661186, EPI_ISL_2661187, EPI_ISL_2661188, EPI_ISL_2661189, EPI_ISL_2661190, EPI_ISL_2661191, EPI_ISL_2661192, EPI_ISL_2661194, EPI_ISL_2661195 | CSIR-Centre for Cellular and Molecular Biology | CSIR-Centre for Cellular and Molecular Biology-INSACOG | B Himasri,Amareshwar Vodapalli,Ara Sreenivas,Shreekant Verma,Valli Nagalakshmi Undamatla,Payel Mukherjee, Lamuk Zaveri, Onkar Kulkarni,Tulasi Nagabandi, Sofia Banu, Vidhyadhari Methuku,Priya Nurkurthy, Sumedha Avadhanula, Archana Bharadwaj Siva, Karthik Bharadwaj Tallapaka,Rakesh K Mishra,Divya Tej Sowpati |
| EPI_ISL_2661196, EPI_ISL_2661197, EPI_ISL_2661198, EPI_ISL_2661199, EPI_ISL_2661200, EPI_ISL_2661201 | CSIR-Centre for Cellular and Molecular Biology | CSIR-Centre for Cellular and Molecular Biology-INSACOG | Lamuk Zaveri,Tulasi Nagabandi,Payel Mukherjee,Onkar Kulkarni, Sofia Banu, Ara Sreenivas,Shreekant Verma, Amareshwar Vodapalli ,B Himasri,Valli Nagalakshmi Undamatla, Sumedha Avadhanula, Priya Nurkurthy, Vidhyadhari Methuku, Archana Bharadwaj Siva, Karthik Bharadwaj Tallapaka,Rakesh K Mishra,Divya Tej Sowpati |
| EPI_ISL_2661203, EPI_ISL_2661204, EPI_ISL_2661205, EPI_ISL_2661206, EPI_ISL_2661207, EPI_ISL_2661208, EPI_ISL_2661209, EPI_ISL_2661210, EPI_ISL_2661211, EPI_ISL_2661213, EPI_ISL_2661214, EPI_ISL_2661215, EPI_ISL_2661216, EPI_ISL_2661217, EPI_ISL_2661218 | | | |
| see above | CSIR-Centre for Cellular and Molecular Biology | CSIR-Centre for Cellular and Molecular Biology-INSACOG | Tulasi Nagabandi,Lamuk Zaveri,Payel Mukherjee,Onkar Kulkarni, Sofia Banu, Ara Sreenivas,Shreekant Verma, Amareshwar Vodapalli ,B Himasri,Valli Nagalakshmi Undamatla, Priya Nurkurthy,Sumedha Avadhanula, Vidhyadhari Methuku, Archana Bharadwaj Siva, Karthik Bharadwaj Tallapaka,Rakesh K Mishra,Divya Tej Sowpati |
| EPI_ISL_2661219, EPI_ISL_2661220, | CSIR-Centre for Cellular and Molecular Biology | CSIR-Centre for Cellular and Molecular Biology-INSACOG | Ara Sreenivas,Shreekant Verma, Amareshwar Vodapalli ,B Himasri,Valli Nagalakshmi Undamatla,Payel Mukherjee, Lamuk Zaveri, Onkar Kulkarni,Tulasi Nagabandi, Sofia Banu, ,Priya Nurkurthy, Vidhyadhari Methuku, Sumedha Avadhanula, Archana Bharadwaj Siva, Karthik Bharadwaj Tallapaka,Rakesh K Mishra,Divya Tej Sowpati |

| | | | |
|---|---|---|---|
| EPI_ISL_2661221 | | | |
| EPI_ISL_2661222, EPI_ISL_2661223, EPI_ISL_2661224 | CSIR-Centre for Cellular and Molecular Biology | CSIR-Centre for Cellular and Molecular Biology-INSACOG | Shreekant Verma, Amareshwar Vodapalli ,Ara Sreenivas,B Himasri,Valli Nagalakshmi Undamatla,Payel Mukherjee, Lamuk Zaveri, Onkar Kulkarni,Tulasi Nagabandi, Sofia Banu, Sumedha Avadhanula, Priya Nurkurthy, Vidhyadhari Methuku, Archana Bharadwaj Siva, Karthik Bharadwaj Tallapaka,Rakesh K Mishra,Divya Tej Sowpati |
| EPI_ISL_2661226, EPI_ISL_2661227 | CSIR-Centre for Cellular and Molecular Biology | CSIR-Centre for Cellular and Molecular Biology-INSACOG | Amareshwar Vodapalli,Ara Sreenivas,Shreekant Verma, B Himasri,Valli Nagalakshmi Undamatla,Payel Mukherjee, Lamuk Zaveri, Onkar Kulkarni,Tulasi Nagabandi, Sofia Banu, ,Priya Nurkurthy, Vidhyadhari Methuku, Sumedha Avadhanula, Archana Bharadwaj Siva, Karthik Bharadwaj Tallapaka,Rakesh K Mishra,Divya Tej Sowpati |
| EPI_ISL_2661228, EPI_ISL_2661229 | CSIR-Centre for Cellular and Molecular Biology | CSIR-Centre for Cellular and Molecular Biology-INSACOG | B Himasri,Amareshwar Vodapalli,Ara Sreenivas,Shreekant Verma,Valli Nagalakshmi Undamatla,Payel Mukherjee, Lamuk Zaveri, Onkar Kulkarni,Tulasi Nagabandi, Sofia Banu, Vidhyadhari Methuku,Priya Nurkurthy, Sumedha Avadhanula, Archana Bharadwaj Siva, Karthik Bharadwaj Tallapaka,Rakesh K Mishra,Divya Tej Sowpati |
| EPI_ISL_2661231, EPI_ISL_2661232, EPI_ISL_2661233, EPI_ISL_2661234, EPI_ISL_2661235, EPI_ISL_2661236, EPI_ISL_2661237 | CSIR-Centre for Cellular and Molecular Biology | CSIR-Centre for Cellular and Molecular Biology-INSACOG | Payel Mukherjee, Lamuk Zaveri, Onkar Kulkarni,Tulasi Nagabandi, Sofia Banu, Ara Sreenivas,Shreekant Verma, Amareshwar Vodapalli ,B Himasri,Valli Nagalakshmi Undamatla, Sumedha Avadhanula, Vidhyadhari Methuku, Priya Nurkurthy, Archana Bharadwaj Siva, Karthik Bharadwaj Tallapaka,Rakesh K Mishra,Divya Tej Sowpati |
| EPI_ISL_2661238, EPI_ISL_2661239, EPI_ISL_2661240, EPI_ISL_2661241, EPI_ISL_2661242, EPI_ISL_2661243, EPI_ISL_2661244, EPI_ISL_2661245, EPI_ISL_2661246, EPI_ISL_2661247, EPI_ISL_2661248, EPI_ISL_2661249, EPI_ISL_2661250 | | | |
| see above | CSIR-Centre for Cellular and Molecular Biology | CSIR-Centre for Cellular and Molecular Biology-INSACOG | Sofia Banu,Payel Mukherjee, Lamuk Zaveri, Onkar Kulkarni,Tulasi Nagabandi, Ara Sreenivas,Shreekant Verma, Amareshwar Vodapalli ,B Himasri,Valli Nagalakshmi Undamatla, Sumedha Avadhanula, Vidhyadhari Methuku, Priya Nurkurthy, Archana Bharadwaj Siva, Karthik Bharadwaj Tallapaka,Rakesh K Mishra,Divya Tej Sowpati |
| EPI_ISL_2661251, EPI_ISL_2661253, EPI_ISL_2661254, EPI_ISL_2661255, EPI_ISL_2661256, EPI_ISL_2661257, EPI_ISL_2661258, EPI_ISL_2661259, EPI_ISL_2661260, EPI_ISL_2661261, EPI_ISL_2661262, EPI_ISL_2661263, EPI_ISL_2661264, EPI_ISL_2661265 | | | |
| see above | CSIR-Centre for Cellular and Molecular Biology | CSIR-Centre for Cellular and Molecular Biology-INSACOG | Lamuk Zaveri,Tulasi Nagabandi,Payel Mukherjee,Onkar Kulkarni, Sofia Banu, Ara Sreenivas,Shreekant Verma, Amareshwar Vodapalli ,B Himasri,Valli Nagalakshmi Undamatla, Sumedha Avadhanula, Priya Nurkurthy, Vidhyadhari Methuku, Archana Bharadwaj Siva, Karthik Bharadwaj Tallapaka,Rakesh K Mishra,Divya Tej Sowpati |
| EPI_ISL_2661267, EPI_ISL_2661268, EPI_ISL_2661269, EPI_ISL_2661270, EPI_ISL_2661271, EPI_ISL_2661272, EPI_ISL_2661273, EPI_ISL_2661274, EPI_ISL_2661275, EPI_ISL_2661276, EPI_ISL_2661279, EPI_ISL_2661281, EPI_ISL_2661282, EPI_ISL_2661283, EPI_ISL_2661284 | | | |
| see above | CSIR-Centre for Cellular and Molecular Biology | CSIR-Centre for Cellular and Molecular Biology-INSACOG | Tulasi Nagabandi,Lamuk Zaveri,Payel Mukherjee,Onkar Kulkarni, Sofia Banu, Ara Sreenivas,Shreekant Verma, Amareshwar Vodapalli ,B Himasri,Valli Nagalakshmi Undamatla, Priya Nurkurthy,Sumedha Avadhanula, Vidhyadhari Methuku, Archana Bharadwaj Siva, Karthik Bharadwaj Tallapaka,Rakesh K Mishra,Divya Tej Sowpati |
| EPI_ISL_2661285, EPI_ISL_2661286, EPI_ISL_2661287, EPI_ISL_2661288, EPI_ISL_2661289, EPI_ISL_2661290, EPI_ISL_2661291, EPI_ISL_2661292, EPI_ISL_2661293, EPI_ISL_2661294, EPI_ISL_2661295, EPI_ISL_2661296, EPI_ISL_2661297, EPI_ISL_2661298, EPI_ISL_2661299, EPI_ISL_2661300, EPI_ISL_2661301 | | | |
| see above | CSIR-Centre for Cellular and Molecular Biology | CSIR-Centre for Cellular and Molecular Biology-INSACOG | Ara Sreenivas,Shreekant Verma, Amareshwar Vodapalli ,B Himasri,Valli Nagalakshmi Undamatla,Payel Mukherjee, Lamuk Zaveri, Onkar Kulkarni,Tulasi Nagabandi, Sofia Banu, ,Priya Nurkurthy, Vidhyadhari Methuku, Sumedha Avadhanula, Archana Bharadwaj Siva, Karthik Bharadwaj Tallapaka,Rakesh K Mishra,Divya Tej Sowpati |
| EPI_ISL_2661302, EPI_ISL_2661303, EPI_ISL_2661304, EPI_ISL_2661305, EPI_ISL_2661306, EPI_ISL_2661307, EPI_ISL_2661308, EPI_ISL_2661310, EPI_ISL_2661311, EPI_ISL_2661312, EPI_ISL_2661313, EPI_ISL_2661314, EPI_ISL_2661315, EPI_ISL_2661316 | | | |
| see above | CSIR-Centre for Cellular and Molecular Biology | CSIR-Centre for Cellular and Molecular Biology-INSACOG | Shreekant Verma, Amareshwar Vodapalli ,Ara Sreenivas,B Himasri,Valli Nagalakshmi Undamatla,Payel Mukherjee, Lamuk Zaveri, Onkar Kulkarni,Tulasi Nagabandi, Sofia Banu, Sumedha Avadhanula, Priya Nurkurthy, Vidhyadhari Methuku, Archana Bharadwaj Siva, Karthik Bharadwaj Tallapaka,Rakesh K Mishra,Divya Tej Sowpati |
| EPI_ISL_2661318, EPI_ISL_2661319, EPI_ISL_2661320, EPI_ISL_2661321 | CSIR-Centre for Cellular and Molecular Biology | CSIR-Centre for Cellular and Molecular Biology-INSACOG | Sumedha Avadhanula, Lamuk Zaveri, Tulasi Nagabandi, Ara Sreenivas,Shreekant Verma, Amareshwar Vodapalli ,B Himasri,Valli Nagalakshmi Undamatla, Vidhyadhari Methuku, Priya Nurkurthy, Payel Mukherjee,Onkar Kulkarni, Sofia Banu, Archana Bharadwaj Siva, Karthik Bharadwaj Tallapaka,Rakesh K Mishra,Divya Tej Sowpati |
| EPI_ISL_2661322, EPI_ISL_2661323, EPI_ISL_2661324, EPI_ISL_2661325, EPI_ISL_2661326, EPI_ISL_2661327, EPI_ISL_2661328, EPI_ISL_2661329, EPI_ISL_2661330, EPI_ISL_2661331, EPI_ISL_2661332, EPI_ISL_2661333, EPI_ISL_2661334, EPI_ISL_2661335, EPI_ISL_2661336, EPI_ISL_2661337, EPI_ISL_2661338, EPI_ISL_2661339, EPI_ISL_2661340, EPI_ISL_2661341, EPI_ISL_2661342 | | | |
| see above | CSIR-Centre for Cellular and Molecular Biology | CSIR-Centre for Cellular and Molecular Biology-INSACOG | Amareshwar Vodapalli,Ara Sreenivas,Shreekant Verma, B Himasri,Valli Nagalakshmi Undamatla,Payel Mukherjee, Lamuk Zaveri, Onkar Kulkarni,Tulasi Nagabandi, Sofia Banu, ,Priya Nurkurthy, Vidhyadhari Methuku, Sumedha Avadhanula, Archana Bharadwaj Siva, Karthik Bharadwaj Tallapaka,Rakesh K Mishra,Divya Tej Sowpati |
| EPI_ISL_2661343, EPI_ISL_2661344, EPI_ISL_2661345, EPI_ISL_2661346, EPI_ISL_2661347, EPI_ISL_2661348, EPI_ISL_2661349, EPI_ISL_2661350, EPI_ISL_2661351, EPI_ISL_2661352, EPI_ISL_2661353, EPI_ISL_2661354 | | | |
| see above | CSIR-Centre for Cellular and Molecular Biology | CSIR-Centre for Cellular and Molecular Biology-INSACOG | B Himasri,Amareshwar Vodapalli,Ara Sreenivas,Shreekant Verma,Valli Nagalakshmi Undamatla,Payel Mukherjee, Lamuk Zaveri, Onkar Kulkarni,Tulasi Nagabandi, Sofia Banu, Vidhyadhari Methuku,Priya Nurkurthy, Sumedha Avadhanula, Archana Bharadwaj Siva, Karthik Bharadwaj Tallapaka,Rakesh K Mishra,Divya Tej Sowpati |
| EPI_ISL_2661355, EPI_ISL_2661356, EPI_ISL_2661357, EPI_ISL_2661358, EPI_ISL_2661359, EPI_ISL_2661360, EPI_ISL_2661361 | CSIR-Centre for Cellular and Molecular Biology | CSIR-Centre for Cellular and Molecular Biology-INSACOG | Lamuk Zaveri,Tulasi Nagabandi,Payel Mukherjee,Onkar Kulkarni, Sofia Banu, Ara Sreenivas,Shreekant Verma, Amareshwar Vodapalli ,B Himasri,Valli Nagalakshmi Undamatla, Sumedha Avadhanula, Priya Nurkurthy, Vidhyadhari Methuku, Archana Bharadwaj Siva, Karthik Bharadwaj Tallapaka,Rakesh K Mishra,Divya Tej Sowpati |
| EPI_ISL_2661362, EPI_ISL_2661363, EPI_ISL_2661364, EPI_ISL_2661365, EPI_ISL_2661366, EPI_ISL_2661367, EPI_ISL_2661368, EPI_ISL_2661369, EPI_ISL_2661370, EPI_ISL_2661371, EPI_ISL_2661372, EPI_ISL_2661373, EPI_ISL_2661375, EPI_ISL_2661376, EPI_ISL_2661377 | | | |
| see above | CSIR-Centre for Cellular and Molecular Biology | CSIR-Centre for Cellular and Molecular Biology-INSACOG | Tulasi Nagabandi,Lamuk Zaveri,Payel Mukherjee,Onkar Kulkarni, Sofia Banu, Ara Sreenivas,Shreekant Verma, Amareshwar Vodapalli ,B Himasri,Valli Nagalakshmi Undamatla, Priya Nurkurthy,Sumedha Avadhanula, Vidhyadhari Methuku, Archana Bharadwaj Siva, Karthik Bharadwaj Tallapaka,Rakesh K Mishra,Divya Tej Sowpati |
| EPI_ISL_2661378 | CSIR-Centre for Cellular and Molecular Biology | CSIR-Centre for Cellular and Molecular Biology-INSACOG | Ara Sreenivas,Shreekant Verma, Amareshwar Vodapalli ,B Himasri,Valli Nagalakshmi Undamatla,Payel Mukherjee, Lamuk Zaveri, Onkar Kulkarni,Tulasi Nagabandi, Sofia Banu, ,Priya Nurkurthy, Vidhyadhari Methuku, Sumedha Avadhanula, Archana Bharadwaj Siva, Karthik Bharadwaj Tallapaka,Rakesh K Mishra,Divya Tej Sowpati |
| EPI_ISL_2661382, EPI_ISL_2661383, EPI_ISL_2661385, EPI_ISL_2661386, EPI_ISL_2661387, EPI_ISL_2661388, EPI_ISL_2661390 | RUHS College of Medical Sciences | CSIR-Centre for Cellular and Molecular Biology - INSACOG | Dr. Gaurav Dalela,Dr. Vaseem Naheed Baig,Dr. Sonali Sharma,Dr. Ramesh Sharma,Dr. Nilofer Khayyam,Dr. Jitendra Panda,Dr. Sudhanshu Kacker,Dr. Raja Babu Panwar,Dr. Rajeev Gupta, Payel Mukherjee, Lamuk Zaveri, Jandhyala Sai Krishna,Onkar Kulkarni,Tulasi Nagabandi, Sofia Banu, Ara Sreenivas,Shreekant Verma, Amareshwar Vodapalli ,B Himasri,Valli Nagalakshmi Undamatla, Sumedha Avadhanula, Vidhyadhari Methuku, Priya Nurkurthy, Archana Bharadwaj Siva, Karthik Bharadwaj Tallapaka,Rakesh K Mishra,Divya Tej Sowpati |
| EPI_ISL_2661392, EPI_ISL_2661393, EPI_ISL_2661394 | RUHS College of Medical Sciences | CSIR-Centre for Cellular and Molecular Biology - INSACOG | Dr. Gaurav Dalela,Dr. Vaseem Naheed Baig,Dr. Sonali Sharma,Dr. Ramesh Sharma,Dr. Nilofer Khayyam,Dr. Jitendra Panda,Dr. Sudhanshu Kacker,Dr. Raja Babu Panwar,Dr. Rajeev Gupta, Lamuk Zaveri,Tulasi Nagabandi,Payel Mukherjee,Onkar Kulkarni,Jandhyala Sai Krishna, Sofia Banu, Ara Sreenivas,Shreekant Verma, Amareshwar Vodapalli ,B Himasri,Valli Nagalakshmi Undamatla, Sumedha Avadhanula, Priya Nurkurthy, Vidhyadhari Methuku, Archana Bharadwaj Siva, Karthik Bharadwaj Tallapaka,Rakesh K Mishra,Divya Tej Sowpati |
| EPI_ISL_2661395 | RUHS College of Medical Sciences | CSIR-Centre for Cellular and Molecular Biology - INSACOG | Dr. Gaurav Dalela,Dr. Vaseem Naheed Baig,Dr. Sonali Sharma,Dr. Ramesh Sharma,Dr. Nilofer Khayyam,Dr. Jitendra Panda,Dr. Sudhanshu Kacker,Dr. Raja Babu Panwar,Dr. Rajeev Gupta, Sofia Banu,Payel Mukherjee, Lamuk Zaveri, Jandhyala Sai Krishna,Onkar Kulkarni,Tulasi Nagabandi, Ara Sreenivas,Shreekant Verma, Amareshwar Vodapalli ,B Himasri,Valli Nagalakshmi Undamatla, Sumedha Avadhanula, Vidhyadhari Methuku, Priya Nurkurthy, Archana Bharadwaj Siva, Karthik Bharadwaj Tallapaka,Rakesh K Mishra,Divya Tej Sowpati |
| EPI_ISL_2661396, EPI_ISL_2661397, EPI_ISL_2661398, EPI_ISL_2661399, EPI_ISL_2661400 | RUHS College of Medical Sciences | CSIR-Centre for Cellular and Molecular Biology - INSACOG | Dr. Gaurav Dalela,Dr. Vaseem Naheed Baig,Dr. Sonali Sharma,Dr. Ramesh Sharma,Dr. Nilofer Khayyam,Dr. Jitendra Panda,Dr. Sudhanshu Kacker,Dr. Raja Babu Panwar,Dr. Rajeev Gupta, Lamuk Zaveri,Tulasi Nagabandi,Payel Mukherjee,Onkar Kulkarni,Jandhyala Sai Krishna, Sofia Banu, Ara Sreenivas,Shreekant Verma, Amareshwar Vodapalli ,B Himasri,Valli Nagalakshmi Undamatla, Sumedha Avadhanula, Priya Nurkurthy, Vidhyadhari Methuku, Archana Bharadwaj Siva, Karthik Bharadwaj Tallapaka,Rakesh K Mishra,Divya Tej Sowpati |
| EPI_ISL_2661401, EPI_ISL_2661402, EPI_ISL_2661403, EPI_ISL_2661404, EPI_ISL_2661405, EPI_ISL_2661406, EPI_ISL_2661407, EPI_ISL_2661408, EPI_ISL_2661409, EPI_ISL_2661410 | RUHS College of Medical Sciences | CSIR-Centre for Cellular and Molecular Biology - INSACOG | Dr. Gaurav Dalela,Dr. Vaseem Naheed Baig,Dr. Sonali Sharma,Dr. Ramesh Sharma,Dr. Nilofer Khayyam,Dr. Jitendra Panda,Dr. Sudhanshu Kacker,Dr. Raja Babu Panwar,Dr. Rajeev Gupta, Tulasi Nagabandi,Lamuk Zaveri,Jandhyala Sai Krishna,Payel Mukherjee,Onkar Kulkarni, Sofia Banu, Ara Sreenivas,Shreekant Verma, Amareshwar Vodapalli ,B Himasri,Valli Nagalakshmi Undamatla, Priya Nurkurthy,Sumedha Avadhanula, Vidhyadhari Methuku, Archana Bharadwaj Siva, Karthik Bharadwaj Tallapaka,Rakesh K Mishra,Divya Tej Sowpati |
| EPI_ISL_2661411 | RUHS College of Medical Sciences | CSIR-Centre for Cellular and Molecular Biology - INSACOG | Dr. Gaurav Dalela,Dr. Vaseem Naheed Baig,Dr. Sonali Sharma,Dr. Ramesh Sharma,Dr. Nilofer Khayyam,Dr. Jitendra Panda,Dr. Sudhanshu Kacker,Dr. Raja Babu Panwar,Dr. Rajeev Gupta, Ara Sreenivas,Shreekant Verma, Amareshwar Vodapalli ,B Himasri,Jandhyala Sai Krishna,Valli Nagalakshmi Undamatla,Payel Mukherjee, Lamuk Zaveri, Onkar Kulkarni,Tulasi Nagabandi, Sofia Banu, Priya Nurkurthy, Vidhyadhari Methuku, Sumedha Avadhanula, Archana Bharadwaj Siva, Karthik Bharadwaj Tallapaka,Rakesh K Mishra,Divya Tej Sowpati |
| EPI_ISL_2661412 | RUHS College of Medical Sciences | CSIR-Centre for Cellular and Molecular Biology - INSACOG | Dr. Gaurav Dalela,Dr. Vaseem Naheed Baig,Dr. Sonali Sharma,Dr. Ramesh Sharma,Dr. Nilofer Khayyam,Dr. Jitendra Panda,Dr. Sudhanshu Kacker,Dr. Raja Babu Panwar,Dr. Rajeev Gupta, Jandhyala Sai Krishna,Onkar Kulkarni, Lamuk Zaveri, Tulasi Nagabandi,Sofia Banu,Payel Mukherjee, Ara Sreenivas,Shreekant Verma, Amareshwar Vodapalli ,B Himasri,Valli Nagalakshmi Undamatla, Sumedha Avadhanula, Vidhyadhari Methuku, Priya Nurkurthy, Archana Bharadwaj Siva, Karthik Bharadwaj Tallapaka,Rakesh K Mishra,Divya Tej Sowpati |
| EPI_ISL_2661414, EPI_ISL_2661415 | RUHS College of Medical Sciences | CSIR-Centre for Cellular and Molecular Biology - INSACOG | Dr. Gaurav Dalela,Dr. Vaseem Naheed Baig,Dr. Sonali Sharma,Dr. Ramesh Sharma,Dr. Nilofer Khayyam,Dr. Jitendra Panda,Dr. Sudhanshu Kacker,Dr. Raja Babu Panwar,Dr. Rajeev Gupta, Ara Sreenivas,Shreekant Verma, Amareshwar Vodapalli ,B Himasri,Jandhyala Sai Krishna,Valli Nagalakshmi Undamatla,Payel Mukherjee, Lamuk Zaveri, Onkar Kulkarni,Tulasi Nagabandi, Sofia Banu, Priya Nurkurthy, Vidhyadhari Methuku, Sumedha Avadhanula, Archana Bharadwaj Siva, Karthik Bharadwaj Tallapaka,Rakesh K Mishra,Divya Tej Sowpati |
| EPI_ISL_2661416 | RUHS College of Medical Sciences | CSIR-Centre for Cellular and Molecular Biology - INSACOG | Dr. Gaurav Dalela,Dr. Vaseem Naheed Baig,Dr. Sonali Sharma,Dr. Ramesh Sharma,Dr. Nilofer Khayyam,Dr. Jitendra Panda,Dr. Sudhanshu Kacker,Dr. Raja Babu Panwar,Dr. Rajeev Gupta, Onkar Kulkarni, Lamuk Zaveri, Jandhyala Sai Krishna,Tulasi Nagabandi,Sofia Banu,Payel Mukherjee, Ara Sreenivas,Shreekant Verma, Amareshwar Vodapalli ,B Himasri,Valli Nagalakshmi Undamatla, Sumedha Avadhanula, Vidhyadhari Methuku, Priya Nurkurthy, Archana Bharadwaj Siva, Karthik Bharadwaj Tallapaka,Rakesh K Mishra,Divya Tej Sowpati |
| EPI_ISL_2661418, EPI_ISL_2661420, EPI_ISL_2661421 | RUHS College of Medical Sciences | CSIR-Centre for Cellular and Molecular Biology - INSACOG | Dr. Gaurav Dalela,Dr. Vaseem Naheed Baig,Dr. Sonali Sharma,Dr. Ramesh Sharma,Dr. Nilofer Khayyam,Dr. Jitendra Panda,Dr. Sudhanshu Kacker,Dr. Raja Babu Panwar,Dr. Rajeev Gupta, Ara Sreenivas,Shreekant Verma, Amareshwar Vodapalli ,B Himasri,Jandhyala Sai Krishna,Valli Nagalakshmi Undamatla,Payel Mukherjee, Lamuk Zaveri, Onkar Kulkarni,Tulasi Nagabandi, Sofia Banu, Priya Nurkurthy, Vidhyadhari Methuku, Sumedha Avadhanula, Archana Bharadwaj Siva, Karthik Bharadwaj Tallapaka,Rakesh K Mishra,Divya Tej Sowpati |

| | | | |
|---|---|---|---|
| EPI_ISL_2661422, EPI_ISL_2661423, EPI_ISL_2661424, EPI_ISL_2661425, EPI_ISL_2661426, EPI_ISL_2661427, EPI_ISL_2661428, EPI_ISL_2661429, EPI_ISL_2661430, EPI_ISL_2661431, EPI_ISL_2661432 | | | |
| see above | RUHS College of Medical Sciences | CSIR-Centre for Cellular and Molecular Biology - INSACOG | Dr. Gaurav Dalela,Dr. Vaseem Naheed Baig,Dr. Sonali Sharma,Dr. Ramesh Sharma,Dr. Nilofer Khayyam,Dr. Jitendra Panda,Dr. Sudhanshu Kacker,Dr. Raja Babu Panwar,Dr. Rajeev Gupta, Shreekant Verma, Amareshwar Vodapalli ,Ara Sreenivas,B Himasri,Valli Nagalakshmi Undamatla,Payel Mukherjee, Lamuk Zaveri, Onkar Kulkarni,Tulasi Nagabandi,Jandhyala Sai Krishna, Sofia Banu, Sumedha Avadhanula, Priya Nurkurthy, Vidhyadhari Methuku, Archana Bharadwaj Siva, Karthik Bharadwaj Tallapaka,Rakesh K Mishra,Divya Tej Sowpati |
| EPI_ISL_2661433, EPI_ISL_2661434 | RUHS College of Medical Sciences | CSIR-Centre for Cellular and Molecular Biology - INSACOG | Dr. Gaurav Dalela,Dr. Vaseem Naheed Baig,Dr. Sonali Sharma,Dr. Ramesh Sharma,Dr. Nilofer Khayyam,Dr. Jitendra Panda,Dr. Sudhanshu Kacker,Dr. Raja Babu Panwar,Dr. Rajeev Gupta, Amareshwar Vodapalli,Ara Sreenivas,Shreekant Verma, B Himasri,Valli Nagalakshmi Undamatla,Payel Mukherjee, Lamuk Zaveri, Onkar Kulkarni,Tulasi Nagabandi, Sofia Banu,Jandhyala Sai Krishna ,Priya Nurkurthy, Vidhyadhari Methuku, Sumedha Avadhanula, Archana Bharadwaj Siva, Karthik Bharadwaj Tallapaka,Rakesh K Mishra,Divya Tej Sowpati |
| EPI_ISL_2661435 | RUHS College of Medical Sciences | CSIR-Centre for Cellular and Molecular Biology - INSACOG | Dr. Gaurav Dalela,Dr. Vaseem Naheed Baig,Dr. Sonali Sharma,Dr. Ramesh Sharma,Dr. Nilofer Khayyam,Dr. Jitendra Panda,Dr. Sudhanshu Kacker,Dr. Raja Babu Panwar,Dr. Rajeev Gupta, Jandhyala Sai Krishna,Onkar Kulkarni, Lamuk Zaveri, Tulasi Nagabandi,Sofia Banu,Payel Mukherjee, Ara Sreenivas,Shreekant Verma, Amareshwar Vodapalli ,B Himasri,Valli Nagalakshmi Undamatla, Sumedha Avadhanula, Vidhyadhari Methuku, Priya Nurkurthy, Archana Bharadwaj Siva, Karthik Bharadwaj Tallapaka,Rakesh K Mishra,Divya Tej Sowpati |
| EPI_ISL_2661440, EPI_ISL_2661441, EPI_ISL_2661443 | RUHS College of Medical Sciences | CSIR-Centre for Cellular and Molecular Biology - INSACOG | Dr. Gaurav Dalela,Dr. Vaseem Naheed Baig,Dr. Sonali Sharma,Dr. Ramesh Sharma,Dr. Nilofer Khayyam,Dr. Jitendra Panda,Dr. Sudhanshu Kacker,Dr. Raja Babu Panwar,Dr. Rajeev Gupta, Amareshwar Vodapalli,Ara Sreenivas,Shreekant Verma, B Himasri,Valli Nagalakshmi Undamatla,Payel Mukherjee, Lamuk Zaveri, Onkar Kulkarni,Tulasi Nagabandi, Sofia Banu,Jandhyala Sai Krishna ,Priya Nurkurthy, Vidhyadhari Methuku, Sumedha Avadhanula, Archana Bharadwaj Siva, Karthik Bharadwaj Tallapaka,Rakesh K Mishra,Divya Tej Sowpati |
| EPI_ISL_2661444 | RUHS College of Medical Sciences | CSIR-Centre for Cellular and Molecular Biology - INSACOG | Dr. Gaurav Dalela,Dr. Vaseem Naheed Baig,Dr. Sonali Sharma,Dr. Ramesh Sharma,Dr. Nilofer Khayyam,Dr. Jitendra Panda,Dr. Sudhanshu Kacker,Dr. Raja Babu Panwar,Dr. Rajeev Gupta, B Himasri,Amareshwar Vodapalli,Ara Sreenivas,Shreekant Verma,Valli Nagalakshmi Undamatla,Jandhyala Sai Krishna,Payel Mukherjee, Lamuk Zaveri, Onkar Kulkarni,Tulasi Nagabandi, Sofia Banu, Vidhyadhari Methuku,Priya Nurkurthy, Sumedha Avadhanula, Archana Bharadwaj Siva, Karthik Bharadwaj Tallapaka,Rakesh K Mishra,Divya Tej Sowpati |
| EPI_ISL_2661446 | RUHS College of Medical Sciences | CSIR-Centre for Cellular and Molecular Biology - INSACOG | Dr. Gaurav Dalela,Dr. Vaseem Naheed Baig,Dr. Sonali Sharma,Dr. Ramesh Sharma,Dr. Nilofer Khayyam,Dr. Jitendra Panda,Dr. Sudhanshu Kacker,Dr. Raja Babu Panwar,Dr. Rajeev Gupta, Onkar Kulkarni, Lamuk Zaveri, Jandhyala Sai Krishna,Tulasi Nagabandi,Sofia Banu,Payel Mukherjee, Ara Sreenivas,Shreekant Verma, Amareshwar Vodapalli ,B Himasri,Valli Nagalakshmi Undamatla, Sumedha Avadhanula, Vidhyadhari Methuku, Priya Nurkurthy, Archana Bharadwaj Siva, Karthik Bharadwaj Tallapaka,Rakesh K Mishra,Divya Tej Sowpati |
| EPI_ISL_2661448, EPI_ISL_2661450, EPI_ISL_2661451, EPI_ISL_2661452, EPI_ISL_2661453 | RUHS College of Medical Sciences | CSIR-Centre for Cellular and Molecular Biology - INSACOG | Dr. Gaurav Dalela,Dr. Vaseem Naheed Baig,Dr. Sonali Sharma,Dr. Ramesh Sharma,Dr. Nilofer Khayyam,Dr. Jitendra Panda,Dr. Sudhanshu Kacker,Dr. Raja Babu Panwar,Dr. Rajeev Gupta, B Himasri,Amareshwar Vodapalli,Ara Sreenivas,Shreekant Verma,Valli Nagalakshmi Undamatla,Jandhyala Sai Krishna,Payel Mukherjee, Lamuk Zaveri, Onkar Kulkarni,Tulasi Nagabandi, Sofia Banu, Vidhyadhari Methuku,Priya Nurkurthy, Sumedha Avadhanula, Archana Bharadwaj Siva, Karthik Bharadwaj Tallapaka,Rakesh K Mishra,Divya Tej Sowpati |
| EPI_ISL_2661454, EPI_ISL_2661455, EPI_ISL_2661456, EPI_ISL_2661457, EPI_ISL_2661458, EPI_ISL_2661460, EPI_ISL_2661461 | RUHS College of Medical Sciences | CSIR-Centre for Cellular and Molecular Biology - INSACOG | Dr. Gaurav Dalela,Dr. Vaseem Naheed Baig,Dr. Sonali Sharma,Dr. Ramesh Sharma,Dr. Nilofer Khayyam,Dr. Jitendra Panda,Dr. Sudhanshu Kacker,Dr. Raja Babu Panwar,Dr. Rajeev Gupta, Jandhyala Sai Krishna,Onkar Kulkarni, Lamuk Zaveri, Tulasi Nagabandi,Sofia Banu,Payel Mukherjee, Ara Sreenivas,Shreekant Verma, Amareshwar Vodapalli ,B Himasri,Valli Nagalakshmi Undamatla, Sumedha Avadhanula, Vidhyadhari Methuku, Priya Nurkurthy, Archana Bharadwaj Siva, Karthik Bharadwaj Tallapaka,Rakesh K Mishra,Divya Tej Sowpati |
| EPI_ISL_2661462, EPI_ISL_2661464 | RUHS College of Medical Sciences | CSIR-Centre for Cellular and Molecular Biology - INSACOG | Dr. Gaurav Dalela,Dr. Vaseem Naheed Baig,Dr. Sonali Sharma,Dr. Ramesh Sharma,Dr. Nilofer Khayyam,Dr. Jitendra Panda,Dr. Sudhanshu Kacker,Dr. Raja Babu Panwar,Dr. Rajeev Gupta, Onkar Kulkarni, Lamuk Zaveri, Jandhyala Sai Krishna,Tulasi Nagabandi,Sofia Banu,Payel Mukherjee, Ara Sreenivas,Shreekant Verma, Amareshwar Vodapalli ,B Himasri,Valli Nagalakshmi Undamatla, Sumedha Avadhanula, Vidhyadhari Methuku, Priya Nurkurthy, Archana Bharadwaj Siva, Karthik Bharadwaj Tallapaka,Rakesh K Mishra,Divya Tej Sowpati |
| EPI_ISL_2661465, EPI_ISL_2661467, EPI_ISL_2661490 | US Air Force School of Aerospace Medicine | US Air Force School of Aerospace Medicine | Anthony Fries, Jennifer Meyer, William Gruner, Amanda Javorina, Carol Garrett, Sarah Purves, Clarise Starr, Elizabeth Macias |
| EPI_ISL_2661499 | US Air Force School of Aerospace Medicine | US Air Force School of Aerospace Medicine | Anthony Fries, Jennifer Meyer, William Gruner, Amanda Javorina, Carol Garrett, Sarah Purves, Clarise Starr, Elizabeth Macias, Fritz Castillo, Cole Anderson |
| EPI_ISL_2661516, EPI_ISL_2661518, EPI_ISL_2661520, EPI_ISL_2661522, EPI_ISL_2661523 | Queensland Health Forensic and Scientific Services | Queensland Health Forensic and Scientific Services | Son Nguyen |
| EPI_ISL_2661539, EPI_ISL_2661540, EPI_ISL_2661541 | UHTL, University Hospitals | UHTL, University Hospitals | Sadri,N., Alouani,D., Song,X. |
| EPI_ISL_2661552, EPI_ISL_2661557, EPI_ISL_2661558, EPI_ISL_2661564, EPI_ISL_2661566, EPI_ISL_2661572, EPI_ISL_2661574, EPI_ISL_2661575, EPI_ISL_2661576, EPI_ISL_2661577, EPI_ISL_2661578, EPI_ISL_2661579, EPI_ISL_2661593 | | | |
| see above | National Institute of Public Health | National Institute of Public Health | Helena Jirincova et al |
| EPI_ISL_2662287 | Reditus Laboratories | Reditus Laboratories | Joshua J. Geltz, Ph.D., Robert M. Sgambelluri, Ph.D., Rex Dyer, Ph.D., Cassy Philips, M.S., Alexa Eichelberger, M.S. |
| EPI_ISL_2662307, EPI_ISL_2662313, EPI_ISL_2662321, EPI_ISL_2662356, EPI_ISL_2662389, EPI_ISL_2662390, EPI_ISL_2662401, EPI_ISL_2662428, EPI_ISL_2662433, EPI_ISL_2662435, EPI_ISL_2662439 | | | |
| see above | Viollier AG | Department of Biosystems Science and Engineering, ETH Zürich | Andrea Patrizia Salzmann, Henriette Kurth, Christiane Beckmann, Ivan Topolsky, Chaoran Chen, Sarah Nadeau, Kim Philipp Jablonski, Lara Fuhrmann, Niko Beerenwinkel, Christoph Noppen, Olivier Kobel, Maurice Redondo, Tanja Stadler |
| EPI_ISL_2662453 | Laboratorio HUB -Azienda Ospedaliero Universitaria - AOU – Cagliari | Laboratorio SPOKE Biologia Molecolare -Azienda Ospedaliero Universitaria - AOU – Cagliari | Sara Fais, Valentina Medda, Alessandra Scano, Germano Orrù, Miriam Loddo, Riccardo Cappai, Ferdinando Coghe |
| EPI_ISL_2662589, EPI_ISL_2662600 | TAMBO MEMORIAL LABORATORY | National Institute for Communicable Diseases of the National Health Laboratory Service | Amoako DG, Everatt J, Scheepers C, Mohale T, Ntuli N, Mahlangu B, Mnguni A, Ismail A, Bhiman JN |
| EPI_ISL_2662623 | CHRIS HANI BARAGWANATH LABORATORY | National Institute for Communicable Diseases of the National Health Laboratory Service | Amoako DG, Everatt J, Scheepers C, Mohale T, Ntuli N, Mahlangu B, Mnguni A, Ismail A, Bhiman JN |
| EPI_ISL_2662730, EPI_ISL_2662752, EPI_ISL_2662759, EPI_ISL_2662801 | DPHL | Delaware Public Health Lab | Rebecca Savage |
| EPI_ISL_2662975 | Kaleida Health Laboratories | University at Buffalo Genomics and Bioinformatics Core | Jonathan Bard, Natalie Lamb, Alyssa Pohlman, Brandon Marzullo, Amanda Boccolucci, Norma Nowak, Donald Yergeau, Jennifer Surtees |
| EPI_ISL_2663000, EPI_ISL_2663001, EPI_ISL_2663002, EPI_ISL_2663003, EPI_ISL_2663004, EPI_ISL_2663005, EPI_ISL_2663006, EPI_ISL_2663007, EPI_ISL_2663008, EPI_ISL_2663009, EPI_ISL_2663010, EPI_ISL_2663011, EPI_ISL_2663012, EPI_ISL_2663013, EPI_ISL_2663015, EPI_ISL_2663016, EPI_ISL_2663017, EPI_ISL_2663018, EPI_ISL_2663019, EPI_ISL_2663020, EPI_ISL_2663021, EPI_ISL_2663022, EPI_ISL_2663023, EPI_ISL_2663024, EPI_ISL_2663025, EPI_ISL_2663026, EPI_ISL_2663027, EPI_ISL_2663028, EPI_ISL_2663029, EPI_ISL_2663030, EPI_ISL_2663031, EPI_ISL_2663032, EPI_ISL_2663033, EPI_ISL_2663034, EPI_ISL_2663035, EPI_ISL_2663036, EPI_ISL_2663037, EPI_ISL_2663038, EPI_ISL_2663039, EPI_ISL_2663040, EPI_ISL_2663041, EPI_ISL_2663042, EPI_ISL_2663043, EPI_ISL_2663044, EPI_ISL_2663046, EPI_ISL_2663047, EPI_ISL_2663048, EPI_ISL_2663049, EPI_ISL_2663051, EPI_ISL_2663052, EPI_ISL_2663053, EPI_ISL_2663055, EPI_ISL_2663056, EPI_ISL_2663057, EPI_ISL_2663059, EPI_ISL_2663060, EPI_ISL_2663061, EPI_ISL_2663062, EPI_ISL_2663063, EPI_ISL_2663064, EPI_ISL_2663065, EPI_ISL_2663066, EPI_ISL_2663067, EPI_ISL_2663068, EPI_ISL_2663069, EPI_ISL_2663070, EPI_ISL_2663072, EPI_ISL_2663073, EPI_ISL_2663074, EPI_ISL_2663075, EPI_ISL_2663076, EPI_ISL_2663077, EPI_ISL_2663078, EPI_ISL_2663079, EPI_ISL_2663081, EPI_ISL_2663082, EPI_ISL_2663083, EPI_ISL_2663084, EPI_ISL_2663085, EPI_ISL_2663086, EPI_ISL_2663087, EPI_ISL_2663088, EPI_ISL_2663089, EPI_ISL_2663090, EPI_ISL_2663091, EPI_ISL_2663092, EPI_ISL_2663093, EPI_ISL_2663094, EPI_ISL_2663095, EPI_ISL_2663096, EPI_ISL_2663097, EPI_ISL_2663098, EPI_ISL_2663099, EPI_ISL_2663101, EPI_ISL_2663102, EPI_ISL_2663103, EPI_ISL_2663105, EPI_ISL_2663106, EPI_ISL_2663109, EPI_ISL_2663110, EPI_ISL_2663111, EPI_ISL_2663113, EPI_ISL_2663114, EPI_ISL_2663116, EPI_ISL_2663119, EPI_ISL_2663122, EPI_ISL_2663125, EPI_ISL_2663130 | | | |
| see above | N.F. Gamaleya Research Center for Epidemiology and Microbiology | WHO National Influenza Centre Russian Federation | Andrey Komissarov, Artem Fadeev, Kseniya Komissarova, Oula Mansour, Kirill Varchenko, Mikhail Bakaev, Tamila Musaeva, Maria Timofeeva, Veronika Eder, Maria Pisareva, Nikita Yolshin, Daria Danilenko, Elizaveta Divisenko, Alina Odintsova, Elena Shidlovskaya, Ludmila Vasilchenko, Nadezhda Kuznetsova, Maria Nikiforova, Alexey Shchetinin, Artem Tkachuk, Andrei Pochtovyi, Andrei Siniavin, Denis Kleymenov, Evgeniia Bykonia, Liubov Popova, Elena Mazunina, Inna Dolzhikova, Anna Kovyrshina, Andrei Botikov, Dmitry Shcheblyakov, Vladimir Gushchin, Denis Logunov, Alexander Gintsburg, Ksenia Safina, Elena Nabieva, Georgii Bazykin, Dmitry Lioznov |
| EPI_ISL_2663131 | Ochsner Health | Bioinfoexperts, LLC | Judy Crabtree, Amy Feehan, Rebecca Rose, David J. Nolan, Tessa LaFleur, Julia-Garcia-Diaz, Lucio Miele, Susanna L. Lamers |
| EPI_ISL_2663133, EPI_ISL_2663136, EPI_ISL_2663138, EPI_ISL_2663141, EPI_ISL_2663144, EPI_ISL_2663147, EPI_ISL_2663150, EPI_ISL_2663153, EPI_ISL_2663156, EPI_ISL_2663159, EPI_ISL_2663161, EPI_ISL_2663164, EPI_ISL_2663167, EPI_ISL_2663175, EPI_ISL_2663178, EPI_ISL_2663183, EPI_ISL_2663186, EPI_ISL_2663189, EPI_ISL_2663191 | | | |
| see above | N.F. Gamaleya Research Center for Epidemiology and Microbiology | WHO National Influenza Centre Russian Federation | Andrey Komissarov, Artem Fadeev, Kseniya Komissarova, Oula Mansour, Kirill Varchenko, Mikhail Bakaev, Tamila Musaeva, Maria Timofeeva, Veronika Eder, Maria Pisareva, Nikita Yolshin, Daria Danilenko, Elizaveta Divisenko, Alina Odintsova, Elena Shidlovskaya, Ludmila Vasilchenko, Nadezhda Kuznetsova, Maria Nikiforova, Alexey Shchetinin, Artem Tkachuk, Andrei Pochtovyi, Andrei Siniavin, Denis Kleymenov, Evgeniia Bykonia, Liubov Popova, Elena Mazunina, Inna Dolzhikova, Anna Kovyrshina, Andrei Botikov, Dmitry Shcheblyakov, Vladimir Gushchin, Denis Logunov, Alexander Gintsburg, Ksenia Safina, Elena Nabieva, Georgii Bazykin, Dmitry Lioznov |
| EPI_ISL_2663197 | Minnesota Department of Health, Public Health Laboratory | Minnesota Department of Health, Public Health Laboratory | Alexandra Lorentz, Jacob Garfin, Matt Plumb, and Xiong Wang |
| EPI_ISL_2663199 | Maple Grove Hospital | Minnesota Department of Health, Public Health Laboratory | Alexandra Lorentz, Jacob Garfin, Matt Plumb, and Xiong Wang |
| EPI_ISL_2663205, EPI_ISL_2663210, EPI_ISL_2663211, EPI_ISL_2663214, EPI_ISL_2663218, EPI_ISL_2663219, EPI_ISL_2663222, EPI_ISL_2663223, EPI_ISL_2663224, EPI_ISL_2663227 | HealthPartners Central Lab | Minnesota Department of Health, Public Health Laboratory | Alexandra Lorentz, Jacob Garfin, Matt Plumb, and Xiong Wang |
| EPI_ISL_2663244, EPI_ISL_2663245, EPI_ISL_2663246, EPI_ISL_2663248, EPI_ISL_2663250, EPI_ISL_2663251 | Fondazione IRCCS Ca' Granda Ospedale Maggiore Policlinico | Fondazione IRCCS Ca' Granda Ospedale Maggiore Policlinico | Ferruccio Ceriotti, Sara Uceda Renteria |
| EPI_ISL_2663318, EPI_ISL_2663319, EPI_ISL_2663320, | Communicable Disease Laboratory, Public Health Directorate | Communicable Disease Laboratory, Public Health Directorate | Alwasti,H., AlHujairi,Z., AlAbbas,Z., Marhoon,A., Almoamen,G. |

| | | | |
|---|---|---|---|
| EPI_ISL_2663321, EPI_ISL_2663322, EPI_ISL_2663323, EPI_ISL_2663324 | | | |
| EPI_ISL_2663483, EPI_ISL_2663489, EPI_ISL_2663492, EPI_ISL_2663508, EPI_ISL_2663510 | Virginia Division of Consolidated Laboratory Services | Virginia Division of Consolidated Laboratory Services | Virginia Division of Consolidated Laboratory Services |
| EPI_ISL_2663544, EPI_ISL_2663546, EPI_ISL_2663553 | NYU Langone Health | Departments of Pathology and Medicine, New York University School of Medicine | Adriana Heguy, Dacia Dimartino, Emily Guzman, Christian Marier, Peter Meyn, Sitharam Ramaswami, Gael Westby, Paul Zappile, Yutong Zhang, Paolo Cotzia, Guiqing Wang |
| EPI_ISL_2663797, EPI_ISL_2663798, EPI_ISL_2663799, EPI_ISL_2663801, EPI_ISL_2663802, EPI_ISL_2663803, EPI_ISL_2663804, EPI_ISL_2663805, EPI_ISL_2663806, EPI_ISL_2663807, EPI_ISL_2663808, EPI_ISL_2663809, EPI_ISL_2663810, EPI_ISL_2663811, EPI_ISL_2663812, EPI_ISL_2663813, EPI_ISL_2663814, EPI_ISL_2663815, EPI_ISL_2663816 | | | |
| see above | Lighthouse Lab in Milton Keynes | Wellcome Sanger Institute for the COVID-19 Genomics UK (COG-UK) Consortium | The Lighthouse Lab in Milton Keynes and Alex Alderton, Roberto Amato, Jeffrey Barrett, Sonia Goncalves, Ewan Harrison, David K. Jackson, Ian Johnston, Dominic Kwiatkowski, Cordelia Langford, John Sillitoe on behalf of the Wellcome Sanger Institute COVID-19 Surveillance Team |
| EPI_ISL_2663817, EPI_ISL_2663818 | Lighthouse Lab in Glasgow | Wellcome Sanger Institute for the COVID-19 Genomics UK (COG-UK) Consortium | Harper VanSteenhouse, Yumi Kasai, David Gray, Carol Clugston, Anna Dominiczak and Alex Alderton, Roberto Amato, Jeffrey Barrett, Sonia Goncalves, Ewan Harrison, David K. Jackson, Ian Johnston, Dominic Kwiatkowski, Cordelia Langford, John Sillitoe on behalf of the Wellcome Sanger Institute COVID-19 Surveillance Team |
| EPI_ISL_2663819, EPI_ISL_2663820, EPI_ISL_2663821, EPI_ISL_2663822 | Lighthouse Lab in Milton Keynes | Wellcome Sanger Institute for the COVID-19 Genomics UK (COG-UK) Consortium | The Lighthouse Lab in Milton Keynes and Alex Alderton, Roberto Amato, Jeffrey Barrett, Sonia Goncalves, Ewan Harrison, David K. Jackson, Ian Johnston, Dominic Kwiatkowski, Cordelia Langford, John Sillitoe on behalf of the Wellcome Sanger Institute COVID-19 Surveillance Team |
| EPI_ISL_2663823 | Lighthouse Lab in Glasgow | Wellcome Sanger Institute for the COVID-19 Genomics UK (COG-UK) Consortium | Harper VanSteenhouse, Yumi Kasai, David Gray, Carol Clugston, Anna Dominiczak and Alex Alderton, Roberto Amato, Jeffrey Barrett, Sonia Goncalves, Ewan Harrison, David K. Jackson, Ian Johnston, Dominic Kwiatkowski, Cordelia Langford, John Sillitoe on behalf of the Wellcome Sanger Institute COVID-19 Surveillance Team |
| EPI_ISL_2663824, EPI_ISL_2663825, EPI_ISL_2663826 | Lighthouse Lab in Milton Keynes | Wellcome Sanger Institute for the COVID-19 Genomics UK (COG-UK) Consortium | The Lighthouse Lab in Milton Keynes and Alex Alderton, Roberto Amato, Jeffrey Barrett, Sonia Goncalves, Ewan Harrison, David K. Jackson, Ian Johnston, Dominic Kwiatkowski, Cordelia Langford, John Sillitoe on behalf of the Wellcome Sanger Institute COVID-19 Surveillance Team |
| EPI_ISL_2663827, EPI_ISL_2663828, EPI_ISL_2663829 | Lighthouse Lab in Glasgow | Wellcome Sanger Institute for the COVID-19 Genomics UK (COG-UK) Consortium | Harper VanSteenhouse, Yumi Kasai, David Gray, Carol Clugston, Anna Dominiczak and Alex Alderton, Roberto Amato, Jeffrey Barrett, Sonia Goncalves, Ewan Harrison, David K. Jackson, Ian Johnston, Dominic Kwiatkowski, Cordelia Langford, John Sillitoe on behalf of the Wellcome Sanger Institute COVID-19 Surveillance Team |
| EPI_ISL_2663830, EPI_ISL_2663831, EPI_ISL_2663832, EPI_ISL_2663833, EPI_ISL_2663834, EPI_ISL_2663835, EPI_ISL_2663836, EPI_ISL_2663837 | Lighthouse Lab in Milton Keynes | Wellcome Sanger Institute for the COVID-19 Genomics UK (COG-UK) Consortium | The Lighthouse Lab in Milton Keynes and Alex Alderton, Roberto Amato, Jeffrey Barrett, Sonia Goncalves, Ewan Harrison, David K. Jackson, Ian Johnston, Dominic Kwiatkowski, Cordelia Langford, John Sillitoe on behalf of the Wellcome Sanger Institute COVID-19 Surveillance Team |
| EPI_ISL_2663838 | Lighthouse Lab in Glasgow | Wellcome Sanger Institute for the COVID-19 Genomics UK (COG-UK) Consortium | Harper VanSteenhouse, Yumi Kasai, David Gray, Carol Clugston, Anna Dominiczak and Alex Alderton, Roberto Amato, Jeffrey Barrett, Sonia Goncalves, Ewan Harrison, David K. Jackson, Ian Johnston, Dominic Kwiatkowski, Cordelia Langford, John Sillitoe on behalf of the Wellcome Sanger Institute COVID-19 Surveillance Team |
| EPI_ISL_2663839, EPI_ISL_2663840, EPI_ISL_2663841, EPI_ISL_2663842, EPI_ISL_2663843, EPI_ISL_2663844, EPI_ISL_2663845, EPI_ISL_2663846, EPI_ISL_2663847, EPI_ISL_2663848 | Lighthouse Lab in Milton Keynes | Wellcome Sanger Institute for the COVID-19 Genomics UK (COG-UK) Consortium | The Lighthouse Lab in Milton Keynes and Alex Alderton, Roberto Amato, Jeffrey Barrett, Sonia Goncalves, Ewan Harrison, David K. Jackson, Ian Johnston, Dominic Kwiatkowski, Cordelia Langford, John Sillitoe on behalf of the Wellcome Sanger Institute COVID-19 Surveillance Team |
| EPI_ISL_2663849 | Lighthouse Lab in Glasgow | Wellcome Sanger Institute for the COVID-19 Genomics UK (COG-UK) Consortium | Harper VanSteenhouse, Yumi Kasai, David Gray, Carol Clugston, Anna Dominiczak and Alex Alderton, Roberto Amato, Jeffrey Barrett, Sonia Goncalves, Ewan Harrison, David K. Jackson, Ian Johnston, Dominic Kwiatkowski, Cordelia Langford, John Sillitoe on behalf of the Wellcome Sanger Institute COVID-19 Surveillance Team |
| EPI_ISL_2663850, EPI_ISL_2663851, EPI_ISL_2663852, EPI_ISL_2663853, EPI_ISL_2663854 | Lighthouse Lab in Milton Keynes | Wellcome Sanger Institute for the COVID-19 Genomics UK (COG-UK) Consortium | The Lighthouse Lab in Milton Keynes and Alex Alderton, Roberto Amato, Jeffrey Barrett, Sonia Goncalves, Ewan Harrison, David K. Jackson, Ian Johnston, Dominic Kwiatkowski, Cordelia Langford, John Sillitoe on behalf of the Wellcome Sanger Institute COVID-19 Surveillance Team |
| EPI_ISL_2663855 | Lighthouse Lab in Glasgow | Wellcome Sanger Institute for the COVID-19 Genomics UK (COG-UK) Consortium | Harper VanSteenhouse, Yumi Kasai, David Gray, Carol Clugston, Anna Dominiczak and Alex Alderton, Roberto Amato, Jeffrey Barrett, Sonia Goncalves, Ewan Harrison, David K. Jackson, Ian Johnston, Dominic Kwiatkowski, Cordelia Langford, John Sillitoe on behalf of the Wellcome Sanger Institute COVID-19 Surveillance Team |
| EPI_ISL_2663856, EPI_ISL_2663857, EPI_ISL_2663858, EPI_ISL_2663859, EPI_ISL_2663860, EPI_ISL_2663861, EPI_ISL_2663862, EPI_ISL_2663863 | Lighthouse Lab in Milton Keynes | Wellcome Sanger Institute for the COVID-19 Genomics UK (COG-UK) Consortium | The Lighthouse Lab in Milton Keynes and Alex Alderton, Roberto Amato, Jeffrey Barrett, Sonia Goncalves, Ewan Harrison, David K. Jackson, Ian Johnston, Dominic Kwiatkowski, Cordelia Langford, John Sillitoe on behalf of the Wellcome Sanger Institute COVID-19 Surveillance Team |
| EPI_ISL_2663865, EPI_ISL_2663866 | Lighthouse Lab in Glasgow | Wellcome Sanger Institute for the COVID-19 Genomics UK (COG-UK) Consortium | Harper VanSteenhouse, Yumi Kasai, David Gray, Carol Clugston, Anna Dominiczak and Alex Alderton, Roberto Amato, Jeffrey Barrett, Sonia Goncalves, Ewan Harrison, David K. Jackson, Ian Johnston, Dominic Kwiatkowski, Cordelia Langford, John Sillitoe on behalf of the Wellcome Sanger Institute COVID-19 Surveillance Team |
| EPI_ISL_2663867, EPI_ISL_2663868, EPI_ISL_2663869, EPI_ISL_2663870 | Lighthouse Lab in Milton Keynes | Wellcome Sanger Institute for the COVID-19 Genomics UK (COG-UK) Consortium | The Lighthouse Lab in Milton Keynes and Alex Alderton, Roberto Amato, Jeffrey Barrett, Sonia Goncalves, Ewan Harrison, David K. Jackson, Ian Johnston, Dominic Kwiatkowski, Cordelia Langford, John Sillitoe on behalf of the Wellcome Sanger Institute COVID-19 Surveillance Team |
| EPI_ISL_2663871 | Lighthouse Lab in Glasgow | Wellcome Sanger Institute for the COVID-19 Genomics UK (COG-UK) Consortium | Harper VanSteenhouse, Yumi Kasai, David Gray, Carol Clugston, Anna Dominiczak and Alex Alderton, Roberto Amato, Jeffrey Barrett, Sonia Goncalves, Ewan Harrison, David K. Jackson, Ian Johnston, Dominic Kwiatkowski, Cordelia Langford, John Sillitoe on behalf of the Wellcome Sanger Institute COVID-19 Surveillance Team |
| EPI_ISL_2663872, EPI_ISL_2663873, EPI_ISL_2663874, EPI_ISL_2663875, EPI_ISL_2663876, EPI_ISL_2663877, EPI_ISL_2663878, EPI_ISL_2663879, EPI_ISL_2663880, EPI_ISL_2663881 | Lighthouse Lab in Milton Keynes | Wellcome Sanger Institute for the COVID-19 Genomics UK (COG-UK) Consortium | The Lighthouse Lab in Milton Keynes and Alex Alderton, Roberto Amato, Jeffrey Barrett, Sonia Goncalves, Ewan Harrison, David K. Jackson, Ian Johnston, Dominic Kwiatkowski, Cordelia Langford, John Sillitoe on behalf of the Wellcome Sanger Institute COVID-19 Surveillance Team |
| EPI_ISL_2663882 | Lighthouse Lab in Glasgow | Wellcome Sanger Institute for the COVID-19 Genomics UK (COG-UK) Consortium | Harper VanSteenhouse, Yumi Kasai, David Gray, Carol Clugston, Anna Dominiczak and Alex Alderton, Roberto Amato, Jeffrey Barrett, Sonia Goncalves, Ewan Harrison, David K. Jackson, Ian Johnston, Dominic Kwiatkowski, Cordelia Langford, John Sillitoe on behalf of the Wellcome Sanger Institute COVID-19 Surveillance Team |
| EPI_ISL_2663883, EPI_ISL_2663884, EPI_ISL_2663885 | Lighthouse Lab in Milton Keynes | Wellcome Sanger Institute for the COVID-19 Genomics UK (COG-UK) Consortium | The Lighthouse Lab in Milton Keynes and Alex Alderton, Roberto Amato, Jeffrey Barrett, Sonia Goncalves, Ewan Harrison, David K. Jackson, Ian Johnston, Dominic Kwiatkowski, Cordelia Langford, John Sillitoe on behalf of the Wellcome Sanger Institute COVID-19 Surveillance Team |
| EPI_ISL_2663886 | Lighthouse Lab in Glasgow | Wellcome Sanger Institute for the COVID-19 Genomics UK (COG-UK) Consortium | Harper VanSteenhouse, Yumi Kasai, David Gray, Carol Clugston, Anna Dominiczak and Alex Alderton, Roberto Amato, Jeffrey Barrett, Sonia Goncalves, Ewan Harrison, David K. Jackson, Ian Johnston, Dominic Kwiatkowski, Cordelia Langford, John Sillitoe on behalf of the Wellcome Sanger Institute COVID-19 Surveillance Team |
| EPI_ISL_2663887, EPI_ISL_2663888, EPI_ISL_2663889, EPI_ISL_2663890, EPI_ISL_2663891, EPI_ISL_2663892, EPI_ISL_2663893, EPI_ISL_2663894, EPI_ISL_2663895, EPI_ISL_2663896, EPI_ISL_2663897, EPI_ISL_2663898, EPI_ISL_2663899, EPI_ISL_2663900, EPI_ISL_2663901 | | | |
| see above | Lighthouse Lab in Milton Keynes | Wellcome Sanger Institute for the COVID-19 Genomics UK (COG-UK) Consortium | The Lighthouse Lab in Milton Keynes and Alex Alderton, Roberto Amato, Jeffrey Barrett, Sonia Goncalves, Ewan Harrison, David K. Jackson, Ian Johnston, Dominic Kwiatkowski, Cordelia Langford, John Sillitoe on behalf of the Wellcome Sanger Institute COVID-19 Surveillance Team |
| EPI_ISL_2663902, EPI_ISL_2663903 | Lighthouse Lab in Glasgow | Wellcome Sanger Institute for the COVID-19 Genomics UK (COG-UK) Consortium | Harper VanSteenhouse, Yumi Kasai, David Gray, Carol Clugston, Anna Dominiczak and Alex Alderton, Roberto Amato, Jeffrey Barrett, Sonia Goncalves, Ewan Harrison, David K. Jackson, Ian Johnston, Dominic Kwiatkowski, Cordelia Langford, John Sillitoe on behalf of the Wellcome Sanger Institute COVID-19 Surveillance Team |
| EPI_ISL_2663904, EPI_ISL_2663905, | Lighthouse Lab in Milton Keynes | Wellcome Sanger Institute for the COVID-19 Genomics UK (COG-UK) Consortium | The Lighthouse Lab in Milton Keynes and Alex Alderton, Roberto Amato, Jeffrey Barrett, Sonia Goncalves, Ewan Harrison, David K. Jackson, Ian Johnston, Dominic Kwiatkowski, Cordelia Langford, John Sillitoe on behalf of the Wellcome Sanger Institute COVID-19 Surveillance Team |

| | | | |
|---|---|---|---|
| EPI_ISL_2663906 | | | |
| EPI_ISL_2663907 | Lighthouse Lab in Glasgow | Wellcome Sanger Institute for the COVID-19 Genomics UK (COG-UK) Consortium | Harper VanSteenhouse, Yumi Kasai, David Gray, Carol Clugston, Anna Dominiczak and Alex Alderton, Roberto Amato, Jeffrey Barrett, Sonia Goncalves, Ewan Harrison, David K. Jackson, Ian Johnston, Dominic Kwiatkowski, Cordelia Langford, John Sillitoe on behalf of the Wellcome Sanger Institute COVID-19 Surveillance Team |
| EPI_ISL_2663908, EPI_ISL_2663909, EPI_ISL_2663910, EPI_ISL_2663911, EPI_ISL_2663912, EPI_ISL_2663913, EPI_ISL_2663914, EPI_ISL_2663915, EPI_ISL_2663916 | Lighthouse Lab in Milton Keynes | Wellcome Sanger Institute for the COVID-19 Genomics UK (COG-UK) Consortium | The Lighthouse Lab in Milton Keynes and Alex Alderton, Roberto Amato, Jeffrey Barrett, Sonia Goncalves, Ewan Harrison, David K. Jackson, Ian Johnston, Dominic Kwiatkowski, Cordelia Langford, John Sillitoe on behalf of the Wellcome Sanger Institute COVID-19 Surveillance Team |
| EPI_ISL_2663917 | Lighthouse Lab in Glasgow | Wellcome Sanger Institute for the COVID-19 Genomics UK (COG-UK) Consortium | Harper VanSteenhouse, Yumi Kasai, David Gray, Carol Clugston, Anna Dominiczak and Alex Alderton, Roberto Amato, Jeffrey Barrett, Sonia Goncalves, Ewan Harrison, David K. Jackson, Ian Johnston, Dominic Kwiatkowski, Cordelia Langford, John Sillitoe on behalf of the Wellcome Sanger Institute COVID-19 Surveillance Team |
| EPI_ISL_2663918, EPI_ISL_2663919, EPI_ISL_2663920, EPI_ISL_2663921, EPI_ISL_2663923, EPI_ISL_2663924, EPI_ISL_2663925, EPI_ISL_2663926, EPI_ISL_2663927, EPI_ISL_2663928, EPI_ISL_2663929 | | | |
| see above | Lighthouse Lab in Milton Keynes | Wellcome Sanger Institute for the COVID-19 Genomics UK (COG-UK) Consortium | The Lighthouse Lab in Milton Keynes and Alex Alderton, Roberto Amato, Jeffrey Barrett, Sonia Goncalves, Ewan Harrison, David K. Jackson, Ian Johnston, Dominic Kwiatkowski, Cordelia Langford, John Sillitoe on behalf of the Wellcome Sanger Institute COVID-19 Surveillance Team |
| EPI_ISL_2663931 | Lighthouse Lab in Glasgow | Wellcome Sanger Institute for the COVID-19 Genomics UK (COG-UK) Consortium | Harper VanSteenhouse, Yumi Kasai, David Gray, Carol Clugston, Anna Dominiczak and Alex Alderton, Roberto Amato, Jeffrey Barrett, Sonia Goncalves, Ewan Harrison, David K. Jackson, Ian Johnston, Dominic Kwiatkowski, Cordelia Langford, John Sillitoe on behalf of the Wellcome Sanger Institute COVID-19 Surveillance Team |
| EPI_ISL_2663932 | Lighthouse Lab in Milton Keynes | Wellcome Sanger Institute for the COVID-19 Genomics UK (COG-UK) Consortium | The Lighthouse Lab in Milton Keynes and Alex Alderton, Roberto Amato, Jeffrey Barrett, Sonia Goncalves, Ewan Harrison, David K. Jackson, Ian Johnston, Dominic Kwiatkowski, Cordelia Langford, John Sillitoe on behalf of the Wellcome Sanger Institute COVID-19 Surveillance Team |
| EPI_ISL_2663933, EPI_ISL_2663934 | Lighthouse Lab in Glasgow | Wellcome Sanger Institute for the COVID-19 Genomics UK (COG-UK) Consortium | Harper VanSteenhouse, Yumi Kasai, David Gray, Carol Clugston, Anna Dominiczak and Alex Alderton, Roberto Amato, Jeffrey Barrett, Sonia Goncalves, Ewan Harrison, David K. Jackson, Ian Johnston, Dominic Kwiatkowski, Cordelia Langford, John Sillitoe on behalf of the Wellcome Sanger Institute COVID-19 Surveillance Team |
| EPI_ISL_2663935, EPI_ISL_2663936, EPI_ISL_2663937, EPI_ISL_2663938, EPI_ISL_2663939, EPI_ISL_2663940 | Lighthouse Lab in Milton Keynes | Wellcome Sanger Institute for the COVID-19 Genomics UK (COG-UK) Consortium | The Lighthouse Lab in Milton Keynes and Alex Alderton, Roberto Amato, Jeffrey Barrett, Sonia Goncalves, Ewan Harrison, David K. Jackson, Ian Johnston, Dominic Kwiatkowski, Cordelia Langford, John Sillitoe on behalf of the Wellcome Sanger Institute COVID-19 Surveillance Team |
| EPI_ISL_2663941 | Lighthouse Lab in Glasgow | Wellcome Sanger Institute for the COVID-19 Genomics UK (COG-UK) Consortium | Harper VanSteenhouse, Yumi Kasai, David Gray, Carol Clugston, Anna Dominiczak and Alex Alderton, Roberto Amato, Jeffrey Barrett, Sonia Goncalves, Ewan Harrison, David K. Jackson, Ian Johnston, Dominic Kwiatkowski, Cordelia Langford, John Sillitoe on behalf of the Wellcome Sanger Institute COVID-19 Surveillance Team |
| EPI_ISL_2663942, EPI_ISL_2663943, EPI_ISL_2663944, EPI_ISL_2663945, EPI_ISL_2663946, EPI_ISL_2663947, EPI_ISL_2663948, EPI_ISL_2663949, EPI_ISL_2663950 | Lighthouse Lab in Milton Keynes | Wellcome Sanger Institute for the COVID-19 Genomics UK (COG-UK) Consortium | The Lighthouse Lab in Milton Keynes and Alex Alderton, Roberto Amato, Jeffrey Barrett, Sonia Goncalves, Ewan Harrison, David K. Jackson, Ian Johnston, Dominic Kwiatkowski, Cordelia Langford, John Sillitoe on behalf of the Wellcome Sanger Institute COVID-19 Surveillance Team |
| EPI_ISL_2663951, EPI_ISL_2663952 | Lighthouse Lab in Glasgow | Wellcome Sanger Institute for the COVID-19 Genomics UK (COG-UK) Consortium | Harper VanSteenhouse, Yumi Kasai, David Gray, Carol Clugston, Anna Dominiczak and Alex Alderton, Roberto Amato, Jeffrey Barrett, Sonia Goncalves, Ewan Harrison, David K. Jackson, Ian Johnston, Dominic Kwiatkowski, Cordelia Langford, John Sillitoe on behalf of the Wellcome Sanger Institute COVID-19 Surveillance Team |
| EPI_ISL_2663953, EPI_ISL_2663954, EPI_ISL_2663955, EPI_ISL_2663956, EPI_ISL_2663957 | Lighthouse Lab in Milton Keynes | Wellcome Sanger Institute for the COVID-19 Genomics UK (COG-UK) Consortium | The Lighthouse Lab in Milton Keynes and Alex Alderton, Roberto Amato, Jeffrey Barrett, Sonia Goncalves, Ewan Harrison, David K. Jackson, Ian Johnston, Dominic Kwiatkowski, Cordelia Langford, John Sillitoe on behalf of the Wellcome Sanger Institute COVID-19 Surveillance Team |
| EPI_ISL_2663958 | Lighthouse Lab in Glasgow | Wellcome Sanger Institute for the COVID-19 Genomics UK (COG-UK) Consortium | Harper VanSteenhouse, Yumi Kasai, David Gray, Carol Clugston, Anna Dominiczak and Alex Alderton, Roberto Amato, Jeffrey Barrett, Sonia Goncalves, Ewan Harrison, David K. Jackson, Ian Johnston, Dominic Kwiatkowski, Cordelia Langford, John Sillitoe on behalf of the Wellcome Sanger Institute COVID-19 Surveillance Team |
| EPI_ISL_2663959, EPI_ISL_2663960, EPI_ISL_2663961, EPI_ISL_2663962 | Lighthouse Lab in Milton Keynes | Wellcome Sanger Institute for the COVID-19 Genomics UK (COG-UK) Consortium | The Lighthouse Lab in Milton Keynes and Alex Alderton, Roberto Amato, Jeffrey Barrett, Sonia Goncalves, Ewan Harrison, David K. Jackson, Ian Johnston, Dominic Kwiatkowski, Cordelia Langford, John Sillitoe on behalf of the Wellcome Sanger Institute COVID-19 Surveillance Team |
| EPI_ISL_2663963 | Lighthouse Lab in Glasgow | Wellcome Sanger Institute for the COVID-19 Genomics UK (COG-UK) Consortium | Harper VanSteenhouse, Yumi Kasai, David Gray, Carol Clugston, Anna Dominiczak and Alex Alderton, Roberto Amato, Jeffrey Barrett, Sonia Goncalves, Ewan Harrison, David K. Jackson, Ian Johnston, Dominic Kwiatkowski, Cordelia Langford, John Sillitoe on behalf of the Wellcome Sanger Institute COVID-19 Surveillance Team |
| EPI_ISL_2663964, EPI_ISL_2663965 | Lighthouse Lab in Milton Keynes | Wellcome Sanger Institute for the COVID-19 Genomics UK (COG-UK) Consortium | The Lighthouse Lab in Milton Keynes and Alex Alderton, Roberto Amato, Jeffrey Barrett, Sonia Goncalves, Ewan Harrison, David K. Jackson, Ian Johnston, Dominic Kwiatkowski, Cordelia Langford, John Sillitoe on behalf of the Wellcome Sanger Institute COVID-19 Surveillance Team |
| EPI_ISL_2663966 | Lighthouse Lab in Glasgow | Wellcome Sanger Institute for the COVID-19 Genomics UK (COG-UK) Consortium | Harper VanSteenhouse, Yumi Kasai, David Gray, Carol Clugston, Anna Dominiczak and Alex Alderton, Roberto Amato, Jeffrey Barrett, Sonia Goncalves, Ewan Harrison, David K. Jackson, Ian Johnston, Dominic Kwiatkowski, Cordelia Langford, John Sillitoe on behalf of the Wellcome Sanger Institute COVID-19 Surveillance Team |
| EPI_ISL_2663967 | Lighthouse Lab in Milton Keynes | Wellcome Sanger Institute for the COVID-19 Genomics UK (COG-UK) Consortium | The Lighthouse Lab in Milton Keynes and Alex Alderton, Roberto Amato, Jeffrey Barrett, Sonia Goncalves, Ewan Harrison, David K. Jackson, Ian Johnston, Dominic Kwiatkowski, Cordelia Langford, John Sillitoe on behalf of the Wellcome Sanger Institute COVID-19 Surveillance Team |
| EPI_ISL_2663968 | Lighthouse Lab in Glasgow | Wellcome Sanger Institute for the COVID-19 Genomics UK (COG-UK) Consortium | Harper VanSteenhouse, Yumi Kasai, David Gray, Carol Clugston, Anna Dominiczak and Alex Alderton, Roberto Amato, Jeffrey Barrett, Sonia Goncalves, Ewan Harrison, David K. Jackson, Ian Johnston, Dominic Kwiatkowski, Cordelia Langford, John Sillitoe on behalf of the Wellcome Sanger Institute COVID-19 Surveillance Team |
| EPI_ISL_2663971, EPI_ISL_2663972, EPI_ISL_2663973, EPI_ISL_2663974 | Lighthouse Lab in Milton Keynes | Wellcome Sanger Institute for the COVID-19 Genomics UK (COG-UK) Consortium | The Lighthouse Lab in Milton Keynes and Alex Alderton, Roberto Amato, Jeffrey Barrett, Sonia Goncalves, Ewan Harrison, David K. Jackson, Ian Johnston, Dominic Kwiatkowski, Cordelia Langford, John Sillitoe on behalf of the Wellcome Sanger Institute COVID-19 Surveillance Team |
| EPI_ISL_2663975 | Lighthouse Lab in Glasgow | Wellcome Sanger Institute for the COVID-19 Genomics UK (COG-UK) Consortium | Harper VanSteenhouse, Yumi Kasai, David Gray, Carol Clugston, Anna Dominiczak and Alex Alderton, Roberto Amato, Jeffrey Barrett, Sonia Goncalves, Ewan Harrison, David K. Jackson, Ian Johnston, Dominic Kwiatkowski, Cordelia Langford, John Sillitoe on behalf of the Wellcome Sanger Institute COVID-19 Surveillance Team |
| EPI_ISL_2663976, EPI_ISL_2663977, EPI_ISL_2663978, EPI_ISL_2663979, EPI_ISL_2663980, EPI_ISL_2663981, EPI_ISL_2663982, EPI_ISL_2663983, EPI_ISL_2663984 | Lighthouse Lab in Milton Keynes | Wellcome Sanger Institute for the COVID-19 Genomics UK (COG-UK) Consortium | The Lighthouse Lab in Milton Keynes and Alex Alderton, Roberto Amato, Jeffrey Barrett, Sonia Goncalves, Ewan Harrison, David K. Jackson, Ian Johnston, Dominic Kwiatkowski, Cordelia Langford, John Sillitoe on behalf of the Wellcome Sanger Institute COVID-19 Surveillance Team |
| EPI_ISL_2663986 | Lighthouse Lab in Glasgow | Wellcome Sanger Institute for the COVID-19 Genomics UK (COG-UK) Consortium | Harper VanSteenhouse, Yumi Kasai, David Gray, Carol Clugston, Anna Dominiczak and Alex Alderton, Roberto Amato, Jeffrey Barrett, Sonia Goncalves, Ewan Harrison, David K. Jackson, Ian Johnston, Dominic Kwiatkowski, Cordelia Langford, John Sillitoe on behalf of the Wellcome Sanger Institute COVID-19 Surveillance Team |
| EPI_ISL_2663987 | Lighthouse Lab in Milton Keynes | Wellcome Sanger Institute for the COVID-19 Genomics UK (COG-UK) Consortium | The Lighthouse Lab in Milton Keynes and Alex Alderton, Roberto Amato, Jeffrey Barrett, Sonia Goncalves, Ewan Harrison, David K. Jackson, Ian Johnston, Dominic Kwiatkowski, Cordelia Langford, John Sillitoe on behalf of the Wellcome Sanger Institute COVID-19 Surveillance Team |
| EPI_ISL_2663988 | Lighthouse Lab in Glasgow | Wellcome Sanger Institute for the COVID-19 Genomics UK (COG-UK) Consortium | Harper VanSteenhouse, Yumi Kasai, David Gray, Carol Clugston, Anna Dominiczak and Alex Alderton, Roberto Amato, Jeffrey Barrett, Sonia Goncalves, Ewan Harrison, David K. Jackson, Ian Johnston, Dominic Kwiatkowski, Cordelia Langford, John Sillitoe on behalf of the Wellcome Sanger Institute COVID-19 Surveillance Team |
| EPI_ISL_2663989, EPI_ISL_2663990, EPI_ISL_2663991, EPI_ISL_2663992, EPI_ISL_2663993, EPI_ISL_2663994, EPI_ISL_2663995, EPI_ISL_2663996, EPI_ISL_2663997, EPI_ISL_2663998, EPI_ISL_2663999, EPI_ISL_2664000, EPI_ISL_2664001, EPI_ISL_2664002, EPI_ISL_2664003, EPI_ISL_2664004, EPI_ISL_2664005 | | | |
| see above | Lighthouse Lab in Milton Keynes | Wellcome Sanger Institute for the COVID-19 Genomics UK (COG-UK) Consortium | The Lighthouse Lab in Milton Keynes and Alex Alderton, Roberto Amato, Jeffrey Barrett, Sonia Goncalves, Ewan Harrison, David K. Jackson, Ian Johnston, Dominic Kwiatkowski, Cordelia Langford, John Sillitoe on behalf of the Wellcome Sanger Institute COVID-19 Surveillance Team |
| EPI_ISL_2664006 | Lighthouse Lab in Glasgow | Wellcome Sanger Institute for the COVID-19 Genomics UK (COG-UK) Consortium | Harper VanSteenhouse, Yumi Kasai, David Gray, Carol Clugston, Anna Dominiczak and Alex Alderton, Roberto Amato, Jeffrey Barrett, Sonia Goncalves, Ewan Harrison, David K. Jackson, Ian Johnston, Dominic Kwiatkowski, Cordelia Langford, John Sillitoe on behalf of the Wellcome Sanger Institute COVID-19 Surveillance Team |
| EPI_ISL_2664007 | Lighthouse Lab in Milton Keynes | Wellcome Sanger Institute for the COVID-19 Genomics UK (COG-UK) Consortium | The Lighthouse Lab in Milton Keynes and Alex Alderton, Roberto Amato, Jeffrey Barrett, Sonia Goncalves, Ewan Harrison, David K. Jackson, Ian Johnston, Dominic Kwiatkowski, Cordelia Langford, John Sillitoe on behalf of the Wellcome Sanger Institute COVID-19 Surveillance Team |
| EPI_ISL_2664008 | Lighthouse Lab in Glasgow | Wellcome Sanger Institute for the COVID-19 Genomics UK (COG-UK) Consortium | Harper VanSteenhouse, Yumi Kasai, David Gray, Carol Clugston, Anna Dominiczak and Alex Alderton, Roberto Amato, Jeffrey Barrett, Sonia Goncalves, Ewan Harrison, David K. Jackson, Ian Johnston, Dominic Kwiatkowski, Cordelia Langford, John Sillitoe on behalf of the Wellcome Sanger Institute COVID-19 Surveillance Team |
| EPI_ISL_2664009 | Lighthouse Lab in Milton Keynes | Wellcome Sanger Institute for the COVID-19 Genomics UK (COG-UK) Consortium | The Lighthouse Lab in Milton Keynes and Alex Alderton, Roberto Amato, Jeffrey Barrett, Sonia Goncalves, Ewan Harrison, David K. Jackson, Ian Johnston, Dominic Kwiatkowski, Cordelia Langford, John Sillitoe on behalf of the Wellcome Sanger Institute COVID-19 Surveillance Team |

| | | | |
|---|---|---|---|
| EPI_ISL_2664010, EPI_ISL_2664011, EPI_ISL_2664012 | Lighthouse Lab in Glasgow | Wellcome Sanger Institute for the COVID-19 Genomics UK (COG-UK) Consortium | Harper VanSteenhouse, Yumi Kasai, David Gray, Carol Clugston, Anna Dominiczak and Alex Alderton, Roberto Amato, Jeffrey Barrett, Sonia Goncalves, Ewan Harrison, David K. Jackson, Ian Johnston, Dominic Kwiatkowski, Cordelia Langford, John Sillitoe on behalf of the Wellcome Sanger Institute COVID-19 Surveillance Team |
| EPI_ISL_2664013, EPI_ISL_2664014, EPI_ISL_2664015 | Lighthouse Lab in Alderley Park | Wellcome Sanger Institute for the COVID-19 Genomics UK (COG-UK) Consortium | Jacquelyn Wynn, Mairead Hyland, The Lighthouse Lab in Alderley Park and Alex Alderton, Roberto Amato, Jeffrey Barrett, Sonia Goncalves, Ewan Harrison, David K. Jackson, Ian Johnston, Dominic Kwiatkowski, Cordelia Langford, John Sillitoe on behalf of the Wellcome Sanger Institute COVID-19 Surveillance Team |
| EPI_ISL_2664016 | Lighthouse Lab in Milton Keynes | Wellcome Sanger Institute for the COVID-19 Genomics UK (COG-UK) Consortium | The Lighthouse Lab in Milton Keynes and Alex Alderton, Roberto Amato, Jeffrey Barrett, Sonia Goncalves, Ewan Harrison, David K. Jackson, Ian Johnston, Dominic Kwiatkowski, Cordelia Langford, John Sillitoe on behalf of the Wellcome Sanger Institute COVID-19 Surveillance Team |
| EPI_ISL_2664017 | Lighthouse Lab in Alderley Park | Wellcome Sanger Institute for the COVID-19 Genomics UK (COG-UK) Consortium | Jacquelyn Wynn, Mairead Hyland, The Lighthouse Lab in Alderley Park and Alex Alderton, Roberto Amato, Jeffrey Barrett, Sonia Goncalves, Ewan Harrison, David K. Jackson, Ian Johnston, Dominic Kwiatkowski, Cordelia Langford, John Sillitoe on behalf of the Wellcome Sanger Institute COVID-19 Surveillance Team |
| EPI_ISL_2664018 | Lighthouse Lab in Glasgow | Wellcome Sanger Institute for the COVID-19 Genomics UK (COG-UK) Consortium | Harper VanSteenhouse, Yumi Kasai, David Gray, Carol Clugston, Anna Dominiczak and Alex Alderton, Roberto Amato, Jeffrey Barrett, Sonia Goncalves, Ewan Harrison, David K. Jackson, Ian Johnston, Dominic Kwiatkowski, Cordelia Langford, John Sillitoe on behalf of the Wellcome Sanger Institute COVID-19 Surveillance Team |
| EPI_ISL_2664019, EPI_ISL_2664020 | Lighthouse Lab in Milton Keynes | Wellcome Sanger Institute for the COVID-19 Genomics UK (COG-UK) Consortium | The Lighthouse Lab in Milton Keynes and Alex Alderton, Roberto Amato, Jeffrey Barrett, Sonia Goncalves, Ewan Harrison, David K. Jackson, Ian Johnston, Dominic Kwiatkowski, Cordelia Langford, John Sillitoe on behalf of the Wellcome Sanger Institute COVID-19 Surveillance Team |
| EPI_ISL_2664021, EPI_ISL_2664022 | Lighthouse Lab in Alderley Park | Wellcome Sanger Institute for the COVID-19 Genomics UK (COG-UK) Consortium | Jacquelyn Wynn, Mairead Hyland, The Lighthouse Lab in Alderley Park and Alex Alderton, Roberto Amato, Jeffrey Barrett, Sonia Goncalves, Ewan Harrison, David K. Jackson, Ian Johnston, Dominic Kwiatkowski, Cordelia Langford, John Sillitoe on behalf of the Wellcome Sanger Institute COVID-19 Surveillance Team |
| EPI_ISL_2664023 | Lighthouse Lab in Milton Keynes | Wellcome Sanger Institute for the COVID-19 Genomics UK (COG-UK) Consortium | The Lighthouse Lab in Milton Keynes and Alex Alderton, Roberto Amato, Jeffrey Barrett, Sonia Goncalves, Ewan Harrison, David K. Jackson, Ian Johnston, Dominic Kwiatkowski, Cordelia Langford, John Sillitoe on behalf of the Wellcome Sanger Institute COVID-19 Surveillance Team |
| EPI_ISL_2664024, EPI_ISL_2664025 | Lighthouse Lab in Alderley Park | Wellcome Sanger Institute for the COVID-19 Genomics UK (COG-UK) Consortium | Jacquelyn Wynn, Mairead Hyland, The Lighthouse Lab in Alderley Park and Alex Alderton, Roberto Amato, Jeffrey Barrett, Sonia Goncalves, Ewan Harrison, David K. Jackson, Ian Johnston, Dominic Kwiatkowski, Cordelia Langford, John Sillitoe on behalf of the Wellcome Sanger Institute COVID-19 Surveillance Team |
| EPI_ISL_2664026 | Lighthouse Lab in Milton Keynes | Wellcome Sanger Institute for the COVID-19 Genomics UK (COG-UK) Consortium | The Lighthouse Lab in Milton Keynes and Alex Alderton, Roberto Amato, Jeffrey Barrett, Sonia Goncalves, Ewan Harrison, David K. Jackson, Ian Johnston, Dominic Kwiatkowski, Cordelia Langford, John Sillitoe on behalf of the Wellcome Sanger Institute COVID-19 Surveillance Team |
| EPI_ISL_2664027 | Lighthouse Lab in Glasgow | Wellcome Sanger Institute for the COVID-19 Genomics UK (COG-UK) Consortium | Harper VanSteenhouse, Yumi Kasai, David Gray, Carol Clugston, Anna Dominiczak and Alex Alderton, Roberto Amato, Jeffrey Barrett, Sonia Goncalves, Ewan Harrison, David K. Jackson, Ian Johnston, Dominic Kwiatkowski, Cordelia Langford, John Sillitoe on behalf of the Wellcome Sanger Institute COVID-19 Surveillance Team |
| EPI_ISL_2664028, EPI_ISL_2664029, EPI_ISL_2664030 | Lighthouse Lab in Milton Keynes | Wellcome Sanger Institute for the COVID-19 Genomics UK (COG-UK) Consortium | The Lighthouse Lab in Milton Keynes and Alex Alderton, Roberto Amato, Jeffrey Barrett, Sonia Goncalves, Ewan Harrison, David K. Jackson, Ian Johnston, Dominic Kwiatkowski, Cordelia Langford, John Sillitoe on behalf of the Wellcome Sanger Institute COVID-19 Surveillance Team |
| EPI_ISL_2664031, EPI_ISL_2664032 | Lighthouse Lab in Alderley Park | Wellcome Sanger Institute for the COVID-19 Genomics UK (COG-UK) Consortium | Jacquelyn Wynn, Mairead Hyland, The Lighthouse Lab in Alderley Park and Alex Alderton, Roberto Amato, Jeffrey Barrett, Sonia Goncalves, Ewan Harrison, David K. Jackson, Ian Johnston, Dominic Kwiatkowski, Cordelia Langford, John Sillitoe on behalf of the Wellcome Sanger Institute COVID-19 Surveillance Team |
| EPI_ISL_2664033, EPI_ISL_2664034 | Lighthouse Lab in Milton Keynes | Wellcome Sanger Institute for the COVID-19 Genomics UK (COG-UK) Consortium | The Lighthouse Lab in Milton Keynes and Alex Alderton, Roberto Amato, Jeffrey Barrett, Sonia Goncalves, Ewan Harrison, David K. Jackson, Ian Johnston, Dominic Kwiatkowski, Cordelia Langford, John Sillitoe on behalf of the Wellcome Sanger Institute COVID-19 Surveillance Team |
| EPI_ISL_2664035, EPI_ISL_2664036 | Lighthouse Lab in Glasgow | Wellcome Sanger Institute for the COVID-19 Genomics UK (COG-UK) Consortium | Harper VanSteenhouse, Yumi Kasai, David Gray, Carol Clugston, Anna Dominiczak and Alex Alderton, Roberto Amato, Jeffrey Barrett, Sonia Goncalves, Ewan Harrison, David K. Jackson, Ian Johnston, Dominic Kwiatkowski, Cordelia Langford, John Sillitoe on behalf of the Wellcome Sanger Institute COVID-19 Surveillance Team |
| EPI_ISL_2664037, EPI_ISL_2664039 | Lighthouse Lab in Alderley Park | Wellcome Sanger Institute for the COVID-19 Genomics UK (COG-UK) Consortium | Jacquelyn Wynn, Mairead Hyland, The Lighthouse Lab in Alderley Park and Alex Alderton, Roberto Amato, Jeffrey Barrett, Sonia Goncalves, Ewan Harrison, David K. Jackson, Ian Johnston, Dominic Kwiatkowski, Cordelia Langford, John Sillitoe on behalf of the Wellcome Sanger Institute COVID-19 Surveillance Team |
| EPI_ISL_2664040 | Lighthouse Lab in Glasgow | Wellcome Sanger Institute for the COVID-19 Genomics UK (COG-UK) Consortium | Harper VanSteenhouse, Yumi Kasai, David Gray, Carol Clugston, Anna Dominiczak and Alex Alderton, Roberto Amato, Jeffrey Barrett, Sonia Goncalves, Ewan Harrison, David K. Jackson, Ian Johnston, Dominic Kwiatkowski, Cordelia Langford, John Sillitoe on behalf of the Wellcome Sanger Institute COVID-19 Surveillance Team |
| EPI_ISL_2664041, EPI_ISL_2664042, EPI_ISL_2664043 | Lighthouse Lab in Alderley Park | Wellcome Sanger Institute for the COVID-19 Genomics UK (COG-UK) Consortium | Jacquelyn Wynn, Mairead Hyland, The Lighthouse Lab in Alderley Park and Alex Alderton, Roberto Amato, Jeffrey Barrett, Sonia Goncalves, Ewan Harrison, David K. Jackson, Ian Johnston, Dominic Kwiatkowski, Cordelia Langford, John Sillitoe on behalf of the Wellcome Sanger Institute COVID-19 Surveillance Team |
| EPI_ISL_2664044, EPI_ISL_2664045 | Lighthouse Lab in Milton Keynes | Wellcome Sanger Institute for the COVID-19 Genomics UK (COG-UK) Consortium | The Lighthouse Lab in Milton Keynes and Alex Alderton, Roberto Amato, Jeffrey Barrett, Sonia Goncalves, Ewan Harrison, David K. Jackson, Ian Johnston, Dominic Kwiatkowski, Cordelia Langford, John Sillitoe on behalf of the Wellcome Sanger Institute COVID-19 Surveillance Team |
| EPI_ISL_2664046, EPI_ISL_2664047 | Lighthouse Lab in Alderley Park | Wellcome Sanger Institute for the COVID-19 Genomics UK (COG-UK) Consortium | Jacquelyn Wynn, Mairead Hyland, The Lighthouse Lab in Alderley Park and Alex Alderton, Roberto Amato, Jeffrey Barrett, Sonia Goncalves, Ewan Harrison, David K. Jackson, Ian Johnston, Dominic Kwiatkowski, Cordelia Langford, John Sillitoe on behalf of the Wellcome Sanger Institute COVID-19 Surveillance Team |
| EPI_ISL_2664048 | Lighthouse Lab in Milton Keynes | Wellcome Sanger Institute for the COVID-19 Genomics UK (COG-UK) Consortium | The Lighthouse Lab in Milton Keynes and Alex Alderton, Roberto Amato, Jeffrey Barrett, Sonia Goncalves, Ewan Harrison, David K. Jackson, Ian Johnston, Dominic Kwiatkowski, Cordelia Langford, John Sillitoe on behalf of the Wellcome Sanger Institute COVID-19 Surveillance Team |
| EPI_ISL_2664049 | Lighthouse Lab in Alderley Park | Wellcome Sanger Institute for the COVID-19 Genomics UK (COG-UK) Consortium | Jacquelyn Wynn, Mairead Hyland, The Lighthouse Lab in Alderley Park and Alex Alderton, Roberto Amato, Jeffrey Barrett, Sonia Goncalves, Ewan Harrison, David K. Jackson, Ian Johnston, Dominic Kwiatkowski, Cordelia Langford, John Sillitoe on behalf of the Wellcome Sanger Institute COVID-19 Surveillance Team |
| EPI_ISL_2664050, EPI_ISL_2664051, EPI_ISL_2664052, EPI_ISL_2664053 | Lighthouse Lab in Milton Keynes | Wellcome Sanger Institute for the COVID-19 Genomics UK (COG-UK) Consortium | The Lighthouse Lab in Milton Keynes and Alex Alderton, Roberto Amato, Jeffrey Barrett, Sonia Goncalves, Ewan Harrison, David K. Jackson, Ian Johnston, Dominic Kwiatkowski, Cordelia Langford, John Sillitoe on behalf of the Wellcome Sanger Institute COVID-19 Surveillance Team |
| EPI_ISL_2664054 | Lighthouse Lab in Glasgow | Wellcome Sanger Institute for the COVID-19 Genomics UK (COG-UK) Consortium | Harper VanSteenhouse, Yumi Kasai, David Gray, Carol Clugston, Anna Dominiczak and Alex Alderton, Roberto Amato, Jeffrey Barrett, Sonia Goncalves, Ewan Harrison, David K. Jackson, Ian Johnston, Dominic Kwiatkowski, Cordelia Langford, John Sillitoe on behalf of the Wellcome Sanger Institute COVID-19 Surveillance Team |
| EPI_ISL_2664055 | Lighthouse Lab in Milton Keynes | Wellcome Sanger Institute for the COVID-19 Genomics UK (COG-UK) Consortium | The Lighthouse Lab in Milton Keynes and Alex Alderton, Roberto Amato, Jeffrey Barrett, Sonia Goncalves, Ewan Harrison, David K. Jackson, Ian Johnston, Dominic Kwiatkowski, Cordelia Langford, John Sillitoe on behalf of the Wellcome Sanger Institute COVID-19 Surveillance Team |
| EPI_ISL_2664056 | Lighthouse Lab in Alderley Park | Wellcome Sanger Institute for the COVID-19 Genomics UK (COG-UK) Consortium | Jacquelyn Wynn, Mairead Hyland, The Lighthouse Lab in Alderley Park and Alex Alderton, Roberto Amato, Jeffrey Barrett, Sonia Goncalves, Ewan Harrison, David K. Jackson, Ian Johnston, Dominic Kwiatkowski, Cordelia Langford, John Sillitoe on behalf of the Wellcome Sanger Institute COVID-19 Surveillance Team |
| EPI_ISL_2664057, EPI_ISL_2664058 | Lighthouse Lab in Milton Keynes | Wellcome Sanger Institute for the COVID-19 Genomics UK (COG-UK) Consortium | The Lighthouse Lab in Milton Keynes and Alex Alderton, Roberto Amato, Jeffrey Barrett, Sonia Goncalves, Ewan Harrison, David K. Jackson, Ian Johnston, Dominic Kwiatkowski, Cordelia Langford, John Sillitoe on behalf of the Wellcome Sanger Institute COVID-19 Surveillance Team |
| EPI_ISL_2664060 | Lighthouse Lab in Glasgow | Wellcome Sanger Institute for the COVID-19 Genomics UK (COG-UK) Consortium | Harper VanSteenhouse, Yumi Kasai, David Gray, Carol Clugston, Anna Dominiczak and Alex Alderton, Roberto Amato, Jeffrey Barrett, Sonia Goncalves, Ewan Harrison, David K. Jackson, Ian Johnston, Dominic Kwiatkowski, Cordelia Langford, John Sillitoe on behalf of the Wellcome Sanger Institute COVID-19 Surveillance Team |
| EPI_ISL_2664061 | Lighthouse Lab in Milton Keynes | Wellcome Sanger Institute for the COVID-19 Genomics UK (COG-UK) Consortium | The Lighthouse Lab in Milton Keynes and Alex Alderton, Roberto Amato, Jeffrey Barrett, Sonia Goncalves, Ewan Harrison, David K. Jackson, Ian Johnston, Dominic Kwiatkowski, Cordelia Langford, John Sillitoe on behalf of the Wellcome Sanger Institute COVID-19 Surveillance Team |
| EPI_ISL_2664062 | Lighthouse Lab in Alderley Park | Wellcome Sanger Institute for the COVID-19 Genomics UK (COG-UK) Consortium | Jacquelyn Wynn, Mairead Hyland, The Lighthouse Lab in Alderley Park and Alex Alderton, Roberto Amato, Jeffrey Barrett, Sonia Goncalves, Ewan Harrison, David K. Jackson, Ian Johnston, Dominic Kwiatkowski, Cordelia Langford, John Sillitoe on behalf of the Wellcome Sanger Institute COVID-19 Surveillance Team |
| EPI_ISL_2664063 | Lighthouse Lab in Glasgow | Wellcome Sanger Institute for the COVID-19 Genomics UK (COG-UK) Consortium | Harper VanSteenhouse, Yumi Kasai, David Gray, Carol Clugston, Anna Dominiczak and Alex Alderton, Roberto Amato, Jeffrey Barrett, Sonia Goncalves, Ewan Harrison, David K. Jackson, Ian Johnston, Dominic Kwiatkowski, Cordelia Langford, John Sillitoe on behalf of the Wellcome Sanger Institute COVID-19 Surveillance Team |
| EPI_ISL_2664064, EPI_ISL_2664065, EPI_ISL_2664066 | Lighthouse Lab in Milton Keynes | Wellcome Sanger Institute for the COVID-19 Genomics UK (COG-UK) Consortium | The Lighthouse Lab in Milton Keynes and Alex Alderton, Roberto Amato, Jeffrey Barrett, Sonia Goncalves, Ewan Harrison, David K. Jackson, Ian Johnston, Dominic Kwiatkowski, Cordelia Langford, John Sillitoe on behalf of the Wellcome Sanger Institute COVID-19 Surveillance Team |
| EPI_ISL_2664067, EPI_ISL_2664068, EPI_ISL_2664069, EPI_ISL_2664070, EPI_ISL_2664071 | Lighthouse Lab in Alderley Park | Wellcome Sanger Institute for the COVID-19 Genomics UK (COG-UK) Consortium | Jacquelyn Wynn, Mairead Hyland, The Lighthouse Lab in Alderley Park and Alex Alderton, Roberto Amato, Jeffrey Barrett, Sonia Goncalves, Ewan Harrison, David K. Jackson, Ian Johnston, Dominic Kwiatkowski, Cordelia Langford, John Sillitoe on behalf of the Wellcome Sanger Institute COVID-19 Surveillance Team |
| EPI_ISL_2664072 | Lighthouse Lab in Milton Keynes | Wellcome Sanger Institute for the COVID-19 Genomics UK (COG-UK) Consortium | The Lighthouse Lab in Milton Keynes and Alex Alderton, Roberto Amato, Jeffrey Barrett, Sonia Goncalves, Ewan Harrison, David K. Jackson, Ian Johnston, Dominic Kwiatkowski, Cordelia Langford, John Sillitoe on behalf of the Wellcome Sanger Institute COVID-19 Surveillance Team |
| EPI_ISL_2664074, EPI_ISL_2664075 | Lighthouse Lab in Alderley Park | Wellcome Sanger Institute for the COVID-19 Genomics UK (COG-UK) Consortium | Jacquelyn Wynn, Mairead Hyland, The Lighthouse Lab in Alderley Park and Alex Alderton, Roberto Amato, Jeffrey Barrett, Sonia Goncalves, Ewan Harrison, David K. Jackson, Ian Johnston, Dominic Kwiatkowski, Cordelia Langford, John Sillitoe on behalf of the Wellcome Sanger Institute COVID-19 Surveillance Team |
| EPI_ISL_2664076, EPI_ISL_2664077 | Lighthouse Lab in Milton Keynes | Wellcome Sanger Institute for the COVID-19 Genomics UK (COG-UK) Consortium | The Lighthouse Lab in Milton Keynes and Alex Alderton, Roberto Amato, Jeffrey Barrett, Sonia Goncalves, Ewan Harrison, David K. Jackson, Ian Johnston, Dominic Kwiatkowski, Cordelia Langford, John Sillitoe on behalf of the Wellcome Sanger Institute COVID-19 Surveillance Team |
| EPI_ISL_2664078 | Lighthouse Lab in Alderley Park | Wellcome Sanger Institute for the COVID-19 Genomics UK (COG-UK) Consortium | Jacquelyn Wynn, Mairead Hyland, The Lighthouse Lab in Alderley Park and Alex Alderton, Roberto Amato, Jeffrey Barrett, Sonia Goncalves, Ewan Harrison, David K. Jackson, Ian Johnston, Dominic Kwiatkowski, Cordelia Langford, John Sillitoe on behalf of the Wellcome Sanger Institute COVID-19 Surveillance Team |
| EPI_ISL_2664079, EPI_ISL_2664080, EPI_ISL_2664081 | Lighthouse Lab in Milton Keynes | Wellcome Sanger Institute for the COVID-19 Genomics UK (COG-UK) Consortium | The Lighthouse Lab in Milton Keynes and Alex Alderton, Roberto Amato, Jeffrey Barrett, Sonia Goncalves, Ewan Harrison, David K. Jackson, Ian Johnston, Dominic Kwiatkowski, Cordelia Langford, John Sillitoe on behalf of the Wellcome Sanger Institute COVID-19 Surveillance Team |
| EPI_ISL_2664082 | Lighthouse Lab in Alderley Park | Wellcome Sanger Institute for the COVID-19 Genomics UK (COG-UK) Consortium | Jacquelyn Wynn, Mairead Hyland, The Lighthouse Lab in Alderley Park and Alex Alderton, Roberto Amato, Jeffrey Barrett, Sonia Goncalves, Ewan Harrison, David K. Jackson, Ian Johnston, Dominic Kwiatkowski, Cordelia Langford, John Sillitoe on behalf of the Wellcome Sanger Institute COVID-19 Surveillance Team |
| EPI_ISL_2664083, EPI_ISL_2664084 | Lighthouse Lab in Milton Keynes | Wellcome Sanger Institute for the COVID-19 Genomics UK (COG-UK) Consortium | The Lighthouse Lab in Milton Keynes and Alex Alderton, Roberto Amato, Jeffrey Barrett, Sonia Goncalves, Ewan Harrison, David K. Jackson, Ian Johnston, Dominic Kwiatkowski, Cordelia Langford, John Sillitoe on behalf of the Wellcome Sanger Institute COVID-19 Surveillance Team |
| EPI_ISL_2664085 | Lighthouse Lab in Alderley Park | Wellcome Sanger Institute for the COVID-19 Genomics UK (COG-UK) | Jacquelyn Wynn, Mairead Hyland, The Lighthouse Lab in Alderley Park and Alex Alderton, Roberto Amato, Jeffrey Barrett, Sonia Goncalves, Ewan Harrison, David K. Jackson, Ian Johnston, Dominic Kwiatkowski, Cordelia Langford, John Sillitoe on |

| | | | |
|---|---|---|---|
| | | Consortium | behalf of the Wellcome Sanger Institute COVID-19 Surveillance Team |
| EPI_ISL_2664086 | Lighthouse Lab in Milton Keynes | Wellcome Sanger Institute for the COVID-19 Genomics UK (COG-UK) Consortium | The Lighthouse Lab in Milton Keynes and Alex Alderton, Roberto Amato, Jeffrey Barrett, Sonia Goncalves, Ewan Harrison, David K. Jackson, Ian Johnston, Dominic Kwiatkowski, Cordelia Langford, John Sillitoe on behalf of the Wellcome Sanger Institute COVID-19 Surveillance Team |
| EPI_ISL_2664087 | Lighthouse Lab in Glasgow | Wellcome Sanger Institute for the COVID-19 Genomics UK (COG-UK) Consortium | Harper VanSteenhouse, Yumi Kasai, David Gray, Carol Clugston, Anna Dominiczak and Alex Alderton, Roberto Amato, Jeffrey Barrett, Sonia Goncalves, Ewan Harrison, David K. Jackson, Ian Johnston, Dominic Kwiatkowski, Cordelia Langford, John Sillitoe on behalf of the Wellcome Sanger Institute COVID-19 Surveillance Team |
| EPI_ISL_2664088, EPI_ISL_2664089, EPI_ISL_2664090 | Lighthouse Lab in Alderley Park | Wellcome Sanger Institute for the COVID-19 Genomics UK (COG-UK) Consortium | Jacquelyn Wynn, Mairead Hyland, The Lighthouse Lab in Alderley Park and Alex Alderton, Roberto Amato, Jeffrey Barrett, Sonia Goncalves, Ewan Harrison, David K. Jackson, Ian Johnston, Dominic Kwiatkowski, Cordelia Langford, John Sillitoe on behalf of the Wellcome Sanger Institute COVID-19 Surveillance Team |
| EPI_ISL_2664091 | Lighthouse Lab in Milton Keynes | Wellcome Sanger Institute for the COVID-19 Genomics UK (COG-UK) Consortium | The Lighthouse Lab in Milton Keynes and Alex Alderton, Roberto Amato, Jeffrey Barrett, Sonia Goncalves, Ewan Harrison, David K. Jackson, Ian Johnston, Dominic Kwiatkowski, Cordelia Langford, John Sillitoe on behalf of the Wellcome Sanger Institute COVID-19 Surveillance Team |
| EPI_ISL_2664092, EPI_ISL_2664093 | Lighthouse Lab in Glasgow | Wellcome Sanger Institute for the COVID-19 Genomics UK (COG-UK) Consortium | Harper VanSteenhouse, Yumi Kasai, David Gray, Carol Clugston, Anna Dominiczak and Alex Alderton, Roberto Amato, Jeffrey Barrett, Sonia Goncalves, Ewan Harrison, David K. Jackson, Ian Johnston, Dominic Kwiatkowski, Cordelia Langford, John Sillitoe on behalf of the Wellcome Sanger Institute COVID-19 Surveillance Team |
| EPI_ISL_2664094 | Lighthouse Lab in Alderley Park | Wellcome Sanger Institute for the COVID-19 Genomics UK (COG-UK) Consortium | Jacquelyn Wynn, Mairead Hyland, The Lighthouse Lab in Alderley Park and Alex Alderton, Roberto Amato, Jeffrey Barrett, Sonia Goncalves, Ewan Harrison, David K. Jackson, Ian Johnston, Dominic Kwiatkowski, Cordelia Langford, John Sillitoe on behalf of the Wellcome Sanger Institute COVID-19 Surveillance Team |
| EPI_ISL_2664095 | Lighthouse Lab in Glasgow | Wellcome Sanger Institute for the COVID-19 Genomics UK (COG-UK) Consortium | Harper VanSteenhouse, Yumi Kasai, David Gray, Carol Clugston, Anna Dominiczak and Alex Alderton, Roberto Amato, Jeffrey Barrett, Sonia Goncalves, Ewan Harrison, David K. Jackson, Ian Johnston, Dominic Kwiatkowski, Cordelia Langford, John Sillitoe on behalf of the Wellcome Sanger Institute COVID-19 Surveillance Team |
| EPI_ISL_2664096, EPI_ISL_2664097, EPI_ISL_2664098 | Lighthouse Lab in Alderley Park | Wellcome Sanger Institute for the COVID-19 Genomics UK (COG-UK) Consortium | Jacquelyn Wynn, Mairead Hyland, The Lighthouse Lab in Alderley Park and Alex Alderton, Roberto Amato, Jeffrey Barrett, Sonia Goncalves, Ewan Harrison, David K. Jackson, Ian Johnston, Dominic Kwiatkowski, Cordelia Langford, John Sillitoe on behalf of the Wellcome Sanger Institute COVID-19 Surveillance Team |
| EPI_ISL_2664099 | Lighthouse Lab in Glasgow | Wellcome Sanger Institute for the COVID-19 Genomics UK (COG-UK) Consortium | Harper VanSteenhouse, Yumi Kasai, David Gray, Carol Clugston, Anna Dominiczak and Alex Alderton, Roberto Amato, Jeffrey Barrett, Sonia Goncalves, Ewan Harrison, David K. Jackson, Ian Johnston, Dominic Kwiatkowski, Cordelia Langford, John Sillitoe on behalf of the Wellcome Sanger Institute COVID-19 Surveillance Team |
| EPI_ISL_2664100 | Lighthouse Lab in Alderley Park | Wellcome Sanger Institute for the COVID-19 Genomics UK (COG-UK) Consortium | Jacquelyn Wynn, Mairead Hyland, The Lighthouse Lab in Alderley Park and Alex Alderton, Roberto Amato, Jeffrey Barrett, Sonia Goncalves, Ewan Harrison, David K. Jackson, Ian Johnston, Dominic Kwiatkowski, Cordelia Langford, John Sillitoe on behalf of the Wellcome Sanger Institute COVID-19 Surveillance Team |
| EPI_ISL_2664101, EPI_ISL_2664102, EPI_ISL_2664103 | Lighthouse Lab in Milton Keynes | Wellcome Sanger Institute for the COVID-19 Genomics UK (COG-UK) Consortium | The Lighthouse Lab in Milton Keynes and Alex Alderton, Roberto Amato, Jeffrey Barrett, Sonia Goncalves, Ewan Harrison, David K. Jackson, Ian Johnston, Dominic Kwiatkowski, Cordelia Langford, John Sillitoe on behalf of the Wellcome Sanger Institute COVID-19 Surveillance Team |
| EPI_ISL_2664104 | Lighthouse Lab in Alderley Park | Wellcome Sanger Institute for the COVID-19 Genomics UK (COG-UK) Consortium | Jacquelyn Wynn, Mairead Hyland, The Lighthouse Lab in Alderley Park and Alex Alderton, Roberto Amato, Jeffrey Barrett, Sonia Goncalves, Ewan Harrison, David K. Jackson, Ian Johnston, Dominic Kwiatkowski, Cordelia Langford, John Sillitoe on behalf of the Wellcome Sanger Institute COVID-19 Surveillance Team |
| EPI_ISL_2664105 | Lighthouse Lab in Glasgow | Wellcome Sanger Institute for the COVID-19 Genomics UK (COG-UK) Consortium | Harper VanSteenhouse, Yumi Kasai, David Gray, Carol Clugston, Anna Dominiczak and Alex Alderton, Roberto Amato, Jeffrey Barrett, Sonia Goncalves, Ewan Harrison, David K. Jackson, Ian Johnston, Dominic Kwiatkowski, Cordelia Langford, John Sillitoe on behalf of the Wellcome Sanger Institute COVID-19 Surveillance Team |
| EPI_ISL_2664106, EPI_ISL_2664107, EPI_ISL_2664108, EPI_ISL_2664109 | Lighthouse Lab in Alderley Park | Wellcome Sanger Institute for the COVID-19 Genomics UK (COG-UK) Consortium | Jacquelyn Wynn, Mairead Hyland, The Lighthouse Lab in Alderley Park and Alex Alderton, Roberto Amato, Jeffrey Barrett, Sonia Goncalves, Ewan Harrison, David K. Jackson, Ian Johnston, Dominic Kwiatkowski, Cordelia Langford, John Sillitoe on behalf of the Wellcome Sanger Institute COVID-19 Surveillance Team |
| EPI_ISL_2664110 | Lighthouse Lab in Glasgow | Wellcome Sanger Institute for the COVID-19 Genomics UK (COG-UK) Consortium | Harper VanSteenhouse, Yumi Kasai, David Gray, Carol Clugston, Anna Dominiczak and Alex Alderton, Roberto Amato, Jeffrey Barrett, Sonia Goncalves, Ewan Harrison, David K. Jackson, Ian Johnston, Dominic Kwiatkowski, Cordelia Langford, John Sillitoe on behalf of the Wellcome Sanger Institute COVID-19 Surveillance Team |
| EPI_ISL_2664111 | Lighthouse Lab in Alderley Park | Wellcome Sanger Institute for the COVID-19 Genomics UK (COG-UK) Consortium | Jacquelyn Wynn, Mairead Hyland, The Lighthouse Lab in Alderley Park and Alex Alderton, Roberto Amato, Jeffrey Barrett, Sonia Goncalves, Ewan Harrison, David K. Jackson, Ian Johnston, Dominic Kwiatkowski, Cordelia Langford, John Sillitoe on behalf of the Wellcome Sanger Institute COVID-19 Surveillance Team |
| EPI_ISL_2664112, EPI_ISL_2664113 | Lighthouse Lab in Milton Keynes | Wellcome Sanger Institute for the COVID-19 Genomics UK (COG-UK) Consortium | The Lighthouse Lab in Milton Keynes and Alex Alderton, Roberto Amato, Jeffrey Barrett, Sonia Goncalves, Ewan Harrison, David K. Jackson, Ian Johnston, Dominic Kwiatkowski, Cordelia Langford, John Sillitoe on behalf of the Wellcome Sanger Institute COVID-19 Surveillance Team |
| EPI_ISL_2664114 | Lighthouse Lab in Glasgow | Wellcome Sanger Institute for the COVID-19 Genomics UK (COG-UK) Consortium | Harper VanSteenhouse, Yumi Kasai, David Gray, Carol Clugston, Anna Dominiczak and Alex Alderton, Roberto Amato, Jeffrey Barrett, Sonia Goncalves, Ewan Harrison, David K. Jackson, Ian Johnston, Dominic Kwiatkowski, Cordelia Langford, John Sillitoe on behalf of the Wellcome Sanger Institute COVID-19 Surveillance Team |
| EPI_ISL_2664115 | Lighthouse Lab in Milton Keynes | Wellcome Sanger Institute for the COVID-19 Genomics UK (COG-UK) Consortium | The Lighthouse Lab in Milton Keynes and Alex Alderton, Roberto Amato, Jeffrey Barrett, Sonia Goncalves, Ewan Harrison, David K. Jackson, Ian Johnston, Dominic Kwiatkowski, Cordelia Langford, John Sillitoe on behalf of the Wellcome Sanger Institute COVID-19 Surveillance Team |
| EPI_ISL_2664116 | Lighthouse Lab in Glasgow | Wellcome Sanger Institute for the COVID-19 Genomics UK (COG-UK) Consortium | Harper VanSteenhouse, Yumi Kasai, David Gray, Carol Clugston, Anna Dominiczak and Alex Alderton, Roberto Amato, Jeffrey Barrett, Sonia Goncalves, Ewan Harrison, David K. Jackson, Ian Johnston, Dominic Kwiatkowski, Cordelia Langford, John Sillitoe on behalf of the Wellcome Sanger Institute COVID-19 Surveillance Team |
| EPI_ISL_2664117, EPI_ISL_2664118 | Lighthouse Lab in Milton Keynes | Wellcome Sanger Institute for the COVID-19 Genomics UK (COG-UK) Consortium | The Lighthouse Lab in Milton Keynes and Alex Alderton, Roberto Amato, Jeffrey Barrett, Sonia Goncalves, Ewan Harrison, David K. Jackson, Ian Johnston, Dominic Kwiatkowski, Cordelia Langford, John Sillitoe on behalf of the Wellcome Sanger Institute COVID-19 Surveillance Team |
| EPI_ISL_2664119 | Lighthouse Lab in Alderley Park | Wellcome Sanger Institute for the COVID-19 Genomics UK (COG-UK) Consortium | Jacquelyn Wynn, Mairead Hyland, The Lighthouse Lab in Alderley Park and Alex Alderton, Roberto Amato, Jeffrey Barrett, Sonia Goncalves, Ewan Harrison, David K. Jackson, Ian Johnston, Dominic Kwiatkowski, Cordelia Langford, John Sillitoe on behalf of the Wellcome Sanger Institute COVID-19 Surveillance Team |
| EPI_ISL_2664120 | Lighthouse Lab in Milton Keynes | Wellcome Sanger Institute for the COVID-19 Genomics UK (COG-UK) Consortium | The Lighthouse Lab in Milton Keynes and Alex Alderton, Roberto Amato, Jeffrey Barrett, Sonia Goncalves, Ewan Harrison, David K. Jackson, Ian Johnston, Dominic Kwiatkowski, Cordelia Langford, John Sillitoe on behalf of the Wellcome Sanger Institute COVID-19 Surveillance Team |
| EPI_ISL_2664121 | Lighthouse Lab in Alderley Park | Wellcome Sanger Institute for the COVID-19 Genomics UK (COG-UK) Consortium | Jacquelyn Wynn, Mairead Hyland, The Lighthouse Lab in Alderley Park and Alex Alderton, Roberto Amato, Jeffrey Barrett, Sonia Goncalves, Ewan Harrison, David K. Jackson, Ian Johnston, Dominic Kwiatkowski, Cordelia Langford, John Sillitoe on behalf of the Wellcome Sanger Institute COVID-19 Surveillance Team |
| EPI_ISL_2664122 | Lighthouse Lab in Glasgow | Wellcome Sanger Institute for the COVID-19 Genomics UK (COG-UK) Consortium | Harper VanSteenhouse, Yumi Kasai, David Gray, Carol Clugston, Anna Dominiczak and Alex Alderton, Roberto Amato, Jeffrey Barrett, Sonia Goncalves, Ewan Harrison, David K. Jackson, Ian Johnston, Dominic Kwiatkowski, Cordelia Langford, John Sillitoe on behalf of the Wellcome Sanger Institute COVID-19 Surveillance Team |
| EPI_ISL_2664123 | Lighthouse Lab in Alderley Park | Wellcome Sanger Institute for the COVID-19 Genomics UK (COG-UK) Consortium | Jacquelyn Wynn, Mairead Hyland, The Lighthouse Lab in Alderley Park and Alex Alderton, Roberto Amato, Jeffrey Barrett, Sonia Goncalves, Ewan Harrison, David K. Jackson, Ian Johnston, Dominic Kwiatkowski, Cordelia Langford, John Sillitoe on behalf of the Wellcome Sanger Institute COVID-19 Surveillance Team |
| EPI_ISL_2664124 | Lighthouse Lab in Glasgow | Wellcome Sanger Institute for the COVID-19 Genomics UK (COG-UK) Consortium | Harper VanSteenhouse, Yumi Kasai, David Gray, Carol Clugston, Anna Dominiczak and Alex Alderton, Roberto Amato, Jeffrey Barrett, Sonia Goncalves, Ewan Harrison, David K. Jackson, Ian Johnston, Dominic Kwiatkowski, Cordelia Langford, John Sillitoe on behalf of the Wellcome Sanger Institute COVID-19 Surveillance Team |
| EPI_ISL_2664125, EPI_ISL_2664126 | Lighthouse Lab in Alderley Park | Wellcome Sanger Institute for the COVID-19 Genomics UK (COG-UK) Consortium | Jacquelyn Wynn, Mairead Hyland, The Lighthouse Lab in Alderley Park and Alex Alderton, Roberto Amato, Jeffrey Barrett, Sonia Goncalves, Ewan Harrison, David K. Jackson, Ian Johnston, Dominic Kwiatkowski, Cordelia Langford, John Sillitoe on behalf of the Wellcome Sanger Institute COVID-19 Surveillance Team |
| EPI_ISL_2664127 | Lighthouse Lab in Glasgow | Wellcome Sanger Institute for the COVID-19 Genomics UK (COG-UK) Consortium | Harper VanSteenhouse, Yumi Kasai, David Gray, Carol Clugston, Anna Dominiczak and Alex Alderton, Roberto Amato, Jeffrey Barrett, Sonia Goncalves, Ewan Harrison, David K. Jackson, Ian Johnston, Dominic Kwiatkowski, Cordelia Langford, John Sillitoe on behalf of the Wellcome Sanger Institute COVID-19 Surveillance Team |
| EPI_ISL_2664128 | Lighthouse Lab in Alderley Park | Wellcome Sanger Institute for the COVID-19 Genomics UK (COG-UK) Consortium | Jacquelyn Wynn, Mairead Hyland, The Lighthouse Lab in Alderley Park and Alex Alderton, Roberto Amato, Jeffrey Barrett, Sonia Goncalves, Ewan Harrison, David K. Jackson, Ian Johnston, Dominic Kwiatkowski, Cordelia Langford, John Sillitoe on behalf of the Wellcome Sanger Institute COVID-19 Surveillance Team |
| EPI_ISL_2664129 | Lighthouse Lab in Glasgow | Wellcome Sanger Institute for the COVID-19 Genomics UK (COG-UK) Consortium | Harper VanSteenhouse, Yumi Kasai, David Gray, Carol Clugston, Anna Dominiczak and Alex Alderton, Roberto Amato, Jeffrey Barrett, Sonia Goncalves, Ewan Harrison, David K. Jackson, Ian Johnston, Dominic Kwiatkowski, Cordelia Langford, John Sillitoe on behalf of the Wellcome Sanger Institute COVID-19 Surveillance Team |
| EPI_ISL_2664130 | Lighthouse Lab in Milton Keynes | Wellcome Sanger Institute for the COVID-19 Genomics UK (COG-UK) Consortium | The Lighthouse Lab in Milton Keynes and Alex Alderton, Roberto Amato, Jeffrey Barrett, Sonia Goncalves, Ewan Harrison, David K. Jackson, Ian Johnston, Dominic Kwiatkowski, Cordelia Langford, John Sillitoe on behalf of the Wellcome Sanger Institute COVID-19 Surveillance Team |
| EPI_ISL_2664131, EPI_ISL_2664132 | Lighthouse Lab in Glasgow | Wellcome Sanger Institute for the COVID-19 Genomics UK (COG-UK) Consortium | Harper VanSteenhouse, Yumi Kasai, David Gray, Carol Clugston, Anna Dominiczak and Alex Alderton, Roberto Amato, Jeffrey Barrett, Sonia Goncalves, Ewan Harrison, David K. Jackson, Ian Johnston, Dominic Kwiatkowski, Cordelia Langford, John Sillitoe on behalf of the Wellcome Sanger Institute COVID-19 Surveillance Team |
| EPI_ISL_2664133 | Lighthouse Lab in Milton Keynes | Wellcome Sanger Institute for the COVID-19 Genomics UK (COG-UK) Consortium | The Lighthouse Lab in Milton Keynes and Alex Alderton, Roberto Amato, Jeffrey Barrett, Sonia Goncalves, Ewan Harrison, David K. Jackson, Ian Johnston, Dominic Kwiatkowski, Cordelia Langford, John Sillitoe on behalf of the Wellcome Sanger Institute COVID-19 Surveillance Team |
| EPI_ISL_2664134 | Lighthouse Lab in Glasgow | Wellcome Sanger Institute for the COVID-19 Genomics UK (COG-UK) Consortium | Harper VanSteenhouse, Yumi Kasai, David Gray, Carol Clugston, Anna Dominiczak and Alex Alderton, Roberto Amato, Jeffrey Barrett, Sonia Goncalves, Ewan Harrison, David K. Jackson, Ian Johnston, Dominic Kwiatkowski, Cordelia Langford, John Sillitoe on behalf of the Wellcome Sanger Institute COVID-19 Surveillance Team |
| EPI_ISL_2664135 | Lighthouse Lab in Alderley Park | Wellcome Sanger Institute for the COVID-19 Genomics UK (COG-UK) Consortium | Jacquelyn Wynn, Mairead Hyland, The Lighthouse Lab in Alderley Park and Alex Alderton, Roberto Amato, Jeffrey Barrett, Sonia Goncalves, Ewan Harrison, David K. Jackson, Ian Johnston, Dominic Kwiatkowski, Cordelia Langford, John Sillitoe on behalf of the Wellcome Sanger Institute COVID-19 Surveillance Team |
| EPI_ISL_2664136 | Lighthouse Lab in Milton Keynes | Wellcome Sanger Institute for the COVID-19 Genomics UK (COG-UK) Consortium | The Lighthouse Lab in Milton Keynes and Alex Alderton, Roberto Amato, Jeffrey Barrett, Sonia Goncalves, Ewan Harrison, David K. Jackson, Ian Johnston, Dominic Kwiatkowski, Cordelia Langford, John Sillitoe on behalf of the Wellcome Sanger Institute COVID-19 Surveillance Team |
| EPI_ISL_2664137 | Lighthouse Lab in Alderley Park | Wellcome Sanger Institute for the COVID-19 Genomics UK (COG-UK) Consortium | Jacquelyn Wynn, Mairead Hyland, The Lighthouse Lab in Alderley Park and Alex Alderton, Roberto Amato, Jeffrey Barrett, Sonia Goncalves, Ewan Harrison, David K. Jackson, Ian Johnston, Dominic Kwiatkowski, Cordelia Langford, John Sillitoe on behalf of the Wellcome Sanger Institute COVID-19 Surveillance Team |
| EPI_ISL_2664138, EPI_ISL_2664139 | Lighthouse Lab in Milton Keynes | Wellcome Sanger Institute for the COVID-19 Genomics UK (COG-UK) Consortium | The Lighthouse Lab in Milton Keynes and Alex Alderton, Roberto Amato, Jeffrey Barrett, Sonia Goncalves, Ewan Harrison, David K. Jackson, Ian Johnston, Dominic Kwiatkowski, Cordelia Langford, John Sillitoe on behalf of the Wellcome Sanger Institute COVID-19 Surveillance Team |
| EPI_ISL_2664140 | Lighthouse Lab in Alderley Park | Wellcome Sanger Institute for the COVID-19 Genomics UK (COG-UK) Consortium | Jacquelyn Wynn, Mairead Hyland, The Lighthouse Lab in Alderley Park and Alex Alderton, Roberto Amato, Jeffrey Barrett, Sonia Goncalves, Ewan Harrison, David K. Jackson, Ian Johnston, Dominic Kwiatkowski, Cordelia Langford, John Sillitoe on behalf of the Wellcome Sanger Institute COVID-19 Surveillance Team |
| EPI_ISL_2664141, EPI_ISL_2664142 | Lighthouse Lab in Milton Keynes | Wellcome Sanger Institute for the COVID-19 Genomics UK (COG-UK) Consortium | The Lighthouse Lab in Milton Keynes and Alex Alderton, Roberto Amato, Jeffrey Barrett, Sonia Goncalves, Ewan Harrison, David K. Jackson, Ian Johnston, Dominic Kwiatkowski, Cordelia Langford, John Sillitoe on behalf of the Wellcome Sanger Institute COVID-19 Surveillance Team |
| EPI_ISL_2664143 | Lighthouse Lab in Glasgow | Wellcome Sanger Institute for the COVID-19 Genomics UK (COG-UK) Consortium | Harper VanSteenhouse, Yumi Kasai, David Gray, Carol Clugston, Anna Dominiczak and Alex Alderton, Roberto Amato, Jeffrey Barrett, Sonia Goncalves, Ewan Harrison, David K. Jackson, Ian Johnston, Dominic Kwiatkowski, Cordelia Langford, John Sillitoe on behalf of the Wellcome Sanger Institute COVID-19 Surveillance Team |
| EPI_ISL_2664144, | Lighthouse Lab in Milton Keynes | Wellcome Sanger Institute for the COVID-19 Genomics UK (COG-UK) | The Lighthouse Lab in Milton Keynes and Alex Alderton, Roberto Amato, Jeffrey Barrett, Sonia Goncalves, Ewan Harrison, David K. Jackson, Ian Johnston, Dominic Kwiatkowski, Cordelia Langford, John Sillitoe on behalf of the Wellcome Sanger |

| | | | |
|---|---|---|---|
| EPI_ISL_2664145 | | Consortium | Institute COVID-19 Surveillance Team |
| EPI_ISL_2664146 | Lighthouse Lab in Glasgow | Wellcome Sanger Institute for the COVID-19 Genomics UK (COG-UK) Consortium | Harper VanSteenhouse, Yumi Kasai, David Gray, Carol Clugston, Anna Dominiczak and Alex Alderton, Roberto Amato, Jeffrey Barrett, Sonia Goncalves, Ewan Harrison, David K. Jackson, Ian Johnston, Dominic Kwiatkowski, Cordelia Langford, John Sillitoe on behalf of the Wellcome Sanger Institute COVID-19 Surveillance Team |
| EPI_ISL_2664147, EPI_ISL_2664148, EPI_ISL_2664149 | Lighthouse Lab in Milton Keynes | Wellcome Sanger Institute for the COVID-19 Genomics UK (COG-UK) Consortium | The Lighthouse Lab in Milton Keynes and Alex Alderton, Roberto Amato, Jeffrey Barrett, Sonia Goncalves, Ewan Harrison, David K. Jackson, Ian Johnston, Dominic Kwiatkowski, Cordelia Langford, John Sillitoe on behalf of the Wellcome Sanger Institute COVID-19 Surveillance Team |
| EPI_ISL_2664150 | Lighthouse Lab in Glasgow | Wellcome Sanger Institute for the COVID-19 Genomics UK (COG-UK) Consortium | Harper VanSteenhouse, Yumi Kasai, David Gray, Carol Clugston, Anna Dominiczak and Alex Alderton, Roberto Amato, Jeffrey Barrett, Sonia Goncalves, Ewan Harrison, David K. Jackson, Ian Johnston, Dominic Kwiatkowski, Cordelia Langford, John Sillitoe on behalf of the Wellcome Sanger Institute COVID-19 Surveillance Team |
| EPI_ISL_2664151, EPI_ISL_2664152 | Lighthouse Lab in Milton Keynes | Wellcome Sanger Institute for the COVID-19 Genomics UK (COG-UK) Consortium | The Lighthouse Lab in Milton Keynes and Alex Alderton, Roberto Amato, Jeffrey Barrett, Sonia Goncalves, Ewan Harrison, David K. Jackson, Ian Johnston, Dominic Kwiatkowski, Cordelia Langford, John Sillitoe on behalf of the Wellcome Sanger Institute COVID-19 Surveillance Team |
| EPI_ISL_2664153 | Lighthouse Lab in Alderley Park | Wellcome Sanger Institute for the COVID-19 Genomics UK (COG-UK) Consortium | Jacquelyn Wynn, Mairead Hyland, The Lighthouse Lab in Alderley Park and Alex Alderton, Roberto Amato, Jeffrey Barrett, Sonia Goncalves, Ewan Harrison, David K. Jackson, Ian Johnston, Dominic Kwiatkowski, Cordelia Langford, John Sillitoe on behalf of the Wellcome Sanger Institute COVID-19 Surveillance Team |
| EPI_ISL_2664154 | Lighthouse Lab in Glasgow | Wellcome Sanger Institute for the COVID-19 Genomics UK (COG-UK) Consortium | Harper VanSteenhouse, Yumi Kasai, David Gray, Carol Clugston, Anna Dominiczak and Alex Alderton, Roberto Amato, Jeffrey Barrett, Sonia Goncalves, Ewan Harrison, David K. Jackson, Ian Johnston, Dominic Kwiatkowski, Cordelia Langford, John Sillitoe on behalf of the Wellcome Sanger Institute COVID-19 Surveillance Team |
| EPI_ISL_2664155 | Lighthouse Lab in Milton Keynes | Wellcome Sanger Institute for the COVID-19 Genomics UK (COG-UK) Consortium | The Lighthouse Lab in Milton Keynes and Alex Alderton, Roberto Amato, Jeffrey Barrett, Sonia Goncalves, Ewan Harrison, David K. Jackson, Ian Johnston, Dominic Kwiatkowski, Cordelia Langford, John Sillitoe on behalf of the Wellcome Sanger Institute COVID-19 Surveillance Team |
| EPI_ISL_2664156 | Lighthouse Lab in Glasgow | Wellcome Sanger Institute for the COVID-19 Genomics UK (COG-UK) Consortium | Harper VanSteenhouse, Yumi Kasai, David Gray, Carol Clugston, Anna Dominiczak and Alex Alderton, Roberto Amato, Jeffrey Barrett, Sonia Goncalves, Ewan Harrison, David K. Jackson, Ian Johnston, Dominic Kwiatkowski, Cordelia Langford, John Sillitoe on behalf of the Wellcome Sanger Institute COVID-19 Surveillance Team |
| EPI_ISL_2664157 | Lighthouse Lab in Alderley Park | Wellcome Sanger Institute for the COVID-19 Genomics UK (COG-UK) Consortium | Jacquelyn Wynn, Mairead Hyland, The Lighthouse Lab in Alderley Park and Alex Alderton, Roberto Amato, Jeffrey Barrett, Sonia Goncalves, Ewan Harrison, David K. Jackson, Ian Johnston, Dominic Kwiatkowski, Cordelia Langford, John Sillitoe on behalf of the Wellcome Sanger Institute COVID-19 Surveillance Team |
| EPI_ISL_2664158 | Lighthouse Lab in Glasgow | Wellcome Sanger Institute for the COVID-19 Genomics UK (COG-UK) Consortium | Harper VanSteenhouse, Yumi Kasai, David Gray, Carol Clugston, Anna Dominiczak and Alex Alderton, Roberto Amato, Jeffrey Barrett, Sonia Goncalves, Ewan Harrison, David K. Jackson, Ian Johnston, Dominic Kwiatkowski, Cordelia Langford, John Sillitoe on behalf of the Wellcome Sanger Institute COVID-19 Surveillance Team |
| EPI_ISL_2664159 | Lighthouse Lab in Alderley Park | Wellcome Sanger Institute for the COVID-19 Genomics UK (COG-UK) Consortium | Jacquelyn Wynn, Mairead Hyland, The Lighthouse Lab in Alderley Park and Alex Alderton, Roberto Amato, Jeffrey Barrett, Sonia Goncalves, Ewan Harrison, David K. Jackson, Ian Johnston, Dominic Kwiatkowski, Cordelia Langford, John Sillitoe on behalf of the Wellcome Sanger Institute COVID-19 Surveillance Team |
| EPI_ISL_2664160 | Lighthouse Lab in Glasgow | Wellcome Sanger Institute for the COVID-19 Genomics UK (COG-UK) Consortium | Harper VanSteenhouse, Yumi Kasai, David Gray, Carol Clugston, Anna Dominiczak and Alex Alderton, Roberto Amato, Jeffrey Barrett, Sonia Goncalves, Ewan Harrison, David K. Jackson, Ian Johnston, Dominic Kwiatkowski, Cordelia Langford, John Sillitoe on behalf of the Wellcome Sanger Institute COVID-19 Surveillance Team |
| EPI_ISL_2664161 | Lighthouse Lab in Milton Keynes | Wellcome Sanger Institute for the COVID-19 Genomics UK (COG-UK) Consortium | The Lighthouse Lab in Milton Keynes and Alex Alderton, Roberto Amato, Jeffrey Barrett, Sonia Goncalves, Ewan Harrison, David K. Jackson, Ian Johnston, Dominic Kwiatkowski, Cordelia Langford, John Sillitoe on behalf of the Wellcome Sanger Institute COVID-19 Surveillance Team |
| EPI_ISL_2664162, EPI_ISL_2664163, EPI_ISL_2664164 | Lighthouse Lab in Alderley Park | Wellcome Sanger Institute for the COVID-19 Genomics UK (COG-UK) Consortium | Jacquelyn Wynn, Mairead Hyland, The Lighthouse Lab in Alderley Park and Alex Alderton, Roberto Amato, Jeffrey Barrett, Sonia Goncalves, Ewan Harrison, David K. Jackson, Ian Johnston, Dominic Kwiatkowski, Cordelia Langford, John Sillitoe on behalf of the Wellcome Sanger Institute COVID-19 Surveillance Team |
| EPI_ISL_2664165 | Lighthouse Lab in Milton Keynes | Wellcome Sanger Institute for the COVID-19 Genomics UK (COG-UK) Consortium | The Lighthouse Lab in Milton Keynes and Alex Alderton, Roberto Amato, Jeffrey Barrett, Sonia Goncalves, Ewan Harrison, David K. Jackson, Ian Johnston, Dominic Kwiatkowski, Cordelia Langford, John Sillitoe on behalf of the Wellcome Sanger Institute COVID-19 Surveillance Team |
| EPI_ISL_2664166, EPI_ISL_2664167, EPI_ISL_2664168 | Lighthouse Lab in Alderley Park | Wellcome Sanger Institute for the COVID-19 Genomics UK (COG-UK) Consortium | Jacquelyn Wynn, Mairead Hyland, The Lighthouse Lab in Alderley Park and Alex Alderton, Roberto Amato, Jeffrey Barrett, Sonia Goncalves, Ewan Harrison, David K. Jackson, Ian Johnston, Dominic Kwiatkowski, Cordelia Langford, John Sillitoe on behalf of the Wellcome Sanger Institute COVID-19 Surveillance Team |
| EPI_ISL_2664169 | Lighthouse Lab in Milton Keynes | Wellcome Sanger Institute for the COVID-19 Genomics UK (COG-UK) Consortium | The Lighthouse Lab in Milton Keynes and Alex Alderton, Roberto Amato, Jeffrey Barrett, Sonia Goncalves, Ewan Harrison, David K. Jackson, Ian Johnston, Dominic Kwiatkowski, Cordelia Langford, John Sillitoe on behalf of the Wellcome Sanger Institute COVID-19 Surveillance Team |
| EPI_ISL_2664170, EPI_ISL_2664171 | Lighthouse Lab in Alderley Park | Wellcome Sanger Institute for the COVID-19 Genomics UK (COG-UK) Consortium | Jacquelyn Wynn, Mairead Hyland, The Lighthouse Lab in Alderley Park and Alex Alderton, Roberto Amato, Jeffrey Barrett, Sonia Goncalves, Ewan Harrison, David K. Jackson, Ian Johnston, Dominic Kwiatkowski, Cordelia Langford, John Sillitoe on behalf of the Wellcome Sanger Institute COVID-19 Surveillance Team |
| EPI_ISL_2664172 | Lighthouse Lab in Glasgow | Wellcome Sanger Institute for the COVID-19 Genomics UK (COG-UK) Consortium | Harper VanSteenhouse, Yumi Kasai, David Gray, Carol Clugston, Anna Dominiczak and Alex Alderton, Roberto Amato, Jeffrey Barrett, Sonia Goncalves, Ewan Harrison, David K. Jackson, Ian Johnston, Dominic Kwiatkowski, Cordelia Langford, John Sillitoe on behalf of the Wellcome Sanger Institute COVID-19 Surveillance Team |
| EPI_ISL_2664173, EPI_ISL_2664174 | Lighthouse Lab in Milton Keynes | Wellcome Sanger Institute for the COVID-19 Genomics UK (COG-UK) Consortium | The Lighthouse Lab in Milton Keynes and Alex Alderton, Roberto Amato, Jeffrey Barrett, Sonia Goncalves, Ewan Harrison, David K. Jackson, Ian Johnston, Dominic Kwiatkowski, Cordelia Langford, John Sillitoe on behalf of the Wellcome Sanger Institute COVID-19 Surveillance Team |
| EPI_ISL_2664175 | Lighthouse Lab in Alderley Park | Wellcome Sanger Institute for the COVID-19 Genomics UK (COG-UK) Consortium | Jacquelyn Wynn, Mairead Hyland, The Lighthouse Lab in Alderley Park and Alex Alderton, Roberto Amato, Jeffrey Barrett, Sonia Goncalves, Ewan Harrison, David K. Jackson, Ian Johnston, Dominic Kwiatkowski, Cordelia Langford, John Sillitoe on behalf of the Wellcome Sanger Institute COVID-19 Surveillance Team |
| EPI_ISL_2664176 | Lighthouse Lab in Glasgow | Wellcome Sanger Institute for the COVID-19 Genomics UK (COG-UK) Consortium | Harper VanSteenhouse, Yumi Kasai, David Gray, Carol Clugston, Anna Dominiczak and Alex Alderton, Roberto Amato, Jeffrey Barrett, Sonia Goncalves, Ewan Harrison, David K. Jackson, Ian Johnston, Dominic Kwiatkowski, Cordelia Langford, John Sillitoe on behalf of the Wellcome Sanger Institute COVID-19 Surveillance Team |
| EPI_ISL_2664177 | Lighthouse Lab in Milton Keynes | Wellcome Sanger Institute for the COVID-19 Genomics UK (COG-UK) Consortium | The Lighthouse Lab in Milton Keynes and Alex Alderton, Roberto Amato, Jeffrey Barrett, Sonia Goncalves, Ewan Harrison, David K. Jackson, Ian Johnston, Dominic Kwiatkowski, Cordelia Langford, John Sillitoe on behalf of the Wellcome Sanger Institute COVID-19 Surveillance Team |
| EPI_ISL_2664178 | Lighthouse Lab in Alderley Park | Wellcome Sanger Institute for the COVID-19 Genomics UK (COG-UK) Consortium | Jacquelyn Wynn, Mairead Hyland, The Lighthouse Lab in Alderley Park and Alex Alderton, Roberto Amato, Jeffrey Barrett, Sonia Goncalves, Ewan Harrison, David K. Jackson, Ian Johnston, Dominic Kwiatkowski, Cordelia Langford, John Sillitoe on behalf of the Wellcome Sanger Institute COVID-19 Surveillance Team |
| EPI_ISL_2664179 | Lighthouse Lab in Milton Keynes | Wellcome Sanger Institute for the COVID-19 Genomics UK (COG-UK) Consortium | The Lighthouse Lab in Milton Keynes and Alex Alderton, Roberto Amato, Jeffrey Barrett, Sonia Goncalves, Ewan Harrison, David K. Jackson, Ian Johnston, Dominic Kwiatkowski, Cordelia Langford, John Sillitoe on behalf of the Wellcome Sanger Institute COVID-19 Surveillance Team |
| EPI_ISL_2664180 | Lighthouse Lab in Glasgow | Wellcome Sanger Institute for the COVID-19 Genomics UK (COG-UK) Consortium | Harper VanSteenhouse, Yumi Kasai, David Gray, Carol Clugston, Anna Dominiczak and Alex Alderton, Roberto Amato, Jeffrey Barrett, Sonia Goncalves, Ewan Harrison, David K. Jackson, Ian Johnston, Dominic Kwiatkowski, Cordelia Langford, John Sillitoe on behalf of the Wellcome Sanger Institute COVID-19 Surveillance Team |
| EPI_ISL_2664181 | Lighthouse Lab in Milton Keynes | Wellcome Sanger Institute for the COVID-19 Genomics UK (COG-UK) Consortium | The Lighthouse Lab in Milton Keynes and Alex Alderton, Roberto Amato, Jeffrey Barrett, Sonia Goncalves, Ewan Harrison, David K. Jackson, Ian Johnston, Dominic Kwiatkowski, Cordelia Langford, John Sillitoe on behalf of the Wellcome Sanger Institute COVID-19 Surveillance Team |
| EPI_ISL_2664182 | Lighthouse Lab in Glasgow | Wellcome Sanger Institute for the COVID-19 Genomics UK (COG-UK) Consortium | Harper VanSteenhouse, Yumi Kasai, David Gray, Carol Clugston, Anna Dominiczak and Alex Alderton, Roberto Amato, Jeffrey Barrett, Sonia Goncalves, Ewan Harrison, David K. Jackson, Ian Johnston, Dominic Kwiatkowski, Cordelia Langford, John Sillitoe on behalf of the Wellcome Sanger Institute COVID-19 Surveillance Team |
| EPI_ISL_2664183 | Lighthouse Lab in Milton Keynes | Wellcome Sanger Institute for the COVID-19 Genomics UK (COG-UK) Consortium | The Lighthouse Lab in Milton Keynes and Alex Alderton, Roberto Amato, Jeffrey Barrett, Sonia Goncalves, Ewan Harrison, David K. Jackson, Ian Johnston, Dominic Kwiatkowski, Cordelia Langford, John Sillitoe on behalf of the Wellcome Sanger Institute COVID-19 Surveillance Team |
| EPI_ISL_2664184 | Lighthouse Lab in Glasgow | Wellcome Sanger Institute for the COVID-19 Genomics UK (COG-UK) Consortium | Harper VanSteenhouse, Yumi Kasai, David Gray, Carol Clugston, Anna Dominiczak and Alex Alderton, Roberto Amato, Jeffrey Barrett, Sonia Goncalves, Ewan Harrison, David K. Jackson, Ian Johnston, Dominic Kwiatkowski, Cordelia Langford, John Sillitoe on behalf of the Wellcome Sanger Institute COVID-19 Surveillance Team |
| EPI_ISL_2664185 | Lighthouse Lab in Milton Keynes | Wellcome Sanger Institute for the COVID-19 Genomics UK (COG-UK) Consortium | The Lighthouse Lab in Milton Keynes and Alex Alderton, Roberto Amato, Jeffrey Barrett, Sonia Goncalves, Ewan Harrison, David K. Jackson, Ian Johnston, Dominic Kwiatkowski, Cordelia Langford, John Sillitoe on behalf of the Wellcome Sanger Institute COVID-19 Surveillance Team |
| EPI_ISL_2664186, EPI_ISL_2664187, EPI_ISL_2664188, EPI_ISL_2664189 | Lighthouse Lab in Alderley Park | Wellcome Sanger Institute for the COVID-19 Genomics UK (COG-UK) Consortium | Jacquelyn Wynn, Mairead Hyland, The Lighthouse Lab in Alderley Park and Alex Alderton, Roberto Amato, Jeffrey Barrett, Sonia Goncalves, Ewan Harrison, David K. Jackson, Ian Johnston, Dominic Kwiatkowski, Cordelia Langford, John Sillitoe on behalf of the Wellcome Sanger Institute COVID-19 Surveillance Team |
| EPI_ISL_2664190 | Lighthouse Lab in Glasgow | Wellcome Sanger Institute for the COVID-19 Genomics UK (COG-UK) Consortium | Harper VanSteenhouse, Yumi Kasai, David Gray, Carol Clugston, Anna Dominiczak and Alex Alderton, Roberto Amato, Jeffrey Barrett, Sonia Goncalves, Ewan Harrison, David K. Jackson, Ian Johnston, Dominic Kwiatkowski, Cordelia Langford, John Sillitoe on behalf of the Wellcome Sanger Institute COVID-19 Surveillance Team |
| EPI_ISL_2664191 | Lighthouse Lab in Alderley Park | Wellcome Sanger Institute for the COVID-19 Genomics UK (COG-UK) Consortium | Jacquelyn Wynn, Mairead Hyland, The Lighthouse Lab in Alderley Park and Alex Alderton, Roberto Amato, Jeffrey Barrett, Sonia Goncalves, Ewan Harrison, David K. Jackson, Ian Johnston, Dominic Kwiatkowski, Cordelia Langford, John Sillitoe on behalf of the Wellcome Sanger Institute COVID-19 Surveillance Team |
| EPI_ISL_2664192 | Lighthouse Lab in Milton Keynes | Wellcome Sanger Institute for the COVID-19 Genomics UK (COG-UK) Consortium | The Lighthouse Lab in Milton Keynes and Alex Alderton, Roberto Amato, Jeffrey Barrett, Sonia Goncalves, Ewan Harrison, David K. Jackson, Ian Johnston, Dominic Kwiatkowski, Cordelia Langford, John Sillitoe on behalf of the Wellcome Sanger Institute COVID-19 Surveillance Team |
| EPI_ISL_2664193 | Lighthouse Lab in Glasgow | Wellcome Sanger Institute for the COVID-19 Genomics UK (COG-UK) Consortium | Harper VanSteenhouse, Yumi Kasai, David Gray, Carol Clugston, Anna Dominiczak and Alex Alderton, Roberto Amato, Jeffrey Barrett, Sonia Goncalves, Ewan Harrison, David K. Jackson, Ian Johnston, Dominic Kwiatkowski, Cordelia Langford, John Sillitoe on behalf of the Wellcome Sanger Institute COVID-19 Surveillance Team |
| EPI_ISL_2664194, EPI_ISL_2664195 | Lighthouse Lab in Alderley Park | Wellcome Sanger Institute for the COVID-19 Genomics UK (COG-UK) Consortium | Jacquelyn Wynn, Mairead Hyland, The Lighthouse Lab in Alderley Park and Alex Alderton, Roberto Amato, Jeffrey Barrett, Sonia Goncalves, Ewan Harrison, David K. Jackson, Ian Johnston, Dominic Kwiatkowski, Cordelia Langford, John Sillitoe on behalf of the Wellcome Sanger Institute COVID-19 Surveillance Team |
| EPI_ISL_2664196, EPI_ISL_2664197 | Lighthouse Lab in Milton Keynes | Wellcome Sanger Institute for the COVID-19 Genomics UK (COG-UK) Consortium | The Lighthouse Lab in Milton Keynes and Alex Alderton, Roberto Amato, Jeffrey Barrett, Sonia Goncalves, Ewan Harrison, David K. Jackson, Ian Johnston, Dominic Kwiatkowski, Cordelia Langford, John Sillitoe on behalf of the Wellcome Sanger Institute COVID-19 Surveillance Team |
| EPI_ISL_2664198 | Lighthouse Lab in Alderley Park | Wellcome Sanger Institute for the COVID-19 Genomics UK (COG-UK) Consortium | Jacquelyn Wynn, Mairead Hyland, The Lighthouse Lab in Alderley Park and Alex Alderton, Roberto Amato, Jeffrey Barrett, Sonia Goncalves, Ewan Harrison, David K. Jackson, Ian Johnston, Dominic Kwiatkowski, Cordelia Langford, John Sillitoe on behalf of the Wellcome Sanger Institute COVID-19 Surveillance Team |
| EPI_ISL_2664199, EPI_ISL_2664200 | Lighthouse Lab in Glasgow | Wellcome Sanger Institute for the COVID-19 Genomics UK (COG-UK) Consortium | Harper VanSteenhouse, Yumi Kasai, David Gray, Carol Clugston, Anna Dominiczak and Alex Alderton, Roberto Amato, Jeffrey Barrett, Sonia Goncalves, Ewan Harrison, David K. Jackson, Ian Johnston, Dominic Kwiatkowski, Cordelia Langford, John Sillitoe on behalf of the Wellcome Sanger Institute COVID-19 Surveillance Team |
| EPI_ISL_2664201, EPI_ISL_2664202, EPI_ISL_2664203 | Lighthouse Lab in Alderley Park | Wellcome Sanger Institute for the COVID-19 Genomics UK (COG-UK) Consortium | Jacquelyn Wynn, Mairead Hyland, The Lighthouse Lab in Alderley Park and Alex Alderton, Roberto Amato, Jeffrey Barrett, Sonia Goncalves, Ewan Harrison, David K. Jackson, Ian Johnston, Dominic Kwiatkowski, Cordelia Langford, John Sillitoe on behalf of the Wellcome Sanger Institute COVID-19 Surveillance Team |
| EPI_ISL_2664204 | Lighthouse Lab in Milton Keynes | Wellcome Sanger Institute for the COVID-19 Genomics UK (COG-UK) Consortium | The Lighthouse Lab in Milton Keynes and Alex Alderton, Roberto Amato, Jeffrey Barrett, Sonia Goncalves, Ewan Harrison, David K. Jackson, Ian Johnston, Dominic Kwiatkowski, Cordelia Langford, John Sillitoe on behalf of the Wellcome Sanger Institute COVID-19 Surveillance Team |

| | | | |
|---|---|---|---|
| EPI_ISL_2664205 | Lighthouse Lab in Alderley Park | Wellcome Sanger Institute for the COVID-19 Genomics UK (COG-UK) Consortium | Jacquelyn Wynn, Mairead Hyland, The Lighthouse Lab in Alderley Park and Alex Alderton, Roberto Amato, Jeffrey Barrett, Sonia Goncalves, Ewan Harrison, David K. Jackson, Ian Johnston, Dominic Kwiatkowski, Cordelia Langford, John Sillitoe on behalf of the Wellcome Sanger Institute COVID-19 Surveillance Team |
| EPI_ISL_2664206 | Lighthouse Lab in Glasgow | Wellcome Sanger Institute for the COVID-19 Genomics UK (COG-UK) Consortium | Harper VanSteenhouse, Yumi Kasai, David Gray, Carol Clugston, Anna Dominiczak and Alex Alderton, Roberto Amato, Jeffrey Barrett, Sonia Goncalves, Ewan Harrison, David K. Jackson, Ian Johnston, Dominic Kwiatkowski, Cordelia Langford, John Sillitoe on behalf of the Wellcome Sanger Institute COVID-19 Surveillance Team |
| EPI_ISL_2664207 | Lighthouse Lab in Alderley Park | Wellcome Sanger Institute for the COVID-19 Genomics UK (COG-UK) Consortium | Jacquelyn Wynn, Mairead Hyland, The Lighthouse Lab in Alderley Park and Alex Alderton, Roberto Amato, Jeffrey Barrett, Sonia Goncalves, Ewan Harrison, David K. Jackson, Ian Johnston, Dominic Kwiatkowski, Cordelia Langford, John Sillitoe on behalf of the Wellcome Sanger Institute COVID-19 Surveillance Team |
| EPI_ISL_2664208 | Lighthouse Lab in Glasgow | Wellcome Sanger Institute for the COVID-19 Genomics UK (COG-UK) Consortium | Harper VanSteenhouse, Yumi Kasai, David Gray, Carol Clugston, Anna Dominiczak and Alex Alderton, Roberto Amato, Jeffrey Barrett, Sonia Goncalves, Ewan Harrison, David K. Jackson, Ian Johnston, Dominic Kwiatkowski, Cordelia Langford, John Sillitoe on behalf of the Wellcome Sanger Institute COVID-19 Surveillance Team |
| EPI_ISL_2664209, EPI_ISL_2664210, EPI_ISL_2664211, EPI_ISL_2664212, EPI_ISL_2664213, EPI_ISL_2664214 | Lighthouse Lab in Alderley Park | Wellcome Sanger Institute for the COVID-19 Genomics UK (COG-UK) Consortium | Jacquelyn Wynn, Mairead Hyland, The Lighthouse Lab in Alderley Park and Alex Alderton, Roberto Amato, Jeffrey Barrett, Sonia Goncalves, Ewan Harrison, David K. Jackson, Ian Johnston, Dominic Kwiatkowski, Cordelia Langford, John Sillitoe on behalf of the Wellcome Sanger Institute COVID-19 Surveillance Team |
| EPI_ISL_2664215 | Lighthouse Lab in Milton Keynes | Wellcome Sanger Institute for the COVID-19 Genomics UK (COG-UK) Consortium | The Lighthouse Lab in Milton Keynes and Alex Alderton, Roberto Amato, Jeffrey Barrett, Sonia Goncalves, Ewan Harrison, David K. Jackson, Ian Johnston, Dominic Kwiatkowski, Cordelia Langford, John Sillitoe on behalf of the Wellcome Sanger Institute COVID-19 Surveillance Team |
| EPI_ISL_2664216, EPI_ISL_2664217 | Lighthouse Lab in Alderley Park | Wellcome Sanger Institute for the COVID-19 Genomics UK (COG-UK) Consortium | Jacquelyn Wynn, Mairead Hyland, The Lighthouse Lab in Alderley Park and Alex Alderton, Roberto Amato, Jeffrey Barrett, Sonia Goncalves, Ewan Harrison, David K. Jackson, Ian Johnston, Dominic Kwiatkowski, Cordelia Langford, John Sillitoe on behalf of the Wellcome Sanger Institute COVID-19 Surveillance Team |
| EPI_ISL_2664218 | Lighthouse Lab in Milton Keynes | Wellcome Sanger Institute for the COVID-19 Genomics UK (COG-UK) Consortium | The Lighthouse Lab in Milton Keynes and Alex Alderton, Roberto Amato, Jeffrey Barrett, Sonia Goncalves, Ewan Harrison, David K. Jackson, Ian Johnston, Dominic Kwiatkowski, Cordelia Langford, John Sillitoe on behalf of the Wellcome Sanger Institute COVID-19 Surveillance Team |
| EPI_ISL_2664219 | Lighthouse Lab in Alderley Park | Wellcome Sanger Institute for the COVID-19 Genomics UK (COG-UK) Consortium | Jacquelyn Wynn, Mairead Hyland, The Lighthouse Lab in Alderley Park and Alex Alderton, Roberto Amato, Jeffrey Barrett, Sonia Goncalves, Ewan Harrison, David K. Jackson, Ian Johnston, Dominic Kwiatkowski, Cordelia Langford, John Sillitoe on behalf of the Wellcome Sanger Institute COVID-19 Surveillance Team |
| EPI_ISL_2664220, EPI_ISL_2664221, EPI_ISL_2664222 | Lighthouse Lab in Milton Keynes | Wellcome Sanger Institute for the COVID-19 Genomics UK (COG-UK) Consortium | The Lighthouse Lab in Milton Keynes and Alex Alderton, Roberto Amato, Jeffrey Barrett, Sonia Goncalves, Ewan Harrison, David K. Jackson, Ian Johnston, Dominic Kwiatkowski, Cordelia Langford, John Sillitoe on behalf of the Wellcome Sanger Institute COVID-19 Surveillance Team |
| EPI_ISL_2664223, EPI_ISL_2664224 | Lighthouse Lab in Glasgow | Wellcome Sanger Institute for the COVID-19 Genomics UK (COG-UK) Consortium | Harper VanSteenhouse, Yumi Kasai, David Gray, Carol Clugston, Anna Dominiczak and Alex Alderton, Roberto Amato, Jeffrey Barrett, Sonia Goncalves, Ewan Harrison, David K. Jackson, Ian Johnston, Dominic Kwiatkowski, Cordelia Langford, John Sillitoe on behalf of the Wellcome Sanger Institute COVID-19 Surveillance Team |
| EPI_ISL_2664225, EPI_ISL_2664226 | Lighthouse Lab in Alderley Park | Wellcome Sanger Institute for the COVID-19 Genomics UK (COG-UK) Consortium | Jacquelyn Wynn, Mairead Hyland, The Lighthouse Lab in Alderley Park and Alex Alderton, Roberto Amato, Jeffrey Barrett, Sonia Goncalves, Ewan Harrison, David K. Jackson, Ian Johnston, Dominic Kwiatkowski, Cordelia Langford, John Sillitoe on behalf of the Wellcome Sanger Institute COVID-19 Surveillance Team |
| EPI_ISL_2664227, EPI_ISL_2664228 | Lighthouse Lab in Milton Keynes | Wellcome Sanger Institute for the COVID-19 Genomics UK (COG-UK) Consortium | The Lighthouse Lab in Milton Keynes and Alex Alderton, Roberto Amato, Jeffrey Barrett, Sonia Goncalves, Ewan Harrison, David K. Jackson, Ian Johnston, Dominic Kwiatkowski, Cordelia Langford, John Sillitoe on behalf of the Wellcome Sanger Institute COVID-19 Surveillance Team |
| EPI_ISL_2664229 | Lighthouse Lab in Alderley Park | Wellcome Sanger Institute for the COVID-19 Genomics UK (COG-UK) Consortium | Jacquelyn Wynn, Mairead Hyland, The Lighthouse Lab in Alderley Park and Alex Alderton, Roberto Amato, Jeffrey Barrett, Sonia Goncalves, Ewan Harrison, David K. Jackson, Ian Johnston, Dominic Kwiatkowski, Cordelia Langford, John Sillitoe on behalf of the Wellcome Sanger Institute COVID-19 Surveillance Team |
| EPI_ISL_2664230, EPI_ISL_2664231 | Lighthouse Lab in Milton Keynes | Wellcome Sanger Institute for the COVID-19 Genomics UK (COG-UK) Consortium | The Lighthouse Lab in Milton Keynes and Alex Alderton, Roberto Amato, Jeffrey Barrett, Sonia Goncalves, Ewan Harrison, David K. Jackson, Ian Johnston, Dominic Kwiatkowski, Cordelia Langford, John Sillitoe on behalf of the Wellcome Sanger Institute COVID-19 Surveillance Team |
| EPI_ISL_2664232 | Lighthouse Lab in Alderley Park | Wellcome Sanger Institute for the COVID-19 Genomics UK (COG-UK) Consortium | Jacquelyn Wynn, Mairead Hyland, The Lighthouse Lab in Alderley Park and Alex Alderton, Roberto Amato, Jeffrey Barrett, Sonia Goncalves, Ewan Harrison, David K. Jackson, Ian Johnston, Dominic Kwiatkowski, Cordelia Langford, John Sillitoe on behalf of the Wellcome Sanger Institute COVID-19 Surveillance Team |
| EPI_ISL_2664234 | Lighthouse Lab in Glasgow | Wellcome Sanger Institute for the COVID-19 Genomics UK (COG-UK) Consortium | Harper VanSteenhouse, Yumi Kasai, David Gray, Carol Clugston, Anna Dominiczak and Alex Alderton, Roberto Amato, Jeffrey Barrett, Sonia Goncalves, Ewan Harrison, David K. Jackson, Ian Johnston, Dominic Kwiatkowski, Cordelia Langford, John Sillitoe on behalf of the Wellcome Sanger Institute COVID-19 Surveillance Team |
| EPI_ISL_2664235 | Lighthouse Lab in Milton Keynes | Wellcome Sanger Institute for the COVID-19 Genomics UK (COG-UK) Consortium | The Lighthouse Lab in Milton Keynes and Alex Alderton, Roberto Amato, Jeffrey Barrett, Sonia Goncalves, Ewan Harrison, David K. Jackson, Ian Johnston, Dominic Kwiatkowski, Cordelia Langford, John Sillitoe on behalf of the Wellcome Sanger Institute COVID-19 Surveillance Team |
| EPI_ISL_2664236 | Lighthouse Lab in Alderley Park | Wellcome Sanger Institute for the COVID-19 Genomics UK (COG-UK) Consortium | Jacquelyn Wynn, Mairead Hyland, The Lighthouse Lab in Alderley Park and Alex Alderton, Roberto Amato, Jeffrey Barrett, Sonia Goncalves, Ewan Harrison, David K. Jackson, Ian Johnston, Dominic Kwiatkowski, Cordelia Langford, John Sillitoe on behalf of the Wellcome Sanger Institute COVID-19 Surveillance Team |
| EPI_ISL_2664237 | Lighthouse Lab in Glasgow | Wellcome Sanger Institute for the COVID-19 Genomics UK (COG-UK) Consortium | Harper VanSteenhouse, Yumi Kasai, David Gray, Carol Clugston, Anna Dominiczak and Alex Alderton, Roberto Amato, Jeffrey Barrett, Sonia Goncalves, Ewan Harrison, David K. Jackson, Ian Johnston, Dominic Kwiatkowski, Cordelia Langford, John Sillitoe on behalf of the Wellcome Sanger Institute COVID-19 Surveillance Team |
| EPI_ISL_2664238, EPI_ISL_2664239 | Lighthouse Lab in Milton Keynes | Wellcome Sanger Institute for the COVID-19 Genomics UK (COG-UK) Consortium | The Lighthouse Lab in Milton Keynes and Alex Alderton, Roberto Amato, Jeffrey Barrett, Sonia Goncalves, Ewan Harrison, David K. Jackson, Ian Johnston, Dominic Kwiatkowski, Cordelia Langford, John Sillitoe on behalf of the Wellcome Sanger Institute COVID-19 Surveillance Team |
| EPI_ISL_2664240 | Lighthouse Lab in Glasgow | Wellcome Sanger Institute for the COVID-19 Genomics UK (COG-UK) Consortium | Harper VanSteenhouse, Yumi Kasai, David Gray, Carol Clugston, Anna Dominiczak and Alex Alderton, Roberto Amato, Jeffrey Barrett, Sonia Goncalves, Ewan Harrison, David K. Jackson, Ian Johnston, Dominic Kwiatkowski, Cordelia Langford, John Sillitoe on behalf of the Wellcome Sanger Institute COVID-19 Surveillance Team |
| EPI_ISL_2664241 | Lighthouse Lab in Milton Keynes | Wellcome Sanger Institute for the COVID-19 Genomics UK (COG-UK) Consortium | The Lighthouse Lab in Milton Keynes and Alex Alderton, Roberto Amato, Jeffrey Barrett, Sonia Goncalves, Ewan Harrison, David K. Jackson, Ian Johnston, Dominic Kwiatkowski, Cordelia Langford, John Sillitoe on behalf of the Wellcome Sanger Institute COVID-19 Surveillance Team |
| EPI_ISL_2664242 | Lighthouse Lab in Alderley Park | Wellcome Sanger Institute for the COVID-19 Genomics UK (COG-UK) Consortium | Jacquelyn Wynn, Mairead Hyland, The Lighthouse Lab in Alderley Park and Alex Alderton, Roberto Amato, Jeffrey Barrett, Sonia Goncalves, Ewan Harrison, David K. Jackson, Ian Johnston, Dominic Kwiatkowski, Cordelia Langford, John Sillitoe on behalf of the Wellcome Sanger Institute COVID-19 Surveillance Team |
| EPI_ISL_2664243, EPI_ISL_2664244, EPI_ISL_2664245 | Lighthouse Lab in Milton Keynes | Wellcome Sanger Institute for the COVID-19 Genomics UK (COG-UK) Consortium | The Lighthouse Lab in Milton Keynes and Alex Alderton, Roberto Amato, Jeffrey Barrett, Sonia Goncalves, Ewan Harrison, David K. Jackson, Ian Johnston, Dominic Kwiatkowski, Cordelia Langford, John Sillitoe on behalf of the Wellcome Sanger Institute COVID-19 Surveillance Team |
| EPI_ISL_2664246 | Lighthouse Lab in Alderley Park | Wellcome Sanger Institute for the COVID-19 Genomics UK (COG-UK) Consortium | Jacquelyn Wynn, Mairead Hyland, The Lighthouse Lab in Alderley Park and Alex Alderton, Roberto Amato, Jeffrey Barrett, Sonia Goncalves, Ewan Harrison, David K. Jackson, Ian Johnston, Dominic Kwiatkowski, Cordelia Langford, John Sillitoe on behalf of the Wellcome Sanger Institute COVID-19 Surveillance Team |
| EPI_ISL_2664247 | Lighthouse Lab in Milton Keynes | Wellcome Sanger Institute for the COVID-19 Genomics UK (COG-UK) Consortium | The Lighthouse Lab in Milton Keynes and Alex Alderton, Roberto Amato, Jeffrey Barrett, Sonia Goncalves, Ewan Harrison, David K. Jackson, Ian Johnston, Dominic Kwiatkowski, Cordelia Langford, John Sillitoe on behalf of the Wellcome Sanger Institute COVID-19 Surveillance Team |
| EPI_ISL_2664248 | Lighthouse Lab in Alderley Park | Wellcome Sanger Institute for the COVID-19 Genomics UK (COG-UK) Consortium | Jacquelyn Wynn, Mairead Hyland, The Lighthouse Lab in Alderley Park and Alex Alderton, Roberto Amato, Jeffrey Barrett, Sonia Goncalves, Ewan Harrison, David K. Jackson, Ian Johnston, Dominic Kwiatkowski, Cordelia Langford, John Sillitoe on behalf of the Wellcome Sanger Institute COVID-19 Surveillance Team |
| EPI_ISL_2664249 | Lighthouse Lab in Milton Keynes | Wellcome Sanger Institute for the COVID-19 Genomics UK (COG-UK) Consortium | The Lighthouse Lab in Milton Keynes and Alex Alderton, Roberto Amato, Jeffrey Barrett, Sonia Goncalves, Ewan Harrison, David K. Jackson, Ian Johnston, Dominic Kwiatkowski, Cordelia Langford, John Sillitoe on behalf of the Wellcome Sanger Institute COVID-19 Surveillance Team |
| EPI_ISL_2664250 | Lighthouse Lab in Glasgow | Wellcome Sanger Institute for the COVID-19 Genomics UK (COG-UK) Consortium | Harper VanSteenhouse, Yumi Kasai, David Gray, Carol Clugston, Anna Dominiczak and Alex Alderton, Roberto Amato, Jeffrey Barrett, Sonia Goncalves, Ewan Harrison, David K. Jackson, Ian Johnston, Dominic Kwiatkowski, Cordelia Langford, John Sillitoe on behalf of the Wellcome Sanger Institute COVID-19 Surveillance Team |
| EPI_ISL_2664251 | Lighthouse Lab in Alderley Park | Wellcome Sanger Institute for the COVID-19 Genomics UK (COG-UK) Consortium | Jacquelyn Wynn, Mairead Hyland, The Lighthouse Lab in Alderley Park and Alex Alderton, Roberto Amato, Jeffrey Barrett, Sonia Goncalves, Ewan Harrison, David K. Jackson, Ian Johnston, Dominic Kwiatkowski, Cordelia Langford, John Sillitoe on behalf of the Wellcome Sanger Institute COVID-19 Surveillance Team |
| EPI_ISL_2664252 | Lighthouse Lab in Milton Keynes | Wellcome Sanger Institute for the COVID-19 Genomics UK (COG-UK) Consortium | The Lighthouse Lab in Milton Keynes and Alex Alderton, Roberto Amato, Jeffrey Barrett, Sonia Goncalves, Ewan Harrison, David K. Jackson, Ian Johnston, Dominic Kwiatkowski, Cordelia Langford, John Sillitoe on behalf of the Wellcome Sanger Institute COVID-19 Surveillance Team |
| EPI_ISL_2664253 | Lighthouse Lab in Alderley Park | Wellcome Sanger Institute for the COVID-19 Genomics UK (COG-UK) Consortium | Jacquelyn Wynn, Mairead Hyland, The Lighthouse Lab in Alderley Park and Alex Alderton, Roberto Amato, Jeffrey Barrett, Sonia Goncalves, Ewan Harrison, David K. Jackson, Ian Johnston, Dominic Kwiatkowski, Cordelia Langford, John Sillitoe on behalf of the Wellcome Sanger Institute COVID-19 Surveillance Team |
| EPI_ISL_2664254 | Lighthouse Lab in Glasgow | Wellcome Sanger Institute for the COVID-19 Genomics UK (COG-UK) Consortium | Harper VanSteenhouse, Yumi Kasai, David Gray, Carol Clugston, Anna Dominiczak and Alex Alderton, Roberto Amato, Jeffrey Barrett, Sonia Goncalves, Ewan Harrison, David K. Jackson, Ian Johnston, Dominic Kwiatkowski, Cordelia Langford, John Sillitoe on behalf of the Wellcome Sanger Institute COVID-19 Surveillance Team |
| EPI_ISL_2664255, EPI_ISL_2664256, EPI_ISL_2664257 | Lighthouse Lab in Milton Keynes | Wellcome Sanger Institute for the COVID-19 Genomics UK (COG-UK) Consortium | The Lighthouse Lab in Milton Keynes and Alex Alderton, Roberto Amato, Jeffrey Barrett, Sonia Goncalves, Ewan Harrison, David K. Jackson, Ian Johnston, Dominic Kwiatkowski, Cordelia Langford, John Sillitoe on behalf of the Wellcome Sanger Institute COVID-19 Surveillance Team |
| EPI_ISL_2664258, EPI_ISL_2664259, EPI_ISL_2664260 | Lighthouse Lab in Alderley Park | Wellcome Sanger Institute for the COVID-19 Genomics UK (COG-UK) Consortium | Jacquelyn Wynn, Mairead Hyland, The Lighthouse Lab in Alderley Park and Alex Alderton, Roberto Amato, Jeffrey Barrett, Sonia Goncalves, Ewan Harrison, David K. Jackson, Ian Johnston, Dominic Kwiatkowski, Cordelia Langford, John Sillitoe on behalf of the Wellcome Sanger Institute COVID-19 Surveillance Team |
| EPI_ISL_2664261, EPI_ISL_2664262 | Lighthouse Lab in Milton Keynes | Wellcome Sanger Institute for the COVID-19 Genomics UK (COG-UK) Consortium | The Lighthouse Lab in Milton Keynes and Alex Alderton, Roberto Amato, Jeffrey Barrett, Sonia Goncalves, Ewan Harrison, David K. Jackson, Ian Johnston, Dominic Kwiatkowski, Cordelia Langford, John Sillitoe on behalf of the Wellcome Sanger Institute COVID-19 Surveillance Team |
| EPI_ISL_2664263, EPI_ISL_2664264, EPI_ISL_2664265 | Lighthouse Lab in Alderley Park | Wellcome Sanger Institute for the COVID-19 Genomics UK (COG-UK) Consortium | Jacquelyn Wynn, Mairead Hyland, The Lighthouse Lab in Alderley Park and Alex Alderton, Roberto Amato, Jeffrey Barrett, Sonia Goncalves, Ewan Harrison, David K. Jackson, Ian Johnston, Dominic Kwiatkowski, Cordelia Langford, John Sillitoe on behalf of the Wellcome Sanger Institute COVID-19 Surveillance Team |
| EPI_ISL_2664266, EPI_ISL_2664267 | Lighthouse Lab in Glasgow | Wellcome Sanger Institute for the COVID-19 Genomics UK (COG-UK) Consortium | Harper VanSteenhouse, Yumi Kasai, David Gray, Carol Clugston, Anna Dominiczak and Alex Alderton, Roberto Amato, Jeffrey Barrett, Sonia Goncalves, Ewan Harrison, David K. Jackson, Ian Johnston, Dominic Kwiatkowski, Cordelia Langford, John Sillitoe on behalf of the Wellcome Sanger Institute COVID-19 Surveillance Team |
| EPI_ISL_2664268 | Lighthouse Lab in Milton Keynes | Wellcome Sanger Institute for the COVID-19 Genomics UK (COG-UK) Consortium | The Lighthouse Lab in Milton Keynes and Alex Alderton, Roberto Amato, Jeffrey Barrett, Sonia Goncalves, Ewan Harrison, David K. Jackson, Ian Johnston, Dominic Kwiatkowski, Cordelia Langford, John Sillitoe on behalf of the Wellcome Sanger Institute COVID-19 Surveillance Team |
| EPI_ISL_2664269 | Lighthouse Lab in Glasgow | Wellcome Sanger Institute for the COVID-19 Genomics UK (COG-UK) Consortium | Harper VanSteenhouse, Yumi Kasai, David Gray, Carol Clugston, Anna Dominiczak and Alex Alderton, Roberto Amato, Jeffrey Barrett, Sonia Goncalves, Ewan Harrison, David K. Jackson, Ian Johnston, Dominic Kwiatkowski, Cordelia Langford, John Sillitoe on behalf of the Wellcome Sanger Institute COVID-19 Surveillance Team |
| EPI_ISL_2664270 | Lighthouse Lab in Alderley Park | Wellcome Sanger Institute for the COVID-19 Genomics UK (COG-UK) | Jacquelyn Wynn, Mairead Hyland, The Lighthouse Lab in Alderley Park and Alex Alderton, Roberto Amato, Jeffrey Barrett, Sonia Goncalves, Ewan Harrison, David K. Jackson, Ian Johnston, Dominic Kwiatkowski, Cordelia Langford, John Sillitoe on |

| | | | |
|---|---|---|---|
| | | Consortium | behalf of the Wellcome Sanger Institute COVID-19 Surveillance Team |
| EPI_ISL_2664271 | Lighthouse Lab in Milton Keynes | Wellcome Sanger Institute for the COVID-19 Genomics UK (COG-UK) Consortium | The Lighthouse Lab in Milton Keynes and Alex Alderton, Roberto Amato, Jeffrey Barrett, Sonia Goncalves, Ewan Harrison, David K. Jackson, Ian Johnston, Dominic Kwiatkowski, Cordelia Langford, John Sillitoe on behalf of the Wellcome Sanger Institute COVID-19 Surveillance Team |
| EPI_ISL_2664272 | Lighthouse Lab in Glasgow | Wellcome Sanger Institute for the COVID-19 Genomics UK (COG-UK) Consortium | Harper VanSteenhouse, Yumi Kasai, David Gray, Carol Clugston, Anna Dominiczak and Alex Alderton, Roberto Amato, Jeffrey Barrett, Sonia Goncalves, Ewan Harrison, David K. Jackson, Ian Johnston, Dominic Kwiatkowski, Cordelia Langford, John Sillitoe on behalf of the Wellcome Sanger Institute COVID-19 Surveillance Team |
| EPI_ISL_2664273 | Lighthouse Lab in Alderley Park | Wellcome Sanger Institute for the COVID-19 Genomics UK (COG-UK) Consortium | Jacquelyn Wynn, Mairead Hyland, The Lighthouse Lab in Alderley Park and Alex Alderton, Roberto Amato, Jeffrey Barrett, Sonia Goncalves, Ewan Harrison, David K. Jackson, Ian Johnston, Dominic Kwiatkowski, Cordelia Langford, John Sillitoe on behalf of the Wellcome Sanger Institute COVID-19 Surveillance Team |
| EPI_ISL_2664274, EPI_ISL_2664275, EPI_ISL_2664276, EPI_ISL_2664277 | Lighthouse Lab in Glasgow | Wellcome Sanger Institute for the COVID-19 Genomics UK (COG-UK) Consortium | Harper VanSteenhouse, Yumi Kasai, David Gray, Carol Clugston, Anna Dominiczak and Alex Alderton, Roberto Amato, Jeffrey Barrett, Sonia Goncalves, Ewan Harrison, David K. Jackson, Ian Johnston, Dominic Kwiatkowski, Cordelia Langford, John Sillitoe on behalf of the Wellcome Sanger Institute COVID-19 Surveillance Team |
| EPI_ISL_2664278, EPI_ISL_2664279 | Lighthouse Lab in Milton Keynes | Wellcome Sanger Institute for the COVID-19 Genomics UK (COG-UK) Consortium | The Lighthouse Lab in Milton Keynes and Alex Alderton, Roberto Amato, Jeffrey Barrett, Sonia Goncalves, Ewan Harrison, David K. Jackson, Ian Johnston, Dominic Kwiatkowski, Cordelia Langford, John Sillitoe on behalf of the Wellcome Sanger Institute COVID-19 Surveillance Team |
| EPI_ISL_2664280, EPI_ISL_2664281 | Lighthouse Lab in Glasgow | Wellcome Sanger Institute for the COVID-19 Genomics UK (COG-UK) Consortium | Harper VanSteenhouse, Yumi Kasai, David Gray, Carol Clugston, Anna Dominiczak and Alex Alderton, Roberto Amato, Jeffrey Barrett, Sonia Goncalves, Ewan Harrison, David K. Jackson, Ian Johnston, Dominic Kwiatkowski, Cordelia Langford, John Sillitoe on behalf of the Wellcome Sanger Institute COVID-19 Surveillance Team |
| EPI_ISL_2664282 | Lighthouse Lab in Alderley Park | Wellcome Sanger Institute for the COVID-19 Genomics UK (COG-UK) Consortium | Jacquelyn Wynn, Mairead Hyland, The Lighthouse Lab in Alderley Park and Alex Alderton, Roberto Amato, Jeffrey Barrett, Sonia Goncalves, Ewan Harrison, David K. Jackson, Ian Johnston, Dominic Kwiatkowski, Cordelia Langford, John Sillitoe on behalf of the Wellcome Sanger Institute COVID-19 Surveillance Team |
| EPI_ISL_2664283 | Lighthouse Lab in Glasgow | Wellcome Sanger Institute for the COVID-19 Genomics UK (COG-UK) Consortium | Harper VanSteenhouse, Yumi Kasai, David Gray, Carol Clugston, Anna Dominiczak and Alex Alderton, Roberto Amato, Jeffrey Barrett, Sonia Goncalves, Ewan Harrison, David K. Jackson, Ian Johnston, Dominic Kwiatkowski, Cordelia Langford, John Sillitoe on behalf of the Wellcome Sanger Institute COVID-19 Surveillance Team |
| EPI_ISL_2664284 | Lighthouse Lab in Milton Keynes | Wellcome Sanger Institute for the COVID-19 Genomics UK (COG-UK) Consortium | The Lighthouse Lab in Milton Keynes and Alex Alderton, Roberto Amato, Jeffrey Barrett, Sonia Goncalves, Ewan Harrison, David K. Jackson, Ian Johnston, Dominic Kwiatkowski, Cordelia Langford, John Sillitoe on behalf of the Wellcome Sanger Institute COVID-19 Surveillance Team |
| EPI_ISL_2664285 | Lighthouse Lab in Glasgow | Wellcome Sanger Institute for the COVID-19 Genomics UK (COG-UK) Consortium | Harper VanSteenhouse, Yumi Kasai, David Gray, Carol Clugston, Anna Dominiczak and Alex Alderton, Roberto Amato, Jeffrey Barrett, Sonia Goncalves, Ewan Harrison, David K. Jackson, Ian Johnston, Dominic Kwiatkowski, Cordelia Langford, John Sillitoe on behalf of the Wellcome Sanger Institute COVID-19 Surveillance Team |
| EPI_ISL_2664286 | Lighthouse Lab in Alderley Park | Wellcome Sanger Institute for the COVID-19 Genomics UK (COG-UK) Consortium | Jacquelyn Wynn, Mairead Hyland, The Lighthouse Lab in Alderley Park and Alex Alderton, Roberto Amato, Jeffrey Barrett, Sonia Goncalves, Ewan Harrison, David K. Jackson, Ian Johnston, Dominic Kwiatkowski, Cordelia Langford, John Sillitoe on behalf of the Wellcome Sanger Institute COVID-19 Surveillance Team |
| EPI_ISL_2664287, EPI_ISL_2664288 | Lighthouse Lab in Milton Keynes | Wellcome Sanger Institute for the COVID-19 Genomics UK (COG-UK) Consortium | The Lighthouse Lab in Milton Keynes and Alex Alderton, Roberto Amato, Jeffrey Barrett, Sonia Goncalves, Ewan Harrison, David K. Jackson, Ian Johnston, Dominic Kwiatkowski, Cordelia Langford, John Sillitoe on behalf of the Wellcome Sanger Institute COVID-19 Surveillance Team |
| EPI_ISL_2664289 | Lighthouse Lab in Glasgow | Wellcome Sanger Institute for the COVID-19 Genomics UK (COG-UK) Consortium | Harper VanSteenhouse, Yumi Kasai, David Gray, Carol Clugston, Anna Dominiczak and Alex Alderton, Roberto Amato, Jeffrey Barrett, Sonia Goncalves, Ewan Harrison, David K. Jackson, Ian Johnston, Dominic Kwiatkowski, Cordelia Langford, John Sillitoe on behalf of the Wellcome Sanger Institute COVID-19 Surveillance Team |
| EPI_ISL_2664290 | Lighthouse Lab in Milton Keynes | Wellcome Sanger Institute for the COVID-19 Genomics UK (COG-UK) Consortium | The Lighthouse Lab in Milton Keynes and Alex Alderton, Roberto Amato, Jeffrey Barrett, Sonia Goncalves, Ewan Harrison, David K. Jackson, Ian Johnston, Dominic Kwiatkowski, Cordelia Langford, John Sillitoe on behalf of the Wellcome Sanger Institute COVID-19 Surveillance Team |
| EPI_ISL_2664291, EPI_ISL_2664292 | Lighthouse Lab in Alderley Park | Wellcome Sanger Institute for the COVID-19 Genomics UK (COG-UK) Consortium | Jacquelyn Wynn, Mairead Hyland, The Lighthouse Lab in Alderley Park and Alex Alderton, Roberto Amato, Jeffrey Barrett, Sonia Goncalves, Ewan Harrison, David K. Jackson, Ian Johnston, Dominic Kwiatkowski, Cordelia Langford, John Sillitoe on behalf of the Wellcome Sanger Institute COVID-19 Surveillance Team |
| EPI_ISL_2664293 | Lighthouse Lab in Glasgow | Wellcome Sanger Institute for the COVID-19 Genomics UK (COG-UK) Consortium | Harper VanSteenhouse, Yumi Kasai, David Gray, Carol Clugston, Anna Dominiczak and Alex Alderton, Roberto Amato, Jeffrey Barrett, Sonia Goncalves, Ewan Harrison, David K. Jackson, Ian Johnston, Dominic Kwiatkowski, Cordelia Langford, John Sillitoe on behalf of the Wellcome Sanger Institute COVID-19 Surveillance Team |
| EPI_ISL_2664294, EPI_ISL_2664295, EPI_ISL_2664296, EPI_ISL_2664297 | Lighthouse Lab in Alderley Park | Wellcome Sanger Institute for the COVID-19 Genomics UK (COG-UK) Consortium | Jacquelyn Wynn, Mairead Hyland, The Lighthouse Lab in Alderley Park and Alex Alderton, Roberto Amato, Jeffrey Barrett, Sonia Goncalves, Ewan Harrison, David K. Jackson, Ian Johnston, Dominic Kwiatkowski, Cordelia Langford, John Sillitoe on behalf of the Wellcome Sanger Institute COVID-19 Surveillance Team |
| EPI_ISL_2664298 | Lighthouse Lab in Milton Keynes | Wellcome Sanger Institute for the COVID-19 Genomics UK (COG-UK) Consortium | The Lighthouse Lab in Milton Keynes and Alex Alderton, Roberto Amato, Jeffrey Barrett, Sonia Goncalves, Ewan Harrison, David K. Jackson, Ian Johnston, Dominic Kwiatkowski, Cordelia Langford, John Sillitoe on behalf of the Wellcome Sanger Institute COVID-19 Surveillance Team |
| EPI_ISL_2664299 | Lighthouse Lab in Glasgow | Wellcome Sanger Institute for the COVID-19 Genomics UK (COG-UK) Consortium | Harper VanSteenhouse, Yumi Kasai, David Gray, Carol Clugston, Anna Dominiczak and Alex Alderton, Roberto Amato, Jeffrey Barrett, Sonia Goncalves, Ewan Harrison, David K. Jackson, Ian Johnston, Dominic Kwiatkowski, Cordelia Langford, John Sillitoe on behalf of the Wellcome Sanger Institute COVID-19 Surveillance Team |
| EPI_ISL_2664300, EPI_ISL_2664301, EPI_ISL_2664302 | Lighthouse Lab in Alderley Park | Wellcome Sanger Institute for the COVID-19 Genomics UK (COG-UK) Consortium | Jacquelyn Wynn, Mairead Hyland, The Lighthouse Lab in Alderley Park and Alex Alderton, Roberto Amato, Jeffrey Barrett, Sonia Goncalves, Ewan Harrison, David K. Jackson, Ian Johnston, Dominic Kwiatkowski, Cordelia Langford, John Sillitoe on behalf of the Wellcome Sanger Institute COVID-19 Surveillance Team |
| EPI_ISL_2664303 | Lighthouse Lab in Milton Keynes | Wellcome Sanger Institute for the COVID-19 Genomics UK (COG-UK) Consortium | The Lighthouse Lab in Milton Keynes and Alex Alderton, Roberto Amato, Jeffrey Barrett, Sonia Goncalves, Ewan Harrison, David K. Jackson, Ian Johnston, Dominic Kwiatkowski, Cordelia Langford, John Sillitoe on behalf of the Wellcome Sanger Institute COVID-19 Surveillance Team |
| EPI_ISL_2664304 | Lighthouse Lab in Alderley Park | Wellcome Sanger Institute for the COVID-19 Genomics UK (COG-UK) Consortium | Jacquelyn Wynn, Mairead Hyland, The Lighthouse Lab in Alderley Park and Alex Alderton, Roberto Amato, Jeffrey Barrett, Sonia Goncalves, Ewan Harrison, David K. Jackson, Ian Johnston, Dominic Kwiatkowski, Cordelia Langford, John Sillitoe on behalf of the Wellcome Sanger Institute COVID-19 Surveillance Team |
| EPI_ISL_2664305, EPI_ISL_2664306, EPI_ISL_2664307 | Lighthouse Lab in Milton Keynes | Wellcome Sanger Institute for the COVID-19 Genomics UK (COG-UK) Consortium | The Lighthouse Lab in Milton Keynes and Alex Alderton, Roberto Amato, Jeffrey Barrett, Sonia Goncalves, Ewan Harrison, David K. Jackson, Ian Johnston, Dominic Kwiatkowski, Cordelia Langford, John Sillitoe on behalf of the Wellcome Sanger Institute COVID-19 Surveillance Team |
| EPI_ISL_2664308 | Lighthouse Lab in Alderley Park | Wellcome Sanger Institute for the COVID-19 Genomics UK (COG-UK) Consortium | Jacquelyn Wynn, Mairead Hyland, The Lighthouse Lab in Alderley Park and Alex Alderton, Roberto Amato, Jeffrey Barrett, Sonia Goncalves, Ewan Harrison, David K. Jackson, Ian Johnston, Dominic Kwiatkowski, Cordelia Langford, John Sillitoe on behalf of the Wellcome Sanger Institute COVID-19 Surveillance Team |
| EPI_ISL_2664309 | Lighthouse Lab in Milton Keynes | Wellcome Sanger Institute for the COVID-19 Genomics UK (COG-UK) Consortium | The Lighthouse Lab in Milton Keynes and Alex Alderton, Roberto Amato, Jeffrey Barrett, Sonia Goncalves, Ewan Harrison, David K. Jackson, Ian Johnston, Dominic Kwiatkowski, Cordelia Langford, John Sillitoe on behalf of the Wellcome Sanger Institute COVID-19 Surveillance Team |
| EPI_ISL_2664310 | Lighthouse Lab in Glasgow | Wellcome Sanger Institute for the COVID-19 Genomics UK (COG-UK) Consortium | Harper VanSteenhouse, Yumi Kasai, David Gray, Carol Clugston, Anna Dominiczak and Alex Alderton, Roberto Amato, Jeffrey Barrett, Sonia Goncalves, Ewan Harrison, David K. Jackson, Ian Johnston, Dominic Kwiatkowski, Cordelia Langford, John Sillitoe on behalf of the Wellcome Sanger Institute COVID-19 Surveillance Team |
| EPI_ISL_2664311 | Lighthouse Lab in Alderley Park | Wellcome Sanger Institute for the COVID-19 Genomics UK (COG-UK) Consortium | Jacquelyn Wynn, Mairead Hyland, The Lighthouse Lab in Alderley Park and Alex Alderton, Roberto Amato, Jeffrey Barrett, Sonia Goncalves, Ewan Harrison, David K. Jackson, Ian Johnston, Dominic Kwiatkowski, Cordelia Langford, John Sillitoe on behalf of the Wellcome Sanger Institute COVID-19 Surveillance Team |
| EPI_ISL_2664312 | Lighthouse Lab in Milton Keynes | Wellcome Sanger Institute for the COVID-19 Genomics UK (COG-UK) Consortium | The Lighthouse Lab in Milton Keynes and Alex Alderton, Roberto Amato, Jeffrey Barrett, Sonia Goncalves, Ewan Harrison, David K. Jackson, Ian Johnston, Dominic Kwiatkowski, Cordelia Langford, John Sillitoe on behalf of the Wellcome Sanger Institute COVID-19 Surveillance Team |
| EPI_ISL_2664313, EPI_ISL_2664314 | Lighthouse Lab in Glasgow | Wellcome Sanger Institute for the COVID-19 Genomics UK (COG-UK) Consortium | Harper VanSteenhouse, Yumi Kasai, David Gray, Carol Clugston, Anna Dominiczak and Alex Alderton, Roberto Amato, Jeffrey Barrett, Sonia Goncalves, Ewan Harrison, David K. Jackson, Ian Johnston, Dominic Kwiatkowski, Cordelia Langford, John Sillitoe on behalf of the Wellcome Sanger Institute COVID-19 Surveillance Team |
| EPI_ISL_2664315 | Lighthouse Lab in Milton Keynes | Wellcome Sanger Institute for the COVID-19 Genomics UK (COG-UK) Consortium | The Lighthouse Lab in Milton Keynes and Alex Alderton, Roberto Amato, Jeffrey Barrett, Sonia Goncalves, Ewan Harrison, David K. Jackson, Ian Johnston, Dominic Kwiatkowski, Cordelia Langford, John Sillitoe on behalf of the Wellcome Sanger Institute COVID-19 Surveillance Team |
| EPI_ISL_2664316, EPI_ISL_2664317, EPI_ISL_2664318, EPI_ISL_2664319 | Lighthouse Lab in Alderley Park | Wellcome Sanger Institute for the COVID-19 Genomics UK (COG-UK) Consortium | Jacquelyn Wynn, Mairead Hyland, The Lighthouse Lab in Alderley Park and Alex Alderton, Roberto Amato, Jeffrey Barrett, Sonia Goncalves, Ewan Harrison, David K. Jackson, Ian Johnston, Dominic Kwiatkowski, Cordelia Langford, John Sillitoe on behalf of the Wellcome Sanger Institute COVID-19 Surveillance Team |
| EPI_ISL_2664320 | Lighthouse Lab in Glasgow | Wellcome Sanger Institute for the COVID-19 Genomics UK (COG-UK) Consortium | Harper VanSteenhouse, Yumi Kasai, David Gray, Carol Clugston, Anna Dominiczak and Alex Alderton, Roberto Amato, Jeffrey Barrett, Sonia Goncalves, Ewan Harrison, David K. Jackson, Ian Johnston, Dominic Kwiatkowski, Cordelia Langford, John Sillitoe on behalf of the Wellcome Sanger Institute COVID-19 Surveillance Team |
| EPI_ISL_2664321, EPI_ISL_2664322 | Lighthouse Lab in Milton Keynes | Wellcome Sanger Institute for the COVID-19 Genomics UK (COG-UK) Consortium | The Lighthouse Lab in Milton Keynes and Alex Alderton, Roberto Amato, Jeffrey Barrett, Sonia Goncalves, Ewan Harrison, David K. Jackson, Ian Johnston, Dominic Kwiatkowski, Cordelia Langford, John Sillitoe on behalf of the Wellcome Sanger Institute COVID-19 Surveillance Team |
| EPI_ISL_2664323 | Lighthouse Lab in Glasgow | Wellcome Sanger Institute for the COVID-19 Genomics UK (COG-UK) Consortium | Harper VanSteenhouse, Yumi Kasai, David Gray, Carol Clugston, Anna Dominiczak and Alex Alderton, Roberto Amato, Jeffrey Barrett, Sonia Goncalves, Ewan Harrison, David K. Jackson, Ian Johnston, Dominic Kwiatkowski, Cordelia Langford, John Sillitoe on behalf of the Wellcome Sanger Institute COVID-19 Surveillance Team |
| EPI_ISL_2664324 | Lighthouse Lab in Alderley Park | Wellcome Sanger Institute for the COVID-19 Genomics UK (COG-UK) Consortium | Jacquelyn Wynn, Mairead Hyland, The Lighthouse Lab in Alderley Park and Alex Alderton, Roberto Amato, Jeffrey Barrett, Sonia Goncalves, Ewan Harrison, David K. Jackson, Ian Johnston, Dominic Kwiatkowski, Cordelia Langford, John Sillitoe on behalf of the Wellcome Sanger Institute COVID-19 Surveillance Team |
| EPI_ISL_2664325, EPI_ISL_2664326, EPI_ISL_2664327, EPI_ISL_2664328 | Lighthouse Lab in Milton Keynes | Wellcome Sanger Institute for the COVID-19 Genomics UK (COG-UK) Consortium | The Lighthouse Lab in Milton Keynes and Alex Alderton, Roberto Amato, Jeffrey Barrett, Sonia Goncalves, Ewan Harrison, David K. Jackson, Ian Johnston, Dominic Kwiatkowski, Cordelia Langford, John Sillitoe on behalf of the Wellcome Sanger Institute COVID-19 Surveillance Team |
| EPI_ISL_2664329 | Lighthouse Lab in Alderley Park | Wellcome Sanger Institute for the COVID-19 Genomics UK (COG-UK) Consortium | Jacquelyn Wynn, Mairead Hyland, The Lighthouse Lab in Alderley Park and Alex Alderton, Roberto Amato, Jeffrey Barrett, Sonia Goncalves, Ewan Harrison, David K. Jackson, Ian Johnston, Dominic Kwiatkowski, Cordelia Langford, John Sillitoe on behalf of the Wellcome Sanger Institute COVID-19 Surveillance Team |
| EPI_ISL_2664330, EPI_ISL_2664331 | Lighthouse Lab in Glasgow | Wellcome Sanger Institute for the COVID-19 Genomics UK (COG-UK) Consortium | Harper VanSteenhouse, Yumi Kasai, David Gray, Carol Clugston, Anna Dominiczak and Alex Alderton, Roberto Amato, Jeffrey Barrett, Sonia Goncalves, Ewan Harrison, David K. Jackson, Ian Johnston, Dominic Kwiatkowski, Cordelia Langford, John Sillitoe on behalf of the Wellcome Sanger Institute COVID-19 Surveillance Team |
| EPI_ISL_2664332, EPI_ISL_2664333, EPI_ISL_2664334 | Lighthouse Lab in Milton Keynes | Wellcome Sanger Institute for the COVID-19 Genomics UK (COG-UK) Consortium | The Lighthouse Lab in Milton Keynes and Alex Alderton, Roberto Amato, Jeffrey Barrett, Sonia Goncalves, Ewan Harrison, David K. Jackson, Ian Johnston, Dominic Kwiatkowski, Cordelia Langford, John Sillitoe on behalf of the Wellcome Sanger Institute COVID-19 Surveillance Team |
| EPI_ISL_2664335, EPI_ISL_2664336, EPI_ISL_2664337 | Lighthouse Lab in Alderley Park | Wellcome Sanger Institute for the COVID-19 Genomics UK (COG-UK) Consortium | Jacquelyn Wynn, Mairead Hyland, The Lighthouse Lab in Alderley Park and Alex Alderton, Roberto Amato, Jeffrey Barrett, Sonia Goncalves, Ewan Harrison, David K. Jackson, Ian Johnston, Dominic Kwiatkowski, Cordelia Langford, John Sillitoe on behalf of the Wellcome Sanger Institute COVID-19 Surveillance Team |

| | | | |
|---|---|---|---|
| EPI_ISL_2664338, EPI_ISL_2664339 | Lighthouse Lab in Glasgow | Wellcome Sanger Institute for the COVID-19 Genomics UK (COG-UK) Consortium | Harper VanSteenhouse, Yumi Kasai, David Gray, Carol Clugston, Anna Dominiczak and Alex Alderton, Roberto Amato, Jeffrey Barrett, Sonia Goncalves, Ewan Harrison, David K. Jackson, Ian Johnston, Dominic Kwiatkowski, Cordelia Langford, John Sillitoe on behalf of the Wellcome Sanger Institute COVID-19 Surveillance Team |
| EPI_ISL_2664340 | Lighthouse Lab in Milton Keynes | Wellcome Sanger Institute for the COVID-19 Genomics UK (COG-UK) Consortium | The Lighthouse Lab in Milton Keynes and Alex Alderton, Roberto Amato, Jeffrey Barrett, Sonia Goncalves, Ewan Harrison, David K. Jackson, Ian Johnston, Dominic Kwiatkowski, Cordelia Langford, John Sillitoe on behalf of the Wellcome Sanger Institute COVID-19 Surveillance Team |
| EPI_ISL_2664341 | Lighthouse Lab in Alderley Park | Wellcome Sanger Institute for the COVID-19 Genomics UK (COG-UK) Consortium | Jacquelyn Wynn, Mairead Hyland, The Lighthouse Lab in Alderley Park and Alex Alderton, Roberto Amato, Jeffrey Barrett, Sonia Goncalves, Ewan Harrison, David K. Jackson, Ian Johnston, Dominic Kwiatkowski, Cordelia Langford, John Sillitoe on behalf of the Wellcome Sanger Institute COVID-19 Surveillance Team |
| EPI_ISL_2664342 | Lighthouse Lab in Milton Keynes | Wellcome Sanger Institute for the COVID-19 Genomics UK (COG-UK) Consortium | The Lighthouse Lab in Milton Keynes and Alex Alderton, Roberto Amato, Jeffrey Barrett, Sonia Goncalves, Ewan Harrison, David K. Jackson, Ian Johnston, Dominic Kwiatkowski, Cordelia Langford, John Sillitoe on behalf of the Wellcome Sanger Institute COVID-19 Surveillance Team |
| EPI_ISL_2664343 | Lighthouse Lab in Glasgow | Wellcome Sanger Institute for the COVID-19 Genomics UK (COG-UK) Consortium | Harper VanSteenhouse, Yumi Kasai, David Gray, Carol Clugston, Anna Dominiczak and Alex Alderton, Roberto Amato, Jeffrey Barrett, Sonia Goncalves, Ewan Harrison, David K. Jackson, Ian Johnston, Dominic Kwiatkowski, Cordelia Langford, John Sillitoe on behalf of the Wellcome Sanger Institute COVID-19 Surveillance Team |
| EPI_ISL_2664344, EPI_ISL_2664345 | Lighthouse Lab in Milton Keynes | Wellcome Sanger Institute for the COVID-19 Genomics UK (COG-UK) Consortium | The Lighthouse Lab in Milton Keynes and Alex Alderton, Roberto Amato, Jeffrey Barrett, Sonia Goncalves, Ewan Harrison, David K. Jackson, Ian Johnston, Dominic Kwiatkowski, Cordelia Langford, John Sillitoe on behalf of the Wellcome Sanger Institute COVID-19 Surveillance Team |
| EPI_ISL_2664346 | Lighthouse Lab in Alderley Park | Wellcome Sanger Institute for the COVID-19 Genomics UK (COG-UK) Consortium | Jacquelyn Wynn, Mairead Hyland, The Lighthouse Lab in Alderley Park and Alex Alderton, Roberto Amato, Jeffrey Barrett, Sonia Goncalves, Ewan Harrison, David K. Jackson, Ian Johnston, Dominic Kwiatkowski, Cordelia Langford, John Sillitoe on behalf of the Wellcome Sanger Institute COVID-19 Surveillance Team |
| EPI_ISL_2664347, EPI_ISL_2664348 | Lighthouse Lab in Milton Keynes | Wellcome Sanger Institute for the COVID-19 Genomics UK (COG-UK) Consortium | The Lighthouse Lab in Milton Keynes and Alex Alderton, Roberto Amato, Jeffrey Barrett, Sonia Goncalves, Ewan Harrison, David K. Jackson, Ian Johnston, Dominic Kwiatkowski, Cordelia Langford, John Sillitoe on behalf of the Wellcome Sanger Institute COVID-19 Surveillance Team |
| EPI_ISL_2664349, EPI_ISL_2664350 | Lighthouse Lab in Alderley Park | Wellcome Sanger Institute for the COVID-19 Genomics UK (COG-UK) Consortium | Jacquelyn Wynn, Mairead Hyland, The Lighthouse Lab in Alderley Park and Alex Alderton, Roberto Amato, Jeffrey Barrett, Sonia Goncalves, Ewan Harrison, David K. Jackson, Ian Johnston, Dominic Kwiatkowski, Cordelia Langford, John Sillitoe on behalf of the Wellcome Sanger Institute COVID-19 Surveillance Team |
| EPI_ISL_2664351, EPI_ISL_2664352 | Lighthouse Lab in Milton Keynes | Wellcome Sanger Institute for the COVID-19 Genomics UK (COG-UK) Consortium | The Lighthouse Lab in Milton Keynes and Alex Alderton, Roberto Amato, Jeffrey Barrett, Sonia Goncalves, Ewan Harrison, David K. Jackson, Ian Johnston, Dominic Kwiatkowski, Cordelia Langford, John Sillitoe on behalf of the Wellcome Sanger Institute COVID-19 Surveillance Team |
| EPI_ISL_2664353 | Lighthouse Lab in Glasgow | Wellcome Sanger Institute for the COVID-19 Genomics UK (COG-UK) Consortium | Harper VanSteenhouse, Yumi Kasai, David Gray, Carol Clugston, Anna Dominiczak and Alex Alderton, Roberto Amato, Jeffrey Barrett, Sonia Goncalves, Ewan Harrison, David K. Jackson, Ian Johnston, Dominic Kwiatkowski, Cordelia Langford, John Sillitoe on behalf of the Wellcome Sanger Institute COVID-19 Surveillance Team |
| EPI_ISL_2664354 | Lighthouse Lab in Milton Keynes | Wellcome Sanger Institute for the COVID-19 Genomics UK (COG-UK) Consortium | The Lighthouse Lab in Milton Keynes and Alex Alderton, Roberto Amato, Jeffrey Barrett, Sonia Goncalves, Ewan Harrison, David K. Jackson, Ian Johnston, Dominic Kwiatkowski, Cordelia Langford, John Sillitoe on behalf of the Wellcome Sanger Institute COVID-19 Surveillance Team |
| EPI_ISL_2664355 | Lighthouse Lab in Glasgow | Wellcome Sanger Institute for the COVID-19 Genomics UK (COG-UK) Consortium | Harper VanSteenhouse, Yumi Kasai, David Gray, Carol Clugston, Anna Dominiczak and Alex Alderton, Roberto Amato, Jeffrey Barrett, Sonia Goncalves, Ewan Harrison, David K. Jackson, Ian Johnston, Dominic Kwiatkowski, Cordelia Langford, John Sillitoe on behalf of the Wellcome Sanger Institute COVID-19 Surveillance Team |
| EPI_ISL_2664356 | Lighthouse Lab in Alderley Park | Wellcome Sanger Institute for the COVID-19 Genomics UK (COG-UK) Consortium | Jacquelyn Wynn, Mairead Hyland, The Lighthouse Lab in Alderley Park and Alex Alderton, Roberto Amato, Jeffrey Barrett, Sonia Goncalves, Ewan Harrison, David K. Jackson, Ian Johnston, Dominic Kwiatkowski, Cordelia Langford, John Sillitoe on behalf of the Wellcome Sanger Institute COVID-19 Surveillance Team |
| EPI_ISL_2664357 | Lighthouse Lab in Milton Keynes | Wellcome Sanger Institute for the COVID-19 Genomics UK (COG-UK) Consortium | The Lighthouse Lab in Milton Keynes and Alex Alderton, Roberto Amato, Jeffrey Barrett, Sonia Goncalves, Ewan Harrison, David K. Jackson, Ian Johnston, Dominic Kwiatkowski, Cordelia Langford, John Sillitoe on behalf of the Wellcome Sanger Institute COVID-19 Surveillance Team |
| EPI_ISL_2664358 | Lighthouse Lab in Alderley Park | Wellcome Sanger Institute for the COVID-19 Genomics UK (COG-UK) Consortium | Jacquelyn Wynn, Mairead Hyland, The Lighthouse Lab in Alderley Park and Alex Alderton, Roberto Amato, Jeffrey Barrett, Sonia Goncalves, Ewan Harrison, David K. Jackson, Ian Johnston, Dominic Kwiatkowski, Cordelia Langford, John Sillitoe on behalf of the Wellcome Sanger Institute COVID-19 Surveillance Team |
| EPI_ISL_2664359 | Lighthouse Lab in Milton Keynes | Wellcome Sanger Institute for the COVID-19 Genomics UK (COG-UK) Consortium | The Lighthouse Lab in Milton Keynes and Alex Alderton, Roberto Amato, Jeffrey Barrett, Sonia Goncalves, Ewan Harrison, David K. Jackson, Ian Johnston, Dominic Kwiatkowski, Cordelia Langford, John Sillitoe on behalf of the Wellcome Sanger Institute COVID-19 Surveillance Team |
| EPI_ISL_2664360 | Lighthouse Lab in Alderley Park | Wellcome Sanger Institute for the COVID-19 Genomics UK (COG-UK) Consortium | Jacquelyn Wynn, Mairead Hyland, The Lighthouse Lab in Alderley Park and Alex Alderton, Roberto Amato, Jeffrey Barrett, Sonia Goncalves, Ewan Harrison, David K. Jackson, Ian Johnston, Dominic Kwiatkowski, Cordelia Langford, John Sillitoe on behalf of the Wellcome Sanger Institute COVID-19 Surveillance Team |
| EPI_ISL_2664361 | Lighthouse Lab in Glasgow | Wellcome Sanger Institute for the COVID-19 Genomics UK (COG-UK) Consortium | Harper VanSteenhouse, Yumi Kasai, David Gray, Carol Clugston, Anna Dominiczak and Alex Alderton, Roberto Amato, Jeffrey Barrett, Sonia Goncalves, Ewan Harrison, David K. Jackson, Ian Johnston, Dominic Kwiatkowski, Cordelia Langford, John Sillitoe on behalf of the Wellcome Sanger Institute COVID-19 Surveillance Team |
| EPI_ISL_2664362 | Lighthouse Lab in Alderley Park | Wellcome Sanger Institute for the COVID-19 Genomics UK (COG-UK) Consortium | Jacquelyn Wynn, Mairead Hyland, The Lighthouse Lab in Alderley Park and Alex Alderton, Roberto Amato, Jeffrey Barrett, Sonia Goncalves, Ewan Harrison, David K. Jackson, Ian Johnston, Dominic Kwiatkowski, Cordelia Langford, John Sillitoe on behalf of the Wellcome Sanger Institute COVID-19 Surveillance Team |
| EPI_ISL_2664363, EPI_ISL_2664364 | Lighthouse Lab in Glasgow | Wellcome Sanger Institute for the COVID-19 Genomics UK (COG-UK) Consortium | Harper VanSteenhouse, Yumi Kasai, David Gray, Carol Clugston, Anna Dominiczak and Alex Alderton, Roberto Amato, Jeffrey Barrett, Sonia Goncalves, Ewan Harrison, David K. Jackson, Ian Johnston, Dominic Kwiatkowski, Cordelia Langford, John Sillitoe on behalf of the Wellcome Sanger Institute COVID-19 Surveillance Team |
| EPI_ISL_2664365, EPI_ISL_2664366 | Lighthouse Lab in Alderley Park | Wellcome Sanger Institute for the COVID-19 Genomics UK (COG-UK) Consortium | Jacquelyn Wynn, Mairead Hyland, The Lighthouse Lab in Alderley Park and Alex Alderton, Roberto Amato, Jeffrey Barrett, Sonia Goncalves, Ewan Harrison, David K. Jackson, Ian Johnston, Dominic Kwiatkowski, Cordelia Langford, John Sillitoe on behalf of the Wellcome Sanger Institute COVID-19 Surveillance Team |
| EPI_ISL_2664367 | Lighthouse Lab in Milton Keynes | Wellcome Sanger Institute for the COVID-19 Genomics UK (COG-UK) Consortium | The Lighthouse Lab in Milton Keynes and Alex Alderton, Roberto Amato, Jeffrey Barrett, Sonia Goncalves, Ewan Harrison, David K. Jackson, Ian Johnston, Dominic Kwiatkowski, Cordelia Langford, John Sillitoe on behalf of the Wellcome Sanger Institute COVID-19 Surveillance Team |
| EPI_ISL_2664368, EPI_ISL_2664369, EPI_ISL_2664370 | Lighthouse Lab in Alderley Park | Wellcome Sanger Institute for the COVID-19 Genomics UK (COG-UK) Consortium | Jacquelyn Wynn, Mairead Hyland, The Lighthouse Lab in Alderley Park and Alex Alderton, Roberto Amato, Jeffrey Barrett, Sonia Goncalves, Ewan Harrison, David K. Jackson, Ian Johnston, Dominic Kwiatkowski, Cordelia Langford, John Sillitoe on behalf of the Wellcome Sanger Institute COVID-19 Surveillance Team |
| EPI_ISL_2664371 | Lighthouse Lab in Milton Keynes | Wellcome Sanger Institute for the COVID-19 Genomics UK (COG-UK) Consortium | The Lighthouse Lab in Milton Keynes and Alex Alderton, Roberto Amato, Jeffrey Barrett, Sonia Goncalves, Ewan Harrison, David K. Jackson, Ian Johnston, Dominic Kwiatkowski, Cordelia Langford, John Sillitoe on behalf of the Wellcome Sanger Institute COVID-19 Surveillance Team |
| EPI_ISL_2664372, EPI_ISL_2664373, EPI_ISL_2664374, EPI_ISL_2664375 | Lighthouse Lab in Alderley Park | Wellcome Sanger Institute for the COVID-19 Genomics UK (COG-UK) Consortium | Jacquelyn Wynn, Mairead Hyland, The Lighthouse Lab in Alderley Park and Alex Alderton, Roberto Amato, Jeffrey Barrett, Sonia Goncalves, Ewan Harrison, David K. Jackson, Ian Johnston, Dominic Kwiatkowski, Cordelia Langford, John Sillitoe on behalf of the Wellcome Sanger Institute COVID-19 Surveillance Team |
| EPI_ISL_2664376 | Lighthouse Lab in Glasgow | Wellcome Sanger Institute for the COVID-19 Genomics UK (COG-UK) Consortium | Harper VanSteenhouse, Yumi Kasai, David Gray, Carol Clugston, Anna Dominiczak and Alex Alderton, Roberto Amato, Jeffrey Barrett, Sonia Goncalves, Ewan Harrison, David K. Jackson, Ian Johnston, Dominic Kwiatkowski, Cordelia Langford, John Sillitoe on behalf of the Wellcome Sanger Institute COVID-19 Surveillance Team |
| EPI_ISL_2664377 | Lighthouse Lab in Alderley Park | Wellcome Sanger Institute for the COVID-19 Genomics UK (COG-UK) Consortium | Jacquelyn Wynn, Mairead Hyland, The Lighthouse Lab in Alderley Park and Alex Alderton, Roberto Amato, Jeffrey Barrett, Sonia Goncalves, Ewan Harrison, David K. Jackson, Ian Johnston, Dominic Kwiatkowski, Cordelia Langford, John Sillitoe on behalf of the Wellcome Sanger Institute COVID-19 Surveillance Team |
| EPI_ISL_2664378, EPI_ISL_2664379 | Lighthouse Lab in Milton Keynes | Wellcome Sanger Institute for the COVID-19 Genomics UK (COG-UK) Consortium | The Lighthouse Lab in Milton Keynes and Alex Alderton, Roberto Amato, Jeffrey Barrett, Sonia Goncalves, Ewan Harrison, David K. Jackson, Ian Johnston, Dominic Kwiatkowski, Cordelia Langford, John Sillitoe on behalf of the Wellcome Sanger Institute COVID-19 Surveillance Team |
| EPI_ISL_2664380 | Lighthouse Lab in Alderley Park | Wellcome Sanger Institute for the COVID-19 Genomics UK (COG-UK) Consortium | Jacquelyn Wynn, Mairead Hyland, The Lighthouse Lab in Alderley Park and Alex Alderton, Roberto Amato, Jeffrey Barrett, Sonia Goncalves, Ewan Harrison, David K. Jackson, Ian Johnston, Dominic Kwiatkowski, Cordelia Langford, John Sillitoe on behalf of the Wellcome Sanger Institute COVID-19 Surveillance Team |
| EPI_ISL_2664381, EPI_ISL_2664382, EPI_ISL_2664383 | Lighthouse Lab in Glasgow | Wellcome Sanger Institute for the COVID-19 Genomics UK (COG-UK) Consortium | Harper VanSteenhouse, Yumi Kasai, David Gray, Carol Clugston, Anna Dominiczak and Alex Alderton, Roberto Amato, Jeffrey Barrett, Sonia Goncalves, Ewan Harrison, David K. Jackson, Ian Johnston, Dominic Kwiatkowski, Cordelia Langford, John Sillitoe on behalf of the Wellcome Sanger Institute COVID-19 Surveillance Team |
| EPI_ISL_2664384 | Lighthouse Lab in Alderley Park | Wellcome Sanger Institute for the COVID-19 Genomics UK (COG-UK) Consortium | Jacquelyn Wynn, Mairead Hyland, The Lighthouse Lab in Alderley Park and Alex Alderton, Roberto Amato, Jeffrey Barrett, Sonia Goncalves, Ewan Harrison, David K. Jackson, Ian Johnston, Dominic Kwiatkowski, Cordelia Langford, John Sillitoe on behalf of the Wellcome Sanger Institute COVID-19 Surveillance Team |
| EPI_ISL_2664385 | Lighthouse Lab in Milton Keynes | Wellcome Sanger Institute for the COVID-19 Genomics UK (COG-UK) Consortium | The Lighthouse Lab in Milton Keynes and Alex Alderton, Roberto Amato, Jeffrey Barrett, Sonia Goncalves, Ewan Harrison, David K. Jackson, Ian Johnston, Dominic Kwiatkowski, Cordelia Langford, John Sillitoe on behalf of the Wellcome Sanger Institute COVID-19 Surveillance Team |
| EPI_ISL_2664386, EPI_ISL_2664387 | Lighthouse Lab in Glasgow | Wellcome Sanger Institute for the COVID-19 Genomics UK (COG-UK) Consortium | Harper VanSteenhouse, Yumi Kasai, David Gray, Carol Clugston, Anna Dominiczak and Alex Alderton, Roberto Amato, Jeffrey Barrett, Sonia Goncalves, Ewan Harrison, David K. Jackson, Ian Johnston, Dominic Kwiatkowski, Cordelia Langford, John Sillitoe on behalf of the Wellcome Sanger Institute COVID-19 Surveillance Team |
| EPI_ISL_2664388 | Lighthouse Lab in Milton Keynes | Wellcome Sanger Institute for the COVID-19 Genomics UK (COG-UK) Consortium | The Lighthouse Lab in Milton Keynes and Alex Alderton, Roberto Amato, Jeffrey Barrett, Sonia Goncalves, Ewan Harrison, David K. Jackson, Ian Johnston, Dominic Kwiatkowski, Cordelia Langford, John Sillitoe on behalf of the Wellcome Sanger Institute COVID-19 Surveillance Team |
| EPI_ISL_2664389, EPI_ISL_2664390 | Lighthouse Lab in Alderley Park | Wellcome Sanger Institute for the COVID-19 Genomics UK (COG-UK) Consortium | Jacquelyn Wynn, Mairead Hyland, The Lighthouse Lab in Alderley Park and Alex Alderton, Roberto Amato, Jeffrey Barrett, Sonia Goncalves, Ewan Harrison, David K. Jackson, Ian Johnston, Dominic Kwiatkowski, Cordelia Langford, John Sillitoe on behalf of the Wellcome Sanger Institute COVID-19 Surveillance Team |
| EPI_ISL_2664391 | Lighthouse Lab in Milton Keynes | Wellcome Sanger Institute for the COVID-19 Genomics UK (COG-UK) Consortium | The Lighthouse Lab in Milton Keynes and Alex Alderton, Roberto Amato, Jeffrey Barrett, Sonia Goncalves, Ewan Harrison, David K. Jackson, Ian Johnston, Dominic Kwiatkowski, Cordelia Langford, John Sillitoe on behalf of the Wellcome Sanger Institute COVID-19 Surveillance Team |
| EPI_ISL_2664392 | Lighthouse Lab in Alderley Park | Wellcome Sanger Institute for the COVID-19 Genomics UK (COG-UK) Consortium | Jacquelyn Wynn, Mairead Hyland, The Lighthouse Lab in Alderley Park and Alex Alderton, Roberto Amato, Jeffrey Barrett, Sonia Goncalves, Ewan Harrison, David K. Jackson, Ian Johnston, Dominic Kwiatkowski, Cordelia Langford, John Sillitoe on behalf of the Wellcome Sanger Institute COVID-19 Surveillance Team |
| EPI_ISL_2664393, EPI_ISL_2664394, EPI_ISL_2664395 | Lighthouse Lab in Milton Keynes | Wellcome Sanger Institute for the COVID-19 Genomics UK (COG-UK) Consortium | The Lighthouse Lab in Milton Keynes and Alex Alderton, Roberto Amato, Jeffrey Barrett, Sonia Goncalves, Ewan Harrison, David K. Jackson, Ian Johnston, Dominic Kwiatkowski, Cordelia Langford, John Sillitoe on behalf of the Wellcome Sanger Institute COVID-19 Surveillance Team |
| EPI_ISL_2664396, EPI_ISL_2664397 | Lighthouse Lab in Alderley Park | Wellcome Sanger Institute for the COVID-19 Genomics UK (COG-UK) Consortium | Jacquelyn Wynn, Mairead Hyland, The Lighthouse Lab in Alderley Park and Alex Alderton, Roberto Amato, Jeffrey Barrett, Sonia Goncalves, Ewan Harrison, David K. Jackson, Ian Johnston, Dominic Kwiatkowski, Cordelia Langford, John Sillitoe on behalf of the Wellcome Sanger Institute COVID-19 Surveillance Team |
| EPI_ISL_2664398 | Lighthouse Lab in Milton Keynes | Wellcome Sanger Institute for the COVID-19 Genomics UK (COG-UK) Consortium | The Lighthouse Lab in Milton Keynes and Alex Alderton, Roberto Amato, Jeffrey Barrett, Sonia Goncalves, Ewan Harrison, David K. Jackson, Ian Johnston, Dominic Kwiatkowski, Cordelia Langford, John Sillitoe on behalf of the Wellcome Sanger Institute COVID-19 Surveillance Team |
| EPI_ISL_2664399, EPI_ISL_2664400, EPI_ISL_2664401, EPI_ISL_2664402, EPI_ISL_2664403 | Lighthouse Lab in Alderley Park | Wellcome Sanger Institute for the COVID-19 Genomics UK (COG-UK) Consortium | Jacquelyn Wynn, Mairead Hyland, The Lighthouse Lab in Alderley Park and Alex Alderton, Roberto Amato, Jeffrey Barrett, Sonia Goncalves, Ewan Harrison, David K. Jackson, Ian Johnston, Dominic Kwiatkowski, Cordelia Langford, John Sillitoe on behalf of the Wellcome Sanger Institute COVID-19 Surveillance Team |

| | | | |
|---|---|---|---|
| EPI_ISL_2664404 | Lighthouse Lab in Glasgow | Wellcome Sanger Institute for the COVID-19 Genomics UK (COG-UK) Consortium | Harper VanSteenhouse, Yumi Kasai, David Gray, Carol Clugston, Anna Dominiczak and Alex Alderton, Roberto Amato, Jeffrey Barrett, Sonia Goncalves, Ewan Harrison, David K. Jackson, Ian Johnston, Dominic Kwiatkowski, Cordelia Langford, John Sillitoe on behalf of the Wellcome Sanger Institute COVID-19 Surveillance Team |
| EPI_ISL_2664405 | Lighthouse Lab in Milton Keynes | Wellcome Sanger Institute for the COVID-19 Genomics UK (COG-UK) Consortium | The Lighthouse Lab in Milton Keynes and Alex Alderton, Roberto Amato, Jeffrey Barrett, Sonia Goncalves, Ewan Harrison, David K. Jackson, Ian Johnston, Dominic Kwiatkowski, Cordelia Langford, John Sillitoe on behalf of the Wellcome Sanger Institute COVID-19 Surveillance Team |
| EPI_ISL_2664406 | Lighthouse Lab in Alderley Park | Wellcome Sanger Institute for the COVID-19 Genomics UK (COG-UK) Consortium | Jacquelyn Wynn, Mairead Hyland, The Lighthouse Lab in Alderley Park and Alex Alderton, Roberto Amato, Jeffrey Barrett, Sonia Goncalves, Ewan Harrison, David K. Jackson, Ian Johnston, Dominic Kwiatkowski, Cordelia Langford, John Sillitoe on behalf of the Wellcome Sanger Institute COVID-19 Surveillance Team |
| EPI_ISL_2664407 | Lighthouse Lab in Glasgow | Wellcome Sanger Institute for the COVID-19 Genomics UK (COG-UK) Consortium | Harper VanSteenhouse, Yumi Kasai, David Gray, Carol Clugston, Anna Dominiczak and Alex Alderton, Roberto Amato, Jeffrey Barrett, Sonia Goncalves, Ewan Harrison, David K. Jackson, Ian Johnston, Dominic Kwiatkowski, Cordelia Langford, John Sillitoe on behalf of the Wellcome Sanger Institute COVID-19 Surveillance Team |
| EPI_ISL_2664408 | Lighthouse Lab in Alderley Park | Wellcome Sanger Institute for the COVID-19 Genomics UK (COG-UK) Consortium | Jacquelyn Wynn, Mairead Hyland, The Lighthouse Lab in Alderley Park and Alex Alderton, Roberto Amato, Jeffrey Barrett, Sonia Goncalves, Ewan Harrison, David K. Jackson, Ian Johnston, Dominic Kwiatkowski, Cordelia Langford, John Sillitoe on behalf of the Wellcome Sanger Institute COVID-19 Surveillance Team |
| EPI_ISL_2664409 | Lighthouse Lab in Milton Keynes | Wellcome Sanger Institute for the COVID-19 Genomics UK (COG-UK) Consortium | The Lighthouse Lab in Milton Keynes and Alex Alderton, Roberto Amato, Jeffrey Barrett, Sonia Goncalves, Ewan Harrison, David K. Jackson, Ian Johnston, Dominic Kwiatkowski, Cordelia Langford, John Sillitoe on behalf of the Wellcome Sanger Institute COVID-19 Surveillance Team |
| EPI_ISL_2664410 | Lighthouse Lab in Glasgow | Wellcome Sanger Institute for the COVID-19 Genomics UK (COG-UK) Consortium | Harper VanSteenhouse, Yumi Kasai, David Gray, Carol Clugston, Anna Dominiczak and Alex Alderton, Roberto Amato, Jeffrey Barrett, Sonia Goncalves, Ewan Harrison, David K. Jackson, Ian Johnston, Dominic Kwiatkowski, Cordelia Langford, John Sillitoe on behalf of the Wellcome Sanger Institute COVID-19 Surveillance Team |
| EPI_ISL_2664411 | Lighthouse Lab in Alderley Park | Wellcome Sanger Institute for the COVID-19 Genomics UK (COG-UK) Consortium | Jacquelyn Wynn, Mairead Hyland, The Lighthouse Lab in Alderley Park and Alex Alderton, Roberto Amato, Jeffrey Barrett, Sonia Goncalves, Ewan Harrison, David K. Jackson, Ian Johnston, Dominic Kwiatkowski, Cordelia Langford, John Sillitoe on behalf of the Wellcome Sanger Institute COVID-19 Surveillance Team |
| EPI_ISL_2664412 | Lighthouse Lab in Milton Keynes | Wellcome Sanger Institute for the COVID-19 Genomics UK (COG-UK) Consortium | The Lighthouse Lab in Milton Keynes and Alex Alderton, Roberto Amato, Jeffrey Barrett, Sonia Goncalves, Ewan Harrison, David K. Jackson, Ian Johnston, Dominic Kwiatkowski, Cordelia Langford, John Sillitoe on behalf of the Wellcome Sanger Institute COVID-19 Surveillance Team |
| EPI_ISL_2664413 | Lighthouse Lab in Alderley Park | Wellcome Sanger Institute for the COVID-19 Genomics UK (COG-UK) Consortium | Jacquelyn Wynn, Mairead Hyland, The Lighthouse Lab in Alderley Park and Alex Alderton, Roberto Amato, Jeffrey Barrett, Sonia Goncalves, Ewan Harrison, David K. Jackson, Ian Johnston, Dominic Kwiatkowski, Cordelia Langford, John Sillitoe on behalf of the Wellcome Sanger Institute COVID-19 Surveillance Team |
| EPI_ISL_2664414, EPI_ISL_2664415 | Lighthouse Lab in Milton Keynes | Wellcome Sanger Institute for the COVID-19 Genomics UK (COG-UK) Consortium | The Lighthouse Lab in Milton Keynes and Alex Alderton, Roberto Amato, Jeffrey Barrett, Sonia Goncalves, Ewan Harrison, David K. Jackson, Ian Johnston, Dominic Kwiatkowski, Cordelia Langford, John Sillitoe on behalf of the Wellcome Sanger Institute COVID-19 Surveillance Team |
| EPI_ISL_2664416 | Lighthouse Lab in Alderley Park | Wellcome Sanger Institute for the COVID-19 Genomics UK (COG-UK) Consortium | Jacquelyn Wynn, Mairead Hyland, The Lighthouse Lab in Alderley Park and Alex Alderton, Roberto Amato, Jeffrey Barrett, Sonia Goncalves, Ewan Harrison, David K. Jackson, Ian Johnston, Dominic Kwiatkowski, Cordelia Langford, John Sillitoe on behalf of the Wellcome Sanger Institute COVID-19 Surveillance Team |
| EPI_ISL_2664417 | Lighthouse Lab in Milton Keynes | Wellcome Sanger Institute for the COVID-19 Genomics UK (COG-UK) Consortium | The Lighthouse Lab in Milton Keynes and Alex Alderton, Roberto Amato, Jeffrey Barrett, Sonia Goncalves, Ewan Harrison, David K. Jackson, Ian Johnston, Dominic Kwiatkowski, Cordelia Langford, John Sillitoe on behalf of the Wellcome Sanger Institute COVID-19 Surveillance Team |
| EPI_ISL_2664418 | Lighthouse Lab in Alderley Park | Wellcome Sanger Institute for the COVID-19 Genomics UK (COG-UK) Consortium | Jacquelyn Wynn, Mairead Hyland, The Lighthouse Lab in Alderley Park and Alex Alderton, Roberto Amato, Jeffrey Barrett, Sonia Goncalves, Ewan Harrison, David K. Jackson, Ian Johnston, Dominic Kwiatkowski, Cordelia Langford, John Sillitoe on behalf of the Wellcome Sanger Institute COVID-19 Surveillance Team |
| EPI_ISL_2664419 | Lighthouse Lab in Glasgow | Wellcome Sanger Institute for the COVID-19 Genomics UK (COG-UK) Consortium | Harper VanSteenhouse, Yumi Kasai, David Gray, Carol Clugston, Anna Dominiczak and Alex Alderton, Roberto Amato, Jeffrey Barrett, Sonia Goncalves, Ewan Harrison, David K. Jackson, Ian Johnston, Dominic Kwiatkowski, Cordelia Langford, John Sillitoe on behalf of the Wellcome Sanger Institute COVID-19 Surveillance Team |
| EPI_ISL_2664420 | Lighthouse Lab in Alderley Park | Wellcome Sanger Institute for the COVID-19 Genomics UK (COG-UK) Consortium | Jacquelyn Wynn, Mairead Hyland, The Lighthouse Lab in Alderley Park and Alex Alderton, Roberto Amato, Jeffrey Barrett, Sonia Goncalves, Ewan Harrison, David K. Jackson, Ian Johnston, Dominic Kwiatkowski, Cordelia Langford, John Sillitoe on behalf of the Wellcome Sanger Institute COVID-19 Surveillance Team |
| EPI_ISL_2664421 | Lighthouse Lab in Milton Keynes | Wellcome Sanger Institute for the COVID-19 Genomics UK (COG-UK) Consortium | The Lighthouse Lab in Milton Keynes and Alex Alderton, Roberto Amato, Jeffrey Barrett, Sonia Goncalves, Ewan Harrison, David K. Jackson, Ian Johnston, Dominic Kwiatkowski, Cordelia Langford, John Sillitoe on behalf of the Wellcome Sanger Institute COVID-19 Surveillance Team |
| EPI_ISL_2664422 | Lighthouse Lab in Alderley Park | Wellcome Sanger Institute for the COVID-19 Genomics UK (COG-UK) Consortium | Jacquelyn Wynn, Mairead Hyland, The Lighthouse Lab in Alderley Park and Alex Alderton, Roberto Amato, Jeffrey Barrett, Sonia Goncalves, Ewan Harrison, David K. Jackson, Ian Johnston, Dominic Kwiatkowski, Cordelia Langford, John Sillitoe on behalf of the Wellcome Sanger Institute COVID-19 Surveillance Team |
| EPI_ISL_2664423 | Lighthouse Lab in Milton Keynes | Wellcome Sanger Institute for the COVID-19 Genomics UK (COG-UK) Consortium | The Lighthouse Lab in Milton Keynes and Alex Alderton, Roberto Amato, Jeffrey Barrett, Sonia Goncalves, Ewan Harrison, David K. Jackson, Ian Johnston, Dominic Kwiatkowski, Cordelia Langford, John Sillitoe on behalf of the Wellcome Sanger Institute COVID-19 Surveillance Team |
| EPI_ISL_2664424 | Lighthouse Lab in Alderley Park | Wellcome Sanger Institute for the COVID-19 Genomics UK (COG-UK) Consortium | Jacquelyn Wynn, Mairead Hyland, The Lighthouse Lab in Alderley Park and Alex Alderton, Roberto Amato, Jeffrey Barrett, Sonia Goncalves, Ewan Harrison, David K. Jackson, Ian Johnston, Dominic Kwiatkowski, Cordelia Langford, John Sillitoe on behalf of the Wellcome Sanger Institute COVID-19 Surveillance Team |
| EPI_ISL_2664425, EPI_ISL_2664426 | Lighthouse Lab in Milton Keynes | Wellcome Sanger Institute for the COVID-19 Genomics UK (COG-UK) Consortium | The Lighthouse Lab in Milton Keynes and Alex Alderton, Roberto Amato, Jeffrey Barrett, Sonia Goncalves, Ewan Harrison, David K. Jackson, Ian Johnston, Dominic Kwiatkowski, Cordelia Langford, John Sillitoe on behalf of the Wellcome Sanger Institute COVID-19 Surveillance Team |
| EPI_ISL_2664427 | Lighthouse Lab in Glasgow | Wellcome Sanger Institute for the COVID-19 Genomics UK (COG-UK) Consortium | Harper VanSteenhouse, Yumi Kasai, David Gray, Carol Clugston, Anna Dominiczak and Alex Alderton, Roberto Amato, Jeffrey Barrett, Sonia Goncalves, Ewan Harrison, David K. Jackson, Ian Johnston, Dominic Kwiatkowski, Cordelia Langford, John Sillitoe on behalf of the Wellcome Sanger Institute COVID-19 Surveillance Team |
| EPI_ISL_2664428, EPI_ISL_2664429 | Lighthouse Lab in Alderley Park | Wellcome Sanger Institute for the COVID-19 Genomics UK (COG-UK) Consortium | Jacquelyn Wynn, Mairead Hyland, The Lighthouse Lab in Alderley Park and Alex Alderton, Roberto Amato, Jeffrey Barrett, Sonia Goncalves, Ewan Harrison, David K. Jackson, Ian Johnston, Dominic Kwiatkowski, Cordelia Langford, John Sillitoe on behalf of the Wellcome Sanger Institute COVID-19 Surveillance Team |
| EPI_ISL_2664430, EPI_ISL_2664431, EPI_ISL_2664432, EPI_ISL_2664433 | Lighthouse Lab in Milton Keynes | Wellcome Sanger Institute for the COVID-19 Genomics UK (COG-UK) Consortium | The Lighthouse Lab in Milton Keynes and Alex Alderton, Roberto Amato, Jeffrey Barrett, Sonia Goncalves, Ewan Harrison, David K. Jackson, Ian Johnston, Dominic Kwiatkowski, Cordelia Langford, John Sillitoe on behalf of the Wellcome Sanger Institute COVID-19 Surveillance Team |
| EPI_ISL_2664434, EPI_ISL_2664435, EPI_ISL_2664436 | Lighthouse Lab in Glasgow | Wellcome Sanger Institute for the COVID-19 Genomics UK (COG-UK) Consortium | Harper VanSteenhouse, Yumi Kasai, David Gray, Carol Clugston, Anna Dominiczak and Alex Alderton, Roberto Amato, Jeffrey Barrett, Sonia Goncalves, Ewan Harrison, David K. Jackson, Ian Johnston, Dominic Kwiatkowski, Cordelia Langford, John Sillitoe on behalf of the Wellcome Sanger Institute COVID-19 Surveillance Team |
| EPI_ISL_2664437 | Lighthouse Lab in Alderley Park | Wellcome Sanger Institute for the COVID-19 Genomics UK (COG-UK) Consortium | Jacquelyn Wynn, Mairead Hyland, The Lighthouse Lab in Alderley Park and Alex Alderton, Roberto Amato, Jeffrey Barrett, Sonia Goncalves, Ewan Harrison, David K. Jackson, Ian Johnston, Dominic Kwiatkowski, Cordelia Langford, John Sillitoe on behalf of the Wellcome Sanger Institute COVID-19 Surveillance Team |
| EPI_ISL_2664438 | Lighthouse Lab in Milton Keynes | Wellcome Sanger Institute for the COVID-19 Genomics UK (COG-UK) Consortium | The Lighthouse Lab in Milton Keynes and Alex Alderton, Roberto Amato, Jeffrey Barrett, Sonia Goncalves, Ewan Harrison, David K. Jackson, Ian Johnston, Dominic Kwiatkowski, Cordelia Langford, John Sillitoe on behalf of the Wellcome Sanger Institute COVID-19 Surveillance Team |
| EPI_ISL_2664439, EPI_ISL_2664440 | Lighthouse Lab in Alderley Park | Wellcome Sanger Institute for the COVID-19 Genomics UK (COG-UK) Consortium | Jacquelyn Wynn, Mairead Hyland, The Lighthouse Lab in Alderley Park and Alex Alderton, Roberto Amato, Jeffrey Barrett, Sonia Goncalves, Ewan Harrison, David K. Jackson, Ian Johnston, Dominic Kwiatkowski, Cordelia Langford, John Sillitoe on behalf of the Wellcome Sanger Institute COVID-19 Surveillance Team |
| EPI_ISL_2664441 | Lighthouse Lab in Glasgow | Wellcome Sanger Institute for the COVID-19 Genomics UK (COG-UK) Consortium | Harper VanSteenhouse, Yumi Kasai, David Gray, Carol Clugston, Anna Dominiczak and Alex Alderton, Roberto Amato, Jeffrey Barrett, Sonia Goncalves, Ewan Harrison, David K. Jackson, Ian Johnston, Dominic Kwiatkowski, Cordelia Langford, John Sillitoe on behalf of the Wellcome Sanger Institute COVID-19 Surveillance Team |
| EPI_ISL_2664442 | Lighthouse Lab in Alderley Park | Wellcome Sanger Institute for the COVID-19 Genomics UK (COG-UK) Consortium | Jacquelyn Wynn, Mairead Hyland, The Lighthouse Lab in Alderley Park and Alex Alderton, Roberto Amato, Jeffrey Barrett, Sonia Goncalves, Ewan Harrison, David K. Jackson, Ian Johnston, Dominic Kwiatkowski, Cordelia Langford, John Sillitoe on behalf of the Wellcome Sanger Institute COVID-19 Surveillance Team |
| EPI_ISL_2664443, EPI_ISL_2664444 | Lighthouse Lab in Milton Keynes | Wellcome Sanger Institute for the COVID-19 Genomics UK (COG-UK) Consortium | The Lighthouse Lab in Milton Keynes and Alex Alderton, Roberto Amato, Jeffrey Barrett, Sonia Goncalves, Ewan Harrison, David K. Jackson, Ian Johnston, Dominic Kwiatkowski, Cordelia Langford, John Sillitoe on behalf of the Wellcome Sanger Institute COVID-19 Surveillance Team |
| EPI_ISL_2664445, EPI_ISL_2664446 | Lighthouse Lab in Glasgow | Wellcome Sanger Institute for the COVID-19 Genomics UK (COG-UK) Consortium | Harper VanSteenhouse, Yumi Kasai, David Gray, Carol Clugston, Anna Dominiczak and Alex Alderton, Roberto Amato, Jeffrey Barrett, Sonia Goncalves, Ewan Harrison, David K. Jackson, Ian Johnston, Dominic Kwiatkowski, Cordelia Langford, John Sillitoe on behalf of the Wellcome Sanger Institute COVID-19 Surveillance Team |
| EPI_ISL_2664447, EPI_ISL_2664448, EPI_ISL_2664449, EPI_ISL_2664450 | Lighthouse Lab in Alderley Park | Wellcome Sanger Institute for the COVID-19 Genomics UK (COG-UK) Consortium | Jacquelyn Wynn, Mairead Hyland, The Lighthouse Lab in Alderley Park and Alex Alderton, Roberto Amato, Jeffrey Barrett, Sonia Goncalves, Ewan Harrison, David K. Jackson, Ian Johnston, Dominic Kwiatkowski, Cordelia Langford, John Sillitoe on behalf of the Wellcome Sanger Institute COVID-19 Surveillance Team |
| EPI_ISL_2664451 | Lighthouse Lab in Glasgow | Wellcome Sanger Institute for the COVID-19 Genomics UK (COG-UK) Consortium | Harper VanSteenhouse, Yumi Kasai, David Gray, Carol Clugston, Anna Dominiczak and Alex Alderton, Roberto Amato, Jeffrey Barrett, Sonia Goncalves, Ewan Harrison, David K. Jackson, Ian Johnston, Dominic Kwiatkowski, Cordelia Langford, John Sillitoe on behalf of the Wellcome Sanger Institute COVID-19 Surveillance Team |
| EPI_ISL_2664452 | Lighthouse Lab in Milton Keynes | Wellcome Sanger Institute for the COVID-19 Genomics UK (COG-UK) Consortium | The Lighthouse Lab in Milton Keynes and Alex Alderton, Roberto Amato, Jeffrey Barrett, Sonia Goncalves, Ewan Harrison, David K. Jackson, Ian Johnston, Dominic Kwiatkowski, Cordelia Langford, John Sillitoe on behalf of the Wellcome Sanger Institute COVID-19 Surveillance Team |
| EPI_ISL_2664453, EPI_ISL_2664454, EPI_ISL_2664455 | Lighthouse Lab in Glasgow | Wellcome Sanger Institute for the COVID-19 Genomics UK (COG-UK) Consortium | Harper VanSteenhouse, Yumi Kasai, David Gray, Carol Clugston, Anna Dominiczak and Alex Alderton, Roberto Amato, Jeffrey Barrett, Sonia Goncalves, Ewan Harrison, David K. Jackson, Ian Johnston, Dominic Kwiatkowski, Cordelia Langford, John Sillitoe on behalf of the Wellcome Sanger Institute COVID-19 Surveillance Team |
| EPI_ISL_2664456, EPI_ISL_2664457, EPI_ISL_2664458 | Lighthouse Lab in Alderley Park | Wellcome Sanger Institute for the COVID-19 Genomics UK (COG-UK) Consortium | Jacquelyn Wynn, Mairead Hyland, The Lighthouse Lab in Alderley Park and Alex Alderton, Roberto Amato, Jeffrey Barrett, Sonia Goncalves, Ewan Harrison, David K. Jackson, Ian Johnston, Dominic Kwiatkowski, Cordelia Langford, John Sillitoe on behalf of the Wellcome Sanger Institute COVID-19 Surveillance Team |
| EPI_ISL_2664459, EPI_ISL_2664460, EPI_ISL_2664461 | Lighthouse Lab in Milton Keynes | Wellcome Sanger Institute for the COVID-19 Genomics UK (COG-UK) Consortium | The Lighthouse Lab in Milton Keynes and Alex Alderton, Roberto Amato, Jeffrey Barrett, Sonia Goncalves, Ewan Harrison, David K. Jackson, Ian Johnston, Dominic Kwiatkowski, Cordelia Langford, John Sillitoe on behalf of the Wellcome Sanger Institute COVID-19 Surveillance Team |
| EPI_ISL_2664462 | Lighthouse Lab in Alderley Park | Wellcome Sanger Institute for the COVID-19 Genomics UK (COG-UK) Consortium | Jacquelyn Wynn, Mairead Hyland, The Lighthouse Lab in Alderley Park and Alex Alderton, Roberto Amato, Jeffrey Barrett, Sonia Goncalves, Ewan Harrison, David K. Jackson, Ian Johnston, Dominic Kwiatkowski, Cordelia Langford, John Sillitoe on behalf of the Wellcome Sanger Institute COVID-19 Surveillance Team |
| EPI_ISL_2664463 | Lighthouse Lab in Milton Keynes | Wellcome Sanger Institute for the COVID-19 Genomics UK (COG-UK) Consortium | The Lighthouse Lab in Milton Keynes and Alex Alderton, Roberto Amato, Jeffrey Barrett, Sonia Goncalves, Ewan Harrison, David K. Jackson, Ian Johnston, Dominic Kwiatkowski, Cordelia Langford, John Sillitoe on behalf of the Wellcome Sanger Institute COVID-19 Surveillance Team |
| EPI_ISL_2664464, EPI_ISL_2664465 | Lighthouse Lab in Alderley Park | Wellcome Sanger Institute for the COVID-19 Genomics UK (COG-UK) Consortium | Jacquelyn Wynn, Mairead Hyland, The Lighthouse Lab in Alderley Park and Alex Alderton, Roberto Amato, Jeffrey Barrett, Sonia Goncalves, Ewan Harrison, David K. Jackson, Ian Johnston, Dominic Kwiatkowski, Cordelia Langford, John Sillitoe on behalf of the Wellcome Sanger Institute COVID-19 Surveillance Team |
| EPI_ISL_2664466 | Lighthouse Lab in Glasgow | Wellcome Sanger Institute for the COVID-19 Genomics UK (COG-UK) Consortium | Harper VanSteenhouse, Yumi Kasai, David Gray, Carol Clugston, Anna Dominiczak and Alex Alderton, Roberto Amato, Jeffrey Barrett, Sonia Goncalves, Ewan Harrison, David K. Jackson, Ian Johnston, Dominic Kwiatkowski, Cordelia Langford, John Sillitoe on behalf of the Wellcome Sanger Institute COVID-19 Surveillance Team |

| | | | |
|---|---|---|---|
| EPI_ISL_2664467 | Lighthouse Lab in Alderley Park | Wellcome Sanger Institute for the COVID-19 Genomics UK (COG-UK) Consortium | Jacquelyn Wynn, Mairead Hyland, The Lighthouse Lab in Alderley Park and Alex Alderton, Roberto Amato, Jeffrey Barrett, Sonia Goncalves, Ewan Harrison, David K. Jackson, Ian Johnston, Dominic Kwiatkowski, Cordelia Langford, John Sillitoe on behalf of the Wellcome Sanger Institute COVID-19 Surveillance Team |
| EPI_ISL_2664468 | Lighthouse Laboratory Plymouth | Wellcome Sanger Institute for the COVID-19 Genomics UK (COG-UK) Consortium | Lighthouse Laboratory Plymouth and Alex Alderton, Roberto Amato, Jeffrey Barrett, Sonia Goncalves, Ewan Harrison, David K. Jackson, Ian Johnston, Dominic Kwiatkowski, Cordelia Langford, John Sillitoe on behalf of the Wellcome Sanger Institute COVID-19 Surveillance Team |
| EPI_ISL_2664469 | Lighthouse Lab in Milton Keynes | Wellcome Sanger Institute for the COVID-19 Genomics UK (COG-UK) Consortium | The Lighthouse Lab in Milton Keynes and Alex Alderton, Roberto Amato, Jeffrey Barrett, Sonia Goncalves, Ewan Harrison, David K. Jackson, Ian Johnston, Dominic Kwiatkowski, Cordelia Langford, John Sillitoe on behalf of the Wellcome Sanger Institute COVID-19 Surveillance Team |
| EPI_ISL_2664470, EPI_ISL_2664471 | Lighthouse Lab in Alderley Park | Wellcome Sanger Institute for the COVID-19 Genomics UK (COG-UK) Consortium | Jacquelyn Wynn, Mairead Hyland, The Lighthouse Lab in Alderley Park and Alex Alderton, Roberto Amato, Jeffrey Barrett, Sonia Goncalves, Ewan Harrison, David K. Jackson, Ian Johnston, Dominic Kwiatkowski, Cordelia Langford, John Sillitoe on behalf of the Wellcome Sanger Institute COVID-19 Surveillance Team |
| EPI_ISL_2664472, EPI_ISL_2664473 | Lighthouse Lab in Glasgow | Wellcome Sanger Institute for the COVID-19 Genomics UK (COG-UK) Consortium | Harper VanSteenhouse, Yumi Kasai, David Gray, Carol Clugston, Anna Dominiczak and Alex Alderton, Roberto Amato, Jeffrey Barrett, Sonia Goncalves, Ewan Harrison, David K. Jackson, Ian Johnston, Dominic Kwiatkowski, Cordelia Langford, John Sillitoe on behalf of the Wellcome Sanger Institute COVID-19 Surveillance Team |
| EPI_ISL_2664474 | Lighthouse Lab in Alderley Park | Wellcome Sanger Institute for the COVID-19 Genomics UK (COG-UK) Consortium | Jacquelyn Wynn, Mairead Hyland, The Lighthouse Lab in Alderley Park and Alex Alderton, Roberto Amato, Jeffrey Barrett, Sonia Goncalves, Ewan Harrison, David K. Jackson, Ian Johnston, Dominic Kwiatkowski, Cordelia Langford, John Sillitoe on behalf of the Wellcome Sanger Institute COVID-19 Surveillance Team |
| EPI_ISL_2664475, EPI_ISL_2664476 | Lighthouse Lab in Milton Keynes | Wellcome Sanger Institute for the COVID-19 Genomics UK (COG-UK) Consortium | The Lighthouse Lab in Milton Keynes and Alex Alderton, Roberto Amato, Jeffrey Barrett, Sonia Goncalves, Ewan Harrison, David K. Jackson, Ian Johnston, Dominic Kwiatkowski, Cordelia Langford, John Sillitoe on behalf of the Wellcome Sanger Institute COVID-19 Surveillance Team |
| EPI_ISL_2664477 | Lighthouse Lab in Alderley Park | Wellcome Sanger Institute for the COVID-19 Genomics UK (COG-UK) Consortium | Jacquelyn Wynn, Mairead Hyland, The Lighthouse Lab in Alderley Park and Alex Alderton, Roberto Amato, Jeffrey Barrett, Sonia Goncalves, Ewan Harrison, David K. Jackson, Ian Johnston, Dominic Kwiatkowski, Cordelia Langford, John Sillitoe on behalf of the Wellcome Sanger Institute COVID-19 Surveillance Team |
| EPI_ISL_2664478, EPI_ISL_2664479 | Lighthouse Laboratory Plymouth | Wellcome Sanger Institute for the COVID-19 Genomics UK (COG-UK) Consortium | Lighthouse Laboratory Plymouth and Alex Alderton, Roberto Amato, Jeffrey Barrett, Sonia Goncalves, Ewan Harrison, David K. Jackson, Ian Johnston, Dominic Kwiatkowski, Cordelia Langford, John Sillitoe on behalf of the Wellcome Sanger Institute COVID-19 Surveillance Team |
| EPI_ISL_2664480, EPI_ISL_2664481 | Lighthouse Lab in Milton Keynes | Wellcome Sanger Institute for the COVID-19 Genomics UK (COG-UK) Consortium | The Lighthouse Lab in Milton Keynes and Alex Alderton, Roberto Amato, Jeffrey Barrett, Sonia Goncalves, Ewan Harrison, David K. Jackson, Ian Johnston, Dominic Kwiatkowski, Cordelia Langford, John Sillitoe on behalf of the Wellcome Sanger Institute COVID-19 Surveillance Team |
| EPI_ISL_2664482 | Lighthouse Lab in Glasgow | Wellcome Sanger Institute for the COVID-19 Genomics UK (COG-UK) Consortium | Harper VanSteenhouse, Yumi Kasai, David Gray, Carol Clugston, Anna Dominiczak and Alex Alderton, Roberto Amato, Jeffrey Barrett, Sonia Goncalves, Ewan Harrison, David K. Jackson, Ian Johnston, Dominic Kwiatkowski, Cordelia Langford, John Sillitoe on behalf of the Wellcome Sanger Institute COVID-19 Surveillance Team |
| EPI_ISL_2664483 | Health Services Laboratories | Wellcome Sanger Institute for the COVID-19 Genomics UK (COG-UK) Consortium | Health Services Laboratories and Alex Alderton, Roberto Amato, Jeffrey Barrett, Sonia Goncalves, Ewan Harrison, David K. Jackson, Ian Johnston, Dominic Kwiatkowski, Cordelia Langford, John Sillitoe on behalf of the Wellcome Sanger Institute COVID-19 Surveillance Team |
| EPI_ISL_2664484, EPI_ISL_2664485 | Lighthouse Laboratory Plymouth | Wellcome Sanger Institute for the COVID-19 Genomics UK (COG-UK) Consortium | Lighthouse Laboratory Plymouth and Alex Alderton, Roberto Amato, Jeffrey Barrett, Sonia Goncalves, Ewan Harrison, David K. Jackson, Ian Johnston, Dominic Kwiatkowski, Cordelia Langford, John Sillitoe on behalf of the Wellcome Sanger Institute COVID-19 Surveillance Team |
| EPI_ISL_2664486 | Lighthouse Lab in Alderley Park | Wellcome Sanger Institute for the COVID-19 Genomics UK (COG-UK) Consortium | Jacquelyn Wynn, Mairead Hyland, The Lighthouse Lab in Alderley Park and Alex Alderton, Roberto Amato, Jeffrey Barrett, Sonia Goncalves, Ewan Harrison, David K. Jackson, Ian Johnston, Dominic Kwiatkowski, Cordelia Langford, John Sillitoe on behalf of the Wellcome Sanger Institute COVID-19 Surveillance Team |
| EPI_ISL_2664487 | Lighthouse Laboratory Plymouth | Wellcome Sanger Institute for the COVID-19 Genomics UK (COG-UK) Consortium | Lighthouse Laboratory Plymouth and Alex Alderton, Roberto Amato, Jeffrey Barrett, Sonia Goncalves, Ewan Harrison, David K. Jackson, Ian Johnston, Dominic Kwiatkowski, Cordelia Langford, John Sillitoe on behalf of the Wellcome Sanger Institute COVID-19 Surveillance Team |
| EPI_ISL_2664488 | Lighthouse Lab in Alderley Park | Wellcome Sanger Institute for the COVID-19 Genomics UK (COG-UK) Consortium | Jacquelyn Wynn, Mairead Hyland, The Lighthouse Lab in Alderley Park and Alex Alderton, Roberto Amato, Jeffrey Barrett, Sonia Goncalves, Ewan Harrison, David K. Jackson, Ian Johnston, Dominic Kwiatkowski, Cordelia Langford, John Sillitoe on behalf of the Wellcome Sanger Institute COVID-19 Surveillance Team |
| EPI_ISL_2664489 | Lighthouse Laboratory Plymouth | Wellcome Sanger Institute for the COVID-19 Genomics UK (COG-UK) Consortium | Lighthouse Laboratory Plymouth and Alex Alderton, Roberto Amato, Jeffrey Barrett, Sonia Goncalves, Ewan Harrison, David K. Jackson, Ian Johnston, Dominic Kwiatkowski, Cordelia Langford, John Sillitoe on behalf of the Wellcome Sanger Institute COVID-19 Surveillance Team |
| EPI_ISL_2664490 | Lighthouse Lab in Milton Keynes | Wellcome Sanger Institute for the COVID-19 Genomics UK (COG-UK) Consortium | The Lighthouse Lab in Milton Keynes and Alex Alderton, Roberto Amato, Jeffrey Barrett, Sonia Goncalves, Ewan Harrison, David K. Jackson, Ian Johnston, Dominic Kwiatkowski, Cordelia Langford, John Sillitoe on behalf of the Wellcome Sanger Institute COVID-19 Surveillance Team |
| EPI_ISL_2664491 | Lighthouse Lab in Alderley Park | Wellcome Sanger Institute for the COVID-19 Genomics UK (COG-UK) Consortium | Jacquelyn Wynn, Mairead Hyland, The Lighthouse Lab in Alderley Park and Alex Alderton, Roberto Amato, Jeffrey Barrett, Sonia Goncalves, Ewan Harrison, David K. Jackson, Ian Johnston, Dominic Kwiatkowski, Cordelia Langford, John Sillitoe on behalf of the Wellcome Sanger Institute COVID-19 Surveillance Team |
| EPI_ISL_2664492 | Lighthouse Lab in Glasgow | Wellcome Sanger Institute for the COVID-19 Genomics UK (COG-UK) Consortium | Harper VanSteenhouse, Yumi Kasai, David Gray, Carol Clugston, Anna Dominiczak and Alex Alderton, Roberto Amato, Jeffrey Barrett, Sonia Goncalves, Ewan Harrison, David K. Jackson, Ian Johnston, Dominic Kwiatkowski, Cordelia Langford, John Sillitoe on behalf of the Wellcome Sanger Institute COVID-19 Surveillance Team |
| EPI_ISL_2664493 | Lighthouse Laboratory Plymouth | Wellcome Sanger Institute for the COVID-19 Genomics UK (COG-UK) Consortium | Lighthouse Laboratory Plymouth and Alex Alderton, Roberto Amato, Jeffrey Barrett, Sonia Goncalves, Ewan Harrison, David K. Jackson, Ian Johnston, Dominic Kwiatkowski, Cordelia Langford, John Sillitoe on behalf of the Wellcome Sanger Institute COVID-19 Surveillance Team |
| EPI_ISL_2664494 | Lighthouse Lab in Milton Keynes | Wellcome Sanger Institute for the COVID-19 Genomics UK (COG-UK) Consortium | The Lighthouse Lab in Milton Keynes and Alex Alderton, Roberto Amato, Jeffrey Barrett, Sonia Goncalves, Ewan Harrison, David K. Jackson, Ian Johnston, Dominic Kwiatkowski, Cordelia Langford, John Sillitoe on behalf of the Wellcome Sanger Institute COVID-19 Surveillance Team |
| EPI_ISL_2664495 | Lighthouse Lab in Alderley Park | Wellcome Sanger Institute for the COVID-19 Genomics UK (COG-UK) Consortium | Jacquelyn Wynn, Mairead Hyland, The Lighthouse Lab in Alderley Park and Alex Alderton, Roberto Amato, Jeffrey Barrett, Sonia Goncalves, Ewan Harrison, David K. Jackson, Ian Johnston, Dominic Kwiatkowski, Cordelia Langford, John Sillitoe on behalf of the Wellcome Sanger Institute COVID-19 Surveillance Team |
| EPI_ISL_2664496 | Lighthouse Laboratory Plymouth | Wellcome Sanger Institute for the COVID-19 Genomics UK (COG-UK) Consortium | Lighthouse Laboratory Plymouth and Alex Alderton, Roberto Amato, Jeffrey Barrett, Sonia Goncalves, Ewan Harrison, David K. Jackson, Ian Johnston, Dominic Kwiatkowski, Cordelia Langford, John Sillitoe on behalf of the Wellcome Sanger Institute COVID-19 Surveillance Team |
| EPI_ISL_2664497, EPI_ISL_2664498 | Lighthouse Lab in Milton Keynes | Wellcome Sanger Institute for the COVID-19 Genomics UK (COG-UK) Consortium | The Lighthouse Lab in Milton Keynes and Alex Alderton, Roberto Amato, Jeffrey Barrett, Sonia Goncalves, Ewan Harrison, David K. Jackson, Ian Johnston, Dominic Kwiatkowski, Cordelia Langford, John Sillitoe on behalf of the Wellcome Sanger Institute COVID-19 Surveillance Team |
| EPI_ISL_2664499 | Lighthouse Laboratory Plymouth | Wellcome Sanger Institute for the COVID-19 Genomics UK (COG-UK) Consortium | Lighthouse Laboratory Plymouth and Alex Alderton, Roberto Amato, Jeffrey Barrett, Sonia Goncalves, Ewan Harrison, David K. Jackson, Ian Johnston, Dominic Kwiatkowski, Cordelia Langford, John Sillitoe on behalf of the Wellcome Sanger Institute COVID-19 Surveillance Team |
| EPI_ISL_2664500 | Lighthouse Lab in Alderley Park | Wellcome Sanger Institute for the COVID-19 Genomics UK (COG-UK) Consortium | Jacquelyn Wynn, Mairead Hyland, The Lighthouse Lab in Alderley Park and Alex Alderton, Roberto Amato, Jeffrey Barrett, Sonia Goncalves, Ewan Harrison, David K. Jackson, Ian Johnston, Dominic Kwiatkowski, Cordelia Langford, John Sillitoe on behalf of the Wellcome Sanger Institute COVID-19 Surveillance Team |
| EPI_ISL_2664501, EPI_ISL_2664502 | Lighthouse Laboratory Plymouth | Wellcome Sanger Institute for the COVID-19 Genomics UK (COG-UK) Consortium | Lighthouse Laboratory Plymouth and Alex Alderton, Roberto Amato, Jeffrey Barrett, Sonia Goncalves, Ewan Harrison, David K. Jackson, Ian Johnston, Dominic Kwiatkowski, Cordelia Langford, John Sillitoe on behalf of the Wellcome Sanger Institute COVID-19 Surveillance Team |
| EPI_ISL_2664503 | Lighthouse Lab in Alderley Park | Wellcome Sanger Institute for the COVID-19 Genomics UK (COG-UK) Consortium | Jacquelyn Wynn, Mairead Hyland, The Lighthouse Lab in Alderley Park and Alex Alderton, Roberto Amato, Jeffrey Barrett, Sonia Goncalves, Ewan Harrison, David K. Jackson, Ian Johnston, Dominic Kwiatkowski, Cordelia Langford, John Sillitoe on behalf of the Wellcome Sanger Institute COVID-19 Surveillance Team |
| EPI_ISL_2664504, EPI_ISL_2664505, EPI_ISL_2664506, EPI_ISL_2664507 | Lighthouse Laboratory Plymouth | Wellcome Sanger Institute for the COVID-19 Genomics UK (COG-UK) Consortium | Lighthouse Laboratory Plymouth and Alex Alderton, Roberto Amato, Jeffrey Barrett, Sonia Goncalves, Ewan Harrison, David K. Jackson, Ian Johnston, Dominic Kwiatkowski, Cordelia Langford, John Sillitoe on behalf of the Wellcome Sanger Institute COVID-19 Surveillance Team |
| EPI_ISL_2664508 | Lighthouse Lab in Alderley Park | Wellcome Sanger Institute for the COVID-19 Genomics UK (COG-UK) Consortium | Jacquelyn Wynn, Mairead Hyland, The Lighthouse Lab in Alderley Park and Alex Alderton, Roberto Amato, Jeffrey Barrett, Sonia Goncalves, Ewan Harrison, David K. Jackson, Ian Johnston, Dominic Kwiatkowski, Cordelia Langford, John Sillitoe on behalf of the Wellcome Sanger Institute COVID-19 Surveillance Team |
| EPI_ISL_2664509, EPI_ISL_2664510 | Lighthouse Lab in Milton Keynes | Wellcome Sanger Institute for the COVID-19 Genomics UK (COG-UK) Consortium | The Lighthouse Lab in Milton Keynes and Alex Alderton, Roberto Amato, Jeffrey Barrett, Sonia Goncalves, Ewan Harrison, David K. Jackson, Ian Johnston, Dominic Kwiatkowski, Cordelia Langford, John Sillitoe on behalf of the Wellcome Sanger Institute COVID-19 Surveillance Team |
| EPI_ISL_2664511 | Lighthouse Laboratory Plymouth | Wellcome Sanger Institute for the COVID-19 Genomics UK (COG-UK) Consortium | Lighthouse Laboratory Plymouth and Alex Alderton, Roberto Amato, Jeffrey Barrett, Sonia Goncalves, Ewan Harrison, David K. Jackson, Ian Johnston, Dominic Kwiatkowski, Cordelia Langford, John Sillitoe on behalf of the Wellcome Sanger Institute COVID-19 Surveillance Team |
| EPI_ISL_2664512, EPI_ISL_2664513 | Lighthouse Lab in Milton Keynes | Wellcome Sanger Institute for the COVID-19 Genomics UK (COG-UK) Consortium | The Lighthouse Lab in Milton Keynes and Alex Alderton, Roberto Amato, Jeffrey Barrett, Sonia Goncalves, Ewan Harrison, David K. Jackson, Ian Johnston, Dominic Kwiatkowski, Cordelia Langford, John Sillitoe on behalf of the Wellcome Sanger Institute COVID-19 Surveillance Team |
| EPI_ISL_2664514 | Lighthouse Lab in Alderley Park | Wellcome Sanger Institute for the COVID-19 Genomics UK (COG-UK) Consortium | Jacquelyn Wynn, Mairead Hyland, The Lighthouse Lab in Alderley Park and Alex Alderton, Roberto Amato, Jeffrey Barrett, Sonia Goncalves, Ewan Harrison, David K. Jackson, Ian Johnston, Dominic Kwiatkowski, Cordelia Langford, John Sillitoe on behalf of the Wellcome Sanger Institute COVID-19 Surveillance Team |
| EPI_ISL_2664515 | Lighthouse Lab in Milton Keynes | Wellcome Sanger Institute for the COVID-19 Genomics UK (COG-UK) Consortium | The Lighthouse Lab in Milton Keynes and Alex Alderton, Roberto Amato, Jeffrey Barrett, Sonia Goncalves, Ewan Harrison, David K. Jackson, Ian Johnston, Dominic Kwiatkowski, Cordelia Langford, John Sillitoe on behalf of the Wellcome Sanger Institute COVID-19 Surveillance Team |
| EPI_ISL_2664516 | Lighthouse Lab in Alderley Park | Wellcome Sanger Institute for the COVID-19 Genomics UK (COG-UK) Consortium | Jacquelyn Wynn, Mairead Hyland, The Lighthouse Lab in Alderley Park and Alex Alderton, Roberto Amato, Jeffrey Barrett, Sonia Goncalves, Ewan Harrison, David K. Jackson, Ian Johnston, Dominic Kwiatkowski, Cordelia Langford, John Sillitoe on behalf of the Wellcome Sanger Institute COVID-19 Surveillance Team |
| EPI_ISL_2664517 | Lighthouse Laboratory Plymouth | Wellcome Sanger Institute for the COVID-19 Genomics UK (COG-UK) Consortium | Lighthouse Laboratory Plymouth and Alex Alderton, Roberto Amato, Jeffrey Barrett, Sonia Goncalves, Ewan Harrison, David K. Jackson, Ian Johnston, Dominic Kwiatkowski, Cordelia Langford, John Sillitoe on behalf of the Wellcome Sanger Institute COVID-19 Surveillance Team |
| EPI_ISL_2664518 | Lighthouse Lab in Milton Keynes | Wellcome Sanger Institute for the COVID-19 Genomics UK (COG-UK) Consortium | The Lighthouse Lab in Milton Keynes and Alex Alderton, Roberto Amato, Jeffrey Barrett, Sonia Goncalves, Ewan Harrison, David K. Jackson, Ian Johnston, Dominic Kwiatkowski, Cordelia Langford, John Sillitoe on behalf of the Wellcome Sanger Institute COVID-19 Surveillance Team |
| EPI_ISL_2664519 | Lighthouse Lab in Alderley Park | Wellcome Sanger Institute for the COVID-19 Genomics UK (COG-UK) Consortium | Jacquelyn Wynn, Mairead Hyland, The Lighthouse Lab in Alderley Park and Alex Alderton, Roberto Amato, Jeffrey Barrett, Sonia Goncalves, Ewan Harrison, David K. Jackson, Ian Johnston, Dominic Kwiatkowski, Cordelia Langford, John Sillitoe on behalf of the Wellcome Sanger Institute COVID-19 Surveillance Team |
| EPI_ISL_2664520 | Lighthouse Laboratory Plymouth | Wellcome Sanger Institute for the COVID-19 Genomics UK (COG-UK) Consortium | Lighthouse Laboratory Plymouth and Alex Alderton, Roberto Amato, Jeffrey Barrett, Sonia Goncalves, Ewan Harrison, David K. Jackson, Ian Johnston, Dominic Kwiatkowski, Cordelia Langford, John Sillitoe on behalf of the Wellcome Sanger Institute COVID-19 Surveillance Team |
| EPI_ISL_2664521, EPI_ISL_2664522 | Lighthouse Lab in Milton Keynes | Wellcome Sanger Institute for the COVID-19 Genomics UK (COG-UK) Consortium | The Lighthouse Lab in Milton Keynes and Alex Alderton, Roberto Amato, Jeffrey Barrett, Sonia Goncalves, Ewan Harrison, David K. Jackson, Ian Johnston, Dominic Kwiatkowski, Cordelia Langford, John Sillitoe on behalf of the Wellcome Sanger Institute COVID-19 Surveillance Team |
| EPI_ISL_2664523 | Lighthouse Laboratory Plymouth | Wellcome Sanger Institute for the COVID-19 Genomics UK (COG-UK) Consortium | Lighthouse Laboratory Plymouth and Alex Alderton, Roberto Amato, Jeffrey Barrett, Sonia Goncalves, Ewan Harrison, David K. Jackson, Ian Johnston, Dominic Kwiatkowski, Cordelia Langford, John Sillitoe on behalf of the Wellcome Sanger Institute COVID-19 Surveillance Team |
| EPI_ISL_2664524 | Lighthouse Lab in Milton Keynes | Wellcome Sanger Institute for the COVID-19 Genomics UK (COG-UK) Consortium | The Lighthouse Lab in Milton Keynes and Alex Alderton, Roberto Amato, Jeffrey Barrett, Sonia Goncalves, Ewan Harrison, David K. Jackson, Ian Johnston, Dominic Kwiatkowski, Cordelia Langford, John Sillitoe on behalf of the Wellcome Sanger Institute COVID-19 Surveillance Team |

| | | | |
|---|---|---|---|
| EPI_ISL_2664525 | Lighthouse Laboratory Plymouth | Wellcome Sanger Institute for the COVID-19 Genomics UK (COG-UK) Consortium | Lighthouse Laboratory Plymouth and Alex Alderton, Roberto Amato, Jeffrey Barrett, Sonia Goncalves, Ewan Harrison, David K. Jackson, Ian Johnston, Dominic Kwiatkowski, Cordelia Langford, John Sillitoe on behalf of the Wellcome Sanger Institute COVID-19 Surveillance Team |
| EPI_ISL_2664526 | Lighthouse Lab in Glasgow | Wellcome Sanger Institute for the COVID-19 Genomics UK (COG-UK) Consortium | Harper VanSteenhouse, Yumi Kasai, David Gray, Carol Clugston, Anna Dominiczak and Alex Alderton, Roberto Amato, Jeffrey Barrett, Sonia Goncalves, Ewan Harrison, David K. Jackson, Ian Johnston, Dominic Kwiatkowski, Cordelia Langford, John Sillitoe on behalf of the Wellcome Sanger Institute COVID-19 Surveillance Team |
| EPI_ISL_2664527 | Lighthouse Laboratory Plymouth | Wellcome Sanger Institute for the COVID-19 Genomics UK (COG-UK) Consortium | Lighthouse Laboratory Plymouth and Alex Alderton, Roberto Amato, Jeffrey Barrett, Sonia Goncalves, Ewan Harrison, David K. Jackson, Ian Johnston, Dominic Kwiatkowski, Cordelia Langford, John Sillitoe on behalf of the Wellcome Sanger Institute COVID-19 Surveillance Team |
| EPI_ISL_2664528 | Lighthouse Lab in Glasgow | Wellcome Sanger Institute for the COVID-19 Genomics UK (COG-UK) Consortium | Harper VanSteenhouse, Yumi Kasai, David Gray, Carol Clugston, Anna Dominiczak and Alex Alderton, Roberto Amato, Jeffrey Barrett, Sonia Goncalves, Ewan Harrison, David K. Jackson, Ian Johnston, Dominic Kwiatkowski, Cordelia Langford, John Sillitoe on behalf of the Wellcome Sanger Institute COVID-19 Surveillance Team |
| EPI_ISL_2664529 | Lighthouse Laboratory Plymouth | Wellcome Sanger Institute for the COVID-19 Genomics UK (COG-UK) Consortium | Lighthouse Laboratory Plymouth and Alex Alderton, Roberto Amato, Jeffrey Barrett, Sonia Goncalves, Ewan Harrison, David K. Jackson, Ian Johnston, Dominic Kwiatkowski, Cordelia Langford, John Sillitoe on behalf of the Wellcome Sanger Institute COVID-19 Surveillance Team |
| EPI_ISL_2664530, EPI_ISL_2664531, EPI_ISL_2664532 | Lighthouse Lab in Milton Keynes | Wellcome Sanger Institute for the COVID-19 Genomics UK (COG-UK) Consortium | The Lighthouse Lab in Milton Keynes and Alex Alderton, Roberto Amato, Jeffrey Barrett, Sonia Goncalves, Ewan Harrison, David K. Jackson, Ian Johnston, Dominic Kwiatkowski, Cordelia Langford, John Sillitoe on behalf of the Wellcome Sanger Institute COVID-19 Surveillance Team |
| EPI_ISL_2664533, EPI_ISL_2664534 | Lighthouse Laboratory Plymouth | Wellcome Sanger Institute for the COVID-19 Genomics UK (COG-UK) Consortium | Lighthouse Laboratory Plymouth and Alex Alderton, Roberto Amato, Jeffrey Barrett, Sonia Goncalves, Ewan Harrison, David K. Jackson, Ian Johnston, Dominic Kwiatkowski, Cordelia Langford, John Sillitoe on behalf of the Wellcome Sanger Institute COVID-19 Surveillance Team |
| EPI_ISL_2664535 | Lighthouse Lab in Milton Keynes | Wellcome Sanger Institute for the COVID-19 Genomics UK (COG-UK) Consortium | The Lighthouse Lab in Milton Keynes and Alex Alderton, Roberto Amato, Jeffrey Barrett, Sonia Goncalves, Ewan Harrison, David K. Jackson, Ian Johnston, Dominic Kwiatkowski, Cordelia Langford, John Sillitoe on behalf of the Wellcome Sanger Institute COVID-19 Surveillance Team |
| EPI_ISL_2664536 | Lighthouse Lab in Alderley Park | Wellcome Sanger Institute for the COVID-19 Genomics UK (COG-UK) Consortium | Jacquelyn Wynn, Mairead Hyland, The Lighthouse Lab in Alderley Park and Alex Alderton, Roberto Amato, Jeffrey Barrett, Sonia Goncalves, Ewan Harrison, David K. Jackson, Ian Johnston, Dominic Kwiatkowski, Cordelia Langford, John Sillitoe on behalf of the Wellcome Sanger Institute COVID-19 Surveillance Team |
| EPI_ISL_2664537 | Lighthouse Laboratory Plymouth | Wellcome Sanger Institute for the COVID-19 Genomics UK (COG-UK) Consortium | Lighthouse Laboratory Plymouth and Alex Alderton, Roberto Amato, Jeffrey Barrett, Sonia Goncalves, Ewan Harrison, David K. Jackson, Ian Johnston, Dominic Kwiatkowski, Cordelia Langford, John Sillitoe on behalf of the Wellcome Sanger Institute COVID-19 Surveillance Team |
| EPI_ISL_2664538 | Lighthouse Lab in Milton Keynes | Wellcome Sanger Institute for the COVID-19 Genomics UK (COG-UK) Consortium | The Lighthouse Lab in Milton Keynes and Alex Alderton, Roberto Amato, Jeffrey Barrett, Sonia Goncalves, Ewan Harrison, David K. Jackson, Ian Johnston, Dominic Kwiatkowski, Cordelia Langford, John Sillitoe on behalf of the Wellcome Sanger Institute COVID-19 Surveillance Team |
| EPI_ISL_2664539 | Lighthouse Lab in Alderley Park | Wellcome Sanger Institute for the COVID-19 Genomics UK (COG-UK) Consortium | Jacquelyn Wynn, Mairead Hyland, The Lighthouse Lab in Alderley Park and Alex Alderton, Roberto Amato, Jeffrey Barrett, Sonia Goncalves, Ewan Harrison, David K. Jackson, Ian Johnston, Dominic Kwiatkowski, Cordelia Langford, John Sillitoe on behalf of the Wellcome Sanger Institute COVID-19 Surveillance Team |
| EPI_ISL_2664540 | Health Services Laboratories | Wellcome Sanger Institute for the COVID-19 Genomics UK (COG-UK) Consortium | Health Services Laboratories and Alex Alderton, Roberto Amato, Jeffrey Barrett, Sonia Goncalves, Ewan Harrison, David K. Jackson, Ian Johnston, Dominic Kwiatkowski, Cordelia Langford, John Sillitoe on behalf of the Wellcome Sanger Institute COVID-19 Surveillance Team |
| EPI_ISL_2664541 | Lighthouse Lab in Milton Keynes | Wellcome Sanger Institute for the COVID-19 Genomics UK (COG-UK) Consortium | The Lighthouse Lab in Milton Keynes and Alex Alderton, Roberto Amato, Jeffrey Barrett, Sonia Goncalves, Ewan Harrison, David K. Jackson, Ian Johnston, Dominic Kwiatkowski, Cordelia Langford, John Sillitoe on behalf of the Wellcome Sanger Institute COVID-19 Surveillance Team |
| EPI_ISL_2664542 | Lighthouse Lab in Alderley Park | Wellcome Sanger Institute for the COVID-19 Genomics UK (COG-UK) Consortium | Jacquelyn Wynn, Mairead Hyland, The Lighthouse Lab in Alderley Park and Alex Alderton, Roberto Amato, Jeffrey Barrett, Sonia Goncalves, Ewan Harrison, David K. Jackson, Ian Johnston, Dominic Kwiatkowski, Cordelia Langford, John Sillitoe on behalf of the Wellcome Sanger Institute COVID-19 Surveillance Team |
| EPI_ISL_2664543 | Lighthouse Lab in Milton Keynes | Wellcome Sanger Institute for the COVID-19 Genomics UK (COG-UK) Consortium | The Lighthouse Lab in Milton Keynes and Alex Alderton, Roberto Amato, Jeffrey Barrett, Sonia Goncalves, Ewan Harrison, David K. Jackson, Ian Johnston, Dominic Kwiatkowski, Cordelia Langford, John Sillitoe on behalf of the Wellcome Sanger Institute COVID-19 Surveillance Team |
| EPI_ISL_2664544 | Lighthouse Laboratory Plymouth | Wellcome Sanger Institute for the COVID-19 Genomics UK (COG-UK) Consortium | Lighthouse Laboratory Plymouth and Alex Alderton, Roberto Amato, Jeffrey Barrett, Sonia Goncalves, Ewan Harrison, David K. Jackson, Ian Johnston, Dominic Kwiatkowski, Cordelia Langford, John Sillitoe on behalf of the Wellcome Sanger Institute COVID-19 Surveillance Team |
| EPI_ISL_2664545 | Lighthouse Lab in Alderley Park | Wellcome Sanger Institute for the COVID-19 Genomics UK (COG-UK) Consortium | Jacquelyn Wynn, Mairead Hyland, The Lighthouse Lab in Alderley Park and Alex Alderton, Roberto Amato, Jeffrey Barrett, Sonia Goncalves, Ewan Harrison, David K. Jackson, Ian Johnston, Dominic Kwiatkowski, Cordelia Langford, John Sillitoe on behalf of the Wellcome Sanger Institute COVID-19 Surveillance Team |
| EPI_ISL_2664546 | Health Services Laboratories | Wellcome Sanger Institute for the COVID-19 Genomics UK (COG-UK) Consortium | Health Services Laboratories and Alex Alderton, Roberto Amato, Jeffrey Barrett, Sonia Goncalves, Ewan Harrison, David K. Jackson, Ian Johnston, Dominic Kwiatkowski, Cordelia Langford, John Sillitoe on behalf of the Wellcome Sanger Institute COVID-19 Surveillance Team |
| EPI_ISL_2664547, EPI_ISL_2664548, EPI_ISL_2664549 | Lighthouse Lab in Alderley Park | Wellcome Sanger Institute for the COVID-19 Genomics UK (COG-UK) Consortium | Jacquelyn Wynn, Mairead Hyland, The Lighthouse Lab in Alderley Park and Alex Alderton, Roberto Amato, Jeffrey Barrett, Sonia Goncalves, Ewan Harrison, David K. Jackson, Ian Johnston, Dominic Kwiatkowski, Cordelia Langford, John Sillitoe on behalf of the Wellcome Sanger Institute COVID-19 Surveillance Team |
| EPI_ISL_2664550, EPI_ISL_2664551 | Lighthouse Laboratory Plymouth | Wellcome Sanger Institute for the COVID-19 Genomics UK (COG-UK) Consortium | Lighthouse Laboratory Plymouth and Alex Alderton, Roberto Amato, Jeffrey Barrett, Sonia Goncalves, Ewan Harrison, David K. Jackson, Ian Johnston, Dominic Kwiatkowski, Cordelia Langford, John Sillitoe on behalf of the Wellcome Sanger Institute COVID-19 Surveillance Team |
| EPI_ISL_2664552 | Lighthouse Lab in Alderley Park | Wellcome Sanger Institute for the COVID-19 Genomics UK (COG-UK) Consortium | Jacquelyn Wynn, Mairead Hyland, The Lighthouse Lab in Alderley Park and Alex Alderton, Roberto Amato, Jeffrey Barrett, Sonia Goncalves, Ewan Harrison, David K. Jackson, Ian Johnston, Dominic Kwiatkowski, Cordelia Langford, John Sillitoe on behalf of the Wellcome Sanger Institute COVID-19 Surveillance Team |
| EPI_ISL_2664553, EPI_ISL_2664554, EPI_ISL_2664555 | Lighthouse Laboratory Plymouth | Wellcome Sanger Institute for the COVID-19 Genomics UK (COG-UK) Consortium | Lighthouse Laboratory Plymouth and Alex Alderton, Roberto Amato, Jeffrey Barrett, Sonia Goncalves, Ewan Harrison, David K. Jackson, Ian Johnston, Dominic Kwiatkowski, Cordelia Langford, John Sillitoe on behalf of the Wellcome Sanger Institute COVID-19 Surveillance Team |
| EPI_ISL_2664556, EPI_ISL_2664557 | Lighthouse Lab in Alderley Park | Wellcome Sanger Institute for the COVID-19 Genomics UK (COG-UK) Consortium | Jacquelyn Wynn, Mairead Hyland, The Lighthouse Lab in Alderley Park and Alex Alderton, Roberto Amato, Jeffrey Barrett, Sonia Goncalves, Ewan Harrison, David K. Jackson, Ian Johnston, Dominic Kwiatkowski, Cordelia Langford, John Sillitoe on behalf of the Wellcome Sanger Institute COVID-19 Surveillance Team |
| EPI_ISL_2664558 | Lighthouse Lab in Glasgow | Wellcome Sanger Institute for the COVID-19 Genomics UK (COG-UK) Consortium | Harper VanSteenhouse, Yumi Kasai, David Gray, Carol Clugston, Anna Dominiczak and Alex Alderton, Roberto Amato, Jeffrey Barrett, Sonia Goncalves, Ewan Harrison, David K. Jackson, Ian Johnston, Dominic Kwiatkowski, Cordelia Langford, John Sillitoe on behalf of the Wellcome Sanger Institute COVID-19 Surveillance Team |
| EPI_ISL_2664559, EPI_ISL_2664560 | Lighthouse Lab in Milton Keynes | Wellcome Sanger Institute for the COVID-19 Genomics UK (COG-UK) Consortium | The Lighthouse Lab in Milton Keynes and Alex Alderton, Roberto Amato, Jeffrey Barrett, Sonia Goncalves, Ewan Harrison, David K. Jackson, Ian Johnston, Dominic Kwiatkowski, Cordelia Langford, John Sillitoe on behalf of the Wellcome Sanger Institute COVID-19 Surveillance Team |
| EPI_ISL_2664561 | Lighthouse Laboratory Plymouth | Wellcome Sanger Institute for the COVID-19 Genomics UK (COG-UK) Consortium | Lighthouse Laboratory Plymouth and Alex Alderton, Roberto Amato, Jeffrey Barrett, Sonia Goncalves, Ewan Harrison, David K. Jackson, Ian Johnston, Dominic Kwiatkowski, Cordelia Langford, John Sillitoe on behalf of the Wellcome Sanger Institute COVID-19 Surveillance Team |
| EPI_ISL_2664562 | Lighthouse Lab in Alderley Park | Wellcome Sanger Institute for the COVID-19 Genomics UK (COG-UK) Consortium | Jacquelyn Wynn, Mairead Hyland, The Lighthouse Lab in Alderley Park and Alex Alderton, Roberto Amato, Jeffrey Barrett, Sonia Goncalves, Ewan Harrison, David K. Jackson, Ian Johnston, Dominic Kwiatkowski, Cordelia Langford, John Sillitoe on behalf of the Wellcome Sanger Institute COVID-19 Surveillance Team |
| EPI_ISL_2664563 | Lighthouse Laboratory Plymouth | Wellcome Sanger Institute for the COVID-19 Genomics UK (COG-UK) Consortium | Lighthouse Laboratory Plymouth and Alex Alderton, Roberto Amato, Jeffrey Barrett, Sonia Goncalves, Ewan Harrison, David K. Jackson, Ian Johnston, Dominic Kwiatkowski, Cordelia Langford, John Sillitoe on behalf of the Wellcome Sanger Institute COVID-19 Surveillance Team |
| EPI_ISL_2664564 | Lighthouse Lab in Alderley Park | Wellcome Sanger Institute for the COVID-19 Genomics UK (COG-UK) Consortium | Jacquelyn Wynn, Mairead Hyland, The Lighthouse Lab in Alderley Park and Alex Alderton, Roberto Amato, Jeffrey Barrett, Sonia Goncalves, Ewan Harrison, David K. Jackson, Ian Johnston, Dominic Kwiatkowski, Cordelia Langford, John Sillitoe on behalf of the Wellcome Sanger Institute COVID-19 Surveillance Team |
| EPI_ISL_2664565 | Lighthouse Laboratory Plymouth | Wellcome Sanger Institute for the COVID-19 Genomics UK (COG-UK) Consortium | Lighthouse Laboratory Plymouth and Alex Alderton, Roberto Amato, Jeffrey Barrett, Sonia Goncalves, Ewan Harrison, David K. Jackson, Ian Johnston, Dominic Kwiatkowski, Cordelia Langford, John Sillitoe on behalf of the Wellcome Sanger Institute COVID-19 Surveillance Team |
| EPI_ISL_2664566, EPI_ISL_2664567 | Lighthouse Lab in Alderley Park | Wellcome Sanger Institute for the COVID-19 Genomics UK (COG-UK) Consortium | Jacquelyn Wynn, Mairead Hyland, The Lighthouse Lab in Alderley Park and Alex Alderton, Roberto Amato, Jeffrey Barrett, Sonia Goncalves, Ewan Harrison, David K. Jackson, Ian Johnston, Dominic Kwiatkowski, Cordelia Langford, John Sillitoe on behalf of the Wellcome Sanger Institute COVID-19 Surveillance Team |
| EPI_ISL_2664568, EPI_ISL_2664569 | Lighthouse Lab in Glasgow | Wellcome Sanger Institute for the COVID-19 Genomics UK (COG-UK) Consortium | Harper VanSteenhouse, Yumi Kasai, David Gray, Carol Clugston, Anna Dominiczak and Alex Alderton, Roberto Amato, Jeffrey Barrett, Sonia Goncalves, Ewan Harrison, David K. Jackson, Ian Johnston, Dominic Kwiatkowski, Cordelia Langford, John Sillitoe on behalf of the Wellcome Sanger Institute COVID-19 Surveillance Team |
| EPI_ISL_2664570 | Lighthouse Lab in Alderley Park | Wellcome Sanger Institute for the COVID-19 Genomics UK (COG-UK) Consortium | Jacquelyn Wynn, Mairead Hyland, The Lighthouse Lab in Alderley Park and Alex Alderton, Roberto Amato, Jeffrey Barrett, Sonia Goncalves, Ewan Harrison, David K. Jackson, Ian Johnston, Dominic Kwiatkowski, Cordelia Langford, John Sillitoe on behalf of the Wellcome Sanger Institute COVID-19 Surveillance Team |
| EPI_ISL_2664571, EPI_ISL_2664572 | Lighthouse Lab in Glasgow | Wellcome Sanger Institute for the COVID-19 Genomics UK (COG-UK) Consortium | Harper VanSteenhouse, Yumi Kasai, David Gray, Carol Clugston, Anna Dominiczak and Alex Alderton, Roberto Amato, Jeffrey Barrett, Sonia Goncalves, Ewan Harrison, David K. Jackson, Ian Johnston, Dominic Kwiatkowski, Cordelia Langford, John Sillitoe on behalf of the Wellcome Sanger Institute COVID-19 Surveillance Team |
| EPI_ISL_2664573, EPI_ISL_2664574 | Lighthouse Lab in Alderley Park | Wellcome Sanger Institute for the COVID-19 Genomics UK (COG-UK) Consortium | Jacquelyn Wynn, Mairead Hyland, The Lighthouse Lab in Alderley Park and Alex Alderton, Roberto Amato, Jeffrey Barrett, Sonia Goncalves, Ewan Harrison, David K. Jackson, Ian Johnston, Dominic Kwiatkowski, Cordelia Langford, John Sillitoe on behalf of the Wellcome Sanger Institute COVID-19 Surveillance Team |
| EPI_ISL_2664575 | Lighthouse Lab in Glasgow | Wellcome Sanger Institute for the COVID-19 Genomics UK (COG-UK) Consortium | Harper VanSteenhouse, Yumi Kasai, David Gray, Carol Clugston, Anna Dominiczak and Alex Alderton, Roberto Amato, Jeffrey Barrett, Sonia Goncalves, Ewan Harrison, David K. Jackson, Ian Johnston, Dominic Kwiatkowski, Cordelia Langford, John Sillitoe on behalf of the Wellcome Sanger Institute COVID-19 Surveillance Team |
| EPI_ISL_2664576, EPI_ISL_2664577 | Lighthouse Lab in Milton Keynes | Wellcome Sanger Institute for the COVID-19 Genomics UK (COG-UK) Consortium | The Lighthouse Lab in Milton Keynes and Alex Alderton, Roberto Amato, Jeffrey Barrett, Sonia Goncalves, Ewan Harrison, David K. Jackson, Ian Johnston, Dominic Kwiatkowski, Cordelia Langford, John Sillitoe on behalf of the Wellcome Sanger Institute COVID-19 Surveillance Team |
| EPI_ISL_2664578 | Lighthouse Laboratory Plymouth | Wellcome Sanger Institute for the COVID-19 Genomics UK (COG-UK) Consortium | Lighthouse Laboratory Plymouth and Alex Alderton, Roberto Amato, Jeffrey Barrett, Sonia Goncalves, Ewan Harrison, David K. Jackson, Ian Johnston, Dominic Kwiatkowski, Cordelia Langford, John Sillitoe on behalf of the Wellcome Sanger Institute COVID-19 Surveillance Team |
| EPI_ISL_2664579, EPI_ISL_2664581, EPI_ISL_2664582 | Lighthouse Lab in Alderley Park | Wellcome Sanger Institute for the COVID-19 Genomics UK (COG-UK) Consortium | Jacquelyn Wynn, Mairead Hyland, The Lighthouse Lab in Alderley Park and Alex Alderton, Roberto Amato, Jeffrey Barrett, Sonia Goncalves, Ewan Harrison, David K. Jackson, Ian Johnston, Dominic Kwiatkowski, Cordelia Langford, John Sillitoe on behalf of the Wellcome Sanger Institute COVID-19 Surveillance Team |
| EPI_ISL_2664583 | Lighthouse Lab in Milton Keynes | Wellcome Sanger Institute for the COVID-19 Genomics UK (COG-UK) Consortium | The Lighthouse Lab in Milton Keynes and Alex Alderton, Roberto Amato, Jeffrey Barrett, Sonia Goncalves, Ewan Harrison, David K. Jackson, Ian Johnston, Dominic Kwiatkowski, Cordelia Langford, John Sillitoe on behalf of the Wellcome Sanger Institute COVID-19 Surveillance Team |
| EPI_ISL_2664584, EPI_ISL_2664585 | Lighthouse Lab in Alderley Park | Wellcome Sanger Institute for the COVID-19 Genomics UK (COG-UK) Consortium | Jacquelyn Wynn, Mairead Hyland, The Lighthouse Lab in Alderley Park and Alex Alderton, Roberto Amato, Jeffrey Barrett, Sonia Goncalves, Ewan Harrison, David K. Jackson, Ian Johnston, Dominic Kwiatkowski, Cordelia Langford, John Sillitoe on behalf of the Wellcome Sanger Institute COVID-19 Surveillance Team |
| EPI_ISL_2664587 | Lighthouse Lab in Glasgow | Wellcome Sanger Institute for the COVID-19 Genomics UK (COG-UK) Consortium | Harper VanSteenhouse, Yumi Kasai, David Gray, Carol Clugston, Anna Dominiczak and Alex Alderton, Roberto Amato, Jeffrey Barrett, Sonia Goncalves, Ewan Harrison, David K. Jackson, Ian Johnston, Dominic Kwiatkowski, Cordelia Langford, John Sillitoe on behalf of the Wellcome Sanger Institute COVID-19 Surveillance Team |
| EPI_ISL_2664588, | Lighthouse Laboratory Plymouth | Wellcome Sanger Institute for the COVID-19 Genomics UK (COG-UK) | Lighthouse Laboratory Plymouth and Alex Alderton, Roberto Amato, Jeffrey Barrett, Sonia Goncalves, Ewan Harrison, David K. Jackson, Ian Johnston, Dominic Kwiatkowski, Cordelia Langford, John Sillitoe on behalf of the Wellcome Sanger |

| | | | |
|---|---|---|---|
| EPI_ISL_2664589, EPI_ISL_2664590 | | Consortium | Institute COVID-19 Surveillance Team |
| EPI_ISL_2664591 | Lighthouse Lab in Milton Keynes | Wellcome Sanger Institute for the COVID-19 Genomics UK (COG-UK) Consortium | The Lighthouse Lab in Milton Keynes and Alex Alderton, Roberto Amato, Jeffrey Barrett, Sonia Goncalves, Ewan Harrison, David K. Jackson, Ian Johnston, Dominic Kwiatkowski, Cordelia Langford, John Sillitoe on behalf of the Wellcome Sanger Institute COVID-19 Surveillance Team |
| EPI_ISL_2664592 | Lighthouse Lab in Glasgow | Wellcome Sanger Institute for the COVID-19 Genomics UK (COG-UK) Consortium | Harper VanSteenhouse, Yumi Kasai, David Gray, Carol Clugston, Anna Dominiczak and Alex Alderton, Roberto Amato, Jeffrey Barrett, Sonia Goncalves, Ewan Harrison, David K. Jackson, Ian Johnston, Dominic Kwiatkowski, Cordelia Langford, John Sillitoe on behalf of the Wellcome Sanger Institute COVID-19 Surveillance Team |
| EPI_ISL_2664596 | Health Services Laboratories | Wellcome Sanger Institute for the COVID-19 Genomics UK (COG-UK) Consortium | Health Services Laboratories and Alex Alderton, Roberto Amato, Jeffrey Barrett, Sonia Goncalves, Ewan Harrison, David K. Jackson, Ian Johnston, Dominic Kwiatkowski, Cordelia Langford, John Sillitoe on behalf of the Wellcome Sanger Institute COVID-19 Surveillance Team |
| EPI_ISL_2664597, EPI_ISL_2664598, EPI_ISL_2664599 | Lighthouse Lab in Alderley Park | Wellcome Sanger Institute for the COVID-19 Genomics UK (COG-UK) Consortium | Jacquelyn Wynn, Mairead Hyland, The Lighthouse Lab in Alderley Park and Alex Alderton, Roberto Amato, Jeffrey Barrett, Sonia Goncalves, Ewan Harrison, David K. Jackson, Ian Johnston, Dominic Kwiatkowski, Cordelia Langford, John Sillitoe on behalf of the Wellcome Sanger Institute COVID-19 Surveillance Team |
| EPI_ISL_2664600 | Lighthouse Lab in Milton Keynes | Wellcome Sanger Institute for the COVID-19 Genomics UK (COG-UK) Consortium | The Lighthouse Lab in Milton Keynes and Alex Alderton, Roberto Amato, Jeffrey Barrett, Sonia Goncalves, Ewan Harrison, David K. Jackson, Ian Johnston, Dominic Kwiatkowski, Cordelia Langford, John Sillitoe on behalf of the Wellcome Sanger Institute COVID-19 Surveillance Team |
| EPI_ISL_2664601 | Lighthouse Lab in Alderley Park | Wellcome Sanger Institute for the COVID-19 Genomics UK (COG-UK) Consortium | Jacquelyn Wynn, Mairead Hyland, The Lighthouse Lab in Alderley Park and Alex Alderton, Roberto Amato, Jeffrey Barrett, Sonia Goncalves, Ewan Harrison, David K. Jackson, Ian Johnston, Dominic Kwiatkowski, Cordelia Langford, John Sillitoe on behalf of the Wellcome Sanger Institute COVID-19 Surveillance Team |
| EPI_ISL_2664602, EPI_ISL_2664603, EPI_ISL_2664604 | Lighthouse Laboratory Plymouth | Wellcome Sanger Institute for the COVID-19 Genomics UK (COG-UK) Consortium | Lighthouse Laboratory Plymouth and Alex Alderton, Roberto Amato, Jeffrey Barrett, Sonia Goncalves, Ewan Harrison, David K. Jackson, Ian Johnston, Dominic Kwiatkowski, Cordelia Langford, John Sillitoe on behalf of the Wellcome Sanger Institute COVID-19 Surveillance Team |
| EPI_ISL_2664605 | Lighthouse Lab in Glasgow | Wellcome Sanger Institute for the COVID-19 Genomics UK (COG-UK) Consortium | Harper VanSteenhouse, Yumi Kasai, David Gray, Carol Clugston, Anna Dominiczak and Alex Alderton, Roberto Amato, Jeffrey Barrett, Sonia Goncalves, Ewan Harrison, David K. Jackson, Ian Johnston, Dominic Kwiatkowski, Cordelia Langford, John Sillitoe on behalf of the Wellcome Sanger Institute COVID-19 Surveillance Team |
| EPI_ISL_2664606, EPI_ISL_2664607 | Lighthouse Laboratory Plymouth | Wellcome Sanger Institute for the COVID-19 Genomics UK (COG-UK) Consortium | Lighthouse Laboratory Plymouth and Alex Alderton, Roberto Amato, Jeffrey Barrett, Sonia Goncalves, Ewan Harrison, David K. Jackson, Ian Johnston, Dominic Kwiatkowski, Cordelia Langford, John Sillitoe on behalf of the Wellcome Sanger Institute COVID-19 Surveillance Team |
| EPI_ISL_2664608, EPI_ISL_2664609 | Lighthouse Lab in Milton Keynes | Wellcome Sanger Institute for the COVID-19 Genomics UK (COG-UK) Consortium | The Lighthouse Lab in Milton Keynes and Alex Alderton, Roberto Amato, Jeffrey Barrett, Sonia Goncalves, Ewan Harrison, David K. Jackson, Ian Johnston, Dominic Kwiatkowski, Cordelia Langford, John Sillitoe on behalf of the Wellcome Sanger Institute COVID-19 Surveillance Team |
| EPI_ISL_2664610 | Lighthouse Lab in Alderley Park | Wellcome Sanger Institute for the COVID-19 Genomics UK (COG-UK) Consortium | Jacquelyn Wynn, Mairead Hyland, The Lighthouse Lab in Alderley Park and Alex Alderton, Roberto Amato, Jeffrey Barrett, Sonia Goncalves, Ewan Harrison, David K. Jackson, Ian Johnston, Dominic Kwiatkowski, Cordelia Langford, John Sillitoe on behalf of the Wellcome Sanger Institute COVID-19 Surveillance Team |
| EPI_ISL_2664611 | Lighthouse Lab in Milton Keynes | Wellcome Sanger Institute for the COVID-19 Genomics UK (COG-UK) Consortium | The Lighthouse Lab in Milton Keynes and Alex Alderton, Roberto Amato, Jeffrey Barrett, Sonia Goncalves, Ewan Harrison, David K. Jackson, Ian Johnston, Dominic Kwiatkowski, Cordelia Langford, John Sillitoe on behalf of the Wellcome Sanger Institute COVID-19 Surveillance Team |
| EPI_ISL_2664612 | Lighthouse Lab in Alderley Park | Wellcome Sanger Institute for the COVID-19 Genomics UK (COG-UK) Consortium | Jacquelyn Wynn, Mairead Hyland, The Lighthouse Lab in Alderley Park and Alex Alderton, Roberto Amato, Jeffrey Barrett, Sonia Goncalves, Ewan Harrison, David K. Jackson, Ian Johnston, Dominic Kwiatkowski, Cordelia Langford, John Sillitoe on behalf of the Wellcome Sanger Institute COVID-19 Surveillance Team |
| EPI_ISL_2664613, EPI_ISL_2664614 | Lighthouse Laboratory Plymouth | Wellcome Sanger Institute for the COVID-19 Genomics UK (COG-UK) Consortium | Lighthouse Laboratory Plymouth and Alex Alderton, Roberto Amato, Jeffrey Barrett, Sonia Goncalves, Ewan Harrison, David K. Jackson, Ian Johnston, Dominic Kwiatkowski, Cordelia Langford, John Sillitoe on behalf of the Wellcome Sanger Institute COVID-19 Surveillance Team |
| EPI_ISL_2664615 | Lighthouse Lab in Alderley Park | Wellcome Sanger Institute for the COVID-19 Genomics UK (COG-UK) Consortium | Jacquelyn Wynn, Mairead Hyland, The Lighthouse Lab in Alderley Park and Alex Alderton, Roberto Amato, Jeffrey Barrett, Sonia Goncalves, Ewan Harrison, David K. Jackson, Ian Johnston, Dominic Kwiatkowski, Cordelia Langford, John Sillitoe on behalf of the Wellcome Sanger Institute COVID-19 Surveillance Team |
| EPI_ISL_2664616 | Lighthouse Lab in Milton Keynes | Wellcome Sanger Institute for the COVID-19 Genomics UK (COG-UK) Consortium | The Lighthouse Lab in Milton Keynes and Alex Alderton, Roberto Amato, Jeffrey Barrett, Sonia Goncalves, Ewan Harrison, David K. Jackson, Ian Johnston, Dominic Kwiatkowski, Cordelia Langford, John Sillitoe on behalf of the Wellcome Sanger Institute COVID-19 Surveillance Team |
| EPI_ISL_2664617 | Lighthouse Lab in Alderley Park | Wellcome Sanger Institute for the COVID-19 Genomics UK (COG-UK) Consortium | Jacquelyn Wynn, Mairead Hyland, The Lighthouse Lab in Alderley Park and Alex Alderton, Roberto Amato, Jeffrey Barrett, Sonia Goncalves, Ewan Harrison, David K. Jackson, Ian Johnston, Dominic Kwiatkowski, Cordelia Langford, John Sillitoe on behalf of the Wellcome Sanger Institute COVID-19 Surveillance Team |
| EPI_ISL_2664618, EPI_ISL_2664619 | Lighthouse Lab in Milton Keynes | Wellcome Sanger Institute for the COVID-19 Genomics UK (COG-UK) Consortium | The Lighthouse Lab in Milton Keynes and Alex Alderton, Roberto Amato, Jeffrey Barrett, Sonia Goncalves, Ewan Harrison, David K. Jackson, Ian Johnston, Dominic Kwiatkowski, Cordelia Langford, John Sillitoe on behalf of the Wellcome Sanger Institute COVID-19 Surveillance Team |
| EPI_ISL_2664620 | Lighthouse Lab in Alderley Park | Wellcome Sanger Institute for the COVID-19 Genomics UK (COG-UK) Consortium | Jacquelyn Wynn, Mairead Hyland, The Lighthouse Lab in Alderley Park and Alex Alderton, Roberto Amato, Jeffrey Barrett, Sonia Goncalves, Ewan Harrison, David K. Jackson, Ian Johnston, Dominic Kwiatkowski, Cordelia Langford, John Sillitoe on behalf of the Wellcome Sanger Institute COVID-19 Surveillance Team |
| EPI_ISL_2664621 | Lighthouse Lab in Milton Keynes | Wellcome Sanger Institute for the COVID-19 Genomics UK (COG-UK) Consortium | The Lighthouse Lab in Milton Keynes and Alex Alderton, Roberto Amato, Jeffrey Barrett, Sonia Goncalves, Ewan Harrison, David K. Jackson, Ian Johnston, Dominic Kwiatkowski, Cordelia Langford, John Sillitoe on behalf of the Wellcome Sanger Institute COVID-19 Surveillance Team |
| EPI_ISL_2664622 | Lighthouse Laboratory Plymouth | Wellcome Sanger Institute for the COVID-19 Genomics UK (COG-UK) Consortium | Lighthouse Laboratory Plymouth and Alex Alderton, Roberto Amato, Jeffrey Barrett, Sonia Goncalves, Ewan Harrison, David K. Jackson, Ian Johnston, Dominic Kwiatkowski, Cordelia Langford, John Sillitoe on behalf of the Wellcome Sanger Institute COVID-19 Surveillance Team |
| EPI_ISL_2664623, EPI_ISL_2664624 | Lighthouse Lab in Milton Keynes | Wellcome Sanger Institute for the COVID-19 Genomics UK (COG-UK) Consortium | The Lighthouse Lab in Milton Keynes and Alex Alderton, Roberto Amato, Jeffrey Barrett, Sonia Goncalves, Ewan Harrison, David K. Jackson, Ian Johnston, Dominic Kwiatkowski, Cordelia Langford, John Sillitoe on behalf of the Wellcome Sanger Institute COVID-19 Surveillance Team |
| EPI_ISL_2664625 | Lighthouse Laboratory Plymouth | Wellcome Sanger Institute for the COVID-19 Genomics UK (COG-UK) Consortium | Lighthouse Laboratory Plymouth and Alex Alderton, Roberto Amato, Jeffrey Barrett, Sonia Goncalves, Ewan Harrison, David K. Jackson, Ian Johnston, Dominic Kwiatkowski, Cordelia Langford, John Sillitoe on behalf of the Wellcome Sanger Institute COVID-19 Surveillance Team |
| EPI_ISL_2664626, EPI_ISL_2664627, EPI_ISL_2664628 | Lighthouse Lab in Milton Keynes | Wellcome Sanger Institute for the COVID-19 Genomics UK (COG-UK) Consortium | The Lighthouse Lab in Milton Keynes and Alex Alderton, Roberto Amato, Jeffrey Barrett, Sonia Goncalves, Ewan Harrison, David K. Jackson, Ian Johnston, Dominic Kwiatkowski, Cordelia Langford, John Sillitoe on behalf of the Wellcome Sanger Institute COVID-19 Surveillance Team |
| EPI_ISL_2664629 | Lighthouse Laboratory Plymouth | Wellcome Sanger Institute for the COVID-19 Genomics UK (COG-UK) Consortium | Lighthouse Laboratory Plymouth and Alex Alderton, Roberto Amato, Jeffrey Barrett, Sonia Goncalves, Ewan Harrison, David K. Jackson, Ian Johnston, Dominic Kwiatkowski, Cordelia Langford, John Sillitoe on behalf of the Wellcome Sanger Institute COVID-19 Surveillance Team |
| EPI_ISL_2664630 | Lighthouse Lab in Milton Keynes | Wellcome Sanger Institute for the COVID-19 Genomics UK (COG-UK) Consortium | The Lighthouse Lab in Milton Keynes and Alex Alderton, Roberto Amato, Jeffrey Barrett, Sonia Goncalves, Ewan Harrison, David K. Jackson, Ian Johnston, Dominic Kwiatkowski, Cordelia Langford, John Sillitoe on behalf of the Wellcome Sanger Institute COVID-19 Surveillance Team |
| EPI_ISL_2664631 | Lighthouse Laboratory Plymouth | Wellcome Sanger Institute for the COVID-19 Genomics UK (COG-UK) Consortium | Lighthouse Laboratory Plymouth and Alex Alderton, Roberto Amato, Jeffrey Barrett, Sonia Goncalves, Ewan Harrison, David K. Jackson, Ian Johnston, Dominic Kwiatkowski, Cordelia Langford, John Sillitoe on behalf of the Wellcome Sanger Institute COVID-19 Surveillance Team |
| EPI_ISL_2664632 | Lighthouse Lab in Glasgow | Wellcome Sanger Institute for the COVID-19 Genomics UK (COG-UK) Consortium | Harper VanSteenhouse, Yumi Kasai, David Gray, Carol Clugston, Anna Dominiczak and Alex Alderton, Roberto Amato, Jeffrey Barrett, Sonia Goncalves, Ewan Harrison, David K. Jackson, Ian Johnston, Dominic Kwiatkowski, Cordelia Langford, John Sillitoe on behalf of the Wellcome Sanger Institute COVID-19 Surveillance Team |
| EPI_ISL_2664633 | Lighthouse Lab in Milton Keynes | Wellcome Sanger Institute for the COVID-19 Genomics UK (COG-UK) Consortium | The Lighthouse Lab in Milton Keynes and Alex Alderton, Roberto Amato, Jeffrey Barrett, Sonia Goncalves, Ewan Harrison, David K. Jackson, Ian Johnston, Dominic Kwiatkowski, Cordelia Langford, John Sillitoe on behalf of the Wellcome Sanger Institute COVID-19 Surveillance Team |
| EPI_ISL_2664634, EPI_ISL_2664635 | Lighthouse Laboratory Plymouth | Wellcome Sanger Institute for the COVID-19 Genomics UK (COG-UK) Consortium | Lighthouse Laboratory Plymouth and Alex Alderton, Roberto Amato, Jeffrey Barrett, Sonia Goncalves, Ewan Harrison, David K. Jackson, Ian Johnston, Dominic Kwiatkowski, Cordelia Langford, John Sillitoe on behalf of the Wellcome Sanger Institute COVID-19 Surveillance Team |
| EPI_ISL_2664636 | Lighthouse Lab in Alderley Park | Wellcome Sanger Institute for the COVID-19 Genomics UK (COG-UK) Consortium | Jacquelyn Wynn, Mairead Hyland, The Lighthouse Lab in Alderley Park and Alex Alderton, Roberto Amato, Jeffrey Barrett, Sonia Goncalves, Ewan Harrison, David K. Jackson, Ian Johnston, Dominic Kwiatkowski, Cordelia Langford, John Sillitoe on behalf of the Wellcome Sanger Institute COVID-19 Surveillance Team |
| EPI_ISL_2664637, EPI_ISL_2664638 | Lighthouse Lab in Milton Keynes | Wellcome Sanger Institute for the COVID-19 Genomics UK (COG-UK) Consortium | The Lighthouse Lab in Milton Keynes and Alex Alderton, Roberto Amato, Jeffrey Barrett, Sonia Goncalves, Ewan Harrison, David K. Jackson, Ian Johnston, Dominic Kwiatkowski, Cordelia Langford, John Sillitoe on behalf of the Wellcome Sanger Institute COVID-19 Surveillance Team |
| EPI_ISL_2664639 | Lighthouse Laboratory Plymouth | Wellcome Sanger Institute for the COVID-19 Genomics UK (COG-UK) Consortium | Lighthouse Laboratory Plymouth and Alex Alderton, Roberto Amato, Jeffrey Barrett, Sonia Goncalves, Ewan Harrison, David K. Jackson, Ian Johnston, Dominic Kwiatkowski, Cordelia Langford, John Sillitoe on behalf of the Wellcome Sanger Institute COVID-19 Surveillance Team |
| EPI_ISL_2664640 | Lighthouse Lab in Alderley Park | Wellcome Sanger Institute for the COVID-19 Genomics UK (COG-UK) Consortium | Jacquelyn Wynn, Mairead Hyland, The Lighthouse Lab in Alderley Park and Alex Alderton, Roberto Amato, Jeffrey Barrett, Sonia Goncalves, Ewan Harrison, David K. Jackson, Ian Johnston, Dominic Kwiatkowski, Cordelia Langford, John Sillitoe on behalf of the Wellcome Sanger Institute COVID-19 Surveillance Team |
| EPI_ISL_2664641 | Lighthouse Lab in Milton Keynes | Wellcome Sanger Institute for the COVID-19 Genomics UK (COG-UK) Consortium | The Lighthouse Lab in Milton Keynes and Alex Alderton, Roberto Amato, Jeffrey Barrett, Sonia Goncalves, Ewan Harrison, David K. Jackson, Ian Johnston, Dominic Kwiatkowski, Cordelia Langford, John Sillitoe on behalf of the Wellcome Sanger Institute COVID-19 Surveillance Team |
| EPI_ISL_2664642 | Lighthouse Laboratory Plymouth | Wellcome Sanger Institute for the COVID-19 Genomics UK (COG-UK) Consortium | Lighthouse Laboratory Plymouth and Alex Alderton, Roberto Amato, Jeffrey Barrett, Sonia Goncalves, Ewan Harrison, David K. Jackson, Ian Johnston, Dominic Kwiatkowski, Cordelia Langford, John Sillitoe on behalf of the Wellcome Sanger Institute COVID-19 Surveillance Team |
| EPI_ISL_2664643, EPI_ISL_2664644 | Lighthouse Lab in Alderley Park | Wellcome Sanger Institute for the COVID-19 Genomics UK (COG-UK) Consortium | Jacquelyn Wynn, Mairead Hyland, The Lighthouse Lab in Alderley Park and Alex Alderton, Roberto Amato, Jeffrey Barrett, Sonia Goncalves, Ewan Harrison, David K. Jackson, Ian Johnston, Dominic Kwiatkowski, Cordelia Langford, John Sillitoe on behalf of the Wellcome Sanger Institute COVID-19 Surveillance Team |
| EPI_ISL_2664645, EPI_ISL_2664646 | Lighthouse Lab in Milton Keynes | Wellcome Sanger Institute for the COVID-19 Genomics UK (COG-UK) Consortium | The Lighthouse Lab in Milton Keynes and Alex Alderton, Roberto Amato, Jeffrey Barrett, Sonia Goncalves, Ewan Harrison, David K. Jackson, Ian Johnston, Dominic Kwiatkowski, Cordelia Langford, John Sillitoe on behalf of the Wellcome Sanger Institute COVID-19 Surveillance Team |
| EPI_ISL_2664647 | Lighthouse Lab in Glasgow | Wellcome Sanger Institute for the COVID-19 Genomics UK (COG-UK) Consortium | Harper VanSteenhouse, Yumi Kasai, David Gray, Carol Clugston, Anna Dominiczak and Alex Alderton, Roberto Amato, Jeffrey Barrett, Sonia Goncalves, Ewan Harrison, David K. Jackson, Ian Johnston, Dominic Kwiatkowski, Cordelia Langford, John Sillitoe on behalf of the Wellcome Sanger Institute COVID-19 Surveillance Team |
| EPI_ISL_2664648 | Lighthouse Lab in Alderley Park | Wellcome Sanger Institute for the COVID-19 Genomics UK (COG-UK) Consortium | Jacquelyn Wynn, Mairead Hyland, The Lighthouse Lab in Alderley Park and Alex Alderton, Roberto Amato, Jeffrey Barrett, Sonia Goncalves, Ewan Harrison, David K. Jackson, Ian Johnston, Dominic Kwiatkowski, Cordelia Langford, John Sillitoe on behalf of the Wellcome Sanger Institute COVID-19 Surveillance Team |
| EPI_ISL_2664649 | Lighthouse Laboratory Plymouth | Wellcome Sanger Institute for the COVID-19 Genomics UK (COG-UK) Consortium | Lighthouse Laboratory Plymouth and Alex Alderton, Roberto Amato, Jeffrey Barrett, Sonia Goncalves, Ewan Harrison, David K. Jackson, Ian Johnston, Dominic Kwiatkowski, Cordelia Langford, John Sillitoe on behalf of the Wellcome Sanger Institute COVID-19 Surveillance Team |
| EPI_ISL_2664650 | Lighthouse Lab in Glasgow | Wellcome Sanger Institute for the COVID-19 Genomics UK (COG-UK) Consortium | Harper VanSteenhouse, Yumi Kasai, David Gray, Carol Clugston, Anna Dominiczak and Alex Alderton, Roberto Amato, Jeffrey Barrett, Sonia Goncalves, Ewan Harrison, David K. Jackson, Ian Johnston, Dominic Kwiatkowski, Cordelia Langford, John Sillitoe on behalf of the Wellcome Sanger Institute COVID-19 Surveillance Team |
| EPI_ISL_2664651, | Lighthouse Lab in Alderley Park | Wellcome Sanger Institute for the COVID-19 Genomics UK (COG-UK) | Jacquelyn Wynn, Mairead Hyland, The Lighthouse Lab in Alderley Park and Alex Alderton, Roberto Amato, Jeffrey Barrett, Sonia Goncalves, Ewan Harrison, David K. Jackson, Ian Johnston, Dominic Kwiatkowski, Cordelia Langford, John Sillitoe on |

| | | | |
|---|---|---|---|
| EPI_ISL_2664652, EPI_ISL_2664653 | | Consortium | behalf of the Wellcome Sanger Institute COVID-19 Surveillance Team |
| EPI_ISL_2664654 | Lighthouse Laboratory Plymouth | Wellcome Sanger Institute for the COVID-19 Genomics UK (COG-UK) Consortium | Lighthouse Laboratory Plymouth and Alex Alderton, Roberto Amato, Jeffrey Barrett, Sonia Goncalves, Ewan Harrison, David K. Jackson, Ian Johnston, Dominic Kwiatkowski, Cordelia Langford, John Sillitoe on behalf of the Wellcome Sanger Institute COVID-19 Surveillance Team |
| EPI_ISL_2664655 | Lighthouse Lab in Milton Keynes | Wellcome Sanger Institute for the COVID-19 Genomics UK (COG-UK) Consortium | The Lighthouse Lab in Milton Keynes and Alex Alderton, Roberto Amato, Jeffrey Barrett, Sonia Goncalves, Ewan Harrison, David K. Jackson, Ian Johnston, Dominic Kwiatkowski, Cordelia Langford, John Sillitoe on behalf of the Wellcome Sanger Institute COVID-19 Surveillance Team |
| EPI_ISL_2664656, EPI_ISL_2664657, EPI_ISL_2664658 | Lighthouse Laboratory Plymouth | Wellcome Sanger Institute for the COVID-19 Genomics UK (COG-UK) Consortium | Lighthouse Laboratory Plymouth and Alex Alderton, Roberto Amato, Jeffrey Barrett, Sonia Goncalves, Ewan Harrison, David K. Jackson, Ian Johnston, Dominic Kwiatkowski, Cordelia Langford, John Sillitoe on behalf of the Wellcome Sanger Institute COVID-19 Surveillance Team |
| EPI_ISL_2664659 | Lighthouse Lab in Alderley Park | Wellcome Sanger Institute for the COVID-19 Genomics UK (COG-UK) Consortium | Jacquelyn Wynn, Mairead Hyland, The Lighthouse Lab in Alderley Park and Alex Alderton, Roberto Amato, Jeffrey Barrett, Sonia Goncalves, Ewan Harrison, David K. Jackson, Ian Johnston, Dominic Kwiatkowski, Cordelia Langford, John Sillitoe on behalf of the Wellcome Sanger Institute COVID-19 Surveillance Team |
| EPI_ISL_2664660 | Lighthouse Laboratory Plymouth | Wellcome Sanger Institute for the COVID-19 Genomics UK (COG-UK) Consortium | Lighthouse Laboratory Plymouth and Alex Alderton, Roberto Amato, Jeffrey Barrett, Sonia Goncalves, Ewan Harrison, David K. Jackson, Ian Johnston, Dominic Kwiatkowski, Cordelia Langford, John Sillitoe on behalf of the Wellcome Sanger Institute COVID-19 Surveillance Team |
| EPI_ISL_2664661 | Lighthouse Lab in Milton Keynes | Wellcome Sanger Institute for the COVID-19 Genomics UK (COG-UK) Consortium | The Lighthouse Lab in Milton Keynes and Alex Alderton, Roberto Amato, Jeffrey Barrett, Sonia Goncalves, Ewan Harrison, David K. Jackson, Ian Johnston, Dominic Kwiatkowski, Cordelia Langford, John Sillitoe on behalf of the Wellcome Sanger Institute COVID-19 Surveillance Team |
| EPI_ISL_2664662 | Lighthouse Lab in Glasgow | Wellcome Sanger Institute for the COVID-19 Genomics UK (COG-UK) Consortium | Harper VanSteenhouse, Yumi Kasai, David Gray, Carol Clugston, Anna Dominiczak and Alex Alderton, Roberto Amato, Jeffrey Barrett, Sonia Goncalves, Ewan Harrison, David K. Jackson, Ian Johnston, Dominic Kwiatkowski, Cordelia Langford, John Sillitoe on behalf of the Wellcome Sanger Institute COVID-19 Surveillance Team |
| EPI_ISL_2664663 | Lighthouse Lab in Milton Keynes | Wellcome Sanger Institute for the COVID-19 Genomics UK (COG-UK) Consortium | The Lighthouse Lab in Milton Keynes and Alex Alderton, Roberto Amato, Jeffrey Barrett, Sonia Goncalves, Ewan Harrison, David K. Jackson, Ian Johnston, Dominic Kwiatkowski, Cordelia Langford, John Sillitoe on behalf of the Wellcome Sanger Institute COVID-19 Surveillance Team |
| EPI_ISL_2664665 | Lighthouse Lab in Alderley Park | Wellcome Sanger Institute for the COVID-19 Genomics UK (COG-UK) Consortium | Jacquelyn Wynn, Mairead Hyland, The Lighthouse Lab in Alderley Park and Alex Alderton, Roberto Amato, Jeffrey Barrett, Sonia Goncalves, Ewan Harrison, David K. Jackson, Ian Johnston, Dominic Kwiatkowski, Cordelia Langford, John Sillitoe on behalf of the Wellcome Sanger Institute COVID-19 Surveillance Team |
| EPI_ISL_2664666, EPI_ISL_2664667 | Lighthouse Lab in Milton Keynes | Wellcome Sanger Institute for the COVID-19 Genomics UK (COG-UK) Consortium | The Lighthouse Lab in Milton Keynes and Alex Alderton, Roberto Amato, Jeffrey Barrett, Sonia Goncalves, Ewan Harrison, David K. Jackson, Ian Johnston, Dominic Kwiatkowski, Cordelia Langford, John Sillitoe on behalf of the Wellcome Sanger Institute COVID-19 Surveillance Team |
| EPI_ISL_2664668 | Lighthouse Lab in Alderley Park | Wellcome Sanger Institute for the COVID-19 Genomics UK (COG-UK) Consortium | Jacquelyn Wynn, Mairead Hyland, The Lighthouse Lab in Alderley Park and Alex Alderton, Roberto Amato, Jeffrey Barrett, Sonia Goncalves, Ewan Harrison, David K. Jackson, Ian Johnston, Dominic Kwiatkowski, Cordelia Langford, John Sillitoe on behalf of the Wellcome Sanger Institute COVID-19 Surveillance Team |
| EPI_ISL_2664669 | Lighthouse Lab in Glasgow | Wellcome Sanger Institute for the COVID-19 Genomics UK (COG-UK) Consortium | Harper VanSteenhouse, Yumi Kasai, David Gray, Carol Clugston, Anna Dominiczak and Alex Alderton, Roberto Amato, Jeffrey Barrett, Sonia Goncalves, Ewan Harrison, David K. Jackson, Ian Johnston, Dominic Kwiatkowski, Cordelia Langford, John Sillitoe on behalf of the Wellcome Sanger Institute COVID-19 Surveillance Team |
| EPI_ISL_2664670 | Lighthouse Laboratory Plymouth | Wellcome Sanger Institute for the COVID-19 Genomics UK (COG-UK) Consortium | Lighthouse Laboratory Plymouth and Alex Alderton, Roberto Amato, Jeffrey Barrett, Sonia Goncalves, Ewan Harrison, David K. Jackson, Ian Johnston, Dominic Kwiatkowski, Cordelia Langford, John Sillitoe on behalf of the Wellcome Sanger Institute COVID-19 Surveillance Team |
| EPI_ISL_2664671 | Lighthouse Lab in Milton Keynes | Wellcome Sanger Institute for the COVID-19 Genomics UK (COG-UK) Consortium | The Lighthouse Lab in Milton Keynes and Alex Alderton, Roberto Amato, Jeffrey Barrett, Sonia Goncalves, Ewan Harrison, David K. Jackson, Ian Johnston, Dominic Kwiatkowski, Cordelia Langford, John Sillitoe on behalf of the Wellcome Sanger Institute COVID-19 Surveillance Team |
| EPI_ISL_2664672 | Lighthouse Laboratory Plymouth | Wellcome Sanger Institute for the COVID-19 Genomics UK (COG-UK) Consortium | Lighthouse Laboratory Plymouth and Alex Alderton, Roberto Amato, Jeffrey Barrett, Sonia Goncalves, Ewan Harrison, David K. Jackson, Ian Johnston, Dominic Kwiatkowski, Cordelia Langford, John Sillitoe on behalf of the Wellcome Sanger Institute COVID-19 Surveillance Team |
| EPI_ISL_2664673, EPI_ISL_2664674, EPI_ISL_2664675 | Lighthouse Lab in Milton Keynes | Wellcome Sanger Institute for the COVID-19 Genomics UK (COG-UK) Consortium | The Lighthouse Lab in Milton Keynes and Alex Alderton, Roberto Amato, Jeffrey Barrett, Sonia Goncalves, Ewan Harrison, David K. Jackson, Ian Johnston, Dominic Kwiatkowski, Cordelia Langford, John Sillitoe on behalf of the Wellcome Sanger Institute COVID-19 Surveillance Team |
| EPI_ISL_2664676, EPI_ISL_2664677, EPI_ISL_2664678 | Lighthouse Laboratory Plymouth | Wellcome Sanger Institute for the COVID-19 Genomics UK (COG-UK) Consortium | Lighthouse Laboratory Plymouth and Alex Alderton, Roberto Amato, Jeffrey Barrett, Sonia Goncalves, Ewan Harrison, David K. Jackson, Ian Johnston, Dominic Kwiatkowski, Cordelia Langford, John Sillitoe on behalf of the Wellcome Sanger Institute COVID-19 Surveillance Team |
| EPI_ISL_2664679 | Lighthouse Lab in Milton Keynes | Wellcome Sanger Institute for the COVID-19 Genomics UK (COG-UK) Consortium | The Lighthouse Lab in Milton Keynes and Alex Alderton, Roberto Amato, Jeffrey Barrett, Sonia Goncalves, Ewan Harrison, David K. Jackson, Ian Johnston, Dominic Kwiatkowski, Cordelia Langford, John Sillitoe on behalf of the Wellcome Sanger Institute COVID-19 Surveillance Team |
| EPI_ISL_2664680 | Lighthouse Lab in Alderley Park | Wellcome Sanger Institute for the COVID-19 Genomics UK (COG-UK) Consortium | Jacquelyn Wynn, Mairead Hyland, The Lighthouse Lab in Alderley Park and Alex Alderton, Roberto Amato, Jeffrey Barrett, Sonia Goncalves, Ewan Harrison, David K. Jackson, Ian Johnston, Dominic Kwiatkowski, Cordelia Langford, John Sillitoe on behalf of the Wellcome Sanger Institute COVID-19 Surveillance Team |
| EPI_ISL_2664681 | Lighthouse Lab in Glasgow | Wellcome Sanger Institute for the COVID-19 Genomics UK (COG-UK) Consortium | Harper VanSteenhouse, Yumi Kasai, David Gray, Carol Clugston, Anna Dominiczak and Alex Alderton, Roberto Amato, Jeffrey Barrett, Sonia Goncalves, Ewan Harrison, David K. Jackson, Ian Johnston, Dominic Kwiatkowski, Cordelia Langford, John Sillitoe on behalf of the Wellcome Sanger Institute COVID-19 Surveillance Team |
| EPI_ISL_2664682 | Lighthouse Laboratory Plymouth | Wellcome Sanger Institute for the COVID-19 Genomics UK (COG-UK) Consortium | Lighthouse Laboratory Plymouth and Alex Alderton, Roberto Amato, Jeffrey Barrett, Sonia Goncalves, Ewan Harrison, David K. Jackson, Ian Johnston, Dominic Kwiatkowski, Cordelia Langford, John Sillitoe on behalf of the Wellcome Sanger Institute COVID-19 Surveillance Team |
| EPI_ISL_2664683, EPI_ISL_2664684 | Lighthouse Lab in Glasgow | Wellcome Sanger Institute for the COVID-19 Genomics UK (COG-UK) Consortium | Harper VanSteenhouse, Yumi Kasai, David Gray, Carol Clugston, Anna Dominiczak and Alex Alderton, Roberto Amato, Jeffrey Barrett, Sonia Goncalves, Ewan Harrison, David K. Jackson, Ian Johnston, Dominic Kwiatkowski, Cordelia Langford, John Sillitoe on behalf of the Wellcome Sanger Institute COVID-19 Surveillance Team |
| EPI_ISL_2664685 | Lighthouse Lab in Alderley Park | Wellcome Sanger Institute for the COVID-19 Genomics UK (COG-UK) Consortium | Jacquelyn Wynn, Mairead Hyland, The Lighthouse Lab in Alderley Park and Alex Alderton, Roberto Amato, Jeffrey Barrett, Sonia Goncalves, Ewan Harrison, David K. Jackson, Ian Johnston, Dominic Kwiatkowski, Cordelia Langford, John Sillitoe on behalf of the Wellcome Sanger Institute COVID-19 Surveillance Team |
| EPI_ISL_2664686 | Lighthouse Lab in Glasgow | Wellcome Sanger Institute for the COVID-19 Genomics UK (COG-UK) Consortium | Harper VanSteenhouse, Yumi Kasai, David Gray, Carol Clugston, Anna Dominiczak and Alex Alderton, Roberto Amato, Jeffrey Barrett, Sonia Goncalves, Ewan Harrison, David K. Jackson, Ian Johnston, Dominic Kwiatkowski, Cordelia Langford, John Sillitoe on behalf of the Wellcome Sanger Institute COVID-19 Surveillance Team |
| EPI_ISL_2664687 | Lighthouse Lab in Milton Keynes | Wellcome Sanger Institute for the COVID-19 Genomics UK (COG-UK) Consortium | The Lighthouse Lab in Milton Keynes and Alex Alderton, Roberto Amato, Jeffrey Barrett, Sonia Goncalves, Ewan Harrison, David K. Jackson, Ian Johnston, Dominic Kwiatkowski, Cordelia Langford, John Sillitoe on behalf of the Wellcome Sanger Institute COVID-19 Surveillance Team |
| EPI_ISL_2664688 | Lighthouse Laboratory Plymouth | Wellcome Sanger Institute for the COVID-19 Genomics UK (COG-UK) Consortium | Lighthouse Laboratory Plymouth and Alex Alderton, Roberto Amato, Jeffrey Barrett, Sonia Goncalves, Ewan Harrison, David K. Jackson, Ian Johnston, Dominic Kwiatkowski, Cordelia Langford, John Sillitoe on behalf of the Wellcome Sanger Institute COVID-19 Surveillance Team |
| EPI_ISL_2664689 | Lighthouse Lab in Glasgow | Wellcome Sanger Institute for the COVID-19 Genomics UK (COG-UK) Consortium | Harper VanSteenhouse, Yumi Kasai, David Gray, Carol Clugston, Anna Dominiczak and Alex Alderton, Roberto Amato, Jeffrey Barrett, Sonia Goncalves, Ewan Harrison, David K. Jackson, Ian Johnston, Dominic Kwiatkowski, Cordelia Langford, John Sillitoe on behalf of the Wellcome Sanger Institute COVID-19 Surveillance Team |
| EPI_ISL_2664690 | Lighthouse Lab in Milton Keynes | Wellcome Sanger Institute for the COVID-19 Genomics UK (COG-UK) Consortium | The Lighthouse Lab in Milton Keynes and Alex Alderton, Roberto Amato, Jeffrey Barrett, Sonia Goncalves, Ewan Harrison, David K. Jackson, Ian Johnston, Dominic Kwiatkowski, Cordelia Langford, John Sillitoe on behalf of the Wellcome Sanger Institute COVID-19 Surveillance Team |
| EPI_ISL_2664691 | Lighthouse Lab in Alderley Park | Wellcome Sanger Institute for the COVID-19 Genomics UK (COG-UK) Consortium | Jacquelyn Wynn, Mairead Hyland, The Lighthouse Lab in Alderley Park and Alex Alderton, Roberto Amato, Jeffrey Barrett, Sonia Goncalves, Ewan Harrison, David K. Jackson, Ian Johnston, Dominic Kwiatkowski, Cordelia Langford, John Sillitoe on behalf of the Wellcome Sanger Institute COVID-19 Surveillance Team |
| EPI_ISL_2664692 | Lighthouse Lab in Milton Keynes | Wellcome Sanger Institute for the COVID-19 Genomics UK (COG-UK) Consortium | The Lighthouse Lab in Milton Keynes and Alex Alderton, Roberto Amato, Jeffrey Barrett, Sonia Goncalves, Ewan Harrison, David K. Jackson, Ian Johnston, Dominic Kwiatkowski, Cordelia Langford, John Sillitoe on behalf of the Wellcome Sanger Institute COVID-19 Surveillance Team |
| EPI_ISL_2664693 | Lighthouse Lab in Alderley Park | Wellcome Sanger Institute for the COVID-19 Genomics UK (COG-UK) Consortium | Jacquelyn Wynn, Mairead Hyland, The Lighthouse Lab in Alderley Park and Alex Alderton, Roberto Amato, Jeffrey Barrett, Sonia Goncalves, Ewan Harrison, David K. Jackson, Ian Johnston, Dominic Kwiatkowski, Cordelia Langford, John Sillitoe on behalf of the Wellcome Sanger Institute COVID-19 Surveillance Team |
| EPI_ISL_2664694 | Lighthouse Lab in Glasgow | Wellcome Sanger Institute for the COVID-19 Genomics UK (COG-UK) Consortium | Harper VanSteenhouse, Yumi Kasai, David Gray, Carol Clugston, Anna Dominiczak and Alex Alderton, Roberto Amato, Jeffrey Barrett, Sonia Goncalves, Ewan Harrison, David K. Jackson, Ian Johnston, Dominic Kwiatkowski, Cordelia Langford, John Sillitoe on behalf of the Wellcome Sanger Institute COVID-19 Surveillance Team |
| EPI_ISL_2664695 | Lighthouse Laboratory Plymouth | Wellcome Sanger Institute for the COVID-19 Genomics UK (COG-UK) Consortium | Lighthouse Laboratory Plymouth and Alex Alderton, Roberto Amato, Jeffrey Barrett, Sonia Goncalves, Ewan Harrison, David K. Jackson, Ian Johnston, Dominic Kwiatkowski, Cordelia Langford, John Sillitoe on behalf of the Wellcome Sanger Institute COVID-19 Surveillance Team |
| EPI_ISL_2664696, EPI_ISL_2664697, EPI_ISL_2664698, EPI_ISL_2664699 | Lighthouse Lab in Milton Keynes | Wellcome Sanger Institute for the COVID-19 Genomics UK (COG-UK) Consortium | The Lighthouse Lab in Milton Keynes and Alex Alderton, Roberto Amato, Jeffrey Barrett, Sonia Goncalves, Ewan Harrison, David K. Jackson, Ian Johnston, Dominic Kwiatkowski, Cordelia Langford, John Sillitoe on behalf of the Wellcome Sanger Institute COVID-19 Surveillance Team |
| EPI_ISL_2664700 | Health Services Laboratories | Wellcome Sanger Institute for the COVID-19 Genomics UK (COG-UK) Consortium | Health Services Laboratories and Alex Alderton, Roberto Amato, Jeffrey Barrett, Sonia Goncalves, Ewan Harrison, David K. Jackson, Ian Johnston, Dominic Kwiatkowski, Cordelia Langford, John Sillitoe on behalf of the Wellcome Sanger Institute COVID-19 Surveillance Team |
| EPI_ISL_2664701, EPI_ISL_2664702 | Lighthouse Laboratory Plymouth | Wellcome Sanger Institute for the COVID-19 Genomics UK (COG-UK) Consortium | Lighthouse Laboratory Plymouth and Alex Alderton, Roberto Amato, Jeffrey Barrett, Sonia Goncalves, Ewan Harrison, David K. Jackson, Ian Johnston, Dominic Kwiatkowski, Cordelia Langford, John Sillitoe on behalf of the Wellcome Sanger Institute COVID-19 Surveillance Team |
| EPI_ISL_2664703 | Lighthouse Lab in Milton Keynes | Wellcome Sanger Institute for the COVID-19 Genomics UK (COG-UK) Consortium | The Lighthouse Lab in Milton Keynes and Alex Alderton, Roberto Amato, Jeffrey Barrett, Sonia Goncalves, Ewan Harrison, David K. Jackson, Ian Johnston, Dominic Kwiatkowski, Cordelia Langford, John Sillitoe on behalf of the Wellcome Sanger Institute COVID-19 Surveillance Team |
| EPI_ISL_2664704 | Lighthouse Laboratory Plymouth | Wellcome Sanger Institute for the COVID-19 Genomics UK (COG-UK) Consortium | Lighthouse Laboratory Plymouth and Alex Alderton, Roberto Amato, Jeffrey Barrett, Sonia Goncalves, Ewan Harrison, David K. Jackson, Ian Johnston, Dominic Kwiatkowski, Cordelia Langford, John Sillitoe on behalf of the Wellcome Sanger Institute COVID-19 Surveillance Team |
| EPI_ISL_2664705, EPI_ISL_2664706 | Lighthouse Lab in Alderley Park | Wellcome Sanger Institute for the COVID-19 Genomics UK (COG-UK) Consortium | Jacquelyn Wynn, Mairead Hyland, The Lighthouse Lab in Alderley Park and Alex Alderton, Roberto Amato, Jeffrey Barrett, Sonia Goncalves, Ewan Harrison, David K. Jackson, Ian Johnston, Dominic Kwiatkowski, Cordelia Langford, John Sillitoe on behalf of the Wellcome Sanger Institute COVID-19 Surveillance Team |
| EPI_ISL_2664707 | Lighthouse Laboratory Plymouth | Wellcome Sanger Institute for the COVID-19 Genomics UK (COG-UK) Consortium | Lighthouse Laboratory Plymouth and Alex Alderton, Roberto Amato, Jeffrey Barrett, Sonia Goncalves, Ewan Harrison, David K. Jackson, Ian Johnston, Dominic Kwiatkowski, Cordelia Langford, John Sillitoe on behalf of the Wellcome Sanger Institute COVID-19 Surveillance Team |
| EPI_ISL_2664708, EPI_ISL_2664709, EPI_ISL_2664710, EPI_ISL_2664711, EPI_ISL_2664712 | Lighthouse Lab in Alderley Park | Wellcome Sanger Institute for the COVID-19 Genomics UK (COG-UK) Consortium | Jacquelyn Wynn, Mairead Hyland, The Lighthouse Lab in Alderley Park and Alex Alderton, Roberto Amato, Jeffrey Barrett, Sonia Goncalves, Ewan Harrison, David K. Jackson, Ian Johnston, Dominic Kwiatkowski, Cordelia Langford, John Sillitoe on behalf of the Wellcome Sanger Institute COVID-19 Surveillance Team |

| | | | |
|---|---|---|---|
| EPI_ISL_2664713, EPI_ISL_2664714 | Lighthouse Lab in Milton Keynes | Wellcome Sanger Institute for the COVID-19 Genomics UK (COG-UK) Consortium | The Lighthouse Lab in Milton Keynes and Alex Alderton, Roberto Amato, Jeffrey Barrett, Sonia Goncalves, Ewan Harrison, David K. Jackson, Ian Johnston, Dominic Kwiatkowski, Cordelia Langford, John Sillitoe on behalf of the Wellcome Sanger Institute COVID-19 Surveillance Team |
| EPI_ISL_2664715 | Lighthouse Lab in Alderley Park | Wellcome Sanger Institute for the COVID-19 Genomics UK (COG-UK) Consortium | Jacquelyn Wynn, Mairead Hyland, The Lighthouse Lab in Alderley Park and Alex Alderton, Roberto Amato, Jeffrey Barrett, Sonia Goncalves, Ewan Harrison, David K. Jackson, Ian Johnston, Dominic Kwiatkowski, Cordelia Langford, John Sillitoe on behalf of the Wellcome Sanger Institute COVID-19 Surveillance Team |
| EPI_ISL_2664716 | Lighthouse Laboratory Plymouth | Wellcome Sanger Institute for the COVID-19 Genomics UK (COG-UK) Consortium | Lighthouse Laboratory Plymouth and Alex Alderton, Roberto Amato, Jeffrey Barrett, Sonia Goncalves, Ewan Harrison, David K. Jackson, Ian Johnston, Dominic Kwiatkowski, Cordelia Langford, John Sillitoe on behalf of the Wellcome Sanger Institute COVID-19 Surveillance Team |
| EPI_ISL_2664717, EPI_ISL_2664718 | Lighthouse Lab in Alderley Park | Wellcome Sanger Institute for the COVID-19 Genomics UK (COG-UK) Consortium | Jacquelyn Wynn, Mairead Hyland, The Lighthouse Lab in Alderley Park and Alex Alderton, Roberto Amato, Jeffrey Barrett, Sonia Goncalves, Ewan Harrison, David K. Jackson, Ian Johnston, Dominic Kwiatkowski, Cordelia Langford, John Sillitoe on behalf of the Wellcome Sanger Institute COVID-19 Surveillance Team |
| EPI_ISL_2664719 | Lighthouse Lab in Glasgow | Wellcome Sanger Institute for the COVID-19 Genomics UK (COG-UK) Consortium | Harper VanSteenhouse, Yumi Kasai, David Gray, Carol Clugston, Anna Dominiczak and Alex Alderton, Roberto Amato, Jeffrey Barrett, Sonia Goncalves, Ewan Harrison, David K. Jackson, Ian Johnston, Dominic Kwiatkowski, Cordelia Langford, John Sillitoe on behalf of the Wellcome Sanger Institute COVID-19 Surveillance Team |
| EPI_ISL_2664720 | Lighthouse Lab in Milton Keynes | Wellcome Sanger Institute for the COVID-19 Genomics UK (COG-UK) Consortium | The Lighthouse Lab in Milton Keynes and Alex Alderton, Roberto Amato, Jeffrey Barrett, Sonia Goncalves, Ewan Harrison, David K. Jackson, Ian Johnston, Dominic Kwiatkowski, Cordelia Langford, John Sillitoe on behalf of the Wellcome Sanger Institute COVID-19 Surveillance Team |
| EPI_ISL_2664721, EPI_ISL_2664722, EPI_ISL_2664723 | Lighthouse Lab in Alderley Park | Wellcome Sanger Institute for the COVID-19 Genomics UK (COG-UK) Consortium | Jacquelyn Wynn, Mairead Hyland, The Lighthouse Lab in Alderley Park and Alex Alderton, Roberto Amato, Jeffrey Barrett, Sonia Goncalves, Ewan Harrison, David K. Jackson, Ian Johnston, Dominic Kwiatkowski, Cordelia Langford, John Sillitoe on behalf of the Wellcome Sanger Institute COVID-19 Surveillance Team |
| EPI_ISL_2664724 | Lighthouse Lab in Glasgow | Wellcome Sanger Institute for the COVID-19 Genomics UK (COG-UK) Consortium | Harper VanSteenhouse, Yumi Kasai, David Gray, Carol Clugston, Anna Dominiczak and Alex Alderton, Roberto Amato, Jeffrey Barrett, Sonia Goncalves, Ewan Harrison, David K. Jackson, Ian Johnston, Dominic Kwiatkowski, Cordelia Langford, John Sillitoe on behalf of the Wellcome Sanger Institute COVID-19 Surveillance Team |
| EPI_ISL_2664725, EPI_ISL_2664726 | Lighthouse Lab in Milton Keynes | Wellcome Sanger Institute for the COVID-19 Genomics UK (COG-UK) Consortium | The Lighthouse Lab in Milton Keynes and Alex Alderton, Roberto Amato, Jeffrey Barrett, Sonia Goncalves, Ewan Harrison, David K. Jackson, Ian Johnston, Dominic Kwiatkowski, Cordelia Langford, John Sillitoe on behalf of the Wellcome Sanger Institute COVID-19 Surveillance Team |
| EPI_ISL_2664727 | Lighthouse Laboratory Plymouth | Wellcome Sanger Institute for the COVID-19 Genomics UK (COG-UK) Consortium | Lighthouse Laboratory Plymouth and Alex Alderton, Roberto Amato, Jeffrey Barrett, Sonia Goncalves, Ewan Harrison, David K. Jackson, Ian Johnston, Dominic Kwiatkowski, Cordelia Langford, John Sillitoe on behalf of the Wellcome Sanger Institute COVID-19 Surveillance Team |
| EPI_ISL_2664728 | Lighthouse Lab in Milton Keynes | Wellcome Sanger Institute for the COVID-19 Genomics UK (COG-UK) Consortium | The Lighthouse Lab in Milton Keynes and Alex Alderton, Roberto Amato, Jeffrey Barrett, Sonia Goncalves, Ewan Harrison, David K. Jackson, Ian Johnston, Dominic Kwiatkowski, Cordelia Langford, John Sillitoe on behalf of the Wellcome Sanger Institute COVID-19 Surveillance Team |
| EPI_ISL_2664729 | Lighthouse Laboratory Plymouth | Wellcome Sanger Institute for the COVID-19 Genomics UK (COG-UK) Consortium | Lighthouse Laboratory Plymouth and Alex Alderton, Roberto Amato, Jeffrey Barrett, Sonia Goncalves, Ewan Harrison, David K. Jackson, Ian Johnston, Dominic Kwiatkowski, Cordelia Langford, John Sillitoe on behalf of the Wellcome Sanger Institute COVID-19 Surveillance Team |
| EPI_ISL_2664730, EPI_ISL_2664731 | Lighthouse Lab in Alderley Park | Wellcome Sanger Institute for the COVID-19 Genomics UK (COG-UK) Consortium | Jacquelyn Wynn, Mairead Hyland, The Lighthouse Lab in Alderley Park and Alex Alderton, Roberto Amato, Jeffrey Barrett, Sonia Goncalves, Ewan Harrison, David K. Jackson, Ian Johnston, Dominic Kwiatkowski, Cordelia Langford, John Sillitoe on behalf of the Wellcome Sanger Institute COVID-19 Surveillance Team |
| EPI_ISL_2664732 | Lighthouse Laboratory Plymouth | Wellcome Sanger Institute for the COVID-19 Genomics UK (COG-UK) Consortium | Lighthouse Laboratory Plymouth and Alex Alderton, Roberto Amato, Jeffrey Barrett, Sonia Goncalves, Ewan Harrison, David K. Jackson, Ian Johnston, Dominic Kwiatkowski, Cordelia Langford, John Sillitoe on behalf of the Wellcome Sanger Institute COVID-19 Surveillance Team |
| EPI_ISL_2664733 | Lighthouse Lab in Glasgow | Wellcome Sanger Institute for the COVID-19 Genomics UK (COG-UK) Consortium | Harper VanSteenhouse, Yumi Kasai, David Gray, Carol Clugston, Anna Dominiczak and Alex Alderton, Roberto Amato, Jeffrey Barrett, Sonia Goncalves, Ewan Harrison, David K. Jackson, Ian Johnston, Dominic Kwiatkowski, Cordelia Langford, John Sillitoe on behalf of the Wellcome Sanger Institute COVID-19 Surveillance Team |
| EPI_ISL_2664734, EPI_ISL_2664735, EPI_ISL_2664736 | Lighthouse Lab in Milton Keynes | Wellcome Sanger Institute for the COVID-19 Genomics UK (COG-UK) Consortium | The Lighthouse Lab in Milton Keynes and Alex Alderton, Roberto Amato, Jeffrey Barrett, Sonia Goncalves, Ewan Harrison, David K. Jackson, Ian Johnston, Dominic Kwiatkowski, Cordelia Langford, John Sillitoe on behalf of the Wellcome Sanger Institute COVID-19 Surveillance Team |
| EPI_ISL_2664737 | Lighthouse Laboratory Plymouth | Wellcome Sanger Institute for the COVID-19 Genomics UK (COG-UK) Consortium | Lighthouse Laboratory Plymouth and Alex Alderton, Roberto Amato, Jeffrey Barrett, Sonia Goncalves, Ewan Harrison, David K. Jackson, Ian Johnston, Dominic Kwiatkowski, Cordelia Langford, John Sillitoe on behalf of the Wellcome Sanger Institute COVID-19 Surveillance Team |
| EPI_ISL_2664738, EPI_ISL_2664739 | Lighthouse Lab in Alderley Park | Wellcome Sanger Institute for the COVID-19 Genomics UK (COG-UK) Consortium | Jacquelyn Wynn, Mairead Hyland, The Lighthouse Lab in Alderley Park and Alex Alderton, Roberto Amato, Jeffrey Barrett, Sonia Goncalves, Ewan Harrison, David K. Jackson, Ian Johnston, Dominic Kwiatkowski, Cordelia Langford, John Sillitoe on behalf of the Wellcome Sanger Institute COVID-19 Surveillance Team |
| EPI_ISL_2664740, EPI_ISL_2664741 | Lighthouse Laboratory Plymouth | Wellcome Sanger Institute for the COVID-19 Genomics UK (COG-UK) Consortium | Lighthouse Laboratory Plymouth and Alex Alderton, Roberto Amato, Jeffrey Barrett, Sonia Goncalves, Ewan Harrison, David K. Jackson, Ian Johnston, Dominic Kwiatkowski, Cordelia Langford, John Sillitoe on behalf of the Wellcome Sanger Institute COVID-19 Surveillance Team |
| EPI_ISL_2664742 | Lighthouse Lab in Alderley Park | Wellcome Sanger Institute for the COVID-19 Genomics UK (COG-UK) Consortium | Jacquelyn Wynn, Mairead Hyland, The Lighthouse Lab in Alderley Park and Alex Alderton, Roberto Amato, Jeffrey Barrett, Sonia Goncalves, Ewan Harrison, David K. Jackson, Ian Johnston, Dominic Kwiatkowski, Cordelia Langford, John Sillitoe on behalf of the Wellcome Sanger Institute COVID-19 Surveillance Team |
| EPI_ISL_2664743 | Lighthouse Laboratory Plymouth | Wellcome Sanger Institute for the COVID-19 Genomics UK (COG-UK) Consortium | Lighthouse Laboratory Plymouth and Alex Alderton, Roberto Amato, Jeffrey Barrett, Sonia Goncalves, Ewan Harrison, David K. Jackson, Ian Johnston, Dominic Kwiatkowski, Cordelia Langford, John Sillitoe on behalf of the Wellcome Sanger Institute COVID-19 Surveillance Team |
| EPI_ISL_2664744 | Lighthouse Lab in Alderley Park | Wellcome Sanger Institute for the COVID-19 Genomics UK (COG-UK) Consortium | Jacquelyn Wynn, Mairead Hyland, The Lighthouse Lab in Alderley Park and Alex Alderton, Roberto Amato, Jeffrey Barrett, Sonia Goncalves, Ewan Harrison, David K. Jackson, Ian Johnston, Dominic Kwiatkowski, Cordelia Langford, John Sillitoe on behalf of the Wellcome Sanger Institute COVID-19 Surveillance Team |
| EPI_ISL_2664746, EPI_ISL_2664747, EPI_ISL_2664748 | Lighthouse Lab in Milton Keynes | Wellcome Sanger Institute for the COVID-19 Genomics UK (COG-UK) Consortium | The Lighthouse Lab in Milton Keynes and Alex Alderton, Roberto Amato, Jeffrey Barrett, Sonia Goncalves, Ewan Harrison, David K. Jackson, Ian Johnston, Dominic Kwiatkowski, Cordelia Langford, John Sillitoe on behalf of the Wellcome Sanger Institute COVID-19 Surveillance Team |
| EPI_ISL_2664749, EPI_ISL_2664751 | Lighthouse Laboratory Plymouth | Wellcome Sanger Institute for the COVID-19 Genomics UK (COG-UK) Consortium | Lighthouse Laboratory Plymouth and Alex Alderton, Roberto Amato, Jeffrey Barrett, Sonia Goncalves, Ewan Harrison, David K. Jackson, Ian Johnston, Dominic Kwiatkowski, Cordelia Langford, John Sillitoe on behalf of the Wellcome Sanger Institute COVID-19 Surveillance Team |
| EPI_ISL_2664752 | Lighthouse Lab in Glasgow | Wellcome Sanger Institute for the COVID-19 Genomics UK (COG-UK) Consortium | Harper VanSteenhouse, Yumi Kasai, David Gray, Carol Clugston, Anna Dominiczak and Alex Alderton, Roberto Amato, Jeffrey Barrett, Sonia Goncalves, Ewan Harrison, David K. Jackson, Ian Johnston, Dominic Kwiatkowski, Cordelia Langford, John Sillitoe on behalf of the Wellcome Sanger Institute COVID-19 Surveillance Team |
| EPI_ISL_2664753 | Lighthouse Lab in Alderley Park | Wellcome Sanger Institute for the COVID-19 Genomics UK (COG-UK) Consortium | Jacquelyn Wynn, Mairead Hyland, The Lighthouse Lab in Alderley Park and Alex Alderton, Roberto Amato, Jeffrey Barrett, Sonia Goncalves, Ewan Harrison, David K. Jackson, Ian Johnston, Dominic Kwiatkowski, Cordelia Langford, John Sillitoe on behalf of the Wellcome Sanger Institute COVID-19 Surveillance Team |
| EPI_ISL_2664754 | Lighthouse Laboratory Plymouth | Wellcome Sanger Institute for the COVID-19 Genomics UK (COG-UK) Consortium | Lighthouse Laboratory Plymouth and Alex Alderton, Roberto Amato, Jeffrey Barrett, Sonia Goncalves, Ewan Harrison, David K. Jackson, Ian Johnston, Dominic Kwiatkowski, Cordelia Langford, John Sillitoe on behalf of the Wellcome Sanger Institute COVID-19 Surveillance Team |
| EPI_ISL_2664755 | Lighthouse Lab in Milton Keynes | Wellcome Sanger Institute for the COVID-19 Genomics UK (COG-UK) Consortium | The Lighthouse Lab in Milton Keynes and Alex Alderton, Roberto Amato, Jeffrey Barrett, Sonia Goncalves, Ewan Harrison, David K. Jackson, Ian Johnston, Dominic Kwiatkowski, Cordelia Langford, John Sillitoe on behalf of the Wellcome Sanger Institute COVID-19 Surveillance Team |
| EPI_ISL_2664756 | Lighthouse Lab in Alderley Park | Wellcome Sanger Institute for the COVID-19 Genomics UK (COG-UK) Consortium | Jacquelyn Wynn, Mairead Hyland, The Lighthouse Lab in Alderley Park and Alex Alderton, Roberto Amato, Jeffrey Barrett, Sonia Goncalves, Ewan Harrison, David K. Jackson, Ian Johnston, Dominic Kwiatkowski, Cordelia Langford, John Sillitoe on behalf of the Wellcome Sanger Institute COVID-19 Surveillance Team |
| EPI_ISL_2664757, EPI_ISL_2664758 | Lighthouse Lab in Milton Keynes | Wellcome Sanger Institute for the COVID-19 Genomics UK (COG-UK) Consortium | The Lighthouse Lab in Milton Keynes and Alex Alderton, Roberto Amato, Jeffrey Barrett, Sonia Goncalves, Ewan Harrison, David K. Jackson, Ian Johnston, Dominic Kwiatkowski, Cordelia Langford, John Sillitoe on behalf of the Wellcome Sanger Institute COVID-19 Surveillance Team |
| EPI_ISL_2664759, EPI_ISL_2664760, EPI_ISL_2664761 | Lighthouse Lab in Alderley Park | Wellcome Sanger Institute for the COVID-19 Genomics UK (COG-UK) Consortium | Jacquelyn Wynn, Mairead Hyland, The Lighthouse Lab in Alderley Park and Alex Alderton, Roberto Amato, Jeffrey Barrett, Sonia Goncalves, Ewan Harrison, David K. Jackson, Ian Johnston, Dominic Kwiatkowski, Cordelia Langford, John Sillitoe on behalf of the Wellcome Sanger Institute COVID-19 Surveillance Team |
| EPI_ISL_2664762 | Lighthouse Lab in Milton Keynes | Wellcome Sanger Institute for the COVID-19 Genomics UK (COG-UK) Consortium | The Lighthouse Lab in Milton Keynes and Alex Alderton, Roberto Amato, Jeffrey Barrett, Sonia Goncalves, Ewan Harrison, David K. Jackson, Ian Johnston, Dominic Kwiatkowski, Cordelia Langford, John Sillitoe on behalf of the Wellcome Sanger Institute COVID-19 Surveillance Team |
| EPI_ISL_2664763 | Lighthouse Lab in Alderley Park | Wellcome Sanger Institute for the COVID-19 Genomics UK (COG-UK) Consortium | Jacquelyn Wynn, Mairead Hyland, The Lighthouse Lab in Alderley Park and Alex Alderton, Roberto Amato, Jeffrey Barrett, Sonia Goncalves, Ewan Harrison, David K. Jackson, Ian Johnston, Dominic Kwiatkowski, Cordelia Langford, John Sillitoe on behalf of the Wellcome Sanger Institute COVID-19 Surveillance Team |
| EPI_ISL_2664764 | Lighthouse Lab in Milton Keynes | Wellcome Sanger Institute for the COVID-19 Genomics UK (COG-UK) Consortium | The Lighthouse Lab in Milton Keynes and Alex Alderton, Roberto Amato, Jeffrey Barrett, Sonia Goncalves, Ewan Harrison, David K. Jackson, Ian Johnston, Dominic Kwiatkowski, Cordelia Langford, John Sillitoe on behalf of the Wellcome Sanger Institute COVID-19 Surveillance Team |
| EPI_ISL_2664765 | Lighthouse Laboratory Plymouth | Wellcome Sanger Institute for the COVID-19 Genomics UK (COG-UK) Consortium | Lighthouse Laboratory Plymouth and Alex Alderton, Roberto Amato, Jeffrey Barrett, Sonia Goncalves, Ewan Harrison, David K. Jackson, Ian Johnston, Dominic Kwiatkowski, Cordelia Langford, John Sillitoe on behalf of the Wellcome Sanger Institute COVID-19 Surveillance Team |
| EPI_ISL_2664766, EPI_ISL_2664767, EPI_ISL_2664768 | Lighthouse Lab in Alderley Park | Wellcome Sanger Institute for the COVID-19 Genomics UK (COG-UK) Consortium | Jacquelyn Wynn, Mairead Hyland, The Lighthouse Lab in Alderley Park and Alex Alderton, Roberto Amato, Jeffrey Barrett, Sonia Goncalves, Ewan Harrison, David K. Jackson, Ian Johnston, Dominic Kwiatkowski, Cordelia Langford, John Sillitoe on behalf of the Wellcome Sanger Institute COVID-19 Surveillance Team |
| EPI_ISL_2664769, EPI_ISL_2664771, EPI_ISL_2664772, EPI_ISL_2664773, EPI_ISL_2664774 | Lighthouse Laboratory Plymouth | Wellcome Sanger Institute for the COVID-19 Genomics UK (COG-UK) Consortium | Lighthouse Laboratory Plymouth and Alex Alderton, Roberto Amato, Jeffrey Barrett, Sonia Goncalves, Ewan Harrison, David K. Jackson, Ian Johnston, Dominic Kwiatkowski, Cordelia Langford, John Sillitoe on behalf of the Wellcome Sanger Institute COVID-19 Surveillance Team |
| EPI_ISL_2664775, EPI_ISL_2664776, EPI_ISL_2664777, EPI_ISL_2664778 | Lighthouse Lab in Alderley Park | Wellcome Sanger Institute for the COVID-19 Genomics UK (COG-UK) Consortium | Jacquelyn Wynn, Mairead Hyland, The Lighthouse Lab in Alderley Park and Alex Alderton, Roberto Amato, Jeffrey Barrett, Sonia Goncalves, Ewan Harrison, David K. Jackson, Ian Johnston, Dominic Kwiatkowski, Cordelia Langford, John Sillitoe on behalf of the Wellcome Sanger Institute COVID-19 Surveillance Team |
| EPI_ISL_2664779 | Lighthouse Laboratory Plymouth | Wellcome Sanger Institute for the COVID-19 Genomics UK (COG-UK) Consortium | Lighthouse Laboratory Plymouth and Alex Alderton, Roberto Amato, Jeffrey Barrett, Sonia Goncalves, Ewan Harrison, David K. Jackson, Ian Johnston, Dominic Kwiatkowski, Cordelia Langford, John Sillitoe on behalf of the Wellcome Sanger Institute COVID-19 Surveillance Team |
| EPI_ISL_2664780, EPI_ISL_2664781 | Lighthouse Lab in Milton Keynes | Wellcome Sanger Institute for the COVID-19 Genomics UK (COG-UK) Consortium | The Lighthouse Lab in Milton Keynes and Alex Alderton, Roberto Amato, Jeffrey Barrett, Sonia Goncalves, Ewan Harrison, David K. Jackson, Ian Johnston, Dominic Kwiatkowski, Cordelia Langford, John Sillitoe on behalf of the Wellcome Sanger Institute COVID-19 Surveillance Team |
| EPI_ISL_2664782 | Lighthouse Lab in Alderley Park | Wellcome Sanger Institute for the COVID-19 Genomics UK (COG-UK) Consortium | Jacquelyn Wynn, Mairead Hyland, The Lighthouse Lab in Alderley Park and Alex Alderton, Roberto Amato, Jeffrey Barrett, Sonia Goncalves, Ewan Harrison, David K. Jackson, Ian Johnston, Dominic Kwiatkowski, Cordelia Langford, John Sillitoe on behalf of the Wellcome Sanger Institute COVID-19 Surveillance Team |

| | | | |
|---|---|---|---|
| EPI_ISL_2664783 | Lighthouse Laboratory Plymouth | Wellcome Sanger Institute for the COVID-19 Genomics UK (COG-UK) Consortium | Lighthouse Laboratory Plymouth and Alex Alderton, Roberto Amato, Jeffrey Barrett, Sonia Goncalves, Ewan Harrison, David K. Jackson, Ian Johnston, Dominic Kwiatkowski, Cordelia Langford, John Sillitoe on behalf of the Wellcome Sanger Institute COVID-19 Surveillance Team |
| EPI_ISL_2664784 | Lighthouse Lab in Milton Keynes | Wellcome Sanger Institute for the COVID-19 Genomics UK (COG-UK) Consortium | The Lighthouse Lab in Milton Keynes and Alex Alderton, Roberto Amato, Jeffrey Barrett, Sonia Goncalves, Ewan Harrison, David K. Jackson, Ian Johnston, Dominic Kwiatkowski, Cordelia Langford, John Sillitoe on behalf of the Wellcome Sanger Institute COVID-19 Surveillance Team |
| EPI_ISL_2664785, EPI_ISL_2664786 | Lighthouse Lab in Alderley Park | Wellcome Sanger Institute for the COVID-19 Genomics UK (COG-UK) Consortium | Jacquelyn Wynn, Mairead Hyland, The Lighthouse Lab in Alderley Park and Alex Alderton, Roberto Amato, Jeffrey Barrett, Sonia Goncalves, Ewan Harrison, David K. Jackson, Ian Johnston, Dominic Kwiatkowski, Cordelia Langford, John Sillitoe on behalf of the Wellcome Sanger Institute COVID-19 Surveillance Team |
| EPI_ISL_2664787 | Lighthouse Laboratory Plymouth | Wellcome Sanger Institute for the COVID-19 Genomics UK (COG-UK) Consortium | Lighthouse Laboratory Plymouth and Alex Alderton, Roberto Amato, Jeffrey Barrett, Sonia Goncalves, Ewan Harrison, David K. Jackson, Ian Johnston, Dominic Kwiatkowski, Cordelia Langford, John Sillitoe on behalf of the Wellcome Sanger Institute COVID-19 Surveillance Team |
| EPI_ISL_2664788 | Lighthouse Lab in Milton Keynes | Wellcome Sanger Institute for the COVID-19 Genomics UK (COG-UK) Consortium | The Lighthouse Lab in Milton Keynes and Alex Alderton, Roberto Amato, Jeffrey Barrett, Sonia Goncalves, Ewan Harrison, David K. Jackson, Ian Johnston, Dominic Kwiatkowski, Cordelia Langford, John Sillitoe on behalf of the Wellcome Sanger Institute COVID-19 Surveillance Team |
| EPI_ISL_2664790 | Lighthouse Laboratory Plymouth | Wellcome Sanger Institute for the COVID-19 Genomics UK (COG-UK) Consortium | Lighthouse Laboratory Plymouth and Alex Alderton, Roberto Amato, Jeffrey Barrett, Sonia Goncalves, Ewan Harrison, David K. Jackson, Ian Johnston, Dominic Kwiatkowski, Cordelia Langford, John Sillitoe on behalf of the Wellcome Sanger Institute COVID-19 Surveillance Team |
| EPI_ISL_2664791 | Lighthouse Lab in Alderley Park | Wellcome Sanger Institute for the COVID-19 Genomics UK (COG-UK) Consortium | Jacquelyn Wynn, Mairead Hyland, The Lighthouse Lab in Alderley Park and Alex Alderton, Roberto Amato, Jeffrey Barrett, Sonia Goncalves, Ewan Harrison, David K. Jackson, Ian Johnston, Dominic Kwiatkowski, Cordelia Langford, John Sillitoe on behalf of the Wellcome Sanger Institute COVID-19 Surveillance Team |
| EPI_ISL_2664792, EPI_ISL_2664793 | Lighthouse Laboratory Plymouth | Wellcome Sanger Institute for the COVID-19 Genomics UK (COG-UK) Consortium | Lighthouse Laboratory Plymouth and Alex Alderton, Roberto Amato, Jeffrey Barrett, Sonia Goncalves, Ewan Harrison, David K. Jackson, Ian Johnston, Dominic Kwiatkowski, Cordelia Langford, John Sillitoe on behalf of the Wellcome Sanger Institute COVID-19 Surveillance Team |
| EPI_ISL_2664794, EPI_ISL_2664795 | Lighthouse Lab in Milton Keynes | Wellcome Sanger Institute for the COVID-19 Genomics UK (COG-UK) Consortium | The Lighthouse Lab in Milton Keynes and Alex Alderton, Roberto Amato, Jeffrey Barrett, Sonia Goncalves, Ewan Harrison, David K. Jackson, Ian Johnston, Dominic Kwiatkowski, Cordelia Langford, John Sillitoe on behalf of the Wellcome Sanger Institute COVID-19 Surveillance Team |
| EPI_ISL_2664796 | Lighthouse Laboratory Plymouth | Wellcome Sanger Institute for the COVID-19 Genomics UK (COG-UK) Consortium | Lighthouse Laboratory Plymouth and Alex Alderton, Roberto Amato, Jeffrey Barrett, Sonia Goncalves, Ewan Harrison, David K. Jackson, Ian Johnston, Dominic Kwiatkowski, Cordelia Langford, John Sillitoe on behalf of the Wellcome Sanger Institute COVID-19 Surveillance Team |
| EPI_ISL_2664797, EPI_ISL_2664798 | Lighthouse Lab in Alderley Park | Wellcome Sanger Institute for the COVID-19 Genomics UK (COG-UK) Consortium | Jacquelyn Wynn, Mairead Hyland, The Lighthouse Lab in Alderley Park and Alex Alderton, Roberto Amato, Jeffrey Barrett, Sonia Goncalves, Ewan Harrison, David K. Jackson, Ian Johnston, Dominic Kwiatkowski, Cordelia Langford, John Sillitoe on behalf of the Wellcome Sanger Institute COVID-19 Surveillance Team |
| EPI_ISL_2664799 | Lighthouse Lab in Milton Keynes | Wellcome Sanger Institute for the COVID-19 Genomics UK (COG-UK) Consortium | The Lighthouse Lab in Milton Keynes and Alex Alderton, Roberto Amato, Jeffrey Barrett, Sonia Goncalves, Ewan Harrison, David K. Jackson, Ian Johnston, Dominic Kwiatkowski, Cordelia Langford, John Sillitoe on behalf of the Wellcome Sanger Institute COVID-19 Surveillance Team |
| EPI_ISL_2664800 | Lighthouse Laboratory Plymouth | Wellcome Sanger Institute for the COVID-19 Genomics UK (COG-UK) Consortium | Lighthouse Laboratory Plymouth and Alex Alderton, Roberto Amato, Jeffrey Barrett, Sonia Goncalves, Ewan Harrison, David K. Jackson, Ian Johnston, Dominic Kwiatkowski, Cordelia Langford, John Sillitoe on behalf of the Wellcome Sanger Institute COVID-19 Surveillance Team |
| EPI_ISL_2664801 | Lighthouse Lab in Milton Keynes | Wellcome Sanger Institute for the COVID-19 Genomics UK (COG-UK) Consortium | The Lighthouse Lab in Milton Keynes and Alex Alderton, Roberto Amato, Jeffrey Barrett, Sonia Goncalves, Ewan Harrison, David K. Jackson, Ian Johnston, Dominic Kwiatkowski, Cordelia Langford, John Sillitoe on behalf of the Wellcome Sanger Institute COVID-19 Surveillance Team |
| EPI_ISL_2664802 | Lighthouse Laboratory Plymouth | Wellcome Sanger Institute for the COVID-19 Genomics UK (COG-UK) Consortium | Lighthouse Laboratory Plymouth and Alex Alderton, Roberto Amato, Jeffrey Barrett, Sonia Goncalves, Ewan Harrison, David K. Jackson, Ian Johnston, Dominic Kwiatkowski, Cordelia Langford, John Sillitoe on behalf of the Wellcome Sanger Institute COVID-19 Surveillance Team |
| EPI_ISL_2664803 | Lighthouse Lab in Milton Keynes | Wellcome Sanger Institute for the COVID-19 Genomics UK (COG-UK) Consortium | The Lighthouse Lab in Milton Keynes and Alex Alderton, Roberto Amato, Jeffrey Barrett, Sonia Goncalves, Ewan Harrison, David K. Jackson, Ian Johnston, Dominic Kwiatkowski, Cordelia Langford, John Sillitoe on behalf of the Wellcome Sanger Institute COVID-19 Surveillance Team |
| EPI_ISL_2664804 | Lighthouse Lab in Glasgow | Wellcome Sanger Institute for the COVID-19 Genomics UK (COG-UK) Consortium | Harper VanSteenhouse, Yumi Kasai, David Gray, Carol Clugston, Anna Dominiczak and Alex Alderton, Roberto Amato, Jeffrey Barrett, Sonia Goncalves, Ewan Harrison, David K. Jackson, Ian Johnston, Dominic Kwiatkowski, Cordelia Langford, John Sillitoe on behalf of the Wellcome Sanger Institute COVID-19 Surveillance Team |
| EPI_ISL_2664805 | Lighthouse Lab in Milton Keynes | Wellcome Sanger Institute for the COVID-19 Genomics UK (COG-UK) Consortium | The Lighthouse Lab in Milton Keynes and Alex Alderton, Roberto Amato, Jeffrey Barrett, Sonia Goncalves, Ewan Harrison, David K. Jackson, Ian Johnston, Dominic Kwiatkowski, Cordelia Langford, John Sillitoe on behalf of the Wellcome Sanger Institute COVID-19 Surveillance Team |
| EPI_ISL_2664806 | Lighthouse Lab in Glasgow | Wellcome Sanger Institute for the COVID-19 Genomics UK (COG-UK) Consortium | Harper VanSteenhouse, Yumi Kasai, David Gray, Carol Clugston, Anna Dominiczak and Alex Alderton, Roberto Amato, Jeffrey Barrett, Sonia Goncalves, Ewan Harrison, David K. Jackson, Ian Johnston, Dominic Kwiatkowski, Cordelia Langford, John Sillitoe on behalf of the Wellcome Sanger Institute COVID-19 Surveillance Team |
| EPI_ISL_2664807, EPI_ISL_2664808 | Lighthouse Lab in Alderley Park | Wellcome Sanger Institute for the COVID-19 Genomics UK (COG-UK) Consortium | Jacquelyn Wynn, Mairead Hyland, The Lighthouse Lab in Alderley Park and Alex Alderton, Roberto Amato, Jeffrey Barrett, Sonia Goncalves, Ewan Harrison, David K. Jackson, Ian Johnston, Dominic Kwiatkowski, Cordelia Langford, John Sillitoe on behalf of the Wellcome Sanger Institute COVID-19 Surveillance Team |
| EPI_ISL_2664809, EPI_ISL_2664810, EPI_ISL_2664811 | Lighthouse Lab in Milton Keynes | Wellcome Sanger Institute for the COVID-19 Genomics UK (COG-UK) Consortium | The Lighthouse Lab in Milton Keynes and Alex Alderton, Roberto Amato, Jeffrey Barrett, Sonia Goncalves, Ewan Harrison, David K. Jackson, Ian Johnston, Dominic Kwiatkowski, Cordelia Langford, John Sillitoe on behalf of the Wellcome Sanger Institute COVID-19 Surveillance Team |
| EPI_ISL_2664812 | Lighthouse Laboratory Plymouth | Wellcome Sanger Institute for the COVID-19 Genomics UK (COG-UK) Consortium | Lighthouse Laboratory Plymouth and Alex Alderton, Roberto Amato, Jeffrey Barrett, Sonia Goncalves, Ewan Harrison, David K. Jackson, Ian Johnston, Dominic Kwiatkowski, Cordelia Langford, John Sillitoe on behalf of the Wellcome Sanger Institute COVID-19 Surveillance Team |
| EPI_ISL_2664813, EPI_ISL_2664814, EPI_ISL_2664815 | Lighthouse Lab in Alderley Park | Wellcome Sanger Institute for the COVID-19 Genomics UK (COG-UK) Consortium | Jacquelyn Wynn, Mairead Hyland, The Lighthouse Lab in Alderley Park and Alex Alderton, Roberto Amato, Jeffrey Barrett, Sonia Goncalves, Ewan Harrison, David K. Jackson, Ian Johnston, Dominic Kwiatkowski, Cordelia Langford, John Sillitoe on behalf of the Wellcome Sanger Institute COVID-19 Surveillance Team |
| EPI_ISL_2664816 | Lighthouse Lab in Milton Keynes | Wellcome Sanger Institute for the COVID-19 Genomics UK (COG-UK) Consortium | The Lighthouse Lab in Milton Keynes and Alex Alderton, Roberto Amato, Jeffrey Barrett, Sonia Goncalves, Ewan Harrison, David K. Jackson, Ian Johnston, Dominic Kwiatkowski, Cordelia Langford, John Sillitoe on behalf of the Wellcome Sanger Institute COVID-19 Surveillance Team |
| EPI_ISL_2664817 | Health Services Laboratories | Wellcome Sanger Institute for the COVID-19 Genomics UK (COG-UK) Consortium | Health Services Laboratories and Alex Alderton, Roberto Amato, Jeffrey Barrett, Sonia Goncalves, Ewan Harrison, David K. Jackson, Ian Johnston, Dominic Kwiatkowski, Cordelia Langford, John Sillitoe on behalf of the Wellcome Sanger Institute COVID-19 Surveillance Team |
| EPI_ISL_2664818, EPI_ISL_2664819 | Lighthouse Laboratory Plymouth | Wellcome Sanger Institute for the COVID-19 Genomics UK (COG-UK) Consortium | Lighthouse Laboratory Plymouth and Alex Alderton, Roberto Amato, Jeffrey Barrett, Sonia Goncalves, Ewan Harrison, David K. Jackson, Ian Johnston, Dominic Kwiatkowski, Cordelia Langford, John Sillitoe on behalf of the Wellcome Sanger Institute COVID-19 Surveillance Team |
| EPI_ISL_2664820 | Lighthouse Lab in Milton Keynes | Wellcome Sanger Institute for the COVID-19 Genomics UK (COG-UK) Consortium | The Lighthouse Lab in Milton Keynes and Alex Alderton, Roberto Amato, Jeffrey Barrett, Sonia Goncalves, Ewan Harrison, David K. Jackson, Ian Johnston, Dominic Kwiatkowski, Cordelia Langford, John Sillitoe on behalf of the Wellcome Sanger Institute COVID-19 Surveillance Team |
| EPI_ISL_2664821 | Lighthouse Lab in Alderley Park | Wellcome Sanger Institute for the COVID-19 Genomics UK (COG-UK) Consortium | Jacquelyn Wynn, Mairead Hyland, The Lighthouse Lab in Alderley Park and Alex Alderton, Roberto Amato, Jeffrey Barrett, Sonia Goncalves, Ewan Harrison, David K. Jackson, Ian Johnston, Dominic Kwiatkowski, Cordelia Langford, John Sillitoe on behalf of the Wellcome Sanger Institute COVID-19 Surveillance Team |
| EPI_ISL_2664822 | Lighthouse Lab in Milton Keynes | Wellcome Sanger Institute for the COVID-19 Genomics UK (COG-UK) Consortium | The Lighthouse Lab in Milton Keynes and Alex Alderton, Roberto Amato, Jeffrey Barrett, Sonia Goncalves, Ewan Harrison, David K. Jackson, Ian Johnston, Dominic Kwiatkowski, Cordelia Langford, John Sillitoe on behalf of the Wellcome Sanger Institute COVID-19 Surveillance Team |
| EPI_ISL_2664823, EPI_ISL_2664824, EPI_ISL_2664825 | Lighthouse Laboratory Plymouth | Wellcome Sanger Institute for the COVID-19 Genomics UK (COG-UK) Consortium | Lighthouse Laboratory Plymouth and Alex Alderton, Roberto Amato, Jeffrey Barrett, Sonia Goncalves, Ewan Harrison, David K. Jackson, Ian Johnston, Dominic Kwiatkowski, Cordelia Langford, John Sillitoe on behalf of the Wellcome Sanger Institute COVID-19 Surveillance Team |
| EPI_ISL_2664826, EPI_ISL_2664827 | Lighthouse Lab in Milton Keynes | Wellcome Sanger Institute for the COVID-19 Genomics UK (COG-UK) Consortium | The Lighthouse Lab in Milton Keynes and Alex Alderton, Roberto Amato, Jeffrey Barrett, Sonia Goncalves, Ewan Harrison, David K. Jackson, Ian Johnston, Dominic Kwiatkowski, Cordelia Langford, John Sillitoe on behalf of the Wellcome Sanger Institute COVID-19 Surveillance Team |
| EPI_ISL_2664828 | Lighthouse Lab in Glasgow | Wellcome Sanger Institute for the COVID-19 Genomics UK (COG-UK) Consortium | Harper VanSteenhouse, Yumi Kasai, David Gray, Carol Clugston, Anna Dominiczak and Alex Alderton, Roberto Amato, Jeffrey Barrett, Sonia Goncalves, Ewan Harrison, David K. Jackson, Ian Johnston, Dominic Kwiatkowski, Cordelia Langford, John Sillitoe on behalf of the Wellcome Sanger Institute COVID-19 Surveillance Team |
| EPI_ISL_2664829 | Lighthouse Lab in Alderley Park | Wellcome Sanger Institute for the COVID-19 Genomics UK (COG-UK) Consortium | Jacquelyn Wynn, Mairead Hyland, The Lighthouse Lab in Alderley Park and Alex Alderton, Roberto Amato, Jeffrey Barrett, Sonia Goncalves, Ewan Harrison, David K. Jackson, Ian Johnston, Dominic Kwiatkowski, Cordelia Langford, John Sillitoe on behalf of the Wellcome Sanger Institute COVID-19 Surveillance Team |
| EPI_ISL_2664830 | Lighthouse Laboratory Plymouth | Wellcome Sanger Institute for the COVID-19 Genomics UK (COG-UK) Consortium | Lighthouse Laboratory Plymouth and Alex Alderton, Roberto Amato, Jeffrey Barrett, Sonia Goncalves, Ewan Harrison, David K. Jackson, Ian Johnston, Dominic Kwiatkowski, Cordelia Langford, John Sillitoe on behalf of the Wellcome Sanger Institute COVID-19 Surveillance Team |
| EPI_ISL_2664831, EPI_ISL_2664832 | Lighthouse Lab in Alderley Park | Wellcome Sanger Institute for the COVID-19 Genomics UK (COG-UK) Consortium | Jacquelyn Wynn, Mairead Hyland, The Lighthouse Lab in Alderley Park and Alex Alderton, Roberto Amato, Jeffrey Barrett, Sonia Goncalves, Ewan Harrison, David K. Jackson, Ian Johnston, Dominic Kwiatkowski, Cordelia Langford, John Sillitoe on behalf of the Wellcome Sanger Institute COVID-19 Surveillance Team |
| EPI_ISL_2664833, EPI_ISL_2664834 | Lighthouse Laboratory Plymouth | Wellcome Sanger Institute for the COVID-19 Genomics UK (COG-UK) Consortium | Lighthouse Laboratory Plymouth and Alex Alderton, Roberto Amato, Jeffrey Barrett, Sonia Goncalves, Ewan Harrison, David K. Jackson, Ian Johnston, Dominic Kwiatkowski, Cordelia Langford, John Sillitoe on behalf of the Wellcome Sanger Institute COVID-19 Surveillance Team |
| EPI_ISL_2664835 | Lighthouse Lab in Alderley Park | Wellcome Sanger Institute for the COVID-19 Genomics UK (COG-UK) Consortium | Jacquelyn Wynn, Mairead Hyland, The Lighthouse Lab in Alderley Park and Alex Alderton, Roberto Amato, Jeffrey Barrett, Sonia Goncalves, Ewan Harrison, David K. Jackson, Ian Johnston, Dominic Kwiatkowski, Cordelia Langford, John Sillitoe on behalf of the Wellcome Sanger Institute COVID-19 Surveillance Team |
| EPI_ISL_2664836 | Lighthouse Laboratory Plymouth | Wellcome Sanger Institute for the COVID-19 Genomics UK (COG-UK) Consortium | Lighthouse Laboratory Plymouth and Alex Alderton, Roberto Amato, Jeffrey Barrett, Sonia Goncalves, Ewan Harrison, David K. Jackson, Ian Johnston, Dominic Kwiatkowski, Cordelia Langford, John Sillitoe on behalf of the Wellcome Sanger Institute COVID-19 Surveillance Team |
| EPI_ISL_2664837 | Lighthouse Lab in Milton Keynes | Wellcome Sanger Institute for the COVID-19 Genomics UK (COG-UK) Consortium | The Lighthouse Lab in Milton Keynes and Alex Alderton, Roberto Amato, Jeffrey Barrett, Sonia Goncalves, Ewan Harrison, David K. Jackson, Ian Johnston, Dominic Kwiatkowski, Cordelia Langford, John Sillitoe on behalf of the Wellcome Sanger Institute COVID-19 Surveillance Team |
| EPI_ISL_2664838 | Health Services Laboratories | Wellcome Sanger Institute for the COVID-19 Genomics UK (COG-UK) Consortium | Health Services Laboratories and Alex Alderton, Roberto Amato, Jeffrey Barrett, Sonia Goncalves, Ewan Harrison, David K. Jackson, Ian Johnston, Dominic Kwiatkowski, Cordelia Langford, John Sillitoe on behalf of the Wellcome Sanger Institute COVID-19 Surveillance Team |
| EPI_ISL_2664839 | Lighthouse Lab in Milton Keynes | Wellcome Sanger Institute for the COVID-19 Genomics UK (COG-UK) Consortium | The Lighthouse Lab in Milton Keynes and Alex Alderton, Roberto Amato, Jeffrey Barrett, Sonia Goncalves, Ewan Harrison, David K. Jackson, Ian Johnston, Dominic Kwiatkowski, Cordelia Langford, John Sillitoe on behalf of the Wellcome Sanger Institute COVID-19 Surveillance Team |
| EPI_ISL_2664840 | Lighthouse Laboratory Plymouth | Wellcome Sanger Institute for the COVID-19 Genomics UK (COG-UK) Consortium | Lighthouse Laboratory Plymouth and Alex Alderton, Roberto Amato, Jeffrey Barrett, Sonia Goncalves, Ewan Harrison, David K. Jackson, Ian Johnston, Dominic Kwiatkowski, Cordelia Langford, John Sillitoe on behalf of the Wellcome Sanger Institute COVID-19 Surveillance Team |
| EPI_ISL_2664841, EPI_ISL_2664842 | Lighthouse Lab in Alderley Park | Wellcome Sanger Institute for the COVID-19 Genomics UK (COG-UK) Consortium | Jacquelyn Wynn, Mairead Hyland, The Lighthouse Lab in Alderley Park and Alex Alderton, Roberto Amato, Jeffrey Barrett, Sonia Goncalves, Ewan Harrison, David K. Jackson, Ian Johnston, Dominic Kwiatkowski, Cordelia Langford, John Sillitoe on behalf of the Wellcome Sanger Institute COVID-19 Surveillance Team |
| EPI_ISL_2664843, EPI_ISL_2664844 | Lighthouse Lab in Milton Keynes | Wellcome Sanger Institute for the COVID-19 Genomics UK (COG-UK) Consortium | The Lighthouse Lab in Milton Keynes and Alex Alderton, Roberto Amato, Jeffrey Barrett, Sonia Goncalves, Ewan Harrison, David K. Jackson, Ian Johnston, Dominic Kwiatkowski, Cordelia Langford, John Sillitoe on behalf of the Wellcome Sanger Institute COVID-19 Surveillance Team |

| | | | |
|---|---|---|---|
| EPI_ISL_2664845 | Lighthouse Laboratory Plymouth | Wellcome Sanger Institute for the COVID-19 Genomics UK (COG-UK) Consortium | Lighthouse Laboratory Plymouth and Alex Alderton, Roberto Amato, Jeffrey Barrett, Sonia Goncalves, Ewan Harrison, David K. Jackson, Ian Johnston, Dominic Kwiatkowski, Cordelia Langford, John Sillitoe on behalf of the Wellcome Sanger Institute COVID-19 Surveillance Team |
| EPI_ISL_2664846, EPI_ISL_2664847, EPI_ISL_2664848 | Lighthouse Lab in Alderley Park | Wellcome Sanger Institute for the COVID-19 Genomics UK (COG-UK) Consortium | Jacquelyn Wynn, Mairead Hyland, The Lighthouse Lab in Alderley Park and Alex Alderton, Roberto Amato, Jeffrey Barrett, Sonia Goncalves, Ewan Harrison, David K. Jackson, Ian Johnston, Dominic Kwiatkowski, Cordelia Langford, John Sillitoe on behalf of the Wellcome Sanger Institute COVID-19 Surveillance Team |
| EPI_ISL_2664849 | Health Services Laboratories | Wellcome Sanger Institute for the COVID-19 Genomics UK (COG-UK) Consortium | Health Services Laboratories and Alex Alderton, Roberto Amato, Jeffrey Barrett, Sonia Goncalves, Ewan Harrison, David K. Jackson, Ian Johnston, Dominic Kwiatkowski, Cordelia Langford, John Sillitoe on behalf of the Wellcome Sanger Institute COVID-19 Surveillance Team |
| EPI_ISL_2664850 | Lighthouse Laboratory Plymouth | Wellcome Sanger Institute for the COVID-19 Genomics UK (COG-UK) Consortium | Lighthouse Laboratory Plymouth and Alex Alderton, Roberto Amato, Jeffrey Barrett, Sonia Goncalves, Ewan Harrison, David K. Jackson, Ian Johnston, Dominic Kwiatkowski, Cordelia Langford, John Sillitoe on behalf of the Wellcome Sanger Institute COVID-19 Surveillance Team |
| EPI_ISL_2664851 | Lighthouse Lab in Milton Keynes | Wellcome Sanger Institute for the COVID-19 Genomics UK (COG-UK) Consortium | The Lighthouse Lab in Milton Keynes and Alex Alderton, Roberto Amato, Jeffrey Barrett, Sonia Goncalves, Ewan Harrison, David K. Jackson, Ian Johnston, Dominic Kwiatkowski, Cordelia Langford, John Sillitoe on behalf of the Wellcome Sanger Institute COVID-19 Surveillance Team |
| EPI_ISL_2664852 | Lighthouse Laboratory Plymouth | Wellcome Sanger Institute for the COVID-19 Genomics UK (COG-UK) Consortium | Lighthouse Laboratory Plymouth and Alex Alderton, Roberto Amato, Jeffrey Barrett, Sonia Goncalves, Ewan Harrison, David K. Jackson, Ian Johnston, Dominic Kwiatkowski, Cordelia Langford, John Sillitoe on behalf of the Wellcome Sanger Institute COVID-19 Surveillance Team |
| EPI_ISL_2664853 | Lighthouse Lab in Milton Keynes | Wellcome Sanger Institute for the COVID-19 Genomics UK (COG-UK) Consortium | The Lighthouse Lab in Milton Keynes and Alex Alderton, Roberto Amato, Jeffrey Barrett, Sonia Goncalves, Ewan Harrison, David K. Jackson, Ian Johnston, Dominic Kwiatkowski, Cordelia Langford, John Sillitoe on behalf of the Wellcome Sanger Institute COVID-19 Surveillance Team |
| EPI_ISL_2664854 | Lighthouse Lab in Alderley Park | Wellcome Sanger Institute for the COVID-19 Genomics UK (COG-UK) Consortium | Jacquelyn Wynn, Mairead Hyland, The Lighthouse Lab in Alderley Park and Alex Alderton, Roberto Amato, Jeffrey Barrett, Sonia Goncalves, Ewan Harrison, David K. Jackson, Ian Johnston, Dominic Kwiatkowski, Cordelia Langford, John Sillitoe on behalf of the Wellcome Sanger Institute COVID-19 Surveillance Team |
| EPI_ISL_2664855 | Lighthouse Laboratory Plymouth | Wellcome Sanger Institute for the COVID-19 Genomics UK (COG-UK) Consortium | Lighthouse Laboratory Plymouth and Alex Alderton, Roberto Amato, Jeffrey Barrett, Sonia Goncalves, Ewan Harrison, David K. Jackson, Ian Johnston, Dominic Kwiatkowski, Cordelia Langford, John Sillitoe on behalf of the Wellcome Sanger Institute COVID-19 Surveillance Team |
| EPI_ISL_2664856, EPI_ISL_2664858 | Lighthouse Lab in Milton Keynes | Wellcome Sanger Institute for the COVID-19 Genomics UK (COG-UK) Consortium | The Lighthouse Lab in Milton Keynes and Alex Alderton, Roberto Amato, Jeffrey Barrett, Sonia Goncalves, Ewan Harrison, David K. Jackson, Ian Johnston, Dominic Kwiatkowski, Cordelia Langford, John Sillitoe on behalf of the Wellcome Sanger Institute COVID-19 Surveillance Team |
| EPI_ISL_2664859, EPI_ISL_2664860, EPI_ISL_2664861, EPI_ISL_2664862, EPI_ISL_2664863, EPI_ISL_2664864 | Lighthouse Lab in Alderley Park | Wellcome Sanger Institute for the COVID-19 Genomics UK (COG-UK) Consortium | Jacquelyn Wynn, Mairead Hyland, The Lighthouse Lab in Alderley Park and Alex Alderton, Roberto Amato, Jeffrey Barrett, Sonia Goncalves, Ewan Harrison, David K. Jackson, Ian Johnston, Dominic Kwiatkowski, Cordelia Langford, John Sillitoe on behalf of the Wellcome Sanger Institute COVID-19 Surveillance Team |
| EPI_ISL_2664865 | Lighthouse Lab in Milton Keynes | Wellcome Sanger Institute for the COVID-19 Genomics UK (COG-UK) Consortium | The Lighthouse Lab in Milton Keynes and Alex Alderton, Roberto Amato, Jeffrey Barrett, Sonia Goncalves, Ewan Harrison, David K. Jackson, Ian Johnston, Dominic Kwiatkowski, Cordelia Langford, John Sillitoe on behalf of the Wellcome Sanger Institute COVID-19 Surveillance Team |
| EPI_ISL_2664866 | Lighthouse Laboratory Plymouth | Wellcome Sanger Institute for the COVID-19 Genomics UK (COG-UK) Consortium | Lighthouse Laboratory Plymouth and Alex Alderton, Roberto Amato, Jeffrey Barrett, Sonia Goncalves, Ewan Harrison, David K. Jackson, Ian Johnston, Dominic Kwiatkowski, Cordelia Langford, John Sillitoe on behalf of the Wellcome Sanger Institute COVID-19 Surveillance Team |
| EPI_ISL_2664867 | Lighthouse Lab in Milton Keynes | Wellcome Sanger Institute for the COVID-19 Genomics UK (COG-UK) Consortium | The Lighthouse Lab in Milton Keynes and Alex Alderton, Roberto Amato, Jeffrey Barrett, Sonia Goncalves, Ewan Harrison, David K. Jackson, Ian Johnston, Dominic Kwiatkowski, Cordelia Langford, John Sillitoe on behalf of the Wellcome Sanger Institute COVID-19 Surveillance Team |
| EPI_ISL_2664868, EPI_ISL_2664869 | Lighthouse Laboratory Plymouth | Wellcome Sanger Institute for the COVID-19 Genomics UK (COG-UK) Consortium | Lighthouse Laboratory Plymouth and Alex Alderton, Roberto Amato, Jeffrey Barrett, Sonia Goncalves, Ewan Harrison, David K. Jackson, Ian Johnston, Dominic Kwiatkowski, Cordelia Langford, John Sillitoe on behalf of the Wellcome Sanger Institute COVID-19 Surveillance Team |
| EPI_ISL_2664870 | Lighthouse Lab in Alderley Park | Wellcome Sanger Institute for the COVID-19 Genomics UK (COG-UK) Consortium | Jacquelyn Wynn, Mairead Hyland, The Lighthouse Lab in Alderley Park and Alex Alderton, Roberto Amato, Jeffrey Barrett, Sonia Goncalves, Ewan Harrison, David K. Jackson, Ian Johnston, Dominic Kwiatkowski, Cordelia Langford, John Sillitoe on behalf of the Wellcome Sanger Institute COVID-19 Surveillance Team |
| EPI_ISL_2664871 | Lighthouse Lab in Glasgow | Wellcome Sanger Institute for the COVID-19 Genomics UK (COG-UK) Consortium | Harper VanSteenhouse, Yumi Kasai, David Gray, Carol Clugston, Anna Dominiczak and Alex Alderton, Roberto Amato, Jeffrey Barrett, Sonia Goncalves, Ewan Harrison, David K. Jackson, Ian Johnston, Dominic Kwiatkowski, Cordelia Langford, John Sillitoe on behalf of the Wellcome Sanger Institute COVID-19 Surveillance Team |
| EPI_ISL_2664872 | Lighthouse Lab in Alderley Park | Wellcome Sanger Institute for the COVID-19 Genomics UK (COG-UK) Consortium | Jacquelyn Wynn, Mairead Hyland, The Lighthouse Lab in Alderley Park and Alex Alderton, Roberto Amato, Jeffrey Barrett, Sonia Goncalves, Ewan Harrison, David K. Jackson, Ian Johnston, Dominic Kwiatkowski, Cordelia Langford, John Sillitoe on behalf of the Wellcome Sanger Institute COVID-19 Surveillance Team |
| EPI_ISL_2664873 | Lighthouse Lab in Milton Keynes | Wellcome Sanger Institute for the COVID-19 Genomics UK (COG-UK) Consortium | The Lighthouse Lab in Milton Keynes and Alex Alderton, Roberto Amato, Jeffrey Barrett, Sonia Goncalves, Ewan Harrison, David K. Jackson, Ian Johnston, Dominic Kwiatkowski, Cordelia Langford, John Sillitoe on behalf of the Wellcome Sanger Institute COVID-19 Surveillance Team |
| EPI_ISL_2664874, EPI_ISL_2664875, EPI_ISL_2664876, EPI_ISL_2664877 | Lighthouse Laboratory Plymouth | Wellcome Sanger Institute for the COVID-19 Genomics UK (COG-UK) Consortium | Lighthouse Laboratory Plymouth and Alex Alderton, Roberto Amato, Jeffrey Barrett, Sonia Goncalves, Ewan Harrison, David K. Jackson, Ian Johnston, Dominic Kwiatkowski, Cordelia Langford, John Sillitoe on behalf of the Wellcome Sanger Institute COVID-19 Surveillance Team |
| EPI_ISL_2664878 | Lighthouse Lab in Milton Keynes | Wellcome Sanger Institute for the COVID-19 Genomics UK (COG-UK) Consortium | The Lighthouse Lab in Milton Keynes and Alex Alderton, Roberto Amato, Jeffrey Barrett, Sonia Goncalves, Ewan Harrison, David K. Jackson, Ian Johnston, Dominic Kwiatkowski, Cordelia Langford, John Sillitoe on behalf of the Wellcome Sanger Institute COVID-19 Surveillance Team |
| EPI_ISL_2664879 | Health Services Laboratories | Wellcome Sanger Institute for the COVID-19 Genomics UK (COG-UK) Consortium | Health Services Laboratories and Alex Alderton, Roberto Amato, Jeffrey Barrett, Sonia Goncalves, Ewan Harrison, David K. Jackson, Ian Johnston, Dominic Kwiatkowski, Cordelia Langford, John Sillitoe on behalf of the Wellcome Sanger Institute COVID-19 Surveillance Team |
| EPI_ISL_2664880 | Lighthouse Laboratory Plymouth | Wellcome Sanger Institute for the COVID-19 Genomics UK (COG-UK) Consortium | Lighthouse Laboratory Plymouth and Alex Alderton, Roberto Amato, Jeffrey Barrett, Sonia Goncalves, Ewan Harrison, David K. Jackson, Ian Johnston, Dominic Kwiatkowski, Cordelia Langford, John Sillitoe on behalf of the Wellcome Sanger Institute COVID-19 Surveillance Team |
| EPI_ISL_2664881 | Lighthouse Lab in Glasgow | Wellcome Sanger Institute for the COVID-19 Genomics UK (COG-UK) Consortium | Harper VanSteenhouse, Yumi Kasai, David Gray, Carol Clugston, Anna Dominiczak and Alex Alderton, Roberto Amato, Jeffrey Barrett, Sonia Goncalves, Ewan Harrison, David K. Jackson, Ian Johnston, Dominic Kwiatkowski, Cordelia Langford, John Sillitoe on behalf of the Wellcome Sanger Institute COVID-19 Surveillance Team |
| EPI_ISL_2664882 | Lighthouse Laboratory Plymouth | Wellcome Sanger Institute for the COVID-19 Genomics UK (COG-UK) Consortium | Lighthouse Laboratory Plymouth and Alex Alderton, Roberto Amato, Jeffrey Barrett, Sonia Goncalves, Ewan Harrison, David K. Jackson, Ian Johnston, Dominic Kwiatkowski, Cordelia Langford, John Sillitoe on behalf of the Wellcome Sanger Institute COVID-19 Surveillance Team |
| EPI_ISL_2664883 | Lighthouse Lab in Alderley Park | Wellcome Sanger Institute for the COVID-19 Genomics UK (COG-UK) Consortium | Jacquelyn Wynn, Mairead Hyland, The Lighthouse Lab in Alderley Park and Alex Alderton, Roberto Amato, Jeffrey Barrett, Sonia Goncalves, Ewan Harrison, David K. Jackson, Ian Johnston, Dominic Kwiatkowski, Cordelia Langford, John Sillitoe on behalf of the Wellcome Sanger Institute COVID-19 Surveillance Team |
| EPI_ISL_2664884, EPI_ISL_2664885 | Lighthouse Lab in Milton Keynes | Wellcome Sanger Institute for the COVID-19 Genomics UK (COG-UK) Consortium | The Lighthouse Lab in Milton Keynes and Alex Alderton, Roberto Amato, Jeffrey Barrett, Sonia Goncalves, Ewan Harrison, David K. Jackson, Ian Johnston, Dominic Kwiatkowski, Cordelia Langford, John Sillitoe on behalf of the Wellcome Sanger Institute COVID-19 Surveillance Team |
| EPI_ISL_2664886 | Lighthouse Laboratory Plymouth | Wellcome Sanger Institute for the COVID-19 Genomics UK (COG-UK) Consortium | Lighthouse Laboratory Plymouth and Alex Alderton, Roberto Amato, Jeffrey Barrett, Sonia Goncalves, Ewan Harrison, David K. Jackson, Ian Johnston, Dominic Kwiatkowski, Cordelia Langford, John Sillitoe on behalf of the Wellcome Sanger Institute COVID-19 Surveillance Team |
| EPI_ISL_2664887, EPI_ISL_2664888 | Lighthouse Lab in Milton Keynes | Wellcome Sanger Institute for the COVID-19 Genomics UK (COG-UK) Consortium | The Lighthouse Lab in Milton Keynes and Alex Alderton, Roberto Amato, Jeffrey Barrett, Sonia Goncalves, Ewan Harrison, David K. Jackson, Ian Johnston, Dominic Kwiatkowski, Cordelia Langford, John Sillitoe on behalf of the Wellcome Sanger Institute COVID-19 Surveillance Team |
| EPI_ISL_2664889 | Lighthouse Lab in Alderley Park | Wellcome Sanger Institute for the COVID-19 Genomics UK (COG-UK) Consortium | Jacquelyn Wynn, Mairead Hyland, The Lighthouse Lab in Alderley Park and Alex Alderton, Roberto Amato, Jeffrey Barrett, Sonia Goncalves, Ewan Harrison, David K. Jackson, Ian Johnston, Dominic Kwiatkowski, Cordelia Langford, John Sillitoe on behalf of the Wellcome Sanger Institute COVID-19 Surveillance Team |
| EPI_ISL_2664890 | Lighthouse Lab in Milton Keynes | Wellcome Sanger Institute for the COVID-19 Genomics UK (COG-UK) Consortium | The Lighthouse Lab in Milton Keynes and Alex Alderton, Roberto Amato, Jeffrey Barrett, Sonia Goncalves, Ewan Harrison, David K. Jackson, Ian Johnston, Dominic Kwiatkowski, Cordelia Langford, John Sillitoe on behalf of the Wellcome Sanger Institute COVID-19 Surveillance Team |
| EPI_ISL_2664891 | Lighthouse Laboratory Plymouth | Wellcome Sanger Institute for the COVID-19 Genomics UK (COG-UK) Consortium | Lighthouse Laboratory Plymouth and Alex Alderton, Roberto Amato, Jeffrey Barrett, Sonia Goncalves, Ewan Harrison, David K. Jackson, Ian Johnston, Dominic Kwiatkowski, Cordelia Langford, John Sillitoe on behalf of the Wellcome Sanger Institute COVID-19 Surveillance Team |
| EPI_ISL_2664892 | Lighthouse Lab in Milton Keynes | Wellcome Sanger Institute for the COVID-19 Genomics UK (COG-UK) Consortium | The Lighthouse Lab in Milton Keynes and Alex Alderton, Roberto Amato, Jeffrey Barrett, Sonia Goncalves, Ewan Harrison, David K. Jackson, Ian Johnston, Dominic Kwiatkowski, Cordelia Langford, John Sillitoe on behalf of the Wellcome Sanger Institute COVID-19 Surveillance Team |
| EPI_ISL_2664893 | Lighthouse Lab in Alderley Park | Wellcome Sanger Institute for the COVID-19 Genomics UK (COG-UK) Consortium | Jacquelyn Wynn, Mairead Hyland, The Lighthouse Lab in Alderley Park and Alex Alderton, Roberto Amato, Jeffrey Barrett, Sonia Goncalves, Ewan Harrison, David K. Jackson, Ian Johnston, Dominic Kwiatkowski, Cordelia Langford, John Sillitoe on behalf of the Wellcome Sanger Institute COVID-19 Surveillance Team |
| EPI_ISL_2664895 | Lighthouse Lab in Milton Keynes | Wellcome Sanger Institute for the COVID-19 Genomics UK (COG-UK) Consortium | The Lighthouse Lab in Milton Keynes and Alex Alderton, Roberto Amato, Jeffrey Barrett, Sonia Goncalves, Ewan Harrison, David K. Jackson, Ian Johnston, Dominic Kwiatkowski, Cordelia Langford, John Sillitoe on behalf of the Wellcome Sanger Institute COVID-19 Surveillance Team |
| EPI_ISL_2664896, EPI_ISL_2664897, EPI_ISL_2664898 | Lighthouse Laboratory Plymouth | Wellcome Sanger Institute for the COVID-19 Genomics UK (COG-UK) Consortium | Lighthouse Laboratory Plymouth and Alex Alderton, Roberto Amato, Jeffrey Barrett, Sonia Goncalves, Ewan Harrison, David K. Jackson, Ian Johnston, Dominic Kwiatkowski, Cordelia Langford, John Sillitoe on behalf of the Wellcome Sanger Institute COVID-19 Surveillance Team |
| EPI_ISL_2664899, EPI_ISL_2664900, EPI_ISL_2664901 | Lighthouse Lab in Milton Keynes | Wellcome Sanger Institute for the COVID-19 Genomics UK (COG-UK) Consortium | The Lighthouse Lab in Milton Keynes and Alex Alderton, Roberto Amato, Jeffrey Barrett, Sonia Goncalves, Ewan Harrison, David K. Jackson, Ian Johnston, Dominic Kwiatkowski, Cordelia Langford, John Sillitoe on behalf of the Wellcome Sanger Institute COVID-19 Surveillance Team |
| EPI_ISL_2664902, EPI_ISL_2664903 | Lighthouse Laboratory Plymouth | Wellcome Sanger Institute for the COVID-19 Genomics UK (COG-UK) Consortium | Lighthouse Laboratory Plymouth and Alex Alderton, Roberto Amato, Jeffrey Barrett, Sonia Goncalves, Ewan Harrison, David K. Jackson, Ian Johnston, Dominic Kwiatkowski, Cordelia Langford, John Sillitoe on behalf of the Wellcome Sanger Institute COVID-19 Surveillance Team |
| EPI_ISL_2664904 | Lighthouse Lab in Alderley Park | Wellcome Sanger Institute for the COVID-19 Genomics UK (COG-UK) Consortium | Jacquelyn Wynn, Mairead Hyland, The Lighthouse Lab in Alderley Park and Alex Alderton, Roberto Amato, Jeffrey Barrett, Sonia Goncalves, Ewan Harrison, David K. Jackson, Ian Johnston, Dominic Kwiatkowski, Cordelia Langford, John Sillitoe on behalf of the Wellcome Sanger Institute COVID-19 Surveillance Team |
| EPI_ISL_2664905 | Lighthouse Laboratory Plymouth | Wellcome Sanger Institute for the COVID-19 Genomics UK (COG-UK) Consortium | Lighthouse Laboratory Plymouth and Alex Alderton, Roberto Amato, Jeffrey Barrett, Sonia Goncalves, Ewan Harrison, David K. Jackson, Ian Johnston, Dominic Kwiatkowski, Cordelia Langford, John Sillitoe on behalf of the Wellcome Sanger Institute COVID-19 Surveillance Team |
| EPI_ISL_2664906 | Lighthouse Lab in Glasgow | Wellcome Sanger Institute for the COVID-19 Genomics UK (COG-UK) Consortium | Harper VanSteenhouse, Yumi Kasai, David Gray, Carol Clugston, Anna Dominiczak and Alex Alderton, Roberto Amato, Jeffrey Barrett, Sonia Goncalves, Ewan Harrison, David K. Jackson, Ian Johnston, Dominic Kwiatkowski, Cordelia Langford, John Sillitoe on behalf of the Wellcome Sanger Institute COVID-19 Surveillance Team |
| EPI_ISL_2664907 | Lighthouse Lab in Alderley Park | Wellcome Sanger Institute for the COVID-19 Genomics UK (COG-UK) | Jacquelyn Wynn, Mairead Hyland, The Lighthouse Lab in Alderley Park and Alex Alderton, Roberto Amato, Jeffrey Barrett, Sonia Goncalves, Ewan Harrison, David K. Jackson, Ian Johnston, Dominic Kwiatkowski, Cordelia Langford, John Sillitoe on |

| | | | |
|---|---|---|---|
| | | Consortium | behalf of the Wellcome Sanger Institute COVID-19 Surveillance Team |
| EPI_ISL_2664908 | Lighthouse Laboratory Plymouth | Wellcome Sanger Institute for the COVID-19 Genomics UK (COG-UK) Consortium | Lighthouse Laboratory Plymouth and Alex Alderton, Roberto Amato, Jeffrey Barrett, Sonia Goncalves, Ewan Harrison, David K. Jackson, Ian Johnston, Dominic Kwiatkowski, Cordelia Langford, John Sillitoe on behalf of the Wellcome Sanger Institute COVID-19 Surveillance Team |
| EPI_ISL_2664909 | Lighthouse Lab in Alderley Park | Wellcome Sanger Institute for the COVID-19 Genomics UK (COG-UK) Consortium | Jacquelyn Wynn, Mairead Hyland, The Lighthouse Lab in Alderley Park and Alex Alderton, Roberto Amato, Jeffrey Barrett, Sonia Goncalves, Ewan Harrison, David K. Jackson, Ian Johnston, Dominic Kwiatkowski, Cordelia Langford, John Sillitoe on behalf of the Wellcome Sanger Institute COVID-19 Surveillance Team |
| EPI_ISL_2664910 | Lighthouse Lab in Glasgow | Wellcome Sanger Institute for the COVID-19 Genomics UK (COG-UK) Consortium | Harper VanSteenhouse, Yumi Kasai, David Gray, Carol Clugston, Anna Dominiczak and Alex Alderton, Roberto Amato, Jeffrey Barrett, Sonia Goncalves, Ewan Harrison, David K. Jackson, Ian Johnston, Dominic Kwiatkowski, Cordelia Langford, John Sillitoe on behalf of the Wellcome Sanger Institute COVID-19 Surveillance Team |
| EPI_ISL_2664911 | Lighthouse Lab in Alderley Park | Wellcome Sanger Institute for the COVID-19 Genomics UK (COG-UK) Consortium | Jacquelyn Wynn, Mairead Hyland, The Lighthouse Lab in Alderley Park and Alex Alderton, Roberto Amato, Jeffrey Barrett, Sonia Goncalves, Ewan Harrison, David K. Jackson, Ian Johnston, Dominic Kwiatkowski, Cordelia Langford, John Sillitoe on behalf of the Wellcome Sanger Institute COVID-19 Surveillance Team |
| EPI_ISL_2664912 | Lighthouse Laboratory Plymouth | Wellcome Sanger Institute for the COVID-19 Genomics UK (COG-UK) Consortium | Lighthouse Laboratory Plymouth and Alex Alderton, Roberto Amato, Jeffrey Barrett, Sonia Goncalves, Ewan Harrison, David K. Jackson, Ian Johnston, Dominic Kwiatkowski, Cordelia Langford, John Sillitoe on behalf of the Wellcome Sanger Institute COVID-19 Surveillance Team |
| EPI_ISL_2664913 | Lighthouse Lab in Alderley Park | Wellcome Sanger Institute for the COVID-19 Genomics UK (COG-UK) Consortium | Jacquelyn Wynn, Mairead Hyland, The Lighthouse Lab in Alderley Park and Alex Alderton, Roberto Amato, Jeffrey Barrett, Sonia Goncalves, Ewan Harrison, David K. Jackson, Ian Johnston, Dominic Kwiatkowski, Cordelia Langford, John Sillitoe on behalf of the Wellcome Sanger Institute COVID-19 Surveillance Team |
| EPI_ISL_2664914 | Lighthouse Laboratory Plymouth | Wellcome Sanger Institute for the COVID-19 Genomics UK (COG-UK) Consortium | Lighthouse Laboratory Plymouth and Alex Alderton, Roberto Amato, Jeffrey Barrett, Sonia Goncalves, Ewan Harrison, David K. Jackson, Ian Johnston, Dominic Kwiatkowski, Cordelia Langford, John Sillitoe on behalf of the Wellcome Sanger Institute COVID-19 Surveillance Team |
| EPI_ISL_2664915 | Lighthouse Lab in Milton Keynes | Wellcome Sanger Institute for the COVID-19 Genomics UK (COG-UK) Consortium | The Lighthouse Lab in Milton Keynes and Alex Alderton, Roberto Amato, Jeffrey Barrett, Sonia Goncalves, Ewan Harrison, David K. Jackson, Ian Johnston, Dominic Kwiatkowski, Cordelia Langford, John Sillitoe on behalf of the Wellcome Sanger Institute COVID-19 Surveillance Team |
| EPI_ISL_2664916 | Lighthouse Lab in Alderley Park | Wellcome Sanger Institute for the COVID-19 Genomics UK (COG-UK) Consortium | Jacquelyn Wynn, Mairead Hyland, The Lighthouse Lab in Alderley Park and Alex Alderton, Roberto Amato, Jeffrey Barrett, Sonia Goncalves, Ewan Harrison, David K. Jackson, Ian Johnston, Dominic Kwiatkowski, Cordelia Langford, John Sillitoe on behalf of the Wellcome Sanger Institute COVID-19 Surveillance Team |
| EPI_ISL_2664917 | Lighthouse Lab in Milton Keynes | Wellcome Sanger Institute for the COVID-19 Genomics UK (COG-UK) Consortium | The Lighthouse Lab in Milton Keynes and Alex Alderton, Roberto Amato, Jeffrey Barrett, Sonia Goncalves, Ewan Harrison, David K. Jackson, Ian Johnston, Dominic Kwiatkowski, Cordelia Langford, John Sillitoe on behalf of the Wellcome Sanger Institute COVID-19 Surveillance Team |
| EPI_ISL_2664918, EPI_ISL_2664919 | Lighthouse Laboratory Plymouth | Wellcome Sanger Institute for the COVID-19 Genomics UK (COG-UK) Consortium | Lighthouse Laboratory Plymouth and Alex Alderton, Roberto Amato, Jeffrey Barrett, Sonia Goncalves, Ewan Harrison, David K. Jackson, Ian Johnston, Dominic Kwiatkowski, Cordelia Langford, John Sillitoe on behalf of the Wellcome Sanger Institute COVID-19 Surveillance Team |
| EPI_ISL_2664920, EPI_ISL_2664921 | Lighthouse Lab in Alderley Park | Wellcome Sanger Institute for the COVID-19 Genomics UK (COG-UK) Consortium | Jacquelyn Wynn, Mairead Hyland, The Lighthouse Lab in Alderley Park and Alex Alderton, Roberto Amato, Jeffrey Barrett, Sonia Goncalves, Ewan Harrison, David K. Jackson, Ian Johnston, Dominic Kwiatkowski, Cordelia Langford, John Sillitoe on behalf of the Wellcome Sanger Institute COVID-19 Surveillance Team |
| EPI_ISL_2664922, EPI_ISL_2664923 | Lighthouse Laboratory Plymouth | Wellcome Sanger Institute for the COVID-19 Genomics UK (COG-UK) Consortium | Lighthouse Laboratory Plymouth and Alex Alderton, Roberto Amato, Jeffrey Barrett, Sonia Goncalves, Ewan Harrison, David K. Jackson, Ian Johnston, Dominic Kwiatkowski, Cordelia Langford, John Sillitoe on behalf of the Wellcome Sanger Institute COVID-19 Surveillance Team |
| EPI_ISL_2664924 | Lighthouse Lab in Glasgow | Wellcome Sanger Institute for the COVID-19 Genomics UK (COG-UK) Consortium | Harper VanSteenhouse, Yumi Kasai, David Gray, Carol Clugston, Anna Dominiczak and Alex Alderton, Roberto Amato, Jeffrey Barrett, Sonia Goncalves, Ewan Harrison, David K. Jackson, Ian Johnston, Dominic Kwiatkowski, Cordelia Langford, John Sillitoe on behalf of the Wellcome Sanger Institute COVID-19 Surveillance Team |
| EPI_ISL_2664925 | Lighthouse Lab in Alderley Park | Wellcome Sanger Institute for the COVID-19 Genomics UK (COG-UK) Consortium | Jacquelyn Wynn, Mairead Hyland, The Lighthouse Lab in Alderley Park and Alex Alderton, Roberto Amato, Jeffrey Barrett, Sonia Goncalves, Ewan Harrison, David K. Jackson, Ian Johnston, Dominic Kwiatkowski, Cordelia Langford, John Sillitoe on behalf of the Wellcome Sanger Institute COVID-19 Surveillance Team |
| EPI_ISL_2664926 | Lighthouse Laboratory Plymouth | Wellcome Sanger Institute for the COVID-19 Genomics UK (COG-UK) Consortium | Lighthouse Laboratory Plymouth and Alex Alderton, Roberto Amato, Jeffrey Barrett, Sonia Goncalves, Ewan Harrison, David K. Jackson, Ian Johnston, Dominic Kwiatkowski, Cordelia Langford, John Sillitoe on behalf of the Wellcome Sanger Institute COVID-19 Surveillance Team |
| EPI_ISL_2664927 | Lighthouse Lab in Alderley Park | Wellcome Sanger Institute for the COVID-19 Genomics UK (COG-UK) Consortium | Jacquelyn Wynn, Mairead Hyland, The Lighthouse Lab in Alderley Park and Alex Alderton, Roberto Amato, Jeffrey Barrett, Sonia Goncalves, Ewan Harrison, David K. Jackson, Ian Johnston, Dominic Kwiatkowski, Cordelia Langford, John Sillitoe on behalf of the Wellcome Sanger Institute COVID-19 Surveillance Team |
| EPI_ISL_2664928 | Lighthouse Laboratory Plymouth | Wellcome Sanger Institute for the COVID-19 Genomics UK (COG-UK) Consortium | Lighthouse Laboratory Plymouth and Alex Alderton, Roberto Amato, Jeffrey Barrett, Sonia Goncalves, Ewan Harrison, David K. Jackson, Ian Johnston, Dominic Kwiatkowski, Cordelia Langford, John Sillitoe on behalf of the Wellcome Sanger Institute COVID-19 Surveillance Team |
| EPI_ISL_2664929 | Lighthouse Lab in Glasgow | Wellcome Sanger Institute for the COVID-19 Genomics UK (COG-UK) Consortium | Harper VanSteenhouse, Yumi Kasai, David Gray, Carol Clugston, Anna Dominiczak and Alex Alderton, Roberto Amato, Jeffrey Barrett, Sonia Goncalves, Ewan Harrison, David K. Jackson, Ian Johnston, Dominic Kwiatkowski, Cordelia Langford, John Sillitoe on behalf of the Wellcome Sanger Institute COVID-19 Surveillance Team |
| EPI_ISL_2664930, EPI_ISL_2664931 | Lighthouse Lab in Milton Keynes | Wellcome Sanger Institute for the COVID-19 Genomics UK (COG-UK) Consortium | The Lighthouse Lab in Milton Keynes and Alex Alderton, Roberto Amato, Jeffrey Barrett, Sonia Goncalves, Ewan Harrison, David K. Jackson, Ian Johnston, Dominic Kwiatkowski, Cordelia Langford, John Sillitoe on behalf of the Wellcome Sanger Institute COVID-19 Surveillance Team |
| EPI_ISL_2664932 | Health Services Laboratories | Wellcome Sanger Institute for the COVID-19 Genomics UK (COG-UK) Consortium | Health Services Laboratories and Alex Alderton, Roberto Amato, Jeffrey Barrett, Sonia Goncalves, Ewan Harrison, David K. Jackson, Ian Johnston, Dominic Kwiatkowski, Cordelia Langford, John Sillitoe on behalf of the Wellcome Sanger Institute COVID-19 Surveillance Team |
| EPI_ISL_2664933 | Lighthouse Lab in Milton Keynes | Wellcome Sanger Institute for the COVID-19 Genomics UK (COG-UK) Consortium | The Lighthouse Lab in Milton Keynes and Alex Alderton, Roberto Amato, Jeffrey Barrett, Sonia Goncalves, Ewan Harrison, David K. Jackson, Ian Johnston, Dominic Kwiatkowski, Cordelia Langford, John Sillitoe on behalf of the Wellcome Sanger Institute COVID-19 Surveillance Team |
| EPI_ISL_2664934 | Lighthouse Lab in Glasgow | Wellcome Sanger Institute for the COVID-19 Genomics UK (COG-UK) Consortium | Harper VanSteenhouse, Yumi Kasai, David Gray, Carol Clugston, Anna Dominiczak and Alex Alderton, Roberto Amato, Jeffrey Barrett, Sonia Goncalves, Ewan Harrison, David K. Jackson, Ian Johnston, Dominic Kwiatkowski, Cordelia Langford, John Sillitoe on behalf of the Wellcome Sanger Institute COVID-19 Surveillance Team |
| EPI_ISL_2664935, EPI_ISL_2664936, EPI_ISL_2664938, EPI_ISL_2664939 | Lighthouse Lab in Alderley Park | Wellcome Sanger Institute for the COVID-19 Genomics UK (COG-UK) Consortium | Jacquelyn Wynn, Mairead Hyland, The Lighthouse Lab in Alderley Park and Alex Alderton, Roberto Amato, Jeffrey Barrett, Sonia Goncalves, Ewan Harrison, David K. Jackson, Ian Johnston, Dominic Kwiatkowski, Cordelia Langford, John Sillitoe on behalf of the Wellcome Sanger Institute COVID-19 Surveillance Team |
| EPI_ISL_2664940 | Lighthouse Lab in Milton Keynes | Wellcome Sanger Institute for the COVID-19 Genomics UK (COG-UK) Consortium | The Lighthouse Lab in Milton Keynes and Alex Alderton, Roberto Amato, Jeffrey Barrett, Sonia Goncalves, Ewan Harrison, David K. Jackson, Ian Johnston, Dominic Kwiatkowski, Cordelia Langford, John Sillitoe on behalf of the Wellcome Sanger Institute COVID-19 Surveillance Team |
| EPI_ISL_2664941 | Lighthouse Lab in Alderley Park | Wellcome Sanger Institute for the COVID-19 Genomics UK (COG-UK) Consortium | Jacquelyn Wynn, Mairead Hyland, The Lighthouse Lab in Alderley Park and Alex Alderton, Roberto Amato, Jeffrey Barrett, Sonia Goncalves, Ewan Harrison, David K. Jackson, Ian Johnston, Dominic Kwiatkowski, Cordelia Langford, John Sillitoe on behalf of the Wellcome Sanger Institute COVID-19 Surveillance Team |
| EPI_ISL_2664942 | Lighthouse Laboratory Plymouth | Wellcome Sanger Institute for the COVID-19 Genomics UK (COG-UK) Consortium | Lighthouse Laboratory Plymouth and Alex Alderton, Roberto Amato, Jeffrey Barrett, Sonia Goncalves, Ewan Harrison, David K. Jackson, Ian Johnston, Dominic Kwiatkowski, Cordelia Langford, John Sillitoe on behalf of the Wellcome Sanger Institute COVID-19 Surveillance Team |
| EPI_ISL_2664943 | Lighthouse Lab in Alderley Park | Wellcome Sanger Institute for the COVID-19 Genomics UK (COG-UK) Consortium | Jacquelyn Wynn, Mairead Hyland, The Lighthouse Lab in Alderley Park and Alex Alderton, Roberto Amato, Jeffrey Barrett, Sonia Goncalves, Ewan Harrison, David K. Jackson, Ian Johnston, Dominic Kwiatkowski, Cordelia Langford, John Sillitoe on behalf of the Wellcome Sanger Institute COVID-19 Surveillance Team |
| EPI_ISL_2664944 | Lighthouse Lab in Milton Keynes | Wellcome Sanger Institute for the COVID-19 Genomics UK (COG-UK) Consortium | The Lighthouse Lab in Milton Keynes and Alex Alderton, Roberto Amato, Jeffrey Barrett, Sonia Goncalves, Ewan Harrison, David K. Jackson, Ian Johnston, Dominic Kwiatkowski, Cordelia Langford, John Sillitoe on behalf of the Wellcome Sanger Institute COVID-19 Surveillance Team |
| EPI_ISL_2664945 | Lighthouse Lab in Glasgow | Wellcome Sanger Institute for the COVID-19 Genomics UK (COG-UK) Consortium | Harper VanSteenhouse, Yumi Kasai, David Gray, Carol Clugston, Anna Dominiczak and Alex Alderton, Roberto Amato, Jeffrey Barrett, Sonia Goncalves, Ewan Harrison, David K. Jackson, Ian Johnston, Dominic Kwiatkowski, Cordelia Langford, John Sillitoe on behalf of the Wellcome Sanger Institute COVID-19 Surveillance Team |
| EPI_ISL_2664946 | Lighthouse Lab in Milton Keynes | Wellcome Sanger Institute for the COVID-19 Genomics UK (COG-UK) Consortium | The Lighthouse Lab in Milton Keynes and Alex Alderton, Roberto Amato, Jeffrey Barrett, Sonia Goncalves, Ewan Harrison, David K. Jackson, Ian Johnston, Dominic Kwiatkowski, Cordelia Langford, John Sillitoe on behalf of the Wellcome Sanger Institute COVID-19 Surveillance Team |
| EPI_ISL_2664947, EPI_ISL_2664948, EPI_ISL_2664949 | Lighthouse Lab in Alderley Park | Wellcome Sanger Institute for the COVID-19 Genomics UK (COG-UK) Consortium | Jacquelyn Wynn, Mairead Hyland, The Lighthouse Lab in Alderley Park and Alex Alderton, Roberto Amato, Jeffrey Barrett, Sonia Goncalves, Ewan Harrison, David K. Jackson, Ian Johnston, Dominic Kwiatkowski, Cordelia Langford, John Sillitoe on behalf of the Wellcome Sanger Institute COVID-19 Surveillance Team |
| EPI_ISL_2664950 | Lighthouse Lab in Milton Keynes | Wellcome Sanger Institute for the COVID-19 Genomics UK (COG-UK) Consortium | The Lighthouse Lab in Milton Keynes and Alex Alderton, Roberto Amato, Jeffrey Barrett, Sonia Goncalves, Ewan Harrison, David K. Jackson, Ian Johnston, Dominic Kwiatkowski, Cordelia Langford, John Sillitoe on behalf of the Wellcome Sanger Institute COVID-19 Surveillance Team |
| EPI_ISL_2664951 | Lighthouse Laboratory Plymouth | Wellcome Sanger Institute for the COVID-19 Genomics UK (COG-UK) Consortium | Lighthouse Laboratory Plymouth and Alex Alderton, Roberto Amato, Jeffrey Barrett, Sonia Goncalves, Ewan Harrison, David K. Jackson, Ian Johnston, Dominic Kwiatkowski, Cordelia Langford, John Sillitoe on behalf of the Wellcome Sanger Institute COVID-19 Surveillance Team |
| EPI_ISL_2664952, EPI_ISL_2664953, EPI_ISL_2664954 | Lighthouse Lab in Alderley Park | Wellcome Sanger Institute for the COVID-19 Genomics UK (COG-UK) Consortium | Jacquelyn Wynn, Mairead Hyland, The Lighthouse Lab in Alderley Park and Alex Alderton, Roberto Amato, Jeffrey Barrett, Sonia Goncalves, Ewan Harrison, David K. Jackson, Ian Johnston, Dominic Kwiatkowski, Cordelia Langford, John Sillitoe on behalf of the Wellcome Sanger Institute COVID-19 Surveillance Team |
| EPI_ISL_2664955 | Lighthouse Laboratory Plymouth | Wellcome Sanger Institute for the COVID-19 Genomics UK (COG-UK) Consortium | Lighthouse Laboratory Plymouth and Alex Alderton, Roberto Amato, Jeffrey Barrett, Sonia Goncalves, Ewan Harrison, David K. Jackson, Ian Johnston, Dominic Kwiatkowski, Cordelia Langford, John Sillitoe on behalf of the Wellcome Sanger Institute COVID-19 Surveillance Team |
| EPI_ISL_2664956 | Health Services Laboratories | Wellcome Sanger Institute for the COVID-19 Genomics UK (COG-UK) Consortium | Health Services Laboratories and Alex Alderton, Roberto Amato, Jeffrey Barrett, Sonia Goncalves, Ewan Harrison, David K. Jackson, Ian Johnston, Dominic Kwiatkowski, Cordelia Langford, John Sillitoe on behalf of the Wellcome Sanger Institute COVID-19 Surveillance Team |
| EPI_ISL_2664957 | Lighthouse Lab in Milton Keynes | Wellcome Sanger Institute for the COVID-19 Genomics UK (COG-UK) Consortium | The Lighthouse Lab in Milton Keynes and Alex Alderton, Roberto Amato, Jeffrey Barrett, Sonia Goncalves, Ewan Harrison, David K. Jackson, Ian Johnston, Dominic Kwiatkowski, Cordelia Langford, John Sillitoe on behalf of the Wellcome Sanger Institute COVID-19 Surveillance Team |
| EPI_ISL_2664958 | Lighthouse Laboratory Plymouth | Wellcome Sanger Institute for the COVID-19 Genomics UK (COG-UK) Consortium | Lighthouse Laboratory Plymouth and Alex Alderton, Roberto Amato, Jeffrey Barrett, Sonia Goncalves, Ewan Harrison, David K. Jackson, Ian Johnston, Dominic Kwiatkowski, Cordelia Langford, John Sillitoe on behalf of the Wellcome Sanger Institute COVID-19 Surveillance Team |
| EPI_ISL_2664959 | Lighthouse Lab in Alderley Park | Wellcome Sanger Institute for the COVID-19 Genomics UK (COG-UK) Consortium | Jacquelyn Wynn, Mairead Hyland, The Lighthouse Lab in Alderley Park and Alex Alderton, Roberto Amato, Jeffrey Barrett, Sonia Goncalves, Ewan Harrison, David K. Jackson, Ian Johnston, Dominic Kwiatkowski, Cordelia Langford, John Sillitoe on behalf of the Wellcome Sanger Institute COVID-19 Surveillance Team |
| EPI_ISL_2664960 | Health Services Laboratories | Wellcome Sanger Institute for the COVID-19 Genomics UK (COG-UK) Consortium | Health Services Laboratories and Alex Alderton, Roberto Amato, Jeffrey Barrett, Sonia Goncalves, Ewan Harrison, David K. Jackson, Ian Johnston, Dominic Kwiatkowski, Cordelia Langford, John Sillitoe on behalf of the Wellcome Sanger Institute COVID-19 Surveillance Team |
| EPI_ISL_2664961 | Lighthouse Laboratory Plymouth | Wellcome Sanger Institute for the COVID-19 Genomics UK (COG-UK) Consortium | Lighthouse Laboratory Plymouth and Alex Alderton, Roberto Amato, Jeffrey Barrett, Sonia Goncalves, Ewan Harrison, David K. Jackson, Ian Johnston, Dominic Kwiatkowski, Cordelia Langford, John Sillitoe on behalf of the Wellcome Sanger Institute COVID-19 Surveillance Team |
| EPI_ISL_2664962, | Lighthouse Lab in Alderley Park | Wellcome Sanger Institute for the COVID-19 Genomics UK (COG-UK) | Jacquelyn Wynn, Mairead Hyland, The Lighthouse Lab in Alderley Park and Alex Alderton, Roberto Amato, Jeffrey Barrett, Sonia Goncalves, Ewan Harrison, David K. Jackson, Ian Johnston, Dominic Kwiatkowski, Cordelia Langford, John Sillitoe on |

| | | | |
|---|---|---|---|
| EPI_ISL_2664963, EPI_ISL_2664964, EPI_ISL_2664965 | | Consortium | behalf of the Wellcome Sanger Institute COVID-19 Surveillance Team |
| EPI_ISL_2664966, EPI_ISL_2664967 | Lighthouse Laboratory Plymouth | Wellcome Sanger Institute for the COVID-19 Genomics UK (COG-UK) Consortium | Lighthouse Laboratory Plymouth and Alex Alderton, Roberto Amato, Jeffrey Barrett, Sonia Goncalves, Ewan Harrison, David K. Jackson, Ian Johnston, Dominic Kwiatkowski, Cordelia Langford, John Sillitoe on behalf of the Wellcome Sanger Institute COVID-19 Surveillance Team |
| EPI_ISL_2664968, EPI_ISL_2664969 | Lighthouse Lab in Alderley Park | Wellcome Sanger Institute for the COVID-19 Genomics UK (COG-UK) Consortium | Jacquelyn Wynn, Mairead Hyland, The Lighthouse Lab in Alderley Park and Alex Alderton, Roberto Amato, Jeffrey Barrett, Sonia Goncalves, Ewan Harrison, David K. Jackson, Ian Johnston, Dominic Kwiatkowski, Cordelia Langford, John Sillitoe on behalf of the Wellcome Sanger Institute COVID-19 Surveillance Team |
| EPI_ISL_2664970 | Lighthouse Laboratory Plymouth | Wellcome Sanger Institute for the COVID-19 Genomics UK (COG-UK) Consortium | Lighthouse Laboratory Plymouth and Alex Alderton, Roberto Amato, Jeffrey Barrett, Sonia Goncalves, Ewan Harrison, David K. Jackson, Ian Johnston, Dominic Kwiatkowski, Cordelia Langford, John Sillitoe on behalf of the Wellcome Sanger Institute COVID-19 Surveillance Team |
| EPI_ISL_2664971 | Lighthouse Lab in Glasgow | Wellcome Sanger Institute for the COVID-19 Genomics UK (COG-UK) Consortium | Harper VanSteenhouse, Yumi Kasai, David Gray, Carol Clugston, Anna Dominiczak and Alex Alderton, Roberto Amato, Jeffrey Barrett, Sonia Goncalves, Ewan Harrison, David K. Jackson, Ian Johnston, Dominic Kwiatkowski, Cordelia Langford, John Sillitoe on behalf of the Wellcome Sanger Institute COVID-19 Surveillance Team |
| EPI_ISL_2664972 | Lighthouse Laboratory Plymouth | Wellcome Sanger Institute for the COVID-19 Genomics UK (COG-UK) Consortium | Lighthouse Laboratory Plymouth and Alex Alderton, Roberto Amato, Jeffrey Barrett, Sonia Goncalves, Ewan Harrison, David K. Jackson, Ian Johnston, Dominic Kwiatkowski, Cordelia Langford, John Sillitoe on behalf of the Wellcome Sanger Institute COVID-19 Surveillance Team |
| EPI_ISL_2664973 | Lighthouse Lab in Alderley Park | Wellcome Sanger Institute for the COVID-19 Genomics UK (COG-UK) Consortium | Jacquelyn Wynn, Mairead Hyland, The Lighthouse Lab in Alderley Park and Alex Alderton, Roberto Amato, Jeffrey Barrett, Sonia Goncalves, Ewan Harrison, David K. Jackson, Ian Johnston, Dominic Kwiatkowski, Cordelia Langford, John Sillitoe on behalf of the Wellcome Sanger Institute COVID-19 Surveillance Team |
| EPI_ISL_2664974 | Lighthouse Laboratory Plymouth | Wellcome Sanger Institute for the COVID-19 Genomics UK (COG-UK) Consortium | Lighthouse Laboratory Plymouth and Alex Alderton, Roberto Amato, Jeffrey Barrett, Sonia Goncalves, Ewan Harrison, David K. Jackson, Ian Johnston, Dominic Kwiatkowski, Cordelia Langford, John Sillitoe on behalf of the Wellcome Sanger Institute COVID-19 Surveillance Team |
| EPI_ISL_2664975 | Lighthouse Lab in Milton Keynes | Wellcome Sanger Institute for the COVID-19 Genomics UK (COG-UK) Consortium | The Lighthouse Lab in Milton Keynes and Alex Alderton, Roberto Amato, Jeffrey Barrett, Sonia Goncalves, Ewan Harrison, David K. Jackson, Ian Johnston, Dominic Kwiatkowski, Cordelia Langford, John Sillitoe on behalf of the Wellcome Sanger Institute COVID-19 Surveillance Team |
| EPI_ISL_2664977 | Lighthouse Lab in Glasgow | Wellcome Sanger Institute for the COVID-19 Genomics UK (COG-UK) Consortium | Harper VanSteenhouse, Yumi Kasai, David Gray, Carol Clugston, Anna Dominiczak and Alex Alderton, Roberto Amato, Jeffrey Barrett, Sonia Goncalves, Ewan Harrison, David K. Jackson, Ian Johnston, Dominic Kwiatkowski, Cordelia Langford, John Sillitoe on behalf of the Wellcome Sanger Institute COVID-19 Surveillance Team |
| EPI_ISL_2664978 | Lighthouse Lab in Alderley Park | Wellcome Sanger Institute for the COVID-19 Genomics UK (COG-UK) Consortium | Jacquelyn Wynn, Mairead Hyland, The Lighthouse Lab in Alderley Park and Alex Alderton, Roberto Amato, Jeffrey Barrett, Sonia Goncalves, Ewan Harrison, David K. Jackson, Ian Johnston, Dominic Kwiatkowski, Cordelia Langford, John Sillitoe on behalf of the Wellcome Sanger Institute COVID-19 Surveillance Team |
| EPI_ISL_2664979 | Lighthouse Laboratory Plymouth | Wellcome Sanger Institute for the COVID-19 Genomics UK (COG-UK) Consortium | Lighthouse Laboratory Plymouth and Alex Alderton, Roberto Amato, Jeffrey Barrett, Sonia Goncalves, Ewan Harrison, David K. Jackson, Ian Johnston, Dominic Kwiatkowski, Cordelia Langford, John Sillitoe on behalf of the Wellcome Sanger Institute COVID-19 Surveillance Team |
| EPI_ISL_2664980 | Lighthouse Lab in Milton Keynes | Wellcome Sanger Institute for the COVID-19 Genomics UK (COG-UK) Consortium | The Lighthouse Lab in Milton Keynes and Alex Alderton, Roberto Amato, Jeffrey Barrett, Sonia Goncalves, Ewan Harrison, David K. Jackson, Ian Johnston, Dominic Kwiatkowski, Cordelia Langford, John Sillitoe on behalf of the Wellcome Sanger Institute COVID-19 Surveillance Team |
| EPI_ISL_2664981 | Lighthouse Lab in Alderley Park | Wellcome Sanger Institute for the COVID-19 Genomics UK (COG-UK) Consortium | Jacquelyn Wynn, Mairead Hyland, The Lighthouse Lab in Alderley Park and Alex Alderton, Roberto Amato, Jeffrey Barrett, Sonia Goncalves, Ewan Harrison, David K. Jackson, Ian Johnston, Dominic Kwiatkowski, Cordelia Langford, John Sillitoe on behalf of the Wellcome Sanger Institute COVID-19 Surveillance Team |
| EPI_ISL_2664982 | Lighthouse Laboratory Plymouth | Wellcome Sanger Institute for the COVID-19 Genomics UK (COG-UK) Consortium | Lighthouse Laboratory Plymouth and Alex Alderton, Roberto Amato, Jeffrey Barrett, Sonia Goncalves, Ewan Harrison, David K. Jackson, Ian Johnston, Dominic Kwiatkowski, Cordelia Langford, John Sillitoe on behalf of the Wellcome Sanger Institute COVID-19 Surveillance Team |
| EPI_ISL_2664983, EPI_ISL_2664984, EPI_ISL_2664985 | Lighthouse Lab in Alderley Park | Wellcome Sanger Institute for the COVID-19 Genomics UK (COG-UK) Consortium | Jacquelyn Wynn, Mairead Hyland, The Lighthouse Lab in Alderley Park and Alex Alderton, Roberto Amato, Jeffrey Barrett, Sonia Goncalves, Ewan Harrison, David K. Jackson, Ian Johnston, Dominic Kwiatkowski, Cordelia Langford, John Sillitoe on behalf of the Wellcome Sanger Institute COVID-19 Surveillance Team |
| EPI_ISL_2664986 | Lighthouse Laboratory Plymouth | Wellcome Sanger Institute for the COVID-19 Genomics UK (COG-UK) Consortium | Lighthouse Laboratory Plymouth and Alex Alderton, Roberto Amato, Jeffrey Barrett, Sonia Goncalves, Ewan Harrison, David K. Jackson, Ian Johnston, Dominic Kwiatkowski, Cordelia Langford, John Sillitoe on behalf of the Wellcome Sanger Institute COVID-19 Surveillance Team |
| EPI_ISL_2664987, EPI_ISL_2664988, EPI_ISL_2664989 | Lighthouse Lab in Milton Keynes | Wellcome Sanger Institute for the COVID-19 Genomics UK (COG-UK) Consortium | The Lighthouse Lab in Milton Keynes and Alex Alderton, Roberto Amato, Jeffrey Barrett, Sonia Goncalves, Ewan Harrison, David K. Jackson, Ian Johnston, Dominic Kwiatkowski, Cordelia Langford, John Sillitoe on behalf of the Wellcome Sanger Institute COVID-19 Surveillance Team |
| EPI_ISL_2664990, EPI_ISL_2664991, EPI_ISL_2664992 | Lighthouse Lab in Alderley Park | Wellcome Sanger Institute for the COVID-19 Genomics UK (COG-UK) Consortium | Jacquelyn Wynn, Mairead Hyland, The Lighthouse Lab in Alderley Park and Alex Alderton, Roberto Amato, Jeffrey Barrett, Sonia Goncalves, Ewan Harrison, David K. Jackson, Ian Johnston, Dominic Kwiatkowski, Cordelia Langford, John Sillitoe on behalf of the Wellcome Sanger Institute COVID-19 Surveillance Team |
| EPI_ISL_2664993, EPI_ISL_2664994 | Lighthouse Laboratory Plymouth | Wellcome Sanger Institute for the COVID-19 Genomics UK (COG-UK) Consortium | Lighthouse Laboratory Plymouth and Alex Alderton, Roberto Amato, Jeffrey Barrett, Sonia Goncalves, Ewan Harrison, David K. Jackson, Ian Johnston, Dominic Kwiatkowski, Cordelia Langford, John Sillitoe on behalf of the Wellcome Sanger Institute COVID-19 Surveillance Team |
| EPI_ISL_2664995 | Lighthouse Lab in Milton Keynes | Wellcome Sanger Institute for the COVID-19 Genomics UK (COG-UK) Consortium | The Lighthouse Lab in Milton Keynes and Alex Alderton, Roberto Amato, Jeffrey Barrett, Sonia Goncalves, Ewan Harrison, David K. Jackson, Ian Johnston, Dominic Kwiatkowski, Cordelia Langford, John Sillitoe on behalf of the Wellcome Sanger Institute COVID-19 Surveillance Team |
| EPI_ISL_2664996 | Lighthouse Lab in Alderley Park | Wellcome Sanger Institute for the COVID-19 Genomics UK (COG-UK) Consortium | Jacquelyn Wynn, Mairead Hyland, The Lighthouse Lab in Alderley Park and Alex Alderton, Roberto Amato, Jeffrey Barrett, Sonia Goncalves, Ewan Harrison, David K. Jackson, Ian Johnston, Dominic Kwiatkowski, Cordelia Langford, John Sillitoe on behalf of the Wellcome Sanger Institute COVID-19 Surveillance Team |
| EPI_ISL_2664997 | Lighthouse Lab in Milton Keynes | Wellcome Sanger Institute for the COVID-19 Genomics UK (COG-UK) Consortium | The Lighthouse Lab in Milton Keynes and Alex Alderton, Roberto Amato, Jeffrey Barrett, Sonia Goncalves, Ewan Harrison, David K. Jackson, Ian Johnston, Dominic Kwiatkowski, Cordelia Langford, John Sillitoe on behalf of the Wellcome Sanger Institute COVID-19 Surveillance Team |
| EPI_ISL_2664998 | Lighthouse Lab in Glasgow | Wellcome Sanger Institute for the COVID-19 Genomics UK (COG-UK) Consortium | Harper VanSteenhouse, Yumi Kasai, David Gray, Carol Clugston, Anna Dominiczak and Alex Alderton, Roberto Amato, Jeffrey Barrett, Sonia Goncalves, Ewan Harrison, David K. Jackson, Ian Johnston, Dominic Kwiatkowski, Cordelia Langford, John Sillitoe on behalf of the Wellcome Sanger Institute COVID-19 Surveillance Team |
| EPI_ISL_2664999, EPI_ISL_2665000 | Lighthouse Lab in Milton Keynes | Wellcome Sanger Institute for the COVID-19 Genomics UK (COG-UK) Consortium | The Lighthouse Lab in Milton Keynes and Alex Alderton, Roberto Amato, Jeffrey Barrett, Sonia Goncalves, Ewan Harrison, David K. Jackson, Ian Johnston, Dominic Kwiatkowski, Cordelia Langford, John Sillitoe on behalf of the Wellcome Sanger Institute COVID-19 Surveillance Team |
| EPI_ISL_2665001 | Lighthouse Lab in Alderley Park | Wellcome Sanger Institute for the COVID-19 Genomics UK (COG-UK) Consortium | Jacquelyn Wynn, Mairead Hyland, The Lighthouse Lab in Alderley Park and Alex Alderton, Roberto Amato, Jeffrey Barrett, Sonia Goncalves, Ewan Harrison, David K. Jackson, Ian Johnston, Dominic Kwiatkowski, Cordelia Langford, John Sillitoe on behalf of the Wellcome Sanger Institute COVID-19 Surveillance Team |
| EPI_ISL_2665002 | Lighthouse Laboratory Plymouth | Wellcome Sanger Institute for the COVID-19 Genomics UK (COG-UK) Consortium | Lighthouse Laboratory Plymouth and Alex Alderton, Roberto Amato, Jeffrey Barrett, Sonia Goncalves, Ewan Harrison, David K. Jackson, Ian Johnston, Dominic Kwiatkowski, Cordelia Langford, John Sillitoe on behalf of the Wellcome Sanger Institute COVID-19 Surveillance Team |
| EPI_ISL_2665003, EPI_ISL_2665004 | Lighthouse Lab in Milton Keynes | Wellcome Sanger Institute for the COVID-19 Genomics UK (COG-UK) Consortium | The Lighthouse Lab in Milton Keynes and Alex Alderton, Roberto Amato, Jeffrey Barrett, Sonia Goncalves, Ewan Harrison, David K. Jackson, Ian Johnston, Dominic Kwiatkowski, Cordelia Langford, John Sillitoe on behalf of the Wellcome Sanger Institute COVID-19 Surveillance Team |
| EPI_ISL_2665005, EPI_ISL_2665006, EPI_ISL_2665007 | Lighthouse Lab in Alderley Park | Wellcome Sanger Institute for the COVID-19 Genomics UK (COG-UK) Consortium | Jacquelyn Wynn, Mairead Hyland, The Lighthouse Lab in Alderley Park and Alex Alderton, Roberto Amato, Jeffrey Barrett, Sonia Goncalves, Ewan Harrison, David K. Jackson, Ian Johnston, Dominic Kwiatkowski, Cordelia Langford, John Sillitoe on behalf of the Wellcome Sanger Institute COVID-19 Surveillance Team |
| EPI_ISL_2665008 | Lighthouse Lab in Glasgow | Wellcome Sanger Institute for the COVID-19 Genomics UK (COG-UK) Consortium | Harper VanSteenhouse, Yumi Kasai, David Gray, Carol Clugston, Anna Dominiczak and Alex Alderton, Roberto Amato, Jeffrey Barrett, Sonia Goncalves, Ewan Harrison, David K. Jackson, Ian Johnston, Dominic Kwiatkowski, Cordelia Langford, John Sillitoe on behalf of the Wellcome Sanger Institute COVID-19 Surveillance Team |
| EPI_ISL_2665009 | Lighthouse Lab in Alderley Park | Wellcome Sanger Institute for the COVID-19 Genomics UK (COG-UK) Consortium | Jacquelyn Wynn, Mairead Hyland, The Lighthouse Lab in Alderley Park and Alex Alderton, Roberto Amato, Jeffrey Barrett, Sonia Goncalves, Ewan Harrison, David K. Jackson, Ian Johnston, Dominic Kwiatkowski, Cordelia Langford, John Sillitoe on behalf of the Wellcome Sanger Institute COVID-19 Surveillance Team |
| EPI_ISL_2665010 | Health Services Laboratories | Wellcome Sanger Institute for the COVID-19 Genomics UK (COG-UK) Consortium | Health Services Laboratories and Alex Alderton, Roberto Amato, Jeffrey Barrett, Sonia Goncalves, Ewan Harrison, David K. Jackson, Ian Johnston, Dominic Kwiatkowski, Cordelia Langford, John Sillitoe on behalf of the Wellcome Sanger Institute COVID-19 Surveillance Team |
| EPI_ISL_2665011 | Lighthouse Lab in Milton Keynes | Wellcome Sanger Institute for the COVID-19 Genomics UK (COG-UK) Consortium | The Lighthouse Lab in Milton Keynes and Alex Alderton, Roberto Amato, Jeffrey Barrett, Sonia Goncalves, Ewan Harrison, David K. Jackson, Ian Johnston, Dominic Kwiatkowski, Cordelia Langford, John Sillitoe on behalf of the Wellcome Sanger Institute COVID-19 Surveillance Team |
| EPI_ISL_2665012 | Lighthouse Laboratory Plymouth | Wellcome Sanger Institute for the COVID-19 Genomics UK (COG-UK) Consortium | Lighthouse Laboratory Plymouth and Alex Alderton, Roberto Amato, Jeffrey Barrett, Sonia Goncalves, Ewan Harrison, David K. Jackson, Ian Johnston, Dominic Kwiatkowski, Cordelia Langford, John Sillitoe on behalf of the Wellcome Sanger Institute COVID-19 Surveillance Team |
| EPI_ISL_2665013 | Lighthouse Lab in Milton Keynes | Wellcome Sanger Institute for the COVID-19 Genomics UK (COG-UK) Consortium | The Lighthouse Lab in Milton Keynes and Alex Alderton, Roberto Amato, Jeffrey Barrett, Sonia Goncalves, Ewan Harrison, David K. Jackson, Ian Johnston, Dominic Kwiatkowski, Cordelia Langford, John Sillitoe on behalf of the Wellcome Sanger Institute COVID-19 Surveillance Team |
| EPI_ISL_2665014 | Lighthouse Laboratory Plymouth | Wellcome Sanger Institute for the COVID-19 Genomics UK (COG-UK) Consortium | Lighthouse Laboratory Plymouth and Alex Alderton, Roberto Amato, Jeffrey Barrett, Sonia Goncalves, Ewan Harrison, David K. Jackson, Ian Johnston, Dominic Kwiatkowski, Cordelia Langford, John Sillitoe on behalf of the Wellcome Sanger Institute COVID-19 Surveillance Team |
| EPI_ISL_2665015 | Lighthouse Lab in Milton Keynes | Wellcome Sanger Institute for the COVID-19 Genomics UK (COG-UK) Consortium | The Lighthouse Lab in Milton Keynes and Alex Alderton, Roberto Amato, Jeffrey Barrett, Sonia Goncalves, Ewan Harrison, David K. Jackson, Ian Johnston, Dominic Kwiatkowski, Cordelia Langford, John Sillitoe on behalf of the Wellcome Sanger Institute COVID-19 Surveillance Team |
| EPI_ISL_2665016 | Lighthouse Lab in Alderley Park | Wellcome Sanger Institute for the COVID-19 Genomics UK (COG-UK) Consortium | Jacquelyn Wynn, Mairead Hyland, The Lighthouse Lab in Alderley Park and Alex Alderton, Roberto Amato, Jeffrey Barrett, Sonia Goncalves, Ewan Harrison, David K. Jackson, Ian Johnston, Dominic Kwiatkowski, Cordelia Langford, John Sillitoe on behalf of the Wellcome Sanger Institute COVID-19 Surveillance Team |
| EPI_ISL_2665017 | Lighthouse Lab in Milton Keynes | Wellcome Sanger Institute for the COVID-19 Genomics UK (COG-UK) Consortium | The Lighthouse Lab in Milton Keynes and Alex Alderton, Roberto Amato, Jeffrey Barrett, Sonia Goncalves, Ewan Harrison, David K. Jackson, Ian Johnston, Dominic Kwiatkowski, Cordelia Langford, John Sillitoe on behalf of the Wellcome Sanger Institute COVID-19 Surveillance Team |
| EPI_ISL_2665018 | Lighthouse Lab in Alderley Park | Wellcome Sanger Institute for the COVID-19 Genomics UK (COG-UK) Consortium | Jacquelyn Wynn, Mairead Hyland, The Lighthouse Lab in Alderley Park and Alex Alderton, Roberto Amato, Jeffrey Barrett, Sonia Goncalves, Ewan Harrison, David K. Jackson, Ian Johnston, Dominic Kwiatkowski, Cordelia Langford, John Sillitoe on behalf of the Wellcome Sanger Institute COVID-19 Surveillance Team |
| EPI_ISL_2665019, EPI_ISL_2665020, EPI_ISL_2665021 | Lighthouse Lab in Milton Keynes | Wellcome Sanger Institute for the COVID-19 Genomics UK (COG-UK) Consortium | The Lighthouse Lab in Milton Keynes and Alex Alderton, Roberto Amato, Jeffrey Barrett, Sonia Goncalves, Ewan Harrison, David K. Jackson, Ian Johnston, Dominic Kwiatkowski, Cordelia Langford, John Sillitoe on behalf of the Wellcome Sanger Institute COVID-19 Surveillance Team |
| EPI_ISL_2665022 | Lighthouse Lab in Alderley Park | Wellcome Sanger Institute for the COVID-19 Genomics UK (COG-UK) Consortium | Jacquelyn Wynn, Mairead Hyland, The Lighthouse Lab in Alderley Park and Alex Alderton, Roberto Amato, Jeffrey Barrett, Sonia Goncalves, Ewan Harrison, David K. Jackson, Ian Johnston, Dominic Kwiatkowski, Cordelia Langford, John Sillitoe on behalf of the Wellcome Sanger Institute COVID-19 Surveillance Team |
| EPI_ISL_2665023 | Lighthouse Lab in Milton Keynes | Wellcome Sanger Institute for the COVID-19 Genomics UK (COG-UK) Consortium | The Lighthouse Lab in Milton Keynes and Alex Alderton, Roberto Amato, Jeffrey Barrett, Sonia Goncalves, Ewan Harrison, David K. Jackson, Ian Johnston, Dominic Kwiatkowski, Cordelia Langford, John Sillitoe on behalf of the Wellcome Sanger Institute COVID-19 Surveillance Team |

| | | | |
|---|---|---|---|
| EPI_ISL_2665024 | Lighthouse Laboratory Plymouth | Wellcome Sanger Institute for the COVID-19 Genomics UK (COG-UK) Consortium | Lighthouse Laboratory Plymouth and Alex Alderton, Roberto Amato, Jeffrey Barrett, Sonia Goncalves, Ewan Harrison, David K. Jackson, Ian Johnston, Dominic Kwiatkowski, Cordelia Langford, John Sillitoe on behalf of the Wellcome Sanger Institute COVID-19 Surveillance Team |
| EPI_ISL_2665025 | Lighthouse Lab in Milton Keynes | Wellcome Sanger Institute for the COVID-19 Genomics UK (COG-UK) Consortium | The Lighthouse Lab in Milton Keynes and Alex Alderton, Roberto Amato, Jeffrey Barrett, Sonia Goncalves, Ewan Harrison, David K. Jackson, Ian Johnston, Dominic Kwiatkowski, Cordelia Langford, John Sillitoe on behalf of the Wellcome Sanger Institute COVID-19 Surveillance Team |
| EPI_ISL_2665026 | Lighthouse Lab in Alderley Park | Wellcome Sanger Institute for the COVID-19 Genomics UK (COG-UK) Consortium | Jacquelyn Wynn, Mairead Hyland, The Lighthouse Lab in Alderley Park and Alex Alderton, Roberto Amato, Jeffrey Barrett, Sonia Goncalves, Ewan Harrison, David K. Jackson, Ian Johnston, Dominic Kwiatkowski, Cordelia Langford, John Sillitoe on behalf of the Wellcome Sanger Institute COVID-19 Surveillance Team |
| EPI_ISL_2665027 | Lighthouse Laboratory Plymouth | Wellcome Sanger Institute for the COVID-19 Genomics UK (COG-UK) Consortium | Lighthouse Laboratory Plymouth and Alex Alderton, Roberto Amato, Jeffrey Barrett, Sonia Goncalves, Ewan Harrison, David K. Jackson, Ian Johnston, Dominic Kwiatkowski, Cordelia Langford, John Sillitoe on behalf of the Wellcome Sanger Institute COVID-19 Surveillance Team |
| EPI_ISL_2665028, EPI_ISL_2665029 | Lighthouse Lab in Milton Keynes | Wellcome Sanger Institute for the COVID-19 Genomics UK (COG-UK) Consortium | The Lighthouse Lab in Milton Keynes and Alex Alderton, Roberto Amato, Jeffrey Barrett, Sonia Goncalves, Ewan Harrison, David K. Jackson, Ian Johnston, Dominic Kwiatkowski, Cordelia Langford, John Sillitoe on behalf of the Wellcome Sanger Institute COVID-19 Surveillance Team |
| EPI_ISL_2665030 | Lighthouse Lab in Alderley Park | Wellcome Sanger Institute for the COVID-19 Genomics UK (COG-UK) Consortium | Jacquelyn Wynn, Mairead Hyland, The Lighthouse Lab in Alderley Park and Alex Alderton, Roberto Amato, Jeffrey Barrett, Sonia Goncalves, Ewan Harrison, David K. Jackson, Ian Johnston, Dominic Kwiatkowski, Cordelia Langford, John Sillitoe on behalf of the Wellcome Sanger Institute COVID-19 Surveillance Team |
| EPI_ISL_2665031, EPI_ISL_2665032 | Lighthouse Lab in Milton Keynes | Wellcome Sanger Institute for the COVID-19 Genomics UK (COG-UK) Consortium | The Lighthouse Lab in Milton Keynes and Alex Alderton, Roberto Amato, Jeffrey Barrett, Sonia Goncalves, Ewan Harrison, David K. Jackson, Ian Johnston, Dominic Kwiatkowski, Cordelia Langford, John Sillitoe on behalf of the Wellcome Sanger Institute COVID-19 Surveillance Team |
| EPI_ISL_2665033 | Lighthouse Lab in Alderley Park | Wellcome Sanger Institute for the COVID-19 Genomics UK (COG-UK) Consortium | Jacquelyn Wynn, Mairead Hyland, The Lighthouse Lab in Alderley Park and Alex Alderton, Roberto Amato, Jeffrey Barrett, Sonia Goncalves, Ewan Harrison, David K. Jackson, Ian Johnston, Dominic Kwiatkowski, Cordelia Langford, John Sillitoe on behalf of the Wellcome Sanger Institute COVID-19 Surveillance Team |
| EPI_ISL_2665034, EPI_ISL_2665035, EPI_ISL_2665037, EPI_ISL_2665038, EPI_ISL_2665039, EPI_ISL_2665040 | Lighthouse Lab in Milton Keynes | Wellcome Sanger Institute for the COVID-19 Genomics UK (COG-UK) Consortium | The Lighthouse Lab in Milton Keynes and Alex Alderton, Roberto Amato, Jeffrey Barrett, Sonia Goncalves, Ewan Harrison, David K. Jackson, Ian Johnston, Dominic Kwiatkowski, Cordelia Langford, John Sillitoe on behalf of the Wellcome Sanger Institute COVID-19 Surveillance Team |
| EPI_ISL_2665041, EPI_ISL_2665042, EPI_ISL_2665043, EPI_ISL_2665044 | Lighthouse Lab in Alderley Park | Wellcome Sanger Institute for the COVID-19 Genomics UK (COG-UK) Consortium | Jacquelyn Wynn, Mairead Hyland, The Lighthouse Lab in Alderley Park and Alex Alderton, Roberto Amato, Jeffrey Barrett, Sonia Goncalves, Ewan Harrison, David K. Jackson, Ian Johnston, Dominic Kwiatkowski, Cordelia Langford, John Sillitoe on behalf of the Wellcome Sanger Institute COVID-19 Surveillance Team |
| EPI_ISL_2665045, EPI_ISL_2665046, EPI_ISL_2665047, EPI_ISL_2665048, EPI_ISL_2665049, EPI_ISL_2665050, EPI_ISL_2665051, EPI_ISL_2665052 | Lighthouse Lab in Milton Keynes | Wellcome Sanger Institute for the COVID-19 Genomics UK (COG-UK) Consortium | The Lighthouse Lab in Milton Keynes and Alex Alderton, Roberto Amato, Jeffrey Barrett, Sonia Goncalves, Ewan Harrison, David K. Jackson, Ian Johnston, Dominic Kwiatkowski, Cordelia Langford, John Sillitoe on behalf of the Wellcome Sanger Institute COVID-19 Surveillance Team |
| EPI_ISL_2665053, EPI_ISL_2665054 | Lighthouse Lab in Alderley Park | Wellcome Sanger Institute for the COVID-19 Genomics UK (COG-UK) Consortium | Jacquelyn Wynn, Mairead Hyland, The Lighthouse Lab in Alderley Park and Alex Alderton, Roberto Amato, Jeffrey Barrett, Sonia Goncalves, Ewan Harrison, David K. Jackson, Ian Johnston, Dominic Kwiatkowski, Cordelia Langford, John Sillitoe on behalf of the Wellcome Sanger Institute COVID-19 Surveillance Team |
| EPI_ISL_2665055 | Lighthouse Lab in Milton Keynes | Wellcome Sanger Institute for the COVID-19 Genomics UK (COG-UK) Consortium | The Lighthouse Lab in Milton Keynes and Alex Alderton, Roberto Amato, Jeffrey Barrett, Sonia Goncalves, Ewan Harrison, David K. Jackson, Ian Johnston, Dominic Kwiatkowski, Cordelia Langford, John Sillitoe on behalf of the Wellcome Sanger Institute COVID-19 Surveillance Team |
| EPI_ISL_2665056 | Lighthouse Lab in Alderley Park | Wellcome Sanger Institute for the COVID-19 Genomics UK (COG-UK) Consortium | Jacquelyn Wynn, Mairead Hyland, The Lighthouse Lab in Alderley Park and Alex Alderton, Roberto Amato, Jeffrey Barrett, Sonia Goncalves, Ewan Harrison, David K. Jackson, Ian Johnston, Dominic Kwiatkowski, Cordelia Langford, John Sillitoe on behalf of the Wellcome Sanger Institute COVID-19 Surveillance Team |
| EPI_ISL_2665057 | Lighthouse Lab in Milton Keynes | Wellcome Sanger Institute for the COVID-19 Genomics UK (COG-UK) Consortium | The Lighthouse Lab in Milton Keynes and Alex Alderton, Roberto Amato, Jeffrey Barrett, Sonia Goncalves, Ewan Harrison, David K. Jackson, Ian Johnston, Dominic Kwiatkowski, Cordelia Langford, John Sillitoe on behalf of the Wellcome Sanger Institute COVID-19 Surveillance Team |
| EPI_ISL_2665058 | Lighthouse Lab in Alderley Park | Wellcome Sanger Institute for the COVID-19 Genomics UK (COG-UK) Consortium | Jacquelyn Wynn, Mairead Hyland, The Lighthouse Lab in Alderley Park and Alex Alderton, Roberto Amato, Jeffrey Barrett, Sonia Goncalves, Ewan Harrison, David K. Jackson, Ian Johnston, Dominic Kwiatkowski, Cordelia Langford, John Sillitoe on behalf of the Wellcome Sanger Institute COVID-19 Surveillance Team |
| EPI_ISL_2665059, EPI_ISL_2665060, EPI_ISL_2665061, EPI_ISL_2665062, EPI_ISL_2665063, EPI_ISL_2665064, EPI_ISL_2665065, EPI_ISL_2665066, EPI_ISL_2665067, EPI_ISL_2665068, EPI_ISL_2665069 | | | |
| see above | Lighthouse Lab in Milton Keynes | Wellcome Sanger Institute for the COVID-19 Genomics UK (COG-UK) Consortium | The Lighthouse Lab in Milton Keynes and Alex Alderton, Roberto Amato, Jeffrey Barrett, Sonia Goncalves, Ewan Harrison, David K. Jackson, Ian Johnston, Dominic Kwiatkowski, Cordelia Langford, John Sillitoe on behalf of the Wellcome Sanger Institute COVID-19 Surveillance Team |
| EPI_ISL_2665070 | Lighthouse Lab in Alderley Park | Wellcome Sanger Institute for the COVID-19 Genomics UK (COG-UK) Consortium | Jacquelyn Wynn, Mairead Hyland, The Lighthouse Lab in Alderley Park and Alex Alderton, Roberto Amato, Jeffrey Barrett, Sonia Goncalves, Ewan Harrison, David K. Jackson, Ian Johnston, Dominic Kwiatkowski, Cordelia Langford, John Sillitoe on behalf of the Wellcome Sanger Institute COVID-19 Surveillance Team |
| EPI_ISL_2665071, EPI_ISL_2665072, EPI_ISL_2665073, EPI_ISL_2665074 | Lighthouse Lab in Milton Keynes | Wellcome Sanger Institute for the COVID-19 Genomics UK (COG-UK) Consortium | The Lighthouse Lab in Milton Keynes and Alex Alderton, Roberto Amato, Jeffrey Barrett, Sonia Goncalves, Ewan Harrison, David K. Jackson, Ian Johnston, Dominic Kwiatkowski, Cordelia Langford, John Sillitoe on behalf of the Wellcome Sanger Institute COVID-19 Surveillance Team |
| EPI_ISL_2665075 | Lighthouse Lab in Alderley Park | Wellcome Sanger Institute for the COVID-19 Genomics UK (COG-UK) Consortium | Jacquelyn Wynn, Mairead Hyland, The Lighthouse Lab in Alderley Park and Alex Alderton, Roberto Amato, Jeffrey Barrett, Sonia Goncalves, Ewan Harrison, David K. Jackson, Ian Johnston, Dominic Kwiatkowski, Cordelia Langford, John Sillitoe on behalf of the Wellcome Sanger Institute COVID-19 Surveillance Team |
| EPI_ISL_2665076 | Lighthouse Lab in Milton Keynes | Wellcome Sanger Institute for the COVID-19 Genomics UK (COG-UK) Consortium | The Lighthouse Lab in Milton Keynes and Alex Alderton, Roberto Amato, Jeffrey Barrett, Sonia Goncalves, Ewan Harrison, David K. Jackson, Ian Johnston, Dominic Kwiatkowski, Cordelia Langford, John Sillitoe on behalf of the Wellcome Sanger Institute COVID-19 Surveillance Team |
| EPI_ISL_2665077 | Lighthouse Lab in Alderley Park | Wellcome Sanger Institute for the COVID-19 Genomics UK (COG-UK) Consortium | Jacquelyn Wynn, Mairead Hyland, The Lighthouse Lab in Alderley Park and Alex Alderton, Roberto Amato, Jeffrey Barrett, Sonia Goncalves, Ewan Harrison, David K. Jackson, Ian Johnston, Dominic Kwiatkowski, Cordelia Langford, John Sillitoe on behalf of the Wellcome Sanger Institute COVID-19 Surveillance Team |
| EPI_ISL_2665078, EPI_ISL_2665079, EPI_ISL_2665080, EPI_ISL_2665081, EPI_ISL_2665082, EPI_ISL_2665083, EPI_ISL_2665084 | Lighthouse Lab in Milton Keynes | Wellcome Sanger Institute for the COVID-19 Genomics UK (COG-UK) Consortium | The Lighthouse Lab in Milton Keynes and Alex Alderton, Roberto Amato, Jeffrey Barrett, Sonia Goncalves, Ewan Harrison, David K. Jackson, Ian Johnston, Dominic Kwiatkowski, Cordelia Langford, John Sillitoe on behalf of the Wellcome Sanger Institute COVID-19 Surveillance Team |
| EPI_ISL_2665085 | Lighthouse Lab in Alderley Park | Wellcome Sanger Institute for the COVID-19 Genomics UK (COG-UK) Consortium | Jacquelyn Wynn, Mairead Hyland, The Lighthouse Lab in Alderley Park and Alex Alderton, Roberto Amato, Jeffrey Barrett, Sonia Goncalves, Ewan Harrison, David K. Jackson, Ian Johnston, Dominic Kwiatkowski, Cordelia Langford, John Sillitoe on behalf of the Wellcome Sanger Institute COVID-19 Surveillance Team |
| EPI_ISL_2665086, EPI_ISL_2665087, EPI_ISL_2665088, EPI_ISL_2665089, EPI_ISL_2665090, EPI_ISL_2665091, EPI_ISL_2665092, EPI_ISL_2665093, EPI_ISL_2665094 | Lighthouse Lab in Milton Keynes | Wellcome Sanger Institute for the COVID-19 Genomics UK (COG-UK) Consortium | The Lighthouse Lab in Milton Keynes and Alex Alderton, Roberto Amato, Jeffrey Barrett, Sonia Goncalves, Ewan Harrison, David K. Jackson, Ian Johnston, Dominic Kwiatkowski, Cordelia Langford, John Sillitoe on behalf of the Wellcome Sanger Institute COVID-19 Surveillance Team |
| EPI_ISL_2665095, EPI_ISL_2665096, EPI_ISL_2665097 | Lighthouse Lab in Alderley Park | Wellcome Sanger Institute for the COVID-19 Genomics UK (COG-UK) Consortium | Jacquelyn Wynn, Mairead Hyland, The Lighthouse Lab in Alderley Park and Alex Alderton, Roberto Amato, Jeffrey Barrett, Sonia Goncalves, Ewan Harrison, David K. Jackson, Ian Johnston, Dominic Kwiatkowski, Cordelia Langford, John Sillitoe on behalf of the Wellcome Sanger Institute COVID-19 Surveillance Team |
| EPI_ISL_2665098, EPI_ISL_2665099, EPI_ISL_2665100, EPI_ISL_2665101, EPI_ISL_2665102, EPI_ISL_2665103, EPI_ISL_2665104, EPI_ISL_2665105 | Lighthouse Lab in Milton Keynes | Wellcome Sanger Institute for the COVID-19 Genomics UK (COG-UK) Consortium | The Lighthouse Lab in Milton Keynes and Alex Alderton, Roberto Amato, Jeffrey Barrett, Sonia Goncalves, Ewan Harrison, David K. Jackson, Ian Johnston, Dominic Kwiatkowski, Cordelia Langford, John Sillitoe on behalf of the Wellcome Sanger Institute COVID-19 Surveillance Team |
| EPI_ISL_2665106 | Lighthouse Lab in Alderley Park | Wellcome Sanger Institute for the COVID-19 Genomics UK (COG-UK) Consortium | Jacquelyn Wynn, Mairead Hyland, The Lighthouse Lab in Alderley Park and Alex Alderton, Roberto Amato, Jeffrey Barrett, Sonia Goncalves, Ewan Harrison, David K. Jackson, Ian Johnston, Dominic Kwiatkowski, Cordelia Langford, John Sillitoe on behalf of the Wellcome Sanger Institute COVID-19 Surveillance Team |
| EPI_ISL_2665107, EPI_ISL_2665108, EPI_ISL_2665109, EPI_ISL_2665110, EPI_ISL_2665111, EPI_ISL_2665112 | Lighthouse Lab in Milton Keynes | Wellcome Sanger Institute for the COVID-19 Genomics UK (COG-UK) Consortium | The Lighthouse Lab in Milton Keynes and Alex Alderton, Roberto Amato, Jeffrey Barrett, Sonia Goncalves, Ewan Harrison, David K. Jackson, Ian Johnston, Dominic Kwiatkowski, Cordelia Langford, John Sillitoe on behalf of the Wellcome Sanger Institute COVID-19 Surveillance Team |

| | | | |
|---|---|---|---|
| EPI_ISL_2665113 | Lighthouse Lab in Alderley Park | Wellcome Sanger Institute for the COVID-19 Genomics UK (COG-UK) Consortium | Jacquelyn Wynn, Mairead Hyland, The Lighthouse Lab in Alderley Park and Alex Alderton, Roberto Amato, Jeffrey Barrett, Sonia Goncalves, Ewan Harrison, David K. Jackson, Ian Johnston, Dominic Kwiatkowski, Cordelia Langford, John Sillitoe on behalf of the Wellcome Sanger Institute COVID-19 Surveillance Team |
| EPI_ISL_2665114, EPI_ISL_2665115 | Lighthouse Lab in Milton Keynes | Wellcome Sanger Institute for the COVID-19 Genomics UK (COG-UK) Consortium | The Lighthouse Lab in Milton Keynes and Alex Alderton, Roberto Amato, Jeffrey Barrett, Sonia Goncalves, Ewan Harrison, David K. Jackson, Ian Johnston, Dominic Kwiatkowski, Cordelia Langford, John Sillitoe on behalf of the Wellcome Sanger Institute COVID-19 Surveillance Team |
| EPI_ISL_2665116 | Lighthouse Lab in Alderley Park | Wellcome Sanger Institute for the COVID-19 Genomics UK (COG-UK) Consortium | Jacquelyn Wynn, Mairead Hyland, The Lighthouse Lab in Alderley Park and Alex Alderton, Roberto Amato, Jeffrey Barrett, Sonia Goncalves, Ewan Harrison, David K. Jackson, Ian Johnston, Dominic Kwiatkowski, Cordelia Langford, John Sillitoe on behalf of the Wellcome Sanger Institute COVID-19 Surveillance Team |
| EPI_ISL_2665117, EPI_ISL_2665118 | Lighthouse Lab in Milton Keynes | Wellcome Sanger Institute for the COVID-19 Genomics UK (COG-UK) Consortium | The Lighthouse Lab in Milton Keynes and Alex Alderton, Roberto Amato, Jeffrey Barrett, Sonia Goncalves, Ewan Harrison, David K. Jackson, Ian Johnston, Dominic Kwiatkowski, Cordelia Langford, John Sillitoe on behalf of the Wellcome Sanger Institute COVID-19 Surveillance Team |
| EPI_ISL_2665119, EPI_ISL_2665120, EPI_ISL_2665121 | Lighthouse Lab in Alderley Park | Wellcome Sanger Institute for the COVID-19 Genomics UK (COG-UK) Consortium | Jacquelyn Wynn, Mairead Hyland, The Lighthouse Lab in Alderley Park and Alex Alderton, Roberto Amato, Jeffrey Barrett, Sonia Goncalves, Ewan Harrison, David K. Jackson, Ian Johnston, Dominic Kwiatkowski, Cordelia Langford, John Sillitoe on behalf of the Wellcome Sanger Institute COVID-19 Surveillance Team |
| EPI_ISL_2665122 | Lighthouse Lab in Milton Keynes | Wellcome Sanger Institute for the COVID-19 Genomics UK (COG-UK) Consortium | The Lighthouse Lab in Milton Keynes and Alex Alderton, Roberto Amato, Jeffrey Barrett, Sonia Goncalves, Ewan Harrison, David K. Jackson, Ian Johnston, Dominic Kwiatkowski, Cordelia Langford, John Sillitoe on behalf of the Wellcome Sanger Institute COVID-19 Surveillance Team |
| EPI_ISL_2665123 | Lighthouse Lab in Alderley Park | Wellcome Sanger Institute for the COVID-19 Genomics UK (COG-UK) Consortium | Jacquelyn Wynn, Mairead Hyland, The Lighthouse Lab in Alderley Park and Alex Alderton, Roberto Amato, Jeffrey Barrett, Sonia Goncalves, Ewan Harrison, David K. Jackson, Ian Johnston, Dominic Kwiatkowski, Cordelia Langford, John Sillitoe on behalf of the Wellcome Sanger Institute COVID-19 Surveillance Team |
| EPI_ISL_2665124, EPI_ISL_2665125, EPI_ISL_2665126 | Lighthouse Lab in Milton Keynes | Wellcome Sanger Institute for the COVID-19 Genomics UK (COG-UK) Consortium | The Lighthouse Lab in Milton Keynes and Alex Alderton, Roberto Amato, Jeffrey Barrett, Sonia Goncalves, Ewan Harrison, David K. Jackson, Ian Johnston, Dominic Kwiatkowski, Cordelia Langford, John Sillitoe on behalf of the Wellcome Sanger Institute COVID-19 Surveillance Team |
| EPI_ISL_2665127 | Lighthouse Lab in Alderley Park | Wellcome Sanger Institute for the COVID-19 Genomics UK (COG-UK) Consortium | Jacquelyn Wynn, Mairead Hyland, The Lighthouse Lab in Alderley Park and Alex Alderton, Roberto Amato, Jeffrey Barrett, Sonia Goncalves, Ewan Harrison, David K. Jackson, Ian Johnston, Dominic Kwiatkowski, Cordelia Langford, John Sillitoe on behalf of the Wellcome Sanger Institute COVID-19 Surveillance Team |
| EPI_ISL_2665129, EPI_ISL_2665130 | Lighthouse Lab in Milton Keynes | Wellcome Sanger Institute for the COVID-19 Genomics UK (COG-UK) Consortium | The Lighthouse Lab in Milton Keynes and Alex Alderton, Roberto Amato, Jeffrey Barrett, Sonia Goncalves, Ewan Harrison, David K. Jackson, Ian Johnston, Dominic Kwiatkowski, Cordelia Langford, John Sillitoe on behalf of the Wellcome Sanger Institute COVID-19 Surveillance Team |
| EPI_ISL_2665131 | Lighthouse Lab in Alderley Park | Wellcome Sanger Institute for the COVID-19 Genomics UK (COG-UK) Consortium | Jacquelyn Wynn, Mairead Hyland, The Lighthouse Lab in Alderley Park and Alex Alderton, Roberto Amato, Jeffrey Barrett, Sonia Goncalves, Ewan Harrison, David K. Jackson, Ian Johnston, Dominic Kwiatkowski, Cordelia Langford, John Sillitoe on behalf of the Wellcome Sanger Institute COVID-19 Surveillance Team |
| EPI_ISL_2665132 | Lighthouse Lab in Milton Keynes | Wellcome Sanger Institute for the COVID-19 Genomics UK (COG-UK) Consortium | The Lighthouse Lab in Milton Keynes and Alex Alderton, Roberto Amato, Jeffrey Barrett, Sonia Goncalves, Ewan Harrison, David K. Jackson, Ian Johnston, Dominic Kwiatkowski, Cordelia Langford, John Sillitoe on behalf of the Wellcome Sanger Institute COVID-19 Surveillance Team |
| EPI_ISL_2665133 | Lighthouse Lab in Alderley Park | Wellcome Sanger Institute for the COVID-19 Genomics UK (COG-UK) Consortium | Jacquelyn Wynn, Mairead Hyland, The Lighthouse Lab in Alderley Park and Alex Alderton, Roberto Amato, Jeffrey Barrett, Sonia Goncalves, Ewan Harrison, David K. Jackson, Ian Johnston, Dominic Kwiatkowski, Cordelia Langford, John Sillitoe on behalf of the Wellcome Sanger Institute COVID-19 Surveillance Team |
| EPI_ISL_2665134, EPI_ISL_2665135, EPI_ISL_2665136, EPI_ISL_2665137, EPI_ISL_2665138 | Lighthouse Lab in Milton Keynes | Wellcome Sanger Institute for the COVID-19 Genomics UK (COG-UK) Consortium | The Lighthouse Lab in Milton Keynes and Alex Alderton, Roberto Amato, Jeffrey Barrett, Sonia Goncalves, Ewan Harrison, David K. Jackson, Ian Johnston, Dominic Kwiatkowski, Cordelia Langford, John Sillitoe on behalf of the Wellcome Sanger Institute COVID-19 Surveillance Team |
| EPI_ISL_2665139, EPI_ISL_2665140, EPI_ISL_2665141 | Lighthouse Lab in Alderley Park | Wellcome Sanger Institute for the COVID-19 Genomics UK (COG-UK) Consortium | Jacquelyn Wynn, Mairead Hyland, The Lighthouse Lab in Alderley Park and Alex Alderton, Roberto Amato, Jeffrey Barrett, Sonia Goncalves, Ewan Harrison, David K. Jackson, Ian Johnston, Dominic Kwiatkowski, Cordelia Langford, John Sillitoe on behalf of the Wellcome Sanger Institute COVID-19 Surveillance Team |
| EPI_ISL_2665142, EPI_ISL_2665143, EPI_ISL_2665144, EPI_ISL_2665145, EPI_ISL_2665146 | Lighthouse Lab in Milton Keynes | Wellcome Sanger Institute for the COVID-19 Genomics UK (COG-UK) Consortium | The Lighthouse Lab in Milton Keynes and Alex Alderton, Roberto Amato, Jeffrey Barrett, Sonia Goncalves, Ewan Harrison, David K. Jackson, Ian Johnston, Dominic Kwiatkowski, Cordelia Langford, John Sillitoe on behalf of the Wellcome Sanger Institute COVID-19 Surveillance Team |
| EPI_ISL_2665147 | Lighthouse Lab in Alderley Park | Wellcome Sanger Institute for the COVID-19 Genomics UK (COG-UK) Consortium | Jacquelyn Wynn, Mairead Hyland, The Lighthouse Lab in Alderley Park and Alex Alderton, Roberto Amato, Jeffrey Barrett, Sonia Goncalves, Ewan Harrison, David K. Jackson, Ian Johnston, Dominic Kwiatkowski, Cordelia Langford, John Sillitoe on behalf of the Wellcome Sanger Institute COVID-19 Surveillance Team |
| EPI_ISL_2665148, EPI_ISL_2665149 | Lighthouse Lab in Milton Keynes | Wellcome Sanger Institute for the COVID-19 Genomics UK (COG-UK) Consortium | The Lighthouse Lab in Milton Keynes and Alex Alderton, Roberto Amato, Jeffrey Barrett, Sonia Goncalves, Ewan Harrison, David K. Jackson, Ian Johnston, Dominic Kwiatkowski, Cordelia Langford, John Sillitoe on behalf of the Wellcome Sanger Institute COVID-19 Surveillance Team |
| EPI_ISL_2665150 | Lighthouse Lab in Alderley Park | Wellcome Sanger Institute for the COVID-19 Genomics UK (COG-UK) Consortium | Jacquelyn Wynn, Mairead Hyland, The Lighthouse Lab in Alderley Park and Alex Alderton, Roberto Amato, Jeffrey Barrett, Sonia Goncalves, Ewan Harrison, David K. Jackson, Ian Johnston, Dominic Kwiatkowski, Cordelia Langford, John Sillitoe on behalf of the Wellcome Sanger Institute COVID-19 Surveillance Team |
| EPI_ISL_2665151, EPI_ISL_2665152, EPI_ISL_2665153, EPI_ISL_2665154, EPI_ISL_2665155, EPI_ISL_2665156 | Lighthouse Lab in Milton Keynes | Wellcome Sanger Institute for the COVID-19 Genomics UK (COG-UK) Consortium | The Lighthouse Lab in Milton Keynes and Alex Alderton, Roberto Amato, Jeffrey Barrett, Sonia Goncalves, Ewan Harrison, David K. Jackson, Ian Johnston, Dominic Kwiatkowski, Cordelia Langford, John Sillitoe on behalf of the Wellcome Sanger Institute COVID-19 Surveillance Team |
| EPI_ISL_2665157, EPI_ISL_2665158 | Lighthouse Lab in Alderley Park | Wellcome Sanger Institute for the COVID-19 Genomics UK (COG-UK) Consortium | Jacquelyn Wynn, Mairead Hyland, The Lighthouse Lab in Alderley Park and Alex Alderton, Roberto Amato, Jeffrey Barrett, Sonia Goncalves, Ewan Harrison, David K. Jackson, Ian Johnston, Dominic Kwiatkowski, Cordelia Langford, John Sillitoe on behalf of the Wellcome Sanger Institute COVID-19 Surveillance Team |
| EPI_ISL_2665159 | Lighthouse Lab in Milton Keynes | Wellcome Sanger Institute for the COVID-19 Genomics UK (COG-UK) Consortium | The Lighthouse Lab in Milton Keynes and Alex Alderton, Roberto Amato, Jeffrey Barrett, Sonia Goncalves, Ewan Harrison, David K. Jackson, Ian Johnston, Dominic Kwiatkowski, Cordelia Langford, John Sillitoe on behalf of the Wellcome Sanger Institute COVID-19 Surveillance Team |
| EPI_ISL_2665160 | Lighthouse Lab in Alderley Park | Wellcome Sanger Institute for the COVID-19 Genomics UK (COG-UK) Consortium | Jacquelyn Wynn, Mairead Hyland, The Lighthouse Lab in Alderley Park and Alex Alderton, Roberto Amato, Jeffrey Barrett, Sonia Goncalves, Ewan Harrison, David K. Jackson, Ian Johnston, Dominic Kwiatkowski, Cordelia Langford, John Sillitoe on behalf of the Wellcome Sanger Institute COVID-19 Surveillance Team |
| EPI_ISL_2665161, EPI_ISL_2665162, EPI_ISL_2665163 | Lighthouse Lab in Milton Keynes | Wellcome Sanger Institute for the COVID-19 Genomics UK (COG-UK) Consortium | The Lighthouse Lab in Milton Keynes and Alex Alderton, Roberto Amato, Jeffrey Barrett, Sonia Goncalves, Ewan Harrison, David K. Jackson, Ian Johnston, Dominic Kwiatkowski, Cordelia Langford, John Sillitoe on behalf of the Wellcome Sanger Institute COVID-19 Surveillance Team |
| EPI_ISL_2665164 | Lighthouse Lab in Alderley Park | Wellcome Sanger Institute for the COVID-19 Genomics UK (COG-UK) Consortium | Jacquelyn Wynn, Mairead Hyland, The Lighthouse Lab in Alderley Park and Alex Alderton, Roberto Amato, Jeffrey Barrett, Sonia Goncalves, Ewan Harrison, David K. Jackson, Ian Johnston, Dominic Kwiatkowski, Cordelia Langford, John Sillitoe on behalf of the Wellcome Sanger Institute COVID-19 Surveillance Team |
| EPI_ISL_2665165, EPI_ISL_2665166 | Lighthouse Lab in Milton Keynes | Wellcome Sanger Institute for the COVID-19 Genomics UK (COG-UK) Consortium | The Lighthouse Lab in Milton Keynes and Alex Alderton, Roberto Amato, Jeffrey Barrett, Sonia Goncalves, Ewan Harrison, David K. Jackson, Ian Johnston, Dominic Kwiatkowski, Cordelia Langford, John Sillitoe on behalf of the Wellcome Sanger Institute COVID-19 Surveillance Team |
| EPI_ISL_2665167, EPI_ISL_2665168 | Lighthouse Lab in Alderley Park | Wellcome Sanger Institute for the COVID-19 Genomics UK (COG-UK) Consortium | Jacquelyn Wynn, Mairead Hyland, The Lighthouse Lab in Alderley Park and Alex Alderton, Roberto Amato, Jeffrey Barrett, Sonia Goncalves, Ewan Harrison, David K. Jackson, Ian Johnston, Dominic Kwiatkowski, Cordelia Langford, John Sillitoe on behalf of the Wellcome Sanger Institute COVID-19 Surveillance Team |
| EPI_ISL_2665169, EPI_ISL_2665170, EPI_ISL_2665171 | Lighthouse Lab in Milton Keynes | Wellcome Sanger Institute for the COVID-19 Genomics UK (COG-UK) Consortium | The Lighthouse Lab in Milton Keynes and Alex Alderton, Roberto Amato, Jeffrey Barrett, Sonia Goncalves, Ewan Harrison, David K. Jackson, Ian Johnston, Dominic Kwiatkowski, Cordelia Langford, John Sillitoe on behalf of the Wellcome Sanger Institute COVID-19 Surveillance Team |
| EPI_ISL_2665172 | Lighthouse Lab in Alderley Park | Wellcome Sanger Institute for the COVID-19 Genomics UK (COG-UK) Consortium | Jacquelyn Wynn, Mairead Hyland, The Lighthouse Lab in Alderley Park and Alex Alderton, Roberto Amato, Jeffrey Barrett, Sonia Goncalves, Ewan Harrison, David K. Jackson, Ian Johnston, Dominic Kwiatkowski, Cordelia Langford, John Sillitoe on behalf of the Wellcome Sanger Institute COVID-19 Surveillance Team |
| EPI_ISL_2665173, EPI_ISL_2665174, EPI_ISL_2665175, EPI_ISL_2665176, EPI_ISL_2665177, EPI_ISL_2665178 | Lighthouse Lab in Milton Keynes | Wellcome Sanger Institute for the COVID-19 Genomics UK (COG-UK) Consortium | The Lighthouse Lab in Milton Keynes and Alex Alderton, Roberto Amato, Jeffrey Barrett, Sonia Goncalves, Ewan Harrison, David K. Jackson, Ian Johnston, Dominic Kwiatkowski, Cordelia Langford, John Sillitoe on behalf of the Wellcome Sanger Institute COVID-19 Surveillance Team |
| EPI_ISL_2665179, EPI_ISL_2665180 | Lighthouse Lab in Alderley Park | Wellcome Sanger Institute for the COVID-19 Genomics UK (COG-UK) Consortium | Jacquelyn Wynn, Mairead Hyland, The Lighthouse Lab in Alderley Park and Alex Alderton, Roberto Amato, Jeffrey Barrett, Sonia Goncalves, Ewan Harrison, David K. Jackson, Ian Johnston, Dominic Kwiatkowski, Cordelia Langford, John Sillitoe on behalf of the Wellcome Sanger Institute COVID-19 Surveillance Team |
| EPI_ISL_2665181, EPI_ISL_2665182, EPI_ISL_2665183 | Lighthouse Lab in Milton Keynes | Wellcome Sanger Institute for the COVID-19 Genomics UK (COG-UK) Consortium | The Lighthouse Lab in Milton Keynes and Alex Alderton, Roberto Amato, Jeffrey Barrett, Sonia Goncalves, Ewan Harrison, David K. Jackson, Ian Johnston, Dominic Kwiatkowski, Cordelia Langford, John Sillitoe on behalf of the Wellcome Sanger Institute COVID-19 Surveillance Team |
| EPI_ISL_2665184 | Lighthouse Lab in Alderley Park | Wellcome Sanger Institute for the COVID-19 Genomics UK (COG-UK) Consortium | Jacquelyn Wynn, Mairead Hyland, The Lighthouse Lab in Alderley Park and Alex Alderton, Roberto Amato, Jeffrey Barrett, Sonia Goncalves, Ewan Harrison, David K. Jackson, Ian Johnston, Dominic Kwiatkowski, Cordelia Langford, John Sillitoe on behalf of the Wellcome Sanger Institute COVID-19 Surveillance Team |
| EPI_ISL_2665185 | Lighthouse Lab in Milton Keynes | Wellcome Sanger Institute for the COVID-19 Genomics UK (COG-UK) Consortium | The Lighthouse Lab in Milton Keynes and Alex Alderton, Roberto Amato, Jeffrey Barrett, Sonia Goncalves, Ewan Harrison, David K. Jackson, Ian Johnston, Dominic Kwiatkowski, Cordelia Langford, John Sillitoe on behalf of the Wellcome Sanger Institute COVID-19 Surveillance Team |
| EPI_ISL_2665186, EPI_ISL_2665187 | Lighthouse Lab in Alderley Park | Wellcome Sanger Institute for the COVID-19 Genomics UK (COG-UK) Consortium | Jacquelyn Wynn, Mairead Hyland, The Lighthouse Lab in Alderley Park and Alex Alderton, Roberto Amato, Jeffrey Barrett, Sonia Goncalves, Ewan Harrison, David K. Jackson, Ian Johnston, Dominic Kwiatkowski, Cordelia Langford, John Sillitoe on behalf of the Wellcome Sanger Institute COVID-19 Surveillance Team |
| EPI_ISL_2665188 | Lighthouse Lab in Milton Keynes | Wellcome Sanger Institute for the COVID-19 Genomics UK (COG-UK) Consortium | The Lighthouse Lab in Milton Keynes and Alex Alderton, Roberto Amato, Jeffrey Barrett, Sonia Goncalves, Ewan Harrison, David K. Jackson, Ian Johnston, Dominic Kwiatkowski, Cordelia Langford, John Sillitoe on behalf of the Wellcome Sanger Institute COVID-19 Surveillance Team |
| EPI_ISL_2665189, EPI_ISL_2665190, | Lighthouse Lab in Alderley Park | Wellcome Sanger Institute for the COVID-19 Genomics UK (COG-UK) Consortium | Jacquelyn Wynn, Mairead Hyland, The Lighthouse Lab in Alderley Park and Alex Alderton, Roberto Amato, Jeffrey Barrett, Sonia Goncalves, Ewan Harrison, David K. Jackson, Ian Johnston, Dominic Kwiatkowski, Cordelia Langford, John Sillitoe on behalf of the Wellcome Sanger Institute COVID-19 Surveillance Team |

| | | | |
|---|---|---|---|
| EPI_ISL_2665191, EPI_ISL_2665192 | | | |
| EPI_ISL_2665193, EPI_ISL_2665194, EPI_ISL_2665195, EPI_ISL_2665196, EPI_ISL_2665197 | Lighthouse Lab in Milton Keynes | Wellcome Sanger Institute for the COVID-19 Genomics UK (COG-UK) Consortium | The Lighthouse Lab in Milton Keynes and Alex Alderton, Roberto Amato, Jeffrey Barrett, Sonia Goncalves, Ewan Harrison, David K. Jackson, Ian Johnston, Dominic Kwiatkowski, Cordelia Langford, John Sillitoe on behalf of the Wellcome Sanger Institute COVID-19 Surveillance Team |
| EPI_ISL_2665198 | Lighthouse Lab in Alderley Park | Wellcome Sanger Institute for the COVID-19 Genomics UK (COG-UK) Consortium | Jacquelyn Wynn, Mairead Hyland, The Lighthouse Lab in Alderley Park and Alex Alderton, Roberto Amato, Jeffrey Barrett, Sonia Goncalves, Ewan Harrison, David K. Jackson, Ian Johnston, Dominic Kwiatkowski, Cordelia Langford, John Sillitoe on behalf of the Wellcome Sanger Institute COVID-19 Surveillance Team |
| EPI_ISL_2665199 | Lighthouse Lab in Milton Keynes | Wellcome Sanger Institute for the COVID-19 Genomics UK (COG-UK) Consortium | The Lighthouse Lab in Milton Keynes and Alex Alderton, Roberto Amato, Jeffrey Barrett, Sonia Goncalves, Ewan Harrison, David K. Jackson, Ian Johnston, Dominic Kwiatkowski, Cordelia Langford, John Sillitoe on behalf of the Wellcome Sanger Institute COVID-19 Surveillance Team |
| EPI_ISL_2665200 | Lighthouse Lab in Alderley Park | Wellcome Sanger Institute for the COVID-19 Genomics UK (COG-UK) Consortium | Jacquelyn Wynn, Mairead Hyland, The Lighthouse Lab in Alderley Park and Alex Alderton, Roberto Amato, Jeffrey Barrett, Sonia Goncalves, Ewan Harrison, David K. Jackson, Ian Johnston, Dominic Kwiatkowski, Cordelia Langford, John Sillitoe on behalf of the Wellcome Sanger Institute COVID-19 Surveillance Team |
| EPI_ISL_2665201, EPI_ISL_2665202, EPI_ISL_2665203, EPI_ISL_2665204, EPI_ISL_2665205, EPI_ISL_2665207 | Lighthouse Lab in Milton Keynes | Wellcome Sanger Institute for the COVID-19 Genomics UK (COG-UK) Consortium | The Lighthouse Lab in Milton Keynes and Alex Alderton, Roberto Amato, Jeffrey Barrett, Sonia Goncalves, Ewan Harrison, David K. Jackson, Ian Johnston, Dominic Kwiatkowski, Cordelia Langford, John Sillitoe on behalf of the Wellcome Sanger Institute COVID-19 Surveillance Team |
| EPI_ISL_2665208 | Lighthouse Lab in Alderley Park | Wellcome Sanger Institute for the COVID-19 Genomics UK (COG-UK) Consortium | Jacquelyn Wynn, Mairead Hyland, The Lighthouse Lab in Alderley Park and Alex Alderton, Roberto Amato, Jeffrey Barrett, Sonia Goncalves, Ewan Harrison, David K. Jackson, Ian Johnston, Dominic Kwiatkowski, Cordelia Langford, John Sillitoe on behalf of the Wellcome Sanger Institute COVID-19 Surveillance Team |
| EPI_ISL_2665209, EPI_ISL_2665210, EPI_ISL_2665211, EPI_ISL_2665212, EPI_ISL_2665213, EPI_ISL_2665214, EPI_ISL_2665216, EPI_ISL_2665217 | Lighthouse Lab in Milton Keynes | Wellcome Sanger Institute for the COVID-19 Genomics UK (COG-UK) Consortium | The Lighthouse Lab in Milton Keynes and Alex Alderton, Roberto Amato, Jeffrey Barrett, Sonia Goncalves, Ewan Harrison, David K. Jackson, Ian Johnston, Dominic Kwiatkowski, Cordelia Langford, John Sillitoe on behalf of the Wellcome Sanger Institute COVID-19 Surveillance Team |
| EPI_ISL_2665218 | Lighthouse Lab in Alderley Park | Wellcome Sanger Institute for the COVID-19 Genomics UK (COG-UK) Consortium | Jacquelyn Wynn, Mairead Hyland, The Lighthouse Lab in Alderley Park and Alex Alderton, Roberto Amato, Jeffrey Barrett, Sonia Goncalves, Ewan Harrison, David K. Jackson, Ian Johnston, Dominic Kwiatkowski, Cordelia Langford, John Sillitoe on behalf of the Wellcome Sanger Institute COVID-19 Surveillance Team |
| EPI_ISL_2665219, EPI_ISL_2665220, EPI_ISL_2665221, EPI_ISL_2665222, EPI_ISL_2665223, EPI_ISL_2665224 | Lighthouse Lab in Milton Keynes | Wellcome Sanger Institute for the COVID-19 Genomics UK (COG-UK) Consortium | The Lighthouse Lab in Milton Keynes and Alex Alderton, Roberto Amato, Jeffrey Barrett, Sonia Goncalves, Ewan Harrison, David K. Jackson, Ian Johnston, Dominic Kwiatkowski, Cordelia Langford, John Sillitoe on behalf of the Wellcome Sanger Institute COVID-19 Surveillance Team |
| EPI_ISL_2665225 | Lighthouse Lab in Alderley Park | Wellcome Sanger Institute for the COVID-19 Genomics UK (COG-UK) Consortium | Jacquelyn Wynn, Mairead Hyland, The Lighthouse Lab in Alderley Park and Alex Alderton, Roberto Amato, Jeffrey Barrett, Sonia Goncalves, Ewan Harrison, David K. Jackson, Ian Johnston, Dominic Kwiatkowski, Cordelia Langford, John Sillitoe on behalf of the Wellcome Sanger Institute COVID-19 Surveillance Team |
| EPI_ISL_2665226, EPI_ISL_2665228, EPI_ISL_2665229 | Lighthouse Lab in Milton Keynes | Wellcome Sanger Institute for the COVID-19 Genomics UK (COG-UK) Consortium | The Lighthouse Lab in Milton Keynes and Alex Alderton, Roberto Amato, Jeffrey Barrett, Sonia Goncalves, Ewan Harrison, David K. Jackson, Ian Johnston, Dominic Kwiatkowski, Cordelia Langford, John Sillitoe on behalf of the Wellcome Sanger Institute COVID-19 Surveillance Team |
| EPI_ISL_2665230 | Lighthouse Lab in Alderley Park | Wellcome Sanger Institute for the COVID-19 Genomics UK (COG-UK) Consortium | Jacquelyn Wynn, Mairead Hyland, The Lighthouse Lab in Alderley Park and Alex Alderton, Roberto Amato, Jeffrey Barrett, Sonia Goncalves, Ewan Harrison, David K. Jackson, Ian Johnston, Dominic Kwiatkowski, Cordelia Langford, John Sillitoe on behalf of the Wellcome Sanger Institute COVID-19 Surveillance Team |
| EPI_ISL_2665231, EPI_ISL_2665232, EPI_ISL_2665233, EPI_ISL_2665234, EPI_ISL_2665235 | Lighthouse Lab in Milton Keynes | Wellcome Sanger Institute for the COVID-19 Genomics UK (COG-UK) Consortium | The Lighthouse Lab in Milton Keynes and Alex Alderton, Roberto Amato, Jeffrey Barrett, Sonia Goncalves, Ewan Harrison, David K. Jackson, Ian Johnston, Dominic Kwiatkowski, Cordelia Langford, John Sillitoe on behalf of the Wellcome Sanger Institute COVID-19 Surveillance Team |
| EPI_ISL_2665236 | Lighthouse Lab in Alderley Park | Wellcome Sanger Institute for the COVID-19 Genomics UK (COG-UK) Consortium | Jacquelyn Wynn, Mairead Hyland, The Lighthouse Lab in Alderley Park and Alex Alderton, Roberto Amato, Jeffrey Barrett, Sonia Goncalves, Ewan Harrison, David K. Jackson, Ian Johnston, Dominic Kwiatkowski, Cordelia Langford, John Sillitoe on behalf of the Wellcome Sanger Institute COVID-19 Surveillance Team |
| EPI_ISL_2665237, EPI_ISL_2665238, EPI_ISL_2665239, EPI_ISL_2665240, EPI_ISL_2665241, EPI_ISL_2665242, EPI_ISL_2665243, EPI_ISL_2665244, EPI_ISL_2665245 | Lighthouse Lab in Milton Keynes | Wellcome Sanger Institute for the COVID-19 Genomics UK (COG-UK) Consortium | The Lighthouse Lab in Milton Keynes and Alex Alderton, Roberto Amato, Jeffrey Barrett, Sonia Goncalves, Ewan Harrison, David K. Jackson, Ian Johnston, Dominic Kwiatkowski, Cordelia Langford, John Sillitoe on behalf of the Wellcome Sanger Institute COVID-19 Surveillance Team |
| EPI_ISL_2665246, EPI_ISL_2665247 | Lighthouse Lab in Alderley Park | Wellcome Sanger Institute for the COVID-19 Genomics UK (COG-UK) Consortium | Jacquelyn Wynn, Mairead Hyland, The Lighthouse Lab in Alderley Park and Alex Alderton, Roberto Amato, Jeffrey Barrett, Sonia Goncalves, Ewan Harrison, David K. Jackson, Ian Johnston, Dominic Kwiatkowski, Cordelia Langford, John Sillitoe on behalf of the Wellcome Sanger Institute COVID-19 Surveillance Team |
| EPI_ISL_2665248, EPI_ISL_2665249, EPI_ISL_2665250 | Lighthouse Lab in Milton Keynes | Wellcome Sanger Institute for the COVID-19 Genomics UK (COG-UK) Consortium | The Lighthouse Lab in Milton Keynes and Alex Alderton, Roberto Amato, Jeffrey Barrett, Sonia Goncalves, Ewan Harrison, David K. Jackson, Ian Johnston, Dominic Kwiatkowski, Cordelia Langford, John Sillitoe on behalf of the Wellcome Sanger Institute COVID-19 Surveillance Team |
| EPI_ISL_2665251 | Lighthouse Lab in Alderley Park | Wellcome Sanger Institute for the COVID-19 Genomics UK (COG-UK) Consortium | Jacquelyn Wynn, Mairead Hyland, The Lighthouse Lab in Alderley Park and Alex Alderton, Roberto Amato, Jeffrey Barrett, Sonia Goncalves, Ewan Harrison, David K. Jackson, Ian Johnston, Dominic Kwiatkowski, Cordelia Langford, John Sillitoe on behalf of the Wellcome Sanger Institute COVID-19 Surveillance Team |
| EPI_ISL_2665252, EPI_ISL_2665253, EPI_ISL_2665254 | Lighthouse Lab in Milton Keynes | Wellcome Sanger Institute for the COVID-19 Genomics UK (COG-UK) Consortium | The Lighthouse Lab in Milton Keynes and Alex Alderton, Roberto Amato, Jeffrey Barrett, Sonia Goncalves, Ewan Harrison, David K. Jackson, Ian Johnston, Dominic Kwiatkowski, Cordelia Langford, John Sillitoe on behalf of the Wellcome Sanger Institute COVID-19 Surveillance Team |
| EPI_ISL_2665255 | Lighthouse Lab in Alderley Park | Wellcome Sanger Institute for the COVID-19 Genomics UK (COG-UK) Consortium | Jacquelyn Wynn, Mairead Hyland, The Lighthouse Lab in Alderley Park and Alex Alderton, Roberto Amato, Jeffrey Barrett, Sonia Goncalves, Ewan Harrison, David K. Jackson, Ian Johnston, Dominic Kwiatkowski, Cordelia Langford, John Sillitoe on behalf of the Wellcome Sanger Institute COVID-19 Surveillance Team |
| EPI_ISL_2665256, EPI_ISL_2665257, EPI_ISL_2665258, EPI_ISL_2665259 | Lighthouse Lab in Milton Keynes | Wellcome Sanger Institute for the COVID-19 Genomics UK (COG-UK) Consortium | The Lighthouse Lab in Milton Keynes and Alex Alderton, Roberto Amato, Jeffrey Barrett, Sonia Goncalves, Ewan Harrison, David K. Jackson, Ian Johnston, Dominic Kwiatkowski, Cordelia Langford, John Sillitoe on behalf of the Wellcome Sanger Institute COVID-19 Surveillance Team |
| EPI_ISL_2665260 | Lighthouse Lab in Alderley Park | Wellcome Sanger Institute for the COVID-19 Genomics UK (COG-UK) Consortium | Jacquelyn Wynn, Mairead Hyland, The Lighthouse Lab in Alderley Park and Alex Alderton, Roberto Amato, Jeffrey Barrett, Sonia Goncalves, Ewan Harrison, David K. Jackson, Ian Johnston, Dominic Kwiatkowski, Cordelia Langford, John Sillitoe on behalf of the Wellcome Sanger Institute COVID-19 Surveillance Team |
| EPI_ISL_2665261 | Lighthouse Lab in Milton Keynes | Wellcome Sanger Institute for the COVID-19 Genomics UK (COG-UK) Consortium | The Lighthouse Lab in Milton Keynes and Alex Alderton, Roberto Amato, Jeffrey Barrett, Sonia Goncalves, Ewan Harrison, David K. Jackson, Ian Johnston, Dominic Kwiatkowski, Cordelia Langford, John Sillitoe on behalf of the Wellcome Sanger Institute COVID-19 Surveillance Team |
| EPI_ISL_2665262 | Lighthouse Lab in Alderley Park | Wellcome Sanger Institute for the COVID-19 Genomics UK (COG-UK) Consortium | Jacquelyn Wynn, Mairead Hyland, The Lighthouse Lab in Alderley Park and Alex Alderton, Roberto Amato, Jeffrey Barrett, Sonia Goncalves, Ewan Harrison, David K. Jackson, Ian Johnston, Dominic Kwiatkowski, Cordelia Langford, John Sillitoe on behalf of the Wellcome Sanger Institute COVID-19 Surveillance Team |
| EPI_ISL_2665263, EPI_ISL_2665264, EPI_ISL_2665265 | Lighthouse Lab in Milton Keynes | Wellcome Sanger Institute for the COVID-19 Genomics UK (COG-UK) Consortium | The Lighthouse Lab in Milton Keynes and Alex Alderton, Roberto Amato, Jeffrey Barrett, Sonia Goncalves, Ewan Harrison, David K. Jackson, Ian Johnston, Dominic Kwiatkowski, Cordelia Langford, John Sillitoe on behalf of the Wellcome Sanger Institute COVID-19 Surveillance Team |
| EPI_ISL_2665266 | Lighthouse Lab in Alderley Park | Wellcome Sanger Institute for the COVID-19 Genomics UK (COG-UK) Consortium | Jacquelyn Wynn, Mairead Hyland, The Lighthouse Lab in Alderley Park and Alex Alderton, Roberto Amato, Jeffrey Barrett, Sonia Goncalves, Ewan Harrison, David K. Jackson, Ian Johnston, Dominic Kwiatkowski, Cordelia Langford, John Sillitoe on behalf of the Wellcome Sanger Institute COVID-19 Surveillance Team |
| EPI_ISL_2665267, EPI_ISL_2665268, EPI_ISL_2665269, EPI_ISL_2665270, EPI_ISL_2665271, EPI_ISL_2665272 | Lighthouse Lab in Milton Keynes | Wellcome Sanger Institute for the COVID-19 Genomics UK (COG-UK) Consortium | The Lighthouse Lab in Milton Keynes and Alex Alderton, Roberto Amato, Jeffrey Barrett, Sonia Goncalves, Ewan Harrison, David K. Jackson, Ian Johnston, Dominic Kwiatkowski, Cordelia Langford, John Sillitoe on behalf of the Wellcome Sanger Institute COVID-19 Surveillance Team |
| EPI_ISL_2665273 | Lighthouse Lab in Alderley Park | Wellcome Sanger Institute for the COVID-19 Genomics UK (COG-UK) Consortium | Jacquelyn Wynn, Mairead Hyland, The Lighthouse Lab in Alderley Park and Alex Alderton, Roberto Amato, Jeffrey Barrett, Sonia Goncalves, Ewan Harrison, David K. Jackson, Ian Johnston, Dominic Kwiatkowski, Cordelia Langford, John Sillitoe on behalf of the Wellcome Sanger Institute COVID-19 Surveillance Team |
| EPI_ISL_2665274 | Lighthouse Lab in Milton Keynes | Wellcome Sanger Institute for the COVID-19 Genomics UK (COG-UK) Consortium | The Lighthouse Lab in Milton Keynes and Alex Alderton, Roberto Amato, Jeffrey Barrett, Sonia Goncalves, Ewan Harrison, David K. Jackson, Ian Johnston, Dominic Kwiatkowski, Cordelia Langford, John Sillitoe on behalf of the Wellcome Sanger Institute COVID-19 Surveillance Team |

| | | | |
|---|---|---|---|
| EPI_ISL_2665275 | Lighthouse Lab in Alderley Park | Wellcome Sanger Institute for the COVID-19 Genomics UK (COG-UK) Consortium | Jacquelyn Wynn, Mairead Hyland, The Lighthouse Lab in Alderley Park and Alex Alderton, Roberto Amato, Jeffrey Barrett, Sonia Goncalves, Ewan Harrison, David K. Jackson, Ian Johnston, Dominic Kwiatkowski, Cordelia Langford, John Sillitoe on behalf of the Wellcome Sanger Institute COVID-19 Surveillance Team |
| EPI_ISL_2665276 | Lighthouse Lab in Milton Keynes | Wellcome Sanger Institute for the COVID-19 Genomics UK (COG-UK) Consortium | The Lighthouse Lab in Milton Keynes and Alex Alderton, Roberto Amato, Jeffrey Barrett, Sonia Goncalves, Ewan Harrison, David K. Jackson, Ian Johnston, Dominic Kwiatkowski, Cordelia Langford, John Sillitoe on behalf of the Wellcome Sanger Institute COVID-19 Surveillance Team |
| EPI_ISL_2665277, EPI_ISL_2665278 | Lighthouse Lab in Alderley Park | Wellcome Sanger Institute for the COVID-19 Genomics UK (COG-UK) Consortium | Jacquelyn Wynn, Mairead Hyland, The Lighthouse Lab in Alderley Park and Alex Alderton, Roberto Amato, Jeffrey Barrett, Sonia Goncalves, Ewan Harrison, David K. Jackson, Ian Johnston, Dominic Kwiatkowski, Cordelia Langford, John Sillitoe on behalf of the Wellcome Sanger Institute COVID-19 Surveillance Team |
| EPI_ISL_2665279, EPI_ISL_2665280 | Lighthouse Lab in Milton Keynes | Wellcome Sanger Institute for the COVID-19 Genomics UK (COG-UK) Consortium | The Lighthouse Lab in Milton Keynes and Alex Alderton, Roberto Amato, Jeffrey Barrett, Sonia Goncalves, Ewan Harrison, David K. Jackson, Ian Johnston, Dominic Kwiatkowski, Cordelia Langford, John Sillitoe on behalf of the Wellcome Sanger Institute COVID-19 Surveillance Team |
| EPI_ISL_2665281, EPI_ISL_2665282 | Lighthouse Lab in Alderley Park | Wellcome Sanger Institute for the COVID-19 Genomics UK (COG-UK) Consortium | Jacquelyn Wynn, Mairead Hyland, The Lighthouse Lab in Alderley Park and Alex Alderton, Roberto Amato, Jeffrey Barrett, Sonia Goncalves, Ewan Harrison, David K. Jackson, Ian Johnston, Dominic Kwiatkowski, Cordelia Langford, John Sillitoe on behalf of the Wellcome Sanger Institute COVID-19 Surveillance Team |
| EPI_ISL_2665283, EPI_ISL_2665284 | Lighthouse Lab in Milton Keynes | Wellcome Sanger Institute for the COVID-19 Genomics UK (COG-UK) Consortium | The Lighthouse Lab in Milton Keynes and Alex Alderton, Roberto Amato, Jeffrey Barrett, Sonia Goncalves, Ewan Harrison, David K. Jackson, Ian Johnston, Dominic Kwiatkowski, Cordelia Langford, John Sillitoe on behalf of the Wellcome Sanger Institute COVID-19 Surveillance Team |
| EPI_ISL_2665285 | Lighthouse Lab in Alderley Park | Wellcome Sanger Institute for the COVID-19 Genomics UK (COG-UK) Consortium | Jacquelyn Wynn, Mairead Hyland, The Lighthouse Lab in Alderley Park and Alex Alderton, Roberto Amato, Jeffrey Barrett, Sonia Goncalves, Ewan Harrison, David K. Jackson, Ian Johnston, Dominic Kwiatkowski, Cordelia Langford, John Sillitoe on behalf of the Wellcome Sanger Institute COVID-19 Surveillance Team |
| EPI_ISL_2665286, EPI_ISL_2665287, EPI_ISL_2665288, EPI_ISL_2665289, EPI_ISL_2665290, EPI_ISL_2665291, EPI_ISL_2665292 | Lighthouse Lab in Milton Keynes | Wellcome Sanger Institute for the COVID-19 Genomics UK (COG-UK) Consortium | The Lighthouse Lab in Milton Keynes and Alex Alderton, Roberto Amato, Jeffrey Barrett, Sonia Goncalves, Ewan Harrison, David K. Jackson, Ian Johnston, Dominic Kwiatkowski, Cordelia Langford, John Sillitoe on behalf of the Wellcome Sanger Institute COVID-19 Surveillance Team |
| EPI_ISL_2665293 | Lighthouse Lab in Alderley Park | Wellcome Sanger Institute for the COVID-19 Genomics UK (COG-UK) Consortium | Jacquelyn Wynn, Mairead Hyland, The Lighthouse Lab in Alderley Park and Alex Alderton, Roberto Amato, Jeffrey Barrett, Sonia Goncalves, Ewan Harrison, David K. Jackson, Ian Johnston, Dominic Kwiatkowski, Cordelia Langford, John Sillitoe on behalf of the Wellcome Sanger Institute COVID-19 Surveillance Team |
| EPI_ISL_2665294, EPI_ISL_2665295 | Lighthouse Lab in Milton Keynes | Wellcome Sanger Institute for the COVID-19 Genomics UK (COG-UK) Consortium | The Lighthouse Lab in Milton Keynes and Alex Alderton, Roberto Amato, Jeffrey Barrett, Sonia Goncalves, Ewan Harrison, David K. Jackson, Ian Johnston, Dominic Kwiatkowski, Cordelia Langford, John Sillitoe on behalf of the Wellcome Sanger Institute COVID-19 Surveillance Team |
| EPI_ISL_2665296, EPI_ISL_2665297 | Lighthouse Lab in Alderley Park | Wellcome Sanger Institute for the COVID-19 Genomics UK (COG-UK) Consortium | Jacquelyn Wynn, Mairead Hyland, The Lighthouse Lab in Alderley Park and Alex Alderton, Roberto Amato, Jeffrey Barrett, Sonia Goncalves, Ewan Harrison, David K. Jackson, Ian Johnston, Dominic Kwiatkowski, Cordelia Langford, John Sillitoe on behalf of the Wellcome Sanger Institute COVID-19 Surveillance Team |
| EPI_ISL_2665298, EPI_ISL_2665300, EPI_ISL_2665301 | Lighthouse Lab in Milton Keynes | Wellcome Sanger Institute for the COVID-19 Genomics UK (COG-UK) Consortium | The Lighthouse Lab in Milton Keynes and Alex Alderton, Roberto Amato, Jeffrey Barrett, Sonia Goncalves, Ewan Harrison, David K. Jackson, Ian Johnston, Dominic Kwiatkowski, Cordelia Langford, John Sillitoe on behalf of the Wellcome Sanger Institute COVID-19 Surveillance Team |
| EPI_ISL_2665302 | Lighthouse Lab in Alderley Park | Wellcome Sanger Institute for the COVID-19 Genomics UK (COG-UK) Consortium | Jacquelyn Wynn, Mairead Hyland, The Lighthouse Lab in Alderley Park and Alex Alderton, Roberto Amato, Jeffrey Barrett, Sonia Goncalves, Ewan Harrison, David K. Jackson, Ian Johnston, Dominic Kwiatkowski, Cordelia Langford, John Sillitoe on behalf of the Wellcome Sanger Institute COVID-19 Surveillance Team |
| EPI_ISL_2665303, EPI_ISL_2665304, EPI_ISL_2665305, EPI_ISL_2665306, EPI_ISL_2665307, EPI_ISL_2665308, EPI_ISL_2665309, EPI_ISL_2665310 | Lighthouse Lab in Milton Keynes | Wellcome Sanger Institute for the COVID-19 Genomics UK (COG-UK) Consortium | The Lighthouse Lab in Milton Keynes and Alex Alderton, Roberto Amato, Jeffrey Barrett, Sonia Goncalves, Ewan Harrison, David K. Jackson, Ian Johnston, Dominic Kwiatkowski, Cordelia Langford, John Sillitoe on behalf of the Wellcome Sanger Institute COVID-19 Surveillance Team |
| EPI_ISL_2665311 | Lighthouse Lab in Alderley Park | Wellcome Sanger Institute for the COVID-19 Genomics UK (COG-UK) Consortium | Jacquelyn Wynn, Mairead Hyland, The Lighthouse Lab in Alderley Park and Alex Alderton, Roberto Amato, Jeffrey Barrett, Sonia Goncalves, Ewan Harrison, David K. Jackson, Ian Johnston, Dominic Kwiatkowski, Cordelia Langford, John Sillitoe on behalf of the Wellcome Sanger Institute COVID-19 Surveillance Team |
| EPI_ISL_2665312, EPI_ISL_2665313, EPI_ISL_2665314, EPI_ISL_2665315, EPI_ISL_2665316, EPI_ISL_2665317, EPI_ISL_2665318, EPI_ISL_2665319, EPI_ISL_2665320, EPI_ISL_2665321, EPI_ISL_2665322, EPI_ISL_2665323, EPI_ISL_2665324, EPI_ISL_2665325, EPI_ISL_2665326, EPI_ISL_2665327, EPI_ISL_2665328, EPI_ISL_2665329 | | | |
| see above | Lighthouse Lab in Milton Keynes | Wellcome Sanger Institute for the COVID-19 Genomics UK (COG-UK) Consortium | The Lighthouse Lab in Milton Keynes and Alex Alderton, Roberto Amato, Jeffrey Barrett, Sonia Goncalves, Ewan Harrison, David K. Jackson, Ian Johnston, Dominic Kwiatkowski, Cordelia Langford, John Sillitoe on behalf of the Wellcome Sanger Institute COVID-19 Surveillance Team |
| EPI_ISL_2665330 | Lighthouse Lab in Alderley Park | Wellcome Sanger Institute for the COVID-19 Genomics UK (COG-UK) Consortium | Jacquelyn Wynn, Mairead Hyland, The Lighthouse Lab in Alderley Park and Alex Alderton, Roberto Amato, Jeffrey Barrett, Sonia Goncalves, Ewan Harrison, David K. Jackson, Ian Johnston, Dominic Kwiatkowski, Cordelia Langford, John Sillitoe on behalf of the Wellcome Sanger Institute COVID-19 Surveillance Team |
| EPI_ISL_2665331 | Lighthouse Lab in Milton Keynes | Wellcome Sanger Institute for the COVID-19 Genomics UK (COG-UK) Consortium | The Lighthouse Lab in Milton Keynes and Alex Alderton, Roberto Amato, Jeffrey Barrett, Sonia Goncalves, Ewan Harrison, David K. Jackson, Ian Johnston, Dominic Kwiatkowski, Cordelia Langford, John Sillitoe on behalf of the Wellcome Sanger Institute COVID-19 Surveillance Team |
| EPI_ISL_2665332 | Lighthouse Lab in Alderley Park | Wellcome Sanger Institute for the COVID-19 Genomics UK (COG-UK) Consortium | Jacquelyn Wynn, Mairead Hyland, The Lighthouse Lab in Alderley Park and Alex Alderton, Roberto Amato, Jeffrey Barrett, Sonia Goncalves, Ewan Harrison, David K. Jackson, Ian Johnston, Dominic Kwiatkowski, Cordelia Langford, John Sillitoe on behalf of the Wellcome Sanger Institute COVID-19 Surveillance Team |
| EPI_ISL_2665333, EPI_ISL_2665334, EPI_ISL_2665335, EPI_ISL_2665336, EPI_ISL_2665337, EPI_ISL_2665338, EPI_ISL_2665339, EPI_ISL_2665341, EPI_ISL_2665342, EPI_ISL_2665343, EPI_ISL_2665344, EPI_ISL_2665345, EPI_ISL_2665346, EPI_ISL_2665348, EPI_ISL_2665349, EPI_ISL_2665350, EPI_ISL_2665351, EPI_ISL_2665352 | | | |
| see above | Lighthouse Lab in Milton Keynes | Wellcome Sanger Institute for the COVID-19 Genomics UK (COG-UK) Consortium | The Lighthouse Lab in Milton Keynes and Alex Alderton, Roberto Amato, Jeffrey Barrett, Sonia Goncalves, Ewan Harrison, David K. Jackson, Ian Johnston, Dominic Kwiatkowski, Cordelia Langford, John Sillitoe on behalf of the Wellcome Sanger Institute COVID-19 Surveillance Team |
| EPI_ISL_2665353 | Lighthouse Lab in Alderley Park | Wellcome Sanger Institute for the COVID-19 Genomics UK (COG-UK) Consortium | Jacquelyn Wynn, Mairead Hyland, The Lighthouse Lab in Alderley Park and Alex Alderton, Roberto Amato, Jeffrey Barrett, Sonia Goncalves, Ewan Harrison, David K. Jackson, Ian Johnston, Dominic Kwiatkowski, Cordelia Langford, John Sillitoe on behalf of the Wellcome Sanger Institute COVID-19 Surveillance Team |
| EPI_ISL_2665354, EPI_ISL_2665355, EPI_ISL_2665356, EPI_ISL_2665357, EPI_ISL_2665358 | Lighthouse Lab in Milton Keynes | Wellcome Sanger Institute for the COVID-19 Genomics UK (COG-UK) Consortium | The Lighthouse Lab in Milton Keynes and Alex Alderton, Roberto Amato, Jeffrey Barrett, Sonia Goncalves, Ewan Harrison, David K. Jackson, Ian Johnston, Dominic Kwiatkowski, Cordelia Langford, John Sillitoe on behalf of the Wellcome Sanger Institute COVID-19 Surveillance Team |
| EPI_ISL_2665359, EPI_ISL_2665360 | Lighthouse Lab in Alderley Park | Wellcome Sanger Institute for the COVID-19 Genomics UK (COG-UK) Consortium | Jacquelyn Wynn, Mairead Hyland, The Lighthouse Lab in Alderley Park and Alex Alderton, Roberto Amato, Jeffrey Barrett, Sonia Goncalves, Ewan Harrison, David K. Jackson, Ian Johnston, Dominic Kwiatkowski, Cordelia Langford, John Sillitoe on behalf of the Wellcome Sanger Institute COVID-19 Surveillance Team |
| EPI_ISL_2665361, EPI_ISL_2665362 | Lighthouse Lab in Milton Keynes | Wellcome Sanger Institute for the COVID-19 Genomics UK (COG-UK) Consortium | The Lighthouse Lab in Milton Keynes and Alex Alderton, Roberto Amato, Jeffrey Barrett, Sonia Goncalves, Ewan Harrison, David K. Jackson, Ian Johnston, Dominic Kwiatkowski, Cordelia Langford, John Sillitoe on behalf of the Wellcome Sanger Institute COVID-19 Surveillance Team |
| EPI_ISL_2665363 | Lighthouse Lab in Alderley Park | Wellcome Sanger Institute for the COVID-19 Genomics UK (COG-UK) Consortium | Jacquelyn Wynn, Mairead Hyland, The Lighthouse Lab in Alderley Park and Alex Alderton, Roberto Amato, Jeffrey Barrett, Sonia Goncalves, Ewan Harrison, David K. Jackson, Ian Johnston, Dominic Kwiatkowski, Cordelia Langford, John Sillitoe on behalf of the Wellcome Sanger Institute COVID-19 Surveillance Team |
| EPI_ISL_2665364, EPI_ISL_2665365, EPI_ISL_2665366, EPI_ISL_2665367, EPI_ISL_2665368, EPI_ISL_2665369 | Lighthouse Lab in Milton Keynes | Wellcome Sanger Institute for the COVID-19 Genomics UK (COG-UK) Consortium | The Lighthouse Lab in Milton Keynes and Alex Alderton, Roberto Amato, Jeffrey Barrett, Sonia Goncalves, Ewan Harrison, David K. Jackson, Ian Johnston, Dominic Kwiatkowski, Cordelia Langford, John Sillitoe on behalf of the Wellcome Sanger Institute COVID-19 Surveillance Team |
| EPI_ISL_2665370 | Lighthouse Lab in Alderley Park | Wellcome Sanger Institute for the COVID-19 Genomics UK (COG-UK) Consortium | Jacquelyn Wynn, Mairead Hyland, The Lighthouse Lab in Alderley Park and Alex Alderton, Roberto Amato, Jeffrey Barrett, Sonia Goncalves, Ewan Harrison, David K. Jackson, Ian Johnston, Dominic Kwiatkowski, Cordelia Langford, John Sillitoe on behalf of the Wellcome Sanger Institute COVID-19 Surveillance Team |
| EPI_ISL_2665371, EPI_ISL_2665372, EPI_ISL_2665373 | Lighthouse Lab in Milton Keynes | Wellcome Sanger Institute for the COVID-19 Genomics UK (COG-UK) Consortium | The Lighthouse Lab in Milton Keynes and Alex Alderton, Roberto Amato, Jeffrey Barrett, Sonia Goncalves, Ewan Harrison, David K. Jackson, Ian Johnston, Dominic Kwiatkowski, Cordelia Langford, John Sillitoe on behalf of the Wellcome Sanger Institute COVID-19 Surveillance Team |
| EPI_ISL_2665374 | Lighthouse Lab in Alderley Park | Wellcome Sanger Institute for the COVID-19 Genomics UK (COG-UK) Consortium | Jacquelyn Wynn, Mairead Hyland, The Lighthouse Lab in Alderley Park and Alex Alderton, Roberto Amato, Jeffrey Barrett, Sonia Goncalves, Ewan Harrison, David K. Jackson, Ian Johnston, Dominic Kwiatkowski, Cordelia Langford, John Sillitoe on behalf of the Wellcome Sanger Institute COVID-19 Surveillance Team |
| EPI_ISL_2665375, EPI_ISL_2665376 | Lighthouse Lab in Milton Keynes | Wellcome Sanger Institute for the COVID-19 Genomics UK (COG-UK) Consortium | The Lighthouse Lab in Milton Keynes and Alex Alderton, Roberto Amato, Jeffrey Barrett, Sonia Goncalves, Ewan Harrison, David K. Jackson, Ian Johnston, Dominic Kwiatkowski, Cordelia Langford, John Sillitoe on behalf of the Wellcome Sanger Institute COVID-19 Surveillance Team |
| EPI_ISL_2665377 | Lighthouse Lab in Alderley Park | Wellcome Sanger Institute for the COVID-19 Genomics UK (COG-UK) Consortium | Jacquelyn Wynn, Mairead Hyland, The Lighthouse Lab in Alderley Park and Alex Alderton, Roberto Amato, Jeffrey Barrett, Sonia Goncalves, Ewan Harrison, David K. Jackson, Ian Johnston, Dominic Kwiatkowski, Cordelia Langford, John Sillitoe on behalf of the Wellcome Sanger Institute COVID-19 Surveillance Team |
| EPI_ISL_2665378 | Lighthouse Lab in Milton Keynes | Wellcome Sanger Institute for the COVID-19 Genomics UK (COG-UK) Consortium | The Lighthouse Lab in Milton Keynes and Alex Alderton, Roberto Amato, Jeffrey Barrett, Sonia Goncalves, Ewan Harrison, David K. Jackson, Ian Johnston, Dominic Kwiatkowski, Cordelia Langford, John Sillitoe on behalf of the Wellcome Sanger Institute COVID-19 Surveillance Team |
| EPI_ISL_2665379, EPI_ISL_2665380 | Lighthouse Lab in Alderley Park | Wellcome Sanger Institute for the COVID-19 Genomics UK (COG-UK) Consortium | Jacquelyn Wynn, Mairead Hyland, The Lighthouse Lab in Alderley Park and Alex Alderton, Roberto Amato, Jeffrey Barrett, Sonia Goncalves, Ewan Harrison, David K. Jackson, Ian Johnston, Dominic Kwiatkowski, Cordelia Langford, John Sillitoe on behalf of the Wellcome Sanger Institute COVID-19 Surveillance Team |
| EPI_ISL_2665381, EPI_ISL_2665382, EPI_ISL_2665383, EPI_ISL_2665384 | Lighthouse Lab in Milton Keynes | Wellcome Sanger Institute for the COVID-19 Genomics UK (COG-UK) Consortium | The Lighthouse Lab in Milton Keynes and Alex Alderton, Roberto Amato, Jeffrey Barrett, Sonia Goncalves, Ewan Harrison, David K. Jackson, Ian Johnston, Dominic Kwiatkowski, Cordelia Langford, John Sillitoe on behalf of the Wellcome Sanger Institute COVID-19 Surveillance Team |
| EPI_ISL_2665385, EPI_ISL_2665386, | Lighthouse Laboratory Plymouth | Wellcome Sanger Institute for the COVID-19 Genomics UK (COG-UK) Consortium | Lighthouse Laboratory Plymouth and Alex Alderton, Roberto Amato, Jeffrey Barrett, Sonia Goncalves, Ewan Harrison, David K. Jackson, Ian Johnston, Dominic Kwiatkowski, Cordelia Langford, John Sillitoe on behalf of the Wellcome Sanger Institute COVID-19 Surveillance Team |

| | | | |
|---|---|---|---|
| EPI_ISL_2665387 | | | |
| EPI_ISL_2665388, EPI_ISL_2665389, EPI_ISL_2665390, EPI_ISL_2665391, EPI_ISL_2665392, EPI_ISL_2665393 | Lighthouse Lab in Milton Keynes | Wellcome Sanger Institute for the COVID-19 Genomics UK (COG-UK) Consortium | The Lighthouse Lab in Milton Keynes and Alex Alderton, Roberto Amato, Jeffrey Barrett, Sonia Goncalves, Ewan Harrison, David K. Jackson, Ian Johnston, Dominic Kwiatkowski, Cordelia Langford, John Sillitoe on behalf of the Wellcome Sanger Institute COVID-19 Surveillance Team |
| EPI_ISL_2665415, EPI_ISL_2665447, EPI_ISL_2665480, EPI_ISL_2665507, EPI_ISL_2665596, EPI_ISL_2665812, EPI_ISL_2665815, EPI_ISL_2665855, EPI_ISL_2665873, EPI_ISL_2665915, EPI_ISL_2665924, EPI_ISL_2666000, EPI_ISL_2666027, EPI_ISL_2666041, EPI_ISL_2666048, EPI_ISL_2666131, EPI_ISL_2666132, EPI_ISL_2666155, EPI_ISL_2666159, EPI_ISL_2666226, EPI_ISL_2666264, EPI_ISL_2666270, EPI_ISL_2666301, EPI_ISL_2666349, EPI_ISL_2666384, EPI_ISL_2666392, EPI_ISL_2666405, EPI_ISL_2666451, EPI_ISL_2666493, EPI_ISL_2666513, EPI_ISL_2666515, EPI_ISL_2666562, EPI_ISL_2666700, EPI_ISL_2666711, EPI_ISL_2666744, EPI_ISL_2666760, EPI_ISL_2666807, EPI_ISL_2666809 | | | |
| see above | Department of Bacteria, Parasites and Fungi, Statens Serum Institut, Copenhagen, Denmark | Statens Serum Institut Bioinformatics and Microbial Genomics | Danish Covid-19 Genome Consortium |
| EPI_ISL_2666813 | Molekylær Medicinsk Afdeling, Aarhus University Hospital, Aarhus, Denmark | Statens Serum Institut Bioinformatics and Microbial Genomics | Danish Covid-19 Genome Consortium |
| EPI_ISL_2666828, EPI_ISL_2666913, EPI_ISL_2666968, EPI_ISL_2666994, EPI_ISL_2667010, EPI_ISL_2667192, EPI_ISL_2667204, EPI_ISL_2667226, EPI_ISL_2667233, EPI_ISL_2667325, EPI_ISL_2667342, EPI_ISL_2667354, EPI_ISL_2667413, EPI_ISL_2667457 | | | |
| see above | Department of Bacteria, Parasites and Fungi, Statens Serum Institut, Copenhagen, Denmark | Statens Serum Institut Bioinformatics and Microbial Genomics | Danish Covid-19 Genome Consortium |
| EPI_ISL_2667495 | Molekylær Medicinsk Afdeling, Aarhus University Hospital, Aarhus, Denmark | Statens Serum Institut Bioinformatics and Microbial Genomics | Danish Covid-19 Genome Consortium |
| EPI_ISL_2667519, EPI_ISL_2667541, EPI_ISL_2667607, EPI_ISL_2667612, EPI_ISL_2667628, EPI_ISL_2667646, EPI_ISL_2667724, EPI_ISL_2667866, EPI_ISL_2667871, EPI_ISL_2667875, EPI_ISL_2667987, EPI_ISL_2668017, EPI_ISL_2668071, EPI_ISL_2668073, EPI_ISL_2668151, EPI_ISL_2668182, EPI_ISL_2668183, EPI_ISL_2668318, EPI_ISL_2668328, EPI_ISL_2668362, EPI_ISL_2668369, EPI_ISL_2668435, EPI_ISL_2668506, EPI_ISL_2668510, EPI_ISL_2668554, EPI_ISL_2668655, EPI_ISL_2668671, EPI_ISL_2668703, EPI_ISL_2668734, EPI_ISL_2668747, EPI_ISL_2668757, EPI_ISL_2668783, EPI_ISL_2668807, EPI_ISL_2668914, EPI_ISL_2668987, EPI_ISL_2669041, EPI_ISL_2669045, EPI_ISL_2669138, EPI_ISL_2669191, EPI_ISL_2669196, EPI_ISL_2669198, EPI_ISL_2669235, EPI_ISL_2669256, EPI_ISL_2669261, EPI_ISL_2669310, EPI_ISL_2669320, EPI_ISL_2669409, EPI_ISL_2669478, EPI_ISL_2669548, EPI_ISL_2669583, EPI_ISL_2669593, EPI_ISL_2669604, EPI_ISL_2669614, EPI_ISL_2669643, EPI_ISL_2669764, EPI_ISL_2669787, EPI_ISL_2669899, EPI_ISL_2669906, EPI_ISL_2669907, EPI_ISL_2669925, EPI_ISL_2669928, EPI_ISL_2670009, EPI_ISL_2670010, EPI_ISL_2670071, EPI_ISL_2670090, EPI_ISL_2670150, EPI_ISL_2670156, EPI_ISL_2670207, EPI_ISL_2670242 | | | |
| see above | Department of Bacteria, Parasites and Fungi, Statens Serum Institut, Copenhagen, Denmark | Statens Serum Institut Bioinformatics and Microbial Genomics | Danish Covid-19 Genome Consortium |
| EPI_ISL_2670318 | Department of Clinical Microbiology, Odense University Hospital, Odense, Denmark | Statens Serum Institut Bioinformatics and Microbial Genomics | Danish Covid-19 Genome Consortium |
| EPI_ISL_2670371, EPI_ISL_2670378, EPI_ISL_2670389, EPI_ISL_2670462, EPI_ISL_2670472, EPI_ISL_2670481, EPI_ISL_2670483, EPI_ISL_2670491, EPI_ISL_2670493, EPI_ISL_2670501, EPI_ISL_2670516, EPI_ISL_2670522, EPI_ISL_2670547, EPI_ISL_2670580, EPI_ISL_2670611, EPI_ISL_2670695, EPI_ISL_2670699, EPI_ISL_2670769, EPI_ISL_2670831, EPI_ISL_2670846, EPI_ISL_2670854, EPI_ISL_2670887, EPI_ISL_2670908, EPI_ISL_2670978, EPI_ISL_2670986, EPI_ISL_2671046, EPI_ISL_2671244 | | | |
| see above | Department of Bacteria, Parasites and Fungi, Statens Serum Institut, Copenhagen, Denmark | Statens Serum Institut Bioinformatics and Microbial Genomics | Danish Covid-19 Genome Consortium |
| EPI_ISL_2671366, EPI_ISL_2671368, EPI_ISL_2671371, EPI_ISL_2671374, EPI_ISL_2671377, EPI_ISL_2671381, EPI_ISL_2671385, EPI_ISL_2671386, EPI_ISL_2671387, EPI_ISL_2671391, EPI_ISL_2671395, EPI_ISL_2671397, EPI_ISL_2671399, EPI_ISL_2671405, EPI_ISL_2671407, EPI_ISL_2671408, EPI_ISL_2671411 | | | |
| see above | Microbiology Department, Laboratori Clínic Metropolitana Nord. Hospital Universitari Germans Trias i Pujol | Can Ruti SARS-CoV-2 Sequencing Hub (HUGTiP/IrsiCaixa/IGTP) | Marc Noguera-Julian, Pilar Armengol, Ignacio Blanco, Antoni E Bordoy, Francesc Catala-Moll, Pere-Joan Cardona, Maria Casadellà, Cristina Casañ, Gemma Clara, Bonaventura Clotet, Cristina Esteban, Montserrat Giménez, Mercedes Guerrero, Anna Not, Roger Paredes, Mariona Parera, Verónica Saludes, Alba Sánchez, and Elisa Martró on behalf of the Can Ruti SARS-CoV-2 Sequencing Hub |
| EPI_ISL_2671413 | Microbiologia CATLAB | Can Ruti SARS-CoV-2 Sequencing Hub (HUGTiP/IrsiCaixa/IGTP) | Marc Noguera-Julian, Pilar Armengol, Ignacio Blanco, Antoni E Bordoy, Francesc Catala-Moll, Pere-Joan Cardona, Maria Casadellà, Cristina Casañ, Gemma Clara, Bonaventura Clotet, Cristina Esteban, Montserrat Giménez, Mercedes Guerrero, Anna Not, Roger Paredes, Mariona Parera, Verónica Saludes, Alba Sánchez, and Elisa Martró on behalf of the Can Ruti SARS-CoV-2 Sequencing Hub |
| EPI_ISL_2671418, EPI_ISL_2671419, EPI_ISL_2671420, EPI_ISL_2671421, EPI_ISL_2671422, EPI_ISL_2671429, EPI_ISL_2671430, EPI_ISL_2671433 | Microbiology Department, Laboratori Clínic Metropolitana Nord. Hospital Universitari Germans Trias i Pujol | Can Ruti SARS-CoV-2 Sequencing Hub (HUGTiP/IrsiCaixa/IGTP) | Marc Noguera-Julian, Pilar Armengol, Ignacio Blanco, Antoni E Bordoy, Francesc Catala-Moll, Pere-Joan Cardona, Maria Casadellà, Cristina Casañ, Gemma Clara, Bonaventura Clotet, Cristina Esteban, Montserrat Giménez, Mercedes Guerrero, Anna Not, Roger Paredes, Mariona Parera, Verónica Saludes, Alba Sánchez, and Elisa Martró on behalf of the Can Ruti SARS-CoV-2 Sequencing Hub |
| EPI_ISL_2671436 | Microbiologia CATLAB | Can Ruti SARS-CoV-2 Sequencing Hub (HUGTiP/IrsiCaixa/IGTP) | Marc Noguera-Julian, Pilar Armengol, Ignacio Blanco, Antoni E Bordoy, Francesc Catala-Moll, Pere-Joan Cardona, Maria Casadellà, Cristina Casañ, Gemma Clara, Bonaventura Clotet, Cristina Esteban, Montserrat Giménez, Mercedes Guerrero, Anna Not, Roger Paredes, Mariona Parera, Verónica Saludes, Alba Sánchez, and Elisa Martró on behalf of the Can Ruti SARS-CoV-2 Sequencing Hub |
| EPI_ISL_2671440, EPI_ISL_2671441, EPI_ISL_2671442, EPI_ISL_2671443, EPI_ISL_2671451, EPI_ISL_2671452, EPI_ISL_2671453, EPI_ISL_2671455 | Microbiology Department, Laboratori Clínic Metropolitana Nord. Hospital Universitari Germans Trias i Pujol | Can Ruti SARS-CoV-2 Sequencing Hub (HUGTiP/IrsiCaixa/IGTP) | Marc Noguera-Julian, Pilar Armengol, Ignacio Blanco, Antoni E Bordoy, Francesc Catala-Moll, Pere-Joan Cardona, Maria Casadellà, Cristina Casañ, Gemma Clara, Bonaventura Clotet, Cristina Esteban, Montserrat Giménez, Mercedes Guerrero, Anna Not, Roger Paredes, Mariona Parera, Verónica Saludes, Alba Sánchez, and Elisa Martró on behalf of the Can Ruti SARS-CoV-2 Sequencing Hub |
| EPI_ISL_2671456, EPI_ISL_2671457, EPI_ISL_2671462, EPI_ISL_2671481 | Center for Laboratory Medicine | Center for Laboratory Medicine | Yannick Gerth |
| EPI_ISL_2671484 | Centro de Investigacion Biomedica de Occidente (CIBO) | Unidad de Genomica Avanzada | Consorcio Mexicano de Vigilancia Genomica (CoViGen-Mex). Authors (in alphabetical order): Julio Elias Alvarado-Yaah, Carlos F. Arias, Santiago avila-Rios, Eduardo Becerril-Vargas, Victor Hugo Borja-Aburto, Celia Boukadida, Cristobal Chaidez-Quiroz, Juan Bautista Chale-Dzul, Ricardo Ciria Merce, Andreu Comas-Garcia, Celida Duque Molina, Julissa Enciso-Ibarra, Jose Antonio Enciso-Moreno, Gloria Elena Espinosa-Ayala, Fernando Fontove-Herrera, Daniel Fregoso-Rueda, Victor Eduardo Garcia-Arias, Alejandra Garcia-Gasca, Bruno Gomez-Gil, Jean Pierre Gonzalez, Irvin Gonzalez-Lopez, Concepcion Grajales-Muñiz, Ricardo Grande, Rosa Maria Gutierrez Rios, Jesus Hernandez, Alejandra Hernandez-Teran, Alfredo Herrera-Estrella, Carla Ivon Herrera-Najera, Pavel Isa, Veronica Mata-Haro, Daniel Lira Morales, Susana Lopez, Antonio Loza Roman, Brenda Irasema Maldonado-Meza, Bernardo Martinez-Miguel, Jose Arturo Martinez-Orozco, Celida Martinez-Rodriguez, Margarita Matias-Florentino, Fidencio Mejia-Nepomuceno, Maria Guadalupe de Jesus Mireles-Rivera, Gloria Maria Molina-Salinas, Hector Montoya-Fuentes, , Mario Mujica-Sanchez, Jose Esteban Muñoz-Medina, Jose de Jesus Nuñez-Contreras, Alicia Ocaña-Mondragon, Luis Alberto Ochoa-Carrera, Hector Esteban Paz-Juarez, Marissa Perez-Garcia, Francisco Pulido, Helen Haydee Fernanda Ramirez-Plascencia, Jorge Salas-Hernandez, Angel Gustavo Salas-Lais, Alejandro Sanchez-Flores, Clara Esperanza Santacruz-Tinoco, Maria Guadalupe Santiago-Mauricio, Selene Zarate, Nelly Selem-Mojica, Blanca Taboada, Gloria Vazquez, Joel Armando Vazquez-Perez. |
| EPI_ISL_2671488, EPI_ISL_2671539, EPI_ISL_2671563 | Laboratorio Central de Epidemiologia (LCE) | Unidad de Genomica Avanzada | Consorcio Mexicano de Vigilancia Genomica (CoViGen-Mex). Authors (in alphabetical order): Julio Elias Alvarado-Yaah, Carlos F. Arias, Santiago avila-Rios, Eduardo Becerril-Vargas, Victor Hugo Borja-Aburto, Celia Boukadida, Cristobal Chaidez-Quiroz, Juan Bautista Chale-Dzul, Ricardo Ciria Merce, Andreu Comas-Garcia, Celida Duque Molina, Julissa Enciso-Ibarra, Jose Antonio Enciso-Moreno, Gloria Elena Espinosa-Ayala, Fernando Fontove-Herrera, Daniel Fregoso-Rueda, Victor Eduardo Garcia-Arias, Alejandra Garcia-Gasca, Bruno Gomez-Gil, Jean Pierre Gonzalez, Irvin Gonzalez-Lopez, Concepcion Grajales-Muñiz, Ricardo Grande, Rosa Maria Gutierrez Rios, Jesus Hernandez, Alejandra Hernandez-Teran, Alfredo Herrera-Estrella, Carla Ivon Herrera-Najera, Pavel Isa, Veronica Mata-Haro, Daniel Lira Morales, Susana Lopez, Antonio Loza Roman, Brenda Irasema Maldonado-Meza, Bernardo Martinez-Miguel, Jose Arturo Martinez-Orozco, Celida Martinez-Rodriguez, Margarita Matias-Florentino, Fidencio Mejia-Nepomuceno, Maria Guadalupe de Jesus Mireles-Rivera, Gloria Maria Molina-Salinas, Hector Montoya-Fuentes, , Mario Mujica-Sanchez, Jose Esteban Muñoz-Medina, Jose de Jesus Nuñez-Contreras, Alicia Ocaña-Mondragon, Luis Alberto Ochoa-Carrera, Hector Esteban Paz-Juarez, Marissa Perez-Garcia, Francisco Pulido, Helen Haydee Fernanda Ramirez-Plascencia, Jorge Salas-Hernandez, Angel Gustavo Salas-Lais, Alejandro Sanchez-Flores, Clara Esperanza Santacruz-Tinoco, Maria Guadalupe Santiago-Mauricio, Selene Zarate, Nelly Selem-Mojica, Blanca Taboada, Gloria Vazquez, Joel Armando Vazquez-Perez. |
| EPI_ISL_2671635, EPI_ISL_2671636 | Centro de Investigacion Biomedica del Noreste (CIBIN) | Unidad de Genomica Avanzada | Consorcio Mexicano de Vigilancia Genomica (CoViGen-Mex). Authors (in alphabetical order): Julio Elias Alvarado-Yaah, Carlos F. Arias, Santiago avila-Rios, Eduardo Becerril-Vargas, Victor Hugo Borja-Aburto, Celia Boukadida, Cristobal Chaidez-Quiroz, Juan Bautista Chale-Dzul, Ricardo Ciria Merce, Andreu Comas-Garcia, Celida Duque Molina, Julissa Enciso-Ibarra, Jose Antonio Enciso-Moreno, Gloria Elena Espinosa-Ayala, Fernando Fontove-Herrera, Daniel Fregoso-Rueda, Victor Eduardo Garcia-Arias, Alejandra Garcia-Gasca, Bruno Gomez-Gil, Jean Pierre Gonzalez, Irvin Gonzalez-Lopez, Concepcion Grajales-Muñiz, Ricardo Grande, Rosa Maria Gutierrez Rios, Jesus Hernandez, Alejandra Hernandez-Teran, Alfredo Herrera-Estrella, Carla Ivon Herrera-Najera, Pavel Isa, Veronica Mata-Haro, Daniel Lira Morales, Susana Lopez, Antonio Loza Roman, Brenda Irasema Maldonado-Meza, Bernardo Martinez-Miguel, Jose Arturo Martinez-Orozco, Celida Martinez-Rodriguez, Margarita Matias-Florentino, Fidencio Mejia-Nepomuceno, Maria Guadalupe de Jesus Mireles-Rivera, Gloria Maria Molina-Salinas, Hector Montoya-Fuentes, , Mario Mujica-Sanchez, Jose Esteban Muñoz-Medina, Jose de Jesus Nuñez-Contreras, Alicia Ocaña-Mondragon, Luis Alberto Ochoa-Carrera, Hector Esteban Paz-Juarez, Marissa Perez-Garcia, Francisco Pulido, Helen Haydee Fernanda Ramirez-Plascencia, Jorge Salas-Hernandez, Angel Gustavo Salas-Lais, Alejandro Sanchez-Flores, Clara Esperanza Santacruz-Tinoco, Maria Guadalupe Santiago-Mauricio, Selene Zarate, Nelly Selem-Mojica, Blanca Taboada, Gloria Vazquez, Joel Armando Vazquez-Perez. |
| EPI_ISL_2671649 | Laboratorio Central de Epidemiologia (LCE) | Unidad de Genomica Avanzada | Consorcio Mexicano de Vigilancia Genomica (CoViGen-Mex). Authors (in alphabetical order): Julio Elias Alvarado-Yaah, Carlos F. Arias, Santiago avila-Rios, Eduardo Becerril-Vargas, Victor Hugo Borja-Aburto, Celia Boukadida, Cristobal Chaidez-Quiroz, Juan Bautista Chale-Dzul, Ricardo Ciria Merce, Andreu Comas-Garcia, Celida Duque Molina, Julissa Enciso-Ibarra, Jose Antonio Enciso-Moreno, Gloria Elena Espinosa-Ayala, Fernando Fontove-Herrera, Daniel Fregoso-Rueda, Victor Eduardo Garcia-Arias, Alejandra Garcia-Gasca, Bruno Gomez-Gil, Jean Pierre Gonzalez, Irvin Gonzalez-Lopez, Concepcion Grajales-Muñiz, Ricardo Grande, Rosa Maria Gutierrez Rios, Jesus Hernandez, Alejandra Hernandez-Teran, Alfredo Herrera-Estrella, Carla Ivon Herrera-Najera, Pavel Isa, Veronica Mata-Haro, Daniel Lira Morales, Susana Lopez, Antonio Loza Roman, Brenda Irasema Maldonado-Meza, Bernardo Martinez-Miguel, Jose Arturo Martinez-Orozco, Celida Martinez-Rodriguez, Margarita Matias-Florentino, Fidencio Mejia-Nepomuceno, Maria Guadalupe de Jesus Mireles-Rivera, Gloria Maria Molina-Salinas, Hector Montoya-Fuentes, , Mario Mujica-Sanchez, Jose Esteban Muñoz-Medina, Jose de Jesus Nuñez-Contreras, Alicia Ocaña-Mondragon, Luis Alberto Ochoa-Carrera, Hector Esteban Paz-Juarez, Marissa Perez-Garcia, Francisco Pulido, Helen Haydee Fernanda Ramirez-Plascencia, Jorge Salas-Hernandez, Angel Gustavo Salas-Lais, Alejandro Sanchez-Flores, Clara Esperanza Santacruz-Tinoco, Maria Guadalupe Santiago-Mauricio, Selene Zarate, Nelly Selem-Mojica, Blanca Taboada, Gloria Vazquez, Joel Armando Vazquez-Perez. |
| EPI_ISL_2671900, EPI_ISL_2671901, EPI_ISL_2671902 | Indian Council of Medical Research-National Institute of Virology, Microbial Containment Complex | Indian Council of Medical Research-National Institute of Virology, Microbial Containment Complex | Pragya D Yadav |
| EPI_ISL_2672018, EPI_ISL_2672078 | TGen North | TGen North | Jolene Bowers, Heather Centner, Chris French, Hayley Yaglom, Ashlyn Pfeiffer, Darrin Lemmer, Dave Engelthaler, The Arizona COVID Genomics Union (ACGU) |
| EPI_ISL_2672092, EPI_ISL_2672106, EPI_ISL_2672110, EPI_ISL_2672115, EPI_ISL_2672117, EPI_ISL_2672125, EPI_ISL_2672127, EPI_ISL_2672129, EPI_ISL_2672130, EPI_ISL_2672132, EPI_ISL_2672137, EPI_ISL_2672144 | | | |
| see above | IN State Department of Health Laboratory Services | IN State Department of Health Laboratory Services | Cassandra Campion, Jamie Yeadon, Brian Pope, Lixia Liu, Kyle Brownlee, Melissa Hindenlang, Mark Glazier |
| EPI_ISL_2672150, EPI_ISL_2672208 | TGen North | TGen North | Jolene Bowers, Heather Centner, Chris French, Hayley Yaglom, Ashlyn Pfeiffer, Darrin Lemmer, Dave Engelthaler, The Arizona COVID Genomics Union (ACGU) |
| EPI_ISL_2672302, EPI_ISL_2672306, EPI_ISL_2672318, EPI_ISL_2672409, EPI_ISL_2672434, EPI_ISL_2672470, EPI_ISL_2672520, EPI_ISL_2672558, EPI_ISL_2672581, EPI_ISL_2672582, EPI_ISL_2672592, EPI_ISL_2672593, EPI_ISL_2672602, EPI_ISL_2672603, EPI_ISL_2672604, EPI_ISL_2672614, EPI_ISL_2672615, EPI_ISL_2672616, EPI_ISL_2672627, EPI_ISL_2672628, EPI_ISL_2672637, EPI_ISL_2672639, EPI_ISL_2672651, EPI_ISL_2672660, EPI_ISL_2672661, EPI_ISL_2672683, EPI_ISL_2672711, EPI_ISL_2672738, EPI_ISL_2672741, EPI_ISL_2672757, EPI_ISL_2672780, EPI_ISL_2672787, EPI_ISL_2672790, EPI_ISL_2672798, EPI_ISL_2672817, EPI_ISL_2672820, EPI_ISL_2672822, EPI_ISL_2672823, EPI_ISL_2672843, EPI_ISL_2672845, EPI_ISL_2672853, EPI_ISL_2672858, EPI_ISL_2672865, EPI_ISL_2672898, EPI_ISL_2672899, EPI_ISL_2672901, EPI_ISL_2672925, EPI_ISL_2672930, EPI_ISL_2672935, EPI_ISL_2672973, EPI_ISL_2672998, EPI_ISL_2673016, EPI_ISL_2673039, EPI_ISL_2673048, EPI_ISL_2673050, EPI_ISL_2673072, EPI_ISL_2673073, EPI_ISL_2673081, EPI_ISL_2673082, EPI_ISL_2673091, EPI_ISL_2673092, EPI_ISL_2673107, EPI_ISL_2673108, EPI_ISL_2673112, EPI_ISL_2673115, EPI_ISL_2673116, | | | |

| | | | |
|---|---|---|---|
| EPI_ISL_2673117, EPI_ISL_2673124, EPI_ISL_2673125, EPI_ISL_2673133, EPI_ISL_2673134, EPI_ISL_2673135, EPI_ISL_2673136, EPI_ISL_2673139, EPI_ISL_2673141, EPI_ISL_2673142, EPI_ISL_2673143, EPI_ISL_2673144, EPI_ISL_2673146, EPI_ISL_2673148, EPI_ISL_2673149, EPI_ISL_2673152, EPI_ISL_2673153, EPI_ISL_2673156, EPI_ISL_2673160, EPI_ISL_2673162, EPI_ISL_2673172, EPI_ISL_2673173, EPI_ISL_2673183, EPI_ISL_2673184, EPI_ISL_2673193, EPI_ISL_2673202, EPI_ISL_2673204, EPI_ISL_2673208, EPI_ISL_2673222, EPI_ISL_2673256, EPI_ISL_2673332, EPI_ISL_2673341, EPI_ISL_2673366, EPI_ISL_2673369, EPI_ISL_2673370, EPI_ISL_2673377, EPI_ISL_2673381, EPI_ISL_2673385, EPI_ISL_2673387, EPI_ISL_2673388, EPI_ISL_2673389, EPI_ISL_2673394, EPI_ISL_2673395 | | | |
| see above | Dutch COVID-19 response team | National Institute for Public Health and the Environment (RIVM) | Adam Meijer, Harry Vennema, Dirk Eggink, Jeroen Cremer, Sharon van den Brink, Bas van der Veer, AnneMarie van den Brandt, Lisa Wijsman, Kim Freriks, Ryanne Jaarsma, Euníce Then, Lynn Aarts, Sanne Bos, Melissa van Tuil, Linda van de Nes, Florian Zwagemaker, Dennis Schmitz, Annelies Kroneman, Karim Hajji, Chantal Reusken, on behalf of the national COVID-19 response team |
| EPI_ISL_2673654, EPI_ISL_2673660 | SIESP CHIETI - DRIVE IN VASTO | Istituto Zooprofilattico Sperimentale dell'Abruzzo e Molise "G. Caporale" | Lorusso A, Marcacci M, Di Domenico M, Ancora M, Curini V, Di Lollo Valeria, Mangone I, Rinaldi A, Delli Compagni E, Scialabba S, Caporale M, Di Pasquale A, Cammà C, Puglia I, Calistri P, Savini G |
| EPI_ISL_2673676, EPI_ISL_2673677, EPI_ISL_2673678 | OSPEDALE CIVILE TERAMO - CENTRO TRASFUSIONALE | Istituto Zooprofilattico Sperimentale dell'Abruzzo e Molise "G. Caporale" | Lorusso A, Marcacci M, Di Domenico M, Ancora M, Curini V, Di Lollo Valeria, Mangone I, Rinaldi A, Delli Compagni E, Scialabba S, Caporale M, Di Pasquale A, Cammà C, Puglia I, Calistri P, Savini G |
| EPI_ISL_2673685, EPI_ISL_2673687, EPI_ISL_2673689 | Plateforme de testing Namuroise | Plateforme de testing Namuroise | Nicolas GILLIARD, Lesly NYINKEU K.; Demars Aurore ; Maschietto Céline ; Nobis Chloé ; Otto Gaetan ; Denis Olivier ; Degosserie Jonathan ; Mullier François |
| EPI_ISL_2673713, EPI_ISL_2673714, EPI_ISL_2673716, EPI_ISL_2673719, EPI_ISL_2673723, EPI_ISL_2673736, EPI_ISL_2673737, EPI_ISL_2673738 | Furst Medical Laboratory | Norwegian Institute of Public Health, Department of Virology | Kathrine Stene-Johansen, Kamilla Heddeland Instefjord, Hilde Elshaug, Garcia Llorente Ignacio, Jon Bråte, Engebretsen Serina Beate,Pedersen Benedikte Nevjen, Line Victoria Moen, Debech Nadia, Atiya R Ali,Marie Paulsen Madsen, Rasmus Riis Kopperud, Hilde Vollan, Karoline Bragstad, Olav Hungnes |
| EPI_ISL_2673739 | Medical Microbiology Unit, Department for Laboratory Medicine, Drammen Hospital, Vestre Viken Health Trust, | Norwegian Institute of Public Health, Department of Virology | Kathrine Stene-Johansen, Kamilla Heddeland Instefjord, Hilde Elshaug, Garcia Llorente Ignacio, Jon Bråte, Engebretsen Serina Beate,Pedersen Benedikte Nevjen, Line Victoria Moen, Debech Nadia, Atiya R Ali,Marie Paulsen Madsen, Rasmus Riis Kopperud, Hilde Vollan, Karoline Bragstad, Olav Hungnes |
| EPI_ISL_2673740, EPI_ISL_2673741, EPI_ISL_2673744, EPI_ISL_2673745, EPI_ISL_2673746, EPI_ISL_2673748 | Furst Medical Laboratory | Norwegian Institute of Public Health, Department of Virology | Kathrine Stene-Johansen, Kamilla Heddeland Instefjord, Hilde Elshaug, Garcia Llorente Ignacio, Jon Bråte, Engebretsen Serina Beate,Pedersen Benedikte Nevjen, Line Victoria Moen, Debech Nadia, Atiya R Ali,Marie Paulsen Madsen, Rasmus Riis Kopperud, Hilde Vollan, Karoline Bragstad, Olav Hungnes |
| EPI_ISL_2673752 | Akershus University Hospital, Department for Microbiology and Infectious Disease Control | Norwegian Institute of Public Health, Department of Virology | Kathrine Stene-Johansen, Kamilla Heddeland Instefjord, Hilde Elshaug, Garcia Llorente Ignacio, Jon Bråte, Engebretsen Serina Beate,Pedersen Benedikte Nevjen, Line Victoria Moen, Debech Nadia, Atiya R Ali,Marie Paulsen Madsen, Rasmus Riis Kopperud, Hilde Vollan, Karoline Bragstad, Olav Hungnes |
| EPI_ISL_2673753, EPI_ISL_2673764, EPI_ISL_2673766 | Furst Medical Laboratory | Norwegian Institute of Public Health, Department of Virology | Kathrine Stene-Johansen, Kamilla Heddeland Instefjord, Hilde Elshaug, Garcia Llorente Ignacio, Jon Bråte, Engebretsen Serina Beate,Pedersen Benedikte Nevjen, Line Victoria Moen, Debech Nadia, Atiya R Ali,Marie Paulsen Madsen, Rasmus Riis Kopperud, Hilde Vollan, Karoline Bragstad, Olav Hungnes |
| EPI_ISL_2673831, EPI_ISL_2673842 | Nastavni zavod za javno zdravstvo Splitsko- Dalmatinske Županije | Hrvatski zavod za javno zdravstvo | Irena Tabain, Ivana Ferenčak |
| EPI_ISL_2674065 | National Public Health Laboratory/Pokhara Academy of Health Sciences | National Public Health Laboratory/CSIR-Institute of Genomics and Integrative Biology | National Public Health Laboratory Team /Pokhara Academy of Health Sciences COVID 19 PCR Team |
| EPI_ISL_2674066, EPI_ISL_2674067 | National Public Health Laboratory/Bharatpur Hospital COVID-19 Diagnostic Lab | National Public Health Laboratory/CSIR-Institute of Genomics and Integrative Biology | National Public Health Laboratory Team /Bharatpur Hospital COVID-19 Diagnostic PCR Team |
| EPI_ISL_2674068, EPI_ISL_2674069 | National Public Health Laboratory/Palpa HospitalCOVID-19 Lab | National Public Health Laboratory/CSIR-Institute of Genomics and Integrative Biology | National Public Health Laboratory Team /Palpa HospitalCOVID-19 PCR Team |
| EPI_ISL_2674070, EPI_ISL_2674071, EPI_ISL_2674072 | National Public Health Laboratory/Provincial Public Health Laboratory -Gandaki | National Public Health Laboratory/CSIR-Institute of Genomics and Integrative Biology | National Public Health Laboratory Team /Provincial Public Health Laboratory -Gandaki Team |
| EPI_ISL_2674073, EPI_ISL_2674074 | National Public Health Laboratory/B.P Koirala Institute of Health Science | National Public Health Laboratory/CSIR-Institute of Genomics and Integrative Biology | National Public Health Laboratory Team /B.P Koirala Institute of Health Science COVID 19 PCR Team |
| EPI_ISL_2674075 | National Public Health Laboratory/B.P Koirala Institute of Health Science | National Public Health Laboratory/CSIR-Institute of Genomics and Integrative Biology | National Public Health Laboratory Team /B.P Koirala Institute of Health Science COVID 19 PCR Team |
| EPI_ISL_2674076 | National Public Health Laboratory/Lumbini Provincial Hospital COVID 19 PCR Lab | National Public Health Laboratory/CSIR-Institute of Genomics and Integrative Biology | National Public Health Laboratory Team /Lumbini Provincial Hospital COVID 19 PCR Lab Team |
| EPI_ISL_2674077, EPI_ISL_2674078 | National Public Health Laboratory/Lumbini Provincial Hospital COVID 19 PCR Lab | National Public Health Laboratory/CSIR-Institute of Genomics and Integrative Biology | National Public Health Laboratory Team /Lumbini Provincial Hospital COVID 19 PCR Lab Team |
| EPI_ISL_2674079 | National Public Health Laboratory/B.P Koirala Memorial Cancer Hospital | National Public Health Laboratory/CSIR-Institute of Genomics and Integrative Biology | National Public Health Laboratory Team /B.P Koirala Memorial Cancer Hospital COVID 19 PCR Team |
| EPI_ISL_2674080, EPI_ISL_2674081 | National Public Health Laboratory/B.P Koirala Memorial Cancer Hospital COVID 19 PCR Lab | National Public Health Laboratory/CSIR-Institute of Genomics and Integrative Biology | National Public Health Laboratory Team /B.P Koirala Memorial Cancer Hospital COVID 19 PCR Lab Team |
| EPI_ISL_2674082, EPI_ISL_2674083, EPI_ISL_2674084 | National Public Health Laboratory/Trishuli Hospital COVID 19 PCR Lab | National Public Health Laboratory/CSIR-Institute of Genomics and Integrative Biology | National Public Health Laboratory Team /Trishuli Hospital COVID 19 PCR Lab Team |
| EPI_ISL_2674085, EPI_ISL_2674086, EPI_ISL_2674087 | National Public Health Laboratory/Provincial Public Health Laboratory -2 | National Public Health Laboratory/CSIR-Institute of Genomics and Integrative Biology | National Public Health Laboratory Team /Provincial Public Health Laboratory -2 Team |
| EPI_ISL_2674089, EPI_ISL_2674090, EPI_ISL_2674091, EPI_ISL_2674092, EPI_ISL_2674093, EPI_ISL_2674094, EPI_ISL_2674095, EPI_ISL_2674096, EPI_ISL_2674097, EPI_ISL_2674099, EPI_ISL_2674100, EPI_ISL_2674101, EPI_ISL_2674102, EPI_ISL_2674103 | | | |
| see above | National Public Health Laboratory | National Public Health Laboratory/CSIR-Institute of Genomics and Integrative Biology | National Public Health Laboratory Team |
| EPI_ISL_2674104, EPI_ISL_2674105 | National Public Health Laboratory/Provincial Public Health Laboratory -Gandaki | National Public Health Laboratory/CSIR-Institute of Genomics and Integrative Biology | National Public Health Laboratory Team /Provincial Public Health Laboratory -Gandaki Team |
| EPI_ISL_2674106 | National Public Health Laboratory/Trishuli Hospital COVID 19 PCR Lab | National Public Health Laboratory/CSIR-Institute of Genomics and Integrative Biology | National Public Health Laboratory Team /Trishuli Hospital COVID 19 PCR Lab Team |
| EPI_ISL_2674107 | National Public Health Laboratory/Provincial Public Health Laboratory -Gandaki | National Public Health Laboratory/CSIR-Institute of Genomics and Integrative Biology | National Public Health Laboratory Team /Provincial Public Health Laboratory -Gandaki Team |
| EPI_ISL_2674108, EPI_ISL_2674109 | National Public Health Laboratory/Provincial Public Health Laboratory -Lumbini | National Public Health Laboratory/CSIR-Institute of Genomics and Integrative Biology | National Public Health Laboratory Team /Provincial Public Health Laboratory -Lumbini Team |
| EPI_ISL_2674110, EPI_ISL_2674111 | National Public Health Laboratory | National Public Health Laboratory/CSIR-Institute of Genomics and Integrative Biology | National Public Health Laboratory Team |
| EPI_ISL_2674112 | National Public Health Laboratory/Nepal Medical College & Teaching Hospital | National Public Health Laboratory/CSIR-Institute of Genomics and Integrative Biology | National Public Health Laboratory Team/Nepal Medical College & Teaching Hospital COVID 19 PCR Team |
| EPI_ISL_2674214, EPI_ISL_2674215, EPI_ISL_2674216, EPI_ISL_2674217, EPI_ISL_2674218, EPI_ISL_2674219, EPI_ISL_2674220, EPI_ISL_2674221, EPI_ISL_2674222, EPI_ISL_2674223, EPI_ISL_2674224, EPI_ISL_2674225, EPI_ISL_2674270, EPI_ISL_2674271, EPI_ISL_2674272, EPI_ISL_2674273, EPI_ISL_2674274, EPI_ISL_2674275, EPI_ISL_2674276, EPI_ISL_2674277, EPI_ISL_2674278, EPI_ISL_2674279 | | | |
| see above | National Institute of Public Health | State Veterinary Institute Prague | Nagy,A;Cernikov,L;Stara,M;Suri,T;Vecerova,J;Jirincova,H |
| EPI_ISL_2674807, EPI_ISL_2674808, EPI_ISL_2674810, EPI_ISL_2674811 | SARS-CoV-2 testing team, National Institute of Infectious Diseases | Pathogen Genomics Center, National Institute of Infectious Diseases | Tsuyoshi Sekizuka, Kentaro Itokawa, Rina Tanaka, Masanori Hashino, Nozomu Hanaoka, Masumichi Saito, Naomi Nojiri, Hazuka Y Furihata, Hiromizu Takahashi, Tsuguto Fujimoto, Makoto Kuroda |
| EPI_ISL_2674901, EPI_ISL_2674902 | LESP Ciudad de Mexico | Instituto de Diagnostico y Referencia Epidemiologicos (INDRE) | Claudia Wong-Arambula, Abril Rodriguez-Maldonado, Vanessa Rivero-Arredondo, Ariadna Medina-Benitez, Joaquin Quiroz-Mercado, Sergio Rangel-Guerrero, Natividad Cruz-Ortiz, Tatiana Nunez-Garcia, Gisela Barrera-Badillo, Lucia Hernandez-Rivas, Irma Lopez-Martinez, Ernesto Ramirez-Gonzalez. |
| EPI_ISL_2674903 | LESP Jalisco | Instituto de Diagnostico y Referencia Epidemiologicos (INDRE) | Claudia Wong-Arambula, Abril Rodriguez-Maldonado, Vanessa Rivero-Arredondo, Ariadna Medina-Benitez, Joaquin Quiroz-Mercado, Sergio Rangel-Guerrero, Natividad Cruz-Ortiz, Tatiana Nunez-Garcia, Gisela Barrera-Badillo, Lucia Hernandez-Rivas, Irma Lopez-Martinez, Ernesto Ramirez-Gonzalez. |
| EPI_ISL_2674904 | LESP Ciudad de Mexico | Instituto de Diagnostico y Referencia Epidemiologicos (INDRE) | Claudia Wong-Arambula, Abril Rodriguez-Maldonado, Vanessa Rivero-Arredondo, Ariadna Medina-Benitez, Joaquin Quiroz-Mercado, Sergio Rangel-Guerrero, Natividad Cruz-Ortiz, Tatiana Nunez-Garcia, Gisela Barrera-Badillo, Lucia Hernandez-Rivas, Irma Lopez-Martinez, Ernesto Ramirez-Gonzalez. |
| EPI_ISL_2674905 | LESP Yucatan | Instituto de Diagnostico y Referencia Epidemiologicos (INDRE) | Claudia Wong-Arambula, Abril Rodriguez-Maldonado, Vanessa Rivero-Arredondo, Ariadna Medina-Benitez, Joaquin Quiroz-Mercado, Sergio Rangel-Guerrero, Natividad Cruz-Ortiz, Tatiana Nunez-Garcia, Gisela Barrera-Badillo, Lucia Hernandez-Rivas, Irma Lopez-Martinez, Ernesto Ramirez-Gonzalez. |
| EPI_ISL_2675078, EPI_ISL_2675080, | Vestfold Hospital, Toensberg Department of Microbiology | Norwegian Institute of Public Health, Department of Virology | Kathrine Stene-Johansen, Kamilla Heddeland Instefjord, Hilde Elshaug, Garcia Llorente Ignacio, Jon Bråte, Engebretsen Serina Beate,Pedersen Benedikte Nevjen, Line Victoria Moen, Debech Nadia, Atiya R Ali,Marie Paulsen Madsen, Rasmus Riis Kopperud, Hilde Vollan, Karoline Bragstad, Olav Hungnes |

| | | | |
|---|---|---|---|
| EPI_ISL_2675093, EPI_ISL_2675094 | | | |
| EPI_ISL_2675096 | Furst Medical Laboratory | Norwegian Institute of Public Health, Department of Virology | Kathrine Stene-Johansen, Kamilla Heddeland Instefjord, Hilde Elshaug, Garcia Llorente Ignacio, Jon Bråte, Engebretsen Serina Beate,Pedersen Benedikte Nevjen, Line Victoria Moen, Debech Nadia, Atiya R Ali,Marie Paulsen Madsen, Rasmus Riis Kopperud, Hilde Vollan, Karoline Bragstad, Olav Hungnes |
| EPI_ISL_2675189, EPI_ISL_2675190, EPI_ISL_2675191, EPI_ISL_2675194, EPI_ISL_2675195, EPI_ISL_2675196, EPI_ISL_2675199, EPI_ISL_2675201 | Wichita State University - Molecular Diagnostics Lab | Kansas Health and Environmental Lab | Mike Grose, Katherine Wiggins, Jonathan Barnell, Ben Olsen, and Phil Adam |
| EPI_ISL_2675204, EPI_ISL_2675208, EPI_ISL_2675209 | Houston Health Dept. | Houston Health Dept. | Ryker Penn, Pamela Brown, Adolpho Lara, Yanlai Lai |
| EPI_ISL_2675216, EPI_ISL_2675217 | South Dakota Public Health Laboratory | South Dakota Public Health Laboratory | Jacob Garfin and Chris Carlson |
| EPI_ISL_2675251 | Queensland Health Forensic and Scientific Services | Queensland Health Forensic and Scientific Services | Son Nguyen |
| EPI_ISL_2675257, EPI_ISL_2675259 | ZOL | Jessa | Severine Berden et al. on behalf of the Jessa_cmdLab |
| EPI_ISL_2675263 | Jessa | Jessa | Severine Berden et al. on behalf of the Jessa_cmdLab |
| EPI_ISL_2675282, EPI_ISL_2675290, EPI_ISL_2675294, EPI_ISL_2675296, EPI_ISL_2675298, EPI_ISL_2675303 | Hospital General Universitario Gregorio Marañón | Hospital General Universitario Gregorio Marañón | Sergio Buenestado Serrano, Pedro Sola Campoy, Laura Pérez-Lago, Cristina Rodriguez-Grande, Marta Herranz Martin, Victor Manuel de la Cueva, Julia Suárez, Pilar Catalán, Patricia Muñoz, Darío García de Viedma |
| EPI_ISL_2675305, EPI_ISL_2675306, EPI_ISL_2675307, EPI_ISL_2675308, EPI_ISL_2675309, EPI_ISL_2675310, EPI_ISL_2675311, EPI_ISL_2675312, EPI_ISL_2675313, EPI_ISL_2675314, EPI_ISL_2675315, EPI_ISL_2675316, EPI_ISL_2675317, EPI_ISL_2675318, EPI_ISL_2675319, EPI_ISL_2675320, EPI_ISL_2675321, EPI_ISL_2675322, EPI_ISL_2675323, EPI_ISL_2675324, EPI_ISL_2675325, EPI_ISL_2675326, EPI_ISL_2675327, EPI_ISL_2675328, EPI_ISL_2675329, EPI_ISL_2675330, EPI_ISL_2675331, EPI_ISL_2675332, EPI_ISL_2675333, EPI_ISL_2675334, EPI_ISL_2675335, EPI_ISL_2675336, EPI_ISL_2675337, EPI_ISL_2675338, EPI_ISL_2675339, EPI_ISL_2675340, EPI_ISL_2675341, EPI_ISL_2675342, EPI_ISL_2675343, EPI_ISL_2675344, EPI_ISL_2675345, EPI_ISL_2675346, EPI_ISL_2675347, EPI_ISL_2675348, EPI_ISL_2675349, EPI_ISL_2675350, EPI_ISL_2675351, EPI_ISL_2675352, EPI_ISL_2675353, EPI_ISL_2675355, EPI_ISL_2675356, EPI_ISL_2675357, EPI_ISL_2675358, EPI_ISL_2675359, EPI_ISL_2675360, EPI_ISL_2675361, EPI_ISL_2675362, EPI_ISL_2675363, EPI_ISL_2675364, EPI_ISL_2675365, EPI_ISL_2675366, EPI_ISL_2675367, EPI_ISL_2675368, EPI_ISL_2675369, EPI_ISL_2675370, EPI_ISL_2675371, EPI_ISL_2675372, EPI_ISL_2675373, EPI_ISL_2675374, EPI_ISL_2675375, EPI_ISL_2675377, EPI_ISL_2675378, EPI_ISL_2675379, EPI_ISL_2675380, EPI_ISL_2675381, EPI_ISL_2675382, EPI_ISL_2675383, EPI_ISL_2675384, EPI_ISL_2675385, EPI_ISL_2675386, EPI_ISL_2675387, EPI_ISL_2675388, EPI_ISL_2675389, EPI_ISL_2675390, EPI_ISL_2675391, EPI_ISL_2675392 | | | |
| see above | INSACOG-NAGALAND | National Institute of Biomedical Genomics – INSACOG | Arindam Maitra, Vinotsole Khamo, Nidhan Kumar Biswas, Chenole Keppen, Sreedhar Chinnaswamy, Wetetsho Kapfo, Saumitra Das |
| EPI_ISL_2675405, EPI_ISL_2675428, EPI_ISL_2675451, EPI_ISL_2675462, EPI_ISL_2675534, EPI_ISL_2675618, EPI_ISL_2675638, EPI_ISL_2675655, EPI_ISL_2675739 | Department of Bacteria, Parasites and Fungi, Statens Serum Institut, Copenhagen, Denmark | Statens Serum Institut Bioinformatics and Microbial Genomics | Danish Covid-19 Genome Consortium |
| EPI_ISL_2675890, EPI_ISL_2675891, EPI_ISL_2675892, EPI_ISL_2675893, EPI_ISL_2675894, EPI_ISL_2675895, EPI_ISL_2675896, EPI_ISL_2675897, EPI_ISL_2675898, EPI_ISL_2675899, EPI_ISL_2675900, EPI_ISL_2675901, EPI_ISL_2675902, EPI_ISL_2675903, EPI_ISL_2675904, EPI_ISL_2675905, EPI_ISL_2675906, EPI_ISL_2675907, EPI_ISL_2675910, EPI_ISL_2675911, EPI_ISL_2675912, EPI_ISL_2675913, EPI_ISL_2675914, EPI_ISL_2675915, EPI_ISL_2675916, EPI_ISL_2675918, EPI_ISL_2675919, EPI_ISL_2675920, EPI_ISL_2675921, EPI_ISL_2675922, EPI_ISL_2675923, EPI_ISL_2675924, EPI_ISL_2675927, EPI_ISL_2675928, EPI_ISL_2675929, EPI_ISL_2675930, EPI_ISL_2675931, EPI_ISL_2675932, EPI_ISL_2675933, EPI_ISL_2675934, EPI_ISL_2675935, EPI_ISL_2675936, EPI_ISL_2675937, EPI_ISL_2675938, EPI_ISL_2675939, EPI_ISL_2675940, EPI_ISL_2675941, EPI_ISL_2675942, EPI_ISL_2675943, EPI_ISL_2675946, EPI_ISL_2675947, EPI_ISL_2675948, EPI_ISL_2675949, EPI_ISL_2675950, EPI_ISL_2675951, EPI_ISL_2675952, EPI_ISL_2675953, EPI_ISL_2675954, EPI_ISL_2675955, EPI_ISL_2675956, EPI_ISL_2675957, EPI_ISL_2675958, EPI_ISL_2675959, EPI_ISL_2675960, EPI_ISL_2675961, EPI_ISL_2675962, EPI_ISL_2675963, EPI_ISL_2675964, EPI_ISL_2675965, EPI_ISL_2675966, EPI_ISL_2675967, EPI_ISL_2675968, EPI_ISL_2675970, EPI_ISL_2675971, EPI_ISL_2675973, EPI_ISL_2675974, EPI_ISL_2675975, EPI_ISL_2675976, EPI_ISL_2675977, EPI_ISL_2675978, EPI_ISL_2675979, EPI_ISL_2675980, EPI_ISL_2675981, EPI_ISL_2675983, EPI_ISL_2675985, EPI_ISL_2675987, EPI_ISL_2675988, EPI_ISL_2675990, EPI_ISL_2675991, EPI_ISL_2675992, EPI_ISL_2675993, EPI_ISL_2675994, EPI_ISL_2675995, EPI_ISL_2675996, EPI_ISL_2675997, EPI_ISL_2675998, EPI_ISL_2675999, EPI_ISL_2676000, EPI_ISL_2676001, EPI_ISL_2676002, EPI_ISL_2676003, EPI_ISL_2676004, EPI_ISL_2676005, EPI_ISL_2676006, EPI_ISL_2676007, EPI_ISL_2676008, EPI_ISL_2676009, EPI_ISL_2676011, EPI_ISL_2676012, EPI_ISL_2676014, EPI_ISL_2676015, EPI_ISL_2676016, EPI_ISL_2676017, EPI_ISL_2676018, EPI_ISL_2676019, EPI_ISL_2676020, EPI_ISL_2676021, EPI_ISL_2676022, EPI_ISL_2676023, EPI_ISL_2676024, EPI_ISL_2676027, EPI_ISL_2676028, EPI_ISL_2676029, EPI_ISL_2676030, EPI_ISL_2676032, EPI_ISL_2676033, EPI_ISL_2676034, EPI_ISL_2676035, EPI_ISL_2676036, EPI_ISL_2676037, EPI_ISL_2676038, EPI_ISL_2676039, EPI_ISL_2676040, EPI_ISL_2676041, EPI_ISL_2676042, EPI_ISL_2676044, EPI_ISL_2676046, EPI_ISL_2676047, EPI_ISL_2676048, EPI_ISL_2676049, EPI_ISL_2676050, EPI_ISL_2676051, EPI_ISL_2676052, EPI_ISL_2676053, EPI_ISL_2676054, EPI_ISL_2676055, EPI_ISL_2676056, EPI_ISL_2676057, EPI_ISL_2676058, EPI_ISL_2676059, EPI_ISL_2676060, EPI_ISL_2676061, EPI_ISL_2676062, EPI_ISL_2676063, EPI_ISL_2676064, EPI_ISL_2676065, EPI_ISL_2676066, EPI_ISL_2676067, EPI_ISL_2676068, EPI_ISL_2676069, EPI_ISL_2676070, EPI_ISL_2676071, EPI_ISL_2676072, EPI_ISL_2676073, EPI_ISL_2676074, EPI_ISL_2676075, EPI_ISL_2676076, EPI_ISL_2676077, EPI_ISL_2676078, EPI_ISL_2676079, EPI_ISL_2676080, EPI_ISL_2676081, EPI_ISL_2676082, EPI_ISL_2676083, EPI_ISL_2676084, EPI_ISL_2676085, EPI_ISL_2676086, EPI_ISL_2676087, EPI_ISL_2676088, EPI_ISL_2676089, EPI_ISL_2676090, EPI_ISL_2676091, EPI_ISL_2676092, EPI_ISL_2676093, EPI_ISL_2676094, EPI_ISL_2676095, EPI_ISL_2676096, EPI_ISL_2676097, EPI_ISL_2676098, EPI_ISL_2676099, EPI_ISL_2676100, EPI_ISL_2676101, EPI_ISL_2676102, EPI_ISL_2676103, EPI_ISL_2676104, EPI_ISL_2676105, EPI_ISL_2676106, EPI_ISL_2676107, EPI_ISL_2676108, EPI_ISL_2676109, EPI_ISL_2676110, EPI_ISL_2676111, EPI_ISL_2676112, EPI_ISL_2676113, EPI_ISL_2676114, EPI_ISL_2676115, EPI_ISL_2676116, EPI_ISL_2676117, EPI_ISL_2676118, EPI_ISL_2676119, EPI_ISL_2676120, EPI_ISL_2676121, EPI_ISL_2676122, EPI_ISL_2676123, EPI_ISL_2676124, EPI_ISL_2676125, EPI_ISL_2676126, EPI_ISL_2676127, EPI_ISL_2676128, EPI_ISL_2676129, EPI_ISL_2676130, EPI_ISL_2676132, EPI_ISL_2676133, EPI_ISL_2676134, EPI_ISL_2676135, EPI_ISL_2676136, EPI_ISL_2676137, EPI_ISL_2676138, EPI_ISL_2676139, EPI_ISL_2676140, EPI_ISL_2676141, EPI_ISL_2676142, EPI_ISL_2676143, EPI_ISL_2676144, EPI_ISL_2676145, EPI_ISL_2676146, EPI_ISL_2676147, EPI_ISL_2676148, EPI_ISL_2676149, EPI_ISL_2676150, EPI_ISL_2676151, EPI_ISL_2676152, EPI_ISL_2676153, EPI_ISL_2676154, EPI_ISL_2676155, EPI_ISL_2676156, EPI_ISL_2676157, EPI_ISL_2676158, EPI_ISL_2676159, EPI_ISL_2676160, EPI_ISL_2676161, EPI_ISL_2676162, EPI_ISL_2676163, EPI_ISL_2676164, EPI_ISL_2676165, EPI_ISL_2676166, EPI_ISL_2676167, EPI_ISL_2676168, EPI_ISL_2676169, EPI_ISL_2676171, EPI_ISL_2676172, EPI_ISL_2676173, EPI_ISL_2676175, EPI_ISL_2676176, EPI_ISL_2676177, EPI_ISL_2676178, EPI_ISL_2676179, EPI_ISL_2676180, EPI_ISL_2676181, EPI_ISL_2676182, EPI_ISL_2676183, EPI_ISL_2676184, EPI_ISL_2676185, EPI_ISL_2676186, EPI_ISL_2676187, EPI_ISL_2676188, EPI_ISL_2676189, EPI_ISL_2676190, EPI_ISL_2676191, EPI_ISL_2676192, EPI_ISL_2676193, EPI_ISL_2676194, EPI_ISL_2676195, EPI_ISL_2676196, EPI_ISL_2676197, EPI_ISL_2676198, EPI_ISL_2676199, EPI_ISL_2676200, EPI_ISL_2676201, EPI_ISL_2676202, EPI_ISL_2676203, EPI_ISL_2676204, EPI_ISL_2676205, EPI_ISL_2676206, EPI_ISL_2676207, EPI_ISL_2676208, EPI_ISL_2676209, EPI_ISL_2676210, EPI_ISL_2676211, EPI_ISL_2676212, EPI_ISL_2676213, EPI_ISL_2676214, EPI_ISL_2676215, EPI_ISL_2676216, EPI_ISL_2676217, EPI_ISL_2676218, EPI_ISL_2676220, EPI_ISL_2676221, EPI_ISL_2676222, EPI_ISL_2676223, EPI_ISL_2676224, EPI_ISL_2676225, EPI_ISL_2676226, EPI_ISL_2676227, EPI_ISL_2676228, EPI_ISL_2676229, EPI_ISL_2676230, EPI_ISL_2676231, EPI_ISL_2676232, EPI_ISL_2676233, EPI_ISL_2676234, EPI_ISL_2676235, EPI_ISL_2676236, EPI_ISL_2676237, EPI_ISL_2676238, EPI_ISL_2676239, EPI_ISL_2676240, EPI_ISL_2676241, EPI_ISL_2676242, EPI_ISL_2676243, EPI_ISL_2676244, EPI_ISL_2676245, EPI_ISL_2676246, EPI_ISL_2676247, EPI_ISL_2676248, EPI_ISL_2676249, EPI_ISL_2676250, EPI_ISL_2676251, EPI_ISL_2676253, EPI_ISL_2676254, EPI_ISL_2676256, EPI_ISL_2676257, EPI_ISL_2676258, EPI_ISL_2676259, EPI_ISL_2676260, EPI_ISL_2676261, EPI_ISL_2676262, EPI_ISL_2676263, EPI_ISL_2676264, EPI_ISL_2676265, EPI_ISL_2676266, EPI_ISL_2676267, EPI_ISL_2676268, EPI_ISL_2676269, EPI_ISL_2676270, EPI_ISL_2676271, EPI_ISL_2676272, EPI_ISL_2676273, EPI_ISL_2676274, EPI_ISL_2676275, EPI_ISL_2676276, EPI_ISL_2676277, EPI_ISL_2676278, EPI_ISL_2676279, EPI_ISL_2676280, EPI_ISL_2676281, EPI_ISL_2676282, EPI_ISL_2676283, EPI_ISL_2676284, EPI_ISL_2676285, EPI_ISL_2676286, EPI_ISL_2676287, EPI_ISL_2676288, EPI_ISL_2676289, EPI_ISL_2676290, EPI_ISL_2676291, EPI_ISL_2676292, EPI_ISL_2676293, EPI_ISL_2676294, EPI_ISL_2676295, EPI_ISL_2676296, EPI_ISL_2676297, EPI_ISL_2676298, EPI_ISL_2676299, EPI_ISL_2676300, EPI_ISL_2676301, EPI_ISL_2676302, EPI_ISL_2676303, EPI_ISL_2676304, EPI_ISL_2676305, EPI_ISL_2676306, EPI_ISL_2676307, EPI_ISL_2676308, EPI_ISL_2676309, EPI_ISL_2676310, EPI_ISL_2676311, EPI_ISL_2676312, EPI_ISL_2676313, EPI_ISL_2676314, EPI_ISL_2676315, EPI_ISL_2676316, EPI_ISL_2676317, EPI_ISL_2676318, EPI_ISL_2676319, EPI_ISL_2676320, EPI_ISL_2676321, EPI_ISL_2676322, EPI_ISL_2676323, EPI_ISL_2676324, EPI_ISL_2676325, EPI_ISL_2676326, EPI_ISL_2676327, EPI_ISL_2676328, EPI_ISL_2676329, EPI_ISL_2676330, EPI_ISL_2676331, EPI_ISL_2676332, EPI_ISL_2676333, EPI_ISL_2676334, EPI_ISL_2676335, EPI_ISL_2676336, EPI_ISL_2676337, EPI_ISL_2676338, EPI_ISL_2676339, EPI_ISL_2676340, EPI_ISL_2676341, EPI_ISL_2676342, EPI_ISL_2676343, EPI_ISL_2676344, EPI_ISL_2676345, EPI_ISL_2676346, EPI_ISL_2676347, EPI_ISL_2676348, EPI_ISL_2676349, EPI_ISL_2676350, EPI_ISL_2676351, EPI_ISL_2676352, EPI_ISL_2676353, EPI_ISL_2676354, EPI_ISL_2676355, EPI_ISL_2676356, EPI_ISL_2676357, EPI_ISL_2676358, EPI_ISL_2676359, EPI_ISL_2676360, EPI_ISL_2676361, EPI_ISL_2676362, EPI_ISL_2676363, EPI_ISL_2676364, EPI_ISL_2676365, EPI_ISL_2676366, EPI_ISL_2676367, EPI_ISL_2676368, EPI_ISL_2676369, EPI_ISL_2676370, EPI_ISL_2676371, EPI_ISL_2676372, EPI_ISL_2676373, EPI_ISL_2676374, EPI_ISL_2676375, EPI_ISL_2676376, EPI_ISL_2676378, EPI_ISL_2676379, EPI_ISL_2676380, EPI_ISL_2676381, EPI_ISL_2676382, EPI_ISL_2676383, EPI_ISL_2676384, EPI_ISL_2676385, EPI_ISL_2676386, EPI_ISL_2676387, EPI_ISL_2676388, EPI_ISL_2676389, EPI_ISL_2676390, EPI_ISL_2676391, EPI_ISL_2676392, EPI_ISL_2676393, EPI_ISL_2676394, EPI_ISL_2676395, EPI_ISL_2676396, EPI_ISL_2676397, EPI_ISL_2676398, EPI_ISL_2676399, EPI_ISL_2676400, EPI_ISL_2676401, EPI_ISL_2676402 | | | |
| see above | INSACOG-WB | National Institute of Biomedical Genomics – INSACOG | Arindam Maitra, Bhaswati Bandyopadhyay, Nidhan Kumar Biswas, Tamal Ghosh, Sreedhar Chinnaswamy, Ajay Chakraborti, Saumitra Das |
| EPI_ISL_2676404, EPI_ISL_2676407, EPI_ISL_2676409 | DC Public Health Lab/ Dept. of Forensic Sciences | DC Public Health Lab/ Dept. of Forensic Sciences | Janis Doss, Scott Nguyen, Elizabeth Zelaya, Sarah Scott, Connie Maza, Monica Mann, Brittany Hamilton, David Payne, Jocelyn Hauser |
| EPI_ISL_2676431, EPI_ISL_2676432, EPI_ISL_2676433, EPI_ISL_2676434 | Wyoming Public Health Laboratory | Wyoming Public Health Laboratory | Jim Mildenberger, Taylor Fearing, Channing Weber, Ashley Norberg, Chayse Rowley, Marley Goetz, Brian Dominguez, Elliot Thomasson, Sam Britz, Cari Sloma, Robert Petit, and Rob Christensen |
| EPI_ISL_2676543, EPI_ISL_2676545, EPI_ISL_2676546, EPI_ISL_2676548, EPI_ISL_2676549, EPI_ISL_2676550 | LABORATOIRE FORTE BIO ST PAUL LES DAX | CNR Virus des Infections Respiratoires - France SUD | Antonin Bal, Gregory Destras, Gwendolyne Burfin, Hadrien Regue, Quentin Semanas, Martine Valette, Bruno Lina, Laurence Josset |
| EPI_ISL_2676552 | AX BIO OCEAN | CNR Virus des Infections Respiratoires - France SUD | Antonin Bal, Gregory Destras, Gwendolyne Burfin, Hadrien Regue, Quentin Semanas, Martine Valette, Bruno Lina, Laurence Josset |
| EPI_ISL_2676565, EPI_ISL_2676568 | CERBALLIANCE - LAMAT | CNR Virus des Infections Respiratoires - France SUD | Antonin Bal, Gregory Destras, Gwendolyne Burfin, Hadrien Regue, Quentin Semanas, Martine Valette, Bruno Lina, Laurence Josset |
| EPI_ISL_2676572 | CNR Virus des Infections Respiratoires - France SUD | CNR Virus des Infections Respiratoires - France SUD | Antonin Bal, Gregory Destras, Gwendolyne Burfin, Hadrien Regue, Quentin Semanas, Martine Valette, Bruno Lina, Laurence Josset |
| EPI_ISL_2676604, EPI_ISL_2676609, EPI_ISL_2676612, EPI_ISL_2676613, EPI_ISL_2676617, EPI_ISL_2676619 | LAM BIOPOLE ORTHEZ | CNR Virus des Infections Respiratoires - France SUD | Antonin Bal, Gregory Destras, Gwendolyne Burfin, Hadrien Regue, Quentin Semanas, Martine Valette, Bruno Lina, Laurence Josset |
| EPI_ISL_2676628 | CNR Virus des Infections Respiratoires - France SUD | CNR Virus des Infections Respiratoires - France SUD | Antonin Bal, Gregory Destras, Gwendolyne Burfin, Hadrien Regue, Quentin Semanas, Martine Valette, Bruno Lina, Laurence Josset |
| EPI_ISL_2676637, EPI_ISL_2676639 | LABORATOIRE CERBALLIANCE PLT VILLON | CNR Virus des Infections Respiratoires - France SUD | Antonin Bal, Gregory Destras, Gwendolyne Burfin, Hadrien Regue, Quentin Semanas, Martine Valette, Bruno Lina, Laurence Josset |
| EPI_ISL_2676683 | LABORATOIRE DE BIOLOGIE MEDICALE | CNR Virus des Infections Respiratoires - France SUD | Antonin Bal, Gregory Destras, Gwendolyne Burfin, Hadrien Regue, Quentin Semanas, Martine Valette, Bruno Lina, Laurence Josset |
| EPI_ISL_2676684 | HOPITAL SAINT ANDRE | CNR Virus des Infections Respiratoires - France SUD | Antonin Bal, Gregory Destras, Gwendolyne Burfin, Hadrien Regue, Quentin Semanas, Martine Valette, Bruno Lina, Laurence Josset |
| EPI_ISL_2676690, EPI_ISL_2676692 | LABORATOIRE d'ANALYSES de BIOLOGIE MEDICALES | CNR Virus des Infections Respiratoires - France SUD | Antonin Bal, Gregory Destras, Gwendolyne Burfin, Hadrien Regue, Quentin Semanas, Martine Valette, Bruno Lina, Laurence Josset |
| EPI_ISL_2676696 | UNIBIO ROMANS GAMBETTA | CNR Virus des Infections Respiratoires - France SUD | Antonin Bal, Gregory Destras, Gwendolyne Burfin, Hadrien Regue, Quentin Semanas, Martine Valette, Bruno Lina, Laurence Josset |
| EPI_ISL_2676711 | LABORATOIRE CERBALLIANCE PLT VILLON | CNR Virus des Infections Respiratoires - France SUD | Antonin Bal, Gregory Destras, Gwendolyne Burfin, Hadrien Regue, Quentin Semanas, Martine Valette, Bruno Lina, Laurence Josset |

| | | | |
|---|---|---|---|
| EPI_ISL_2676750 | MIRIALIS CLUSES BECHET | CNR Virus des Infections Respiratoires - France SUD | Antonin Bal, Gregory Destras, Gwendolyne Burfin, Hadrien Regue, Quentin Semanas, Martine Valette, Bruno Lina, Laurence Josset |
| EPI_ISL_2676752, EPI_ISL_2676754, EPI_ISL_2676757 | LBM UNIBIO VALENTIN | CNR Virus des Infections Respiratoires - France SUD | Antonin Bal, Gregory Destras, Gwendolyne Burfin, Hadrien Regue, Quentin Semanas, Martine Valette, Bruno Lina, Laurence Josset |
| EPI_ISL_2676766, EPI_ISL_2676767, EPI_ISL_2676768 | MIRIALIS CLUSES BECHET | CNR Virus des Infections Respiratoires - France SUD | Antonin Bal, Gregory Destras, Gwendolyne Burfin, Hadrien Regue, Quentin Semanas, Martine Valette, Bruno Lina, Laurence Josset |
| EPI_ISL_2676774 | ASTRALAB TASSIGNY | CNR Virus des Infections Respiratoires - France SUD | Antonin Bal, Gregory Destras, Gwendolyne Burfin, Hadrien Regue, Quentin Semanas, Martine Valette, Bruno Lina, Laurence Josset |
| EPI_ISL_2676777 | LABORATOIRE BIOESTEREL | CNR Virus des Infections Respiratoires - France SUD | Antonin Bal, Gregory Destras, Gwendolyne Burfin, Hadrien Regue, Quentin Semanas, Martine Valette, Bruno Lina, Laurence Josset |
| EPI_ISL_2676779 | ORIAPOLE | CNR Virus des Infections Respiratoires - France SUD | Antonin Bal, Gregory Destras, Gwendolyne Burfin, Hadrien Regue, Quentin Semanas, Martine Valette, Bruno Lina, Laurence Josset |
| EPI_ISL_2676780 | ASTRALAB TASSIGNY | CNR Virus des Infections Respiratoires - France SUD | Antonin Bal, Gregory Destras, Gwendolyne Burfin, Hadrien Regue, Quentin Semanas, Martine Valette, Bruno Lina, Laurence Josset |
| EPI_ISL_2676781, EPI_ISL_2676782, EPI_ISL_2676784, EPI_ISL_2676786 | MIRIALIS CLUSES BECHET | CNR Virus des Infections Respiratoires - France SUD | Antonin Bal, Gregory Destras, Gwendolyne Burfin, Hadrien Regue, Quentin Semanas, Martine Valette, Bruno Lina, Laurence Josset |
| EPI_ISL_2676790 | LABORATOIRE BIOESTEREL | CNR Virus des Infections Respiratoires - France SUD | Antonin Bal, Gregory Destras, Gwendolyne Burfin, Hadrien Regue, Quentin Semanas, Martine Valette, Bruno Lina, Laurence Josset |
| EPI_ISL_2676799, EPI_ISL_2676800, EPI_ISL_2676801, EPI_ISL_2676802 | CNR Virus des Infections Respiratoires - France SUD | CNR Virus des Infections Respiratoires - France SUD | Antonin Bal, Gregory Destras, Gwendolyne Burfin, Hadrien Regue, Quentin Semanas, Martine Valette, Bruno Lina, Laurence Josset |
| EPI_ISL_2677318 | Labortorio Central de Saude Publica do Estado do Rio de Janeiro (LACEN/RJ) | Laboratory of Respiratory Viruses and Measles, Oswaldo Cruz Institute, FIOCRUZ | Paola Resende, Alex Pauvolid-Corrêa, Mia Ferreira de Araujo, Ana Beatriz Machado Lima,Luciana Appolinario, Fernando Motta, Anna Carolina Paixao, Ana Carolina Mendonca, Alice Sampaio Rocha, Taina Venas, Elisa Cavalcante Pereira, Renata Serrano Lopes, Andrea Cony Cavalcanti, Marilda Siqueira on behalf of the Fiocruz COVID-19 Genomic Surveillance Network |
| EPI_ISL_2677426, EPI_ISL_2677427, EPI_ISL_2677569, EPI_ISL_2677611 | Synlab Haut de France | UMR 8199/1283 EGID | Derhourhi Mehdi |
| EPI_ISL_2677633, EPI_ISL_2677634, EPI_ISL_2677635, EPI_ISL_2677636, EPI_ISL_2677637, EPI_ISL_2677638, EPI_ISL_2677639, EPI_ISL_2677642, EPI_ISL_2677644 | Wichita State University - Molecular Diagnostics Lab | Kansas Health and Environmental Lab | Katherine Wiggins, Mike Grose, Jonathan Barnell, Ben Olsen, and Phil Adam |
| EPI_ISL_2678055 | Indian Council of Medical Research-National Institute of Virology, Microbial Containment Complex | Indian Council of Medical Research-National Institute of Virology, Microbial Containment Complex | Pragya D Yadav |
| EPI_ISL_2678093, EPI_ISL_2678096, EPI_ISL_2678097 | Institute for Urban Disease Control and Prevention | COVID-19 Network Investigations (CONI) Alliance | Elizabeth Batty, Wasun Chantratita, Thanat Chookajorn, Stefan Fernandez, Angkana Huang, Anthony R. Jones, Khajohn Joonlasak, Chonticha Klungtong, Theerarat Kochakarn, Namfon Kotanan, Krittikorn Kumpornsin, Duangkamon Loesbanluechai, Wudtichai Manasatienkij, Bhakbhoom Panthan, Ekawat Pasomsub, Kingkan Rakmanee, Insee Sensorn, Janjira Thaipadungpanit, Arporn Wangwiwatsin, Treewat Watthanachockchai, Kamolthip Atsawawaranunt, Prayuth Kaewmalang, Amornmas Kongklieng, Pukkaporn Parnwijitkul, Vichan Pawun |
| EPI_ISL_2678103, EPI_ISL_2678104, EPI_ISL_2678105 | Ramathibodi Hospital | COVID-19 Network Investigations (CONI) Alliance | Elizabeth Batty, Wasun Chantratita, Thanat Chookajorn, Stefan Fernandez, Angkana Huang, Anthony R. Jones, Khajohn Joonlasak, Chonticha Klungtong, Theerarat Kochakarn, Namfon Kotanan, Krittikorn Kumpornsin, Duangkamon Loesbanluechai, Wudtichai Manasatienkij, Bhakbhoom Panthan, Ekawat Pasomsub, Kingkan Rakmanee, Insee Sensorn, Janjira Thaipadungpanit, Arporn Wangwiwatsin, Treewat Watthanachockchai |
| EPI_ISL_2678164 | The Caribbean Public Health Agency | Carrington Lab, Department of Preclinical Sciences, Faculty of Medical Sciences, The University of the West Indies, St Augustine Campus | Nikita S. D. Sahadeo, Arianne Brown-Jordan, Sarah Hill, Vernie Ramkissoon, Anushka Ramjag, Kenneth George, Avery Hinds, Risha Singh, SueMin Nathaniel, Nuno Faria, Oliver Pybus, Gabriel Escobar, Christine V. F. Carrington |
| EPI_ISL_2678383, EPI_ISL_2678385 | Virologisches Institut des Universitätsklinikums Erlangen | Robert Koch Institute | unknown |
| EPI_ISL_2678387, EPI_ISL_2678388, EPI_ISL_2678389, EPI_ISL_2678396, EPI_ISL_2678401, EPI_ISL_2678407, EPI_ISL_2678410, EPI_ISL_2678417, EPI_ISL_2678421, EPI_ISL_2678426, EPI_ISL_2678435 | | | |
| see above | Labor Becker & Kollegen (Standort München) | Robert Koch Institute | unknown |
| EPI_ISL_2678452 | Uniklinikum Carl Gustav Carus an der TU Dresden; Institut für Virologie | Robert Koch Institute | unknown |
| EPI_ISL_2678453, EPI_ISL_2678456, EPI_ISL_2678464, EPI_ISL_2678466, EPI_ISL_2678469 | Städtisches Klinikum Dresden Institut für Klinische Chemie und Laboratoriumsmedizin | Robert Koch Institute | unknown |
| EPI_ISL_2678472, EPI_ISL_2678473, EPI_ISL_2678474, EPI_ISL_2678475, EPI_ISL_2678478, EPI_ISL_2678480, EPI_ISL_2678482, EPI_ISL_2678484, EPI_ISL_2678485, EPI_ISL_2678486, EPI_ISL_2678487, EPI_ISL_2678492, EPI_ISL_2678494, EPI_ISL_2678496, EPI_ISL_2678497, EPI_ISL_2678498 | | | |
| see above | Eurofins LifeCodexx GmbH | Robert Koch Institute | unknown |
| EPI_ISL_2678501 | MVZ Labor Dr. Quade & Kollegen GmbH | Robert Koch Institute | unknown |
| EPI_ISL_2678544, EPI_ISL_2678553, EPI_ISL_2678576, EPI_ISL_2678600, EPI_ISL_2678602, EPI_ISL_2678607 | Labor Blackholm MVZ | Robert Koch Institute | unknown |
| EPI_ISL_2678608 | MVZ Labor Dr. Quade & Kollegen GmbH | Robert Koch Institute | unknown |
| EPI_ISL_2678611, EPI_ISL_2678614, EPI_ISL_2678643, EPI_ISL_2678644, EPI_ISL_2678647 | Labor Blackholm MVZ | Robert Koch Institute | unknown |
| EPI_ISL_2678652 | Labor Dr. Wisplinghoff - Köln | Robert Koch Institute | unknown |
| EPI_ISL_2678657, EPI_ISL_2678658, EPI_ISL_2678662, EPI_ISL_2678663, EPI_ISL_2678667, EPI_ISL_2678668 | MDI Limbach Berlin GmbH; MVZ Labor Berlin | Robert Koch Institute | unknown |
| EPI_ISL_2678669, EPI_ISL_2678670 | Labor Prof. Dr. G. Enders MVZ GbR | Robert Koch Institute | unknown |
| EPI_ISL_2678671 | MDI Limbach Berlin GmbH; MVZ Labor Berlin | Robert Koch Institute | unknown |
| EPI_ISL_2678675 | Labor Prof. Dr. G. Enders MVZ GbR | Robert Koch Institute | unknown |
| EPI_ISL_2678677, EPI_ISL_2678679, EPI_ISL_2678689, EPI_ISL_2678690, EPI_ISL_2678700, EPI_ISL_2678701, EPI_ISL_2678703, EPI_ISL_2678704, EPI_ISL_2678705, EPI_ISL_2678706 | MDI Limbach Berlin GmbH; MVZ Labor Berlin | Robert Koch Institute | unknown |
| EPI_ISL_2678708, EPI_ISL_2678712 | MVZ Medizinisches Labor Hannover GmbH | Robert Koch Institute | unknown |

| | | | |
|---|---|---|---|
| EPI_ISL_2678716, EPI_ISL_2678717, EPI_ISL_2678718 | MVZ Labor Dr. Reising-Ackermann und Kollegen | Robert Koch Institute | unknown |
| EPI_ISL_2678729 | MVZ Medizinisches Labor Hannover GmbH | Robert Koch Institute | unknown |
| EPI_ISL_2678730 | Limbach - MVZ Labor Eveld & Kollegen Essen | Robert Koch Institute | unknown |
| EPI_ISL_2678743 | Klinikum Ernst von Bergmann gemeinnützige GmbH - stationärer Bereich | Robert Koch Institute | unknown |
| EPI_ISL_2678749, EPI_ISL_2678751, EPI_ISL_2678755, EPI_ISL_2678756, EPI_ISL_2678760, EPI_ISL_2678766, EPI_ISL_2678768, EPI_ISL_2678787, EPI_ISL_2678792, EPI_ISL_2678793, EPI_ISL_2678794, EPI_ISL_2678796, EPI_ISL_2678802 | | | |
| see above | Labor ZOTZ\|KLIMAS; MVZ Düsseldorf-Centrum | Robert Koch Institute | unknown |
| EPI_ISL_2678936, EPI_ISL_2678939, EPI_ISL_2678941, EPI_ISL_2678942, EPI_ISL_2678946, EPI_ISL_2678950, EPI_ISL_2678955, EPI_ISL_2678958, EPI_ISL_2678960, EPI_ISL_2678961, EPI_ISL_2678963, EPI_ISL_2678967, EPI_ISL_2678969, EPI_ISL_2678970, EPI_ISL_2678973, EPI_ISL_2678975, EPI_ISL_2678979, EPI_ISL_2678980, EPI_ISL_2678982, EPI_ISL_2679237, EPI_ISL_2679247, EPI_ISL_2679255, EPI_ISL_2679257, EPI_ISL_2679258, EPI_ISL_2679271, EPI_ISL_2679273, EPI_ISL_2679279, EPI_ISL_2679288, EPI_ISL_2679293, EPI_ISL_2679305, EPI_ISL_2679309, EPI_ISL_2679313, EPI_ISL_2679315, EPI_ISL_2679317, EPI_ISL_2679324, EPI_ISL_2679328, EPI_ISL_2679332, EPI_ISL_2679334, EPI_ISL_2679337, EPI_ISL_2679341, EPI_ISL_2679437, EPI_ISL_2679439, EPI_ISL_2679446, EPI_ISL_2679450, EPI_ISL_2679494, EPI_ISL_2679514, EPI_ISL_2679564, EPI_ISL_2679613, EPI_ISL_2679628, EPI_ISL_2679634 | | | |
| see above | Utah Public Health Laboratory | Utah Public Health Laboratory | Erin L. Young, Kelly F. Oakeson, Tara Gallagher |
| EPI_ISL_2680325 | Department of Bacteria, Parasites and Fungi, Statens Serum Institut, Copenhagen, Denmark | Statens Serum Institut Bioinformatics and Microbial Genomics | Danish Covid-19 Genome Consortium |
| EPI_ISL_2680471, EPI_ISL_2680478, EPI_ISL_2680480, EPI_ISL_2680487, EPI_ISL_2680490 | Humboldt County Public Health Laboratory | Humboldt County Public Health Laboratory | Jeremy Corrigan |
| EPI_ISL_2680526, EPI_ISL_2680722 | Department of Bacteria, Parasites and Fungi, Statens Serum Institut, Copenhagen, Denmark | Statens Serum Institut Bioinformatics and Microbial Genomics | Danish Covid-19 Genome Consortium |
| EPI_ISL_2680919, EPI_ISL_2680921, EPI_ISL_2680926 | Laboratorios Delia Barraza | Laboratorio nacional-LANIIA-CIAD | Consorcio Mexicano de Vigilancia Genómica (CoViGen-Mex). Authors (in alphabetical order): Julio Elias Alvarado-Yaah, Carlos F. Arias, Santiago Ávila-Ríos, Eduardo Becerril-Vargas, Víctor Hugo Borja-Aburto, Celia Boukadida, Cristóbal Cháidez-Quiróz, Juan Bautista Chale-Dzul, Ricardo Ciria Merce, Andreu Comas-García, Célida Duque Molina, Julissa Enciso-Ibarra, José Antonio Enciso-Moreno, Gloria Elena Espinosa-Ayala, Fernando Fontove-Herrera, Daniel Fregoso-Rueda, Victor Eduardo García-Arias, Alejandra García-Gasca, Bruno Gómez-Gil, Jean Pierre González, Irvin González-López, Concepción Grajales-Muñiz, Ricardo Grande, Rosa María Gutiérrez Rios, Jesús Hernández, Alejandra Hernández-Terán, Alfredo Herrera-Estrella, Carla Ivón Herrera-Najera, Pavel Isa, Verónica Mata-Haro, Daniel Lira Morales, Susana Lopez, Antonio Loza Román, Brenda Irasema Maldonado-Meza, Bernardo Martínez-Miguel, José Arturo Martínez-Orozco, Célida Martínez-Rodríguez, Margarita Matías-Florentino, Fidencio Mejía-Nepomuceno, María Guadalupe de Jesús Mireles-Rivera, Gloria María Molina-Salinas, Hector Montoya-Fuentes, , Mario Mújica-Sánchez, José Esteban Muñoz-Medina, José de Jesús Nuñez-Contreras, Alicia Ocaña-Mondragón, Luis Alberto Ochoa-Carrera, Hector Esteban Paz-Juárez, Marissa Perez-Garcia, Francisco Pulido, Helen Haydee Fernanda Ramírez-Plascencia, Jorge Salas-Hernández, Angel Gustavo Salas-Lais, Alejandro Sánchez-Flores, Clara Esperanza Santacruz-Tinoco, María Guadalupe Santiago-Mauricio, Selene Zárate, Nelly Sélem-Mojica, Blanca Taboada, Gloria Vazquez, Joel Armando Vázquez-Pérez. Laboratorios Delia Barraza: María Delia Barraza Sámano, Claudia Soto Félix, Melissa García Angulo. |
| EPI_ISL_2680929 | Hospital Fátima S.A. de C.V. | Laboratorio nacional-LANIIA-CIAD | Consorcio Mexicano de Vigilancia Genómica (CoViGen-Mex). Authors (in alphabetical order): Julio Elias Alvarado-Yaah, Carlos F. Arias, Santiago Ávila-Ríos, Eduardo Becerril-Vargas, Víctor Hugo Borja-Aburto, Celia Boukadida, Cristóbal Cháidez-Quiróz, Juan Bautista Chale-Dzul, Ricardo Ciria Merce, Andreu Comas-García, Célida Duque Molina, Julissa Enciso-Ibarra, José Antonio Enciso-Moreno, Gloria Elena Espinosa-Ayala, Fernando Fontove-Herrera, Daniel Fregoso-Rueda, Victor Eduardo García-Arias, Alejandra García-Gasca, Bruno Gómez-Gil, Jean Pierre González, Irvin González-López, Concepción Grajales-Muñiz, Ricardo Grande, Rosa María Gutiérrez Rios, Jesús Hernández, Alejandra Hernández-Terán, Alfredo Herrera-Estrella, Carla Ivón Herrera-Najera, Pavel Isa, Verónica Mata-Haro, Daniel Lira Morales, Susana Lopez, Antonio Loza Román, Brenda Irasema Maldonado-Meza, Bernardo Martínez-Miguel, José Arturo Martínez-Orozco, Célida Martínez-Rodríguez, Margarita Matías-Florentino, Fidencio Mejía-Nepomuceno, María Guadalupe de Jesús Mireles-Rivera, Gloria María Molina-Salinas, Hector Montoya-Fuentes, , Mario Mújica-Sánchez, José Esteban Muñoz-Medina, José de Jesús Nuñez-Contreras, Alicia Ocaña-Mondragón, Luis Alberto Ochoa-Carrera, Hector Esteban Paz-Juárez, Marissa Perez-Garcia, Francisco Pulido, Helen Haydee Fernanda Ramírez-Plascencia, Jorge Salas-Hernández, Angel Gustavo Salas-Lais, Alejandro Sánchez-Flores, Clara Esperanza Santacruz-Tinoco, María Guadalupe Santiago-Mauricio, Selene Zárate, Nelly Sélem-Mojica, Blanca Taboada, Gloria Vazquez, Joel Armando Vázquez-Pérez. |
| EPI_ISL_2680931, EPI_ISL_2680937 | Hospital Margarita Maza de Juárez | Microbial Genomics Laboratory | Consorcio Mexicano de Vigilancia Genómica (CoViGen-Mex). Authors (in alphabetical order): Julio Elias Alvarado-Yaah, Carlos F. Arias, Santiago Ávila-Ríos, Eduardo Becerril-Vargas, Víctor Hugo Borja-Aburto, Celia Boukadida, Cristóbal Cháidez-Quiróz, Juan Bautista Chale-Dzul, Ricardo Ciria Merce, Andreu Comas-García, Célida Duque Molina, Julissa Enciso-Ibarra, José Antonio Enciso-Moreno, Gloria Elena Espinosa-Ayala, Fernando Fontove-Herrera, Daniel Fregoso-Rueda, Victor Eduardo García-Arias, Alejandra García-Gasca, Bruno Gómez-Gil, Jean Pierre González, Irvin González-López, Concepción Grajales-Muñiz, Ricardo Grande, Rosa María Gutiérrez Rios, Jesús Hernández, Alejandra Hernández-Terán, Alfredo Herrera-Estrella, Carla Ivón Herrera-Najera, Pavel Isa, Verónica Mata-Haro, Daniel Lira Morales, Susana Lopez, Antonio Loza Román, Brenda Irasema Maldonado-Meza, Bernardo Martínez-Miguel, José Arturo Martínez-Orozco, Célida Martínez-Rodríguez, Margarita Matías-Florentino, Fidencio Mejía-Nepomuceno, María Guadalupe de Jesús Mireles-Rivera, Gloria María Molina-Salinas, Hector Montoya-Fuentes, , Mario Mújica-Sánchez, José Esteban Muñoz-Medina, José de Jesús Nuñez-Contreras, Alicia Ocaña-Mondragón, Luis Alberto Ochoa-Carrera, Hector Esteban Paz-Juárez, Marissa Perez-Garcia, Francisco Pulido, Helen Haydee Fernanda Ramírez-Plascencia, Jorge Salas-Hernández, Angel Gustavo Salas-Lais, Alejandro Sánchez-Flores, Clara Esperanza Santacruz-Tinoco, María Guadalupe Santiago-Mauricio, Selene Zárate, Nelly Sélem-Mojica, Blanca Taboada, Gloria Vazquez, Joel Armando Vázquez-Pérez. |
| EPI_ISL_2680954, EPI_ISL_2680956, EPI_ISL_2680963, EPI_ISL_2680966, EPI_ISL_2680969, EPI_ISL_2680970, EPI_ISL_2680971 | Azienda Sanitaria dell'Alto Adige - Laboratorio Aziendale di Microbiologia e Virologia | Azienda Sanitaria dell'Alto Adige | Irene Bianconi |
| EPI_ISL_2680981 | Australian Clinical Labs | NSW Health Pathology - Institute of Clinical Pathology and Medical Research; Westmead Hospital; University of Sydney | CIDM-PH et al. |
| EPI_ISL_2680982, EPI_ISL_2680983 | St Vincent's Pathology (SydPath) | NSW Health Pathology - Institute of Clinical Pathology and Medical Research; Westmead Hospital; University of Sydney | CIDM-PH et al. |
| EPI_ISL_2680984, EPI_ISL_2680985 | Laverty Pathology | NSW Health Pathology - Institute of Clinical Pathology and Medical Research; Westmead Hospital; University of Sydney | CIDM-PH et al. |
| EPI_ISL_2680986 | Ministry of Health, Jaber Al-Ahmad Hospital | Virology Unit, Department of Microbiology, Faculty of Medicine, Kuwait University | Nada Madi, Hussain Safar, Ebaa Al-Awadhi, Anfal Al-Adwani |
| EPI_ISL_2681042 | UM im. Karola Marcinkowskiego w Poznaniu Laboratorium UCA_Covid-19 Centrum Biologii Medycznej | 1. Tricity SARS-CoV-2 sequencing consortium: University of Gdansk, Medical University of Gdansk, Vaxican Ltd., Invicta Ltd. 2. National Institute of Public Health - National Institute of Hygiene, Warsaw, Poland | Maciej Kosinski, Celina Cybulska, Krystyna Bienkowska Szewczyk, Maciej Grzybek, Karolina Gackowska, Marcin Lubocki, Katarzyna Groth, Lukasz Rabalski, Katarzyna Zacharczuk, Magdalena Nowakowska, Małgorzata Sadkowska-Todys, Tomasz Wołkowicz |
| EPI_ISL_2681059, EPI_ISL_2681064, EPI_ISL_2681065, EPI_ISL_2681071, EPI_ISL_2681073, EPI_ISL_2681113 | Laboratorio Central de Epidemiología (LCE) | Instituto de Biotecnología de la UNAM | Consorcio Mexicano de Vigilancia Genómica (CoViGen-Mex). Authors (in alphabetical order): Julio Elias Alvarado-Yaah, Carlos F. Arias, Santiago Ávila-Ríos, Eduardo Becerril-Vargas, Víctor Hugo Borja-Aburto, Celia Boukadida, Cristóbal Cháidez-Quiróz, Juan Bautista Chale-Dzul, Ricardo Ciria Merce, Andreu Comas-García, Célida Duque Molina, Julissa Enciso-Ibarra, José Antonio Enciso-Moreno, Gloria Elena Espinosa-Ayala, Fernando Fontove-Herrera, Daniel Fregoso-Rueda, Victor Eduardo García-Arias, Alejandra García-Gasca, Bruno Gómez-Gil, Jean Pierre González, Irvin González-López, Concepción Grajales-Muñiz, Ricardo Grande, Rosa María Gutiérrez Rios, Jesús Hernández, Alejandra Hernández-Terán, Alfredo Herrera-Estrella, Carla Ivón Herrera-Najera, Pavel Isa, Verónica Mata-Haro, Daniel Lira Morales, Susana Lopez, Antonio Loza Román, Brenda Irasema Maldonado-Meza, Bernardo Martínez-Miguel, José Arturo Martínez-Orozco, Célida Martínez-Rodríguez, Margarita Matías-Florentino, Fidencio Mejía-Nepomuceno, María Guadalupe de Jesús Mireles-Rivera, Gloria María Molina-Salinas, Hector Montoya-Fuentes, , Mario Mújica-Sánchez, José Esteban Muñoz-Medina, José de Jesús Nuñez-Contreras, Alicia Ocaña-Mondragón, Luis Alberto Ochoa-Carrera, Hector Esteban Paz-Juárez, Marissa Perez-Garcia, Francisco Pulido, Helen Haydee Fernanda Ramírez-Plascencia, Jorge Salas-Hernández, Angel Gustavo Salas-Lais, Alejandro Sánchez-Flores, Clara Esperanza Santacruz-Tinoco, María Guadalupe Santiago-Mauricio, Selene Zárate, Nelly Sélem-Mojica, Blanca Taboada, Gloria Vazquez, Joel Armando Vázquez-Pérez. |
| EPI_ISL_2681130 | Unidad de Investigación Médica de Yucatán (UIMY) | Instituto de Biotecnología de la UNAM | Consorcio Mexicano de Vigilancia Genómica (CoViGen-Mex). Authors (in alphabetical order): Julio Elias Alvarado-Yaah, Carlos F. Arias, Santiago Ávila-Ríos, Eduardo Becerril-Vargas, Víctor Hugo Borja-Aburto, Celia Boukadida, Cristóbal Cháidez-Quiróz, Juan Bautista Chale-Dzul, Ricardo Ciria Merce, Andreu Comas-García, Célida Duque Molina, Julissa Enciso-Ibarra, José Antonio Enciso-Moreno, Gloria Elena Espinosa-Ayala, Fernando Fontove-Herrera, Daniel Fregoso-Rueda, Victor Eduardo García-Arias, Alejandra García-Gasca, Bruno Gómez-Gil, Jean Pierre González, Irvin González-López, Concepción Grajales-Muñiz, Ricardo Grande, Rosa María Gutiérrez Rios, Jesús Hernández, Alejandra Hernández-Terán, Alfredo Herrera-Estrella, Carla Ivón Herrera-Najera, Pavel Isa, Verónica Mata-Haro, Daniel Lira Morales, Susana Lopez, Antonio Loza Román, Brenda Irasema Maldonado-Meza, Bernardo Martínez-Miguel, José Arturo Martínez-Orozco, Célida Martínez-Rodríguez, Margarita Matías-Florentino, Fidencio Mejía-Nepomuceno, María Guadalupe de Jesús Mireles-Rivera, Gloria María Molina-Salinas, Hector Montoya-Fuentes, , Mario Mújica-Sánchez, José Esteban Muñoz-Medina, José de Jesús Nuñez-Contreras, Alicia Ocaña-Mondragón, Luis Alberto Ochoa-Carrera, Hector Esteban Paz-Juárez, Marissa Perez-Garcia, Francisco Pulido, Helen Haydee Fernanda Ramírez-Plascencia, Jorge Salas-Hernández, Angel Gustavo Salas-Lais, Alejandro Sánchez-Flores, Clara Esperanza Santacruz-Tinoco, María Guadalupe Santiago-Mauricio, Selene Zárate, Nelly Sélem-Mojica, Blanca Taboada, Gloria Vazquez, Joel Armando Vázquez-Pérez. |
| EPI_ISL_2681140, EPI_ISL_2681151, EPI_ISL_2681155, EPI_ISL_2681156, EPI_ISL_2681175, EPI_ISL_2681181, EPI_ISL_2681184, EPI_ISL_2681188, EPI_ISL_2681189, EPI_ISL_2681193, EPI_ISL_2681194, EPI_ISL_2681196, EPI_ISL_2681199, EPI_ISL_2681205, EPI_ISL_2681214 | | | |
| see above | Laboratorio Central de Epidemiología (LCE) | Instituto de Biotecnología de la UNAM | Consorcio Mexicano de Vigilancia Genómica (CoViGen-Mex). Authors (in alphabetical order): Julio Elias Alvarado-Yaah, Carlos F. Arias, Santiago Ávila-Ríos, Eduardo Becerril-Vargas, Víctor Hugo Borja-Aburto, Celia Boukadida, Cristóbal Cháidez-Quiróz, Juan Bautista Chale-Dzul, Ricardo Ciria Merce, Andreu Comas-García, Célida Duque Molina, Julissa Enciso-Ibarra, José Antonio Enciso-Moreno, Gloria Elena Espinosa-Ayala, Fernando Fontove-Herrera, Daniel Fregoso-Rueda, Victor Eduardo García-Arias, Alejandra García-Gasca, Bruno Gómez-Gil, Jean Pierre González, Irvin González-López, Concepción Grajales-Muñiz, Ricardo Grande, Rosa María Gutiérrez Rios, Jesús Hernández, Alejandra Hernández-Terán, Alfredo Herrera-Estrella, Carla Ivón Herrera-Najera, Pavel Isa, Verónica Mata-Haro, Daniel Lira Morales, Susana Lopez, Antonio Loza Román, Brenda Irasema Maldonado-Meza, Bernardo Martínez-Miguel, José Arturo Martínez-Orozco, Célida Martínez-Rodríguez, Margarita Matías-Florentino, Fidencio Mejía-Nepomuceno, María Guadalupe de Jesús Mireles-Rivera, Gloria María Molina-Salinas, Hector Montoya-Fuentes, , Mario Mújica-Sánchez, José Esteban Muñoz-Medina, José de Jesús Nuñez-Contreras, Alicia Ocaña-Mondragón, Luis Alberto Ochoa-Carrera, Hector Esteban Paz-Juárez, Marissa Perez-Garcia, Francisco Pulido, Helen Haydee Fernanda Ramírez-Plascencia, Jorge Salas-Hernández, Angel Gustavo Salas-Lais, Alejandro Sánchez-Flores, Clara Esperanza Santacruz-Tinoco, María Guadalupe Santiago-Mauricio, Selene Zárate, Nelly Sélem-Mojica, Blanca Taboada, Gloria Vazquez, Joel Armando Vázquez-Pérez. |
| EPI_ISL_2681218 | Unidad de Investigación Médica de Yucatán (UIMY) | Instituto de Biotecnología de la UNAM | Consorcio Mexicano de Vigilancia Genómica (CoViGen-Mex). Authors (in alphabetical order): Julio Elias Alvarado-Yaah, Carlos F. Arias, Santiago Ávila-Ríos, Eduardo Becerril-Vargas, Víctor Hugo Borja-Aburto, Celia Boukadida, Cristóbal Cháidez-Quiróz, Juan Bautista Chale-Dzul, Ricardo Ciria Merce, Andreu Comas-García, Célida Duque Molina, Julissa Enciso-Ibarra, José Antonio Enciso-Moreno, Gloria Elena Espinosa-Ayala, Fernando Fontove-Herrera, Daniel Fregoso-Rueda, Victor Eduardo García-Arias, Alejandra García-Gasca, Bruno Gómez-Gil, Jean Pierre González, Irvin González-López, Concepción Grajales-Muñiz, Ricardo Grande, Rosa María Gutiérrez Rios, Jesús Hernández, Alejandra Hernández-Terán, Alfredo Herrera-Estrella, Carla Ivón Herrera-Najera, Pavel Isa, Verónica Mata-Haro, Daniel Lira Morales, Susana Lopez, Antonio Loza Román, Brenda Irasema Maldonado-Meza, Bernardo Martínez-Miguel, José Arturo Martínez-Orozco, Célida Martínez-Rodríguez, Margarita Matías-Florentino, Fidencio Mejía-Nepomuceno, María Guadalupe de Jesús Mireles-Rivera, Gloria María Molina-Salinas, Hector Montoya-Fuentes, , Mario Mújica-Sánchez, José Esteban Muñoz-Medina, José de Jesús Nuñez-Contreras, Alicia Ocaña-Mondragón, Luis Alberto Ochoa-Carrera, Hector Esteban Paz-Juárez, Marissa Perez-Garcia, Francisco Pulido, Helen Haydee Fernanda Ramírez-Plascencia, Jorge Salas-Hernández, Angel Gustavo Salas-Lais, Alejandro Sánchez-Flores, Clara Esperanza Santacruz-Tinoco, María Guadalupe Santiago-Mauricio, Selene Zárate, Nelly Sélem-Mojica, Blanca Taboada, Gloria Vazquez, Joel Armando Vázquez-Pérez. |
| EPI_ISL_2681229, EPI_ISL_2681231, EPI_ISL_2681251, EPI_ISL_2681266, EPI_ISL_2681267, EPI_ISL_2681306 | Laboratorio Central de Epidemiología (LCE) | Instituto de Biotecnología de la UNAM | Consorcio Mexicano de Vigilancia Genómica (CoViGen-Mex). Authors (in alphabetical order): Julio Elias Alvarado-Yaah, Carlos F. Arias, Santiago Ávila-Ríos, Eduardo Becerril-Vargas, Víctor Hugo Borja-Aburto, Celia Boukadida, Cristóbal Cháidez-Quiróz, Juan Bautista Chale-Dzul, Ricardo Ciria Merce, Andreu Comas-García, Célida Duque Molina, Julissa Enciso-Ibarra, José Antonio Enciso-Moreno, Gloria Elena Espinosa-Ayala, Fernando Fontove-Herrera, Daniel Fregoso-Rueda, Victor Eduardo García-Arias, Alejandra García-Gasca, Bruno Gómez-Gil, Jean Pierre González, Irvin González-López, Concepción Grajales-Muñiz, Ricardo Grande, Rosa María Gutiérrez Rios, Jesús Hernández, Alejandra Hernández-Terán, Alfredo Herrera-Estrella, Carla Ivón Herrera-Najera, Pavel Isa, Verónica Mata-Haro, Daniel Lira Morales, Susana Lopez, Antonio Loza Román, Brenda Irasema Maldonado-Meza, Bernardo Martínez-Miguel, José Arturo Martínez-Orozco, Célida Martínez-Rodríguez, Margarita Matías-Florentino, Fidencio Mejía-Nepomuceno, María Guadalupe de Jesús Mireles-Rivera, Gloria María Molina-Salinas, Hector Montoya-Fuentes, , Mario Mújica-Sánchez, José Esteban Muñoz-Medina, José de Jesús Nuñez-Contreras, Alicia Ocaña-Mondragón, Luis Alberto Ochoa-Carrera, Hector Esteban Paz-Juárez, Marissa Perez-Garcia, Francisco Pulido, Helen Haydee Fernanda Ramírez-Plascencia, Jorge Salas-Hernández, Angel Gustavo Salas-Lais, Alejandro |

| | | | |
|---|---|---|---|
| | | | Sánchez-Flores, Clara Esperanza Santacruz-Tinoco, María Guadalupe Santiago-Mauricio, Selene Zárate, Nelly Sélem-Mojica, Blanca Taboada, Gloria Vazquez, Joel Armando Vázquez-Pérez. |
| EPI_ISL_2681326 | Unidad de Investigación Médica de Yucatán (UIMY) | Instituto de Biotecnología de la UNAM | Consorcio Mexicano de Vigilancia Genómica (CoViGen-Mex). Authors (in alphabetical order): Julio Elias Alvarado-Yaah, Carlos F. Arias, Santiago Ávila-Ríos, Eduardo Becerril-Vargas, Víctor Hugo Borja-Aburto, Celia Boukadida, Cristóbal Cháidez-Quiróz, Juan Bautista Chale-Dzul, Ricardo Ciria Merce, Andreu Comas-García, Célida Duque Molina, Julissa Enciso-Ibarra, José Antonio Enciso-Moreno, Gloria Elena Espinosa-Ayala, Fernando Fontove-Herrera, Daniel Fregoso-Rueda, Victor Eduardo García-Arias, Alejandra García-Gasca, Bruno Gómez-Gil, Jean Pierre González, Irvin González-López, Concepción Grajales-Muñiz, Ricardo Grande, Rosa María Gutiérrez Rios, Jesús Hernández, Alejandra Hernández-Terán, Alfredo Herrera-Estrella, Carla Ivón Herrera-Najera, Pavel Isa, Verónica Mata-Haro, Daniel Lira Morales, Susana Lopez, Antonio Loza Román, Brenda Irasema Maldonado-Meza, Bernardo Martínez-Miguel, José Arturo Martínez-Orozco, Célida Martínez-Rodríguez, Margarita Matías-Florentino, Fidencio Mejía-Nepomuceno, María Guadalupe de Jesús Mireles-Rivera, Gloria María Molina-Salinas, Hector Montoya-Fuentes, , Mario Mújica-Sánchez, José Esteban Muñoz-Medina, José de Jesús Nuñez-Contreras, Alicia Ocaña-Mondragón, Luis Alberto Ochoa-Carrera, Hector Esteban Paz-Juárez, Marissa Perez-Garcia, Francisco Pulido, Helen Haydee Fernanda Ramírez-Plascencia, Jorge Salas-Hernández, Angel Gustavo Salas-Lais, Alejandro Sánchez-Flores, Clara Esperanza Santacruz-Tinoco, María Guadalupe Santiago-Mauricio, Selene Zárate, Nelly Sélem-Mojica, Blanca Taboada, Gloria Vazquez, Joel Armando Vázquez-Pérez. |
| EPI_ISL_2681371 | Laboratorio Central de Epidemiología (LCE) | Instituto de Biotecnología de la UNAM | Consorcio Mexicano de Vigilancia Genómica (CoViGen-Mex). Authors (in alphabetical order): Julio Elias Alvarado-Yaah, Carlos F. Arias, Santiago Ávila-Ríos, Eduardo Becerril-Vargas, Víctor Hugo Borja-Aburto, Celia Boukadida, Cristóbal Cháidez-Quiróz, Juan Bautista Chale-Dzul, Ricardo Ciria Merce, Andreu Comas-García, Célida Duque Molina, Julissa Enciso-Ibarra, José Antonio Enciso-Moreno, Gloria Elena Espinosa-Ayala, Fernando Fontove-Herrera, Daniel Fregoso-Rueda, Victor Eduardo García-Arias, Alejandra García-Gasca, Bruno Gómez-Gil, Jean Pierre González, Irvin González-López, Concepción Grajales-Muñiz, Ricardo Grande, Rosa María Gutiérrez Rios, Jesús Hernández, Alejandra Hernández-Terán, Alfredo Herrera-Estrella, Carla Ivón Herrera-Najera, Pavel Isa, Verónica Mata-Haro, Daniel Lira Morales, Susana Lopez, Antonio Loza Román, Brenda Irasema Maldonado-Meza, Bernardo Martínez-Miguel, José Arturo Martínez-Orozco, Célida Martínez-Rodríguez, Margarita Matías-Florentino, Fidencio Mejía-Nepomuceno, María Guadalupe de Jesús Mireles-Rivera, Gloria María Molina-Salinas, Hector Montoya-Fuentes, , Mario Mújica-Sánchez, José Esteban Muñoz-Medina, José de Jesús Nuñez-Contreras, Alicia Ocaña-Mondragón, Luis Alberto Ochoa-Carrera, Hector Esteban Paz-Juárez, Marissa Perez-Garcia, Francisco Pulido, Helen Haydee Fernanda Ramírez-Plascencia, Jorge Salas-Hernández, Angel Gustavo Salas-Lais, Alejandro Sánchez-Flores, Clara Esperanza Santacruz-Tinoco, María Guadalupe Santiago-Mauricio, Selene Zárate, Nelly Sélem-Mojica, Blanca Taboada, Gloria Vazquez, Joel Armando Vázquez-Pérez. |
| EPI_ISL_2681389, EPI_ISL_2681391, EPI_ISL_2681392, EPI_ISL_2681393 | National Institute of Public Health | State Veterinary Institute Prague | Nagy,A;Cernikov,L;Stara,M;Suri,T;Vecerova,J;Jirincova,H |
| EPI_ISL_2681450 | Althaia. Xarxa Assistencial Universitària de Manresa | IrsiCaixa | Marc Noguera-Julian, Mariona Parera, Maria Casadellà, Pilar Armengol, Francesc Catala-Moll, Roger Paredes, Bonaventura Clotet Gloria Trujillo, Rafel Perez Vidal, Jaume Trapé, Carolina Gonzalez-Fernandez, Miquel Micó, Antonia Flor, Roger Paredes, Eulalia Grau, Bonaventura Clotet |
| EPI_ISL_2681451, EPI_ISL_2681452 | ZOL | Jessa | Berden et al. on behalf of the Jessa_cmdLab |
| EPI_ISL_2681456, EPI_ISL_2681459, EPI_ISL_2681460, EPI_ISL_2681461, EPI_ISL_2681463, EPI_ISL_2681464, EPI_ISL_2681465, EPI_ISL_2681466, EPI_ISL_2681471, EPI_ISL_2681473, EPI_ISL_2681474, EPI_ISL_2681476, EPI_ISL_2681479, EPI_ISL_2681482, EPI_ISL_2681483 | | | |
| see above | University Hospitals of Geneva, Laboratory of Virology | HUG, Laboratory of Virology and the Health2030 Genome Center | Samuel Cordey, Ana Rita Goncalves, Laurent Kaiser, Lorenzo Cerutti, Henri Pegeot, Melyssa Elies, Deborah Penet, Keith Harshman, Ioannis Xenarios, Emmanouil Dermitzakis |
| EPI_ISL_2681552, EPI_ISL_2681556, EPI_ISL_2681567, EPI_ISL_2681568, EPI_ISL_2681576, EPI_ISL_2681578, EPI_ISL_2681585, EPI_ISL_2681593, EPI_ISL_2681594, EPI_ISL_2681598, EPI_ISL_2681599, EPI_ISL_2681600, EPI_ISL_2681601, EPI_ISL_2681606, EPI_ISL_2681607, EPI_ISL_2681609, EPI_ISL_2681610, EPI_ISL_2681615, EPI_ISL_2681621, EPI_ISL_2681625, EPI_ISL_2681633, EPI_ISL_2681639 | | | |
| see above | Microbiology Department, Laboratori Clínic Metropolitana Nord. Hospital Universitari Germans Trias i Pujol | Can Ruti SARS-CoV-2 Sequencing Hub (HUGTiP/IrsiCaixa/IGTP) | Marc Noguera-Julian, Pilar Armengol, Ignacio Blanco, Antoni E Bordoy, Francesc Catala-Moll, Pere-Joan Cardona, Maria Casadellà, Cristina Casañ, Gemma Clara, Bonaventura Clotet, Cristina Esteban, Montserrat Giménez, Mercedes Guerrero, Anna Not, Roger Paredes, Mariona Parera, Verónica Saludes, Alba Sánchez, and Elisa Martró on behalf of the Can Ruti SARS-CoV-2 Sequencing Hub |
| EPI_ISL_2681641 | Lighthouse Lab in Milton Keynes | Wellcome Sanger Institute for the COVID-19 Genomics UK (COG-UK) Consortium | The Lighthouse Lab in Milton Keynes and Alex Alderton, Roberto Amato, Jeffrey Barrett, Sonia Goncalves, Ewan Harrison, David K. Jackson, Ian Johnston, Dominic Kwiatkowski, Cordelia Langford, John Sillitoe on behalf of the Wellcome Sanger Institute COVID-19 Surveillance Team |
| EPI_ISL_2681642, EPI_ISL_2681643, EPI_ISL_2681644, EPI_ISL_2681645 | Lighthouse Lab in Alderley Park | Wellcome Sanger Institute for the COVID-19 Genomics UK (COG-UK) Consortium | Jacquelyn Wynn, Mairead Hyland, The Lighthouse Lab in Alderley Park and Alex Alderton, Roberto Amato, Jeffrey Barrett, Sonia Goncalves, Ewan Harrison, David K. Jackson, Ian Johnston, Dominic Kwiatkowski, Cordelia Langford, John Sillitoe on behalf of the Wellcome Sanger Institute COVID-19 Surveillance Team |
| EPI_ISL_2681646 | Lighthouse Lab in Milton Keynes | Wellcome Sanger Institute for the COVID-19 Genomics UK (COG-UK) Consortium | The Lighthouse Lab in Milton Keynes and Alex Alderton, Roberto Amato, Jeffrey Barrett, Sonia Goncalves, Ewan Harrison, David K. Jackson, Ian Johnston, Dominic Kwiatkowski, Cordelia Langford, John Sillitoe on behalf of the Wellcome Sanger Institute COVID-19 Surveillance Team |
| EPI_ISL_2681647 | Lighthouse Lab in Alderley Park | Wellcome Sanger Institute for the COVID-19 Genomics UK (COG-UK) Consortium | Jacquelyn Wynn, Mairead Hyland, The Lighthouse Lab in Alderley Park and Alex Alderton, Roberto Amato, Jeffrey Barrett, Sonia Goncalves, Ewan Harrison, David K. Jackson, Ian Johnston, Dominic Kwiatkowski, Cordelia Langford, John Sillitoe on behalf of the Wellcome Sanger Institute COVID-19 Surveillance Team |
| EPI_ISL_2681648, EPI_ISL_2681649, EPI_ISL_2681650 | Lighthouse Lab in Milton Keynes | Wellcome Sanger Institute for the COVID-19 Genomics UK (COG-UK) Consortium | The Lighthouse Lab in Milton Keynes and Alex Alderton, Roberto Amato, Jeffrey Barrett, Sonia Goncalves, Ewan Harrison, David K. Jackson, Ian Johnston, Dominic Kwiatkowski, Cordelia Langford, John Sillitoe on behalf of the Wellcome Sanger Institute COVID-19 Surveillance Team |
| EPI_ISL_2681651, EPI_ISL_2681652, EPI_ISL_2681653, EPI_ISL_2681654, EPI_ISL_2681655 | Lighthouse Lab in Alderley Park | Wellcome Sanger Institute for the COVID-19 Genomics UK (COG-UK) Consortium | Jacquelyn Wynn, Mairead Hyland, The Lighthouse Lab in Alderley Park and Alex Alderton, Roberto Amato, Jeffrey Barrett, Sonia Goncalves, Ewan Harrison, David K. Jackson, Ian Johnston, Dominic Kwiatkowski, Cordelia Langford, John Sillitoe on behalf of the Wellcome Sanger Institute COVID-19 Surveillance Team |
| EPI_ISL_2681656, EPI_ISL_2681657 | Lighthouse Lab in Glasgow | Wellcome Sanger Institute for the COVID-19 Genomics UK (COG-UK) Consortium | Harper VanSteenhouse, Yumi Kasai, David Gray, Carol Clugston, Anna Dominiczak and Alex Alderton, Roberto Amato, Jeffrey Barrett, Sonia Goncalves, Ewan Harrison, David K. Jackson, Ian Johnston, Dominic Kwiatkowski, Cordelia Langford, John Sillitoe on behalf of the Wellcome Sanger Institute COVID-19 Surveillance Team |
| EPI_ISL_2681658, EPI_ISL_2681659 | Lighthouse Lab in Alderley Park | Wellcome Sanger Institute for the COVID-19 Genomics UK (COG-UK) Consortium | Jacquelyn Wynn, Mairead Hyland, The Lighthouse Lab in Alderley Park and Alex Alderton, Roberto Amato, Jeffrey Barrett, Sonia Goncalves, Ewan Harrison, David K. Jackson, Ian Johnston, Dominic Kwiatkowski, Cordelia Langford, John Sillitoe on behalf of the Wellcome Sanger Institute COVID-19 Surveillance Team |
| EPI_ISL_2681660 | Lighthouse Lab in Glasgow | HUG Sanger Institute for the COVID-19 Genomics UK (COG-UK) Consortium | Harper VanSteenhouse, Yumi Kasai, David Gray, Carol Clugston, Anna Dominiczak and Alex Alderton, Roberto Amato, Jeffrey Barrett, Sonia Goncalves, Ewan Harrison, David K. Jackson, Ian Johnston, Dominic Kwiatkowski, Cordelia Langford, John Sillitoe on behalf of the Wellcome Sanger Institute COVID-19 Surveillance Team |
| EPI_ISL_2681662, EPI_ISL_2681663 | Lighthouse Lab in Alderley Park | Wellcome Sanger Institute for the COVID-19 Genomics UK (COG-UK) Consortium | Jacquelyn Wynn, Mairead Hyland, The Lighthouse Lab in Alderley Park and Alex Alderton, Roberto Amato, Jeffrey Barrett, Sonia Goncalves, Ewan Harrison, David K. Jackson, Ian Johnston, Dominic Kwiatkowski, Cordelia Langford, John Sillitoe on behalf of the Wellcome Sanger Institute COVID-19 Surveillance Team |
| EPI_ISL_2681664 | Lighthouse Lab in Milton Keynes | Wellcome Sanger Institute for the COVID-19 Genomics UK (COG-UK) Consortium | The Lighthouse Lab in Milton Keynes and Alex Alderton, Roberto Amato, Jeffrey Barrett, Sonia Goncalves, Ewan Harrison, David K. Jackson, Ian Johnston, Dominic Kwiatkowski, Cordelia Langford, John Sillitoe on behalf of the Wellcome Sanger Institute COVID-19 Surveillance Team |
| EPI_ISL_2681665, EPI_ISL_2681666 | Lighthouse Lab in Glasgow | Wellcome Sanger Institute for the COVID-19 Genomics UK (COG-UK) Consortium | Harper VanSteenhouse, Yumi Kasai, David Gray, Carol Clugston, Anna Dominiczak and Alex Alderton, Roberto Amato, Jeffrey Barrett, Sonia Goncalves, Ewan Harrison, David K. Jackson, Ian Johnston, Dominic Kwiatkowski, Cordelia Langford, John Sillitoe on behalf of the Wellcome Sanger Institute COVID-19 Surveillance Team |
| EPI_ISL_2681667 | Lighthouse Lab in Milton Keynes | Wellcome Sanger Institute for the COVID-19 Genomics UK (COG-UK) Consortium | The Lighthouse Lab in Milton Keynes and Alex Alderton, Roberto Amato, Jeffrey Barrett, Sonia Goncalves, Ewan Harrison, David K. Jackson, Ian Johnston, Dominic Kwiatkowski, Cordelia Langford, John Sillitoe on behalf of the Wellcome Sanger Institute COVID-19 Surveillance Team |
| EPI_ISL_2681668, EPI_ISL_2681669 | Lighthouse Lab in Alderley Park | Wellcome Sanger Institute for the COVID-19 Genomics UK (COG-UK) Consortium | Jacquelyn Wynn, Mairead Hyland, The Lighthouse Lab in Alderley Park and Alex Alderton, Roberto Amato, Jeffrey Barrett, Sonia Goncalves, Ewan Harrison, David K. Jackson, Ian Johnston, Dominic Kwiatkowski, Cordelia Langford, John Sillitoe on behalf of the Wellcome Sanger Institute COVID-19 Surveillance Team |
| EPI_ISL_2681670 | Lighthouse Lab in Milton Keynes | Wellcome Sanger Institute for the COVID-19 Genomics UK (COG-UK) Consortium | The Lighthouse Lab in Milton Keynes and Alex Alderton, Roberto Amato, Jeffrey Barrett, Sonia Goncalves, Ewan Harrison, David K. Jackson, Ian Johnston, Dominic Kwiatkowski, Cordelia Langford, John Sillitoe on behalf of the Wellcome Sanger Institute COVID-19 Surveillance Team |
| EPI_ISL_2681671 | Lighthouse Lab in Alderley Park | Wellcome Sanger Institute for the COVID-19 Genomics UK (COG-UK) Consortium | Jacquelyn Wynn, Mairead Hyland, The Lighthouse Lab in Alderley Park and Alex Alderton, Roberto Amato, Jeffrey Barrett, Sonia Goncalves, Ewan Harrison, David K. Jackson, Ian Johnston, Dominic Kwiatkowski, Cordelia Langford, John Sillitoe on behalf of the Wellcome Sanger Institute COVID-19 Surveillance Team |
| EPI_ISL_2681672 | Lighthouse Lab in Milton Keynes | Wellcome Sanger Institute for the COVID-19 Genomics UK (COG-UK) Consortium | The Lighthouse Lab in Milton Keynes and Alex Alderton, Roberto Amato, Jeffrey Barrett, Sonia Goncalves, Ewan Harrison, David K. Jackson, Ian Johnston, Dominic Kwiatkowski, Cordelia Langford, John Sillitoe on behalf of the Wellcome Sanger Institute COVID-19 Surveillance Team |
| EPI_ISL_2681673 | Lighthouse Lab in Glasgow | Wellcome Sanger Institute for the COVID-19 Genomics UK (COG-UK) Consortium | Harper VanSteenhouse, Yumi Kasai, David Gray, Carol Clugston, Anna Dominiczak and Alex Alderton, Roberto Amato, Jeffrey Barrett, Sonia Goncalves, Ewan Harrison, David K. Jackson, Ian Johnston, Dominic Kwiatkowski, Cordelia Langford, John Sillitoe on behalf of the Wellcome Sanger Institute COVID-19 Surveillance Team |
| EPI_ISL_2681674 | Lighthouse Lab in Alderley Park | Wellcome Sanger Institute for the COVID-19 Genomics UK (COG-UK) Consortium | Jacquelyn Wynn, Mairead Hyland, The Lighthouse Lab in Alderley Park and Alex Alderton, Roberto Amato, Jeffrey Barrett, Sonia Goncalves, Ewan Harrison, David K. Jackson, Ian Johnston, Dominic Kwiatkowski, Cordelia Langford, John Sillitoe on behalf of the Wellcome Sanger Institute COVID-19 Surveillance Team |
| EPI_ISL_2681675 | Lighthouse Lab in Milton Keynes | Wellcome Sanger Institute for the COVID-19 Genomics UK (COG-UK) Consortium | The Lighthouse Lab in Milton Keynes and Alex Alderton, Roberto Amato, Jeffrey Barrett, Sonia Goncalves, Ewan Harrison, David K. Jackson, Ian Johnston, Dominic Kwiatkowski, Cordelia Langford, John Sillitoe on behalf of the Wellcome Sanger Institute COVID-19 Surveillance Team |
| EPI_ISL_2681676, EPI_ISL_2681677, EPI_ISL_2681678 | Lighthouse Lab in Alderley Park | Wellcome Sanger Institute for the COVID-19 Genomics UK (COG-UK) Consortium | Jacquelyn Wynn, Mairead Hyland, The Lighthouse Lab in Alderley Park and Alex Alderton, Roberto Amato, Jeffrey Barrett, Sonia Goncalves, Ewan Harrison, David K. Jackson, Ian Johnston, Dominic Kwiatkowski, Cordelia Langford, John Sillitoe on behalf of the Wellcome Sanger Institute COVID-19 Surveillance Team |
| EPI_ISL_2681679 | Lighthouse Lab in Milton Keynes | Wellcome Sanger Institute for the COVID-19 Genomics UK (COG-UK) Consortium | The Lighthouse Lab in Milton Keynes and Alex Alderton, Roberto Amato, Jeffrey Barrett, Sonia Goncalves, Ewan Harrison, David K. Jackson, Ian Johnston, Dominic Kwiatkowski, Cordelia Langford, John Sillitoe on behalf of the Wellcome Sanger Institute COVID-19 Surveillance Team |
| EPI_ISL_2681680 | Lighthouse Lab in Alderley Park | Wellcome Sanger Institute for the COVID-19 Genomics UK (COG-UK) Consortium | Jacquelyn Wynn, Mairead Hyland, The Lighthouse Lab in Alderley Park and Alex Alderton, Roberto Amato, Jeffrey Barrett, Sonia Goncalves, Ewan Harrison, David K. Jackson, Ian Johnston, Dominic Kwiatkowski, Cordelia Langford, John Sillitoe on behalf of the Wellcome Sanger Institute COVID-19 Surveillance Team |
| EPI_ISL_2681681, EPI_ISL_2681682 | Lighthouse Lab in Milton Keynes | Wellcome Sanger Institute for the COVID-19 Genomics UK (COG-UK) Consortium | The Lighthouse Lab in Milton Keynes and Alex Alderton, Roberto Amato, Jeffrey Barrett, Sonia Goncalves, Ewan Harrison, David K. Jackson, Ian Johnston, Dominic Kwiatkowski, Cordelia Langford, John Sillitoe on behalf of the Wellcome Sanger Institute COVID-19 Surveillance Team |
| EPI_ISL_2681683, EPI_ISL_2681684 | Lighthouse Lab in Alderley Park | Wellcome Sanger Institute for the COVID-19 Genomics UK (COG-UK) Consortium | Jacquelyn Wynn, Mairead Hyland, The Lighthouse Lab in Alderley Park and Alex Alderton, Roberto Amato, Jeffrey Barrett, Sonia Goncalves, Ewan Harrison, David K. Jackson, Ian Johnston, Dominic Kwiatkowski, Cordelia Langford, John Sillitoe on behalf of the Wellcome Sanger Institute COVID-19 Surveillance Team |
| EPI_ISL_2681685 | Lighthouse Lab in Milton Keynes | Wellcome Sanger Institute for the COVID-19 Genomics UK (COG-UK) Consortium | The Lighthouse Lab in Milton Keynes and Alex Alderton, Roberto Amato, Jeffrey Barrett, Sonia Goncalves, Ewan Harrison, David K. Jackson, Ian Johnston, Dominic Kwiatkowski, Cordelia Langford, John Sillitoe on behalf of the Wellcome Sanger Institute COVID-19 Surveillance Team |
| EPI_ISL_2681686, EPI_ISL_2681687 | Lighthouse Lab in Alderley Park | Wellcome Sanger Institute for the COVID-19 Genomics UK (COG-UK) Consortium | Jacquelyn Wynn, Mairead Hyland, The Lighthouse Lab in Alderley Park and Alex Alderton, Roberto Amato, Jeffrey Barrett, Sonia Goncalves, Ewan Harrison, David K. Jackson, Ian Johnston, Dominic Kwiatkowski, Cordelia Langford, John Sillitoe on behalf of the Wellcome Sanger Institute COVID-19 Surveillance Team |
| EPI_ISL_2681688, EPI_ISL_2681689, EPI_ISL_2681690, EPI_ISL_2681691 | Lighthouse Lab in Milton Keynes | Wellcome Sanger Institute for the COVID-19 Genomics UK (COG-UK) Consortium | The Lighthouse Lab in Milton Keynes and Alex Alderton, Roberto Amato, Jeffrey Barrett, Sonia Goncalves, Ewan Harrison, David K. Jackson, Ian Johnston, Dominic Kwiatkowski, Cordelia Langford, John Sillitoe on behalf of the Wellcome Sanger Institute COVID-19 Surveillance Team |

| | | | |
|---|---|---|---|
| EPI_ISL_2681692, EPI_ISL_2681693, EPI_ISL_2681694, EPI_ISL_2681695 | Lighthouse Lab in Alderley Park | Wellcome Sanger Institute for the COVID-19 Genomics UK (COG-UK) Consortium | Jacquelyn Wynn, Mairead Hyland, The Lighthouse Lab in Alderley Park and Alex Alderton, Roberto Amato, Jeffrey Barrett, Sonia Goncalves, Ewan Harrison, David K. Jackson, Ian Johnston, Dominic Kwiatkowski, Cordelia Langford, John Sillitoe on behalf of the Wellcome Sanger Institute COVID-19 Surveillance Team |
| EPI_ISL_2681696 | Lighthouse Lab in Milton Keynes | Wellcome Sanger Institute for the COVID-19 Genomics UK (COG-UK) Consortium | The Lighthouse Lab in Milton Keynes and Alex Alderton, Roberto Amato, Jeffrey Barrett, Sonia Goncalves, Ewan Harrison, David K. Jackson, Ian Johnston, Dominic Kwiatkowski, Cordelia Langford, John Sillitoe on behalf of the Wellcome Sanger Institute COVID-19 Surveillance Team |
| EPI_ISL_2681697, EPI_ISL_2681698, EPI_ISL_2681699, EPI_ISL_2681700 | Lighthouse Lab in Alderley Park | Wellcome Sanger Institute for the COVID-19 Genomics UK (COG-UK) Consortium | Jacquelyn Wynn, Mairead Hyland, The Lighthouse Lab in Alderley Park and Alex Alderton, Roberto Amato, Jeffrey Barrett, Sonia Goncalves, Ewan Harrison, David K. Jackson, Ian Johnston, Dominic Kwiatkowski, Cordelia Langford, John Sillitoe on behalf of the Wellcome Sanger Institute COVID-19 Surveillance Team |
| EPI_ISL_2681701 | Lighthouse Lab in Milton Keynes | Wellcome Sanger Institute for the COVID-19 Genomics UK (COG-UK) Consortium | The Lighthouse Lab in Milton Keynes and Alex Alderton, Roberto Amato, Jeffrey Barrett, Sonia Goncalves, Ewan Harrison, David K. Jackson, Ian Johnston, Dominic Kwiatkowski, Cordelia Langford, John Sillitoe on behalf of the Wellcome Sanger Institute COVID-19 Surveillance Team |
| EPI_ISL_2681702, EPI_ISL_2681703, EPI_ISL_2681704 | Lighthouse Lab in Alderley Park | Wellcome Sanger Institute for the COVID-19 Genomics UK (COG-UK) Consortium | Jacquelyn Wynn, Mairead Hyland, The Lighthouse Lab in Alderley Park and Alex Alderton, Roberto Amato, Jeffrey Barrett, Sonia Goncalves, Ewan Harrison, David K. Jackson, Ian Johnston, Dominic Kwiatkowski, Cordelia Langford, John Sillitoe on behalf of the Wellcome Sanger Institute COVID-19 Surveillance Team |
| EPI_ISL_2681705, EPI_ISL_2681706 | Lighthouse Lab in Milton Keynes | Wellcome Sanger Institute for the COVID-19 Genomics UK (COG-UK) Consortium | The Lighthouse Lab in Milton Keynes and Alex Alderton, Roberto Amato, Jeffrey Barrett, Sonia Goncalves, Ewan Harrison, David K. Jackson, Ian Johnston, Dominic Kwiatkowski, Cordelia Langford, John Sillitoe on behalf of the Wellcome Sanger Institute COVID-19 Surveillance Team |
| EPI_ISL_2681707, EPI_ISL_2681709, EPI_ISL_2681710 | Lighthouse Lab in Alderley Park | Wellcome Sanger Institute for the COVID-19 Genomics UK (COG-UK) Consortium | Jacquelyn Wynn, Mairead Hyland, The Lighthouse Lab in Alderley Park and Alex Alderton, Roberto Amato, Jeffrey Barrett, Sonia Goncalves, Ewan Harrison, David K. Jackson, Ian Johnston, Dominic Kwiatkowski, Cordelia Langford, John Sillitoe on behalf of the Wellcome Sanger Institute COVID-19 Surveillance Team |
| EPI_ISL_2681711 | Lighthouse Lab in Glasgow | Wellcome Sanger Institute for the COVID-19 Genomics UK (COG-UK) Consortium | Harper VanSteenhouse, Yumi Kasai, David Gray, Carol Clugston, Anna Dominiczak and Alex Alderton, Roberto Amato, Jeffrey Barrett, Sonia Goncalves, Ewan Harrison, David K. Jackson, Ian Johnston, Dominic Kwiatkowski, Cordelia Langford, John Sillitoe on behalf of the Wellcome Sanger Institute COVID-19 Surveillance Team |
| EPI_ISL_2681712 | Lighthouse Lab in Alderley Park | Wellcome Sanger Institute for the COVID-19 Genomics UK (COG-UK) Consortium | Jacquelyn Wynn, Mairead Hyland, The Lighthouse Lab in Alderley Park and Alex Alderton, Roberto Amato, Jeffrey Barrett, Sonia Goncalves, Ewan Harrison, David K. Jackson, Ian Johnston, Dominic Kwiatkowski, Cordelia Langford, John Sillitoe on behalf of the Wellcome Sanger Institute COVID-19 Surveillance Team |
| EPI_ISL_2681713 | Lighthouse Lab in Milton Keynes | Wellcome Sanger Institute for the COVID-19 Genomics UK (COG-UK) Consortium | The Lighthouse Lab in Milton Keynes and Alex Alderton, Roberto Amato, Jeffrey Barrett, Sonia Goncalves, Ewan Harrison, David K. Jackson, Ian Johnston, Dominic Kwiatkowski, Cordelia Langford, John Sillitoe on behalf of the Wellcome Sanger Institute COVID-19 Surveillance Team |
| EPI_ISL_2681714 | Lighthouse Lab in Alderley Park | Wellcome Sanger Institute for the COVID-19 Genomics UK (COG-UK) Consortium | Jacquelyn Wynn, Mairead Hyland, The Lighthouse Lab in Alderley Park and Alex Alderton, Roberto Amato, Jeffrey Barrett, Sonia Goncalves, Ewan Harrison, David K. Jackson, Ian Johnston, Dominic Kwiatkowski, Cordelia Langford, John Sillitoe on behalf of the Wellcome Sanger Institute COVID-19 Surveillance Team |
| EPI_ISL_2681715, EPI_ISL_2681716 | Lighthouse Lab in Milton Keynes | Wellcome Sanger Institute for the COVID-19 Genomics UK (COG-UK) Consortium | The Lighthouse Lab in Milton Keynes and Alex Alderton, Roberto Amato, Jeffrey Barrett, Sonia Goncalves, Ewan Harrison, David K. Jackson, Ian Johnston, Dominic Kwiatkowski, Cordelia Langford, John Sillitoe on behalf of the Wellcome Sanger Institute COVID-19 Surveillance Team |
| EPI_ISL_2681717 | Lighthouse Lab in Alderley Park | Wellcome Sanger Institute for the COVID-19 Genomics UK (COG-UK) Consortium | Jacquelyn Wynn, Mairead Hyland, The Lighthouse Lab in Alderley Park and Alex Alderton, Roberto Amato, Jeffrey Barrett, Sonia Goncalves, Ewan Harrison, David K. Jackson, Ian Johnston, Dominic Kwiatkowski, Cordelia Langford, John Sillitoe on behalf of the Wellcome Sanger Institute COVID-19 Surveillance Team |
| EPI_ISL_2681718 | Lighthouse Lab in Glasgow | Wellcome Sanger Institute for the COVID-19 Genomics UK (COG-UK) Consortium | Harper VanSteenhouse, Yumi Kasai, David Gray, Carol Clugston, Anna Dominiczak and Alex Alderton, Roberto Amato, Jeffrey Barrett, Sonia Goncalves, Ewan Harrison, David K. Jackson, Ian Johnston, Dominic Kwiatkowski, Cordelia Langford, John Sillitoe on behalf of the Wellcome Sanger Institute COVID-19 Surveillance Team |
| EPI_ISL_2681719 | Lighthouse Lab in Milton Keynes | Wellcome Sanger Institute for the COVID-19 Genomics UK (COG-UK) Consortium | The Lighthouse Lab in Milton Keynes and Alex Alderton, Roberto Amato, Jeffrey Barrett, Sonia Goncalves, Ewan Harrison, David K. Jackson, Ian Johnston, Dominic Kwiatkowski, Cordelia Langford, John Sillitoe on behalf of the Wellcome Sanger Institute COVID-19 Surveillance Team |
| EPI_ISL_2681720, EPI_ISL_2681721 | Lighthouse Lab in Glasgow | Wellcome Sanger Institute for the COVID-19 Genomics UK (COG-UK) Consortium | Harper VanSteenhouse, Yumi Kasai, David Gray, Carol Clugston, Anna Dominiczak and Alex Alderton, Roberto Amato, Jeffrey Barrett, Sonia Goncalves, Ewan Harrison, David K. Jackson, Ian Johnston, Dominic Kwiatkowski, Cordelia Langford, John Sillitoe on behalf of the Wellcome Sanger Institute COVID-19 Surveillance Team |
| EPI_ISL_2681722, EPI_ISL_2681723, EPI_ISL_2681724, EPI_ISL_2681725, EPI_ISL_2681726 | Lighthouse Lab in Alderley Park | Wellcome Sanger Institute for the COVID-19 Genomics UK (COG-UK) Consortium | Jacquelyn Wynn, Mairead Hyland, The Lighthouse Lab in Alderley Park and Alex Alderton, Roberto Amato, Jeffrey Barrett, Sonia Goncalves, Ewan Harrison, David K. Jackson, Ian Johnston, Dominic Kwiatkowski, Cordelia Langford, John Sillitoe on behalf of the Wellcome Sanger Institute COVID-19 Surveillance Team |
| EPI_ISL_2681727, EPI_ISL_2681728 | Lighthouse Lab in Glasgow | Wellcome Sanger Institute for the COVID-19 Genomics UK (COG-UK) Consortium | Harper VanSteenhouse, Yumi Kasai, David Gray, Carol Clugston, Anna Dominiczak and Alex Alderton, Roberto Amato, Jeffrey Barrett, Sonia Goncalves, Ewan Harrison, David K. Jackson, Ian Johnston, Dominic Kwiatkowski, Cordelia Langford, John Sillitoe on behalf of the Wellcome Sanger Institute COVID-19 Surveillance Team |
| EPI_ISL_2681729 | Lighthouse Lab in Alderley Park | Wellcome Sanger Institute for the COVID-19 Genomics UK (COG-UK) Consortium | Jacquelyn Wynn, Mairead Hyland, The Lighthouse Lab in Alderley Park and Alex Alderton, Roberto Amato, Jeffrey Barrett, Sonia Goncalves, Ewan Harrison, David K. Jackson, Ian Johnston, Dominic Kwiatkowski, Cordelia Langford, John Sillitoe on behalf of the Wellcome Sanger Institute COVID-19 Surveillance Team |
| EPI_ISL_2681730, EPI_ISL_2681731 | Lighthouse Lab in Milton Keynes | Wellcome Sanger Institute for the COVID-19 Genomics UK (COG-UK) Consortium | The Lighthouse Lab in Milton Keynes and Alex Alderton, Roberto Amato, Jeffrey Barrett, Sonia Goncalves, Ewan Harrison, David K. Jackson, Ian Johnston, Dominic Kwiatkowski, Cordelia Langford, John Sillitoe on behalf of the Wellcome Sanger Institute COVID-19 Surveillance Team |
| EPI_ISL_2681732 | Lighthouse Lab in Alderley Park | Wellcome Sanger Institute for the COVID-19 Genomics UK (COG-UK) Consortium | Jacquelyn Wynn, Mairead Hyland, The Lighthouse Lab in Alderley Park and Alex Alderton, Roberto Amato, Jeffrey Barrett, Sonia Goncalves, Ewan Harrison, David K. Jackson, Ian Johnston, Dominic Kwiatkowski, Cordelia Langford, John Sillitoe on behalf of the Wellcome Sanger Institute COVID-19 Surveillance Team |
| EPI_ISL_2681733 | Lighthouse Lab in Milton Keynes | Wellcome Sanger Institute for the COVID-19 Genomics UK (COG-UK) Consortium | The Lighthouse Lab in Milton Keynes and Alex Alderton, Roberto Amato, Jeffrey Barrett, Sonia Goncalves, Ewan Harrison, David K. Jackson, Ian Johnston, Dominic Kwiatkowski, Cordelia Langford, John Sillitoe on behalf of the Wellcome Sanger Institute COVID-19 Surveillance Team |
| EPI_ISL_2681734 | Lighthouse Lab in Alderley Park | Wellcome Sanger Institute for the COVID-19 Genomics UK (COG-UK) Consortium | Jacquelyn Wynn, Mairead Hyland, The Lighthouse Lab in Alderley Park and Alex Alderton, Roberto Amato, Jeffrey Barrett, Sonia Goncalves, Ewan Harrison, David K. Jackson, Ian Johnston, Dominic Kwiatkowski, Cordelia Langford, John Sillitoe on behalf of the Wellcome Sanger Institute COVID-19 Surveillance Team |
| EPI_ISL_2681735, EPI_ISL_2681736 | Lighthouse Lab in Milton Keynes | Wellcome Sanger Institute for the COVID-19 Genomics UK (COG-UK) Consortium | The Lighthouse Lab in Milton Keynes and Alex Alderton, Roberto Amato, Jeffrey Barrett, Sonia Goncalves, Ewan Harrison, David K. Jackson, Ian Johnston, Dominic Kwiatkowski, Cordelia Langford, John Sillitoe on behalf of the Wellcome Sanger Institute COVID-19 Surveillance Team |
| EPI_ISL_2681737 | Lighthouse Lab in Alderley Park | Wellcome Sanger Institute for the COVID-19 Genomics UK (COG-UK) Consortium | Jacquelyn Wynn, Mairead Hyland, The Lighthouse Lab in Alderley Park and Alex Alderton, Roberto Amato, Jeffrey Barrett, Sonia Goncalves, Ewan Harrison, David K. Jackson, Ian Johnston, Dominic Kwiatkowski, Cordelia Langford, John Sillitoe on behalf of the Wellcome Sanger Institute COVID-19 Surveillance Team |
| EPI_ISL_2681738 | Lighthouse Lab in Milton Keynes | Wellcome Sanger Institute for the COVID-19 Genomics UK (COG-UK) Consortium | The Lighthouse Lab in Milton Keynes and Alex Alderton, Roberto Amato, Jeffrey Barrett, Sonia Goncalves, Ewan Harrison, David K. Jackson, Ian Johnston, Dominic Kwiatkowski, Cordelia Langford, John Sillitoe on behalf of the Wellcome Sanger Institute COVID-19 Surveillance Team |
| EPI_ISL_2681739, EPI_ISL_2681740 | Lighthouse Lab in Alderley Park | Wellcome Sanger Institute for the COVID-19 Genomics UK (COG-UK) Consortium | Jacquelyn Wynn, Mairead Hyland, The Lighthouse Lab in Alderley Park and Alex Alderton, Roberto Amato, Jeffrey Barrett, Sonia Goncalves, Ewan Harrison, David K. Jackson, Ian Johnston, Dominic Kwiatkowski, Cordelia Langford, John Sillitoe on behalf of the Wellcome Sanger Institute COVID-19 Surveillance Team |
| EPI_ISL_2681741 | Lighthouse Lab in Milton Keynes | Wellcome Sanger Institute for the COVID-19 Genomics UK (COG-UK) Consortium | The Lighthouse Lab in Milton Keynes and Alex Alderton, Roberto Amato, Jeffrey Barrett, Sonia Goncalves, Ewan Harrison, David K. Jackson, Ian Johnston, Dominic Kwiatkowski, Cordelia Langford, John Sillitoe on behalf of the Wellcome Sanger Institute COVID-19 Surveillance Team |
| EPI_ISL_2681742 | Lighthouse Lab in Alderley Park | Wellcome Sanger Institute for the COVID-19 Genomics UK (COG-UK) Consortium | Jacquelyn Wynn, Mairead Hyland, The Lighthouse Lab in Alderley Park and Alex Alderton, Roberto Amato, Jeffrey Barrett, Sonia Goncalves, Ewan Harrison, David K. Jackson, Ian Johnston, Dominic Kwiatkowski, Cordelia Langford, John Sillitoe on behalf of the Wellcome Sanger Institute COVID-19 Surveillance Team |
| EPI_ISL_2681743 | Lighthouse Lab in Milton Keynes | Wellcome Sanger Institute for the COVID-19 Genomics UK (COG-UK) Consortium | The Lighthouse Lab in Milton Keynes and Alex Alderton, Roberto Amato, Jeffrey Barrett, Sonia Goncalves, Ewan Harrison, David K. Jackson, Ian Johnston, Dominic Kwiatkowski, Cordelia Langford, John Sillitoe on behalf of the Wellcome Sanger Institute COVID-19 Surveillance Team |
| EPI_ISL_2681744, EPI_ISL_2681745 | Lighthouse Lab in Alderley Park | Wellcome Sanger Institute for the COVID-19 Genomics UK (COG-UK) Consortium | Jacquelyn Wynn, Mairead Hyland, The Lighthouse Lab in Alderley Park and Alex Alderton, Roberto Amato, Jeffrey Barrett, Sonia Goncalves, Ewan Harrison, David K. Jackson, Ian Johnston, Dominic Kwiatkowski, Cordelia Langford, John Sillitoe on behalf of the Wellcome Sanger Institute COVID-19 Surveillance Team |
| EPI_ISL_2681746 | Lighthouse Lab in Milton Keynes | Wellcome Sanger Institute for the COVID-19 Genomics UK (COG-UK) Consortium | The Lighthouse Lab in Milton Keynes and Alex Alderton, Roberto Amato, Jeffrey Barrett, Sonia Goncalves, Ewan Harrison, David K. Jackson, Ian Johnston, Dominic Kwiatkowski, Cordelia Langford, John Sillitoe on behalf of the Wellcome Sanger Institute COVID-19 Surveillance Team |
| EPI_ISL_2681747 | Lighthouse Lab in Alderley Park | Wellcome Sanger Institute for the COVID-19 Genomics UK (COG-UK) Consortium | Jacquelyn Wynn, Mairead Hyland, The Lighthouse Lab in Alderley Park and Alex Alderton, Roberto Amato, Jeffrey Barrett, Sonia Goncalves, Ewan Harrison, David K. Jackson, Ian Johnston, Dominic Kwiatkowski, Cordelia Langford, John Sillitoe on behalf of the Wellcome Sanger Institute COVID-19 Surveillance Team |
| EPI_ISL_2681748 | Lighthouse Lab in Milton Keynes | Wellcome Sanger Institute for the COVID-19 Genomics UK (COG-UK) Consortium | The Lighthouse Lab in Milton Keynes and Alex Alderton, Roberto Amato, Jeffrey Barrett, Sonia Goncalves, Ewan Harrison, David K. Jackson, Ian Johnston, Dominic Kwiatkowski, Cordelia Langford, John Sillitoe on behalf of the Wellcome Sanger Institute COVID-19 Surveillance Team |
| EPI_ISL_2681749, EPI_ISL_2681750, EPI_ISL_2681751, EPI_ISL_2681752 | Lighthouse Lab in Alderley Park | Wellcome Sanger Institute for the COVID-19 Genomics UK (COG-UK) Consortium | Jacquelyn Wynn, Mairead Hyland, The Lighthouse Lab in Alderley Park and Alex Alderton, Roberto Amato, Jeffrey Barrett, Sonia Goncalves, Ewan Harrison, David K. Jackson, Ian Johnston, Dominic Kwiatkowski, Cordelia Langford, John Sillitoe on behalf of the Wellcome Sanger Institute COVID-19 Surveillance Team |
| EPI_ISL_2681753 | Lighthouse Lab in Milton Keynes | Wellcome Sanger Institute for the COVID-19 Genomics UK (COG-UK) Consortium | The Lighthouse Lab in Milton Keynes and Alex Alderton, Roberto Amato, Jeffrey Barrett, Sonia Goncalves, Ewan Harrison, David K. Jackson, Ian Johnston, Dominic Kwiatkowski, Cordelia Langford, John Sillitoe on behalf of the Wellcome Sanger Institute COVID-19 Surveillance Team |
| EPI_ISL_2681754, EPI_ISL_2681755 | Lighthouse Lab in Alderley Park | Wellcome Sanger Institute for the COVID-19 Genomics UK (COG-UK) Consortium | Jacquelyn Wynn, Mairead Hyland, The Lighthouse Lab in Alderley Park and Alex Alderton, Roberto Amato, Jeffrey Barrett, Sonia Goncalves, Ewan Harrison, David K. Jackson, Ian Johnston, Dominic Kwiatkowski, Cordelia Langford, John Sillitoe on behalf of the Wellcome Sanger Institute COVID-19 Surveillance Team |
| EPI_ISL_2681756 | Lighthouse Lab in Milton Keynes | Wellcome Sanger Institute for the COVID-19 Genomics UK (COG-UK) Consortium | The Lighthouse Lab in Milton Keynes and Alex Alderton, Roberto Amato, Jeffrey Barrett, Sonia Goncalves, Ewan Harrison, David K. Jackson, Ian Johnston, Dominic Kwiatkowski, Cordelia Langford, John Sillitoe on behalf of the Wellcome Sanger Institute COVID-19 Surveillance Team |
| EPI_ISL_2681757, EPI_ISL_2681758, EPI_ISL_2681759, EPI_ISL_2681760 | Lighthouse Lab in Alderley Park | Wellcome Sanger Institute for the COVID-19 Genomics UK (COG-UK) Consortium | Jacquelyn Wynn, Mairead Hyland, The Lighthouse Lab in Alderley Park and Alex Alderton, Roberto Amato, Jeffrey Barrett, Sonia Goncalves, Ewan Harrison, David K. Jackson, Ian Johnston, Dominic Kwiatkowski, Cordelia Langford, John Sillitoe on behalf of the Wellcome Sanger Institute COVID-19 Surveillance Team |
| EPI_ISL_2681761, | Lighthouse Lab in Milton Keynes | Wellcome Sanger Institute for the COVID-19 Genomics UK (COG-UK) | The Lighthouse Lab in Milton Keynes and Alex Alderton, Roberto Amato, Jeffrey Barrett, Sonia Goncalves, Ewan Harrison, David K. Jackson, Ian Johnston, Dominic Kwiatkowski, Cordelia Langford, John Sillitoe on behalf of the Wellcome Sanger |

| | | | |
|---|---|---|---|
| EPI_ISL_2681762 | | Consortium | Institute COVID-19 Surveillance Team |
| EPI_ISL_2681763 | Lighthouse Lab in Alderley Park | Wellcome Sanger Institute for the COVID-19 Genomics UK (COG-UK) Consortium | Jacquelyn Wynn, Mairead Hyland, The Lighthouse Lab in Alderley Park and Alex Alderton, Roberto Amato, Jeffrey Barrett, Sonia Goncalves, Ewan Harrison, David K. Jackson, Ian Johnston, Dominic Kwiatkowski, Cordelia Langford, John Sillitoe on behalf of the Wellcome Sanger Institute COVID-19 Surveillance Team |
| EPI_ISL_2681764 | Lighthouse Lab in Milton Keynes | Wellcome Sanger Institute for the COVID-19 Genomics UK (COG-UK) Consortium | The Lighthouse Lab in Milton Keynes and Alex Alderton, Roberto Amato, Jeffrey Barrett, Sonia Goncalves, Ewan Harrison, David K. Jackson, Ian Johnston, Dominic Kwiatkowski, Cordelia Langford, John Sillitoe on behalf of the Wellcome Sanger Institute COVID-19 Surveillance Team |
| EPI_ISL_2681765, EPI_ISL_2681766, EPI_ISL_2681767 | Lighthouse Lab in Alderley Park | Wellcome Sanger Institute for the COVID-19 Genomics UK (COG-UK) Consortium | Jacquelyn Wynn, Mairead Hyland, The Lighthouse Lab in Alderley Park and Alex Alderton, Roberto Amato, Jeffrey Barrett, Sonia Goncalves, Ewan Harrison, David K. Jackson, Ian Johnston, Dominic Kwiatkowski, Cordelia Langford, John Sillitoe on behalf of the Wellcome Sanger Institute COVID-19 Surveillance Team |
| EPI_ISL_2681768 | Lighthouse Lab in Glasgow | Wellcome Sanger Institute for the COVID-19 Genomics UK (COG-UK) Consortium | Harper VanSteenhouse, Yumi Kasai, David Gray, Carol Clugston, Anna Dominiczak and Alex Alderton, Roberto Amato, Jeffrey Barrett, Sonia Goncalves, Ewan Harrison, David K. Jackson, Ian Johnston, Dominic Kwiatkowski, Cordelia Langford, John Sillitoe on behalf of the Wellcome Sanger Institute COVID-19 Surveillance Team |
| EPI_ISL_2681769, EPI_ISL_2681770, EPI_ISL_2681771 | Lighthouse Lab in Milton Keynes | Wellcome Sanger Institute for the COVID-19 Genomics UK (COG-UK) Consortium | The Lighthouse Lab in Milton Keynes and Alex Alderton, Roberto Amato, Jeffrey Barrett, Sonia Goncalves, Ewan Harrison, David K. Jackson, Ian Johnston, Dominic Kwiatkowski, Cordelia Langford, John Sillitoe on behalf of the Wellcome Sanger Institute COVID-19 Surveillance Team |
| EPI_ISL_2681772 | Lighthouse Lab in Alderley Park | Wellcome Sanger Institute for the COVID-19 Genomics UK (COG-UK) Consortium | Jacquelyn Wynn, Mairead Hyland, The Lighthouse Lab in Alderley Park and Alex Alderton, Roberto Amato, Jeffrey Barrett, Sonia Goncalves, Ewan Harrison, David K. Jackson, Ian Johnston, Dominic Kwiatkowski, Cordelia Langford, John Sillitoe on behalf of the Wellcome Sanger Institute COVID-19 Surveillance Team |
| EPI_ISL_2681773 | Lighthouse Lab in Milton Keynes | Wellcome Sanger Institute for the COVID-19 Genomics UK (COG-UK) Consortium | The Lighthouse Lab in Milton Keynes and Alex Alderton, Roberto Amato, Jeffrey Barrett, Sonia Goncalves, Ewan Harrison, David K. Jackson, Ian Johnston, Dominic Kwiatkowski, Cordelia Langford, John Sillitoe on behalf of the Wellcome Sanger Institute COVID-19 Surveillance Team |
| EPI_ISL_2681774, EPI_ISL_2681775, EPI_ISL_2681776, EPI_ISL_2681777 | Lighthouse Lab in Alderley Park | Wellcome Sanger Institute for the COVID-19 Genomics UK (COG-UK) Consortium | Jacquelyn Wynn, Mairead Hyland, The Lighthouse Lab in Alderley Park and Alex Alderton, Roberto Amato, Jeffrey Barrett, Sonia Goncalves, Ewan Harrison, David K. Jackson, Ian Johnston, Dominic Kwiatkowski, Cordelia Langford, John Sillitoe on behalf of the Wellcome Sanger Institute COVID-19 Surveillance Team |
| EPI_ISL_2681778 | Lighthouse Lab in Milton Keynes | Wellcome Sanger Institute for the COVID-19 Genomics UK (COG-UK) Consortium | The Lighthouse Lab in Milton Keynes and Alex Alderton, Roberto Amato, Jeffrey Barrett, Sonia Goncalves, Ewan Harrison, David K. Jackson, Ian Johnston, Dominic Kwiatkowski, Cordelia Langford, John Sillitoe on behalf of the Wellcome Sanger Institute COVID-19 Surveillance Team |
| EPI_ISL_2681779 | Lighthouse Lab in Glasgow | Wellcome Sanger Institute for the COVID-19 Genomics UK (COG-UK) Consortium | Harper VanSteenhouse, Yumi Kasai, David Gray, Carol Clugston, Anna Dominiczak and Alex Alderton, Roberto Amato, Jeffrey Barrett, Sonia Goncalves, Ewan Harrison, David K. Jackson, Ian Johnston, Dominic Kwiatkowski, Cordelia Langford, John Sillitoe on behalf of the Wellcome Sanger Institute COVID-19 Surveillance Team |
| EPI_ISL_2681780, EPI_ISL_2681781, EPI_ISL_2681782, EPI_ISL_2681783, EPI_ISL_2681784, EPI_ISL_2681785, EPI_ISL_2681786, EPI_ISL_2681787 | Lighthouse Lab in Alderley Park | Wellcome Sanger Institute for the COVID-19 Genomics UK (COG-UK) Consortium | Jacquelyn Wynn, Mairead Hyland, The Lighthouse Lab in Alderley Park and Alex Alderton, Roberto Amato, Jeffrey Barrett, Sonia Goncalves, Ewan Harrison, David K. Jackson, Ian Johnston, Dominic Kwiatkowski, Cordelia Langford, John Sillitoe on behalf of the Wellcome Sanger Institute COVID-19 Surveillance Team |
| EPI_ISL_2681788 | Lighthouse Lab in Milton Keynes | Wellcome Sanger Institute for the COVID-19 Genomics UK (COG-UK) Consortium | The Lighthouse Lab in Milton Keynes and Alex Alderton, Roberto Amato, Jeffrey Barrett, Sonia Goncalves, Ewan Harrison, David K. Jackson, Ian Johnston, Dominic Kwiatkowski, Cordelia Langford, John Sillitoe on behalf of the Wellcome Sanger Institute COVID-19 Surveillance Team |
| EPI_ISL_2681789, EPI_ISL_2681790, EPI_ISL_2681791 | Lighthouse Lab in Alderley Park | Wellcome Sanger Institute for the COVID-19 Genomics UK (COG-UK) Consortium | Jacquelyn Wynn, Mairead Hyland, The Lighthouse Lab in Alderley Park and Alex Alderton, Roberto Amato, Jeffrey Barrett, Sonia Goncalves, Ewan Harrison, David K. Jackson, Ian Johnston, Dominic Kwiatkowski, Cordelia Langford, John Sillitoe on behalf of the Wellcome Sanger Institute COVID-19 Surveillance Team |
| EPI_ISL_2681792 | Lighthouse Lab in Milton Keynes | Wellcome Sanger Institute for the COVID-19 Genomics UK (COG-UK) Consortium | The Lighthouse Lab in Milton Keynes and Alex Alderton, Roberto Amato, Jeffrey Barrett, Sonia Goncalves, Ewan Harrison, David K. Jackson, Ian Johnston, Dominic Kwiatkowski, Cordelia Langford, John Sillitoe on behalf of the Wellcome Sanger Institute COVID-19 Surveillance Team |
| EPI_ISL_2681793, EPI_ISL_2681794 | Lighthouse Lab in Alderley Park | Wellcome Sanger Institute for the COVID-19 Genomics UK (COG-UK) Consortium | Jacquelyn Wynn, Mairead Hyland, The Lighthouse Lab in Alderley Park and Alex Alderton, Roberto Amato, Jeffrey Barrett, Sonia Goncalves, Ewan Harrison, David K. Jackson, Ian Johnston, Dominic Kwiatkowski, Cordelia Langford, John Sillitoe on behalf of the Wellcome Sanger Institute COVID-19 Surveillance Team |
| EPI_ISL_2681795 | Lighthouse Lab in Milton Keynes | Wellcome Sanger Institute for the COVID-19 Genomics UK (COG-UK) Consortium | The Lighthouse Lab in Milton Keynes and Alex Alderton, Roberto Amato, Jeffrey Barrett, Sonia Goncalves, Ewan Harrison, David K. Jackson, Ian Johnston, Dominic Kwiatkowski, Cordelia Langford, John Sillitoe on behalf of the Wellcome Sanger Institute COVID-19 Surveillance Team |
| EPI_ISL_2681796 | Lighthouse Lab in Alderley Park | Wellcome Sanger Institute for the COVID-19 Genomics UK (COG-UK) Consortium | Jacquelyn Wynn, Mairead Hyland, The Lighthouse Lab in Alderley Park and Alex Alderton, Roberto Amato, Jeffrey Barrett, Sonia Goncalves, Ewan Harrison, David K. Jackson, Ian Johnston, Dominic Kwiatkowski, Cordelia Langford, John Sillitoe on behalf of the Wellcome Sanger Institute COVID-19 Surveillance Team |
| EPI_ISL_2681797, EPI_ISL_2681798 | Lighthouse Lab in Milton Keynes | Wellcome Sanger Institute for the COVID-19 Genomics UK (COG-UK) Consortium | The Lighthouse Lab in Milton Keynes and Alex Alderton, Roberto Amato, Jeffrey Barrett, Sonia Goncalves, Ewan Harrison, David K. Jackson, Ian Johnston, Dominic Kwiatkowski, Cordelia Langford, John Sillitoe on behalf of the Wellcome Sanger Institute COVID-19 Surveillance Team |
| EPI_ISL_2681799, EPI_ISL_2681800, EPI_ISL_2681802, EPI_ISL_2681803, EPI_ISL_2681804, EPI_ISL_2681805 | Lighthouse Lab in Alderley Park | Wellcome Sanger Institute for the COVID-19 Genomics UK (COG-UK) Consortium | Jacquelyn Wynn, Mairead Hyland, The Lighthouse Lab in Alderley Park and Alex Alderton, Roberto Amato, Jeffrey Barrett, Sonia Goncalves, Ewan Harrison, David K. Jackson, Ian Johnston, Dominic Kwiatkowski, Cordelia Langford, John Sillitoe on behalf of the Wellcome Sanger Institute COVID-19 Surveillance Team |
| EPI_ISL_2681806 | Lighthouse Lab in Milton Keynes | Wellcome Sanger Institute for the COVID-19 Genomics UK (COG-UK) Consortium | The Lighthouse Lab in Milton Keynes and Alex Alderton, Roberto Amato, Jeffrey Barrett, Sonia Goncalves, Ewan Harrison, David K. Jackson, Ian Johnston, Dominic Kwiatkowski, Cordelia Langford, John Sillitoe on behalf of the Wellcome Sanger Institute COVID-19 Surveillance Team |
| EPI_ISL_2681807, EPI_ISL_2681808, EPI_ISL_2681809 | Lighthouse Lab in Alderley Park | Wellcome Sanger Institute for the COVID-19 Genomics UK (COG-UK) Consortium | Jacquelyn Wynn, Mairead Hyland, The Lighthouse Lab in Alderley Park and Alex Alderton, Roberto Amato, Jeffrey Barrett, Sonia Goncalves, Ewan Harrison, David K. Jackson, Ian Johnston, Dominic Kwiatkowski, Cordelia Langford, John Sillitoe on behalf of the Wellcome Sanger Institute COVID-19 Surveillance Team |
| EPI_ISL_2681810 | Lighthouse Lab in Milton Keynes | Wellcome Sanger Institute for the COVID-19 Genomics UK (COG-UK) Consortium | The Lighthouse Lab in Milton Keynes and Alex Alderton, Roberto Amato, Jeffrey Barrett, Sonia Goncalves, Ewan Harrison, David K. Jackson, Ian Johnston, Dominic Kwiatkowski, Cordelia Langford, John Sillitoe on behalf of the Wellcome Sanger Institute COVID-19 Surveillance Team |
| EPI_ISL_2681811, EPI_ISL_2681812, EPI_ISL_2681813, EPI_ISL_2681814, EPI_ISL_2681815, EPI_ISL_2681816, EPI_ISL_2681817, EPI_ISL_2681818 | Lighthouse Lab in Alderley Park | Wellcome Sanger Institute for the COVID-19 Genomics UK (COG-UK) Consortium | Jacquelyn Wynn, Mairead Hyland, The Lighthouse Lab in Alderley Park and Alex Alderton, Roberto Amato, Jeffrey Barrett, Sonia Goncalves, Ewan Harrison, David K. Jackson, Ian Johnston, Dominic Kwiatkowski, Cordelia Langford, John Sillitoe on behalf of the Wellcome Sanger Institute COVID-19 Surveillance Team |
| EPI_ISL_2681819 | Lighthouse Lab in Milton Keynes | Wellcome Sanger Institute for the COVID-19 Genomics UK (COG-UK) Consortium | The Lighthouse Lab in Milton Keynes and Alex Alderton, Roberto Amato, Jeffrey Barrett, Sonia Goncalves, Ewan Harrison, David K. Jackson, Ian Johnston, Dominic Kwiatkowski, Cordelia Langford, John Sillitoe on behalf of the Wellcome Sanger Institute COVID-19 Surveillance Team |
| EPI_ISL_2681820 | Lighthouse Lab in Alderley Park | Wellcome Sanger Institute for the COVID-19 Genomics UK (COG-UK) Consortium | Jacquelyn Wynn, Mairead Hyland, The Lighthouse Lab in Alderley Park and Alex Alderton, Roberto Amato, Jeffrey Barrett, Sonia Goncalves, Ewan Harrison, David K. Jackson, Ian Johnston, Dominic Kwiatkowski, Cordelia Langford, John Sillitoe on behalf of the Wellcome Sanger Institute COVID-19 Surveillance Team |
| EPI_ISL_2681821 | Lighthouse Lab in Milton Keynes | Wellcome Sanger Institute for the COVID-19 Genomics UK (COG-UK) Consortium | The Lighthouse Lab in Milton Keynes and Alex Alderton, Roberto Amato, Jeffrey Barrett, Sonia Goncalves, Ewan Harrison, David K. Jackson, Ian Johnston, Dominic Kwiatkowski, Cordelia Langford, John Sillitoe on behalf of the Wellcome Sanger Institute COVID-19 Surveillance Team |
| EPI_ISL_2681822, EPI_ISL_2681823, EPI_ISL_2681824, EPI_ISL_2681825, EPI_ISL_2681826, EPI_ISL_2681827 | Lighthouse Lab in Alderley Park | Wellcome Sanger Institute for the COVID-19 Genomics UK (COG-UK) Consortium | Jacquelyn Wynn, Mairead Hyland, The Lighthouse Lab in Alderley Park and Alex Alderton, Roberto Amato, Jeffrey Barrett, Sonia Goncalves, Ewan Harrison, David K. Jackson, Ian Johnston, Dominic Kwiatkowski, Cordelia Langford, John Sillitoe on behalf of the Wellcome Sanger Institute COVID-19 Surveillance Team |
| EPI_ISL_2681828 | Lighthouse Lab in Milton Keynes | Wellcome Sanger Institute for the COVID-19 Genomics UK (COG-UK) Consortium | The Lighthouse Lab in Milton Keynes and Alex Alderton, Roberto Amato, Jeffrey Barrett, Sonia Goncalves, Ewan Harrison, David K. Jackson, Ian Johnston, Dominic Kwiatkowski, Cordelia Langford, John Sillitoe on behalf of the Wellcome Sanger Institute COVID-19 Surveillance Team |
| EPI_ISL_2681829 | Lighthouse Lab in Glasgow | Wellcome Sanger Institute for the COVID-19 Genomics UK (COG-UK) Consortium | Harper VanSteenhouse, Yumi Kasai, David Gray, Carol Clugston, Anna Dominiczak and Alex Alderton, Roberto Amato, Jeffrey Barrett, Sonia Goncalves, Ewan Harrison, David K. Jackson, Ian Johnston, Dominic Kwiatkowski, Cordelia Langford, John Sillitoe on behalf of the Wellcome Sanger Institute COVID-19 Surveillance Team |
| EPI_ISL_2681830 | Lighthouse Lab in Milton Keynes | Wellcome Sanger Institute for the COVID-19 Genomics UK (COG-UK) Consortium | The Lighthouse Lab in Milton Keynes and Alex Alderton, Roberto Amato, Jeffrey Barrett, Sonia Goncalves, Ewan Harrison, David K. Jackson, Ian Johnston, Dominic Kwiatkowski, Cordelia Langford, John Sillitoe on behalf of the Wellcome Sanger Institute COVID-19 Surveillance Team |
| EPI_ISL_2681831, EPI_ISL_2681832, EPI_ISL_2681833, EPI_ISL_2681834, EPI_ISL_2681835 | Lighthouse Lab in Alderley Park | Wellcome Sanger Institute for the COVID-19 Genomics UK (COG-UK) Consortium | Jacquelyn Wynn, Mairead Hyland, The Lighthouse Lab in Alderley Park and Alex Alderton, Roberto Amato, Jeffrey Barrett, Sonia Goncalves, Ewan Harrison, David K. Jackson, Ian Johnston, Dominic Kwiatkowski, Cordelia Langford, John Sillitoe on behalf of the Wellcome Sanger Institute COVID-19 Surveillance Team |
| EPI_ISL_2681836 | Lighthouse Lab in Milton Keynes | Wellcome Sanger Institute for the COVID-19 Genomics UK (COG-UK) Consortium | The Lighthouse Lab in Milton Keynes and Alex Alderton, Roberto Amato, Jeffrey Barrett, Sonia Goncalves, Ewan Harrison, David K. Jackson, Ian Johnston, Dominic Kwiatkowski, Cordelia Langford, John Sillitoe on behalf of the Wellcome Sanger Institute COVID-19 Surveillance Team |
| EPI_ISL_2681837 | Lighthouse Lab in Alderley Park | Wellcome Sanger Institute for the COVID-19 Genomics UK (COG-UK) | Jacquelyn Wynn, Mairead Hyland, The Lighthouse Lab in Alderley Park and Alex Alderton, Roberto Amato, Jeffrey Barrett, Sonia Goncalves, Ewan Harrison, David K. Jackson, Ian Johnston, Dominic Kwiatkowski, Cordelia Langford, John Sillitoe on |

| | | | |
|---|---|---|---|
| | | Consortium | behalf of the Wellcome Sanger Institute COVID-19 Surveillance Team |
| EPI_ISL_2681838 | Lighthouse Lab in Milton Keynes | Wellcome Sanger Institute for the COVID-19 Genomics UK (COG-UK) Consortium | The Lighthouse Lab in Milton Keynes and Alex Alderton, Roberto Amato, Jeffrey Barrett, Sonia Goncalves, Ewan Harrison, David K. Jackson, Ian Johnston, Dominic Kwiatkowski, Cordelia Langford, John Sillitoe on behalf of the Wellcome Sanger Institute COVID-19 Surveillance Team |
| EPI_ISL_2681839 | Lighthouse Lab in Alderley Park | Wellcome Sanger Institute for the COVID-19 Genomics UK (COG-UK) Consortium | Jacquelyn Wynn, Mairead Hyland, The Lighthouse Lab in Alderley Park and Alex Alderton, Roberto Amato, Jeffrey Barrett, Sonia Goncalves, Ewan Harrison, David K. Jackson, Ian Johnston, Dominic Kwiatkowski, Cordelia Langford, John Sillitoe on behalf of the Wellcome Sanger Institute COVID-19 Surveillance Team |
| EPI_ISL_2681840 | Lighthouse Lab in Milton Keynes | Wellcome Sanger Institute for the COVID-19 Genomics UK (COG-UK) Consortium | The Lighthouse Lab in Milton Keynes and Alex Alderton, Roberto Amato, Jeffrey Barrett, Sonia Goncalves, Ewan Harrison, David K. Jackson, Ian Johnston, Dominic Kwiatkowski, Cordelia Langford, John Sillitoe on behalf of the Wellcome Sanger Institute COVID-19 Surveillance Team |
| EPI_ISL_2681841 | Lighthouse Lab in Alderley Park | Wellcome Sanger Institute for the COVID-19 Genomics UK (COG-UK) Consortium | Jacquelyn Wynn, Mairead Hyland, The Lighthouse Lab in Alderley Park and Alex Alderton, Roberto Amato, Jeffrey Barrett, Sonia Goncalves, Ewan Harrison, David K. Jackson, Ian Johnston, Dominic Kwiatkowski, Cordelia Langford, John Sillitoe on behalf of the Wellcome Sanger Institute COVID-19 Surveillance Team |
| EPI_ISL_2681842, EPI_ISL_2681843 | Lighthouse Lab in Milton Keynes | Wellcome Sanger Institute for the COVID-19 Genomics UK (COG-UK) Consortium | The Lighthouse Lab in Milton Keynes and Alex Alderton, Roberto Amato, Jeffrey Barrett, Sonia Goncalves, Ewan Harrison, David K. Jackson, Ian Johnston, Dominic Kwiatkowski, Cordelia Langford, John Sillitoe on behalf of the Wellcome Sanger Institute COVID-19 Surveillance Team |
| EPI_ISL_2681844 | Lighthouse Lab in Glasgow | Wellcome Sanger Institute for the COVID-19 Genomics UK (COG-UK) Consortium | Harper VanSteenhouse, Yumi Kasai, David Gray, Carol Clugston, Anna Dominiczak and Alex Alderton, Roberto Amato, Jeffrey Barrett, Sonia Goncalves, Ewan Harrison, David K. Jackson, Ian Johnston, Dominic Kwiatkowski, Cordelia Langford, John Sillitoe on behalf of the Wellcome Sanger Institute COVID-19 Surveillance Team |
| EPI_ISL_2681845, EPI_ISL_2681846 | Lighthouse Lab in Alderley Park | Wellcome Sanger Institute for the COVID-19 Genomics UK (COG-UK) Consortium | Jacquelyn Wynn, Mairead Hyland, The Lighthouse Lab in Alderley Park and Alex Alderton, Roberto Amato, Jeffrey Barrett, Sonia Goncalves, Ewan Harrison, David K. Jackson, Ian Johnston, Dominic Kwiatkowski, Cordelia Langford, John Sillitoe on behalf of the Wellcome Sanger Institute COVID-19 Surveillance Team |
| EPI_ISL_2681847 | Lighthouse Lab in Glasgow | Wellcome Sanger Institute for the COVID-19 Genomics UK (COG-UK) Consortium | Harper VanSteenhouse, Yumi Kasai, David Gray, Carol Clugston, Anna Dominiczak and Alex Alderton, Roberto Amato, Jeffrey Barrett, Sonia Goncalves, Ewan Harrison, David K. Jackson, Ian Johnston, Dominic Kwiatkowski, Cordelia Langford, John Sillitoe on behalf of the Wellcome Sanger Institute COVID-19 Surveillance Team |
| EPI_ISL_2681848, EPI_ISL_2681849 | Lighthouse Lab in Milton Keynes | Wellcome Sanger Institute for the COVID-19 Genomics UK (COG-UK) Consortium | The Lighthouse Lab in Milton Keynes and Alex Alderton, Roberto Amato, Jeffrey Barrett, Sonia Goncalves, Ewan Harrison, David K. Jackson, Ian Johnston, Dominic Kwiatkowski, Cordelia Langford, John Sillitoe on behalf of the Wellcome Sanger Institute COVID-19 Surveillance Team |
| EPI_ISL_2681850, EPI_ISL_2681851, EPI_ISL_2681852, EPI_ISL_2681853 | Lighthouse Lab in Alderley Park | Wellcome Sanger Institute for the COVID-19 Genomics UK (COG-UK) Consortium | Jacquelyn Wynn, Mairead Hyland, The Lighthouse Lab in Alderley Park and Alex Alderton, Roberto Amato, Jeffrey Barrett, Sonia Goncalves, Ewan Harrison, David K. Jackson, Ian Johnston, Dominic Kwiatkowski, Cordelia Langford, John Sillitoe on behalf of the Wellcome Sanger Institute COVID-19 Surveillance Team |
| EPI_ISL_2681854 | Lighthouse Lab in Milton Keynes | Wellcome Sanger Institute for the COVID-19 Genomics UK (COG-UK) Consortium | The Lighthouse Lab in Milton Keynes and Alex Alderton, Roberto Amato, Jeffrey Barrett, Sonia Goncalves, Ewan Harrison, David K. Jackson, Ian Johnston, Dominic Kwiatkowski, Cordelia Langford, John Sillitoe on behalf of the Wellcome Sanger Institute COVID-19 Surveillance Team |
| EPI_ISL_2681855, EPI_ISL_2681856, EPI_ISL_2681857, EPI_ISL_2681858, EPI_ISL_2681859, EPI_ISL_2681860, EPI_ISL_2681861 | Lighthouse Lab in Alderley Park | Wellcome Sanger Institute for the COVID-19 Genomics UK (COG-UK) Consortium | Jacquelyn Wynn, Mairead Hyland, The Lighthouse Lab in Alderley Park and Alex Alderton, Roberto Amato, Jeffrey Barrett, Sonia Goncalves, Ewan Harrison, David K. Jackson, Ian Johnston, Dominic Kwiatkowski, Cordelia Langford, John Sillitoe on behalf of the Wellcome Sanger Institute COVID-19 Surveillance Team |
| EPI_ISL_2681862 | Lighthouse Lab in Glasgow | Wellcome Sanger Institute for the COVID-19 Genomics UK (COG-UK) Consortium | Harper VanSteenhouse, Yumi Kasai, David Gray, Carol Clugston, Anna Dominiczak and Alex Alderton, Roberto Amato, Jeffrey Barrett, Sonia Goncalves, Ewan Harrison, David K. Jackson, Ian Johnston, Dominic Kwiatkowski, Cordelia Langford, John Sillitoe on behalf of the Wellcome Sanger Institute COVID-19 Surveillance Team |
| EPI_ISL_2681863 | Lighthouse Lab in Milton Keynes | Wellcome Sanger Institute for the COVID-19 Genomics UK (COG-UK) Consortium | The Lighthouse Lab in Milton Keynes and Alex Alderton, Roberto Amato, Jeffrey Barrett, Sonia Goncalves, Ewan Harrison, David K. Jackson, Ian Johnston, Dominic Kwiatkowski, Cordelia Langford, John Sillitoe on behalf of the Wellcome Sanger Institute COVID-19 Surveillance Team |
| EPI_ISL_2681864 | Lighthouse Lab in Alderley Park | Wellcome Sanger Institute for the COVID-19 Genomics UK (COG-UK) Consortium | Jacquelyn Wynn, Mairead Hyland, The Lighthouse Lab in Alderley Park and Alex Alderton, Roberto Amato, Jeffrey Barrett, Sonia Goncalves, Ewan Harrison, David K. Jackson, Ian Johnston, Dominic Kwiatkowski, Cordelia Langford, John Sillitoe on behalf of the Wellcome Sanger Institute COVID-19 Surveillance Team |
| EPI_ISL_2681865 | Lighthouse Lab in Glasgow | Wellcome Sanger Institute for the COVID-19 Genomics UK (COG-UK) Consortium | Harper VanSteenhouse, Yumi Kasai, David Gray, Carol Clugston, Anna Dominiczak and Alex Alderton, Roberto Amato, Jeffrey Barrett, Sonia Goncalves, Ewan Harrison, David K. Jackson, Ian Johnston, Dominic Kwiatkowski, Cordelia Langford, John Sillitoe on behalf of the Wellcome Sanger Institute COVID-19 Surveillance Team |
| EPI_ISL_2681866 | Lighthouse Lab in Alderley Park | Wellcome Sanger Institute for the COVID-19 Genomics UK (COG-UK) Consortium | Jacquelyn Wynn, Mairead Hyland, The Lighthouse Lab in Alderley Park and Alex Alderton, Roberto Amato, Jeffrey Barrett, Sonia Goncalves, Ewan Harrison, David K. Jackson, Ian Johnston, Dominic Kwiatkowski, Cordelia Langford, John Sillitoe on behalf of the Wellcome Sanger Institute COVID-19 Surveillance Team |
| EPI_ISL_2681867 | Lighthouse Lab in Glasgow | Wellcome Sanger Institute for the COVID-19 Genomics UK (COG-UK) Consortium | Harper VanSteenhouse, Yumi Kasai, David Gray, Carol Clugston, Anna Dominiczak and Alex Alderton, Roberto Amato, Jeffrey Barrett, Sonia Goncalves, Ewan Harrison, David K. Jackson, Ian Johnston, Dominic Kwiatkowski, Cordelia Langford, John Sillitoe on behalf of the Wellcome Sanger Institute COVID-19 Surveillance Team |
| EPI_ISL_2681868 | Lighthouse Lab in Milton Keynes | Wellcome Sanger Institute for the COVID-19 Genomics UK (COG-UK) Consortium | The Lighthouse Lab in Milton Keynes and Alex Alderton, Roberto Amato, Jeffrey Barrett, Sonia Goncalves, Ewan Harrison, David K. Jackson, Ian Johnston, Dominic Kwiatkowski, Cordelia Langford, John Sillitoe on behalf of the Wellcome Sanger Institute COVID-19 Surveillance Team |
| EPI_ISL_2681869, EPI_ISL_2681870, EPI_ISL_2681871 | Lighthouse Lab in Alderley Park | Wellcome Sanger Institute for the COVID-19 Genomics UK (COG-UK) Consortium | Jacquelyn Wynn, Mairead Hyland, The Lighthouse Lab in Alderley Park and Alex Alderton, Roberto Amato, Jeffrey Barrett, Sonia Goncalves, Ewan Harrison, David K. Jackson, Ian Johnston, Dominic Kwiatkowski, Cordelia Langford, John Sillitoe on behalf of the Wellcome Sanger Institute COVID-19 Surveillance Team |
| EPI_ISL_2681872 | Lighthouse Lab in Milton Keynes | Wellcome Sanger Institute for the COVID-19 Genomics UK (COG-UK) Consortium | The Lighthouse Lab in Milton Keynes and Alex Alderton, Roberto Amato, Jeffrey Barrett, Sonia Goncalves, Ewan Harrison, David K. Jackson, Ian Johnston, Dominic Kwiatkowski, Cordelia Langford, John Sillitoe on behalf of the Wellcome Sanger Institute COVID-19 Surveillance Team |
| EPI_ISL_2681873 | Lighthouse Lab in Alderley Park | Wellcome Sanger Institute for the COVID-19 Genomics UK (COG-UK) Consortium | Jacquelyn Wynn, Mairead Hyland, The Lighthouse Lab in Alderley Park and Alex Alderton, Roberto Amato, Jeffrey Barrett, Sonia Goncalves, Ewan Harrison, David K. Jackson, Ian Johnston, Dominic Kwiatkowski, Cordelia Langford, John Sillitoe on behalf of the Wellcome Sanger Institute COVID-19 Surveillance Team |
| EPI_ISL_2681874, EPI_ISL_2681875 | Lighthouse Lab in Milton Keynes | Wellcome Sanger Institute for the COVID-19 Genomics UK (COG-UK) Consortium | The Lighthouse Lab in Milton Keynes and Alex Alderton, Roberto Amato, Jeffrey Barrett, Sonia Goncalves, Ewan Harrison, David K. Jackson, Ian Johnston, Dominic Kwiatkowski, Cordelia Langford, John Sillitoe on behalf of the Wellcome Sanger Institute COVID-19 Surveillance Team |
| EPI_ISL_2681876 | Lighthouse Lab in Glasgow | Wellcome Sanger Institute for the COVID-19 Genomics UK (COG-UK) Consortium | Harper VanSteenhouse, Yumi Kasai, David Gray, Carol Clugston, Anna Dominiczak and Alex Alderton, Roberto Amato, Jeffrey Barrett, Sonia Goncalves, Ewan Harrison, David K. Jackson, Ian Johnston, Dominic Kwiatkowski, Cordelia Langford, John Sillitoe on behalf of the Wellcome Sanger Institute COVID-19 Surveillance Team |
| EPI_ISL_2681877 | Lighthouse Lab in Alderley Park | Wellcome Sanger Institute for the COVID-19 Genomics UK (COG-UK) Consortium | Jacquelyn Wynn, Mairead Hyland, The Lighthouse Lab in Alderley Park and Alex Alderton, Roberto Amato, Jeffrey Barrett, Sonia Goncalves, Ewan Harrison, David K. Jackson, Ian Johnston, Dominic Kwiatkowski, Cordelia Langford, John Sillitoe on behalf of the Wellcome Sanger Institute COVID-19 Surveillance Team |
| EPI_ISL_2681878 | Lighthouse Lab in Glasgow | Wellcome Sanger Institute for the COVID-19 Genomics UK (COG-UK) Consortium | Harper VanSteenhouse, Yumi Kasai, David Gray, Carol Clugston, Anna Dominiczak and Alex Alderton, Roberto Amato, Jeffrey Barrett, Sonia Goncalves, Ewan Harrison, David K. Jackson, Ian Johnston, Dominic Kwiatkowski, Cordelia Langford, John Sillitoe on behalf of the Wellcome Sanger Institute COVID-19 Surveillance Team |
| EPI_ISL_2681879 | Lighthouse Lab in Milton Keynes | Wellcome Sanger Institute for the COVID-19 Genomics UK (COG-UK) Consortium | The Lighthouse Lab in Milton Keynes and Alex Alderton, Roberto Amato, Jeffrey Barrett, Sonia Goncalves, Ewan Harrison, David K. Jackson, Ian Johnston, Dominic Kwiatkowski, Cordelia Langford, John Sillitoe on behalf of the Wellcome Sanger Institute COVID-19 Surveillance Team |
| EPI_ISL_2681880, EPI_ISL_2681881, EPI_ISL_2681882 | Lighthouse Lab in Alderley Park | Wellcome Sanger Institute for the COVID-19 Genomics UK (COG-UK) Consortium | Jacquelyn Wynn, Mairead Hyland, The Lighthouse Lab in Alderley Park and Alex Alderton, Roberto Amato, Jeffrey Barrett, Sonia Goncalves, Ewan Harrison, David K. Jackson, Ian Johnston, Dominic Kwiatkowski, Cordelia Langford, John Sillitoe on behalf of the Wellcome Sanger Institute COVID-19 Surveillance Team |
| EPI_ISL_2681883 | Lighthouse Lab in Milton Keynes | Wellcome Sanger Institute for the COVID-19 Genomics UK (COG-UK) Consortium | The Lighthouse Lab in Milton Keynes and Alex Alderton, Roberto Amato, Jeffrey Barrett, Sonia Goncalves, Ewan Harrison, David K. Jackson, Ian Johnston, Dominic Kwiatkowski, Cordelia Langford, John Sillitoe on behalf of the Wellcome Sanger Institute COVID-19 Surveillance Team |
| EPI_ISL_2681885 | Lighthouse Lab in Alderley Park | Wellcome Sanger Institute for the COVID-19 Genomics UK (COG-UK) Consortium | Jacquelyn Wynn, Mairead Hyland, The Lighthouse Lab in Alderley Park and Alex Alderton, Roberto Amato, Jeffrey Barrett, Sonia Goncalves, Ewan Harrison, David K. Jackson, Ian Johnston, Dominic Kwiatkowski, Cordelia Langford, John Sillitoe on behalf of the Wellcome Sanger Institute COVID-19 Surveillance Team |
| EPI_ISL_2681886 | Lighthouse Lab in Milton Keynes | Wellcome Sanger Institute for the COVID-19 Genomics UK (COG-UK) Consortium | The Lighthouse Lab in Milton Keynes and Alex Alderton, Roberto Amato, Jeffrey Barrett, Sonia Goncalves, Ewan Harrison, David K. Jackson, Ian Johnston, Dominic Kwiatkowski, Cordelia Langford, John Sillitoe on behalf of the Wellcome Sanger Institute COVID-19 Surveillance Team |
| EPI_ISL_2681887 | Lighthouse Lab in Alderley Park | Wellcome Sanger Institute for the COVID-19 Genomics UK (COG-UK) Consortium | Jacquelyn Wynn, Mairead Hyland, The Lighthouse Lab in Alderley Park and Alex Alderton, Roberto Amato, Jeffrey Barrett, Sonia Goncalves, Ewan Harrison, David K. Jackson, Ian Johnston, Dominic Kwiatkowski, Cordelia Langford, John Sillitoe on behalf of the Wellcome Sanger Institute COVID-19 Surveillance Team |
| EPI_ISL_2681888 | Lighthouse Lab in Milton Keynes | Wellcome Sanger Institute for the COVID-19 Genomics UK (COG-UK) Consortium | The Lighthouse Lab in Milton Keynes and Alex Alderton, Roberto Amato, Jeffrey Barrett, Sonia Goncalves, Ewan Harrison, David K. Jackson, Ian Johnston, Dominic Kwiatkowski, Cordelia Langford, John Sillitoe on behalf of the Wellcome Sanger Institute COVID-19 Surveillance Team |
| EPI_ISL_2681889 | Lighthouse Lab in Glasgow | Wellcome Sanger Institute for the COVID-19 Genomics UK (COG-UK) Consortium | Harper VanSteenhouse, Yumi Kasai, David Gray, Carol Clugston, Anna Dominiczak and Alex Alderton, Roberto Amato, Jeffrey Barrett, Sonia Goncalves, Ewan Harrison, David K. Jackson, Ian Johnston, Dominic Kwiatkowski, Cordelia Langford, John Sillitoe on behalf of the Wellcome Sanger Institute COVID-19 Surveillance Team |
| EPI_ISL_2681890 | Lighthouse Lab in Alderley Park | Wellcome Sanger Institute for the COVID-19 Genomics UK (COG-UK) Consortium | Jacquelyn Wynn, Mairead Hyland, The Lighthouse Lab in Alderley Park and Alex Alderton, Roberto Amato, Jeffrey Barrett, Sonia Goncalves, Ewan Harrison, David K. Jackson, Ian Johnston, Dominic Kwiatkowski, Cordelia Langford, John Sillitoe on behalf of the Wellcome Sanger Institute COVID-19 Surveillance Team |
| EPI_ISL_2681891 | Lighthouse Lab in Milton Keynes | Wellcome Sanger Institute for the COVID-19 Genomics UK (COG-UK) Consortium | The Lighthouse Lab in Milton Keynes and Alex Alderton, Roberto Amato, Jeffrey Barrett, Sonia Goncalves, Ewan Harrison, David K. Jackson, Ian Johnston, Dominic Kwiatkowski, Cordelia Langford, John Sillitoe on behalf of the Wellcome Sanger Institute COVID-19 Surveillance Team |
| EPI_ISL_2681892, EPI_ISL_2681894, EPI_ISL_2681895 | Lighthouse Lab in Alderley Park | Wellcome Sanger Institute for the COVID-19 Genomics UK (COG-UK) Consortium | Jacquelyn Wynn, Mairead Hyland, The Lighthouse Lab in Alderley Park and Alex Alderton, Roberto Amato, Jeffrey Barrett, Sonia Goncalves, Ewan Harrison, David K. Jackson, Ian Johnston, Dominic Kwiatkowski, Cordelia Langford, John Sillitoe on behalf of the Wellcome Sanger Institute COVID-19 Surveillance Team |
| EPI_ISL_2681896, EPI_ISL_2681897 | Lighthouse Lab in Milton Keynes | Wellcome Sanger Institute for the COVID-19 Genomics UK (COG-UK) Consortium | The Lighthouse Lab in Milton Keynes and Alex Alderton, Roberto Amato, Jeffrey Barrett, Sonia Goncalves, Ewan Harrison, David K. Jackson, Ian Johnston, Dominic Kwiatkowski, Cordelia Langford, John Sillitoe on behalf of the Wellcome Sanger Institute COVID-19 Surveillance Team |
| EPI_ISL_2681898, EPI_ISL_2681899 | Lighthouse Lab in Alderley Park | Wellcome Sanger Institute for the COVID-19 Genomics UK (COG-UK) Consortium | Jacquelyn Wynn, Mairead Hyland, The Lighthouse Lab in Alderley Park and Alex Alderton, Roberto Amato, Jeffrey Barrett, Sonia Goncalves, Ewan Harrison, David K. Jackson, Ian Johnston, Dominic Kwiatkowski, Cordelia Langford, John Sillitoe on behalf of the Wellcome Sanger Institute COVID-19 Surveillance Team |
| EPI_ISL_2681900 | Lighthouse Lab in Milton Keynes | Wellcome Sanger Institute for the COVID-19 Genomics UK (COG-UK) Consortium | The Lighthouse Lab in Milton Keynes and Alex Alderton, Roberto Amato, Jeffrey Barrett, Sonia Goncalves, Ewan Harrison, David K. Jackson, Ian Johnston, Dominic Kwiatkowski, Cordelia Langford, John Sillitoe on behalf of the Wellcome Sanger Institute COVID-19 Surveillance Team |
| EPI_ISL_2681901 | Lighthouse Lab in Alderley Park | Wellcome Sanger Institute for the COVID-19 Genomics UK (COG-UK) | Jacquelyn Wynn, Mairead Hyland, The Lighthouse Lab in Alderley Park and Alex Alderton, Roberto Amato, Jeffrey Barrett, Sonia Goncalves, Ewan Harrison, David K. Jackson, Ian Johnston, Dominic Kwiatkowski, Cordelia Langford, John Sillitoe on |

| | | | |
|---|---|---|---|
| | | Consortium | behalf of the Wellcome Sanger Institute COVID-19 Surveillance Team |
| EPI_ISL_2681902 | Lighthouse Lab in Glasgow | Wellcome Sanger Institute for the COVID-19 Genomics UK (COG-UK) Consortium | Harper VanSteenhouse, Yumi Kasai, David Gray, Carol Clugston, Anna Dominiczak and Alex Alderton, Roberto Amato, Jeffrey Barrett, Sonia Goncalves, Ewan Harrison, David K. Jackson, Ian Johnston, Dominic Kwiatkowski, Cordelia Langford, John Sillitoe on behalf of the Wellcome Sanger Institute COVID-19 Surveillance Team |
| EPI_ISL_2681903, EPI_ISL_2681904, EPI_ISL_2681905, EPI_ISL_2681906, EPI_ISL_2681907, EPI_ISL_2681908, EPI_ISL_2681909, EPI_ISL_2681910, EPI_ISL_2681911 | Lighthouse Lab in Alderley Park | Wellcome Sanger Institute for the COVID-19 Genomics UK (COG-UK) Consortium | Jacquelyn Wynn, Mairead Hyland, The Lighthouse Lab in Alderley Park and Alex Alderton, Roberto Amato, Jeffrey Barrett, Sonia Goncalves, Ewan Harrison, David K. Jackson, Ian Johnston, Dominic Kwiatkowski, Cordelia Langford, John Sillitoe on behalf of the Wellcome Sanger Institute COVID-19 Surveillance Team |
| EPI_ISL_2681912, EPI_ISL_2681913 | Lighthouse Lab in Milton Keynes | Wellcome Sanger Institute for the COVID-19 Genomics UK (COG-UK) Consortium | The Lighthouse Lab in Milton Keynes and Alex Alderton, Roberto Amato, Jeffrey Barrett, Sonia Goncalves, Ewan Harrison, David K. Jackson, Ian Johnston, Dominic Kwiatkowski, Cordelia Langford, John Sillitoe on behalf of the Wellcome Sanger Institute COVID-19 Surveillance Team |
| EPI_ISL_2681914 | Lighthouse Lab in Glasgow | Wellcome Sanger Institute for the COVID-19 Genomics UK (COG-UK) Consortium | Harper VanSteenhouse, Yumi Kasai, David Gray, Carol Clugston, Anna Dominiczak and Alex Alderton, Roberto Amato, Jeffrey Barrett, Sonia Goncalves, Ewan Harrison, David K. Jackson, Ian Johnston, Dominic Kwiatkowski, Cordelia Langford, John Sillitoe on behalf of the Wellcome Sanger Institute COVID-19 Surveillance Team |
| EPI_ISL_2681915 | Lighthouse Lab in Alderley Park | Wellcome Sanger Institute for the COVID-19 Genomics UK (COG-UK) Consortium | Jacquelyn Wynn, Mairead Hyland, The Lighthouse Lab in Alderley Park and Alex Alderton, Roberto Amato, Jeffrey Barrett, Sonia Goncalves, Ewan Harrison, David K. Jackson, Ian Johnston, Dominic Kwiatkowski, Cordelia Langford, John Sillitoe on behalf of the Wellcome Sanger Institute COVID-19 Surveillance Team |
| EPI_ISL_2681916 | Lighthouse Lab in Milton Keynes | Wellcome Sanger Institute for the COVID-19 Genomics UK (COG-UK) Consortium | The Lighthouse Lab in Milton Keynes and Alex Alderton, Roberto Amato, Jeffrey Barrett, Sonia Goncalves, Ewan Harrison, David K. Jackson, Ian Johnston, Dominic Kwiatkowski, Cordelia Langford, John Sillitoe on behalf of the Wellcome Sanger Institute COVID-19 Surveillance Team |
| EPI_ISL_2681917 | Lighthouse Lab in Alderley Park | Wellcome Sanger Institute for the COVID-19 Genomics UK (COG-UK) Consortium | Jacquelyn Wynn, Mairead Hyland, The Lighthouse Lab in Alderley Park and Alex Alderton, Roberto Amato, Jeffrey Barrett, Sonia Goncalves, Ewan Harrison, David K. Jackson, Ian Johnston, Dominic Kwiatkowski, Cordelia Langford, John Sillitoe on behalf of the Wellcome Sanger Institute COVID-19 Surveillance Team |
| EPI_ISL_2681918 | Lighthouse Lab in Milton Keynes | Wellcome Sanger Institute for the COVID-19 Genomics UK (COG-UK) Consortium | The Lighthouse Lab in Milton Keynes and Alex Alderton, Roberto Amato, Jeffrey Barrett, Sonia Goncalves, Ewan Harrison, David K. Jackson, Ian Johnston, Dominic Kwiatkowski, Cordelia Langford, John Sillitoe on behalf of the Wellcome Sanger Institute COVID-19 Surveillance Team |
| EPI_ISL_2681919 | Lighthouse Lab in Alderley Park | Wellcome Sanger Institute for the COVID-19 Genomics UK (COG-UK) Consortium | Jacquelyn Wynn, Mairead Hyland, The Lighthouse Lab in Alderley Park and Alex Alderton, Roberto Amato, Jeffrey Barrett, Sonia Goncalves, Ewan Harrison, David K. Jackson, Ian Johnston, Dominic Kwiatkowski, Cordelia Langford, John Sillitoe on behalf of the Wellcome Sanger Institute COVID-19 Surveillance Team |
| EPI_ISL_2681920, EPI_ISL_2681921 | Lighthouse Lab in Milton Keynes | Wellcome Sanger Institute for the COVID-19 Genomics UK (COG-UK) Consortium | The Lighthouse Lab in Milton Keynes and Alex Alderton, Roberto Amato, Jeffrey Barrett, Sonia Goncalves, Ewan Harrison, David K. Jackson, Ian Johnston, Dominic Kwiatkowski, Cordelia Langford, John Sillitoe on behalf of the Wellcome Sanger Institute COVID-19 Surveillance Team |
| EPI_ISL_2681922 | Lighthouse Lab in Alderley Park | Wellcome Sanger Institute for the COVID-19 Genomics UK (COG-UK) Consortium | Jacquelyn Wynn, Mairead Hyland, The Lighthouse Lab in Alderley Park and Alex Alderton, Roberto Amato, Jeffrey Barrett, Sonia Goncalves, Ewan Harrison, David K. Jackson, Ian Johnston, Dominic Kwiatkowski, Cordelia Langford, John Sillitoe on behalf of the Wellcome Sanger Institute COVID-19 Surveillance Team |
| EPI_ISL_2681923 | Lighthouse Lab in Glasgow | Wellcome Sanger Institute for the COVID-19 Genomics UK (COG-UK) Consortium | Harper VanSteenhouse, Yumi Kasai, David Gray, Carol Clugston, Anna Dominiczak and Alex Alderton, Roberto Amato, Jeffrey Barrett, Sonia Goncalves, Ewan Harrison, David K. Jackson, Ian Johnston, Dominic Kwiatkowski, Cordelia Langford, John Sillitoe on behalf of the Wellcome Sanger Institute COVID-19 Surveillance Team |
| EPI_ISL_2681924, EPI_ISL_2681925 | Lighthouse Lab in Alderley Park | Wellcome Sanger Institute for the COVID-19 Genomics UK (COG-UK) Consortium | Jacquelyn Wynn, Mairead Hyland, The Lighthouse Lab in Alderley Park and Alex Alderton, Roberto Amato, Jeffrey Barrett, Sonia Goncalves, Ewan Harrison, David K. Jackson, Ian Johnston, Dominic Kwiatkowski, Cordelia Langford, John Sillitoe on behalf of the Wellcome Sanger Institute COVID-19 Surveillance Team |
| EPI_ISL_2681926 | Lighthouse Lab in Glasgow | Wellcome Sanger Institute for the COVID-19 Genomics UK (COG-UK) Consortium | Harper VanSteenhouse, Yumi Kasai, David Gray, Carol Clugston, Anna Dominiczak and Alex Alderton, Roberto Amato, Jeffrey Barrett, Sonia Goncalves, Ewan Harrison, David K. Jackson, Ian Johnston, Dominic Kwiatkowski, Cordelia Langford, John Sillitoe on behalf of the Wellcome Sanger Institute COVID-19 Surveillance Team |
| EPI_ISL_2681927 | Lighthouse Lab in Alderley Park | Wellcome Sanger Institute for the COVID-19 Genomics UK (COG-UK) Consortium | Jacquelyn Wynn, Mairead Hyland, The Lighthouse Lab in Alderley Park and Alex Alderton, Roberto Amato, Jeffrey Barrett, Sonia Goncalves, Ewan Harrison, David K. Jackson, Ian Johnston, Dominic Kwiatkowski, Cordelia Langford, John Sillitoe on behalf of the Wellcome Sanger Institute COVID-19 Surveillance Team |
| EPI_ISL_2681929, EPI_ISL_2681930, EPI_ISL_2681931, EPI_ISL_2681932, EPI_ISL_2681934, EPI_ISL_2681935, EPI_ISL_2681937, EPI_ISL_2681938 | Lighthouse Lab in Glasgow | Wellcome Sanger Institute for the COVID-19 Genomics UK (COG-UK) Consortium | Harper VanSteenhouse, Yumi Kasai, David Gray, Carol Clugston, Anna Dominiczak and Alex Alderton, Roberto Amato, Jeffrey Barrett, Sonia Goncalves, Ewan Harrison, David K. Jackson, Ian Johnston, Dominic Kwiatkowski, Cordelia Langford, John Sillitoe on behalf of the Wellcome Sanger Institute COVID-19 Surveillance Team |
| EPI_ISL_2681939 | Lighthouse Lab in Milton Keynes | Wellcome Sanger Institute for the COVID-19 Genomics UK (COG-UK) Consortium | The Lighthouse Lab in Milton Keynes and Alex Alderton, Roberto Amato, Jeffrey Barrett, Sonia Goncalves, Ewan Harrison, David K. Jackson, Ian Johnston, Dominic Kwiatkowski, Cordelia Langford, John Sillitoe on behalf of the Wellcome Sanger Institute COVID-19 Surveillance Team |
| EPI_ISL_2681940, EPI_ISL_2681941, EPI_ISL_2681942, EPI_ISL_2681943, EPI_ISL_2681944, EPI_ISL_2681945, EPI_ISL_2681946, EPI_ISL_2681948, EPI_ISL_2681950, EPI_ISL_2681951, EPI_ISL_2681952, EPI_ISL_2681954, EPI_ISL_2681955, EPI_ISL_2681956, EPI_ISL_2681958, EPI_ISL_2681959, EPI_ISL_2681960, EPI_ISL_2681961, EPI_ISL_2681962, EPI_ISL_2681963 | | | |
| see above | Lighthouse Lab in Glasgow | Wellcome Sanger Institute for the COVID-19 Genomics UK (COG-UK) Consortium | Harper VanSteenhouse, Yumi Kasai, David Gray, Carol Clugston, Anna Dominiczak and Alex Alderton, Roberto Amato, Jeffrey Barrett, Sonia Goncalves, Ewan Harrison, David K. Jackson, Ian Johnston, Dominic Kwiatkowski, Cordelia Langford, John Sillitoe on behalf of the Wellcome Sanger Institute COVID-19 Surveillance Team |
| EPI_ISL_2681964 | Lighthouse Lab in Milton Keynes | Wellcome Sanger Institute for the COVID-19 Genomics UK (COG-UK) Consortium | The Lighthouse Lab in Milton Keynes and Alex Alderton, Roberto Amato, Jeffrey Barrett, Sonia Goncalves, Ewan Harrison, David K. Jackson, Ian Johnston, Dominic Kwiatkowski, Cordelia Langford, John Sillitoe on behalf of the Wellcome Sanger Institute COVID-19 Surveillance Team |
| EPI_ISL_2681965, EPI_ISL_2681966, EPI_ISL_2681967, EPI_ISL_2681968, EPI_ISL_2681969, EPI_ISL_2681970, EPI_ISL_2681971, EPI_ISL_2681972, EPI_ISL_2681974, EPI_ISL_2681975, EPI_ISL_2681976, EPI_ISL_2681977, EPI_ISL_2681978, EPI_ISL_2681979 | | | |
| see above | Lighthouse Lab in Glasgow | Wellcome Sanger Institute for the COVID-19 Genomics UK (COG-UK) Consortium | Harper VanSteenhouse, Yumi Kasai, David Gray, Carol Clugston, Anna Dominiczak and Alex Alderton, Roberto Amato, Jeffrey Barrett, Sonia Goncalves, Ewan Harrison, David K. Jackson, Ian Johnston, Dominic Kwiatkowski, Cordelia Langford, John Sillitoe on behalf of the Wellcome Sanger Institute COVID-19 Surveillance Team |
| EPI_ISL_2681980 | Lighthouse Lab in Milton Keynes | Wellcome Sanger Institute for the COVID-19 Genomics UK (COG-UK) Consortium | The Lighthouse Lab in Milton Keynes and Alex Alderton, Roberto Amato, Jeffrey Barrett, Sonia Goncalves, Ewan Harrison, David K. Jackson, Ian Johnston, Dominic Kwiatkowski, Cordelia Langford, John Sillitoe on behalf of the Wellcome Sanger Institute COVID-19 Surveillance Team |
| EPI_ISL_2681981, EPI_ISL_2681982, EPI_ISL_2681983, EPI_ISL_2681984, EPI_ISL_2681985, EPI_ISL_2681986, EPI_ISL_2681987, EPI_ISL_2681988, EPI_ISL_2681989, EPI_ISL_2681990, EPI_ISL_2681991, EPI_ISL_2681992, EPI_ISL_2681993, EPI_ISL_2681994, EPI_ISL_2681995, EPI_ISL_2681997, EPI_ISL_2681998, EPI_ISL_2681999, EPI_ISL_2682000, EPI_ISL_2682001, EPI_ISL_2682002, EPI_ISL_2682003, EPI_ISL_2682004, EPI_ISL_2682005, EPI_ISL_2682006, EPI_ISL_2682007, EPI_ISL_2682008, EPI_ISL_2682009, EPI_ISL_2682010, EPI_ISL_2682011, EPI_ISL_2682012, EPI_ISL_2682013, EPI_ISL_2682014, EPI_ISL_2682015, EPI_ISL_2682016, EPI_ISL_2682017, EPI_ISL_2682018, EPI_ISL_2682019, EPI_ISL_2682020 | | | |
| see above | Lighthouse Lab in Glasgow | Wellcome Sanger Institute for the COVID-19 Genomics UK (COG-UK) Consortium | Harper VanSteenhouse, Yumi Kasai, David Gray, Carol Clugston, Anna Dominiczak and Alex Alderton, Roberto Amato, Jeffrey Barrett, Sonia Goncalves, Ewan Harrison, David K. Jackson, Ian Johnston, Dominic Kwiatkowski, Cordelia Langford, John Sillitoe on behalf of the Wellcome Sanger Institute COVID-19 Surveillance Team |
| EPI_ISL_2682021 | Lighthouse Lab in Milton Keynes | Wellcome Sanger Institute for the COVID-19 Genomics UK (COG-UK) Consortium | The Lighthouse Lab in Milton Keynes and Alex Alderton, Roberto Amato, Jeffrey Barrett, Sonia Goncalves, Ewan Harrison, David K. Jackson, Ian Johnston, Dominic Kwiatkowski, Cordelia Langford, John Sillitoe on behalf of the Wellcome Sanger Institute COVID-19 Surveillance Team |
| EPI_ISL_2682023, EPI_ISL_2682025, EPI_ISL_2682026, EPI_ISL_2682027, EPI_ISL_2682028, EPI_ISL_2682029, EPI_ISL_2682030, EPI_ISL_2682031, EPI_ISL_2682032, EPI_ISL_2682033, EPI_ISL_2682034, EPI_ISL_2682035, EPI_ISL_2682036, EPI_ISL_2682037, EPI_ISL_2682038, EPI_ISL_2682039, EPI_ISL_2682040, EPI_ISL_2682042, EPI_ISL_2682043, EPI_ISL_2682044, EPI_ISL_2682045, EPI_ISL_2682047, EPI_ISL_2682048, EPI_ISL_2682049, EPI_ISL_2682050, EPI_ISL_2682051, EPI_ISL_2682053, EPI_ISL_2682054, EPI_ISL_2682055, EPI_ISL_2682056, EPI_ISL_2682057, EPI_ISL_2682058 | | | |
| see above | Lighthouse Lab in Glasgow | Wellcome Sanger Institute for the COVID-19 Genomics UK (COG-UK) Consortium | Harper VanSteenhouse, Yumi Kasai, David Gray, Carol Clugston, Anna Dominiczak and Alex Alderton, Roberto Amato, Jeffrey Barrett, Sonia Goncalves, Ewan Harrison, David K. Jackson, Ian Johnston, Dominic Kwiatkowski, Cordelia Langford, John Sillitoe on behalf of the Wellcome Sanger Institute COVID-19 Surveillance Team |
| EPI_ISL_2682059 | Health Services Laboratories | Wellcome Sanger Institute for the COVID-19 Genomics UK (COG-UK) Consortium | Health Services Laboratories and Alex Alderton, Roberto Amato, Jeffrey Barrett, Sonia Goncalves, Ewan Harrison, David K. Jackson, Ian Johnston, Dominic Kwiatkowski, Cordelia Langford, John Sillitoe on behalf of the Wellcome Sanger Institute COVID-19 Surveillance Team |
| EPI_ISL_2682060, EPI_ISL_2682061, EPI_ISL_2682062 | Lighthouse Lab in Milton Keynes | Wellcome Sanger Institute for the COVID-19 Genomics UK (COG-UK) Consortium | The Lighthouse Lab in Milton Keynes and Alex Alderton, Roberto Amato, Jeffrey Barrett, Sonia Goncalves, Ewan Harrison, David K. Jackson, Ian Johnston, Dominic Kwiatkowski, Cordelia Langford, John Sillitoe on behalf of the Wellcome Sanger Institute COVID-19 Surveillance Team |
| EPI_ISL_2682063 | Lighthouse Lab in Glasgow | Wellcome Sanger Institute for the COVID-19 Genomics UK (COG-UK) Consortium | Harper VanSteenhouse, Yumi Kasai, David Gray, Carol Clugston, Anna Dominiczak and Alex Alderton, Roberto Amato, Jeffrey Barrett, Sonia Goncalves, Ewan Harrison, David K. Jackson, Ian Johnston, Dominic Kwiatkowski, Cordelia Langford, John Sillitoe on behalf of the Wellcome Sanger Institute COVID-19 Surveillance Team |
| EPI_ISL_2682064, EPI_ISL_2682065 | Health Services Laboratories | Wellcome Sanger Institute for the COVID-19 Genomics UK (COG-UK) Consortium | Health Services Laboratories and Alex Alderton, Roberto Amato, Jeffrey Barrett, Sonia Goncalves, Ewan Harrison, David K. Jackson, Ian Johnston, Dominic Kwiatkowski, Cordelia Langford, John Sillitoe on behalf of the Wellcome Sanger Institute COVID-19 Surveillance Team |
| EPI_ISL_2682066, EPI_ISL_2682067 | Lighthouse Lab in Milton Keynes | Wellcome Sanger Institute for the COVID-19 Genomics UK (COG-UK) Consortium | The Lighthouse Lab in Milton Keynes and Alex Alderton, Roberto Amato, Jeffrey Barrett, Sonia Goncalves, Ewan Harrison, David K. Jackson, Ian Johnston, Dominic Kwiatkowski, Cordelia Langford, John Sillitoe on behalf of the Wellcome Sanger Institute COVID-19 Surveillance Team |
| EPI_ISL_2682068, EPI_ISL_2682069 | Lighthouse Lab in Glasgow | Wellcome Sanger Institute for the COVID-19 Genomics UK (COG-UK) Consortium | Harper VanSteenhouse, Yumi Kasai, David Gray, Carol Clugston, Anna Dominiczak and Alex Alderton, Roberto Amato, Jeffrey Barrett, Sonia Goncalves, Ewan Harrison, David K. Jackson, Ian Johnston, Dominic Kwiatkowski, Cordelia Langford, John Sillitoe on behalf of the Wellcome Sanger Institute COVID-19 Surveillance Team |
| EPI_ISL_2682070 | Lighthouse Lab in Milton Keynes | Wellcome Sanger Institute for the COVID-19 Genomics UK (COG-UK) Consortium | The Lighthouse Lab in Milton Keynes and Alex Alderton, Roberto Amato, Jeffrey Barrett, Sonia Goncalves, Ewan Harrison, David K. Jackson, Ian Johnston, Dominic Kwiatkowski, Cordelia Langford, John Sillitoe on behalf of the Wellcome Sanger Institute COVID-19 Surveillance Team |
| EPI_ISL_2682071 | Lighthouse Lab in Glasgow | Wellcome Sanger Institute for the COVID-19 Genomics UK (COG-UK) Consortium | Harper VanSteenhouse, Yumi Kasai, David Gray, Carol Clugston, Anna Dominiczak and Alex Alderton, Roberto Amato, Jeffrey Barrett, Sonia Goncalves, Ewan Harrison, David K. Jackson, Ian Johnston, Dominic Kwiatkowski, Cordelia Langford, John Sillitoe on behalf of the Wellcome Sanger Institute COVID-19 Surveillance Team |
| EPI_ISL_2682072 | Lighthouse Lab in Milton Keynes | Wellcome Sanger Institute for the COVID-19 Genomics UK (COG-UK) Consortium | The Lighthouse Lab in Milton Keynes and Alex Alderton, Roberto Amato, Jeffrey Barrett, Sonia Goncalves, Ewan Harrison, David K. Jackson, Ian Johnston, Dominic Kwiatkowski, Cordelia Langford, John Sillitoe on behalf of the Wellcome Sanger Institute COVID-19 Surveillance Team |
| EPI_ISL_2682073 | Lighthouse Lab in Glasgow | Wellcome Sanger Institute for the COVID-19 Genomics UK (COG-UK) Consortium | Harper VanSteenhouse, Yumi Kasai, David Gray, Carol Clugston, Anna Dominiczak and Alex Alderton, Roberto Amato, Jeffrey Barrett, Sonia Goncalves, Ewan Harrison, David K. Jackson, Ian Johnston, Dominic Kwiatkowski, Cordelia Langford, John Sillitoe on behalf of the Wellcome Sanger Institute COVID-19 Surveillance Team |
| EPI_ISL_2682074, EPI_ISL_2682075, EPI_ISL_2682076, | Health Services Laboratories | Wellcome Sanger Institute for the COVID-19 Genomics UK (COG-UK) Consortium | Health Services Laboratories and Alex Alderton, Roberto Amato, Jeffrey Barrett, Sonia Goncalves, Ewan Harrison, David K. Jackson, Ian Johnston, Dominic Kwiatkowski, Cordelia Langford, John Sillitoe on behalf of the Wellcome Sanger Institute COVID-19 Surveillance Team |

| | | | |
|---|---|---|---|
| EPI_ISL_2682077, EPI_ISL_2682078, EPI_ISL_2682079, EPI_ISL_2682080 | | | |
| EPI_ISL_2682081 | Lighthouse Lab in Milton Keynes | Wellcome Sanger Institute for the COVID-19 Genomics UK (COG-UK) Consortium | The Lighthouse Lab in Milton Keynes and Alex Alderton, Roberto Amato, Jeffrey Barrett, Sonia Goncalves, Ewan Harrison, David K. Jackson, Ian Johnston, Dominic Kwiatkowski, Cordelia Langford, John Sillitoe on behalf of the Wellcome Sanger Institute COVID-19 Surveillance Team |
| EPI_ISL_2682082, EPI_ISL_2682083 | Lighthouse Lab in Glasgow | Wellcome Sanger Institute for the COVID-19 Genomics UK (COG-UK) Consortium | Harper VanSteenhouse, Yumi Kasai, David Gray, Carol Clugston, Anna Dominiczak and Alex Alderton, Roberto Amato, Jeffrey Barrett, Sonia Goncalves, Ewan Harrison, David K. Jackson, Ian Johnston, Dominic Kwiatkowski, Cordelia Langford, John Sillitoe on behalf of the Wellcome Sanger Institute COVID-19 Surveillance Team |
| EPI_ISL_2682084 | Lighthouse Lab in Milton Keynes | Wellcome Sanger Institute for the COVID-19 Genomics UK (COG-UK) Consortium | The Lighthouse Lab in Milton Keynes and Alex Alderton, Roberto Amato, Jeffrey Barrett, Sonia Goncalves, Ewan Harrison, David K. Jackson, Ian Johnston, Dominic Kwiatkowski, Cordelia Langford, John Sillitoe on behalf of the Wellcome Sanger Institute COVID-19 Surveillance Team |
| EPI_ISL_2682085 | Lighthouse Lab in Glasgow | Wellcome Sanger Institute for the COVID-19 Genomics UK (COG-UK) Consortium | Harper VanSteenhouse, Yumi Kasai, David Gray, Carol Clugston, Anna Dominiczak and Alex Alderton, Roberto Amato, Jeffrey Barrett, Sonia Goncalves, Ewan Harrison, David K. Jackson, Ian Johnston, Dominic Kwiatkowski, Cordelia Langford, John Sillitoe on behalf of the Wellcome Sanger Institute COVID-19 Surveillance Team |
| EPI_ISL_2682086 | Lighthouse Lab in Milton Keynes | Wellcome Sanger Institute for the COVID-19 Genomics UK (COG-UK) Consortium | The Lighthouse Lab in Milton Keynes and Alex Alderton, Roberto Amato, Jeffrey Barrett, Sonia Goncalves, Ewan Harrison, David K. Jackson, Ian Johnston, Dominic Kwiatkowski, Cordelia Langford, John Sillitoe on behalf of the Wellcome Sanger Institute COVID-19 Surveillance Team |
| EPI_ISL_2682087, EPI_ISL_2682088, EPI_ISL_2682089, EPI_ISL_2682090 | Health Services Laboratories | Wellcome Sanger Institute for the COVID-19 Genomics UK (COG-UK) Consortium | Health Services Laboratories and Alex Alderton, Roberto Amato, Jeffrey Barrett, Sonia Goncalves, Ewan Harrison, David K. Jackson, Ian Johnston, Dominic Kwiatkowski, Cordelia Langford, John Sillitoe on behalf of the Wellcome Sanger Institute COVID-19 Surveillance Team |
| EPI_ISL_2682091 | Lighthouse Lab in Milton Keynes | Wellcome Sanger Institute for the COVID-19 Genomics UK (COG-UK) Consortium | The Lighthouse Lab in Milton Keynes and Alex Alderton, Roberto Amato, Jeffrey Barrett, Sonia Goncalves, Ewan Harrison, David K. Jackson, Ian Johnston, Dominic Kwiatkowski, Cordelia Langford, John Sillitoe on behalf of the Wellcome Sanger Institute COVID-19 Surveillance Team |
| EPI_ISL_2682092 | Health Services Laboratories | Wellcome Sanger Institute for the COVID-19 Genomics UK (COG-UK) Consortium | Health Services Laboratories and Alex Alderton, Roberto Amato, Jeffrey Barrett, Sonia Goncalves, Ewan Harrison, David K. Jackson, Ian Johnston, Dominic Kwiatkowski, Cordelia Langford, John Sillitoe on behalf of the Wellcome Sanger Institute COVID-19 Surveillance Team |
| EPI_ISL_2682093 | Lighthouse Lab in Glasgow | Wellcome Sanger Institute for the COVID-19 Genomics UK (COG-UK) Consortium | Harper VanSteenhouse, Yumi Kasai, David Gray, Carol Clugston, Anna Dominiczak and Alex Alderton, Roberto Amato, Jeffrey Barrett, Sonia Goncalves, Ewan Harrison, David K. Jackson, Ian Johnston, Dominic Kwiatkowski, Cordelia Langford, John Sillitoe on behalf of the Wellcome Sanger Institute COVID-19 Surveillance Team |
| EPI_ISL_2682094 | Lighthouse Lab in Milton Keynes | Wellcome Sanger Institute for the COVID-19 Genomics UK (COG-UK) Consortium | The Lighthouse Lab in Milton Keynes and Alex Alderton, Roberto Amato, Jeffrey Barrett, Sonia Goncalves, Ewan Harrison, David K. Jackson, Ian Johnston, Dominic Kwiatkowski, Cordelia Langford, John Sillitoe on behalf of the Wellcome Sanger Institute COVID-19 Surveillance Team |
| EPI_ISL_2682095, EPI_ISL_2682096 | Health Services Laboratories | Wellcome Sanger Institute for the COVID-19 Genomics UK (COG-UK) Consortium | Health Services Laboratories and Alex Alderton, Roberto Amato, Jeffrey Barrett, Sonia Goncalves, Ewan Harrison, David K. Jackson, Ian Johnston, Dominic Kwiatkowski, Cordelia Langford, John Sillitoe on behalf of the Wellcome Sanger Institute COVID-19 Surveillance Team |
| EPI_ISL_2682097 | Lighthouse Lab in Glasgow | Wellcome Sanger Institute for the COVID-19 Genomics UK (COG-UK) Consortium | Harper VanSteenhouse, Yumi Kasai, David Gray, Carol Clugston, Anna Dominiczak and Alex Alderton, Roberto Amato, Jeffrey Barrett, Sonia Goncalves, Ewan Harrison, David K. Jackson, Ian Johnston, Dominic Kwiatkowski, Cordelia Langford, John Sillitoe on behalf of the Wellcome Sanger Institute COVID-19 Surveillance Team |
| EPI_ISL_2682098 | Health Services Laboratories | Wellcome Sanger Institute for the COVID-19 Genomics UK (COG-UK) Consortium | Health Services Laboratories and Alex Alderton, Roberto Amato, Jeffrey Barrett, Sonia Goncalves, Ewan Harrison, David K. Jackson, Ian Johnston, Dominic Kwiatkowski, Cordelia Langford, John Sillitoe on behalf of the Wellcome Sanger Institute COVID-19 Surveillance Team |
| EPI_ISL_2682099 | Lighthouse Lab in Milton Keynes | Wellcome Sanger Institute for the COVID-19 Genomics UK (COG-UK) Consortium | The Lighthouse Lab in Milton Keynes and Alex Alderton, Roberto Amato, Jeffrey Barrett, Sonia Goncalves, Ewan Harrison, David K. Jackson, Ian Johnston, Dominic Kwiatkowski, Cordelia Langford, John Sillitoe on behalf of the Wellcome Sanger Institute COVID-19 Surveillance Team |
| EPI_ISL_2682100 | Lighthouse Lab in Glasgow | Wellcome Sanger Institute for the COVID-19 Genomics UK (COG-UK) Consortium | Harper VanSteenhouse, Yumi Kasai, David Gray, Carol Clugston, Anna Dominiczak and Alex Alderton, Roberto Amato, Jeffrey Barrett, Sonia Goncalves, Ewan Harrison, David K. Jackson, Ian Johnston, Dominic Kwiatkowski, Cordelia Langford, John Sillitoe on behalf of the Wellcome Sanger Institute COVID-19 Surveillance Team |
| EPI_ISL_2682101, EPI_ISL_2682102, EPI_ISL_2682103, EPI_ISL_2682104 | Health Services Laboratories | Wellcome Sanger Institute for the COVID-19 Genomics UK (COG-UK) Consortium | Health Services Laboratories and Alex Alderton, Roberto Amato, Jeffrey Barrett, Sonia Goncalves, Ewan Harrison, David K. Jackson, Ian Johnston, Dominic Kwiatkowski, Cordelia Langford, John Sillitoe on behalf of the Wellcome Sanger Institute COVID-19 Surveillance Team |
| EPI_ISL_2682105 | Lighthouse Lab in Glasgow | Wellcome Sanger Institute for the COVID-19 Genomics UK (COG-UK) Consortium | Harper VanSteenhouse, Yumi Kasai, David Gray, Carol Clugston, Anna Dominiczak and Alex Alderton, Roberto Amato, Jeffrey Barrett, Sonia Goncalves, Ewan Harrison, David K. Jackson, Ian Johnston, Dominic Kwiatkowski, Cordelia Langford, John Sillitoe on behalf of the Wellcome Sanger Institute COVID-19 Surveillance Team |
| EPI_ISL_2682106 | Health Services Laboratories | Wellcome Sanger Institute for the COVID-19 Genomics UK (COG-UK) Consortium | Health Services Laboratories and Alex Alderton, Roberto Amato, Jeffrey Barrett, Sonia Goncalves, Ewan Harrison, David K. Jackson, Ian Johnston, Dominic Kwiatkowski, Cordelia Langford, John Sillitoe on behalf of the Wellcome Sanger Institute COVID-19 Surveillance Team |
| EPI_ISL_2682107 | Lighthouse Lab in Glasgow | Wellcome Sanger Institute for the COVID-19 Genomics UK (COG-UK) Consortium | Harper VanSteenhouse, Yumi Kasai, David Gray, Carol Clugston, Anna Dominiczak and Alex Alderton, Roberto Amato, Jeffrey Barrett, Sonia Goncalves, Ewan Harrison, David K. Jackson, Ian Johnston, Dominic Kwiatkowski, Cordelia Langford, John Sillitoe on behalf of the Wellcome Sanger Institute COVID-19 Surveillance Team |
| EPI_ISL_2682108, EPI_ISL_2682109, EPI_ISL_2682110 | Lighthouse Lab in Milton Keynes | Wellcome Sanger Institute for the COVID-19 Genomics UK (COG-UK) Consortium | The Lighthouse Lab in Milton Keynes and Alex Alderton, Roberto Amato, Jeffrey Barrett, Sonia Goncalves, Ewan Harrison, David K. Jackson, Ian Johnston, Dominic Kwiatkowski, Cordelia Langford, John Sillitoe on behalf of the Wellcome Sanger Institute COVID-19 Surveillance Team |
| EPI_ISL_2682111 | Health Services Laboratories | Wellcome Sanger Institute for the COVID-19 Genomics UK (COG-UK) Consortium | Health Services Laboratories and Alex Alderton, Roberto Amato, Jeffrey Barrett, Sonia Goncalves, Ewan Harrison, David K. Jackson, Ian Johnston, Dominic Kwiatkowski, Cordelia Langford, John Sillitoe on behalf of the Wellcome Sanger Institute COVID-19 Surveillance Team |
| EPI_ISL_2682112 | Lighthouse Lab in Milton Keynes | Wellcome Sanger Institute for the COVID-19 Genomics UK (COG-UK) Consortium | The Lighthouse Lab in Milton Keynes and Alex Alderton, Roberto Amato, Jeffrey Barrett, Sonia Goncalves, Ewan Harrison, David K. Jackson, Ian Johnston, Dominic Kwiatkowski, Cordelia Langford, John Sillitoe on behalf of the Wellcome Sanger Institute COVID-19 Surveillance Team |
| EPI_ISL_2682113 | Health Services Laboratories | Wellcome Sanger Institute for the COVID-19 Genomics UK (COG-UK) Consortium | Health Services Laboratories and Alex Alderton, Roberto Amato, Jeffrey Barrett, Sonia Goncalves, Ewan Harrison, David K. Jackson, Ian Johnston, Dominic Kwiatkowski, Cordelia Langford, John Sillitoe on behalf of the Wellcome Sanger Institute COVID-19 Surveillance Team |
| EPI_ISL_2682114 | Lighthouse Lab in Glasgow | Wellcome Sanger Institute for the COVID-19 Genomics UK (COG-UK) Consortium | Harper VanSteenhouse, Yumi Kasai, David Gray, Carol Clugston, Anna Dominiczak and Alex Alderton, Roberto Amato, Jeffrey Barrett, Sonia Goncalves, Ewan Harrison, David K. Jackson, Ian Johnston, Dominic Kwiatkowski, Cordelia Langford, John Sillitoe on behalf of the Wellcome Sanger Institute COVID-19 Surveillance Team |
| EPI_ISL_2682115 | Health Services Laboratories | Wellcome Sanger Institute for the COVID-19 Genomics UK (COG-UK) Consortium | Health Services Laboratories and Alex Alderton, Roberto Amato, Jeffrey Barrett, Sonia Goncalves, Ewan Harrison, David K. Jackson, Ian Johnston, Dominic Kwiatkowski, Cordelia Langford, John Sillitoe on behalf of the Wellcome Sanger Institute COVID-19 Surveillance Team |
| EPI_ISL_2682116 | Lighthouse Lab in Milton Keynes | Wellcome Sanger Institute for the COVID-19 Genomics UK (COG-UK) Consortium | The Lighthouse Lab in Milton Keynes and Alex Alderton, Roberto Amato, Jeffrey Barrett, Sonia Goncalves, Ewan Harrison, David K. Jackson, Ian Johnston, Dominic Kwiatkowski, Cordelia Langford, John Sillitoe on behalf of the Wellcome Sanger Institute COVID-19 Surveillance Team |
| EPI_ISL_2682117 | Health Services Laboratories | Wellcome Sanger Institute for the COVID-19 Genomics UK (COG-UK) Consortium | Health Services Laboratories and Alex Alderton, Roberto Amato, Jeffrey Barrett, Sonia Goncalves, Ewan Harrison, David K. Jackson, Ian Johnston, Dominic Kwiatkowski, Cordelia Langford, John Sillitoe on behalf of the Wellcome Sanger Institute COVID-19 Surveillance Team |
| EPI_ISL_2682118, EPI_ISL_2682119 | Lighthouse Lab in Milton Keynes | Wellcome Sanger Institute for the COVID-19 Genomics UK (COG-UK) Consortium | The Lighthouse Lab in Milton Keynes and Alex Alderton, Roberto Amato, Jeffrey Barrett, Sonia Goncalves, Ewan Harrison, David K. Jackson, Ian Johnston, Dominic Kwiatkowski, Cordelia Langford, John Sillitoe on behalf of the Wellcome Sanger Institute COVID-19 Surveillance Team |
| EPI_ISL_2682120 | Lighthouse Lab in Glasgow | Wellcome Sanger Institute for the COVID-19 Genomics UK (COG-UK) Consortium | Harper VanSteenhouse, Yumi Kasai, David Gray, Carol Clugston, Anna Dominiczak and Alex Alderton, Roberto Amato, Jeffrey Barrett, Sonia Goncalves, Ewan Harrison, David K. Jackson, Ian Johnston, Dominic Kwiatkowski, Cordelia Langford, John Sillitoe on behalf of the Wellcome Sanger Institute COVID-19 Surveillance Team |
| EPI_ISL_2682121 | Health Services Laboratories | Wellcome Sanger Institute for the COVID-19 Genomics UK (COG-UK) Consortium | Health Services Laboratories and Alex Alderton, Roberto Amato, Jeffrey Barrett, Sonia Goncalves, Ewan Harrison, David K. Jackson, Ian Johnston, Dominic Kwiatkowski, Cordelia Langford, John Sillitoe on behalf of the Wellcome Sanger Institute COVID-19 Surveillance Team |
| EPI_ISL_2682122 | Lighthouse Lab in Milton Keynes | Wellcome Sanger Institute for the COVID-19 Genomics UK (COG-UK) Consortium | The Lighthouse Lab in Milton Keynes and Alex Alderton, Roberto Amato, Jeffrey Barrett, Sonia Goncalves, Ewan Harrison, David K. Jackson, Ian Johnston, Dominic Kwiatkowski, Cordelia Langford, John Sillitoe on behalf of the Wellcome Sanger Institute COVID-19 Surveillance Team |
| EPI_ISL_2682123, EPI_ISL_2682124 | Lighthouse Lab in Glasgow | Wellcome Sanger Institute for the COVID-19 Genomics UK (COG-UK) Consortium | Harper VanSteenhouse, Yumi Kasai, David Gray, Carol Clugston, Anna Dominiczak and Alex Alderton, Roberto Amato, Jeffrey Barrett, Sonia Goncalves, Ewan Harrison, David K. Jackson, Ian Johnston, Dominic Kwiatkowski, Cordelia Langford, John Sillitoe on behalf of the Wellcome Sanger Institute COVID-19 Surveillance Team |
| EPI_ISL_2682125, EPI_ISL_2682126, EPI_ISL_2682127, EPI_ISL_2682128, EPI_ISL_2682129 | Lighthouse Lab in Milton Keynes | Wellcome Sanger Institute for the COVID-19 Genomics UK (COG-UK) Consortium | The Lighthouse Lab in Milton Keynes and Alex Alderton, Roberto Amato, Jeffrey Barrett, Sonia Goncalves, Ewan Harrison, David K. Jackson, Ian Johnston, Dominic Kwiatkowski, Cordelia Langford, John Sillitoe on behalf of the Wellcome Sanger Institute COVID-19 Surveillance Team |
| EPI_ISL_2682130 | Lighthouse Lab in Glasgow | Wellcome Sanger Institute for the COVID-19 Genomics UK (COG-UK) Consortium | Harper VanSteenhouse, Yumi Kasai, David Gray, Carol Clugston, Anna Dominiczak and Alex Alderton, Roberto Amato, Jeffrey Barrett, Sonia Goncalves, Ewan Harrison, David K. Jackson, Ian Johnston, Dominic Kwiatkowski, Cordelia Langford, John Sillitoe on behalf of the Wellcome Sanger Institute COVID-19 Surveillance Team |
| EPI_ISL_2682131, EPI_ISL_2682132 | Health Services Laboratories | Wellcome Sanger Institute for the COVID-19 Genomics UK (COG-UK) Consortium | Health Services Laboratories and Alex Alderton, Roberto Amato, Jeffrey Barrett, Sonia Goncalves, Ewan Harrison, David K. Jackson, Ian Johnston, Dominic Kwiatkowski, Cordelia Langford, John Sillitoe on behalf of the Wellcome Sanger Institute COVID-19 Surveillance Team |
| EPI_ISL_2682133 | Lighthouse Lab in Glasgow | Wellcome Sanger Institute for the COVID-19 Genomics UK (COG-UK) Consortium | Harper VanSteenhouse, Yumi Kasai, David Gray, Carol Clugston, Anna Dominiczak and Alex Alderton, Roberto Amato, Jeffrey Barrett, Sonia Goncalves, Ewan Harrison, David K. Jackson, Ian Johnston, Dominic Kwiatkowski, Cordelia Langford, John Sillitoe on behalf of the Wellcome Sanger Institute COVID-19 Surveillance Team |
| EPI_ISL_2682134 | Health Services Laboratories | Wellcome Sanger Institute for the COVID-19 Genomics UK (COG-UK) Consortium | Health Services Laboratories and Alex Alderton, Roberto Amato, Jeffrey Barrett, Sonia Goncalves, Ewan Harrison, David K. Jackson, Ian Johnston, Dominic Kwiatkowski, Cordelia Langford, John Sillitoe on behalf of the Wellcome Sanger Institute COVID-19 Surveillance Team |
| EPI_ISL_2682135 | Lighthouse Lab in Milton Keynes | Wellcome Sanger Institute for the COVID-19 Genomics UK (COG-UK) Consortium | The Lighthouse Lab in Milton Keynes and Alex Alderton, Roberto Amato, Jeffrey Barrett, Sonia Goncalves, Ewan Harrison, David K. Jackson, Ian Johnston, Dominic Kwiatkowski, Cordelia Langford, John Sillitoe on behalf of the Wellcome Sanger Institute COVID-19 Surveillance Team |
| EPI_ISL_2682136, EPI_ISL_2682137 | Health Services Laboratories | Wellcome Sanger Institute for the COVID-19 Genomics UK (COG-UK) Consortium | Health Services Laboratories and Alex Alderton, Roberto Amato, Jeffrey Barrett, Sonia Goncalves, Ewan Harrison, David K. Jackson, Ian Johnston, Dominic Kwiatkowski, Cordelia Langford, John Sillitoe on behalf of the Wellcome Sanger Institute COVID-19 Surveillance Team |
| EPI_ISL_2682138 | Lighthouse Lab in Glasgow | Wellcome Sanger Institute for the COVID-19 Genomics UK (COG-UK) Consortium | Harper VanSteenhouse, Yumi Kasai, David Gray, Carol Clugston, Anna Dominiczak and Alex Alderton, Roberto Amato, Jeffrey Barrett, Sonia Goncalves, Ewan Harrison, David K. Jackson, Ian Johnston, Dominic Kwiatkowski, Cordelia Langford, John Sillitoe on behalf of the Wellcome Sanger Institute COVID-19 Surveillance Team |

| | | | |
|---|---|---|---|
| EPI_ISL_2682139 | Health Services Laboratories | Wellcome Sanger Institute for the COVID-19 Genomics UK (COG-UK) Consortium | Health Services Laboratories and Alex Alderton, Roberto Amato, Jeffrey Barrett, Sonia Goncalves, Ewan Harrison, David K. Jackson, Ian Johnston, Dominic Kwiatkowski, Cordelia Langford, John Sillitoe on behalf of the Wellcome Sanger Institute COVID-19 Surveillance Team |
| EPI_ISL_2682140 | Lighthouse Lab in Glasgow | Wellcome Sanger Institute for the COVID-19 Genomics UK (COG-UK) Consortium | Harper VanSteenhouse, Yumi Kasai, David Gray, Carol Clugston, Anna Dominiczak and Alex Alderton, Roberto Amato, Jeffrey Barrett, Sonia Goncalves, Ewan Harrison, David K. Jackson, Ian Johnston, Dominic Kwiatkowski, Cordelia Langford, John Sillitoe on behalf of the Wellcome Sanger Institute COVID-19 Surveillance Team |
| EPI_ISL_2682141 | Lighthouse Lab in Milton Keynes | Wellcome Sanger Institute for the COVID-19 Genomics UK (COG-UK) Consortium | The Lighthouse Lab in Milton Keynes and Alex Alderton, Roberto Amato, Jeffrey Barrett, Sonia Goncalves, Ewan Harrison, David K. Jackson, Ian Johnston, Dominic Kwiatkowski, Cordelia Langford, John Sillitoe on behalf of the Wellcome Sanger Institute COVID-19 Surveillance Team |
| EPI_ISL_2682142 | Health Services Laboratories | Wellcome Sanger Institute for the COVID-19 Genomics UK (COG-UK) Consortium | Health Services Laboratories and Alex Alderton, Roberto Amato, Jeffrey Barrett, Sonia Goncalves, Ewan Harrison, David K. Jackson, Ian Johnston, Dominic Kwiatkowski, Cordelia Langford, John Sillitoe on behalf of the Wellcome Sanger Institute COVID-19 Surveillance Team |
| EPI_ISL_2682143 | Lighthouse Lab in Milton Keynes | Wellcome Sanger Institute for the COVID-19 Genomics UK (COG-UK) Consortium | The Lighthouse Lab in Milton Keynes and Alex Alderton, Roberto Amato, Jeffrey Barrett, Sonia Goncalves, Ewan Harrison, David K. Jackson, Ian Johnston, Dominic Kwiatkowski, Cordelia Langford, John Sillitoe on behalf of the Wellcome Sanger Institute COVID-19 Surveillance Team |
| EPI_ISL_2682144, EPI_ISL_2682145 | Lighthouse Lab in Glasgow | Wellcome Sanger Institute for the COVID-19 Genomics UK (COG-UK) Consortium | Harper VanSteenhouse, Yumi Kasai, David Gray, Carol Clugston, Anna Dominiczak and Alex Alderton, Roberto Amato, Jeffrey Barrett, Sonia Goncalves, Ewan Harrison, David K. Jackson, Ian Johnston, Dominic Kwiatkowski, Cordelia Langford, John Sillitoe on behalf of the Wellcome Sanger Institute COVID-19 Surveillance Team |
| EPI_ISL_2682146 | Health Services Laboratories | Wellcome Sanger Institute for the COVID-19 Genomics UK (COG-UK) Consortium | Health Services Laboratories and Alex Alderton, Roberto Amato, Jeffrey Barrett, Sonia Goncalves, Ewan Harrison, David K. Jackson, Ian Johnston, Dominic Kwiatkowski, Cordelia Langford, John Sillitoe on behalf of the Wellcome Sanger Institute COVID-19 Surveillance Team |
| EPI_ISL_2682147, EPI_ISL_2682148 | Lighthouse Lab in Milton Keynes | Wellcome Sanger Institute for the COVID-19 Genomics UK (COG-UK) Consortium | The Lighthouse Lab in Milton Keynes and Alex Alderton, Roberto Amato, Jeffrey Barrett, Sonia Goncalves, Ewan Harrison, David K. Jackson, Ian Johnston, Dominic Kwiatkowski, Cordelia Langford, John Sillitoe on behalf of the Wellcome Sanger Institute COVID-19 Surveillance Team |
| EPI_ISL_2682149 | Lighthouse Lab in Glasgow | Wellcome Sanger Institute for the COVID-19 Genomics UK (COG-UK) Consortium | Harper VanSteenhouse, Yumi Kasai, David Gray, Carol Clugston, Anna Dominiczak and Alex Alderton, Roberto Amato, Jeffrey Barrett, Sonia Goncalves, Ewan Harrison, David K. Jackson, Ian Johnston, Dominic Kwiatkowski, Cordelia Langford, John Sillitoe on behalf of the Wellcome Sanger Institute COVID-19 Surveillance Team |
| EPI_ISL_2682150 | Lighthouse Lab in Milton Keynes | Wellcome Sanger Institute for the COVID-19 Genomics UK (COG-UK) Consortium | The Lighthouse Lab in Milton Keynes and Alex Alderton, Roberto Amato, Jeffrey Barrett, Sonia Goncalves, Ewan Harrison, David K. Jackson, Ian Johnston, Dominic Kwiatkowski, Cordelia Langford, John Sillitoe on behalf of the Wellcome Sanger Institute COVID-19 Surveillance Team |
| EPI_ISL_2682151 | Health Services Laboratories | Wellcome Sanger Institute for the COVID-19 Genomics UK (COG-UK) Consortium | Health Services Laboratories and Alex Alderton, Roberto Amato, Jeffrey Barrett, Sonia Goncalves, Ewan Harrison, David K. Jackson, Ian Johnston, Dominic Kwiatkowski, Cordelia Langford, John Sillitoe on behalf of the Wellcome Sanger Institute COVID-19 Surveillance Team |
| EPI_ISL_2682152 | Lighthouse Lab in Milton Keynes | Wellcome Sanger Institute for the COVID-19 Genomics UK (COG-UK) Consortium | The Lighthouse Lab in Milton Keynes and Alex Alderton, Roberto Amato, Jeffrey Barrett, Sonia Goncalves, Ewan Harrison, David K. Jackson, Ian Johnston, Dominic Kwiatkowski, Cordelia Langford, John Sillitoe on behalf of the Wellcome Sanger Institute COVID-19 Surveillance Team |
| EPI_ISL_2682153 | Health Services Laboratories | Wellcome Sanger Institute for the COVID-19 Genomics UK (COG-UK) Consortium | Health Services Laboratories and Alex Alderton, Roberto Amato, Jeffrey Barrett, Sonia Goncalves, Ewan Harrison, David K. Jackson, Ian Johnston, Dominic Kwiatkowski, Cordelia Langford, John Sillitoe on behalf of the Wellcome Sanger Institute COVID-19 Surveillance Team |
| EPI_ISL_2682154, EPI_ISL_2682155, EPI_ISL_2682156 | Lighthouse Lab in Milton Keynes | Wellcome Sanger Institute for the COVID-19 Genomics UK (COG-UK) Consortium | The Lighthouse Lab in Milton Keynes and Alex Alderton, Roberto Amato, Jeffrey Barrett, Sonia Goncalves, Ewan Harrison, David K. Jackson, Ian Johnston, Dominic Kwiatkowski, Cordelia Langford, John Sillitoe on behalf of the Wellcome Sanger Institute COVID-19 Surveillance Team |
| EPI_ISL_2682157 | Lighthouse Lab in Glasgow | Wellcome Sanger Institute for the COVID-19 Genomics UK (COG-UK) Consortium | Harper VanSteenhouse, Yumi Kasai, David Gray, Carol Clugston, Anna Dominiczak and Alex Alderton, Roberto Amato, Jeffrey Barrett, Sonia Goncalves, Ewan Harrison, David K. Jackson, Ian Johnston, Dominic Kwiatkowski, Cordelia Langford, John Sillitoe on behalf of the Wellcome Sanger Institute COVID-19 Surveillance Team |
| EPI_ISL_2682158 | Lighthouse Lab in Milton Keynes | Wellcome Sanger Institute for the COVID-19 Genomics UK (COG-UK) Consortium | The Lighthouse Lab in Milton Keynes and Alex Alderton, Roberto Amato, Jeffrey Barrett, Sonia Goncalves, Ewan Harrison, David K. Jackson, Ian Johnston, Dominic Kwiatkowski, Cordelia Langford, John Sillitoe on behalf of the Wellcome Sanger Institute COVID-19 Surveillance Team |
| EPI_ISL_2682159 | Health Services Laboratories | Wellcome Sanger Institute for the COVID-19 Genomics UK (COG-UK) Consortium | Health Services Laboratories and Alex Alderton, Roberto Amato, Jeffrey Barrett, Sonia Goncalves, Ewan Harrison, David K. Jackson, Ian Johnston, Dominic Kwiatkowski, Cordelia Langford, John Sillitoe on behalf of the Wellcome Sanger Institute COVID-19 Surveillance Team |
| EPI_ISL_2682160, EPI_ISL_2682161 | Lighthouse Lab in Glasgow | Wellcome Sanger Institute for the COVID-19 Genomics UK (COG-UK) Consortium | Harper VanSteenhouse, Yumi Kasai, David Gray, Carol Clugston, Anna Dominiczak and Alex Alderton, Roberto Amato, Jeffrey Barrett, Sonia Goncalves, Ewan Harrison, David K. Jackson, Ian Johnston, Dominic Kwiatkowski, Cordelia Langford, John Sillitoe on behalf of the Wellcome Sanger Institute COVID-19 Surveillance Team |
| EPI_ISL_2682162, EPI_ISL_2682163 | Lighthouse Lab in Milton Keynes | Wellcome Sanger Institute for the COVID-19 Genomics UK (COG-UK) Consortium | The Lighthouse Lab in Milton Keynes and Alex Alderton, Roberto Amato, Jeffrey Barrett, Sonia Goncalves, Ewan Harrison, David K. Jackson, Ian Johnston, Dominic Kwiatkowski, Cordelia Langford, John Sillitoe on behalf of the Wellcome Sanger Institute COVID-19 Surveillance Team |
| EPI_ISL_2682164, EPI_ISL_2682165 | Health Services Laboratories | Wellcome Sanger Institute for the COVID-19 Genomics UK (COG-UK) Consortium | Health Services Laboratories and Alex Alderton, Roberto Amato, Jeffrey Barrett, Sonia Goncalves, Ewan Harrison, David K. Jackson, Ian Johnston, Dominic Kwiatkowski, Cordelia Langford, John Sillitoe on behalf of the Wellcome Sanger Institute COVID-19 Surveillance Team |
| EPI_ISL_2682166 | Lighthouse Lab in Glasgow | Wellcome Sanger Institute for the COVID-19 Genomics UK (COG-UK) Consortium | Harper VanSteenhouse, Yumi Kasai, David Gray, Carol Clugston, Anna Dominiczak and Alex Alderton, Roberto Amato, Jeffrey Barrett, Sonia Goncalves, Ewan Harrison, David K. Jackson, Ian Johnston, Dominic Kwiatkowski, Cordelia Langford, John Sillitoe on behalf of the Wellcome Sanger Institute COVID-19 Surveillance Team |
| EPI_ISL_2682167, EPI_ISL_2682168, EPI_ISL_2682169 | Health Services Laboratories | Wellcome Sanger Institute for the COVID-19 Genomics UK (COG-UK) Consortium | Health Services Laboratories and Alex Alderton, Roberto Amato, Jeffrey Barrett, Sonia Goncalves, Ewan Harrison, David K. Jackson, Ian Johnston, Dominic Kwiatkowski, Cordelia Langford, John Sillitoe on behalf of the Wellcome Sanger Institute COVID-19 Surveillance Team |
| EPI_ISL_2682170 | Lighthouse Lab in Milton Keynes | Wellcome Sanger Institute for the COVID-19 Genomics UK (COG-UK) Consortium | The Lighthouse Lab in Milton Keynes and Alex Alderton, Roberto Amato, Jeffrey Barrett, Sonia Goncalves, Ewan Harrison, David K. Jackson, Ian Johnston, Dominic Kwiatkowski, Cordelia Langford, John Sillitoe on behalf of the Wellcome Sanger Institute COVID-19 Surveillance Team |
| EPI_ISL_2682171 | Lighthouse Lab in Glasgow | Wellcome Sanger Institute for the COVID-19 Genomics UK (COG-UK) Consortium | Harper VanSteenhouse, Yumi Kasai, David Gray, Carol Clugston, Anna Dominiczak and Alex Alderton, Roberto Amato, Jeffrey Barrett, Sonia Goncalves, Ewan Harrison, David K. Jackson, Ian Johnston, Dominic Kwiatkowski, Cordelia Langford, John Sillitoe on behalf of the Wellcome Sanger Institute COVID-19 Surveillance Team |
| EPI_ISL_2682172 | Lighthouse Lab in Milton Keynes | Wellcome Sanger Institute for the COVID-19 Genomics UK (COG-UK) Consortium | The Lighthouse Lab in Milton Keynes and Alex Alderton, Roberto Amato, Jeffrey Barrett, Sonia Goncalves, Ewan Harrison, David K. Jackson, Ian Johnston, Dominic Kwiatkowski, Cordelia Langford, John Sillitoe on behalf of the Wellcome Sanger Institute COVID-19 Surveillance Team |
| EPI_ISL_2682173 | Lighthouse Lab in Glasgow | Wellcome Sanger Institute for the COVID-19 Genomics UK (COG-UK) Consortium | Harper VanSteenhouse, Yumi Kasai, David Gray, Carol Clugston, Anna Dominiczak and Alex Alderton, Roberto Amato, Jeffrey Barrett, Sonia Goncalves, Ewan Harrison, David K. Jackson, Ian Johnston, Dominic Kwiatkowski, Cordelia Langford, John Sillitoe on behalf of the Wellcome Sanger Institute COVID-19 Surveillance Team |
| EPI_ISL_2682174, EPI_ISL_2682175 | Health Services Laboratories | Wellcome Sanger Institute for the COVID-19 Genomics UK (COG-UK) Consortium | Health Services Laboratories and Alex Alderton, Roberto Amato, Jeffrey Barrett, Sonia Goncalves, Ewan Harrison, David K. Jackson, Ian Johnston, Dominic Kwiatkowski, Cordelia Langford, John Sillitoe on behalf of the Wellcome Sanger Institute COVID-19 Surveillance Team |
| EPI_ISL_2682176 | Lighthouse Lab in Glasgow | Wellcome Sanger Institute for the COVID-19 Genomics UK (COG-UK) Consortium | Harper VanSteenhouse, Yumi Kasai, David Gray, Carol Clugston, Anna Dominiczak and Alex Alderton, Roberto Amato, Jeffrey Barrett, Sonia Goncalves, Ewan Harrison, David K. Jackson, Ian Johnston, Dominic Kwiatkowski, Cordelia Langford, John Sillitoe on behalf of the Wellcome Sanger Institute COVID-19 Surveillance Team |
| EPI_ISL_2682177, EPI_ISL_2682178, EPI_ISL_2682179 | Lighthouse Lab in Milton Keynes | Wellcome Sanger Institute for the COVID-19 Genomics UK (COG-UK) Consortium | The Lighthouse Lab in Milton Keynes and Alex Alderton, Roberto Amato, Jeffrey Barrett, Sonia Goncalves, Ewan Harrison, David K. Jackson, Ian Johnston, Dominic Kwiatkowski, Cordelia Langford, John Sillitoe on behalf of the Wellcome Sanger Institute COVID-19 Surveillance Team |
| EPI_ISL_2682180 | Lighthouse Lab in Glasgow | Wellcome Sanger Institute for the COVID-19 Genomics UK (COG-UK) Consortium | Harper VanSteenhouse, Yumi Kasai, David Gray, Carol Clugston, Anna Dominiczak and Alex Alderton, Roberto Amato, Jeffrey Barrett, Sonia Goncalves, Ewan Harrison, David K. Jackson, Ian Johnston, Dominic Kwiatkowski, Cordelia Langford, John Sillitoe on behalf of the Wellcome Sanger Institute COVID-19 Surveillance Team |
| EPI_ISL_2682181, EPI_ISL_2682182 | Lighthouse Lab in Milton Keynes | Wellcome Sanger Institute for the COVID-19 Genomics UK (COG-UK) Consortium | The Lighthouse Lab in Milton Keynes and Alex Alderton, Roberto Amato, Jeffrey Barrett, Sonia Goncalves, Ewan Harrison, David K. Jackson, Ian Johnston, Dominic Kwiatkowski, Cordelia Langford, John Sillitoe on behalf of the Wellcome Sanger Institute COVID-19 Surveillance Team |
| EPI_ISL_2682183 | Health Services Laboratories | Wellcome Sanger Institute for the COVID-19 Genomics UK (COG-UK) Consortium | Health Services Laboratories and Alex Alderton, Roberto Amato, Jeffrey Barrett, Sonia Goncalves, Ewan Harrison, David K. Jackson, Ian Johnston, Dominic Kwiatkowski, Cordelia Langford, John Sillitoe on behalf of the Wellcome Sanger Institute COVID-19 Surveillance Team |
| EPI_ISL_2682184 | Lighthouse Lab in Milton Keynes | Wellcome Sanger Institute for the COVID-19 Genomics UK (COG-UK) Consortium | The Lighthouse Lab in Milton Keynes and Alex Alderton, Roberto Amato, Jeffrey Barrett, Sonia Goncalves, Ewan Harrison, David K. Jackson, Ian Johnston, Dominic Kwiatkowski, Cordelia Langford, John Sillitoe on behalf of the Wellcome Sanger Institute COVID-19 Surveillance Team |
| EPI_ISL_2682185, EPI_ISL_2682186 | Health Services Laboratories | Wellcome Sanger Institute for the COVID-19 Genomics UK (COG-UK) Consortium | Health Services Laboratories and Alex Alderton, Roberto Amato, Jeffrey Barrett, Sonia Goncalves, Ewan Harrison, David K. Jackson, Ian Johnston, Dominic Kwiatkowski, Cordelia Langford, John Sillitoe on behalf of the Wellcome Sanger Institute COVID-19 Surveillance Team |
| EPI_ISL_2682187, EPI_ISL_2682188 | Lighthouse Lab in Milton Keynes | Wellcome Sanger Institute for the COVID-19 Genomics UK (COG-UK) Consortium | The Lighthouse Lab in Milton Keynes and Alex Alderton, Roberto Amato, Jeffrey Barrett, Sonia Goncalves, Ewan Harrison, David K. Jackson, Ian Johnston, Dominic Kwiatkowski, Cordelia Langford, John Sillitoe on behalf of the Wellcome Sanger Institute COVID-19 Surveillance Team |
| EPI_ISL_2682189 | Health Services Laboratories | Wellcome Sanger Institute for the COVID-19 Genomics UK (COG-UK) Consortium | Health Services Laboratories and Alex Alderton, Roberto Amato, Jeffrey Barrett, Sonia Goncalves, Ewan Harrison, David K. Jackson, Ian Johnston, Dominic Kwiatkowski, Cordelia Langford, John Sillitoe on behalf of the Wellcome Sanger Institute COVID-19 Surveillance Team |
| EPI_ISL_2682190 | Lighthouse Lab in Glasgow | Wellcome Sanger Institute for the COVID-19 Genomics UK (COG-UK) Consortium | Harper VanSteenhouse, Yumi Kasai, David Gray, Carol Clugston, Anna Dominiczak and Alex Alderton, Roberto Amato, Jeffrey Barrett, Sonia Goncalves, Ewan Harrison, David K. Jackson, Ian Johnston, Dominic Kwiatkowski, Cordelia Langford, John Sillitoe on behalf of the Wellcome Sanger Institute COVID-19 Surveillance Team |
| EPI_ISL_2682191 | Health Services Laboratories | Wellcome Sanger Institute for the COVID-19 Genomics UK (COG-UK) Consortium | Health Services Laboratories and Alex Alderton, Roberto Amato, Jeffrey Barrett, Sonia Goncalves, Ewan Harrison, David K. Jackson, Ian Johnston, Dominic Kwiatkowski, Cordelia Langford, John Sillitoe on behalf of the Wellcome Sanger Institute COVID-19 Surveillance Team |
| EPI_ISL_2682192 | Lighthouse Lab in Glasgow | Wellcome Sanger Institute for the COVID-19 Genomics UK (COG-UK) Consortium | Harper VanSteenhouse, Yumi Kasai, David Gray, Carol Clugston, Anna Dominiczak and Alex Alderton, Roberto Amato, Jeffrey Barrett, Sonia Goncalves, Ewan Harrison, David K. Jackson, Ian Johnston, Dominic Kwiatkowski, Cordelia Langford, John Sillitoe on behalf of the Wellcome Sanger Institute COVID-19 Surveillance Team |
| EPI_ISL_2682193 | Lighthouse Lab in Milton Keynes | Wellcome Sanger Institute for the COVID-19 Genomics UK (COG-UK) Consortium | The Lighthouse Lab in Milton Keynes and Alex Alderton, Roberto Amato, Jeffrey Barrett, Sonia Goncalves, Ewan Harrison, David K. Jackson, Ian Johnston, Dominic Kwiatkowski, Cordelia Langford, John Sillitoe on behalf of the Wellcome Sanger Institute COVID-19 Surveillance Team |
| EPI_ISL_2682194, EPI_ISL_2682195 | Health Services Laboratories | Wellcome Sanger Institute for the COVID-19 Genomics UK (COG-UK) Consortium | Health Services Laboratories and Alex Alderton, Roberto Amato, Jeffrey Barrett, Sonia Goncalves, Ewan Harrison, David K. Jackson, Ian Johnston, Dominic Kwiatkowski, Cordelia Langford, John Sillitoe on behalf of the Wellcome Sanger Institute COVID-19 Surveillance Team |
| EPI_ISL_2682196, EPI_ISL_2682197 | Lighthouse Lab in Milton Keynes | Wellcome Sanger Institute for the COVID-19 Genomics UK (COG-UK) Consortium | The Lighthouse Lab in Milton Keynes and Alex Alderton, Roberto Amato, Jeffrey Barrett, Sonia Goncalves, Ewan Harrison, David K. Jackson, Ian Johnston, Dominic Kwiatkowski, Cordelia Langford, John Sillitoe on behalf of the Wellcome Sanger Institute COVID-19 Surveillance Team |
| EPI_ISL_2682198 | Lighthouse Lab in Glasgow | Wellcome Sanger Institute for the COVID-19 Genomics UK (COG-UK) Consortium | Harper VanSteenhouse, Yumi Kasai, David Gray, Carol Clugston, Anna Dominiczak and Alex Alderton, Roberto Amato, Jeffrey Barrett, Sonia Goncalves, Ewan Harrison, David K. Jackson, Ian Johnston, Dominic Kwiatkowski, Cordelia Langford, John Sillitoe on behalf of the Wellcome Sanger Institute COVID-19 Surveillance Team |
| EPI_ISL_2682199, EPI_ISL_2682200 | Health Services Laboratories | Wellcome Sanger Institute for the COVID-19 Genomics UK (COG-UK) Consortium | Health Services Laboratories and Alex Alderton, Roberto Amato, Jeffrey Barrett, Sonia Goncalves, Ewan Harrison, David K. Jackson, Ian Johnston, Dominic Kwiatkowski, Cordelia Langford, John Sillitoe on behalf of the Wellcome Sanger Institute COVID-19 Surveillance Team |

| | | | |
|---|---|---|---|
| EPI_ISL_2682201 | Lighthouse Lab in Milton Keynes | Wellcome Sanger Institute for the COVID-19 Genomics UK (COG-UK) Consortium | The Lighthouse Lab in Milton Keynes and Alex Alderton, Roberto Amato, Jeffrey Barrett, Sonia Goncalves, Ewan Harrison, David K. Jackson, Ian Johnston, Dominic Kwiatkowski, Cordelia Langford, John Sillitoe on behalf of the Wellcome Sanger Institute COVID-19 Surveillance Team |
| EPI_ISL_2682202 | Health Services Laboratories | Wellcome Sanger Institute for the COVID-19 Genomics UK (COG-UK) Consortium | Health Services Laboratories and Alex Alderton, Roberto Amato, Jeffrey Barrett, Sonia Goncalves, Ewan Harrison, David K. Jackson, Ian Johnston, Dominic Kwiatkowski, Cordelia Langford, John Sillitoe on behalf of the Wellcome Sanger Institute COVID-19 Surveillance Team |
| EPI_ISL_2682203 | Lighthouse Lab in Milton Keynes | Wellcome Sanger Institute for the COVID-19 Genomics UK (COG-UK) Consortium | The Lighthouse Lab in Milton Keynes and Alex Alderton, Roberto Amato, Jeffrey Barrett, Sonia Goncalves, Ewan Harrison, David K. Jackson, Ian Johnston, Dominic Kwiatkowski, Cordelia Langford, John Sillitoe on behalf of the Wellcome Sanger Institute COVID-19 Surveillance Team |
| EPI_ISL_2682204 | Lighthouse Lab in Glasgow | Wellcome Sanger Institute for the COVID-19 Genomics UK (COG-UK) Consortium | Harper VanSteenhouse, Yumi Kasai, David Gray, Carol Clugston, Anna Dominiczak and Alex Alderton, Roberto Amato, Jeffrey Barrett, Sonia Goncalves, Ewan Harrison, David K. Jackson, Ian Johnston, Dominic Kwiatkowski, Cordelia Langford, John Sillitoe on behalf of the Wellcome Sanger Institute COVID-19 Surveillance Team |
| EPI_ISL_2682205, EPI_ISL_2682206 | Lighthouse Lab in Milton Keynes | Wellcome Sanger Institute for the COVID-19 Genomics UK (COG-UK) Consortium | The Lighthouse Lab in Milton Keynes and Alex Alderton, Roberto Amato, Jeffrey Barrett, Sonia Goncalves, Ewan Harrison, David K. Jackson, Ian Johnston, Dominic Kwiatkowski, Cordelia Langford, John Sillitoe on behalf of the Wellcome Sanger Institute COVID-19 Surveillance Team |
| EPI_ISL_2682207, EPI_ISL_2682208 | Health Services Laboratories | Wellcome Sanger Institute for the COVID-19 Genomics UK (COG-UK) Consortium | Health Services Laboratories and Alex Alderton, Roberto Amato, Jeffrey Barrett, Sonia Goncalves, Ewan Harrison, David K. Jackson, Ian Johnston, Dominic Kwiatkowski, Cordelia Langford, John Sillitoe on behalf of the Wellcome Sanger Institute COVID-19 Surveillance Team |
| EPI_ISL_2682209 | Lighthouse Lab in Milton Keynes | Wellcome Sanger Institute for the COVID-19 Genomics UK (COG-UK) Consortium | The Lighthouse Lab in Milton Keynes and Alex Alderton, Roberto Amato, Jeffrey Barrett, Sonia Goncalves, Ewan Harrison, David K. Jackson, Ian Johnston, Dominic Kwiatkowski, Cordelia Langford, John Sillitoe on behalf of the Wellcome Sanger Institute COVID-19 Surveillance Team |
| EPI_ISL_2682210, EPI_ISL_2682211 | Health Services Laboratories | Wellcome Sanger Institute for the COVID-19 Genomics UK (COG-UK) Consortium | Health Services Laboratories and Alex Alderton, Roberto Amato, Jeffrey Barrett, Sonia Goncalves, Ewan Harrison, David K. Jackson, Ian Johnston, Dominic Kwiatkowski, Cordelia Langford, John Sillitoe on behalf of the Wellcome Sanger Institute COVID-19 Surveillance Team |
| EPI_ISL_2682212, EPI_ISL_2682213 | Lighthouse Lab in Glasgow | Wellcome Sanger Institute for the COVID-19 Genomics UK (COG-UK) Consortium | Harper VanSteenhouse, Yumi Kasai, David Gray, Carol Clugston, Anna Dominiczak and Alex Alderton, Roberto Amato, Jeffrey Barrett, Sonia Goncalves, Ewan Harrison, David K. Jackson, Ian Johnston, Dominic Kwiatkowski, Cordelia Langford, John Sillitoe on behalf of the Wellcome Sanger Institute COVID-19 Surveillance Team |
| EPI_ISL_2682214 | Lighthouse Lab in Milton Keynes | Wellcome Sanger Institute for the COVID-19 Genomics UK (COG-UK) Consortium | The Lighthouse Lab in Milton Keynes and Alex Alderton, Roberto Amato, Jeffrey Barrett, Sonia Goncalves, Ewan Harrison, David K. Jackson, Ian Johnston, Dominic Kwiatkowski, Cordelia Langford, John Sillitoe on behalf of the Wellcome Sanger Institute COVID-19 Surveillance Team |
| EPI_ISL_2682215, EPI_ISL_2682216 | Health Services Laboratories | Wellcome Sanger Institute for the COVID-19 Genomics UK (COG-UK) Consortium | Health Services Laboratories and Alex Alderton, Roberto Amato, Jeffrey Barrett, Sonia Goncalves, Ewan Harrison, David K. Jackson, Ian Johnston, Dominic Kwiatkowski, Cordelia Langford, John Sillitoe on behalf of the Wellcome Sanger Institute COVID-19 Surveillance Team |
| EPI_ISL_2682217 | Lighthouse Lab in Milton Keynes | Wellcome Sanger Institute for the COVID-19 Genomics UK (COG-UK) Consortium | The Lighthouse Lab in Milton Keynes and Alex Alderton, Roberto Amato, Jeffrey Barrett, Sonia Goncalves, Ewan Harrison, David K. Jackson, Ian Johnston, Dominic Kwiatkowski, Cordelia Langford, John Sillitoe on behalf of the Wellcome Sanger Institute COVID-19 Surveillance Team |
| EPI_ISL_2682218 | Health Services Laboratories | Wellcome Sanger Institute for the COVID-19 Genomics UK (COG-UK) Consortium | Health Services Laboratories and Alex Alderton, Roberto Amato, Jeffrey Barrett, Sonia Goncalves, Ewan Harrison, David K. Jackson, Ian Johnston, Dominic Kwiatkowski, Cordelia Langford, John Sillitoe on behalf of the Wellcome Sanger Institute COVID-19 Surveillance Team |
| EPI_ISL_2682219, EPI_ISL_2682220 | Lighthouse Lab in Milton Keynes | Wellcome Sanger Institute for the COVID-19 Genomics UK (COG-UK) Consortium | The Lighthouse Lab in Milton Keynes and Alex Alderton, Roberto Amato, Jeffrey Barrett, Sonia Goncalves, Ewan Harrison, David K. Jackson, Ian Johnston, Dominic Kwiatkowski, Cordelia Langford, John Sillitoe on behalf of the Wellcome Sanger Institute COVID-19 Surveillance Team |
| EPI_ISL_2682221, EPI_ISL_2682222 | Health Services Laboratories | Wellcome Sanger Institute for the COVID-19 Genomics UK (COG-UK) Consortium | Health Services Laboratories and Alex Alderton, Roberto Amato, Jeffrey Barrett, Sonia Goncalves, Ewan Harrison, David K. Jackson, Ian Johnston, Dominic Kwiatkowski, Cordelia Langford, John Sillitoe on behalf of the Wellcome Sanger Institute COVID-19 Surveillance Team |
| EPI_ISL_2682223, EPI_ISL_2682224 | Lighthouse Lab in Milton Keynes | Wellcome Sanger Institute for the COVID-19 Genomics UK (COG-UK) Consortium | The Lighthouse Lab in Milton Keynes and Alex Alderton, Roberto Amato, Jeffrey Barrett, Sonia Goncalves, Ewan Harrison, David K. Jackson, Ian Johnston, Dominic Kwiatkowski, Cordelia Langford, John Sillitoe on behalf of the Wellcome Sanger Institute COVID-19 Surveillance Team |
| EPI_ISL_2682225 | Health Services Laboratories | Wellcome Sanger Institute for the COVID-19 Genomics UK (COG-UK) Consortium | Health Services Laboratories and Alex Alderton, Roberto Amato, Jeffrey Barrett, Sonia Goncalves, Ewan Harrison, David K. Jackson, Ian Johnston, Dominic Kwiatkowski, Cordelia Langford, John Sillitoe on behalf of the Wellcome Sanger Institute COVID-19 Surveillance Team |
| EPI_ISL_2682226 | Lighthouse Lab in Glasgow | Wellcome Sanger Institute for the COVID-19 Genomics UK (COG-UK) Consortium | Harper VanSteenhouse, Yumi Kasai, David Gray, Carol Clugston, Anna Dominiczak and Alex Alderton, Roberto Amato, Jeffrey Barrett, Sonia Goncalves, Ewan Harrison, David K. Jackson, Ian Johnston, Dominic Kwiatkowski, Cordelia Langford, John Sillitoe on behalf of the Wellcome Sanger Institute COVID-19 Surveillance Team |
| EPI_ISL_2682227, EPI_ISL_2682228 | Health Services Laboratories | Wellcome Sanger Institute for the COVID-19 Genomics UK (COG-UK) Consortium | Health Services Laboratories and Alex Alderton, Roberto Amato, Jeffrey Barrett, Sonia Goncalves, Ewan Harrison, David K. Jackson, Ian Johnston, Dominic Kwiatkowski, Cordelia Langford, John Sillitoe on behalf of the Wellcome Sanger Institute COVID-19 Surveillance Team |
| EPI_ISL_2682229, EPI_ISL_2682230 | Lighthouse Lab in Glasgow | Wellcome Sanger Institute for the COVID-19 Genomics UK (COG-UK) Consortium | Harper VanSteenhouse, Yumi Kasai, David Gray, Carol Clugston, Anna Dominiczak and Alex Alderton, Roberto Amato, Jeffrey Barrett, Sonia Goncalves, Ewan Harrison, David K. Jackson, Ian Johnston, Dominic Kwiatkowski, Cordelia Langford, John Sillitoe on behalf of the Wellcome Sanger Institute COVID-19 Surveillance Team |
| EPI_ISL_2682231, EPI_ISL_2682232 | Health Services Laboratories | Wellcome Sanger Institute for the COVID-19 Genomics UK (COG-UK) Consortium | Health Services Laboratories and Alex Alderton, Roberto Amato, Jeffrey Barrett, Sonia Goncalves, Ewan Harrison, David K. Jackson, Ian Johnston, Dominic Kwiatkowski, Cordelia Langford, John Sillitoe on behalf of the Wellcome Sanger Institute COVID-19 Surveillance Team |
| EPI_ISL_2682233 | Lighthouse Lab in Glasgow | Wellcome Sanger Institute for the COVID-19 Genomics UK (COG-UK) Consortium | Harper VanSteenhouse, Yumi Kasai, David Gray, Carol Clugston, Anna Dominiczak and Alex Alderton, Roberto Amato, Jeffrey Barrett, Sonia Goncalves, Ewan Harrison, David K. Jackson, Ian Johnston, Dominic Kwiatkowski, Cordelia Langford, John Sillitoe on behalf of the Wellcome Sanger Institute COVID-19 Surveillance Team |
| EPI_ISL_2682234, EPI_ISL_2682235, EPI_ISL_2682236 | Lighthouse Lab in Milton Keynes | Wellcome Sanger Institute for the COVID-19 Genomics UK (COG-UK) Consortium | The Lighthouse Lab in Milton Keynes and Alex Alderton, Roberto Amato, Jeffrey Barrett, Sonia Goncalves, Ewan Harrison, David K. Jackson, Ian Johnston, Dominic Kwiatkowski, Cordelia Langford, John Sillitoe on behalf of the Wellcome Sanger Institute COVID-19 Surveillance Team |
| EPI_ISL_2682237, EPI_ISL_2682238 | Health Services Laboratories | Wellcome Sanger Institute for the COVID-19 Genomics UK (COG-UK) Consortium | Health Services Laboratories and Alex Alderton, Roberto Amato, Jeffrey Barrett, Sonia Goncalves, Ewan Harrison, David K. Jackson, Ian Johnston, Dominic Kwiatkowski, Cordelia Langford, John Sillitoe on behalf of the Wellcome Sanger Institute COVID-19 Surveillance Team |
| EPI_ISL_2682239 | Lighthouse Lab in Milton Keynes | Wellcome Sanger Institute for the COVID-19 Genomics UK (COG-UK) Consortium | The Lighthouse Lab in Milton Keynes and Alex Alderton, Roberto Amato, Jeffrey Barrett, Sonia Goncalves, Ewan Harrison, David K. Jackson, Ian Johnston, Dominic Kwiatkowski, Cordelia Langford, John Sillitoe on behalf of the Wellcome Sanger Institute COVID-19 Surveillance Team |
| EPI_ISL_2682240, EPI_ISL_2682241, EPI_ISL_2682242 | Health Services Laboratories | Wellcome Sanger Institute for the COVID-19 Genomics UK (COG-UK) Consortium | Health Services Laboratories and Alex Alderton, Roberto Amato, Jeffrey Barrett, Sonia Goncalves, Ewan Harrison, David K. Jackson, Ian Johnston, Dominic Kwiatkowski, Cordelia Langford, John Sillitoe on behalf of the Wellcome Sanger Institute COVID-19 Surveillance Team |
| EPI_ISL_2682243 | Lighthouse Lab in Milton Keynes | Wellcome Sanger Institute for the COVID-19 Genomics UK (COG-UK) Consortium | The Lighthouse Lab in Milton Keynes and Alex Alderton, Roberto Amato, Jeffrey Barrett, Sonia Goncalves, Ewan Harrison, David K. Jackson, Ian Johnston, Dominic Kwiatkowski, Cordelia Langford, John Sillitoe on behalf of the Wellcome Sanger Institute COVID-19 Surveillance Team |
| EPI_ISL_2682244, EPI_ISL_2682245 | Health Services Laboratories | Wellcome Sanger Institute for the COVID-19 Genomics UK (COG-UK) Consortium | Health Services Laboratories and Alex Alderton, Roberto Amato, Jeffrey Barrett, Sonia Goncalves, Ewan Harrison, David K. Jackson, Ian Johnston, Dominic Kwiatkowski, Cordelia Langford, John Sillitoe on behalf of the Wellcome Sanger Institute COVID-19 Surveillance Team |
| EPI_ISL_2682246, EPI_ISL_2682247 | Lighthouse Lab in Milton Keynes | Wellcome Sanger Institute for the COVID-19 Genomics UK (COG-UK) Consortium | The Lighthouse Lab in Milton Keynes and Alex Alderton, Roberto Amato, Jeffrey Barrett, Sonia Goncalves, Ewan Harrison, David K. Jackson, Ian Johnston, Dominic Kwiatkowski, Cordelia Langford, John Sillitoe on behalf of the Wellcome Sanger Institute COVID-19 Surveillance Team |
| EPI_ISL_2682248, EPI_ISL_2682249 | Health Services Laboratories | Wellcome Sanger Institute for the COVID-19 Genomics UK (COG-UK) Consortium | Health Services Laboratories and Alex Alderton, Roberto Amato, Jeffrey Barrett, Sonia Goncalves, Ewan Harrison, David K. Jackson, Ian Johnston, Dominic Kwiatkowski, Cordelia Langford, John Sillitoe on behalf of the Wellcome Sanger Institute COVID-19 Surveillance Team |
| EPI_ISL_2682250 | Lighthouse Lab in Milton Keynes | Wellcome Sanger Institute for the COVID-19 Genomics UK (COG-UK) Consortium | The Lighthouse Lab in Milton Keynes and Alex Alderton, Roberto Amato, Jeffrey Barrett, Sonia Goncalves, Ewan Harrison, David K. Jackson, Ian Johnston, Dominic Kwiatkowski, Cordelia Langford, John Sillitoe on behalf of the Wellcome Sanger Institute COVID-19 Surveillance Team |
| EPI_ISL_2682251 | Health Services Laboratories | Wellcome Sanger Institute for the COVID-19 Genomics UK (COG-UK) Consortium | Health Services Laboratories and Alex Alderton, Roberto Amato, Jeffrey Barrett, Sonia Goncalves, Ewan Harrison, David K. Jackson, Ian Johnston, Dominic Kwiatkowski, Cordelia Langford, John Sillitoe on behalf of the Wellcome Sanger Institute COVID-19 Surveillance Team |
| EPI_ISL_2682252 | Lighthouse Lab in Glasgow | Wellcome Sanger Institute for the COVID-19 Genomics UK (COG-UK) Consortium | Harper VanSteenhouse, Yumi Kasai, David Gray, Carol Clugston, Anna Dominiczak and Alex Alderton, Roberto Amato, Jeffrey Barrett, Sonia Goncalves, Ewan Harrison, David K. Jackson, Ian Johnston, Dominic Kwiatkowski, Cordelia Langford, John Sillitoe on behalf of the Wellcome Sanger Institute COVID-19 Surveillance Team |
| EPI_ISL_2682253 | Lighthouse Lab in Milton Keynes | Wellcome Sanger Institute for the COVID-19 Genomics UK (COG-UK) Consortium | The Lighthouse Lab in Milton Keynes and Alex Alderton, Roberto Amato, Jeffrey Barrett, Sonia Goncalves, Ewan Harrison, David K. Jackson, Ian Johnston, Dominic Kwiatkowski, Cordelia Langford, John Sillitoe on behalf of the Wellcome Sanger Institute COVID-19 Surveillance Team |
| EPI_ISL_2682254 | Health Services Laboratories | Wellcome Sanger Institute for the COVID-19 Genomics UK (COG-UK) Consortium | Health Services Laboratories and Alex Alderton, Roberto Amato, Jeffrey Barrett, Sonia Goncalves, Ewan Harrison, David K. Jackson, Ian Johnston, Dominic Kwiatkowski, Cordelia Langford, John Sillitoe on behalf of the Wellcome Sanger Institute COVID-19 Surveillance Team |
| EPI_ISL_2682255 | Lighthouse Lab in Milton Keynes | Wellcome Sanger Institute for the COVID-19 Genomics UK (COG-UK) Consortium | The Lighthouse Lab in Milton Keynes and Alex Alderton, Roberto Amato, Jeffrey Barrett, Sonia Goncalves, Ewan Harrison, David K. Jackson, Ian Johnston, Dominic Kwiatkowski, Cordelia Langford, John Sillitoe on behalf of the Wellcome Sanger Institute COVID-19 Surveillance Team |
| EPI_ISL_2682256 | Lighthouse Lab in Glasgow | Wellcome Sanger Institute for the COVID-19 Genomics UK (COG-UK) Consortium | Harper VanSteenhouse, Yumi Kasai, David Gray, Carol Clugston, Anna Dominiczak and Alex Alderton, Roberto Amato, Jeffrey Barrett, Sonia Goncalves, Ewan Harrison, David K. Jackson, Ian Johnston, Dominic Kwiatkowski, Cordelia Langford, John Sillitoe on behalf of the Wellcome Sanger Institute COVID-19 Surveillance Team |
| EPI_ISL_2682257 | Lighthouse Lab in Milton Keynes | Wellcome Sanger Institute for the COVID-19 Genomics UK (COG-UK) Consortium | The Lighthouse Lab in Milton Keynes and Alex Alderton, Roberto Amato, Jeffrey Barrett, Sonia Goncalves, Ewan Harrison, David K. Jackson, Ian Johnston, Dominic Kwiatkowski, Cordelia Langford, John Sillitoe on behalf of the Wellcome Sanger Institute COVID-19 Surveillance Team |
| EPI_ISL_2682258 | Health Services Laboratories | Wellcome Sanger Institute for the COVID-19 Genomics UK (COG-UK) Consortium | Health Services Laboratories and Alex Alderton, Roberto Amato, Jeffrey Barrett, Sonia Goncalves, Ewan Harrison, David K. Jackson, Ian Johnston, Dominic Kwiatkowski, Cordelia Langford, John Sillitoe on behalf of the Wellcome Sanger Institute COVID-19 Surveillance Team |
| EPI_ISL_2682259 | Lighthouse Lab in Milton Keynes | Wellcome Sanger Institute for the COVID-19 Genomics UK (COG-UK) Consortium | The Lighthouse Lab in Milton Keynes and Alex Alderton, Roberto Amato, Jeffrey Barrett, Sonia Goncalves, Ewan Harrison, David K. Jackson, Ian Johnston, Dominic Kwiatkowski, Cordelia Langford, John Sillitoe on behalf of the Wellcome Sanger Institute COVID-19 Surveillance Team |
| EPI_ISL_2682260 | Lighthouse Lab in Glasgow | Wellcome Sanger Institute for the COVID-19 Genomics UK (COG-UK) Consortium | Harper VanSteenhouse, Yumi Kasai, David Gray, Carol Clugston, Anna Dominiczak and Alex Alderton, Roberto Amato, Jeffrey Barrett, Sonia Goncalves, Ewan Harrison, David K. Jackson, Ian Johnston, Dominic Kwiatkowski, Cordelia Langford, John Sillitoe on behalf of the Wellcome Sanger Institute COVID-19 Surveillance Team |
| EPI_ISL_2682261 | Health Services Laboratories | Wellcome Sanger Institute for the COVID-19 Genomics UK (COG-UK) Consortium | Health Services Laboratories and Alex Alderton, Roberto Amato, Jeffrey Barrett, Sonia Goncalves, Ewan Harrison, David K. Jackson, Ian Johnston, Dominic Kwiatkowski, Cordelia Langford, John Sillitoe on behalf of the Wellcome Sanger Institute COVID-19 Surveillance Team |
| EPI_ISL_2682262 | Lighthouse Lab in Glasgow | Wellcome Sanger Institute for the COVID-19 Genomics UK (COG-UK) Consortium | Harper VanSteenhouse, Yumi Kasai, David Gray, Carol Clugston, Anna Dominiczak and Alex Alderton, Roberto Amato, Jeffrey Barrett, Sonia Goncalves, Ewan Harrison, David K. Jackson, Ian Johnston, Dominic Kwiatkowski, Cordelia Langford, John Sillitoe on behalf of the Wellcome Sanger Institute COVID-19 Surveillance Team |
| EPI_ISL_2682263, EPI_ISL_2682264 | Lighthouse Lab in Milton Keynes | Wellcome Sanger Institute for the COVID-19 Genomics UK (COG-UK) Consortium | The Lighthouse Lab in Milton Keynes and Alex Alderton, Roberto Amato, Jeffrey Barrett, Sonia Goncalves, Ewan Harrison, David K. Jackson, Ian Johnston, Dominic Kwiatkowski, Cordelia Langford, John Sillitoe on behalf of the Wellcome Sanger Institute COVID-19 Surveillance Team |

| | | | |
|---|---|---|---|
| EPI_ISL_2682265 | Lighthouse Lab in Glasgow | Wellcome Sanger Institute for the COVID-19 Genomics UK (COG-UK) Consortium | Harper VanSteenhouse, Yumi Kasai, David Gray, Carol Clugston, Anna Dominiczak and Alex Alderton, Roberto Amato, Jeffrey Barrett, Sonia Goncalves, Ewan Harrison, David K. Jackson, Ian Johnston, Dominic Kwiatkowski, Cordelia Langford, John Sillitoe on behalf of the Wellcome Sanger Institute COVID-19 Surveillance Team |
| EPI_ISL_2682266 | Health Services Laboratories | Wellcome Sanger Institute for the COVID-19 Genomics UK (COG-UK) Consortium | Health Services Laboratories and Alex Alderton, Roberto Amato, Jeffrey Barrett, Sonia Goncalves, Ewan Harrison, David K. Jackson, Ian Johnston, Dominic Kwiatkowski, Cordelia Langford, John Sillitoe on behalf of the Wellcome Sanger Institute COVID-19 Surveillance Team |
| EPI_ISL_2682267 | Lighthouse Lab in Milton Keynes | Wellcome Sanger Institute for the COVID-19 Genomics UK (COG-UK) Consortium | The Lighthouse Lab in Milton Keynes and Alex Alderton, Roberto Amato, Jeffrey Barrett, Sonia Goncalves, Ewan Harrison, David K. Jackson, Ian Johnston, Dominic Kwiatkowski, Cordelia Langford, John Sillitoe on behalf of the Wellcome Sanger Institute COVID-19 Surveillance Team |
| EPI_ISL_2682268, EPI_ISL_2682269, EPI_ISL_2682270, EPI_ISL_2682271 | Health Services Laboratories | Wellcome Sanger Institute for the COVID-19 Genomics UK (COG-UK) Consortium | Health Services Laboratories and Alex Alderton, Roberto Amato, Jeffrey Barrett, Sonia Goncalves, Ewan Harrison, David K. Jackson, Ian Johnston, Dominic Kwiatkowski, Cordelia Langford, John Sillitoe on behalf of the Wellcome Sanger Institute COVID-19 Surveillance Team |
| EPI_ISL_2682272 | Lighthouse Lab in Milton Keynes | Wellcome Sanger Institute for the COVID-19 Genomics UK (COG-UK) Consortium | The Lighthouse Lab in Milton Keynes and Alex Alderton, Roberto Amato, Jeffrey Barrett, Sonia Goncalves, Ewan Harrison, David K. Jackson, Ian Johnston, Dominic Kwiatkowski, Cordelia Langford, John Sillitoe on behalf of the Wellcome Sanger Institute COVID-19 Surveillance Team |
| EPI_ISL_2682273 | Health Services Laboratories | Wellcome Sanger Institute for the COVID-19 Genomics UK (COG-UK) Consortium | Health Services Laboratories and Alex Alderton, Roberto Amato, Jeffrey Barrett, Sonia Goncalves, Ewan Harrison, David K. Jackson, Ian Johnston, Dominic Kwiatkowski, Cordelia Langford, John Sillitoe on behalf of the Wellcome Sanger Institute COVID-19 Surveillance Team |
| EPI_ISL_2682274 | Lighthouse Lab in Glasgow | Wellcome Sanger Institute for the COVID-19 Genomics UK (COG-UK) Consortium | Harper VanSteenhouse, Yumi Kasai, David Gray, Carol Clugston, Anna Dominiczak and Alex Alderton, Roberto Amato, Jeffrey Barrett, Sonia Goncalves, Ewan Harrison, David K. Jackson, Ian Johnston, Dominic Kwiatkowski, Cordelia Langford, John Sillitoe on behalf of the Wellcome Sanger Institute COVID-19 Surveillance Team |
| EPI_ISL_2682275 | Lighthouse Lab in Milton Keynes | Wellcome Sanger Institute for the COVID-19 Genomics UK (COG-UK) Consortium | The Lighthouse Lab in Milton Keynes and Alex Alderton, Roberto Amato, Jeffrey Barrett, Sonia Goncalves, Ewan Harrison, David K. Jackson, Ian Johnston, Dominic Kwiatkowski, Cordelia Langford, John Sillitoe on behalf of the Wellcome Sanger Institute COVID-19 Surveillance Team |
| EPI_ISL_2682276 | Lighthouse Lab in Glasgow | Wellcome Sanger Institute for the COVID-19 Genomics UK (COG-UK) Consortium | Harper VanSteenhouse, Yumi Kasai, David Gray, Carol Clugston, Anna Dominiczak and Alex Alderton, Roberto Amato, Jeffrey Barrett, Sonia Goncalves, Ewan Harrison, David K. Jackson, Ian Johnston, Dominic Kwiatkowski, Cordelia Langford, John Sillitoe on behalf of the Wellcome Sanger Institute COVID-19 Surveillance Team |
| EPI_ISL_2682277 | Lighthouse Lab in Milton Keynes | Wellcome Sanger Institute for the COVID-19 Genomics UK (COG-UK) Consortium | The Lighthouse Lab in Milton Keynes and Alex Alderton, Roberto Amato, Jeffrey Barrett, Sonia Goncalves, Ewan Harrison, David K. Jackson, Ian Johnston, Dominic Kwiatkowski, Cordelia Langford, John Sillitoe on behalf of the Wellcome Sanger Institute COVID-19 Surveillance Team |
| EPI_ISL_2682278, EPI_ISL_2682279 | Health Services Laboratories | Wellcome Sanger Institute for the COVID-19 Genomics UK (COG-UK) Consortium | Health Services Laboratories and Alex Alderton, Roberto Amato, Jeffrey Barrett, Sonia Goncalves, Ewan Harrison, David K. Jackson, Ian Johnston, Dominic Kwiatkowski, Cordelia Langford, John Sillitoe on behalf of the Wellcome Sanger Institute COVID-19 Surveillance Team |
| EPI_ISL_2682280, EPI_ISL_2682281 | Lighthouse Lab in Milton Keynes | Wellcome Sanger Institute for the COVID-19 Genomics UK (COG-UK) Consortium | The Lighthouse Lab in Milton Keynes and Alex Alderton, Roberto Amato, Jeffrey Barrett, Sonia Goncalves, Ewan Harrison, David K. Jackson, Ian Johnston, Dominic Kwiatkowski, Cordelia Langford, John Sillitoe on behalf of the Wellcome Sanger Institute COVID-19 Surveillance Team |
| EPI_ISL_2682282, EPI_ISL_2682283 | Health Services Laboratories | Wellcome Sanger Institute for the COVID-19 Genomics UK (COG-UK) Consortium | Health Services Laboratories and Alex Alderton, Roberto Amato, Jeffrey Barrett, Sonia Goncalves, Ewan Harrison, David K. Jackson, Ian Johnston, Dominic Kwiatkowski, Cordelia Langford, John Sillitoe on behalf of the Wellcome Sanger Institute COVID-19 Surveillance Team |
| EPI_ISL_2682284 | Lighthouse Lab in Glasgow | Wellcome Sanger Institute for the COVID-19 Genomics UK (COG-UK) Consortium | Harper VanSteenhouse, Yumi Kasai, David Gray, Carol Clugston, Anna Dominiczak and Alex Alderton, Roberto Amato, Jeffrey Barrett, Sonia Goncalves, Ewan Harrison, David K. Jackson, Ian Johnston, Dominic Kwiatkowski, Cordelia Langford, John Sillitoe on behalf of the Wellcome Sanger Institute COVID-19 Surveillance Team |
| EPI_ISL_2682285, EPI_ISL_2682286, EPI_ISL_2682287 | Lighthouse Lab in Alderley Park | Wellcome Sanger Institute for the COVID-19 Genomics UK (COG-UK) Consortium | Jacquelyn Wynn, Mairead Hyland, The Lighthouse Lab in Alderley Park and Alex Alderton, Roberto Amato, Jeffrey Barrett, Sonia Goncalves, Ewan Harrison, David K. Jackson, Ian Johnston, Dominic Kwiatkowski, Cordelia Langford, John Sillitoe on behalf of the Wellcome Sanger Institute COVID-19 Surveillance Team |
| EPI_ISL_2682288 | Health Services Laboratories | Wellcome Sanger Institute for the COVID-19 Genomics UK (COG-UK) Consortium | Health Services Laboratories and Alex Alderton, Roberto Amato, Jeffrey Barrett, Sonia Goncalves, Ewan Harrison, David K. Jackson, Ian Johnston, Dominic Kwiatkowski, Cordelia Langford, John Sillitoe on behalf of the Wellcome Sanger Institute COVID-19 Surveillance Team |
| EPI_ISL_2682289, EPI_ISL_2682290 | Lighthouse Lab in Alderley Park | Wellcome Sanger Institute for the COVID-19 Genomics UK (COG-UK) Consortium | Jacquelyn Wynn, Mairead Hyland, The Lighthouse Lab in Alderley Park and Alex Alderton, Roberto Amato, Jeffrey Barrett, Sonia Goncalves, Ewan Harrison, David K. Jackson, Ian Johnston, Dominic Kwiatkowski, Cordelia Langford, John Sillitoe on behalf of the Wellcome Sanger Institute COVID-19 Surveillance Team |
| EPI_ISL_2682291, EPI_ISL_2682292 | Health Services Laboratories | Wellcome Sanger Institute for the COVID-19 Genomics UK (COG-UK) Consortium | Health Services Laboratories and Alex Alderton, Roberto Amato, Jeffrey Barrett, Sonia Goncalves, Ewan Harrison, David K. Jackson, Ian Johnston, Dominic Kwiatkowski, Cordelia Langford, John Sillitoe on behalf of the Wellcome Sanger Institute COVID-19 Surveillance Team |
| EPI_ISL_2682293, EPI_ISL_2682294, EPI_ISL_2682295 | Lighthouse Lab in Alderley Park | Wellcome Sanger Institute for the COVID-19 Genomics UK (COG-UK) Consortium | Jacquelyn Wynn, Mairead Hyland, The Lighthouse Lab in Alderley Park and Alex Alderton, Roberto Amato, Jeffrey Barrett, Sonia Goncalves, Ewan Harrison, David K. Jackson, Ian Johnston, Dominic Kwiatkowski, Cordelia Langford, John Sillitoe on behalf of the Wellcome Sanger Institute COVID-19 Surveillance Team |
| EPI_ISL_2682296 | Health Services Laboratories | Wellcome Sanger Institute for the COVID-19 Genomics UK (COG-UK) Consortium | Health Services Laboratories and Alex Alderton, Roberto Amato, Jeffrey Barrett, Sonia Goncalves, Ewan Harrison, David K. Jackson, Ian Johnston, Dominic Kwiatkowski, Cordelia Langford, John Sillitoe on behalf of the Wellcome Sanger Institute COVID-19 Surveillance Team |
| EPI_ISL_2682297, EPI_ISL_2682298, EPI_ISL_2682299, EPI_ISL_2682300, EPI_ISL_2682301, EPI_ISL_2682302, EPI_ISL_2682303, EPI_ISL_2682304, EPI_ISL_2682305, EPI_ISL_2682306, EPI_ISL_2682307, EPI_ISL_2682308, EPI_ISL_2682309, EPI_ISL_2682310, EPI_ISL_2682311, EPI_ISL_2682312 | | | |
| see above | Lighthouse Lab in Alderley Park | Wellcome Sanger Institute for the COVID-19 Genomics UK (COG-UK) Consortium | Jacquelyn Wynn, Mairead Hyland, The Lighthouse Lab in Alderley Park and Alex Alderton, Roberto Amato, Jeffrey Barrett, Sonia Goncalves, Ewan Harrison, David K. Jackson, Ian Johnston, Dominic Kwiatkowski, Cordelia Langford, John Sillitoe on behalf of the Wellcome Sanger Institute COVID-19 Surveillance Team |
| EPI_ISL_2682313 | Health Services Laboratories | Wellcome Sanger Institute for the COVID-19 Genomics UK (COG-UK) Consortium | Health Services Laboratories and Alex Alderton, Roberto Amato, Jeffrey Barrett, Sonia Goncalves, Ewan Harrison, David K. Jackson, Ian Johnston, Dominic Kwiatkowski, Cordelia Langford, John Sillitoe on behalf of the Wellcome Sanger Institute COVID-19 Surveillance Team |
| EPI_ISL_2682314, EPI_ISL_2682315, EPI_ISL_2682316, EPI_ISL_2682317, EPI_ISL_2682318, EPI_ISL_2682319 | Lighthouse Lab in Alderley Park | Wellcome Sanger Institute for the COVID-19 Genomics UK (COG-UK) Consortium | Jacquelyn Wynn, Mairead Hyland, The Lighthouse Lab in Alderley Park and Alex Alderton, Roberto Amato, Jeffrey Barrett, Sonia Goncalves, Ewan Harrison, David K. Jackson, Ian Johnston, Dominic Kwiatkowski, Cordelia Langford, John Sillitoe on behalf of the Wellcome Sanger Institute COVID-19 Surveillance Team |
| EPI_ISL_2682320 | Health Services Laboratories | Wellcome Sanger Institute for the COVID-19 Genomics UK (COG-UK) Consortium | Health Services Laboratories and Alex Alderton, Roberto Amato, Jeffrey Barrett, Sonia Goncalves, Ewan Harrison, David K. Jackson, Ian Johnston, Dominic Kwiatkowski, Cordelia Langford, John Sillitoe on behalf of the Wellcome Sanger Institute COVID-19 Surveillance Team |
| EPI_ISL_2682321, EPI_ISL_2682322, EPI_ISL_2682323 | Lighthouse Lab in Alderley Park | Wellcome Sanger Institute for the COVID-19 Genomics UK (COG-UK) Consortium | Jacquelyn Wynn, Mairead Hyland, The Lighthouse Lab in Alderley Park and Alex Alderton, Roberto Amato, Jeffrey Barrett, Sonia Goncalves, Ewan Harrison, David K. Jackson, Ian Johnston, Dominic Kwiatkowski, Cordelia Langford, John Sillitoe on behalf of the Wellcome Sanger Institute COVID-19 Surveillance Team |
| EPI_ISL_2682325 | Health Services Laboratories | Wellcome Sanger Institute for the COVID-19 Genomics UK (COG-UK) Consortium | Health Services Laboratories and Alex Alderton, Roberto Amato, Jeffrey Barrett, Sonia Goncalves, Ewan Harrison, David K. Jackson, Ian Johnston, Dominic Kwiatkowski, Cordelia Langford, John Sillitoe on behalf of the Wellcome Sanger Institute COVID-19 Surveillance Team |
| EPI_ISL_2682327, EPI_ISL_2682328, EPI_ISL_2682329, EPI_ISL_2682330, EPI_ISL_2682331, EPI_ISL_2682332 | Lighthouse Lab in Alderley Park | Wellcome Sanger Institute for the COVID-19 Genomics UK (COG-UK) Consortium | Jacquelyn Wynn, Mairead Hyland, The Lighthouse Lab in Alderley Park and Alex Alderton, Roberto Amato, Jeffrey Barrett, Sonia Goncalves, Ewan Harrison, David K. Jackson, Ian Johnston, Dominic Kwiatkowski, Cordelia Langford, John Sillitoe on behalf of the Wellcome Sanger Institute COVID-19 Surveillance Team |
| EPI_ISL_2682333, EPI_ISL_2682334 | Health Services Laboratories | Wellcome Sanger Institute for the COVID-19 Genomics UK (COG-UK) Consortium | Health Services Laboratories and Alex Alderton, Roberto Amato, Jeffrey Barrett, Sonia Goncalves, Ewan Harrison, David K. Jackson, Ian Johnston, Dominic Kwiatkowski, Cordelia Langford, John Sillitoe on behalf of the Wellcome Sanger Institute COVID-19 Surveillance Team |
| EPI_ISL_2682335, EPI_ISL_2682336, EPI_ISL_2682337 | Lighthouse Lab in Alderley Park | Wellcome Sanger Institute for the COVID-19 Genomics UK (COG-UK) Consortium | Jacquelyn Wynn, Mairead Hyland, The Lighthouse Lab in Alderley Park and Alex Alderton, Roberto Amato, Jeffrey Barrett, Sonia Goncalves, Ewan Harrison, David K. Jackson, Ian Johnston, Dominic Kwiatkowski, Cordelia Langford, John Sillitoe on behalf of the Wellcome Sanger Institute COVID-19 Surveillance Team |
| EPI_ISL_2682338 | Health Services Laboratories | Wellcome Sanger Institute for the COVID-19 Genomics UK (COG-UK) Consortium | Health Services Laboratories and Alex Alderton, Roberto Amato, Jeffrey Barrett, Sonia Goncalves, Ewan Harrison, David K. Jackson, Ian Johnston, Dominic Kwiatkowski, Cordelia Langford, John Sillitoe on behalf of the Wellcome Sanger Institute COVID-19 Surveillance Team |
| EPI_ISL_2682339, EPI_ISL_2682340, EPI_ISL_2682341, EPI_ISL_2682342, EPI_ISL_2682343, EPI_ISL_2682344, EPI_ISL_2682345, EPI_ISL_2682346, EPI_ISL_2682347, EPI_ISL_2682348, EPI_ISL_2682349 | | | |
| see above | Lighthouse Lab in Alderley Park | Wellcome Sanger Institute for the COVID-19 Genomics UK (COG-UK) Consortium | Jacquelyn Wynn, Mairead Hyland, The Lighthouse Lab in Alderley Park and Alex Alderton, Roberto Amato, Jeffrey Barrett, Sonia Goncalves, Ewan Harrison, David K. Jackson, Ian Johnston, Dominic Kwiatkowski, Cordelia Langford, John Sillitoe on behalf of the Wellcome Sanger Institute COVID-19 Surveillance Team |
| EPI_ISL_2682350, EPI_ISL_2682351 | Health Services Laboratories | Wellcome Sanger Institute for the COVID-19 Genomics UK (COG-UK) Consortium | Health Services Laboratories and Alex Alderton, Roberto Amato, Jeffrey Barrett, Sonia Goncalves, Ewan Harrison, David K. Jackson, Ian Johnston, Dominic Kwiatkowski, Cordelia Langford, John Sillitoe on behalf of the Wellcome Sanger Institute COVID-19 Surveillance Team |
| EPI_ISL_2682352, EPI_ISL_2682353 | Lighthouse Lab in Alderley Park | Wellcome Sanger Institute for the COVID-19 Genomics UK (COG-UK) Consortium | Jacquelyn Wynn, Mairead Hyland, The Lighthouse Lab in Alderley Park and Alex Alderton, Roberto Amato, Jeffrey Barrett, Sonia Goncalves, Ewan Harrison, David K. Jackson, Ian Johnston, Dominic Kwiatkowski, Cordelia Langford, John Sillitoe on behalf of the Wellcome Sanger Institute COVID-19 Surveillance Team |
| EPI_ISL_2682354 | Health Services Laboratories | Wellcome Sanger Institute for the COVID-19 Genomics UK (COG-UK) Consortium | Health Services Laboratories and Alex Alderton, Roberto Amato, Jeffrey Barrett, Sonia Goncalves, Ewan Harrison, David K. Jackson, Ian Johnston, Dominic Kwiatkowski, Cordelia Langford, John Sillitoe on behalf of the Wellcome Sanger Institute COVID-19 Surveillance Team |
| EPI_ISL_2682355, EPI_ISL_2682356 | Lighthouse Lab in Alderley Park | Wellcome Sanger Institute for the COVID-19 Genomics UK (COG-UK) Consortium | Jacquelyn Wynn, Mairead Hyland, The Lighthouse Lab in Alderley Park and Alex Alderton, Roberto Amato, Jeffrey Barrett, Sonia Goncalves, Ewan Harrison, David K. Jackson, Ian Johnston, Dominic Kwiatkowski, Cordelia Langford, John Sillitoe on behalf of the Wellcome Sanger Institute COVID-19 Surveillance Team |
| EPI_ISL_2682357 | Health Services Laboratories | Wellcome Sanger Institute for the COVID-19 Genomics UK (COG-UK) Consortium | Health Services Laboratories and Alex Alderton, Roberto Amato, Jeffrey Barrett, Sonia Goncalves, Ewan Harrison, David K. Jackson, Ian Johnston, Dominic Kwiatkowski, Cordelia Langford, John Sillitoe on behalf of the Wellcome Sanger Institute COVID-19 Surveillance Team |
| EPI_ISL_2682358, EPI_ISL_2682359, EPI_ISL_2682360, EPI_ISL_2682361, EPI_ISL_2682362, EPI_ISL_2682363, EPI_ISL_2682364, EPI_ISL_2682365, EPI_ISL_2682366, EPI_ISL_2682367, EPI_ISL_2682368, EPI_ISL_2682369, EPI_ISL_2682370 | | | |
| see above | Lighthouse Lab in Alderley Park | Wellcome Sanger Institute for the COVID-19 Genomics UK (COG-UK) Consortium | Jacquelyn Wynn, Mairead Hyland, The Lighthouse Lab in Alderley Park and Alex Alderton, Roberto Amato, Jeffrey Barrett, Sonia Goncalves, Ewan Harrison, David K. Jackson, Ian Johnston, Dominic Kwiatkowski, Cordelia Langford, John Sillitoe on behalf of the Wellcome Sanger Institute COVID-19 Surveillance Team |
| EPI_ISL_2682371 | Health Services Laboratories | Wellcome Sanger Institute for the COVID-19 Genomics UK (COG-UK) | Health Services Laboratories and Alex Alderton, Roberto Amato, Jeffrey Barrett, Sonia Goncalves, Ewan Harrison, David K. Jackson, Ian Johnston, Dominic Kwiatkowski, Cordelia Langford, John Sillitoe on behalf of the Wellcome Sanger Institute |

| | | | |
|---|---|---|---|
| | | Consortium | COVID-19 Surveillance Team |
| EPI_ISL_2682372, EPI_ISL_2682373, EPI_ISL_2682374, EPI_ISL_2682375, EPI_ISL_2682376, EPI_ISL_2682377, EPI_ISL_2682378, EPI_ISL_2682379, EPI_ISL_2682380, EPI_ISL_2682381, EPI_ISL_2682382, EPI_ISL_2682383, EPI_ISL_2682384, EPI_ISL_2682385, EPI_ISL_2682386, EPI_ISL_2682387, EPI_ISL_2682388, EPI_ISL_2682389, EPI_ISL_2682390, EPI_ISL_2682391, EPI_ISL_2682392, EPI_ISL_2682393, EPI_ISL_2682394 | | | |
| see above | Lighthouse Lab in Alderley Park | Wellcome Sanger Institute for the COVID-19 Genomics UK (COG-UK) Consortium | Jacquelyn Wynn, Mairead Hyland, The Lighthouse Lab in Alderley Park and Alex Alderton, Roberto Amato, Jeffrey Barrett, Sonia Goncalves, Ewan Harrison, David K. Jackson, Ian Johnston, Dominic Kwiatkowski, Cordelia Langford, John Sillitoe on behalf of the Wellcome Sanger Institute COVID-19 Surveillance Team |
| EPI_ISL_2682395 | Health Services Laboratories | Wellcome Sanger Institute for the COVID-19 Genomics UK (COG-UK) Consortium | Health Services Laboratories and Alex Alderton, Roberto Amato, Jeffrey Barrett, Sonia Goncalves, Ewan Harrison, David K. Jackson, Ian Johnston, Dominic Kwiatkowski, Cordelia Langford, John Sillitoe on behalf of the Wellcome Sanger Institute COVID-19 Surveillance Team |
| EPI_ISL_2682396, EPI_ISL_2682397, EPI_ISL_2682398, EPI_ISL_2682399 | Lighthouse Lab in Alderley Park | Wellcome Sanger Institute for the COVID-19 Genomics UK (COG-UK) Consortium | Jacquelyn Wynn, Mairead Hyland, The Lighthouse Lab in Alderley Park and Alex Alderton, Roberto Amato, Jeffrey Barrett, Sonia Goncalves, Ewan Harrison, David K. Jackson, Ian Johnston, Dominic Kwiatkowski, Cordelia Langford, John Sillitoe on behalf of the Wellcome Sanger Institute COVID-19 Surveillance Team |
| EPI_ISL_2682400 | Health Services Laboratories | Wellcome Sanger Institute for the COVID-19 Genomics UK (COG-UK) Consortium | Health Services Laboratories and Alex Alderton, Roberto Amato, Jeffrey Barrett, Sonia Goncalves, Ewan Harrison, David K. Jackson, Ian Johnston, Dominic Kwiatkowski, Cordelia Langford, John Sillitoe on behalf of the Wellcome Sanger Institute COVID-19 Surveillance Team |
| EPI_ISL_2682401, EPI_ISL_2682402, EPI_ISL_2682403, EPI_ISL_2682404, EPI_ISL_2682405, EPI_ISL_2682406, EPI_ISL_2682407, EPI_ISL_2682408, EPI_ISL_2682409, EPI_ISL_2682410, EPI_ISL_2682411, EPI_ISL_2682412, EPI_ISL_2682413, EPI_ISL_2682414, EPI_ISL_2682415 | | | |
| see above | Lighthouse Lab in Alderley Park | Wellcome Sanger Institute for the COVID-19 Genomics UK (COG-UK) Consortium | Jacquelyn Wynn, Mairead Hyland, The Lighthouse Lab in Alderley Park and Alex Alderton, Roberto Amato, Jeffrey Barrett, Sonia Goncalves, Ewan Harrison, David K. Jackson, Ian Johnston, Dominic Kwiatkowski, Cordelia Langford, John Sillitoe on behalf of the Wellcome Sanger Institute COVID-19 Surveillance Team |
| EPI_ISL_2682416 | Health Services Laboratories | Wellcome Sanger Institute for the COVID-19 Genomics UK (COG-UK) Consortium | Health Services Laboratories and Alex Alderton, Roberto Amato, Jeffrey Barrett, Sonia Goncalves, Ewan Harrison, David K. Jackson, Ian Johnston, Dominic Kwiatkowski, Cordelia Langford, John Sillitoe on behalf of the Wellcome Sanger Institute COVID-19 Surveillance Team |
| EPI_ISL_2682417, EPI_ISL_2682418, EPI_ISL_2682419, EPI_ISL_2682420, EPI_ISL_2682421, EPI_ISL_2682422, EPI_ISL_2682423, EPI_ISL_2682424, EPI_ISL_2682425, EPI_ISL_2682426, EPI_ISL_2682427, EPI_ISL_2682428, EPI_ISL_2682429, EPI_ISL_2682430, EPI_ISL_2682431, EPI_ISL_2682432, EPI_ISL_2682433, EPI_ISL_2682434, EPI_ISL_2682435, EPI_ISL_2682436, EPI_ISL_2682437, EPI_ISL_2682438, EPI_ISL_2682439, EPI_ISL_2682440, EPI_ISL_2682441 | | | |
| see above | Lighthouse Lab in Alderley Park | Wellcome Sanger Institute for the COVID-19 Genomics UK (COG-UK) Consortium | Jacquelyn Wynn, Mairead Hyland, The Lighthouse Lab in Alderley Park and Alex Alderton, Roberto Amato, Jeffrey Barrett, Sonia Goncalves, Ewan Harrison, David K. Jackson, Ian Johnston, Dominic Kwiatkowski, Cordelia Langford, John Sillitoe on behalf of the Wellcome Sanger Institute COVID-19 Surveillance Team |
| EPI_ISL_2682442 | Health Services Laboratories | Wellcome Sanger Institute for the COVID-19 Genomics UK (COG-UK) Consortium | Health Services Laboratories and Alex Alderton, Roberto Amato, Jeffrey Barrett, Sonia Goncalves, Ewan Harrison, David K. Jackson, Ian Johnston, Dominic Kwiatkowski, Cordelia Langford, John Sillitoe on behalf of the Wellcome Sanger Institute COVID-19 Surveillance Team |
| EPI_ISL_2682443, EPI_ISL_2682444 | Lighthouse Lab in Alderley Park | Wellcome Sanger Institute for the COVID-19 Genomics UK (COG-UK) Consortium | Jacquelyn Wynn, Mairead Hyland, The Lighthouse Lab in Alderley Park and Alex Alderton, Roberto Amato, Jeffrey Barrett, Sonia Goncalves, Ewan Harrison, David K. Jackson, Ian Johnston, Dominic Kwiatkowski, Cordelia Langford, John Sillitoe on behalf of the Wellcome Sanger Institute COVID-19 Surveillance Team |
| EPI_ISL_2682445 | Health Services Laboratories | Wellcome Sanger Institute for the COVID-19 Genomics UK (COG-UK) Consortium | Health Services Laboratories and Alex Alderton, Roberto Amato, Jeffrey Barrett, Sonia Goncalves, Ewan Harrison, David K. Jackson, Ian Johnston, Dominic Kwiatkowski, Cordelia Langford, John Sillitoe on behalf of the Wellcome Sanger Institute COVID-19 Surveillance Team |
| EPI_ISL_2682446, EPI_ISL_2682447, EPI_ISL_2682448, EPI_ISL_2682449, EPI_ISL_2682450, EPI_ISL_2682451, EPI_ISL_2682452, EPI_ISL_2682453, EPI_ISL_2682454 | Lighthouse Lab in Alderley Park | Wellcome Sanger Institute for the COVID-19 Genomics UK (COG-UK) Consortium | Jacquelyn Wynn, Mairead Hyland, The Lighthouse Lab in Alderley Park and Alex Alderton, Roberto Amato, Jeffrey Barrett, Sonia Goncalves, Ewan Harrison, David K. Jackson, Ian Johnston, Dominic Kwiatkowski, Cordelia Langford, John Sillitoe on behalf of the Wellcome Sanger Institute COVID-19 Surveillance Team |
| EPI_ISL_2682455 | Health Services Laboratories | Wellcome Sanger Institute for the COVID-19 Genomics UK (COG-UK) Consortium | Health Services Laboratories and Alex Alderton, Roberto Amato, Jeffrey Barrett, Sonia Goncalves, Ewan Harrison, David K. Jackson, Ian Johnston, Dominic Kwiatkowski, Cordelia Langford, John Sillitoe on behalf of the Wellcome Sanger Institute COVID-19 Surveillance Team |
| EPI_ISL_2682456 | Lighthouse Lab in Alderley Park | Wellcome Sanger Institute for the COVID-19 Genomics UK (COG-UK) Consortium | Jacquelyn Wynn, Mairead Hyland, The Lighthouse Lab in Alderley Park and Alex Alderton, Roberto Amato, Jeffrey Barrett, Sonia Goncalves, Ewan Harrison, David K. Jackson, Ian Johnston, Dominic Kwiatkowski, Cordelia Langford, John Sillitoe on behalf of the Wellcome Sanger Institute COVID-19 Surveillance Team |
| EPI_ISL_2682457 | Health Services Laboratories | Wellcome Sanger Institute for the COVID-19 Genomics UK (COG-UK) Consortium | Health Services Laboratories and Alex Alderton, Roberto Amato, Jeffrey Barrett, Sonia Goncalves, Ewan Harrison, David K. Jackson, Ian Johnston, Dominic Kwiatkowski, Cordelia Langford, John Sillitoe on behalf of the Wellcome Sanger Institute COVID-19 Surveillance Team |
| EPI_ISL_2682458, EPI_ISL_2682459, EPI_ISL_2682460, EPI_ISL_2682461, EPI_ISL_2682462, EPI_ISL_2682463, EPI_ISL_2682464, EPI_ISL_2682465, EPI_ISL_2682466 | Lighthouse Lab in Alderley Park | Wellcome Sanger Institute for the COVID-19 Genomics UK (COG-UK) Consortium | Jacquelyn Wynn, Mairead Hyland, The Lighthouse Lab in Alderley Park and Alex Alderton, Roberto Amato, Jeffrey Barrett, Sonia Goncalves, Ewan Harrison, David K. Jackson, Ian Johnston, Dominic Kwiatkowski, Cordelia Langford, John Sillitoe on behalf of the Wellcome Sanger Institute COVID-19 Surveillance Team |
| EPI_ISL_2682467 | Health Services Laboratories | Wellcome Sanger Institute for the COVID-19 Genomics UK (COG-UK) Consortium | Health Services Laboratories and Alex Alderton, Roberto Amato, Jeffrey Barrett, Sonia Goncalves, Ewan Harrison, David K. Jackson, Ian Johnston, Dominic Kwiatkowski, Cordelia Langford, John Sillitoe on behalf of the Wellcome Sanger Institute COVID-19 Surveillance Team |
| EPI_ISL_2682468, EPI_ISL_2682469, EPI_ISL_2682470 | Lighthouse Lab in Alderley Park | Wellcome Sanger Institute for the COVID-19 Genomics UK (COG-UK) Consortium | Jacquelyn Wynn, Mairead Hyland, The Lighthouse Lab in Alderley Park and Alex Alderton, Roberto Amato, Jeffrey Barrett, Sonia Goncalves, Ewan Harrison, David K. Jackson, Ian Johnston, Dominic Kwiatkowski, Cordelia Langford, John Sillitoe on behalf of the Wellcome Sanger Institute COVID-19 Surveillance Team |
| EPI_ISL_2682471 | Health Services Laboratories | Wellcome Sanger Institute for the COVID-19 Genomics UK (COG-UK) Consortium | Health Services Laboratories and Alex Alderton, Roberto Amato, Jeffrey Barrett, Sonia Goncalves, Ewan Harrison, David K. Jackson, Ian Johnston, Dominic Kwiatkowski, Cordelia Langford, John Sillitoe on behalf of the Wellcome Sanger Institute COVID-19 Surveillance Team |
| EPI_ISL_2682472 | Lighthouse Lab in Alderley Park | Wellcome Sanger Institute for the COVID-19 Genomics UK (COG-UK) Consortium | Jacquelyn Wynn, Mairead Hyland, The Lighthouse Lab in Alderley Park and Alex Alderton, Roberto Amato, Jeffrey Barrett, Sonia Goncalves, Ewan Harrison, David K. Jackson, Ian Johnston, Dominic Kwiatkowski, Cordelia Langford, John Sillitoe on behalf of the Wellcome Sanger Institute COVID-19 Surveillance Team |
| EPI_ISL_2682473 | Health Services Laboratories | Wellcome Sanger Institute for the COVID-19 Genomics UK (COG-UK) Consortium | Health Services Laboratories and Alex Alderton, Roberto Amato, Jeffrey Barrett, Sonia Goncalves, Ewan Harrison, David K. Jackson, Ian Johnston, Dominic Kwiatkowski, Cordelia Langford, John Sillitoe on behalf of the Wellcome Sanger Institute COVID-19 Surveillance Team |
| EPI_ISL_2682474, EPI_ISL_2682475, EPI_ISL_2682476, EPI_ISL_2682477, EPI_ISL_2682478, EPI_ISL_2682479, EPI_ISL_2682480, EPI_ISL_2682481, EPI_ISL_2682482, EPI_ISL_2682483, EPI_ISL_2682484, EPI_ISL_2682485, EPI_ISL_2682486, EPI_ISL_2682487, EPI_ISL_2682488, EPI_ISL_2682489 | | | |
| see above | Lighthouse Lab in Alderley Park | Wellcome Sanger Institute for the COVID-19 Genomics UK (COG-UK) Consortium | Jacquelyn Wynn, Mairead Hyland, The Lighthouse Lab in Alderley Park and Alex Alderton, Roberto Amato, Jeffrey Barrett, Sonia Goncalves, Ewan Harrison, David K. Jackson, Ian Johnston, Dominic Kwiatkowski, Cordelia Langford, John Sillitoe on behalf of the Wellcome Sanger Institute COVID-19 Surveillance Team |
| EPI_ISL_2682490 | Health Services Laboratories | Wellcome Sanger Institute for the COVID-19 Genomics UK (COG-UK) Consortium | Health Services Laboratories and Alex Alderton, Roberto Amato, Jeffrey Barrett, Sonia Goncalves, Ewan Harrison, David K. Jackson, Ian Johnston, Dominic Kwiatkowski, Cordelia Langford, John Sillitoe on behalf of the Wellcome Sanger Institute COVID-19 Surveillance Team |
| EPI_ISL_2682491, EPI_ISL_2682492, EPI_ISL_2682493, EPI_ISL_2682494, EPI_ISL_2682495, EPI_ISL_2682496, EPI_ISL_2682497, EPI_ISL_2682498, EPI_ISL_2682499, EPI_ISL_2682500, EPI_ISL_2682501, EPI_ISL_2682502 | | | |
| see above | Lighthouse Lab in Alderley Park | Wellcome Sanger Institute for the COVID-19 Genomics UK (COG-UK) Consortium | Jacquelyn Wynn, Mairead Hyland, The Lighthouse Lab in Alderley Park and Alex Alderton, Roberto Amato, Jeffrey Barrett, Sonia Goncalves, Ewan Harrison, David K. Jackson, Ian Johnston, Dominic Kwiatkowski, Cordelia Langford, John Sillitoe on behalf of the Wellcome Sanger Institute COVID-19 Surveillance Team |
| EPI_ISL_2682503 | Health Services Laboratories | Wellcome Sanger Institute for the COVID-19 Genomics UK (COG-UK) Consortium | Health Services Laboratories and Alex Alderton, Roberto Amato, Jeffrey Barrett, Sonia Goncalves, Ewan Harrison, David K. Jackson, Ian Johnston, Dominic Kwiatkowski, Cordelia Langford, John Sillitoe on behalf of the Wellcome Sanger Institute COVID-19 Surveillance Team |
| EPI_ISL_2682504, EPI_ISL_2682505, EPI_ISL_2682506, EPI_ISL_2682507, EPI_ISL_2682508, EPI_ISL_2682509, EPI_ISL_2682510, EPI_ISL_2682511, EPI_ISL_2682512, EPI_ISL_2682513, EPI_ISL_2682514, EPI_ISL_2682515, EPI_ISL_2682516, EPI_ISL_2682517, EPI_ISL_2682518, EPI_ISL_2682519, EPI_ISL_2682520, EPI_ISL_2682521, EPI_ISL_2682522 | | | |
| see above | Lighthouse Lab in Alderley Park | Wellcome Sanger Institute for the COVID-19 Genomics UK (COG-UK) Consortium | Jacquelyn Wynn, Mairead Hyland, The Lighthouse Lab in Alderley Park and Alex Alderton, Roberto Amato, Jeffrey Barrett, Sonia Goncalves, Ewan Harrison, David K. Jackson, Ian Johnston, Dominic Kwiatkowski, Cordelia Langford, John Sillitoe on behalf of the Wellcome Sanger Institute COVID-19 Surveillance Team |
| EPI_ISL_2682523, EPI_ISL_2682524 | Health Services Laboratories | Wellcome Sanger Institute for the COVID-19 Genomics UK (COG-UK) Consortium | Health Services Laboratories and Alex Alderton, Roberto Amato, Jeffrey Barrett, Sonia Goncalves, Ewan Harrison, David K. Jackson, Ian Johnston, Dominic Kwiatkowski, Cordelia Langford, John Sillitoe on behalf of the Wellcome Sanger Institute COVID-19 Surveillance Team |
| EPI_ISL_2682525 | Lighthouse Lab in Alderley Park | Wellcome Sanger Institute for the COVID-19 Genomics UK (COG-UK) Consortium | Jacquelyn Wynn, Mairead Hyland, The Lighthouse Lab in Alderley Park and Alex Alderton, Roberto Amato, Jeffrey Barrett, Sonia Goncalves, Ewan Harrison, David K. Jackson, Ian Johnston, Dominic Kwiatkowski, Cordelia Langford, John Sillitoe on behalf of the Wellcome Sanger Institute COVID-19 Surveillance Team |
| EPI_ISL_2682526 | Health Services Laboratories | Wellcome Sanger Institute for the COVID-19 Genomics UK (COG-UK) Consortium | Health Services Laboratories and Alex Alderton, Roberto Amato, Jeffrey Barrett, Sonia Goncalves, Ewan Harrison, David K. Jackson, Ian Johnston, Dominic Kwiatkowski, Cordelia Langford, John Sillitoe on behalf of the Wellcome Sanger Institute COVID-19 Surveillance Team |
| EPI_ISL_2682527, EPI_ISL_2682528, EPI_ISL_2682529, EPI_ISL_2682530, EPI_ISL_2682531, EPI_ISL_2682532, EPI_ISL_2682533, EPI_ISL_2682534, EPI_ISL_2682535, EPI_ISL_2682536 | Lighthouse Lab in Alderley Park | Wellcome Sanger Institute for the COVID-19 Genomics UK (COG-UK) Consortium | Jacquelyn Wynn, Mairead Hyland, The Lighthouse Lab in Alderley Park and Alex Alderton, Roberto Amato, Jeffrey Barrett, Sonia Goncalves, Ewan Harrison, David K. Jackson, Ian Johnston, Dominic Kwiatkowski, Cordelia Langford, John Sillitoe on behalf of the Wellcome Sanger Institute COVID-19 Surveillance Team |
| EPI_ISL_2682537, EPI_ISL_2682538 | Health Services Laboratories | Wellcome Sanger Institute for the COVID-19 Genomics UK (COG-UK) Consortium | Health Services Laboratories and Alex Alderton, Roberto Amato, Jeffrey Barrett, Sonia Goncalves, Ewan Harrison, David K. Jackson, Ian Johnston, Dominic Kwiatkowski, Cordelia Langford, John Sillitoe on behalf of the Wellcome Sanger Institute COVID-19 Surveillance Team |

| | | | |
|---|---|---|---|
| EPI_ISL_2682539, EPI_ISL_2682540, EPI_ISL_2682541, EPI_ISL_2682542, EPI_ISL_2682543, EPI_ISL_2682544 | Lighthouse Lab in Alderley Park | Wellcome Sanger Institute for the COVID-19 Genomics UK (COG-UK) Consortium | Jacquelyn Wynn, Mairead Hyland, The Lighthouse Lab in Alderley Park and Alex Alderton, Roberto Amato, Jeffrey Barrett, Sonia Goncalves, Ewan Harrison, David K. Jackson, Ian Johnston, Dominic Kwiatkowski, Cordelia Langford, John Sillitoe on behalf of the Wellcome Sanger Institute COVID-19 Surveillance Team |
| EPI_ISL_2682545, EPI_ISL_2682546 | Health Services Laboratories | Wellcome Sanger Institute for the COVID-19 Genomics UK (COG-UK) Consortium | Health Services Laboratories and Alex Alderton, Roberto Amato, Jeffrey Barrett, Sonia Goncalves, Ewan Harrison, David K. Jackson, Ian Johnston, Dominic Kwiatkowski, Cordelia Langford, John Sillitoe on behalf of the Wellcome Sanger Institute COVID-19 Surveillance Team |
| EPI_ISL_2682547, EPI_ISL_2682548 | Lighthouse Lab in Alderley Park | Wellcome Sanger Institute for the COVID-19 Genomics UK (COG-UK) Consortium | Jacquelyn Wynn, Mairead Hyland, The Lighthouse Lab in Alderley Park and Alex Alderton, Roberto Amato, Jeffrey Barrett, Sonia Goncalves, Ewan Harrison, David K. Jackson, Ian Johnston, Dominic Kwiatkowski, Cordelia Langford, John Sillitoe on behalf of the Wellcome Sanger Institute COVID-19 Surveillance Team |
| EPI_ISL_2682549 | Health Services Laboratories | Wellcome Sanger Institute for the COVID-19 Genomics UK (COG-UK) Consortium | Health Services Laboratories and Alex Alderton, Roberto Amato, Jeffrey Barrett, Sonia Goncalves, Ewan Harrison, David K. Jackson, Ian Johnston, Dominic Kwiatkowski, Cordelia Langford, John Sillitoe on behalf of the Wellcome Sanger Institute COVID-19 Surveillance Team |
| EPI_ISL_2682550, EPI_ISL_2682551, EPI_ISL_2682552, EPI_ISL_2682553, EPI_ISL_2682554, EPI_ISL_2682555, EPI_ISL_2682556, EPI_ISL_2682557 | Lighthouse Lab in Alderley Park | Wellcome Sanger Institute for the COVID-19 Genomics UK (COG-UK) Consortium | Jacquelyn Wynn, Mairead Hyland, The Lighthouse Lab in Alderley Park and Alex Alderton, Roberto Amato, Jeffrey Barrett, Sonia Goncalves, Ewan Harrison, David K. Jackson, Ian Johnston, Dominic Kwiatkowski, Cordelia Langford, John Sillitoe on behalf of the Wellcome Sanger Institute COVID-19 Surveillance Team |
| EPI_ISL_2682558 | Health Services Laboratories | Wellcome Sanger Institute for the COVID-19 Genomics UK (COG-UK) Consortium | Health Services Laboratories and Alex Alderton, Roberto Amato, Jeffrey Barrett, Sonia Goncalves, Ewan Harrison, David K. Jackson, Ian Johnston, Dominic Kwiatkowski, Cordelia Langford, John Sillitoe on behalf of the Wellcome Sanger Institute COVID-19 Surveillance Team |
| EPI_ISL_2682559, EPI_ISL_2682560, EPI_ISL_2682561, EPI_ISL_2682562, EPI_ISL_2682563, EPI_ISL_2682564, EPI_ISL_2682565, EPI_ISL_2682566, EPI_ISL_2682567, EPI_ISL_2682568 | Lighthouse Lab in Alderley Park | Wellcome Sanger Institute for the COVID-19 Genomics UK (COG-UK) Consortium | Jacquelyn Wynn, Mairead Hyland, The Lighthouse Lab in Alderley Park and Alex Alderton, Roberto Amato, Jeffrey Barrett, Sonia Goncalves, Ewan Harrison, David K. Jackson, Ian Johnston, Dominic Kwiatkowski, Cordelia Langford, John Sillitoe on behalf of the Wellcome Sanger Institute COVID-19 Surveillance Team |
| EPI_ISL_2682569 | Health Services Laboratories | Wellcome Sanger Institute for the COVID-19 Genomics UK (COG-UK) Consortium | Health Services Laboratories and Alex Alderton, Roberto Amato, Jeffrey Barrett, Sonia Goncalves, Ewan Harrison, David K. Jackson, Ian Johnston, Dominic Kwiatkowski, Cordelia Langford, John Sillitoe on behalf of the Wellcome Sanger Institute COVID-19 Surveillance Team |
| EPI_ISL_2682570, EPI_ISL_2682571, EPI_ISL_2682572, EPI_ISL_2682573, EPI_ISL_2682574, EPI_ISL_2682575, EPI_ISL_2682576, EPI_ISL_2682577, EPI_ISL_2682578, EPI_ISL_2682579, EPI_ISL_2682580, EPI_ISL_2682581, EPI_ISL_2682582, EPI_ISL_2682583, EPI_ISL_2682584, EPI_ISL_2682585, EPI_ISL_2682586, EPI_ISL_2682587, EPI_ISL_2682588, EPI_ISL_2682589, EPI_ISL_2682590, EPI_ISL_2682591, EPI_ISL_2682592, EPI_ISL_2682593 | | | |
| see above | Lighthouse Lab in Alderley Park | Wellcome Sanger Institute for the COVID-19 Genomics UK (COG-UK) Consortium | Jacquelyn Wynn, Mairead Hyland, The Lighthouse Lab in Alderley Park and Alex Alderton, Roberto Amato, Jeffrey Barrett, Sonia Goncalves, Ewan Harrison, David K. Jackson, Ian Johnston, Dominic Kwiatkowski, Cordelia Langford, John Sillitoe on behalf of the Wellcome Sanger Institute COVID-19 Surveillance Team |
| EPI_ISL_2682594, EPI_ISL_2682595 | Health Services Laboratories | Wellcome Sanger Institute for the COVID-19 Genomics UK (COG-UK) Consortium | Health Services Laboratories and Alex Alderton, Roberto Amato, Jeffrey Barrett, Sonia Goncalves, Ewan Harrison, David K. Jackson, Ian Johnston, Dominic Kwiatkowski, Cordelia Langford, John Sillitoe on behalf of the Wellcome Sanger Institute COVID-19 Surveillance Team |
| EPI_ISL_2682596, EPI_ISL_2682598, EPI_ISL_2682599, EPI_ISL_2682600, EPI_ISL_2682601, EPI_ISL_2682602, EPI_ISL_2682603, EPI_ISL_2682604, EPI_ISL_2682605, EPI_ISL_2682606, EPI_ISL_2682607, EPI_ISL_2682608, EPI_ISL_2682609, EPI_ISL_2682610, EPI_ISL_2682611, EPI_ISL_2682612, EPI_ISL_2682613, EPI_ISL_2682614, EPI_ISL_2682615, EPI_ISL_2682616, EPI_ISL_2682617, EPI_ISL_2682618, EPI_ISL_2682619, EPI_ISL_2682620, EPI_ISL_2682621, EPI_ISL_2682622 | | | |
| see above | Lighthouse Lab in Alderley Park | Wellcome Sanger Institute for the COVID-19 Genomics UK (COG-UK) Consortium | Jacquelyn Wynn, Mairead Hyland, The Lighthouse Lab in Alderley Park and Alex Alderton, Roberto Amato, Jeffrey Barrett, Sonia Goncalves, Ewan Harrison, David K. Jackson, Ian Johnston, Dominic Kwiatkowski, Cordelia Langford, John Sillitoe on behalf of the Wellcome Sanger Institute COVID-19 Surveillance Team |
| EPI_ISL_2682623 | Health Services Laboratories | Wellcome Sanger Institute for the COVID-19 Genomics UK (COG-UK) Consortium | Health Services Laboratories and Alex Alderton, Roberto Amato, Jeffrey Barrett, Sonia Goncalves, Ewan Harrison, David K. Jackson, Ian Johnston, Dominic Kwiatkowski, Cordelia Langford, John Sillitoe on behalf of the Wellcome Sanger Institute COVID-19 Surveillance Team |
| EPI_ISL_2682624, EPI_ISL_2682625, EPI_ISL_2682626, EPI_ISL_2682627, EPI_ISL_2682628, EPI_ISL_2682629 | Lighthouse Lab in Alderley Park | Wellcome Sanger Institute for the COVID-19 Genomics UK (COG-UK) Consortium | Jacquelyn Wynn, Mairead Hyland, The Lighthouse Lab in Alderley Park and Alex Alderton, Roberto Amato, Jeffrey Barrett, Sonia Goncalves, Ewan Harrison, David K. Jackson, Ian Johnston, Dominic Kwiatkowski, Cordelia Langford, John Sillitoe on behalf of the Wellcome Sanger Institute COVID-19 Surveillance Team |
| EPI_ISL_2682630 | Health Services Laboratories | Wellcome Sanger Institute for the COVID-19 Genomics UK (COG-UK) Consortium | Health Services Laboratories and Alex Alderton, Roberto Amato, Jeffrey Barrett, Sonia Goncalves, Ewan Harrison, David K. Jackson, Ian Johnston, Dominic Kwiatkowski, Cordelia Langford, John Sillitoe on behalf of the Wellcome Sanger Institute COVID-19 Surveillance Team |
| EPI_ISL_2682631, EPI_ISL_2682632, EPI_ISL_2682633, EPI_ISL_2682634, EPI_ISL_2682635, EPI_ISL_2682636 | Lighthouse Lab in Alderley Park | Wellcome Sanger Institute for the COVID-19 Genomics UK (COG-UK) Consortium | Jacquelyn Wynn, Mairead Hyland, The Lighthouse Lab in Alderley Park and Alex Alderton, Roberto Amato, Jeffrey Barrett, Sonia Goncalves, Ewan Harrison, David K. Jackson, Ian Johnston, Dominic Kwiatkowski, Cordelia Langford, John Sillitoe on behalf of the Wellcome Sanger Institute COVID-19 Surveillance Team |
| EPI_ISL_2682637 | Health Services Laboratories | Wellcome Sanger Institute for the COVID-19 Genomics UK (COG-UK) Consortium | Health Services Laboratories and Alex Alderton, Roberto Amato, Jeffrey Barrett, Sonia Goncalves, Ewan Harrison, David K. Jackson, Ian Johnston, Dominic Kwiatkowski, Cordelia Langford, John Sillitoe on behalf of the Wellcome Sanger Institute COVID-19 Surveillance Team |
| EPI_ISL_2682638, EPI_ISL_2682639, EPI_ISL_2682640, EPI_ISL_2682641, EPI_ISL_2682642, EPI_ISL_2682643, EPI_ISL_2682644, EPI_ISL_2682645 | Lighthouse Lab in Alderley Park | Wellcome Sanger Institute for the COVID-19 Genomics UK (COG-UK) Consortium | Jacquelyn Wynn, Mairead Hyland, The Lighthouse Lab in Alderley Park and Alex Alderton, Roberto Amato, Jeffrey Barrett, Sonia Goncalves, Ewan Harrison, David K. Jackson, Ian Johnston, Dominic Kwiatkowski, Cordelia Langford, John Sillitoe on behalf of the Wellcome Sanger Institute COVID-19 Surveillance Team |
| EPI_ISL_2682646 | Health Services Laboratories | Wellcome Sanger Institute for the COVID-19 Genomics UK (COG-UK) Consortium | Health Services Laboratories and Alex Alderton, Roberto Amato, Jeffrey Barrett, Sonia Goncalves, Ewan Harrison, David K. Jackson, Ian Johnston, Dominic Kwiatkowski, Cordelia Langford, John Sillitoe on behalf of the Wellcome Sanger Institute COVID-19 Surveillance Team |
| EPI_ISL_2682647 | Lighthouse Lab in Alderley Park | Wellcome Sanger Institute for the COVID-19 Genomics UK (COG-UK) Consortium | Jacquelyn Wynn, Mairead Hyland, The Lighthouse Lab in Alderley Park and Alex Alderton, Roberto Amato, Jeffrey Barrett, Sonia Goncalves, Ewan Harrison, David K. Jackson, Ian Johnston, Dominic Kwiatkowski, Cordelia Langford, John Sillitoe on behalf of the Wellcome Sanger Institute COVID-19 Surveillance Team |
| EPI_ISL_2682648 | Health Services Laboratories | Wellcome Sanger Institute for the COVID-19 Genomics UK (COG-UK) Consortium | Health Services Laboratories and Alex Alderton, Roberto Amato, Jeffrey Barrett, Sonia Goncalves, Ewan Harrison, David K. Jackson, Ian Johnston, Dominic Kwiatkowski, Cordelia Langford, John Sillitoe on behalf of the Wellcome Sanger Institute COVID-19 Surveillance Team |
| EPI_ISL_2682649, EPI_ISL_2682650 | Lighthouse Lab in Alderley Park | Wellcome Sanger Institute for the COVID-19 Genomics UK (COG-UK) Consortium | Jacquelyn Wynn, Mairead Hyland, The Lighthouse Lab in Alderley Park and Alex Alderton, Roberto Amato, Jeffrey Barrett, Sonia Goncalves, Ewan Harrison, David K. Jackson, Ian Johnston, Dominic Kwiatkowski, Cordelia Langford, John Sillitoe on behalf of the Wellcome Sanger Institute COVID-19 Surveillance Team |
| EPI_ISL_2682651 | Health Services Laboratories | Wellcome Sanger Institute for the COVID-19 Genomics UK (COG-UK) Consortium | Health Services Laboratories and Alex Alderton, Roberto Amato, Jeffrey Barrett, Sonia Goncalves, Ewan Harrison, David K. Jackson, Ian Johnston, Dominic Kwiatkowski, Cordelia Langford, John Sillitoe on behalf of the Wellcome Sanger Institute COVID-19 Surveillance Team |
| EPI_ISL_2682652 | Lighthouse Lab in Alderley Park | Wellcome Sanger Institute for the COVID-19 Genomics UK (COG-UK) Consortium | Jacquelyn Wynn, Mairead Hyland, The Lighthouse Lab in Alderley Park and Alex Alderton, Roberto Amato, Jeffrey Barrett, Sonia Goncalves, Ewan Harrison, David K. Jackson, Ian Johnston, Dominic Kwiatkowski, Cordelia Langford, John Sillitoe on behalf of the Wellcome Sanger Institute COVID-19 Surveillance Team |
| EPI_ISL_2682653 | Lighthouse Lab in Glasgow | Wellcome Sanger Institute for the COVID-19 Genomics UK (COG-UK) Consortium | Harper VanSteenhouse, Yumi Kasai, David Gray, Carol Clugston, Anna Dominiczak and Alex Alderton, Roberto Amato, Jeffrey Barrett, Sonia Goncalves, Ewan Harrison, David K. Jackson, Ian Johnston, Dominic Kwiatkowski, Cordelia Langford, John Sillitoe on behalf of the Wellcome Sanger Institute COVID-19 Surveillance Team |
| EPI_ISL_2682654 | Lighthouse Lab in Alderley Park | Wellcome Sanger Institute for the COVID-19 Genomics UK (COG-UK) Consortium | Jacquelyn Wynn, Mairead Hyland, The Lighthouse Lab in Alderley Park and Alex Alderton, Roberto Amato, Jeffrey Barrett, Sonia Goncalves, Ewan Harrison, David K. Jackson, Ian Johnston, Dominic Kwiatkowski, Cordelia Langford, John Sillitoe on behalf of the Wellcome Sanger Institute COVID-19 Surveillance Team |
| EPI_ISL_2682655, EPI_ISL_2682656, EPI_ISL_2682657 | Lighthouse Lab in Glasgow | Wellcome Sanger Institute for the COVID-19 Genomics UK (COG-UK) Consortium | Harper VanSteenhouse, Yumi Kasai, David Gray, Carol Clugston, Anna Dominiczak and Alex Alderton, Roberto Amato, Jeffrey Barrett, Sonia Goncalves, Ewan Harrison, David K. Jackson, Ian Johnston, Dominic Kwiatkowski, Cordelia Langford, John Sillitoe on behalf of the Wellcome Sanger Institute COVID-19 Surveillance Team |
| EPI_ISL_2682658 | Lighthouse Lab in Alderley Park | Wellcome Sanger Institute for the COVID-19 Genomics UK (COG-UK) Consortium | Jacquelyn Wynn, Mairead Hyland, The Lighthouse Lab in Alderley Park and Alex Alderton, Roberto Amato, Jeffrey Barrett, Sonia Goncalves, Ewan Harrison, David K. Jackson, Ian Johnston, Dominic Kwiatkowski, Cordelia Langford, John Sillitoe on behalf of the Wellcome Sanger Institute COVID-19 Surveillance Team |
| EPI_ISL_2682659, EPI_ISL_2682660, EPI_ISL_2682661 | Lighthouse Lab in Glasgow | Wellcome Sanger Institute for the COVID-19 Genomics UK (COG-UK) Consortium | Harper VanSteenhouse, Yumi Kasai, David Gray, Carol Clugston, Anna Dominiczak and Alex Alderton, Roberto Amato, Jeffrey Barrett, Sonia Goncalves, Ewan Harrison, David K. Jackson, Ian Johnston, Dominic Kwiatkowski, Cordelia Langford, John Sillitoe on behalf of the Wellcome Sanger Institute COVID-19 Surveillance Team |
| EPI_ISL_2682662 | Lighthouse Lab in Alderley Park | Wellcome Sanger Institute for the COVID-19 Genomics UK (COG-UK) | Jacquelyn Wynn, Mairead Hyland, The Lighthouse Lab in Alderley Park and Alex Alderton, Roberto Amato, Jeffrey Barrett, Sonia Goncalves, Ewan Harrison, David K. Jackson, Ian Johnston, Dominic Kwiatkowski, Cordelia Langford, John Sillitoe on |

| | | | |
|---|---|---|---|
| | | Consortium | behalf of the Wellcome Sanger Institute COVID-19 Surveillance Team |
| EPI_ISL_2682663 | Lighthouse Lab in Glasgow | Wellcome Sanger Institute for the COVID-19 Genomics UK (COG-UK) Consortium | Harper VanSteenhouse, Yumi Kasai, David Gray, Carol Clugston, Anna Dominiczak and Alex Alderton, Roberto Amato, Jeffrey Barrett, Sonia Goncalves, Ewan Harrison, David K. Jackson, Ian Johnston, Dominic Kwiatkowski, Cordelia Langford, John Sillitoe on behalf of the Wellcome Sanger Institute COVID-19 Surveillance Team |
| EPI_ISL_2682664 | Lighthouse Lab in Alderley Park | Wellcome Sanger Institute for the COVID-19 Genomics UK (COG-UK) Consortium | Jacquelyn Wynn, Mairead Hyland, The Lighthouse Lab in Alderley Park and Alex Alderton, Roberto Amato, Jeffrey Barrett, Sonia Goncalves, Ewan Harrison, David K. Jackson, Ian Johnston, Dominic Kwiatkowski, Cordelia Langford, John Sillitoe on behalf of the Wellcome Sanger Institute COVID-19 Surveillance Team |
| EPI_ISL_2682665 | Lighthouse Lab in Glasgow | Wellcome Sanger Institute for the COVID-19 Genomics UK (COG-UK) Consortium | Harper VanSteenhouse, Yumi Kasai, David Gray, Carol Clugston, Anna Dominiczak and Alex Alderton, Roberto Amato, Jeffrey Barrett, Sonia Goncalves, Ewan Harrison, David K. Jackson, Ian Johnston, Dominic Kwiatkowski, Cordelia Langford, John Sillitoe on behalf of the Wellcome Sanger Institute COVID-19 Surveillance Team |
| EPI_ISL_2682666, EPI_ISL_2682667, EPI_ISL_2682668 | Lighthouse Lab in Alderley Park | Wellcome Sanger Institute for the COVID-19 Genomics UK (COG-UK) Consortium | Jacquelyn Wynn, Mairead Hyland, The Lighthouse Lab in Alderley Park and Alex Alderton, Roberto Amato, Jeffrey Barrett, Sonia Goncalves, Ewan Harrison, David K. Jackson, Ian Johnston, Dominic Kwiatkowski, Cordelia Langford, John Sillitoe on behalf of the Wellcome Sanger Institute COVID-19 Surveillance Team |
| EPI_ISL_2682669, EPI_ISL_2682670 | Lighthouse Lab in Glasgow | Wellcome Sanger Institute for the COVID-19 Genomics UK (COG-UK) Consortium | Harper VanSteenhouse, Yumi Kasai, David Gray, Carol Clugston, Anna Dominiczak and Alex Alderton, Roberto Amato, Jeffrey Barrett, Sonia Goncalves, Ewan Harrison, David K. Jackson, Ian Johnston, Dominic Kwiatkowski, Cordelia Langford, John Sillitoe on behalf of the Wellcome Sanger Institute COVID-19 Surveillance Team |
| EPI_ISL_2682671 | Lighthouse Lab in Alderley Park | Wellcome Sanger Institute for the COVID-19 Genomics UK (COG-UK) Consortium | Jacquelyn Wynn, Mairead Hyland, The Lighthouse Lab in Alderley Park and Alex Alderton, Roberto Amato, Jeffrey Barrett, Sonia Goncalves, Ewan Harrison, David K. Jackson, Ian Johnston, Dominic Kwiatkowski, Cordelia Langford, John Sillitoe on behalf of the Wellcome Sanger Institute COVID-19 Surveillance Team |
| EPI_ISL_2682672 | Lighthouse Lab in Glasgow | Wellcome Sanger Institute for the COVID-19 Genomics UK (COG-UK) Consortium | Harper VanSteenhouse, Yumi Kasai, David Gray, Carol Clugston, Anna Dominiczak and Alex Alderton, Roberto Amato, Jeffrey Barrett, Sonia Goncalves, Ewan Harrison, David K. Jackson, Ian Johnston, Dominic Kwiatkowski, Cordelia Langford, John Sillitoe on behalf of the Wellcome Sanger Institute COVID-19 Surveillance Team |
| EPI_ISL_2682673, EPI_ISL_2682674, EPI_ISL_2682675 | Lighthouse Lab in Alderley Park | Wellcome Sanger Institute for the COVID-19 Genomics UK (COG-UK) Consortium | Jacquelyn Wynn, Mairead Hyland, The Lighthouse Lab in Alderley Park and Alex Alderton, Roberto Amato, Jeffrey Barrett, Sonia Goncalves, Ewan Harrison, David K. Jackson, Ian Johnston, Dominic Kwiatkowski, Cordelia Langford, John Sillitoe on behalf of the Wellcome Sanger Institute COVID-19 Surveillance Team |
| EPI_ISL_2682676, EPI_ISL_2682677 | Lighthouse Lab in Glasgow | Wellcome Sanger Institute for the COVID-19 Genomics UK (COG-UK) Consortium | Harper VanSteenhouse, Yumi Kasai, David Gray, Carol Clugston, Anna Dominiczak and Alex Alderton, Roberto Amato, Jeffrey Barrett, Sonia Goncalves, Ewan Harrison, David K. Jackson, Ian Johnston, Dominic Kwiatkowski, Cordelia Langford, John Sillitoe on behalf of the Wellcome Sanger Institute COVID-19 Surveillance Team |
| EPI_ISL_2682678 | Lighthouse Lab in Alderley Park | Wellcome Sanger Institute for the COVID-19 Genomics UK (COG-UK) Consortium | Jacquelyn Wynn, Mairead Hyland, The Lighthouse Lab in Alderley Park and Alex Alderton, Roberto Amato, Jeffrey Barrett, Sonia Goncalves, Ewan Harrison, David K. Jackson, Ian Johnston, Dominic Kwiatkowski, Cordelia Langford, John Sillitoe on behalf of the Wellcome Sanger Institute COVID-19 Surveillance Team |
| EPI_ISL_2682679 | Lighthouse Lab in Glasgow | Wellcome Sanger Institute for the COVID-19 Genomics UK (COG-UK) Consortium | Harper VanSteenhouse, Yumi Kasai, David Gray, Carol Clugston, Anna Dominiczak and Alex Alderton, Roberto Amato, Jeffrey Barrett, Sonia Goncalves, Ewan Harrison, David K. Jackson, Ian Johnston, Dominic Kwiatkowski, Cordelia Langford, John Sillitoe on behalf of the Wellcome Sanger Institute COVID-19 Surveillance Team |
| EPI_ISL_2682681 | Lighthouse Lab in Alderley Park | Wellcome Sanger Institute for the COVID-19 Genomics UK (COG-UK) Consortium | Jacquelyn Wynn, Mairead Hyland, The Lighthouse Lab in Alderley Park and Alex Alderton, Roberto Amato, Jeffrey Barrett, Sonia Goncalves, Ewan Harrison, David K. Jackson, Ian Johnston, Dominic Kwiatkowski, Cordelia Langford, John Sillitoe on behalf of the Wellcome Sanger Institute COVID-19 Surveillance Team |
| EPI_ISL_2682682 | Lighthouse Lab in Glasgow | Wellcome Sanger Institute for the COVID-19 Genomics UK (COG-UK) Consortium | Harper VanSteenhouse, Yumi Kasai, David Gray, Carol Clugston, Anna Dominiczak and Alex Alderton, Roberto Amato, Jeffrey Barrett, Sonia Goncalves, Ewan Harrison, David K. Jackson, Ian Johnston, Dominic Kwiatkowski, Cordelia Langford, John Sillitoe on behalf of the Wellcome Sanger Institute COVID-19 Surveillance Team |
| EPI_ISL_2682683 | Lighthouse Lab in Alderley Park | Wellcome Sanger Institute for the COVID-19 Genomics UK (COG-UK) Consortium | Jacquelyn Wynn, Mairead Hyland, The Lighthouse Lab in Alderley Park and Alex Alderton, Roberto Amato, Jeffrey Barrett, Sonia Goncalves, Ewan Harrison, David K. Jackson, Ian Johnston, Dominic Kwiatkowski, Cordelia Langford, John Sillitoe on behalf of the Wellcome Sanger Institute COVID-19 Surveillance Team |
| EPI_ISL_2682684, EPI_ISL_2682685, EPI_ISL_2682686 | Lighthouse Lab in Glasgow | Wellcome Sanger Institute for the COVID-19 Genomics UK (COG-UK) Consortium | Harper VanSteenhouse, Yumi Kasai, David Gray, Carol Clugston, Anna Dominiczak and Alex Alderton, Roberto Amato, Jeffrey Barrett, Sonia Goncalves, Ewan Harrison, David K. Jackson, Ian Johnston, Dominic Kwiatkowski, Cordelia Langford, John Sillitoe on behalf of the Wellcome Sanger Institute COVID-19 Surveillance Team |
| EPI_ISL_2682687, EPI_ISL_2682688 | Lighthouse Lab in Alderley Park | Wellcome Sanger Institute for the COVID-19 Genomics UK (COG-UK) Consortium | Jacquelyn Wynn, Mairead Hyland, The Lighthouse Lab in Alderley Park and Alex Alderton, Roberto Amato, Jeffrey Barrett, Sonia Goncalves, Ewan Harrison, David K. Jackson, Ian Johnston, Dominic Kwiatkowski, Cordelia Langford, John Sillitoe on behalf of the Wellcome Sanger Institute COVID-19 Surveillance Team |
| EPI_ISL_2682689 | Lighthouse Lab in Glasgow | Wellcome Sanger Institute for the COVID-19 Genomics UK (COG-UK) Consortium | Harper VanSteenhouse, Yumi Kasai, David Gray, Carol Clugston, Anna Dominiczak and Alex Alderton, Roberto Amato, Jeffrey Barrett, Sonia Goncalves, Ewan Harrison, David K. Jackson, Ian Johnston, Dominic Kwiatkowski, Cordelia Langford, John Sillitoe on behalf of the Wellcome Sanger Institute COVID-19 Surveillance Team |
| EPI_ISL_2682690 | Lighthouse Lab in Alderley Park | Wellcome Sanger Institute for the COVID-19 Genomics UK (COG-UK) Consortium | Jacquelyn Wynn, Mairead Hyland, The Lighthouse Lab in Alderley Park and Alex Alderton, Roberto Amato, Jeffrey Barrett, Sonia Goncalves, Ewan Harrison, David K. Jackson, Ian Johnston, Dominic Kwiatkowski, Cordelia Langford, John Sillitoe on behalf of the Wellcome Sanger Institute COVID-19 Surveillance Team |
| EPI_ISL_2682691, EPI_ISL_2682692, EPI_ISL_2682693, EPI_ISL_2682694 | Lighthouse Lab in Glasgow | Wellcome Sanger Institute for the COVID-19 Genomics UK (COG-UK) Consortium | Harper VanSteenhouse, Yumi Kasai, David Gray, Carol Clugston, Anna Dominiczak and Alex Alderton, Roberto Amato, Jeffrey Barrett, Sonia Goncalves, Ewan Harrison, David K. Jackson, Ian Johnston, Dominic Kwiatkowski, Cordelia Langford, John Sillitoe on behalf of the Wellcome Sanger Institute COVID-19 Surveillance Team |
| EPI_ISL_2682695 | Lighthouse Lab in Alderley Park | Wellcome Sanger Institute for the COVID-19 Genomics UK (COG-UK) Consortium | Jacquelyn Wynn, Mairead Hyland, The Lighthouse Lab in Alderley Park and Alex Alderton, Roberto Amato, Jeffrey Barrett, Sonia Goncalves, Ewan Harrison, David K. Jackson, Ian Johnston, Dominic Kwiatkowski, Cordelia Langford, John Sillitoe on behalf of the Wellcome Sanger Institute COVID-19 Surveillance Team |
| EPI_ISL_2682696, EPI_ISL_2682698, EPI_ISL_2682699 | Lighthouse Lab in Glasgow | Wellcome Sanger Institute for the COVID-19 Genomics UK (COG-UK) Consortium | Harper VanSteenhouse, Yumi Kasai, David Gray, Carol Clugston, Anna Dominiczak and Alex Alderton, Roberto Amato, Jeffrey Barrett, Sonia Goncalves, Ewan Harrison, David K. Jackson, Ian Johnston, Dominic Kwiatkowski, Cordelia Langford, John Sillitoe on behalf of the Wellcome Sanger Institute COVID-19 Surveillance Team |
| EPI_ISL_2682700 | Lighthouse Lab in Alderley Park | Wellcome Sanger Institute for the COVID-19 Genomics UK (COG-UK) Consortium | Jacquelyn Wynn, Mairead Hyland, The Lighthouse Lab in Alderley Park and Alex Alderton, Roberto Amato, Jeffrey Barrett, Sonia Goncalves, Ewan Harrison, David K. Jackson, Ian Johnston, Dominic Kwiatkowski, Cordelia Langford, John Sillitoe on behalf of the Wellcome Sanger Institute COVID-19 Surveillance Team |
| EPI_ISL_2682701 | Lighthouse Lab in Glasgow | Wellcome Sanger Institute for the COVID-19 Genomics UK (COG-UK) Consortium | Harper VanSteenhouse, Yumi Kasai, David Gray, Carol Clugston, Anna Dominiczak and Alex Alderton, Roberto Amato, Jeffrey Barrett, Sonia Goncalves, Ewan Harrison, David K. Jackson, Ian Johnston, Dominic Kwiatkowski, Cordelia Langford, John Sillitoe on behalf of the Wellcome Sanger Institute COVID-19 Surveillance Team |
| EPI_ISL_2682702 | Lighthouse Lab in Alderley Park | Wellcome Sanger Institute for the COVID-19 Genomics UK (COG-UK) Consortium | Jacquelyn Wynn, Mairead Hyland, The Lighthouse Lab in Alderley Park and Alex Alderton, Roberto Amato, Jeffrey Barrett, Sonia Goncalves, Ewan Harrison, David K. Jackson, Ian Johnston, Dominic Kwiatkowski, Cordelia Langford, John Sillitoe on behalf of the Wellcome Sanger Institute COVID-19 Surveillance Team |
| EPI_ISL_2682703, EPI_ISL_2682704 | Lighthouse Lab in Glasgow | Wellcome Sanger Institute for the COVID-19 Genomics UK (COG-UK) Consortium | Harper VanSteenhouse, Yumi Kasai, David Gray, Carol Clugston, Anna Dominiczak and Alex Alderton, Roberto Amato, Jeffrey Barrett, Sonia Goncalves, Ewan Harrison, David K. Jackson, Ian Johnston, Dominic Kwiatkowski, Cordelia Langford, John Sillitoe on behalf of the Wellcome Sanger Institute COVID-19 Surveillance Team |
| EPI_ISL_2682705, EPI_ISL_2682706, EPI_ISL_2682707 | Lighthouse Lab in Alderley Park | Wellcome Sanger Institute for the COVID-19 Genomics UK (COG-UK) Consortium | Jacquelyn Wynn, Mairead Hyland, The Lighthouse Lab in Alderley Park and Alex Alderton, Roberto Amato, Jeffrey Barrett, Sonia Goncalves, Ewan Harrison, David K. Jackson, Ian Johnston, Dominic Kwiatkowski, Cordelia Langford, John Sillitoe on behalf of the Wellcome Sanger Institute COVID-19 Surveillance Team |
| EPI_ISL_2682708 | Lighthouse Lab in Glasgow | Wellcome Sanger Institute for the COVID-19 Genomics UK (COG-UK) Consortium | Harper VanSteenhouse, Yumi Kasai, David Gray, Carol Clugston, Anna Dominiczak and Alex Alderton, Roberto Amato, Jeffrey Barrett, Sonia Goncalves, Ewan Harrison, David K. Jackson, Ian Johnston, Dominic Kwiatkowski, Cordelia Langford, John Sillitoe on behalf of the Wellcome Sanger Institute COVID-19 Surveillance Team |
| EPI_ISL_2682709 | Lighthouse Lab in Alderley Park | Wellcome Sanger Institute for the COVID-19 Genomics UK (COG-UK) Consortium | Jacquelyn Wynn, Mairead Hyland, The Lighthouse Lab in Alderley Park and Alex Alderton, Roberto Amato, Jeffrey Barrett, Sonia Goncalves, Ewan Harrison, David K. Jackson, Ian Johnston, Dominic Kwiatkowski, Cordelia Langford, John Sillitoe on behalf of the Wellcome Sanger Institute COVID-19 Surveillance Team |
| EPI_ISL_2682710, EPI_ISL_2682711, EPI_ISL_2682712 | Lighthouse Lab in Glasgow | Wellcome Sanger Institute for the COVID-19 Genomics UK (COG-UK) Consortium | Harper VanSteenhouse, Yumi Kasai, David Gray, Carol Clugston, Anna Dominiczak and Alex Alderton, Roberto Amato, Jeffrey Barrett, Sonia Goncalves, Ewan Harrison, David K. Jackson, Ian Johnston, Dominic Kwiatkowski, Cordelia Langford, John Sillitoe on behalf of the Wellcome Sanger Institute COVID-19 Surveillance Team |
| EPI_ISL_2682713, EPI_ISL_2682714 | Lighthouse Lab in Alderley Park | Wellcome Sanger Institute for the COVID-19 Genomics UK (COG-UK) Consortium | Jacquelyn Wynn, Mairead Hyland, The Lighthouse Lab in Alderley Park and Alex Alderton, Roberto Amato, Jeffrey Barrett, Sonia Goncalves, Ewan Harrison, David K. Jackson, Ian Johnston, Dominic Kwiatkowski, Cordelia Langford, John Sillitoe on behalf of the Wellcome Sanger Institute COVID-19 Surveillance Team |
| EPI_ISL_2682715, EPI_ISL_2682716 | Lighthouse Lab in Glasgow | Wellcome Sanger Institute for the COVID-19 Genomics UK (COG-UK) Consortium | Harper VanSteenhouse, Yumi Kasai, David Gray, Carol Clugston, Anna Dominiczak and Alex Alderton, Roberto Amato, Jeffrey Barrett, Sonia Goncalves, Ewan Harrison, David K. Jackson, Ian Johnston, Dominic Kwiatkowski, Cordelia Langford, John Sillitoe on behalf of the Wellcome Sanger Institute COVID-19 Surveillance Team |
| EPI_ISL_2682717 | Lighthouse Lab in Alderley Park | Wellcome Sanger Institute for the COVID-19 Genomics UK (COG-UK) Consortium | Jacquelyn Wynn, Mairead Hyland, The Lighthouse Lab in Alderley Park and Alex Alderton, Roberto Amato, Jeffrey Barrett, Sonia Goncalves, Ewan Harrison, David K. Jackson, Ian Johnston, Dominic Kwiatkowski, Cordelia Langford, John Sillitoe on behalf of the Wellcome Sanger Institute COVID-19 Surveillance Team |
| EPI_ISL_2682718 | Lighthouse Lab in Glasgow | Wellcome Sanger Institute for the COVID-19 Genomics UK (COG-UK) Consortium | Harper VanSteenhouse, Yumi Kasai, David Gray, Carol Clugston, Anna Dominiczak and Alex Alderton, Roberto Amato, Jeffrey Barrett, Sonia Goncalves, Ewan Harrison, David K. Jackson, Ian Johnston, Dominic Kwiatkowski, Cordelia Langford, John Sillitoe on behalf of the Wellcome Sanger Institute COVID-19 Surveillance Team |
| EPI_ISL_2682719, EPI_ISL_2682720, EPI_ISL_2682721, EPI_ISL_2682722, EPI_ISL_2682723 | Lighthouse Lab in Alderley Park | Wellcome Sanger Institute for the COVID-19 Genomics UK (COG-UK) Consortium | Jacquelyn Wynn, Mairead Hyland, The Lighthouse Lab in Alderley Park and Alex Alderton, Roberto Amato, Jeffrey Barrett, Sonia Goncalves, Ewan Harrison, David K. Jackson, Ian Johnston, Dominic Kwiatkowski, Cordelia Langford, John Sillitoe on behalf of the Wellcome Sanger Institute COVID-19 Surveillance Team |
| EPI_ISL_2682724 | Lighthouse Lab in Glasgow | Wellcome Sanger Institute for the COVID-19 Genomics UK (COG-UK) Consortium | Harper VanSteenhouse, Yumi Kasai, David Gray, Carol Clugston, Anna Dominiczak and Alex Alderton, Roberto Amato, Jeffrey Barrett, Sonia Goncalves, Ewan Harrison, David K. Jackson, Ian Johnston, Dominic Kwiatkowski, Cordelia Langford, John Sillitoe on behalf of the Wellcome Sanger Institute COVID-19 Surveillance Team |
| EPI_ISL_2682725, EPI_ISL_2682726 | Lighthouse Lab in Alderley Park | Wellcome Sanger Institute for the COVID-19 Genomics UK (COG-UK) Consortium | Jacquelyn Wynn, Mairead Hyland, The Lighthouse Lab in Alderley Park and Alex Alderton, Roberto Amato, Jeffrey Barrett, Sonia Goncalves, Ewan Harrison, David K. Jackson, Ian Johnston, Dominic Kwiatkowski, Cordelia Langford, John Sillitoe on behalf of the Wellcome Sanger Institute COVID-19 Surveillance Team |
| EPI_ISL_2682727, EPI_ISL_2682728, EPI_ISL_2682729, EPI_ISL_2682730, EPI_ISL_2682731 | Lighthouse Lab in Glasgow | Wellcome Sanger Institute for the COVID-19 Genomics UK (COG-UK) Consortium | Harper VanSteenhouse, Yumi Kasai, David Gray, Carol Clugston, Anna Dominiczak and Alex Alderton, Roberto Amato, Jeffrey Barrett, Sonia Goncalves, Ewan Harrison, David K. Jackson, Ian Johnston, Dominic Kwiatkowski, Cordelia Langford, John Sillitoe on behalf of the Wellcome Sanger Institute COVID-19 Surveillance Team |
| EPI_ISL_2682732, EPI_ISL_2682733, EPI_ISL_2682734, EPI_ISL_2682735, | Lighthouse Lab in Alderley Park | Wellcome Sanger Institute for the COVID-19 Genomics UK (COG-UK) Consortium | Jacquelyn Wynn, Mairead Hyland, The Lighthouse Lab in Alderley Park and Alex Alderton, Roberto Amato, Jeffrey Barrett, Sonia Goncalves, Ewan Harrison, David K. Jackson, Ian Johnston, Dominic Kwiatkowski, Cordelia Langford, John Sillitoe on behalf of the Wellcome Sanger Institute COVID-19 Surveillance Team |

| | | | |
|---|---|---|---|
| EPI_ISL_2682736 | | | |
| EPI_ISL_2682737 | Lighthouse Lab in Glasgow | Wellcome Sanger Institute for the COVID-19 Genomics UK (COG-UK) Consortium | Harper VanSteenhouse, Yumi Kasai, David Gray, Carol Clugston, Anna Dominiczak and Alex Alderton, Roberto Amato, Jeffrey Barrett, Sonia Goncalves, Ewan Harrison, David K. Jackson, Ian Johnston, Dominic Kwiatkowski, Cordelia Langford, John Sillitoe on behalf of the Wellcome Sanger Institute COVID-19 Surveillance Team |
| EPI_ISL_2682738, EPI_ISL_2682740, EPI_ISL_2682741 | Lighthouse Lab in Alderley Park | Wellcome Sanger Institute for the COVID-19 Genomics UK (COG-UK) Consortium | Jacquelyn Wynn, Mairead Hyland, The Lighthouse Lab in Alderley Park and Alex Alderton, Roberto Amato, Jeffrey Barrett, Sonia Goncalves, Ewan Harrison, David K. Jackson, Ian Johnston, Dominic Kwiatkowski, Cordelia Langford, John Sillitoe on behalf of the Wellcome Sanger Institute COVID-19 Surveillance Team |
| EPI_ISL_2682742 | Lighthouse Lab in Glasgow | Wellcome Sanger Institute for the COVID-19 Genomics UK (COG-UK) Consortium | Harper VanSteenhouse, Yumi Kasai, David Gray, Carol Clugston, Anna Dominiczak and Alex Alderton, Roberto Amato, Jeffrey Barrett, Sonia Goncalves, Ewan Harrison, David K. Jackson, Ian Johnston, Dominic Kwiatkowski, Cordelia Langford, John Sillitoe on behalf of the Wellcome Sanger Institute COVID-19 Surveillance Team |
| EPI_ISL_2682743 | Lighthouse Lab in Alderley Park | Wellcome Sanger Institute for the COVID-19 Genomics UK (COG-UK) Consortium | Jacquelyn Wynn, Mairead Hyland, The Lighthouse Lab in Alderley Park and Alex Alderton, Roberto Amato, Jeffrey Barrett, Sonia Goncalves, Ewan Harrison, David K. Jackson, Ian Johnston, Dominic Kwiatkowski, Cordelia Langford, John Sillitoe on behalf of the Wellcome Sanger Institute COVID-19 Surveillance Team |
| EPI_ISL_2682744, EPI_ISL_2682745 | Lighthouse Lab in Glasgow | Wellcome Sanger Institute for the COVID-19 Genomics UK (COG-UK) Consortium | Harper VanSteenhouse, Yumi Kasai, David Gray, Carol Clugston, Anna Dominiczak and Alex Alderton, Roberto Amato, Jeffrey Barrett, Sonia Goncalves, Ewan Harrison, David K. Jackson, Ian Johnston, Dominic Kwiatkowski, Cordelia Langford, John Sillitoe on behalf of the Wellcome Sanger Institute COVID-19 Surveillance Team |
| EPI_ISL_2682746 | Lighthouse Lab in Alderley Park | Wellcome Sanger Institute for the COVID-19 Genomics UK (COG-UK) Consortium | Jacquelyn Wynn, Mairead Hyland, The Lighthouse Lab in Alderley Park and Alex Alderton, Roberto Amato, Jeffrey Barrett, Sonia Goncalves, Ewan Harrison, David K. Jackson, Ian Johnston, Dominic Kwiatkowski, Cordelia Langford, John Sillitoe on behalf of the Wellcome Sanger Institute COVID-19 Surveillance Team |
| EPI_ISL_2682747 | Lighthouse Lab in Glasgow | Wellcome Sanger Institute for the COVID-19 Genomics UK (COG-UK) Consortium | Harper VanSteenhouse, Yumi Kasai, David Gray, Carol Clugston, Anna Dominiczak and Alex Alderton, Roberto Amato, Jeffrey Barrett, Sonia Goncalves, Ewan Harrison, David K. Jackson, Ian Johnston, Dominic Kwiatkowski, Cordelia Langford, John Sillitoe on behalf of the Wellcome Sanger Institute COVID-19 Surveillance Team |
| EPI_ISL_2682748, EPI_ISL_2682749 | Lighthouse Lab in Alderley Park | Wellcome Sanger Institute for the COVID-19 Genomics UK (COG-UK) Consortium | Jacquelyn Wynn, Mairead Hyland, The Lighthouse Lab in Alderley Park and Alex Alderton, Roberto Amato, Jeffrey Barrett, Sonia Goncalves, Ewan Harrison, David K. Jackson, Ian Johnston, Dominic Kwiatkowski, Cordelia Langford, John Sillitoe on behalf of the Wellcome Sanger Institute COVID-19 Surveillance Team |
| EPI_ISL_2682751, EPI_ISL_2682752 | Lighthouse Lab in Glasgow | Wellcome Sanger Institute for the COVID-19 Genomics UK (COG-UK) Consortium | Harper VanSteenhouse, Yumi Kasai, David Gray, Carol Clugston, Anna Dominiczak and Alex Alderton, Roberto Amato, Jeffrey Barrett, Sonia Goncalves, Ewan Harrison, David K. Jackson, Ian Johnston, Dominic Kwiatkowski, Cordelia Langford, John Sillitoe on behalf of the Wellcome Sanger Institute COVID-19 Surveillance Team |
| EPI_ISL_2682753 | Lighthouse Lab in Alderley Park | Wellcome Sanger Institute for the COVID-19 Genomics UK (COG-UK) Consortium | Jacquelyn Wynn, Mairead Hyland, The Lighthouse Lab in Alderley Park and Alex Alderton, Roberto Amato, Jeffrey Barrett, Sonia Goncalves, Ewan Harrison, David K. Jackson, Ian Johnston, Dominic Kwiatkowski, Cordelia Langford, John Sillitoe on behalf of the Wellcome Sanger Institute COVID-19 Surveillance Team |
| EPI_ISL_2682754 | Lighthouse Lab in Glasgow | Wellcome Sanger Institute for the COVID-19 Genomics UK (COG-UK) Consortium | Harper VanSteenhouse, Yumi Kasai, David Gray, Carol Clugston, Anna Dominiczak and Alex Alderton, Roberto Amato, Jeffrey Barrett, Sonia Goncalves, Ewan Harrison, David K. Jackson, Ian Johnston, Dominic Kwiatkowski, Cordelia Langford, John Sillitoe on behalf of the Wellcome Sanger Institute COVID-19 Surveillance Team |
| EPI_ISL_2682755, EPI_ISL_2682756, EPI_ISL_2682757 | Lighthouse Lab in Alderley Park | Wellcome Sanger Institute for the COVID-19 Genomics UK (COG-UK) Consortium | Jacquelyn Wynn, Mairead Hyland, The Lighthouse Lab in Alderley Park and Alex Alderton, Roberto Amato, Jeffrey Barrett, Sonia Goncalves, Ewan Harrison, David K. Jackson, Ian Johnston, Dominic Kwiatkowski, Cordelia Langford, John Sillitoe on behalf of the Wellcome Sanger Institute COVID-19 Surveillance Team |
| EPI_ISL_2682758 | Lighthouse Lab in Glasgow | Wellcome Sanger Institute for the COVID-19 Genomics UK (COG-UK) Consortium | Harper VanSteenhouse, Yumi Kasai, David Gray, Carol Clugston, Anna Dominiczak and Alex Alderton, Roberto Amato, Jeffrey Barrett, Sonia Goncalves, Ewan Harrison, David K. Jackson, Ian Johnston, Dominic Kwiatkowski, Cordelia Langford, John Sillitoe on behalf of the Wellcome Sanger Institute COVID-19 Surveillance Team |
| EPI_ISL_2682759 | Lighthouse Lab in Alderley Park | Wellcome Sanger Institute for the COVID-19 Genomics UK (COG-UK) Consortium | Jacquelyn Wynn, Mairead Hyland, The Lighthouse Lab in Alderley Park and Alex Alderton, Roberto Amato, Jeffrey Barrett, Sonia Goncalves, Ewan Harrison, David K. Jackson, Ian Johnston, Dominic Kwiatkowski, Cordelia Langford, John Sillitoe on behalf of the Wellcome Sanger Institute COVID-19 Surveillance Team |
| EPI_ISL_2682760, EPI_ISL_2682761 | Lighthouse Lab in Glasgow | Wellcome Sanger Institute for the COVID-19 Genomics UK (COG-UK) Consortium | Harper VanSteenhouse, Yumi Kasai, David Gray, Carol Clugston, Anna Dominiczak and Alex Alderton, Roberto Amato, Jeffrey Barrett, Sonia Goncalves, Ewan Harrison, David K. Jackson, Ian Johnston, Dominic Kwiatkowski, Cordelia Langford, John Sillitoe on behalf of the Wellcome Sanger Institute COVID-19 Surveillance Team |
| EPI_ISL_2682762, EPI_ISL_2682763, EPI_ISL_2682764, EPI_ISL_2682765 | Lighthouse Lab in Alderley Park | Wellcome Sanger Institute for the COVID-19 Genomics UK (COG-UK) Consortium | Jacquelyn Wynn, Mairead Hyland, The Lighthouse Lab in Alderley Park and Alex Alderton, Roberto Amato, Jeffrey Barrett, Sonia Goncalves, Ewan Harrison, David K. Jackson, Ian Johnston, Dominic Kwiatkowski, Cordelia Langford, John Sillitoe on behalf of the Wellcome Sanger Institute COVID-19 Surveillance Team |
| EPI_ISL_2682766, EPI_ISL_2682767, EPI_ISL_2682768, EPI_ISL_2682769 | Lighthouse Lab in Glasgow | Wellcome Sanger Institute for the COVID-19 Genomics UK (COG-UK) Consortium | Harper VanSteenhouse, Yumi Kasai, David Gray, Carol Clugston, Anna Dominiczak and Alex Alderton, Roberto Amato, Jeffrey Barrett, Sonia Goncalves, Ewan Harrison, David K. Jackson, Ian Johnston, Dominic Kwiatkowski, Cordelia Langford, John Sillitoe on behalf of the Wellcome Sanger Institute COVID-19 Surveillance Team |
| EPI_ISL_2682770 | Lighthouse Lab in Alderley Park | Wellcome Sanger Institute for the COVID-19 Genomics UK (COG-UK) Consortium | Jacquelyn Wynn, Mairead Hyland, The Lighthouse Lab in Alderley Park and Alex Alderton, Roberto Amato, Jeffrey Barrett, Sonia Goncalves, Ewan Harrison, David K. Jackson, Ian Johnston, Dominic Kwiatkowski, Cordelia Langford, John Sillitoe on behalf of the Wellcome Sanger Institute COVID-19 Surveillance Team |
| EPI_ISL_2682771, EPI_ISL_2682772 | Lighthouse Lab in Glasgow | Wellcome Sanger Institute for the COVID-19 Genomics UK (COG-UK) Consortium | Harper VanSteenhouse, Yumi Kasai, David Gray, Carol Clugston, Anna Dominiczak and Alex Alderton, Roberto Amato, Jeffrey Barrett, Sonia Goncalves, Ewan Harrison, David K. Jackson, Ian Johnston, Dominic Kwiatkowski, Cordelia Langford, John Sillitoe on behalf of the Wellcome Sanger Institute COVID-19 Surveillance Team |
| EPI_ISL_2682773, EPI_ISL_2682775, EPI_ISL_2682776 | Lighthouse Lab in Alderley Park | Wellcome Sanger Institute for the COVID-19 Genomics UK (COG-UK) Consortium | Jacquelyn Wynn, Mairead Hyland, The Lighthouse Lab in Alderley Park and Alex Alderton, Roberto Amato, Jeffrey Barrett, Sonia Goncalves, Ewan Harrison, David K. Jackson, Ian Johnston, Dominic Kwiatkowski, Cordelia Langford, John Sillitoe on behalf of the Wellcome Sanger Institute COVID-19 Surveillance Team |
| EPI_ISL_2682777, EPI_ISL_2682778 | Lighthouse Lab in Glasgow | Wellcome Sanger Institute for the COVID-19 Genomics UK (COG-UK) Consortium | Harper VanSteenhouse, Yumi Kasai, David Gray, Carol Clugston, Anna Dominiczak and Alex Alderton, Roberto Amato, Jeffrey Barrett, Sonia Goncalves, Ewan Harrison, David K. Jackson, Ian Johnston, Dominic Kwiatkowski, Cordelia Langford, John Sillitoe on behalf of the Wellcome Sanger Institute COVID-19 Surveillance Team |
| EPI_ISL_2682779 | Lighthouse Lab in Alderley Park | Wellcome Sanger Institute for the COVID-19 Genomics UK (COG-UK) Consortium | Jacquelyn Wynn, Mairead Hyland, The Lighthouse Lab in Alderley Park and Alex Alderton, Roberto Amato, Jeffrey Barrett, Sonia Goncalves, Ewan Harrison, David K. Jackson, Ian Johnston, Dominic Kwiatkowski, Cordelia Langford, John Sillitoe on behalf of the Wellcome Sanger Institute COVID-19 Surveillance Team |
| EPI_ISL_2682780 | Lighthouse Lab in Glasgow | Wellcome Sanger Institute for the COVID-19 Genomics UK (COG-UK) Consortium | Harper VanSteenhouse, Yumi Kasai, David Gray, Carol Clugston, Anna Dominiczak and Alex Alderton, Roberto Amato, Jeffrey Barrett, Sonia Goncalves, Ewan Harrison, David K. Jackson, Ian Johnston, Dominic Kwiatkowski, Cordelia Langford, John Sillitoe on behalf of the Wellcome Sanger Institute COVID-19 Surveillance Team |
| EPI_ISL_2682781, EPI_ISL_2682782 | Lighthouse Lab in Alderley Park | Wellcome Sanger Institute for the COVID-19 Genomics UK (COG-UK) Consortium | Jacquelyn Wynn, Mairead Hyland, The Lighthouse Lab in Alderley Park and Alex Alderton, Roberto Amato, Jeffrey Barrett, Sonia Goncalves, Ewan Harrison, David K. Jackson, Ian Johnston, Dominic Kwiatkowski, Cordelia Langford, John Sillitoe on behalf of the Wellcome Sanger Institute COVID-19 Surveillance Team |
| EPI_ISL_2682783 | Lighthouse Lab in Glasgow | Wellcome Sanger Institute for the COVID-19 Genomics UK (COG-UK) Consortium | Harper VanSteenhouse, Yumi Kasai, David Gray, Carol Clugston, Anna Dominiczak and Alex Alderton, Roberto Amato, Jeffrey Barrett, Sonia Goncalves, Ewan Harrison, David K. Jackson, Ian Johnston, Dominic Kwiatkowski, Cordelia Langford, John Sillitoe on behalf of the Wellcome Sanger Institute COVID-19 Surveillance Team |
| EPI_ISL_2682784 | Lighthouse Lab in Alderley Park | Wellcome Sanger Institute for the COVID-19 Genomics UK (COG-UK) Consortium | Jacquelyn Wynn, Mairead Hyland, The Lighthouse Lab in Alderley Park and Alex Alderton, Roberto Amato, Jeffrey Barrett, Sonia Goncalves, Ewan Harrison, David K. Jackson, Ian Johnston, Dominic Kwiatkowski, Cordelia Langford, John Sillitoe on behalf of the Wellcome Sanger Institute COVID-19 Surveillance Team |
| EPI_ISL_2682785 | Lighthouse Lab in Glasgow | Wellcome Sanger Institute for the COVID-19 Genomics UK (COG-UK) Consortium | Harper VanSteenhouse, Yumi Kasai, David Gray, Carol Clugston, Anna Dominiczak and Alex Alderton, Roberto Amato, Jeffrey Barrett, Sonia Goncalves, Ewan Harrison, David K. Jackson, Ian Johnston, Dominic Kwiatkowski, Cordelia Langford, John Sillitoe on behalf of the Wellcome Sanger Institute COVID-19 Surveillance Team |
| EPI_ISL_2682786, EPI_ISL_2682787, EPI_ISL_2682788 | Lighthouse Lab in Alderley Park | Wellcome Sanger Institute for the COVID-19 Genomics UK (COG-UK) Consortium | Jacquelyn Wynn, Mairead Hyland, The Lighthouse Lab in Alderley Park and Alex Alderton, Roberto Amato, Jeffrey Barrett, Sonia Goncalves, Ewan Harrison, David K. Jackson, Ian Johnston, Dominic Kwiatkowski, Cordelia Langford, John Sillitoe on behalf of the Wellcome Sanger Institute COVID-19 Surveillance Team |
| EPI_ISL_2682789 | Lighthouse Lab in Glasgow | Wellcome Sanger Institute for the COVID-19 Genomics UK (COG-UK) Consortium | Harper VanSteenhouse, Yumi Kasai, David Gray, Carol Clugston, Anna Dominiczak and Alex Alderton, Roberto Amato, Jeffrey Barrett, Sonia Goncalves, Ewan Harrison, David K. Jackson, Ian Johnston, Dominic Kwiatkowski, Cordelia Langford, John Sillitoe on behalf of the Wellcome Sanger Institute COVID-19 Surveillance Team |
| EPI_ISL_2682790 | Lighthouse Lab in Alderley Park | Wellcome Sanger Institute for the COVID-19 Genomics UK (COG-UK) Consortium | Jacquelyn Wynn, Mairead Hyland, The Lighthouse Lab in Alderley Park and Alex Alderton, Roberto Amato, Jeffrey Barrett, Sonia Goncalves, Ewan Harrison, David K. Jackson, Ian Johnston, Dominic Kwiatkowski, Cordelia Langford, John Sillitoe on behalf of the Wellcome Sanger Institute COVID-19 Surveillance Team |
| EPI_ISL_2682791 | Lighthouse Lab in Glasgow | Wellcome Sanger Institute for the COVID-19 Genomics UK (COG-UK) Consortium | Harper VanSteenhouse, Yumi Kasai, David Gray, Carol Clugston, Anna Dominiczak and Alex Alderton, Roberto Amato, Jeffrey Barrett, Sonia Goncalves, Ewan Harrison, David K. Jackson, Ian Johnston, Dominic Kwiatkowski, Cordelia Langford, John Sillitoe on behalf of the Wellcome Sanger Institute COVID-19 Surveillance Team |
| EPI_ISL_2682792 | Lighthouse Lab in Alderley Park | Wellcome Sanger Institute for the COVID-19 Genomics UK (COG-UK) Consortium | Jacquelyn Wynn, Mairead Hyland, The Lighthouse Lab in Alderley Park and Alex Alderton, Roberto Amato, Jeffrey Barrett, Sonia Goncalves, Ewan Harrison, David K. Jackson, Ian Johnston, Dominic Kwiatkowski, Cordelia Langford, John Sillitoe on behalf of the Wellcome Sanger Institute COVID-19 Surveillance Team |
| EPI_ISL_2682793 | Lighthouse Lab in Glasgow | Wellcome Sanger Institute for the COVID-19 Genomics UK (COG-UK) Consortium | Harper VanSteenhouse, Yumi Kasai, David Gray, Carol Clugston, Anna Dominiczak and Alex Alderton, Roberto Amato, Jeffrey Barrett, Sonia Goncalves, Ewan Harrison, David K. Jackson, Ian Johnston, Dominic Kwiatkowski, Cordelia Langford, John Sillitoe on behalf of the Wellcome Sanger Institute COVID-19 Surveillance Team |
| EPI_ISL_2682794 | Lighthouse Lab in Alderley Park | Wellcome Sanger Institute for the COVID-19 Genomics UK (COG-UK) Consortium | Jacquelyn Wynn, Mairead Hyland, The Lighthouse Lab in Alderley Park and Alex Alderton, Roberto Amato, Jeffrey Barrett, Sonia Goncalves, Ewan Harrison, David K. Jackson, Ian Johnston, Dominic Kwiatkowski, Cordelia Langford, John Sillitoe on behalf of the Wellcome Sanger Institute COVID-19 Surveillance Team |
| EPI_ISL_2682795 | Lighthouse Lab in Glasgow | Wellcome Sanger Institute for the COVID-19 Genomics UK (COG-UK) Consortium | Harper VanSteenhouse, Yumi Kasai, David Gray, Carol Clugston, Anna Dominiczak and Alex Alderton, Roberto Amato, Jeffrey Barrett, Sonia Goncalves, Ewan Harrison, David K. Jackson, Ian Johnston, Dominic Kwiatkowski, Cordelia Langford, John Sillitoe on behalf of the Wellcome Sanger Institute COVID-19 Surveillance Team |
| EPI_ISL_2682796 | Lighthouse Lab in Alderley Park | Wellcome Sanger Institute for the COVID-19 Genomics UK (COG-UK) Consortium | Jacquelyn Wynn, Mairead Hyland, The Lighthouse Lab in Alderley Park and Alex Alderton, Roberto Amato, Jeffrey Barrett, Sonia Goncalves, Ewan Harrison, David K. Jackson, Ian Johnston, Dominic Kwiatkowski, Cordelia Langford, John Sillitoe on behalf of the Wellcome Sanger Institute COVID-19 Surveillance Team |
| EPI_ISL_2682797 | Lighthouse Lab in Glasgow | Wellcome Sanger Institute for the COVID-19 Genomics UK (COG-UK) Consortium | Harper VanSteenhouse, Yumi Kasai, David Gray, Carol Clugston, Anna Dominiczak and Alex Alderton, Roberto Amato, Jeffrey Barrett, Sonia Goncalves, Ewan Harrison, David K. Jackson, Ian Johnston, Dominic Kwiatkowski, Cordelia Langford, John Sillitoe on behalf of the Wellcome Sanger Institute COVID-19 Surveillance Team |
| EPI_ISL_2682798 | Lighthouse Lab in Alderley Park | Wellcome Sanger Institute for the COVID-19 Genomics UK (COG-UK) Consortium | Jacquelyn Wynn, Mairead Hyland, The Lighthouse Lab in Alderley Park and Alex Alderton, Roberto Amato, Jeffrey Barrett, Sonia Goncalves, Ewan Harrison, David K. Jackson, Ian Johnston, Dominic Kwiatkowski, Cordelia Langford, John Sillitoe on behalf of the Wellcome Sanger Institute COVID-19 Surveillance Team |
| EPI_ISL_2682799, EPI_ISL_2682800 | Lighthouse Lab in Glasgow | Wellcome Sanger Institute for the COVID-19 Genomics UK (COG-UK) Consortium | Harper VanSteenhouse, Yumi Kasai, David Gray, Carol Clugston, Anna Dominiczak and Alex Alderton, Roberto Amato, Jeffrey Barrett, Sonia Goncalves, Ewan Harrison, David K. Jackson, Ian Johnston, Dominic Kwiatkowski, Cordelia Langford, John Sillitoe on behalf of the Wellcome Sanger Institute COVID-19 Surveillance Team |
| EPI_ISL_2682802 | Lighthouse Lab in Alderley Park | Wellcome Sanger Institute for the COVID-19 Genomics UK (COG-UK) Consortium | Jacquelyn Wynn, Mairead Hyland, The Lighthouse Lab in Alderley Park and Alex Alderton, Roberto Amato, Jeffrey Barrett, Sonia Goncalves, Ewan Harrison, David K. Jackson, Ian Johnston, Dominic Kwiatkowski, Cordelia Langford, John Sillitoe on behalf of the Wellcome Sanger Institute COVID-19 Surveillance Team |
| EPI_ISL_2682803 | Lighthouse Lab in Glasgow | Wellcome Sanger Institute for the COVID-19 Genomics UK (COG-UK) Consortium | Harper VanSteenhouse, Yumi Kasai, David Gray, Carol Clugston, Anna Dominiczak and Alex Alderton, Roberto Amato, Jeffrey Barrett, Sonia Goncalves, Ewan Harrison, David K. Jackson, Ian Johnston, Dominic Kwiatkowski, Cordelia Langford, John Sillitoe on behalf of the Wellcome Sanger Institute COVID-19 Surveillance Team |
| EPI_ISL_2682804, | Lighthouse Lab in Alderley Park | Wellcome Sanger Institute for the COVID-19 Genomics UK (COG-UK) | Jacquelyn Wynn, Mairead Hyland, The Lighthouse Lab in Alderley Park and Alex Alderton, Roberto Amato, Jeffrey Barrett, Sonia Goncalves, Ewan Harrison, David K. Jackson, Ian Johnston, Dominic Kwiatkowski, Cordelia Langford, John Sillitoe on |

| | | | |
|---|---|---|---|
| EPI_ISL_2682805 | | Consortium | behalf of the Wellcome Sanger Institute COVID-19 Surveillance Team |
| EPI_ISL_2682806 | Lighthouse Lab in Glasgow | Wellcome Sanger Institute for the COVID-19 Genomics UK (COG-UK) Consortium | Harper VanSteenhouse, Yumi Kasai, David Gray, Carol Clugston, Anna Dominiczak and Alex Alderton, Roberto Amato, Jeffrey Barrett, Sonia Goncalves, Ewan Harrison, David K. Jackson, Ian Johnston, Dominic Kwiatkowski, Cordelia Langford, John Sillitoe on behalf of the Wellcome Sanger Institute COVID-19 Surveillance Team |
| EPI_ISL_2682807, EPI_ISL_2682808 | Lighthouse Lab in Alderley Park | Wellcome Sanger Institute for the COVID-19 Genomics UK (COG-UK) Consortium | Jacquelyn Wynn, Mairead Hyland, The Lighthouse Lab in Alderley Park and Alex Alderton, Roberto Amato, Jeffrey Barrett, Sonia Goncalves, Ewan Harrison, David K. Jackson, Ian Johnston, Dominic Kwiatkowski, Cordelia Langford, John Sillitoe on behalf of the Wellcome Sanger Institute COVID-19 Surveillance Team |
| EPI_ISL_2682809, EPI_ISL_2682810 | Lighthouse Lab in Glasgow | Wellcome Sanger Institute for the COVID-19 Genomics UK (COG-UK) Consortium | Harper VanSteenhouse, Yumi Kasai, David Gray, Carol Clugston, Anna Dominiczak and Alex Alderton, Roberto Amato, Jeffrey Barrett, Sonia Goncalves, Ewan Harrison, David K. Jackson, Ian Johnston, Dominic Kwiatkowski, Cordelia Langford, John Sillitoe on behalf of the Wellcome Sanger Institute COVID-19 Surveillance Team |
| EPI_ISL_2682811, EPI_ISL_2682812 | Lighthouse Lab in Alderley Park | Wellcome Sanger Institute for the COVID-19 Genomics UK (COG-UK) Consortium | Jacquelyn Wynn, Mairead Hyland, The Lighthouse Lab in Alderley Park and Alex Alderton, Roberto Amato, Jeffrey Barrett, Sonia Goncalves, Ewan Harrison, David K. Jackson, Ian Johnston, Dominic Kwiatkowski, Cordelia Langford, John Sillitoe on behalf of the Wellcome Sanger Institute COVID-19 Surveillance Team |
| EPI_ISL_2682813, EPI_ISL_2682814 | Lighthouse Lab in Glasgow | Wellcome Sanger Institute for the COVID-19 Genomics UK (COG-UK) Consortium | Harper VanSteenhouse, Yumi Kasai, David Gray, Carol Clugston, Anna Dominiczak and Alex Alderton, Roberto Amato, Jeffrey Barrett, Sonia Goncalves, Ewan Harrison, David K. Jackson, Ian Johnston, Dominic Kwiatkowski, Cordelia Langford, John Sillitoe on behalf of the Wellcome Sanger Institute COVID-19 Surveillance Team |
| EPI_ISL_2682815, EPI_ISL_2682816, EPI_ISL_2682817, EPI_ISL_2682818, EPI_ISL_2682819 | Lighthouse Lab in Alderley Park | Wellcome Sanger Institute for the COVID-19 Genomics UK (COG-UK) Consortium | Jacquelyn Wynn, Mairead Hyland, The Lighthouse Lab in Alderley Park and Alex Alderton, Roberto Amato, Jeffrey Barrett, Sonia Goncalves, Ewan Harrison, David K. Jackson, Ian Johnston, Dominic Kwiatkowski, Cordelia Langford, John Sillitoe on behalf of the Wellcome Sanger Institute COVID-19 Surveillance Team |
| EPI_ISL_2682820 | Lighthouse Lab in Glasgow | Wellcome Sanger Institute for the COVID-19 Genomics UK (COG-UK) Consortium | Harper VanSteenhouse, Yumi Kasai, David Gray, Carol Clugston, Anna Dominiczak and Alex Alderton, Roberto Amato, Jeffrey Barrett, Sonia Goncalves, Ewan Harrison, David K. Jackson, Ian Johnston, Dominic Kwiatkowski, Cordelia Langford, John Sillitoe on behalf of the Wellcome Sanger Institute COVID-19 Surveillance Team |
| EPI_ISL_2682821, EPI_ISL_2682822, EPI_ISL_2682823, EPI_ISL_2682824, EPI_ISL_2682825, EPI_ISL_2682826 | Lighthouse Lab in Alderley Park | Wellcome Sanger Institute for the COVID-19 Genomics UK (COG-UK) Consortium | Jacquelyn Wynn, Mairead Hyland, The Lighthouse Lab in Alderley Park and Alex Alderton, Roberto Amato, Jeffrey Barrett, Sonia Goncalves, Ewan Harrison, David K. Jackson, Ian Johnston, Dominic Kwiatkowski, Cordelia Langford, John Sillitoe on behalf of the Wellcome Sanger Institute COVID-19 Surveillance Team |
| EPI_ISL_2682827, EPI_ISL_2682828 | Lighthouse Lab in Glasgow | Wellcome Sanger Institute for the COVID-19 Genomics UK (COG-UK) Consortium | Harper VanSteenhouse, Yumi Kasai, David Gray, Carol Clugston, Anna Dominiczak and Alex Alderton, Roberto Amato, Jeffrey Barrett, Sonia Goncalves, Ewan Harrison, David K. Jackson, Ian Johnston, Dominic Kwiatkowski, Cordelia Langford, John Sillitoe on behalf of the Wellcome Sanger Institute COVID-19 Surveillance Team |
| EPI_ISL_2682829, EPI_ISL_2682830 | Lighthouse Lab in Alderley Park | Wellcome Sanger Institute for the COVID-19 Genomics UK (COG-UK) Consortium | Jacquelyn Wynn, Mairead Hyland, The Lighthouse Lab in Alderley Park and Alex Alderton, Roberto Amato, Jeffrey Barrett, Sonia Goncalves, Ewan Harrison, David K. Jackson, Ian Johnston, Dominic Kwiatkowski, Cordelia Langford, John Sillitoe on behalf of the Wellcome Sanger Institute COVID-19 Surveillance Team |
| EPI_ISL_2682831 | Lighthouse Lab in Glasgow | Wellcome Sanger Institute for the COVID-19 Genomics UK (COG-UK) Consortium | Harper VanSteenhouse, Yumi Kasai, David Gray, Carol Clugston, Anna Dominiczak and Alex Alderton, Roberto Amato, Jeffrey Barrett, Sonia Goncalves, Ewan Harrison, David K. Jackson, Ian Johnston, Dominic Kwiatkowski, Cordelia Langford, John Sillitoe on behalf of the Wellcome Sanger Institute COVID-19 Surveillance Team |
| EPI_ISL_2682832, EPI_ISL_2682833 | Lighthouse Lab in Alderley Park | Wellcome Sanger Institute for the COVID-19 Genomics UK (COG-UK) Consortium | Jacquelyn Wynn, Mairead Hyland, The Lighthouse Lab in Alderley Park and Alex Alderton, Roberto Amato, Jeffrey Barrett, Sonia Goncalves, Ewan Harrison, David K. Jackson, Ian Johnston, Dominic Kwiatkowski, Cordelia Langford, John Sillitoe on behalf of the Wellcome Sanger Institute COVID-19 Surveillance Team |
| EPI_ISL_2682834, EPI_ISL_2682835 | Lighthouse Lab in Glasgow | Wellcome Sanger Institute for the COVID-19 Genomics UK (COG-UK) Consortium | Harper VanSteenhouse, Yumi Kasai, David Gray, Carol Clugston, Anna Dominiczak and Alex Alderton, Roberto Amato, Jeffrey Barrett, Sonia Goncalves, Ewan Harrison, David K. Jackson, Ian Johnston, Dominic Kwiatkowski, Cordelia Langford, John Sillitoe on behalf of the Wellcome Sanger Institute COVID-19 Surveillance Team |
| EPI_ISL_2682836, EPI_ISL_2682837, EPI_ISL_2682838 | Lighthouse Lab in Alderley Park | Wellcome Sanger Institute for the COVID-19 Genomics UK (COG-UK) Consortium | Jacquelyn Wynn, Mairead Hyland, The Lighthouse Lab in Alderley Park and Alex Alderton, Roberto Amato, Jeffrey Barrett, Sonia Goncalves, Ewan Harrison, David K. Jackson, Ian Johnston, Dominic Kwiatkowski, Cordelia Langford, John Sillitoe on behalf of the Wellcome Sanger Institute COVID-19 Surveillance Team |
| EPI_ISL_2682840, EPI_ISL_2682841 | Lighthouse Lab in Glasgow | Wellcome Sanger Institute for the COVID-19 Genomics UK (COG-UK) Consortium | Harper VanSteenhouse, Yumi Kasai, David Gray, Carol Clugston, Anna Dominiczak and Alex Alderton, Roberto Amato, Jeffrey Barrett, Sonia Goncalves, Ewan Harrison, David K. Jackson, Ian Johnston, Dominic Kwiatkowski, Cordelia Langford, John Sillitoe on behalf of the Wellcome Sanger Institute COVID-19 Surveillance Team |
| EPI_ISL_2682842, EPI_ISL_2682843, EPI_ISL_2682844, EPI_ISL_2682845 | Lighthouse Lab in Alderley Park | Wellcome Sanger Institute for the COVID-19 Genomics UK (COG-UK) Consortium | Jacquelyn Wynn, Mairead Hyland, The Lighthouse Lab in Alderley Park and Alex Alderton, Roberto Amato, Jeffrey Barrett, Sonia Goncalves, Ewan Harrison, David K. Jackson, Ian Johnston, Dominic Kwiatkowski, Cordelia Langford, John Sillitoe on behalf of the Wellcome Sanger Institute COVID-19 Surveillance Team |
| EPI_ISL_2682846, EPI_ISL_2682847, EPI_ISL_2682848, EPI_ISL_2682849, EPI_ISL_2682850, EPI_ISL_2682851, EPI_ISL_2682852 | Lighthouse Lab in Glasgow | Wellcome Sanger Institute for the COVID-19 Genomics UK (COG-UK) Consortium | Harper VanSteenhouse, Yumi Kasai, David Gray, Carol Clugston, Anna Dominiczak and Alex Alderton, Roberto Amato, Jeffrey Barrett, Sonia Goncalves, Ewan Harrison, David K. Jackson, Ian Johnston, Dominic Kwiatkowski, Cordelia Langford, John Sillitoe on behalf of the Wellcome Sanger Institute COVID-19 Surveillance Team |
| EPI_ISL_2682853, EPI_ISL_2682854 | Lighthouse Lab in Alderley Park | Wellcome Sanger Institute for the COVID-19 Genomics UK (COG-UK) Consortium | Jacquelyn Wynn, Mairead Hyland, The Lighthouse Lab in Alderley Park and Alex Alderton, Roberto Amato, Jeffrey Barrett, Sonia Goncalves, Ewan Harrison, David K. Jackson, Ian Johnston, Dominic Kwiatkowski, Cordelia Langford, John Sillitoe on behalf of the Wellcome Sanger Institute COVID-19 Surveillance Team |
| EPI_ISL_2682855 | Lighthouse Lab in Glasgow | Wellcome Sanger Institute for the COVID-19 Genomics UK (COG-UK) Consortium | Harper VanSteenhouse, Yumi Kasai, David Gray, Carol Clugston, Anna Dominiczak and Alex Alderton, Roberto Amato, Jeffrey Barrett, Sonia Goncalves, Ewan Harrison, David K. Jackson, Ian Johnston, Dominic Kwiatkowski, Cordelia Langford, John Sillitoe on behalf of the Wellcome Sanger Institute COVID-19 Surveillance Team |
| EPI_ISL_2682856, EPI_ISL_2682857, EPI_ISL_2682858 | Lighthouse Lab in Alderley Park | Wellcome Sanger Institute for the COVID-19 Genomics UK (COG-UK) Consortium | Jacquelyn Wynn, Mairead Hyland, The Lighthouse Lab in Alderley Park and Alex Alderton, Roberto Amato, Jeffrey Barrett, Sonia Goncalves, Ewan Harrison, David K. Jackson, Ian Johnston, Dominic Kwiatkowski, Cordelia Langford, John Sillitoe on behalf of the Wellcome Sanger Institute COVID-19 Surveillance Team |
| EPI_ISL_2682859, EPI_ISL_2682860, EPI_ISL_2682861 | Lighthouse Lab in Glasgow | Wellcome Sanger Institute for the COVID-19 Genomics UK (COG-UK) Consortium | Harper VanSteenhouse, Yumi Kasai, David Gray, Carol Clugston, Anna Dominiczak and Alex Alderton, Roberto Amato, Jeffrey Barrett, Sonia Goncalves, Ewan Harrison, David K. Jackson, Ian Johnston, Dominic Kwiatkowski, Cordelia Langford, John Sillitoe on behalf of the Wellcome Sanger Institute COVID-19 Surveillance Team |
| EPI_ISL_2682862, EPI_ISL_2682864, EPI_ISL_2682865, EPI_ISL_2682866 | Lighthouse Lab in Alderley Park | Wellcome Sanger Institute for the COVID-19 Genomics UK (COG-UK) Consortium | Jacquelyn Wynn, Mairead Hyland, The Lighthouse Lab in Alderley Park and Alex Alderton, Roberto Amato, Jeffrey Barrett, Sonia Goncalves, Ewan Harrison, David K. Jackson, Ian Johnston, Dominic Kwiatkowski, Cordelia Langford, John Sillitoe on behalf of the Wellcome Sanger Institute COVID-19 Surveillance Team |
| EPI_ISL_2682867, EPI_ISL_2682868, EPI_ISL_2682869, EPI_ISL_2682870, EPI_ISL_2682871, EPI_ISL_2682872 | Lighthouse Lab in Glasgow | Wellcome Sanger Institute for the COVID-19 Genomics UK (COG-UK) Consortium | Harper VanSteenhouse, Yumi Kasai, David Gray, Carol Clugston, Anna Dominiczak and Alex Alderton, Roberto Amato, Jeffrey Barrett, Sonia Goncalves, Ewan Harrison, David K. Jackson, Ian Johnston, Dominic Kwiatkowski, Cordelia Langford, John Sillitoe on behalf of the Wellcome Sanger Institute COVID-19 Surveillance Team |
| EPI_ISL_2682873 | Lighthouse Lab in Alderley Park | Wellcome Sanger Institute for the COVID-19 Genomics UK (COG-UK) Consortium | Jacquelyn Wynn, Mairead Hyland, The Lighthouse Lab in Alderley Park and Alex Alderton, Roberto Amato, Jeffrey Barrett, Sonia Goncalves, Ewan Harrison, David K. Jackson, Ian Johnston, Dominic Kwiatkowski, Cordelia Langford, John Sillitoe on behalf of the Wellcome Sanger Institute COVID-19 Surveillance Team |
| EPI_ISL_2682874 | Lighthouse Lab in Glasgow | Wellcome Sanger Institute for the COVID-19 Genomics UK (COG-UK) Consortium | Harper VanSteenhouse, Yumi Kasai, David Gray, Carol Clugston, Anna Dominiczak and Alex Alderton, Roberto Amato, Jeffrey Barrett, Sonia Goncalves, Ewan Harrison, David K. Jackson, Ian Johnston, Dominic Kwiatkowski, Cordelia Langford, John Sillitoe on behalf of the Wellcome Sanger Institute COVID-19 Surveillance Team |
| EPI_ISL_2682875, EPI_ISL_2682876 | Lighthouse Lab in Alderley Park | Wellcome Sanger Institute for the COVID-19 Genomics UK (COG-UK) Consortium | Jacquelyn Wynn, Mairead Hyland, The Lighthouse Lab in Alderley Park and Alex Alderton, Roberto Amato, Jeffrey Barrett, Sonia Goncalves, Ewan Harrison, David K. Jackson, Ian Johnston, Dominic Kwiatkowski, Cordelia Langford, John Sillitoe on behalf of the Wellcome Sanger Institute COVID-19 Surveillance Team |
| EPI_ISL_2682877 | Lighthouse Lab in Glasgow | Wellcome Sanger Institute for the COVID-19 Genomics UK (COG-UK) Consortium | Harper VanSteenhouse, Yumi Kasai, David Gray, Carol Clugston, Anna Dominiczak and Alex Alderton, Roberto Amato, Jeffrey Barrett, Sonia Goncalves, Ewan Harrison, David K. Jackson, Ian Johnston, Dominic Kwiatkowski, Cordelia Langford, John Sillitoe on behalf of the Wellcome Sanger Institute COVID-19 Surveillance Team |
| EPI_ISL_2682878 | Lighthouse Lab in Alderley Park | Wellcome Sanger Institute for the COVID-19 Genomics UK (COG-UK) Consortium | Jacquelyn Wynn, Mairead Hyland, The Lighthouse Lab in Alderley Park and Alex Alderton, Roberto Amato, Jeffrey Barrett, Sonia Goncalves, Ewan Harrison, David K. Jackson, Ian Johnston, Dominic Kwiatkowski, Cordelia Langford, John Sillitoe on behalf of the Wellcome Sanger Institute COVID-19 Surveillance Team |
| EPI_ISL_2682879 | Lighthouse Lab in Glasgow | Wellcome Sanger Institute for the COVID-19 Genomics UK (COG-UK) Consortium | Harper VanSteenhouse, Yumi Kasai, David Gray, Carol Clugston, Anna Dominiczak and Alex Alderton, Roberto Amato, Jeffrey Barrett, Sonia Goncalves, Ewan Harrison, David K. Jackson, Ian Johnston, Dominic Kwiatkowski, Cordelia Langford, John Sillitoe on behalf of the Wellcome Sanger Institute COVID-19 Surveillance Team |
| EPI_ISL_2682881, EPI_ISL_2682882 | Lighthouse Lab in Alderley Park | Wellcome Sanger Institute for the COVID-19 Genomics UK (COG-UK) Consortium | Jacquelyn Wynn, Mairead Hyland, The Lighthouse Lab in Alderley Park and Alex Alderton, Roberto Amato, Jeffrey Barrett, Sonia Goncalves, Ewan Harrison, David K. Jackson, Ian Johnston, Dominic Kwiatkowski, Cordelia Langford, John Sillitoe on behalf of the Wellcome Sanger Institute COVID-19 Surveillance Team |
| EPI_ISL_2682883 | Lighthouse Lab in Glasgow | Wellcome Sanger Institute for the COVID-19 Genomics UK (COG-UK) Consortium | Harper VanSteenhouse, Yumi Kasai, David Gray, Carol Clugston, Anna Dominiczak and Alex Alderton, Roberto Amato, Jeffrey Barrett, Sonia Goncalves, Ewan Harrison, David K. Jackson, Ian Johnston, Dominic Kwiatkowski, Cordelia Langford, John Sillitoe on behalf of the Wellcome Sanger Institute COVID-19 Surveillance Team |
| EPI_ISL_2682884 | Lighthouse Lab in Alderley Park | Wellcome Sanger Institute for the COVID-19 Genomics UK (COG-UK) Consortium | Jacquelyn Wynn, Mairead Hyland, The Lighthouse Lab in Alderley Park and Alex Alderton, Roberto Amato, Jeffrey Barrett, Sonia Goncalves, Ewan Harrison, David K. Jackson, Ian Johnston, Dominic Kwiatkowski, Cordelia Langford, John Sillitoe on behalf of the Wellcome Sanger Institute COVID-19 Surveillance Team |
| EPI_ISL_2682885 | Lighthouse Lab in Glasgow | Wellcome Sanger Institute for the COVID-19 Genomics UK (COG-UK) Consortium | Harper VanSteenhouse, Yumi Kasai, David Gray, Carol Clugston, Anna Dominiczak and Alex Alderton, Roberto Amato, Jeffrey Barrett, Sonia Goncalves, Ewan Harrison, David K. Jackson, Ian Johnston, Dominic Kwiatkowski, Cordelia Langford, John Sillitoe on behalf of the Wellcome Sanger Institute COVID-19 Surveillance Team |
| EPI_ISL_2682886, EPI_ISL_2682888 | Lighthouse Lab in Alderley Park | Wellcome Sanger Institute for the COVID-19 Genomics UK (COG-UK) Consortium | Jacquelyn Wynn, Mairead Hyland, The Lighthouse Lab in Alderley Park and Alex Alderton, Roberto Amato, Jeffrey Barrett, Sonia Goncalves, Ewan Harrison, David K. Jackson, Ian Johnston, Dominic Kwiatkowski, Cordelia Langford, John Sillitoe on behalf of the Wellcome Sanger Institute COVID-19 Surveillance Team |
| EPI_ISL_2682889 | Lighthouse Lab in Glasgow | Wellcome Sanger Institute for the COVID-19 Genomics UK (COG-UK) Consortium | Harper VanSteenhouse, Yumi Kasai, David Gray, Carol Clugston, Anna Dominiczak and Alex Alderton, Roberto Amato, Jeffrey Barrett, Sonia Goncalves, Ewan Harrison, David K. Jackson, Ian Johnston, Dominic Kwiatkowski, Cordelia Langford, John Sillitoe on behalf of the Wellcome Sanger Institute COVID-19 Surveillance Team |

| | | | |
|---|---|---|---|
| EPI_ISL_2682890 | Lighthouse Lab in Alderley Park | Wellcome Sanger Institute for the COVID-19 Genomics UK (COG-UK) Consortium | Jacquelyn Wynn, Mairead Hyland, The Lighthouse Lab in Alderley Park and Alex Alderton, Roberto Amato, Jeffrey Barrett, Sonia Goncalves, Ewan Harrison, David K. Jackson, Ian Johnston, Dominic Kwiatkowski, Cordelia Langford, John Sillitoe on behalf of the Wellcome Sanger Institute COVID-19 Surveillance Team |
| EPI_ISL_2682891, EPI_ISL_2682892 | Lighthouse Lab in Glasgow | Wellcome Sanger Institute for the COVID-19 Genomics UK (COG-UK) Consortium | Harper VanSteenhouse, Yumi Kasai, David Gray, Carol Clugston, Anna Dominiczak and Alex Alderton, Roberto Amato, Jeffrey Barrett, Sonia Goncalves, Ewan Harrison, David K. Jackson, Ian Johnston, Dominic Kwiatkowski, Cordelia Langford, John Sillitoe on behalf of the Wellcome Sanger Institute COVID-19 Surveillance Team |
| EPI_ISL_2682893 | Lighthouse Lab in Alderley Park | Wellcome Sanger Institute for the COVID-19 Genomics UK (COG-UK) Consortium | Jacquelyn Wynn, Mairead Hyland, The Lighthouse Lab in Alderley Park and Alex Alderton, Roberto Amato, Jeffrey Barrett, Sonia Goncalves, Ewan Harrison, David K. Jackson, Ian Johnston, Dominic Kwiatkowski, Cordelia Langford, John Sillitoe on behalf of the Wellcome Sanger Institute COVID-19 Surveillance Team |
| EPI_ISL_2682894, EPI_ISL_2682895 | Lighthouse Lab in Glasgow | Wellcome Sanger Institute for the COVID-19 Genomics UK (COG-UK) Consortium | Harper VanSteenhouse, Yumi Kasai, David Gray, Carol Clugston, Anna Dominiczak and Alex Alderton, Roberto Amato, Jeffrey Barrett, Sonia Goncalves, Ewan Harrison, David K. Jackson, Ian Johnston, Dominic Kwiatkowski, Cordelia Langford, John Sillitoe on behalf of the Wellcome Sanger Institute COVID-19 Surveillance Team |
| EPI_ISL_2682896, EPI_ISL_2682897, EPI_ISL_2682898 | Lighthouse Lab in Alderley Park | Wellcome Sanger Institute for the COVID-19 Genomics UK (COG-UK) Consortium | Jacquelyn Wynn, Mairead Hyland, The Lighthouse Lab in Alderley Park and Alex Alderton, Roberto Amato, Jeffrey Barrett, Sonia Goncalves, Ewan Harrison, David K. Jackson, Ian Johnston, Dominic Kwiatkowski, Cordelia Langford, John Sillitoe on behalf of the Wellcome Sanger Institute COVID-19 Surveillance Team |
| EPI_ISL_2682899, EPI_ISL_2682900, EPI_ISL_2682901 | Lighthouse Lab in Glasgow | Wellcome Sanger Institute for the COVID-19 Genomics UK (COG-UK) Consortium | Harper VanSteenhouse, Yumi Kasai, David Gray, Carol Clugston, Anna Dominiczak and Alex Alderton, Roberto Amato, Jeffrey Barrett, Sonia Goncalves, Ewan Harrison, David K. Jackson, Ian Johnston, Dominic Kwiatkowski, Cordelia Langford, John Sillitoe on behalf of the Wellcome Sanger Institute COVID-19 Surveillance Team |
| EPI_ISL_2682902, EPI_ISL_2682903 | Lighthouse Lab in Alderley Park | Wellcome Sanger Institute for the COVID-19 Genomics UK (COG-UK) Consortium | Jacquelyn Wynn, Mairead Hyland, The Lighthouse Lab in Alderley Park and Alex Alderton, Roberto Amato, Jeffrey Barrett, Sonia Goncalves, Ewan Harrison, David K. Jackson, Ian Johnston, Dominic Kwiatkowski, Cordelia Langford, John Sillitoe on behalf of the Wellcome Sanger Institute COVID-19 Surveillance Team |
| EPI_ISL_2682904, EPI_ISL_2682905, EPI_ISL_2682906 | Lighthouse Lab in Glasgow | Wellcome Sanger Institute for the COVID-19 Genomics UK (COG-UK) Consortium | Harper VanSteenhouse, Yumi Kasai, David Gray, Carol Clugston, Anna Dominiczak and Alex Alderton, Roberto Amato, Jeffrey Barrett, Sonia Goncalves, Ewan Harrison, David K. Jackson, Ian Johnston, Dominic Kwiatkowski, Cordelia Langford, John Sillitoe on behalf of the Wellcome Sanger Institute COVID-19 Surveillance Team |
| EPI_ISL_2682907 | Lighthouse Lab in Alderley Park | Wellcome Sanger Institute for the COVID-19 Genomics UK (COG-UK) Consortium | Jacquelyn Wynn, Mairead Hyland, The Lighthouse Lab in Alderley Park and Alex Alderton, Roberto Amato, Jeffrey Barrett, Sonia Goncalves, Ewan Harrison, David K. Jackson, Ian Johnston, Dominic Kwiatkowski, Cordelia Langford, John Sillitoe on behalf of the Wellcome Sanger Institute COVID-19 Surveillance Team |
| EPI_ISL_2682908, EPI_ISL_2682909 | Lighthouse Lab in Glasgow | Wellcome Sanger Institute for the COVID-19 Genomics UK (COG-UK) Consortium | Harper VanSteenhouse, Yumi Kasai, David Gray, Carol Clugston, Anna Dominiczak and Alex Alderton, Roberto Amato, Jeffrey Barrett, Sonia Goncalves, Ewan Harrison, David K. Jackson, Ian Johnston, Dominic Kwiatkowski, Cordelia Langford, John Sillitoe on behalf of the Wellcome Sanger Institute COVID-19 Surveillance Team |
| EPI_ISL_2682910 | Lighthouse Lab in Alderley Park | Wellcome Sanger Institute for the COVID-19 Genomics UK (COG-UK) Consortium | Jacquelyn Wynn, Mairead Hyland, The Lighthouse Lab in Alderley Park and Alex Alderton, Roberto Amato, Jeffrey Barrett, Sonia Goncalves, Ewan Harrison, David K. Jackson, Ian Johnston, Dominic Kwiatkowski, Cordelia Langford, John Sillitoe on behalf of the Wellcome Sanger Institute COVID-19 Surveillance Team |
| EPI_ISL_2682911, EPI_ISL_2682912 | Lighthouse Lab in Glasgow | Wellcome Sanger Institute for the COVID-19 Genomics UK (COG-UK) Consortium | Harper VanSteenhouse, Yumi Kasai, David Gray, Carol Clugston, Anna Dominiczak and Alex Alderton, Roberto Amato, Jeffrey Barrett, Sonia Goncalves, Ewan Harrison, David K. Jackson, Ian Johnston, Dominic Kwiatkowski, Cordelia Langford, John Sillitoe on behalf of the Wellcome Sanger Institute COVID-19 Surveillance Team |
| EPI_ISL_2682913, EPI_ISL_2682914 | Lighthouse Lab in Alderley Park | Wellcome Sanger Institute for the COVID-19 Genomics UK (COG-UK) Consortium | Jacquelyn Wynn, Mairead Hyland, The Lighthouse Lab in Alderley Park and Alex Alderton, Roberto Amato, Jeffrey Barrett, Sonia Goncalves, Ewan Harrison, David K. Jackson, Ian Johnston, Dominic Kwiatkowski, Cordelia Langford, John Sillitoe on behalf of the Wellcome Sanger Institute COVID-19 Surveillance Team |
| EPI_ISL_2682915, EPI_ISL_2682916, EPI_ISL_2682917, EPI_ISL_2682918 | Lighthouse Lab in Glasgow | Wellcome Sanger Institute for the COVID-19 Genomics UK (COG-UK) Consortium | Harper VanSteenhouse, Yumi Kasai, David Gray, Carol Clugston, Anna Dominiczak and Alex Alderton, Roberto Amato, Jeffrey Barrett, Sonia Goncalves, Ewan Harrison, David K. Jackson, Ian Johnston, Dominic Kwiatkowski, Cordelia Langford, John Sillitoe on behalf of the Wellcome Sanger Institute COVID-19 Surveillance Team |
| EPI_ISL_2682919, EPI_ISL_2682920 | Lighthouse Lab in Alderley Park | Wellcome Sanger Institute for the COVID-19 Genomics UK (COG-UK) Consortium | Jacquelyn Wynn, Mairead Hyland, The Lighthouse Lab in Alderley Park and Alex Alderton, Roberto Amato, Jeffrey Barrett, Sonia Goncalves, Ewan Harrison, David K. Jackson, Ian Johnston, Dominic Kwiatkowski, Cordelia Langford, John Sillitoe on behalf of the Wellcome Sanger Institute COVID-19 Surveillance Team |
| EPI_ISL_2682921, EPI_ISL_2682922, EPI_ISL_2682923 | Lighthouse Lab in Glasgow | Wellcome Sanger Institute for the COVID-19 Genomics UK (COG-UK) Consortium | Harper VanSteenhouse, Yumi Kasai, David Gray, Carol Clugston, Anna Dominiczak and Alex Alderton, Roberto Amato, Jeffrey Barrett, Sonia Goncalves, Ewan Harrison, David K. Jackson, Ian Johnston, Dominic Kwiatkowski, Cordelia Langford, John Sillitoe on behalf of the Wellcome Sanger Institute COVID-19 Surveillance Team |
| EPI_ISL_2682924, EPI_ISL_2682925 | Lighthouse Lab in Alderley Park | Wellcome Sanger Institute for the COVID-19 Genomics UK (COG-UK) Consortium | Jacquelyn Wynn, Mairead Hyland, The Lighthouse Lab in Alderley Park and Alex Alderton, Roberto Amato, Jeffrey Barrett, Sonia Goncalves, Ewan Harrison, David K. Jackson, Ian Johnston, Dominic Kwiatkowski, Cordelia Langford, John Sillitoe on behalf of the Wellcome Sanger Institute COVID-19 Surveillance Team |
| EPI_ISL_2682926, EPI_ISL_2682927 | Lighthouse Lab in Glasgow | Wellcome Sanger Institute for the COVID-19 Genomics UK (COG-UK) Consortium | Harper VanSteenhouse, Yumi Kasai, David Gray, Carol Clugston, Anna Dominiczak and Alex Alderton, Roberto Amato, Jeffrey Barrett, Sonia Goncalves, Ewan Harrison, David K. Jackson, Ian Johnston, Dominic Kwiatkowski, Cordelia Langford, John Sillitoe on behalf of the Wellcome Sanger Institute COVID-19 Surveillance Team |
| EPI_ISL_2682928, EPI_ISL_2682929 | Lighthouse Lab in Alderley Park | Wellcome Sanger Institute for the COVID-19 Genomics UK (COG-UK) Consortium | Jacquelyn Wynn, Mairead Hyland, The Lighthouse Lab in Alderley Park and Alex Alderton, Roberto Amato, Jeffrey Barrett, Sonia Goncalves, Ewan Harrison, David K. Jackson, Ian Johnston, Dominic Kwiatkowski, Cordelia Langford, John Sillitoe on behalf of the Wellcome Sanger Institute COVID-19 Surveillance Team |
| EPI_ISL_2682930, EPI_ISL_2682931 | Lighthouse Lab in Glasgow | Wellcome Sanger Institute for the COVID-19 Genomics UK (COG-UK) Consortium | Harper VanSteenhouse, Yumi Kasai, David Gray, Carol Clugston, Anna Dominiczak and Alex Alderton, Roberto Amato, Jeffrey Barrett, Sonia Goncalves, Ewan Harrison, David K. Jackson, Ian Johnston, Dominic Kwiatkowski, Cordelia Langford, John Sillitoe on behalf of the Wellcome Sanger Institute COVID-19 Surveillance Team |
| EPI_ISL_2682932, EPI_ISL_2682933 | Lighthouse Lab in Alderley Park | Wellcome Sanger Institute for the COVID-19 Genomics UK (COG-UK) Consortium | Jacquelyn Wynn, Mairead Hyland, The Lighthouse Lab in Alderley Park and Alex Alderton, Roberto Amato, Jeffrey Barrett, Sonia Goncalves, Ewan Harrison, David K. Jackson, Ian Johnston, Dominic Kwiatkowski, Cordelia Langford, John Sillitoe on behalf of the Wellcome Sanger Institute COVID-19 Surveillance Team |
| EPI_ISL_2682934 | Lighthouse Lab in Glasgow | Wellcome Sanger Institute for the COVID-19 Genomics UK (COG-UK) Consortium | Harper VanSteenhouse, Yumi Kasai, David Gray, Carol Clugston, Anna Dominiczak and Alex Alderton, Roberto Amato, Jeffrey Barrett, Sonia Goncalves, Ewan Harrison, David K. Jackson, Ian Johnston, Dominic Kwiatkowski, Cordelia Langford, John Sillitoe on behalf of the Wellcome Sanger Institute COVID-19 Surveillance Team |
| EPI_ISL_2682935, EPI_ISL_2682936, EPI_ISL_2682937, EPI_ISL_2682938, EPI_ISL_2682939 | Lighthouse Lab in Alderley Park | Wellcome Sanger Institute for the COVID-19 Genomics UK (COG-UK) Consortium | Jacquelyn Wynn, Mairead Hyland, The Lighthouse Lab in Alderley Park and Alex Alderton, Roberto Amato, Jeffrey Barrett, Sonia Goncalves, Ewan Harrison, David K. Jackson, Ian Johnston, Dominic Kwiatkowski, Cordelia Langford, John Sillitoe on behalf of the Wellcome Sanger Institute COVID-19 Surveillance Team |
| EPI_ISL_2682940 | Lighthouse Lab in Glasgow | Wellcome Sanger Institute for the COVID-19 Genomics UK (COG-UK) Consortium | Harper VanSteenhouse, Yumi Kasai, David Gray, Carol Clugston, Anna Dominiczak and Alex Alderton, Roberto Amato, Jeffrey Barrett, Sonia Goncalves, Ewan Harrison, David K. Jackson, Ian Johnston, Dominic Kwiatkowski, Cordelia Langford, John Sillitoe on behalf of the Wellcome Sanger Institute COVID-19 Surveillance Team |
| EPI_ISL_2682941, EPI_ISL_2682942, EPI_ISL_2682943, EPI_ISL_2682944, EPI_ISL_2682945 | Lighthouse Lab in Alderley Park | Wellcome Sanger Institute for the COVID-19 Genomics UK (COG-UK) Consortium | Jacquelyn Wynn, Mairead Hyland, The Lighthouse Lab in Alderley Park and Alex Alderton, Roberto Amato, Jeffrey Barrett, Sonia Goncalves, Ewan Harrison, David K. Jackson, Ian Johnston, Dominic Kwiatkowski, Cordelia Langford, John Sillitoe on behalf of the Wellcome Sanger Institute COVID-19 Surveillance Team |
| EPI_ISL_2682946, EPI_ISL_2682947, EPI_ISL_2682948 | Lighthouse Lab in Glasgow | Wellcome Sanger Institute for the COVID-19 Genomics UK (COG-UK) Consortium | Harper VanSteenhouse, Yumi Kasai, David Gray, Carol Clugston, Anna Dominiczak and Alex Alderton, Roberto Amato, Jeffrey Barrett, Sonia Goncalves, Ewan Harrison, David K. Jackson, Ian Johnston, Dominic Kwiatkowski, Cordelia Langford, John Sillitoe on behalf of the Wellcome Sanger Institute COVID-19 Surveillance Team |
| EPI_ISL_2682949, EPI_ISL_2682950 | Lighthouse Lab in Alderley Park | Wellcome Sanger Institute for the COVID-19 Genomics UK (COG-UK) Consortium | Jacquelyn Wynn, Mairead Hyland, The Lighthouse Lab in Alderley Park and Alex Alderton, Roberto Amato, Jeffrey Barrett, Sonia Goncalves, Ewan Harrison, David K. Jackson, Ian Johnston, Dominic Kwiatkowski, Cordelia Langford, John Sillitoe on behalf of the Wellcome Sanger Institute COVID-19 Surveillance Team |
| EPI_ISL_2682951 | Lighthouse Lab in Glasgow | Wellcome Sanger Institute for the COVID-19 Genomics UK (COG-UK) Consortium | Harper VanSteenhouse, Yumi Kasai, David Gray, Carol Clugston, Anna Dominiczak and Alex Alderton, Roberto Amato, Jeffrey Barrett, Sonia Goncalves, Ewan Harrison, David K. Jackson, Ian Johnston, Dominic Kwiatkowski, Cordelia Langford, John Sillitoe on behalf of the Wellcome Sanger Institute COVID-19 Surveillance Team |
| EPI_ISL_2682952 | Lighthouse Lab in Alderley Park | Wellcome Sanger Institute for the COVID-19 Genomics UK (COG-UK) Consortium | Jacquelyn Wynn, Mairead Hyland, The Lighthouse Lab in Alderley Park and Alex Alderton, Roberto Amato, Jeffrey Barrett, Sonia Goncalves, Ewan Harrison, David K. Jackson, Ian Johnston, Dominic Kwiatkowski, Cordelia Langford, John Sillitoe on behalf of the Wellcome Sanger Institute COVID-19 Surveillance Team |
| EPI_ISL_2682953 | Lighthouse Lab in Glasgow | Wellcome Sanger Institute for the COVID-19 Genomics UK (COG-UK) Consortium | Harper VanSteenhouse, Yumi Kasai, David Gray, Carol Clugston, Anna Dominiczak and Alex Alderton, Roberto Amato, Jeffrey Barrett, Sonia Goncalves, Ewan Harrison, David K. Jackson, Ian Johnston, Dominic Kwiatkowski, Cordelia Langford, John Sillitoe on behalf of the Wellcome Sanger Institute COVID-19 Surveillance Team |
| EPI_ISL_2682954, EPI_ISL_2682955 | Lighthouse Lab in Alderley Park | Wellcome Sanger Institute for the COVID-19 Genomics UK (COG-UK) Consortium | Jacquelyn Wynn, Mairead Hyland, The Lighthouse Lab in Alderley Park and Alex Alderton, Roberto Amato, Jeffrey Barrett, Sonia Goncalves, Ewan Harrison, David K. Jackson, Ian Johnston, Dominic Kwiatkowski, Cordelia Langford, John Sillitoe on behalf of the Wellcome Sanger Institute COVID-19 Surveillance Team |
| EPI_ISL_2682956 | Lighthouse Lab in Glasgow | Wellcome Sanger Institute for the COVID-19 Genomics UK (COG-UK) Consortium | Harper VanSteenhouse, Yumi Kasai, David Gray, Carol Clugston, Anna Dominiczak and Alex Alderton, Roberto Amato, Jeffrey Barrett, Sonia Goncalves, Ewan Harrison, David K. Jackson, Ian Johnston, Dominic Kwiatkowski, Cordelia Langford, John Sillitoe on behalf of the Wellcome Sanger Institute COVID-19 Surveillance Team |
| EPI_ISL_2682957, EPI_ISL_2682958, EPI_ISL_2682960, EPI_ISL_2682961 | Lighthouse Lab in Alderley Park | Wellcome Sanger Institute for the COVID-19 Genomics UK (COG-UK) Consortium | Jacquelyn Wynn, Mairead Hyland, The Lighthouse Lab in Alderley Park and Alex Alderton, Roberto Amato, Jeffrey Barrett, Sonia Goncalves, Ewan Harrison, David K. Jackson, Ian Johnston, Dominic Kwiatkowski, Cordelia Langford, John Sillitoe on behalf of the Wellcome Sanger Institute COVID-19 Surveillance Team |
| EPI_ISL_2682962, EPI_ISL_2682963, EPI_ISL_2682964, EPI_ISL_2682965 | Lighthouse Lab in Glasgow | Wellcome Sanger Institute for the COVID-19 Genomics UK (COG-UK) Consortium | Harper VanSteenhouse, Yumi Kasai, David Gray, Carol Clugston, Anna Dominiczak and Alex Alderton, Roberto Amato, Jeffrey Barrett, Sonia Goncalves, Ewan Harrison, David K. Jackson, Ian Johnston, Dominic Kwiatkowski, Cordelia Langford, John Sillitoe on behalf of the Wellcome Sanger Institute COVID-19 Surveillance Team |
| EPI_ISL_2682966 | Lighthouse Lab in Alderley Park | Wellcome Sanger Institute for the COVID-19 Genomics UK (COG-UK) Consortium | Jacquelyn Wynn, Mairead Hyland, The Lighthouse Lab in Alderley Park and Alex Alderton, Roberto Amato, Jeffrey Barrett, Sonia Goncalves, Ewan Harrison, David K. Jackson, Ian Johnston, Dominic Kwiatkowski, Cordelia Langford, John Sillitoe on behalf of the Wellcome Sanger Institute COVID-19 Surveillance Team |
| EPI_ISL_2682967, EPI_ISL_2682968 | Lighthouse Lab in Glasgow | Wellcome Sanger Institute for the COVID-19 Genomics UK (COG-UK) Consortium | Harper VanSteenhouse, Yumi Kasai, David Gray, Carol Clugston, Anna Dominiczak and Alex Alderton, Roberto Amato, Jeffrey Barrett, Sonia Goncalves, Ewan Harrison, David K. Jackson, Ian Johnston, Dominic Kwiatkowski, Cordelia Langford, John Sillitoe on behalf of the Wellcome Sanger Institute COVID-19 Surveillance Team |
| EPI_ISL_2682969 | Lighthouse Lab in Alderley Park | Wellcome Sanger Institute for the COVID-19 Genomics UK (COG-UK) Consortium | Jacquelyn Wynn, Mairead Hyland, The Lighthouse Lab in Alderley Park and Alex Alderton, Roberto Amato, Jeffrey Barrett, Sonia Goncalves, Ewan Harrison, David K. Jackson, Ian Johnston, Dominic Kwiatkowski, Cordelia Langford, John Sillitoe on behalf of the Wellcome Sanger Institute COVID-19 Surveillance Team |
| EPI_ISL_2682970 | Lighthouse Lab in Glasgow | Wellcome Sanger Institute for the COVID-19 Genomics UK (COG-UK) Consortium | Harper VanSteenhouse, Yumi Kasai, David Gray, Carol Clugston, Anna Dominiczak and Alex Alderton, Roberto Amato, Jeffrey Barrett, Sonia Goncalves, Ewan Harrison, David K. Jackson, Ian Johnston, Dominic Kwiatkowski, Cordelia Langford, John Sillitoe on behalf of the Wellcome Sanger Institute COVID-19 Surveillance Team |

| | | | |
|---|---|---|---|
| EPI_ISL_2682971 | Lighthouse Lab in Alderley Park | Wellcome Sanger Institute for the COVID-19 Genomics UK (COG-UK) Consortium | Jacquelyn Wynn, Mairead Hyland, The Lighthouse Lab in Alderley Park and Alex Alderton, Roberto Amato, Jeffrey Barrett, Sonia Goncalves, Ewan Harrison, David K. Jackson, Ian Johnston, Dominic Kwiatkowski, Cordelia Langford, John Sillitoe on behalf of the Wellcome Sanger Institute COVID-19 Surveillance Team |
| EPI_ISL_2682972, EPI_ISL_2682973, EPI_ISL_2682975, EPI_ISL_2682976, EPI_ISL_2682977, EPI_ISL_2682978, EPI_ISL_2682979, EPI_ISL_2682980, EPI_ISL_2682981, EPI_ISL_2682982, EPI_ISL_2682983, EPI_ISL_2682984, EPI_ISL_2682985, EPI_ISL_2682986, EPI_ISL_2682987, EPI_ISL_2682988, EPI_ISL_2682990, EPI_ISL_2682991, EPI_ISL_2682992, EPI_ISL_2682993, EPI_ISL_2682994, EPI_ISL_2682995, EPI_ISL_2682996, EPI_ISL_2682997, EPI_ISL_2682998, EPI_ISL_2682999, EPI_ISL_2683000, EPI_ISL_2683001, EPI_ISL_2683003, EPI_ISL_2683004 | | | |
| see above | Lighthouse Lab in Glasgow | Wellcome Sanger Institute for the COVID-19 Genomics UK (COG-UK) Consortium | Harper VanSteenhouse, Yumi Kasai, David Gray, Carol Clugston, Anna Dominiczak and Alex Alderton, Roberto Amato, Jeffrey Barrett, Sonia Goncalves, Ewan Harrison, David K. Jackson, Ian Johnston, Dominic Kwiatkowski, Cordelia Langford, John Sillitoe on behalf of the Wellcome Sanger Institute COVID-19 Surveillance Team |
| EPI_ISL_2683005 | Lighthouse Lab in Alderley Park | Wellcome Sanger Institute for the COVID-19 Genomics UK (COG-UK) Consortium | Jacquelyn Wynn, Mairead Hyland, The Lighthouse Lab in Alderley Park and Alex Alderton, Roberto Amato, Jeffrey Barrett, Sonia Goncalves, Ewan Harrison, David K. Jackson, Ian Johnston, Dominic Kwiatkowski, Cordelia Langford, John Sillitoe on behalf of the Wellcome Sanger Institute COVID-19 Surveillance Team |
| EPI_ISL_2683006, EPI_ISL_2683007, EPI_ISL_2683008, EPI_ISL_2683009, EPI_ISL_2683010, EPI_ISL_2683011, EPI_ISL_2683013, EPI_ISL_2683015 | Lighthouse Lab in Glasgow | Wellcome Sanger Institute for the COVID-19 Genomics UK (COG-UK) Consortium | Harper VanSteenhouse, Yumi Kasai, David Gray, Carol Clugston, Anna Dominiczak and Alex Alderton, Roberto Amato, Jeffrey Barrett, Sonia Goncalves, Ewan Harrison, David K. Jackson, Ian Johnston, Dominic Kwiatkowski, Cordelia Langford, John Sillitoe on behalf of the Wellcome Sanger Institute COVID-19 Surveillance Team |
| EPI_ISL_2683016 | Lighthouse Lab in Alderley Park | Wellcome Sanger Institute for the COVID-19 Genomics UK (COG-UK) Consortium | Jacquelyn Wynn, Mairead Hyland, The Lighthouse Lab in Alderley Park and Alex Alderton, Roberto Amato, Jeffrey Barrett, Sonia Goncalves, Ewan Harrison, David K. Jackson, Ian Johnston, Dominic Kwiatkowski, Cordelia Langford, John Sillitoe on behalf of the Wellcome Sanger Institute COVID-19 Surveillance Team |
| EPI_ISL_2683017, EPI_ISL_2683018, EPI_ISL_2683019, EPI_ISL_2683020, EPI_ISL_2683021, EPI_ISL_2683022, EPI_ISL_2683023, EPI_ISL_2683024, EPI_ISL_2683025, EPI_ISL_2683026, EPI_ISL_2683027, EPI_ISL_2683028, EPI_ISL_2683029, EPI_ISL_2683030, EPI_ISL_2683031, EPI_ISL_2683032, EPI_ISL_2683033, EPI_ISL_2683034, EPI_ISL_2683035, EPI_ISL_2683036, EPI_ISL_2683037, EPI_ISL_2683038, EPI_ISL_2683039, EPI_ISL_2683040, EPI_ISL_2683041, EPI_ISL_2683042, EPI_ISL_2683043, EPI_ISL_2683044, EPI_ISL_2683045, EPI_ISL_2683046, EPI_ISL_2683047, EPI_ISL_2683048, EPI_ISL_2683049, EPI_ISL_2683050, EPI_ISL_2683051, EPI_ISL_2683052, EPI_ISL_2683053, EPI_ISL_2683054, EPI_ISL_2683055, EPI_ISL_2683056, EPI_ISL_2683058, EPI_ISL_2683059, EPI_ISL_2683060, EPI_ISL_2683061, EPI_ISL_2683062, EPI_ISL_2683063, EPI_ISL_2683064, EPI_ISL_2683065, EPI_ISL_2683066, EPI_ISL_2683067, EPI_ISL_2683068, EPI_ISL_2683069, EPI_ISL_2683070, EPI_ISL_2683071, EPI_ISL_2683072, EPI_ISL_2683073, EPI_ISL_2683074, EPI_ISL_2683075, EPI_ISL_2683076, EPI_ISL_2683077, EPI_ISL_2683078, EPI_ISL_2683079, EPI_ISL_2683080, EPI_ISL_2683082, EPI_ISL_2683083, EPI_ISL_2683085, EPI_ISL_2683086, EPI_ISL_2683087, EPI_ISL_2683088, EPI_ISL_2683089, EPI_ISL_2683090, EPI_ISL_2683091, EPI_ISL_2683092, EPI_ISL_2683093, EPI_ISL_2683094, EPI_ISL_2683095, EPI_ISL_2683096, EPI_ISL_2683097, EPI_ISL_2683098, EPI_ISL_2683099, EPI_ISL_2683100, EPI_ISL_2683101, EPI_ISL_2683102, EPI_ISL_2683103, EPI_ISL_2683105, EPI_ISL_2683106, EPI_ISL_2683107, EPI_ISL_2683108, EPI_ISL_2683109, EPI_ISL_2683110 | | | |
| see above | Lighthouse Lab in Glasgow | Wellcome Sanger Institute for the COVID-19 Genomics UK (COG-UK) Consortium | Harper VanSteenhouse, Yumi Kasai, David Gray, Carol Clugston, Anna Dominiczak and Alex Alderton, Roberto Amato, Jeffrey Barrett, Sonia Goncalves, Ewan Harrison, David K. Jackson, Ian Johnston, Dominic Kwiatkowski, Cordelia Langford, John Sillitoe on behalf of the Wellcome Sanger Institute COVID-19 Surveillance Team |
| EPI_ISL_2683111, EPI_ISL_2683112 | Lighthouse Lab in Alderley Park | Wellcome Sanger Institute for the COVID-19 Genomics UK (COG-UK) Consortium | Jacquelyn Wynn, Mairead Hyland, The Lighthouse Lab in Alderley Park and Alex Alderton, Roberto Amato, Jeffrey Barrett, Sonia Goncalves, Ewan Harrison, David K. Jackson, Ian Johnston, Dominic Kwiatkowski, Cordelia Langford, John Sillitoe on behalf of the Wellcome Sanger Institute COVID-19 Surveillance Team |
| EPI_ISL_2683113 | Lighthouse Lab in Milton Keynes | Wellcome Sanger Institute for the COVID-19 Genomics UK (COG-UK) Consortium | The Lighthouse Lab in Milton Keynes and Alex Alderton, Roberto Amato, Jeffrey Barrett, Sonia Goncalves, Ewan Harrison, David K. Jackson, Ian Johnston, Dominic Kwiatkowski, Cordelia Langford, John Sillitoe on behalf of the Wellcome Sanger Institute COVID-19 Surveillance Team |
| EPI_ISL_2683114 | Lighthouse Lab in Alderley Park | Wellcome Sanger Institute for the COVID-19 Genomics UK (COG-UK) Consortium | Jacquelyn Wynn, Mairead Hyland, The Lighthouse Lab in Alderley Park and Alex Alderton, Roberto Amato, Jeffrey Barrett, Sonia Goncalves, Ewan Harrison, David K. Jackson, Ian Johnston, Dominic Kwiatkowski, Cordelia Langford, John Sillitoe on behalf of the Wellcome Sanger Institute COVID-19 Surveillance Team |
| EPI_ISL_2683115, EPI_ISL_2683116, EPI_ISL_2683117, EPI_ISL_2683118, EPI_ISL_2683119, EPI_ISL_2683120, EPI_ISL_2683121, EPI_ISL_2683122, EPI_ISL_2683123, EPI_ISL_2683124, EPI_ISL_2683125, EPI_ISL_2683126, EPI_ISL_2683127, EPI_ISL_2683128, EPI_ISL_2683129, EPI_ISL_2683130, EPI_ISL_2683131, EPI_ISL_2683132, EPI_ISL_2683133, EPI_ISL_2683134, EPI_ISL_2683135, EPI_ISL_2683136, EPI_ISL_2683137, EPI_ISL_2683138, EPI_ISL_2683139, EPI_ISL_2683140, EPI_ISL_2683141, EPI_ISL_2683142, EPI_ISL_2683143, EPI_ISL_2683144, EPI_ISL_2683145, EPI_ISL_2683146, EPI_ISL_2683147, EPI_ISL_2683148, EPI_ISL_2683149, EPI_ISL_2683150, EPI_ISL_2683151, EPI_ISL_2683152, EPI_ISL_2683153, EPI_ISL_2683154, EPI_ISL_2683155, EPI_ISL_2683156, EPI_ISL_2683157, EPI_ISL_2683158, EPI_ISL_2683159, EPI_ISL_2683161, EPI_ISL_2683162, EPI_ISL_2683163, EPI_ISL_2683164, EPI_ISL_2683165, EPI_ISL_2683166, EPI_ISL_2683167, EPI_ISL_2683168, EPI_ISL_2683169, EPI_ISL_2683170, EPI_ISL_2683171, EPI_ISL_2683172, EPI_ISL_2683173, EPI_ISL_2683174, EPI_ISL_2683175, EPI_ISL_2683177, EPI_ISL_2683178, EPI_ISL_2683179, EPI_ISL_2683180, EPI_ISL_2683181, EPI_ISL_2683182, EPI_ISL_2683183, EPI_ISL_2683184, EPI_ISL_2683185, EPI_ISL_2683186, EPI_ISL_2683187, EPI_ISL_2683188, EPI_ISL_2683189, EPI_ISL_2683190, EPI_ISL_2683191, EPI_ISL_2683192, EPI_ISL_2683193, EPI_ISL_2683194, EPI_ISL_2683195, EPI_ISL_2683196, EPI_ISL_2683197, EPI_ISL_2683198, EPI_ISL_2683200, EPI_ISL_2683201, EPI_ISL_2683202, EPI_ISL_2683204, EPI_ISL_2683205, EPI_ISL_2683206, EPI_ISL_2683207, EPI_ISL_2683208, EPI_ISL_2683209, EPI_ISL_2683210, EPI_ISL_2683211, EPI_ISL_2683213, EPI_ISL_2683214, EPI_ISL_2683215, EPI_ISL_2683216, EPI_ISL_2683217, EPI_ISL_2683218, EPI_ISL_2683219, EPI_ISL_2683220, EPI_ISL_2683221, EPI_ISL_2683222, EPI_ISL_2683223, EPI_ISL_2683224, EPI_ISL_2683225, EPI_ISL_2683226, EPI_ISL_2683227, EPI_ISL_2683228, EPI_ISL_2683229, EPI_ISL_2683230, EPI_ISL_2683231, EPI_ISL_2683232, EPI_ISL_2683233, EPI_ISL_2683234, EPI_ISL_2683235, EPI_ISL_2683236, EPI_ISL_2683237, EPI_ISL_2683238, EPI_ISL_2683239, EPI_ISL_2683241, EPI_ISL_2683242, EPI_ISL_2683243, EPI_ISL_2683244, EPI_ISL_2683245, EPI_ISL_2683246, EPI_ISL_2683248, EPI_ISL_2683249, EPI_ISL_2683250, EPI_ISL_2683251, EPI_ISL_2683252, EPI_ISL_2683253, EPI_ISL_2683254, EPI_ISL_2683255, EPI_ISL_2683256, EPI_ISL_2683257, EPI_ISL_2683258, EPI_ISL_2683259, EPI_ISL_2683261, EPI_ISL_2683262, EPI_ISL_2683263, EPI_ISL_2683264, EPI_ISL_2683265, EPI_ISL_2683266, EPI_ISL_2683267, EPI_ISL_2683268, EPI_ISL_2683269, EPI_ISL_2683270, EPI_ISL_2683271, EPI_ISL_2683272, EPI_ISL_2683273, EPI_ISL_2683274, EPI_ISL_2683275, EPI_ISL_2683277, EPI_ISL_2683278, EPI_ISL_2683279, EPI_ISL_2683281, EPI_ISL_2683282, EPI_ISL_2683283, EPI_ISL_2683284, EPI_ISL_2683285, EPI_ISL_2683286, EPI_ISL_2683287, EPI_ISL_2683288, EPI_ISL_2683289, EPI_ISL_2683290, EPI_ISL_2683291, EPI_ISL_2683292, EPI_ISL_2683293, EPI_ISL_2683294, EPI_ISL_2683295, EPI_ISL_2683296, EPI_ISL_2683297, EPI_ISL_2683298, EPI_ISL_2683299, EPI_ISL_2683300, EPI_ISL_2683301, EPI_ISL_2683302, EPI_ISL_2683303, EPI_ISL_2683304, EPI_ISL_2683305, EPI_ISL_2683306, EPI_ISL_2683307, EPI_ISL_2683308, EPI_ISL_2683309, EPI_ISL_2683310, EPI_ISL_2683311, EPI_ISL_2683312, EPI_ISL_2683313, EPI_ISL_2683314, EPI_ISL_2683315, EPI_ISL_2683316, EPI_ISL_2683317, EPI_ISL_2683318, EPI_ISL_2683319, EPI_ISL_2683321, EPI_ISL_2683322, EPI_ISL_2683323, EPI_ISL_2683324, EPI_ISL_2683325, EPI_ISL_2683326, EPI_ISL_2683327, EPI_ISL_2683328, EPI_ISL_2683329, EPI_ISL_2683330, EPI_ISL_2683332, EPI_ISL_2683333, EPI_ISL_2683334, EPI_ISL_2683335, EPI_ISL_2683336, EPI_ISL_2683337, EPI_ISL_2683338, EPI_ISL_2683339, EPI_ISL_2683340, EPI_ISL_2683341, EPI_ISL_2683343, EPI_ISL_2683344, EPI_ISL_2683345, EPI_ISL_2683347, EPI_ISL_2683349, EPI_ISL_2683350, EPI_ISL_2683351, EPI_ISL_2683352, EPI_ISL_2683353, EPI_ISL_2683354, EPI_ISL_2683355, EPI_ISL_2683356, EPI_ISL_2683357, EPI_ISL_2683358, EPI_ISL_2683359, EPI_ISL_2683360, EPI_ISL_2683361, EPI_ISL_2683362, EPI_ISL_2683363, EPI_ISL_2683364, EPI_ISL_2683365, EPI_ISL_2683366, EPI_ISL_2683367, EPI_ISL_2683368, EPI_ISL_2683369, EPI_ISL_2683370, EPI_ISL_2683371, EPI_ISL_2683372, EPI_ISL_2683373, EPI_ISL_2683374, EPI_ISL_2683375, EPI_ISL_2683376, EPI_ISL_2683377, EPI_ISL_2683378, EPI_ISL_2683379, EPI_ISL_2683380, EPI_ISL_2683381, EPI_ISL_2683382, EPI_ISL_2683383, EPI_ISL_2683384, EPI_ISL_2683385, EPI_ISL_2683386, EPI_ISL_2683387, EPI_ISL_2683388, EPI_ISL_2683389, EPI_ISL_2683390, EPI_ISL_2683391, EPI_ISL_2683392, EPI_ISL_2683393, EPI_ISL_2683394, EPI_ISL_2683395, EPI_ISL_2683396, EPI_ISL_2683398, EPI_ISL_2683400, EPI_ISL_2683401, EPI_ISL_2683402, EPI_ISL_2683403, EPI_ISL_2683404, EPI_ISL_2683405, EPI_ISL_2683406, EPI_ISL_2683407, EPI_ISL_2683408, EPI_ISL_2683409, EPI_ISL_2683410, EPI_ISL_2683411, EPI_ISL_2683412, EPI_ISL_2683413, EPI_ISL_2683414, EPI_ISL_2683415, EPI_ISL_2683416, EPI_ISL_2683417, EPI_ISL_2683418, EPI_ISL_2683419, EPI_ISL_2683420, EPI_ISL_2683421, EPI_ISL_2683422, EPI_ISL_2683423, EPI_ISL_2683424, EPI_ISL_2683425, EPI_ISL_2683427, EPI_ISL_2683429, EPI_ISL_2683430, EPI_ISL_2683431, EPI_ISL_2683432, EPI_ISL_2683433, EPI_ISL_2683435, EPI_ISL_2683436, EPI_ISL_2683437, EPI_ISL_2683438, EPI_ISL_2683439, EPI_ISL_2683440, EPI_ISL_2683441, EPI_ISL_2683443, EPI_ISL_2683444, EPI_ISL_2683445, EPI_ISL_2683446, EPI_ISL_2683447, EPI_ISL_2683448, EPI_ISL_2683449, EPI_ISL_2683450, EPI_ISL_2683451, EPI_ISL_2683452, EPI_ISL_2683453, EPI_ISL_2683454, EPI_ISL_2683455, EPI_ISL_2683456, EPI_ISL_2683457, EPI_ISL_2683458, EPI_ISL_2683459, EPI_ISL_2683461, EPI_ISL_2683462, EPI_ISL_2683463, EPI_ISL_2683464, EPI_ISL_2683465, EPI_ISL_2683466, EPI_ISL_2683467, EPI_ISL_2683468, EPI_ISL_2683469, EPI_ISL_2683470, EPI_ISL_2683471, EPI_ISL_2683472, EPI_ISL_2683473, EPI_ISL_2683474, EPI_ISL_2683475, EPI_ISL_2683476, EPI_ISL_2683477, EPI_ISL_2683478, EPI_ISL_2683479, EPI_ISL_2683480, EPI_ISL_2683481, EPI_ISL_2683482, EPI_ISL_2683483, EPI_ISL_2683484, EPI_ISL_2683485, EPI_ISL_2683486, EPI_ISL_2683488, EPI_ISL_2683489, EPI_ISL_2683490 | | | |
| see above | Lighthouse Lab in Glasgow | Wellcome Sanger Institute for the COVID-19 Genomics UK (COG-UK) Consortium | Harper VanSteenhouse, Yumi Kasai, David Gray, Carol Clugston, Anna Dominiczak and Alex Alderton, Roberto Amato, Jeffrey Barrett, Sonia Goncalves, Ewan Harrison, David K. Jackson, Ian Johnston, Dominic Kwiatkowski, Cordelia Langford, John Sillitoe on behalf of the Wellcome Sanger Institute COVID-19 Surveillance Team |
| EPI_ISL_2684111 | Department of Bacteria, Parasites and Fungi, Statens Serum Institut, Copenhagen, Denmark | Statens Serum Institut Bioinformatics and Microbial Genomics | Danish Covid-19 Genome Consortium |
| EPI_ISL_2684247, EPI_ISL_2684307, EPI_ISL_2684314, EPI_ISL_2684640, EPI_ISL_2684814, EPI_ISL_2684859, EPI_ISL_2684941, EPI_ISL_2684963, EPI_ISL_2685126, EPI_ISL_2685164, EPI_ISL_2685428, EPI_ISL_2685467, EPI_ISL_2685473, EPI_ISL_2685474, EPI_ISL_2685521, EPI_ISL_2685602, EPI_ISL_2685650, EPI_ISL_2685736, EPI_ISL_2685763 | | | |
| see above | Statens Serum Institut Bioinformatics and Microbial Genomics | Statens Serum Institut Bioinformatics and Microbial Genomics | Danish Covid-19 Genome Consortium |
| EPI_ISL_2685852, EPI_ISL_2685853, EPI_ISL_2685856, EPI_ISL_2685858, EPI_ISL_2685859, EPI_ISL_2685870, EPI_ISL_2685873, EPI_ISL_2685882, EPI_ISL_2685884, EPI_ISL_2685885 | SYNLAB | GIGA Medical Genomics | Keith Durkin, Maria Artesi, Sébastien Bontems, Raphaël Boreux, Bouchra Boujemla, Nathalie Renotte, Cécile Meex, Pierrette Melin, Marie-Pierre Hayette, Vincent Bours |
| EPI_ISL_2685887, EPI_ISL_2685889 | Department of Clinical Microbiology | GIGA Medical Genomics | Keith Durkin, Maria Artesi, Sébastien Bontems, Raphaël Boreux, Bouchra Boujemla, Nathalie Renotte, Cécile Meex, Pierrette Melin, Marie-Pierre Hayette, Vincent Bours |
| EPI_ISL_2685898 | SYNLAB | GIGA Medical Genomics | Keith Durkin, Maria Artesi, Sébastien Bontems, Raphaël Boreux, Bouchra Boujemla, Nathalie Renotte, Cécile Meex, Pierrette Melin, Marie-Pierre Hayette, Vincent Bours |
| EPI_ISL_2686029 | Università degli Studi di Perugia | Istituto Zooprofilattico Sperimentale dell'Abruzzo e Molise "G. Caporale" | Mencacci A, Camilloni B, Biagetti M, Giammarioli M, Lorusso A, Marcacci M, Di Domenico M, Ancora M, Curini V, Mangone I, Rinaldi A, Delli Compagni E, Scialabba S, Di Pasquale A, Cammà C, Puglia I, Calistri P, Savini G |
| EPI_ISL_2686054, EPI_ISL_2686055 | Laboratorio Analisi Osp. Città di Castello - Azienda USL Umbria1 | Istituto Zooprofilattico Sperimentale dell'Abruzzo e Molise "G. Caporale" | Malagigi V, Taccani P,Lorusso A, Marcacci M, Di Domenico M, Ancora M, Curini V, Mangone I, Rinaldi A, Delli Compagni E, Scialabba S, Di Pasquale A, Cammà C, Puglia I, Calistri P, Savini G |
| EPI_ISL_2686067 | Università degli Studi di Perugia | Istituto Zooprofilattico Sperimentale dell'Abruzzo e Molise "G. Caporale" | Mencacci A, Camilloni B, Biagetti M, Giammarioli M, Lorusso A, Marcacci M, Di Domenico M, Ancora M, Curini V, Mangone I, Rinaldi A, Delli Compagni E, Scialabba S, Di Pasquale A, Cammà C, Puglia I, Calistri P, Savini G |
| EPI_ISL_2686070, EPI_ISL_2686071, EPI_ISL_2686072, EPI_ISL_2686073 | Laboratorio Analisi Osp. Città di Castello - Azienda USL Umbria1 | Istituto Zooprofilattico Sperimentale dell'Abruzzo e Molise "G. Caporale" | Malagigi V, Taccani P,Lorusso A, Marcacci M, Di Domenico M, Ancora M, Curini V, Mangone I, Rinaldi A, Delli Compagni E, Scialabba S, Di Pasquale A, Cammà C, Puglia I, Calistri P, Savini G |
| EPI_ISL_2686077 | Azienda USL Umbria 2 | Istituto Zooprofilattico Sperimentale dell'Abruzzo e Molise "G. Caporale" | Proietti A, Pistoni E, Lorusso A, Marcacci M, Di Domenico M, Ancora M, Curini V, Mangone I, Rinaldi A, Delli Compagni E, Scialabba S, Di Pasquale A, Cammà C, Puglia I, Calistri P, Savini G |
| EPI_ISL_2686086 | SELAS LABOSUD | Inovie AS GenBio | VERCRUYSSEN Florian LAURENT Pauline |
| EPI_ISL_2686087 | Inovie AS LABOSUD PROVENCE(IA) | Inovie AS GenBio | VERCRUYSSEN Florian LAURENT Pauline |
| EPI_ISL_2686088, EPI_ISL_2686089, EPI_ISL_2686091, EPI_ISL_2686092, EPI_ISL_2686097, EPI_ISL_2686101, EPI_ISL_2686103, EPI_ISL_2686105, EPI_ISL_2686108, EPI_ISL_2686109, EPI_ISL_2686110, EPI_ISL_2686112, EPI_ISL_2686113, EPI_ISL_2686115, EPI_ISL_2686117, EPI_ISL_2686119, EPI_ISL_2686121, EPI_ISL_2686122, EPI_ISL_2686123, EPI_ISL_2686125, EPI_ISL_2686127, EPI_ISL_2686128, EPI_ISL_2686129, EPI_ISL_2686131, EPI_ISL_2686132, EPI_ISL_2686133, EPI_ISL_2686135, EPI_ISL_2686137, EPI_ISL_2686138, EPI_ISL_2686140, EPI_ISL_2686141, EPI_ISL_2686142, EPI_ISL_2686144, EPI_ISL_2686146, EPI_ISL_2686148, EPI_ISL_2686149, EPI_ISL_2686151, EPI_ISL_2686152, EPI_ISL_2686154, EPI_ISL_2686155, EPI_ISL_2686156, EPI_ISL_2686157, EPI_ISL_2686158, EPI_ISL_2686159, EPI_ISL_2686161, EPI_ISL_2686162, EPI_ISL_2686163, EPI_ISL_2686164, EPI_ISL_2686165, EPI_ISL_2686166, EPI_ISL_2686167, EPI_ISL_2686168, EPI_ISL_2686169, EPI_ISL_2686170, EPI_ISL_2686171, EPI_ISL_2686172, EPI_ISL_2686173, EPI_ISL_2686175, EPI_ISL_2686176, EPI_ISL_2686177, EPI_ISL_2686179, EPI_ISL_2686180, EPI_ISL_2686181, EPI_ISL_2686182, EPI_ISL_2686183, EPI_ISL_2686184, EPI_ISL_2686185, EPI_ISL_2686186, EPI_ISL_2686187, EPI_ISL_2686188, EPI_ISL_2686189, EPI_ISL_2686190, EPI_ISL_2686191, EPI_ISL_2686192, EPI_ISL_2686193, EPI_ISL_2686194, EPI_ISL_2686195, EPI_ISL_2686196, EPI_ISL_2686197, EPI_ISL_2686198, EPI_ISL_2686199, EPI_ISL_2686200, EPI_ISL_2686201, EPI_ISL_2686202 | | | |
| see above | Amazon Dx, AUV2 Laboratory | Amazon Dx, AUV2 Laboratory | Luke Meredith |
| EPI_ISL_2686217, EPI_ISL_2686218, EPI_ISL_2686219, EPI_ISL_2686220, EPI_ISL_2686226, EPI_ISL_2686227 | MKK Mühlenkreiskliniken, Johannes Wesling Klinikum Minden | Hannover Medical School, Institute of Virology | Lars Steinbrück |
| EPI_ISL_2686276 | National Institute of Public Health | National Institute of Public Health | Helena Jirincova, Jaromira Vecerova, Timotej Suri, Dusan Trnka, Alexander Nagy |
| EPI_ISL_2686393 | AZ Sint-Jan Brugge-Oostende AV | AZ Sint Jan | Marijke Reynders, Patrick Descheemaeker, Laurien Hoornaert, Thomas Van Landschoot, Marie Madeleine Chabert-Consen, Jorn Hellemans |
| EPI_ISL_2686394, EPI_ISL_2686396, EPI_ISL_2686397, EPI_ISL_2686398, EPI_ISL_2686399, EPI_ISL_2686400, EPI_ISL_2686401, EPI_ISL_2686403, EPI_ISL_2686404, EPI_ISL_2686405, EPI_ISL_2686406, EPI_ISL_2686413, EPI_ISL_2686416, EPI_ISL_2686417, EPI_ISL_2686418, EPI_ISL_2686423, EPI_ISL_2686428, EPI_ISL_2686429, EPI_ISL_2686432, EPI_ISL_2686433, EPI_ISL_2686434, EPI_ISL_2686435, EPI_ISL_2686437, EPI_ISL_2686438, EPI_ISL_2686440, EPI_ISL_2686446, EPI_ISL_2686447, EPI_ISL_2686458, EPI_ISL_2686462, EPI_ISL_2686464, EPI_ISL_2686466, EPI_ISL_2686467, EPI_ISL_2686471, EPI_ISL_2686485, EPI_ISL_2686504, EPI_ISL_2686533, EPI_ISL_2686534, EPI_ISL_2686535, EPI_ISL_2686536, EPI_ISL_2686537, EPI_ISL_2686538, EPI_ISL_2686542, EPI_ISL_2686543, EPI_ISL_2686544, | | | |

| | | | |
|---|---|---|---|
| EPI_ISL_2686545, EPI_ISL_2686546, EPI_ISL_2686556, EPI_ISL_2686560, EPI_ISL_2686563, EPI_ISL_2686565, EPI_ISL_2686566, EPI_ISL_2686567, EPI_ISL_2686570, EPI_ISL_2686571, EPI_ISL_2686572, EPI_ISL_2686575, EPI_ISL_2686576, EPI_ISL_2686577, EPI_ISL_2686579, EPI_ISL_2686580, EPI_ISL_2686582, EPI_ISL_2686583, EPI_ISL_2686588, EPI_ISL_2686590, EPI_ISL_2686591, EPI_ISL_2686594, EPI_ISL_2686596, EPI_ISL_2686598, EPI_ISL_2686599, EPI_ISL_2686600, EPI_ISL_2686601, EPI_ISL_2686605, EPI_ISL_2686608, EPI_ISL_2686612, EPI_ISL_2686613, EPI_ISL_2686614, EPI_ISL_2686615, EPI_ISL_2686616, EPI_ISL_2686617, EPI_ISL_2686620, EPI_ISL_2686621, EPI_ISL_2686622, EPI_ISL_2686624, EPI_ISL_2686625, EPI_ISL_2686626, EPI_ISL_2686627, EPI_ISL_2686628, EPI_ISL_2686633, EPI_ISL_2686635, EPI_ISL_2686636, EPI_ISL_2686638, EPI_ISL_2686639, EPI_ISL_2686641, EPI_ISL_2686642, EPI_ISL_2686643, EPI_ISL_2686645, EPI_ISL_2686652, EPI_ISL_2686653, EPI_ISL_2686654, EPI_ISL_2686660, EPI_ISL_2686661, EPI_ISL_2686663, EPI_ISL_2686666, EPI_ISL_2686669, EPI_ISL_2686670, EPI_ISL_2686671, EPI_ISL_2686672, EPI_ISL_2686673, EPI_ISL_2686674, EPI_ISL_2686675, EPI_ISL_2686676, EPI_ISL_2686678, EPI_ISL_2686679, EPI_ISL_2686680, EPI_ISL_2686681, EPI_ISL_2686682, EPI_ISL_2686683, EPI_ISL_2686685, EPI_ISL_2686686, EPI_ISL_2686687, EPI_ISL_2686688, EPI_ISL_2686689, EPI_ISL_2686692, EPI_ISL_2686693, EPI_ISL_2686695, EPI_ISL_2686696, EPI_ISL_2686697, EPI_ISL_2686698, EPI_ISL_2686699, EPI_ISL_2686702, EPI_ISL_2686705, EPI_ISL_2686706, EPI_ISL_2686708, EPI_ISL_2686716, EPI_ISL_2686737, EPI_ISL_2686741, EPI_ISL_2686742, EPI_ISL_2686746, EPI_ISL_2686749, EPI_ISL_2686753, EPI_ISL_2686755, EPI_ISL_2686756, EPI_ISL_2686758, EPI_ISL_2686759, EPI_ISL_2686761, EPI_ISL_2686770, EPI_ISL_2686802, EPI_ISL_2686803, EPI_ISL_2686804, EPI_ISL_2686805, EPI_ISL_2686806, EPI_ISL_2686807, EPI_ISL_2686808, EPI_ISL_2686811 | | | |
| see above | Fulgent Genetics | Centers for Disease Control and Prevention Division of Viral Diseases, Pathogen Discovery | Dakota Howard, Dhwani Batra, Peter W. Cook, Kara Moser, Adrian Paskey, Jason Caravas, Benjamin Rambo-Martin, Shatavia Morrison, Christopher Gulvick, Scott Sammons, Yvette Unoarumhi, Darlene Wagner, Matthew Schmerer, Harry Gao, Mickey Li, John Gao, Joseph Fierro, Benafsh Sapra, Becky Tsai, Yan Meng, Doreen Ng, James Xie, Clinton R. Paden, Duncan MacCannell |
| EPI_ISL_2686816, EPI_ISL_2686817, EPI_ISL_2686820, EPI_ISL_2686821, EPI_ISL_2686822 | CHUV | Laboratory of genomics and metagenomics | Trestan Pillonel, Damien Jacot, Sébastien Aeby, Gilbert Greub, Claire Bertelli |
| EPI_ISL_2686827 | EHNV | Laboratory of genomics and metagenomics | Trestan Pillonel, Damien Jacot, Sébastien Aeby, Gilbert Greub, Claire Bertelli |
| EPI_ISL_2686838 | CHUV | Laboratory of genomics and metagenomics | Trestan Pillonel, Damien Jacot, Sébastien Aeby, Gilbert Greub, Claire Bertelli |
| EPI_ISL_2686840, EPI_ISL_2686866, EPI_ISL_2686872 | Infinity Biologix | Centers for Disease Control and Prevention Division of Viral Diseases, Pathogen Discovery | Dakota Howard, Dhwani Batra, Peter W. Cook, Kara Moser, Adrian Paskey, Jason Caravas, Benjamin Rambo-Martin, Shatavia Morrison, Christopher Gulvick, Scott Sammons, Yvette Unoarumhi, Darlene Wagner, Matthew Schmerer, Christian Bixby, Yihe Wang, Jonathan Schultz, Chirayu Goswami, Russ Hager, Robin Grimwood, Clinton R. Paden, Duncan MacCannell |
| EPI_ISL_2686905, EPI_ISL_2686907, EPI_ISL_2686910, EPI_ISL_2686912, EPI_ISL_2686913, EPI_ISL_2686914, EPI_ISL_2686916, EPI_ISL_2686918, EPI_ISL_2686936, EPI_ISL_2686939, EPI_ISL_2686946, EPI_ISL_2686948, EPI_ISL_2686952, EPI_ISL_2686963, EPI_ISL_2686965, EPI_ISL_2686966, EPI_ISL_2686972, EPI_ISL_2686973, EPI_ISL_2686987, EPI_ISL_2686991, EPI_ISL_2687001, EPI_ISL_2687014, EPI_ISL_2687023, EPI_ISL_2687025, EPI_ISL_2687026, EPI_ISL_2687037, EPI_ISL_2687056, EPI_ISL_2687066, EPI_ISL_2687067, EPI_ISL_2687091, EPI_ISL_2687092, EPI_ISL_2687101, EPI_ISL_2687102, EPI_ISL_2687103, EPI_ISL_2687104, EPI_ISL_2687107, EPI_ISL_2687109, EPI_ISL_2687110, EPI_ISL_2687118, EPI_ISL_2687128, EPI_ISL_2687129, EPI_ISL_2687138, EPI_ISL_2687139, EPI_ISL_2687146, EPI_ISL_2687148, EPI_ISL_2687152, EPI_ISL_2687153, EPI_ISL_2687156, EPI_ISL_2687157, EPI_ISL_2687161, EPI_ISL_2687164, EPI_ISL_2687166, EPI_ISL_2687181, EPI_ISL_2687182, EPI_ISL_2687192, EPI_ISL_2687201, EPI_ISL_2687204, EPI_ISL_2687206, EPI_ISL_2687208, EPI_ISL_2687228, EPI_ISL_2687237, EPI_ISL_2687241, EPI_ISL_2687250, EPI_ISL_2687252, EPI_ISL_2687261, EPI_ISL_2687264, EPI_ISL_2687272, EPI_ISL_2687273, EPI_ISL_2687275, EPI_ISL_2687294, EPI_ISL_2687295, EPI_ISL_2687311, EPI_ISL_2687312, EPI_ISL_2687313, EPI_ISL_2687317, EPI_ISL_2687318, EPI_ISL_2687320, EPI_ISL_2687322, EPI_ISL_2687323, EPI_ISL_2687324, EPI_ISL_2687327, EPI_ISL_2687331, EPI_ISL_2687339, EPI_ISL_2687347, EPI_ISL_2687353, EPI_ISL_2687354, EPI_ISL_2687358, EPI_ISL_2687368, EPI_ISL_2687369, EPI_ISL_2687370, EPI_ISL_2687371, EPI_ISL_2687372, EPI_ISL_2687377, EPI_ISL_2687389, EPI_ISL_2687395, EPI_ISL_2687400, EPI_ISL_2687406, EPI_ISL_2687412, EPI_ISL_2687415, EPI_ISL_2687417, EPI_ISL_2687420, EPI_ISL_2687421, EPI_ISL_2687424, EPI_ISL_2687426, EPI_ISL_2687427, EPI_ISL_2687434, EPI_ISL_2687438, EPI_ISL_2687439, EPI_ISL_2687448, EPI_ISL_2687449, EPI_ISL_2687451, EPI_ISL_2687453, EPI_ISL_2687456, EPI_ISL_2687458, EPI_ISL_2687460, EPI_ISL_2687467, EPI_ISL_2687468, EPI_ISL_2687470, EPI_ISL_2687471, EPI_ISL_2687475, EPI_ISL_2687476, EPI_ISL_2687477, EPI_ISL_2687478, EPI_ISL_2687482, EPI_ISL_2687485, EPI_ISL_2687486, EPI_ISL_2687496, EPI_ISL_2687497, EPI_ISL_2687498, EPI_ISL_2687500, EPI_ISL_2687502, EPI_ISL_2687513, EPI_ISL_2687515, EPI_ISL_2687518, EPI_ISL_2687520, EPI_ISL_2687523, EPI_ISL_2687524, EPI_ISL_2687532, EPI_ISL_2687540, EPI_ISL_2687547, EPI_ISL_2687557, EPI_ISL_2687560, EPI_ISL_2687561, EPI_ISL_2687566, EPI_ISL_2687567, EPI_ISL_2687570, EPI_ISL_2687575, EPI_ISL_2687579, EPI_ISL_2687580 | | | |
| see above | Aegis Sciences Corporation | Centers for Disease Control and Prevention Division of Viral Diseases, Pathogen Discovery | Dakota Howard, Dhwani Batra, Peter W. Cook, Kara Moser, Adrian Paskey, Jason Caravas, Benjamin Rambo-Martin, Shatavia Morrison, Christopher Gulvick, Scott Sammons, Yvette Unoarumhi, Darlene Wagner, Matthew Schmerer, Cyndi Clark, Patrick Campbell, Rob Case, Vikramsinha Ghorpade, Holly Houdeshell, Ola Kvalvaag, Dillon Nall, Ethan Sanders, Alec Vest, Shaun Westlund, Matthew Hardison, Clinton R. Paden, Duncan MacCannell |
| EPI_ISL_2687944, EPI_ISL_2687960, EPI_ISL_2687968, EPI_ISL_2687977 | National Platform bis UMONS/Jolimont | National Platform bis UMONS/Jolimont | François Dufrasne, Guillaume Bayon-Vicente, Florian Juszczak, Gautier Detry, Ruddy Wattiez |
| EPI_ISL_2687993 | National Institute of Public Health | National Institute of Infectious Diseases-Prof. Dr. Matei Bals Molecular Diagnostics Laboratory | Corina Casangiu, Leontina Banica, Marius Surleac, Petre Milu, Robert Hohan, Simona Paraschiv, Dan Otelea |
| EPI_ISL_2687994 | Department of Public Health Bucharest | National Institute of Infectious Diseases-Prof. Dr. Matei Bals Molecular Diagnostics Laboratory | Corina Casangiu, Leontina Banica, Marius Surleac, Petre Milu, Robert Hohan, Simona Paraschiv, Dan Otelea |
| EPI_ISL_2687995, EPI_ISL_2687996 | National Institute of Infectious Diseases-Prof. Dr. Matei Bals Molecular Diagnostics Laboratory | National Institute of Infectious Diseases-Prof. Dr. Matei Bals Molecular Diagnostics Laboratory | Corina Casangiu, Leontina Banica, Marius Surleac, Petre Milu, Robert Hohan, Simona Paraschiv, Dan Otelea |
| EPI_ISL_2687997 | Department of Public Health Bucharest | National Institute of Infectious Diseases-Prof. Dr. Matei Bals Molecular Diagnostics Laboratory | Corina Casangiu, Leontina Banica, Marius Surleac, Petre Milu, Robert Hohan, Simona Paraschiv, Dan Otelea |
| EPI_ISL_2688190, EPI_ISL_2688192, EPI_ISL_2688193 | Kansas Health and Environmental Lab | Kansas Health and Environmental Lab | Mike Grose, Katherine Wiggins, Jonathan Barnell, Ben Olsen, and Phil Adam |
| EPI_ISL_2688194 | IN State Department of Health Laboratory Services | IN State Department of Health Laboratory Services | Cassandra Campion, Jamie Yeadon, Brian Pope, Lixia Liu, Kyle Brownlee, Melissa Hindenlang, Mark Glazier |
| EPI_ISL_2688195 | Kansas Health and Environmental Lab | Kansas Health and Environmental Lab | Mike Grose, Katherine Wiggins, Jonathan Barnell, Ben Olsen, and Phil Adam |
| EPI_ISL_2688197 | IN State Department of Health Laboratory Services | IN State Department of Health Laboratory Services | Cassandra Campion, Jamie Yeadon, Brian Pope, Lixia Liu, Kyle Brownlee, Melissa Hindenlang, Mark Glazier |
| EPI_ISL_2688198, EPI_ISL_2688199 | Kansas Health and Environmental Lab | Kansas Health and Environmental Lab | Mike Grose, Katherine Wiggins, Jonathan Barnell, Ben Olsen, and Phil Adam |
| EPI_ISL_2688200 | IN State Department of Health Laboratory Services | IN State Department of Health Laboratory Services | Cassandra Campion, Jamie Yeadon, Brian Pope, Lixia Liu, Kyle Brownlee, Melissa Hindenlang, Mark Glazier |
| EPI_ISL_2688201 | Kansas Health and Environmental Lab | Kansas Health and Environmental Lab | Mike Grose, Katherine Wiggins, Jonathan Barnell, Ben Olsen, and Phil Adam |
| EPI_ISL_2688203 | IN State Department of Health Laboratory Services | IN State Department of Health Laboratory Services | Cassandra Campion, Jamie Yeadon, Brian Pope, Lixia Liu, Kyle Brownlee, Melissa Hindenlang, Mark Glazier |
| EPI_ISL_2688204, EPI_ISL_2688205 | Kansas Health and Environmental Lab | Kansas Health and Environmental Lab | Mike Grose, Katherine Wiggins, Jonathan Barnell, Ben Olsen, and Phil Adam |
| EPI_ISL_2688206 | IN State Department of Health Laboratory Services | IN State Department of Health Laboratory Services | Cassandra Campion, Jamie Yeadon, Brian Pope, Lixia Liu, Kyle Brownlee, Melissa Hindenlang, Mark Glazier |
| EPI_ISL_2688207, EPI_ISL_2688208 | Kansas Health and Environmental Lab | Kansas Health and Environmental Lab | Mike Grose, Katherine Wiggins, Jonathan Barnell, Ben Olsen, and Phil Adam |
| EPI_ISL_2688209 | IN State Department of Health Laboratory Services | IN State Department of Health Laboratory Services | Cassandra Campion, Jamie Yeadon, Brian Pope, Lixia Liu, Kyle Brownlee, Melissa Hindenlang, Mark Glazier |
| EPI_ISL_2688211 | Kansas Health and Environmental Lab | Kansas Health and Environmental Lab | Mike Grose, Katherine Wiggins, Jonathan Barnell, Ben Olsen, and Phil Adam |
| EPI_ISL_2688212 | IN State Department of Health Laboratory Services | IN State Department of Health Laboratory Services | Cassandra Campion, Jamie Yeadon, Brian Pope, Lixia Liu, Kyle Brownlee, Melissa Hindenlang, Mark Glazier |
| EPI_ISL_2688213, EPI_ISL_2688214 | Kansas Health and Environmental Lab | Kansas Health and Environmental Lab | Mike Grose, Katherine Wiggins, Jonathan Barnell, Ben Olsen, and Phil Adam |
| EPI_ISL_2688215 | IN State Department of Health Laboratory Services | IN State Department of Health Laboratory Services | Cassandra Campion, Jamie Yeadon, Brian Pope, Lixia Liu, Kyle Brownlee, Melissa Hindenlang, Mark Glazier |
| EPI_ISL_2688216, EPI_ISL_2688217 | Kansas Health and Environmental Lab | Kansas Health and Environmental Lab | Mike Grose, Katherine Wiggins, Jonathan Barnell, Ben Olsen, and Phil Adam |
| EPI_ISL_2688218 | IN State Department of Health Laboratory Services | IN State Department of Health Laboratory Services | Cassandra Campion, Jamie Yeadon, Brian Pope, Lixia Liu, Kyle Brownlee, Melissa Hindenlang, Mark Glazier |
| EPI_ISL_2688219, EPI_ISL_2688222 | Kansas Health and Environmental Lab | Kansas Health and Environmental Lab | Mike Grose, Katherine Wiggins, Jonathan Barnell, Ben Olsen, and Phil Adam |
| EPI_ISL_2688223 | IN State Department of Health Laboratory Services | IN State Department of Health Laboratory Services | Cassandra Campion, Jamie Yeadon, Brian Pope, Lixia Liu, Kyle Brownlee, Melissa Hindenlang, Mark Glazier |
| EPI_ISL_2688224, EPI_ISL_2688226 | Kansas Health and Environmental Lab | Kansas Health and Environmental Lab | Mike Grose, Katherine Wiggins, Jonathan Barnell, Ben Olsen, and Phil Adam |
| EPI_ISL_2688227 | IN State Department of Health Laboratory Services | IN State Department of Health Laboratory Services | Cassandra Campion, Jamie Yeadon, Brian Pope, Lixia Liu, Kyle Brownlee, Melissa Hindenlang, Mark Glazier |
| EPI_ISL_2688228, EPI_ISL_2688229, EPI_ISL_2688231, EPI_ISL_2688232 | Kansas Health and Environmental Lab | Kansas Health and Environmental Lab | Mike Grose, Katherine Wiggins, Jonathan Barnell, Ben Olsen, and Phil Adam |
| EPI_ISL_2688233 | IN State Department of Health Laboratory Services | IN State Department of Health Laboratory Services | Cassandra Campion, Jamie Yeadon, Brian Pope, Lixia Liu, Kyle Brownlee, Melissa Hindenlang, Mark Glazier |
| EPI_ISL_2688234, EPI_ISL_2688235 | Kansas Health and Environmental Lab | Kansas Health and Environmental Lab | Mike Grose, Katherine Wiggins, Jonathan Barnell, Ben Olsen, and Phil Adam |
| EPI_ISL_2688236 | IN State Department of Health Laboratory Services | IN State Department of Health Laboratory Services | Cassandra Campion, Jamie Yeadon, Brian Pope, Lixia Liu, Kyle Brownlee, Melissa Hindenlang, Mark Glazier |
| EPI_ISL_2688237 | Kansas Health and Environmental Lab | Kansas Health and Environmental Lab | Mike Grose, Katherine Wiggins, Jonathan Barnell, Ben Olsen, and Phil Adam |
| EPI_ISL_2688240, EPI_ISL_2688242, EPI_ISL_2688244, EPI_ISL_2688246, EPI_ISL_2688247 | IN State Department of Health Laboratory Services | IN State Department of Health Laboratory Services | Cassandra Campion, Jamie Yeadon, Brian Pope, Lixia Liu, Kyle Brownlee, Melissa Hindenlang, Mark Glazier |
| EPI_ISL_2688250, EPI_ISL_2688251 | AP SSO | CSIR-Centre for Cellular and Molecular Biology-INSACOG | Shreekant Verma, Amareshwar Vodapalli ,Ara Sreenivas,B Himasri,Valli Nagalakshmi Undamatla,Payel Mukherjee, Lamuk Zaveri, Onkar Kulkarni,Tulasi Nagabandi, Sofia Banu, Sumedha Avadhanula, Priya Nurkurthy, Vidhyadhari Methuku, Archana Bharadwaj Siva, Karthik Bharadwaj Tallapaka,Rakesh K Mishra,Divya Tej Sowpati |
| EPI_ISL_2688252, EPI_ISL_2688253 | AP SSO | CSIR-Centre for Cellular and Molecular Biology-INSACOG | Amareshwar Vodapalli,Ara Sreenivas,Shreekant Verma, B Himasri,Valli Nagalakshmi Undamatla,Payel Mukherjee, Lamuk Zaveri, Onkar Kulkarni,Tulasi Nagabandi, Sofia Banu, ,Priya Nurkurthy, Vidhyadhari Methuku, Sumedha Avadhanula, Archana Bharadwaj Siva, Karthik Bharadwaj Tallapaka,Rakesh K Mishra,Divya Tej Sowpati |
| EPI_ISL_2688254 | PASTEUR MOLECULAR LABORATORY - INSACOG MEGHALAYA | CSIR-Centre for Cellular and Molecular Biology-INSACOG | R. Lyngdoh, E Shadap , E Marak,Lamuk Zaveri,Tulasi Nagabandi,Payel Mukherjee,Onkar Kulkarni, Sofia Banu, Ara Sreenivas,Shreekant Verma, Amareshwar Vodapalli ,B Himasri,Valli Nagalakshmi Undamatla, Sumedha Avadhanula, Priya Nurkurthy, Vidhyadhari Methuku, Archana Bharadwaj Siva, Karthik Bharadwaj Tallapaka,Rakesh K Mishra,Divya Tej Sowpati |
| EPI_ISL_2688257 | CSIR-Centre for Cellular and Molecular Biology | CSIR-Centre for Cellular and Molecular Biology-INSACOG | Lamuk Zaveri,Tulasi Nagabandi,Payel Mukherjee,Onkar Kulkarni, Sofia Banu, Ara Sreenivas,Shreekant Verma, Amareshwar Vodapalli ,B Himasri,Valli Nagalakshmi Undamatla, Sumedha Avadhanula, Priya Nurkurthy, Vidhyadhari Methuku, |

| | | | |
|---|---|---|---|
| | | | Archana Bharadwaj Siva, Karthik Bharadwaj Tallapaka,Rakesh K Mishra,Divya Tej Sowpati |
| EPI_ISL_2688258 | CSIR-Centre for Cellular and Molecular Biology | CSIR-Centre for Cellular and Molecular Biology-INSACOG | Ara Sreenivas,Shreekant Verma, Amareshwar Vodapalli ,B Himasri,Valli Nagalakshmi Undamatla,Payel Mukherjee, Lamuk Zaveri, Onkar Kulkarni,Tulasi Nagabandi, Sofia Banu, ,Priya Nurkurthy, Vidhyadhari Methuku, Sumedha Avadhanula, Archana Bharadwaj Siva, Karthik Bharadwaj Tallapaka,Rakesh K Mishra,Divya Tej Sowpati |
| EPI_ISL_2688325, EPI_ISL_2688357, EPI_ISL_2688365 | Multiplex DX | Multiplex DX | Diana Drobna, Veronika Mancikova, Jakub Kovac, Pavol Cekan |
| EPI_ISL_2688533, EPI_ISL_2688550, EPI_ISL_2688554, EPI_ISL_2688555, EPI_ISL_2688556, EPI_ISL_2688558 | National Health Laboratory Service, South Africa | KRISP, KZn Research Innovation and Sequencing Platform | Giandhari Jennifer, Pillay Sureshnee, Yajna Ramphal, Naidoo Yeshnee, Tshabuila Derek,Tegally Houriiyah, San James, Wilkinson Eduan, de Oliveira Tulio |
| EPI_ISL_2688593, EPI_ISL_2688598 | Furst Medical Laboratory | Norwegian Institute of Public Health, Department of Virology | Kathrine Stene-Johansen, Kamilla Heddeland Instefjord, Hilde Elshaug, Garcia Llorente Ignacio, Jon Bråte, Engebretsen Serina Beate,Pedersen Benedikte Nevjen, Line Victoria Moen, Debech Nadia, Atiya R Ali,Marie Paulsen Madsen, Rasmus Riis Kopperud, Hilde Vollan, Karoline Bragstad, Olav Hungnes |
| EPI_ISL_2688682 | Hôpital Jacques Monod | Centre Hospitalier Universitaire de Rouen Laboratoire de Virologie | Alice Moisan, Fabienne De Oliveira, Marie Leoz |
| EPI_ISL_2688806, EPI_ISL_2688812, EPI_ISL_2688813, EPI_ISL_2688817, EPI_ISL_2688818 | Massachusetts State Public Health Laboratory | Massachusetts State Public Health Laboratory | Andrew Lang, Timelia Fink, Glen Gallagher, Sandra Smole |
| EPI_ISL_2689021 | Minnesota Department of Health, Public Health Laboratory | Minnesota Department of Health, Public Health Laboratory | Alexandra Lorentz, Jacob Garfin, Matt Plumb, and Xiong Wang |
| EPI_ISL_2689022 | Allina Health Laboratory | Minnesota Department of Health, Public Health Laboratory | Alexandra Lorentz, Jacob Garfin, Matt Plumb, and Xiong Wang |
| EPI_ISL_2689024 | Minnesota Department of Health, Public Health Laboratory | Minnesota Department of Health, Public Health Laboratory | Alexandra Lorentz, Jacob Garfin, Matt Plumb, and Xiong Wang |
| EPI_ISL_2689049, EPI_ISL_2689051, EPI_ISL_2689052, EPI_ISL_2689053, EPI_ISL_2689108, EPI_ISL_2689109 | MD PHL | MD PHL | Maryland Department of Health Laboratories Administration |
| EPI_ISL_2689116, EPI_ISL_2689117, EPI_ISL_2689119, EPI_ISL_2689120, EPI_ISL_2689121, EPI_ISL_2689122 | Nebraska Public Health Laboratory | NPHL COVID-19 Response Team | NPHL COVID-19 Response Team |
| EPI_ISL_2689168, EPI_ISL_2689556, EPI_ISL_2689576 | Labor Dr. Fenner und Kollegen | Heinrich Pette Institute, Leibniz Institute for Experimental Virology | Alexis Robitaille, Thomas Günther, Johannes Knobloch, Martin Aepfelbacher, Nicole Fischer, Adam Grundhoff |
| EPI_ISL_2689602 | NM Dept. Health, Scientific Laboratory Division | Genomics and Discovery, Respiratory Viruses Branch, Division of Viral Diseases, Centers for Disease Control and Prevention | Jing Zhang, Anna Kelleher, Ying Tao, Yan Li, Brian Lynch, Krista Queen, Anna Uehara, Peter Cook, Han Jia Justin Ng, Rachel Marine, Clinton R. Paden, Dhwani Batra, Haibin Wang, Tara Coalter, Jasmine Padilla, Morgan Davis, Mili Sheth, Sarah Nobles, Mark Burroughs, Justin Lee, Adam Retchless, Suxiang Tong |
| EPI_ISL_2689604 | AR Dept. of Health-PHL, Molecular Diagnostics | Genomics and Discovery, Respiratory Viruses Branch, Division of Viral Diseases, Centers for Disease Control and Prevention | Jing Zhang, Anna Kelleher, Ying Tao, Yan Li, Brian Lynch, Krista Queen, Anna Uehara, Peter Cook, Han Jia Justin Ng, Rachel Marine, Clinton R. Paden, Dhwani Batra, Haibin Wang, Tara Coalter, Jasmine Padilla, Morgan Davis, Mili Sheth, Sarah Nobles, Mark Burroughs, Justin Lee, Adam Retchless, Suxiang Tong |
| EPI_ISL_2689637 | MLD Szpital Św. Elżbiety | National Institute of Public Health - National Institute of Hygiene | Wołkowicz Tomasz, Zacharczuk Katarzyna, Sadkowska-Todys Małgorzata, Gierczyńki Rafał |
| EPI_ISL_2689658 | National Institute of Public Health - National Institute of Hygiene | National Institute of Public Health - National Institute of Hygiene | Wołkowicz Tomasz, Zacharczuk Katarzyna, Sadkowska-Todys Małgorzata, Gierczyńki Rafał |
| EPI_ISL_2689669, EPI_ISL_2689671, EPI_ISL_2689681, EPI_ISL_2689685 | Nebraska Public Health Laboratory | NPHL COVID-19 Response Team | NPHL COVID-19 Response Team |
| EPI_ISL_2689692, EPI_ISL_2689695, EPI_ISL_2689701, EPI_ISL_2689708, EPI_ISL_2689730, EPI_ISL_2689734 | Servicio de Microbiologia Hospital Ramon y Cajal | Servicio de Microbiologia Hospital Ramon y Cajal | Ponce M, Galan JC, Martinez L. Abreu M, y Gonzalez-Alba JM |
| EPI_ISL_2689755, EPI_ISL_2689756, EPI_ISL_2689757 | Kansas Health and Environmental Lab | Kansas Health and Environmental Lab | Katherine Wiggins, Mike Grose, Jonathan Barnell, Ben Olsen, and Phil Adam |
| EPI_ISL_2689758, EPI_ISL_2689759, EPI_ISL_2689764, EPI_ISL_2689765, EPI_ISL_2689767, EPI_ISL_2689771, EPI_ISL_2689772, EPI_ISL_2689773, EPI_ISL_2689774, EPI_ISL_2689775, EPI_ISL_2689776, EPI_ISL_2689778, EPI_ISL_2689779, EPI_ISL_2689780, EPI_ISL_2689781, EPI_ISL_2689782 | | | |
| see above | Wichita State University - Molecular Diagnostics Lab | Kansas Health and Environmental Lab | Katherine Wiggins, Mike Grose, Jonathan Barnell, Ben Olsen, and Phil Adam |
| EPI_ISL_2689867 | MO State Public Health Laboratory | Centers for Disease Control and Prevention Division of Viral Diseases, Pathogen Discovery | Mili Sheth, Sarah Nobles, Jasmine Padilla, Mark Burroughs, Shoshona Le, Katie Dillon, Peter Cook, Clinton R. Paden, Dhwani Batra, Krista Queen, Kristen Knipe, Dakota Howard, Yvette Unoarumhi, Darlene Wagner, Matthew Schmerer, Ben L. Rambo-Martin, Kristine Lacek, Sam Shepard, Alison Laufer Halpin, Dave Wentworth, Vivien Dugan, Suxiang Tong, Justin Lee |
| EPI_ISL_2689888 | IA State Hygienic Laboratory | Centers for Disease Control and Prevention Division of Viral Diseases, Pathogen Discovery | Mili Sheth, Sarah Nobles, Jasmine Padilla, Mark Burroughs, Shoshona Le, Katie Dillon, Peter Cook, Clinton R. Paden, Dhwani Batra, Krista Queen, Kristen Knipe, Dakota Howard, Yvette Unoarumhi, Darlene Wagner, Matthew Schmerer, Ben L. Rambo-Martin, Kristine Lacek, Sam Shepard, Alison Laufer Halpin, Dave Wentworth, Vivien Dugan, Suxiang Tong, Justin Lee |
| EPI_ISL_2689895, EPI_ISL_2689898, EPI_ISL_2689899, EPI_ISL_2689900, EPI_ISL_2689903, EPI_ISL_2689904, EPI_ISL_2689906 | WY Public Health Laboratory | Centers for Disease Control and Prevention Division of Viral Diseases, Pathogen Discovery | Mili Sheth, Sarah Nobles, Jasmine Padilla, Mark Burroughs, Shoshona Le, Katie Dillon, Peter Cook, Clinton R. Paden, Dhwani Batra, Krista Queen, Kristen Knipe, Dakota Howard, Yvette Unoarumhi, Darlene Wagner, Matthew Schmerer, Ben L. Rambo-Martin, Kristine Lacek, Sam Shepard, Alison Laufer Halpin, Dave Wentworth, Vivien Dugan, Suxiang Tong, Justin Lee |
| EPI_ISL_2689936 | NJ Public Health and Environmental Laboratories | Centers for Disease Control and Prevention Division of Viral Diseases, Pathogen Discovery | Mili Sheth, Sarah Nobles, Jasmine Padilla, Mark Burroughs, Shoshona Le, Katie Dillon, Peter Cook, Clinton R. Paden, Dhwani Batra, Krista Queen, Kristen Knipe, Dakota Howard, Yvette Unoarumhi, Darlene Wagner, Matthew Schmerer, Ben L. Rambo-Martin, Kristine Lacek, Sam Shepard, Alison Laufer Halpin, Dave Wentworth, Vivien Dugan, Suxiang Tong, Justin Lee |
| EPI_ISL_2689941, EPI_ISL_2689944 | MI - Michigan Department of Health and Human Services - Bureau of Laboratories | Centers for Disease Control and Prevention Division of Viral Diseases, Pathogen Discovery | Mili Sheth, Sarah Nobles, Jasmine Padilla, Mark Burroughs, Shoshona Le, Katie Dillon, Peter Cook, Clinton R. Paden, Dhwani Batra, Krista Queen, Kristen Knipe, Dakota Howard, Yvette Unoarumhi, Darlene Wagner, Matthew Schmerer, Ben L. Rambo-Martin, Kristine Lacek, Sam Shepard, Alison Laufer Halpin, Dave Wentworth, Vivien Dugan, Suxiang Tong, Justin Lee |
| EPI_ISL_2689970, EPI_ISL_2689972 | "AR Dept. of Health-PHL, Molecular Diagnostics" | Centers for Disease Control and Prevention Division of Viral Diseases, Pathogen Discovery | Mili Sheth, Sarah Nobles, Jasmine Padilla, Mark Burroughs, Shoshona Le, Katie Dillon, Peter Cook, Clinton R. Paden, Dhwani Batra, Krista Queen, Kristen Knipe, Dakota Howard, Yvette Unoarumhi, Darlene Wagner, Matthew Schmerer, Ben L. Rambo-Martin, Kristine Lacek, Sam Shepard, Alison Laufer Halpin, Dave Wentworth, Vivien Dugan, Suxiang Tong, Justin Lee |
| EPI_ISL_2689989, EPI_ISL_2689992, EPI_ISL_2689993, EPI_ISL_2689996, EPI_ISL_2690000, EPI_ISL_2690004, EPI_ISL_2690007, EPI_ISL_2690008 | IN State Department of Health Laboratory Services | Centers for Disease Control and Prevention Division of Viral Diseases, Pathogen Discovery | Mili Sheth, Sarah Nobles, Jasmine Padilla, Mark Burroughs, Shoshona Le, Katie Dillon, Peter Cook, Clinton R. Paden, Dhwani Batra, Krista Queen, Kristen Knipe, Dakota Howard, Yvette Unoarumhi, Darlene Wagner, Matthew Schmerer, Ben L. Rambo-Martin, Kristine Lacek, Sam Shepard, Alison Laufer Halpin, Dave Wentworth, Vivien Dugan, Suxiang Tong, Justin Lee |
| EPI_ISL_2690022, EPI_ISL_2690024, EPI_ISL_2690025, EPI_ISL_2690026 | UT-Unified State Labs: Public Health Utah DOH | Centers for Disease Control and Prevention Division of Viral Diseases, Pathogen Discovery | Mili Sheth, Sarah Nobles, Jasmine Padilla, Mark Burroughs, Shoshona Le, Katie Dillon, Peter Cook, Clinton R. Paden, Dhwani Batra, Krista Queen, Kristen Knipe, Dakota Howard, Yvette Unoarumhi, Darlene Wagner, Matthew Schmerer, Ben L. Rambo-Martin, Kristine Lacek, Sam Shepard, Alison Laufer Halpin, Dave Wentworth, Vivien Dugan, Suxiang Tong, Justin Lee |
| EPI_ISL_2690031 | GA Department of Public Health Laboratory | Centers for Disease Control and Prevention Division of Viral Diseases, Pathogen Discovery | Mili Sheth, Sarah Nobles, Jasmine Padilla, Mark Burroughs, Shoshona Le, Katie Dillon, Peter Cook, Clinton R. Paden, Dhwani Batra, Krista Queen, Kristen Knipe, Dakota Howard, Yvette Unoarumhi, Darlene Wagner, Matthew Schmerer, Ben L. Rambo-Martin, Kristine Lacek, Sam Shepard, Alison Laufer Halpin, Dave Wentworth, Vivien Dugan, Suxiang Tong, Justin Lee |
| EPI_ISL_2690048, EPI_ISL_2690049, EPI_ISL_2690050, EPI_ISL_2690052 | KS Health and Environmental Laboratories | Centers for Disease Control and Prevention Division of Viral Diseases, Pathogen Discovery | Mili Sheth, Sarah Nobles, Jasmine Padilla, Mark Burroughs, Shoshona Le, Katie Dillon, Peter Cook, Clinton R. Paden, Dhwani Batra, Krista Queen, Kristen Knipe, Dakota Howard, Yvette Unoarumhi, Darlene Wagner, Matthew Schmerer, Ben L. Rambo-Martin, Kristine Lacek, Sam Shepard, Alison Laufer Halpin, Dave Wentworth, Vivien Dugan, Suxiang Tong, Justin Lee |
| EPI_ISL_2690101 | WA State Department of Health | Centers for Disease Control and Prevention Division of Viral Diseases, Pathogen Discovery | Mili Sheth, Sarah Nobles, Jasmine Padilla, Mark Burroughs, Shoshona Le, Katie Dillon, Peter Cook, Clinton R. Paden, Dhwani Batra, Krista Queen, Kristen Knipe, Dakota Howard, Yvette Unoarumhi, Darlene Wagner, Matthew Schmerer, Ben L. Rambo-Martin, Kristine Lacek, Sam Shepard, Alison Laufer Halpin, Dave Wentworth, Vivien Dugan, Suxiang Tong, Justin Lee |

| | | | |
|---|---|---|---|
| EPI_ISL_2690124, EPI_ISL_2690125 | MT Public Health Laboratory | Centers for Disease Control and Prevention Division of Viral Diseases, Pathogen Discovery | Mili Sheth, Sarah Nobles, Jasmine Padilla, Mark Burroughs, Shoshona Le, Katie Dillon, Peter Cook, Clinton R. Paden, Dhwani Batra, Krista Queen, Kristen Knipe, Dakota Howard, Yvette Unoarumhi, Darlene Wagner, Matthew Schmerer, Ben L. Rambo-Martin, Kristine Lacek, Sam Shepard, Alison Laufer Halpin, Dave Wentworth, Vivien Dugan, Suxiang Tong, Justin Lee |
| EPI_ISL_2690248, EPI_ISL_2690249, EPI_ISL_2690250, EPI_ISL_2690254, EPI_ISL_2690255 | University of Mississippi Medical Center, Department of Pathology | University of Mississippi Medical Center, Molecular and Genomics Core Facility | Ashley C. Johnson, Wenjie Wu, Ithiel J. Frame, Krishna K. Ayyalasomayajula, Michael R. Garrett, D. Ashley Robinson |
| EPI_ISL_2690260, EPI_ISL_2690261, EPI_ISL_2690262, EPI_ISL_2690263, EPI_ISL_2690264, EPI_ISL_2690265, EPI_ISL_2690266, EPI_ISL_2690267, EPI_ISL_2690268, EPI_ISL_2690269 | MS Public Health Laboratory | University of Mississippi Medical Center, Molecular and Genomics Core Facility | Ashley C. Johnson, Wenjie Wu, Ithiel J. Frame, Krishna K. Ayyalasomayajula, Michael R. Garrett, D. Ashley Robinson |
| EPI_ISL_2690271, EPI_ISL_2690273, EPI_ISL_2690274, EPI_ISL_2690275, EPI_ISL_2690276, EPI_ISL_2690277, EPI_ISL_2690278, EPI_ISL_2690279, EPI_ISL_2690280, EPI_ISL_2690281, EPI_ISL_2690282, EPI_ISL_2690283, EPI_ISL_2690285, EPI_ISL_2690286, EPI_ISL_2690287, EPI_ISL_2690288, EPI_ISL_2690289, EPI_ISL_2690290, EPI_ISL_2690294, EPI_ISL_2690295 | | | |
| see above | Nevada State Public Health Laboratory | Nevada State Public Health Laboratory | Andrew Gorzalski, Mark Pandori |
| EPI_ISL_2690345, EPI_ISL_2690349, EPI_ISL_2690358, EPI_ISL_2690362, EPI_ISL_2690384, EPI_ISL_2690395, EPI_ISL_2690399 | Arizona State University | Arizona State University | Peter T. Skidmore, LaRinda A. Holland, Matthew F. Smith, Rabia Maqsood, Nicholas J. Mellor, Joy M. Blain, Valerie Harris, Joshua LaBaer, Vel Murugan, Efrem S. Lim |
| EPI_ISL_2690479, EPI_ISL_2690480, EPI_ISL_2690481, EPI_ISL_2690482, EPI_ISL_2690483, EPI_ISL_2690484, EPI_ISL_2690485, EPI_ISL_2690486, EPI_ISL_2690487, EPI_ISL_2690488, EPI_ISL_2690489, EPI_ISL_2690490, EPI_ISL_2690491, EPI_ISL_2690492, EPI_ISL_2690493, EPI_ISL_2690494, EPI_ISL_2690495, EPI_ISL_2690496, EPI_ISL_2690497, EPI_ISL_2690498, EPI_ISL_2690499, EPI_ISL_2690500, EPI_ISL_2690502, EPI_ISL_2690503, EPI_ISL_2690504, EPI_ISL_2690505, EPI_ISL_2690506, EPI_ISL_2690507, EPI_ISL_2690508, EPI_ISL_2690509, EPI_ISL_2690510, EPI_ISL_2690511, EPI_ISL_2690512, EPI_ISL_2690513 | | | |
| see above | MRC/UVRI & LSHTM Uganda Research Unit, Central Public Health Laboratories, Rakai Health Sciences Program | MRC/UVRI & LSHTM Uganda Research Unit, Central Public Health Laboratories | Matthew Cotten, Dan Lule Bugembe, My V.T. Phan, Pontiano Kaleebu,Isaac Sseeewanyana, Patrick Semanda, Susan Nabadda, Hellen Nansumba |
| EPI_ISL_2691231, EPI_ISL_2691232, EPI_ISL_2691239, EPI_ISL_2691249, EPI_ISL_2691268, EPI_ISL_2691273, EPI_ISL_2691282 | Santa Clara Valley Medical Center | Santa Clara County Public Health Laboratory | Santa Clara County Public Health Department |
| EPI_ISL_2691301 | PAMF Palo Alto | Santa Clara County Public Health Laboratory | Santa Clara County Public Health Department |
| EPI_ISL_2691310 | Santa Clara Valley Medical Center | Santa Clara County Public Health Laboratory | Santa Clara County Public Health Department |
| EPI_ISL_2691707 | PathWest Laboratory Medicine WA Microbial Surveillance Unit | PathWest Laboratory Medicine WA Microbial Surveillance Unit | QEII Medical Centre, Hospital Ave, Nedlands WA 6009 |
| EPI_ISL_2691905, EPI_ISL_2691910 | Johns Hopkins Hospital Department of Pathology | Johns Hopkins Hospital Department of Pathology | C. Paul Morris, Chun Huai Luo, Adannaya Amadi, Matthew Schwartz, Nicholas Gallagher, Heba H. Mostafa |
| EPI_ISL_2691947 | Rush University Medical Center | RIPHL at Rush University Medical Center | Stefan Green, Kevin Kunstman, Max Kolton, Marieta Hyde, Laura Furtado, Felix Araujo Perez, Mary Hayden, Joyce Houlihan, Diane Springer |
| EPI_ISL_2692117, EPI_ISL_2692119, EPI_ISL_2692127 | University of Illinois at Chicago | RIPHL at Rush University Medical Center | Stefan Green, Kevin Kunstman, Max Kolton, Marieta Hyde, Laura Furtado, Felix Araujo Perez, Julie Bentea, Omar Perez |
| EPI_ISL_2692145, EPI_ISL_2692146, EPI_ISL_2692148, EPI_ISL_2692155, EPI_ISL_2692156, EPI_ISL_2692163 | Montana Public Health Laboratory | Montana Public Health Laboratory | Joy Ritter, Michelle Mozer, Carrie Biskupiak, Deborah Gibson, Michael Dills |
| EPI_ISL_2692230 | Seattle Flu Study | Seattle Flu Study | Deborah A. Nickerson, Chris D. Frazar, Jover Lee, Benjamin Pelle, Erica Ryke, Matthew Richardson, Amanda Adler, Elisabeth Brandstetter, Peter D. Han, Kairsten Fay, Misja Ilcisin, Kirsten Lacombe, Thomas R. Sibley, Melissa Truong, Caitlin R. Wolf, Karen Cowgill, Stephanie Schrag, Jeff Duchin, Michael Boeckh, Janet A. Englund, Michael Famulare, Barry R. Lutz, Mark J. Rieder, Lea M. Starita, Matthew Thompson, Helen Y. Chu, Trevor Bedford, Jay Shendure |
| EPI_ISL_2692454, EPI_ISL_2692468 | Atlas Genomics | Seattle Flu Study | Deborah A. Nickerson, Chris D. Frazar, Jover Lee, Benjamin Pelle, Erica Ryke, Matthew Richardson, Amanda Adler, Elisabeth Brandstetter, Peter D. Han, Kairsten Fay, Misja Ilcisin, Kirsten Lacombe, Thomas R. Sibley, Melissa Truong, Caitlin R. Wolf, Romesh Gautom, Geoff Melly, Brian Hiatt, Philip Dykema, Scott Lindquist, Michael Boeckh, Janet A. Englund, Michael Famulare, Barry R. Lutz, Mark J. Rieder, Lea M. Starita, Matthew Thompson, Helen Y. Chu, Jay Shendure, Trevor Bedford |
| EPI_ISL_2692535 | GA Department of Public Health Laboratory | Genomics and Discovery, Respiratory Viruses Branch, Division of Viral Diseases, Centers for Disease Control and Prevention | Jing Zhang, Anna Kelleher, Ying Tao, Yan Li, Brian Lynch, Krista Queen, Anna Uehara, Peter Cook, Han Jia Justin Ng, Rachel Marine, Clinton R. Paden, Dhwani Batra, Haibin Wang, Tara Coalter, Jasmine Padilla, Morgan Davis, Mili Sheth, Sarah Nobles, Mark Burroughs, Justin Lee, Adam Retchless, Suxiang Tong |
| EPI_ISL_2692536, EPI_ISL_2692537 | OK Public Health Laboratory, Oklahoma State DOH | Genomics and Discovery, Respiratory Viruses Branch, Division of Viral Diseases, Centers for Disease Control and Prevention | Jing Zhang, Anna Kelleher, Ying Tao, Yan Li, Brian Lynch, Krista Queen, Anna Uehara, Peter Cook, Han Jia Justin Ng, Rachel Marine, Clinton R. Paden, Dhwani Batra, Haibin Wang, Tara Coalter, Jasmine Padilla, Morgan Davis, Mili Sheth, Sarah Nobles, Mark Burroughs, Justin Lee, Adam Retchless, Suxiang Tong |
| EPI_ISL_2692592, EPI_ISL_2692635, EPI_ISL_2692642, EPI_ISL_2692702, EPI_ISL_2692705, EPI_ISL_2692707, EPI_ISL_2692708, EPI_ISL_2692709, EPI_ISL_2692710, EPI_ISL_2692711, EPI_ISL_2692713, EPI_ISL_2692714, EPI_ISL_2692716, EPI_ISL_2692717, EPI_ISL_2692719, EPI_ISL_2692720, EPI_ISL_2692721, EPI_ISL_2692722, EPI_ISL_2692723, EPI_ISL_2692724, EPI_ISL_2692725, EPI_ISL_2692726, EPI_ISL_2692728, EPI_ISL_2692729, EPI_ISL_2692730, EPI_ISL_2692731, EPI_ISL_2692732, EPI_ISL_2692733, EPI_ISL_2692735, EPI_ISL_2692736, EPI_ISL_2692737, EPI_ISL_2692738, EPI_ISL_2692740, EPI_ISL_2692741, EPI_ISL_2692742, EPI_ISL_2692743, EPI_ISL_2692746, EPI_ISL_2692747, EPI_ISL_2692748, EPI_ISL_2692749, EPI_ISL_2692751, EPI_ISL_2692752, EPI_ISL_2692753, EPI_ISL_2692756, EPI_ISL_2692757, EPI_ISL_2692758, EPI_ISL_2692759, EPI_ISL_2692760, EPI_ISL_2692761, EPI_ISL_2692763, EPI_ISL_2692765, EPI_ISL_2692766, EPI_ISL_2692767, EPI_ISL_2692768, EPI_ISL_2692770, EPI_ISL_2692771, EPI_ISL_2692773, EPI_ISL_2692775, EPI_ISL_2692776, EPI_ISL_2692777, EPI_ISL_2692778, EPI_ISL_2692784, EPI_ISL_2692791, EPI_ISL_2692826, EPI_ISL_2692827, EPI_ISL_2692829, EPI_ISL_2692832 | | | |
| see above | Salud Digna | Instituto Nacional de Medicina Genomica | Cedro-Tanda A, Hidalgo-Miranda A, Mendoza-Vargas A, Reyes-Grajeda JP, Abraham Campos-Romero, Moreno-Camacho José Luis, Rodriguez-Gallegos Jorge, Luna-Ruiz Marco, Gonzalez-Barrera D, Rangel-DeLeon D, Munguia-Garza P, Ramirez-Vega O, Escobar-Arrazola, M, Herrera-Montalvo LA. |
| EPI_ISL_2692834, EPI_ISL_2692835, EPI_ISL_2692836, EPI_ISL_2692837, EPI_ISL_2692838, EPI_ISL_2692839, EPI_ISL_2692840, EPI_ISL_2692842, EPI_ISL_2692843, EPI_ISL_2692845, EPI_ISL_2692846, EPI_ISL_2692847 | | | |
| see above | Instituto Nacional de Medicina Genomica | Instituto Nacional de Medicina Genomica | Cedro-Tanda A, Hidalgo-Miranda A, Mendoza-Vargas A, Reyes-Grajeda JP, Abraham Campos-Romero, Moreno-Camacho José Luis, Rodriguez-Gallegos Jorge, Luna-Ruiz Marco, Gonzalez-Barrera D, Rangel-DeLeon D, Munguia-Garza P, Ramirez-Vega O, Escobar-Arrazola, M, Herrera-Montalvo LA. |
| EPI_ISL_2692858, EPI_ISL_2692859, EPI_ISL_2692860, EPI_ISL_2692861, EPI_ISL_2692862, EPI_ISL_2692863 | Child Health Research Foundation | Child Health Research Foundation | CHRF Bangladesh Genomics Team |
| EPI_ISL_2692924, EPI_ISL_2692925, EPI_ISL_2692946, EPI_ISL_2692972 | Wojewodzka Stacja Sanitarno-Epidemiologiczna w Olsztynie, Laboratorium Badan Epidemiologiczno-Klinicznych | Wojewodzka Stacja Sanitarno-Epidemiologiczna w Olsztynie, Laboratorium Badan Epidemiologiczno-Klinicznych | Sylwia Krzetowska, Monika Czerminska, Ewa Liszewska, Tomasz Jakubczak, Marta Lukian, Patryk Bielecki, Aleksandra Kobiatko, Emilia Tarabasz, Paulina Rozycka, Barbara Dolinska |
| EPI_ISL_2693033, EPI_ISL_2693034, EPI_ISL_2693035 | Oregon State Public Health Laboratory | Oregon State Public Health Laboratory | Rafia Razzaque, Eugene Yeboah, Vanda Makris, Laura Tsaknaridis, John Fontana and Shane Sevey |
| EPI_ISL_2693036, EPI_ISL_2693038, EPI_ISL_2693039, EPI_ISL_2693041, EPI_ISL_2693042, EPI_ISL_2693043, EPI_ISL_2693046, EPI_ISL_2693047, EPI_ISL_2693050, EPI_ISL_2693053, EPI_ISL_2693055, EPI_ISL_2693056, EPI_ISL_2693058, EPI_ISL_2693059, EPI_ISL_2693061, EPI_ISL_2693063, EPI_ISL_2693064, EPI_ISL_2693067, EPI_ISL_2693068, EPI_ISL_2693069, EPI_ISL_2693071, EPI_ISL_2693073, EPI_ISL_2693076, EPI_ISL_2693077, EPI_ISL_2693078, EPI_ISL_2693079, EPI_ISL_2693080, EPI_ISL_2693081, EPI_ISL_2693082, EPI_ISL_2693083, EPI_ISL_2693084, EPI_ISL_2693085, EPI_ISL_2693086, EPI_ISL_2693087 | | | |
| see above | NHLS_VIRO | KRISP, KZN Research Innovation and Sequencing Platform | Giandhari Jennifer, Pillay Sureshnee, Yajna Ramphal, Naidoo Yeshnee, Tshabuila Derek,Tegally Houriiyah, San James, Wilkinson Eduan, de Oliveira Tulio |
| EPI_ISL_2693088 | Ampath laboratories- Sharks | KRISP, KZN Research Innovation and Sequencing Platform | Giandhari Jennifer, Pillay Sureshnee, Yajna Ramphal, Naidoo Yeshnee, Tshabuila Derek,Tegally Houriiyah, San James, Wilkinson Eduan, de Oliveira Tulio |
| EPI_ISL_2693089, EPI_ISL_2693090, EPI_ISL_2693091, EPI_ISL_2693092, EPI_ISL_2693093, EPI_ISL_2693094, EPI_ISL_2693095 | NHLS_VIRO | KRISP, KZN Research Innovation and Sequencing Platform | Giandhari Jennifer, Pillay Sureshnee, Yajna Ramphal, Naidoo Yeshnee, Tshabuila Derek,Tegally Houriiyah, San James, Wilkinson Eduan, de Oliveira Tulio |
| EPI_ISL_2693109, EPI_ISL_2693110 | St James's Hospital, Dublin 8, Ireland. | ELDA biotech | Deirdre M. Broderick, Peter R. Flanagan, Brendan Crowley |
| EPI_ISL_2693147, EPI_ISL_2693150, EPI_ISL_2693156, EPI_ISL_2693158, EPI_ISL_2693161, EPI_ISL_2693162, EPI_ISL_2693163, EPI_ISL_2693166, EPI_ISL_2693167, EPI_ISL_2693168, EPI_ISL_2693169, EPI_ISL_2693172, EPI_ISL_2693173, EPI_ISL_2693175, EPI_ISL_2693176, EPI_ISL_2693177, EPI_ISL_2693182, EPI_ISL_2693184, EPI_ISL_2693185, EPI_ISL_2693186, EPI_ISL_2693189, EPI_ISL_2693192, | | | |

| | | | |
|---|---|---|---|
| EPI_ISL_2693198, EPI_ISL_2693201, EPI_ISL_2693212, EPI_ISL_2693214, EPI_ISL_2693215, EPI_ISL_2693219, EPI_ISL_2693221, EPI_ISL_2693222, EPI_ISL_2693224, EPI_ISL_2693228, EPI_ISL_2693231, EPI_ISL_2693232, EPI_ISL_2693233, EPI_ISL_2693234, EPI_ISL_2693243, EPI_ISL_2693246, EPI_ISL_2693248, EPI_ISL_2693249, EPI_ISL_2693251, EPI_ISL_2693255, EPI_ISL_2693257, EPI_ISL_2693261, EPI_ISL_2693271, EPI_ISL_2693272, EPI_ISL_2693274, EPI_ISL_2693283, EPI_ISL_2693284, EPI_ISL_2693285, EPI_ISL_2693286, EPI_ISL_2693291, EPI_ISL_2693293, EPI_ISL_2693296, EPI_ISL_2693302, EPI_ISL_2693304, EPI_ISL_2693315 | | | |
| see above | UW Virology Lab | UW Virology Lab | Pavitra Roychoudhury, Hong Xie, Lasata Shrestha, Tien V. Nguyen, Shah Mohamed Bakhash, Michelle Lin, Noah R. Baker, Ricardo Perez, Sean Ellis, Nathan Breit, Robert J. Livingston, Meei-Li Huang, Keith R Jerome, Patrick Mathias, Alexander Greninger |
| EPI_ISL_2693479, EPI_ISL_2693481, EPI_ISL_2693484, EPI_ISL_2693485, EPI_ISL_2693487, EPI_ISL_2693488 | Zavod za javno zdravstvo Međimurske Županije | Hrvatski zavod za javno zdravstvo | Irena Tabain, Ivana Ferenčak |
| EPI_ISL_2693525, EPI_ISL_2693526, EPI_ISL_2693528, EPI_ISL_2693531, EPI_ISL_2693532, EPI_ISL_2693533, EPI_ISL_2693534, EPI_ISL_2693535, EPI_ISL_2693536, EPI_ISL_2693537, EPI_ISL_2693538, EPI_ISL_2693539, EPI_ISL_2693540, EPI_ISL_2693541, EPI_ISL_2693542, EPI_ISL_2693543, EPI_ISL_2693544, EPI_ISL_2693545 | | | |
| see above | Zavod za javno zdravstvo Šibensko- Kninske Županije | Hrvatski zavod za javno zdravstvo | Irena Tabain, Ivana Ferenčak |
| EPI_ISL_2693549 | Klinika za infektivne bolesti "Dr. Fran Mihaljević" | Hrvatski zavod za javno zdravstvo | Irena Tabain, Ivana Ferenčak |
| EPI_ISL_2693577 | Zavod za javno zdravstvo Šibensko- Kninske Županije | Hrvatski zavod za javno zdravstvo | Irena Tabain, Ivana Ferenčak |
| EPI_ISL_2693669 | Surabaya Health Office | Institute of Tropical Disease, Universitas Airlangga | Kazufumi Shimizu, Krisnoadi Rahardjo, Aldise M Nastri, Jezzy R Dewantari, Rima R Prasetya, Febria Rachmanita, Herlin Ferliana, Gatot Soegiarto, Laksmi Wulandari, Resti Yudhawati, Yasuko Mori, Soetjipto, Maria I Lusida |
| EPI_ISL_2693670 | Surabaya Health Office | Institute of Tropical Disease, Universitas Airlangga | Rima R Prasetya, Krisnoadi Rahardjo, Aldise M Nastri, Jezzy R Dewantari, Febria Rachmanita, Herlin Ferliana, Gatot Soegiarto, Laksmi Wulandari, Resti Yudhawati, Yasuko Mori, Soetjipto, Kazufumi Shimizu, Maria I Lusida |
| EPI_ISL_2693671 | Mitra Keluarga Hospital Waru | Institute of Tropical Disease, Universitas Airlangga | Krisnoadi Rahardjo, Aldise M Nastri, Jezzy R Dewantari, Rima R Prasetya, Febria Rachmanita, Herlin Ferliana, Gatot Soegiarto, Laksmi Wulandari, Resti Yudhawati, Yasuko Mori, Soetjipto, Kazufumi Shimizu, Maria I Lusida |
| EPI_ISL_2693679, EPI_ISL_2693687, EPI_ISL_2693688, EPI_ISL_2693690, EPI_ISL_2693692, EPI_ISL_2693694, EPI_ISL_2693696, EPI_ISL_2693697, EPI_ISL_2693698, EPI_ISL_2693703, EPI_ISL_2693710, EPI_ISL_2693713, EPI_ISL_2693715, EPI_ISL_2693716, EPI_ISL_2693719, EPI_ISL_2693720, EPI_ISL_2693729, EPI_ISL_2693731, EPI_ISL_2693732, EPI_ISL_2693737, EPI_ISL_2693742, EPI_ISL_2693747, EPI_ISL_2693748, EPI_ISL_2693757, EPI_ISL_2693766, EPI_ISL_2693769 | | | |
| see above | UW Virology Lab | UW Virology Lab | Pavitra Roychoudhury, Hong Xie, Lasata Shrestha, Shah Mohamed Bakhash, Tien V. Nguyen, Noah R. Baker, Sean Ellis, Meei-Li Huang, Keith R Jerome, Alexander Greninger |
| EPI_ISL_2693825, EPI_ISL_2693827, EPI_ISL_2693830, EPI_ISL_2693831, EPI_ISL_2693832, EPI_ISL_2693833, EPI_ISL_2693835, EPI_ISL_2693836, EPI_ISL_2693842, EPI_ISL_2693845, EPI_ISL_2693851, EPI_ISL_2693864, EPI_ISL_2693865, EPI_ISL_2693875, EPI_ISL_2693876, EPI_ISL_2693877, EPI_ISL_2693878, EPI_ISL_2693879, EPI_ISL_2693880, EPI_ISL_2693885, EPI_ISL_2693886, EPI_ISL_2693887, EPI_ISL_2693888, EPI_ISL_2693891, EPI_ISL_2693894, EPI_ISL_2693895, EPI_ISL_2693896, EPI_ISL_2693897, EPI_ISL_2693898, EPI_ISL_2693899, EPI_ISL_2693900, EPI_ISL_2693901 | | | |
| see above | NHLS_VIRO Eastern Cape | KRISP, KZN Research Innovation and Sequencing Platform | Giandhari Jennifer, Pillay Sureshnee, Yajna Ramphal, Naidoo Yeshnee, Tshabuila Derek,Tegally Houriiyah, San James, Wilkinson Eduan, de Oliveira Tulio |
| EPI_ISL_2693902, EPI_ISL_2693903, EPI_ISL_2693904 | A. Krumbholz, Labor Dr. Krause und Kollegen MVZ GmbH, Kiel | Charité Universitätsmedizin Berlin, Institute of Virology | Victor M Corman, Barbara Mühlemann, Julia Schneider, Jörn Beheim-Schwarzbach, Tobias Bleicker, Julia Tesch, Talitha Veith, Terry Jones, Christian Drosten |
| EPI_ISL_2693911, EPI_ISL_2693913, EPI_ISL_2693914, EPI_ISL_2693915, EPI_ISL_2693919, EPI_ISL_2693920, EPI_ISL_2693921, EPI_ISL_2693926, EPI_ISL_2693936, EPI_ISL_2693937, EPI_ISL_2693938, EPI_ISL_2693942 | | | |
| see above | Virology Unit, Institut Pasteur du Cambodge | Virology Unit, Institut Pasteur du Cambodge | Cecile Troupin, Leakhena Pum, Jurre Y Siegers, Ly Sovann, Kraing Sidonn, Yi Sengdoeurn, Chin Savuth, Chau Darapheak, Veasna Duong, Erik A Karlsson |
| EPI_ISL_2693959, EPI_ISL_2693960, EPI_ISL_2693961, EPI_ISL_2693964 | laboratorio Microbiologia PO Cardarelli | laboratorio Microbiologia PO Cardarelli | Felice V., Niro G. Scutella' M. |
| EPI_ISL_2694525, EPI_ISL_2694580, EPI_ISL_2694581 | Nacionalinis maisto ir veterinarijos rizikos vertinimo institutas | National Public Health Surveillance Laboratory | Lukas Zemaitis, Migle Gabrielaite, Jelena Razmuk, Svajune Muralyte, Ana Steponkiene, Lukas Vasionis, Danas Baksa |
| EPI_ISL_2694582, EPI_ISL_2694583, EPI_ISL_2694584 | UAB InMedica | National Public Health Surveillance Laboratory | Lukas Zemaitis, Migle Gabrielaite, Jelena Razmuk, Svajune Muralyte, Ana Steponkiene, Lukas Vasionis, Danas Baksa |
| EPI_ISL_2694594 | PathCare, Cape Town | Division of Medical Virology, National Health Laboratory Service (NHLS), Tygerberg Hospital / Stellenbosch University | Sisonke, Susan Engelbrecht, Tongai Maponga, Bronwyn Kleinhans, Tania Stander, Jean Maritz, Petra Raimond, Houriiyah Tegally, San Emmanuel James, Gert van Zyl, Tulio de Oliveira, Wolfgang Preiser |
| EPI_ISL_2694595, EPI_ISL_2694596, EPI_ISL_2694597, EPI_ISL_2694599, EPI_ISL_2694600, EPI_ISL_2694602 | PathCare, Cape Town | Division of Medical Virology, National Health Laboratory Service (NHLS), Tygerberg Hospital / Stellenbosch University | Susan Engelbrecht, Tongai Maponga, Bronwyn Kleinhans, Tania Stander, Jean Maritz, Petra Raimond, Houriiyah Tegally, San Emmanuel James, Gert van Zyl, Tulio de Oliveira, Wolfgang Preiser |
| EPI_ISL_2694605 | PathCare, Cape Town | Division of Medical Virology, National Health Laboratory Service (NHLS), Tygerberg Hospital / Stellenbosch University | Sisonke, Susan Engelbrecht, Tongai Maponga, Bronwyn Kleinhans, Tania Stander, Jean Maritz, Petra Raimond, Houriiyah Tegally, San Emmanuel James, Gert van Zyl, Tulio de Oliveira, Wolfgang Preiser |
| EPI_ISL_2694609, EPI_ISL_2694610 | PathCare, Cape Town | Division of Medical Virology, National Health Laboratory Service (NHLS), Tygerberg Hospital / Stellenbosch University | Susan Engelbrecht, Tongai Maponga, Bronwyn Kleinhans, Tania Stander, Jean Maritz, Petra Raimond, Houriiyah Tegally, San Emmanuel James, Gert van Zyl, Tulio de Oliveira, Wolfgang Preiser |
| EPI_ISL_2694613, EPI_ISL_2694616, EPI_ISL_2694619, EPI_ISL_2694623 | TAMBO MEMORIAL | National Institute for Communicable Diseases of the National Health Laboratory Service | Amoako DG, Everatt J, Scheepers C, Mohale T, Ntuli N, Mahlangu B, Mnguni A, Ismail A, Bhiman JN |
| EPI_ISL_2694627, EPI_ISL_2694636, EPI_ISL_2694637, EPI_ISL_2694638, EPI_ISL_2694640, EPI_ISL_2694642, EPI_ISL_2694644, EPI_ISL_2694645, EPI_ISL_2694657, EPI_ISL_2694658, EPI_ISL_2694659, EPI_ISL_2694663, EPI_ISL_2694664, EPI_ISL_2694665, EPI_ISL_2694666, EPI_ISL_2694667, EPI_ISL_2694670, EPI_ISL_2694671 | | | |
| see above | CHRIS HANI BARAGWANATH | National Institute for Communicable Diseases of the National Health Laboratory Service | Amoako DG, Everatt J, Scheepers C, Mohale T, Ntuli N, Mahlangu B, Mnguni A, Ismail A, Bhiman JN |
| EPI_ISL_2694678, EPI_ISL_2694680, EPI_ISL_2694687, EPI_ISL_2694689, EPI_ISL_2694690, EPI_ISL_2694691, EPI_ISL_2694692, EPI_ISL_2694693, EPI_ISL_2694698, EPI_ISL_2694699, EPI_ISL_2694700, EPI_ISL_2694701, EPI_ISL_2694702, EPI_ISL_2694709, EPI_ISL_2694715, EPI_ISL_2694716, EPI_ISL_2694718, EPI_ISL_2694719, EPI_ISL_2694723, EPI_ISL_2694724, EPI_ISL_2694725 | | | |
| see above | TAMBO MEMORIAL | National Institute for Communicable Diseases of the National Health Laboratory Service | Amoako DG, Everatt J, Scheepers C, Mohale T, Ntuli N, Mahlangu B, Mnguni A, Ismail A, Bhiman JN |
| EPI_ISL_2694759, EPI_ISL_2694775, EPI_ISL_2694787 | Diagnostyka. Laboratoria Medyczne. | 1. ViroGenetics - BSL3 Laboratory of Virology, Małopolska Centre of Biotechnology, Jagiellonian University; 2. Diagtron Laboratoria Łukasz Rąbalski | Rabalski L., Gromowski,T., Rogalska-Kupiec M., Swadzba J., Wydmanski,W., Mazur-Panasiuk,N., Kowalski,M., Maciej Kosinski, Natalia Derewonko, Szulc,P., Sylwia Januszczak, Labaj,P.P., Pyrc,K. |
| EPI_ISL_2694801, EPI_ISL_2694804, EPI_ISL_2694807, EPI_ISL_2694808, EPI_ISL_2694810, EPI_ISL_2694815, EPI_ISL_2694816, EPI_ISL_2694821, EPI_ISL_2694826, EPI_ISL_2694827, EPI_ISL_2694828, EPI_ISL_2694829, EPI_ISL_2694832, EPI_ISL_2694834, EPI_ISL_2694839, EPI_ISL_2694841, EPI_ISL_2694842, EPI_ISL_2694847, EPI_ISL_2694850, EPI_ISL_2694856, EPI_ISL_2694859 | | | |
| see above | Labor Berlin Charité Vivantes GmbH / Institut für Virologie | Charité Universitätsmedizin Berlin, Institut für Virologie/Labor Berlin | Peter Menzel, Christine Stephan, Rolf Schwarzer, Victor M Corman, Barbara Mühlemann, Terry Jones, Christian Drosten |
| EPI_ISL_2694865 | Laboratorio di Riferimento Regionale della Sicilia Occidentale per l'Emergenza COVID-19 | Laboratorio di Riferimento Regionale della Sicilia Occidentale per l'Emergenza COVID-19 | Fabio Tramuto, Carmelo Massimo Maida, Daniela Di Naro, Giulia Randazzo, Walter Mazzucco, Giorgio Graziano, Vincenzo Restivo, Claudio Costantino, Francesco Vitale |
| EPI_ISL_2694881, EPI_ISL_2694882, EPI_ISL_2694883, EPI_ISL_2694884, EPI_ISL_2694885, EPI_ISL_2694886 | Affidea | Instituto Nacional de Saude (INSA) | Borges et al |
| EPI_ISL_2694888, EPI_ISL_2694889, EPI_ISL_2694890, EPI_ISL_2694892, EPI_ISL_2694893, EPI_ISL_2694894, EPI_ISL_2694895, EPI_ISL_2694896, EPI_ISL_2694897, EPI_ISL_2694898, EPI_ISL_2694899, EPI_ISL_2694900 | | | |
| see above | H Santarem | Instituto Nacional de Saude (INSA) | Borges et al |
| EPI_ISL_2694902, EPI_ISL_2694903, EPI_ISL_2694904, EPI_ISL_2694905, EPI_ISL_2694906, EPI_ISL_2694907, EPI_ISL_2694908, EPI_ISL_2694909, EPI_ISL_2694910, EPI_ISL_2694911, EPI_ISL_2694912, EPI_ISL_2694915, EPI_ISL_2694916, EPI_ISL_2694917, EPI_ISL_2694918, EPI_ISL_2694920, EPI_ISL_2694921, EPI_ISL_2694922, EPI_ISL_2694923, EPI_ISL_2694925 | | | |
| see above | H Garcia de Orta | Instituto Nacional de Saude (INSA) | Borges et al |
| EPI_ISL_2694927, EPI_ISL_2694928, EPI_ISL_2694929, EPI_ISL_2694930, EPI_ISL_2694931, EPI_ISL_2694932, EPI_ISL_2694933, EPI_ISL_2694934, EPI_ISL_2694935, EPI_ISL_2694936, EPI_ISL_2694938, EPI_ISL_2694939, EPI_ISL_2694940, EPI_ISL_2694941, EPI_ISL_2694942, EPI_ISL_2694943, EPI_ISL_2694944, EPI_ISL_2694945, EPI_ISL_2694946 | | | |
| see above | CHLC | Instituto Nacional de Saude (INSA) | Borges et al |
| EPI_ISL_2694947, EPI_ISL_2694948, EPI_ISL_2694949, EPI_ISL_2694950, EPI_ISL_2694952, EPI_ISL_2694953, EPI_ISL_2694954, EPI_ISL_2694956, EPI_ISL_2694957, EPI_ISL_2694958, EPI_ISL_2694960, EPI_ISL_2694961, EPI_ISL_2694962, EPI_ISL_2694963, EPI_ISL_2694964, EPI_ISL_2694965, EPI_ISL_2694966 | | | |
| see above | H Fernando Fonseca | Instituto Nacional de Saude (INSA) | Borges et al |
| EPI_ISL_2695015, EPI_ISL_2695057, EPI_ISL_2695068, EPI_ISL_2695071, EPI_ISL_2695072 | H Divino Espirito Santo - Ponta Delgada | Instituto Nacional de Saude (INSA) | Borges et al |
| EPI_ISL_2695073, EPI_ISL_2695074, EPI_ISL_2695075, EPI_ISL_2695076, EPI_ISL_2695077, EPI_ISL_2695078, EPI_ISL_2695079, EPI_ISL_2695080, EPI_ISL_2695081, EPI_ISL_2695082, EPI_ISL_2695083, EPI_ISL_2695084, EPI_ISL_2695085, EPI_ISL_2695086, EPI_ISL_2695088, EPI_ISL_2695089, EPI_ISL_2695090, EPI_ISL_2695091, EPI_ISL_2695092 | | | |
| see above | H Vila Franca Xira | Instituto Nacional de Saude (INSA) | Borges et al |
| EPI_ISL_2695093, EPI_ISL_2695094, EPI_ISL_2695096, EPI_ISL_2695097, EPI_ISL_2695099, EPI_ISL_2695103, EPI_ISL_2695104, EPI_ISL_2695105, EPI_ISL_2695106, EPI_ISL_2695107, EPI_ISL_2695108, EPI_ISL_2695109, EPI_ISL_2695110, EPI_ISL_2695111, EPI_ISL_2695112, EPI_ISL_2695113, EPI_ISL_2695114, EPI_ISL_2695115, EPI_ISL_2695116, EPI_ISL_2695117, EPI_ISL_2695119, EPI_ISL_2695120, EPI_ISL_2695121, EPI_ISL_2695122, EPI_ISL_2695123 | | | |

| | | | |
|---|---|---|---|
| see above | CHULN - H Santa Maria | Instituto Nacional de Saude (INSA) | Borges et al |
| EPI_ISL_2695124, EPI_ISL_2695125, EPI_ISL_2695126, EPI_ISL_2695127, EPI_ISL_2695128, EPI_ISL_2695129, EPI_ISL_2695130, EPI_ISL_2695131, EPI_ISL_2695132, EPI_ISL_2695133, EPI_ISL_2695134, EPI_ISL_2695135, EPI_ISL_2695136, EPI_ISL_2695137, EPI_ISL_2695138, EPI_ISL_2695139, EPI_ISL_2695140, EPI_ISL_2695141, EPI_ISL_2695142, EPI_ISL_2695143, EPI_ISL_2695144, EPI_ISL_2695145, EPI_ISL_2695146, EPI_ISL_2695147, EPI_ISL_2695148, EPI_ISL_2695149, EPI_ISL_2695150, EPI_ISL_2695151, EPI_ISL_2695152, EPI_ISL_2695153, EPI_ISL_2695154, EPI_ISL_2695155, EPI_ISL_2695156, EPI_ISL_2695157, EPI_ISL_2695158 | | | |
| see above | ULS Castelo Branco | Instituto Nacional de Saude (INSA) | Borges et al |
| EPI_ISL_2695159, EPI_ISL_2695160, EPI_ISL_2695161, EPI_ISL_2695162, EPI_ISL_2695163 | CH Setubal | Instituto Nacional de Saude (INSA) | Borges et al |
| EPI_ISL_2695164, EPI_ISL_2695165, EPI_ISL_2695166, EPI_ISL_2695167, EPI_ISL_2695168, EPI_ISL_2695169, EPI_ISL_2695170, EPI_ISL_2695171, EPI_ISL_2695172, EPI_ISL_2695173, EPI_ISL_2695174, EPI_ISL_2695175, EPI_ISL_2695176, EPI_ISL_2695177, EPI_ISL_2695178, EPI_ISL_2695179, EPI_ISL_2695180, EPI_ISL_2695181, EPI_ISL_2695182, EPI_ISL_2695183, EPI_ISL_2695184, EPI_ISL_2695185, EPI_ISL_2695186, EPI_ISL_2695187, EPI_ISL_2695188, EPI_ISL_2695189, EPI_ISL_2695190, EPI_ISL_2695191, EPI_ISL_2695192, EPI_ISL_2695193, EPI_ISL_2695194, EPI_ISL_2695195, EPI_ISL_2695196, EPI_ISL_2695197, EPI_ISL_2695198, EPI_ISL_2695199, EPI_ISL_2695200, EPI_ISL_2695201, EPI_ISL_2695202, EPI_ISL_2695203, EPI_ISL_2695204, EPI_ISL_2695205, EPI_ISL_2695206, EPI_ISL_2695207, EPI_ISL_2695208, EPI_ISL_2695209, EPI_ISL_2695210, EPI_ISL_2695211, EPI_ISL_2695212, EPI_ISL_2695213, EPI_ISL_2695214, EPI_ISL_2695215, EPI_ISL_2695217, EPI_ISL_2695218, EPI_ISL_2695219, EPI_ISL_2695220, EPI_ISL_2695221 | | | |
| see above | ULS Litoral Alentejano | Instituto Nacional de Saude (INSA) and Institute of Biomedicine (iBiMed), Universidade de Aveiro | Borges et al |
| EPI_ISL_2695225 | ABC Algarve | Instituto Nacional de Saude (INSA) and Institute of Biomedicine (iBiMed), Universidade de Aveiro | Borges et al |
| EPI_ISL_2695234, EPI_ISL_2695238, EPI_ISL_2695239, EPI_ISL_2695241, EPI_ISL_2695243, EPI_ISL_2695247, EPI_ISL_2695257 | SESARAM | Instituto Nacional de Saude (INSA) | Borges et al |
| EPI_ISL_2695258 | ABC Algarve | Instituto Nacional de Saude (INSA) | Borges et al |
| EPI_ISL_2695260, EPI_ISL_2695261, EPI_ISL_2695262, EPI_ISL_2695263, EPI_ISL_2695264, EPI_ISL_2695265, EPI_ISL_2695266, EPI_ISL_2695268 | CH Porto - H Sto Antonio | Instituto Nacional de Saude (INSA) | Borges et al |
| EPI_ISL_2695269 | LAC Arunce | Instituto Nacional de Saude (INSA) | Borges et al |
| EPI_ISL_2695271, EPI_ISL_2695285, EPI_ISL_2695287 | H Braga | Instituto Nacional de Saude (INSA) | Borges et al |
| EPI_ISL_2695290, EPI_ISL_2695291, EPI_ISL_2695293, EPI_ISL_2695295, EPI_ISL_2695296, EPI_ISL_2695297, EPI_ISL_2695298, EPI_ISL_2695299, EPI_ISL_2695300, EPI_ISL_2695301, EPI_ISL_2695302, EPI_ISL_2695303, EPI_ISL_2695305 | | | |
| see above | CHTMAD | Instituto Nacional de Saude (INSA) | Borges et al |
| EPI_ISL_2695306, EPI_ISL_2695307, EPI_ISL_2695308, EPI_ISL_2695309, EPI_ISL_2695310, EPI_ISL_2695311, EPI_ISL_2695312, EPI_ISL_2695313, EPI_ISL_2695314, EPI_ISL_2695315 | ULS Guarda | Instituto Nacional de Saude (INSA) | Borges et al |
| EPI_ISL_2695316, EPI_ISL_2695317, EPI_ISL_2695318, EPI_ISL_2695319, EPI_ISL_2695320, EPI_ISL_2695321 | FMUP | Instituto Nacional de Saude (INSA) | Borges et al |
| EPI_ISL_2695322, EPI_ISL_2695328, EPI_ISL_2695330, EPI_ISL_2695331, EPI_ISL_2695332, EPI_ISL_2695333, EPI_ISL_2695334, EPI_ISL_2695337, EPI_ISL_2695338 | ULSM - Matosinhos | Instituto Nacional de Saude (INSA) | Borges et al |
| EPI_ISL_2695341, EPI_ISL_2695347, EPI_ISL_2695348 | CHEDV | Instituto Nacional de Saude (INSA) | Borges et al |
| EPI_ISL_2695349, EPI_ISL_2695350, EPI_ISL_2695351, EPI_ISL_2695353, EPI_ISL_2695354, EPI_ISL_2695355, EPI_ISL_2695356, EPI_ISL_2695357, EPI_ISL_2695358, EPI_ISL_2695359, EPI_ISL_2695360, EPI_ISL_2695361, EPI_ISL_2695362 | | | |
| see above | Lab Dra Celeste Formosinho | Instituto Nacional de Saude (INSA) | Borges et al |
| EPI_ISL_2695363, EPI_ISL_2695364, EPI_ISL_2695365, EPI_ISL_2695367, EPI_ISL_2695368, EPI_ISL_2695369, EPI_ISL_2695372, EPI_ISL_2695373, EPI_ISL_2695374, EPI_ISL_2695375, EPI_ISL_2695376, EPI_ISL_2695377, EPI_ISL_2695378, EPI_ISL_2695379, EPI_ISL_2695380, EPI_ISL_2695381, EPI_ISL_2695383, EPI_ISL_2695384, EPI_ISL_2695385, EPI_ISL_2695387 | | | |
| see above | Labeto - CAB - Leiria | Instituto Nacional de Saude (INSA) | Borges et al |
| EPI_ISL_2695388 | ARS Algarve - Laboratorio Laura Ayres | Instituto Nacional de Saude (INSA) | Borges et al |
| EPI_ISL_2695389 | CH Tondela Viseu | Instituto Nacional de Saude (INSA) | Borges et al |
| EPI_ISL_2695396, EPI_ISL_2695397, EPI_ISL_2695402, EPI_ISL_2695403, EPI_ISL_2695404, EPI_ISL_2695405, EPI_ISL_2695408 | Lab La Salete Robles - VN Famalicao | Instituto Nacional de Saude (INSA) | Borges et al |
| EPI_ISL_2695410 | ULS Alto Minho | Instituto Nacional de Saude (INSA) | Borges et al |
| EPI_ISL_2695412, EPI_ISL_2695416 | H Guimaraes | Instituto Nacional de Saude (INSA) | Borges et al |
| EPI_ISL_2695418, EPI_ISL_2695419, EPI_ISL_2695420, EPI_ISL_2695421, EPI_ISL_2695422, EPI_ISL_2695423, EPI_ISL_2695425, EPI_ISL_2695426, EPI_ISL_2695428, EPI_ISL_2695429, EPI_ISL_2695430, EPI_ISL_2695431, EPI_ISL_2695432, EPI_ISL_2695433, EPI_ISL_2695435, EPI_ISL_2695436 | | | |
| see above | UTAD | Instituto Nacional de Saude (INSA) | Borges et al |
| EPI_ISL_2695437, EPI_ISL_2695438, EPI_ISL_2695440, EPI_ISL_2695442, EPI_ISL_2695445, EPI_ISL_2695446, EPI_ISL_2695448, EPI_ISL_2695449, EPI_ISL_2695450, EPI_ISL_2695451, EPI_ISL_2695452, EPI_ISL_2695453, EPI_ISL_2695454, EPI_ISL_2695455, EPI_ISL_2695456, EPI_ISL_2695457, EPI_ISL_2695458, EPI_ISL_2695459, EPI_ISL_2695460, EPI_ISL_2695461, EPI_ISL_2695462, EPI_ISL_2695464, EPI_ISL_2695465, EPI_ISL_2695466, EPI_ISL_2695467, EPI_ISL_2695468, EPI_ISL_2695469, EPI_ISL_2695471 | | | |
| see above | ARS Algarve - Laboratorio Laura Ayres | Instituto Nacional de Saude (INSA) | Borges et al |
| EPI_ISL_2695472, EPI_ISL_2695473 | LAC Mesquita e Damiao - Aves | Instituto Nacional de Saude (INSA) | Borges et al |
| EPI_ISL_2695474, EPI_ISL_2695476, EPI_ISL_2695479, EPI_ISL_2695480, EPI_ISL_2695481, EPI_ISL_2695484, EPI_ISL_2695486, EPI_ISL_2695487, EPI_ISL_2695488, EPI_ISL_2695489, EPI_ISL_2695490, EPI_ISL_2695491, EPI_ISL_2695495, EPI_ISL_2695496, EPI_ISL_2695497, EPI_ISL_2695498, EPI_ISL_2695499, EPI_ISL_2695500, EPI_ISL_2695501, EPI_ISL_2695502, EPI_ISL_2695503 | | | |
| see above | ARS Algarve - Laboratorio Laura Ayres | Instituto Nacional de Saude (INSA) | Borges et al |
| EPI_ISL_2695505 | TAMBO MEMORIAL | National Institute for Communicable Diseases of the National Health Laboratory Service | Amoako DG, Everatt J, Scheepers C, Mohale T, Ntuli N, Mahlangu B, Mnguni A, Ismail A, Bhiman JN |
| EPI_ISL_2695513 | KOPANONG | National Institute for Communicable Diseases of the National Health | Amoako DG, Everatt J, Scheepers C, Mohale T, Ntuli N, Mahlangu B, Mnguni A, Ismail A, Bhiman JN |

| | | | |
|---|---|---|---|
| | | Laboratory Service | |
| EPI_ISL_2695514, EPI_ISL_2695515, EPI_ISL_2695516, EPI_ISL_2695520, EPI_ISL_2695522, EPI_ISL_2695526 | TAMBO MEMORIAL | National Institute for Communicable Diseases of the National Health Laboratory Service | Amoako DG, Everatt J, Scheepers C, Mohale T, Ntuli N, Mahlangu B, Mnguni A, Ismail A, Bhiman JN |
| EPI_ISL_2695530, EPI_ISL_2695531, EPI_ISL_2695532, EPI_ISL_2695533, EPI_ISL_2695538, EPI_ISL_2695550, EPI_ISL_2695551, EPI_ISL_2695556, EPI_ISL_2695562, EPI_ISL_2695569, EPI_ISL_2695573, EPI_ISL_2695574, EPI_ISL_2695576 | | | |
| see above | POLOKWANE MANKWENG HOSPITAL | National Institute for Communicable Diseases of the National Health Laboratory Service | Amoako DG, Everatt J, Scheepers C, Mohale T, Ntuli N, Mahlangu B, Mnguni A, Ismail A, Bhiman JN |
| EPI_ISL_2695583, EPI_ISL_2695586 | PATHCARE LABORATORY | National Institute for Communicable Diseases of the National Health Laboratory Service | Amoako DG, Everatt J, Scheepers C, Mohale T, Ntuli N, Mahlangu B, Mnguni A, Ismail A, Bhiman JN |
| EPI_ISL_2695588, EPI_ISL_2695589, EPI_ISL_2695594, EPI_ISL_2695598, EPI_ISL_2695600 | HELEN JOSEPH | National Institute for Communicable Diseases of the National Health Laboratory Service | Amoako DG, Everatt J, Scheepers C, Mohale T, Ntuli N, Mahlangu B, Mnguni A, Ismail A, Bhiman JN |
| EPI_ISL_2695625, EPI_ISL_2695633 | ROB FERREIRA LABORATORY | National Institute for Communicable Diseases of the National Health Laboratory Service | Amoako DG, Everatt J, Scheepers C, Mohale T, Ntuli N, Mahlangu B, Mnguni A, Ismail A, Bhiman JN |
| EPI_ISL_2695635, EPI_ISL_2695636, EPI_ISL_2695637, EPI_ISL_2695639, EPI_ISL_2695640, EPI_ISL_2695641, EPI_ISL_2695653, EPI_ISL_2695655, EPI_ISL_2695656, EPI_ISL_2695657, EPI_ISL_2695658, EPI_ISL_2695662, EPI_ISL_2695663, EPI_ISL_2695664, EPI_ISL_2695666, EPI_ISL_2695668, EPI_ISL_2695669, EPI_ISL_2695672, EPI_ISL_2695673, EPI_ISL_2695674, EPI_ISL_2695675, EPI_ISL_2695676, EPI_ISL_2695679, EPI_ISL_2695681, EPI_ISL_2695682, EPI_ISL_2695683, EPI_ISL_2695688, EPI_ISL_2695689, EPI_ISL_2695697, EPI_ISL_2695700 | | | |
| see above | KIMBERLEY LABORATORY | National Institute for Communicable Diseases of the National Health Laboratory Service | Amoako DG, Everatt J, Scheepers C, Mohale T, Ntuli N, Mahlangu B, Mnguni A, Ismail A, Bhiman JN |
| EPI_ISL_2695726 | ROB FERREIRA LABORATORY | National Institute for Communicable Diseases of the National Health Laboratory Service | Amoako DG, Everatt J, Scheepers C, Mohale T, Ntuli N, Mahlangu B, Mnguni A, Ismail A, Bhiman JN |
| EPI_ISL_2695729, EPI_ISL_2695731, EPI_ISL_2695740, EPI_ISL_2695741, EPI_ISL_2695743, EPI_ISL_2695745, EPI_ISL_2695747, EPI_ISL_2695748, EPI_ISL_2695750, EPI_ISL_2695751 | HELEN JOSEPH | National Institute for Communicable Diseases of the National Health Laboratory Service | Amoako DG, Everatt J, Scheepers C, Mohale T, Ntuli N, Mahlangu B, Mnguni A, Ismail A, Bhiman JN |
| EPI_ISL_2695756, EPI_ISL_2695758 | ROB FERREIRA LABORATORY | National Institute for Communicable Diseases of the National Health Laboratory Service | Amoako DG, Everatt J, Scheepers C, Mohale T, Ntuli N, Mahlangu B, Mnguni A, Ismail A, Bhiman JN |
| EPI_ISL_2695777, EPI_ISL_2695780, EPI_ISL_2695781, EPI_ISL_2695783 | TAMBO MEMORIAL LABORATORY | National Institute for Communicable Diseases of the National Health Laboratory Service | Amoako DG, Everatt J, Scheepers C, Mohale T, Ntuli N, Mahlangu B, Mnguni A, Ismail A, Bhiman JN |
| EPI_ISL_2695790 | Queensland Health Forensic and Scientific Services | Queensland Health Forensic and Scientific Services | Son Nguyen |
| EPI_ISL_2695791, EPI_ISL_2695792, EPI_ISL_2695793, EPI_ISL_2695794, EPI_ISL_2695795, EPI_ISL_2695796 | Area of Virology, Serology and Virology Division (SAViD), New South Wales Health Pathology Randwick | Virology Research Laboratory; Area of Virology, Serology and Virology Division (SAViD), New South Wales Health Pathology Randwick | Foster, C.; Au, J.; Ruiz Silva, M.; Deveson, I.; Bull, R.; Van Hal, S.; Rawlinson, W. |
| EPI_ISL_2695797, EPI_ISL_2695798, EPI_ISL_2695799, EPI_ISL_2695800 | New South Wales Health Pathology Royal Prince Alfred Hospital | Microbiology RPAH | Foster, C.; Au, J.; Ruiz Silva, M.; Deveson, I.; Bull, R.; Van Hal, S.; Rawlinson, W. |
| EPI_ISL_2695817, EPI_ISL_2695818, EPI_ISL_2695820, EPI_ISL_2695821, EPI_ISL_2695822, EPI_ISL_2695829, EPI_ISL_2695841, EPI_ISL_2695842, EPI_ISL_2695843, EPI_ISL_2695844, EPI_ISL_2695845, EPI_ISL_2695847 | | | |
| see above | PathCare, Cape Town | Division of Medical Virology, National Health Laboratory Service (NHLS), Tygerberg Hospital / Stellenbosch University | Susan Engelbrecht, Tongai Maponga, Bronwyn Kleinhans, Tania Stander, Jean Maritz, Petra Raimond, San Emmanual James, Tulio de Oliveira, Gert van Zyl, Wolfgang Preiser |
| EPI_ISL_2696274, EPI_ISL_2696275, EPI_ISL_2696276, EPI_ISL_2696277, EPI_ISL_2696278, EPI_ISL_2696279, EPI_ISL_2696280 | New South Wales Health Pathology Royal Prince Alfred Hospital | Microbiology RPAH | Foster, C.; Au, J.; Ruiz Silva, M.; Deveson, I.; Bull, R.; Van Hal, S.; Rawlinson, W. |
| EPI_ISL_2696720, EPI_ISL_2696880 | Laboratory, Gravity Diagnostics | Laboratory, Gravity Diagnostics | Diagnostics,G. |
| EPI_ISL_2698239 | Public Health Laboratory, Minnesota Department of Health | University of Minnesota Genomics Center | Daryl M. Gohl, John Garbe, Jaquelyn Kuriger-Laber, Corbin Dirkx, and Sean Wang |
| EPI_ISL_2698321 | AULSS 6 Euganea | Istituto Zooprofilattico Sperimentale delle Venezie | Adelaide Milani, Alessia Schivo, Annalisa Salviato, Elisa Palumbo, Erika Giorgia Quaranta, Luca Tassoni, Ambra Pastori, Edoardo Giussani, Alice Fusaro, Isabella Monne, Calogero Terregino, Antonia Ricci |
| EPI_ISL_2698425, EPI_ISL_2698433, EPI_ISL_2698442, EPI_ISL_2698444, EPI_ISL_2698447, EPI_ISL_2698450, EPI_ISL_2698455, EPI_ISL_2698458, EPI_ISL_2698461, EPI_ISL_2698467, EPI_ISL_2698470, EPI_ISL_2698473, EPI_ISL_2698476, EPI_ISL_2698478, EPI_ISL_2698484, EPI_ISL_2698487 | | | |
| see above | WHO National Influenza Centre Russian Federation | WHO National Influenza Centre Russian Federation | Andrey Komissarov, Artem Fadeev, Kseniya Komissarova, Oula Mansour, Kirill Varchenko, Mikhail Bakaev, Tamila Musaeva, Veronika Eder, Maria Pisareva, Nikita Yolshin, Daria Danilenko, Ksenia Safina, Elena Nabieva, Georgii Bazykin, Dmitry Lioznov |
| EPI_ISL_2698489 | Public Health Laboratory, Minnesota Department of Health | University of Minnesota Genomics Center | Daryl M. Gohl, John Garbe, Jaquelyn Kuriger-Laber, Corbin Dirkx, and Sean Wang |
| EPI_ISL_2698490, EPI_ISL_2698502, EPI_ISL_2698504, EPI_ISL_2698507, EPI_ISL_2698512, EPI_ISL_2698520, EPI_ISL_2698522, EPI_ISL_2698530, EPI_ISL_2698532, EPI_ISL_2698535, EPI_ISL_2698537, EPI_ISL_2698539, EPI_ISL_2698542, EPI_ISL_2698545, EPI_ISL_2698547, EPI_ISL_2698549, EPI_ISL_2698552, EPI_ISL_2698554, EPI_ISL_2698557, EPI_ISL_2698559, EPI_ISL_2698562, EPI_ISL_2698564, EPI_ISL_2698567, EPI_ISL_2698570, EPI_ISL_2698572, EPI_ISL_2698575, EPI_ISL_2698577, EPI_ISL_2698579, EPI_ISL_2698585, EPI_ISL_2698586, EPI_ISL_2698588, EPI_ISL_2698590, EPI_ISL_2698591, EPI_ISL_2698593, EPI_ISL_2698595, EPI_ISL_2698598, EPI_ISL_2698601, EPI_ISL_2698604, EPI_ISL_2698610, EPI_ISL_2698612, EPI_ISL_2698615, EPI_ISL_2698620, EPI_ISL_2698622, EPI_ISL_2698625, EPI_ISL_2698628, EPI_ISL_2698631, EPI_ISL_2698634, EPI_ISL_2698636, EPI_ISL_2698639, EPI_ISL_2698641, EPI_ISL_2698644, EPI_ISL_2698646, EPI_ISL_2698648, EPI_ISL_2698651, EPI_ISL_2698653, EPI_ISL_2698655, EPI_ISL_2698658, EPI_ISL_2698661, EPI_ISL_2698664, EPI_ISL_2698666, EPI_ISL_2698669, EPI_ISL_2698672, EPI_ISL_2698675, EPI_ISL_2698678, EPI_ISL_2698680 | | | |
| see above | WHO National Influenza Centre Russian Federation | WHO National Influenza Centre Russian Federation | Andrey Komissarov, Artem Fadeev, Kseniya Komissarova, Oula Mansour, Kirill Varchenko, Mikhail Bakaev, Tamila Musaeva, Veronika Eder, Maria Pisareva, Nikita Yolshin, Daria Danilenko, Ksenia Safina, Elena Nabieva, Georgii Bazykin, Dmitry Lioznov |
| EPI_ISL_2699510 | Public Health Laboratory, Minnesota Department of Health | University of Minnesota Genomics Center | Daryl M. Gohl, John Garbe, Jaquelyn Kuriger-Laber, Corbin Dirkx, and Sean Wang |
| EPI_ISL_2699520 | Lighthouse Lab in Milton Keynes | Wellcome Sanger Institute for the COVID-19 Genomics UK (COG-UK) Consortium | The Lighthouse Lab in Milton Keynes and Alex Alderton, Roberto Amato, Jeffrey Barrett, Sonia Goncalves, Ewan Harrison, David K. Jackson, Ian Johnston, Dominic Kwiatkowski, Cordelia Langford, John Sillitoe on behalf of the Wellcome Sanger Institute COVID-19 Surveillance Team |
| EPI_ISL_2699521 | Lighthouse Lab in Glasgow | Wellcome Sanger Institute for the COVID-19 Genomics UK (COG-UK) Consortium | Harper VanSteenhouse, Yumi Kasai, David Gray, Carol Clugston, Anna Dominiczak and Alex Alderton, Roberto Amato, Jeffrey Barrett, Sonia Goncalves, Ewan Harrison, David K. Jackson, Ian Johnston, Dominic Kwiatkowski, Cordelia Langford, John Sillitoe on behalf of the Wellcome Sanger Institute COVID-19 Surveillance Team |
| EPI_ISL_2699522, EPI_ISL_2699523, EPI_ISL_2699524 | Lighthouse Lab in Milton Keynes | Wellcome Sanger Institute for the COVID-19 Genomics UK (COG-UK) Consortium | The Lighthouse Lab in Milton Keynes and Alex Alderton, Roberto Amato, Jeffrey Barrett, Sonia Goncalves, Ewan Harrison, David K. Jackson, Ian Johnston, Dominic Kwiatkowski, Cordelia Langford, John Sillitoe on behalf of the Wellcome Sanger Institute COVID-19 Surveillance Team |
| EPI_ISL_2699525, EPI_ISL_2699526 | Lighthouse Lab in Glasgow | Wellcome Sanger Institute for the COVID-19 Genomics UK (COG-UK) Consortium | Harper VanSteenhouse, Yumi Kasai, David Gray, Carol Clugston, Anna Dominiczak and Alex Alderton, Roberto Amato, Jeffrey Barrett, Sonia Goncalves, Ewan Harrison, David K. Jackson, Ian Johnston, Dominic Kwiatkowski, Cordelia Langford, John Sillitoe on behalf of the Wellcome Sanger Institute COVID-19 Surveillance Team |
| EPI_ISL_2699527 | Lighthouse Laboratory Plymouth | Wellcome Sanger Institute for the COVID-19 Genomics UK (COG-UK) Consortium | Lighthouse Laboratory Plymouth and Alex Alderton, Roberto Amato, Jeffrey Barrett, Sonia Goncalves, Ewan Harrison, David K. Jackson, Ian Johnston, Dominic Kwiatkowski, Cordelia Langford, John Sillitoe on behalf of the Wellcome Sanger Institute COVID-19 Surveillance Team |
| EPI_ISL_2699528, EPI_ISL_2699529, EPI_ISL_2699530 | Lighthouse Lab in Glasgow | Wellcome Sanger Institute for the COVID-19 Genomics UK (COG-UK) Consortium | Harper VanSteenhouse, Yumi Kasai, David Gray, Carol Clugston, Anna Dominiczak and Alex Alderton, Roberto Amato, Jeffrey Barrett, Sonia Goncalves, Ewan Harrison, David K. Jackson, Ian Johnston, Dominic Kwiatkowski, Cordelia Langford, John Sillitoe on behalf of the Wellcome Sanger Institute COVID-19 Surveillance Team |
| EPI_ISL_2699531 | Lighthouse Laboratory Plymouth | Wellcome Sanger Institute for the COVID-19 Genomics UK (COG-UK) Consortium | Lighthouse Laboratory Plymouth and Alex Alderton, Roberto Amato, Jeffrey Barrett, Sonia Goncalves, Ewan Harrison, David K. Jackson, Ian Johnston, Dominic Kwiatkowski, Cordelia Langford, John Sillitoe on behalf of the Wellcome Sanger Institute COVID-19 Surveillance Team |
| EPI_ISL_2699532, EPI_ISL_2699533, | Lighthouse Lab in Glasgow | Wellcome Sanger Institute for the COVID-19 Genomics UK (COG-UK) Consortium | Harper VanSteenhouse, Yumi Kasai, David Gray, Carol Clugston, Anna Dominiczak and Alex Alderton, Roberto Amato, Jeffrey Barrett, Sonia Goncalves, Ewan Harrison, David K. Jackson, Ian Johnston, Dominic Kwiatkowski, Cordelia Langford, John Sillitoe on behalf of the Wellcome Sanger Institute COVID-19 Surveillance Team |

| | | | |
|---|---|---|---|
| EPI_ISL_2699534 | | | |
| EPI_ISL_2699535, EPI_ISL_2699536, EPI_ISL_2699537, EPI_ISL_2699538 | Lighthouse Lab in Milton Keynes | Wellcome Sanger Institute for the COVID-19 Genomics UK (COG-UK) Consortium | The Lighthouse Lab in Milton Keynes and Alex Alderton, Roberto Amato, Jeffrey Barrett, Sonia Goncalves, Ewan Harrison, David K. Jackson, Ian Johnston, Dominic Kwiatkowski, Cordelia Langford, John Sillitoe on behalf of the Wellcome Sanger Institute COVID-19 Surveillance Team |
| EPI_ISL_2699539, EPI_ISL_2699540, EPI_ISL_2699541 | Lighthouse Lab in Glasgow | Wellcome Sanger Institute for the COVID-19 Genomics UK (COG-UK) Consortium | Harper VanSteenhouse, Yumi Kasai, David Gray, Carol Clugston, Anna Dominiczak and Alex Alderton, Roberto Amato, Jeffrey Barrett, Sonia Goncalves, Ewan Harrison, David K. Jackson, Ian Johnston, Dominic Kwiatkowski, Cordelia Langford, John Sillitoe on behalf of the Wellcome Sanger Institute COVID-19 Surveillance Team |
| EPI_ISL_2699542 | Lighthouse Lab in Milton Keynes | Wellcome Sanger Institute for the COVID-19 Genomics UK (COG-UK) Consortium | The Lighthouse Lab in Milton Keynes and Alex Alderton, Roberto Amato, Jeffrey Barrett, Sonia Goncalves, Ewan Harrison, David K. Jackson, Ian Johnston, Dominic Kwiatkowski, Cordelia Langford, John Sillitoe on behalf of the Wellcome Sanger Institute COVID-19 Surveillance Team |
| EPI_ISL_2699543 | Lighthouse Laboratory Plymouth | Wellcome Sanger Institute for the COVID-19 Genomics UK (COG-UK) Consortium | Lighthouse Laboratory Plymouth and Alex Alderton, Roberto Amato, Jeffrey Barrett, Sonia Goncalves, Ewan Harrison, David K. Jackson, Ian Johnston, Dominic Kwiatkowski, Cordelia Langford, John Sillitoe on behalf of the Wellcome Sanger Institute COVID-19 Surveillance Team |
| EPI_ISL_2699544 | Lighthouse Lab in Glasgow | Wellcome Sanger Institute for the COVID-19 Genomics UK (COG-UK) Consortium | Harper VanSteenhouse, Yumi Kasai, David Gray, Carol Clugston, Anna Dominiczak and Alex Alderton, Roberto Amato, Jeffrey Barrett, Sonia Goncalves, Ewan Harrison, David K. Jackson, Ian Johnston, Dominic Kwiatkowski, Cordelia Langford, John Sillitoe on behalf of the Wellcome Sanger Institute COVID-19 Surveillance Team |
| EPI_ISL_2699545, EPI_ISL_2699546 | Lighthouse Laboratory Plymouth | Wellcome Sanger Institute for the COVID-19 Genomics UK (COG-UK) Consortium | Lighthouse Laboratory Plymouth and Alex Alderton, Roberto Amato, Jeffrey Barrett, Sonia Goncalves, Ewan Harrison, David K. Jackson, Ian Johnston, Dominic Kwiatkowski, Cordelia Langford, John Sillitoe on behalf of the Wellcome Sanger Institute COVID-19 Surveillance Team |
| EPI_ISL_2699547 | Lighthouse Lab in Milton Keynes | Wellcome Sanger Institute for the COVID-19 Genomics UK (COG-UK) Consortium | The Lighthouse Lab in Milton Keynes and Alex Alderton, Roberto Amato, Jeffrey Barrett, Sonia Goncalves, Ewan Harrison, David K. Jackson, Ian Johnston, Dominic Kwiatkowski, Cordelia Langford, John Sillitoe on behalf of the Wellcome Sanger Institute COVID-19 Surveillance Team |
| EPI_ISL_2699548, EPI_ISL_2699549 | Lighthouse Lab in Glasgow | Wellcome Sanger Institute for the COVID-19 Genomics UK (COG-UK) Consortium | Harper VanSteenhouse, Yumi Kasai, David Gray, Carol Clugston, Anna Dominiczak and Alex Alderton, Roberto Amato, Jeffrey Barrett, Sonia Goncalves, Ewan Harrison, David K. Jackson, Ian Johnston, Dominic Kwiatkowski, Cordelia Langford, John Sillitoe on behalf of the Wellcome Sanger Institute COVID-19 Surveillance Team |
| EPI_ISL_2699550, EPI_ISL_2699551, EPI_ISL_2699552 | Lighthouse Lab in Milton Keynes | Wellcome Sanger Institute for the COVID-19 Genomics UK (COG-UK) Consortium | The Lighthouse Lab in Milton Keynes and Alex Alderton, Roberto Amato, Jeffrey Barrett, Sonia Goncalves, Ewan Harrison, David K. Jackson, Ian Johnston, Dominic Kwiatkowski, Cordelia Langford, John Sillitoe on behalf of the Wellcome Sanger Institute COVID-19 Surveillance Team |
| EPI_ISL_2699553, EPI_ISL_2699554 | Lighthouse Lab in Glasgow | Wellcome Sanger Institute for the COVID-19 Genomics UK (COG-UK) Consortium | Harper VanSteenhouse, Yumi Kasai, David Gray, Carol Clugston, Anna Dominiczak and Alex Alderton, Roberto Amato, Jeffrey Barrett, Sonia Goncalves, Ewan Harrison, David K. Jackson, Ian Johnston, Dominic Kwiatkowski, Cordelia Langford, John Sillitoe on behalf of the Wellcome Sanger Institute COVID-19 Surveillance Team |
| EPI_ISL_2699555 | Lighthouse Lab in Milton Keynes | Wellcome Sanger Institute for the COVID-19 Genomics UK (COG-UK) Consortium | The Lighthouse Lab in Milton Keynes and Alex Alderton, Roberto Amato, Jeffrey Barrett, Sonia Goncalves, Ewan Harrison, David K. Jackson, Ian Johnston, Dominic Kwiatkowski, Cordelia Langford, John Sillitoe on behalf of the Wellcome Sanger Institute COVID-19 Surveillance Team |
| EPI_ISL_2699556, EPI_ISL_2699557, EPI_ISL_2699558, EPI_ISL_2699559, EPI_ISL_2699560 | Lighthouse Lab in Glasgow | Wellcome Sanger Institute for the COVID-19 Genomics UK (COG-UK) Consortium | Harper VanSteenhouse, Yumi Kasai, David Gray, Carol Clugston, Anna Dominiczak and Alex Alderton, Roberto Amato, Jeffrey Barrett, Sonia Goncalves, Ewan Harrison, David K. Jackson, Ian Johnston, Dominic Kwiatkowski, Cordelia Langford, John Sillitoe on behalf of the Wellcome Sanger Institute COVID-19 Surveillance Team |
| EPI_ISL_2699561, EPI_ISL_2699562 | Lighthouse Lab in Milton Keynes | Wellcome Sanger Institute for the COVID-19 Genomics UK (COG-UK) Consortium | The Lighthouse Lab in Milton Keynes and Alex Alderton, Roberto Amato, Jeffrey Barrett, Sonia Goncalves, Ewan Harrison, David K. Jackson, Ian Johnston, Dominic Kwiatkowski, Cordelia Langford, John Sillitoe on behalf of the Wellcome Sanger Institute COVID-19 Surveillance Team |
| EPI_ISL_2699563, EPI_ISL_2699564, EPI_ISL_2699565 | Lighthouse Lab in Glasgow | Wellcome Sanger Institute for the COVID-19 Genomics UK (COG-UK) Consortium | Harper VanSteenhouse, Yumi Kasai, David Gray, Carol Clugston, Anna Dominiczak and Alex Alderton, Roberto Amato, Jeffrey Barrett, Sonia Goncalves, Ewan Harrison, David K. Jackson, Ian Johnston, Dominic Kwiatkowski, Cordelia Langford, John Sillitoe on behalf of the Wellcome Sanger Institute COVID-19 Surveillance Team |
| EPI_ISL_2699566, EPI_ISL_2699567, EPI_ISL_2699568, EPI_ISL_2699569, EPI_ISL_2699570, EPI_ISL_2699571 | Lighthouse Lab in Milton Keynes | Wellcome Sanger Institute for the COVID-19 Genomics UK (COG-UK) Consortium | The Lighthouse Lab in Milton Keynes and Alex Alderton, Roberto Amato, Jeffrey Barrett, Sonia Goncalves, Ewan Harrison, David K. Jackson, Ian Johnston, Dominic Kwiatkowski, Cordelia Langford, John Sillitoe on behalf of the Wellcome Sanger Institute COVID-19 Surveillance Team |
| EPI_ISL_2699572, EPI_ISL_2699573 | Lighthouse Lab in Glasgow | Wellcome Sanger Institute for the COVID-19 Genomics UK (COG-UK) Consortium | Harper VanSteenhouse, Yumi Kasai, David Gray, Carol Clugston, Anna Dominiczak and Alex Alderton, Roberto Amato, Jeffrey Barrett, Sonia Goncalves, Ewan Harrison, David K. Jackson, Ian Johnston, Dominic Kwiatkowski, Cordelia Langford, John Sillitoe on behalf of the Wellcome Sanger Institute COVID-19 Surveillance Team |
| EPI_ISL_2699574, EPI_ISL_2699575 | Lighthouse Lab in Milton Keynes | Wellcome Sanger Institute for the COVID-19 Genomics UK (COG-UK) Consortium | The Lighthouse Lab in Milton Keynes and Alex Alderton, Roberto Amato, Jeffrey Barrett, Sonia Goncalves, Ewan Harrison, David K. Jackson, Ian Johnston, Dominic Kwiatkowski, Cordelia Langford, John Sillitoe on behalf of the Wellcome Sanger Institute COVID-19 Surveillance Team |
| EPI_ISL_2699576, EPI_ISL_2699577 | Lighthouse Lab in Glasgow | Wellcome Sanger Institute for the COVID-19 Genomics UK (COG-UK) Consortium | Harper VanSteenhouse, Yumi Kasai, David Gray, Carol Clugston, Anna Dominiczak and Alex Alderton, Roberto Amato, Jeffrey Barrett, Sonia Goncalves, Ewan Harrison, David K. Jackson, Ian Johnston, Dominic Kwiatkowski, Cordelia Langford, John Sillitoe on behalf of the Wellcome Sanger Institute COVID-19 Surveillance Team |
| EPI_ISL_2699578 | Lighthouse Lab in Milton Keynes | Wellcome Sanger Institute for the COVID-19 Genomics UK (COG-UK) Consortium | The Lighthouse Lab in Milton Keynes and Alex Alderton, Roberto Amato, Jeffrey Barrett, Sonia Goncalves, Ewan Harrison, David K. Jackson, Ian Johnston, Dominic Kwiatkowski, Cordelia Langford, John Sillitoe on behalf of the Wellcome Sanger Institute COVID-19 Surveillance Team |
| EPI_ISL_2699579, EPI_ISL_2699580 | Lighthouse Lab in Glasgow | Wellcome Sanger Institute for the COVID-19 Genomics UK (COG-UK) Consortium | Harper VanSteenhouse, Yumi Kasai, David Gray, Carol Clugston, Anna Dominiczak and Alex Alderton, Roberto Amato, Jeffrey Barrett, Sonia Goncalves, Ewan Harrison, David K. Jackson, Ian Johnston, Dominic Kwiatkowski, Cordelia Langford, John Sillitoe on behalf of the Wellcome Sanger Institute COVID-19 Surveillance Team |
| EPI_ISL_2699581 | Lighthouse Lab in Milton Keynes | Wellcome Sanger Institute for the COVID-19 Genomics UK (COG-UK) Consortium | The Lighthouse Lab in Milton Keynes and Alex Alderton, Roberto Amato, Jeffrey Barrett, Sonia Goncalves, Ewan Harrison, David K. Jackson, Ian Johnston, Dominic Kwiatkowski, Cordelia Langford, John Sillitoe on behalf of the Wellcome Sanger Institute COVID-19 Surveillance Team |
| EPI_ISL_2699582, EPI_ISL_2699583, EPI_ISL_2699584, EPI_ISL_2699585 | Lighthouse Lab in Glasgow | Wellcome Sanger Institute for the COVID-19 Genomics UK (COG-UK) Consortium | Harper VanSteenhouse, Yumi Kasai, David Gray, Carol Clugston, Anna Dominiczak and Alex Alderton, Roberto Amato, Jeffrey Barrett, Sonia Goncalves, Ewan Harrison, David K. Jackson, Ian Johnston, Dominic Kwiatkowski, Cordelia Langford, John Sillitoe on behalf of the Wellcome Sanger Institute COVID-19 Surveillance Team |
| EPI_ISL_2699586, EPI_ISL_2699587 | Lighthouse Lab in Milton Keynes | Wellcome Sanger Institute for the COVID-19 Genomics UK (COG-UK) Consortium | The Lighthouse Lab in Milton Keynes and Alex Alderton, Roberto Amato, Jeffrey Barrett, Sonia Goncalves, Ewan Harrison, David K. Jackson, Ian Johnston, Dominic Kwiatkowski, Cordelia Langford, John Sillitoe on behalf of the Wellcome Sanger Institute COVID-19 Surveillance Team |
| EPI_ISL_2699588 | Lighthouse Lab in Glasgow | Wellcome Sanger Institute for the COVID-19 Genomics UK (COG-UK) Consortium | Harper VanSteenhouse, Yumi Kasai, David Gray, Carol Clugston, Anna Dominiczak and Alex Alderton, Roberto Amato, Jeffrey Barrett, Sonia Goncalves, Ewan Harrison, David K. Jackson, Ian Johnston, Dominic Kwiatkowski, Cordelia Langford, John Sillitoe on behalf of the Wellcome Sanger Institute COVID-19 Surveillance Team |
| EPI_ISL_2699589 | Lighthouse Lab in Milton Keynes | Wellcome Sanger Institute for the COVID-19 Genomics UK (COG-UK) Consortium | The Lighthouse Lab in Milton Keynes and Alex Alderton, Roberto Amato, Jeffrey Barrett, Sonia Goncalves, Ewan Harrison, David K. Jackson, Ian Johnston, Dominic Kwiatkowski, Cordelia Langford, John Sillitoe on behalf of the Wellcome Sanger Institute COVID-19 Surveillance Team |
| EPI_ISL_2699590, EPI_ISL_2699591, EPI_ISL_2699592, EPI_ISL_2699593 | Lighthouse Lab in Glasgow | Wellcome Sanger Institute for the COVID-19 Genomics UK (COG-UK) Consortium | Harper VanSteenhouse, Yumi Kasai, David Gray, Carol Clugston, Anna Dominiczak and Alex Alderton, Roberto Amato, Jeffrey Barrett, Sonia Goncalves, Ewan Harrison, David K. Jackson, Ian Johnston, Dominic Kwiatkowski, Cordelia Langford, John Sillitoe on behalf of the Wellcome Sanger Institute COVID-19 Surveillance Team |
| EPI_ISL_2699594 | Lighthouse Lab in Milton Keynes | Wellcome Sanger Institute for the COVID-19 Genomics UK (COG-UK) Consortium | The Lighthouse Lab in Milton Keynes and Alex Alderton, Roberto Amato, Jeffrey Barrett, Sonia Goncalves, Ewan Harrison, David K. Jackson, Ian Johnston, Dominic Kwiatkowski, Cordelia Langford, John Sillitoe on behalf of the Wellcome Sanger Institute COVID-19 Surveillance Team |
| EPI_ISL_2699595 | Lighthouse Laboratory Plymouth | Wellcome Sanger Institute for the COVID-19 Genomics UK (COG-UK) Consortium | Lighthouse Laboratory Plymouth and Alex Alderton, Roberto Amato, Jeffrey Barrett, Sonia Goncalves, Ewan Harrison, David K. Jackson, Ian Johnston, Dominic Kwiatkowski, Cordelia Langford, John Sillitoe on behalf of the Wellcome Sanger Institute COVID-19 Surveillance Team |
| EPI_ISL_2699596 | Lighthouse Lab in Milton Keynes | Wellcome Sanger Institute for the COVID-19 Genomics UK (COG-UK) Consortium | The Lighthouse Lab in Milton Keynes and Alex Alderton, Roberto Amato, Jeffrey Barrett, Sonia Goncalves, Ewan Harrison, David K. Jackson, Ian Johnston, Dominic Kwiatkowski, Cordelia Langford, John Sillitoe on behalf of the Wellcome Sanger Institute COVID-19 Surveillance Team |
| EPI_ISL_2699597 | Lighthouse Laboratory Plymouth | Wellcome Sanger Institute for the COVID-19 Genomics UK (COG-UK) Consortium | Lighthouse Laboratory Plymouth and Alex Alderton, Roberto Amato, Jeffrey Barrett, Sonia Goncalves, Ewan Harrison, David K. Jackson, Ian Johnston, Dominic Kwiatkowski, Cordelia Langford, John Sillitoe on behalf of the Wellcome Sanger Institute COVID-19 Surveillance Team |
| EPI_ISL_2699598 | Lighthouse Lab in Glasgow | Wellcome Sanger Institute for the COVID-19 Genomics UK (COG-UK) Consortium | Harper VanSteenhouse, Yumi Kasai, David Gray, Carol Clugston, Anna Dominiczak and Alex Alderton, Roberto Amato, Jeffrey Barrett, Sonia Goncalves, Ewan Harrison, David K. Jackson, Ian Johnston, Dominic Kwiatkowski, Cordelia Langford, John Sillitoe on behalf of the Wellcome Sanger Institute COVID-19 Surveillance Team |
| EPI_ISL_2699599 | Lighthouse Laboratory Plymouth | Wellcome Sanger Institute for the COVID-19 Genomics UK (COG-UK) Consortium | Lighthouse Laboratory Plymouth and Alex Alderton, Roberto Amato, Jeffrey Barrett, Sonia Goncalves, Ewan Harrison, David K. Jackson, Ian Johnston, Dominic Kwiatkowski, Cordelia Langford, John Sillitoe on behalf of the Wellcome Sanger Institute COVID-19 Surveillance Team |
| EPI_ISL_2699600, EPI_ISL_2699601, EPI_ISL_2699602, EPI_ISL_2699603 | Lighthouse Lab in Glasgow | Wellcome Sanger Institute for the COVID-19 Genomics UK (COG-UK) Consortium | Harper VanSteenhouse, Yumi Kasai, David Gray, Carol Clugston, Anna Dominiczak and Alex Alderton, Roberto Amato, Jeffrey Barrett, Sonia Goncalves, Ewan Harrison, David K. Jackson, Ian Johnston, Dominic Kwiatkowski, Cordelia Langford, John Sillitoe on behalf of the Wellcome Sanger Institute COVID-19 Surveillance Team |
| EPI_ISL_2699604, EPI_ISL_2699605, EPI_ISL_2699606, EPI_ISL_2699607 | Lighthouse Lab in Milton Keynes | Wellcome Sanger Institute for the COVID-19 Genomics UK (COG-UK) Consortium | The Lighthouse Lab in Milton Keynes and Alex Alderton, Roberto Amato, Jeffrey Barrett, Sonia Goncalves, Ewan Harrison, David K. Jackson, Ian Johnston, Dominic Kwiatkowski, Cordelia Langford, John Sillitoe on behalf of the Wellcome Sanger Institute COVID-19 Surveillance Team |
| EPI_ISL_2699608, EPI_ISL_2699609 | Lighthouse Laboratory Plymouth | Wellcome Sanger Institute for the COVID-19 Genomics UK (COG-UK) Consortium | Lighthouse Laboratory Plymouth and Alex Alderton, Roberto Amato, Jeffrey Barrett, Sonia Goncalves, Ewan Harrison, David K. Jackson, Ian Johnston, Dominic Kwiatkowski, Cordelia Langford, John Sillitoe on behalf of the Wellcome Sanger Institute COVID-19 Surveillance Team |
| EPI_ISL_2699610, EPI_ISL_2699611, EPI_ISL_2699612 | Lighthouse Lab in Milton Keynes | Wellcome Sanger Institute for the COVID-19 Genomics UK (COG-UK) Consortium | The Lighthouse Lab in Milton Keynes and Alex Alderton, Roberto Amato, Jeffrey Barrett, Sonia Goncalves, Ewan Harrison, David K. Jackson, Ian Johnston, Dominic Kwiatkowski, Cordelia Langford, John Sillitoe on behalf of the Wellcome Sanger Institute COVID-19 Surveillance Team |

| | | | |
|---|---|---|---|
| EPI_ISL_2699613 | Lighthouse Lab in Glasgow | Wellcome Sanger Institute for the COVID-19 Genomics UK (COG-UK) Consortium | Harper VanSteenhouse, Yumi Kasai, David Gray, Carol Clugston, Anna Dominiczak and Alex Alderton, Roberto Amato, Jeffrey Barrett, Sonia Goncalves, Ewan Harrison, David K. Jackson, Ian Johnston, Dominic Kwiatkowski, Cordelia Langford, John Sillitoe on behalf of the Wellcome Sanger Institute COVID-19 Surveillance Team |
| EPI_ISL_2699614, EPI_ISL_2699615, EPI_ISL_2699616 | Lighthouse Lab in Milton Keynes | Wellcome Sanger Institute for the COVID-19 Genomics UK (COG-UK) Consortium | The Lighthouse Lab in Milton Keynes and Alex Alderton, Roberto Amato, Jeffrey Barrett, Sonia Goncalves, Ewan Harrison, David K. Jackson, Ian Johnston, Dominic Kwiatkowski, Cordelia Langford, John Sillitoe on behalf of the Wellcome Sanger Institute COVID-19 Surveillance Team |
| EPI_ISL_2699618, EPI_ISL_2699619 | Lighthouse Lab in Glasgow | Wellcome Sanger Institute for the COVID-19 Genomics UK (COG-UK) Consortium | Harper VanSteenhouse, Yumi Kasai, David Gray, Carol Clugston, Anna Dominiczak and Alex Alderton, Roberto Amato, Jeffrey Barrett, Sonia Goncalves, Ewan Harrison, David K. Jackson, Ian Johnston, Dominic Kwiatkowski, Cordelia Langford, John Sillitoe on behalf of the Wellcome Sanger Institute COVID-19 Surveillance Team |
| EPI_ISL_2699620 | Lighthouse Lab in Milton Keynes | Wellcome Sanger Institute for the COVID-19 Genomics UK (COG-UK) Consortium | The Lighthouse Lab in Milton Keynes and Alex Alderton, Roberto Amato, Jeffrey Barrett, Sonia Goncalves, Ewan Harrison, David K. Jackson, Ian Johnston, Dominic Kwiatkowski, Cordelia Langford, John Sillitoe on behalf of the Wellcome Sanger Institute COVID-19 Surveillance Team |
| EPI_ISL_2699621 | Lighthouse Lab in Glasgow | Wellcome Sanger Institute for the COVID-19 Genomics UK (COG-UK) Consortium | Harper VanSteenhouse, Yumi Kasai, David Gray, Carol Clugston, Anna Dominiczak and Alex Alderton, Roberto Amato, Jeffrey Barrett, Sonia Goncalves, Ewan Harrison, David K. Jackson, Ian Johnston, Dominic Kwiatkowski, Cordelia Langford, John Sillitoe on behalf of the Wellcome Sanger Institute COVID-19 Surveillance Team |
| EPI_ISL_2699622, EPI_ISL_2699623, EPI_ISL_2699624, EPI_ISL_2699625 | Lighthouse Lab in Milton Keynes | Wellcome Sanger Institute for the COVID-19 Genomics UK (COG-UK) Consortium | The Lighthouse Lab in Milton Keynes and Alex Alderton, Roberto Amato, Jeffrey Barrett, Sonia Goncalves, Ewan Harrison, David K. Jackson, Ian Johnston, Dominic Kwiatkowski, Cordelia Langford, John Sillitoe on behalf of the Wellcome Sanger Institute COVID-19 Surveillance Team |
| EPI_ISL_2699626 | Lighthouse Lab in Glasgow | Wellcome Sanger Institute for the COVID-19 Genomics UK (COG-UK) Consortium | Harper VanSteenhouse, Yumi Kasai, David Gray, Carol Clugston, Anna Dominiczak and Alex Alderton, Roberto Amato, Jeffrey Barrett, Sonia Goncalves, Ewan Harrison, David K. Jackson, Ian Johnston, Dominic Kwiatkowski, Cordelia Langford, John Sillitoe on behalf of the Wellcome Sanger Institute COVID-19 Surveillance Team |
| EPI_ISL_2699628, EPI_ISL_2699629, EPI_ISL_2699630 | Lighthouse Lab in Milton Keynes | Wellcome Sanger Institute for the COVID-19 Genomics UK (COG-UK) Consortium | The Lighthouse Lab in Milton Keynes and Alex Alderton, Roberto Amato, Jeffrey Barrett, Sonia Goncalves, Ewan Harrison, David K. Jackson, Ian Johnston, Dominic Kwiatkowski, Cordelia Langford, John Sillitoe on behalf of the Wellcome Sanger Institute COVID-19 Surveillance Team |
| EPI_ISL_2699631 | Lighthouse Lab in Glasgow | Wellcome Sanger Institute for the COVID-19 Genomics UK (COG-UK) Consortium | Harper VanSteenhouse, Yumi Kasai, David Gray, Carol Clugston, Anna Dominiczak and Alex Alderton, Roberto Amato, Jeffrey Barrett, Sonia Goncalves, Ewan Harrison, David K. Jackson, Ian Johnston, Dominic Kwiatkowski, Cordelia Langford, John Sillitoe on behalf of the Wellcome Sanger Institute COVID-19 Surveillance Team |
| EPI_ISL_2699632 | Lighthouse Laboratory Plymouth | Wellcome Sanger Institute for the COVID-19 Genomics UK (COG-UK) Consortium | Lighthouse Laboratory Plymouth and Alex Alderton, Roberto Amato, Jeffrey Barrett, Sonia Goncalves, Ewan Harrison, David K. Jackson, Ian Johnston, Dominic Kwiatkowski, Cordelia Langford, John Sillitoe on behalf of the Wellcome Sanger Institute COVID-19 Surveillance Team |
| EPI_ISL_2699633, EPI_ISL_2699634, EPI_ISL_2699635, EPI_ISL_2699636, EPI_ISL_2699637, EPI_ISL_2699638, EPI_ISL_2699639 | Lighthouse Lab in Milton Keynes | Wellcome Sanger Institute for the COVID-19 Genomics UK (COG-UK) Consortium | The Lighthouse Lab in Milton Keynes and Alex Alderton, Roberto Amato, Jeffrey Barrett, Sonia Goncalves, Ewan Harrison, David K. Jackson, Ian Johnston, Dominic Kwiatkowski, Cordelia Langford, John Sillitoe on behalf of the Wellcome Sanger Institute COVID-19 Surveillance Team |
| EPI_ISL_2699640, EPI_ISL_2699642 | Lighthouse Lab in Glasgow | Wellcome Sanger Institute for the COVID-19 Genomics UK (COG-UK) Consortium | Harper VanSteenhouse, Yumi Kasai, David Gray, Carol Clugston, Anna Dominiczak and Alex Alderton, Roberto Amato, Jeffrey Barrett, Sonia Goncalves, Ewan Harrison, David K. Jackson, Ian Johnston, Dominic Kwiatkowski, Cordelia Langford, John Sillitoe on behalf of the Wellcome Sanger Institute COVID-19 Surveillance Team |
| EPI_ISL_2699643 | Lighthouse Lab in Milton Keynes | Wellcome Sanger Institute for the COVID-19 Genomics UK (COG-UK) Consortium | The Lighthouse Lab in Milton Keynes and Alex Alderton, Roberto Amato, Jeffrey Barrett, Sonia Goncalves, Ewan Harrison, David K. Jackson, Ian Johnston, Dominic Kwiatkowski, Cordelia Langford, John Sillitoe on behalf of the Wellcome Sanger Institute COVID-19 Surveillance Team |
| EPI_ISL_2699644, EPI_ISL_2699645 | Lighthouse Lab in Glasgow | Wellcome Sanger Institute for the COVID-19 Genomics UK (COG-UK) Consortium | Harper VanSteenhouse, Yumi Kasai, David Gray, Carol Clugston, Anna Dominiczak and Alex Alderton, Roberto Amato, Jeffrey Barrett, Sonia Goncalves, Ewan Harrison, David K. Jackson, Ian Johnston, Dominic Kwiatkowski, Cordelia Langford, John Sillitoe on behalf of the Wellcome Sanger Institute COVID-19 Surveillance Team |
| EPI_ISL_2699646 | Lighthouse Lab in Milton Keynes | Wellcome Sanger Institute for the COVID-19 Genomics UK (COG-UK) Consortium | The Lighthouse Lab in Milton Keynes and Alex Alderton, Roberto Amato, Jeffrey Barrett, Sonia Goncalves, Ewan Harrison, David K. Jackson, Ian Johnston, Dominic Kwiatkowski, Cordelia Langford, John Sillitoe on behalf of the Wellcome Sanger Institute COVID-19 Surveillance Team |
| EPI_ISL_2699647, EPI_ISL_2699649 | Lighthouse Laboratory Plymouth | Wellcome Sanger Institute for the COVID-19 Genomics UK (COG-UK) Consortium | Lighthouse Laboratory Plymouth and Alex Alderton, Roberto Amato, Jeffrey Barrett, Sonia Goncalves, Ewan Harrison, David K. Jackson, Ian Johnston, Dominic Kwiatkowski, Cordelia Langford, John Sillitoe on behalf of the Wellcome Sanger Institute COVID-19 Surveillance Team |
| EPI_ISL_2699650, EPI_ISL_2699651 | Lighthouse Lab in Milton Keynes | Wellcome Sanger Institute for the COVID-19 Genomics UK (COG-UK) Consortium | The Lighthouse Lab in Milton Keynes and Alex Alderton, Roberto Amato, Jeffrey Barrett, Sonia Goncalves, Ewan Harrison, David K. Jackson, Ian Johnston, Dominic Kwiatkowski, Cordelia Langford, John Sillitoe on behalf of the Wellcome Sanger Institute COVID-19 Surveillance Team |
| EPI_ISL_2699652 | Lighthouse Lab in Glasgow | Wellcome Sanger Institute for the COVID-19 Genomics UK (COG-UK) Consortium | Harper VanSteenhouse, Yumi Kasai, David Gray, Carol Clugston, Anna Dominiczak and Alex Alderton, Roberto Amato, Jeffrey Barrett, Sonia Goncalves, Ewan Harrison, David K. Jackson, Ian Johnston, Dominic Kwiatkowski, Cordelia Langford, John Sillitoe on behalf of the Wellcome Sanger Institute COVID-19 Surveillance Team |
| EPI_ISL_2699653, EPI_ISL_2699654 | Lighthouse Lab in Milton Keynes | Wellcome Sanger Institute for the COVID-19 Genomics UK (COG-UK) Consortium | The Lighthouse Lab in Milton Keynes and Alex Alderton, Roberto Amato, Jeffrey Barrett, Sonia Goncalves, Ewan Harrison, David K. Jackson, Ian Johnston, Dominic Kwiatkowski, Cordelia Langford, John Sillitoe on behalf of the Wellcome Sanger Institute COVID-19 Surveillance Team |
| EPI_ISL_2699655, EPI_ISL_2699656 | Lighthouse Lab in Glasgow | Wellcome Sanger Institute for the COVID-19 Genomics UK (COG-UK) Consortium | Harper VanSteenhouse, Yumi Kasai, David Gray, Carol Clugston, Anna Dominiczak and Alex Alderton, Roberto Amato, Jeffrey Barrett, Sonia Goncalves, Ewan Harrison, David K. Jackson, Ian Johnston, Dominic Kwiatkowski, Cordelia Langford, John Sillitoe on behalf of the Wellcome Sanger Institute COVID-19 Surveillance Team |
| EPI_ISL_2699657 | Lighthouse Lab in Milton Keynes | Wellcome Sanger Institute for the COVID-19 Genomics UK (COG-UK) Consortium | The Lighthouse Lab in Milton Keynes and Alex Alderton, Roberto Amato, Jeffrey Barrett, Sonia Goncalves, Ewan Harrison, David K. Jackson, Ian Johnston, Dominic Kwiatkowski, Cordelia Langford, John Sillitoe on behalf of the Wellcome Sanger Institute COVID-19 Surveillance Team |
| EPI_ISL_2699658, EPI_ISL_2699659 | Lighthouse Lab in Glasgow | Wellcome Sanger Institute for the COVID-19 Genomics UK (COG-UK) Consortium | Harper VanSteenhouse, Yumi Kasai, David Gray, Carol Clugston, Anna Dominiczak and Alex Alderton, Roberto Amato, Jeffrey Barrett, Sonia Goncalves, Ewan Harrison, David K. Jackson, Ian Johnston, Dominic Kwiatkowski, Cordelia Langford, John Sillitoe on behalf of the Wellcome Sanger Institute COVID-19 Surveillance Team |
| EPI_ISL_2699660 | Lighthouse Lab in Milton Keynes | Wellcome Sanger Institute for the COVID-19 Genomics UK (COG-UK) Consortium | The Lighthouse Lab in Milton Keynes and Alex Alderton, Roberto Amato, Jeffrey Barrett, Sonia Goncalves, Ewan Harrison, David K. Jackson, Ian Johnston, Dominic Kwiatkowski, Cordelia Langford, John Sillitoe on behalf of the Wellcome Sanger Institute COVID-19 Surveillance Team |
| EPI_ISL_2699661 | Lighthouse Lab in Glasgow | Wellcome Sanger Institute for the COVID-19 Genomics UK (COG-UK) Consortium | Harper VanSteenhouse, Yumi Kasai, David Gray, Carol Clugston, Anna Dominiczak and Alex Alderton, Roberto Amato, Jeffrey Barrett, Sonia Goncalves, Ewan Harrison, David K. Jackson, Ian Johnston, Dominic Kwiatkowski, Cordelia Langford, John Sillitoe on behalf of the Wellcome Sanger Institute COVID-19 Surveillance Team |
| EPI_ISL_2699662, EPI_ISL_2699663 | Lighthouse Lab in Milton Keynes | Wellcome Sanger Institute for the COVID-19 Genomics UK (COG-UK) Consortium | The Lighthouse Lab in Milton Keynes and Alex Alderton, Roberto Amato, Jeffrey Barrett, Sonia Goncalves, Ewan Harrison, David K. Jackson, Ian Johnston, Dominic Kwiatkowski, Cordelia Langford, John Sillitoe on behalf of the Wellcome Sanger Institute COVID-19 Surveillance Team |
| EPI_ISL_2699664 | Lighthouse Laboratory Plymouth | Wellcome Sanger Institute for the COVID-19 Genomics UK (COG-UK) Consortium | Lighthouse Laboratory Plymouth and Alex Alderton, Roberto Amato, Jeffrey Barrett, Sonia Goncalves, Ewan Harrison, David K. Jackson, Ian Johnston, Dominic Kwiatkowski, Cordelia Langford, John Sillitoe on behalf of the Wellcome Sanger Institute COVID-19 Surveillance Team |
| EPI_ISL_2699665 | Lighthouse Lab in Milton Keynes | Wellcome Sanger Institute for the COVID-19 Genomics UK (COG-UK) Consortium | The Lighthouse Lab in Milton Keynes and Alex Alderton, Roberto Amato, Jeffrey Barrett, Sonia Goncalves, Ewan Harrison, David K. Jackson, Ian Johnston, Dominic Kwiatkowski, Cordelia Langford, John Sillitoe on behalf of the Wellcome Sanger Institute COVID-19 Surveillance Team |
| EPI_ISL_2699666 | Lighthouse Laboratory Plymouth | Wellcome Sanger Institute for the COVID-19 Genomics UK (COG-UK) Consortium | Lighthouse Laboratory Plymouth and Alex Alderton, Roberto Amato, Jeffrey Barrett, Sonia Goncalves, Ewan Harrison, David K. Jackson, Ian Johnston, Dominic Kwiatkowski, Cordelia Langford, John Sillitoe on behalf of the Wellcome Sanger Institute COVID-19 Surveillance Team |
| EPI_ISL_2699667 | Lighthouse Lab in Glasgow | Wellcome Sanger Institute for the COVID-19 Genomics UK (COG-UK) Consortium | Harper VanSteenhouse, Yumi Kasai, David Gray, Carol Clugston, Anna Dominiczak and Alex Alderton, Roberto Amato, Jeffrey Barrett, Sonia Goncalves, Ewan Harrison, David K. Jackson, Ian Johnston, Dominic Kwiatkowski, Cordelia Langford, John Sillitoe on behalf of the Wellcome Sanger Institute COVID-19 Surveillance Team |
| EPI_ISL_2699668, EPI_ISL_2699669, EPI_ISL_2699670 | Lighthouse Lab in Milton Keynes | Wellcome Sanger Institute for the COVID-19 Genomics UK (COG-UK) Consortium | The Lighthouse Lab in Milton Keynes and Alex Alderton, Roberto Amato, Jeffrey Barrett, Sonia Goncalves, Ewan Harrison, David K. Jackson, Ian Johnston, Dominic Kwiatkowski, Cordelia Langford, John Sillitoe on behalf of the Wellcome Sanger Institute COVID-19 Surveillance Team |
| EPI_ISL_2699671, EPI_ISL_2699672, EPI_ISL_2699673 | Lighthouse Lab in Glasgow | Wellcome Sanger Institute for the COVID-19 Genomics UK (COG-UK) Consortium | Harper VanSteenhouse, Yumi Kasai, David Gray, Carol Clugston, Anna Dominiczak and Alex Alderton, Roberto Amato, Jeffrey Barrett, Sonia Goncalves, Ewan Harrison, David K. Jackson, Ian Johnston, Dominic Kwiatkowski, Cordelia Langford, John Sillitoe on behalf of the Wellcome Sanger Institute COVID-19 Surveillance Team |
| EPI_ISL_2699674 | Lighthouse Laboratory Plymouth | Wellcome Sanger Institute for the COVID-19 Genomics UK (COG-UK) Consortium | Lighthouse Laboratory Plymouth and Alex Alderton, Roberto Amato, Jeffrey Barrett, Sonia Goncalves, Ewan Harrison, David K. Jackson, Ian Johnston, Dominic Kwiatkowski, Cordelia Langford, John Sillitoe on behalf of the Wellcome Sanger Institute COVID-19 Surveillance Team |
| EPI_ISL_2699675, EPI_ISL_2699676 | Lighthouse Lab in Milton Keynes | Wellcome Sanger Institute for the COVID-19 Genomics UK (COG-UK) Consortium | The Lighthouse Lab in Milton Keynes and Alex Alderton, Roberto Amato, Jeffrey Barrett, Sonia Goncalves, Ewan Harrison, David K. Jackson, Ian Johnston, Dominic Kwiatkowski, Cordelia Langford, John Sillitoe on behalf of the Wellcome Sanger Institute COVID-19 Surveillance Team |
| EPI_ISL_2699677 | Lighthouse Lab in Glasgow | Wellcome Sanger Institute for the COVID-19 Genomics UK (COG-UK) Consortium | Harper VanSteenhouse, Yumi Kasai, David Gray, Carol Clugston, Anna Dominiczak and Alex Alderton, Roberto Amato, Jeffrey Barrett, Sonia Goncalves, Ewan Harrison, David K. Jackson, Ian Johnston, Dominic Kwiatkowski, Cordelia Langford, John Sillitoe on behalf of the Wellcome Sanger Institute COVID-19 Surveillance Team |
| EPI_ISL_2699678 | Lighthouse Lab in Milton Keynes | Wellcome Sanger Institute for the COVID-19 Genomics UK (COG-UK) Consortium | The Lighthouse Lab in Milton Keynes and Alex Alderton, Roberto Amato, Jeffrey Barrett, Sonia Goncalves, Ewan Harrison, David K. Jackson, Ian Johnston, Dominic Kwiatkowski, Cordelia Langford, John Sillitoe on behalf of the Wellcome Sanger Institute COVID-19 Surveillance Team |
| EPI_ISL_2699679 | Lighthouse Lab in Glasgow | Wellcome Sanger Institute for the COVID-19 Genomics UK (COG-UK) Consortium | Harper VanSteenhouse, Yumi Kasai, David Gray, Carol Clugston, Anna Dominiczak and Alex Alderton, Roberto Amato, Jeffrey Barrett, Sonia Goncalves, Ewan Harrison, David K. Jackson, Ian Johnston, Dominic Kwiatkowski, Cordelia Langford, John Sillitoe on behalf of the Wellcome Sanger Institute COVID-19 Surveillance Team |
| EPI_ISL_2699680, EPI_ISL_2699681, EPI_ISL_2699682 | Lighthouse Lab in Milton Keynes | Wellcome Sanger Institute for the COVID-19 Genomics UK (COG-UK) Consortium | The Lighthouse Lab in Milton Keynes and Alex Alderton, Roberto Amato, Jeffrey Barrett, Sonia Goncalves, Ewan Harrison, David K. Jackson, Ian Johnston, Dominic Kwiatkowski, Cordelia Langford, John Sillitoe on behalf of the Wellcome Sanger Institute COVID-19 Surveillance Team |
| EPI_ISL_2699683, EPI_ISL_2699684 | Lighthouse Lab in Glasgow | Wellcome Sanger Institute for the COVID-19 Genomics UK (COG-UK) Consortium | Harper VanSteenhouse, Yumi Kasai, David Gray, Carol Clugston, Anna Dominiczak and Alex Alderton, Roberto Amato, Jeffrey Barrett, Sonia Goncalves, Ewan Harrison, David K. Jackson, Ian Johnston, Dominic Kwiatkowski, Cordelia Langford, John Sillitoe on behalf of the Wellcome Sanger Institute COVID-19 Surveillance Team |
| EPI_ISL_2699685 | Lighthouse Laboratory Plymouth | Wellcome Sanger Institute for the COVID-19 Genomics UK (COG-UK) Consortium | Lighthouse Laboratory Plymouth and Alex Alderton, Roberto Amato, Jeffrey Barrett, Sonia Goncalves, Ewan Harrison, David K. Jackson, Ian Johnston, Dominic Kwiatkowski, Cordelia Langford, John Sillitoe on behalf of the Wellcome Sanger Institute COVID-19 Surveillance Team |
| EPI_ISL_2699686 | Lighthouse Lab in Milton Keynes | Wellcome Sanger Institute for the COVID-19 Genomics UK (COG-UK) Consortium | The Lighthouse Lab in Milton Keynes and Alex Alderton, Roberto Amato, Jeffrey Barrett, Sonia Goncalves, Ewan Harrison, David K. Jackson, Ian Johnston, Dominic Kwiatkowski, Cordelia Langford, John Sillitoe on behalf of the Wellcome Sanger Institute COVID-19 Surveillance Team |
| EPI_ISL_2699687 | Lighthouse Lab in Glasgow | Wellcome Sanger Institute for the COVID-19 Genomics UK (COG-UK) | Harper VanSteenhouse, Yumi Kasai, David Gray, Carol Clugston, Anna Dominiczak and Alex Alderton, Roberto Amato, Jeffrey Barrett, Sonia Goncalves, Ewan Harrison, David K. Jackson, Ian Johnston, Dominic Kwiatkowski, Cordelia Langford, |

| | | | |
|---|---|---|---|
| | | Consortium | John Sillitoe on behalf of the Wellcome Sanger Institute COVID-19 Surveillance Team |
| EPI_ISL_2699688, EPI_ISL_2699689 | Lighthouse Lab in Milton Keynes | Wellcome Sanger Institute for the COVID-19 Genomics UK (COG-UK) Consortium | The Lighthouse Lab in Milton Keynes and Alex Alderton, Roberto Amato, Jeffrey Barrett, Sonia Goncalves, Ewan Harrison, David K. Jackson, Ian Johnston, Dominic Kwiatkowski, Cordelia Langford, John Sillitoe on behalf of the Wellcome Sanger Institute COVID-19 Surveillance Team |
| EPI_ISL_2699690 | Lighthouse Laboratory Plymouth | Wellcome Sanger Institute for the COVID-19 Genomics UK (COG-UK) Consortium | Lighthouse Laboratory Plymouth and Alex Alderton, Roberto Amato, Jeffrey Barrett, Sonia Goncalves, Ewan Harrison, David K. Jackson, Ian Johnston, Dominic Kwiatkowski, Cordelia Langford, John Sillitoe on behalf of the Wellcome Sanger Institute COVID-19 Surveillance Team |
| EPI_ISL_2699691 | Lighthouse Lab in Milton Keynes | Wellcome Sanger Institute for the COVID-19 Genomics UK (COG-UK) Consortium | The Lighthouse Lab in Milton Keynes and Alex Alderton, Roberto Amato, Jeffrey Barrett, Sonia Goncalves, Ewan Harrison, David K. Jackson, Ian Johnston, Dominic Kwiatkowski, Cordelia Langford, John Sillitoe on behalf of the Wellcome Sanger Institute COVID-19 Surveillance Team |
| EPI_ISL_2699692 | Lighthouse Lab in Glasgow | Wellcome Sanger Institute for the COVID-19 Genomics UK (COG-UK) Consortium | Harper VanSteenhouse, Yumi Kasai, David Gray, Carol Clugston, Anna Dominiczak and Alex Alderton, Roberto Amato, Jeffrey Barrett, Sonia Goncalves, Ewan Harrison, David K. Jackson, Ian Johnston, Dominic Kwiatkowski, Cordelia Langford, John Sillitoe on behalf of the Wellcome Sanger Institute COVID-19 Surveillance Team |
| EPI_ISL_2699693, EPI_ISL_2699694 | Lighthouse Lab in Milton Keynes | Wellcome Sanger Institute for the COVID-19 Genomics UK (COG-UK) Consortium | The Lighthouse Lab in Milton Keynes and Alex Alderton, Roberto Amato, Jeffrey Barrett, Sonia Goncalves, Ewan Harrison, David K. Jackson, Ian Johnston, Dominic Kwiatkowski, Cordelia Langford, John Sillitoe on behalf of the Wellcome Sanger Institute COVID-19 Surveillance Team |
| EPI_ISL_2699695 | Lighthouse Lab in Glasgow | Wellcome Sanger Institute for the COVID-19 Genomics UK (COG-UK) Consortium | Harper VanSteenhouse, Yumi Kasai, David Gray, Carol Clugston, Anna Dominiczak and Alex Alderton, Roberto Amato, Jeffrey Barrett, Sonia Goncalves, Ewan Harrison, David K. Jackson, Ian Johnston, Dominic Kwiatkowski, Cordelia Langford, John Sillitoe on behalf of the Wellcome Sanger Institute COVID-19 Surveillance Team |
| EPI_ISL_2699696, EPI_ISL_2699697 | Lighthouse Lab in Milton Keynes | Wellcome Sanger Institute for the COVID-19 Genomics UK (COG-UK) Consortium | The Lighthouse Lab in Milton Keynes and Alex Alderton, Roberto Amato, Jeffrey Barrett, Sonia Goncalves, Ewan Harrison, David K. Jackson, Ian Johnston, Dominic Kwiatkowski, Cordelia Langford, John Sillitoe on behalf of the Wellcome Sanger Institute COVID-19 Surveillance Team |
| EPI_ISL_2699698 | Lighthouse Lab in Glasgow | Wellcome Sanger Institute for the COVID-19 Genomics UK (COG-UK) Consortium | Harper VanSteenhouse, Yumi Kasai, David Gray, Carol Clugston, Anna Dominiczak and Alex Alderton, Roberto Amato, Jeffrey Barrett, Sonia Goncalves, Ewan Harrison, David K. Jackson, Ian Johnston, Dominic Kwiatkowski, Cordelia Langford, John Sillitoe on behalf of the Wellcome Sanger Institute COVID-19 Surveillance Team |
| EPI_ISL_2699699 | Lighthouse Lab in Milton Keynes | Wellcome Sanger Institute for the COVID-19 Genomics UK (COG-UK) Consortium | The Lighthouse Lab in Milton Keynes and Alex Alderton, Roberto Amato, Jeffrey Barrett, Sonia Goncalves, Ewan Harrison, David K. Jackson, Ian Johnston, Dominic Kwiatkowski, Cordelia Langford, John Sillitoe on behalf of the Wellcome Sanger Institute COVID-19 Surveillance Team |
| EPI_ISL_2699700 | Lighthouse Lab in Glasgow | Wellcome Sanger Institute for the COVID-19 Genomics UK (COG-UK) Consortium | Harper VanSteenhouse, Yumi Kasai, David Gray, Carol Clugston, Anna Dominiczak and Alex Alderton, Roberto Amato, Jeffrey Barrett, Sonia Goncalves, Ewan Harrison, David K. Jackson, Ian Johnston, Dominic Kwiatkowski, Cordelia Langford, John Sillitoe on behalf of the Wellcome Sanger Institute COVID-19 Surveillance Team |
| EPI_ISL_2699701 | Lighthouse Lab in Milton Keynes | Wellcome Sanger Institute for the COVID-19 Genomics UK (COG-UK) Consortium | The Lighthouse Lab in Milton Keynes and Alex Alderton, Roberto Amato, Jeffrey Barrett, Sonia Goncalves, Ewan Harrison, David K. Jackson, Ian Johnston, Dominic Kwiatkowski, Cordelia Langford, John Sillitoe on behalf of the Wellcome Sanger Institute COVID-19 Surveillance Team |
| EPI_ISL_2699702 | Lighthouse Laboratory Plymouth | Wellcome Sanger Institute for the COVID-19 Genomics UK (COG-UK) Consortium | Lighthouse Laboratory Plymouth and Alex Alderton, Roberto Amato, Jeffrey Barrett, Sonia Goncalves, Ewan Harrison, David K. Jackson, Ian Johnston, Dominic Kwiatkowski, Cordelia Langford, John Sillitoe on behalf of the Wellcome Sanger Institute COVID-19 Surveillance Team |
| EPI_ISL_2699703 | Lighthouse Lab in Milton Keynes | Wellcome Sanger Institute for the COVID-19 Genomics UK (COG-UK) Consortium | The Lighthouse Lab in Milton Keynes and Alex Alderton, Roberto Amato, Jeffrey Barrett, Sonia Goncalves, Ewan Harrison, David K. Jackson, Ian Johnston, Dominic Kwiatkowski, Cordelia Langford, John Sillitoe on behalf of the Wellcome Sanger Institute COVID-19 Surveillance Team |
| EPI_ISL_2699704, EPI_ISL_2699705, EPI_ISL_2699706, EPI_ISL_2699707, EPI_ISL_2699709 | Lighthouse Lab in Glasgow | Wellcome Sanger Institute for the COVID-19 Genomics UK (COG-UK) Consortium | Harper VanSteenhouse, Yumi Kasai, David Gray, Carol Clugston, Anna Dominiczak and Alex Alderton, Roberto Amato, Jeffrey Barrett, Sonia Goncalves, Ewan Harrison, David K. Jackson, Ian Johnston, Dominic Kwiatkowski, Cordelia Langford, John Sillitoe on behalf of the Wellcome Sanger Institute COVID-19 Surveillance Team |
| EPI_ISL_2699710, EPI_ISL_2699711, EPI_ISL_2699712 | Lighthouse Lab in Milton Keynes | Wellcome Sanger Institute for the COVID-19 Genomics UK (COG-UK) Consortium | The Lighthouse Lab in Milton Keynes and Alex Alderton, Roberto Amato, Jeffrey Barrett, Sonia Goncalves, Ewan Harrison, David K. Jackson, Ian Johnston, Dominic Kwiatkowski, Cordelia Langford, John Sillitoe on behalf of the Wellcome Sanger Institute COVID-19 Surveillance Team |
| EPI_ISL_2699713 | Lighthouse Lab in Glasgow | Wellcome Sanger Institute for the COVID-19 Genomics UK (COG-UK) Consortium | Harper VanSteenhouse, Yumi Kasai, David Gray, Carol Clugston, Anna Dominiczak and Alex Alderton, Roberto Amato, Jeffrey Barrett, Sonia Goncalves, Ewan Harrison, David K. Jackson, Ian Johnston, Dominic Kwiatkowski, Cordelia Langford, John Sillitoe on behalf of the Wellcome Sanger Institute COVID-19 Surveillance Team |
| EPI_ISL_2699714 | Lighthouse Lab in Milton Keynes | Wellcome Sanger Institute for the COVID-19 Genomics UK (COG-UK) Consortium | The Lighthouse Lab in Milton Keynes and Alex Alderton, Roberto Amato, Jeffrey Barrett, Sonia Goncalves, Ewan Harrison, David K. Jackson, Ian Johnston, Dominic Kwiatkowski, Cordelia Langford, John Sillitoe on behalf of the Wellcome Sanger Institute COVID-19 Surveillance Team |
| EPI_ISL_2699715 | Lighthouse Lab in Glasgow | Wellcome Sanger Institute for the COVID-19 Genomics UK (COG-UK) Consortium | Harper VanSteenhouse, Yumi Kasai, David Gray, Carol Clugston, Anna Dominiczak and Alex Alderton, Roberto Amato, Jeffrey Barrett, Sonia Goncalves, Ewan Harrison, David K. Jackson, Ian Johnston, Dominic Kwiatkowski, Cordelia Langford, John Sillitoe on behalf of the Wellcome Sanger Institute COVID-19 Surveillance Team |
| EPI_ISL_2699716, EPI_ISL_2699717 | Lighthouse Lab in Milton Keynes | Wellcome Sanger Institute for the COVID-19 Genomics UK (COG-UK) Consortium | The Lighthouse Lab in Milton Keynes and Alex Alderton, Roberto Amato, Jeffrey Barrett, Sonia Goncalves, Ewan Harrison, David K. Jackson, Ian Johnston, Dominic Kwiatkowski, Cordelia Langford, John Sillitoe on behalf of the Wellcome Sanger Institute COVID-19 Surveillance Team |
| EPI_ISL_2699718 | Lighthouse Lab in Glasgow | Wellcome Sanger Institute for the COVID-19 Genomics UK (COG-UK) Consortium | Harper VanSteenhouse, Yumi Kasai, David Gray, Carol Clugston, Anna Dominiczak and Alex Alderton, Roberto Amato, Jeffrey Barrett, Sonia Goncalves, Ewan Harrison, David K. Jackson, Ian Johnston, Dominic Kwiatkowski, Cordelia Langford, John Sillitoe on behalf of the Wellcome Sanger Institute COVID-19 Surveillance Team |
| EPI_ISL_2699720, EPI_ISL_2699721, EPI_ISL_2699722, EPI_ISL_2699723, EPI_ISL_2699724 | Lighthouse Lab in Milton Keynes | Wellcome Sanger Institute for the COVID-19 Genomics UK (COG-UK) Consortium | The Lighthouse Lab in Milton Keynes and Alex Alderton, Roberto Amato, Jeffrey Barrett, Sonia Goncalves, Ewan Harrison, David K. Jackson, Ian Johnston, Dominic Kwiatkowski, Cordelia Langford, John Sillitoe on behalf of the Wellcome Sanger Institute COVID-19 Surveillance Team |
| EPI_ISL_2699725 | Lighthouse Lab in Glasgow | Wellcome Sanger Institute for the COVID-19 Genomics UK (COG-UK) Consortium | Harper VanSteenhouse, Yumi Kasai, David Gray, Carol Clugston, Anna Dominiczak and Alex Alderton, Roberto Amato, Jeffrey Barrett, Sonia Goncalves, Ewan Harrison, David K. Jackson, Ian Johnston, Dominic Kwiatkowski, Cordelia Langford, John Sillitoe on behalf of the Wellcome Sanger Institute COVID-19 Surveillance Team |
| EPI_ISL_2699726 | Lighthouse Lab in Milton Keynes | Wellcome Sanger Institute for the COVID-19 Genomics UK (COG-UK) Consortium | The Lighthouse Lab in Milton Keynes and Alex Alderton, Roberto Amato, Jeffrey Barrett, Sonia Goncalves, Ewan Harrison, David K. Jackson, Ian Johnston, Dominic Kwiatkowski, Cordelia Langford, John Sillitoe on behalf of the Wellcome Sanger Institute COVID-19 Surveillance Team |
| EPI_ISL_2699727, EPI_ISL_2699728 | Lighthouse Lab in Glasgow | Wellcome Sanger Institute for the COVID-19 Genomics UK (COG-UK) Consortium | Harper VanSteenhouse, Yumi Kasai, David Gray, Carol Clugston, Anna Dominiczak and Alex Alderton, Roberto Amato, Jeffrey Barrett, Sonia Goncalves, Ewan Harrison, David K. Jackson, Ian Johnston, Dominic Kwiatkowski, Cordelia Langford, John Sillitoe on behalf of the Wellcome Sanger Institute COVID-19 Surveillance Team |
| EPI_ISL_2699729, EPI_ISL_2699730 | Lighthouse Lab in Milton Keynes | Wellcome Sanger Institute for the COVID-19 Genomics UK (COG-UK) Consortium | The Lighthouse Lab in Milton Keynes and Alex Alderton, Roberto Amato, Jeffrey Barrett, Sonia Goncalves, Ewan Harrison, David K. Jackson, Ian Johnston, Dominic Kwiatkowski, Cordelia Langford, John Sillitoe on behalf of the Wellcome Sanger Institute COVID-19 Surveillance Team |
| EPI_ISL_2699731 | Lighthouse Lab in Glasgow | Wellcome Sanger Institute for the COVID-19 Genomics UK (COG-UK) Consortium | Harper VanSteenhouse, Yumi Kasai, David Gray, Carol Clugston, Anna Dominiczak and Alex Alderton, Roberto Amato, Jeffrey Barrett, Sonia Goncalves, Ewan Harrison, David K. Jackson, Ian Johnston, Dominic Kwiatkowski, Cordelia Langford, John Sillitoe on behalf of the Wellcome Sanger Institute COVID-19 Surveillance Team |
| EPI_ISL_2699732 | Lighthouse Lab in Milton Keynes | Wellcome Sanger Institute for the COVID-19 Genomics UK (COG-UK) Consortium | The Lighthouse Lab in Milton Keynes and Alex Alderton, Roberto Amato, Jeffrey Barrett, Sonia Goncalves, Ewan Harrison, David K. Jackson, Ian Johnston, Dominic Kwiatkowski, Cordelia Langford, John Sillitoe on behalf of the Wellcome Sanger Institute COVID-19 Surveillance Team |
| EPI_ISL_2699733 | Lighthouse Lab in Glasgow | Wellcome Sanger Institute for the COVID-19 Genomics UK (COG-UK) Consortium | Harper VanSteenhouse, Yumi Kasai, David Gray, Carol Clugston, Anna Dominiczak and Alex Alderton, Roberto Amato, Jeffrey Barrett, Sonia Goncalves, Ewan Harrison, David K. Jackson, Ian Johnston, Dominic Kwiatkowski, Cordelia Langford, John Sillitoe on behalf of the Wellcome Sanger Institute COVID-19 Surveillance Team |
| EPI_ISL_2699734, EPI_ISL_2699735, EPI_ISL_2699736, EPI_ISL_2699737, EPI_ISL_2699738, EPI_ISL_2699740 | Lighthouse Lab in Milton Keynes | Wellcome Sanger Institute for the COVID-19 Genomics UK (COG-UK) Consortium | The Lighthouse Lab in Milton Keynes and Alex Alderton, Roberto Amato, Jeffrey Barrett, Sonia Goncalves, Ewan Harrison, David K. Jackson, Ian Johnston, Dominic Kwiatkowski, Cordelia Langford, John Sillitoe on behalf of the Wellcome Sanger Institute COVID-19 Surveillance Team |
| EPI_ISL_2699741 | Lighthouse Lab in Glasgow | Wellcome Sanger Institute for the COVID-19 Genomics UK (COG-UK) Consortium | Harper VanSteenhouse, Yumi Kasai, David Gray, Carol Clugston, Anna Dominiczak and Alex Alderton, Roberto Amato, Jeffrey Barrett, Sonia Goncalves, Ewan Harrison, David K. Jackson, Ian Johnston, Dominic Kwiatkowski, Cordelia Langford, John Sillitoe on behalf of the Wellcome Sanger Institute COVID-19 Surveillance Team |
| EPI_ISL_2699742, EPI_ISL_2699744 | Lighthouse Lab in Milton Keynes | Wellcome Sanger Institute for the COVID-19 Genomics UK (COG-UK) Consortium | The Lighthouse Lab in Milton Keynes and Alex Alderton, Roberto Amato, Jeffrey Barrett, Sonia Goncalves, Ewan Harrison, David K. Jackson, Ian Johnston, Dominic Kwiatkowski, Cordelia Langford, John Sillitoe on behalf of the Wellcome Sanger Institute COVID-19 Surveillance Team |
| EPI_ISL_2699745 | Lighthouse Lab in Glasgow | Wellcome Sanger Institute for the COVID-19 Genomics UK (COG-UK) Consortium | Harper VanSteenhouse, Yumi Kasai, David Gray, Carol Clugston, Anna Dominiczak and Alex Alderton, Roberto Amato, Jeffrey Barrett, Sonia Goncalves, Ewan Harrison, David K. Jackson, Ian Johnston, Dominic Kwiatkowski, Cordelia Langford, John Sillitoe on behalf of the Wellcome Sanger Institute COVID-19 Surveillance Team |
| EPI_ISL_2699746 | Lighthouse Laboratory Plymouth | Wellcome Sanger Institute for the COVID-19 Genomics UK (COG-UK) Consortium | Lighthouse Laboratory Plymouth and Alex Alderton, Roberto Amato, Jeffrey Barrett, Sonia Goncalves, Ewan Harrison, David K. Jackson, Ian Johnston, Dominic Kwiatkowski, Cordelia Langford, John Sillitoe on behalf of the Wellcome Sanger Institute COVID-19 Surveillance Team |
| EPI_ISL_2699747 | Lighthouse Lab in Glasgow | Wellcome Sanger Institute for the COVID-19 Genomics UK (COG-UK) Consortium | Harper VanSteenhouse, Yumi Kasai, David Gray, Carol Clugston, Anna Dominiczak and Alex Alderton, Roberto Amato, Jeffrey Barrett, Sonia Goncalves, Ewan Harrison, David K. Jackson, Ian Johnston, Dominic Kwiatkowski, Cordelia Langford, John Sillitoe on behalf of the Wellcome Sanger Institute COVID-19 Surveillance Team |
| EPI_ISL_2699748, EPI_ISL_2699749 | Lighthouse Lab in Milton Keynes | Wellcome Sanger Institute for the COVID-19 Genomics UK (COG-UK) Consortium | The Lighthouse Lab in Milton Keynes and Alex Alderton, Roberto Amato, Jeffrey Barrett, Sonia Goncalves, Ewan Harrison, David K. Jackson, Ian Johnston, Dominic Kwiatkowski, Cordelia Langford, John Sillitoe on behalf of the Wellcome Sanger Institute COVID-19 Surveillance Team |
| EPI_ISL_2699750 | Lighthouse Laboratory Plymouth | Wellcome Sanger Institute for the COVID-19 Genomics UK (COG-UK) Consortium | Lighthouse Laboratory Plymouth and Alex Alderton, Roberto Amato, Jeffrey Barrett, Sonia Goncalves, Ewan Harrison, David K. Jackson, Ian Johnston, Dominic Kwiatkowski, Cordelia Langford, John Sillitoe on behalf of the Wellcome Sanger Institute COVID-19 Surveillance Team |
| EPI_ISL_2699751 | Lighthouse Lab in Glasgow | Wellcome Sanger Institute for the COVID-19 Genomics UK (COG-UK) Consortium | Harper VanSteenhouse, Yumi Kasai, David Gray, Carol Clugston, Anna Dominiczak and Alex Alderton, Roberto Amato, Jeffrey Barrett, Sonia Goncalves, Ewan Harrison, David K. Jackson, Ian Johnston, Dominic Kwiatkowski, Cordelia Langford, John Sillitoe on behalf of the Wellcome Sanger Institute COVID-19 Surveillance Team |
| EPI_ISL_2699752 | Lighthouse Lab in Milton Keynes | Wellcome Sanger Institute for the COVID-19 Genomics UK (COG-UK) Consortium | The Lighthouse Lab in Milton Keynes and Alex Alderton, Roberto Amato, Jeffrey Barrett, Sonia Goncalves, Ewan Harrison, David K. Jackson, Ian Johnston, Dominic Kwiatkowski, Cordelia Langford, John Sillitoe on behalf of the Wellcome Sanger Institute COVID-19 Surveillance Team |
| EPI_ISL_2699753, EPI_ISL_2699754 | Lighthouse Lab in Glasgow | Wellcome Sanger Institute for the COVID-19 Genomics UK (COG-UK) Consortium | Harper VanSteenhouse, Yumi Kasai, David Gray, Carol Clugston, Anna Dominiczak and Alex Alderton, Roberto Amato, Jeffrey Barrett, Sonia Goncalves, Ewan Harrison, David K. Jackson, Ian Johnston, Dominic Kwiatkowski, Cordelia Langford, John Sillitoe on behalf of the Wellcome Sanger Institute COVID-19 Surveillance Team |
| EPI_ISL_2699755, EPI_ISL_2699756, EPI_ISL_2699757, | Lighthouse Lab in Milton Keynes | Wellcome Sanger Institute for the COVID-19 Genomics UK (COG-UK) Consortium | The Lighthouse Lab in Milton Keynes and Alex Alderton, Roberto Amato, Jeffrey Barrett, Sonia Goncalves, Ewan Harrison, David K. Jackson, Ian Johnston, Dominic Kwiatkowski, Cordelia Langford, John Sillitoe on behalf of the Wellcome Sanger Institute COVID-19 Surveillance Team |

| | | | |
|---|---|---|---|
| EPI_ISL_2699758, EPI_ISL_2699759, EPI_ISL_2699760 | | | |
| EPI_ISL_2699761 | Lighthouse Lab in Glasgow | Wellcome Sanger Institute for the COVID-19 Genomics UK (COG-UK) Consortium | Harper VanSteenhouse, Yumi Kasai, David Gray, Carol Clugston, Anna Dominiczak and Alex Alderton, Roberto Amato, Jeffrey Barrett, Sonia Goncalves, Ewan Harrison, David K. Jackson, Ian Johnston, Dominic Kwiatkowski, Cordelia Langford, John Sillitoe on behalf of the Wellcome Sanger Institute COVID-19 Surveillance Team |
| EPI_ISL_2699762 | Lighthouse Laboratory Plymouth | Wellcome Sanger Institute for the COVID-19 Genomics UK (COG-UK) Consortium | Lighthouse Laboratory Plymouth and Alex Alderton, Roberto Amato, Jeffrey Barrett, Sonia Goncalves, Ewan Harrison, David K. Jackson, Ian Johnston, Dominic Kwiatkowski, Cordelia Langford, John Sillitoe on behalf of the Wellcome Sanger Institute COVID-19 Surveillance Team |
| EPI_ISL_2699763 | Lighthouse Lab in Milton Keynes | Wellcome Sanger Institute for the COVID-19 Genomics UK (COG-UK) Consortium | The Lighthouse Lab in Milton Keynes and Alex Alderton, Roberto Amato, Jeffrey Barrett, Sonia Goncalves, Ewan Harrison, David K. Jackson, Ian Johnston, Dominic Kwiatkowski, Cordelia Langford, John Sillitoe on behalf of the Wellcome Sanger Institute COVID-19 Surveillance Team |
| EPI_ISL_2699764 | Lighthouse Lab in Glasgow | Wellcome Sanger Institute for the COVID-19 Genomics UK (COG-UK) Consortium | Harper VanSteenhouse, Yumi Kasai, David Gray, Carol Clugston, Anna Dominiczak and Alex Alderton, Roberto Amato, Jeffrey Barrett, Sonia Goncalves, Ewan Harrison, David K. Jackson, Ian Johnston, Dominic Kwiatkowski, Cordelia Langford, John Sillitoe on behalf of the Wellcome Sanger Institute COVID-19 Surveillance Team |
| EPI_ISL_2699765, EPI_ISL_2699766, EPI_ISL_2699767 | Lighthouse Lab in Milton Keynes | Wellcome Sanger Institute for the COVID-19 Genomics UK (COG-UK) Consortium | The Lighthouse Lab in Milton Keynes and Alex Alderton, Roberto Amato, Jeffrey Barrett, Sonia Goncalves, Ewan Harrison, David K. Jackson, Ian Johnston, Dominic Kwiatkowski, Cordelia Langford, John Sillitoe on behalf of the Wellcome Sanger Institute COVID-19 Surveillance Team |
| EPI_ISL_2699768, EPI_ISL_2699769, EPI_ISL_2699770 | Lighthouse Lab in Glasgow | Wellcome Sanger Institute for the COVID-19 Genomics UK (COG-UK) Consortium | Harper VanSteenhouse, Yumi Kasai, David Gray, Carol Clugston, Anna Dominiczak and Alex Alderton, Roberto Amato, Jeffrey Barrett, Sonia Goncalves, Ewan Harrison, David K. Jackson, Ian Johnston, Dominic Kwiatkowski, Cordelia Langford, John Sillitoe on behalf of the Wellcome Sanger Institute COVID-19 Surveillance Team |
| EPI_ISL_2699771 | Lighthouse Lab in Milton Keynes | Wellcome Sanger Institute for the COVID-19 Genomics UK (COG-UK) Consortium | The Lighthouse Lab in Milton Keynes and Alex Alderton, Roberto Amato, Jeffrey Barrett, Sonia Goncalves, Ewan Harrison, David K. Jackson, Ian Johnston, Dominic Kwiatkowski, Cordelia Langford, John Sillitoe on behalf of the Wellcome Sanger Institute COVID-19 Surveillance Team |
| EPI_ISL_2699772 | Lighthouse Lab in Glasgow | Wellcome Sanger Institute for the COVID-19 Genomics UK (COG-UK) Consortium | Harper VanSteenhouse, Yumi Kasai, David Gray, Carol Clugston, Anna Dominiczak and Alex Alderton, Roberto Amato, Jeffrey Barrett, Sonia Goncalves, Ewan Harrison, David K. Jackson, Ian Johnston, Dominic Kwiatkowski, Cordelia Langford, John Sillitoe on behalf of the Wellcome Sanger Institute COVID-19 Surveillance Team |
| EPI_ISL_2699773, EPI_ISL_2699774, EPI_ISL_2699775 | Lighthouse Lab in Milton Keynes | Wellcome Sanger Institute for the COVID-19 Genomics UK (COG-UK) Consortium | The Lighthouse Lab in Milton Keynes and Alex Alderton, Roberto Amato, Jeffrey Barrett, Sonia Goncalves, Ewan Harrison, David K. Jackson, Ian Johnston, Dominic Kwiatkowski, Cordelia Langford, John Sillitoe on behalf of the Wellcome Sanger Institute COVID-19 Surveillance Team |
| EPI_ISL_2699776, EPI_ISL_2699777 | Lighthouse Lab in Glasgow | Wellcome Sanger Institute for the COVID-19 Genomics UK (COG-UK) Consortium | Harper VanSteenhouse, Yumi Kasai, David Gray, Carol Clugston, Anna Dominiczak and Alex Alderton, Roberto Amato, Jeffrey Barrett, Sonia Goncalves, Ewan Harrison, David K. Jackson, Ian Johnston, Dominic Kwiatkowski, Cordelia Langford, John Sillitoe on behalf of the Wellcome Sanger Institute COVID-19 Surveillance Team |
| EPI_ISL_2699778, EPI_ISL_2699779 | Lighthouse Lab in Milton Keynes | Wellcome Sanger Institute for the COVID-19 Genomics UK (COG-UK) Consortium | The Lighthouse Lab in Milton Keynes and Alex Alderton, Roberto Amato, Jeffrey Barrett, Sonia Goncalves, Ewan Harrison, David K. Jackson, Ian Johnston, Dominic Kwiatkowski, Cordelia Langford, John Sillitoe on behalf of the Wellcome Sanger Institute COVID-19 Surveillance Team |
| EPI_ISL_2699780, EPI_ISL_2699781 | Lighthouse Lab in Glasgow | Wellcome Sanger Institute for the COVID-19 Genomics UK (COG-UK) Consortium | Harper VanSteenhouse, Yumi Kasai, David Gray, Carol Clugston, Anna Dominiczak and Alex Alderton, Roberto Amato, Jeffrey Barrett, Sonia Goncalves, Ewan Harrison, David K. Jackson, Ian Johnston, Dominic Kwiatkowski, Cordelia Langford, John Sillitoe on behalf of the Wellcome Sanger Institute COVID-19 Surveillance Team |
| EPI_ISL_2699782, EPI_ISL_2699783, EPI_ISL_2699785 | Lighthouse Lab in Milton Keynes | Wellcome Sanger Institute for the COVID-19 Genomics UK (COG-UK) Consortium | The Lighthouse Lab in Milton Keynes and Alex Alderton, Roberto Amato, Jeffrey Barrett, Sonia Goncalves, Ewan Harrison, David K. Jackson, Ian Johnston, Dominic Kwiatkowski, Cordelia Langford, John Sillitoe on behalf of the Wellcome Sanger Institute COVID-19 Surveillance Team |
| EPI_ISL_2699786, EPI_ISL_2699788, EPI_ISL_2699789, EPI_ISL_2699790, EPI_ISL_2699791 | Lighthouse Lab in Glasgow | Wellcome Sanger Institute for the COVID-19 Genomics UK (COG-UK) Consortium | Harper VanSteenhouse, Yumi Kasai, David Gray, Carol Clugston, Anna Dominiczak and Alex Alderton, Roberto Amato, Jeffrey Barrett, Sonia Goncalves, Ewan Harrison, David K. Jackson, Ian Johnston, Dominic Kwiatkowski, Cordelia Langford, John Sillitoe on behalf of the Wellcome Sanger Institute COVID-19 Surveillance Team |
| EPI_ISL_2699792 | Lighthouse Laboratory Plymouth | Wellcome Sanger Institute for the COVID-19 Genomics UK (COG-UK) Consortium | Lighthouse Laboratory Plymouth and Alex Alderton, Roberto Amato, Jeffrey Barrett, Sonia Goncalves, Ewan Harrison, David K. Jackson, Ian Johnston, Dominic Kwiatkowski, Cordelia Langford, John Sillitoe on behalf of the Wellcome Sanger Institute COVID-19 Surveillance Team |
| EPI_ISL_2699793 | Lighthouse Lab in Glasgow | Wellcome Sanger Institute for the COVID-19 Genomics UK (COG-UK) Consortium | Harper VanSteenhouse, Yumi Kasai, David Gray, Carol Clugston, Anna Dominiczak and Alex Alderton, Roberto Amato, Jeffrey Barrett, Sonia Goncalves, Ewan Harrison, David K. Jackson, Ian Johnston, Dominic Kwiatkowski, Cordelia Langford, John Sillitoe on behalf of the Wellcome Sanger Institute COVID-19 Surveillance Team |
| EPI_ISL_2699794 | Lighthouse Laboratory Plymouth | Wellcome Sanger Institute for the COVID-19 Genomics UK (COG-UK) Consortium | Lighthouse Laboratory Plymouth and Alex Alderton, Roberto Amato, Jeffrey Barrett, Sonia Goncalves, Ewan Harrison, David K. Jackson, Ian Johnston, Dominic Kwiatkowski, Cordelia Langford, John Sillitoe on behalf of the Wellcome Sanger Institute COVID-19 Surveillance Team |
| EPI_ISL_2699795 | Lighthouse Lab in Glasgow | Wellcome Sanger Institute for the COVID-19 Genomics UK (COG-UK) Consortium | Harper VanSteenhouse, Yumi Kasai, David Gray, Carol Clugston, Anna Dominiczak and Alex Alderton, Roberto Amato, Jeffrey Barrett, Sonia Goncalves, Ewan Harrison, David K. Jackson, Ian Johnston, Dominic Kwiatkowski, Cordelia Langford, John Sillitoe on behalf of the Wellcome Sanger Institute COVID-19 Surveillance Team |
| EPI_ISL_2699796 | Lighthouse Lab in Milton Keynes | Wellcome Sanger Institute for the COVID-19 Genomics UK (COG-UK) Consortium | The Lighthouse Lab in Milton Keynes and Alex Alderton, Roberto Amato, Jeffrey Barrett, Sonia Goncalves, Ewan Harrison, David K. Jackson, Ian Johnston, Dominic Kwiatkowski, Cordelia Langford, John Sillitoe on behalf of the Wellcome Sanger Institute COVID-19 Surveillance Team |
| EPI_ISL_2699797, EPI_ISL_2699798 | Lighthouse Lab in Glasgow | Wellcome Sanger Institute for the COVID-19 Genomics UK (COG-UK) Consortium | Harper VanSteenhouse, Yumi Kasai, David Gray, Carol Clugston, Anna Dominiczak and Alex Alderton, Roberto Amato, Jeffrey Barrett, Sonia Goncalves, Ewan Harrison, David K. Jackson, Ian Johnston, Dominic Kwiatkowski, Cordelia Langford, John Sillitoe on behalf of the Wellcome Sanger Institute COVID-19 Surveillance Team |
| EPI_ISL_2699799, EPI_ISL_2699800, EPI_ISL_2699801, EPI_ISL_2699802, EPI_ISL_2699803 | Lighthouse Lab in Milton Keynes | Wellcome Sanger Institute for the COVID-19 Genomics UK (COG-UK) Consortium | The Lighthouse Lab in Milton Keynes and Alex Alderton, Roberto Amato, Jeffrey Barrett, Sonia Goncalves, Ewan Harrison, David K. Jackson, Ian Johnston, Dominic Kwiatkowski, Cordelia Langford, John Sillitoe on behalf of the Wellcome Sanger Institute COVID-19 Surveillance Team |
| EPI_ISL_2699804 | Lighthouse Lab in Glasgow | Wellcome Sanger Institute for the COVID-19 Genomics UK (COG-UK) Consortium | Harper VanSteenhouse, Yumi Kasai, David Gray, Carol Clugston, Anna Dominiczak and Alex Alderton, Roberto Amato, Jeffrey Barrett, Sonia Goncalves, Ewan Harrison, David K. Jackson, Ian Johnston, Dominic Kwiatkowski, Cordelia Langford, John Sillitoe on behalf of the Wellcome Sanger Institute COVID-19 Surveillance Team |
| EPI_ISL_2699805 | Lighthouse Lab in Milton Keynes | Wellcome Sanger Institute for the COVID-19 Genomics UK (COG-UK) Consortium | The Lighthouse Lab in Milton Keynes and Alex Alderton, Roberto Amato, Jeffrey Barrett, Sonia Goncalves, Ewan Harrison, David K. Jackson, Ian Johnston, Dominic Kwiatkowski, Cordelia Langford, John Sillitoe on behalf of the Wellcome Sanger Institute COVID-19 Surveillance Team |
| EPI_ISL_2699806 | Lighthouse Laboratory Plymouth | Wellcome Sanger Institute for the COVID-19 Genomics UK (COG-UK) Consortium | Lighthouse Laboratory Plymouth and Alex Alderton, Roberto Amato, Jeffrey Barrett, Sonia Goncalves, Ewan Harrison, David K. Jackson, Ian Johnston, Dominic Kwiatkowski, Cordelia Langford, John Sillitoe on behalf of the Wellcome Sanger Institute COVID-19 Surveillance Team |
| EPI_ISL_2699807, EPI_ISL_2699808 | Lighthouse Lab in Glasgow | Wellcome Sanger Institute for the COVID-19 Genomics UK (COG-UK) Consortium | Harper VanSteenhouse, Yumi Kasai, David Gray, Carol Clugston, Anna Dominiczak and Alex Alderton, Roberto Amato, Jeffrey Barrett, Sonia Goncalves, Ewan Harrison, David K. Jackson, Ian Johnston, Dominic Kwiatkowski, Cordelia Langford, John Sillitoe on behalf of the Wellcome Sanger Institute COVID-19 Surveillance Team |
| EPI_ISL_2699809, EPI_ISL_2699810 | Lighthouse Lab in Milton Keynes | Wellcome Sanger Institute for the COVID-19 Genomics UK (COG-UK) Consortium | The Lighthouse Lab in Milton Keynes and Alex Alderton, Roberto Amato, Jeffrey Barrett, Sonia Goncalves, Ewan Harrison, David K. Jackson, Ian Johnston, Dominic Kwiatkowski, Cordelia Langford, John Sillitoe on behalf of the Wellcome Sanger Institute COVID-19 Surveillance Team |
| EPI_ISL_2699811, EPI_ISL_2699812, EPI_ISL_2699813 | Lighthouse Lab in Glasgow | Wellcome Sanger Institute for the COVID-19 Genomics UK (COG-UK) Consortium | Harper VanSteenhouse, Yumi Kasai, David Gray, Carol Clugston, Anna Dominiczak and Alex Alderton, Roberto Amato, Jeffrey Barrett, Sonia Goncalves, Ewan Harrison, David K. Jackson, Ian Johnston, Dominic Kwiatkowski, Cordelia Langford, John Sillitoe on behalf of the Wellcome Sanger Institute COVID-19 Surveillance Team |
| EPI_ISL_2699814, EPI_ISL_2699815, EPI_ISL_2699816 | Lighthouse Lab in Milton Keynes | Wellcome Sanger Institute for the COVID-19 Genomics UK (COG-UK) Consortium | The Lighthouse Lab in Milton Keynes and Alex Alderton, Roberto Amato, Jeffrey Barrett, Sonia Goncalves, Ewan Harrison, David K. Jackson, Ian Johnston, Dominic Kwiatkowski, Cordelia Langford, John Sillitoe on behalf of the Wellcome Sanger Institute COVID-19 Surveillance Team |
| EPI_ISL_2699817 | Lighthouse Lab in Glasgow | Wellcome Sanger Institute for the COVID-19 Genomics UK (COG-UK) Consortium | Harper VanSteenhouse, Yumi Kasai, David Gray, Carol Clugston, Anna Dominiczak and Alex Alderton, Roberto Amato, Jeffrey Barrett, Sonia Goncalves, Ewan Harrison, David K. Jackson, Ian Johnston, Dominic Kwiatkowski, Cordelia Langford, John Sillitoe on behalf of the Wellcome Sanger Institute COVID-19 Surveillance Team |
| EPI_ISL_2699818, EPI_ISL_2699819 | Lighthouse Lab in Milton Keynes | Wellcome Sanger Institute for the COVID-19 Genomics UK (COG-UK) Consortium | The Lighthouse Lab in Milton Keynes and Alex Alderton, Roberto Amato, Jeffrey Barrett, Sonia Goncalves, Ewan Harrison, David K. Jackson, Ian Johnston, Dominic Kwiatkowski, Cordelia Langford, John Sillitoe on behalf of the Wellcome Sanger Institute COVID-19 Surveillance Team |
| EPI_ISL_2699820 | Lighthouse Lab in Glasgow | Wellcome Sanger Institute for the COVID-19 Genomics UK (COG-UK) Consortium | Harper VanSteenhouse, Yumi Kasai, David Gray, Carol Clugston, Anna Dominiczak and Alex Alderton, Roberto Amato, Jeffrey Barrett, Sonia Goncalves, Ewan Harrison, David K. Jackson, Ian Johnston, Dominic Kwiatkowski, Cordelia Langford, John Sillitoe on behalf of the Wellcome Sanger Institute COVID-19 Surveillance Team |
| EPI_ISL_2699821 | Lighthouse Laboratory Plymouth | Wellcome Sanger Institute for the COVID-19 Genomics UK (COG-UK) Consortium | Lighthouse Laboratory Plymouth and Alex Alderton, Roberto Amato, Jeffrey Barrett, Sonia Goncalves, Ewan Harrison, David K. Jackson, Ian Johnston, Dominic Kwiatkowski, Cordelia Langford, John Sillitoe on behalf of the Wellcome Sanger Institute COVID-19 Surveillance Team |
| EPI_ISL_2699822, EPI_ISL_2699823 | Lighthouse Lab in Milton Keynes | Wellcome Sanger Institute for the COVID-19 Genomics UK (COG-UK) Consortium | The Lighthouse Lab in Milton Keynes and Alex Alderton, Roberto Amato, Jeffrey Barrett, Sonia Goncalves, Ewan Harrison, David K. Jackson, Ian Johnston, Dominic Kwiatkowski, Cordelia Langford, John Sillitoe on behalf of the Wellcome Sanger Institute COVID-19 Surveillance Team |
| EPI_ISL_2699824 | Lighthouse Lab in Glasgow | Wellcome Sanger Institute for the COVID-19 Genomics UK (COG-UK) Consortium | Harper VanSteenhouse, Yumi Kasai, David Gray, Carol Clugston, Anna Dominiczak and Alex Alderton, Roberto Amato, Jeffrey Barrett, Sonia Goncalves, Ewan Harrison, David K. Jackson, Ian Johnston, Dominic Kwiatkowski, Cordelia Langford, John Sillitoe on behalf of the Wellcome Sanger Institute COVID-19 Surveillance Team |
| EPI_ISL_2699825, EPI_ISL_2699826 | Lighthouse Laboratory Plymouth | Wellcome Sanger Institute for the COVID-19 Genomics UK (COG-UK) Consortium | Lighthouse Laboratory Plymouth and Alex Alderton, Roberto Amato, Jeffrey Barrett, Sonia Goncalves, Ewan Harrison, David K. Jackson, Ian Johnston, Dominic Kwiatkowski, Cordelia Langford, John Sillitoe on behalf of the Wellcome Sanger Institute COVID-19 Surveillance Team |
| EPI_ISL_2699827, EPI_ISL_2699828 | Lighthouse Lab in Milton Keynes | Wellcome Sanger Institute for the COVID-19 Genomics UK (COG-UK) Consortium | The Lighthouse Lab in Milton Keynes and Alex Alderton, Roberto Amato, Jeffrey Barrett, Sonia Goncalves, Ewan Harrison, David K. Jackson, Ian Johnston, Dominic Kwiatkowski, Cordelia Langford, John Sillitoe on behalf of the Wellcome Sanger Institute COVID-19 Surveillance Team |
| EPI_ISL_2699829 | Lighthouse Lab in Glasgow | Wellcome Sanger Institute for the COVID-19 Genomics UK (COG-UK) Consortium | Harper VanSteenhouse, Yumi Kasai, David Gray, Carol Clugston, Anna Dominiczak and Alex Alderton, Roberto Amato, Jeffrey Barrett, Sonia Goncalves, Ewan Harrison, David K. Jackson, Ian Johnston, Dominic Kwiatkowski, Cordelia Langford, John Sillitoe on behalf of the Wellcome Sanger Institute COVID-19 Surveillance Team |
| EPI_ISL_2699830 | Lighthouse Lab in Milton Keynes | Wellcome Sanger Institute for the COVID-19 Genomics UK (COG-UK) Consortium | The Lighthouse Lab in Milton Keynes and Alex Alderton, Roberto Amato, Jeffrey Barrett, Sonia Goncalves, Ewan Harrison, David K. Jackson, Ian Johnston, Dominic Kwiatkowski, Cordelia Langford, John Sillitoe on behalf of the Wellcome Sanger Institute COVID-19 Surveillance Team |
| EPI_ISL_2699831, EPI_ISL_2699832 | Lighthouse Lab in Glasgow | Wellcome Sanger Institute for the COVID-19 Genomics UK (COG-UK) Consortium | Harper VanSteenhouse, Yumi Kasai, David Gray, Carol Clugston, Anna Dominiczak and Alex Alderton, Roberto Amato, Jeffrey Barrett, Sonia Goncalves, Ewan Harrison, David K. Jackson, Ian Johnston, Dominic Kwiatkowski, Cordelia Langford, John Sillitoe on behalf of the Wellcome Sanger Institute COVID-19 Surveillance Team |

| | | | |
|---|---|---|---|
| EPI_ISL_2699834, EPI_ISL_2699835 | Lighthouse Lab in Milton Keynes | Wellcome Sanger Institute for the COVID-19 Genomics UK (COG-UK) Consortium | The Lighthouse Lab in Milton Keynes and Alex Alderton, Roberto Amato, Jeffrey Barrett, Sonia Goncalves, Ewan Harrison, David K. Jackson, Ian Johnston, Dominic Kwiatkowski, Cordelia Langford, John Sillitoe on behalf of the Wellcome Sanger Institute COVID-19 Surveillance Team |
| EPI_ISL_2699836 | Lighthouse Lab in Glasgow | Wellcome Sanger Institute for the COVID-19 Genomics UK (COG-UK) Consortium | Harper VanSteenhouse, Yumi Kasai, David Gray, Carol Clugston, Anna Dominiczak and Alex Alderton, Roberto Amato, Jeffrey Barrett, Sonia Goncalves, Ewan Harrison, David K. Jackson, Ian Johnston, Dominic Kwiatkowski, Cordelia Langford, John Sillitoe on behalf of the Wellcome Sanger Institute COVID-19 Surveillance Team |
| EPI_ISL_2699837 | Lighthouse Lab in Milton Keynes | Wellcome Sanger Institute for the COVID-19 Genomics UK (COG-UK) Consortium | The Lighthouse Lab in Milton Keynes and Alex Alderton, Roberto Amato, Jeffrey Barrett, Sonia Goncalves, Ewan Harrison, David K. Jackson, Ian Johnston, Dominic Kwiatkowski, Cordelia Langford, John Sillitoe on behalf of the Wellcome Sanger Institute COVID-19 Surveillance Team |
| EPI_ISL_2699838 | Lighthouse Lab in Glasgow | Wellcome Sanger Institute for the COVID-19 Genomics UK (COG-UK) Consortium | Harper VanSteenhouse, Yumi Kasai, David Gray, Carol Clugston, Anna Dominiczak and Alex Alderton, Roberto Amato, Jeffrey Barrett, Sonia Goncalves, Ewan Harrison, David K. Jackson, Ian Johnston, Dominic Kwiatkowski, Cordelia Langford, John Sillitoe on behalf of the Wellcome Sanger Institute COVID-19 Surveillance Team |
| EPI_ISL_2699839, EPI_ISL_2699840 | Lighthouse Lab in Milton Keynes | Wellcome Sanger Institute for the COVID-19 Genomics UK (COG-UK) Consortium | The Lighthouse Lab in Milton Keynes and Alex Alderton, Roberto Amato, Jeffrey Barrett, Sonia Goncalves, Ewan Harrison, David K. Jackson, Ian Johnston, Dominic Kwiatkowski, Cordelia Langford, John Sillitoe on behalf of the Wellcome Sanger Institute COVID-19 Surveillance Team |
| EPI_ISL_2699841 | Lighthouse Lab in Glasgow | Wellcome Sanger Institute for the COVID-19 Genomics UK (COG-UK) Consortium | Harper VanSteenhouse, Yumi Kasai, David Gray, Carol Clugston, Anna Dominiczak and Alex Alderton, Roberto Amato, Jeffrey Barrett, Sonia Goncalves, Ewan Harrison, David K. Jackson, Ian Johnston, Dominic Kwiatkowski, Cordelia Langford, John Sillitoe on behalf of the Wellcome Sanger Institute COVID-19 Surveillance Team |
| EPI_ISL_2699842, EPI_ISL_2699843, EPI_ISL_2699844, EPI_ISL_2699845, EPI_ISL_2699846 | Lighthouse Lab in Milton Keynes | Wellcome Sanger Institute for the COVID-19 Genomics UK (COG-UK) Consortium | The Lighthouse Lab in Milton Keynes and Alex Alderton, Roberto Amato, Jeffrey Barrett, Sonia Goncalves, Ewan Harrison, David K. Jackson, Ian Johnston, Dominic Kwiatkowski, Cordelia Langford, John Sillitoe on behalf of the Wellcome Sanger Institute COVID-19 Surveillance Team |
| EPI_ISL_2699847 | Lighthouse Lab in Glasgow | Wellcome Sanger Institute for the COVID-19 Genomics UK (COG-UK) Consortium | Harper VanSteenhouse, Yumi Kasai, David Gray, Carol Clugston, Anna Dominiczak and Alex Alderton, Roberto Amato, Jeffrey Barrett, Sonia Goncalves, Ewan Harrison, David K. Jackson, Ian Johnston, Dominic Kwiatkowski, Cordelia Langford, John Sillitoe on behalf of the Wellcome Sanger Institute COVID-19 Surveillance Team |
| EPI_ISL_2699848, EPI_ISL_2699849 | Lighthouse Lab in Milton Keynes | Wellcome Sanger Institute for the COVID-19 Genomics UK (COG-UK) Consortium | The Lighthouse Lab in Milton Keynes and Alex Alderton, Roberto Amato, Jeffrey Barrett, Sonia Goncalves, Ewan Harrison, David K. Jackson, Ian Johnston, Dominic Kwiatkowski, Cordelia Langford, John Sillitoe on behalf of the Wellcome Sanger Institute COVID-19 Surveillance Team |
| EPI_ISL_2699850 | Lighthouse Laboratory Plymouth | Wellcome Sanger Institute for the COVID-19 Genomics UK (COG-UK) Consortium | Lighthouse Laboratory Plymouth and Alex Alderton, Roberto Amato, Jeffrey Barrett, Sonia Goncalves, Ewan Harrison, David K. Jackson, Ian Johnston, Dominic Kwiatkowski, Cordelia Langford, John Sillitoe on behalf of the Wellcome Sanger Institute COVID-19 Surveillance Team |
| EPI_ISL_2699851, EPI_ISL_2699852 | Lighthouse Lab in Glasgow | Wellcome Sanger Institute for the COVID-19 Genomics UK (COG-UK) Consortium | Harper VanSteenhouse, Yumi Kasai, David Gray, Carol Clugston, Anna Dominiczak and Alex Alderton, Roberto Amato, Jeffrey Barrett, Sonia Goncalves, Ewan Harrison, David K. Jackson, Ian Johnston, Dominic Kwiatkowski, Cordelia Langford, John Sillitoe on behalf of the Wellcome Sanger Institute COVID-19 Surveillance Team |
| EPI_ISL_2699853 | Lighthouse Lab in Milton Keynes | Wellcome Sanger Institute for the COVID-19 Genomics UK (COG-UK) Consortium | The Lighthouse Lab in Milton Keynes and Alex Alderton, Roberto Amato, Jeffrey Barrett, Sonia Goncalves, Ewan Harrison, David K. Jackson, Ian Johnston, Dominic Kwiatkowski, Cordelia Langford, John Sillitoe on behalf of the Wellcome Sanger Institute COVID-19 Surveillance Team |
| EPI_ISL_2699854 | Lighthouse Lab in Glasgow | Wellcome Sanger Institute for the COVID-19 Genomics UK (COG-UK) Consortium | Harper VanSteenhouse, Yumi Kasai, David Gray, Carol Clugston, Anna Dominiczak and Alex Alderton, Roberto Amato, Jeffrey Barrett, Sonia Goncalves, Ewan Harrison, David K. Jackson, Ian Johnston, Dominic Kwiatkowski, Cordelia Langford, John Sillitoe on behalf of the Wellcome Sanger Institute COVID-19 Surveillance Team |
| EPI_ISL_2699855 | Lighthouse Lab in Milton Keynes | Wellcome Sanger Institute for the COVID-19 Genomics UK (COG-UK) Consortium | The Lighthouse Lab in Milton Keynes and Alex Alderton, Roberto Amato, Jeffrey Barrett, Sonia Goncalves, Ewan Harrison, David K. Jackson, Ian Johnston, Dominic Kwiatkowski, Cordelia Langford, John Sillitoe on behalf of the Wellcome Sanger Institute COVID-19 Surveillance Team |
| EPI_ISL_2699856 | Lighthouse Lab in Glasgow | Wellcome Sanger Institute for the COVID-19 Genomics UK (COG-UK) Consortium | Harper VanSteenhouse, Yumi Kasai, David Gray, Carol Clugston, Anna Dominiczak and Alex Alderton, Roberto Amato, Jeffrey Barrett, Sonia Goncalves, Ewan Harrison, David K. Jackson, Ian Johnston, Dominic Kwiatkowski, Cordelia Langford, John Sillitoe on behalf of the Wellcome Sanger Institute COVID-19 Surveillance Team |
| EPI_ISL_2699857 | Lighthouse Laboratory Plymouth | Wellcome Sanger Institute for the COVID-19 Genomics UK (COG-UK) Consortium | Lighthouse Laboratory Plymouth and Alex Alderton, Roberto Amato, Jeffrey Barrett, Sonia Goncalves, Ewan Harrison, David K. Jackson, Ian Johnston, Dominic Kwiatkowski, Cordelia Langford, John Sillitoe on behalf of the Wellcome Sanger Institute COVID-19 Surveillance Team |
| EPI_ISL_2699858, EPI_ISL_2699859, EPI_ISL_2699860 | Lighthouse Lab in Glasgow | Wellcome Sanger Institute for the COVID-19 Genomics UK (COG-UK) Consortium | Harper VanSteenhouse, Yumi Kasai, David Gray, Carol Clugston, Anna Dominiczak and Alex Alderton, Roberto Amato, Jeffrey Barrett, Sonia Goncalves, Ewan Harrison, David K. Jackson, Ian Johnston, Dominic Kwiatkowski, Cordelia Langford, John Sillitoe on behalf of the Wellcome Sanger Institute COVID-19 Surveillance Team |
| EPI_ISL_2699861 | Lighthouse Lab in Milton Keynes | Wellcome Sanger Institute for the COVID-19 Genomics UK (COG-UK) Consortium | The Lighthouse Lab in Milton Keynes and Alex Alderton, Roberto Amato, Jeffrey Barrett, Sonia Goncalves, Ewan Harrison, David K. Jackson, Ian Johnston, Dominic Kwiatkowski, Cordelia Langford, John Sillitoe on behalf of the Wellcome Sanger Institute COVID-19 Surveillance Team |
| EPI_ISL_2699862 | Lighthouse Lab in Glasgow | Wellcome Sanger Institute for the COVID-19 Genomics UK (COG-UK) Consortium | Harper VanSteenhouse, Yumi Kasai, David Gray, Carol Clugston, Anna Dominiczak and Alex Alderton, Roberto Amato, Jeffrey Barrett, Sonia Goncalves, Ewan Harrison, David K. Jackson, Ian Johnston, Dominic Kwiatkowski, Cordelia Langford, John Sillitoe on behalf of the Wellcome Sanger Institute COVID-19 Surveillance Team |
| EPI_ISL_2699863, EPI_ISL_2699864 | Lighthouse Lab in Milton Keynes | Wellcome Sanger Institute for the COVID-19 Genomics UK (COG-UK) Consortium | The Lighthouse Lab in Milton Keynes and Alex Alderton, Roberto Amato, Jeffrey Barrett, Sonia Goncalves, Ewan Harrison, David K. Jackson, Ian Johnston, Dominic Kwiatkowski, Cordelia Langford, John Sillitoe on behalf of the Wellcome Sanger Institute COVID-19 Surveillance Team |
| EPI_ISL_2699865 | Lighthouse Lab in Glasgow | Wellcome Sanger Institute for the COVID-19 Genomics UK (COG-UK) Consortium | Harper VanSteenhouse, Yumi Kasai, David Gray, Carol Clugston, Anna Dominiczak and Alex Alderton, Roberto Amato, Jeffrey Barrett, Sonia Goncalves, Ewan Harrison, David K. Jackson, Ian Johnston, Dominic Kwiatkowski, Cordelia Langford, John Sillitoe on behalf of the Wellcome Sanger Institute COVID-19 Surveillance Team |
| EPI_ISL_2699866, EPI_ISL_2699867 | Lighthouse Lab in Milton Keynes | Wellcome Sanger Institute for the COVID-19 Genomics UK (COG-UK) Consortium | The Lighthouse Lab in Milton Keynes and Alex Alderton, Roberto Amato, Jeffrey Barrett, Sonia Goncalves, Ewan Harrison, David K. Jackson, Ian Johnston, Dominic Kwiatkowski, Cordelia Langford, John Sillitoe on behalf of the Wellcome Sanger Institute COVID-19 Surveillance Team |
| EPI_ISL_2699868 | Lighthouse Lab in Alderley Park | Wellcome Sanger Institute for the COVID-19 Genomics UK (COG-UK) Consortium | Jacquelyn Wynn, Mairead Hyland, The Lighthouse Lab in Alderley Park and Alex Alderton, Roberto Amato, Jeffrey Barrett, Sonia Goncalves, Ewan Harrison, David K. Jackson, Ian Johnston, Dominic Kwiatkowski, Cordelia Langford, John Sillitoe on behalf of the Wellcome Sanger Institute COVID-19 Surveillance Team |
| EPI_ISL_2699869 | Lighthouse Lab in Milton Keynes | Wellcome Sanger Institute for the COVID-19 Genomics UK (COG-UK) Consortium | The Lighthouse Lab in Milton Keynes and Alex Alderton, Roberto Amato, Jeffrey Barrett, Sonia Goncalves, Ewan Harrison, David K. Jackson, Ian Johnston, Dominic Kwiatkowski, Cordelia Langford, John Sillitoe on behalf of the Wellcome Sanger Institute COVID-19 Surveillance Team |
| EPI_ISL_2699870, EPI_ISL_2699871 | Lighthouse Lab in Alderley Park | Wellcome Sanger Institute for the COVID-19 Genomics UK (COG-UK) Consortium | Jacquelyn Wynn, Mairead Hyland, The Lighthouse Lab in Alderley Park and Alex Alderton, Roberto Amato, Jeffrey Barrett, Sonia Goncalves, Ewan Harrison, David K. Jackson, Ian Johnston, Dominic Kwiatkowski, Cordelia Langford, John Sillitoe on behalf of the Wellcome Sanger Institute COVID-19 Surveillance Team |
| EPI_ISL_2699872 | Lighthouse Lab in Milton Keynes | Wellcome Sanger Institute for the COVID-19 Genomics UK (COG-UK) Consortium | The Lighthouse Lab in Milton Keynes and Alex Alderton, Roberto Amato, Jeffrey Barrett, Sonia Goncalves, Ewan Harrison, David K. Jackson, Ian Johnston, Dominic Kwiatkowski, Cordelia Langford, John Sillitoe on behalf of the Wellcome Sanger Institute COVID-19 Surveillance Team |
| EPI_ISL_2699873 | Lighthouse Lab in Alderley Park | Wellcome Sanger Institute for the COVID-19 Genomics UK (COG-UK) Consortium | Jacquelyn Wynn, Mairead Hyland, The Lighthouse Lab in Alderley Park and Alex Alderton, Roberto Amato, Jeffrey Barrett, Sonia Goncalves, Ewan Harrison, David K. Jackson, Ian Johnston, Dominic Kwiatkowski, Cordelia Langford, John Sillitoe on behalf of the Wellcome Sanger Institute COVID-19 Surveillance Team |
| EPI_ISL_2699874, EPI_ISL_2699875, EPI_ISL_2699876, EPI_ISL_2699877, EPI_ISL_2699878, EPI_ISL_2699879, EPI_ISL_2699880, EPI_ISL_2699881, EPI_ISL_2699882, EPI_ISL_2699883, EPI_ISL_2699884, EPI_ISL_2699885 | | | |
| see above | Lighthouse Lab in Milton Keynes | Wellcome Sanger Institute for the COVID-19 Genomics UK (COG-UK) Consortium | The Lighthouse Lab in Milton Keynes and Alex Alderton, Roberto Amato, Jeffrey Barrett, Sonia Goncalves, Ewan Harrison, David K. Jackson, Ian Johnston, Dominic Kwiatkowski, Cordelia Langford, John Sillitoe on behalf of the Wellcome Sanger Institute COVID-19 Surveillance Team |
| EPI_ISL_2699886, EPI_ISL_2699887 | Lighthouse Lab in Alderley Park | Wellcome Sanger Institute for the COVID-19 Genomics UK (COG-UK) Consortium | Jacquelyn Wynn, Mairead Hyland, The Lighthouse Lab in Alderley Park and Alex Alderton, Roberto Amato, Jeffrey Barrett, Sonia Goncalves, Ewan Harrison, David K. Jackson, Ian Johnston, Dominic Kwiatkowski, Cordelia Langford, John Sillitoe on behalf of the Wellcome Sanger Institute COVID-19 Surveillance Team |
| EPI_ISL_2699888 | Lighthouse Lab in Milton Keynes | Wellcome Sanger Institute for the COVID-19 Genomics UK (COG-UK) Consortium | The Lighthouse Lab in Milton Keynes and Alex Alderton, Roberto Amato, Jeffrey Barrett, Sonia Goncalves, Ewan Harrison, David K. Jackson, Ian Johnston, Dominic Kwiatkowski, Cordelia Langford, John Sillitoe on behalf of the Wellcome Sanger Institute COVID-19 Surveillance Team |
| EPI_ISL_2699889 | Lighthouse Lab in Alderley Park | Wellcome Sanger Institute for the COVID-19 Genomics UK (COG-UK) Consortium | Jacquelyn Wynn, Mairead Hyland, The Lighthouse Lab in Alderley Park and Alex Alderton, Roberto Amato, Jeffrey Barrett, Sonia Goncalves, Ewan Harrison, David K. Jackson, Ian Johnston, Dominic Kwiatkowski, Cordelia Langford, John Sillitoe on behalf of the Wellcome Sanger Institute COVID-19 Surveillance Team |
| EPI_ISL_2699890, EPI_ISL_2699891 | Lighthouse Lab in Milton Keynes | Wellcome Sanger Institute for the COVID-19 Genomics UK (COG-UK) Consortium | The Lighthouse Lab in Milton Keynes and Alex Alderton, Roberto Amato, Jeffrey Barrett, Sonia Goncalves, Ewan Harrison, David K. Jackson, Ian Johnston, Dominic Kwiatkowski, Cordelia Langford, John Sillitoe on behalf of the Wellcome Sanger Institute COVID-19 Surveillance Team |
| EPI_ISL_2699892 | Lighthouse Lab in Alderley Park | Wellcome Sanger Institute for the COVID-19 Genomics UK (COG-UK) Consortium | Jacquelyn Wynn, Mairead Hyland, The Lighthouse Lab in Alderley Park and Alex Alderton, Roberto Amato, Jeffrey Barrett, Sonia Goncalves, Ewan Harrison, David K. Jackson, Ian Johnston, Dominic Kwiatkowski, Cordelia Langford, John Sillitoe on behalf of the Wellcome Sanger Institute COVID-19 Surveillance Team |
| EPI_ISL_2699893, EPI_ISL_2699894, EPI_ISL_2699895 | Lighthouse Lab in Milton Keynes | Wellcome Sanger Institute for the COVID-19 Genomics UK (COG-UK) Consortium | The Lighthouse Lab in Milton Keynes and Alex Alderton, Roberto Amato, Jeffrey Barrett, Sonia Goncalves, Ewan Harrison, David K. Jackson, Ian Johnston, Dominic Kwiatkowski, Cordelia Langford, John Sillitoe on behalf of the Wellcome Sanger Institute COVID-19 Surveillance Team |
| EPI_ISL_2699896 | Lighthouse Lab in Alderley Park | Wellcome Sanger Institute for the COVID-19 Genomics UK (COG-UK) Consortium | Jacquelyn Wynn, Mairead Hyland, The Lighthouse Lab in Alderley Park and Alex Alderton, Roberto Amato, Jeffrey Barrett, Sonia Goncalves, Ewan Harrison, David K. Jackson, Ian Johnston, Dominic Kwiatkowski, Cordelia Langford, John Sillitoe on behalf of the Wellcome Sanger Institute COVID-19 Surveillance Team |
| EPI_ISL_2699897, EPI_ISL_2699898 | Lighthouse Lab in Milton Keynes | Wellcome Sanger Institute for the COVID-19 Genomics UK (COG-UK) Consortium | The Lighthouse Lab in Milton Keynes and Alex Alderton, Roberto Amato, Jeffrey Barrett, Sonia Goncalves, Ewan Harrison, David K. Jackson, Ian Johnston, Dominic Kwiatkowski, Cordelia Langford, John Sillitoe on behalf of the Wellcome Sanger Institute COVID-19 Surveillance Team |
| EPI_ISL_2699899 | Lighthouse Lab in Alderley Park | Wellcome Sanger Institute for the COVID-19 Genomics UK (COG-UK) Consortium | Jacquelyn Wynn, Mairead Hyland, The Lighthouse Lab in Alderley Park and Alex Alderton, Roberto Amato, Jeffrey Barrett, Sonia Goncalves, Ewan Harrison, David K. Jackson, Ian Johnston, Dominic Kwiatkowski, Cordelia Langford, John Sillitoe on behalf of the Wellcome Sanger Institute COVID-19 Surveillance Team |
| EPI_ISL_2699900, EPI_ISL_2699901, EPI_ISL_2699902, EPI_ISL_2699904, EPI_ISL_2699905, EPI_ISL_2699906 | Lighthouse Lab in Milton Keynes | Wellcome Sanger Institute for the COVID-19 Genomics UK (COG-UK) Consortium | The Lighthouse Lab in Milton Keynes and Alex Alderton, Roberto Amato, Jeffrey Barrett, Sonia Goncalves, Ewan Harrison, David K. Jackson, Ian Johnston, Dominic Kwiatkowski, Cordelia Langford, John Sillitoe on behalf of the Wellcome Sanger Institute COVID-19 Surveillance Team |
| EPI_ISL_2699907, EPI_ISL_2699908, EPI_ISL_2699909, EPI_ISL_2699910 | Lighthouse Lab in Alderley Park | Wellcome Sanger Institute for the COVID-19 Genomics UK (COG-UK) Consortium | Jacquelyn Wynn, Mairead Hyland, The Lighthouse Lab in Alderley Park and Alex Alderton, Roberto Amato, Jeffrey Barrett, Sonia Goncalves, Ewan Harrison, David K. Jackson, Ian Johnston, Dominic Kwiatkowski, Cordelia Langford, John Sillitoe on behalf of the Wellcome Sanger Institute COVID-19 Surveillance Team |
| EPI_ISL_2699911, EPI_ISL_2699912 | Lighthouse Lab in Milton Keynes | Wellcome Sanger Institute for the COVID-19 Genomics UK (COG-UK) Consortium | The Lighthouse Lab in Milton Keynes and Alex Alderton, Roberto Amato, Jeffrey Barrett, Sonia Goncalves, Ewan Harrison, David K. Jackson, Ian Johnston, Dominic Kwiatkowski, Cordelia Langford, John Sillitoe on behalf of the Wellcome Sanger Institute COVID-19 Surveillance Team |

| | | | |
|---|---|---|---|
| EPI_ISL_2699913, EPI_ISL_2699914 | Lighthouse Lab in Alderley Park | Wellcome Sanger Institute for the COVID-19 Genomics UK (COG-UK) Consortium | Jacquelyn Wynn, Mairead Hyland, The Lighthouse Lab in Alderley Park and Alex Alderton, Roberto Amato, Jeffrey Barrett, Sonia Goncalves, Ewan Harrison, David K. Jackson, Ian Johnston, Dominic Kwiatkowski, Cordelia Langford, John Sillitoe on behalf of the Wellcome Sanger Institute COVID-19 Surveillance Team |
| EPI_ISL_2699915, EPI_ISL_2699916, EPI_ISL_2699917, EPI_ISL_2699918 | Lighthouse Lab in Milton Keynes | Wellcome Sanger Institute for the COVID-19 Genomics UK (COG-UK) Consortium | The Lighthouse Lab in Milton Keynes and Alex Alderton, Roberto Amato, Jeffrey Barrett, Sonia Goncalves, Ewan Harrison, David K. Jackson, Ian Johnston, Dominic Kwiatkowski, Cordelia Langford, John Sillitoe on behalf of the Wellcome Sanger Institute COVID-19 Surveillance Team |
| EPI_ISL_2699919 | Lighthouse Lab in Alderley Park | Wellcome Sanger Institute for the COVID-19 Genomics UK (COG-UK) Consortium | Jacquelyn Wynn, Mairead Hyland, The Lighthouse Lab in Alderley Park and Alex Alderton, Roberto Amato, Jeffrey Barrett, Sonia Goncalves, Ewan Harrison, David K. Jackson, Ian Johnston, Dominic Kwiatkowski, Cordelia Langford, John Sillitoe on behalf of the Wellcome Sanger Institute COVID-19 Surveillance Team |
| EPI_ISL_2699921, EPI_ISL_2699922, EPI_ISL_2699923, EPI_ISL_2699924, EPI_ISL_2699925, EPI_ISL_2699926, EPI_ISL_2699927 | Lighthouse Lab in Milton Keynes | Wellcome Sanger Institute for the COVID-19 Genomics UK (COG-UK) Consortium | The Lighthouse Lab in Milton Keynes and Alex Alderton, Roberto Amato, Jeffrey Barrett, Sonia Goncalves, Ewan Harrison, David K. Jackson, Ian Johnston, Dominic Kwiatkowski, Cordelia Langford, John Sillitoe on behalf of the Wellcome Sanger Institute COVID-19 Surveillance Team |
| EPI_ISL_2699928 | Lighthouse Lab in Alderley Park | Wellcome Sanger Institute for the COVID-19 Genomics UK (COG-UK) Consortium | Jacquelyn Wynn, Mairead Hyland, The Lighthouse Lab in Alderley Park and Alex Alderton, Roberto Amato, Jeffrey Barrett, Sonia Goncalves, Ewan Harrison, David K. Jackson, Ian Johnston, Dominic Kwiatkowski, Cordelia Langford, John Sillitoe on behalf of the Wellcome Sanger Institute COVID-19 Surveillance Team |
| EPI_ISL_2699929, EPI_ISL_2699930 | Lighthouse Lab in Milton Keynes | Wellcome Sanger Institute for the COVID-19 Genomics UK (COG-UK) Consortium | The Lighthouse Lab in Milton Keynes and Alex Alderton, Roberto Amato, Jeffrey Barrett, Sonia Goncalves, Ewan Harrison, David K. Jackson, Ian Johnston, Dominic Kwiatkowski, Cordelia Langford, John Sillitoe on behalf of the Wellcome Sanger Institute COVID-19 Surveillance Team |
| EPI_ISL_2699931, EPI_ISL_2699932, EPI_ISL_2699933, EPI_ISL_2699934 | Lighthouse Lab in Alderley Park | Wellcome Sanger Institute for the COVID-19 Genomics UK (COG-UK) Consortium | Jacquelyn Wynn, Mairead Hyland, The Lighthouse Lab in Alderley Park and Alex Alderton, Roberto Amato, Jeffrey Barrett, Sonia Goncalves, Ewan Harrison, David K. Jackson, Ian Johnston, Dominic Kwiatkowski, Cordelia Langford, John Sillitoe on behalf of the Wellcome Sanger Institute COVID-19 Surveillance Team |
| EPI_ISL_2699935 | Lighthouse Lab in Milton Keynes | Wellcome Sanger Institute for the COVID-19 Genomics UK (COG-UK) Consortium | The Lighthouse Lab in Milton Keynes and Alex Alderton, Roberto Amato, Jeffrey Barrett, Sonia Goncalves, Ewan Harrison, David K. Jackson, Ian Johnston, Dominic Kwiatkowski, Cordelia Langford, John Sillitoe on behalf of the Wellcome Sanger Institute COVID-19 Surveillance Team |
| EPI_ISL_2699936 | Lighthouse Lab in Alderley Park | Wellcome Sanger Institute for the COVID-19 Genomics UK (COG-UK) Consortium | Jacquelyn Wynn, Mairead Hyland, The Lighthouse Lab in Alderley Park and Alex Alderton, Roberto Amato, Jeffrey Barrett, Sonia Goncalves, Ewan Harrison, David K. Jackson, Ian Johnston, Dominic Kwiatkowski, Cordelia Langford, John Sillitoe on behalf of the Wellcome Sanger Institute COVID-19 Surveillance Team |
| EPI_ISL_2699937, EPI_ISL_2699938, EPI_ISL_2699939, EPI_ISL_2699940, EPI_ISL_2699941, EPI_ISL_2699942, EPI_ISL_2699943, EPI_ISL_2699944, EPI_ISL_2699945 | Lighthouse Lab in Milton Keynes | Wellcome Sanger Institute for the COVID-19 Genomics UK (COG-UK) Consortium | The Lighthouse Lab in Milton Keynes and Alex Alderton, Roberto Amato, Jeffrey Barrett, Sonia Goncalves, Ewan Harrison, David K. Jackson, Ian Johnston, Dominic Kwiatkowski, Cordelia Langford, John Sillitoe on behalf of the Wellcome Sanger Institute COVID-19 Surveillance Team |
| EPI_ISL_2699946, EPI_ISL_2699947 | Lighthouse Lab in Alderley Park | Wellcome Sanger Institute for the COVID-19 Genomics UK (COG-UK) Consortium | Jacquelyn Wynn, Mairead Hyland, The Lighthouse Lab in Alderley Park and Alex Alderton, Roberto Amato, Jeffrey Barrett, Sonia Goncalves, Ewan Harrison, David K. Jackson, Ian Johnston, Dominic Kwiatkowski, Cordelia Langford, John Sillitoe on behalf of the Wellcome Sanger Institute COVID-19 Surveillance Team |
| EPI_ISL_2699948, EPI_ISL_2699949, EPI_ISL_2699950, EPI_ISL_2699951 | Lighthouse Lab in Milton Keynes | Wellcome Sanger Institute for the COVID-19 Genomics UK (COG-UK) Consortium | The Lighthouse Lab in Milton Keynes and Alex Alderton, Roberto Amato, Jeffrey Barrett, Sonia Goncalves, Ewan Harrison, David K. Jackson, Ian Johnston, Dominic Kwiatkowski, Cordelia Langford, John Sillitoe on behalf of the Wellcome Sanger Institute COVID-19 Surveillance Team |
| EPI_ISL_2699952, EPI_ISL_2699953 | Lighthouse Lab in Alderley Park | Wellcome Sanger Institute for the COVID-19 Genomics UK (COG-UK) Consortium | Jacquelyn Wynn, Mairead Hyland, The Lighthouse Lab in Alderley Park and Alex Alderton, Roberto Amato, Jeffrey Barrett, Sonia Goncalves, Ewan Harrison, David K. Jackson, Ian Johnston, Dominic Kwiatkowski, Cordelia Langford, John Sillitoe on behalf of the Wellcome Sanger Institute COVID-19 Surveillance Team |
| EPI_ISL_2699954 | Lighthouse Lab in Milton Keynes | Wellcome Sanger Institute for the COVID-19 Genomics UK (COG-UK) Consortium | The Lighthouse Lab in Milton Keynes and Alex Alderton, Roberto Amato, Jeffrey Barrett, Sonia Goncalves, Ewan Harrison, David K. Jackson, Ian Johnston, Dominic Kwiatkowski, Cordelia Langford, John Sillitoe on behalf of the Wellcome Sanger Institute COVID-19 Surveillance Team |
| EPI_ISL_2699955 | Lighthouse Lab in Alderley Park | Wellcome Sanger Institute for the COVID-19 Genomics UK (COG-UK) Consortium | Jacquelyn Wynn, Mairead Hyland, The Lighthouse Lab in Alderley Park and Alex Alderton, Roberto Amato, Jeffrey Barrett, Sonia Goncalves, Ewan Harrison, David K. Jackson, Ian Johnston, Dominic Kwiatkowski, Cordelia Langford, John Sillitoe on behalf of the Wellcome Sanger Institute COVID-19 Surveillance Team |
| EPI_ISL_2699956, EPI_ISL_2699957, EPI_ISL_2699958, EPI_ISL_2699959, EPI_ISL_2699960, EPI_ISL_2699961, EPI_ISL_2699962, EPI_ISL_2699963, EPI_ISL_2699964 | Lighthouse Lab in Milton Keynes | Wellcome Sanger Institute for the COVID-19 Genomics UK (COG-UK) Consortium | The Lighthouse Lab in Milton Keynes and Alex Alderton, Roberto Amato, Jeffrey Barrett, Sonia Goncalves, Ewan Harrison, David K. Jackson, Ian Johnston, Dominic Kwiatkowski, Cordelia Langford, John Sillitoe on behalf of the Wellcome Sanger Institute COVID-19 Surveillance Team |
| EPI_ISL_2699965 | Lighthouse Lab in Alderley Park | Wellcome Sanger Institute for the COVID-19 Genomics UK (COG-UK) Consortium | Jacquelyn Wynn, Mairead Hyland, The Lighthouse Lab in Alderley Park and Alex Alderton, Roberto Amato, Jeffrey Barrett, Sonia Goncalves, Ewan Harrison, David K. Jackson, Ian Johnston, Dominic Kwiatkowski, Cordelia Langford, John Sillitoe on behalf of the Wellcome Sanger Institute COVID-19 Surveillance Team |
| EPI_ISL_2699966, EPI_ISL_2699967 | Lighthouse Lab in Milton Keynes | Wellcome Sanger Institute for the COVID-19 Genomics UK (COG-UK) Consortium | The Lighthouse Lab in Milton Keynes and Alex Alderton, Roberto Amato, Jeffrey Barrett, Sonia Goncalves, Ewan Harrison, David K. Jackson, Ian Johnston, Dominic Kwiatkowski, Cordelia Langford, John Sillitoe on behalf of the Wellcome Sanger Institute COVID-19 Surveillance Team |
| EPI_ISL_2699968 | Lighthouse Lab in Alderley Park | Wellcome Sanger Institute for the COVID-19 Genomics UK (COG-UK) Consortium | Jacquelyn Wynn, Mairead Hyland, The Lighthouse Lab in Alderley Park and Alex Alderton, Roberto Amato, Jeffrey Barrett, Sonia Goncalves, Ewan Harrison, David K. Jackson, Ian Johnston, Dominic Kwiatkowski, Cordelia Langford, John Sillitoe on behalf of the Wellcome Sanger Institute COVID-19 Surveillance Team |
| EPI_ISL_2699969 | Lighthouse Lab in Milton Keynes | Wellcome Sanger Institute for the COVID-19 Genomics UK (COG-UK) Consortium | The Lighthouse Lab in Milton Keynes and Alex Alderton, Roberto Amato, Jeffrey Barrett, Sonia Goncalves, Ewan Harrison, David K. Jackson, Ian Johnston, Dominic Kwiatkowski, Cordelia Langford, John Sillitoe on behalf of the Wellcome Sanger Institute COVID-19 Surveillance Team |
| EPI_ISL_2699970 | Lighthouse Lab in Alderley Park | Wellcome Sanger Institute for the COVID-19 Genomics UK (COG-UK) Consortium | Jacquelyn Wynn, Mairead Hyland, The Lighthouse Lab in Alderley Park and Alex Alderton, Roberto Amato, Jeffrey Barrett, Sonia Goncalves, Ewan Harrison, David K. Jackson, Ian Johnston, Dominic Kwiatkowski, Cordelia Langford, John Sillitoe on behalf of the Wellcome Sanger Institute COVID-19 Surveillance Team |
| EPI_ISL_2699971 | Lighthouse Lab in Milton Keynes | Wellcome Sanger Institute for the COVID-19 Genomics UK (COG-UK) Consortium | The Lighthouse Lab in Milton Keynes and Alex Alderton, Roberto Amato, Jeffrey Barrett, Sonia Goncalves, Ewan Harrison, David K. Jackson, Ian Johnston, Dominic Kwiatkowski, Cordelia Langford, John Sillitoe on behalf of the Wellcome Sanger Institute COVID-19 Surveillance Team |
| EPI_ISL_2699972, EPI_ISL_2699973, EPI_ISL_2699974 | Lighthouse Lab in Alderley Park | Wellcome Sanger Institute for the COVID-19 Genomics UK (COG-UK) Consortium | Jacquelyn Wynn, Mairead Hyland, The Lighthouse Lab in Alderley Park and Alex Alderton, Roberto Amato, Jeffrey Barrett, Sonia Goncalves, Ewan Harrison, David K. Jackson, Ian Johnston, Dominic Kwiatkowski, Cordelia Langford, John Sillitoe on behalf of the Wellcome Sanger Institute COVID-19 Surveillance Team |
| EPI_ISL_2699975, EPI_ISL_2699976, EPI_ISL_2699977 | Lighthouse Lab in Milton Keynes | Wellcome Sanger Institute for the COVID-19 Genomics UK (COG-UK) Consortium | The Lighthouse Lab in Milton Keynes and Alex Alderton, Roberto Amato, Jeffrey Barrett, Sonia Goncalves, Ewan Harrison, David K. Jackson, Ian Johnston, Dominic Kwiatkowski, Cordelia Langford, John Sillitoe on behalf of the Wellcome Sanger Institute COVID-19 Surveillance Team |
| EPI_ISL_2699978 | Lighthouse Lab in Alderley Park | Wellcome Sanger Institute for the COVID-19 Genomics UK (COG-UK) Consortium | Jacquelyn Wynn, Mairead Hyland, The Lighthouse Lab in Alderley Park and Alex Alderton, Roberto Amato, Jeffrey Barrett, Sonia Goncalves, Ewan Harrison, David K. Jackson, Ian Johnston, Dominic Kwiatkowski, Cordelia Langford, John Sillitoe on behalf of the Wellcome Sanger Institute COVID-19 Surveillance Team |
| EPI_ISL_2699979, EPI_ISL_2699980, EPI_ISL_2699981 | Lighthouse Lab in Milton Keynes | Wellcome Sanger Institute for the COVID-19 Genomics UK (COG-UK) Consortium | The Lighthouse Lab in Milton Keynes and Alex Alderton, Roberto Amato, Jeffrey Barrett, Sonia Goncalves, Ewan Harrison, David K. Jackson, Ian Johnston, Dominic Kwiatkowski, Cordelia Langford, John Sillitoe on behalf of the Wellcome Sanger Institute COVID-19 Surveillance Team |
| EPI_ISL_2699982 | Lighthouse Lab in Alderley Park | Wellcome Sanger Institute for the COVID-19 Genomics UK (COG-UK) Consortium | Jacquelyn Wynn, Mairead Hyland, The Lighthouse Lab in Alderley Park and Alex Alderton, Roberto Amato, Jeffrey Barrett, Sonia Goncalves, Ewan Harrison, David K. Jackson, Ian Johnston, Dominic Kwiatkowski, Cordelia Langford, John Sillitoe on behalf of the Wellcome Sanger Institute COVID-19 Surveillance Team |
| EPI_ISL_2699983 | Lighthouse Lab in Milton Keynes | Wellcome Sanger Institute for the COVID-19 Genomics UK (COG-UK) Consortium | The Lighthouse Lab in Milton Keynes and Alex Alderton, Roberto Amato, Jeffrey Barrett, Sonia Goncalves, Ewan Harrison, David K. Jackson, Ian Johnston, Dominic Kwiatkowski, Cordelia Langford, John Sillitoe on behalf of the Wellcome Sanger Institute COVID-19 Surveillance Team |
| EPI_ISL_2699984 | Lighthouse Lab in Alderley Park | Wellcome Sanger Institute for the COVID-19 Genomics UK (COG-UK) Consortium | Jacquelyn Wynn, Mairead Hyland, The Lighthouse Lab in Alderley Park and Alex Alderton, Roberto Amato, Jeffrey Barrett, Sonia Goncalves, Ewan Harrison, David K. Jackson, Ian Johnston, Dominic Kwiatkowski, Cordelia Langford, John Sillitoe on behalf of the Wellcome Sanger Institute COVID-19 Surveillance Team |
| EPI_ISL_2699985, EPI_ISL_2699986, EPI_ISL_2699987, EPI_ISL_2699988, EPI_ISL_2699989, EPI_ISL_2699990, EPI_ISL_2699991 | Lighthouse Lab in Milton Keynes | Wellcome Sanger Institute for the COVID-19 Genomics UK (COG-UK) Consortium | The Lighthouse Lab in Milton Keynes and Alex Alderton, Roberto Amato, Jeffrey Barrett, Sonia Goncalves, Ewan Harrison, David K. Jackson, Ian Johnston, Dominic Kwiatkowski, Cordelia Langford, John Sillitoe on behalf of the Wellcome Sanger Institute COVID-19 Surveillance Team |
| EPI_ISL_2699992 | Lighthouse Lab in Alderley Park | Wellcome Sanger Institute for the COVID-19 Genomics UK (COG-UK) Consortium | Jacquelyn Wynn, Mairead Hyland, The Lighthouse Lab in Alderley Park and Alex Alderton, Roberto Amato, Jeffrey Barrett, Sonia Goncalves, Ewan Harrison, David K. Jackson, Ian Johnston, Dominic Kwiatkowski, Cordelia Langford, John Sillitoe on behalf of the Wellcome Sanger Institute COVID-19 Surveillance Team |

| | | | |
|---|---|---|---|
| EPI_ISL_2699993 | Lighthouse Lab in Milton Keynes | Wellcome Sanger Institute for the COVID-19 Genomics UK (COG-UK) Consortium | The Lighthouse Lab in Milton Keynes and Alex Alderton, Roberto Amato, Jeffrey Barrett, Sonia Goncalves, Ewan Harrison, David K. Jackson, Ian Johnston, Dominic Kwiatkowski, Cordelia Langford, John Sillitoe on behalf of the Wellcome Sanger Institute COVID-19 Surveillance Team |
| EPI_ISL_2699994 | Lighthouse Lab in Alderley Park | Wellcome Sanger Institute for the COVID-19 Genomics UK (COG-UK) Consortium | Jacquelyn Wynn, Mairead Hyland, The Lighthouse Lab in Alderley Park and Alex Alderton, Roberto Amato, Jeffrey Barrett, Sonia Goncalves, Ewan Harrison, David K. Jackson, Ian Johnston, Dominic Kwiatkowski, Cordelia Langford, John Sillitoe on behalf of the Wellcome Sanger Institute COVID-19 Surveillance Team |
| EPI_ISL_2699995, EPI_ISL_2699996, EPI_ISL_2699997, EPI_ISL_2699999 | Lighthouse Lab in Milton Keynes | Wellcome Sanger Institute for the COVID-19 Genomics UK (COG-UK) Consortium | The Lighthouse Lab in Milton Keynes and Alex Alderton, Roberto Amato, Jeffrey Barrett, Sonia Goncalves, Ewan Harrison, David K. Jackson, Ian Johnston, Dominic Kwiatkowski, Cordelia Langford, John Sillitoe on behalf of the Wellcome Sanger Institute COVID-19 Surveillance Team |
| EPI_ISL_2700000 | Lighthouse Lab in Alderley Park | Wellcome Sanger Institute for the COVID-19 Genomics UK (COG-UK) Consortium | Jacquelyn Wynn, Mairead Hyland, The Lighthouse Lab in Alderley Park and Alex Alderton, Roberto Amato, Jeffrey Barrett, Sonia Goncalves, Ewan Harrison, David K. Jackson, Ian Johnston, Dominic Kwiatkowski, Cordelia Langford, John Sillitoe on behalf of the Wellcome Sanger Institute COVID-19 Surveillance Team |
| EPI_ISL_2700001, EPI_ISL_2700002, EPI_ISL_2700003, EPI_ISL_2700004, EPI_ISL_2700005, EPI_ISL_2700006, EPI_ISL_2700007, EPI_ISL_2700008, EPI_ISL_2700009, EPI_ISL_2700010, EPI_ISL_2700011 | | | |
| see above | Lighthouse Lab in Milton Keynes | Wellcome Sanger Institute for the COVID-19 Genomics UK (COG-UK) Consortium | The Lighthouse Lab in Milton Keynes and Alex Alderton, Roberto Amato, Jeffrey Barrett, Sonia Goncalves, Ewan Harrison, David K. Jackson, Ian Johnston, Dominic Kwiatkowski, Cordelia Langford, John Sillitoe on behalf of the Wellcome Sanger Institute COVID-19 Surveillance Team |
| EPI_ISL_2700012 | Lighthouse Lab in Alderley Park | Wellcome Sanger Institute for the COVID-19 Genomics UK (COG-UK) Consortium | Jacquelyn Wynn, Mairead Hyland, The Lighthouse Lab in Alderley Park and Alex Alderton, Roberto Amato, Jeffrey Barrett, Sonia Goncalves, Ewan Harrison, David K. Jackson, Ian Johnston, Dominic Kwiatkowski, Cordelia Langford, John Sillitoe on behalf of the Wellcome Sanger Institute COVID-19 Surveillance Team |
| EPI_ISL_2700013, EPI_ISL_2700014, EPI_ISL_2700015, EPI_ISL_2700016 | Lighthouse Lab in Milton Keynes | Wellcome Sanger Institute for the COVID-19 Genomics UK (COG-UK) Consortium | The Lighthouse Lab in Milton Keynes and Alex Alderton, Roberto Amato, Jeffrey Barrett, Sonia Goncalves, Ewan Harrison, David K. Jackson, Ian Johnston, Dominic Kwiatkowski, Cordelia Langford, John Sillitoe on behalf of the Wellcome Sanger Institute COVID-19 Surveillance Team |
| EPI_ISL_2700017, EPI_ISL_2700018 | Lighthouse Lab in Alderley Park | Wellcome Sanger Institute for the COVID-19 Genomics UK (COG-UK) Consortium | Jacquelyn Wynn, Mairead Hyland, The Lighthouse Lab in Alderley Park and Alex Alderton, Roberto Amato, Jeffrey Barrett, Sonia Goncalves, Ewan Harrison, David K. Jackson, Ian Johnston, Dominic Kwiatkowski, Cordelia Langford, John Sillitoe on behalf of the Wellcome Sanger Institute COVID-19 Surveillance Team |
| EPI_ISL_2700019, EPI_ISL_2700020, EPI_ISL_2700021, EPI_ISL_2700022 | Lighthouse Lab in Milton Keynes | Wellcome Sanger Institute for the COVID-19 Genomics UK (COG-UK) Consortium | The Lighthouse Lab in Milton Keynes and Alex Alderton, Roberto Amato, Jeffrey Barrett, Sonia Goncalves, Ewan Harrison, David K. Jackson, Ian Johnston, Dominic Kwiatkowski, Cordelia Langford, John Sillitoe on behalf of the Wellcome Sanger Institute COVID-19 Surveillance Team |
| EPI_ISL_2700023, EPI_ISL_2700024 | Lighthouse Lab in Alderley Park | Wellcome Sanger Institute for the COVID-19 Genomics UK (COG-UK) Consortium | Jacquelyn Wynn, Mairead Hyland, The Lighthouse Lab in Alderley Park and Alex Alderton, Roberto Amato, Jeffrey Barrett, Sonia Goncalves, Ewan Harrison, David K. Jackson, Ian Johnston, Dominic Kwiatkowski, Cordelia Langford, John Sillitoe on behalf of the Wellcome Sanger Institute COVID-19 Surveillance Team |
| EPI_ISL_2700025 | Lighthouse Lab in Milton Keynes | Wellcome Sanger Institute for the COVID-19 Genomics UK (COG-UK) Consortium | The Lighthouse Lab in Milton Keynes and Alex Alderton, Roberto Amato, Jeffrey Barrett, Sonia Goncalves, Ewan Harrison, David K. Jackson, Ian Johnston, Dominic Kwiatkowski, Cordelia Langford, John Sillitoe on behalf of the Wellcome Sanger Institute COVID-19 Surveillance Team |
| EPI_ISL_2700026 | Lighthouse Lab in Alderley Park | Wellcome Sanger Institute for the COVID-19 Genomics UK (COG-UK) Consortium | Jacquelyn Wynn, Mairead Hyland, The Lighthouse Lab in Alderley Park and Alex Alderton, Roberto Amato, Jeffrey Barrett, Sonia Goncalves, Ewan Harrison, David K. Jackson, Ian Johnston, Dominic Kwiatkowski, Cordelia Langford, John Sillitoe on behalf of the Wellcome Sanger Institute COVID-19 Surveillance Team |
| EPI_ISL_2700027, EPI_ISL_2700028, EPI_ISL_2700029, EPI_ISL_2700030, EPI_ISL_2700031, EPI_ISL_2700032, EPI_ISL_2700033, EPI_ISL_2700034, EPI_ISL_2700035, EPI_ISL_2700036, EPI_ISL_2700037, EPI_ISL_2700038, EPI_ISL_2700039, EPI_ISL_2700040, EPI_ISL_2700041, EPI_ISL_2700042, EPI_ISL_2700043, EPI_ISL_2700044 | | | |
| see above | Lighthouse Lab in Milton Keynes | Wellcome Sanger Institute for the COVID-19 Genomics UK (COG-UK) Consortium | The Lighthouse Lab in Milton Keynes and Alex Alderton, Roberto Amato, Jeffrey Barrett, Sonia Goncalves, Ewan Harrison, David K. Jackson, Ian Johnston, Dominic Kwiatkowski, Cordelia Langford, John Sillitoe on behalf of the Wellcome Sanger Institute COVID-19 Surveillance Team |
| EPI_ISL_2700045 | Lighthouse Lab in Alderley Park | Wellcome Sanger Institute for the COVID-19 Genomics UK (COG-UK) Consortium | Jacquelyn Wynn, Mairead Hyland, The Lighthouse Lab in Alderley Park and Alex Alderton, Roberto Amato, Jeffrey Barrett, Sonia Goncalves, Ewan Harrison, David K. Jackson, Ian Johnston, Dominic Kwiatkowski, Cordelia Langford, John Sillitoe on behalf of the Wellcome Sanger Institute COVID-19 Surveillance Team |
| EPI_ISL_2700046 | Lighthouse Lab in Milton Keynes | Wellcome Sanger Institute for the COVID-19 Genomics UK (COG-UK) Consortium | The Lighthouse Lab in Milton Keynes and Alex Alderton, Roberto Amato, Jeffrey Barrett, Sonia Goncalves, Ewan Harrison, David K. Jackson, Ian Johnston, Dominic Kwiatkowski, Cordelia Langford, John Sillitoe on behalf of the Wellcome Sanger Institute COVID-19 Surveillance Team |
| EPI_ISL_2700047 | Lighthouse Lab in Alderley Park | Wellcome Sanger Institute for the COVID-19 Genomics UK (COG-UK) Consortium | Jacquelyn Wynn, Mairead Hyland, The Lighthouse Lab in Alderley Park and Alex Alderton, Roberto Amato, Jeffrey Barrett, Sonia Goncalves, Ewan Harrison, David K. Jackson, Ian Johnston, Dominic Kwiatkowski, Cordelia Langford, John Sillitoe on behalf of the Wellcome Sanger Institute COVID-19 Surveillance Team |
| EPI_ISL_2700048, EPI_ISL_2700049 | Lighthouse Lab in Milton Keynes | Wellcome Sanger Institute for the COVID-19 Genomics UK (COG-UK) Consortium | The Lighthouse Lab in Milton Keynes and Alex Alderton, Roberto Amato, Jeffrey Barrett, Sonia Goncalves, Ewan Harrison, David K. Jackson, Ian Johnston, Dominic Kwiatkowski, Cordelia Langford, John Sillitoe on behalf of the Wellcome Sanger Institute COVID-19 Surveillance Team |
| EPI_ISL_2700050 | Lighthouse Lab in Alderley Park | Wellcome Sanger Institute for the COVID-19 Genomics UK (COG-UK) Consortium | Jacquelyn Wynn, Mairead Hyland, The Lighthouse Lab in Alderley Park and Alex Alderton, Roberto Amato, Jeffrey Barrett, Sonia Goncalves, Ewan Harrison, David K. Jackson, Ian Johnston, Dominic Kwiatkowski, Cordelia Langford, John Sillitoe on behalf of the Wellcome Sanger Institute COVID-19 Surveillance Team |
| EPI_ISL_2700051, EPI_ISL_2700052, EPI_ISL_2700053, EPI_ISL_2700054, EPI_ISL_2700055, EPI_ISL_2700056 | Lighthouse Lab in Milton Keynes | Wellcome Sanger Institute for the COVID-19 Genomics UK (COG-UK) Consortium | The Lighthouse Lab in Milton Keynes and Alex Alderton, Roberto Amato, Jeffrey Barrett, Sonia Goncalves, Ewan Harrison, David K. Jackson, Ian Johnston, Dominic Kwiatkowski, Cordelia Langford, John Sillitoe on behalf of the Wellcome Sanger Institute COVID-19 Surveillance Team |
| EPI_ISL_2700057 | Lighthouse Lab in Alderley Park | Wellcome Sanger Institute for the COVID-19 Genomics UK (COG-UK) Consortium | Jacquelyn Wynn, Mairead Hyland, The Lighthouse Lab in Alderley Park and Alex Alderton, Roberto Amato, Jeffrey Barrett, Sonia Goncalves, Ewan Harrison, David K. Jackson, Ian Johnston, Dominic Kwiatkowski, Cordelia Langford, John Sillitoe on behalf of the Wellcome Sanger Institute COVID-19 Surveillance Team |
| EPI_ISL_2700058, EPI_ISL_2700059, EPI_ISL_2700060, EPI_ISL_2700061, EPI_ISL_2700062, EPI_ISL_2700063, EPI_ISL_2700064, EPI_ISL_2700065, EPI_ISL_2700066, EPI_ISL_2700067, EPI_ISL_2700068 | | | |
| see above | Lighthouse Lab in Milton Keynes | Wellcome Sanger Institute for the COVID-19 Genomics UK (COG-UK) Consortium | The Lighthouse Lab in Milton Keynes and Alex Alderton, Roberto Amato, Jeffrey Barrett, Sonia Goncalves, Ewan Harrison, David K. Jackson, Ian Johnston, Dominic Kwiatkowski, Cordelia Langford, John Sillitoe on behalf of the Wellcome Sanger Institute COVID-19 Surveillance Team |
| EPI_ISL_2700069 | Lighthouse Lab in Alderley Park | Wellcome Sanger Institute for the COVID-19 Genomics UK (COG-UK) Consortium | Jacquelyn Wynn, Mairead Hyland, The Lighthouse Lab in Alderley Park and Alex Alderton, Roberto Amato, Jeffrey Barrett, Sonia Goncalves, Ewan Harrison, David K. Jackson, Ian Johnston, Dominic Kwiatkowski, Cordelia Langford, John Sillitoe on behalf of the Wellcome Sanger Institute COVID-19 Surveillance Team |
| EPI_ISL_2700070, EPI_ISL_2700071, EPI_ISL_2700072, EPI_ISL_2700073, EPI_ISL_2700074, EPI_ISL_2700075, EPI_ISL_2700076 | Lighthouse Lab in Milton Keynes | Wellcome Sanger Institute for the COVID-19 Genomics UK (COG-UK) Consortium | The Lighthouse Lab in Milton Keynes and Alex Alderton, Roberto Amato, Jeffrey Barrett, Sonia Goncalves, Ewan Harrison, David K. Jackson, Ian Johnston, Dominic Kwiatkowski, Cordelia Langford, John Sillitoe on behalf of the Wellcome Sanger Institute COVID-19 Surveillance Team |
| EPI_ISL_2700077, EPI_ISL_2700078, EPI_ISL_2700079 | Lighthouse Lab in Alderley Park | Wellcome Sanger Institute for the COVID-19 Genomics UK (COG-UK) Consortium | Jacquelyn Wynn, Mairead Hyland, The Lighthouse Lab in Alderley Park and Alex Alderton, Roberto Amato, Jeffrey Barrett, Sonia Goncalves, Ewan Harrison, David K. Jackson, Ian Johnston, Dominic Kwiatkowski, Cordelia Langford, John Sillitoe on behalf of the Wellcome Sanger Institute COVID-19 Surveillance Team |
| EPI_ISL_2700080 | Lighthouse Lab in Milton Keynes | Wellcome Sanger Institute for the COVID-19 Genomics UK (COG-UK) Consortium | The Lighthouse Lab in Milton Keynes and Alex Alderton, Roberto Amato, Jeffrey Barrett, Sonia Goncalves, Ewan Harrison, David K. Jackson, Ian Johnston, Dominic Kwiatkowski, Cordelia Langford, John Sillitoe on behalf of the Wellcome Sanger Institute COVID-19 Surveillance Team |
| EPI_ISL_2700081, EPI_ISL_2700082, EPI_ISL_2700083 | Lighthouse Lab in Alderley Park | Wellcome Sanger Institute for the COVID-19 Genomics UK (COG-UK) Consortium | Jacquelyn Wynn, Mairead Hyland, The Lighthouse Lab in Alderley Park and Alex Alderton, Roberto Amato, Jeffrey Barrett, Sonia Goncalves, Ewan Harrison, David K. Jackson, Ian Johnston, Dominic Kwiatkowski, Cordelia Langford, John Sillitoe on behalf of the Wellcome Sanger Institute COVID-19 Surveillance Team |
| EPI_ISL_2700084, EPI_ISL_2700085, EPI_ISL_2700086, EPI_ISL_2700087, EPI_ISL_2700088 | Lighthouse Lab in Milton Keynes | Wellcome Sanger Institute for the COVID-19 Genomics UK (COG-UK) Consortium | The Lighthouse Lab in Milton Keynes and Alex Alderton, Roberto Amato, Jeffrey Barrett, Sonia Goncalves, Ewan Harrison, David K. Jackson, Ian Johnston, Dominic Kwiatkowski, Cordelia Langford, John Sillitoe on behalf of the Wellcome Sanger Institute COVID-19 Surveillance Team |
| EPI_ISL_2700089 | Lighthouse Lab in Alderley Park | Wellcome Sanger Institute for the COVID-19 Genomics UK (COG-UK) Consortium | Jacquelyn Wynn, Mairead Hyland, The Lighthouse Lab in Alderley Park and Alex Alderton, Roberto Amato, Jeffrey Barrett, Sonia Goncalves, Ewan Harrison, David K. Jackson, Ian Johnston, Dominic Kwiatkowski, Cordelia Langford, John Sillitoe on behalf of the Wellcome Sanger Institute COVID-19 Surveillance Team |
| EPI_ISL_2700090, EPI_ISL_2700091, EPI_ISL_2700092, EPI_ISL_2700093, EPI_ISL_2700094, EPI_ISL_2700095, EPI_ISL_2700096 | Lighthouse Lab in Milton Keynes | Wellcome Sanger Institute for the COVID-19 Genomics UK (COG-UK) Consortium | The Lighthouse Lab in Milton Keynes and Alex Alderton, Roberto Amato, Jeffrey Barrett, Sonia Goncalves, Ewan Harrison, David K. Jackson, Ian Johnston, Dominic Kwiatkowski, Cordelia Langford, John Sillitoe on behalf of the Wellcome Sanger Institute COVID-19 Surveillance Team |
| EPI_ISL_2700097, EPI_ISL_2700098, EPI_ISL_2700099 | Lighthouse Lab in Alderley Park | Wellcome Sanger Institute for the COVID-19 Genomics UK (COG-UK) Consortium | Jacquelyn Wynn, Mairead Hyland, The Lighthouse Lab in Alderley Park and Alex Alderton, Roberto Amato, Jeffrey Barrett, Sonia Goncalves, Ewan Harrison, David K. Jackson, Ian Johnston, Dominic Kwiatkowski, Cordelia Langford, John Sillitoe on behalf of the Wellcome Sanger Institute COVID-19 Surveillance Team |
| EPI_ISL_2700100 | Lighthouse Lab in Milton Keynes | Wellcome Sanger Institute for the COVID-19 Genomics UK (COG-UK) Consortium | The Lighthouse Lab in Milton Keynes and Alex Alderton, Roberto Amato, Jeffrey Barrett, Sonia Goncalves, Ewan Harrison, David K. Jackson, Ian Johnston, Dominic Kwiatkowski, Cordelia Langford, John Sillitoe on behalf of the Wellcome Sanger Institute COVID-19 Surveillance Team |
| EPI_ISL_2700101 | Lighthouse Lab in Alderley Park | Wellcome Sanger Institute for the COVID-19 Genomics UK (COG-UK) Consortium | Jacquelyn Wynn, Mairead Hyland, The Lighthouse Lab in Alderley Park and Alex Alderton, Roberto Amato, Jeffrey Barrett, Sonia Goncalves, Ewan Harrison, David K. Jackson, Ian Johnston, Dominic Kwiatkowski, Cordelia Langford, John Sillitoe on behalf of the Wellcome Sanger Institute COVID-19 Surveillance Team |
| EPI_ISL_2700102, | Lighthouse Lab in Milton Keynes | Wellcome Sanger Institute for the COVID-19 Genomics UK (COG-UK) | The Lighthouse Lab in Milton Keynes and Alex Alderton, Roberto Amato, Jeffrey Barrett, Sonia Goncalves, Ewan Harrison, David K. Jackson, Ian Johnston, Dominic Kwiatkowski, Cordelia Langford, John Sillitoe on behalf of the Wellcome Sanger |

| | | | |
|---|---|---|---|
| EPI_ISL_2700103 | | Consortium | Institute COVID-19 Surveillance Team |
| EPI_ISL_2700104 | Lighthouse Lab in Alderley Park | Wellcome Sanger Institute for the COVID-19 Genomics UK (COG-UK) Consortium | Jacquelyn Wynn, Mairead Hyland, The Lighthouse Lab in Alderley Park and Alex Alderton, Roberto Amato, Jeffrey Barrett, Sonia Goncalves, Ewan Harrison, David K. Jackson, Ian Johnston, Dominic Kwiatkowski, Cordelia Langford, John Sillitoe on behalf of the Wellcome Sanger Institute COVID-19 Surveillance Team |
| EPI_ISL_2700105, EPI_ISL_2700106, EPI_ISL_2700107, EPI_ISL_2700108, EPI_ISL_2700109, EPI_ISL_2700110, EPI_ISL_2700111, EPI_ISL_2700112, EPI_ISL_2700113 | Lighthouse Lab in Milton Keynes | Wellcome Sanger Institute for the COVID-19 Genomics UK (COG-UK) Consortium | The Lighthouse Lab in Milton Keynes and Alex Alderton, Roberto Amato, Jeffrey Barrett, Sonia Goncalves, Ewan Harrison, David K. Jackson, Ian Johnston, Dominic Kwiatkowski, Cordelia Langford, John Sillitoe on behalf of the Wellcome Sanger Institute COVID-19 Surveillance Team |
| EPI_ISL_2700114 | Lighthouse Lab in Alderley Park | Wellcome Sanger Institute for the COVID-19 Genomics UK (COG-UK) Consortium | Jacquelyn Wynn, Mairead Hyland, The Lighthouse Lab in Alderley Park and Alex Alderton, Roberto Amato, Jeffrey Barrett, Sonia Goncalves, Ewan Harrison, David K. Jackson, Ian Johnston, Dominic Kwiatkowski, Cordelia Langford, John Sillitoe on behalf of the Wellcome Sanger Institute COVID-19 Surveillance Team |
| EPI_ISL_2700115, EPI_ISL_2700116, EPI_ISL_2700117, EPI_ISL_2700118, EPI_ISL_2700119, EPI_ISL_2700120 | Lighthouse Lab in Milton Keynes | Wellcome Sanger Institute for the COVID-19 Genomics UK (COG-UK) Consortium | The Lighthouse Lab in Milton Keynes and Alex Alderton, Roberto Amato, Jeffrey Barrett, Sonia Goncalves, Ewan Harrison, David K. Jackson, Ian Johnston, Dominic Kwiatkowski, Cordelia Langford, John Sillitoe on behalf of the Wellcome Sanger Institute COVID-19 Surveillance Team |
| EPI_ISL_2700121, EPI_ISL_2700122 | Lighthouse Lab in Alderley Park | Wellcome Sanger Institute for the COVID-19 Genomics UK (COG-UK) Consortium | Jacquelyn Wynn, Mairead Hyland, The Lighthouse Lab in Alderley Park and Alex Alderton, Roberto Amato, Jeffrey Barrett, Sonia Goncalves, Ewan Harrison, David K. Jackson, Ian Johnston, Dominic Kwiatkowski, Cordelia Langford, John Sillitoe on behalf of the Wellcome Sanger Institute COVID-19 Surveillance Team |
| EPI_ISL_2700123 | Lighthouse Lab in Milton Keynes | Wellcome Sanger Institute for the COVID-19 Genomics UK (COG-UK) Consortium | The Lighthouse Lab in Milton Keynes and Alex Alderton, Roberto Amato, Jeffrey Barrett, Sonia Goncalves, Ewan Harrison, David K. Jackson, Ian Johnston, Dominic Kwiatkowski, Cordelia Langford, John Sillitoe on behalf of the Wellcome Sanger Institute COVID-19 Surveillance Team |
| EPI_ISL_2700124 | Lighthouse Lab in Alderley Park | Wellcome Sanger Institute for the COVID-19 Genomics UK (COG-UK) Consortium | Jacquelyn Wynn, Mairead Hyland, The Lighthouse Lab in Alderley Park and Alex Alderton, Roberto Amato, Jeffrey Barrett, Sonia Goncalves, Ewan Harrison, David K. Jackson, Ian Johnston, Dominic Kwiatkowski, Cordelia Langford, John Sillitoe on behalf of the Wellcome Sanger Institute COVID-19 Surveillance Team |
| EPI_ISL_2700125, EPI_ISL_2700126, EPI_ISL_2700127 | Lighthouse Lab in Milton Keynes | Wellcome Sanger Institute for the COVID-19 Genomics UK (COG-UK) Consortium | The Lighthouse Lab in Milton Keynes and Alex Alderton, Roberto Amato, Jeffrey Barrett, Sonia Goncalves, Ewan Harrison, David K. Jackson, Ian Johnston, Dominic Kwiatkowski, Cordelia Langford, John Sillitoe on behalf of the Wellcome Sanger Institute COVID-19 Surveillance Team |
| EPI_ISL_2700128 | Lighthouse Lab in Alderley Park | Wellcome Sanger Institute for the COVID-19 Genomics UK (COG-UK) Consortium | Jacquelyn Wynn, Mairead Hyland, The Lighthouse Lab in Alderley Park and Alex Alderton, Roberto Amato, Jeffrey Barrett, Sonia Goncalves, Ewan Harrison, David K. Jackson, Ian Johnston, Dominic Kwiatkowski, Cordelia Langford, John Sillitoe on behalf of the Wellcome Sanger Institute COVID-19 Surveillance Team |
| EPI_ISL_2700129, EPI_ISL_2700130, EPI_ISL_2700131, EPI_ISL_2700132, EPI_ISL_2700133, EPI_ISL_2700134, EPI_ISL_2700135, EPI_ISL_2700136, EPI_ISL_2700137, EPI_ISL_2700138, EPI_ISL_2700139, EPI_ISL_2700140 | | | |
| see above | Lighthouse Lab in Milton Keynes | Wellcome Sanger Institute for the COVID-19 Genomics UK (COG-UK) Consortium | The Lighthouse Lab in Milton Keynes and Alex Alderton, Roberto Amato, Jeffrey Barrett, Sonia Goncalves, Ewan Harrison, David K. Jackson, Ian Johnston, Dominic Kwiatkowski, Cordelia Langford, John Sillitoe on behalf of the Wellcome Sanger Institute COVID-19 Surveillance Team |
| EPI_ISL_2700141, EPI_ISL_2700142, EPI_ISL_2700143 | Lighthouse Lab in Alderley Park | Wellcome Sanger Institute for the COVID-19 Genomics UK (COG-UK) Consortium | Jacquelyn Wynn, Mairead Hyland, The Lighthouse Lab in Alderley Park and Alex Alderton, Roberto Amato, Jeffrey Barrett, Sonia Goncalves, Ewan Harrison, David K. Jackson, Ian Johnston, Dominic Kwiatkowski, Cordelia Langford, John Sillitoe on behalf of the Wellcome Sanger Institute COVID-19 Surveillance Team |
| EPI_ISL_2700144, EPI_ISL_2700145, EPI_ISL_2700146, EPI_ISL_2700147, EPI_ISL_2700148, EPI_ISL_2700149, EPI_ISL_2700150, EPI_ISL_2700151 | Lighthouse Lab in Milton Keynes | Wellcome Sanger Institute for the COVID-19 Genomics UK (COG-UK) Consortium | The Lighthouse Lab in Milton Keynes and Alex Alderton, Roberto Amato, Jeffrey Barrett, Sonia Goncalves, Ewan Harrison, David K. Jackson, Ian Johnston, Dominic Kwiatkowski, Cordelia Langford, John Sillitoe on behalf of the Wellcome Sanger Institute COVID-19 Surveillance Team |
| EPI_ISL_2700152 | Lighthouse Lab in Alderley Park | Wellcome Sanger Institute for the COVID-19 Genomics UK (COG-UK) Consortium | Jacquelyn Wynn, Mairead Hyland, The Lighthouse Lab in Alderley Park and Alex Alderton, Roberto Amato, Jeffrey Barrett, Sonia Goncalves, Ewan Harrison, David K. Jackson, Ian Johnston, Dominic Kwiatkowski, Cordelia Langford, John Sillitoe on behalf of the Wellcome Sanger Institute COVID-19 Surveillance Team |
| EPI_ISL_2700153, EPI_ISL_2700154, EPI_ISL_2700155, EPI_ISL_2700156, EPI_ISL_2700157, EPI_ISL_2700158 | Lighthouse Lab in Milton Keynes | Wellcome Sanger Institute for the COVID-19 Genomics UK (COG-UK) Consortium | The Lighthouse Lab in Milton Keynes and Alex Alderton, Roberto Amato, Jeffrey Barrett, Sonia Goncalves, Ewan Harrison, David K. Jackson, Ian Johnston, Dominic Kwiatkowski, Cordelia Langford, John Sillitoe on behalf of the Wellcome Sanger Institute COVID-19 Surveillance Team |
| EPI_ISL_2700159 | Lighthouse Lab in Alderley Park | Wellcome Sanger Institute for the COVID-19 Genomics UK (COG-UK) Consortium | Jacquelyn Wynn, Mairead Hyland, The Lighthouse Lab in Alderley Park and Alex Alderton, Roberto Amato, Jeffrey Barrett, Sonia Goncalves, Ewan Harrison, David K. Jackson, Ian Johnston, Dominic Kwiatkowski, Cordelia Langford, John Sillitoe on behalf of the Wellcome Sanger Institute COVID-19 Surveillance Team |
| EPI_ISL_2700160, EPI_ISL_2700161, EPI_ISL_2700162, EPI_ISL_2700163 | Lighthouse Lab in Milton Keynes | Wellcome Sanger Institute for the COVID-19 Genomics UK (COG-UK) Consortium | The Lighthouse Lab in Milton Keynes and Alex Alderton, Roberto Amato, Jeffrey Barrett, Sonia Goncalves, Ewan Harrison, David K. Jackson, Ian Johnston, Dominic Kwiatkowski, Cordelia Langford, John Sillitoe on behalf of the Wellcome Sanger Institute COVID-19 Surveillance Team |
| EPI_ISL_2700164, EPI_ISL_2700165, EPI_ISL_2700166 | Lighthouse Lab in Alderley Park | Wellcome Sanger Institute for the COVID-19 Genomics UK (COG-UK) Consortium | Jacquelyn Wynn, Mairead Hyland, The Lighthouse Lab in Alderley Park and Alex Alderton, Roberto Amato, Jeffrey Barrett, Sonia Goncalves, Ewan Harrison, David K. Jackson, Ian Johnston, Dominic Kwiatkowski, Cordelia Langford, John Sillitoe on behalf of the Wellcome Sanger Institute COVID-19 Surveillance Team |
| EPI_ISL_2700167, EPI_ISL_2700168, EPI_ISL_2700169, EPI_ISL_2700170 | Lighthouse Lab in Milton Keynes | Wellcome Sanger Institute for the COVID-19 Genomics UK (COG-UK) Consortium | The Lighthouse Lab in Milton Keynes and Alex Alderton, Roberto Amato, Jeffrey Barrett, Sonia Goncalves, Ewan Harrison, David K. Jackson, Ian Johnston, Dominic Kwiatkowski, Cordelia Langford, John Sillitoe on behalf of the Wellcome Sanger Institute COVID-19 Surveillance Team |
| EPI_ISL_2700171 | Lighthouse Lab in Alderley Park | Wellcome Sanger Institute for the COVID-19 Genomics UK (COG-UK) Consortium | Jacquelyn Wynn, Mairead Hyland, The Lighthouse Lab in Alderley Park and Alex Alderton, Roberto Amato, Jeffrey Barrett, Sonia Goncalves, Ewan Harrison, David K. Jackson, Ian Johnston, Dominic Kwiatkowski, Cordelia Langford, John Sillitoe on behalf of the Wellcome Sanger Institute COVID-19 Surveillance Team |
| EPI_ISL_2700172, EPI_ISL_2700173, EPI_ISL_2700174, EPI_ISL_2700175, EPI_ISL_2700176, EPI_ISL_2700177 | Lighthouse Lab in Milton Keynes | Wellcome Sanger Institute for the COVID-19 Genomics UK (COG-UK) Consortium | The Lighthouse Lab in Milton Keynes and Alex Alderton, Roberto Amato, Jeffrey Barrett, Sonia Goncalves, Ewan Harrison, David K. Jackson, Ian Johnston, Dominic Kwiatkowski, Cordelia Langford, John Sillitoe on behalf of the Wellcome Sanger Institute COVID-19 Surveillance Team |
| EPI_ISL_2700178 | Lighthouse Lab in Alderley Park | Wellcome Sanger Institute for the COVID-19 Genomics UK (COG-UK) Consortium | Jacquelyn Wynn, Mairead Hyland, The Lighthouse Lab in Alderley Park and Alex Alderton, Roberto Amato, Jeffrey Barrett, Sonia Goncalves, Ewan Harrison, David K. Jackson, Ian Johnston, Dominic Kwiatkowski, Cordelia Langford, John Sillitoe on behalf of the Wellcome Sanger Institute COVID-19 Surveillance Team |
| EPI_ISL_2700179, EPI_ISL_2700180, EPI_ISL_2700181, EPI_ISL_2700182 | Lighthouse Lab in Milton Keynes | Wellcome Sanger Institute for the COVID-19 Genomics UK (COG-UK) Consortium | The Lighthouse Lab in Milton Keynes and Alex Alderton, Roberto Amato, Jeffrey Barrett, Sonia Goncalves, Ewan Harrison, David K. Jackson, Ian Johnston, Dominic Kwiatkowski, Cordelia Langford, John Sillitoe on behalf of the Wellcome Sanger Institute COVID-19 Surveillance Team |
| EPI_ISL_2700183, EPI_ISL_2700184, EPI_ISL_2700185 | Lighthouse Lab in Alderley Park | Wellcome Sanger Institute for the COVID-19 Genomics UK (COG-UK) Consortium | Jacquelyn Wynn, Mairead Hyland, The Lighthouse Lab in Alderley Park and Alex Alderton, Roberto Amato, Jeffrey Barrett, Sonia Goncalves, Ewan Harrison, David K. Jackson, Ian Johnston, Dominic Kwiatkowski, Cordelia Langford, John Sillitoe on behalf of the Wellcome Sanger Institute COVID-19 Surveillance Team |
| EPI_ISL_2700186, EPI_ISL_2700187 | Lighthouse Lab in Milton Keynes | Wellcome Sanger Institute for the COVID-19 Genomics UK (COG-UK) Consortium | The Lighthouse Lab in Milton Keynes and Alex Alderton, Roberto Amato, Jeffrey Barrett, Sonia Goncalves, Ewan Harrison, David K. Jackson, Ian Johnston, Dominic Kwiatkowski, Cordelia Langford, John Sillitoe on behalf of the Wellcome Sanger Institute COVID-19 Surveillance Team |
| EPI_ISL_2700188 | Lighthouse Lab in Alderley Park | Wellcome Sanger Institute for the COVID-19 Genomics UK (COG-UK) Consortium | Jacquelyn Wynn, Mairead Hyland, The Lighthouse Lab in Alderley Park and Alex Alderton, Roberto Amato, Jeffrey Barrett, Sonia Goncalves, Ewan Harrison, David K. Jackson, Ian Johnston, Dominic Kwiatkowski, Cordelia Langford, John Sillitoe on behalf of the Wellcome Sanger Institute COVID-19 Surveillance Team |
| EPI_ISL_2700189, EPI_ISL_2700190 | Lighthouse Lab in Milton Keynes | Wellcome Sanger Institute for the COVID-19 Genomics UK (COG-UK) Consortium | The Lighthouse Lab in Milton Keynes and Alex Alderton, Roberto Amato, Jeffrey Barrett, Sonia Goncalves, Ewan Harrison, David K. Jackson, Ian Johnston, Dominic Kwiatkowski, Cordelia Langford, John Sillitoe on behalf of the Wellcome Sanger Institute COVID-19 Surveillance Team |
| EPI_ISL_2700191 | Lighthouse Lab in Alderley Park | Wellcome Sanger Institute for the COVID-19 Genomics UK (COG-UK) Consortium | Jacquelyn Wynn, Mairead Hyland, The Lighthouse Lab in Alderley Park and Alex Alderton, Roberto Amato, Jeffrey Barrett, Sonia Goncalves, Ewan Harrison, David K. Jackson, Ian Johnston, Dominic Kwiatkowski, Cordelia Langford, John Sillitoe on behalf of the Wellcome Sanger Institute COVID-19 Surveillance Team |
| EPI_ISL_2700192 | Lighthouse Lab in Milton Keynes | Wellcome Sanger Institute for the COVID-19 Genomics UK (COG-UK) Consortium | The Lighthouse Lab in Milton Keynes and Alex Alderton, Roberto Amato, Jeffrey Barrett, Sonia Goncalves, Ewan Harrison, David K. Jackson, Ian Johnston, Dominic Kwiatkowski, Cordelia Langford, John Sillitoe on behalf of the Wellcome Sanger Institute COVID-19 Surveillance Team |
| EPI_ISL_2700193 | Lighthouse Lab in Alderley Park | Wellcome Sanger Institute for the COVID-19 Genomics UK (COG-UK) Consortium | Jacquelyn Wynn, Mairead Hyland, The Lighthouse Lab in Alderley Park and Alex Alderton, Roberto Amato, Jeffrey Barrett, Sonia Goncalves, Ewan Harrison, David K. Jackson, Ian Johnston, Dominic Kwiatkowski, Cordelia Langford, John Sillitoe on behalf of the Wellcome Sanger Institute COVID-19 Surveillance Team |
| EPI_ISL_2700194 | Lighthouse Lab in Milton Keynes | Wellcome Sanger Institute for the COVID-19 Genomics UK (COG-UK) | The Lighthouse Lab in Milton Keynes and Alex Alderton, Roberto Amato, Jeffrey Barrett, Sonia Goncalves, Ewan Harrison, David K. Jackson, Ian Johnston, Dominic Kwiatkowski, Cordelia Langford, John Sillitoe on behalf of the Wellcome Sanger |

| | | | |
|---|---|---|---|
| | | Consortium | Institute COVID-19 Surveillance Team |
| EPI_ISL_2700195, EPI_ISL_2700196 | Lighthouse Lab in Alderley Park | Wellcome Sanger Institute for the COVID-19 Genomics UK (COG-UK) Consortium | Jacquelyn Wynn, Mairead Hyland, The Lighthouse Lab in Alderley Park and Alex Alderton, Roberto Amato, Jeffrey Barrett, Sonia Goncalves, Ewan Harrison, David K. Jackson, Ian Johnston, Dominic Kwiatkowski, Cordelia Langford, John Sillitoe on behalf of the Wellcome Sanger Institute COVID-19 Surveillance Team |
| EPI_ISL_2700197 | Lighthouse Lab in Milton Keynes | Wellcome Sanger Institute for the COVID-19 Genomics UK (COG-UK) Consortium | The Lighthouse Lab in Milton Keynes and Alex Alderton, Roberto Amato, Jeffrey Barrett, Sonia Goncalves, Ewan Harrison, David K. Jackson, Ian Johnston, Dominic Kwiatkowski, Cordelia Langford, John Sillitoe on behalf of the Wellcome Sanger Institute COVID-19 Surveillance Team |
| EPI_ISL_2700198 | Lighthouse Lab in Alderley Park | Wellcome Sanger Institute for the COVID-19 Genomics UK (COG-UK) Consortium | Jacquelyn Wynn, Mairead Hyland, The Lighthouse Lab in Alderley Park and Alex Alderton, Roberto Amato, Jeffrey Barrett, Sonia Goncalves, Ewan Harrison, David K. Jackson, Ian Johnston, Dominic Kwiatkowski, Cordelia Langford, John Sillitoe on behalf of the Wellcome Sanger Institute COVID-19 Surveillance Team |
| EPI_ISL_2700199 | Lighthouse Lab in Milton Keynes | Wellcome Sanger Institute for the COVID-19 Genomics UK (COG-UK) Consortium | The Lighthouse Lab in Milton Keynes and Alex Alderton, Roberto Amato, Jeffrey Barrett, Sonia Goncalves, Ewan Harrison, David K. Jackson, Ian Johnston, Dominic Kwiatkowski, Cordelia Langford, John Sillitoe on behalf of the Wellcome Sanger Institute COVID-19 Surveillance Team |
| EPI_ISL_2700200, EPI_ISL_2700201 | Lighthouse Lab in Alderley Park | Wellcome Sanger Institute for the COVID-19 Genomics UK (COG-UK) Consortium | Jacquelyn Wynn, Mairead Hyland, The Lighthouse Lab in Alderley Park and Alex Alderton, Roberto Amato, Jeffrey Barrett, Sonia Goncalves, Ewan Harrison, David K. Jackson, Ian Johnston, Dominic Kwiatkowski, Cordelia Langford, John Sillitoe on behalf of the Wellcome Sanger Institute COVID-19 Surveillance Team |
| EPI_ISL_2700202, EPI_ISL_2700203, EPI_ISL_2700204, EPI_ISL_2700205, EPI_ISL_2700206, EPI_ISL_2700207 | Lighthouse Lab in Milton Keynes | Wellcome Sanger Institute for the COVID-19 Genomics UK (COG-UK) Consortium | The Lighthouse Lab in Milton Keynes and Alex Alderton, Roberto Amato, Jeffrey Barrett, Sonia Goncalves, Ewan Harrison, David K. Jackson, Ian Johnston, Dominic Kwiatkowski, Cordelia Langford, John Sillitoe on behalf of the Wellcome Sanger Institute COVID-19 Surveillance Team |
| EPI_ISL_2700208, EPI_ISL_2700209 | Lighthouse Lab in Alderley Park | Wellcome Sanger Institute for the COVID-19 Genomics UK (COG-UK) Consortium | Jacquelyn Wynn, Mairead Hyland, The Lighthouse Lab in Alderley Park and Alex Alderton, Roberto Amato, Jeffrey Barrett, Sonia Goncalves, Ewan Harrison, David K. Jackson, Ian Johnston, Dominic Kwiatkowski, Cordelia Langford, John Sillitoe on behalf of the Wellcome Sanger Institute COVID-19 Surveillance Team |
| EPI_ISL_2700210, EPI_ISL_2700211 | Lighthouse Lab in Milton Keynes | Wellcome Sanger Institute for the COVID-19 Genomics UK (COG-UK) Consortium | The Lighthouse Lab in Milton Keynes and Alex Alderton, Roberto Amato, Jeffrey Barrett, Sonia Goncalves, Ewan Harrison, David K. Jackson, Ian Johnston, Dominic Kwiatkowski, Cordelia Langford, John Sillitoe on behalf of the Wellcome Sanger Institute COVID-19 Surveillance Team |
| EPI_ISL_2700212 | Lighthouse Lab in Alderley Park | Wellcome Sanger Institute for the COVID-19 Genomics UK (COG-UK) Consortium | Jacquelyn Wynn, Mairead Hyland, The Lighthouse Lab in Alderley Park and Alex Alderton, Roberto Amato, Jeffrey Barrett, Sonia Goncalves, Ewan Harrison, David K. Jackson, Ian Johnston, Dominic Kwiatkowski, Cordelia Langford, John Sillitoe on behalf of the Wellcome Sanger Institute COVID-19 Surveillance Team |
| EPI_ISL_2700213 | Lighthouse Lab in Milton Keynes | Wellcome Sanger Institute for the COVID-19 Genomics UK (COG-UK) Consortium | The Lighthouse Lab in Milton Keynes and Alex Alderton, Roberto Amato, Jeffrey Barrett, Sonia Goncalves, Ewan Harrison, David K. Jackson, Ian Johnston, Dominic Kwiatkowski, Cordelia Langford, John Sillitoe on behalf of the Wellcome Sanger Institute COVID-19 Surveillance Team |
| EPI_ISL_2700214 | Lighthouse Lab in Glasgow | Wellcome Sanger Institute for the COVID-19 Genomics UK (COG-UK) Consortium | Harper VanSteenhouse, Yumi Kasai, David Gray, Carol Clugston, Anna Dominiczak and Alex Alderton, Roberto Amato, Jeffrey Barrett, Sonia Goncalves, Ewan Harrison, David K. Jackson, Ian Johnston, Dominic Kwiatkowski, Cordelia Langford, John Sillitoe on behalf of the Wellcome Sanger Institute COVID-19 Surveillance Team |
| EPI_ISL_2700215 | Lighthouse Lab in Milton Keynes | Wellcome Sanger Institute for the COVID-19 Genomics UK (COG-UK) Consortium | The Lighthouse Lab in Milton Keynes and Alex Alderton, Roberto Amato, Jeffrey Barrett, Sonia Goncalves, Ewan Harrison, David K. Jackson, Ian Johnston, Dominic Kwiatkowski, Cordelia Langford, John Sillitoe on behalf of the Wellcome Sanger Institute COVID-19 Surveillance Team |
| EPI_ISL_2700216, EPI_ISL_2700217, EPI_ISL_2700218, EPI_ISL_2700219, EPI_ISL_2700220, EPI_ISL_2700221, EPI_ISL_2700222, EPI_ISL_2700223, EPI_ISL_2700224, EPI_ISL_2700226 | Lighthouse Lab in Glasgow | Wellcome Sanger Institute for the COVID-19 Genomics UK (COG-UK) Consortium | Harper VanSteenhouse, Yumi Kasai, David Gray, Carol Clugston, Anna Dominiczak and Alex Alderton, Roberto Amato, Jeffrey Barrett, Sonia Goncalves, Ewan Harrison, David K. Jackson, Ian Johnston, Dominic Kwiatkowski, Cordelia Langford, John Sillitoe on behalf of the Wellcome Sanger Institute COVID-19 Surveillance Team |
| EPI_ISL_2700227 | Lighthouse Lab in Milton Keynes | Wellcome Sanger Institute for the COVID-19 Genomics UK (COG-UK) Consortium | The Lighthouse Lab in Milton Keynes and Alex Alderton, Roberto Amato, Jeffrey Barrett, Sonia Goncalves, Ewan Harrison, David K. Jackson, Ian Johnston, Dominic Kwiatkowski, Cordelia Langford, John Sillitoe on behalf of the Wellcome Sanger Institute COVID-19 Surveillance Team |
| EPI_ISL_2700228, EPI_ISL_2700229 | Lighthouse Lab in Glasgow | Wellcome Sanger Institute for the COVID-19 Genomics UK (COG-UK) Consortium | Harper VanSteenhouse, Yumi Kasai, David Gray, Carol Clugston, Anna Dominiczak and Alex Alderton, Roberto Amato, Jeffrey Barrett, Sonia Goncalves, Ewan Harrison, David K. Jackson, Ian Johnston, Dominic Kwiatkowski, Cordelia Langford, John Sillitoe on behalf of the Wellcome Sanger Institute COVID-19 Surveillance Team |
| EPI_ISL_2700230, EPI_ISL_2700231, EPI_ISL_2700232, EPI_ISL_2700233 | Lighthouse Lab in Milton Keynes | Wellcome Sanger Institute for the COVID-19 Genomics UK (COG-UK) Consortium | The Lighthouse Lab in Milton Keynes and Alex Alderton, Roberto Amato, Jeffrey Barrett, Sonia Goncalves, Ewan Harrison, David K. Jackson, Ian Johnston, Dominic Kwiatkowski, Cordelia Langford, John Sillitoe on behalf of the Wellcome Sanger Institute COVID-19 Surveillance Team |
| EPI_ISL_2700234 | Lighthouse Lab in Glasgow | Wellcome Sanger Institute for the COVID-19 Genomics UK (COG-UK) Consortium | Harper VanSteenhouse, Yumi Kasai, David Gray, Carol Clugston, Anna Dominiczak and Alex Alderton, Roberto Amato, Jeffrey Barrett, Sonia Goncalves, Ewan Harrison, David K. Jackson, Ian Johnston, Dominic Kwiatkowski, Cordelia Langford, John Sillitoe on behalf of the Wellcome Sanger Institute COVID-19 Surveillance Team |
| EPI_ISL_2700235, EPI_ISL_2700236, EPI_ISL_2700237, EPI_ISL_2700238, EPI_ISL_2700239, EPI_ISL_2700240 | Lighthouse Lab in Milton Keynes | Wellcome Sanger Institute for the COVID-19 Genomics UK (COG-UK) Consortium | The Lighthouse Lab in Milton Keynes and Alex Alderton, Roberto Amato, Jeffrey Barrett, Sonia Goncalves, Ewan Harrison, David K. Jackson, Ian Johnston, Dominic Kwiatkowski, Cordelia Langford, John Sillitoe on behalf of the Wellcome Sanger Institute COVID-19 Surveillance Team |
| EPI_ISL_2700242 | Lighthouse Lab in Glasgow | Wellcome Sanger Institute for the COVID-19 Genomics UK (COG-UK) Consortium | Harper VanSteenhouse, Yumi Kasai, David Gray, Carol Clugston, Anna Dominiczak and Alex Alderton, Roberto Amato, Jeffrey Barrett, Sonia Goncalves, Ewan Harrison, David K. Jackson, Ian Johnston, Dominic Kwiatkowski, Cordelia Langford, John Sillitoe on behalf of the Wellcome Sanger Institute COVID-19 Surveillance Team |
| EPI_ISL_2700243, EPI_ISL_2700244 | Lighthouse Lab in Milton Keynes | Wellcome Sanger Institute for the COVID-19 Genomics UK (COG-UK) Consortium | The Lighthouse Lab in Milton Keynes and Alex Alderton, Roberto Amato, Jeffrey Barrett, Sonia Goncalves, Ewan Harrison, David K. Jackson, Ian Johnston, Dominic Kwiatkowski, Cordelia Langford, John Sillitoe on behalf of the Wellcome Sanger Institute COVID-19 Surveillance Team |
| EPI_ISL_2700245 | Lighthouse Lab in Glasgow | Wellcome Sanger Institute for the COVID-19 Genomics UK (COG-UK) Consortium | Harper VanSteenhouse, Yumi Kasai, David Gray, Carol Clugston, Anna Dominiczak and Alex Alderton, Roberto Amato, Jeffrey Barrett, Sonia Goncalves, Ewan Harrison, David K. Jackson, Ian Johnston, Dominic Kwiatkowski, Cordelia Langford, John Sillitoe on behalf of the Wellcome Sanger Institute COVID-19 Surveillance Team |
| EPI_ISL_2700246 | Lighthouse Lab in Milton Keynes | Wellcome Sanger Institute for the COVID-19 Genomics UK (COG-UK) Consortium | The Lighthouse Lab in Milton Keynes and Alex Alderton, Roberto Amato, Jeffrey Barrett, Sonia Goncalves, Ewan Harrison, David K. Jackson, Ian Johnston, Dominic Kwiatkowski, Cordelia Langford, John Sillitoe on behalf of the Wellcome Sanger Institute COVID-19 Surveillance Team |
| EPI_ISL_2700247 | Lighthouse Lab in Glasgow | Wellcome Sanger Institute for the COVID-19 Genomics UK (COG-UK) Consortium | Harper VanSteenhouse, Yumi Kasai, David Gray, Carol Clugston, Anna Dominiczak and Alex Alderton, Roberto Amato, Jeffrey Barrett, Sonia Goncalves, Ewan Harrison, David K. Jackson, Ian Johnston, Dominic Kwiatkowski, Cordelia Langford, John Sillitoe on behalf of the Wellcome Sanger Institute COVID-19 Surveillance Team |
| EPI_ISL_2700248, EPI_ISL_2700249, EPI_ISL_2700250, EPI_ISL_2700251, EPI_ISL_2700252, EPI_ISL_2700253, EPI_ISL_2700255 | Lighthouse Lab in Milton Keynes | Wellcome Sanger Institute for the COVID-19 Genomics UK (COG-UK) Consortium | The Lighthouse Lab in Milton Keynes and Alex Alderton, Roberto Amato, Jeffrey Barrett, Sonia Goncalves, Ewan Harrison, David K. Jackson, Ian Johnston, Dominic Kwiatkowski, Cordelia Langford, John Sillitoe on behalf of the Wellcome Sanger Institute COVID-19 Surveillance Team |
| EPI_ISL_2700256, EPI_ISL_2700257, EPI_ISL_2700258, EPI_ISL_2700259, EPI_ISL_2700260 | Lighthouse Lab in Glasgow | Wellcome Sanger Institute for the COVID-19 Genomics UK (COG-UK) Consortium | Harper VanSteenhouse, Yumi Kasai, David Gray, Carol Clugston, Anna Dominiczak and Alex Alderton, Roberto Amato, Jeffrey Barrett, Sonia Goncalves, Ewan Harrison, David K. Jackson, Ian Johnston, Dominic Kwiatkowski, Cordelia Langford, John Sillitoe on behalf of the Wellcome Sanger Institute COVID-19 Surveillance Team |
| EPI_ISL_2700261, EPI_ISL_2700262, EPI_ISL_2700263 | Lighthouse Lab in Milton Keynes | Wellcome Sanger Institute for the COVID-19 Genomics UK (COG-UK) Consortium | The Lighthouse Lab in Milton Keynes and Alex Alderton, Roberto Amato, Jeffrey Barrett, Sonia Goncalves, Ewan Harrison, David K. Jackson, Ian Johnston, Dominic Kwiatkowski, Cordelia Langford, John Sillitoe on behalf of the Wellcome Sanger Institute COVID-19 Surveillance Team |
| EPI_ISL_2700264, EPI_ISL_2700265, EPI_ISL_2700266, EPI_ISL_2700267, EPI_ISL_2700268, EPI_ISL_2700269, EPI_ISL_2700270 | Lighthouse Lab in Glasgow | Wellcome Sanger Institute for the COVID-19 Genomics UK (COG-UK) Consortium | Harper VanSteenhouse, Yumi Kasai, David Gray, Carol Clugston, Anna Dominiczak and Alex Alderton, Roberto Amato, Jeffrey Barrett, Sonia Goncalves, Ewan Harrison, David K. Jackson, Ian Johnston, Dominic Kwiatkowski, Cordelia Langford, John Sillitoe on behalf of the Wellcome Sanger Institute COVID-19 Surveillance Team |
| EPI_ISL_2700271, EPI_ISL_2700272 | Lighthouse Lab in Milton Keynes | Wellcome Sanger Institute for the COVID-19 Genomics UK (COG-UK) Consortium | The Lighthouse Lab in Milton Keynes and Alex Alderton, Roberto Amato, Jeffrey Barrett, Sonia Goncalves, Ewan Harrison, David K. Jackson, Ian Johnston, Dominic Kwiatkowski, Cordelia Langford, John Sillitoe on behalf of the Wellcome Sanger Institute COVID-19 Surveillance Team |
| EPI_ISL_2700273, EPI_ISL_2700274 | Lighthouse Lab in Glasgow | Wellcome Sanger Institute for the COVID-19 Genomics UK (COG-UK) Consortium | Harper VanSteenhouse, Yumi Kasai, David Gray, Carol Clugston, Anna Dominiczak and Alex Alderton, Roberto Amato, Jeffrey Barrett, Sonia Goncalves, Ewan Harrison, David K. Jackson, Ian Johnston, Dominic Kwiatkowski, Cordelia Langford, John Sillitoe on behalf of the Wellcome Sanger Institute COVID-19 Surveillance Team |
| EPI_ISL_2700275, EPI_ISL_2700276, EPI_ISL_2700278, EPI_ISL_2700279, EPI_ISL_2700280, EPI_ISL_2700281, | Lighthouse Lab in Milton Keynes | Wellcome Sanger Institute for the COVID-19 Genomics UK (COG-UK) Consortium | The Lighthouse Lab in Milton Keynes and Alex Alderton, Roberto Amato, Jeffrey Barrett, Sonia Goncalves, Ewan Harrison, David K. Jackson, Ian Johnston, Dominic Kwiatkowski, Cordelia Langford, John Sillitoe on behalf of the Wellcome Sanger Institute COVID-19 Surveillance Team |

| | | | |
|---|---|---|---|
| EPI_ISL_2700282, EPI_ISL_2700283, EPI_ISL_2700284 | | | |
| EPI_ISL_2700285, EPI_ISL_2700286, EPI_ISL_2700287, EPI_ISL_2700288, EPI_ISL_2700289, EPI_ISL_2700290 | Lighthouse Lab in Glasgow | Wellcome Sanger Institute for the COVID-19 Genomics UK (COG-UK) Consortium | Harper VanSteenhouse, Yumi Kasai, David Gray, Carol Clugston, Anna Dominiczak and Alex Alderton, Roberto Amato, Jeffrey Barrett, Sonia Goncalves, Ewan Harrison, David K. Jackson, Ian Johnston, Dominic Kwiatkowski, Cordelia Langford, John Sillitoe on behalf of the Wellcome Sanger Institute COVID-19 Surveillance Team |
| EPI_ISL_2700291, EPI_ISL_2700292, EPI_ISL_2700293 | Lighthouse Lab in Milton Keynes | Wellcome Sanger Institute for the COVID-19 Genomics UK (COG-UK) Consortium | The Lighthouse Lab in Milton Keynes and Alex Alderton, Roberto Amato, Jeffrey Barrett, Sonia Goncalves, Ewan Harrison, David K. Jackson, Ian Johnston, Dominic Kwiatkowski, Cordelia Langford, John Sillitoe on behalf of the Wellcome Sanger Institute COVID-19 Surveillance Team |
| EPI_ISL_2700294, EPI_ISL_2700295 | Lighthouse Lab in Glasgow | Wellcome Sanger Institute for the COVID-19 Genomics UK (COG-UK) Consortium | Harper VanSteenhouse, Yumi Kasai, David Gray, Carol Clugston, Anna Dominiczak and Alex Alderton, Roberto Amato, Jeffrey Barrett, Sonia Goncalves, Ewan Harrison, David K. Jackson, Ian Johnston, Dominic Kwiatkowski, Cordelia Langford, John Sillitoe on behalf of the Wellcome Sanger Institute COVID-19 Surveillance Team |
| EPI_ISL_2700296 | Lighthouse Lab in Milton Keynes | Wellcome Sanger Institute for the COVID-19 Genomics UK (COG-UK) Consortium | The Lighthouse Lab in Milton Keynes and Alex Alderton, Roberto Amato, Jeffrey Barrett, Sonia Goncalves, Ewan Harrison, David K. Jackson, Ian Johnston, Dominic Kwiatkowski, Cordelia Langford, John Sillitoe on behalf of the Wellcome Sanger Institute COVID-19 Surveillance Team |
| EPI_ISL_2700297 | Lighthouse Lab in Glasgow | Wellcome Sanger Institute for the COVID-19 Genomics UK (COG-UK) Consortium | Harper VanSteenhouse, Yumi Kasai, David Gray, Carol Clugston, Anna Dominiczak and Alex Alderton, Roberto Amato, Jeffrey Barrett, Sonia Goncalves, Ewan Harrison, David K. Jackson, Ian Johnston, Dominic Kwiatkowski, Cordelia Langford, John Sillitoe on behalf of the Wellcome Sanger Institute COVID-19 Surveillance Team |
| EPI_ISL_2700298 | Lighthouse Lab in Milton Keynes | Wellcome Sanger Institute for the COVID-19 Genomics UK (COG-UK) Consortium | The Lighthouse Lab in Milton Keynes and Alex Alderton, Roberto Amato, Jeffrey Barrett, Sonia Goncalves, Ewan Harrison, David K. Jackson, Ian Johnston, Dominic Kwiatkowski, Cordelia Langford, John Sillitoe on behalf of the Wellcome Sanger Institute COVID-19 Surveillance Team |
| EPI_ISL_2700299 | Lighthouse Lab in Glasgow | Wellcome Sanger Institute for the COVID-19 Genomics UK (COG-UK) Consortium | Harper VanSteenhouse, Yumi Kasai, David Gray, Carol Clugston, Anna Dominiczak and Alex Alderton, Roberto Amato, Jeffrey Barrett, Sonia Goncalves, Ewan Harrison, David K. Jackson, Ian Johnston, Dominic Kwiatkowski, Cordelia Langford, John Sillitoe on behalf of the Wellcome Sanger Institute COVID-19 Surveillance Team |
| EPI_ISL_2700300 | Lighthouse Lab in Milton Keynes | Wellcome Sanger Institute for the COVID-19 Genomics UK (COG-UK) Consortium | The Lighthouse Lab in Milton Keynes and Alex Alderton, Roberto Amato, Jeffrey Barrett, Sonia Goncalves, Ewan Harrison, David K. Jackson, Ian Johnston, Dominic Kwiatkowski, Cordelia Langford, John Sillitoe on behalf of the Wellcome Sanger Institute COVID-19 Surveillance Team |
| EPI_ISL_2700301 | Lighthouse Lab in Glasgow | Wellcome Sanger Institute for the COVID-19 Genomics UK (COG-UK) Consortium | Harper VanSteenhouse, Yumi Kasai, David Gray, Carol Clugston, Anna Dominiczak and Alex Alderton, Roberto Amato, Jeffrey Barrett, Sonia Goncalves, Ewan Harrison, David K. Jackson, Ian Johnston, Dominic Kwiatkowski, Cordelia Langford, John Sillitoe on behalf of the Wellcome Sanger Institute COVID-19 Surveillance Team |
| EPI_ISL_2700302, EPI_ISL_2700303 | Lighthouse Lab in Milton Keynes | Wellcome Sanger Institute for the COVID-19 Genomics UK (COG-UK) Consortium | The Lighthouse Lab in Milton Keynes and Alex Alderton, Roberto Amato, Jeffrey Barrett, Sonia Goncalves, Ewan Harrison, David K. Jackson, Ian Johnston, Dominic Kwiatkowski, Cordelia Langford, John Sillitoe on behalf of the Wellcome Sanger Institute COVID-19 Surveillance Team |
| EPI_ISL_2700304 | Lighthouse Lab in Glasgow | Wellcome Sanger Institute for the COVID-19 Genomics UK (COG-UK) Consortium | Harper VanSteenhouse, Yumi Kasai, David Gray, Carol Clugston, Anna Dominiczak and Alex Alderton, Roberto Amato, Jeffrey Barrett, Sonia Goncalves, Ewan Harrison, David K. Jackson, Ian Johnston, Dominic Kwiatkowski, Cordelia Langford, John Sillitoe on behalf of the Wellcome Sanger Institute COVID-19 Surveillance Team |
| EPI_ISL_2700305, EPI_ISL_2700306, EPI_ISL_2700307 | Lighthouse Lab in Milton Keynes | Wellcome Sanger Institute for the COVID-19 Genomics UK (COG-UK) Consortium | The Lighthouse Lab in Milton Keynes and Alex Alderton, Roberto Amato, Jeffrey Barrett, Sonia Goncalves, Ewan Harrison, David K. Jackson, Ian Johnston, Dominic Kwiatkowski, Cordelia Langford, John Sillitoe on behalf of the Wellcome Sanger Institute COVID-19 Surveillance Team |
| EPI_ISL_2700308, EPI_ISL_2700309 | Lighthouse Lab in Glasgow | Wellcome Sanger Institute for the COVID-19 Genomics UK (COG-UK) Consortium | Harper VanSteenhouse, Yumi Kasai, David Gray, Carol Clugston, Anna Dominiczak and Alex Alderton, Roberto Amato, Jeffrey Barrett, Sonia Goncalves, Ewan Harrison, David K. Jackson, Ian Johnston, Dominic Kwiatkowski, Cordelia Langford, John Sillitoe on behalf of the Wellcome Sanger Institute COVID-19 Surveillance Team |
| EPI_ISL_2700310 | Lighthouse Lab in Milton Keynes | Wellcome Sanger Institute for the COVID-19 Genomics UK (COG-UK) Consortium | The Lighthouse Lab in Milton Keynes and Alex Alderton, Roberto Amato, Jeffrey Barrett, Sonia Goncalves, Ewan Harrison, David K. Jackson, Ian Johnston, Dominic Kwiatkowski, Cordelia Langford, John Sillitoe on behalf of the Wellcome Sanger Institute COVID-19 Surveillance Team |
| EPI_ISL_2700311, EPI_ISL_2700312, EPI_ISL_2700313, EPI_ISL_2700314, EPI_ISL_2700315 | Lighthouse Lab in Glasgow | Wellcome Sanger Institute for the COVID-19 Genomics UK (COG-UK) Consortium | Harper VanSteenhouse, Yumi Kasai, David Gray, Carol Clugston, Anna Dominiczak and Alex Alderton, Roberto Amato, Jeffrey Barrett, Sonia Goncalves, Ewan Harrison, David K. Jackson, Ian Johnston, Dominic Kwiatkowski, Cordelia Langford, John Sillitoe on behalf of the Wellcome Sanger Institute COVID-19 Surveillance Team |
| EPI_ISL_2700316 | Lighthouse Lab in Milton Keynes | Wellcome Sanger Institute for the COVID-19 Genomics UK (COG-UK) Consortium | The Lighthouse Lab in Milton Keynes and Alex Alderton, Roberto Amato, Jeffrey Barrett, Sonia Goncalves, Ewan Harrison, David K. Jackson, Ian Johnston, Dominic Kwiatkowski, Cordelia Langford, John Sillitoe on behalf of the Wellcome Sanger Institute COVID-19 Surveillance Team |
| EPI_ISL_2700317 | Lighthouse Lab in Glasgow | Wellcome Sanger Institute for the COVID-19 Genomics UK (COG-UK) Consortium | Harper VanSteenhouse, Yumi Kasai, David Gray, Carol Clugston, Anna Dominiczak and Alex Alderton, Roberto Amato, Jeffrey Barrett, Sonia Goncalves, Ewan Harrison, David K. Jackson, Ian Johnston, Dominic Kwiatkowski, Cordelia Langford, John Sillitoe on behalf of the Wellcome Sanger Institute COVID-19 Surveillance Team |
| EPI_ISL_2700318 | Lighthouse Lab in Milton Keynes | Wellcome Sanger Institute for the COVID-19 Genomics UK (COG-UK) Consortium | The Lighthouse Lab in Milton Keynes and Alex Alderton, Roberto Amato, Jeffrey Barrett, Sonia Goncalves, Ewan Harrison, David K. Jackson, Ian Johnston, Dominic Kwiatkowski, Cordelia Langford, John Sillitoe on behalf of the Wellcome Sanger Institute COVID-19 Surveillance Team |
| EPI_ISL_2700319 | Lighthouse Lab in Glasgow | Wellcome Sanger Institute for the COVID-19 Genomics UK (COG-UK) Consortium | Harper VanSteenhouse, Yumi Kasai, David Gray, Carol Clugston, Anna Dominiczak and Alex Alderton, Roberto Amato, Jeffrey Barrett, Sonia Goncalves, Ewan Harrison, David K. Jackson, Ian Johnston, Dominic Kwiatkowski, Cordelia Langford, John Sillitoe on behalf of the Wellcome Sanger Institute COVID-19 Surveillance Team |
| EPI_ISL_2700320, EPI_ISL_2700321, EPI_ISL_2700322 | Lighthouse Lab in Milton Keynes | Wellcome Sanger Institute for the COVID-19 Genomics UK (COG-UK) Consortium | The Lighthouse Lab in Milton Keynes and Alex Alderton, Roberto Amato, Jeffrey Barrett, Sonia Goncalves, Ewan Harrison, David K. Jackson, Ian Johnston, Dominic Kwiatkowski, Cordelia Langford, John Sillitoe on behalf of the Wellcome Sanger Institute COVID-19 Surveillance Team |
| EPI_ISL_2700323 | Lighthouse Lab in Glasgow | Wellcome Sanger Institute for the COVID-19 Genomics UK (COG-UK) Consortium | Harper VanSteenhouse, Yumi Kasai, David Gray, Carol Clugston, Anna Dominiczak and Alex Alderton, Roberto Amato, Jeffrey Barrett, Sonia Goncalves, Ewan Harrison, David K. Jackson, Ian Johnston, Dominic Kwiatkowski, Cordelia Langford, John Sillitoe on behalf of the Wellcome Sanger Institute COVID-19 Surveillance Team |
| EPI_ISL_2700324, EPI_ISL_2700325, EPI_ISL_2700326 | Lighthouse Lab in Milton Keynes | Wellcome Sanger Institute for the COVID-19 Genomics UK (COG-UK) Consortium | The Lighthouse Lab in Milton Keynes and Alex Alderton, Roberto Amato, Jeffrey Barrett, Sonia Goncalves, Ewan Harrison, David K. Jackson, Ian Johnston, Dominic Kwiatkowski, Cordelia Langford, John Sillitoe on behalf of the Wellcome Sanger Institute COVID-19 Surveillance Team |
| EPI_ISL_2700327 | Lighthouse Lab in Glasgow | Wellcome Sanger Institute for the COVID-19 Genomics UK (COG-UK) Consortium | Harper VanSteenhouse, Yumi Kasai, David Gray, Carol Clugston, Anna Dominiczak and Alex Alderton, Roberto Amato, Jeffrey Barrett, Sonia Goncalves, Ewan Harrison, David K. Jackson, Ian Johnston, Dominic Kwiatkowski, Cordelia Langford, John Sillitoe on behalf of the Wellcome Sanger Institute COVID-19 Surveillance Team |
| EPI_ISL_2700328, EPI_ISL_2700329 | Lighthouse Lab in Milton Keynes | Wellcome Sanger Institute for the COVID-19 Genomics UK (COG-UK) Consortium | The Lighthouse Lab in Milton Keynes and Alex Alderton, Roberto Amato, Jeffrey Barrett, Sonia Goncalves, Ewan Harrison, David K. Jackson, Ian Johnston, Dominic Kwiatkowski, Cordelia Langford, John Sillitoe on behalf of the Wellcome Sanger Institute COVID-19 Surveillance Team |
| EPI_ISL_2700330, EPI_ISL_2700331, EPI_ISL_2700332 | Lighthouse Lab in Glasgow | Wellcome Sanger Institute for the COVID-19 Genomics UK (COG-UK) Consortium | Harper VanSteenhouse, Yumi Kasai, David Gray, Carol Clugston, Anna Dominiczak and Alex Alderton, Roberto Amato, Jeffrey Barrett, Sonia Goncalves, Ewan Harrison, David K. Jackson, Ian Johnston, Dominic Kwiatkowski, Cordelia Langford, John Sillitoe on behalf of the Wellcome Sanger Institute COVID-19 Surveillance Team |
| EPI_ISL_2700333 | Lighthouse Lab in Milton Keynes | Wellcome Sanger Institute for the COVID-19 Genomics UK (COG-UK) Consortium | The Lighthouse Lab in Milton Keynes and Alex Alderton, Roberto Amato, Jeffrey Barrett, Sonia Goncalves, Ewan Harrison, David K. Jackson, Ian Johnston, Dominic Kwiatkowski, Cordelia Langford, John Sillitoe on behalf of the Wellcome Sanger Institute COVID-19 Surveillance Team |
| EPI_ISL_2700334, EPI_ISL_2700335, EPI_ISL_2700336 | Lighthouse Lab in Glasgow | Wellcome Sanger Institute for the COVID-19 Genomics UK (COG-UK) Consortium | Harper VanSteenhouse, Yumi Kasai, David Gray, Carol Clugston, Anna Dominiczak and Alex Alderton, Roberto Amato, Jeffrey Barrett, Sonia Goncalves, Ewan Harrison, David K. Jackson, Ian Johnston, Dominic Kwiatkowski, Cordelia Langford, John Sillitoe on behalf of the Wellcome Sanger Institute COVID-19 Surveillance Team |
| EPI_ISL_2700337, EPI_ISL_2700338 | Lighthouse Lab in Milton Keynes | Wellcome Sanger Institute for the COVID-19 Genomics UK (COG-UK) Consortium | The Lighthouse Lab in Milton Keynes and Alex Alderton, Roberto Amato, Jeffrey Barrett, Sonia Goncalves, Ewan Harrison, David K. Jackson, Ian Johnston, Dominic Kwiatkowski, Cordelia Langford, John Sillitoe on behalf of the Wellcome Sanger Institute COVID-19 Surveillance Team |
| EPI_ISL_2700339, EPI_ISL_2700340, EPI_ISL_2700341, EPI_ISL_2700342, EPI_ISL_2700343 | Lighthouse Lab in Glasgow | Wellcome Sanger Institute for the COVID-19 Genomics UK (COG-UK) Consortium | Harper VanSteenhouse, Yumi Kasai, David Gray, Carol Clugston, Anna Dominiczak and Alex Alderton, Roberto Amato, Jeffrey Barrett, Sonia Goncalves, Ewan Harrison, David K. Jackson, Ian Johnston, Dominic Kwiatkowski, Cordelia Langford, John Sillitoe on behalf of the Wellcome Sanger Institute COVID-19 Surveillance Team |
| EPI_ISL_2700344 | Lighthouse Lab in Milton Keynes | Wellcome Sanger Institute for the COVID-19 Genomics UK (COG-UK) Consortium | The Lighthouse Lab in Milton Keynes and Alex Alderton, Roberto Amato, Jeffrey Barrett, Sonia Goncalves, Ewan Harrison, David K. Jackson, Ian Johnston, Dominic Kwiatkowski, Cordelia Langford, John Sillitoe on behalf of the Wellcome Sanger Institute COVID-19 Surveillance Team |
| EPI_ISL_2700345 | Lighthouse Lab in Glasgow | Wellcome Sanger Institute for the COVID-19 Genomics UK (COG-UK) Consortium | Harper VanSteenhouse, Yumi Kasai, David Gray, Carol Clugston, Anna Dominiczak and Alex Alderton, Roberto Amato, Jeffrey Barrett, Sonia Goncalves, Ewan Harrison, David K. Jackson, Ian Johnston, Dominic Kwiatkowski, Cordelia Langford, John Sillitoe on behalf of the Wellcome Sanger Institute COVID-19 Surveillance Team |
| EPI_ISL_2700346, EPI_ISL_2700347 | Lighthouse Lab in Milton Keynes | Wellcome Sanger Institute for the COVID-19 Genomics UK (COG-UK) Consortium | The Lighthouse Lab in Milton Keynes and Alex Alderton, Roberto Amato, Jeffrey Barrett, Sonia Goncalves, Ewan Harrison, David K. Jackson, Ian Johnston, Dominic Kwiatkowski, Cordelia Langford, John Sillitoe on behalf of the Wellcome Sanger Institute COVID-19 Surveillance Team |
| EPI_ISL_2700350 | Lighthouse Lab in Glasgow | Wellcome Sanger Institute for the COVID-19 Genomics UK (COG-UK) Consortium | Harper VanSteenhouse, Yumi Kasai, David Gray, Carol Clugston, Anna Dominiczak and Alex Alderton, Roberto Amato, Jeffrey Barrett, Sonia Goncalves, Ewan Harrison, David K. Jackson, Ian Johnston, Dominic Kwiatkowski, Cordelia Langford, John Sillitoe on behalf of the Wellcome Sanger Institute COVID-19 Surveillance Team |
| EPI_ISL_2700351, EPI_ISL_2700352 | Lighthouse Lab in Milton Keynes | Wellcome Sanger Institute for the COVID-19 Genomics UK (COG-UK) Consortium | The Lighthouse Lab in Milton Keynes and Alex Alderton, Roberto Amato, Jeffrey Barrett, Sonia Goncalves, Ewan Harrison, David K. Jackson, Ian Johnston, Dominic Kwiatkowski, Cordelia Langford, John Sillitoe on behalf of the Wellcome Sanger Institute COVID-19 Surveillance Team |
| EPI_ISL_2700353, EPI_ISL_2700354 | Lighthouse Lab in Glasgow | Wellcome Sanger Institute for the COVID-19 Genomics UK (COG-UK) Consortium | Harper VanSteenhouse, Yumi Kasai, David Gray, Carol Clugston, Anna Dominiczak and Alex Alderton, Roberto Amato, Jeffrey Barrett, Sonia Goncalves, Ewan Harrison, David K. Jackson, Ian Johnston, Dominic Kwiatkowski, Cordelia Langford, John Sillitoe on behalf of the Wellcome Sanger Institute COVID-19 Surveillance Team |
| EPI_ISL_2700355, EPI_ISL_2700356 | Lighthouse Lab in Milton Keynes | Wellcome Sanger Institute for the COVID-19 Genomics UK (COG-UK) Consortium | The Lighthouse Lab in Milton Keynes and Alex Alderton, Roberto Amato, Jeffrey Barrett, Sonia Goncalves, Ewan Harrison, David K. Jackson, Ian Johnston, Dominic Kwiatkowski, Cordelia Langford, John Sillitoe on behalf of the Wellcome Sanger Institute COVID-19 Surveillance Team |
| EPI_ISL_2700357, EPI_ISL_2700358 | Lighthouse Lab in Glasgow | Wellcome Sanger Institute for the COVID-19 Genomics UK (COG-UK) Consortium | Harper VanSteenhouse, Yumi Kasai, David Gray, Carol Clugston, Anna Dominiczak and Alex Alderton, Roberto Amato, Jeffrey Barrett, Sonia Goncalves, Ewan Harrison, David K. Jackson, Ian Johnston, Dominic Kwiatkowski, Cordelia Langford, John Sillitoe on behalf of the Wellcome Sanger Institute COVID-19 Surveillance Team |

| | | | |
|---|---|---|---|
| EPI_ISL_2700359, EPI_ISL_2700360, EPI_ISL_2700361, EPI_ISL_2700362, EPI_ISL_2700363, EPI_ISL_2700364, EPI_ISL_2700365 | Lighthouse Lab in Milton Keynes | Wellcome Sanger Institute for the COVID-19 Genomics UK (COG-UK) Consortium | The Lighthouse Lab in Milton Keynes and Alex Alderton, Roberto Amato, Jeffrey Barrett, Sonia Goncalves, Ewan Harrison, David K. Jackson, Ian Johnston, Dominic Kwiatkowski, Cordelia Langford, John Sillitoe on behalf of the Wellcome Sanger Institute COVID-19 Surveillance Team |
| EPI_ISL_2700366, EPI_ISL_2700367, EPI_ISL_2700368, EPI_ISL_2700369, EPI_ISL_2700370 | Lighthouse Lab in Glasgow | Wellcome Sanger Institute for the COVID-19 Genomics UK (COG-UK) Consortium | Harper VanSteenhouse, Yumi Kasai, David Gray, Carol Clugston, Anna Dominiczak and Alex Alderton, Roberto Amato, Jeffrey Barrett, Sonia Goncalves, Ewan Harrison, David K. Jackson, Ian Johnston, Dominic Kwiatkowski, Cordelia Langford, John Sillitoe on behalf of the Wellcome Sanger Institute COVID-19 Surveillance Team |
| EPI_ISL_2700371 | Lighthouse Lab in Milton Keynes | Wellcome Sanger Institute for the COVID-19 Genomics UK (COG-UK) Consortium | The Lighthouse Lab in Milton Keynes and Alex Alderton, Roberto Amato, Jeffrey Barrett, Sonia Goncalves, Ewan Harrison, David K. Jackson, Ian Johnston, Dominic Kwiatkowski, Cordelia Langford, John Sillitoe on behalf of the Wellcome Sanger Institute COVID-19 Surveillance Team |
| EPI_ISL_2700372, EPI_ISL_2700373, EPI_ISL_2700374, EPI_ISL_2700375 | Lighthouse Lab in Glasgow | Wellcome Sanger Institute for the COVID-19 Genomics UK (COG-UK) Consortium | Harper VanSteenhouse, Yumi Kasai, David Gray, Carol Clugston, Anna Dominiczak and Alex Alderton, Roberto Amato, Jeffrey Barrett, Sonia Goncalves, Ewan Harrison, David K. Jackson, Ian Johnston, Dominic Kwiatkowski, Cordelia Langford, John Sillitoe on behalf of the Wellcome Sanger Institute COVID-19 Surveillance Team |
| EPI_ISL_2700376 | Lighthouse Lab in Milton Keynes | Wellcome Sanger Institute for the COVID-19 Genomics UK (COG-UK) Consortium | The Lighthouse Lab in Milton Keynes and Alex Alderton, Roberto Amato, Jeffrey Barrett, Sonia Goncalves, Ewan Harrison, David K. Jackson, Ian Johnston, Dominic Kwiatkowski, Cordelia Langford, John Sillitoe on behalf of the Wellcome Sanger Institute COVID-19 Surveillance Team |
| EPI_ISL_2700377 | Lighthouse Lab in Glasgow | Wellcome Sanger Institute for the COVID-19 Genomics UK (COG-UK) Consortium | Harper VanSteenhouse, Yumi Kasai, David Gray, Carol Clugston, Anna Dominiczak and Alex Alderton, Roberto Amato, Jeffrey Barrett, Sonia Goncalves, Ewan Harrison, David K. Jackson, Ian Johnston, Dominic Kwiatkowski, Cordelia Langford, John Sillitoe on behalf of the Wellcome Sanger Institute COVID-19 Surveillance Team |
| EPI_ISL_2700378, EPI_ISL_2700379, EPI_ISL_2700380, EPI_ISL_2700381 | Lighthouse Lab in Milton Keynes | Wellcome Sanger Institute for the COVID-19 Genomics UK (COG-UK) Consortium | The Lighthouse Lab in Milton Keynes and Alex Alderton, Roberto Amato, Jeffrey Barrett, Sonia Goncalves, Ewan Harrison, David K. Jackson, Ian Johnston, Dominic Kwiatkowski, Cordelia Langford, John Sillitoe on behalf of the Wellcome Sanger Institute COVID-19 Surveillance Team |
| EPI_ISL_2700382, EPI_ISL_2700383 | Lighthouse Lab in Glasgow | Wellcome Sanger Institute for the COVID-19 Genomics UK (COG-UK) Consortium | Harper VanSteenhouse, Yumi Kasai, David Gray, Carol Clugston, Anna Dominiczak and Alex Alderton, Roberto Amato, Jeffrey Barrett, Sonia Goncalves, Ewan Harrison, David K. Jackson, Ian Johnston, Dominic Kwiatkowski, Cordelia Langford, John Sillitoe on behalf of the Wellcome Sanger Institute COVID-19 Surveillance Team |
| EPI_ISL_2700385 | Lighthouse Lab in Milton Keynes | Wellcome Sanger Institute for the COVID-19 Genomics UK (COG-UK) Consortium | The Lighthouse Lab in Milton Keynes and Alex Alderton, Roberto Amato, Jeffrey Barrett, Sonia Goncalves, Ewan Harrison, David K. Jackson, Ian Johnston, Dominic Kwiatkowski, Cordelia Langford, John Sillitoe on behalf of the Wellcome Sanger Institute COVID-19 Surveillance Team |
| EPI_ISL_2700386, EPI_ISL_2700387 | Lighthouse Lab in Glasgow | Wellcome Sanger Institute for the COVID-19 Genomics UK (COG-UK) Consortium | Harper VanSteenhouse, Yumi Kasai, David Gray, Carol Clugston, Anna Dominiczak and Alex Alderton, Roberto Amato, Jeffrey Barrett, Sonia Goncalves, Ewan Harrison, David K. Jackson, Ian Johnston, Dominic Kwiatkowski, Cordelia Langford, John Sillitoe on behalf of the Wellcome Sanger Institute COVID-19 Surveillance Team |
| EPI_ISL_2700388, EPI_ISL_2700389 | Lighthouse Lab in Milton Keynes | Wellcome Sanger Institute for the COVID-19 Genomics UK (COG-UK) Consortium | The Lighthouse Lab in Milton Keynes and Alex Alderton, Roberto Amato, Jeffrey Barrett, Sonia Goncalves, Ewan Harrison, David K. Jackson, Ian Johnston, Dominic Kwiatkowski, Cordelia Langford, John Sillitoe on behalf of the Wellcome Sanger Institute COVID-19 Surveillance Team |
| EPI_ISL_2700390, EPI_ISL_2700391, EPI_ISL_2700392, EPI_ISL_2700393, EPI_ISL_2700394 | Lighthouse Lab in Glasgow | Wellcome Sanger Institute for the COVID-19 Genomics UK (COG-UK) Consortium | Harper VanSteenhouse, Yumi Kasai, David Gray, Carol Clugston, Anna Dominiczak and Alex Alderton, Roberto Amato, Jeffrey Barrett, Sonia Goncalves, Ewan Harrison, David K. Jackson, Ian Johnston, Dominic Kwiatkowski, Cordelia Langford, John Sillitoe on behalf of the Wellcome Sanger Institute COVID-19 Surveillance Team |
| EPI_ISL_2700395, EPI_ISL_2700396 | Lighthouse Lab in Milton Keynes | Wellcome Sanger Institute for the COVID-19 Genomics UK (COG-UK) Consortium | The Lighthouse Lab in Milton Keynes and Alex Alderton, Roberto Amato, Jeffrey Barrett, Sonia Goncalves, Ewan Harrison, David K. Jackson, Ian Johnston, Dominic Kwiatkowski, Cordelia Langford, John Sillitoe on behalf of the Wellcome Sanger Institute COVID-19 Surveillance Team |
| EPI_ISL_2700397, EPI_ISL_2700398, EPI_ISL_2700399, EPI_ISL_2700400, EPI_ISL_2700401 | Lighthouse Lab in Glasgow | Wellcome Sanger Institute for the COVID-19 Genomics UK (COG-UK) Consortium | Harper VanSteenhouse, Yumi Kasai, David Gray, Carol Clugston, Anna Dominiczak and Alex Alderton, Roberto Amato, Jeffrey Barrett, Sonia Goncalves, Ewan Harrison, David K. Jackson, Ian Johnston, Dominic Kwiatkowski, Cordelia Langford, John Sillitoe on behalf of the Wellcome Sanger Institute COVID-19 Surveillance Team |
| EPI_ISL_2700402 | Lighthouse Lab in Milton Keynes | Wellcome Sanger Institute for the COVID-19 Genomics UK (COG-UK) Consortium | The Lighthouse Lab in Milton Keynes and Alex Alderton, Roberto Amato, Jeffrey Barrett, Sonia Goncalves, Ewan Harrison, David K. Jackson, Ian Johnston, Dominic Kwiatkowski, Cordelia Langford, John Sillitoe on behalf of the Wellcome Sanger Institute COVID-19 Surveillance Team |
| EPI_ISL_2700403 | Lighthouse Lab in Glasgow | Wellcome Sanger Institute for the COVID-19 Genomics UK (COG-UK) Consortium | Harper VanSteenhouse, Yumi Kasai, David Gray, Carol Clugston, Anna Dominiczak and Alex Alderton, Roberto Amato, Jeffrey Barrett, Sonia Goncalves, Ewan Harrison, David K. Jackson, Ian Johnston, Dominic Kwiatkowski, Cordelia Langford, John Sillitoe on behalf of the Wellcome Sanger Institute COVID-19 Surveillance Team |
| EPI_ISL_2700404 | Lighthouse Lab in Milton Keynes | Wellcome Sanger Institute for the COVID-19 Genomics UK (COG-UK) Consortium | The Lighthouse Lab in Milton Keynes and Alex Alderton, Roberto Amato, Jeffrey Barrett, Sonia Goncalves, Ewan Harrison, David K. Jackson, Ian Johnston, Dominic Kwiatkowski, Cordelia Langford, John Sillitoe on behalf of the Wellcome Sanger Institute COVID-19 Surveillance Team |
| EPI_ISL_2700405 | Lighthouse Lab in Glasgow | Wellcome Sanger Institute for the COVID-19 Genomics UK (COG-UK) Consortium | Harper VanSteenhouse, Yumi Kasai, David Gray, Carol Clugston, Anna Dominiczak and Alex Alderton, Roberto Amato, Jeffrey Barrett, Sonia Goncalves, Ewan Harrison, David K. Jackson, Ian Johnston, Dominic Kwiatkowski, Cordelia Langford, John Sillitoe on behalf of the Wellcome Sanger Institute COVID-19 Surveillance Team |
| EPI_ISL_2700406, EPI_ISL_2700407, EPI_ISL_2700408 | Lighthouse Lab in Milton Keynes | Wellcome Sanger Institute for the COVID-19 Genomics UK (COG-UK) Consortium | The Lighthouse Lab in Milton Keynes and Alex Alderton, Roberto Amato, Jeffrey Barrett, Sonia Goncalves, Ewan Harrison, David K. Jackson, Ian Johnston, Dominic Kwiatkowski, Cordelia Langford, John Sillitoe on behalf of the Wellcome Sanger Institute COVID-19 Surveillance Team |
| EPI_ISL_2700409 | Lighthouse Lab in Glasgow | Wellcome Sanger Institute for the COVID-19 Genomics UK (COG-UK) Consortium | Harper VanSteenhouse, Yumi Kasai, David Gray, Carol Clugston, Anna Dominiczak and Alex Alderton, Roberto Amato, Jeffrey Barrett, Sonia Goncalves, Ewan Harrison, David K. Jackson, Ian Johnston, Dominic Kwiatkowski, Cordelia Langford, John Sillitoe on behalf of the Wellcome Sanger Institute COVID-19 Surveillance Team |
| EPI_ISL_2700410 | Lighthouse Lab in Milton Keynes | Wellcome Sanger Institute for the COVID-19 Genomics UK (COG-UK) Consortium | The Lighthouse Lab in Milton Keynes and Alex Alderton, Roberto Amato, Jeffrey Barrett, Sonia Goncalves, Ewan Harrison, David K. Jackson, Ian Johnston, Dominic Kwiatkowski, Cordelia Langford, John Sillitoe on behalf of the Wellcome Sanger Institute COVID-19 Surveillance Team |
| EPI_ISL_2700411 | Lighthouse Lab in Glasgow | Wellcome Sanger Institute for the COVID-19 Genomics UK (COG-UK) Consortium | Harper VanSteenhouse, Yumi Kasai, David Gray, Carol Clugston, Anna Dominiczak and Alex Alderton, Roberto Amato, Jeffrey Barrett, Sonia Goncalves, Ewan Harrison, David K. Jackson, Ian Johnston, Dominic Kwiatkowski, Cordelia Langford, John Sillitoe on behalf of the Wellcome Sanger Institute COVID-19 Surveillance Team |
| EPI_ISL_2700412, EPI_ISL_2700413 | Lighthouse Lab in Milton Keynes | Wellcome Sanger Institute for the COVID-19 Genomics UK (COG-UK) Consortium | The Lighthouse Lab in Milton Keynes and Alex Alderton, Roberto Amato, Jeffrey Barrett, Sonia Goncalves, Ewan Harrison, David K. Jackson, Ian Johnston, Dominic Kwiatkowski, Cordelia Langford, John Sillitoe on behalf of the Wellcome Sanger Institute COVID-19 Surveillance Team |
| EPI_ISL_2700414 | Lighthouse Lab in Glasgow | Wellcome Sanger Institute for the COVID-19 Genomics UK (COG-UK) Consortium | Harper VanSteenhouse, Yumi Kasai, David Gray, Carol Clugston, Anna Dominiczak and Alex Alderton, Roberto Amato, Jeffrey Barrett, Sonia Goncalves, Ewan Harrison, David K. Jackson, Ian Johnston, Dominic Kwiatkowski, Cordelia Langford, John Sillitoe on behalf of the Wellcome Sanger Institute COVID-19 Surveillance Team |
| EPI_ISL_2700415 | Lighthouse Lab in Milton Keynes | Wellcome Sanger Institute for the COVID-19 Genomics UK (COG-UK) Consortium | The Lighthouse Lab in Milton Keynes and Alex Alderton, Roberto Amato, Jeffrey Barrett, Sonia Goncalves, Ewan Harrison, David K. Jackson, Ian Johnston, Dominic Kwiatkowski, Cordelia Langford, John Sillitoe on behalf of the Wellcome Sanger Institute COVID-19 Surveillance Team |
| EPI_ISL_2700416, EPI_ISL_2700417 | Lighthouse Lab in Glasgow | Wellcome Sanger Institute for the COVID-19 Genomics UK (COG-UK) Consortium | Harper VanSteenhouse, Yumi Kasai, David Gray, Carol Clugston, Anna Dominiczak and Alex Alderton, Roberto Amato, Jeffrey Barrett, Sonia Goncalves, Ewan Harrison, David K. Jackson, Ian Johnston, Dominic Kwiatkowski, Cordelia Langford, John Sillitoe on behalf of the Wellcome Sanger Institute COVID-19 Surveillance Team |
| EPI_ISL_2700418, EPI_ISL_2700419, EPI_ISL_2700420, EPI_ISL_2700421, EPI_ISL_2700422, EPI_ISL_2700423, EPI_ISL_2700424, EPI_ISL_2700425 | Lighthouse Lab in Milton Keynes | Wellcome Sanger Institute for the COVID-19 Genomics UK (COG-UK) Consortium | The Lighthouse Lab in Milton Keynes and Alex Alderton, Roberto Amato, Jeffrey Barrett, Sonia Goncalves, Ewan Harrison, David K. Jackson, Ian Johnston, Dominic Kwiatkowski, Cordelia Langford, John Sillitoe on behalf of the Wellcome Sanger Institute COVID-19 Surveillance Team |
| EPI_ISL_2700426 | Lighthouse Lab in Glasgow | Wellcome Sanger Institute for the COVID-19 Genomics UK (COG-UK) Consortium | Harper VanSteenhouse, Yumi Kasai, David Gray, Carol Clugston, Anna Dominiczak and Alex Alderton, Roberto Amato, Jeffrey Barrett, Sonia Goncalves, Ewan Harrison, David K. Jackson, Ian Johnston, Dominic Kwiatkowski, Cordelia Langford, John Sillitoe on behalf of the Wellcome Sanger Institute COVID-19 Surveillance Team |
| EPI_ISL_2700427 | Lighthouse Lab in Milton Keynes | Wellcome Sanger Institute for the COVID-19 Genomics UK (COG-UK) Consortium | The Lighthouse Lab in Milton Keynes and Alex Alderton, Roberto Amato, Jeffrey Barrett, Sonia Goncalves, Ewan Harrison, David K. Jackson, Ian Johnston, Dominic Kwiatkowski, Cordelia Langford, John Sillitoe on behalf of the Wellcome Sanger Institute COVID-19 Surveillance Team |
| EPI_ISL_2700428 | Lighthouse Lab in Glasgow | Wellcome Sanger Institute for the COVID-19 Genomics UK (COG-UK) Consortium | Harper VanSteenhouse, Yumi Kasai, David Gray, Carol Clugston, Anna Dominiczak and Alex Alderton, Roberto Amato, Jeffrey Barrett, Sonia Goncalves, Ewan Harrison, David K. Jackson, Ian Johnston, Dominic Kwiatkowski, Cordelia Langford, John Sillitoe on behalf of the Wellcome Sanger Institute COVID-19 Surveillance Team |
| EPI_ISL_2700429, EPI_ISL_2700430, EPI_ISL_2700431 | Lighthouse Lab in Milton Keynes | Wellcome Sanger Institute for the COVID-19 Genomics UK (COG-UK) Consortium | The Lighthouse Lab in Milton Keynes and Alex Alderton, Roberto Amato, Jeffrey Barrett, Sonia Goncalves, Ewan Harrison, David K. Jackson, Ian Johnston, Dominic Kwiatkowski, Cordelia Langford, John Sillitoe on behalf of the Wellcome Sanger Institute COVID-19 Surveillance Team |
| EPI_ISL_2700432 | Lighthouse Lab in Glasgow | Wellcome Sanger Institute for the COVID-19 Genomics UK (COG-UK) Consortium | Harper VanSteenhouse, Yumi Kasai, David Gray, Carol Clugston, Anna Dominiczak and Alex Alderton, Roberto Amato, Jeffrey Barrett, Sonia Goncalves, Ewan Harrison, David K. Jackson, Ian Johnston, Dominic Kwiatkowski, Cordelia Langford, John Sillitoe on behalf of the Wellcome Sanger Institute COVID-19 Surveillance Team |
| EPI_ISL_2700433, EPI_ISL_2700434, EPI_ISL_2700435, EPI_ISL_2700436 | Lighthouse Lab in Milton Keynes | Wellcome Sanger Institute for the COVID-19 Genomics UK (COG-UK) Consortium | The Lighthouse Lab in Milton Keynes and Alex Alderton, Roberto Amato, Jeffrey Barrett, Sonia Goncalves, Ewan Harrison, David K. Jackson, Ian Johnston, Dominic Kwiatkowski, Cordelia Langford, John Sillitoe on behalf of the Wellcome Sanger Institute COVID-19 Surveillance Team |
| EPI_ISL_2700437, EPI_ISL_2700438, | Lighthouse Lab in Glasgow | Wellcome Sanger Institute for the COVID-19 Genomics UK (COG-UK) Consortium | Harper VanSteenhouse, Yumi Kasai, David Gray, Carol Clugston, Anna Dominiczak and Alex Alderton, Roberto Amato, Jeffrey Barrett, Sonia Goncalves, Ewan Harrison, David K. Jackson, Ian Johnston, Dominic Kwiatkowski, Cordelia Langford, John Sillitoe on behalf of the Wellcome Sanger Institute COVID-19 Surveillance Team |

| | | | |
|---|---|---|---|
| EPI_ISL_2700439, EPI_ISL_2700440 | | | |
| EPI_ISL_2700441, EPI_ISL_2700442, EPI_ISL_2700443, EPI_ISL_2700444, EPI_ISL_2700445, EPI_ISL_2700446 | Lighthouse Lab in Milton Keynes | Wellcome Sanger Institute for the COVID-19 Genomics UK (COG-UK) Consortium | The Lighthouse Lab in Milton Keynes and Alex Alderton, Roberto Amato, Jeffrey Barrett, Sonia Goncalves, Ewan Harrison, David K. Jackson, Ian Johnston, Dominic Kwiatkowski, Cordelia Langford, John Sillitoe on behalf of the Wellcome Sanger Institute COVID-19 Surveillance Team |
| EPI_ISL_2700447 | Lighthouse Lab in Glasgow | Wellcome Sanger Institute for the COVID-19 Genomics UK (COG-UK) Consortium | Harper VanSteenhouse, Yumi Kasai, David Gray, Carol Clugston, Anna Dominiczak and Alex Alderton, Roberto Amato, Jeffrey Barrett, Sonia Goncalves, Ewan Harrison, David K. Jackson, Ian Johnston, Dominic Kwiatkowski, Cordelia Langford, John Sillitoe on behalf of the Wellcome Sanger Institute COVID-19 Surveillance Team |
| EPI_ISL_2700448 | Lighthouse Lab in Milton Keynes | Wellcome Sanger Institute for the COVID-19 Genomics UK (COG-UK) Consortium | The Lighthouse Lab in Milton Keynes and Alex Alderton, Roberto Amato, Jeffrey Barrett, Sonia Goncalves, Ewan Harrison, David K. Jackson, Ian Johnston, Dominic Kwiatkowski, Cordelia Langford, John Sillitoe on behalf of the Wellcome Sanger Institute COVID-19 Surveillance Team |
| EPI_ISL_2700449, EPI_ISL_2700450, EPI_ISL_2700451, EPI_ISL_2700452, EPI_ISL_2700453 | Lighthouse Lab in Glasgow | Wellcome Sanger Institute for the COVID-19 Genomics UK (COG-UK) Consortium | Harper VanSteenhouse, Yumi Kasai, David Gray, Carol Clugston, Anna Dominiczak and Alex Alderton, Roberto Amato, Jeffrey Barrett, Sonia Goncalves, Ewan Harrison, David K. Jackson, Ian Johnston, Dominic Kwiatkowski, Cordelia Langford, John Sillitoe on behalf of the Wellcome Sanger Institute COVID-19 Surveillance Team |
| EPI_ISL_2700454, EPI_ISL_2700455, EPI_ISL_2700456 | Lighthouse Lab in Milton Keynes | Wellcome Sanger Institute for the COVID-19 Genomics UK (COG-UK) Consortium | The Lighthouse Lab in Milton Keynes and Alex Alderton, Roberto Amato, Jeffrey Barrett, Sonia Goncalves, Ewan Harrison, David K. Jackson, Ian Johnston, Dominic Kwiatkowski, Cordelia Langford, John Sillitoe on behalf of the Wellcome Sanger Institute COVID-19 Surveillance Team |
| EPI_ISL_2700457, EPI_ISL_2700458, EPI_ISL_2700459 | Lighthouse Lab in Glasgow | Wellcome Sanger Institute for the COVID-19 Genomics UK (COG-UK) Consortium | Harper VanSteenhouse, Yumi Kasai, David Gray, Carol Clugston, Anna Dominiczak and Alex Alderton, Roberto Amato, Jeffrey Barrett, Sonia Goncalves, Ewan Harrison, David K. Jackson, Ian Johnston, Dominic Kwiatkowski, Cordelia Langford, John Sillitoe on behalf of the Wellcome Sanger Institute COVID-19 Surveillance Team |
| EPI_ISL_2700460, EPI_ISL_2700461, EPI_ISL_2700463 | Lighthouse Lab in Milton Keynes | Wellcome Sanger Institute for the COVID-19 Genomics UK (COG-UK) Consortium | The Lighthouse Lab in Milton Keynes and Alex Alderton, Roberto Amato, Jeffrey Barrett, Sonia Goncalves, Ewan Harrison, David K. Jackson, Ian Johnston, Dominic Kwiatkowski, Cordelia Langford, John Sillitoe on behalf of the Wellcome Sanger Institute COVID-19 Surveillance Team |
| EPI_ISL_2700464 | Lighthouse Lab in Glasgow | Wellcome Sanger Institute for the COVID-19 Genomics UK (COG-UK) Consortium | Harper VanSteenhouse, Yumi Kasai, David Gray, Carol Clugston, Anna Dominiczak and Alex Alderton, Roberto Amato, Jeffrey Barrett, Sonia Goncalves, Ewan Harrison, David K. Jackson, Ian Johnston, Dominic Kwiatkowski, Cordelia Langford, John Sillitoe on behalf of the Wellcome Sanger Institute COVID-19 Surveillance Team |
| EPI_ISL_2700465 | Lighthouse Lab in Milton Keynes | Wellcome Sanger Institute for the COVID-19 Genomics UK (COG-UK) Consortium | The Lighthouse Lab in Milton Keynes and Alex Alderton, Roberto Amato, Jeffrey Barrett, Sonia Goncalves, Ewan Harrison, David K. Jackson, Ian Johnston, Dominic Kwiatkowski, Cordelia Langford, John Sillitoe on behalf of the Wellcome Sanger Institute COVID-19 Surveillance Team |
| EPI_ISL_2700466 | Lighthouse Lab in Glasgow | Wellcome Sanger Institute for the COVID-19 Genomics UK (COG-UK) Consortium | Harper VanSteenhouse, Yumi Kasai, David Gray, Carol Clugston, Anna Dominiczak and Alex Alderton, Roberto Amato, Jeffrey Barrett, Sonia Goncalves, Ewan Harrison, David K. Jackson, Ian Johnston, Dominic Kwiatkowski, Cordelia Langford, John Sillitoe on behalf of the Wellcome Sanger Institute COVID-19 Surveillance Team |
| EPI_ISL_2700467 | Lighthouse Lab in Milton Keynes | Wellcome Sanger Institute for the COVID-19 Genomics UK (COG-UK) Consortium | The Lighthouse Lab in Milton Keynes and Alex Alderton, Roberto Amato, Jeffrey Barrett, Sonia Goncalves, Ewan Harrison, David K. Jackson, Ian Johnston, Dominic Kwiatkowski, Cordelia Langford, John Sillitoe on behalf of the Wellcome Sanger Institute COVID-19 Surveillance Team |
| EPI_ISL_2700469, EPI_ISL_2700470 | Lighthouse Lab in Glasgow | Wellcome Sanger Institute for the COVID-19 Genomics UK (COG-UK) Consortium | Harper VanSteenhouse, Yumi Kasai, David Gray, Carol Clugston, Anna Dominiczak and Alex Alderton, Roberto Amato, Jeffrey Barrett, Sonia Goncalves, Ewan Harrison, David K. Jackson, Ian Johnston, Dominic Kwiatkowski, Cordelia Langford, John Sillitoe on behalf of the Wellcome Sanger Institute COVID-19 Surveillance Team |
| EPI_ISL_2700471, EPI_ISL_2700472 | Lighthouse Lab in Milton Keynes | Wellcome Sanger Institute for the COVID-19 Genomics UK (COG-UK) Consortium | The Lighthouse Lab in Milton Keynes and Alex Alderton, Roberto Amato, Jeffrey Barrett, Sonia Goncalves, Ewan Harrison, David K. Jackson, Ian Johnston, Dominic Kwiatkowski, Cordelia Langford, John Sillitoe on behalf of the Wellcome Sanger Institute COVID-19 Surveillance Team |
| EPI_ISL_2700473 | Lighthouse Lab in Glasgow | Wellcome Sanger Institute for the COVID-19 Genomics UK (COG-UK) Consortium | Harper VanSteenhouse, Yumi Kasai, David Gray, Carol Clugston, Anna Dominiczak and Alex Alderton, Roberto Amato, Jeffrey Barrett, Sonia Goncalves, Ewan Harrison, David K. Jackson, Ian Johnston, Dominic Kwiatkowski, Cordelia Langford, John Sillitoe on behalf of the Wellcome Sanger Institute COVID-19 Surveillance Team |
| EPI_ISL_2700474, EPI_ISL_2700475, EPI_ISL_2700476, EPI_ISL_2700477 | Lighthouse Lab in Milton Keynes | Wellcome Sanger Institute for the COVID-19 Genomics UK (COG-UK) Consortium | The Lighthouse Lab in Milton Keynes and Alex Alderton, Roberto Amato, Jeffrey Barrett, Sonia Goncalves, Ewan Harrison, David K. Jackson, Ian Johnston, Dominic Kwiatkowski, Cordelia Langford, John Sillitoe on behalf of the Wellcome Sanger Institute COVID-19 Surveillance Team |
| EPI_ISL_2700478, EPI_ISL_2700479, EPI_ISL_2700480, EPI_ISL_2700481, EPI_ISL_2700482, EPI_ISL_2700483, EPI_ISL_2700484, EPI_ISL_2700485 | Lighthouse Lab in Glasgow | Wellcome Sanger Institute for the COVID-19 Genomics UK (COG-UK) Consortium | Harper VanSteenhouse, Yumi Kasai, David Gray, Carol Clugston, Anna Dominiczak and Alex Alderton, Roberto Amato, Jeffrey Barrett, Sonia Goncalves, Ewan Harrison, David K. Jackson, Ian Johnston, Dominic Kwiatkowski, Cordelia Langford, John Sillitoe on behalf of the Wellcome Sanger Institute COVID-19 Surveillance Team |
| EPI_ISL_2700486 | Lighthouse Lab in Milton Keynes | Wellcome Sanger Institute for the COVID-19 Genomics UK (COG-UK) Consortium | The Lighthouse Lab in Milton Keynes and Alex Alderton, Roberto Amato, Jeffrey Barrett, Sonia Goncalves, Ewan Harrison, David K. Jackson, Ian Johnston, Dominic Kwiatkowski, Cordelia Langford, John Sillitoe on behalf of the Wellcome Sanger Institute COVID-19 Surveillance Team |
| EPI_ISL_2700487 | Lighthouse Lab in Glasgow | Wellcome Sanger Institute for the COVID-19 Genomics UK (COG-UK) Consortium | Harper VanSteenhouse, Yumi Kasai, David Gray, Carol Clugston, Anna Dominiczak and Alex Alderton, Roberto Amato, Jeffrey Barrett, Sonia Goncalves, Ewan Harrison, David K. Jackson, Ian Johnston, Dominic Kwiatkowski, Cordelia Langford, John Sillitoe on behalf of the Wellcome Sanger Institute COVID-19 Surveillance Team |
| EPI_ISL_2700488, EPI_ISL_2700489 | Lighthouse Lab in Milton Keynes | Wellcome Sanger Institute for the COVID-19 Genomics UK (COG-UK) Consortium | The Lighthouse Lab in Milton Keynes and Alex Alderton, Roberto Amato, Jeffrey Barrett, Sonia Goncalves, Ewan Harrison, David K. Jackson, Ian Johnston, Dominic Kwiatkowski, Cordelia Langford, John Sillitoe on behalf of the Wellcome Sanger Institute COVID-19 Surveillance Team |
| EPI_ISL_2700490 | Lighthouse Lab in Glasgow | Wellcome Sanger Institute for the COVID-19 Genomics UK (COG-UK) Consortium | Harper VanSteenhouse, Yumi Kasai, David Gray, Carol Clugston, Anna Dominiczak and Alex Alderton, Roberto Amato, Jeffrey Barrett, Sonia Goncalves, Ewan Harrison, David K. Jackson, Ian Johnston, Dominic Kwiatkowski, Cordelia Langford, John Sillitoe on behalf of the Wellcome Sanger Institute COVID-19 Surveillance Team |
| EPI_ISL_2700491 | Lighthouse Lab in Milton Keynes | Wellcome Sanger Institute for the COVID-19 Genomics UK (COG-UK) Consortium | The Lighthouse Lab in Milton Keynes and Alex Alderton, Roberto Amato, Jeffrey Barrett, Sonia Goncalves, Ewan Harrison, David K. Jackson, Ian Johnston, Dominic Kwiatkowski, Cordelia Langford, John Sillitoe on behalf of the Wellcome Sanger Institute COVID-19 Surveillance Team |
| EPI_ISL_2700492, EPI_ISL_2700493 | Lighthouse Lab in Glasgow | Wellcome Sanger Institute for the COVID-19 Genomics UK (COG-UK) Consortium | Harper VanSteenhouse, Yumi Kasai, David Gray, Carol Clugston, Anna Dominiczak and Alex Alderton, Roberto Amato, Jeffrey Barrett, Sonia Goncalves, Ewan Harrison, David K. Jackson, Ian Johnston, Dominic Kwiatkowski, Cordelia Langford, John Sillitoe on behalf of the Wellcome Sanger Institute COVID-19 Surveillance Team |
| EPI_ISL_2700494 | Lighthouse Lab in Milton Keynes | Wellcome Sanger Institute for the COVID-19 Genomics UK (COG-UK) Consortium | The Lighthouse Lab in Milton Keynes and Alex Alderton, Roberto Amato, Jeffrey Barrett, Sonia Goncalves, Ewan Harrison, David K. Jackson, Ian Johnston, Dominic Kwiatkowski, Cordelia Langford, John Sillitoe on behalf of the Wellcome Sanger Institute COVID-19 Surveillance Team |
| EPI_ISL_2700495 | Lighthouse Lab in Glasgow | Wellcome Sanger Institute for the COVID-19 Genomics UK (COG-UK) Consortium | Harper VanSteenhouse, Yumi Kasai, David Gray, Carol Clugston, Anna Dominiczak and Alex Alderton, Roberto Amato, Jeffrey Barrett, Sonia Goncalves, Ewan Harrison, David K. Jackson, Ian Johnston, Dominic Kwiatkowski, Cordelia Langford, John Sillitoe on behalf of the Wellcome Sanger Institute COVID-19 Surveillance Team |
| EPI_ISL_2700496, EPI_ISL_2700497, EPI_ISL_2700498 | Lighthouse Lab in Milton Keynes | Wellcome Sanger Institute for the COVID-19 Genomics UK (COG-UK) Consortium | The Lighthouse Lab in Milton Keynes and Alex Alderton, Roberto Amato, Jeffrey Barrett, Sonia Goncalves, Ewan Harrison, David K. Jackson, Ian Johnston, Dominic Kwiatkowski, Cordelia Langford, John Sillitoe on behalf of the Wellcome Sanger Institute COVID-19 Surveillance Team |
| EPI_ISL_2700499 | Lighthouse Lab in Glasgow | Wellcome Sanger Institute for the COVID-19 Genomics UK (COG-UK) Consortium | Harper VanSteenhouse, Yumi Kasai, David Gray, Carol Clugston, Anna Dominiczak and Alex Alderton, Roberto Amato, Jeffrey Barrett, Sonia Goncalves, Ewan Harrison, David K. Jackson, Ian Johnston, Dominic Kwiatkowski, Cordelia Langford, John Sillitoe on behalf of the Wellcome Sanger Institute COVID-19 Surveillance Team |
| EPI_ISL_2700500 | Lighthouse Lab in Milton Keynes | Wellcome Sanger Institute for the COVID-19 Genomics UK (COG-UK) Consortium | The Lighthouse Lab in Milton Keynes and Alex Alderton, Roberto Amato, Jeffrey Barrett, Sonia Goncalves, Ewan Harrison, David K. Jackson, Ian Johnston, Dominic Kwiatkowski, Cordelia Langford, John Sillitoe on behalf of the Wellcome Sanger Institute COVID-19 Surveillance Team |
| EPI_ISL_2700501 | Lighthouse Lab in Glasgow | Wellcome Sanger Institute for the COVID-19 Genomics UK (COG-UK) Consortium | Harper VanSteenhouse, Yumi Kasai, David Gray, Carol Clugston, Anna Dominiczak and Alex Alderton, Roberto Amato, Jeffrey Barrett, Sonia Goncalves, Ewan Harrison, David K. Jackson, Ian Johnston, Dominic Kwiatkowski, Cordelia Langford, John Sillitoe on behalf of the Wellcome Sanger Institute COVID-19 Surveillance Team |
| EPI_ISL_2700502 | Lighthouse Lab in Milton Keynes | Wellcome Sanger Institute for the COVID-19 Genomics UK (COG-UK) Consortium | The Lighthouse Lab in Milton Keynes and Alex Alderton, Roberto Amato, Jeffrey Barrett, Sonia Goncalves, Ewan Harrison, David K. Jackson, Ian Johnston, Dominic Kwiatkowski, Cordelia Langford, John Sillitoe on behalf of the Wellcome Sanger Institute COVID-19 Surveillance Team |
| EPI_ISL_2700503 | Lighthouse Lab in Glasgow | Wellcome Sanger Institute for the COVID-19 Genomics UK (COG-UK) Consortium | Harper VanSteenhouse, Yumi Kasai, David Gray, Carol Clugston, Anna Dominiczak and Alex Alderton, Roberto Amato, Jeffrey Barrett, Sonia Goncalves, Ewan Harrison, David K. Jackson, Ian Johnston, Dominic Kwiatkowski, Cordelia Langford, John Sillitoe on behalf of the Wellcome Sanger Institute COVID-19 Surveillance Team |
| EPI_ISL_2700504, EPI_ISL_2700505 | Lighthouse Lab in Milton Keynes | Wellcome Sanger Institute for the COVID-19 Genomics UK (COG-UK) Consortium | The Lighthouse Lab in Milton Keynes and Alex Alderton, Roberto Amato, Jeffrey Barrett, Sonia Goncalves, Ewan Harrison, David K. Jackson, Ian Johnston, Dominic Kwiatkowski, Cordelia Langford, John Sillitoe on behalf of the Wellcome Sanger Institute COVID-19 Surveillance Team |
| EPI_ISL_2700506 | Lighthouse Lab in Glasgow | Wellcome Sanger Institute for the COVID-19 Genomics UK (COG-UK) Consortium | Harper VanSteenhouse, Yumi Kasai, David Gray, Carol Clugston, Anna Dominiczak and Alex Alderton, Roberto Amato, Jeffrey Barrett, Sonia Goncalves, Ewan Harrison, David K. Jackson, Ian Johnston, Dominic Kwiatkowski, Cordelia Langford, John Sillitoe on behalf of the Wellcome Sanger Institute COVID-19 Surveillance Team |
| EPI_ISL_2700507 | Lighthouse Lab in Milton Keynes | Wellcome Sanger Institute for the COVID-19 Genomics UK (COG-UK) Consortium | The Lighthouse Lab in Milton Keynes and Alex Alderton, Roberto Amato, Jeffrey Barrett, Sonia Goncalves, Ewan Harrison, David K. Jackson, Ian Johnston, Dominic Kwiatkowski, Cordelia Langford, John Sillitoe on behalf of the Wellcome Sanger Institute COVID-19 Surveillance Team |
| EPI_ISL_2700508 | Lighthouse Lab in Glasgow | Wellcome Sanger Institute for the COVID-19 Genomics UK (COG-UK) Consortium | Harper VanSteenhouse, Yumi Kasai, David Gray, Carol Clugston, Anna Dominiczak and Alex Alderton, Roberto Amato, Jeffrey Barrett, Sonia Goncalves, Ewan Harrison, David K. Jackson, Ian Johnston, Dominic Kwiatkowski, Cordelia Langford, John Sillitoe on behalf of the Wellcome Sanger Institute COVID-19 Surveillance Team |
| EPI_ISL_2700509 | Lighthouse Lab in Milton Keynes | Wellcome Sanger Institute for the COVID-19 Genomics UK (COG-UK) Consortium | The Lighthouse Lab in Milton Keynes and Alex Alderton, Roberto Amato, Jeffrey Barrett, Sonia Goncalves, Ewan Harrison, David K. Jackson, Ian Johnston, Dominic Kwiatkowski, Cordelia Langford, John Sillitoe on behalf of the Wellcome Sanger Institute COVID-19 Surveillance Team |
| EPI_ISL_2700510, EPI_ISL_2700511, EPI_ISL_2700512 | Lighthouse Lab in Glasgow | Wellcome Sanger Institute for the COVID-19 Genomics UK (COG-UK) Consortium | Harper VanSteenhouse, Yumi Kasai, David Gray, Carol Clugston, Anna Dominiczak and Alex Alderton, Roberto Amato, Jeffrey Barrett, Sonia Goncalves, Ewan Harrison, David K. Jackson, Ian Johnston, Dominic Kwiatkowski, Cordelia Langford, John Sillitoe on behalf of the Wellcome Sanger Institute COVID-19 Surveillance Team |

| | | | |
|---|---|---|---|
| EPI_ISL_2700513 | Lighthouse Lab in Milton Keynes | Wellcome Sanger Institute for the COVID-19 Genomics UK (COG-UK) Consortium | The Lighthouse Lab in Milton Keynes and Alex Alderton, Roberto Amato, Jeffrey Barrett, Sonia Goncalves, Ewan Harrison, David K. Jackson, Ian Johnston, Dominic Kwiatkowski, Cordelia Langford, John Sillitoe on behalf of the Wellcome Sanger Institute COVID-19 Surveillance Team |
| EPI_ISL_2700514 | Lighthouse Lab in Glasgow | Wellcome Sanger Institute for the COVID-19 Genomics UK (COG-UK) Consortium | Harper VanSteenhouse, Yumi Kasai, David Gray, Carol Clugston, Anna Dominiczak and Alex Alderton, Roberto Amato, Jeffrey Barrett, Sonia Goncalves, Ewan Harrison, David K. Jackson, Ian Johnston, Dominic Kwiatkowski, Cordelia Langford, John Sillitoe on behalf of the Wellcome Sanger Institute COVID-19 Surveillance Team |
| EPI_ISL_2700515 | Lighthouse Lab in Milton Keynes | Wellcome Sanger Institute for the COVID-19 Genomics UK (COG-UK) Consortium | The Lighthouse Lab in Milton Keynes and Alex Alderton, Roberto Amato, Jeffrey Barrett, Sonia Goncalves, Ewan Harrison, David K. Jackson, Ian Johnston, Dominic Kwiatkowski, Cordelia Langford, John Sillitoe on behalf of the Wellcome Sanger Institute COVID-19 Surveillance Team |
| EPI_ISL_2700516 | Lighthouse Lab in Glasgow | Wellcome Sanger Institute for the COVID-19 Genomics UK (COG-UK) Consortium | Harper VanSteenhouse, Yumi Kasai, David Gray, Carol Clugston, Anna Dominiczak and Alex Alderton, Roberto Amato, Jeffrey Barrett, Sonia Goncalves, Ewan Harrison, David K. Jackson, Ian Johnston, Dominic Kwiatkowski, Cordelia Langford, John Sillitoe on behalf of the Wellcome Sanger Institute COVID-19 Surveillance Team |
| EPI_ISL_2700517 | Lighthouse Lab in Milton Keynes | Wellcome Sanger Institute for the COVID-19 Genomics UK (COG-UK) Consortium | The Lighthouse Lab in Milton Keynes and Alex Alderton, Roberto Amato, Jeffrey Barrett, Sonia Goncalves, Ewan Harrison, David K. Jackson, Ian Johnston, Dominic Kwiatkowski, Cordelia Langford, John Sillitoe on behalf of the Wellcome Sanger Institute COVID-19 Surveillance Team |
| EPI_ISL_2700518 | Lighthouse Lab in Glasgow | Wellcome Sanger Institute for the COVID-19 Genomics UK (COG-UK) Consortium | Harper VanSteenhouse, Yumi Kasai, David Gray, Carol Clugston, Anna Dominiczak and Alex Alderton, Roberto Amato, Jeffrey Barrett, Sonia Goncalves, Ewan Harrison, David K. Jackson, Ian Johnston, Dominic Kwiatkowski, Cordelia Langford, John Sillitoe on behalf of the Wellcome Sanger Institute COVID-19 Surveillance Team |
| EPI_ISL_2700519 | Lighthouse Lab in Milton Keynes | Wellcome Sanger Institute for the COVID-19 Genomics UK (COG-UK) Consortium | The Lighthouse Lab in Milton Keynes and Alex Alderton, Roberto Amato, Jeffrey Barrett, Sonia Goncalves, Ewan Harrison, David K. Jackson, Ian Johnston, Dominic Kwiatkowski, Cordelia Langford, John Sillitoe on behalf of the Wellcome Sanger Institute COVID-19 Surveillance Team |
| EPI_ISL_2700520, EPI_ISL_2700521 | Lighthouse Lab in Glasgow | Wellcome Sanger Institute for the COVID-19 Genomics UK (COG-UK) Consortium | Harper VanSteenhouse, Yumi Kasai, David Gray, Carol Clugston, Anna Dominiczak and Alex Alderton, Roberto Amato, Jeffrey Barrett, Sonia Goncalves, Ewan Harrison, David K. Jackson, Ian Johnston, Dominic Kwiatkowski, Cordelia Langford, John Sillitoe on behalf of the Wellcome Sanger Institute COVID-19 Surveillance Team |
| EPI_ISL_2700522 | Lighthouse Lab in Milton Keynes | Wellcome Sanger Institute for the COVID-19 Genomics UK (COG-UK) Consortium | The Lighthouse Lab in Milton Keynes and Alex Alderton, Roberto Amato, Jeffrey Barrett, Sonia Goncalves, Ewan Harrison, David K. Jackson, Ian Johnston, Dominic Kwiatkowski, Cordelia Langford, John Sillitoe on behalf of the Wellcome Sanger Institute COVID-19 Surveillance Team |
| EPI_ISL_2700523 | Lighthouse Lab in Glasgow | Wellcome Sanger Institute for the COVID-19 Genomics UK (COG-UK) Consortium | Harper VanSteenhouse, Yumi Kasai, David Gray, Carol Clugston, Anna Dominiczak and Alex Alderton, Roberto Amato, Jeffrey Barrett, Sonia Goncalves, Ewan Harrison, David K. Jackson, Ian Johnston, Dominic Kwiatkowski, Cordelia Langford, John Sillitoe on behalf of the Wellcome Sanger Institute COVID-19 Surveillance Team |
| EPI_ISL_2700524 | Lighthouse Lab in Milton Keynes | Wellcome Sanger Institute for the COVID-19 Genomics UK (COG-UK) Consortium | The Lighthouse Lab in Milton Keynes and Alex Alderton, Roberto Amato, Jeffrey Barrett, Sonia Goncalves, Ewan Harrison, David K. Jackson, Ian Johnston, Dominic Kwiatkowski, Cordelia Langford, John Sillitoe on behalf of the Wellcome Sanger Institute COVID-19 Surveillance Team |
| EPI_ISL_2700525 | Lighthouse Lab in Glasgow | Wellcome Sanger Institute for the COVID-19 Genomics UK (COG-UK) Consortium | Harper VanSteenhouse, Yumi Kasai, David Gray, Carol Clugston, Anna Dominiczak and Alex Alderton, Roberto Amato, Jeffrey Barrett, Sonia Goncalves, Ewan Harrison, David K. Jackson, Ian Johnston, Dominic Kwiatkowski, Cordelia Langford, John Sillitoe on behalf of the Wellcome Sanger Institute COVID-19 Surveillance Team |
| EPI_ISL_2700526 | Lighthouse Lab in Milton Keynes | Wellcome Sanger Institute for the COVID-19 Genomics UK (COG-UK) Consortium | The Lighthouse Lab in Milton Keynes and Alex Alderton, Roberto Amato, Jeffrey Barrett, Sonia Goncalves, Ewan Harrison, David K. Jackson, Ian Johnston, Dominic Kwiatkowski, Cordelia Langford, John Sillitoe on behalf of the Wellcome Sanger Institute COVID-19 Surveillance Team |
| EPI_ISL_2700527 | Lighthouse Lab in Glasgow | Wellcome Sanger Institute for the COVID-19 Genomics UK (COG-UK) Consortium | Harper VanSteenhouse, Yumi Kasai, David Gray, Carol Clugston, Anna Dominiczak and Alex Alderton, Roberto Amato, Jeffrey Barrett, Sonia Goncalves, Ewan Harrison, David K. Jackson, Ian Johnston, Dominic Kwiatkowski, Cordelia Langford, John Sillitoe on behalf of the Wellcome Sanger Institute COVID-19 Surveillance Team |
| EPI_ISL_2700528, EPI_ISL_2700529, EPI_ISL_2700530 | Lighthouse Lab in Milton Keynes | Wellcome Sanger Institute for the COVID-19 Genomics UK (COG-UK) Consortium | The Lighthouse Lab in Milton Keynes and Alex Alderton, Roberto Amato, Jeffrey Barrett, Sonia Goncalves, Ewan Harrison, David K. Jackson, Ian Johnston, Dominic Kwiatkowski, Cordelia Langford, John Sillitoe on behalf of the Wellcome Sanger Institute COVID-19 Surveillance Team |
| EPI_ISL_2700531, EPI_ISL_2700532, EPI_ISL_2700533 | Lighthouse Lab in Glasgow | Wellcome Sanger Institute for the COVID-19 Genomics UK (COG-UK) Consortium | Harper VanSteenhouse, Yumi Kasai, David Gray, Carol Clugston, Anna Dominiczak and Alex Alderton, Roberto Amato, Jeffrey Barrett, Sonia Goncalves, Ewan Harrison, David K. Jackson, Ian Johnston, Dominic Kwiatkowski, Cordelia Langford, John Sillitoe on behalf of the Wellcome Sanger Institute COVID-19 Surveillance Team |
| EPI_ISL_2700534, EPI_ISL_2700535 | Lighthouse Lab in Milton Keynes | Wellcome Sanger Institute for the COVID-19 Genomics UK (COG-UK) Consortium | The Lighthouse Lab in Milton Keynes and Alex Alderton, Roberto Amato, Jeffrey Barrett, Sonia Goncalves, Ewan Harrison, David K. Jackson, Ian Johnston, Dominic Kwiatkowski, Cordelia Langford, John Sillitoe on behalf of the Wellcome Sanger Institute COVID-19 Surveillance Team |
| EPI_ISL_2700536 | Lighthouse Lab in Glasgow | Wellcome Sanger Institute for the COVID-19 Genomics UK (COG-UK) Consortium | Harper VanSteenhouse, Yumi Kasai, David Gray, Carol Clugston, Anna Dominiczak and Alex Alderton, Roberto Amato, Jeffrey Barrett, Sonia Goncalves, Ewan Harrison, David K. Jackson, Ian Johnston, Dominic Kwiatkowski, Cordelia Langford, John Sillitoe on behalf of the Wellcome Sanger Institute COVID-19 Surveillance Team |
| EPI_ISL_2700537, EPI_ISL_2700538 | Lighthouse Lab in Milton Keynes | Wellcome Sanger Institute for the COVID-19 Genomics UK (COG-UK) Consortium | The Lighthouse Lab in Milton Keynes and Alex Alderton, Roberto Amato, Jeffrey Barrett, Sonia Goncalves, Ewan Harrison, David K. Jackson, Ian Johnston, Dominic Kwiatkowski, Cordelia Langford, John Sillitoe on behalf of the Wellcome Sanger Institute COVID-19 Surveillance Team |
| EPI_ISL_2700539, EPI_ISL_2700541, EPI_ISL_2700542, EPI_ISL_2700543 | Lighthouse Lab in Glasgow | Wellcome Sanger Institute for the COVID-19 Genomics UK (COG-UK) Consortium | Harper VanSteenhouse, Yumi Kasai, David Gray, Carol Clugston, Anna Dominiczak and Alex Alderton, Roberto Amato, Jeffrey Barrett, Sonia Goncalves, Ewan Harrison, David K. Jackson, Ian Johnston, Dominic Kwiatkowski, Cordelia Langford, John Sillitoe on behalf of the Wellcome Sanger Institute COVID-19 Surveillance Team |
| EPI_ISL_2700544 | Lighthouse Lab in Milton Keynes | Wellcome Sanger Institute for the COVID-19 Genomics UK (COG-UK) Consortium | The Lighthouse Lab in Milton Keynes and Alex Alderton, Roberto Amato, Jeffrey Barrett, Sonia Goncalves, Ewan Harrison, David K. Jackson, Ian Johnston, Dominic Kwiatkowski, Cordelia Langford, John Sillitoe on behalf of the Wellcome Sanger Institute COVID-19 Surveillance Team |
| EPI_ISL_2700545, EPI_ISL_2700546 | Lighthouse Lab in Glasgow | Wellcome Sanger Institute for the COVID-19 Genomics UK (COG-UK) Consortium | Harper VanSteenhouse, Yumi Kasai, David Gray, Carol Clugston, Anna Dominiczak and Alex Alderton, Roberto Amato, Jeffrey Barrett, Sonia Goncalves, Ewan Harrison, David K. Jackson, Ian Johnston, Dominic Kwiatkowski, Cordelia Langford, John Sillitoe on behalf of the Wellcome Sanger Institute COVID-19 Surveillance Team |
| EPI_ISL_2700547, EPI_ISL_2700548 | Lighthouse Lab in Milton Keynes | Wellcome Sanger Institute for the COVID-19 Genomics UK (COG-UK) Consortium | The Lighthouse Lab in Milton Keynes and Alex Alderton, Roberto Amato, Jeffrey Barrett, Sonia Goncalves, Ewan Harrison, David K. Jackson, Ian Johnston, Dominic Kwiatkowski, Cordelia Langford, John Sillitoe on behalf of the Wellcome Sanger Institute COVID-19 Surveillance Team |
| EPI_ISL_2700549 | Lighthouse Lab in Glasgow | Wellcome Sanger Institute for the COVID-19 Genomics UK (COG-UK) Consortium | Harper VanSteenhouse, Yumi Kasai, David Gray, Carol Clugston, Anna Dominiczak and Alex Alderton, Roberto Amato, Jeffrey Barrett, Sonia Goncalves, Ewan Harrison, David K. Jackson, Ian Johnston, Dominic Kwiatkowski, Cordelia Langford, John Sillitoe on behalf of the Wellcome Sanger Institute COVID-19 Surveillance Team |
| EPI_ISL_2700550, EPI_ISL_2700551, EPI_ISL_2700552, EPI_ISL_2700553 | Lighthouse Lab in Milton Keynes | Wellcome Sanger Institute for the COVID-19 Genomics UK (COG-UK) Consortium | The Lighthouse Lab in Milton Keynes and Alex Alderton, Roberto Amato, Jeffrey Barrett, Sonia Goncalves, Ewan Harrison, David K. Jackson, Ian Johnston, Dominic Kwiatkowski, Cordelia Langford, John Sillitoe on behalf of the Wellcome Sanger Institute COVID-19 Surveillance Team |
| EPI_ISL_2700554 | Lighthouse Lab in Glasgow | Wellcome Sanger Institute for the COVID-19 Genomics UK (COG-UK) Consortium | Harper VanSteenhouse, Yumi Kasai, David Gray, Carol Clugston, Anna Dominiczak and Alex Alderton, Roberto Amato, Jeffrey Barrett, Sonia Goncalves, Ewan Harrison, David K. Jackson, Ian Johnston, Dominic Kwiatkowski, Cordelia Langford, John Sillitoe on behalf of the Wellcome Sanger Institute COVID-19 Surveillance Team |
| EPI_ISL_2700555, EPI_ISL_2700556 | Lighthouse Lab in Milton Keynes | Wellcome Sanger Institute for the COVID-19 Genomics UK (COG-UK) Consortium | The Lighthouse Lab in Milton Keynes and Alex Alderton, Roberto Amato, Jeffrey Barrett, Sonia Goncalves, Ewan Harrison, David K. Jackson, Ian Johnston, Dominic Kwiatkowski, Cordelia Langford, John Sillitoe on behalf of the Wellcome Sanger Institute COVID-19 Surveillance Team |
| EPI_ISL_2700557 | Lighthouse Lab in Glasgow | Wellcome Sanger Institute for the COVID-19 Genomics UK (COG-UK) Consortium | Harper VanSteenhouse, Yumi Kasai, David Gray, Carol Clugston, Anna Dominiczak and Alex Alderton, Roberto Amato, Jeffrey Barrett, Sonia Goncalves, Ewan Harrison, David K. Jackson, Ian Johnston, Dominic Kwiatkowski, Cordelia Langford, John Sillitoe on behalf of the Wellcome Sanger Institute COVID-19 Surveillance Team |
| EPI_ISL_2700558, EPI_ISL_2700559 | Lighthouse Lab in Milton Keynes | Wellcome Sanger Institute for the COVID-19 Genomics UK (COG-UK) Consortium | The Lighthouse Lab in Milton Keynes and Alex Alderton, Roberto Amato, Jeffrey Barrett, Sonia Goncalves, Ewan Harrison, David K. Jackson, Ian Johnston, Dominic Kwiatkowski, Cordelia Langford, John Sillitoe on behalf of the Wellcome Sanger Institute COVID-19 Surveillance Team |
| EPI_ISL_2700560, EPI_ISL_2700561, EPI_ISL_2700562, EPI_ISL_2700563, EPI_ISL_2700564 | Lighthouse Lab in Glasgow | Wellcome Sanger Institute for the COVID-19 Genomics UK (COG-UK) Consortium | Harper VanSteenhouse, Yumi Kasai, David Gray, Carol Clugston, Anna Dominiczak and Alex Alderton, Roberto Amato, Jeffrey Barrett, Sonia Goncalves, Ewan Harrison, David K. Jackson, Ian Johnston, Dominic Kwiatkowski, Cordelia Langford, John Sillitoe on behalf of the Wellcome Sanger Institute COVID-19 Surveillance Team |
| EPI_ISL_2700565 | Lighthouse Lab in Milton Keynes | Wellcome Sanger Institute for the COVID-19 Genomics UK (COG-UK) Consortium | The Lighthouse Lab in Milton Keynes and Alex Alderton, Roberto Amato, Jeffrey Barrett, Sonia Goncalves, Ewan Harrison, David K. Jackson, Ian Johnston, Dominic Kwiatkowski, Cordelia Langford, John Sillitoe on behalf of the Wellcome Sanger Institute COVID-19 Surveillance Team |
| EPI_ISL_2700566, EPI_ISL_2700567 | Lighthouse Lab in Glasgow | Wellcome Sanger Institute for the COVID-19 Genomics UK (COG-UK) Consortium | Harper VanSteenhouse, Yumi Kasai, David Gray, Carol Clugston, Anna Dominiczak and Alex Alderton, Roberto Amato, Jeffrey Barrett, Sonia Goncalves, Ewan Harrison, David K. Jackson, Ian Johnston, Dominic Kwiatkowski, Cordelia Langford, John Sillitoe on behalf of the Wellcome Sanger Institute COVID-19 Surveillance Team |
| EPI_ISL_2700568 | Lighthouse Lab in Milton Keynes | Wellcome Sanger Institute for the COVID-19 Genomics UK (COG-UK) Consortium | The Lighthouse Lab in Milton Keynes and Alex Alderton, Roberto Amato, Jeffrey Barrett, Sonia Goncalves, Ewan Harrison, David K. Jackson, Ian Johnston, Dominic Kwiatkowski, Cordelia Langford, John Sillitoe on behalf of the Wellcome Sanger Institute COVID-19 Surveillance Team |
| EPI_ISL_2700569 | Lighthouse Lab in Glasgow | Wellcome Sanger Institute for the COVID-19 Genomics UK (COG-UK) Consortium | Harper VanSteenhouse, Yumi Kasai, David Gray, Carol Clugston, Anna Dominiczak and Alex Alderton, Roberto Amato, Jeffrey Barrett, Sonia Goncalves, Ewan Harrison, David K. Jackson, Ian Johnston, Dominic Kwiatkowski, Cordelia Langford, John Sillitoe on behalf of the Wellcome Sanger Institute COVID-19 Surveillance Team |
| EPI_ISL_2700570 | Lighthouse Lab in Milton Keynes | Wellcome Sanger Institute for the COVID-19 Genomics UK (COG-UK) Consortium | The Lighthouse Lab in Milton Keynes and Alex Alderton, Roberto Amato, Jeffrey Barrett, Sonia Goncalves, Ewan Harrison, David K. Jackson, Ian Johnston, Dominic Kwiatkowski, Cordelia Langford, John Sillitoe on behalf of the Wellcome Sanger Institute COVID-19 Surveillance Team |
| EPI_ISL_2700571 | Lighthouse Lab in Glasgow | Wellcome Sanger Institute for the COVID-19 Genomics UK (COG-UK) Consortium | Harper VanSteenhouse, Yumi Kasai, David Gray, Carol Clugston, Anna Dominiczak and Alex Alderton, Roberto Amato, Jeffrey Barrett, Sonia Goncalves, Ewan Harrison, David K. Jackson, Ian Johnston, Dominic Kwiatkowski, Cordelia Langford, John Sillitoe on behalf of the Wellcome Sanger Institute COVID-19 Surveillance Team |
| EPI_ISL_2700572 | Lighthouse Lab in Milton Keynes | Wellcome Sanger Institute for the COVID-19 Genomics UK (COG-UK) Consortium | The Lighthouse Lab in Milton Keynes and Alex Alderton, Roberto Amato, Jeffrey Barrett, Sonia Goncalves, Ewan Harrison, David K. Jackson, Ian Johnston, Dominic Kwiatkowski, Cordelia Langford, John Sillitoe on behalf of the Wellcome Sanger Institute COVID-19 Surveillance Team |
| EPI_ISL_2700573 | Lighthouse Lab in Glasgow | Wellcome Sanger Institute for the COVID-19 Genomics UK (COG-UK) Consortium | Harper VanSteenhouse, Yumi Kasai, David Gray, Carol Clugston, Anna Dominiczak and Alex Alderton, Roberto Amato, Jeffrey Barrett, Sonia Goncalves, Ewan Harrison, David K. Jackson, Ian Johnston, Dominic Kwiatkowski, Cordelia Langford, John Sillitoe on behalf of the Wellcome Sanger Institute COVID-19 Surveillance Team |
| EPI_ISL_2700574 | Lighthouse Lab in Milton Keynes | Wellcome Sanger Institute for the COVID-19 Genomics UK (COG-UK) Consortium | The Lighthouse Lab in Milton Keynes and Alex Alderton, Roberto Amato, Jeffrey Barrett, Sonia Goncalves, Ewan Harrison, David K. Jackson, Ian Johnston, Dominic Kwiatkowski, Cordelia Langford, John Sillitoe on behalf of the Wellcome Sanger Institute COVID-19 Surveillance Team |
| EPI_ISL_2700575 | Lighthouse Lab in Glasgow | Wellcome Sanger Institute for the COVID-19 Genomics UK (COG-UK) Consortium | Harper VanSteenhouse, Yumi Kasai, David Gray, Carol Clugston, Anna Dominiczak and Alex Alderton, Roberto Amato, Jeffrey Barrett, Sonia Goncalves, Ewan Harrison, David K. Jackson, Ian Johnston, Dominic Kwiatkowski, Cordelia Langford, John Sillitoe on behalf of the Wellcome Sanger Institute COVID-19 Surveillance Team |
| EPI_ISL_2700576 | Lighthouse Lab in Milton Keynes | Wellcome Sanger Institute for the COVID-19 Genomics UK (COG-UK) Consortium | The Lighthouse Lab in Milton Keynes and Alex Alderton, Roberto Amato, Jeffrey Barrett, Sonia Goncalves, Ewan Harrison, David K. Jackson, Ian Johnston, Dominic Kwiatkowski, Cordelia Langford, John Sillitoe on behalf of the Wellcome Sanger Institute COVID-19 Surveillance Team |

| | | | |
|---|---|---|---|
| EPI_ISL_2700577, EPI_ISL_2700578, EPI_ISL_2700579, EPI_ISL_2700580, EPI_ISL_2700581, EPI_ISL_2700582, EPI_ISL_2700583, EPI_ISL_2700584, EPI_ISL_2700585, EPI_ISL_2700586, EPI_ISL_2700587, EPI_ISL_2700588, EPI_ISL_2700589, EPI_ISL_2700590, EPI_ISL_2700591, EPI_ISL_2700592, EPI_ISL_2700593, EPI_ISL_2700594, EPI_ISL_2700595, EPI_ISL_2700596, EPI_ISL_2700597, EPI_ISL_2700598, EPI_ISL_2700599, EPI_ISL_2700600, EPI_ISL_2700601, EPI_ISL_2700602, EPI_ISL_2700603, EPI_ISL_2700604, EPI_ISL_2700605, EPI_ISL_2700606, EPI_ISL_2700607, EPI_ISL_2700608, EPI_ISL_2700609, EPI_ISL_2700610, EPI_ISL_2700611, EPI_ISL_2700612, EPI_ISL_2700613, EPI_ISL_2700614, EPI_ISL_2700615, EPI_ISL_2700616, EPI_ISL_2700617, EPI_ISL_2700618, EPI_ISL_2700619, EPI_ISL_2700620, EPI_ISL_2700621, EPI_ISL_2700623, EPI_ISL_2700624, EPI_ISL_2700625, EPI_ISL_2700626, EPI_ISL_2700627, EPI_ISL_2700628, EPI_ISL_2700629, EPI_ISL_2700630, EPI_ISL_2700631, EPI_ISL_2700632, EPI_ISL_2700633, EPI_ISL_2700634, EPI_ISL_2700635, EPI_ISL_2700636, EPI_ISL_2700637, EPI_ISL_2700638, EPI_ISL_2700639, EPI_ISL_2700640, EPI_ISL_2700641, EPI_ISL_2700642, EPI_ISL_2700643, EPI_ISL_2700644, EPI_ISL_2700645, EPI_ISL_2700646, EPI_ISL_2700647, EPI_ISL_2700648, EPI_ISL_2700649, EPI_ISL_2700650, EPI_ISL_2700651, EPI_ISL_2700652, EPI_ISL_2700653, EPI_ISL_2700654, EPI_ISL_2700655, EPI_ISL_2700656, EPI_ISL_2700657, EPI_ISL_2700658, EPI_ISL_2700659, EPI_ISL_2700660, EPI_ISL_2700661, EPI_ISL_2700662, EPI_ISL_2700663, EPI_ISL_2700664, EPI_ISL_2700665, EPI_ISL_2700666, EPI_ISL_2700667, EPI_ISL_2700668, EPI_ISL_2700669, EPI_ISL_2700670, EPI_ISL_2700671, EPI_ISL_2700672, EPI_ISL_2700673, EPI_ISL_2700674, EPI_ISL_2700675, EPI_ISL_2700676, EPI_ISL_2700677, EPI_ISL_2700678, EPI_ISL_2700679, EPI_ISL_2700680, EPI_ISL_2700681, EPI_ISL_2700682, EPI_ISL_2700683, EPI_ISL_2700684, EPI_ISL_2700685, EPI_ISL_2700686, EPI_ISL_2700687, EPI_ISL_2700688, EPI_ISL_2700689, EPI_ISL_2700690, EPI_ISL_2700691, EPI_ISL_2700692, EPI_ISL_2700693, EPI_ISL_2700694, EPI_ISL_2700695, EPI_ISL_2700696, EPI_ISL_2700697, EPI_ISL_2700698, EPI_ISL_2700699, EPI_ISL_2700701, EPI_ISL_2700702, EPI_ISL_2700703, EPI_ISL_2700704, EPI_ISL_2700705, EPI_ISL_2700706, EPI_ISL_2700707, EPI_ISL_2700708, EPI_ISL_2700709, EPI_ISL_2700710, EPI_ISL_2700711, EPI_ISL_2700712, EPI_ISL_2700713, EPI_ISL_2700714, EPI_ISL_2700715, EPI_ISL_2700716, EPI_ISL_2700717, EPI_ISL_2700718, EPI_ISL_2700719, EPI_ISL_2700720, EPI_ISL_2700721, EPI_ISL_2700722, EPI_ISL_2700723, EPI_ISL_2700724, EPI_ISL_2700725, EPI_ISL_2700726, EPI_ISL_2700727, EPI_ISL_2700728, EPI_ISL_2700729, EPI_ISL_2700730, EPI_ISL_2700731, EPI_ISL_2700732, EPI_ISL_2700733, EPI_ISL_2700734, EPI_ISL_2700735, EPI_ISL_2700736, EPI_ISL_2700737, EPI_ISL_2700738, EPI_ISL_2700739, EPI_ISL_2700740, EPI_ISL_2700741, EPI_ISL_2700742, EPI_ISL_2700743, EPI_ISL_2700744, EPI_ISL_2700745, EPI_ISL_2700746, EPI_ISL_2700747, EPI_ISL_2700748, EPI_ISL_2700749, EPI_ISL_2700750, EPI_ISL_2700751, EPI_ISL_2700752, EPI_ISL_2700753, EPI_ISL_2700754, EPI_ISL_2700755, EPI_ISL_2700756, EPI_ISL_2700757, EPI_ISL_2700758, EPI_ISL_2700759, EPI_ISL_2700760, EPI_ISL_2700761, EPI_ISL_2700762, EPI_ISL_2700763, EPI_ISL_2700764, EPI_ISL_2700765, EPI_ISL_2700766, EPI_ISL_2700767, EPI_ISL_2700768, EPI_ISL_2700769, EPI_ISL_2700770, EPI_ISL_2700771, EPI_ISL_2700772, EPI_ISL_2700773, EPI_ISL_2700774, EPI_ISL_2700775, EPI_ISL_2700776, EPI_ISL_2700777, EPI_ISL_2700778, EPI_ISL_2700779, EPI_ISL_2700780, EPI_ISL_2700781, EPI_ISL_2700782, EPI_ISL_2700783, EPI_ISL_2700784, EPI_ISL_2700785, EPI_ISL_2700786, EPI_ISL_2700787, EPI_ISL_2700788, EPI_ISL_2700789, EPI_ISL_2700790, EPI_ISL_2700791, EPI_ISL_2700792, EPI_ISL_2700793, EPI_ISL_2700794, EPI_ISL_2700795, EPI_ISL_2700796, EPI_ISL_2700797, EPI_ISL_2700798, EPI_ISL_2700799, EPI_ISL_2700801, EPI_ISL_2700802, EPI_ISL_2700803, EPI_ISL_2700804, EPI_ISL_2700805, EPI_ISL_2700806, EPI_ISL_2700807, EPI_ISL_2700808, EPI_ISL_2700809, EPI_ISL_2700810, EPI_ISL_2700811, EPI_ISL_2700812, EPI_ISL_2700813, EPI_ISL_2700814, EPI_ISL_2700815, EPI_ISL_2700816, EPI_ISL_2700817, EPI_ISL_2700818, EPI_ISL_2700819, EPI_ISL_2700820, EPI_ISL_2700821, EPI_ISL_2700822, EPI_ISL_2700823, EPI_ISL_2700824, EPI_ISL_2700825, EPI_ISL_2700826, EPI_ISL_2700827, EPI_ISL_2700828, EPI_ISL_2700829, EPI_ISL_2700830, EPI_ISL_2700831, EPI_ISL_2700832, EPI_ISL_2700833, EPI_ISL_2700834, EPI_ISL_2700835, EPI_ISL_2700836, EPI_ISL_2700837, EPI_ISL_2700838, EPI_ISL_2700839, EPI_ISL_2700840, EPI_ISL_2700841, EPI_ISL_2700842, EPI_ISL_2700843, EPI_ISL_2700844, EPI_ISL_2700845, EPI_ISL_2700846, EPI_ISL_2700847, EPI_ISL_2700848, EPI_ISL_2700849, EPI_ISL_2700850, EPI_ISL_2700851, EPI_ISL_2700852, EPI_ISL_2700853, EPI_ISL_2700854, EPI_ISL_2700855, EPI_ISL_2700856, EPI_ISL_2700857, EPI_ISL_2700858, EPI_ISL_2700859, EPI_ISL_2700860, EPI_ISL_2700861, EPI_ISL_2700862, EPI_ISL_2700863, EPI_ISL_2700864, EPI_ISL_2700865, EPI_ISL_2700866, EPI_ISL_2700867, EPI_ISL_2700868, EPI_ISL_2700869, EPI_ISL_2700870, EPI_ISL_2700871, EPI_ISL_2700872, EPI_ISL_2700873, EPI_ISL_2700874, EPI_ISL_2700875, EPI_ISL_2700876, EPI_ISL_2700877, EPI_ISL_2700878, EPI_ISL_2700879, EPI_ISL_2700880, EPI_ISL_2700881, EPI_ISL_2700882, EPI_ISL_2700883, EPI_ISL_2700884, EPI_ISL_2700885, EPI_ISL_2700886, EPI_ISL_2700887, EPI_ISL_2700888, EPI_ISL_2700889, EPI_ISL_2700890, EPI_ISL_2700891, EPI_ISL_2700892, EPI_ISL_2700893, EPI_ISL_2700894, EPI_ISL_2700895, EPI_ISL_2700896, EPI_ISL_2700897, EPI_ISL_2700898, EPI_ISL_2700899, EPI_ISL_2700900, EPI_ISL_2700901, EPI_ISL_2700902, EPI_ISL_2700903, EPI_ISL_2700904, EPI_ISL_2700905, EPI_ISL_2700906, EPI_ISL_2700907, EPI_ISL_2700908, EPI_ISL_2700909, EPI_ISL_2700910, EPI_ISL_2700911, EPI_ISL_2700912, EPI_ISL_2700913, EPI_ISL_2700914, EPI_ISL_2700915, EPI_ISL_2700916, EPI_ISL_2700917, EPI_ISL_2700918, EPI_ISL_2700919, EPI_ISL_2700920, EPI_ISL_2700921, EPI_ISL_2700922, EPI_ISL_2700923, EPI_ISL_2700924, EPI_ISL_2700925, EPI_ISL_2700926, EPI_ISL_2700927, EPI_ISL_2700928, EPI_ISL_2700929, EPI_ISL_2700930, EPI_ISL_2700931, EPI_ISL_2700932, EPI_ISL_2700933, EPI_ISL_2700934, EPI_ISL_2700935, EPI_ISL_2700936, EPI_ISL_2700937, EPI_ISL_2700938 | | | |
| see above | Lighthouse Lab in Alderley Park | Wellcome Sanger Institute for the COVID-19 Genomics UK (COG-UK) Consortium | Jacquelyn Wynn, Mairead Hyland, The Lighthouse Lab in Alderley Park and Alex Alderton, Roberto Amato, Jeffrey Barrett, Sonia Goncalves, Ewan Harrison, David K. Jackson, Ian Johnston, Dominic Kwiatkowski, Cordelia Langford, John Sillitoe on behalf of the Wellcome Sanger Institute COVID-19 Surveillance Team |
| EPI_ISL_2700939 | Lighthouse Lab in Glasgow | Wellcome Sanger Institute for the COVID-19 Genomics UK (COG-UK) Consortium | Harper VanSteenhouse, Yumi Kasai, David Gray, Carol Clugston, Anna Dominiczak and Alex Alderton, Roberto Amato, Jeffrey Barrett, Sonia Goncalves, Ewan Harrison, David K. Jackson, Ian Johnston, Dominic Kwiatkowski, Cordelia Langford, John Sillitoe on behalf of the Wellcome Sanger Institute COVID-19 Surveillance Team |
| EPI_ISL_2700940, EPI_ISL_2700941, EPI_ISL_2700942, EPI_ISL_2700943, EPI_ISL_2700944 | Lighthouse Lab in Alderley Park | Wellcome Sanger Institute for the COVID-19 Genomics UK (COG-UK) Consortium | Jacquelyn Wynn, Mairead Hyland, The Lighthouse Lab in Alderley Park and Alex Alderton, Roberto Amato, Jeffrey Barrett, Sonia Goncalves, Ewan Harrison, David K. Jackson, Ian Johnston, Dominic Kwiatkowski, Cordelia Langford, John Sillitoe on behalf of the Wellcome Sanger Institute COVID-19 Surveillance Team |
| EPI_ISL_2700945, EPI_ISL_2700946, EPI_ISL_2700947, EPI_ISL_2700948, EPI_ISL_2700949, EPI_ISL_2700950 | Lighthouse Lab in Glasgow | Wellcome Sanger Institute for the COVID-19 Genomics UK (COG-UK) Consortium | Harper VanSteenhouse, Yumi Kasai, David Gray, Carol Clugston, Anna Dominiczak and Alex Alderton, Roberto Amato, Jeffrey Barrett, Sonia Goncalves, Ewan Harrison, David K. Jackson, Ian Johnston, Dominic Kwiatkowski, Cordelia Langford, John Sillitoe on behalf of the Wellcome Sanger Institute COVID-19 Surveillance Team |
| EPI_ISL_2700951, EPI_ISL_2700952 | Lighthouse Lab in Alderley Park | Wellcome Sanger Institute for the COVID-19 Genomics UK (COG-UK) Consortium | Jacquelyn Wynn, Mairead Hyland, The Lighthouse Lab in Alderley Park and Alex Alderton, Roberto Amato, Jeffrey Barrett, Sonia Goncalves, Ewan Harrison, David K. Jackson, Ian Johnston, Dominic Kwiatkowski, Cordelia Langford, John Sillitoe on behalf of the Wellcome Sanger Institute COVID-19 Surveillance Team |
| EPI_ISL_2700953, EPI_ISL_2700954 | Lighthouse Lab in Glasgow | Wellcome Sanger Institute for the COVID-19 Genomics UK (COG-UK) Consortium | Harper VanSteenhouse, Yumi Kasai, David Gray, Carol Clugston, Anna Dominiczak and Alex Alderton, Roberto Amato, Jeffrey Barrett, Sonia Goncalves, Ewan Harrison, David K. Jackson, Ian Johnston, Dominic Kwiatkowski, Cordelia Langford, John Sillitoe on behalf of the Wellcome Sanger Institute COVID-19 Surveillance Team |
| EPI_ISL_2700955 | Lighthouse Lab in Alderley Park | Wellcome Sanger Institute for the COVID-19 Genomics UK (COG-UK) Consortium | Jacquelyn Wynn, Mairead Hyland, The Lighthouse Lab in Alderley Park and Alex Alderton, Roberto Amato, Jeffrey Barrett, Sonia Goncalves, Ewan Harrison, David K. Jackson, Ian Johnston, Dominic Kwiatkowski, Cordelia Langford, John Sillitoe on behalf of the Wellcome Sanger Institute COVID-19 Surveillance Team |
| EPI_ISL_2700956, EPI_ISL_2700957, EPI_ISL_2700958 | Lighthouse Lab in Glasgow | Wellcome Sanger Institute for the COVID-19 Genomics UK (COG-UK) Consortium | Harper VanSteenhouse, Yumi Kasai, David Gray, Carol Clugston, Anna Dominiczak and Alex Alderton, Roberto Amato, Jeffrey Barrett, Sonia Goncalves, Ewan Harrison, David K. Jackson, Ian Johnston, Dominic Kwiatkowski, Cordelia Langford, John Sillitoe on behalf of the Wellcome Sanger Institute COVID-19 Surveillance Team |
| EPI_ISL_2700959 | Lighthouse Lab in Alderley Park | Wellcome Sanger Institute for the COVID-19 Genomics UK (COG-UK) Consortium | Jacquelyn Wynn, Mairead Hyland, The Lighthouse Lab in Alderley Park and Alex Alderton, Roberto Amato, Jeffrey Barrett, Sonia Goncalves, Ewan Harrison, David K. Jackson, Ian Johnston, Dominic Kwiatkowski, Cordelia Langford, John Sillitoe on behalf of the Wellcome Sanger Institute COVID-19 Surveillance Team |
| EPI_ISL_2700960, EPI_ISL_2700961, EPI_ISL_2700962, EPI_ISL_2700964, EPI_ISL_2700965, EPI_ISL_2700966 | Lighthouse Lab in Glasgow | Wellcome Sanger Institute for the COVID-19 Genomics UK (COG-UK) Consortium | Harper VanSteenhouse, Yumi Kasai, David Gray, Carol Clugston, Anna Dominiczak and Alex Alderton, Roberto Amato, Jeffrey Barrett, Sonia Goncalves, Ewan Harrison, David K. Jackson, Ian Johnston, Dominic Kwiatkowski, Cordelia Langford, John Sillitoe on behalf of the Wellcome Sanger Institute COVID-19 Surveillance Team |
| EPI_ISL_2700967 | Lighthouse Lab in Alderley Park | Wellcome Sanger Institute for the COVID-19 Genomics UK (COG-UK) Consortium | Jacquelyn Wynn, Mairead Hyland, The Lighthouse Lab in Alderley Park and Alex Alderton, Roberto Amato, Jeffrey Barrett, Sonia Goncalves, Ewan Harrison, David K. Jackson, Ian Johnston, Dominic Kwiatkowski, Cordelia Langford, John Sillitoe on behalf of the Wellcome Sanger Institute COVID-19 Surveillance Team |
| EPI_ISL_2700968 | Lighthouse Lab in Glasgow | Wellcome Sanger Institute for the COVID-19 Genomics UK (COG-UK) Consortium | Harper VanSteenhouse, Yumi Kasai, David Gray, Carol Clugston, Anna Dominiczak and Alex Alderton, Roberto Amato, Jeffrey Barrett, Sonia Goncalves, Ewan Harrison, David K. Jackson, Ian Johnston, Dominic Kwiatkowski, Cordelia Langford, John Sillitoe on behalf of the Wellcome Sanger Institute COVID-19 Surveillance Team |
| EPI_ISL_2700969 | Lighthouse Lab in Alderley Park | Wellcome Sanger Institute for the COVID-19 Genomics UK (COG-UK) Consortium | Jacquelyn Wynn, Mairead Hyland, The Lighthouse Lab in Alderley Park and Alex Alderton, Roberto Amato, Jeffrey Barrett, Sonia Goncalves, Ewan Harrison, David K. Jackson, Ian Johnston, Dominic Kwiatkowski, Cordelia Langford, John Sillitoe on behalf of the Wellcome Sanger Institute COVID-19 Surveillance Team |
| EPI_ISL_2700970 | Lighthouse Lab in Glasgow | Wellcome Sanger Institute for the COVID-19 Genomics UK (COG-UK) Consortium | Harper VanSteenhouse, Yumi Kasai, David Gray, Carol Clugston, Anna Dominiczak and Alex Alderton, Roberto Amato, Jeffrey Barrett, Sonia Goncalves, Ewan Harrison, David K. Jackson, Ian Johnston, Dominic Kwiatkowski, Cordelia Langford, John Sillitoe on behalf of the Wellcome Sanger Institute COVID-19 Surveillance Team |
| EPI_ISL_2700971 | Lighthouse Lab in Alderley Park | Wellcome Sanger Institute for the COVID-19 Genomics UK (COG-UK) Consortium | Jacquelyn Wynn, Mairead Hyland, The Lighthouse Lab in Alderley Park and Alex Alderton, Roberto Amato, Jeffrey Barrett, Sonia Goncalves, Ewan Harrison, David K. Jackson, Ian Johnston, Dominic Kwiatkowski, Cordelia Langford, John Sillitoe on behalf of the Wellcome Sanger Institute COVID-19 Surveillance Team |
| EPI_ISL_2700972, EPI_ISL_2700973, EPI_ISL_2700974 | Lighthouse Lab in Glasgow | Wellcome Sanger Institute for the COVID-19 Genomics UK (COG-UK) Consortium | Harper VanSteenhouse, Yumi Kasai, David Gray, Carol Clugston, Anna Dominiczak and Alex Alderton, Roberto Amato, Jeffrey Barrett, Sonia Goncalves, Ewan Harrison, David K. Jackson, Ian Johnston, Dominic Kwiatkowski, Cordelia Langford, John Sillitoe on behalf of the Wellcome Sanger Institute COVID-19 Surveillance Team |
| EPI_ISL_2700975 | Lighthouse Lab in Alderley Park | Wellcome Sanger Institute for the COVID-19 Genomics UK (COG-UK) Consortium | Jacquelyn Wynn, Mairead Hyland, The Lighthouse Lab in Alderley Park and Alex Alderton, Roberto Amato, Jeffrey Barrett, Sonia Goncalves, Ewan Harrison, David K. Jackson, Ian Johnston, Dominic Kwiatkowski, Cordelia Langford, John Sillitoe on behalf of the Wellcome Sanger Institute COVID-19 Surveillance Team |
| EPI_ISL_2700976, EPI_ISL_2700977 | Lighthouse Lab in Glasgow | Wellcome Sanger Institute for the COVID-19 Genomics UK (COG-UK) Consortium | Harper VanSteenhouse, Yumi Kasai, David Gray, Carol Clugston, Anna Dominiczak and Alex Alderton, Roberto Amato, Jeffrey Barrett, Sonia Goncalves, Ewan Harrison, David K. Jackson, Ian Johnston, Dominic Kwiatkowski, Cordelia Langford, John Sillitoe on behalf of the Wellcome Sanger Institute COVID-19 Surveillance Team |
| EPI_ISL_2700978 | Lighthouse Lab in Alderley Park | Wellcome Sanger Institute for the COVID-19 Genomics UK (COG-UK) Consortium | Jacquelyn Wynn, Mairead Hyland, The Lighthouse Lab in Alderley Park and Alex Alderton, Roberto Amato, Jeffrey Barrett, Sonia Goncalves, Ewan Harrison, David K. Jackson, Ian Johnston, Dominic Kwiatkowski, Cordelia Langford, John Sillitoe on behalf of the Wellcome Sanger Institute COVID-19 Surveillance Team |
| EPI_ISL_2700979 | Lighthouse Lab in Glasgow | Wellcome Sanger Institute for the COVID-19 Genomics UK (COG-UK) Consortium | Harper VanSteenhouse, Yumi Kasai, David Gray, Carol Clugston, Anna Dominiczak and Alex Alderton, Roberto Amato, Jeffrey Barrett, Sonia Goncalves, Ewan Harrison, David K. Jackson, Ian Johnston, Dominic Kwiatkowski, Cordelia Langford, John Sillitoe on behalf of the Wellcome Sanger Institute COVID-19 Surveillance Team |
| EPI_ISL_2700980, EPI_ISL_2700981, EPI_ISL_2700982 | Lighthouse Lab in Alderley Park | Wellcome Sanger Institute for the COVID-19 Genomics UK (COG-UK) Consortium | Jacquelyn Wynn, Mairead Hyland, The Lighthouse Lab in Alderley Park and Alex Alderton, Roberto Amato, Jeffrey Barrett, Sonia Goncalves, Ewan Harrison, David K. Jackson, Ian Johnston, Dominic Kwiatkowski, Cordelia Langford, John Sillitoe on behalf of the Wellcome Sanger Institute COVID-19 Surveillance Team |
| EPI_ISL_2700983 | Lighthouse Lab in Glasgow | Wellcome Sanger Institute for the COVID-19 Genomics UK (COG-UK) Consortium | Harper VanSteenhouse, Yumi Kasai, David Gray, Carol Clugston, Anna Dominiczak and Alex Alderton, Roberto Amato, Jeffrey Barrett, Sonia Goncalves, Ewan Harrison, David K. Jackson, Ian Johnston, Dominic Kwiatkowski, Cordelia Langford, John Sillitoe on behalf of the Wellcome Sanger Institute COVID-19 Surveillance Team |
| EPI_ISL_2700984 | Lighthouse Lab in Alderley Park | Wellcome Sanger Institute for the COVID-19 Genomics UK (COG-UK) Consortium | Jacquelyn Wynn, Mairead Hyland, The Lighthouse Lab in Alderley Park and Alex Alderton, Roberto Amato, Jeffrey Barrett, Sonia Goncalves, Ewan Harrison, David K. Jackson, Ian Johnston, Dominic Kwiatkowski, Cordelia Langford, John Sillitoe on behalf of the Wellcome Sanger Institute COVID-19 Surveillance Team |
| EPI_ISL_2700985, EPI_ISL_2700986, EPI_ISL_2700987 | Lighthouse Lab in Glasgow | Wellcome Sanger Institute for the COVID-19 Genomics UK (COG-UK) Consortium | Harper VanSteenhouse, Yumi Kasai, David Gray, Carol Clugston, Anna Dominiczak and Alex Alderton, Roberto Amato, Jeffrey Barrett, Sonia Goncalves, Ewan Harrison, David K. Jackson, Ian Johnston, Dominic Kwiatkowski, Cordelia Langford, John Sillitoe on behalf of the Wellcome Sanger Institute COVID-19 Surveillance Team |
| EPI_ISL_2700988 | Lighthouse Lab in Alderley Park | Wellcome Sanger Institute for the COVID-19 Genomics UK (COG-UK) Consortium | Jacquelyn Wynn, Mairead Hyland, The Lighthouse Lab in Alderley Park and Alex Alderton, Roberto Amato, Jeffrey Barrett, Sonia Goncalves, Ewan Harrison, David K. Jackson, Ian Johnston, Dominic Kwiatkowski, Cordelia Langford, John Sillitoe on behalf of the Wellcome Sanger Institute COVID-19 Surveillance Team |
| EPI_ISL_2700989 | Lighthouse Lab in Glasgow | Wellcome Sanger Institute for the COVID-19 Genomics UK (COG-UK) Consortium | Harper VanSteenhouse, Yumi Kasai, David Gray, Carol Clugston, Anna Dominiczak and Alex Alderton, Roberto Amato, Jeffrey Barrett, Sonia Goncalves, Ewan Harrison, David K. Jackson, Ian Johnston, Dominic Kwiatkowski, Cordelia Langford, John Sillitoe on behalf of the Wellcome Sanger Institute COVID-19 Surveillance Team |
| EPI_ISL_2700990, EPI_ISL_2700991, EPI_ISL_2700992, EPI_ISL_2700993, EPI_ISL_2700994, EPI_ISL_2700995 | Lighthouse Lab in Alderley Park | Wellcome Sanger Institute for the COVID-19 Genomics UK (COG-UK) Consortium | Jacquelyn Wynn, Mairead Hyland, The Lighthouse Lab in Alderley Park and Alex Alderton, Roberto Amato, Jeffrey Barrett, Sonia Goncalves, Ewan Harrison, David K. Jackson, Ian Johnston, Dominic Kwiatkowski, Cordelia Langford, John Sillitoe on behalf of the Wellcome Sanger Institute COVID-19 Surveillance Team |
| EPI_ISL_2700996 | Lighthouse Lab in Glasgow | Wellcome Sanger Institute for the COVID-19 Genomics UK (COG-UK) Consortium | Harper VanSteenhouse, Yumi Kasai, David Gray, Carol Clugston, Anna Dominiczak and Alex Alderton, Roberto Amato, Jeffrey Barrett, Sonia Goncalves, Ewan Harrison, David K. Jackson, Ian Johnston, Dominic Kwiatkowski, Cordelia Langford, John Sillitoe on behalf of the Wellcome Sanger Institute COVID-19 Surveillance Team |
| EPI_ISL_2700997, EPI_ISL_2700998, EPI_ISL_2700999, EPI_ISL_2701000 | Lighthouse Lab in Alderley Park | Wellcome Sanger Institute for the COVID-19 Genomics UK (COG-UK) Consortium | Jacquelyn Wynn, Mairead Hyland, The Lighthouse Lab in Alderley Park and Alex Alderton, Roberto Amato, Jeffrey Barrett, Sonia Goncalves, Ewan Harrison, David K. Jackson, Ian Johnston, Dominic Kwiatkowski, Cordelia Langford, John Sillitoe on behalf of the Wellcome Sanger Institute COVID-19 Surveillance Team |
| EPI_ISL_2701001, | Lighthouse Lab in Glasgow | Wellcome Sanger Institute for the COVID-19 Genomics UK (COG-UK) | Harper VanSteenhouse, Yumi Kasai, David Gray, Carol Clugston, Anna Dominiczak and Alex Alderton, Roberto Amato, Jeffrey Barrett, Sonia Goncalves, Ewan Harrison, David K. Jackson, Ian Johnston, Dominic Kwiatkowski, Cordelia Langford, |

| | | | |
|---|---|---|---|
| EPI_ISL_2701002 | | Consortium | John Sillitoe on behalf of the Wellcome Sanger Institute COVID-19 Surveillance Team |
| EPI_ISL_2701003 | Lighthouse Lab in Alderley Park | Wellcome Sanger Institute for the COVID-19 Genomics UK (COG-UK) Consortium | Jacquelyn Wynn, Mairead Hyland, The Lighthouse Lab in Alderley Park and Alex Alderton, Roberto Amato, Jeffrey Barrett, Sonia Goncalves, Ewan Harrison, David K. Jackson, Ian Johnston, Dominic Kwiatkowski, Cordelia Langford, John Sillitoe on behalf of the Wellcome Sanger Institute COVID-19 Surveillance Team |
| EPI_ISL_2701004, EPI_ISL_2701005, EPI_ISL_2701006 | Lighthouse Lab in Glasgow | Wellcome Sanger Institute for the COVID-19 Genomics UK (COG-UK) Consortium | Harper VanSteenhouse, Yumi Kasai, David Gray, Carol Clugston, Anna Dominiczak and Alex Alderton, Roberto Amato, Jeffrey Barrett, Sonia Goncalves, Ewan Harrison, David K. Jackson, Ian Johnston, Dominic Kwiatkowski, Cordelia Langford, John Sillitoe on behalf of the Wellcome Sanger Institute COVID-19 Surveillance Team |
| EPI_ISL_2701007 | Lighthouse Lab in Alderley Park | Wellcome Sanger Institute for the COVID-19 Genomics UK (COG-UK) Consortium | Jacquelyn Wynn, Mairead Hyland, The Lighthouse Lab in Alderley Park and Alex Alderton, Roberto Amato, Jeffrey Barrett, Sonia Goncalves, Ewan Harrison, David K. Jackson, Ian Johnston, Dominic Kwiatkowski, Cordelia Langford, John Sillitoe on behalf of the Wellcome Sanger Institute COVID-19 Surveillance Team |
| EPI_ISL_2701008, EPI_ISL_2701009, EPI_ISL_2701010, EPI_ISL_2701011 | Lighthouse Lab in Glasgow | Wellcome Sanger Institute for the COVID-19 Genomics UK (COG-UK) Consortium | Harper VanSteenhouse, Yumi Kasai, David Gray, Carol Clugston, Anna Dominiczak and Alex Alderton, Roberto Amato, Jeffrey Barrett, Sonia Goncalves, Ewan Harrison, David K. Jackson, Ian Johnston, Dominic Kwiatkowski, Cordelia Langford, John Sillitoe on behalf of the Wellcome Sanger Institute COVID-19 Surveillance Team |
| EPI_ISL_2701012, EPI_ISL_2701013, EPI_ISL_2701014 | Lighthouse Lab in Alderley Park | Wellcome Sanger Institute for the COVID-19 Genomics UK (COG-UK) Consortium | Jacquelyn Wynn, Mairead Hyland, The Lighthouse Lab in Alderley Park and Alex Alderton, Roberto Amato, Jeffrey Barrett, Sonia Goncalves, Ewan Harrison, David K. Jackson, Ian Johnston, Dominic Kwiatkowski, Cordelia Langford, John Sillitoe on behalf of the Wellcome Sanger Institute COVID-19 Surveillance Team |
| EPI_ISL_2701015 | Lighthouse Lab in Glasgow | Wellcome Sanger Institute for the COVID-19 Genomics UK (COG-UK) Consortium | Harper VanSteenhouse, Yumi Kasai, David Gray, Carol Clugston, Anna Dominiczak and Alex Alderton, Roberto Amato, Jeffrey Barrett, Sonia Goncalves, Ewan Harrison, David K. Jackson, Ian Johnston, Dominic Kwiatkowski, Cordelia Langford, John Sillitoe on behalf of the Wellcome Sanger Institute COVID-19 Surveillance Team |
| EPI_ISL_2701016 | Lighthouse Lab in Alderley Park | Wellcome Sanger Institute for the COVID-19 Genomics UK (COG-UK) Consortium | Jacquelyn Wynn, Mairead Hyland, The Lighthouse Lab in Alderley Park and Alex Alderton, Roberto Amato, Jeffrey Barrett, Sonia Goncalves, Ewan Harrison, David K. Jackson, Ian Johnston, Dominic Kwiatkowski, Cordelia Langford, John Sillitoe on behalf of the Wellcome Sanger Institute COVID-19 Surveillance Team |
| EPI_ISL_2701017 | Lighthouse Lab in Glasgow | Wellcome Sanger Institute for the COVID-19 Genomics UK (COG-UK) Consortium | Harper VanSteenhouse, Yumi Kasai, David Gray, Carol Clugston, Anna Dominiczak and Alex Alderton, Roberto Amato, Jeffrey Barrett, Sonia Goncalves, Ewan Harrison, David K. Jackson, Ian Johnston, Dominic Kwiatkowski, Cordelia Langford, John Sillitoe on behalf of the Wellcome Sanger Institute COVID-19 Surveillance Team |
| EPI_ISL_2701018 | Lighthouse Lab in Alderley Park | Wellcome Sanger Institute for the COVID-19 Genomics UK (COG-UK) Consortium | Jacquelyn Wynn, Mairead Hyland, The Lighthouse Lab in Alderley Park and Alex Alderton, Roberto Amato, Jeffrey Barrett, Sonia Goncalves, Ewan Harrison, David K. Jackson, Ian Johnston, Dominic Kwiatkowski, Cordelia Langford, John Sillitoe on behalf of the Wellcome Sanger Institute COVID-19 Surveillance Team |
| EPI_ISL_2701019 | Lighthouse Lab in Glasgow | Wellcome Sanger Institute for the COVID-19 Genomics UK (COG-UK) Consortium | Harper VanSteenhouse, Yumi Kasai, David Gray, Carol Clugston, Anna Dominiczak and Alex Alderton, Roberto Amato, Jeffrey Barrett, Sonia Goncalves, Ewan Harrison, David K. Jackson, Ian Johnston, Dominic Kwiatkowski, Cordelia Langford, John Sillitoe on behalf of the Wellcome Sanger Institute COVID-19 Surveillance Team |
| EPI_ISL_2701020, EPI_ISL_2701021 | Lighthouse Lab in Alderley Park | Wellcome Sanger Institute for the COVID-19 Genomics UK (COG-UK) Consortium | Jacquelyn Wynn, Mairead Hyland, The Lighthouse Lab in Alderley Park and Alex Alderton, Roberto Amato, Jeffrey Barrett, Sonia Goncalves, Ewan Harrison, David K. Jackson, Ian Johnston, Dominic Kwiatkowski, Cordelia Langford, John Sillitoe on behalf of the Wellcome Sanger Institute COVID-19 Surveillance Team |
| EPI_ISL_2701022 | Lighthouse Lab in Glasgow | Wellcome Sanger Institute for the COVID-19 Genomics UK (COG-UK) Consortium | Harper VanSteenhouse, Yumi Kasai, David Gray, Carol Clugston, Anna Dominiczak and Alex Alderton, Roberto Amato, Jeffrey Barrett, Sonia Goncalves, Ewan Harrison, David K. Jackson, Ian Johnston, Dominic Kwiatkowski, Cordelia Langford, John Sillitoe on behalf of the Wellcome Sanger Institute COVID-19 Surveillance Team |
| EPI_ISL_2701023 | Lighthouse Lab in Alderley Park | Wellcome Sanger Institute for the COVID-19 Genomics UK (COG-UK) Consortium | Jacquelyn Wynn, Mairead Hyland, The Lighthouse Lab in Alderley Park and Alex Alderton, Roberto Amato, Jeffrey Barrett, Sonia Goncalves, Ewan Harrison, David K. Jackson, Ian Johnston, Dominic Kwiatkowski, Cordelia Langford, John Sillitoe on behalf of the Wellcome Sanger Institute COVID-19 Surveillance Team |
| EPI_ISL_2701024, EPI_ISL_2701025 | Lighthouse Lab in Glasgow | Wellcome Sanger Institute for the COVID-19 Genomics UK (COG-UK) Consortium | Harper VanSteenhouse, Yumi Kasai, David Gray, Carol Clugston, Anna Dominiczak and Alex Alderton, Roberto Amato, Jeffrey Barrett, Sonia Goncalves, Ewan Harrison, David K. Jackson, Ian Johnston, Dominic Kwiatkowski, Cordelia Langford, John Sillitoe on behalf of the Wellcome Sanger Institute COVID-19 Surveillance Team |
| EPI_ISL_2701026, EPI_ISL_2701027, EPI_ISL_2701028, EPI_ISL_2701029, EPI_ISL_2701030, EPI_ISL_2701031 | Lighthouse Lab in Alderley Park | Wellcome Sanger Institute for the COVID-19 Genomics UK (COG-UK) Consortium | Jacquelyn Wynn, Mairead Hyland, The Lighthouse Lab in Alderley Park and Alex Alderton, Roberto Amato, Jeffrey Barrett, Sonia Goncalves, Ewan Harrison, David K. Jackson, Ian Johnston, Dominic Kwiatkowski, Cordelia Langford, John Sillitoe on behalf of the Wellcome Sanger Institute COVID-19 Surveillance Team |
| EPI_ISL_2701032, EPI_ISL_2701033, EPI_ISL_2701034 | Lighthouse Lab in Glasgow | Wellcome Sanger Institute for the COVID-19 Genomics UK (COG-UK) Consortium | Harper VanSteenhouse, Yumi Kasai, David Gray, Carol Clugston, Anna Dominiczak and Alex Alderton, Roberto Amato, Jeffrey Barrett, Sonia Goncalves, Ewan Harrison, David K. Jackson, Ian Johnston, Dominic Kwiatkowski, Cordelia Langford, John Sillitoe on behalf of the Wellcome Sanger Institute COVID-19 Surveillance Team |
| EPI_ISL_2701035 | Lighthouse Lab in Alderley Park | Wellcome Sanger Institute for the COVID-19 Genomics UK (COG-UK) Consortium | Jacquelyn Wynn, Mairead Hyland, The Lighthouse Lab in Alderley Park and Alex Alderton, Roberto Amato, Jeffrey Barrett, Sonia Goncalves, Ewan Harrison, David K. Jackson, Ian Johnston, Dominic Kwiatkowski, Cordelia Langford, John Sillitoe on behalf of the Wellcome Sanger Institute COVID-19 Surveillance Team |
| EPI_ISL_2701036 | Lighthouse Lab in Glasgow | Wellcome Sanger Institute for the COVID-19 Genomics UK (COG-UK) Consortium | Harper VanSteenhouse, Yumi Kasai, David Gray, Carol Clugston, Anna Dominiczak and Alex Alderton, Roberto Amato, Jeffrey Barrett, Sonia Goncalves, Ewan Harrison, David K. Jackson, Ian Johnston, Dominic Kwiatkowski, Cordelia Langford, John Sillitoe on behalf of the Wellcome Sanger Institute COVID-19 Surveillance Team |
| EPI_ISL_2701037 | Lighthouse Lab in Alderley Park | Wellcome Sanger Institute for the COVID-19 Genomics UK (COG-UK) Consortium | Jacquelyn Wynn, Mairead Hyland, The Lighthouse Lab in Alderley Park and Alex Alderton, Roberto Amato, Jeffrey Barrett, Sonia Goncalves, Ewan Harrison, David K. Jackson, Ian Johnston, Dominic Kwiatkowski, Cordelia Langford, John Sillitoe on behalf of the Wellcome Sanger Institute COVID-19 Surveillance Team |
| EPI_ISL_2701038 | Lighthouse Lab in Glasgow | Wellcome Sanger Institute for the COVID-19 Genomics UK (COG-UK) Consortium | Harper VanSteenhouse, Yumi Kasai, David Gray, Carol Clugston, Anna Dominiczak and Alex Alderton, Roberto Amato, Jeffrey Barrett, Sonia Goncalves, Ewan Harrison, David K. Jackson, Ian Johnston, Dominic Kwiatkowski, Cordelia Langford, John Sillitoe on behalf of the Wellcome Sanger Institute COVID-19 Surveillance Team |
| EPI_ISL_2701039, EPI_ISL_2701041 | Lighthouse Lab in Alderley Park | Wellcome Sanger Institute for the COVID-19 Genomics UK (COG-UK) Consortium | Jacquelyn Wynn, Mairead Hyland, The Lighthouse Lab in Alderley Park and Alex Alderton, Roberto Amato, Jeffrey Barrett, Sonia Goncalves, Ewan Harrison, David K. Jackson, Ian Johnston, Dominic Kwiatkowski, Cordelia Langford, John Sillitoe on behalf of the Wellcome Sanger Institute COVID-19 Surveillance Team |
| EPI_ISL_2701042, EPI_ISL_2701043, EPI_ISL_2701044 | Lighthouse Lab in Glasgow | Wellcome Sanger Institute for the COVID-19 Genomics UK (COG-UK) Consortium | Harper VanSteenhouse, Yumi Kasai, David Gray, Carol Clugston, Anna Dominiczak and Alex Alderton, Roberto Amato, Jeffrey Barrett, Sonia Goncalves, Ewan Harrison, David K. Jackson, Ian Johnston, Dominic Kwiatkowski, Cordelia Langford, John Sillitoe on behalf of the Wellcome Sanger Institute COVID-19 Surveillance Team |
| EPI_ISL_2701045, EPI_ISL_2701046 | Lighthouse Lab in Alderley Park | Wellcome Sanger Institute for the COVID-19 Genomics UK (COG-UK) Consortium | Jacquelyn Wynn, Mairead Hyland, The Lighthouse Lab in Alderley Park and Alex Alderton, Roberto Amato, Jeffrey Barrett, Sonia Goncalves, Ewan Harrison, David K. Jackson, Ian Johnston, Dominic Kwiatkowski, Cordelia Langford, John Sillitoe on behalf of the Wellcome Sanger Institute COVID-19 Surveillance Team |
| EPI_ISL_2701047 | Lighthouse Lab in Glasgow | Wellcome Sanger Institute for the COVID-19 Genomics UK (COG-UK) Consortium | Harper VanSteenhouse, Yumi Kasai, David Gray, Carol Clugston, Anna Dominiczak and Alex Alderton, Roberto Amato, Jeffrey Barrett, Sonia Goncalves, Ewan Harrison, David K. Jackson, Ian Johnston, Dominic Kwiatkowski, Cordelia Langford, John Sillitoe on behalf of the Wellcome Sanger Institute COVID-19 Surveillance Team |
| EPI_ISL_2701048 | Lighthouse Lab in Alderley Park | Wellcome Sanger Institute for the COVID-19 Genomics UK (COG-UK) Consortium | Jacquelyn Wynn, Mairead Hyland, The Lighthouse Lab in Alderley Park and Alex Alderton, Roberto Amato, Jeffrey Barrett, Sonia Goncalves, Ewan Harrison, David K. Jackson, Ian Johnston, Dominic Kwiatkowski, Cordelia Langford, John Sillitoe on behalf of the Wellcome Sanger Institute COVID-19 Surveillance Team |
| EPI_ISL_2701049, EPI_ISL_2701050 | Lighthouse Lab in Glasgow | Wellcome Sanger Institute for the COVID-19 Genomics UK (COG-UK) Consortium | Harper VanSteenhouse, Yumi Kasai, David Gray, Carol Clugston, Anna Dominiczak and Alex Alderton, Roberto Amato, Jeffrey Barrett, Sonia Goncalves, Ewan Harrison, David K. Jackson, Ian Johnston, Dominic Kwiatkowski, Cordelia Langford, John Sillitoe on behalf of the Wellcome Sanger Institute COVID-19 Surveillance Team |
| EPI_ISL_2701051, EPI_ISL_2701052 | Lighthouse Lab in Alderley Park | Wellcome Sanger Institute for the COVID-19 Genomics UK (COG-UK) Consortium | Jacquelyn Wynn, Mairead Hyland, The Lighthouse Lab in Alderley Park and Alex Alderton, Roberto Amato, Jeffrey Barrett, Sonia Goncalves, Ewan Harrison, David K. Jackson, Ian Johnston, Dominic Kwiatkowski, Cordelia Langford, John Sillitoe on behalf of the Wellcome Sanger Institute COVID-19 Surveillance Team |
| EPI_ISL_2701053 | Lighthouse Lab in Glasgow | Wellcome Sanger Institute for the COVID-19 Genomics UK (COG-UK) Consortium | Harper VanSteenhouse, Yumi Kasai, David Gray, Carol Clugston, Anna Dominiczak and Alex Alderton, Roberto Amato, Jeffrey Barrett, Sonia Goncalves, Ewan Harrison, David K. Jackson, Ian Johnston, Dominic Kwiatkowski, Cordelia Langford, John Sillitoe on behalf of the Wellcome Sanger Institute COVID-19 Surveillance Team |
| EPI_ISL_2701054 | Lighthouse Lab in Alderley Park | Wellcome Sanger Institute for the COVID-19 Genomics UK (COG-UK) Consortium | Jacquelyn Wynn, Mairead Hyland, The Lighthouse Lab in Alderley Park and Alex Alderton, Roberto Amato, Jeffrey Barrett, Sonia Goncalves, Ewan Harrison, David K. Jackson, Ian Johnston, Dominic Kwiatkowski, Cordelia Langford, John Sillitoe on behalf of the Wellcome Sanger Institute COVID-19 Surveillance Team |
| EPI_ISL_2701055 | Lighthouse Lab in Glasgow | Wellcome Sanger Institute for the COVID-19 Genomics UK (COG-UK) Consortium | Harper VanSteenhouse, Yumi Kasai, David Gray, Carol Clugston, Anna Dominiczak and Alex Alderton, Roberto Amato, Jeffrey Barrett, Sonia Goncalves, Ewan Harrison, David K. Jackson, Ian Johnston, Dominic Kwiatkowski, Cordelia Langford, John Sillitoe on behalf of the Wellcome Sanger Institute COVID-19 Surveillance Team |
| EPI_ISL_2701056, EPI_ISL_2701057, EPI_ISL_2701058 | Lighthouse Lab in Alderley Park | Wellcome Sanger Institute for the COVID-19 Genomics UK (COG-UK) Consortium | Jacquelyn Wynn, Mairead Hyland, The Lighthouse Lab in Alderley Park and Alex Alderton, Roberto Amato, Jeffrey Barrett, Sonia Goncalves, Ewan Harrison, David K. Jackson, Ian Johnston, Dominic Kwiatkowski, Cordelia Langford, John Sillitoe on behalf of the Wellcome Sanger Institute COVID-19 Surveillance Team |
| EPI_ISL_2701059 | Lighthouse Lab in Glasgow | Wellcome Sanger Institute for the COVID-19 Genomics UK (COG-UK) Consortium | Harper VanSteenhouse, Yumi Kasai, David Gray, Carol Clugston, Anna Dominiczak and Alex Alderton, Roberto Amato, Jeffrey Barrett, Sonia Goncalves, Ewan Harrison, David K. Jackson, Ian Johnston, Dominic Kwiatkowski, Cordelia Langford, John Sillitoe on behalf of the Wellcome Sanger Institute COVID-19 Surveillance Team |
| EPI_ISL_2701060 | Lighthouse Lab in Alderley Park | Wellcome Sanger Institute for the COVID-19 Genomics UK (COG-UK) Consortium | Jacquelyn Wynn, Mairead Hyland, The Lighthouse Lab in Alderley Park and Alex Alderton, Roberto Amato, Jeffrey Barrett, Sonia Goncalves, Ewan Harrison, David K. Jackson, Ian Johnston, Dominic Kwiatkowski, Cordelia Langford, John Sillitoe on behalf of the Wellcome Sanger Institute COVID-19 Surveillance Team |
| EPI_ISL_2701061 | Lighthouse Lab in Glasgow | Wellcome Sanger Institute for the COVID-19 Genomics UK (COG-UK) Consortium | Harper VanSteenhouse, Yumi Kasai, David Gray, Carol Clugston, Anna Dominiczak and Alex Alderton, Roberto Amato, Jeffrey Barrett, Sonia Goncalves, Ewan Harrison, David K. Jackson, Ian Johnston, Dominic Kwiatkowski, Cordelia Langford, John Sillitoe on behalf of the Wellcome Sanger Institute COVID-19 Surveillance Team |
| EPI_ISL_2701062, EPI_ISL_2701063, EPI_ISL_2701064 | Lighthouse Lab in Alderley Park | Wellcome Sanger Institute for the COVID-19 Genomics UK (COG-UK) Consortium | Jacquelyn Wynn, Mairead Hyland, The Lighthouse Lab in Alderley Park and Alex Alderton, Roberto Amato, Jeffrey Barrett, Sonia Goncalves, Ewan Harrison, David K. Jackson, Ian Johnston, Dominic Kwiatkowski, Cordelia Langford, John Sillitoe on behalf of the Wellcome Sanger Institute COVID-19 Surveillance Team |
| EPI_ISL_2701065 | Lighthouse Lab in Glasgow | Wellcome Sanger Institute for the COVID-19 Genomics UK (COG-UK) Consortium | Harper VanSteenhouse, Yumi Kasai, David Gray, Carol Clugston, Anna Dominiczak and Alex Alderton, Roberto Amato, Jeffrey Barrett, Sonia Goncalves, Ewan Harrison, David K. Jackson, Ian Johnston, Dominic Kwiatkowski, Cordelia Langford, John Sillitoe on behalf of the Wellcome Sanger Institute COVID-19 Surveillance Team |
| EPI_ISL_2701066, EPI_ISL_2701067 | Lighthouse Lab in Alderley Park | Wellcome Sanger Institute for the COVID-19 Genomics UK (COG-UK) Consortium | Jacquelyn Wynn, Mairead Hyland, The Lighthouse Lab in Alderley Park and Alex Alderton, Roberto Amato, Jeffrey Barrett, Sonia Goncalves, Ewan Harrison, David K. Jackson, Ian Johnston, Dominic Kwiatkowski, Cordelia Langford, John Sillitoe on behalf of the Wellcome Sanger Institute COVID-19 Surveillance Team |
| EPI_ISL_2701068 | Lighthouse Lab in Glasgow | Wellcome Sanger Institute for the COVID-19 Genomics UK (COG-UK) Consortium | Harper VanSteenhouse, Yumi Kasai, David Gray, Carol Clugston, Anna Dominiczak and Alex Alderton, Roberto Amato, Jeffrey Barrett, Sonia Goncalves, Ewan Harrison, David K. Jackson, Ian Johnston, Dominic Kwiatkowski, Cordelia Langford, John Sillitoe on behalf of the Wellcome Sanger Institute COVID-19 Surveillance Team |
| EPI_ISL_2701069 | Lighthouse Lab in Alderley Park | Wellcome Sanger Institute for the COVID-19 Genomics UK (COG-UK) Consortium | Jacquelyn Wynn, Mairead Hyland, The Lighthouse Lab in Alderley Park and Alex Alderton, Roberto Amato, Jeffrey Barrett, Sonia Goncalves, Ewan Harrison, David K. Jackson, Ian Johnston, Dominic Kwiatkowski, Cordelia Langford, John Sillitoe on behalf of the Wellcome Sanger Institute COVID-19 Surveillance Team |
| EPI_ISL_2701070 | Lighthouse Lab in Glasgow | Wellcome Sanger Institute for the COVID-19 Genomics UK (COG-UK) Consortium | Harper VanSteenhouse, Yumi Kasai, David Gray, Carol Clugston, Anna Dominiczak and Alex Alderton, Roberto Amato, Jeffrey Barrett, Sonia Goncalves, Ewan Harrison, David K. Jackson, Ian Johnston, Dominic Kwiatkowski, Cordelia Langford, John Sillitoe on behalf of the Wellcome Sanger Institute COVID-19 Surveillance Team |

| | | | |
|---|---|---|---|
| EPI_ISL_2701071 | Lighthouse Lab in Alderley Park | Wellcome Sanger Institute for the COVID-19 Genomics UK (COG-UK) Consortium | Jacquelyn Wynn, Mairead Hyland, The Lighthouse Lab in Alderley Park and Alex Alderton, Roberto Amato, Jeffrey Barrett, Sonia Goncalves, Ewan Harrison, David K. Jackson, Ian Johnston, Dominic Kwiatkowski, Cordelia Langford, John Sillitoe on behalf of the Wellcome Sanger Institute COVID-19 Surveillance Team |
| EPI_ISL_2701072, EPI_ISL_2701073 | Lighthouse Lab in Glasgow | Wellcome Sanger Institute for the COVID-19 Genomics UK (COG-UK) Consortium | Harper VanSteenhouse, Yumi Kasai, David Gray, Carol Clugston, Anna Dominiczak and Alex Alderton, Roberto Amato, Jeffrey Barrett, Sonia Goncalves, Ewan Harrison, David K. Jackson, Ian Johnston, Dominic Kwiatkowski, Cordelia Langford, John Sillitoe on behalf of the Wellcome Sanger Institute COVID-19 Surveillance Team |
| EPI_ISL_2701074, EPI_ISL_2701075, EPI_ISL_2701076, EPI_ISL_2701077, EPI_ISL_2701078, EPI_ISL_2701079 | Lighthouse Lab in Alderley Park | Wellcome Sanger Institute for the COVID-19 Genomics UK (COG-UK) Consortium | Jacquelyn Wynn, Mairead Hyland, The Lighthouse Lab in Alderley Park and Alex Alderton, Roberto Amato, Jeffrey Barrett, Sonia Goncalves, Ewan Harrison, David K. Jackson, Ian Johnston, Dominic Kwiatkowski, Cordelia Langford, John Sillitoe on behalf of the Wellcome Sanger Institute COVID-19 Surveillance Team |
| EPI_ISL_2701080, EPI_ISL_2701081, EPI_ISL_2701082, EPI_ISL_2701083, EPI_ISL_2701084 | Lighthouse Lab in Glasgow | Wellcome Sanger Institute for the COVID-19 Genomics UK (COG-UK) Consortium | Harper VanSteenhouse, Yumi Kasai, David Gray, Carol Clugston, Anna Dominiczak and Alex Alderton, Roberto Amato, Jeffrey Barrett, Sonia Goncalves, Ewan Harrison, David K. Jackson, Ian Johnston, Dominic Kwiatkowski, Cordelia Langford, John Sillitoe on behalf of the Wellcome Sanger Institute COVID-19 Surveillance Team |
| EPI_ISL_2701085, EPI_ISL_2701087 | Lighthouse Lab in Alderley Park | Wellcome Sanger Institute for the COVID-19 Genomics UK (COG-UK) Consortium | Jacquelyn Wynn, Mairead Hyland, The Lighthouse Lab in Alderley Park and Alex Alderton, Roberto Amato, Jeffrey Barrett, Sonia Goncalves, Ewan Harrison, David K. Jackson, Ian Johnston, Dominic Kwiatkowski, Cordelia Langford, John Sillitoe on behalf of the Wellcome Sanger Institute COVID-19 Surveillance Team |
| EPI_ISL_2701088, EPI_ISL_2701089, EPI_ISL_2701090, EPI_ISL_2701091, EPI_ISL_2701092, EPI_ISL_2701093, EPI_ISL_2701094, EPI_ISL_2701095 | Lighthouse Lab in Glasgow | Wellcome Sanger Institute for the COVID-19 Genomics UK (COG-UK) Consortium | Harper VanSteenhouse, Yumi Kasai, David Gray, Carol Clugston, Anna Dominiczak and Alex Alderton, Roberto Amato, Jeffrey Barrett, Sonia Goncalves, Ewan Harrison, David K. Jackson, Ian Johnston, Dominic Kwiatkowski, Cordelia Langford, John Sillitoe on behalf of the Wellcome Sanger Institute COVID-19 Surveillance Team |
| EPI_ISL_2701096, EPI_ISL_2701097, EPI_ISL_2701098 | Lighthouse Lab in Alderley Park | Wellcome Sanger Institute for the COVID-19 Genomics UK (COG-UK) Consortium | Jacquelyn Wynn, Mairead Hyland, The Lighthouse Lab in Alderley Park and Alex Alderton, Roberto Amato, Jeffrey Barrett, Sonia Goncalves, Ewan Harrison, David K. Jackson, Ian Johnston, Dominic Kwiatkowski, Cordelia Langford, John Sillitoe on behalf of the Wellcome Sanger Institute COVID-19 Surveillance Team |
| EPI_ISL_2701099 | Lighthouse Lab in Glasgow | Wellcome Sanger Institute for the COVID-19 Genomics UK (COG-UK) Consortium | Harper VanSteenhouse, Yumi Kasai, David Gray, Carol Clugston, Anna Dominiczak and Alex Alderton, Roberto Amato, Jeffrey Barrett, Sonia Goncalves, Ewan Harrison, David K. Jackson, Ian Johnston, Dominic Kwiatkowski, Cordelia Langford, John Sillitoe on behalf of the Wellcome Sanger Institute COVID-19 Surveillance Team |
| EPI_ISL_2701100, EPI_ISL_2701101, EPI_ISL_2701102 | Lighthouse Lab in Alderley Park | Wellcome Sanger Institute for the COVID-19 Genomics UK (COG-UK) Consortium | Jacquelyn Wynn, Mairead Hyland, The Lighthouse Lab in Alderley Park and Alex Alderton, Roberto Amato, Jeffrey Barrett, Sonia Goncalves, Ewan Harrison, David K. Jackson, Ian Johnston, Dominic Kwiatkowski, Cordelia Langford, John Sillitoe on behalf of the Wellcome Sanger Institute COVID-19 Surveillance Team |
| EPI_ISL_2701103 | Lighthouse Lab in Glasgow | Wellcome Sanger Institute for the COVID-19 Genomics UK (COG-UK) Consortium | Harper VanSteenhouse, Yumi Kasai, David Gray, Carol Clugston, Anna Dominiczak and Alex Alderton, Roberto Amato, Jeffrey Barrett, Sonia Goncalves, Ewan Harrison, David K. Jackson, Ian Johnston, Dominic Kwiatkowski, Cordelia Langford, John Sillitoe on behalf of the Wellcome Sanger Institute COVID-19 Surveillance Team |
| EPI_ISL_2701104 | Lighthouse Lab in Alderley Park | Wellcome Sanger Institute for the COVID-19 Genomics UK (COG-UK) Consortium | Jacquelyn Wynn, Mairead Hyland, The Lighthouse Lab in Alderley Park and Alex Alderton, Roberto Amato, Jeffrey Barrett, Sonia Goncalves, Ewan Harrison, David K. Jackson, Ian Johnston, Dominic Kwiatkowski, Cordelia Langford, John Sillitoe on behalf of the Wellcome Sanger Institute COVID-19 Surveillance Team |
| EPI_ISL_2701105 | Lighthouse Lab in Glasgow | Wellcome Sanger Institute for the COVID-19 Genomics UK (COG-UK) Consortium | Harper VanSteenhouse, Yumi Kasai, David Gray, Carol Clugston, Anna Dominiczak and Alex Alderton, Roberto Amato, Jeffrey Barrett, Sonia Goncalves, Ewan Harrison, David K. Jackson, Ian Johnston, Dominic Kwiatkowski, Cordelia Langford, John Sillitoe on behalf of the Wellcome Sanger Institute COVID-19 Surveillance Team |
| EPI_ISL_2701106 | Lighthouse Lab in Alderley Park | Wellcome Sanger Institute for the COVID-19 Genomics UK (COG-UK) Consortium | Jacquelyn Wynn, Mairead Hyland, The Lighthouse Lab in Alderley Park and Alex Alderton, Roberto Amato, Jeffrey Barrett, Sonia Goncalves, Ewan Harrison, David K. Jackson, Ian Johnston, Dominic Kwiatkowski, Cordelia Langford, John Sillitoe on behalf of the Wellcome Sanger Institute COVID-19 Surveillance Team |
| EPI_ISL_2701107 | Lighthouse Lab in Glasgow | Wellcome Sanger Institute for the COVID-19 Genomics UK (COG-UK) Consortium | Harper VanSteenhouse, Yumi Kasai, David Gray, Carol Clugston, Anna Dominiczak and Alex Alderton, Roberto Amato, Jeffrey Barrett, Sonia Goncalves, Ewan Harrison, David K. Jackson, Ian Johnston, Dominic Kwiatkowski, Cordelia Langford, John Sillitoe on behalf of the Wellcome Sanger Institute COVID-19 Surveillance Team |
| EPI_ISL_2701108, EPI_ISL_2701109, EPI_ISL_2701110 | Lighthouse Lab in Alderley Park | Wellcome Sanger Institute for the COVID-19 Genomics UK (COG-UK) Consortium | Jacquelyn Wynn, Mairead Hyland, The Lighthouse Lab in Alderley Park and Alex Alderton, Roberto Amato, Jeffrey Barrett, Sonia Goncalves, Ewan Harrison, David K. Jackson, Ian Johnston, Dominic Kwiatkowski, Cordelia Langford, John Sillitoe on behalf of the Wellcome Sanger Institute COVID-19 Surveillance Team |
| EPI_ISL_2701111, EPI_ISL_2701112, EPI_ISL_2701113, EPI_ISL_2701114, EPI_ISL_2701115 | Lighthouse Lab in Glasgow | Wellcome Sanger Institute for the COVID-19 Genomics UK (COG-UK) Consortium | Harper VanSteenhouse, Yumi Kasai, David Gray, Carol Clugston, Anna Dominiczak and Alex Alderton, Roberto Amato, Jeffrey Barrett, Sonia Goncalves, Ewan Harrison, David K. Jackson, Ian Johnston, Dominic Kwiatkowski, Cordelia Langford, John Sillitoe on behalf of the Wellcome Sanger Institute COVID-19 Surveillance Team |
| EPI_ISL_2701116, EPI_ISL_2701117 | Lighthouse Lab in Alderley Park | Wellcome Sanger Institute for the COVID-19 Genomics UK (COG-UK) Consortium | Jacquelyn Wynn, Mairead Hyland, The Lighthouse Lab in Alderley Park and Alex Alderton, Roberto Amato, Jeffrey Barrett, Sonia Goncalves, Ewan Harrison, David K. Jackson, Ian Johnston, Dominic Kwiatkowski, Cordelia Langford, John Sillitoe on behalf of the Wellcome Sanger Institute COVID-19 Surveillance Team |
| EPI_ISL_2701118 | Lighthouse Lab in Glasgow | Wellcome Sanger Institute for the COVID-19 Genomics UK (COG-UK) Consortium | Harper VanSteenhouse, Yumi Kasai, David Gray, Carol Clugston, Anna Dominiczak and Alex Alderton, Roberto Amato, Jeffrey Barrett, Sonia Goncalves, Ewan Harrison, David K. Jackson, Ian Johnston, Dominic Kwiatkowski, Cordelia Langford, John Sillitoe on behalf of the Wellcome Sanger Institute COVID-19 Surveillance Team |
| EPI_ISL_2701119 | Lighthouse Lab in Alderley Park | Wellcome Sanger Institute for the COVID-19 Genomics UK (COG-UK) Consortium | Jacquelyn Wynn, Mairead Hyland, The Lighthouse Lab in Alderley Park and Alex Alderton, Roberto Amato, Jeffrey Barrett, Sonia Goncalves, Ewan Harrison, David K. Jackson, Ian Johnston, Dominic Kwiatkowski, Cordelia Langford, John Sillitoe on behalf of the Wellcome Sanger Institute COVID-19 Surveillance Team |
| EPI_ISL_2701120 | Lighthouse Lab in Glasgow | Wellcome Sanger Institute for the COVID-19 Genomics UK (COG-UK) Consortium | Harper VanSteenhouse, Yumi Kasai, David Gray, Carol Clugston, Anna Dominiczak and Alex Alderton, Roberto Amato, Jeffrey Barrett, Sonia Goncalves, Ewan Harrison, David K. Jackson, Ian Johnston, Dominic Kwiatkowski, Cordelia Langford, John Sillitoe on behalf of the Wellcome Sanger Institute COVID-19 Surveillance Team |
| EPI_ISL_2701121, EPI_ISL_2701122 | Lighthouse Lab in Alderley Park | Wellcome Sanger Institute for the COVID-19 Genomics UK (COG-UK) Consortium | Jacquelyn Wynn, Mairead Hyland, The Lighthouse Lab in Alderley Park and Alex Alderton, Roberto Amato, Jeffrey Barrett, Sonia Goncalves, Ewan Harrison, David K. Jackson, Ian Johnston, Dominic Kwiatkowski, Cordelia Langford, John Sillitoe on behalf of the Wellcome Sanger Institute COVID-19 Surveillance Team |
| EPI_ISL_2701123 | Lighthouse Lab in Glasgow | Wellcome Sanger Institute for the COVID-19 Genomics UK (COG-UK) Consortium | Harper VanSteenhouse, Yumi Kasai, David Gray, Carol Clugston, Anna Dominiczak and Alex Alderton, Roberto Amato, Jeffrey Barrett, Sonia Goncalves, Ewan Harrison, David K. Jackson, Ian Johnston, Dominic Kwiatkowski, Cordelia Langford, John Sillitoe on behalf of the Wellcome Sanger Institute COVID-19 Surveillance Team |
| EPI_ISL_2701124 | Lighthouse Lab in Alderley Park | Wellcome Sanger Institute for the COVID-19 Genomics UK (COG-UK) Consortium | Jacquelyn Wynn, Mairead Hyland, The Lighthouse Lab in Alderley Park and Alex Alderton, Roberto Amato, Jeffrey Barrett, Sonia Goncalves, Ewan Harrison, David K. Jackson, Ian Johnston, Dominic Kwiatkowski, Cordelia Langford, John Sillitoe on behalf of the Wellcome Sanger Institute COVID-19 Surveillance Team |
| EPI_ISL_2701125, EPI_ISL_2701126 | Lighthouse Lab in Glasgow | Wellcome Sanger Institute for the COVID-19 Genomics UK (COG-UK) Consortium | Harper VanSteenhouse, Yumi Kasai, David Gray, Carol Clugston, Anna Dominiczak and Alex Alderton, Roberto Amato, Jeffrey Barrett, Sonia Goncalves, Ewan Harrison, David K. Jackson, Ian Johnston, Dominic Kwiatkowski, Cordelia Langford, John Sillitoe on behalf of the Wellcome Sanger Institute COVID-19 Surveillance Team |
| EPI_ISL_2701127 | Lighthouse Lab in Alderley Park | Wellcome Sanger Institute for the COVID-19 Genomics UK (COG-UK) Consortium | Jacquelyn Wynn, Mairead Hyland, The Lighthouse Lab in Alderley Park and Alex Alderton, Roberto Amato, Jeffrey Barrett, Sonia Goncalves, Ewan Harrison, David K. Jackson, Ian Johnston, Dominic Kwiatkowski, Cordelia Langford, John Sillitoe on behalf of the Wellcome Sanger Institute COVID-19 Surveillance Team |
| EPI_ISL_2701128 | Lighthouse Lab in Glasgow | Wellcome Sanger Institute for the COVID-19 Genomics UK (COG-UK) Consortium | Harper VanSteenhouse, Yumi Kasai, David Gray, Carol Clugston, Anna Dominiczak and Alex Alderton, Roberto Amato, Jeffrey Barrett, Sonia Goncalves, Ewan Harrison, David K. Jackson, Ian Johnston, Dominic Kwiatkowski, Cordelia Langford, John Sillitoe on behalf of the Wellcome Sanger Institute COVID-19 Surveillance Team |
| EPI_ISL_2701129 | Lighthouse Lab in Alderley Park | Wellcome Sanger Institute for the COVID-19 Genomics UK (COG-UK) Consortium | Jacquelyn Wynn, Mairead Hyland, The Lighthouse Lab in Alderley Park and Alex Alderton, Roberto Amato, Jeffrey Barrett, Sonia Goncalves, Ewan Harrison, David K. Jackson, Ian Johnston, Dominic Kwiatkowski, Cordelia Langford, John Sillitoe on behalf of the Wellcome Sanger Institute COVID-19 Surveillance Team |
| EPI_ISL_2701130 | Lighthouse Lab in Glasgow | Wellcome Sanger Institute for the COVID-19 Genomics UK (COG-UK) Consortium | Harper VanSteenhouse, Yumi Kasai, David Gray, Carol Clugston, Anna Dominiczak and Alex Alderton, Roberto Amato, Jeffrey Barrett, Sonia Goncalves, Ewan Harrison, David K. Jackson, Ian Johnston, Dominic Kwiatkowski, Cordelia Langford, John Sillitoe on behalf of the Wellcome Sanger Institute COVID-19 Surveillance Team |
| EPI_ISL_2701131 | Lighthouse Lab in Alderley Park | Wellcome Sanger Institute for the COVID-19 Genomics UK (COG-UK) Consortium | Jacquelyn Wynn, Mairead Hyland, The Lighthouse Lab in Alderley Park and Alex Alderton, Roberto Amato, Jeffrey Barrett, Sonia Goncalves, Ewan Harrison, David K. Jackson, Ian Johnston, Dominic Kwiatkowski, Cordelia Langford, John Sillitoe on behalf of the Wellcome Sanger Institute COVID-19 Surveillance Team |
| EPI_ISL_2701132 | Lighthouse Lab in Glasgow | Wellcome Sanger Institute for the COVID-19 Genomics UK (COG-UK) Consortium | Harper VanSteenhouse, Yumi Kasai, David Gray, Carol Clugston, Anna Dominiczak and Alex Alderton, Roberto Amato, Jeffrey Barrett, Sonia Goncalves, Ewan Harrison, David K. Jackson, Ian Johnston, Dominic Kwiatkowski, Cordelia Langford, John Sillitoe on behalf of the Wellcome Sanger Institute COVID-19 Surveillance Team |
| EPI_ISL_2701133 | Lighthouse Lab in Alderley Park | Wellcome Sanger Institute for the COVID-19 Genomics UK (COG-UK) Consortium | Jacquelyn Wynn, Mairead Hyland, The Lighthouse Lab in Alderley Park and Alex Alderton, Roberto Amato, Jeffrey Barrett, Sonia Goncalves, Ewan Harrison, David K. Jackson, Ian Johnston, Dominic Kwiatkowski, Cordelia Langford, John Sillitoe on behalf of the Wellcome Sanger Institute COVID-19 Surveillance Team |
| EPI_ISL_2701134 | Lighthouse Lab in Glasgow | Wellcome Sanger Institute for the COVID-19 Genomics UK (COG-UK) Consortium | Harper VanSteenhouse, Yumi Kasai, David Gray, Carol Clugston, Anna Dominiczak and Alex Alderton, Roberto Amato, Jeffrey Barrett, Sonia Goncalves, Ewan Harrison, David K. Jackson, Ian Johnston, Dominic Kwiatkowski, Cordelia Langford, John Sillitoe on behalf of the Wellcome Sanger Institute COVID-19 Surveillance Team |
| EPI_ISL_2701135, EPI_ISL_2701137, EPI_ISL_2701138 | Lighthouse Lab in Alderley Park | Wellcome Sanger Institute for the COVID-19 Genomics UK (COG-UK) Consortium | Jacquelyn Wynn, Mairead Hyland, The Lighthouse Lab in Alderley Park and Alex Alderton, Roberto Amato, Jeffrey Barrett, Sonia Goncalves, Ewan Harrison, David K. Jackson, Ian Johnston, Dominic Kwiatkowski, Cordelia Langford, John Sillitoe on behalf of the Wellcome Sanger Institute COVID-19 Surveillance Team |
| EPI_ISL_2701139, EPI_ISL_2701140 | Lighthouse Lab in Glasgow | Wellcome Sanger Institute for the COVID-19 Genomics UK (COG-UK) Consortium | Harper VanSteenhouse, Yumi Kasai, David Gray, Carol Clugston, Anna Dominiczak and Alex Alderton, Roberto Amato, Jeffrey Barrett, Sonia Goncalves, Ewan Harrison, David K. Jackson, Ian Johnston, Dominic Kwiatkowski, Cordelia Langford, John Sillitoe on behalf of the Wellcome Sanger Institute COVID-19 Surveillance Team |
| EPI_ISL_2701141 | Lighthouse Lab in Alderley Park | Wellcome Sanger Institute for the COVID-19 Genomics UK (COG-UK) Consortium | Jacquelyn Wynn, Mairead Hyland, The Lighthouse Lab in Alderley Park and Alex Alderton, Roberto Amato, Jeffrey Barrett, Sonia Goncalves, Ewan Harrison, David K. Jackson, Ian Johnston, Dominic Kwiatkowski, Cordelia Langford, John Sillitoe on behalf of the Wellcome Sanger Institute COVID-19 Surveillance Team |
| EPI_ISL_2701142 | Lighthouse Lab in Glasgow | Wellcome Sanger Institute for the COVID-19 Genomics UK (COG-UK) Consortium | Harper VanSteenhouse, Yumi Kasai, David Gray, Carol Clugston, Anna Dominiczak and Alex Alderton, Roberto Amato, Jeffrey Barrett, Sonia Goncalves, Ewan Harrison, David K. Jackson, Ian Johnston, Dominic Kwiatkowski, Cordelia Langford, John Sillitoe on behalf of the Wellcome Sanger Institute COVID-19 Surveillance Team |
| EPI_ISL_2701143 | Lighthouse Lab in Alderley Park | Wellcome Sanger Institute for the COVID-19 Genomics UK (COG-UK) Consortium | Jacquelyn Wynn, Mairead Hyland, The Lighthouse Lab in Alderley Park and Alex Alderton, Roberto Amato, Jeffrey Barrett, Sonia Goncalves, Ewan Harrison, David K. Jackson, Ian Johnston, Dominic Kwiatkowski, Cordelia Langford, John Sillitoe on behalf of the Wellcome Sanger Institute COVID-19 Surveillance Team |

| | | | |
|---|---|---|---|
| EPI_ISL_2701144 | Lighthouse Lab in Glasgow | Wellcome Sanger Institute for the COVID-19 Genomics UK (COG-UK) Consortium | Harper VanSteenhouse, Yumi Kasai, David Gray, Carol Clugston, Anna Dominiczak and Alex Alderton, Roberto Amato, Jeffrey Barrett, Sonia Goncalves, Ewan Harrison, David K. Jackson, Ian Johnston, Dominic Kwiatkowski, Cordelia Langford, John Sillitoe on behalf of the Wellcome Sanger Institute COVID-19 Surveillance Team |
| EPI_ISL_2701145, EPI_ISL_2701146 | Lighthouse Lab in Alderley Park | Wellcome Sanger Institute for the COVID-19 Genomics UK (COG-UK) Consortium | Jacquelyn Wynn, Mairead Hyland, The Lighthouse Lab in Alderley Park and Alex Alderton, Roberto Amato, Jeffrey Barrett, Sonia Goncalves, Ewan Harrison, David K. Jackson, Ian Johnston, Dominic Kwiatkowski, Cordelia Langford, John Sillitoe on behalf of the Wellcome Sanger Institute COVID-19 Surveillance Team |
| EPI_ISL_2701147 | Lighthouse Lab in Glasgow | Wellcome Sanger Institute for the COVID-19 Genomics UK (COG-UK) Consortium | Harper VanSteenhouse, Yumi Kasai, David Gray, Carol Clugston, Anna Dominiczak and Alex Alderton, Roberto Amato, Jeffrey Barrett, Sonia Goncalves, Ewan Harrison, David K. Jackson, Ian Johnston, Dominic Kwiatkowski, Cordelia Langford, John Sillitoe on behalf of the Wellcome Sanger Institute COVID-19 Surveillance Team |
| EPI_ISL_2701148 | Lighthouse Lab in Alderley Park | Wellcome Sanger Institute for the COVID-19 Genomics UK (COG-UK) Consortium | Jacquelyn Wynn, Mairead Hyland, The Lighthouse Lab in Alderley Park and Alex Alderton, Roberto Amato, Jeffrey Barrett, Sonia Goncalves, Ewan Harrison, David K. Jackson, Ian Johnston, Dominic Kwiatkowski, Cordelia Langford, John Sillitoe on behalf of the Wellcome Sanger Institute COVID-19 Surveillance Team |
| EPI_ISL_2701149, EPI_ISL_2701150, EPI_ISL_2701151 | Lighthouse Lab in Glasgow | Wellcome Sanger Institute for the COVID-19 Genomics UK (COG-UK) Consortium | Harper VanSteenhouse, Yumi Kasai, David Gray, Carol Clugston, Anna Dominiczak and Alex Alderton, Roberto Amato, Jeffrey Barrett, Sonia Goncalves, Ewan Harrison, David K. Jackson, Ian Johnston, Dominic Kwiatkowski, Cordelia Langford, John Sillitoe on behalf of the Wellcome Sanger Institute COVID-19 Surveillance Team |
| EPI_ISL_2701152 | Lighthouse Lab in Alderley Park | Wellcome Sanger Institute for the COVID-19 Genomics UK (COG-UK) Consortium | Jacquelyn Wynn, Mairead Hyland, The Lighthouse Lab in Alderley Park and Alex Alderton, Roberto Amato, Jeffrey Barrett, Sonia Goncalves, Ewan Harrison, David K. Jackson, Ian Johnston, Dominic Kwiatkowski, Cordelia Langford, John Sillitoe on behalf of the Wellcome Sanger Institute COVID-19 Surveillance Team |
| EPI_ISL_2701153 | Lighthouse Lab in Glasgow | Wellcome Sanger Institute for the COVID-19 Genomics UK (COG-UK) Consortium | Harper VanSteenhouse, Yumi Kasai, David Gray, Carol Clugston, Anna Dominiczak and Alex Alderton, Roberto Amato, Jeffrey Barrett, Sonia Goncalves, Ewan Harrison, David K. Jackson, Ian Johnston, Dominic Kwiatkowski, Cordelia Langford, John Sillitoe on behalf of the Wellcome Sanger Institute COVID-19 Surveillance Team |
| EPI_ISL_2701154, EPI_ISL_2701155 | Lighthouse Lab in Alderley Park | Wellcome Sanger Institute for the COVID-19 Genomics UK (COG-UK) Consortium | Jacquelyn Wynn, Mairead Hyland, The Lighthouse Lab in Alderley Park and Alex Alderton, Roberto Amato, Jeffrey Barrett, Sonia Goncalves, Ewan Harrison, David K. Jackson, Ian Johnston, Dominic Kwiatkowski, Cordelia Langford, John Sillitoe on behalf of the Wellcome Sanger Institute COVID-19 Surveillance Team |
| EPI_ISL_2701156, EPI_ISL_2701157, EPI_ISL_2701158 | Lighthouse Lab in Glasgow | Wellcome Sanger Institute for the COVID-19 Genomics UK (COG-UK) Consortium | Harper VanSteenhouse, Yumi Kasai, David Gray, Carol Clugston, Anna Dominiczak and Alex Alderton, Roberto Amato, Jeffrey Barrett, Sonia Goncalves, Ewan Harrison, David K. Jackson, Ian Johnston, Dominic Kwiatkowski, Cordelia Langford, John Sillitoe on behalf of the Wellcome Sanger Institute COVID-19 Surveillance Team |
| EPI_ISL_2701159, EPI_ISL_2701160, EPI_ISL_2701161, EPI_ISL_2701162, EPI_ISL_2701163, EPI_ISL_2701164 | Lighthouse Lab in Alderley Park | Wellcome Sanger Institute for the COVID-19 Genomics UK (COG-UK) Consortium | Jacquelyn Wynn, Mairead Hyland, The Lighthouse Lab in Alderley Park and Alex Alderton, Roberto Amato, Jeffrey Barrett, Sonia Goncalves, Ewan Harrison, David K. Jackson, Ian Johnston, Dominic Kwiatkowski, Cordelia Langford, John Sillitoe on behalf of the Wellcome Sanger Institute COVID-19 Surveillance Team |
| EPI_ISL_2701165 | Lighthouse Lab in Glasgow | Wellcome Sanger Institute for the COVID-19 Genomics UK (COG-UK) Consortium | Harper VanSteenhouse, Yumi Kasai, David Gray, Carol Clugston, Anna Dominiczak and Alex Alderton, Roberto Amato, Jeffrey Barrett, Sonia Goncalves, Ewan Harrison, David K. Jackson, Ian Johnston, Dominic Kwiatkowski, Cordelia Langford, John Sillitoe on behalf of the Wellcome Sanger Institute COVID-19 Surveillance Team |
| EPI_ISL_2701166, EPI_ISL_2701167, EPI_ISL_2701168, EPI_ISL_2701169, EPI_ISL_2701170 | Lighthouse Lab in Alderley Park | Wellcome Sanger Institute for the COVID-19 Genomics UK (COG-UK) Consortium | Jacquelyn Wynn, Mairead Hyland, The Lighthouse Lab in Alderley Park and Alex Alderton, Roberto Amato, Jeffrey Barrett, Sonia Goncalves, Ewan Harrison, David K. Jackson, Ian Johnston, Dominic Kwiatkowski, Cordelia Langford, John Sillitoe on behalf of the Wellcome Sanger Institute COVID-19 Surveillance Team |
| EPI_ISL_2701171, EPI_ISL_2701172, EPI_ISL_2701173, EPI_ISL_2701174 | Lighthouse Lab in Glasgow | Wellcome Sanger Institute for the COVID-19 Genomics UK (COG-UK) Consortium | Harper VanSteenhouse, Yumi Kasai, David Gray, Carol Clugston, Anna Dominiczak and Alex Alderton, Roberto Amato, Jeffrey Barrett, Sonia Goncalves, Ewan Harrison, David K. Jackson, Ian Johnston, Dominic Kwiatkowski, Cordelia Langford, John Sillitoe on behalf of the Wellcome Sanger Institute COVID-19 Surveillance Team |
| EPI_ISL_2701175 | Lighthouse Lab in Alderley Park | Wellcome Sanger Institute for the COVID-19 Genomics UK (COG-UK) Consortium | Jacquelyn Wynn, Mairead Hyland, The Lighthouse Lab in Alderley Park and Alex Alderton, Roberto Amato, Jeffrey Barrett, Sonia Goncalves, Ewan Harrison, David K. Jackson, Ian Johnston, Dominic Kwiatkowski, Cordelia Langford, John Sillitoe on behalf of the Wellcome Sanger Institute COVID-19 Surveillance Team |
| EPI_ISL_2701176 | Lighthouse Lab in Glasgow | Wellcome Sanger Institute for the COVID-19 Genomics UK (COG-UK) Consortium | Harper VanSteenhouse, Yumi Kasai, David Gray, Carol Clugston, Anna Dominiczak and Alex Alderton, Roberto Amato, Jeffrey Barrett, Sonia Goncalves, Ewan Harrison, David K. Jackson, Ian Johnston, Dominic Kwiatkowski, Cordelia Langford, John Sillitoe on behalf of the Wellcome Sanger Institute COVID-19 Surveillance Team |
| EPI_ISL_2701177 | Lighthouse Lab in Alderley Park | Wellcome Sanger Institute for the COVID-19 Genomics UK (COG-UK) Consortium | Jacquelyn Wynn, Mairead Hyland, The Lighthouse Lab in Alderley Park and Alex Alderton, Roberto Amato, Jeffrey Barrett, Sonia Goncalves, Ewan Harrison, David K. Jackson, Ian Johnston, Dominic Kwiatkowski, Cordelia Langford, John Sillitoe on behalf of the Wellcome Sanger Institute COVID-19 Surveillance Team |
| EPI_ISL_2701178, EPI_ISL_2701179 | Lighthouse Lab in Glasgow | Wellcome Sanger Institute for the COVID-19 Genomics UK (COG-UK) Consortium | Harper VanSteenhouse, Yumi Kasai, David Gray, Carol Clugston, Anna Dominiczak and Alex Alderton, Roberto Amato, Jeffrey Barrett, Sonia Goncalves, Ewan Harrison, David K. Jackson, Ian Johnston, Dominic Kwiatkowski, Cordelia Langford, John Sillitoe on behalf of the Wellcome Sanger Institute COVID-19 Surveillance Team |
| EPI_ISL_2701180 | Lighthouse Lab in Alderley Park | Wellcome Sanger Institute for the COVID-19 Genomics UK (COG-UK) Consortium | Jacquelyn Wynn, Mairead Hyland, The Lighthouse Lab in Alderley Park and Alex Alderton, Roberto Amato, Jeffrey Barrett, Sonia Goncalves, Ewan Harrison, David K. Jackson, Ian Johnston, Dominic Kwiatkowski, Cordelia Langford, John Sillitoe on behalf of the Wellcome Sanger Institute COVID-19 Surveillance Team |
| EPI_ISL_2701181 | Lighthouse Lab in Glasgow | Wellcome Sanger Institute for the COVID-19 Genomics UK (COG-UK) Consortium | Harper VanSteenhouse, Yumi Kasai, David Gray, Carol Clugston, Anna Dominiczak and Alex Alderton, Roberto Amato, Jeffrey Barrett, Sonia Goncalves, Ewan Harrison, David K. Jackson, Ian Johnston, Dominic Kwiatkowski, Cordelia Langford, John Sillitoe on behalf of the Wellcome Sanger Institute COVID-19 Surveillance Team |
| EPI_ISL_2701182 | Lighthouse Lab in Alderley Park | Wellcome Sanger Institute for the COVID-19 Genomics UK (COG-UK) Consortium | Jacquelyn Wynn, Mairead Hyland, The Lighthouse Lab in Alderley Park and Alex Alderton, Roberto Amato, Jeffrey Barrett, Sonia Goncalves, Ewan Harrison, David K. Jackson, Ian Johnston, Dominic Kwiatkowski, Cordelia Langford, John Sillitoe on behalf of the Wellcome Sanger Institute COVID-19 Surveillance Team |
| EPI_ISL_2701183 | Lighthouse Lab in Glasgow | Wellcome Sanger Institute for the COVID-19 Genomics UK (COG-UK) Consortium | Harper VanSteenhouse, Yumi Kasai, David Gray, Carol Clugston, Anna Dominiczak and Alex Alderton, Roberto Amato, Jeffrey Barrett, Sonia Goncalves, Ewan Harrison, David K. Jackson, Ian Johnston, Dominic Kwiatkowski, Cordelia Langford, John Sillitoe on behalf of the Wellcome Sanger Institute COVID-19 Surveillance Team |
| EPI_ISL_2701184, EPI_ISL_2701185 | Lighthouse Lab in Alderley Park | Wellcome Sanger Institute for the COVID-19 Genomics UK (COG-UK) Consortium | Jacquelyn Wynn, Mairead Hyland, The Lighthouse Lab in Alderley Park and Alex Alderton, Roberto Amato, Jeffrey Barrett, Sonia Goncalves, Ewan Harrison, David K. Jackson, Ian Johnston, Dominic Kwiatkowski, Cordelia Langford, John Sillitoe on behalf of the Wellcome Sanger Institute COVID-19 Surveillance Team |
| EPI_ISL_2701186 | Lighthouse Lab in Glasgow | Wellcome Sanger Institute for the COVID-19 Genomics UK (COG-UK) Consortium | Harper VanSteenhouse, Yumi Kasai, David Gray, Carol Clugston, Anna Dominiczak and Alex Alderton, Roberto Amato, Jeffrey Barrett, Sonia Goncalves, Ewan Harrison, David K. Jackson, Ian Johnston, Dominic Kwiatkowski, Cordelia Langford, John Sillitoe on behalf of the Wellcome Sanger Institute COVID-19 Surveillance Team |
| EPI_ISL_2701187, EPI_ISL_2701188, EPI_ISL_2701189, EPI_ISL_2701190 | Lighthouse Lab in Alderley Park | Wellcome Sanger Institute for the COVID-19 Genomics UK (COG-UK) Consortium | Jacquelyn Wynn, Mairead Hyland, The Lighthouse Lab in Alderley Park and Alex Alderton, Roberto Amato, Jeffrey Barrett, Sonia Goncalves, Ewan Harrison, David K. Jackson, Ian Johnston, Dominic Kwiatkowski, Cordelia Langford, John Sillitoe on behalf of the Wellcome Sanger Institute COVID-19 Surveillance Team |
| EPI_ISL_2701191, EPI_ISL_2701192 | Lighthouse Lab in Glasgow | Wellcome Sanger Institute for the COVID-19 Genomics UK (COG-UK) Consortium | Harper VanSteenhouse, Yumi Kasai, David Gray, Carol Clugston, Anna Dominiczak and Alex Alderton, Roberto Amato, Jeffrey Barrett, Sonia Goncalves, Ewan Harrison, David K. Jackson, Ian Johnston, Dominic Kwiatkowski, Cordelia Langford, John Sillitoe on behalf of the Wellcome Sanger Institute COVID-19 Surveillance Team |
| EPI_ISL_2701193 | Lighthouse Lab in Alderley Park | Wellcome Sanger Institute for the COVID-19 Genomics UK (COG-UK) Consortium | Jacquelyn Wynn, Mairead Hyland, The Lighthouse Lab in Alderley Park and Alex Alderton, Roberto Amato, Jeffrey Barrett, Sonia Goncalves, Ewan Harrison, David K. Jackson, Ian Johnston, Dominic Kwiatkowski, Cordelia Langford, John Sillitoe on behalf of the Wellcome Sanger Institute COVID-19 Surveillance Team |
| EPI_ISL_2701194 | Lighthouse Lab in Glasgow | Wellcome Sanger Institute for the COVID-19 Genomics UK (COG-UK) Consortium | Harper VanSteenhouse, Yumi Kasai, David Gray, Carol Clugston, Anna Dominiczak and Alex Alderton, Roberto Amato, Jeffrey Barrett, Sonia Goncalves, Ewan Harrison, David K. Jackson, Ian Johnston, Dominic Kwiatkowski, Cordelia Langford, John Sillitoe on behalf of the Wellcome Sanger Institute COVID-19 Surveillance Team |
| EPI_ISL_2701195, EPI_ISL_2701196, EPI_ISL_2701197 | Lighthouse Lab in Alderley Park | Wellcome Sanger Institute for the COVID-19 Genomics UK (COG-UK) Consortium | Jacquelyn Wynn, Mairead Hyland, The Lighthouse Lab in Alderley Park and Alex Alderton, Roberto Amato, Jeffrey Barrett, Sonia Goncalves, Ewan Harrison, David K. Jackson, Ian Johnston, Dominic Kwiatkowski, Cordelia Langford, John Sillitoe on behalf of the Wellcome Sanger Institute COVID-19 Surveillance Team |
| EPI_ISL_2701198 | Lighthouse Lab in Glasgow | Wellcome Sanger Institute for the COVID-19 Genomics UK (COG-UK) Consortium | Harper VanSteenhouse, Yumi Kasai, David Gray, Carol Clugston, Anna Dominiczak and Alex Alderton, Roberto Amato, Jeffrey Barrett, Sonia Goncalves, Ewan Harrison, David K. Jackson, Ian Johnston, Dominic Kwiatkowski, Cordelia Langford, John Sillitoe on behalf of the Wellcome Sanger Institute COVID-19 Surveillance Team |
| EPI_ISL_2701199 | Lighthouse Lab in Alderley Park | Wellcome Sanger Institute for the COVID-19 Genomics UK (COG-UK) Consortium | Jacquelyn Wynn, Mairead Hyland, The Lighthouse Lab in Alderley Park and Alex Alderton, Roberto Amato, Jeffrey Barrett, Sonia Goncalves, Ewan Harrison, David K. Jackson, Ian Johnston, Dominic Kwiatkowski, Cordelia Langford, John Sillitoe on behalf of the Wellcome Sanger Institute COVID-19 Surveillance Team |
| EPI_ISL_2701200 | Lighthouse Lab in Glasgow | Wellcome Sanger Institute for the COVID-19 Genomics UK (COG-UK) Consortium | Harper VanSteenhouse, Yumi Kasai, David Gray, Carol Clugston, Anna Dominiczak and Alex Alderton, Roberto Amato, Jeffrey Barrett, Sonia Goncalves, Ewan Harrison, David K. Jackson, Ian Johnston, Dominic Kwiatkowski, Cordelia Langford, John Sillitoe on behalf of the Wellcome Sanger Institute COVID-19 Surveillance Team |
| EPI_ISL_2701201 | Lighthouse Lab in Alderley Park | Wellcome Sanger Institute for the COVID-19 Genomics UK (COG-UK) Consortium | Jacquelyn Wynn, Mairead Hyland, The Lighthouse Lab in Alderley Park and Alex Alderton, Roberto Amato, Jeffrey Barrett, Sonia Goncalves, Ewan Harrison, David K. Jackson, Ian Johnston, Dominic Kwiatkowski, Cordelia Langford, John Sillitoe on behalf of the Wellcome Sanger Institute COVID-19 Surveillance Team |
| EPI_ISL_2701202, EPI_ISL_2701203, EPI_ISL_2701204, EPI_ISL_2701205 | Lighthouse Lab in Glasgow | Wellcome Sanger Institute for the COVID-19 Genomics UK (COG-UK) Consortium | Harper VanSteenhouse, Yumi Kasai, David Gray, Carol Clugston, Anna Dominiczak and Alex Alderton, Roberto Amato, Jeffrey Barrett, Sonia Goncalves, Ewan Harrison, David K. Jackson, Ian Johnston, Dominic Kwiatkowski, Cordelia Langford, John Sillitoe on behalf of the Wellcome Sanger Institute COVID-19 Surveillance Team |
| EPI_ISL_2701206, EPI_ISL_2701207 | Lighthouse Lab in Alderley Park | Wellcome Sanger Institute for the COVID-19 Genomics UK (COG-UK) Consortium | Jacquelyn Wynn, Mairead Hyland, The Lighthouse Lab in Alderley Park and Alex Alderton, Roberto Amato, Jeffrey Barrett, Sonia Goncalves, Ewan Harrison, David K. Jackson, Ian Johnston, Dominic Kwiatkowski, Cordelia Langford, John Sillitoe on behalf of the Wellcome Sanger Institute COVID-19 Surveillance Team |
| EPI_ISL_2701208 | Lighthouse Lab in Glasgow | Wellcome Sanger Institute for the COVID-19 Genomics UK (COG-UK) Consortium | Harper VanSteenhouse, Yumi Kasai, David Gray, Carol Clugston, Anna Dominiczak and Alex Alderton, Roberto Amato, Jeffrey Barrett, Sonia Goncalves, Ewan Harrison, David K. Jackson, Ian Johnston, Dominic Kwiatkowski, Cordelia Langford, John Sillitoe on behalf of the Wellcome Sanger Institute COVID-19 Surveillance Team |
| EPI_ISL_2701209 | Lighthouse Lab in Alderley Park | Wellcome Sanger Institute for the COVID-19 Genomics UK (COG-UK) Consortium | Jacquelyn Wynn, Mairead Hyland, The Lighthouse Lab in Alderley Park and Alex Alderton, Roberto Amato, Jeffrey Barrett, Sonia Goncalves, Ewan Harrison, David K. Jackson, Ian Johnston, Dominic Kwiatkowski, Cordelia Langford, John Sillitoe on behalf of the Wellcome Sanger Institute COVID-19 Surveillance Team |
| EPI_ISL_2701210, EPI_ISL_2701211 | Lighthouse Lab in Glasgow | Wellcome Sanger Institute for the COVID-19 Genomics UK (COG-UK) Consortium | Harper VanSteenhouse, Yumi Kasai, David Gray, Carol Clugston, Anna Dominiczak and Alex Alderton, Roberto Amato, Jeffrey Barrett, Sonia Goncalves, Ewan Harrison, David K. Jackson, Ian Johnston, Dominic Kwiatkowski, Cordelia Langford, John Sillitoe on behalf of the Wellcome Sanger Institute COVID-19 Surveillance Team |
| EPI_ISL_2701212, EPI_ISL_2701213 | Lighthouse Lab in Alderley Park | Wellcome Sanger Institute for the COVID-19 Genomics UK (COG-UK) Consortium | Jacquelyn Wynn, Mairead Hyland, The Lighthouse Lab in Alderley Park and Alex Alderton, Roberto Amato, Jeffrey Barrett, Sonia Goncalves, Ewan Harrison, David K. Jackson, Ian Johnston, Dominic Kwiatkowski, Cordelia Langford, John Sillitoe on behalf of the Wellcome Sanger Institute COVID-19 Surveillance Team |
| EPI_ISL_2701214 | Lighthouse Lab in Glasgow | Wellcome Sanger Institute for the COVID-19 Genomics UK (COG-UK) | Harper VanSteenhouse, Yumi Kasai, David Gray, Carol Clugston, Anna Dominiczak and Alex Alderton, Roberto Amato, Jeffrey Barrett, Sonia Goncalves, Ewan Harrison, David K. Jackson, Ian Johnston, Dominic Kwiatkowski, Cordelia Langford, |

| | | | |
|---|---|---|---|
| | | Consortium | John Sillitoe on behalf of the Wellcome Sanger Institute COVID-19 Surveillance Team |
| EPI_ISL_2701215, EPI_ISL_2701216 | Lighthouse Lab in Alderley Park | Wellcome Sanger Institute for the COVID-19 Genomics UK (COG-UK) Consortium | Jacquelyn Wynn, Mairead Hyland, The Lighthouse Lab in Alderley Park and Alex Alderton, Roberto Amato, Jeffrey Barrett, Sonia Goncalves, Ewan Harrison, David K. Jackson, Ian Johnston, Dominic Kwiatkowski, Cordelia Langford, John Sillitoe on behalf of the Wellcome Sanger Institute COVID-19 Surveillance Team |
| EPI_ISL_2701217 | Lighthouse Lab in Glasgow | Wellcome Sanger Institute for the COVID-19 Genomics UK (COG-UK) Consortium | Harper VanSteenhouse, Yumi Kasai, David Gray, Carol Clugston, Anna Dominiczak and Alex Alderton, Roberto Amato, Jeffrey Barrett, Sonia Goncalves, Ewan Harrison, David K. Jackson, Ian Johnston, Dominic Kwiatkowski, Cordelia Langford, John Sillitoe on behalf of the Wellcome Sanger Institute COVID-19 Surveillance Team |
| EPI_ISL_2701218 | Lighthouse Lab in Alderley Park | Wellcome Sanger Institute for the COVID-19 Genomics UK (COG-UK) Consortium | Jacquelyn Wynn, Mairead Hyland, The Lighthouse Lab in Alderley Park and Alex Alderton, Roberto Amato, Jeffrey Barrett, Sonia Goncalves, Ewan Harrison, David K. Jackson, Ian Johnston, Dominic Kwiatkowski, Cordelia Langford, John Sillitoe on behalf of the Wellcome Sanger Institute COVID-19 Surveillance Team |
| EPI_ISL_2701219, EPI_ISL_2701220, EPI_ISL_2701221, EPI_ISL_2701222, EPI_ISL_2701223, EPI_ISL_2701224 | Lighthouse Lab in Glasgow | Wellcome Sanger Institute for the COVID-19 Genomics UK (COG-UK) Consortium | Harper VanSteenhouse, Yumi Kasai, David Gray, Carol Clugston, Anna Dominiczak and Alex Alderton, Roberto Amato, Jeffrey Barrett, Sonia Goncalves, Ewan Harrison, David K. Jackson, Ian Johnston, Dominic Kwiatkowski, Cordelia Langford, John Sillitoe on behalf of the Wellcome Sanger Institute COVID-19 Surveillance Team |
| EPI_ISL_2701225, EPI_ISL_2701226, EPI_ISL_2701227 | Lighthouse Lab in Alderley Park | Wellcome Sanger Institute for the COVID-19 Genomics UK (COG-UK) Consortium | Jacquelyn Wynn, Mairead Hyland, The Lighthouse Lab in Alderley Park and Alex Alderton, Roberto Amato, Jeffrey Barrett, Sonia Goncalves, Ewan Harrison, David K. Jackson, Ian Johnston, Dominic Kwiatkowski, Cordelia Langford, John Sillitoe on behalf of the Wellcome Sanger Institute COVID-19 Surveillance Team |
| EPI_ISL_2701228 | Lighthouse Lab in Glasgow | Wellcome Sanger Institute for the COVID-19 Genomics UK (COG-UK) Consortium | Harper VanSteenhouse, Yumi Kasai, David Gray, Carol Clugston, Anna Dominiczak and Alex Alderton, Roberto Amato, Jeffrey Barrett, Sonia Goncalves, Ewan Harrison, David K. Jackson, Ian Johnston, Dominic Kwiatkowski, Cordelia Langford, John Sillitoe on behalf of the Wellcome Sanger Institute COVID-19 Surveillance Team |
| EPI_ISL_2701229, EPI_ISL_2701230, EPI_ISL_2701231 | Lighthouse Lab in Alderley Park | Wellcome Sanger Institute for the COVID-19 Genomics UK (COG-UK) Consortium | Jacquelyn Wynn, Mairead Hyland, The Lighthouse Lab in Alderley Park and Alex Alderton, Roberto Amato, Jeffrey Barrett, Sonia Goncalves, Ewan Harrison, David K. Jackson, Ian Johnston, Dominic Kwiatkowski, Cordelia Langford, John Sillitoe on behalf of the Wellcome Sanger Institute COVID-19 Surveillance Team |
| EPI_ISL_2701232, EPI_ISL_2701234 | Lighthouse Lab in Glasgow | Wellcome Sanger Institute for the COVID-19 Genomics UK (COG-UK) Consortium | Harper VanSteenhouse, Yumi Kasai, David Gray, Carol Clugston, Anna Dominiczak and Alex Alderton, Roberto Amato, Jeffrey Barrett, Sonia Goncalves, Ewan Harrison, David K. Jackson, Ian Johnston, Dominic Kwiatkowski, Cordelia Langford, John Sillitoe on behalf of the Wellcome Sanger Institute COVID-19 Surveillance Team |
| EPI_ISL_2701235 | Lighthouse Lab in Alderley Park | Wellcome Sanger Institute for the COVID-19 Genomics UK (COG-UK) Consortium | Jacquelyn Wynn, Mairead Hyland, The Lighthouse Lab in Alderley Park and Alex Alderton, Roberto Amato, Jeffrey Barrett, Sonia Goncalves, Ewan Harrison, David K. Jackson, Ian Johnston, Dominic Kwiatkowski, Cordelia Langford, John Sillitoe on behalf of the Wellcome Sanger Institute COVID-19 Surveillance Team |
| EPI_ISL_2701236 | Lighthouse Lab in Glasgow | Wellcome Sanger Institute for the COVID-19 Genomics UK (COG-UK) Consortium | Harper VanSteenhouse, Yumi Kasai, David Gray, Carol Clugston, Anna Dominiczak and Alex Alderton, Roberto Amato, Jeffrey Barrett, Sonia Goncalves, Ewan Harrison, David K. Jackson, Ian Johnston, Dominic Kwiatkowski, Cordelia Langford, John Sillitoe on behalf of the Wellcome Sanger Institute COVID-19 Surveillance Team |
| EPI_ISL_2701237, EPI_ISL_2701238 | Lighthouse Lab in Alderley Park | Wellcome Sanger Institute for the COVID-19 Genomics UK (COG-UK) Consortium | Jacquelyn Wynn, Mairead Hyland, The Lighthouse Lab in Alderley Park and Alex Alderton, Roberto Amato, Jeffrey Barrett, Sonia Goncalves, Ewan Harrison, David K. Jackson, Ian Johnston, Dominic Kwiatkowski, Cordelia Langford, John Sillitoe on behalf of the Wellcome Sanger Institute COVID-19 Surveillance Team |
| EPI_ISL_2701239 | Lighthouse Lab in Glasgow | Wellcome Sanger Institute for the COVID-19 Genomics UK (COG-UK) Consortium | Harper VanSteenhouse, Yumi Kasai, David Gray, Carol Clugston, Anna Dominiczak and Alex Alderton, Roberto Amato, Jeffrey Barrett, Sonia Goncalves, Ewan Harrison, David K. Jackson, Ian Johnston, Dominic Kwiatkowski, Cordelia Langford, John Sillitoe on behalf of the Wellcome Sanger Institute COVID-19 Surveillance Team |
| EPI_ISL_2701240 | Lighthouse Lab in Alderley Park | Wellcome Sanger Institute for the COVID-19 Genomics UK (COG-UK) Consortium | Jacquelyn Wynn, Mairead Hyland, The Lighthouse Lab in Alderley Park and Alex Alderton, Roberto Amato, Jeffrey Barrett, Sonia Goncalves, Ewan Harrison, David K. Jackson, Ian Johnston, Dominic Kwiatkowski, Cordelia Langford, John Sillitoe on behalf of the Wellcome Sanger Institute COVID-19 Surveillance Team |
| EPI_ISL_2701241, EPI_ISL_2701242 | Lighthouse Lab in Glasgow | Wellcome Sanger Institute for the COVID-19 Genomics UK (COG-UK) Consortium | Harper VanSteenhouse, Yumi Kasai, David Gray, Carol Clugston, Anna Dominiczak and Alex Alderton, Roberto Amato, Jeffrey Barrett, Sonia Goncalves, Ewan Harrison, David K. Jackson, Ian Johnston, Dominic Kwiatkowski, Cordelia Langford, John Sillitoe on behalf of the Wellcome Sanger Institute COVID-19 Surveillance Team |
| EPI_ISL_2701243, EPI_ISL_2701244 | Lighthouse Lab in Alderley Park | Wellcome Sanger Institute for the COVID-19 Genomics UK (COG-UK) Consortium | Jacquelyn Wynn, Mairead Hyland, The Lighthouse Lab in Alderley Park and Alex Alderton, Roberto Amato, Jeffrey Barrett, Sonia Goncalves, Ewan Harrison, David K. Jackson, Ian Johnston, Dominic Kwiatkowski, Cordelia Langford, John Sillitoe on behalf of the Wellcome Sanger Institute COVID-19 Surveillance Team |
| EPI_ISL_2701245 | Lighthouse Lab in Glasgow | Wellcome Sanger Institute for the COVID-19 Genomics UK (COG-UK) Consortium | Harper VanSteenhouse, Yumi Kasai, David Gray, Carol Clugston, Anna Dominiczak and Alex Alderton, Roberto Amato, Jeffrey Barrett, Sonia Goncalves, Ewan Harrison, David K. Jackson, Ian Johnston, Dominic Kwiatkowski, Cordelia Langford, John Sillitoe on behalf of the Wellcome Sanger Institute COVID-19 Surveillance Team |
| EPI_ISL_2701246, EPI_ISL_2701247, EPI_ISL_2701248, EPI_ISL_2701249, EPI_ISL_2701250 | Lighthouse Lab in Alderley Park | Wellcome Sanger Institute for the COVID-19 Genomics UK (COG-UK) Consortium | Jacquelyn Wynn, Mairead Hyland, The Lighthouse Lab in Alderley Park and Alex Alderton, Roberto Amato, Jeffrey Barrett, Sonia Goncalves, Ewan Harrison, David K. Jackson, Ian Johnston, Dominic Kwiatkowski, Cordelia Langford, John Sillitoe on behalf of the Wellcome Sanger Institute COVID-19 Surveillance Team |
| EPI_ISL_2701251, EPI_ISL_2701252 | Lighthouse Lab in Glasgow | Wellcome Sanger Institute for the COVID-19 Genomics UK (COG-UK) Consortium | Harper VanSteenhouse, Yumi Kasai, David Gray, Carol Clugston, Anna Dominiczak and Alex Alderton, Roberto Amato, Jeffrey Barrett, Sonia Goncalves, Ewan Harrison, David K. Jackson, Ian Johnston, Dominic Kwiatkowski, Cordelia Langford, John Sillitoe on behalf of the Wellcome Sanger Institute COVID-19 Surveillance Team |
| EPI_ISL_2701253 | Lighthouse Lab in Alderley Park | Wellcome Sanger Institute for the COVID-19 Genomics UK (COG-UK) Consortium | Jacquelyn Wynn, Mairead Hyland, The Lighthouse Lab in Alderley Park and Alex Alderton, Roberto Amato, Jeffrey Barrett, Sonia Goncalves, Ewan Harrison, David K. Jackson, Ian Johnston, Dominic Kwiatkowski, Cordelia Langford, John Sillitoe on behalf of the Wellcome Sanger Institute COVID-19 Surveillance Team |
| EPI_ISL_2701254 | Lighthouse Lab in Glasgow | Wellcome Sanger Institute for the COVID-19 Genomics UK (COG-UK) Consortium | Harper VanSteenhouse, Yumi Kasai, David Gray, Carol Clugston, Anna Dominiczak and Alex Alderton, Roberto Amato, Jeffrey Barrett, Sonia Goncalves, Ewan Harrison, David K. Jackson, Ian Johnston, Dominic Kwiatkowski, Cordelia Langford, John Sillitoe on behalf of the Wellcome Sanger Institute COVID-19 Surveillance Team |
| EPI_ISL_2701255 | Lighthouse Lab in Alderley Park | Wellcome Sanger Institute for the COVID-19 Genomics UK (COG-UK) Consortium | Jacquelyn Wynn, Mairead Hyland, The Lighthouse Lab in Alderley Park and Alex Alderton, Roberto Amato, Jeffrey Barrett, Sonia Goncalves, Ewan Harrison, David K. Jackson, Ian Johnston, Dominic Kwiatkowski, Cordelia Langford, John Sillitoe on behalf of the Wellcome Sanger Institute COVID-19 Surveillance Team |
| EPI_ISL_2701256, EPI_ISL_2701257, EPI_ISL_2701258, EPI_ISL_2701259, EPI_ISL_2701260, EPI_ISL_2701261 | Lighthouse Lab in Glasgow | Wellcome Sanger Institute for the COVID-19 Genomics UK (COG-UK) Consortium | Harper VanSteenhouse, Yumi Kasai, David Gray, Carol Clugston, Anna Dominiczak and Alex Alderton, Roberto Amato, Jeffrey Barrett, Sonia Goncalves, Ewan Harrison, David K. Jackson, Ian Johnston, Dominic Kwiatkowski, Cordelia Langford, John Sillitoe on behalf of the Wellcome Sanger Institute COVID-19 Surveillance Team |
| EPI_ISL_2701262, EPI_ISL_2701263 | Lighthouse Lab in Alderley Park | Wellcome Sanger Institute for the COVID-19 Genomics UK (COG-UK) Consortium | Jacquelyn Wynn, Mairead Hyland, The Lighthouse Lab in Alderley Park and Alex Alderton, Roberto Amato, Jeffrey Barrett, Sonia Goncalves, Ewan Harrison, David K. Jackson, Ian Johnston, Dominic Kwiatkowski, Cordelia Langford, John Sillitoe on behalf of the Wellcome Sanger Institute COVID-19 Surveillance Team |
| EPI_ISL_2701264 | Lighthouse Lab in Glasgow | Wellcome Sanger Institute for the COVID-19 Genomics UK (COG-UK) Consortium | Harper VanSteenhouse, Yumi Kasai, David Gray, Carol Clugston, Anna Dominiczak and Alex Alderton, Roberto Amato, Jeffrey Barrett, Sonia Goncalves, Ewan Harrison, David K. Jackson, Ian Johnston, Dominic Kwiatkowski, Cordelia Langford, John Sillitoe on behalf of the Wellcome Sanger Institute COVID-19 Surveillance Team |
| EPI_ISL_2701265 | Lighthouse Lab in Alderley Park | Wellcome Sanger Institute for the COVID-19 Genomics UK (COG-UK) Consortium | Jacquelyn Wynn, Mairead Hyland, The Lighthouse Lab in Alderley Park and Alex Alderton, Roberto Amato, Jeffrey Barrett, Sonia Goncalves, Ewan Harrison, David K. Jackson, Ian Johnston, Dominic Kwiatkowski, Cordelia Langford, John Sillitoe on behalf of the Wellcome Sanger Institute COVID-19 Surveillance Team |
| EPI_ISL_2701266 | Lighthouse Lab in Glasgow | Wellcome Sanger Institute for the COVID-19 Genomics UK (COG-UK) Consortium | Harper VanSteenhouse, Yumi Kasai, David Gray, Carol Clugston, Anna Dominiczak and Alex Alderton, Roberto Amato, Jeffrey Barrett, Sonia Goncalves, Ewan Harrison, David K. Jackson, Ian Johnston, Dominic Kwiatkowski, Cordelia Langford, John Sillitoe on behalf of the Wellcome Sanger Institute COVID-19 Surveillance Team |
| EPI_ISL_2701267, EPI_ISL_2701268 | Lighthouse Lab in Alderley Park | Wellcome Sanger Institute for the COVID-19 Genomics UK (COG-UK) Consortium | Jacquelyn Wynn, Mairead Hyland, The Lighthouse Lab in Alderley Park and Alex Alderton, Roberto Amato, Jeffrey Barrett, Sonia Goncalves, Ewan Harrison, David K. Jackson, Ian Johnston, Dominic Kwiatkowski, Cordelia Langford, John Sillitoe on behalf of the Wellcome Sanger Institute COVID-19 Surveillance Team |
| EPI_ISL_2701269 | Lighthouse Lab in Glasgow | Wellcome Sanger Institute for the COVID-19 Genomics UK (COG-UK) Consortium | Harper VanSteenhouse, Yumi Kasai, David Gray, Carol Clugston, Anna Dominiczak and Alex Alderton, Roberto Amato, Jeffrey Barrett, Sonia Goncalves, Ewan Harrison, David K. Jackson, Ian Johnston, Dominic Kwiatkowski, Cordelia Langford, John Sillitoe on behalf of the Wellcome Sanger Institute COVID-19 Surveillance Team |
| EPI_ISL_2701270, EPI_ISL_2701271, EPI_ISL_2701272 | Lighthouse Lab in Alderley Park | Wellcome Sanger Institute for the COVID-19 Genomics UK (COG-UK) Consortium | Jacquelyn Wynn, Mairead Hyland, The Lighthouse Lab in Alderley Park and Alex Alderton, Roberto Amato, Jeffrey Barrett, Sonia Goncalves, Ewan Harrison, David K. Jackson, Ian Johnston, Dominic Kwiatkowski, Cordelia Langford, John Sillitoe on behalf of the Wellcome Sanger Institute COVID-19 Surveillance Team |
| EPI_ISL_2701273, EPI_ISL_2701274, EPI_ISL_2701275, EPI_ISL_2701277, EPI_ISL_2701278, EPI_ISL_2701279, EPI_ISL_2701280 | Lighthouse Lab in Glasgow | Wellcome Sanger Institute for the COVID-19 Genomics UK (COG-UK) Consortium | Harper VanSteenhouse, Yumi Kasai, David Gray, Carol Clugston, Anna Dominiczak and Alex Alderton, Roberto Amato, Jeffrey Barrett, Sonia Goncalves, Ewan Harrison, David K. Jackson, Ian Johnston, Dominic Kwiatkowski, Cordelia Langford, John Sillitoe on behalf of the Wellcome Sanger Institute COVID-19 Surveillance Team |
| EPI_ISL_2701281 | Lighthouse Lab in Alderley Park | Wellcome Sanger Institute for the COVID-19 Genomics UK (COG-UK) Consortium | Jacquelyn Wynn, Mairead Hyland, The Lighthouse Lab in Alderley Park and Alex Alderton, Roberto Amato, Jeffrey Barrett, Sonia Goncalves, Ewan Harrison, David K. Jackson, Ian Johnston, Dominic Kwiatkowski, Cordelia Langford, John Sillitoe on behalf of the Wellcome Sanger Institute COVID-19 Surveillance Team |
| EPI_ISL_2701282, EPI_ISL_2701283 | Lighthouse Lab in Glasgow | Wellcome Sanger Institute for the COVID-19 Genomics UK (COG-UK) Consortium | Harper VanSteenhouse, Yumi Kasai, David Gray, Carol Clugston, Anna Dominiczak and Alex Alderton, Roberto Amato, Jeffrey Barrett, Sonia Goncalves, Ewan Harrison, David K. Jackson, Ian Johnston, Dominic Kwiatkowski, Cordelia Langford, John Sillitoe on behalf of the Wellcome Sanger Institute COVID-19 Surveillance Team |
| EPI_ISL_2701284, EPI_ISL_2701285, EPI_ISL_2701286, EPI_ISL_2701287 | Lighthouse Lab in Alderley Park | Wellcome Sanger Institute for the COVID-19 Genomics UK (COG-UK) Consortium | Jacquelyn Wynn, Mairead Hyland, The Lighthouse Lab in Alderley Park and Alex Alderton, Roberto Amato, Jeffrey Barrett, Sonia Goncalves, Ewan Harrison, David K. Jackson, Ian Johnston, Dominic Kwiatkowski, Cordelia Langford, John Sillitoe on behalf of the Wellcome Sanger Institute COVID-19 Surveillance Team |
| EPI_ISL_2701288 | Lighthouse Lab in Glasgow | Wellcome Sanger Institute for the COVID-19 Genomics UK (COG-UK) Consortium | Harper VanSteenhouse, Yumi Kasai, David Gray, Carol Clugston, Anna Dominiczak and Alex Alderton, Roberto Amato, Jeffrey Barrett, Sonia Goncalves, Ewan Harrison, David K. Jackson, Ian Johnston, Dominic Kwiatkowski, Cordelia Langford, John Sillitoe on behalf of the Wellcome Sanger Institute COVID-19 Surveillance Team |
| EPI_ISL_2701289, EPI_ISL_2701290, EPI_ISL_2701291, EPI_ISL_2701292, EPI_ISL_2701293, | Lighthouse Lab in Alderley Park | Wellcome Sanger Institute for the COVID-19 Genomics UK (COG-UK) Consortium | Jacquelyn Wynn, Mairead Hyland, The Lighthouse Lab in Alderley Park and Alex Alderton, Roberto Amato, Jeffrey Barrett, Sonia Goncalves, Ewan Harrison, David K. Jackson, Ian Johnston, Dominic Kwiatkowski, Cordelia Langford, John Sillitoe on behalf of the Wellcome Sanger Institute COVID-19 Surveillance Team |

| | | | |
|---|---|---|---|
| EPI_ISL_2701294 | | | |
| EPI_ISL_2701295, EPI_ISL_2701296, EPI_ISL_2701297, EPI_ISL_2701298, EPI_ISL_2701299, EPI_ISL_2701300 | Lighthouse Lab in Glasgow | Wellcome Sanger Institute for the COVID-19 Genomics UK (COG-UK) Consortium | Harper VanSteenhouse, Yumi Kasai, David Gray, Carol Clugston, Anna Dominiczak and Alex Alderton, Roberto Amato, Jeffrey Barrett, Sonia Goncalves, Ewan Harrison, David K. Jackson, Ian Johnston, Dominic Kwiatkowski, Cordelia Langford, John Sillitoe on behalf of the Wellcome Sanger Institute COVID-19 Surveillance Team |
| EPI_ISL_2701301, EPI_ISL_2701302 | Lighthouse Lab in Alderley Park | Wellcome Sanger Institute for the COVID-19 Genomics UK (COG-UK) Consortium | Jacquelyn Wynn, Mairead Hyland, The Lighthouse Lab in Alderley Park and Alex Alderton, Roberto Amato, Jeffrey Barrett, Sonia Goncalves, Ewan Harrison, David K. Jackson, Ian Johnston, Dominic Kwiatkowski, Cordelia Langford, John Sillitoe on behalf of the Wellcome Sanger Institute COVID-19 Surveillance Team |
| EPI_ISL_2701303 | Lighthouse Lab in Glasgow | Wellcome Sanger Institute for the COVID-19 Genomics UK (COG-UK) Consortium | Harper VanSteenhouse, Yumi Kasai, David Gray, Carol Clugston, Anna Dominiczak and Alex Alderton, Roberto Amato, Jeffrey Barrett, Sonia Goncalves, Ewan Harrison, David K. Jackson, Ian Johnston, Dominic Kwiatkowski, Cordelia Langford, John Sillitoe on behalf of the Wellcome Sanger Institute COVID-19 Surveillance Team |
| EPI_ISL_2701304 | Lighthouse Lab in Alderley Park | Wellcome Sanger Institute for the COVID-19 Genomics UK (COG-UK) Consortium | Jacquelyn Wynn, Mairead Hyland, The Lighthouse Lab in Alderley Park and Alex Alderton, Roberto Amato, Jeffrey Barrett, Sonia Goncalves, Ewan Harrison, David K. Jackson, Ian Johnston, Dominic Kwiatkowski, Cordelia Langford, John Sillitoe on behalf of the Wellcome Sanger Institute COVID-19 Surveillance Team |
| EPI_ISL_2701305 | Lighthouse Lab in Glasgow | Wellcome Sanger Institute for the COVID-19 Genomics UK (COG-UK) Consortium | Harper VanSteenhouse, Yumi Kasai, David Gray, Carol Clugston, Anna Dominiczak and Alex Alderton, Roberto Amato, Jeffrey Barrett, Sonia Goncalves, Ewan Harrison, David K. Jackson, Ian Johnston, Dominic Kwiatkowski, Cordelia Langford, John Sillitoe on behalf of the Wellcome Sanger Institute COVID-19 Surveillance Team |
| EPI_ISL_2701306 | Lighthouse Lab in Alderley Park | Wellcome Sanger Institute for the COVID-19 Genomics UK (COG-UK) Consortium | Jacquelyn Wynn, Mairead Hyland, The Lighthouse Lab in Alderley Park and Alex Alderton, Roberto Amato, Jeffrey Barrett, Sonia Goncalves, Ewan Harrison, David K. Jackson, Ian Johnston, Dominic Kwiatkowski, Cordelia Langford, John Sillitoe on behalf of the Wellcome Sanger Institute COVID-19 Surveillance Team |
| EPI_ISL_2701307, EPI_ISL_2701308 | Lighthouse Lab in Glasgow | Wellcome Sanger Institute for the COVID-19 Genomics UK (COG-UK) Consortium | Harper VanSteenhouse, Yumi Kasai, David Gray, Carol Clugston, Anna Dominiczak and Alex Alderton, Roberto Amato, Jeffrey Barrett, Sonia Goncalves, Ewan Harrison, David K. Jackson, Ian Johnston, Dominic Kwiatkowski, Cordelia Langford, John Sillitoe on behalf of the Wellcome Sanger Institute COVID-19 Surveillance Team |
| EPI_ISL_2701309 | Lighthouse Lab in Milton Keynes | Wellcome Sanger Institute for the COVID-19 Genomics UK (COG-UK) Consortium | The Lighthouse Lab in Milton Keynes and Alex Alderton, Roberto Amato, Jeffrey Barrett, Sonia Goncalves, Ewan Harrison, David K. Jackson, Ian Johnston, Dominic Kwiatkowski, Cordelia Langford, John Sillitoe on behalf of the Wellcome Sanger Institute COVID-19 Surveillance Team |
| EPI_ISL_2701310 | Lighthouse Lab in Alderley Park | Wellcome Sanger Institute for the COVID-19 Genomics UK (COG-UK) Consortium | Jacquelyn Wynn, Mairead Hyland, The Lighthouse Lab in Alderley Park and Alex Alderton, Roberto Amato, Jeffrey Barrett, Sonia Goncalves, Ewan Harrison, David K. Jackson, Ian Johnston, Dominic Kwiatkowski, Cordelia Langford, John Sillitoe on behalf of the Wellcome Sanger Institute COVID-19 Surveillance Team |
| EPI_ISL_2701311 | Lighthouse Lab in Milton Keynes | Wellcome Sanger Institute for the COVID-19 Genomics UK (COG-UK) Consortium | The Lighthouse Lab in Milton Keynes and Alex Alderton, Roberto Amato, Jeffrey Barrett, Sonia Goncalves, Ewan Harrison, David K. Jackson, Ian Johnston, Dominic Kwiatkowski, Cordelia Langford, John Sillitoe on behalf of the Wellcome Sanger Institute COVID-19 Surveillance Team |
| EPI_ISL_2701312, EPI_ISL_2701313 | Lighthouse Lab in Alderley Park | Wellcome Sanger Institute for the COVID-19 Genomics UK (COG-UK) Consortium | Jacquelyn Wynn, Mairead Hyland, The Lighthouse Lab in Alderley Park and Alex Alderton, Roberto Amato, Jeffrey Barrett, Sonia Goncalves, Ewan Harrison, David K. Jackson, Ian Johnston, Dominic Kwiatkowski, Cordelia Langford, John Sillitoe on behalf of the Wellcome Sanger Institute COVID-19 Surveillance Team |
| EPI_ISL_2701314 | Lighthouse Lab in Glasgow | Wellcome Sanger Institute for the COVID-19 Genomics UK (COG-UK) Consortium | Harper VanSteenhouse, Yumi Kasai, David Gray, Carol Clugston, Anna Dominiczak and Alex Alderton, Roberto Amato, Jeffrey Barrett, Sonia Goncalves, Ewan Harrison, David K. Jackson, Ian Johnston, Dominic Kwiatkowski, Cordelia Langford, John Sillitoe on behalf of the Wellcome Sanger Institute COVID-19 Surveillance Team |
| EPI_ISL_2701315 | Lighthouse Lab in Milton Keynes | Wellcome Sanger Institute for the COVID-19 Genomics UK (COG-UK) Consortium | The Lighthouse Lab in Milton Keynes and Alex Alderton, Roberto Amato, Jeffrey Barrett, Sonia Goncalves, Ewan Harrison, David K. Jackson, Ian Johnston, Dominic Kwiatkowski, Cordelia Langford, John Sillitoe on behalf of the Wellcome Sanger Institute COVID-19 Surveillance Team |
| EPI_ISL_2701316, EPI_ISL_2701317 | Lighthouse Lab in Glasgow | Wellcome Sanger Institute for the COVID-19 Genomics UK (COG-UK) Consortium | Harper VanSteenhouse, Yumi Kasai, David Gray, Carol Clugston, Anna Dominiczak and Alex Alderton, Roberto Amato, Jeffrey Barrett, Sonia Goncalves, Ewan Harrison, David K. Jackson, Ian Johnston, Dominic Kwiatkowski, Cordelia Langford, John Sillitoe on behalf of the Wellcome Sanger Institute COVID-19 Surveillance Team |
| EPI_ISL_2701318, EPI_ISL_2701319 | Lighthouse Lab in Milton Keynes | Wellcome Sanger Institute for the COVID-19 Genomics UK (COG-UK) Consortium | The Lighthouse Lab in Milton Keynes and Alex Alderton, Roberto Amato, Jeffrey Barrett, Sonia Goncalves, Ewan Harrison, David K. Jackson, Ian Johnston, Dominic Kwiatkowski, Cordelia Langford, John Sillitoe on behalf of the Wellcome Sanger Institute COVID-19 Surveillance Team |
| EPI_ISL_2701320 | Lighthouse Lab in Alderley Park | Wellcome Sanger Institute for the COVID-19 Genomics UK (COG-UK) Consortium | Jacquelyn Wynn, Mairead Hyland, The Lighthouse Lab in Alderley Park and Alex Alderton, Roberto Amato, Jeffrey Barrett, Sonia Goncalves, Ewan Harrison, David K. Jackson, Ian Johnston, Dominic Kwiatkowski, Cordelia Langford, John Sillitoe on behalf of the Wellcome Sanger Institute COVID-19 Surveillance Team |
| EPI_ISL_2701321, EPI_ISL_2701322 | Lighthouse Lab in Glasgow | Wellcome Sanger Institute for the COVID-19 Genomics UK (COG-UK) Consortium | Harper VanSteenhouse, Yumi Kasai, David Gray, Carol Clugston, Anna Dominiczak and Alex Alderton, Roberto Amato, Jeffrey Barrett, Sonia Goncalves, Ewan Harrison, David K. Jackson, Ian Johnston, Dominic Kwiatkowski, Cordelia Langford, John Sillitoe on behalf of the Wellcome Sanger Institute COVID-19 Surveillance Team |
| EPI_ISL_2701323 | Lighthouse Lab in Milton Keynes | Wellcome Sanger Institute for the COVID-19 Genomics UK (COG-UK) Consortium | The Lighthouse Lab in Milton Keynes and Alex Alderton, Roberto Amato, Jeffrey Barrett, Sonia Goncalves, Ewan Harrison, David K. Jackson, Ian Johnston, Dominic Kwiatkowski, Cordelia Langford, John Sillitoe on behalf of the Wellcome Sanger Institute COVID-19 Surveillance Team |
| EPI_ISL_2701324 | Lighthouse Lab in Glasgow | Wellcome Sanger Institute for the COVID-19 Genomics UK (COG-UK) Consortium | Harper VanSteenhouse, Yumi Kasai, David Gray, Carol Clugston, Anna Dominiczak and Alex Alderton, Roberto Amato, Jeffrey Barrett, Sonia Goncalves, Ewan Harrison, David K. Jackson, Ian Johnston, Dominic Kwiatkowski, Cordelia Langford, John Sillitoe on behalf of the Wellcome Sanger Institute COVID-19 Surveillance Team |
| EPI_ISL_2701325, EPI_ISL_2701326 | Lighthouse Lab in Milton Keynes | Wellcome Sanger Institute for the COVID-19 Genomics UK (COG-UK) Consortium | The Lighthouse Lab in Milton Keynes and Alex Alderton, Roberto Amato, Jeffrey Barrett, Sonia Goncalves, Ewan Harrison, David K. Jackson, Ian Johnston, Dominic Kwiatkowski, Cordelia Langford, John Sillitoe on behalf of the Wellcome Sanger Institute COVID-19 Surveillance Team |
| EPI_ISL_2701327 | Lighthouse Lab in Glasgow | Wellcome Sanger Institute for the COVID-19 Genomics UK (COG-UK) Consortium | Harper VanSteenhouse, Yumi Kasai, David Gray, Carol Clugston, Anna Dominiczak and Alex Alderton, Roberto Amato, Jeffrey Barrett, Sonia Goncalves, Ewan Harrison, David K. Jackson, Ian Johnston, Dominic Kwiatkowski, Cordelia Langford, John Sillitoe on behalf of the Wellcome Sanger Institute COVID-19 Surveillance Team |
| EPI_ISL_2701328 | Lighthouse Lab in Alderley Park | Wellcome Sanger Institute for the COVID-19 Genomics UK (COG-UK) Consortium | Jacquelyn Wynn, Mairead Hyland, The Lighthouse Lab in Alderley Park and Alex Alderton, Roberto Amato, Jeffrey Barrett, Sonia Goncalves, Ewan Harrison, David K. Jackson, Ian Johnston, Dominic Kwiatkowski, Cordelia Langford, John Sillitoe on behalf of the Wellcome Sanger Institute COVID-19 Surveillance Team |
| EPI_ISL_2701329, EPI_ISL_2701330 | Lighthouse Lab in Glasgow | Wellcome Sanger Institute for the COVID-19 Genomics UK (COG-UK) Consortium | Harper VanSteenhouse, Yumi Kasai, David Gray, Carol Clugston, Anna Dominiczak and Alex Alderton, Roberto Amato, Jeffrey Barrett, Sonia Goncalves, Ewan Harrison, David K. Jackson, Ian Johnston, Dominic Kwiatkowski, Cordelia Langford, John Sillitoe on behalf of the Wellcome Sanger Institute COVID-19 Surveillance Team |
| EPI_ISL_2701331 | Lighthouse Lab in Milton Keynes | Wellcome Sanger Institute for the COVID-19 Genomics UK (COG-UK) Consortium | The Lighthouse Lab in Milton Keynes and Alex Alderton, Roberto Amato, Jeffrey Barrett, Sonia Goncalves, Ewan Harrison, David K. Jackson, Ian Johnston, Dominic Kwiatkowski, Cordelia Langford, John Sillitoe on behalf of the Wellcome Sanger Institute COVID-19 Surveillance Team |
| EPI_ISL_2701332 | Lighthouse Lab in Alderley Park | Wellcome Sanger Institute for the COVID-19 Genomics UK (COG-UK) Consortium | Jacquelyn Wynn, Mairead Hyland, The Lighthouse Lab in Alderley Park and Alex Alderton, Roberto Amato, Jeffrey Barrett, Sonia Goncalves, Ewan Harrison, David K. Jackson, Ian Johnston, Dominic Kwiatkowski, Cordelia Langford, John Sillitoe on behalf of the Wellcome Sanger Institute COVID-19 Surveillance Team |
| EPI_ISL_2701333 | Lighthouse Lab in Glasgow | Wellcome Sanger Institute for the COVID-19 Genomics UK (COG-UK) Consortium | Harper VanSteenhouse, Yumi Kasai, David Gray, Carol Clugston, Anna Dominiczak and Alex Alderton, Roberto Amato, Jeffrey Barrett, Sonia Goncalves, Ewan Harrison, David K. Jackson, Ian Johnston, Dominic Kwiatkowski, Cordelia Langford, John Sillitoe on behalf of the Wellcome Sanger Institute COVID-19 Surveillance Team |
| EPI_ISL_2701334 | Lighthouse Lab in Alderley Park | Wellcome Sanger Institute for the COVID-19 Genomics UK (COG-UK) Consortium | Jacquelyn Wynn, Mairead Hyland, The Lighthouse Lab in Alderley Park and Alex Alderton, Roberto Amato, Jeffrey Barrett, Sonia Goncalves, Ewan Harrison, David K. Jackson, Ian Johnston, Dominic Kwiatkowski, Cordelia Langford, John Sillitoe on behalf of the Wellcome Sanger Institute COVID-19 Surveillance Team |
| EPI_ISL_2701335, EPI_ISL_2701336 | Lighthouse Lab in Milton Keynes | Wellcome Sanger Institute for the COVID-19 Genomics UK (COG-UK) Consortium | The Lighthouse Lab in Milton Keynes and Alex Alderton, Roberto Amato, Jeffrey Barrett, Sonia Goncalves, Ewan Harrison, David K. Jackson, Ian Johnston, Dominic Kwiatkowski, Cordelia Langford, John Sillitoe on behalf of the Wellcome Sanger Institute COVID-19 Surveillance Team |
| EPI_ISL_2701337, EPI_ISL_2701338, EPI_ISL_2701339, EPI_ISL_2701340, EPI_ISL_2701341 | Lighthouse Lab in Glasgow | Wellcome Sanger Institute for the COVID-19 Genomics UK (COG-UK) Consortium | Harper VanSteenhouse, Yumi Kasai, David Gray, Carol Clugston, Anna Dominiczak and Alex Alderton, Roberto Amato, Jeffrey Barrett, Sonia Goncalves, Ewan Harrison, David K. Jackson, Ian Johnston, Dominic Kwiatkowski, Cordelia Langford, John Sillitoe on behalf of the Wellcome Sanger Institute COVID-19 Surveillance Team |
| EPI_ISL_2701342 | Lighthouse Lab in Alderley Park | Wellcome Sanger Institute for the COVID-19 Genomics UK (COG-UK) Consortium | Jacquelyn Wynn, Mairead Hyland, The Lighthouse Lab in Alderley Park and Alex Alderton, Roberto Amato, Jeffrey Barrett, Sonia Goncalves, Ewan Harrison, David K. Jackson, Ian Johnston, Dominic Kwiatkowski, Cordelia Langford, John Sillitoe on behalf of the Wellcome Sanger Institute COVID-19 Surveillance Team |
| EPI_ISL_2701343 | Lighthouse Lab in Milton Keynes | Wellcome Sanger Institute for the COVID-19 Genomics UK (COG-UK) Consortium | The Lighthouse Lab in Milton Keynes and Alex Alderton, Roberto Amato, Jeffrey Barrett, Sonia Goncalves, Ewan Harrison, David K. Jackson, Ian Johnston, Dominic Kwiatkowski, Cordelia Langford, John Sillitoe on behalf of the Wellcome Sanger Institute COVID-19 Surveillance Team |
| EPI_ISL_2701344 | Lighthouse Lab in Alderley Park | Wellcome Sanger Institute for the COVID-19 Genomics UK (COG-UK) Consortium | Jacquelyn Wynn, Mairead Hyland, The Lighthouse Lab in Alderley Park and Alex Alderton, Roberto Amato, Jeffrey Barrett, Sonia Goncalves, Ewan Harrison, David K. Jackson, Ian Johnston, Dominic Kwiatkowski, Cordelia Langford, John Sillitoe on behalf of the Wellcome Sanger Institute COVID-19 Surveillance Team |
| EPI_ISL_2701345, EPI_ISL_2701346, EPI_ISL_2701347, EPI_ISL_2701348, EPI_ISL_2701349, EPI_ISL_2701350, EPI_ISL_2701351 | Lighthouse Lab in Glasgow | Wellcome Sanger Institute for the COVID-19 Genomics UK (COG-UK) Consortium | Harper VanSteenhouse, Yumi Kasai, David Gray, Carol Clugston, Anna Dominiczak and Alex Alderton, Roberto Amato, Jeffrey Barrett, Sonia Goncalves, Ewan Harrison, David K. Jackson, Ian Johnston, Dominic Kwiatkowski, Cordelia Langford, John Sillitoe on behalf of the Wellcome Sanger Institute COVID-19 Surveillance Team |
| EPI_ISL_2701352, EPI_ISL_2701353 | Lighthouse Lab in Alderley Park | Wellcome Sanger Institute for the COVID-19 Genomics UK (COG-UK) Consortium | Jacquelyn Wynn, Mairead Hyland, The Lighthouse Lab in Alderley Park and Alex Alderton, Roberto Amato, Jeffrey Barrett, Sonia Goncalves, Ewan Harrison, David K. Jackson, Ian Johnston, Dominic Kwiatkowski, Cordelia Langford, John Sillitoe on behalf of the Wellcome Sanger Institute COVID-19 Surveillance Team |
| EPI_ISL_2701354 | Lighthouse Lab in Glasgow | Wellcome Sanger Institute for the COVID-19 Genomics UK (COG-UK) Consortium | Harper VanSteenhouse, Yumi Kasai, David Gray, Carol Clugston, Anna Dominiczak and Alex Alderton, Roberto Amato, Jeffrey Barrett, Sonia Goncalves, Ewan Harrison, David K. Jackson, Ian Johnston, Dominic Kwiatkowski, Cordelia Langford, John Sillitoe on behalf of the Wellcome Sanger Institute COVID-19 Surveillance Team |
| EPI_ISL_2701355 | Lighthouse Lab in Alderley Park | Wellcome Sanger Institute for the COVID-19 Genomics UK (COG-UK) Consortium | Jacquelyn Wynn, Mairead Hyland, The Lighthouse Lab in Alderley Park and Alex Alderton, Roberto Amato, Jeffrey Barrett, Sonia Goncalves, Ewan Harrison, David K. Jackson, Ian Johnston, Dominic Kwiatkowski, Cordelia Langford, John Sillitoe on behalf of the Wellcome Sanger Institute COVID-19 Surveillance Team |
| EPI_ISL_2701356 | Lighthouse Lab in Glasgow | Wellcome Sanger Institute for the COVID-19 Genomics UK (COG-UK) Consortium | Harper VanSteenhouse, Yumi Kasai, David Gray, Carol Clugston, Anna Dominiczak and Alex Alderton, Roberto Amato, Jeffrey Barrett, Sonia Goncalves, Ewan Harrison, David K. Jackson, Ian Johnston, Dominic Kwiatkowski, Cordelia Langford, John Sillitoe on behalf of the Wellcome Sanger Institute COVID-19 Surveillance Team |
| EPI_ISL_2701357 | Lighthouse Lab in Alderley Park | Wellcome Sanger Institute for the COVID-19 Genomics UK (COG-UK) Consortium | Jacquelyn Wynn, Mairead Hyland, The Lighthouse Lab in Alderley Park and Alex Alderton, Roberto Amato, Jeffrey Barrett, Sonia Goncalves, Ewan Harrison, David K. Jackson, Ian Johnston, Dominic Kwiatkowski, Cordelia Langford, John Sillitoe on behalf of the Wellcome Sanger Institute COVID-19 Surveillance Team |
| EPI_ISL_2701358 | Lighthouse Lab in Milton Keynes | Wellcome Sanger Institute for the COVID-19 Genomics UK (COG-UK) Consortium | The Lighthouse Lab in Milton Keynes and Alex Alderton, Roberto Amato, Jeffrey Barrett, Sonia Goncalves, Ewan Harrison, David K. Jackson, Ian Johnston, Dominic Kwiatkowski, Cordelia Langford, John Sillitoe on behalf of the Wellcome Sanger Institute COVID-19 Surveillance Team |

| | | | |
|---|---|---|---|
| EPI_ISL_2701359 | Lighthouse Lab in Alderley Park | Wellcome Sanger Institute for the COVID-19 Genomics UK (COG-UK) Consortium | Jacquelyn Wynn, Mairead Hyland, The Lighthouse Lab in Alderley Park and Alex Alderton, Roberto Amato, Jeffrey Barrett, Sonia Goncalves, Ewan Harrison, David K. Jackson, Ian Johnston, Dominic Kwiatkowski, Cordelia Langford, John Sillitoe on behalf of the Wellcome Sanger Institute COVID-19 Surveillance Team |
| EPI_ISL_2701360 | Lighthouse Lab in Glasgow | Wellcome Sanger Institute for the COVID-19 Genomics UK (COG-UK) Consortium | Harper VanSteenhouse, Yumi Kasai, David Gray, Carol Clugston, Anna Dominiczak and Alex Alderton, Roberto Amato, Jeffrey Barrett, Sonia Goncalves, Ewan Harrison, David K. Jackson, Ian Johnston, Dominic Kwiatkowski, Cordelia Langford, John Sillitoe on behalf of the Wellcome Sanger Institute COVID-19 Surveillance Team |
| EPI_ISL_2701361, EPI_ISL_2701362, EPI_ISL_2701363 | Lighthouse Lab in Milton Keynes | Wellcome Sanger Institute for the COVID-19 Genomics UK (COG-UK) Consortium | The Lighthouse Lab in Milton Keynes and Alex Alderton, Roberto Amato, Jeffrey Barrett, Sonia Goncalves, Ewan Harrison, David K. Jackson, Ian Johnston, Dominic Kwiatkowski, Cordelia Langford, John Sillitoe on behalf of the Wellcome Sanger Institute COVID-19 Surveillance Team |
| EPI_ISL_2701364 | Lighthouse Lab in Glasgow | Wellcome Sanger Institute for the COVID-19 Genomics UK (COG-UK) Consortium | Harper VanSteenhouse, Yumi Kasai, David Gray, Carol Clugston, Anna Dominiczak and Alex Alderton, Roberto Amato, Jeffrey Barrett, Sonia Goncalves, Ewan Harrison, David K. Jackson, Ian Johnston, Dominic Kwiatkowski, Cordelia Langford, John Sillitoe on behalf of the Wellcome Sanger Institute COVID-19 Surveillance Team |
| EPI_ISL_2701365 | Lighthouse Lab in Milton Keynes | Wellcome Sanger Institute for the COVID-19 Genomics UK (COG-UK) Consortium | The Lighthouse Lab in Milton Keynes and Alex Alderton, Roberto Amato, Jeffrey Barrett, Sonia Goncalves, Ewan Harrison, David K. Jackson, Ian Johnston, Dominic Kwiatkowski, Cordelia Langford, John Sillitoe on behalf of the Wellcome Sanger Institute COVID-19 Surveillance Team |
| EPI_ISL_2701366 | Lighthouse Lab in Glasgow | Wellcome Sanger Institute for the COVID-19 Genomics UK (COG-UK) Consortium | Harper VanSteenhouse, Yumi Kasai, David Gray, Carol Clugston, Anna Dominiczak and Alex Alderton, Roberto Amato, Jeffrey Barrett, Sonia Goncalves, Ewan Harrison, David K. Jackson, Ian Johnston, Dominic Kwiatkowski, Cordelia Langford, John Sillitoe on behalf of the Wellcome Sanger Institute COVID-19 Surveillance Team |
| EPI_ISL_2701367, EPI_ISL_2701368 | Lighthouse Lab in Alderley Park | Wellcome Sanger Institute for the COVID-19 Genomics UK (COG-UK) Consortium | Jacquelyn Wynn, Mairead Hyland, The Lighthouse Lab in Alderley Park and Alex Alderton, Roberto Amato, Jeffrey Barrett, Sonia Goncalves, Ewan Harrison, David K. Jackson, Ian Johnston, Dominic Kwiatkowski, Cordelia Langford, John Sillitoe on behalf of the Wellcome Sanger Institute COVID-19 Surveillance Team |
| EPI_ISL_2701369, EPI_ISL_2701370 | Lighthouse Lab in Glasgow | Wellcome Sanger Institute for the COVID-19 Genomics UK (COG-UK) Consortium | Harper VanSteenhouse, Yumi Kasai, David Gray, Carol Clugston, Anna Dominiczak and Alex Alderton, Roberto Amato, Jeffrey Barrett, Sonia Goncalves, Ewan Harrison, David K. Jackson, Ian Johnston, Dominic Kwiatkowski, Cordelia Langford, John Sillitoe on behalf of the Wellcome Sanger Institute COVID-19 Surveillance Team |
| EPI_ISL_2701371 | Lighthouse Lab in Milton Keynes | Wellcome Sanger Institute for the COVID-19 Genomics UK (COG-UK) Consortium | The Lighthouse Lab in Milton Keynes and Alex Alderton, Roberto Amato, Jeffrey Barrett, Sonia Goncalves, Ewan Harrison, David K. Jackson, Ian Johnston, Dominic Kwiatkowski, Cordelia Langford, John Sillitoe on behalf of the Wellcome Sanger Institute COVID-19 Surveillance Team |
| EPI_ISL_2701372 | Lighthouse Lab in Glasgow | Wellcome Sanger Institute for the COVID-19 Genomics UK (COG-UK) Consortium | Harper VanSteenhouse, Yumi Kasai, David Gray, Carol Clugston, Anna Dominiczak and Alex Alderton, Roberto Amato, Jeffrey Barrett, Sonia Goncalves, Ewan Harrison, David K. Jackson, Ian Johnston, Dominic Kwiatkowski, Cordelia Langford, John Sillitoe on behalf of the Wellcome Sanger Institute COVID-19 Surveillance Team |
| EPI_ISL_2701373 | Lighthouse Lab in Milton Keynes | Wellcome Sanger Institute for the COVID-19 Genomics UK (COG-UK) Consortium | The Lighthouse Lab in Milton Keynes and Alex Alderton, Roberto Amato, Jeffrey Barrett, Sonia Goncalves, Ewan Harrison, David K. Jackson, Ian Johnston, Dominic Kwiatkowski, Cordelia Langford, John Sillitoe on behalf of the Wellcome Sanger Institute COVID-19 Surveillance Team |
| EPI_ISL_2701374, EPI_ISL_2701375 | Lighthouse Lab in Glasgow | Wellcome Sanger Institute for the COVID-19 Genomics UK (COG-UK) Consortium | Harper VanSteenhouse, Yumi Kasai, David Gray, Carol Clugston, Anna Dominiczak and Alex Alderton, Roberto Amato, Jeffrey Barrett, Sonia Goncalves, Ewan Harrison, David K. Jackson, Ian Johnston, Dominic Kwiatkowski, Cordelia Langford, John Sillitoe on behalf of the Wellcome Sanger Institute COVID-19 Surveillance Team |
| EPI_ISL_2701376, EPI_ISL_2701377, EPI_ISL_2701378 | Lighthouse Lab in Alderley Park | Wellcome Sanger Institute for the COVID-19 Genomics UK (COG-UK) Consortium | Jacquelyn Wynn, Mairead Hyland, The Lighthouse Lab in Alderley Park and Alex Alderton, Roberto Amato, Jeffrey Barrett, Sonia Goncalves, Ewan Harrison, David K. Jackson, Ian Johnston, Dominic Kwiatkowski, Cordelia Langford, John Sillitoe on behalf of the Wellcome Sanger Institute COVID-19 Surveillance Team |
| EPI_ISL_2701379, EPI_ISL_2701380, EPI_ISL_2701381 | Lighthouse Lab in Glasgow | Wellcome Sanger Institute for the COVID-19 Genomics UK (COG-UK) Consortium | Harper VanSteenhouse, Yumi Kasai, David Gray, Carol Clugston, Anna Dominiczak and Alex Alderton, Roberto Amato, Jeffrey Barrett, Sonia Goncalves, Ewan Harrison, David K. Jackson, Ian Johnston, Dominic Kwiatkowski, Cordelia Langford, John Sillitoe on behalf of the Wellcome Sanger Institute COVID-19 Surveillance Team |
| EPI_ISL_2701382 | Lighthouse Lab in Milton Keynes | Wellcome Sanger Institute for the COVID-19 Genomics UK (COG-UK) Consortium | The Lighthouse Lab in Milton Keynes and Alex Alderton, Roberto Amato, Jeffrey Barrett, Sonia Goncalves, Ewan Harrison, David K. Jackson, Ian Johnston, Dominic Kwiatkowski, Cordelia Langford, John Sillitoe on behalf of the Wellcome Sanger Institute COVID-19 Surveillance Team |
| EPI_ISL_2701383 | Lighthouse Lab in Glasgow | Wellcome Sanger Institute for the COVID-19 Genomics UK (COG-UK) Consortium | Harper VanSteenhouse, Yumi Kasai, David Gray, Carol Clugston, Anna Dominiczak and Alex Alderton, Roberto Amato, Jeffrey Barrett, Sonia Goncalves, Ewan Harrison, David K. Jackson, Ian Johnston, Dominic Kwiatkowski, Cordelia Langford, John Sillitoe on behalf of the Wellcome Sanger Institute COVID-19 Surveillance Team |
| EPI_ISL_2701384 | Lighthouse Lab in Alderley Park | Wellcome Sanger Institute for the COVID-19 Genomics UK (COG-UK) Consortium | Jacquelyn Wynn, Mairead Hyland, The Lighthouse Lab in Alderley Park and Alex Alderton, Roberto Amato, Jeffrey Barrett, Sonia Goncalves, Ewan Harrison, David K. Jackson, Ian Johnston, Dominic Kwiatkowski, Cordelia Langford, John Sillitoe on behalf of the Wellcome Sanger Institute COVID-19 Surveillance Team |
| EPI_ISL_2701385 | Lighthouse Lab in Glasgow | Wellcome Sanger Institute for the COVID-19 Genomics UK (COG-UK) Consortium | Harper VanSteenhouse, Yumi Kasai, David Gray, Carol Clugston, Anna Dominiczak and Alex Alderton, Roberto Amato, Jeffrey Barrett, Sonia Goncalves, Ewan Harrison, David K. Jackson, Ian Johnston, Dominic Kwiatkowski, Cordelia Langford, John Sillitoe on behalf of the Wellcome Sanger Institute COVID-19 Surveillance Team |
| EPI_ISL_2701386, EPI_ISL_2701387 | Lighthouse Lab in Alderley Park | Wellcome Sanger Institute for the COVID-19 Genomics UK (COG-UK) Consortium | Jacquelyn Wynn, Mairead Hyland, The Lighthouse Lab in Alderley Park and Alex Alderton, Roberto Amato, Jeffrey Barrett, Sonia Goncalves, Ewan Harrison, David K. Jackson, Ian Johnston, Dominic Kwiatkowski, Cordelia Langford, John Sillitoe on behalf of the Wellcome Sanger Institute COVID-19 Surveillance Team |
| EPI_ISL_2701388 | Lighthouse Lab in Milton Keynes | Wellcome Sanger Institute for the COVID-19 Genomics UK (COG-UK) Consortium | The Lighthouse Lab in Milton Keynes and Alex Alderton, Roberto Amato, Jeffrey Barrett, Sonia Goncalves, Ewan Harrison, David K. Jackson, Ian Johnston, Dominic Kwiatkowski, Cordelia Langford, John Sillitoe on behalf of the Wellcome Sanger Institute COVID-19 Surveillance Team |
| EPI_ISL_2701389, EPI_ISL_2701390, EPI_ISL_2701391 | Lighthouse Lab in Glasgow | Wellcome Sanger Institute for the COVID-19 Genomics UK (COG-UK) Consortium | Harper VanSteenhouse, Yumi Kasai, David Gray, Carol Clugston, Anna Dominiczak and Alex Alderton, Roberto Amato, Jeffrey Barrett, Sonia Goncalves, Ewan Harrison, David K. Jackson, Ian Johnston, Dominic Kwiatkowski, Cordelia Langford, John Sillitoe on behalf of the Wellcome Sanger Institute COVID-19 Surveillance Team |
| EPI_ISL_2701392, EPI_ISL_2701393 | Lighthouse Lab in Alderley Park | Wellcome Sanger Institute for the COVID-19 Genomics UK (COG-UK) Consortium | Jacquelyn Wynn, Mairead Hyland, The Lighthouse Lab in Alderley Park and Alex Alderton, Roberto Amato, Jeffrey Barrett, Sonia Goncalves, Ewan Harrison, David K. Jackson, Ian Johnston, Dominic Kwiatkowski, Cordelia Langford, John Sillitoe on behalf of the Wellcome Sanger Institute COVID-19 Surveillance Team |
| EPI_ISL_2701394 | Lighthouse Lab in Milton Keynes | Wellcome Sanger Institute for the COVID-19 Genomics UK (COG-UK) Consortium | The Lighthouse Lab in Milton Keynes and Alex Alderton, Roberto Amato, Jeffrey Barrett, Sonia Goncalves, Ewan Harrison, David K. Jackson, Ian Johnston, Dominic Kwiatkowski, Cordelia Langford, John Sillitoe on behalf of the Wellcome Sanger Institute COVID-19 Surveillance Team |
| EPI_ISL_2701395, EPI_ISL_2701396 | Lighthouse Lab in Alderley Park | Wellcome Sanger Institute for the COVID-19 Genomics UK (COG-UK) Consortium | Jacquelyn Wynn, Mairead Hyland, The Lighthouse Lab in Alderley Park and Alex Alderton, Roberto Amato, Jeffrey Barrett, Sonia Goncalves, Ewan Harrison, David K. Jackson, Ian Johnston, Dominic Kwiatkowski, Cordelia Langford, John Sillitoe on behalf of the Wellcome Sanger Institute COVID-19 Surveillance Team |
| EPI_ISL_2701397 | Lighthouse Lab in Glasgow | Wellcome Sanger Institute for the COVID-19 Genomics UK (COG-UK) Consortium | Harper VanSteenhouse, Yumi Kasai, David Gray, Carol Clugston, Anna Dominiczak and Alex Alderton, Roberto Amato, Jeffrey Barrett, Sonia Goncalves, Ewan Harrison, David K. Jackson, Ian Johnston, Dominic Kwiatkowski, Cordelia Langford, John Sillitoe on behalf of the Wellcome Sanger Institute COVID-19 Surveillance Team |
| EPI_ISL_2701398 | Lighthouse Lab in Alderley Park | Wellcome Sanger Institute for the COVID-19 Genomics UK (COG-UK) Consortium | Jacquelyn Wynn, Mairead Hyland, The Lighthouse Lab in Alderley Park and Alex Alderton, Roberto Amato, Jeffrey Barrett, Sonia Goncalves, Ewan Harrison, David K. Jackson, Ian Johnston, Dominic Kwiatkowski, Cordelia Langford, John Sillitoe on behalf of the Wellcome Sanger Institute COVID-19 Surveillance Team |
| EPI_ISL_2701399, EPI_ISL_2701400, EPI_ISL_2701401 | Lighthouse Lab in Glasgow | Wellcome Sanger Institute for the COVID-19 Genomics UK (COG-UK) Consortium | Harper VanSteenhouse, Yumi Kasai, David Gray, Carol Clugston, Anna Dominiczak and Alex Alderton, Roberto Amato, Jeffrey Barrett, Sonia Goncalves, Ewan Harrison, David K. Jackson, Ian Johnston, Dominic Kwiatkowski, Cordelia Langford, John Sillitoe on behalf of the Wellcome Sanger Institute COVID-19 Surveillance Team |
| EPI_ISL_2701402, EPI_ISL_2701403 | Lighthouse Lab in Milton Keynes | Wellcome Sanger Institute for the COVID-19 Genomics UK (COG-UK) Consortium | The Lighthouse Lab in Milton Keynes and Alex Alderton, Roberto Amato, Jeffrey Barrett, Sonia Goncalves, Ewan Harrison, David K. Jackson, Ian Johnston, Dominic Kwiatkowski, Cordelia Langford, John Sillitoe on behalf of the Wellcome Sanger Institute COVID-19 Surveillance Team |
| EPI_ISL_2701404, EPI_ISL_2701405 | Lighthouse Lab in Glasgow | Wellcome Sanger Institute for the COVID-19 Genomics UK (COG-UK) Consortium | Harper VanSteenhouse, Yumi Kasai, David Gray, Carol Clugston, Anna Dominiczak and Alex Alderton, Roberto Amato, Jeffrey Barrett, Sonia Goncalves, Ewan Harrison, David K. Jackson, Ian Johnston, Dominic Kwiatkowski, Cordelia Langford, John Sillitoe on behalf of the Wellcome Sanger Institute COVID-19 Surveillance Team |
| EPI_ISL_2701406 | Lighthouse Lab in Alderley Park | Wellcome Sanger Institute for the COVID-19 Genomics UK (COG-UK) Consortium | Jacquelyn Wynn, Mairead Hyland, The Lighthouse Lab in Alderley Park and Alex Alderton, Roberto Amato, Jeffrey Barrett, Sonia Goncalves, Ewan Harrison, David K. Jackson, Ian Johnston, Dominic Kwiatkowski, Cordelia Langford, John Sillitoe on behalf of the Wellcome Sanger Institute COVID-19 Surveillance Team |
| EPI_ISL_2701407 | Lighthouse Lab in Glasgow | Wellcome Sanger Institute for the COVID-19 Genomics UK (COG-UK) Consortium | Harper VanSteenhouse, Yumi Kasai, David Gray, Carol Clugston, Anna Dominiczak and Alex Alderton, Roberto Amato, Jeffrey Barrett, Sonia Goncalves, Ewan Harrison, David K. Jackson, Ian Johnston, Dominic Kwiatkowski, Cordelia Langford, John Sillitoe on behalf of the Wellcome Sanger Institute COVID-19 Surveillance Team |
| EPI_ISL_2701408 | Lighthouse Lab in Milton Keynes | Wellcome Sanger Institute for the COVID-19 Genomics UK (COG-UK) Consortium | The Lighthouse Lab in Milton Keynes and Alex Alderton, Roberto Amato, Jeffrey Barrett, Sonia Goncalves, Ewan Harrison, David K. Jackson, Ian Johnston, Dominic Kwiatkowski, Cordelia Langford, John Sillitoe on behalf of the Wellcome Sanger Institute COVID-19 Surveillance Team |
| EPI_ISL_2701409 | Lighthouse Lab in Glasgow | Wellcome Sanger Institute for the COVID-19 Genomics UK (COG-UK) Consortium | Harper VanSteenhouse, Yumi Kasai, David Gray, Carol Clugston, Anna Dominiczak and Alex Alderton, Roberto Amato, Jeffrey Barrett, Sonia Goncalves, Ewan Harrison, David K. Jackson, Ian Johnston, Dominic Kwiatkowski, Cordelia Langford, John Sillitoe on behalf of the Wellcome Sanger Institute COVID-19 Surveillance Team |
| EPI_ISL_2701410 | Lighthouse Lab in Alderley Park | Wellcome Sanger Institute for the COVID-19 Genomics UK (COG-UK) Consortium | Jacquelyn Wynn, Mairead Hyland, The Lighthouse Lab in Alderley Park and Alex Alderton, Roberto Amato, Jeffrey Barrett, Sonia Goncalves, Ewan Harrison, David K. Jackson, Ian Johnston, Dominic Kwiatkowski, Cordelia Langford, John Sillitoe on behalf of the Wellcome Sanger Institute COVID-19 Surveillance Team |
| EPI_ISL_2701411 | Lighthouse Lab in Glasgow | Wellcome Sanger Institute for the COVID-19 Genomics UK (COG-UK) Consortium | Harper VanSteenhouse, Yumi Kasai, David Gray, Carol Clugston, Anna Dominiczak and Alex Alderton, Roberto Amato, Jeffrey Barrett, Sonia Goncalves, Ewan Harrison, David K. Jackson, Ian Johnston, Dominic Kwiatkowski, Cordelia Langford, John Sillitoe on behalf of the Wellcome Sanger Institute COVID-19 Surveillance Team |
| EPI_ISL_2701412, EPI_ISL_2701413 | Lighthouse Lab in Alderley Park | Wellcome Sanger Institute for the COVID-19 Genomics UK (COG-UK) Consortium | Jacquelyn Wynn, Mairead Hyland, The Lighthouse Lab in Alderley Park and Alex Alderton, Roberto Amato, Jeffrey Barrett, Sonia Goncalves, Ewan Harrison, David K. Jackson, Ian Johnston, Dominic Kwiatkowski, Cordelia Langford, John Sillitoe on behalf of the Wellcome Sanger Institute COVID-19 Surveillance Team |
| EPI_ISL_2701414 | Lighthouse Lab in Milton Keynes | Wellcome Sanger Institute for the COVID-19 Genomics UK (COG-UK) Consortium | The Lighthouse Lab in Milton Keynes and Alex Alderton, Roberto Amato, Jeffrey Barrett, Sonia Goncalves, Ewan Harrison, David K. Jackson, Ian Johnston, Dominic Kwiatkowski, Cordelia Langford, John Sillitoe on behalf of the Wellcome Sanger Institute COVID-19 Surveillance Team |
| EPI_ISL_2701415 | Lighthouse Lab in Glasgow | Wellcome Sanger Institute for the COVID-19 Genomics UK (COG-UK) Consortium | Harper VanSteenhouse, Yumi Kasai, David Gray, Carol Clugston, Anna Dominiczak and Alex Alderton, Roberto Amato, Jeffrey Barrett, Sonia Goncalves, Ewan Harrison, David K. Jackson, Ian Johnston, Dominic Kwiatkowski, Cordelia Langford, John Sillitoe on behalf of the Wellcome Sanger Institute COVID-19 Surveillance Team |
| EPI_ISL_2701416 | Lighthouse Lab in Alderley Park | Wellcome Sanger Institute for the COVID-19 Genomics UK (COG-UK) Consortium | Jacquelyn Wynn, Mairead Hyland, The Lighthouse Lab in Alderley Park and Alex Alderton, Roberto Amato, Jeffrey Barrett, Sonia Goncalves, Ewan Harrison, David K. Jackson, Ian Johnston, Dominic Kwiatkowski, Cordelia Langford, John Sillitoe on behalf of the Wellcome Sanger Institute COVID-19 Surveillance Team |
| EPI_ISL_2701417 | Lighthouse Lab in Milton Keynes | Wellcome Sanger Institute for the COVID-19 Genomics UK (COG-UK) Consortium | The Lighthouse Lab in Milton Keynes and Alex Alderton, Roberto Amato, Jeffrey Barrett, Sonia Goncalves, Ewan Harrison, David K. Jackson, Ian Johnston, Dominic Kwiatkowski, Cordelia Langford, John Sillitoe on behalf of the Wellcome Sanger Institute COVID-19 Surveillance Team |
| EPI_ISL_2701418, EPI_ISL_2701419 | Lighthouse Lab in Alderley Park | Wellcome Sanger Institute for the COVID-19 Genomics UK (COG-UK) Consortium | Jacquelyn Wynn, Mairead Hyland, The Lighthouse Lab in Alderley Park and Alex Alderton, Roberto Amato, Jeffrey Barrett, Sonia Goncalves, Ewan Harrison, David K. Jackson, Ian Johnston, Dominic Kwiatkowski, Cordelia Langford, John Sillitoe on behalf of the Wellcome Sanger Institute COVID-19 Surveillance Team |
| EPI_ISL_2701420 | Lighthouse Lab in Milton Keynes | Wellcome Sanger Institute for the COVID-19 Genomics UK (COG-UK) Consortium | The Lighthouse Lab in Milton Keynes and Alex Alderton, Roberto Amato, Jeffrey Barrett, Sonia Goncalves, Ewan Harrison, David K. Jackson, Ian Johnston, Dominic Kwiatkowski, Cordelia Langford, John Sillitoe on behalf of the Wellcome Sanger Institute COVID-19 Surveillance Team |
| EPI_ISL_2701421 | Lighthouse Lab in Alderley Park | Wellcome Sanger Institute for the COVID-19 Genomics UK (COG-UK) Consortium | Jacquelyn Wynn, Mairead Hyland, The Lighthouse Lab in Alderley Park and Alex Alderton, Roberto Amato, Jeffrey Barrett, Sonia Goncalves, Ewan Harrison, David K. Jackson, Ian Johnston, Dominic Kwiatkowski, Cordelia Langford, John Sillitoe on behalf of the Wellcome Sanger Institute COVID-19 Surveillance Team |

| | | | |
|---|---|---|---|
| EPI_ISL_2701422 | Lighthouse Lab in Milton Keynes | Wellcome Sanger Institute for the COVID-19 Genomics UK (COG-UK) Consortium | The Lighthouse Lab in Milton Keynes and Alex Alderton, Roberto Amato, Jeffrey Barrett, Sonia Goncalves, Ewan Harrison, David K. Jackson, Ian Johnston, Dominic Kwiatkowski, Cordelia Langford, John Sillitoe on behalf of the Wellcome Sanger Institute COVID-19 Surveillance Team |
| EPI_ISL_2701423 | Lighthouse Lab in Glasgow | Wellcome Sanger Institute for the COVID-19 Genomics UK (COG-UK) Consortium | Harper VanSteenhouse, Yumi Kasai, David Gray, Carol Clugston, Anna Dominiczak and Alex Alderton, Roberto Amato, Jeffrey Barrett, Sonia Goncalves, Ewan Harrison, David K. Jackson, Ian Johnston, Dominic Kwiatkowski, Cordelia Langford, John Sillitoe on behalf of the Wellcome Sanger Institute COVID-19 Surveillance Team |
| EPI_ISL_2701424 | Lighthouse Lab in Milton Keynes | Wellcome Sanger Institute for the COVID-19 Genomics UK (COG-UK) Consortium | The Lighthouse Lab in Milton Keynes and Alex Alderton, Roberto Amato, Jeffrey Barrett, Sonia Goncalves, Ewan Harrison, David K. Jackson, Ian Johnston, Dominic Kwiatkowski, Cordelia Langford, John Sillitoe on behalf of the Wellcome Sanger Institute COVID-19 Surveillance Team |
| EPI_ISL_2701425, EPI_ISL_2701426 | Lighthouse Lab in Glasgow | Wellcome Sanger Institute for the COVID-19 Genomics UK (COG-UK) Consortium | Harper VanSteenhouse, Yumi Kasai, David Gray, Carol Clugston, Anna Dominiczak and Alex Alderton, Roberto Amato, Jeffrey Barrett, Sonia Goncalves, Ewan Harrison, David K. Jackson, Ian Johnston, Dominic Kwiatkowski, Cordelia Langford, John Sillitoe on behalf of the Wellcome Sanger Institute COVID-19 Surveillance Team |
| EPI_ISL_2701427 | Lighthouse Lab in Milton Keynes | Wellcome Sanger Institute for the COVID-19 Genomics UK (COG-UK) Consortium | The Lighthouse Lab in Milton Keynes and Alex Alderton, Roberto Amato, Jeffrey Barrett, Sonia Goncalves, Ewan Harrison, David K. Jackson, Ian Johnston, Dominic Kwiatkowski, Cordelia Langford, John Sillitoe on behalf of the Wellcome Sanger Institute COVID-19 Surveillance Team |
| EPI_ISL_2701428 | Lighthouse Lab in Glasgow | Wellcome Sanger Institute for the COVID-19 Genomics UK (COG-UK) Consortium | Harper VanSteenhouse, Yumi Kasai, David Gray, Carol Clugston, Anna Dominiczak and Alex Alderton, Roberto Amato, Jeffrey Barrett, Sonia Goncalves, Ewan Harrison, David K. Jackson, Ian Johnston, Dominic Kwiatkowski, Cordelia Langford, John Sillitoe on behalf of the Wellcome Sanger Institute COVID-19 Surveillance Team |
| EPI_ISL_2701429 | Lighthouse Lab in Alderley Park | Wellcome Sanger Institute for the COVID-19 Genomics UK (COG-UK) Consortium | Jacquelyn Wynn, Mairead Hyland, The Lighthouse Lab in Alderley Park and Alex Alderton, Roberto Amato, Jeffrey Barrett, Sonia Goncalves, Ewan Harrison, David K. Jackson, Ian Johnston, Dominic Kwiatkowski, Cordelia Langford, John Sillitoe on behalf of the Wellcome Sanger Institute COVID-19 Surveillance Team |
| EPI_ISL_2701430, EPI_ISL_2701431 | Lighthouse Lab in Glasgow | Wellcome Sanger Institute for the COVID-19 Genomics UK (COG-UK) Consortium | Harper VanSteenhouse, Yumi Kasai, David Gray, Carol Clugston, Anna Dominiczak and Alex Alderton, Roberto Amato, Jeffrey Barrett, Sonia Goncalves, Ewan Harrison, David K. Jackson, Ian Johnston, Dominic Kwiatkowski, Cordelia Langford, John Sillitoe on behalf of the Wellcome Sanger Institute COVID-19 Surveillance Team |
| EPI_ISL_2701432 | Lighthouse Lab in Milton Keynes | Wellcome Sanger Institute for the COVID-19 Genomics UK (COG-UK) Consortium | The Lighthouse Lab in Milton Keynes and Alex Alderton, Roberto Amato, Jeffrey Barrett, Sonia Goncalves, Ewan Harrison, David K. Jackson, Ian Johnston, Dominic Kwiatkowski, Cordelia Langford, John Sillitoe on behalf of the Wellcome Sanger Institute COVID-19 Surveillance Team |
| EPI_ISL_2701433 | Lighthouse Lab in Alderley Park | Wellcome Sanger Institute for the COVID-19 Genomics UK (COG-UK) Consortium | Jacquelyn Wynn, Mairead Hyland, The Lighthouse Lab in Alderley Park and Alex Alderton, Roberto Amato, Jeffrey Barrett, Sonia Goncalves, Ewan Harrison, David K. Jackson, Ian Johnston, Dominic Kwiatkowski, Cordelia Langford, John Sillitoe on behalf of the Wellcome Sanger Institute COVID-19 Surveillance Team |
| EPI_ISL_2701434 | Lighthouse Lab in Glasgow | Wellcome Sanger Institute for the COVID-19 Genomics UK (COG-UK) Consortium | Harper VanSteenhouse, Yumi Kasai, David Gray, Carol Clugston, Anna Dominiczak and Alex Alderton, Roberto Amato, Jeffrey Barrett, Sonia Goncalves, Ewan Harrison, David K. Jackson, Ian Johnston, Dominic Kwiatkowski, Cordelia Langford, John Sillitoe on behalf of the Wellcome Sanger Institute COVID-19 Surveillance Team |
| EPI_ISL_2701435 | Lighthouse Lab in Alderley Park | Wellcome Sanger Institute for the COVID-19 Genomics UK (COG-UK) Consortium | Jacquelyn Wynn, Mairead Hyland, The Lighthouse Lab in Alderley Park and Alex Alderton, Roberto Amato, Jeffrey Barrett, Sonia Goncalves, Ewan Harrison, David K. Jackson, Ian Johnston, Dominic Kwiatkowski, Cordelia Langford, John Sillitoe on behalf of the Wellcome Sanger Institute COVID-19 Surveillance Team |
| EPI_ISL_2701436, EPI_ISL_2701437 | Lighthouse Lab in Milton Keynes | Wellcome Sanger Institute for the COVID-19 Genomics UK (COG-UK) Consortium | The Lighthouse Lab in Milton Keynes and Alex Alderton, Roberto Amato, Jeffrey Barrett, Sonia Goncalves, Ewan Harrison, David K. Jackson, Ian Johnston, Dominic Kwiatkowski, Cordelia Langford, John Sillitoe on behalf of the Wellcome Sanger Institute COVID-19 Surveillance Team |
| EPI_ISL_2701438, EPI_ISL_2701439 | Lighthouse Lab in Glasgow | Wellcome Sanger Institute for the COVID-19 Genomics UK (COG-UK) Consortium | Harper VanSteenhouse, Yumi Kasai, David Gray, Carol Clugston, Anna Dominiczak and Alex Alderton, Roberto Amato, Jeffrey Barrett, Sonia Goncalves, Ewan Harrison, David K. Jackson, Ian Johnston, Dominic Kwiatkowski, Cordelia Langford, John Sillitoe on behalf of the Wellcome Sanger Institute COVID-19 Surveillance Team |
| EPI_ISL_2701440 | Lighthouse Lab in Alderley Park | Wellcome Sanger Institute for the COVID-19 Genomics UK (COG-UK) Consortium | Jacquelyn Wynn, Mairead Hyland, The Lighthouse Lab in Alderley Park and Alex Alderton, Roberto Amato, Jeffrey Barrett, Sonia Goncalves, Ewan Harrison, David K. Jackson, Ian Johnston, Dominic Kwiatkowski, Cordelia Langford, John Sillitoe on behalf of the Wellcome Sanger Institute COVID-19 Surveillance Team |
| EPI_ISL_2701441 | Lighthouse Lab in Glasgow | Wellcome Sanger Institute for the COVID-19 Genomics UK (COG-UK) Consortium | Harper VanSteenhouse, Yumi Kasai, David Gray, Carol Clugston, Anna Dominiczak and Alex Alderton, Roberto Amato, Jeffrey Barrett, Sonia Goncalves, Ewan Harrison, David K. Jackson, Ian Johnston, Dominic Kwiatkowski, Cordelia Langford, John Sillitoe on behalf of the Wellcome Sanger Institute COVID-19 Surveillance Team |
| EPI_ISL_2701442 | Lighthouse Lab in Alderley Park | Wellcome Sanger Institute for the COVID-19 Genomics UK (COG-UK) Consortium | Jacquelyn Wynn, Mairead Hyland, The Lighthouse Lab in Alderley Park and Alex Alderton, Roberto Amato, Jeffrey Barrett, Sonia Goncalves, Ewan Harrison, David K. Jackson, Ian Johnston, Dominic Kwiatkowski, Cordelia Langford, John Sillitoe on behalf of the Wellcome Sanger Institute COVID-19 Surveillance Team |
| EPI_ISL_2701443, EPI_ISL_2701444 | Lighthouse Lab in Glasgow | Wellcome Sanger Institute for the COVID-19 Genomics UK (COG-UK) Consortium | Harper VanSteenhouse, Yumi Kasai, David Gray, Carol Clugston, Anna Dominiczak and Alex Alderton, Roberto Amato, Jeffrey Barrett, Sonia Goncalves, Ewan Harrison, David K. Jackson, Ian Johnston, Dominic Kwiatkowski, Cordelia Langford, John Sillitoe on behalf of the Wellcome Sanger Institute COVID-19 Surveillance Team |
| EPI_ISL_2701445 | Lighthouse Lab in Milton Keynes | Wellcome Sanger Institute for the COVID-19 Genomics UK (COG-UK) Consortium | The Lighthouse Lab in Milton Keynes and Alex Alderton, Roberto Amato, Jeffrey Barrett, Sonia Goncalves, Ewan Harrison, David K. Jackson, Ian Johnston, Dominic Kwiatkowski, Cordelia Langford, John Sillitoe on behalf of the Wellcome Sanger Institute COVID-19 Surveillance Team |
| EPI_ISL_2701446, EPI_ISL_2701447 | Lighthouse Lab in Alderley Park | Wellcome Sanger Institute for the COVID-19 Genomics UK (COG-UK) Consortium | Jacquelyn Wynn, Mairead Hyland, The Lighthouse Lab in Alderley Park and Alex Alderton, Roberto Amato, Jeffrey Barrett, Sonia Goncalves, Ewan Harrison, David K. Jackson, Ian Johnston, Dominic Kwiatkowski, Cordelia Langford, John Sillitoe on behalf of the Wellcome Sanger Institute COVID-19 Surveillance Team |
| EPI_ISL_2701448 | Lighthouse Lab in Glasgow | Wellcome Sanger Institute for the COVID-19 Genomics UK (COG-UK) Consortium | Harper VanSteenhouse, Yumi Kasai, David Gray, Carol Clugston, Anna Dominiczak and Alex Alderton, Roberto Amato, Jeffrey Barrett, Sonia Goncalves, Ewan Harrison, David K. Jackson, Ian Johnston, Dominic Kwiatkowski, Cordelia Langford, John Sillitoe on behalf of the Wellcome Sanger Institute COVID-19 Surveillance Team |
| EPI_ISL_2701449 | Lighthouse Lab in Milton Keynes | Wellcome Sanger Institute for the COVID-19 Genomics UK (COG-UK) Consortium | The Lighthouse Lab in Milton Keynes and Alex Alderton, Roberto Amato, Jeffrey Barrett, Sonia Goncalves, Ewan Harrison, David K. Jackson, Ian Johnston, Dominic Kwiatkowski, Cordelia Langford, John Sillitoe on behalf of the Wellcome Sanger Institute COVID-19 Surveillance Team |
| EPI_ISL_2701450, EPI_ISL_2701451 | Lighthouse Lab in Alderley Park | Wellcome Sanger Institute for the COVID-19 Genomics UK (COG-UK) Consortium | Jacquelyn Wynn, Mairead Hyland, The Lighthouse Lab in Alderley Park and Alex Alderton, Roberto Amato, Jeffrey Barrett, Sonia Goncalves, Ewan Harrison, David K. Jackson, Ian Johnston, Dominic Kwiatkowski, Cordelia Langford, John Sillitoe on behalf of the Wellcome Sanger Institute COVID-19 Surveillance Team |
| EPI_ISL_2701452 | Lighthouse Lab in Glasgow | Wellcome Sanger Institute for the COVID-19 Genomics UK (COG-UK) Consortium | Harper VanSteenhouse, Yumi Kasai, David Gray, Carol Clugston, Anna Dominiczak and Alex Alderton, Roberto Amato, Jeffrey Barrett, Sonia Goncalves, Ewan Harrison, David K. Jackson, Ian Johnston, Dominic Kwiatkowski, Cordelia Langford, John Sillitoe on behalf of the Wellcome Sanger Institute COVID-19 Surveillance Team |
| EPI_ISL_2701453 | Lighthouse Lab in Milton Keynes | Wellcome Sanger Institute for the COVID-19 Genomics UK (COG-UK) Consortium | The Lighthouse Lab in Milton Keynes and Alex Alderton, Roberto Amato, Jeffrey Barrett, Sonia Goncalves, Ewan Harrison, David K. Jackson, Ian Johnston, Dominic Kwiatkowski, Cordelia Langford, John Sillitoe on behalf of the Wellcome Sanger Institute COVID-19 Surveillance Team |
| EPI_ISL_2701454, EPI_ISL_2701455, EPI_ISL_2701456, EPI_ISL_2701457, EPI_ISL_2701458, EPI_ISL_2701459, EPI_ISL_2701460 | Lighthouse Lab in Glasgow | Wellcome Sanger Institute for the COVID-19 Genomics UK (COG-UK) Consortium | Harper VanSteenhouse, Yumi Kasai, David Gray, Carol Clugston, Anna Dominiczak and Alex Alderton, Roberto Amato, Jeffrey Barrett, Sonia Goncalves, Ewan Harrison, David K. Jackson, Ian Johnston, Dominic Kwiatkowski, Cordelia Langford, John Sillitoe on behalf of the Wellcome Sanger Institute COVID-19 Surveillance Team |
| EPI_ISL_2701461 | Lighthouse Lab in Alderley Park | Wellcome Sanger Institute for the COVID-19 Genomics UK (COG-UK) Consortium | Jacquelyn Wynn, Mairead Hyland, The Lighthouse Lab in Alderley Park and Alex Alderton, Roberto Amato, Jeffrey Barrett, Sonia Goncalves, Ewan Harrison, David K. Jackson, Ian Johnston, Dominic Kwiatkowski, Cordelia Langford, John Sillitoe on behalf of the Wellcome Sanger Institute COVID-19 Surveillance Team |
| EPI_ISL_2701462 | Lighthouse Lab in Glasgow | Wellcome Sanger Institute for the COVID-19 Genomics UK (COG-UK) Consortium | Harper VanSteenhouse, Yumi Kasai, David Gray, Carol Clugston, Anna Dominiczak and Alex Alderton, Roberto Amato, Jeffrey Barrett, Sonia Goncalves, Ewan Harrison, David K. Jackson, Ian Johnston, Dominic Kwiatkowski, Cordelia Langford, John Sillitoe on behalf of the Wellcome Sanger Institute COVID-19 Surveillance Team |
| EPI_ISL_2701463 | Lighthouse Lab in Alderley Park | Wellcome Sanger Institute for the COVID-19 Genomics UK (COG-UK) Consortium | Jacquelyn Wynn, Mairead Hyland, The Lighthouse Lab in Alderley Park and Alex Alderton, Roberto Amato, Jeffrey Barrett, Sonia Goncalves, Ewan Harrison, David K. Jackson, Ian Johnston, Dominic Kwiatkowski, Cordelia Langford, John Sillitoe on behalf of the Wellcome Sanger Institute COVID-19 Surveillance Team |
| EPI_ISL_2701464 | Lighthouse Lab in Glasgow | Wellcome Sanger Institute for the COVID-19 Genomics UK (COG-UK) Consortium | Harper VanSteenhouse, Yumi Kasai, David Gray, Carol Clugston, Anna Dominiczak and Alex Alderton, Roberto Amato, Jeffrey Barrett, Sonia Goncalves, Ewan Harrison, David K. Jackson, Ian Johnston, Dominic Kwiatkowski, Cordelia Langford, John Sillitoe on behalf of the Wellcome Sanger Institute COVID-19 Surveillance Team |
| EPI_ISL_2701465 | Lighthouse Lab in Alderley Park | Wellcome Sanger Institute for the COVID-19 Genomics UK (COG-UK) Consortium | Jacquelyn Wynn, Mairead Hyland, The Lighthouse Lab in Alderley Park and Alex Alderton, Roberto Amato, Jeffrey Barrett, Sonia Goncalves, Ewan Harrison, David K. Jackson, Ian Johnston, Dominic Kwiatkowski, Cordelia Langford, John Sillitoe on behalf of the Wellcome Sanger Institute COVID-19 Surveillance Team |
| EPI_ISL_2701466 | Lighthouse Lab in Milton Keynes | Wellcome Sanger Institute for the COVID-19 Genomics UK (COG-UK) Consortium | The Lighthouse Lab in Milton Keynes and Alex Alderton, Roberto Amato, Jeffrey Barrett, Sonia Goncalves, Ewan Harrison, David K. Jackson, Ian Johnston, Dominic Kwiatkowski, Cordelia Langford, John Sillitoe on behalf of the Wellcome Sanger Institute COVID-19 Surveillance Team |
| EPI_ISL_2701467 | Lighthouse Lab in Alderley Park | Wellcome Sanger Institute for the COVID-19 Genomics UK (COG-UK) Consortium | Jacquelyn Wynn, Mairead Hyland, The Lighthouse Lab in Alderley Park and Alex Alderton, Roberto Amato, Jeffrey Barrett, Sonia Goncalves, Ewan Harrison, David K. Jackson, Ian Johnston, Dominic Kwiatkowski, Cordelia Langford, John Sillitoe on behalf of the Wellcome Sanger Institute COVID-19 Surveillance Team |
| EPI_ISL_2701468, EPI_ISL_2701469, EPI_ISL_2701470, EPI_ISL_2701471 | Lighthouse Lab in Glasgow | Wellcome Sanger Institute for the COVID-19 Genomics UK (COG-UK) Consortium | Harper VanSteenhouse, Yumi Kasai, David Gray, Carol Clugston, Anna Dominiczak and Alex Alderton, Roberto Amato, Jeffrey Barrett, Sonia Goncalves, Ewan Harrison, David K. Jackson, Ian Johnston, Dominic Kwiatkowski, Cordelia Langford, John Sillitoe on behalf of the Wellcome Sanger Institute COVID-19 Surveillance Team |
| EPI_ISL_2701472 | Lighthouse Lab in Alderley Park | Wellcome Sanger Institute for the COVID-19 Genomics UK (COG-UK) Consortium | Jacquelyn Wynn, Mairead Hyland, The Lighthouse Lab in Alderley Park and Alex Alderton, Roberto Amato, Jeffrey Barrett, Sonia Goncalves, Ewan Harrison, David K. Jackson, Ian Johnston, Dominic Kwiatkowski, Cordelia Langford, John Sillitoe on behalf of the Wellcome Sanger Institute COVID-19 Surveillance Team |
| EPI_ISL_2701473 | Lighthouse Lab in Milton Keynes | Wellcome Sanger Institute for the COVID-19 Genomics UK (COG-UK) Consortium | The Lighthouse Lab in Milton Keynes and Alex Alderton, Roberto Amato, Jeffrey Barrett, Sonia Goncalves, Ewan Harrison, David K. Jackson, Ian Johnston, Dominic Kwiatkowski, Cordelia Langford, John Sillitoe on behalf of the Wellcome Sanger Institute COVID-19 Surveillance Team |
| EPI_ISL_2701474, EPI_ISL_2701475, EPI_ISL_2701476 | Lighthouse Lab in Glasgow | Wellcome Sanger Institute for the COVID-19 Genomics UK (COG-UK) Consortium | Harper VanSteenhouse, Yumi Kasai, David Gray, Carol Clugston, Anna Dominiczak and Alex Alderton, Roberto Amato, Jeffrey Barrett, Sonia Goncalves, Ewan Harrison, David K. Jackson, Ian Johnston, Dominic Kwiatkowski, Cordelia Langford, John Sillitoe on behalf of the Wellcome Sanger Institute COVID-19 Surveillance Team |
| EPI_ISL_2701477 | Lighthouse Lab in Alderley Park | Wellcome Sanger Institute for the COVID-19 Genomics UK (COG-UK) Consortium | Jacquelyn Wynn, Mairead Hyland, The Lighthouse Lab in Alderley Park and Alex Alderton, Roberto Amato, Jeffrey Barrett, Sonia Goncalves, Ewan Harrison, David K. Jackson, Ian Johnston, Dominic Kwiatkowski, Cordelia Langford, John Sillitoe on behalf of the Wellcome Sanger Institute COVID-19 Surveillance Team |
| EPI_ISL_2701478, EPI_ISL_2701479, EPI_ISL_2701480 | Lighthouse Lab in Glasgow | Wellcome Sanger Institute for the COVID-19 Genomics UK (COG-UK) Consortium | Harper VanSteenhouse, Yumi Kasai, David Gray, Carol Clugston, Anna Dominiczak and Alex Alderton, Roberto Amato, Jeffrey Barrett, Sonia Goncalves, Ewan Harrison, David K. Jackson, Ian Johnston, Dominic Kwiatkowski, Cordelia Langford, John Sillitoe on behalf of the Wellcome Sanger Institute COVID-19 Surveillance Team |
| EPI_ISL_2701481 | Lighthouse Lab in Milton Keynes | Wellcome Sanger Institute for the COVID-19 Genomics UK (COG-UK) Consortium | The Lighthouse Lab in Milton Keynes and Alex Alderton, Roberto Amato, Jeffrey Barrett, Sonia Goncalves, Ewan Harrison, David K. Jackson, Ian Johnston, Dominic Kwiatkowski, Cordelia Langford, John Sillitoe on behalf of the Wellcome Sanger Institute COVID-19 Surveillance Team |
| EPI_ISL_2701482 | Lighthouse Lab in Glasgow | Wellcome Sanger Institute for the COVID-19 Genomics UK (COG-UK) Consortium | Harper VanSteenhouse, Yumi Kasai, David Gray, Carol Clugston, Anna Dominiczak and Alex Alderton, Roberto Amato, Jeffrey Barrett, Sonia Goncalves, Ewan Harrison, David K. Jackson, Ian Johnston, Dominic Kwiatkowski, Cordelia Langford, John Sillitoe on behalf of the Wellcome Sanger Institute COVID-19 Surveillance Team |
| EPI_ISL_2701483, | Lighthouse Lab in Alderley Park | Wellcome Sanger Institute for the COVID-19 Genomics UK (COG-UK) | Jacquelyn Wynn, Mairead Hyland, The Lighthouse Lab in Alderley Park and Alex Alderton, Roberto Amato, Jeffrey Barrett, Sonia Goncalves, Ewan Harrison, David K. Jackson, Ian Johnston, Dominic Kwiatkowski, Cordelia Langford, John Sillitoe on |

| | | | |
|---|---|---|---|
| EPI_ISL_2701484 | | Consortium | behalf of the Wellcome Sanger Institute COVID-19 Surveillance Team |
| EPI_ISL_2701485 | Lighthouse Lab in Milton Keynes | Wellcome Sanger Institute for the COVID-19 Genomics UK (COG-UK) Consortium | The Lighthouse Lab in Milton Keynes and Alex Alderton, Roberto Amato, Jeffrey Barrett, Sonia Goncalves, Ewan Harrison, David K. Jackson, Ian Johnston, Dominic Kwiatkowski, Cordelia Langford, John Sillitoe on behalf of the Wellcome Sanger Institute COVID-19 Surveillance Team |
| EPI_ISL_2701486 | Lighthouse Lab in Glasgow | Wellcome Sanger Institute for the COVID-19 Genomics UK (COG-UK) Consortium | Harper VanSteenhouse, Yumi Kasai, David Gray, Carol Clugston, Anna Dominiczak and Alex Alderton, Roberto Amato, Jeffrey Barrett, Sonia Goncalves, Ewan Harrison, David K. Jackson, Ian Johnston, Dominic Kwiatkowski, Cordelia Langford, John Sillitoe on behalf of the Wellcome Sanger Institute COVID-19 Surveillance Team |
| EPI_ISL_2701487 | Lighthouse Lab in Milton Keynes | Wellcome Sanger Institute for the COVID-19 Genomics UK (COG-UK) Consortium | The Lighthouse Lab in Milton Keynes and Alex Alderton, Roberto Amato, Jeffrey Barrett, Sonia Goncalves, Ewan Harrison, David K. Jackson, Ian Johnston, Dominic Kwiatkowski, Cordelia Langford, John Sillitoe on behalf of the Wellcome Sanger Institute COVID-19 Surveillance Team |
| EPI_ISL_2701488, EPI_ISL_2701489, EPI_ISL_2701490, EPI_ISL_2701491, EPI_ISL_2701492 | Lighthouse Lab in Glasgow | Wellcome Sanger Institute for the COVID-19 Genomics UK (COG-UK) Consortium | Harper VanSteenhouse, Yumi Kasai, David Gray, Carol Clugston, Anna Dominiczak and Alex Alderton, Roberto Amato, Jeffrey Barrett, Sonia Goncalves, Ewan Harrison, David K. Jackson, Ian Johnston, Dominic Kwiatkowski, Cordelia Langford, John Sillitoe on behalf of the Wellcome Sanger Institute COVID-19 Surveillance Team |
| EPI_ISL_2701493 | Lighthouse Lab in Milton Keynes | Wellcome Sanger Institute for the COVID-19 Genomics UK (COG-UK) Consortium | The Lighthouse Lab in Milton Keynes and Alex Alderton, Roberto Amato, Jeffrey Barrett, Sonia Goncalves, Ewan Harrison, David K. Jackson, Ian Johnston, Dominic Kwiatkowski, Cordelia Langford, John Sillitoe on behalf of the Wellcome Sanger Institute COVID-19 Surveillance Team |
| EPI_ISL_2701494 | Lighthouse Lab in Alderley Park | Wellcome Sanger Institute for the COVID-19 Genomics UK (COG-UK) Consortium | Jacquelyn Wynn, Mairead Hyland, The Lighthouse Lab in Alderley Park and Alex Alderton, Roberto Amato, Jeffrey Barrett, Sonia Goncalves, Ewan Harrison, David K. Jackson, Ian Johnston, Dominic Kwiatkowski, Cordelia Langford, John Sillitoe on behalf of the Wellcome Sanger Institute COVID-19 Surveillance Team |
| EPI_ISL_2701495, EPI_ISL_2701496, EPI_ISL_2701497 | Lighthouse Lab in Milton Keynes | Wellcome Sanger Institute for the COVID-19 Genomics UK (COG-UK) Consortium | The Lighthouse Lab in Milton Keynes and Alex Alderton, Roberto Amato, Jeffrey Barrett, Sonia Goncalves, Ewan Harrison, David K. Jackson, Ian Johnston, Dominic Kwiatkowski, Cordelia Langford, John Sillitoe on behalf of the Wellcome Sanger Institute COVID-19 Surveillance Team |
| EPI_ISL_2701498 | Lighthouse Lab in Glasgow | Wellcome Sanger Institute for the COVID-19 Genomics UK (COG-UK) Consortium | Harper VanSteenhouse, Yumi Kasai, David Gray, Carol Clugston, Anna Dominiczak and Alex Alderton, Roberto Amato, Jeffrey Barrett, Sonia Goncalves, Ewan Harrison, David K. Jackson, Ian Johnston, Dominic Kwiatkowski, Cordelia Langford, John Sillitoe on behalf of the Wellcome Sanger Institute COVID-19 Surveillance Team |
| EPI_ISL_2701499, EPI_ISL_2701500 | Lighthouse Lab in Milton Keynes | Wellcome Sanger Institute for the COVID-19 Genomics UK (COG-UK) Consortium | The Lighthouse Lab in Milton Keynes and Alex Alderton, Roberto Amato, Jeffrey Barrett, Sonia Goncalves, Ewan Harrison, David K. Jackson, Ian Johnston, Dominic Kwiatkowski, Cordelia Langford, John Sillitoe on behalf of the Wellcome Sanger Institute COVID-19 Surveillance Team |
| EPI_ISL_2701501, EPI_ISL_2701502, EPI_ISL_2701503 | Lighthouse Lab in Alderley Park | Wellcome Sanger Institute for the COVID-19 Genomics UK (COG-UK) Consortium | Jacquelyn Wynn, Mairead Hyland, The Lighthouse Lab in Alderley Park and Alex Alderton, Roberto Amato, Jeffrey Barrett, Sonia Goncalves, Ewan Harrison, David K. Jackson, Ian Johnston, Dominic Kwiatkowski, Cordelia Langford, John Sillitoe on behalf of the Wellcome Sanger Institute COVID-19 Surveillance Team |
| EPI_ISL_2701504 | Lighthouse Lab in Milton Keynes | Wellcome Sanger Institute for the COVID-19 Genomics UK (COG-UK) Consortium | The Lighthouse Lab in Milton Keynes and Alex Alderton, Roberto Amato, Jeffrey Barrett, Sonia Goncalves, Ewan Harrison, David K. Jackson, Ian Johnston, Dominic Kwiatkowski, Cordelia Langford, John Sillitoe on behalf of the Wellcome Sanger Institute COVID-19 Surveillance Team |
| EPI_ISL_2701505, EPI_ISL_2701508 | Lighthouse Lab in Glasgow | Wellcome Sanger Institute for the COVID-19 Genomics UK (COG-UK) Consortium | Harper VanSteenhouse, Yumi Kasai, David Gray, Carol Clugston, Anna Dominiczak and Alex Alderton, Roberto Amato, Jeffrey Barrett, Sonia Goncalves, Ewan Harrison, David K. Jackson, Ian Johnston, Dominic Kwiatkowski, Cordelia Langford, John Sillitoe on behalf of the Wellcome Sanger Institute COVID-19 Surveillance Team |
| EPI_ISL_2701509 | Lighthouse Lab in Milton Keynes | Wellcome Sanger Institute for the COVID-19 Genomics UK (COG-UK) Consortium | The Lighthouse Lab in Milton Keynes and Alex Alderton, Roberto Amato, Jeffrey Barrett, Sonia Goncalves, Ewan Harrison, David K. Jackson, Ian Johnston, Dominic Kwiatkowski, Cordelia Langford, John Sillitoe on behalf of the Wellcome Sanger Institute COVID-19 Surveillance Team |
| EPI_ISL_2701510, EPI_ISL_2701511 | Lighthouse Lab in Alderley Park | Wellcome Sanger Institute for the COVID-19 Genomics UK (COG-UK) Consortium | Jacquelyn Wynn, Mairead Hyland, The Lighthouse Lab in Alderley Park and Alex Alderton, Roberto Amato, Jeffrey Barrett, Sonia Goncalves, Ewan Harrison, David K. Jackson, Ian Johnston, Dominic Kwiatkowski, Cordelia Langford, John Sillitoe on behalf of the Wellcome Sanger Institute COVID-19 Surveillance Team |
| EPI_ISL_2701512, EPI_ISL_2701513, EPI_ISL_2701514 | Lighthouse Lab in Glasgow | Wellcome Sanger Institute for the COVID-19 Genomics UK (COG-UK) Consortium | Harper VanSteenhouse, Yumi Kasai, David Gray, Carol Clugston, Anna Dominiczak and Alex Alderton, Roberto Amato, Jeffrey Barrett, Sonia Goncalves, Ewan Harrison, David K. Jackson, Ian Johnston, Dominic Kwiatkowski, Cordelia Langford, John Sillitoe on behalf of the Wellcome Sanger Institute COVID-19 Surveillance Team |
| EPI_ISL_2701515 | Lighthouse Lab in Milton Keynes | Wellcome Sanger Institute for the COVID-19 Genomics UK (COG-UK) Consortium | The Lighthouse Lab in Milton Keynes and Alex Alderton, Roberto Amato, Jeffrey Barrett, Sonia Goncalves, Ewan Harrison, David K. Jackson, Ian Johnston, Dominic Kwiatkowski, Cordelia Langford, John Sillitoe on behalf of the Wellcome Sanger Institute COVID-19 Surveillance Team |
| EPI_ISL_2701516, EPI_ISL_2701517, EPI_ISL_2701518, EPI_ISL_2701519 | Lighthouse Lab in Glasgow | Wellcome Sanger Institute for the COVID-19 Genomics UK (COG-UK) Consortium | Harper VanSteenhouse, Yumi Kasai, David Gray, Carol Clugston, Anna Dominiczak and Alex Alderton, Roberto Amato, Jeffrey Barrett, Sonia Goncalves, Ewan Harrison, David K. Jackson, Ian Johnston, Dominic Kwiatkowski, Cordelia Langford, John Sillitoe on behalf of the Wellcome Sanger Institute COVID-19 Surveillance Team |
| EPI_ISL_2701520, EPI_ISL_2701521 | Lighthouse Lab in Milton Keynes | Wellcome Sanger Institute for the COVID-19 Genomics UK (COG-UK) Consortium | The Lighthouse Lab in Milton Keynes and Alex Alderton, Roberto Amato, Jeffrey Barrett, Sonia Goncalves, Ewan Harrison, David K. Jackson, Ian Johnston, Dominic Kwiatkowski, Cordelia Langford, John Sillitoe on behalf of the Wellcome Sanger Institute COVID-19 Surveillance Team |
| EPI_ISL_2701522, EPI_ISL_2701523 | Lighthouse Lab in Glasgow | Wellcome Sanger Institute for the COVID-19 Genomics UK (COG-UK) Consortium | Harper VanSteenhouse, Yumi Kasai, David Gray, Carol Clugston, Anna Dominiczak and Alex Alderton, Roberto Amato, Jeffrey Barrett, Sonia Goncalves, Ewan Harrison, David K. Jackson, Ian Johnston, Dominic Kwiatkowski, Cordelia Langford, John Sillitoe on behalf of the Wellcome Sanger Institute COVID-19 Surveillance Team |
| EPI_ISL_2701524 | Lighthouse Lab in Alderley Park | Wellcome Sanger Institute for the COVID-19 Genomics UK (COG-UK) Consortium | Jacquelyn Wynn, Mairead Hyland, The Lighthouse Lab in Alderley Park and Alex Alderton, Roberto Amato, Jeffrey Barrett, Sonia Goncalves, Ewan Harrison, David K. Jackson, Ian Johnston, Dominic Kwiatkowski, Cordelia Langford, John Sillitoe on behalf of the Wellcome Sanger Institute COVID-19 Surveillance Team |
| EPI_ISL_2701525 | Lighthouse Lab in Glasgow | Wellcome Sanger Institute for the COVID-19 Genomics UK (COG-UK) Consortium | Harper VanSteenhouse, Yumi Kasai, David Gray, Carol Clugston, Anna Dominiczak and Alex Alderton, Roberto Amato, Jeffrey Barrett, Sonia Goncalves, Ewan Harrison, David K. Jackson, Ian Johnston, Dominic Kwiatkowski, Cordelia Langford, John Sillitoe on behalf of the Wellcome Sanger Institute COVID-19 Surveillance Team |
| EPI_ISL_2701526 | Lighthouse Lab in Milton Keynes | Wellcome Sanger Institute for the COVID-19 Genomics UK (COG-UK) Consortium | The Lighthouse Lab in Milton Keynes and Alex Alderton, Roberto Amato, Jeffrey Barrett, Sonia Goncalves, Ewan Harrison, David K. Jackson, Ian Johnston, Dominic Kwiatkowski, Cordelia Langford, John Sillitoe on behalf of the Wellcome Sanger Institute COVID-19 Surveillance Team |
| EPI_ISL_2701527, EPI_ISL_2701528, EPI_ISL_2701529, EPI_ISL_2701530 | Lighthouse Lab in Glasgow | Wellcome Sanger Institute for the COVID-19 Genomics UK (COG-UK) Consortium | Harper VanSteenhouse, Yumi Kasai, David Gray, Carol Clugston, Anna Dominiczak and Alex Alderton, Roberto Amato, Jeffrey Barrett, Sonia Goncalves, Ewan Harrison, David K. Jackson, Ian Johnston, Dominic Kwiatkowski, Cordelia Langford, John Sillitoe on behalf of the Wellcome Sanger Institute COVID-19 Surveillance Team |
| EPI_ISL_2701531 | Lighthouse Lab in Milton Keynes | Wellcome Sanger Institute for the COVID-19 Genomics UK (COG-UK) Consortium | The Lighthouse Lab in Milton Keynes and Alex Alderton, Roberto Amato, Jeffrey Barrett, Sonia Goncalves, Ewan Harrison, David K. Jackson, Ian Johnston, Dominic Kwiatkowski, Cordelia Langford, John Sillitoe on behalf of the Wellcome Sanger Institute COVID-19 Surveillance Team |
| EPI_ISL_2701532 | Lighthouse Lab in Alderley Park | Wellcome Sanger Institute for the COVID-19 Genomics UK (COG-UK) Consortium | Jacquelyn Wynn, Mairead Hyland, The Lighthouse Lab in Alderley Park and Alex Alderton, Roberto Amato, Jeffrey Barrett, Sonia Goncalves, Ewan Harrison, David K. Jackson, Ian Johnston, Dominic Kwiatkowski, Cordelia Langford, John Sillitoe on behalf of the Wellcome Sanger Institute COVID-19 Surveillance Team |
| EPI_ISL_2701533 | Lighthouse Lab in Milton Keynes | Wellcome Sanger Institute for the COVID-19 Genomics UK (COG-UK) Consortium | The Lighthouse Lab in Milton Keynes and Alex Alderton, Roberto Amato, Jeffrey Barrett, Sonia Goncalves, Ewan Harrison, David K. Jackson, Ian Johnston, Dominic Kwiatkowski, Cordelia Langford, John Sillitoe on behalf of the Wellcome Sanger Institute COVID-19 Surveillance Team |
| EPI_ISL_2701534 | Lighthouse Lab in Glasgow | Wellcome Sanger Institute for the COVID-19 Genomics UK (COG-UK) Consortium | Harper VanSteenhouse, Yumi Kasai, David Gray, Carol Clugston, Anna Dominiczak and Alex Alderton, Roberto Amato, Jeffrey Barrett, Sonia Goncalves, Ewan Harrison, David K. Jackson, Ian Johnston, Dominic Kwiatkowski, Cordelia Langford, John Sillitoe on behalf of the Wellcome Sanger Institute COVID-19 Surveillance Team |
| EPI_ISL_2701535 | Lighthouse Lab in Alderley Park | Wellcome Sanger Institute for the COVID-19 Genomics UK (COG-UK) Consortium | Jacquelyn Wynn, Mairead Hyland, The Lighthouse Lab in Alderley Park and Alex Alderton, Roberto Amato, Jeffrey Barrett, Sonia Goncalves, Ewan Harrison, David K. Jackson, Ian Johnston, Dominic Kwiatkowski, Cordelia Langford, John Sillitoe on behalf of the Wellcome Sanger Institute COVID-19 Surveillance Team |
| EPI_ISL_2701536, EPI_ISL_2701537 | Lighthouse Lab in Glasgow | Wellcome Sanger Institute for the COVID-19 Genomics UK (COG-UK) Consortium | Harper VanSteenhouse, Yumi Kasai, David Gray, Carol Clugston, Anna Dominiczak and Alex Alderton, Roberto Amato, Jeffrey Barrett, Sonia Goncalves, Ewan Harrison, David K. Jackson, Ian Johnston, Dominic Kwiatkowski, Cordelia Langford, John Sillitoe on behalf of the Wellcome Sanger Institute COVID-19 Surveillance Team |
| EPI_ISL_2701538 | Lighthouse Lab in Milton Keynes | Wellcome Sanger Institute for the COVID-19 Genomics UK (COG-UK) Consortium | The Lighthouse Lab in Milton Keynes and Alex Alderton, Roberto Amato, Jeffrey Barrett, Sonia Goncalves, Ewan Harrison, David K. Jackson, Ian Johnston, Dominic Kwiatkowski, Cordelia Langford, John Sillitoe on behalf of the Wellcome Sanger Institute COVID-19 Surveillance Team |
| EPI_ISL_2701539, EPI_ISL_2701540, EPI_ISL_2701541, EPI_ISL_2701543 | Lighthouse Lab in Glasgow | Wellcome Sanger Institute for the COVID-19 Genomics UK (COG-UK) Consortium | Harper VanSteenhouse, Yumi Kasai, David Gray, Carol Clugston, Anna Dominiczak and Alex Alderton, Roberto Amato, Jeffrey Barrett, Sonia Goncalves, Ewan Harrison, David K. Jackson, Ian Johnston, Dominic Kwiatkowski, Cordelia Langford, John Sillitoe on behalf of the Wellcome Sanger Institute COVID-19 Surveillance Team |
| EPI_ISL_2701544 | Lighthouse Lab in Alderley Park | Wellcome Sanger Institute for the COVID-19 Genomics UK (COG-UK) Consortium | Jacquelyn Wynn, Mairead Hyland, The Lighthouse Lab in Alderley Park and Alex Alderton, Roberto Amato, Jeffrey Barrett, Sonia Goncalves, Ewan Harrison, David K. Jackson, Ian Johnston, Dominic Kwiatkowski, Cordelia Langford, John Sillitoe on behalf of the Wellcome Sanger Institute COVID-19 Surveillance Team |
| EPI_ISL_2701545, EPI_ISL_2701546, EPI_ISL_2701547, EPI_ISL_2701548 | Lighthouse Lab in Glasgow | Wellcome Sanger Institute for the COVID-19 Genomics UK (COG-UK) Consortium | Harper VanSteenhouse, Yumi Kasai, David Gray, Carol Clugston, Anna Dominiczak and Alex Alderton, Roberto Amato, Jeffrey Barrett, Sonia Goncalves, Ewan Harrison, David K. Jackson, Ian Johnston, Dominic Kwiatkowski, Cordelia Langford, John Sillitoe on behalf of the Wellcome Sanger Institute COVID-19 Surveillance Team |
| EPI_ISL_2701549 | Lighthouse Lab in Milton Keynes | Wellcome Sanger Institute for the COVID-19 Genomics UK (COG-UK) Consortium | The Lighthouse Lab in Milton Keynes and Alex Alderton, Roberto Amato, Jeffrey Barrett, Sonia Goncalves, Ewan Harrison, David K. Jackson, Ian Johnston, Dominic Kwiatkowski, Cordelia Langford, John Sillitoe on behalf of the Wellcome Sanger Institute COVID-19 Surveillance Team |
| EPI_ISL_2701550 | Lighthouse Lab in Alderley Park | Wellcome Sanger Institute for the COVID-19 Genomics UK (COG-UK) Consortium | Jacquelyn Wynn, Mairead Hyland, The Lighthouse Lab in Alderley Park and Alex Alderton, Roberto Amato, Jeffrey Barrett, Sonia Goncalves, Ewan Harrison, David K. Jackson, Ian Johnston, Dominic Kwiatkowski, Cordelia Langford, John Sillitoe on behalf of the Wellcome Sanger Institute COVID-19 Surveillance Team |
| EPI_ISL_2701551, EPI_ISL_2701552 | Lighthouse Lab in Milton Keynes | Wellcome Sanger Institute for the COVID-19 Genomics UK (COG-UK) Consortium | The Lighthouse Lab in Milton Keynes and Alex Alderton, Roberto Amato, Jeffrey Barrett, Sonia Goncalves, Ewan Harrison, David K. Jackson, Ian Johnston, Dominic Kwiatkowski, Cordelia Langford, John Sillitoe on behalf of the Wellcome Sanger Institute COVID-19 Surveillance Team |
| EPI_ISL_2701553, EPI_ISL_2701554, EPI_ISL_2701555 | Lighthouse Lab in Alderley Park | Wellcome Sanger Institute for the COVID-19 Genomics UK (COG-UK) Consortium | Jacquelyn Wynn, Mairead Hyland, The Lighthouse Lab in Alderley Park and Alex Alderton, Roberto Amato, Jeffrey Barrett, Sonia Goncalves, Ewan Harrison, David K. Jackson, Ian Johnston, Dominic Kwiatkowski, Cordelia Langford, John Sillitoe on behalf of the Wellcome Sanger Institute COVID-19 Surveillance Team |
| EPI_ISL_2701556 | Lighthouse Lab in Glasgow | Wellcome Sanger Institute for the COVID-19 Genomics UK (COG-UK) Consortium | Harper VanSteenhouse, Yumi Kasai, David Gray, Carol Clugston, Anna Dominiczak and Alex Alderton, Roberto Amato, Jeffrey Barrett, Sonia Goncalves, Ewan Harrison, David K. Jackson, Ian Johnston, Dominic Kwiatkowski, Cordelia Langford, John Sillitoe on behalf of the Wellcome Sanger Institute COVID-19 Surveillance Team |

| | | | |
|---|---|---|---|
| EPI_ISL_2701557 | Lighthouse Lab in Milton Keynes | Wellcome Sanger Institute for the COVID-19 Genomics UK (COG-UK) Consortium | The Lighthouse Lab in Milton Keynes and Alex Alderton, Roberto Amato, Jeffrey Barrett, Sonia Goncalves, Ewan Harrison, David K. Jackson, Ian Johnston, Dominic Kwiatkowski, Cordelia Langford, John Sillitoe on behalf of the Wellcome Sanger Institute COVID-19 Surveillance Team |
| EPI_ISL_2701558 | Lighthouse Lab in Glasgow | Wellcome Sanger Institute for the COVID-19 Genomics UK (COG-UK) Consortium | Harper VanSteenhouse, Yumi Kasai, David Gray, Carol Clugston, Anna Dominiczak and Alex Alderton, Roberto Amato, Jeffrey Barrett, Sonia Goncalves, Ewan Harrison, David K. Jackson, Ian Johnston, Dominic Kwiatkowski, Cordelia Langford, John Sillitoe on behalf of the Wellcome Sanger Institute COVID-19 Surveillance Team |
| EPI_ISL_2701559 | Lighthouse Lab in Milton Keynes | Wellcome Sanger Institute for the COVID-19 Genomics UK (COG-UK) Consortium | The Lighthouse Lab in Milton Keynes and Alex Alderton, Roberto Amato, Jeffrey Barrett, Sonia Goncalves, Ewan Harrison, David K. Jackson, Ian Johnston, Dominic Kwiatkowski, Cordelia Langford, John Sillitoe on behalf of the Wellcome Sanger Institute COVID-19 Surveillance Team |
| EPI_ISL_2701560 | Lighthouse Lab in Alderley Park | Wellcome Sanger Institute for the COVID-19 Genomics UK (COG-UK) Consortium | Jacquelyn Wynn, Mairead Hyland, The Lighthouse Lab in Alderley Park and Alex Alderton, Roberto Amato, Jeffrey Barrett, Sonia Goncalves, Ewan Harrison, David K. Jackson, Ian Johnston, Dominic Kwiatkowski, Cordelia Langford, John Sillitoe on behalf of the Wellcome Sanger Institute COVID-19 Surveillance Team |
| EPI_ISL_2701562, EPI_ISL_2701563, EPI_ISL_2701564 | Lighthouse Lab in Glasgow | Wellcome Sanger Institute for the COVID-19 Genomics UK (COG-UK) Consortium | Harper VanSteenhouse, Yumi Kasai, David Gray, Carol Clugston, Anna Dominiczak and Alex Alderton, Roberto Amato, Jeffrey Barrett, Sonia Goncalves, Ewan Harrison, David K. Jackson, Ian Johnston, Dominic Kwiatkowski, Cordelia Langford, John Sillitoe on behalf of the Wellcome Sanger Institute COVID-19 Surveillance Team |
| EPI_ISL_2701566 | Lighthouse Lab in Alderley Park | Wellcome Sanger Institute for the COVID-19 Genomics UK (COG-UK) Consortium | Jacquelyn Wynn, Mairead Hyland, The Lighthouse Lab in Alderley Park and Alex Alderton, Roberto Amato, Jeffrey Barrett, Sonia Goncalves, Ewan Harrison, David K. Jackson, Ian Johnston, Dominic Kwiatkowski, Cordelia Langford, John Sillitoe on behalf of the Wellcome Sanger Institute COVID-19 Surveillance Team |
| EPI_ISL_2701567, EPI_ISL_2701568, EPI_ISL_2701569 | Lighthouse Lab in Glasgow | Wellcome Sanger Institute for the COVID-19 Genomics UK (COG-UK) Consortium | Harper VanSteenhouse, Yumi Kasai, David Gray, Carol Clugston, Anna Dominiczak and Alex Alderton, Roberto Amato, Jeffrey Barrett, Sonia Goncalves, Ewan Harrison, David K. Jackson, Ian Johnston, Dominic Kwiatkowski, Cordelia Langford, John Sillitoe on behalf of the Wellcome Sanger Institute COVID-19 Surveillance Team |
| EPI_ISL_2701570 | Lighthouse Lab in Milton Keynes | Wellcome Sanger Institute for the COVID-19 Genomics UK (COG-UK) Consortium | The Lighthouse Lab in Milton Keynes and Alex Alderton, Roberto Amato, Jeffrey Barrett, Sonia Goncalves, Ewan Harrison, David K. Jackson, Ian Johnston, Dominic Kwiatkowski, Cordelia Langford, John Sillitoe on behalf of the Wellcome Sanger Institute COVID-19 Surveillance Team |
| EPI_ISL_2701571 | Lighthouse Lab in Alderley Park | Wellcome Sanger Institute for the COVID-19 Genomics UK (COG-UK) Consortium | Jacquelyn Wynn, Mairead Hyland, The Lighthouse Lab in Alderley Park and Alex Alderton, Roberto Amato, Jeffrey Barrett, Sonia Goncalves, Ewan Harrison, David K. Jackson, Ian Johnston, Dominic Kwiatkowski, Cordelia Langford, John Sillitoe on behalf of the Wellcome Sanger Institute COVID-19 Surveillance Team |
| EPI_ISL_2701572 | Lighthouse Lab in Milton Keynes | Wellcome Sanger Institute for the COVID-19 Genomics UK (COG-UK) Consortium | The Lighthouse Lab in Milton Keynes and Alex Alderton, Roberto Amato, Jeffrey Barrett, Sonia Goncalves, Ewan Harrison, David K. Jackson, Ian Johnston, Dominic Kwiatkowski, Cordelia Langford, John Sillitoe on behalf of the Wellcome Sanger Institute COVID-19 Surveillance Team |
| EPI_ISL_2701573 | Lighthouse Lab in Alderley Park | Wellcome Sanger Institute for the COVID-19 Genomics UK (COG-UK) Consortium | Jacquelyn Wynn, Mairead Hyland, The Lighthouse Lab in Alderley Park and Alex Alderton, Roberto Amato, Jeffrey Barrett, Sonia Goncalves, Ewan Harrison, David K. Jackson, Ian Johnston, Dominic Kwiatkowski, Cordelia Langford, John Sillitoe on behalf of the Wellcome Sanger Institute COVID-19 Surveillance Team |
| EPI_ISL_2701574 | Lighthouse Lab in Glasgow | Wellcome Sanger Institute for the COVID-19 Genomics UK (COG-UK) Consortium | Harper VanSteenhouse, Yumi Kasai, David Gray, Carol Clugston, Anna Dominiczak and Alex Alderton, Roberto Amato, Jeffrey Barrett, Sonia Goncalves, Ewan Harrison, David K. Jackson, Ian Johnston, Dominic Kwiatkowski, Cordelia Langford, John Sillitoe on behalf of the Wellcome Sanger Institute COVID-19 Surveillance Team |
| EPI_ISL_2701575 | Lighthouse Lab in Alderley Park | Wellcome Sanger Institute for the COVID-19 Genomics UK (COG-UK) Consortium | Jacquelyn Wynn, Mairead Hyland, The Lighthouse Lab in Alderley Park and Alex Alderton, Roberto Amato, Jeffrey Barrett, Sonia Goncalves, Ewan Harrison, David K. Jackson, Ian Johnston, Dominic Kwiatkowski, Cordelia Langford, John Sillitoe on behalf of the Wellcome Sanger Institute COVID-19 Surveillance Team |
| EPI_ISL_2701576 | Lighthouse Lab in Glasgow | Wellcome Sanger Institute for the COVID-19 Genomics UK (COG-UK) Consortium | Harper VanSteenhouse, Yumi Kasai, David Gray, Carol Clugston, Anna Dominiczak and Alex Alderton, Roberto Amato, Jeffrey Barrett, Sonia Goncalves, Ewan Harrison, David K. Jackson, Ian Johnston, Dominic Kwiatkowski, Cordelia Langford, John Sillitoe on behalf of the Wellcome Sanger Institute COVID-19 Surveillance Team |
| EPI_ISL_2701577, EPI_ISL_2701578 | Lighthouse Lab in Milton Keynes | Wellcome Sanger Institute for the COVID-19 Genomics UK (COG-UK) Consortium | The Lighthouse Lab in Milton Keynes and Alex Alderton, Roberto Amato, Jeffrey Barrett, Sonia Goncalves, Ewan Harrison, David K. Jackson, Ian Johnston, Dominic Kwiatkowski, Cordelia Langford, John Sillitoe on behalf of the Wellcome Sanger Institute COVID-19 Surveillance Team |
| EPI_ISL_2701579, EPI_ISL_2701580 | Lighthouse Lab in Glasgow | Wellcome Sanger Institute for the COVID-19 Genomics UK (COG-UK) Consortium | Harper VanSteenhouse, Yumi Kasai, David Gray, Carol Clugston, Anna Dominiczak and Alex Alderton, Roberto Amato, Jeffrey Barrett, Sonia Goncalves, Ewan Harrison, David K. Jackson, Ian Johnston, Dominic Kwiatkowski, Cordelia Langford, John Sillitoe on behalf of the Wellcome Sanger Institute COVID-19 Surveillance Team |
| EPI_ISL_2701581 | Lighthouse Lab in Alderley Park | Wellcome Sanger Institute for the COVID-19 Genomics UK (COG-UK) Consortium | Jacquelyn Wynn, Mairead Hyland, The Lighthouse Lab in Alderley Park and Alex Alderton, Roberto Amato, Jeffrey Barrett, Sonia Goncalves, Ewan Harrison, David K. Jackson, Ian Johnston, Dominic Kwiatkowski, Cordelia Langford, John Sillitoe on behalf of the Wellcome Sanger Institute COVID-19 Surveillance Team |
| EPI_ISL_2701582 | Lighthouse Lab in Glasgow | Wellcome Sanger Institute for the COVID-19 Genomics UK (COG-UK) Consortium | Harper VanSteenhouse, Yumi Kasai, David Gray, Carol Clugston, Anna Dominiczak and Alex Alderton, Roberto Amato, Jeffrey Barrett, Sonia Goncalves, Ewan Harrison, David K. Jackson, Ian Johnston, Dominic Kwiatkowski, Cordelia Langford, John Sillitoe on behalf of the Wellcome Sanger Institute COVID-19 Surveillance Team |
| EPI_ISL_2701583 | Lighthouse Lab in Milton Keynes | Wellcome Sanger Institute for the COVID-19 Genomics UK (COG-UK) Consortium | The Lighthouse Lab in Milton Keynes and Alex Alderton, Roberto Amato, Jeffrey Barrett, Sonia Goncalves, Ewan Harrison, David K. Jackson, Ian Johnston, Dominic Kwiatkowski, Cordelia Langford, John Sillitoe on behalf of the Wellcome Sanger Institute COVID-19 Surveillance Team |
| EPI_ISL_2701584 | Lighthouse Lab in Alderley Park | Wellcome Sanger Institute for the COVID-19 Genomics UK (COG-UK) Consortium | Jacquelyn Wynn, Mairead Hyland, The Lighthouse Lab in Alderley Park and Alex Alderton, Roberto Amato, Jeffrey Barrett, Sonia Goncalves, Ewan Harrison, David K. Jackson, Ian Johnston, Dominic Kwiatkowski, Cordelia Langford, John Sillitoe on behalf of the Wellcome Sanger Institute COVID-19 Surveillance Team |
| EPI_ISL_2701585 | Lighthouse Lab in Glasgow | Wellcome Sanger Institute for the COVID-19 Genomics UK (COG-UK) Consortium | Harper VanSteenhouse, Yumi Kasai, David Gray, Carol Clugston, Anna Dominiczak and Alex Alderton, Roberto Amato, Jeffrey Barrett, Sonia Goncalves, Ewan Harrison, David K. Jackson, Ian Johnston, Dominic Kwiatkowski, Cordelia Langford, John Sillitoe on behalf of the Wellcome Sanger Institute COVID-19 Surveillance Team |
| EPI_ISL_2701586, EPI_ISL_2701587 | Lighthouse Lab in Milton Keynes | Wellcome Sanger Institute for the COVID-19 Genomics UK (COG-UK) Consortium | The Lighthouse Lab in Milton Keynes and Alex Alderton, Roberto Amato, Jeffrey Barrett, Sonia Goncalves, Ewan Harrison, David K. Jackson, Ian Johnston, Dominic Kwiatkowski, Cordelia Langford, John Sillitoe on behalf of the Wellcome Sanger Institute COVID-19 Surveillance Team |
| EPI_ISL_2701588 | Lighthouse Lab in Alderley Park | Wellcome Sanger Institute for the COVID-19 Genomics UK (COG-UK) Consortium | Jacquelyn Wynn, Mairead Hyland, The Lighthouse Lab in Alderley Park and Alex Alderton, Roberto Amato, Jeffrey Barrett, Sonia Goncalves, Ewan Harrison, David K. Jackson, Ian Johnston, Dominic Kwiatkowski, Cordelia Langford, John Sillitoe on behalf of the Wellcome Sanger Institute COVID-19 Surveillance Team |
| EPI_ISL_2701589, EPI_ISL_2701590, EPI_ISL_2701591 | Lighthouse Lab in Glasgow | Wellcome Sanger Institute for the COVID-19 Genomics UK (COG-UK) Consortium | Harper VanSteenhouse, Yumi Kasai, David Gray, Carol Clugston, Anna Dominiczak and Alex Alderton, Roberto Amato, Jeffrey Barrett, Sonia Goncalves, Ewan Harrison, David K. Jackson, Ian Johnston, Dominic Kwiatkowski, Cordelia Langford, John Sillitoe on behalf of the Wellcome Sanger Institute COVID-19 Surveillance Team |
| EPI_ISL_2701592, EPI_ISL_2701593, EPI_ISL_2701594 | Lighthouse Lab in Alderley Park | Wellcome Sanger Institute for the COVID-19 Genomics UK (COG-UK) Consortium | Jacquelyn Wynn, Mairead Hyland, The Lighthouse Lab in Alderley Park and Alex Alderton, Roberto Amato, Jeffrey Barrett, Sonia Goncalves, Ewan Harrison, David K. Jackson, Ian Johnston, Dominic Kwiatkowski, Cordelia Langford, John Sillitoe on behalf of the Wellcome Sanger Institute COVID-19 Surveillance Team |
| EPI_ISL_2701595, EPI_ISL_2701596 | Lighthouse Lab in Milton Keynes | Wellcome Sanger Institute for the COVID-19 Genomics UK (COG-UK) Consortium | The Lighthouse Lab in Milton Keynes and Alex Alderton, Roberto Amato, Jeffrey Barrett, Sonia Goncalves, Ewan Harrison, David K. Jackson, Ian Johnston, Dominic Kwiatkowski, Cordelia Langford, John Sillitoe on behalf of the Wellcome Sanger Institute COVID-19 Surveillance Team |
| EPI_ISL_2701597 | Lighthouse Lab in Glasgow | Wellcome Sanger Institute for the COVID-19 Genomics UK (COG-UK) Consortium | Harper VanSteenhouse, Yumi Kasai, David Gray, Carol Clugston, Anna Dominiczak and Alex Alderton, Roberto Amato, Jeffrey Barrett, Sonia Goncalves, Ewan Harrison, David K. Jackson, Ian Johnston, Dominic Kwiatkowski, Cordelia Langford, John Sillitoe on behalf of the Wellcome Sanger Institute COVID-19 Surveillance Team |
| EPI_ISL_2701598 | Lighthouse Lab in Alderley Park | Wellcome Sanger Institute for the COVID-19 Genomics UK (COG-UK) Consortium | Jacquelyn Wynn, Mairead Hyland, The Lighthouse Lab in Alderley Park and Alex Alderton, Roberto Amato, Jeffrey Barrett, Sonia Goncalves, Ewan Harrison, David K. Jackson, Ian Johnston, Dominic Kwiatkowski, Cordelia Langford, John Sillitoe on behalf of the Wellcome Sanger Institute COVID-19 Surveillance Team |
| EPI_ISL_2701599, EPI_ISL_2701600 | Lighthouse Lab in Glasgow | Wellcome Sanger Institute for the COVID-19 Genomics UK (COG-UK) Consortium | Harper VanSteenhouse, Yumi Kasai, David Gray, Carol Clugston, Anna Dominiczak and Alex Alderton, Roberto Amato, Jeffrey Barrett, Sonia Goncalves, Ewan Harrison, David K. Jackson, Ian Johnston, Dominic Kwiatkowski, Cordelia Langford, John Sillitoe on behalf of the Wellcome Sanger Institute COVID-19 Surveillance Team |
| EPI_ISL_2701601 | Lighthouse Lab in Alderley Park | Wellcome Sanger Institute for the COVID-19 Genomics UK (COG-UK) Consortium | Jacquelyn Wynn, Mairead Hyland, The Lighthouse Lab in Alderley Park and Alex Alderton, Roberto Amato, Jeffrey Barrett, Sonia Goncalves, Ewan Harrison, David K. Jackson, Ian Johnston, Dominic Kwiatkowski, Cordelia Langford, John Sillitoe on behalf of the Wellcome Sanger Institute COVID-19 Surveillance Team |
| EPI_ISL_2701602 | Lighthouse Lab in Glasgow | Wellcome Sanger Institute for the COVID-19 Genomics UK (COG-UK) Consortium | Harper VanSteenhouse, Yumi Kasai, David Gray, Carol Clugston, Anna Dominiczak and Alex Alderton, Roberto Amato, Jeffrey Barrett, Sonia Goncalves, Ewan Harrison, David K. Jackson, Ian Johnston, Dominic Kwiatkowski, Cordelia Langford, John Sillitoe on behalf of the Wellcome Sanger Institute COVID-19 Surveillance Team |
| EPI_ISL_2701603 | Lighthouse Lab in Alderley Park | Wellcome Sanger Institute for the COVID-19 Genomics UK (COG-UK) Consortium | Jacquelyn Wynn, Mairead Hyland, The Lighthouse Lab in Alderley Park and Alex Alderton, Roberto Amato, Jeffrey Barrett, Sonia Goncalves, Ewan Harrison, David K. Jackson, Ian Johnston, Dominic Kwiatkowski, Cordelia Langford, John Sillitoe on behalf of the Wellcome Sanger Institute COVID-19 Surveillance Team |
| EPI_ISL_2701604, EPI_ISL_2701605 | Lighthouse Lab in Glasgow | Wellcome Sanger Institute for the COVID-19 Genomics UK (COG-UK) Consortium | Harper VanSteenhouse, Yumi Kasai, David Gray, Carol Clugston, Anna Dominiczak and Alex Alderton, Roberto Amato, Jeffrey Barrett, Sonia Goncalves, Ewan Harrison, David K. Jackson, Ian Johnston, Dominic Kwiatkowski, Cordelia Langford, John Sillitoe on behalf of the Wellcome Sanger Institute COVID-19 Surveillance Team |
| EPI_ISL_2701606, EPI_ISL_2701607, EPI_ISL_2701608 | Lighthouse Lab in Alderley Park | Wellcome Sanger Institute for the COVID-19 Genomics UK (COG-UK) Consortium | Jacquelyn Wynn, Mairead Hyland, The Lighthouse Lab in Alderley Park and Alex Alderton, Roberto Amato, Jeffrey Barrett, Sonia Goncalves, Ewan Harrison, David K. Jackson, Ian Johnston, Dominic Kwiatkowski, Cordelia Langford, John Sillitoe on behalf of the Wellcome Sanger Institute COVID-19 Surveillance Team |
| EPI_ISL_2701609, EPI_ISL_2701610 | Lighthouse Lab in Milton Keynes | Wellcome Sanger Institute for the COVID-19 Genomics UK (COG-UK) Consortium | The Lighthouse Lab in Milton Keynes and Alex Alderton, Roberto Amato, Jeffrey Barrett, Sonia Goncalves, Ewan Harrison, David K. Jackson, Ian Johnston, Dominic Kwiatkowski, Cordelia Langford, John Sillitoe on behalf of the Wellcome Sanger Institute COVID-19 Surveillance Team |
| EPI_ISL_2701611 | Lighthouse Lab in Alderley Park | Wellcome Sanger Institute for the COVID-19 Genomics UK (COG-UK) Consortium | Jacquelyn Wynn, Mairead Hyland, The Lighthouse Lab in Alderley Park and Alex Alderton, Roberto Amato, Jeffrey Barrett, Sonia Goncalves, Ewan Harrison, David K. Jackson, Ian Johnston, Dominic Kwiatkowski, Cordelia Langford, John Sillitoe on behalf of the Wellcome Sanger Institute COVID-19 Surveillance Team |
| EPI_ISL_2701612 | Lighthouse Lab in Milton Keynes | Wellcome Sanger Institute for the COVID-19 Genomics UK (COG-UK) Consortium | The Lighthouse Lab in Milton Keynes and Alex Alderton, Roberto Amato, Jeffrey Barrett, Sonia Goncalves, Ewan Harrison, David K. Jackson, Ian Johnston, Dominic Kwiatkowski, Cordelia Langford, John Sillitoe on behalf of the Wellcome Sanger Institute COVID-19 Surveillance Team |
| EPI_ISL_2701613 | Lighthouse Lab in Alderley Park | Wellcome Sanger Institute for the COVID-19 Genomics UK (COG-UK) Consortium | Jacquelyn Wynn, Mairead Hyland, The Lighthouse Lab in Alderley Park and Alex Alderton, Roberto Amato, Jeffrey Barrett, Sonia Goncalves, Ewan Harrison, David K. Jackson, Ian Johnston, Dominic Kwiatkowski, Cordelia Langford, John Sillitoe on behalf of the Wellcome Sanger Institute COVID-19 Surveillance Team |
| EPI_ISL_2701614, EPI_ISL_2701615 | Lighthouse Lab in Glasgow | Wellcome Sanger Institute for the COVID-19 Genomics UK (COG-UK) Consortium | Harper VanSteenhouse, Yumi Kasai, David Gray, Carol Clugston, Anna Dominiczak and Alex Alderton, Roberto Amato, Jeffrey Barrett, Sonia Goncalves, Ewan Harrison, David K. Jackson, Ian Johnston, Dominic Kwiatkowski, Cordelia Langford, John Sillitoe on behalf of the Wellcome Sanger Institute COVID-19 Surveillance Team |
| EPI_ISL_2701616, EPI_ISL_2701617, EPI_ISL_2701618, EPI_ISL_2701619 | Lighthouse Lab in Alderley Park | Wellcome Sanger Institute for the COVID-19 Genomics UK (COG-UK) Consortium | Jacquelyn Wynn, Mairead Hyland, The Lighthouse Lab in Alderley Park and Alex Alderton, Roberto Amato, Jeffrey Barrett, Sonia Goncalves, Ewan Harrison, David K. Jackson, Ian Johnston, Dominic Kwiatkowski, Cordelia Langford, John Sillitoe on behalf of the Wellcome Sanger Institute COVID-19 Surveillance Team |
| EPI_ISL_2701620 | Lighthouse Lab in Milton Keynes | Wellcome Sanger Institute for the COVID-19 Genomics UK (COG-UK) Consortium | The Lighthouse Lab in Milton Keynes and Alex Alderton, Roberto Amato, Jeffrey Barrett, Sonia Goncalves, Ewan Harrison, David K. Jackson, Ian Johnston, Dominic Kwiatkowski, Cordelia Langford, John Sillitoe on behalf of the Wellcome Sanger Institute COVID-19 Surveillance Team |
| EPI_ISL_2701621 | Lighthouse Lab in Glasgow | Wellcome Sanger Institute for the COVID-19 Genomics UK (COG-UK) Consortium | Harper VanSteenhouse, Yumi Kasai, David Gray, Carol Clugston, Anna Dominiczak and Alex Alderton, Roberto Amato, Jeffrey Barrett, Sonia Goncalves, Ewan Harrison, David K. Jackson, Ian Johnston, Dominic Kwiatkowski, Cordelia Langford, John Sillitoe on behalf of the Wellcome Sanger Institute COVID-19 Surveillance Team |

| | | | |
|---|---|---|---|
| EPI_ISL_2701622 | Lighthouse Lab in Alderley Park | Wellcome Sanger Institute for the COVID-19 Genomics UK (COG-UK) Consortium | Jacquelyn Wynn, Mairead Hyland, The Lighthouse Lab in Alderley Park and Alex Alderton, Roberto Amato, Jeffrey Barrett, Sonia Goncalves, Ewan Harrison, David K. Jackson, Ian Johnston, Dominic Kwiatkowski, Cordelia Langford, John Sillitoe on behalf of the Wellcome Sanger Institute COVID-19 Surveillance Team |
| EPI_ISL_2701623 | Lighthouse Lab in Milton Keynes | Wellcome Sanger Institute for the COVID-19 Genomics UK (COG-UK) Consortium | The Lighthouse Lab in Milton Keynes and Alex Alderton, Roberto Amato, Jeffrey Barrett, Sonia Goncalves, Ewan Harrison, David K. Jackson, Ian Johnston, Dominic Kwiatkowski, Cordelia Langford, John Sillitoe on behalf of the Wellcome Sanger Institute COVID-19 Surveillance Team |
| EPI_ISL_2701625 | Lighthouse Lab in Alderley Park | Wellcome Sanger Institute for the COVID-19 Genomics UK (COG-UK) Consortium | Jacquelyn Wynn, Mairead Hyland, The Lighthouse Lab in Alderley Park and Alex Alderton, Roberto Amato, Jeffrey Barrett, Sonia Goncalves, Ewan Harrison, David K. Jackson, Ian Johnston, Dominic Kwiatkowski, Cordelia Langford, John Sillitoe on behalf of the Wellcome Sanger Institute COVID-19 Surveillance Team |
| EPI_ISL_2701626 | Lighthouse Lab in Milton Keynes | Wellcome Sanger Institute for the COVID-19 Genomics UK (COG-UK) Consortium | The Lighthouse Lab in Milton Keynes and Alex Alderton, Roberto Amato, Jeffrey Barrett, Sonia Goncalves, Ewan Harrison, David K. Jackson, Ian Johnston, Dominic Kwiatkowski, Cordelia Langford, John Sillitoe on behalf of the Wellcome Sanger Institute COVID-19 Surveillance Team |
| EPI_ISL_2701627 | Lighthouse Lab in Glasgow | Wellcome Sanger Institute for the COVID-19 Genomics UK (COG-UK) Consortium | Harper VanSteenhouse, Yumi Kasai, David Gray, Carol Clugston, Anna Dominiczak and Alex Alderton, Roberto Amato, Jeffrey Barrett, Sonia Goncalves, Ewan Harrison, David K. Jackson, Ian Johnston, Dominic Kwiatkowski, Cordelia Langford, John Sillitoe on behalf of the Wellcome Sanger Institute COVID-19 Surveillance Team |
| EPI_ISL_2701628 | Lighthouse Lab in Alderley Park | Wellcome Sanger Institute for the COVID-19 Genomics UK (COG-UK) Consortium | Jacquelyn Wynn, Mairead Hyland, The Lighthouse Lab in Alderley Park and Alex Alderton, Roberto Amato, Jeffrey Barrett, Sonia Goncalves, Ewan Harrison, David K. Jackson, Ian Johnston, Dominic Kwiatkowski, Cordelia Langford, John Sillitoe on behalf of the Wellcome Sanger Institute COVID-19 Surveillance Team |
| EPI_ISL_2701629, EPI_ISL_2701630 | Lighthouse Lab in Glasgow | Wellcome Sanger Institute for the COVID-19 Genomics UK (COG-UK) Consortium | Harper VanSteenhouse, Yumi Kasai, David Gray, Carol Clugston, Anna Dominiczak and Alex Alderton, Roberto Amato, Jeffrey Barrett, Sonia Goncalves, Ewan Harrison, David K. Jackson, Ian Johnston, Dominic Kwiatkowski, Cordelia Langford, John Sillitoe on behalf of the Wellcome Sanger Institute COVID-19 Surveillance Team |
| EPI_ISL_2701631, EPI_ISL_2701632 | Lighthouse Lab in Alderley Park | Wellcome Sanger Institute for the COVID-19 Genomics UK (COG-UK) Consortium | Jacquelyn Wynn, Mairead Hyland, The Lighthouse Lab in Alderley Park and Alex Alderton, Roberto Amato, Jeffrey Barrett, Sonia Goncalves, Ewan Harrison, David K. Jackson, Ian Johnston, Dominic Kwiatkowski, Cordelia Langford, John Sillitoe on behalf of the Wellcome Sanger Institute COVID-19 Surveillance Team |
| EPI_ISL_2701633 | Lighthouse Lab in Glasgow | Wellcome Sanger Institute for the COVID-19 Genomics UK (COG-UK) Consortium | Harper VanSteenhouse, Yumi Kasai, David Gray, Carol Clugston, Anna Dominiczak and Alex Alderton, Roberto Amato, Jeffrey Barrett, Sonia Goncalves, Ewan Harrison, David K. Jackson, Ian Johnston, Dominic Kwiatkowski, Cordelia Langford, John Sillitoe on behalf of the Wellcome Sanger Institute COVID-19 Surveillance Team |
| EPI_ISL_2701634 | Lighthouse Lab in Alderley Park | Wellcome Sanger Institute for the COVID-19 Genomics UK (COG-UK) Consortium | Jacquelyn Wynn, Mairead Hyland, The Lighthouse Lab in Alderley Park and Alex Alderton, Roberto Amato, Jeffrey Barrett, Sonia Goncalves, Ewan Harrison, David K. Jackson, Ian Johnston, Dominic Kwiatkowski, Cordelia Langford, John Sillitoe on behalf of the Wellcome Sanger Institute COVID-19 Surveillance Team |
| EPI_ISL_2701635 | Lighthouse Lab in Glasgow | Wellcome Sanger Institute for the COVID-19 Genomics UK (COG-UK) Consortium | Harper VanSteenhouse, Yumi Kasai, David Gray, Carol Clugston, Anna Dominiczak and Alex Alderton, Roberto Amato, Jeffrey Barrett, Sonia Goncalves, Ewan Harrison, David K. Jackson, Ian Johnston, Dominic Kwiatkowski, Cordelia Langford, John Sillitoe on behalf of the Wellcome Sanger Institute COVID-19 Surveillance Team |
| EPI_ISL_2701637 | Lighthouse Lab in Milton Keynes | Wellcome Sanger Institute for the COVID-19 Genomics UK (COG-UK) Consortium | The Lighthouse Lab in Milton Keynes and Alex Alderton, Roberto Amato, Jeffrey Barrett, Sonia Goncalves, Ewan Harrison, David K. Jackson, Ian Johnston, Dominic Kwiatkowski, Cordelia Langford, John Sillitoe on behalf of the Wellcome Sanger Institute COVID-19 Surveillance Team |
| EPI_ISL_2701638 | Lighthouse Lab in Glasgow | Wellcome Sanger Institute for the COVID-19 Genomics UK (COG-UK) Consortium | Harper VanSteenhouse, Yumi Kasai, David Gray, Carol Clugston, Anna Dominiczak and Alex Alderton, Roberto Amato, Jeffrey Barrett, Sonia Goncalves, Ewan Harrison, David K. Jackson, Ian Johnston, Dominic Kwiatkowski, Cordelia Langford, John Sillitoe on behalf of the Wellcome Sanger Institute COVID-19 Surveillance Team |
| EPI_ISL_2701639 | Lighthouse Lab in Alderley Park | Wellcome Sanger Institute for the COVID-19 Genomics UK (COG-UK) Consortium | Jacquelyn Wynn, Mairead Hyland, The Lighthouse Lab in Alderley Park and Alex Alderton, Roberto Amato, Jeffrey Barrett, Sonia Goncalves, Ewan Harrison, David K. Jackson, Ian Johnston, Dominic Kwiatkowski, Cordelia Langford, John Sillitoe on behalf of the Wellcome Sanger Institute COVID-19 Surveillance Team |
| EPI_ISL_2701641, EPI_ISL_2701642 | Lighthouse Lab in Milton Keynes | Wellcome Sanger Institute for the COVID-19 Genomics UK (COG-UK) Consortium | The Lighthouse Lab in Milton Keynes and Alex Alderton, Roberto Amato, Jeffrey Barrett, Sonia Goncalves, Ewan Harrison, David K. Jackson, Ian Johnston, Dominic Kwiatkowski, Cordelia Langford, John Sillitoe on behalf of the Wellcome Sanger Institute COVID-19 Surveillance Team |
| EPI_ISL_2701643 | Lighthouse Lab in Glasgow | Wellcome Sanger Institute for the COVID-19 Genomics UK (COG-UK) Consortium | Harper VanSteenhouse, Yumi Kasai, David Gray, Carol Clugston, Anna Dominiczak and Alex Alderton, Roberto Amato, Jeffrey Barrett, Sonia Goncalves, Ewan Harrison, David K. Jackson, Ian Johnston, Dominic Kwiatkowski, Cordelia Langford, John Sillitoe on behalf of the Wellcome Sanger Institute COVID-19 Surveillance Team |
| EPI_ISL_2701644, EPI_ISL_2701645 | Lighthouse Lab in Alderley Park | Wellcome Sanger Institute for the COVID-19 Genomics UK (COG-UK) Consortium | Jacquelyn Wynn, Mairead Hyland, The Lighthouse Lab in Alderley Park and Alex Alderton, Roberto Amato, Jeffrey Barrett, Sonia Goncalves, Ewan Harrison, David K. Jackson, Ian Johnston, Dominic Kwiatkowski, Cordelia Langford, John Sillitoe on behalf of the Wellcome Sanger Institute COVID-19 Surveillance Team |
| EPI_ISL_2701646, EPI_ISL_2701647 | Lighthouse Lab in Glasgow | Wellcome Sanger Institute for the COVID-19 Genomics UK (COG-UK) Consortium | Harper VanSteenhouse, Yumi Kasai, David Gray, Carol Clugston, Anna Dominiczak and Alex Alderton, Roberto Amato, Jeffrey Barrett, Sonia Goncalves, Ewan Harrison, David K. Jackson, Ian Johnston, Dominic Kwiatkowski, Cordelia Langford, John Sillitoe on behalf of the Wellcome Sanger Institute COVID-19 Surveillance Team |
| EPI_ISL_2701648 | Lighthouse Lab in Milton Keynes | Wellcome Sanger Institute for the COVID-19 Genomics UK (COG-UK) Consortium | The Lighthouse Lab in Milton Keynes and Alex Alderton, Roberto Amato, Jeffrey Barrett, Sonia Goncalves, Ewan Harrison, David K. Jackson, Ian Johnston, Dominic Kwiatkowski, Cordelia Langford, John Sillitoe on behalf of the Wellcome Sanger Institute COVID-19 Surveillance Team |
| EPI_ISL_2701649 | Lighthouse Lab in Alderley Park | Wellcome Sanger Institute for the COVID-19 Genomics UK (COG-UK) Consortium | Jacquelyn Wynn, Mairead Hyland, The Lighthouse Lab in Alderley Park and Alex Alderton, Roberto Amato, Jeffrey Barrett, Sonia Goncalves, Ewan Harrison, David K. Jackson, Ian Johnston, Dominic Kwiatkowski, Cordelia Langford, John Sillitoe on behalf of the Wellcome Sanger Institute COVID-19 Surveillance Team |
| EPI_ISL_2701650 | Lighthouse Lab in Milton Keynes | Wellcome Sanger Institute for the COVID-19 Genomics UK (COG-UK) Consortium | The Lighthouse Lab in Milton Keynes and Alex Alderton, Roberto Amato, Jeffrey Barrett, Sonia Goncalves, Ewan Harrison, David K. Jackson, Ian Johnston, Dominic Kwiatkowski, Cordelia Langford, John Sillitoe on behalf of the Wellcome Sanger Institute COVID-19 Surveillance Team |
| EPI_ISL_2701651 | Lighthouse Lab in Glasgow | Wellcome Sanger Institute for the COVID-19 Genomics UK (COG-UK) Consortium | Harper VanSteenhouse, Yumi Kasai, David Gray, Carol Clugston, Anna Dominiczak and Alex Alderton, Roberto Amato, Jeffrey Barrett, Sonia Goncalves, Ewan Harrison, David K. Jackson, Ian Johnston, Dominic Kwiatkowski, Cordelia Langford, John Sillitoe on behalf of the Wellcome Sanger Institute COVID-19 Surveillance Team |
| EPI_ISL_2701652, EPI_ISL_2701654, EPI_ISL_2701655, EPI_ISL_2701656, EPI_ISL_2701657, EPI_ISL_2701658 | Lighthouse Lab in Milton Keynes | Wellcome Sanger Institute for the COVID-19 Genomics UK (COG-UK) Consortium | The Lighthouse Lab in Milton Keynes and Alex Alderton, Roberto Amato, Jeffrey Barrett, Sonia Goncalves, Ewan Harrison, David K. Jackson, Ian Johnston, Dominic Kwiatkowski, Cordelia Langford, John Sillitoe on behalf of the Wellcome Sanger Institute COVID-19 Surveillance Team |
| EPI_ISL_2701659 | Lighthouse Lab in Glasgow | Wellcome Sanger Institute for the COVID-19 Genomics UK (COG-UK) Consortium | Harper VanSteenhouse, Yumi Kasai, David Gray, Carol Clugston, Anna Dominiczak and Alex Alderton, Roberto Amato, Jeffrey Barrett, Sonia Goncalves, Ewan Harrison, David K. Jackson, Ian Johnston, Dominic Kwiatkowski, Cordelia Langford, John Sillitoe on behalf of the Wellcome Sanger Institute COVID-19 Surveillance Team |
| EPI_ISL_2701660 | Lighthouse Lab in Alderley Park | Wellcome Sanger Institute for the COVID-19 Genomics UK (COG-UK) Consortium | Jacquelyn Wynn, Mairead Hyland, The Lighthouse Lab in Alderley Park and Alex Alderton, Roberto Amato, Jeffrey Barrett, Sonia Goncalves, Ewan Harrison, David K. Jackson, Ian Johnston, Dominic Kwiatkowski, Cordelia Langford, John Sillitoe on behalf of the Wellcome Sanger Institute COVID-19 Surveillance Team |
| EPI_ISL_2701661, EPI_ISL_2701662, EPI_ISL_2701663 | Lighthouse Lab in Glasgow | Wellcome Sanger Institute for the COVID-19 Genomics UK (COG-UK) Consortium | Harper VanSteenhouse, Yumi Kasai, David Gray, Carol Clugston, Anna Dominiczak and Alex Alderton, Roberto Amato, Jeffrey Barrett, Sonia Goncalves, Ewan Harrison, David K. Jackson, Ian Johnston, Dominic Kwiatkowski, Cordelia Langford, John Sillitoe on behalf of the Wellcome Sanger Institute COVID-19 Surveillance Team |
| EPI_ISL_2701664, EPI_ISL_2701665, EPI_ISL_2701666 | Lighthouse Lab in Alderley Park | Wellcome Sanger Institute for the COVID-19 Genomics UK (COG-UK) Consortium | Jacquelyn Wynn, Mairead Hyland, The Lighthouse Lab in Alderley Park and Alex Alderton, Roberto Amato, Jeffrey Barrett, Sonia Goncalves, Ewan Harrison, David K. Jackson, Ian Johnston, Dominic Kwiatkowski, Cordelia Langford, John Sillitoe on behalf of the Wellcome Sanger Institute COVID-19 Surveillance Team |
| EPI_ISL_2701667 | Lighthouse Lab in Glasgow | Wellcome Sanger Institute for the COVID-19 Genomics UK (COG-UK) Consortium | Harper VanSteenhouse, Yumi Kasai, David Gray, Carol Clugston, Anna Dominiczak and Alex Alderton, Roberto Amato, Jeffrey Barrett, Sonia Goncalves, Ewan Harrison, David K. Jackson, Ian Johnston, Dominic Kwiatkowski, Cordelia Langford, John Sillitoe on behalf of the Wellcome Sanger Institute COVID-19 Surveillance Team |
| EPI_ISL_2701668 | Lighthouse Lab in Milton Keynes | Wellcome Sanger Institute for the COVID-19 Genomics UK (COG-UK) Consortium | The Lighthouse Lab in Milton Keynes and Alex Alderton, Roberto Amato, Jeffrey Barrett, Sonia Goncalves, Ewan Harrison, David K. Jackson, Ian Johnston, Dominic Kwiatkowski, Cordelia Langford, John Sillitoe on behalf of the Wellcome Sanger Institute COVID-19 Surveillance Team |
| EPI_ISL_2701669 | Lighthouse Lab in Alderley Park | Wellcome Sanger Institute for the COVID-19 Genomics UK (COG-UK) Consortium | Jacquelyn Wynn, Mairead Hyland, The Lighthouse Lab in Alderley Park and Alex Alderton, Roberto Amato, Jeffrey Barrett, Sonia Goncalves, Ewan Harrison, David K. Jackson, Ian Johnston, Dominic Kwiatkowski, Cordelia Langford, John Sillitoe on behalf of the Wellcome Sanger Institute COVID-19 Surveillance Team |
| EPI_ISL_2701670, EPI_ISL_2701671, EPI_ISL_2701672 | Lighthouse Lab in Glasgow | Wellcome Sanger Institute for the COVID-19 Genomics UK (COG-UK) Consortium | Harper VanSteenhouse, Yumi Kasai, David Gray, Carol Clugston, Anna Dominiczak and Alex Alderton, Roberto Amato, Jeffrey Barrett, Sonia Goncalves, Ewan Harrison, David K. Jackson, Ian Johnston, Dominic Kwiatkowski, Cordelia Langford, John Sillitoe on behalf of the Wellcome Sanger Institute COVID-19 Surveillance Team |
| EPI_ISL_2701673 | Lighthouse Lab in Alderley Park | Wellcome Sanger Institute for the COVID-19 Genomics UK (COG-UK) Consortium | Jacquelyn Wynn, Mairead Hyland, The Lighthouse Lab in Alderley Park and Alex Alderton, Roberto Amato, Jeffrey Barrett, Sonia Goncalves, Ewan Harrison, David K. Jackson, Ian Johnston, Dominic Kwiatkowski, Cordelia Langford, John Sillitoe on behalf of the Wellcome Sanger Institute COVID-19 Surveillance Team |
| EPI_ISL_2701674 | Lighthouse Lab in Glasgow | Wellcome Sanger Institute for the COVID-19 Genomics UK (COG-UK) Consortium | Harper VanSteenhouse, Yumi Kasai, David Gray, Carol Clugston, Anna Dominiczak and Alex Alderton, Roberto Amato, Jeffrey Barrett, Sonia Goncalves, Ewan Harrison, David K. Jackson, Ian Johnston, Dominic Kwiatkowski, Cordelia Langford, John Sillitoe on behalf of the Wellcome Sanger Institute COVID-19 Surveillance Team |
| EPI_ISL_2701675 | Lighthouse Lab in Alderley Park | Wellcome Sanger Institute for the COVID-19 Genomics UK (COG-UK) Consortium | Jacquelyn Wynn, Mairead Hyland, The Lighthouse Lab in Alderley Park and Alex Alderton, Roberto Amato, Jeffrey Barrett, Sonia Goncalves, Ewan Harrison, David K. Jackson, Ian Johnston, Dominic Kwiatkowski, Cordelia Langford, John Sillitoe on behalf of the Wellcome Sanger Institute COVID-19 Surveillance Team |
| EPI_ISL_2701676, EPI_ISL_2701677 | Lighthouse Lab in Milton Keynes | Wellcome Sanger Institute for the COVID-19 Genomics UK (COG-UK) Consortium | The Lighthouse Lab in Milton Keynes and Alex Alderton, Roberto Amato, Jeffrey Barrett, Sonia Goncalves, Ewan Harrison, David K. Jackson, Ian Johnston, Dominic Kwiatkowski, Cordelia Langford, John Sillitoe on behalf of the Wellcome Sanger Institute COVID-19 Surveillance Team |
| EPI_ISL_2701678, EPI_ISL_2701679, EPI_ISL_2701680 | Lighthouse Lab in Glasgow | Wellcome Sanger Institute for the COVID-19 Genomics UK (COG-UK) Consortium | Harper VanSteenhouse, Yumi Kasai, David Gray, Carol Clugston, Anna Dominiczak and Alex Alderton, Roberto Amato, Jeffrey Barrett, Sonia Goncalves, Ewan Harrison, David K. Jackson, Ian Johnston, Dominic Kwiatkowski, Cordelia Langford, John Sillitoe on behalf of the Wellcome Sanger Institute COVID-19 Surveillance Team |
| EPI_ISL_2701681 | Lighthouse Lab in Milton Keynes | Wellcome Sanger Institute for the COVID-19 Genomics UK (COG-UK) Consortium | The Lighthouse Lab in Milton Keynes and Alex Alderton, Roberto Amato, Jeffrey Barrett, Sonia Goncalves, Ewan Harrison, David K. Jackson, Ian Johnston, Dominic Kwiatkowski, Cordelia Langford, John Sillitoe on behalf of the Wellcome Sanger Institute COVID-19 Surveillance Team |
| EPI_ISL_2701682, EPI_ISL_2701683 | Lighthouse Lab in Glasgow | Wellcome Sanger Institute for the COVID-19 Genomics UK (COG-UK) Consortium | Harper VanSteenhouse, Yumi Kasai, David Gray, Carol Clugston, Anna Dominiczak and Alex Alderton, Roberto Amato, Jeffrey Barrett, Sonia Goncalves, Ewan Harrison, David K. Jackson, Ian Johnston, Dominic Kwiatkowski, Cordelia Langford, John Sillitoe on behalf of the Wellcome Sanger Institute COVID-19 Surveillance Team |
| EPI_ISL_2701684, EPI_ISL_2701685, EPI_ISL_2701686 | Lighthouse Lab in Alderley Park | Wellcome Sanger Institute for the COVID-19 Genomics UK (COG-UK) Consortium | Jacquelyn Wynn, Mairead Hyland, The Lighthouse Lab in Alderley Park and Alex Alderton, Roberto Amato, Jeffrey Barrett, Sonia Goncalves, Ewan Harrison, David K. Jackson, Ian Johnston, Dominic Kwiatkowski, Cordelia Langford, John Sillitoe on behalf of the Wellcome Sanger Institute COVID-19 Surveillance Team |
| EPI_ISL_2701687, EPI_ISL_2701688 | Lighthouse Lab in Glasgow | Wellcome Sanger Institute for the COVID-19 Genomics UK (COG-UK) Consortium | Harper VanSteenhouse, Yumi Kasai, David Gray, Carol Clugston, Anna Dominiczak and Alex Alderton, Roberto Amato, Jeffrey Barrett, Sonia Goncalves, Ewan Harrison, David K. Jackson, Ian Johnston, Dominic Kwiatkowski, Cordelia Langford, John Sillitoe on behalf of the Wellcome Sanger Institute COVID-19 Surveillance Team |
| EPI_ISL_2701689 | Lighthouse Lab in Milton Keynes | Wellcome Sanger Institute for the COVID-19 Genomics UK (COG-UK) Consortium | The Lighthouse Lab in Milton Keynes and Alex Alderton, Roberto Amato, Jeffrey Barrett, Sonia Goncalves, Ewan Harrison, David K. Jackson, Ian Johnston, Dominic Kwiatkowski, Cordelia Langford, John Sillitoe on behalf of the Wellcome Sanger Institute COVID-19 Surveillance Team |
| EPI_ISL_2701690 | Lighthouse Lab in Alderley Park | Wellcome Sanger Institute for the COVID-19 Genomics UK (COG-UK) | Jacquelyn Wynn, Mairead Hyland, The Lighthouse Lab in Alderley Park and Alex Alderton, Roberto Amato, Jeffrey Barrett, Sonia Goncalves, Ewan Harrison, David K. Jackson, Ian Johnston, Dominic Kwiatkowski, Cordelia Langford, John Sillitoe on |

| | | | |
|---|---|---|---|
| | | Consortium | behalf of the Wellcome Sanger Institute COVID-19 Surveillance Team |
| EPI_ISL_2701691 | Lighthouse Lab in Glasgow | Wellcome Sanger Institute for the COVID-19 Genomics UK (COG-UK) Consortium | Harper VanSteenhouse, Yumi Kasai, David Gray, Carol Clugston, Anna Dominiczak and Alex Alderton, Roberto Amato, Jeffrey Barrett, Sonia Goncalves, Ewan Harrison, David K. Jackson, Ian Johnston, Dominic Kwiatkowski, Cordelia Langford, John Sillitoe on behalf of the Wellcome Sanger Institute COVID-19 Surveillance Team |
| EPI_ISL_2701692 | Lighthouse Lab in Milton Keynes | Wellcome Sanger Institute for the COVID-19 Genomics UK (COG-UK) Consortium | The Lighthouse Lab in Milton Keynes and Alex Alderton, Roberto Amato, Jeffrey Barrett, Sonia Goncalves, Ewan Harrison, David K. Jackson, Ian Johnston, Dominic Kwiatkowski, Cordelia Langford, John Sillitoe on behalf of the Wellcome Sanger Institute COVID-19 Surveillance Team |
| EPI_ISL_2701693 | Lighthouse Lab in Alderley Park | Wellcome Sanger Institute for the COVID-19 Genomics UK (COG-UK) Consortium | Jacquelyn Wynn, Mairead Hyland, The Lighthouse Lab in Alderley Park and Alex Alderton, Roberto Amato, Jeffrey Barrett, Sonia Goncalves, Ewan Harrison, David K. Jackson, Ian Johnston, Dominic Kwiatkowski, Cordelia Langford, John Sillitoe on behalf of the Wellcome Sanger Institute COVID-19 Surveillance Team |
| EPI_ISL_2701694 | Lighthouse Lab in Glasgow | Wellcome Sanger Institute for the COVID-19 Genomics UK (COG-UK) Consortium | Harper VanSteenhouse, Yumi Kasai, David Gray, Carol Clugston, Anna Dominiczak and Alex Alderton, Roberto Amato, Jeffrey Barrett, Sonia Goncalves, Ewan Harrison, David K. Jackson, Ian Johnston, Dominic Kwiatkowski, Cordelia Langford, John Sillitoe on behalf of the Wellcome Sanger Institute COVID-19 Surveillance Team |
| EPI_ISL_2701695, EPI_ISL_2701696 | Lighthouse Lab in Alderley Park | Wellcome Sanger Institute for the COVID-19 Genomics UK (COG-UK) Consortium | Jacquelyn Wynn, Mairead Hyland, The Lighthouse Lab in Alderley Park and Alex Alderton, Roberto Amato, Jeffrey Barrett, Sonia Goncalves, Ewan Harrison, David K. Jackson, Ian Johnston, Dominic Kwiatkowski, Cordelia Langford, John Sillitoe on behalf of the Wellcome Sanger Institute COVID-19 Surveillance Team |
| EPI_ISL_2701697 | Lighthouse Lab in Glasgow | Wellcome Sanger Institute for the COVID-19 Genomics UK (COG-UK) Consortium | Harper VanSteenhouse, Yumi Kasai, David Gray, Carol Clugston, Anna Dominiczak and Alex Alderton, Roberto Amato, Jeffrey Barrett, Sonia Goncalves, Ewan Harrison, David K. Jackson, Ian Johnston, Dominic Kwiatkowski, Cordelia Langford, John Sillitoe on behalf of the Wellcome Sanger Institute COVID-19 Surveillance Team |
| EPI_ISL_2701698 | Lighthouse Lab in Milton Keynes | Wellcome Sanger Institute for the COVID-19 Genomics UK (COG-UK) Consortium | The Lighthouse Lab in Milton Keynes and Alex Alderton, Roberto Amato, Jeffrey Barrett, Sonia Goncalves, Ewan Harrison, David K. Jackson, Ian Johnston, Dominic Kwiatkowski, Cordelia Langford, John Sillitoe on behalf of the Wellcome Sanger Institute COVID-19 Surveillance Team |
| EPI_ISL_2701699 | Lighthouse Lab in Glasgow | Wellcome Sanger Institute for the COVID-19 Genomics UK (COG-UK) Consortium | Harper VanSteenhouse, Yumi Kasai, David Gray, Carol Clugston, Anna Dominiczak and Alex Alderton, Roberto Amato, Jeffrey Barrett, Sonia Goncalves, Ewan Harrison, David K. Jackson, Ian Johnston, Dominic Kwiatkowski, Cordelia Langford, John Sillitoe on behalf of the Wellcome Sanger Institute COVID-19 Surveillance Team |
| EPI_ISL_2701700 | Lighthouse Lab in Milton Keynes | Wellcome Sanger Institute for the COVID-19 Genomics UK (COG-UK) Consortium | The Lighthouse Lab in Milton Keynes and Alex Alderton, Roberto Amato, Jeffrey Barrett, Sonia Goncalves, Ewan Harrison, David K. Jackson, Ian Johnston, Dominic Kwiatkowski, Cordelia Langford, John Sillitoe on behalf of the Wellcome Sanger Institute COVID-19 Surveillance Team |
| EPI_ISL_2701701 | Lighthouse Lab in Glasgow | Wellcome Sanger Institute for the COVID-19 Genomics UK (COG-UK) Consortium | Harper VanSteenhouse, Yumi Kasai, David Gray, Carol Clugston, Anna Dominiczak and Alex Alderton, Roberto Amato, Jeffrey Barrett, Sonia Goncalves, Ewan Harrison, David K. Jackson, Ian Johnston, Dominic Kwiatkowski, Cordelia Langford, John Sillitoe on behalf of the Wellcome Sanger Institute COVID-19 Surveillance Team |
| EPI_ISL_2701702, EPI_ISL_2701703, EPI_ISL_2701704 | Lighthouse Lab in Milton Keynes | Wellcome Sanger Institute for the COVID-19 Genomics UK (COG-UK) Consortium | The Lighthouse Lab in Milton Keynes and Alex Alderton, Roberto Amato, Jeffrey Barrett, Sonia Goncalves, Ewan Harrison, David K. Jackson, Ian Johnston, Dominic Kwiatkowski, Cordelia Langford, John Sillitoe on behalf of the Wellcome Sanger Institute COVID-19 Surveillance Team |
| EPI_ISL_2701705 | Lighthouse Lab in Alderley Park | Wellcome Sanger Institute for the COVID-19 Genomics UK (COG-UK) Consortium | Jacquelyn Wynn, Mairead Hyland, The Lighthouse Lab in Alderley Park and Alex Alderton, Roberto Amato, Jeffrey Barrett, Sonia Goncalves, Ewan Harrison, David K. Jackson, Ian Johnston, Dominic Kwiatkowski, Cordelia Langford, John Sillitoe on behalf of the Wellcome Sanger Institute COVID-19 Surveillance Team |
| EPI_ISL_2701706 | Lighthouse Lab in Milton Keynes | Wellcome Sanger Institute for the COVID-19 Genomics UK (COG-UK) Consortium | The Lighthouse Lab in Milton Keynes and Alex Alderton, Roberto Amato, Jeffrey Barrett, Sonia Goncalves, Ewan Harrison, David K. Jackson, Ian Johnston, Dominic Kwiatkowski, Cordelia Langford, John Sillitoe on behalf of the Wellcome Sanger Institute COVID-19 Surveillance Team |
| EPI_ISL_2701707 | Lighthouse Lab in Alderley Park | Wellcome Sanger Institute for the COVID-19 Genomics UK (COG-UK) Consortium | Jacquelyn Wynn, Mairead Hyland, The Lighthouse Lab in Alderley Park and Alex Alderton, Roberto Amato, Jeffrey Barrett, Sonia Goncalves, Ewan Harrison, David K. Jackson, Ian Johnston, Dominic Kwiatkowski, Cordelia Langford, John Sillitoe on behalf of the Wellcome Sanger Institute COVID-19 Surveillance Team |
| EPI_ISL_2701708 | Lighthouse Lab in Glasgow | Wellcome Sanger Institute for the COVID-19 Genomics UK (COG-UK) Consortium | Harper VanSteenhouse, Yumi Kasai, David Gray, Carol Clugston, Anna Dominiczak and Alex Alderton, Roberto Amato, Jeffrey Barrett, Sonia Goncalves, Ewan Harrison, David K. Jackson, Ian Johnston, Dominic Kwiatkowski, Cordelia Langford, John Sillitoe on behalf of the Wellcome Sanger Institute COVID-19 Surveillance Team |
| EPI_ISL_2701709 | Lighthouse Lab in Alderley Park | Wellcome Sanger Institute for the COVID-19 Genomics UK (COG-UK) Consortium | Jacquelyn Wynn, Mairead Hyland, The Lighthouse Lab in Alderley Park and Alex Alderton, Roberto Amato, Jeffrey Barrett, Sonia Goncalves, Ewan Harrison, David K. Jackson, Ian Johnston, Dominic Kwiatkowski, Cordelia Langford, John Sillitoe on behalf of the Wellcome Sanger Institute COVID-19 Surveillance Team |
| EPI_ISL_2701710, EPI_ISL_2701711 | Lighthouse Lab in Glasgow | Wellcome Sanger Institute for the COVID-19 Genomics UK (COG-UK) Consortium | Harper VanSteenhouse, Yumi Kasai, David Gray, Carol Clugston, Anna Dominiczak and Alex Alderton, Roberto Amato, Jeffrey Barrett, Sonia Goncalves, Ewan Harrison, David K. Jackson, Ian Johnston, Dominic Kwiatkowski, Cordelia Langford, John Sillitoe on behalf of the Wellcome Sanger Institute COVID-19 Surveillance Team |
| EPI_ISL_2701712 | Lighthouse Lab in Milton Keynes | Wellcome Sanger Institute for the COVID-19 Genomics UK (COG-UK) Consortium | The Lighthouse Lab in Milton Keynes and Alex Alderton, Roberto Amato, Jeffrey Barrett, Sonia Goncalves, Ewan Harrison, David K. Jackson, Ian Johnston, Dominic Kwiatkowski, Cordelia Langford, John Sillitoe on behalf of the Wellcome Sanger Institute COVID-19 Surveillance Team |
| EPI_ISL_2701713, EPI_ISL_2701714, EPI_ISL_2701715 | Lighthouse Lab in Glasgow | Wellcome Sanger Institute for the COVID-19 Genomics UK (COG-UK) Consortium | Harper VanSteenhouse, Yumi Kasai, David Gray, Carol Clugston, Anna Dominiczak and Alex Alderton, Roberto Amato, Jeffrey Barrett, Sonia Goncalves, Ewan Harrison, David K. Jackson, Ian Johnston, Dominic Kwiatkowski, Cordelia Langford, John Sillitoe on behalf of the Wellcome Sanger Institute COVID-19 Surveillance Team |
| EPI_ISL_2701716, EPI_ISL_2701717 | Lighthouse Lab in Alderley Park | Wellcome Sanger Institute for the COVID-19 Genomics UK (COG-UK) Consortium | Jacquelyn Wynn, Mairead Hyland, The Lighthouse Lab in Alderley Park and Alex Alderton, Roberto Amato, Jeffrey Barrett, Sonia Goncalves, Ewan Harrison, David K. Jackson, Ian Johnston, Dominic Kwiatkowski, Cordelia Langford, John Sillitoe on behalf of the Wellcome Sanger Institute COVID-19 Surveillance Team |
| EPI_ISL_2701718, EPI_ISL_2701719 | Lighthouse Lab in Glasgow | Wellcome Sanger Institute for the COVID-19 Genomics UK (COG-UK) Consortium | Harper VanSteenhouse, Yumi Kasai, David Gray, Carol Clugston, Anna Dominiczak and Alex Alderton, Roberto Amato, Jeffrey Barrett, Sonia Goncalves, Ewan Harrison, David K. Jackson, Ian Johnston, Dominic Kwiatkowski, Cordelia Langford, John Sillitoe on behalf of the Wellcome Sanger Institute COVID-19 Surveillance Team |
| EPI_ISL_2701720 | Lighthouse Lab in Alderley Park | Wellcome Sanger Institute for the COVID-19 Genomics UK (COG-UK) Consortium | Jacquelyn Wynn, Mairead Hyland, The Lighthouse Lab in Alderley Park and Alex Alderton, Roberto Amato, Jeffrey Barrett, Sonia Goncalves, Ewan Harrison, David K. Jackson, Ian Johnston, Dominic Kwiatkowski, Cordelia Langford, John Sillitoe on behalf of the Wellcome Sanger Institute COVID-19 Surveillance Team |
| EPI_ISL_2701721, EPI_ISL_2701722, EPI_ISL_2701723 | Lighthouse Lab in Milton Keynes | Wellcome Sanger Institute for the COVID-19 Genomics UK (COG-UK) Consortium | The Lighthouse Lab in Milton Keynes and Alex Alderton, Roberto Amato, Jeffrey Barrett, Sonia Goncalves, Ewan Harrison, David K. Jackson, Ian Johnston, Dominic Kwiatkowski, Cordelia Langford, John Sillitoe on behalf of the Wellcome Sanger Institute COVID-19 Surveillance Team |
| EPI_ISL_2701724, EPI_ISL_2701725 | Lighthouse Lab in Glasgow | Wellcome Sanger Institute for the COVID-19 Genomics UK (COG-UK) Consortium | Harper VanSteenhouse, Yumi Kasai, David Gray, Carol Clugston, Anna Dominiczak and Alex Alderton, Roberto Amato, Jeffrey Barrett, Sonia Goncalves, Ewan Harrison, David K. Jackson, Ian Johnston, Dominic Kwiatkowski, Cordelia Langford, John Sillitoe on behalf of the Wellcome Sanger Institute COVID-19 Surveillance Team |
| EPI_ISL_2701726 | Lighthouse Lab in Alderley Park | Wellcome Sanger Institute for the COVID-19 Genomics UK (COG-UK) Consortium | Jacquelyn Wynn, Mairead Hyland, The Lighthouse Lab in Alderley Park and Alex Alderton, Roberto Amato, Jeffrey Barrett, Sonia Goncalves, Ewan Harrison, David K. Jackson, Ian Johnston, Dominic Kwiatkowski, Cordelia Langford, John Sillitoe on behalf of the Wellcome Sanger Institute COVID-19 Surveillance Team |
| EPI_ISL_2701727, EPI_ISL_2701728, EPI_ISL_2701729, EPI_ISL_2701730 | Lighthouse Lab in Glasgow | Wellcome Sanger Institute for the COVID-19 Genomics UK (COG-UK) Consortium | Harper VanSteenhouse, Yumi Kasai, David Gray, Carol Clugston, Anna Dominiczak and Alex Alderton, Roberto Amato, Jeffrey Barrett, Sonia Goncalves, Ewan Harrison, David K. Jackson, Ian Johnston, Dominic Kwiatkowski, Cordelia Langford, John Sillitoe on behalf of the Wellcome Sanger Institute COVID-19 Surveillance Team |
| EPI_ISL_2701731 | Lighthouse Lab in Milton Keynes | Wellcome Sanger Institute for the COVID-19 Genomics UK (COG-UK) Consortium | The Lighthouse Lab in Milton Keynes and Alex Alderton, Roberto Amato, Jeffrey Barrett, Sonia Goncalves, Ewan Harrison, David K. Jackson, Ian Johnston, Dominic Kwiatkowski, Cordelia Langford, John Sillitoe on behalf of the Wellcome Sanger Institute COVID-19 Surveillance Team |
| EPI_ISL_2701732 | Lighthouse Lab in Glasgow | Wellcome Sanger Institute for the COVID-19 Genomics UK (COG-UK) Consortium | Harper VanSteenhouse, Yumi Kasai, David Gray, Carol Clugston, Anna Dominiczak and Alex Alderton, Roberto Amato, Jeffrey Barrett, Sonia Goncalves, Ewan Harrison, David K. Jackson, Ian Johnston, Dominic Kwiatkowski, Cordelia Langford, John Sillitoe on behalf of the Wellcome Sanger Institute COVID-19 Surveillance Team |
| EPI_ISL_2701733 | Lighthouse Lab in Milton Keynes | Wellcome Sanger Institute for the COVID-19 Genomics UK (COG-UK) Consortium | The Lighthouse Lab in Milton Keynes and Alex Alderton, Roberto Amato, Jeffrey Barrett, Sonia Goncalves, Ewan Harrison, David K. Jackson, Ian Johnston, Dominic Kwiatkowski, Cordelia Langford, John Sillitoe on behalf of the Wellcome Sanger Institute COVID-19 Surveillance Team |
| EPI_ISL_2701734 | Lighthouse Lab in Glasgow | Wellcome Sanger Institute for the COVID-19 Genomics UK (COG-UK) Consortium | Harper VanSteenhouse, Yumi Kasai, David Gray, Carol Clugston, Anna Dominiczak and Alex Alderton, Roberto Amato, Jeffrey Barrett, Sonia Goncalves, Ewan Harrison, David K. Jackson, Ian Johnston, Dominic Kwiatkowski, Cordelia Langford, John Sillitoe on behalf of the Wellcome Sanger Institute COVID-19 Surveillance Team |
| EPI_ISL_2701735 | Lighthouse Lab in Alderley Park | Wellcome Sanger Institute for the COVID-19 Genomics UK (COG-UK) Consortium | Jacquelyn Wynn, Mairead Hyland, The Lighthouse Lab in Alderley Park and Alex Alderton, Roberto Amato, Jeffrey Barrett, Sonia Goncalves, Ewan Harrison, David K. Jackson, Ian Johnston, Dominic Kwiatkowski, Cordelia Langford, John Sillitoe on behalf of the Wellcome Sanger Institute COVID-19 Surveillance Team |
| EPI_ISL_2701736 | Lighthouse Lab in Milton Keynes | Wellcome Sanger Institute for the COVID-19 Genomics UK (COG-UK) Consortium | The Lighthouse Lab in Milton Keynes and Alex Alderton, Roberto Amato, Jeffrey Barrett, Sonia Goncalves, Ewan Harrison, David K. Jackson, Ian Johnston, Dominic Kwiatkowski, Cordelia Langford, John Sillitoe on behalf of the Wellcome Sanger Institute COVID-19 Surveillance Team |
| EPI_ISL_2701737, EPI_ISL_2701738 | Lighthouse Lab in Glasgow | Wellcome Sanger Institute for the COVID-19 Genomics UK (COG-UK) Consortium | Harper VanSteenhouse, Yumi Kasai, David Gray, Carol Clugston, Anna Dominiczak and Alex Alderton, Roberto Amato, Jeffrey Barrett, Sonia Goncalves, Ewan Harrison, David K. Jackson, Ian Johnston, Dominic Kwiatkowski, Cordelia Langford, John Sillitoe on behalf of the Wellcome Sanger Institute COVID-19 Surveillance Team |
| EPI_ISL_2701739 | Lighthouse Lab in Alderley Park | Wellcome Sanger Institute for the COVID-19 Genomics UK (COG-UK) Consortium | Jacquelyn Wynn, Mairead Hyland, The Lighthouse Lab in Alderley Park and Alex Alderton, Roberto Amato, Jeffrey Barrett, Sonia Goncalves, Ewan Harrison, David K. Jackson, Ian Johnston, Dominic Kwiatkowski, Cordelia Langford, John Sillitoe on behalf of the Wellcome Sanger Institute COVID-19 Surveillance Team |
| EPI_ISL_2701740 | Lighthouse Lab in Glasgow | Wellcome Sanger Institute for the COVID-19 Genomics UK (COG-UK) Consortium | Harper VanSteenhouse, Yumi Kasai, David Gray, Carol Clugston, Anna Dominiczak and Alex Alderton, Roberto Amato, Jeffrey Barrett, Sonia Goncalves, Ewan Harrison, David K. Jackson, Ian Johnston, Dominic Kwiatkowski, Cordelia Langford, John Sillitoe on behalf of the Wellcome Sanger Institute COVID-19 Surveillance Team |
| EPI_ISL_2701741 | Lighthouse Lab in Milton Keynes | Wellcome Sanger Institute for the COVID-19 Genomics UK (COG-UK) Consortium | The Lighthouse Lab in Milton Keynes and Alex Alderton, Roberto Amato, Jeffrey Barrett, Sonia Goncalves, Ewan Harrison, David K. Jackson, Ian Johnston, Dominic Kwiatkowski, Cordelia Langford, John Sillitoe on behalf of the Wellcome Sanger Institute COVID-19 Surveillance Team |
| EPI_ISL_2701742 | Lighthouse Lab in Glasgow | Wellcome Sanger Institute for the COVID-19 Genomics UK (COG-UK) Consortium | Harper VanSteenhouse, Yumi Kasai, David Gray, Carol Clugston, Anna Dominiczak and Alex Alderton, Roberto Amato, Jeffrey Barrett, Sonia Goncalves, Ewan Harrison, David K. Jackson, Ian Johnston, Dominic Kwiatkowski, Cordelia Langford, John Sillitoe on behalf of the Wellcome Sanger Institute COVID-19 Surveillance Team |
| EPI_ISL_2701743 | Lighthouse Lab in Milton Keynes | Wellcome Sanger Institute for the COVID-19 Genomics UK (COG-UK) Consortium | The Lighthouse Lab in Milton Keynes and Alex Alderton, Roberto Amato, Jeffrey Barrett, Sonia Goncalves, Ewan Harrison, David K. Jackson, Ian Johnston, Dominic Kwiatkowski, Cordelia Langford, John Sillitoe on behalf of the Wellcome Sanger Institute COVID-19 Surveillance Team |
| EPI_ISL_2701744 | Lighthouse Lab in Alderley Park | Wellcome Sanger Institute for the COVID-19 Genomics UK (COG-UK) Consortium | Jacquelyn Wynn, Mairead Hyland, The Lighthouse Lab in Alderley Park and Alex Alderton, Roberto Amato, Jeffrey Barrett, Sonia Goncalves, Ewan Harrison, David K. Jackson, Ian Johnston, Dominic Kwiatkowski, Cordelia Langford, John Sillitoe on behalf of the Wellcome Sanger Institute COVID-19 Surveillance Team |
| EPI_ISL_2701745 | Lighthouse Lab in Milton Keynes | Wellcome Sanger Institute for the COVID-19 Genomics UK (COG-UK) Consortium | The Lighthouse Lab in Milton Keynes and Alex Alderton, Roberto Amato, Jeffrey Barrett, Sonia Goncalves, Ewan Harrison, David K. Jackson, Ian Johnston, Dominic Kwiatkowski, Cordelia Langford, John Sillitoe on behalf of the Wellcome Sanger Institute COVID-19 Surveillance Team |
| EPI_ISL_2701746 | Lighthouse Lab in Alderley Park | Wellcome Sanger Institute for the COVID-19 Genomics UK (COG-UK) Consortium | Jacquelyn Wynn, Mairead Hyland, The Lighthouse Lab in Alderley Park and Alex Alderton, Roberto Amato, Jeffrey Barrett, Sonia Goncalves, Ewan Harrison, David K. Jackson, Ian Johnston, Dominic Kwiatkowski, Cordelia Langford, John Sillitoe on behalf of the Wellcome Sanger Institute COVID-19 Surveillance Team |
| EPI_ISL_2701747 | Lighthouse Lab in Glasgow | Wellcome Sanger Institute for the COVID-19 Genomics UK (COG-UK) Consortium | Harper VanSteenhouse, Yumi Kasai, David Gray, Carol Clugston, Anna Dominiczak and Alex Alderton, Roberto Amato, Jeffrey Barrett, Sonia Goncalves, Ewan Harrison, David K. Jackson, Ian Johnston, Dominic Kwiatkowski, Cordelia Langford, John Sillitoe on behalf of the Wellcome Sanger Institute COVID-19 Surveillance Team |
| EPI_ISL_2701748 | Lighthouse Lab in Milton Keynes | Wellcome Sanger Institute for the COVID-19 Genomics UK (COG-UK) | The Lighthouse Lab in Milton Keynes and Alex Alderton, Roberto Amato, Jeffrey Barrett, Sonia Goncalves, Ewan Harrison, David K. Jackson, Ian Johnston, Dominic Kwiatkowski, Cordelia Langford, John Sillitoe on behalf of the Wellcome Sanger |

| | | | |
|---|---|---|---|
| | | Consortium | Institute COVID-19 Surveillance Team |
| EPI_ISL_2701749 | Lighthouse Lab in Alderley Park | Wellcome Sanger Institute for the COVID-19 Genomics UK (COG-UK) Consortium | Jacquelyn Wynn, Mairead Hyland, The Lighthouse Lab in Alderley Park and Alex Alderton, Roberto Amato, Jeffrey Barrett, Sonia Goncalves, Ewan Harrison, David K. Jackson, Ian Johnston, Dominic Kwiatkowski, Cordelia Langford, John Sillitoe on behalf of the Wellcome Sanger Institute COVID-19 Surveillance Team |
| EPI_ISL_2701750 | Lighthouse Lab in Glasgow | Wellcome Sanger Institute for the COVID-19 Genomics UK (COG-UK) Consortium | Harper VanSteenhouse, Yumi Kasai, David Gray, Carol Clugston, Anna Dominiczak and Alex Alderton, Roberto Amato, Jeffrey Barrett, Sonia Goncalves, Ewan Harrison, David K. Jackson, Ian Johnston, Dominic Kwiatkowski, Cordelia Langford, John Sillitoe on behalf of the Wellcome Sanger Institute COVID-19 Surveillance Team |
| EPI_ISL_2701751 | Lighthouse Lab in Milton Keynes | Wellcome Sanger Institute for the COVID-19 Genomics UK (COG-UK) Consortium | The Lighthouse Lab in Milton Keynes and Alex Alderton, Roberto Amato, Jeffrey Barrett, Sonia Goncalves, Ewan Harrison, David K. Jackson, Ian Johnston, Dominic Kwiatkowski, Cordelia Langford, John Sillitoe on behalf of the Wellcome Sanger Institute COVID-19 Surveillance Team |
| EPI_ISL_2701752 | Lighthouse Lab in Alderley Park | Wellcome Sanger Institute for the COVID-19 Genomics UK (COG-UK) Consortium | Jacquelyn Wynn, Mairead Hyland, The Lighthouse Lab in Alderley Park and Alex Alderton, Roberto Amato, Jeffrey Barrett, Sonia Goncalves, Ewan Harrison, David K. Jackson, Ian Johnston, Dominic Kwiatkowski, Cordelia Langford, John Sillitoe on behalf of the Wellcome Sanger Institute COVID-19 Surveillance Team |
| EPI_ISL_2701753 | Lighthouse Lab in Milton Keynes | Wellcome Sanger Institute for the COVID-19 Genomics UK (COG-UK) Consortium | The Lighthouse Lab in Milton Keynes and Alex Alderton, Roberto Amato, Jeffrey Barrett, Sonia Goncalves, Ewan Harrison, David K. Jackson, Ian Johnston, Dominic Kwiatkowski, Cordelia Langford, John Sillitoe on behalf of the Wellcome Sanger Institute COVID-19 Surveillance Team |
| EPI_ISL_2701754 | Lighthouse Lab in Alderley Park | Wellcome Sanger Institute for the COVID-19 Genomics UK (COG-UK) Consortium | Jacquelyn Wynn, Mairead Hyland, The Lighthouse Lab in Alderley Park and Alex Alderton, Roberto Amato, Jeffrey Barrett, Sonia Goncalves, Ewan Harrison, David K. Jackson, Ian Johnston, Dominic Kwiatkowski, Cordelia Langford, John Sillitoe on behalf of the Wellcome Sanger Institute COVID-19 Surveillance Team |
| EPI_ISL_2701755 | Lighthouse Lab in Milton Keynes | Wellcome Sanger Institute for the COVID-19 Genomics UK (COG-UK) Consortium | The Lighthouse Lab in Milton Keynes and Alex Alderton, Roberto Amato, Jeffrey Barrett, Sonia Goncalves, Ewan Harrison, David K. Jackson, Ian Johnston, Dominic Kwiatkowski, Cordelia Langford, John Sillitoe on behalf of the Wellcome Sanger Institute COVID-19 Surveillance Team |
| EPI_ISL_2701756, EPI_ISL_2701757, EPI_ISL_2701758, EPI_ISL_2701759, EPI_ISL_2701760, EPI_ISL_2701761, EPI_ISL_2701762, EPI_ISL_2701763, EPI_ISL_2701764 | Lighthouse Lab in Glasgow | Wellcome Sanger Institute for the COVID-19 Genomics UK (COG-UK) Consortium | Harper VanSteenhouse, Yumi Kasai, David Gray, Carol Clugston, Anna Dominiczak and Alex Alderton, Roberto Amato, Jeffrey Barrett, Sonia Goncalves, Ewan Harrison, David K. Jackson, Ian Johnston, Dominic Kwiatkowski, Cordelia Langford, John Sillitoe on behalf of the Wellcome Sanger Institute COVID-19 Surveillance Team |
| EPI_ISL_2701765 | Lighthouse Lab in Alderley Park | Wellcome Sanger Institute for the COVID-19 Genomics UK (COG-UK) Consortium | Jacquelyn Wynn, Mairead Hyland, The Lighthouse Lab in Alderley Park and Alex Alderton, Roberto Amato, Jeffrey Barrett, Sonia Goncalves, Ewan Harrison, David K. Jackson, Ian Johnston, Dominic Kwiatkowski, Cordelia Langford, John Sillitoe on behalf of the Wellcome Sanger Institute COVID-19 Surveillance Team |
| EPI_ISL_2701766, EPI_ISL_2701767 | Lighthouse Lab in Glasgow | Wellcome Sanger Institute for the COVID-19 Genomics UK (COG-UK) Consortium | Harper VanSteenhouse, Yumi Kasai, David Gray, Carol Clugston, Anna Dominiczak and Alex Alderton, Roberto Amato, Jeffrey Barrett, Sonia Goncalves, Ewan Harrison, David K. Jackson, Ian Johnston, Dominic Kwiatkowski, Cordelia Langford, John Sillitoe on behalf of the Wellcome Sanger Institute COVID-19 Surveillance Team |
| EPI_ISL_2701768, EPI_ISL_2701769, EPI_ISL_2701770, EPI_ISL_2701771, EPI_ISL_2701772 | Lighthouse Lab in Milton Keynes | Wellcome Sanger Institute for the COVID-19 Genomics UK (COG-UK) Consortium | The Lighthouse Lab in Milton Keynes and Alex Alderton, Roberto Amato, Jeffrey Barrett, Sonia Goncalves, Ewan Harrison, David K. Jackson, Ian Johnston, Dominic Kwiatkowski, Cordelia Langford, John Sillitoe on behalf of the Wellcome Sanger Institute COVID-19 Surveillance Team |
| EPI_ISL_2701773 | Lighthouse Lab in Alderley Park | Wellcome Sanger Institute for the COVID-19 Genomics UK (COG-UK) Consortium | Jacquelyn Wynn, Mairead Hyland, The Lighthouse Lab in Alderley Park and Alex Alderton, Roberto Amato, Jeffrey Barrett, Sonia Goncalves, Ewan Harrison, David K. Jackson, Ian Johnston, Dominic Kwiatkowski, Cordelia Langford, John Sillitoe on behalf of the Wellcome Sanger Institute COVID-19 Surveillance Team |
| EPI_ISL_2701774, EPI_ISL_2701775, EPI_ISL_2701776 | Lighthouse Lab in Glasgow | Wellcome Sanger Institute for the COVID-19 Genomics UK (COG-UK) Consortium | Harper VanSteenhouse, Yumi Kasai, David Gray, Carol Clugston, Anna Dominiczak and Alex Alderton, Roberto Amato, Jeffrey Barrett, Sonia Goncalves, Ewan Harrison, David K. Jackson, Ian Johnston, Dominic Kwiatkowski, Cordelia Langford, John Sillitoe on behalf of the Wellcome Sanger Institute COVID-19 Surveillance Team |
| EPI_ISL_2701777, EPI_ISL_2701778, EPI_ISL_2701779, EPI_ISL_2701780, EPI_ISL_2701782 | Lighthouse Lab in Alderley Park | Wellcome Sanger Institute for the COVID-19 Genomics UK (COG-UK) Consortium | Jacquelyn Wynn, Mairead Hyland, The Lighthouse Lab in Alderley Park and Alex Alderton, Roberto Amato, Jeffrey Barrett, Sonia Goncalves, Ewan Harrison, David K. Jackson, Ian Johnston, Dominic Kwiatkowski, Cordelia Langford, John Sillitoe on behalf of the Wellcome Sanger Institute COVID-19 Surveillance Team |
| EPI_ISL_2701784 | Lighthouse Lab in Glasgow | Wellcome Sanger Institute for the COVID-19 Genomics UK (COG-UK) Consortium | Harper VanSteenhouse, Yumi Kasai, David Gray, Carol Clugston, Anna Dominiczak and Alex Alderton, Roberto Amato, Jeffrey Barrett, Sonia Goncalves, Ewan Harrison, David K. Jackson, Ian Johnston, Dominic Kwiatkowski, Cordelia Langford, John Sillitoe on behalf of the Wellcome Sanger Institute COVID-19 Surveillance Team |
| EPI_ISL_2701785 | Lighthouse Lab in Milton Keynes | Wellcome Sanger Institute for the COVID-19 Genomics UK (COG-UK) Consortium | The Lighthouse Lab in Milton Keynes and Alex Alderton, Roberto Amato, Jeffrey Barrett, Sonia Goncalves, Ewan Harrison, David K. Jackson, Ian Johnston, Dominic Kwiatkowski, Cordelia Langford, John Sillitoe on behalf of the Wellcome Sanger Institute COVID-19 Surveillance Team |
| EPI_ISL_2701786 | Lighthouse Lab in Alderley Park | Wellcome Sanger Institute for the COVID-19 Genomics UK (COG-UK) Consortium | Jacquelyn Wynn, Mairead Hyland, The Lighthouse Lab in Alderley Park and Alex Alderton, Roberto Amato, Jeffrey Barrett, Sonia Goncalves, Ewan Harrison, David K. Jackson, Ian Johnston, Dominic Kwiatkowski, Cordelia Langford, John Sillitoe on behalf of the Wellcome Sanger Institute COVID-19 Surveillance Team |
| EPI_ISL_2701787, EPI_ISL_2701788, EPI_ISL_2701789 | Lighthouse Lab in Glasgow | Wellcome Sanger Institute for the COVID-19 Genomics UK (COG-UK) Consortium | Harper VanSteenhouse, Yumi Kasai, David Gray, Carol Clugston, Anna Dominiczak and Alex Alderton, Roberto Amato, Jeffrey Barrett, Sonia Goncalves, Ewan Harrison, David K. Jackson, Ian Johnston, Dominic Kwiatkowski, Cordelia Langford, John Sillitoe on behalf of the Wellcome Sanger Institute COVID-19 Surveillance Team |
| EPI_ISL_2701790 | Lighthouse Lab in Milton Keynes | Wellcome Sanger Institute for the COVID-19 Genomics UK (COG-UK) Consortium | The Lighthouse Lab in Milton Keynes and Alex Alderton, Roberto Amato, Jeffrey Barrett, Sonia Goncalves, Ewan Harrison, David K. Jackson, Ian Johnston, Dominic Kwiatkowski, Cordelia Langford, John Sillitoe on behalf of the Wellcome Sanger Institute COVID-19 Surveillance Team |
| EPI_ISL_2701791, EPI_ISL_2701792, EPI_ISL_2701793, EPI_ISL_2701794 | Lighthouse Lab in Glasgow | Wellcome Sanger Institute for the COVID-19 Genomics UK (COG-UK) Consortium | Harper VanSteenhouse, Yumi Kasai, David Gray, Carol Clugston, Anna Dominiczak and Alex Alderton, Roberto Amato, Jeffrey Barrett, Sonia Goncalves, Ewan Harrison, David K. Jackson, Ian Johnston, Dominic Kwiatkowski, Cordelia Langford, John Sillitoe on behalf of the Wellcome Sanger Institute COVID-19 Surveillance Team |
| EPI_ISL_2701795, EPI_ISL_2701796 | Lighthouse Lab in Alderley Park | Wellcome Sanger Institute for the COVID-19 Genomics UK (COG-UK) Consortium | Jacquelyn Wynn, Mairead Hyland, The Lighthouse Lab in Alderley Park and Alex Alderton, Roberto Amato, Jeffrey Barrett, Sonia Goncalves, Ewan Harrison, David K. Jackson, Ian Johnston, Dominic Kwiatkowski, Cordelia Langford, John Sillitoe on behalf of the Wellcome Sanger Institute COVID-19 Surveillance Team |
| EPI_ISL_2701797, EPI_ISL_2701798, EPI_ISL_2701799, EPI_ISL_2701800 | Lighthouse Lab in Glasgow | Wellcome Sanger Institute for the COVID-19 Genomics UK (COG-UK) Consortium | Harper VanSteenhouse, Yumi Kasai, David Gray, Carol Clugston, Anna Dominiczak and Alex Alderton, Roberto Amato, Jeffrey Barrett, Sonia Goncalves, Ewan Harrison, David K. Jackson, Ian Johnston, Dominic Kwiatkowski, Cordelia Langford, John Sillitoe on behalf of the Wellcome Sanger Institute COVID-19 Surveillance Team |
| EPI_ISL_2701801, EPI_ISL_2701802 | Lighthouse Lab in Alderley Park | Wellcome Sanger Institute for the COVID-19 Genomics UK (COG-UK) Consortium | Jacquelyn Wynn, Mairead Hyland, The Lighthouse Lab in Alderley Park and Alex Alderton, Roberto Amato, Jeffrey Barrett, Sonia Goncalves, Ewan Harrison, David K. Jackson, Ian Johnston, Dominic Kwiatkowski, Cordelia Langford, John Sillitoe on behalf of the Wellcome Sanger Institute COVID-19 Surveillance Team |
| EPI_ISL_2701803 | Lighthouse Lab in Milton Keynes | Wellcome Sanger Institute for the COVID-19 Genomics UK (COG-UK) Consortium | The Lighthouse Lab in Milton Keynes and Alex Alderton, Roberto Amato, Jeffrey Barrett, Sonia Goncalves, Ewan Harrison, David K. Jackson, Ian Johnston, Dominic Kwiatkowski, Cordelia Langford, John Sillitoe on behalf of the Wellcome Sanger Institute COVID-19 Surveillance Team |
| EPI_ISL_2701804 | Lighthouse Lab in Glasgow | Wellcome Sanger Institute for the COVID-19 Genomics UK (COG-UK) Consortium | Harper VanSteenhouse, Yumi Kasai, David Gray, Carol Clugston, Anna Dominiczak and Alex Alderton, Roberto Amato, Jeffrey Barrett, Sonia Goncalves, Ewan Harrison, David K. Jackson, Ian Johnston, Dominic Kwiatkowski, Cordelia Langford, John Sillitoe on behalf of the Wellcome Sanger Institute COVID-19 Surveillance Team |
| EPI_ISL_2701805 | Lighthouse Lab in Milton Keynes | Wellcome Sanger Institute for the COVID-19 Genomics UK (COG-UK) Consortium | The Lighthouse Lab in Milton Keynes and Alex Alderton, Roberto Amato, Jeffrey Barrett, Sonia Goncalves, Ewan Harrison, David K. Jackson, Ian Johnston, Dominic Kwiatkowski, Cordelia Langford, John Sillitoe on behalf of the Wellcome Sanger Institute COVID-19 Surveillance Team |
| EPI_ISL_2701806, EPI_ISL_2701807 | Lighthouse Lab in Glasgow | Wellcome Sanger Institute for the COVID-19 Genomics UK (COG-UK) Consortium | Harper VanSteenhouse, Yumi Kasai, David Gray, Carol Clugston, Anna Dominiczak and Alex Alderton, Roberto Amato, Jeffrey Barrett, Sonia Goncalves, Ewan Harrison, David K. Jackson, Ian Johnston, Dominic Kwiatkowski, Cordelia Langford, John Sillitoe on behalf of the Wellcome Sanger Institute COVID-19 Surveillance Team |
| EPI_ISL_2701808, EPI_ISL_2701809 | Lighthouse Lab in Alderley Park | Wellcome Sanger Institute for the COVID-19 Genomics UK (COG-UK) Consortium | Jacquelyn Wynn, Mairead Hyland, The Lighthouse Lab in Alderley Park and Alex Alderton, Roberto Amato, Jeffrey Barrett, Sonia Goncalves, Ewan Harrison, David K. Jackson, Ian Johnston, Dominic Kwiatkowski, Cordelia Langford, John Sillitoe on behalf of the Wellcome Sanger Institute COVID-19 Surveillance Team |
| EPI_ISL_2701810, EPI_ISL_2701811 | Lighthouse Lab in Milton Keynes | Wellcome Sanger Institute for the COVID-19 Genomics UK (COG-UK) Consortium | The Lighthouse Lab in Milton Keynes and Alex Alderton, Roberto Amato, Jeffrey Barrett, Sonia Goncalves, Ewan Harrison, David K. Jackson, Ian Johnston, Dominic Kwiatkowski, Cordelia Langford, John Sillitoe on behalf of the Wellcome Sanger Institute COVID-19 Surveillance Team |
| EPI_ISL_2701812, EPI_ISL_2701813 | Lighthouse Lab in Alderley Park | Wellcome Sanger Institute for the COVID-19 Genomics UK (COG-UK) Consortium | Jacquelyn Wynn, Mairead Hyland, The Lighthouse Lab in Alderley Park and Alex Alderton, Roberto Amato, Jeffrey Barrett, Sonia Goncalves, Ewan Harrison, David K. Jackson, Ian Johnston, Dominic Kwiatkowski, Cordelia Langford, John Sillitoe on behalf of the Wellcome Sanger Institute COVID-19 Surveillance Team |
| EPI_ISL_2701814 | Lighthouse Lab in Glasgow | Wellcome Sanger Institute for the COVID-19 Genomics UK (COG-UK) Consortium | Harper VanSteenhouse, Yumi Kasai, David Gray, Carol Clugston, Anna Dominiczak and Alex Alderton, Roberto Amato, Jeffrey Barrett, Sonia Goncalves, Ewan Harrison, David K. Jackson, Ian Johnston, Dominic Kwiatkowski, Cordelia Langford, John Sillitoe on behalf of the Wellcome Sanger Institute COVID-19 Surveillance Team |
| EPI_ISL_2701815 | Lighthouse Lab in Alderley Park | Wellcome Sanger Institute for the COVID-19 Genomics UK (COG-UK) Consortium | Jacquelyn Wynn, Mairead Hyland, The Lighthouse Lab in Alderley Park and Alex Alderton, Roberto Amato, Jeffrey Barrett, Sonia Goncalves, Ewan Harrison, David K. Jackson, Ian Johnston, Dominic Kwiatkowski, Cordelia Langford, John Sillitoe on behalf of the Wellcome Sanger Institute COVID-19 Surveillance Team |
| EPI_ISL_2701816 | Lighthouse Lab in Milton Keynes | Wellcome Sanger Institute for the COVID-19 Genomics UK (COG-UK) Consortium | The Lighthouse Lab in Milton Keynes and Alex Alderton, Roberto Amato, Jeffrey Barrett, Sonia Goncalves, Ewan Harrison, David K. Jackson, Ian Johnston, Dominic Kwiatkowski, Cordelia Langford, John Sillitoe on behalf of the Wellcome Sanger Institute COVID-19 Surveillance Team |
| EPI_ISL_2701817 | Lighthouse Lab in Alderley Park | Wellcome Sanger Institute for the COVID-19 Genomics UK (COG-UK) Consortium | Jacquelyn Wynn, Mairead Hyland, The Lighthouse Lab in Alderley Park and Alex Alderton, Roberto Amato, Jeffrey Barrett, Sonia Goncalves, Ewan Harrison, David K. Jackson, Ian Johnston, Dominic Kwiatkowski, Cordelia Langford, John Sillitoe on behalf of the Wellcome Sanger Institute COVID-19 Surveillance Team |
| EPI_ISL_2701818 | Lighthouse Lab in Milton Keynes | Wellcome Sanger Institute for the COVID-19 Genomics UK (COG-UK) Consortium | The Lighthouse Lab in Milton Keynes and Alex Alderton, Roberto Amato, Jeffrey Barrett, Sonia Goncalves, Ewan Harrison, David K. Jackson, Ian Johnston, Dominic Kwiatkowski, Cordelia Langford, John Sillitoe on behalf of the Wellcome Sanger Institute COVID-19 Surveillance Team |
| EPI_ISL_2701819, EPI_ISL_2701820, EPI_ISL_2701821 | Lighthouse Lab in Alderley Park | Wellcome Sanger Institute for the COVID-19 Genomics UK (COG-UK) Consortium | Jacquelyn Wynn, Mairead Hyland, The Lighthouse Lab in Alderley Park and Alex Alderton, Roberto Amato, Jeffrey Barrett, Sonia Goncalves, Ewan Harrison, David K. Jackson, Ian Johnston, Dominic Kwiatkowski, Cordelia Langford, John Sillitoe on behalf of the Wellcome Sanger Institute COVID-19 Surveillance Team |
| EPI_ISL_2701822, | Lighthouse Lab in Milton Keynes | Wellcome Sanger Institute for the COVID-19 Genomics UK (COG-UK) | The Lighthouse Lab in Milton Keynes and Alex Alderton, Roberto Amato, Jeffrey Barrett, Sonia Goncalves, Ewan Harrison, David K. Jackson, Ian Johnston, Dominic Kwiatkowski, Cordelia Langford, John Sillitoe on behalf of the Wellcome Sanger |

| | | | |
|---|---|---|---|
| EPI_ISL_2701824 | | Consortium | Institute COVID-19 Surveillance Team |
| EPI_ISL_2701825 | Lighthouse Lab in Glasgow | Wellcome Sanger Institute for the COVID-19 Genomics UK (COG-UK) Consortium | Harper VanSteenhouse, Yumi Kasai, David Gray, Carol Clugston, Anna Dominiczak and Alex Alderton, Roberto Amato, Jeffrey Barrett, Sonia Goncalves, Ewan Harrison, David K. Jackson, Ian Johnston, Dominic Kwiatkowski, Cordelia Langford, John Sillitoe on behalf of the Wellcome Sanger Institute COVID-19 Surveillance Team |
| EPI_ISL_2701826, EPI_ISL_2701827 | Lighthouse Lab in Alderley Park | Wellcome Sanger Institute for the COVID-19 Genomics UK (COG-UK) Consortium | Jacquelyn Wynn, Mairead Hyland, The Lighthouse Lab in Alderley Park and Alex Alderton, Roberto Amato, Jeffrey Barrett, Sonia Goncalves, Ewan Harrison, David K. Jackson, Ian Johnston, Dominic Kwiatkowski, Cordelia Langford, John Sillitoe on behalf of the Wellcome Sanger Institute COVID-19 Surveillance Team |
| EPI_ISL_2701828 | Lighthouse Lab in Glasgow | Wellcome Sanger Institute for the COVID-19 Genomics UK (COG-UK) Consortium | Harper VanSteenhouse, Yumi Kasai, David Gray, Carol Clugston, Anna Dominiczak and Alex Alderton, Roberto Amato, Jeffrey Barrett, Sonia Goncalves, Ewan Harrison, David K. Jackson, Ian Johnston, Dominic Kwiatkowski, Cordelia Langford, John Sillitoe on behalf of the Wellcome Sanger Institute COVID-19 Surveillance Team |
| EPI_ISL_2701829 | Lighthouse Lab in Alderley Park | Wellcome Sanger Institute for the COVID-19 Genomics UK (COG-UK) Consortium | Jacquelyn Wynn, Mairead Hyland, The Lighthouse Lab in Alderley Park and Alex Alderton, Roberto Amato, Jeffrey Barrett, Sonia Goncalves, Ewan Harrison, David K. Jackson, Ian Johnston, Dominic Kwiatkowski, Cordelia Langford, John Sillitoe on behalf of the Wellcome Sanger Institute COVID-19 Surveillance Team |
| EPI_ISL_2701830 | Lighthouse Lab in Milton Keynes | Wellcome Sanger Institute for the COVID-19 Genomics UK (COG-UK) Consortium | The Lighthouse Lab in Milton Keynes and Alex Alderton, Roberto Amato, Jeffrey Barrett, Sonia Goncalves, Ewan Harrison, David K. Jackson, Ian Johnston, Dominic Kwiatkowski, Cordelia Langford, John Sillitoe on behalf of the Wellcome Sanger Institute COVID-19 Surveillance Team |
| EPI_ISL_2701831 | Lighthouse Lab in Alderley Park | Wellcome Sanger Institute for the COVID-19 Genomics UK (COG-UK) Consortium | Jacquelyn Wynn, Mairead Hyland, The Lighthouse Lab in Alderley Park and Alex Alderton, Roberto Amato, Jeffrey Barrett, Sonia Goncalves, Ewan Harrison, David K. Jackson, Ian Johnston, Dominic Kwiatkowski, Cordelia Langford, John Sillitoe on behalf of the Wellcome Sanger Institute COVID-19 Surveillance Team |
| EPI_ISL_2701832, EPI_ISL_2701833 | Lighthouse Lab in Glasgow | Wellcome Sanger Institute for the COVID-19 Genomics UK (COG-UK) Consortium | Harper VanSteenhouse, Yumi Kasai, David Gray, Carol Clugston, Anna Dominiczak and Alex Alderton, Roberto Amato, Jeffrey Barrett, Sonia Goncalves, Ewan Harrison, David K. Jackson, Ian Johnston, Dominic Kwiatkowski, Cordelia Langford, John Sillitoe on behalf of the Wellcome Sanger Institute COVID-19 Surveillance Team |
| EPI_ISL_2701834 | Lighthouse Lab in Alderley Park | Wellcome Sanger Institute for the COVID-19 Genomics UK (COG-UK) Consortium | Jacquelyn Wynn, Mairead Hyland, The Lighthouse Lab in Alderley Park and Alex Alderton, Roberto Amato, Jeffrey Barrett, Sonia Goncalves, Ewan Harrison, David K. Jackson, Ian Johnston, Dominic Kwiatkowski, Cordelia Langford, John Sillitoe on behalf of the Wellcome Sanger Institute COVID-19 Surveillance Team |
| EPI_ISL_2701835 | Lighthouse Lab in Glasgow | Wellcome Sanger Institute for the COVID-19 Genomics UK (COG-UK) Consortium | Harper VanSteenhouse, Yumi Kasai, David Gray, Carol Clugston, Anna Dominiczak and Alex Alderton, Roberto Amato, Jeffrey Barrett, Sonia Goncalves, Ewan Harrison, David K. Jackson, Ian Johnston, Dominic Kwiatkowski, Cordelia Langford, John Sillitoe on behalf of the Wellcome Sanger Institute COVID-19 Surveillance Team |
| EPI_ISL_2701836, EPI_ISL_2701837 | Lighthouse Lab in Alderley Park | Wellcome Sanger Institute for the COVID-19 Genomics UK (COG-UK) Consortium | Jacquelyn Wynn, Mairead Hyland, The Lighthouse Lab in Alderley Park and Alex Alderton, Roberto Amato, Jeffrey Barrett, Sonia Goncalves, Ewan Harrison, David K. Jackson, Ian Johnston, Dominic Kwiatkowski, Cordelia Langford, John Sillitoe on behalf of the Wellcome Sanger Institute COVID-19 Surveillance Team |
| EPI_ISL_2701838, EPI_ISL_2701839, EPI_ISL_2701840, EPI_ISL_2701841, EPI_ISL_2701842, EPI_ISL_2701843, EPI_ISL_2701844, EPI_ISL_2701845 | Lighthouse Lab in Glasgow | Wellcome Sanger Institute for the COVID-19 Genomics UK (COG-UK) Consortium | Harper VanSteenhouse, Yumi Kasai, David Gray, Carol Clugston, Anna Dominiczak and Alex Alderton, Roberto Amato, Jeffrey Barrett, Sonia Goncalves, Ewan Harrison, David K. Jackson, Ian Johnston, Dominic Kwiatkowski, Cordelia Langford, John Sillitoe on behalf of the Wellcome Sanger Institute COVID-19 Surveillance Team |
| EPI_ISL_2701846 | Lighthouse Lab in Alderley Park | Wellcome Sanger Institute for the COVID-19 Genomics UK (COG-UK) Consortium | Jacquelyn Wynn, Mairead Hyland, The Lighthouse Lab in Alderley Park and Alex Alderton, Roberto Amato, Jeffrey Barrett, Sonia Goncalves, Ewan Harrison, David K. Jackson, Ian Johnston, Dominic Kwiatkowski, Cordelia Langford, John Sillitoe on behalf of the Wellcome Sanger Institute COVID-19 Surveillance Team |
| EPI_ISL_2701847 | Lighthouse Lab in Milton Keynes | Wellcome Sanger Institute for the COVID-19 Genomics UK (COG-UK) Consortium | The Lighthouse Lab in Milton Keynes and Alex Alderton, Roberto Amato, Jeffrey Barrett, Sonia Goncalves, Ewan Harrison, David K. Jackson, Ian Johnston, Dominic Kwiatkowski, Cordelia Langford, John Sillitoe on behalf of the Wellcome Sanger Institute COVID-19 Surveillance Team |
| EPI_ISL_2701848, EPI_ISL_2701849 | Lighthouse Lab in Glasgow | Wellcome Sanger Institute for the COVID-19 Genomics UK (COG-UK) Consortium | Harper VanSteenhouse, Yumi Kasai, David Gray, Carol Clugston, Anna Dominiczak and Alex Alderton, Roberto Amato, Jeffrey Barrett, Sonia Goncalves, Ewan Harrison, David K. Jackson, Ian Johnston, Dominic Kwiatkowski, Cordelia Langford, John Sillitoe on behalf of the Wellcome Sanger Institute COVID-19 Surveillance Team |
| EPI_ISL_2701850, EPI_ISL_2701851 | Lighthouse Lab in Milton Keynes | Wellcome Sanger Institute for the COVID-19 Genomics UK (COG-UK) Consortium | The Lighthouse Lab in Milton Keynes and Alex Alderton, Roberto Amato, Jeffrey Barrett, Sonia Goncalves, Ewan Harrison, David K. Jackson, Ian Johnston, Dominic Kwiatkowski, Cordelia Langford, John Sillitoe on behalf of the Wellcome Sanger Institute COVID-19 Surveillance Team |
| EPI_ISL_2701852 | Lighthouse Lab in Glasgow | Wellcome Sanger Institute for the COVID-19 Genomics UK (COG-UK) Consortium | Harper VanSteenhouse, Yumi Kasai, David Gray, Carol Clugston, Anna Dominiczak and Alex Alderton, Roberto Amato, Jeffrey Barrett, Sonia Goncalves, Ewan Harrison, David K. Jackson, Ian Johnston, Dominic Kwiatkowski, Cordelia Langford, John Sillitoe on behalf of the Wellcome Sanger Institute COVID-19 Surveillance Team |
| EPI_ISL_2701853, EPI_ISL_2701854 | Lighthouse Lab in Alderley Park | Wellcome Sanger Institute for the COVID-19 Genomics UK (COG-UK) Consortium | Jacquelyn Wynn, Mairead Hyland, The Lighthouse Lab in Alderley Park and Alex Alderton, Roberto Amato, Jeffrey Barrett, Sonia Goncalves, Ewan Harrison, David K. Jackson, Ian Johnston, Dominic Kwiatkowski, Cordelia Langford, John Sillitoe on behalf of the Wellcome Sanger Institute COVID-19 Surveillance Team |
| EPI_ISL_2701855 | Lighthouse Lab in Glasgow | Wellcome Sanger Institute for the COVID-19 Genomics UK (COG-UK) Consortium | Harper VanSteenhouse, Yumi Kasai, David Gray, Carol Clugston, Anna Dominiczak and Alex Alderton, Roberto Amato, Jeffrey Barrett, Sonia Goncalves, Ewan Harrison, David K. Jackson, Ian Johnston, Dominic Kwiatkowski, Cordelia Langford, John Sillitoe on behalf of the Wellcome Sanger Institute COVID-19 Surveillance Team |
| EPI_ISL_2701856 | Lighthouse Lab in Alderley Park | Wellcome Sanger Institute for the COVID-19 Genomics UK (COG-UK) Consortium | Jacquelyn Wynn, Mairead Hyland, The Lighthouse Lab in Alderley Park and Alex Alderton, Roberto Amato, Jeffrey Barrett, Sonia Goncalves, Ewan Harrison, David K. Jackson, Ian Johnston, Dominic Kwiatkowski, Cordelia Langford, John Sillitoe on behalf of the Wellcome Sanger Institute COVID-19 Surveillance Team |
| EPI_ISL_2701857, EPI_ISL_2701858, EPI_ISL_2701859 | Lighthouse Lab in Glasgow | Wellcome Sanger Institute for the COVID-19 Genomics UK (COG-UK) Consortium | Harper VanSteenhouse, Yumi Kasai, David Gray, Carol Clugston, Anna Dominiczak and Alex Alderton, Roberto Amato, Jeffrey Barrett, Sonia Goncalves, Ewan Harrison, David K. Jackson, Ian Johnston, Dominic Kwiatkowski, Cordelia Langford, John Sillitoe on behalf of the Wellcome Sanger Institute COVID-19 Surveillance Team |
| EPI_ISL_2701860 | Lighthouse Lab in Alderley Park | Wellcome Sanger Institute for the COVID-19 Genomics UK (COG-UK) Consortium | Jacquelyn Wynn, Mairead Hyland, The Lighthouse Lab in Alderley Park and Alex Alderton, Roberto Amato, Jeffrey Barrett, Sonia Goncalves, Ewan Harrison, David K. Jackson, Ian Johnston, Dominic Kwiatkowski, Cordelia Langford, John Sillitoe on behalf of the Wellcome Sanger Institute COVID-19 Surveillance Team |
| EPI_ISL_2701861 | Lighthouse Lab in Milton Keynes | Wellcome Sanger Institute for the COVID-19 Genomics UK (COG-UK) Consortium | The Lighthouse Lab in Milton Keynes and Alex Alderton, Roberto Amato, Jeffrey Barrett, Sonia Goncalves, Ewan Harrison, David K. Jackson, Ian Johnston, Dominic Kwiatkowski, Cordelia Langford, John Sillitoe on behalf of the Wellcome Sanger Institute COVID-19 Surveillance Team |
| EPI_ISL_2701862 | Lighthouse Lab in Alderley Park | Wellcome Sanger Institute for the COVID-19 Genomics UK (COG-UK) Consortium | Jacquelyn Wynn, Mairead Hyland, The Lighthouse Lab in Alderley Park and Alex Alderton, Roberto Amato, Jeffrey Barrett, Sonia Goncalves, Ewan Harrison, David K. Jackson, Ian Johnston, Dominic Kwiatkowski, Cordelia Langford, John Sillitoe on behalf of the Wellcome Sanger Institute COVID-19 Surveillance Team |
| EPI_ISL_2701863, EPI_ISL_2701864 | Lighthouse Lab in Milton Keynes | Wellcome Sanger Institute for the COVID-19 Genomics UK (COG-UK) Consortium | The Lighthouse Lab in Milton Keynes and Alex Alderton, Roberto Amato, Jeffrey Barrett, Sonia Goncalves, Ewan Harrison, David K. Jackson, Ian Johnston, Dominic Kwiatkowski, Cordelia Langford, John Sillitoe on behalf of the Wellcome Sanger Institute COVID-19 Surveillance Team |
| EPI_ISL_2701865 | Lighthouse Lab in Glasgow | Wellcome Sanger Institute for the COVID-19 Genomics UK (COG-UK) Consortium | Harper VanSteenhouse, Yumi Kasai, David Gray, Carol Clugston, Anna Dominiczak and Alex Alderton, Roberto Amato, Jeffrey Barrett, Sonia Goncalves, Ewan Harrison, David K. Jackson, Ian Johnston, Dominic Kwiatkowski, Cordelia Langford, John Sillitoe on behalf of the Wellcome Sanger Institute COVID-19 Surveillance Team |
| EPI_ISL_2701866, EPI_ISL_2701867 | Lighthouse Lab in Milton Keynes | Wellcome Sanger Institute for the COVID-19 Genomics UK (COG-UK) Consortium | The Lighthouse Lab in Milton Keynes and Alex Alderton, Roberto Amato, Jeffrey Barrett, Sonia Goncalves, Ewan Harrison, David K. Jackson, Ian Johnston, Dominic Kwiatkowski, Cordelia Langford, John Sillitoe on behalf of the Wellcome Sanger Institute COVID-19 Surveillance Team |
| EPI_ISL_2701868 | Lighthouse Lab in Alderley Park | Wellcome Sanger Institute for the COVID-19 Genomics UK (COG-UK) Consortium | Jacquelyn Wynn, Mairead Hyland, The Lighthouse Lab in Alderley Park and Alex Alderton, Roberto Amato, Jeffrey Barrett, Sonia Goncalves, Ewan Harrison, David K. Jackson, Ian Johnston, Dominic Kwiatkowski, Cordelia Langford, John Sillitoe on behalf of the Wellcome Sanger Institute COVID-19 Surveillance Team |
| EPI_ISL_2701869 | Lighthouse Lab in Glasgow | Wellcome Sanger Institute for the COVID-19 Genomics UK (COG-UK) Consortium | Harper VanSteenhouse, Yumi Kasai, David Gray, Carol Clugston, Anna Dominiczak and Alex Alderton, Roberto Amato, Jeffrey Barrett, Sonia Goncalves, Ewan Harrison, David K. Jackson, Ian Johnston, Dominic Kwiatkowski, Cordelia Langford, John Sillitoe on behalf of the Wellcome Sanger Institute COVID-19 Surveillance Team |
| EPI_ISL_2701870 | Lighthouse Lab in Alderley Park | Wellcome Sanger Institute for the COVID-19 Genomics UK (COG-UK) Consortium | Jacquelyn Wynn, Mairead Hyland, The Lighthouse Lab in Alderley Park and Alex Alderton, Roberto Amato, Jeffrey Barrett, Sonia Goncalves, Ewan Harrison, David K. Jackson, Ian Johnston, Dominic Kwiatkowski, Cordelia Langford, John Sillitoe on behalf of the Wellcome Sanger Institute COVID-19 Surveillance Team |
| EPI_ISL_2701871, EPI_ISL_2701872, EPI_ISL_2701873, EPI_ISL_2701874, EPI_ISL_2701875, EPI_ISL_2701876 | Lighthouse Lab in Glasgow | Wellcome Sanger Institute for the COVID-19 Genomics UK (COG-UK) Consortium | Harper VanSteenhouse, Yumi Kasai, David Gray, Carol Clugston, Anna Dominiczak and Alex Alderton, Roberto Amato, Jeffrey Barrett, Sonia Goncalves, Ewan Harrison, David K. Jackson, Ian Johnston, Dominic Kwiatkowski, Cordelia Langford, John Sillitoe on behalf of the Wellcome Sanger Institute COVID-19 Surveillance Team |
| EPI_ISL_2701877 | Lighthouse Lab in Milton Keynes | Wellcome Sanger Institute for the COVID-19 Genomics UK (COG-UK) Consortium | The Lighthouse Lab in Milton Keynes and Alex Alderton, Roberto Amato, Jeffrey Barrett, Sonia Goncalves, Ewan Harrison, David K. Jackson, Ian Johnston, Dominic Kwiatkowski, Cordelia Langford, John Sillitoe on behalf of the Wellcome Sanger Institute COVID-19 Surveillance Team |
| EPI_ISL_2701878 | Lighthouse Lab in Alderley Park | Wellcome Sanger Institute for the COVID-19 Genomics UK (COG-UK) Consortium | Jacquelyn Wynn, Mairead Hyland, The Lighthouse Lab in Alderley Park and Alex Alderton, Roberto Amato, Jeffrey Barrett, Sonia Goncalves, Ewan Harrison, David K. Jackson, Ian Johnston, Dominic Kwiatkowski, Cordelia Langford, John Sillitoe on behalf of the Wellcome Sanger Institute COVID-19 Surveillance Team |
| EPI_ISL_2701879 | Lighthouse Lab in Milton Keynes | Wellcome Sanger Institute for the COVID-19 Genomics UK (COG-UK) Consortium | The Lighthouse Lab in Milton Keynes and Alex Alderton, Roberto Amato, Jeffrey Barrett, Sonia Goncalves, Ewan Harrison, David K. Jackson, Ian Johnston, Dominic Kwiatkowski, Cordelia Langford, John Sillitoe on behalf of the Wellcome Sanger Institute COVID-19 Surveillance Team |
| EPI_ISL_2701880 | Lighthouse Lab in Alderley Park | Wellcome Sanger Institute for the COVID-19 Genomics UK (COG-UK) Consortium | Jacquelyn Wynn, Mairead Hyland, The Lighthouse Lab in Alderley Park and Alex Alderton, Roberto Amato, Jeffrey Barrett, Sonia Goncalves, Ewan Harrison, David K. Jackson, Ian Johnston, Dominic Kwiatkowski, Cordelia Langford, John Sillitoe on behalf of the Wellcome Sanger Institute COVID-19 Surveillance Team |
| EPI_ISL_2701881 | Lighthouse Lab in Milton Keynes | Wellcome Sanger Institute for the COVID-19 Genomics UK (COG-UK) Consortium | The Lighthouse Lab in Milton Keynes and Alex Alderton, Roberto Amato, Jeffrey Barrett, Sonia Goncalves, Ewan Harrison, David K. Jackson, Ian Johnston, Dominic Kwiatkowski, Cordelia Langford, John Sillitoe on behalf of the Wellcome Sanger Institute COVID-19 Surveillance Team |
| EPI_ISL_2701882 | Lighthouse Lab in Glasgow | Wellcome Sanger Institute for the COVID-19 Genomics UK (COG-UK) Consortium | Harper VanSteenhouse, Yumi Kasai, David Gray, Carol Clugston, Anna Dominiczak and Alex Alderton, Roberto Amato, Jeffrey Barrett, Sonia Goncalves, Ewan Harrison, David K. Jackson, Ian Johnston, Dominic Kwiatkowski, Cordelia Langford, John Sillitoe on behalf of the Wellcome Sanger Institute COVID-19 Surveillance Team |
| EPI_ISL_2701883, EPI_ISL_2701884 | Lighthouse Lab in Alderley Park | Wellcome Sanger Institute for the COVID-19 Genomics UK (COG-UK) Consortium | Jacquelyn Wynn, Mairead Hyland, The Lighthouse Lab in Alderley Park and Alex Alderton, Roberto Amato, Jeffrey Barrett, Sonia Goncalves, Ewan Harrison, David K. Jackson, Ian Johnston, Dominic Kwiatkowski, Cordelia Langford, John Sillitoe on behalf of the Wellcome Sanger Institute COVID-19 Surveillance Team |
| EPI_ISL_2701885 | Lighthouse Lab in Milton Keynes | Wellcome Sanger Institute for the COVID-19 Genomics UK (COG-UK) Consortium | The Lighthouse Lab in Milton Keynes and Alex Alderton, Roberto Amato, Jeffrey Barrett, Sonia Goncalves, Ewan Harrison, David K. Jackson, Ian Johnston, Dominic Kwiatkowski, Cordelia Langford, John Sillitoe on behalf of the Wellcome Sanger Institute COVID-19 Surveillance Team |
| EPI_ISL_2701886, EPI_ISL_2701887 | Lighthouse Lab in Glasgow | Wellcome Sanger Institute for the COVID-19 Genomics UK (COG-UK) Consortium | Harper VanSteenhouse, Yumi Kasai, David Gray, Carol Clugston, Anna Dominiczak and Alex Alderton, Roberto Amato, Jeffrey Barrett, Sonia Goncalves, Ewan Harrison, David K. Jackson, Ian Johnston, Dominic Kwiatkowski, Cordelia Langford, John Sillitoe on behalf of the Wellcome Sanger Institute COVID-19 Surveillance Team |
| EPI_ISL_2701888 | Lighthouse Lab in Alderley Park | Wellcome Sanger Institute for the COVID-19 Genomics UK (COG-UK) Consortium | Jacquelyn Wynn, Mairead Hyland, The Lighthouse Lab in Alderley Park and Alex Alderton, Roberto Amato, Jeffrey Barrett, Sonia Goncalves, Ewan Harrison, David K. Jackson, Ian Johnston, Dominic Kwiatkowski, Cordelia Langford, John Sillitoe on behalf of the Wellcome Sanger Institute COVID-19 Surveillance Team |

| | | | |
|---|---|---|---|
| EPI_ISL_2701889 | Lighthouse Lab in Glasgow | Wellcome Sanger Institute for the COVID-19 Genomics UK (COG-UK) Consortium | Harper VanSteenhouse, Yumi Kasai, David Gray, Carol Clugston, Anna Dominiczak and Alex Alderton, Roberto Amato, Jeffrey Barrett, Sonia Goncalves, Ewan Harrison, David K. Jackson, Ian Johnston, Dominic Kwiatkowski, Cordelia Langford, John Sillitoe on behalf of the Wellcome Sanger Institute COVID-19 Surveillance Team |
| EPI_ISL_2701890 | Lighthouse Lab in Alderley Park | Wellcome Sanger Institute for the COVID-19 Genomics UK (COG-UK) Consortium | Jacquelyn Wynn, Mairead Hyland, The Lighthouse Lab in Alderley Park and Alex Alderton, Roberto Amato, Jeffrey Barrett, Sonia Goncalves, Ewan Harrison, David K. Jackson, Ian Johnston, Dominic Kwiatkowski, Cordelia Langford, John Sillitoe on behalf of the Wellcome Sanger Institute COVID-19 Surveillance Team |
| EPI_ISL_2701891 | Lighthouse Lab in Glasgow | Wellcome Sanger Institute for the COVID-19 Genomics UK (COG-UK) Consortium | Harper VanSteenhouse, Yumi Kasai, David Gray, Carol Clugston, Anna Dominiczak and Alex Alderton, Roberto Amato, Jeffrey Barrett, Sonia Goncalves, Ewan Harrison, David K. Jackson, Ian Johnston, Dominic Kwiatkowski, Cordelia Langford, John Sillitoe on behalf of the Wellcome Sanger Institute COVID-19 Surveillance Team |
| EPI_ISL_2701892 | Lighthouse Lab in Milton Keynes | Wellcome Sanger Institute for the COVID-19 Genomics UK (COG-UK) Consortium | The Lighthouse Lab in Milton Keynes and Alex Alderton, Roberto Amato, Jeffrey Barrett, Sonia Goncalves, Ewan Harrison, David K. Jackson, Ian Johnston, Dominic Kwiatkowski, Cordelia Langford, John Sillitoe on behalf of the Wellcome Sanger Institute COVID-19 Surveillance Team |
| EPI_ISL_2701893 | Lighthouse Lab in Glasgow | Wellcome Sanger Institute for the COVID-19 Genomics UK (COG-UK) Consortium | Harper VanSteenhouse, Yumi Kasai, David Gray, Carol Clugston, Anna Dominiczak and Alex Alderton, Roberto Amato, Jeffrey Barrett, Sonia Goncalves, Ewan Harrison, David K. Jackson, Ian Johnston, Dominic Kwiatkowski, Cordelia Langford, John Sillitoe on behalf of the Wellcome Sanger Institute COVID-19 Surveillance Team |
| EPI_ISL_2701894, EPI_ISL_2701895 | Lighthouse Lab in Milton Keynes | Wellcome Sanger Institute for the COVID-19 Genomics UK (COG-UK) Consortium | The Lighthouse Lab in Milton Keynes and Alex Alderton, Roberto Amato, Jeffrey Barrett, Sonia Goncalves, Ewan Harrison, David K. Jackson, Ian Johnston, Dominic Kwiatkowski, Cordelia Langford, John Sillitoe on behalf of the Wellcome Sanger Institute COVID-19 Surveillance Team |
| EPI_ISL_2701896, EPI_ISL_2701898, EPI_ISL_2701899, EPI_ISL_2701900 | Lighthouse Lab in Glasgow | Wellcome Sanger Institute for the COVID-19 Genomics UK (COG-UK) Consortium | Harper VanSteenhouse, Yumi Kasai, David Gray, Carol Clugston, Anna Dominiczak and Alex Alderton, Roberto Amato, Jeffrey Barrett, Sonia Goncalves, Ewan Harrison, David K. Jackson, Ian Johnston, Dominic Kwiatkowski, Cordelia Langford, John Sillitoe on behalf of the Wellcome Sanger Institute COVID-19 Surveillance Team |
| EPI_ISL_2701901 | Lighthouse Lab in Alderley Park | Wellcome Sanger Institute for the COVID-19 Genomics UK (COG-UK) Consortium | Jacquelyn Wynn, Mairead Hyland, The Lighthouse Lab in Alderley Park and Alex Alderton, Roberto Amato, Jeffrey Barrett, Sonia Goncalves, Ewan Harrison, David K. Jackson, Ian Johnston, Dominic Kwiatkowski, Cordelia Langford, John Sillitoe on behalf of the Wellcome Sanger Institute COVID-19 Surveillance Team |
| EPI_ISL_2701902 | Lighthouse Lab in Milton Keynes | Wellcome Sanger Institute for the COVID-19 Genomics UK (COG-UK) Consortium | The Lighthouse Lab in Milton Keynes and Alex Alderton, Roberto Amato, Jeffrey Barrett, Sonia Goncalves, Ewan Harrison, David K. Jackson, Ian Johnston, Dominic Kwiatkowski, Cordelia Langford, John Sillitoe on behalf of the Wellcome Sanger Institute COVID-19 Surveillance Team |
| EPI_ISL_2701903 | Lighthouse Lab in Glasgow | Wellcome Sanger Institute for the COVID-19 Genomics UK (COG-UK) Consortium | Harper VanSteenhouse, Yumi Kasai, David Gray, Carol Clugston, Anna Dominiczak and Alex Alderton, Roberto Amato, Jeffrey Barrett, Sonia Goncalves, Ewan Harrison, David K. Jackson, Ian Johnston, Dominic Kwiatkowski, Cordelia Langford, John Sillitoe on behalf of the Wellcome Sanger Institute COVID-19 Surveillance Team |
| EPI_ISL_2701904 | Lighthouse Lab in Alderley Park | Wellcome Sanger Institute for the COVID-19 Genomics UK (COG-UK) Consortium | Jacquelyn Wynn, Mairead Hyland, The Lighthouse Lab in Alderley Park and Alex Alderton, Roberto Amato, Jeffrey Barrett, Sonia Goncalves, Ewan Harrison, David K. Jackson, Ian Johnston, Dominic Kwiatkowski, Cordelia Langford, John Sillitoe on behalf of the Wellcome Sanger Institute COVID-19 Surveillance Team |
| EPI_ISL_2701905 | Lighthouse Lab in Glasgow | Wellcome Sanger Institute for the COVID-19 Genomics UK (COG-UK) Consortium | Harper VanSteenhouse, Yumi Kasai, David Gray, Carol Clugston, Anna Dominiczak and Alex Alderton, Roberto Amato, Jeffrey Barrett, Sonia Goncalves, Ewan Harrison, David K. Jackson, Ian Johnston, Dominic Kwiatkowski, Cordelia Langford, John Sillitoe on behalf of the Wellcome Sanger Institute COVID-19 Surveillance Team |
| EPI_ISL_2701906 | Lighthouse Lab in Alderley Park | Wellcome Sanger Institute for the COVID-19 Genomics UK (COG-UK) Consortium | Jacquelyn Wynn, Mairead Hyland, The Lighthouse Lab in Alderley Park and Alex Alderton, Roberto Amato, Jeffrey Barrett, Sonia Goncalves, Ewan Harrison, David K. Jackson, Ian Johnston, Dominic Kwiatkowski, Cordelia Langford, John Sillitoe on behalf of the Wellcome Sanger Institute COVID-19 Surveillance Team |
| EPI_ISL_2701907, EPI_ISL_2701908 | Lighthouse Lab in Glasgow | Wellcome Sanger Institute for the COVID-19 Genomics UK (COG-UK) Consortium | Harper VanSteenhouse, Yumi Kasai, David Gray, Carol Clugston, Anna Dominiczak and Alex Alderton, Roberto Amato, Jeffrey Barrett, Sonia Goncalves, Ewan Harrison, David K. Jackson, Ian Johnston, Dominic Kwiatkowski, Cordelia Langford, John Sillitoe on behalf of the Wellcome Sanger Institute COVID-19 Surveillance Team |
| EPI_ISL_2701910 | Lighthouse Lab in Alderley Park | Wellcome Sanger Institute for the COVID-19 Genomics UK (COG-UK) Consortium | Jacquelyn Wynn, Mairead Hyland, The Lighthouse Lab in Alderley Park and Alex Alderton, Roberto Amato, Jeffrey Barrett, Sonia Goncalves, Ewan Harrison, David K. Jackson, Ian Johnston, Dominic Kwiatkowski, Cordelia Langford, John Sillitoe on behalf of the Wellcome Sanger Institute COVID-19 Surveillance Team |
| EPI_ISL_2701911, EPI_ISL_2701912 | Lighthouse Lab in Milton Keynes | Wellcome Sanger Institute for the COVID-19 Genomics UK (COG-UK) Consortium | The Lighthouse Lab in Milton Keynes and Alex Alderton, Roberto Amato, Jeffrey Barrett, Sonia Goncalves, Ewan Harrison, David K. Jackson, Ian Johnston, Dominic Kwiatkowski, Cordelia Langford, John Sillitoe on behalf of the Wellcome Sanger Institute COVID-19 Surveillance Team |
| EPI_ISL_2701913 | Lighthouse Lab in Alderley Park | Wellcome Sanger Institute for the COVID-19 Genomics UK (COG-UK) Consortium | Jacquelyn Wynn, Mairead Hyland, The Lighthouse Lab in Alderley Park and Alex Alderton, Roberto Amato, Jeffrey Barrett, Sonia Goncalves, Ewan Harrison, David K. Jackson, Ian Johnston, Dominic Kwiatkowski, Cordelia Langford, John Sillitoe on behalf of the Wellcome Sanger Institute COVID-19 Surveillance Team |
| EPI_ISL_2701914 | Lighthouse Lab in Glasgow | Wellcome Sanger Institute for the COVID-19 Genomics UK (COG-UK) Consortium | Harper VanSteenhouse, Yumi Kasai, David Gray, Carol Clugston, Anna Dominiczak and Alex Alderton, Roberto Amato, Jeffrey Barrett, Sonia Goncalves, Ewan Harrison, David K. Jackson, Ian Johnston, Dominic Kwiatkowski, Cordelia Langford, John Sillitoe on behalf of the Wellcome Sanger Institute COVID-19 Surveillance Team |
| EPI_ISL_2701915, EPI_ISL_2701916 | Lighthouse Lab in Milton Keynes | Wellcome Sanger Institute for the COVID-19 Genomics UK (COG-UK) Consortium | The Lighthouse Lab in Milton Keynes and Alex Alderton, Roberto Amato, Jeffrey Barrett, Sonia Goncalves, Ewan Harrison, David K. Jackson, Ian Johnston, Dominic Kwiatkowski, Cordelia Langford, John Sillitoe on behalf of the Wellcome Sanger Institute COVID-19 Surveillance Team |
| EPI_ISL_2701917, EPI_ISL_2701918 | Lighthouse Lab in Glasgow | Wellcome Sanger Institute for the COVID-19 Genomics UK (COG-UK) Consortium | Harper VanSteenhouse, Yumi Kasai, David Gray, Carol Clugston, Anna Dominiczak and Alex Alderton, Roberto Amato, Jeffrey Barrett, Sonia Goncalves, Ewan Harrison, David K. Jackson, Ian Johnston, Dominic Kwiatkowski, Cordelia Langford, John Sillitoe on behalf of the Wellcome Sanger Institute COVID-19 Surveillance Team |
| EPI_ISL_2701919 | Lighthouse Lab in Alderley Park | Wellcome Sanger Institute for the COVID-19 Genomics UK (COG-UK) Consortium | Jacquelyn Wynn, Mairead Hyland, The Lighthouse Lab in Alderley Park and Alex Alderton, Roberto Amato, Jeffrey Barrett, Sonia Goncalves, Ewan Harrison, David K. Jackson, Ian Johnston, Dominic Kwiatkowski, Cordelia Langford, John Sillitoe on behalf of the Wellcome Sanger Institute COVID-19 Surveillance Team |
| EPI_ISL_2701920, EPI_ISL_2701921, EPI_ISL_2701922, EPI_ISL_2701924, EPI_ISL_2701925 | Lighthouse Lab in Glasgow | Wellcome Sanger Institute for the COVID-19 Genomics UK (COG-UK) Consortium | Harper VanSteenhouse, Yumi Kasai, David Gray, Carol Clugston, Anna Dominiczak and Alex Alderton, Roberto Amato, Jeffrey Barrett, Sonia Goncalves, Ewan Harrison, David K. Jackson, Ian Johnston, Dominic Kwiatkowski, Cordelia Langford, John Sillitoe on behalf of the Wellcome Sanger Institute COVID-19 Surveillance Team |
| EPI_ISL_2701926 | Lighthouse Lab in Milton Keynes | Wellcome Sanger Institute for the COVID-19 Genomics UK (COG-UK) Consortium | The Lighthouse Lab in Milton Keynes and Alex Alderton, Roberto Amato, Jeffrey Barrett, Sonia Goncalves, Ewan Harrison, David K. Jackson, Ian Johnston, Dominic Kwiatkowski, Cordelia Langford, John Sillitoe on behalf of the Wellcome Sanger Institute COVID-19 Surveillance Team |
| EPI_ISL_2701927, EPI_ISL_2701928 | Lighthouse Lab in Glasgow | Wellcome Sanger Institute for the COVID-19 Genomics UK (COG-UK) Consortium | Harper VanSteenhouse, Yumi Kasai, David Gray, Carol Clugston, Anna Dominiczak and Alex Alderton, Roberto Amato, Jeffrey Barrett, Sonia Goncalves, Ewan Harrison, David K. Jackson, Ian Johnston, Dominic Kwiatkowski, Cordelia Langford, John Sillitoe on behalf of the Wellcome Sanger Institute COVID-19 Surveillance Team |
| EPI_ISL_2701929 | Lighthouse Lab in Milton Keynes | Wellcome Sanger Institute for the COVID-19 Genomics UK (COG-UK) Consortium | The Lighthouse Lab in Milton Keynes and Alex Alderton, Roberto Amato, Jeffrey Barrett, Sonia Goncalves, Ewan Harrison, David K. Jackson, Ian Johnston, Dominic Kwiatkowski, Cordelia Langford, John Sillitoe on behalf of the Wellcome Sanger Institute COVID-19 Surveillance Team |
| EPI_ISL_2701930 | Lighthouse Lab in Glasgow | Wellcome Sanger Institute for the COVID-19 Genomics UK (COG-UK) Consortium | Harper VanSteenhouse, Yumi Kasai, David Gray, Carol Clugston, Anna Dominiczak and Alex Alderton, Roberto Amato, Jeffrey Barrett, Sonia Goncalves, Ewan Harrison, David K. Jackson, Ian Johnston, Dominic Kwiatkowski, Cordelia Langford, John Sillitoe on behalf of the Wellcome Sanger Institute COVID-19 Surveillance Team |
| EPI_ISL_2701931, EPI_ISL_2701932 | Lighthouse Lab in Milton Keynes | Wellcome Sanger Institute for the COVID-19 Genomics UK (COG-UK) Consortium | The Lighthouse Lab in Milton Keynes and Alex Alderton, Roberto Amato, Jeffrey Barrett, Sonia Goncalves, Ewan Harrison, David K. Jackson, Ian Johnston, Dominic Kwiatkowski, Cordelia Langford, John Sillitoe on behalf of the Wellcome Sanger Institute COVID-19 Surveillance Team |
| EPI_ISL_2701933 | Lighthouse Lab in Alderley Park | Wellcome Sanger Institute for the COVID-19 Genomics UK (COG-UK) Consortium | Jacquelyn Wynn, Mairead Hyland, The Lighthouse Lab in Alderley Park and Alex Alderton, Roberto Amato, Jeffrey Barrett, Sonia Goncalves, Ewan Harrison, David K. Jackson, Ian Johnston, Dominic Kwiatkowski, Cordelia Langford, John Sillitoe on behalf of the Wellcome Sanger Institute COVID-19 Surveillance Team |
| EPI_ISL_2701934, EPI_ISL_2701935 | Lighthouse Lab in Glasgow | Wellcome Sanger Institute for the COVID-19 Genomics UK (COG-UK) Consortium | Harper VanSteenhouse, Yumi Kasai, David Gray, Carol Clugston, Anna Dominiczak and Alex Alderton, Roberto Amato, Jeffrey Barrett, Sonia Goncalves, Ewan Harrison, David K. Jackson, Ian Johnston, Dominic Kwiatkowski, Cordelia Langford, John Sillitoe on behalf of the Wellcome Sanger Institute COVID-19 Surveillance Team |
| EPI_ISL_2701936 | Lighthouse Lab in Milton Keynes | Wellcome Sanger Institute for the COVID-19 Genomics UK (COG-UK) Consortium | The Lighthouse Lab in Milton Keynes and Alex Alderton, Roberto Amato, Jeffrey Barrett, Sonia Goncalves, Ewan Harrison, David K. Jackson, Ian Johnston, Dominic Kwiatkowski, Cordelia Langford, John Sillitoe on behalf of the Wellcome Sanger Institute COVID-19 Surveillance Team |
| EPI_ISL_2701937, EPI_ISL_2701938, EPI_ISL_2701939 | Lighthouse Lab in Alderley Park | Wellcome Sanger Institute for the COVID-19 Genomics UK (COG-UK) Consortium | Jacquelyn Wynn, Mairead Hyland, The Lighthouse Lab in Alderley Park and Alex Alderton, Roberto Amato, Jeffrey Barrett, Sonia Goncalves, Ewan Harrison, David K. Jackson, Ian Johnston, Dominic Kwiatkowski, Cordelia Langford, John Sillitoe on behalf of the Wellcome Sanger Institute COVID-19 Surveillance Team |
| EPI_ISL_2701940 | Lighthouse Lab in Milton Keynes | Wellcome Sanger Institute for the COVID-19 Genomics UK (COG-UK) Consortium | The Lighthouse Lab in Milton Keynes and Alex Alderton, Roberto Amato, Jeffrey Barrett, Sonia Goncalves, Ewan Harrison, David K. Jackson, Ian Johnston, Dominic Kwiatkowski, Cordelia Langford, John Sillitoe on behalf of the Wellcome Sanger Institute COVID-19 Surveillance Team |
| EPI_ISL_2701941 | Lighthouse Lab in Alderley Park | Wellcome Sanger Institute for the COVID-19 Genomics UK (COG-UK) Consortium | Jacquelyn Wynn, Mairead Hyland, The Lighthouse Lab in Alderley Park and Alex Alderton, Roberto Amato, Jeffrey Barrett, Sonia Goncalves, Ewan Harrison, David K. Jackson, Ian Johnston, Dominic Kwiatkowski, Cordelia Langford, John Sillitoe on behalf of the Wellcome Sanger Institute COVID-19 Surveillance Team |
| EPI_ISL_2701943, EPI_ISL_2701944 | Lighthouse Lab in Milton Keynes | Wellcome Sanger Institute for the COVID-19 Genomics UK (COG-UK) Consortium | The Lighthouse Lab in Milton Keynes and Alex Alderton, Roberto Amato, Jeffrey Barrett, Sonia Goncalves, Ewan Harrison, David K. Jackson, Ian Johnston, Dominic Kwiatkowski, Cordelia Langford, John Sillitoe on behalf of the Wellcome Sanger Institute COVID-19 Surveillance Team |
| EPI_ISL_2701945 | Lighthouse Lab in Alderley Park | Wellcome Sanger Institute for the COVID-19 Genomics UK (COG-UK) Consortium | Jacquelyn Wynn, Mairead Hyland, The Lighthouse Lab in Alderley Park and Alex Alderton, Roberto Amato, Jeffrey Barrett, Sonia Goncalves, Ewan Harrison, David K. Jackson, Ian Johnston, Dominic Kwiatkowski, Cordelia Langford, John Sillitoe on behalf of the Wellcome Sanger Institute COVID-19 Surveillance Team |
| EPI_ISL_2701946 | Lighthouse Lab in Milton Keynes | Wellcome Sanger Institute for the COVID-19 Genomics UK (COG-UK) Consortium | The Lighthouse Lab in Milton Keynes and Alex Alderton, Roberto Amato, Jeffrey Barrett, Sonia Goncalves, Ewan Harrison, David K. Jackson, Ian Johnston, Dominic Kwiatkowski, Cordelia Langford, John Sillitoe on behalf of the Wellcome Sanger Institute COVID-19 Surveillance Team |
| EPI_ISL_2701947, EPI_ISL_2701948, EPI_ISL_2701949 | Lighthouse Lab in Glasgow | Wellcome Sanger Institute for the COVID-19 Genomics UK (COG-UK) Consortium | Harper VanSteenhouse, Yumi Kasai, David Gray, Carol Clugston, Anna Dominiczak and Alex Alderton, Roberto Amato, Jeffrey Barrett, Sonia Goncalves, Ewan Harrison, David K. Jackson, Ian Johnston, Dominic Kwiatkowski, Cordelia Langford, John Sillitoe on behalf of the Wellcome Sanger Institute COVID-19 Surveillance Team |
| EPI_ISL_2701950 | Lighthouse Lab in Alderley Park | Wellcome Sanger Institute for the COVID-19 Genomics UK (COG-UK) Consortium | Jacquelyn Wynn, Mairead Hyland, The Lighthouse Lab in Alderley Park and Alex Alderton, Roberto Amato, Jeffrey Barrett, Sonia Goncalves, Ewan Harrison, David K. Jackson, Ian Johnston, Dominic Kwiatkowski, Cordelia Langford, John Sillitoe on behalf of the Wellcome Sanger Institute COVID-19 Surveillance Team |
| EPI_ISL_2701951, EPI_ISL_2701952 | Lighthouse Lab in Milton Keynes | Wellcome Sanger Institute for the COVID-19 Genomics UK (COG-UK) Consortium | The Lighthouse Lab in Milton Keynes and Alex Alderton, Roberto Amato, Jeffrey Barrett, Sonia Goncalves, Ewan Harrison, David K. Jackson, Ian Johnston, Dominic Kwiatkowski, Cordelia Langford, John Sillitoe on behalf of the Wellcome Sanger Institute COVID-19 Surveillance Team |
| EPI_ISL_2701954 | Lighthouse Lab in Glasgow | Wellcome Sanger Institute for the COVID-19 Genomics UK (COG-UK) Consortium | Harper VanSteenhouse, Yumi Kasai, David Gray, Carol Clugston, Anna Dominiczak and Alex Alderton, Roberto Amato, Jeffrey Barrett, Sonia Goncalves, Ewan Harrison, David K. Jackson, Ian Johnston, Dominic Kwiatkowski, Cordelia Langford, John Sillitoe on behalf of the Wellcome Sanger Institute COVID-19 Surveillance Team |
| EPI_ISL_2701955 | Lighthouse Lab in Milton Keynes | Wellcome Sanger Institute for the COVID-19 Genomics UK (COG-UK) Consortium | The Lighthouse Lab in Milton Keynes and Alex Alderton, Roberto Amato, Jeffrey Barrett, Sonia Goncalves, Ewan Harrison, David K. Jackson, Ian Johnston, Dominic Kwiatkowski, Cordelia Langford, John Sillitoe on behalf of the Wellcome Sanger Institute COVID-19 Surveillance Team |
| EPI_ISL_2701956 | Lighthouse Lab in Alderley Park | Wellcome Sanger Institute for the COVID-19 Genomics UK (COG-UK) Consortium | Jacquelyn Wynn, Mairead Hyland, The Lighthouse Lab in Alderley Park and Alex Alderton, Roberto Amato, Jeffrey Barrett, Sonia Goncalves, Ewan Harrison, David K. Jackson, Ian Johnston, Dominic Kwiatkowski, Cordelia Langford, John Sillitoe on behalf of the Wellcome Sanger Institute COVID-19 Surveillance Team |
| EPI_ISL_2701957 | Lighthouse Lab in Milton Keynes | Wellcome Sanger Institute for the COVID-19 Genomics UK (COG-UK) | The Lighthouse Lab in Milton Keynes and Alex Alderton, Roberto Amato, Jeffrey Barrett, Sonia Goncalves, Ewan Harrison, David K. Jackson, Ian Johnston, Dominic Kwiatkowski, Cordelia Langford, John Sillitoe on behalf of the Wellcome Sanger |

| | | | |
|---|---|---|---|
| | | Consortium | Institute COVID-19 Surveillance Team |
| EPI_ISL_2701958, EPI_ISL_2701959 | Lighthouse Lab in Glasgow | Wellcome Sanger Institute for the COVID-19 Genomics UK (COG-UK) Consortium | Harper VanSteenhouse, Yumi Kasai, David Gray, Carol Clugston, Anna Dominiczak and Alex Alderton, Roberto Amato, Jeffrey Barrett, Sonia Goncalves, Ewan Harrison, David K. Jackson, Ian Johnston, Dominic Kwiatkowski, Cordelia Langford, John Sillitoe on behalf of the Wellcome Sanger Institute COVID-19 Surveillance Team |
| EPI_ISL_2701960 | Lighthouse Lab in Milton Keynes | Wellcome Sanger Institute for the COVID-19 Genomics UK (COG-UK) Consortium | The Lighthouse Lab in Milton Keynes and Alex Alderton, Roberto Amato, Jeffrey Barrett, Sonia Goncalves, Ewan Harrison, David K. Jackson, Ian Johnston, Dominic Kwiatkowski, Cordelia Langford, John Sillitoe on behalf of the Wellcome Sanger Institute COVID-19 Surveillance Team |
| EPI_ISL_2701962 | Lighthouse Lab in Glasgow | Wellcome Sanger Institute for the COVID-19 Genomics UK (COG-UK) Consortium | Harper VanSteenhouse, Yumi Kasai, David Gray, Carol Clugston, Anna Dominiczak and Alex Alderton, Roberto Amato, Jeffrey Barrett, Sonia Goncalves, Ewan Harrison, David K. Jackson, Ian Johnston, Dominic Kwiatkowski, Cordelia Langford, John Sillitoe on behalf of the Wellcome Sanger Institute COVID-19 Surveillance Team |
| EPI_ISL_2701963 | Lighthouse Lab in Alderley Park | Wellcome Sanger Institute for the COVID-19 Genomics UK (COG-UK) Consortium | Jacquelyn Wynn, Mairead Hyland, The Lighthouse Lab in Alderley Park and Alex Alderton, Roberto Amato, Jeffrey Barrett, Sonia Goncalves, Ewan Harrison, David K. Jackson, Ian Johnston, Dominic Kwiatkowski, Cordelia Langford, John Sillitoe on behalf of the Wellcome Sanger Institute COVID-19 Surveillance Team |
| EPI_ISL_2701964 | Lighthouse Lab in Milton Keynes | Wellcome Sanger Institute for the COVID-19 Genomics UK (COG-UK) Consortium | The Lighthouse Lab in Milton Keynes and Alex Alderton, Roberto Amato, Jeffrey Barrett, Sonia Goncalves, Ewan Harrison, David K. Jackson, Ian Johnston, Dominic Kwiatkowski, Cordelia Langford, John Sillitoe on behalf of the Wellcome Sanger Institute COVID-19 Surveillance Team |
| EPI_ISL_2701965, EPI_ISL_2701966 | Lighthouse Lab in Alderley Park | Wellcome Sanger Institute for the COVID-19 Genomics UK (COG-UK) Consortium | Jacquelyn Wynn, Mairead Hyland, The Lighthouse Lab in Alderley Park and Alex Alderton, Roberto Amato, Jeffrey Barrett, Sonia Goncalves, Ewan Harrison, David K. Jackson, Ian Johnston, Dominic Kwiatkowski, Cordelia Langford, John Sillitoe on behalf of the Wellcome Sanger Institute COVID-19 Surveillance Team |
| EPI_ISL_2701967 | Lighthouse Lab in Milton Keynes | Wellcome Sanger Institute for the COVID-19 Genomics UK (COG-UK) Consortium | The Lighthouse Lab in Milton Keynes and Alex Alderton, Roberto Amato, Jeffrey Barrett, Sonia Goncalves, Ewan Harrison, David K. Jackson, Ian Johnston, Dominic Kwiatkowski, Cordelia Langford, John Sillitoe on behalf of the Wellcome Sanger Institute COVID-19 Surveillance Team |
| EPI_ISL_2701968 | Lighthouse Lab in Alderley Park | Wellcome Sanger Institute for the COVID-19 Genomics UK (COG-UK) Consortium | Jacquelyn Wynn, Mairead Hyland, The Lighthouse Lab in Alderley Park and Alex Alderton, Roberto Amato, Jeffrey Barrett, Sonia Goncalves, Ewan Harrison, David K. Jackson, Ian Johnston, Dominic Kwiatkowski, Cordelia Langford, John Sillitoe on behalf of the Wellcome Sanger Institute COVID-19 Surveillance Team |
| EPI_ISL_2701969, EPI_ISL_2701970 | Lighthouse Lab in Milton Keynes | Wellcome Sanger Institute for the COVID-19 Genomics UK (COG-UK) Consortium | The Lighthouse Lab in Milton Keynes and Alex Alderton, Roberto Amato, Jeffrey Barrett, Sonia Goncalves, Ewan Harrison, David K. Jackson, Ian Johnston, Dominic Kwiatkowski, Cordelia Langford, John Sillitoe on behalf of the Wellcome Sanger Institute COVID-19 Surveillance Team |
| EPI_ISL_2701971, EPI_ISL_2701972 | Lighthouse Lab in Glasgow | Wellcome Sanger Institute for the COVID-19 Genomics UK (COG-UK) Consortium | Harper VanSteenhouse, Yumi Kasai, David Gray, Carol Clugston, Anna Dominiczak and Alex Alderton, Roberto Amato, Jeffrey Barrett, Sonia Goncalves, Ewan Harrison, David K. Jackson, Ian Johnston, Dominic Kwiatkowski, Cordelia Langford, John Sillitoe on behalf of the Wellcome Sanger Institute COVID-19 Surveillance Team |
| EPI_ISL_2701973 | Lighthouse Lab in Alderley Park | Wellcome Sanger Institute for the COVID-19 Genomics UK (COG-UK) Consortium | Jacquelyn Wynn, Mairead Hyland, The Lighthouse Lab in Alderley Park and Alex Alderton, Roberto Amato, Jeffrey Barrett, Sonia Goncalves, Ewan Harrison, David K. Jackson, Ian Johnston, Dominic Kwiatkowski, Cordelia Langford, John Sillitoe on behalf of the Wellcome Sanger Institute COVID-19 Surveillance Team |
| EPI_ISL_2701974, EPI_ISL_2701975, EPI_ISL_2701976, EPI_ISL_2701977 | Lighthouse Lab in Glasgow | Wellcome Sanger Institute for the COVID-19 Genomics UK (COG-UK) Consortium | Harper VanSteenhouse, Yumi Kasai, David Gray, Carol Clugston, Anna Dominiczak and Alex Alderton, Roberto Amato, Jeffrey Barrett, Sonia Goncalves, Ewan Harrison, David K. Jackson, Ian Johnston, Dominic Kwiatkowski, Cordelia Langford, John Sillitoe on behalf of the Wellcome Sanger Institute COVID-19 Surveillance Team |
| EPI_ISL_2701978, EPI_ISL_2701979 | Lighthouse Lab in Alderley Park | Wellcome Sanger Institute for the COVID-19 Genomics UK (COG-UK) Consortium | Jacquelyn Wynn, Mairead Hyland, The Lighthouse Lab in Alderley Park and Alex Alderton, Roberto Amato, Jeffrey Barrett, Sonia Goncalves, Ewan Harrison, David K. Jackson, Ian Johnston, Dominic Kwiatkowski, Cordelia Langford, John Sillitoe on behalf of the Wellcome Sanger Institute COVID-19 Surveillance Team |
| EPI_ISL_2701980 | Lighthouse Lab in Glasgow | Wellcome Sanger Institute for the COVID-19 Genomics UK (COG-UK) Consortium | Harper VanSteenhouse, Yumi Kasai, David Gray, Carol Clugston, Anna Dominiczak and Alex Alderton, Roberto Amato, Jeffrey Barrett, Sonia Goncalves, Ewan Harrison, David K. Jackson, Ian Johnston, Dominic Kwiatkowski, Cordelia Langford, John Sillitoe on behalf of the Wellcome Sanger Institute COVID-19 Surveillance Team |
| EPI_ISL_2701981 | Lighthouse Lab in Milton Keynes | Wellcome Sanger Institute for the COVID-19 Genomics UK (COG-UK) Consortium | The Lighthouse Lab in Milton Keynes and Alex Alderton, Roberto Amato, Jeffrey Barrett, Sonia Goncalves, Ewan Harrison, David K. Jackson, Ian Johnston, Dominic Kwiatkowski, Cordelia Langford, John Sillitoe on behalf of the Wellcome Sanger Institute COVID-19 Surveillance Team |
| EPI_ISL_2701982 | Lighthouse Lab in Alderley Park | Wellcome Sanger Institute for the COVID-19 Genomics UK (COG-UK) Consortium | Jacquelyn Wynn, Mairead Hyland, The Lighthouse Lab in Alderley Park and Alex Alderton, Roberto Amato, Jeffrey Barrett, Sonia Goncalves, Ewan Harrison, David K. Jackson, Ian Johnston, Dominic Kwiatkowski, Cordelia Langford, John Sillitoe on behalf of the Wellcome Sanger Institute COVID-19 Surveillance Team |
| EPI_ISL_2701983 | Lighthouse Lab in Glasgow | Wellcome Sanger Institute for the COVID-19 Genomics UK (COG-UK) Consortium | Harper VanSteenhouse, Yumi Kasai, David Gray, Carol Clugston, Anna Dominiczak and Alex Alderton, Roberto Amato, Jeffrey Barrett, Sonia Goncalves, Ewan Harrison, David K. Jackson, Ian Johnston, Dominic Kwiatkowski, Cordelia Langford, John Sillitoe on behalf of the Wellcome Sanger Institute COVID-19 Surveillance Team |
| EPI_ISL_2701984 | Lighthouse Lab in Alderley Park | Wellcome Sanger Institute for the COVID-19 Genomics UK (COG-UK) Consortium | Jacquelyn Wynn, Mairead Hyland, The Lighthouse Lab in Alderley Park and Alex Alderton, Roberto Amato, Jeffrey Barrett, Sonia Goncalves, Ewan Harrison, David K. Jackson, Ian Johnston, Dominic Kwiatkowski, Cordelia Langford, John Sillitoe on behalf of the Wellcome Sanger Institute COVID-19 Surveillance Team |
| EPI_ISL_2701985, EPI_ISL_2701986 | Lighthouse Lab in Milton Keynes | Wellcome Sanger Institute for the COVID-19 Genomics UK (COG-UK) Consortium | The Lighthouse Lab in Milton Keynes and Alex Alderton, Roberto Amato, Jeffrey Barrett, Sonia Goncalves, Ewan Harrison, David K. Jackson, Ian Johnston, Dominic Kwiatkowski, Cordelia Langford, John Sillitoe on behalf of the Wellcome Sanger Institute COVID-19 Surveillance Team |
| EPI_ISL_2701987 | Lighthouse Lab in Alderley Park | Wellcome Sanger Institute for the COVID-19 Genomics UK (COG-UK) Consortium | Jacquelyn Wynn, Mairead Hyland, The Lighthouse Lab in Alderley Park and Alex Alderton, Roberto Amato, Jeffrey Barrett, Sonia Goncalves, Ewan Harrison, David K. Jackson, Ian Johnston, Dominic Kwiatkowski, Cordelia Langford, John Sillitoe on behalf of the Wellcome Sanger Institute COVID-19 Surveillance Team |
| EPI_ISL_2701988 | Lighthouse Lab in Milton Keynes | Wellcome Sanger Institute for the COVID-19 Genomics UK (COG-UK) Consortium | The Lighthouse Lab in Milton Keynes and Alex Alderton, Roberto Amato, Jeffrey Barrett, Sonia Goncalves, Ewan Harrison, David K. Jackson, Ian Johnston, Dominic Kwiatkowski, Cordelia Langford, John Sillitoe on behalf of the Wellcome Sanger Institute COVID-19 Surveillance Team |
| EPI_ISL_2701989 | Lighthouse Lab in Glasgow | Wellcome Sanger Institute for the COVID-19 Genomics UK (COG-UK) Consortium | Harper VanSteenhouse, Yumi Kasai, David Gray, Carol Clugston, Anna Dominiczak and Alex Alderton, Roberto Amato, Jeffrey Barrett, Sonia Goncalves, Ewan Harrison, David K. Jackson, Ian Johnston, Dominic Kwiatkowski, Cordelia Langford, John Sillitoe on behalf of the Wellcome Sanger Institute COVID-19 Surveillance Team |
| EPI_ISL_2701990 | Lighthouse Lab in Alderley Park | Wellcome Sanger Institute for the COVID-19 Genomics UK (COG-UK) Consortium | Jacquelyn Wynn, Mairead Hyland, The Lighthouse Lab in Alderley Park and Alex Alderton, Roberto Amato, Jeffrey Barrett, Sonia Goncalves, Ewan Harrison, David K. Jackson, Ian Johnston, Dominic Kwiatkowski, Cordelia Langford, John Sillitoe on behalf of the Wellcome Sanger Institute COVID-19 Surveillance Team |
| EPI_ISL_2701991, EPI_ISL_2701992 | Lighthouse Lab in Glasgow | Wellcome Sanger Institute for the COVID-19 Genomics UK (COG-UK) Consortium | Harper VanSteenhouse, Yumi Kasai, David Gray, Carol Clugston, Anna Dominiczak and Alex Alderton, Roberto Amato, Jeffrey Barrett, Sonia Goncalves, Ewan Harrison, David K. Jackson, Ian Johnston, Dominic Kwiatkowski, Cordelia Langford, John Sillitoe on behalf of the Wellcome Sanger Institute COVID-19 Surveillance Team |
| EPI_ISL_2701993 | Lighthouse Lab in Alderley Park | Wellcome Sanger Institute for the COVID-19 Genomics UK (COG-UK) Consortium | Jacquelyn Wynn, Mairead Hyland, The Lighthouse Lab in Alderley Park and Alex Alderton, Roberto Amato, Jeffrey Barrett, Sonia Goncalves, Ewan Harrison, David K. Jackson, Ian Johnston, Dominic Kwiatkowski, Cordelia Langford, John Sillitoe on behalf of the Wellcome Sanger Institute COVID-19 Surveillance Team |
| EPI_ISL_2701994 | Lighthouse Lab in Glasgow | Wellcome Sanger Institute for the COVID-19 Genomics UK (COG-UK) Consortium | Harper VanSteenhouse, Yumi Kasai, David Gray, Carol Clugston, Anna Dominiczak and Alex Alderton, Roberto Amato, Jeffrey Barrett, Sonia Goncalves, Ewan Harrison, David K. Jackson, Ian Johnston, Dominic Kwiatkowski, Cordelia Langford, John Sillitoe on behalf of the Wellcome Sanger Institute COVID-19 Surveillance Team |
| EPI_ISL_2701995, EPI_ISL_2701996, EPI_ISL_2701997 | Lighthouse Lab in Alderley Park | Wellcome Sanger Institute for the COVID-19 Genomics UK (COG-UK) Consortium | Jacquelyn Wynn, Mairead Hyland, The Lighthouse Lab in Alderley Park and Alex Alderton, Roberto Amato, Jeffrey Barrett, Sonia Goncalves, Ewan Harrison, David K. Jackson, Ian Johnston, Dominic Kwiatkowski, Cordelia Langford, John Sillitoe on behalf of the Wellcome Sanger Institute COVID-19 Surveillance Team |
| EPI_ISL_2701998, EPI_ISL_2701999, EPI_ISL_2702000, EPI_ISL_2702001, EPI_ISL_2702002 | Lighthouse Lab in Glasgow | Wellcome Sanger Institute for the COVID-19 Genomics UK (COG-UK) Consortium | Harper VanSteenhouse, Yumi Kasai, David Gray, Carol Clugston, Anna Dominiczak and Alex Alderton, Roberto Amato, Jeffrey Barrett, Sonia Goncalves, Ewan Harrison, David K. Jackson, Ian Johnston, Dominic Kwiatkowski, Cordelia Langford, John Sillitoe on behalf of the Wellcome Sanger Institute COVID-19 Surveillance Team |
| EPI_ISL_2702003 | Lighthouse Lab in Milton Keynes | Wellcome Sanger Institute for the COVID-19 Genomics UK (COG-UK) Consortium | The Lighthouse Lab in Milton Keynes and Alex Alderton, Roberto Amato, Jeffrey Barrett, Sonia Goncalves, Ewan Harrison, David K. Jackson, Ian Johnston, Dominic Kwiatkowski, Cordelia Langford, John Sillitoe on behalf of the Wellcome Sanger Institute COVID-19 Surveillance Team |
| EPI_ISL_2702004 | Lighthouse Lab in Alderley Park | Wellcome Sanger Institute for the COVID-19 Genomics UK (COG-UK) Consortium | Jacquelyn Wynn, Mairead Hyland, The Lighthouse Lab in Alderley Park and Alex Alderton, Roberto Amato, Jeffrey Barrett, Sonia Goncalves, Ewan Harrison, David K. Jackson, Ian Johnston, Dominic Kwiatkowski, Cordelia Langford, John Sillitoe on behalf of the Wellcome Sanger Institute COVID-19 Surveillance Team |
| EPI_ISL_2702005, EPI_ISL_2702006, EPI_ISL_2702007, EPI_ISL_2702009, EPI_ISL_2702010 | Lighthouse Lab in Glasgow | Wellcome Sanger Institute for the COVID-19 Genomics UK (COG-UK) Consortium | Harper VanSteenhouse, Yumi Kasai, David Gray, Carol Clugston, Anna Dominiczak and Alex Alderton, Roberto Amato, Jeffrey Barrett, Sonia Goncalves, Ewan Harrison, David K. Jackson, Ian Johnston, Dominic Kwiatkowski, Cordelia Langford, John Sillitoe on behalf of the Wellcome Sanger Institute COVID-19 Surveillance Team |
| EPI_ISL_2702011 | Lighthouse Lab in Alderley Park | Wellcome Sanger Institute for the COVID-19 Genomics UK (COG-UK) Consortium | Jacquelyn Wynn, Mairead Hyland, The Lighthouse Lab in Alderley Park and Alex Alderton, Roberto Amato, Jeffrey Barrett, Sonia Goncalves, Ewan Harrison, David K. Jackson, Ian Johnston, Dominic Kwiatkowski, Cordelia Langford, John Sillitoe on behalf of the Wellcome Sanger Institute COVID-19 Surveillance Team |
| EPI_ISL_2702012, EPI_ISL_2702013 | Lighthouse Lab in Glasgow | Wellcome Sanger Institute for the COVID-19 Genomics UK (COG-UK) Consortium | Harper VanSteenhouse, Yumi Kasai, David Gray, Carol Clugston, Anna Dominiczak and Alex Alderton, Roberto Amato, Jeffrey Barrett, Sonia Goncalves, Ewan Harrison, David K. Jackson, Ian Johnston, Dominic Kwiatkowski, Cordelia Langford, John Sillitoe on behalf of the Wellcome Sanger Institute COVID-19 Surveillance Team |
| EPI_ISL_2702014 | Lighthouse Lab in Alderley Park | Wellcome Sanger Institute for the COVID-19 Genomics UK (COG-UK) Consortium | Jacquelyn Wynn, Mairead Hyland, The Lighthouse Lab in Alderley Park and Alex Alderton, Roberto Amato, Jeffrey Barrett, Sonia Goncalves, Ewan Harrison, David K. Jackson, Ian Johnston, Dominic Kwiatkowski, Cordelia Langford, John Sillitoe on behalf of the Wellcome Sanger Institute COVID-19 Surveillance Team |
| EPI_ISL_2702015 | Lighthouse Lab in Milton Keynes | Wellcome Sanger Institute for the COVID-19 Genomics UK (COG-UK) Consortium | The Lighthouse Lab in Milton Keynes and Alex Alderton, Roberto Amato, Jeffrey Barrett, Sonia Goncalves, Ewan Harrison, David K. Jackson, Ian Johnston, Dominic Kwiatkowski, Cordelia Langford, John Sillitoe on behalf of the Wellcome Sanger Institute COVID-19 Surveillance Team |
| EPI_ISL_2702016 | Lighthouse Lab in Glasgow | Wellcome Sanger Institute for the COVID-19 Genomics UK (COG-UK) Consortium | Harper VanSteenhouse, Yumi Kasai, David Gray, Carol Clugston, Anna Dominiczak and Alex Alderton, Roberto Amato, Jeffrey Barrett, Sonia Goncalves, Ewan Harrison, David K. Jackson, Ian Johnston, Dominic Kwiatkowski, Cordelia Langford, John Sillitoe on behalf of the Wellcome Sanger Institute COVID-19 Surveillance Team |
| EPI_ISL_2702017 | Lighthouse Lab in Milton Keynes | Wellcome Sanger Institute for the COVID-19 Genomics UK (COG-UK) Consortium | The Lighthouse Lab in Milton Keynes and Alex Alderton, Roberto Amato, Jeffrey Barrett, Sonia Goncalves, Ewan Harrison, David K. Jackson, Ian Johnston, Dominic Kwiatkowski, Cordelia Langford, John Sillitoe on behalf of the Wellcome Sanger Institute COVID-19 Surveillance Team |
| EPI_ISL_2702018, EPI_ISL_2702019 | Lighthouse Lab in Glasgow | Wellcome Sanger Institute for the COVID-19 Genomics UK (COG-UK) Consortium | Harper VanSteenhouse, Yumi Kasai, David Gray, Carol Clugston, Anna Dominiczak and Alex Alderton, Roberto Amato, Jeffrey Barrett, Sonia Goncalves, Ewan Harrison, David K. Jackson, Ian Johnston, Dominic Kwiatkowski, Cordelia Langford, John Sillitoe on behalf of the Wellcome Sanger Institute COVID-19 Surveillance Team |
| EPI_ISL_2702020 | Lighthouse Lab in Alderley Park | Wellcome Sanger Institute for the COVID-19 Genomics UK (COG-UK) Consortium | Jacquelyn Wynn, Mairead Hyland, The Lighthouse Lab in Alderley Park and Alex Alderton, Roberto Amato, Jeffrey Barrett, Sonia Goncalves, Ewan Harrison, David K. Jackson, Ian Johnston, Dominic Kwiatkowski, Cordelia Langford, John Sillitoe on behalf of the Wellcome Sanger Institute COVID-19 Surveillance Team |
| EPI_ISL_2702021 | Lighthouse Lab in Glasgow | Wellcome Sanger Institute for the COVID-19 Genomics UK (COG-UK) Consortium | Harper VanSteenhouse, Yumi Kasai, David Gray, Carol Clugston, Anna Dominiczak and Alex Alderton, Roberto Amato, Jeffrey Barrett, Sonia Goncalves, Ewan Harrison, David K. Jackson, Ian Johnston, Dominic Kwiatkowski, Cordelia Langford, John Sillitoe on behalf of the Wellcome Sanger Institute COVID-19 Surveillance Team |
| EPI_ISL_2702022 | Lighthouse Lab in Alderley Park | Wellcome Sanger Institute for the COVID-19 Genomics UK (COG-UK) Consortium | Jacquelyn Wynn, Mairead Hyland, The Lighthouse Lab in Alderley Park and Alex Alderton, Roberto Amato, Jeffrey Barrett, Sonia Goncalves, Ewan Harrison, David K. Jackson, Ian Johnston, Dominic Kwiatkowski, Cordelia Langford, John Sillitoe on behalf of the Wellcome Sanger Institute COVID-19 Surveillance Team |

| | | | |
|---|---|---|---|
| EPI_ISL_2702023 | Lighthouse Lab in Glasgow | Wellcome Sanger Institute for the COVID-19 Genomics UK (COG-UK) Consortium | Harper VanSteenhouse, Yumi Kasai, David Gray, Carol Clugston, Anna Dominiczak and Alex Alderton, Roberto Amato, Jeffrey Barrett, Sonia Goncalves, Ewan Harrison, David K. Jackson, Ian Johnston, Dominic Kwiatkowski, Cordelia Langford, John Sillitoe on behalf of the Wellcome Sanger Institute COVID-19 Surveillance Team |
| EPI_ISL_2702024 | Lighthouse Lab in Alderley Park | Wellcome Sanger Institute for the COVID-19 Genomics UK (COG-UK) Consortium | Jacquelyn Wynn, Mairead Hyland, The Lighthouse Lab in Alderley Park and Alex Alderton, Roberto Amato, Jeffrey Barrett, Sonia Goncalves, Ewan Harrison, David K. Jackson, Ian Johnston, Dominic Kwiatkowski, Cordelia Langford, John Sillitoe on behalf of the Wellcome Sanger Institute COVID-19 Surveillance Team |
| EPI_ISL_2702026 | Lighthouse Lab in Glasgow | Wellcome Sanger Institute for the COVID-19 Genomics UK (COG-UK) Consortium | Harper VanSteenhouse, Yumi Kasai, David Gray, Carol Clugston, Anna Dominiczak and Alex Alderton, Roberto Amato, Jeffrey Barrett, Sonia Goncalves, Ewan Harrison, David K. Jackson, Ian Johnston, Dominic Kwiatkowski, Cordelia Langford, John Sillitoe on behalf of the Wellcome Sanger Institute COVID-19 Surveillance Team |
| EPI_ISL_2702027, EPI_ISL_2702028 | Lighthouse Lab in Milton Keynes | Wellcome Sanger Institute for the COVID-19 Genomics UK (COG-UK) Consortium | The Lighthouse Lab in Milton Keynes and Alex Alderton, Roberto Amato, Jeffrey Barrett, Sonia Goncalves, Ewan Harrison, David K. Jackson, Ian Johnston, Dominic Kwiatkowski, Cordelia Langford, John Sillitoe on behalf of the Wellcome Sanger Institute COVID-19 Surveillance Team |
| EPI_ISL_2702029 | Lighthouse Lab in Glasgow | Wellcome Sanger Institute for the COVID-19 Genomics UK (COG-UK) Consortium | Harper VanSteenhouse, Yumi Kasai, David Gray, Carol Clugston, Anna Dominiczak and Alex Alderton, Roberto Amato, Jeffrey Barrett, Sonia Goncalves, Ewan Harrison, David K. Jackson, Ian Johnston, Dominic Kwiatkowski, Cordelia Langford, John Sillitoe on behalf of the Wellcome Sanger Institute COVID-19 Surveillance Team |
| EPI_ISL_2702030 | Lighthouse Lab in Alderley Park | Wellcome Sanger Institute for the COVID-19 Genomics UK (COG-UK) Consortium | Jacquelyn Wynn, Mairead Hyland, The Lighthouse Lab in Alderley Park and Alex Alderton, Roberto Amato, Jeffrey Barrett, Sonia Goncalves, Ewan Harrison, David K. Jackson, Ian Johnston, Dominic Kwiatkowski, Cordelia Langford, John Sillitoe on behalf of the Wellcome Sanger Institute COVID-19 Surveillance Team |
| EPI_ISL_2702032 | Lighthouse Lab in Milton Keynes | Wellcome Sanger Institute for the COVID-19 Genomics UK (COG-UK) Consortium | The Lighthouse Lab in Milton Keynes and Alex Alderton, Roberto Amato, Jeffrey Barrett, Sonia Goncalves, Ewan Harrison, David K. Jackson, Ian Johnston, Dominic Kwiatkowski, Cordelia Langford, John Sillitoe on behalf of the Wellcome Sanger Institute COVID-19 Surveillance Team |
| EPI_ISL_2702033 | Lighthouse Lab in Glasgow | Wellcome Sanger Institute for the COVID-19 Genomics UK (COG-UK) Consortium | Harper VanSteenhouse, Yumi Kasai, David Gray, Carol Clugston, Anna Dominiczak and Alex Alderton, Roberto Amato, Jeffrey Barrett, Sonia Goncalves, Ewan Harrison, David K. Jackson, Ian Johnston, Dominic Kwiatkowski, Cordelia Langford, John Sillitoe on behalf of the Wellcome Sanger Institute COVID-19 Surveillance Team |
| EPI_ISL_2702034, EPI_ISL_2702035, EPI_ISL_2702037 | Lighthouse Lab in Alderley Park | Wellcome Sanger Institute for the COVID-19 Genomics UK (COG-UK) Consortium | Jacquelyn Wynn, Mairead Hyland, The Lighthouse Lab in Alderley Park and Alex Alderton, Roberto Amato, Jeffrey Barrett, Sonia Goncalves, Ewan Harrison, David K. Jackson, Ian Johnston, Dominic Kwiatkowski, Cordelia Langford, John Sillitoe on behalf of the Wellcome Sanger Institute COVID-19 Surveillance Team |
| EPI_ISL_2702038 | Lighthouse Lab in Glasgow | Wellcome Sanger Institute for the COVID-19 Genomics UK (COG-UK) Consortium | Harper VanSteenhouse, Yumi Kasai, David Gray, Carol Clugston, Anna Dominiczak and Alex Alderton, Roberto Amato, Jeffrey Barrett, Sonia Goncalves, Ewan Harrison, David K. Jackson, Ian Johnston, Dominic Kwiatkowski, Cordelia Langford, John Sillitoe on behalf of the Wellcome Sanger Institute COVID-19 Surveillance Team |
| EPI_ISL_2702039, EPI_ISL_2702040 | Lighthouse Lab in Alderley Park | Wellcome Sanger Institute for the COVID-19 Genomics UK (COG-UK) Consortium | Jacquelyn Wynn, Mairead Hyland, The Lighthouse Lab in Alderley Park and Alex Alderton, Roberto Amato, Jeffrey Barrett, Sonia Goncalves, Ewan Harrison, David K. Jackson, Ian Johnston, Dominic Kwiatkowski, Cordelia Langford, John Sillitoe on behalf of the Wellcome Sanger Institute COVID-19 Surveillance Team |
| EPI_ISL_2702041 | Lighthouse Lab in Milton Keynes | Wellcome Sanger Institute for the COVID-19 Genomics UK (COG-UK) Consortium | The Lighthouse Lab in Milton Keynes and Alex Alderton, Roberto Amato, Jeffrey Barrett, Sonia Goncalves, Ewan Harrison, David K. Jackson, Ian Johnston, Dominic Kwiatkowski, Cordelia Langford, John Sillitoe on behalf of the Wellcome Sanger Institute COVID-19 Surveillance Team |
| EPI_ISL_2702042, EPI_ISL_2702043, EPI_ISL_2702044 | Lighthouse Lab in Glasgow | Wellcome Sanger Institute for the COVID-19 Genomics UK (COG-UK) Consortium | Harper VanSteenhouse, Yumi Kasai, David Gray, Carol Clugston, Anna Dominiczak and Alex Alderton, Roberto Amato, Jeffrey Barrett, Sonia Goncalves, Ewan Harrison, David K. Jackson, Ian Johnston, Dominic Kwiatkowski, Cordelia Langford, John Sillitoe on behalf of the Wellcome Sanger Institute COVID-19 Surveillance Team |
| EPI_ISL_2702045 | Lighthouse Lab in Alderley Park | Wellcome Sanger Institute for the COVID-19 Genomics UK (COG-UK) Consortium | Jacquelyn Wynn, Mairead Hyland, The Lighthouse Lab in Alderley Park and Alex Alderton, Roberto Amato, Jeffrey Barrett, Sonia Goncalves, Ewan Harrison, David K. Jackson, Ian Johnston, Dominic Kwiatkowski, Cordelia Langford, John Sillitoe on behalf of the Wellcome Sanger Institute COVID-19 Surveillance Team |
| EPI_ISL_2702046, EPI_ISL_2702047 | Lighthouse Lab in Glasgow | Wellcome Sanger Institute for the COVID-19 Genomics UK (COG-UK) Consortium | Harper VanSteenhouse, Yumi Kasai, David Gray, Carol Clugston, Anna Dominiczak and Alex Alderton, Roberto Amato, Jeffrey Barrett, Sonia Goncalves, Ewan Harrison, David K. Jackson, Ian Johnston, Dominic Kwiatkowski, Cordelia Langford, John Sillitoe on behalf of the Wellcome Sanger Institute COVID-19 Surveillance Team |
| EPI_ISL_2702048 | Lighthouse Lab in Alderley Park | Wellcome Sanger Institute for the COVID-19 Genomics UK (COG-UK) Consortium | Jacquelyn Wynn, Mairead Hyland, The Lighthouse Lab in Alderley Park and Alex Alderton, Roberto Amato, Jeffrey Barrett, Sonia Goncalves, Ewan Harrison, David K. Jackson, Ian Johnston, Dominic Kwiatkowski, Cordelia Langford, John Sillitoe on behalf of the Wellcome Sanger Institute COVID-19 Surveillance Team |
| EPI_ISL_2702049, EPI_ISL_2702050 | Lighthouse Lab in Glasgow | Wellcome Sanger Institute for the COVID-19 Genomics UK (COG-UK) Consortium | Harper VanSteenhouse, Yumi Kasai, David Gray, Carol Clugston, Anna Dominiczak and Alex Alderton, Roberto Amato, Jeffrey Barrett, Sonia Goncalves, Ewan Harrison, David K. Jackson, Ian Johnston, Dominic Kwiatkowski, Cordelia Langford, John Sillitoe on behalf of the Wellcome Sanger Institute COVID-19 Surveillance Team |
| EPI_ISL_2702051 | Lighthouse Lab in Milton Keynes | Wellcome Sanger Institute for the COVID-19 Genomics UK (COG-UK) Consortium | The Lighthouse Lab in Milton Keynes and Alex Alderton, Roberto Amato, Jeffrey Barrett, Sonia Goncalves, Ewan Harrison, David K. Jackson, Ian Johnston, Dominic Kwiatkowski, Cordelia Langford, John Sillitoe on behalf of the Wellcome Sanger Institute COVID-19 Surveillance Team |
| EPI_ISL_2702052, EPI_ISL_2702053 | Lighthouse Lab in Glasgow | Wellcome Sanger Institute for the COVID-19 Genomics UK (COG-UK) Consortium | Harper VanSteenhouse, Yumi Kasai, David Gray, Carol Clugston, Anna Dominiczak and Alex Alderton, Roberto Amato, Jeffrey Barrett, Sonia Goncalves, Ewan Harrison, David K. Jackson, Ian Johnston, Dominic Kwiatkowski, Cordelia Langford, John Sillitoe on behalf of the Wellcome Sanger Institute COVID-19 Surveillance Team |
| EPI_ISL_2702054, EPI_ISL_2702055 | Lighthouse Lab in Alderley Park | Wellcome Sanger Institute for the COVID-19 Genomics UK (COG-UK) Consortium | Jacquelyn Wynn, Mairead Hyland, The Lighthouse Lab in Alderley Park and Alex Alderton, Roberto Amato, Jeffrey Barrett, Sonia Goncalves, Ewan Harrison, David K. Jackson, Ian Johnston, Dominic Kwiatkowski, Cordelia Langford, John Sillitoe on behalf of the Wellcome Sanger Institute COVID-19 Surveillance Team |
| EPI_ISL_2702056 | Lighthouse Lab in Milton Keynes | Wellcome Sanger Institute for the COVID-19 Genomics UK (COG-UK) Consortium | The Lighthouse Lab in Milton Keynes and Alex Alderton, Roberto Amato, Jeffrey Barrett, Sonia Goncalves, Ewan Harrison, David K. Jackson, Ian Johnston, Dominic Kwiatkowski, Cordelia Langford, John Sillitoe on behalf of the Wellcome Sanger Institute COVID-19 Surveillance Team |
| EPI_ISL_2702057 | Lighthouse Lab in Glasgow | Wellcome Sanger Institute for the COVID-19 Genomics UK (COG-UK) Consortium | Harper VanSteenhouse, Yumi Kasai, David Gray, Carol Clugston, Anna Dominiczak and Alex Alderton, Roberto Amato, Jeffrey Barrett, Sonia Goncalves, Ewan Harrison, David K. Jackson, Ian Johnston, Dominic Kwiatkowski, Cordelia Langford, John Sillitoe on behalf of the Wellcome Sanger Institute COVID-19 Surveillance Team |
| EPI_ISL_2702058 | Lighthouse Lab in Alderley Park | Wellcome Sanger Institute for the COVID-19 Genomics UK (COG-UK) Consortium | Jacquelyn Wynn, Mairead Hyland, The Lighthouse Lab in Alderley Park and Alex Alderton, Roberto Amato, Jeffrey Barrett, Sonia Goncalves, Ewan Harrison, David K. Jackson, Ian Johnston, Dominic Kwiatkowski, Cordelia Langford, John Sillitoe on behalf of the Wellcome Sanger Institute COVID-19 Surveillance Team |
| EPI_ISL_2702059, EPI_ISL_2702060 | Lighthouse Lab in Glasgow | Wellcome Sanger Institute for the COVID-19 Genomics UK (COG-UK) Consortium | Harper VanSteenhouse, Yumi Kasai, David Gray, Carol Clugston, Anna Dominiczak and Alex Alderton, Roberto Amato, Jeffrey Barrett, Sonia Goncalves, Ewan Harrison, David K. Jackson, Ian Johnston, Dominic Kwiatkowski, Cordelia Langford, John Sillitoe on behalf of the Wellcome Sanger Institute COVID-19 Surveillance Team |
| EPI_ISL_2702061, EPI_ISL_2702062 | Lighthouse Lab in Milton Keynes | Wellcome Sanger Institute for the COVID-19 Genomics UK (COG-UK) Consortium | The Lighthouse Lab in Milton Keynes and Alex Alderton, Roberto Amato, Jeffrey Barrett, Sonia Goncalves, Ewan Harrison, David K. Jackson, Ian Johnston, Dominic Kwiatkowski, Cordelia Langford, John Sillitoe on behalf of the Wellcome Sanger Institute COVID-19 Surveillance Team |
| EPI_ISL_2702063 | Lighthouse Lab in Glasgow | Wellcome Sanger Institute for the COVID-19 Genomics UK (COG-UK) Consortium | Harper VanSteenhouse, Yumi Kasai, David Gray, Carol Clugston, Anna Dominiczak and Alex Alderton, Roberto Amato, Jeffrey Barrett, Sonia Goncalves, Ewan Harrison, David K. Jackson, Ian Johnston, Dominic Kwiatkowski, Cordelia Langford, John Sillitoe on behalf of the Wellcome Sanger Institute COVID-19 Surveillance Team |
| EPI_ISL_2702064 | Lighthouse Lab in Milton Keynes | Wellcome Sanger Institute for the COVID-19 Genomics UK (COG-UK) Consortium | The Lighthouse Lab in Milton Keynes and Alex Alderton, Roberto Amato, Jeffrey Barrett, Sonia Goncalves, Ewan Harrison, David K. Jackson, Ian Johnston, Dominic Kwiatkowski, Cordelia Langford, John Sillitoe on behalf of the Wellcome Sanger Institute COVID-19 Surveillance Team |
| EPI_ISL_2702065, EPI_ISL_2702066 | Lighthouse Lab in Glasgow | Wellcome Sanger Institute for the COVID-19 Genomics UK (COG-UK) Consortium | Harper VanSteenhouse, Yumi Kasai, David Gray, Carol Clugston, Anna Dominiczak and Alex Alderton, Roberto Amato, Jeffrey Barrett, Sonia Goncalves, Ewan Harrison, David K. Jackson, Ian Johnston, Dominic Kwiatkowski, Cordelia Langford, John Sillitoe on behalf of the Wellcome Sanger Institute COVID-19 Surveillance Team |
| EPI_ISL_2702067 | Lighthouse Lab in Alderley Park | Wellcome Sanger Institute for the COVID-19 Genomics UK (COG-UK) Consortium | Jacquelyn Wynn, Mairead Hyland, The Lighthouse Lab in Alderley Park and Alex Alderton, Roberto Amato, Jeffrey Barrett, Sonia Goncalves, Ewan Harrison, David K. Jackson, Ian Johnston, Dominic Kwiatkowski, Cordelia Langford, John Sillitoe on behalf of the Wellcome Sanger Institute COVID-19 Surveillance Team |
| EPI_ISL_2702068 | Lighthouse Lab in Glasgow | Wellcome Sanger Institute for the COVID-19 Genomics UK (COG-UK) Consortium | Harper VanSteenhouse, Yumi Kasai, David Gray, Carol Clugston, Anna Dominiczak and Alex Alderton, Roberto Amato, Jeffrey Barrett, Sonia Goncalves, Ewan Harrison, David K. Jackson, Ian Johnston, Dominic Kwiatkowski, Cordelia Langford, John Sillitoe on behalf of the Wellcome Sanger Institute COVID-19 Surveillance Team |
| EPI_ISL_2702069 | Lighthouse Lab in Milton Keynes | Wellcome Sanger Institute for the COVID-19 Genomics UK (COG-UK) Consortium | The Lighthouse Lab in Milton Keynes and Alex Alderton, Roberto Amato, Jeffrey Barrett, Sonia Goncalves, Ewan Harrison, David K. Jackson, Ian Johnston, Dominic Kwiatkowski, Cordelia Langford, John Sillitoe on behalf of the Wellcome Sanger Institute COVID-19 Surveillance Team |
| EPI_ISL_2702070 | Lighthouse Lab in Glasgow | Wellcome Sanger Institute for the COVID-19 Genomics UK (COG-UK) Consortium | Harper VanSteenhouse, Yumi Kasai, David Gray, Carol Clugston, Anna Dominiczak and Alex Alderton, Roberto Amato, Jeffrey Barrett, Sonia Goncalves, Ewan Harrison, David K. Jackson, Ian Johnston, Dominic Kwiatkowski, Cordelia Langford, John Sillitoe on behalf of the Wellcome Sanger Institute COVID-19 Surveillance Team |
| EPI_ISL_2702071, EPI_ISL_2702072 | Lighthouse Lab in Alderley Park | Wellcome Sanger Institute for the COVID-19 Genomics UK (COG-UK) Consortium | Jacquelyn Wynn, Mairead Hyland, The Lighthouse Lab in Alderley Park and Alex Alderton, Roberto Amato, Jeffrey Barrett, Sonia Goncalves, Ewan Harrison, David K. Jackson, Ian Johnston, Dominic Kwiatkowski, Cordelia Langford, John Sillitoe on behalf of the Wellcome Sanger Institute COVID-19 Surveillance Team |
| EPI_ISL_2702073, EPI_ISL_2702074 | Lighthouse Lab in Milton Keynes | Wellcome Sanger Institute for the COVID-19 Genomics UK (COG-UK) Consortium | The Lighthouse Lab in Milton Keynes and Alex Alderton, Roberto Amato, Jeffrey Barrett, Sonia Goncalves, Ewan Harrison, David K. Jackson, Ian Johnston, Dominic Kwiatkowski, Cordelia Langford, John Sillitoe on behalf of the Wellcome Sanger Institute COVID-19 Surveillance Team |
| EPI_ISL_2702075, EPI_ISL_2702076 | Lighthouse Lab in Glasgow | Wellcome Sanger Institute for the COVID-19 Genomics UK (COG-UK) Consortium | Harper VanSteenhouse, Yumi Kasai, David Gray, Carol Clugston, Anna Dominiczak and Alex Alderton, Roberto Amato, Jeffrey Barrett, Sonia Goncalves, Ewan Harrison, David K. Jackson, Ian Johnston, Dominic Kwiatkowski, Cordelia Langford, John Sillitoe on behalf of the Wellcome Sanger Institute COVID-19 Surveillance Team |
| EPI_ISL_2702077 | Lighthouse Lab in Milton Keynes | Wellcome Sanger Institute for the COVID-19 Genomics UK (COG-UK) Consortium | The Lighthouse Lab in Milton Keynes and Alex Alderton, Roberto Amato, Jeffrey Barrett, Sonia Goncalves, Ewan Harrison, David K. Jackson, Ian Johnston, Dominic Kwiatkowski, Cordelia Langford, John Sillitoe on behalf of the Wellcome Sanger Institute COVID-19 Surveillance Team |
| EPI_ISL_2702078 | Lighthouse Lab in Alderley Park | Wellcome Sanger Institute for the COVID-19 Genomics UK (COG-UK) Consortium | Jacquelyn Wynn, Mairead Hyland, The Lighthouse Lab in Alderley Park and Alex Alderton, Roberto Amato, Jeffrey Barrett, Sonia Goncalves, Ewan Harrison, David K. Jackson, Ian Johnston, Dominic Kwiatkowski, Cordelia Langford, John Sillitoe on behalf of the Wellcome Sanger Institute COVID-19 Surveillance Team |
| EPI_ISL_2702079, EPI_ISL_2702080 | Lighthouse Lab in Glasgow | Wellcome Sanger Institute for the COVID-19 Genomics UK (COG-UK) Consortium | Harper VanSteenhouse, Yumi Kasai, David Gray, Carol Clugston, Anna Dominiczak and Alex Alderton, Roberto Amato, Jeffrey Barrett, Sonia Goncalves, Ewan Harrison, David K. Jackson, Ian Johnston, Dominic Kwiatkowski, Cordelia Langford, John Sillitoe on behalf of the Wellcome Sanger Institute COVID-19 Surveillance Team |
| EPI_ISL_2702081 | Lighthouse Lab in Milton Keynes | Wellcome Sanger Institute for the COVID-19 Genomics UK (COG-UK) Consortium | The Lighthouse Lab in Milton Keynes and Alex Alderton, Roberto Amato, Jeffrey Barrett, Sonia Goncalves, Ewan Harrison, David K. Jackson, Ian Johnston, Dominic Kwiatkowski, Cordelia Langford, John Sillitoe on behalf of the Wellcome Sanger Institute COVID-19 Surveillance Team |
| EPI_ISL_2702083 | Lighthouse Lab in Glasgow | Wellcome Sanger Institute for the COVID-19 Genomics UK (COG-UK) Consortium | Harper VanSteenhouse, Yumi Kasai, David Gray, Carol Clugston, Anna Dominiczak and Alex Alderton, Roberto Amato, Jeffrey Barrett, Sonia Goncalves, Ewan Harrison, David K. Jackson, Ian Johnston, Dominic Kwiatkowski, Cordelia Langford, John Sillitoe on behalf of the Wellcome Sanger Institute COVID-19 Surveillance Team |
| EPI_ISL_2702084, EPI_ISL_2702085 | Lighthouse Lab in Alderley Park | Wellcome Sanger Institute for the COVID-19 Genomics UK (COG-UK) Consortium | Jacquelyn Wynn, Mairead Hyland, The Lighthouse Lab in Alderley Park and Alex Alderton, Roberto Amato, Jeffrey Barrett, Sonia Goncalves, Ewan Harrison, David K. Jackson, Ian Johnston, Dominic Kwiatkowski, Cordelia Langford, John Sillitoe on behalf of the Wellcome Sanger Institute COVID-19 Surveillance Team |
| EPI_ISL_2702086 | Lighthouse Lab in Milton Keynes | Wellcome Sanger Institute for the COVID-19 Genomics UK (COG-UK) Consortium | The Lighthouse Lab in Milton Keynes and Alex Alderton, Roberto Amato, Jeffrey Barrett, Sonia Goncalves, Ewan Harrison, David K. Jackson, Ian Johnston, Dominic Kwiatkowski, Cordelia Langford, John Sillitoe on behalf of the Wellcome Sanger Institute COVID-19 Surveillance Team |
| EPI_ISL_2702087 | Lighthouse Lab in Alderley Park | Wellcome Sanger Institute for the COVID-19 Genomics UK (COG-UK) Consortium | Jacquelyn Wynn, Mairead Hyland, The Lighthouse Lab in Alderley Park and Alex Alderton, Roberto Amato, Jeffrey Barrett, Sonia Goncalves, Ewan Harrison, David K. Jackson, Ian Johnston, Dominic Kwiatkowski, Cordelia Langford, John Sillitoe on behalf of the Wellcome Sanger Institute COVID-19 Surveillance Team |
| EPI_ISL_2702088 | Lighthouse Lab in Milton Keynes | Wellcome Sanger Institute for the COVID-19 Genomics UK (COG-UK) Consortium | The Lighthouse Lab in Milton Keynes and Alex Alderton, Roberto Amato, Jeffrey Barrett, Sonia Goncalves, Ewan Harrison, David K. Jackson, Ian Johnston, Dominic Kwiatkowski, Cordelia Langford, John Sillitoe on behalf of the Wellcome Sanger Institute COVID-19 Surveillance Team |
| EPI_ISL_2702089 | Lighthouse Lab in Alderley Park | Wellcome Sanger Institute for the COVID-19 Genomics UK (COG-UK) | Jacquelyn Wynn, Mairead Hyland, The Lighthouse Lab in Alderley Park and Alex Alderton, Roberto Amato, Jeffrey Barrett, Sonia Goncalves, Ewan Harrison, David K. Jackson, Ian Johnston, Dominic Kwiatkowski, Cordelia Langford, John Sillitoe on |

| | | | |
|---|---|---|---|
| | | Consortium | behalf of the Wellcome Sanger Institute COVID-19 Surveillance Team |
| EPI_ISL_2702090 | Lighthouse Lab in Milton Keynes | Wellcome Sanger Institute for the COVID-19 Genomics UK (COG-UK) Consortium | The Lighthouse Lab in Milton Keynes and Alex Alderton, Roberto Amato, Jeffrey Barrett, Sonia Goncalves, Ewan Harrison, David K. Jackson, Ian Johnston, Dominic Kwiatkowski, Cordelia Langford, John Sillitoe on behalf of the Wellcome Sanger Institute COVID-19 Surveillance Team |
| EPI_ISL_2702091 | Lighthouse Lab in Alderley Park | Wellcome Sanger Institute for the COVID-19 Genomics UK (COG-UK) Consortium | Jacquelyn Wynn, Mairead Hyland, The Lighthouse Lab in Alderley Park and Alex Alderton, Roberto Amato, Jeffrey Barrett, Sonia Goncalves, Ewan Harrison, David K. Jackson, Ian Johnston, Dominic Kwiatkowski, Cordelia Langford, John Sillitoe on behalf of the Wellcome Sanger Institute COVID-19 Surveillance Team |
| EPI_ISL_2702092, EPI_ISL_2702093, EPI_ISL_2702094 | Lighthouse Lab in Glasgow | Wellcome Sanger Institute for the COVID-19 Genomics UK (COG-UK) Consortium | Harper VanSteenhouse, Yumi Kasai, David Gray, Carol Clugston, Anna Dominiczak and Alex Alderton, Roberto Amato, Jeffrey Barrett, Sonia Goncalves, Ewan Harrison, David K. Jackson, Ian Johnston, Dominic Kwiatkowski, Cordelia Langford, John Sillitoe on behalf of the Wellcome Sanger Institute COVID-19 Surveillance Team |
| EPI_ISL_2702095 | Lighthouse Lab in Alderley Park | Wellcome Sanger Institute for the COVID-19 Genomics UK (COG-UK) Consortium | Jacquelyn Wynn, Mairead Hyland, The Lighthouse Lab in Alderley Park and Alex Alderton, Roberto Amato, Jeffrey Barrett, Sonia Goncalves, Ewan Harrison, David K. Jackson, Ian Johnston, Dominic Kwiatkowski, Cordelia Langford, John Sillitoe on behalf of the Wellcome Sanger Institute COVID-19 Surveillance Team |
| EPI_ISL_2702096, EPI_ISL_2702097 | Lighthouse Lab in Glasgow | Wellcome Sanger Institute for the COVID-19 Genomics UK (COG-UK) Consortium | Harper VanSteenhouse, Yumi Kasai, David Gray, Carol Clugston, Anna Dominiczak and Alex Alderton, Roberto Amato, Jeffrey Barrett, Sonia Goncalves, Ewan Harrison, David K. Jackson, Ian Johnston, Dominic Kwiatkowski, Cordelia Langford, John Sillitoe on behalf of the Wellcome Sanger Institute COVID-19 Surveillance Team |
| EPI_ISL_2702098 | Lighthouse Lab in Alderley Park | Wellcome Sanger Institute for the COVID-19 Genomics UK (COG-UK) Consortium | Jacquelyn Wynn, Mairead Hyland, The Lighthouse Lab in Alderley Park and Alex Alderton, Roberto Amato, Jeffrey Barrett, Sonia Goncalves, Ewan Harrison, David K. Jackson, Ian Johnston, Dominic Kwiatkowski, Cordelia Langford, John Sillitoe on behalf of the Wellcome Sanger Institute COVID-19 Surveillance Team |
| EPI_ISL_2702099 | Lighthouse Lab in Milton Keynes | Wellcome Sanger Institute for the COVID-19 Genomics UK (COG-UK) Consortium | The Lighthouse Lab in Milton Keynes and Alex Alderton, Roberto Amato, Jeffrey Barrett, Sonia Goncalves, Ewan Harrison, David K. Jackson, Ian Johnston, Dominic Kwiatkowski, Cordelia Langford, John Sillitoe on behalf of the Wellcome Sanger Institute COVID-19 Surveillance Team |
| EPI_ISL_2702100 | Lighthouse Lab in Alderley Park | Wellcome Sanger Institute for the COVID-19 Genomics UK (COG-UK) Consortium | Jacquelyn Wynn, Mairead Hyland, The Lighthouse Lab in Alderley Park and Alex Alderton, Roberto Amato, Jeffrey Barrett, Sonia Goncalves, Ewan Harrison, David K. Jackson, Ian Johnston, Dominic Kwiatkowski, Cordelia Langford, John Sillitoe on behalf of the Wellcome Sanger Institute COVID-19 Surveillance Team |
| EPI_ISL_2702101 | Lighthouse Lab in Milton Keynes | Wellcome Sanger Institute for the COVID-19 Genomics UK (COG-UK) Consortium | The Lighthouse Lab in Milton Keynes and Alex Alderton, Roberto Amato, Jeffrey Barrett, Sonia Goncalves, Ewan Harrison, David K. Jackson, Ian Johnston, Dominic Kwiatkowski, Cordelia Langford, John Sillitoe on behalf of the Wellcome Sanger Institute COVID-19 Surveillance Team |
| EPI_ISL_2702102, EPI_ISL_2702103, EPI_ISL_2702104, EPI_ISL_2702105, EPI_ISL_2702106 | Lighthouse Lab in Glasgow | Wellcome Sanger Institute for the COVID-19 Genomics UK (COG-UK) Consortium | Harper VanSteenhouse, Yumi Kasai, David Gray, Carol Clugston, Anna Dominiczak and Alex Alderton, Roberto Amato, Jeffrey Barrett, Sonia Goncalves, Ewan Harrison, David K. Jackson, Ian Johnston, Dominic Kwiatkowski, Cordelia Langford, John Sillitoe on behalf of the Wellcome Sanger Institute COVID-19 Surveillance Team |
| EPI_ISL_2702107 | Lighthouse Lab in Alderley Park | Wellcome Sanger Institute for the COVID-19 Genomics UK (COG-UK) Consortium | Jacquelyn Wynn, Mairead Hyland, The Lighthouse Lab in Alderley Park and Alex Alderton, Roberto Amato, Jeffrey Barrett, Sonia Goncalves, Ewan Harrison, David K. Jackson, Ian Johnston, Dominic Kwiatkowski, Cordelia Langford, John Sillitoe on behalf of the Wellcome Sanger Institute COVID-19 Surveillance Team |
| EPI_ISL_2702108 | Lighthouse Lab in Glasgow | Wellcome Sanger Institute for the COVID-19 Genomics UK (COG-UK) Consortium | Harper VanSteenhouse, Yumi Kasai, David Gray, Carol Clugston, Anna Dominiczak and Alex Alderton, Roberto Amato, Jeffrey Barrett, Sonia Goncalves, Ewan Harrison, David K. Jackson, Ian Johnston, Dominic Kwiatkowski, Cordelia Langford, John Sillitoe on behalf of the Wellcome Sanger Institute COVID-19 Surveillance Team |
| EPI_ISL_2702109 | Lighthouse Lab in Alderley Park | Wellcome Sanger Institute for the COVID-19 Genomics UK (COG-UK) Consortium | Jacquelyn Wynn, Mairead Hyland, The Lighthouse Lab in Alderley Park and Alex Alderton, Roberto Amato, Jeffrey Barrett, Sonia Goncalves, Ewan Harrison, David K. Jackson, Ian Johnston, Dominic Kwiatkowski, Cordelia Langford, John Sillitoe on behalf of the Wellcome Sanger Institute COVID-19 Surveillance Team |
| EPI_ISL_2702110 | Lighthouse Lab in Glasgow | Wellcome Sanger Institute for the COVID-19 Genomics UK (COG-UK) Consortium | Harper VanSteenhouse, Yumi Kasai, David Gray, Carol Clugston, Anna Dominiczak and Alex Alderton, Roberto Amato, Jeffrey Barrett, Sonia Goncalves, Ewan Harrison, David K. Jackson, Ian Johnston, Dominic Kwiatkowski, Cordelia Langford, John Sillitoe on behalf of the Wellcome Sanger Institute COVID-19 Surveillance Team |
| EPI_ISL_2702111, EPI_ISL_2702112, EPI_ISL_2702113 | Lighthouse Lab in Alderley Park | Wellcome Sanger Institute for the COVID-19 Genomics UK (COG-UK) Consortium | Jacquelyn Wynn, Mairead Hyland, The Lighthouse Lab in Alderley Park and Alex Alderton, Roberto Amato, Jeffrey Barrett, Sonia Goncalves, Ewan Harrison, David K. Jackson, Ian Johnston, Dominic Kwiatkowski, Cordelia Langford, John Sillitoe on behalf of the Wellcome Sanger Institute COVID-19 Surveillance Team |
| EPI_ISL_2702114 | Lighthouse Lab in Milton Keynes | Wellcome Sanger Institute for the COVID-19 Genomics UK (COG-UK) Consortium | The Lighthouse Lab in Milton Keynes and Alex Alderton, Roberto Amato, Jeffrey Barrett, Sonia Goncalves, Ewan Harrison, David K. Jackson, Ian Johnston, Dominic Kwiatkowski, Cordelia Langford, John Sillitoe on behalf of the Wellcome Sanger Institute COVID-19 Surveillance Team |
| EPI_ISL_2702115, EPI_ISL_2702116 | Lighthouse Lab in Glasgow | Wellcome Sanger Institute for the COVID-19 Genomics UK (COG-UK) Consortium | Harper VanSteenhouse, Yumi Kasai, David Gray, Carol Clugston, Anna Dominiczak and Alex Alderton, Roberto Amato, Jeffrey Barrett, Sonia Goncalves, Ewan Harrison, David K. Jackson, Ian Johnston, Dominic Kwiatkowski, Cordelia Langford, John Sillitoe on behalf of the Wellcome Sanger Institute COVID-19 Surveillance Team |
| EPI_ISL_2702117 | Lighthouse Lab in Alderley Park | Wellcome Sanger Institute for the COVID-19 Genomics UK (COG-UK) Consortium | Jacquelyn Wynn, Mairead Hyland, The Lighthouse Lab in Alderley Park and Alex Alderton, Roberto Amato, Jeffrey Barrett, Sonia Goncalves, Ewan Harrison, David K. Jackson, Ian Johnston, Dominic Kwiatkowski, Cordelia Langford, John Sillitoe on behalf of the Wellcome Sanger Institute COVID-19 Surveillance Team |
| EPI_ISL_2702118 | Lighthouse Lab in Glasgow | Wellcome Sanger Institute for the COVID-19 Genomics UK (COG-UK) Consortium | Harper VanSteenhouse, Yumi Kasai, David Gray, Carol Clugston, Anna Dominiczak and Alex Alderton, Roberto Amato, Jeffrey Barrett, Sonia Goncalves, Ewan Harrison, David K. Jackson, Ian Johnston, Dominic Kwiatkowski, Cordelia Langford, John Sillitoe on behalf of the Wellcome Sanger Institute COVID-19 Surveillance Team |
| EPI_ISL_2702119 | Lighthouse Lab in Milton Keynes | Wellcome Sanger Institute for the COVID-19 Genomics UK (COG-UK) Consortium | The Lighthouse Lab in Milton Keynes and Alex Alderton, Roberto Amato, Jeffrey Barrett, Sonia Goncalves, Ewan Harrison, David K. Jackson, Ian Johnston, Dominic Kwiatkowski, Cordelia Langford, John Sillitoe on behalf of the Wellcome Sanger Institute COVID-19 Surveillance Team |
| EPI_ISL_2702120 | Lighthouse Lab in Alderley Park | Wellcome Sanger Institute for the COVID-19 Genomics UK (COG-UK) Consortium | Jacquelyn Wynn, Mairead Hyland, The Lighthouse Lab in Alderley Park and Alex Alderton, Roberto Amato, Jeffrey Barrett, Sonia Goncalves, Ewan Harrison, David K. Jackson, Ian Johnston, Dominic Kwiatkowski, Cordelia Langford, John Sillitoe on behalf of the Wellcome Sanger Institute COVID-19 Surveillance Team |
| EPI_ISL_2702121, EPI_ISL_2702122 | Lighthouse Lab in Glasgow | Wellcome Sanger Institute for the COVID-19 Genomics UK (COG-UK) Consortium | Harper VanSteenhouse, Yumi Kasai, David Gray, Carol Clugston, Anna Dominiczak and Alex Alderton, Roberto Amato, Jeffrey Barrett, Sonia Goncalves, Ewan Harrison, David K. Jackson, Ian Johnston, Dominic Kwiatkowski, Cordelia Langford, John Sillitoe on behalf of the Wellcome Sanger Institute COVID-19 Surveillance Team |
| EPI_ISL_2702123 | Lighthouse Lab in Milton Keynes | Wellcome Sanger Institute for the COVID-19 Genomics UK (COG-UK) Consortium | The Lighthouse Lab in Milton Keynes and Alex Alderton, Roberto Amato, Jeffrey Barrett, Sonia Goncalves, Ewan Harrison, David K. Jackson, Ian Johnston, Dominic Kwiatkowski, Cordelia Langford, John Sillitoe on behalf of the Wellcome Sanger Institute COVID-19 Surveillance Team |
| EPI_ISL_2702124 | Lighthouse Lab in Glasgow | Wellcome Sanger Institute for the COVID-19 Genomics UK (COG-UK) Consortium | Harper VanSteenhouse, Yumi Kasai, David Gray, Carol Clugston, Anna Dominiczak and Alex Alderton, Roberto Amato, Jeffrey Barrett, Sonia Goncalves, Ewan Harrison, David K. Jackson, Ian Johnston, Dominic Kwiatkowski, Cordelia Langford, John Sillitoe on behalf of the Wellcome Sanger Institute COVID-19 Surveillance Team |
| EPI_ISL_2702125 | Lighthouse Lab in Milton Keynes | Wellcome Sanger Institute for the COVID-19 Genomics UK (COG-UK) Consortium | The Lighthouse Lab in Milton Keynes and Alex Alderton, Roberto Amato, Jeffrey Barrett, Sonia Goncalves, Ewan Harrison, David K. Jackson, Ian Johnston, Dominic Kwiatkowski, Cordelia Langford, John Sillitoe on behalf of the Wellcome Sanger Institute COVID-19 Surveillance Team |
| EPI_ISL_2702126 | Lighthouse Lab in Glasgow | Wellcome Sanger Institute for the COVID-19 Genomics UK (COG-UK) Consortium | Harper VanSteenhouse, Yumi Kasai, David Gray, Carol Clugston, Anna Dominiczak and Alex Alderton, Roberto Amato, Jeffrey Barrett, Sonia Goncalves, Ewan Harrison, David K. Jackson, Ian Johnston, Dominic Kwiatkowski, Cordelia Langford, John Sillitoe on behalf of the Wellcome Sanger Institute COVID-19 Surveillance Team |
| EPI_ISL_2702127 | Lighthouse Lab in Milton Keynes | Wellcome Sanger Institute for the COVID-19 Genomics UK (COG-UK) Consortium | The Lighthouse Lab in Milton Keynes and Alex Alderton, Roberto Amato, Jeffrey Barrett, Sonia Goncalves, Ewan Harrison, David K. Jackson, Ian Johnston, Dominic Kwiatkowski, Cordelia Langford, John Sillitoe on behalf of the Wellcome Sanger Institute COVID-19 Surveillance Team |
| EPI_ISL_2702128 | Lighthouse Lab in Glasgow | Wellcome Sanger Institute for the COVID-19 Genomics UK (COG-UK) Consortium | Harper VanSteenhouse, Yumi Kasai, David Gray, Carol Clugston, Anna Dominiczak and Alex Alderton, Roberto Amato, Jeffrey Barrett, Sonia Goncalves, Ewan Harrison, David K. Jackson, Ian Johnston, Dominic Kwiatkowski, Cordelia Langford, John Sillitoe on behalf of the Wellcome Sanger Institute COVID-19 Surveillance Team |
| EPI_ISL_2702129, EPI_ISL_2702130 | Lighthouse Lab in Alderley Park | Wellcome Sanger Institute for the COVID-19 Genomics UK (COG-UK) Consortium | Jacquelyn Wynn, Mairead Hyland, The Lighthouse Lab in Alderley Park and Alex Alderton, Roberto Amato, Jeffrey Barrett, Sonia Goncalves, Ewan Harrison, David K. Jackson, Ian Johnston, Dominic Kwiatkowski, Cordelia Langford, John Sillitoe on behalf of the Wellcome Sanger Institute COVID-19 Surveillance Team |
| EPI_ISL_2702132, EPI_ISL_2702133 | Lighthouse Lab in Glasgow | Wellcome Sanger Institute for the COVID-19 Genomics UK (COG-UK) Consortium | Harper VanSteenhouse, Yumi Kasai, David Gray, Carol Clugston, Anna Dominiczak and Alex Alderton, Roberto Amato, Jeffrey Barrett, Sonia Goncalves, Ewan Harrison, David K. Jackson, Ian Johnston, Dominic Kwiatkowski, Cordelia Langford, John Sillitoe on behalf of the Wellcome Sanger Institute COVID-19 Surveillance Team |
| EPI_ISL_2702134 | Lighthouse Lab in Alderley Park | Wellcome Sanger Institute for the COVID-19 Genomics UK (COG-UK) Consortium | Jacquelyn Wynn, Mairead Hyland, The Lighthouse Lab in Alderley Park and Alex Alderton, Roberto Amato, Jeffrey Barrett, Sonia Goncalves, Ewan Harrison, David K. Jackson, Ian Johnston, Dominic Kwiatkowski, Cordelia Langford, John Sillitoe on behalf of the Wellcome Sanger Institute COVID-19 Surveillance Team |
| EPI_ISL_2702135 | Lighthouse Lab in Glasgow | Wellcome Sanger Institute for the COVID-19 Genomics UK (COG-UK) Consortium | Harper VanSteenhouse, Yumi Kasai, David Gray, Carol Clugston, Anna Dominiczak and Alex Alderton, Roberto Amato, Jeffrey Barrett, Sonia Goncalves, Ewan Harrison, David K. Jackson, Ian Johnston, Dominic Kwiatkowski, Cordelia Langford, John Sillitoe on behalf of the Wellcome Sanger Institute COVID-19 Surveillance Team |
| EPI_ISL_2702136 | Lighthouse Lab in Alderley Park | Wellcome Sanger Institute for the COVID-19 Genomics UK (COG-UK) Consortium | Jacquelyn Wynn, Mairead Hyland, The Lighthouse Lab in Alderley Park and Alex Alderton, Roberto Amato, Jeffrey Barrett, Sonia Goncalves, Ewan Harrison, David K. Jackson, Ian Johnston, Dominic Kwiatkowski, Cordelia Langford, John Sillitoe on behalf of the Wellcome Sanger Institute COVID-19 Surveillance Team |
| EPI_ISL_2702137, EPI_ISL_2702138 | Lighthouse Lab in Glasgow | Wellcome Sanger Institute for the COVID-19 Genomics UK (COG-UK) Consortium | Harper VanSteenhouse, Yumi Kasai, David Gray, Carol Clugston, Anna Dominiczak and Alex Alderton, Roberto Amato, Jeffrey Barrett, Sonia Goncalves, Ewan Harrison, David K. Jackson, Ian Johnston, Dominic Kwiatkowski, Cordelia Langford, John Sillitoe on behalf of the Wellcome Sanger Institute COVID-19 Surveillance Team |
| EPI_ISL_2702139, EPI_ISL_2702140 | Lighthouse Lab in Alderley Park | Wellcome Sanger Institute for the COVID-19 Genomics UK (COG-UK) Consortium | Jacquelyn Wynn, Mairead Hyland, The Lighthouse Lab in Alderley Park and Alex Alderton, Roberto Amato, Jeffrey Barrett, Sonia Goncalves, Ewan Harrison, David K. Jackson, Ian Johnston, Dominic Kwiatkowski, Cordelia Langford, John Sillitoe on behalf of the Wellcome Sanger Institute COVID-19 Surveillance Team |
| EPI_ISL_2702141, EPI_ISL_2702142 | Lighthouse Lab in Glasgow | Wellcome Sanger Institute for the COVID-19 Genomics UK (COG-UK) Consortium | Harper VanSteenhouse, Yumi Kasai, David Gray, Carol Clugston, Anna Dominiczak and Alex Alderton, Roberto Amato, Jeffrey Barrett, Sonia Goncalves, Ewan Harrison, David K. Jackson, Ian Johnston, Dominic Kwiatkowski, Cordelia Langford, John Sillitoe on behalf of the Wellcome Sanger Institute COVID-19 Surveillance Team |
| EPI_ISL_2702143, EPI_ISL_2702144, EPI_ISL_2702145 | Lighthouse Lab in Milton Keynes | Wellcome Sanger Institute for the COVID-19 Genomics UK (COG-UK) Consortium | The Lighthouse Lab in Milton Keynes and Alex Alderton, Roberto Amato, Jeffrey Barrett, Sonia Goncalves, Ewan Harrison, David K. Jackson, Ian Johnston, Dominic Kwiatkowski, Cordelia Langford, John Sillitoe on behalf of the Wellcome Sanger Institute COVID-19 Surveillance Team |
| EPI_ISL_2702147 | Lighthouse Lab in Glasgow | Wellcome Sanger Institute for the COVID-19 Genomics UK (COG-UK) Consortium | Harper VanSteenhouse, Yumi Kasai, David Gray, Carol Clugston, Anna Dominiczak and Alex Alderton, Roberto Amato, Jeffrey Barrett, Sonia Goncalves, Ewan Harrison, David K. Jackson, Ian Johnston, Dominic Kwiatkowski, Cordelia Langford, John Sillitoe on behalf of the Wellcome Sanger Institute COVID-19 Surveillance Team |
| EPI_ISL_2702148 | Lighthouse Lab in Alderley Park | Wellcome Sanger Institute for the COVID-19 Genomics UK (COG-UK) Consortium | Jacquelyn Wynn, Mairead Hyland, The Lighthouse Lab in Alderley Park and Alex Alderton, Roberto Amato, Jeffrey Barrett, Sonia Goncalves, Ewan Harrison, David K. Jackson, Ian Johnston, Dominic Kwiatkowski, Cordelia Langford, John Sillitoe on behalf of the Wellcome Sanger Institute COVID-19 Surveillance Team |
| EPI_ISL_2702149 | Lighthouse Lab in Milton Keynes | Wellcome Sanger Institute for the COVID-19 Genomics UK (COG-UK) Consortium | The Lighthouse Lab in Milton Keynes and Alex Alderton, Roberto Amato, Jeffrey Barrett, Sonia Goncalves, Ewan Harrison, David K. Jackson, Ian Johnston, Dominic Kwiatkowski, Cordelia Langford, John Sillitoe on behalf of the Wellcome Sanger Institute COVID-19 Surveillance Team |
| EPI_ISL_2702151 | Lighthouse Lab in Alderley Park | Wellcome Sanger Institute for the COVID-19 Genomics UK (COG-UK) Consortium | Jacquelyn Wynn, Mairead Hyland, The Lighthouse Lab in Alderley Park and Alex Alderton, Roberto Amato, Jeffrey Barrett, Sonia Goncalves, Ewan Harrison, David K. Jackson, Ian Johnston, Dominic Kwiatkowski, Cordelia Langford, John Sillitoe on behalf of the Wellcome Sanger Institute COVID-19 Surveillance Team |
| EPI_ISL_2702152, EPI_ISL_2702153 | Lighthouse Lab in Glasgow | Wellcome Sanger Institute for the COVID-19 Genomics UK (COG-UK) Consortium | Harper VanSteenhouse, Yumi Kasai, David Gray, Carol Clugston, Anna Dominiczak and Alex Alderton, Roberto Amato, Jeffrey Barrett, Sonia Goncalves, Ewan Harrison, David K. Jackson, Ian Johnston, Dominic Kwiatkowski, Cordelia Langford, John Sillitoe on behalf of the Wellcome Sanger Institute COVID-19 Surveillance Team |

| | | | |
|---|---|---|---|
| EPI_ISL_2702154 | Lighthouse Lab in Alderley Park | Wellcome Sanger Institute for the COVID-19 Genomics UK (COG-UK) Consortium | Jacquelyn Wynn, Mairead Hyland, The Lighthouse Lab in Alderley Park and Alex Alderton, Roberto Amato, Jeffrey Barrett, Sonia Goncalves, Ewan Harrison, David K. Jackson, Ian Johnston, Dominic Kwiatkowski, Cordelia Langford, John Sillitoe on behalf of the Wellcome Sanger Institute COVID-19 Surveillance Team |
| EPI_ISL_2702155 | Lighthouse Lab in Glasgow | Wellcome Sanger Institute for the COVID-19 Genomics UK (COG-UK) Consortium | Harper VanSteenhouse, Yumi Kasai, David Gray, Carol Clugston, Anna Dominiczak and Alex Alderton, Roberto Amato, Jeffrey Barrett, Sonia Goncalves, Ewan Harrison, David K. Jackson, Ian Johnston, Dominic Kwiatkowski, Cordelia Langford, John Sillitoe on behalf of the Wellcome Sanger Institute COVID-19 Surveillance Team |
| EPI_ISL_2702156 | Lighthouse Lab in Alderley Park | Wellcome Sanger Institute for the COVID-19 Genomics UK (COG-UK) Consortium | Jacquelyn Wynn, Mairead Hyland, The Lighthouse Lab in Alderley Park and Alex Alderton, Roberto Amato, Jeffrey Barrett, Sonia Goncalves, Ewan Harrison, David K. Jackson, Ian Johnston, Dominic Kwiatkowski, Cordelia Langford, John Sillitoe on behalf of the Wellcome Sanger Institute COVID-19 Surveillance Team |
| EPI_ISL_2702157, EPI_ISL_2702158 | Lighthouse Lab in Glasgow | Wellcome Sanger Institute for the COVID-19 Genomics UK (COG-UK) Consortium | Harper VanSteenhouse, Yumi Kasai, David Gray, Carol Clugston, Anna Dominiczak and Alex Alderton, Roberto Amato, Jeffrey Barrett, Sonia Goncalves, Ewan Harrison, David K. Jackson, Ian Johnston, Dominic Kwiatkowski, Cordelia Langford, John Sillitoe on behalf of the Wellcome Sanger Institute COVID-19 Surveillance Team |
| EPI_ISL_2702159 | Lighthouse Lab in Milton Keynes | Wellcome Sanger Institute for the COVID-19 Genomics UK (COG-UK) Consortium | The Lighthouse Lab in Milton Keynes and Alex Alderton, Roberto Amato, Jeffrey Barrett, Sonia Goncalves, Ewan Harrison, David K. Jackson, Ian Johnston, Dominic Kwiatkowski, Cordelia Langford, John Sillitoe on behalf of the Wellcome Sanger Institute COVID-19 Surveillance Team |
| EPI_ISL_2702160 | Lighthouse Lab in Alderley Park | Wellcome Sanger Institute for the COVID-19 Genomics UK (COG-UK) Consortium | Jacquelyn Wynn, Mairead Hyland, The Lighthouse Lab in Alderley Park and Alex Alderton, Roberto Amato, Jeffrey Barrett, Sonia Goncalves, Ewan Harrison, David K. Jackson, Ian Johnston, Dominic Kwiatkowski, Cordelia Langford, John Sillitoe on behalf of the Wellcome Sanger Institute COVID-19 Surveillance Team |
| EPI_ISL_2702161, EPI_ISL_2702162 | Lighthouse Lab in Glasgow | Wellcome Sanger Institute for the COVID-19 Genomics UK (COG-UK) Consortium | Harper VanSteenhouse, Yumi Kasai, David Gray, Carol Clugston, Anna Dominiczak and Alex Alderton, Roberto Amato, Jeffrey Barrett, Sonia Goncalves, Ewan Harrison, David K. Jackson, Ian Johnston, Dominic Kwiatkowski, Cordelia Langford, John Sillitoe on behalf of the Wellcome Sanger Institute COVID-19 Surveillance Team |
| EPI_ISL_2702163 | Lighthouse Lab in Milton Keynes | Wellcome Sanger Institute for the COVID-19 Genomics UK (COG-UK) Consortium | The Lighthouse Lab in Milton Keynes and Alex Alderton, Roberto Amato, Jeffrey Barrett, Sonia Goncalves, Ewan Harrison, David K. Jackson, Ian Johnston, Dominic Kwiatkowski, Cordelia Langford, John Sillitoe on behalf of the Wellcome Sanger Institute COVID-19 Surveillance Team |
| EPI_ISL_2702164, EPI_ISL_2702165 | Lighthouse Lab in Glasgow | Wellcome Sanger Institute for the COVID-19 Genomics UK (COG-UK) Consortium | Harper VanSteenhouse, Yumi Kasai, David Gray, Carol Clugston, Anna Dominiczak and Alex Alderton, Roberto Amato, Jeffrey Barrett, Sonia Goncalves, Ewan Harrison, David K. Jackson, Ian Johnston, Dominic Kwiatkowski, Cordelia Langford, John Sillitoe on behalf of the Wellcome Sanger Institute COVID-19 Surveillance Team |
| EPI_ISL_2702166 | Lighthouse Lab in Milton Keynes | Wellcome Sanger Institute for the COVID-19 Genomics UK (COG-UK) Consortium | The Lighthouse Lab in Milton Keynes and Alex Alderton, Roberto Amato, Jeffrey Barrett, Sonia Goncalves, Ewan Harrison, David K. Jackson, Ian Johnston, Dominic Kwiatkowski, Cordelia Langford, John Sillitoe on behalf of the Wellcome Sanger Institute COVID-19 Surveillance Team |
| EPI_ISL_2702167 | Lighthouse Lab in Glasgow | Wellcome Sanger Institute for the COVID-19 Genomics UK (COG-UK) Consortium | Harper VanSteenhouse, Yumi Kasai, David Gray, Carol Clugston, Anna Dominiczak and Alex Alderton, Roberto Amato, Jeffrey Barrett, Sonia Goncalves, Ewan Harrison, David K. Jackson, Ian Johnston, Dominic Kwiatkowski, Cordelia Langford, John Sillitoe on behalf of the Wellcome Sanger Institute COVID-19 Surveillance Team |
| EPI_ISL_2702168 | Lighthouse Lab in Milton Keynes | Wellcome Sanger Institute for the COVID-19 Genomics UK (COG-UK) Consortium | The Lighthouse Lab in Milton Keynes and Alex Alderton, Roberto Amato, Jeffrey Barrett, Sonia Goncalves, Ewan Harrison, David K. Jackson, Ian Johnston, Dominic Kwiatkowski, Cordelia Langford, John Sillitoe on behalf of the Wellcome Sanger Institute COVID-19 Surveillance Team |
| EPI_ISL_2702169 | Lighthouse Lab in Glasgow | Wellcome Sanger Institute for the COVID-19 Genomics UK (COG-UK) Consortium | Harper VanSteenhouse, Yumi Kasai, David Gray, Carol Clugston, Anna Dominiczak and Alex Alderton, Roberto Amato, Jeffrey Barrett, Sonia Goncalves, Ewan Harrison, David K. Jackson, Ian Johnston, Dominic Kwiatkowski, Cordelia Langford, John Sillitoe on behalf of the Wellcome Sanger Institute COVID-19 Surveillance Team |
| EPI_ISL_2702170 | Lighthouse Lab in Alderley Park | Wellcome Sanger Institute for the COVID-19 Genomics UK (COG-UK) Consortium | Jacquelyn Wynn, Mairead Hyland, The Lighthouse Lab in Alderley Park and Alex Alderton, Roberto Amato, Jeffrey Barrett, Sonia Goncalves, Ewan Harrison, David K. Jackson, Ian Johnston, Dominic Kwiatkowski, Cordelia Langford, John Sillitoe on behalf of the Wellcome Sanger Institute COVID-19 Surveillance Team |
| EPI_ISL_2702171 | Lighthouse Lab in Glasgow | Wellcome Sanger Institute for the COVID-19 Genomics UK (COG-UK) Consortium | Harper VanSteenhouse, Yumi Kasai, David Gray, Carol Clugston, Anna Dominiczak and Alex Alderton, Roberto Amato, Jeffrey Barrett, Sonia Goncalves, Ewan Harrison, David K. Jackson, Ian Johnston, Dominic Kwiatkowski, Cordelia Langford, John Sillitoe on behalf of the Wellcome Sanger Institute COVID-19 Surveillance Team |
| EPI_ISL_2702172 | Lighthouse Lab in Milton Keynes | Wellcome Sanger Institute for the COVID-19 Genomics UK (COG-UK) Consortium | The Lighthouse Lab in Milton Keynes and Alex Alderton, Roberto Amato, Jeffrey Barrett, Sonia Goncalves, Ewan Harrison, David K. Jackson, Ian Johnston, Dominic Kwiatkowski, Cordelia Langford, John Sillitoe on behalf of the Wellcome Sanger Institute COVID-19 Surveillance Team |
| EPI_ISL_2702173, EPI_ISL_2702174, EPI_ISL_2702175 | Lighthouse Lab in Glasgow | Wellcome Sanger Institute for the COVID-19 Genomics UK (COG-UK) Consortium | Harper VanSteenhouse, Yumi Kasai, David Gray, Carol Clugston, Anna Dominiczak and Alex Alderton, Roberto Amato, Jeffrey Barrett, Sonia Goncalves, Ewan Harrison, David K. Jackson, Ian Johnston, Dominic Kwiatkowski, Cordelia Langford, John Sillitoe on behalf of the Wellcome Sanger Institute COVID-19 Surveillance Team |
| EPI_ISL_2702176 | Lighthouse Lab in Alderley Park | Wellcome Sanger Institute for the COVID-19 Genomics UK (COG-UK) Consortium | Jacquelyn Wynn, Mairead Hyland, The Lighthouse Lab in Alderley Park and Alex Alderton, Roberto Amato, Jeffrey Barrett, Sonia Goncalves, Ewan Harrison, David K. Jackson, Ian Johnston, Dominic Kwiatkowski, Cordelia Langford, John Sillitoe on behalf of the Wellcome Sanger Institute COVID-19 Surveillance Team |
| EPI_ISL_2702177, EPI_ISL_2702178, EPI_ISL_2702179 | Lighthouse Lab in Glasgow | Wellcome Sanger Institute for the COVID-19 Genomics UK (COG-UK) Consortium | Harper VanSteenhouse, Yumi Kasai, David Gray, Carol Clugston, Anna Dominiczak and Alex Alderton, Roberto Amato, Jeffrey Barrett, Sonia Goncalves, Ewan Harrison, David K. Jackson, Ian Johnston, Dominic Kwiatkowski, Cordelia Langford, John Sillitoe on behalf of the Wellcome Sanger Institute COVID-19 Surveillance Team |
| EPI_ISL_2702180 | Lighthouse Lab in Alderley Park | Wellcome Sanger Institute for the COVID-19 Genomics UK (COG-UK) Consortium | Jacquelyn Wynn, Mairead Hyland, The Lighthouse Lab in Alderley Park and Alex Alderton, Roberto Amato, Jeffrey Barrett, Sonia Goncalves, Ewan Harrison, David K. Jackson, Ian Johnston, Dominic Kwiatkowski, Cordelia Langford, John Sillitoe on behalf of the Wellcome Sanger Institute COVID-19 Surveillance Team |
| EPI_ISL_2702181 | Lighthouse Lab in Glasgow | Wellcome Sanger Institute for the COVID-19 Genomics UK (COG-UK) Consortium | Harper VanSteenhouse, Yumi Kasai, David Gray, Carol Clugston, Anna Dominiczak and Alex Alderton, Roberto Amato, Jeffrey Barrett, Sonia Goncalves, Ewan Harrison, David K. Jackson, Ian Johnston, Dominic Kwiatkowski, Cordelia Langford, John Sillitoe on behalf of the Wellcome Sanger Institute COVID-19 Surveillance Team |
| EPI_ISL_2702182 | Lighthouse Lab in Milton Keynes | Wellcome Sanger Institute for the COVID-19 Genomics UK (COG-UK) Consortium | The Lighthouse Lab in Milton Keynes and Alex Alderton, Roberto Amato, Jeffrey Barrett, Sonia Goncalves, Ewan Harrison, David K. Jackson, Ian Johnston, Dominic Kwiatkowski, Cordelia Langford, John Sillitoe on behalf of the Wellcome Sanger Institute COVID-19 Surveillance Team |
| EPI_ISL_2702183, EPI_ISL_2702184 | Lighthouse Lab in Alderley Park | Wellcome Sanger Institute for the COVID-19 Genomics UK (COG-UK) Consortium | Jacquelyn Wynn, Mairead Hyland, The Lighthouse Lab in Alderley Park and Alex Alderton, Roberto Amato, Jeffrey Barrett, Sonia Goncalves, Ewan Harrison, David K. Jackson, Ian Johnston, Dominic Kwiatkowski, Cordelia Langford, John Sillitoe on behalf of the Wellcome Sanger Institute COVID-19 Surveillance Team |
| EPI_ISL_2702185 | Lighthouse Lab in Glasgow | Wellcome Sanger Institute for the COVID-19 Genomics UK (COG-UK) Consortium | Harper VanSteenhouse, Yumi Kasai, David Gray, Carol Clugston, Anna Dominiczak and Alex Alderton, Roberto Amato, Jeffrey Barrett, Sonia Goncalves, Ewan Harrison, David K. Jackson, Ian Johnston, Dominic Kwiatkowski, Cordelia Langford, John Sillitoe on behalf of the Wellcome Sanger Institute COVID-19 Surveillance Team |
| EPI_ISL_2702186 | Lighthouse Lab in Milton Keynes | Wellcome Sanger Institute for the COVID-19 Genomics UK (COG-UK) Consortium | The Lighthouse Lab in Milton Keynes and Alex Alderton, Roberto Amato, Jeffrey Barrett, Sonia Goncalves, Ewan Harrison, David K. Jackson, Ian Johnston, Dominic Kwiatkowski, Cordelia Langford, John Sillitoe on behalf of the Wellcome Sanger Institute COVID-19 Surveillance Team |
| EPI_ISL_2702187, EPI_ISL_2702188 | Lighthouse Lab in Glasgow | Wellcome Sanger Institute for the COVID-19 Genomics UK (COG-UK) Consortium | Harper VanSteenhouse, Yumi Kasai, David Gray, Carol Clugston, Anna Dominiczak and Alex Alderton, Roberto Amato, Jeffrey Barrett, Sonia Goncalves, Ewan Harrison, David K. Jackson, Ian Johnston, Dominic Kwiatkowski, Cordelia Langford, John Sillitoe on behalf of the Wellcome Sanger Institute COVID-19 Surveillance Team |
| EPI_ISL_2702189 | Lighthouse Lab in Milton Keynes | Wellcome Sanger Institute for the COVID-19 Genomics UK (COG-UK) Consortium | The Lighthouse Lab in Milton Keynes and Alex Alderton, Roberto Amato, Jeffrey Barrett, Sonia Goncalves, Ewan Harrison, David K. Jackson, Ian Johnston, Dominic Kwiatkowski, Cordelia Langford, John Sillitoe on behalf of the Wellcome Sanger Institute COVID-19 Surveillance Team |
| EPI_ISL_2702190, EPI_ISL_2702191 | Lighthouse Lab in Glasgow | Wellcome Sanger Institute for the COVID-19 Genomics UK (COG-UK) Consortium | Harper VanSteenhouse, Yumi Kasai, David Gray, Carol Clugston, Anna Dominiczak and Alex Alderton, Roberto Amato, Jeffrey Barrett, Sonia Goncalves, Ewan Harrison, David K. Jackson, Ian Johnston, Dominic Kwiatkowski, Cordelia Langford, John Sillitoe on behalf of the Wellcome Sanger Institute COVID-19 Surveillance Team |
| EPI_ISL_2702192, EPI_ISL_2702193 | Lighthouse Lab in Milton Keynes | Wellcome Sanger Institute for the COVID-19 Genomics UK (COG-UK) Consortium | The Lighthouse Lab in Milton Keynes and Alex Alderton, Roberto Amato, Jeffrey Barrett, Sonia Goncalves, Ewan Harrison, David K. Jackson, Ian Johnston, Dominic Kwiatkowski, Cordelia Langford, John Sillitoe on behalf of the Wellcome Sanger Institute COVID-19 Surveillance Team |
| EPI_ISL_2702194 | Lighthouse Lab in Alderley Park | Wellcome Sanger Institute for the COVID-19 Genomics UK (COG-UK) Consortium | Jacquelyn Wynn, Mairead Hyland, The Lighthouse Lab in Alderley Park and Alex Alderton, Roberto Amato, Jeffrey Barrett, Sonia Goncalves, Ewan Harrison, David K. Jackson, Ian Johnston, Dominic Kwiatkowski, Cordelia Langford, John Sillitoe on behalf of the Wellcome Sanger Institute COVID-19 Surveillance Team |
| EPI_ISL_2702195 | Lighthouse Lab in Glasgow | Wellcome Sanger Institute for the COVID-19 Genomics UK (COG-UK) Consortium | Harper VanSteenhouse, Yumi Kasai, David Gray, Carol Clugston, Anna Dominiczak and Alex Alderton, Roberto Amato, Jeffrey Barrett, Sonia Goncalves, Ewan Harrison, David K. Jackson, Ian Johnston, Dominic Kwiatkowski, Cordelia Langford, John Sillitoe on behalf of the Wellcome Sanger Institute COVID-19 Surveillance Team |
| EPI_ISL_2702196 | Lighthouse Lab in Milton Keynes | Wellcome Sanger Institute for the COVID-19 Genomics UK (COG-UK) Consortium | The Lighthouse Lab in Milton Keynes and Alex Alderton, Roberto Amato, Jeffrey Barrett, Sonia Goncalves, Ewan Harrison, David K. Jackson, Ian Johnston, Dominic Kwiatkowski, Cordelia Langford, John Sillitoe on behalf of the Wellcome Sanger Institute COVID-19 Surveillance Team |
| EPI_ISL_2702197, EPI_ISL_2702198 | Lighthouse Lab in Alderley Park | Wellcome Sanger Institute for the COVID-19 Genomics UK (COG-UK) Consortium | Jacquelyn Wynn, Mairead Hyland, The Lighthouse Lab in Alderley Park and Alex Alderton, Roberto Amato, Jeffrey Barrett, Sonia Goncalves, Ewan Harrison, David K. Jackson, Ian Johnston, Dominic Kwiatkowski, Cordelia Langford, John Sillitoe on behalf of the Wellcome Sanger Institute COVID-19 Surveillance Team |
| EPI_ISL_2702199 | Lighthouse Lab in Glasgow | Wellcome Sanger Institute for the COVID-19 Genomics UK (COG-UK) Consortium | Harper VanSteenhouse, Yumi Kasai, David Gray, Carol Clugston, Anna Dominiczak and Alex Alderton, Roberto Amato, Jeffrey Barrett, Sonia Goncalves, Ewan Harrison, David K. Jackson, Ian Johnston, Dominic Kwiatkowski, Cordelia Langford, John Sillitoe on behalf of the Wellcome Sanger Institute COVID-19 Surveillance Team |
| EPI_ISL_2702200 | Lighthouse Lab in Milton Keynes | Wellcome Sanger Institute for the COVID-19 Genomics UK (COG-UK) Consortium | The Lighthouse Lab in Milton Keynes and Alex Alderton, Roberto Amato, Jeffrey Barrett, Sonia Goncalves, Ewan Harrison, David K. Jackson, Ian Johnston, Dominic Kwiatkowski, Cordelia Langford, John Sillitoe on behalf of the Wellcome Sanger Institute COVID-19 Surveillance Team |
| EPI_ISL_2702201 | Lighthouse Lab in Alderley Park | Wellcome Sanger Institute for the COVID-19 Genomics UK (COG-UK) Consortium | Jacquelyn Wynn, Mairead Hyland, The Lighthouse Lab in Alderley Park and Alex Alderton, Roberto Amato, Jeffrey Barrett, Sonia Goncalves, Ewan Harrison, David K. Jackson, Ian Johnston, Dominic Kwiatkowski, Cordelia Langford, John Sillitoe on behalf of the Wellcome Sanger Institute COVID-19 Surveillance Team |
| EPI_ISL_2702202, EPI_ISL_2702203, EPI_ISL_2702204 | Lighthouse Lab in Glasgow | Wellcome Sanger Institute for the COVID-19 Genomics UK (COG-UK) Consortium | Harper VanSteenhouse, Yumi Kasai, David Gray, Carol Clugston, Anna Dominiczak and Alex Alderton, Roberto Amato, Jeffrey Barrett, Sonia Goncalves, Ewan Harrison, David K. Jackson, Ian Johnston, Dominic Kwiatkowski, Cordelia Langford, John Sillitoe on behalf of the Wellcome Sanger Institute COVID-19 Surveillance Team |
| EPI_ISL_2702205 | Lighthouse Lab in Milton Keynes | Wellcome Sanger Institute for the COVID-19 Genomics UK (COG-UK) Consortium | The Lighthouse Lab in Milton Keynes and Alex Alderton, Roberto Amato, Jeffrey Barrett, Sonia Goncalves, Ewan Harrison, David K. Jackson, Ian Johnston, Dominic Kwiatkowski, Cordelia Langford, John Sillitoe on behalf of the Wellcome Sanger Institute COVID-19 Surveillance Team |
| EPI_ISL_2702206 | Lighthouse Lab in Glasgow | Wellcome Sanger Institute for the COVID-19 Genomics UK (COG-UK) Consortium | Harper VanSteenhouse, Yumi Kasai, David Gray, Carol Clugston, Anna Dominiczak and Alex Alderton, Roberto Amato, Jeffrey Barrett, Sonia Goncalves, Ewan Harrison, David K. Jackson, Ian Johnston, Dominic Kwiatkowski, Cordelia Langford, John Sillitoe on behalf of the Wellcome Sanger Institute COVID-19 Surveillance Team |
| EPI_ISL_2702207 | Lighthouse Lab in Milton Keynes | Wellcome Sanger Institute for the COVID-19 Genomics UK (COG-UK) Consortium | The Lighthouse Lab in Milton Keynes and Alex Alderton, Roberto Amato, Jeffrey Barrett, Sonia Goncalves, Ewan Harrison, David K. Jackson, Ian Johnston, Dominic Kwiatkowski, Cordelia Langford, John Sillitoe on behalf of the Wellcome Sanger Institute COVID-19 Surveillance Team |
| EPI_ISL_2702208, EPI_ISL_2702210 | Lighthouse Lab in Glasgow | Wellcome Sanger Institute for the COVID-19 Genomics UK (COG-UK) Consortium | Harper VanSteenhouse, Yumi Kasai, David Gray, Carol Clugston, Anna Dominiczak and Alex Alderton, Roberto Amato, Jeffrey Barrett, Sonia Goncalves, Ewan Harrison, David K. Jackson, Ian Johnston, Dominic Kwiatkowski, Cordelia Langford, John Sillitoe on behalf of the Wellcome Sanger Institute COVID-19 Surveillance Team |
| EPI_ISL_2702211 | Lighthouse Lab in Alderley Park | Wellcome Sanger Institute for the COVID-19 Genomics UK (COG-UK) Consortium | Jacquelyn Wynn, Mairead Hyland, The Lighthouse Lab in Alderley Park and Alex Alderton, Roberto Amato, Jeffrey Barrett, Sonia Goncalves, Ewan Harrison, David K. Jackson, Ian Johnston, Dominic Kwiatkowski, Cordelia Langford, John Sillitoe on behalf of the Wellcome Sanger Institute COVID-19 Surveillance Team |
| EPI_ISL_2702212 | Lighthouse Lab in Glasgow | Wellcome Sanger Institute for the COVID-19 Genomics UK (COG-UK) Consortium | Harper VanSteenhouse, Yumi Kasai, David Gray, Carol Clugston, Anna Dominiczak and Alex Alderton, Roberto Amato, Jeffrey Barrett, Sonia Goncalves, Ewan Harrison, David K. Jackson, Ian Johnston, Dominic Kwiatkowski, Cordelia Langford, John Sillitoe on behalf of the Wellcome Sanger Institute COVID-19 Surveillance Team |
| EPI_ISL_2702213 | Lighthouse Lab in Alderley Park | Wellcome Sanger Institute for the COVID-19 Genomics UK (COG-UK) Consortium | Jacquelyn Wynn, Mairead Hyland, The Lighthouse Lab in Alderley Park and Alex Alderton, Roberto Amato, Jeffrey Barrett, Sonia Goncalves, Ewan Harrison, David K. Jackson, Ian Johnston, Dominic Kwiatkowski, Cordelia Langford, John Sillitoe on behalf of the Wellcome Sanger Institute COVID-19 Surveillance Team |

| | | | |
|---|---|---|---|
| EPI_ISL_2702214, EPI_ISL_2702215 | Lighthouse Lab in Milton Keynes | Wellcome Sanger Institute for the COVID-19 Genomics UK (COG-UK) Consortium | The Lighthouse Lab in Milton Keynes and Alex Alderton, Roberto Amato, Jeffrey Barrett, Sonia Goncalves, Ewan Harrison, David K. Jackson, Ian Johnston, Dominic Kwiatkowski, Cordelia Langford, John Sillitoe on behalf of the Wellcome Sanger Institute COVID-19 Surveillance Team |
| EPI_ISL_2702216 | Lighthouse Lab in Alderley Park | Wellcome Sanger Institute for the COVID-19 Genomics UK (COG-UK) Consortium | Jacquelyn Wynn, Mairead Hyland, The Lighthouse Lab in Alderley Park and Alex Alderton, Roberto Amato, Jeffrey Barrett, Sonia Goncalves, Ewan Harrison, David K. Jackson, Ian Johnston, Dominic Kwiatkowski, Cordelia Langford, John Sillitoe on behalf of the Wellcome Sanger Institute COVID-19 Surveillance Team |
| EPI_ISL_2702217 | Lighthouse Lab in Milton Keynes | Wellcome Sanger Institute for the COVID-19 Genomics UK (COG-UK) Consortium | The Lighthouse Lab in Milton Keynes and Alex Alderton, Roberto Amato, Jeffrey Barrett, Sonia Goncalves, Ewan Harrison, David K. Jackson, Ian Johnston, Dominic Kwiatkowski, Cordelia Langford, John Sillitoe on behalf of the Wellcome Sanger Institute COVID-19 Surveillance Team |
| EPI_ISL_2702218 | Lighthouse Lab in Glasgow | Wellcome Sanger Institute for the COVID-19 Genomics UK (COG-UK) Consortium | Harper VanSteenhouse, Yumi Kasai, David Gray, Carol Clugston, Anna Dominiczak and Alex Alderton, Roberto Amato, Jeffrey Barrett, Sonia Goncalves, Ewan Harrison, David K. Jackson, Ian Johnston, Dominic Kwiatkowski, Cordelia Langford, John Sillitoe on behalf of the Wellcome Sanger Institute COVID-19 Surveillance Team |
| EPI_ISL_2702219 | Lighthouse Lab in Milton Keynes | Wellcome Sanger Institute for the COVID-19 Genomics UK (COG-UK) Consortium | The Lighthouse Lab in Milton Keynes and Alex Alderton, Roberto Amato, Jeffrey Barrett, Sonia Goncalves, Ewan Harrison, David K. Jackson, Ian Johnston, Dominic Kwiatkowski, Cordelia Langford, John Sillitoe on behalf of the Wellcome Sanger Institute COVID-19 Surveillance Team |
| EPI_ISL_2702220, EPI_ISL_2702221 | Lighthouse Lab in Glasgow | Wellcome Sanger Institute for the COVID-19 Genomics UK (COG-UK) Consortium | Harper VanSteenhouse, Yumi Kasai, David Gray, Carol Clugston, Anna Dominiczak and Alex Alderton, Roberto Amato, Jeffrey Barrett, Sonia Goncalves, Ewan Harrison, David K. Jackson, Ian Johnston, Dominic Kwiatkowski, Cordelia Langford, John Sillitoe on behalf of the Wellcome Sanger Institute COVID-19 Surveillance Team |
| EPI_ISL_2702222 | Lighthouse Lab in Alderley Park | Wellcome Sanger Institute for the COVID-19 Genomics UK (COG-UK) Consortium | Jacquelyn Wynn, Mairead Hyland, The Lighthouse Lab in Alderley Park and Alex Alderton, Roberto Amato, Jeffrey Barrett, Sonia Goncalves, Ewan Harrison, David K. Jackson, Ian Johnston, Dominic Kwiatkowski, Cordelia Langford, John Sillitoe on behalf of the Wellcome Sanger Institute COVID-19 Surveillance Team |
| EPI_ISL_2702223 | Lighthouse Lab in Milton Keynes | Wellcome Sanger Institute for the COVID-19 Genomics UK (COG-UK) Consortium | The Lighthouse Lab in Milton Keynes and Alex Alderton, Roberto Amato, Jeffrey Barrett, Sonia Goncalves, Ewan Harrison, David K. Jackson, Ian Johnston, Dominic Kwiatkowski, Cordelia Langford, John Sillitoe on behalf of the Wellcome Sanger Institute COVID-19 Surveillance Team |
| EPI_ISL_2702224 | Lighthouse Lab in Alderley Park | Wellcome Sanger Institute for the COVID-19 Genomics UK (COG-UK) Consortium | Jacquelyn Wynn, Mairead Hyland, The Lighthouse Lab in Alderley Park and Alex Alderton, Roberto Amato, Jeffrey Barrett, Sonia Goncalves, Ewan Harrison, David K. Jackson, Ian Johnston, Dominic Kwiatkowski, Cordelia Langford, John Sillitoe on behalf of the Wellcome Sanger Institute COVID-19 Surveillance Team |
| EPI_ISL_2702225 | Lighthouse Lab in Milton Keynes | Wellcome Sanger Institute for the COVID-19 Genomics UK (COG-UK) Consortium | The Lighthouse Lab in Milton Keynes and Alex Alderton, Roberto Amato, Jeffrey Barrett, Sonia Goncalves, Ewan Harrison, David K. Jackson, Ian Johnston, Dominic Kwiatkowski, Cordelia Langford, John Sillitoe on behalf of the Wellcome Sanger Institute COVID-19 Surveillance Team |
| EPI_ISL_2702226 | Lighthouse Lab in Alderley Park | Wellcome Sanger Institute for the COVID-19 Genomics UK (COG-UK) Consortium | Jacquelyn Wynn, Mairead Hyland, The Lighthouse Lab in Alderley Park and Alex Alderton, Roberto Amato, Jeffrey Barrett, Sonia Goncalves, Ewan Harrison, David K. Jackson, Ian Johnston, Dominic Kwiatkowski, Cordelia Langford, John Sillitoe on behalf of the Wellcome Sanger Institute COVID-19 Surveillance Team |
| EPI_ISL_2702227 | Lighthouse Lab in Glasgow | Wellcome Sanger Institute for the COVID-19 Genomics UK (COG-UK) Consortium | Harper VanSteenhouse, Yumi Kasai, David Gray, Carol Clugston, Anna Dominiczak and Alex Alderton, Roberto Amato, Jeffrey Barrett, Sonia Goncalves, Ewan Harrison, David K. Jackson, Ian Johnston, Dominic Kwiatkowski, Cordelia Langford, John Sillitoe on behalf of the Wellcome Sanger Institute COVID-19 Surveillance Team |
| EPI_ISL_2702228, EPI_ISL_2702229 | Lighthouse Lab in Alderley Park | Wellcome Sanger Institute for the COVID-19 Genomics UK (COG-UK) Consortium | Jacquelyn Wynn, Mairead Hyland, The Lighthouse Lab in Alderley Park and Alex Alderton, Roberto Amato, Jeffrey Barrett, Sonia Goncalves, Ewan Harrison, David K. Jackson, Ian Johnston, Dominic Kwiatkowski, Cordelia Langford, John Sillitoe on behalf of the Wellcome Sanger Institute COVID-19 Surveillance Team |
| EPI_ISL_2702230 | Lighthouse Lab in Glasgow | Wellcome Sanger Institute for the COVID-19 Genomics UK (COG-UK) Consortium | Harper VanSteenhouse, Yumi Kasai, David Gray, Carol Clugston, Anna Dominiczak and Alex Alderton, Roberto Amato, Jeffrey Barrett, Sonia Goncalves, Ewan Harrison, David K. Jackson, Ian Johnston, Dominic Kwiatkowski, Cordelia Langford, John Sillitoe on behalf of the Wellcome Sanger Institute COVID-19 Surveillance Team |
| EPI_ISL_2702231 | Lighthouse Lab in Alderley Park | Wellcome Sanger Institute for the COVID-19 Genomics UK (COG-UK) Consortium | Jacquelyn Wynn, Mairead Hyland, The Lighthouse Lab in Alderley Park and Alex Alderton, Roberto Amato, Jeffrey Barrett, Sonia Goncalves, Ewan Harrison, David K. Jackson, Ian Johnston, Dominic Kwiatkowski, Cordelia Langford, John Sillitoe on behalf of the Wellcome Sanger Institute COVID-19 Surveillance Team |
| EPI_ISL_2702232 | Lighthouse Lab in Glasgow | Wellcome Sanger Institute for the COVID-19 Genomics UK (COG-UK) Consortium | Harper VanSteenhouse, Yumi Kasai, David Gray, Carol Clugston, Anna Dominiczak and Alex Alderton, Roberto Amato, Jeffrey Barrett, Sonia Goncalves, Ewan Harrison, David K. Jackson, Ian Johnston, Dominic Kwiatkowski, Cordelia Langford, John Sillitoe on behalf of the Wellcome Sanger Institute COVID-19 Surveillance Team |
| EPI_ISL_2702233 | Lighthouse Lab in Milton Keynes | Wellcome Sanger Institute for the COVID-19 Genomics UK (COG-UK) Consortium | The Lighthouse Lab in Milton Keynes and Alex Alderton, Roberto Amato, Jeffrey Barrett, Sonia Goncalves, Ewan Harrison, David K. Jackson, Ian Johnston, Dominic Kwiatkowski, Cordelia Langford, John Sillitoe on behalf of the Wellcome Sanger Institute COVID-19 Surveillance Team |
| EPI_ISL_2702234, EPI_ISL_2702235 | Lighthouse Lab in Glasgow | Wellcome Sanger Institute for the COVID-19 Genomics UK (COG-UK) Consortium | Harper VanSteenhouse, Yumi Kasai, David Gray, Carol Clugston, Anna Dominiczak and Alex Alderton, Roberto Amato, Jeffrey Barrett, Sonia Goncalves, Ewan Harrison, David K. Jackson, Ian Johnston, Dominic Kwiatkowski, Cordelia Langford, John Sillitoe on behalf of the Wellcome Sanger Institute COVID-19 Surveillance Team |
| EPI_ISL_2702236, EPI_ISL_2702237 | Lighthouse Lab in Alderley Park | Wellcome Sanger Institute for the COVID-19 Genomics UK (COG-UK) Consortium | Jacquelyn Wynn, Mairead Hyland, The Lighthouse Lab in Alderley Park and Alex Alderton, Roberto Amato, Jeffrey Barrett, Sonia Goncalves, Ewan Harrison, David K. Jackson, Ian Johnston, Dominic Kwiatkowski, Cordelia Langford, John Sillitoe on behalf of the Wellcome Sanger Institute COVID-19 Surveillance Team |
| EPI_ISL_2702238, EPI_ISL_2702239, EPI_ISL_2702240 | Lighthouse Lab in Glasgow | Wellcome Sanger Institute for the COVID-19 Genomics UK (COG-UK) Consortium | Harper VanSteenhouse, Yumi Kasai, David Gray, Carol Clugston, Anna Dominiczak and Alex Alderton, Roberto Amato, Jeffrey Barrett, Sonia Goncalves, Ewan Harrison, David K. Jackson, Ian Johnston, Dominic Kwiatkowski, Cordelia Langford, John Sillitoe on behalf of the Wellcome Sanger Institute COVID-19 Surveillance Team |
| EPI_ISL_2702241, EPI_ISL_2702242 | Lighthouse Lab in Alderley Park | Wellcome Sanger Institute for the COVID-19 Genomics UK (COG-UK) Consortium | Jacquelyn Wynn, Mairead Hyland, The Lighthouse Lab in Alderley Park and Alex Alderton, Roberto Amato, Jeffrey Barrett, Sonia Goncalves, Ewan Harrison, David K. Jackson, Ian Johnston, Dominic Kwiatkowski, Cordelia Langford, John Sillitoe on behalf of the Wellcome Sanger Institute COVID-19 Surveillance Team |
| EPI_ISL_2702243, EPI_ISL_2702244, EPI_ISL_2702245, EPI_ISL_2702246, EPI_ISL_2702247, EPI_ISL_2702248, EPI_ISL_2702249, EPI_ISL_2702250, EPI_ISL_2702251, EPI_ISL_2702252, EPI_ISL_2702253, EPI_ISL_2702254 | | | |
| see above | Lighthouse Lab in Glasgow | Wellcome Sanger Institute for the COVID-19 Genomics UK (COG-UK) Consortium | Harper VanSteenhouse, Yumi Kasai, David Gray, Carol Clugston, Anna Dominiczak and Alex Alderton, Roberto Amato, Jeffrey Barrett, Sonia Goncalves, Ewan Harrison, David K. Jackson, Ian Johnston, Dominic Kwiatkowski, Cordelia Langford, John Sillitoe on behalf of the Wellcome Sanger Institute COVID-19 Surveillance Team |
| EPI_ISL_2702255 | Lighthouse Lab in Milton Keynes | Wellcome Sanger Institute for the COVID-19 Genomics UK (COG-UK) Consortium | The Lighthouse Lab in Milton Keynes and Alex Alderton, Roberto Amato, Jeffrey Barrett, Sonia Goncalves, Ewan Harrison, David K. Jackson, Ian Johnston, Dominic Kwiatkowski, Cordelia Langford, John Sillitoe on behalf of the Wellcome Sanger Institute COVID-19 Surveillance Team |
| EPI_ISL_2702256, EPI_ISL_2702257, EPI_ISL_2702258 | Lighthouse Lab in Alderley Park | Wellcome Sanger Institute for the COVID-19 Genomics UK (COG-UK) Consortium | Jacquelyn Wynn, Mairead Hyland, The Lighthouse Lab in Alderley Park and Alex Alderton, Roberto Amato, Jeffrey Barrett, Sonia Goncalves, Ewan Harrison, David K. Jackson, Ian Johnston, Dominic Kwiatkowski, Cordelia Langford, John Sillitoe on behalf of the Wellcome Sanger Institute COVID-19 Surveillance Team |
| EPI_ISL_2702259 | Lighthouse Lab in Glasgow | Wellcome Sanger Institute for the COVID-19 Genomics UK (COG-UK) Consortium | Harper VanSteenhouse, Yumi Kasai, David Gray, Carol Clugston, Anna Dominiczak and Alex Alderton, Roberto Amato, Jeffrey Barrett, Sonia Goncalves, Ewan Harrison, David K. Jackson, Ian Johnston, Dominic Kwiatkowski, Cordelia Langford, John Sillitoe on behalf of the Wellcome Sanger Institute COVID-19 Surveillance Team |
| EPI_ISL_2702260, EPI_ISL_2702261 | Lighthouse Lab in Milton Keynes | Wellcome Sanger Institute for the COVID-19 Genomics UK (COG-UK) Consortium | The Lighthouse Lab in Milton Keynes and Alex Alderton, Roberto Amato, Jeffrey Barrett, Sonia Goncalves, Ewan Harrison, David K. Jackson, Ian Johnston, Dominic Kwiatkowski, Cordelia Langford, John Sillitoe on behalf of the Wellcome Sanger Institute COVID-19 Surveillance Team |
| EPI_ISL_2702262, EPI_ISL_2702263, EPI_ISL_2702264, EPI_ISL_2702265, EPI_ISL_2702266, EPI_ISL_2702267, EPI_ISL_2702268, EPI_ISL_2702269, EPI_ISL_2702270, EPI_ISL_2702271, EPI_ISL_2702272, EPI_ISL_2702273 | | | |
| see above | Lighthouse Lab in Glasgow | Wellcome Sanger Institute for the COVID-19 Genomics UK (COG-UK) Consortium | Harper VanSteenhouse, Yumi Kasai, David Gray, Carol Clugston, Anna Dominiczak and Alex Alderton, Roberto Amato, Jeffrey Barrett, Sonia Goncalves, Ewan Harrison, David K. Jackson, Ian Johnston, Dominic Kwiatkowski, Cordelia Langford, John Sillitoe on behalf of the Wellcome Sanger Institute COVID-19 Surveillance Team |
| EPI_ISL_2702274 | Lighthouse Lab in Alderley Park | Wellcome Sanger Institute for the COVID-19 Genomics UK (COG-UK) Consortium | Jacquelyn Wynn, Mairead Hyland, The Lighthouse Lab in Alderley Park and Alex Alderton, Roberto Amato, Jeffrey Barrett, Sonia Goncalves, Ewan Harrison, David K. Jackson, Ian Johnston, Dominic Kwiatkowski, Cordelia Langford, John Sillitoe on behalf of the Wellcome Sanger Institute COVID-19 Surveillance Team |
| EPI_ISL_2702275 | Lighthouse Lab in Milton Keynes | Wellcome Sanger Institute for the COVID-19 Genomics UK (COG-UK) Consortium | The Lighthouse Lab in Milton Keynes and Alex Alderton, Roberto Amato, Jeffrey Barrett, Sonia Goncalves, Ewan Harrison, David K. Jackson, Ian Johnston, Dominic Kwiatkowski, Cordelia Langford, John Sillitoe on behalf of the Wellcome Sanger Institute COVID-19 Surveillance Team |
| EPI_ISL_2702276 | Lighthouse Lab in Glasgow | Wellcome Sanger Institute for the COVID-19 Genomics UK (COG-UK) Consortium | Harper VanSteenhouse, Yumi Kasai, David Gray, Carol Clugston, Anna Dominiczak and Alex Alderton, Roberto Amato, Jeffrey Barrett, Sonia Goncalves, Ewan Harrison, David K. Jackson, Ian Johnston, Dominic Kwiatkowski, Cordelia Langford, John Sillitoe on behalf of the Wellcome Sanger Institute COVID-19 Surveillance Team |
| EPI_ISL_2702277 | Lighthouse Lab in Milton Keynes | Wellcome Sanger Institute for the COVID-19 Genomics UK (COG-UK) Consortium | The Lighthouse Lab in Milton Keynes and Alex Alderton, Roberto Amato, Jeffrey Barrett, Sonia Goncalves, Ewan Harrison, David K. Jackson, Ian Johnston, Dominic Kwiatkowski, Cordelia Langford, John Sillitoe on behalf of the Wellcome Sanger Institute COVID-19 Surveillance Team |
| EPI_ISL_2702278, EPI_ISL_2702279, EPI_ISL_2702280 | Lighthouse Lab in Glasgow | Wellcome Sanger Institute for the COVID-19 Genomics UK (COG-UK) Consortium | Harper VanSteenhouse, Yumi Kasai, David Gray, Carol Clugston, Anna Dominiczak and Alex Alderton, Roberto Amato, Jeffrey Barrett, Sonia Goncalves, Ewan Harrison, David K. Jackson, Ian Johnston, Dominic Kwiatkowski, Cordelia Langford, John Sillitoe on behalf of the Wellcome Sanger Institute COVID-19 Surveillance Team |
| EPI_ISL_2702281 | Lighthouse Lab in Alderley Park | Wellcome Sanger Institute for the COVID-19 Genomics UK (COG-UK) Consortium | Jacquelyn Wynn, Mairead Hyland, The Lighthouse Lab in Alderley Park and Alex Alderton, Roberto Amato, Jeffrey Barrett, Sonia Goncalves, Ewan Harrison, David K. Jackson, Ian Johnston, Dominic Kwiatkowski, Cordelia Langford, John Sillitoe on behalf of the Wellcome Sanger Institute COVID-19 Surveillance Team |
| EPI_ISL_2702282, EPI_ISL_2702283 | Lighthouse Lab in Milton Keynes | Wellcome Sanger Institute for the COVID-19 Genomics UK (COG-UK) Consortium | The Lighthouse Lab in Milton Keynes and Alex Alderton, Roberto Amato, Jeffrey Barrett, Sonia Goncalves, Ewan Harrison, David K. Jackson, Ian Johnston, Dominic Kwiatkowski, Cordelia Langford, John Sillitoe on behalf of the Wellcome Sanger Institute COVID-19 Surveillance Team |
| EPI_ISL_2702284 | Lighthouse Lab in Glasgow | Wellcome Sanger Institute for the COVID-19 Genomics UK (COG-UK) Consortium | Harper VanSteenhouse, Yumi Kasai, David Gray, Carol Clugston, Anna Dominiczak and Alex Alderton, Roberto Amato, Jeffrey Barrett, Sonia Goncalves, Ewan Harrison, David K. Jackson, Ian Johnston, Dominic Kwiatkowski, Cordelia Langford, John Sillitoe on behalf of the Wellcome Sanger Institute COVID-19 Surveillance Team |
| EPI_ISL_2702285 | Lighthouse Lab in Milton Keynes | Wellcome Sanger Institute for the COVID-19 Genomics UK (COG-UK) Consortium | The Lighthouse Lab in Milton Keynes and Alex Alderton, Roberto Amato, Jeffrey Barrett, Sonia Goncalves, Ewan Harrison, David K. Jackson, Ian Johnston, Dominic Kwiatkowski, Cordelia Langford, John Sillitoe on behalf of the Wellcome Sanger Institute COVID-19 Surveillance Team |
| EPI_ISL_2702286 | Lighthouse Lab in Glasgow | Wellcome Sanger Institute for the COVID-19 Genomics UK (COG-UK) Consortium | Harper VanSteenhouse, Yumi Kasai, David Gray, Carol Clugston, Anna Dominiczak and Alex Alderton, Roberto Amato, Jeffrey Barrett, Sonia Goncalves, Ewan Harrison, David K. Jackson, Ian Johnston, Dominic Kwiatkowski, Cordelia Langford, John Sillitoe on behalf of the Wellcome Sanger Institute COVID-19 Surveillance Team |
| EPI_ISL_2702287 | Lighthouse Lab in Milton Keynes | Wellcome Sanger Institute for the COVID-19 Genomics UK (COG-UK) Consortium | The Lighthouse Lab in Milton Keynes and Alex Alderton, Roberto Amato, Jeffrey Barrett, Sonia Goncalves, Ewan Harrison, David K. Jackson, Ian Johnston, Dominic Kwiatkowski, Cordelia Langford, John Sillitoe on behalf of the Wellcome Sanger Institute COVID-19 Surveillance Team |
| EPI_ISL_2702288 | Lighthouse Lab in Alderley Park | Wellcome Sanger Institute for the COVID-19 Genomics UK (COG-UK) Consortium | Jacquelyn Wynn, Mairead Hyland, The Lighthouse Lab in Alderley Park and Alex Alderton, Roberto Amato, Jeffrey Barrett, Sonia Goncalves, Ewan Harrison, David K. Jackson, Ian Johnston, Dominic Kwiatkowski, Cordelia Langford, John Sillitoe on behalf of the Wellcome Sanger Institute COVID-19 Surveillance Team |
| EPI_ISL_2702289, EPI_ISL_2702290, EPI_ISL_2702291, EPI_ISL_2702292, EPI_ISL_2702293, EPI_ISL_2702294 | Lighthouse Lab in Glasgow | Wellcome Sanger Institute for the COVID-19 Genomics UK (COG-UK) Consortium | Harper VanSteenhouse, Yumi Kasai, David Gray, Carol Clugston, Anna Dominiczak and Alex Alderton, Roberto Amato, Jeffrey Barrett, Sonia Goncalves, Ewan Harrison, David K. Jackson, Ian Johnston, Dominic Kwiatkowski, Cordelia Langford, John Sillitoe on behalf of the Wellcome Sanger Institute COVID-19 Surveillance Team |
| EPI_ISL_2702295 | Lighthouse Lab in Milton Keynes | Wellcome Sanger Institute for the COVID-19 Genomics UK (COG-UK) Consortium | The Lighthouse Lab in Milton Keynes and Alex Alderton, Roberto Amato, Jeffrey Barrett, Sonia Goncalves, Ewan Harrison, David K. Jackson, Ian Johnston, Dominic Kwiatkowski, Cordelia Langford, John Sillitoe on behalf of the Wellcome Sanger Institute COVID-19 Surveillance Team |

| | | | |
|---|---|---|---|
| EPI_ISL_2702296, EPI_ISL_2702297 | Lighthouse Lab in Glasgow | Wellcome Sanger Institute for the COVID-19 Genomics UK (COG-UK) Consortium | Harper VanSteenhouse, Yumi Kasai, David Gray, Carol Clugston, Anna Dominiczak and Alex Alderton, Roberto Amato, Jeffrey Barrett, Sonia Goncalves, Ewan Harrison, David K. Jackson, Ian Johnston, Dominic Kwiatkowski, Cordelia Langford, John Sillitoe on behalf of the Wellcome Sanger Institute COVID-19 Surveillance Team |
| EPI_ISL_2702298 | Lighthouse Lab in Alderley Park | Wellcome Sanger Institute for the COVID-19 Genomics UK (COG-UK) Consortium | Jacquelyn Wynn, Mairead Hyland, The Lighthouse Lab in Alderley Park and Alex Alderton, Roberto Amato, Jeffrey Barrett, Sonia Goncalves, Ewan Harrison, David K. Jackson, Ian Johnston, Dominic Kwiatkowski, Cordelia Langford, John Sillitoe on behalf of the Wellcome Sanger Institute COVID-19 Surveillance Team |
| EPI_ISL_2702299 | Lighthouse Lab in Milton Keynes | Wellcome Sanger Institute for the COVID-19 Genomics UK (COG-UK) Consortium | The Lighthouse Lab in Milton Keynes and Alex Alderton, Roberto Amato, Jeffrey Barrett, Sonia Goncalves, Ewan Harrison, David K. Jackson, Ian Johnston, Dominic Kwiatkowski, Cordelia Langford, John Sillitoe on behalf of the Wellcome Sanger Institute COVID-19 Surveillance Team |
| EPI_ISL_2702300, EPI_ISL_2702301 | Lighthouse Lab in Alderley Park | Wellcome Sanger Institute for the COVID-19 Genomics UK (COG-UK) Consortium | Jacquelyn Wynn, Mairead Hyland, The Lighthouse Lab in Alderley Park and Alex Alderton, Roberto Amato, Jeffrey Barrett, Sonia Goncalves, Ewan Harrison, David K. Jackson, Ian Johnston, Dominic Kwiatkowski, Cordelia Langford, John Sillitoe on behalf of the Wellcome Sanger Institute COVID-19 Surveillance Team |
| EPI_ISL_2702302 | Lighthouse Lab in Glasgow | Wellcome Sanger Institute for the COVID-19 Genomics UK (COG-UK) Consortium | Harper VanSteenhouse, Yumi Kasai, David Gray, Carol Clugston, Anna Dominiczak and Alex Alderton, Roberto Amato, Jeffrey Barrett, Sonia Goncalves, Ewan Harrison, David K. Jackson, Ian Johnston, Dominic Kwiatkowski, Cordelia Langford, John Sillitoe on behalf of the Wellcome Sanger Institute COVID-19 Surveillance Team |
| EPI_ISL_2702303, EPI_ISL_2702304 | Lighthouse Lab in Milton Keynes | Wellcome Sanger Institute for the COVID-19 Genomics UK (COG-UK) Consortium | The Lighthouse Lab in Milton Keynes and Alex Alderton, Roberto Amato, Jeffrey Barrett, Sonia Goncalves, Ewan Harrison, David K. Jackson, Ian Johnston, Dominic Kwiatkowski, Cordelia Langford, John Sillitoe on behalf of the Wellcome Sanger Institute COVID-19 Surveillance Team |
| EPI_ISL_2702305 | Lighthouse Lab in Alderley Park | Wellcome Sanger Institute for the COVID-19 Genomics UK (COG-UK) Consortium | Jacquelyn Wynn, Mairead Hyland, The Lighthouse Lab in Alderley Park and Alex Alderton, Roberto Amato, Jeffrey Barrett, Sonia Goncalves, Ewan Harrison, David K. Jackson, Ian Johnston, Dominic Kwiatkowski, Cordelia Langford, John Sillitoe on behalf of the Wellcome Sanger Institute COVID-19 Surveillance Team |
| EPI_ISL_2702306 | Lighthouse Lab in Milton Keynes | Wellcome Sanger Institute for the COVID-19 Genomics UK (COG-UK) Consortium | The Lighthouse Lab in Milton Keynes and Alex Alderton, Roberto Amato, Jeffrey Barrett, Sonia Goncalves, Ewan Harrison, David K. Jackson, Ian Johnston, Dominic Kwiatkowski, Cordelia Langford, John Sillitoe on behalf of the Wellcome Sanger Institute COVID-19 Surveillance Team |
| EPI_ISL_2702307 | Lighthouse Lab in Glasgow | Wellcome Sanger Institute for the COVID-19 Genomics UK (COG-UK) Consortium | Harper VanSteenhouse, Yumi Kasai, David Gray, Carol Clugston, Anna Dominiczak and Alex Alderton, Roberto Amato, Jeffrey Barrett, Sonia Goncalves, Ewan Harrison, David K. Jackson, Ian Johnston, Dominic Kwiatkowski, Cordelia Langford, John Sillitoe on behalf of the Wellcome Sanger Institute COVID-19 Surveillance Team |
| EPI_ISL_2702308, EPI_ISL_2702309 | Lighthouse Lab in Milton Keynes | Wellcome Sanger Institute for the COVID-19 Genomics UK (COG-UK) Consortium | The Lighthouse Lab in Milton Keynes and Alex Alderton, Roberto Amato, Jeffrey Barrett, Sonia Goncalves, Ewan Harrison, David K. Jackson, Ian Johnston, Dominic Kwiatkowski, Cordelia Langford, John Sillitoe on behalf of the Wellcome Sanger Institute COVID-19 Surveillance Team |
| EPI_ISL_2702310 | Lighthouse Lab in Glasgow | Wellcome Sanger Institute for the COVID-19 Genomics UK (COG-UK) Consortium | Harper VanSteenhouse, Yumi Kasai, David Gray, Carol Clugston, Anna Dominiczak and Alex Alderton, Roberto Amato, Jeffrey Barrett, Sonia Goncalves, Ewan Harrison, David K. Jackson, Ian Johnston, Dominic Kwiatkowski, Cordelia Langford, John Sillitoe on behalf of the Wellcome Sanger Institute COVID-19 Surveillance Team |
| EPI_ISL_2702311, EPI_ISL_2702312, EPI_ISL_2702313, EPI_ISL_2702315, EPI_ISL_2702316 | Lighthouse Lab in Milton Keynes | Wellcome Sanger Institute for the COVID-19 Genomics UK (COG-UK) Consortium | The Lighthouse Lab in Milton Keynes and Alex Alderton, Roberto Amato, Jeffrey Barrett, Sonia Goncalves, Ewan Harrison, David K. Jackson, Ian Johnston, Dominic Kwiatkowski, Cordelia Langford, John Sillitoe on behalf of the Wellcome Sanger Institute COVID-19 Surveillance Team |
| EPI_ISL_2702317, EPI_ISL_2702318 | Lighthouse Lab in Alderley Park | Wellcome Sanger Institute for the COVID-19 Genomics UK (COG-UK) Consortium | Jacquelyn Wynn, Mairead Hyland, The Lighthouse Lab in Alderley Park and Alex Alderton, Roberto Amato, Jeffrey Barrett, Sonia Goncalves, Ewan Harrison, David K. Jackson, Ian Johnston, Dominic Kwiatkowski, Cordelia Langford, John Sillitoe on behalf of the Wellcome Sanger Institute COVID-19 Surveillance Team |
| EPI_ISL_2702319 | Lighthouse Lab in Glasgow | Wellcome Sanger Institute for the COVID-19 Genomics UK (COG-UK) Consortium | Harper VanSteenhouse, Yumi Kasai, David Gray, Carol Clugston, Anna Dominiczak and Alex Alderton, Roberto Amato, Jeffrey Barrett, Sonia Goncalves, Ewan Harrison, David K. Jackson, Ian Johnston, Dominic Kwiatkowski, Cordelia Langford, John Sillitoe on behalf of the Wellcome Sanger Institute COVID-19 Surveillance Team |
| EPI_ISL_2702320 | Lighthouse Lab in Milton Keynes | Wellcome Sanger Institute for the COVID-19 Genomics UK (COG-UK) Consortium | The Lighthouse Lab in Milton Keynes and Alex Alderton, Roberto Amato, Jeffrey Barrett, Sonia Goncalves, Ewan Harrison, David K. Jackson, Ian Johnston, Dominic Kwiatkowski, Cordelia Langford, John Sillitoe on behalf of the Wellcome Sanger Institute COVID-19 Surveillance Team |
| EPI_ISL_2702321, EPI_ISL_2702322, EPI_ISL_2702323, EPI_ISL_2702324, EPI_ISL_2702325 | Lighthouse Lab in Glasgow | Wellcome Sanger Institute for the COVID-19 Genomics UK (COG-UK) Consortium | Harper VanSteenhouse, Yumi Kasai, David Gray, Carol Clugston, Anna Dominiczak and Alex Alderton, Roberto Amato, Jeffrey Barrett, Sonia Goncalves, Ewan Harrison, David K. Jackson, Ian Johnston, Dominic Kwiatkowski, Cordelia Langford, John Sillitoe on behalf of the Wellcome Sanger Institute COVID-19 Surveillance Team |
| EPI_ISL_2702326 | Lighthouse Lab in Milton Keynes | Wellcome Sanger Institute for the COVID-19 Genomics UK (COG-UK) Consortium | The Lighthouse Lab in Milton Keynes and Alex Alderton, Roberto Amato, Jeffrey Barrett, Sonia Goncalves, Ewan Harrison, David K. Jackson, Ian Johnston, Dominic Kwiatkowski, Cordelia Langford, John Sillitoe on behalf of the Wellcome Sanger Institute COVID-19 Surveillance Team |
| EPI_ISL_2702327, EPI_ISL_2702328 | Lighthouse Lab in Glasgow | Wellcome Sanger Institute for the COVID-19 Genomics UK (COG-UK) Consortium | Harper VanSteenhouse, Yumi Kasai, David Gray, Carol Clugston, Anna Dominiczak and Alex Alderton, Roberto Amato, Jeffrey Barrett, Sonia Goncalves, Ewan Harrison, David K. Jackson, Ian Johnston, Dominic Kwiatkowski, Cordelia Langford, John Sillitoe on behalf of the Wellcome Sanger Institute COVID-19 Surveillance Team |
| EPI_ISL_2702329, EPI_ISL_2702330 | Lighthouse Lab in Milton Keynes | Wellcome Sanger Institute for the COVID-19 Genomics UK (COG-UK) Consortium | The Lighthouse Lab in Milton Keynes and Alex Alderton, Roberto Amato, Jeffrey Barrett, Sonia Goncalves, Ewan Harrison, David K. Jackson, Ian Johnston, Dominic Kwiatkowski, Cordelia Langford, John Sillitoe on behalf of the Wellcome Sanger Institute COVID-19 Surveillance Team |
| EPI_ISL_2702331 | Lighthouse Lab in Alderley Park | Wellcome Sanger Institute for the COVID-19 Genomics UK (COG-UK) Consortium | Jacquelyn Wynn, Mairead Hyland, The Lighthouse Lab in Alderley Park and Alex Alderton, Roberto Amato, Jeffrey Barrett, Sonia Goncalves, Ewan Harrison, David K. Jackson, Ian Johnston, Dominic Kwiatkowski, Cordelia Langford, John Sillitoe on behalf of the Wellcome Sanger Institute COVID-19 Surveillance Team |
| EPI_ISL_2702332, EPI_ISL_2702333 | Lighthouse Lab in Glasgow | Wellcome Sanger Institute for the COVID-19 Genomics UK (COG-UK) Consortium | Harper VanSteenhouse, Yumi Kasai, David Gray, Carol Clugston, Anna Dominiczak and Alex Alderton, Roberto Amato, Jeffrey Barrett, Sonia Goncalves, Ewan Harrison, David K. Jackson, Ian Johnston, Dominic Kwiatkowski, Cordelia Langford, John Sillitoe on behalf of the Wellcome Sanger Institute COVID-19 Surveillance Team |
| EPI_ISL_2702334 | Lighthouse Lab in Alderley Park | Wellcome Sanger Institute for the COVID-19 Genomics UK (COG-UK) Consortium | Jacquelyn Wynn, Mairead Hyland, The Lighthouse Lab in Alderley Park and Alex Alderton, Roberto Amato, Jeffrey Barrett, Sonia Goncalves, Ewan Harrison, David K. Jackson, Ian Johnston, Dominic Kwiatkowski, Cordelia Langford, John Sillitoe on behalf of the Wellcome Sanger Institute COVID-19 Surveillance Team |
| EPI_ISL_2702335, EPI_ISL_2702336, EPI_ISL_2702337 | Lighthouse Lab in Glasgow | Wellcome Sanger Institute for the COVID-19 Genomics UK (COG-UK) Consortium | Harper VanSteenhouse, Yumi Kasai, David Gray, Carol Clugston, Anna Dominiczak and Alex Alderton, Roberto Amato, Jeffrey Barrett, Sonia Goncalves, Ewan Harrison, David K. Jackson, Ian Johnston, Dominic Kwiatkowski, Cordelia Langford, John Sillitoe on behalf of the Wellcome Sanger Institute COVID-19 Surveillance Team |
| EPI_ISL_2702339, EPI_ISL_2702340 | Lighthouse Lab in Milton Keynes | Wellcome Sanger Institute for the COVID-19 Genomics UK (COG-UK) Consortium | The Lighthouse Lab in Milton Keynes and Alex Alderton, Roberto Amato, Jeffrey Barrett, Sonia Goncalves, Ewan Harrison, David K. Jackson, Ian Johnston, Dominic Kwiatkowski, Cordelia Langford, John Sillitoe on behalf of the Wellcome Sanger Institute COVID-19 Surveillance Team |
| EPI_ISL_2702341 | Lighthouse Lab in Alderley Park | Wellcome Sanger Institute for the COVID-19 Genomics UK (COG-UK) Consortium | Jacquelyn Wynn, Mairead Hyland, The Lighthouse Lab in Alderley Park and Alex Alderton, Roberto Amato, Jeffrey Barrett, Sonia Goncalves, Ewan Harrison, David K. Jackson, Ian Johnston, Dominic Kwiatkowski, Cordelia Langford, John Sillitoe on behalf of the Wellcome Sanger Institute COVID-19 Surveillance Team |
| EPI_ISL_2702342, EPI_ISL_2702343, EPI_ISL_2702344 | Lighthouse Lab in Glasgow | Wellcome Sanger Institute for the COVID-19 Genomics UK (COG-UK) Consortium | Harper VanSteenhouse, Yumi Kasai, David Gray, Carol Clugston, Anna Dominiczak and Alex Alderton, Roberto Amato, Jeffrey Barrett, Sonia Goncalves, Ewan Harrison, David K. Jackson, Ian Johnston, Dominic Kwiatkowski, Cordelia Langford, John Sillitoe on behalf of the Wellcome Sanger Institute COVID-19 Surveillance Team |
| EPI_ISL_2702345 | Lighthouse Lab in Alderley Park | Wellcome Sanger Institute for the COVID-19 Genomics UK (COG-UK) Consortium | Jacquelyn Wynn, Mairead Hyland, The Lighthouse Lab in Alderley Park and Alex Alderton, Roberto Amato, Jeffrey Barrett, Sonia Goncalves, Ewan Harrison, David K. Jackson, Ian Johnston, Dominic Kwiatkowski, Cordelia Langford, John Sillitoe on behalf of the Wellcome Sanger Institute COVID-19 Surveillance Team |
| EPI_ISL_2702346 | Lighthouse Lab in Glasgow | Wellcome Sanger Institute for the COVID-19 Genomics UK (COG-UK) Consortium | Harper VanSteenhouse, Yumi Kasai, David Gray, Carol Clugston, Anna Dominiczak and Alex Alderton, Roberto Amato, Jeffrey Barrett, Sonia Goncalves, Ewan Harrison, David K. Jackson, Ian Johnston, Dominic Kwiatkowski, Cordelia Langford, John Sillitoe on behalf of the Wellcome Sanger Institute COVID-19 Surveillance Team |
| EPI_ISL_2702347 | Lighthouse Lab in Alderley Park | Wellcome Sanger Institute for the COVID-19 Genomics UK (COG-UK) Consortium | Jacquelyn Wynn, Mairead Hyland, The Lighthouse Lab in Alderley Park and Alex Alderton, Roberto Amato, Jeffrey Barrett, Sonia Goncalves, Ewan Harrison, David K. Jackson, Ian Johnston, Dominic Kwiatkowski, Cordelia Langford, John Sillitoe on behalf of the Wellcome Sanger Institute COVID-19 Surveillance Team |
| EPI_ISL_2702348 | Lighthouse Lab in Glasgow | Wellcome Sanger Institute for the COVID-19 Genomics UK (COG-UK) Consortium | Harper VanSteenhouse, Yumi Kasai, David Gray, Carol Clugston, Anna Dominiczak and Alex Alderton, Roberto Amato, Jeffrey Barrett, Sonia Goncalves, Ewan Harrison, David K. Jackson, Ian Johnston, Dominic Kwiatkowski, Cordelia Langford, John Sillitoe on behalf of the Wellcome Sanger Institute COVID-19 Surveillance Team |
| EPI_ISL_2702349 | Lighthouse Lab in Milton Keynes | Wellcome Sanger Institute for the COVID-19 Genomics UK (COG-UK) Consortium | The Lighthouse Lab in Milton Keynes and Alex Alderton, Roberto Amato, Jeffrey Barrett, Sonia Goncalves, Ewan Harrison, David K. Jackson, Ian Johnston, Dominic Kwiatkowski, Cordelia Langford, John Sillitoe on behalf of the Wellcome Sanger Institute COVID-19 Surveillance Team |
| EPI_ISL_2702350, EPI_ISL_2702351 | Lighthouse Lab in Glasgow | Wellcome Sanger Institute for the COVID-19 Genomics UK (COG-UK) Consortium | Harper VanSteenhouse, Yumi Kasai, David Gray, Carol Clugston, Anna Dominiczak and Alex Alderton, Roberto Amato, Jeffrey Barrett, Sonia Goncalves, Ewan Harrison, David K. Jackson, Ian Johnston, Dominic Kwiatkowski, Cordelia Langford, John Sillitoe on behalf of the Wellcome Sanger Institute COVID-19 Surveillance Team |
| EPI_ISL_2702352 | Lighthouse Lab in Alderley Park | Wellcome Sanger Institute for the COVID-19 Genomics UK (COG-UK) Consortium | Jacquelyn Wynn, Mairead Hyland, The Lighthouse Lab in Alderley Park and Alex Alderton, Roberto Amato, Jeffrey Barrett, Sonia Goncalves, Ewan Harrison, David K. Jackson, Ian Johnston, Dominic Kwiatkowski, Cordelia Langford, John Sillitoe on behalf of the Wellcome Sanger Institute COVID-19 Surveillance Team |
| EPI_ISL_2702353, EPI_ISL_2702355 | Lighthouse Lab in Glasgow | Wellcome Sanger Institute for the COVID-19 Genomics UK (COG-UK) Consortium | Harper VanSteenhouse, Yumi Kasai, David Gray, Carol Clugston, Anna Dominiczak and Alex Alderton, Roberto Amato, Jeffrey Barrett, Sonia Goncalves, Ewan Harrison, David K. Jackson, Ian Johnston, Dominic Kwiatkowski, Cordelia Langford, John Sillitoe on behalf of the Wellcome Sanger Institute COVID-19 Surveillance Team |
| EPI_ISL_2702357, EPI_ISL_2702359 | Lighthouse Lab in Alderley Park | Wellcome Sanger Institute for the COVID-19 Genomics UK (COG-UK) Consortium | Jacquelyn Wynn, Mairead Hyland, The Lighthouse Lab in Alderley Park and Alex Alderton, Roberto Amato, Jeffrey Barrett, Sonia Goncalves, Ewan Harrison, David K. Jackson, Ian Johnston, Dominic Kwiatkowski, Cordelia Langford, John Sillitoe on behalf of the Wellcome Sanger Institute COVID-19 Surveillance Team |
| EPI_ISL_2702360 | Lighthouse Lab in Glasgow | Wellcome Sanger Institute for the COVID-19 Genomics UK (COG-UK) Consortium | Harper VanSteenhouse, Yumi Kasai, David Gray, Carol Clugston, Anna Dominiczak and Alex Alderton, Roberto Amato, Jeffrey Barrett, Sonia Goncalves, Ewan Harrison, David K. Jackson, Ian Johnston, Dominic Kwiatkowski, Cordelia Langford, John Sillitoe on behalf of the Wellcome Sanger Institute COVID-19 Surveillance Team |
| EPI_ISL_2702361 | Lighthouse Lab in Milton Keynes | Wellcome Sanger Institute for the COVID-19 Genomics UK (COG-UK) Consortium | The Lighthouse Lab in Milton Keynes and Alex Alderton, Roberto Amato, Jeffrey Barrett, Sonia Goncalves, Ewan Harrison, David K. Jackson, Ian Johnston, Dominic Kwiatkowski, Cordelia Langford, John Sillitoe on behalf of the Wellcome Sanger Institute COVID-19 Surveillance Team |
| EPI_ISL_2702362 | Lighthouse Lab in Alderley Park | Wellcome Sanger Institute for the COVID-19 Genomics UK (COG-UK) Consortium | Jacquelyn Wynn, Mairead Hyland, The Lighthouse Lab in Alderley Park and Alex Alderton, Roberto Amato, Jeffrey Barrett, Sonia Goncalves, Ewan Harrison, David K. Jackson, Ian Johnston, Dominic Kwiatkowski, Cordelia Langford, John Sillitoe on behalf of the Wellcome Sanger Institute COVID-19 Surveillance Team |
| EPI_ISL_2702363, EPI_ISL_2702364 | Lighthouse Lab in Glasgow | Wellcome Sanger Institute for the COVID-19 Genomics UK (COG-UK) Consortium | Harper VanSteenhouse, Yumi Kasai, David Gray, Carol Clugston, Anna Dominiczak and Alex Alderton, Roberto Amato, Jeffrey Barrett, Sonia Goncalves, Ewan Harrison, David K. Jackson, Ian Johnston, Dominic Kwiatkowski, Cordelia Langford, John Sillitoe on behalf of the Wellcome Sanger Institute COVID-19 Surveillance Team |
| EPI_ISL_2702366 | Lighthouse Lab in Alderley Park | Wellcome Sanger Institute for the COVID-19 Genomics UK (COG-UK) Consortium | Jacquelyn Wynn, Mairead Hyland, The Lighthouse Lab in Alderley Park and Alex Alderton, Roberto Amato, Jeffrey Barrett, Sonia Goncalves, Ewan Harrison, David K. Jackson, Ian Johnston, Dominic Kwiatkowski, Cordelia Langford, John Sillitoe on behalf of the Wellcome Sanger Institute COVID-19 Surveillance Team |
| EPI_ISL_2702367 | Lighthouse Lab in Glasgow | Wellcome Sanger Institute for the COVID-19 Genomics UK (COG-UK) Consortium | Harper VanSteenhouse, Yumi Kasai, David Gray, Carol Clugston, Anna Dominiczak and Alex Alderton, Roberto Amato, Jeffrey Barrett, Sonia Goncalves, Ewan Harrison, David K. Jackson, Ian Johnston, Dominic Kwiatkowski, Cordelia Langford, John Sillitoe on behalf of the Wellcome Sanger Institute COVID-19 Surveillance Team |
| EPI_ISL_2702368 | Lighthouse Lab in Alderley Park | Wellcome Sanger Institute for the COVID-19 Genomics UK (COG-UK) Consortium | Jacquelyn Wynn, Mairead Hyland, The Lighthouse Lab in Alderley Park and Alex Alderton, Roberto Amato, Jeffrey Barrett, Sonia Goncalves, Ewan Harrison, David K. Jackson, Ian Johnston, Dominic Kwiatkowski, Cordelia Langford, John Sillitoe on behalf of the Wellcome Sanger Institute COVID-19 Surveillance Team |

| | | | |
|---|---|---|---|
| EPI_ISL_2702369, EPI_ISL_2702370 | Lighthouse Lab in Glasgow | Wellcome Sanger Institute for the COVID-19 Genomics UK (COG-UK) Consortium | Harper VanSteenhouse, Yumi Kasai, David Gray, Carol Clugston, Anna Dominiczak and Alex Alderton, Roberto Amato, Jeffrey Barrett, Sonia Goncalves, Ewan Harrison, David K. Jackson, Ian Johnston, Dominic Kwiatkowski, Cordelia Langford, John Sillitoe on behalf of the Wellcome Sanger Institute COVID-19 Surveillance Team |
| EPI_ISL_2702371 | Lighthouse Lab in Alderley Park | Wellcome Sanger Institute for the COVID-19 Genomics UK (COG-UK) Consortium | Jacquelyn Wynn, Mairead Hyland, The Lighthouse Lab in Alderley Park and Alex Alderton, Roberto Amato, Jeffrey Barrett, Sonia Goncalves, Ewan Harrison, David K. Jackson, Ian Johnston, Dominic Kwiatkowski, Cordelia Langford, John Sillitoe on behalf of the Wellcome Sanger Institute COVID-19 Surveillance Team |
| EPI_ISL_2702372 | Lighthouse Lab in Milton Keynes | Wellcome Sanger Institute for the COVID-19 Genomics UK (COG-UK) Consortium | The Lighthouse Lab in Milton Keynes and Alex Alderton, Roberto Amato, Jeffrey Barrett, Sonia Goncalves, Ewan Harrison, David K. Jackson, Ian Johnston, Dominic Kwiatkowski, Cordelia Langford, John Sillitoe on behalf of the Wellcome Sanger Institute COVID-19 Surveillance Team |
| EPI_ISL_2702373 | Lighthouse Lab in Alderley Park | Wellcome Sanger Institute for the COVID-19 Genomics UK (COG-UK) Consortium | Jacquelyn Wynn, Mairead Hyland, The Lighthouse Lab in Alderley Park and Alex Alderton, Roberto Amato, Jeffrey Barrett, Sonia Goncalves, Ewan Harrison, David K. Jackson, Ian Johnston, Dominic Kwiatkowski, Cordelia Langford, John Sillitoe on behalf of the Wellcome Sanger Institute COVID-19 Surveillance Team |
| EPI_ISL_2702374 | Lighthouse Lab in Glasgow | Wellcome Sanger Institute for the COVID-19 Genomics UK (COG-UK) Consortium | Harper VanSteenhouse, Yumi Kasai, David Gray, Carol Clugston, Anna Dominiczak and Alex Alderton, Roberto Amato, Jeffrey Barrett, Sonia Goncalves, Ewan Harrison, David K. Jackson, Ian Johnston, Dominic Kwiatkowski, Cordelia Langford, John Sillitoe on behalf of the Wellcome Sanger Institute COVID-19 Surveillance Team |
| EPI_ISL_2702375 | Lighthouse Lab in Alderley Park | Wellcome Sanger Institute for the COVID-19 Genomics UK (COG-UK) Consortium | Jacquelyn Wynn, Mairead Hyland, The Lighthouse Lab in Alderley Park and Alex Alderton, Roberto Amato, Jeffrey Barrett, Sonia Goncalves, Ewan Harrison, David K. Jackson, Ian Johnston, Dominic Kwiatkowski, Cordelia Langford, John Sillitoe on behalf of the Wellcome Sanger Institute COVID-19 Surveillance Team |
| EPI_ISL_2702376 | Lighthouse Lab in Milton Keynes | Wellcome Sanger Institute for the COVID-19 Genomics UK (COG-UK) Consortium | The Lighthouse Lab in Milton Keynes and Alex Alderton, Roberto Amato, Jeffrey Barrett, Sonia Goncalves, Ewan Harrison, David K. Jackson, Ian Johnston, Dominic Kwiatkowski, Cordelia Langford, John Sillitoe on behalf of the Wellcome Sanger Institute COVID-19 Surveillance Team |
| EPI_ISL_2702377, EPI_ISL_2702378 | Lighthouse Lab in Glasgow | Wellcome Sanger Institute for the COVID-19 Genomics UK (COG-UK) Consortium | Harper VanSteenhouse, Yumi Kasai, David Gray, Carol Clugston, Anna Dominiczak and Alex Alderton, Roberto Amato, Jeffrey Barrett, Sonia Goncalves, Ewan Harrison, David K. Jackson, Ian Johnston, Dominic Kwiatkowski, Cordelia Langford, John Sillitoe on behalf of the Wellcome Sanger Institute COVID-19 Surveillance Team |
| EPI_ISL_2702379 | Lighthouse Lab in Alderley Park | Wellcome Sanger Institute for the COVID-19 Genomics UK (COG-UK) Consortium | Jacquelyn Wynn, Mairead Hyland, The Lighthouse Lab in Alderley Park and Alex Alderton, Roberto Amato, Jeffrey Barrett, Sonia Goncalves, Ewan Harrison, David K. Jackson, Ian Johnston, Dominic Kwiatkowski, Cordelia Langford, John Sillitoe on behalf of the Wellcome Sanger Institute COVID-19 Surveillance Team |
| EPI_ISL_2702380 | Lighthouse Lab in Glasgow | Wellcome Sanger Institute for the COVID-19 Genomics UK (COG-UK) Consortium | Harper VanSteenhouse, Yumi Kasai, David Gray, Carol Clugston, Anna Dominiczak and Alex Alderton, Roberto Amato, Jeffrey Barrett, Sonia Goncalves, Ewan Harrison, David K. Jackson, Ian Johnston, Dominic Kwiatkowski, Cordelia Langford, John Sillitoe on behalf of the Wellcome Sanger Institute COVID-19 Surveillance Team |
| EPI_ISL_2702381 | Lighthouse Lab in Milton Keynes | Wellcome Sanger Institute for the COVID-19 Genomics UK (COG-UK) Consortium | The Lighthouse Lab in Milton Keynes and Alex Alderton, Roberto Amato, Jeffrey Barrett, Sonia Goncalves, Ewan Harrison, David K. Jackson, Ian Johnston, Dominic Kwiatkowski, Cordelia Langford, John Sillitoe on behalf of the Wellcome Sanger Institute COVID-19 Surveillance Team |
| EPI_ISL_2702382, EPI_ISL_2702383 | Lighthouse Lab in Glasgow | Wellcome Sanger Institute for the COVID-19 Genomics UK (COG-UK) Consortium | Harper VanSteenhouse, Yumi Kasai, David Gray, Carol Clugston, Anna Dominiczak and Alex Alderton, Roberto Amato, Jeffrey Barrett, Sonia Goncalves, Ewan Harrison, David K. Jackson, Ian Johnston, Dominic Kwiatkowski, Cordelia Langford, John Sillitoe on behalf of the Wellcome Sanger Institute COVID-19 Surveillance Team |
| EPI_ISL_2702384 | Lighthouse Lab in Alderley Park | Wellcome Sanger Institute for the COVID-19 Genomics UK (COG-UK) Consortium | Jacquelyn Wynn, Mairead Hyland, The Lighthouse Lab in Alderley Park and Alex Alderton, Roberto Amato, Jeffrey Barrett, Sonia Goncalves, Ewan Harrison, David K. Jackson, Ian Johnston, Dominic Kwiatkowski, Cordelia Langford, John Sillitoe on behalf of the Wellcome Sanger Institute COVID-19 Surveillance Team |
| EPI_ISL_2702385, EPI_ISL_2702386 | Lighthouse Lab in Milton Keynes | Wellcome Sanger Institute for the COVID-19 Genomics UK (COG-UK) Consortium | The Lighthouse Lab in Milton Keynes and Alex Alderton, Roberto Amato, Jeffrey Barrett, Sonia Goncalves, Ewan Harrison, David K. Jackson, Ian Johnston, Dominic Kwiatkowski, Cordelia Langford, John Sillitoe on behalf of the Wellcome Sanger Institute COVID-19 Surveillance Team |
| EPI_ISL_2702387, EPI_ISL_2702388, EPI_ISL_2702389, EPI_ISL_2702390 | Lighthouse Lab in Glasgow | Wellcome Sanger Institute for the COVID-19 Genomics UK (COG-UK) Consortium | Harper VanSteenhouse, Yumi Kasai, David Gray, Carol Clugston, Anna Dominiczak and Alex Alderton, Roberto Amato, Jeffrey Barrett, Sonia Goncalves, Ewan Harrison, David K. Jackson, Ian Johnston, Dominic Kwiatkowski, Cordelia Langford, John Sillitoe on behalf of the Wellcome Sanger Institute COVID-19 Surveillance Team |
| EPI_ISL_2702391 | Lighthouse Lab in Alderley Park | Wellcome Sanger Institute for the COVID-19 Genomics UK (COG-UK) Consortium | Jacquelyn Wynn, Mairead Hyland, The Lighthouse Lab in Alderley Park and Alex Alderton, Roberto Amato, Jeffrey Barrett, Sonia Goncalves, Ewan Harrison, David K. Jackson, Ian Johnston, Dominic Kwiatkowski, Cordelia Langford, John Sillitoe on behalf of the Wellcome Sanger Institute COVID-19 Surveillance Team |
| EPI_ISL_2702392, EPI_ISL_2702393, EPI_ISL_2702394, EPI_ISL_2702395 | Lighthouse Lab in Glasgow | Wellcome Sanger Institute for the COVID-19 Genomics UK (COG-UK) Consortium | Harper VanSteenhouse, Yumi Kasai, David Gray, Carol Clugston, Anna Dominiczak and Alex Alderton, Roberto Amato, Jeffrey Barrett, Sonia Goncalves, Ewan Harrison, David K. Jackson, Ian Johnston, Dominic Kwiatkowski, Cordelia Langford, John Sillitoe on behalf of the Wellcome Sanger Institute COVID-19 Surveillance Team |
| EPI_ISL_2702396 | Lighthouse Lab in Milton Keynes | Wellcome Sanger Institute for the COVID-19 Genomics UK (COG-UK) Consortium | The Lighthouse Lab in Milton Keynes and Alex Alderton, Roberto Amato, Jeffrey Barrett, Sonia Goncalves, Ewan Harrison, David K. Jackson, Ian Johnston, Dominic Kwiatkowski, Cordelia Langford, John Sillitoe on behalf of the Wellcome Sanger Institute COVID-19 Surveillance Team |
| EPI_ISL_2702397, EPI_ISL_2702398, EPI_ISL_2702399, EPI_ISL_2702400, EPI_ISL_2702401, EPI_ISL_2702402, EPI_ISL_2702403, EPI_ISL_2702404, EPI_ISL_2702405, EPI_ISL_2702406, EPI_ISL_2702407 | | | |
| see above | Lighthouse Lab in Glasgow | Wellcome Sanger Institute for the COVID-19 Genomics UK (COG-UK) Consortium | Harper VanSteenhouse, Yumi Kasai, David Gray, Carol Clugston, Anna Dominiczak and Alex Alderton, Roberto Amato, Jeffrey Barrett, Sonia Goncalves, Ewan Harrison, David K. Jackson, Ian Johnston, Dominic Kwiatkowski, Cordelia Langford, John Sillitoe on behalf of the Wellcome Sanger Institute COVID-19 Surveillance Team |
| EPI_ISL_2702408 | Lighthouse Lab in Alderley Park | Wellcome Sanger Institute for the COVID-19 Genomics UK (COG-UK) Consortium | Jacquelyn Wynn, Mairead Hyland, The Lighthouse Lab in Alderley Park and Alex Alderton, Roberto Amato, Jeffrey Barrett, Sonia Goncalves, Ewan Harrison, David K. Jackson, Ian Johnston, Dominic Kwiatkowski, Cordelia Langford, John Sillitoe on behalf of the Wellcome Sanger Institute COVID-19 Surveillance Team |
| EPI_ISL_2702409, EPI_ISL_2702410, EPI_ISL_2702411, EPI_ISL_2702412, EPI_ISL_2702413, EPI_ISL_2702415, EPI_ISL_2702417, EPI_ISL_2702418, EPI_ISL_2702419, EPI_ISL_2702420 | Lighthouse Lab in Glasgow | Wellcome Sanger Institute for the COVID-19 Genomics UK (COG-UK) Consortium | Harper VanSteenhouse, Yumi Kasai, David Gray, Carol Clugston, Anna Dominiczak and Alex Alderton, Roberto Amato, Jeffrey Barrett, Sonia Goncalves, Ewan Harrison, David K. Jackson, Ian Johnston, Dominic Kwiatkowski, Cordelia Langford, John Sillitoe on behalf of the Wellcome Sanger Institute COVID-19 Surveillance Team |
| EPI_ISL_2702421, EPI_ISL_2702422 | Lighthouse Lab in Alderley Park | Wellcome Sanger Institute for the COVID-19 Genomics UK (COG-UK) Consortium | Jacquelyn Wynn, Mairead Hyland, The Lighthouse Lab in Alderley Park and Alex Alderton, Roberto Amato, Jeffrey Barrett, Sonia Goncalves, Ewan Harrison, David K. Jackson, Ian Johnston, Dominic Kwiatkowski, Cordelia Langford, John Sillitoe on behalf of the Wellcome Sanger Institute COVID-19 Surveillance Team |
| EPI_ISL_2702423, EPI_ISL_2702425, EPI_ISL_2702426, EPI_ISL_2702427, EPI_ISL_2702428, EPI_ISL_2702430, EPI_ISL_2702431, EPI_ISL_2702432, EPI_ISL_2702433, EPI_ISL_2702434, EPI_ISL_2702435, EPI_ISL_2702436, EPI_ISL_2702437, EPI_ISL_2702438, EPI_ISL_2702439, EPI_ISL_2702440, EPI_ISL_2702441, EPI_ISL_2702442, EPI_ISL_2702443, EPI_ISL_2702444, EPI_ISL_2702445, EPI_ISL_2702446, EPI_ISL_2702447, EPI_ISL_2702448, EPI_ISL_2702449, EPI_ISL_2702450, EPI_ISL_2702451, EPI_ISL_2702452, EPI_ISL_2702453, EPI_ISL_2702454, EPI_ISL_2702455, EPI_ISL_2702456, EPI_ISL_2702457 | | | |
| see above | Lighthouse Lab in Glasgow | Wellcome Sanger Institute for the COVID-19 Genomics UK (COG-UK) Consortium | Harper VanSteenhouse, Yumi Kasai, David Gray, Carol Clugston, Anna Dominiczak and Alex Alderton, Roberto Amato, Jeffrey Barrett, Sonia Goncalves, Ewan Harrison, David K. Jackson, Ian Johnston, Dominic Kwiatkowski, Cordelia Langford, John Sillitoe on behalf of the Wellcome Sanger Institute COVID-19 Surveillance Team |
| EPI_ISL_2702458, EPI_ISL_2702459 | Lighthouse Lab in Alderley Park | Wellcome Sanger Institute for the COVID-19 Genomics UK (COG-UK) Consortium | Jacquelyn Wynn, Mairead Hyland, The Lighthouse Lab in Alderley Park and Alex Alderton, Roberto Amato, Jeffrey Barrett, Sonia Goncalves, Ewan Harrison, David K. Jackson, Ian Johnston, Dominic Kwiatkowski, Cordelia Langford, John Sillitoe on behalf of the Wellcome Sanger Institute COVID-19 Surveillance Team |
| EPI_ISL_2702460 | Lighthouse Lab in Glasgow | Wellcome Sanger Institute for the COVID-19 Genomics UK (COG-UK) Consortium | Harper VanSteenhouse, Yumi Kasai, David Gray, Carol Clugston, Anna Dominiczak and Alex Alderton, Roberto Amato, Jeffrey Barrett, Sonia Goncalves, Ewan Harrison, David K. Jackson, Ian Johnston, Dominic Kwiatkowski, Cordelia Langford, John Sillitoe on behalf of the Wellcome Sanger Institute COVID-19 Surveillance Team |
| EPI_ISL_2702461, EPI_ISL_2702462, EPI_ISL_2702464, EPI_ISL_2702465, EPI_ISL_2702466 | Lighthouse Lab in Alderley Park | Wellcome Sanger Institute for the COVID-19 Genomics UK (COG-UK) Consortium | Jacquelyn Wynn, Mairead Hyland, The Lighthouse Lab in Alderley Park and Alex Alderton, Roberto Amato, Jeffrey Barrett, Sonia Goncalves, Ewan Harrison, David K. Jackson, Ian Johnston, Dominic Kwiatkowski, Cordelia Langford, John Sillitoe on behalf of the Wellcome Sanger Institute COVID-19 Surveillance Team |
| EPI_ISL_2702467 | Lighthouse Lab in Glasgow | Wellcome Sanger Institute for the COVID-19 Genomics UK (COG-UK) Consortium | Harper VanSteenhouse, Yumi Kasai, David Gray, Carol Clugston, Anna Dominiczak and Alex Alderton, Roberto Amato, Jeffrey Barrett, Sonia Goncalves, Ewan Harrison, David K. Jackson, Ian Johnston, Dominic Kwiatkowski, Cordelia Langford, John Sillitoe on behalf of the Wellcome Sanger Institute COVID-19 Surveillance Team |
| EPI_ISL_2702468, EPI_ISL_2702469 | Lighthouse Lab in Alderley Park | Wellcome Sanger Institute for the COVID-19 Genomics UK (COG-UK) Consortium | Jacquelyn Wynn, Mairead Hyland, The Lighthouse Lab in Alderley Park and Alex Alderton, Roberto Amato, Jeffrey Barrett, Sonia Goncalves, Ewan Harrison, David K. Jackson, Ian Johnston, Dominic Kwiatkowski, Cordelia Langford, John Sillitoe on behalf of the Wellcome Sanger Institute COVID-19 Surveillance Team |
| EPI_ISL_2702470 | Lighthouse Lab in Glasgow | Wellcome Sanger Institute for the COVID-19 Genomics UK (COG-UK) Consortium | Harper VanSteenhouse, Yumi Kasai, David Gray, Carol Clugston, Anna Dominiczak and Alex Alderton, Roberto Amato, Jeffrey Barrett, Sonia Goncalves, Ewan Harrison, David K. Jackson, Ian Johnston, Dominic Kwiatkowski, Cordelia Langford, John Sillitoe on behalf of the Wellcome Sanger Institute COVID-19 Surveillance Team |
| EPI_ISL_2702471 | Lighthouse Lab in Alderley Park | Wellcome Sanger Institute for the COVID-19 Genomics UK (COG-UK) Consortium | Jacquelyn Wynn, Mairead Hyland, The Lighthouse Lab in Alderley Park and Alex Alderton, Roberto Amato, Jeffrey Barrett, Sonia Goncalves, Ewan Harrison, David K. Jackson, Ian Johnston, Dominic Kwiatkowski, Cordelia Langford, John Sillitoe on behalf of the Wellcome Sanger Institute COVID-19 Surveillance Team |
| EPI_ISL_2702472 | Lighthouse Lab in Glasgow | Wellcome Sanger Institute for the COVID-19 Genomics UK (COG-UK) Consortium | Harper VanSteenhouse, Yumi Kasai, David Gray, Carol Clugston, Anna Dominiczak and Alex Alderton, Roberto Amato, Jeffrey Barrett, Sonia Goncalves, Ewan Harrison, David K. Jackson, Ian Johnston, Dominic Kwiatkowski, Cordelia Langford, John Sillitoe on behalf of the Wellcome Sanger Institute COVID-19 Surveillance Team |
| EPI_ISL_2702473 | Lighthouse Lab in Alderley Park | Wellcome Sanger Institute for the COVID-19 Genomics UK (COG-UK) Consortium | Jacquelyn Wynn, Mairead Hyland, The Lighthouse Lab in Alderley Park and Alex Alderton, Roberto Amato, Jeffrey Barrett, Sonia Goncalves, Ewan Harrison, David K. Jackson, Ian Johnston, Dominic Kwiatkowski, Cordelia Langford, John Sillitoe on behalf of the Wellcome Sanger Institute COVID-19 Surveillance Team |
| EPI_ISL_2702474, EPI_ISL_2702475 | Lighthouse Lab in Glasgow | Wellcome Sanger Institute for the COVID-19 Genomics UK (COG-UK) Consortium | Harper VanSteenhouse, Yumi Kasai, David Gray, Carol Clugston, Anna Dominiczak and Alex Alderton, Roberto Amato, Jeffrey Barrett, Sonia Goncalves, Ewan Harrison, David K. Jackson, Ian Johnston, Dominic Kwiatkowski, Cordelia Langford, John Sillitoe on behalf of the Wellcome Sanger Institute COVID-19 Surveillance Team |
| EPI_ISL_2702476, EPI_ISL_2702478 | Lighthouse Lab in Alderley Park | Wellcome Sanger Institute for the COVID-19 Genomics UK (COG-UK) Consortium | Jacquelyn Wynn, Mairead Hyland, The Lighthouse Lab in Alderley Park and Alex Alderton, Roberto Amato, Jeffrey Barrett, Sonia Goncalves, Ewan Harrison, David K. Jackson, Ian Johnston, Dominic Kwiatkowski, Cordelia Langford, John Sillitoe on behalf of the Wellcome Sanger Institute COVID-19 Surveillance Team |
| EPI_ISL_2702479 | Lighthouse Lab in Glasgow | Wellcome Sanger Institute for the COVID-19 Genomics UK (COG-UK) Consortium | Harper VanSteenhouse, Yumi Kasai, David Gray, Carol Clugston, Anna Dominiczak and Alex Alderton, Roberto Amato, Jeffrey Barrett, Sonia Goncalves, Ewan Harrison, David K. Jackson, Ian Johnston, Dominic Kwiatkowski, Cordelia Langford, John Sillitoe on behalf of the Wellcome Sanger Institute COVID-19 Surveillance Team |
| EPI_ISL_2702480, EPI_ISL_2702481, EPI_ISL_2702482 | Lighthouse Lab in Alderley Park | Wellcome Sanger Institute for the COVID-19 Genomics UK (COG-UK) Consortium | Jacquelyn Wynn, Mairead Hyland, The Lighthouse Lab in Alderley Park and Alex Alderton, Roberto Amato, Jeffrey Barrett, Sonia Goncalves, Ewan Harrison, David K. Jackson, Ian Johnston, Dominic Kwiatkowski, Cordelia Langford, John Sillitoe on behalf of the Wellcome Sanger Institute COVID-19 Surveillance Team |
| EPI_ISL_2702483 | Lighthouse Lab in Glasgow | Wellcome Sanger Institute for the COVID-19 Genomics UK (COG-UK) Consortium | Harper VanSteenhouse, Yumi Kasai, David Gray, Carol Clugston, Anna Dominiczak and Alex Alderton, Roberto Amato, Jeffrey Barrett, Sonia Goncalves, Ewan Harrison, David K. Jackson, Ian Johnston, Dominic Kwiatkowski, Cordelia Langford, John Sillitoe on behalf of the Wellcome Sanger Institute COVID-19 Surveillance Team |

| | | | |
|---|---|---|---|
| EPI_ISL_2702484, EPI_ISL_2702485, EPI_ISL_2702486, EPI_ISL_2702487, EPI_ISL_2702488 | Lighthouse Lab in Alderley Park | Wellcome Sanger Institute for the COVID-19 Genomics UK (COG-UK) Consortium | Jacquelyn Wynn, Mairead Hyland, The Lighthouse Lab in Alderley Park and Alex Alderton, Roberto Amato, Jeffrey Barrett, Sonia Goncalves, Ewan Harrison, David K. Jackson, Ian Johnston, Dominic Kwiatkowski, Cordelia Langford, John Sillitoe on behalf of the Wellcome Sanger Institute COVID-19 Surveillance Team |
| EPI_ISL_2702489 | Lighthouse Lab in Glasgow | Wellcome Sanger Institute for the COVID-19 Genomics UK (COG-UK) Consortium | Harper VanSteenhouse, Yumi Kasai, David Gray, Carol Clugston, Anna Dominiczak and Alex Alderton, Roberto Amato, Jeffrey Barrett, Sonia Goncalves, Ewan Harrison, David K. Jackson, Ian Johnston, Dominic Kwiatkowski, Cordelia Langford, John Sillitoe on behalf of the Wellcome Sanger Institute COVID-19 Surveillance Team |
| EPI_ISL_2702490, EPI_ISL_2702491, EPI_ISL_2702492, EPI_ISL_2702493, EPI_ISL_2702494, EPI_ISL_2702495, EPI_ISL_2702496, EPI_ISL_2702497, EPI_ISL_2702498, EPI_ISL_2702499, EPI_ISL_2702500 | | | |
| see above | Lighthouse Lab in Alderley Park | Wellcome Sanger Institute for the COVID-19 Genomics UK (COG-UK) Consortium | Jacquelyn Wynn, Mairead Hyland, The Lighthouse Lab in Alderley Park and Alex Alderton, Roberto Amato, Jeffrey Barrett, Sonia Goncalves, Ewan Harrison, David K. Jackson, Ian Johnston, Dominic Kwiatkowski, Cordelia Langford, John Sillitoe on behalf of the Wellcome Sanger Institute COVID-19 Surveillance Team |
| EPI_ISL_2702501 | Lighthouse Lab in Glasgow | Wellcome Sanger Institute for the COVID-19 Genomics UK (COG-UK) Consortium | Harper VanSteenhouse, Yumi Kasai, David Gray, Carol Clugston, Anna Dominiczak and Alex Alderton, Roberto Amato, Jeffrey Barrett, Sonia Goncalves, Ewan Harrison, David K. Jackson, Ian Johnston, Dominic Kwiatkowski, Cordelia Langford, John Sillitoe on behalf of the Wellcome Sanger Institute COVID-19 Surveillance Team |
| EPI_ISL_2702502, EPI_ISL_2702503, EPI_ISL_2702504 | Lighthouse Lab in Alderley Park | Wellcome Sanger Institute for the COVID-19 Genomics UK (COG-UK) Consortium | Jacquelyn Wynn, Mairead Hyland, The Lighthouse Lab in Alderley Park and Alex Alderton, Roberto Amato, Jeffrey Barrett, Sonia Goncalves, Ewan Harrison, David K. Jackson, Ian Johnston, Dominic Kwiatkowski, Cordelia Langford, John Sillitoe on behalf of the Wellcome Sanger Institute COVID-19 Surveillance Team |
| EPI_ISL_2702505 | Lighthouse Lab in Glasgow | Wellcome Sanger Institute for the COVID-19 Genomics UK (COG-UK) Consortium | Harper VanSteenhouse, Yumi Kasai, David Gray, Carol Clugston, Anna Dominiczak and Alex Alderton, Roberto Amato, Jeffrey Barrett, Sonia Goncalves, Ewan Harrison, David K. Jackson, Ian Johnston, Dominic Kwiatkowski, Cordelia Langford, John Sillitoe on behalf of the Wellcome Sanger Institute COVID-19 Surveillance Team |
| EPI_ISL_2702506, EPI_ISL_2702507 | Lighthouse Lab in Alderley Park | Wellcome Sanger Institute for the COVID-19 Genomics UK (COG-UK) Consortium | Jacquelyn Wynn, Mairead Hyland, The Lighthouse Lab in Alderley Park and Alex Alderton, Roberto Amato, Jeffrey Barrett, Sonia Goncalves, Ewan Harrison, David K. Jackson, Ian Johnston, Dominic Kwiatkowski, Cordelia Langford, John Sillitoe on behalf of the Wellcome Sanger Institute COVID-19 Surveillance Team |
| EPI_ISL_2702508 | Lighthouse Lab in Glasgow | Wellcome Sanger Institute for the COVID-19 Genomics UK (COG-UK) Consortium | Harper VanSteenhouse, Yumi Kasai, David Gray, Carol Clugston, Anna Dominiczak and Alex Alderton, Roberto Amato, Jeffrey Barrett, Sonia Goncalves, Ewan Harrison, David K. Jackson, Ian Johnston, Dominic Kwiatkowski, Cordelia Langford, John Sillitoe on behalf of the Wellcome Sanger Institute COVID-19 Surveillance Team |
| EPI_ISL_2702509, EPI_ISL_2702510, EPI_ISL_2702511, EPI_ISL_2702512, EPI_ISL_2702513, EPI_ISL_2702514, EPI_ISL_2702515 | Lighthouse Lab in Alderley Park | Wellcome Sanger Institute for the COVID-19 Genomics UK (COG-UK) Consortium | Jacquelyn Wynn, Mairead Hyland, The Lighthouse Lab in Alderley Park and Alex Alderton, Roberto Amato, Jeffrey Barrett, Sonia Goncalves, Ewan Harrison, David K. Jackson, Ian Johnston, Dominic Kwiatkowski, Cordelia Langford, John Sillitoe on behalf of the Wellcome Sanger Institute COVID-19 Surveillance Team |
| EPI_ISL_2702516 | Lighthouse Lab in Glasgow | Wellcome Sanger Institute for the COVID-19 Genomics UK (COG-UK) Consortium | Harper VanSteenhouse, Yumi Kasai, David Gray, Carol Clugston, Anna Dominiczak and Alex Alderton, Roberto Amato, Jeffrey Barrett, Sonia Goncalves, Ewan Harrison, David K. Jackson, Ian Johnston, Dominic Kwiatkowski, Cordelia Langford, John Sillitoe on behalf of the Wellcome Sanger Institute COVID-19 Surveillance Team |
| EPI_ISL_2702517, EPI_ISL_2702518, EPI_ISL_2702519 | Lighthouse Lab in Alderley Park | Wellcome Sanger Institute for the COVID-19 Genomics UK (COG-UK) Consortium | Jacquelyn Wynn, Mairead Hyland, The Lighthouse Lab in Alderley Park and Alex Alderton, Roberto Amato, Jeffrey Barrett, Sonia Goncalves, Ewan Harrison, David K. Jackson, Ian Johnston, Dominic Kwiatkowski, Cordelia Langford, John Sillitoe on behalf of the Wellcome Sanger Institute COVID-19 Surveillance Team |
| EPI_ISL_2702520, EPI_ISL_2702521 | Lighthouse Lab in Glasgow | Wellcome Sanger Institute for the COVID-19 Genomics UK (COG-UK) Consortium | Harper VanSteenhouse, Yumi Kasai, David Gray, Carol Clugston, Anna Dominiczak and Alex Alderton, Roberto Amato, Jeffrey Barrett, Sonia Goncalves, Ewan Harrison, David K. Jackson, Ian Johnston, Dominic Kwiatkowski, Cordelia Langford, John Sillitoe on behalf of the Wellcome Sanger Institute COVID-19 Surveillance Team |
| EPI_ISL_2702522, EPI_ISL_2702523 | Lighthouse Lab in Alderley Park | Wellcome Sanger Institute for the COVID-19 Genomics UK (COG-UK) Consortium | Jacquelyn Wynn, Mairead Hyland, The Lighthouse Lab in Alderley Park and Alex Alderton, Roberto Amato, Jeffrey Barrett, Sonia Goncalves, Ewan Harrison, David K. Jackson, Ian Johnston, Dominic Kwiatkowski, Cordelia Langford, John Sillitoe on behalf of the Wellcome Sanger Institute COVID-19 Surveillance Team |
| EPI_ISL_2702524 | Lighthouse Lab in Glasgow | Wellcome Sanger Institute for the COVID-19 Genomics UK (COG-UK) Consortium | Harper VanSteenhouse, Yumi Kasai, David Gray, Carol Clugston, Anna Dominiczak and Alex Alderton, Roberto Amato, Jeffrey Barrett, Sonia Goncalves, Ewan Harrison, David K. Jackson, Ian Johnston, Dominic Kwiatkowski, Cordelia Langford, John Sillitoe on behalf of the Wellcome Sanger Institute COVID-19 Surveillance Team |
| EPI_ISL_2702525, EPI_ISL_2702526, EPI_ISL_2702527, EPI_ISL_2702528, EPI_ISL_2702529, EPI_ISL_2702530, EPI_ISL_2702531, EPI_ISL_2702532, EPI_ISL_2702533 | Lighthouse Lab in Alderley Park | Wellcome Sanger Institute for the COVID-19 Genomics UK (COG-UK) Consortium | Jacquelyn Wynn, Mairead Hyland, The Lighthouse Lab in Alderley Park and Alex Alderton, Roberto Amato, Jeffrey Barrett, Sonia Goncalves, Ewan Harrison, David K. Jackson, Ian Johnston, Dominic Kwiatkowski, Cordelia Langford, John Sillitoe on behalf of the Wellcome Sanger Institute COVID-19 Surveillance Team |
| EPI_ISL_2702534 | Lighthouse Lab in Glasgow | Wellcome Sanger Institute for the COVID-19 Genomics UK (COG-UK) Consortium | Harper VanSteenhouse, Yumi Kasai, David Gray, Carol Clugston, Anna Dominiczak and Alex Alderton, Roberto Amato, Jeffrey Barrett, Sonia Goncalves, Ewan Harrison, David K. Jackson, Ian Johnston, Dominic Kwiatkowski, Cordelia Langford, John Sillitoe on behalf of the Wellcome Sanger Institute COVID-19 Surveillance Team |
| EPI_ISL_2702535, EPI_ISL_2702536, EPI_ISL_2702537, EPI_ISL_2702538, EPI_ISL_2702539, EPI_ISL_2702540, EPI_ISL_2702541, EPI_ISL_2702542, EPI_ISL_2702543, EPI_ISL_2702544, EPI_ISL_2702545, EPI_ISL_2702546, EPI_ISL_2702547, EPI_ISL_2702548, EPI_ISL_2702549, EPI_ISL_2702550, EPI_ISL_2702551, EPI_ISL_2702552 | | | |
| see above | Lighthouse Lab in Alderley Park | Wellcome Sanger Institute for the COVID-19 Genomics UK (COG-UK) Consortium | Jacquelyn Wynn, Mairead Hyland, The Lighthouse Lab in Alderley Park and Alex Alderton, Roberto Amato, Jeffrey Barrett, Sonia Goncalves, Ewan Harrison, David K. Jackson, Ian Johnston, Dominic Kwiatkowski, Cordelia Langford, John Sillitoe on behalf of the Wellcome Sanger Institute COVID-19 Surveillance Team |
| EPI_ISL_2702553 | Lighthouse Lab in Glasgow | Wellcome Sanger Institute for the COVID-19 Genomics UK (COG-UK) Consortium | Harper VanSteenhouse, Yumi Kasai, David Gray, Carol Clugston, Anna Dominiczak and Alex Alderton, Roberto Amato, Jeffrey Barrett, Sonia Goncalves, Ewan Harrison, David K. Jackson, Ian Johnston, Dominic Kwiatkowski, Cordelia Langford, John Sillitoe on behalf of the Wellcome Sanger Institute COVID-19 Surveillance Team |
| EPI_ISL_2702554, EPI_ISL_2702555, EPI_ISL_2702556, EPI_ISL_2702557, EPI_ISL_2702558, EPI_ISL_2702559, EPI_ISL_2702560 | Lighthouse Lab in Alderley Park | Wellcome Sanger Institute for the COVID-19 Genomics UK (COG-UK) Consortium | Jacquelyn Wynn, Mairead Hyland, The Lighthouse Lab in Alderley Park and Alex Alderton, Roberto Amato, Jeffrey Barrett, Sonia Goncalves, Ewan Harrison, David K. Jackson, Ian Johnston, Dominic Kwiatkowski, Cordelia Langford, John Sillitoe on behalf of the Wellcome Sanger Institute COVID-19 Surveillance Team |
| EPI_ISL_2702561 | Lighthouse Lab in Glasgow | Wellcome Sanger Institute for the COVID-19 Genomics UK (COG-UK) Consortium | Harper VanSteenhouse, Yumi Kasai, David Gray, Carol Clugston, Anna Dominiczak and Alex Alderton, Roberto Amato, Jeffrey Barrett, Sonia Goncalves, Ewan Harrison, David K. Jackson, Ian Johnston, Dominic Kwiatkowski, Cordelia Langford, John Sillitoe on behalf of the Wellcome Sanger Institute COVID-19 Surveillance Team |
| EPI_ISL_2702562, EPI_ISL_2702563, EPI_ISL_2702564, EPI_ISL_2702565, EPI_ISL_2702566, EPI_ISL_2702567, EPI_ISL_2702568, EPI_ISL_2702569, EPI_ISL_2702570, EPI_ISL_2702571, EPI_ISL_2702572, EPI_ISL_2702573 | | | |
| see above | Lighthouse Lab in Alderley Park | Wellcome Sanger Institute for the COVID-19 Genomics UK (COG-UK) Consortium | Jacquelyn Wynn, Mairead Hyland, The Lighthouse Lab in Alderley Park and Alex Alderton, Roberto Amato, Jeffrey Barrett, Sonia Goncalves, Ewan Harrison, David K. Jackson, Ian Johnston, Dominic Kwiatkowski, Cordelia Langford, John Sillitoe on behalf of the Wellcome Sanger Institute COVID-19 Surveillance Team |
| EPI_ISL_2702574 | Lighthouse Lab in Glasgow | Wellcome Sanger Institute for the COVID-19 Genomics UK (COG-UK) Consortium | Harper VanSteenhouse, Yumi Kasai, David Gray, Carol Clugston, Anna Dominiczak and Alex Alderton, Roberto Amato, Jeffrey Barrett, Sonia Goncalves, Ewan Harrison, David K. Jackson, Ian Johnston, Dominic Kwiatkowski, Cordelia Langford, John Sillitoe on behalf of the Wellcome Sanger Institute COVID-19 Surveillance Team |
| EPI_ISL_2702575 | Lighthouse Lab in Alderley Park | Wellcome Sanger Institute for the COVID-19 Genomics UK (COG-UK) Consortium | Jacquelyn Wynn, Mairead Hyland, The Lighthouse Lab in Alderley Park and Alex Alderton, Roberto Amato, Jeffrey Barrett, Sonia Goncalves, Ewan Harrison, David K. Jackson, Ian Johnston, Dominic Kwiatkowski, Cordelia Langford, John Sillitoe on behalf of the Wellcome Sanger Institute COVID-19 Surveillance Team |
| EPI_ISL_2702576 | Lighthouse Lab in Glasgow | Wellcome Sanger Institute for the COVID-19 Genomics UK (COG-UK) Consortium | Harper VanSteenhouse, Yumi Kasai, David Gray, Carol Clugston, Anna Dominiczak and Alex Alderton, Roberto Amato, Jeffrey Barrett, Sonia Goncalves, Ewan Harrison, David K. Jackson, Ian Johnston, Dominic Kwiatkowski, Cordelia Langford, John Sillitoe on behalf of the Wellcome Sanger Institute COVID-19 Surveillance Team |
| EPI_ISL_2702577, EPI_ISL_2702578, EPI_ISL_2702579 | Lighthouse Lab in Alderley Park | Wellcome Sanger Institute for the COVID-19 Genomics UK (COG-UK) Consortium | Jacquelyn Wynn, Mairead Hyland, The Lighthouse Lab in Alderley Park and Alex Alderton, Roberto Amato, Jeffrey Barrett, Sonia Goncalves, Ewan Harrison, David K. Jackson, Ian Johnston, Dominic Kwiatkowski, Cordelia Langford, John Sillitoe on behalf of the Wellcome Sanger Institute COVID-19 Surveillance Team |
| EPI_ISL_2702580, EPI_ISL_2702581 | Lighthouse Lab in Glasgow | Wellcome Sanger Institute for the COVID-19 Genomics UK (COG-UK) Consortium | Harper VanSteenhouse, Yumi Kasai, David Gray, Carol Clugston, Anna Dominiczak and Alex Alderton, Roberto Amato, Jeffrey Barrett, Sonia Goncalves, Ewan Harrison, David K. Jackson, Ian Johnston, Dominic Kwiatkowski, Cordelia Langford, John Sillitoe on behalf of the Wellcome Sanger Institute COVID-19 Surveillance Team |
| EPI_ISL_2702582, EPI_ISL_2702583, EPI_ISL_2702584, EPI_ISL_2702585, EPI_ISL_2702586 | Lighthouse Lab in Alderley Park | Wellcome Sanger Institute for the COVID-19 Genomics UK (COG-UK) Consortium | Jacquelyn Wynn, Mairead Hyland, The Lighthouse Lab in Alderley Park and Alex Alderton, Roberto Amato, Jeffrey Barrett, Sonia Goncalves, Ewan Harrison, David K. Jackson, Ian Johnston, Dominic Kwiatkowski, Cordelia Langford, John Sillitoe on behalf of the Wellcome Sanger Institute COVID-19 Surveillance Team |
| EPI_ISL_2702587 | Lighthouse Lab in Glasgow | Wellcome Sanger Institute for the COVID-19 Genomics UK (COG-UK) Consortium | Harper VanSteenhouse, Yumi Kasai, David Gray, Carol Clugston, Anna Dominiczak and Alex Alderton, Roberto Amato, Jeffrey Barrett, Sonia Goncalves, Ewan Harrison, David K. Jackson, Ian Johnston, Dominic Kwiatkowski, Cordelia Langford, John Sillitoe on behalf of the Wellcome Sanger Institute COVID-19 Surveillance Team |
| EPI_ISL_2702588, EPI_ISL_2702589, EPI_ISL_2702590, EPI_ISL_2702591 | Lighthouse Lab in Alderley Park | Wellcome Sanger Institute for the COVID-19 Genomics UK (COG-UK) Consortium | Jacquelyn Wynn, Mairead Hyland, The Lighthouse Lab in Alderley Park and Alex Alderton, Roberto Amato, Jeffrey Barrett, Sonia Goncalves, Ewan Harrison, David K. Jackson, Ian Johnston, Dominic Kwiatkowski, Cordelia Langford, John Sillitoe on behalf of the Wellcome Sanger Institute COVID-19 Surveillance Team |
| EPI_ISL_2702592, EPI_ISL_2702593 | Lighthouse Lab in Glasgow | Wellcome Sanger Institute for the COVID-19 Genomics UK (COG-UK) Consortium | Harper VanSteenhouse, Yumi Kasai, David Gray, Carol Clugston, Anna Dominiczak and Alex Alderton, Roberto Amato, Jeffrey Barrett, Sonia Goncalves, Ewan Harrison, David K. Jackson, Ian Johnston, Dominic Kwiatkowski, Cordelia Langford, John Sillitoe on behalf of the Wellcome Sanger Institute COVID-19 Surveillance Team |
| EPI_ISL_2702594, EPI_ISL_2702595 | Lighthouse Lab in Alderley Park | Wellcome Sanger Institute for the COVID-19 Genomics UK (COG-UK) Consortium | Jacquelyn Wynn, Mairead Hyland, The Lighthouse Lab in Alderley Park and Alex Alderton, Roberto Amato, Jeffrey Barrett, Sonia Goncalves, Ewan Harrison, David K. Jackson, Ian Johnston, Dominic Kwiatkowski, Cordelia Langford, John Sillitoe on behalf of the Wellcome Sanger Institute COVID-19 Surveillance Team |
| EPI_ISL_2702596 | Lighthouse Lab in Glasgow | Wellcome Sanger Institute for the COVID-19 Genomics UK (COG-UK) Consortium | Harper VanSteenhouse, Yumi Kasai, David Gray, Carol Clugston, Anna Dominiczak and Alex Alderton, Roberto Amato, Jeffrey Barrett, Sonia Goncalves, Ewan Harrison, David K. Jackson, Ian Johnston, Dominic Kwiatkowski, Cordelia Langford, John Sillitoe on behalf of the Wellcome Sanger Institute COVID-19 Surveillance Team |

| | | | |
|---|---|---|---|
| EPI_ISL_2702597, EPI_ISL_2702598, EPI_ISL_2702599, EPI_ISL_2702600, EPI_ISL_2702601, EPI_ISL_2702602, EPI_ISL_2702603 | Lighthouse Lab in Alderley Park | Wellcome Sanger Institute for the COVID-19 Genomics UK (COG-UK) Consortium | Jacquelyn Wynn, Mairead Hyland, The Lighthouse Lab in Alderley Park and Alex Alderton, Roberto Amato, Jeffrey Barrett, Sonia Goncalves, Ewan Harrison, David K. Jackson, Ian Johnston, Dominic Kwiatkowski, Cordelia Langford, John Sillitoe on behalf of the Wellcome Sanger Institute COVID-19 Surveillance Team |
| EPI_ISL_2702604 | Lighthouse Lab in Glasgow | Wellcome Sanger Institute for the COVID-19 Genomics UK (COG-UK) Consortium | Harper VanSteenhouse, Yumi Kasai, David Gray, Carol Clugston, Anna Dominiczak and Alex Alderton, Roberto Amato, Jeffrey Barrett, Sonia Goncalves, Ewan Harrison, David K. Jackson, Ian Johnston, Dominic Kwiatkowski, Cordelia Langford, John Sillitoe on behalf of the Wellcome Sanger Institute COVID-19 Surveillance Team |
| EPI_ISL_2702605 | Lighthouse Lab in Alderley Park | Wellcome Sanger Institute for the COVID-19 Genomics UK (COG-UK) Consortium | Jacquelyn Wynn, Mairead Hyland, The Lighthouse Lab in Alderley Park and Alex Alderton, Roberto Amato, Jeffrey Barrett, Sonia Goncalves, Ewan Harrison, David K. Jackson, Ian Johnston, Dominic Kwiatkowski, Cordelia Langford, John Sillitoe on behalf of the Wellcome Sanger Institute COVID-19 Surveillance Team |
| EPI_ISL_2702606 | Lighthouse Lab in Glasgow | Wellcome Sanger Institute for the COVID-19 Genomics UK (COG-UK) Consortium | Harper VanSteenhouse, Yumi Kasai, David Gray, Carol Clugston, Anna Dominiczak and Alex Alderton, Roberto Amato, Jeffrey Barrett, Sonia Goncalves, Ewan Harrison, David K. Jackson, Ian Johnston, Dominic Kwiatkowski, Cordelia Langford, John Sillitoe on behalf of the Wellcome Sanger Institute COVID-19 Surveillance Team |
| EPI_ISL_2702607 | Lighthouse Lab in Alderley Park | Wellcome Sanger Institute for the COVID-19 Genomics UK (COG-UK) Consortium | Jacquelyn Wynn, Mairead Hyland, The Lighthouse Lab in Alderley Park and Alex Alderton, Roberto Amato, Jeffrey Barrett, Sonia Goncalves, Ewan Harrison, David K. Jackson, Ian Johnston, Dominic Kwiatkowski, Cordelia Langford, John Sillitoe on behalf of the Wellcome Sanger Institute COVID-19 Surveillance Team |
| EPI_ISL_2702608, EPI_ISL_2702609 | Lighthouse Lab in Glasgow | Wellcome Sanger Institute for the COVID-19 Genomics UK (COG-UK) Consortium | Harper VanSteenhouse, Yumi Kasai, David Gray, Carol Clugston, Anna Dominiczak and Alex Alderton, Roberto Amato, Jeffrey Barrett, Sonia Goncalves, Ewan Harrison, David K. Jackson, Ian Johnston, Dominic Kwiatkowski, Cordelia Langford, John Sillitoe on behalf of the Wellcome Sanger Institute COVID-19 Surveillance Team |
| EPI_ISL_2702610, EPI_ISL_2702611, EPI_ISL_2702612, EPI_ISL_2702613, EPI_ISL_2702614, EPI_ISL_2702615, EPI_ISL_2702616, EPI_ISL_2702617, EPI_ISL_2702618, EPI_ISL_2702620, EPI_ISL_2702621 | | | |
| see above | Lighthouse Lab in Alderley Park | Wellcome Sanger Institute for the COVID-19 Genomics UK (COG-UK) Consortium | Jacquelyn Wynn, Mairead Hyland, The Lighthouse Lab in Alderley Park and Alex Alderton, Roberto Amato, Jeffrey Barrett, Sonia Goncalves, Ewan Harrison, David K. Jackson, Ian Johnston, Dominic Kwiatkowski, Cordelia Langford, John Sillitoe on behalf of the Wellcome Sanger Institute COVID-19 Surveillance Team |
| EPI_ISL_2702622 | Lighthouse Lab in Glasgow | Wellcome Sanger Institute for the COVID-19 Genomics UK (COG-UK) Consortium | Harper VanSteenhouse, Yumi Kasai, David Gray, Carol Clugston, Anna Dominiczak and Alex Alderton, Roberto Amato, Jeffrey Barrett, Sonia Goncalves, Ewan Harrison, David K. Jackson, Ian Johnston, Dominic Kwiatkowski, Cordelia Langford, John Sillitoe on behalf of the Wellcome Sanger Institute COVID-19 Surveillance Team |
| EPI_ISL_2702623, EPI_ISL_2702624, EPI_ISL_2702625, EPI_ISL_2702626, EPI_ISL_2702627, EPI_ISL_2702628 | Lighthouse Lab in Alderley Park | Wellcome Sanger Institute for the COVID-19 Genomics UK (COG-UK) Consortium | Jacquelyn Wynn, Mairead Hyland, The Lighthouse Lab in Alderley Park and Alex Alderton, Roberto Amato, Jeffrey Barrett, Sonia Goncalves, Ewan Harrison, David K. Jackson, Ian Johnston, Dominic Kwiatkowski, Cordelia Langford, John Sillitoe on behalf of the Wellcome Sanger Institute COVID-19 Surveillance Team |
| EPI_ISL_2702629 | Lighthouse Lab in Glasgow | Wellcome Sanger Institute for the COVID-19 Genomics UK (COG-UK) Consortium | Harper VanSteenhouse, Yumi Kasai, David Gray, Carol Clugston, Anna Dominiczak and Alex Alderton, Roberto Amato, Jeffrey Barrett, Sonia Goncalves, Ewan Harrison, David K. Jackson, Ian Johnston, Dominic Kwiatkowski, Cordelia Langford, John Sillitoe on behalf of the Wellcome Sanger Institute COVID-19 Surveillance Team |
| EPI_ISL_2702630, EPI_ISL_2702631, EPI_ISL_2702632, EPI_ISL_2702633 | Lighthouse Lab in Alderley Park | Wellcome Sanger Institute for the COVID-19 Genomics UK (COG-UK) Consortium | Jacquelyn Wynn, Mairead Hyland, The Lighthouse Lab in Alderley Park and Alex Alderton, Roberto Amato, Jeffrey Barrett, Sonia Goncalves, Ewan Harrison, David K. Jackson, Ian Johnston, Dominic Kwiatkowski, Cordelia Langford, John Sillitoe on behalf of the Wellcome Sanger Institute COVID-19 Surveillance Team |
| EPI_ISL_2702634, EPI_ISL_2702635 | Lighthouse Lab in Glasgow | Wellcome Sanger Institute for the COVID-19 Genomics UK (COG-UK) Consortium | Harper VanSteenhouse, Yumi Kasai, David Gray, Carol Clugston, Anna Dominiczak and Alex Alderton, Roberto Amato, Jeffrey Barrett, Sonia Goncalves, Ewan Harrison, David K. Jackson, Ian Johnston, Dominic Kwiatkowski, Cordelia Langford, John Sillitoe on behalf of the Wellcome Sanger Institute COVID-19 Surveillance Team |
| EPI_ISL_2702636, EPI_ISL_2702637, EPI_ISL_2702638 | Lighthouse Lab in Alderley Park | Wellcome Sanger Institute for the COVID-19 Genomics UK (COG-UK) Consortium | Jacquelyn Wynn, Mairead Hyland, The Lighthouse Lab in Alderley Park and Alex Alderton, Roberto Amato, Jeffrey Barrett, Sonia Goncalves, Ewan Harrison, David K. Jackson, Ian Johnston, Dominic Kwiatkowski, Cordelia Langford, John Sillitoe on behalf of the Wellcome Sanger Institute COVID-19 Surveillance Team |
| EPI_ISL_2702639 | Lighthouse Lab in Glasgow | Wellcome Sanger Institute for the COVID-19 Genomics UK (COG-UK) Consortium | Harper VanSteenhouse, Yumi Kasai, David Gray, Carol Clugston, Anna Dominiczak and Alex Alderton, Roberto Amato, Jeffrey Barrett, Sonia Goncalves, Ewan Harrison, David K. Jackson, Ian Johnston, Dominic Kwiatkowski, Cordelia Langford, John Sillitoe on behalf of the Wellcome Sanger Institute COVID-19 Surveillance Team |
| EPI_ISL_2702640, EPI_ISL_2702641 | Lighthouse Lab in Alderley Park | Wellcome Sanger Institute for the COVID-19 Genomics UK (COG-UK) Consortium | Jacquelyn Wynn, Mairead Hyland, The Lighthouse Lab in Alderley Park and Alex Alderton, Roberto Amato, Jeffrey Barrett, Sonia Goncalves, Ewan Harrison, David K. Jackson, Ian Johnston, Dominic Kwiatkowski, Cordelia Langford, John Sillitoe on behalf of the Wellcome Sanger Institute COVID-19 Surveillance Team |
| EPI_ISL_2702642 | Lighthouse Lab in Glasgow | Wellcome Sanger Institute for the COVID-19 Genomics UK (COG-UK) Consortium | Harper VanSteenhouse, Yumi Kasai, David Gray, Carol Clugston, Anna Dominiczak and Alex Alderton, Roberto Amato, Jeffrey Barrett, Sonia Goncalves, Ewan Harrison, David K. Jackson, Ian Johnston, Dominic Kwiatkowski, Cordelia Langford, John Sillitoe on behalf of the Wellcome Sanger Institute COVID-19 Surveillance Team |
| EPI_ISL_2702643, EPI_ISL_2702644 | Lighthouse Lab in Alderley Park | Wellcome Sanger Institute for the COVID-19 Genomics UK (COG-UK) Consortium | Jacquelyn Wynn, Mairead Hyland, The Lighthouse Lab in Alderley Park and Alex Alderton, Roberto Amato, Jeffrey Barrett, Sonia Goncalves, Ewan Harrison, David K. Jackson, Ian Johnston, Dominic Kwiatkowski, Cordelia Langford, John Sillitoe on behalf of the Wellcome Sanger Institute COVID-19 Surveillance Team |
| EPI_ISL_2702645 | Lighthouse Lab in Glasgow | Wellcome Sanger Institute for the COVID-19 Genomics UK (COG-UK) Consortium | Harper VanSteenhouse, Yumi Kasai, David Gray, Carol Clugston, Anna Dominiczak and Alex Alderton, Roberto Amato, Jeffrey Barrett, Sonia Goncalves, Ewan Harrison, David K. Jackson, Ian Johnston, Dominic Kwiatkowski, Cordelia Langford, John Sillitoe on behalf of the Wellcome Sanger Institute COVID-19 Surveillance Team |
| EPI_ISL_2702646, EPI_ISL_2702647 | Lighthouse Lab in Alderley Park | Wellcome Sanger Institute for the COVID-19 Genomics UK (COG-UK) Consortium | Jacquelyn Wynn, Mairead Hyland, The Lighthouse Lab in Alderley Park and Alex Alderton, Roberto Amato, Jeffrey Barrett, Sonia Goncalves, Ewan Harrison, David K. Jackson, Ian Johnston, Dominic Kwiatkowski, Cordelia Langford, John Sillitoe on behalf of the Wellcome Sanger Institute COVID-19 Surveillance Team |
| EPI_ISL_2702648, EPI_ISL_2702649 | Lighthouse Lab in Glasgow | Wellcome Sanger Institute for the COVID-19 Genomics UK (COG-UK) Consortium | Harper VanSteenhouse, Yumi Kasai, David Gray, Carol Clugston, Anna Dominiczak and Alex Alderton, Roberto Amato, Jeffrey Barrett, Sonia Goncalves, Ewan Harrison, David K. Jackson, Ian Johnston, Dominic Kwiatkowski, Cordelia Langford, John Sillitoe on behalf of the Wellcome Sanger Institute COVID-19 Surveillance Team |
| EPI_ISL_2702650, EPI_ISL_2702651, EPI_ISL_2702652, EPI_ISL_2702653, EPI_ISL_2702654, EPI_ISL_2702655, EPI_ISL_2702656, EPI_ISL_2702657, EPI_ISL_2702658 | Lighthouse Lab in Alderley Park | Wellcome Sanger Institute for the COVID-19 Genomics UK (COG-UK) Consortium | Jacquelyn Wynn, Mairead Hyland, The Lighthouse Lab in Alderley Park and Alex Alderton, Roberto Amato, Jeffrey Barrett, Sonia Goncalves, Ewan Harrison, David K. Jackson, Ian Johnston, Dominic Kwiatkowski, Cordelia Langford, John Sillitoe on behalf of the Wellcome Sanger Institute COVID-19 Surveillance Team |
| EPI_ISL_2702659 | Lighthouse Lab in Glasgow | Wellcome Sanger Institute for the COVID-19 Genomics UK (COG-UK) Consortium | Harper VanSteenhouse, Yumi Kasai, David Gray, Carol Clugston, Anna Dominiczak and Alex Alderton, Roberto Amato, Jeffrey Barrett, Sonia Goncalves, Ewan Harrison, David K. Jackson, Ian Johnston, Dominic Kwiatkowski, Cordelia Langford, John Sillitoe on behalf of the Wellcome Sanger Institute COVID-19 Surveillance Team |
| EPI_ISL_2702660, EPI_ISL_2702661 | Lighthouse Lab in Alderley Park | Wellcome Sanger Institute for the COVID-19 Genomics UK (COG-UK) Consortium | Jacquelyn Wynn, Mairead Hyland, The Lighthouse Lab in Alderley Park and Alex Alderton, Roberto Amato, Jeffrey Barrett, Sonia Goncalves, Ewan Harrison, David K. Jackson, Ian Johnston, Dominic Kwiatkowski, Cordelia Langford, John Sillitoe on behalf of the Wellcome Sanger Institute COVID-19 Surveillance Team |
| EPI_ISL_2702662, EPI_ISL_2702663 | Lighthouse Lab in Glasgow | Wellcome Sanger Institute for the COVID-19 Genomics UK (COG-UK) Consortium | Harper VanSteenhouse, Yumi Kasai, David Gray, Carol Clugston, Anna Dominiczak and Alex Alderton, Roberto Amato, Jeffrey Barrett, Sonia Goncalves, Ewan Harrison, David K. Jackson, Ian Johnston, Dominic Kwiatkowski, Cordelia Langford, John Sillitoe on behalf of the Wellcome Sanger Institute COVID-19 Surveillance Team |
| EPI_ISL_2702664, EPI_ISL_2702665 | Lighthouse Lab in Alderley Park | Wellcome Sanger Institute for the COVID-19 Genomics UK (COG-UK) Consortium | Jacquelyn Wynn, Mairead Hyland, The Lighthouse Lab in Alderley Park and Alex Alderton, Roberto Amato, Jeffrey Barrett, Sonia Goncalves, Ewan Harrison, David K. Jackson, Ian Johnston, Dominic Kwiatkowski, Cordelia Langford, John Sillitoe on behalf of the Wellcome Sanger Institute COVID-19 Surveillance Team |
| EPI_ISL_2702666 | Lighthouse Lab in Glasgow | Wellcome Sanger Institute for the COVID-19 Genomics UK (COG-UK) Consortium | Harper VanSteenhouse, Yumi Kasai, David Gray, Carol Clugston, Anna Dominiczak and Alex Alderton, Roberto Amato, Jeffrey Barrett, Sonia Goncalves, Ewan Harrison, David K. Jackson, Ian Johnston, Dominic Kwiatkowski, Cordelia Langford, John Sillitoe on behalf of the Wellcome Sanger Institute COVID-19 Surveillance Team |
| EPI_ISL_2702667, EPI_ISL_2702668, EPI_ISL_2702669 | Lighthouse Lab in Alderley Park | Wellcome Sanger Institute for the COVID-19 Genomics UK (COG-UK) Consortium | Jacquelyn Wynn, Mairead Hyland, The Lighthouse Lab in Alderley Park and Alex Alderton, Roberto Amato, Jeffrey Barrett, Sonia Goncalves, Ewan Harrison, David K. Jackson, Ian Johnston, Dominic Kwiatkowski, Cordelia Langford, John Sillitoe on behalf of the Wellcome Sanger Institute COVID-19 Surveillance Team |
| EPI_ISL_2702670 | Lighthouse Lab in Glasgow | Wellcome Sanger Institute for the COVID-19 Genomics UK (COG-UK) Consortium | Harper VanSteenhouse, Yumi Kasai, David Gray, Carol Clugston, Anna Dominiczak and Alex Alderton, Roberto Amato, Jeffrey Barrett, Sonia Goncalves, Ewan Harrison, David K. Jackson, Ian Johnston, Dominic Kwiatkowski, Cordelia Langford, John Sillitoe on behalf of the Wellcome Sanger Institute COVID-19 Surveillance Team |
| EPI_ISL_2702671 | Lighthouse Lab in Alderley Park | Wellcome Sanger Institute for the COVID-19 Genomics UK (COG-UK) Consortium | Jacquelyn Wynn, Mairead Hyland, The Lighthouse Lab in Alderley Park and Alex Alderton, Roberto Amato, Jeffrey Barrett, Sonia Goncalves, Ewan Harrison, David K. Jackson, Ian Johnston, Dominic Kwiatkowski, Cordelia Langford, John Sillitoe on behalf of the Wellcome Sanger Institute COVID-19 Surveillance Team |
| EPI_ISL_2702672 | Lighthouse Lab in Glasgow | Wellcome Sanger Institute for the COVID-19 Genomics UK (COG-UK) Consortium | Harper VanSteenhouse, Yumi Kasai, David Gray, Carol Clugston, Anna Dominiczak and Alex Alderton, Roberto Amato, Jeffrey Barrett, Sonia Goncalves, Ewan Harrison, David K. Jackson, Ian Johnston, Dominic Kwiatkowski, Cordelia Langford, John Sillitoe on behalf of the Wellcome Sanger Institute COVID-19 Surveillance Team |
| EPI_ISL_2702673, EPI_ISL_2702674, EPI_ISL_2702675 | Lighthouse Lab in Alderley Park | Wellcome Sanger Institute for the COVID-19 Genomics UK (COG-UK) Consortium | Jacquelyn Wynn, Mairead Hyland, The Lighthouse Lab in Alderley Park and Alex Alderton, Roberto Amato, Jeffrey Barrett, Sonia Goncalves, Ewan Harrison, David K. Jackson, Ian Johnston, Dominic Kwiatkowski, Cordelia Langford, John Sillitoe on behalf of the Wellcome Sanger Institute COVID-19 Surveillance Team |
| EPI_ISL_2702676 | Lighthouse Lab in Glasgow | Wellcome Sanger Institute for the COVID-19 Genomics UK (COG-UK) Consortium | Harper VanSteenhouse, Yumi Kasai, David Gray, Carol Clugston, Anna Dominiczak and Alex Alderton, Roberto Amato, Jeffrey Barrett, Sonia Goncalves, Ewan Harrison, David K. Jackson, Ian Johnston, Dominic Kwiatkowski, Cordelia Langford, John Sillitoe on behalf of the Wellcome Sanger Institute COVID-19 Surveillance Team |
| EPI_ISL_2702677, EPI_ISL_2702678, EPI_ISL_2702679, EPI_ISL_2702680, EPI_ISL_2702681, EPI_ISL_2702682 | Lighthouse Lab in Alderley Park | Wellcome Sanger Institute for the COVID-19 Genomics UK (COG-UK) Consortium | Jacquelyn Wynn, Mairead Hyland, The Lighthouse Lab in Alderley Park and Alex Alderton, Roberto Amato, Jeffrey Barrett, Sonia Goncalves, Ewan Harrison, David K. Jackson, Ian Johnston, Dominic Kwiatkowski, Cordelia Langford, John Sillitoe on behalf of the Wellcome Sanger Institute COVID-19 Surveillance Team |
| EPI_ISL_2702683 | Lighthouse Lab in Glasgow | Wellcome Sanger Institute for the COVID-19 Genomics UK (COG-UK) Consortium | Harper VanSteenhouse, Yumi Kasai, David Gray, Carol Clugston, Anna Dominiczak and Alex Alderton, Roberto Amato, Jeffrey Barrett, Sonia Goncalves, Ewan Harrison, David K. Jackson, Ian Johnston, Dominic Kwiatkowski, Cordelia Langford, John Sillitoe on behalf of the Wellcome Sanger Institute COVID-19 Surveillance Team |
| EPI_ISL_2702684, | Lighthouse Lab in Alderley Park | Wellcome Sanger Institute for the COVID-19 Genomics UK (COG-UK) | Jacquelyn Wynn, Mairead Hyland, The Lighthouse Lab in Alderley Park and Alex Alderton, Roberto Amato, Jeffrey Barrett, Sonia Goncalves, Ewan Harrison, David K. Jackson, Ian Johnston, Dominic Kwiatkowski, Cordelia Langford, John Sillitoe on |

| | | | |
|---|---|---|---|
| EPI_ISL_2702685, EPI_ISL_2702686 | | Consortium | behalf of the Wellcome Sanger Institute COVID-19 Surveillance Team |
| EPI_ISL_2702687, EPI_ISL_2702688 | Lighthouse Lab in Glasgow | Wellcome Sanger Institute for the COVID-19 Genomics UK (COG-UK) Consortium | Harper VanSteenhouse, Yumi Kasai, David Gray, Carol Clugston, Anna Dominiczak and Alex Alderton, Roberto Amato, Jeffrey Barrett, Sonia Goncalves, Ewan Harrison, David K. Jackson, Ian Johnston, Dominic Kwiatkowski, Cordelia Langford, John Sillitoe on behalf of the Wellcome Sanger Institute COVID-19 Surveillance Team |
| EPI_ISL_2702689 | Lighthouse Lab in Alderley Park | Wellcome Sanger Institute for the COVID-19 Genomics UK (COG-UK) Consortium | Jacquelyn Wynn, Mairead Hyland, The Lighthouse Lab in Alderley Park and Alex Alderton, Roberto Amato, Jeffrey Barrett, Sonia Goncalves, Ewan Harrison, David K. Jackson, Ian Johnston, Dominic Kwiatkowski, Cordelia Langford, John Sillitoe on behalf of the Wellcome Sanger Institute COVID-19 Surveillance Team |
| EPI_ISL_2702690, EPI_ISL_2702691 | Lighthouse Lab in Glasgow | Wellcome Sanger Institute for the COVID-19 Genomics UK (COG-UK) Consortium | Harper VanSteenhouse, Yumi Kasai, David Gray, Carol Clugston, Anna Dominiczak and Alex Alderton, Roberto Amato, Jeffrey Barrett, Sonia Goncalves, Ewan Harrison, David K. Jackson, Ian Johnston, Dominic Kwiatkowski, Cordelia Langford, John Sillitoe on behalf of the Wellcome Sanger Institute COVID-19 Surveillance Team |
| EPI_ISL_2702692, EPI_ISL_2702693 | Lighthouse Lab in Alderley Park | Wellcome Sanger Institute for the COVID-19 Genomics UK (COG-UK) Consortium | Jacquelyn Wynn, Mairead Hyland, The Lighthouse Lab in Alderley Park and Alex Alderton, Roberto Amato, Jeffrey Barrett, Sonia Goncalves, Ewan Harrison, David K. Jackson, Ian Johnston, Dominic Kwiatkowski, Cordelia Langford, John Sillitoe on behalf of the Wellcome Sanger Institute COVID-19 Surveillance Team |
| EPI_ISL_2702694 | Lighthouse Lab in Glasgow | Wellcome Sanger Institute for the COVID-19 Genomics UK (COG-UK) Consortium | Harper VanSteenhouse, Yumi Kasai, David Gray, Carol Clugston, Anna Dominiczak and Alex Alderton, Roberto Amato, Jeffrey Barrett, Sonia Goncalves, Ewan Harrison, David K. Jackson, Ian Johnston, Dominic Kwiatkowski, Cordelia Langford, John Sillitoe on behalf of the Wellcome Sanger Institute COVID-19 Surveillance Team |
| EPI_ISL_2702695 | Lighthouse Lab in Alderley Park | Wellcome Sanger Institute for the COVID-19 Genomics UK (COG-UK) Consortium | Jacquelyn Wynn, Mairead Hyland, The Lighthouse Lab in Alderley Park and Alex Alderton, Roberto Amato, Jeffrey Barrett, Sonia Goncalves, Ewan Harrison, David K. Jackson, Ian Johnston, Dominic Kwiatkowski, Cordelia Langford, John Sillitoe on behalf of the Wellcome Sanger Institute COVID-19 Surveillance Team |
| EPI_ISL_2702696 | Lighthouse Lab in Glasgow | Wellcome Sanger Institute for the COVID-19 Genomics UK (COG-UK) Consortium | Harper VanSteenhouse, Yumi Kasai, David Gray, Carol Clugston, Anna Dominiczak and Alex Alderton, Roberto Amato, Jeffrey Barrett, Sonia Goncalves, Ewan Harrison, David K. Jackson, Ian Johnston, Dominic Kwiatkowski, Cordelia Langford, John Sillitoe on behalf of the Wellcome Sanger Institute COVID-19 Surveillance Team |
| EPI_ISL_2702697 | Lighthouse Lab in Alderley Park | Wellcome Sanger Institute for the COVID-19 Genomics UK (COG-UK) Consortium | Jacquelyn Wynn, Mairead Hyland, The Lighthouse Lab in Alderley Park and Alex Alderton, Roberto Amato, Jeffrey Barrett, Sonia Goncalves, Ewan Harrison, David K. Jackson, Ian Johnston, Dominic Kwiatkowski, Cordelia Langford, John Sillitoe on behalf of the Wellcome Sanger Institute COVID-19 Surveillance Team |
| EPI_ISL_2702698 | Lighthouse Lab in Glasgow | Wellcome Sanger Institute for the COVID-19 Genomics UK (COG-UK) Consortium | Harper VanSteenhouse, Yumi Kasai, David Gray, Carol Clugston, Anna Dominiczak and Alex Alderton, Roberto Amato, Jeffrey Barrett, Sonia Goncalves, Ewan Harrison, David K. Jackson, Ian Johnston, Dominic Kwiatkowski, Cordelia Langford, John Sillitoe on behalf of the Wellcome Sanger Institute COVID-19 Surveillance Team |
| EPI_ISL_2702699 | Lighthouse Lab in Alderley Park | Wellcome Sanger Institute for the COVID-19 Genomics UK (COG-UK) Consortium | Jacquelyn Wynn, Mairead Hyland, The Lighthouse Lab in Alderley Park and Alex Alderton, Roberto Amato, Jeffrey Barrett, Sonia Goncalves, Ewan Harrison, David K. Jackson, Ian Johnston, Dominic Kwiatkowski, Cordelia Langford, John Sillitoe on behalf of the Wellcome Sanger Institute COVID-19 Surveillance Team |
| EPI_ISL_2702700 | Lighthouse Lab in Glasgow | Wellcome Sanger Institute for the COVID-19 Genomics UK (COG-UK) Consortium | Harper VanSteenhouse, Yumi Kasai, David Gray, Carol Clugston, Anna Dominiczak and Alex Alderton, Roberto Amato, Jeffrey Barrett, Sonia Goncalves, Ewan Harrison, David K. Jackson, Ian Johnston, Dominic Kwiatkowski, Cordelia Langford, John Sillitoe on behalf of the Wellcome Sanger Institute COVID-19 Surveillance Team |
| EPI_ISL_2702701, EPI_ISL_2702702, EPI_ISL_2702703, EPI_ISL_2702704 | Lighthouse Lab in Alderley Park | Wellcome Sanger Institute for the COVID-19 Genomics UK (COG-UK) Consortium | Jacquelyn Wynn, Mairead Hyland, The Lighthouse Lab in Alderley Park and Alex Alderton, Roberto Amato, Jeffrey Barrett, Sonia Goncalves, Ewan Harrison, David K. Jackson, Ian Johnston, Dominic Kwiatkowski, Cordelia Langford, John Sillitoe on behalf of the Wellcome Sanger Institute COVID-19 Surveillance Team |
| EPI_ISL_2702705 | Lighthouse Lab in Glasgow | Wellcome Sanger Institute for the COVID-19 Genomics UK (COG-UK) Consortium | Harper VanSteenhouse, Yumi Kasai, David Gray, Carol Clugston, Anna Dominiczak and Alex Alderton, Roberto Amato, Jeffrey Barrett, Sonia Goncalves, Ewan Harrison, David K. Jackson, Ian Johnston, Dominic Kwiatkowski, Cordelia Langford, John Sillitoe on behalf of the Wellcome Sanger Institute COVID-19 Surveillance Team |
| EPI_ISL_2702706, EPI_ISL_2702707, EPI_ISL_2702708, EPI_ISL_2702709, EPI_ISL_2702710, EPI_ISL_2702711 | Lighthouse Lab in Alderley Park | Wellcome Sanger Institute for the COVID-19 Genomics UK (COG-UK) Consortium | Jacquelyn Wynn, Mairead Hyland, The Lighthouse Lab in Alderley Park and Alex Alderton, Roberto Amato, Jeffrey Barrett, Sonia Goncalves, Ewan Harrison, David K. Jackson, Ian Johnston, Dominic Kwiatkowski, Cordelia Langford, John Sillitoe on behalf of the Wellcome Sanger Institute COVID-19 Surveillance Team |
| EPI_ISL_2702712 | Lighthouse Lab in Glasgow | Wellcome Sanger Institute for the COVID-19 Genomics UK (COG-UK) Consortium | Harper VanSteenhouse, Yumi Kasai, David Gray, Carol Clugston, Anna Dominiczak and Alex Alderton, Roberto Amato, Jeffrey Barrett, Sonia Goncalves, Ewan Harrison, David K. Jackson, Ian Johnston, Dominic Kwiatkowski, Cordelia Langford, John Sillitoe on behalf of the Wellcome Sanger Institute COVID-19 Surveillance Team |
| EPI_ISL_2702713, EPI_ISL_2702714, EPI_ISL_2702715, EPI_ISL_2702716, EPI_ISL_2702717, EPI_ISL_2702718 | Lighthouse Lab in Alderley Park | Wellcome Sanger Institute for the COVID-19 Genomics UK (COG-UK) Consortium | Jacquelyn Wynn, Mairead Hyland, The Lighthouse Lab in Alderley Park and Alex Alderton, Roberto Amato, Jeffrey Barrett, Sonia Goncalves, Ewan Harrison, David K. Jackson, Ian Johnston, Dominic Kwiatkowski, Cordelia Langford, John Sillitoe on behalf of the Wellcome Sanger Institute COVID-19 Surveillance Team |
| EPI_ISL_2702719 | Lighthouse Lab in Glasgow | Wellcome Sanger Institute for the COVID-19 Genomics UK (COG-UK) Consortium | Harper VanSteenhouse, Yumi Kasai, David Gray, Carol Clugston, Anna Dominiczak and Alex Alderton, Roberto Amato, Jeffrey Barrett, Sonia Goncalves, Ewan Harrison, David K. Jackson, Ian Johnston, Dominic Kwiatkowski, Cordelia Langford, John Sillitoe on behalf of the Wellcome Sanger Institute COVID-19 Surveillance Team |
| EPI_ISL_2702720, EPI_ISL_2702721, EPI_ISL_2702722 | Lighthouse Lab in Alderley Park | Wellcome Sanger Institute for the COVID-19 Genomics UK (COG-UK) Consortium | Jacquelyn Wynn, Mairead Hyland, The Lighthouse Lab in Alderley Park and Alex Alderton, Roberto Amato, Jeffrey Barrett, Sonia Goncalves, Ewan Harrison, David K. Jackson, Ian Johnston, Dominic Kwiatkowski, Cordelia Langford, John Sillitoe on behalf of the Wellcome Sanger Institute COVID-19 Surveillance Team |
| EPI_ISL_2702723 | Lighthouse Lab in Glasgow | Wellcome Sanger Institute for the COVID-19 Genomics UK (COG-UK) Consortium | Harper VanSteenhouse, Yumi Kasai, David Gray, Carol Clugston, Anna Dominiczak and Alex Alderton, Roberto Amato, Jeffrey Barrett, Sonia Goncalves, Ewan Harrison, David K. Jackson, Ian Johnston, Dominic Kwiatkowski, Cordelia Langford, John Sillitoe on behalf of the Wellcome Sanger Institute COVID-19 Surveillance Team |
| EPI_ISL_2702724 | Lighthouse Lab in Alderley Park | Wellcome Sanger Institute for the COVID-19 Genomics UK (COG-UK) Consortium | Jacquelyn Wynn, Mairead Hyland, The Lighthouse Lab in Alderley Park and Alex Alderton, Roberto Amato, Jeffrey Barrett, Sonia Goncalves, Ewan Harrison, David K. Jackson, Ian Johnston, Dominic Kwiatkowski, Cordelia Langford, John Sillitoe on behalf of the Wellcome Sanger Institute COVID-19 Surveillance Team |
| EPI_ISL_2702725 | Lighthouse Lab in Glasgow | Wellcome Sanger Institute for the COVID-19 Genomics UK (COG-UK) Consortium | Harper VanSteenhouse, Yumi Kasai, David Gray, Carol Clugston, Anna Dominiczak and Alex Alderton, Roberto Amato, Jeffrey Barrett, Sonia Goncalves, Ewan Harrison, David K. Jackson, Ian Johnston, Dominic Kwiatkowski, Cordelia Langford, John Sillitoe on behalf of the Wellcome Sanger Institute COVID-19 Surveillance Team |
| EPI_ISL_2702726, EPI_ISL_2702727, EPI_ISL_2702728 | Lighthouse Lab in Alderley Park | Wellcome Sanger Institute for the COVID-19 Genomics UK (COG-UK) Consortium | Jacquelyn Wynn, Mairead Hyland, The Lighthouse Lab in Alderley Park and Alex Alderton, Roberto Amato, Jeffrey Barrett, Sonia Goncalves, Ewan Harrison, David K. Jackson, Ian Johnston, Dominic Kwiatkowski, Cordelia Langford, John Sillitoe on behalf of the Wellcome Sanger Institute COVID-19 Surveillance Team |
| EPI_ISL_2702729 | Lighthouse Lab in Glasgow | Wellcome Sanger Institute for the COVID-19 Genomics UK (COG-UK) Consortium | Harper VanSteenhouse, Yumi Kasai, David Gray, Carol Clugston, Anna Dominiczak and Alex Alderton, Roberto Amato, Jeffrey Barrett, Sonia Goncalves, Ewan Harrison, David K. Jackson, Ian Johnston, Dominic Kwiatkowski, Cordelia Langford, John Sillitoe on behalf of the Wellcome Sanger Institute COVID-19 Surveillance Team |
| EPI_ISL_2702730, EPI_ISL_2702731, EPI_ISL_2702732 | Lighthouse Lab in Alderley Park | Wellcome Sanger Institute for the COVID-19 Genomics UK (COG-UK) Consortium | Jacquelyn Wynn, Mairead Hyland, The Lighthouse Lab in Alderley Park and Alex Alderton, Roberto Amato, Jeffrey Barrett, Sonia Goncalves, Ewan Harrison, David K. Jackson, Ian Johnston, Dominic Kwiatkowski, Cordelia Langford, John Sillitoe on behalf of the Wellcome Sanger Institute COVID-19 Surveillance Team |
| EPI_ISL_2702733 | Lighthouse Lab in Glasgow | Wellcome Sanger Institute for the COVID-19 Genomics UK (COG-UK) Consortium | Harper VanSteenhouse, Yumi Kasai, David Gray, Carol Clugston, Anna Dominiczak and Alex Alderton, Roberto Amato, Jeffrey Barrett, Sonia Goncalves, Ewan Harrison, David K. Jackson, Ian Johnston, Dominic Kwiatkowski, Cordelia Langford, John Sillitoe on behalf of the Wellcome Sanger Institute COVID-19 Surveillance Team |
| EPI_ISL_2702734, EPI_ISL_2702735 | Lighthouse Lab in Alderley Park | Wellcome Sanger Institute for the COVID-19 Genomics UK (COG-UK) Consortium | Jacquelyn Wynn, Mairead Hyland, The Lighthouse Lab in Alderley Park and Alex Alderton, Roberto Amato, Jeffrey Barrett, Sonia Goncalves, Ewan Harrison, David K. Jackson, Ian Johnston, Dominic Kwiatkowski, Cordelia Langford, John Sillitoe on behalf of the Wellcome Sanger Institute COVID-19 Surveillance Team |
| EPI_ISL_2702736 | Lighthouse Lab in Glasgow | Wellcome Sanger Institute for the COVID-19 Genomics UK (COG-UK) Consortium | Harper VanSteenhouse, Yumi Kasai, David Gray, Carol Clugston, Anna Dominiczak and Alex Alderton, Roberto Amato, Jeffrey Barrett, Sonia Goncalves, Ewan Harrison, David K. Jackson, Ian Johnston, Dominic Kwiatkowski, Cordelia Langford, John Sillitoe on behalf of the Wellcome Sanger Institute COVID-19 Surveillance Team |
| EPI_ISL_2702737, EPI_ISL_2702738, EPI_ISL_2702739, EPI_ISL_2702740 | Lighthouse Lab in Alderley Park | Wellcome Sanger Institute for the COVID-19 Genomics UK (COG-UK) Consortium | Jacquelyn Wynn, Mairead Hyland, The Lighthouse Lab in Alderley Park and Alex Alderton, Roberto Amato, Jeffrey Barrett, Sonia Goncalves, Ewan Harrison, David K. Jackson, Ian Johnston, Dominic Kwiatkowski, Cordelia Langford, John Sillitoe on behalf of the Wellcome Sanger Institute COVID-19 Surveillance Team |
| EPI_ISL_2702741 | Lighthouse Lab in Glasgow | Wellcome Sanger Institute for the COVID-19 Genomics UK (COG-UK) Consortium | Harper VanSteenhouse, Yumi Kasai, David Gray, Carol Clugston, Anna Dominiczak and Alex Alderton, Roberto Amato, Jeffrey Barrett, Sonia Goncalves, Ewan Harrison, David K. Jackson, Ian Johnston, Dominic Kwiatkowski, Cordelia Langford, John Sillitoe on behalf of the Wellcome Sanger Institute COVID-19 Surveillance Team |
| EPI_ISL_2702742, EPI_ISL_2702743, EPI_ISL_2702744, EPI_ISL_2702745 | Lighthouse Lab in Alderley Park | Wellcome Sanger Institute for the COVID-19 Genomics UK (COG-UK) Consortium | Jacquelyn Wynn, Mairead Hyland, The Lighthouse Lab in Alderley Park and Alex Alderton, Roberto Amato, Jeffrey Barrett, Sonia Goncalves, Ewan Harrison, David K. Jackson, Ian Johnston, Dominic Kwiatkowski, Cordelia Langford, John Sillitoe on behalf of the Wellcome Sanger Institute COVID-19 Surveillance Team |
| EPI_ISL_2702746 | Lighthouse Lab in Glasgow | Wellcome Sanger Institute for the COVID-19 Genomics UK (COG-UK) Consortium | Harper VanSteenhouse, Yumi Kasai, David Gray, Carol Clugston, Anna Dominiczak and Alex Alderton, Roberto Amato, Jeffrey Barrett, Sonia Goncalves, Ewan Harrison, David K. Jackson, Ian Johnston, Dominic Kwiatkowski, Cordelia Langford, John Sillitoe on behalf of the Wellcome Sanger Institute COVID-19 Surveillance Team |
| EPI_ISL_2702747 | Lighthouse Lab in Alderley Park | Wellcome Sanger Institute for the COVID-19 Genomics UK (COG-UK) Consortium | Jacquelyn Wynn, Mairead Hyland, The Lighthouse Lab in Alderley Park and Alex Alderton, Roberto Amato, Jeffrey Barrett, Sonia Goncalves, Ewan Harrison, David K. Jackson, Ian Johnston, Dominic Kwiatkowski, Cordelia Langford, John Sillitoe on behalf of the Wellcome Sanger Institute COVID-19 Surveillance Team |
| EPI_ISL_2702748 | Lighthouse Lab in Glasgow | Wellcome Sanger Institute for the COVID-19 Genomics UK (COG-UK) Consortium | Harper VanSteenhouse, Yumi Kasai, David Gray, Carol Clugston, Anna Dominiczak and Alex Alderton, Roberto Amato, Jeffrey Barrett, Sonia Goncalves, Ewan Harrison, David K. Jackson, Ian Johnston, Dominic Kwiatkowski, Cordelia Langford, John Sillitoe on behalf of the Wellcome Sanger Institute COVID-19 Surveillance Team |
| EPI_ISL_2702749, EPI_ISL_2702750, EPI_ISL_2702751, EPI_ISL_2702752 | Lighthouse Lab in Alderley Park | Wellcome Sanger Institute for the COVID-19 Genomics UK (COG-UK) Consortium | Jacquelyn Wynn, Mairead Hyland, The Lighthouse Lab in Alderley Park and Alex Alderton, Roberto Amato, Jeffrey Barrett, Sonia Goncalves, Ewan Harrison, David K. Jackson, Ian Johnston, Dominic Kwiatkowski, Cordelia Langford, John Sillitoe on behalf of the Wellcome Sanger Institute COVID-19 Surveillance Team |
| EPI_ISL_2702753, EPI_ISL_2702754, EPI_ISL_2702755 | Lighthouse Lab in Glasgow | Wellcome Sanger Institute for the COVID-19 Genomics UK (COG-UK) Consortium | Harper VanSteenhouse, Yumi Kasai, David Gray, Carol Clugston, Anna Dominiczak and Alex Alderton, Roberto Amato, Jeffrey Barrett, Sonia Goncalves, Ewan Harrison, David K. Jackson, Ian Johnston, Dominic Kwiatkowski, Cordelia Langford, John Sillitoe on behalf of the Wellcome Sanger Institute COVID-19 Surveillance Team |
| EPI_ISL_2702756, EPI_ISL_2702757, | Lighthouse Lab in Alderley Park | Wellcome Sanger Institute for the COVID-19 Genomics UK (COG-UK) Consortium | Jacquelyn Wynn, Mairead Hyland, The Lighthouse Lab in Alderley Park and Alex Alderton, Roberto Amato, Jeffrey Barrett, Sonia Goncalves, Ewan Harrison, David K. Jackson, Ian Johnston, Dominic Kwiatkowski, Cordelia Langford, John Sillitoe on behalf of the Wellcome Sanger Institute COVID-19 Surveillance Team |

| | | | |
|---|---|---|---|
| EPI_ISL_2702758, EPI_ISL_2702759, EPI_ISL_2702760, EPI_ISL_2702761, EPI_ISL_2702762, EPI_ISL_2702763, EPI_ISL_2702764 | | | |
| EPI_ISL_2702765 | Lighthouse Lab in Glasgow | Wellcome Sanger Institute for the COVID-19 Genomics UK (COG-UK) Consortium | Harper VanSteenhouse, Yumi Kasai, David Gray, Carol Clugston, Anna Dominiczak and Alex Alderton, Roberto Amato, Jeffrey Barrett, Sonia Goncalves, Ewan Harrison, David K. Jackson, Ian Johnston, Dominic Kwiatkowski, Cordelia Langford, John Sillitoe on behalf of the Wellcome Sanger Institute COVID-19 Surveillance Team |
| EPI_ISL_2702766, EPI_ISL_2702767, EPI_ISL_2702768, EPI_ISL_2702769 | Lighthouse Lab in Alderley Park | Wellcome Sanger Institute for the COVID-19 Genomics UK (COG-UK) Consortium | Jacquelyn Wynn, Mairead Hyland, The Lighthouse Lab in Alderley Park and Alex Alderton, Roberto Amato, Jeffrey Barrett, Sonia Goncalves, Ewan Harrison, David K. Jackson, Ian Johnston, Dominic Kwiatkowski, Cordelia Langford, John Sillitoe on behalf of the Wellcome Sanger Institute COVID-19 Surveillance Team |
| EPI_ISL_2702770 | Lighthouse Lab in Glasgow | Wellcome Sanger Institute for the COVID-19 Genomics UK (COG-UK) Consortium | Harper VanSteenhouse, Yumi Kasai, David Gray, Carol Clugston, Anna Dominiczak and Alex Alderton, Roberto Amato, Jeffrey Barrett, Sonia Goncalves, Ewan Harrison, David K. Jackson, Ian Johnston, Dominic Kwiatkowski, Cordelia Langford, John Sillitoe on behalf of the Wellcome Sanger Institute COVID-19 Surveillance Team |
| EPI_ISL_2702771, EPI_ISL_2702772 | Lighthouse Lab in Alderley Park | Wellcome Sanger Institute for the COVID-19 Genomics UK (COG-UK) Consortium | Jacquelyn Wynn, Mairead Hyland, The Lighthouse Lab in Alderley Park and Alex Alderton, Roberto Amato, Jeffrey Barrett, Sonia Goncalves, Ewan Harrison, David K. Jackson, Ian Johnston, Dominic Kwiatkowski, Cordelia Langford, John Sillitoe on behalf of the Wellcome Sanger Institute COVID-19 Surveillance Team |
| EPI_ISL_2702773 | Lighthouse Lab in Glasgow | Wellcome Sanger Institute for the COVID-19 Genomics UK (COG-UK) Consortium | Harper VanSteenhouse, Yumi Kasai, David Gray, Carol Clugston, Anna Dominiczak and Alex Alderton, Roberto Amato, Jeffrey Barrett, Sonia Goncalves, Ewan Harrison, David K. Jackson, Ian Johnston, Dominic Kwiatkowski, Cordelia Langford, John Sillitoe on behalf of the Wellcome Sanger Institute COVID-19 Surveillance Team |
| EPI_ISL_2702774 | Lighthouse Lab in Alderley Park | Wellcome Sanger Institute for the COVID-19 Genomics UK (COG-UK) Consortium | Jacquelyn Wynn, Mairead Hyland, The Lighthouse Lab in Alderley Park and Alex Alderton, Roberto Amato, Jeffrey Barrett, Sonia Goncalves, Ewan Harrison, David K. Jackson, Ian Johnston, Dominic Kwiatkowski, Cordelia Langford, John Sillitoe on behalf of the Wellcome Sanger Institute COVID-19 Surveillance Team |
| EPI_ISL_2702775 | Lighthouse Lab in Glasgow | Wellcome Sanger Institute for the COVID-19 Genomics UK (COG-UK) Consortium | Harper VanSteenhouse, Yumi Kasai, David Gray, Carol Clugston, Anna Dominiczak and Alex Alderton, Roberto Amato, Jeffrey Barrett, Sonia Goncalves, Ewan Harrison, David K. Jackson, Ian Johnston, Dominic Kwiatkowski, Cordelia Langford, John Sillitoe on behalf of the Wellcome Sanger Institute COVID-19 Surveillance Team |
| EPI_ISL_2702776, EPI_ISL_2702777, EPI_ISL_2702778, EPI_ISL_2702779 | Lighthouse Lab in Alderley Park | Wellcome Sanger Institute for the COVID-19 Genomics UK (COG-UK) Consortium | Jacquelyn Wynn, Mairead Hyland, The Lighthouse Lab in Alderley Park and Alex Alderton, Roberto Amato, Jeffrey Barrett, Sonia Goncalves, Ewan Harrison, David K. Jackson, Ian Johnston, Dominic Kwiatkowski, Cordelia Langford, John Sillitoe on behalf of the Wellcome Sanger Institute COVID-19 Surveillance Team |
| EPI_ISL_2702780 | Lighthouse Lab in Glasgow | Wellcome Sanger Institute for the COVID-19 Genomics UK (COG-UK) Consortium | Harper VanSteenhouse, Yumi Kasai, David Gray, Carol Clugston, Anna Dominiczak and Alex Alderton, Roberto Amato, Jeffrey Barrett, Sonia Goncalves, Ewan Harrison, David K. Jackson, Ian Johnston, Dominic Kwiatkowski, Cordelia Langford, John Sillitoe on behalf of the Wellcome Sanger Institute COVID-19 Surveillance Team |
| EPI_ISL_2702781 | Lighthouse Lab in Alderley Park | Wellcome Sanger Institute for the COVID-19 Genomics UK (COG-UK) Consortium | Jacquelyn Wynn, Mairead Hyland, The Lighthouse Lab in Alderley Park and Alex Alderton, Roberto Amato, Jeffrey Barrett, Sonia Goncalves, Ewan Harrison, David K. Jackson, Ian Johnston, Dominic Kwiatkowski, Cordelia Langford, John Sillitoe on behalf of the Wellcome Sanger Institute COVID-19 Surveillance Team |
| EPI_ISL_2702782, EPI_ISL_2702783 | Lighthouse Lab in Glasgow | Wellcome Sanger Institute for the COVID-19 Genomics UK (COG-UK) Consortium | Harper VanSteenhouse, Yumi Kasai, David Gray, Carol Clugston, Anna Dominiczak and Alex Alderton, Roberto Amato, Jeffrey Barrett, Sonia Goncalves, Ewan Harrison, David K. Jackson, Ian Johnston, Dominic Kwiatkowski, Cordelia Langford, John Sillitoe on behalf of the Wellcome Sanger Institute COVID-19 Surveillance Team |
| EPI_ISL_2702784, EPI_ISL_2702785 | Lighthouse Lab in Alderley Park | Wellcome Sanger Institute for the COVID-19 Genomics UK (COG-UK) Consortium | Jacquelyn Wynn, Mairead Hyland, The Lighthouse Lab in Alderley Park and Alex Alderton, Roberto Amato, Jeffrey Barrett, Sonia Goncalves, Ewan Harrison, David K. Jackson, Ian Johnston, Dominic Kwiatkowski, Cordelia Langford, John Sillitoe on behalf of the Wellcome Sanger Institute COVID-19 Surveillance Team |
| EPI_ISL_2702786, EPI_ISL_2702787 | Lighthouse Lab in Glasgow | Wellcome Sanger Institute for the COVID-19 Genomics UK (COG-UK) Consortium | Harper VanSteenhouse, Yumi Kasai, David Gray, Carol Clugston, Anna Dominiczak and Alex Alderton, Roberto Amato, Jeffrey Barrett, Sonia Goncalves, Ewan Harrison, David K. Jackson, Ian Johnston, Dominic Kwiatkowski, Cordelia Langford, John Sillitoe on behalf of the Wellcome Sanger Institute COVID-19 Surveillance Team |
| EPI_ISL_2702788, EPI_ISL_2702789, EPI_ISL_2702790, EPI_ISL_2702791, EPI_ISL_2702792 | Lighthouse Lab in Alderley Park | Wellcome Sanger Institute for the COVID-19 Genomics UK (COG-UK) Consortium | Jacquelyn Wynn, Mairead Hyland, The Lighthouse Lab in Alderley Park and Alex Alderton, Roberto Amato, Jeffrey Barrett, Sonia Goncalves, Ewan Harrison, David K. Jackson, Ian Johnston, Dominic Kwiatkowski, Cordelia Langford, John Sillitoe on behalf of the Wellcome Sanger Institute COVID-19 Surveillance Team |
| EPI_ISL_2702793, EPI_ISL_2702794 | Lighthouse Lab in Glasgow | Wellcome Sanger Institute for the COVID-19 Genomics UK (COG-UK) Consortium | Harper VanSteenhouse, Yumi Kasai, David Gray, Carol Clugston, Anna Dominiczak and Alex Alderton, Roberto Amato, Jeffrey Barrett, Sonia Goncalves, Ewan Harrison, David K. Jackson, Ian Johnston, Dominic Kwiatkowski, Cordelia Langford, John Sillitoe on behalf of the Wellcome Sanger Institute COVID-19 Surveillance Team |
| EPI_ISL_2702795 | Lighthouse Lab in Alderley Park | Wellcome Sanger Institute for the COVID-19 Genomics UK (COG-UK) Consortium | Jacquelyn Wynn, Mairead Hyland, The Lighthouse Lab in Alderley Park and Alex Alderton, Roberto Amato, Jeffrey Barrett, Sonia Goncalves, Ewan Harrison, David K. Jackson, Ian Johnston, Dominic Kwiatkowski, Cordelia Langford, John Sillitoe on behalf of the Wellcome Sanger Institute COVID-19 Surveillance Team |
| EPI_ISL_2702796 | Lighthouse Lab in Glasgow | Wellcome Sanger Institute for the COVID-19 Genomics UK (COG-UK) Consortium | Harper VanSteenhouse, Yumi Kasai, David Gray, Carol Clugston, Anna Dominiczak and Alex Alderton, Roberto Amato, Jeffrey Barrett, Sonia Goncalves, Ewan Harrison, David K. Jackson, Ian Johnston, Dominic Kwiatkowski, Cordelia Langford, John Sillitoe on behalf of the Wellcome Sanger Institute COVID-19 Surveillance Team |
| EPI_ISL_2702797, EPI_ISL_2702798, EPI_ISL_2702799 | Lighthouse Lab in Alderley Park | Wellcome Sanger Institute for the COVID-19 Genomics UK (COG-UK) Consortium | Jacquelyn Wynn, Mairead Hyland, The Lighthouse Lab in Alderley Park and Alex Alderton, Roberto Amato, Jeffrey Barrett, Sonia Goncalves, Ewan Harrison, David K. Jackson, Ian Johnston, Dominic Kwiatkowski, Cordelia Langford, John Sillitoe on behalf of the Wellcome Sanger Institute COVID-19 Surveillance Team |
| EPI_ISL_2702800 | Lighthouse Lab in Glasgow | Wellcome Sanger Institute for the COVID-19 Genomics UK (COG-UK) Consortium | Harper VanSteenhouse, Yumi Kasai, David Gray, Carol Clugston, Anna Dominiczak and Alex Alderton, Roberto Amato, Jeffrey Barrett, Sonia Goncalves, Ewan Harrison, David K. Jackson, Ian Johnston, Dominic Kwiatkowski, Cordelia Langford, John Sillitoe on behalf of the Wellcome Sanger Institute COVID-19 Surveillance Team |
| EPI_ISL_2702801, EPI_ISL_2702802, EPI_ISL_2702803 | Lighthouse Lab in Alderley Park | Wellcome Sanger Institute for the COVID-19 Genomics UK (COG-UK) Consortium | Jacquelyn Wynn, Mairead Hyland, The Lighthouse Lab in Alderley Park and Alex Alderton, Roberto Amato, Jeffrey Barrett, Sonia Goncalves, Ewan Harrison, David K. Jackson, Ian Johnston, Dominic Kwiatkowski, Cordelia Langford, John Sillitoe on behalf of the Wellcome Sanger Institute COVID-19 Surveillance Team |
| EPI_ISL_2702804, EPI_ISL_2702805 | Lighthouse Lab in Glasgow | Wellcome Sanger Institute for the COVID-19 Genomics UK (COG-UK) Consortium | Harper VanSteenhouse, Yumi Kasai, David Gray, Carol Clugston, Anna Dominiczak and Alex Alderton, Roberto Amato, Jeffrey Barrett, Sonia Goncalves, Ewan Harrison, David K. Jackson, Ian Johnston, Dominic Kwiatkowski, Cordelia Langford, John Sillitoe on behalf of the Wellcome Sanger Institute COVID-19 Surveillance Team |
| EPI_ISL_2702806, EPI_ISL_2702807 | Lighthouse Lab in Alderley Park | Wellcome Sanger Institute for the COVID-19 Genomics UK (COG-UK) Consortium | Jacquelyn Wynn, Mairead Hyland, The Lighthouse Lab in Alderley Park and Alex Alderton, Roberto Amato, Jeffrey Barrett, Sonia Goncalves, Ewan Harrison, David K. Jackson, Ian Johnston, Dominic Kwiatkowski, Cordelia Langford, John Sillitoe on behalf of the Wellcome Sanger Institute COVID-19 Surveillance Team |
| EPI_ISL_2702808 | Lighthouse Lab in Glasgow | Wellcome Sanger Institute for the COVID-19 Genomics UK (COG-UK) Consortium | Harper VanSteenhouse, Yumi Kasai, David Gray, Carol Clugston, Anna Dominiczak and Alex Alderton, Roberto Amato, Jeffrey Barrett, Sonia Goncalves, Ewan Harrison, David K. Jackson, Ian Johnston, Dominic Kwiatkowski, Cordelia Langford, John Sillitoe on behalf of the Wellcome Sanger Institute COVID-19 Surveillance Team |
| EPI_ISL_2702809 | Lighthouse Lab in Alderley Park | Wellcome Sanger Institute for the COVID-19 Genomics UK (COG-UK) Consortium | Jacquelyn Wynn, Mairead Hyland, The Lighthouse Lab in Alderley Park and Alex Alderton, Roberto Amato, Jeffrey Barrett, Sonia Goncalves, Ewan Harrison, David K. Jackson, Ian Johnston, Dominic Kwiatkowski, Cordelia Langford, John Sillitoe on behalf of the Wellcome Sanger Institute COVID-19 Surveillance Team |
| EPI_ISL_2702810 | Lighthouse Lab in Glasgow | Wellcome Sanger Institute for the COVID-19 Genomics UK (COG-UK) Consortium | Harper VanSteenhouse, Yumi Kasai, David Gray, Carol Clugston, Anna Dominiczak and Alex Alderton, Roberto Amato, Jeffrey Barrett, Sonia Goncalves, Ewan Harrison, David K. Jackson, Ian Johnston, Dominic Kwiatkowski, Cordelia Langford, John Sillitoe on behalf of the Wellcome Sanger Institute COVID-19 Surveillance Team |
| EPI_ISL_2702811, EPI_ISL_2702812 | Lighthouse Lab in Alderley Park | Wellcome Sanger Institute for the COVID-19 Genomics UK (COG-UK) Consortium | Jacquelyn Wynn, Mairead Hyland, The Lighthouse Lab in Alderley Park and Alex Alderton, Roberto Amato, Jeffrey Barrett, Sonia Goncalves, Ewan Harrison, David K. Jackson, Ian Johnston, Dominic Kwiatkowski, Cordelia Langford, John Sillitoe on behalf of the Wellcome Sanger Institute COVID-19 Surveillance Team |
| EPI_ISL_2702813, EPI_ISL_2702814 | Lighthouse Lab in Milton Keynes | Wellcome Sanger Institute for the COVID-19 Genomics UK (COG-UK) Consortium | The Lighthouse Lab in Milton Keynes and Alex Alderton, Roberto Amato, Jeffrey Barrett, Sonia Goncalves, Ewan Harrison, David K. Jackson, Ian Johnston, Dominic Kwiatkowski, Cordelia Langford, John Sillitoe on behalf of the Wellcome Sanger Institute COVID-19 Surveillance Team |
| EPI_ISL_2702815 | Lighthouse Lab in Alderley Park | Wellcome Sanger Institute for the COVID-19 Genomics UK (COG-UK) Consortium | Jacquelyn Wynn, Mairead Hyland, The Lighthouse Lab in Alderley Park and Alex Alderton, Roberto Amato, Jeffrey Barrett, Sonia Goncalves, Ewan Harrison, David K. Jackson, Ian Johnston, Dominic Kwiatkowski, Cordelia Langford, John Sillitoe on behalf of the Wellcome Sanger Institute COVID-19 Surveillance Team |
| EPI_ISL_2702816 | Lighthouse Lab in Milton Keynes | Wellcome Sanger Institute for the COVID-19 Genomics UK (COG-UK) Consortium | The Lighthouse Lab in Milton Keynes and Alex Alderton, Roberto Amato, Jeffrey Barrett, Sonia Goncalves, Ewan Harrison, David K. Jackson, Ian Johnston, Dominic Kwiatkowski, Cordelia Langford, John Sillitoe on behalf of the Wellcome Sanger Institute COVID-19 Surveillance Team |
| EPI_ISL_2702817 | Lighthouse Lab in Alderley Park | Wellcome Sanger Institute for the COVID-19 Genomics UK (COG-UK) Consortium | Jacquelyn Wynn, Mairead Hyland, The Lighthouse Lab in Alderley Park and Alex Alderton, Roberto Amato, Jeffrey Barrett, Sonia Goncalves, Ewan Harrison, David K. Jackson, Ian Johnston, Dominic Kwiatkowski, Cordelia Langford, John Sillitoe on behalf of the Wellcome Sanger Institute COVID-19 Surveillance Team |
| EPI_ISL_2702818, EPI_ISL_2702819, EPI_ISL_2702820, EPI_ISL_2702821 | Lighthouse Lab in Milton Keynes | Wellcome Sanger Institute for the COVID-19 Genomics UK (COG-UK) Consortium | The Lighthouse Lab in Milton Keynes and Alex Alderton, Roberto Amato, Jeffrey Barrett, Sonia Goncalves, Ewan Harrison, David K. Jackson, Ian Johnston, Dominic Kwiatkowski, Cordelia Langford, John Sillitoe on behalf of the Wellcome Sanger Institute COVID-19 Surveillance Team |
| EPI_ISL_2702822, EPI_ISL_2702823 | Lighthouse Lab in Alderley Park | Wellcome Sanger Institute for the COVID-19 Genomics UK (COG-UK) Consortium | Jacquelyn Wynn, Mairead Hyland, The Lighthouse Lab in Alderley Park and Alex Alderton, Roberto Amato, Jeffrey Barrett, Sonia Goncalves, Ewan Harrison, David K. Jackson, Ian Johnston, Dominic Kwiatkowski, Cordelia Langford, John Sillitoe on behalf of the Wellcome Sanger Institute COVID-19 Surveillance Team |
| EPI_ISL_2702825, EPI_ISL_2702826, EPI_ISL_2702827, EPI_ISL_2702828 | Lighthouse Lab in Milton Keynes | Wellcome Sanger Institute for the COVID-19 Genomics UK (COG-UK) Consortium | The Lighthouse Lab in Milton Keynes and Alex Alderton, Roberto Amato, Jeffrey Barrett, Sonia Goncalves, Ewan Harrison, David K. Jackson, Ian Johnston, Dominic Kwiatkowski, Cordelia Langford, John Sillitoe on behalf of the Wellcome Sanger Institute COVID-19 Surveillance Team |
| EPI_ISL_2702829 | Lighthouse Lab in Alderley Park | Wellcome Sanger Institute for the COVID-19 Genomics UK (COG-UK) Consortium | Jacquelyn Wynn, Mairead Hyland, The Lighthouse Lab in Alderley Park and Alex Alderton, Roberto Amato, Jeffrey Barrett, Sonia Goncalves, Ewan Harrison, David K. Jackson, Ian Johnston, Dominic Kwiatkowski, Cordelia Langford, John Sillitoe on behalf of the Wellcome Sanger Institute COVID-19 Surveillance Team |
| EPI_ISL_2702830, EPI_ISL_2702831, EPI_ISL_2702832, EPI_ISL_2702833, EPI_ISL_2702834, EPI_ISL_2702835, EPI_ISL_2702836, | Lighthouse Lab in Milton Keynes | Wellcome Sanger Institute for the COVID-19 Genomics UK (COG-UK) Consortium | The Lighthouse Lab in Milton Keynes and Alex Alderton, Roberto Amato, Jeffrey Barrett, Sonia Goncalves, Ewan Harrison, David K. Jackson, Ian Johnston, Dominic Kwiatkowski, Cordelia Langford, John Sillitoe on behalf of the Wellcome Sanger Institute COVID-19 Surveillance Team |

| | | | |
|---|---|---|---|
| EPI_ISL_2702837, EPI_ISL_2702838, EPI_ISL_2702839 | | | |
| EPI_ISL_2702840 | Lighthouse Lab in Alderley Park | Wellcome Sanger Institute for the COVID-19 Genomics UK (COG-UK) Consortium | Jacquelyn Wynn, Mairead Hyland, The Lighthouse Lab in Alderley Park and Alex Alderton, Roberto Amato, Jeffrey Barrett, Sonia Goncalves, Ewan Harrison, David K. Jackson, Ian Johnston, Dominic Kwiatkowski, Cordelia Langford, John Sillitoe on behalf of the Wellcome Sanger Institute COVID-19 Surveillance Team |
| EPI_ISL_2702841, EPI_ISL_2702842 | Lighthouse Lab in Milton Keynes | Wellcome Sanger Institute for the COVID-19 Genomics UK (COG-UK) Consortium | The Lighthouse Lab in Milton Keynes and Alex Alderton, Roberto Amato, Jeffrey Barrett, Sonia Goncalves, Ewan Harrison, David K. Jackson, Ian Johnston, Dominic Kwiatkowski, Cordelia Langford, John Sillitoe on behalf of the Wellcome Sanger Institute COVID-19 Surveillance Team |
| EPI_ISL_2702843 | Lighthouse Lab in Alderley Park | Wellcome Sanger Institute for the COVID-19 Genomics UK (COG-UK) Consortium | Jacquelyn Wynn, Mairead Hyland, The Lighthouse Lab in Alderley Park and Alex Alderton, Roberto Amato, Jeffrey Barrett, Sonia Goncalves, Ewan Harrison, David K. Jackson, Ian Johnston, Dominic Kwiatkowski, Cordelia Langford, John Sillitoe on behalf of the Wellcome Sanger Institute COVID-19 Surveillance Team |
| EPI_ISL_2702845, EPI_ISL_2702846 | Lighthouse Lab in Milton Keynes | Wellcome Sanger Institute for the COVID-19 Genomics UK (COG-UK) Consortium | The Lighthouse Lab in Milton Keynes and Alex Alderton, Roberto Amato, Jeffrey Barrett, Sonia Goncalves, Ewan Harrison, David K. Jackson, Ian Johnston, Dominic Kwiatkowski, Cordelia Langford, John Sillitoe on behalf of the Wellcome Sanger Institute COVID-19 Surveillance Team |
| EPI_ISL_2702847 | Lighthouse Lab in Alderley Park | Wellcome Sanger Institute for the COVID-19 Genomics UK (COG-UK) Consortium | Jacquelyn Wynn, Mairead Hyland, The Lighthouse Lab in Alderley Park and Alex Alderton, Roberto Amato, Jeffrey Barrett, Sonia Goncalves, Ewan Harrison, David K. Jackson, Ian Johnston, Dominic Kwiatkowski, Cordelia Langford, John Sillitoe on behalf of the Wellcome Sanger Institute COVID-19 Surveillance Team |
| EPI_ISL_2702848, EPI_ISL_2702849 | Lighthouse Lab in Milton Keynes | Wellcome Sanger Institute for the COVID-19 Genomics UK (COG-UK) Consortium | The Lighthouse Lab in Milton Keynes and Alex Alderton, Roberto Amato, Jeffrey Barrett, Sonia Goncalves, Ewan Harrison, David K. Jackson, Ian Johnston, Dominic Kwiatkowski, Cordelia Langford, John Sillitoe on behalf of the Wellcome Sanger Institute COVID-19 Surveillance Team |
| EPI_ISL_2702850 | Lighthouse Lab in Alderley Park | Wellcome Sanger Institute for the COVID-19 Genomics UK (COG-UK) Consortium | Jacquelyn Wynn, Mairead Hyland, The Lighthouse Lab in Alderley Park and Alex Alderton, Roberto Amato, Jeffrey Barrett, Sonia Goncalves, Ewan Harrison, David K. Jackson, Ian Johnston, Dominic Kwiatkowski, Cordelia Langford, John Sillitoe on behalf of the Wellcome Sanger Institute COVID-19 Surveillance Team |
| EPI_ISL_2702851, EPI_ISL_2702852, EPI_ISL_2702853 | Lighthouse Lab in Milton Keynes | Wellcome Sanger Institute for the COVID-19 Genomics UK (COG-UK) Consortium | The Lighthouse Lab in Milton Keynes and Alex Alderton, Roberto Amato, Jeffrey Barrett, Sonia Goncalves, Ewan Harrison, David K. Jackson, Ian Johnston, Dominic Kwiatkowski, Cordelia Langford, John Sillitoe on behalf of the Wellcome Sanger Institute COVID-19 Surveillance Team |
| EPI_ISL_2702854 | Lighthouse Lab in Alderley Park | Wellcome Sanger Institute for the COVID-19 Genomics UK (COG-UK) Consortium | Jacquelyn Wynn, Mairead Hyland, The Lighthouse Lab in Alderley Park and Alex Alderton, Roberto Amato, Jeffrey Barrett, Sonia Goncalves, Ewan Harrison, David K. Jackson, Ian Johnston, Dominic Kwiatkowski, Cordelia Langford, John Sillitoe on behalf of the Wellcome Sanger Institute COVID-19 Surveillance Team |
| EPI_ISL_2702855 | Lighthouse Lab in Milton Keynes | Wellcome Sanger Institute for the COVID-19 Genomics UK (COG-UK) Consortium | The Lighthouse Lab in Milton Keynes and Alex Alderton, Roberto Amato, Jeffrey Barrett, Sonia Goncalves, Ewan Harrison, David K. Jackson, Ian Johnston, Dominic Kwiatkowski, Cordelia Langford, John Sillitoe on behalf of the Wellcome Sanger Institute COVID-19 Surveillance Team |
| EPI_ISL_2702857, EPI_ISL_2702858, EPI_ISL_2702859, EPI_ISL_2702860 | Lighthouse Lab in Alderley Park | Wellcome Sanger Institute for the COVID-19 Genomics UK (COG-UK) Consortium | Jacquelyn Wynn, Mairead Hyland, The Lighthouse Lab in Alderley Park and Alex Alderton, Roberto Amato, Jeffrey Barrett, Sonia Goncalves, Ewan Harrison, David K. Jackson, Ian Johnston, Dominic Kwiatkowski, Cordelia Langford, John Sillitoe on behalf of the Wellcome Sanger Institute COVID-19 Surveillance Team |
| EPI_ISL_2702861, EPI_ISL_2702862, EPI_ISL_2702863 | Lighthouse Lab in Milton Keynes | Wellcome Sanger Institute for the COVID-19 Genomics UK (COG-UK) Consortium | The Lighthouse Lab in Milton Keynes and Alex Alderton, Roberto Amato, Jeffrey Barrett, Sonia Goncalves, Ewan Harrison, David K. Jackson, Ian Johnston, Dominic Kwiatkowski, Cordelia Langford, John Sillitoe on behalf of the Wellcome Sanger Institute COVID-19 Surveillance Team |
| EPI_ISL_2702864, EPI_ISL_2702865, EPI_ISL_2702866 | Lighthouse Lab in Alderley Park | Wellcome Sanger Institute for the COVID-19 Genomics UK (COG-UK) Consortium | Jacquelyn Wynn, Mairead Hyland, The Lighthouse Lab in Alderley Park and Alex Alderton, Roberto Amato, Jeffrey Barrett, Sonia Goncalves, Ewan Harrison, David K. Jackson, Ian Johnston, Dominic Kwiatkowski, Cordelia Langford, John Sillitoe on behalf of the Wellcome Sanger Institute COVID-19 Surveillance Team |
| EPI_ISL_2702867 | Lighthouse Lab in Milton Keynes | Wellcome Sanger Institute for the COVID-19 Genomics UK (COG-UK) Consortium | The Lighthouse Lab in Milton Keynes and Alex Alderton, Roberto Amato, Jeffrey Barrett, Sonia Goncalves, Ewan Harrison, David K. Jackson, Ian Johnston, Dominic Kwiatkowski, Cordelia Langford, John Sillitoe on behalf of the Wellcome Sanger Institute COVID-19 Surveillance Team |
| EPI_ISL_2702868 | Lighthouse Lab in Alderley Park | Wellcome Sanger Institute for the COVID-19 Genomics UK (COG-UK) Consortium | Jacquelyn Wynn, Mairead Hyland, The Lighthouse Lab in Alderley Park and Alex Alderton, Roberto Amato, Jeffrey Barrett, Sonia Goncalves, Ewan Harrison, David K. Jackson, Ian Johnston, Dominic Kwiatkowski, Cordelia Langford, John Sillitoe on behalf of the Wellcome Sanger Institute COVID-19 Surveillance Team |
| EPI_ISL_2702869, EPI_ISL_2702870, EPI_ISL_2702871, EPI_ISL_2702872, EPI_ISL_2702873, EPI_ISL_2702874, EPI_ISL_2702875 | Lighthouse Lab in Milton Keynes | Wellcome Sanger Institute for the COVID-19 Genomics UK (COG-UK) Consortium | The Lighthouse Lab in Milton Keynes and Alex Alderton, Roberto Amato, Jeffrey Barrett, Sonia Goncalves, Ewan Harrison, David K. Jackson, Ian Johnston, Dominic Kwiatkowski, Cordelia Langford, John Sillitoe on behalf of the Wellcome Sanger Institute COVID-19 Surveillance Team |
| EPI_ISL_2702876 | Lighthouse Lab in Alderley Park | Wellcome Sanger Institute for the COVID-19 Genomics UK (COG-UK) Consortium | Jacquelyn Wynn, Mairead Hyland, The Lighthouse Lab in Alderley Park and Alex Alderton, Roberto Amato, Jeffrey Barrett, Sonia Goncalves, Ewan Harrison, David K. Jackson, Ian Johnston, Dominic Kwiatkowski, Cordelia Langford, John Sillitoe on behalf of the Wellcome Sanger Institute COVID-19 Surveillance Team |
| EPI_ISL_2702877, EPI_ISL_2702878, EPI_ISL_2702879, EPI_ISL_2702880, EPI_ISL_2702881, EPI_ISL_2702882, EPI_ISL_2702883 | Lighthouse Lab in Milton Keynes | Wellcome Sanger Institute for the COVID-19 Genomics UK (COG-UK) Consortium | The Lighthouse Lab in Milton Keynes and Alex Alderton, Roberto Amato, Jeffrey Barrett, Sonia Goncalves, Ewan Harrison, David K. Jackson, Ian Johnston, Dominic Kwiatkowski, Cordelia Langford, John Sillitoe on behalf of the Wellcome Sanger Institute COVID-19 Surveillance Team |
| EPI_ISL_2702884 | Lighthouse Lab in Alderley Park | Wellcome Sanger Institute for the COVID-19 Genomics UK (COG-UK) Consortium | Jacquelyn Wynn, Mairead Hyland, The Lighthouse Lab in Alderley Park and Alex Alderton, Roberto Amato, Jeffrey Barrett, Sonia Goncalves, Ewan Harrison, David K. Jackson, Ian Johnston, Dominic Kwiatkowski, Cordelia Langford, John Sillitoe on behalf of the Wellcome Sanger Institute COVID-19 Surveillance Team |
| EPI_ISL_2702885, EPI_ISL_2702886, EPI_ISL_2702887, EPI_ISL_2702888, EPI_ISL_2702889 | Lighthouse Lab in Milton Keynes | Wellcome Sanger Institute for the COVID-19 Genomics UK (COG-UK) Consortium | The Lighthouse Lab in Milton Keynes and Alex Alderton, Roberto Amato, Jeffrey Barrett, Sonia Goncalves, Ewan Harrison, David K. Jackson, Ian Johnston, Dominic Kwiatkowski, Cordelia Langford, John Sillitoe on behalf of the Wellcome Sanger Institute COVID-19 Surveillance Team |
| EPI_ISL_2702890 | Lighthouse Lab in Alderley Park | Wellcome Sanger Institute for the COVID-19 Genomics UK (COG-UK) Consortium | Jacquelyn Wynn, Mairead Hyland, The Lighthouse Lab in Alderley Park and Alex Alderton, Roberto Amato, Jeffrey Barrett, Sonia Goncalves, Ewan Harrison, David K. Jackson, Ian Johnston, Dominic Kwiatkowski, Cordelia Langford, John Sillitoe on behalf of the Wellcome Sanger Institute COVID-19 Surveillance Team |
| EPI_ISL_2702891, EPI_ISL_2702892, EPI_ISL_2702893, EPI_ISL_2702894, EPI_ISL_2702895, EPI_ISL_2702896 | Lighthouse Lab in Milton Keynes | Wellcome Sanger Institute for the COVID-19 Genomics UK (COG-UK) Consortium | The Lighthouse Lab in Milton Keynes and Alex Alderton, Roberto Amato, Jeffrey Barrett, Sonia Goncalves, Ewan Harrison, David K. Jackson, Ian Johnston, Dominic Kwiatkowski, Cordelia Langford, John Sillitoe on behalf of the Wellcome Sanger Institute COVID-19 Surveillance Team |
| EPI_ISL_2702897 | Lighthouse Lab in Alderley Park | Wellcome Sanger Institute for the COVID-19 Genomics UK (COG-UK) Consortium | Jacquelyn Wynn, Mairead Hyland, The Lighthouse Lab in Alderley Park and Alex Alderton, Roberto Amato, Jeffrey Barrett, Sonia Goncalves, Ewan Harrison, David K. Jackson, Ian Johnston, Dominic Kwiatkowski, Cordelia Langford, John Sillitoe on behalf of the Wellcome Sanger Institute COVID-19 Surveillance Team |
| EPI_ISL_2702898, EPI_ISL_2702899, EPI_ISL_2702900, EPI_ISL_2702901, EPI_ISL_2702902, EPI_ISL_2702903, EPI_ISL_2702904 | Lighthouse Lab in Milton Keynes | Wellcome Sanger Institute for the COVID-19 Genomics UK (COG-UK) Consortium | The Lighthouse Lab in Milton Keynes and Alex Alderton, Roberto Amato, Jeffrey Barrett, Sonia Goncalves, Ewan Harrison, David K. Jackson, Ian Johnston, Dominic Kwiatkowski, Cordelia Langford, John Sillitoe on behalf of the Wellcome Sanger Institute COVID-19 Surveillance Team |
| EPI_ISL_2702905 | Lighthouse Lab in Alderley Park | Wellcome Sanger Institute for the COVID-19 Genomics UK (COG-UK) Consortium | Jacquelyn Wynn, Mairead Hyland, The Lighthouse Lab in Alderley Park and Alex Alderton, Roberto Amato, Jeffrey Barrett, Sonia Goncalves, Ewan Harrison, David K. Jackson, Ian Johnston, Dominic Kwiatkowski, Cordelia Langford, John Sillitoe on behalf of the Wellcome Sanger Institute COVID-19 Surveillance Team |
| EPI_ISL_2702906, EPI_ISL_2702907, EPI_ISL_2702908, EPI_ISL_2702909 | Lighthouse Lab in Milton Keynes | Wellcome Sanger Institute for the COVID-19 Genomics UK (COG-UK) Consortium | The Lighthouse Lab in Milton Keynes and Alex Alderton, Roberto Amato, Jeffrey Barrett, Sonia Goncalves, Ewan Harrison, David K. Jackson, Ian Johnston, Dominic Kwiatkowski, Cordelia Langford, John Sillitoe on behalf of the Wellcome Sanger Institute COVID-19 Surveillance Team |
| EPI_ISL_2702910 | Lighthouse Lab in Alderley Park | Wellcome Sanger Institute for the COVID-19 Genomics UK (COG-UK) Consortium | Jacquelyn Wynn, Mairead Hyland, The Lighthouse Lab in Alderley Park and Alex Alderton, Roberto Amato, Jeffrey Barrett, Sonia Goncalves, Ewan Harrison, David K. Jackson, Ian Johnston, Dominic Kwiatkowski, Cordelia Langford, John Sillitoe on behalf of the Wellcome Sanger Institute COVID-19 Surveillance Team |
| EPI_ISL_2702911, EPI_ISL_2702912, EPI_ISL_2702913, EPI_ISL_2702914, EPI_ISL_2702915 | Lighthouse Lab in Milton Keynes | Wellcome Sanger Institute for the COVID-19 Genomics UK (COG-UK) Consortium | The Lighthouse Lab in Milton Keynes and Alex Alderton, Roberto Amato, Jeffrey Barrett, Sonia Goncalves, Ewan Harrison, David K. Jackson, Ian Johnston, Dominic Kwiatkowski, Cordelia Langford, John Sillitoe on behalf of the Wellcome Sanger Institute COVID-19 Surveillance Team |
| EPI_ISL_2702916 | Lighthouse Lab in Alderley Park | Wellcome Sanger Institute for the COVID-19 Genomics UK (COG-UK) Consortium | Jacquelyn Wynn, Mairead Hyland, The Lighthouse Lab in Alderley Park and Alex Alderton, Roberto Amato, Jeffrey Barrett, Sonia Goncalves, Ewan Harrison, David K. Jackson, Ian Johnston, Dominic Kwiatkowski, Cordelia Langford, John Sillitoe on behalf of the Wellcome Sanger Institute COVID-19 Surveillance Team |
| EPI_ISL_2702917, EPI_ISL_2702918, EPI_ISL_2702919, EPI_ISL_2702920 | Lighthouse Lab in Milton Keynes | Wellcome Sanger Institute for the COVID-19 Genomics UK (COG-UK) Consortium | The Lighthouse Lab in Milton Keynes and Alex Alderton, Roberto Amato, Jeffrey Barrett, Sonia Goncalves, Ewan Harrison, David K. Jackson, Ian Johnston, Dominic Kwiatkowski, Cordelia Langford, John Sillitoe on behalf of the Wellcome Sanger Institute COVID-19 Surveillance Team |

| | | | |
|---|---|---|---|
| EPI_ISL_2702921 | Lighthouse Lab in Alderley Park | Wellcome Sanger Institute for the COVID-19 Genomics UK (COG-UK) Consortium | Jacquelyn Wynn, Mairead Hyland, The Lighthouse Lab in Alderley Park and Alex Alderton, Roberto Amato, Jeffrey Barrett, Sonia Goncalves, Ewan Harrison, David K. Jackson, Ian Johnston, Dominic Kwiatkowski, Cordelia Langford, John Sillitoe on behalf of the Wellcome Sanger Institute COVID-19 Surveillance Team |
| EPI_ISL_2702922 | Lighthouse Lab in Milton Keynes | Wellcome Sanger Institute for the COVID-19 Genomics UK (COG-UK) Consortium | The Lighthouse Lab in Milton Keynes and Alex Alderton, Roberto Amato, Jeffrey Barrett, Sonia Goncalves, Ewan Harrison, David K. Jackson, Ian Johnston, Dominic Kwiatkowski, Cordelia Langford, John Sillitoe on behalf of the Wellcome Sanger Institute COVID-19 Surveillance Team |
| EPI_ISL_2702923, EPI_ISL_2702924 | Lighthouse Lab in Alderley Park | Wellcome Sanger Institute for the COVID-19 Genomics UK (COG-UK) Consortium | Jacquelyn Wynn, Mairead Hyland, The Lighthouse Lab in Alderley Park and Alex Alderton, Roberto Amato, Jeffrey Barrett, Sonia Goncalves, Ewan Harrison, David K. Jackson, Ian Johnston, Dominic Kwiatkowski, Cordelia Langford, John Sillitoe on behalf of the Wellcome Sanger Institute COVID-19 Surveillance Team |
| EPI_ISL_2702925 | Lighthouse Lab in Milton Keynes | Wellcome Sanger Institute for the COVID-19 Genomics UK (COG-UK) Consortium | The Lighthouse Lab in Milton Keynes and Alex Alderton, Roberto Amato, Jeffrey Barrett, Sonia Goncalves, Ewan Harrison, David K. Jackson, Ian Johnston, Dominic Kwiatkowski, Cordelia Langford, John Sillitoe on behalf of the Wellcome Sanger Institute COVID-19 Surveillance Team |
| EPI_ISL_2702926, EPI_ISL_2702927 | Lighthouse Lab in Alderley Park | Wellcome Sanger Institute for the COVID-19 Genomics UK (COG-UK) Consortium | Jacquelyn Wynn, Mairead Hyland, The Lighthouse Lab in Alderley Park and Alex Alderton, Roberto Amato, Jeffrey Barrett, Sonia Goncalves, Ewan Harrison, David K. Jackson, Ian Johnston, Dominic Kwiatkowski, Cordelia Langford, John Sillitoe on behalf of the Wellcome Sanger Institute COVID-19 Surveillance Team |
| EPI_ISL_2702928, EPI_ISL_2702929, EPI_ISL_2702930, EPI_ISL_2702931, EPI_ISL_2702932, EPI_ISL_2702933 | Lighthouse Lab in Milton Keynes | Wellcome Sanger Institute for the COVID-19 Genomics UK (COG-UK) Consortium | The Lighthouse Lab in Milton Keynes and Alex Alderton, Roberto Amato, Jeffrey Barrett, Sonia Goncalves, Ewan Harrison, David K. Jackson, Ian Johnston, Dominic Kwiatkowski, Cordelia Langford, John Sillitoe on behalf of the Wellcome Sanger Institute COVID-19 Surveillance Team |
| EPI_ISL_2702934 | Lighthouse Lab in Alderley Park | Wellcome Sanger Institute for the COVID-19 Genomics UK (COG-UK) Consortium | Jacquelyn Wynn, Mairead Hyland, The Lighthouse Lab in Alderley Park and Alex Alderton, Roberto Amato, Jeffrey Barrett, Sonia Goncalves, Ewan Harrison, David K. Jackson, Ian Johnston, Dominic Kwiatkowski, Cordelia Langford, John Sillitoe on behalf of the Wellcome Sanger Institute COVID-19 Surveillance Team |
| EPI_ISL_2702935, EPI_ISL_2702936, EPI_ISL_2702937, EPI_ISL_2702938, EPI_ISL_2702939, EPI_ISL_2702940, EPI_ISL_2702941, EPI_ISL_2702942 | Lighthouse Lab in Milton Keynes | Wellcome Sanger Institute for the COVID-19 Genomics UK (COG-UK) Consortium | The Lighthouse Lab in Milton Keynes and Alex Alderton, Roberto Amato, Jeffrey Barrett, Sonia Goncalves, Ewan Harrison, David K. Jackson, Ian Johnston, Dominic Kwiatkowski, Cordelia Langford, John Sillitoe on behalf of the Wellcome Sanger Institute COVID-19 Surveillance Team |
| EPI_ISL_2702943 | Lighthouse Lab in Alderley Park | Wellcome Sanger Institute for the COVID-19 Genomics UK (COG-UK) Consortium | Jacquelyn Wynn, Mairead Hyland, The Lighthouse Lab in Alderley Park and Alex Alderton, Roberto Amato, Jeffrey Barrett, Sonia Goncalves, Ewan Harrison, David K. Jackson, Ian Johnston, Dominic Kwiatkowski, Cordelia Langford, John Sillitoe on behalf of the Wellcome Sanger Institute COVID-19 Surveillance Team |
| EPI_ISL_2702944 | Lighthouse Lab in Milton Keynes | Wellcome Sanger Institute for the COVID-19 Genomics UK (COG-UK) Consortium | The Lighthouse Lab in Milton Keynes and Alex Alderton, Roberto Amato, Jeffrey Barrett, Sonia Goncalves, Ewan Harrison, David K. Jackson, Ian Johnston, Dominic Kwiatkowski, Cordelia Langford, John Sillitoe on behalf of the Wellcome Sanger Institute COVID-19 Surveillance Team |
| EPI_ISL_2702945 | Lighthouse Lab in Alderley Park | Wellcome Sanger Institute for the COVID-19 Genomics UK (COG-UK) Consortium | Jacquelyn Wynn, Mairead Hyland, The Lighthouse Lab in Alderley Park and Alex Alderton, Roberto Amato, Jeffrey Barrett, Sonia Goncalves, Ewan Harrison, David K. Jackson, Ian Johnston, Dominic Kwiatkowski, Cordelia Langford, John Sillitoe on behalf of the Wellcome Sanger Institute COVID-19 Surveillance Team |
| EPI_ISL_2702946, EPI_ISL_2702947, EPI_ISL_2702948, EPI_ISL_2702949 | Lighthouse Lab in Milton Keynes | Wellcome Sanger Institute for the COVID-19 Genomics UK (COG-UK) Consortium | The Lighthouse Lab in Milton Keynes and Alex Alderton, Roberto Amato, Jeffrey Barrett, Sonia Goncalves, Ewan Harrison, David K. Jackson, Ian Johnston, Dominic Kwiatkowski, Cordelia Langford, John Sillitoe on behalf of the Wellcome Sanger Institute COVID-19 Surveillance Team |
| EPI_ISL_2702950 | Lighthouse Lab in Alderley Park | Wellcome Sanger Institute for the COVID-19 Genomics UK (COG-UK) Consortium | Jacquelyn Wynn, Mairead Hyland, The Lighthouse Lab in Alderley Park and Alex Alderton, Roberto Amato, Jeffrey Barrett, Sonia Goncalves, Ewan Harrison, David K. Jackson, Ian Johnston, Dominic Kwiatkowski, Cordelia Langford, John Sillitoe on behalf of the Wellcome Sanger Institute COVID-19 Surveillance Team |
| EPI_ISL_2702951, EPI_ISL_2702952, EPI_ISL_2702953, EPI_ISL_2702954, EPI_ISL_2702955, EPI_ISL_2702956, EPI_ISL_2702957 | Lighthouse Lab in Milton Keynes | Wellcome Sanger Institute for the COVID-19 Genomics UK (COG-UK) Consortium | The Lighthouse Lab in Milton Keynes and Alex Alderton, Roberto Amato, Jeffrey Barrett, Sonia Goncalves, Ewan Harrison, David K. Jackson, Ian Johnston, Dominic Kwiatkowski, Cordelia Langford, John Sillitoe on behalf of the Wellcome Sanger Institute COVID-19 Surveillance Team |
| EPI_ISL_2702958 | Lighthouse Lab in Alderley Park | Wellcome Sanger Institute for the COVID-19 Genomics UK (COG-UK) Consortium | Jacquelyn Wynn, Mairead Hyland, The Lighthouse Lab in Alderley Park and Alex Alderton, Roberto Amato, Jeffrey Barrett, Sonia Goncalves, Ewan Harrison, David K. Jackson, Ian Johnston, Dominic Kwiatkowski, Cordelia Langford, John Sillitoe on behalf of the Wellcome Sanger Institute COVID-19 Surveillance Team |
| EPI_ISL_2702959, EPI_ISL_2702960, EPI_ISL_2702961 | Lighthouse Lab in Milton Keynes | Wellcome Sanger Institute for the COVID-19 Genomics UK (COG-UK) Consortium | The Lighthouse Lab in Milton Keynes and Alex Alderton, Roberto Amato, Jeffrey Barrett, Sonia Goncalves, Ewan Harrison, David K. Jackson, Ian Johnston, Dominic Kwiatkowski, Cordelia Langford, John Sillitoe on behalf of the Wellcome Sanger Institute COVID-19 Surveillance Team |
| EPI_ISL_2702962 | Lighthouse Lab in Alderley Park | Wellcome Sanger Institute for the COVID-19 Genomics UK (COG-UK) Consortium | Jacquelyn Wynn, Mairead Hyland, The Lighthouse Lab in Alderley Park and Alex Alderton, Roberto Amato, Jeffrey Barrett, Sonia Goncalves, Ewan Harrison, David K. Jackson, Ian Johnston, Dominic Kwiatkowski, Cordelia Langford, John Sillitoe on behalf of the Wellcome Sanger Institute COVID-19 Surveillance Team |
| EPI_ISL_2702963 | Lighthouse Lab in Milton Keynes | Wellcome Sanger Institute for the COVID-19 Genomics UK (COG-UK) Consortium | The Lighthouse Lab in Milton Keynes and Alex Alderton, Roberto Amato, Jeffrey Barrett, Sonia Goncalves, Ewan Harrison, David K. Jackson, Ian Johnston, Dominic Kwiatkowski, Cordelia Langford, John Sillitoe on behalf of the Wellcome Sanger Institute COVID-19 Surveillance Team |
| EPI_ISL_2702964 | Lighthouse Lab in Alderley Park | Wellcome Sanger Institute for the COVID-19 Genomics UK (COG-UK) Consortium | Jacquelyn Wynn, Mairead Hyland, The Lighthouse Lab in Alderley Park and Alex Alderton, Roberto Amato, Jeffrey Barrett, Sonia Goncalves, Ewan Harrison, David K. Jackson, Ian Johnston, Dominic Kwiatkowski, Cordelia Langford, John Sillitoe on behalf of the Wellcome Sanger Institute COVID-19 Surveillance Team |
| EPI_ISL_2702965, EPI_ISL_2702966, EPI_ISL_2702967 | Lighthouse Lab in Milton Keynes | Wellcome Sanger Institute for the COVID-19 Genomics UK (COG-UK) Consortium | The Lighthouse Lab in Milton Keynes and Alex Alderton, Roberto Amato, Jeffrey Barrett, Sonia Goncalves, Ewan Harrison, David K. Jackson, Ian Johnston, Dominic Kwiatkowski, Cordelia Langford, John Sillitoe on behalf of the Wellcome Sanger Institute COVID-19 Surveillance Team |
| EPI_ISL_2702968 | Lighthouse Lab in Alderley Park | Wellcome Sanger Institute for the COVID-19 Genomics UK (COG-UK) Consortium | Jacquelyn Wynn, Mairead Hyland, The Lighthouse Lab in Alderley Park and Alex Alderton, Roberto Amato, Jeffrey Barrett, Sonia Goncalves, Ewan Harrison, David K. Jackson, Ian Johnston, Dominic Kwiatkowski, Cordelia Langford, John Sillitoe on behalf of the Wellcome Sanger Institute COVID-19 Surveillance Team |
| EPI_ISL_2702969, EPI_ISL_2702970, EPI_ISL_2702971, EPI_ISL_2702972, EPI_ISL_2702973, EPI_ISL_2702974, EPI_ISL_2702975 | Lighthouse Lab in Milton Keynes | Wellcome Sanger Institute for the COVID-19 Genomics UK (COG-UK) Consortium | The Lighthouse Lab in Milton Keynes and Alex Alderton, Roberto Amato, Jeffrey Barrett, Sonia Goncalves, Ewan Harrison, David K. Jackson, Ian Johnston, Dominic Kwiatkowski, Cordelia Langford, John Sillitoe on behalf of the Wellcome Sanger Institute COVID-19 Surveillance Team |
| EPI_ISL_2702977 | Lighthouse Lab in Alderley Park | Wellcome Sanger Institute for the COVID-19 Genomics UK (COG-UK) Consortium | Jacquelyn Wynn, Mairead Hyland, The Lighthouse Lab in Alderley Park and Alex Alderton, Roberto Amato, Jeffrey Barrett, Sonia Goncalves, Ewan Harrison, David K. Jackson, Ian Johnston, Dominic Kwiatkowski, Cordelia Langford, John Sillitoe on behalf of the Wellcome Sanger Institute COVID-19 Surveillance Team |
| EPI_ISL_2702979, EPI_ISL_2702980, EPI_ISL_2702981 | Lighthouse Lab in Milton Keynes | Wellcome Sanger Institute for the COVID-19 Genomics UK (COG-UK) Consortium | The Lighthouse Lab in Milton Keynes and Alex Alderton, Roberto Amato, Jeffrey Barrett, Sonia Goncalves, Ewan Harrison, David K. Jackson, Ian Johnston, Dominic Kwiatkowski, Cordelia Langford, John Sillitoe on behalf of the Wellcome Sanger Institute COVID-19 Surveillance Team |
| EPI_ISL_2702982 | Lighthouse Lab in Alderley Park | Wellcome Sanger Institute for the COVID-19 Genomics UK (COG-UK) Consortium | Jacquelyn Wynn, Mairead Hyland, The Lighthouse Lab in Alderley Park and Alex Alderton, Roberto Amato, Jeffrey Barrett, Sonia Goncalves, Ewan Harrison, David K. Jackson, Ian Johnston, Dominic Kwiatkowski, Cordelia Langford, John Sillitoe on behalf of the Wellcome Sanger Institute COVID-19 Surveillance Team |
| EPI_ISL_2702983 | Lighthouse Lab in Milton Keynes | Wellcome Sanger Institute for the COVID-19 Genomics UK (COG-UK) Consortium | The Lighthouse Lab in Milton Keynes and Alex Alderton, Roberto Amato, Jeffrey Barrett, Sonia Goncalves, Ewan Harrison, David K. Jackson, Ian Johnston, Dominic Kwiatkowski, Cordelia Langford, John Sillitoe on behalf of the Wellcome Sanger Institute COVID-19 Surveillance Team |
| EPI_ISL_2702984 | Lighthouse Lab in Alderley Park | Wellcome Sanger Institute for the COVID-19 Genomics UK (COG-UK) Consortium | Jacquelyn Wynn, Mairead Hyland, The Lighthouse Lab in Alderley Park and Alex Alderton, Roberto Amato, Jeffrey Barrett, Sonia Goncalves, Ewan Harrison, David K. Jackson, Ian Johnston, Dominic Kwiatkowski, Cordelia Langford, John Sillitoe on behalf of the Wellcome Sanger Institute COVID-19 Surveillance Team |
| EPI_ISL_2702985, EPI_ISL_2702986, EPI_ISL_2702987, EPI_ISL_2702988 | Lighthouse Lab in Milton Keynes | Wellcome Sanger Institute for the COVID-19 Genomics UK (COG-UK) Consortium | The Lighthouse Lab in Milton Keynes and Alex Alderton, Roberto Amato, Jeffrey Barrett, Sonia Goncalves, Ewan Harrison, David K. Jackson, Ian Johnston, Dominic Kwiatkowski, Cordelia Langford, John Sillitoe on behalf of the Wellcome Sanger Institute COVID-19 Surveillance Team |
| EPI_ISL_2702989 | Lighthouse Lab in Alderley Park | Wellcome Sanger Institute for the COVID-19 Genomics UK (COG-UK) Consortium | Jacquelyn Wynn, Mairead Hyland, The Lighthouse Lab in Alderley Park and Alex Alderton, Roberto Amato, Jeffrey Barrett, Sonia Goncalves, Ewan Harrison, David K. Jackson, Ian Johnston, Dominic Kwiatkowski, Cordelia Langford, John Sillitoe on behalf of the Wellcome Sanger Institute COVID-19 Surveillance Team |
| EPI_ISL_2702990 | Lighthouse Lab in Milton Keynes | Wellcome Sanger Institute for the COVID-19 Genomics UK (COG-UK) Consortium | The Lighthouse Lab in Milton Keynes and Alex Alderton, Roberto Amato, Jeffrey Barrett, Sonia Goncalves, Ewan Harrison, David K. Jackson, Ian Johnston, Dominic Kwiatkowski, Cordelia Langford, John Sillitoe on behalf of the Wellcome Sanger Institute COVID-19 Surveillance Team |
| EPI_ISL_2702991, EPI_ISL_2702992, EPI_ISL_2702993 | Lighthouse Lab in Alderley Park | Wellcome Sanger Institute for the COVID-19 Genomics UK (COG-UK) Consortium | Jacquelyn Wynn, Mairead Hyland, The Lighthouse Lab in Alderley Park and Alex Alderton, Roberto Amato, Jeffrey Barrett, Sonia Goncalves, Ewan Harrison, David K. Jackson, Ian Johnston, Dominic Kwiatkowski, Cordelia Langford, John Sillitoe on behalf of the Wellcome Sanger Institute COVID-19 Surveillance Team |
| EPI_ISL_2702994, EPI_ISL_2702995, EPI_ISL_2702996 | Lighthouse Lab in Milton Keynes | Wellcome Sanger Institute for the COVID-19 Genomics UK (COG-UK) Consortium | The Lighthouse Lab in Milton Keynes and Alex Alderton, Roberto Amato, Jeffrey Barrett, Sonia Goncalves, Ewan Harrison, David K. Jackson, Ian Johnston, Dominic Kwiatkowski, Cordelia Langford, John Sillitoe on behalf of the Wellcome Sanger Institute COVID-19 Surveillance Team |
| EPI_ISL_2702997 | Lighthouse Lab in Alderley Park | Wellcome Sanger Institute for the COVID-19 Genomics UK (COG-UK) Consortium | Jacquelyn Wynn, Mairead Hyland, The Lighthouse Lab in Alderley Park and Alex Alderton, Roberto Amato, Jeffrey Barrett, Sonia Goncalves, Ewan Harrison, David K. Jackson, Ian Johnston, Dominic Kwiatkowski, Cordelia Langford, John Sillitoe on behalf of the Wellcome Sanger Institute COVID-19 Surveillance Team |

| | | | |
|---|---|---|---|
| EPI_ISL_2702998, EPI_ISL_2702999, EPI_ISL_2703000, EPI_ISL_2703001, EPI_ISL_2703002 | Lighthouse Lab in Milton Keynes | Wellcome Sanger Institute for the COVID-19 Genomics UK (COG-UK) Consortium | The Lighthouse Lab in Milton Keynes and Alex Alderton, Roberto Amato, Jeffrey Barrett, Sonia Goncalves, Ewan Harrison, David K. Jackson, Ian Johnston, Dominic Kwiatkowski, Cordelia Langford, John Sillitoe on behalf of the Wellcome Sanger Institute COVID-19 Surveillance Team |
| EPI_ISL_2703003, EPI_ISL_2703004, EPI_ISL_2703005, EPI_ISL_2703006, EPI_ISL_2703007, EPI_ISL_2703008 | Lighthouse Lab in Alderley Park | Wellcome Sanger Institute for the COVID-19 Genomics UK (COG-UK) Consortium | Jacquelyn Wynn, Mairead Hyland, The Lighthouse Lab in Alderley Park and Alex Alderton, Roberto Amato, Jeffrey Barrett, Sonia Goncalves, Ewan Harrison, David K. Jackson, Ian Johnston, Dominic Kwiatkowski, Cordelia Langford, John Sillitoe on behalf of the Wellcome Sanger Institute COVID-19 Surveillance Team |
| EPI_ISL_2703009, EPI_ISL_2703010, EPI_ISL_2703011, EPI_ISL_2703012, EPI_ISL_2703013, EPI_ISL_2703014, EPI_ISL_2703015, EPI_ISL_2703016 | Lighthouse Lab in Milton Keynes | Wellcome Sanger Institute for the COVID-19 Genomics UK (COG-UK) Consortium | The Lighthouse Lab in Milton Keynes and Alex Alderton, Roberto Amato, Jeffrey Barrett, Sonia Goncalves, Ewan Harrison, David K. Jackson, Ian Johnston, Dominic Kwiatkowski, Cordelia Langford, John Sillitoe on behalf of the Wellcome Sanger Institute COVID-19 Surveillance Team |
| EPI_ISL_2703017 | Lighthouse Lab in Alderley Park | Wellcome Sanger Institute for the COVID-19 Genomics UK (COG-UK) Consortium | Jacquelyn Wynn, Mairead Hyland, The Lighthouse Lab in Alderley Park and Alex Alderton, Roberto Amato, Jeffrey Barrett, Sonia Goncalves, Ewan Harrison, David K. Jackson, Ian Johnston, Dominic Kwiatkowski, Cordelia Langford, John Sillitoe on behalf of the Wellcome Sanger Institute COVID-19 Surveillance Team |
| EPI_ISL_2703018, EPI_ISL_2703019, EPI_ISL_2703020, EPI_ISL_2703021 | Lighthouse Lab in Milton Keynes | Wellcome Sanger Institute for the COVID-19 Genomics UK (COG-UK) Consortium | The Lighthouse Lab in Milton Keynes and Alex Alderton, Roberto Amato, Jeffrey Barrett, Sonia Goncalves, Ewan Harrison, David K. Jackson, Ian Johnston, Dominic Kwiatkowski, Cordelia Langford, John Sillitoe on behalf of the Wellcome Sanger Institute COVID-19 Surveillance Team |
| EPI_ISL_2703022, EPI_ISL_2703023, EPI_ISL_2703024 | Lighthouse Lab in Alderley Park | Wellcome Sanger Institute for the COVID-19 Genomics UK (COG-UK) Consortium | Jacquelyn Wynn, Mairead Hyland, The Lighthouse Lab in Alderley Park and Alex Alderton, Roberto Amato, Jeffrey Barrett, Sonia Goncalves, Ewan Harrison, David K. Jackson, Ian Johnston, Dominic Kwiatkowski, Cordelia Langford, John Sillitoe on behalf of the Wellcome Sanger Institute COVID-19 Surveillance Team |
| EPI_ISL_2703025, EPI_ISL_2703026, EPI_ISL_2703027 | Lighthouse Lab in Milton Keynes | Wellcome Sanger Institute for the COVID-19 Genomics UK (COG-UK) Consortium | The Lighthouse Lab in Milton Keynes and Alex Alderton, Roberto Amato, Jeffrey Barrett, Sonia Goncalves, Ewan Harrison, David K. Jackson, Ian Johnston, Dominic Kwiatkowski, Cordelia Langford, John Sillitoe on behalf of the Wellcome Sanger Institute COVID-19 Surveillance Team |
| EPI_ISL_2703028, EPI_ISL_2703029, EPI_ISL_2703030, EPI_ISL_2703031 | Lighthouse Lab in Alderley Park | Wellcome Sanger Institute for the COVID-19 Genomics UK (COG-UK) Consortium | Jacquelyn Wynn, Mairead Hyland, The Lighthouse Lab in Alderley Park and Alex Alderton, Roberto Amato, Jeffrey Barrett, Sonia Goncalves, Ewan Harrison, David K. Jackson, Ian Johnston, Dominic Kwiatkowski, Cordelia Langford, John Sillitoe on behalf of the Wellcome Sanger Institute COVID-19 Surveillance Team |
| EPI_ISL_2703032 | Lighthouse Lab in Milton Keynes | Wellcome Sanger Institute for the COVID-19 Genomics UK (COG-UK) Consortium | The Lighthouse Lab in Milton Keynes and Alex Alderton, Roberto Amato, Jeffrey Barrett, Sonia Goncalves, Ewan Harrison, David K. Jackson, Ian Johnston, Dominic Kwiatkowski, Cordelia Langford, John Sillitoe on behalf of the Wellcome Sanger Institute COVID-19 Surveillance Team |
| EPI_ISL_2703033 | Lighthouse Lab in Alderley Park | Wellcome Sanger Institute for the COVID-19 Genomics UK (COG-UK) Consortium | Jacquelyn Wynn, Mairead Hyland, The Lighthouse Lab in Alderley Park and Alex Alderton, Roberto Amato, Jeffrey Barrett, Sonia Goncalves, Ewan Harrison, David K. Jackson, Ian Johnston, Dominic Kwiatkowski, Cordelia Langford, John Sillitoe on behalf of the Wellcome Sanger Institute COVID-19 Surveillance Team |
| EPI_ISL_2703034 | Lighthouse Lab in Milton Keynes | Wellcome Sanger Institute for the COVID-19 Genomics UK (COG-UK) Consortium | The Lighthouse Lab in Milton Keynes and Alex Alderton, Roberto Amato, Jeffrey Barrett, Sonia Goncalves, Ewan Harrison, David K. Jackson, Ian Johnston, Dominic Kwiatkowski, Cordelia Langford, John Sillitoe on behalf of the Wellcome Sanger Institute COVID-19 Surveillance Team |
| EPI_ISL_2703035 | Lighthouse Lab in Alderley Park | Wellcome Sanger Institute for the COVID-19 Genomics UK (COG-UK) Consortium | Jacquelyn Wynn, Mairead Hyland, The Lighthouse Lab in Alderley Park and Alex Alderton, Roberto Amato, Jeffrey Barrett, Sonia Goncalves, Ewan Harrison, David K. Jackson, Ian Johnston, Dominic Kwiatkowski, Cordelia Langford, John Sillitoe on behalf of the Wellcome Sanger Institute COVID-19 Surveillance Team |
| EPI_ISL_2703036, EPI_ISL_2703037, EPI_ISL_2703038, EPI_ISL_2703039, EPI_ISL_2703040, EPI_ISL_2703041, EPI_ISL_2703042, EPI_ISL_2703043, EPI_ISL_2703044 | Lighthouse Lab in Milton Keynes | Wellcome Sanger Institute for the COVID-19 Genomics UK (COG-UK) Consortium | The Lighthouse Lab in Milton Keynes and Alex Alderton, Roberto Amato, Jeffrey Barrett, Sonia Goncalves, Ewan Harrison, David K. Jackson, Ian Johnston, Dominic Kwiatkowski, Cordelia Langford, John Sillitoe on behalf of the Wellcome Sanger Institute COVID-19 Surveillance Team |
| EPI_ISL_2703045 | Lighthouse Lab in Alderley Park | Wellcome Sanger Institute for the COVID-19 Genomics UK (COG-UK) Consortium | Jacquelyn Wynn, Mairead Hyland, The Lighthouse Lab in Alderley Park and Alex Alderton, Roberto Amato, Jeffrey Barrett, Sonia Goncalves, Ewan Harrison, David K. Jackson, Ian Johnston, Dominic Kwiatkowski, Cordelia Langford, John Sillitoe on behalf of the Wellcome Sanger Institute COVID-19 Surveillance Team |
| EPI_ISL_2703046, EPI_ISL_2703047, EPI_ISL_2703048, EPI_ISL_2703049 | Lighthouse Lab in Milton Keynes | Wellcome Sanger Institute for the COVID-19 Genomics UK (COG-UK) Consortium | The Lighthouse Lab in Milton Keynes and Alex Alderton, Roberto Amato, Jeffrey Barrett, Sonia Goncalves, Ewan Harrison, David K. Jackson, Ian Johnston, Dominic Kwiatkowski, Cordelia Langford, John Sillitoe on behalf of the Wellcome Sanger Institute COVID-19 Surveillance Team |
| EPI_ISL_2703050 | Lighthouse Lab in Alderley Park | Wellcome Sanger Institute for the COVID-19 Genomics UK (COG-UK) Consortium | Jacquelyn Wynn, Mairead Hyland, The Lighthouse Lab in Alderley Park and Alex Alderton, Roberto Amato, Jeffrey Barrett, Sonia Goncalves, Ewan Harrison, David K. Jackson, Ian Johnston, Dominic Kwiatkowski, Cordelia Langford, John Sillitoe on behalf of the Wellcome Sanger Institute COVID-19 Surveillance Team |
| EPI_ISL_2703051, EPI_ISL_2703052, EPI_ISL_2703053, EPI_ISL_2703055, EPI_ISL_2703056, EPI_ISL_2703057, EPI_ISL_2703058, EPI_ISL_2703059, EPI_ISL_2703060, EPI_ISL_2703061, EPI_ISL_2703062 | | | |
| see above | Lighthouse Lab in Milton Keynes | Wellcome Sanger Institute for the COVID-19 Genomics UK (COG-UK) Consortium | The Lighthouse Lab in Milton Keynes and Alex Alderton, Roberto Amato, Jeffrey Barrett, Sonia Goncalves, Ewan Harrison, David K. Jackson, Ian Johnston, Dominic Kwiatkowski, Cordelia Langford, John Sillitoe on behalf of the Wellcome Sanger Institute COVID-19 Surveillance Team |
| EPI_ISL_2703063 | Lighthouse Lab in Alderley Park | Wellcome Sanger Institute for the COVID-19 Genomics UK (COG-UK) Consortium | Jacquelyn Wynn, Mairead Hyland, The Lighthouse Lab in Alderley Park and Alex Alderton, Roberto Amato, Jeffrey Barrett, Sonia Goncalves, Ewan Harrison, David K. Jackson, Ian Johnston, Dominic Kwiatkowski, Cordelia Langford, John Sillitoe on behalf of the Wellcome Sanger Institute COVID-19 Surveillance Team |
| EPI_ISL_2703064, EPI_ISL_2703065, EPI_ISL_2703066, EPI_ISL_2703067, EPI_ISL_2703069, EPI_ISL_2703070 | Lighthouse Lab in Milton Keynes | Wellcome Sanger Institute for the COVID-19 Genomics UK (COG-UK) Consortium | The Lighthouse Lab in Milton Keynes and Alex Alderton, Roberto Amato, Jeffrey Barrett, Sonia Goncalves, Ewan Harrison, David K. Jackson, Ian Johnston, Dominic Kwiatkowski, Cordelia Langford, John Sillitoe on behalf of the Wellcome Sanger Institute COVID-19 Surveillance Team |
| EPI_ISL_2703071 | Lighthouse Lab in Alderley Park | Wellcome Sanger Institute for the COVID-19 Genomics UK (COG-UK) Consortium | Jacquelyn Wynn, Mairead Hyland, The Lighthouse Lab in Alderley Park and Alex Alderton, Roberto Amato, Jeffrey Barrett, Sonia Goncalves, Ewan Harrison, David K. Jackson, Ian Johnston, Dominic Kwiatkowski, Cordelia Langford, John Sillitoe on behalf of the Wellcome Sanger Institute COVID-19 Surveillance Team |
| EPI_ISL_2703072, EPI_ISL_2703073, EPI_ISL_2703074, EPI_ISL_2703075, EPI_ISL_2703076, EPI_ISL_2703077, EPI_ISL_2703078 | Lighthouse Lab in Milton Keynes | Wellcome Sanger Institute for the COVID-19 Genomics UK (COG-UK) Consortium | The Lighthouse Lab in Milton Keynes and Alex Alderton, Roberto Amato, Jeffrey Barrett, Sonia Goncalves, Ewan Harrison, David K. Jackson, Ian Johnston, Dominic Kwiatkowski, Cordelia Langford, John Sillitoe on behalf of the Wellcome Sanger Institute COVID-19 Surveillance Team |
| EPI_ISL_2703079 | Lighthouse Lab in Alderley Park | Wellcome Sanger Institute for the COVID-19 Genomics UK (COG-UK) Consortium | Jacquelyn Wynn, Mairead Hyland, The Lighthouse Lab in Alderley Park and Alex Alderton, Roberto Amato, Jeffrey Barrett, Sonia Goncalves, Ewan Harrison, David K. Jackson, Ian Johnston, Dominic Kwiatkowski, Cordelia Langford, John Sillitoe on behalf of the Wellcome Sanger Institute COVID-19 Surveillance Team |
| EPI_ISL_2703080, EPI_ISL_2703081, EPI_ISL_2703082, EPI_ISL_2703083 | Lighthouse Lab in Milton Keynes | Wellcome Sanger Institute for the COVID-19 Genomics UK (COG-UK) Consortium | The Lighthouse Lab in Milton Keynes and Alex Alderton, Roberto Amato, Jeffrey Barrett, Sonia Goncalves, Ewan Harrison, David K. Jackson, Ian Johnston, Dominic Kwiatkowski, Cordelia Langford, John Sillitoe on behalf of the Wellcome Sanger Institute COVID-19 Surveillance Team |
| EPI_ISL_2703084 | Lighthouse Lab in Alderley Park | Wellcome Sanger Institute for the COVID-19 Genomics UK (COG-UK) Consortium | Jacquelyn Wynn, Mairead Hyland, The Lighthouse Lab in Alderley Park and Alex Alderton, Roberto Amato, Jeffrey Barrett, Sonia Goncalves, Ewan Harrison, David K. Jackson, Ian Johnston, Dominic Kwiatkowski, Cordelia Langford, John Sillitoe on behalf of the Wellcome Sanger Institute COVID-19 Surveillance Team |
| EPI_ISL_2703085 | Lighthouse Lab in Milton Keynes | Wellcome Sanger Institute for the COVID-19 Genomics UK (COG-UK) Consortium | The Lighthouse Lab in Milton Keynes and Alex Alderton, Roberto Amato, Jeffrey Barrett, Sonia Goncalves, Ewan Harrison, David K. Jackson, Ian Johnston, Dominic Kwiatkowski, Cordelia Langford, John Sillitoe on behalf of the Wellcome Sanger Institute COVID-19 Surveillance Team |
| EPI_ISL_2703086 | Lighthouse Lab in Alderley Park | Wellcome Sanger Institute for the COVID-19 Genomics UK (COG-UK) Consortium | Jacquelyn Wynn, Mairead Hyland, The Lighthouse Lab in Alderley Park and Alex Alderton, Roberto Amato, Jeffrey Barrett, Sonia Goncalves, Ewan Harrison, David K. Jackson, Ian Johnston, Dominic Kwiatkowski, Cordelia Langford, John Sillitoe on behalf of the Wellcome Sanger Institute COVID-19 Surveillance Team |
| EPI_ISL_2703087, EPI_ISL_2703088, EPI_ISL_2703089 | Lighthouse Lab in Milton Keynes | Wellcome Sanger Institute for the COVID-19 Genomics UK (COG-UK) Consortium | The Lighthouse Lab in Milton Keynes and Alex Alderton, Roberto Amato, Jeffrey Barrett, Sonia Goncalves, Ewan Harrison, David K. Jackson, Ian Johnston, Dominic Kwiatkowski, Cordelia Langford, John Sillitoe on behalf of the Wellcome Sanger Institute COVID-19 Surveillance Team |
| EPI_ISL_2703090 | Lighthouse Lab in Alderley Park | Wellcome Sanger Institute for the COVID-19 Genomics UK (COG-UK) Consortium | Jacquelyn Wynn, Mairead Hyland, The Lighthouse Lab in Alderley Park and Alex Alderton, Roberto Amato, Jeffrey Barrett, Sonia Goncalves, Ewan Harrison, David K. Jackson, Ian Johnston, Dominic Kwiatkowski, Cordelia Langford, John Sillitoe on behalf of the Wellcome Sanger Institute COVID-19 Surveillance Team |

| | | | |
|---|---|---|---|
| EPI_ISL_2703091, EPI_ISL_2703092 | Lighthouse Lab in Milton Keynes | Wellcome Sanger Institute for the COVID-19 Genomics UK (COG-UK) Consortium | The Lighthouse Lab in Milton Keynes and Alex Alderton, Roberto Amato, Jeffrey Barrett, Sonia Goncalves, Ewan Harrison, David K. Jackson, Ian Johnston, Dominic Kwiatkowski, Cordelia Langford, John Sillitoe on behalf of the Wellcome Sanger Institute COVID-19 Surveillance Team |
| EPI_ISL_2703093 | Lighthouse Lab in Alderley Park | Wellcome Sanger Institute for the COVID-19 Genomics UK (COG-UK) Consortium | Jacquelyn Wynn, Mairead Hyland, The Lighthouse Lab in Alderley Park and Alex Alderton, Roberto Amato, Jeffrey Barrett, Sonia Goncalves, Ewan Harrison, David K. Jackson, Ian Johnston, Dominic Kwiatkowski, Cordelia Langford, John Sillitoe on behalf of the Wellcome Sanger Institute COVID-19 Surveillance Team |
| EPI_ISL_2703094 | Lighthouse Lab in Milton Keynes | Wellcome Sanger Institute for the COVID-19 Genomics UK (COG-UK) Consortium | The Lighthouse Lab in Milton Keynes and Alex Alderton, Roberto Amato, Jeffrey Barrett, Sonia Goncalves, Ewan Harrison, David K. Jackson, Ian Johnston, Dominic Kwiatkowski, Cordelia Langford, John Sillitoe on behalf of the Wellcome Sanger Institute COVID-19 Surveillance Team |
| EPI_ISL_2703095, EPI_ISL_2703096 | Lighthouse Lab in Alderley Park | Wellcome Sanger Institute for the COVID-19 Genomics UK (COG-UK) Consortium | Jacquelyn Wynn, Mairead Hyland, The Lighthouse Lab in Alderley Park and Alex Alderton, Roberto Amato, Jeffrey Barrett, Sonia Goncalves, Ewan Harrison, David K. Jackson, Ian Johnston, Dominic Kwiatkowski, Cordelia Langford, John Sillitoe on behalf of the Wellcome Sanger Institute COVID-19 Surveillance Team |
| EPI_ISL_2703097, EPI_ISL_2703098, EPI_ISL_2703099, EPI_ISL_2703100 | Lighthouse Lab in Milton Keynes | Wellcome Sanger Institute for the COVID-19 Genomics UK (COG-UK) Consortium | The Lighthouse Lab in Milton Keynes and Alex Alderton, Roberto Amato, Jeffrey Barrett, Sonia Goncalves, Ewan Harrison, David K. Jackson, Ian Johnston, Dominic Kwiatkowski, Cordelia Langford, John Sillitoe on behalf of the Wellcome Sanger Institute COVID-19 Surveillance Team |
| EPI_ISL_2703101 | Lighthouse Lab in Alderley Park | Wellcome Sanger Institute for the COVID-19 Genomics UK (COG-UK) Consortium | Jacquelyn Wynn, Mairead Hyland, The Lighthouse Lab in Alderley Park and Alex Alderton, Roberto Amato, Jeffrey Barrett, Sonia Goncalves, Ewan Harrison, David K. Jackson, Ian Johnston, Dominic Kwiatkowski, Cordelia Langford, John Sillitoe on behalf of the Wellcome Sanger Institute COVID-19 Surveillance Team |
| EPI_ISL_2703102 | Lighthouse Lab in Milton Keynes | Wellcome Sanger Institute for the COVID-19 Genomics UK (COG-UK) Consortium | The Lighthouse Lab in Milton Keynes and Alex Alderton, Roberto Amato, Jeffrey Barrett, Sonia Goncalves, Ewan Harrison, David K. Jackson, Ian Johnston, Dominic Kwiatkowski, Cordelia Langford, John Sillitoe on behalf of the Wellcome Sanger Institute COVID-19 Surveillance Team |
| EPI_ISL_2703103 | Lighthouse Lab in Alderley Park | Wellcome Sanger Institute for the COVID-19 Genomics UK (COG-UK) Consortium | Jacquelyn Wynn, Mairead Hyland, The Lighthouse Lab in Alderley Park and Alex Alderton, Roberto Amato, Jeffrey Barrett, Sonia Goncalves, Ewan Harrison, David K. Jackson, Ian Johnston, Dominic Kwiatkowski, Cordelia Langford, John Sillitoe on behalf of the Wellcome Sanger Institute COVID-19 Surveillance Team |
| EPI_ISL_2703104, EPI_ISL_2703105 | Lighthouse Lab in Milton Keynes | Wellcome Sanger Institute for the COVID-19 Genomics UK (COG-UK) Consortium | The Lighthouse Lab in Milton Keynes and Alex Alderton, Roberto Amato, Jeffrey Barrett, Sonia Goncalves, Ewan Harrison, David K. Jackson, Ian Johnston, Dominic Kwiatkowski, Cordelia Langford, John Sillitoe on behalf of the Wellcome Sanger Institute COVID-19 Surveillance Team |
| EPI_ISL_2703106 | Lighthouse Lab in Alderley Park | Wellcome Sanger Institute for the COVID-19 Genomics UK (COG-UK) Consortium | Jacquelyn Wynn, Mairead Hyland, The Lighthouse Lab in Alderley Park and Alex Alderton, Roberto Amato, Jeffrey Barrett, Sonia Goncalves, Ewan Harrison, David K. Jackson, Ian Johnston, Dominic Kwiatkowski, Cordelia Langford, John Sillitoe on behalf of the Wellcome Sanger Institute COVID-19 Surveillance Team |
| EPI_ISL_2703107, EPI_ISL_2703108, EPI_ISL_2703109, EPI_ISL_2703110, EPI_ISL_2703111, EPI_ISL_2703113, EPI_ISL_2703114 | Lighthouse Lab in Milton Keynes | Wellcome Sanger Institute for the COVID-19 Genomics UK (COG-UK) Consortium | The Lighthouse Lab in Milton Keynes and Alex Alderton, Roberto Amato, Jeffrey Barrett, Sonia Goncalves, Ewan Harrison, David K. Jackson, Ian Johnston, Dominic Kwiatkowski, Cordelia Langford, John Sillitoe on behalf of the Wellcome Sanger Institute COVID-19 Surveillance Team |
| EPI_ISL_2703115 | Lighthouse Lab in Alderley Park | Wellcome Sanger Institute for the COVID-19 Genomics UK (COG-UK) Consortium | Jacquelyn Wynn, Mairead Hyland, The Lighthouse Lab in Alderley Park and Alex Alderton, Roberto Amato, Jeffrey Barrett, Sonia Goncalves, Ewan Harrison, David K. Jackson, Ian Johnston, Dominic Kwiatkowski, Cordelia Langford, John Sillitoe on behalf of the Wellcome Sanger Institute COVID-19 Surveillance Team |
| EPI_ISL_2703116, EPI_ISL_2703117, EPI_ISL_2703118, EPI_ISL_2703119 | Lighthouse Lab in Milton Keynes | Wellcome Sanger Institute for the COVID-19 Genomics UK (COG-UK) Consortium | The Lighthouse Lab in Milton Keynes and Alex Alderton, Roberto Amato, Jeffrey Barrett, Sonia Goncalves, Ewan Harrison, David K. Jackson, Ian Johnston, Dominic Kwiatkowski, Cordelia Langford, John Sillitoe on behalf of the Wellcome Sanger Institute COVID-19 Surveillance Team |
| EPI_ISL_2703120 | Lighthouse Lab in Alderley Park | Wellcome Sanger Institute for the COVID-19 Genomics UK (COG-UK) Consortium | Jacquelyn Wynn, Mairead Hyland, The Lighthouse Lab in Alderley Park and Alex Alderton, Roberto Amato, Jeffrey Barrett, Sonia Goncalves, Ewan Harrison, David K. Jackson, Ian Johnston, Dominic Kwiatkowski, Cordelia Langford, John Sillitoe on behalf of the Wellcome Sanger Institute COVID-19 Surveillance Team |
| EPI_ISL_2703121 | Lighthouse Lab in Milton Keynes | Wellcome Sanger Institute for the COVID-19 Genomics UK (COG-UK) Consortium | The Lighthouse Lab in Milton Keynes and Alex Alderton, Roberto Amato, Jeffrey Barrett, Sonia Goncalves, Ewan Harrison, David K. Jackson, Ian Johnston, Dominic Kwiatkowski, Cordelia Langford, John Sillitoe on behalf of the Wellcome Sanger Institute COVID-19 Surveillance Team |
| EPI_ISL_2703122 | Lighthouse Lab in Alderley Park | Wellcome Sanger Institute for the COVID-19 Genomics UK (COG-UK) Consortium | Jacquelyn Wynn, Mairead Hyland, The Lighthouse Lab in Alderley Park and Alex Alderton, Roberto Amato, Jeffrey Barrett, Sonia Goncalves, Ewan Harrison, David K. Jackson, Ian Johnston, Dominic Kwiatkowski, Cordelia Langford, John Sillitoe on behalf of the Wellcome Sanger Institute COVID-19 Surveillance Team |
| EPI_ISL_2703123, EPI_ISL_2703124, EPI_ISL_2703125, EPI_ISL_2703126, EPI_ISL_2703127 | Lighthouse Lab in Milton Keynes | Wellcome Sanger Institute for the COVID-19 Genomics UK (COG-UK) Consortium | The Lighthouse Lab in Milton Keynes and Alex Alderton, Roberto Amato, Jeffrey Barrett, Sonia Goncalves, Ewan Harrison, David K. Jackson, Ian Johnston, Dominic Kwiatkowski, Cordelia Langford, John Sillitoe on behalf of the Wellcome Sanger Institute COVID-19 Surveillance Team |
| EPI_ISL_2703128 | Lighthouse Lab in Alderley Park | Wellcome Sanger Institute for the COVID-19 Genomics UK (COG-UK) Consortium | Jacquelyn Wynn, Mairead Hyland, The Lighthouse Lab in Alderley Park and Alex Alderton, Roberto Amato, Jeffrey Barrett, Sonia Goncalves, Ewan Harrison, David K. Jackson, Ian Johnston, Dominic Kwiatkowski, Cordelia Langford, John Sillitoe on behalf of the Wellcome Sanger Institute COVID-19 Surveillance Team |
| EPI_ISL_2703129 | Lighthouse Lab in Milton Keynes | Wellcome Sanger Institute for the COVID-19 Genomics UK (COG-UK) Consortium | The Lighthouse Lab in Milton Keynes and Alex Alderton, Roberto Amato, Jeffrey Barrett, Sonia Goncalves, Ewan Harrison, David K. Jackson, Ian Johnston, Dominic Kwiatkowski, Cordelia Langford, John Sillitoe on behalf of the Wellcome Sanger Institute COVID-19 Surveillance Team |
| EPI_ISL_2703130 | Lighthouse Lab in Alderley Park | Wellcome Sanger Institute for the COVID-19 Genomics UK (COG-UK) Consortium | Jacquelyn Wynn, Mairead Hyland, The Lighthouse Lab in Alderley Park and Alex Alderton, Roberto Amato, Jeffrey Barrett, Sonia Goncalves, Ewan Harrison, David K. Jackson, Ian Johnston, Dominic Kwiatkowski, Cordelia Langford, John Sillitoe on behalf of the Wellcome Sanger Institute COVID-19 Surveillance Team |
| EPI_ISL_2703131, EPI_ISL_2703132 | Lighthouse Lab in Milton Keynes | Wellcome Sanger Institute for the COVID-19 Genomics UK (COG-UK) Consortium | The Lighthouse Lab in Milton Keynes and Alex Alderton, Roberto Amato, Jeffrey Barrett, Sonia Goncalves, Ewan Harrison, David K. Jackson, Ian Johnston, Dominic Kwiatkowski, Cordelia Langford, John Sillitoe on behalf of the Wellcome Sanger Institute COVID-19 Surveillance Team |
| EPI_ISL_2703133, EPI_ISL_2703134 | Lighthouse Lab in Alderley Park | Wellcome Sanger Institute for the COVID-19 Genomics UK (COG-UK) Consortium | Jacquelyn Wynn, Mairead Hyland, The Lighthouse Lab in Alderley Park and Alex Alderton, Roberto Amato, Jeffrey Barrett, Sonia Goncalves, Ewan Harrison, David K. Jackson, Ian Johnston, Dominic Kwiatkowski, Cordelia Langford, John Sillitoe on behalf of the Wellcome Sanger Institute COVID-19 Surveillance Team |
| EPI_ISL_2703135 | Lighthouse Lab in Milton Keynes | Wellcome Sanger Institute for the COVID-19 Genomics UK (COG-UK) Consortium | The Lighthouse Lab in Milton Keynes and Alex Alderton, Roberto Amato, Jeffrey Barrett, Sonia Goncalves, Ewan Harrison, David K. Jackson, Ian Johnston, Dominic Kwiatkowski, Cordelia Langford, John Sillitoe on behalf of the Wellcome Sanger Institute COVID-19 Surveillance Team |
| EPI_ISL_2703136, EPI_ISL_2703137 | Lighthouse Lab in Alderley Park | Wellcome Sanger Institute for the COVID-19 Genomics UK (COG-UK) Consortium | Jacquelyn Wynn, Mairead Hyland, The Lighthouse Lab in Alderley Park and Alex Alderton, Roberto Amato, Jeffrey Barrett, Sonia Goncalves, Ewan Harrison, David K. Jackson, Ian Johnston, Dominic Kwiatkowski, Cordelia Langford, John Sillitoe on behalf of the Wellcome Sanger Institute COVID-19 Surveillance Team |
| EPI_ISL_2703138 | Lighthouse Lab in Milton Keynes | Wellcome Sanger Institute for the COVID-19 Genomics UK (COG-UK) Consortium | The Lighthouse Lab in Milton Keynes and Alex Alderton, Roberto Amato, Jeffrey Barrett, Sonia Goncalves, Ewan Harrison, David K. Jackson, Ian Johnston, Dominic Kwiatkowski, Cordelia Langford, John Sillitoe on behalf of the Wellcome Sanger Institute COVID-19 Surveillance Team |
| EPI_ISL_2703139 | Lighthouse Lab in Alderley Park | Wellcome Sanger Institute for the COVID-19 Genomics UK (COG-UK) Consortium | Jacquelyn Wynn, Mairead Hyland, The Lighthouse Lab in Alderley Park and Alex Alderton, Roberto Amato, Jeffrey Barrett, Sonia Goncalves, Ewan Harrison, David K. Jackson, Ian Johnston, Dominic Kwiatkowski, Cordelia Langford, John Sillitoe on behalf of the Wellcome Sanger Institute COVID-19 Surveillance Team |
| EPI_ISL_2703140, EPI_ISL_2703141, EPI_ISL_2703142, EPI_ISL_2703143, EPI_ISL_2703144, EPI_ISL_2703145, EPI_ISL_2703146, EPI_ISL_2703147, EPI_ISL_2703148, EPI_ISL_2703150, EPI_ISL_2703151, EPI_ISL_2703152, EPI_ISL_2703153, EPI_ISL_2703154 | | | |
| see above | Lighthouse Lab in Milton Keynes | Wellcome Sanger Institute for the COVID-19 Genomics UK (COG-UK) Consortium | The Lighthouse Lab in Milton Keynes and Alex Alderton, Roberto Amato, Jeffrey Barrett, Sonia Goncalves, Ewan Harrison, David K. Jackson, Ian Johnston, Dominic Kwiatkowski, Cordelia Langford, John Sillitoe on behalf of the Wellcome Sanger Institute COVID-19 Surveillance Team |
| EPI_ISL_2703155, EPI_ISL_2703156, EPI_ISL_2703157, EPI_ISL_2703158 | Lighthouse Lab in Alderley Park | Wellcome Sanger Institute for the COVID-19 Genomics UK (COG-UK) Consortium | Jacquelyn Wynn, Mairead Hyland, The Lighthouse Lab in Alderley Park and Alex Alderton, Roberto Amato, Jeffrey Barrett, Sonia Goncalves, Ewan Harrison, David K. Jackson, Ian Johnston, Dominic Kwiatkowski, Cordelia Langford, John Sillitoe on behalf of the Wellcome Sanger Institute COVID-19 Surveillance Team |
| EPI_ISL_2703159 | Lighthouse Lab in Milton Keynes | Wellcome Sanger Institute for the COVID-19 Genomics UK (COG-UK) Consortium | The Lighthouse Lab in Milton Keynes and Alex Alderton, Roberto Amato, Jeffrey Barrett, Sonia Goncalves, Ewan Harrison, David K. Jackson, Ian Johnston, Dominic Kwiatkowski, Cordelia Langford, John Sillitoe on behalf of the Wellcome Sanger Institute COVID-19 Surveillance Team |
| EPI_ISL_2703160, EPI_ISL_2703161, EPI_ISL_2703162 | Lighthouse Lab in Alderley Park | Wellcome Sanger Institute for the COVID-19 Genomics UK (COG-UK) Consortium | Jacquelyn Wynn, Mairead Hyland, The Lighthouse Lab in Alderley Park and Alex Alderton, Roberto Amato, Jeffrey Barrett, Sonia Goncalves, Ewan Harrison, David K. Jackson, Ian Johnston, Dominic Kwiatkowski, Cordelia Langford, John Sillitoe on behalf of the Wellcome Sanger Institute COVID-19 Surveillance Team |
| EPI_ISL_2703163 | Lighthouse Lab in Milton Keynes | Wellcome Sanger Institute for the COVID-19 Genomics UK (COG-UK) Consortium | The Lighthouse Lab in Milton Keynes and Alex Alderton, Roberto Amato, Jeffrey Barrett, Sonia Goncalves, Ewan Harrison, David K. Jackson, Ian Johnston, Dominic Kwiatkowski, Cordelia Langford, John Sillitoe on behalf of the Wellcome Sanger Institute COVID-19 Surveillance Team |
| EPI_ISL_2703164, EPI_ISL_2703165, EPI_ISL_2703166 | Lighthouse Lab in Alderley Park | Wellcome Sanger Institute for the COVID-19 Genomics UK (COG-UK) Consortium | Jacquelyn Wynn, Mairead Hyland, The Lighthouse Lab in Alderley Park and Alex Alderton, Roberto Amato, Jeffrey Barrett, Sonia Goncalves, Ewan Harrison, David K. Jackson, Ian Johnston, Dominic Kwiatkowski, Cordelia Langford, John Sillitoe on behalf of the Wellcome Sanger Institute COVID-19 Surveillance Team |
| EPI_ISL_2703167 | Lighthouse Lab in Milton Keynes | Wellcome Sanger Institute for the COVID-19 Genomics UK (COG-UK) Consortium | The Lighthouse Lab in Milton Keynes and Alex Alderton, Roberto Amato, Jeffrey Barrett, Sonia Goncalves, Ewan Harrison, David K. Jackson, Ian Johnston, Dominic Kwiatkowski, Cordelia Langford, John Sillitoe on behalf of the Wellcome Sanger Institute COVID-19 Surveillance Team |
| EPI_ISL_2703168 | Lighthouse Lab in Alderley Park | Wellcome Sanger Institute for the COVID-19 Genomics UK (COG-UK) Consortium | Jacquelyn Wynn, Mairead Hyland, The Lighthouse Lab in Alderley Park and Alex Alderton, Roberto Amato, Jeffrey Barrett, Sonia Goncalves, Ewan Harrison, David K. Jackson, Ian Johnston, Dominic Kwiatkowski, Cordelia Langford, John Sillitoe on behalf of the Wellcome Sanger Institute COVID-19 Surveillance Team |
| EPI_ISL_2703169 | Lighthouse Lab in Milton Keynes | Wellcome Sanger Institute for the COVID-19 Genomics UK (COG-UK) Consortium | The Lighthouse Lab in Milton Keynes and Alex Alderton, Roberto Amato, Jeffrey Barrett, Sonia Goncalves, Ewan Harrison, David K. Jackson, Ian Johnston, Dominic Kwiatkowski, Cordelia Langford, John Sillitoe on behalf of the Wellcome Sanger Institute COVID-19 Surveillance Team |
| EPI_ISL_2703170, EPI_ISL_2703171, EPI_ISL_2703172, EPI_ISL_2703173, EPI_ISL_2703174, EPI_ISL_2703175, EPI_ISL_2703176, | Lighthouse Lab in Alderley Park | Wellcome Sanger Institute for the COVID-19 Genomics UK (COG-UK) Consortium | Jacquelyn Wynn, Mairead Hyland, The Lighthouse Lab in Alderley Park and Alex Alderton, Roberto Amato, Jeffrey Barrett, Sonia Goncalves, Ewan Harrison, David K. Jackson, Ian Johnston, Dominic Kwiatkowski, Cordelia Langford, John Sillitoe on behalf of the Wellcome Sanger Institute COVID-19 Surveillance Team |

| | | | |
|---|---|---|---|
| EPI_ISL_2703177, EPI_ISL_2703178, EPI_ISL_2703179 | | | |
| EPI_ISL_2703180, EPI_ISL_2703181, EPI_ISL_2703182 | Lighthouse Lab in Milton Keynes | Wellcome Sanger Institute for the COVID-19 Genomics UK (COG-UK) Consortium | The Lighthouse Lab in Milton Keynes and Alex Alderton, Roberto Amato, Jeffrey Barrett, Sonia Goncalves, Ewan Harrison, David K. Jackson, Ian Johnston, Dominic Kwiatkowski, Cordelia Langford, John Sillitoe on behalf of the Wellcome Sanger Institute COVID-19 Surveillance Team |
| EPI_ISL_2703183 | Lighthouse Lab in Alderley Park | Wellcome Sanger Institute for the COVID-19 Genomics UK (COG-UK) Consortium | Jacquelyn Wynn, Mairead Hyland, The Lighthouse Lab in Alderley Park and Alex Alderton, Roberto Amato, Jeffrey Barrett, Sonia Goncalves, Ewan Harrison, David K. Jackson, Ian Johnston, Dominic Kwiatkowski, Cordelia Langford, John Sillitoe on behalf of the Wellcome Sanger Institute COVID-19 Surveillance Team |
| EPI_ISL_2703184, EPI_ISL_2703185 | Lighthouse Lab in Milton Keynes | Wellcome Sanger Institute for the COVID-19 Genomics UK (COG-UK) Consortium | The Lighthouse Lab in Milton Keynes and Alex Alderton, Roberto Amato, Jeffrey Barrett, Sonia Goncalves, Ewan Harrison, David K. Jackson, Ian Johnston, Dominic Kwiatkowski, Cordelia Langford, John Sillitoe on behalf of the Wellcome Sanger Institute COVID-19 Surveillance Team |
| EPI_ISL_2703186 | Lighthouse Lab in Alderley Park | Wellcome Sanger Institute for the COVID-19 Genomics UK (COG-UK) Consortium | Jacquelyn Wynn, Mairead Hyland, The Lighthouse Lab in Alderley Park and Alex Alderton, Roberto Amato, Jeffrey Barrett, Sonia Goncalves, Ewan Harrison, David K. Jackson, Ian Johnston, Dominic Kwiatkowski, Cordelia Langford, John Sillitoe on behalf of the Wellcome Sanger Institute COVID-19 Surveillance Team |
| EPI_ISL_2703187 | Lighthouse Lab in Milton Keynes | Wellcome Sanger Institute for the COVID-19 Genomics UK (COG-UK) Consortium | The Lighthouse Lab in Milton Keynes and Alex Alderton, Roberto Amato, Jeffrey Barrett, Sonia Goncalves, Ewan Harrison, David K. Jackson, Ian Johnston, Dominic Kwiatkowski, Cordelia Langford, John Sillitoe on behalf of the Wellcome Sanger Institute COVID-19 Surveillance Team |
| EPI_ISL_2703188 | Lighthouse Lab in Alderley Park | Wellcome Sanger Institute for the COVID-19 Genomics UK (COG-UK) Consortium | Jacquelyn Wynn, Mairead Hyland, The Lighthouse Lab in Alderley Park and Alex Alderton, Roberto Amato, Jeffrey Barrett, Sonia Goncalves, Ewan Harrison, David K. Jackson, Ian Johnston, Dominic Kwiatkowski, Cordelia Langford, John Sillitoe on behalf of the Wellcome Sanger Institute COVID-19 Surveillance Team |
| EPI_ISL_2703189 | Lighthouse Lab in Milton Keynes | Wellcome Sanger Institute for the COVID-19 Genomics UK (COG-UK) Consortium | The Lighthouse Lab in Milton Keynes and Alex Alderton, Roberto Amato, Jeffrey Barrett, Sonia Goncalves, Ewan Harrison, David K. Jackson, Ian Johnston, Dominic Kwiatkowski, Cordelia Langford, John Sillitoe on behalf of the Wellcome Sanger Institute COVID-19 Surveillance Team |
| EPI_ISL_2703190, EPI_ISL_2703191, EPI_ISL_2703192, EPI_ISL_2703193, EPI_ISL_2703194 | Lighthouse Lab in Alderley Park | Wellcome Sanger Institute for the COVID-19 Genomics UK (COG-UK) Consortium | Jacquelyn Wynn, Mairead Hyland, The Lighthouse Lab in Alderley Park and Alex Alderton, Roberto Amato, Jeffrey Barrett, Sonia Goncalves, Ewan Harrison, David K. Jackson, Ian Johnston, Dominic Kwiatkowski, Cordelia Langford, John Sillitoe on behalf of the Wellcome Sanger Institute COVID-19 Surveillance Team |
| EPI_ISL_2703195 | Lighthouse Lab in Milton Keynes | Wellcome Sanger Institute for the COVID-19 Genomics UK (COG-UK) Consortium | The Lighthouse Lab in Milton Keynes and Alex Alderton, Roberto Amato, Jeffrey Barrett, Sonia Goncalves, Ewan Harrison, David K. Jackson, Ian Johnston, Dominic Kwiatkowski, Cordelia Langford, John Sillitoe on behalf of the Wellcome Sanger Institute COVID-19 Surveillance Team |
| EPI_ISL_2703196, EPI_ISL_2703197, EPI_ISL_2703198, EPI_ISL_2703199 | Lighthouse Lab in Alderley Park | Wellcome Sanger Institute for the COVID-19 Genomics UK (COG-UK) Consortium | Jacquelyn Wynn, Mairead Hyland, The Lighthouse Lab in Alderley Park and Alex Alderton, Roberto Amato, Jeffrey Barrett, Sonia Goncalves, Ewan Harrison, David K. Jackson, Ian Johnston, Dominic Kwiatkowski, Cordelia Langford, John Sillitoe on behalf of the Wellcome Sanger Institute COVID-19 Surveillance Team |
| EPI_ISL_2703200 | Lighthouse Lab in Milton Keynes | Wellcome Sanger Institute for the COVID-19 Genomics UK (COG-UK) Consortium | The Lighthouse Lab in Milton Keynes and Alex Alderton, Roberto Amato, Jeffrey Barrett, Sonia Goncalves, Ewan Harrison, David K. Jackson, Ian Johnston, Dominic Kwiatkowski, Cordelia Langford, John Sillitoe on behalf of the Wellcome Sanger Institute COVID-19 Surveillance Team |
| EPI_ISL_2703201, EPI_ISL_2703202, EPI_ISL_2703203, EPI_ISL_2703204, EPI_ISL_2703205, EPI_ISL_2703206, EPI_ISL_2703207, EPI_ISL_2703208, EPI_ISL_2703209 | Lighthouse Lab in Alderley Park | Wellcome Sanger Institute for the COVID-19 Genomics UK (COG-UK) Consortium | Jacquelyn Wynn, Mairead Hyland, The Lighthouse Lab in Alderley Park and Alex Alderton, Roberto Amato, Jeffrey Barrett, Sonia Goncalves, Ewan Harrison, David K. Jackson, Ian Johnston, Dominic Kwiatkowski, Cordelia Langford, John Sillitoe on behalf of the Wellcome Sanger Institute COVID-19 Surveillance Team |
| EPI_ISL_2703210 | Lighthouse Lab in Milton Keynes | Wellcome Sanger Institute for the COVID-19 Genomics UK (COG-UK) Consortium | The Lighthouse Lab in Milton Keynes and Alex Alderton, Roberto Amato, Jeffrey Barrett, Sonia Goncalves, Ewan Harrison, David K. Jackson, Ian Johnston, Dominic Kwiatkowski, Cordelia Langford, John Sillitoe on behalf of the Wellcome Sanger Institute COVID-19 Surveillance Team |
| EPI_ISL_2703211 | Lighthouse Lab in Alderley Park | Wellcome Sanger Institute for the COVID-19 Genomics UK (COG-UK) Consortium | Jacquelyn Wynn, Mairead Hyland, The Lighthouse Lab in Alderley Park and Alex Alderton, Roberto Amato, Jeffrey Barrett, Sonia Goncalves, Ewan Harrison, David K. Jackson, Ian Johnston, Dominic Kwiatkowski, Cordelia Langford, John Sillitoe on behalf of the Wellcome Sanger Institute COVID-19 Surveillance Team |
| EPI_ISL_2703212 | Lighthouse Lab in Milton Keynes | Wellcome Sanger Institute for the COVID-19 Genomics UK (COG-UK) Consortium | The Lighthouse Lab in Milton Keynes and Alex Alderton, Roberto Amato, Jeffrey Barrett, Sonia Goncalves, Ewan Harrison, David K. Jackson, Ian Johnston, Dominic Kwiatkowski, Cordelia Langford, John Sillitoe on behalf of the Wellcome Sanger Institute COVID-19 Surveillance Team |
| EPI_ISL_2703213, EPI_ISL_2703214 | Lighthouse Lab in Alderley Park | Wellcome Sanger Institute for the COVID-19 Genomics UK (COG-UK) Consortium | Jacquelyn Wynn, Mairead Hyland, The Lighthouse Lab in Alderley Park and Alex Alderton, Roberto Amato, Jeffrey Barrett, Sonia Goncalves, Ewan Harrison, David K. Jackson, Ian Johnston, Dominic Kwiatkowski, Cordelia Langford, John Sillitoe on behalf of the Wellcome Sanger Institute COVID-19 Surveillance Team |
| EPI_ISL_2703215 | Lighthouse Lab in Milton Keynes | Wellcome Sanger Institute for the COVID-19 Genomics UK (COG-UK) Consortium | The Lighthouse Lab in Milton Keynes and Alex Alderton, Roberto Amato, Jeffrey Barrett, Sonia Goncalves, Ewan Harrison, David K. Jackson, Ian Johnston, Dominic Kwiatkowski, Cordelia Langford, John Sillitoe on behalf of the Wellcome Sanger Institute COVID-19 Surveillance Team |
| EPI_ISL_2703216, EPI_ISL_2703217, EPI_ISL_2703218, EPI_ISL_2703219, EPI_ISL_2703220, EPI_ISL_2703222, EPI_ISL_2703223, EPI_ISL_2703224, EPI_ISL_2703225, EPI_ISL_2703226, EPI_ISL_2703227, EPI_ISL_2703228, EPI_ISL_2703229, EPI_ISL_2703230, EPI_ISL_2703231, EPI_ISL_2703232, EPI_ISL_2703233, EPI_ISL_2703234, EPI_ISL_2703235, EPI_ISL_2703236 | | | |
| see above | Lighthouse Lab in Alderley Park | Wellcome Sanger Institute for the COVID-19 Genomics UK (COG-UK) Consortium | Jacquelyn Wynn, Mairead Hyland, The Lighthouse Lab in Alderley Park and Alex Alderton, Roberto Amato, Jeffrey Barrett, Sonia Goncalves, Ewan Harrison, David K. Jackson, Ian Johnston, Dominic Kwiatkowski, Cordelia Langford, John Sillitoe on behalf of the Wellcome Sanger Institute COVID-19 Surveillance Team |
| EPI_ISL_2703237 | Lighthouse Lab in Milton Keynes | Wellcome Sanger Institute for the COVID-19 Genomics UK (COG-UK) Consortium | The Lighthouse Lab in Milton Keynes and Alex Alderton, Roberto Amato, Jeffrey Barrett, Sonia Goncalves, Ewan Harrison, David K. Jackson, Ian Johnston, Dominic Kwiatkowski, Cordelia Langford, John Sillitoe on behalf of the Wellcome Sanger Institute COVID-19 Surveillance Team |
| EPI_ISL_2703238, EPI_ISL_2703239, EPI_ISL_2703240, EPI_ISL_2703241, EPI_ISL_2703242 | Lighthouse Lab in Alderley Park | Wellcome Sanger Institute for the COVID-19 Genomics UK (COG-UK) Consortium | Jacquelyn Wynn, Mairead Hyland, The Lighthouse Lab in Alderley Park and Alex Alderton, Roberto Amato, Jeffrey Barrett, Sonia Goncalves, Ewan Harrison, David K. Jackson, Ian Johnston, Dominic Kwiatkowski, Cordelia Langford, John Sillitoe on behalf of the Wellcome Sanger Institute COVID-19 Surveillance Team |
| EPI_ISL_2703243 | Lighthouse Lab in Milton Keynes | Wellcome Sanger Institute for the COVID-19 Genomics UK (COG-UK) Consortium | The Lighthouse Lab in Milton Keynes and Alex Alderton, Roberto Amato, Jeffrey Barrett, Sonia Goncalves, Ewan Harrison, David K. Jackson, Ian Johnston, Dominic Kwiatkowski, Cordelia Langford, John Sillitoe on behalf of the Wellcome Sanger Institute COVID-19 Surveillance Team |
| EPI_ISL_2703244, EPI_ISL_2703245, EPI_ISL_2703246, EPI_ISL_2703247 | Lighthouse Lab in Alderley Park | Wellcome Sanger Institute for the COVID-19 Genomics UK (COG-UK) Consortium | Jacquelyn Wynn, Mairead Hyland, The Lighthouse Lab in Alderley Park and Alex Alderton, Roberto Amato, Jeffrey Barrett, Sonia Goncalves, Ewan Harrison, David K. Jackson, Ian Johnston, Dominic Kwiatkowski, Cordelia Langford, John Sillitoe on behalf of the Wellcome Sanger Institute COVID-19 Surveillance Team |
| EPI_ISL_2703248 | Lighthouse Lab in Milton Keynes | Wellcome Sanger Institute for the COVID-19 Genomics UK (COG-UK) Consortium | The Lighthouse Lab in Milton Keynes and Alex Alderton, Roberto Amato, Jeffrey Barrett, Sonia Goncalves, Ewan Harrison, David K. Jackson, Ian Johnston, Dominic Kwiatkowski, Cordelia Langford, John Sillitoe on behalf of the Wellcome Sanger Institute COVID-19 Surveillance Team |
| EPI_ISL_2703249, EPI_ISL_2703250, EPI_ISL_2703251, EPI_ISL_2703252, EPI_ISL_2703253, EPI_ISL_2703254 | Lighthouse Lab in Alderley Park | Wellcome Sanger Institute for the COVID-19 Genomics UK (COG-UK) Consortium | Jacquelyn Wynn, Mairead Hyland, The Lighthouse Lab in Alderley Park and Alex Alderton, Roberto Amato, Jeffrey Barrett, Sonia Goncalves, Ewan Harrison, David K. Jackson, Ian Johnston, Dominic Kwiatkowski, Cordelia Langford, John Sillitoe on behalf of the Wellcome Sanger Institute COVID-19 Surveillance Team |
| EPI_ISL_2703255 | Lighthouse Lab in Milton Keynes | Wellcome Sanger Institute for the COVID-19 Genomics UK (COG-UK) Consortium | The Lighthouse Lab in Milton Keynes and Alex Alderton, Roberto Amato, Jeffrey Barrett, Sonia Goncalves, Ewan Harrison, David K. Jackson, Ian Johnston, Dominic Kwiatkowski, Cordelia Langford, John Sillitoe on behalf of the Wellcome Sanger Institute COVID-19 Surveillance Team |
| EPI_ISL_2703256, EPI_ISL_2703257, EPI_ISL_2703258, EPI_ISL_2703259, EPI_ISL_2703260 | Lighthouse Lab in Alderley Park | Wellcome Sanger Institute for the COVID-19 Genomics UK (COG-UK) Consortium | Jacquelyn Wynn, Mairead Hyland, The Lighthouse Lab in Alderley Park and Alex Alderton, Roberto Amato, Jeffrey Barrett, Sonia Goncalves, Ewan Harrison, David K. Jackson, Ian Johnston, Dominic Kwiatkowski, Cordelia Langford, John Sillitoe on behalf of the Wellcome Sanger Institute COVID-19 Surveillance Team |
| EPI_ISL_2703261 | Lighthouse Lab in Milton Keynes | Wellcome Sanger Institute for the COVID-19 Genomics UK (COG-UK) Consortium | The Lighthouse Lab in Milton Keynes and Alex Alderton, Roberto Amato, Jeffrey Barrett, Sonia Goncalves, Ewan Harrison, David K. Jackson, Ian Johnston, Dominic Kwiatkowski, Cordelia Langford, John Sillitoe on behalf of the Wellcome Sanger Institute COVID-19 Surveillance Team |
| EPI_ISL_2703262, EPI_ISL_2703263, EPI_ISL_2703264, EPI_ISL_2703265, EPI_ISL_2703266 | Lighthouse Lab in Alderley Park | Wellcome Sanger Institute for the COVID-19 Genomics UK (COG-UK) Consortium | Jacquelyn Wynn, Mairead Hyland, The Lighthouse Lab in Alderley Park and Alex Alderton, Roberto Amato, Jeffrey Barrett, Sonia Goncalves, Ewan Harrison, David K. Jackson, Ian Johnston, Dominic Kwiatkowski, Cordelia Langford, John Sillitoe on behalf of the Wellcome Sanger Institute COVID-19 Surveillance Team |
| EPI_ISL_2703267 | Lighthouse Lab in Milton Keynes | Wellcome Sanger Institute for the COVID-19 Genomics UK (COG-UK) Consortium | The Lighthouse Lab in Milton Keynes and Alex Alderton, Roberto Amato, Jeffrey Barrett, Sonia Goncalves, Ewan Harrison, David K. Jackson, Ian Johnston, Dominic Kwiatkowski, Cordelia Langford, John Sillitoe on behalf of the Wellcome Sanger Institute COVID-19 Surveillance Team |
| EPI_ISL_2703268 | Lighthouse Lab in Alderley Park | Wellcome Sanger Institute for the COVID-19 Genomics UK (COG-UK) Consortium | Jacquelyn Wynn, Mairead Hyland, The Lighthouse Lab in Alderley Park and Alex Alderton, Roberto Amato, Jeffrey Barrett, Sonia Goncalves, Ewan Harrison, David K. Jackson, Ian Johnston, Dominic Kwiatkowski, Cordelia Langford, John Sillitoe on behalf of the Wellcome Sanger Institute COVID-19 Surveillance Team |
| EPI_ISL_2703269, EPI_ISL_2703270, EPI_ISL_2703271 | Lighthouse Lab in Milton Keynes | Wellcome Sanger Institute for the COVID-19 Genomics UK (COG-UK) Consortium | The Lighthouse Lab in Milton Keynes and Alex Alderton, Roberto Amato, Jeffrey Barrett, Sonia Goncalves, Ewan Harrison, David K. Jackson, Ian Johnston, Dominic Kwiatkowski, Cordelia Langford, John Sillitoe on behalf of the Wellcome Sanger Institute COVID-19 Surveillance Team |
| EPI_ISL_2703272 | Lighthouse Lab in Alderley Park | Wellcome Sanger Institute for the COVID-19 Genomics UK (COG-UK) Consortium | Jacquelyn Wynn, Mairead Hyland, The Lighthouse Lab in Alderley Park and Alex Alderton, Roberto Amato, Jeffrey Barrett, Sonia Goncalves, Ewan Harrison, David K. Jackson, Ian Johnston, Dominic Kwiatkowski, Cordelia Langford, John Sillitoe on |

| | | | |
|---|---|---|---|
| | | Consortium | behalf of the Wellcome Sanger Institute COVID-19 Surveillance Team |
| EPI_ISL_2703273 | Lighthouse Lab in Milton Keynes | Wellcome Sanger Institute for the COVID-19 Genomics UK (COG-UK) Consortium | The Lighthouse Lab in Milton Keynes and Alex Alderton, Roberto Amato, Jeffrey Barrett, Sonia Goncalves, Ewan Harrison, David K. Jackson, Ian Johnston, Dominic Kwiatkowski, Cordelia Langford, John Sillitoe on behalf of the Wellcome Sanger Institute COVID-19 Surveillance Team |
| EPI_ISL_2703274 | Lighthouse Lab in Alderley Park | Wellcome Sanger Institute for the COVID-19 Genomics UK (COG-UK) Consortium | Jacquelyn Wynn, Mairead Hyland, The Lighthouse Lab in Alderley Park and Alex Alderton, Roberto Amato, Jeffrey Barrett, Sonia Goncalves, Ewan Harrison, David K. Jackson, Ian Johnston, Dominic Kwiatkowski, Cordelia Langford, John Sillitoe on behalf of the Wellcome Sanger Institute COVID-19 Surveillance Team |
| EPI_ISL_2703275, EPI_ISL_2703276 | Lighthouse Lab in Milton Keynes | Wellcome Sanger Institute for the COVID-19 Genomics UK (COG-UK) Consortium | The Lighthouse Lab in Milton Keynes and Alex Alderton, Roberto Amato, Jeffrey Barrett, Sonia Goncalves, Ewan Harrison, David K. Jackson, Ian Johnston, Dominic Kwiatkowski, Cordelia Langford, John Sillitoe on behalf of the Wellcome Sanger Institute COVID-19 Surveillance Team |
| EPI_ISL_2703277, EPI_ISL_2703278 | Lighthouse Lab in Alderley Park | Wellcome Sanger Institute for the COVID-19 Genomics UK (COG-UK) Consortium | Jacquelyn Wynn, Mairead Hyland, The Lighthouse Lab in Alderley Park and Alex Alderton, Roberto Amato, Jeffrey Barrett, Sonia Goncalves, Ewan Harrison, David K. Jackson, Ian Johnston, Dominic Kwiatkowski, Cordelia Langford, John Sillitoe on behalf of the Wellcome Sanger Institute COVID-19 Surveillance Team |
| EPI_ISL_2703279 | Lighthouse Lab in Milton Keynes | Wellcome Sanger Institute for the COVID-19 Genomics UK (COG-UK) Consortium | The Lighthouse Lab in Milton Keynes and Alex Alderton, Roberto Amato, Jeffrey Barrett, Sonia Goncalves, Ewan Harrison, David K. Jackson, Ian Johnston, Dominic Kwiatkowski, Cordelia Langford, John Sillitoe on behalf of the Wellcome Sanger Institute COVID-19 Surveillance Team |
| EPI_ISL_2703280, EPI_ISL_2703281, EPI_ISL_2703282, EPI_ISL_2703283, EPI_ISL_2703284 | Lighthouse Lab in Alderley Park | Wellcome Sanger Institute for the COVID-19 Genomics UK (COG-UK) Consortium | Jacquelyn Wynn, Mairead Hyland, The Lighthouse Lab in Alderley Park and Alex Alderton, Roberto Amato, Jeffrey Barrett, Sonia Goncalves, Ewan Harrison, David K. Jackson, Ian Johnston, Dominic Kwiatkowski, Cordelia Langford, John Sillitoe on behalf of the Wellcome Sanger Institute COVID-19 Surveillance Team |
| EPI_ISL_2703285, EPI_ISL_2703286 | Lighthouse Lab in Milton Keynes | Wellcome Sanger Institute for the COVID-19 Genomics UK (COG-UK) Consortium | The Lighthouse Lab in Milton Keynes and Alex Alderton, Roberto Amato, Jeffrey Barrett, Sonia Goncalves, Ewan Harrison, David K. Jackson, Ian Johnston, Dominic Kwiatkowski, Cordelia Langford, John Sillitoe on behalf of the Wellcome Sanger Institute COVID-19 Surveillance Team |
| EPI_ISL_2703287, EPI_ISL_2703288, EPI_ISL_2703289, EPI_ISL_2703290 | Lighthouse Lab in Alderley Park | Wellcome Sanger Institute for the COVID-19 Genomics UK (COG-UK) Consortium | Jacquelyn Wynn, Mairead Hyland, The Lighthouse Lab in Alderley Park and Alex Alderton, Roberto Amato, Jeffrey Barrett, Sonia Goncalves, Ewan Harrison, David K. Jackson, Ian Johnston, Dominic Kwiatkowski, Cordelia Langford, John Sillitoe on behalf of the Wellcome Sanger Institute COVID-19 Surveillance Team |
| EPI_ISL_2703291, EPI_ISL_2703292 | Lighthouse Lab in Milton Keynes | Wellcome Sanger Institute for the COVID-19 Genomics UK (COG-UK) Consortium | The Lighthouse Lab in Milton Keynes and Alex Alderton, Roberto Amato, Jeffrey Barrett, Sonia Goncalves, Ewan Harrison, David K. Jackson, Ian Johnston, Dominic Kwiatkowski, Cordelia Langford, John Sillitoe on behalf of the Wellcome Sanger Institute COVID-19 Surveillance Team |
| EPI_ISL_2703293, EPI_ISL_2703294, EPI_ISL_2703295, EPI_ISL_2703296, EPI_ISL_2703297, EPI_ISL_2703298, EPI_ISL_2703299, EPI_ISL_2703300, EPI_ISL_2703301, EPI_ISL_2703302, EPI_ISL_2703303, EPI_ISL_2703304, EPI_ISL_2703305 | | | |
| see above | Lighthouse Lab in Alderley Park | Wellcome Sanger Institute for the COVID-19 Genomics UK (COG-UK) Consortium | Jacquelyn Wynn, Mairead Hyland, The Lighthouse Lab in Alderley Park and Alex Alderton, Roberto Amato, Jeffrey Barrett, Sonia Goncalves, Ewan Harrison, David K. Jackson, Ian Johnston, Dominic Kwiatkowski, Cordelia Langford, John Sillitoe on behalf of the Wellcome Sanger Institute COVID-19 Surveillance Team |
| EPI_ISL_2703306 | Lighthouse Lab in Milton Keynes | Wellcome Sanger Institute for the COVID-19 Genomics UK (COG-UK) Consortium | The Lighthouse Lab in Milton Keynes and Alex Alderton, Roberto Amato, Jeffrey Barrett, Sonia Goncalves, Ewan Harrison, David K. Jackson, Ian Johnston, Dominic Kwiatkowski, Cordelia Langford, John Sillitoe on behalf of the Wellcome Sanger Institute COVID-19 Surveillance Team |
| EPI_ISL_2703307 | Lighthouse Lab in Alderley Park | Wellcome Sanger Institute for the COVID-19 Genomics UK (COG-UK) Consortium | Jacquelyn Wynn, Mairead Hyland, The Lighthouse Lab in Alderley Park and Alex Alderton, Roberto Amato, Jeffrey Barrett, Sonia Goncalves, Ewan Harrison, David K. Jackson, Ian Johnston, Dominic Kwiatkowski, Cordelia Langford, John Sillitoe on behalf of the Wellcome Sanger Institute COVID-19 Surveillance Team |
| EPI_ISL_2703308 | Lighthouse Lab in Milton Keynes | Wellcome Sanger Institute for the COVID-19 Genomics UK (COG-UK) Consortium | The Lighthouse Lab in Milton Keynes and Alex Alderton, Roberto Amato, Jeffrey Barrett, Sonia Goncalves, Ewan Harrison, David K. Jackson, Ian Johnston, Dominic Kwiatkowski, Cordelia Langford, John Sillitoe on behalf of the Wellcome Sanger Institute COVID-19 Surveillance Team |
| EPI_ISL_2703309, EPI_ISL_2703310, EPI_ISL_2703311 | Lighthouse Lab in Alderley Park | Wellcome Sanger Institute for the COVID-19 Genomics UK (COG-UK) Consortium | Jacquelyn Wynn, Mairead Hyland, The Lighthouse Lab in Alderley Park and Alex Alderton, Roberto Amato, Jeffrey Barrett, Sonia Goncalves, Ewan Harrison, David K. Jackson, Ian Johnston, Dominic Kwiatkowski, Cordelia Langford, John Sillitoe on behalf of the Wellcome Sanger Institute COVID-19 Surveillance Team |
| EPI_ISL_2703312 | Lighthouse Lab in Milton Keynes | Wellcome Sanger Institute for the COVID-19 Genomics UK (COG-UK) Consortium | The Lighthouse Lab in Milton Keynes and Alex Alderton, Roberto Amato, Jeffrey Barrett, Sonia Goncalves, Ewan Harrison, David K. Jackson, Ian Johnston, Dominic Kwiatkowski, Cordelia Langford, John Sillitoe on behalf of the Wellcome Sanger Institute COVID-19 Surveillance Team |
| EPI_ISL_2703313, EPI_ISL_2703314, EPI_ISL_2703315, EPI_ISL_2703316 | Lighthouse Lab in Alderley Park | Wellcome Sanger Institute for the COVID-19 Genomics UK (COG-UK) Consortium | Jacquelyn Wynn, Mairead Hyland, The Lighthouse Lab in Alderley Park and Alex Alderton, Roberto Amato, Jeffrey Barrett, Sonia Goncalves, Ewan Harrison, David K. Jackson, Ian Johnston, Dominic Kwiatkowski, Cordelia Langford, John Sillitoe on behalf of the Wellcome Sanger Institute COVID-19 Surveillance Team |
| EPI_ISL_2703317, EPI_ISL_2703318 | Lighthouse Lab in Milton Keynes | Wellcome Sanger Institute for the COVID-19 Genomics UK (COG-UK) Consortium | The Lighthouse Lab in Milton Keynes and Alex Alderton, Roberto Amato, Jeffrey Barrett, Sonia Goncalves, Ewan Harrison, David K. Jackson, Ian Johnston, Dominic Kwiatkowski, Cordelia Langford, John Sillitoe on behalf of the Wellcome Sanger Institute COVID-19 Surveillance Team |
| EPI_ISL_2703319 | Lighthouse Lab in Alderley Park | Wellcome Sanger Institute for the COVID-19 Genomics UK (COG-UK) Consortium | Jacquelyn Wynn, Mairead Hyland, The Lighthouse Lab in Alderley Park and Alex Alderton, Roberto Amato, Jeffrey Barrett, Sonia Goncalves, Ewan Harrison, David K. Jackson, Ian Johnston, Dominic Kwiatkowski, Cordelia Langford, John Sillitoe on behalf of the Wellcome Sanger Institute COVID-19 Surveillance Team |
| EPI_ISL_2703320 | Lighthouse Lab in Milton Keynes | Wellcome Sanger Institute for the COVID-19 Genomics UK (COG-UK) Consortium | The Lighthouse Lab in Milton Keynes and Alex Alderton, Roberto Amato, Jeffrey Barrett, Sonia Goncalves, Ewan Harrison, David K. Jackson, Ian Johnston, Dominic Kwiatkowski, Cordelia Langford, John Sillitoe on behalf of the Wellcome Sanger Institute COVID-19 Surveillance Team |
| EPI_ISL_2703321 | Lighthouse Lab in Alderley Park | Wellcome Sanger Institute for the COVID-19 Genomics UK (COG-UK) Consortium | Jacquelyn Wynn, Mairead Hyland, The Lighthouse Lab in Alderley Park and Alex Alderton, Roberto Amato, Jeffrey Barrett, Sonia Goncalves, Ewan Harrison, David K. Jackson, Ian Johnston, Dominic Kwiatkowski, Cordelia Langford, John Sillitoe on behalf of the Wellcome Sanger Institute COVID-19 Surveillance Team |
| EPI_ISL_2703322, EPI_ISL_2703323 | Lighthouse Lab in Milton Keynes | Wellcome Sanger Institute for the COVID-19 Genomics UK (COG-UK) Consortium | The Lighthouse Lab in Milton Keynes and Alex Alderton, Roberto Amato, Jeffrey Barrett, Sonia Goncalves, Ewan Harrison, David K. Jackson, Ian Johnston, Dominic Kwiatkowski, Cordelia Langford, John Sillitoe on behalf of the Wellcome Sanger Institute COVID-19 Surveillance Team |
| EPI_ISL_2703324, EPI_ISL_2703325, EPI_ISL_2703326 | Lighthouse Lab in Alderley Park | Wellcome Sanger Institute for the COVID-19 Genomics UK (COG-UK) Consortium | Jacquelyn Wynn, Mairead Hyland, The Lighthouse Lab in Alderley Park and Alex Alderton, Roberto Amato, Jeffrey Barrett, Sonia Goncalves, Ewan Harrison, David K. Jackson, Ian Johnston, Dominic Kwiatkowski, Cordelia Langford, John Sillitoe on behalf of the Wellcome Sanger Institute COVID-19 Surveillance Team |
| EPI_ISL_2703327 | Lighthouse Lab in Milton Keynes | Wellcome Sanger Institute for the COVID-19 Genomics UK (COG-UK) Consortium | The Lighthouse Lab in Milton Keynes and Alex Alderton, Roberto Amato, Jeffrey Barrett, Sonia Goncalves, Ewan Harrison, David K. Jackson, Ian Johnston, Dominic Kwiatkowski, Cordelia Langford, John Sillitoe on behalf of the Wellcome Sanger Institute COVID-19 Surveillance Team |
| EPI_ISL_2703328 | Lighthouse Lab in Alderley Park | Wellcome Sanger Institute for the COVID-19 Genomics UK (COG-UK) Consortium | Jacquelyn Wynn, Mairead Hyland, The Lighthouse Lab in Alderley Park and Alex Alderton, Roberto Amato, Jeffrey Barrett, Sonia Goncalves, Ewan Harrison, David K. Jackson, Ian Johnston, Dominic Kwiatkowski, Cordelia Langford, John Sillitoe on behalf of the Wellcome Sanger Institute COVID-19 Surveillance Team |
| EPI_ISL_2703329, EPI_ISL_2703330 | Lighthouse Lab in Milton Keynes | Wellcome Sanger Institute for the COVID-19 Genomics UK (COG-UK) Consortium | The Lighthouse Lab in Milton Keynes and Alex Alderton, Roberto Amato, Jeffrey Barrett, Sonia Goncalves, Ewan Harrison, David K. Jackson, Ian Johnston, Dominic Kwiatkowski, Cordelia Langford, John Sillitoe on behalf of the Wellcome Sanger Institute COVID-19 Surveillance Team |
| EPI_ISL_2703331, EPI_ISL_2703332, EPI_ISL_2703333, EPI_ISL_2703334, EPI_ISL_2703335, EPI_ISL_2703336 | Lighthouse Lab in Alderley Park | Wellcome Sanger Institute for the COVID-19 Genomics UK (COG-UK) Consortium | Jacquelyn Wynn, Mairead Hyland, The Lighthouse Lab in Alderley Park and Alex Alderton, Roberto Amato, Jeffrey Barrett, Sonia Goncalves, Ewan Harrison, David K. Jackson, Ian Johnston, Dominic Kwiatkowski, Cordelia Langford, John Sillitoe on behalf of the Wellcome Sanger Institute COVID-19 Surveillance Team |
| EPI_ISL_2703337 | Lighthouse Lab in Milton Keynes | Wellcome Sanger Institute for the COVID-19 Genomics UK (COG-UK) Consortium | The Lighthouse Lab in Milton Keynes and Alex Alderton, Roberto Amato, Jeffrey Barrett, Sonia Goncalves, Ewan Harrison, David K. Jackson, Ian Johnston, Dominic Kwiatkowski, Cordelia Langford, John Sillitoe on behalf of the Wellcome Sanger Institute COVID-19 Surveillance Team |
| EPI_ISL_2703338, EPI_ISL_2703339, EPI_ISL_2703340 | Lighthouse Lab in Alderley Park | Wellcome Sanger Institute for the COVID-19 Genomics UK (COG-UK) Consortium | Jacquelyn Wynn, Mairead Hyland, The Lighthouse Lab in Alderley Park and Alex Alderton, Roberto Amato, Jeffrey Barrett, Sonia Goncalves, Ewan Harrison, David K. Jackson, Ian Johnston, Dominic Kwiatkowski, Cordelia Langford, John Sillitoe on behalf of the Wellcome Sanger Institute COVID-19 Surveillance Team |
| EPI_ISL_2703341 | Lighthouse Lab in Milton Keynes | Wellcome Sanger Institute for the COVID-19 Genomics UK (COG-UK) Consortium | The Lighthouse Lab in Milton Keynes and Alex Alderton, Roberto Amato, Jeffrey Barrett, Sonia Goncalves, Ewan Harrison, David K. Jackson, Ian Johnston, Dominic Kwiatkowski, Cordelia Langford, John Sillitoe on behalf of the Wellcome Sanger Institute COVID-19 Surveillance Team |
| EPI_ISL_2703342, EPI_ISL_2703343, EPI_ISL_2703344, EPI_ISL_2703345, EPI_ISL_2703346, EPI_ISL_2703347 | Lighthouse Lab in Alderley Park | Wellcome Sanger Institute for the COVID-19 Genomics UK (COG-UK) Consortium | Jacquelyn Wynn, Mairead Hyland, The Lighthouse Lab in Alderley Park and Alex Alderton, Roberto Amato, Jeffrey Barrett, Sonia Goncalves, Ewan Harrison, David K. Jackson, Ian Johnston, Dominic Kwiatkowski, Cordelia Langford, John Sillitoe on behalf of the Wellcome Sanger Institute COVID-19 Surveillance Team |
| EPI_ISL_2703348, EPI_ISL_2703349 | Lighthouse Lab in Milton Keynes | Wellcome Sanger Institute for the COVID-19 Genomics UK (COG-UK) Consortium | The Lighthouse Lab in Milton Keynes and Alex Alderton, Roberto Amato, Jeffrey Barrett, Sonia Goncalves, Ewan Harrison, David K. Jackson, Ian Johnston, Dominic Kwiatkowski, Cordelia Langford, John Sillitoe on behalf of the Wellcome Sanger Institute COVID-19 Surveillance Team |
| EPI_ISL_2703350, EPI_ISL_2703351, EPI_ISL_2703352, EPI_ISL_2703353, EPI_ISL_2703354, EPI_ISL_2703355, EPI_ISL_2703356, EPI_ISL_2703357, EPI_ISL_2703358, EPI_ISL_2703359, EPI_ISL_2703360, EPI_ISL_2703361 | | | |
| see above | Lighthouse Lab in Alderley Park | Wellcome Sanger Institute for the COVID-19 Genomics UK (COG-UK) Consortium | Jacquelyn Wynn, Mairead Hyland, The Lighthouse Lab in Alderley Park and Alex Alderton, Roberto Amato, Jeffrey Barrett, Sonia Goncalves, Ewan Harrison, David K. Jackson, Ian Johnston, Dominic Kwiatkowski, Cordelia Langford, John Sillitoe on behalf of the Wellcome Sanger Institute COVID-19 Surveillance Team |
| EPI_ISL_2703362 | Lighthouse Lab in Milton Keynes | Wellcome Sanger Institute for the COVID-19 Genomics UK (COG-UK) Consortium | The Lighthouse Lab in Milton Keynes and Alex Alderton, Roberto Amato, Jeffrey Barrett, Sonia Goncalves, Ewan Harrison, David K. Jackson, Ian Johnston, Dominic Kwiatkowski, Cordelia Langford, John Sillitoe on behalf of the Wellcome Sanger Institute COVID-19 Surveillance Team |
| EPI_ISL_2703363, EPI_ISL_2703364, EPI_ISL_2703365 | Lighthouse Lab in Alderley Park | Wellcome Sanger Institute for the COVID-19 Genomics UK (COG-UK) Consortium | Jacquelyn Wynn, Mairead Hyland, The Lighthouse Lab in Alderley Park and Alex Alderton, Roberto Amato, Jeffrey Barrett, Sonia Goncalves, Ewan Harrison, David K. Jackson, Ian Johnston, Dominic Kwiatkowski, Cordelia Langford, John Sillitoe on behalf of the Wellcome Sanger Institute COVID-19 Surveillance Team |
| EPI_ISL_2703366 | Lighthouse Lab in Milton Keynes | Wellcome Sanger Institute for the COVID-19 Genomics UK (COG-UK) Consortium | The Lighthouse Lab in Milton Keynes and Alex Alderton, Roberto Amato, Jeffrey Barrett, Sonia Goncalves, Ewan Harrison, David K. Jackson, Ian Johnston, Dominic Kwiatkowski, Cordelia Langford, John Sillitoe on behalf of the Wellcome Sanger Institute COVID-19 Surveillance Team |
| EPI_ISL_2703367 | Lighthouse Lab in Alderley Park | Wellcome Sanger Institute for the COVID-19 Genomics UK (COG-UK) | Jacquelyn Wynn, Mairead Hyland, The Lighthouse Lab in Alderley Park and Alex Alderton, Roberto Amato, Jeffrey Barrett, Sonia Goncalves, Ewan Harrison, David K. Jackson, Ian Johnston, Dominic Kwiatkowski, Cordelia Langford, John Sillitoe on |

| | | | |
|---|---|---|---|
| | | Consortium | behalf of the Wellcome Sanger Institute COVID-19 Surveillance Team |
| EPI_ISL_2703368 | Lighthouse Lab in Milton Keynes | Wellcome Sanger Institute for the COVID-19 Genomics UK (COG-UK) Consortium | The Lighthouse Lab in Milton Keynes and Alex Alderton, Roberto Amato, Jeffrey Barrett, Sonia Goncalves, Ewan Harrison, David K. Jackson, Ian Johnston, Dominic Kwiatkowski, Cordelia Langford, John Sillitoe on behalf of the Wellcome Sanger Institute COVID-19 Surveillance Team |
| EPI_ISL_2703369, EPI_ISL_2703370, EPI_ISL_2703371, EPI_ISL_2703372 | Lighthouse Lab in Alderley Park | Wellcome Sanger Institute for the COVID-19 Genomics UK (COG-UK) Consortium | Jacquelyn Wynn, Mairead Hyland, The Lighthouse Lab in Alderley Park and Alex Alderton, Roberto Amato, Jeffrey Barrett, Sonia Goncalves, Ewan Harrison, David K. Jackson, Ian Johnston, Dominic Kwiatkowski, Cordelia Langford, John Sillitoe on behalf of the Wellcome Sanger Institute COVID-19 Surveillance Team |
| EPI_ISL_2703373, EPI_ISL_2703374 | Lighthouse Lab in Milton Keynes | Wellcome Sanger Institute for the COVID-19 Genomics UK (COG-UK) Consortium | The Lighthouse Lab in Milton Keynes and Alex Alderton, Roberto Amato, Jeffrey Barrett, Sonia Goncalves, Ewan Harrison, David K. Jackson, Ian Johnston, Dominic Kwiatkowski, Cordelia Langford, John Sillitoe on behalf of the Wellcome Sanger Institute COVID-19 Surveillance Team |
| EPI_ISL_2703375, EPI_ISL_2703376, EPI_ISL_2703377, EPI_ISL_2703378, EPI_ISL_2703380 | Lighthouse Lab in Alderley Park | Wellcome Sanger Institute for the COVID-19 Genomics UK (COG-UK) Consortium | Jacquelyn Wynn, Mairead Hyland, The Lighthouse Lab in Alderley Park and Alex Alderton, Roberto Amato, Jeffrey Barrett, Sonia Goncalves, Ewan Harrison, David K. Jackson, Ian Johnston, Dominic Kwiatkowski, Cordelia Langford, John Sillitoe on behalf of the Wellcome Sanger Institute COVID-19 Surveillance Team |
| EPI_ISL_2703381, EPI_ISL_2703382, EPI_ISL_2703384 | Lighthouse Lab in Milton Keynes | Wellcome Sanger Institute for the COVID-19 Genomics UK (COG-UK) Consortium | The Lighthouse Lab in Milton Keynes and Alex Alderton, Roberto Amato, Jeffrey Barrett, Sonia Goncalves, Ewan Harrison, David K. Jackson, Ian Johnston, Dominic Kwiatkowski, Cordelia Langford, John Sillitoe on behalf of the Wellcome Sanger Institute COVID-19 Surveillance Team |
| EPI_ISL_2703385, EPI_ISL_2703386, EPI_ISL_2703387, EPI_ISL_2703388, EPI_ISL_2703389, EPI_ISL_2703390, EPI_ISL_2703391, EPI_ISL_2703392, EPI_ISL_2703393, EPI_ISL_2703394, EPI_ISL_2703395, EPI_ISL_2703396, EPI_ISL_2703397 | | | |
| see above | Lighthouse Lab in Alderley Park | Wellcome Sanger Institute for the COVID-19 Genomics UK (COG-UK) Consortium | Jacquelyn Wynn, Mairead Hyland, The Lighthouse Lab in Alderley Park and Alex Alderton, Roberto Amato, Jeffrey Barrett, Sonia Goncalves, Ewan Harrison, David K. Jackson, Ian Johnston, Dominic Kwiatkowski, Cordelia Langford, John Sillitoe on behalf of the Wellcome Sanger Institute COVID-19 Surveillance Team |
| EPI_ISL_2703398 | Lighthouse Lab in Milton Keynes | Wellcome Sanger Institute for the COVID-19 Genomics UK (COG-UK) Consortium | The Lighthouse Lab in Milton Keynes and Alex Alderton, Roberto Amato, Jeffrey Barrett, Sonia Goncalves, Ewan Harrison, David K. Jackson, Ian Johnston, Dominic Kwiatkowski, Cordelia Langford, John Sillitoe on behalf of the Wellcome Sanger Institute COVID-19 Surveillance Team |
| EPI_ISL_2703399, EPI_ISL_2703400, EPI_ISL_2703401, EPI_ISL_2703402, EPI_ISL_2703403 | Lighthouse Lab in Alderley Park | Wellcome Sanger Institute for the COVID-19 Genomics UK (COG-UK) Consortium | Jacquelyn Wynn, Mairead Hyland, The Lighthouse Lab in Alderley Park and Alex Alderton, Roberto Amato, Jeffrey Barrett, Sonia Goncalves, Ewan Harrison, David K. Jackson, Ian Johnston, Dominic Kwiatkowski, Cordelia Langford, John Sillitoe on behalf of the Wellcome Sanger Institute COVID-19 Surveillance Team |
| EPI_ISL_2703404, EPI_ISL_2703405 | Lighthouse Lab in Milton Keynes | Wellcome Sanger Institute for the COVID-19 Genomics UK (COG-UK) Consortium | The Lighthouse Lab in Milton Keynes and Alex Alderton, Roberto Amato, Jeffrey Barrett, Sonia Goncalves, Ewan Harrison, David K. Jackson, Ian Johnston, Dominic Kwiatkowski, Cordelia Langford, John Sillitoe on behalf of the Wellcome Sanger Institute COVID-19 Surveillance Team |
| EPI_ISL_2703406, EPI_ISL_2703407, EPI_ISL_2703408, EPI_ISL_2703409, EPI_ISL_2703410, EPI_ISL_2703411 | Lighthouse Lab in Alderley Park | Wellcome Sanger Institute for the COVID-19 Genomics UK (COG-UK) Consortium | Jacquelyn Wynn, Mairead Hyland, The Lighthouse Lab in Alderley Park and Alex Alderton, Roberto Amato, Jeffrey Barrett, Sonia Goncalves, Ewan Harrison, David K. Jackson, Ian Johnston, Dominic Kwiatkowski, Cordelia Langford, John Sillitoe on behalf of the Wellcome Sanger Institute COVID-19 Surveillance Team |
| EPI_ISL_2703412 | Lighthouse Lab in Milton Keynes | Wellcome Sanger Institute for the COVID-19 Genomics UK (COG-UK) Consortium | The Lighthouse Lab in Milton Keynes and Alex Alderton, Roberto Amato, Jeffrey Barrett, Sonia Goncalves, Ewan Harrison, David K. Jackson, Ian Johnston, Dominic Kwiatkowski, Cordelia Langford, John Sillitoe on behalf of the Wellcome Sanger Institute COVID-19 Surveillance Team |
| EPI_ISL_2703413, EPI_ISL_2703414, EPI_ISL_2703415, EPI_ISL_2703416, EPI_ISL_2703417, EPI_ISL_2703418, EPI_ISL_2703419 | Lighthouse Lab in Alderley Park | Wellcome Sanger Institute for the COVID-19 Genomics UK (COG-UK) Consortium | Jacquelyn Wynn, Mairead Hyland, The Lighthouse Lab in Alderley Park and Alex Alderton, Roberto Amato, Jeffrey Barrett, Sonia Goncalves, Ewan Harrison, David K. Jackson, Ian Johnston, Dominic Kwiatkowski, Cordelia Langford, John Sillitoe on behalf of the Wellcome Sanger Institute COVID-19 Surveillance Team |
| EPI_ISL_2703420 | Lighthouse Lab in Milton Keynes | Wellcome Sanger Institute for the COVID-19 Genomics UK (COG-UK) Consortium | The Lighthouse Lab in Milton Keynes and Alex Alderton, Roberto Amato, Jeffrey Barrett, Sonia Goncalves, Ewan Harrison, David K. Jackson, Ian Johnston, Dominic Kwiatkowski, Cordelia Langford, John Sillitoe on behalf of the Wellcome Sanger Institute COVID-19 Surveillance Team |
| EPI_ISL_2703421, EPI_ISL_2703422 | Lighthouse Lab in Alderley Park | Wellcome Sanger Institute for the COVID-19 Genomics UK (COG-UK) Consortium | Jacquelyn Wynn, Mairead Hyland, The Lighthouse Lab in Alderley Park and Alex Alderton, Roberto Amato, Jeffrey Barrett, Sonia Goncalves, Ewan Harrison, David K. Jackson, Ian Johnston, Dominic Kwiatkowski, Cordelia Langford, John Sillitoe on behalf of the Wellcome Sanger Institute COVID-19 Surveillance Team |
| EPI_ISL_2703423, EPI_ISL_2703424 | Lighthouse Lab in Milton Keynes | Wellcome Sanger Institute for the COVID-19 Genomics UK (COG-UK) Consortium | The Lighthouse Lab in Milton Keynes and Alex Alderton, Roberto Amato, Jeffrey Barrett, Sonia Goncalves, Ewan Harrison, David K. Jackson, Ian Johnston, Dominic Kwiatkowski, Cordelia Langford, John Sillitoe on behalf of the Wellcome Sanger Institute COVID-19 Surveillance Team |
| EPI_ISL_2703425, EPI_ISL_2703426, EPI_ISL_2703427, EPI_ISL_2703428, EPI_ISL_2703429, EPI_ISL_2703430, EPI_ISL_2703431, EPI_ISL_2703432, EPI_ISL_2703433, EPI_ISL_2703434, EPI_ISL_2703435, EPI_ISL_2703436, EPI_ISL_2703437 | | | |
| see above | Lighthouse Lab in Alderley Park | Wellcome Sanger Institute for the COVID-19 Genomics UK (COG-UK) Consortium | Jacquelyn Wynn, Mairead Hyland, The Lighthouse Lab in Alderley Park and Alex Alderton, Roberto Amato, Jeffrey Barrett, Sonia Goncalves, Ewan Harrison, David K. Jackson, Ian Johnston, Dominic Kwiatkowski, Cordelia Langford, John Sillitoe on behalf of the Wellcome Sanger Institute COVID-19 Surveillance Team |
| EPI_ISL_2703438 | Lighthouse Lab in Milton Keynes | Wellcome Sanger Institute for the COVID-19 Genomics UK (COG-UK) Consortium | The Lighthouse Lab in Milton Keynes and Alex Alderton, Roberto Amato, Jeffrey Barrett, Sonia Goncalves, Ewan Harrison, David K. Jackson, Ian Johnston, Dominic Kwiatkowski, Cordelia Langford, John Sillitoe on behalf of the Wellcome Sanger Institute COVID-19 Surveillance Team |
| EPI_ISL_2703439, EPI_ISL_2703440, EPI_ISL_2703442 | Lighthouse Lab in Alderley Park | Wellcome Sanger Institute for the COVID-19 Genomics UK (COG-UK) Consortium | Jacquelyn Wynn, Mairead Hyland, The Lighthouse Lab in Alderley Park and Alex Alderton, Roberto Amato, Jeffrey Barrett, Sonia Goncalves, Ewan Harrison, David K. Jackson, Ian Johnston, Dominic Kwiatkowski, Cordelia Langford, John Sillitoe on behalf of the Wellcome Sanger Institute COVID-19 Surveillance Team |
| EPI_ISL_2703443 | Lighthouse Lab in Milton Keynes | Wellcome Sanger Institute for the COVID-19 Genomics UK (COG-UK) Consortium | The Lighthouse Lab in Milton Keynes and Alex Alderton, Roberto Amato, Jeffrey Barrett, Sonia Goncalves, Ewan Harrison, David K. Jackson, Ian Johnston, Dominic Kwiatkowski, Cordelia Langford, John Sillitoe on behalf of the Wellcome Sanger Institute COVID-19 Surveillance Team |
| EPI_ISL_2703444 | Lighthouse Lab in Alderley Park | Wellcome Sanger Institute for the COVID-19 Genomics UK (COG-UK) Consortium | Jacquelyn Wynn, Mairead Hyland, The Lighthouse Lab in Alderley Park and Alex Alderton, Roberto Amato, Jeffrey Barrett, Sonia Goncalves, Ewan Harrison, David K. Jackson, Ian Johnston, Dominic Kwiatkowski, Cordelia Langford, John Sillitoe on behalf of the Wellcome Sanger Institute COVID-19 Surveillance Team |
| EPI_ISL_2703445, EPI_ISL_2703446 | Lighthouse Lab in Milton Keynes | Wellcome Sanger Institute for the COVID-19 Genomics UK (COG-UK) Consortium | The Lighthouse Lab in Milton Keynes and Alex Alderton, Roberto Amato, Jeffrey Barrett, Sonia Goncalves, Ewan Harrison, David K. Jackson, Ian Johnston, Dominic Kwiatkowski, Cordelia Langford, John Sillitoe on behalf of the Wellcome Sanger Institute COVID-19 Surveillance Team |
| EPI_ISL_2703447, EPI_ISL_2703448, EPI_ISL_2703450, EPI_ISL_2703451, EPI_ISL_2703452, EPI_ISL_2703453 | Lighthouse Lab in Alderley Park | Wellcome Sanger Institute for the COVID-19 Genomics UK (COG-UK) Consortium | Jacquelyn Wynn, Mairead Hyland, The Lighthouse Lab in Alderley Park and Alex Alderton, Roberto Amato, Jeffrey Barrett, Sonia Goncalves, Ewan Harrison, David K. Jackson, Ian Johnston, Dominic Kwiatkowski, Cordelia Langford, John Sillitoe on behalf of the Wellcome Sanger Institute COVID-19 Surveillance Team |
| EPI_ISL_2703454 | Lighthouse Lab in Milton Keynes | Wellcome Sanger Institute for the COVID-19 Genomics UK (COG-UK) Consortium | The Lighthouse Lab in Milton Keynes and Alex Alderton, Roberto Amato, Jeffrey Barrett, Sonia Goncalves, Ewan Harrison, David K. Jackson, Ian Johnston, Dominic Kwiatkowski, Cordelia Langford, John Sillitoe on behalf of the Wellcome Sanger Institute COVID-19 Surveillance Team |
| EPI_ISL_2703455, EPI_ISL_2703456 | Lighthouse Lab in Alderley Park | Wellcome Sanger Institute for the COVID-19 Genomics UK (COG-UK) Consortium | Jacquelyn Wynn, Mairead Hyland, The Lighthouse Lab in Alderley Park and Alex Alderton, Roberto Amato, Jeffrey Barrett, Sonia Goncalves, Ewan Harrison, David K. Jackson, Ian Johnston, Dominic Kwiatkowski, Cordelia Langford, John Sillitoe on behalf of the Wellcome Sanger Institute COVID-19 Surveillance Team |
| EPI_ISL_2703457 | Lighthouse Lab in Milton Keynes | Wellcome Sanger Institute for the COVID-19 Genomics UK (COG-UK) Consortium | The Lighthouse Lab in Milton Keynes and Alex Alderton, Roberto Amato, Jeffrey Barrett, Sonia Goncalves, Ewan Harrison, David K. Jackson, Ian Johnston, Dominic Kwiatkowski, Cordelia Langford, John Sillitoe on behalf of the Wellcome Sanger Institute COVID-19 Surveillance Team |
| EPI_ISL_2703458, EPI_ISL_2703459, EPI_ISL_2703460, EPI_ISL_2703461, EPI_ISL_2703462, EPI_ISL_2703463, EPI_ISL_2703464, EPI_ISL_2703465, EPI_ISL_2703466 | Lighthouse Lab in Alderley Park | Wellcome Sanger Institute for the COVID-19 Genomics UK (COG-UK) Consortium | Jacquelyn Wynn, Mairead Hyland, The Lighthouse Lab in Alderley Park and Alex Alderton, Roberto Amato, Jeffrey Barrett, Sonia Goncalves, Ewan Harrison, David K. Jackson, Ian Johnston, Dominic Kwiatkowski, Cordelia Langford, John Sillitoe on behalf of the Wellcome Sanger Institute COVID-19 Surveillance Team |
| EPI_ISL_2703467 | Lighthouse Lab in Milton Keynes | Wellcome Sanger Institute for the COVID-19 Genomics UK (COG-UK) Consortium | The Lighthouse Lab in Milton Keynes and Alex Alderton, Roberto Amato, Jeffrey Barrett, Sonia Goncalves, Ewan Harrison, David K. Jackson, Ian Johnston, Dominic Kwiatkowski, Cordelia Langford, John Sillitoe on behalf of the Wellcome Sanger Institute COVID-19 Surveillance Team |
| EPI_ISL_2703468 | Lighthouse Lab in Alderley Park | Wellcome Sanger Institute for the COVID-19 Genomics UK (COG-UK) Consortium | Jacquelyn Wynn, Mairead Hyland, The Lighthouse Lab in Alderley Park and Alex Alderton, Roberto Amato, Jeffrey Barrett, Sonia Goncalves, Ewan Harrison, David K. Jackson, Ian Johnston, Dominic Kwiatkowski, Cordelia Langford, John Sillitoe on behalf of the Wellcome Sanger Institute COVID-19 Surveillance Team |
| EPI_ISL_2703469 | Lighthouse Lab in Milton Keynes | Wellcome Sanger Institute for the COVID-19 Genomics UK (COG-UK) Consortium | The Lighthouse Lab in Milton Keynes and Alex Alderton, Roberto Amato, Jeffrey Barrett, Sonia Goncalves, Ewan Harrison, David K. Jackson, Ian Johnston, Dominic Kwiatkowski, Cordelia Langford, John Sillitoe on behalf of the Wellcome Sanger Institute COVID-19 Surveillance Team |
| EPI_ISL_2703470, EPI_ISL_2703471, EPI_ISL_2703472 | Lighthouse Lab in Alderley Park | Wellcome Sanger Institute for the COVID-19 Genomics UK (COG-UK) Consortium | Jacquelyn Wynn, Mairead Hyland, The Lighthouse Lab in Alderley Park and Alex Alderton, Roberto Amato, Jeffrey Barrett, Sonia Goncalves, Ewan Harrison, David K. Jackson, Ian Johnston, Dominic Kwiatkowski, Cordelia Langford, John Sillitoe on behalf of the Wellcome Sanger Institute COVID-19 Surveillance Team |
| EPI_ISL_2703473 | Lighthouse Lab in Milton Keynes | Wellcome Sanger Institute for the COVID-19 Genomics UK (COG-UK) Consortium | The Lighthouse Lab in Milton Keynes and Alex Alderton, Roberto Amato, Jeffrey Barrett, Sonia Goncalves, Ewan Harrison, David K. Jackson, Ian Johnston, Dominic Kwiatkowski, Cordelia Langford, John Sillitoe on behalf of the Wellcome Sanger Institute COVID-19 Surveillance Team |

| | | | |
|---|---|---|---|
| EPI_ISL_2703474 | Lighthouse Lab in Alderley Park | Wellcome Sanger Institute for the COVID-19 Genomics UK (COG-UK) Consortium | Jacquelyn Wynn, Mairead Hyland, The Lighthouse Lab in Alderley Park and Alex Alderton, Roberto Amato, Jeffrey Barrett, Sonia Goncalves, Ewan Harrison, David K. Jackson, Ian Johnston, Dominic Kwiatkowski, Cordelia Langford, John Sillitoe on behalf of the Wellcome Sanger Institute COVID-19 Surveillance Team |
| EPI_ISL_2703475 | Lighthouse Lab in Milton Keynes | Wellcome Sanger Institute for the COVID-19 Genomics UK (COG-UK) Consortium | The Lighthouse Lab in Milton Keynes and Alex Alderton, Roberto Amato, Jeffrey Barrett, Sonia Goncalves, Ewan Harrison, David K. Jackson, Ian Johnston, Dominic Kwiatkowski, Cordelia Langford, John Sillitoe on behalf of the Wellcome Sanger Institute COVID-19 Surveillance Team |
| EPI_ISL_2703476, EPI_ISL_2703477, EPI_ISL_2703478, EPI_ISL_2703479, EPI_ISL_2703480, EPI_ISL_2703481, EPI_ISL_2703482, EPI_ISL_2703483 | Lighthouse Lab in Alderley Park | Wellcome Sanger Institute for the COVID-19 Genomics UK (COG-UK) Consortium | Jacquelyn Wynn, Mairead Hyland, The Lighthouse Lab in Alderley Park and Alex Alderton, Roberto Amato, Jeffrey Barrett, Sonia Goncalves, Ewan Harrison, David K. Jackson, Ian Johnston, Dominic Kwiatkowski, Cordelia Langford, John Sillitoe on behalf of the Wellcome Sanger Institute COVID-19 Surveillance Team |
| EPI_ISL_2703484 | Lighthouse Lab in Milton Keynes | Wellcome Sanger Institute for the COVID-19 Genomics UK (COG-UK) Consortium | The Lighthouse Lab in Milton Keynes and Alex Alderton, Roberto Amato, Jeffrey Barrett, Sonia Goncalves, Ewan Harrison, David K. Jackson, Ian Johnston, Dominic Kwiatkowski, Cordelia Langford, John Sillitoe on behalf of the Wellcome Sanger Institute COVID-19 Surveillance Team |
| EPI_ISL_2703485, EPI_ISL_2703486, EPI_ISL_2703487, EPI_ISL_2703488 | Lighthouse Lab in Alderley Park | Wellcome Sanger Institute for the COVID-19 Genomics UK (COG-UK) Consortium | Jacquelyn Wynn, Mairead Hyland, The Lighthouse Lab in Alderley Park and Alex Alderton, Roberto Amato, Jeffrey Barrett, Sonia Goncalves, Ewan Harrison, David K. Jackson, Ian Johnston, Dominic Kwiatkowski, Cordelia Langford, John Sillitoe on behalf of the Wellcome Sanger Institute COVID-19 Surveillance Team |
| EPI_ISL_2703489 | Lighthouse Lab in Milton Keynes | Wellcome Sanger Institute for the COVID-19 Genomics UK (COG-UK) Consortium | The Lighthouse Lab in Milton Keynes and Alex Alderton, Roberto Amato, Jeffrey Barrett, Sonia Goncalves, Ewan Harrison, David K. Jackson, Ian Johnston, Dominic Kwiatkowski, Cordelia Langford, John Sillitoe on behalf of the Wellcome Sanger Institute COVID-19 Surveillance Team |
| EPI_ISL_2703490, EPI_ISL_2703491, EPI_ISL_2703492 | Lighthouse Lab in Alderley Park | Wellcome Sanger Institute for the COVID-19 Genomics UK (COG-UK) Consortium | Jacquelyn Wynn, Mairead Hyland, The Lighthouse Lab in Alderley Park and Alex Alderton, Roberto Amato, Jeffrey Barrett, Sonia Goncalves, Ewan Harrison, David K. Jackson, Ian Johnston, Dominic Kwiatkowski, Cordelia Langford, John Sillitoe on behalf of the Wellcome Sanger Institute COVID-19 Surveillance Team |
| EPI_ISL_2703493 | Lighthouse Lab in Milton Keynes | Wellcome Sanger Institute for the COVID-19 Genomics UK (COG-UK) Consortium | The Lighthouse Lab in Milton Keynes and Alex Alderton, Roberto Amato, Jeffrey Barrett, Sonia Goncalves, Ewan Harrison, David K. Jackson, Ian Johnston, Dominic Kwiatkowski, Cordelia Langford, John Sillitoe on behalf of the Wellcome Sanger Institute COVID-19 Surveillance Team |
| EPI_ISL_2703494, EPI_ISL_2703495, EPI_ISL_2703496, EPI_ISL_2703497, EPI_ISL_2703498, EPI_ISL_2703499, EPI_ISL_2703500, EPI_ISL_2703501 | Lighthouse Lab in Alderley Park | Wellcome Sanger Institute for the COVID-19 Genomics UK (COG-UK) Consortium | Jacquelyn Wynn, Mairead Hyland, The Lighthouse Lab in Alderley Park and Alex Alderton, Roberto Amato, Jeffrey Barrett, Sonia Goncalves, Ewan Harrison, David K. Jackson, Ian Johnston, Dominic Kwiatkowski, Cordelia Langford, John Sillitoe on behalf of the Wellcome Sanger Institute COVID-19 Surveillance Team |
| EPI_ISL_2703502 | Lighthouse Lab in Milton Keynes | Wellcome Sanger Institute for the COVID-19 Genomics UK (COG-UK) Consortium | The Lighthouse Lab in Milton Keynes and Alex Alderton, Roberto Amato, Jeffrey Barrett, Sonia Goncalves, Ewan Harrison, David K. Jackson, Ian Johnston, Dominic Kwiatkowski, Cordelia Langford, John Sillitoe on behalf of the Wellcome Sanger Institute COVID-19 Surveillance Team |
| EPI_ISL_2703503 | Lighthouse Lab in Alderley Park | Wellcome Sanger Institute for the COVID-19 Genomics UK (COG-UK) Consortium | Jacquelyn Wynn, Mairead Hyland, The Lighthouse Lab in Alderley Park and Alex Alderton, Roberto Amato, Jeffrey Barrett, Sonia Goncalves, Ewan Harrison, David K. Jackson, Ian Johnston, Dominic Kwiatkowski, Cordelia Langford, John Sillitoe on behalf of the Wellcome Sanger Institute COVID-19 Surveillance Team |
| EPI_ISL_2703504 | Lighthouse Lab in Glasgow | Wellcome Sanger Institute for the COVID-19 Genomics UK (COG-UK) Consortium | Harper VanSteenhouse, Yumi Kasai, David Gray, Carol Clugston, Anna Dominiczak and Alex Alderton, Roberto Amato, Jeffrey Barrett, Sonia Goncalves, Ewan Harrison, David K. Jackson, Ian Johnston, Dominic Kwiatkowski, Cordelia Langford, John Sillitoe on behalf of the Wellcome Sanger Institute COVID-19 Surveillance Team |
| EPI_ISL_2703505 | Lighthouse Lab in Milton Keynes | Wellcome Sanger Institute for the COVID-19 Genomics UK (COG-UK) Consortium | The Lighthouse Lab in Milton Keynes and Alex Alderton, Roberto Amato, Jeffrey Barrett, Sonia Goncalves, Ewan Harrison, David K. Jackson, Ian Johnston, Dominic Kwiatkowski, Cordelia Langford, John Sillitoe on behalf of the Wellcome Sanger Institute COVID-19 Surveillance Team |
| EPI_ISL_2703506 | Lighthouse Lab in Glasgow | Wellcome Sanger Institute for the COVID-19 Genomics UK (COG-UK) Consortium | Harper VanSteenhouse, Yumi Kasai, David Gray, Carol Clugston, Anna Dominiczak and Alex Alderton, Roberto Amato, Jeffrey Barrett, Sonia Goncalves, Ewan Harrison, David K. Jackson, Ian Johnston, Dominic Kwiatkowski, Cordelia Langford, John Sillitoe on behalf of the Wellcome Sanger Institute COVID-19 Surveillance Team |
| EPI_ISL_2703507 | Lighthouse Laboratory Plymouth | Wellcome Sanger Institute for the COVID-19 Genomics UK (COG-UK) Consortium | Lighthouse Laboratory Plymouth and Alex Alderton, Roberto Amato, Jeffrey Barrett, Sonia Goncalves, Ewan Harrison, David K. Jackson, Ian Johnston, Dominic Kwiatkowski, Cordelia Langford, John Sillitoe on behalf of the Wellcome Sanger Institute COVID-19 Surveillance Team |
| EPI_ISL_2703508 | Lighthouse Lab in Milton Keynes | Wellcome Sanger Institute for the COVID-19 Genomics UK (COG-UK) Consortium | The Lighthouse Lab in Milton Keynes and Alex Alderton, Roberto Amato, Jeffrey Barrett, Sonia Goncalves, Ewan Harrison, David K. Jackson, Ian Johnston, Dominic Kwiatkowski, Cordelia Langford, John Sillitoe on behalf of the Wellcome Sanger Institute COVID-19 Surveillance Team |
| EPI_ISL_2703509 | Lighthouse Lab in Glasgow | Wellcome Sanger Institute for the COVID-19 Genomics UK (COG-UK) Consortium | Harper VanSteenhouse, Yumi Kasai, David Gray, Carol Clugston, Anna Dominiczak and Alex Alderton, Roberto Amato, Jeffrey Barrett, Sonia Goncalves, Ewan Harrison, David K. Jackson, Ian Johnston, Dominic Kwiatkowski, Cordelia Langford, John Sillitoe on behalf of the Wellcome Sanger Institute COVID-19 Surveillance Team |
| EPI_ISL_2703510, EPI_ISL_2703511, EPI_ISL_2703512 | Lighthouse Lab in Alderley Park | Wellcome Sanger Institute for the COVID-19 Genomics UK (COG-UK) Consortium | Jacquelyn Wynn, Mairead Hyland, The Lighthouse Lab in Alderley Park and Alex Alderton, Roberto Amato, Jeffrey Barrett, Sonia Goncalves, Ewan Harrison, David K. Jackson, Ian Johnston, Dominic Kwiatkowski, Cordelia Langford, John Sillitoe on behalf of the Wellcome Sanger Institute COVID-19 Surveillance Team |
| EPI_ISL_2703513 | Lighthouse Lab in Glasgow | Wellcome Sanger Institute for the COVID-19 Genomics UK (COG-UK) Consortium | Harper VanSteenhouse, Yumi Kasai, David Gray, Carol Clugston, Anna Dominiczak and Alex Alderton, Roberto Amato, Jeffrey Barrett, Sonia Goncalves, Ewan Harrison, David K. Jackson, Ian Johnston, Dominic Kwiatkowski, Cordelia Langford, John Sillitoe on behalf of the Wellcome Sanger Institute COVID-19 Surveillance Team |
| EPI_ISL_2703514 | Lighthouse Lab in Alderley Park | Wellcome Sanger Institute for the COVID-19 Genomics UK (COG-UK) Consortium | Jacquelyn Wynn, Mairead Hyland, The Lighthouse Lab in Alderley Park and Alex Alderton, Roberto Amato, Jeffrey Barrett, Sonia Goncalves, Ewan Harrison, David K. Jackson, Ian Johnston, Dominic Kwiatkowski, Cordelia Langford, John Sillitoe on behalf of the Wellcome Sanger Institute COVID-19 Surveillance Team |
| EPI_ISL_2703515, EPI_ISL_2703516 | Lighthouse Lab in Milton Keynes | Wellcome Sanger Institute for the COVID-19 Genomics UK (COG-UK) Consortium | The Lighthouse Lab in Milton Keynes and Alex Alderton, Roberto Amato, Jeffrey Barrett, Sonia Goncalves, Ewan Harrison, David K. Jackson, Ian Johnston, Dominic Kwiatkowski, Cordelia Langford, John Sillitoe on behalf of the Wellcome Sanger Institute COVID-19 Surveillance Team |
| EPI_ISL_2703517 | Lighthouse Lab in Glasgow | Wellcome Sanger Institute for the COVID-19 Genomics UK (COG-UK) Consortium | Harper VanSteenhouse, Yumi Kasai, David Gray, Carol Clugston, Anna Dominiczak and Alex Alderton, Roberto Amato, Jeffrey Barrett, Sonia Goncalves, Ewan Harrison, David K. Jackson, Ian Johnston, Dominic Kwiatkowski, Cordelia Langford, John Sillitoe on behalf of the Wellcome Sanger Institute COVID-19 Surveillance Team |
| EPI_ISL_2703518 | Lighthouse Lab in Alderley Park | Wellcome Sanger Institute for the COVID-19 Genomics UK (COG-UK) Consortium | Jacquelyn Wynn, Mairead Hyland, The Lighthouse Lab in Alderley Park and Alex Alderton, Roberto Amato, Jeffrey Barrett, Sonia Goncalves, Ewan Harrison, David K. Jackson, Ian Johnston, Dominic Kwiatkowski, Cordelia Langford, John Sillitoe on behalf of the Wellcome Sanger Institute COVID-19 Surveillance Team |
| EPI_ISL_2703519 | Lighthouse Lab in Milton Keynes | Wellcome Sanger Institute for the COVID-19 Genomics UK (COG-UK) Consortium | The Lighthouse Lab in Milton Keynes and Alex Alderton, Roberto Amato, Jeffrey Barrett, Sonia Goncalves, Ewan Harrison, David K. Jackson, Ian Johnston, Dominic Kwiatkowski, Cordelia Langford, John Sillitoe on behalf of the Wellcome Sanger Institute COVID-19 Surveillance Team |
| EPI_ISL_2703520, EPI_ISL_2703521 | Lighthouse Lab in Alderley Park | Wellcome Sanger Institute for the COVID-19 Genomics UK (COG-UK) Consortium | Jacquelyn Wynn, Mairead Hyland, The Lighthouse Lab in Alderley Park and Alex Alderton, Roberto Amato, Jeffrey Barrett, Sonia Goncalves, Ewan Harrison, David K. Jackson, Ian Johnston, Dominic Kwiatkowski, Cordelia Langford, John Sillitoe on behalf of the Wellcome Sanger Institute COVID-19 Surveillance Team |
| EPI_ISL_2703522 | Lighthouse Lab in Milton Keynes | Wellcome Sanger Institute for the COVID-19 Genomics UK (COG-UK) Consortium | The Lighthouse Lab in Milton Keynes and Alex Alderton, Roberto Amato, Jeffrey Barrett, Sonia Goncalves, Ewan Harrison, David K. Jackson, Ian Johnston, Dominic Kwiatkowski, Cordelia Langford, John Sillitoe on behalf of the Wellcome Sanger Institute COVID-19 Surveillance Team |
| EPI_ISL_2703523 | Lighthouse Lab in Alderley Park | Wellcome Sanger Institute for the COVID-19 Genomics UK (COG-UK) Consortium | Jacquelyn Wynn, Mairead Hyland, The Lighthouse Lab in Alderley Park and Alex Alderton, Roberto Amato, Jeffrey Barrett, Sonia Goncalves, Ewan Harrison, David K. Jackson, Ian Johnston, Dominic Kwiatkowski, Cordelia Langford, John Sillitoe on behalf of the Wellcome Sanger Institute COVID-19 Surveillance Team |
| EPI_ISL_2703524, EPI_ISL_2703525 | Lighthouse Lab in Glasgow | Wellcome Sanger Institute for the COVID-19 Genomics UK (COG-UK) Consortium | Harper VanSteenhouse, Yumi Kasai, David Gray, Carol Clugston, Anna Dominiczak and Alex Alderton, Roberto Amato, Jeffrey Barrett, Sonia Goncalves, Ewan Harrison, David K. Jackson, Ian Johnston, Dominic Kwiatkowski, Cordelia Langford, John Sillitoe on behalf of the Wellcome Sanger Institute COVID-19 Surveillance Team |
| EPI_ISL_2703526 | Lighthouse Lab in Alderley Park | Wellcome Sanger Institute for the COVID-19 Genomics UK (COG-UK) Consortium | Jacquelyn Wynn, Mairead Hyland, The Lighthouse Lab in Alderley Park and Alex Alderton, Roberto Amato, Jeffrey Barrett, Sonia Goncalves, Ewan Harrison, David K. Jackson, Ian Johnston, Dominic Kwiatkowski, Cordelia Langford, John Sillitoe on behalf of the Wellcome Sanger Institute COVID-19 Surveillance Team |
| EPI_ISL_2703527 | Lighthouse Lab in Milton Keynes | Wellcome Sanger Institute for the COVID-19 Genomics UK (COG-UK) Consortium | The Lighthouse Lab in Milton Keynes and Alex Alderton, Roberto Amato, Jeffrey Barrett, Sonia Goncalves, Ewan Harrison, David K. Jackson, Ian Johnston, Dominic Kwiatkowski, Cordelia Langford, John Sillitoe on behalf of the Wellcome Sanger Institute COVID-19 Surveillance Team |
| EPI_ISL_2703528 | Lighthouse Lab in Glasgow | Wellcome Sanger Institute for the COVID-19 Genomics UK (COG-UK) Consortium | Harper VanSteenhouse, Yumi Kasai, David Gray, Carol Clugston, Anna Dominiczak and Alex Alderton, Roberto Amato, Jeffrey Barrett, Sonia Goncalves, Ewan Harrison, David K. Jackson, Ian Johnston, Dominic Kwiatkowski, Cordelia Langford, John Sillitoe on behalf of the Wellcome Sanger Institute COVID-19 Surveillance Team |
| EPI_ISL_2703529 | Lighthouse Lab in Alderley Park | Wellcome Sanger Institute for the COVID-19 Genomics UK (COG-UK) Consortium | Jacquelyn Wynn, Mairead Hyland, The Lighthouse Lab in Alderley Park and Alex Alderton, Roberto Amato, Jeffrey Barrett, Sonia Goncalves, Ewan Harrison, David K. Jackson, Ian Johnston, Dominic Kwiatkowski, Cordelia Langford, John Sillitoe on behalf of the Wellcome Sanger Institute COVID-19 Surveillance Team |
| EPI_ISL_2703530 | Lighthouse Lab in Glasgow | Wellcome Sanger Institute for the COVID-19 Genomics UK (COG-UK) Consortium | Harper VanSteenhouse, Yumi Kasai, David Gray, Carol Clugston, Anna Dominiczak and Alex Alderton, Roberto Amato, Jeffrey Barrett, Sonia Goncalves, Ewan Harrison, David K. Jackson, Ian Johnston, Dominic Kwiatkowski, Cordelia Langford, John Sillitoe on behalf of the Wellcome Sanger Institute COVID-19 Surveillance Team |
| EPI_ISL_2703531, EPI_ISL_2703532, EPI_ISL_2703533 | Lighthouse Lab in Alderley Park | Wellcome Sanger Institute for the COVID-19 Genomics UK (COG-UK) Consortium | Jacquelyn Wynn, Mairead Hyland, The Lighthouse Lab in Alderley Park and Alex Alderton, Roberto Amato, Jeffrey Barrett, Sonia Goncalves, Ewan Harrison, David K. Jackson, Ian Johnston, Dominic Kwiatkowski, Cordelia Langford, John Sillitoe on behalf of the Wellcome Sanger Institute COVID-19 Surveillance Team |
| EPI_ISL_2703534 | Lighthouse Lab in Milton Keynes | Wellcome Sanger Institute for the COVID-19 Genomics UK (COG-UK) Consortium | The Lighthouse Lab in Milton Keynes and Alex Alderton, Roberto Amato, Jeffrey Barrett, Sonia Goncalves, Ewan Harrison, David K. Jackson, Ian Johnston, Dominic Kwiatkowski, Cordelia Langford, John Sillitoe on behalf of the Wellcome Sanger Institute COVID-19 Surveillance Team |
| EPI_ISL_2703535 | Lighthouse Lab in Alderley Park | Wellcome Sanger Institute for the COVID-19 Genomics UK (COG-UK) Consortium | Jacquelyn Wynn, Mairead Hyland, The Lighthouse Lab in Alderley Park and Alex Alderton, Roberto Amato, Jeffrey Barrett, Sonia Goncalves, Ewan Harrison, David K. Jackson, Ian Johnston, Dominic Kwiatkowski, Cordelia Langford, John Sillitoe on behalf of the Wellcome Sanger Institute COVID-19 Surveillance Team |
| EPI_ISL_2703536, EPI_ISL_2703537 | Lighthouse Lab in Glasgow | Wellcome Sanger Institute for the COVID-19 Genomics UK (COG-UK) Consortium | Harper VanSteenhouse, Yumi Kasai, David Gray, Carol Clugston, Anna Dominiczak and Alex Alderton, Roberto Amato, Jeffrey Barrett, Sonia Goncalves, Ewan Harrison, David K. Jackson, Ian Johnston, Dominic Kwiatkowski, Cordelia Langford, John Sillitoe on behalf of the Wellcome Sanger Institute COVID-19 Surveillance Team |
| EPI_ISL_2703538 | Lighthouse Lab in Milton Keynes | Wellcome Sanger Institute for the COVID-19 Genomics UK (COG-UK) Consortium | The Lighthouse Lab in Milton Keynes and Alex Alderton, Roberto Amato, Jeffrey Barrett, Sonia Goncalves, Ewan Harrison, David K. Jackson, Ian Johnston, Dominic Kwiatkowski, Cordelia Langford, John Sillitoe on behalf of the Wellcome Sanger Institute COVID-19 Surveillance Team |

| | | | |
|---|---|---|---|
| EPI_ISL_2703539, EPI_ISL_2703540, EPI_ISL_2703542, EPI_ISL_2703543, EPI_ISL_2703544 | Lighthouse Lab in Alderley Park | Wellcome Sanger Institute for the COVID-19 Genomics UK (COG-UK) Consortium | Jacquelyn Wynn, Mairead Hyland, The Lighthouse Lab in Alderley Park and Alex Alderton, Roberto Amato, Jeffrey Barrett, Sonia Goncalves, Ewan Harrison, David K. Jackson, Ian Johnston, Dominic Kwiatkowski, Cordelia Langford, John Sillitoe on behalf of the Wellcome Sanger Institute COVID-19 Surveillance Team |
| EPI_ISL_2703545 | Lighthouse Lab in Milton Keynes | Wellcome Sanger Institute for the COVID-19 Genomics UK (COG-UK) Consortium | The Lighthouse Lab in Milton Keynes and Alex Alderton, Roberto Amato, Jeffrey Barrett, Sonia Goncalves, Ewan Harrison, David K. Jackson, Ian Johnston, Dominic Kwiatkowski, Cordelia Langford, John Sillitoe on behalf of the Wellcome Sanger Institute COVID-19 Surveillance Team |
| EPI_ISL_2703546, EPI_ISL_2703547 | Lighthouse Lab in Alderley Park | Wellcome Sanger Institute for the COVID-19 Genomics UK (COG-UK) Consortium | Jacquelyn Wynn, Mairead Hyland, The Lighthouse Lab in Alderley Park and Alex Alderton, Roberto Amato, Jeffrey Barrett, Sonia Goncalves, Ewan Harrison, David K. Jackson, Ian Johnston, Dominic Kwiatkowski, Cordelia Langford, John Sillitoe on behalf of the Wellcome Sanger Institute COVID-19 Surveillance Team |
| EPI_ISL_2703548, EPI_ISL_2703549 | Lighthouse Lab in Milton Keynes | Wellcome Sanger Institute for the COVID-19 Genomics UK (COG-UK) Consortium | The Lighthouse Lab in Milton Keynes and Alex Alderton, Roberto Amato, Jeffrey Barrett, Sonia Goncalves, Ewan Harrison, David K. Jackson, Ian Johnston, Dominic Kwiatkowski, Cordelia Langford, John Sillitoe on behalf of the Wellcome Sanger Institute COVID-19 Surveillance Team |
| EPI_ISL_2703550 | Lighthouse Lab in Glasgow | Wellcome Sanger Institute for the COVID-19 Genomics UK (COG-UK) Consortium | Harper VanSteenhouse, Yumi Kasai, David Gray, Carol Clugston, Anna Dominiczak and Alex Alderton, Roberto Amato, Jeffrey Barrett, Sonia Goncalves, Ewan Harrison, David K. Jackson, Ian Johnston, Dominic Kwiatkowski, Cordelia Langford, John Sillitoe on behalf of the Wellcome Sanger Institute COVID-19 Surveillance Team |
| EPI_ISL_2703551 | Lighthouse Lab in Alderley Park | Wellcome Sanger Institute for the COVID-19 Genomics UK (COG-UK) Consortium | Jacquelyn Wynn, Mairead Hyland, The Lighthouse Lab in Alderley Park and Alex Alderton, Roberto Amato, Jeffrey Barrett, Sonia Goncalves, Ewan Harrison, David K. Jackson, Ian Johnston, Dominic Kwiatkowski, Cordelia Langford, John Sillitoe on behalf of the Wellcome Sanger Institute COVID-19 Surveillance Team |
| EPI_ISL_2703552 | Lighthouse Lab in Milton Keynes | Wellcome Sanger Institute for the COVID-19 Genomics UK (COG-UK) Consortium | The Lighthouse Lab in Milton Keynes and Alex Alderton, Roberto Amato, Jeffrey Barrett, Sonia Goncalves, Ewan Harrison, David K. Jackson, Ian Johnston, Dominic Kwiatkowski, Cordelia Langford, John Sillitoe on behalf of the Wellcome Sanger Institute COVID-19 Surveillance Team |
| EPI_ISL_2703553 | Lighthouse Lab in Alderley Park | Wellcome Sanger Institute for the COVID-19 Genomics UK (COG-UK) Consortium | Jacquelyn Wynn, Mairead Hyland, The Lighthouse Lab in Alderley Park and Alex Alderton, Roberto Amato, Jeffrey Barrett, Sonia Goncalves, Ewan Harrison, David K. Jackson, Ian Johnston, Dominic Kwiatkowski, Cordelia Langford, John Sillitoe on behalf of the Wellcome Sanger Institute COVID-19 Surveillance Team |
| EPI_ISL_2703554 | Lighthouse Lab in Glasgow | Wellcome Sanger Institute for the COVID-19 Genomics UK (COG-UK) Consortium | Harper VanSteenhouse, Yumi Kasai, David Gray, Carol Clugston, Anna Dominiczak and Alex Alderton, Roberto Amato, Jeffrey Barrett, Sonia Goncalves, Ewan Harrison, David K. Jackson, Ian Johnston, Dominic Kwiatkowski, Cordelia Langford, John Sillitoe on behalf of the Wellcome Sanger Institute COVID-19 Surveillance Team |
| EPI_ISL_2703555, EPI_ISL_2703556 | Lighthouse Lab in Milton Keynes | Wellcome Sanger Institute for the COVID-19 Genomics UK (COG-UK) Consortium | The Lighthouse Lab in Milton Keynes and Alex Alderton, Roberto Amato, Jeffrey Barrett, Sonia Goncalves, Ewan Harrison, David K. Jackson, Ian Johnston, Dominic Kwiatkowski, Cordelia Langford, John Sillitoe on behalf of the Wellcome Sanger Institute COVID-19 Surveillance Team |
| EPI_ISL_2703557 | Lighthouse Lab in Alderley Park | Wellcome Sanger Institute for the COVID-19 Genomics UK (COG-UK) Consortium | Jacquelyn Wynn, Mairead Hyland, The Lighthouse Lab in Alderley Park and Alex Alderton, Roberto Amato, Jeffrey Barrett, Sonia Goncalves, Ewan Harrison, David K. Jackson, Ian Johnston, Dominic Kwiatkowski, Cordelia Langford, John Sillitoe on behalf of the Wellcome Sanger Institute COVID-19 Surveillance Team |
| EPI_ISL_2703558, EPI_ISL_2703559, EPI_ISL_2703560 | Lighthouse Lab in Milton Keynes | Wellcome Sanger Institute for the COVID-19 Genomics UK (COG-UK) Consortium | The Lighthouse Lab in Milton Keynes and Alex Alderton, Roberto Amato, Jeffrey Barrett, Sonia Goncalves, Ewan Harrison, David K. Jackson, Ian Johnston, Dominic Kwiatkowski, Cordelia Langford, John Sillitoe on behalf of the Wellcome Sanger Institute COVID-19 Surveillance Team |
| EPI_ISL_2703561, EPI_ISL_2703562 | Lighthouse Lab in Glasgow | Wellcome Sanger Institute for the COVID-19 Genomics UK (COG-UK) Consortium | Harper VanSteenhouse, Yumi Kasai, David Gray, Carol Clugston, Anna Dominiczak and Alex Alderton, Roberto Amato, Jeffrey Barrett, Sonia Goncalves, Ewan Harrison, David K. Jackson, Ian Johnston, Dominic Kwiatkowski, Cordelia Langford, John Sillitoe on behalf of the Wellcome Sanger Institute COVID-19 Surveillance Team |
| EPI_ISL_2703563 | Lighthouse Lab in Alderley Park | Wellcome Sanger Institute for the COVID-19 Genomics UK (COG-UK) Consortium | Jacquelyn Wynn, Mairead Hyland, The Lighthouse Lab in Alderley Park and Alex Alderton, Roberto Amato, Jeffrey Barrett, Sonia Goncalves, Ewan Harrison, David K. Jackson, Ian Johnston, Dominic Kwiatkowski, Cordelia Langford, John Sillitoe on behalf of the Wellcome Sanger Institute COVID-19 Surveillance Team |
| EPI_ISL_2703564 | Lighthouse Lab in Milton Keynes | Wellcome Sanger Institute for the COVID-19 Genomics UK (COG-UK) Consortium | The Lighthouse Lab in Milton Keynes and Alex Alderton, Roberto Amato, Jeffrey Barrett, Sonia Goncalves, Ewan Harrison, David K. Jackson, Ian Johnston, Dominic Kwiatkowski, Cordelia Langford, John Sillitoe on behalf of the Wellcome Sanger Institute COVID-19 Surveillance Team |
| EPI_ISL_2703565, EPI_ISL_2703566, EPI_ISL_2703567, EPI_ISL_2703568, EPI_ISL_2703569 | Lighthouse Lab in Alderley Park | Wellcome Sanger Institute for the COVID-19 Genomics UK (COG-UK) Consortium | Jacquelyn Wynn, Mairead Hyland, The Lighthouse Lab in Alderley Park and Alex Alderton, Roberto Amato, Jeffrey Barrett, Sonia Goncalves, Ewan Harrison, David K. Jackson, Ian Johnston, Dominic Kwiatkowski, Cordelia Langford, John Sillitoe on behalf of the Wellcome Sanger Institute COVID-19 Surveillance Team |
| EPI_ISL_2703570, EPI_ISL_2703571, EPI_ISL_2703572 | Lighthouse Lab in Milton Keynes | Wellcome Sanger Institute for the COVID-19 Genomics UK (COG-UK) Consortium | The Lighthouse Lab in Milton Keynes and Alex Alderton, Roberto Amato, Jeffrey Barrett, Sonia Goncalves, Ewan Harrison, David K. Jackson, Ian Johnston, Dominic Kwiatkowski, Cordelia Langford, John Sillitoe on behalf of the Wellcome Sanger Institute COVID-19 Surveillance Team |
| EPI_ISL_2703573 | Lighthouse Laboratory Plymouth | Wellcome Sanger Institute for the COVID-19 Genomics UK (COG-UK) Consortium | Lighthouse Laboratory Plymouth and Alex Alderton, Roberto Amato, Jeffrey Barrett, Sonia Goncalves, Ewan Harrison, David K. Jackson, Ian Johnston, Dominic Kwiatkowski, Cordelia Langford, John Sillitoe on behalf of the Wellcome Sanger Institute COVID-19 Surveillance Team |
| EPI_ISL_2703574, EPI_ISL_2703575 | Lighthouse Lab in Alderley Park | Wellcome Sanger Institute for the COVID-19 Genomics UK (COG-UK) Consortium | Jacquelyn Wynn, Mairead Hyland, The Lighthouse Lab in Alderley Park and Alex Alderton, Roberto Amato, Jeffrey Barrett, Sonia Goncalves, Ewan Harrison, David K. Jackson, Ian Johnston, Dominic Kwiatkowski, Cordelia Langford, John Sillitoe on behalf of the Wellcome Sanger Institute COVID-19 Surveillance Team |
| EPI_ISL_2703576, EPI_ISL_2703577 | Lighthouse Lab in Milton Keynes | Wellcome Sanger Institute for the COVID-19 Genomics UK (COG-UK) Consortium | The Lighthouse Lab in Milton Keynes and Alex Alderton, Roberto Amato, Jeffrey Barrett, Sonia Goncalves, Ewan Harrison, David K. Jackson, Ian Johnston, Dominic Kwiatkowski, Cordelia Langford, John Sillitoe on behalf of the Wellcome Sanger Institute COVID-19 Surveillance Team |
| EPI_ISL_2703578 | Lighthouse Lab in Glasgow | Wellcome Sanger Institute for the COVID-19 Genomics UK (COG-UK) Consortium | Harper VanSteenhouse, Yumi Kasai, David Gray, Carol Clugston, Anna Dominiczak and Alex Alderton, Roberto Amato, Jeffrey Barrett, Sonia Goncalves, Ewan Harrison, David K. Jackson, Ian Johnston, Dominic Kwiatkowski, Cordelia Langford, John Sillitoe on behalf of the Wellcome Sanger Institute COVID-19 Surveillance Team |
| EPI_ISL_2703579, EPI_ISL_2703580 | Lighthouse Lab in Milton Keynes | Wellcome Sanger Institute for the COVID-19 Genomics UK (COG-UK) Consortium | The Lighthouse Lab in Milton Keynes and Alex Alderton, Roberto Amato, Jeffrey Barrett, Sonia Goncalves, Ewan Harrison, David K. Jackson, Ian Johnston, Dominic Kwiatkowski, Cordelia Langford, John Sillitoe on behalf of the Wellcome Sanger Institute COVID-19 Surveillance Team |
| EPI_ISL_2703581, EPI_ISL_2703582 | Lighthouse Lab in Glasgow | Wellcome Sanger Institute for the COVID-19 Genomics UK (COG-UK) Consortium | Harper VanSteenhouse, Yumi Kasai, David Gray, Carol Clugston, Anna Dominiczak and Alex Alderton, Roberto Amato, Jeffrey Barrett, Sonia Goncalves, Ewan Harrison, David K. Jackson, Ian Johnston, Dominic Kwiatkowski, Cordelia Langford, John Sillitoe on behalf of the Wellcome Sanger Institute COVID-19 Surveillance Team |
| EPI_ISL_2703583 | Lighthouse Lab in Milton Keynes | Wellcome Sanger Institute for the COVID-19 Genomics UK (COG-UK) Consortium | The Lighthouse Lab in Milton Keynes and Alex Alderton, Roberto Amato, Jeffrey Barrett, Sonia Goncalves, Ewan Harrison, David K. Jackson, Ian Johnston, Dominic Kwiatkowski, Cordelia Langford, John Sillitoe on behalf of the Wellcome Sanger Institute COVID-19 Surveillance Team |
| EPI_ISL_2703584, EPI_ISL_2703585 | Lighthouse Lab in Alderley Park | Wellcome Sanger Institute for the COVID-19 Genomics UK (COG-UK) Consortium | Jacquelyn Wynn, Mairead Hyland, The Lighthouse Lab in Alderley Park and Alex Alderton, Roberto Amato, Jeffrey Barrett, Sonia Goncalves, Ewan Harrison, David K. Jackson, Ian Johnston, Dominic Kwiatkowski, Cordelia Langford, John Sillitoe on behalf of the Wellcome Sanger Institute COVID-19 Surveillance Team |
| EPI_ISL_2703586 | Lighthouse Laboratory Plymouth | Wellcome Sanger Institute for the COVID-19 Genomics UK (COG-UK) Consortium | Lighthouse Laboratory Plymouth and Alex Alderton, Roberto Amato, Jeffrey Barrett, Sonia Goncalves, Ewan Harrison, David K. Jackson, Ian Johnston, Dominic Kwiatkowski, Cordelia Langford, John Sillitoe on behalf of the Wellcome Sanger Institute COVID-19 Surveillance Team |
| EPI_ISL_2703587 | Lighthouse Lab in Milton Keynes | Wellcome Sanger Institute for the COVID-19 Genomics UK (COG-UK) Consortium | The Lighthouse Lab in Milton Keynes and Alex Alderton, Roberto Amato, Jeffrey Barrett, Sonia Goncalves, Ewan Harrison, David K. Jackson, Ian Johnston, Dominic Kwiatkowski, Cordelia Langford, John Sillitoe on behalf of the Wellcome Sanger Institute COVID-19 Surveillance Team |
| EPI_ISL_2703588 | Lighthouse Lab in Glasgow | Wellcome Sanger Institute for the COVID-19 Genomics UK (COG-UK) Consortium | Harper VanSteenhouse, Yumi Kasai, David Gray, Carol Clugston, Anna Dominiczak and Alex Alderton, Roberto Amato, Jeffrey Barrett, Sonia Goncalves, Ewan Harrison, David K. Jackson, Ian Johnston, Dominic Kwiatkowski, Cordelia Langford, John Sillitoe on behalf of the Wellcome Sanger Institute COVID-19 Surveillance Team |
| EPI_ISL_2703589, EPI_ISL_2703590 | Lighthouse Lab in Alderley Park | Wellcome Sanger Institute for the COVID-19 Genomics UK (COG-UK) Consortium | Jacquelyn Wynn, Mairead Hyland, The Lighthouse Lab in Alderley Park and Alex Alderton, Roberto Amato, Jeffrey Barrett, Sonia Goncalves, Ewan Harrison, David K. Jackson, Ian Johnston, Dominic Kwiatkowski, Cordelia Langford, John Sillitoe on behalf of the Wellcome Sanger Institute COVID-19 Surveillance Team |
| EPI_ISL_2703591 | Lighthouse Lab in Glasgow | Wellcome Sanger Institute for the COVID-19 Genomics UK (COG-UK) Consortium | Harper VanSteenhouse, Yumi Kasai, David Gray, Carol Clugston, Anna Dominiczak and Alex Alderton, Roberto Amato, Jeffrey Barrett, Sonia Goncalves, Ewan Harrison, David K. Jackson, Ian Johnston, Dominic Kwiatkowski, Cordelia Langford, John Sillitoe on behalf of the Wellcome Sanger Institute COVID-19 Surveillance Team |
| EPI_ISL_2703592 | Lighthouse Lab in Milton Keynes | Wellcome Sanger Institute for the COVID-19 Genomics UK (COG-UK) Consortium | The Lighthouse Lab in Milton Keynes and Alex Alderton, Roberto Amato, Jeffrey Barrett, Sonia Goncalves, Ewan Harrison, David K. Jackson, Ian Johnston, Dominic Kwiatkowski, Cordelia Langford, John Sillitoe on behalf of the Wellcome Sanger Institute COVID-19 Surveillance Team |
| EPI_ISL_2703593 | Lighthouse Lab in Alderley Park | Wellcome Sanger Institute for the COVID-19 Genomics UK (COG-UK) Consortium | Jacquelyn Wynn, Mairead Hyland, The Lighthouse Lab in Alderley Park and Alex Alderton, Roberto Amato, Jeffrey Barrett, Sonia Goncalves, Ewan Harrison, David K. Jackson, Ian Johnston, Dominic Kwiatkowski, Cordelia Langford, John Sillitoe on behalf of the Wellcome Sanger Institute COVID-19 Surveillance Team |
| EPI_ISL_2703594 | Lighthouse Lab in Glasgow | Wellcome Sanger Institute for the COVID-19 Genomics UK (COG-UK) Consortium | Harper VanSteenhouse, Yumi Kasai, David Gray, Carol Clugston, Anna Dominiczak and Alex Alderton, Roberto Amato, Jeffrey Barrett, Sonia Goncalves, Ewan Harrison, David K. Jackson, Ian Johnston, Dominic Kwiatkowski, Cordelia Langford, John Sillitoe on behalf of the Wellcome Sanger Institute COVID-19 Surveillance Team |
| EPI_ISL_2703595, EPI_ISL_2703596, EPI_ISL_2703597 | Lighthouse Lab in Alderley Park | Wellcome Sanger Institute for the COVID-19 Genomics UK (COG-UK) Consortium | Jacquelyn Wynn, Mairead Hyland, The Lighthouse Lab in Alderley Park and Alex Alderton, Roberto Amato, Jeffrey Barrett, Sonia Goncalves, Ewan Harrison, David K. Jackson, Ian Johnston, Dominic Kwiatkowski, Cordelia Langford, John Sillitoe on behalf of the Wellcome Sanger Institute COVID-19 Surveillance Team |
| EPI_ISL_2703598 | Lighthouse Lab in Glasgow | Wellcome Sanger Institute for the COVID-19 Genomics UK (COG-UK) Consortium | Harper VanSteenhouse, Yumi Kasai, David Gray, Carol Clugston, Anna Dominiczak and Alex Alderton, Roberto Amato, Jeffrey Barrett, Sonia Goncalves, Ewan Harrison, David K. Jackson, Ian Johnston, Dominic Kwiatkowski, Cordelia Langford, John Sillitoe on behalf of the Wellcome Sanger Institute COVID-19 Surveillance Team |
| EPI_ISL_2703599, EPI_ISL_2703600 | Lighthouse Lab in Milton Keynes | Wellcome Sanger Institute for the COVID-19 Genomics UK (COG-UK) Consortium | The Lighthouse Lab in Milton Keynes and Alex Alderton, Roberto Amato, Jeffrey Barrett, Sonia Goncalves, Ewan Harrison, David K. Jackson, Ian Johnston, Dominic Kwiatkowski, Cordelia Langford, John Sillitoe on behalf of the Wellcome Sanger Institute COVID-19 Surveillance Team |
| EPI_ISL_2703602, EPI_ISL_2703603 | Lighthouse Lab in Alderley Park | Wellcome Sanger Institute for the COVID-19 Genomics UK (COG-UK) Consortium | Jacquelyn Wynn, Mairead Hyland, The Lighthouse Lab in Alderley Park and Alex Alderton, Roberto Amato, Jeffrey Barrett, Sonia Goncalves, Ewan Harrison, David K. Jackson, Ian Johnston, Dominic Kwiatkowski, Cordelia Langford, John Sillitoe on behalf of the Wellcome Sanger Institute COVID-19 Surveillance Team |
| EPI_ISL_2703604 | Lighthouse Lab in Glasgow | Wellcome Sanger Institute for the COVID-19 Genomics UK (COG-UK) Consortium | Harper VanSteenhouse, Yumi Kasai, David Gray, Carol Clugston, Anna Dominiczak and Alex Alderton, Roberto Amato, Jeffrey Barrett, Sonia Goncalves, Ewan Harrison, David K. Jackson, Ian Johnston, Dominic Kwiatkowski, Cordelia Langford, John Sillitoe on behalf of the Wellcome Sanger Institute COVID-19 Surveillance Team |
| EPI_ISL_2703605 | Lighthouse Lab in Milton Keynes | Wellcome Sanger Institute for the COVID-19 Genomics UK (COG-UK) Consortium | The Lighthouse Lab in Milton Keynes and Alex Alderton, Roberto Amato, Jeffrey Barrett, Sonia Goncalves, Ewan Harrison, David K. Jackson, Ian Johnston, Dominic Kwiatkowski, Cordelia Langford, John Sillitoe on behalf of the Wellcome Sanger Institute COVID-19 Surveillance Team |
| EPI_ISL_2703606 | Lighthouse Lab in Glasgow | Wellcome Sanger Institute for the COVID-19 Genomics UK (COG-UK) Consortium | Harper VanSteenhouse, Yumi Kasai, David Gray, Carol Clugston, Anna Dominiczak and Alex Alderton, Roberto Amato, Jeffrey Barrett, Sonia Goncalves, Ewan Harrison, David K. Jackson, Ian Johnston, Dominic Kwiatkowski, Cordelia Langford, John Sillitoe on behalf of the Wellcome Sanger Institute COVID-19 Surveillance Team |
| EPI_ISL_2703607 | Lighthouse Lab in Milton Keynes | Wellcome Sanger Institute for the COVID-19 Genomics UK (COG-UK) Consortium | The Lighthouse Lab in Milton Keynes and Alex Alderton, Roberto Amato, Jeffrey Barrett, Sonia Goncalves, Ewan Harrison, David K. Jackson, Ian Johnston, Dominic Kwiatkowski, Cordelia Langford, John Sillitoe on behalf of the Wellcome Sanger Institute COVID-19 Surveillance Team |

| | | | |
|---|---|---|---|
| EPI_ISL_2703609, EPI_ISL_2703610, EPI_ISL_2703611, EPI_ISL_2703612 | Lighthouse Lab in Alderley Park | Wellcome Sanger Institute for the COVID-19 Genomics UK (COG-UK) Consortium | Jacquelyn Wynn, Mairead Hyland, The Lighthouse Lab in Alderley Park and Alex Alderton, Roberto Amato, Jeffrey Barrett, Sonia Goncalves, Ewan Harrison, David K. Jackson, Ian Johnston, Dominic Kwiatkowski, Cordelia Langford, John Sillitoe on behalf of the Wellcome Sanger Institute COVID-19 Surveillance Team |
| EPI_ISL_2703613 | Lighthouse Lab in Glasgow | Wellcome Sanger Institute for the COVID-19 Genomics UK (COG-UK) Consortium | Harper VanSteenhouse, Yumi Kasai, David Gray, Carol Clugston, Anna Dominiczak and Alex Alderton, Roberto Amato, Jeffrey Barrett, Sonia Goncalves, Ewan Harrison, David K. Jackson, Ian Johnston, Dominic Kwiatkowski, Cordelia Langford, John Sillitoe on behalf of the Wellcome Sanger Institute COVID-19 Surveillance Team |
| EPI_ISL_2703614, EPI_ISL_2703615, EPI_ISL_2703616, EPI_ISL_2703617 | Lighthouse Lab in Alderley Park | Wellcome Sanger Institute for the COVID-19 Genomics UK (COG-UK) Consortium | Jacquelyn Wynn, Mairead Hyland, The Lighthouse Lab in Alderley Park and Alex Alderton, Roberto Amato, Jeffrey Barrett, Sonia Goncalves, Ewan Harrison, David K. Jackson, Ian Johnston, Dominic Kwiatkowski, Cordelia Langford, John Sillitoe on behalf of the Wellcome Sanger Institute COVID-19 Surveillance Team |
| EPI_ISL_2703618, EPI_ISL_2703619 | Lighthouse Lab in Milton Keynes | Wellcome Sanger Institute for the COVID-19 Genomics UK (COG-UK) Consortium | The Lighthouse Lab in Milton Keynes and Alex Alderton, Roberto Amato, Jeffrey Barrett, Sonia Goncalves, Ewan Harrison, David K. Jackson, Ian Johnston, Dominic Kwiatkowski, Cordelia Langford, John Sillitoe on behalf of the Wellcome Sanger Institute COVID-19 Surveillance Team |
| EPI_ISL_2703620 | Lighthouse Lab in Alderley Park | Wellcome Sanger Institute for the COVID-19 Genomics UK (COG-UK) Consortium | Jacquelyn Wynn, Mairead Hyland, The Lighthouse Lab in Alderley Park and Alex Alderton, Roberto Amato, Jeffrey Barrett, Sonia Goncalves, Ewan Harrison, David K. Jackson, Ian Johnston, Dominic Kwiatkowski, Cordelia Langford, John Sillitoe on behalf of the Wellcome Sanger Institute COVID-19 Surveillance Team |
| EPI_ISL_2703621 | Lighthouse Lab in Milton Keynes | Wellcome Sanger Institute for the COVID-19 Genomics UK (COG-UK) Consortium | The Lighthouse Lab in Milton Keynes and Alex Alderton, Roberto Amato, Jeffrey Barrett, Sonia Goncalves, Ewan Harrison, David K. Jackson, Ian Johnston, Dominic Kwiatkowski, Cordelia Langford, John Sillitoe on behalf of the Wellcome Sanger Institute COVID-19 Surveillance Team |
| EPI_ISL_2703622 | Lighthouse Lab in Glasgow | Wellcome Sanger Institute for the COVID-19 Genomics UK (COG-UK) Consortium | Harper VanSteenhouse, Yumi Kasai, David Gray, Carol Clugston, Anna Dominiczak and Alex Alderton, Roberto Amato, Jeffrey Barrett, Sonia Goncalves, Ewan Harrison, David K. Jackson, Ian Johnston, Dominic Kwiatkowski, Cordelia Langford, John Sillitoe on behalf of the Wellcome Sanger Institute COVID-19 Surveillance Team |
| EPI_ISL_2703623 | Lighthouse Lab in Milton Keynes | Wellcome Sanger Institute for the COVID-19 Genomics UK (COG-UK) Consortium | The Lighthouse Lab in Milton Keynes and Alex Alderton, Roberto Amato, Jeffrey Barrett, Sonia Goncalves, Ewan Harrison, David K. Jackson, Ian Johnston, Dominic Kwiatkowski, Cordelia Langford, John Sillitoe on behalf of the Wellcome Sanger Institute COVID-19 Surveillance Team |
| EPI_ISL_2703624, EPI_ISL_2703625, EPI_ISL_2703626, EPI_ISL_2703627 | Lighthouse Lab in Alderley Park | Wellcome Sanger Institute for the COVID-19 Genomics UK (COG-UK) Consortium | Jacquelyn Wynn, Mairead Hyland, The Lighthouse Lab in Alderley Park and Alex Alderton, Roberto Amato, Jeffrey Barrett, Sonia Goncalves, Ewan Harrison, David K. Jackson, Ian Johnston, Dominic Kwiatkowski, Cordelia Langford, John Sillitoe on behalf of the Wellcome Sanger Institute COVID-19 Surveillance Team |
| EPI_ISL_2703628 | Lighthouse Lab in Milton Keynes | Wellcome Sanger Institute for the COVID-19 Genomics UK (COG-UK) Consortium | The Lighthouse Lab in Milton Keynes and Alex Alderton, Roberto Amato, Jeffrey Barrett, Sonia Goncalves, Ewan Harrison, David K. Jackson, Ian Johnston, Dominic Kwiatkowski, Cordelia Langford, John Sillitoe on behalf of the Wellcome Sanger Institute COVID-19 Surveillance Team |
| EPI_ISL_2703629, EPI_ISL_2703630, EPI_ISL_2703631 | Lighthouse Lab in Alderley Park | Wellcome Sanger Institute for the COVID-19 Genomics UK (COG-UK) Consortium | Jacquelyn Wynn, Mairead Hyland, The Lighthouse Lab in Alderley Park and Alex Alderton, Roberto Amato, Jeffrey Barrett, Sonia Goncalves, Ewan Harrison, David K. Jackson, Ian Johnston, Dominic Kwiatkowski, Cordelia Langford, John Sillitoe on behalf of the Wellcome Sanger Institute COVID-19 Surveillance Team |
| EPI_ISL_2703632, EPI_ISL_2703633, EPI_ISL_2703635, EPI_ISL_2703636 | Lighthouse Lab in Glasgow | Wellcome Sanger Institute for the COVID-19 Genomics UK (COG-UK) Consortium | Harper VanSteenhouse, Yumi Kasai, David Gray, Carol Clugston, Anna Dominiczak and Alex Alderton, Roberto Amato, Jeffrey Barrett, Sonia Goncalves, Ewan Harrison, David K. Jackson, Ian Johnston, Dominic Kwiatkowski, Cordelia Langford, John Sillitoe on behalf of the Wellcome Sanger Institute COVID-19 Surveillance Team |
| EPI_ISL_2703637, EPI_ISL_2703638 | Lighthouse Lab in Milton Keynes | Wellcome Sanger Institute for the COVID-19 Genomics UK (COG-UK) Consortium | The Lighthouse Lab in Milton Keynes and Alex Alderton, Roberto Amato, Jeffrey Barrett, Sonia Goncalves, Ewan Harrison, David K. Jackson, Ian Johnston, Dominic Kwiatkowski, Cordelia Langford, John Sillitoe on behalf of the Wellcome Sanger Institute COVID-19 Surveillance Team |
| EPI_ISL_2703639 | Lighthouse Lab in Glasgow | Wellcome Sanger Institute for the COVID-19 Genomics UK (COG-UK) Consortium | Harper VanSteenhouse, Yumi Kasai, David Gray, Carol Clugston, Anna Dominiczak and Alex Alderton, Roberto Amato, Jeffrey Barrett, Sonia Goncalves, Ewan Harrison, David K. Jackson, Ian Johnston, Dominic Kwiatkowski, Cordelia Langford, John Sillitoe on behalf of the Wellcome Sanger Institute COVID-19 Surveillance Team |
| EPI_ISL_2703640, EPI_ISL_2703641 | Lighthouse Lab in Milton Keynes | Wellcome Sanger Institute for the COVID-19 Genomics UK (COG-UK) Consortium | The Lighthouse Lab in Milton Keynes and Alex Alderton, Roberto Amato, Jeffrey Barrett, Sonia Goncalves, Ewan Harrison, David K. Jackson, Ian Johnston, Dominic Kwiatkowski, Cordelia Langford, John Sillitoe on behalf of the Wellcome Sanger Institute COVID-19 Surveillance Team |
| EPI_ISL_2703642 | Lighthouse Lab in Glasgow | Wellcome Sanger Institute for the COVID-19 Genomics UK (COG-UK) Consortium | Harper VanSteenhouse, Yumi Kasai, David Gray, Carol Clugston, Anna Dominiczak and Alex Alderton, Roberto Amato, Jeffrey Barrett, Sonia Goncalves, Ewan Harrison, David K. Jackson, Ian Johnston, Dominic Kwiatkowski, Cordelia Langford, John Sillitoe on behalf of the Wellcome Sanger Institute COVID-19 Surveillance Team |
| EPI_ISL_2703643 | Lighthouse Lab in Milton Keynes | Wellcome Sanger Institute for the COVID-19 Genomics UK (COG-UK) Consortium | The Lighthouse Lab in Milton Keynes and Alex Alderton, Roberto Amato, Jeffrey Barrett, Sonia Goncalves, Ewan Harrison, David K. Jackson, Ian Johnston, Dominic Kwiatkowski, Cordelia Langford, John Sillitoe on behalf of the Wellcome Sanger Institute COVID-19 Surveillance Team |
| EPI_ISL_2703644, EPI_ISL_2703645, EPI_ISL_2703646 | Lighthouse Lab in Alderley Park | Wellcome Sanger Institute for the COVID-19 Genomics UK (COG-UK) Consortium | Jacquelyn Wynn, Mairead Hyland, The Lighthouse Lab in Alderley Park and Alex Alderton, Roberto Amato, Jeffrey Barrett, Sonia Goncalves, Ewan Harrison, David K. Jackson, Ian Johnston, Dominic Kwiatkowski, Cordelia Langford, John Sillitoe on behalf of the Wellcome Sanger Institute COVID-19 Surveillance Team |
| EPI_ISL_2703647 | Lighthouse Lab in Glasgow | Wellcome Sanger Institute for the COVID-19 Genomics UK (COG-UK) Consortium | Harper VanSteenhouse, Yumi Kasai, David Gray, Carol Clugston, Anna Dominiczak and Alex Alderton, Roberto Amato, Jeffrey Barrett, Sonia Goncalves, Ewan Harrison, David K. Jackson, Ian Johnston, Dominic Kwiatkowski, Cordelia Langford, John Sillitoe on behalf of the Wellcome Sanger Institute COVID-19 Surveillance Team |
| EPI_ISL_2703648, EPI_ISL_2703649 | Lighthouse Lab in Milton Keynes | Wellcome Sanger Institute for the COVID-19 Genomics UK (COG-UK) Consortium | The Lighthouse Lab in Milton Keynes and Alex Alderton, Roberto Amato, Jeffrey Barrett, Sonia Goncalves, Ewan Harrison, David K. Jackson, Ian Johnston, Dominic Kwiatkowski, Cordelia Langford, John Sillitoe on behalf of the Wellcome Sanger Institute COVID-19 Surveillance Team |
| EPI_ISL_2703650, EPI_ISL_2703651 | Lighthouse Lab in Alderley Park | Wellcome Sanger Institute for the COVID-19 Genomics UK (COG-UK) Consortium | Jacquelyn Wynn, Mairead Hyland, The Lighthouse Lab in Alderley Park and Alex Alderton, Roberto Amato, Jeffrey Barrett, Sonia Goncalves, Ewan Harrison, David K. Jackson, Ian Johnston, Dominic Kwiatkowski, Cordelia Langford, John Sillitoe on behalf of the Wellcome Sanger Institute COVID-19 Surveillance Team |
| EPI_ISL_2703652 | Lighthouse Lab in Milton Keynes | Wellcome Sanger Institute for the COVID-19 Genomics UK (COG-UK) Consortium | The Lighthouse Lab in Milton Keynes and Alex Alderton, Roberto Amato, Jeffrey Barrett, Sonia Goncalves, Ewan Harrison, David K. Jackson, Ian Johnston, Dominic Kwiatkowski, Cordelia Langford, John Sillitoe on behalf of the Wellcome Sanger Institute COVID-19 Surveillance Team |
| EPI_ISL_2703653 | Lighthouse Lab in Alderley Park | Wellcome Sanger Institute for the COVID-19 Genomics UK (COG-UK) Consortium | Jacquelyn Wynn, Mairead Hyland, The Lighthouse Lab in Alderley Park and Alex Alderton, Roberto Amato, Jeffrey Barrett, Sonia Goncalves, Ewan Harrison, David K. Jackson, Ian Johnston, Dominic Kwiatkowski, Cordelia Langford, John Sillitoe on behalf of the Wellcome Sanger Institute COVID-19 Surveillance Team |
| EPI_ISL_2703654 | Lighthouse Lab in Milton Keynes | Wellcome Sanger Institute for the COVID-19 Genomics UK (COG-UK) Consortium | The Lighthouse Lab in Milton Keynes and Alex Alderton, Roberto Amato, Jeffrey Barrett, Sonia Goncalves, Ewan Harrison, David K. Jackson, Ian Johnston, Dominic Kwiatkowski, Cordelia Langford, John Sillitoe on behalf of the Wellcome Sanger Institute COVID-19 Surveillance Team |
| EPI_ISL_2703655, EPI_ISL_2703656 | Lighthouse Lab in Alderley Park | Wellcome Sanger Institute for the COVID-19 Genomics UK (COG-UK) Consortium | Jacquelyn Wynn, Mairead Hyland, The Lighthouse Lab in Alderley Park and Alex Alderton, Roberto Amato, Jeffrey Barrett, Sonia Goncalves, Ewan Harrison, David K. Jackson, Ian Johnston, Dominic Kwiatkowski, Cordelia Langford, John Sillitoe on behalf of the Wellcome Sanger Institute COVID-19 Surveillance Team |
| EPI_ISL_2703657 | Lighthouse Lab in Glasgow | Wellcome Sanger Institute for the COVID-19 Genomics UK (COG-UK) Consortium | Harper VanSteenhouse, Yumi Kasai, David Gray, Carol Clugston, Anna Dominiczak and Alex Alderton, Roberto Amato, Jeffrey Barrett, Sonia Goncalves, Ewan Harrison, David K. Jackson, Ian Johnston, Dominic Kwiatkowski, Cordelia Langford, John Sillitoe on behalf of the Wellcome Sanger Institute COVID-19 Surveillance Team |
| EPI_ISL_2703658 | Lighthouse Lab in Milton Keynes | Wellcome Sanger Institute for the COVID-19 Genomics UK (COG-UK) Consortium | The Lighthouse Lab in Milton Keynes and Alex Alderton, Roberto Amato, Jeffrey Barrett, Sonia Goncalves, Ewan Harrison, David K. Jackson, Ian Johnston, Dominic Kwiatkowski, Cordelia Langford, John Sillitoe on behalf of the Wellcome Sanger Institute COVID-19 Surveillance Team |
| EPI_ISL_2703659 | Lighthouse Lab in Glasgow | Wellcome Sanger Institute for the COVID-19 Genomics UK (COG-UK) Consortium | Harper VanSteenhouse, Yumi Kasai, David Gray, Carol Clugston, Anna Dominiczak and Alex Alderton, Roberto Amato, Jeffrey Barrett, Sonia Goncalves, Ewan Harrison, David K. Jackson, Ian Johnston, Dominic Kwiatkowski, Cordelia Langford, John Sillitoe on behalf of the Wellcome Sanger Institute COVID-19 Surveillance Team |
| EPI_ISL_2703660, EPI_ISL_2703661 | Lighthouse Lab in Alderley Park | Wellcome Sanger Institute for the COVID-19 Genomics UK (COG-UK) Consortium | Jacquelyn Wynn, Mairead Hyland, The Lighthouse Lab in Alderley Park and Alex Alderton, Roberto Amato, Jeffrey Barrett, Sonia Goncalves, Ewan Harrison, David K. Jackson, Ian Johnston, Dominic Kwiatkowski, Cordelia Langford, John Sillitoe on behalf of the Wellcome Sanger Institute COVID-19 Surveillance Team |
| EPI_ISL_2703662 | Lighthouse Lab in Milton Keynes | Wellcome Sanger Institute for the COVID-19 Genomics UK (COG-UK) Consortium | The Lighthouse Lab in Milton Keynes and Alex Alderton, Roberto Amato, Jeffrey Barrett, Sonia Goncalves, Ewan Harrison, David K. Jackson, Ian Johnston, Dominic Kwiatkowski, Cordelia Langford, John Sillitoe on behalf of the Wellcome Sanger Institute COVID-19 Surveillance Team |
| EPI_ISL_2703663 | Lighthouse Lab in Glasgow | Wellcome Sanger Institute for the COVID-19 Genomics UK (COG-UK) Consortium | Harper VanSteenhouse, Yumi Kasai, David Gray, Carol Clugston, Anna Dominiczak and Alex Alderton, Roberto Amato, Jeffrey Barrett, Sonia Goncalves, Ewan Harrison, David K. Jackson, Ian Johnston, Dominic Kwiatkowski, Cordelia Langford, John Sillitoe on behalf of the Wellcome Sanger Institute COVID-19 Surveillance Team |
| EPI_ISL_2703664 | Lighthouse Lab in Alderley Park | Wellcome Sanger Institute for the COVID-19 Genomics UK (COG-UK) Consortium | Jacquelyn Wynn, Mairead Hyland, The Lighthouse Lab in Alderley Park and Alex Alderton, Roberto Amato, Jeffrey Barrett, Sonia Goncalves, Ewan Harrison, David K. Jackson, Ian Johnston, Dominic Kwiatkowski, Cordelia Langford, John Sillitoe on behalf of the Wellcome Sanger Institute COVID-19 Surveillance Team |
| EPI_ISL_2703665 | Lighthouse Lab in Milton Keynes | Wellcome Sanger Institute for the COVID-19 Genomics UK (COG-UK) Consortium | The Lighthouse Lab in Milton Keynes and Alex Alderton, Roberto Amato, Jeffrey Barrett, Sonia Goncalves, Ewan Harrison, David K. Jackson, Ian Johnston, Dominic Kwiatkowski, Cordelia Langford, John Sillitoe on behalf of the Wellcome Sanger Institute COVID-19 Surveillance Team |
| EPI_ISL_2703666, EPI_ISL_2703667 | Lighthouse Lab in Glasgow | Wellcome Sanger Institute for the COVID-19 Genomics UK (COG-UK) Consortium | Harper VanSteenhouse, Yumi Kasai, David Gray, Carol Clugston, Anna Dominiczak and Alex Alderton, Roberto Amato, Jeffrey Barrett, Sonia Goncalves, Ewan Harrison, David K. Jackson, Ian Johnston, Dominic Kwiatkowski, Cordelia Langford, John Sillitoe on behalf of the Wellcome Sanger Institute COVID-19 Surveillance Team |
| EPI_ISL_2703668, EPI_ISL_2703669, EPI_ISL_2703670, EPI_ISL_2703671, EPI_ISL_2703672 | Lighthouse Lab in Alderley Park | Wellcome Sanger Institute for the COVID-19 Genomics UK (COG-UK) Consortium | Jacquelyn Wynn, Mairead Hyland, The Lighthouse Lab in Alderley Park and Alex Alderton, Roberto Amato, Jeffrey Barrett, Sonia Goncalves, Ewan Harrison, David K. Jackson, Ian Johnston, Dominic Kwiatkowski, Cordelia Langford, John Sillitoe on behalf of the Wellcome Sanger Institute COVID-19 Surveillance Team |
| EPI_ISL_2703673 | Lighthouse Lab in Glasgow | Wellcome Sanger Institute for the COVID-19 Genomics UK (COG-UK) Consortium | Harper VanSteenhouse, Yumi Kasai, David Gray, Carol Clugston, Anna Dominiczak and Alex Alderton, Roberto Amato, Jeffrey Barrett, Sonia Goncalves, Ewan Harrison, David K. Jackson, Ian Johnston, Dominic Kwiatkowski, Cordelia Langford, John Sillitoe on behalf of the Wellcome Sanger Institute COVID-19 Surveillance Team |
| EPI_ISL_2703674, EPI_ISL_2703676, EPI_ISL_2703678 | Lighthouse Lab in Alderley Park | Wellcome Sanger Institute for the COVID-19 Genomics UK (COG-UK) Consortium | Jacquelyn Wynn, Mairead Hyland, The Lighthouse Lab in Alderley Park and Alex Alderton, Roberto Amato, Jeffrey Barrett, Sonia Goncalves, Ewan Harrison, David K. Jackson, Ian Johnston, Dominic Kwiatkowski, Cordelia Langford, John Sillitoe on behalf of the Wellcome Sanger Institute COVID-19 Surveillance Team |
| EPI_ISL_2703679 | Lighthouse Lab in Milton Keynes | Wellcome Sanger Institute for the COVID-19 Genomics UK (COG-UK) Consortium | The Lighthouse Lab in Milton Keynes and Alex Alderton, Roberto Amato, Jeffrey Barrett, Sonia Goncalves, Ewan Harrison, David K. Jackson, Ian Johnston, Dominic Kwiatkowski, Cordelia Langford, John Sillitoe on behalf of the Wellcome Sanger Institute COVID-19 Surveillance Team |
| EPI_ISL_2703680 | Lighthouse Lab in Glasgow | Wellcome Sanger Institute for the COVID-19 Genomics UK (COG-UK) Consortium | Harper VanSteenhouse, Yumi Kasai, David Gray, Carol Clugston, Anna Dominiczak and Alex Alderton, Roberto Amato, Jeffrey Barrett, Sonia Goncalves, Ewan Harrison, David K. Jackson, Ian Johnston, Dominic Kwiatkowski, Cordelia Langford, John Sillitoe on behalf of the Wellcome Sanger Institute COVID-19 Surveillance Team |
| EPI_ISL_2703681 | Lighthouse Lab in Alderley Park | Wellcome Sanger Institute for the COVID-19 Genomics UK (COG-UK) | Jacquelyn Wynn, Mairead Hyland, The Lighthouse Lab in Alderley Park and Alex Alderton, Roberto Amato, Jeffrey Barrett, Sonia Goncalves, Ewan Harrison, David K. Jackson, Ian Johnston, Dominic Kwiatkowski, Cordelia Langford, John Sillitoe on |

| | | | |
|---|---|---|---|
| | | Consortium | behalf of the Wellcome Sanger Institute COVID-19 Surveillance Team |
| EPI_ISL_2703682 | Lighthouse Lab in Milton Keynes | Wellcome Sanger Institute for the COVID-19 Genomics UK (COG-UK) Consortium | The Lighthouse Lab in Milton Keynes and Alex Alderton, Roberto Amato, Jeffrey Barrett, Sonia Goncalves, Ewan Harrison, David K. Jackson, Ian Johnston, Dominic Kwiatkowski, Cordelia Langford, John Sillitoe on behalf of the Wellcome Sanger Institute COVID-19 Surveillance Team |
| EPI_ISL_2703684 | Lighthouse Lab in Glasgow | Wellcome Sanger Institute for the COVID-19 Genomics UK (COG-UK) Consortium | Harper VanSteenhouse, Yumi Kasai, David Gray, Carol Clugston, Anna Dominiczak and Alex Alderton, Roberto Amato, Jeffrey Barrett, Sonia Goncalves, Ewan Harrison, David K. Jackson, Ian Johnston, Dominic Kwiatkowski, Cordelia Langford, John Sillitoe on behalf of the Wellcome Sanger Institute COVID-19 Surveillance Team |
| EPI_ISL_2703685, EPI_ISL_2703686, EPI_ISL_2703687, EPI_ISL_2703688, EPI_ISL_2703689 | Lighthouse Lab in Alderley Park | Wellcome Sanger Institute for the COVID-19 Genomics UK (COG-UK) Consortium | Jacquelyn Wynn, Mairead Hyland, The Lighthouse Lab in Alderley Park and Alex Alderton, Roberto Amato, Jeffrey Barrett, Sonia Goncalves, Ewan Harrison, David K. Jackson, Ian Johnston, Dominic Kwiatkowski, Cordelia Langford, John Sillitoe on behalf of the Wellcome Sanger Institute COVID-19 Surveillance Team |
| EPI_ISL_2703690, EPI_ISL_2703691, EPI_ISL_2703692 | Lighthouse Lab in Glasgow | Wellcome Sanger Institute for the COVID-19 Genomics UK (COG-UK) Consortium | Harper VanSteenhouse, Yumi Kasai, David Gray, Carol Clugston, Anna Dominiczak and Alex Alderton, Roberto Amato, Jeffrey Barrett, Sonia Goncalves, Ewan Harrison, David K. Jackson, Ian Johnston, Dominic Kwiatkowski, Cordelia Langford, John Sillitoe on behalf of the Wellcome Sanger Institute COVID-19 Surveillance Team |
| EPI_ISL_2703693 | Lighthouse Lab in Alderley Park | Wellcome Sanger Institute for the COVID-19 Genomics UK (COG-UK) Consortium | Jacquelyn Wynn, Mairead Hyland, The Lighthouse Lab in Alderley Park and Alex Alderton, Roberto Amato, Jeffrey Barrett, Sonia Goncalves, Ewan Harrison, David K. Jackson, Ian Johnston, Dominic Kwiatkowski, Cordelia Langford, John Sillitoe on behalf of the Wellcome Sanger Institute COVID-19 Surveillance Team |
| EPI_ISL_2703694 | Lighthouse Laboratory Plymouth | Wellcome Sanger Institute for the COVID-19 Genomics UK (COG-UK) Consortium | Lighthouse Laboratory Plymouth and Alex Alderton, Roberto Amato, Jeffrey Barrett, Sonia Goncalves, Ewan Harrison, David K. Jackson, Ian Johnston, Dominic Kwiatkowski, Cordelia Langford, John Sillitoe on behalf of the Wellcome Sanger Institute COVID-19 Surveillance Team |
| EPI_ISL_2703695 | Lighthouse Lab in Milton Keynes | Wellcome Sanger Institute for the COVID-19 Genomics UK (COG-UK) Consortium | The Lighthouse Lab in Milton Keynes and Alex Alderton, Roberto Amato, Jeffrey Barrett, Sonia Goncalves, Ewan Harrison, David K. Jackson, Ian Johnston, Dominic Kwiatkowski, Cordelia Langford, John Sillitoe on behalf of the Wellcome Sanger Institute COVID-19 Surveillance Team |
| EPI_ISL_2703696, EPI_ISL_2703697 | Lighthouse Lab in Glasgow | Wellcome Sanger Institute for the COVID-19 Genomics UK (COG-UK) Consortium | Harper VanSteenhouse, Yumi Kasai, David Gray, Carol Clugston, Anna Dominiczak and Alex Alderton, Roberto Amato, Jeffrey Barrett, Sonia Goncalves, Ewan Harrison, David K. Jackson, Ian Johnston, Dominic Kwiatkowski, Cordelia Langford, John Sillitoe on behalf of the Wellcome Sanger Institute COVID-19 Surveillance Team |
| EPI_ISL_2703698, EPI_ISL_2703699 | Lighthouse Lab in Alderley Park | Wellcome Sanger Institute for the COVID-19 Genomics UK (COG-UK) Consortium | Jacquelyn Wynn, Mairead Hyland, The Lighthouse Lab in Alderley Park and Alex Alderton, Roberto Amato, Jeffrey Barrett, Sonia Goncalves, Ewan Harrison, David K. Jackson, Ian Johnston, Dominic Kwiatkowski, Cordelia Langford, John Sillitoe on behalf of the Wellcome Sanger Institute COVID-19 Surveillance Team |
| EPI_ISL_2703700, EPI_ISL_2703701 | Lighthouse Lab in Milton Keynes | Wellcome Sanger Institute for the COVID-19 Genomics UK (COG-UK) Consortium | The Lighthouse Lab in Milton Keynes and Alex Alderton, Roberto Amato, Jeffrey Barrett, Sonia Goncalves, Ewan Harrison, David K. Jackson, Ian Johnston, Dominic Kwiatkowski, Cordelia Langford, John Sillitoe on behalf of the Wellcome Sanger Institute COVID-19 Surveillance Team |
| EPI_ISL_2703702, EPI_ISL_2703704, EPI_ISL_2703705, EPI_ISL_2703706 | Lighthouse Lab in Alderley Park | Wellcome Sanger Institute for the COVID-19 Genomics UK (COG-UK) Consortium | Jacquelyn Wynn, Mairead Hyland, The Lighthouse Lab in Alderley Park and Alex Alderton, Roberto Amato, Jeffrey Barrett, Sonia Goncalves, Ewan Harrison, David K. Jackson, Ian Johnston, Dominic Kwiatkowski, Cordelia Langford, John Sillitoe on behalf of the Wellcome Sanger Institute COVID-19 Surveillance Team |
| EPI_ISL_2703707 | Lighthouse Lab in Milton Keynes | Wellcome Sanger Institute for the COVID-19 Genomics UK (COG-UK) Consortium | The Lighthouse Lab in Milton Keynes and Alex Alderton, Roberto Amato, Jeffrey Barrett, Sonia Goncalves, Ewan Harrison, David K. Jackson, Ian Johnston, Dominic Kwiatkowski, Cordelia Langford, John Sillitoe on behalf of the Wellcome Sanger Institute COVID-19 Surveillance Team |
| EPI_ISL_2703708 | Lighthouse Lab in Glasgow | Wellcome Sanger Institute for the COVID-19 Genomics UK (COG-UK) Consortium | Harper VanSteenhouse, Yumi Kasai, David Gray, Carol Clugston, Anna Dominiczak and Alex Alderton, Roberto Amato, Jeffrey Barrett, Sonia Goncalves, Ewan Harrison, David K. Jackson, Ian Johnston, Dominic Kwiatkowski, Cordelia Langford, John Sillitoe on behalf of the Wellcome Sanger Institute COVID-19 Surveillance Team |
| EPI_ISL_2703709, EPI_ISL_2703710, EPI_ISL_2703711, EPI_ISL_2703712 | Lighthouse Lab in Milton Keynes | Wellcome Sanger Institute for the COVID-19 Genomics UK (COG-UK) Consortium | The Lighthouse Lab in Milton Keynes and Alex Alderton, Roberto Amato, Jeffrey Barrett, Sonia Goncalves, Ewan Harrison, David K. Jackson, Ian Johnston, Dominic Kwiatkowski, Cordelia Langford, John Sillitoe on behalf of the Wellcome Sanger Institute COVID-19 Surveillance Team |
| EPI_ISL_2703713, EPI_ISL_2703714 | Lighthouse Lab in Glasgow | Wellcome Sanger Institute for the COVID-19 Genomics UK (COG-UK) Consortium | Harper VanSteenhouse, Yumi Kasai, David Gray, Carol Clugston, Anna Dominiczak and Alex Alderton, Roberto Amato, Jeffrey Barrett, Sonia Goncalves, Ewan Harrison, David K. Jackson, Ian Johnston, Dominic Kwiatkowski, Cordelia Langford, John Sillitoe on behalf of the Wellcome Sanger Institute COVID-19 Surveillance Team |
| EPI_ISL_2703715 | Lighthouse Lab in Alderley Park | Wellcome Sanger Institute for the COVID-19 Genomics UK (COG-UK) Consortium | Jacquelyn Wynn, Mairead Hyland, The Lighthouse Lab in Alderley Park and Alex Alderton, Roberto Amato, Jeffrey Barrett, Sonia Goncalves, Ewan Harrison, David K. Jackson, Ian Johnston, Dominic Kwiatkowski, Cordelia Langford, John Sillitoe on behalf of the Wellcome Sanger Institute COVID-19 Surveillance Team |
| EPI_ISL_2703716, EPI_ISL_2703717 | Lighthouse Lab in Milton Keynes | Wellcome Sanger Institute for the COVID-19 Genomics UK (COG-UK) Consortium | The Lighthouse Lab in Milton Keynes and Alex Alderton, Roberto Amato, Jeffrey Barrett, Sonia Goncalves, Ewan Harrison, David K. Jackson, Ian Johnston, Dominic Kwiatkowski, Cordelia Langford, John Sillitoe on behalf of the Wellcome Sanger Institute COVID-19 Surveillance Team |
| EPI_ISL_2703718 | Lighthouse Lab in Alderley Park | Wellcome Sanger Institute for the COVID-19 Genomics UK (COG-UK) Consortium | Jacquelyn Wynn, Mairead Hyland, The Lighthouse Lab in Alderley Park and Alex Alderton, Roberto Amato, Jeffrey Barrett, Sonia Goncalves, Ewan Harrison, David K. Jackson, Ian Johnston, Dominic Kwiatkowski, Cordelia Langford, John Sillitoe on behalf of the Wellcome Sanger Institute COVID-19 Surveillance Team |
| EPI_ISL_2703719, EPI_ISL_2703721 | Lighthouse Lab in Milton Keynes | Wellcome Sanger Institute for the COVID-19 Genomics UK (COG-UK) Consortium | The Lighthouse Lab in Milton Keynes and Alex Alderton, Roberto Amato, Jeffrey Barrett, Sonia Goncalves, Ewan Harrison, David K. Jackson, Ian Johnston, Dominic Kwiatkowski, Cordelia Langford, John Sillitoe on behalf of the Wellcome Sanger Institute COVID-19 Surveillance Team |
| EPI_ISL_2703722 | Lighthouse Lab in Glasgow | Wellcome Sanger Institute for the COVID-19 Genomics UK (COG-UK) Consortium | Harper VanSteenhouse, Yumi Kasai, David Gray, Carol Clugston, Anna Dominiczak and Alex Alderton, Roberto Amato, Jeffrey Barrett, Sonia Goncalves, Ewan Harrison, David K. Jackson, Ian Johnston, Dominic Kwiatkowski, Cordelia Langford, John Sillitoe on behalf of the Wellcome Sanger Institute COVID-19 Surveillance Team |
| EPI_ISL_2703723, EPI_ISL_2703724 | Lighthouse Lab in Alderley Park | Wellcome Sanger Institute for the COVID-19 Genomics UK (COG-UK) Consortium | Jacquelyn Wynn, Mairead Hyland, The Lighthouse Lab in Alderley Park and Alex Alderton, Roberto Amato, Jeffrey Barrett, Sonia Goncalves, Ewan Harrison, David K. Jackson, Ian Johnston, Dominic Kwiatkowski, Cordelia Langford, John Sillitoe on behalf of the Wellcome Sanger Institute COVID-19 Surveillance Team |
| EPI_ISL_2703725 | Lighthouse Lab in Glasgow | Wellcome Sanger Institute for the COVID-19 Genomics UK (COG-UK) Consortium | Harper VanSteenhouse, Yumi Kasai, David Gray, Carol Clugston, Anna Dominiczak and Alex Alderton, Roberto Amato, Jeffrey Barrett, Sonia Goncalves, Ewan Harrison, David K. Jackson, Ian Johnston, Dominic Kwiatkowski, Cordelia Langford, John Sillitoe on behalf of the Wellcome Sanger Institute COVID-19 Surveillance Team |
| EPI_ISL_2703726, EPI_ISL_2703727 | Lighthouse Lab in Alderley Park | Wellcome Sanger Institute for the COVID-19 Genomics UK (COG-UK) Consortium | Jacquelyn Wynn, Mairead Hyland, The Lighthouse Lab in Alderley Park and Alex Alderton, Roberto Amato, Jeffrey Barrett, Sonia Goncalves, Ewan Harrison, David K. Jackson, Ian Johnston, Dominic Kwiatkowski, Cordelia Langford, John Sillitoe on behalf of the Wellcome Sanger Institute COVID-19 Surveillance Team |
| EPI_ISL_2703728 | Lighthouse Lab in Milton Keynes | Wellcome Sanger Institute for the COVID-19 Genomics UK (COG-UK) Consortium | The Lighthouse Lab in Milton Keynes and Alex Alderton, Roberto Amato, Jeffrey Barrett, Sonia Goncalves, Ewan Harrison, David K. Jackson, Ian Johnston, Dominic Kwiatkowski, Cordelia Langford, John Sillitoe on behalf of the Wellcome Sanger Institute COVID-19 Surveillance Team |
| EPI_ISL_2703729 | Lighthouse Lab in Glasgow | Wellcome Sanger Institute for the COVID-19 Genomics UK (COG-UK) Consortium | Harper VanSteenhouse, Yumi Kasai, David Gray, Carol Clugston, Anna Dominiczak and Alex Alderton, Roberto Amato, Jeffrey Barrett, Sonia Goncalves, Ewan Harrison, David K. Jackson, Ian Johnston, Dominic Kwiatkowski, Cordelia Langford, John Sillitoe on behalf of the Wellcome Sanger Institute COVID-19 Surveillance Team |
| EPI_ISL_2703730 | Lighthouse Lab in Alderley Park | Wellcome Sanger Institute for the COVID-19 Genomics UK (COG-UK) Consortium | Jacquelyn Wynn, Mairead Hyland, The Lighthouse Lab in Alderley Park and Alex Alderton, Roberto Amato, Jeffrey Barrett, Sonia Goncalves, Ewan Harrison, David K. Jackson, Ian Johnston, Dominic Kwiatkowski, Cordelia Langford, John Sillitoe on behalf of the Wellcome Sanger Institute COVID-19 Surveillance Team |
| EPI_ISL_2703731 | Lighthouse Lab in Glasgow | Wellcome Sanger Institute for the COVID-19 Genomics UK (COG-UK) Consortium | Harper VanSteenhouse, Yumi Kasai, David Gray, Carol Clugston, Anna Dominiczak and Alex Alderton, Roberto Amato, Jeffrey Barrett, Sonia Goncalves, Ewan Harrison, David K. Jackson, Ian Johnston, Dominic Kwiatkowski, Cordelia Langford, John Sillitoe on behalf of the Wellcome Sanger Institute COVID-19 Surveillance Team |
| EPI_ISL_2703732, EPI_ISL_2703733, EPI_ISL_2703734, EPI_ISL_2703735, EPI_ISL_2703736 | Lighthouse Lab in Alderley Park | Wellcome Sanger Institute for the COVID-19 Genomics UK (COG-UK) Consortium | Jacquelyn Wynn, Mairead Hyland, The Lighthouse Lab in Alderley Park and Alex Alderton, Roberto Amato, Jeffrey Barrett, Sonia Goncalves, Ewan Harrison, David K. Jackson, Ian Johnston, Dominic Kwiatkowski, Cordelia Langford, John Sillitoe on behalf of the Wellcome Sanger Institute COVID-19 Surveillance Team |
| EPI_ISL_2703737 | Lighthouse Lab in Glasgow | Wellcome Sanger Institute for the COVID-19 Genomics UK (COG-UK) Consortium | Harper VanSteenhouse, Yumi Kasai, David Gray, Carol Clugston, Anna Dominiczak and Alex Alderton, Roberto Amato, Jeffrey Barrett, Sonia Goncalves, Ewan Harrison, David K. Jackson, Ian Johnston, Dominic Kwiatkowski, Cordelia Langford, John Sillitoe on behalf of the Wellcome Sanger Institute COVID-19 Surveillance Team |
| EPI_ISL_2703738 | Lighthouse Lab in Alderley Park | Wellcome Sanger Institute for the COVID-19 Genomics UK (COG-UK) Consortium | Jacquelyn Wynn, Mairead Hyland, The Lighthouse Lab in Alderley Park and Alex Alderton, Roberto Amato, Jeffrey Barrett, Sonia Goncalves, Ewan Harrison, David K. Jackson, Ian Johnston, Dominic Kwiatkowski, Cordelia Langford, John Sillitoe on behalf of the Wellcome Sanger Institute COVID-19 Surveillance Team |
| EPI_ISL_2703739, EPI_ISL_2703740 | Lighthouse Lab in Glasgow | Wellcome Sanger Institute for the COVID-19 Genomics UK (COG-UK) Consortium | Harper VanSteenhouse, Yumi Kasai, David Gray, Carol Clugston, Anna Dominiczak and Alex Alderton, Roberto Amato, Jeffrey Barrett, Sonia Goncalves, Ewan Harrison, David K. Jackson, Ian Johnston, Dominic Kwiatkowski, Cordelia Langford, John Sillitoe on behalf of the Wellcome Sanger Institute COVID-19 Surveillance Team |
| EPI_ISL_2703742, EPI_ISL_2703743, EPI_ISL_2703744 | Lighthouse Lab in Alderley Park | Wellcome Sanger Institute for the COVID-19 Genomics UK (COG-UK) Consortium | Jacquelyn Wynn, Mairead Hyland, The Lighthouse Lab in Alderley Park and Alex Alderton, Roberto Amato, Jeffrey Barrett, Sonia Goncalves, Ewan Harrison, David K. Jackson, Ian Johnston, Dominic Kwiatkowski, Cordelia Langford, John Sillitoe on behalf of the Wellcome Sanger Institute COVID-19 Surveillance Team |
| EPI_ISL_2703745, EPI_ISL_2703746 | Lighthouse Lab in Milton Keynes | Wellcome Sanger Institute for the COVID-19 Genomics UK (COG-UK) Consortium | The Lighthouse Lab in Milton Keynes and Alex Alderton, Roberto Amato, Jeffrey Barrett, Sonia Goncalves, Ewan Harrison, David K. Jackson, Ian Johnston, Dominic Kwiatkowski, Cordelia Langford, John Sillitoe on behalf of the Wellcome Sanger Institute COVID-19 Surveillance Team |
| EPI_ISL_2703747 | Lighthouse Lab in Alderley Park | Wellcome Sanger Institute for the COVID-19 Genomics UK (COG-UK) Consortium | Jacquelyn Wynn, Mairead Hyland, The Lighthouse Lab in Alderley Park and Alex Alderton, Roberto Amato, Jeffrey Barrett, Sonia Goncalves, Ewan Harrison, David K. Jackson, Ian Johnston, Dominic Kwiatkowski, Cordelia Langford, John Sillitoe on behalf of the Wellcome Sanger Institute COVID-19 Surveillance Team |
| EPI_ISL_2703748 | Lighthouse Lab in Glasgow | Wellcome Sanger Institute for the COVID-19 Genomics UK (COG-UK) Consortium | Harper VanSteenhouse, Yumi Kasai, David Gray, Carol Clugston, Anna Dominiczak and Alex Alderton, Roberto Amato, Jeffrey Barrett, Sonia Goncalves, Ewan Harrison, David K. Jackson, Ian Johnston, Dominic Kwiatkowski, Cordelia Langford, John Sillitoe on behalf of the Wellcome Sanger Institute COVID-19 Surveillance Team |
| EPI_ISL_2703749, EPI_ISL_2703750 | Lighthouse Lab in Alderley Park | Wellcome Sanger Institute for the COVID-19 Genomics UK (COG-UK) Consortium | Jacquelyn Wynn, Mairead Hyland, The Lighthouse Lab in Alderley Park and Alex Alderton, Roberto Amato, Jeffrey Barrett, Sonia Goncalves, Ewan Harrison, David K. Jackson, Ian Johnston, Dominic Kwiatkowski, Cordelia Langford, John Sillitoe on behalf of the Wellcome Sanger Institute COVID-19 Surveillance Team |
| EPI_ISL_2703751 | Lighthouse Lab in Glasgow | Wellcome Sanger Institute for the COVID-19 Genomics UK (COG-UK) Consortium | Harper VanSteenhouse, Yumi Kasai, David Gray, Carol Clugston, Anna Dominiczak and Alex Alderton, Roberto Amato, Jeffrey Barrett, Sonia Goncalves, Ewan Harrison, David K. Jackson, Ian Johnston, Dominic Kwiatkowski, Cordelia Langford, John Sillitoe on behalf of the Wellcome Sanger Institute COVID-19 Surveillance Team |
| EPI_ISL_2703752 | Lighthouse Lab in Alderley Park | Wellcome Sanger Institute for the COVID-19 Genomics UK (COG-UK) Consortium | Jacquelyn Wynn, Mairead Hyland, The Lighthouse Lab in Alderley Park and Alex Alderton, Roberto Amato, Jeffrey Barrett, Sonia Goncalves, Ewan Harrison, David K. Jackson, Ian Johnston, Dominic Kwiatkowski, Cordelia Langford, John Sillitoe on behalf of the Wellcome Sanger Institute COVID-19 Surveillance Team |
| EPI_ISL_2703753 | Lighthouse Lab in Milton Keynes | Wellcome Sanger Institute for the COVID-19 Genomics UK (COG-UK) Consortium | The Lighthouse Lab in Milton Keynes and Alex Alderton, Roberto Amato, Jeffrey Barrett, Sonia Goncalves, Ewan Harrison, David K. Jackson, Ian Johnston, Dominic Kwiatkowski, Cordelia Langford, John Sillitoe on behalf of the Wellcome Sanger Institute COVID-19 Surveillance Team |
| EPI_ISL_2703754, | Lighthouse Lab in Alderley Park | Wellcome Sanger Institute for the COVID-19 Genomics UK (COG-UK) | Jacquelyn Wynn, Mairead Hyland, The Lighthouse Lab in Alderley Park and Alex Alderton, Roberto Amato, Jeffrey Barrett, Sonia Goncalves, Ewan Harrison, David K. Jackson, Ian Johnston, Dominic Kwiatkowski, Cordelia Langford, John Sillitoe on |

| | | | |
|---|---|---|---|
| EPI_ISL_2703755, EPI_ISL_2703756 | | Consortium | behalf of the Wellcome Sanger Institute COVID-19 Surveillance Team |
| EPI_ISL_2703757, EPI_ISL_2703758 | Lighthouse Lab in Glasgow | Wellcome Sanger Institute for the COVID-19 Genomics UK (COG-UK) Consortium | Harper VanSteenhouse, Yumi Kasai, David Gray, Carol Clugston, Anna Dominiczak and Alex Alderton, Roberto Amato, Jeffrey Barrett, Sonia Goncalves, Ewan Harrison, David K. Jackson, Ian Johnston, Dominic Kwiatkowski, Cordelia Langford, John Sillitoe on behalf of the Wellcome Sanger Institute COVID-19 Surveillance Team |
| EPI_ISL_2703759 | Lighthouse Lab in Milton Keynes | Wellcome Sanger Institute for the COVID-19 Genomics UK (COG-UK) Consortium | The Lighthouse Lab in Milton Keynes and Alex Alderton, Roberto Amato, Jeffrey Barrett, Sonia Goncalves, Ewan Harrison, David K. Jackson, Ian Johnston, Dominic Kwiatkowski, Cordelia Langford, John Sillitoe on behalf of the Wellcome Sanger Institute COVID-19 Surveillance Team |
| EPI_ISL_2703760, EPI_ISL_2703761 | Lighthouse Lab in Alderley Park | Wellcome Sanger Institute for the COVID-19 Genomics UK (COG-UK) Consortium | Jacquelyn Wynn, Mairead Hyland, The Lighthouse Lab in Alderley Park and Alex Alderton, Roberto Amato, Jeffrey Barrett, Sonia Goncalves, Ewan Harrison, David K. Jackson, Ian Johnston, Dominic Kwiatkowski, Cordelia Langford, John Sillitoe on behalf of the Wellcome Sanger Institute COVID-19 Surveillance Team |
| EPI_ISL_2703762 | Lighthouse Lab in Milton Keynes | Wellcome Sanger Institute for the COVID-19 Genomics UK (COG-UK) Consortium | The Lighthouse Lab in Milton Keynes and Alex Alderton, Roberto Amato, Jeffrey Barrett, Sonia Goncalves, Ewan Harrison, David K. Jackson, Ian Johnston, Dominic Kwiatkowski, Cordelia Langford, John Sillitoe on behalf of the Wellcome Sanger Institute COVID-19 Surveillance Team |
| EPI_ISL_2703763 | Lighthouse Lab in Alderley Park | Wellcome Sanger Institute for the COVID-19 Genomics UK (COG-UK) Consortium | Jacquelyn Wynn, Mairead Hyland, The Lighthouse Lab in Alderley Park and Alex Alderton, Roberto Amato, Jeffrey Barrett, Sonia Goncalves, Ewan Harrison, David K. Jackson, Ian Johnston, Dominic Kwiatkowski, Cordelia Langford, John Sillitoe on behalf of the Wellcome Sanger Institute COVID-19 Surveillance Team |
| EPI_ISL_2703764 | Lighthouse Lab in Milton Keynes | Wellcome Sanger Institute for the COVID-19 Genomics UK (COG-UK) Consortium | The Lighthouse Lab in Milton Keynes and Alex Alderton, Roberto Amato, Jeffrey Barrett, Sonia Goncalves, Ewan Harrison, David K. Jackson, Ian Johnston, Dominic Kwiatkowski, Cordelia Langford, John Sillitoe on behalf of the Wellcome Sanger Institute COVID-19 Surveillance Team |
| EPI_ISL_2703765 | Lighthouse Lab in Alderley Park | Wellcome Sanger Institute for the COVID-19 Genomics UK (COG-UK) Consortium | Jacquelyn Wynn, Mairead Hyland, The Lighthouse Lab in Alderley Park and Alex Alderton, Roberto Amato, Jeffrey Barrett, Sonia Goncalves, Ewan Harrison, David K. Jackson, Ian Johnston, Dominic Kwiatkowski, Cordelia Langford, John Sillitoe on behalf of the Wellcome Sanger Institute COVID-19 Surveillance Team |
| EPI_ISL_2703766 | Lighthouse Lab in Milton Keynes | Wellcome Sanger Institute for the COVID-19 Genomics UK (COG-UK) Consortium | The Lighthouse Lab in Milton Keynes and Alex Alderton, Roberto Amato, Jeffrey Barrett, Sonia Goncalves, Ewan Harrison, David K. Jackson, Ian Johnston, Dominic Kwiatkowski, Cordelia Langford, John Sillitoe on behalf of the Wellcome Sanger Institute COVID-19 Surveillance Team |
| EPI_ISL_2703767 | Lighthouse Lab in Glasgow | Wellcome Sanger Institute for the COVID-19 Genomics UK (COG-UK) Consortium | Harper VanSteenhouse, Yumi Kasai, David Gray, Carol Clugston, Anna Dominiczak and Alex Alderton, Roberto Amato, Jeffrey Barrett, Sonia Goncalves, Ewan Harrison, David K. Jackson, Ian Johnston, Dominic Kwiatkowski, Cordelia Langford, John Sillitoe on behalf of the Wellcome Sanger Institute COVID-19 Surveillance Team |
| EPI_ISL_2703768 | Lighthouse Lab in Milton Keynes | Wellcome Sanger Institute for the COVID-19 Genomics UK (COG-UK) Consortium | The Lighthouse Lab in Milton Keynes and Alex Alderton, Roberto Amato, Jeffrey Barrett, Sonia Goncalves, Ewan Harrison, David K. Jackson, Ian Johnston, Dominic Kwiatkowski, Cordelia Langford, John Sillitoe on behalf of the Wellcome Sanger Institute COVID-19 Surveillance Team |
| EPI_ISL_2703769, EPI_ISL_2703770 | Lighthouse Lab in Glasgow | Wellcome Sanger Institute for the COVID-19 Genomics UK (COG-UK) Consortium | Harper VanSteenhouse, Yumi Kasai, David Gray, Carol Clugston, Anna Dominiczak and Alex Alderton, Roberto Amato, Jeffrey Barrett, Sonia Goncalves, Ewan Harrison, David K. Jackson, Ian Johnston, Dominic Kwiatkowski, Cordelia Langford, John Sillitoe on behalf of the Wellcome Sanger Institute COVID-19 Surveillance Team |
| EPI_ISL_2703772, EPI_ISL_2703773, EPI_ISL_2703774, EPI_ISL_2703775, EPI_ISL_2703776 | Lighthouse Lab in Alderley Park | Wellcome Sanger Institute for the COVID-19 Genomics UK (COG-UK) Consortium | Jacquelyn Wynn, Mairead Hyland, The Lighthouse Lab in Alderley Park and Alex Alderton, Roberto Amato, Jeffrey Barrett, Sonia Goncalves, Ewan Harrison, David K. Jackson, Ian Johnston, Dominic Kwiatkowski, Cordelia Langford, John Sillitoe on behalf of the Wellcome Sanger Institute COVID-19 Surveillance Team |
| EPI_ISL_2703777 | Lighthouse Lab in Glasgow | Wellcome Sanger Institute for the COVID-19 Genomics UK (COG-UK) Consortium | Harper VanSteenhouse, Yumi Kasai, David Gray, Carol Clugston, Anna Dominiczak and Alex Alderton, Roberto Amato, Jeffrey Barrett, Sonia Goncalves, Ewan Harrison, David K. Jackson, Ian Johnston, Dominic Kwiatkowski, Cordelia Langford, John Sillitoe on behalf of the Wellcome Sanger Institute COVID-19 Surveillance Team |
| EPI_ISL_2703778 | Lighthouse Lab in Alderley Park | Wellcome Sanger Institute for the COVID-19 Genomics UK (COG-UK) Consortium | Jacquelyn Wynn, Mairead Hyland, The Lighthouse Lab in Alderley Park and Alex Alderton, Roberto Amato, Jeffrey Barrett, Sonia Goncalves, Ewan Harrison, David K. Jackson, Ian Johnston, Dominic Kwiatkowski, Cordelia Langford, John Sillitoe on behalf of the Wellcome Sanger Institute COVID-19 Surveillance Team |
| EPI_ISL_2703779, EPI_ISL_2703780 | Lighthouse Lab in Glasgow | Wellcome Sanger Institute for the COVID-19 Genomics UK (COG-UK) Consortium | Harper VanSteenhouse, Yumi Kasai, David Gray, Carol Clugston, Anna Dominiczak and Alex Alderton, Roberto Amato, Jeffrey Barrett, Sonia Goncalves, Ewan Harrison, David K. Jackson, Ian Johnston, Dominic Kwiatkowski, Cordelia Langford, John Sillitoe on behalf of the Wellcome Sanger Institute COVID-19 Surveillance Team |
| EPI_ISL_2703781 | Lighthouse Lab in Alderley Park | Wellcome Sanger Institute for the COVID-19 Genomics UK (COG-UK) Consortium | Jacquelyn Wynn, Mairead Hyland, The Lighthouse Lab in Alderley Park and Alex Alderton, Roberto Amato, Jeffrey Barrett, Sonia Goncalves, Ewan Harrison, David K. Jackson, Ian Johnston, Dominic Kwiatkowski, Cordelia Langford, John Sillitoe on behalf of the Wellcome Sanger Institute COVID-19 Surveillance Team |
| EPI_ISL_2703782 | Lighthouse Lab in Milton Keynes | Wellcome Sanger Institute for the COVID-19 Genomics UK (COG-UK) Consortium | The Lighthouse Lab in Milton Keynes and Alex Alderton, Roberto Amato, Jeffrey Barrett, Sonia Goncalves, Ewan Harrison, David K. Jackson, Ian Johnston, Dominic Kwiatkowski, Cordelia Langford, John Sillitoe on behalf of the Wellcome Sanger Institute COVID-19 Surveillance Team |
| EPI_ISL_2703783 | Lighthouse Lab in Glasgow | Wellcome Sanger Institute for the COVID-19 Genomics UK (COG-UK) Consortium | Harper VanSteenhouse, Yumi Kasai, David Gray, Carol Clugston, Anna Dominiczak and Alex Alderton, Roberto Amato, Jeffrey Barrett, Sonia Goncalves, Ewan Harrison, David K. Jackson, Ian Johnston, Dominic Kwiatkowski, Cordelia Langford, John Sillitoe on behalf of the Wellcome Sanger Institute COVID-19 Surveillance Team |
| EPI_ISL_2703784, EPI_ISL_2703785, EPI_ISL_2703786, EPI_ISL_2703787 | Lighthouse Lab in Alderley Park | Wellcome Sanger Institute for the COVID-19 Genomics UK (COG-UK) Consortium | Jacquelyn Wynn, Mairead Hyland, The Lighthouse Lab in Alderley Park and Alex Alderton, Roberto Amato, Jeffrey Barrett, Sonia Goncalves, Ewan Harrison, David K. Jackson, Ian Johnston, Dominic Kwiatkowski, Cordelia Langford, John Sillitoe on behalf of the Wellcome Sanger Institute COVID-19 Surveillance Team |
| EPI_ISL_2703789, EPI_ISL_2703790, EPI_ISL_2703791 | Lighthouse Lab in Glasgow | Wellcome Sanger Institute for the COVID-19 Genomics UK (COG-UK) Consortium | Harper VanSteenhouse, Yumi Kasai, David Gray, Carol Clugston, Anna Dominiczak and Alex Alderton, Roberto Amato, Jeffrey Barrett, Sonia Goncalves, Ewan Harrison, David K. Jackson, Ian Johnston, Dominic Kwiatkowski, Cordelia Langford, John Sillitoe on behalf of the Wellcome Sanger Institute COVID-19 Surveillance Team |
| EPI_ISL_2703792, EPI_ISL_2703793, EPI_ISL_2703794, EPI_ISL_2703795 | Lighthouse Lab in Alderley Park | Wellcome Sanger Institute for the COVID-19 Genomics UK (COG-UK) Consortium | Jacquelyn Wynn, Mairead Hyland, The Lighthouse Lab in Alderley Park and Alex Alderton, Roberto Amato, Jeffrey Barrett, Sonia Goncalves, Ewan Harrison, David K. Jackson, Ian Johnston, Dominic Kwiatkowski, Cordelia Langford, John Sillitoe on behalf of the Wellcome Sanger Institute COVID-19 Surveillance Team |
| EPI_ISL_2703796, EPI_ISL_2703797 | Lighthouse Lab in Glasgow | Wellcome Sanger Institute for the COVID-19 Genomics UK (COG-UK) Consortium | Harper VanSteenhouse, Yumi Kasai, David Gray, Carol Clugston, Anna Dominiczak and Alex Alderton, Roberto Amato, Jeffrey Barrett, Sonia Goncalves, Ewan Harrison, David K. Jackson, Ian Johnston, Dominic Kwiatkowski, Cordelia Langford, John Sillitoe on behalf of the Wellcome Sanger Institute COVID-19 Surveillance Team |
| EPI_ISL_2703798, EPI_ISL_2703799, EPI_ISL_2703800, EPI_ISL_2703801, EPI_ISL_2703802 | Lighthouse Lab in Milton Keynes | Wellcome Sanger Institute for the COVID-19 Genomics UK (COG-UK) Consortium | The Lighthouse Lab in Milton Keynes and Alex Alderton, Roberto Amato, Jeffrey Barrett, Sonia Goncalves, Ewan Harrison, David K. Jackson, Ian Johnston, Dominic Kwiatkowski, Cordelia Langford, John Sillitoe on behalf of the Wellcome Sanger Institute COVID-19 Surveillance Team |
| EPI_ISL_2703803 | Lighthouse Lab in Alderley Park | Wellcome Sanger Institute for the COVID-19 Genomics UK (COG-UK) Consortium | Jacquelyn Wynn, Mairead Hyland, The Lighthouse Lab in Alderley Park and Alex Alderton, Roberto Amato, Jeffrey Barrett, Sonia Goncalves, Ewan Harrison, David K. Jackson, Ian Johnston, Dominic Kwiatkowski, Cordelia Langford, John Sillitoe on behalf of the Wellcome Sanger Institute COVID-19 Surveillance Team |
| EPI_ISL_2703804 | Lighthouse Lab in Milton Keynes | Wellcome Sanger Institute for the COVID-19 Genomics UK (COG-UK) Consortium | The Lighthouse Lab in Milton Keynes and Alex Alderton, Roberto Amato, Jeffrey Barrett, Sonia Goncalves, Ewan Harrison, David K. Jackson, Ian Johnston, Dominic Kwiatkowski, Cordelia Langford, John Sillitoe on behalf of the Wellcome Sanger Institute COVID-19 Surveillance Team |
| EPI_ISL_2703805 | Lighthouse Laboratory Plymouth | Wellcome Sanger Institute for the COVID-19 Genomics UK (COG-UK) Consortium | Lighthouse Laboratory Plymouth and Alex Alderton, Roberto Amato, Jeffrey Barrett, Sonia Goncalves, Ewan Harrison, David K. Jackson, Ian Johnston, Dominic Kwiatkowski, Cordelia Langford, John Sillitoe on behalf of the Wellcome Sanger Institute COVID-19 Surveillance Team |
| EPI_ISL_2703806, EPI_ISL_2703807 | Lighthouse Lab in Glasgow | Wellcome Sanger Institute for the COVID-19 Genomics UK (COG-UK) Consortium | Harper VanSteenhouse, Yumi Kasai, David Gray, Carol Clugston, Anna Dominiczak and Alex Alderton, Roberto Amato, Jeffrey Barrett, Sonia Goncalves, Ewan Harrison, David K. Jackson, Ian Johnston, Dominic Kwiatkowski, Cordelia Langford, John Sillitoe on behalf of the Wellcome Sanger Institute COVID-19 Surveillance Team |
| EPI_ISL_2703808 | Lighthouse Laboratory Plymouth | Wellcome Sanger Institute for the COVID-19 Genomics UK (COG-UK) Consortium | Lighthouse Laboratory Plymouth and Alex Alderton, Roberto Amato, Jeffrey Barrett, Sonia Goncalves, Ewan Harrison, David K. Jackson, Ian Johnston, Dominic Kwiatkowski, Cordelia Langford, John Sillitoe on behalf of the Wellcome Sanger Institute COVID-19 Surveillance Team |
| EPI_ISL_2703809 | Lighthouse Lab in Glasgow | Wellcome Sanger Institute for the COVID-19 Genomics UK (COG-UK) Consortium | Harper VanSteenhouse, Yumi Kasai, David Gray, Carol Clugston, Anna Dominiczak and Alex Alderton, Roberto Amato, Jeffrey Barrett, Sonia Goncalves, Ewan Harrison, David K. Jackson, Ian Johnston, Dominic Kwiatkowski, Cordelia Langford, John Sillitoe on behalf of the Wellcome Sanger Institute COVID-19 Surveillance Team |
| EPI_ISL_2703810 | Lighthouse Lab in Alderley Park | Wellcome Sanger Institute for the COVID-19 Genomics UK (COG-UK) Consortium | Jacquelyn Wynn, Mairead Hyland, The Lighthouse Lab in Alderley Park and Alex Alderton, Roberto Amato, Jeffrey Barrett, Sonia Goncalves, Ewan Harrison, David K. Jackson, Ian Johnston, Dominic Kwiatkowski, Cordelia Langford, John Sillitoe on behalf of the Wellcome Sanger Institute COVID-19 Surveillance Team |
| EPI_ISL_2703811 | Lighthouse Lab in Glasgow | Wellcome Sanger Institute for the COVID-19 Genomics UK (COG-UK) Consortium | Harper VanSteenhouse, Yumi Kasai, David Gray, Carol Clugston, Anna Dominiczak and Alex Alderton, Roberto Amato, Jeffrey Barrett, Sonia Goncalves, Ewan Harrison, David K. Jackson, Ian Johnston, Dominic Kwiatkowski, Cordelia Langford, John Sillitoe on behalf of the Wellcome Sanger Institute COVID-19 Surveillance Team |
| EPI_ISL_2703812 | Lighthouse Lab in Alderley Park | Wellcome Sanger Institute for the COVID-19 Genomics UK (COG-UK) Consortium | Jacquelyn Wynn, Mairead Hyland, The Lighthouse Lab in Alderley Park and Alex Alderton, Roberto Amato, Jeffrey Barrett, Sonia Goncalves, Ewan Harrison, David K. Jackson, Ian Johnston, Dominic Kwiatkowski, Cordelia Langford, John Sillitoe on behalf of the Wellcome Sanger Institute COVID-19 Surveillance Team |
| EPI_ISL_2703813, EPI_ISL_2703814 | Lighthouse Lab in Glasgow | Wellcome Sanger Institute for the COVID-19 Genomics UK (COG-UK) Consortium | Harper VanSteenhouse, Yumi Kasai, David Gray, Carol Clugston, Anna Dominiczak and Alex Alderton, Roberto Amato, Jeffrey Barrett, Sonia Goncalves, Ewan Harrison, David K. Jackson, Ian Johnston, Dominic Kwiatkowski, Cordelia Langford, John Sillitoe on behalf of the Wellcome Sanger Institute COVID-19 Surveillance Team |
| EPI_ISL_2703815, EPI_ISL_2703816, EPI_ISL_2703817 | Lighthouse Lab in Alderley Park | Wellcome Sanger Institute for the COVID-19 Genomics UK (COG-UK) Consortium | Jacquelyn Wynn, Mairead Hyland, The Lighthouse Lab in Alderley Park and Alex Alderton, Roberto Amato, Jeffrey Barrett, Sonia Goncalves, Ewan Harrison, David K. Jackson, Ian Johnston, Dominic Kwiatkowski, Cordelia Langford, John Sillitoe on behalf of the Wellcome Sanger Institute COVID-19 Surveillance Team |
| EPI_ISL_2703818 | Lighthouse Lab in Milton Keynes | Wellcome Sanger Institute for the COVID-19 Genomics UK (COG-UK) Consortium | The Lighthouse Lab in Milton Keynes and Alex Alderton, Roberto Amato, Jeffrey Barrett, Sonia Goncalves, Ewan Harrison, David K. Jackson, Ian Johnston, Dominic Kwiatkowski, Cordelia Langford, John Sillitoe on behalf of the Wellcome Sanger Institute COVID-19 Surveillance Team |
| EPI_ISL_2703819, EPI_ISL_2703820, EPI_ISL_2703821, EPI_ISL_2703822, EPI_ISL_2703823, EPI_ISL_2703824, EPI_ISL_2703825 | Lighthouse Lab in Alderley Park | Wellcome Sanger Institute for the COVID-19 Genomics UK (COG-UK) Consortium | Jacquelyn Wynn, Mairead Hyland, The Lighthouse Lab in Alderley Park and Alex Alderton, Roberto Amato, Jeffrey Barrett, Sonia Goncalves, Ewan Harrison, David K. Jackson, Ian Johnston, Dominic Kwiatkowski, Cordelia Langford, John Sillitoe on behalf of the Wellcome Sanger Institute COVID-19 Surveillance Team |
| EPI_ISL_2703826 | Lighthouse Laboratory Plymouth | Wellcome Sanger Institute for the COVID-19 Genomics UK (COG-UK) Consortium | Lighthouse Laboratory Plymouth and Alex Alderton, Roberto Amato, Jeffrey Barrett, Sonia Goncalves, Ewan Harrison, David K. Jackson, Ian Johnston, Dominic Kwiatkowski, Cordelia Langford, John Sillitoe on behalf of the Wellcome Sanger Institute COVID-19 Surveillance Team |

| | | | |
|---|---|---|---|
| EPI_ISL_2703827 | Lighthouse Lab in Milton Keynes | Wellcome Sanger Institute for the COVID-19 Genomics UK (COG-UK) Consortium | The Lighthouse Lab in Milton Keynes and Alex Alderton, Roberto Amato, Jeffrey Barrett, Sonia Goncalves, Ewan Harrison, David K. Jackson, Ian Johnston, Dominic Kwiatkowski, Cordelia Langford, John Sillitoe on behalf of the Wellcome Sanger Institute COVID-19 Surveillance Team |
| EPI_ISL_2703828 | Lighthouse Lab in Alderley Park | Wellcome Sanger Institute for the COVID-19 Genomics UK (COG-UK) Consortium | Jacquelyn Wynn, Mairead Hyland, The Lighthouse Lab in Alderley Park and Alex Alderton, Roberto Amato, Jeffrey Barrett, Sonia Goncalves, Ewan Harrison, David K. Jackson, Ian Johnston, Dominic Kwiatkowski, Cordelia Langford, John Sillitoe on behalf of the Wellcome Sanger Institute COVID-19 Surveillance Team |
| EPI_ISL_2703829, EPI_ISL_2703830, EPI_ISL_2703831 | Lighthouse Lab in Milton Keynes | Wellcome Sanger Institute for the COVID-19 Genomics UK (COG-UK) Consortium | The Lighthouse Lab in Milton Keynes and Alex Alderton, Roberto Amato, Jeffrey Barrett, Sonia Goncalves, Ewan Harrison, David K. Jackson, Ian Johnston, Dominic Kwiatkowski, Cordelia Langford, John Sillitoe on behalf of the Wellcome Sanger Institute COVID-19 Surveillance Team |
| EPI_ISL_2703832, EPI_ISL_2703833 | Lighthouse Lab in Glasgow | Wellcome Sanger Institute for the COVID-19 Genomics UK (COG-UK) Consortium | Harper VanSteenhouse, Yumi Kasai, David Gray, Carol Clugston, Anna Dominiczak and Alex Alderton, Roberto Amato, Jeffrey Barrett, Sonia Goncalves, Ewan Harrison, David K. Jackson, Ian Johnston, Dominic Kwiatkowski, Cordelia Langford, John Sillitoe on behalf of the Wellcome Sanger Institute COVID-19 Surveillance Team |
| EPI_ISL_2703834 | Lighthouse Lab in Milton Keynes | Wellcome Sanger Institute for the COVID-19 Genomics UK (COG-UK) Consortium | The Lighthouse Lab in Milton Keynes and Alex Alderton, Roberto Amato, Jeffrey Barrett, Sonia Goncalves, Ewan Harrison, David K. Jackson, Ian Johnston, Dominic Kwiatkowski, Cordelia Langford, John Sillitoe on behalf of the Wellcome Sanger Institute COVID-19 Surveillance Team |
| EPI_ISL_2703835, EPI_ISL_2703836, EPI_ISL_2703837, EPI_ISL_2703838, EPI_ISL_2703839, EPI_ISL_2703840, EPI_ISL_2703841, EPI_ISL_2703842, EPI_ISL_2703843, EPI_ISL_2703844, EPI_ISL_2703845, EPI_ISL_2703846, EPI_ISL_2703847, EPI_ISL_2703848, EPI_ISL_2703849, EPI_ISL_2703850, EPI_ISL_2703851, EPI_ISL_2703852, EPI_ISL_2703853, EPI_ISL_2703854, EPI_ISL_2703855, EPI_ISL_2703856, EPI_ISL_2703857, EPI_ISL_2703858, EPI_ISL_2703859, EPI_ISL_2703860, EPI_ISL_2703861, EPI_ISL_2703862, EPI_ISL_2703863, EPI_ISL_2703864, EPI_ISL_2703865, EPI_ISL_2703866, EPI_ISL_2703867, EPI_ISL_2703868, EPI_ISL_2703869, EPI_ISL_2703870, EPI_ISL_2703871, EPI_ISL_2703872, EPI_ISL_2703873, EPI_ISL_2703874, EPI_ISL_2703875, EPI_ISL_2703876, EPI_ISL_2703877, EPI_ISL_2703878, EPI_ISL_2703879, EPI_ISL_2703880, EPI_ISL_2703881, EPI_ISL_2703882, EPI_ISL_2703883, EPI_ISL_2703884, EPI_ISL_2703885, EPI_ISL_2703886, EPI_ISL_2703887, EPI_ISL_2703888, EPI_ISL_2703889, EPI_ISL_2703890, EPI_ISL_2703892, EPI_ISL_2703893, EPI_ISL_2703894, EPI_ISL_2703895, EPI_ISL_2703896, EPI_ISL_2703897, EPI_ISL_2703898, EPI_ISL_2703899, EPI_ISL_2703900, EPI_ISL_2703901, EPI_ISL_2703902, EPI_ISL_2703903, EPI_ISL_2703904, EPI_ISL_2703905, EPI_ISL_2703906, EPI_ISL_2703907, EPI_ISL_2703908, EPI_ISL_2703909, EPI_ISL_2703910, EPI_ISL_2703911, EPI_ISL_2703912, EPI_ISL_2703913, EPI_ISL_2703914, EPI_ISL_2703915, EPI_ISL_2703916, EPI_ISL_2703917, EPI_ISL_2703918, EPI_ISL_2703919, EPI_ISL_2703920, EPI_ISL_2703921, EPI_ISL_2703922, EPI_ISL_2703923, EPI_ISL_2703924, EPI_ISL_2703925, EPI_ISL_2703926, EPI_ISL_2703927, EPI_ISL_2703928, EPI_ISL_2703929, EPI_ISL_2703930, EPI_ISL_2703931, EPI_ISL_2703932, EPI_ISL_2703933, EPI_ISL_2703934, EPI_ISL_2703935, EPI_ISL_2703936, EPI_ISL_2703937, EPI_ISL_2703938, EPI_ISL_2703939, EPI_ISL_2703940, EPI_ISL_2703941, EPI_ISL_2703942, EPI_ISL_2703943, EPI_ISL_2703944, EPI_ISL_2703945, EPI_ISL_2703946, EPI_ISL_2703947, EPI_ISL_2703948, EPI_ISL_2703949, EPI_ISL_2703950, EPI_ISL_2703951, EPI_ISL_2703952, EPI_ISL_2703953, EPI_ISL_2703954, EPI_ISL_2703955, EPI_ISL_2703956, EPI_ISL_2703957, EPI_ISL_2703958, EPI_ISL_2703959, EPI_ISL_2703960, EPI_ISL_2703961, EPI_ISL_2703962, EPI_ISL_2703963, EPI_ISL_2703964, EPI_ISL_2703965, EPI_ISL_2703966, EPI_ISL_2703967, EPI_ISL_2703968, EPI_ISL_2703969, EPI_ISL_2703970, EPI_ISL_2703971, EPI_ISL_2703972, EPI_ISL_2703973, EPI_ISL_2703974, EPI_ISL_2703975, EPI_ISL_2703976, EPI_ISL_2703977, EPI_ISL_2703978, EPI_ISL_2703979, EPI_ISL_2703980, EPI_ISL_2703981, EPI_ISL_2703982, EPI_ISL_2703983, EPI_ISL_2703984, EPI_ISL_2703985, EPI_ISL_2703986, EPI_ISL_2703987, EPI_ISL_2703988, EPI_ISL_2703989, EPI_ISL_2703990, EPI_ISL_2703991, EPI_ISL_2703992, EPI_ISL_2703993, EPI_ISL_2703994, EPI_ISL_2703995, EPI_ISL_2703996, EPI_ISL_2703997, EPI_ISL_2703998, EPI_ISL_2703999, EPI_ISL_2704000, EPI_ISL_2704001, EPI_ISL_2704002, EPI_ISL_2704003, EPI_ISL_2704004, EPI_ISL_2704005, EPI_ISL_2704006, EPI_ISL_2704007, EPI_ISL_2704008, EPI_ISL_2704009, EPI_ISL_2704010, EPI_ISL_2704011, EPI_ISL_2704012, EPI_ISL_2704013, EPI_ISL_2704014, EPI_ISL_2704015, EPI_ISL_2704016, EPI_ISL_2704017, EPI_ISL_2704018, EPI_ISL_2704019, EPI_ISL_2704020, EPI_ISL_2704021, EPI_ISL_2704022, EPI_ISL_2704023, EPI_ISL_2704024, EPI_ISL_2704025, EPI_ISL_2704026, EPI_ISL_2704027, EPI_ISL_2704028, EPI_ISL_2704029, EPI_ISL_2704030, EPI_ISL_2704031, EPI_ISL_2704032, EPI_ISL_2704033, EPI_ISL_2704034, EPI_ISL_2704036, EPI_ISL_2704037, EPI_ISL_2704038, EPI_ISL_2704039, EPI_ISL_2704040, EPI_ISL_2704041, EPI_ISL_2704042, EPI_ISL_2704043, EPI_ISL_2704044, EPI_ISL_2704045, EPI_ISL_2704046, EPI_ISL_2704047, EPI_ISL_2704048, EPI_ISL_2704049, EPI_ISL_2704050, EPI_ISL_2704051, EPI_ISL_2704052, EPI_ISL_2704053, EPI_ISL_2704054, EPI_ISL_2704055, EPI_ISL_2704056, EPI_ISL_2704057, EPI_ISL_2704058, EPI_ISL_2704059, EPI_ISL_2704060, EPI_ISL_2704061, EPI_ISL_2704062, EPI_ISL_2704063, EPI_ISL_2704064, EPI_ISL_2704065, EPI_ISL_2704066, EPI_ISL_2704067, EPI_ISL_2704068, EPI_ISL_2704070, EPI_ISL_2704071, EPI_ISL_2704072, EPI_ISL_2704073, EPI_ISL_2704074, EPI_ISL_2704075, EPI_ISL_2704076, EPI_ISL_2704077, EPI_ISL_2704078, EPI_ISL_2704079, EPI_ISL_2704080, EPI_ISL_2704081, EPI_ISL_2704082, EPI_ISL_2704083, EPI_ISL_2704084, EPI_ISL_2704085, EPI_ISL_2704086, EPI_ISL_2704087, EPI_ISL_2704088, EPI_ISL_2704089, EPI_ISL_2704090, EPI_ISL_2704091, EPI_ISL_2704092, EPI_ISL_2704093, EPI_ISL_2704094, EPI_ISL_2704095, EPI_ISL_2704096, EPI_ISL_2704097, EPI_ISL_2704098, EPI_ISL_2704099, EPI_ISL_2704100, EPI_ISL_2704101, EPI_ISL_2704102, EPI_ISL_2704103, EPI_ISL_2704104, EPI_ISL_2704105, EPI_ISL_2704106, EPI_ISL_2704107, EPI_ISL_2704108, EPI_ISL_2704109, EPI_ISL_2704110, EPI_ISL_2704111, EPI_ISL_2704112, EPI_ISL_2704113, EPI_ISL_2704114, EPI_ISL_2704115, EPI_ISL_2704116, EPI_ISL_2704117, EPI_ISL_2704119, EPI_ISL_2704120, EPI_ISL_2704121, EPI_ISL_2704122, EPI_ISL_2704123, EPI_ISL_2704124, EPI_ISL_2704125, EPI_ISL_2704126, EPI_ISL_2704127, EPI_ISL_2704128, EPI_ISL_2704129, EPI_ISL_2704130, EPI_ISL_2704131, EPI_ISL_2704132, EPI_ISL_2704133, EPI_ISL_2704134, EPI_ISL_2704135, EPI_ISL_2704136, EPI_ISL_2704137, EPI_ISL_2704138, EPI_ISL_2704139, EPI_ISL_2704140, EPI_ISL_2704141, EPI_ISL_2704142, EPI_ISL_2704143, EPI_ISL_2704144, EPI_ISL_2704145, EPI_ISL_2704146, EPI_ISL_2704147, EPI_ISL_2704148, EPI_ISL_2704149, EPI_ISL_2704150, EPI_ISL_2704151, EPI_ISL_2704152, EPI_ISL_2704153, EPI_ISL_2704154, EPI_ISL_2704155, EPI_ISL_2704156, EPI_ISL_2704157, EPI_ISL_2704158, EPI_ISL_2704159, EPI_ISL_2704160, EPI_ISL_2704161, EPI_ISL_2704162, EPI_ISL_2704163, EPI_ISL_2704164, EPI_ISL_2704165, EPI_ISL_2704166, EPI_ISL_2704167, EPI_ISL_2704168, EPI_ISL_2704169, EPI_ISL_2704170, EPI_ISL_2704172, EPI_ISL_2704173, EPI_ISL_2704174, EPI_ISL_2704175, EPI_ISL_2704176, EPI_ISL_2704177, EPI_ISL_2704178, EPI_ISL_2704179, EPI_ISL_2704180, EPI_ISL_2704181, EPI_ISL_2704182, EPI_ISL_2704183, EPI_ISL_2704184 | | | |
| see above | Lighthouse Laboratory Plymouth | Wellcome Sanger Institute for the COVID-19 Genomics UK (COG-UK) Consortium | Lighthouse Laboratory Plymouth and Alex Alderton, Roberto Amato, Jeffrey Barrett, Sonia Goncalves, Ewan Harrison, David K. Jackson, Ian Johnston, Dominic Kwiatkowski, Cordelia Langford, John Sillitoe on behalf of the Wellcome Sanger Institute COVID-19 Surveillance Team |
| EPI_ISL_2704185, EPI_ISL_2704186, EPI_ISL_2704187 | Lighthouse Lab in Alderley Park | Wellcome Sanger Institute for the COVID-19 Genomics UK (COG-UK) Consortium | Jacquelyn Wynn, Mairead Hyland, The Lighthouse Lab in Alderley Park and Alex Alderton, Roberto Amato, Jeffrey Barrett, Sonia Goncalves, Ewan Harrison, David K. Jackson, Ian Johnston, Dominic Kwiatkowski, Cordelia Langford, John Sillitoe on behalf of the Wellcome Sanger Institute COVID-19 Surveillance Team |
| EPI_ISL_2704188 | Health Services Laboratories | Wellcome Sanger Institute for the COVID-19 Genomics UK (COG-UK) Consortium | Health Services Laboratories and Alex Alderton, Roberto Amato, Jeffrey Barrett, Sonia Goncalves, Ewan Harrison, David K. Jackson, Ian Johnston, Dominic Kwiatkowski, Cordelia Langford, John Sillitoe on behalf of the Wellcome Sanger Institute COVID-19 Surveillance Team |
| EPI_ISL_2704189 | Lighthouse Lab in Alderley Park | Wellcome Sanger Institute for the COVID-19 Genomics UK (COG-UK) Consortium | Jacquelyn Wynn, Mairead Hyland, The Lighthouse Lab in Alderley Park and Alex Alderton, Roberto Amato, Jeffrey Barrett, Sonia Goncalves, Ewan Harrison, David K. Jackson, Ian Johnston, Dominic Kwiatkowski, Cordelia Langford, John Sillitoe on behalf of the Wellcome Sanger Institute COVID-19 Surveillance Team |
| EPI_ISL_2704190, EPI_ISL_2704191 | Health Services Laboratories | Wellcome Sanger Institute for the COVID-19 Genomics UK (COG-UK) Consortium | Health Services Laboratories and Alex Alderton, Roberto Amato, Jeffrey Barrett, Sonia Goncalves, Ewan Harrison, David K. Jackson, Ian Johnston, Dominic Kwiatkowski, Cordelia Langford, John Sillitoe on behalf of the Wellcome Sanger Institute COVID-19 Surveillance Team |
| EPI_ISL_2704192, EPI_ISL_2704193 | Lighthouse Lab in Alderley Park | Wellcome Sanger Institute for the COVID-19 Genomics UK (COG-UK) Consortium | Jacquelyn Wynn, Mairead Hyland, The Lighthouse Lab in Alderley Park and Alex Alderton, Roberto Amato, Jeffrey Barrett, Sonia Goncalves, Ewan Harrison, David K. Jackson, Ian Johnston, Dominic Kwiatkowski, Cordelia Langford, John Sillitoe on behalf of the Wellcome Sanger Institute COVID-19 Surveillance Team |
| EPI_ISL_2704194 | Health Services Laboratories | Wellcome Sanger Institute for the COVID-19 Genomics UK (COG-UK) Consortium | Health Services Laboratories and Alex Alderton, Roberto Amato, Jeffrey Barrett, Sonia Goncalves, Ewan Harrison, David K. Jackson, Ian Johnston, Dominic Kwiatkowski, Cordelia Langford, John Sillitoe on behalf of the Wellcome Sanger Institute COVID-19 Surveillance Team |
| EPI_ISL_2704195 | Lighthouse Lab in Alderley Park | Wellcome Sanger Institute for the COVID-19 Genomics UK (COG-UK) Consortium | Jacquelyn Wynn, Mairead Hyland, The Lighthouse Lab in Alderley Park and Alex Alderton, Roberto Amato, Jeffrey Barrett, Sonia Goncalves, Ewan Harrison, David K. Jackson, Ian Johnston, Dominic Kwiatkowski, Cordelia Langford, John Sillitoe on behalf of the Wellcome Sanger Institute COVID-19 Surveillance Team |
| EPI_ISL_2704196 | Health Services Laboratories | Wellcome Sanger Institute for the COVID-19 Genomics UK (COG-UK) Consortium | Health Services Laboratories and Alex Alderton, Roberto Amato, Jeffrey Barrett, Sonia Goncalves, Ewan Harrison, David K. Jackson, Ian Johnston, Dominic Kwiatkowski, Cordelia Langford, John Sillitoe on behalf of the Wellcome Sanger Institute COVID-19 Surveillance Team |
| EPI_ISL_2704197 | Lighthouse Lab in Alderley Park | Wellcome Sanger Institute for the COVID-19 Genomics UK (COG-UK) Consortium | Jacquelyn Wynn, Mairead Hyland, The Lighthouse Lab in Alderley Park and Alex Alderton, Roberto Amato, Jeffrey Barrett, Sonia Goncalves, Ewan Harrison, David K. Jackson, Ian Johnston, Dominic Kwiatkowski, Cordelia Langford, John Sillitoe on behalf of the Wellcome Sanger Institute COVID-19 Surveillance Team |
| EPI_ISL_2704198 | Health Services Laboratories | Wellcome Sanger Institute for the COVID-19 Genomics UK (COG-UK) Consortium | Health Services Laboratories and Alex Alderton, Roberto Amato, Jeffrey Barrett, Sonia Goncalves, Ewan Harrison, David K. Jackson, Ian Johnston, Dominic Kwiatkowski, Cordelia Langford, John Sillitoe on behalf of the Wellcome Sanger Institute COVID-19 Surveillance Team |
| EPI_ISL_2704199 | Lighthouse Lab in Alderley Park | Wellcome Sanger Institute for the COVID-19 Genomics UK (COG-UK) Consortium | Jacquelyn Wynn, Mairead Hyland, The Lighthouse Lab in Alderley Park and Alex Alderton, Roberto Amato, Jeffrey Barrett, Sonia Goncalves, Ewan Harrison, David K. Jackson, Ian Johnston, Dominic Kwiatkowski, Cordelia Langford, John Sillitoe on behalf of the Wellcome Sanger Institute COVID-19 Surveillance Team |
| EPI_ISL_2704200 | Health Services Laboratories | Wellcome Sanger Institute for the COVID-19 Genomics UK (COG-UK) Consortium | Health Services Laboratories and Alex Alderton, Roberto Amato, Jeffrey Barrett, Sonia Goncalves, Ewan Harrison, David K. Jackson, Ian Johnston, Dominic Kwiatkowski, Cordelia Langford, John Sillitoe on behalf of the Wellcome Sanger Institute COVID-19 Surveillance Team |
| EPI_ISL_2704201, EPI_ISL_2704202, EPI_ISL_2704203 | Lighthouse Lab in Alderley Park | Wellcome Sanger Institute for the COVID-19 Genomics UK (COG-UK) Consortium | Jacquelyn Wynn, Mairead Hyland, The Lighthouse Lab in Alderley Park and Alex Alderton, Roberto Amato, Jeffrey Barrett, Sonia Goncalves, Ewan Harrison, David K. Jackson, Ian Johnston, Dominic Kwiatkowski, Cordelia Langford, John Sillitoe on behalf of the Wellcome Sanger Institute COVID-19 Surveillance Team |
| EPI_ISL_2704204 | Health Services Laboratories | Wellcome Sanger Institute for the COVID-19 Genomics UK (COG-UK) Consortium | Health Services Laboratories and Alex Alderton, Roberto Amato, Jeffrey Barrett, Sonia Goncalves, Ewan Harrison, David K. Jackson, Ian Johnston, Dominic Kwiatkowski, Cordelia Langford, John Sillitoe on behalf of the Wellcome Sanger Institute COVID-19 Surveillance Team |
| EPI_ISL_2704205, EPI_ISL_2704206, EPI_ISL_2704207, EPI_ISL_2704208, EPI_ISL_2704209, EPI_ISL_2704210, EPI_ISL_2704211, EPI_ISL_2704212, EPI_ISL_2704213 | Lighthouse Lab in Alderley Park | Wellcome Sanger Institute for the COVID-19 Genomics UK (COG-UK) Consortium | Jacquelyn Wynn, Mairead Hyland, The Lighthouse Lab in Alderley Park and Alex Alderton, Roberto Amato, Jeffrey Barrett, Sonia Goncalves, Ewan Harrison, David K. Jackson, Ian Johnston, Dominic Kwiatkowski, Cordelia Langford, John Sillitoe on behalf of the Wellcome Sanger Institute COVID-19 Surveillance Team |
| EPI_ISL_2704214, EPI_ISL_2704215 | Health Services Laboratories | Wellcome Sanger Institute for the COVID-19 Genomics UK (COG-UK) Consortium | Health Services Laboratories and Alex Alderton, Roberto Amato, Jeffrey Barrett, Sonia Goncalves, Ewan Harrison, David K. Jackson, Ian Johnston, Dominic Kwiatkowski, Cordelia Langford, John Sillitoe on behalf of the Wellcome Sanger Institute COVID-19 Surveillance Team |
| EPI_ISL_2704216, EPI_ISL_2704217 | Lighthouse Lab in Alderley Park | Wellcome Sanger Institute for the COVID-19 Genomics UK (COG-UK) Consortium | Jacquelyn Wynn, Mairead Hyland, The Lighthouse Lab in Alderley Park and Alex Alderton, Roberto Amato, Jeffrey Barrett, Sonia Goncalves, Ewan Harrison, David K. Jackson, Ian Johnston, Dominic Kwiatkowski, Cordelia Langford, John Sillitoe on behalf of the Wellcome Sanger Institute COVID-19 Surveillance Team |
| EPI_ISL_2704218 | Health Services Laboratories | Wellcome Sanger Institute for the COVID-19 Genomics UK (COG-UK) Consortium | Health Services Laboratories and Alex Alderton, Roberto Amato, Jeffrey Barrett, Sonia Goncalves, Ewan Harrison, David K. Jackson, Ian Johnston, Dominic Kwiatkowski, Cordelia Langford, John Sillitoe on behalf of the Wellcome Sanger Institute COVID-19 Surveillance Team |
| EPI_ISL_2704219, EPI_ISL_2704220, EPI_ISL_2704221 | Lighthouse Lab in Alderley Park | Wellcome Sanger Institute for the COVID-19 Genomics UK (COG-UK) Consortium | Jacquelyn Wynn, Mairead Hyland, The Lighthouse Lab in Alderley Park and Alex Alderton, Roberto Amato, Jeffrey Barrett, Sonia Goncalves, Ewan Harrison, David K. Jackson, Ian Johnston, Dominic Kwiatkowski, Cordelia Langford, John Sillitoe on behalf of the Wellcome Sanger Institute COVID-19 Surveillance Team |
| EPI_ISL_2704222 | Health Services Laboratories | Wellcome Sanger Institute for the COVID-19 Genomics UK (COG-UK) Consortium | Health Services Laboratories and Alex Alderton, Roberto Amato, Jeffrey Barrett, Sonia Goncalves, Ewan Harrison, David K. Jackson, Ian Johnston, Dominic Kwiatkowski, Cordelia Langford, John Sillitoe on behalf of the Wellcome Sanger Institute COVID-19 Surveillance Team |
| EPI_ISL_2704223 | Lighthouse Lab in Alderley Park | Wellcome Sanger Institute for the COVID-19 Genomics UK (COG-UK) Consortium | Jacquelyn Wynn, Mairead Hyland, The Lighthouse Lab in Alderley Park and Alex Alderton, Roberto Amato, Jeffrey Barrett, Sonia Goncalves, Ewan Harrison, David K. Jackson, Ian Johnston, Dominic Kwiatkowski, Cordelia Langford, John Sillitoe on behalf of the Wellcome Sanger Institute COVID-19 Surveillance Team |
| EPI_ISL_2704224, EPI_ISL_2704225 | Health Services Laboratories | Wellcome Sanger Institute for the COVID-19 Genomics UK (COG-UK) Consortium | Health Services Laboratories and Alex Alderton, Roberto Amato, Jeffrey Barrett, Sonia Goncalves, Ewan Harrison, David K. Jackson, Ian Johnston, Dominic Kwiatkowski, Cordelia Langford, John Sillitoe on behalf of the Wellcome Sanger Institute COVID-19 Surveillance Team |
| EPI_ISL_2704226 | Lighthouse Lab in Alderley Park | Wellcome Sanger Institute for the COVID-19 Genomics UK (COG-UK) Consortium | Jacquelyn Wynn, Mairead Hyland, The Lighthouse Lab in Alderley Park and Alex Alderton, Roberto Amato, Jeffrey Barrett, Sonia Goncalves, Ewan Harrison, David K. Jackson, Ian Johnston, Dominic Kwiatkowski, Cordelia Langford, John Sillitoe on behalf of the Wellcome Sanger Institute COVID-19 Surveillance Team |
| EPI_ISL_2704227 | Health Services Laboratories | Wellcome Sanger Institute for the COVID-19 Genomics UK (COG-UK) Consortium | Health Services Laboratories and Alex Alderton, Roberto Amato, Jeffrey Barrett, Sonia Goncalves, Ewan Harrison, David K. Jackson, Ian Johnston, Dominic Kwiatkowski, Cordelia Langford, John Sillitoe on behalf of the Wellcome Sanger Institute COVID-19 Surveillance Team |
| EPI_ISL_2704228 | Lighthouse Lab in Alderley Park | Wellcome Sanger Institute for the COVID-19 Genomics UK (COG-UK) Consortium | Jacquelyn Wynn, Mairead Hyland, The Lighthouse Lab in Alderley Park and Alex Alderton, Roberto Amato, Jeffrey Barrett, Sonia Goncalves, Ewan Harrison, David K. Jackson, Ian Johnston, Dominic Kwiatkowski, Cordelia Langford, John Sillitoe on behalf of the Wellcome Sanger Institute COVID-19 Surveillance Team |
| EPI_ISL_2704229 | Health Services Laboratories | Wellcome Sanger Institute for the COVID-19 Genomics UK (COG-UK) Consortium | Health Services Laboratories and Alex Alderton, Roberto Amato, Jeffrey Barrett, Sonia Goncalves, Ewan Harrison, David K. Jackson, Ian Johnston, Dominic Kwiatkowski, Cordelia Langford, John Sillitoe on behalf of the Wellcome Sanger Institute COVID-19 Surveillance Team |
| EPI_ISL_2704230, EPI_ISL_2704231 | Lighthouse Lab in Alderley Park | Wellcome Sanger Institute for the COVID-19 Genomics UK (COG-UK) Consortium | Jacquelyn Wynn, Mairead Hyland, The Lighthouse Lab in Alderley Park and Alex Alderton, Roberto Amato, Jeffrey Barrett, Sonia Goncalves, Ewan Harrison, David K. Jackson, Ian Johnston, Dominic Kwiatkowski, Cordelia Langford, John Sillitoe on behalf of the Wellcome Sanger Institute COVID-19 Surveillance Team |
| EPI_ISL_2704232 | Health Services Laboratories | Wellcome Sanger Institute for the COVID-19 Genomics UK (COG-UK) Consortium | Health Services Laboratories and Alex Alderton, Roberto Amato, Jeffrey Barrett, Sonia Goncalves, Ewan Harrison, David K. Jackson, Ian Johnston, Dominic Kwiatkowski, Cordelia Langford, John Sillitoe on behalf of the Wellcome Sanger Institute COVID-19 Surveillance Team |

| | | | |
|---|---|---|---|
| EPI_ISL_2704233 | Lighthouse Lab in Alderley Park | Wellcome Sanger Institute for the COVID-19 Genomics UK (COG-UK) Consortium | Jacquelyn Wynn, Mairead Hyland, The Lighthouse Lab in Alderley Park and Alex Alderton, Roberto Amato, Jeffrey Barrett, Sonia Goncalves, Ewan Harrison, David K. Jackson, Ian Johnston, Dominic Kwiatkowski, Cordelia Langford, John Sillitoe on behalf of the Wellcome Sanger Institute COVID-19 Surveillance Team |
| EPI_ISL_2704234 | Health Services Laboratories | Wellcome Sanger Institute for the COVID-19 Genomics UK (COG-UK) Consortium | Health Services Laboratories and Alex Alderton, Roberto Amato, Jeffrey Barrett, Sonia Goncalves, Ewan Harrison, David K. Jackson, Ian Johnston, Dominic Kwiatkowski, Cordelia Langford, John Sillitoe on behalf of the Wellcome Sanger Institute COVID-19 Surveillance Team |
| EPI_ISL_2704235, EPI_ISL_2704236 | Lighthouse Lab in Alderley Park | Wellcome Sanger Institute for the COVID-19 Genomics UK (COG-UK) Consortium | Jacquelyn Wynn, Mairead Hyland, The Lighthouse Lab in Alderley Park and Alex Alderton, Roberto Amato, Jeffrey Barrett, Sonia Goncalves, Ewan Harrison, David K. Jackson, Ian Johnston, Dominic Kwiatkowski, Cordelia Langford, John Sillitoe on behalf of the Wellcome Sanger Institute COVID-19 Surveillance Team |
| EPI_ISL_2704237 | Health Services Laboratories | Wellcome Sanger Institute for the COVID-19 Genomics UK (COG-UK) Consortium | Health Services Laboratories and Alex Alderton, Roberto Amato, Jeffrey Barrett, Sonia Goncalves, Ewan Harrison, David K. Jackson, Ian Johnston, Dominic Kwiatkowski, Cordelia Langford, John Sillitoe on behalf of the Wellcome Sanger Institute COVID-19 Surveillance Team |
| EPI_ISL_2704238, EPI_ISL_2704239, EPI_ISL_2704240, EPI_ISL_2704241 | Lighthouse Lab in Alderley Park | Wellcome Sanger Institute for the COVID-19 Genomics UK (COG-UK) Consortium | Jacquelyn Wynn, Mairead Hyland, The Lighthouse Lab in Alderley Park and Alex Alderton, Roberto Amato, Jeffrey Barrett, Sonia Goncalves, Ewan Harrison, David K. Jackson, Ian Johnston, Dominic Kwiatkowski, Cordelia Langford, John Sillitoe on behalf of the Wellcome Sanger Institute COVID-19 Surveillance Team |
| EPI_ISL_2704242 | Health Services Laboratories | Wellcome Sanger Institute for the COVID-19 Genomics UK (COG-UK) Consortium | Health Services Laboratories and Alex Alderton, Roberto Amato, Jeffrey Barrett, Sonia Goncalves, Ewan Harrison, David K. Jackson, Ian Johnston, Dominic Kwiatkowski, Cordelia Langford, John Sillitoe on behalf of the Wellcome Sanger Institute COVID-19 Surveillance Team |
| EPI_ISL_2704243 | Lighthouse Lab in Alderley Park | Wellcome Sanger Institute for the COVID-19 Genomics UK (COG-UK) Consortium | Jacquelyn Wynn, Mairead Hyland, The Lighthouse Lab in Alderley Park and Alex Alderton, Roberto Amato, Jeffrey Barrett, Sonia Goncalves, Ewan Harrison, David K. Jackson, Ian Johnston, Dominic Kwiatkowski, Cordelia Langford, John Sillitoe on behalf of the Wellcome Sanger Institute COVID-19 Surveillance Team |
| EPI_ISL_2704244 | Health Services Laboratories | Wellcome Sanger Institute for the COVID-19 Genomics UK (COG-UK) Consortium | Health Services Laboratories and Alex Alderton, Roberto Amato, Jeffrey Barrett, Sonia Goncalves, Ewan Harrison, David K. Jackson, Ian Johnston, Dominic Kwiatkowski, Cordelia Langford, John Sillitoe on behalf of the Wellcome Sanger Institute COVID-19 Surveillance Team |
| EPI_ISL_2704245 | Lighthouse Lab in Alderley Park | Wellcome Sanger Institute for the COVID-19 Genomics UK (COG-UK) Consortium | Jacquelyn Wynn, Mairead Hyland, The Lighthouse Lab in Alderley Park and Alex Alderton, Roberto Amato, Jeffrey Barrett, Sonia Goncalves, Ewan Harrison, David K. Jackson, Ian Johnston, Dominic Kwiatkowski, Cordelia Langford, John Sillitoe on behalf of the Wellcome Sanger Institute COVID-19 Surveillance Team |
| EPI_ISL_2704246 | Health Services Laboratories | Wellcome Sanger Institute for the COVID-19 Genomics UK (COG-UK) Consortium | Health Services Laboratories and Alex Alderton, Roberto Amato, Jeffrey Barrett, Sonia Goncalves, Ewan Harrison, David K. Jackson, Ian Johnston, Dominic Kwiatkowski, Cordelia Langford, John Sillitoe on behalf of the Wellcome Sanger Institute COVID-19 Surveillance Team |
| EPI_ISL_2704247, EPI_ISL_2704248 | Lighthouse Lab in Alderley Park | Wellcome Sanger Institute for the COVID-19 Genomics UK (COG-UK) Consortium | Jacquelyn Wynn, Mairead Hyland, The Lighthouse Lab in Alderley Park and Alex Alderton, Roberto Amato, Jeffrey Barrett, Sonia Goncalves, Ewan Harrison, David K. Jackson, Ian Johnston, Dominic Kwiatkowski, Cordelia Langford, John Sillitoe on behalf of the Wellcome Sanger Institute COVID-19 Surveillance Team |
| EPI_ISL_2704249 | Health Services Laboratories | Wellcome Sanger Institute for the COVID-19 Genomics UK (COG-UK) Consortium | Health Services Laboratories and Alex Alderton, Roberto Amato, Jeffrey Barrett, Sonia Goncalves, Ewan Harrison, David K. Jackson, Ian Johnston, Dominic Kwiatkowski, Cordelia Langford, John Sillitoe on behalf of the Wellcome Sanger Institute COVID-19 Surveillance Team |
| EPI_ISL_2704250, EPI_ISL_2704251, EPI_ISL_2704252, EPI_ISL_2704253, EPI_ISL_2704254 | Lighthouse Lab in Alderley Park | Wellcome Sanger Institute for the COVID-19 Genomics UK (COG-UK) Consortium | Jacquelyn Wynn, Mairead Hyland, The Lighthouse Lab in Alderley Park and Alex Alderton, Roberto Amato, Jeffrey Barrett, Sonia Goncalves, Ewan Harrison, David K. Jackson, Ian Johnston, Dominic Kwiatkowski, Cordelia Langford, John Sillitoe on behalf of the Wellcome Sanger Institute COVID-19 Surveillance Team |
| EPI_ISL_2704255 | Health Services Laboratories | Wellcome Sanger Institute for the COVID-19 Genomics UK (COG-UK) Consortium | Health Services Laboratories and Alex Alderton, Roberto Amato, Jeffrey Barrett, Sonia Goncalves, Ewan Harrison, David K. Jackson, Ian Johnston, Dominic Kwiatkowski, Cordelia Langford, John Sillitoe on behalf of the Wellcome Sanger Institute COVID-19 Surveillance Team |
| EPI_ISL_2704256 | Lighthouse Lab in Alderley Park | Wellcome Sanger Institute for the COVID-19 Genomics UK (COG-UK) Consortium | Jacquelyn Wynn, Mairead Hyland, The Lighthouse Lab in Alderley Park and Alex Alderton, Roberto Amato, Jeffrey Barrett, Sonia Goncalves, Ewan Harrison, David K. Jackson, Ian Johnston, Dominic Kwiatkowski, Cordelia Langford, John Sillitoe on behalf of the Wellcome Sanger Institute COVID-19 Surveillance Team |
| EPI_ISL_2704257 | Health Services Laboratories | Wellcome Sanger Institute for the COVID-19 Genomics UK (COG-UK) Consortium | Health Services Laboratories and Alex Alderton, Roberto Amato, Jeffrey Barrett, Sonia Goncalves, Ewan Harrison, David K. Jackson, Ian Johnston, Dominic Kwiatkowski, Cordelia Langford, John Sillitoe on behalf of the Wellcome Sanger Institute COVID-19 Surveillance Team |
| EPI_ISL_2704258 | Lighthouse Lab in Alderley Park | Wellcome Sanger Institute for the COVID-19 Genomics UK (COG-UK) Consortium | Jacquelyn Wynn, Mairead Hyland, The Lighthouse Lab in Alderley Park and Alex Alderton, Roberto Amato, Jeffrey Barrett, Sonia Goncalves, Ewan Harrison, David K. Jackson, Ian Johnston, Dominic Kwiatkowski, Cordelia Langford, John Sillitoe on behalf of the Wellcome Sanger Institute COVID-19 Surveillance Team |
| EPI_ISL_2704259 | Health Services Laboratories | Wellcome Sanger Institute for the COVID-19 Genomics UK (COG-UK) Consortium | Health Services Laboratories and Alex Alderton, Roberto Amato, Jeffrey Barrett, Sonia Goncalves, Ewan Harrison, David K. Jackson, Ian Johnston, Dominic Kwiatkowski, Cordelia Langford, John Sillitoe on behalf of the Wellcome Sanger Institute COVID-19 Surveillance Team |
| EPI_ISL_2704260, EPI_ISL_2704261, EPI_ISL_2704262, EPI_ISL_2704263, EPI_ISL_2704264, EPI_ISL_2704265 | Lighthouse Lab in Alderley Park | Wellcome Sanger Institute for the COVID-19 Genomics UK (COG-UK) Consortium | Jacquelyn Wynn, Mairead Hyland, The Lighthouse Lab in Alderley Park and Alex Alderton, Roberto Amato, Jeffrey Barrett, Sonia Goncalves, Ewan Harrison, David K. Jackson, Ian Johnston, Dominic Kwiatkowski, Cordelia Langford, John Sillitoe on behalf of the Wellcome Sanger Institute COVID-19 Surveillance Team |
| EPI_ISL_2704266 | Health Services Laboratories | Wellcome Sanger Institute for the COVID-19 Genomics UK (COG-UK) Consortium | Health Services Laboratories and Alex Alderton, Roberto Amato, Jeffrey Barrett, Sonia Goncalves, Ewan Harrison, David K. Jackson, Ian Johnston, Dominic Kwiatkowski, Cordelia Langford, John Sillitoe on behalf of the Wellcome Sanger Institute COVID-19 Surveillance Team |
| EPI_ISL_2704267, EPI_ISL_2704268, EPI_ISL_2704269 | Lighthouse Lab in Alderley Park | Wellcome Sanger Institute for the COVID-19 Genomics UK (COG-UK) Consortium | Jacquelyn Wynn, Mairead Hyland, The Lighthouse Lab in Alderley Park and Alex Alderton, Roberto Amato, Jeffrey Barrett, Sonia Goncalves, Ewan Harrison, David K. Jackson, Ian Johnston, Dominic Kwiatkowski, Cordelia Langford, John Sillitoe on behalf of the Wellcome Sanger Institute COVID-19 Surveillance Team |
| EPI_ISL_2704270, EPI_ISL_2704271 | Health Services Laboratories | Wellcome Sanger Institute for the COVID-19 Genomics UK (COG-UK) Consortium | Health Services Laboratories and Alex Alderton, Roberto Amato, Jeffrey Barrett, Sonia Goncalves, Ewan Harrison, David K. Jackson, Ian Johnston, Dominic Kwiatkowski, Cordelia Langford, John Sillitoe on behalf of the Wellcome Sanger Institute COVID-19 Surveillance Team |
| EPI_ISL_2704272, EPI_ISL_2704273 | Lighthouse Lab in Alderley Park | Wellcome Sanger Institute for the COVID-19 Genomics UK (COG-UK) Consortium | Jacquelyn Wynn, Mairead Hyland, The Lighthouse Lab in Alderley Park and Alex Alderton, Roberto Amato, Jeffrey Barrett, Sonia Goncalves, Ewan Harrison, David K. Jackson, Ian Johnston, Dominic Kwiatkowski, Cordelia Langford, John Sillitoe on behalf of the Wellcome Sanger Institute COVID-19 Surveillance Team |
| EPI_ISL_2704274, EPI_ISL_2704275 | Health Services Laboratories | Wellcome Sanger Institute for the COVID-19 Genomics UK (COG-UK) Consortium | Health Services Laboratories and Alex Alderton, Roberto Amato, Jeffrey Barrett, Sonia Goncalves, Ewan Harrison, David K. Jackson, Ian Johnston, Dominic Kwiatkowski, Cordelia Langford, John Sillitoe on behalf of the Wellcome Sanger Institute COVID-19 Surveillance Team |
| EPI_ISL_2704276 | Lighthouse Lab in Alderley Park | Wellcome Sanger Institute for the COVID-19 Genomics UK (COG-UK) Consortium | Jacquelyn Wynn, Mairead Hyland, The Lighthouse Lab in Alderley Park and Alex Alderton, Roberto Amato, Jeffrey Barrett, Sonia Goncalves, Ewan Harrison, David K. Jackson, Ian Johnston, Dominic Kwiatkowski, Cordelia Langford, John Sillitoe on behalf of the Wellcome Sanger Institute COVID-19 Surveillance Team |
| EPI_ISL_2704277 | Health Services Laboratories | Wellcome Sanger Institute for the COVID-19 Genomics UK (COG-UK) Consortium | Health Services Laboratories and Alex Alderton, Roberto Amato, Jeffrey Barrett, Sonia Goncalves, Ewan Harrison, David K. Jackson, Ian Johnston, Dominic Kwiatkowski, Cordelia Langford, John Sillitoe on behalf of the Wellcome Sanger Institute COVID-19 Surveillance Team |
| EPI_ISL_2704278, EPI_ISL_2704279, EPI_ISL_2704280, EPI_ISL_2704281, EPI_ISL_2704282, EPI_ISL_2704283, EPI_ISL_2704284, EPI_ISL_2704285, EPI_ISL_2704286, EPI_ISL_2704287 | Lighthouse Lab in Alderley Park | Wellcome Sanger Institute for the COVID-19 Genomics UK (COG-UK) Consortium | Jacquelyn Wynn, Mairead Hyland, The Lighthouse Lab in Alderley Park and Alex Alderton, Roberto Amato, Jeffrey Barrett, Sonia Goncalves, Ewan Harrison, David K. Jackson, Ian Johnston, Dominic Kwiatkowski, Cordelia Langford, John Sillitoe on behalf of the Wellcome Sanger Institute COVID-19 Surveillance Team |
| EPI_ISL_2704288, EPI_ISL_2704289, EPI_ISL_2704291, EPI_ISL_2704292 | Health Services Laboratories | Wellcome Sanger Institute for the COVID-19 Genomics UK (COG-UK) Consortium | Health Services Laboratories and Alex Alderton, Roberto Amato, Jeffrey Barrett, Sonia Goncalves, Ewan Harrison, David K. Jackson, Ian Johnston, Dominic Kwiatkowski, Cordelia Langford, John Sillitoe on behalf of the Wellcome Sanger Institute COVID-19 Surveillance Team |
| EPI_ISL_2704293, EPI_ISL_2704294, EPI_ISL_2704295, EPI_ISL_2704296, EPI_ISL_2704297, EPI_ISL_2704298, EPI_ISL_2704299, EPI_ISL_2704300 | Lighthouse Lab in Alderley Park | Wellcome Sanger Institute for the COVID-19 Genomics UK (COG-UK) Consortium | Jacquelyn Wynn, Mairead Hyland, The Lighthouse Lab in Alderley Park and Alex Alderton, Roberto Amato, Jeffrey Barrett, Sonia Goncalves, Ewan Harrison, David K. Jackson, Ian Johnston, Dominic Kwiatkowski, Cordelia Langford, John Sillitoe on behalf of the Wellcome Sanger Institute COVID-19 Surveillance Team |
| EPI_ISL_2704301 | Health Services Laboratories | Wellcome Sanger Institute for the COVID-19 Genomics UK (COG-UK) Consortium | Health Services Laboratories and Alex Alderton, Roberto Amato, Jeffrey Barrett, Sonia Goncalves, Ewan Harrison, David K. Jackson, Ian Johnston, Dominic Kwiatkowski, Cordelia Langford, John Sillitoe on behalf of the Wellcome Sanger Institute COVID-19 Surveillance Team |
| EPI_ISL_2704302, EPI_ISL_2704303, EPI_ISL_2704304, EPI_ISL_2704305, EPI_ISL_2704306 | Lighthouse Lab in Alderley Park | Wellcome Sanger Institute for the COVID-19 Genomics UK (COG-UK) Consortium | Jacquelyn Wynn, Mairead Hyland, The Lighthouse Lab in Alderley Park and Alex Alderton, Roberto Amato, Jeffrey Barrett, Sonia Goncalves, Ewan Harrison, David K. Jackson, Ian Johnston, Dominic Kwiatkowski, Cordelia Langford, John Sillitoe on behalf of the Wellcome Sanger Institute COVID-19 Surveillance Team |
| EPI_ISL_2704307 | Health Services Laboratories | Wellcome Sanger Institute for the COVID-19 Genomics UK (COG-UK) Consortium | Health Services Laboratories and Alex Alderton, Roberto Amato, Jeffrey Barrett, Sonia Goncalves, Ewan Harrison, David K. Jackson, Ian Johnston, Dominic Kwiatkowski, Cordelia Langford, John Sillitoe on behalf of the Wellcome Sanger Institute COVID-19 Surveillance Team |
| EPI_ISL_2704308, EPI_ISL_2704309 | Lighthouse Lab in Alderley Park | Wellcome Sanger Institute for the COVID-19 Genomics UK (COG-UK) Consortium | Jacquelyn Wynn, Mairead Hyland, The Lighthouse Lab in Alderley Park and Alex Alderton, Roberto Amato, Jeffrey Barrett, Sonia Goncalves, Ewan Harrison, David K. Jackson, Ian Johnston, Dominic Kwiatkowski, Cordelia Langford, John Sillitoe on behalf of the Wellcome Sanger Institute COVID-19 Surveillance Team |
| EPI_ISL_2704310 | Health Services Laboratories | Wellcome Sanger Institute for the COVID-19 Genomics UK (COG-UK) | Health Services Laboratories and Alex Alderton, Roberto Amato, Jeffrey Barrett, Sonia Goncalves, Ewan Harrison, David K. Jackson, Ian Johnston, Dominic Kwiatkowski, Cordelia Langford, John Sillitoe on behalf of the Wellcome Sanger Institute |

| | | | |
|---|---|---|---|
| | | Consortium | COVID-19 Surveillance Team |
| EPI_ISL_2704311 | Lighthouse Lab in Alderley Park | Wellcome Sanger Institute for the COVID-19 Genomics UK (COG-UK) Consortium | Jacquelyn Wynn, Mairead Hyland, The Lighthouse Lab in Alderley Park and Alex Alderton, Roberto Amato, Jeffrey Barrett, Sonia Goncalves, Ewan Harrison, David K. Jackson, Ian Johnston, Dominic Kwiatkowski, Cordelia Langford, John Sillitoe on behalf of the Wellcome Sanger Institute COVID-19 Surveillance Team |
| EPI_ISL_2704312 | Health Services Laboratories | Wellcome Sanger Institute for the COVID-19 Genomics UK (COG-UK) Consortium | Health Services Laboratories and Alex Alderton, Roberto Amato, Jeffrey Barrett, Sonia Goncalves, Ewan Harrison, David K. Jackson, Ian Johnston, Dominic Kwiatkowski, Cordelia Langford, John Sillitoe on behalf of the Wellcome Sanger Institute COVID-19 Surveillance Team |
| EPI_ISL_2704313 | Lighthouse Lab in Alderley Park | Wellcome Sanger Institute for the COVID-19 Genomics UK (COG-UK) Consortium | Jacquelyn Wynn, Mairead Hyland, The Lighthouse Lab in Alderley Park and Alex Alderton, Roberto Amato, Jeffrey Barrett, Sonia Goncalves, Ewan Harrison, David K. Jackson, Ian Johnston, Dominic Kwiatkowski, Cordelia Langford, John Sillitoe on behalf of the Wellcome Sanger Institute COVID-19 Surveillance Team |
| EPI_ISL_2704314 | Health Services Laboratories | Wellcome Sanger Institute for the COVID-19 Genomics UK (COG-UK) Consortium | Health Services Laboratories and Alex Alderton, Roberto Amato, Jeffrey Barrett, Sonia Goncalves, Ewan Harrison, David K. Jackson, Ian Johnston, Dominic Kwiatkowski, Cordelia Langford, John Sillitoe on behalf of the Wellcome Sanger Institute COVID-19 Surveillance Team |
| EPI_ISL_2704315 | Lighthouse Lab in Alderley Park | Wellcome Sanger Institute for the COVID-19 Genomics UK (COG-UK) Consortium | Jacquelyn Wynn, Mairead Hyland, The Lighthouse Lab in Alderley Park and Alex Alderton, Roberto Amato, Jeffrey Barrett, Sonia Goncalves, Ewan Harrison, David K. Jackson, Ian Johnston, Dominic Kwiatkowski, Cordelia Langford, John Sillitoe on behalf of the Wellcome Sanger Institute COVID-19 Surveillance Team |
| EPI_ISL_2704316, EPI_ISL_2704317, EPI_ISL_2704318, EPI_ISL_2704319 | Health Services Laboratories | Wellcome Sanger Institute for the COVID-19 Genomics UK (COG-UK) Consortium | Health Services Laboratories and Alex Alderton, Roberto Amato, Jeffrey Barrett, Sonia Goncalves, Ewan Harrison, David K. Jackson, Ian Johnston, Dominic Kwiatkowski, Cordelia Langford, John Sillitoe on behalf of the Wellcome Sanger Institute COVID-19 Surveillance Team |
| EPI_ISL_2704320, EPI_ISL_2704321, EPI_ISL_2704322, EPI_ISL_2704323, EPI_ISL_2704324, EPI_ISL_2704325, EPI_ISL_2704326, EPI_ISL_2704327, EPI_ISL_2704328, EPI_ISL_2704329, EPI_ISL_2704330, EPI_ISL_2704331, EPI_ISL_2704332 | | | |
| see above | Lighthouse Lab in Alderley Park | Wellcome Sanger Institute for the COVID-19 Genomics UK (COG-UK) Consortium | Jacquelyn Wynn, Mairead Hyland, The Lighthouse Lab in Alderley Park and Alex Alderton, Roberto Amato, Jeffrey Barrett, Sonia Goncalves, Ewan Harrison, David K. Jackson, Ian Johnston, Dominic Kwiatkowski, Cordelia Langford, John Sillitoe on behalf of the Wellcome Sanger Institute COVID-19 Surveillance Team |
| EPI_ISL_2704333 | Health Services Laboratories | Wellcome Sanger Institute for the COVID-19 Genomics UK (COG-UK) Consortium | Health Services Laboratories and Alex Alderton, Roberto Amato, Jeffrey Barrett, Sonia Goncalves, Ewan Harrison, David K. Jackson, Ian Johnston, Dominic Kwiatkowski, Cordelia Langford, John Sillitoe on behalf of the Wellcome Sanger Institute COVID-19 Surveillance Team |
| EPI_ISL_2704334, EPI_ISL_2704335 | Lighthouse Lab in Alderley Park | Wellcome Sanger Institute for the COVID-19 Genomics UK (COG-UK) Consortium | Jacquelyn Wynn, Mairead Hyland, The Lighthouse Lab in Alderley Park and Alex Alderton, Roberto Amato, Jeffrey Barrett, Sonia Goncalves, Ewan Harrison, David K. Jackson, Ian Johnston, Dominic Kwiatkowski, Cordelia Langford, John Sillitoe on behalf of the Wellcome Sanger Institute COVID-19 Surveillance Team |
| EPI_ISL_2704336, EPI_ISL_2704337, EPI_ISL_2704338 | Health Services Laboratories | Wellcome Sanger Institute for the COVID-19 Genomics UK (COG-UK) Consortium | Health Services Laboratories and Alex Alderton, Roberto Amato, Jeffrey Barrett, Sonia Goncalves, Ewan Harrison, David K. Jackson, Ian Johnston, Dominic Kwiatkowski, Cordelia Langford, John Sillitoe on behalf of the Wellcome Sanger Institute COVID-19 Surveillance Team |
| EPI_ISL_2704339 | Lighthouse Lab in Alderley Park | Wellcome Sanger Institute for the COVID-19 Genomics UK (COG-UK) Consortium | Jacquelyn Wynn, Mairead Hyland, The Lighthouse Lab in Alderley Park and Alex Alderton, Roberto Amato, Jeffrey Barrett, Sonia Goncalves, Ewan Harrison, David K. Jackson, Ian Johnston, Dominic Kwiatkowski, Cordelia Langford, John Sillitoe on behalf of the Wellcome Sanger Institute COVID-19 Surveillance Team |
| EPI_ISL_2704340 | Health Services Laboratories | Wellcome Sanger Institute for the COVID-19 Genomics UK (COG-UK) Consortium | Health Services Laboratories and Alex Alderton, Roberto Amato, Jeffrey Barrett, Sonia Goncalves, Ewan Harrison, David K. Jackson, Ian Johnston, Dominic Kwiatkowski, Cordelia Langford, John Sillitoe on behalf of the Wellcome Sanger Institute COVID-19 Surveillance Team |
| EPI_ISL_2704341, EPI_ISL_2704342, EPI_ISL_2704343, EPI_ISL_2704344, EPI_ISL_2704345 | Lighthouse Lab in Alderley Park | Wellcome Sanger Institute for the COVID-19 Genomics UK (COG-UK) Consortium | Jacquelyn Wynn, Mairead Hyland, The Lighthouse Lab in Alderley Park and Alex Alderton, Roberto Amato, Jeffrey Barrett, Sonia Goncalves, Ewan Harrison, David K. Jackson, Ian Johnston, Dominic Kwiatkowski, Cordelia Langford, John Sillitoe on behalf of the Wellcome Sanger Institute COVID-19 Surveillance Team |
| EPI_ISL_2704346 | Health Services Laboratories | Wellcome Sanger Institute for the COVID-19 Genomics UK (COG-UK) Consortium | Health Services Laboratories and Alex Alderton, Roberto Amato, Jeffrey Barrett, Sonia Goncalves, Ewan Harrison, David K. Jackson, Ian Johnston, Dominic Kwiatkowski, Cordelia Langford, John Sillitoe on behalf of the Wellcome Sanger Institute COVID-19 Surveillance Team |
| EPI_ISL_2704347, EPI_ISL_2704348 | Lighthouse Lab in Alderley Park | Wellcome Sanger Institute for the COVID-19 Genomics UK (COG-UK) Consortium | Jacquelyn Wynn, Mairead Hyland, The Lighthouse Lab in Alderley Park and Alex Alderton, Roberto Amato, Jeffrey Barrett, Sonia Goncalves, Ewan Harrison, David K. Jackson, Ian Johnston, Dominic Kwiatkowski, Cordelia Langford, John Sillitoe on behalf of the Wellcome Sanger Institute COVID-19 Surveillance Team |
| EPI_ISL_2704349 | Health Services Laboratories | Wellcome Sanger Institute for the COVID-19 Genomics UK (COG-UK) Consortium | Health Services Laboratories and Alex Alderton, Roberto Amato, Jeffrey Barrett, Sonia Goncalves, Ewan Harrison, David K. Jackson, Ian Johnston, Dominic Kwiatkowski, Cordelia Langford, John Sillitoe on behalf of the Wellcome Sanger Institute COVID-19 Surveillance Team |
| EPI_ISL_2704350, EPI_ISL_2704351, EPI_ISL_2704352, EPI_ISL_2704353, EPI_ISL_2704354, EPI_ISL_2704355 | Lighthouse Lab in Alderley Park | Wellcome Sanger Institute for the COVID-19 Genomics UK (COG-UK) Consortium | Jacquelyn Wynn, Mairead Hyland, The Lighthouse Lab in Alderley Park and Alex Alderton, Roberto Amato, Jeffrey Barrett, Sonia Goncalves, Ewan Harrison, David K. Jackson, Ian Johnston, Dominic Kwiatkowski, Cordelia Langford, John Sillitoe on behalf of the Wellcome Sanger Institute COVID-19 Surveillance Team |
| EPI_ISL_2704356 | Health Services Laboratories | Wellcome Sanger Institute for the COVID-19 Genomics UK (COG-UK) Consortium | Health Services Laboratories and Alex Alderton, Roberto Amato, Jeffrey Barrett, Sonia Goncalves, Ewan Harrison, David K. Jackson, Ian Johnston, Dominic Kwiatkowski, Cordelia Langford, John Sillitoe on behalf of the Wellcome Sanger Institute COVID-19 Surveillance Team |
| EPI_ISL_2704357, EPI_ISL_2704358 | Lighthouse Lab in Alderley Park | Wellcome Sanger Institute for the COVID-19 Genomics UK (COG-UK) Consortium | Jacquelyn Wynn, Mairead Hyland, The Lighthouse Lab in Alderley Park and Alex Alderton, Roberto Amato, Jeffrey Barrett, Sonia Goncalves, Ewan Harrison, David K. Jackson, Ian Johnston, Dominic Kwiatkowski, Cordelia Langford, John Sillitoe on behalf of the Wellcome Sanger Institute COVID-19 Surveillance Team |
| EPI_ISL_2704359 | Health Services Laboratories | Wellcome Sanger Institute for the COVID-19 Genomics UK (COG-UK) Consortium | Health Services Laboratories and Alex Alderton, Roberto Amato, Jeffrey Barrett, Sonia Goncalves, Ewan Harrison, David K. Jackson, Ian Johnston, Dominic Kwiatkowski, Cordelia Langford, John Sillitoe on behalf of the Wellcome Sanger Institute COVID-19 Surveillance Team |
| EPI_ISL_2704360 | Lighthouse Lab in Alderley Park | Wellcome Sanger Institute for the COVID-19 Genomics UK (COG-UK) Consortium | Jacquelyn Wynn, Mairead Hyland, The Lighthouse Lab in Alderley Park and Alex Alderton, Roberto Amato, Jeffrey Barrett, Sonia Goncalves, Ewan Harrison, David K. Jackson, Ian Johnston, Dominic Kwiatkowski, Cordelia Langford, John Sillitoe on behalf of the Wellcome Sanger Institute COVID-19 Surveillance Team |
| EPI_ISL_2704361 | Health Services Laboratories | Wellcome Sanger Institute for the COVID-19 Genomics UK (COG-UK) Consortium | Health Services Laboratories and Alex Alderton, Roberto Amato, Jeffrey Barrett, Sonia Goncalves, Ewan Harrison, David K. Jackson, Ian Johnston, Dominic Kwiatkowski, Cordelia Langford, John Sillitoe on behalf of the Wellcome Sanger Institute COVID-19 Surveillance Team |
| EPI_ISL_2704362, EPI_ISL_2704363, EPI_ISL_2704364, EPI_ISL_2704365, EPI_ISL_2704366, EPI_ISL_2704367, EPI_ISL_2704368, EPI_ISL_2704369, EPI_ISL_2704370 | Lighthouse Lab in Alderley Park | Wellcome Sanger Institute for the COVID-19 Genomics UK (COG-UK) Consortium | Jacquelyn Wynn, Mairead Hyland, The Lighthouse Lab in Alderley Park and Alex Alderton, Roberto Amato, Jeffrey Barrett, Sonia Goncalves, Ewan Harrison, David K. Jackson, Ian Johnston, Dominic Kwiatkowski, Cordelia Langford, John Sillitoe on behalf of the Wellcome Sanger Institute COVID-19 Surveillance Team |
| EPI_ISL_2704371 | Health Services Laboratories | Wellcome Sanger Institute for the COVID-19 Genomics UK (COG-UK) Consortium | Health Services Laboratories and Alex Alderton, Roberto Amato, Jeffrey Barrett, Sonia Goncalves, Ewan Harrison, David K. Jackson, Ian Johnston, Dominic Kwiatkowski, Cordelia Langford, John Sillitoe on behalf of the Wellcome Sanger Institute COVID-19 Surveillance Team |
| EPI_ISL_2704372, EPI_ISL_2704373, EPI_ISL_2704374, EPI_ISL_2704375, EPI_ISL_2704376, EPI_ISL_2704377 | Lighthouse Lab in Alderley Park | Wellcome Sanger Institute for the COVID-19 Genomics UK (COG-UK) Consortium | Jacquelyn Wynn, Mairead Hyland, The Lighthouse Lab in Alderley Park and Alex Alderton, Roberto Amato, Jeffrey Barrett, Sonia Goncalves, Ewan Harrison, David K. Jackson, Ian Johnston, Dominic Kwiatkowski, Cordelia Langford, John Sillitoe on behalf of the Wellcome Sanger Institute COVID-19 Surveillance Team |
| EPI_ISL_2704378, EPI_ISL_2704379 | Health Services Laboratories | Wellcome Sanger Institute for the COVID-19 Genomics UK (COG-UK) Consortium | Health Services Laboratories and Alex Alderton, Roberto Amato, Jeffrey Barrett, Sonia Goncalves, Ewan Harrison, David K. Jackson, Ian Johnston, Dominic Kwiatkowski, Cordelia Langford, John Sillitoe on behalf of the Wellcome Sanger Institute COVID-19 Surveillance Team |
| EPI_ISL_2704380, EPI_ISL_2704381, EPI_ISL_2704382, EPI_ISL_2704383, EPI_ISL_2704384 | Lighthouse Lab in Alderley Park | Wellcome Sanger Institute for the COVID-19 Genomics UK (COG-UK) Consortium | Jacquelyn Wynn, Mairead Hyland, The Lighthouse Lab in Alderley Park and Alex Alderton, Roberto Amato, Jeffrey Barrett, Sonia Goncalves, Ewan Harrison, David K. Jackson, Ian Johnston, Dominic Kwiatkowski, Cordelia Langford, John Sillitoe on behalf of the Wellcome Sanger Institute COVID-19 Surveillance Team |
| EPI_ISL_2704385 | Health Services Laboratories | Wellcome Sanger Institute for the COVID-19 Genomics UK (COG-UK) Consortium | Health Services Laboratories and Alex Alderton, Roberto Amato, Jeffrey Barrett, Sonia Goncalves, Ewan Harrison, David K. Jackson, Ian Johnston, Dominic Kwiatkowski, Cordelia Langford, John Sillitoe on behalf of the Wellcome Sanger Institute COVID-19 Surveillance Team |
| EPI_ISL_2704386, EPI_ISL_2704388 | Lighthouse Lab in Alderley Park | Wellcome Sanger Institute for the COVID-19 Genomics UK (COG-UK) Consortium | Jacquelyn Wynn, Mairead Hyland, The Lighthouse Lab in Alderley Park and Alex Alderton, Roberto Amato, Jeffrey Barrett, Sonia Goncalves, Ewan Harrison, David K. Jackson, Ian Johnston, Dominic Kwiatkowski, Cordelia Langford, John Sillitoe on behalf of the Wellcome Sanger Institute COVID-19 Surveillance Team |
| EPI_ISL_2704389, EPI_ISL_2704390 | Health Services Laboratories | Wellcome Sanger Institute for the COVID-19 Genomics UK (COG-UK) Consortium | Health Services Laboratories and Alex Alderton, Roberto Amato, Jeffrey Barrett, Sonia Goncalves, Ewan Harrison, David K. Jackson, Ian Johnston, Dominic Kwiatkowski, Cordelia Langford, John Sillitoe on behalf of the Wellcome Sanger Institute COVID-19 Surveillance Team |
| EPI_ISL_2704391 | Lighthouse Lab in Alderley Park | Wellcome Sanger Institute for the COVID-19 Genomics UK (COG-UK) Consortium | Jacquelyn Wynn, Mairead Hyland, The Lighthouse Lab in Alderley Park and Alex Alderton, Roberto Amato, Jeffrey Barrett, Sonia Goncalves, Ewan Harrison, David K. Jackson, Ian Johnston, Dominic Kwiatkowski, Cordelia Langford, John Sillitoe on behalf of the Wellcome Sanger Institute COVID-19 Surveillance Team |
| EPI_ISL_2704392 | Health Services Laboratories | Wellcome Sanger Institute for the COVID-19 Genomics UK (COG-UK) Consortium | Health Services Laboratories and Alex Alderton, Roberto Amato, Jeffrey Barrett, Sonia Goncalves, Ewan Harrison, David K. Jackson, Ian Johnston, Dominic Kwiatkowski, Cordelia Langford, John Sillitoe on behalf of the Wellcome Sanger Institute COVID-19 Surveillance Team |
| EPI_ISL_2704393, EPI_ISL_2704394, EPI_ISL_2704395 | Lighthouse Lab in Alderley Park | Wellcome Sanger Institute for the COVID-19 Genomics UK (COG-UK) Consortium | Jacquelyn Wynn, Mairead Hyland, The Lighthouse Lab in Alderley Park and Alex Alderton, Roberto Amato, Jeffrey Barrett, Sonia Goncalves, Ewan Harrison, David K. Jackson, Ian Johnston, Dominic Kwiatkowski, Cordelia Langford, John Sillitoe on behalf of the Wellcome Sanger Institute COVID-19 Surveillance Team |
| EPI_ISL_2704396 | Health Services Laboratories | Wellcome Sanger Institute for the COVID-19 Genomics UK (COG-UK) | Health Services Laboratories and Alex Alderton, Roberto Amato, Jeffrey Barrett, Sonia Goncalves, Ewan Harrison, David K. Jackson, Ian Johnston, Dominic Kwiatkowski, Cordelia Langford, John Sillitoe on behalf of the Wellcome Sanger Institute |

| | | | |
|---|---|---|---|
| | | Consortium | COVID-19 Surveillance Team |
| EPI_ISL_2704397, EPI_ISL_2704398, EPI_ISL_2704399 | Lighthouse Lab in Alderley Park | Wellcome Sanger Institute for the COVID-19 Genomics UK (COG-UK) Consortium | Jacquelyn Wynn, Mairead Hyland, The Lighthouse Lab in Alderley Park and Alex Alderton, Roberto Amato, Jeffrey Barrett, Sonia Goncalves, Ewan Harrison, David K. Jackson, Ian Johnston, Dominic Kwiatkowski, Cordelia Langford, John Sillitoe on behalf of the Wellcome Sanger Institute COVID-19 Surveillance Team |
| EPI_ISL_2704400 | Health Services Laboratories | Wellcome Sanger Institute for the COVID-19 Genomics UK (COG-UK) Consortium | Health Services Laboratories and Alex Alderton, Roberto Amato, Jeffrey Barrett, Sonia Goncalves, Ewan Harrison, David K. Jackson, Ian Johnston, Dominic Kwiatkowski, Cordelia Langford, John Sillitoe on behalf of the Wellcome Sanger Institute COVID-19 Surveillance Team |
| EPI_ISL_2704401 | Lighthouse Lab in Alderley Park | Wellcome Sanger Institute for the COVID-19 Genomics UK (COG-UK) Consortium | Jacquelyn Wynn, Mairead Hyland, The Lighthouse Lab in Alderley Park and Alex Alderton, Roberto Amato, Jeffrey Barrett, Sonia Goncalves, Ewan Harrison, David K. Jackson, Ian Johnston, Dominic Kwiatkowski, Cordelia Langford, John Sillitoe on behalf of the Wellcome Sanger Institute COVID-19 Surveillance Team |
| EPI_ISL_2704402 | Health Services Laboratories | Wellcome Sanger Institute for the COVID-19 Genomics UK (COG-UK) Consortium | Health Services Laboratories and Alex Alderton, Roberto Amato, Jeffrey Barrett, Sonia Goncalves, Ewan Harrison, David K. Jackson, Ian Johnston, Dominic Kwiatkowski, Cordelia Langford, John Sillitoe on behalf of the Wellcome Sanger Institute COVID-19 Surveillance Team |
| EPI_ISL_2704403 | Lighthouse Lab in Alderley Park | Wellcome Sanger Institute for the COVID-19 Genomics UK (COG-UK) Consortium | Jacquelyn Wynn, Mairead Hyland, The Lighthouse Lab in Alderley Park and Alex Alderton, Roberto Amato, Jeffrey Barrett, Sonia Goncalves, Ewan Harrison, David K. Jackson, Ian Johnston, Dominic Kwiatkowski, Cordelia Langford, John Sillitoe on behalf of the Wellcome Sanger Institute COVID-19 Surveillance Team |
| EPI_ISL_2704404 | Health Services Laboratories | Wellcome Sanger Institute for the COVID-19 Genomics UK (COG-UK) Consortium | Health Services Laboratories and Alex Alderton, Roberto Amato, Jeffrey Barrett, Sonia Goncalves, Ewan Harrison, David K. Jackson, Ian Johnston, Dominic Kwiatkowski, Cordelia Langford, John Sillitoe on behalf of the Wellcome Sanger Institute COVID-19 Surveillance Team |
| EPI_ISL_2704405, EPI_ISL_2704406, EPI_ISL_2704407, EPI_ISL_2704408, EPI_ISL_2704409, EPI_ISL_2704410, EPI_ISL_2704411 | Lighthouse Lab in Alderley Park | Wellcome Sanger Institute for the COVID-19 Genomics UK (COG-UK) Consortium | Jacquelyn Wynn, Mairead Hyland, The Lighthouse Lab in Alderley Park and Alex Alderton, Roberto Amato, Jeffrey Barrett, Sonia Goncalves, Ewan Harrison, David K. Jackson, Ian Johnston, Dominic Kwiatkowski, Cordelia Langford, John Sillitoe on behalf of the Wellcome Sanger Institute COVID-19 Surveillance Team |
| EPI_ISL_2704412, EPI_ISL_2704413, EPI_ISL_2704414, EPI_ISL_2704415 | Health Services Laboratories | Wellcome Sanger Institute for the COVID-19 Genomics UK (COG-UK) Consortium | Health Services Laboratories and Alex Alderton, Roberto Amato, Jeffrey Barrett, Sonia Goncalves, Ewan Harrison, David K. Jackson, Ian Johnston, Dominic Kwiatkowski, Cordelia Langford, John Sillitoe on behalf of the Wellcome Sanger Institute COVID-19 Surveillance Team |
| EPI_ISL_2704416, EPI_ISL_2704417, EPI_ISL_2704418 | Lighthouse Lab in Alderley Park | Wellcome Sanger Institute for the COVID-19 Genomics UK (COG-UK) Consortium | Jacquelyn Wynn, Mairead Hyland, The Lighthouse Lab in Alderley Park and Alex Alderton, Roberto Amato, Jeffrey Barrett, Sonia Goncalves, Ewan Harrison, David K. Jackson, Ian Johnston, Dominic Kwiatkowski, Cordelia Langford, John Sillitoe on behalf of the Wellcome Sanger Institute COVID-19 Surveillance Team |
| EPI_ISL_2704419 | Health Services Laboratories | Wellcome Sanger Institute for the COVID-19 Genomics UK (COG-UK) Consortium | Health Services Laboratories and Alex Alderton, Roberto Amato, Jeffrey Barrett, Sonia Goncalves, Ewan Harrison, David K. Jackson, Ian Johnston, Dominic Kwiatkowski, Cordelia Langford, John Sillitoe on behalf of the Wellcome Sanger Institute COVID-19 Surveillance Team |
| EPI_ISL_2704420 | Lighthouse Lab in Alderley Park | Wellcome Sanger Institute for the COVID-19 Genomics UK (COG-UK) Consortium | Jacquelyn Wynn, Mairead Hyland, The Lighthouse Lab in Alderley Park and Alex Alderton, Roberto Amato, Jeffrey Barrett, Sonia Goncalves, Ewan Harrison, David K. Jackson, Ian Johnston, Dominic Kwiatkowski, Cordelia Langford, John Sillitoe on behalf of the Wellcome Sanger Institute COVID-19 Surveillance Team |
| EPI_ISL_2704421 | Health Services Laboratories | Wellcome Sanger Institute for the COVID-19 Genomics UK (COG-UK) Consortium | Health Services Laboratories and Alex Alderton, Roberto Amato, Jeffrey Barrett, Sonia Goncalves, Ewan Harrison, David K. Jackson, Ian Johnston, Dominic Kwiatkowski, Cordelia Langford, John Sillitoe on behalf of the Wellcome Sanger Institute COVID-19 Surveillance Team |
| EPI_ISL_2704422, EPI_ISL_2704423, EPI_ISL_2704424, EPI_ISL_2704425, EPI_ISL_2704426, EPI_ISL_2704427, EPI_ISL_2704428, EPI_ISL_2704429, EPI_ISL_2704430, EPI_ISL_2704431, EPI_ISL_2704432, EPI_ISL_2704433, EPI_ISL_2704434, EPI_ISL_2704435, EPI_ISL_2704436, EPI_ISL_2704437, EPI_ISL_2704438, EPI_ISL_2704439, EPI_ISL_2704440, EPI_ISL_2704441, EPI_ISL_2704442, EPI_ISL_2704443, EPI_ISL_2704444, EPI_ISL_2704446, EPI_ISL_2704447, EPI_ISL_2704448 | | | |
| see above | Lighthouse Lab in Alderley Park | Wellcome Sanger Institute for the COVID-19 Genomics UK (COG-UK) Consortium | Jacquelyn Wynn, Mairead Hyland, The Lighthouse Lab in Alderley Park and Alex Alderton, Roberto Amato, Jeffrey Barrett, Sonia Goncalves, Ewan Harrison, David K. Jackson, Ian Johnston, Dominic Kwiatkowski, Cordelia Langford, John Sillitoe on behalf of the Wellcome Sanger Institute COVID-19 Surveillance Team |
| EPI_ISL_2704449 | Health Services Laboratories | Wellcome Sanger Institute for the COVID-19 Genomics UK (COG-UK) Consortium | Health Services Laboratories and Alex Alderton, Roberto Amato, Jeffrey Barrett, Sonia Goncalves, Ewan Harrison, David K. Jackson, Ian Johnston, Dominic Kwiatkowski, Cordelia Langford, John Sillitoe on behalf of the Wellcome Sanger Institute COVID-19 Surveillance Team |
| EPI_ISL_2704450 | Lighthouse Lab in Alderley Park | Wellcome Sanger Institute for the COVID-19 Genomics UK (COG-UK) Consortium | Jacquelyn Wynn, Mairead Hyland, The Lighthouse Lab in Alderley Park and Alex Alderton, Roberto Amato, Jeffrey Barrett, Sonia Goncalves, Ewan Harrison, David K. Jackson, Ian Johnston, Dominic Kwiatkowski, Cordelia Langford, John Sillitoe on behalf of the Wellcome Sanger Institute COVID-19 Surveillance Team |
| EPI_ISL_2704451 | Health Services Laboratories | Wellcome Sanger Institute for the COVID-19 Genomics UK (COG-UK) Consortium | Health Services Laboratories and Alex Alderton, Roberto Amato, Jeffrey Barrett, Sonia Goncalves, Ewan Harrison, David K. Jackson, Ian Johnston, Dominic Kwiatkowski, Cordelia Langford, John Sillitoe on behalf of the Wellcome Sanger Institute COVID-19 Surveillance Team |
| EPI_ISL_2704452, EPI_ISL_2704453, EPI_ISL_2704454, EPI_ISL_2704455, EPI_ISL_2704456 | Lighthouse Lab in Alderley Park | Wellcome Sanger Institute for the COVID-19 Genomics UK (COG-UK) Consortium | Jacquelyn Wynn, Mairead Hyland, The Lighthouse Lab in Alderley Park and Alex Alderton, Roberto Amato, Jeffrey Barrett, Sonia Goncalves, Ewan Harrison, David K. Jackson, Ian Johnston, Dominic Kwiatkowski, Cordelia Langford, John Sillitoe on behalf of the Wellcome Sanger Institute COVID-19 Surveillance Team |
| EPI_ISL_2704457 | Health Services Laboratories | Wellcome Sanger Institute for the COVID-19 Genomics UK (COG-UK) Consortium | Health Services Laboratories and Alex Alderton, Roberto Amato, Jeffrey Barrett, Sonia Goncalves, Ewan Harrison, David K. Jackson, Ian Johnston, Dominic Kwiatkowski, Cordelia Langford, John Sillitoe on behalf of the Wellcome Sanger Institute COVID-19 Surveillance Team |
| EPI_ISL_2704458, EPI_ISL_2704459, EPI_ISL_2704460, EPI_ISL_2704461, EPI_ISL_2704462, EPI_ISL_2704463, EPI_ISL_2704464, EPI_ISL_2704465 | Lighthouse Lab in Alderley Park | Wellcome Sanger Institute for the COVID-19 Genomics UK (COG-UK) Consortium | Jacquelyn Wynn, Mairead Hyland, The Lighthouse Lab in Alderley Park and Alex Alderton, Roberto Amato, Jeffrey Barrett, Sonia Goncalves, Ewan Harrison, David K. Jackson, Ian Johnston, Dominic Kwiatkowski, Cordelia Langford, John Sillitoe on behalf of the Wellcome Sanger Institute COVID-19 Surveillance Team |
| EPI_ISL_2704466, EPI_ISL_2704467 | Health Services Laboratories | Wellcome Sanger Institute for the COVID-19 Genomics UK (COG-UK) Consortium | Health Services Laboratories and Alex Alderton, Roberto Amato, Jeffrey Barrett, Sonia Goncalves, Ewan Harrison, David K. Jackson, Ian Johnston, Dominic Kwiatkowski, Cordelia Langford, John Sillitoe on behalf of the Wellcome Sanger Institute COVID-19 Surveillance Team |
| EPI_ISL_2704468 | Lighthouse Lab in Alderley Park | Wellcome Sanger Institute for the COVID-19 Genomics UK (COG-UK) Consortium | Jacquelyn Wynn, Mairead Hyland, The Lighthouse Lab in Alderley Park and Alex Alderton, Roberto Amato, Jeffrey Barrett, Sonia Goncalves, Ewan Harrison, David K. Jackson, Ian Johnston, Dominic Kwiatkowski, Cordelia Langford, John Sillitoe on behalf of the Wellcome Sanger Institute COVID-19 Surveillance Team |
| EPI_ISL_2704469, EPI_ISL_2704470 | Health Services Laboratories | Wellcome Sanger Institute for the COVID-19 Genomics UK (COG-UK) Consortium | Health Services Laboratories and Alex Alderton, Roberto Amato, Jeffrey Barrett, Sonia Goncalves, Ewan Harrison, David K. Jackson, Ian Johnston, Dominic Kwiatkowski, Cordelia Langford, John Sillitoe on behalf of the Wellcome Sanger Institute COVID-19 Surveillance Team |
| EPI_ISL_2704471, EPI_ISL_2704472, EPI_ISL_2704473, EPI_ISL_2704474, EPI_ISL_2704475, EPI_ISL_2704476, EPI_ISL_2704477 | Lighthouse Lab in Alderley Park | Wellcome Sanger Institute for the COVID-19 Genomics UK (COG-UK) Consortium | Jacquelyn Wynn, Mairead Hyland, The Lighthouse Lab in Alderley Park and Alex Alderton, Roberto Amato, Jeffrey Barrett, Sonia Goncalves, Ewan Harrison, David K. Jackson, Ian Johnston, Dominic Kwiatkowski, Cordelia Langford, John Sillitoe on behalf of the Wellcome Sanger Institute COVID-19 Surveillance Team |
| EPI_ISL_2704478 | Health Services Laboratories | Wellcome Sanger Institute for the COVID-19 Genomics UK (COG-UK) Consortium | Health Services Laboratories and Alex Alderton, Roberto Amato, Jeffrey Barrett, Sonia Goncalves, Ewan Harrison, David K. Jackson, Ian Johnston, Dominic Kwiatkowski, Cordelia Langford, John Sillitoe on behalf of the Wellcome Sanger Institute COVID-19 Surveillance Team |
| EPI_ISL_2704479 | Lighthouse Lab in Alderley Park | Wellcome Sanger Institute for the COVID-19 Genomics UK (COG-UK) Consortium | Jacquelyn Wynn, Mairead Hyland, The Lighthouse Lab in Alderley Park and Alex Alderton, Roberto Amato, Jeffrey Barrett, Sonia Goncalves, Ewan Harrison, David K. Jackson, Ian Johnston, Dominic Kwiatkowski, Cordelia Langford, John Sillitoe on behalf of the Wellcome Sanger Institute COVID-19 Surveillance Team |
| EPI_ISL_2704480, EPI_ISL_2704481, EPI_ISL_2704482 | Health Services Laboratories | Wellcome Sanger Institute for the COVID-19 Genomics UK (COG-UK) Consortium | Health Services Laboratories and Alex Alderton, Roberto Amato, Jeffrey Barrett, Sonia Goncalves, Ewan Harrison, David K. Jackson, Ian Johnston, Dominic Kwiatkowski, Cordelia Langford, John Sillitoe on behalf of the Wellcome Sanger Institute COVID-19 Surveillance Team |
| EPI_ISL_2704483 | Lighthouse Lab in Alderley Park | Wellcome Sanger Institute for the COVID-19 Genomics UK (COG-UK) Consortium | Jacquelyn Wynn, Mairead Hyland, The Lighthouse Lab in Alderley Park and Alex Alderton, Roberto Amato, Jeffrey Barrett, Sonia Goncalves, Ewan Harrison, David K. Jackson, Ian Johnston, Dominic Kwiatkowski, Cordelia Langford, John Sillitoe on behalf of the Wellcome Sanger Institute COVID-19 Surveillance Team |
| EPI_ISL_2704484 | Health Services Laboratories | Wellcome Sanger Institute for the COVID-19 Genomics UK (COG-UK) Consortium | Health Services Laboratories and Alex Alderton, Roberto Amato, Jeffrey Barrett, Sonia Goncalves, Ewan Harrison, David K. Jackson, Ian Johnston, Dominic Kwiatkowski, Cordelia Langford, John Sillitoe on behalf of the Wellcome Sanger Institute COVID-19 Surveillance Team |
| EPI_ISL_2704485, EPI_ISL_2704486, EPI_ISL_2704488, EPI_ISL_2704489, EPI_ISL_2704490, EPI_ISL_2704491 | Lighthouse Lab in Alderley Park | Wellcome Sanger Institute for the COVID-19 Genomics UK (COG-UK) Consortium | Jacquelyn Wynn, Mairead Hyland, The Lighthouse Lab in Alderley Park and Alex Alderton, Roberto Amato, Jeffrey Barrett, Sonia Goncalves, Ewan Harrison, David K. Jackson, Ian Johnston, Dominic Kwiatkowski, Cordelia Langford, John Sillitoe on behalf of the Wellcome Sanger Institute COVID-19 Surveillance Team |
| EPI_ISL_2704492 | Health Services Laboratories | Wellcome Sanger Institute for the COVID-19 Genomics UK (COG-UK) Consortium | Health Services Laboratories and Alex Alderton, Roberto Amato, Jeffrey Barrett, Sonia Goncalves, Ewan Harrison, David K. Jackson, Ian Johnston, Dominic Kwiatkowski, Cordelia Langford, John Sillitoe on behalf of the Wellcome Sanger Institute COVID-19 Surveillance Team |
| EPI_ISL_2704493 | Lighthouse Lab in Alderley Park | Wellcome Sanger Institute for the COVID-19 Genomics UK (COG-UK) Consortium | Jacquelyn Wynn, Mairead Hyland, The Lighthouse Lab in Alderley Park and Alex Alderton, Roberto Amato, Jeffrey Barrett, Sonia Goncalves, Ewan Harrison, David K. Jackson, Ian Johnston, Dominic Kwiatkowski, Cordelia Langford, John Sillitoe on behalf of the Wellcome Sanger Institute COVID-19 Surveillance Team |
| EPI_ISL_2704494 | Health Services Laboratories | Wellcome Sanger Institute for the COVID-19 Genomics UK (COG-UK) Consortium | Health Services Laboratories and Alex Alderton, Roberto Amato, Jeffrey Barrett, Sonia Goncalves, Ewan Harrison, David K. Jackson, Ian Johnston, Dominic Kwiatkowski, Cordelia Langford, John Sillitoe on behalf of the Wellcome Sanger Institute COVID-19 Surveillance Team |

| | | | |
|---|---|---|---|
| EPI_ISL_2704495 | Lighthouse Lab in Alderley Park | Wellcome Sanger Institute for the COVID-19 Genomics UK (COG-UK) Consortium | Jacquelyn Wynn, Mairead Hyland, The Lighthouse Lab in Alderley Park and Alex Alderton, Roberto Amato, Jeffrey Barrett, Sonia Goncalves, Ewan Harrison, David K. Jackson, Ian Johnston, Dominic Kwiatkowski, Cordelia Langford, John Sillitoe on behalf of the Wellcome Sanger Institute COVID-19 Surveillance Team |
| EPI_ISL_2704496 | Health Services Laboratories | Wellcome Sanger Institute for the COVID-19 Genomics UK (COG-UK) Consortium | Health Services Laboratories and Alex Alderton, Roberto Amato, Jeffrey Barrett, Sonia Goncalves, Ewan Harrison, David K. Jackson, Ian Johnston, Dominic Kwiatkowski, Cordelia Langford, John Sillitoe on behalf of the Wellcome Sanger Institute COVID-19 Surveillance Team |
| EPI_ISL_2704497 | Lighthouse Lab in Alderley Park | Wellcome Sanger Institute for the COVID-19 Genomics UK (COG-UK) Consortium | Jacquelyn Wynn, Mairead Hyland, The Lighthouse Lab in Alderley Park and Alex Alderton, Roberto Amato, Jeffrey Barrett, Sonia Goncalves, Ewan Harrison, David K. Jackson, Ian Johnston, Dominic Kwiatkowski, Cordelia Langford, John Sillitoe on behalf of the Wellcome Sanger Institute COVID-19 Surveillance Team |
| EPI_ISL_2704498 | Health Services Laboratories | Wellcome Sanger Institute for the COVID-19 Genomics UK (COG-UK) Consortium | Health Services Laboratories and Alex Alderton, Roberto Amato, Jeffrey Barrett, Sonia Goncalves, Ewan Harrison, David K. Jackson, Ian Johnston, Dominic Kwiatkowski, Cordelia Langford, John Sillitoe on behalf of the Wellcome Sanger Institute COVID-19 Surveillance Team |
| EPI_ISL_2704499, EPI_ISL_2704500, EPI_ISL_2704501, EPI_ISL_2704502, EPI_ISL_2704503, EPI_ISL_2704504, EPI_ISL_2704505, EPI_ISL_2704506, EPI_ISL_2704507, EPI_ISL_2704508, EPI_ISL_2704509, EPI_ISL_2704510 | | | |
| see above | Lighthouse Lab in Alderley Park | Wellcome Sanger Institute for the COVID-19 Genomics UK (COG-UK) Consortium | Jacquelyn Wynn, Mairead Hyland, The Lighthouse Lab in Alderley Park and Alex Alderton, Roberto Amato, Jeffrey Barrett, Sonia Goncalves, Ewan Harrison, David K. Jackson, Ian Johnston, Dominic Kwiatkowski, Cordelia Langford, John Sillitoe on behalf of the Wellcome Sanger Institute COVID-19 Surveillance Team |
| EPI_ISL_2704511 | Health Services Laboratories | Wellcome Sanger Institute for the COVID-19 Genomics UK (COG-UK) Consortium | Health Services Laboratories and Alex Alderton, Roberto Amato, Jeffrey Barrett, Sonia Goncalves, Ewan Harrison, David K. Jackson, Ian Johnston, Dominic Kwiatkowski, Cordelia Langford, John Sillitoe on behalf of the Wellcome Sanger Institute COVID-19 Surveillance Team |
| EPI_ISL_2704512 | Lighthouse Lab in Alderley Park | Wellcome Sanger Institute for the COVID-19 Genomics UK (COG-UK) Consortium | Jacquelyn Wynn, Mairead Hyland, The Lighthouse Lab in Alderley Park and Alex Alderton, Roberto Amato, Jeffrey Barrett, Sonia Goncalves, Ewan Harrison, David K. Jackson, Ian Johnston, Dominic Kwiatkowski, Cordelia Langford, John Sillitoe on behalf of the Wellcome Sanger Institute COVID-19 Surveillance Team |
| EPI_ISL_2704513 | Health Services Laboratories | Wellcome Sanger Institute for the COVID-19 Genomics UK (COG-UK) Consortium | Health Services Laboratories and Alex Alderton, Roberto Amato, Jeffrey Barrett, Sonia Goncalves, Ewan Harrison, David K. Jackson, Ian Johnston, Dominic Kwiatkowski, Cordelia Langford, John Sillitoe on behalf of the Wellcome Sanger Institute COVID-19 Surveillance Team |
| EPI_ISL_2704514, EPI_ISL_2704515, EPI_ISL_2704516, EPI_ISL_2704517, EPI_ISL_2704518, EPI_ISL_2704519, EPI_ISL_2704520, EPI_ISL_2704521, EPI_ISL_2704522, EPI_ISL_2704523, EPI_ISL_2704524, EPI_ISL_2704525, EPI_ISL_2704526 | | | |
| see above | Lighthouse Lab in Alderley Park | Wellcome Sanger Institute for the COVID-19 Genomics UK (COG-UK) Consortium | Jacquelyn Wynn, Mairead Hyland, The Lighthouse Lab in Alderley Park and Alex Alderton, Roberto Amato, Jeffrey Barrett, Sonia Goncalves, Ewan Harrison, David K. Jackson, Ian Johnston, Dominic Kwiatkowski, Cordelia Langford, John Sillitoe on behalf of the Wellcome Sanger Institute COVID-19 Surveillance Team |
| EPI_ISL_2704527 | Health Services Laboratories | Wellcome Sanger Institute for the COVID-19 Genomics UK (COG-UK) Consortium | Health Services Laboratories and Alex Alderton, Roberto Amato, Jeffrey Barrett, Sonia Goncalves, Ewan Harrison, David K. Jackson, Ian Johnston, Dominic Kwiatkowski, Cordelia Langford, John Sillitoe on behalf of the Wellcome Sanger Institute COVID-19 Surveillance Team |
| EPI_ISL_2704528, EPI_ISL_2704529, EPI_ISL_2704530, EPI_ISL_2704531 | Lighthouse Lab in Alderley Park | Wellcome Sanger Institute for the COVID-19 Genomics UK (COG-UK) Consortium | Jacquelyn Wynn, Mairead Hyland, The Lighthouse Lab in Alderley Park and Alex Alderton, Roberto Amato, Jeffrey Barrett, Sonia Goncalves, Ewan Harrison, David K. Jackson, Ian Johnston, Dominic Kwiatkowski, Cordelia Langford, John Sillitoe on behalf of the Wellcome Sanger Institute COVID-19 Surveillance Team |
| EPI_ISL_2704532 | Health Services Laboratories | Wellcome Sanger Institute for the COVID-19 Genomics UK (COG-UK) Consortium | Health Services Laboratories and Alex Alderton, Roberto Amato, Jeffrey Barrett, Sonia Goncalves, Ewan Harrison, David K. Jackson, Ian Johnston, Dominic Kwiatkowski, Cordelia Langford, John Sillitoe on behalf of the Wellcome Sanger Institute COVID-19 Surveillance Team |
| EPI_ISL_2704533 | Lighthouse Lab in Glasgow | Wellcome Sanger Institute for the COVID-19 Genomics UK (COG-UK) Consortium | Harper VanSteenhouse, Yumi Kasai, David Gray, Carol Clugston, Anna Dominiczak and Alex Alderton, Roberto Amato, Jeffrey Barrett, Sonia Goncalves, Ewan Harrison, David K. Jackson, Ian Johnston, Dominic Kwiatkowski, Cordelia Langford, John Sillitoe on behalf of the Wellcome Sanger Institute COVID-19 Surveillance Team |
| EPI_ISL_2704535 | Lighthouse Lab in Milton Keynes | Wellcome Sanger Institute for the COVID-19 Genomics UK (COG-UK) Consortium | The Lighthouse Lab in Milton Keynes and Alex Alderton, Roberto Amato, Jeffrey Barrett, Sonia Goncalves, Ewan Harrison, David K. Jackson, Ian Johnston, Dominic Kwiatkowski, Cordelia Langford, John Sillitoe on behalf of the Wellcome Sanger Institute COVID-19 Surveillance Team |
| EPI_ISL_2704536, EPI_ISL_2704537, EPI_ISL_2704538, EPI_ISL_2704539, EPI_ISL_2704540, EPI_ISL_2704541, EPI_ISL_2704542, EPI_ISL_2704543, EPI_ISL_2704544, EPI_ISL_2704545, EPI_ISL_2704546, EPI_ISL_2704548, EPI_ISL_2704549, EPI_ISL_2704550, EPI_ISL_2704551, EPI_ISL_2704552, EPI_ISL_2704553, EPI_ISL_2704554, EPI_ISL_2704555, EPI_ISL_2704556, EPI_ISL_2704557, EPI_ISL_2704558, EPI_ISL_2704559, EPI_ISL_2704560, EPI_ISL_2704561, EPI_ISL_2704563, EPI_ISL_2704565, EPI_ISL_2704567, EPI_ISL_2704570, EPI_ISL_2704572, EPI_ISL_2704573, EPI_ISL_2704574, EPI_ISL_2704575, EPI_ISL_2704577, EPI_ISL_2704578, EPI_ISL_2704580, EPI_ISL_2704581, EPI_ISL_2704582, EPI_ISL_2704583, EPI_ISL_2704584, EPI_ISL_2704585, EPI_ISL_2704587, EPI_ISL_2704588, EPI_ISL_2704589, EPI_ISL_2704590, EPI_ISL_2704591, EPI_ISL_2704592, EPI_ISL_2704593, EPI_ISL_2704594, EPI_ISL_2704595, EPI_ISL_2704596, EPI_ISL_2704598, EPI_ISL_2704599, EPI_ISL_2704600, EPI_ISL_2704601, EPI_ISL_2704602, EPI_ISL_2704603, EPI_ISL_2704604, EPI_ISL_2704605, EPI_ISL_2704606, EPI_ISL_2704607, EPI_ISL_2704608, EPI_ISL_2704609, EPI_ISL_2704610, EPI_ISL_2704611, EPI_ISL_2704612, EPI_ISL_2704613, EPI_ISL_2704615, EPI_ISL_2704616, EPI_ISL_2704617, EPI_ISL_2704618, EPI_ISL_2704620, EPI_ISL_2704621, EPI_ISL_2704622, EPI_ISL_2704623, EPI_ISL_2704624, EPI_ISL_2704625, EPI_ISL_2704626, EPI_ISL_2704627, EPI_ISL_2704628, EPI_ISL_2704629, EPI_ISL_2704630, EPI_ISL_2704631, EPI_ISL_2704632, EPI_ISL_2704633, EPI_ISL_2704635, EPI_ISL_2704636, EPI_ISL_2704637, EPI_ISL_2704638, EPI_ISL_2704639, EPI_ISL_2704640, EPI_ISL_2704641, EPI_ISL_2704643, EPI_ISL_2704644, EPI_ISL_2704645, EPI_ISL_2704646, EPI_ISL_2704647, EPI_ISL_2704648, EPI_ISL_2704649, EPI_ISL_2704650, EPI_ISL_2704651, EPI_ISL_2704652, EPI_ISL_2704653, EPI_ISL_2704654, EPI_ISL_2704655, EPI_ISL_2704656, EPI_ISL_2704657, EPI_ISL_2704658, EPI_ISL_2704659, EPI_ISL_2704660, EPI_ISL_2704661, EPI_ISL_2704662, EPI_ISL_2704663, EPI_ISL_2704664, EPI_ISL_2704665 | | | |
| see above | Lighthouse Lab in Glasgow | Wellcome Sanger Institute for the COVID-19 Genomics UK (COG-UK) Consortium | Harper VanSteenhouse, Yumi Kasai, David Gray, Carol Clugston, Anna Dominiczak and Alex Alderton, Roberto Amato, Jeffrey Barrett, Sonia Goncalves, Ewan Harrison, David K. Jackson, Ian Johnston, Dominic Kwiatkowski, Cordelia Langford, John Sillitoe on behalf of the Wellcome Sanger Institute COVID-19 Surveillance Team |
| EPI_ISL_2704666 | Lighthouse Lab in Milton Keynes | Wellcome Sanger Institute for the COVID-19 Genomics UK (COG-UK) Consortium | The Lighthouse Lab in Milton Keynes and Alex Alderton, Roberto Amato, Jeffrey Barrett, Sonia Goncalves, Ewan Harrison, David K. Jackson, Ian Johnston, Dominic Kwiatkowski, Cordelia Langford, John Sillitoe on behalf of the Wellcome Sanger Institute COVID-19 Surveillance Team |
| EPI_ISL_2704667 | Lighthouse Lab in Glasgow | Wellcome Sanger Institute for the COVID-19 Genomics UK (COG-UK) Consortium | Harper VanSteenhouse, Yumi Kasai, David Gray, Carol Clugston, Anna Dominiczak and Alex Alderton, Roberto Amato, Jeffrey Barrett, Sonia Goncalves, Ewan Harrison, David K. Jackson, Ian Johnston, Dominic Kwiatkowski, Cordelia Langford, John Sillitoe on behalf of the Wellcome Sanger Institute COVID-19 Surveillance Team |
| EPI_ISL_2704668, EPI_ISL_2704669, EPI_ISL_2704670, EPI_ISL_2704672, EPI_ISL_2704673, EPI_ISL_2704674 | Lighthouse Lab in Alderley Park | Wellcome Sanger Institute for the COVID-19 Genomics UK (COG-UK) Consortium | Jacquelyn Wynn, Mairead Hyland, The Lighthouse Lab in Alderley Park and Alex Alderton, Roberto Amato, Jeffrey Barrett, Sonia Goncalves, Ewan Harrison, David K. Jackson, Ian Johnston, Dominic Kwiatkowski, Cordelia Langford, John Sillitoe on behalf of the Wellcome Sanger Institute COVID-19 Surveillance Team |
| EPI_ISL_2704675 | Lighthouse Lab in Milton Keynes | Wellcome Sanger Institute for the COVID-19 Genomics UK (COG-UK) Consortium | The Lighthouse Lab in Milton Keynes and Alex Alderton, Roberto Amato, Jeffrey Barrett, Sonia Goncalves, Ewan Harrison, David K. Jackson, Ian Johnston, Dominic Kwiatkowski, Cordelia Langford, John Sillitoe on behalf of the Wellcome Sanger Institute COVID-19 Surveillance Team |
| EPI_ISL_2704676 | Lighthouse Lab in Glasgow | Wellcome Sanger Institute for the COVID-19 Genomics UK (COG-UK) Consortium | Harper VanSteenhouse, Yumi Kasai, David Gray, Carol Clugston, Anna Dominiczak and Alex Alderton, Roberto Amato, Jeffrey Barrett, Sonia Goncalves, Ewan Harrison, David K. Jackson, Ian Johnston, Dominic Kwiatkowski, Cordelia Langford, John Sillitoe on behalf of the Wellcome Sanger Institute COVID-19 Surveillance Team |
| EPI_ISL_2704678, EPI_ISL_2704679, EPI_ISL_2704680, EPI_ISL_2704681, EPI_ISL_2704682, EPI_ISL_2704683, EPI_ISL_2704684, EPI_ISL_2704685, EPI_ISL_2704686, EPI_ISL_2704687, EPI_ISL_2704688, EPI_ISL_2704689, EPI_ISL_2704690, EPI_ISL_2704691, EPI_ISL_2704692, EPI_ISL_2704693, EPI_ISL_2704694, EPI_ISL_2704695, EPI_ISL_2704696, EPI_ISL_2704698, EPI_ISL_2704699, EPI_ISL_2704700, EPI_ISL_2704701, EPI_ISL_2704702, EPI_ISL_2704704, EPI_ISL_2704705, EPI_ISL_2704706, EPI_ISL_2704707, EPI_ISL_2704708, EPI_ISL_2704709, EPI_ISL_2704710, EPI_ISL_2704711, EPI_ISL_2704712, EPI_ISL_2704713, EPI_ISL_2704714, EPI_ISL_2704715, EPI_ISL_2704716, EPI_ISL_2704717, EPI_ISL_2704718, EPI_ISL_2704719, EPI_ISL_2704720, EPI_ISL_2704721, EPI_ISL_2704722, EPI_ISL_2704723, EPI_ISL_2704724, EPI_ISL_2704725, EPI_ISL_2704726, EPI_ISL_2704727, EPI_ISL_2704728, EPI_ISL_2704729, EPI_ISL_2704730, EPI_ISL_2704731, EPI_ISL_2704732, EPI_ISL_2704733, EPI_ISL_2704734, EPI_ISL_2704735, EPI_ISL_2704736, EPI_ISL_2704737, EPI_ISL_2704738, EPI_ISL_2704739, EPI_ISL_2704740, EPI_ISL_2704741, EPI_ISL_2704742, EPI_ISL_2704743 | | | |
| see above | Department of Pathology, University of Cambridge | COVID-19 Genomics UK (COG-UK) Consortium | Aminu S. Jahun, Yasmin Chaudhry, Iliana Georgana, Myra Hosmillo, Rhys Izuagbe, William L. Hamilton, Martin D. Curran, Surendra Parmar, Ian Goodfellow |
| EPI_ISL_2704823, EPI_ISL_2704824, EPI_ISL_2704826, EPI_ISL_2704827, EPI_ISL_2704828, EPI_ISL_2704829, EPI_ISL_2704830, EPI_ISL_2704831, EPI_ISL_2704832, EPI_ISL_2704834, EPI_ISL_2704835, EPI_ISL_2704836, EPI_ISL_2704837, EPI_ISL_2704838, EPI_ISL_2704839, EPI_ISL_2704840, EPI_ISL_2704841, EPI_ISL_2704842, EPI_ISL_2704843, EPI_ISL_2704844, EPI_ISL_2704845, EPI_ISL_2704846, EPI_ISL_2704847, EPI_ISL_2704848, EPI_ISL_2704850, EPI_ISL_2704851, EPI_ISL_2704852, EPI_ISL_2704853, EPI_ISL_2704854, EPI_ISL_2704855, EPI_ISL_2704856, EPI_ISL_2704857, EPI_ISL_2704858, EPI_ISL_2704860, EPI_ISL_2704861, EPI_ISL_2704862, EPI_ISL_2704864, EPI_ISL_2704865, EPI_ISL_2704866, EPI_ISL_2704867, EPI_ISL_2704868, EPI_ISL_2704870, EPI_ISL_2704871, EPI_ISL_2704872, EPI_ISL_2704873, EPI_ISL_2704879, EPI_ISL_2704882, EPI_ISL_2704883, EPI_ISL_2704885, EPI_ISL_2704886, EPI_ISL_2704887, EPI_ISL_2704888, EPI_ISL_2704889, EPI_ISL_2704890, EPI_ISL_2704891, EPI_ISL_2704892, EPI_ISL_2704893, EPI_ISL_2704894, EPI_ISL_2704895, EPI_ISL_2704896, EPI_ISL_2704897, EPI_ISL_2704898, EPI_ISL_2704899, EPI_ISL_2704900, EPI_ISL_2704901, EPI_ISL_2704902, EPI_ISL_2704903, EPI_ISL_2704904, EPI_ISL_2704905, EPI_ISL_2704906, EPI_ISL_2704907, EPI_ISL_2704908, EPI_ISL_2704909, EPI_ISL_2704910, EPI_ISL_2704911, EPI_ISL_2704912, EPI_ISL_2704913, EPI_ISL_2704914, EPI_ISL_2704915, EPI_ISL_2704916, EPI_ISL_2704917, EPI_ISL_2704918, EPI_ISL_2704919, EPI_ISL_2704920, EPI_ISL_2704921, EPI_ISL_2704923, EPI_ISL_2704924, EPI_ISL_2704926, EPI_ISL_2704927, EPI_ISL_2704928, EPI_ISL_2704929, EPI_ISL_2704930, EPI_ISL_2704931, EPI_ISL_2704932, EPI_ISL_2704933, EPI_ISL_2704934, EPI_ISL_2704935, EPI_ISL_2704936, EPI_ISL_2704937, EPI_ISL_2704938, EPI_ISL_2704939, EPI_ISL_2704940 | | | |
| see above | Virology Department, Royal Infirmary of Edinburgh, NHS Lothian / School of Biological Sciences, University of Edinburgh | COVID-19 Genomics UK (COG-UK) Consortium | McHugh M, Dewar R, Cotton S, Gallagher A, Maloney D, Rooke S, O'Toole Á, Scher E, Hill V, McCrone JT, Colquhoun R, Yu X, Jackson B, Rambaut A, Templeton K |
| EPI_ISL_2705472, EPI_ISL_2705473, EPI_ISL_2705474, EPI_ISL_2705475, EPI_ISL_2705476, EPI_ISL_2705477, EPI_ISL_2705478, EPI_ISL_2705479, EPI_ISL_2705480, EPI_ISL_2705481, EPI_ISL_2705482 | | | |
| see above | Liverpool Clinical Laboratories | COVID-19 Genomics UK (COG-UK) Consortium | Sam Haldenby, Alistair Darby, Steve Paterson, Anita Lucaci, Julian Hiscox, M Almsaud, A Alrezaihi, Muhannad Alruwaili, Stuart D Armstrong, Jones Benjamin, Eleanor G Bentley, Anu Chawla, Jordan J Clark, Angela Cowell, Richard Eccles, Isabel García-Dorival, Matthew Gemmell, Alessandro Gerada, PKF Gilmore, Richard Gregory, Ximeng Han, Catherine Hartley, Margaret Hughes, Miren Iturriza-Gomara, James Johnson, L Luu, Jenifer Manson, Charlotte Nelson, Elaine O'Toole, Cassie Olateju, Rebekah Penrice-Randal , Lucille Rainbow, N.P Randle, Trevor Ian Robinson, Parul Sharma, Ghada T Shawli, James P Stewart, Neil Swainston, Ecaterina Vamos, Joanne Watts, Mark Whitehead, Hermione Webster |
| EPI_ISL_2705483, EPI_ISL_2705485, EPI_ISL_2705486, EPI_ISL_2705487, EPI_ISL_2705488, EPI_ISL_2705489, EPI_ISL_2705490, EPI_ISL_2705491, EPI_ISL_2705492, EPI_ISL_2705493, EPI_ISL_2705494, EPI_ISL_2705495 | | | |
| see above | Barts Health NHS Trust | COVID-19 Genomics UK (COG-UK) Consortium | CUTINO-MOGUEL, Maria-Teresa; HARRINGTON, David; OWOYEMI, Dola; KULASEGARAN-SHYLINI, Raghavendran; BROAD, Claire; KELE, Beatrix |
| EPI_ISL_2705496, EPI_ISL_2705498, EPI_ISL_2705499, EPI_ISL_2705500, EPI_ISL_2705501, EPI_ISL_2705502, EPI_ISL_2705503, EPI_ISL_2705504, EPI_ISL_2705505, EPI_ISL_2705506, EPI_ISL_2705507, EPI_ISL_2705508, EPI_ISL_2705509, EPI_ISL_2705510, EPI_ISL_2705511, EPI_ISL_2705512, EPI_ISL_2705513, EPI_ISL_2705514, EPI_ISL_2705515, EPI_ISL_2705516, EPI_ISL_2705517, EPI_ISL_2705518, EPI_ISL_2705519, EPI_ISL_2705520, EPI_ISL_2705521, EPI_ISL_2705522, EPI_ISL_2705523, EPI_ISL_2705524, EPI_ISL_2705525, EPI_ISL_2705526, EPI_ISL_2705527, EPI_ISL_2705528, EPI_ISL_2705529, EPI_ISL_2705530, EPI_ISL_2705531, EPI_ISL_2705532, EPI_ISL_2705533, EPI_ISL_2705534, EPI_ISL_2705536, EPI_ISL_2705537, EPI_ISL_2705538, EPI_ISL_2705539, EPI_ISL_2705540, EPI_ISL_2705541, EPI_ISL_2705542, EPI_ISL_2705543, EPI_ISL_2705544, EPI_ISL_2705545, EPI_ISL_2705546, EPI_ISL_2705547, EPI_ISL_2705548, EPI_ISL_2705549, EPI_ISL_2705550, EPI_ISL_2705551, EPI_ISL_2705552, EPI_ISL_2705553, EPI_ISL_2705554, EPI_ISL_2705555, EPI_ISL_2705556, EPI_ISL_2705557, EPI_ISL_2705558, EPI_ISL_2705559, EPI_ISL_2705560, EPI_ISL_2705561, EPI_ISL_2705562, EPI_ISL_2705563, EPI_ISL_2705564, EPI_ISL_2705567, EPI_ISL_2705568, EPI_ISL_2705569, EPI_ISL_2705570, EPI_ISL_2705571, EPI_ISL_2705572, EPI_ISL_2705573, EPI_ISL_2705574 | | | |
| see above | University College London, Great Ormond Street Hospital for Children NHS Foundation Trust, Imperial College Healthcare NHS Trust | COVID-19 Genomics UK (COG-UK) Consortium | Sergi Castellano, Rachel Williams, Mark Kristiansen, Paola Resende Silva, Sunando Roy, Tony Brooks, Helena Tutill, Paola Niola, Patricia Dyal, Charlotte Williams, Leysa Forrest, Yasmin Panchbhaya, Jacqueline Findlay, Samuel Weeks, Julianne Brown, Kathryn Harris, Paul Randell, James Price, Alison Holmes, Judith Breuer |
| EPI_ISL_2705575 | Centers for Disease Control, R.O.C. (Taiwan) | Centers for Disease Control, R.O.C. (Taiwan) | Ji-Rong Yang, Yu-Chi-Lin, Jung-Jung Mu, Ming-Tsan-Liu |
| EPI_ISL_2705576, EPI_ISL_2705577, EPI_ISL_2705578, EPI_ISL_2705579, EPI_ISL_2705580, EPI_ISL_2705581, EPI_ISL_2705582, EPI_ISL_2705584, EPI_ISL_2705585, EPI_ISL_2705586, EPI_ISL_2705587, EPI_ISL_2705588, EPI_ISL_2705589, EPI_ISL_2705590, EPI_ISL_2705591, EPI_ISL_2705592, EPI_ISL_2705593, EPI_ISL_2705594, EPI_ISL_2705595, EPI_ISL_2705596, EPI_ISL_2705597, EPI_ISL_2705598, EPI_ISL_2705599, EPI_ISL_2705600, EPI_ISL_2705601, EPI_ISL_2705603, EPI_ISL_2705604, EPI_ISL_2705605, EPI_ISL_2705606, EPI_ISL_2705607, EPI_ISL_2705608, EPI_ISL_2705609, EPI_ISL_2705610, EPI_ISL_2705611, EPI_ISL_2705612, EPI_ISL_2705613, EPI_ISL_2705614, EPI_ISL_2705615, EPI_ISL_2705616, EPI_ISL_2705617, EPI_ISL_2705618, EPI_ISL_2705619, EPI_ISL_2705620, EPI_ISL_2705621, EPI_ISL_2705622, EPI_ISL_2705624, EPI_ISL_2705625, EPI_ISL_2705626, EPI_ISL_2705627, EPI_ISL_2705628, EPI_ISL_2705629, EPI_ISL_2705630, EPI_ISL_2705631, EPI_ISL_2705632, EPI_ISL_2705633, EPI_ISL_2705634, EPI_ISL_2705635, EPI_ISL_2705636, EPI_ISL_2705637, EPI_ISL_2705638, EPI_ISL_2705639, EPI_ISL_2705640, EPI_ISL_2705641, EPI_ISL_2705642, EPI_ISL_2705643, EPI_ISL_2705644, EPI_ISL_2705645, EPI_ISL_2705646, EPI_ISL_2705647, EPI_ISL_2705648, EPI_ISL_2705649, EPI_ISL_2705650, EPI_ISL_2705651, EPI_ISL_2705652, EPI_ISL_2705653, EPI_ISL_2705655, EPI_ISL_2705656, EPI_ISL_2705657, EPI_ISL_2705658, EPI_ISL_2705659, EPI_ISL_2705660, EPI_ISL_2705661, EPI_ISL_2705662, EPI_ISL_2705663, EPI_ISL_2705664, EPI_ISL_2705665, EPI_ISL_2705666, EPI_ISL_2705667, EPI_ISL_2705668, EPI_ISL_2705669, EPI_ISL_2705670, EPI_ISL_2705671, EPI_ISL_2705672, EPI_ISL_2705673, EPI_ISL_2705674, EPI_ISL_2705675, EPI_ISL_2705676, EPI_ISL_2705677, EPI_ISL_2705678, EPI_ISL_2705679, EPI_ISL_2705680, EPI_ISL_2705681, EPI_ISL_2705682, EPI_ISL_2705683, EPI_ISL_2705684, EPI_ISL_2705685, EPI_ISL_2705686, EPI_ISL_2705687, EPI_ISL_2705688, EPI_ISL_2705689, EPI_ISL_2705690, EPI_ISL_2705691, EPI_ISL_2705692, EPI_ISL_2705693, EPI_ISL_2705694, EPI_ISL_2705695, EPI_ISL_2705696, EPI_ISL_2705697, EPI_ISL_2705698, EPI_ISL_2705699, EPI_ISL_2705700, EPI_ISL_2705701, EPI_ISL_2705702, EPI_ISL_2705703, EPI_ISL_2705704, EPI_ISL_2705705, EPI_ISL_2705706, EPI_ISL_2705707, EPI_ISL_2705708, EPI_ISL_2705709, EPI_ISL_2705710, EPI_ISL_2705711, EPI_ISL_2705712, EPI_ISL_2705713, EPI_ISL_2705714, EPI_ISL_2705715, EPI_ISL_2705716, EPI_ISL_2705717, EPI_ISL_2705718, EPI_ISL_2705719, EPI_ISL_2705720, EPI_ISL_2705721, EPI_ISL_2705722, EPI_ISL_2705723, EPI_ISL_2705724, EPI_ISL_2705725, EPI_ISL_2705726, EPI_ISL_2705727, EPI_ISL_2705728, EPI_ISL_2705729, EPI_ISL_2705730, EPI_ISL_2705732, EPI_ISL_2705733, EPI_ISL_2705734, EPI_ISL_2705735, EPI_ISL_2705736, EPI_ISL_2705737, EPI_ISL_2705738, EPI_ISL_2705739, EPI_ISL_2705740, EPI_ISL_2705741, EPI_ISL_2705742, EPI_ISL_2705743, EPI_ISL_2705744, EPI_ISL_2705745, EPI_ISL_2705746, EPI_ISL_2705747, EPI_ISL_2705748, EPI_ISL_2705749, EPI_ISL_2705750, EPI_ISL_2705751, EPI_ISL_2705752, EPI_ISL_2705753, EPI_ISL_2705754, EPI_ISL_2705755, EPI_ISL_2705756, EPI_ISL_2705757, EPI_ISL_2705758, EPI_ISL_2705759, EPI_ISL_2705760, EPI_ISL_2705761, EPI_ISL_2705762, EPI_ISL_2705763, EPI_ISL_2705764, EPI_ISL_2705765, EPI_ISL_2705766, EPI_ISL_2705767, EPI_ISL_2705768, EPI_ISL_2705769, EPI_ISL_2705770, EPI_ISL_2705771, EPI_ISL_2705772, EPI_ISL_2705773, EPI_ISL_2705774, EPI_ISL_2705775, EPI_ISL_2705776, EPI_ISL_2705778, EPI_ISL_2705779, EPI_ISL_2705780, EPI_ISL_2705781, EPI_ISL_2705782, EPI_ISL_2705783, EPI_ISL_2705784, EPI_ISL_2705785, EPI_ISL_2705786, EPI_ISL_2705787, EPI_ISL_2705788, EPI_ISL_2705789, EPI_ISL_2705790, EPI_ISL_2705791, EPI_ISL_2705792, EPI_ISL_2705793, EPI_ISL_2705794, EPI_ISL_2705795, EPI_ISL_2705796, EPI_ISL_2705797, EPI_ISL_2705798, EPI_ISL_2705799, EPI_ISL_2705800, EPI_ISL_2705801, EPI_ISL_2705802, EPI_ISL_2705803, EPI_ISL_2705804, EPI_ISL_2705805, EPI_ISL_2705806, EPI_ISL_2705807, EPI_ISL_2705809, EPI_ISL_2705810, EPI_ISL_2705811, EPI_ISL_2705812, EPI_ISL_2705813, EPI_ISL_2705814, EPI_ISL_2705815, EPI_ISL_2705816, EPI_ISL_2705817, EPI_ISL_2705818, EPI_ISL_2705819, EPI_ISL_2705820, EPI_ISL_2705821, EPI_ISL_2705822, EPI_ISL_2705823, EPI_ISL_2705824, EPI_ISL_2705826, EPI_ISL_2705827, EPI_ISL_2705828, EPI_ISL_2705829, EPI_ISL_2705830, EPI_ISL_2705831, EPI_ISL_2705832, EPI_ISL_2705833, EPI_ISL_2705834, EPI_ISL_2705835, EPI_ISL_2705836, EPI_ISL_2705838, EPI_ISL_2705839, EPI_ISL_2705840, EPI_ISL_2705841, EPI_ISL_2705842, EPI_ISL_2705843, EPI_ISL_2705844, EPI_ISL_2705845, EPI_ISL_2705846, EPI_ISL_2705847, EPI_ISL_2705848, EPI_ISL_2705849, EPI_ISL_2705850, EPI_ISL_2705851, EPI_ISL_2705852, EPI_ISL_2705853, EPI_ISL_2705854, EPI_ISL_2705855, EPI_ISL_2705856, EPI_ISL_2705857, EPI_ISL_2705858, EPI_ISL_2705859, EPI_ISL_2705860, EPI_ISL_2705861, EPI_ISL_2705862, EPI_ISL_2705863, EPI_ISL_2705864, EPI_ISL_2705865, EPI_ISL_2705866, EPI_ISL_2705867, EPI_ISL_2705868, EPI_ISL_2705869, EPI_ISL_2705870, EPI_ISL_2705871, EPI_ISL_2705872, EPI_ISL_2705873, EPI_ISL_2705874, EPI_ISL_2705875, EPI_ISL_2705876, EPI_ISL_2705877, EPI_ISL_2705878, EPI_ISL_2705879, EPI_ISL_2705880, EPI_ISL_2705881, EPI_ISL_2705882, EPI_ISL_2705883, EPI_ISL_2705884, EPI_ISL_2705885, EPI_ISL_2705886, EPI_ISL_2705887, EPI_ISL_2705888, EPI_ISL_2705889, EPI_ISL_2705890, EPI_ISL_2705891, EPI_ISL_2705892, EPI_ISL_2705893, EPI_ISL_2705894, EPI_ISL_2705895, EPI_ISL_2705896, EPI_ISL_2705897, EPI_ISL_2705899, EPI_ISL_2705900, EPI_ISL_2705901, EPI_ISL_2705902, EPI_ISL_2705903, EPI_ISL_2705904, EPI_ISL_2705905, EPI_ISL_2705906, EPI_ISL_2705907, EPI_ISL_2705908, EPI_ISL_2705909, EPI_ISL_2705910, EPI_ISL_2705911, EPI_ISL_2705912, EPI_ISL_2705913, EPI_ISL_2705914, EPI_ISL_2705915, EPI_ISL_2705916, EPI_ISL_2705917, EPI_ISL_2705918, EPI_ISL_2705919, EPI_ISL_2705920, EPI_ISL_2705921, EPI_ISL_2705922, EPI_ISL_2705923, EPI_ISL_2705924, EPI_ISL_2705925, EPI_ISL_2705926, EPI_ISL_2705927, EPI_ISL_2705928, EPI_ISL_2705929 | | | |

see above | University College London, Great Ormond Street Hospital for Children NHS Foundation Trust, Imperial College Healthcare NHS Trust | COVID-19 Genomics UK (COG-UK) Consortium | Sergi Castellano, Rachel Williams, Mark Kristiansen, Paola Resende Silva, Sunando Roy, Tony Brooks, Helena Tutill, Paola Niola, Patricia Dyal, Charlotte Williams, Leysa Forrest, Yasmin Panchbhaya, Jacqueline Findlay, Samuel Weeks, Julianne Brown, Kathryn Harris, Paul Randell, James Price, Alison Holmes, Judith Breuer

EPI_ISL_2705930, EPI_ISL_2705931, EPI_ISL_2705932, EPI_ISL_2705933, EPI_ISL_2705934, EPI_ISL_2705935, EPI_ISL_2705936, EPI_ISL_2705937, EPI_ISL_2705938, EPI_ISL_2705940, EPI_ISL_2705941, EPI_ISL_2705942, EPI_ISL_2705944, EPI_ISL_2705945, EPI_ISL_2705946, EPI_ISL_2705947, EPI_ISL_2705948, EPI_ISL_2705949, EPI_ISL_2705950, EPI_ISL_2705951, EPI_ISL_2705952, EPI_ISL_2705953, EPI_ISL_2705954, EPI_ISL_2705955, EPI_ISL_2705956, EPI_ISL_2705957, EPI_ISL_2705958, EPI_ISL_2705960, EPI_ISL_2705961, EPI_ISL_2705962, EPI_ISL_2705963, EPI_ISL_2705964, EPI_ISL_2705965, EPI_ISL_2705966, EPI_ISL_2705967, EPI_ISL_2705968, EPI_ISL_2705969, EPI_ISL_2705970, EPI_ISL_2705971, EPI_ISL_2705972, EPI_ISL_2705973, EPI_ISL_2705974, EPI_ISL_2705975, EPI_ISL_2705976, EPI_ISL_2705977, EPI_ISL_2705978, EPI_ISL_2705979, EPI_ISL_2705981, EPI_ISL_2705982, EPI_ISL_2705983, EPI_ISL_2705984, EPI_ISL_2705985, EPI_ISL_2705986, EPI_ISL_2705988, EPI_ISL_2705989, EPI_ISL_2705990, EPI_ISL_2705991, EPI_ISL_2705993, EPI_ISL_2705994, EPI_ISL_2705995, EPI_ISL_2705996, EPI_ISL_2705997, EPI_ISL_2705998, EPI_ISL_2705999, EPI_ISL_2706000, EPI_ISL_2706001, EPI_ISL_2706002, EPI_ISL_2706003, EPI_ISL_2706004, EPI_ISL_2706005, EPI_ISL_2706006, EPI_ISL_2706008, EPI_ISL_2706010, EPI_ISL_2706011, EPI_ISL_2706012, EPI_ISL_2706013, EPI_ISL_2706014, EPI_ISL_2706015, EPI_ISL_2706016, EPI_ISL_2706017, EPI_ISL_2706019, EPI_ISL_2706020, EPI_ISL_2706021, EPI_ISL_2706023, EPI_ISL_2706024, EPI_ISL_2706025, EPI_ISL_2706026, EPI_ISL_2706027, EPI_ISL_2706028, EPI_ISL_2706029, EPI_ISL_2706030, EPI_ISL_2706031, EPI_ISL_2706032, EPI_ISL_2706033, EPI_ISL_2706034, EPI_ISL_2706035, EPI_ISL_2706036, EPI_ISL_2706037, EPI_ISL_2706038, EPI_ISL_2706039, EPI_ISL_2706040, EPI_ISL_2706041, EPI_ISL_2706042, EPI_ISL_2706043, EPI_ISL_2706044, EPI_ISL_2706045, EPI_ISL_2706046, EPI_ISL_2706047, EPI_ISL_2706048, EPI_ISL_2706049, EPI_ISL_2706050, EPI_ISL_2706052, EPI_ISL_2706053, EPI_ISL_2706054, EPI_ISL_2706056, EPI_ISL_2706057, EPI_ISL_2706058, EPI_ISL_2706059, EPI_ISL_2706060, EPI_ISL_2706061, EPI_ISL_2706062, EPI_ISL_2706063, EPI_ISL_2706064, EPI_ISL_2706065, EPI_ISL_2706066, EPI_ISL_2706067, EPI_ISL_2706068, EPI_ISL_2706069, EPI_ISL_2706070, EPI_ISL_2706071, EPI_ISL_2706072, EPI_ISL_2706073, EPI_ISL_2706074, EPI_ISL_2706075, EPI_ISL_2706076, EPI_ISL_2706077, EPI_ISL_2706078, EPI_ISL_2706079, EPI_ISL_2706080, EPI_ISL_2706081, EPI_ISL_2706082, EPI_ISL_2706084, EPI_ISL_2706085, EPI_ISL_2706086, EPI_ISL_2706087, EPI_ISL_2706088, EPI_ISL_2706089, EPI_ISL_2706090, EPI_ISL_2706091, EPI_ISL_2706092, EPI_ISL_2706093, EPI_ISL_2706094, EPI_ISL_2706095, EPI_ISL_2706096, EPI_ISL_2706097, EPI_ISL_2706098, EPI_ISL_2706099, EPI_ISL_2706100, EPI_ISL_2706101, EPI_ISL_2706102, EPI_ISL_2706103, EPI_ISL_2706104, EPI_ISL_2706105, EPI_ISL_2706107, EPI_ISL_2706108, EPI_ISL_2706109, EPI_ISL_2706110, EPI_ISL_2706111, EPI_ISL_2706112, EPI_ISL_2706113, EPI_ISL_2706114, EPI_ISL_2706116, EPI_ISL_2706117, EPI_ISL_2706118, EPI_ISL_2706119, EPI_ISL_2706120, EPI_ISL_2706121, EPI_ISL_2706122, EPI_ISL_2706123, EPI_ISL_2706124, EPI_ISL_2706125, EPI_ISL_2706126, EPI_ISL_2706127, EPI_ISL_2706128, EPI_ISL_2706129, EPI_ISL_2706130, EPI_ISL_2706131, EPI_ISL_2706135, EPI_ISL_2706136, EPI_ISL_2706137, EPI_ISL_2706138, EPI_ISL_2706139, EPI_ISL_2706141, EPI_ISL_2706142, EPI_ISL_2706143, EPI_ISL_2706144, EPI_ISL_2706145, EPI_ISL_2706146, EPI_ISL_2706147, EPI_ISL_2706148, EPI_ISL_2706149, EPI_ISL_2706150, EPI_ISL_2706151, EPI_ISL_2706152, EPI_ISL_2706153, EPI_ISL_2706154, EPI_ISL_2706155, EPI_ISL_2706157, EPI_ISL_2706158, EPI_ISL_2706159, EPI_ISL_2706160, EPI_ISL_2706161, EPI_ISL_2706162, EPI_ISL_2706163, EPI_ISL_2706164, EPI_ISL_2706165, EPI_ISL_2706166, EPI_ISL_2706167, EPI_ISL_2706168, EPI_ISL_2706169, EPI_ISL_2706170, EPI_ISL_2706171, EPI_ISL_2706172, EPI_ISL_2706173, EPI_ISL_2706174, EPI_ISL_2706175, EPI_ISL_2706176, EPI_ISL_2706177, EPI_ISL_2706178, EPI_ISL_2706179, EPI_ISL_2706180, EPI_ISL_2706181, EPI_ISL_2706182, EPI_ISL_2706183, EPI_ISL_2706184, EPI_ISL_2706185, EPI_ISL_2706186, EPI_ISL_2706187, EPI_ISL_2706188, EPI_ISL_2706189, EPI_ISL_2706190, EPI_ISL_2706191, EPI_ISL_2706192, EPI_ISL_2706193, EPI_ISL_2706194, EPI_ISL_2706195, EPI_ISL_2706196, EPI_ISL_2706197, EPI_ISL_2706198, EPI_ISL_2706199, EPI_ISL_2706200, EPI_ISL_2706201, EPI_ISL_2706202, EPI_ISL_2706203, EPI_ISL_2706204, EPI_ISL_2706205, EPI_ISL_2706206, EPI_ISL_2706207, EPI_ISL_2706208, EPI_ISL_2706209, EPI_ISL_2706210, EPI_ISL_2706211, EPI_ISL_2706212, EPI_ISL_2706213, EPI_ISL_2706214, EPI_ISL_2706215, EPI_ISL_2706216, EPI_ISL_2706217, EPI_ISL_2706218, EPI_ISL_2706219, EPI_ISL_2706220, EPI_ISL_2706221, EPI_ISL_2706222, EPI_ISL_2706223, EPI_ISL_2706224, EPI_ISL_2706225, EPI_ISL_2706226, EPI_ISL_2706227, EPI_ISL_2706228, EPI_ISL_2706229, EPI_ISL_2706230, EPI_ISL_2706231, EPI_ISL_2706232, EPI_ISL_2706233, EPI_ISL_2706234, EPI_ISL_2706235, EPI_ISL_2706236, EPI_ISL_2706237, EPI_ISL_2706238, EPI_ISL_2706240, EPI_ISL_2706241, EPI_ISL_2706242, EPI_ISL_2706243, EPI_ISL_2706244, EPI_ISL_2706245, EPI_ISL_2706246, EPI_ISL_2706247, EPI_ISL_2706248, EPI_ISL_2706249, EPI_ISL_2706250, EPI_ISL_2706251, EPI_ISL_2706252, EPI_ISL_2706253, EPI_ISL_2706254, EPI_ISL_2706255, EPI_ISL_2706256, EPI_ISL_2706257, EPI_ISL_2706258, EPI_ISL_2706259, EPI_ISL_2706260, EPI_ISL_2706261, EPI_ISL_2706262, EPI_ISL_2706263, EPI_ISL_2706264, EPI_ISL_2706265, EPI_ISL_2706266, EPI_ISL_2706267, EPI_ISL_2706268, EPI_ISL_2706270, EPI_ISL_2706271, EPI_ISL_2706272, EPI_ISL_2706273, EPI_ISL_2706274, EPI_ISL_2706275, EPI_ISL_2706276, EPI_ISL_2706277, EPI_ISL_2706278, EPI_ISL_2706279, EPI_ISL_2706280, EPI_ISL_2706281, EPI_ISL_2706282, EPI_ISL_2706283, EPI_ISL_2706284, EPI_ISL_2706285, EPI_ISL_2706286, EPI_ISL_2706287, EPI_ISL_2706288, EPI_ISL_2706289, EPI_ISL_2706290, EPI_ISL_2706291, EPI_ISL_2706292, EPI_ISL_2706293, EPI_ISL_2706294, EPI_ISL_2706295, EPI_ISL_2706296, EPI_ISL_2706297, EPI_ISL_2706298, EPI_ISL_2706299, EPI_ISL_2706300, EPI_ISL_2706301, EPI_ISL_2706302, EPI_ISL_2706303, EPI_ISL_2706304, EPI_ISL_2706305, EPI_ISL_2706306, EPI_ISL_2706307, EPI_ISL_2706308, EPI_ISL_2706309, EPI_ISL_2706310, EPI_ISL_2706311, EPI_ISL_2706312, EPI_ISL_2706313, EPI_ISL_2706314, EPI_ISL_2706315, EPI_ISL_2706316, EPI_ISL_2706317, EPI_ISL_2706318, EPI_ISL_2706319, EPI_ISL_2706320, EPI_ISL_2706321, EPI_ISL_2706322, EPI_ISL_2706323, EPI_ISL_2706324, EPI_ISL_2706325, EPI_ISL_2706326, EPI_ISL_2706327, EPI_ISL_2706328, EPI_ISL_2706329, EPI_ISL_2706330, EPI_ISL_2706331, EPI_ISL_2706332, EPI_ISL_2706333, EPI_ISL_2706334, EPI_ISL_2706335, EPI_ISL_2706336, EPI_ISL_2706337, EPI_ISL_2706338, EPI_ISL_2706339, EPI_ISL_2706341, EPI_ISL_2706342, EPI_ISL_2706343, EPI_ISL_2706344, EPI_ISL_2706345, EPI_ISL_2706346, EPI_ISL_2706347, EPI_ISL_2706348, EPI_ISL_2706349, EPI_ISL_2706350, EPI_ISL_2706351, EPI_ISL_2706352, EPI_ISL_2706353, EPI_ISL_2706354, EPI_ISL_2706355, EPI_ISL_2706356, EPI_ISL_2706357, EPI_ISL_2706358, EPI_ISL_2706359, EPI_ISL_2706360, EPI_ISL_2706361, EPI_ISL_2706362, EPI_ISL_2706364, EPI_ISL_2706365, EPI_ISL_2706366, EPI_ISL_2706367, EPI_ISL_2706368, EPI_ISL_2706369, EPI_ISL_2706370, EPI_ISL_2706371, EPI_ISL_2706372, EPI_ISL_2706373, EPI_ISL_2706374, EPI_ISL_2706375, EPI_ISL_2706376, EPI_ISL_2706377, EPI_ISL_2706378, EPI_ISL_2706379, EPI_ISL_2706380, EPI_ISL_2706381, EPI_ISL_2706382, EPI_ISL_2706383, EPI_ISL_2706384, EPI_ISL_2706385, EPI_ISL_2706386, EPI_ISL_2706387, EPI_ISL_2706388, EPI_ISL_2706389, EPI_ISL_2706390, EPI_ISL_2706391, EPI_ISL_2706392, EPI_ISL_2706393, EPI_ISL_2706394, EPI_ISL_2706395, EPI_ISL_2706396, EPI_ISL_2706397, EPI_ISL_2706398, EPI_ISL_2706399, EPI_ISL_2706400, EPI_ISL_2706401, EPI_ISL_2706402, EPI_ISL_2706403, EPI_ISL_2706404, EPI_ISL_2706405, EPI_ISL_2706406, EPI_ISL_2706407, EPI_ISL_2706408, EPI_ISL_2706409, EPI_ISL_2706410, EPI_ISL_2706411, EPI_ISL_2706412, EPI_ISL_2706413, EPI_ISL_2706414, EPI_ISL_2706415, EPI_ISL_2706416, EPI_ISL_2706417, EPI_ISL_2706418, EPI_ISL_2706419, EPI_ISL_2706420, EPI_ISL_2706422, EPI_ISL_2706423, EPI_ISL_2706424, EPI_ISL_2706425, EPI_ISL_2706426, EPI_ISL_2706427, EPI_ISL_2706428, EPI_ISL_2706429, EPI_ISL_2706430, EPI_ISL_2706431, EPI_ISL_2706432, EPI_ISL_2706433, EPI_ISL_2706434, EPI_ISL_2706435, EPI_ISL_2706436, EPI_ISL_2706437, EPI_ISL_2706438, EPI_ISL_2706439, EPI_ISL_2706440, EPI_ISL_2706441, EPI_ISL_2706442, EPI_ISL_2706443, EPI_ISL_2706444, EPI_ISL_2706445, EPI_ISL_2706446, EPI_ISL_2706447, EPI_ISL_2706448, EPI_ISL_2706449, EPI_ISL_2706450, EPI_ISL_2706451, EPI_ISL_2706452, EPI_ISL_2706453, EPI_ISL_2706454, EPI_ISL_2706455, EPI_ISL_2706456, EPI_ISL_2706457, EPI_ISL_2706458, EPI_ISL_2706459, EPI_ISL_2706460, EPI_ISL_2706461, EPI_ISL_2706462, EPI_ISL_2706463, EPI_ISL_2706464, EPI_ISL_2706465, EPI_ISL_2706466, EPI_ISL_2706467, EPI_ISL_2706468, EPI_ISL_2706469, EPI_ISL_2706470, EPI_ISL_2706471, EPI_ISL_2706472, EPI_ISL_2706473, EPI_ISL_2706474, EPI_ISL_2706475, EPI_ISL_2706476, EPI_ISL_2706477, EPI_ISL_2706479, EPI_ISL_2706480, EPI_ISL_2706481, EPI_ISL_2706482, EPI_ISL_2706483, EPI_ISL_2706484, EPI_ISL_2706485, EPI_ISL_2706486, EPI_ISL_2706487, EPI_ISL_2706488, EPI_ISL_2706489, EPI_ISL_2706490, EPI_ISL_2706491, EPI_ISL_2706492, EPI_ISL_2706493, EPI_ISL_2706494, EPI_ISL_2706495, EPI_ISL_2706496, EPI_ISL_2706497, EPI_ISL_2706498, EPI_ISL_2706499, EPI_ISL_2706500, EPI_ISL_2706501, EPI_ISL_2706502, EPI_ISL_2706503, EPI_ISL_2706504, EPI_ISL_2706505, EPI_ISL_2706506, EPI_ISL_2706507, EPI_ISL_2706508, EPI_ISL_2706509, EPI_ISL_2706510, EPI_ISL_2706511, EPI_ISL_2706512, EPI_ISL_2706513, EPI_ISL_2706514, EPI_ISL_2706515, EPI_ISL_2706516, EPI_ISL_2706517, EPI_ISL_2706518, EPI_ISL_2706519, EPI_ISL_2706520, EPI_ISL_2706521, EPI_ISL_2706522, EPI_ISL_2706523, EPI_ISL_2706524, EPI_ISL_2706525, EPI_ISL_2706526, EPI_ISL_2706527, EPI_ISL_2706528, EPI_ISL_2706529, EPI_ISL_2706530, EPI_ISL_2706531, EPI_ISL_2706532, EPI_ISL_2706533, EPI_ISL_2706534, EPI_ISL_2706535, EPI_ISL_2706536, EPI_ISL_2706537, EPI_ISL_2706538, EPI_ISL_2706539, EPI_ISL_2706540, EPI_ISL_2706541, EPI_ISL_2706542, EPI_ISL_2706543, EPI_ISL_2706544, EPI_ISL_2706545, EPI_ISL_2706546, EPI_ISL_2706547, EPI_ISL_2706548, EPI_ISL_2706549, EPI_ISL_2706550, EPI_ISL_2706551, EPI_ISL_2706552, EPI_ISL_2706553, EPI_ISL_2706554, EPI_ISL_2706555, EPI_ISL_2706556, EPI_ISL_2706557, EPI_ISL_2706558, EPI_ISL_2706559, EPI_ISL_2706560, EPI_ISL_2706561, EPI_ISL_2706562, EPI_ISL_2706563, EPI_ISL_2706564, EPI_ISL_2706565, EPI_ISL_2706566, EPI_ISL_2706567, EPI_ISL_2706568, EPI_ISL_2706569, EPI_ISL_2706570, EPI_ISL_2706571, EPI_ISL_2706572, EPI_ISL_2706573, EPI_ISL_2706574, EPI_ISL_2706575, EPI_ISL_2706576, EPI_ISL_2706577, EPI_ISL_2706578, EPI_ISL_2706579, EPI_ISL_2706580, EPI_ISL_2706581, EPI_ISL_2706582, EPI_ISL_2706583, EPI_ISL_2706584, EPI_ISL_2706585, EPI_ISL_2706586, EPI_ISL_2706587, EPI_ISL_2706588, EPI_ISL_2706589, EPI_ISL_2706590, EPI_ISL_2706591, EPI_ISL_2706592, EPI_ISL_2706593, EPI_ISL_2706594, EPI_ISL_2706595, EPI_ISL_2706596, EPI_ISL_2706597, EPI_ISL_2706598, EPI_ISL_2706599, EPI_ISL_2706600, EPI_ISL_2706601, EPI_ISL_2706602, EPI_ISL_2706603, EPI_ISL_2706604, EPI_ISL_2706605, EPI_ISL_2706606, EPI_ISL_2706607, EPI_ISL_2706608, EPI_ISL_2706609, EPI_ISL_2706610, EPI_ISL_2706611, EPI_ISL_2706612, EPI_ISL_2706613, EPI_ISL_2706614, EPI_ISL_2706615, EPI_ISL_2706616, EPI_ISL_2706617, EPI_ISL_2706618, EPI_ISL_2706619, EPI_ISL_2706620, EPI_ISL_2706621, EPI_ISL_2706622, EPI_ISL_2706623, EPI_ISL_2706624, EPI_ISL_2706625, EPI_ISL_2706626, EPI_ISL_2706627, EPI_ISL_2706628, EPI_ISL_2706629, EPI_ISL_2706630, EPI_ISL_2706632, EPI_ISL_2706633, EPI_ISL_2706634, EPI_ISL_2706635, EPI_ISL_2706636, EPI_ISL_2706637, EPI_ISL_2706638, EPI_ISL_2706639, EPI_ISL_2706640, EPI_ISL_2706641, EPI_ISL_2706642, EPI_ISL_2706643

see above | Northumbria University / South Tees Hospitals NHS Foundation Trust / North Cumbria Integrated Care NHS Foundation Trust / North Tees and Hartlepool NHS Foundation Trust / Newcastle Hospitals NHS Foundation Trust | COVID-19 Genomics UK (COG-UK) Consortium | Darren L Smith,Andrew Nelson,Matthew Bashton,Greg R Young,Joshua Loh,John Allan,Mohammad A Tariq,Giles S Holt,Gary Black,Wen C Yew,Lynn Dover,Paul Baker,Steve Liggett,Sarah Essex,Jane Greenaway,Debra Padgett,Clive Graham,Garren Scott,Edward Barton,Emma Swindells,Brendan Payne,Jennifer Collins,Yusri Taha,Gary Eltringham

EPI_ISL_2706644, EPI_ISL_2706645, EPI_ISL_2706646, EPI_ISL_2706647, EPI_ISL_2706648, EPI_ISL_2706649, EPI_ISL_2706650, EPI_ISL_2706651, EPI_ISL_2706652, EPI_ISL_2706653, EPI_ISL_2706654, EPI_ISL_2706655, EPI_ISL_2706656, EPI_ISL_2706657, EPI_ISL_2706659, EPI_ISL_2706660, EPI_ISL_2706662, EPI_ISL_2706663, EPI_ISL_2706664, EPI_ISL_2706665, EPI_ISL_2706666, EPI_ISL_2706667, EPI_ISL_2706668, EPI_ISL_2706669, EPI_ISL_2706670, EPI_ISL_2706671, EPI_ISL_2706672, EPI_ISL_2706673, EPI_ISL_2706674, EPI_ISL_2706676, EPI_ISL_2706677, EPI_ISL_2706679, EPI_ISL_2706680, EPI_ISL_2706681, EPI_ISL_2706682, EPI_ISL_2706683, EPI_ISL_2706684, EPI_ISL_2706685, EPI_ISL_2706687, EPI_ISL_2706688, EPI_ISL_2706689, EPI_ISL_2706690, EPI_ISL_2706691, EPI_ISL_2706692, EPI_ISL_2706693, EPI_ISL_2706694, EPI_ISL_2706695, EPI_ISL_2706696, EPI_ISL_2706697, EPI_ISL_2706698, EPI_ISL_2706699, EPI_ISL_2706700, EPI_ISL_2706701, EPI_ISL_2706706, EPI_ISL_2706707, EPI_ISL_2706708, EPI_ISL_2706709, EPI_ISL_2706713, EPI_ISL_2706715, EPI_ISL_2706716, EPI_ISL_2706717, EPI_ISL_2706718, EPI_ISL_2706719, EPI_ISL_2706721, EPI_ISL_2706722, EPI_ISL_2706723, EPI_ISL_2706724, EPI_ISL_2706725, EPI_ISL_2706727, EPI_ISL_2706728, EPI_ISL_2706729, EPI_ISL_2706730, EPI_ISL_2706731, EPI_ISL_2706732, EPI_ISL_2706733, EPI_ISL_2706734, EPI_ISL_2706736, EPI_ISL_2706737, EPI_ISL_2706738, EPI_ISL_2706739, EPI_ISL_2706740, EPI_ISL_2706741, EPI_ISL_2706742, EPI_ISL_2706743, EPI_ISL_2706744, EPI_ISL_2706745, EPI_ISL_2706746, EPI_ISL_2706747, EPI_ISL_2706748, EPI_ISL_2706749, EPI_ISL_2706750, EPI_ISL_2706751, EPI_ISL_2706752, EPI_ISL_2706753, EPI_ISL_2706754, EPI_ISL_2706755, EPI_ISL_2706756, EPI_ISL_2706757, EPI_ISL_2706758, EPI_ISL_2706759, EPI_ISL_2706760, EPI_ISL_2706761, EPI_ISL_2706762, EPI_ISL_2706763, EPI_ISL_2706764, EPI_ISL_2706766, EPI_ISL_2706767, EPI_ISL_2706768, EPI_ISL_2706769, EPI_ISL_2706770, EPI_ISL_2706772, EPI_ISL_2706773, EPI_ISL_2706774, EPI_ISL_2706776, EPI_ISL_2706779, EPI_ISL_2706780, EPI_ISL_2706781, EPI_ISL_2706783, EPI_ISL_2706785, EPI_ISL_2706786, EPI_ISL_2706787, EPI_ISL_2706788, EPI_ISL_2706789, EPI_ISL_2706791, EPI_ISL_2706792, EPI_ISL_2706793, EPI_ISL_2706794, EPI_ISL_2706795, EPI_ISL_2706796, EPI_ISL_2706798, EPI_ISL_2706799, EPI_ISL_2706800, EPI_ISL_2706801, EPI_ISL_2706802, EPI_ISL_2706803, EPI_ISL_2706804, EPI_ISL_2706805, EPI_ISL_2706806, EPI_ISL_2706807, EPI_ISL_2706808, EPI_ISL_2706809, EPI_ISL_2706810, EPI_ISL_2706811, EPI_ISL_2706812, EPI_ISL_2706816, EPI_ISL_2706817, EPI_ISL_2706818, EPI_ISL_2706820, EPI_ISL_2706821, EPI_ISL_2706822, EPI_ISL_2706823, EPI_ISL_2706824, EPI_ISL_2706825, EPI_ISL_2706826, EPI_ISL_2706827, EPI_ISL_2706828, EPI_ISL_2706829, EPI_ISL_2706830, EPI_ISL_2706831, EPI_ISL_2706832, EPI_ISL_2706833, EPI_ISL_2706835, EPI_ISL_2706836, EPI_ISL_2706837, EPI_ISL_2706838, EPI_ISL_2706839, EPI_ISL_2706840, EPI_ISL_2706843, EPI_ISL_2706845, EPI_ISL_2706846, EPI_ISL_2706847, EPI_ISL_2706849, EPI_ISL_2706851, EPI_ISL_2706852, EPI_ISL_2706855, EPI_ISL_2706857, EPI_ISL_2706858, EPI_ISL_2706859, EPI_ISL_2706862, EPI_ISL_2706863, EPI_ISL_2706866, EPI_ISL_2706867, EPI_ISL_2706868, EPI_ISL_2706869, EPI_ISL_2706870, EPI_ISL_2706871, EPI_ISL_2706873, EPI_ISL_2706874, EPI_ISL_2706875, EPI_ISL_2706876, EPI_ISL_2706877, EPI_ISL_2706879, EPI_ISL_2706880, EPI_ISL_2706881, EPI_ISL_2706883, EPI_ISL_2706885, EPI_ISL_2706886, EPI_ISL_2706887, EPI_ISL_2706888, EPI_ISL_2706889, EPI_ISL_2706890, EPI_ISL_2706891, EPI_ISL_2706892, EPI_ISL_2706893, EPI_ISL_2706894, EPI_ISL_2706897, EPI_ISL_2706898, EPI_ISL_2706899, EPI_ISL_2706900, EPI_ISL_2706903, EPI_ISL_2706904, EPI_ISL_2706905, EPI_ISL_2706906, EPI_ISL_2706907, EPI_ISL_2706908, EPI_ISL_2706910, EPI_ISL_2706911, EPI_ISL_2706913, EPI_ISL_2706914, EPI_ISL_2706915, EPI_ISL_2706916, EPI_ISL_2706919, EPI_ISL_2706920, EPI_ISL_2706922, EPI_ISL_2706923, EPI_ISL_2706924, EPI_ISL_2706925, EPI_ISL_2706926, EPI_ISL_2706927, EPI_ISL_2706928, EPI_ISL_2706929, EPI_ISL_2706930, EPI_ISL_2706931, EPI_ISL_2706932, EPI_ISL_2706933, EPI_ISL_2706934, EPI_ISL_2706935, EPI_ISL_2706936, EPI_ISL_2706937, EPI_ISL_2706938, EPI_ISL_2706939, EPI_ISL_2706940, EPI_ISL_2706941, EPI_ISL_2706942, EPI_ISL_2706943, EPI_ISL_2706944, EPI_ISL_2706945, EPI_ISL_2706946, EPI_ISL_2706947, EPI_ISL_2706948, EPI_ISL_2706949, EPI_ISL_2706950, EPI_ISL_2706951, EPI_ISL_2706952, EPI_ISL_2706953, EPI_ISL_2706954, EPI_ISL_2706955, EPI_ISL_2706957, EPI_ISL_2706958, EPI_ISL_2706959, EPI_ISL_2706960, EPI_ISL_2706961, EPI_ISL_2706962, EPI_ISL_2706963, EPI_ISL_2706964, EPI_ISL_2706965, EPI_ISL_2706966, EPI_ISL_2706967, EPI_ISL_2706968, EPI_ISL_2706969, EPI_ISL_2706970, EPI_ISL_2706971, EPI_ISL_2706972, EPI_ISL_2706973, EPI_ISL_2706974, EPI_ISL_2706975, EPI_ISL_2706976, EPI_ISL_2706977, EPI_ISL_2706978, EPI_ISL_2706979, EPI_ISL_2706981, EPI_ISL_2706983, EPI_ISL_2706984, EPI_ISL_2706985, EPI_ISL_2706986, EPI_ISL_2706987, EPI_ISL_2706988, EPI_ISL_2706990, EPI_ISL_2706991, EPI_ISL_2706992, EPI_ISL_2706993, EPI_ISL_2706994, EPI_ISL_2706995, EPI_ISL_2706996, EPI_ISL_2706997, EPI_ISL_2706998, EPI_ISL_2706999, EPI_ISL_2707000, EPI_ISL_2707001, EPI_ISL_2707002, EPI_ISL_2707003, EPI_ISL_2707004, EPI_ISL_2707005, EPI_ISL_2707006, EPI_ISL_2707007, EPI_ISL_2707008, EPI_ISL_2707009, EPI_ISL_2707010, EPI_ISL_2707011, EPI_ISL_2707012, EPI_ISL_2707013, EPI_ISL_2707014, EPI_ISL_2707015, EPI_ISL_2707016, EPI_ISL_2707017, EPI_ISL_2707018, EPI_ISL_2707019, EPI_ISL_2707020, EPI_ISL_2707021, EPI_ISL_2707022, EPI_ISL_2707023, EPI_ISL_2707024, EPI_ISL_2707025, EPI_ISL_2707026, EPI_ISL_2707027, EPI_ISL_2707028, EPI_ISL_2707029, EPI_ISL_2707030, EPI_ISL_2707031, EPI_ISL_2707032, EPI_ISL_2707033, EPI_ISL_2707034, EPI_ISL_2707035, EPI_ISL_2707036, EPI_ISL_2707037, EPI_ISL_2707038, EPI_ISL_2707039, EPI_ISL_2707040, EPI_ISL_2707041, EPI_ISL_2707042, EPI_ISL_2707043, EPI_ISL_2707044, EPI_ISL_2707045, EPI_ISL_2707046, EPI_ISL_2707047, EPI_ISL_2707048, EPI_ISL_2707049, EPI_ISL_2707050, EPI_ISL_2707051, EPI_ISL_2707052, EPI_ISL_2707053, EPI_ISL_2707054, EPI_ISL_2707055, EPI_ISL_2707056, EPI_ISL_2707057, EPI_ISL_2707058, EPI_ISL_2707059, EPI_ISL_2707060, EPI_ISL_2707061, EPI_ISL_2707062, EPI_ISL_2707063, EPI_ISL_2707064, EPI_ISL_2707065, EPI_ISL_2707066, EPI_ISL_2707067, EPI_ISL_2707068, EPI_ISL_2707069, EPI_ISL_2707070, EPI_ISL_2707071, EPI_ISL_2707072, EPI_ISL_2707073, EPI_ISL_2707076, EPI_ISL_2707077, EPI_ISL_2707078, EPI_ISL_2707079, EPI_ISL_2707080, EPI_ISL_2707081, EPI_ISL_2707082, EPI_ISL_2707083, EPI_ISL_2707084, EPI_ISL_2707085, EPI_ISL_2707087, EPI_ISL_2707088, EPI_ISL_2707089, EPI_ISL_2707090, EPI_ISL_2707091, EPI_ISL_2707092, EPI_ISL_2707093, EPI_ISL_2707094, EPI_ISL_2707095, EPI_ISL_2707096, EPI_ISL_2707097, EPI_ISL_2707098, EPI_ISL_2707099, EPI_ISL_2707100, EPI_ISL_2707101, EPI_ISL_2707102, EPI_ISL_2707103, EPI_ISL_2707104, EPI_ISL_2707105, EPI_ISL_2707106, EPI_ISL_2707107, EPI_ISL_2707109, EPI_ISL_2707110, EPI_ISL_2707111, EPI_ISL_2707112, EPI_ISL_2707113, EPI_ISL_2707114, EPI_ISL_2707115, EPI_ISL_2707116, EPI_ISL_2707117, EPI_ISL_2707118, EPI_ISL_2707119, EPI_ISL_2707120, EPI_ISL_2707121, EPI_ISL_2707122, EPI_ISL_2707123, EPI_ISL_2707124, EPI_ISL_2707125, EPI_ISL_2707126, EPI_ISL_2707127, EPI_ISL_2707128, EPI_ISL_2707129, EPI_ISL_2707130, EPI_ISL_2707131, EPI_ISL_2707132, EPI_ISL_2707133, EPI_ISL_2707134, EPI_ISL_2707135, EPI_ISL_2707136, EPI_ISL_2707137, EPI_ISL_2707138, EPI_ISL_2707139, EPI_ISL_2707140, EPI_ISL_2707141, EPI_ISL_2707142, EPI_ISL_2707143, EPI_ISL_2707144, EPI_ISL_2707145, EPI_ISL_2707146, EPI_ISL_2707147, EPI_ISL_2707148, EPI_ISL_2707149, EPI_ISL_2707150, EPI_ISL_2707151, EPI_ISL_2707152, EPI_ISL_2707153, EPI_ISL_2707154, EPI_ISL_2707155, EPI_ISL_2707156, EPI_ISL_2707157, EPI_ISL_2707158, EPI_ISL_2707159, EPI_ISL_2707160, EPI_ISL_2707161, EPI_ISL_2707162, EPI_ISL_2707163, EPI_ISL_2707164, EPI_ISL_2707165, EPI_ISL_2707166, EPI_ISL_2707167, EPI_ISL_2707168, EPI_ISL_2707169, EPI_ISL_2707170, EPI_ISL_2707171, EPI_ISL_2707172, EPI_ISL_2707173, EPI_ISL_2707174, EPI_ISL_2707175, EPI_ISL_2707176, EPI_ISL_2707177, EPI_ISL_2707178, EPI_ISL_2707179, EPI_ISL_2707180, EPI_ISL_2707181, EPI_ISL_2707182, EPI_ISL_2707183, EPI_ISL_2707184, EPI_ISL_2707185, EPI_ISL_2707186, EPI_ISL_2707187, EPI_ISL_2707189, EPI_ISL_2707190, EPI_ISL_2707191, EPI_ISL_2707192, EPI_ISL_2707193, EPI_ISL_2707194, EPI_ISL_2707195, EPI_ISL_2707196, EPI_ISL_2707197, EPI_ISL_2707198, EPI_ISL_2707199, EPI_ISL_2707200, EPI_ISL_2707201, EPI_ISL_2707202, EPI_ISL_2707203, EPI_ISL_2707204, EPI_ISL_2707205, EPI_ISL_2707206, EPI_ISL_2707207, EPI_ISL_2707208, EPI_ISL_2707209, EPI_ISL_2707210, EPI_ISL_2707211, EPI_ISL_2707212, EPI_ISL_2707213, EPI_ISL_2707214, EPI_ISL_2707215, EPI_ISL_2707216, EPI_ISL_2707217, EPI_ISL_2707218, EPI_ISL_2707219, EPI_ISL_2707220, EPI_ISL_2707221, EPI_ISL_2707222, EPI_ISL_2707223, EPI_ISL_2707224, EPI_ISL_2707225, EPI_ISL_2707226, EPI_ISL_2707227, EPI_ISL_2707228, EPI_ISL_2707229, EPI_ISL_2707230, EPI_ISL_2707231, EPI_ISL_2707232, EPI_ISL_2707233, EPI_ISL_2707234, EPI_ISL_2707236, EPI_ISL_2707237, EPI_ISL_2707238, EPI_ISL_2707239, EPI_ISL_2707240, EPI_ISL_2707241, EPI_ISL_2707242, EPI_ISL_2707243, EPI_ISL_2707244, EPI_ISL_2707245, EPI_ISL_2707246, EPI_ISL_2707247, EPI_ISL_2707248, EPI_ISL_2707249, EPI_ISL_2707251, EPI_ISL_2707252, EPI_ISL_2707253, EPI_ISL_2707254, EPI_ISL_2707255, EPI_ISL_2707256, EPI_ISL_2707257, EPI_ISL_2707258, EPI_ISL_2707259, EPI_ISL_2707260, EPI_ISL_2707261, EPI_ISL_2707262, EPI_ISL_2707263, EPI_ISL_2707264, EPI_ISL_2707265, EPI_ISL_2707266, EPI_ISL_2707267, EPI_ISL_2707268, EPI_ISL_2707269, EPI_ISL_2707270, EPI_ISL_2707271, EPI_ISL_2707272, EPI_ISL_2707273, EPI_ISL_2707274, EPI_ISL_2707275, EPI_ISL_2707276, EPI_ISL_2707277, EPI_ISL_2707278, EPI_ISL_2707279, EPI_ISL_2707280, EPI_ISL_2707281, EPI_ISL_2707282, EPI_ISL_2707283, EPI_ISL_2707284, EPI_ISL_2707285, EPI_ISL_2707286, EPI_ISL_2707287, EPI_ISL_2707288, EPI_ISL_2707289, EPI_ISL_2707290, EPI_ISL_2707291, EPI_ISL_2707292, EPI_ISL_2707293, EPI_ISL_2707294, EPI_ISL_2707295, EPI_ISL_2707296, EPI_ISL_2707297, EPI_ISL_2707298, EPI_ISL_2707299, EPI_ISL_2707300, EPI_ISL_2707301, EPI_ISL_2707302, EPI_ISL_2707303, EPI_ISL_2707304, EPI_ISL_2707305, EPI_ISL_2707306, EPI_ISL_2707307, EPI_ISL_2707308, EPI_ISL_2707310

see above | Quadram Institute Bioscience | COVID-19 Genomics UK (COG-UK) Consortium | Dave J. Baker, Gemma L. Kay, Alp Aydin, Thanh Le-Viet, Steven Rudder, Ana P. Tedim, Anastasia Kolyva, Maria Diaz, Leonardo de Oliveira Martins, Nabil-Fareed Alikhan, Lizzie Meadows, Rachael Stanley, Ngozi Elumogo, Muhammed Yasir, Nicholas M. Thomson, Alexander J Trotter, Rachel Gilroy, Samuel Bloomfield, Claire Stuart, Andrew Bell, Reenesh Prakash, Samir Dervisevic, Alison E. Mather, John Wain, Mark Webber, Andrew J. Page, Justin O'Grady

EPI_ISL_2707311, EPI_ISL_2707312, EPI_ISL_2707313, EPI_ISL_2707315, EPI_ISL_2707317, EPI_ISL_2707318, EPI_ISL_2707319, EPI_ISL_2707320, EPI_ISL_2707321, EPI_ISL_2707322, EPI_ISL_2707323, EPI_ISL_2707324, EPI_ISL_2707325, EPI_ISL_2707326, EPI_ISL_2707327, EPI_ISL_2707328, EPI_ISL_2707329, EPI_ISL_2707331, EPI_ISL_2707332, EPI_ISL_2707333, EPI_ISL_2707335, EPI_ISL_2707336, EPI_ISL_2707337, EPI_ISL_2707338, EPI_ISL_2707341, EPI_ISL_2707342, EPI_ISL_2707343, EPI_ISL_2707346, EPI_ISL_2707347, EPI_ISL_2707350, EPI_ISL_2707351, EPI_ISL_2707352, EPI_ISL_2707353, EPI_ISL_2707354, EPI_ISL_2707355, EPI_ISL_2707356, EPI_ISL_2707357, EPI_ISL_2707358

see above | Oxford Viromics, NDM, University of Oxford; Oxford University Hospitals; Basingstoke and North Hampshire Hospital | COVID-19 Genomics UK (COG-UK) Consortium | Tanya Golubchik, David Bonsall, George Macintyre, Amy Trebes, Mariateresa de Cesare, Catrin Moore, Alex Mobbs, Anita Justice, Robert Shaw, Monique Andersson, Timothy Peto, Emma Wise, Nathan Moore, Jessica Lynch, Nick Cortes, Matilde Mori, Stephen Kidd, David Buck, John Todd, Christophe Fraser

EPI_ISL_2707359, EPI_ISL_2707360, EPI_ISL_2707361, EPI_ISL_2707362, EPI_ISL_2707363, EPI_ISL_2707364, EPI_ISL_2707365, EPI_ISL_2707366, EPI_ISL_2707367, EPI_ISL_2707368, EPI_ISL_2707369, EPI_ISL_2707370, EPI_ISL_2707371, EPI_ISL_2707372, EPI_ISL_2707373, EPI_ISL_2707374, EPI_ISL_2707375, EPI_ISL_2707376, EPI_ISL_2707377, EPI_ISL_2707378, EPI_ISL_2707379, EPI_ISL_2707380, EPI_ISL_2707381, EPI_ISL_2707382, EPI_ISL_2707383, EPI_ISL_2707384, EPI_ISL_2707385, EPI_ISL_2707386, EPI_ISL_2707387, EPI_ISL_2707388, EPI_ISL_2707389, EPI_ISL_2707390, EPI_ISL_2707391, EPI_ISL_2707392, EPI_ISL_2707393, EPI_ISL_2707394, EPI_ISL_2707395, EPI_ISL_2707396, EPI_ISL_2707397, EPI_ISL_2707398, EPI_ISL_2707399, EPI_ISL_2707400, EPI_ISL_2707401, EPI_ISL_2707404, EPI_ISL_2707405, EPI_ISL_2707406, EPI_ISL_2707407, EPI_ISL_2707408, EPI_ISL_2707409, EPI_ISL_2707410, EPI_ISL_2707411, EPI_ISL_2707412, EPI_ISL_2707413, EPI_ISL_2707415, EPI_ISL_2707417, EPI_ISL_2707418, EPI_ISL_2707419, EPI_ISL_2707420, EPI_ISL_2707421, EPI_ISL_2707422, EPI_ISL_2707423, EPI_ISL_2707424, EPI_ISL_2707426, EPI_ISL_2707427, EPI_ISL_2707428, EPI_ISL_2707429, EPI_ISL_2707431, EPI_ISL_2707432, EPI_ISL_2707433, EPI_ISL_2707434, EPI_ISL_2707435, EPI_ISL_2707436, EPI_ISL_2707437, EPI_ISL_2707438, EPI_ISL_2707439, EPI_ISL_2707440, EPI_ISL_2707441, EPI_ISL_2707442, EPI_ISL_2707443, EPI_ISL_2707444, EPI_ISL_2707445, EPI_ISL_2707446, EPI_ISL_2707448, EPI_ISL_2707449, EPI_ISL_2707450, EPI_ISL_2707451, EPI_ISL_2707452, EPI_ISL_2707453, EPI_ISL_2707454, EPI_ISL_2707455, EPI_ISL_2707456, EPI_ISL_2707457, EPI_ISL_2707458, EPI_ISL_2707459, EPI_ISL_2707460, EPI_ISL_2707461, EPI_ISL_2707462, EPI_ISL_2707463, EPI_ISL_2707464, EPI_ISL_2707465, EPI_ISL_2707467, EPI_ISL_2707468, EPI_ISL_2707469, EPI_ISL_2707470, EPI_ISL_2707471, EPI_ISL_2707472, EPI_ISL_2707473, EPI_ISL_2707474, EPI_ISL_2707475, EPI_ISL_2707476, EPI_ISL_2707477, EPI_ISL_2707478, EPI_ISL_2707479, EPI_ISL_2707480, EPI_ISL_2707481, EPI_ISL_2707482, EPI_ISL_2707483, EPI_ISL_2707484, EPI_ISL_2707485, EPI_ISL_2707486, EPI_ISL_2707487, EPI_ISL_2707489, EPI_ISL_2707490, EPI_ISL_2707492, EPI_ISL_2707493, EPI_ISL_2707494, EPI_ISL_2707495, EPI_ISL_2707496, EPI_ISL_2707497, EPI_ISL_2707498, EPI_ISL_2707499, EPI_ISL_2707500, EPI_ISL_2707501, EPI_ISL_2707502, EPI_ISL_2707503, EPI_ISL_2707504, EPI_ISL_2707505, EPI_ISL_2707506, EPI_ISL_2707507, EPI_ISL_2707508, EPI_ISL_2707509, EPI_ISL_2707510, EPI_ISL_2707511, EPI_ISL_2707513, EPI_ISL_2707515, EPI_ISL_2707516, EPI_ISL_2707517, EPI_ISL_2707518, EPI_ISL_2707519, EPI_ISL_2707520, EPI_ISL_2707522, EPI_ISL_2707523, EPI_ISL_2707526, EPI_ISL_2707528, EPI_ISL_2707529, EPI_ISL_2707532, EPI_ISL_2707533, EPI_ISL_2707534, EPI_ISL_2707535, EPI_ISL_2707536, EPI_ISL_2707537, EPI_ISL_2707538, EPI_ISL_2707539, EPI_ISL_2707541, EPI_ISL_2707542, EPI_ISL_2707543, EPI_ISL_2707544, EPI_ISL_2707545, EPI_ISL_2707546, EPI_ISL_2707547, EPI_ISL_2707548, EPI_ISL_2707549, EPI_ISL_2707550, EPI_ISL_2707551, EPI_ISL_2707552, EPI_ISL_2707553, EPI_ISL_2707554, EPI_ISL_2707555, EPI_ISL_2707556, EPI_ISL_2707557, EPI_ISL_2707558, EPI_ISL_2707559, EPI_ISL_2707560, EPI_ISL_2707561, EPI_ISL_2707562, EPI_ISL_2707563, EPI_ISL_2707564, EPI_ISL_2707565, EPI_ISL_2707566, EPI_ISL_2707567, EPI_ISL_2707568, EPI_ISL_2707569, EPI_ISL_2707570, EPI_ISL_2707571, EPI_ISL_2707572, EPI_ISL_2707573, EPI_ISL_2707574, EPI_ISL_2707575, EPI_ISL_2707576, EPI_ISL_2707577, EPI_ISL_2707578, EPI_ISL_2707579, EPI_ISL_2707580, EPI_ISL_2707581, EPI_ISL_2707582, EPI_ISL_2707583, EPI_ISL_2707584, EPI_ISL_2707585, EPI_ISL_2707586, EPI_ISL_2707587, EPI_ISL_2707588, EPI_ISL_2707589, EPI_ISL_2707590, EPI_ISL_2707591, EPI_ISL_2707592, EPI_ISL_2707593, EPI_ISL_2707594, EPI_ISL_2707595, EPI_ISL_2707596, EPI_ISL_2707597, EPI_ISL_2707598, EPI_ISL_2707599, EPI_ISL_2707600, EPI_ISL_2707601, EPI_ISL_2707602, EPI_ISL_2707603, EPI_ISL_2707604, EPI_ISL_2707605, EPI_ISL_2707606, EPI_ISL_2707607, EPI_ISL_2707608, EPI_ISL_2707610, EPI_ISL_2707611, EPI_ISL_2707612, EPI_ISL_2707613, EPI_ISL_2707614, EPI_ISL_2707616, EPI_ISL_2707617, EPI_ISL_2707618, EPI_ISL_2707619, EPI_ISL_2707620, EPI_ISL_2707621, EPI_ISL_2707622, EPI_ISL_2707623, EPI_ISL_2707624, EPI_ISL_2707625, EPI_ISL_2707626, EPI_ISL_2707627, EPI_ISL_2707629, EPI_ISL_2707630, EPI_ISL_2707631, EPI_ISL_2707633, EPI_ISL_2707635, EPI_ISL_2707636, EPI_ISL_2707637, EPI_ISL_2707638, EPI_ISL_2707639, EPI_ISL_2707640, EPI_ISL_2707641, EPI_ISL_2707642, EPI_ISL_2707643, EPI_ISL_2707644, EPI_ISL_2707645, EPI_ISL_2707646, EPI_ISL_2707647, EPI_ISL_2707648, EPI_ISL_2707649, EPI_ISL_2707650, EPI_ISL_2707651, EPI_ISL_2707652, EPI_ISL_2707653, EPI_ISL_2707654, EPI_ISL_2707655, EPI_ISL_2707656, EPI_ISL_2707657, EPI_ISL_2707658, EPI_ISL_2707659, EPI_ISL_2707660, EPI_ISL_2707661, EPI_ISL_2707662, EPI_ISL_2707663, EPI_ISL_2707664, EPI_ISL_2707665, EPI_ISL_2707666, EPI_ISL_2707667, EPI_ISL_2707668, EPI_ISL_2707669, EPI_ISL_2707670, EPI_ISL_2707671, EPI_ISL_2707672, EPI_ISL_2707673, EPI_ISL_2707674, EPI_ISL_2707676, EPI_ISL_2707677, EPI_ISL_2707678, EPI_ISL_2707679, EPI_ISL_2707680, EPI_ISL_2707682, EPI_ISL_2707683, EPI_ISL_2707684, EPI_ISL_2707685, EPI_ISL_2707687, EPI_ISL_2707688, EPI_ISL_2707689, EPI_ISL_2707690, EPI_ISL_2707692, EPI_ISL_2707693, EPI_ISL_2707695, EPI_ISL_2707696, EPI_ISL_2707697, EPI_ISL_2707698, EPI_ISL_2707699, EPI_ISL_2707700, EPI_ISL_2707703, EPI_ISL_2707704, EPI_ISL_2707707, EPI_ISL_2707708, EPI_ISL_2707709, EPI_ISL_2707710, EPI_ISL_2707711, EPI_ISL_2707712, EPI_ISL_2707713, EPI_ISL_2707714, EPI_ISL_2707715, EPI_ISL_2707716, EPI_ISL_2707717, EPI_ISL_2707718, EPI_ISL_2707719, EPI_ISL_2707720, EPI_ISL_2707721, EPI_ISL_2707722, EPI_ISL_2707723, EPI_ISL_2707724, EPI_ISL_2707725, EPI_ISL_2707726, EPI_ISL_2707727, EPI_ISL_2707728, EPI_ISL_2707729, EPI_ISL_2707730, EPI_ISL_2707731, EPI_ISL_2707732, EPI_ISL_2707733, EPI_ISL_2707734, EPI_ISL_2707735, EPI_ISL_2707736, EPI_ISL_2707737, EPI_ISL_2707738, EPI_ISL_2707739, EPI_ISL_2707740, EPI_ISL_2707741, EPI_ISL_2707742, EPI_ISL_2707743, EPI_ISL_2707744, EPI_ISL_2707745, EPI_ISL_2707746, EPI_ISL_2707747, EPI_ISL_2707748, EPI_ISL_2707749, EPI_ISL_2707750, EPI_ISL_2707751, EPI_ISL_2707752, EPI_ISL_2707753, EPI_ISL_2707754, EPI_ISL_2707755, EPI_ISL_2707756, EPI_ISL_2707757, EPI_ISL_2707758, EPI_ISL_2707759, EPI_ISL_2707760, EPI_ISL_2707761, EPI_ISL_2707762, EPI_ISL_2707763, EPI_ISL_2707764, EPI_ISL_2707765, EPI_ISL_2707766, EPI_ISL_2707767, EPI_ISL_2707769, EPI_ISL_2707770, EPI_ISL_2707771, EPI_ISL_2707772, EPI_ISL_2707773, EPI_ISL_2707774, EPI_ISL_2707775, EPI_ISL_2707776, EPI_ISL_2707777, EPI_ISL_2707778, EPI_ISL_2707779, EPI_ISL_2707780, EPI_ISL_2707781, EPI_ISL_2707782, EPI_ISL_2707783, EPI_ISL_2707784, EPI_ISL_2707786, EPI_ISL_2707787, EPI_ISL_2707788, EPI_ISL_2707789, EPI_ISL_2707790, EPI_ISL_2707791, EPI_ISL_2707792, EPI_ISL_2707793, EPI_ISL_2707794, EPI_ISL_2707795, EPI_ISL_2707797, EPI_ISL_2707798, EPI_ISL_2707799, EPI_ISL_2707800, EPI_ISL_2707801, EPI_ISL_2707802, EPI_ISL_2707803, EPI_ISL_2707804, EPI_ISL_2707805, EPI_ISL_2707807, EPI_ISL_2707808, EPI_ISL_2707809, EPI_ISL_2707810, EPI_ISL_2707811, EPI_ISL_2707813, EPI_ISL_2707814, EPI_ISL_2707815, EPI_ISL_2707816, EPI_ISL_2707817, EPI_ISL_2707819, EPI_ISL_2707820, EPI_ISL_2707821, EPI_ISL_2707822, EPI_ISL_2707823, EPI_ISL_2707824, EPI_ISL_2707825, EPI_ISL_2707826, EPI_ISL_2707827, EPI_ISL_2707828, EPI_ISL_2707829, EPI_ISL_2707830, EPI_ISL_2707831, EPI_ISL_2707832, EPI_ISL_2707833, EPI_ISL_2707834, EPI_ISL_2707835, EPI_ISL_2707836, EPI_ISL_2707837, EPI_ISL_2707838, EPI_ISL_2707839, EPI_ISL_2707840, EPI_ISL_2707841, EPI_ISL_2707842, EPI_ISL_2707843, EPI_ISL_2707844, EPI_ISL_2707845, EPI_ISL_2707846, EPI_ISL_2707847, EPI_ISL_2707848, EPI_ISL_2707849, EPI_ISL_2707850, EPI_ISL_2707851, EPI_ISL_2707852, EPI_ISL_2707853, EPI_ISL_2707854, EPI_ISL_2707855, EPI_ISL_2707856, EPI_ISL_2707857, EPI_ISL_2707858, EPI_ISL_2707859, EPI_ISL_2707860, EPI_ISL_2707861, EPI_ISL_2707862, EPI_ISL_2707863, EPI_ISL_2707864, EPI_ISL_2707865, EPI_ISL_2707866, EPI_ISL_2707867, EPI_ISL_2707868, EPI_ISL_2707869, EPI_ISL_2707870, EPI_ISL_2707871, EPI_ISL_2707872, EPI_ISL_2707873, EPI_ISL_2707874, EPI_ISL_2707875, EPI_ISL_2707876, EPI_ISL_2707877, EPI_ISL_2707878, EPI_ISL_2707879, EPI_ISL_2707881, EPI_ISL_2707882, EPI_ISL_2707883, EPI_ISL_2707884, EPI_ISL_2707885, EPI_ISL_2707886, EPI_ISL_2707887, EPI_ISL_2707888, EPI_ISL_2707889, EPI_ISL_2707890, EPI_ISL_2707891, EPI_ISL_2707892, EPI_ISL_2707893, EPI_ISL_2707894, EPI_ISL_2707895, EPI_ISL_2707896, EPI_ISL_2707897, EPI_ISL_2707898, EPI_ISL_2707899, EPI_ISL_2707900, EPI_ISL_2707901, EPI_ISL_2707902, EPI_ISL_2707903, EPI_ISL_2707904, EPI_ISL_2707905, EPI_ISL_2707906, EPI_ISL_2707907, EPI_ISL_2707908, EPI_ISL_2707909, EPI_ISL_2707910, EPI_ISL_2707911, EPI_ISL_2707912, EPI_ISL_2707913, EPI_ISL_2707914, EPI_ISL_2707915, EPI_ISL_2707916, EPI_ISL_2707917, EPI_ISL_2707918, EPI_ISL_2707919, EPI_ISL_2707920, EPI_ISL_2707921, EPI_ISL_2707922, EPI_ISL_2707923, EPI_ISL_2707924, EPI_ISL_2707925, EPI_ISL_2707926, EPI_ISL_2707927, EPI_ISL_2707928, EPI_ISL_2707929, EPI_ISL_2707930, EPI_ISL_2707931, EPI_ISL_2707932, EPI_ISL_2707933, EPI_ISL_2707934, EPI_ISL_2707935, EPI_ISL_2707936, EPI_ISL_2707937, EPI_ISL_2707938, EPI_ISL_2707939, EPI_ISL_2707940, EPI_ISL_2707941, EPI_ISL_2707942, EPI_ISL_2707943, EPI_ISL_2707944, EPI_ISL_2707945, EPI_ISL_2707946, EPI_ISL_2707947, EPI_ISL_2707948, EPI_ISL_2707949, EPI_ISL_2707950, EPI_ISL_2707951, EPI_ISL_2707952, EPI_ISL_2707953, EPI_ISL_2707954, EPI_ISL_2707955, EPI_ISL_2707956, EPI_ISL_2707957, EPI_ISL_2707958, EPI_ISL_2707959, EPI_ISL_2707960, EPI_ISL_2707961, EPI_ISL_2707962, EPI_ISL_2707963, EPI_ISL_2707964, EPI_ISL_2707965, EPI_ISL_2707966, EPI_ISL_2707967, EPI_ISL_2707968, EPI_ISL_2707969, EPI_ISL_2707970, EPI_ISL_2707971, EPI_ISL_2707972, EPI_ISL_2707973, EPI_ISL_2707974, EPI_ISL_2707977, EPI_ISL_2707979, EPI_ISL_2707980, EPI_ISL_2707981, EPI_ISL_2707982, EPI_ISL_2707983, EPI_ISL_2707984, EPI_ISL_2707985, EPI_ISL_2707986, EPI_ISL_2707987, EPI_ISL_2707988, EPI_ISL_2707990, EPI_ISL_2707991, EPI_ISL_2707992, EPI_ISL_2707993, EPI_ISL_2707994, EPI_ISL_2707995, EPI_ISL_2707996, EPI_ISL_2707997, EPI_ISL_2707998, EPI_ISL_2707999, EPI_ISL_2708000, EPI_ISL_2708001, EPI_ISL_2708002, EPI_ISL_2708003, EPI_ISL_2708004, EPI_ISL_2708005, EPI_ISL_2708006, EPI_ISL_2708007, EPI_ISL_2708008, EPI_ISL_2708009, EPI_ISL_2708010, EPI_ISL_2708011, EPI_ISL_2708012, EPI_ISL_2708013, EPI_ISL_2708014, EPI_ISL_2708015, EPI_ISL_2708016, EPI_ISL_2708017, EPI_ISL_2708018, EPI_ISL_2708019, EPI_ISL_2708020, EPI_ISL_2708021, EPI_ISL_2708022, EPI_ISL_2708023, EPI_ISL_2708024, EPI_ISL_2708025, EPI_ISL_2708026, EPI_ISL_2708027, EPI_ISL_2708028, EPI_ISL_2708029, EPI_ISL_2708030, EPI_ISL_2708031, EPI_ISL_2708032, EPI_ISL_2708033, EPI_ISL_2708035, EPI_ISL_2708036, EPI_ISL_2708037, EPI_ISL_2708038, EPI_ISL_2708039, EPI_ISL_2708040, EPI_ISL_2708041, EPI_ISL_2708042, EPI_ISL_2708043, EPI_ISL_2708044, EPI_ISL_2708045, EPI_ISL_2708046, EPI_ISL_2708047, EPI_ISL_2708048, EPI_ISL_2708049, EPI_ISL_2708050, EPI_ISL_2708051, EPI_ISL_2708052, EPI_ISL_2708053, EPI_ISL_2708054, EPI_ISL_2708055, EPI_ISL_2708056, EPI_ISL_2708057, EPI_ISL_2708059, EPI_ISL_2708060, EPI_ISL_2708061, EPI_ISL_2708062, EPI_ISL_2708063, EPI_ISL_2708064, EPI_ISL_2708065, EPI_ISL_2708066, EPI_ISL_2708067, EPI_ISL_2708068, EPI_ISL_2708069, EPI_ISL_2708070, EPI_ISL_2708071, EPI_ISL_2708073, EPI_ISL_2708074, EPI_ISL_2708075, EPI_ISL_2708077, EPI_ISL_2708078, EPI_ISL_2708079, EPI_ISL_2708080, EPI_ISL_2708081, EPI_ISL_2708082, EPI_ISL_2708083, EPI_ISL_2708084, EPI_ISL_2708087, EPI_ISL_2708088, EPI_ISL_2708089, EPI_ISL_2708090, EPI_ISL_2708091, EPI_ISL_2708092, EPI_ISL_2708093, EPI_ISL_2708094, EPI_ISL_2708095, EPI_ISL_2708096, EPI_ISL_2708097, EPI_ISL_2708098, EPI_ISL_2708099, EPI_ISL_2708100, EPI_ISL_2708101, EPI_ISL_2708102, EPI_ISL_2708103, EPI_ISL_2708104, EPI_ISL_2708105, EPI_ISL_2708106, EPI_ISL_2708107, EPI_ISL_2708108, EPI_ISL_2708109, EPI_ISL_2708110, EPI_ISL_2708111, EPI_ISL_2708112, EPI_ISL_2708113, EPI_ISL_2708114, EPI_ISL_2708115, EPI_ISL_2708116, EPI_ISL_2708117, EPI_ISL_2708118, EPI_ISL_2708119, EPI_ISL_2708120, EPI_ISL_2708121, EPI_ISL_2708122, EPI_ISL_2708123, EPI_ISL_2708124, EPI_ISL_2708125, EPI_ISL_2708126, EPI_ISL_2708127, EPI_ISL_2708128, EPI_ISL_2708129, EPI_ISL_2708130, EPI_ISL_2708131, EPI_ISL_2708132, EPI_ISL_2708133, EPI_ISL_2708134, EPI_ISL_2708135, EPI_ISL_2708136, EPI_ISL_2708137, EPI_ISL_2708138, EPI_ISL_2708139, EPI_ISL_2708140, EPI_ISL_2708141, EPI_ISL_2708142, EPI_ISL_2708143, EPI_ISL_2708144

| | | | |
|---|---|---|---|
| see above | Originating lab: Wales Specialist Virology Centre Sequencing lab: Pathogen Genomics Unit | Public Health Wales Microbiology Cardiff Wales Specialist Virology Centre | Catherine Moore, Johnathan Evans, Laura Gifford, Malorie Perry, Simon Cottrell, Angela Marchbank, Alec Birchley, Alexander Adams, Amy Gaskin, Bree Gatica-Wilcox, Jason Coombes, Joel Southgate, Lauren Gilbert, Lee Graham, Nicole Pacchiarini, Sara Kumziene-Summerhayes, Sarah Taylor, Sophie Jones, Sara Rey, Matthew Bull, Joanne Watkins, Sally Corden, Tom Connor |
| EPI_ISL_2708145, EPI_ISL_2708146, EPI_ISL_2708147, EPI_ISL_2708148, EPI_ISL_2708149, EPI_ISL_2708150, EPI_ISL_2708151, EPI_ISL_2708152, EPI_ISL_2708153, EPI_ISL_2708154, EPI_ISL_2708155, EPI_ISL_2708156, EPI_ISL_2708157, EPI_ISL_2708158, EPI_ISL_2708159, EPI_ISL_2708160, EPI_ISL_2708161, EPI_ISL_2708162, EPI_ISL_2708163, EPI_ISL_2708164, EPI_ISL_2708165, EPI_ISL_2708166, EPI_ISL_2708167, EPI_ISL_2708168, EPI_ISL_2708169, EPI_ISL_2708170, EPI_ISL_2708171, EPI_ISL_2708172, EPI_ISL_2708173, EPI_ISL_2708174, EPI_ISL_2708175, EPI_ISL_2708176, EPI_ISL_2708177, EPI_ISL_2708178, EPI_ISL_2708179, EPI_ISL_2708180, EPI_ISL_2708181, EPI_ISL_2708182, EPI_ISL_2708183, EPI_ISL_2708184, EPI_ISL_2708185, EPI_ISL_2708186, EPI_ISL_2708187, EPI_ISL_2708188, EPI_ISL_2708189, EPI_ISL_2708190, EPI_ISL_2708191, EPI_ISL_2708192, EPI_ISL_2708193, EPI_ISL_2708194, EPI_ISL_2708195, EPI_ISL_2708196, EPI_ISL_2708197, EPI_ISL_2708198, EPI_ISL_2708199, EPI_ISL_2708200, EPI_ISL_2708201, EPI_ISL_2708202, EPI_ISL_2708203, EPI_ISL_2708204, EPI_ISL_2708205, EPI_ISL_2708206, EPI_ISL_2708207, EPI_ISL_2708208, EPI_ISL_2708209, EPI_ISL_2708210, EPI_ISL_2708211, EPI_ISL_2708212, EPI_ISL_2708213, EPI_ISL_2708214, EPI_ISL_2708215, EPI_ISL_2708216, EPI_ISL_2708217, EPI_ISL_2708218, EPI_ISL_2708219, EPI_ISL_2708220, EPI_ISL_2708221, EPI_ISL_2708222, EPI_ISL_2708223, EPI_ISL_2708224, EPI_ISL_2708225, EPI_ISL_2708226, EPI_ISL_2708227, EPI_ISL_2708228, EPI_ISL_2708229, EPI_ISL_2708230, EPI_ISL_2708231, EPI_ISL_2708232, EPI_ISL_2708233, EPI_ISL_2708234, EPI_ISL_2708235, EPI_ISL_2708236, EPI_ISL_2708237, EPI_ISL_2708238, EPI_ISL_2708239, EPI_ISL_2708240, EPI_ISL_2708241, EPI_ISL_2708242, EPI_ISL_2708243, EPI_ISL_2708244, EPI_ISL_2708245, EPI_ISL_2708246, EPI_ISL_2708247, EPI_ISL_2708248, EPI_ISL_2708249, EPI_ISL_2708250, EPI_ISL_2708251, EPI_ISL_2708252, EPI_ISL_2708253, EPI_ISL_2708254, EPI_ISL_2708255, EPI_ISL_2708256, EPI_ISL_2708257, EPI_ISL_2708258, EPI_ISL_2708259, EPI_ISL_2708260, EPI_ISL_2708261, EPI_ISL_2708262, EPI_ISL_2708263, EPI_ISL_2708264, EPI_ISL_2708265, EPI_ISL_2708266, EPI_ISL_2708267, EPI_ISL_2708268, EPI_ISL_2708269, EPI_ISL_2708270, EPI_ISL_2708271, EPI_ISL_2708272, EPI_ISL_2708273, EPI_ISL_2708274, EPI_ISL_2708275, EPI_ISL_2708276, EPI_ISL_2708277, EPI_ISL_2708278, EPI_ISL_2708279, EPI_ISL_2708280, EPI_ISL_2708281, EPI_ISL_2708282, EPI_ISL_2708283, EPI_ISL_2708284, EPI_ISL_2708285, EPI_ISL_2708286, EPI_ISL_2708287, EPI_ISL_2708288, EPI_ISL_2708289, EPI_ISL_2708290, EPI_ISL_2708291, EPI_ISL_2708292, EPI_ISL_2708293, EPI_ISL_2708294, EPI_ISL_2708295, EPI_ISL_2708296, EPI_ISL_2708297, EPI_ISL_2708298, EPI_ISL_2708299, EPI_ISL_2708300, EPI_ISL_2708301, EPI_ISL_2708302, EPI_ISL_2708303, EPI_ISL_2708304, EPI_ISL_2708305, EPI_ISL_2708306, EPI_ISL_2708307, EPI_ISL_2708308, EPI_ISL_2708309, EPI_ISL_2708310, EPI_ISL_2708311, EPI_ISL_2708312, EPI_ISL_2708313, EPI_ISL_2708314, EPI_ISL_2708315, EPI_ISL_2708316, EPI_ISL_2708317, EPI_ISL_2708318, EPI_ISL_2708319, EPI_ISL_2708320, EPI_ISL_2708321, EPI_ISL_2708322, EPI_ISL_2708323, EPI_ISL_2708324, EPI_ISL_2708325, EPI_ISL_2708326, EPI_ISL_2708327, EPI_ISL_2708328, EPI_ISL_2708329, EPI_ISL_2708330, EPI_ISL_2708331, EPI_ISL_2708332, EPI_ISL_2708333, EPI_ISL_2708334, EPI_ISL_2708335, EPI_ISL_2708336, EPI_ISL_2708337, EPI_ISL_2708338, EPI_ISL_2708340, EPI_ISL_2708341, EPI_ISL_2708342, EPI_ISL_2708343, EPI_ISL_2708344, EPI_ISL_2708345, EPI_ISL_2708346, EPI_ISL_2708347, EPI_ISL_2708348, EPI_ISL_2708349, EPI_ISL_2708350, EPI_ISL_2708351, EPI_ISL_2708352, EPI_ISL_2708353, EPI_ISL_2708354, EPI_ISL_2708355, EPI_ISL_2708356, EPI_ISL_2708357, EPI_ISL_2708358, EPI_ISL_2708359, EPI_ISL_2708360, EPI_ISL_2708361, EPI_ISL_2708362, EPI_ISL_2708363, EPI_ISL_2708364, EPI_ISL_2708365, EPI_ISL_2708366, EPI_ISL_2708367, EPI_ISL_2708368, EPI_ISL_2708369, EPI_ISL_2708370, EPI_ISL_2708371, EPI_ISL_2708372, EPI_ISL_2708373, EPI_ISL_2708374, EPI_ISL_2708375, EPI_ISL_2708376, EPI_ISL_2708377, EPI_ISL_2708378, EPI_ISL_2708379, EPI_ISL_2708380, EPI_ISL_2708381, EPI_ISL_2708382, EPI_ISL_2708383, EPI_ISL_2708384, EPI_ISL_2708385, EPI_ISL_2708386, EPI_ISL_2708387, EPI_ISL_2708388, EPI_ISL_2708389, EPI_ISL_2708390, EPI_ISL_2708391, EPI_ISL_2708392, EPI_ISL_2708393, EPI_ISL_2708394, EPI_ISL_2708395, EPI_ISL_2708396, EPI_ISL_2708397, EPI_ISL_2708398, EPI_ISL_2708399, EPI_ISL_2708400, EPI_ISL_2708401, EPI_ISL_2708402, EPI_ISL_2708403, EPI_ISL_2708404, EPI_ISL_2708405, EPI_ISL_2708406, EPI_ISL_2708407, EPI_ISL_2708408, EPI_ISL_2708409, EPI_ISL_2708410, EPI_ISL_2708411, EPI_ISL_2708412, EPI_ISL_2708413, EPI_ISL_2708414, EPI_ISL_2708415, EPI_ISL_2708416 | | | |
| see above | Centre for Enzyme Innovation, University of Portsmouth / Translational Research Laboratory, Portsmouth Hospitals NHS Trust / Hampshire Hospitals NHS Foundation Trust / Bournemouth University / University Hospitals Dorset NHS Foundation Trust / Brighton and Sussex University Hospitals NHS Trust / Dartford and Gravesham NHS Trust / Isle of Wight NHS Trust / Maidstone and Tunbridge Wells NHS Trust / University Hospital Southampton NHS Foundation Trust | COVID-19 Genomics UK (COG-UK) Consortium | Samuel Robson, Angela Beckett, Salman Goudarzi, Christopher Fearn, Kate Cook, Katie Loveson, Sharon Glaysher, Scott Elliott, Kelly Bicknell, Sarah Wyllie, Allyson Lloyd, Robert Impey, Anoop Chauhan, Stephen Kidd, Nathan Moore, Nick Cortes, Claire Thomas, Anna Mantzaouratou, Sarah Buchan, Magdalena Barrow, Andrew Butt, Liz Sheridan, Jonnie Seymour, Dorian Crudgington, Ben Macklin, Mohammed Hassan-Ibrahim, Cassandra Malone, Benjamin Cogger, Kevin Tucker, Samirakhon Raupova, Rachael Jeremiah, Anibolina Castigador, Emily Macnaughton, Karen Withell, Kordo Saeed, Jacqui Prieto, Adhyana Mahanama, Buddhini Samaraweera, Siona Silvieri, Emanuela Pelosi, Eleri Wilson-Davies, Sarah Jeremiah, Helen Wheeler, Matthew Harvey, Thea Sass, Helen Umpleby, Stephen Aplin |
| EPI_ISL_2708437, EPI_ISL_2708438, EPI_ISL_2708439, EPI_ISL_2708440, EPI_ISL_2708441, EPI_ISL_2708442, EPI_ISL_2708443, EPI_ISL_2708445, EPI_ISL_2708446, EPI_ISL_2708447, EPI_ISL_2708448, EPI_ISL_2708449, EPI_ISL_2708450, EPI_ISL_2708451, EPI_ISL_2708452, EPI_ISL_2708453, EPI_ISL_2708454, EPI_ISL_2708455, EPI_ISL_2708456, EPI_ISL_2708457, EPI_ISL_2708458, EPI_ISL_2708459, EPI_ISL_2708460, EPI_ISL_2708461, EPI_ISL_2708462, EPI_ISL_2708463, EPI_ISL_2708464, EPI_ISL_2708466, EPI_ISL_2708467, EPI_ISL_2708468, EPI_ISL_2708469, EPI_ISL_2708470, EPI_ISL_2708471, EPI_ISL_2708472, EPI_ISL_2708473, EPI_ISL_2708474, EPI_ISL_2708475, EPI_ISL_2708476, EPI_ISL_2708477, EPI_ISL_2708479, EPI_ISL_2708480, EPI_ISL_2708481, EPI_ISL_2708482, EPI_ISL_2708483, EPI_ISL_2708484, EPI_ISL_2708485, EPI_ISL_2708486, EPI_ISL_2708487, EPI_ISL_2708488, EPI_ISL_2708489, EPI_ISL_2708490, EPI_ISL_2708491, EPI_ISL_2708492, EPI_ISL_2708493, EPI_ISL_2708494, EPI_ISL_2708495, EPI_ISL_2708496, EPI_ISL_2708497, EPI_ISL_2708498, EPI_ISL_2708499, EPI_ISL_2708500, EPI_ISL_2708501, EPI_ISL_2708502, EPI_ISL_2708503, EPI_ISL_2708504, EPI_ISL_2708505, EPI_ISL_2708506, EPI_ISL_2708507, EPI_ISL_2708508, EPI_ISL_2708509, EPI_ISL_2708511, EPI_ISL_2708512, EPI_ISL_2708513, EPI_ISL_2708514, EPI_ISL_2708515, EPI_ISL_2708516, EPI_ISL_2708517, EPI_ISL_2708518, EPI_ISL_2708519, EPI_ISL_2708520, EPI_ISL_2708521, EPI_ISL_2708522, EPI_ISL_2708523, EPI_ISL_2708524, EPI_ISL_2708525, EPI_ISL_2708526, EPI_ISL_2708527, EPI_ISL_2708528, EPI_ISL_2708529, EPI_ISL_2708530, EPI_ISL_2708531, EPI_ISL_2708532, EPI_ISL_2708533, EPI_ISL_2708534, EPI_ISL_2708535, EPI_ISL_2708536, EPI_ISL_2708537, EPI_ISL_2708538, EPI_ISL_2708539, EPI_ISL_2708540, EPI_ISL_2708541, EPI_ISL_2708543, EPI_ISL_2708544, EPI_ISL_2708545, EPI_ISL_2708546, EPI_ISL_2708548, EPI_ISL_2708549, EPI_ISL_2708550, EPI_ISL_2708551, EPI_ISL_2708552, EPI_ISL_2708553, EPI_ISL_2708554, EPI_ISL_2708555, EPI_ISL_2708557, EPI_ISL_2708558, EPI_ISL_2708559, EPI_ISL_2708560, EPI_ISL_2708561, EPI_ISL_2708562, EPI_ISL_2708564, EPI_ISL_2708565, EPI_ISL_2708566, EPI_ISL_2708567, EPI_ISL_2708568, EPI_ISL_2708569, EPI_ISL_2708570, EPI_ISL_2708571, EPI_ISL_2708572, EPI_ISL_2708573, EPI_ISL_2708574, EPI_ISL_2708575, EPI_ISL_2708576, EPI_ISL_2708577, EPI_ISL_2708578, EPI_ISL_2708579, EPI_ISL_2708580, EPI_ISL_2708581, EPI_ISL_2708582, EPI_ISL_2708583, EPI_ISL_2708584, EPI_ISL_2708585, EPI_ISL_2708586, EPI_ISL_2708587, EPI_ISL_2708588, EPI_ISL_2708589, EPI_ISL_2708590, EPI_ISL_2708591, EPI_ISL_2708592, EPI_ISL_2708593, EPI_ISL_2708594, EPI_ISL_2708595, EPI_ISL_2708596, EPI_ISL_2708597, EPI_ISL_2708598, EPI_ISL_2708599, EPI_ISL_2708600, EPI_ISL_2708601, EPI_ISL_2708602, EPI_ISL_2708603, EPI_ISL_2708604, EPI_ISL_2708605, EPI_ISL_2708606, EPI_ISL_2708607, EPI_ISL_2708608, EPI_ISL_2708609, EPI_ISL_2708610, EPI_ISL_2708611, EPI_ISL_2708612, EPI_ISL_2708613, EPI_ISL_2708614, EPI_ISL_2708615, EPI_ISL_2708616, EPI_ISL_2708617, EPI_ISL_2708618, EPI_ISL_2708619, EPI_ISL_2708620, EPI_ISL_2708621, EPI_ISL_2708622, EPI_ISL_2708625, EPI_ISL_2708626, EPI_ISL_2708627, EPI_ISL_2708628, EPI_ISL_2708629, EPI_ISL_2708630, EPI_ISL_2708631, EPI_ISL_2708632, EPI_ISL_2708633, EPI_ISL_2708634, EPI_ISL_2708635, EPI_ISL_2708636, EPI_ISL_2708637, EPI_ISL_2708638, EPI_ISL_2708639, EPI_ISL_2708640, EPI_ISL_2708641, EPI_ISL_2708642, EPI_ISL_2708643, EPI_ISL_2708644, EPI_ISL_2708645, EPI_ISL_2708646, EPI_ISL_2708647, EPI_ISL_2708648, EPI_ISL_2708650, EPI_ISL_2708651, EPI_ISL_2708652, EPI_ISL_2708653, EPI_ISL_2708654, EPI_ISL_2708655, EPI_ISL_2708656, EPI_ISL_2708657, EPI_ISL_2708658, EPI_ISL_2708659, EPI_ISL_2708660, EPI_ISL_2708661, EPI_ISL_2708662, EPI_ISL_2708663, EPI_ISL_2708664, EPI_ISL_2708665, EPI_ISL_2708666, EPI_ISL_2708667, EPI_ISL_2708668, EPI_ISL_2708669, EPI_ISL_2708670, EPI_ISL_2708671, EPI_ISL_2708672, EPI_ISL_2708673, EPI_ISL_2708674, EPI_ISL_2708675, EPI_ISL_2708676, EPI_ISL_2708677, EPI_ISL_2708678, EPI_ISL_2708679, EPI_ISL_2708680, EPI_ISL_2708681, EPI_ISL_2708682, EPI_ISL_2708683, EPI_ISL_2708684, EPI_ISL_2708685, EPI_ISL_2708686, EPI_ISL_2708687, EPI_ISL_2708688, EPI_ISL_2708689, EPI_ISL_2708690, EPI_ISL_2708691, EPI_ISL_2708693, EPI_ISL_2708694, EPI_ISL_2708695, EPI_ISL_2708696, EPI_ISL_2708697, EPI_ISL_2708698, EPI_ISL_2708699, EPI_ISL_2708700, EPI_ISL_2708701, EPI_ISL_2708702, EPI_ISL_2708703, EPI_ISL_2708704, EPI_ISL_2708705, EPI_ISL_2708706, EPI_ISL_2708707, EPI_ISL_2708708, EPI_ISL_2708709, EPI_ISL_2708710, EPI_ISL_2708711, EPI_ISL_2708712, EPI_ISL_2708713, EPI_ISL_2708714, EPI_ISL_2708715, EPI_ISL_2708716, EPI_ISL_2708717, EPI_ISL_2708718, EPI_ISL_2708719, EPI_ISL_2708720, EPI_ISL_2708721, EPI_ISL_2708722, EPI_ISL_2708723, EPI_ISL_2708724, EPI_ISL_2708725, EPI_ISL_2708726, EPI_ISL_2708727, EPI_ISL_2708728, EPI_ISL_2708729, EPI_ISL_2708730, EPI_ISL_2708731, EPI_ISL_2708732, EPI_ISL_2708733, EPI_ISL_2708734, EPI_ISL_2708735, EPI_ISL_2708736, EPI_ISL_2708737, EPI_ISL_2708738, EPI_ISL_2708739, EPI_ISL_2708740, EPI_ISL_2708741, EPI_ISL_2708742, EPI_ISL_2708743, EPI_ISL_2708744, EPI_ISL_2708745, EPI_ISL_2708746, EPI_ISL_2708747, EPI_ISL_2708748, EPI_ISL_2708749, EPI_ISL_2708750, EPI_ISL_2708751, EPI_ISL_2708752, EPI_ISL_2708753, EPI_ISL_2708754, EPI_ISL_2708755, EPI_ISL_2708756, EPI_ISL_2708757, EPI_ISL_2708758, EPI_ISL_2708759, EPI_ISL_2708760, EPI_ISL_2708761, EPI_ISL_2708762, EPI_ISL_2708763, EPI_ISL_2708764, EPI_ISL_2708765, EPI_ISL_2708766, EPI_ISL_2708767, EPI_ISL_2708768, EPI_ISL_2708769, EPI_ISL_2708770, EPI_ISL_2708771, EPI_ISL_2708772, EPI_ISL_2708773, EPI_ISL_2708774, EPI_ISL_2708775, EPI_ISL_2708776, EPI_ISL_2708777, EPI_ISL_2708778, EPI_ISL_2708779, EPI_ISL_2708780, EPI_ISL_2708781, EPI_ISL_2708782, EPI_ISL_2708783, EPI_ISL_2708784, EPI_ISL_2708785, EPI_ISL_2708786, EPI_ISL_2708787, EPI_ISL_2708788, EPI_ISL_2708789, EPI_ISL_2708790, EPI_ISL_2708791, EPI_ISL_2708792, EPI_ISL_2708793, EPI_ISL_2708794, EPI_ISL_2708795, EPI_ISL_2708796, EPI_ISL_2708797, EPI_ISL_2708798, EPI_ISL_2708799, EPI_ISL_2708800, EPI_ISL_2708801, EPI_ISL_2708802, EPI_ISL_2708803, EPI_ISL_2708804, EPI_ISL_2708805, EPI_ISL_2708806, EPI_ISL_2708807, EPI_ISL_2708808, EPI_ISL_2708809, EPI_ISL_2708810, EPI_ISL_2708811, EPI_ISL_2708812, EPI_ISL_2708813, EPI_ISL_2708814, EPI_ISL_2708815, EPI_ISL_2708816, EPI_ISL_2708817, EPI_ISL_2708818, EPI_ISL_2708820, EPI_ISL_2708821, EPI_ISL_2708822, EPI_ISL_2708823, EPI_ISL_2708824, EPI_ISL_2708825, EPI_ISL_2708826, EPI_ISL_2708827, EPI_ISL_2708828, EPI_ISL_2708829, EPI_ISL_2708830, EPI_ISL_2708831, EPI_ISL_2708832, EPI_ISL_2708833, EPI_ISL_2708834, EPI_ISL_2708835, EPI_ISL_2708836, EPI_ISL_2708837, EPI_ISL_2708838, EPI_ISL_2708839, EPI_ISL_2708840, EPI_ISL_2708841 | | | |
| see above | Virology Department, Sheffield Teaching Hospitals NHS Foundation Trust/Department of Infection, Immunity and Cardiovascular Disease, The Medical School, University of Sheffield | COVID-19 Genomics UK (COG-UK) Consortium | Thushan de Silva, Matthew Parker, Nikki Smith, Adri Angyal, Rebecca Brown, Luke Green, Rachel Tucker, Paul Parsons, Danielle Groves, Katie Johnson, Laura Carrilero, Alex Keeley, Dave Partridge, Matthew Wyles, Benjamin Lindsey, Mehmet Yavuz, Mohammad Raza, Cariad Evans |
| EPI_ISL_2708963, EPI_ISL_2708966, EPI_ISL_2708969, EPI_ISL_2708970, EPI_ISL_2708971 | Laboratorio di Microbiologia | Laboratorio di Microbiologia | Martinetti Lucchini Gladys, Valeria Spina |
| EPI_ISL_2709235 | Public Health Laboratory, Minnesota Department of Health | University of Minnesota Genomics Center | Daryl M. Gohl, John Garbe, Jaquelyn Kuriger-Laber, Corbin Dirkx, and Sean Wang |
| EPI_ISL_2709249, EPI_ISL_2709256 | CHU Pontchaillou | CHU Pontchaillou | GROLHIER Claire, DENOUAL Florent, ETCHEVERRY Amandine, SASSI Mohamed, JAGLINE Steven, FEBREAU Christine, PRONIER Charlotte, GALIBERT Marie Dominique, DE TAYRAC Marie, THIBAULT Vincent |
| EPI_ISL_2709259 | Hospital | National Reference Center for Viruses of Respiratory Infections, Institut Pasteur, Paris | Marion Barbet, Sylvie Behillil, Méline Bizard, Angela Brisebarre, Camille Capel, Vincent Enouf, Louise Lefrançois, Frédéric Lemoine, Christophe Malabat, Corinne Maufrais, Damien Mornico, Etienne Simon-Lorière, Maud Vanpeene, Sylvie Van der Werf ,Laurent Bret |
| EPI_ISL_2709262 | Labo Analyses Med | National Reference Center for Viruses of Respiratory Infections, Institut Pasteur, Paris | Marion Barbet, Sylvie Behillil, Méline Bizard, Angela Brisebarre, Camille Capel, Vincent Enouf, Louise Lefrançois, Frédéric Lemoine, Christophe Malabat, Corinne Maufrais, Etienne Simon-Lorière, Maud Vanpeene, Sylvie Van der Werf ,Vicky Le |
| EPI_ISL_2709265 | Labo Analyses Med | National Reference Center for Viruses of Respiratory Infections, Institut Pasteur, Paris | Marion Barbet, Sylvie Behillil, Méline Bizard, Angela Brisebarre, Camille Capel, Vincent Enouf, Louise Lefrançois, Frédéric Lemoine, Christophe Malabat, Corinne Maufrais, Etienne Simon-Lorière, Maud Vanpeene, Sylvie Van der Werf ,Hugues Leroy |
| EPI_ISL_2709277 | Labo Analyses Med | National Reference Center for Viruses of Respiratory Infections, Institut Pasteur, Paris | Marion Barbet, Sylvie Behillil, Méline Bizard, Angela Brisebarre, Camille Capel, Vincent Enouf, Louise Lefrançois, Frédéric Lemoine, Christophe Malabat, Corinne Maufrais, Damien Mornico, Etienne Simon-Lorière, Maud Vanpeene, Sylvie Van der Werf ,Hugues Leroy |
| EPI_ISL_2709282, EPI_ISL_2709283, EPI_ISL_2709288 | Labo Analyses med | National Reference Center for Viruses of Respiratory Infections, Institut Pasteur, Paris | Marion Barbet, Sylvie Behillil, Méline Bizard, Angela Brisebarre, Camille Capel, Vincent Enouf, Louise Lefrançois, Frédéric Lemoine, Christophe Malabat, Corinne Maufrais, Etienne Simon-Lorière, Maud Vanpeene, Sylvie Van der Werf ,OphéLie Said-Delattre |
| EPI_ISL_2709296 | Labo Analyses med | National Reference Center for Viruses of Respiratory Infections, Institut Pasteur, Paris | Marion Barbet, Sylvie Behillil, Méline Bizard, Angela Brisebarre, Camille Capel, Vincent Enouf, Louise Lefrançois, Frédéric Lemoine, Christophe Malabat, Corinne Maufrais, Damien Mornico, Etienne Simon-Lorière, Maud Vanpeene, Sylvie Van der Werf ,OphéLie Said-Delattre |
| EPI_ISL_2709313, EPI_ISL_2709315, EPI_ISL_2709321, EPI_ISL_2709336, EPI_ISL_2709337, EPI_ISL_2709338, EPI_ISL_2709357 | Servicio Microbiología Hospital La Paz | Servicio Microbiología Hospital La Paz | Fernando Lázaro, Rubén Cáceres, Jesús Mingorance Cruz, Elie Dahdouh |
| EPI_ISL_2709358 | RS Pondok Indah Pondok Indah | Eijkman Institute for Molecular Biology, National Research and Innovation Agency | Frilasita A Yudhaputri, Muhammad Rezki Rasyak, Willy Agustine, Hidayat Trimarsanto, Lydia V. Panggalo, Iskandar Adnan, Sukma Oktavianthi, Lidwina Priliani, Edison Johar, Safarina G Malik, Khin Saw Myint, Amin Soebandrio |
| EPI_ISL_2709359 | RS OMNI Pulomas | Eijkman Institute for Molecular Biology, National Research and Innovation Agency | Edison Johar, Frilasita A Yudhaputri, Muhammad Rezki Rasyak, Willy Agustine, Hidayat Trimarsanto, Lydia V. Panggalo, Iskandar Adnan, Sukma Oktavianthi, Lidwina Priliani, Safarina G Malik, Khin Saw Myint, Amin Soebandrio |
| EPI_ISL_2709360 | RS OMNI Pulomas | Eijkman Institute for Molecular Biology, National Research and Innovation Agency | Iskandar Adnan, Sukma Oktavianthi, Lidwina Priliani, Edison Johar, Frilasita A Yudhaputri, Muhammad Rezki Rasyak, Willy Agustine, Hidayat Trimarsanto, Lydia V. Panggalo, Safarina G Malik, Khin Saw Myint, Amin Soebandrio |
| EPI_ISL_2709362 | RS Pusat Pertamina | Eijkman Institute for Molecular Biology, National Research and Innovation Agency | Muhammad Rezki Rasyak, Willy Agustine, Hidayat Trimarsanto, Lydia V. Panggalo, Iskandar Adnan, Sukma Oktavianthi, Lidwina Priliani, Edison Johar, Frilasita A Yudhaputri, Safarina G Malik, Khin Saw Myint, Amin Soebandrio |
| EPI_ISL_2709363 | RS Pusat Pertamina | Eijkman Institute for Molecular Biology, National Research and Innovation Agency | Lydia V. Panggalo, Iskandar Adnan, Sukma Oktavianthi, Lidwina Priliani, Edison Johar, Frilasita A Yudhaputri, Muhammad Rezki Rasyak, Willy Agustine, Hidayat Trimarsanto, Safarina G Malik, Khin Saw Myint, Amin Soebandrio |
| EPI_ISL_2709364 | RS Pusat Pertamina | Eijkman Institute for Molecular Biology, National Research and Innovation Agency | Edison Johar, Frilasita A Yudhaputri, Muhammad Rezki Rasyak, Willy Agustine, Hidayat Trimarsanto, Lydia V. Panggalo, Iskandar Adnan, Sukma Oktavianthi, Lidwina Priliani, Safarina G Malik, Khin Saw Myint, Amin Soebandrio |
| EPI_ISL_2709365 | RS Pusat Pertamina | Eijkman Institute for Molecular Biology, National Research and Innovation Agency | Hidayat Trimarsanto, Lydia V. Panggalo, Iskandar Adnan, Sukma Oktavianthi, Lidwina Priliani, Edison Johar, Frilasita A Yudhaputri, Muhammad Rezki Rasyak, Willy Agustine, Safarina G Malik, Khin Saw Myint, Amin Soebandrio |
| EPI_ISL_2709366 | RS Pelni | Eijkman Institute for Molecular Biology, National Research and Innovation Agency | Willy Agustine, Hidayat Trimarsanto, Lydia V. Panggalo, Iskandar Adnan, Sukma Oktavianthi, Lidwina Priliani, Edison Johar, Frilasita A Yudhaputri, Muhammad Rezki Rasyak, Safarina G Malik, Khin Saw Myint, Amin Soebandrio |
| EPI_ISL_2709367 | RS Pelni | Eijkman Institute for Molecular Biology, National Research and Innovation Agency | Muhammad Rezki Rasyak, Willy Agustine, Hidayat Trimarsanto, Lydia V. Panggalo, Iskandar Adnan, Sukma Oktavianthi, Lidwina Priliani, Edison Johar, Frilasita A Yudhaputri, Safarina G Malik, Khin Saw Myint, Amin Soebandrio |
| EPI_ISL_2709368 | RS Pelni | Eijkman Institute for Molecular Biology, National Research and Innovation Agency | Frilasita A Yudhaputri, Muhammad Rezki Rasyak, Willy Agustine, Hidayat Trimarsanto, Lydia V. Panggalo, Iskandar Adnan, Sukma Oktavianthi, Lidwina Priliani, Edison Johar, Safarina G Malik, Khin Saw Myint, Amin Soebandrio |
| EPI_ISL_2709370 | RS Pusat Pertamina | Eijkman Institute for Molecular Biology, National Research and Innovation Agency | Iskandar Adnan, Sukma Oktavianthi, Lidwina Priliani, Edison Johar, Frilasita A Yudhaputri, Muhammad Rezki Rasyak, Willy Agustine, Hidayat Trimarsanto, Lydia V. Panggalo, Safarina G Malik, Khin Saw Myint, Amin Soebandrio |
| EPI_ISL_2709371 | RS Pusat Pertamina | Eijkman Institute for Molecular Biology, National Research and Innovation Agency | Sukma Oktavianthi, Lidwina Priliani, Edison Johar, Frilasita A Yudhaputri, Muhammad Rezki Rasyak, Willy Agustine, Hidayat Trimarsanto, Lydia V. Panggalo, Iskandar Adnan, Safarina G Malik, Khin Saw Myint, Amin Soebandrio |
| EPI_ISL_2709372 | RS Pusat Pertamina | Eijkman Institute for Molecular Biology, National Research and | Muhammad Rezki Rasyak, Willy Agustine, Hidayat Trimarsanto, Lydia V. Panggalo, Iskandar Adnan, Sukma Oktavianthi, Lidwina Priliani, Edison Johar, Frilasita A Yudhaputri, Safarina G Malik, Khin Saw Myint, Amin Soebandrio |

| | | | |
|---|---|---|---|
| | | Innovation Agency | |
| EPI_ISL_2709373 | RS Pondok Indah Pondok Indah | Eijkman Institute for Molecular Biology, National Research and Innovation Agency | Lydia V. Panggalo, Iskandar Adnan, Sukma Oktavianthi, Lidwina Priliani, Edison Johar, Frilasita A Yudhaputri, Muhammad Rezki Rasyak, Willy Agustine, Hidayat Trimarsanto, Safarina G Malik, Khin Saw Myint, Amin Soebandrio |
| EPI_ISL_2709374 | RS Pondok Indah Pondok Indah | Eijkman Institute for Molecular Biology, National Research and Innovation Agency | Edison Johar, Frilasita A Yudhaputri, Muhammad Rezki Rasyak, Willy Agustine, Hidayat Trimarsanto, Lydia V. Panggalo, Iskandar Adnan, Sukma Oktavianthi, Lidwina Priliani, Safarina G Malik, Khin Saw Myint, Amin Soebandrio |
| EPI_ISL_2709376 | RS Pondok Indah Pondok Indah | Eijkman Institute for Molecular Biology, National Research and Innovation Agency | Willy Agustine, Hidayat Trimarsanto, Lydia V. Panggalo, Iskandar Adnan, Sukma Oktavianthi, Lidwina Priliani, Edison Johar, Frilasita A Yudhaputri, Muhammad Rezki Rasyak, Safarina G Malik, Khin Saw Myint, Amin Soebandrio |
| EPI_ISL_2709378 | RS OMNI Pulomas | Eijkman Institute for Molecular Biology, National Research and Innovation Agency | Frilasita A Yudhaputri, Muhammad Rezki Rasyak, Willy Agustine, Hidayat Trimarsanto, Lydia V. Panggalo, Iskandar Adnan, Sukma Oktavianthi, Lidwina Priliani, Edison Johar, Safarina G Malik, Khin Saw Myint, Amin Soebandrio |
| EPI_ISL_2709379 | RS Pusat Pertamina | Eijkman Institute for Molecular Biology, National Research and Innovation Agency | Edison Johar, Frilasita A Yudhaputri, Muhammad Rezki Rasyak, Willy Agustine, Hidayat Trimarsanto, Lydia V. Panggalo, Iskandar Adnan, Sukma Oktavianthi, Lidwina Priliani, Safarina G Malik, Khin Saw Myint, Amin Soebandrio |
| EPI_ISL_2709380 | RS Pusat Pertamina | Eijkman Institute for Molecular Biology, National Research and Innovation Agency | Iskandar Adnan, Sukma Oktavianthi, Lidwina Priliani, Edison Johar, Frilasita A Yudhaputri, Muhammad Rezki Rasyak, Willy Agustine, Hidayat Trimarsanto, Lydia V. Panggalo, Safarina G Malik, Khin Saw Myint, Amin Soebandrio |
| EPI_ISL_2709381 | RS Pondok Indah Pondok Indah | Eijkman Institute for Molecular Biology, National Research and Innovation Agency | Sukma Oktavianthi, Lidwina Priliani, Edison Johar, Frilasita A Yudhaputri, Muhammad Rezki Rasyak, Willy Agustine, Hidayat Trimarsanto, Lydia V. Panggalo, Iskandar Adnan, Safarina G Malik, Khin Saw Myint, Amin Soebandrio |
| EPI_ISL_2709383 | RS OMNI Pulomas | Eijkman Institute for Molecular Biology, National Research and Innovation Agency | Lydia V. Panggalo, Iskandar Adnan, Sukma Oktavianthi, Lidwina Priliani, Edison Johar, Frilasita A Yudhaputri, Muhammad Rezki Rasyak, Willy Agustine, Hidayat Trimarsanto, Safarina G Malik, Khin Saw Myint, Amin Soebandrio |
| EPI_ISL_2709384 | RSCM | Eijkman Institute for Molecular Biology, National Research and Innovation Agency | Edison Johar, Frilasita A Yudhaputri, Muhammad Rezki Rasyak, Willy Agustine, Hidayat Trimarsanto, Lydia V. Panggalo, Iskandar Adnan, Sukma Oktavianthi, Lidwina Priliani, Safarina G Malik, Khin Saw Myint, Amin Soebandrio |
| EPI_ISL_2709385 | RS Sari Asih Cipondoh | Eijkman Institute for Molecular Biology, National Research and Innovation Agency | Hidayat Trimarsanto, Lydia V. Panggalo, Iskandar Adnan, Sukma Oktavianthi, Lidwina Priliani, Edison Johar, Frilasita A Yudhaputri, Muhammad Rezki Rasyak, Willy Agustine, Safarina G Malik, Khin Saw Myint, Amin Soebandrio |
| EPI_ISL_2709388 | RSCM | Eijkman Institute for Molecular Biology, National Research and Innovation Agency | Frilasita A Yudhaputri, Muhammad Rezki Rasyak, Willy Agustine, Hidayat Trimarsanto, Lydia V. Panggalo, Iskandar Adnan, Sukma Oktavianthi, Lidwina Priliani, Edison Johar, Safarina G Malik, Khin Saw Myint, Amin Soebandrio |
| EPI_ISL_2709389 | RS OMNI Pulomas | Eijkman Institute for Molecular Biology, National Research and Innovation Agency | Edison Johar, Frilasita A Yudhaputri, Muhammad Rezki Rasyak, Willy Agustine, Hidayat Trimarsanto, Lydia V. Panggalo, Iskandar Adnan, Sukma Oktavianthi, Lidwina Priliani, Safarina G Malik, Khin Saw Myint, Amin Soebandrio |
| EPI_ISL_2709390 | RS OMNI Pulomas | Eijkman Institute for Molecular Biology, National Research and Innovation Agency | Iskandar Adnan, Sukma Oktavianthi, Lidwina Priliani, Edison Johar, Frilasita A Yudhaputri, Muhammad Rezki Rasyak, Willy Agustine, Hidayat Trimarsanto, Lydia V. Panggalo, Safarina G Malik, Khin Saw Myint, Amin Soebandrio |
| EPI_ISL_2709391 | RS Pondok Indah Pondok Indah | Eijkman Institute for Molecular Biology, National Research and Innovation Agency | Sukma Oktavianthi, Lidwina Priliani, Edison Johar, Frilasita A Yudhaputri, Muhammad Rezki Rasyak, Willy Agustine, Hidayat Trimarsanto, Lydia V. Panggalo, Iskandar Adnan, Safarina G Malik, Khin Saw Myint, Amin Soebandrio |
| EPI_ISL_2709392 | RS Sentra Medika Cibinong | Eijkman Institute for Molecular Biology, National Research and Innovation Agency | Muhammad Rezki Rasyak, Willy Agustine, Hidayat Trimarsanto, Lydia V. Panggalo, Iskandar Adnan, Sukma Oktavianthi, Lidwina Priliani, Edison Johar, Frilasita A Yudhaputri, Safarina G Malik, Khin Saw Myint, Amin Soebandrio |
| EPI_ISL_2709393 | RSU Adhyaksa | Eijkman Institute for Molecular Biology, National Research and Innovation Agency | Lydia V. Panggalo, Iskandar Adnan, Sukma Oktavianthi, Lidwina Priliani, Edison Johar, Frilasita A Yudhaputri, Muhammad Rezki Rasyak, Willy Agustine, Hidayat Trimarsanto, Safarina G Malik, Khin Saw Myint, Amin Soebandrio |
| EPI_ISL_2709394 | RS Permata Cibubur | Eijkman Institute for Molecular Biology, National Research and Innovation Agency | Edison Johar, Frilasita A Yudhaputri, Muhammad Rezki Rasyak, Willy Agustine, Hidayat Trimarsanto, Lydia V. Panggalo, Iskandar Adnan, Sukma Oktavianthi, Lidwina Priliani, Safarina G Malik, Khin Saw Myint, Amin Soebandrio |
| EPI_ISL_2709395 | RS OMNI Pulomas | Eijkman Institute for Molecular Biology, National Research and Innovation Agency | Hidayat Trimarsanto, Lydia V. Panggalo, Iskandar Adnan, Sukma Oktavianthi, Lidwina Priliani, Edison Johar, Frilasita A Yudhaputri, Muhammad Rezki Rasyak, Willy Agustine, Safarina G Malik, Khin Saw Myint, Amin Soebandrio |
| EPI_ISL_2709396 | RSU HGA Depok | Eijkman Institute for Molecular Biology, National Research and Innovation Agency | Willy Agustine, Hidayat Trimarsanto, Lydia V. Panggalo, Iskandar Adnan, Sukma Oktavianthi, Lidwina Priliani, Edison Johar, Frilasita A Yudhaputri, Muhammad Rezki Rasyak, Safarina G Malik, Khin Saw Myint, Amin Soebandrio |
| EPI_ISL_2709397 | RSU HGA Depok | Eijkman Institute for Molecular Biology, National Research and Innovation Agency | Muhammad Rezki Rasyak, Willy Agustine, Hidayat Trimarsanto, Lydia V. Panggalo, Iskandar Adnan, Sukma Oktavianthi, Lidwina Priliani, Edison Johar, Frilasita A Yudhaputri, Safarina G Malik, Khin Saw Myint, Amin Soebandrio |
| EPI_ISL_2709398 | RSU Adhyaksa | Eijkman Institute for Molecular Biology, National Research and Innovation Agency | Frilasita A Yudhaputri, Muhammad Rezki Rasyak, Willy Agustine, Hidayat Trimarsanto, Lydia V. Panggalo, Iskandar Adnan, Sukma Oktavianthi, Lidwina Priliani, Edison Johar, Safarina G Malik, Khin Saw Myint, Amin Soebandrio |
| EPI_ISL_2709399 | RSU Adhyaksa | Eijkman Institute for Molecular Biology, National Research and Innovation Agency | Edison Johar, Frilasita A Yudhaputri, Muhammad Rezki Rasyak, Willy Agustine, Hidayat Trimarsanto, Lydia V. Panggalo, Iskandar Adnan, Sukma Oktavianthi, Lidwina Priliani, Safarina G Malik, Khin Saw Myint, Amin Soebandrio |
| EPI_ISL_2709400 | RSUD Cileungsi | Eijkman Institute for Molecular Biology, National Research and Innovation Agency | Iskandar Adnan, Sukma Oktavianthi, Lidwina Priliani, Edison Johar, Frilasita A Yudhaputri, Muhammad Rezki Rasyak, Willy Agustine, Hidayat Trimarsanto, Lydia V. Panggalo, Safarina G Malik, Khin Saw Myint, Amin Soebandrio |
| EPI_ISL_2709401 | RS Pondok Indah Pondok Indah | Eijkman Institute for Molecular Biology, National Research and Innovation Agency | Sukma Oktavianthi, Lidwina Priliani, Edison Johar, Frilasita A Yudhaputri, Muhammad Rezki Rasyak, Willy Agustine, Hidayat Trimarsanto, Lydia V. Panggalo, Iskandar Adnan, Safarina G Malik, Khin Saw Myint, Amin Soebandrio |
| EPI_ISL_2709402 | RS Pondok Indah Pondok Indah | Eijkman Institute for Molecular Biology, National Research and Innovation Agency | Muhammad Rezki Rasyak, Willy Agustine, Hidayat Trimarsanto, Lydia V. Panggalo, Iskandar Adnan, Sukma Oktavianthi, Lidwina Priliani, Edison Johar, Frilasita A Yudhaputri, Safarina G Malik, Khin Saw Myint, Amin Soebandrio |
| EPI_ISL_2709403 | RS Premier Bintaro | Eijkman Institute for Molecular Biology, National Research and Innovation Agency | Lydia V. Panggalo, Iskandar Adnan, Sukma Oktavianthi, Lidwina Priliani, Edison Johar, Frilasita A Yudhaputri, Muhammad Rezki Rasyak, Willy Agustine, Hidayat Trimarsanto, Safarina G Malik, Khin Saw Myint, Amin Soebandrio |
| EPI_ISL_2709404 | West Java Health Laboratory | Eijkman Institute for Molecular Biology, National Research and Innovation Agency; West Java Health Laboratory | Edison Johar, Frilasita A Yudhaputri, Muhammad Rezki Rasyak, Willy Agustine, Hidayat Trimarsanto, Lydia V. Panggalo, Iskandar Adnan, Sukma Oktavianthi, Lidwina Priliani, Azzania Fibriani, Ema Rahmawati, Ryan Bayusantika Ristandi, Rifky Waluyajati Rachman, Cut Nur Cinthia Alamanda, Safarina G Malik, Khin Saw Myint, Amin Soebandrio |
| EPI_ISL_2709405 | West Java Health Laboratory | Eijkman Institute for Molecular Biology, National Research and Innovation Agency; West Java Health Laboratory | Frilasita A Yudhaputri, Muhammad Rezki Rasyak, Willy Agustine, Hidayat Trimarsanto, Lydia V. Panggalo, Iskandar Adnan, Sukma Oktavianthi, Lidwina Priliani, Edison Johar, Azzania Fibriani, Ema Rahmawati, Ryan Bayusantika Ristandi, Rifky Waluyajati Rachman, Cut Nur Cinthia Alamanda, Safarina G Malik, Khin Saw Myint, Amin Soebandrio |
| EPI_ISL_2709408 | West Java Health Laboratory | Eijkman Institute for Molecular Biology, National Research and Innovation Agency; West Java Health Laboratory | Iskandar Adnan, Sukma Oktavianthi, Lidwina Priliani, Edison Johar, Frilasita A Yudhaputri, Muhammad Rezki Rasyak, Willy Agustine, Hidayat Trimarsanto, Lydia V. Panggalo, Azzania Fibriani, Ema Rahmawati, Ryan Bayusantika Ristandi, Rifky Waluyajati Rachman, Cut Nur Cinthia Alamanda, Safarina G Malik, Khin Saw Myint, Amin Soebandrio |
| EPI_ISL_2709409 | West Java Health Laboratory | Eijkman Institute for Molecular Biology, National Research and Innovation Agency; West Java Health Laboratory | Sukma Oktavianthi, Lidwina Priliani, Edison Johar, Frilasita A Yudhaputri, Muhammad Rezki Rasyak, Willy Agustine, Hidayat Trimarsanto, Lydia V. Panggalo, Iskandar Adnan, Azzania Fibriani, Ema Rahmawati, Ryan Bayusantika Ristandi, Rifky Waluyajati Rachman, Cut Nur Cinthia Alamanda, Safarina G Malik, Khin Saw Myint, Amin Soebandrio |
| EPI_ISL_2709410 | West Java Health Laboratory | Eijkman Institute for Molecular Biology, National Research and Innovation Agency; West Java Health Laboratory | Lidwina Priliani, Edison Johar, Frilasita A Yudhaputri, Muhammad Rezki Rasyak, Willy Agustine, Hidayat Trimarsanto, Lydia V. Panggalo, Iskandar Adnan, Sukma Oktavianthi, Azzania Fibriani, Ema Rahmawati, Ryan Bayusantika Ristandi, Rifky Waluyajati Rachman, Cut Nur Cinthia Alamanda, Safarina G Malik, Khin Saw Myint, Amin Soebandrio |
| EPI_ISL_2709411 | West Java Health Laboratory | Eijkman Institute for Molecular Biology, National Research and Innovation Agency; West Java Health Laboratory | Edison Johar, Frilasita A Yudhaputri, Muhammad Rezki Rasyak, Willy Agustine, Hidayat Trimarsanto, Lydia V. Panggalo, Iskandar Adnan, Sukma Oktavianthi, Lidwina Priliani, Azzania Fibriani, Ema Rahmawati, Ryan Bayusantika Ristandi, Rifky Waluyajati Rachman, Cut Nur Cinthia Alamanda, Safarina G Malik, Khin Saw Myint, Amin Soebandrio |
| EPI_ISL_2709412 | West Java Health Laboratory | Eijkman Institute for Molecular Biology, National Research and Innovation Agency; West Java Health Laboratory | Frilasita A Yudhaputri, Muhammad Rezki Rasyak, Willy Agustine, Hidayat Trimarsanto, Lydia V. Panggalo, Iskandar Adnan, Sukma Oktavianthi, Lidwina Priliani, Edison Johar, Azzania Fibriani, Ema Rahmawati, Ryan Bayusantika Ristandi, Rifky Waluyajati Rachman, Cut Nur Cinthia Alamanda, Safarina G Malik, Khin Saw Myint, Amin Soebandrio |
| EPI_ISL_2709413 | West Java Health Laboratory | Eijkman Institute for Molecular Biology, National Research and Innovation Agency; West Java Health Laboratory | Muhammad Rezki Rasyak, Willy Agustine, Hidayat Trimarsanto, Lydia V. Panggalo, Iskandar Adnan, Sukma Oktavianthi, Lidwina Priliani, Edison Johar, Frilasita A Yudhaputri, Azzania Fibriani, Ema Rahmawati, Ryan Bayusantika Ristandi, Rifky Waluyajati Rachman, Cut Nur Cinthia Alamanda, Safarina G Malik, Khin Saw Myint, Amin Soebandrio |
| EPI_ISL_2709414 | West Java Health Laboratory | Eijkman Institute for Molecular Biology, National Research and Innovation Agency; West Java Health Laboratory | Willy Agustine, Hidayat Trimarsanto, Lydia V. Panggalo, Iskandar Adnan, Sukma Oktavianthi, Lidwina Priliani, Edison Johar, Frilasita A Yudhaputri, Muhammad Rezki Rasyak, Azzania Fibriani, Ema Rahmawati, Ryan Bayusantika Ristandi, Rifky Waluyajati Rachman, Cut Nur Cinthia Alamanda, Safarina G Malik, Khin Saw Myint, Amin Soebandrio |
| EPI_ISL_2709415 | West Java Health Laboratory | Eijkman Institute for Molecular Biology, National Research and Innovation Agency; West Java Health Laboratory | Hidayat Trimarsanto, Lydia V. Panggalo, Iskandar Adnan, Sukma Oktavianthi, Lidwina Priliani, Edison Johar, Frilasita A Yudhaputri, Muhammad Rezki Rasyak, Willy Agustine, Azzania Fibriani, Ema Rahmawati, Ryan Bayusantika Ristandi, Rifky Waluyajati Rachman, Cut Nur Cinthia Alamanda, Safarina G Malik, Khin Saw Myint, Amin Soebandrio |
| EPI_ISL_2709416 | West Java Health Laboratory | Eijkman Institute for Molecular Biology, National Research and Innovation Agency; West Java Health Laboratory | Lydia V. Panggalo, Iskandar Adnan, Sukma Oktavianthi, Lidwina Priliani, Edison Johar, Frilasita A Yudhaputri, Muhammad Rezki Rasyak, Willy Agustine, Hidayat Trimarsanto, Azzania Fibriani, Ema Rahmawati, Ryan Bayusantika Ristandi, Rifky Waluyajati Rachman, Cut Nur Cinthia Alamanda, Safarina G Malik, Khin Saw Myint, Amin Soebandrio |
| EPI_ISL_2709418 | West Java Health Laboratory | Eijkman Institute for Molecular Biology, National Research and Innovation Agency; West Java Health Laboratory | Sukma Oktavianthi, Lidwina Priliani, Edison Johar, Frilasita A Yudhaputri, Muhammad Rezki Rasyak, Willy Agustine, Hidayat Trimarsanto, Lydia V. Panggalo, Iskandar Adnan, Azzania Fibriani, Ema Rahmawati, Ryan Bayusantika Ristandi, Rifky Waluyajati Rachman, Cut Nur Cinthia Alamanda, Safarina G Malik, Khin Saw Myint, Amin Soebandrio |
| EPI_ISL_2709420 | West Java Health Laboratory | Eijkman Institute for Molecular Biology, National Research and Innovation Agency; West Java Health Laboratory | Edison Johar, Frilasita A Yudhaputri, Muhammad Rezki Rasyak, Willy Agustine, Hidayat Trimarsanto, Lydia V. Panggalo, Iskandar Adnan, Sukma Oktavianthi, Lidwina Priliani, Azzania Fibriani, Ema Rahmawati, Ryan Bayusantika Ristandi, Rifky Waluyajati Rachman, Cut Nur Cinthia Alamanda, Safarina G Malik, Khin Saw Myint, Amin Soebandrio |
| EPI_ISL_2709421 | West Java Health Laboratory | Eijkman Institute for Molecular Biology, National Research and Innovation Agency; West Java Health Laboratory | Frilasita A Yudhaputri, Muhammad Rezki Rasyak, Willy Agustine, Hidayat Trimarsanto, Lydia V. Panggalo, Iskandar Adnan, Sukma Oktavianthi, Lidwina Priliani, Edison Johar, Azzania Fibriani, Ema Rahmawati, Ryan Bayusantika Ristandi, Rifky Waluyajati Rachman, Cut Nur Cinthia Alamanda, Safarina G Malik, Khin Saw Myint, Amin Soebandrio |
| EPI_ISL_2709422 | West Java Health Laboratory | Eijkman Institute for Molecular Biology, National Research and Innovation Agency; West Java Health Laboratory | Muhammad Rezki Rasyak, Willy Agustine, Hidayat Trimarsanto, Lydia V. Panggalo, Iskandar Adnan, Sukma Oktavianthi, Lidwina Priliani, Edison Johar, Frilasita A Yudhaputri, Azzania Fibriani, Ema Rahmawati, Ryan Bayusantika Ristandi, Rifky Waluyajati Rachman, Cut Nur Cinthia Alamanda, Safarina G Malik, Khin Saw Myint, Amin Soebandrio |
| EPI_ISL_2709423 | West Java Health Laboratory | Eijkman Institute for Molecular Biology, National Research and Innovation Agency; West Java Health Laboratory | Willy Agustine, Hidayat Trimarsanto, Lydia V. Panggalo, Iskandar Adnan, Sukma Oktavianthi, Lidwina Priliani, Edison Johar, Frilasita A Yudhaputri, Muhammad Rezki Rasyak, Azzania Fibriani, Ema Rahmawati, Ryan Bayusantika Ristandi, Rifky Waluyajati Rachman, Cut Nur Cinthia Alamanda, Safarina G Malik, Khin Saw Myint, Amin Soebandrio |
| EPI_ISL_2709424 | West Java Health Laboratory | Eijkman Institute for Molecular Biology, National Research and Innovation Agency; West Java Health Laboratory | Hidayat Trimarsanto, Lydia V. Panggalo, Iskandar Adnan, Sukma Oktavianthi, Lidwina Priliani, Edison Johar, Frilasita A Yudhaputri, Muhammad Rezki Rasyak, Willy Agustine, Azzania Fibriani, Ema Rahmawati, Ryan Bayusantika Ristandi, Rifky Waluyajati Rachman, Cut Nur Cinthia Alamanda, Safarina G Malik, Khin Saw Myint, Amin Soebandrio |
| EPI_ISL_2709425 | West Java Health Laboratory | Eijkman Institute for Molecular Biology, National Research and Innovation Agency; West Java Health Laboratory | Lydia V. Panggalo, Iskandar Adnan, Sukma Oktavianthi, Lidwina Priliani, Edison Johar, Frilasita A Yudhaputri, Muhammad Rezki Rasyak, Willy Agustine, Hidayat Trimarsanto, Azzania Fibriani, Ema Rahmawati, Ryan Bayusantika Ristandi, Rifky Waluyajati Rachman, Cut Nur Cinthia Alamanda, Safarina G Malik, Khin Saw Myint, Amin Soebandrio |
| EPI_ISL_2709426 | West Java Health Laboratory | Eijkman Institute for Molecular Biology, National Research and | Iskandar Adnan, Sukma Oktavianthi, Lidwina Priliani, Edison Johar, Frilasita A Yudhaputri, Muhammad Rezki Rasyak, Willy Agustine, Hidayat Trimarsanto, Lydia V. Panggalo, Azzania Fibriani, Ema Rahmawati, Ryan Bayusantika Ristandi, Rifky |

| | | | |
|---|---|---|---|
| | | Innovation Agency; West Java Health Laboratory | Waluyajati Rachman, Cut Nur Cinthia Alamanda, Safarina G Malik, Khin Saw Myint, Amin Soebandrio |
| EPI_ISL_2709427 | West Java Health Laboratory | Eijkman Institute for Molecular Biology, National Research and Innovation Agency; West Java Health Laboratory | Lidwina Priliani, Edison Johar, Frilasita A Yudhaputri, Muhammad Rezki Rasyak, Willy Agustine, Hidayat Trimarsanto, Lydia V. Panggalo, Iskandar Adnan, Sukma Oktavianthi, Azzania Fibriani, Ema Rahmawati, Ryan Bayusantika Ristandi, Rifky Waluyajati Rachman, Cut Nur Cinthia Alamanda, Safarina G Malik, Khin Saw Myint, Amin Soebandrio |
| EPI_ISL_2709430 | West Java Health Laboratory | Eijkman Institute for Molecular Biology, National Research and Innovation Agency; West Java Health Laboratory | Muhammad Rezki Rasyak, Willy Agustine, Hidayat Trimarsanto, Lydia V. Panggalo, Iskandar Adnan, Sukma Oktavianthi, Lidwina Priliani, Edison Johar, Frilasita A Yudhaputri, Azzania Fibriani, Ema Rahmawati, Ryan Bayusantika Ristandi, Rifky Waluyajati Rachman, Cut Nur Cinthia Alamanda, Safarina G Malik, Khin Saw Myint, Amin Soebandrio |
| EPI_ISL_2709431 | West Java Health Laboratory | Eijkman Institute for Molecular Biology, National Research and Innovation Agency; West Java Health Laboratory | Willy Agustine, Hidayat Trimarsanto, Lydia V. Panggalo, Iskandar Adnan, Sukma Oktavianthi, Lidwina Priliani, Edison Johar, Frilasita A Yudhaputri, Muhammad Rezki Rasyak, Azzania Fibriani, Ema Rahmawati, Ryan Bayusantika Ristandi, Rifky Waluyajati Rachman, Cut Nur Cinthia Alamanda, Safarina G Malik, Khin Saw Myint, Amin Soebandrio |
| EPI_ISL_2709432 | West Java Health Laboratory | Eijkman Institute for Molecular Biology, National Research and Innovation Agency; West Java Health Laboratory | Hidayat Trimarsanto, Lydia V. Panggalo, Iskandar Adnan, Sukma Oktavianthi, Lidwina Priliani, Edison Johar, Frilasita A Yudhaputri, Muhammad Rezki Rasyak, Willy Agustine, Azzania Fibriani, Ema Rahmawati, Ryan Bayusantika Ristandi, Rifky Waluyajati Rachman, Cut Nur Cinthia Alamanda, Safarina G Malik, Khin Saw Myint, Amin Soebandrio |
| EPI_ISL_2709433 | West Java Health Laboratory | Eijkman Institute for Molecular Biology, National Research and Innovation Agency; West Java Health Laboratory | Lydia V. Panggalo, Iskandar Adnan, Sukma Oktavianthi, Lidwina Priliani, Edison Johar, Frilasita A Yudhaputri, Muhammad Rezki Rasyak, Willy Agustine, Hidayat Trimarsanto, Azzania Fibriani, Ema Rahmawati, Ryan Bayusantika Ristandi, Rifky Waluyajati Rachman, Cut Nur Cinthia Alamanda, Safarina G Malik, Khin Saw Myint, Amin Soebandrio |
| EPI_ISL_2709434 | West Java Health Laboratory | Eijkman Institute for Molecular Biology, National Research and Innovation Agency; West Java Health Laboratory | Sukma Oktavianthi, Lidwina Priliani, Edison Johar, Frilasita A Yudhaputri, Muhammad Rezki Rasyak, Willy Agustine, Hidayat Trimarsanto, Lydia V. Panggalo, Iskandar Adnan, Azzania Fibriani, Ema Rahmawati, Ryan Bayusantika Ristandi, Rifky Waluyajati Rachman, Cut Nur Cinthia Alamanda, Safarina G Malik, Khin Saw Myint, Amin Soebandrio |
| EPI_ISL_2709435 | West Java Health Laboratory | Eijkman Institute for Molecular Biology, National Research and Innovation Agency; West Java Health Laboratory | Lidwina Priliani, Edison Johar, Frilasita A Yudhaputri, Muhammad Rezki Rasyak, Willy Agustine, Hidayat Trimarsanto, Lydia V. Panggalo, Iskandar Adnan, Sukma Oktavianthi, Azzania Fibriani, Ema Rahmawati, Ryan Bayusantika Ristandi, Rifky Waluyajati Rachman, Cut Nur Cinthia Alamanda, Safarina G Malik, Khin Saw Myint, Amin Soebandrio |
| EPI_ISL_2709436 | West Java Health Laboratory | Eijkman Institute for Molecular Biology, National Research and Innovation Agency; West Java Health Laboratory | Edison Johar, Frilasita A Yudhaputri, Muhammad Rezki Rasyak, Willy Agustine, Hidayat Trimarsanto, Lydia V. Panggalo, Iskandar Adnan, Sukma Oktavianthi, Lidwina Priliani, Azzania Fibriani, Ema Rahmawati, Ryan Bayusantika Ristandi, Rifky Waluyajati Rachman, Cut Nur Cinthia Alamanda, Safarina G Malik, Khin Saw Myint, Amin Soebandrio |
| EPI_ISL_2709437 | West Java Health Laboratory | Eijkman Institute for Molecular Biology, National Research and Innovation Agency; West Java Health Laboratory | Frilasita A Yudhaputri, Muhammad Rezki Rasyak, Willy Agustine, Hidayat Trimarsanto, Lydia V. Panggalo, Iskandar Adnan, Sukma Oktavianthi, Lidwina Priliani, Edison Johar, Azzania Fibriani, Ema Rahmawati, Ryan Bayusantika Ristandi, Rifky Waluyajati Rachman, Cut Nur Cinthia Alamanda, Safarina G Malik, Khin Saw Myint, Amin Soebandrio |
| EPI_ISL_2709438 | West Java Health Laboratory | Eijkman Institute for Molecular Biology, National Research and Innovation Agency; West Java Health Laboratory | Muhammad Rezki Rasyak, Willy Agustine, Hidayat Trimarsanto, Lydia V. Panggalo, Iskandar Adnan, Sukma Oktavianthi, Lidwina Priliani, Edison Johar, Frilasita A Yudhaputri, Azzania Fibriani, Ema Rahmawati, Ryan Bayusantika Ristandi, Rifky Waluyajati Rachman, Cut Nur Cinthia Alamanda, Safarina G Malik, Khin Saw Myint, Amin Soebandrio |
| EPI_ISL_2709439 | West Java Health Laboratory | Eijkman Institute for Molecular Biology, National Research and Innovation Agency; West Java Health Laboratory | Willy Agustine, Hidayat Trimarsanto, Lydia V. Panggalo, Iskandar Adnan, Sukma Oktavianthi, Lidwina Priliani, Edison Johar, Frilasita A Yudhaputri, Muhammad Rezki Rasyak, Azzania Fibriani, Ema Rahmawati, Ryan Bayusantika Ristandi, Rifky Waluyajati Rachman, Cut Nur Cinthia Alamanda, Safarina G Malik, Khin Saw Myint, Amin Soebandrio |
| EPI_ISL_2709440 | West Java Health Laboratory | Eijkman Institute for Molecular Biology, National Research and Innovation Agency; West Java Health Laboratory | Hidayat Trimarsanto, Lydia V. Panggalo, Iskandar Adnan, Sukma Oktavianthi, Lidwina Priliani, Edison Johar, Frilasita A Yudhaputri, Muhammad Rezki Rasyak, Willy Agustine, Azzania Fibriani, Ema Rahmawati, Ryan Bayusantika Ristandi, Rifky Waluyajati Rachman, Cut Nur Cinthia Alamanda, Safarina G Malik, Khin Saw Myint, Amin Soebandrio |
| EPI_ISL_2709441 | West Java Health Laboratory | Eijkman Institute for Molecular Biology, National Research and Innovation Agency; West Java Health Laboratory | Lydia V. Panggalo, Iskandar Adnan, Sukma Oktavianthi, Lidwina Priliani, Edison Johar, Frilasita A Yudhaputri, Muhammad Rezki Rasyak, Willy Agustine, Hidayat Trimarsanto, Azzania Fibriani, Ema Rahmawati, Ryan Bayusantika Ristandi, Rifky Waluyajati Rachman, Cut Nur Cinthia Alamanda, Safarina G Malik, Khin Saw Myint, Amin Soebandrio |
| EPI_ISL_2709442 | West Java Health Laboratory | Eijkman Institute for Molecular Biology, National Research and Innovation Agency; West Java Health Laboratory | Iskandar Adnan, Sukma Oktavianthi, Lidwina Priliani, Edison Johar, Frilasita A Yudhaputri, Muhammad Rezki Rasyak, Willy Agustine, Hidayat Trimarsanto, Lydia V. Panggalo, Azzania Fibriani, Ema Rahmawati, Ryan Bayusantika Ristandi, Rifky Waluyajati Rachman, Cut Nur Cinthia Alamanda, Safarina G Malik, Khin Saw Myint, Amin Soebandrio |
| EPI_ISL_2709443 | West Java Health Laboratory | Eijkman Institute for Molecular Biology, National Research and Innovation Agency; West Java Health Laboratory | Sukma Oktavianthi, Lidwina Priliani, Edison Johar, Frilasita A Yudhaputri, Muhammad Rezki Rasyak, Willy Agustine, Hidayat Trimarsanto, Lydia V. Panggalo, Iskandar Adnan, Azzania Fibriani, Ema Rahmawati, Ryan Bayusantika Ristandi, Rifky Waluyajati Rachman, Cut Nur Cinthia Alamanda, Safarina G Malik, Khin Saw Myint, Amin Soebandrio |
| EPI_ISL_2709444 | West Java Health Laboratory | Eijkman Institute for Molecular Biology, National Research and Innovation Agency; West Java Health Laboratory | Lidwina Priliani, Edison Johar, Frilasita A Yudhaputri, Muhammad Rezki Rasyak, Willy Agustine, Hidayat Trimarsanto, Lydia V. Panggalo, Iskandar Adnan, Sukma Oktavianthi, Azzania Fibriani, Ema Rahmawati, Ryan Bayusantika Ristandi, Rifky Waluyajati Rachman, Cut Nur Cinthia Alamanda, Safarina G Malik, Khin Saw Myint, Amin Soebandrio |
| EPI_ISL_2709445 | West Java Health Laboratory | Eijkman Institute for Molecular Biology, National Research and Innovation Agency; West Java Health Laboratory | Edison Johar, Frilasita A Yudhaputri, Muhammad Rezki Rasyak, Willy Agustine, Hidayat Trimarsanto, Lydia V. Panggalo, Iskandar Adnan, Sukma Oktavianthi, Lidwina Priliani, Azzania Fibriani, Ema Rahmawati, Ryan Bayusantika Ristandi, Rifky Waluyajati Rachman, Cut Nur Cinthia Alamanda, Safarina G Malik, Khin Saw Myint, Amin Soebandrio |
| EPI_ISL_2709446 | West Java Health Laboratory | Eijkman Institute for Molecular Biology, National Research and Innovation Agency; West Java Health Laboratory | Frilasita A Yudhaputri, Muhammad Rezki Rasyak, Willy Agustine, Hidayat Trimarsanto, Lydia V. Panggalo, Iskandar Adnan, Sukma Oktavianthi, Lidwina Priliani, Edison Johar, Azzania Fibriani, Ema Rahmawati, Ryan Bayusantika Ristandi, Rifky Waluyajati Rachman, Cut Nur Cinthia Alamanda, Safarina G Malik, Khin Saw Myint, Amin Soebandrio |
| EPI_ISL_2709449, EPI_ISL_2709450, EPI_ISL_2709451, EPI_ISL_2709452, EPI_ISL_2709453, EPI_ISL_2709454, EPI_ISL_2709456, EPI_ISL_2709459, EPI_ISL_2709460, EPI_ISL_2709468, EPI_ISL_2709469, EPI_ISL_2709471, EPI_ISL_2709472, EPI_ISL_2709473, EPI_ISL_2709475 | | | |
| see above | IN State Department of Health Laboratory Services | IN State Department of Health Laboratory Services | Cassandra Campion, Jamie Yeadon, Brian Pope, Lixia Liu, Kyle Brownlee, Melissa Hindenlang, Mark Glazier |
| EPI_ISL_2709497, EPI_ISL_2709515, EPI_ISL_2709523, EPI_ISL_2709532, EPI_ISL_2709535, EPI_ISL_2709536, EPI_ISL_2709540, EPI_ISL_2709551, EPI_ISL_2709552 | Aegis Sciences Corporation | Centers for Disease Control and Prevention Division of Viral Diseases, Pathogen Discovery | Dakota Howard, Dhwani Batra, Peter W. Cook, Kara Moser, Adrian Paskey, Jason Caravas, Benjamin Rambo-Martin, Shatavia Morrison, Christopher Gulvick, Scott Sammons, Yvette Unoarumhi, Darlene Wagner, Matthew Schmerer, Cyndi Clark, Patrick Campbell, Rob Case, Vikramsinha Ghorpade, Holly Houdeshell, Ola Kvalvaag, Dillon Nall, Ethan Sanders, Alec Vest, Shaun Westlund, Matthew Hardison, Clinton R. Paden, Duncan MacCannell |
| EPI_ISL_2709589 | Tokyo Metropolitan Institute of Public Health | Tokyo Metropolitan Institute of Public Health | Hirofumi Miyake, Kenshirou Kuroki, Maya Isogai, Masaki Hayashi, Takako Yamazaki, Akane Negishi,Takushi Fujiwara, Mami Nagashima, Jun Suzuki, Kenji Sadamasu |
| EPI_ISL_2709603 | AZ Sint-Lucas Gent | AZ Sint-Lucas Gent | Jos Van Acker, Charlotte Verfaillie, Elke Vanlaere, Klara De Rauw |
| EPI_ISL_2709604, EPI_ISL_2709605 | ASL Napoli 1 Centro | AMES Centro Polidiagnostico Strumentale S.r.l. | "Giovanni Savarese, Raffaella Ruggiero, Eloisa Evangelista, Antonella Di Carlo, Luisa Circelli, Luigi D'Amore, Roberto Sirica, Antonio Fico" |
| EPI_ISL_2709736, EPI_ISL_2709737, EPI_ISL_2709742, EPI_ISL_2709749 | National Institute of Public Health | State Veterinary Institute Prague | Nagy,A;Cernikov,L;Stara,M;Suri,T;Vecerova,J;Jirincova,H |
| EPI_ISL_2709752, EPI_ISL_2709753 | Department of Virology, Istituto Zooprofilattico Sperimentale del Lazio e della Toscana (IZSLT) | Department of General Diagnostics; Department of Virology; Istituto Zooprofilattico Sperimentale del Lazio e della Toscana (IZSLT) | Patricia Alba, Giuseppe Manna, Elena L. Diaconu, Fabiola Feltrin, Raffaela Conti, Teresa Scicluna, Virginia Carfora, Antonella Cersini, Alessia Franco, Antonio Battisti. |
| EPI_ISL_2709762, EPI_ISL_2709766 | Laboratorio HUB -Azienda Ospedaliero Universitaria - AOU – Cagliari | Laboratorio SPOKE Biologia Molecolare -Azienda Ospedaliero Universitaria - AOU – Cagliari | Germano Orrù, Sara Fais, Valentina Medda, Alessandra Scano, Miriam Loddo, Riccardo Cappai, Ferdinando Coghe |
| EPI_ISL_2709790, EPI_ISL_2709794 | Institute of Molecular and Translational Medicine / Laboratory of Experimental Medicine, Faculty of Medicine and Dentistry, Palacky University and University Hospital Olomouc | Institute of Molecular and Translational Medicine / Laboratory of Experimental Medicine, Faculty of Medicine and Dentistry, Palacky University | Rastislav Slavkovský, Hana Dřímalová, Vladimíra Koudeláková, Barbora Blumová, Marián Hajdúch |
| EPI_ISL_2709800, EPI_ISL_2709802, EPI_ISL_2709805, EPI_ISL_2709817, EPI_ISL_2709823 | MD PHL | MD PHL | Maryland Department of Health Laboratories Administration |
| EPI_ISL_2709848 | MANKWENG LABORATORY | National Institute for Communicable Diseases of the National Health Laboratory Service | Amoako DG, Everatt J, Scheepers C, Mohale T, Ntuli N, Mahlangu B, Mnguni A, Ismail A, Bhiman JN |
| EPI_ISL_2709853, EPI_ISL_2709854, EPI_ISL_2709855, EPI_ISL_2709856, EPI_ISL_2709857, EPI_ISL_2709858, EPI_ISL_2709865, EPI_ISL_2709867, EPI_ISL_2709869 | POLOKWANE MANKWENG HOSPITAL | National Institute for Communicable Diseases of the National Health Laboratory Service | Amoako DG, Everatt J, Scheepers C, Mohale T, Ntuli N, Mahlangu B, Mnguni A, Ismail A, Bhiman JN |
| EPI_ISL_2709873, EPI_ISL_2709874 | HELEN JOSEPH | National Institute for Communicable Diseases of the National Health Laboratory Service | Amoako DG, Everatt J, Scheepers C, Mohale T, Ntuli N, Mahlangu B, Mnguni A, Ismail A, Bhiman JN |
| EPI_ISL_2709881, EPI_ISL_2709889, EPI_ISL_2709890, EPI_ISL_2709895, EPI_ISL_2709896, EPI_ISL_2709903, EPI_ISL_2709904, EPI_ISL_2709912, EPI_ISL_2709913, EPI_ISL_2709914, EPI_ISL_2709915, EPI_ISL_2709920, EPI_ISL_2709923, EPI_ISL_2709925, EPI_ISL_2709928, EPI_ISL_2709932, EPI_ISL_2709935, EPI_ISL_2709939, EPI_ISL_2709941 | | | |
| see above | CHRIS HANI BARAGWANATH | National Institute for Communicable Diseases of the National Health Laboratory Service | Amoako DG, Everatt J, Scheepers C, Mohale T, Ntuli N, Mahlangu B, Mnguni A, Ismail A, Bhiman JN |
| EPI_ISL_2709944, EPI_ISL_2709947, EPI_ISL_2709948, EPI_ISL_2709950, EPI_ISL_2709951, EPI_ISL_2709954, EPI_ISL_2709955, EPI_ISL_2709958, EPI_ISL_2709967, EPI_ISL_2709970, EPI_ISL_2709973, EPI_ISL_2709975, EPI_ISL_2709977 | | | |
| see above | TAMBO MEMORIAL LABORATORY | National Institute for Communicable Diseases of the National Health Laboratory Service | Amoako DG, Everatt J, Scheepers C, Mohale T, Ntuli N, Mahlangu B, Mnguni A, Ismail A, Bhiman JN |
| EPI_ISL_2709978 | PHOLOSONG LABORATORY | National Institute for Communicable Diseases of the National Health Laboratory Service | Amoako DG, Everatt J, Scheepers C, Mohale T, Ntuli N, Mahlangu B, Mnguni A, Ismail A, Bhiman JN |
| EPI_ISL_2709980, EPI_ISL_2709990, EPI_ISL_2709991, EPI_ISL_2709994 | TAMBO MEMORIAL LABORATORY | National Institute for Communicable Diseases of the National Health Laboratory Service | Amoako DG, Everatt J, Scheepers C, Mohale T, Ntuli N, Mahlangu B, Mnguni A, Ismail A, Bhiman JN |

| | | | |
|---|---|---|---|
| EPI_ISL_2709998 | Public Health Authority of the Slovak Republic | Laboratory of Genomics and Bioinformatics, Comenius University Science Park | Tatiana Sedláčková, Diana Rusňáková, Miroslav Böhmer, Anna Gičová, Jaroslav Budiš, Tomáš Szemes, Jakub Styk |
| EPI_ISL_2710003, EPI_ISL_2710004, EPI_ISL_2710005, EPI_ISL_2710006, EPI_ISL_2710008, EPI_ISL_2710009, EPI_ISL_2710010, EPI_ISL_2710011 | Regional VRDL,Dibrugarh | Regional VRDL,Dibrugarh | B Borkakoty, Mandakini Das Sarma, R Hazarika, C Bhattacharya, A Jakharia, D Biswas, K Narain |
| EPI_ISL_2710084, EPI_ISL_2710085, EPI_ISL_2710086, EPI_ISL_2710087, EPI_ISL_2710088, EPI_ISL_2710089, EPI_ISL_2710090, EPI_ISL_2710091, EPI_ISL_2710092, EPI_ISL_2710093, EPI_ISL_2710094, EPI_ISL_2710095, EPI_ISL_2710096, EPI_ISL_2710097, EPI_ISL_2710098, EPI_ISL_2710099, EPI_ISL_2710100, EPI_ISL_2710101, EPI_ISL_2710102, EPI_ISL_2710103, EPI_ISL_2710104, EPI_ISL_2710105, EPI_ISL_2710106, EPI_ISL_2710107, EPI_ISL_2710108, EPI_ISL_2710109, EPI_ISL_2710110, EPI_ISL_2710111, EPI_ISL_2710112, EPI_ISL_2710113, EPI_ISL_2710114, EPI_ISL_2710115, EPI_ISL_2710116, EPI_ISL_2710117, EPI_ISL_2710118, EPI_ISL_2710119, EPI_ISL_2710120, EPI_ISL_2710121, EPI_ISL_2710122, EPI_ISL_2710123, EPI_ISL_2710124, EPI_ISL_2710125, EPI_ISL_2710126, EPI_ISL_2710127, EPI_ISL_2710128, EPI_ISL_2710129, EPI_ISL_2710130, EPI_ISL_2710131, EPI_ISL_2710132, EPI_ISL_2710133, EPI_ISL_2710134, EPI_ISL_2710135, EPI_ISL_2710136, EPI_ISL_2710137, EPI_ISL_2710138, EPI_ISL_2710139, EPI_ISL_2710140, EPI_ISL_2710141, EPI_ISL_2710142, EPI_ISL_2710143, EPI_ISL_2710144, EPI_ISL_2710145, EPI_ISL_2710146, EPI_ISL_2710147, EPI_ISL_2710148, EPI_ISL_2710149, EPI_ISL_2710150, EPI_ISL_2710151, EPI_ISL_2710152, EPI_ISL_2710153, EPI_ISL_2710154, EPI_ISL_2710155, EPI_ISL_2710156, EPI_ISL_2710157, EPI_ISL_2710158, EPI_ISL_2710159, EPI_ISL_2710160, EPI_ISL_2710161, EPI_ISL_2710162, EPI_ISL_2710163, EPI_ISL_2710164, EPI_ISL_2710165, EPI_ISL_2710166, EPI_ISL_2710167, EPI_ISL_2710168, EPI_ISL_2710169, EPI_ISL_2710170, EPI_ISL_2710171 | | | |
| see above | National Public Health Laboratory, National Centre for Infectious Diseases | National Public Health Laboratory, National Centre for Infectious Diseases | Tze Minn Mak, Katherine Ching, Zhenyang Zhou, Royce Ang, Lin Cui, Raymond Tzer Pin Lin |
| EPI_ISL_2710174, EPI_ISL_2710175, EPI_ISL_2710176 | HOSPITAL UNIVERSITARIO SON ESPASES | HOSPITAL UNIVERSITARIO SON ESPASES | Servicio de Microbiología, Hospital Universitario Son Espases |
| EPI_ISL_2710177 | HOSPITAL UNIVERSITARIO SON LLATZER | HOSPITAL UNIVERSITARIO SON ESPASES | Servicio de Microbiología, Hospital Universitario Son Espases |
| EPI_ISL_2710179, EPI_ISL_2710180, EPI_ISL_2710182, EPI_ISL_2710189, EPI_ISL_2710194, EPI_ISL_2710198, EPI_ISL_2710200 | HOSPITAL UNIVERSITARIO SON ESPASES | HOSPITAL UNIVERSITARIO SON ESPASES | Servicio de Microbiología, Hospital Universitario Son Espases |
| EPI_ISL_2710203 | HOSPITAL CAN MISSES | HOSPITAL UNIVERSITARIO SON ESPASES | Servicio de Microbiología, Hospital Universitario Son Espases |
| EPI_ISL_2710206, EPI_ISL_2710209, EPI_ISL_2710211 | HOSPITAL UNIVERSITARIO SON ESPASES | HOSPITAL UNIVERSITARIO SON ESPASES | Servicio de Microbiología, Hospital Universitario Son Espases |
| EPI_ISL_2710216 | HOSPITAL CAN MISSES | HOSPITAL UNIVERSITARIO SON ESPASES | Servicio de Microbiología, Hospital Universitario Son Espases |
| EPI_ISL_2710230, EPI_ISL_2710232, EPI_ISL_2710234, EPI_ISL_2710235, EPI_ISL_2710236, EPI_ISL_2710240 | HOSPITAL UNIVERSITARIO SON ESPASES | HOSPITAL UNIVERSITARIO SON ESPASES | Servicio de Microbiología, Hospital Universitario Son Espases |
| EPI_ISL_2710249, EPI_ISL_2710250, EPI_ISL_2710252, EPI_ISL_2710253, EPI_ISL_2710255, EPI_ISL_2710256, EPI_ISL_2710257, EPI_ISL_2710258, EPI_ISL_2710259, EPI_ISL_2710260, EPI_ISL_2710261, EPI_ISL_2710262, EPI_ISL_2710264, EPI_ISL_2710265, EPI_ISL_2710266, EPI_ISL_2710267, EPI_ISL_2710268, EPI_ISL_2710269, EPI_ISL_2710270, EPI_ISL_2710271, EPI_ISL_2710272, EPI_ISL_2710273, EPI_ISL_2710274, EPI_ISL_2710275, EPI_ISL_2710277, EPI_ISL_2710278, EPI_ISL_2710280, EPI_ISL_2710284, EPI_ISL_2710286, EPI_ISL_2710287, EPI_ISL_2710288, EPI_ISL_2710290, EPI_ISL_2710292, EPI_ISL_2710294, EPI_ISL_2710295, EPI_ISL_2710297, EPI_ISL_2710298, EPI_ISL_2710299, EPI_ISL_2710300, EPI_ISL_2710301, EPI_ISL_2710302, EPI_ISL_2710303 | | | |
| see above | NHLS Charlotte Maxeke Johannesburg Academic Hospital and the University of the Witwatersrand | KRISP, KZN Research Innovation and Sequencing Platform | Florette Treurnicht, Bulelani Manene, Kathleen Subramoney, Giandhari Jennifer, Pillay Sureshnee, Yajna Ramphal, Naidoo Yeshnee, Tshabuila Derek,Tegally Houriiyah, San James, Wilkinson Eduan, de Oliveira Tulio |
| EPI_ISL_2710308, EPI_ISL_2710312, EPI_ISL_2710316, EPI_ISL_2710318, EPI_ISL_2710319, EPI_ISL_2710320 | BARC / Lancet | KRISP, KZN Research Innovation and Sequencing Platform | Sisonke, Giandhari Jennifer, Pillay Sureshnee, Yajna Ramphal, Naidoo Yeshnee, Tshabuila Derek,Tegally Houriiyah, San James, Wilkinson Eduan, de Oliveira Tulio |
| EPI_ISL_2710346 | Montana Public Health Laboratory | Montana Public Health Laboratory | Joy Ritter, Michelle Mozer, Carrie Biskupiak, Deborah Gibson, Michael Dills |
| EPI_ISL_2710362, EPI_ISL_2710363, EPI_ISL_2710364, EPI_ISL_2710370, EPI_ISL_2710371, EPI_ISL_2710372, EPI_ISL_2710373, EPI_ISL_2710374, EPI_ISL_2710375, EPI_ISL_2710376, EPI_ISL_2710377 | | | |
| see above | Kansas Health and Environmental Lab | Kansas Health and Environmental Lab | Mike Grose, Katherine Wiggins, Jonathan Barnell, Ben Olsen, and Phil Adam |
| EPI_ISL_2710382 | CT Department of Public Health | CT Department of Public Health | Claire Pearson, Tu N. Nguyen |
| EPI_ISL_2710625, EPI_ISL_2710640, EPI_ISL_2710644, EPI_ISL_2710652, EPI_ISL_2710707, EPI_ISL_2710708, EPI_ISL_2710709, EPI_ISL_2710711, EPI_ISL_2710712, EPI_ISL_2710713 | National Virus Reference Laboratory | National Virus Reference Laboratory | Zoe Yandle, Charlene Bennett, Gabriel Gonzalez, Michael Carr, Jonathan Dean, Cillian F De Gascun |
| EPI_ISL_2710798 | Orebro University Hospital, Dept Laboratory Medicine, Clinical Microbiology | Orebro University Hospital | Sundqvist M et al |
| EPI_ISL_2712544, EPI_ISL_2712545, EPI_ISL_2712546, EPI_ISL_2712548, EPI_ISL_2712551 | Sharp HealthCare Laboratory | Andersen lab at Scripps Research | SEARCH Alliance San Diego with Aaron Harding, Jacquelyn Berumen, Cathy Woerle, Liam McGinnis, Art Mendoza, Omid Bakhtar |
| EPI_ISL_2712846, EPI_ISL_2712849, EPI_ISL_2712851, EPI_ISL_2712854, EPI_ISL_2712856, EPI_ISL_2712858, EPI_ISL_2712859, EPI_ISL_2712862, EPI_ISL_2712864, EPI_ISL_2712865, EPI_ISL_2712869, EPI_ISL_2712874, EPI_ISL_2712875, EPI_ISL_2712878, EPI_ISL_2712885, EPI_ISL_2712887, EPI_ISL_2712893, EPI_ISL_2712894 | | | |
| see above | Hospital Universitari Vall d'Hebron - Vall d'Hebron Institut de Recerca | Hospital Universitari Vall d'Hebron - Vall d'Hebron Institut de Recerca | Cristina Andrés, Maria Piñana, Alejandra González-Sánchez, Damir Garcia-Cehic, Ariadna Rando, Juliana Esperalba, Maria Gema Codina, Carla Castillo, Maria Carmen Martin, Tomàs Pumarola, Josep Quer, Andrés Antón |
| EPI_ISL_2712922, EPI_ISL_2712925, EPI_ISL_2712928, EPI_ISL_2712929, EPI_ISL_2712930, EPI_ISL_2712932, EPI_ISL_2712933, EPI_ISL_2712935, EPI_ISL_2712939, EPI_ISL_2712941, EPI_ISL_2712942, EPI_ISL_2712947, EPI_ISL_2712948, EPI_ISL_2712949, EPI_ISL_2712950, EPI_ISL_2712951, EPI_ISL_2712952, EPI_ISL_2712954, EPI_ISL_2712957, EPI_ISL_2712958, EPI_ISL_2712959, EPI_ISL_2712960, EPI_ISL_2712961, EPI_ISL_2712962, EPI_ISL_2712963, EPI_ISL_2712964, EPI_ISL_2712965, EPI_ISL_2712966, EPI_ISL_2712967 | | | |
| see above | Wyoming Public Health Laboratory | Wyoming Public Health Laboratory | Jim Mildenberger, Taylor Fearing, Channing Weber, Ashley Norberg, Chayse Rowley, Marley Goetz, Brian Dominguez, Elliot Thomasson, Sam Britz, Cari Sloma, Robert Petit, and Rob Christensen |
| EPI_ISL_2712987 | ISDH | Quantigen Biosciences | Paul J. Childress |
| EPI_ISL_2712994, EPI_ISL_2712995, EPI_ISL_2712998 | TXDSHS | TXDSHS | Rashmi Tuladhar, Bonnie Oh, Jenny Zhang, Maliha Rahman, Mayela Pedrueza, Anita Pokharel, Karen Bobier, Lorraine Rodriguez, Myong Koag, Chun Wang, Rachel Lee, Grace Kubin |
| EPI_ISL_2713007 | Wyoming Public Health Laboratory | Wyoming Public Health Laboratory | Jim Mildenberger, Taylor Fearing, Channing Weber, Ashley Norberg, Chayse Rowley, Marley Goetz, Brian Dominguez, Elliot Thomasson, Sam Britz, Cari Sloma, Robert Petit, and Rob Christensen |
| EPI_ISL_2713050, EPI_ISL_2713058 | University of Wisconsin-Madison AIDS Vaccine Research Laboratories | University of Wisconsin-Madison AIDS Vaccine Research Laboratories | Gage Moreno, Katarina Braun, et al. AIDS Vaccine Research Laboratories |
| EPI_ISL_2713066 | Oregon State Public Health Laboratory | Oregon State Public Health Laboratory | Rafia Razzaque, Eugene Yeboah, Vanda Makris, Laura Tsaknaridis, John Fontana and Shane Sevey |
| EPI_ISL_2713075 | PathWest Laboratory Medicine WA | PathWest Laboratory Medicine WA | PathWest Laboratory Medicine WA Microbial Surveillance Unit |
| EPI_ISL_2713259, EPI_ISL_2713266, EPI_ISL_2713267, EPI_ISL_2713303, EPI_ISL_2713304, EPI_ISL_2713305, EPI_ISL_2713306, EPI_ISL_2713307, EPI_ISL_2713308, EPI_ISL_2713309, EPI_ISL_2713310, EPI_ISL_2713311, EPI_ISL_2713312, EPI_ISL_2713313 | | | |
| see above | Department of Health Technology and Informatics, The Hong Kong Polytechnic University | Department of Health Technology and Informatics, The Hong Kong Polytechnic University | Gilman Kit-Hang Siu, Lam-Kwong Lee, Kenneth Siu-Sing Leung, Jake Siu-Lun Leung, Wing-Hei Lo, Timothy Ting-Leung Ng, Chloe Toi-Mei Chan, Hoi-Ching Jim, Ivan Tak-Fai Wong, Kingsley King-Gee Tam, Hiu-Yin Lao, Denise Sze-Hang Wong, Alan Ka-Lun Wu, Miranda Chong-Yee Yau, Yvette Wai-Man Lai, Kitty Sau-Chun Fung, Sandy Ka-Yee Chau, Barry Kin-Chung Wong, Wing-Kin To, Kristine Luk, Alex Yat-Man Ho, Tak-Lun Que, Kam-Tong Yip, Wing Cheong Yam, David Ho-Keung Shum, Shea Ping Yip |
| EPI_ISL_2713403, EPI_ISL_2713408, EPI_ISL_2713418, EPI_ISL_2713423, EPI_ISL_2713427, EPI_ISL_2713431, EPI_ISL_2713432, EPI_ISL_2713435, EPI_ISL_2713441, EPI_ISL_2713453, EPI_ISL_2713462, EPI_ISL_2713467, EPI_ISL_2713468, EPI_ISL_2713532, EPI_ISL_2713533, EPI_ISL_2713544, EPI_ISL_2713549, EPI_ISL_2713551, EPI_ISL_2713552, EPI_ISL_2713554, EPI_ISL_2713556, EPI_ISL_2713557, EPI_ISL_2713558, EPI_ISL_2713559, EPI_ISL_2713560, EPI_ISL_2713562, EPI_ISL_2713563, EPI_ISL_2713564, EPI_ISL_2713565, EPI_ISL_2713568, EPI_ISL_2713570, EPI_ISL_2713571, EPI_ISL_2713572, EPI_ISL_2713573, EPI_ISL_2713574, EPI_ISL_2713577, EPI_ISL_2713578, EPI_ISL_2713579, EPI_ISL_2713580, EPI_ISL_2713581, EPI_ISL_2713582, EPI_ISL_2713583, EPI_ISL_2713584, EPI_ISL_2713585, EPI_ISL_2713586, EPI_ISL_2713587, EPI_ISL_2713589, EPI_ISL_2713590, EPI_ISL_2713592, EPI_ISL_2713594, EPI_ISL_2713595, EPI_ISL_2713597, EPI_ISL_2713598, EPI_ISL_2713599, EPI_ISL_2713600, EPI_ISL_2713604, EPI_ISL_2713605, EPI_ISL_2713607, EPI_ISL_2713616, EPI_ISL_2713619, EPI_ISL_2713625, EPI_ISL_2713640, EPI_ISL_2713644, EPI_ISL_2713645, EPI_ISL_2713652, EPI_ISL_2713655, EPI_ISL_2713664, EPI_ISL_2713667, EPI_ISL_2713674, EPI_ISL_2713675, EPI_ISL_2713676, EPI_ISL_2713677, EPI_ISL_2713685, EPI_ISL_2713686, EPI_ISL_2713687, EPI_ISL_2713688, EPI_ISL_2713699, EPI_ISL_2713700, EPI_ISL_2713702, EPI_ISL_2713720, EPI_ISL_2713721, EPI_ISL_2713723, EPI_ISL_2713727, EPI_ISL_2713729, EPI_ISL_2713736, EPI_ISL_2713760, EPI_ISL_2713763, EPI_ISL_2713771, EPI_ISL_2713772, EPI_ISL_2713779, EPI_ISL_2713780, EPI_ISL_2713786, EPI_ISL_2713789, EPI_ISL_2713794, EPI_ISL_2713797, EPI_ISL_2713807, EPI_ISL_2713819, EPI_ISL_2713820, EPI_ISL_2713821, EPI_ISL_2713838, EPI_ISL_2713850, EPI_ISL_2713853, EPI_ISL_2713855, EPI_ISL_2713859, EPI_ISL_2713861, EPI_ISL_2713862, EPI_ISL_2713863, EPI_ISL_2713864, EPI_ISL_2713865, EPI_ISL_2713874, EPI_ISL_2713876, EPI_ISL_2713878, EPI_ISL_2713879, EPI_ISL_2713882, EPI_ISL_2713885, EPI_ISL_2713887, EPI_ISL_2713896, EPI_ISL_2713901, EPI_ISL_2713903, EPI_ISL_2713905, EPI_ISL_2713907, EPI_ISL_2713910, EPI_ISL_2713915, EPI_ISL_2713916, EPI_ISL_2713919, EPI_ISL_2713924, EPI_ISL_2713927, EPI_ISL_2713931, EPI_ISL_2713935, EPI_ISL_2713937, EPI_ISL_2713938, EPI_ISL_2713946, EPI_ISL_2713947, EPI_ISL_2713950, EPI_ISL_2713951, EPI_ISL_2713952, EPI_ISL_2713953, EPI_ISL_2713954, EPI_ISL_2713960, EPI_ISL_2713965, EPI_ISL_2713966, EPI_ISL_2713967, EPI_ISL_2713981, EPI_ISL_2713992, EPI_ISL_2713993, EPI_ISL_2714003, EPI_ISL_2714004, EPI_ISL_2714006, EPI_ISL_2714010, EPI_ISL_2714012, EPI_ISL_2714013, EPI_ISL_2714017, EPI_ISL_2714020, EPI_ISL_2714021, EPI_ISL_2714029, EPI_ISL_2714030, EPI_ISL_2714031, EPI_ISL_2714032, EPI_ISL_2714034, EPI_ISL_2714035, EPI_ISL_2714038, EPI_ISL_2714040, EPI_ISL_2714042, EPI_ISL_2714044, EPI_ISL_2714045, EPI_ISL_2714053, EPI_ISL_2714057, EPI_ISL_2714065, EPI_ISL_2714066, EPI_ISL_2714068, EPI_ISL_2714085, EPI_ISL_2714086, EPI_ISL_2714091, EPI_ISL_2714093, EPI_ISL_2714096, EPI_ISL_2714097, | | | |

| | | | |
|---|---|---|---|
| EPI_ISL_2714099, EPI_ISL_2714101, EPI_ISL_2714108, EPI_ISL_2714116, EPI_ISL_2714117, EPI_ISL_2714118, EPI_ISL_2714119, EPI_ISL_2714120, EPI_ISL_2714121, EPI_ISL_2714122, EPI_ISL_2714127, EPI_ISL_2714129, EPI_ISL_2714131, EPI_ISL_2714133, EPI_ISL_2714136, EPI_ISL_2714138, EPI_ISL_2714140, EPI_ISL_2714141, EPI_ISL_2714148, EPI_ISL_2714152, EPI_ISL_2714155, EPI_ISL_2714156, EPI_ISL_2714158, EPI_ISL_2714161, EPI_ISL_2714163, EPI_ISL_2714165, EPI_ISL_2714166, EPI_ISL_2714171, EPI_ISL_2714173, EPI_ISL_2714175, EPI_ISL_2714177, EPI_ISL_2714178, EPI_ISL_2714181, EPI_ISL_2714182, EPI_ISL_2714191, EPI_ISL_2714192, EPI_ISL_2714193, EPI_ISL_2714194, EPI_ISL_2714195, EPI_ISL_2714196, EPI_ISL_2714197, EPI_ISL_2714198, EPI_ISL_2714199, EPI_ISL_2714200, EPI_ISL_2714203, EPI_ISL_2714204, EPI_ISL_2714205, EPI_ISL_2714212, EPI_ISL_2714213, EPI_ISL_2714215, EPI_ISL_2714216, EPI_ISL_2714217, EPI_ISL_2714220, EPI_ISL_2714221, EPI_ISL_2714222, EPI_ISL_2714224, EPI_ISL_2714226, EPI_ISL_2714227, EPI_ISL_2714239, EPI_ISL_2714242, EPI_ISL_2714246, EPI_ISL_2714251, EPI_ISL_2714255, EPI_ISL_2714256, EPI_ISL_2714258, EPI_ISL_2714260, EPI_ISL_2714261, EPI_ISL_2714263, EPI_ISL_2714264, EPI_ISL_2714266, EPI_ISL_2714275, EPI_ISL_2714276, EPI_ISL_2714278, EPI_ISL_2714286, EPI_ISL_2714288, EPI_ISL_2714289, EPI_ISL_2714296, EPI_ISL_2714297, EPI_ISL_2714299, EPI_ISL_2714305, EPI_ISL_2714306, EPI_ISL_2714307, EPI_ISL_2714308, EPI_ISL_2714310, EPI_ISL_2714311, EPI_ISL_2714314, EPI_ISL_2714315, EPI_ISL_2714316, EPI_ISL_2714319, EPI_ISL_2714323, EPI_ISL_2714331, EPI_ISL_2714333, EPI_ISL_2714334, EPI_ISL_2714341, EPI_ISL_2714343, EPI_ISL_2714346, EPI_ISL_2714347, EPI_ISL_2714348, EPI_ISL_2714349, EPI_ISL_2714353, EPI_ISL_2714354, EPI_ISL_2714355, EPI_ISL_2714384, EPI_ISL_2714385, EPI_ISL_2714388, EPI_ISL_2714392, EPI_ISL_2714410, EPI_ISL_2714411, EPI_ISL_2714460, EPI_ISL_2714475, EPI_ISL_2714490, EPI_ISL_2714491, EPI_ISL_2714492, EPI_ISL_2714493, EPI_ISL_2714494, EPI_ISL_2714495, EPI_ISL_2714496, EPI_ISL_2714498, EPI_ISL_2714501, EPI_ISL_2714502, EPI_ISL_2714503, EPI_ISL_2714504, EPI_ISL_2714505, EPI_ISL_2714516, EPI_ISL_2714518, EPI_ISL_2714530, EPI_ISL_2714532, EPI_ISL_2714533, EPI_ISL_2714537, EPI_ISL_2714542, EPI_ISL_2714543, EPI_ISL_2714544, EPI_ISL_2714545, EPI_ISL_2714562 | | | |
| see above | Public Health Ontario Laboratory | Public Health Ontario Laboratory | Vanessa G Allen, Philip Banh, Yao Chen, Richard de Borja, Alireza Eshaghi, Nahuel Fittipaldi, Christine Frantz, Jonathan B Gubbay, Jennifer L Guthrie, Lawrence Heisler, Esha Joshi, Michael Laszloffy, Aimin Li, Michael CY Li, Dean Maxwell, Sandeep Nagra, Samir N Patel, Jared Simpson, Karthikeyan Sivaraman, Ashleigh Sullivan, Yogi Sundaravadanam, Sarah Teatero, Andre Villegas, Matthew Watson, Sandra Zittermann |
| EPI_ISL_2714772, EPI_ISL_2714773, EPI_ISL_2714775 | Platform BIS UZA/UAntwerpen | Labo Klinische Biologie, UZA | Marie Le Mercier, Jasmine Coppens, Kathleen Holemans, Ines Verbesselt, Basil Britto Xavier, Christine Lammens, Veerle Matheeussen, Herman Goossens |
| EPI_ISL_2714776 | CENTRUM VOOR MEDISCHE ANALYSE | Labo Klinische Biologie, UZA | Marie Le Mercier, Jasmine Coppens, Kathleen Holemans, Ines Verbesselt, Basil Britto Xavier, Christine Lammens, Veerle Matheeussen, Herman Goossens |
| EPI_ISL_2714779, EPI_ISL_2714787, EPI_ISL_2714789, EPI_ISL_2714791, EPI_ISL_2714807, EPI_ISL_2714808, EPI_ISL_2714811, EPI_ISL_2714816, EPI_ISL_2714818, EPI_ISL_2714823, EPI_ISL_2714825, EPI_ISL_2714831, EPI_ISL_2714832, EPI_ISL_2714836 | | | |
| see above | Platform BIS UZA/UAntwerpen | Labo Klinische Biologie, UZA | Marie Le Mercier, Jasmine Coppens, Kathleen Holemans, Ines Verbesselt, Basil Britto Xavier, Christine Lammens, Veerle Matheeussen, Herman Goossens |
| EPI_ISL_2714839, EPI_ISL_2714842, EPI_ISL_2714846 | Algemeen Medisch Labo | Labo Klinische Biologie, UZA | Marie Le Mercier, Jasmine Coppens, Kathleen Holemans, Ines Verbesselt, Basil Britto Xavier, Christine Lammens, Veerle Matheeussen, Herman Goossens |
| EPI_ISL_2714866, EPI_ISL_2714870, EPI_ISL_2714877, EPI_ISL_2714878, EPI_ISL_2714888, EPI_ISL_2714889 | Platform BIS UZA/UAntwerpen | Labo Klinische Biologie, UZA | Marie Le Mercier, Jasmine Coppens, Kathleen Holemans, Ines Verbesselt, Basil Britto Xavier, Christine Lammens, Veerle Matheeussen, Herman Goossens |
| EPI_ISL_2714890, EPI_ISL_2714893, EPI_ISL_2714896, EPI_ISL_2714897, EPI_ISL_2714898, EPI_ISL_2714899, EPI_ISL_2714900, EPI_ISL_2714901 | LHUB-ULB | Labo Klinische Biologie, UZA | Marie Le Mercier, Jasmine Coppens, Kathleen Holemans, Ines Verbesselt, Basil Britto Xavier, Christine Lammens, Veerle Matheeussen, Herman Goossens |
| EPI_ISL_2714949 | Interpath Laboratory | Seattle Flu Study | Deborah A. Nickerson, Chris D. Frazar, Jover Lee, Benjamin Pelle, Erica Ryke, Matthew Richardson, Amanda Adler, Elisabeth Brandstetter, Peter D. Han, Kairsten Fay, Misja Ilcisin, Kirsten Lacombe, Thomas R. Sibley, Melissa Truong, Caitlin R. Wolf, Romesh Gautom, Geoff Melly, Brian Hiatt, Philip Dykema, Scott Lindquist, Michael Boeckh, Janet A. Englund, Michael Famulare, Barry R. Lutz, Mark J. Rieder, Lea M. Starita, Matthew Thompson, Helen Y. Chu, Jay Shendure, Trevor Bedford |
| EPI_ISL_2715015, EPI_ISL_2715020, EPI_ISL_2715046, EPI_ISL_2715053, EPI_ISL_2715057, EPI_ISL_2715061, EPI_ISL_2715062, EPI_ISL_2715065 | Atlas Genomics | Seattle Flu Study | Deborah A. Nickerson, Chris D. Frazar, Jover Lee, Benjamin Pelle, Erica Ryke, Matthew Richardson, Amanda Adler, Elisabeth Brandstetter, Peter D. Han, Kairsten Fay, Misja Ilcisin, Kirsten Lacombe, Thomas R. Sibley, Melissa Truong, Caitlin R. Wolf, Romesh Gautom, Geoff Melly, Brian Hiatt, Philip Dykema, Scott Lindquist, Michael Boeckh, Janet A. Englund, Michael Famulare, Barry R. Lutz, Mark J. Rieder, Lea M. Starita, Matthew Thompson, Helen Y. Chu, Jay Shendure, Trevor Bedford |
| EPI_ISL_2715136, EPI_ISL_2715137 | Northwest Laboratory | Seattle Flu Study | Deborah A. Nickerson, Chris D. Frazar, Jover Lee, Benjamin Pelle, Erica Ryke, Matthew Richardson, Amanda Adler, Elisabeth Brandstetter, Peter D. Han, Kairsten Fay, Misja Ilcisin, Kirsten Lacombe, Thomas R. Sibley, Melissa Truong, Caitlin R. Wolf, Romesh Gautom, Geoff Melly, Brian Hiatt, Philip Dykema, Scott Lindquist, Michael Boeckh, Janet A. Englund, Michael Famulare, Barry R. Lutz, Mark J. Rieder, Lea M. Starita, Matthew Thompson, Helen Y. Chu, Jay Shendure, Trevor Bedford |
| EPI_ISL_2715201, EPI_ISL_2715210, EPI_ISL_2715212, EPI_ISL_2715214 | Atlas Genomics | Seattle Flu Study | Deborah A. Nickerson, Chris D. Frazar, Jover Lee, Benjamin Pelle, Erica Ryke, Matthew Richardson, Amanda Adler, Elisabeth Brandstetter, Peter D. Han, Kairsten Fay, Misja Ilcisin, Kirsten Lacombe, Thomas R. Sibley, Melissa Truong, Caitlin R. Wolf, Romesh Gautom, Geoff Melly, Brian Hiatt, Philip Dykema, Scott Lindquist, Michael Boeckh, Janet A. Englund, Michael Famulare, Barry R. Lutz, Mark J. Rieder, Lea M. Starita, Matthew Thompson, Helen Y. Chu, Jay Shendure, Trevor Bedford |
| EPI_ISL_2715293, EPI_ISL_2715326, EPI_ISL_2715328, EPI_ISL_2715333, EPI_ISL_2715334, EPI_ISL_2715341, EPI_ISL_2715342, EPI_ISL_2715347, EPI_ISL_2715348, EPI_ISL_2715349, EPI_ISL_2715352 | | | |
| see above | US Air Force School of Aerospace Medicine | US Air Force School of Aerospace Medicine | Anthony Fries, Jennifer Meyer, William Gruner, Amanda Javorina, Carol Garrett, Sarah Purves, Clarise Starr, Elizabeth Macias |
| EPI_ISL_2715358 | US Air Force School of Aerospace Medicine | US Air Force School of Aerospace Medicine | Anthony Fries, Jennifer Meyer, William Gruner, Amanda Javorina, Carol Garrett, Sarah Purves, Clarise Starr, Elizabeth Macias, Fritz Castillo, Cole Anderson |
| EPI_ISL_2715369 | Respiratory Viruses Branch, Centers for Disease Control and Prevention | Respiratory Viruses Branch, Centers for Disease Control and Prevention | Sheth,M., Nobles,S., Padilla,J., Burroughs,M., Le,S., Dillon,K., Cook,P., Paden,C.R., Batra,D., Queen,K., Knipe,K., Howard,D., Unoarumhi,Y., Wagner,D., Schmerer,M., Rambo-Martin,B.L., Lacek,K., Shepard,S., Halpin,A.L., Wentworth,D., Dugan,V., Tong,S., Lee,J. |
| EPI_ISL_2715652, EPI_ISL_2715653, EPI_ISL_2715654, EPI_ISL_2715657, EPI_ISL_2715658, EPI_ISL_2715659, EPI_ISL_2715660, EPI_ISL_2715663, EPI_ISL_2715664, EPI_ISL_2715665, EPI_ISL_2715668, EPI_ISL_2715670, EPI_ISL_2715671, EPI_ISL_2715672, EPI_ISL_2715677, EPI_ISL_2715678, EPI_ISL_2715679 | | | |
| see above | Pandemic Response Lab - NYC | Pandemic Response Lab, R&D | Henry Lee, Michael Hammerling, Melissa Hopkins, Cybill del Castillo, Shinyoung Clair Kang, William Ward, Pradeep Bugga, Sol Rey, Dylan Law, Katharine Nelson, Haiping Hao, Jon Laurent |
| EPI_ISL_2716210, EPI_ISL_2716215, EPI_ISL_2716220 | The Jackson Laboratory | The Jackson Laboratory | Long J, Renzette N, Adams M, Omerza G, Kelly K, Li L |
| EPI_ISL_2716234 | Yale Clinical Virology Lab | Grubaugh Lab - Yale School of Public Health | Joseph Fauver, Mallery Breban, Isabel Ott, Tara Alpert, Mary Petrone, Anderson Brito, Chantal Vogels, Annie Watkins, Chaney Kalinich, Jessica Rothman, Marie L. Landry, Nathan Grubaugh |
| EPI_ISL_2716284, EPI_ISL_2716285, EPI_ISL_2716287, EPI_ISL_2716289, EPI_ISL_2716290, EPI_ISL_2716291, EPI_ISL_2716292, EPI_ISL_2716296, EPI_ISL_2716297, EPI_ISL_2716298 | Kansas Health and Environmental Lab | Kansas Health and Environmental Lab | Katherine Wiggins, Mike Grose, Jonathan Barnell, Ben Olsen, and Phil Adam |
| EPI_ISL_2716404 | Arizona State Public Health Laboratory | Arizona State Public Health Laboratory | Trung Huynh, Jessica Escobar, Katherine Fullerton, Nobuko Fukushima, Matthew Contursi, Stacy White, Linda Getsinger, Victor Waddell |
| EPI_ISL_2716413, EPI_ISL_2716414, EPI_ISL_2716415, EPI_ISL_2716416, EPI_ISL_2716417, EPI_ISL_2716418, EPI_ISL_2716419, EPI_ISL_2716420, EPI_ISL_2716421, EPI_ISL_2716422 | Area of Virology, Serology and Virology Division (SAViD), New South Wales Health Pathology Randwick | Virology Research Laboratory; Area of Virology, Serology and Virology Division (SAViD), New South Wales Health Pathology Randwick | Foster, C.; Au, J.; Ruiz Silva, M.; Deveson, I.; Bull, R.; Van Hal, S.; Rawlinson, W. |
| EPI_ISL_2716453, EPI_ISL_2716456, EPI_ISL_2716468 | Washington State Department of Health Public Health Laboratories | Washington State Department of Health Public Health Laboratories | Drew MacKellar, Philip Dykema, Denny Russell, Joenice Gonzalez, Hannah Gray, Geoff Melly, Vanessa De Los Santos, Darren Lucas, JohnAric Peterson, Avi Singh, Rebecca Cao |
| EPI_ISL_2716496 | UW Virology Lab | UW Virology Lab | Pavitra Roychoudhury, Hong Xie, Lasata Shrestha, Shah Mohamed Bakhash, Tien V. Nguyen, Noah R. Baker, Sean Ellis, Meei-Li Huang, Keith R Jerome, Alexander Greninger |
| EPI_ISL_2716535, EPI_ISL_2716536, EPI_ISL_2716537, EPI_ISL_2716539, EPI_ISL_2716540, EPI_ISL_2716541, EPI_ISL_2716549, EPI_ISL_2716552, EPI_ISL_2716554, EPI_ISL_2716555 | ASL Napoli 1 Centro | AMES Centro Polidiagnostico Strumentale S.r.l. | "Giovanni Savarese, Raffaella Ruggiero, Eloisa Evangelista, Antonella Di Carlo, Luisa Circelli, Luigi D'Amore, Roberto Sirica,Maurizio D'Amora Antonio Fico" |
| EPI_ISL_2716727, EPI_ISL_2716728, EPI_ISL_2716730, EPI_ISL_2716731, | UW Virology Lab | UW Virology Lab | Pavitra Roychoudhury, Hong Xie, Lasata Shrestha, Shah Mohamed Bakhash, Tien V. Nguyen, Noah R. Baker, Sean Ellis, Meei-Li Huang, Keith R Jerome, Alexander Greninger |

| | | | |
|---|---|---|---|
| EPI_ISL_2716732, EPI_ISL_2716733, EPI_ISL_2716737, EPI_ISL_2716738, EPI_ISL_2716739 | | | |
| EPI_ISL_2716754 | IRCCS San Gallicano Dermatological Institute | IRCCS Regina Elena National Cancer Institute | Frauke Goeman, Giulia Orlandi, Alice Massacci, Martina Diano, Fabrizio Ensoli, Fulvia Pimpinelli, Maurizio Fanciulli, Giovanni Blandino, Aldo Morrone, Gennaro Ciliberto |
| EPI_ISL_2716755 | IRCCS San Gallicano Dermatological Institute | IRCCS Regina Elena National Cancer Institute | Sabrina Strano, Sara Donzelli, Matteo Pallocca, Antonio Federico, Fabrizio Ensoli, Fulvia Pimpinelli, Maurizio Fanciulli, Giovanni Blandino, Aldo Morrone, Gennaro Ciliberto |
| EPI_ISL_2716756 | IRCCS San Gallicano Dermatological Institute | IRCCS Regina Elena National Cancer Institute | Francesca De Nicola, Giulia Orlandi, Eleonora Sperandio, Fulvia Fraticelli, Fabrizio Ensoli, Fulvia Pimpinelli, Maurizio Fanciulli, Giovanni Blandino, Aldo Morrone, Gennaro Ciliberto |
| EPI_ISL_2716759 | IRCCS San Gallicano Dermatological Institute | IRCCS Regina Elena National Cancer Institute | Sabrina Strano, Sara Donzelli, Eleonora Sperandio, Francesca Maione, Fabrizio Ensoli, Fulvia Pimpinelli, Maurizio Fanciulli, Giovanni Blandino, Aldo Morrone, Gennaro Ciliberto |
| EPI_ISL_2716760 | IRCCS San Gallicano Dermatological Institute | IRCCS Regina Elena National Cancer Institute | Francesca De Nicola, Giulia Orlandi, Alice Massacci, Francisco Obregon, Fabrizio Ensoli, Fulvia Pimpinelli, Maurizio Fanciulli, Giovanni Blandino, Aldo Morrone, Gennaro Ciliberto |
| EPI_ISL_2716761 | IRCCS San Gallicano Dermatological Institute | IRCCS Regina Elena National Cancer Institute | Ludovica Ciuffreda, Sara Donzelli, Matteo Pallocca, Silvia Paluzzi, Fabrizio Ensoli, Fulvia Pimpinelli, Maurizio Fanciulli, Giovanni Blandino, Aldo Morrone, Gennaro Ciliberto |
| EPI_ISL_2716762 | IRCCS San Gallicano Dermatological Institute | IRCCS Regina Elena National Cancer Institute | Frauke Goeman, Giulia Orlandi, Eleonora Sperandio, Sara Petrolo, Fabrizio Ensoli, Fulvia Pimpinelli, Maurizio Fanciulli, Giovanni Blandino, Aldo Morrone, Gennaro Ciliberto |
| EPI_ISL_2716763 | IRCCS San Gallicano Dermatological Institute | IRCCS Regina Elena National Cancer Institute | Sabrina Strano, Sara Donzelli, Alice Massacci, Valentina Ricca, Fabrizio Ensoli, Fulvia Pimpinelli, Maurizio Fanciulli, Giovanni Blandino, Aldo Morrone, Gennaro Ciliberto |
| EPI_ISL_2716764 | IRCCS San Gallicano Dermatological Institute | IRCCS Regina Elena National Cancer Institute | Francesca De Nicola, Giulia Orlandi, Matteo Pallocca, Serena Salvo, Fabrizio Ensoli, Fulvia Pimpinelli, Maurizio Fanciulli, Giovanni Blandino, Aldo Morrone, Gennaro Ciliberto |
| EPI_ISL_2716765 | IRCCS San Gallicano Dermatological Institute | IRCCS Regina Elena National Cancer Institute | Ludovica Ciuffreda, Sara Donzelli, Eleonora Sperandio, Grazia Prignano, Fabrizio Ensoli, Fulvia Pimpinelli, Maurizio Fanciulli, Giovanni Blandino, Aldo Morrone, Gennaro Ciliberto |
| EPI_ISL_2716766 | IRCCS San Gallicano Dermatological Institute | IRCCS Regina Elena National Cancer Institute | Frauke Goeman, Giulia Orlandi, Alice Massacci, Elisabetta Trento, Fabrizio Ensoli, Fulvia Pimpinelli, Maurizio Fanciulli, Giovanni Blandino, Aldo Morrone, Gennaro Ciliberto |
| EPI_ISL_2716767 | IRCCS San Gallicano Dermatological Institute | IRCCS Regina Elena National Cancer Institute | Sabrina Strano, Sara Donzelli, Matteo Pallocca, Arianna Mastrofrancesco, Fabrizio Ensoli, Fulvia Pimpinelli, Maurizio Fanciulli, Giovanni Blandino, Aldo Morrone, Gennaro Ciliberto |
| EPI_ISL_2716768 | IRCCS San Gallicano Dermatological Institute | IRCCS Regina Elena National Cancer Institute | Francesca De Nicola, Giulia Orlandi, Eleonora Sperandio, Giovanna D'agosto, Fabrizio Ensoli, Fulvia Pimpinelli, Maurizio Fanciulli, Giovanni Blandino, Aldo Morrone, Gennaro Ciliberto |
| EPI_ISL_2716769 | IRCCS San Gallicano Dermatological Institute | IRCCS Regina Elena National Cancer Institute | Ludovica Ciuffreda, Sara Donzelli, Alice Massacci, Carla Mottini, Fabrizio Ensoli, Fulvia Pimpinelli, Maurizio Fanciulli, Giovanni Blandino, Aldo Morrone, Gennaro Ciliberto |
| EPI_ISL_2716770 | IRCCS San Gallicano Dermatological Institute | IRCCS Regina Elena National Cancer Institute | Frauke Goeman, Giulia Orlandi, Matteo Pallocca, Martina Pontone, Fabrizio Ensoli, Fulvia Pimpinelli, Maurizio Fanciulli, Giovanni Blandino, Aldo Morrone, Gennaro Ciliberto |
| EPI_ISL_2716771 | IRCCS San Gallicano Dermatological Institute | IRCCS Regina Elena National Cancer Institute | Sabrina Strano, Sara Donzelli, Eleonora Sperandio, di Domenico Enea Gino, Fabrizio Ensoli, Fulvia Pimpinelli, Maurizio Fanciulli, Giovanni Blandino, Aldo Morrone, Gennaro Ciliberto |
| EPI_ISL_2716772 | IRCCS San Gallicano Dermatological Institute | IRCCS Regina Elena National Cancer Institute | Francesca De Nicola, Giulia Orlandi, Alice Massacci, Ilaria Cavallo, Fabrizio Ensoli, Fulvia Pimpinelli, Maurizio Fanciulli, Giovanni Blandino, Aldo Morrone, Gennaro Ciliberto |
| EPI_ISL_2716773 | IRCCS San Gallicano Dermatological Institute | IRCCS Regina Elena National Cancer Institute | Ludovica Ciuffreda, Sara Donzelli, Matteo Pallocca, Francesca Sivori, Fabrizio Ensoli, Fulvia Pimpinelli, Maurizio Fanciulli, Giovanni Blandino, Aldo Morrone, Gennaro Ciliberto |
| EPI_ISL_2716774 | IRCCS San Gallicano Dermatological Institute | IRCCS Regina Elena National Cancer Institute | Frauke Goeman, Giulia Orlandi, Eleonora Sperandio, Andrea Cazzani, Fabrizio Ensoli, Fulvia Pimpinelli, Maurizio Fanciulli, Giovanni Blandino, Aldo Morrone, Gennaro Ciliberto |
| EPI_ISL_2716775 | IRCCS San Gallicano Dermatological Institute | IRCCS Regina Elena National Cancer Institute | Sabrina Strano, Sara Donzelli, Alice Massacci, Ilaria Celesti, Fabrizio Ensoli, Fulvia Pimpinelli, Maurizio Fanciulli, Giovanni Blandino, Aldo Morrone, Gennaro Ciliberto |
| EPI_ISL_2716776 | IRCCS San Gallicano Dermatological Institute | IRCCS Regina Elena National Cancer Institute | Francesca De Nicola, Giulia Orlandi, Matteo Pallocca, Martina Diano, Fabrizio Ensoli, Fulvia Pimpinelli, Maurizio Fanciulli, Giovanni Blandino, Aldo Morrone, Gennaro Ciliberto |
| EPI_ISL_2716777 | IRCCS San Gallicano Dermatological Institute | IRCCS Regina Elena National Cancer Institute | Ludovica Ciuffreda, Sara Donzelli, Eleonora Sperandio, Antonio Federico, Fabrizio Ensoli, Fulvia Pimpinelli, Maurizio Fanciulli, Giovanni Blandino, Aldo Morrone, Gennaro Ciliberto |
| EPI_ISL_2716778 | IRCCS San Gallicano Dermatological Institute | IRCCS Regina Elena National Cancer Institute | Frauke Goeman, Giulia Orlandi, Alice Massacci, Fulvia Fraticelli, Fabrizio Ensoli, Fulvia Pimpinelli, Maurizio Fanciulli, Giovanni Blandino, Aldo Morrone, Gennaro Ciliberto |
| EPI_ISL_2716779 | IRCCS San Gallicano Dermatological Institute | IRCCS Regina Elena National Cancer Institute | Sabrina Strano, Sara Donzelli, Matteo Pallocca, Lorenzo Furzi, Fabrizio Ensoli, Fulvia Pimpinelli, Maurizio Fanciulli, Giovanni Blandino, Aldo Morrone, Gennaro Ciliberto |
| EPI_ISL_2716780 | IRCCS San Gallicano Dermatological Institute | IRCCS Regina Elena National Cancer Institute | Francesca De Nicola, Giulia Orlandi, Eleonora Sperandio, Alessia Lauretti, Fabrizio Ensoli, Fulvia Pimpinelli, Maurizio Fanciulli, Giovanni Blandino, Aldo Morrone, Gennaro Ciliberto |
| EPI_ISL_2716892, EPI_ISL_2716897, EPI_ISL_2716900, EPI_ISL_2716904, EPI_ISL_2716913, EPI_ISL_2716915, EPI_ISL_2716918, EPI_ISL_2716920, EPI_ISL_2716921, EPI_ISL_2716923, EPI_ISL_2716927, EPI_ISL_2716928, EPI_ISL_2716931, EPI_ISL_2716932, EPI_ISL_2716933, EPI_ISL_2716937, EPI_ISL_2716939, EPI_ISL_2716941, EPI_ISL_2716942, EPI_ISL_2716943, EPI_ISL_2716944, EPI_ISL_2716945, EPI_ISL_2716946, EPI_ISL_2716947, EPI_ISL_2716948, EPI_ISL_2716949, EPI_ISL_2716950, EPI_ISL_2716952, EPI_ISL_2716953 | | | |
| see above | Southern Nevada Public Health Laboratory | Southern Nevada Public Health Laboratory | Michael Picker |
| EPI_ISL_2716974 | Lighthouse Lab in Glasgow | Wellcome Sanger Institute for the COVID-19 Genomics UK (COG-UK) Consortium | Harper VanSteenhouse, Yumi Kasai, David Gray, Carol Clugston, Anna Dominiczak and Alex Alderton, Roberto Amato, Jeffrey Barrett, Sonia Goncalves, Ewan Harrison, David K. Jackson, Ian Johnston, Dominic Kwiatkowski, Cordelia Langford, John Sillitoe on behalf of the Wellcome Sanger Institute COVID-19 Surveillance Team |
| EPI_ISL_2716975, EPI_ISL_2716976 | Lighthouse Lab in Milton Keynes | Wellcome Sanger Institute for the COVID-19 Genomics UK (COG-UK) Consortium | The Lighthouse Lab in Milton Keynes and Alex Alderton, Roberto Amato, Jeffrey Barrett, Sonia Goncalves, Ewan Harrison, David K. Jackson, Ian Johnston, Dominic Kwiatkowski, Cordelia Langford, John Sillitoe on behalf of the Wellcome Sanger Institute COVID-19 Surveillance Team |
| EPI_ISL_2716977 | Lighthouse Lab in Alderley Park | Wellcome Sanger Institute for the COVID-19 Genomics UK (COG-UK) Consortium | Jacquelyn Wynn, Mairead Hyland, The Lighthouse Lab in Alderley Park and Alex Alderton, Roberto Amato, Jeffrey Barrett, Sonia Goncalves, Ewan Harrison, David K. Jackson, Ian Johnston, Dominic Kwiatkowski, Cordelia Langford, John Sillitoe on behalf of the Wellcome Sanger Institute COVID-19 Surveillance Team |
| EPI_ISL_2716978 | Health Services Laboratories | Wellcome Sanger Institute for the COVID-19 Genomics UK (COG-UK) Consortium | Health Services Laboratories and Alex Alderton, Roberto Amato, Jeffrey Barrett, Sonia Goncalves, Ewan Harrison, David K. Jackson, Ian Johnston, Dominic Kwiatkowski, Cordelia Langford, John Sillitoe on behalf of the Wellcome Sanger Institute COVID-19 Surveillance Team |
| EPI_ISL_2716979 | Lighthouse Lab in Alderley Park | Wellcome Sanger Institute for the COVID-19 Genomics UK (COG-UK) Consortium | Jacquelyn Wynn, Mairead Hyland, The Lighthouse Lab in Alderley Park and Alex Alderton, Roberto Amato, Jeffrey Barrett, Sonia Goncalves, Ewan Harrison, David K. Jackson, Ian Johnston, Dominic Kwiatkowski, Cordelia Langford, John Sillitoe on behalf of the Wellcome Sanger Institute COVID-19 Surveillance Team |
| EPI_ISL_2716980 | Lighthouse Lab in Glasgow | Wellcome Sanger Institute for the COVID-19 Genomics UK (COG-UK) Consortium | Harper VanSteenhouse, Yumi Kasai, David Gray, Carol Clugston, Anna Dominiczak and Alex Alderton, Roberto Amato, Jeffrey Barrett, Sonia Goncalves, Ewan Harrison, David K. Jackson, Ian Johnston, Dominic Kwiatkowski, Cordelia Langford, John Sillitoe on behalf of the Wellcome Sanger Institute COVID-19 Surveillance Team |
| EPI_ISL_2716981 | Lighthouse Lab in Alderley Park | Wellcome Sanger Institute for the COVID-19 Genomics UK (COG-UK) Consortium | Jacquelyn Wynn, Mairead Hyland, The Lighthouse Lab in Alderley Park and Alex Alderton, Roberto Amato, Jeffrey Barrett, Sonia Goncalves, Ewan Harrison, David K. Jackson, Ian Johnston, Dominic Kwiatkowski, Cordelia Langford, John Sillitoe on behalf of the Wellcome Sanger Institute COVID-19 Surveillance Team |
| EPI_ISL_2716982 | Lighthouse Lab in Milton Keynes | Wellcome Sanger Institute for the COVID-19 Genomics UK (COG-UK) Consortium | The Lighthouse Lab in Milton Keynes and Alex Alderton, Roberto Amato, Jeffrey Barrett, Sonia Goncalves, Ewan Harrison, David K. Jackson, Ian Johnston, Dominic Kwiatkowski, Cordelia Langford, John Sillitoe on behalf of the Wellcome Sanger Institute COVID-19 Surveillance Team |
| EPI_ISL_2716983 | Lighthouse Lab in Glasgow | Wellcome Sanger Institute for the COVID-19 Genomics UK (COG-UK) Consortium | Harper VanSteenhouse, Yumi Kasai, David Gray, Carol Clugston, Anna Dominiczak and Alex Alderton, Roberto Amato, Jeffrey Barrett, Sonia Goncalves, Ewan Harrison, David K. Jackson, Ian Johnston, Dominic Kwiatkowski, Cordelia Langford, John Sillitoe on behalf of the Wellcome Sanger Institute COVID-19 Surveillance Team |
| EPI_ISL_2716984, EPI_ISL_2716985 | Lighthouse Lab in Alderley Park | Wellcome Sanger Institute for the COVID-19 Genomics UK (COG-UK) Consortium | Jacquelyn Wynn, Mairead Hyland, The Lighthouse Lab in Alderley Park and Alex Alderton, Roberto Amato, Jeffrey Barrett, Sonia Goncalves, Ewan Harrison, David K. Jackson, Ian Johnston, Dominic Kwiatkowski, Cordelia Langford, John Sillitoe on behalf of the Wellcome Sanger Institute COVID-19 Surveillance Team |
| EPI_ISL_2716986 | Lighthouse Lab in Milton Keynes | Wellcome Sanger Institute for the COVID-19 Genomics UK (COG-UK) Consortium | The Lighthouse Lab in Milton Keynes and Alex Alderton, Roberto Amato, Jeffrey Barrett, Sonia Goncalves, Ewan Harrison, David K. Jackson, Ian Johnston, Dominic Kwiatkowski, Cordelia Langford, John Sillitoe on behalf of the Wellcome Sanger Institute COVID-19 Surveillance Team |
| EPI_ISL_2716987 | Health Services Laboratories | Wellcome Sanger Institute for the COVID-19 Genomics UK (COG-UK) Consortium | Health Services Laboratories and Alex Alderton, Roberto Amato, Jeffrey Barrett, Sonia Goncalves, Ewan Harrison, David K. Jackson, Ian Johnston, Dominic Kwiatkowski, Cordelia Langford, John Sillitoe on behalf of the Wellcome Sanger Institute COVID-19 Surveillance Team |
| EPI_ISL_2716988 | Lighthouse Lab in Milton Keynes | Wellcome Sanger Institute for the COVID-19 Genomics UK (COG-UK) Consortium | The Lighthouse Lab in Milton Keynes and Alex Alderton, Roberto Amato, Jeffrey Barrett, Sonia Goncalves, Ewan Harrison, David K. Jackson, Ian Johnston, Dominic Kwiatkowski, Cordelia Langford, John Sillitoe on behalf of the Wellcome Sanger Institute COVID-19 Surveillance Team |
| EPI_ISL_2716989, EPI_ISL_2716990, EPI_ISL_2716991 | Lighthouse Lab in Alderley Park | Wellcome Sanger Institute for the COVID-19 Genomics UK (COG-UK) Consortium | Jacquelyn Wynn, Mairead Hyland, The Lighthouse Lab in Alderley Park and Alex Alderton, Roberto Amato, Jeffrey Barrett, Sonia Goncalves, Ewan Harrison, David K. Jackson, Ian Johnston, Dominic Kwiatkowski, Cordelia Langford, John Sillitoe on behalf of the Wellcome Sanger Institute COVID-19 Surveillance Team |
| EPI_ISL_2716992 | Lighthouse Lab in Milton Keynes | Wellcome Sanger Institute for the COVID-19 Genomics UK (COG-UK) Consortium | The Lighthouse Lab in Milton Keynes and Alex Alderton, Roberto Amato, Jeffrey Barrett, Sonia Goncalves, Ewan Harrison, David K. Jackson, Ian Johnston, Dominic Kwiatkowski, Cordelia Langford, John Sillitoe on behalf of the Wellcome Sanger Institute COVID-19 Surveillance Team |
| EPI_ISL_2716993, EPI_ISL_2716994 | Lighthouse Lab in Alderley Park | Wellcome Sanger Institute for the COVID-19 Genomics UK (COG-UK) Consortium | Jacquelyn Wynn, Mairead Hyland, The Lighthouse Lab in Alderley Park and Alex Alderton, Roberto Amato, Jeffrey Barrett, Sonia Goncalves, Ewan Harrison, David K. Jackson, Ian Johnston, Dominic Kwiatkowski, Cordelia Langford, John Sillitoe on behalf of the Wellcome Sanger Institute COVID-19 Surveillance Team |
| EPI_ISL_2716995, EPI_ISL_2716996 | Lighthouse Lab in Milton Keynes | Wellcome Sanger Institute for the COVID-19 Genomics UK (COG-UK) Consortium | The Lighthouse Lab in Milton Keynes and Alex Alderton, Roberto Amato, Jeffrey Barrett, Sonia Goncalves, Ewan Harrison, David K. Jackson, Ian Johnston, Dominic Kwiatkowski, Cordelia Langford, John Sillitoe on behalf of the Wellcome Sanger Institute COVID-19 Surveillance Team |
| EPI_ISL_2716997 | Lighthouse Lab in Alderley Park | Wellcome Sanger Institute for the COVID-19 Genomics UK (COG-UK) Consortium | Jacquelyn Wynn, Mairead Hyland, The Lighthouse Lab in Alderley Park and Alex Alderton, Roberto Amato, Jeffrey Barrett, Sonia Goncalves, Ewan Harrison, David K. Jackson, Ian Johnston, Dominic Kwiatkowski, Cordelia Langford, John Sillitoe on behalf of the Wellcome Sanger Institute COVID-19 Surveillance Team |
| EPI_ISL_2716998, EPI_ISL_2716999 | Lighthouse Lab in Milton Keynes | Wellcome Sanger Institute for the COVID-19 Genomics UK (COG-UK) Consortium | The Lighthouse Lab in Milton Keynes and Alex Alderton, Roberto Amato, Jeffrey Barrett, Sonia Goncalves, Ewan Harrison, David K. Jackson, Ian Johnston, Dominic Kwiatkowski, Cordelia Langford, John Sillitoe on behalf of the Wellcome Sanger Institute COVID-19 Surveillance Team |
| EPI_ISL_2717000 | Health Services Laboratories | Wellcome Sanger Institute for the COVID-19 Genomics UK (COG-UK) Consortium | Health Services Laboratories and Alex Alderton, Roberto Amato, Jeffrey Barrett, Sonia Goncalves, Ewan Harrison, David K. Jackson, Ian Johnston, Dominic Kwiatkowski, Cordelia Langford, John Sillitoe on behalf of the Wellcome Sanger Institute COVID-19 Surveillance Team |
| EPI_ISL_2717001 | Lighthouse Lab in Glasgow | Wellcome Sanger Institute for the COVID-19 Genomics UK (COG-UK) Consortium | Harper VanSteenhouse, Yumi Kasai, David Gray, Carol Clugston, Anna Dominiczak and Alex Alderton, Roberto Amato, Jeffrey Barrett, Sonia Goncalves, Ewan Harrison, David K. Jackson, Ian Johnston, Dominic Kwiatkowski, Cordelia Langford, John Sillitoe on behalf of the Wellcome Sanger Institute COVID-19 Surveillance Team |
| EPI_ISL_2717002, EPI_ISL_2717003 | Lighthouse Lab in Milton Keynes | Wellcome Sanger Institute for the COVID-19 Genomics UK (COG-UK) Consortium | The Lighthouse Lab in Milton Keynes and Alex Alderton, Roberto Amato, Jeffrey Barrett, Sonia Goncalves, Ewan Harrison, David K. Jackson, Ian Johnston, Dominic Kwiatkowski, Cordelia Langford, John Sillitoe on behalf of the Wellcome Sanger Institute COVID-19 Surveillance Team |
| EPI_ISL_2717004 | Lighthouse Lab in Alderley Park | Wellcome Sanger Institute for the COVID-19 Genomics UK (COG-UK) Consortium | Jacquelyn Wynn, Mairead Hyland, The Lighthouse Lab in Alderley Park and Alex Alderton, Roberto Amato, Jeffrey Barrett, Sonia Goncalves, Ewan Harrison, David K. Jackson, Ian Johnston, Dominic Kwiatkowski, Cordelia Langford, John Sillitoe on behalf of the Wellcome Sanger Institute COVID-19 Surveillance Team |
| EPI_ISL_2717005 | Lighthouse Lab in Milton Keynes | Wellcome Sanger Institute for the COVID-19 Genomics UK (COG-UK) Consortium | The Lighthouse Lab in Milton Keynes and Alex Alderton, Roberto Amato, Jeffrey Barrett, Sonia Goncalves, Ewan Harrison, David K. Jackson, Ian Johnston, Dominic Kwiatkowski, Cordelia Langford, John Sillitoe on behalf of the Wellcome Sanger Institute COVID-19 Surveillance Team |
| EPI_ISL_2717006 | Health Services Laboratories | Wellcome Sanger Institute for the COVID-19 Genomics UK (COG-UK) Consortium | Health Services Laboratories and Alex Alderton, Roberto Amato, Jeffrey Barrett, Sonia Goncalves, Ewan Harrison, David K. Jackson, Ian Johnston, Dominic Kwiatkowski, Cordelia Langford, John Sillitoe on behalf of the Wellcome Sanger Institute COVID-19 Surveillance Team |
| EPI_ISL_2717007 | Lighthouse Lab in Milton Keynes | Wellcome Sanger Institute for the COVID-19 Genomics UK (COG-UK) Consortium | The Lighthouse Lab in Milton Keynes and Alex Alderton, Roberto Amato, Jeffrey Barrett, Sonia Goncalves, Ewan Harrison, David K. Jackson, Ian Johnston, Dominic Kwiatkowski, Cordelia Langford, John Sillitoe on behalf of the Wellcome Sanger Institute COVID-19 Surveillance Team |

| | | | |
|---|---|---|---|
| EPI_ISL_2717008 | Health Services Laboratories | Wellcome Sanger Institute for the COVID-19 Genomics UK (COG-UK) Consortium | Health Services Laboratories and Alex Alderton, Roberto Amato, Jeffrey Barrett, Sonia Goncalves, Ewan Harrison, David K. Jackson, Ian Johnston, Dominic Kwiatkowski, Cordelia Langford, John Sillitoe on behalf of the Wellcome Sanger Institute COVID-19 Surveillance Team |
| EPI_ISL_2717009, EPI_ISL_2717010 | Lighthouse Lab in Milton Keynes | Wellcome Sanger Institute for the COVID-19 Genomics UK (COG-UK) Consortium | The Lighthouse Lab in Milton Keynes and Alex Alderton, Roberto Amato, Jeffrey Barrett, Sonia Goncalves, Ewan Harrison, David K. Jackson, Ian Johnston, Dominic Kwiatkowski, Cordelia Langford, John Sillitoe on behalf of the Wellcome Sanger Institute COVID-19 Surveillance Team |
| EPI_ISL_2717011 | Lighthouse Lab in Alderley Park | Wellcome Sanger Institute for the COVID-19 Genomics UK (COG-UK) Consortium | Jacquelyn Wynn, Mairead Hyland, The Lighthouse Lab in Alderley Park and Alex Alderton, Roberto Amato, Jeffrey Barrett, Sonia Goncalves, Ewan Harrison, David K. Jackson, Ian Johnston, Dominic Kwiatkowski, Cordelia Langford, John Sillitoe on behalf of the Wellcome Sanger Institute COVID-19 Surveillance Team |
| EPI_ISL_2717012 | Lighthouse Lab in Milton Keynes | Wellcome Sanger Institute for the COVID-19 Genomics UK (COG-UK) Consortium | The Lighthouse Lab in Milton Keynes and Alex Alderton, Roberto Amato, Jeffrey Barrett, Sonia Goncalves, Ewan Harrison, David K. Jackson, Ian Johnston, Dominic Kwiatkowski, Cordelia Langford, John Sillitoe on behalf of the Wellcome Sanger Institute COVID-19 Surveillance Team |
| EPI_ISL_2717013 | Lighthouse Lab in Alderley Park | Wellcome Sanger Institute for the COVID-19 Genomics UK (COG-UK) Consortium | Jacquelyn Wynn, Mairead Hyland, The Lighthouse Lab in Alderley Park and Alex Alderton, Roberto Amato, Jeffrey Barrett, Sonia Goncalves, Ewan Harrison, David K. Jackson, Ian Johnston, Dominic Kwiatkowski, Cordelia Langford, John Sillitoe on behalf of the Wellcome Sanger Institute COVID-19 Surveillance Team |
| EPI_ISL_2717014 | Lighthouse Lab in Milton Keynes | Wellcome Sanger Institute for the COVID-19 Genomics UK (COG-UK) Consortium | The Lighthouse Lab in Milton Keynes and Alex Alderton, Roberto Amato, Jeffrey Barrett, Sonia Goncalves, Ewan Harrison, David K. Jackson, Ian Johnston, Dominic Kwiatkowski, Cordelia Langford, John Sillitoe on behalf of the Wellcome Sanger Institute COVID-19 Surveillance Team |
| EPI_ISL_2717015 | Lighthouse Lab in Glasgow | Wellcome Sanger Institute for the COVID-19 Genomics UK (COG-UK) Consortium | Harper VanSteenhouse, Yumi Kasai, David Gray, Carol Clugston, Anna Dominiczak and Alex Alderton, Roberto Amato, Jeffrey Barrett, Sonia Goncalves, Ewan Harrison, David K. Jackson, Ian Johnston, Dominic Kwiatkowski, Cordelia Langford, John Sillitoe on behalf of the Wellcome Sanger Institute COVID-19 Surveillance Team |
| EPI_ISL_2717016, EPI_ISL_2717017 | Health Services Laboratories | Wellcome Sanger Institute for the COVID-19 Genomics UK (COG-UK) Consortium | Health Services Laboratories and Alex Alderton, Roberto Amato, Jeffrey Barrett, Sonia Goncalves, Ewan Harrison, David K. Jackson, Ian Johnston, Dominic Kwiatkowski, Cordelia Langford, John Sillitoe on behalf of the Wellcome Sanger Institute COVID-19 Surveillance Team |
| EPI_ISL_2717018 | Lighthouse Lab in Milton Keynes | Wellcome Sanger Institute for the COVID-19 Genomics UK (COG-UK) Consortium | The Lighthouse Lab in Milton Keynes and Alex Alderton, Roberto Amato, Jeffrey Barrett, Sonia Goncalves, Ewan Harrison, David K. Jackson, Ian Johnston, Dominic Kwiatkowski, Cordelia Langford, John Sillitoe on behalf of the Wellcome Sanger Institute COVID-19 Surveillance Team |
| EPI_ISL_2717019, EPI_ISL_2717020, EPI_ISL_2717021 | Lighthouse Lab in Alderley Park | Wellcome Sanger Institute for the COVID-19 Genomics UK (COG-UK) Consortium | Jacquelyn Wynn, Mairead Hyland, The Lighthouse Lab in Alderley Park and Alex Alderton, Roberto Amato, Jeffrey Barrett, Sonia Goncalves, Ewan Harrison, David K. Jackson, Ian Johnston, Dominic Kwiatkowski, Cordelia Langford, John Sillitoe on behalf of the Wellcome Sanger Institute COVID-19 Surveillance Team |
| EPI_ISL_2717022, EPI_ISL_2717023 | Health Services Laboratories | Wellcome Sanger Institute for the COVID-19 Genomics UK (COG-UK) Consortium | Health Services Laboratories and Alex Alderton, Roberto Amato, Jeffrey Barrett, Sonia Goncalves, Ewan Harrison, David K. Jackson, Ian Johnston, Dominic Kwiatkowski, Cordelia Langford, John Sillitoe on behalf of the Wellcome Sanger Institute COVID-19 Surveillance Team |
| EPI_ISL_2717024 | Lighthouse Lab in Milton Keynes | Wellcome Sanger Institute for the COVID-19 Genomics UK (COG-UK) Consortium | The Lighthouse Lab in Milton Keynes and Alex Alderton, Roberto Amato, Jeffrey Barrett, Sonia Goncalves, Ewan Harrison, David K. Jackson, Ian Johnston, Dominic Kwiatkowski, Cordelia Langford, John Sillitoe on behalf of the Wellcome Sanger Institute COVID-19 Surveillance Team |
| EPI_ISL_2717025 | Lighthouse Lab in Glasgow | Wellcome Sanger Institute for the COVID-19 Genomics UK (COG-UK) Consortium | Harper VanSteenhouse, Yumi Kasai, David Gray, Carol Clugston, Anna Dominiczak and Alex Alderton, Roberto Amato, Jeffrey Barrett, Sonia Goncalves, Ewan Harrison, David K. Jackson, Ian Johnston, Dominic Kwiatkowski, Cordelia Langford, John Sillitoe on behalf of the Wellcome Sanger Institute COVID-19 Surveillance Team |
| EPI_ISL_2717026 | Lighthouse Lab in Alderley Park | Wellcome Sanger Institute for the COVID-19 Genomics UK (COG-UK) Consortium | Jacquelyn Wynn, Mairead Hyland, The Lighthouse Lab in Alderley Park and Alex Alderton, Roberto Amato, Jeffrey Barrett, Sonia Goncalves, Ewan Harrison, David K. Jackson, Ian Johnston, Dominic Kwiatkowski, Cordelia Langford, John Sillitoe on behalf of the Wellcome Sanger Institute COVID-19 Surveillance Team |
| EPI_ISL_2717027, EPI_ISL_2717028, EPI_ISL_2717029 | Lighthouse Lab in Milton Keynes | Wellcome Sanger Institute for the COVID-19 Genomics UK (COG-UK) Consortium | The Lighthouse Lab in Milton Keynes and Alex Alderton, Roberto Amato, Jeffrey Barrett, Sonia Goncalves, Ewan Harrison, David K. Jackson, Ian Johnston, Dominic Kwiatkowski, Cordelia Langford, John Sillitoe on behalf of the Wellcome Sanger Institute COVID-19 Surveillance Team |
| EPI_ISL_2717030 | Health Services Laboratories | Wellcome Sanger Institute for the COVID-19 Genomics UK (COG-UK) Consortium | Health Services Laboratories and Alex Alderton, Roberto Amato, Jeffrey Barrett, Sonia Goncalves, Ewan Harrison, David K. Jackson, Ian Johnston, Dominic Kwiatkowski, Cordelia Langford, John Sillitoe on behalf of the Wellcome Sanger Institute COVID-19 Surveillance Team |
| EPI_ISL_2717031 | Lighthouse Lab in Glasgow | Wellcome Sanger Institute for the COVID-19 Genomics UK (COG-UK) Consortium | Harper VanSteenhouse, Yumi Kasai, David Gray, Carol Clugston, Anna Dominiczak and Alex Alderton, Roberto Amato, Jeffrey Barrett, Sonia Goncalves, Ewan Harrison, David K. Jackson, Ian Johnston, Dominic Kwiatkowski, Cordelia Langford, John Sillitoe on behalf of the Wellcome Sanger Institute COVID-19 Surveillance Team |
| EPI_ISL_2717032 | Health Services Laboratories | Wellcome Sanger Institute for the COVID-19 Genomics UK (COG-UK) Consortium | Health Services Laboratories and Alex Alderton, Roberto Amato, Jeffrey Barrett, Sonia Goncalves, Ewan Harrison, David K. Jackson, Ian Johnston, Dominic Kwiatkowski, Cordelia Langford, John Sillitoe on behalf of the Wellcome Sanger Institute COVID-19 Surveillance Team |
| EPI_ISL_2717033, EPI_ISL_2717034, EPI_ISL_2717035, EPI_ISL_2717036 | Lighthouse Lab in Milton Keynes | Wellcome Sanger Institute for the COVID-19 Genomics UK (COG-UK) Consortium | The Lighthouse Lab in Milton Keynes and Alex Alderton, Roberto Amato, Jeffrey Barrett, Sonia Goncalves, Ewan Harrison, David K. Jackson, Ian Johnston, Dominic Kwiatkowski, Cordelia Langford, John Sillitoe on behalf of the Wellcome Sanger Institute COVID-19 Surveillance Team |
| EPI_ISL_2717037 | Lighthouse Lab in Alderley Park | Wellcome Sanger Institute for the COVID-19 Genomics UK (COG-UK) Consortium | Jacquelyn Wynn, Mairead Hyland, The Lighthouse Lab in Alderley Park and Alex Alderton, Roberto Amato, Jeffrey Barrett, Sonia Goncalves, Ewan Harrison, David K. Jackson, Ian Johnston, Dominic Kwiatkowski, Cordelia Langford, John Sillitoe on behalf of the Wellcome Sanger Institute COVID-19 Surveillance Team |
| EPI_ISL_2717038 | Lighthouse Lab in Glasgow | Wellcome Sanger Institute for the COVID-19 Genomics UK (COG-UK) Consortium | Harper VanSteenhouse, Yumi Kasai, David Gray, Carol Clugston, Anna Dominiczak and Alex Alderton, Roberto Amato, Jeffrey Barrett, Sonia Goncalves, Ewan Harrison, David K. Jackson, Ian Johnston, Dominic Kwiatkowski, Cordelia Langford, John Sillitoe on behalf of the Wellcome Sanger Institute COVID-19 Surveillance Team |
| EPI_ISL_2717039 | Lighthouse Lab in Milton Keynes | Wellcome Sanger Institute for the COVID-19 Genomics UK (COG-UK) Consortium | The Lighthouse Lab in Milton Keynes and Alex Alderton, Roberto Amato, Jeffrey Barrett, Sonia Goncalves, Ewan Harrison, David K. Jackson, Ian Johnston, Dominic Kwiatkowski, Cordelia Langford, John Sillitoe on behalf of the Wellcome Sanger Institute COVID-19 Surveillance Team |
| EPI_ISL_2717040 | Lighthouse Lab in Alderley Park | Wellcome Sanger Institute for the COVID-19 Genomics UK (COG-UK) Consortium | Jacquelyn Wynn, Mairead Hyland, The Lighthouse Lab in Alderley Park and Alex Alderton, Roberto Amato, Jeffrey Barrett, Sonia Goncalves, Ewan Harrison, David K. Jackson, Ian Johnston, Dominic Kwiatkowski, Cordelia Langford, John Sillitoe on behalf of the Wellcome Sanger Institute COVID-19 Surveillance Team |
| EPI_ISL_2717041 | Lighthouse Lab in Milton Keynes | Wellcome Sanger Institute for the COVID-19 Genomics UK (COG-UK) Consortium | The Lighthouse Lab in Milton Keynes and Alex Alderton, Roberto Amato, Jeffrey Barrett, Sonia Goncalves, Ewan Harrison, David K. Jackson, Ian Johnston, Dominic Kwiatkowski, Cordelia Langford, John Sillitoe on behalf of the Wellcome Sanger Institute COVID-19 Surveillance Team |
| EPI_ISL_2717042 | Health Services Laboratories | Wellcome Sanger Institute for the COVID-19 Genomics UK (COG-UK) Consortium | Health Services Laboratories and Alex Alderton, Roberto Amato, Jeffrey Barrett, Sonia Goncalves, Ewan Harrison, David K. Jackson, Ian Johnston, Dominic Kwiatkowski, Cordelia Langford, John Sillitoe on behalf of the Wellcome Sanger Institute COVID-19 Surveillance Team |
| EPI_ISL_2717043, EPI_ISL_2717044, EPI_ISL_2717045, EPI_ISL_2717046 | Lighthouse Lab in Milton Keynes | Wellcome Sanger Institute for the COVID-19 Genomics UK (COG-UK) Consortium | The Lighthouse Lab in Milton Keynes and Alex Alderton, Roberto Amato, Jeffrey Barrett, Sonia Goncalves, Ewan Harrison, David K. Jackson, Ian Johnston, Dominic Kwiatkowski, Cordelia Langford, John Sillitoe on behalf of the Wellcome Sanger Institute COVID-19 Surveillance Team |
| EPI_ISL_2717047 | Health Services Laboratories | Wellcome Sanger Institute for the COVID-19 Genomics UK (COG-UK) Consortium | Health Services Laboratories and Alex Alderton, Roberto Amato, Jeffrey Barrett, Sonia Goncalves, Ewan Harrison, David K. Jackson, Ian Johnston, Dominic Kwiatkowski, Cordelia Langford, John Sillitoe on behalf of the Wellcome Sanger Institute COVID-19 Surveillance Team |
| EPI_ISL_2717048 | Lighthouse Lab in Milton Keynes | Wellcome Sanger Institute for the COVID-19 Genomics UK (COG-UK) Consortium | The Lighthouse Lab in Milton Keynes and Alex Alderton, Roberto Amato, Jeffrey Barrett, Sonia Goncalves, Ewan Harrison, David K. Jackson, Ian Johnston, Dominic Kwiatkowski, Cordelia Langford, John Sillitoe on behalf of the Wellcome Sanger Institute COVID-19 Surveillance Team |
| EPI_ISL_2717049 | Lighthouse Lab in Alderley Park | Wellcome Sanger Institute for the COVID-19 Genomics UK (COG-UK) Consortium | Jacquelyn Wynn, Mairead Hyland, The Lighthouse Lab in Alderley Park and Alex Alderton, Roberto Amato, Jeffrey Barrett, Sonia Goncalves, Ewan Harrison, David K. Jackson, Ian Johnston, Dominic Kwiatkowski, Cordelia Langford, John Sillitoe on behalf of the Wellcome Sanger Institute COVID-19 Surveillance Team |
| EPI_ISL_2717050, EPI_ISL_2717051 | Lighthouse Lab in Milton Keynes | Wellcome Sanger Institute for the COVID-19 Genomics UK (COG-UK) Consortium | The Lighthouse Lab in Milton Keynes and Alex Alderton, Roberto Amato, Jeffrey Barrett, Sonia Goncalves, Ewan Harrison, David K. Jackson, Ian Johnston, Dominic Kwiatkowski, Cordelia Langford, John Sillitoe on behalf of the Wellcome Sanger Institute COVID-19 Surveillance Team |
| EPI_ISL_2717052, EPI_ISL_2717053 | Lighthouse Lab in Alderley Park | Wellcome Sanger Institute for the COVID-19 Genomics UK (COG-UK) Consortium | Jacquelyn Wynn, Mairead Hyland, The Lighthouse Lab in Alderley Park and Alex Alderton, Roberto Amato, Jeffrey Barrett, Sonia Goncalves, Ewan Harrison, David K. Jackson, Ian Johnston, Dominic Kwiatkowski, Cordelia Langford, John Sillitoe on behalf of the Wellcome Sanger Institute COVID-19 Surveillance Team |
| EPI_ISL_2717054 | Health Services Laboratories | Wellcome Sanger Institute for the COVID-19 Genomics UK (COG-UK) Consortium | Health Services Laboratories and Alex Alderton, Roberto Amato, Jeffrey Barrett, Sonia Goncalves, Ewan Harrison, David K. Jackson, Ian Johnston, Dominic Kwiatkowski, Cordelia Langford, John Sillitoe on behalf of the Wellcome Sanger Institute COVID-19 Surveillance Team |
| EPI_ISL_2717055, EPI_ISL_2717056 | Lighthouse Lab in Alderley Park | Wellcome Sanger Institute for the COVID-19 Genomics UK (COG-UK) Consortium | Jacquelyn Wynn, Mairead Hyland, The Lighthouse Lab in Alderley Park and Alex Alderton, Roberto Amato, Jeffrey Barrett, Sonia Goncalves, Ewan Harrison, David K. Jackson, Ian Johnston, Dominic Kwiatkowski, Cordelia Langford, John Sillitoe on behalf of the Wellcome Sanger Institute COVID-19 Surveillance Team |
| EPI_ISL_2717057 | Lighthouse Lab in Milton Keynes | Wellcome Sanger Institute for the COVID-19 Genomics UK (COG-UK) Consortium | The Lighthouse Lab in Milton Keynes and Alex Alderton, Roberto Amato, Jeffrey Barrett, Sonia Goncalves, Ewan Harrison, David K. Jackson, Ian Johnston, Dominic Kwiatkowski, Cordelia Langford, John Sillitoe on behalf of the Wellcome Sanger Institute COVID-19 Surveillance Team |
| EPI_ISL_2717058, EPI_ISL_2717059 | Health Services Laboratories | Wellcome Sanger Institute for the COVID-19 Genomics UK (COG-UK) Consortium | Health Services Laboratories and Alex Alderton, Roberto Amato, Jeffrey Barrett, Sonia Goncalves, Ewan Harrison, David K. Jackson, Ian Johnston, Dominic Kwiatkowski, Cordelia Langford, John Sillitoe on behalf of the Wellcome Sanger Institute COVID-19 Surveillance Team |
| EPI_ISL_2717060 | Lighthouse Lab in Milton Keynes | Wellcome Sanger Institute for the COVID-19 Genomics UK (COG-UK) Consortium | The Lighthouse Lab in Milton Keynes and Alex Alderton, Roberto Amato, Jeffrey Barrett, Sonia Goncalves, Ewan Harrison, David K. Jackson, Ian Johnston, Dominic Kwiatkowski, Cordelia Langford, John Sillitoe on behalf of the Wellcome Sanger Institute COVID-19 Surveillance Team |
| EPI_ISL_2717061 | Health Services Laboratories | Wellcome Sanger Institute for the COVID-19 Genomics UK (COG-UK) Consortium | Health Services Laboratories and Alex Alderton, Roberto Amato, Jeffrey Barrett, Sonia Goncalves, Ewan Harrison, David K. Jackson, Ian Johnston, Dominic Kwiatkowski, Cordelia Langford, John Sillitoe on behalf of the Wellcome Sanger Institute COVID-19 Surveillance Team |
| EPI_ISL_2717062 | Lighthouse Lab in Milton Keynes | Wellcome Sanger Institute for the COVID-19 Genomics UK (COG-UK) Consortium | The Lighthouse Lab in Milton Keynes and Alex Alderton, Roberto Amato, Jeffrey Barrett, Sonia Goncalves, Ewan Harrison, David K. Jackson, Ian Johnston, Dominic Kwiatkowski, Cordelia Langford, John Sillitoe on behalf of the Wellcome Sanger Institute COVID-19 Surveillance Team |
| EPI_ISL_2717063 | Lighthouse Lab in Glasgow | Wellcome Sanger Institute for the COVID-19 Genomics UK (COG-UK) Consortium | Harper VanSteenhouse, Yumi Kasai, David Gray, Carol Clugston, Anna Dominiczak and Alex Alderton, Roberto Amato, Jeffrey Barrett, Sonia Goncalves, Ewan Harrison, David K. Jackson, Ian Johnston, Dominic Kwiatkowski, Cordelia Langford, John Sillitoe on behalf of the Wellcome Sanger Institute COVID-19 Surveillance Team |
| EPI_ISL_2717064, EPI_ISL_2717066, EPI_ISL_2717067 | Lighthouse Lab in Milton Keynes | Wellcome Sanger Institute for the COVID-19 Genomics UK (COG-UK) Consortium | The Lighthouse Lab in Milton Keynes and Alex Alderton, Roberto Amato, Jeffrey Barrett, Sonia Goncalves, Ewan Harrison, David K. Jackson, Ian Johnston, Dominic Kwiatkowski, Cordelia Langford, John Sillitoe on behalf of the Wellcome Sanger Institute COVID-19 Surveillance Team |
| EPI_ISL_2717068 | Lighthouse Lab in Alderley Park | Wellcome Sanger Institute for the COVID-19 Genomics UK (COG-UK) Consortium | Jacquelyn Wynn, Mairead Hyland, The Lighthouse Lab in Alderley Park and Alex Alderton, Roberto Amato, Jeffrey Barrett, Sonia Goncalves, Ewan Harrison, David K. Jackson, Ian Johnston, Dominic Kwiatkowski, Cordelia Langford, John Sillitoe on behalf of the Wellcome Sanger Institute COVID-19 Surveillance Team |
| EPI_ISL_2717069 | Lighthouse Lab in Milton Keynes | Wellcome Sanger Institute for the COVID-19 Genomics UK (COG-UK) Consortium | The Lighthouse Lab in Milton Keynes and Alex Alderton, Roberto Amato, Jeffrey Barrett, Sonia Goncalves, Ewan Harrison, David K. Jackson, Ian Johnston, Dominic Kwiatkowski, Cordelia Langford, John Sillitoe on behalf of the Wellcome Sanger Institute COVID-19 Surveillance Team |

| | | | |
|---|---|---|---|
| EPI_ISL_2717070 | Lighthouse Lab in Alderley Park | Wellcome Sanger Institute for the COVID-19 Genomics UK (COG-UK) Consortium | Jacquelyn Wynn, Mairead Hyland, The Lighthouse Lab in Alderley Park and Alex Alderton, Roberto Amato, Jeffrey Barrett, Sonia Goncalves, Ewan Harrison, David K. Jackson, Ian Johnston, Dominic Kwiatkowski, Cordelia Langford, John Sillitoe on behalf of the Wellcome Sanger Institute COVID-19 Surveillance Team |
| EPI_ISL_2717071 | Health Services Laboratories | Wellcome Sanger Institute for the COVID-19 Genomics UK (COG-UK) Consortium | Health Services Laboratories and Alex Alderton, Roberto Amato, Jeffrey Barrett, Sonia Goncalves, Ewan Harrison, David K. Jackson, Ian Johnston, Dominic Kwiatkowski, Cordelia Langford, John Sillitoe on behalf of the Wellcome Sanger Institute COVID-19 Surveillance Team |
| EPI_ISL_2717072 | Lighthouse Lab in Milton Keynes | Wellcome Sanger Institute for the COVID-19 Genomics UK (COG-UK) Consortium | The Lighthouse Lab in Milton Keynes and Alex Alderton, Roberto Amato, Jeffrey Barrett, Sonia Goncalves, Ewan Harrison, David K. Jackson, Ian Johnston, Dominic Kwiatkowski, Cordelia Langford, John Sillitoe on behalf of the Wellcome Sanger Institute COVID-19 Surveillance Team |
| EPI_ISL_2717073, EPI_ISL_2717074 | Lighthouse Lab in Alderley Park | Wellcome Sanger Institute for the COVID-19 Genomics UK (COG-UK) Consortium | Jacquelyn Wynn, Mairead Hyland, The Lighthouse Lab in Alderley Park and Alex Alderton, Roberto Amato, Jeffrey Barrett, Sonia Goncalves, Ewan Harrison, David K. Jackson, Ian Johnston, Dominic Kwiatkowski, Cordelia Langford, John Sillitoe on behalf of the Wellcome Sanger Institute COVID-19 Surveillance Team |
| EPI_ISL_2717075 | Lighthouse Lab in Milton Keynes | Wellcome Sanger Institute for the COVID-19 Genomics UK (COG-UK) Consortium | The Lighthouse Lab in Milton Keynes and Alex Alderton, Roberto Amato, Jeffrey Barrett, Sonia Goncalves, Ewan Harrison, David K. Jackson, Ian Johnston, Dominic Kwiatkowski, Cordelia Langford, John Sillitoe on behalf of the Wellcome Sanger Institute COVID-19 Surveillance Team |
| EPI_ISL_2717076 | Lighthouse Lab in Alderley Park | Wellcome Sanger Institute for the COVID-19 Genomics UK (COG-UK) Consortium | Jacquelyn Wynn, Mairead Hyland, The Lighthouse Lab in Alderley Park and Alex Alderton, Roberto Amato, Jeffrey Barrett, Sonia Goncalves, Ewan Harrison, David K. Jackson, Ian Johnston, Dominic Kwiatkowski, Cordelia Langford, John Sillitoe on behalf of the Wellcome Sanger Institute COVID-19 Surveillance Team |
| EPI_ISL_2717077, EPI_ISL_2717078, EPI_ISL_2717079 | Lighthouse Lab in Milton Keynes | Wellcome Sanger Institute for the COVID-19 Genomics UK (COG-UK) Consortium | The Lighthouse Lab in Milton Keynes and Alex Alderton, Roberto Amato, Jeffrey Barrett, Sonia Goncalves, Ewan Harrison, David K. Jackson, Ian Johnston, Dominic Kwiatkowski, Cordelia Langford, John Sillitoe on behalf of the Wellcome Sanger Institute COVID-19 Surveillance Team |
| EPI_ISL_2717080, EPI_ISL_2717081 | Health Services Laboratories | Wellcome Sanger Institute for the COVID-19 Genomics UK (COG-UK) Consortium | Health Services Laboratories and Alex Alderton, Roberto Amato, Jeffrey Barrett, Sonia Goncalves, Ewan Harrison, David K. Jackson, Ian Johnston, Dominic Kwiatkowski, Cordelia Langford, John Sillitoe on behalf of the Wellcome Sanger Institute COVID-19 Surveillance Team |
| EPI_ISL_2717082, EPI_ISL_2717083, EPI_ISL_2717084 | Lighthouse Lab in Milton Keynes | Wellcome Sanger Institute for the COVID-19 Genomics UK (COG-UK) Consortium | The Lighthouse Lab in Milton Keynes and Alex Alderton, Roberto Amato, Jeffrey Barrett, Sonia Goncalves, Ewan Harrison, David K. Jackson, Ian Johnston, Dominic Kwiatkowski, Cordelia Langford, John Sillitoe on behalf of the Wellcome Sanger Institute COVID-19 Surveillance Team |
| EPI_ISL_2717085 | Lighthouse Lab in Alderley Park | Wellcome Sanger Institute for the COVID-19 Genomics UK (COG-UK) Consortium | Jacquelyn Wynn, Mairead Hyland, The Lighthouse Lab in Alderley Park and Alex Alderton, Roberto Amato, Jeffrey Barrett, Sonia Goncalves, Ewan Harrison, David K. Jackson, Ian Johnston, Dominic Kwiatkowski, Cordelia Langford, John Sillitoe on behalf of the Wellcome Sanger Institute COVID-19 Surveillance Team |
| EPI_ISL_2717086 | Lighthouse Lab in Milton Keynes | Wellcome Sanger Institute for the COVID-19 Genomics UK (COG-UK) Consortium | The Lighthouse Lab in Milton Keynes and Alex Alderton, Roberto Amato, Jeffrey Barrett, Sonia Goncalves, Ewan Harrison, David K. Jackson, Ian Johnston, Dominic Kwiatkowski, Cordelia Langford, John Sillitoe on behalf of the Wellcome Sanger Institute COVID-19 Surveillance Team |
| EPI_ISL_2717087 | Lighthouse Lab in Alderley Park | Wellcome Sanger Institute for the COVID-19 Genomics UK (COG-UK) Consortium | Jacquelyn Wynn, Mairead Hyland, The Lighthouse Lab in Alderley Park and Alex Alderton, Roberto Amato, Jeffrey Barrett, Sonia Goncalves, Ewan Harrison, David K. Jackson, Ian Johnston, Dominic Kwiatkowski, Cordelia Langford, John Sillitoe on behalf of the Wellcome Sanger Institute COVID-19 Surveillance Team |
| EPI_ISL_2717088 | Lighthouse Lab in Milton Keynes | Wellcome Sanger Institute for the COVID-19 Genomics UK (COG-UK) Consortium | The Lighthouse Lab in Milton Keynes and Alex Alderton, Roberto Amato, Jeffrey Barrett, Sonia Goncalves, Ewan Harrison, David K. Jackson, Ian Johnston, Dominic Kwiatkowski, Cordelia Langford, John Sillitoe on behalf of the Wellcome Sanger Institute COVID-19 Surveillance Team |
| EPI_ISL_2717089, EPI_ISL_2717090 | Health Services Laboratories | Wellcome Sanger Institute for the COVID-19 Genomics UK (COG-UK) Consortium | Health Services Laboratories and Alex Alderton, Roberto Amato, Jeffrey Barrett, Sonia Goncalves, Ewan Harrison, David K. Jackson, Ian Johnston, Dominic Kwiatkowski, Cordelia Langford, John Sillitoe on behalf of the Wellcome Sanger Institute COVID-19 Surveillance Team |
| EPI_ISL_2717091 | Lighthouse Lab in Alderley Park | Wellcome Sanger Institute for the COVID-19 Genomics UK (COG-UK) Consortium | Jacquelyn Wynn, Mairead Hyland, The Lighthouse Lab in Alderley Park and Alex Alderton, Roberto Amato, Jeffrey Barrett, Sonia Goncalves, Ewan Harrison, David K. Jackson, Ian Johnston, Dominic Kwiatkowski, Cordelia Langford, John Sillitoe on behalf of the Wellcome Sanger Institute COVID-19 Surveillance Team |
| EPI_ISL_2717092, EPI_ISL_2717093, EPI_ISL_2717094 | Lighthouse Lab in Milton Keynes | Wellcome Sanger Institute for the COVID-19 Genomics UK (COG-UK) Consortium | The Lighthouse Lab in Milton Keynes and Alex Alderton, Roberto Amato, Jeffrey Barrett, Sonia Goncalves, Ewan Harrison, David K. Jackson, Ian Johnston, Dominic Kwiatkowski, Cordelia Langford, John Sillitoe on behalf of the Wellcome Sanger Institute COVID-19 Surveillance Team |
| EPI_ISL_2717095 | Lighthouse Lab in Alderley Park | Wellcome Sanger Institute for the COVID-19 Genomics UK (COG-UK) Consortium | Jacquelyn Wynn, Mairead Hyland, The Lighthouse Lab in Alderley Park and Alex Alderton, Roberto Amato, Jeffrey Barrett, Sonia Goncalves, Ewan Harrison, David K. Jackson, Ian Johnston, Dominic Kwiatkowski, Cordelia Langford, John Sillitoe on behalf of the Wellcome Sanger Institute COVID-19 Surveillance Team |
| EPI_ISL_2717096, EPI_ISL_2717097, EPI_ISL_2717098, EPI_ISL_2717099, EPI_ISL_2717100 | Lighthouse Lab in Milton Keynes | Wellcome Sanger Institute for the COVID-19 Genomics UK (COG-UK) Consortium | The Lighthouse Lab in Milton Keynes and Alex Alderton, Roberto Amato, Jeffrey Barrett, Sonia Goncalves, Ewan Harrison, David K. Jackson, Ian Johnston, Dominic Kwiatkowski, Cordelia Langford, John Sillitoe on behalf of the Wellcome Sanger Institute COVID-19 Surveillance Team |
| EPI_ISL_2717101 | Lighthouse Lab in Alderley Park | Wellcome Sanger Institute for the COVID-19 Genomics UK (COG-UK) Consortium | Jacquelyn Wynn, Mairead Hyland, The Lighthouse Lab in Alderley Park and Alex Alderton, Roberto Amato, Jeffrey Barrett, Sonia Goncalves, Ewan Harrison, David K. Jackson, Ian Johnston, Dominic Kwiatkowski, Cordelia Langford, John Sillitoe on behalf of the Wellcome Sanger Institute COVID-19 Surveillance Team |
| EPI_ISL_2717102, EPI_ISL_2717103 | Lighthouse Lab in Milton Keynes | Wellcome Sanger Institute for the COVID-19 Genomics UK (COG-UK) Consortium | The Lighthouse Lab in Milton Keynes and Alex Alderton, Roberto Amato, Jeffrey Barrett, Sonia Goncalves, Ewan Harrison, David K. Jackson, Ian Johnston, Dominic Kwiatkowski, Cordelia Langford, John Sillitoe on behalf of the Wellcome Sanger Institute COVID-19 Surveillance Team |
| EPI_ISL_2717104 | Health Services Laboratories | Wellcome Sanger Institute for the COVID-19 Genomics UK (COG-UK) Consortium | Health Services Laboratories and Alex Alderton, Roberto Amato, Jeffrey Barrett, Sonia Goncalves, Ewan Harrison, David K. Jackson, Ian Johnston, Dominic Kwiatkowski, Cordelia Langford, John Sillitoe on behalf of the Wellcome Sanger Institute COVID-19 Surveillance Team |
| EPI_ISL_2717105 | Lighthouse Lab in Alderley Park | Wellcome Sanger Institute for the COVID-19 Genomics UK (COG-UK) Consortium | Jacquelyn Wynn, Mairead Hyland, The Lighthouse Lab in Alderley Park and Alex Alderton, Roberto Amato, Jeffrey Barrett, Sonia Goncalves, Ewan Harrison, David K. Jackson, Ian Johnston, Dominic Kwiatkowski, Cordelia Langford, John Sillitoe on behalf of the Wellcome Sanger Institute COVID-19 Surveillance Team |
| EPI_ISL_2717106, EPI_ISL_2717107 | Health Services Laboratories | Wellcome Sanger Institute for the COVID-19 Genomics UK (COG-UK) Consortium | Health Services Laboratories and Alex Alderton, Roberto Amato, Jeffrey Barrett, Sonia Goncalves, Ewan Harrison, David K. Jackson, Ian Johnston, Dominic Kwiatkowski, Cordelia Langford, John Sillitoe on behalf of the Wellcome Sanger Institute COVID-19 Surveillance Team |
| EPI_ISL_2717108 | Lighthouse Lab in Alderley Park | Wellcome Sanger Institute for the COVID-19 Genomics UK (COG-UK) Consortium | Jacquelyn Wynn, Mairead Hyland, The Lighthouse Lab in Alderley Park and Alex Alderton, Roberto Amato, Jeffrey Barrett, Sonia Goncalves, Ewan Harrison, David K. Jackson, Ian Johnston, Dominic Kwiatkowski, Cordelia Langford, John Sillitoe on behalf of the Wellcome Sanger Institute COVID-19 Surveillance Team |
| EPI_ISL_2717109, EPI_ISL_2717110 | Health Services Laboratories | Wellcome Sanger Institute for the COVID-19 Genomics UK (COG-UK) Consortium | Health Services Laboratories and Alex Alderton, Roberto Amato, Jeffrey Barrett, Sonia Goncalves, Ewan Harrison, David K. Jackson, Ian Johnston, Dominic Kwiatkowski, Cordelia Langford, John Sillitoe on behalf of the Wellcome Sanger Institute COVID-19 Surveillance Team |
| EPI_ISL_2717111 | Lighthouse Lab in Glasgow | Wellcome Sanger Institute for the COVID-19 Genomics UK (COG-UK) Consortium | Harper VanSteenhouse, Yumi Kasai, David Gray, Carol Clugston, Anna Dominiczak and Alex Alderton, Roberto Amato, Jeffrey Barrett, Sonia Goncalves, Ewan Harrison, David K. Jackson, Ian Johnston, Dominic Kwiatkowski, Cordelia Langford, John Sillitoe on behalf of the Wellcome Sanger Institute COVID-19 Surveillance Team |
| EPI_ISL_2717112, EPI_ISL_2717113, EPI_ISL_2717114 | Lighthouse Lab in Alderley Park | Wellcome Sanger Institute for the COVID-19 Genomics UK (COG-UK) Consortium | Jacquelyn Wynn, Mairead Hyland, The Lighthouse Lab in Alderley Park and Alex Alderton, Roberto Amato, Jeffrey Barrett, Sonia Goncalves, Ewan Harrison, David K. Jackson, Ian Johnston, Dominic Kwiatkowski, Cordelia Langford, John Sillitoe on behalf of the Wellcome Sanger Institute COVID-19 Surveillance Team |
| EPI_ISL_2717115, EPI_ISL_2717116 | Lighthouse Lab in Milton Keynes | Wellcome Sanger Institute for the COVID-19 Genomics UK (COG-UK) Consortium | The Lighthouse Lab in Milton Keynes and Alex Alderton, Roberto Amato, Jeffrey Barrett, Sonia Goncalves, Ewan Harrison, David K. Jackson, Ian Johnston, Dominic Kwiatkowski, Cordelia Langford, John Sillitoe on behalf of the Wellcome Sanger Institute COVID-19 Surveillance Team |
| EPI_ISL_2717117, EPI_ISL_2717118 | Lighthouse Lab in Alderley Park | Wellcome Sanger Institute for the COVID-19 Genomics UK (COG-UK) Consortium | Jacquelyn Wynn, Mairead Hyland, The Lighthouse Lab in Alderley Park and Alex Alderton, Roberto Amato, Jeffrey Barrett, Sonia Goncalves, Ewan Harrison, David K. Jackson, Ian Johnston, Dominic Kwiatkowski, Cordelia Langford, John Sillitoe on behalf of the Wellcome Sanger Institute COVID-19 Surveillance Team |
| EPI_ISL_2717119 | Lighthouse Lab in Milton Keynes | Wellcome Sanger Institute for the COVID-19 Genomics UK (COG-UK) Consortium | The Lighthouse Lab in Milton Keynes and Alex Alderton, Roberto Amato, Jeffrey Barrett, Sonia Goncalves, Ewan Harrison, David K. Jackson, Ian Johnston, Dominic Kwiatkowski, Cordelia Langford, John Sillitoe on behalf of the Wellcome Sanger Institute COVID-19 Surveillance Team |
| EPI_ISL_2717120 | Health Services Laboratories | Wellcome Sanger Institute for the COVID-19 Genomics UK (COG-UK) Consortium | Health Services Laboratories and Alex Alderton, Roberto Amato, Jeffrey Barrett, Sonia Goncalves, Ewan Harrison, David K. Jackson, Ian Johnston, Dominic Kwiatkowski, Cordelia Langford, John Sillitoe on behalf of the Wellcome Sanger Institute COVID-19 Surveillance Team |
| EPI_ISL_2717121, EPI_ISL_2717122 | Lighthouse Lab in Milton Keynes | Wellcome Sanger Institute for the COVID-19 Genomics UK (COG-UK) Consortium | The Lighthouse Lab in Milton Keynes and Alex Alderton, Roberto Amato, Jeffrey Barrett, Sonia Goncalves, Ewan Harrison, David K. Jackson, Ian Johnston, Dominic Kwiatkowski, Cordelia Langford, John Sillitoe on behalf of the Wellcome Sanger Institute COVID-19 Surveillance Team |
| EPI_ISL_2717123 | Lighthouse Lab in Alderley Park | Wellcome Sanger Institute for the COVID-19 Genomics UK (COG-UK) Consortium | Jacquelyn Wynn, Mairead Hyland, The Lighthouse Lab in Alderley Park and Alex Alderton, Roberto Amato, Jeffrey Barrett, Sonia Goncalves, Ewan Harrison, David K. Jackson, Ian Johnston, Dominic Kwiatkowski, Cordelia Langford, John Sillitoe on behalf of the Wellcome Sanger Institute COVID-19 Surveillance Team |
| EPI_ISL_2717124 | Lighthouse Lab in Milton Keynes | Wellcome Sanger Institute for the COVID-19 Genomics UK (COG-UK) Consortium | The Lighthouse Lab in Milton Keynes and Alex Alderton, Roberto Amato, Jeffrey Barrett, Sonia Goncalves, Ewan Harrison, David K. Jackson, Ian Johnston, Dominic Kwiatkowski, Cordelia Langford, John Sillitoe on behalf of the Wellcome Sanger Institute COVID-19 Surveillance Team |
| EPI_ISL_2717125, EPI_ISL_2717126 | Lighthouse Lab in Alderley Park | Wellcome Sanger Institute for the COVID-19 Genomics UK (COG-UK) Consortium | Jacquelyn Wynn, Mairead Hyland, The Lighthouse Lab in Alderley Park and Alex Alderton, Roberto Amato, Jeffrey Barrett, Sonia Goncalves, Ewan Harrison, David K. Jackson, Ian Johnston, Dominic Kwiatkowski, Cordelia Langford, John Sillitoe on behalf of the Wellcome Sanger Institute COVID-19 Surveillance Team |
| EPI_ISL_2717127, EPI_ISL_2717128 | Lighthouse Lab in Milton Keynes | Wellcome Sanger Institute for the COVID-19 Genomics UK (COG-UK) Consortium | The Lighthouse Lab in Milton Keynes and Alex Alderton, Roberto Amato, Jeffrey Barrett, Sonia Goncalves, Ewan Harrison, David K. Jackson, Ian Johnston, Dominic Kwiatkowski, Cordelia Langford, John Sillitoe on behalf of the Wellcome Sanger Institute COVID-19 Surveillance Team |
| EPI_ISL_2717129 | Health Services Laboratories | Wellcome Sanger Institute for the COVID-19 Genomics UK (COG-UK) Consortium | Health Services Laboratories and Alex Alderton, Roberto Amato, Jeffrey Barrett, Sonia Goncalves, Ewan Harrison, David K. Jackson, Ian Johnston, Dominic Kwiatkowski, Cordelia Langford, John Sillitoe on behalf of the Wellcome Sanger Institute COVID-19 Surveillance Team |
| EPI_ISL_2717130 | Lighthouse Lab in Alderley Park | Wellcome Sanger Institute for the COVID-19 Genomics UK (COG-UK) Consortium | Jacquelyn Wynn, Mairead Hyland, The Lighthouse Lab in Alderley Park and Alex Alderton, Roberto Amato, Jeffrey Barrett, Sonia Goncalves, Ewan Harrison, David K. Jackson, Ian Johnston, Dominic Kwiatkowski, Cordelia Langford, John Sillitoe on behalf of the Wellcome Sanger Institute COVID-19 Surveillance Team |
| EPI_ISL_2717131, EPI_ISL_2717132, EPI_ISL_2717133 | Lighthouse Lab in Milton Keynes | Wellcome Sanger Institute for the COVID-19 Genomics UK (COG-UK) Consortium | The Lighthouse Lab in Milton Keynes and Alex Alderton, Roberto Amato, Jeffrey Barrett, Sonia Goncalves, Ewan Harrison, David K. Jackson, Ian Johnston, Dominic Kwiatkowski, Cordelia Langford, John Sillitoe on behalf of the Wellcome Sanger Institute COVID-19 Surveillance Team |
| EPI_ISL_2717134, EPI_ISL_2717135, EPI_ISL_2717136 | Lighthouse Lab in Alderley Park | Wellcome Sanger Institute for the COVID-19 Genomics UK (COG-UK) Consortium | Jacquelyn Wynn, Mairead Hyland, The Lighthouse Lab in Alderley Park and Alex Alderton, Roberto Amato, Jeffrey Barrett, Sonia Goncalves, Ewan Harrison, David K. Jackson, Ian Johnston, Dominic Kwiatkowski, Cordelia Langford, John Sillitoe on behalf of the Wellcome Sanger Institute COVID-19 Surveillance Team |
| EPI_ISL_2717137, EPI_ISL_2717138, EPI_ISL_2717139, EPI_ISL_2717140, | Lighthouse Lab in Milton Keynes | Wellcome Sanger Institute for the COVID-19 Genomics UK (COG-UK) Consortium | The Lighthouse Lab in Milton Keynes and Alex Alderton, Roberto Amato, Jeffrey Barrett, Sonia Goncalves, Ewan Harrison, David K. Jackson, Ian Johnston, Dominic Kwiatkowski, Cordelia Langford, John Sillitoe on behalf of the Wellcome Sanger Institute COVID-19 Surveillance Team |

| | | | |
|---|---|---|---|
| EPI_ISL_2717142 | | | |
| EPI_ISL_2717143 | Lighthouse Lab in Alderley Park | Wellcome Sanger Institute for the COVID-19 Genomics UK (COG-UK) Consortium | Jacquelyn Wynn, Mairead Hyland, The Lighthouse Lab in Alderley Park and Alex Alderton, Roberto Amato, Jeffrey Barrett, Sonia Goncalves, Ewan Harrison, David K. Jackson, Ian Johnston, Dominic Kwiatkowski, Cordelia Langford, John Sillitoe on behalf of the Wellcome Sanger Institute COVID-19 Surveillance Team |
| EPI_ISL_2717144, EPI_ISL_2717145, EPI_ISL_2717146 | Lighthouse Lab in Milton Keynes | Wellcome Sanger Institute for the COVID-19 Genomics UK (COG-UK) Consortium | The Lighthouse Lab in Milton Keynes and Alex Alderton, Roberto Amato, Jeffrey Barrett, Sonia Goncalves, Ewan Harrison, David K. Jackson, Ian Johnston, Dominic Kwiatkowski, Cordelia Langford, John Sillitoe on behalf of the Wellcome Sanger Institute COVID-19 Surveillance Team |
| EPI_ISL_2717147 | Health Services Laboratories | Wellcome Sanger Institute for the COVID-19 Genomics UK (COG-UK) Consortium | Health Services Laboratories and Alex Alderton, Roberto Amato, Jeffrey Barrett, Sonia Goncalves, Ewan Harrison, David K. Jackson, Ian Johnston, Dominic Kwiatkowski, Cordelia Langford, John Sillitoe on behalf of the Wellcome Sanger Institute COVID-19 Surveillance Team |
| EPI_ISL_2717148, EPI_ISL_2717149, EPI_ISL_2717150, EPI_ISL_2717151, EPI_ISL_2717152 | Lighthouse Lab in Milton Keynes | Wellcome Sanger Institute for the COVID-19 Genomics UK (COG-UK) Consortium | The Lighthouse Lab in Milton Keynes and Alex Alderton, Roberto Amato, Jeffrey Barrett, Sonia Goncalves, Ewan Harrison, David K. Jackson, Ian Johnston, Dominic Kwiatkowski, Cordelia Langford, John Sillitoe on behalf of the Wellcome Sanger Institute COVID-19 Surveillance Team |
| EPI_ISL_2717153, EPI_ISL_2717154 | Lighthouse Lab in Alderley Park | Wellcome Sanger Institute for the COVID-19 Genomics UK (COG-UK) Consortium | Jacquelyn Wynn, Mairead Hyland, The Lighthouse Lab in Alderley Park and Alex Alderton, Roberto Amato, Jeffrey Barrett, Sonia Goncalves, Ewan Harrison, David K. Jackson, Ian Johnston, Dominic Kwiatkowski, Cordelia Langford, John Sillitoe on behalf of the Wellcome Sanger Institute COVID-19 Surveillance Team |
| EPI_ISL_2717155 | Health Services Laboratories | Wellcome Sanger Institute for the COVID-19 Genomics UK (COG-UK) Consortium | Health Services Laboratories and Alex Alderton, Roberto Amato, Jeffrey Barrett, Sonia Goncalves, Ewan Harrison, David K. Jackson, Ian Johnston, Dominic Kwiatkowski, Cordelia Langford, John Sillitoe on behalf of the Wellcome Sanger Institute COVID-19 Surveillance Team |
| EPI_ISL_2717156, EPI_ISL_2717157 | Lighthouse Lab in Milton Keynes | Wellcome Sanger Institute for the COVID-19 Genomics UK (COG-UK) Consortium | The Lighthouse Lab in Milton Keynes and Alex Alderton, Roberto Amato, Jeffrey Barrett, Sonia Goncalves, Ewan Harrison, David K. Jackson, Ian Johnston, Dominic Kwiatkowski, Cordelia Langford, John Sillitoe on behalf of the Wellcome Sanger Institute COVID-19 Surveillance Team |
| EPI_ISL_2717158 | Health Services Laboratories | Wellcome Sanger Institute for the COVID-19 Genomics UK (COG-UK) Consortium | Health Services Laboratories and Alex Alderton, Roberto Amato, Jeffrey Barrett, Sonia Goncalves, Ewan Harrison, David K. Jackson, Ian Johnston, Dominic Kwiatkowski, Cordelia Langford, John Sillitoe on behalf of the Wellcome Sanger Institute COVID-19 Surveillance Team |
| EPI_ISL_2717159 | Lighthouse Lab in Milton Keynes | Wellcome Sanger Institute for the COVID-19 Genomics UK (COG-UK) Consortium | The Lighthouse Lab in Milton Keynes and Alex Alderton, Roberto Amato, Jeffrey Barrett, Sonia Goncalves, Ewan Harrison, David K. Jackson, Ian Johnston, Dominic Kwiatkowski, Cordelia Langford, John Sillitoe on behalf of the Wellcome Sanger Institute COVID-19 Surveillance Team |
| EPI_ISL_2717160, EPI_ISL_2717161 | Health Services Laboratories | Wellcome Sanger Institute for the COVID-19 Genomics UK (COG-UK) Consortium | Health Services Laboratories and Alex Alderton, Roberto Amato, Jeffrey Barrett, Sonia Goncalves, Ewan Harrison, David K. Jackson, Ian Johnston, Dominic Kwiatkowski, Cordelia Langford, John Sillitoe on behalf of the Wellcome Sanger Institute COVID-19 Surveillance Team |
| EPI_ISL_2717162, EPI_ISL_2717163 | Lighthouse Lab in Alderley Park | Wellcome Sanger Institute for the COVID-19 Genomics UK (COG-UK) Consortium | Jacquelyn Wynn, Mairead Hyland, The Lighthouse Lab in Alderley Park and Alex Alderton, Roberto Amato, Jeffrey Barrett, Sonia Goncalves, Ewan Harrison, David K. Jackson, Ian Johnston, Dominic Kwiatkowski, Cordelia Langford, John Sillitoe on behalf of the Wellcome Sanger Institute COVID-19 Surveillance Team |
| EPI_ISL_2717164 | Health Services Laboratories | Wellcome Sanger Institute for the COVID-19 Genomics UK (COG-UK) Consortium | Health Services Laboratories and Alex Alderton, Roberto Amato, Jeffrey Barrett, Sonia Goncalves, Ewan Harrison, David K. Jackson, Ian Johnston, Dominic Kwiatkowski, Cordelia Langford, John Sillitoe on behalf of the Wellcome Sanger Institute COVID-19 Surveillance Team |
| EPI_ISL_2717165 | Lighthouse Lab in Alderley Park | Wellcome Sanger Institute for the COVID-19 Genomics UK (COG-UK) Consortium | Jacquelyn Wynn, Mairead Hyland, The Lighthouse Lab in Alderley Park and Alex Alderton, Roberto Amato, Jeffrey Barrett, Sonia Goncalves, Ewan Harrison, David K. Jackson, Ian Johnston, Dominic Kwiatkowski, Cordelia Langford, John Sillitoe on behalf of the Wellcome Sanger Institute COVID-19 Surveillance Team |
| EPI_ISL_2717166 | Health Services Laboratories | Wellcome Sanger Institute for the COVID-19 Genomics UK (COG-UK) Consortium | Health Services Laboratories and Alex Alderton, Roberto Amato, Jeffrey Barrett, Sonia Goncalves, Ewan Harrison, David K. Jackson, Ian Johnston, Dominic Kwiatkowski, Cordelia Langford, John Sillitoe on behalf of the Wellcome Sanger Institute COVID-19 Surveillance Team |
| EPI_ISL_2717167, EPI_ISL_2717168, EPI_ISL_2717169 | Lighthouse Lab in Alderley Park | Wellcome Sanger Institute for the COVID-19 Genomics UK (COG-UK) Consortium | Jacquelyn Wynn, Mairead Hyland, The Lighthouse Lab in Alderley Park and Alex Alderton, Roberto Amato, Jeffrey Barrett, Sonia Goncalves, Ewan Harrison, David K. Jackson, Ian Johnston, Dominic Kwiatkowski, Cordelia Langford, John Sillitoe on behalf of the Wellcome Sanger Institute COVID-19 Surveillance Team |
| EPI_ISL_2717170 | Lighthouse Lab in Milton Keynes | Wellcome Sanger Institute for the COVID-19 Genomics UK (COG-UK) Consortium | The Lighthouse Lab in Milton Keynes and Alex Alderton, Roberto Amato, Jeffrey Barrett, Sonia Goncalves, Ewan Harrison, David K. Jackson, Ian Johnston, Dominic Kwiatkowski, Cordelia Langford, John Sillitoe on behalf of the Wellcome Sanger Institute COVID-19 Surveillance Team |
| EPI_ISL_2717171 | Lighthouse Lab in Glasgow | Wellcome Sanger Institute for the COVID-19 Genomics UK (COG-UK) Consortium | Harper VanSteenhouse, Yumi Kasai, David Gray, Carol Clugston, Anna Dominiczak and Alex Alderton, Roberto Amato, Jeffrey Barrett, Sonia Goncalves, Ewan Harrison, David K. Jackson, Ian Johnston, Dominic Kwiatkowski, Cordelia Langford, John Sillitoe on behalf of the Wellcome Sanger Institute COVID-19 Surveillance Team |
| EPI_ISL_2717172 | Lighthouse Lab in Alderley Park | Wellcome Sanger Institute for the COVID-19 Genomics UK (COG-UK) Consortium | Jacquelyn Wynn, Mairead Hyland, The Lighthouse Lab in Alderley Park and Alex Alderton, Roberto Amato, Jeffrey Barrett, Sonia Goncalves, Ewan Harrison, David K. Jackson, Ian Johnston, Dominic Kwiatkowski, Cordelia Langford, John Sillitoe on behalf of the Wellcome Sanger Institute COVID-19 Surveillance Team |
| EPI_ISL_2717173 | Lighthouse Lab in Milton Keynes | Wellcome Sanger Institute for the COVID-19 Genomics UK (COG-UK) Consortium | The Lighthouse Lab in Milton Keynes and Alex Alderton, Roberto Amato, Jeffrey Barrett, Sonia Goncalves, Ewan Harrison, David K. Jackson, Ian Johnston, Dominic Kwiatkowski, Cordelia Langford, John Sillitoe on behalf of the Wellcome Sanger Institute COVID-19 Surveillance Team |
| EPI_ISL_2717174, EPI_ISL_2717175 | Lighthouse Lab in Alderley Park | Wellcome Sanger Institute for the COVID-19 Genomics UK (COG-UK) Consortium | Jacquelyn Wynn, Mairead Hyland, The Lighthouse Lab in Alderley Park and Alex Alderton, Roberto Amato, Jeffrey Barrett, Sonia Goncalves, Ewan Harrison, David K. Jackson, Ian Johnston, Dominic Kwiatkowski, Cordelia Langford, John Sillitoe on behalf of the Wellcome Sanger Institute COVID-19 Surveillance Team |
| EPI_ISL_2717176 | Health Services Laboratories | Wellcome Sanger Institute for the COVID-19 Genomics UK (COG-UK) Consortium | Health Services Laboratories and Alex Alderton, Roberto Amato, Jeffrey Barrett, Sonia Goncalves, Ewan Harrison, David K. Jackson, Ian Johnston, Dominic Kwiatkowski, Cordelia Langford, John Sillitoe on behalf of the Wellcome Sanger Institute COVID-19 Surveillance Team |
| EPI_ISL_2717177, EPI_ISL_2717178, EPI_ISL_2717179, EPI_ISL_2717180 | Lighthouse Lab in Milton Keynes | Wellcome Sanger Institute for the COVID-19 Genomics UK (COG-UK) Consortium | The Lighthouse Lab in Milton Keynes and Alex Alderton, Roberto Amato, Jeffrey Barrett, Sonia Goncalves, Ewan Harrison, David K. Jackson, Ian Johnston, Dominic Kwiatkowski, Cordelia Langford, John Sillitoe on behalf of the Wellcome Sanger Institute COVID-19 Surveillance Team |
| EPI_ISL_2717181 | Health Services Laboratories | Wellcome Sanger Institute for the COVID-19 Genomics UK (COG-UK) Consortium | Health Services Laboratories and Alex Alderton, Roberto Amato, Jeffrey Barrett, Sonia Goncalves, Ewan Harrison, David K. Jackson, Ian Johnston, Dominic Kwiatkowski, Cordelia Langford, John Sillitoe on behalf of the Wellcome Sanger Institute COVID-19 Surveillance Team |
| EPI_ISL_2717182 | Lighthouse Lab in Alderley Park | Wellcome Sanger Institute for the COVID-19 Genomics UK (COG-UK) Consortium | Jacquelyn Wynn, Mairead Hyland, The Lighthouse Lab in Alderley Park and Alex Alderton, Roberto Amato, Jeffrey Barrett, Sonia Goncalves, Ewan Harrison, David K. Jackson, Ian Johnston, Dominic Kwiatkowski, Cordelia Langford, John Sillitoe on behalf of the Wellcome Sanger Institute COVID-19 Surveillance Team |
| EPI_ISL_2717183 | Lighthouse Lab in Milton Keynes | Wellcome Sanger Institute for the COVID-19 Genomics UK (COG-UK) Consortium | The Lighthouse Lab in Milton Keynes and Alex Alderton, Roberto Amato, Jeffrey Barrett, Sonia Goncalves, Ewan Harrison, David K. Jackson, Ian Johnston, Dominic Kwiatkowski, Cordelia Langford, John Sillitoe on behalf of the Wellcome Sanger Institute COVID-19 Surveillance Team |
| EPI_ISL_2717184 | Lighthouse Lab in Alderley Park | Wellcome Sanger Institute for the COVID-19 Genomics UK (COG-UK) Consortium | Jacquelyn Wynn, Mairead Hyland, The Lighthouse Lab in Alderley Park and Alex Alderton, Roberto Amato, Jeffrey Barrett, Sonia Goncalves, Ewan Harrison, David K. Jackson, Ian Johnston, Dominic Kwiatkowski, Cordelia Langford, John Sillitoe on behalf of the Wellcome Sanger Institute COVID-19 Surveillance Team |
| EPI_ISL_2717185, EPI_ISL_2717186, EPI_ISL_2717187, EPI_ISL_2717188 | Lighthouse Lab in Milton Keynes | Wellcome Sanger Institute for the COVID-19 Genomics UK (COG-UK) Consortium | The Lighthouse Lab in Milton Keynes and Alex Alderton, Roberto Amato, Jeffrey Barrett, Sonia Goncalves, Ewan Harrison, David K. Jackson, Ian Johnston, Dominic Kwiatkowski, Cordelia Langford, John Sillitoe on behalf of the Wellcome Sanger Institute COVID-19 Surveillance Team |
| EPI_ISL_2717189 | Lighthouse Lab in Glasgow | Wellcome Sanger Institute for the COVID-19 Genomics UK (COG-UK) Consortium | Harper VanSteenhouse, Yumi Kasai, David Gray, Carol Clugston, Anna Dominiczak and Alex Alderton, Roberto Amato, Jeffrey Barrett, Sonia Goncalves, Ewan Harrison, David K. Jackson, Ian Johnston, Dominic Kwiatkowski, Cordelia Langford, John Sillitoe on behalf of the Wellcome Sanger Institute COVID-19 Surveillance Team |
| EPI_ISL_2717190 | Lighthouse Lab in Milton Keynes | Wellcome Sanger Institute for the COVID-19 Genomics UK (COG-UK) Consortium | The Lighthouse Lab in Milton Keynes and Alex Alderton, Roberto Amato, Jeffrey Barrett, Sonia Goncalves, Ewan Harrison, David K. Jackson, Ian Johnston, Dominic Kwiatkowski, Cordelia Langford, John Sillitoe on behalf of the Wellcome Sanger Institute COVID-19 Surveillance Team |
| EPI_ISL_2717191 | Lighthouse Lab in Alderley Park | Wellcome Sanger Institute for the COVID-19 Genomics UK (COG-UK) Consortium | Jacquelyn Wynn, Mairead Hyland, The Lighthouse Lab in Alderley Park and Alex Alderton, Roberto Amato, Jeffrey Barrett, Sonia Goncalves, Ewan Harrison, David K. Jackson, Ian Johnston, Dominic Kwiatkowski, Cordelia Langford, John Sillitoe on behalf of the Wellcome Sanger Institute COVID-19 Surveillance Team |
| EPI_ISL_2717192 | Lighthouse Lab in Milton Keynes | Wellcome Sanger Institute for the COVID-19 Genomics UK (COG-UK) Consortium | The Lighthouse Lab in Milton Keynes and Alex Alderton, Roberto Amato, Jeffrey Barrett, Sonia Goncalves, Ewan Harrison, David K. Jackson, Ian Johnston, Dominic Kwiatkowski, Cordelia Langford, John Sillitoe on behalf of the Wellcome Sanger Institute COVID-19 Surveillance Team |
| EPI_ISL_2717193 | Lighthouse Lab in Glasgow | Wellcome Sanger Institute for the COVID-19 Genomics UK (COG-UK) Consortium | Harper VanSteenhouse, Yumi Kasai, David Gray, Carol Clugston, Anna Dominiczak and Alex Alderton, Roberto Amato, Jeffrey Barrett, Sonia Goncalves, Ewan Harrison, David K. Jackson, Ian Johnston, Dominic Kwiatkowski, Cordelia Langford, John Sillitoe on behalf of the Wellcome Sanger Institute COVID-19 Surveillance Team |
| EPI_ISL_2717194, EPI_ISL_2717195, EPI_ISL_2717196 | Health Services Laboratories | Wellcome Sanger Institute for the COVID-19 Genomics UK (COG-UK) Consortium | Health Services Laboratories and Alex Alderton, Roberto Amato, Jeffrey Barrett, Sonia Goncalves, Ewan Harrison, David K. Jackson, Ian Johnston, Dominic Kwiatkowski, Cordelia Langford, John Sillitoe on behalf of the Wellcome Sanger Institute COVID-19 Surveillance Team |
| EPI_ISL_2717197 | Lighthouse Lab in Glasgow | Wellcome Sanger Institute for the COVID-19 Genomics UK (COG-UK) Consortium | Harper VanSteenhouse, Yumi Kasai, David Gray, Carol Clugston, Anna Dominiczak and Alex Alderton, Roberto Amato, Jeffrey Barrett, Sonia Goncalves, Ewan Harrison, David K. Jackson, Ian Johnston, Dominic Kwiatkowski, Cordelia Langford, John Sillitoe on behalf of the Wellcome Sanger Institute COVID-19 Surveillance Team |
| EPI_ISL_2717198 | Lighthouse Lab in Milton Keynes | Wellcome Sanger Institute for the COVID-19 Genomics UK (COG-UK) Consortium | The Lighthouse Lab in Milton Keynes and Alex Alderton, Roberto Amato, Jeffrey Barrett, Sonia Goncalves, Ewan Harrison, David K. Jackson, Ian Johnston, Dominic Kwiatkowski, Cordelia Langford, John Sillitoe on behalf of the Wellcome Sanger Institute COVID-19 Surveillance Team |
| EPI_ISL_2717199 | Lighthouse Lab in Alderley Park | Wellcome Sanger Institute for the COVID-19 Genomics UK (COG-UK) Consortium | Jacquelyn Wynn, Mairead Hyland, The Lighthouse Lab in Alderley Park and Alex Alderton, Roberto Amato, Jeffrey Barrett, Sonia Goncalves, Ewan Harrison, David K. Jackson, Ian Johnston, Dominic Kwiatkowski, Cordelia Langford, John Sillitoe on behalf of the Wellcome Sanger Institute COVID-19 Surveillance Team |
| EPI_ISL_2717200, EPI_ISL_2717201 | Lighthouse Lab in Milton Keynes | Wellcome Sanger Institute for the COVID-19 Genomics UK (COG-UK) Consortium | The Lighthouse Lab in Milton Keynes and Alex Alderton, Roberto Amato, Jeffrey Barrett, Sonia Goncalves, Ewan Harrison, David K. Jackson, Ian Johnston, Dominic Kwiatkowski, Cordelia Langford, John Sillitoe on behalf of the Wellcome Sanger Institute COVID-19 Surveillance Team |
| EPI_ISL_2717202 | Health Services Laboratories | Wellcome Sanger Institute for the COVID-19 Genomics UK (COG-UK) Consortium | Health Services Laboratories and Alex Alderton, Roberto Amato, Jeffrey Barrett, Sonia Goncalves, Ewan Harrison, David K. Jackson, Ian Johnston, Dominic Kwiatkowski, Cordelia Langford, John Sillitoe on behalf of the Wellcome Sanger Institute COVID-19 Surveillance Team |
| EPI_ISL_2717203, EPI_ISL_2717204 | Lighthouse Lab in Alderley Park | Wellcome Sanger Institute for the COVID-19 Genomics UK (COG-UK) Consortium | Jacquelyn Wynn, Mairead Hyland, The Lighthouse Lab in Alderley Park and Alex Alderton, Roberto Amato, Jeffrey Barrett, Sonia Goncalves, Ewan Harrison, David K. Jackson, Ian Johnston, Dominic Kwiatkowski, Cordelia Langford, John Sillitoe on behalf of the Wellcome Sanger Institute COVID-19 Surveillance Team |
| EPI_ISL_2717205, EPI_ISL_2717206, EPI_ISL_2717207 | Lighthouse Lab in Milton Keynes | Wellcome Sanger Institute for the COVID-19 Genomics UK (COG-UK) Consortium | The Lighthouse Lab in Milton Keynes and Alex Alderton, Roberto Amato, Jeffrey Barrett, Sonia Goncalves, Ewan Harrison, David K. Jackson, Ian Johnston, Dominic Kwiatkowski, Cordelia Langford, John Sillitoe on behalf of the Wellcome Sanger Institute COVID-19 Surveillance Team |

| | | | |
|---|---|---|---|
| EPI_ISL_2717208, EPI_ISL_2717209 | Lighthouse Lab in Alderley Park | Wellcome Sanger Institute for the COVID-19 Genomics UK (COG-UK) Consortium | Jacquelyn Wynn, Mairead Hyland, The Lighthouse Lab in Alderley Park and Alex Alderton, Roberto Amato, Jeffrey Barrett, Sonia Goncalves, Ewan Harrison, David K. Jackson, Ian Johnston, Dominic Kwiatkowski, Cordelia Langford, John Sillitoe on behalf of the Wellcome Sanger Institute COVID-19 Surveillance Team |
| EPI_ISL_2717210 | Health Services Laboratories | Wellcome Sanger Institute for the COVID-19 Genomics UK (COG-UK) Consortium | Health Services Laboratories and Alex Alderton, Roberto Amato, Jeffrey Barrett, Sonia Goncalves, Ewan Harrison, David K. Jackson, Ian Johnston, Dominic Kwiatkowski, Cordelia Langford, John Sillitoe on behalf of the Wellcome Sanger Institute COVID-19 Surveillance Team |
| EPI_ISL_2717211 | Lighthouse Lab in Milton Keynes | Wellcome Sanger Institute for the COVID-19 Genomics UK (COG-UK) Consortium | The Lighthouse Lab in Milton Keynes and Alex Alderton, Roberto Amato, Jeffrey Barrett, Sonia Goncalves, Ewan Harrison, David K. Jackson, Ian Johnston, Dominic Kwiatkowski, Cordelia Langford, John Sillitoe on behalf of the Wellcome Sanger Institute COVID-19 Surveillance Team |
| EPI_ISL_2717212 | Lighthouse Lab in Alderley Park | Wellcome Sanger Institute for the COVID-19 Genomics UK (COG-UK) Consortium | Jacquelyn Wynn, Mairead Hyland, The Lighthouse Lab in Alderley Park and Alex Alderton, Roberto Amato, Jeffrey Barrett, Sonia Goncalves, Ewan Harrison, David K. Jackson, Ian Johnston, Dominic Kwiatkowski, Cordelia Langford, John Sillitoe on behalf of the Wellcome Sanger Institute COVID-19 Surveillance Team |
| EPI_ISL_2717213 | Lighthouse Lab in Milton Keynes | Wellcome Sanger Institute for the COVID-19 Genomics UK (COG-UK) Consortium | The Lighthouse Lab in Milton Keynes and Alex Alderton, Roberto Amato, Jeffrey Barrett, Sonia Goncalves, Ewan Harrison, David K. Jackson, Ian Johnston, Dominic Kwiatkowski, Cordelia Langford, John Sillitoe on behalf of the Wellcome Sanger Institute COVID-19 Surveillance Team |
| EPI_ISL_2717214 | Health Services Laboratories | Wellcome Sanger Institute for the COVID-19 Genomics UK (COG-UK) Consortium | Health Services Laboratories and Alex Alderton, Roberto Amato, Jeffrey Barrett, Sonia Goncalves, Ewan Harrison, David K. Jackson, Ian Johnston, Dominic Kwiatkowski, Cordelia Langford, John Sillitoe on behalf of the Wellcome Sanger Institute COVID-19 Surveillance Team |
| EPI_ISL_2717215 | Lighthouse Lab in Milton Keynes | Wellcome Sanger Institute for the COVID-19 Genomics UK (COG-UK) Consortium | The Lighthouse Lab in Milton Keynes and Alex Alderton, Roberto Amato, Jeffrey Barrett, Sonia Goncalves, Ewan Harrison, David K. Jackson, Ian Johnston, Dominic Kwiatkowski, Cordelia Langford, John Sillitoe on behalf of the Wellcome Sanger Institute COVID-19 Surveillance Team |
| EPI_ISL_2717216 | Lighthouse Lab in Alderley Park | Wellcome Sanger Institute for the COVID-19 Genomics UK (COG-UK) Consortium | Jacquelyn Wynn, Mairead Hyland, The Lighthouse Lab in Alderley Park and Alex Alderton, Roberto Amato, Jeffrey Barrett, Sonia Goncalves, Ewan Harrison, David K. Jackson, Ian Johnston, Dominic Kwiatkowski, Cordelia Langford, John Sillitoe on behalf of the Wellcome Sanger Institute COVID-19 Surveillance Team |
| EPI_ISL_2717217 | Health Services Laboratories | Wellcome Sanger Institute for the COVID-19 Genomics UK (COG-UK) Consortium | Health Services Laboratories and Alex Alderton, Roberto Amato, Jeffrey Barrett, Sonia Goncalves, Ewan Harrison, David K. Jackson, Ian Johnston, Dominic Kwiatkowski, Cordelia Langford, John Sillitoe on behalf of the Wellcome Sanger Institute COVID-19 Surveillance Team |
| EPI_ISL_2717219, EPI_ISL_2717220, EPI_ISL_2717221, EPI_ISL_2717222, EPI_ISL_2717223 | Lighthouse Lab in Alderley Park | Wellcome Sanger Institute for the COVID-19 Genomics UK (COG-UK) Consortium | Jacquelyn Wynn, Mairead Hyland, The Lighthouse Lab in Alderley Park and Alex Alderton, Roberto Amato, Jeffrey Barrett, Sonia Goncalves, Ewan Harrison, David K. Jackson, Ian Johnston, Dominic Kwiatkowski, Cordelia Langford, John Sillitoe on behalf of the Wellcome Sanger Institute COVID-19 Surveillance Team |
| EPI_ISL_2717224, EPI_ISL_2717225, EPI_ISL_2717226 | Lighthouse Lab in Milton Keynes | Wellcome Sanger Institute for the COVID-19 Genomics UK (COG-UK) Consortium | The Lighthouse Lab in Milton Keynes and Alex Alderton, Roberto Amato, Jeffrey Barrett, Sonia Goncalves, Ewan Harrison, David K. Jackson, Ian Johnston, Dominic Kwiatkowski, Cordelia Langford, John Sillitoe on behalf of the Wellcome Sanger Institute COVID-19 Surveillance Team |
| EPI_ISL_2717227, EPI_ISL_2717228 | Lighthouse Lab in Alderley Park | Wellcome Sanger Institute for the COVID-19 Genomics UK (COG-UK) Consortium | Jacquelyn Wynn, Mairead Hyland, The Lighthouse Lab in Alderley Park and Alex Alderton, Roberto Amato, Jeffrey Barrett, Sonia Goncalves, Ewan Harrison, David K. Jackson, Ian Johnston, Dominic Kwiatkowski, Cordelia Langford, John Sillitoe on behalf of the Wellcome Sanger Institute COVID-19 Surveillance Team |
| EPI_ISL_2717229 | Lighthouse Lab in Glasgow | Wellcome Sanger Institute for the COVID-19 Genomics UK (COG-UK) Consortium | Harper VanSteenhouse, Yumi Kasai, David Gray, Carol Clugston, Anna Dominiczak and Alex Alderton, Roberto Amato, Jeffrey Barrett, Sonia Goncalves, Ewan Harrison, David K. Jackson, Ian Johnston, Dominic Kwiatkowski, Cordelia Langford, John Sillitoe on behalf of the Wellcome Sanger Institute COVID-19 Surveillance Team |
| EPI_ISL_2717230 | Lighthouse Lab in Milton Keynes | Wellcome Sanger Institute for the COVID-19 Genomics UK (COG-UK) Consortium | The Lighthouse Lab in Milton Keynes and Alex Alderton, Roberto Amato, Jeffrey Barrett, Sonia Goncalves, Ewan Harrison, David K. Jackson, Ian Johnston, Dominic Kwiatkowski, Cordelia Langford, John Sillitoe on behalf of the Wellcome Sanger Institute COVID-19 Surveillance Team |
| EPI_ISL_2717231 | Lighthouse Lab in Alderley Park | Wellcome Sanger Institute for the COVID-19 Genomics UK (COG-UK) Consortium | Jacquelyn Wynn, Mairead Hyland, The Lighthouse Lab in Alderley Park and Alex Alderton, Roberto Amato, Jeffrey Barrett, Sonia Goncalves, Ewan Harrison, David K. Jackson, Ian Johnston, Dominic Kwiatkowski, Cordelia Langford, John Sillitoe on behalf of the Wellcome Sanger Institute COVID-19 Surveillance Team |
| EPI_ISL_2717232, EPI_ISL_2717233, EPI_ISL_2717234, EPI_ISL_2717235, EPI_ISL_2717236, EPI_ISL_2717237 | Lighthouse Lab in Milton Keynes | Wellcome Sanger Institute for the COVID-19 Genomics UK (COG-UK) Consortium | The Lighthouse Lab in Milton Keynes and Alex Alderton, Roberto Amato, Jeffrey Barrett, Sonia Goncalves, Ewan Harrison, David K. Jackson, Ian Johnston, Dominic Kwiatkowski, Cordelia Langford, John Sillitoe on behalf of the Wellcome Sanger Institute COVID-19 Surveillance Team |
| EPI_ISL_2717238 | Health Services Laboratories | Wellcome Sanger Institute for the COVID-19 Genomics UK (COG-UK) Consortium | Health Services Laboratories and Alex Alderton, Roberto Amato, Jeffrey Barrett, Sonia Goncalves, Ewan Harrison, David K. Jackson, Ian Johnston, Dominic Kwiatkowski, Cordelia Langford, John Sillitoe on behalf of the Wellcome Sanger Institute COVID-19 Surveillance Team |
| EPI_ISL_2717239 | Lighthouse Lab in Alderley Park | Wellcome Sanger Institute for the COVID-19 Genomics UK (COG-UK) Consortium | Jacquelyn Wynn, Mairead Hyland, The Lighthouse Lab in Alderley Park and Alex Alderton, Roberto Amato, Jeffrey Barrett, Sonia Goncalves, Ewan Harrison, David K. Jackson, Ian Johnston, Dominic Kwiatkowski, Cordelia Langford, John Sillitoe on behalf of the Wellcome Sanger Institute COVID-19 Surveillance Team |
| EPI_ISL_2717240 | Lighthouse Lab in Glasgow | Wellcome Sanger Institute for the COVID-19 Genomics UK (COG-UK) Consortium | Harper VanSteenhouse, Yumi Kasai, David Gray, Carol Clugston, Anna Dominiczak and Alex Alderton, Roberto Amato, Jeffrey Barrett, Sonia Goncalves, Ewan Harrison, David K. Jackson, Ian Johnston, Dominic Kwiatkowski, Cordelia Langford, John Sillitoe on behalf of the Wellcome Sanger Institute COVID-19 Surveillance Team |
| EPI_ISL_2717241, EPI_ISL_2717242, EPI_ISL_2717243 | Lighthouse Lab in Milton Keynes | Wellcome Sanger Institute for the COVID-19 Genomics UK (COG-UK) Consortium | The Lighthouse Lab in Milton Keynes and Alex Alderton, Roberto Amato, Jeffrey Barrett, Sonia Goncalves, Ewan Harrison, David K. Jackson, Ian Johnston, Dominic Kwiatkowski, Cordelia Langford, John Sillitoe on behalf of the Wellcome Sanger Institute COVID-19 Surveillance Team |
| EPI_ISL_2717244, EPI_ISL_2717245 | Lighthouse Lab in Alderley Park | Wellcome Sanger Institute for the COVID-19 Genomics UK (COG-UK) Consortium | Jacquelyn Wynn, Mairead Hyland, The Lighthouse Lab in Alderley Park and Alex Alderton, Roberto Amato, Jeffrey Barrett, Sonia Goncalves, Ewan Harrison, David K. Jackson, Ian Johnston, Dominic Kwiatkowski, Cordelia Langford, John Sillitoe on behalf of the Wellcome Sanger Institute COVID-19 Surveillance Team |
| EPI_ISL_2717246, EPI_ISL_2717247, EPI_ISL_2717248, EPI_ISL_2717249, EPI_ISL_2717250 | Lighthouse Lab in Milton Keynes | Wellcome Sanger Institute for the COVID-19 Genomics UK (COG-UK) Consortium | The Lighthouse Lab in Milton Keynes and Alex Alderton, Roberto Amato, Jeffrey Barrett, Sonia Goncalves, Ewan Harrison, David K. Jackson, Ian Johnston, Dominic Kwiatkowski, Cordelia Langford, John Sillitoe on behalf of the Wellcome Sanger Institute COVID-19 Surveillance Team |
| EPI_ISL_2717251 | Lighthouse Lab in Alderley Park | Wellcome Sanger Institute for the COVID-19 Genomics UK (COG-UK) Consortium | Jacquelyn Wynn, Mairead Hyland, The Lighthouse Lab in Alderley Park and Alex Alderton, Roberto Amato, Jeffrey Barrett, Sonia Goncalves, Ewan Harrison, David K. Jackson, Ian Johnston, Dominic Kwiatkowski, Cordelia Langford, John Sillitoe on behalf of the Wellcome Sanger Institute COVID-19 Surveillance Team |
| EPI_ISL_2717252, EPI_ISL_2717253, EPI_ISL_2717254 | Lighthouse Lab in Milton Keynes | Wellcome Sanger Institute for the COVID-19 Genomics UK (COG-UK) Consortium | The Lighthouse Lab in Milton Keynes and Alex Alderton, Roberto Amato, Jeffrey Barrett, Sonia Goncalves, Ewan Harrison, David K. Jackson, Ian Johnston, Dominic Kwiatkowski, Cordelia Langford, John Sillitoe on behalf of the Wellcome Sanger Institute COVID-19 Surveillance Team |
| EPI_ISL_2717255 | Lighthouse Lab in Glasgow | Wellcome Sanger Institute for the COVID-19 Genomics UK (COG-UK) Consortium | Harper VanSteenhouse, Yumi Kasai, David Gray, Carol Clugston, Anna Dominiczak and Alex Alderton, Roberto Amato, Jeffrey Barrett, Sonia Goncalves, Ewan Harrison, David K. Jackson, Ian Johnston, Dominic Kwiatkowski, Cordelia Langford, John Sillitoe on behalf of the Wellcome Sanger Institute COVID-19 Surveillance Team |
| EPI_ISL_2717256 | Lighthouse Lab in Alderley Park | Wellcome Sanger Institute for the COVID-19 Genomics UK (COG-UK) Consortium | Jacquelyn Wynn, Mairead Hyland, The Lighthouse Lab in Alderley Park and Alex Alderton, Roberto Amato, Jeffrey Barrett, Sonia Goncalves, Ewan Harrison, David K. Jackson, Ian Johnston, Dominic Kwiatkowski, Cordelia Langford, John Sillitoe on behalf of the Wellcome Sanger Institute COVID-19 Surveillance Team |
| EPI_ISL_2717257 | Lighthouse Lab in Glasgow | Wellcome Sanger Institute for the COVID-19 Genomics UK (COG-UK) Consortium | Harper VanSteenhouse, Yumi Kasai, David Gray, Carol Clugston, Anna Dominiczak and Alex Alderton, Roberto Amato, Jeffrey Barrett, Sonia Goncalves, Ewan Harrison, David K. Jackson, Ian Johnston, Dominic Kwiatkowski, Cordelia Langford, John Sillitoe on behalf of the Wellcome Sanger Institute COVID-19 Surveillance Team |
| EPI_ISL_2717258 | Lighthouse Lab in Milton Keynes | Wellcome Sanger Institute for the COVID-19 Genomics UK (COG-UK) Consortium | The Lighthouse Lab in Milton Keynes and Alex Alderton, Roberto Amato, Jeffrey Barrett, Sonia Goncalves, Ewan Harrison, David K. Jackson, Ian Johnston, Dominic Kwiatkowski, Cordelia Langford, John Sillitoe on behalf of the Wellcome Sanger Institute COVID-19 Surveillance Team |
| EPI_ISL_2717259 | Health Services Laboratories | Wellcome Sanger Institute for the COVID-19 Genomics UK (COG-UK) Consortium | Health Services Laboratories and Alex Alderton, Roberto Amato, Jeffrey Barrett, Sonia Goncalves, Ewan Harrison, David K. Jackson, Ian Johnston, Dominic Kwiatkowski, Cordelia Langford, John Sillitoe on behalf of the Wellcome Sanger Institute COVID-19 Surveillance Team |
| EPI_ISL_2717260 | Lighthouse Lab in Milton Keynes | Wellcome Sanger Institute for the COVID-19 Genomics UK (COG-UK) Consortium | The Lighthouse Lab in Milton Keynes and Alex Alderton, Roberto Amato, Jeffrey Barrett, Sonia Goncalves, Ewan Harrison, David K. Jackson, Ian Johnston, Dominic Kwiatkowski, Cordelia Langford, John Sillitoe on behalf of the Wellcome Sanger Institute COVID-19 Surveillance Team |
| EPI_ISL_2717261 | Health Services Laboratories | Wellcome Sanger Institute for the COVID-19 Genomics UK (COG-UK) Consortium | Health Services Laboratories and Alex Alderton, Roberto Amato, Jeffrey Barrett, Sonia Goncalves, Ewan Harrison, David K. Jackson, Ian Johnston, Dominic Kwiatkowski, Cordelia Langford, John Sillitoe on behalf of the Wellcome Sanger Institute COVID-19 Surveillance Team |
| EPI_ISL_2717262 | Lighthouse Lab in Milton Keynes | Wellcome Sanger Institute for the COVID-19 Genomics UK (COG-UK) Consortium | The Lighthouse Lab in Milton Keynes and Alex Alderton, Roberto Amato, Jeffrey Barrett, Sonia Goncalves, Ewan Harrison, David K. Jackson, Ian Johnston, Dominic Kwiatkowski, Cordelia Langford, John Sillitoe on behalf of the Wellcome Sanger Institute COVID-19 Surveillance Team |
| EPI_ISL_2717263, EPI_ISL_2717264 | Health Services Laboratories | Wellcome Sanger Institute for the COVID-19 Genomics UK (COG-UK) Consortium | Health Services Laboratories and Alex Alderton, Roberto Amato, Jeffrey Barrett, Sonia Goncalves, Ewan Harrison, David K. Jackson, Ian Johnston, Dominic Kwiatkowski, Cordelia Langford, John Sillitoe on behalf of the Wellcome Sanger Institute COVID-19 Surveillance Team |
| EPI_ISL_2717265 | Lighthouse Lab in Alderley Park | Wellcome Sanger Institute for the COVID-19 Genomics UK (COG-UK) Consortium | Jacquelyn Wynn, Mairead Hyland, The Lighthouse Lab in Alderley Park and Alex Alderton, Roberto Amato, Jeffrey Barrett, Sonia Goncalves, Ewan Harrison, David K. Jackson, Ian Johnston, Dominic Kwiatkowski, Cordelia Langford, John Sillitoe on behalf of the Wellcome Sanger Institute COVID-19 Surveillance Team |
| EPI_ISL_2717266 | Health Services Laboratories | Wellcome Sanger Institute for the COVID-19 Genomics UK (COG-UK) Consortium | Health Services Laboratories and Alex Alderton, Roberto Amato, Jeffrey Barrett, Sonia Goncalves, Ewan Harrison, David K. Jackson, Ian Johnston, Dominic Kwiatkowski, Cordelia Langford, John Sillitoe on behalf of the Wellcome Sanger Institute COVID-19 Surveillance Team |
| EPI_ISL_2717267 | Lighthouse Lab in Milton Keynes | Wellcome Sanger Institute for the COVID-19 Genomics UK (COG-UK) Consortium | The Lighthouse Lab in Milton Keynes and Alex Alderton, Roberto Amato, Jeffrey Barrett, Sonia Goncalves, Ewan Harrison, David K. Jackson, Ian Johnston, Dominic Kwiatkowski, Cordelia Langford, John Sillitoe on behalf of the Wellcome Sanger Institute COVID-19 Surveillance Team |
| EPI_ISL_2717268 | Health Services Laboratories | Wellcome Sanger Institute for the COVID-19 Genomics UK (COG-UK) Consortium | Health Services Laboratories and Alex Alderton, Roberto Amato, Jeffrey Barrett, Sonia Goncalves, Ewan Harrison, David K. Jackson, Ian Johnston, Dominic Kwiatkowski, Cordelia Langford, John Sillitoe on behalf of the Wellcome Sanger Institute COVID-19 Surveillance Team |
| EPI_ISL_2717269 | Lighthouse Lab in Alderley Park | Wellcome Sanger Institute for the COVID-19 Genomics UK (COG-UK) Consortium | Jacquelyn Wynn, Mairead Hyland, The Lighthouse Lab in Alderley Park and Alex Alderton, Roberto Amato, Jeffrey Barrett, Sonia Goncalves, Ewan Harrison, David K. Jackson, Ian Johnston, Dominic Kwiatkowski, Cordelia Langford, John Sillitoe on behalf of the Wellcome Sanger Institute COVID-19 Surveillance Team |
| EPI_ISL_2717270, EPI_ISL_2717271 | Health Services Laboratories | Wellcome Sanger Institute for the COVID-19 Genomics UK (COG-UK) Consortium | Health Services Laboratories and Alex Alderton, Roberto Amato, Jeffrey Barrett, Sonia Goncalves, Ewan Harrison, David K. Jackson, Ian Johnston, Dominic Kwiatkowski, Cordelia Langford, John Sillitoe on behalf of the Wellcome Sanger Institute COVID-19 Surveillance Team |
| EPI_ISL_2717272, EPI_ISL_2717273 | Lighthouse Lab in Milton Keynes | Wellcome Sanger Institute for the COVID-19 Genomics UK (COG-UK) Consortium | The Lighthouse Lab in Milton Keynes and Alex Alderton, Roberto Amato, Jeffrey Barrett, Sonia Goncalves, Ewan Harrison, David K. Jackson, Ian Johnston, Dominic Kwiatkowski, Cordelia Langford, John Sillitoe on behalf of the Wellcome Sanger Institute COVID-19 Surveillance Team |

| | | | |
|---|---|---|---|
| EPI_ISL_2717274 | Lighthouse Lab in Glasgow | Wellcome Sanger Institute for the COVID-19 Genomics UK (COG-UK) Consortium | Harper VanSteenhouse, Yumi Kasai, David Gray, Carol Clugston, Anna Dominiczak and Alex Alderton, Roberto Amato, Jeffrey Barrett, Sonia Goncalves, Ewan Harrison, David K. Jackson, Ian Johnston, Dominic Kwiatkowski, Cordelia Langford, John Sillitoe on behalf of the Wellcome Sanger Institute COVID-19 Surveillance Team |
| EPI_ISL_2717275, EPI_ISL_2717276, EPI_ISL_2717277 | Lighthouse Lab in Milton Keynes | Wellcome Sanger Institute for the COVID-19 Genomics UK (COG-UK) Consortium | The Lighthouse Lab in Milton Keynes and Alex Alderton, Roberto Amato, Jeffrey Barrett, Sonia Goncalves, Ewan Harrison, David K. Jackson, Ian Johnston, Dominic Kwiatkowski, Cordelia Langford, John Sillitoe on behalf of the Wellcome Sanger Institute COVID-19 Surveillance Team |
| EPI_ISL_2717278, EPI_ISL_2717279 | Lighthouse Lab in Alderley Park | Wellcome Sanger Institute for the COVID-19 Genomics UK (COG-UK) Consortium | Jacquelyn Wynn, Mairead Hyland, The Lighthouse Lab in Alderley Park and Alex Alderton, Roberto Amato, Jeffrey Barrett, Sonia Goncalves, Ewan Harrison, David K. Jackson, Ian Johnston, Dominic Kwiatkowski, Cordelia Langford, John Sillitoe on behalf of the Wellcome Sanger Institute COVID-19 Surveillance Team |
| EPI_ISL_2717280 | Health Services Laboratories | Wellcome Sanger Institute for the COVID-19 Genomics UK (COG-UK) Consortium | Health Services Laboratories and Alex Alderton, Roberto Amato, Jeffrey Barrett, Sonia Goncalves, Ewan Harrison, David K. Jackson, Ian Johnston, Dominic Kwiatkowski, Cordelia Langford, John Sillitoe on behalf of the Wellcome Sanger Institute COVID-19 Surveillance Team |
| EPI_ISL_2717281 | Lighthouse Lab in Alderley Park | Wellcome Sanger Institute for the COVID-19 Genomics UK (COG-UK) Consortium | Jacquelyn Wynn, Mairead Hyland, The Lighthouse Lab in Alderley Park and Alex Alderton, Roberto Amato, Jeffrey Barrett, Sonia Goncalves, Ewan Harrison, David K. Jackson, Ian Johnston, Dominic Kwiatkowski, Cordelia Langford, John Sillitoe on behalf of the Wellcome Sanger Institute COVID-19 Surveillance Team |
| EPI_ISL_2717282 | Lighthouse Lab in Milton Keynes | Wellcome Sanger Institute for the COVID-19 Genomics UK (COG-UK) Consortium | The Lighthouse Lab in Milton Keynes and Alex Alderton, Roberto Amato, Jeffrey Barrett, Sonia Goncalves, Ewan Harrison, David K. Jackson, Ian Johnston, Dominic Kwiatkowski, Cordelia Langford, John Sillitoe on behalf of the Wellcome Sanger Institute COVID-19 Surveillance Team |
| EPI_ISL_2717283 | Lighthouse Lab in Alderley Park | Wellcome Sanger Institute for the COVID-19 Genomics UK (COG-UK) Consortium | Jacquelyn Wynn, Mairead Hyland, The Lighthouse Lab in Alderley Park and Alex Alderton, Roberto Amato, Jeffrey Barrett, Sonia Goncalves, Ewan Harrison, David K. Jackson, Ian Johnston, Dominic Kwiatkowski, Cordelia Langford, John Sillitoe on behalf of the Wellcome Sanger Institute COVID-19 Surveillance Team |
| EPI_ISL_2717284 | Lighthouse Lab in Glasgow | Wellcome Sanger Institute for the COVID-19 Genomics UK (COG-UK) Consortium | Harper VanSteenhouse, Yumi Kasai, David Gray, Carol Clugston, Anna Dominiczak and Alex Alderton, Roberto Amato, Jeffrey Barrett, Sonia Goncalves, Ewan Harrison, David K. Jackson, Ian Johnston, Dominic Kwiatkowski, Cordelia Langford, John Sillitoe on behalf of the Wellcome Sanger Institute COVID-19 Surveillance Team |
| EPI_ISL_2717285, EPI_ISL_2717286 | Lighthouse Lab in Milton Keynes | Wellcome Sanger Institute for the COVID-19 Genomics UK (COG-UK) Consortium | The Lighthouse Lab in Milton Keynes and Alex Alderton, Roberto Amato, Jeffrey Barrett, Sonia Goncalves, Ewan Harrison, David K. Jackson, Ian Johnston, Dominic Kwiatkowski, Cordelia Langford, John Sillitoe on behalf of the Wellcome Sanger Institute COVID-19 Surveillance Team |
| EPI_ISL_2717287 | Lighthouse Lab in Alderley Park | Wellcome Sanger Institute for the COVID-19 Genomics UK (COG-UK) Consortium | Jacquelyn Wynn, Mairead Hyland, The Lighthouse Lab in Alderley Park and Alex Alderton, Roberto Amato, Jeffrey Barrett, Sonia Goncalves, Ewan Harrison, David K. Jackson, Ian Johnston, Dominic Kwiatkowski, Cordelia Langford, John Sillitoe on behalf of the Wellcome Sanger Institute COVID-19 Surveillance Team |
| EPI_ISL_2717288 | Lighthouse Lab in Milton Keynes | Wellcome Sanger Institute for the COVID-19 Genomics UK (COG-UK) Consortium | The Lighthouse Lab in Milton Keynes and Alex Alderton, Roberto Amato, Jeffrey Barrett, Sonia Goncalves, Ewan Harrison, David K. Jackson, Ian Johnston, Dominic Kwiatkowski, Cordelia Langford, John Sillitoe on behalf of the Wellcome Sanger Institute COVID-19 Surveillance Team |
| EPI_ISL_2717289 | Health Services Laboratories | Wellcome Sanger Institute for the COVID-19 Genomics UK (COG-UK) Consortium | Health Services Laboratories and Alex Alderton, Roberto Amato, Jeffrey Barrett, Sonia Goncalves, Ewan Harrison, David K. Jackson, Ian Johnston, Dominic Kwiatkowski, Cordelia Langford, John Sillitoe on behalf of the Wellcome Sanger Institute COVID-19 Surveillance Team |
| EPI_ISL_2717290 | Lighthouse Lab in Milton Keynes | Wellcome Sanger Institute for the COVID-19 Genomics UK (COG-UK) Consortium | The Lighthouse Lab in Milton Keynes and Alex Alderton, Roberto Amato, Jeffrey Barrett, Sonia Goncalves, Ewan Harrison, David K. Jackson, Ian Johnston, Dominic Kwiatkowski, Cordelia Langford, John Sillitoe on behalf of the Wellcome Sanger Institute COVID-19 Surveillance Team |
| EPI_ISL_2717291 | Lighthouse Lab in Alderley Park | Wellcome Sanger Institute for the COVID-19 Genomics UK (COG-UK) Consortium | Jacquelyn Wynn, Mairead Hyland, The Lighthouse Lab in Alderley Park and Alex Alderton, Roberto Amato, Jeffrey Barrett, Sonia Goncalves, Ewan Harrison, David K. Jackson, Ian Johnston, Dominic Kwiatkowski, Cordelia Langford, John Sillitoe on behalf of the Wellcome Sanger Institute COVID-19 Surveillance Team |
| EPI_ISL_2717292, EPI_ISL_2717293, EPI_ISL_2717294, EPI_ISL_2717295, EPI_ISL_2717296, EPI_ISL_2717297, EPI_ISL_2717298, EPI_ISL_2717299 | Lighthouse Lab in Milton Keynes | Wellcome Sanger Institute for the COVID-19 Genomics UK (COG-UK) Consortium | The Lighthouse Lab in Milton Keynes and Alex Alderton, Roberto Amato, Jeffrey Barrett, Sonia Goncalves, Ewan Harrison, David K. Jackson, Ian Johnston, Dominic Kwiatkowski, Cordelia Langford, John Sillitoe on behalf of the Wellcome Sanger Institute COVID-19 Surveillance Team |
| EPI_ISL_2717300 | Health Services Laboratories | Wellcome Sanger Institute for the COVID-19 Genomics UK (COG-UK) Consortium | Health Services Laboratories and Alex Alderton, Roberto Amato, Jeffrey Barrett, Sonia Goncalves, Ewan Harrison, David K. Jackson, Ian Johnston, Dominic Kwiatkowski, Cordelia Langford, John Sillitoe on behalf of the Wellcome Sanger Institute COVID-19 Surveillance Team |
| EPI_ISL_2717301, EPI_ISL_2717302 | Lighthouse Lab in Alderley Park | Wellcome Sanger Institute for the COVID-19 Genomics UK (COG-UK) Consortium | Jacquelyn Wynn, Mairead Hyland, The Lighthouse Lab in Alderley Park and Alex Alderton, Roberto Amato, Jeffrey Barrett, Sonia Goncalves, Ewan Harrison, David K. Jackson, Ian Johnston, Dominic Kwiatkowski, Cordelia Langford, John Sillitoe on behalf of the Wellcome Sanger Institute COVID-19 Surveillance Team |
| EPI_ISL_2717303 | Health Services Laboratories | Wellcome Sanger Institute for the COVID-19 Genomics UK (COG-UK) Consortium | Health Services Laboratories and Alex Alderton, Roberto Amato, Jeffrey Barrett, Sonia Goncalves, Ewan Harrison, David K. Jackson, Ian Johnston, Dominic Kwiatkowski, Cordelia Langford, John Sillitoe on behalf of the Wellcome Sanger Institute COVID-19 Surveillance Team |
| EPI_ISL_2717304, EPI_ISL_2717305 | Lighthouse Lab in Milton Keynes | Wellcome Sanger Institute for the COVID-19 Genomics UK (COG-UK) Consortium | The Lighthouse Lab in Milton Keynes and Alex Alderton, Roberto Amato, Jeffrey Barrett, Sonia Goncalves, Ewan Harrison, David K. Jackson, Ian Johnston, Dominic Kwiatkowski, Cordelia Langford, John Sillitoe on behalf of the Wellcome Sanger Institute COVID-19 Surveillance Team |
| EPI_ISL_2717306 | Health Services Laboratories | Wellcome Sanger Institute for the COVID-19 Genomics UK (COG-UK) Consortium | Health Services Laboratories and Alex Alderton, Roberto Amato, Jeffrey Barrett, Sonia Goncalves, Ewan Harrison, David K. Jackson, Ian Johnston, Dominic Kwiatkowski, Cordelia Langford, John Sillitoe on behalf of the Wellcome Sanger Institute COVID-19 Surveillance Team |
| EPI_ISL_2717307 | Lighthouse Lab in Alderley Park | Wellcome Sanger Institute for the COVID-19 Genomics UK (COG-UK) Consortium | Jacquelyn Wynn, Mairead Hyland, The Lighthouse Lab in Alderley Park and Alex Alderton, Roberto Amato, Jeffrey Barrett, Sonia Goncalves, Ewan Harrison, David K. Jackson, Ian Johnston, Dominic Kwiatkowski, Cordelia Langford, John Sillitoe on behalf of the Wellcome Sanger Institute COVID-19 Surveillance Team |
| EPI_ISL_2717308, EPI_ISL_2717309 | Lighthouse Lab in Milton Keynes | Wellcome Sanger Institute for the COVID-19 Genomics UK (COG-UK) Consortium | The Lighthouse Lab in Milton Keynes and Alex Alderton, Roberto Amato, Jeffrey Barrett, Sonia Goncalves, Ewan Harrison, David K. Jackson, Ian Johnston, Dominic Kwiatkowski, Cordelia Langford, John Sillitoe on behalf of the Wellcome Sanger Institute COVID-19 Surveillance Team |
| EPI_ISL_2717310, EPI_ISL_2717311 | Lighthouse Lab in Alderley Park | Wellcome Sanger Institute for the COVID-19 Genomics UK (COG-UK) Consortium | Jacquelyn Wynn, Mairead Hyland, The Lighthouse Lab in Alderley Park and Alex Alderton, Roberto Amato, Jeffrey Barrett, Sonia Goncalves, Ewan Harrison, David K. Jackson, Ian Johnston, Dominic Kwiatkowski, Cordelia Langford, John Sillitoe on behalf of the Wellcome Sanger Institute COVID-19 Surveillance Team |
| EPI_ISL_2717312 | Lighthouse Lab in Milton Keynes | Wellcome Sanger Institute for the COVID-19 Genomics UK (COG-UK) Consortium | The Lighthouse Lab in Milton Keynes and Alex Alderton, Roberto Amato, Jeffrey Barrett, Sonia Goncalves, Ewan Harrison, David K. Jackson, Ian Johnston, Dominic Kwiatkowski, Cordelia Langford, John Sillitoe on behalf of the Wellcome Sanger Institute COVID-19 Surveillance Team |
| EPI_ISL_2717313, EPI_ISL_2717314 | Lighthouse Lab in Alderley Park | Wellcome Sanger Institute for the COVID-19 Genomics UK (COG-UK) Consortium | Jacquelyn Wynn, Mairead Hyland, The Lighthouse Lab in Alderley Park and Alex Alderton, Roberto Amato, Jeffrey Barrett, Sonia Goncalves, Ewan Harrison, David K. Jackson, Ian Johnston, Dominic Kwiatkowski, Cordelia Langford, John Sillitoe on behalf of the Wellcome Sanger Institute COVID-19 Surveillance Team |
| EPI_ISL_2717315 | Lighthouse Lab in Milton Keynes | Wellcome Sanger Institute for the COVID-19 Genomics UK (COG-UK) Consortium | The Lighthouse Lab in Milton Keynes and Alex Alderton, Roberto Amato, Jeffrey Barrett, Sonia Goncalves, Ewan Harrison, David K. Jackson, Ian Johnston, Dominic Kwiatkowski, Cordelia Langford, John Sillitoe on behalf of the Wellcome Sanger Institute COVID-19 Surveillance Team |
| EPI_ISL_2717316, EPI_ISL_2717317 | Lighthouse Lab in Alderley Park | Wellcome Sanger Institute for the COVID-19 Genomics UK (COG-UK) Consortium | Jacquelyn Wynn, Mairead Hyland, The Lighthouse Lab in Alderley Park and Alex Alderton, Roberto Amato, Jeffrey Barrett, Sonia Goncalves, Ewan Harrison, David K. Jackson, Ian Johnston, Dominic Kwiatkowski, Cordelia Langford, John Sillitoe on behalf of the Wellcome Sanger Institute COVID-19 Surveillance Team |
| EPI_ISL_2717318, EPI_ISL_2717319 | Lighthouse Lab in Glasgow | Wellcome Sanger Institute for the COVID-19 Genomics UK (COG-UK) Consortium | Harper VanSteenhouse, Yumi Kasai, David Gray, Carol Clugston, Anna Dominiczak and Alex Alderton, Roberto Amato, Jeffrey Barrett, Sonia Goncalves, Ewan Harrison, David K. Jackson, Ian Johnston, Dominic Kwiatkowski, Cordelia Langford, John Sillitoe on behalf of the Wellcome Sanger Institute COVID-19 Surveillance Team |
| EPI_ISL_2717320 | Lighthouse Lab in Milton Keynes | Wellcome Sanger Institute for the COVID-19 Genomics UK (COG-UK) Consortium | The Lighthouse Lab in Milton Keynes and Alex Alderton, Roberto Amato, Jeffrey Barrett, Sonia Goncalves, Ewan Harrison, David K. Jackson, Ian Johnston, Dominic Kwiatkowski, Cordelia Langford, John Sillitoe on behalf of the Wellcome Sanger Institute COVID-19 Surveillance Team |
| EPI_ISL_2717321 | Lighthouse Lab in Glasgow | Wellcome Sanger Institute for the COVID-19 Genomics UK (COG-UK) Consortium | Harper VanSteenhouse, Yumi Kasai, David Gray, Carol Clugston, Anna Dominiczak and Alex Alderton, Roberto Amato, Jeffrey Barrett, Sonia Goncalves, Ewan Harrison, David K. Jackson, Ian Johnston, Dominic Kwiatkowski, Cordelia Langford, John Sillitoe on behalf of the Wellcome Sanger Institute COVID-19 Surveillance Team |
| EPI_ISL_2717322 | Lighthouse Lab in Alderley Park | Wellcome Sanger Institute for the COVID-19 Genomics UK (COG-UK) Consortium | Jacquelyn Wynn, Mairead Hyland, The Lighthouse Lab in Alderley Park and Alex Alderton, Roberto Amato, Jeffrey Barrett, Sonia Goncalves, Ewan Harrison, David K. Jackson, Ian Johnston, Dominic Kwiatkowski, Cordelia Langford, John Sillitoe on behalf of the Wellcome Sanger Institute COVID-19 Surveillance Team |
| EPI_ISL_2717323 | Lighthouse Lab in Glasgow | Wellcome Sanger Institute for the COVID-19 Genomics UK (COG-UK) Consortium | Harper VanSteenhouse, Yumi Kasai, David Gray, Carol Clugston, Anna Dominiczak and Alex Alderton, Roberto Amato, Jeffrey Barrett, Sonia Goncalves, Ewan Harrison, David K. Jackson, Ian Johnston, Dominic Kwiatkowski, Cordelia Langford, John Sillitoe on behalf of the Wellcome Sanger Institute COVID-19 Surveillance Team |
| EPI_ISL_2717324 | Lighthouse Lab in Milton Keynes | Wellcome Sanger Institute for the COVID-19 Genomics UK (COG-UK) Consortium | The Lighthouse Lab in Milton Keynes and Alex Alderton, Roberto Amato, Jeffrey Barrett, Sonia Goncalves, Ewan Harrison, David K. Jackson, Ian Johnston, Dominic Kwiatkowski, Cordelia Langford, John Sillitoe on behalf of the Wellcome Sanger Institute COVID-19 Surveillance Team |
| EPI_ISL_2717325, EPI_ISL_2717326, EPI_ISL_2717327, EPI_ISL_2717328, EPI_ISL_2717329, EPI_ISL_2717330 | Lighthouse Lab in Alderley Park | Wellcome Sanger Institute for the COVID-19 Genomics UK (COG-UK) Consortium | Jacquelyn Wynn, Mairead Hyland, The Lighthouse Lab in Alderley Park and Alex Alderton, Roberto Amato, Jeffrey Barrett, Sonia Goncalves, Ewan Harrison, David K. Jackson, Ian Johnston, Dominic Kwiatkowski, Cordelia Langford, John Sillitoe on behalf of the Wellcome Sanger Institute COVID-19 Surveillance Team |
| EPI_ISL_2717331 | Lighthouse Lab in Milton Keynes | Wellcome Sanger Institute for the COVID-19 Genomics UK (COG-UK) Consortium | The Lighthouse Lab in Milton Keynes and Alex Alderton, Roberto Amato, Jeffrey Barrett, Sonia Goncalves, Ewan Harrison, David K. Jackson, Ian Johnston, Dominic Kwiatkowski, Cordelia Langford, John Sillitoe on behalf of the Wellcome Sanger Institute COVID-19 Surveillance Team |
| EPI_ISL_2717332, EPI_ISL_2717333, EPI_ISL_2717334 | Lighthouse Lab in Alderley Park | Wellcome Sanger Institute for the COVID-19 Genomics UK (COG-UK) Consortium | Jacquelyn Wynn, Mairead Hyland, The Lighthouse Lab in Alderley Park and Alex Alderton, Roberto Amato, Jeffrey Barrett, Sonia Goncalves, Ewan Harrison, David K. Jackson, Ian Johnston, Dominic Kwiatkowski, Cordelia Langford, John Sillitoe on behalf of the Wellcome Sanger Institute COVID-19 Surveillance Team |
| EPI_ISL_2717335 | Health Services Laboratories | Wellcome Sanger Institute for the COVID-19 Genomics UK (COG-UK) Consortium | Health Services Laboratories and Alex Alderton, Roberto Amato, Jeffrey Barrett, Sonia Goncalves, Ewan Harrison, David K. Jackson, Ian Johnston, Dominic Kwiatkowski, Cordelia Langford, John Sillitoe on behalf of the Wellcome Sanger Institute COVID-19 Surveillance Team |
| EPI_ISL_2717336, EPI_ISL_2717337, EPI_ISL_2717338, EPI_ISL_2717339, EPI_ISL_2717340, EPI_ISL_2717341, EPI_ISL_2717342 | Lighthouse Lab in Milton Keynes | Wellcome Sanger Institute for the COVID-19 Genomics UK (COG-UK) Consortium | The Lighthouse Lab in Milton Keynes and Alex Alderton, Roberto Amato, Jeffrey Barrett, Sonia Goncalves, Ewan Harrison, David K. Jackson, Ian Johnston, Dominic Kwiatkowski, Cordelia Langford, John Sillitoe on behalf of the Wellcome Sanger Institute COVID-19 Surveillance Team |

| | | | |
|---|---|---|---|
| EPI_ISL_2717343 | Lighthouse Lab in Alderley Park | Wellcome Sanger Institute for the COVID-19 Genomics UK (COG-UK) Consortium | Jacquelyn Wynn, Mairead Hyland, The Lighthouse Lab in Alderley Park and Alex Alderton, Roberto Amato, Jeffrey Barrett, Sonia Goncalves, Ewan Harrison, David K. Jackson, Ian Johnston, Dominic Kwiatkowski, Cordelia Langford, John Sillitoe on behalf of the Wellcome Sanger Institute COVID-19 Surveillance Team |
| EPI_ISL_2717344, EPI_ISL_2717345 | Lighthouse Lab in Milton Keynes | Wellcome Sanger Institute for the COVID-19 Genomics UK (COG-UK) Consortium | The Lighthouse Lab in Milton Keynes and Alex Alderton, Roberto Amato, Jeffrey Barrett, Sonia Goncalves, Ewan Harrison, David K. Jackson, Ian Johnston, Dominic Kwiatkowski, Cordelia Langford, John Sillitoe on behalf of the Wellcome Sanger Institute COVID-19 Surveillance Team |
| EPI_ISL_2717346 | Lighthouse Lab in Glasgow | Wellcome Sanger Institute for the COVID-19 Genomics UK (COG-UK) Consortium | Harper VanSteenhouse, Yumi Kasai, David Gray, Carol Clugston, Anna Dominiczak and Alex Alderton, Roberto Amato, Jeffrey Barrett, Sonia Goncalves, Ewan Harrison, David K. Jackson, Ian Johnston, Dominic Kwiatkowski, Cordelia Langford, John Sillitoe on behalf of the Wellcome Sanger Institute COVID-19 Surveillance Team |
| EPI_ISL_2717347, EPI_ISL_2717348, EPI_ISL_2717349 | Lighthouse Lab in Milton Keynes | Wellcome Sanger Institute for the COVID-19 Genomics UK (COG-UK) Consortium | The Lighthouse Lab in Milton Keynes and Alex Alderton, Roberto Amato, Jeffrey Barrett, Sonia Goncalves, Ewan Harrison, David K. Jackson, Ian Johnston, Dominic Kwiatkowski, Cordelia Langford, John Sillitoe on behalf of the Wellcome Sanger Institute COVID-19 Surveillance Team |
| EPI_ISL_2717350 | Lighthouse Lab in Alderley Park | Wellcome Sanger Institute for the COVID-19 Genomics UK (COG-UK) Consortium | Jacquelyn Wynn, Mairead Hyland, The Lighthouse Lab in Alderley Park and Alex Alderton, Roberto Amato, Jeffrey Barrett, Sonia Goncalves, Ewan Harrison, David K. Jackson, Ian Johnston, Dominic Kwiatkowski, Cordelia Langford, John Sillitoe on behalf of the Wellcome Sanger Institute COVID-19 Surveillance Team |
| EPI_ISL_2717351 | Lighthouse Lab in Milton Keynes | Wellcome Sanger Institute for the COVID-19 Genomics UK (COG-UK) Consortium | The Lighthouse Lab in Milton Keynes and Alex Alderton, Roberto Amato, Jeffrey Barrett, Sonia Goncalves, Ewan Harrison, David K. Jackson, Ian Johnston, Dominic Kwiatkowski, Cordelia Langford, John Sillitoe on behalf of the Wellcome Sanger Institute COVID-19 Surveillance Team |
| EPI_ISL_2717352, EPI_ISL_2717353 | Lighthouse Lab in Glasgow | Wellcome Sanger Institute for the COVID-19 Genomics UK (COG-UK) Consortium | Harper VanSteenhouse, Yumi Kasai, David Gray, Carol Clugston, Anna Dominiczak and Alex Alderton, Roberto Amato, Jeffrey Barrett, Sonia Goncalves, Ewan Harrison, David K. Jackson, Ian Johnston, Dominic Kwiatkowski, Cordelia Langford, John Sillitoe on behalf of the Wellcome Sanger Institute COVID-19 Surveillance Team |
| EPI_ISL_2717354 | Health Services Laboratories | Wellcome Sanger Institute for the COVID-19 Genomics UK (COG-UK) Consortium | Health Services Laboratories and Alex Alderton, Roberto Amato, Jeffrey Barrett, Sonia Goncalves, Ewan Harrison, David K. Jackson, Ian Johnston, Dominic Kwiatkowski, Cordelia Langford, John Sillitoe on behalf of the Wellcome Sanger Institute COVID-19 Surveillance Team |
| EPI_ISL_2717355 | Lighthouse Lab in Glasgow | Wellcome Sanger Institute for the COVID-19 Genomics UK (COG-UK) Consortium | Harper VanSteenhouse, Yumi Kasai, David Gray, Carol Clugston, Anna Dominiczak and Alex Alderton, Roberto Amato, Jeffrey Barrett, Sonia Goncalves, Ewan Harrison, David K. Jackson, Ian Johnston, Dominic Kwiatkowski, Cordelia Langford, John Sillitoe on behalf of the Wellcome Sanger Institute COVID-19 Surveillance Team |
| EPI_ISL_2717356, EPI_ISL_2717357 | Lighthouse Lab in Milton Keynes | Wellcome Sanger Institute for the COVID-19 Genomics UK (COG-UK) Consortium | The Lighthouse Lab in Milton Keynes and Alex Alderton, Roberto Amato, Jeffrey Barrett, Sonia Goncalves, Ewan Harrison, David K. Jackson, Ian Johnston, Dominic Kwiatkowski, Cordelia Langford, John Sillitoe on behalf of the Wellcome Sanger Institute COVID-19 Surveillance Team |
| EPI_ISL_2717358, EPI_ISL_2717359 | Lighthouse Lab in Alderley Park | Wellcome Sanger Institute for the COVID-19 Genomics UK (COG-UK) Consortium | Jacquelyn Wynn, Mairead Hyland, The Lighthouse Lab in Alderley Park and Alex Alderton, Roberto Amato, Jeffrey Barrett, Sonia Goncalves, Ewan Harrison, David K. Jackson, Ian Johnston, Dominic Kwiatkowski, Cordelia Langford, John Sillitoe on behalf of the Wellcome Sanger Institute COVID-19 Surveillance Team |
| EPI_ISL_2717360 | Health Services Laboratories | Wellcome Sanger Institute for the COVID-19 Genomics UK (COG-UK) Consortium | Health Services Laboratories and Alex Alderton, Roberto Amato, Jeffrey Barrett, Sonia Goncalves, Ewan Harrison, David K. Jackson, Ian Johnston, Dominic Kwiatkowski, Cordelia Langford, John Sillitoe on behalf of the Wellcome Sanger Institute COVID-19 Surveillance Team |
| EPI_ISL_2717361 | Lighthouse Lab in Alderley Park | Wellcome Sanger Institute for the COVID-19 Genomics UK (COG-UK) Consortium | Jacquelyn Wynn, Mairead Hyland, The Lighthouse Lab in Alderley Park and Alex Alderton, Roberto Amato, Jeffrey Barrett, Sonia Goncalves, Ewan Harrison, David K. Jackson, Ian Johnston, Dominic Kwiatkowski, Cordelia Langford, John Sillitoe on behalf of the Wellcome Sanger Institute COVID-19 Surveillance Team |
| EPI_ISL_2717362 | Lighthouse Lab in Glasgow | Wellcome Sanger Institute for the COVID-19 Genomics UK (COG-UK) Consortium | Harper VanSteenhouse, Yumi Kasai, David Gray, Carol Clugston, Anna Dominiczak and Alex Alderton, Roberto Amato, Jeffrey Barrett, Sonia Goncalves, Ewan Harrison, David K. Jackson, Ian Johnston, Dominic Kwiatkowski, Cordelia Langford, John Sillitoe on behalf of the Wellcome Sanger Institute COVID-19 Surveillance Team |
| EPI_ISL_2717363 | Lighthouse Lab in Alderley Park | Wellcome Sanger Institute for the COVID-19 Genomics UK (COG-UK) Consortium | Jacquelyn Wynn, Mairead Hyland, The Lighthouse Lab in Alderley Park and Alex Alderton, Roberto Amato, Jeffrey Barrett, Sonia Goncalves, Ewan Harrison, David K. Jackson, Ian Johnston, Dominic Kwiatkowski, Cordelia Langford, John Sillitoe on behalf of the Wellcome Sanger Institute COVID-19 Surveillance Team |
| EPI_ISL_2717364 | Lighthouse Lab in Glasgow | Wellcome Sanger Institute for the COVID-19 Genomics UK (COG-UK) Consortium | Harper VanSteenhouse, Yumi Kasai, David Gray, Carol Clugston, Anna Dominiczak and Alex Alderton, Roberto Amato, Jeffrey Barrett, Sonia Goncalves, Ewan Harrison, David K. Jackson, Ian Johnston, Dominic Kwiatkowski, Cordelia Langford, John Sillitoe on behalf of the Wellcome Sanger Institute COVID-19 Surveillance Team |
| EPI_ISL_2717365, EPI_ISL_2717366, EPI_ISL_2717367 | Lighthouse Lab in Milton Keynes | Wellcome Sanger Institute for the COVID-19 Genomics UK (COG-UK) Consortium | The Lighthouse Lab in Milton Keynes and Alex Alderton, Roberto Amato, Jeffrey Barrett, Sonia Goncalves, Ewan Harrison, David K. Jackson, Ian Johnston, Dominic Kwiatkowski, Cordelia Langford, John Sillitoe on behalf of the Wellcome Sanger Institute COVID-19 Surveillance Team |
| EPI_ISL_2717368 | Lighthouse Lab in Alderley Park | Wellcome Sanger Institute for the COVID-19 Genomics UK (COG-UK) Consortium | Jacquelyn Wynn, Mairead Hyland, The Lighthouse Lab in Alderley Park and Alex Alderton, Roberto Amato, Jeffrey Barrett, Sonia Goncalves, Ewan Harrison, David K. Jackson, Ian Johnston, Dominic Kwiatkowski, Cordelia Langford, John Sillitoe on behalf of the Wellcome Sanger Institute COVID-19 Surveillance Team |
| EPI_ISL_2717369 | Health Services Laboratories | Wellcome Sanger Institute for the COVID-19 Genomics UK (COG-UK) Consortium | Health Services Laboratories and Alex Alderton, Roberto Amato, Jeffrey Barrett, Sonia Goncalves, Ewan Harrison, David K. Jackson, Ian Johnston, Dominic Kwiatkowski, Cordelia Langford, John Sillitoe on behalf of the Wellcome Sanger Institute COVID-19 Surveillance Team |
| EPI_ISL_2717370, EPI_ISL_2717371, EPI_ISL_2717372 | Lighthouse Lab in Glasgow | Wellcome Sanger Institute for the COVID-19 Genomics UK (COG-UK) Consortium | Harper VanSteenhouse, Yumi Kasai, David Gray, Carol Clugston, Anna Dominiczak and Alex Alderton, Roberto Amato, Jeffrey Barrett, Sonia Goncalves, Ewan Harrison, David K. Jackson, Ian Johnston, Dominic Kwiatkowski, Cordelia Langford, John Sillitoe on behalf of the Wellcome Sanger Institute COVID-19 Surveillance Team |
| EPI_ISL_2717373, EPI_ISL_2717374 | Lighthouse Lab in Alderley Park | Wellcome Sanger Institute for the COVID-19 Genomics UK (COG-UK) Consortium | Jacquelyn Wynn, Mairead Hyland, The Lighthouse Lab in Alderley Park and Alex Alderton, Roberto Amato, Jeffrey Barrett, Sonia Goncalves, Ewan Harrison, David K. Jackson, Ian Johnston, Dominic Kwiatkowski, Cordelia Langford, John Sillitoe on behalf of the Wellcome Sanger Institute COVID-19 Surveillance Team |
| EPI_ISL_2717375, EPI_ISL_2717376 | Lighthouse Lab in Milton Keynes | Wellcome Sanger Institute for the COVID-19 Genomics UK (COG-UK) Consortium | The Lighthouse Lab in Milton Keynes and Alex Alderton, Roberto Amato, Jeffrey Barrett, Sonia Goncalves, Ewan Harrison, David K. Jackson, Ian Johnston, Dominic Kwiatkowski, Cordelia Langford, John Sillitoe on behalf of the Wellcome Sanger Institute COVID-19 Surveillance Team |
| EPI_ISL_2717377 | Lighthouse Lab in Glasgow | Wellcome Sanger Institute for the COVID-19 Genomics UK (COG-UK) Consortium | Harper VanSteenhouse, Yumi Kasai, David Gray, Carol Clugston, Anna Dominiczak and Alex Alderton, Roberto Amato, Jeffrey Barrett, Sonia Goncalves, Ewan Harrison, David K. Jackson, Ian Johnston, Dominic Kwiatkowski, Cordelia Langford, John Sillitoe on behalf of the Wellcome Sanger Institute COVID-19 Surveillance Team |
| EPI_ISL_2717378, EPI_ISL_2717379 | Lighthouse Lab in Milton Keynes | Wellcome Sanger Institute for the COVID-19 Genomics UK (COG-UK) Consortium | The Lighthouse Lab in Milton Keynes and Alex Alderton, Roberto Amato, Jeffrey Barrett, Sonia Goncalves, Ewan Harrison, David K. Jackson, Ian Johnston, Dominic Kwiatkowski, Cordelia Langford, John Sillitoe on behalf of the Wellcome Sanger Institute COVID-19 Surveillance Team |
| EPI_ISL_2717380 | Health Services Laboratories | Wellcome Sanger Institute for the COVID-19 Genomics UK (COG-UK) Consortium | Health Services Laboratories and Alex Alderton, Roberto Amato, Jeffrey Barrett, Sonia Goncalves, Ewan Harrison, David K. Jackson, Ian Johnston, Dominic Kwiatkowski, Cordelia Langford, John Sillitoe on behalf of the Wellcome Sanger Institute COVID-19 Surveillance Team |
| EPI_ISL_2717381, EPI_ISL_2717382 | Lighthouse Lab in Milton Keynes | Wellcome Sanger Institute for the COVID-19 Genomics UK (COG-UK) Consortium | The Lighthouse Lab in Milton Keynes and Alex Alderton, Roberto Amato, Jeffrey Barrett, Sonia Goncalves, Ewan Harrison, David K. Jackson, Ian Johnston, Dominic Kwiatkowski, Cordelia Langford, John Sillitoe on behalf of the Wellcome Sanger Institute COVID-19 Surveillance Team |
| EPI_ISL_2717383, EPI_ISL_2717384 | Lighthouse Lab in Alderley Park | Wellcome Sanger Institute for the COVID-19 Genomics UK (COG-UK) Consortium | Jacquelyn Wynn, Mairead Hyland, The Lighthouse Lab in Alderley Park and Alex Alderton, Roberto Amato, Jeffrey Barrett, Sonia Goncalves, Ewan Harrison, David K. Jackson, Ian Johnston, Dominic Kwiatkowski, Cordelia Langford, John Sillitoe on behalf of the Wellcome Sanger Institute COVID-19 Surveillance Team |
| EPI_ISL_2717385 | Lighthouse Lab in Milton Keynes | Wellcome Sanger Institute for the COVID-19 Genomics UK (COG-UK) Consortium | The Lighthouse Lab in Milton Keynes and Alex Alderton, Roberto Amato, Jeffrey Barrett, Sonia Goncalves, Ewan Harrison, David K. Jackson, Ian Johnston, Dominic Kwiatkowski, Cordelia Langford, John Sillitoe on behalf of the Wellcome Sanger Institute COVID-19 Surveillance Team |
| EPI_ISL_2717386 | Health Services Laboratories | Wellcome Sanger Institute for the COVID-19 Genomics UK (COG-UK) Consortium | Health Services Laboratories and Alex Alderton, Roberto Amato, Jeffrey Barrett, Sonia Goncalves, Ewan Harrison, David K. Jackson, Ian Johnston, Dominic Kwiatkowski, Cordelia Langford, John Sillitoe on behalf of the Wellcome Sanger Institute COVID-19 Surveillance Team |
| EPI_ISL_2717387, EPI_ISL_2717388 | Lighthouse Lab in Milton Keynes | Wellcome Sanger Institute for the COVID-19 Genomics UK (COG-UK) Consortium | The Lighthouse Lab in Milton Keynes and Alex Alderton, Roberto Amato, Jeffrey Barrett, Sonia Goncalves, Ewan Harrison, David K. Jackson, Ian Johnston, Dominic Kwiatkowski, Cordelia Langford, John Sillitoe on behalf of the Wellcome Sanger Institute COVID-19 Surveillance Team |
| EPI_ISL_2717389 | Lighthouse Lab in Glasgow | Wellcome Sanger Institute for the COVID-19 Genomics UK (COG-UK) Consortium | Harper VanSteenhouse, Yumi Kasai, David Gray, Carol Clugston, Anna Dominiczak and Alex Alderton, Roberto Amato, Jeffrey Barrett, Sonia Goncalves, Ewan Harrison, David K. Jackson, Ian Johnston, Dominic Kwiatkowski, Cordelia Langford, John Sillitoe on behalf of the Wellcome Sanger Institute COVID-19 Surveillance Team |
| EPI_ISL_2717390 | Lighthouse Lab in Milton Keynes | Wellcome Sanger Institute for the COVID-19 Genomics UK (COG-UK) Consortium | The Lighthouse Lab in Milton Keynes and Alex Alderton, Roberto Amato, Jeffrey Barrett, Sonia Goncalves, Ewan Harrison, David K. Jackson, Ian Johnston, Dominic Kwiatkowski, Cordelia Langford, John Sillitoe on behalf of the Wellcome Sanger Institute COVID-19 Surveillance Team |
| EPI_ISL_2717391 | Health Services Laboratories | Wellcome Sanger Institute for the COVID-19 Genomics UK (COG-UK) Consortium | Health Services Laboratories and Alex Alderton, Roberto Amato, Jeffrey Barrett, Sonia Goncalves, Ewan Harrison, David K. Jackson, Ian Johnston, Dominic Kwiatkowski, Cordelia Langford, John Sillitoe on behalf of the Wellcome Sanger Institute COVID-19 Surveillance Team |
| EPI_ISL_2717392 | Lighthouse Lab in Alderley Park | Wellcome Sanger Institute for the COVID-19 Genomics UK (COG-UK) Consortium | Jacquelyn Wynn, Mairead Hyland, The Lighthouse Lab in Alderley Park and Alex Alderton, Roberto Amato, Jeffrey Barrett, Sonia Goncalves, Ewan Harrison, David K. Jackson, Ian Johnston, Dominic Kwiatkowski, Cordelia Langford, John Sillitoe on behalf of the Wellcome Sanger Institute COVID-19 Surveillance Team |
| EPI_ISL_2717393, EPI_ISL_2717394 | Lighthouse Lab in Glasgow | Wellcome Sanger Institute for the COVID-19 Genomics UK (COG-UK) Consortium | Harper VanSteenhouse, Yumi Kasai, David Gray, Carol Clugston, Anna Dominiczak and Alex Alderton, Roberto Amato, Jeffrey Barrett, Sonia Goncalves, Ewan Harrison, David K. Jackson, Ian Johnston, Dominic Kwiatkowski, Cordelia Langford, John Sillitoe on behalf of the Wellcome Sanger Institute COVID-19 Surveillance Team |
| EPI_ISL_2717395 | Lighthouse Lab in Alderley Park | Wellcome Sanger Institute for the COVID-19 Genomics UK (COG-UK) Consortium | Jacquelyn Wynn, Mairead Hyland, The Lighthouse Lab in Alderley Park and Alex Alderton, Roberto Amato, Jeffrey Barrett, Sonia Goncalves, Ewan Harrison, David K. Jackson, Ian Johnston, Dominic Kwiatkowski, Cordelia Langford, John Sillitoe on behalf of the Wellcome Sanger Institute COVID-19 Surveillance Team |
| EPI_ISL_2717396, EPI_ISL_2717397, EPI_ISL_2717398, EPI_ISL_2717399 | Lighthouse Lab in Milton Keynes | Wellcome Sanger Institute for the COVID-19 Genomics UK (COG-UK) Consortium | The Lighthouse Lab in Milton Keynes and Alex Alderton, Roberto Amato, Jeffrey Barrett, Sonia Goncalves, Ewan Harrison, David K. Jackson, Ian Johnston, Dominic Kwiatkowski, Cordelia Langford, John Sillitoe on behalf of the Wellcome Sanger Institute COVID-19 Surveillance Team |
| EPI_ISL_2717400 | Health Services Laboratories | Wellcome Sanger Institute for the COVID-19 Genomics UK (COG-UK) Consortium | Health Services Laboratories and Alex Alderton, Roberto Amato, Jeffrey Barrett, Sonia Goncalves, Ewan Harrison, David K. Jackson, Ian Johnston, Dominic Kwiatkowski, Cordelia Langford, John Sillitoe on behalf of the Wellcome Sanger Institute COVID-19 Surveillance Team |
| EPI_ISL_2717401, EPI_ISL_2717402, EPI_ISL_2717403, EPI_ISL_2717404, EPI_ISL_2717405 | Lighthouse Lab in Alderley Park | Wellcome Sanger Institute for the COVID-19 Genomics UK (COG-UK) Consortium | Jacquelyn Wynn, Mairead Hyland, The Lighthouse Lab in Alderley Park and Alex Alderton, Roberto Amato, Jeffrey Barrett, Sonia Goncalves, Ewan Harrison, David K. Jackson, Ian Johnston, Dominic Kwiatkowski, Cordelia Langford, John Sillitoe on behalf of the Wellcome Sanger Institute COVID-19 Surveillance Team |
| EPI_ISL_2717406, EPI_ISL_2717407 | Lighthouse Lab in Milton Keynes | Wellcome Sanger Institute for the COVID-19 Genomics UK (COG-UK) Consortium | The Lighthouse Lab in Milton Keynes and Alex Alderton, Roberto Amato, Jeffrey Barrett, Sonia Goncalves, Ewan Harrison, David K. Jackson, Ian Johnston, Dominic Kwiatkowski, Cordelia Langford, John Sillitoe on behalf of the Wellcome Sanger Institute COVID-19 Surveillance Team |
| EPI_ISL_2717408 | Health Services Laboratories | Wellcome Sanger Institute for the COVID-19 Genomics UK (COG-UK) Consortium | Health Services Laboratories and Alex Alderton, Roberto Amato, Jeffrey Barrett, Sonia Goncalves, Ewan Harrison, David K. Jackson, Ian Johnston, Dominic Kwiatkowski, Cordelia Langford, John Sillitoe on behalf of the Wellcome Sanger Institute COVID-19 Surveillance Team |
| EPI_ISL_2717409 | Lighthouse Lab in Glasgow | Wellcome Sanger Institute for the COVID-19 Genomics UK (COG-UK) Consortium | Harper VanSteenhouse, Yumi Kasai, David Gray, Carol Clugston, Anna Dominiczak and Alex Alderton, Roberto Amato, Jeffrey Barrett, Sonia Goncalves, Ewan Harrison, David K. Jackson, Ian Johnston, Dominic Kwiatkowski, Cordelia Langford, John Sillitoe on behalf of the Wellcome Sanger Institute COVID-19 Surveillance Team |

| | | | |
|---|---|---|---|
| EPI_ISL_2717410 | Lighthouse Lab in Milton Keynes | Wellcome Sanger Institute for the COVID-19 Genomics UK (COG-UK) Consortium | The Lighthouse Lab in Milton Keynes and Alex Alderton, Roberto Amato, Jeffrey Barrett, Sonia Goncalves, Ewan Harrison, David K. Jackson, Ian Johnston, Dominic Kwiatkowski, Cordelia Langford, John Sillitoe on behalf of the Wellcome Sanger Institute COVID-19 Surveillance Team |
| EPI_ISL_2717411 | Lighthouse Lab in Alderley Park | Wellcome Sanger Institute for the COVID-19 Genomics UK (COG-UK) Consortium | Jacquelyn Wynn, Mairead Hyland, The Lighthouse Lab in Alderley Park and Alex Alderton, Roberto Amato, Jeffrey Barrett, Sonia Goncalves, Ewan Harrison, David K. Jackson, Ian Johnston, Dominic Kwiatkowski, Cordelia Langford, John Sillitoe on behalf of the Wellcome Sanger Institute COVID-19 Surveillance Team |
| EPI_ISL_2717412 | Lighthouse Lab in Milton Keynes | Wellcome Sanger Institute for the COVID-19 Genomics UK (COG-UK) Consortium | The Lighthouse Lab in Milton Keynes and Alex Alderton, Roberto Amato, Jeffrey Barrett, Sonia Goncalves, Ewan Harrison, David K. Jackson, Ian Johnston, Dominic Kwiatkowski, Cordelia Langford, John Sillitoe on behalf of the Wellcome Sanger Institute COVID-19 Surveillance Team |
| EPI_ISL_2717413 | Lighthouse Lab in Alderley Park | Wellcome Sanger Institute for the COVID-19 Genomics UK (COG-UK) Consortium | Jacquelyn Wynn, Mairead Hyland, The Lighthouse Lab in Alderley Park and Alex Alderton, Roberto Amato, Jeffrey Barrett, Sonia Goncalves, Ewan Harrison, David K. Jackson, Ian Johnston, Dominic Kwiatkowski, Cordelia Langford, John Sillitoe on behalf of the Wellcome Sanger Institute COVID-19 Surveillance Team |
| EPI_ISL_2717414 | Lighthouse Lab in Milton Keynes | Wellcome Sanger Institute for the COVID-19 Genomics UK (COG-UK) Consortium | The Lighthouse Lab in Milton Keynes and Alex Alderton, Roberto Amato, Jeffrey Barrett, Sonia Goncalves, Ewan Harrison, David K. Jackson, Ian Johnston, Dominic Kwiatkowski, Cordelia Langford, John Sillitoe on behalf of the Wellcome Sanger Institute COVID-19 Surveillance Team |
| EPI_ISL_2717415, EPI_ISL_2717416 | Lighthouse Lab in Alderley Park | Wellcome Sanger Institute for the COVID-19 Genomics UK (COG-UK) Consortium | Jacquelyn Wynn, Mairead Hyland, The Lighthouse Lab in Alderley Park and Alex Alderton, Roberto Amato, Jeffrey Barrett, Sonia Goncalves, Ewan Harrison, David K. Jackson, Ian Johnston, Dominic Kwiatkowski, Cordelia Langford, John Sillitoe on behalf of the Wellcome Sanger Institute COVID-19 Surveillance Team |
| EPI_ISL_2717417 | Lighthouse Lab in Glasgow | Wellcome Sanger Institute for the COVID-19 Genomics UK (COG-UK) Consortium | Harper VanSteenhouse, Yumi Kasai, David Gray, Carol Clugston, Anna Dominiczak and Alex Alderton, Roberto Amato, Jeffrey Barrett, Sonia Goncalves, Ewan Harrison, David K. Jackson, Ian Johnston, Dominic Kwiatkowski, Cordelia Langford, John Sillitoe on behalf of the Wellcome Sanger Institute COVID-19 Surveillance Team |
| EPI_ISL_2717418 | Health Services Laboratories | Wellcome Sanger Institute for the COVID-19 Genomics UK (COG-UK) Consortium | Health Services Laboratories and Alex Alderton, Roberto Amato, Jeffrey Barrett, Sonia Goncalves, Ewan Harrison, David K. Jackson, Ian Johnston, Dominic Kwiatkowski, Cordelia Langford, John Sillitoe on behalf of the Wellcome Sanger Institute COVID-19 Surveillance Team |
| EPI_ISL_2717419 | Lighthouse Lab in Milton Keynes | Wellcome Sanger Institute for the COVID-19 Genomics UK (COG-UK) Consortium | The Lighthouse Lab in Milton Keynes and Alex Alderton, Roberto Amato, Jeffrey Barrett, Sonia Goncalves, Ewan Harrison, David K. Jackson, Ian Johnston, Dominic Kwiatkowski, Cordelia Langford, John Sillitoe on behalf of the Wellcome Sanger Institute COVID-19 Surveillance Team |
| EPI_ISL_2717420, EPI_ISL_2717421, EPI_ISL_2717422 | Lighthouse Lab in Alderley Park | Wellcome Sanger Institute for the COVID-19 Genomics UK (COG-UK) Consortium | Jacquelyn Wynn, Mairead Hyland, The Lighthouse Lab in Alderley Park and Alex Alderton, Roberto Amato, Jeffrey Barrett, Sonia Goncalves, Ewan Harrison, David K. Jackson, Ian Johnston, Dominic Kwiatkowski, Cordelia Langford, John Sillitoe on behalf of the Wellcome Sanger Institute COVID-19 Surveillance Team |
| EPI_ISL_2717423 | Lighthouse Lab in Glasgow | Wellcome Sanger Institute for the COVID-19 Genomics UK (COG-UK) Consortium | Harper VanSteenhouse, Yumi Kasai, David Gray, Carol Clugston, Anna Dominiczak and Alex Alderton, Roberto Amato, Jeffrey Barrett, Sonia Goncalves, Ewan Harrison, David K. Jackson, Ian Johnston, Dominic Kwiatkowski, Cordelia Langford, John Sillitoe on behalf of the Wellcome Sanger Institute COVID-19 Surveillance Team |
| EPI_ISL_2717424 | Lighthouse Lab in Milton Keynes | Wellcome Sanger Institute for the COVID-19 Genomics UK (COG-UK) Consortium | The Lighthouse Lab in Milton Keynes and Alex Alderton, Roberto Amato, Jeffrey Barrett, Sonia Goncalves, Ewan Harrison, David K. Jackson, Ian Johnston, Dominic Kwiatkowski, Cordelia Langford, John Sillitoe on behalf of the Wellcome Sanger Institute COVID-19 Surveillance Team |
| EPI_ISL_2717425, EPI_ISL_2717426 | Lighthouse Lab in Alderley Park | Wellcome Sanger Institute for the COVID-19 Genomics UK (COG-UK) Consortium | Jacquelyn Wynn, Mairead Hyland, The Lighthouse Lab in Alderley Park and Alex Alderton, Roberto Amato, Jeffrey Barrett, Sonia Goncalves, Ewan Harrison, David K. Jackson, Ian Johnston, Dominic Kwiatkowski, Cordelia Langford, John Sillitoe on behalf of the Wellcome Sanger Institute COVID-19 Surveillance Team |
| EPI_ISL_2717427 | Lighthouse Lab in Glasgow | Wellcome Sanger Institute for the COVID-19 Genomics UK (COG-UK) Consortium | Harper VanSteenhouse, Yumi Kasai, David Gray, Carol Clugston, Anna Dominiczak and Alex Alderton, Roberto Amato, Jeffrey Barrett, Sonia Goncalves, Ewan Harrison, David K. Jackson, Ian Johnston, Dominic Kwiatkowski, Cordelia Langford, John Sillitoe on behalf of the Wellcome Sanger Institute COVID-19 Surveillance Team |
| EPI_ISL_2717428, EPI_ISL_2717429 | Lighthouse Lab in Milton Keynes | Wellcome Sanger Institute for the COVID-19 Genomics UK (COG-UK) Consortium | The Lighthouse Lab in Milton Keynes and Alex Alderton, Roberto Amato, Jeffrey Barrett, Sonia Goncalves, Ewan Harrison, David K. Jackson, Ian Johnston, Dominic Kwiatkowski, Cordelia Langford, John Sillitoe on behalf of the Wellcome Sanger Institute COVID-19 Surveillance Team |
| EPI_ISL_2717430 | Lighthouse Lab in Glasgow | Wellcome Sanger Institute for the COVID-19 Genomics UK (COG-UK) Consortium | Harper VanSteenhouse, Yumi Kasai, David Gray, Carol Clugston, Anna Dominiczak and Alex Alderton, Roberto Amato, Jeffrey Barrett, Sonia Goncalves, Ewan Harrison, David K. Jackson, Ian Johnston, Dominic Kwiatkowski, Cordelia Langford, John Sillitoe on behalf of the Wellcome Sanger Institute COVID-19 Surveillance Team |
| EPI_ISL_2717431 | Lighthouse Lab in Milton Keynes | Wellcome Sanger Institute for the COVID-19 Genomics UK (COG-UK) Consortium | The Lighthouse Lab in Milton Keynes and Alex Alderton, Roberto Amato, Jeffrey Barrett, Sonia Goncalves, Ewan Harrison, David K. Jackson, Ian Johnston, Dominic Kwiatkowski, Cordelia Langford, John Sillitoe on behalf of the Wellcome Sanger Institute COVID-19 Surveillance Team |
| EPI_ISL_2717432 | Lighthouse Lab in Glasgow | Wellcome Sanger Institute for the COVID-19 Genomics UK (COG-UK) Consortium | Harper VanSteenhouse, Yumi Kasai, David Gray, Carol Clugston, Anna Dominiczak and Alex Alderton, Roberto Amato, Jeffrey Barrett, Sonia Goncalves, Ewan Harrison, David K. Jackson, Ian Johnston, Dominic Kwiatkowski, Cordelia Langford, John Sillitoe on behalf of the Wellcome Sanger Institute COVID-19 Surveillance Team |
| EPI_ISL_2717433 | Lighthouse Lab in Milton Keynes | Wellcome Sanger Institute for the COVID-19 Genomics UK (COG-UK) Consortium | The Lighthouse Lab in Milton Keynes and Alex Alderton, Roberto Amato, Jeffrey Barrett, Sonia Goncalves, Ewan Harrison, David K. Jackson, Ian Johnston, Dominic Kwiatkowski, Cordelia Langford, John Sillitoe on behalf of the Wellcome Sanger Institute COVID-19 Surveillance Team |
| EPI_ISL_2717434, EPI_ISL_2717435 | Lighthouse Lab in Alderley Park | Wellcome Sanger Institute for the COVID-19 Genomics UK (COG-UK) Consortium | Jacquelyn Wynn, Mairead Hyland, The Lighthouse Lab in Alderley Park and Alex Alderton, Roberto Amato, Jeffrey Barrett, Sonia Goncalves, Ewan Harrison, David K. Jackson, Ian Johnston, Dominic Kwiatkowski, Cordelia Langford, John Sillitoe on behalf of the Wellcome Sanger Institute COVID-19 Surveillance Team |
| EPI_ISL_2717436, EPI_ISL_2717437, EPI_ISL_2717438 | Lighthouse Lab in Milton Keynes | Wellcome Sanger Institute for the COVID-19 Genomics UK (COG-UK) Consortium | The Lighthouse Lab in Milton Keynes and Alex Alderton, Roberto Amato, Jeffrey Barrett, Sonia Goncalves, Ewan Harrison, David K. Jackson, Ian Johnston, Dominic Kwiatkowski, Cordelia Langford, John Sillitoe on behalf of the Wellcome Sanger Institute COVID-19 Surveillance Team |
| EPI_ISL_2717439 | Health Services Laboratories | Wellcome Sanger Institute for the COVID-19 Genomics UK (COG-UK) Consortium | Health Services Laboratories and Alex Alderton, Roberto Amato, Jeffrey Barrett, Sonia Goncalves, Ewan Harrison, David K. Jackson, Ian Johnston, Dominic Kwiatkowski, Cordelia Langford, John Sillitoe on behalf of the Wellcome Sanger Institute COVID-19 Surveillance Team |
| EPI_ISL_2717440, EPI_ISL_2717441 | Lighthouse Lab in Milton Keynes | Wellcome Sanger Institute for the COVID-19 Genomics UK (COG-UK) Consortium | The Lighthouse Lab in Milton Keynes and Alex Alderton, Roberto Amato, Jeffrey Barrett, Sonia Goncalves, Ewan Harrison, David K. Jackson, Ian Johnston, Dominic Kwiatkowski, Cordelia Langford, John Sillitoe on behalf of the Wellcome Sanger Institute COVID-19 Surveillance Team |
| EPI_ISL_2717442, EPI_ISL_2717443 | Lighthouse Lab in Glasgow | Wellcome Sanger Institute for the COVID-19 Genomics UK (COG-UK) Consortium | Harper VanSteenhouse, Yumi Kasai, David Gray, Carol Clugston, Anna Dominiczak and Alex Alderton, Roberto Amato, Jeffrey Barrett, Sonia Goncalves, Ewan Harrison, David K. Jackson, Ian Johnston, Dominic Kwiatkowski, Cordelia Langford, John Sillitoe on behalf of the Wellcome Sanger Institute COVID-19 Surveillance Team |
| EPI_ISL_2717444, EPI_ISL_2717445, EPI_ISL_2717446 | Lighthouse Lab in Alderley Park | Wellcome Sanger Institute for the COVID-19 Genomics UK (COG-UK) Consortium | Jacquelyn Wynn, Mairead Hyland, The Lighthouse Lab in Alderley Park and Alex Alderton, Roberto Amato, Jeffrey Barrett, Sonia Goncalves, Ewan Harrison, David K. Jackson, Ian Johnston, Dominic Kwiatkowski, Cordelia Langford, John Sillitoe on behalf of the Wellcome Sanger Institute COVID-19 Surveillance Team |
| EPI_ISL_2717447 | Lighthouse Lab in Glasgow | Wellcome Sanger Institute for the COVID-19 Genomics UK (COG-UK) Consortium | Harper VanSteenhouse, Yumi Kasai, David Gray, Carol Clugston, Anna Dominiczak and Alex Alderton, Roberto Amato, Jeffrey Barrett, Sonia Goncalves, Ewan Harrison, David K. Jackson, Ian Johnston, Dominic Kwiatkowski, Cordelia Langford, John Sillitoe on behalf of the Wellcome Sanger Institute COVID-19 Surveillance Team |
| EPI_ISL_2717448 | Lighthouse Lab in Milton Keynes | Wellcome Sanger Institute for the COVID-19 Genomics UK (COG-UK) Consortium | The Lighthouse Lab in Milton Keynes and Alex Alderton, Roberto Amato, Jeffrey Barrett, Sonia Goncalves, Ewan Harrison, David K. Jackson, Ian Johnston, Dominic Kwiatkowski, Cordelia Langford, John Sillitoe on behalf of the Wellcome Sanger Institute COVID-19 Surveillance Team |
| EPI_ISL_2717449, EPI_ISL_2717450 | Lighthouse Lab in Glasgow | Wellcome Sanger Institute for the COVID-19 Genomics UK (COG-UK) Consortium | Harper VanSteenhouse, Yumi Kasai, David Gray, Carol Clugston, Anna Dominiczak and Alex Alderton, Roberto Amato, Jeffrey Barrett, Sonia Goncalves, Ewan Harrison, David K. Jackson, Ian Johnston, Dominic Kwiatkowski, Cordelia Langford, John Sillitoe on behalf of the Wellcome Sanger Institute COVID-19 Surveillance Team |
| EPI_ISL_2717451 | Lighthouse Lab in Alderley Park | Wellcome Sanger Institute for the COVID-19 Genomics UK (COG-UK) Consortium | Jacquelyn Wynn, Mairead Hyland, The Lighthouse Lab in Alderley Park and Alex Alderton, Roberto Amato, Jeffrey Barrett, Sonia Goncalves, Ewan Harrison, David K. Jackson, Ian Johnston, Dominic Kwiatkowski, Cordelia Langford, John Sillitoe on behalf of the Wellcome Sanger Institute COVID-19 Surveillance Team |
| EPI_ISL_2717452 | Lighthouse Lab in Milton Keynes | Wellcome Sanger Institute for the COVID-19 Genomics UK (COG-UK) Consortium | The Lighthouse Lab in Milton Keynes and Alex Alderton, Roberto Amato, Jeffrey Barrett, Sonia Goncalves, Ewan Harrison, David K. Jackson, Ian Johnston, Dominic Kwiatkowski, Cordelia Langford, John Sillitoe on behalf of the Wellcome Sanger Institute COVID-19 Surveillance Team |
| EPI_ISL_2717453 | Lighthouse Lab in Glasgow | Wellcome Sanger Institute for the COVID-19 Genomics UK (COG-UK) Consortium | Harper VanSteenhouse, Yumi Kasai, David Gray, Carol Clugston, Anna Dominiczak and Alex Alderton, Roberto Amato, Jeffrey Barrett, Sonia Goncalves, Ewan Harrison, David K. Jackson, Ian Johnston, Dominic Kwiatkowski, Cordelia Langford, John Sillitoe on behalf of the Wellcome Sanger Institute COVID-19 Surveillance Team |
| EPI_ISL_2717454 | Health Services Laboratories | Wellcome Sanger Institute for the COVID-19 Genomics UK (COG-UK) Consortium | Health Services Laboratories and Alex Alderton, Roberto Amato, Jeffrey Barrett, Sonia Goncalves, Ewan Harrison, David K. Jackson, Ian Johnston, Dominic Kwiatkowski, Cordelia Langford, John Sillitoe on behalf of the Wellcome Sanger Institute COVID-19 Surveillance Team |
| EPI_ISL_2717455 | Lighthouse Lab in Milton Keynes | Wellcome Sanger Institute for the COVID-19 Genomics UK (COG-UK) Consortium | The Lighthouse Lab in Milton Keynes and Alex Alderton, Roberto Amato, Jeffrey Barrett, Sonia Goncalves, Ewan Harrison, David K. Jackson, Ian Johnston, Dominic Kwiatkowski, Cordelia Langford, John Sillitoe on behalf of the Wellcome Sanger Institute COVID-19 Surveillance Team |
| EPI_ISL_2717456 | Health Services Laboratories | Wellcome Sanger Institute for the COVID-19 Genomics UK (COG-UK) Consortium | Health Services Laboratories and Alex Alderton, Roberto Amato, Jeffrey Barrett, Sonia Goncalves, Ewan Harrison, David K. Jackson, Ian Johnston, Dominic Kwiatkowski, Cordelia Langford, John Sillitoe on behalf of the Wellcome Sanger Institute COVID-19 Surveillance Team |
| EPI_ISL_2717457, EPI_ISL_2717458 | Lighthouse Lab in Alderley Park | Wellcome Sanger Institute for the COVID-19 Genomics UK (COG-UK) Consortium | Jacquelyn Wynn, Mairead Hyland, The Lighthouse Lab in Alderley Park and Alex Alderton, Roberto Amato, Jeffrey Barrett, Sonia Goncalves, Ewan Harrison, David K. Jackson, Ian Johnston, Dominic Kwiatkowski, Cordelia Langford, John Sillitoe on behalf of the Wellcome Sanger Institute COVID-19 Surveillance Team |
| EPI_ISL_2717459 | Lighthouse Lab in Glasgow | Wellcome Sanger Institute for the COVID-19 Genomics UK (COG-UK) Consortium | Harper VanSteenhouse, Yumi Kasai, David Gray, Carol Clugston, Anna Dominiczak and Alex Alderton, Roberto Amato, Jeffrey Barrett, Sonia Goncalves, Ewan Harrison, David K. Jackson, Ian Johnston, Dominic Kwiatkowski, Cordelia Langford, John Sillitoe on behalf of the Wellcome Sanger Institute COVID-19 Surveillance Team |
| EPI_ISL_2717460 | Health Services Laboratories | Wellcome Sanger Institute for the COVID-19 Genomics UK (COG-UK) Consortium | Health Services Laboratories and Alex Alderton, Roberto Amato, Jeffrey Barrett, Sonia Goncalves, Ewan Harrison, David K. Jackson, Ian Johnston, Dominic Kwiatkowski, Cordelia Langford, John Sillitoe on behalf of the Wellcome Sanger Institute COVID-19 Surveillance Team |
| EPI_ISL_2717462 | Lighthouse Lab in Alderley Park | Wellcome Sanger Institute for the COVID-19 Genomics UK (COG-UK) Consortium | Jacquelyn Wynn, Mairead Hyland, The Lighthouse Lab in Alderley Park and Alex Alderton, Roberto Amato, Jeffrey Barrett, Sonia Goncalves, Ewan Harrison, David K. Jackson, Ian Johnston, Dominic Kwiatkowski, Cordelia Langford, John Sillitoe on behalf of the Wellcome Sanger Institute COVID-19 Surveillance Team |
| EPI_ISL_2717463 | Lighthouse Lab in Milton Keynes | Wellcome Sanger Institute for the COVID-19 Genomics UK (COG-UK) Consortium | The Lighthouse Lab in Milton Keynes and Alex Alderton, Roberto Amato, Jeffrey Barrett, Sonia Goncalves, Ewan Harrison, David K. Jackson, Ian Johnston, Dominic Kwiatkowski, Cordelia Langford, John Sillitoe on behalf of the Wellcome Sanger Institute COVID-19 Surveillance Team |
| EPI_ISL_2717464 | Health Services Laboratories | Wellcome Sanger Institute for the COVID-19 Genomics UK (COG-UK) Consortium | Health Services Laboratories and Alex Alderton, Roberto Amato, Jeffrey Barrett, Sonia Goncalves, Ewan Harrison, David K. Jackson, Ian Johnston, Dominic Kwiatkowski, Cordelia Langford, John Sillitoe on behalf of the Wellcome Sanger Institute COVID-19 Surveillance Team |
| EPI_ISL_2717465, EPI_ISL_2717466 | Lighthouse Lab in Milton Keynes | Wellcome Sanger Institute for the COVID-19 Genomics UK (COG-UK) Consortium | The Lighthouse Lab in Milton Keynes and Alex Alderton, Roberto Amato, Jeffrey Barrett, Sonia Goncalves, Ewan Harrison, David K. Jackson, Ian Johnston, Dominic Kwiatkowski, Cordelia Langford, John Sillitoe on behalf of the Wellcome Sanger Institute COVID-19 Surveillance Team |
| EPI_ISL_2717467 | Lighthouse Lab in Alderley Park | Wellcome Sanger Institute for the COVID-19 Genomics UK (COG-UK) Consortium | Jacquelyn Wynn, Mairead Hyland, The Lighthouse Lab in Alderley Park and Alex Alderton, Roberto Amato, Jeffrey Barrett, Sonia Goncalves, Ewan Harrison, David K. Jackson, Ian Johnston, Dominic Kwiatkowski, Cordelia Langford, John Sillitoe on behalf of the Wellcome Sanger Institute COVID-19 Surveillance Team |
| EPI_ISL_2717468, EPI_ISL_2717469 | Health Services Laboratories | Wellcome Sanger Institute for the COVID-19 Genomics UK (COG-UK) Consortium | Health Services Laboratories and Alex Alderton, Roberto Amato, Jeffrey Barrett, Sonia Goncalves, Ewan Harrison, David K. Jackson, Ian Johnston, Dominic Kwiatkowski, Cordelia Langford, John Sillitoe on behalf of the Wellcome Sanger Institute COVID-19 Surveillance Team |
| EPI_ISL_2717470, EPI_ISL_2717471 | Lighthouse Lab in Milton Keynes | Wellcome Sanger Institute for the COVID-19 Genomics UK (COG-UK) Consortium | The Lighthouse Lab in Milton Keynes and Alex Alderton, Roberto Amato, Jeffrey Barrett, Sonia Goncalves, Ewan Harrison, David K. Jackson, Ian Johnston, Dominic Kwiatkowski, Cordelia Langford, John Sillitoe on behalf of the Wellcome Sanger Institute COVID-19 Surveillance Team |

| | | | |
|---|---|---|---|
| EPI_ISL_2717472 | Lighthouse Lab in Alderley Park | Wellcome Sanger Institute for the COVID-19 Genomics UK (COG-UK) Consortium | Jacquelyn Wynn, Mairead Hyland, The Lighthouse Lab in Alderley Park and Alex Alderton, Roberto Amato, Jeffrey Barrett, Sonia Goncalves, Ewan Harrison, David K. Jackson, Ian Johnston, Dominic Kwiatkowski, Cordelia Langford, John Sillitoe on behalf of the Wellcome Sanger Institute COVID-19 Surveillance Team |
| EPI_ISL_2717473 | Lighthouse Lab in Milton Keynes | Wellcome Sanger Institute for the COVID-19 Genomics UK (COG-UK) Consortium | The Lighthouse Lab in Milton Keynes and Alex Alderton, Roberto Amato, Jeffrey Barrett, Sonia Goncalves, Ewan Harrison, David K. Jackson, Ian Johnston, Dominic Kwiatkowski, Cordelia Langford, John Sillitoe on behalf of the Wellcome Sanger Institute COVID-19 Surveillance Team |
| EPI_ISL_2717475 | Lighthouse Lab in Alderley Park | Wellcome Sanger Institute for the COVID-19 Genomics UK (COG-UK) Consortium | Jacquelyn Wynn, Mairead Hyland, The Lighthouse Lab in Alderley Park and Alex Alderton, Roberto Amato, Jeffrey Barrett, Sonia Goncalves, Ewan Harrison, David K. Jackson, Ian Johnston, Dominic Kwiatkowski, Cordelia Langford, John Sillitoe on behalf of the Wellcome Sanger Institute COVID-19 Surveillance Team |
| EPI_ISL_2717476 | Lighthouse Lab in Glasgow | Wellcome Sanger Institute for the COVID-19 Genomics UK (COG-UK) Consortium | Harper VanSteenhouse, Yumi Kasai, David Gray, Carol Clugston, Anna Dominiczak and Alex Alderton, Roberto Amato, Jeffrey Barrett, Sonia Goncalves, Ewan Harrison, David K. Jackson, Ian Johnston, Dominic Kwiatkowski, Cordelia Langford, John Sillitoe on behalf of the Wellcome Sanger Institute COVID-19 Surveillance Team |
| EPI_ISL_2717477 | Lighthouse Lab in Milton Keynes | Wellcome Sanger Institute for the COVID-19 Genomics UK (COG-UK) Consortium | The Lighthouse Lab in Milton Keynes and Alex Alderton, Roberto Amato, Jeffrey Barrett, Sonia Goncalves, Ewan Harrison, David K. Jackson, Ian Johnston, Dominic Kwiatkowski, Cordelia Langford, John Sillitoe on behalf of the Wellcome Sanger Institute COVID-19 Surveillance Team |
| EPI_ISL_2717478 | Lighthouse Lab in Alderley Park | Wellcome Sanger Institute for the COVID-19 Genomics UK (COG-UK) Consortium | Jacquelyn Wynn, Mairead Hyland, The Lighthouse Lab in Alderley Park and Alex Alderton, Roberto Amato, Jeffrey Barrett, Sonia Goncalves, Ewan Harrison, David K. Jackson, Ian Johnston, Dominic Kwiatkowski, Cordelia Langford, John Sillitoe on behalf of the Wellcome Sanger Institute COVID-19 Surveillance Team |
| EPI_ISL_2717479 | Health Services Laboratories | Wellcome Sanger Institute for the COVID-19 Genomics UK (COG-UK) Consortium | Health Services Laboratories and Alex Alderton, Roberto Amato, Jeffrey Barrett, Sonia Goncalves, Ewan Harrison, David K. Jackson, Ian Johnston, Dominic Kwiatkowski, Cordelia Langford, John Sillitoe on behalf of the Wellcome Sanger Institute COVID-19 Surveillance Team |
| EPI_ISL_2717480 | Lighthouse Lab in Alderley Park | Wellcome Sanger Institute for the COVID-19 Genomics UK (COG-UK) Consortium | Jacquelyn Wynn, Mairead Hyland, The Lighthouse Lab in Alderley Park and Alex Alderton, Roberto Amato, Jeffrey Barrett, Sonia Goncalves, Ewan Harrison, David K. Jackson, Ian Johnston, Dominic Kwiatkowski, Cordelia Langford, John Sillitoe on behalf of the Wellcome Sanger Institute COVID-19 Surveillance Team |
| EPI_ISL_2717481 | Lighthouse Lab in Milton Keynes | Wellcome Sanger Institute for the COVID-19 Genomics UK (COG-UK) Consortium | The Lighthouse Lab in Milton Keynes and Alex Alderton, Roberto Amato, Jeffrey Barrett, Sonia Goncalves, Ewan Harrison, David K. Jackson, Ian Johnston, Dominic Kwiatkowski, Cordelia Langford, John Sillitoe on behalf of the Wellcome Sanger Institute COVID-19 Surveillance Team |
| EPI_ISL_2717482 | Lighthouse Lab in Glasgow | Wellcome Sanger Institute for the COVID-19 Genomics UK (COG-UK) Consortium | Harper VanSteenhouse, Yumi Kasai, David Gray, Carol Clugston, Anna Dominiczak and Alex Alderton, Roberto Amato, Jeffrey Barrett, Sonia Goncalves, Ewan Harrison, David K. Jackson, Ian Johnston, Dominic Kwiatkowski, Cordelia Langford, John Sillitoe on behalf of the Wellcome Sanger Institute COVID-19 Surveillance Team |
| EPI_ISL_2717483 | Health Services Laboratories | Wellcome Sanger Institute for the COVID-19 Genomics UK (COG-UK) Consortium | Health Services Laboratories and Alex Alderton, Roberto Amato, Jeffrey Barrett, Sonia Goncalves, Ewan Harrison, David K. Jackson, Ian Johnston, Dominic Kwiatkowski, Cordelia Langford, John Sillitoe on behalf of the Wellcome Sanger Institute COVID-19 Surveillance Team |
| EPI_ISL_2717484 | Lighthouse Lab in Glasgow | Wellcome Sanger Institute for the COVID-19 Genomics UK (COG-UK) Consortium | Harper VanSteenhouse, Yumi Kasai, David Gray, Carol Clugston, Anna Dominiczak and Alex Alderton, Roberto Amato, Jeffrey Barrett, Sonia Goncalves, Ewan Harrison, David K. Jackson, Ian Johnston, Dominic Kwiatkowski, Cordelia Langford, John Sillitoe on behalf of the Wellcome Sanger Institute COVID-19 Surveillance Team |
| EPI_ISL_2717485 | Health Services Laboratories | Wellcome Sanger Institute for the COVID-19 Genomics UK (COG-UK) Consortium | Health Services Laboratories and Alex Alderton, Roberto Amato, Jeffrey Barrett, Sonia Goncalves, Ewan Harrison, David K. Jackson, Ian Johnston, Dominic Kwiatkowski, Cordelia Langford, John Sillitoe on behalf of the Wellcome Sanger Institute COVID-19 Surveillance Team |
| EPI_ISL_2717486 | Lighthouse Lab in Alderley Park | Wellcome Sanger Institute for the COVID-19 Genomics UK (COG-UK) Consortium | Jacquelyn Wynn, Mairead Hyland, The Lighthouse Lab in Alderley Park and Alex Alderton, Roberto Amato, Jeffrey Barrett, Sonia Goncalves, Ewan Harrison, David K. Jackson, Ian Johnston, Dominic Kwiatkowski, Cordelia Langford, John Sillitoe on behalf of the Wellcome Sanger Institute COVID-19 Surveillance Team |
| EPI_ISL_2717487 | Lighthouse Lab in Milton Keynes | Wellcome Sanger Institute for the COVID-19 Genomics UK (COG-UK) Consortium | The Lighthouse Lab in Milton Keynes and Alex Alderton, Roberto Amato, Jeffrey Barrett, Sonia Goncalves, Ewan Harrison, David K. Jackson, Ian Johnston, Dominic Kwiatkowski, Cordelia Langford, John Sillitoe on behalf of the Wellcome Sanger Institute COVID-19 Surveillance Team |
| EPI_ISL_2717488 | Lighthouse Lab in Alderley Park | Wellcome Sanger Institute for the COVID-19 Genomics UK (COG-UK) Consortium | Jacquelyn Wynn, Mairead Hyland, The Lighthouse Lab in Alderley Park and Alex Alderton, Roberto Amato, Jeffrey Barrett, Sonia Goncalves, Ewan Harrison, David K. Jackson, Ian Johnston, Dominic Kwiatkowski, Cordelia Langford, John Sillitoe on behalf of the Wellcome Sanger Institute COVID-19 Surveillance Team |
| EPI_ISL_2717489 | Lighthouse Lab in Milton Keynes | Wellcome Sanger Institute for the COVID-19 Genomics UK (COG-UK) Consortium | The Lighthouse Lab in Milton Keynes and Alex Alderton, Roberto Amato, Jeffrey Barrett, Sonia Goncalves, Ewan Harrison, David K. Jackson, Ian Johnston, Dominic Kwiatkowski, Cordelia Langford, John Sillitoe on behalf of the Wellcome Sanger Institute COVID-19 Surveillance Team |
| EPI_ISL_2717490 | Health Services Laboratories | Wellcome Sanger Institute for the COVID-19 Genomics UK (COG-UK) Consortium | Health Services Laboratories and Alex Alderton, Roberto Amato, Jeffrey Barrett, Sonia Goncalves, Ewan Harrison, David K. Jackson, Ian Johnston, Dominic Kwiatkowski, Cordelia Langford, John Sillitoe on behalf of the Wellcome Sanger Institute COVID-19 Surveillance Team |
| EPI_ISL_2717491 | Lighthouse Lab in Milton Keynes | Wellcome Sanger Institute for the COVID-19 Genomics UK (COG-UK) Consortium | The Lighthouse Lab in Milton Keynes and Alex Alderton, Roberto Amato, Jeffrey Barrett, Sonia Goncalves, Ewan Harrison, David K. Jackson, Ian Johnston, Dominic Kwiatkowski, Cordelia Langford, John Sillitoe on behalf of the Wellcome Sanger Institute COVID-19 Surveillance Team |
| EPI_ISL_2717492, EPI_ISL_2717493 | Health Services Laboratories | Wellcome Sanger Institute for the COVID-19 Genomics UK (COG-UK) Consortium | Health Services Laboratories and Alex Alderton, Roberto Amato, Jeffrey Barrett, Sonia Goncalves, Ewan Harrison, David K. Jackson, Ian Johnston, Dominic Kwiatkowski, Cordelia Langford, John Sillitoe on behalf of the Wellcome Sanger Institute COVID-19 Surveillance Team |
| EPI_ISL_2717494, EPI_ISL_2717495 | Lighthouse Lab in Alderley Park | Wellcome Sanger Institute for the COVID-19 Genomics UK (COG-UK) Consortium | Jacquelyn Wynn, Mairead Hyland, The Lighthouse Lab in Alderley Park and Alex Alderton, Roberto Amato, Jeffrey Barrett, Sonia Goncalves, Ewan Harrison, David K. Jackson, Ian Johnston, Dominic Kwiatkowski, Cordelia Langford, John Sillitoe on behalf of the Wellcome Sanger Institute COVID-19 Surveillance Team |
| EPI_ISL_2717496 | Lighthouse Lab in Milton Keynes | Wellcome Sanger Institute for the COVID-19 Genomics UK (COG-UK) Consortium | The Lighthouse Lab in Milton Keynes and Alex Alderton, Roberto Amato, Jeffrey Barrett, Sonia Goncalves, Ewan Harrison, David K. Jackson, Ian Johnston, Dominic Kwiatkowski, Cordelia Langford, John Sillitoe on behalf of the Wellcome Sanger Institute COVID-19 Surveillance Team |
| EPI_ISL_2717497 | Lighthouse Lab in Glasgow | Wellcome Sanger Institute for the COVID-19 Genomics UK (COG-UK) Consortium | Harper VanSteenhouse, Yumi Kasai, David Gray, Carol Clugston, Anna Dominiczak and Alex Alderton, Roberto Amato, Jeffrey Barrett, Sonia Goncalves, Ewan Harrison, David K. Jackson, Ian Johnston, Dominic Kwiatkowski, Cordelia Langford, John Sillitoe on behalf of the Wellcome Sanger Institute COVID-19 Surveillance Team |
| EPI_ISL_2717498 | Lighthouse Lab in Alderley Park | Wellcome Sanger Institute for the COVID-19 Genomics UK (COG-UK) Consortium | Jacquelyn Wynn, Mairead Hyland, The Lighthouse Lab in Alderley Park and Alex Alderton, Roberto Amato, Jeffrey Barrett, Sonia Goncalves, Ewan Harrison, David K. Jackson, Ian Johnston, Dominic Kwiatkowski, Cordelia Langford, John Sillitoe on behalf of the Wellcome Sanger Institute COVID-19 Surveillance Team |
| EPI_ISL_2717499 | Lighthouse Lab in Glasgow | Wellcome Sanger Institute for the COVID-19 Genomics UK (COG-UK) Consortium | Harper VanSteenhouse, Yumi Kasai, David Gray, Carol Clugston, Anna Dominiczak and Alex Alderton, Roberto Amato, Jeffrey Barrett, Sonia Goncalves, Ewan Harrison, David K. Jackson, Ian Johnston, Dominic Kwiatkowski, Cordelia Langford, John Sillitoe on behalf of the Wellcome Sanger Institute COVID-19 Surveillance Team |
| EPI_ISL_2717500 | Lighthouse Lab in Alderley Park | Wellcome Sanger Institute for the COVID-19 Genomics UK (COG-UK) Consortium | Jacquelyn Wynn, Mairead Hyland, The Lighthouse Lab in Alderley Park and Alex Alderton, Roberto Amato, Jeffrey Barrett, Sonia Goncalves, Ewan Harrison, David K. Jackson, Ian Johnston, Dominic Kwiatkowski, Cordelia Langford, John Sillitoe on behalf of the Wellcome Sanger Institute COVID-19 Surveillance Team |
| EPI_ISL_2717501 | Lighthouse Lab in Milton Keynes | Wellcome Sanger Institute for the COVID-19 Genomics UK (COG-UK) Consortium | The Lighthouse Lab in Milton Keynes and Alex Alderton, Roberto Amato, Jeffrey Barrett, Sonia Goncalves, Ewan Harrison, David K. Jackson, Ian Johnston, Dominic Kwiatkowski, Cordelia Langford, John Sillitoe on behalf of the Wellcome Sanger Institute COVID-19 Surveillance Team |
| EPI_ISL_2717502 | Lighthouse Lab in Alderley Park | Wellcome Sanger Institute for the COVID-19 Genomics UK (COG-UK) Consortium | Jacquelyn Wynn, Mairead Hyland, The Lighthouse Lab in Alderley Park and Alex Alderton, Roberto Amato, Jeffrey Barrett, Sonia Goncalves, Ewan Harrison, David K. Jackson, Ian Johnston, Dominic Kwiatkowski, Cordelia Langford, John Sillitoe on behalf of the Wellcome Sanger Institute COVID-19 Surveillance Team |
| EPI_ISL_2717503 | Lighthouse Lab in Milton Keynes | Wellcome Sanger Institute for the COVID-19 Genomics UK (COG-UK) Consortium | The Lighthouse Lab in Milton Keynes and Alex Alderton, Roberto Amato, Jeffrey Barrett, Sonia Goncalves, Ewan Harrison, David K. Jackson, Ian Johnston, Dominic Kwiatkowski, Cordelia Langford, John Sillitoe on behalf of the Wellcome Sanger Institute COVID-19 Surveillance Team |
| EPI_ISL_2717504, EPI_ISL_2717505 | Lighthouse Lab in Alderley Park | Wellcome Sanger Institute for the COVID-19 Genomics UK (COG-UK) Consortium | Jacquelyn Wynn, Mairead Hyland, The Lighthouse Lab in Alderley Park and Alex Alderton, Roberto Amato, Jeffrey Barrett, Sonia Goncalves, Ewan Harrison, David K. Jackson, Ian Johnston, Dominic Kwiatkowski, Cordelia Langford, John Sillitoe on behalf of the Wellcome Sanger Institute COVID-19 Surveillance Team |
| EPI_ISL_2717506, EPI_ISL_2717507 | Lighthouse Lab in Milton Keynes | Wellcome Sanger Institute for the COVID-19 Genomics UK (COG-UK) Consortium | The Lighthouse Lab in Milton Keynes and Alex Alderton, Roberto Amato, Jeffrey Barrett, Sonia Goncalves, Ewan Harrison, David K. Jackson, Ian Johnston, Dominic Kwiatkowski, Cordelia Langford, John Sillitoe on behalf of the Wellcome Sanger Institute COVID-19 Surveillance Team |
| EPI_ISL_2717508 | Lighthouse Lab in Alderley Park | Wellcome Sanger Institute for the COVID-19 Genomics UK (COG-UK) Consortium | Jacquelyn Wynn, Mairead Hyland, The Lighthouse Lab in Alderley Park and Alex Alderton, Roberto Amato, Jeffrey Barrett, Sonia Goncalves, Ewan Harrison, David K. Jackson, Ian Johnston, Dominic Kwiatkowski, Cordelia Langford, John Sillitoe on behalf of the Wellcome Sanger Institute COVID-19 Surveillance Team |
| EPI_ISL_2717509 | Lighthouse Lab in Milton Keynes | Wellcome Sanger Institute for the COVID-19 Genomics UK (COG-UK) Consortium | The Lighthouse Lab in Milton Keynes and Alex Alderton, Roberto Amato, Jeffrey Barrett, Sonia Goncalves, Ewan Harrison, David K. Jackson, Ian Johnston, Dominic Kwiatkowski, Cordelia Langford, John Sillitoe on behalf of the Wellcome Sanger Institute COVID-19 Surveillance Team |
| EPI_ISL_2717510 | Lighthouse Lab in Alderley Park | Wellcome Sanger Institute for the COVID-19 Genomics UK (COG-UK) Consortium | Jacquelyn Wynn, Mairead Hyland, The Lighthouse Lab in Alderley Park and Alex Alderton, Roberto Amato, Jeffrey Barrett, Sonia Goncalves, Ewan Harrison, David K. Jackson, Ian Johnston, Dominic Kwiatkowski, Cordelia Langford, John Sillitoe on behalf of the Wellcome Sanger Institute COVID-19 Surveillance Team |
| EPI_ISL_2717511 | Health Services Laboratories | Wellcome Sanger Institute for the COVID-19 Genomics UK (COG-UK) Consortium | Health Services Laboratories and Alex Alderton, Roberto Amato, Jeffrey Barrett, Sonia Goncalves, Ewan Harrison, David K. Jackson, Ian Johnston, Dominic Kwiatkowski, Cordelia Langford, John Sillitoe on behalf of the Wellcome Sanger Institute COVID-19 Surveillance Team |
| EPI_ISL_2717512, EPI_ISL_2717513, EPI_ISL_2717514 | Lighthouse Lab in Milton Keynes | Wellcome Sanger Institute for the COVID-19 Genomics UK (COG-UK) Consortium | The Lighthouse Lab in Milton Keynes and Alex Alderton, Roberto Amato, Jeffrey Barrett, Sonia Goncalves, Ewan Harrison, David K. Jackson, Ian Johnston, Dominic Kwiatkowski, Cordelia Langford, John Sillitoe on behalf of the Wellcome Sanger Institute COVID-19 Surveillance Team |
| EPI_ISL_2717515 | Lighthouse Lab in Glasgow | Wellcome Sanger Institute for the COVID-19 Genomics UK (COG-UK) Consortium | Harper VanSteenhouse, Yumi Kasai, David Gray, Carol Clugston, Anna Dominiczak and Alex Alderton, Roberto Amato, Jeffrey Barrett, Sonia Goncalves, Ewan Harrison, David K. Jackson, Ian Johnston, Dominic Kwiatkowski, Cordelia Langford, John Sillitoe on behalf of the Wellcome Sanger Institute COVID-19 Surveillance Team |
| EPI_ISL_2717516 | Lighthouse Lab in Milton Keynes | Wellcome Sanger Institute for the COVID-19 Genomics UK (COG-UK) Consortium | The Lighthouse Lab in Milton Keynes and Alex Alderton, Roberto Amato, Jeffrey Barrett, Sonia Goncalves, Ewan Harrison, David K. Jackson, Ian Johnston, Dominic Kwiatkowski, Cordelia Langford, John Sillitoe on behalf of the Wellcome Sanger Institute COVID-19 Surveillance Team |
| EPI_ISL_2717517 | Lighthouse Lab in Alderley Park | Wellcome Sanger Institute for the COVID-19 Genomics UK (COG-UK) Consortium | Jacquelyn Wynn, Mairead Hyland, The Lighthouse Lab in Alderley Park and Alex Alderton, Roberto Amato, Jeffrey Barrett, Sonia Goncalves, Ewan Harrison, David K. Jackson, Ian Johnston, Dominic Kwiatkowski, Cordelia Langford, John Sillitoe on behalf of the Wellcome Sanger Institute COVID-19 Surveillance Team |
| EPI_ISL_2717518 | Lighthouse Lab in Glasgow | Wellcome Sanger Institute for the COVID-19 Genomics UK (COG-UK) Consortium | Harper VanSteenhouse, Yumi Kasai, David Gray, Carol Clugston, Anna Dominiczak and Alex Alderton, Roberto Amato, Jeffrey Barrett, Sonia Goncalves, Ewan Harrison, David K. Jackson, Ian Johnston, Dominic Kwiatkowski, Cordelia Langford, John Sillitoe on behalf of the Wellcome Sanger Institute COVID-19 Surveillance Team |
| EPI_ISL_2717519 | Lighthouse Lab in Milton Keynes | Wellcome Sanger Institute for the COVID-19 Genomics UK (COG-UK) Consortium | The Lighthouse Lab in Milton Keynes and Alex Alderton, Roberto Amato, Jeffrey Barrett, Sonia Goncalves, Ewan Harrison, David K. Jackson, Ian Johnston, Dominic Kwiatkowski, Cordelia Langford, John Sillitoe on behalf of the Wellcome Sanger Institute COVID-19 Surveillance Team |
| EPI_ISL_2717520 | Health Services Laboratories | Wellcome Sanger Institute for the COVID-19 Genomics UK (COG-UK) Consortium | Health Services Laboratories and Alex Alderton, Roberto Amato, Jeffrey Barrett, Sonia Goncalves, Ewan Harrison, David K. Jackson, Ian Johnston, Dominic Kwiatkowski, Cordelia Langford, John Sillitoe on behalf of the Wellcome Sanger Institute COVID-19 Surveillance Team |
| EPI_ISL_2717521 | Lighthouse Lab in Milton Keynes | Wellcome Sanger Institute for the COVID-19 Genomics UK (COG-UK) Consortium | The Lighthouse Lab in Milton Keynes and Alex Alderton, Roberto Amato, Jeffrey Barrett, Sonia Goncalves, Ewan Harrison, David K. Jackson, Ian Johnston, Dominic Kwiatkowski, Cordelia Langford, John Sillitoe on behalf of the Wellcome Sanger Institute COVID-19 Surveillance Team |
| EPI_ISL_2717522, EPI_ISL_2717523 | Health Services Laboratories | Wellcome Sanger Institute for the COVID-19 Genomics UK (COG-UK) Consortium | Health Services Laboratories and Alex Alderton, Roberto Amato, Jeffrey Barrett, Sonia Goncalves, Ewan Harrison, David K. Jackson, Ian Johnston, Dominic Kwiatkowski, Cordelia Langford, John Sillitoe on behalf of the Wellcome Sanger Institute COVID-19 Surveillance Team |
| EPI_ISL_2717524 | Lighthouse Lab in Alderley Park | Wellcome Sanger Institute for the COVID-19 Genomics UK (COG-UK) Consortium | Jacquelyn Wynn, Mairead Hyland, The Lighthouse Lab in Alderley Park and Alex Alderton, Roberto Amato, Jeffrey Barrett, Sonia Goncalves, Ewan Harrison, David K. Jackson, Ian Johnston, Dominic Kwiatkowski, Cordelia Langford, John Sillitoe on behalf of the Wellcome Sanger Institute COVID-19 Surveillance Team |

| | | | |
|---|---|---|---|
| EPI_ISL_2717525 | Lighthouse Lab in Glasgow | Wellcome Sanger Institute for the COVID-19 Genomics UK (COG-UK) Consortium | Harper VanSteenhouse, Yumi Kasai, David Gray, Carol Clugston, Anna Dominiczak and Alex Alderton, Roberto Amato, Jeffrey Barrett, Sonia Goncalves, Ewan Harrison, David K. Jackson, Ian Johnston, Dominic Kwiatkowski, Cordelia Langford, John Sillitoe on behalf of the Wellcome Sanger Institute COVID-19 Surveillance Team |
| EPI_ISL_2717526 | Lighthouse Lab in Milton Keynes | Wellcome Sanger Institute for the COVID-19 Genomics UK (COG-UK) Consortium | The Lighthouse Lab in Milton Keynes and Alex Alderton, Roberto Amato, Jeffrey Barrett, Sonia Goncalves, Ewan Harrison, David K. Jackson, Ian Johnston, Dominic Kwiatkowski, Cordelia Langford, John Sillitoe on behalf of the Wellcome Sanger Institute COVID-19 Surveillance Team |
| EPI_ISL_2717527 | Lighthouse Lab in Alderley Park | Wellcome Sanger Institute for the COVID-19 Genomics UK (COG-UK) Consortium | Jacquelyn Wynn, Mairead Hyland, The Lighthouse Lab in Alderley Park and Alex Alderton, Roberto Amato, Jeffrey Barrett, Sonia Goncalves, Ewan Harrison, David K. Jackson, Ian Johnston, Dominic Kwiatkowski, Cordelia Langford, John Sillitoe on behalf of the Wellcome Sanger Institute COVID-19 Surveillance Team |
| EPI_ISL_2717528, EPI_ISL_2717529, EPI_ISL_2717530, EPI_ISL_2717531 | Lighthouse Lab in Milton Keynes | Wellcome Sanger Institute for the COVID-19 Genomics UK (COG-UK) Consortium | The Lighthouse Lab in Milton Keynes and Alex Alderton, Roberto Amato, Jeffrey Barrett, Sonia Goncalves, Ewan Harrison, David K. Jackson, Ian Johnston, Dominic Kwiatkowski, Cordelia Langford, John Sillitoe on behalf of the Wellcome Sanger Institute COVID-19 Surveillance Team |
| EPI_ISL_2717532, EPI_ISL_2717533 | Lighthouse Lab in Alderley Park | Wellcome Sanger Institute for the COVID-19 Genomics UK (COG-UK) Consortium | Jacquelyn Wynn, Mairead Hyland, The Lighthouse Lab in Alderley Park and Alex Alderton, Roberto Amato, Jeffrey Barrett, Sonia Goncalves, Ewan Harrison, David K. Jackson, Ian Johnston, Dominic Kwiatkowski, Cordelia Langford, John Sillitoe on behalf of the Wellcome Sanger Institute COVID-19 Surveillance Team |
| EPI_ISL_2717534 | Health Services Laboratories | Wellcome Sanger Institute for the COVID-19 Genomics UK (COG-UK) Consortium | Health Services Laboratories and Alex Alderton, Roberto Amato, Jeffrey Barrett, Sonia Goncalves, Ewan Harrison, David K. Jackson, Ian Johnston, Dominic Kwiatkowski, Cordelia Langford, John Sillitoe on behalf of the Wellcome Sanger Institute COVID-19 Surveillance Team |
| EPI_ISL_2717535 | Lighthouse Lab in Alderley Park | Wellcome Sanger Institute for the COVID-19 Genomics UK (COG-UK) Consortium | Jacquelyn Wynn, Mairead Hyland, The Lighthouse Lab in Alderley Park and Alex Alderton, Roberto Amato, Jeffrey Barrett, Sonia Goncalves, Ewan Harrison, David K. Jackson, Ian Johnston, Dominic Kwiatkowski, Cordelia Langford, John Sillitoe on behalf of the Wellcome Sanger Institute COVID-19 Surveillance Team |
| EPI_ISL_2717536 | Health Services Laboratories | Wellcome Sanger Institute for the COVID-19 Genomics UK (COG-UK) Consortium | Health Services Laboratories and Alex Alderton, Roberto Amato, Jeffrey Barrett, Sonia Goncalves, Ewan Harrison, David K. Jackson, Ian Johnston, Dominic Kwiatkowski, Cordelia Langford, John Sillitoe on behalf of the Wellcome Sanger Institute COVID-19 Surveillance Team |
| EPI_ISL_2717537, EPI_ISL_2717538 | Lighthouse Lab in Milton Keynes | Wellcome Sanger Institute for the COVID-19 Genomics UK (COG-UK) Consortium | The Lighthouse Lab in Milton Keynes and Alex Alderton, Roberto Amato, Jeffrey Barrett, Sonia Goncalves, Ewan Harrison, David K. Jackson, Ian Johnston, Dominic Kwiatkowski, Cordelia Langford, John Sillitoe on behalf of the Wellcome Sanger Institute COVID-19 Surveillance Team |
| EPI_ISL_2717539 | Lighthouse Lab in Alderley Park | Wellcome Sanger Institute for the COVID-19 Genomics UK (COG-UK) Consortium | Jacquelyn Wynn, Mairead Hyland, The Lighthouse Lab in Alderley Park and Alex Alderton, Roberto Amato, Jeffrey Barrett, Sonia Goncalves, Ewan Harrison, David K. Jackson, Ian Johnston, Dominic Kwiatkowski, Cordelia Langford, John Sillitoe on behalf of the Wellcome Sanger Institute COVID-19 Surveillance Team |
| EPI_ISL_2717540 | Lighthouse Lab in Milton Keynes | Wellcome Sanger Institute for the COVID-19 Genomics UK (COG-UK) Consortium | The Lighthouse Lab in Milton Keynes and Alex Alderton, Roberto Amato, Jeffrey Barrett, Sonia Goncalves, Ewan Harrison, David K. Jackson, Ian Johnston, Dominic Kwiatkowski, Cordelia Langford, John Sillitoe on behalf of the Wellcome Sanger Institute COVID-19 Surveillance Team |
| EPI_ISL_2717541 | Lighthouse Lab in Alderley Park | Wellcome Sanger Institute for the COVID-19 Genomics UK (COG-UK) Consortium | Jacquelyn Wynn, Mairead Hyland, The Lighthouse Lab in Alderley Park and Alex Alderton, Roberto Amato, Jeffrey Barrett, Sonia Goncalves, Ewan Harrison, David K. Jackson, Ian Johnston, Dominic Kwiatkowski, Cordelia Langford, John Sillitoe on behalf of the Wellcome Sanger Institute COVID-19 Surveillance Team |
| EPI_ISL_2717542 | Lighthouse Lab in Glasgow | Wellcome Sanger Institute for the COVID-19 Genomics UK (COG-UK) Consortium | Harper VanSteenhouse, Yumi Kasai, David Gray, Carol Clugston, Anna Dominiczak and Alex Alderton, Roberto Amato, Jeffrey Barrett, Sonia Goncalves, Ewan Harrison, David K. Jackson, Ian Johnston, Dominic Kwiatkowski, Cordelia Langford, John Sillitoe on behalf of the Wellcome Sanger Institute COVID-19 Surveillance Team |
| EPI_ISL_2717543, EPI_ISL_2717544 | Lighthouse Lab in Milton Keynes | Wellcome Sanger Institute for the COVID-19 Genomics UK (COG-UK) Consortium | The Lighthouse Lab in Milton Keynes and Alex Alderton, Roberto Amato, Jeffrey Barrett, Sonia Goncalves, Ewan Harrison, David K. Jackson, Ian Johnston, Dominic Kwiatkowski, Cordelia Langford, John Sillitoe on behalf of the Wellcome Sanger Institute COVID-19 Surveillance Team |
| EPI_ISL_2717545 | Lighthouse Lab in Glasgow | Wellcome Sanger Institute for the COVID-19 Genomics UK (COG-UK) Consortium | Harper VanSteenhouse, Yumi Kasai, David Gray, Carol Clugston, Anna Dominiczak and Alex Alderton, Roberto Amato, Jeffrey Barrett, Sonia Goncalves, Ewan Harrison, David K. Jackson, Ian Johnston, Dominic Kwiatkowski, Cordelia Langford, John Sillitoe on behalf of the Wellcome Sanger Institute COVID-19 Surveillance Team |
| EPI_ISL_2717546, EPI_ISL_2717547 | Lighthouse Lab in Milton Keynes | Wellcome Sanger Institute for the COVID-19 Genomics UK (COG-UK) Consortium | The Lighthouse Lab in Milton Keynes and Alex Alderton, Roberto Amato, Jeffrey Barrett, Sonia Goncalves, Ewan Harrison, David K. Jackson, Ian Johnston, Dominic Kwiatkowski, Cordelia Langford, John Sillitoe on behalf of the Wellcome Sanger Institute COVID-19 Surveillance Team |
| EPI_ISL_2717548, EPI_ISL_2717549 | Lighthouse Lab in Glasgow | Wellcome Sanger Institute for the COVID-19 Genomics UK (COG-UK) Consortium | Harper VanSteenhouse, Yumi Kasai, David Gray, Carol Clugston, Anna Dominiczak and Alex Alderton, Roberto Amato, Jeffrey Barrett, Sonia Goncalves, Ewan Harrison, David K. Jackson, Ian Johnston, Dominic Kwiatkowski, Cordelia Langford, John Sillitoe on behalf of the Wellcome Sanger Institute COVID-19 Surveillance Team |
| EPI_ISL_2717550 | Lighthouse Lab in Alderley Park | Wellcome Sanger Institute for the COVID-19 Genomics UK (COG-UK) Consortium | Jacquelyn Wynn, Mairead Hyland, The Lighthouse Lab in Alderley Park and Alex Alderton, Roberto Amato, Jeffrey Barrett, Sonia Goncalves, Ewan Harrison, David K. Jackson, Ian Johnston, Dominic Kwiatkowski, Cordelia Langford, John Sillitoe on behalf of the Wellcome Sanger Institute COVID-19 Surveillance Team |
| EPI_ISL_2717551, EPI_ISL_2717552, EPI_ISL_2717553 | Lighthouse Lab in Milton Keynes | Wellcome Sanger Institute for the COVID-19 Genomics UK (COG-UK) Consortium | The Lighthouse Lab in Milton Keynes and Alex Alderton, Roberto Amato, Jeffrey Barrett, Sonia Goncalves, Ewan Harrison, David K. Jackson, Ian Johnston, Dominic Kwiatkowski, Cordelia Langford, John Sillitoe on behalf of the Wellcome Sanger Institute COVID-19 Surveillance Team |
| EPI_ISL_2717554, EPI_ISL_2717555 | Lighthouse Lab in Glasgow | Wellcome Sanger Institute for the COVID-19 Genomics UK (COG-UK) Consortium | Harper VanSteenhouse, Yumi Kasai, David Gray, Carol Clugston, Anna Dominiczak and Alex Alderton, Roberto Amato, Jeffrey Barrett, Sonia Goncalves, Ewan Harrison, David K. Jackson, Ian Johnston, Dominic Kwiatkowski, Cordelia Langford, John Sillitoe on behalf of the Wellcome Sanger Institute COVID-19 Surveillance Team |
| EPI_ISL_2717556 | Lighthouse Lab in Milton Keynes | Wellcome Sanger Institute for the COVID-19 Genomics UK (COG-UK) Consortium | The Lighthouse Lab in Milton Keynes and Alex Alderton, Roberto Amato, Jeffrey Barrett, Sonia Goncalves, Ewan Harrison, David K. Jackson, Ian Johnston, Dominic Kwiatkowski, Cordelia Langford, John Sillitoe on behalf of the Wellcome Sanger Institute COVID-19 Surveillance Team |
| EPI_ISL_2717557, EPI_ISL_2717558 | Lighthouse Lab in Glasgow | Wellcome Sanger Institute for the COVID-19 Genomics UK (COG-UK) Consortium | Harper VanSteenhouse, Yumi Kasai, David Gray, Carol Clugston, Anna Dominiczak and Alex Alderton, Roberto Amato, Jeffrey Barrett, Sonia Goncalves, Ewan Harrison, David K. Jackson, Ian Johnston, Dominic Kwiatkowski, Cordelia Langford, John Sillitoe on behalf of the Wellcome Sanger Institute COVID-19 Surveillance Team |
| EPI_ISL_2717559 | Lighthouse Lab in Alderley Park | Wellcome Sanger Institute for the COVID-19 Genomics UK (COG-UK) Consortium | Jacquelyn Wynn, Mairead Hyland, The Lighthouse Lab in Alderley Park and Alex Alderton, Roberto Amato, Jeffrey Barrett, Sonia Goncalves, Ewan Harrison, David K. Jackson, Ian Johnston, Dominic Kwiatkowski, Cordelia Langford, John Sillitoe on behalf of the Wellcome Sanger Institute COVID-19 Surveillance Team |
| EPI_ISL_2717560 | Lighthouse Lab in Milton Keynes | Wellcome Sanger Institute for the COVID-19 Genomics UK (COG-UK) Consortium | The Lighthouse Lab in Milton Keynes and Alex Alderton, Roberto Amato, Jeffrey Barrett, Sonia Goncalves, Ewan Harrison, David K. Jackson, Ian Johnston, Dominic Kwiatkowski, Cordelia Langford, John Sillitoe on behalf of the Wellcome Sanger Institute COVID-19 Surveillance Team |
| EPI_ISL_2717561 | Lighthouse Lab in Glasgow | Wellcome Sanger Institute for the COVID-19 Genomics UK (COG-UK) Consortium | Harper VanSteenhouse, Yumi Kasai, David Gray, Carol Clugston, Anna Dominiczak and Alex Alderton, Roberto Amato, Jeffrey Barrett, Sonia Goncalves, Ewan Harrison, David K. Jackson, Ian Johnston, Dominic Kwiatkowski, Cordelia Langford, John Sillitoe on behalf of the Wellcome Sanger Institute COVID-19 Surveillance Team |
| EPI_ISL_2717562, EPI_ISL_2717563 | Lighthouse Lab in Milton Keynes | Wellcome Sanger Institute for the COVID-19 Genomics UK (COG-UK) Consortium | The Lighthouse Lab in Milton Keynes and Alex Alderton, Roberto Amato, Jeffrey Barrett, Sonia Goncalves, Ewan Harrison, David K. Jackson, Ian Johnston, Dominic Kwiatkowski, Cordelia Langford, John Sillitoe on behalf of the Wellcome Sanger Institute COVID-19 Surveillance Team |
| EPI_ISL_2717564 | Health Services Laboratories | Wellcome Sanger Institute for the COVID-19 Genomics UK (COG-UK) Consortium | Health Services Laboratories and Alex Alderton, Roberto Amato, Jeffrey Barrett, Sonia Goncalves, Ewan Harrison, David K. Jackson, Ian Johnston, Dominic Kwiatkowski, Cordelia Langford, John Sillitoe on behalf of the Wellcome Sanger Institute COVID-19 Surveillance Team |
| EPI_ISL_2717565 | Lighthouse Lab in Alderley Park | Wellcome Sanger Institute for the COVID-19 Genomics UK (COG-UK) Consortium | Jacquelyn Wynn, Mairead Hyland, The Lighthouse Lab in Alderley Park and Alex Alderton, Roberto Amato, Jeffrey Barrett, Sonia Goncalves, Ewan Harrison, David K. Jackson, Ian Johnston, Dominic Kwiatkowski, Cordelia Langford, John Sillitoe on behalf of the Wellcome Sanger Institute COVID-19 Surveillance Team |
| EPI_ISL_2717566, EPI_ISL_2717567, EPI_ISL_2717568 | Lighthouse Lab in Milton Keynes | Wellcome Sanger Institute for the COVID-19 Genomics UK (COG-UK) Consortium | The Lighthouse Lab in Milton Keynes and Alex Alderton, Roberto Amato, Jeffrey Barrett, Sonia Goncalves, Ewan Harrison, David K. Jackson, Ian Johnston, Dominic Kwiatkowski, Cordelia Langford, John Sillitoe on behalf of the Wellcome Sanger Institute COVID-19 Surveillance Team |
| EPI_ISL_2717569 | Lighthouse Lab in Alderley Park | Wellcome Sanger Institute for the COVID-19 Genomics UK (COG-UK) Consortium | Jacquelyn Wynn, Mairead Hyland, The Lighthouse Lab in Alderley Park and Alex Alderton, Roberto Amato, Jeffrey Barrett, Sonia Goncalves, Ewan Harrison, David K. Jackson, Ian Johnston, Dominic Kwiatkowski, Cordelia Langford, John Sillitoe on behalf of the Wellcome Sanger Institute COVID-19 Surveillance Team |
| EPI_ISL_2717570, EPI_ISL_2717571, EPI_ISL_2717572, EPI_ISL_2717573, EPI_ISL_2717574, EPI_ISL_2717575 | Lighthouse Lab in Milton Keynes | Wellcome Sanger Institute for the COVID-19 Genomics UK (COG-UK) Consortium | The Lighthouse Lab in Milton Keynes and Alex Alderton, Roberto Amato, Jeffrey Barrett, Sonia Goncalves, Ewan Harrison, David K. Jackson, Ian Johnston, Dominic Kwiatkowski, Cordelia Langford, John Sillitoe on behalf of the Wellcome Sanger Institute COVID-19 Surveillance Team |
| EPI_ISL_2717576 | Health Services Laboratories | Wellcome Sanger Institute for the COVID-19 Genomics UK (COG-UK) Consortium | Health Services Laboratories and Alex Alderton, Roberto Amato, Jeffrey Barrett, Sonia Goncalves, Ewan Harrison, David K. Jackson, Ian Johnston, Dominic Kwiatkowski, Cordelia Langford, John Sillitoe on behalf of the Wellcome Sanger Institute COVID-19 Surveillance Team |
| EPI_ISL_2717577, EPI_ISL_2717578, EPI_ISL_2717579, EPI_ISL_2717580, EPI_ISL_2717581, EPI_ISL_2717582, EPI_ISL_2717583 | Lighthouse Lab in Alderley Park | Wellcome Sanger Institute for the COVID-19 Genomics UK (COG-UK) Consortium | Jacquelyn Wynn, Mairead Hyland, The Lighthouse Lab in Alderley Park and Alex Alderton, Roberto Amato, Jeffrey Barrett, Sonia Goncalves, Ewan Harrison, David K. Jackson, Ian Johnston, Dominic Kwiatkowski, Cordelia Langford, John Sillitoe on behalf of the Wellcome Sanger Institute COVID-19 Surveillance Team |
| EPI_ISL_2717584 | Lighthouse Lab in Glasgow | Wellcome Sanger Institute for the COVID-19 Genomics UK (COG-UK) Consortium | Harper VanSteenhouse, Yumi Kasai, David Gray, Carol Clugston, Anna Dominiczak and Alex Alderton, Roberto Amato, Jeffrey Barrett, Sonia Goncalves, Ewan Harrison, David K. Jackson, Ian Johnston, Dominic Kwiatkowski, Cordelia Langford, John Sillitoe on behalf of the Wellcome Sanger Institute COVID-19 Surveillance Team |
| EPI_ISL_2717585 | Lighthouse Lab in Alderley Park | Wellcome Sanger Institute for the COVID-19 Genomics UK (COG-UK) Consortium | Jacquelyn Wynn, Mairead Hyland, The Lighthouse Lab in Alderley Park and Alex Alderton, Roberto Amato, Jeffrey Barrett, Sonia Goncalves, Ewan Harrison, David K. Jackson, Ian Johnston, Dominic Kwiatkowski, Cordelia Langford, John Sillitoe on behalf of the Wellcome Sanger Institute COVID-19 Surveillance Team |
| EPI_ISL_2717586 | Lighthouse Lab in Glasgow | Wellcome Sanger Institute for the COVID-19 Genomics UK (COG-UK) Consortium | Harper VanSteenhouse, Yumi Kasai, David Gray, Carol Clugston, Anna Dominiczak and Alex Alderton, Roberto Amato, Jeffrey Barrett, Sonia Goncalves, Ewan Harrison, David K. Jackson, Ian Johnston, Dominic Kwiatkowski, Cordelia Langford, John Sillitoe on behalf of the Wellcome Sanger Institute COVID-19 Surveillance Team |
| EPI_ISL_2717587, EPI_ISL_2717588, EPI_ISL_2717589, EPI_ISL_2717590 | Lighthouse Lab in Alderley Park | Wellcome Sanger Institute for the COVID-19 Genomics UK (COG-UK) Consortium | Jacquelyn Wynn, Mairead Hyland, The Lighthouse Lab in Alderley Park and Alex Alderton, Roberto Amato, Jeffrey Barrett, Sonia Goncalves, Ewan Harrison, David K. Jackson, Ian Johnston, Dominic Kwiatkowski, Cordelia Langford, John Sillitoe on behalf of the Wellcome Sanger Institute COVID-19 Surveillance Team |
| EPI_ISL_2717591 | Lighthouse Lab in Glasgow | Wellcome Sanger Institute for the COVID-19 Genomics UK (COG-UK) Consortium | Harper VanSteenhouse, Yumi Kasai, David Gray, Carol Clugston, Anna Dominiczak and Alex Alderton, Roberto Amato, Jeffrey Barrett, Sonia Goncalves, Ewan Harrison, David K. Jackson, Ian Johnston, Dominic Kwiatkowski, Cordelia Langford, John Sillitoe on behalf of the Wellcome Sanger Institute COVID-19 Surveillance Team |
| EPI_ISL_2717592 | Lighthouse Lab in Alderley Park | Wellcome Sanger Institute for the COVID-19 Genomics UK (COG-UK) Consortium | Jacquelyn Wynn, Mairead Hyland, The Lighthouse Lab in Alderley Park and Alex Alderton, Roberto Amato, Jeffrey Barrett, Sonia Goncalves, Ewan Harrison, David K. Jackson, Ian Johnston, Dominic Kwiatkowski, Cordelia Langford, John Sillitoe on behalf of the Wellcome Sanger Institute COVID-19 Surveillance Team |

| | | | |
|---|---|---|---|
| EPI_ISL_2717593 | Lighthouse Lab in Glasgow | Wellcome Sanger Institute for the COVID-19 Genomics UK (COG-UK) Consortium | Harper VanSteenhouse, Yumi Kasai, David Gray, Carol Clugston, Anna Dominiczak and Alex Alderton, Roberto Amato, Jeffrey Barrett, Sonia Goncalves, Ewan Harrison, David K. Jackson, Ian Johnston, Dominic Kwiatkowski, Cordelia Langford, John Sillitoe on behalf of the Wellcome Sanger Institute COVID-19 Surveillance Team |
| EPI_ISL_2717594 | Lighthouse Lab in Alderley Park | Wellcome Sanger Institute for the COVID-19 Genomics UK (COG-UK) Consortium | Jacquelyn Wynn, Mairead Hyland, The Lighthouse Lab in Alderley Park and Alex Alderton, Roberto Amato, Jeffrey Barrett, Sonia Goncalves, Ewan Harrison, David K. Jackson, Ian Johnston, Dominic Kwiatkowski, Cordelia Langford, John Sillitoe on behalf of the Wellcome Sanger Institute COVID-19 Surveillance Team |
| EPI_ISL_2717595 | Lighthouse Lab in Glasgow | Wellcome Sanger Institute for the COVID-19 Genomics UK (COG-UK) Consortium | Harper VanSteenhouse, Yumi Kasai, David Gray, Carol Clugston, Anna Dominiczak and Alex Alderton, Roberto Amato, Jeffrey Barrett, Sonia Goncalves, Ewan Harrison, David K. Jackson, Ian Johnston, Dominic Kwiatkowski, Cordelia Langford, John Sillitoe on behalf of the Wellcome Sanger Institute COVID-19 Surveillance Team |
| EPI_ISL_2717596, EPI_ISL_2717597 | Lighthouse Lab in Alderley Park | Wellcome Sanger Institute for the COVID-19 Genomics UK (COG-UK) Consortium | Jacquelyn Wynn, Mairead Hyland, The Lighthouse Lab in Alderley Park and Alex Alderton, Roberto Amato, Jeffrey Barrett, Sonia Goncalves, Ewan Harrison, David K. Jackson, Ian Johnston, Dominic Kwiatkowski, Cordelia Langford, John Sillitoe on behalf of the Wellcome Sanger Institute COVID-19 Surveillance Team |
| EPI_ISL_2717598 | Lighthouse Lab in Glasgow | Wellcome Sanger Institute for the COVID-19 Genomics UK (COG-UK) Consortium | Harper VanSteenhouse, Yumi Kasai, David Gray, Carol Clugston, Anna Dominiczak and Alex Alderton, Roberto Amato, Jeffrey Barrett, Sonia Goncalves, Ewan Harrison, David K. Jackson, Ian Johnston, Dominic Kwiatkowski, Cordelia Langford, John Sillitoe on behalf of the Wellcome Sanger Institute COVID-19 Surveillance Team |
| EPI_ISL_2717599, EPI_ISL_2717600, EPI_ISL_2717601, EPI_ISL_2717602, EPI_ISL_2717603 | Lighthouse Lab in Alderley Park | Wellcome Sanger Institute for the COVID-19 Genomics UK (COG-UK) Consortium | Jacquelyn Wynn, Mairead Hyland, The Lighthouse Lab in Alderley Park and Alex Alderton, Roberto Amato, Jeffrey Barrett, Sonia Goncalves, Ewan Harrison, David K. Jackson, Ian Johnston, Dominic Kwiatkowski, Cordelia Langford, John Sillitoe on behalf of the Wellcome Sanger Institute COVID-19 Surveillance Team |
| EPI_ISL_2717604, EPI_ISL_2717605 | Lighthouse Lab in Glasgow | Wellcome Sanger Institute for the COVID-19 Genomics UK (COG-UK) Consortium | Harper VanSteenhouse, Yumi Kasai, David Gray, Carol Clugston, Anna Dominiczak and Alex Alderton, Roberto Amato, Jeffrey Barrett, Sonia Goncalves, Ewan Harrison, David K. Jackson, Ian Johnston, Dominic Kwiatkowski, Cordelia Langford, John Sillitoe on behalf of the Wellcome Sanger Institute COVID-19 Surveillance Team |
| EPI_ISL_2717606 | Lighthouse Lab in Alderley Park | Wellcome Sanger Institute for the COVID-19 Genomics UK (COG-UK) Consortium | Jacquelyn Wynn, Mairead Hyland, The Lighthouse Lab in Alderley Park and Alex Alderton, Roberto Amato, Jeffrey Barrett, Sonia Goncalves, Ewan Harrison, David K. Jackson, Ian Johnston, Dominic Kwiatkowski, Cordelia Langford, John Sillitoe on behalf of the Wellcome Sanger Institute COVID-19 Surveillance Team |
| EPI_ISL_2717607 | Lighthouse Lab in Glasgow | Wellcome Sanger Institute for the COVID-19 Genomics UK (COG-UK) Consortium | Harper VanSteenhouse, Yumi Kasai, David Gray, Carol Clugston, Anna Dominiczak and Alex Alderton, Roberto Amato, Jeffrey Barrett, Sonia Goncalves, Ewan Harrison, David K. Jackson, Ian Johnston, Dominic Kwiatkowski, Cordelia Langford, John Sillitoe on behalf of the Wellcome Sanger Institute COVID-19 Surveillance Team |
| EPI_ISL_2717608, EPI_ISL_2717609, EPI_ISL_2717610, EPI_ISL_2717611, EPI_ISL_2717612, EPI_ISL_2717613 | Lighthouse Lab in Alderley Park | Wellcome Sanger Institute for the COVID-19 Genomics UK (COG-UK) Consortium | Jacquelyn Wynn, Mairead Hyland, The Lighthouse Lab in Alderley Park and Alex Alderton, Roberto Amato, Jeffrey Barrett, Sonia Goncalves, Ewan Harrison, David K. Jackson, Ian Johnston, Dominic Kwiatkowski, Cordelia Langford, John Sillitoe on behalf of the Wellcome Sanger Institute COVID-19 Surveillance Team |
| EPI_ISL_2717614 | Lighthouse Lab in Glasgow | Wellcome Sanger Institute for the COVID-19 Genomics UK (COG-UK) Consortium | Harper VanSteenhouse, Yumi Kasai, David Gray, Carol Clugston, Anna Dominiczak and Alex Alderton, Roberto Amato, Jeffrey Barrett, Sonia Goncalves, Ewan Harrison, David K. Jackson, Ian Johnston, Dominic Kwiatkowski, Cordelia Langford, John Sillitoe on behalf of the Wellcome Sanger Institute COVID-19 Surveillance Team |
| EPI_ISL_2717615, EPI_ISL_2717616, EPI_ISL_2717617 | Lighthouse Lab in Alderley Park | Wellcome Sanger Institute for the COVID-19 Genomics UK (COG-UK) Consortium | Jacquelyn Wynn, Mairead Hyland, The Lighthouse Lab in Alderley Park and Alex Alderton, Roberto Amato, Jeffrey Barrett, Sonia Goncalves, Ewan Harrison, David K. Jackson, Ian Johnston, Dominic Kwiatkowski, Cordelia Langford, John Sillitoe on behalf of the Wellcome Sanger Institute COVID-19 Surveillance Team |
| EPI_ISL_2717618, EPI_ISL_2717619 | Lighthouse Lab in Glasgow | Wellcome Sanger Institute for the COVID-19 Genomics UK (COG-UK) Consortium | Harper VanSteenhouse, Yumi Kasai, David Gray, Carol Clugston, Anna Dominiczak and Alex Alderton, Roberto Amato, Jeffrey Barrett, Sonia Goncalves, Ewan Harrison, David K. Jackson, Ian Johnston, Dominic Kwiatkowski, Cordelia Langford, John Sillitoe on behalf of the Wellcome Sanger Institute COVID-19 Surveillance Team |
| EPI_ISL_2717620 | Lighthouse Lab in Alderley Park | Wellcome Sanger Institute for the COVID-19 Genomics UK (COG-UK) Consortium | Jacquelyn Wynn, Mairead Hyland, The Lighthouse Lab in Alderley Park and Alex Alderton, Roberto Amato, Jeffrey Barrett, Sonia Goncalves, Ewan Harrison, David K. Jackson, Ian Johnston, Dominic Kwiatkowski, Cordelia Langford, John Sillitoe on behalf of the Wellcome Sanger Institute COVID-19 Surveillance Team |
| EPI_ISL_2717621 | Lighthouse Lab in Glasgow | Wellcome Sanger Institute for the COVID-19 Genomics UK (COG-UK) Consortium | Harper VanSteenhouse, Yumi Kasai, David Gray, Carol Clugston, Anna Dominiczak and Alex Alderton, Roberto Amato, Jeffrey Barrett, Sonia Goncalves, Ewan Harrison, David K. Jackson, Ian Johnston, Dominic Kwiatkowski, Cordelia Langford, John Sillitoe on behalf of the Wellcome Sanger Institute COVID-19 Surveillance Team |
| EPI_ISL_2717622, EPI_ISL_2717623, EPI_ISL_2717624 | Lighthouse Lab in Alderley Park | Wellcome Sanger Institute for the COVID-19 Genomics UK (COG-UK) Consortium | Jacquelyn Wynn, Mairead Hyland, The Lighthouse Lab in Alderley Park and Alex Alderton, Roberto Amato, Jeffrey Barrett, Sonia Goncalves, Ewan Harrison, David K. Jackson, Ian Johnston, Dominic Kwiatkowski, Cordelia Langford, John Sillitoe on behalf of the Wellcome Sanger Institute COVID-19 Surveillance Team |
| EPI_ISL_2717625 | Lighthouse Lab in Glasgow | Wellcome Sanger Institute for the COVID-19 Genomics UK (COG-UK) Consortium | Harper VanSteenhouse, Yumi Kasai, David Gray, Carol Clugston, Anna Dominiczak and Alex Alderton, Roberto Amato, Jeffrey Barrett, Sonia Goncalves, Ewan Harrison, David K. Jackson, Ian Johnston, Dominic Kwiatkowski, Cordelia Langford, John Sillitoe on behalf of the Wellcome Sanger Institute COVID-19 Surveillance Team |
| EPI_ISL_2717626, EPI_ISL_2717627, EPI_ISL_2717628, EPI_ISL_2717629, EPI_ISL_2717630 | Lighthouse Lab in Alderley Park | Wellcome Sanger Institute for the COVID-19 Genomics UK (COG-UK) Consortium | Jacquelyn Wynn, Mairead Hyland, The Lighthouse Lab in Alderley Park and Alex Alderton, Roberto Amato, Jeffrey Barrett, Sonia Goncalves, Ewan Harrison, David K. Jackson, Ian Johnston, Dominic Kwiatkowski, Cordelia Langford, John Sillitoe on behalf of the Wellcome Sanger Institute COVID-19 Surveillance Team |
| EPI_ISL_2717631, EPI_ISL_2717632 | Lighthouse Lab in Glasgow | Wellcome Sanger Institute for the COVID-19 Genomics UK (COG-UK) Consortium | Harper VanSteenhouse, Yumi Kasai, David Gray, Carol Clugston, Anna Dominiczak and Alex Alderton, Roberto Amato, Jeffrey Barrett, Sonia Goncalves, Ewan Harrison, David K. Jackson, Ian Johnston, Dominic Kwiatkowski, Cordelia Langford, John Sillitoe on behalf of the Wellcome Sanger Institute COVID-19 Surveillance Team |
| EPI_ISL_2717633, EPI_ISL_2717634, EPI_ISL_2717635, EPI_ISL_2717636, EPI_ISL_2717637, EPI_ISL_2717638, EPI_ISL_2717639, EPI_ISL_2717640, EPI_ISL_2717641 | Lighthouse Lab in Alderley Park | Wellcome Sanger Institute for the COVID-19 Genomics UK (COG-UK) Consortium | Jacquelyn Wynn, Mairead Hyland, The Lighthouse Lab in Alderley Park and Alex Alderton, Roberto Amato, Jeffrey Barrett, Sonia Goncalves, Ewan Harrison, David K. Jackson, Ian Johnston, Dominic Kwiatkowski, Cordelia Langford, John Sillitoe on behalf of the Wellcome Sanger Institute COVID-19 Surveillance Team |
| EPI_ISL_2717642 | Lighthouse Lab in Glasgow | Wellcome Sanger Institute for the COVID-19 Genomics UK (COG-UK) Consortium | Harper VanSteenhouse, Yumi Kasai, David Gray, Carol Clugston, Anna Dominiczak and Alex Alderton, Roberto Amato, Jeffrey Barrett, Sonia Goncalves, Ewan Harrison, David K. Jackson, Ian Johnston, Dominic Kwiatkowski, Cordelia Langford, John Sillitoe on behalf of the Wellcome Sanger Institute COVID-19 Surveillance Team |
| EPI_ISL_2717643, EPI_ISL_2717644, EPI_ISL_2717645, EPI_ISL_2717646, EPI_ISL_2717647, EPI_ISL_2717648, EPI_ISL_2717649, EPI_ISL_2717650, EPI_ISL_2717651 | Lighthouse Lab in Alderley Park | Wellcome Sanger Institute for the COVID-19 Genomics UK (COG-UK) Consortium | Jacquelyn Wynn, Mairead Hyland, The Lighthouse Lab in Alderley Park and Alex Alderton, Roberto Amato, Jeffrey Barrett, Sonia Goncalves, Ewan Harrison, David K. Jackson, Ian Johnston, Dominic Kwiatkowski, Cordelia Langford, John Sillitoe on behalf of the Wellcome Sanger Institute COVID-19 Surveillance Team |
| EPI_ISL_2717652 | Lighthouse Lab in Glasgow | Wellcome Sanger Institute for the COVID-19 Genomics UK (COG-UK) Consortium | Harper VanSteenhouse, Yumi Kasai, David Gray, Carol Clugston, Anna Dominiczak and Alex Alderton, Roberto Amato, Jeffrey Barrett, Sonia Goncalves, Ewan Harrison, David K. Jackson, Ian Johnston, Dominic Kwiatkowski, Cordelia Langford, John Sillitoe on behalf of the Wellcome Sanger Institute COVID-19 Surveillance Team |
| EPI_ISL_2717653 | Lighthouse Lab in Alderley Park | Wellcome Sanger Institute for the COVID-19 Genomics UK (COG-UK) Consortium | Jacquelyn Wynn, Mairead Hyland, The Lighthouse Lab in Alderley Park and Alex Alderton, Roberto Amato, Jeffrey Barrett, Sonia Goncalves, Ewan Harrison, David K. Jackson, Ian Johnston, Dominic Kwiatkowski, Cordelia Langford, John Sillitoe on behalf of the Wellcome Sanger Institute COVID-19 Surveillance Team |
| EPI_ISL_2717654, EPI_ISL_2717655 | Lighthouse Lab in Glasgow | Wellcome Sanger Institute for the COVID-19 Genomics UK (COG-UK) Consortium | Harper VanSteenhouse, Yumi Kasai, David Gray, Carol Clugston, Anna Dominiczak and Alex Alderton, Roberto Amato, Jeffrey Barrett, Sonia Goncalves, Ewan Harrison, David K. Jackson, Ian Johnston, Dominic Kwiatkowski, Cordelia Langford, John Sillitoe on behalf of the Wellcome Sanger Institute COVID-19 Surveillance Team |
| EPI_ISL_2717656, EPI_ISL_2717657, EPI_ISL_2717658, EPI_ISL_2717659, EPI_ISL_2717660, EPI_ISL_2717661, EPI_ISL_2717662, EPI_ISL_2717663, EPI_ISL_2717664, EPI_ISL_2717665, EPI_ISL_2717666 | | | |
| see above | Lighthouse Lab in Alderley Park | Wellcome Sanger Institute for the COVID-19 Genomics UK (COG-UK) Consortium | Jacquelyn Wynn, Mairead Hyland, The Lighthouse Lab in Alderley Park and Alex Alderton, Roberto Amato, Jeffrey Barrett, Sonia Goncalves, Ewan Harrison, David K. Jackson, Ian Johnston, Dominic Kwiatkowski, Cordelia Langford, John Sillitoe on behalf of the Wellcome Sanger Institute COVID-19 Surveillance Team |
| EPI_ISL_2717667 | Lighthouse Lab in Glasgow | Wellcome Sanger Institute for the COVID-19 Genomics UK (COG-UK) Consortium | Harper VanSteenhouse, Yumi Kasai, David Gray, Carol Clugston, Anna Dominiczak and Alex Alderton, Roberto Amato, Jeffrey Barrett, Sonia Goncalves, Ewan Harrison, David K. Jackson, Ian Johnston, Dominic Kwiatkowski, Cordelia Langford, John Sillitoe on behalf of the Wellcome Sanger Institute COVID-19 Surveillance Team |
| EPI_ISL_2717668, EPI_ISL_2717669, EPI_ISL_2717670, EPI_ISL_2717671, EPI_ISL_2717673, EPI_ISL_2717674, EPI_ISL_2717675, EPI_ISL_2717677, EPI_ISL_2717678, EPI_ISL_2717679, EPI_ISL_2717680, EPI_ISL_2717681, EPI_ISL_2717682 | | | |
| see above | Lighthouse Lab in Alderley Park | Wellcome Sanger Institute for the COVID-19 Genomics UK (COG-UK) Consortium | Jacquelyn Wynn, Mairead Hyland, The Lighthouse Lab in Alderley Park and Alex Alderton, Roberto Amato, Jeffrey Barrett, Sonia Goncalves, Ewan Harrison, David K. Jackson, Ian Johnston, Dominic Kwiatkowski, Cordelia Langford, John Sillitoe on behalf of the Wellcome Sanger Institute COVID-19 Surveillance Team |
| EPI_ISL_2717683 | Lighthouse Lab in Glasgow | Wellcome Sanger Institute for the COVID-19 Genomics UK (COG-UK) Consortium | Harper VanSteenhouse, Yumi Kasai, David Gray, Carol Clugston, Anna Dominiczak and Alex Alderton, Roberto Amato, Jeffrey Barrett, Sonia Goncalves, Ewan Harrison, David K. Jackson, Ian Johnston, Dominic Kwiatkowski, Cordelia Langford, John Sillitoe on behalf of the Wellcome Sanger Institute COVID-19 Surveillance Team |
| EPI_ISL_2717684, EPI_ISL_2717685 | Lighthouse Lab in Alderley Park | Wellcome Sanger Institute for the COVID-19 Genomics UK (COG-UK) Consortium | Jacquelyn Wynn, Mairead Hyland, The Lighthouse Lab in Alderley Park and Alex Alderton, Roberto Amato, Jeffrey Barrett, Sonia Goncalves, Ewan Harrison, David K. Jackson, Ian Johnston, Dominic Kwiatkowski, Cordelia Langford, John Sillitoe on behalf of the Wellcome Sanger Institute COVID-19 Surveillance Team |
| EPI_ISL_2717686 | Lighthouse Lab in Glasgow | Wellcome Sanger Institute for the COVID-19 Genomics UK (COG-UK) Consortium | Harper VanSteenhouse, Yumi Kasai, David Gray, Carol Clugston, Anna Dominiczak and Alex Alderton, Roberto Amato, Jeffrey Barrett, Sonia Goncalves, Ewan Harrison, David K. Jackson, Ian Johnston, Dominic Kwiatkowski, Cordelia Langford, John Sillitoe on behalf of the Wellcome Sanger Institute COVID-19 Surveillance Team |
| EPI_ISL_2717687, EPI_ISL_2717688, EPI_ISL_2717689, EPI_ISL_2717690, EPI_ISL_2717691 | Lighthouse Lab in Alderley Park | Wellcome Sanger Institute for the COVID-19 Genomics UK (COG-UK) Consortium | Jacquelyn Wynn, Mairead Hyland, The Lighthouse Lab in Alderley Park and Alex Alderton, Roberto Amato, Jeffrey Barrett, Sonia Goncalves, Ewan Harrison, David K. Jackson, Ian Johnston, Dominic Kwiatkowski, Cordelia Langford, John Sillitoe on behalf of the Wellcome Sanger Institute COVID-19 Surveillance Team |

| | | | |
|---|---|---|---|
| EPI_ISL_2717692, EPI_ISL_2717693, EPI_ISL_2717694 | Lighthouse Lab in Glasgow | Wellcome Sanger Institute for the COVID-19 Genomics UK (COG-UK) Consortium | Harper VanSteenhouse, Yumi Kasai, David Gray, Carol Clugston, Anna Dominiczak and Alex Alderton, Roberto Amato, Jeffrey Barrett, Sonia Goncalves, Ewan Harrison, David K. Jackson, Ian Johnston, Dominic Kwiatkowski, Cordelia Langford, John Sillitoe on behalf of the Wellcome Sanger Institute COVID-19 Surveillance Team |
| EPI_ISL_2717695, EPI_ISL_2717696, EPI_ISL_2717697, EPI_ISL_2717698, EPI_ISL_2717699, EPI_ISL_2717700, EPI_ISL_2717701, EPI_ISL_2717702 | Lighthouse Lab in Alderley Park | Wellcome Sanger Institute for the COVID-19 Genomics UK (COG-UK) Consortium | Jacquelyn Wynn, Mairead Hyland, The Lighthouse Lab in Alderley Park and Alex Alderton, Roberto Amato, Jeffrey Barrett, Sonia Goncalves, Ewan Harrison, David K. Jackson, Ian Johnston, Dominic Kwiatkowski, Cordelia Langford, John Sillitoe on behalf of the Wellcome Sanger Institute COVID-19 Surveillance Team |
| EPI_ISL_2717703 | Lighthouse Lab in Glasgow | Wellcome Sanger Institute for the COVID-19 Genomics UK (COG-UK) Consortium | Harper VanSteenhouse, Yumi Kasai, David Gray, Carol Clugston, Anna Dominiczak and Alex Alderton, Roberto Amato, Jeffrey Barrett, Sonia Goncalves, Ewan Harrison, David K. Jackson, Ian Johnston, Dominic Kwiatkowski, Cordelia Langford, John Sillitoe on behalf of the Wellcome Sanger Institute COVID-19 Surveillance Team |
| EPI_ISL_2717704, EPI_ISL_2717705, EPI_ISL_2717706, EPI_ISL_2717707, EPI_ISL_2717708, EPI_ISL_2717709, EPI_ISL_2717710 | Lighthouse Lab in Alderley Park | Wellcome Sanger Institute for the COVID-19 Genomics UK (COG-UK) Consortium | Jacquelyn Wynn, Mairead Hyland, The Lighthouse Lab in Alderley Park and Alex Alderton, Roberto Amato, Jeffrey Barrett, Sonia Goncalves, Ewan Harrison, David K. Jackson, Ian Johnston, Dominic Kwiatkowski, Cordelia Langford, John Sillitoe on behalf of the Wellcome Sanger Institute COVID-19 Surveillance Team |
| EPI_ISL_2717711 | Lighthouse Lab in Glasgow | Wellcome Sanger Institute for the COVID-19 Genomics UK (COG-UK) Consortium | Harper VanSteenhouse, Yumi Kasai, David Gray, Carol Clugston, Anna Dominiczak and Alex Alderton, Roberto Amato, Jeffrey Barrett, Sonia Goncalves, Ewan Harrison, David K. Jackson, Ian Johnston, Dominic Kwiatkowski, Cordelia Langford, John Sillitoe on behalf of the Wellcome Sanger Institute COVID-19 Surveillance Team |
| EPI_ISL_2717712, EPI_ISL_2717713, EPI_ISL_2717714, EPI_ISL_2717715, EPI_ISL_2717716, EPI_ISL_2717717, EPI_ISL_2717719, EPI_ISL_2717720, EPI_ISL_2717721 | Lighthouse Lab in Alderley Park | Wellcome Sanger Institute for the COVID-19 Genomics UK (COG-UK) Consortium | Jacquelyn Wynn, Mairead Hyland, The Lighthouse Lab in Alderley Park and Alex Alderton, Roberto Amato, Jeffrey Barrett, Sonia Goncalves, Ewan Harrison, David K. Jackson, Ian Johnston, Dominic Kwiatkowski, Cordelia Langford, John Sillitoe on behalf of the Wellcome Sanger Institute COVID-19 Surveillance Team |
| EPI_ISL_2717722 | Lighthouse Lab in Glasgow | Wellcome Sanger Institute for the COVID-19 Genomics UK (COG-UK) Consortium | Harper VanSteenhouse, Yumi Kasai, David Gray, Carol Clugston, Anna Dominiczak and Alex Alderton, Roberto Amato, Jeffrey Barrett, Sonia Goncalves, Ewan Harrison, David K. Jackson, Ian Johnston, Dominic Kwiatkowski, Cordelia Langford, John Sillitoe on behalf of the Wellcome Sanger Institute COVID-19 Surveillance Team |
| EPI_ISL_2717723, EPI_ISL_2717724, EPI_ISL_2717725, EPI_ISL_2717726 | Lighthouse Lab in Alderley Park | Wellcome Sanger Institute for the COVID-19 Genomics UK (COG-UK) Consortium | Jacquelyn Wynn, Mairead Hyland, The Lighthouse Lab in Alderley Park and Alex Alderton, Roberto Amato, Jeffrey Barrett, Sonia Goncalves, Ewan Harrison, David K. Jackson, Ian Johnston, Dominic Kwiatkowski, Cordelia Langford, John Sillitoe on behalf of the Wellcome Sanger Institute COVID-19 Surveillance Team |
| EPI_ISL_2717727 | Lighthouse Lab in Glasgow | Wellcome Sanger Institute for the COVID-19 Genomics UK (COG-UK) Consortium | Harper VanSteenhouse, Yumi Kasai, David Gray, Carol Clugston, Anna Dominiczak and Alex Alderton, Roberto Amato, Jeffrey Barrett, Sonia Goncalves, Ewan Harrison, David K. Jackson, Ian Johnston, Dominic Kwiatkowski, Cordelia Langford, John Sillitoe on behalf of the Wellcome Sanger Institute COVID-19 Surveillance Team |
| EPI_ISL_2717728, EPI_ISL_2717729, EPI_ISL_2717730, EPI_ISL_2717731, EPI_ISL_2717732, EPI_ISL_2717733, EPI_ISL_2717734, EPI_ISL_2717735, EPI_ISL_2717736 | Lighthouse Lab in Alderley Park | Wellcome Sanger Institute for the COVID-19 Genomics UK (COG-UK) Consortium | Jacquelyn Wynn, Mairead Hyland, The Lighthouse Lab in Alderley Park and Alex Alderton, Roberto Amato, Jeffrey Barrett, Sonia Goncalves, Ewan Harrison, David K. Jackson, Ian Johnston, Dominic Kwiatkowski, Cordelia Langford, John Sillitoe on behalf of the Wellcome Sanger Institute COVID-19 Surveillance Team |
| EPI_ISL_2717737, EPI_ISL_2717738 | Lighthouse Lab in Glasgow | Wellcome Sanger Institute for the COVID-19 Genomics UK (COG-UK) Consortium | Harper VanSteenhouse, Yumi Kasai, David Gray, Carol Clugston, Anna Dominiczak and Alex Alderton, Roberto Amato, Jeffrey Barrett, Sonia Goncalves, Ewan Harrison, David K. Jackson, Ian Johnston, Dominic Kwiatkowski, Cordelia Langford, John Sillitoe on behalf of the Wellcome Sanger Institute COVID-19 Surveillance Team |
| EPI_ISL_2717739, EPI_ISL_2717740, EPI_ISL_2717741, EPI_ISL_2717742, EPI_ISL_2717743, EPI_ISL_2717744, EPI_ISL_2717745, EPI_ISL_2717746, EPI_ISL_2717747, EPI_ISL_2717748, EPI_ISL_2717749, EPI_ISL_2717750, EPI_ISL_2717751, EPI_ISL_2717752, EPI_ISL_2717753 | | | |
| see above | Lighthouse Lab in Alderley Park | Wellcome Sanger Institute for the COVID-19 Genomics UK (COG-UK) Consortium | Jacquelyn Wynn, Mairead Hyland, The Lighthouse Lab in Alderley Park and Alex Alderton, Roberto Amato, Jeffrey Barrett, Sonia Goncalves, Ewan Harrison, David K. Jackson, Ian Johnston, Dominic Kwiatkowski, Cordelia Langford, John Sillitoe on behalf of the Wellcome Sanger Institute COVID-19 Surveillance Team |
| EPI_ISL_2717754 | Lighthouse Lab in Glasgow | Wellcome Sanger Institute for the COVID-19 Genomics UK (COG-UK) Consortium | Harper VanSteenhouse, Yumi Kasai, David Gray, Carol Clugston, Anna Dominiczak and Alex Alderton, Roberto Amato, Jeffrey Barrett, Sonia Goncalves, Ewan Harrison, David K. Jackson, Ian Johnston, Dominic Kwiatkowski, Cordelia Langford, John Sillitoe on behalf of the Wellcome Sanger Institute COVID-19 Surveillance Team |
| EPI_ISL_2717755, EPI_ISL_2717756, EPI_ISL_2717757, EPI_ISL_2717758, EPI_ISL_2717759, EPI_ISL_2717760, EPI_ISL_2717761, EPI_ISL_2717762, EPI_ISL_2717763, EPI_ISL_2717764, EPI_ISL_2717765, EPI_ISL_2717766 | | | |
| see above | Lighthouse Lab in Alderley Park | Wellcome Sanger Institute for the COVID-19 Genomics UK (COG-UK) Consortium | Jacquelyn Wynn, Mairead Hyland, The Lighthouse Lab in Alderley Park and Alex Alderton, Roberto Amato, Jeffrey Barrett, Sonia Goncalves, Ewan Harrison, David K. Jackson, Ian Johnston, Dominic Kwiatkowski, Cordelia Langford, John Sillitoe on behalf of the Wellcome Sanger Institute COVID-19 Surveillance Team |
| EPI_ISL_2717767 | Lighthouse Lab in Glasgow | Wellcome Sanger Institute for the COVID-19 Genomics UK (COG-UK) Consortium | Harper VanSteenhouse, Yumi Kasai, David Gray, Carol Clugston, Anna Dominiczak and Alex Alderton, Roberto Amato, Jeffrey Barrett, Sonia Goncalves, Ewan Harrison, David K. Jackson, Ian Johnston, Dominic Kwiatkowski, Cordelia Langford, John Sillitoe on behalf of the Wellcome Sanger Institute COVID-19 Surveillance Team |
| EPI_ISL_2717769, EPI_ISL_2717770, EPI_ISL_2717771, EPI_ISL_2717772, EPI_ISL_2717773, EPI_ISL_2717774, EPI_ISL_2717775, EPI_ISL_2717776, EPI_ISL_2717777 | Lighthouse Lab in Alderley Park | Wellcome Sanger Institute for the COVID-19 Genomics UK (COG-UK) Consortium | Jacquelyn Wynn, Mairead Hyland, The Lighthouse Lab in Alderley Park and Alex Alderton, Roberto Amato, Jeffrey Barrett, Sonia Goncalves, Ewan Harrison, David K. Jackson, Ian Johnston, Dominic Kwiatkowski, Cordelia Langford, John Sillitoe on behalf of the Wellcome Sanger Institute COVID-19 Surveillance Team |
| EPI_ISL_2717779 | Lighthouse Lab in Glasgow | Wellcome Sanger Institute for the COVID-19 Genomics UK (COG-UK) Consortium | Harper VanSteenhouse, Yumi Kasai, David Gray, Carol Clugston, Anna Dominiczak and Alex Alderton, Roberto Amato, Jeffrey Barrett, Sonia Goncalves, Ewan Harrison, David K. Jackson, Ian Johnston, Dominic Kwiatkowski, Cordelia Langford, John Sillitoe on behalf of the Wellcome Sanger Institute COVID-19 Surveillance Team |
| EPI_ISL_2717780, EPI_ISL_2717781, EPI_ISL_2717782, EPI_ISL_2717783 | Lighthouse Lab in Alderley Park | Wellcome Sanger Institute for the COVID-19 Genomics UK (COG-UK) Consortium | Jacquelyn Wynn, Mairead Hyland, The Lighthouse Lab in Alderley Park and Alex Alderton, Roberto Amato, Jeffrey Barrett, Sonia Goncalves, Ewan Harrison, David K. Jackson, Ian Johnston, Dominic Kwiatkowski, Cordelia Langford, John Sillitoe on behalf of the Wellcome Sanger Institute COVID-19 Surveillance Team |
| EPI_ISL_2717784, EPI_ISL_2717785 | Lighthouse Lab in Glasgow | Wellcome Sanger Institute for the COVID-19 Genomics UK (COG-UK) Consortium | Harper VanSteenhouse, Yumi Kasai, David Gray, Carol Clugston, Anna Dominiczak and Alex Alderton, Roberto Amato, Jeffrey Barrett, Sonia Goncalves, Ewan Harrison, David K. Jackson, Ian Johnston, Dominic Kwiatkowski, Cordelia Langford, John Sillitoe on behalf of the Wellcome Sanger Institute COVID-19 Surveillance Team |
| EPI_ISL_2717786, EPI_ISL_2717787, EPI_ISL_2717788 | Lighthouse Lab in Alderley Park | Wellcome Sanger Institute for the COVID-19 Genomics UK (COG-UK) Consortium | Jacquelyn Wynn, Mairead Hyland, The Lighthouse Lab in Alderley Park and Alex Alderton, Roberto Amato, Jeffrey Barrett, Sonia Goncalves, Ewan Harrison, David K. Jackson, Ian Johnston, Dominic Kwiatkowski, Cordelia Langford, John Sillitoe on behalf of the Wellcome Sanger Institute COVID-19 Surveillance Team |
| EPI_ISL_2717789 | Lighthouse Lab in Glasgow | Wellcome Sanger Institute for the COVID-19 Genomics UK (COG-UK) Consortium | Harper VanSteenhouse, Yumi Kasai, David Gray, Carol Clugston, Anna Dominiczak and Alex Alderton, Roberto Amato, Jeffrey Barrett, Sonia Goncalves, Ewan Harrison, David K. Jackson, Ian Johnston, Dominic Kwiatkowski, Cordelia Langford, John Sillitoe on behalf of the Wellcome Sanger Institute COVID-19 Surveillance Team |
| EPI_ISL_2717790, EPI_ISL_2717791, EPI_ISL_2717792, EPI_ISL_2717793 | Lighthouse Lab in Alderley Park | Wellcome Sanger Institute for the COVID-19 Genomics UK (COG-UK) Consortium | Jacquelyn Wynn, Mairead Hyland, The Lighthouse Lab in Alderley Park and Alex Alderton, Roberto Amato, Jeffrey Barrett, Sonia Goncalves, Ewan Harrison, David K. Jackson, Ian Johnston, Dominic Kwiatkowski, Cordelia Langford, John Sillitoe on behalf of the Wellcome Sanger Institute COVID-19 Surveillance Team |
| EPI_ISL_2717794 | Lighthouse Lab in Glasgow | Wellcome Sanger Institute for the COVID-19 Genomics UK (COG-UK) Consortium | Harper VanSteenhouse, Yumi Kasai, David Gray, Carol Clugston, Anna Dominiczak and Alex Alderton, Roberto Amato, Jeffrey Barrett, Sonia Goncalves, Ewan Harrison, David K. Jackson, Ian Johnston, Dominic Kwiatkowski, Cordelia Langford, John Sillitoe on behalf of the Wellcome Sanger Institute COVID-19 Surveillance Team |
| EPI_ISL_2717795 | Lighthouse Lab in Alderley Park | Wellcome Sanger Institute for the COVID-19 Genomics UK (COG-UK) Consortium | Jacquelyn Wynn, Mairead Hyland, The Lighthouse Lab in Alderley Park and Alex Alderton, Roberto Amato, Jeffrey Barrett, Sonia Goncalves, Ewan Harrison, David K. Jackson, Ian Johnston, Dominic Kwiatkowski, Cordelia Langford, John Sillitoe on behalf of the Wellcome Sanger Institute COVID-19 Surveillance Team |
| EPI_ISL_2717796 | Lighthouse Lab in Glasgow | Wellcome Sanger Institute for the COVID-19 Genomics UK (COG-UK) Consortium | Harper VanSteenhouse, Yumi Kasai, David Gray, Carol Clugston, Anna Dominiczak and Alex Alderton, Roberto Amato, Jeffrey Barrett, Sonia Goncalves, Ewan Harrison, David K. Jackson, Ian Johnston, Dominic Kwiatkowski, Cordelia Langford, John Sillitoe on behalf of the Wellcome Sanger Institute COVID-19 Surveillance Team |
| EPI_ISL_2717797, EPI_ISL_2717798, EPI_ISL_2717799 | Lighthouse Lab in Alderley Park | Wellcome Sanger Institute for the COVID-19 Genomics UK (COG-UK) Consortium | Jacquelyn Wynn, Mairead Hyland, The Lighthouse Lab in Alderley Park and Alex Alderton, Roberto Amato, Jeffrey Barrett, Sonia Goncalves, Ewan Harrison, David K. Jackson, Ian Johnston, Dominic Kwiatkowski, Cordelia Langford, John Sillitoe on behalf of the Wellcome Sanger Institute COVID-19 Surveillance Team |
| EPI_ISL_2717800, EPI_ISL_2717801 | Lighthouse Lab in Glasgow | Wellcome Sanger Institute for the COVID-19 Genomics UK (COG-UK) Consortium | Harper VanSteenhouse, Yumi Kasai, David Gray, Carol Clugston, Anna Dominiczak and Alex Alderton, Roberto Amato, Jeffrey Barrett, Sonia Goncalves, Ewan Harrison, David K. Jackson, Ian Johnston, Dominic Kwiatkowski, Cordelia Langford, John Sillitoe on behalf of the Wellcome Sanger Institute COVID-19 Surveillance Team |

| | | | |
|---|---|---|---|
| EPI_ISL_2717802, EPI_ISL_2717803, EPI_ISL_2717804, EPI_ISL_2717805, EPI_ISL_2717806, EPI_ISL_2717807, EPI_ISL_2717808 | Lighthouse Lab in Alderley Park | Wellcome Sanger Institute for the COVID-19 Genomics UK (COG-UK) Consortium | Jacquelyn Wynn, Mairead Hyland, The Lighthouse Lab in Alderley Park and Alex Alderton, Roberto Amato, Jeffrey Barrett, Sonia Goncalves, Ewan Harrison, David K. Jackson, Ian Johnston, Dominic Kwiatkowski, Cordelia Langford, John Sillitoe on behalf of the Wellcome Sanger Institute COVID-19 Surveillance Team |
| EPI_ISL_2717809 | Lighthouse Lab in Milton Keynes | Wellcome Sanger Institute for the COVID-19 Genomics UK (COG-UK) Consortium | The Lighthouse Lab in Milton Keynes and Alex Alderton, Roberto Amato, Jeffrey Barrett, Sonia Goncalves, Ewan Harrison, David K. Jackson, Ian Johnston, Dominic Kwiatkowski, Cordelia Langford, John Sillitoe on behalf of the Wellcome Sanger Institute COVID-19 Surveillance Team |
| EPI_ISL_2717810, EPI_ISL_2717811 | Lighthouse Lab in Alderley Park | Wellcome Sanger Institute for the COVID-19 Genomics UK (COG-UK) Consortium | Jacquelyn Wynn, Mairead Hyland, The Lighthouse Lab in Alderley Park and Alex Alderton, Roberto Amato, Jeffrey Barrett, Sonia Goncalves, Ewan Harrison, David K. Jackson, Ian Johnston, Dominic Kwiatkowski, Cordelia Langford, John Sillitoe on behalf of the Wellcome Sanger Institute COVID-19 Surveillance Team |
| EPI_ISL_2717812, EPI_ISL_2717813, EPI_ISL_2717814 | Lighthouse Lab in Milton Keynes | Wellcome Sanger Institute for the COVID-19 Genomics UK (COG-UK) Consortium | The Lighthouse Lab in Milton Keynes and Alex Alderton, Roberto Amato, Jeffrey Barrett, Sonia Goncalves, Ewan Harrison, David K. Jackson, Ian Johnston, Dominic Kwiatkowski, Cordelia Langford, John Sillitoe on behalf of the Wellcome Sanger Institute COVID-19 Surveillance Team |
| EPI_ISL_2717815, EPI_ISL_2717816 | Lighthouse Lab in Alderley Park | Wellcome Sanger Institute for the COVID-19 Genomics UK (COG-UK) Consortium | Jacquelyn Wynn, Mairead Hyland, The Lighthouse Lab in Alderley Park and Alex Alderton, Roberto Amato, Jeffrey Barrett, Sonia Goncalves, Ewan Harrison, David K. Jackson, Ian Johnston, Dominic Kwiatkowski, Cordelia Langford, John Sillitoe on behalf of the Wellcome Sanger Institute COVID-19 Surveillance Team |
| EPI_ISL_2717817 | Lighthouse Lab in Milton Keynes | Wellcome Sanger Institute for the COVID-19 Genomics UK (COG-UK) Consortium | The Lighthouse Lab in Milton Keynes and Alex Alderton, Roberto Amato, Jeffrey Barrett, Sonia Goncalves, Ewan Harrison, David K. Jackson, Ian Johnston, Dominic Kwiatkowski, Cordelia Langford, John Sillitoe on behalf of the Wellcome Sanger Institute COVID-19 Surveillance Team |
| EPI_ISL_2717818, EPI_ISL_2717819, EPI_ISL_2717820 | Lighthouse Lab in Alderley Park | Wellcome Sanger Institute for the COVID-19 Genomics UK (COG-UK) Consortium | Jacquelyn Wynn, Mairead Hyland, The Lighthouse Lab in Alderley Park and Alex Alderton, Roberto Amato, Jeffrey Barrett, Sonia Goncalves, Ewan Harrison, David K. Jackson, Ian Johnston, Dominic Kwiatkowski, Cordelia Langford, John Sillitoe on behalf of the Wellcome Sanger Institute COVID-19 Surveillance Team |
| EPI_ISL_2717821 | Lighthouse Lab in Milton Keynes | Wellcome Sanger Institute for the COVID-19 Genomics UK (COG-UK) Consortium | The Lighthouse Lab in Milton Keynes and Alex Alderton, Roberto Amato, Jeffrey Barrett, Sonia Goncalves, Ewan Harrison, David K. Jackson, Ian Johnston, Dominic Kwiatkowski, Cordelia Langford, John Sillitoe on behalf of the Wellcome Sanger Institute COVID-19 Surveillance Team |
| EPI_ISL_2717822, EPI_ISL_2717823, EPI_ISL_2717824 | Lighthouse Lab in Alderley Park | Wellcome Sanger Institute for the COVID-19 Genomics UK (COG-UK) Consortium | Jacquelyn Wynn, Mairead Hyland, The Lighthouse Lab in Alderley Park and Alex Alderton, Roberto Amato, Jeffrey Barrett, Sonia Goncalves, Ewan Harrison, David K. Jackson, Ian Johnston, Dominic Kwiatkowski, Cordelia Langford, John Sillitoe on behalf of the Wellcome Sanger Institute COVID-19 Surveillance Team |
| EPI_ISL_2717825 | Lighthouse Lab in Milton Keynes | Wellcome Sanger Institute for the COVID-19 Genomics UK (COG-UK) Consortium | The Lighthouse Lab in Milton Keynes and Alex Alderton, Roberto Amato, Jeffrey Barrett, Sonia Goncalves, Ewan Harrison, David K. Jackson, Ian Johnston, Dominic Kwiatkowski, Cordelia Langford, John Sillitoe on behalf of the Wellcome Sanger Institute COVID-19 Surveillance Team |
| EPI_ISL_2717826 | Lighthouse Lab in Alderley Park | Wellcome Sanger Institute for the COVID-19 Genomics UK (COG-UK) Consortium | Jacquelyn Wynn, Mairead Hyland, The Lighthouse Lab in Alderley Park and Alex Alderton, Roberto Amato, Jeffrey Barrett, Sonia Goncalves, Ewan Harrison, David K. Jackson, Ian Johnston, Dominic Kwiatkowski, Cordelia Langford, John Sillitoe on behalf of the Wellcome Sanger Institute COVID-19 Surveillance Team |
| EPI_ISL_2717827, EPI_ISL_2717828 | Lighthouse Lab in Milton Keynes | Wellcome Sanger Institute for the COVID-19 Genomics UK (COG-UK) Consortium | The Lighthouse Lab in Milton Keynes and Alex Alderton, Roberto Amato, Jeffrey Barrett, Sonia Goncalves, Ewan Harrison, David K. Jackson, Ian Johnston, Dominic Kwiatkowski, Cordelia Langford, John Sillitoe on behalf of the Wellcome Sanger Institute COVID-19 Surveillance Team |
| EPI_ISL_2717829 | Lighthouse Lab in Alderley Park | Wellcome Sanger Institute for the COVID-19 Genomics UK (COG-UK) Consortium | Jacquelyn Wynn, Mairead Hyland, The Lighthouse Lab in Alderley Park and Alex Alderton, Roberto Amato, Jeffrey Barrett, Sonia Goncalves, Ewan Harrison, David K. Jackson, Ian Johnston, Dominic Kwiatkowski, Cordelia Langford, John Sillitoe on behalf of the Wellcome Sanger Institute COVID-19 Surveillance Team |
| EPI_ISL_2717830 | Lighthouse Lab in Milton Keynes | Wellcome Sanger Institute for the COVID-19 Genomics UK (COG-UK) Consortium | The Lighthouse Lab in Milton Keynes and Alex Alderton, Roberto Amato, Jeffrey Barrett, Sonia Goncalves, Ewan Harrison, David K. Jackson, Ian Johnston, Dominic Kwiatkowski, Cordelia Langford, John Sillitoe on behalf of the Wellcome Sanger Institute COVID-19 Surveillance Team |
| EPI_ISL_2717831 | Lighthouse Lab in Alderley Park | Wellcome Sanger Institute for the COVID-19 Genomics UK (COG-UK) Consortium | Jacquelyn Wynn, Mairead Hyland, The Lighthouse Lab in Alderley Park and Alex Alderton, Roberto Amato, Jeffrey Barrett, Sonia Goncalves, Ewan Harrison, David K. Jackson, Ian Johnston, Dominic Kwiatkowski, Cordelia Langford, John Sillitoe on behalf of the Wellcome Sanger Institute COVID-19 Surveillance Team |
| EPI_ISL_2717832, EPI_ISL_2717833, EPI_ISL_2717834, EPI_ISL_2717835, EPI_ISL_2717836 | Lighthouse Lab in Milton Keynes | Wellcome Sanger Institute for the COVID-19 Genomics UK (COG-UK) Consortium | The Lighthouse Lab in Milton Keynes and Alex Alderton, Roberto Amato, Jeffrey Barrett, Sonia Goncalves, Ewan Harrison, David K. Jackson, Ian Johnston, Dominic Kwiatkowski, Cordelia Langford, John Sillitoe on behalf of the Wellcome Sanger Institute COVID-19 Surveillance Team |
| EPI_ISL_2717837, EPI_ISL_2717838, EPI_ISL_2717839, EPI_ISL_2717840 | Lighthouse Lab in Alderley Park | Wellcome Sanger Institute for the COVID-19 Genomics UK (COG-UK) Consortium | Jacquelyn Wynn, Mairead Hyland, The Lighthouse Lab in Alderley Park and Alex Alderton, Roberto Amato, Jeffrey Barrett, Sonia Goncalves, Ewan Harrison, David K. Jackson, Ian Johnston, Dominic Kwiatkowski, Cordelia Langford, John Sillitoe on behalf of the Wellcome Sanger Institute COVID-19 Surveillance Team |
| EPI_ISL_2717841 | Lighthouse Lab in Milton Keynes | Wellcome Sanger Institute for the COVID-19 Genomics UK (COG-UK) Consortium | The Lighthouse Lab in Milton Keynes and Alex Alderton, Roberto Amato, Jeffrey Barrett, Sonia Goncalves, Ewan Harrison, David K. Jackson, Ian Johnston, Dominic Kwiatkowski, Cordelia Langford, John Sillitoe on behalf of the Wellcome Sanger Institute COVID-19 Surveillance Team |
| EPI_ISL_2717842 | Lighthouse Lab in Alderley Park | Wellcome Sanger Institute for the COVID-19 Genomics UK (COG-UK) Consortium | Jacquelyn Wynn, Mairead Hyland, The Lighthouse Lab in Alderley Park and Alex Alderton, Roberto Amato, Jeffrey Barrett, Sonia Goncalves, Ewan Harrison, David K. Jackson, Ian Johnston, Dominic Kwiatkowski, Cordelia Langford, John Sillitoe on behalf of the Wellcome Sanger Institute COVID-19 Surveillance Team |
| EPI_ISL_2717843, EPI_ISL_2717844 | Lighthouse Lab in Milton Keynes | Wellcome Sanger Institute for the COVID-19 Genomics UK (COG-UK) Consortium | The Lighthouse Lab in Milton Keynes and Alex Alderton, Roberto Amato, Jeffrey Barrett, Sonia Goncalves, Ewan Harrison, David K. Jackson, Ian Johnston, Dominic Kwiatkowski, Cordelia Langford, John Sillitoe on behalf of the Wellcome Sanger Institute COVID-19 Surveillance Team |
| EPI_ISL_2717845 | Lighthouse Lab in Alderley Park | Wellcome Sanger Institute for the COVID-19 Genomics UK (COG-UK) Consortium | Jacquelyn Wynn, Mairead Hyland, The Lighthouse Lab in Alderley Park and Alex Alderton, Roberto Amato, Jeffrey Barrett, Sonia Goncalves, Ewan Harrison, David K. Jackson, Ian Johnston, Dominic Kwiatkowski, Cordelia Langford, John Sillitoe on behalf of the Wellcome Sanger Institute COVID-19 Surveillance Team |
| EPI_ISL_2717846 | Lighthouse Lab in Milton Keynes | Wellcome Sanger Institute for the COVID-19 Genomics UK (COG-UK) Consortium | The Lighthouse Lab in Milton Keynes and Alex Alderton, Roberto Amato, Jeffrey Barrett, Sonia Goncalves, Ewan Harrison, David K. Jackson, Ian Johnston, Dominic Kwiatkowski, Cordelia Langford, John Sillitoe on behalf of the Wellcome Sanger Institute COVID-19 Surveillance Team |
| EPI_ISL_2717847, EPI_ISL_2717848, EPI_ISL_2717849, EPI_ISL_2717850, EPI_ISL_2717851 | Lighthouse Lab in Alderley Park | Wellcome Sanger Institute for the COVID-19 Genomics UK (COG-UK) Consortium | Jacquelyn Wynn, Mairead Hyland, The Lighthouse Lab in Alderley Park and Alex Alderton, Roberto Amato, Jeffrey Barrett, Sonia Goncalves, Ewan Harrison, David K. Jackson, Ian Johnston, Dominic Kwiatkowski, Cordelia Langford, John Sillitoe on behalf of the Wellcome Sanger Institute COVID-19 Surveillance Team |
| EPI_ISL_2717852, EPI_ISL_2717853 | Lighthouse Lab in Milton Keynes | Wellcome Sanger Institute for the COVID-19 Genomics UK (COG-UK) Consortium | The Lighthouse Lab in Milton Keynes and Alex Alderton, Roberto Amato, Jeffrey Barrett, Sonia Goncalves, Ewan Harrison, David K. Jackson, Ian Johnston, Dominic Kwiatkowski, Cordelia Langford, John Sillitoe on behalf of the Wellcome Sanger Institute COVID-19 Surveillance Team |
| EPI_ISL_2717854 | Lighthouse Lab in Alderley Park | Wellcome Sanger Institute for the COVID-19 Genomics UK (COG-UK) Consortium | Jacquelyn Wynn, Mairead Hyland, The Lighthouse Lab in Alderley Park and Alex Alderton, Roberto Amato, Jeffrey Barrett, Sonia Goncalves, Ewan Harrison, David K. Jackson, Ian Johnston, Dominic Kwiatkowski, Cordelia Langford, John Sillitoe on behalf of the Wellcome Sanger Institute COVID-19 Surveillance Team |
| EPI_ISL_2717855, EPI_ISL_2717856 | Lighthouse Lab in Milton Keynes | Wellcome Sanger Institute for the COVID-19 Genomics UK (COG-UK) Consortium | The Lighthouse Lab in Milton Keynes and Alex Alderton, Roberto Amato, Jeffrey Barrett, Sonia Goncalves, Ewan Harrison, David K. Jackson, Ian Johnston, Dominic Kwiatkowski, Cordelia Langford, John Sillitoe on behalf of the Wellcome Sanger Institute COVID-19 Surveillance Team |
| EPI_ISL_2717857 | Lighthouse Lab in Alderley Park | Wellcome Sanger Institute for the COVID-19 Genomics UK (COG-UK) Consortium | Jacquelyn Wynn, Mairead Hyland, The Lighthouse Lab in Alderley Park and Alex Alderton, Roberto Amato, Jeffrey Barrett, Sonia Goncalves, Ewan Harrison, David K. Jackson, Ian Johnston, Dominic Kwiatkowski, Cordelia Langford, John Sillitoe on behalf of the Wellcome Sanger Institute COVID-19 Surveillance Team |
| EPI_ISL_2717858 | Lighthouse Lab in Milton Keynes | Wellcome Sanger Institute for the COVID-19 Genomics UK (COG-UK) Consortium | The Lighthouse Lab in Milton Keynes and Alex Alderton, Roberto Amato, Jeffrey Barrett, Sonia Goncalves, Ewan Harrison, David K. Jackson, Ian Johnston, Dominic Kwiatkowski, Cordelia Langford, John Sillitoe on behalf of the Wellcome Sanger Institute COVID-19 Surveillance Team |
| EPI_ISL_2717859 | Lighthouse Lab in Alderley Park | Wellcome Sanger Institute for the COVID-19 Genomics UK (COG-UK) Consortium | Jacquelyn Wynn, Mairead Hyland, The Lighthouse Lab in Alderley Park and Alex Alderton, Roberto Amato, Jeffrey Barrett, Sonia Goncalves, Ewan Harrison, David K. Jackson, Ian Johnston, Dominic Kwiatkowski, Cordelia Langford, John Sillitoe on behalf of the Wellcome Sanger Institute COVID-19 Surveillance Team |
| EPI_ISL_2717860 | Lighthouse Lab in Milton Keynes | Wellcome Sanger Institute for the COVID-19 Genomics UK (COG-UK) Consortium | The Lighthouse Lab in Milton Keynes and Alex Alderton, Roberto Amato, Jeffrey Barrett, Sonia Goncalves, Ewan Harrison, David K. Jackson, Ian Johnston, Dominic Kwiatkowski, Cordelia Langford, John Sillitoe on behalf of the Wellcome Sanger Institute COVID-19 Surveillance Team |
| EPI_ISL_2717861, EPI_ISL_2717862, EPI_ISL_2717863, EPI_ISL_2717864 | Lighthouse Lab in Alderley Park | Wellcome Sanger Institute for the COVID-19 Genomics UK (COG-UK) Consortium | Jacquelyn Wynn, Mairead Hyland, The Lighthouse Lab in Alderley Park and Alex Alderton, Roberto Amato, Jeffrey Barrett, Sonia Goncalves, Ewan Harrison, David K. Jackson, Ian Johnston, Dominic Kwiatkowski, Cordelia Langford, John Sillitoe on behalf of the Wellcome Sanger Institute COVID-19 Surveillance Team |
| EPI_ISL_2717865 | Lighthouse Lab in Milton Keynes | Wellcome Sanger Institute for the COVID-19 Genomics UK (COG-UK) Consortium | The Lighthouse Lab in Milton Keynes and Alex Alderton, Roberto Amato, Jeffrey Barrett, Sonia Goncalves, Ewan Harrison, David K. Jackson, Ian Johnston, Dominic Kwiatkowski, Cordelia Langford, John Sillitoe on behalf of the Wellcome Sanger Institute COVID-19 Surveillance Team |
| EPI_ISL_2717866, EPI_ISL_2717867, EPI_ISL_2717868, EPI_ISL_2717869, EPI_ISL_2717870 | Lighthouse Lab in Alderley Park | Wellcome Sanger Institute for the COVID-19 Genomics UK (COG-UK) Consortium | Jacquelyn Wynn, Mairead Hyland, The Lighthouse Lab in Alderley Park and Alex Alderton, Roberto Amato, Jeffrey Barrett, Sonia Goncalves, Ewan Harrison, David K. Jackson, Ian Johnston, Dominic Kwiatkowski, Cordelia Langford, John Sillitoe on behalf of the Wellcome Sanger Institute COVID-19 Surveillance Team |
| EPI_ISL_2717871, EPI_ISL_2717872, EPI_ISL_2717873, EPI_ISL_2717874, EPI_ISL_2717875, EPI_ISL_2717876 | Lighthouse Lab in Milton Keynes | Wellcome Sanger Institute for the COVID-19 Genomics UK (COG-UK) Consortium | The Lighthouse Lab in Milton Keynes and Alex Alderton, Roberto Amato, Jeffrey Barrett, Sonia Goncalves, Ewan Harrison, David K. Jackson, Ian Johnston, Dominic Kwiatkowski, Cordelia Langford, John Sillitoe on behalf of the Wellcome Sanger Institute COVID-19 Surveillance Team |
| EPI_ISL_2717877, EPI_ISL_2717878 | Lighthouse Lab in Alderley Park | Wellcome Sanger Institute for the COVID-19 Genomics UK (COG-UK) Consortium | Jacquelyn Wynn, Mairead Hyland, The Lighthouse Lab in Alderley Park and Alex Alderton, Roberto Amato, Jeffrey Barrett, Sonia Goncalves, Ewan Harrison, David K. Jackson, Ian Johnston, Dominic Kwiatkowski, Cordelia Langford, John Sillitoe on behalf of the Wellcome Sanger Institute COVID-19 Surveillance Team |

| | | | |
|---|---|---|---|
| EPI_ISL_2717879 | Lighthouse Lab in Milton Keynes | Wellcome Sanger Institute for the COVID-19 Genomics UK (COG-UK) Consortium | The Lighthouse Lab in Milton Keynes and Alex Alderton, Roberto Amato, Jeffrey Barrett, Sonia Goncalves, Ewan Harrison, David K. Jackson, Ian Johnston, Dominic Kwiatkowski, Cordelia Langford, John Sillitoe on behalf of the Wellcome Sanger Institute COVID-19 Surveillance Team |
| EPI_ISL_2717880, EPI_ISL_2717881, EPI_ISL_2717882, EPI_ISL_2717883 | Lighthouse Lab in Alderley Park | Wellcome Sanger Institute for the COVID-19 Genomics UK (COG-UK) Consortium | Jacquelyn Wynn, Mairead Hyland, The Lighthouse Lab in Alderley Park and Alex Alderton, Roberto Amato, Jeffrey Barrett, Sonia Goncalves, Ewan Harrison, David K. Jackson, Ian Johnston, Dominic Kwiatkowski, Cordelia Langford, John Sillitoe on behalf of the Wellcome Sanger Institute COVID-19 Surveillance Team |
| EPI_ISL_2717884, EPI_ISL_2717885, EPI_ISL_2717886 | Lighthouse Lab in Milton Keynes | Wellcome Sanger Institute for the COVID-19 Genomics UK (COG-UK) Consortium | The Lighthouse Lab in Milton Keynes and Alex Alderton, Roberto Amato, Jeffrey Barrett, Sonia Goncalves, Ewan Harrison, David K. Jackson, Ian Johnston, Dominic Kwiatkowski, Cordelia Langford, John Sillitoe on behalf of the Wellcome Sanger Institute COVID-19 Surveillance Team |
| EPI_ISL_2717887, EPI_ISL_2717888 | Lighthouse Lab in Alderley Park | Wellcome Sanger Institute for the COVID-19 Genomics UK (COG-UK) Consortium | Jacquelyn Wynn, Mairead Hyland, The Lighthouse Lab in Alderley Park and Alex Alderton, Roberto Amato, Jeffrey Barrett, Sonia Goncalves, Ewan Harrison, David K. Jackson, Ian Johnston, Dominic Kwiatkowski, Cordelia Langford, John Sillitoe on behalf of the Wellcome Sanger Institute COVID-19 Surveillance Team |
| EPI_ISL_2717889, EPI_ISL_2717890, EPI_ISL_2717891 | Lighthouse Lab in Milton Keynes | Wellcome Sanger Institute for the COVID-19 Genomics UK (COG-UK) Consortium | The Lighthouse Lab in Milton Keynes and Alex Alderton, Roberto Amato, Jeffrey Barrett, Sonia Goncalves, Ewan Harrison, David K. Jackson, Ian Johnston, Dominic Kwiatkowski, Cordelia Langford, John Sillitoe on behalf of the Wellcome Sanger Institute COVID-19 Surveillance Team |
| EPI_ISL_2717892 | Lighthouse Lab in Alderley Park | Wellcome Sanger Institute for the COVID-19 Genomics UK (COG-UK) Consortium | Jacquelyn Wynn, Mairead Hyland, The Lighthouse Lab in Alderley Park and Alex Alderton, Roberto Amato, Jeffrey Barrett, Sonia Goncalves, Ewan Harrison, David K. Jackson, Ian Johnston, Dominic Kwiatkowski, Cordelia Langford, John Sillitoe on behalf of the Wellcome Sanger Institute COVID-19 Surveillance Team |
| EPI_ISL_2717893, EPI_ISL_2717894, EPI_ISL_2717896 | Lighthouse Lab in Milton Keynes | Wellcome Sanger Institute for the COVID-19 Genomics UK (COG-UK) Consortium | The Lighthouse Lab in Milton Keynes and Alex Alderton, Roberto Amato, Jeffrey Barrett, Sonia Goncalves, Ewan Harrison, David K. Jackson, Ian Johnston, Dominic Kwiatkowski, Cordelia Langford, John Sillitoe on behalf of the Wellcome Sanger Institute COVID-19 Surveillance Team |
| EPI_ISL_2717897 | Lighthouse Lab in Alderley Park | Wellcome Sanger Institute for the COVID-19 Genomics UK (COG-UK) Consortium | Jacquelyn Wynn, Mairead Hyland, The Lighthouse Lab in Alderley Park and Alex Alderton, Roberto Amato, Jeffrey Barrett, Sonia Goncalves, Ewan Harrison, David K. Jackson, Ian Johnston, Dominic Kwiatkowski, Cordelia Langford, John Sillitoe on behalf of the Wellcome Sanger Institute COVID-19 Surveillance Team |
| EPI_ISL_2717898 | Lighthouse Lab in Milton Keynes | Wellcome Sanger Institute for the COVID-19 Genomics UK (COG-UK) Consortium | The Lighthouse Lab in Milton Keynes and Alex Alderton, Roberto Amato, Jeffrey Barrett, Sonia Goncalves, Ewan Harrison, David K. Jackson, Ian Johnston, Dominic Kwiatkowski, Cordelia Langford, John Sillitoe on behalf of the Wellcome Sanger Institute COVID-19 Surveillance Team |
| EPI_ISL_2717900, EPI_ISL_2717901 | Lighthouse Lab in Alderley Park | Wellcome Sanger Institute for the COVID-19 Genomics UK (COG-UK) Consortium | Jacquelyn Wynn, Mairead Hyland, The Lighthouse Lab in Alderley Park and Alex Alderton, Roberto Amato, Jeffrey Barrett, Sonia Goncalves, Ewan Harrison, David K. Jackson, Ian Johnston, Dominic Kwiatkowski, Cordelia Langford, John Sillitoe on behalf of the Wellcome Sanger Institute COVID-19 Surveillance Team |
| EPI_ISL_2717902, EPI_ISL_2717903 | Lighthouse Lab in Milton Keynes | Wellcome Sanger Institute for the COVID-19 Genomics UK (COG-UK) Consortium | The Lighthouse Lab in Milton Keynes and Alex Alderton, Roberto Amato, Jeffrey Barrett, Sonia Goncalves, Ewan Harrison, David K. Jackson, Ian Johnston, Dominic Kwiatkowski, Cordelia Langford, John Sillitoe on behalf of the Wellcome Sanger Institute COVID-19 Surveillance Team |
| EPI_ISL_2717904, EPI_ISL_2717905, EPI_ISL_2717906 | Lighthouse Lab in Alderley Park | Wellcome Sanger Institute for the COVID-19 Genomics UK (COG-UK) Consortium | Jacquelyn Wynn, Mairead Hyland, The Lighthouse Lab in Alderley Park and Alex Alderton, Roberto Amato, Jeffrey Barrett, Sonia Goncalves, Ewan Harrison, David K. Jackson, Ian Johnston, Dominic Kwiatkowski, Cordelia Langford, John Sillitoe on behalf of the Wellcome Sanger Institute COVID-19 Surveillance Team |
| EPI_ISL_2717908 | Lighthouse Lab in Milton Keynes | Wellcome Sanger Institute for the COVID-19 Genomics UK (COG-UK) Consortium | The Lighthouse Lab in Milton Keynes and Alex Alderton, Roberto Amato, Jeffrey Barrett, Sonia Goncalves, Ewan Harrison, David K. Jackson, Ian Johnston, Dominic Kwiatkowski, Cordelia Langford, John Sillitoe on behalf of the Wellcome Sanger Institute COVID-19 Surveillance Team |
| EPI_ISL_2717909 | Lighthouse Lab in Alderley Park | Wellcome Sanger Institute for the COVID-19 Genomics UK (COG-UK) Consortium | Jacquelyn Wynn, Mairead Hyland, The Lighthouse Lab in Alderley Park and Alex Alderton, Roberto Amato, Jeffrey Barrett, Sonia Goncalves, Ewan Harrison, David K. Jackson, Ian Johnston, Dominic Kwiatkowski, Cordelia Langford, John Sillitoe on behalf of the Wellcome Sanger Institute COVID-19 Surveillance Team |
| EPI_ISL_2717910 | Lighthouse Lab in Milton Keynes | Wellcome Sanger Institute for the COVID-19 Genomics UK (COG-UK) Consortium | The Lighthouse Lab in Milton Keynes and Alex Alderton, Roberto Amato, Jeffrey Barrett, Sonia Goncalves, Ewan Harrison, David K. Jackson, Ian Johnston, Dominic Kwiatkowski, Cordelia Langford, John Sillitoe on behalf of the Wellcome Sanger Institute COVID-19 Surveillance Team |
| EPI_ISL_2717911, EPI_ISL_2717912, EPI_ISL_2717913, EPI_ISL_2717914 | Lighthouse Lab in Alderley Park | Wellcome Sanger Institute for the COVID-19 Genomics UK (COG-UK) Consortium | Jacquelyn Wynn, Mairead Hyland, The Lighthouse Lab in Alderley Park and Alex Alderton, Roberto Amato, Jeffrey Barrett, Sonia Goncalves, Ewan Harrison, David K. Jackson, Ian Johnston, Dominic Kwiatkowski, Cordelia Langford, John Sillitoe on behalf of the Wellcome Sanger Institute COVID-19 Surveillance Team |
| EPI_ISL_2717915, EPI_ISL_2717916, EPI_ISL_2717918, EPI_ISL_2717919, EPI_ISL_2717920, EPI_ISL_2717921, EPI_ISL_2717922, EPI_ISL_2717923, EPI_ISL_2717924, EPI_ISL_2717925 | Lighthouse Lab in Milton Keynes | Wellcome Sanger Institute for the COVID-19 Genomics UK (COG-UK) Consortium | The Lighthouse Lab in Milton Keynes and Alex Alderton, Roberto Amato, Jeffrey Barrett, Sonia Goncalves, Ewan Harrison, David K. Jackson, Ian Johnston, Dominic Kwiatkowski, Cordelia Langford, John Sillitoe on behalf of the Wellcome Sanger Institute COVID-19 Surveillance Team |
| EPI_ISL_2717926 | Lighthouse Lab in Alderley Park | Wellcome Sanger Institute for the COVID-19 Genomics UK (COG-UK) Consortium | Jacquelyn Wynn, Mairead Hyland, The Lighthouse Lab in Alderley Park and Alex Alderton, Roberto Amato, Jeffrey Barrett, Sonia Goncalves, Ewan Harrison, David K. Jackson, Ian Johnston, Dominic Kwiatkowski, Cordelia Langford, John Sillitoe on behalf of the Wellcome Sanger Institute COVID-19 Surveillance Team |
| EPI_ISL_2717927, EPI_ISL_2717928 | Lighthouse Lab in Milton Keynes | Wellcome Sanger Institute for the COVID-19 Genomics UK (COG-UK) Consortium | The Lighthouse Lab in Milton Keynes and Alex Alderton, Roberto Amato, Jeffrey Barrett, Sonia Goncalves, Ewan Harrison, David K. Jackson, Ian Johnston, Dominic Kwiatkowski, Cordelia Langford, John Sillitoe on behalf of the Wellcome Sanger Institute COVID-19 Surveillance Team |
| EPI_ISL_2717929 | Lighthouse Lab in Alderley Park | Wellcome Sanger Institute for the COVID-19 Genomics UK (COG-UK) Consortium | Jacquelyn Wynn, Mairead Hyland, The Lighthouse Lab in Alderley Park and Alex Alderton, Roberto Amato, Jeffrey Barrett, Sonia Goncalves, Ewan Harrison, David K. Jackson, Ian Johnston, Dominic Kwiatkowski, Cordelia Langford, John Sillitoe on behalf of the Wellcome Sanger Institute COVID-19 Surveillance Team |
| EPI_ISL_2717930 | Lighthouse Lab in Milton Keynes | Wellcome Sanger Institute for the COVID-19 Genomics UK (COG-UK) Consortium | The Lighthouse Lab in Milton Keynes and Alex Alderton, Roberto Amato, Jeffrey Barrett, Sonia Goncalves, Ewan Harrison, David K. Jackson, Ian Johnston, Dominic Kwiatkowski, Cordelia Langford, John Sillitoe on behalf of the Wellcome Sanger Institute COVID-19 Surveillance Team |
| EPI_ISL_2717931, EPI_ISL_2717932, EPI_ISL_2717933, EPI_ISL_2717934, EPI_ISL_2717935, EPI_ISL_2717936 | Lighthouse Lab in Alderley Park | Wellcome Sanger Institute for the COVID-19 Genomics UK (COG-UK) Consortium | Jacquelyn Wynn, Mairead Hyland, The Lighthouse Lab in Alderley Park and Alex Alderton, Roberto Amato, Jeffrey Barrett, Sonia Goncalves, Ewan Harrison, David K. Jackson, Ian Johnston, Dominic Kwiatkowski, Cordelia Langford, John Sillitoe on behalf of the Wellcome Sanger Institute COVID-19 Surveillance Team |
| EPI_ISL_2717937 | Lighthouse Lab in Milton Keynes | Wellcome Sanger Institute for the COVID-19 Genomics UK (COG-UK) Consortium | The Lighthouse Lab in Milton Keynes and Alex Alderton, Roberto Amato, Jeffrey Barrett, Sonia Goncalves, Ewan Harrison, David K. Jackson, Ian Johnston, Dominic Kwiatkowski, Cordelia Langford, John Sillitoe on behalf of the Wellcome Sanger Institute COVID-19 Surveillance Team |
| EPI_ISL_2717938 | Lighthouse Lab in Alderley Park | Wellcome Sanger Institute for the COVID-19 Genomics UK (COG-UK) Consortium | Jacquelyn Wynn, Mairead Hyland, The Lighthouse Lab in Alderley Park and Alex Alderton, Roberto Amato, Jeffrey Barrett, Sonia Goncalves, Ewan Harrison, David K. Jackson, Ian Johnston, Dominic Kwiatkowski, Cordelia Langford, John Sillitoe on behalf of the Wellcome Sanger Institute COVID-19 Surveillance Team |
| EPI_ISL_2717939 | Lighthouse Lab in Milton Keynes | Wellcome Sanger Institute for the COVID-19 Genomics UK (COG-UK) Consortium | The Lighthouse Lab in Milton Keynes and Alex Alderton, Roberto Amato, Jeffrey Barrett, Sonia Goncalves, Ewan Harrison, David K. Jackson, Ian Johnston, Dominic Kwiatkowski, Cordelia Langford, John Sillitoe on behalf of the Wellcome Sanger Institute COVID-19 Surveillance Team |
| EPI_ISL_2717940, EPI_ISL_2717941 | Lighthouse Lab in Alderley Park | Wellcome Sanger Institute for the COVID-19 Genomics UK (COG-UK) Consortium | Jacquelyn Wynn, Mairead Hyland, The Lighthouse Lab in Alderley Park and Alex Alderton, Roberto Amato, Jeffrey Barrett, Sonia Goncalves, Ewan Harrison, David K. Jackson, Ian Johnston, Dominic Kwiatkowski, Cordelia Langford, John Sillitoe on behalf of the Wellcome Sanger Institute COVID-19 Surveillance Team |
| EPI_ISL_2717942, EPI_ISL_2717943, EPI_ISL_2717944 | Lighthouse Lab in Milton Keynes | Wellcome Sanger Institute for the COVID-19 Genomics UK (COG-UK) Consortium | The Lighthouse Lab in Milton Keynes and Alex Alderton, Roberto Amato, Jeffrey Barrett, Sonia Goncalves, Ewan Harrison, David K. Jackson, Ian Johnston, Dominic Kwiatkowski, Cordelia Langford, John Sillitoe on behalf of the Wellcome Sanger Institute COVID-19 Surveillance Team |
| EPI_ISL_2717945 | Lighthouse Lab in Alderley Park | Wellcome Sanger Institute for the COVID-19 Genomics UK (COG-UK) Consortium | Jacquelyn Wynn, Mairead Hyland, The Lighthouse Lab in Alderley Park and Alex Alderton, Roberto Amato, Jeffrey Barrett, Sonia Goncalves, Ewan Harrison, David K. Jackson, Ian Johnston, Dominic Kwiatkowski, Cordelia Langford, John Sillitoe on behalf of the Wellcome Sanger Institute COVID-19 Surveillance Team |
| EPI_ISL_2717946, EPI_ISL_2717947, EPI_ISL_2717948 | Lighthouse Lab in Milton Keynes | Wellcome Sanger Institute for the COVID-19 Genomics UK (COG-UK) Consortium | The Lighthouse Lab in Milton Keynes and Alex Alderton, Roberto Amato, Jeffrey Barrett, Sonia Goncalves, Ewan Harrison, David K. Jackson, Ian Johnston, Dominic Kwiatkowski, Cordelia Langford, John Sillitoe on behalf of the Wellcome Sanger Institute COVID-19 Surveillance Team |
| EPI_ISL_2717949, EPI_ISL_2717950, EPI_ISL_2717951, EPI_ISL_2717952 | Lighthouse Lab in Alderley Park | Wellcome Sanger Institute for the COVID-19 Genomics UK (COG-UK) Consortium | Jacquelyn Wynn, Mairead Hyland, The Lighthouse Lab in Alderley Park and Alex Alderton, Roberto Amato, Jeffrey Barrett, Sonia Goncalves, Ewan Harrison, David K. Jackson, Ian Johnston, Dominic Kwiatkowski, Cordelia Langford, John Sillitoe on behalf of the Wellcome Sanger Institute COVID-19 Surveillance Team |
| EPI_ISL_2717953, EPI_ISL_2717954, EPI_ISL_2717955 | Lighthouse Lab in Milton Keynes | Wellcome Sanger Institute for the COVID-19 Genomics UK (COG-UK) Consortium | The Lighthouse Lab in Milton Keynes and Alex Alderton, Roberto Amato, Jeffrey Barrett, Sonia Goncalves, Ewan Harrison, David K. Jackson, Ian Johnston, Dominic Kwiatkowski, Cordelia Langford, John Sillitoe on behalf of the Wellcome Sanger Institute COVID-19 Surveillance Team |
| EPI_ISL_2717956 | Lighthouse Lab in Alderley Park | Wellcome Sanger Institute for the COVID-19 Genomics UK (COG-UK) Consortium | Jacquelyn Wynn, Mairead Hyland, The Lighthouse Lab in Alderley Park and Alex Alderton, Roberto Amato, Jeffrey Barrett, Sonia Goncalves, Ewan Harrison, David K. Jackson, Ian Johnston, Dominic Kwiatkowski, Cordelia Langford, John Sillitoe on behalf of the Wellcome Sanger Institute COVID-19 Surveillance Team |
| EPI_ISL_2717957 | Lighthouse Lab in Milton Keynes | Wellcome Sanger Institute for the COVID-19 Genomics UK (COG-UK) Consortium | The Lighthouse Lab in Milton Keynes and Alex Alderton, Roberto Amato, Jeffrey Barrett, Sonia Goncalves, Ewan Harrison, David K. Jackson, Ian Johnston, Dominic Kwiatkowski, Cordelia Langford, John Sillitoe on behalf of the Wellcome Sanger Institute COVID-19 Surveillance Team |
| EPI_ISL_2717958, EPI_ISL_2717959, EPI_ISL_2717960, EPI_ISL_2717961 | Lighthouse Lab in Alderley Park | Wellcome Sanger Institute for the COVID-19 Genomics UK (COG-UK) Consortium | Jacquelyn Wynn, Mairead Hyland, The Lighthouse Lab in Alderley Park and Alex Alderton, Roberto Amato, Jeffrey Barrett, Sonia Goncalves, Ewan Harrison, David K. Jackson, Ian Johnston, Dominic Kwiatkowski, Cordelia Langford, John Sillitoe on behalf of the Wellcome Sanger Institute COVID-19 Surveillance Team |

| | | | |
|---|---|---|---|
| EPI_ISL_2717962, EPI_ISL_2717963 | Lighthouse Lab in Milton Keynes | Wellcome Sanger Institute for the COVID-19 Genomics UK (COG-UK) Consortium | The Lighthouse Lab in Milton Keynes and Alex Alderton, Roberto Amato, Jeffrey Barrett, Sonia Goncalves, Ewan Harrison, David K. Jackson, Ian Johnston, Dominic Kwiatkowski, Cordelia Langford, John Sillitoe on behalf of the Wellcome Sanger Institute COVID-19 Surveillance Team |
| EPI_ISL_2717964, EPI_ISL_2717965 | Lighthouse Lab in Alderley Park | Wellcome Sanger Institute for the COVID-19 Genomics UK (COG-UK) Consortium | Jacquelyn Wynn, Mairead Hyland, The Lighthouse Lab in Alderley Park and Alex Alderton, Roberto Amato, Jeffrey Barrett, Sonia Goncalves, Ewan Harrison, David K. Jackson, Ian Johnston, Dominic Kwiatkowski, Cordelia Langford, John Sillitoe on behalf of the Wellcome Sanger Institute COVID-19 Surveillance Team |
| EPI_ISL_2717966 | Lighthouse Lab in Milton Keynes | Wellcome Sanger Institute for the COVID-19 Genomics UK (COG-UK) Consortium | The Lighthouse Lab in Milton Keynes and Alex Alderton, Roberto Amato, Jeffrey Barrett, Sonia Goncalves, Ewan Harrison, David K. Jackson, Ian Johnston, Dominic Kwiatkowski, Cordelia Langford, John Sillitoe on behalf of the Wellcome Sanger Institute COVID-19 Surveillance Team |
| EPI_ISL_2717967, EPI_ISL_2717968 | Lighthouse Lab in Alderley Park | Wellcome Sanger Institute for the COVID-19 Genomics UK (COG-UK) Consortium | Jacquelyn Wynn, Mairead Hyland, The Lighthouse Lab in Alderley Park and Alex Alderton, Roberto Amato, Jeffrey Barrett, Sonia Goncalves, Ewan Harrison, David K. Jackson, Ian Johnston, Dominic Kwiatkowski, Cordelia Langford, John Sillitoe on behalf of the Wellcome Sanger Institute COVID-19 Surveillance Team |
| EPI_ISL_2717969 | Lighthouse Lab in Milton Keynes | Wellcome Sanger Institute for the COVID-19 Genomics UK (COG-UK) Consortium | The Lighthouse Lab in Milton Keynes and Alex Alderton, Roberto Amato, Jeffrey Barrett, Sonia Goncalves, Ewan Harrison, David K. Jackson, Ian Johnston, Dominic Kwiatkowski, Cordelia Langford, John Sillitoe on behalf of the Wellcome Sanger Institute COVID-19 Surveillance Team |
| EPI_ISL_2717970 | Lighthouse Lab in Alderley Park | Wellcome Sanger Institute for the COVID-19 Genomics UK (COG-UK) Consortium | Jacquelyn Wynn, Mairead Hyland, The Lighthouse Lab in Alderley Park and Alex Alderton, Roberto Amato, Jeffrey Barrett, Sonia Goncalves, Ewan Harrison, David K. Jackson, Ian Johnston, Dominic Kwiatkowski, Cordelia Langford, John Sillitoe on behalf of the Wellcome Sanger Institute COVID-19 Surveillance Team |
| EPI_ISL_2717971 | Lighthouse Lab in Milton Keynes | Wellcome Sanger Institute for the COVID-19 Genomics UK (COG-UK) Consortium | The Lighthouse Lab in Milton Keynes and Alex Alderton, Roberto Amato, Jeffrey Barrett, Sonia Goncalves, Ewan Harrison, David K. Jackson, Ian Johnston, Dominic Kwiatkowski, Cordelia Langford, John Sillitoe on behalf of the Wellcome Sanger Institute COVID-19 Surveillance Team |
| EPI_ISL_2717972 | Lighthouse Lab in Alderley Park | Wellcome Sanger Institute for the COVID-19 Genomics UK (COG-UK) Consortium | Jacquelyn Wynn, Mairead Hyland, The Lighthouse Lab in Alderley Park and Alex Alderton, Roberto Amato, Jeffrey Barrett, Sonia Goncalves, Ewan Harrison, David K. Jackson, Ian Johnston, Dominic Kwiatkowski, Cordelia Langford, John Sillitoe on behalf of the Wellcome Sanger Institute COVID-19 Surveillance Team |
| EPI_ISL_2717973 | Lighthouse Lab in Milton Keynes | Wellcome Sanger Institute for the COVID-19 Genomics UK (COG-UK) Consortium | The Lighthouse Lab in Milton Keynes and Alex Alderton, Roberto Amato, Jeffrey Barrett, Sonia Goncalves, Ewan Harrison, David K. Jackson, Ian Johnston, Dominic Kwiatkowski, Cordelia Langford, John Sillitoe on behalf of the Wellcome Sanger Institute COVID-19 Surveillance Team |
| EPI_ISL_2717974 | Lighthouse Lab in Alderley Park | Wellcome Sanger Institute for the COVID-19 Genomics UK (COG-UK) Consortium | Jacquelyn Wynn, Mairead Hyland, The Lighthouse Lab in Alderley Park and Alex Alderton, Roberto Amato, Jeffrey Barrett, Sonia Goncalves, Ewan Harrison, David K. Jackson, Ian Johnston, Dominic Kwiatkowski, Cordelia Langford, John Sillitoe on behalf of the Wellcome Sanger Institute COVID-19 Surveillance Team |
| EPI_ISL_2717975 | Lighthouse Lab in Milton Keynes | Wellcome Sanger Institute for the COVID-19 Genomics UK (COG-UK) Consortium | The Lighthouse Lab in Milton Keynes and Alex Alderton, Roberto Amato, Jeffrey Barrett, Sonia Goncalves, Ewan Harrison, David K. Jackson, Ian Johnston, Dominic Kwiatkowski, Cordelia Langford, John Sillitoe on behalf of the Wellcome Sanger Institute COVID-19 Surveillance Team |
| EPI_ISL_2717976 | Lighthouse Lab in Alderley Park | Wellcome Sanger Institute for the COVID-19 Genomics UK (COG-UK) Consortium | Jacquelyn Wynn, Mairead Hyland, The Lighthouse Lab in Alderley Park and Alex Alderton, Roberto Amato, Jeffrey Barrett, Sonia Goncalves, Ewan Harrison, David K. Jackson, Ian Johnston, Dominic Kwiatkowski, Cordelia Langford, John Sillitoe on behalf of the Wellcome Sanger Institute COVID-19 Surveillance Team |
| EPI_ISL_2717977 | Lighthouse Lab in Milton Keynes | Wellcome Sanger Institute for the COVID-19 Genomics UK (COG-UK) Consortium | The Lighthouse Lab in Milton Keynes and Alex Alderton, Roberto Amato, Jeffrey Barrett, Sonia Goncalves, Ewan Harrison, David K. Jackson, Ian Johnston, Dominic Kwiatkowski, Cordelia Langford, John Sillitoe on behalf of the Wellcome Sanger Institute COVID-19 Surveillance Team |
| EPI_ISL_2717978, EPI_ISL_2717979 | Lighthouse Lab in Alderley Park | Wellcome Sanger Institute for the COVID-19 Genomics UK (COG-UK) Consortium | Jacquelyn Wynn, Mairead Hyland, The Lighthouse Lab in Alderley Park and Alex Alderton, Roberto Amato, Jeffrey Barrett, Sonia Goncalves, Ewan Harrison, David K. Jackson, Ian Johnston, Dominic Kwiatkowski, Cordelia Langford, John Sillitoe on behalf of the Wellcome Sanger Institute COVID-19 Surveillance Team |
| EPI_ISL_2717980 | Lighthouse Lab in Milton Keynes | Wellcome Sanger Institute for the COVID-19 Genomics UK (COG-UK) Consortium | The Lighthouse Lab in Milton Keynes and Alex Alderton, Roberto Amato, Jeffrey Barrett, Sonia Goncalves, Ewan Harrison, David K. Jackson, Ian Johnston, Dominic Kwiatkowski, Cordelia Langford, John Sillitoe on behalf of the Wellcome Sanger Institute COVID-19 Surveillance Team |
| EPI_ISL_2717981 | Lighthouse Lab in Alderley Park | Wellcome Sanger Institute for the COVID-19 Genomics UK (COG-UK) Consortium | Jacquelyn Wynn, Mairead Hyland, The Lighthouse Lab in Alderley Park and Alex Alderton, Roberto Amato, Jeffrey Barrett, Sonia Goncalves, Ewan Harrison, David K. Jackson, Ian Johnston, Dominic Kwiatkowski, Cordelia Langford, John Sillitoe on behalf of the Wellcome Sanger Institute COVID-19 Surveillance Team |
| EPI_ISL_2717982, EPI_ISL_2717983 | Lighthouse Lab in Milton Keynes | Wellcome Sanger Institute for the COVID-19 Genomics UK (COG-UK) Consortium | The Lighthouse Lab in Milton Keynes and Alex Alderton, Roberto Amato, Jeffrey Barrett, Sonia Goncalves, Ewan Harrison, David K. Jackson, Ian Johnston, Dominic Kwiatkowski, Cordelia Langford, John Sillitoe on behalf of the Wellcome Sanger Institute COVID-19 Surveillance Team |
| EPI_ISL_2717984 | Lighthouse Lab in Alderley Park | Wellcome Sanger Institute for the COVID-19 Genomics UK (COG-UK) Consortium | Jacquelyn Wynn, Mairead Hyland, The Lighthouse Lab in Alderley Park and Alex Alderton, Roberto Amato, Jeffrey Barrett, Sonia Goncalves, Ewan Harrison, David K. Jackson, Ian Johnston, Dominic Kwiatkowski, Cordelia Langford, John Sillitoe on behalf of the Wellcome Sanger Institute COVID-19 Surveillance Team |
| EPI_ISL_2717985 | Lighthouse Lab in Milton Keynes | Wellcome Sanger Institute for the COVID-19 Genomics UK (COG-UK) Consortium | The Lighthouse Lab in Milton Keynes and Alex Alderton, Roberto Amato, Jeffrey Barrett, Sonia Goncalves, Ewan Harrison, David K. Jackson, Ian Johnston, Dominic Kwiatkowski, Cordelia Langford, John Sillitoe on behalf of the Wellcome Sanger Institute COVID-19 Surveillance Team |
| EPI_ISL_2717986, EPI_ISL_2717987 | Lighthouse Lab in Alderley Park | Wellcome Sanger Institute for the COVID-19 Genomics UK (COG-UK) Consortium | Jacquelyn Wynn, Mairead Hyland, The Lighthouse Lab in Alderley Park and Alex Alderton, Roberto Amato, Jeffrey Barrett, Sonia Goncalves, Ewan Harrison, David K. Jackson, Ian Johnston, Dominic Kwiatkowski, Cordelia Langford, John Sillitoe on behalf of the Wellcome Sanger Institute COVID-19 Surveillance Team |
| EPI_ISL_2717988 | Lighthouse Lab in Milton Keynes | Wellcome Sanger Institute for the COVID-19 Genomics UK (COG-UK) Consortium | The Lighthouse Lab in Milton Keynes and Alex Alderton, Roberto Amato, Jeffrey Barrett, Sonia Goncalves, Ewan Harrison, David K. Jackson, Ian Johnston, Dominic Kwiatkowski, Cordelia Langford, John Sillitoe on behalf of the Wellcome Sanger Institute COVID-19 Surveillance Team |
| EPI_ISL_2717989, EPI_ISL_2717990 | Lighthouse Lab in Alderley Park | Wellcome Sanger Institute for the COVID-19 Genomics UK (COG-UK) Consortium | Jacquelyn Wynn, Mairead Hyland, The Lighthouse Lab in Alderley Park and Alex Alderton, Roberto Amato, Jeffrey Barrett, Sonia Goncalves, Ewan Harrison, David K. Jackson, Ian Johnston, Dominic Kwiatkowski, Cordelia Langford, John Sillitoe on behalf of the Wellcome Sanger Institute COVID-19 Surveillance Team |
| EPI_ISL_2717991, EPI_ISL_2717992, EPI_ISL_2717993 | Lighthouse Lab in Milton Keynes | Wellcome Sanger Institute for the COVID-19 Genomics UK (COG-UK) Consortium | The Lighthouse Lab in Milton Keynes and Alex Alderton, Roberto Amato, Jeffrey Barrett, Sonia Goncalves, Ewan Harrison, David K. Jackson, Ian Johnston, Dominic Kwiatkowski, Cordelia Langford, John Sillitoe on behalf of the Wellcome Sanger Institute COVID-19 Surveillance Team |
| EPI_ISL_2717994, EPI_ISL_2717995, EPI_ISL_2717996, EPI_ISL_2717997 | Lighthouse Lab in Alderley Park | Wellcome Sanger Institute for the COVID-19 Genomics UK (COG-UK) Consortium | Jacquelyn Wynn, Mairead Hyland, The Lighthouse Lab in Alderley Park and Alex Alderton, Roberto Amato, Jeffrey Barrett, Sonia Goncalves, Ewan Harrison, David K. Jackson, Ian Johnston, Dominic Kwiatkowski, Cordelia Langford, John Sillitoe on behalf of the Wellcome Sanger Institute COVID-19 Surveillance Team |
| EPI_ISL_2717998, EPI_ISL_2717999, EPI_ISL_2718000 | Lighthouse Lab in Milton Keynes | Wellcome Sanger Institute for the COVID-19 Genomics UK (COG-UK) Consortium | The Lighthouse Lab in Milton Keynes and Alex Alderton, Roberto Amato, Jeffrey Barrett, Sonia Goncalves, Ewan Harrison, David K. Jackson, Ian Johnston, Dominic Kwiatkowski, Cordelia Langford, John Sillitoe on behalf of the Wellcome Sanger Institute COVID-19 Surveillance Team |
| EPI_ISL_2718001, EPI_ISL_2718002 | Lighthouse Lab in Alderley Park | Wellcome Sanger Institute for the COVID-19 Genomics UK (COG-UK) Consortium | Jacquelyn Wynn, Mairead Hyland, The Lighthouse Lab in Alderley Park and Alex Alderton, Roberto Amato, Jeffrey Barrett, Sonia Goncalves, Ewan Harrison, David K. Jackson, Ian Johnston, Dominic Kwiatkowski, Cordelia Langford, John Sillitoe on behalf of the Wellcome Sanger Institute COVID-19 Surveillance Team |
| EPI_ISL_2718003 | Lighthouse Lab in Milton Keynes | Wellcome Sanger Institute for the COVID-19 Genomics UK (COG-UK) Consortium | The Lighthouse Lab in Milton Keynes and Alex Alderton, Roberto Amato, Jeffrey Barrett, Sonia Goncalves, Ewan Harrison, David K. Jackson, Ian Johnston, Dominic Kwiatkowski, Cordelia Langford, John Sillitoe on behalf of the Wellcome Sanger Institute COVID-19 Surveillance Team |
| EPI_ISL_2718004 | Lighthouse Lab in Alderley Park | Wellcome Sanger Institute for the COVID-19 Genomics UK (COG-UK) Consortium | Jacquelyn Wynn, Mairead Hyland, The Lighthouse Lab in Alderley Park and Alex Alderton, Roberto Amato, Jeffrey Barrett, Sonia Goncalves, Ewan Harrison, David K. Jackson, Ian Johnston, Dominic Kwiatkowski, Cordelia Langford, John Sillitoe on behalf of the Wellcome Sanger Institute COVID-19 Surveillance Team |
| EPI_ISL_2718005, EPI_ISL_2718006 | Lighthouse Lab in Milton Keynes | Wellcome Sanger Institute for the COVID-19 Genomics UK (COG-UK) Consortium | The Lighthouse Lab in Milton Keynes and Alex Alderton, Roberto Amato, Jeffrey Barrett, Sonia Goncalves, Ewan Harrison, David K. Jackson, Ian Johnston, Dominic Kwiatkowski, Cordelia Langford, John Sillitoe on behalf of the Wellcome Sanger Institute COVID-19 Surveillance Team |
| EPI_ISL_2718007, EPI_ISL_2718008, EPI_ISL_2718009, EPI_ISL_2718010, EPI_ISL_2718011 | Lighthouse Lab in Alderley Park | Wellcome Sanger Institute for the COVID-19 Genomics UK (COG-UK) Consortium | Jacquelyn Wynn, Mairead Hyland, The Lighthouse Lab in Alderley Park and Alex Alderton, Roberto Amato, Jeffrey Barrett, Sonia Goncalves, Ewan Harrison, David K. Jackson, Ian Johnston, Dominic Kwiatkowski, Cordelia Langford, John Sillitoe on behalf of the Wellcome Sanger Institute COVID-19 Surveillance Team |
| EPI_ISL_2718012 | Lighthouse Lab in Milton Keynes | Wellcome Sanger Institute for the COVID-19 Genomics UK (COG-UK) Consortium | The Lighthouse Lab in Milton Keynes and Alex Alderton, Roberto Amato, Jeffrey Barrett, Sonia Goncalves, Ewan Harrison, David K. Jackson, Ian Johnston, Dominic Kwiatkowski, Cordelia Langford, John Sillitoe on behalf of the Wellcome Sanger Institute COVID-19 Surveillance Team |
| EPI_ISL_2718013, EPI_ISL_2718014 | Lighthouse Lab in Alderley Park | Wellcome Sanger Institute for the COVID-19 Genomics UK (COG-UK) Consortium | Jacquelyn Wynn, Mairead Hyland, The Lighthouse Lab in Alderley Park and Alex Alderton, Roberto Amato, Jeffrey Barrett, Sonia Goncalves, Ewan Harrison, David K. Jackson, Ian Johnston, Dominic Kwiatkowski, Cordelia Langford, John Sillitoe on behalf of the Wellcome Sanger Institute COVID-19 Surveillance Team |
| EPI_ISL_2718015, EPI_ISL_2718016 | Lighthouse Lab in Milton Keynes | Wellcome Sanger Institute for the COVID-19 Genomics UK (COG-UK) Consortium | The Lighthouse Lab in Milton Keynes and Alex Alderton, Roberto Amato, Jeffrey Barrett, Sonia Goncalves, Ewan Harrison, David K. Jackson, Ian Johnston, Dominic Kwiatkowski, Cordelia Langford, John Sillitoe on behalf of the Wellcome Sanger Institute COVID-19 Surveillance Team |
| EPI_ISL_2718017, EPI_ISL_2718018 | Lighthouse Lab in Alderley Park | Wellcome Sanger Institute for the COVID-19 Genomics UK (COG-UK) Consortium | Jacquelyn Wynn, Mairead Hyland, The Lighthouse Lab in Alderley Park and Alex Alderton, Roberto Amato, Jeffrey Barrett, Sonia Goncalves, Ewan Harrison, David K. Jackson, Ian Johnston, Dominic Kwiatkowski, Cordelia Langford, John Sillitoe on behalf of the Wellcome Sanger Institute COVID-19 Surveillance Team |
| EPI_ISL_2718019, EPI_ISL_2718020, EPI_ISL_2718021 | Lighthouse Lab in Milton Keynes | Wellcome Sanger Institute for the COVID-19 Genomics UK (COG-UK) Consortium | The Lighthouse Lab in Milton Keynes and Alex Alderton, Roberto Amato, Jeffrey Barrett, Sonia Goncalves, Ewan Harrison, David K. Jackson, Ian Johnston, Dominic Kwiatkowski, Cordelia Langford, John Sillitoe on behalf of the Wellcome Sanger Institute COVID-19 Surveillance Team |
| EPI_ISL_2718022, EPI_ISL_2718023 | Lighthouse Lab in Alderley Park | Wellcome Sanger Institute for the COVID-19 Genomics UK (COG-UK) Consortium | Jacquelyn Wynn, Mairead Hyland, The Lighthouse Lab in Alderley Park and Alex Alderton, Roberto Amato, Jeffrey Barrett, Sonia Goncalves, Ewan Harrison, David K. Jackson, Ian Johnston, Dominic Kwiatkowski, Cordelia Langford, John Sillitoe on behalf of the Wellcome Sanger Institute COVID-19 Surveillance Team |
| EPI_ISL_2718024, EPI_ISL_2718025 | Lighthouse Lab in Milton Keynes | Wellcome Sanger Institute for the COVID-19 Genomics UK (COG-UK) Consortium | The Lighthouse Lab in Milton Keynes and Alex Alderton, Roberto Amato, Jeffrey Barrett, Sonia Goncalves, Ewan Harrison, David K. Jackson, Ian Johnston, Dominic Kwiatkowski, Cordelia Langford, John Sillitoe on behalf of the Wellcome Sanger Institute COVID-19 Surveillance Team |
| EPI_ISL_2718026, EPI_ISL_2718027, EPI_ISL_2718028 | Lighthouse Lab in Alderley Park | Wellcome Sanger Institute for the COVID-19 Genomics UK (COG-UK) Consortium | Jacquelyn Wynn, Mairead Hyland, The Lighthouse Lab in Alderley Park and Alex Alderton, Roberto Amato, Jeffrey Barrett, Sonia Goncalves, Ewan Harrison, David K. Jackson, Ian Johnston, Dominic Kwiatkowski, Cordelia Langford, John Sillitoe on behalf of the Wellcome Sanger Institute COVID-19 Surveillance Team |
| EPI_ISL_2718029 | Lighthouse Lab in Milton Keynes | Wellcome Sanger Institute for the COVID-19 Genomics UK (COG-UK) Consortium | The Lighthouse Lab in Milton Keynes and Alex Alderton, Roberto Amato, Jeffrey Barrett, Sonia Goncalves, Ewan Harrison, David K. Jackson, Ian Johnston, Dominic Kwiatkowski, Cordelia Langford, John Sillitoe on behalf of the Wellcome Sanger Institute COVID-19 Surveillance Team |
| EPI_ISL_2718030, EPI_ISL_2718031 | Lighthouse Lab in Alderley Park | Wellcome Sanger Institute for the COVID-19 Genomics UK (COG-UK) Consortium | Jacquelyn Wynn, Mairead Hyland, The Lighthouse Lab in Alderley Park and Alex Alderton, Roberto Amato, Jeffrey Barrett, Sonia Goncalves, Ewan Harrison, David K. Jackson, Ian Johnston, Dominic Kwiatkowski, Cordelia Langford, John Sillitoe on behalf of the Wellcome Sanger Institute COVID-19 Surveillance Team |
| EPI_ISL_2718032, EPI_ISL_2718033 | Lighthouse Lab in Milton Keynes | Wellcome Sanger Institute for the COVID-19 Genomics UK (COG-UK) Consortium | The Lighthouse Lab in Milton Keynes and Alex Alderton, Roberto Amato, Jeffrey Barrett, Sonia Goncalves, Ewan Harrison, David K. Jackson, Ian Johnston, Dominic Kwiatkowski, Cordelia Langford, John Sillitoe on behalf of the Wellcome Sanger Institute COVID-19 Surveillance Team |
| EPI_ISL_2718034, | Lighthouse Lab in Alderley Park | Wellcome Sanger Institute for the COVID-19 Genomics UK (COG-UK) | Jacquelyn Wynn, Mairead Hyland, The Lighthouse Lab in Alderley Park and Alex Alderton, Roberto Amato, Jeffrey Barrett, Sonia Goncalves, Ewan Harrison, David K. Jackson, Ian Johnston, Dominic Kwiatkowski, Cordelia Langford, John Sillitoe on |

| | | | |
|---|---|---|---|
| EPI_ISL_2718035, EPI_ISL_2718036, EPI_ISL_2718037 | | Consortium | behalf of the Wellcome Sanger Institute COVID-19 Surveillance Team |
| EPI_ISL_2718038 | Lighthouse Lab in Milton Keynes | Wellcome Sanger Institute for the COVID-19 Genomics UK (COG-UK) Consortium | The Lighthouse Lab in Milton Keynes and Alex Alderton, Roberto Amato, Jeffrey Barrett, Sonia Goncalves, Ewan Harrison, David K. Jackson, Ian Johnston, Dominic Kwiatkowski, Cordelia Langford, John Sillitoe on behalf of the Wellcome Sanger Institute COVID-19 Surveillance Team |
| EPI_ISL_2718039, EPI_ISL_2718040 | Lighthouse Lab in Alderley Park | Wellcome Sanger Institute for the COVID-19 Genomics UK (COG-UK) Consortium | Jacquelyn Wynn, Mairead Hyland, The Lighthouse Lab in Alderley Park and Alex Alderton, Roberto Amato, Jeffrey Barrett, Sonia Goncalves, Ewan Harrison, David K. Jackson, Ian Johnston, Dominic Kwiatkowski, Cordelia Langford, John Sillitoe on behalf of the Wellcome Sanger Institute COVID-19 Surveillance Team |
| EPI_ISL_2718041, EPI_ISL_2718042, EPI_ISL_2718043, EPI_ISL_2718044 | Lighthouse Lab in Milton Keynes | Wellcome Sanger Institute for the COVID-19 Genomics UK (COG-UK) Consortium | The Lighthouse Lab in Milton Keynes and Alex Alderton, Roberto Amato, Jeffrey Barrett, Sonia Goncalves, Ewan Harrison, David K. Jackson, Ian Johnston, Dominic Kwiatkowski, Cordelia Langford, John Sillitoe on behalf of the Wellcome Sanger Institute COVID-19 Surveillance Team |
| EPI_ISL_2718045, EPI_ISL_2718046, EPI_ISL_2718047 | Lighthouse Lab in Alderley Park | Wellcome Sanger Institute for the COVID-19 Genomics UK (COG-UK) Consortium | Jacquelyn Wynn, Mairead Hyland, The Lighthouse Lab in Alderley Park and Alex Alderton, Roberto Amato, Jeffrey Barrett, Sonia Goncalves, Ewan Harrison, David K. Jackson, Ian Johnston, Dominic Kwiatkowski, Cordelia Langford, John Sillitoe on behalf of the Wellcome Sanger Institute COVID-19 Surveillance Team |
| EPI_ISL_2718048, EPI_ISL_2718049, EPI_ISL_2718050 | Lighthouse Lab in Milton Keynes | Wellcome Sanger Institute for the COVID-19 Genomics UK (COG-UK) Consortium | The Lighthouse Lab in Milton Keynes and Alex Alderton, Roberto Amato, Jeffrey Barrett, Sonia Goncalves, Ewan Harrison, David K. Jackson, Ian Johnston, Dominic Kwiatkowski, Cordelia Langford, John Sillitoe on behalf of the Wellcome Sanger Institute COVID-19 Surveillance Team |
| EPI_ISL_2718051 | Lighthouse Lab in Alderley Park | Wellcome Sanger Institute for the COVID-19 Genomics UK (COG-UK) Consortium | Jacquelyn Wynn, Mairead Hyland, The Lighthouse Lab in Alderley Park and Alex Alderton, Roberto Amato, Jeffrey Barrett, Sonia Goncalves, Ewan Harrison, David K. Jackson, Ian Johnston, Dominic Kwiatkowski, Cordelia Langford, John Sillitoe on behalf of the Wellcome Sanger Institute COVID-19 Surveillance Team |
| EPI_ISL_2718052 | Lighthouse Lab in Milton Keynes | Wellcome Sanger Institute for the COVID-19 Genomics UK (COG-UK) Consortium | The Lighthouse Lab in Milton Keynes and Alex Alderton, Roberto Amato, Jeffrey Barrett, Sonia Goncalves, Ewan Harrison, David K. Jackson, Ian Johnston, Dominic Kwiatkowski, Cordelia Langford, John Sillitoe on behalf of the Wellcome Sanger Institute COVID-19 Surveillance Team |
| EPI_ISL_2718053, EPI_ISL_2718054 | Lighthouse Lab in Alderley Park | Wellcome Sanger Institute for the COVID-19 Genomics UK (COG-UK) Consortium | Jacquelyn Wynn, Mairead Hyland, The Lighthouse Lab in Alderley Park and Alex Alderton, Roberto Amato, Jeffrey Barrett, Sonia Goncalves, Ewan Harrison, David K. Jackson, Ian Johnston, Dominic Kwiatkowski, Cordelia Langford, John Sillitoe on behalf of the Wellcome Sanger Institute COVID-19 Surveillance Team |
| EPI_ISL_2718055 | Lighthouse Lab in Milton Keynes | Wellcome Sanger Institute for the COVID-19 Genomics UK (COG-UK) Consortium | The Lighthouse Lab in Milton Keynes and Alex Alderton, Roberto Amato, Jeffrey Barrett, Sonia Goncalves, Ewan Harrison, David K. Jackson, Ian Johnston, Dominic Kwiatkowski, Cordelia Langford, John Sillitoe on behalf of the Wellcome Sanger Institute COVID-19 Surveillance Team |
| EPI_ISL_2718056, EPI_ISL_2718057, EPI_ISL_2718058 | Lighthouse Lab in Alderley Park | Wellcome Sanger Institute for the COVID-19 Genomics UK (COG-UK) Consortium | Jacquelyn Wynn, Mairead Hyland, The Lighthouse Lab in Alderley Park and Alex Alderton, Roberto Amato, Jeffrey Barrett, Sonia Goncalves, Ewan Harrison, David K. Jackson, Ian Johnston, Dominic Kwiatkowski, Cordelia Langford, John Sillitoe on behalf of the Wellcome Sanger Institute COVID-19 Surveillance Team |
| EPI_ISL_2718059, EPI_ISL_2718060 | Lighthouse Lab in Milton Keynes | Wellcome Sanger Institute for the COVID-19 Genomics UK (COG-UK) Consortium | The Lighthouse Lab in Milton Keynes and Alex Alderton, Roberto Amato, Jeffrey Barrett, Sonia Goncalves, Ewan Harrison, David K. Jackson, Ian Johnston, Dominic Kwiatkowski, Cordelia Langford, John Sillitoe on behalf of the Wellcome Sanger Institute COVID-19 Surveillance Team |
| EPI_ISL_2718061 | Lighthouse Lab in Alderley Park | Wellcome Sanger Institute for the COVID-19 Genomics UK (COG-UK) Consortium | Jacquelyn Wynn, Mairead Hyland, The Lighthouse Lab in Alderley Park and Alex Alderton, Roberto Amato, Jeffrey Barrett, Sonia Goncalves, Ewan Harrison, David K. Jackson, Ian Johnston, Dominic Kwiatkowski, Cordelia Langford, John Sillitoe on behalf of the Wellcome Sanger Institute COVID-19 Surveillance Team |
| EPI_ISL_2718063 | Lighthouse Lab in Milton Keynes | Wellcome Sanger Institute for the COVID-19 Genomics UK (COG-UK) Consortium | The Lighthouse Lab in Milton Keynes and Alex Alderton, Roberto Amato, Jeffrey Barrett, Sonia Goncalves, Ewan Harrison, David K. Jackson, Ian Johnston, Dominic Kwiatkowski, Cordelia Langford, John Sillitoe on behalf of the Wellcome Sanger Institute COVID-19 Surveillance Team |
| EPI_ISL_2718064 | Lighthouse Lab in Alderley Park | Wellcome Sanger Institute for the COVID-19 Genomics UK (COG-UK) Consortium | Jacquelyn Wynn, Mairead Hyland, The Lighthouse Lab in Alderley Park and Alex Alderton, Roberto Amato, Jeffrey Barrett, Sonia Goncalves, Ewan Harrison, David K. Jackson, Ian Johnston, Dominic Kwiatkowski, Cordelia Langford, John Sillitoe on behalf of the Wellcome Sanger Institute COVID-19 Surveillance Team |
| EPI_ISL_2718065, EPI_ISL_2718066, EPI_ISL_2718067 | Lighthouse Lab in Milton Keynes | Wellcome Sanger Institute for the COVID-19 Genomics UK (COG-UK) Consortium | The Lighthouse Lab in Milton Keynes and Alex Alderton, Roberto Amato, Jeffrey Barrett, Sonia Goncalves, Ewan Harrison, David K. Jackson, Ian Johnston, Dominic Kwiatkowski, Cordelia Langford, John Sillitoe on behalf of the Wellcome Sanger Institute COVID-19 Surveillance Team |
| EPI_ISL_2718068 | Lighthouse Lab in Alderley Park | Wellcome Sanger Institute for the COVID-19 Genomics UK (COG-UK) Consortium | Jacquelyn Wynn, Mairead Hyland, The Lighthouse Lab in Alderley Park and Alex Alderton, Roberto Amato, Jeffrey Barrett, Sonia Goncalves, Ewan Harrison, David K. Jackson, Ian Johnston, Dominic Kwiatkowski, Cordelia Langford, John Sillitoe on behalf of the Wellcome Sanger Institute COVID-19 Surveillance Team |
| EPI_ISL_2718069, EPI_ISL_2718070 | Lighthouse Lab in Milton Keynes | Wellcome Sanger Institute for the COVID-19 Genomics UK (COG-UK) Consortium | The Lighthouse Lab in Milton Keynes and Alex Alderton, Roberto Amato, Jeffrey Barrett, Sonia Goncalves, Ewan Harrison, David K. Jackson, Ian Johnston, Dominic Kwiatkowski, Cordelia Langford, John Sillitoe on behalf of the Wellcome Sanger Institute COVID-19 Surveillance Team |
| EPI_ISL_2718071, EPI_ISL_2718072, EPI_ISL_2718073 | Lighthouse Lab in Alderley Park | Wellcome Sanger Institute for the COVID-19 Genomics UK (COG-UK) Consortium | Jacquelyn Wynn, Mairead Hyland, The Lighthouse Lab in Alderley Park and Alex Alderton, Roberto Amato, Jeffrey Barrett, Sonia Goncalves, Ewan Harrison, David K. Jackson, Ian Johnston, Dominic Kwiatkowski, Cordelia Langford, John Sillitoe on behalf of the Wellcome Sanger Institute COVID-19 Surveillance Team |
| EPI_ISL_2718074, EPI_ISL_2718075 | Lighthouse Lab in Milton Keynes | Wellcome Sanger Institute for the COVID-19 Genomics UK (COG-UK) Consortium | The Lighthouse Lab in Milton Keynes and Alex Alderton, Roberto Amato, Jeffrey Barrett, Sonia Goncalves, Ewan Harrison, David K. Jackson, Ian Johnston, Dominic Kwiatkowski, Cordelia Langford, John Sillitoe on behalf of the Wellcome Sanger Institute COVID-19 Surveillance Team |
| EPI_ISL_2718076 | Lighthouse Lab in Alderley Park | Wellcome Sanger Institute for the COVID-19 Genomics UK (COG-UK) Consortium | Jacquelyn Wynn, Mairead Hyland, The Lighthouse Lab in Alderley Park and Alex Alderton, Roberto Amato, Jeffrey Barrett, Sonia Goncalves, Ewan Harrison, David K. Jackson, Ian Johnston, Dominic Kwiatkowski, Cordelia Langford, John Sillitoe on behalf of the Wellcome Sanger Institute COVID-19 Surveillance Team |
| EPI_ISL_2718077, EPI_ISL_2718078, EPI_ISL_2718079 | Lighthouse Lab in Milton Keynes | Wellcome Sanger Institute for the COVID-19 Genomics UK (COG-UK) Consortium | The Lighthouse Lab in Milton Keynes and Alex Alderton, Roberto Amato, Jeffrey Barrett, Sonia Goncalves, Ewan Harrison, David K. Jackson, Ian Johnston, Dominic Kwiatkowski, Cordelia Langford, John Sillitoe on behalf of the Wellcome Sanger Institute COVID-19 Surveillance Team |
| EPI_ISL_2718080, EPI_ISL_2718081 | Lighthouse Lab in Alderley Park | Wellcome Sanger Institute for the COVID-19 Genomics UK (COG-UK) Consortium | Jacquelyn Wynn, Mairead Hyland, The Lighthouse Lab in Alderley Park and Alex Alderton, Roberto Amato, Jeffrey Barrett, Sonia Goncalves, Ewan Harrison, David K. Jackson, Ian Johnston, Dominic Kwiatkowski, Cordelia Langford, John Sillitoe on behalf of the Wellcome Sanger Institute COVID-19 Surveillance Team |
| EPI_ISL_2718082 | Lighthouse Lab in Milton Keynes | Wellcome Sanger Institute for the COVID-19 Genomics UK (COG-UK) Consortium | The Lighthouse Lab in Milton Keynes and Alex Alderton, Roberto Amato, Jeffrey Barrett, Sonia Goncalves, Ewan Harrison, David K. Jackson, Ian Johnston, Dominic Kwiatkowski, Cordelia Langford, John Sillitoe on behalf of the Wellcome Sanger Institute COVID-19 Surveillance Team |
| EPI_ISL_2718083, EPI_ISL_2718084, EPI_ISL_2718085, EPI_ISL_2718086, EPI_ISL_2718087, EPI_ISL_2718088 | Lighthouse Lab in Alderley Park | Wellcome Sanger Institute for the COVID-19 Genomics UK (COG-UK) Consortium | Jacquelyn Wynn, Mairead Hyland, The Lighthouse Lab in Alderley Park and Alex Alderton, Roberto Amato, Jeffrey Barrett, Sonia Goncalves, Ewan Harrison, David K. Jackson, Ian Johnston, Dominic Kwiatkowski, Cordelia Langford, John Sillitoe on behalf of the Wellcome Sanger Institute COVID-19 Surveillance Team |
| EPI_ISL_2718089 | Lighthouse Lab in Milton Keynes | Wellcome Sanger Institute for the COVID-19 Genomics UK (COG-UK) Consortium | The Lighthouse Lab in Milton Keynes and Alex Alderton, Roberto Amato, Jeffrey Barrett, Sonia Goncalves, Ewan Harrison, David K. Jackson, Ian Johnston, Dominic Kwiatkowski, Cordelia Langford, John Sillitoe on behalf of the Wellcome Sanger Institute COVID-19 Surveillance Team |
| EPI_ISL_2718090 | Lighthouse Lab in Alderley Park | Wellcome Sanger Institute for the COVID-19 Genomics UK (COG-UK) Consortium | Jacquelyn Wynn, Mairead Hyland, The Lighthouse Lab in Alderley Park and Alex Alderton, Roberto Amato, Jeffrey Barrett, Sonia Goncalves, Ewan Harrison, David K. Jackson, Ian Johnston, Dominic Kwiatkowski, Cordelia Langford, John Sillitoe on behalf of the Wellcome Sanger Institute COVID-19 Surveillance Team |
| EPI_ISL_2718091, EPI_ISL_2718092, EPI_ISL_2718093, EPI_ISL_2718094 | Lighthouse Lab in Milton Keynes | Wellcome Sanger Institute for the COVID-19 Genomics UK (COG-UK) Consortium | The Lighthouse Lab in Milton Keynes and Alex Alderton, Roberto Amato, Jeffrey Barrett, Sonia Goncalves, Ewan Harrison, David K. Jackson, Ian Johnston, Dominic Kwiatkowski, Cordelia Langford, John Sillitoe on behalf of the Wellcome Sanger Institute COVID-19 Surveillance Team |
| EPI_ISL_2718095, EPI_ISL_2718096 | Lighthouse Lab in Alderley Park | Wellcome Sanger Institute for the COVID-19 Genomics UK (COG-UK) Consortium | Jacquelyn Wynn, Mairead Hyland, The Lighthouse Lab in Alderley Park and Alex Alderton, Roberto Amato, Jeffrey Barrett, Sonia Goncalves, Ewan Harrison, David K. Jackson, Ian Johnston, Dominic Kwiatkowski, Cordelia Langford, John Sillitoe on behalf of the Wellcome Sanger Institute COVID-19 Surveillance Team |
| EPI_ISL_2718097 | Lighthouse Lab in Milton Keynes | Wellcome Sanger Institute for the COVID-19 Genomics UK (COG-UK) Consortium | The Lighthouse Lab in Milton Keynes and Alex Alderton, Roberto Amato, Jeffrey Barrett, Sonia Goncalves, Ewan Harrison, David K. Jackson, Ian Johnston, Dominic Kwiatkowski, Cordelia Langford, John Sillitoe on behalf of the Wellcome Sanger Institute COVID-19 Surveillance Team |
| EPI_ISL_2718098 | Lighthouse Lab in Alderley Park | Wellcome Sanger Institute for the COVID-19 Genomics UK (COG-UK) Consortium | Jacquelyn Wynn, Mairead Hyland, The Lighthouse Lab in Alderley Park and Alex Alderton, Roberto Amato, Jeffrey Barrett, Sonia Goncalves, Ewan Harrison, David K. Jackson, Ian Johnston, Dominic Kwiatkowski, Cordelia Langford, John Sillitoe on behalf of the Wellcome Sanger Institute COVID-19 Surveillance Team |
| EPI_ISL_2718099, EPI_ISL_2718100 | Lighthouse Lab in Milton Keynes | Wellcome Sanger Institute for the COVID-19 Genomics UK (COG-UK) Consortium | The Lighthouse Lab in Milton Keynes and Alex Alderton, Roberto Amato, Jeffrey Barrett, Sonia Goncalves, Ewan Harrison, David K. Jackson, Ian Johnston, Dominic Kwiatkowski, Cordelia Langford, John Sillitoe on behalf of the Wellcome Sanger Institute COVID-19 Surveillance Team |
| EPI_ISL_2718101 | Lighthouse Lab in Alderley Park | Wellcome Sanger Institute for the COVID-19 Genomics UK (COG-UK) Consortium | Jacquelyn Wynn, Mairead Hyland, The Lighthouse Lab in Alderley Park and Alex Alderton, Roberto Amato, Jeffrey Barrett, Sonia Goncalves, Ewan Harrison, David K. Jackson, Ian Johnston, Dominic Kwiatkowski, Cordelia Langford, John Sillitoe on behalf of the Wellcome Sanger Institute COVID-19 Surveillance Team |
| EPI_ISL_2718102, EPI_ISL_2718103 | Lighthouse Lab in Milton Keynes | Wellcome Sanger Institute for the COVID-19 Genomics UK (COG-UK) Consortium | The Lighthouse Lab in Milton Keynes and Alex Alderton, Roberto Amato, Jeffrey Barrett, Sonia Goncalves, Ewan Harrison, David K. Jackson, Ian Johnston, Dominic Kwiatkowski, Cordelia Langford, John Sillitoe on behalf of the Wellcome Sanger Institute COVID-19 Surveillance Team |
| EPI_ISL_2718104 | Lighthouse Lab in Alderley Park | Wellcome Sanger Institute for the COVID-19 Genomics UK (COG-UK) Consortium | Jacquelyn Wynn, Mairead Hyland, The Lighthouse Lab in Alderley Park and Alex Alderton, Roberto Amato, Jeffrey Barrett, Sonia Goncalves, Ewan Harrison, David K. Jackson, Ian Johnston, Dominic Kwiatkowski, Cordelia Langford, John Sillitoe on behalf of the Wellcome Sanger Institute COVID-19 Surveillance Team |
| EPI_ISL_2718105 | Lighthouse Lab in Milton Keynes | Wellcome Sanger Institute for the COVID-19 Genomics UK (COG-UK) Consortium | The Lighthouse Lab in Milton Keynes and Alex Alderton, Roberto Amato, Jeffrey Barrett, Sonia Goncalves, Ewan Harrison, David K. Jackson, Ian Johnston, Dominic Kwiatkowski, Cordelia Langford, John Sillitoe on behalf of the Wellcome Sanger Institute COVID-19 Surveillance Team |
| EPI_ISL_2718106 | Lighthouse Lab in Alderley Park | Wellcome Sanger Institute for the COVID-19 Genomics UK (COG-UK) Consortium | Jacquelyn Wynn, Mairead Hyland, The Lighthouse Lab in Alderley Park and Alex Alderton, Roberto Amato, Jeffrey Barrett, Sonia Goncalves, Ewan Harrison, David K. Jackson, Ian Johnston, Dominic Kwiatkowski, Cordelia Langford, John Sillitoe on behalf of the Wellcome Sanger Institute COVID-19 Surveillance Team |
| EPI_ISL_2718107, EPI_ISL_2718108, EPI_ISL_2718109, EPI_ISL_2718110, EPI_ISL_2718111 | Lighthouse Lab in Milton Keynes | Wellcome Sanger Institute for the COVID-19 Genomics UK (COG-UK) Consortium | The Lighthouse Lab in Milton Keynes and Alex Alderton, Roberto Amato, Jeffrey Barrett, Sonia Goncalves, Ewan Harrison, David K. Jackson, Ian Johnston, Dominic Kwiatkowski, Cordelia Langford, John Sillitoe on behalf of the Wellcome Sanger Institute COVID-19 Surveillance Team |

| | | | |
|---|---|---|---|
| EPI_ISL_2718112, EPI_ISL_2718113 | Lighthouse Lab in Alderley Park | Wellcome Sanger Institute for the COVID-19 Genomics UK (COG-UK) Consortium | Jacquelyn Wynn, Mairead Hyland, The Lighthouse Lab in Alderley Park and Alex Alderton, Roberto Amato, Jeffrey Barrett, Sonia Goncalves, Ewan Harrison, David K. Jackson, Ian Johnston, Dominic Kwiatkowski, Cordelia Langford, John Sillitoe on behalf of the Wellcome Sanger Institute COVID-19 Surveillance Team |
| EPI_ISL_2718114, EPI_ISL_2718115, EPI_ISL_2718116 | Lighthouse Lab in Milton Keynes | Wellcome Sanger Institute for the COVID-19 Genomics UK (COG-UK) Consortium | The Lighthouse Lab in Milton Keynes and Alex Alderton, Roberto Amato, Jeffrey Barrett, Sonia Goncalves, Ewan Harrison, David K. Jackson, Ian Johnston, Dominic Kwiatkowski, Cordelia Langford, John Sillitoe on behalf of the Wellcome Sanger Institute COVID-19 Surveillance Team |
| EPI_ISL_2718117 | Lighthouse Lab in Alderley Park | Wellcome Sanger Institute for the COVID-19 Genomics UK (COG-UK) Consortium | Jacquelyn Wynn, Mairead Hyland, The Lighthouse Lab in Alderley Park and Alex Alderton, Roberto Amato, Jeffrey Barrett, Sonia Goncalves, Ewan Harrison, David K. Jackson, Ian Johnston, Dominic Kwiatkowski, Cordelia Langford, John Sillitoe on behalf of the Wellcome Sanger Institute COVID-19 Surveillance Team |
| EPI_ISL_2718118, EPI_ISL_2718119 | Lighthouse Lab in Milton Keynes | Wellcome Sanger Institute for the COVID-19 Genomics UK (COG-UK) Consortium | The Lighthouse Lab in Milton Keynes and Alex Alderton, Roberto Amato, Jeffrey Barrett, Sonia Goncalves, Ewan Harrison, David K. Jackson, Ian Johnston, Dominic Kwiatkowski, Cordelia Langford, John Sillitoe on behalf of the Wellcome Sanger Institute COVID-19 Surveillance Team |
| EPI_ISL_2718120, EPI_ISL_2718121, EPI_ISL_2718122 | Lighthouse Lab in Alderley Park | Wellcome Sanger Institute for the COVID-19 Genomics UK (COG-UK) Consortium | Jacquelyn Wynn, Mairead Hyland, The Lighthouse Lab in Alderley Park and Alex Alderton, Roberto Amato, Jeffrey Barrett, Sonia Goncalves, Ewan Harrison, David K. Jackson, Ian Johnston, Dominic Kwiatkowski, Cordelia Langford, John Sillitoe on behalf of the Wellcome Sanger Institute COVID-19 Surveillance Team |
| EPI_ISL_2718123 | Lighthouse Lab in Milton Keynes | Wellcome Sanger Institute for the COVID-19 Genomics UK (COG-UK) Consortium | The Lighthouse Lab in Milton Keynes and Alex Alderton, Roberto Amato, Jeffrey Barrett, Sonia Goncalves, Ewan Harrison, David K. Jackson, Ian Johnston, Dominic Kwiatkowski, Cordelia Langford, John Sillitoe on behalf of the Wellcome Sanger Institute COVID-19 Surveillance Team |
| EPI_ISL_2718124, EPI_ISL_2718125 | Lighthouse Lab in Alderley Park | Wellcome Sanger Institute for the COVID-19 Genomics UK (COG-UK) Consortium | Jacquelyn Wynn, Mairead Hyland, The Lighthouse Lab in Alderley Park and Alex Alderton, Roberto Amato, Jeffrey Barrett, Sonia Goncalves, Ewan Harrison, David K. Jackson, Ian Johnston, Dominic Kwiatkowski, Cordelia Langford, John Sillitoe on behalf of the Wellcome Sanger Institute COVID-19 Surveillance Team |
| EPI_ISL_2718126 | Lighthouse Lab in Milton Keynes | Wellcome Sanger Institute for the COVID-19 Genomics UK (COG-UK) Consortium | The Lighthouse Lab in Milton Keynes and Alex Alderton, Roberto Amato, Jeffrey Barrett, Sonia Goncalves, Ewan Harrison, David K. Jackson, Ian Johnston, Dominic Kwiatkowski, Cordelia Langford, John Sillitoe on behalf of the Wellcome Sanger Institute COVID-19 Surveillance Team |
| EPI_ISL_2718127, EPI_ISL_2718128 | Lighthouse Lab in Alderley Park | Wellcome Sanger Institute for the COVID-19 Genomics UK (COG-UK) Consortium | Jacquelyn Wynn, Mairead Hyland, The Lighthouse Lab in Alderley Park and Alex Alderton, Roberto Amato, Jeffrey Barrett, Sonia Goncalves, Ewan Harrison, David K. Jackson, Ian Johnston, Dominic Kwiatkowski, Cordelia Langford, John Sillitoe on behalf of the Wellcome Sanger Institute COVID-19 Surveillance Team |
| EPI_ISL_2718129 | Lighthouse Lab in Milton Keynes | Wellcome Sanger Institute for the COVID-19 Genomics UK (COG-UK) Consortium | The Lighthouse Lab in Milton Keynes and Alex Alderton, Roberto Amato, Jeffrey Barrett, Sonia Goncalves, Ewan Harrison, David K. Jackson, Ian Johnston, Dominic Kwiatkowski, Cordelia Langford, John Sillitoe on behalf of the Wellcome Sanger Institute COVID-19 Surveillance Team |
| EPI_ISL_2718130 | Lighthouse Lab in Alderley Park | Wellcome Sanger Institute for the COVID-19 Genomics UK (COG-UK) Consortium | Jacquelyn Wynn, Mairead Hyland, The Lighthouse Lab in Alderley Park and Alex Alderton, Roberto Amato, Jeffrey Barrett, Sonia Goncalves, Ewan Harrison, David K. Jackson, Ian Johnston, Dominic Kwiatkowski, Cordelia Langford, John Sillitoe on behalf of the Wellcome Sanger Institute COVID-19 Surveillance Team |
| EPI_ISL_2718131 | Lighthouse Lab in Milton Keynes | Wellcome Sanger Institute for the COVID-19 Genomics UK (COG-UK) Consortium | The Lighthouse Lab in Milton Keynes and Alex Alderton, Roberto Amato, Jeffrey Barrett, Sonia Goncalves, Ewan Harrison, David K. Jackson, Ian Johnston, Dominic Kwiatkowski, Cordelia Langford, John Sillitoe on behalf of the Wellcome Sanger Institute COVID-19 Surveillance Team |
| EPI_ISL_2718132, EPI_ISL_2718133 | Lighthouse Lab in Alderley Park | Wellcome Sanger Institute for the COVID-19 Genomics UK (COG-UK) Consortium | Jacquelyn Wynn, Mairead Hyland, The Lighthouse Lab in Alderley Park and Alex Alderton, Roberto Amato, Jeffrey Barrett, Sonia Goncalves, Ewan Harrison, David K. Jackson, Ian Johnston, Dominic Kwiatkowski, Cordelia Langford, John Sillitoe on behalf of the Wellcome Sanger Institute COVID-19 Surveillance Team |
| EPI_ISL_2718134, EPI_ISL_2718135 | Lighthouse Lab in Milton Keynes | Wellcome Sanger Institute for the COVID-19 Genomics UK (COG-UK) Consortium | The Lighthouse Lab in Milton Keynes and Alex Alderton, Roberto Amato, Jeffrey Barrett, Sonia Goncalves, Ewan Harrison, David K. Jackson, Ian Johnston, Dominic Kwiatkowski, Cordelia Langford, John Sillitoe on behalf of the Wellcome Sanger Institute COVID-19 Surveillance Team |
| EPI_ISL_2718136, EPI_ISL_2718137 | Lighthouse Lab in Alderley Park | Wellcome Sanger Institute for the COVID-19 Genomics UK (COG-UK) Consortium | Jacquelyn Wynn, Mairead Hyland, The Lighthouse Lab in Alderley Park and Alex Alderton, Roberto Amato, Jeffrey Barrett, Sonia Goncalves, Ewan Harrison, David K. Jackson, Ian Johnston, Dominic Kwiatkowski, Cordelia Langford, John Sillitoe on behalf of the Wellcome Sanger Institute COVID-19 Surveillance Team |
| EPI_ISL_2718138, EPI_ISL_2718139 | Lighthouse Lab in Milton Keynes | Wellcome Sanger Institute for the COVID-19 Genomics UK (COG-UK) Consortium | The Lighthouse Lab in Milton Keynes and Alex Alderton, Roberto Amato, Jeffrey Barrett, Sonia Goncalves, Ewan Harrison, David K. Jackson, Ian Johnston, Dominic Kwiatkowski, Cordelia Langford, John Sillitoe on behalf of the Wellcome Sanger Institute COVID-19 Surveillance Team |
| EPI_ISL_2718140, EPI_ISL_2718141 | Lighthouse Lab in Alderley Park | Wellcome Sanger Institute for the COVID-19 Genomics UK (COG-UK) Consortium | Jacquelyn Wynn, Mairead Hyland, The Lighthouse Lab in Alderley Park and Alex Alderton, Roberto Amato, Jeffrey Barrett, Sonia Goncalves, Ewan Harrison, David K. Jackson, Ian Johnston, Dominic Kwiatkowski, Cordelia Langford, John Sillitoe on behalf of the Wellcome Sanger Institute COVID-19 Surveillance Team |
| EPI_ISL_2718142, EPI_ISL_2718143 | Lighthouse Lab in Milton Keynes | Wellcome Sanger Institute for the COVID-19 Genomics UK (COG-UK) Consortium | The Lighthouse Lab in Milton Keynes and Alex Alderton, Roberto Amato, Jeffrey Barrett, Sonia Goncalves, Ewan Harrison, David K. Jackson, Ian Johnston, Dominic Kwiatkowski, Cordelia Langford, John Sillitoe on behalf of the Wellcome Sanger Institute COVID-19 Surveillance Team |
| EPI_ISL_2718144, EPI_ISL_2718145 | Lighthouse Lab in Alderley Park | Wellcome Sanger Institute for the COVID-19 Genomics UK (COG-UK) Consortium | Jacquelyn Wynn, Mairead Hyland, The Lighthouse Lab in Alderley Park and Alex Alderton, Roberto Amato, Jeffrey Barrett, Sonia Goncalves, Ewan Harrison, David K. Jackson, Ian Johnston, Dominic Kwiatkowski, Cordelia Langford, John Sillitoe on behalf of the Wellcome Sanger Institute COVID-19 Surveillance Team |
| EPI_ISL_2718146, EPI_ISL_2718147, EPI_ISL_2718148, EPI_ISL_2718149 | Lighthouse Lab in Milton Keynes | Wellcome Sanger Institute for the COVID-19 Genomics UK (COG-UK) Consortium | The Lighthouse Lab in Milton Keynes and Alex Alderton, Roberto Amato, Jeffrey Barrett, Sonia Goncalves, Ewan Harrison, David K. Jackson, Ian Johnston, Dominic Kwiatkowski, Cordelia Langford, John Sillitoe on behalf of the Wellcome Sanger Institute COVID-19 Surveillance Team |
| EPI_ISL_2718150, EPI_ISL_2718151 | Lighthouse Lab in Alderley Park | Wellcome Sanger Institute for the COVID-19 Genomics UK (COG-UK) Consortium | Jacquelyn Wynn, Mairead Hyland, The Lighthouse Lab in Alderley Park and Alex Alderton, Roberto Amato, Jeffrey Barrett, Sonia Goncalves, Ewan Harrison, David K. Jackson, Ian Johnston, Dominic Kwiatkowski, Cordelia Langford, John Sillitoe on behalf of the Wellcome Sanger Institute COVID-19 Surveillance Team |
| EPI_ISL_2718152 | Lighthouse Lab in Milton Keynes | Wellcome Sanger Institute for the COVID-19 Genomics UK (COG-UK) Consortium | The Lighthouse Lab in Milton Keynes and Alex Alderton, Roberto Amato, Jeffrey Barrett, Sonia Goncalves, Ewan Harrison, David K. Jackson, Ian Johnston, Dominic Kwiatkowski, Cordelia Langford, John Sillitoe on behalf of the Wellcome Sanger Institute COVID-19 Surveillance Team |
| EPI_ISL_2718153, EPI_ISL_2718154, EPI_ISL_2718155 | Lighthouse Lab in Alderley Park | Wellcome Sanger Institute for the COVID-19 Genomics UK (COG-UK) Consortium | Jacquelyn Wynn, Mairead Hyland, The Lighthouse Lab in Alderley Park and Alex Alderton, Roberto Amato, Jeffrey Barrett, Sonia Goncalves, Ewan Harrison, David K. Jackson, Ian Johnston, Dominic Kwiatkowski, Cordelia Langford, John Sillitoe on behalf of the Wellcome Sanger Institute COVID-19 Surveillance Team |
| EPI_ISL_2718156 | Lighthouse Lab in Milton Keynes | Wellcome Sanger Institute for the COVID-19 Genomics UK (COG-UK) Consortium | The Lighthouse Lab in Milton Keynes and Alex Alderton, Roberto Amato, Jeffrey Barrett, Sonia Goncalves, Ewan Harrison, David K. Jackson, Ian Johnston, Dominic Kwiatkowski, Cordelia Langford, John Sillitoe on behalf of the Wellcome Sanger Institute COVID-19 Surveillance Team |
| EPI_ISL_2718157 | Lighthouse Lab in Alderley Park | Wellcome Sanger Institute for the COVID-19 Genomics UK (COG-UK) Consortium | Jacquelyn Wynn, Mairead Hyland, The Lighthouse Lab in Alderley Park and Alex Alderton, Roberto Amato, Jeffrey Barrett, Sonia Goncalves, Ewan Harrison, David K. Jackson, Ian Johnston, Dominic Kwiatkowski, Cordelia Langford, John Sillitoe on behalf of the Wellcome Sanger Institute COVID-19 Surveillance Team |
| EPI_ISL_2718158 | Lighthouse Lab in Milton Keynes | Wellcome Sanger Institute for the COVID-19 Genomics UK (COG-UK) Consortium | The Lighthouse Lab in Milton Keynes and Alex Alderton, Roberto Amato, Jeffrey Barrett, Sonia Goncalves, Ewan Harrison, David K. Jackson, Ian Johnston, Dominic Kwiatkowski, Cordelia Langford, John Sillitoe on behalf of the Wellcome Sanger Institute COVID-19 Surveillance Team |
| EPI_ISL_2718159 | Lighthouse Lab in Alderley Park | Wellcome Sanger Institute for the COVID-19 Genomics UK (COG-UK) Consortium | Jacquelyn Wynn, Mairead Hyland, The Lighthouse Lab in Alderley Park and Alex Alderton, Roberto Amato, Jeffrey Barrett, Sonia Goncalves, Ewan Harrison, David K. Jackson, Ian Johnston, Dominic Kwiatkowski, Cordelia Langford, John Sillitoe on behalf of the Wellcome Sanger Institute COVID-19 Surveillance Team |
| EPI_ISL_2718160 | Lighthouse Lab in Milton Keynes | Wellcome Sanger Institute for the COVID-19 Genomics UK (COG-UK) Consortium | The Lighthouse Lab in Milton Keynes and Alex Alderton, Roberto Amato, Jeffrey Barrett, Sonia Goncalves, Ewan Harrison, David K. Jackson, Ian Johnston, Dominic Kwiatkowski, Cordelia Langford, John Sillitoe on behalf of the Wellcome Sanger Institute COVID-19 Surveillance Team |
| EPI_ISL_2718161 | Lighthouse Lab in Alderley Park | Wellcome Sanger Institute for the COVID-19 Genomics UK (COG-UK) Consortium | Jacquelyn Wynn, Mairead Hyland, The Lighthouse Lab in Alderley Park and Alex Alderton, Roberto Amato, Jeffrey Barrett, Sonia Goncalves, Ewan Harrison, David K. Jackson, Ian Johnston, Dominic Kwiatkowski, Cordelia Langford, John Sillitoe on behalf of the Wellcome Sanger Institute COVID-19 Surveillance Team |
| EPI_ISL_2718162, EPI_ISL_2718163, EPI_ISL_2718164, EPI_ISL_2718165, EPI_ISL_2718166 | Lighthouse Lab in Milton Keynes | Wellcome Sanger Institute for the COVID-19 Genomics UK (COG-UK) Consortium | The Lighthouse Lab in Milton Keynes and Alex Alderton, Roberto Amato, Jeffrey Barrett, Sonia Goncalves, Ewan Harrison, David K. Jackson, Ian Johnston, Dominic Kwiatkowski, Cordelia Langford, John Sillitoe on behalf of the Wellcome Sanger Institute COVID-19 Surveillance Team |
| EPI_ISL_2718167 | Lighthouse Lab in Alderley Park | Wellcome Sanger Institute for the COVID-19 Genomics UK (COG-UK) Consortium | Jacquelyn Wynn, Mairead Hyland, The Lighthouse Lab in Alderley Park and Alex Alderton, Roberto Amato, Jeffrey Barrett, Sonia Goncalves, Ewan Harrison, David K. Jackson, Ian Johnston, Dominic Kwiatkowski, Cordelia Langford, John Sillitoe on behalf of the Wellcome Sanger Institute COVID-19 Surveillance Team |
| EPI_ISL_2718168, EPI_ISL_2718169 | Lighthouse Lab in Milton Keynes | Wellcome Sanger Institute for the COVID-19 Genomics UK (COG-UK) Consortium | The Lighthouse Lab in Milton Keynes and Alex Alderton, Roberto Amato, Jeffrey Barrett, Sonia Goncalves, Ewan Harrison, David K. Jackson, Ian Johnston, Dominic Kwiatkowski, Cordelia Langford, John Sillitoe on behalf of the Wellcome Sanger Institute COVID-19 Surveillance Team |
| EPI_ISL_2718170, EPI_ISL_2718171, EPI_ISL_2718172, EPI_ISL_2718173, EPI_ISL_2718174, EPI_ISL_2718175, EPI_ISL_2718176, EPI_ISL_2718177, EPI_ISL_2718178, EPI_ISL_2718179, EPI_ISL_2718180, EPI_ISL_2718181, EPI_ISL_2718182, EPI_ISL_2718183, EPI_ISL_2718184, EPI_ISL_2718185, EPI_ISL_2718186, EPI_ISL_2718187, EPI_ISL_2718188, EPI_ISL_2718189, EPI_ISL_2718190, EPI_ISL_2718191, EPI_ISL_2718192, EPI_ISL_2718193, EPI_ISL_2718195, EPI_ISL_2718196, EPI_ISL_2718197, EPI_ISL_2718198, EPI_ISL_2718199, EPI_ISL_2718200, EPI_ISL_2718201, EPI_ISL_2718202, EPI_ISL_2718203, EPI_ISL_2718204, EPI_ISL_2718205, EPI_ISL_2718206, EPI_ISL_2718208, EPI_ISL_2718209, EPI_ISL_2718210, EPI_ISL_2718211, EPI_ISL_2718212, EPI_ISL_2718213, EPI_ISL_2718214, EPI_ISL_2718215, EPI_ISL_2718216, EPI_ISL_2718217, EPI_ISL_2718218, EPI_ISL_2718219, EPI_ISL_2718220, EPI_ISL_2718221, EPI_ISL_2718222, EPI_ISL_2718223, EPI_ISL_2718225, EPI_ISL_2718226, EPI_ISL_2718227, EPI_ISL_2718228, EPI_ISL_2718229, EPI_ISL_2718230, EPI_ISL_2718231, EPI_ISL_2718232, EPI_ISL_2718233, EPI_ISL_2718234, EPI_ISL_2718235, EPI_ISL_2718236, EPI_ISL_2718237, EPI_ISL_2718238, EPI_ISL_2718239, EPI_ISL_2718240, EPI_ISL_2718241, EPI_ISL_2718243, EPI_ISL_2718244, EPI_ISL_2718246, EPI_ISL_2718247, EPI_ISL_2718248, EPI_ISL_2718249, EPI_ISL_2718250, EPI_ISL_2718251, EPI_ISL_2718252, EPI_ISL_2718253, EPI_ISL_2718254, EPI_ISL_2718255, EPI_ISL_2718256, EPI_ISL_2718257, EPI_ISL_2718258, EPI_ISL_2718259, EPI_ISL_2718260, EPI_ISL_2718261, EPI_ISL_2718263, EPI_ISL_2718265, EPI_ISL_2718266, EPI_ISL_2718267, EPI_ISL_2718268, EPI_ISL_2718269, EPI_ISL_2718270, EPI_ISL_2718271, EPI_ISL_2718272, EPI_ISL_2718273, EPI_ISL_2718274, EPI_ISL_2718275, EPI_ISL_2718276, EPI_ISL_2718277, EPI_ISL_2718278, EPI_ISL_2718279, EPI_ISL_2718280 | | | |
| see above | Lighthouse Lab in Glasgow | Wellcome Sanger Institute for the COVID-19 Genomics UK (COG-UK) Consortium | Harper VanSteenhouse, Yumi Kasai, David Gray, Carol Clugston, Anna Dominiczak and Alex Alderton, Roberto Amato, Jeffrey Barrett, Sonia Goncalves, Ewan Harrison, David K. Jackson, Ian Johnston, Dominic Kwiatkowski, Cordelia Langford, John Sillitoe on behalf of the Wellcome Sanger Institute COVID-19 Surveillance Team |
| EPI_ISL_2718281, EPI_ISL_2718282, EPI_ISL_2718283, EPI_ISL_2718284 | Lighthouse Lab in Alderley Park | Wellcome Sanger Institute for the COVID-19 Genomics UK (COG-UK) Consortium | Jacquelyn Wynn, Mairead Hyland, The Lighthouse Lab in Alderley Park and Alex Alderton, Roberto Amato, Jeffrey Barrett, Sonia Goncalves, Ewan Harrison, David K. Jackson, Ian Johnston, Dominic Kwiatkowski, Cordelia Langford, John Sillitoe on behalf of the Wellcome Sanger Institute COVID-19 Surveillance Team |
| EPI_ISL_2718285, EPI_ISL_2718286, EPI_ISL_2718287 | Lighthouse Laboratory Plymouth | Wellcome Sanger Institute for the COVID-19 Genomics UK (COG-UK) Consortium | Lighthouse Laboratory Plymouth and Alex Alderton, Roberto Amato, Jeffrey Barrett, Sonia Goncalves, Ewan Harrison, David K. Jackson, Ian Johnston, Dominic Kwiatkowski, Cordelia Langford, John Sillitoe on behalf of the Wellcome Sanger Institute COVID-19 Surveillance Team |
| EPI_ISL_2718288, EPI_ISL_2718289, EPI_ISL_2718290, EPI_ISL_2718291, EPI_ISL_2718292, EPI_ISL_2718293 | Lighthouse Lab in Alderley Park | Wellcome Sanger Institute for the COVID-19 Genomics UK (COG-UK) Consortium | Jacquelyn Wynn, Mairead Hyland, The Lighthouse Lab in Alderley Park and Alex Alderton, Roberto Amato, Jeffrey Barrett, Sonia Goncalves, Ewan Harrison, David K. Jackson, Ian Johnston, Dominic Kwiatkowski, Cordelia Langford, John Sillitoe on behalf of the Wellcome Sanger Institute COVID-19 Surveillance Team |
| EPI_ISL_2718294 | Lighthouse Lab in Milton Keynes | Wellcome Sanger Institute for the COVID-19 Genomics UK (COG-UK) | The Lighthouse Lab in Milton Keynes and Alex Alderton, Roberto Amato, Jeffrey Barrett, Sonia Goncalves, Ewan Harrison, David K. Jackson, Ian Johnston, Dominic Kwiatkowski, Cordelia Langford, John Sillitoe on behalf of the Wellcome Sanger |

| | | | |
|---|---|---|---|
| | | Consortium | Institute COVID-19 Surveillance Team |
| EPI_ISL_2718295 | Lighthouse Lab in Glasgow | Wellcome Sanger Institute for the COVID-19 Genomics UK (COG-UK) Consortium | Harper VanSteenhouse, Yumi Kasai, David Gray, Carol Clugston, Anna Dominiczak and Alex Alderton, Roberto Amato, Jeffrey Barrett, Sonia Goncalves, Ewan Harrison, David K. Jackson, Ian Johnston, Dominic Kwiatkowski, Cordelia Langford, John Sillitoe on behalf of the Wellcome Sanger Institute COVID-19 Surveillance Team |
| EPI_ISL_2718296 | Lighthouse Lab in Alderley Park | Wellcome Sanger Institute for the COVID-19 Genomics UK (COG-UK) Consortium | Jacquelyn Wynn, Mairead Hyland, The Lighthouse Lab in Alderley Park and Alex Alderton, Roberto Amato, Jeffrey Barrett, Sonia Goncalves, Ewan Harrison, David K. Jackson, Ian Johnston, Dominic Kwiatkowski, Cordelia Langford, John Sillitoe on behalf of the Wellcome Sanger Institute COVID-19 Surveillance Team |
| EPI_ISL_2718297 | Lighthouse Lab in Glasgow | Wellcome Sanger Institute for the COVID-19 Genomics UK (COG-UK) Consortium | Harper VanSteenhouse, Yumi Kasai, David Gray, Carol Clugston, Anna Dominiczak and Alex Alderton, Roberto Amato, Jeffrey Barrett, Sonia Goncalves, Ewan Harrison, David K. Jackson, Ian Johnston, Dominic Kwiatkowski, Cordelia Langford, John Sillitoe on behalf of the Wellcome Sanger Institute COVID-19 Surveillance Team |
| EPI_ISL_2718298, EPI_ISL_2718299, EPI_ISL_2718300, EPI_ISL_2718301, EPI_ISL_2718302 | Lighthouse Laboratory Plymouth | Wellcome Sanger Institute for the COVID-19 Genomics UK (COG-UK) Consortium | Lighthouse Laboratory Plymouth and Alex Alderton, Roberto Amato, Jeffrey Barrett, Sonia Goncalves, Ewan Harrison, David K. Jackson, Ian Johnston, Dominic Kwiatkowski, Cordelia Langford, John Sillitoe on behalf of the Wellcome Sanger Institute COVID-19 Surveillance Team |
| EPI_ISL_2718303, EPI_ISL_2718304 | Lighthouse Lab in Alderley Park | Wellcome Sanger Institute for the COVID-19 Genomics UK (COG-UK) Consortium | Jacquelyn Wynn, Mairead Hyland, The Lighthouse Lab in Alderley Park and Alex Alderton, Roberto Amato, Jeffrey Barrett, Sonia Goncalves, Ewan Harrison, David K. Jackson, Ian Johnston, Dominic Kwiatkowski, Cordelia Langford, John Sillitoe on behalf of the Wellcome Sanger Institute COVID-19 Surveillance Team |
| EPI_ISL_2718305 | Lighthouse Lab in Glasgow | Wellcome Sanger Institute for the COVID-19 Genomics UK (COG-UK) Consortium | Harper VanSteenhouse, Yumi Kasai, David Gray, Carol Clugston, Anna Dominiczak and Alex Alderton, Roberto Amato, Jeffrey Barrett, Sonia Goncalves, Ewan Harrison, David K. Jackson, Ian Johnston, Dominic Kwiatkowski, Cordelia Langford, John Sillitoe on behalf of the Wellcome Sanger Institute COVID-19 Surveillance Team |
| EPI_ISL_2718306 | Lighthouse Lab in Alderley Park | Wellcome Sanger Institute for the COVID-19 Genomics UK (COG-UK) Consortium | Jacquelyn Wynn, Mairead Hyland, The Lighthouse Lab in Alderley Park and Alex Alderton, Roberto Amato, Jeffrey Barrett, Sonia Goncalves, Ewan Harrison, David K. Jackson, Ian Johnston, Dominic Kwiatkowski, Cordelia Langford, John Sillitoe on behalf of the Wellcome Sanger Institute COVID-19 Surveillance Team |
| EPI_ISL_2718307 | Lighthouse Laboratory Plymouth | Wellcome Sanger Institute for the COVID-19 Genomics UK (COG-UK) Consortium | Lighthouse Laboratory Plymouth and Alex Alderton, Roberto Amato, Jeffrey Barrett, Sonia Goncalves, Ewan Harrison, David K. Jackson, Ian Johnston, Dominic Kwiatkowski, Cordelia Langford, John Sillitoe on behalf of the Wellcome Sanger Institute COVID-19 Surveillance Team |
| EPI_ISL_2718308 | Lighthouse Lab in Alderley Park | Wellcome Sanger Institute for the COVID-19 Genomics UK (COG-UK) Consortium | Jacquelyn Wynn, Mairead Hyland, The Lighthouse Lab in Alderley Park and Alex Alderton, Roberto Amato, Jeffrey Barrett, Sonia Goncalves, Ewan Harrison, David K. Jackson, Ian Johnston, Dominic Kwiatkowski, Cordelia Langford, John Sillitoe on behalf of the Wellcome Sanger Institute COVID-19 Surveillance Team |
| EPI_ISL_2718309 | Lighthouse Lab in Glasgow | Wellcome Sanger Institute for the COVID-19 Genomics UK (COG-UK) Consortium | Harper VanSteenhouse, Yumi Kasai, David Gray, Carol Clugston, Anna Dominiczak and Alex Alderton, Roberto Amato, Jeffrey Barrett, Sonia Goncalves, Ewan Harrison, David K. Jackson, Ian Johnston, Dominic Kwiatkowski, Cordelia Langford, John Sillitoe on behalf of the Wellcome Sanger Institute COVID-19 Surveillance Team |
| EPI_ISL_2718310 | Lighthouse Laboratory Plymouth | Wellcome Sanger Institute for the COVID-19 Genomics UK (COG-UK) Consortium | Lighthouse Laboratory Plymouth and Alex Alderton, Roberto Amato, Jeffrey Barrett, Sonia Goncalves, Ewan Harrison, David K. Jackson, Ian Johnston, Dominic Kwiatkowski, Cordelia Langford, John Sillitoe on behalf of the Wellcome Sanger Institute COVID-19 Surveillance Team |
| EPI_ISL_2718311, EPI_ISL_2718312 | Lighthouse Lab in Alderley Park | Wellcome Sanger Institute for the COVID-19 Genomics UK (COG-UK) Consortium | Jacquelyn Wynn, Mairead Hyland, The Lighthouse Lab in Alderley Park and Alex Alderton, Roberto Amato, Jeffrey Barrett, Sonia Goncalves, Ewan Harrison, David K. Jackson, Ian Johnston, Dominic Kwiatkowski, Cordelia Langford, John Sillitoe on behalf of the Wellcome Sanger Institute COVID-19 Surveillance Team |
| EPI_ISL_2718313 | Lighthouse Laboratory Plymouth | Wellcome Sanger Institute for the COVID-19 Genomics UK (COG-UK) Consortium | Lighthouse Laboratory Plymouth and Alex Alderton, Roberto Amato, Jeffrey Barrett, Sonia Goncalves, Ewan Harrison, David K. Jackson, Ian Johnston, Dominic Kwiatkowski, Cordelia Langford, John Sillitoe on behalf of the Wellcome Sanger Institute COVID-19 Surveillance Team |
| EPI_ISL_2718314 | Lighthouse Lab in Alderley Park | Wellcome Sanger Institute for the COVID-19 Genomics UK (COG-UK) Consortium | Jacquelyn Wynn, Mairead Hyland, The Lighthouse Lab in Alderley Park and Alex Alderton, Roberto Amato, Jeffrey Barrett, Sonia Goncalves, Ewan Harrison, David K. Jackson, Ian Johnston, Dominic Kwiatkowski, Cordelia Langford, John Sillitoe on behalf of the Wellcome Sanger Institute COVID-19 Surveillance Team |
| EPI_ISL_2718315 | Lighthouse Lab in Glasgow | Wellcome Sanger Institute for the COVID-19 Genomics UK (COG-UK) Consortium | Harper VanSteenhouse, Yumi Kasai, David Gray, Carol Clugston, Anna Dominiczak and Alex Alderton, Roberto Amato, Jeffrey Barrett, Sonia Goncalves, Ewan Harrison, David K. Jackson, Ian Johnston, Dominic Kwiatkowski, Cordelia Langford, John Sillitoe on behalf of the Wellcome Sanger Institute COVID-19 Surveillance Team |
| EPI_ISL_2718316 | Lighthouse Lab in Alderley Park | Wellcome Sanger Institute for the COVID-19 Genomics UK (COG-UK) Consortium | Jacquelyn Wynn, Mairead Hyland, The Lighthouse Lab in Alderley Park and Alex Alderton, Roberto Amato, Jeffrey Barrett, Sonia Goncalves, Ewan Harrison, David K. Jackson, Ian Johnston, Dominic Kwiatkowski, Cordelia Langford, John Sillitoe on behalf of the Wellcome Sanger Institute COVID-19 Surveillance Team |
| EPI_ISL_2718317 | Lighthouse Laboratory Plymouth | Wellcome Sanger Institute for the COVID-19 Genomics UK (COG-UK) Consortium | Lighthouse Laboratory Plymouth and Alex Alderton, Roberto Amato, Jeffrey Barrett, Sonia Goncalves, Ewan Harrison, David K. Jackson, Ian Johnston, Dominic Kwiatkowski, Cordelia Langford, John Sillitoe on behalf of the Wellcome Sanger Institute COVID-19 Surveillance Team |
| EPI_ISL_2718318 | Lighthouse Lab in Alderley Park | Wellcome Sanger Institute for the COVID-19 Genomics UK (COG-UK) Consortium | Jacquelyn Wynn, Mairead Hyland, The Lighthouse Lab in Alderley Park and Alex Alderton, Roberto Amato, Jeffrey Barrett, Sonia Goncalves, Ewan Harrison, David K. Jackson, Ian Johnston, Dominic Kwiatkowski, Cordelia Langford, John Sillitoe on behalf of the Wellcome Sanger Institute COVID-19 Surveillance Team |
| EPI_ISL_2718319 | Lighthouse Laboratory Plymouth | Wellcome Sanger Institute for the COVID-19 Genomics UK (COG-UK) Consortium | Lighthouse Laboratory Plymouth and Alex Alderton, Roberto Amato, Jeffrey Barrett, Sonia Goncalves, Ewan Harrison, David K. Jackson, Ian Johnston, Dominic Kwiatkowski, Cordelia Langford, John Sillitoe on behalf of the Wellcome Sanger Institute COVID-19 Surveillance Team |
| EPI_ISL_2718320 | Lighthouse Lab in Alderley Park | Wellcome Sanger Institute for the COVID-19 Genomics UK (COG-UK) Consortium | Jacquelyn Wynn, Mairead Hyland, The Lighthouse Lab in Alderley Park and Alex Alderton, Roberto Amato, Jeffrey Barrett, Sonia Goncalves, Ewan Harrison, David K. Jackson, Ian Johnston, Dominic Kwiatkowski, Cordelia Langford, John Sillitoe on behalf of the Wellcome Sanger Institute COVID-19 Surveillance Team |
| EPI_ISL_2718321 | Lighthouse Laboratory Plymouth | Wellcome Sanger Institute for the COVID-19 Genomics UK (COG-UK) Consortium | Lighthouse Laboratory Plymouth and Alex Alderton, Roberto Amato, Jeffrey Barrett, Sonia Goncalves, Ewan Harrison, David K. Jackson, Ian Johnston, Dominic Kwiatkowski, Cordelia Langford, John Sillitoe on behalf of the Wellcome Sanger Institute COVID-19 Surveillance Team |
| EPI_ISL_2718322 | Lighthouse Lab in Alderley Park | Wellcome Sanger Institute for the COVID-19 Genomics UK (COG-UK) Consortium | Jacquelyn Wynn, Mairead Hyland, The Lighthouse Lab in Alderley Park and Alex Alderton, Roberto Amato, Jeffrey Barrett, Sonia Goncalves, Ewan Harrison, David K. Jackson, Ian Johnston, Dominic Kwiatkowski, Cordelia Langford, John Sillitoe on behalf of the Wellcome Sanger Institute COVID-19 Surveillance Team |
| EPI_ISL_2718323 | Lighthouse Lab in Glasgow | Wellcome Sanger Institute for the COVID-19 Genomics UK (COG-UK) Consortium | Harper VanSteenhouse, Yumi Kasai, David Gray, Carol Clugston, Anna Dominiczak and Alex Alderton, Roberto Amato, Jeffrey Barrett, Sonia Goncalves, Ewan Harrison, David K. Jackson, Ian Johnston, Dominic Kwiatkowski, Cordelia Langford, John Sillitoe on behalf of the Wellcome Sanger Institute COVID-19 Surveillance Team |
| EPI_ISL_2718324 | Lighthouse Laboratory Plymouth | Wellcome Sanger Institute for the COVID-19 Genomics UK (COG-UK) Consortium | Lighthouse Laboratory Plymouth and Alex Alderton, Roberto Amato, Jeffrey Barrett, Sonia Goncalves, Ewan Harrison, David K. Jackson, Ian Johnston, Dominic Kwiatkowski, Cordelia Langford, John Sillitoe on behalf of the Wellcome Sanger Institute COVID-19 Surveillance Team |
| EPI_ISL_2718325 | Lighthouse Lab in Glasgow | Wellcome Sanger Institute for the COVID-19 Genomics UK (COG-UK) Consortium | Harper VanSteenhouse, Yumi Kasai, David Gray, Carol Clugston, Anna Dominiczak and Alex Alderton, Roberto Amato, Jeffrey Barrett, Sonia Goncalves, Ewan Harrison, David K. Jackson, Ian Johnston, Dominic Kwiatkowski, Cordelia Langford, John Sillitoe on behalf of the Wellcome Sanger Institute COVID-19 Surveillance Team |
| EPI_ISL_2718326 | Lighthouse Lab in Alderley Park | Wellcome Sanger Institute for the COVID-19 Genomics UK (COG-UK) Consortium | Jacquelyn Wynn, Mairead Hyland, The Lighthouse Lab in Alderley Park and Alex Alderton, Roberto Amato, Jeffrey Barrett, Sonia Goncalves, Ewan Harrison, David K. Jackson, Ian Johnston, Dominic Kwiatkowski, Cordelia Langford, John Sillitoe on behalf of the Wellcome Sanger Institute COVID-19 Surveillance Team |
| EPI_ISL_2718327 | Lighthouse Laboratory Plymouth | Wellcome Sanger Institute for the COVID-19 Genomics UK (COG-UK) Consortium | Lighthouse Laboratory Plymouth and Alex Alderton, Roberto Amato, Jeffrey Barrett, Sonia Goncalves, Ewan Harrison, David K. Jackson, Ian Johnston, Dominic Kwiatkowski, Cordelia Langford, John Sillitoe on behalf of the Wellcome Sanger Institute COVID-19 Surveillance Team |
| EPI_ISL_2718328 | Lighthouse Lab in Glasgow | Wellcome Sanger Institute for the COVID-19 Genomics UK (COG-UK) Consortium | Harper VanSteenhouse, Yumi Kasai, David Gray, Carol Clugston, Anna Dominiczak and Alex Alderton, Roberto Amato, Jeffrey Barrett, Sonia Goncalves, Ewan Harrison, David K. Jackson, Ian Johnston, Dominic Kwiatkowski, Cordelia Langford, John Sillitoe on behalf of the Wellcome Sanger Institute COVID-19 Surveillance Team |
| EPI_ISL_2718329 | Lighthouse Laboratory Plymouth | Wellcome Sanger Institute for the COVID-19 Genomics UK (COG-UK) Consortium | Lighthouse Laboratory Plymouth and Alex Alderton, Roberto Amato, Jeffrey Barrett, Sonia Goncalves, Ewan Harrison, David K. Jackson, Ian Johnston, Dominic Kwiatkowski, Cordelia Langford, John Sillitoe on behalf of the Wellcome Sanger Institute COVID-19 Surveillance Team |
| EPI_ISL_2718330 | Lighthouse Lab in Alderley Park | Wellcome Sanger Institute for the COVID-19 Genomics UK (COG-UK) Consortium | Jacquelyn Wynn, Mairead Hyland, The Lighthouse Lab in Alderley Park and Alex Alderton, Roberto Amato, Jeffrey Barrett, Sonia Goncalves, Ewan Harrison, David K. Jackson, Ian Johnston, Dominic Kwiatkowski, Cordelia Langford, John Sillitoe on behalf of the Wellcome Sanger Institute COVID-19 Surveillance Team |
| EPI_ISL_2718331, EPI_ISL_2718332, EPI_ISL_2718333, EPI_ISL_2718334 | Lighthouse Laboratory Plymouth | Wellcome Sanger Institute for the COVID-19 Genomics UK (COG-UK) Consortium | Lighthouse Laboratory Plymouth and Alex Alderton, Roberto Amato, Jeffrey Barrett, Sonia Goncalves, Ewan Harrison, David K. Jackson, Ian Johnston, Dominic Kwiatkowski, Cordelia Langford, John Sillitoe on behalf of the Wellcome Sanger Institute COVID-19 Surveillance Team |
| EPI_ISL_2718335 | Lighthouse Lab in Alderley Park | Wellcome Sanger Institute for the COVID-19 Genomics UK (COG-UK) Consortium | Jacquelyn Wynn, Mairead Hyland, The Lighthouse Lab in Alderley Park and Alex Alderton, Roberto Amato, Jeffrey Barrett, Sonia Goncalves, Ewan Harrison, David K. Jackson, Ian Johnston, Dominic Kwiatkowski, Cordelia Langford, John Sillitoe on behalf of the Wellcome Sanger Institute COVID-19 Surveillance Team |
| EPI_ISL_2718336 | Lighthouse Lab in Glasgow | Wellcome Sanger Institute for the COVID-19 Genomics UK (COG-UK) Consortium | Harper VanSteenhouse, Yumi Kasai, David Gray, Carol Clugston, Anna Dominiczak and Alex Alderton, Roberto Amato, Jeffrey Barrett, Sonia Goncalves, Ewan Harrison, David K. Jackson, Ian Johnston, Dominic Kwiatkowski, Cordelia Langford, John Sillitoe on behalf of the Wellcome Sanger Institute COVID-19 Surveillance Team |
| EPI_ISL_2718337 | Lighthouse Lab in Alderley Park | Wellcome Sanger Institute for the COVID-19 Genomics UK (COG-UK) Consortium | Jacquelyn Wynn, Mairead Hyland, The Lighthouse Lab in Alderley Park and Alex Alderton, Roberto Amato, Jeffrey Barrett, Sonia Goncalves, Ewan Harrison, David K. Jackson, Ian Johnston, Dominic Kwiatkowski, Cordelia Langford, John Sillitoe on behalf of the Wellcome Sanger Institute COVID-19 Surveillance Team |
| EPI_ISL_2718338, EPI_ISL_2718339 | Lighthouse Laboratory Plymouth | Wellcome Sanger Institute for the COVID-19 Genomics UK (COG-UK) Consortium | Lighthouse Laboratory Plymouth and Alex Alderton, Roberto Amato, Jeffrey Barrett, Sonia Goncalves, Ewan Harrison, David K. Jackson, Ian Johnston, Dominic Kwiatkowski, Cordelia Langford, John Sillitoe on behalf of the Wellcome Sanger Institute COVID-19 Surveillance Team |
| EPI_ISL_2718340 | Lighthouse Lab in Glasgow | Wellcome Sanger Institute for the COVID-19 Genomics UK (COG-UK) Consortium | Harper VanSteenhouse, Yumi Kasai, David Gray, Carol Clugston, Anna Dominiczak and Alex Alderton, Roberto Amato, Jeffrey Barrett, Sonia Goncalves, Ewan Harrison, David K. Jackson, Ian Johnston, Dominic Kwiatkowski, Cordelia Langford, John Sillitoe on behalf of the Wellcome Sanger Institute COVID-19 Surveillance Team |
| EPI_ISL_2718341 | Lighthouse Laboratory Plymouth | Wellcome Sanger Institute for the COVID-19 Genomics UK (COG-UK) Consortium | Lighthouse Laboratory Plymouth and Alex Alderton, Roberto Amato, Jeffrey Barrett, Sonia Goncalves, Ewan Harrison, David K. Jackson, Ian Johnston, Dominic Kwiatkowski, Cordelia Langford, John Sillitoe on behalf of the Wellcome Sanger Institute COVID-19 Surveillance Team |
| EPI_ISL_2718342, EPI_ISL_2718343 | Lighthouse Lab in Glasgow | Wellcome Sanger Institute for the COVID-19 Genomics UK (COG-UK) Consortium | Harper VanSteenhouse, Yumi Kasai, David Gray, Carol Clugston, Anna Dominiczak and Alex Alderton, Roberto Amato, Jeffrey Barrett, Sonia Goncalves, Ewan Harrison, David K. Jackson, Ian Johnston, Dominic Kwiatkowski, Cordelia Langford, John Sillitoe on behalf of the Wellcome Sanger Institute COVID-19 Surveillance Team |
| EPI_ISL_2718344 | Lighthouse Lab in Alderley Park | Wellcome Sanger Institute for the COVID-19 Genomics UK (COG-UK) Consortium | Jacquelyn Wynn, Mairead Hyland, The Lighthouse Lab in Alderley Park and Alex Alderton, Roberto Amato, Jeffrey Barrett, Sonia Goncalves, Ewan Harrison, David K. Jackson, Ian Johnston, Dominic Kwiatkowski, Cordelia Langford, John Sillitoe on behalf of the Wellcome Sanger Institute COVID-19 Surveillance Team |
| EPI_ISL_2718345 | Lighthouse Laboratory Plymouth | Wellcome Sanger Institute for the COVID-19 Genomics UK (COG-UK) Consortium | Lighthouse Laboratory Plymouth and Alex Alderton, Roberto Amato, Jeffrey Barrett, Sonia Goncalves, Ewan Harrison, David K. Jackson, Ian Johnston, Dominic Kwiatkowski, Cordelia Langford, John Sillitoe on behalf of the Wellcome Sanger Institute COVID-19 Surveillance Team |
| EPI_ISL_2718346, EPI_ISL_2718347, EPI_ISL_2718348 | Lighthouse Lab in Alderley Park | Wellcome Sanger Institute for the COVID-19 Genomics UK (COG-UK) Consortium | Jacquelyn Wynn, Mairead Hyland, The Lighthouse Lab in Alderley Park and Alex Alderton, Roberto Amato, Jeffrey Barrett, Sonia Goncalves, Ewan Harrison, David K. Jackson, Ian Johnston, Dominic Kwiatkowski, Cordelia Langford, John Sillitoe on behalf of the Wellcome Sanger Institute COVID-19 Surveillance Team |
| EPI_ISL_2718349, EPI_ISL_2718350 | Lighthouse Laboratory Plymouth | Wellcome Sanger Institute for the COVID-19 Genomics UK (COG-UK) Consortium | Lighthouse Laboratory Plymouth and Alex Alderton, Roberto Amato, Jeffrey Barrett, Sonia Goncalves, Ewan Harrison, David K. Jackson, Ian Johnston, Dominic Kwiatkowski, Cordelia Langford, John Sillitoe on behalf of the Wellcome Sanger Institute COVID-19 Surveillance Team |

| | | | |
|---|---|---|---|
| EPI_ISL_2718351, EPI_ISL_2718352 | Lighthouse Lab in Alderley Park | Wellcome Sanger Institute for the COVID-19 Genomics UK (COG-UK) Consortium | Jacquelyn Wynn, Mairead Hyland, The Lighthouse Lab in Alderley Park and Alex Alderton, Roberto Amato, Jeffrey Barrett, Sonia Goncalves, Ewan Harrison, David K. Jackson, Ian Johnston, Dominic Kwiatkowski, Cordelia Langford, John Sillitoe on behalf of the Wellcome Sanger Institute COVID-19 Surveillance Team |
| EPI_ISL_2718353 | Lighthouse Lab in Glasgow | Wellcome Sanger Institute for the COVID-19 Genomics UK (COG-UK) Consortium | Harper VanSteenhouse, Yumi Kasai, David Gray, Carol Clugston, Anna Dominiczak and Alex Alderton, Roberto Amato, Jeffrey Barrett, Sonia Goncalves, Ewan Harrison, David K. Jackson, Ian Johnston, Dominic Kwiatkowski, Cordelia Langford, John Sillitoe on behalf of the Wellcome Sanger Institute COVID-19 Surveillance Team |
| EPI_ISL_2718354 | Lighthouse Lab in Alderley Park | Wellcome Sanger Institute for the COVID-19 Genomics UK (COG-UK) Consortium | Jacquelyn Wynn, Mairead Hyland, The Lighthouse Lab in Alderley Park and Alex Alderton, Roberto Amato, Jeffrey Barrett, Sonia Goncalves, Ewan Harrison, David K. Jackson, Ian Johnston, Dominic Kwiatkowski, Cordelia Langford, John Sillitoe on behalf of the Wellcome Sanger Institute COVID-19 Surveillance Team |
| EPI_ISL_2718355 | Lighthouse Laboratory Plymouth | Wellcome Sanger Institute for the COVID-19 Genomics UK (COG-UK) Consortium | Lighthouse Laboratory Plymouth and Alex Alderton, Roberto Amato, Jeffrey Barrett, Sonia Goncalves, Ewan Harrison, David K. Jackson, Ian Johnston, Dominic Kwiatkowski, Cordelia Langford, John Sillitoe on behalf of the Wellcome Sanger Institute COVID-19 Surveillance Team |
| EPI_ISL_2718356 | Lighthouse Lab in Alderley Park | Wellcome Sanger Institute for the COVID-19 Genomics UK (COG-UK) Consortium | Jacquelyn Wynn, Mairead Hyland, The Lighthouse Lab in Alderley Park and Alex Alderton, Roberto Amato, Jeffrey Barrett, Sonia Goncalves, Ewan Harrison, David K. Jackson, Ian Johnston, Dominic Kwiatkowski, Cordelia Langford, John Sillitoe on behalf of the Wellcome Sanger Institute COVID-19 Surveillance Team |
| EPI_ISL_2718357, EPI_ISL_2718358 | Lighthouse Lab in Glasgow | Wellcome Sanger Institute for the COVID-19 Genomics UK (COG-UK) Consortium | Harper VanSteenhouse, Yumi Kasai, David Gray, Carol Clugston, Anna Dominiczak and Alex Alderton, Roberto Amato, Jeffrey Barrett, Sonia Goncalves, Ewan Harrison, David K. Jackson, Ian Johnston, Dominic Kwiatkowski, Cordelia Langford, John Sillitoe on behalf of the Wellcome Sanger Institute COVID-19 Surveillance Team |
| EPI_ISL_2718359, EPI_ISL_2718360, EPI_ISL_2718361 | Lighthouse Laboratory Plymouth | Wellcome Sanger Institute for the COVID-19 Genomics UK (COG-UK) Consortium | Lighthouse Laboratory Plymouth and Alex Alderton, Roberto Amato, Jeffrey Barrett, Sonia Goncalves, Ewan Harrison, David K. Jackson, Ian Johnston, Dominic Kwiatkowski, Cordelia Langford, John Sillitoe on behalf of the Wellcome Sanger Institute COVID-19 Surveillance Team |
| EPI_ISL_2718362, EPI_ISL_2718363 | Lighthouse Lab in Alderley Park | Wellcome Sanger Institute for the COVID-19 Genomics UK (COG-UK) Consortium | Jacquelyn Wynn, Mairead Hyland, The Lighthouse Lab in Alderley Park and Alex Alderton, Roberto Amato, Jeffrey Barrett, Sonia Goncalves, Ewan Harrison, David K. Jackson, Ian Johnston, Dominic Kwiatkowski, Cordelia Langford, John Sillitoe on behalf of the Wellcome Sanger Institute COVID-19 Surveillance Team |
| EPI_ISL_2718364 | Lighthouse Laboratory Plymouth | Wellcome Sanger Institute for the COVID-19 Genomics UK (COG-UK) Consortium | Lighthouse Laboratory Plymouth and Alex Alderton, Roberto Amato, Jeffrey Barrett, Sonia Goncalves, Ewan Harrison, David K. Jackson, Ian Johnston, Dominic Kwiatkowski, Cordelia Langford, John Sillitoe on behalf of the Wellcome Sanger Institute COVID-19 Surveillance Team |
| EPI_ISL_2718365 | Lighthouse Lab in Alderley Park | Wellcome Sanger Institute for the COVID-19 Genomics UK (COG-UK) Consortium | Jacquelyn Wynn, Mairead Hyland, The Lighthouse Lab in Alderley Park and Alex Alderton, Roberto Amato, Jeffrey Barrett, Sonia Goncalves, Ewan Harrison, David K. Jackson, Ian Johnston, Dominic Kwiatkowski, Cordelia Langford, John Sillitoe on behalf of the Wellcome Sanger Institute COVID-19 Surveillance Team |
| EPI_ISL_2718366 | Lighthouse Lab in Glasgow | Wellcome Sanger Institute for the COVID-19 Genomics UK (COG-UK) Consortium | Harper VanSteenhouse, Yumi Kasai, David Gray, Carol Clugston, Anna Dominiczak and Alex Alderton, Roberto Amato, Jeffrey Barrett, Sonia Goncalves, Ewan Harrison, David K. Jackson, Ian Johnston, Dominic Kwiatkowski, Cordelia Langford, John Sillitoe on behalf of the Wellcome Sanger Institute COVID-19 Surveillance Team |
| EPI_ISL_2718367 | Lighthouse Laboratory Plymouth | Wellcome Sanger Institute for the COVID-19 Genomics UK (COG-UK) Consortium | Lighthouse Laboratory Plymouth and Alex Alderton, Roberto Amato, Jeffrey Barrett, Sonia Goncalves, Ewan Harrison, David K. Jackson, Ian Johnston, Dominic Kwiatkowski, Cordelia Langford, John Sillitoe on behalf of the Wellcome Sanger Institute COVID-19 Surveillance Team |
| EPI_ISL_2718368 | Lighthouse Lab in Glasgow | Wellcome Sanger Institute for the COVID-19 Genomics UK (COG-UK) Consortium | Harper VanSteenhouse, Yumi Kasai, David Gray, Carol Clugston, Anna Dominiczak and Alex Alderton, Roberto Amato, Jeffrey Barrett, Sonia Goncalves, Ewan Harrison, David K. Jackson, Ian Johnston, Dominic Kwiatkowski, Cordelia Langford, John Sillitoe on behalf of the Wellcome Sanger Institute COVID-19 Surveillance Team |
| EPI_ISL_2718369 | Lighthouse Laboratory Plymouth | Wellcome Sanger Institute for the COVID-19 Genomics UK (COG-UK) Consortium | Lighthouse Laboratory Plymouth and Alex Alderton, Roberto Amato, Jeffrey Barrett, Sonia Goncalves, Ewan Harrison, David K. Jackson, Ian Johnston, Dominic Kwiatkowski, Cordelia Langford, John Sillitoe on behalf of the Wellcome Sanger Institute COVID-19 Surveillance Team |
| EPI_ISL_2718370, EPI_ISL_2718371, EPI_ISL_2718372, EPI_ISL_2718373 | Lighthouse Lab in Alderley Park | Wellcome Sanger Institute for the COVID-19 Genomics UK (COG-UK) Consortium | Jacquelyn Wynn, Mairead Hyland, The Lighthouse Lab in Alderley Park and Alex Alderton, Roberto Amato, Jeffrey Barrett, Sonia Goncalves, Ewan Harrison, David K. Jackson, Ian Johnston, Dominic Kwiatkowski, Cordelia Langford, John Sillitoe on behalf of the Wellcome Sanger Institute COVID-19 Surveillance Team |
| EPI_ISL_2718374, EPI_ISL_2718375 | Lighthouse Laboratory Plymouth | Wellcome Sanger Institute for the COVID-19 Genomics UK (COG-UK) Consortium | Lighthouse Laboratory Plymouth and Alex Alderton, Roberto Amato, Jeffrey Barrett, Sonia Goncalves, Ewan Harrison, David K. Jackson, Ian Johnston, Dominic Kwiatkowski, Cordelia Langford, John Sillitoe on behalf of the Wellcome Sanger Institute COVID-19 Surveillance Team |
| EPI_ISL_2718376, EPI_ISL_2718377 | Lighthouse Lab in Glasgow | Wellcome Sanger Institute for the COVID-19 Genomics UK (COG-UK) Consortium | Harper VanSteenhouse, Yumi Kasai, David Gray, Carol Clugston, Anna Dominiczak and Alex Alderton, Roberto Amato, Jeffrey Barrett, Sonia Goncalves, Ewan Harrison, David K. Jackson, Ian Johnston, Dominic Kwiatkowski, Cordelia Langford, John Sillitoe on behalf of the Wellcome Sanger Institute COVID-19 Surveillance Team |
| EPI_ISL_2718378, EPI_ISL_2718379 | Lighthouse Lab in Alderley Park | Wellcome Sanger Institute for the COVID-19 Genomics UK (COG-UK) Consortium | Jacquelyn Wynn, Mairead Hyland, The Lighthouse Lab in Alderley Park and Alex Alderton, Roberto Amato, Jeffrey Barrett, Sonia Goncalves, Ewan Harrison, David K. Jackson, Ian Johnston, Dominic Kwiatkowski, Cordelia Langford, John Sillitoe on behalf of the Wellcome Sanger Institute COVID-19 Surveillance Team |
| EPI_ISL_2718380 | Lighthouse Lab in Glasgow | Wellcome Sanger Institute for the COVID-19 Genomics UK (COG-UK) Consortium | Harper VanSteenhouse, Yumi Kasai, David Gray, Carol Clugston, Anna Dominiczak and Alex Alderton, Roberto Amato, Jeffrey Barrett, Sonia Goncalves, Ewan Harrison, David K. Jackson, Ian Johnston, Dominic Kwiatkowski, Cordelia Langford, John Sillitoe on behalf of the Wellcome Sanger Institute COVID-19 Surveillance Team |
| EPI_ISL_2718381, EPI_ISL_2718382, EPI_ISL_2718383 | Lighthouse Lab in Alderley Park | Wellcome Sanger Institute for the COVID-19 Genomics UK (COG-UK) Consortium | Jacquelyn Wynn, Mairead Hyland, The Lighthouse Lab in Alderley Park and Alex Alderton, Roberto Amato, Jeffrey Barrett, Sonia Goncalves, Ewan Harrison, David K. Jackson, Ian Johnston, Dominic Kwiatkowski, Cordelia Langford, John Sillitoe on behalf of the Wellcome Sanger Institute COVID-19 Surveillance Team |
| EPI_ISL_2718384 | Lighthouse Laboratory Plymouth | Wellcome Sanger Institute for the COVID-19 Genomics UK (COG-UK) Consortium | Lighthouse Laboratory Plymouth and Alex Alderton, Roberto Amato, Jeffrey Barrett, Sonia Goncalves, Ewan Harrison, David K. Jackson, Ian Johnston, Dominic Kwiatkowski, Cordelia Langford, John Sillitoe on behalf of the Wellcome Sanger Institute COVID-19 Surveillance Team |
| EPI_ISL_2718385 | Lighthouse Lab in Alderley Park | Wellcome Sanger Institute for the COVID-19 Genomics UK (COG-UK) Consortium | Jacquelyn Wynn, Mairead Hyland, The Lighthouse Lab in Alderley Park and Alex Alderton, Roberto Amato, Jeffrey Barrett, Sonia Goncalves, Ewan Harrison, David K. Jackson, Ian Johnston, Dominic Kwiatkowski, Cordelia Langford, John Sillitoe on behalf of the Wellcome Sanger Institute COVID-19 Surveillance Team |
| EPI_ISL_2718386, EPI_ISL_2718387 | Lighthouse Laboratory Plymouth | Wellcome Sanger Institute for the COVID-19 Genomics UK (COG-UK) Consortium | Lighthouse Laboratory Plymouth and Alex Alderton, Roberto Amato, Jeffrey Barrett, Sonia Goncalves, Ewan Harrison, David K. Jackson, Ian Johnston, Dominic Kwiatkowski, Cordelia Langford, John Sillitoe on behalf of the Wellcome Sanger Institute COVID-19 Surveillance Team |
| EPI_ISL_2718388 | Lighthouse Lab in Alderley Park | Wellcome Sanger Institute for the COVID-19 Genomics UK (COG-UK) Consortium | Jacquelyn Wynn, Mairead Hyland, The Lighthouse Lab in Alderley Park and Alex Alderton, Roberto Amato, Jeffrey Barrett, Sonia Goncalves, Ewan Harrison, David K. Jackson, Ian Johnston, Dominic Kwiatkowski, Cordelia Langford, John Sillitoe on behalf of the Wellcome Sanger Institute COVID-19 Surveillance Team |
| EPI_ISL_2718389 | Lighthouse Lab in Glasgow | Wellcome Sanger Institute for the COVID-19 Genomics UK (COG-UK) Consortium | Harper VanSteenhouse, Yumi Kasai, David Gray, Carol Clugston, Anna Dominiczak and Alex Alderton, Roberto Amato, Jeffrey Barrett, Sonia Goncalves, Ewan Harrison, David K. Jackson, Ian Johnston, Dominic Kwiatkowski, Cordelia Langford, John Sillitoe on behalf of the Wellcome Sanger Institute COVID-19 Surveillance Team |
| EPI_ISL_2718390, EPI_ISL_2718391 | Lighthouse Lab in Alderley Park | Wellcome Sanger Institute for the COVID-19 Genomics UK (COG-UK) Consortium | Jacquelyn Wynn, Mairead Hyland, The Lighthouse Lab in Alderley Park and Alex Alderton, Roberto Amato, Jeffrey Barrett, Sonia Goncalves, Ewan Harrison, David K. Jackson, Ian Johnston, Dominic Kwiatkowski, Cordelia Langford, John Sillitoe on behalf of the Wellcome Sanger Institute COVID-19 Surveillance Team |
| EPI_ISL_2718392 | Lighthouse Laboratory Plymouth | Wellcome Sanger Institute for the COVID-19 Genomics UK (COG-UK) Consortium | Lighthouse Laboratory Plymouth and Alex Alderton, Roberto Amato, Jeffrey Barrett, Sonia Goncalves, Ewan Harrison, David K. Jackson, Ian Johnston, Dominic Kwiatkowski, Cordelia Langford, John Sillitoe on behalf of the Wellcome Sanger Institute COVID-19 Surveillance Team |
| EPI_ISL_2718393 | Lighthouse Lab in Alderley Park | Wellcome Sanger Institute for the COVID-19 Genomics UK (COG-UK) Consortium | Jacquelyn Wynn, Mairead Hyland, The Lighthouse Lab in Alderley Park and Alex Alderton, Roberto Amato, Jeffrey Barrett, Sonia Goncalves, Ewan Harrison, David K. Jackson, Ian Johnston, Dominic Kwiatkowski, Cordelia Langford, John Sillitoe on behalf of the Wellcome Sanger Institute COVID-19 Surveillance Team |
| EPI_ISL_2718394, EPI_ISL_2718395 | Lighthouse Lab in Glasgow | Wellcome Sanger Institute for the COVID-19 Genomics UK (COG-UK) Consortium | Harper VanSteenhouse, Yumi Kasai, David Gray, Carol Clugston, Anna Dominiczak and Alex Alderton, Roberto Amato, Jeffrey Barrett, Sonia Goncalves, Ewan Harrison, David K. Jackson, Ian Johnston, Dominic Kwiatkowski, Cordelia Langford, John Sillitoe on behalf of the Wellcome Sanger Institute COVID-19 Surveillance Team |
| EPI_ISL_2718396 | Lighthouse Laboratory Plymouth | Wellcome Sanger Institute for the COVID-19 Genomics UK (COG-UK) Consortium | Lighthouse Laboratory Plymouth and Alex Alderton, Roberto Amato, Jeffrey Barrett, Sonia Goncalves, Ewan Harrison, David K. Jackson, Ian Johnston, Dominic Kwiatkowski, Cordelia Langford, John Sillitoe on behalf of the Wellcome Sanger Institute COVID-19 Surveillance Team |
| EPI_ISL_2718397, EPI_ISL_2718398 | Lighthouse Lab in Glasgow | Wellcome Sanger Institute for the COVID-19 Genomics UK (COG-UK) Consortium | Harper VanSteenhouse, Yumi Kasai, David Gray, Carol Clugston, Anna Dominiczak and Alex Alderton, Roberto Amato, Jeffrey Barrett, Sonia Goncalves, Ewan Harrison, David K. Jackson, Ian Johnston, Dominic Kwiatkowski, Cordelia Langford, John Sillitoe on behalf of the Wellcome Sanger Institute COVID-19 Surveillance Team |
| EPI_ISL_2718399 | Lighthouse Laboratory Plymouth | Wellcome Sanger Institute for the COVID-19 Genomics UK (COG-UK) Consortium | Lighthouse Laboratory Plymouth and Alex Alderton, Roberto Amato, Jeffrey Barrett, Sonia Goncalves, Ewan Harrison, David K. Jackson, Ian Johnston, Dominic Kwiatkowski, Cordelia Langford, John Sillitoe on behalf of the Wellcome Sanger Institute COVID-19 Surveillance Team |
| EPI_ISL_2718400, EPI_ISL_2718401 | Lighthouse Lab in Alderley Park | Wellcome Sanger Institute for the COVID-19 Genomics UK (COG-UK) Consortium | Jacquelyn Wynn, Mairead Hyland, The Lighthouse Lab in Alderley Park and Alex Alderton, Roberto Amato, Jeffrey Barrett, Sonia Goncalves, Ewan Harrison, David K. Jackson, Ian Johnston, Dominic Kwiatkowski, Cordelia Langford, John Sillitoe on behalf of the Wellcome Sanger Institute COVID-19 Surveillance Team |
| EPI_ISL_2718402 | Lighthouse Laboratory Plymouth | Wellcome Sanger Institute for the COVID-19 Genomics UK (COG-UK) Consortium | Lighthouse Laboratory Plymouth and Alex Alderton, Roberto Amato, Jeffrey Barrett, Sonia Goncalves, Ewan Harrison, David K. Jackson, Ian Johnston, Dominic Kwiatkowski, Cordelia Langford, John Sillitoe on behalf of the Wellcome Sanger Institute COVID-19 Surveillance Team |
| EPI_ISL_2718403, EPI_ISL_2718404 | Lighthouse Lab in Alderley Park | Wellcome Sanger Institute for the COVID-19 Genomics UK (COG-UK) Consortium | Jacquelyn Wynn, Mairead Hyland, The Lighthouse Lab in Alderley Park and Alex Alderton, Roberto Amato, Jeffrey Barrett, Sonia Goncalves, Ewan Harrison, David K. Jackson, Ian Johnston, Dominic Kwiatkowski, Cordelia Langford, John Sillitoe on behalf of the Wellcome Sanger Institute COVID-19 Surveillance Team |
| EPI_ISL_2718405 | Lighthouse Laboratory Plymouth | Wellcome Sanger Institute for the COVID-19 Genomics UK (COG-UK) Consortium | Lighthouse Laboratory Plymouth and Alex Alderton, Roberto Amato, Jeffrey Barrett, Sonia Goncalves, Ewan Harrison, David K. Jackson, Ian Johnston, Dominic Kwiatkowski, Cordelia Langford, John Sillitoe on behalf of the Wellcome Sanger Institute COVID-19 Surveillance Team |
| EPI_ISL_2718406 | Lighthouse Lab in Alderley Park | Wellcome Sanger Institute for the COVID-19 Genomics UK (COG-UK) Consortium | Jacquelyn Wynn, Mairead Hyland, The Lighthouse Lab in Alderley Park and Alex Alderton, Roberto Amato, Jeffrey Barrett, Sonia Goncalves, Ewan Harrison, David K. Jackson, Ian Johnston, Dominic Kwiatkowski, Cordelia Langford, John Sillitoe on behalf of the Wellcome Sanger Institute COVID-19 Surveillance Team |
| EPI_ISL_2718407, EPI_ISL_2718408, EPI_ISL_2718409 | Lighthouse Laboratory Plymouth | Wellcome Sanger Institute for the COVID-19 Genomics UK (COG-UK) Consortium | Lighthouse Laboratory Plymouth and Alex Alderton, Roberto Amato, Jeffrey Barrett, Sonia Goncalves, Ewan Harrison, David K. Jackson, Ian Johnston, Dominic Kwiatkowski, Cordelia Langford, John Sillitoe on behalf of the Wellcome Sanger Institute COVID-19 Surveillance Team |
| EPI_ISL_2718410 | Lighthouse Lab in Glasgow | Wellcome Sanger Institute for the COVID-19 Genomics UK (COG-UK) Consortium | Harper VanSteenhouse, Yumi Kasai, David Gray, Carol Clugston, Anna Dominiczak and Alex Alderton, Roberto Amato, Jeffrey Barrett, Sonia Goncalves, Ewan Harrison, David K. Jackson, Ian Johnston, Dominic Kwiatkowski, Cordelia Langford, John Sillitoe on behalf of the Wellcome Sanger Institute COVID-19 Surveillance Team |
| EPI_ISL_2718412 | Lighthouse Lab in Alderley Park | Wellcome Sanger Institute for the COVID-19 Genomics UK (COG-UK) Consortium | Jacquelyn Wynn, Mairead Hyland, The Lighthouse Lab in Alderley Park and Alex Alderton, Roberto Amato, Jeffrey Barrett, Sonia Goncalves, Ewan Harrison, David K. Jackson, Ian Johnston, Dominic Kwiatkowski, Cordelia Langford, John Sillitoe on behalf of the Wellcome Sanger Institute COVID-19 Surveillance Team |
| EPI_ISL_2718413 | Lighthouse Laboratory Plymouth | Wellcome Sanger Institute for the COVID-19 Genomics UK (COG-UK) Consortium | Lighthouse Laboratory Plymouth and Alex Alderton, Roberto Amato, Jeffrey Barrett, Sonia Goncalves, Ewan Harrison, David K. Jackson, Ian Johnston, Dominic Kwiatkowski, Cordelia Langford, John Sillitoe on behalf of the Wellcome Sanger Institute COVID-19 Surveillance Team |
| EPI_ISL_2718414, EPI_ISL_2718415 | Lighthouse Lab in Alderley Park | Wellcome Sanger Institute for the COVID-19 Genomics UK (COG-UK) Consortium | Jacquelyn Wynn, Mairead Hyland, The Lighthouse Lab in Alderley Park and Alex Alderton, Roberto Amato, Jeffrey Barrett, Sonia Goncalves, Ewan Harrison, David K. Jackson, Ian Johnston, Dominic Kwiatkowski, Cordelia Langford, John Sillitoe on behalf of the Wellcome Sanger Institute COVID-19 Surveillance Team |
| EPI_ISL_2718416 | Lighthouse Lab in Glasgow | Wellcome Sanger Institute for the COVID-19 Genomics UK (COG-UK) | Harper VanSteenhouse, Yumi Kasai, David Gray, Carol Clugston, Anna Dominiczak and Alex Alderton, Roberto Amato, Jeffrey Barrett, Sonia Goncalves, Ewan Harrison, David K. Jackson, Ian Johnston, Dominic Kwiatkowski, Cordelia Langford, |

| | | | |
|---|---|---|---|
| | | Consortium | John Sillitoe on behalf of the Wellcome Sanger Institute COVID-19 Surveillance Team |
| EPI_ISL_2718417, EPI_ISL_2718418 | Lighthouse Lab in Alderley Park | Wellcome Sanger Institute for the COVID-19 Genomics UK (COG-UK) Consortium | Jacquelyn Wynn, Mairead Hyland, The Lighthouse Lab in Alderley Park and Alex Alderton, Roberto Amato, Jeffrey Barrett, Sonia Goncalves, Ewan Harrison, David K. Jackson, Ian Johnston, Dominic Kwiatkowski, Cordelia Langford, John Sillitoe on behalf of the Wellcome Sanger Institute COVID-19 Surveillance Team |
| EPI_ISL_2718419, EPI_ISL_2718420 | Lighthouse Laboratory Plymouth | Wellcome Sanger Institute for the COVID-19 Genomics UK (COG-UK) Consortium | Lighthouse Laboratory Plymouth and Alex Alderton, Roberto Amato, Jeffrey Barrett, Sonia Goncalves, Ewan Harrison, David K. Jackson, Ian Johnston, Dominic Kwiatkowski, Cordelia Langford, John Sillitoe on behalf of the Wellcome Sanger Institute COVID-19 Surveillance Team |
| EPI_ISL_2718421 | Lighthouse Lab in Glasgow | Wellcome Sanger Institute for the COVID-19 Genomics UK (COG-UK) Consortium | Harper VanSteenhouse, Yumi Kasai, David Gray, Carol Clugston, Anna Dominiczak and Alex Alderton, Roberto Amato, Jeffrey Barrett, Sonia Goncalves, Ewan Harrison, David K. Jackson, Ian Johnston, Dominic Kwiatkowski, Cordelia Langford, John Sillitoe on behalf of the Wellcome Sanger Institute COVID-19 Surveillance Team |
| EPI_ISL_2718422 | Lighthouse Laboratory Plymouth | Wellcome Sanger Institute for the COVID-19 Genomics UK (COG-UK) Consortium | Lighthouse Laboratory Plymouth and Alex Alderton, Roberto Amato, Jeffrey Barrett, Sonia Goncalves, Ewan Harrison, David K. Jackson, Ian Johnston, Dominic Kwiatkowski, Cordelia Langford, John Sillitoe on behalf of the Wellcome Sanger Institute COVID-19 Surveillance Team |
| EPI_ISL_2718423, EPI_ISL_2718424 | Lighthouse Lab in Alderley Park | Wellcome Sanger Institute for the COVID-19 Genomics UK (COG-UK) Consortium | Jacquelyn Wynn, Mairead Hyland, The Lighthouse Lab in Alderley Park and Alex Alderton, Roberto Amato, Jeffrey Barrett, Sonia Goncalves, Ewan Harrison, David K. Jackson, Ian Johnston, Dominic Kwiatkowski, Cordelia Langford, John Sillitoe on behalf of the Wellcome Sanger Institute COVID-19 Surveillance Team |
| EPI_ISL_2718425 | Lighthouse Lab in Glasgow | Wellcome Sanger Institute for the COVID-19 Genomics UK (COG-UK) Consortium | Harper VanSteenhouse, Yumi Kasai, David Gray, Carol Clugston, Anna Dominiczak and Alex Alderton, Roberto Amato, Jeffrey Barrett, Sonia Goncalves, Ewan Harrison, David K. Jackson, Ian Johnston, Dominic Kwiatkowski, Cordelia Langford, John Sillitoe on behalf of the Wellcome Sanger Institute COVID-19 Surveillance Team |
| EPI_ISL_2718426 | Lighthouse Lab in Alderley Park | Wellcome Sanger Institute for the COVID-19 Genomics UK (COG-UK) Consortium | Jacquelyn Wynn, Mairead Hyland, The Lighthouse Lab in Alderley Park and Alex Alderton, Roberto Amato, Jeffrey Barrett, Sonia Goncalves, Ewan Harrison, David K. Jackson, Ian Johnston, Dominic Kwiatkowski, Cordelia Langford, John Sillitoe on behalf of the Wellcome Sanger Institute COVID-19 Surveillance Team |
| EPI_ISL_2718427, EPI_ISL_2718428 | Lighthouse Laboratory Plymouth | Wellcome Sanger Institute for the COVID-19 Genomics UK (COG-UK) Consortium | Lighthouse Laboratory Plymouth and Alex Alderton, Roberto Amato, Jeffrey Barrett, Sonia Goncalves, Ewan Harrison, David K. Jackson, Ian Johnston, Dominic Kwiatkowski, Cordelia Langford, John Sillitoe on behalf of the Wellcome Sanger Institute COVID-19 Surveillance Team |
| EPI_ISL_2718429 | Lighthouse Lab in Glasgow | Wellcome Sanger Institute for the COVID-19 Genomics UK (COG-UK) Consortium | Harper VanSteenhouse, Yumi Kasai, David Gray, Carol Clugston, Anna Dominiczak and Alex Alderton, Roberto Amato, Jeffrey Barrett, Sonia Goncalves, Ewan Harrison, David K. Jackson, Ian Johnston, Dominic Kwiatkowski, Cordelia Langford, John Sillitoe on behalf of the Wellcome Sanger Institute COVID-19 Surveillance Team |
| EPI_ISL_2718430, EPI_ISL_2718431, EPI_ISL_2718432, EPI_ISL_2718433 | Lighthouse Laboratory Plymouth | Wellcome Sanger Institute for the COVID-19 Genomics UK (COG-UK) Consortium | Lighthouse Laboratory Plymouth and Alex Alderton, Roberto Amato, Jeffrey Barrett, Sonia Goncalves, Ewan Harrison, David K. Jackson, Ian Johnston, Dominic Kwiatkowski, Cordelia Langford, John Sillitoe on behalf of the Wellcome Sanger Institute COVID-19 Surveillance Team |
| EPI_ISL_2718434 | Lighthouse Lab in Alderley Park | Wellcome Sanger Institute for the COVID-19 Genomics UK (COG-UK) Consortium | Jacquelyn Wynn, Mairead Hyland, The Lighthouse Lab in Alderley Park and Alex Alderton, Roberto Amato, Jeffrey Barrett, Sonia Goncalves, Ewan Harrison, David K. Jackson, Ian Johnston, Dominic Kwiatkowski, Cordelia Langford, John Sillitoe on behalf of the Wellcome Sanger Institute COVID-19 Surveillance Team |
| EPI_ISL_2718435 | Lighthouse Lab in Glasgow | Wellcome Sanger Institute for the COVID-19 Genomics UK (COG-UK) Consortium | Harper VanSteenhouse, Yumi Kasai, David Gray, Carol Clugston, Anna Dominiczak and Alex Alderton, Roberto Amato, Jeffrey Barrett, Sonia Goncalves, Ewan Harrison, David K. Jackson, Ian Johnston, Dominic Kwiatkowski, Cordelia Langford, John Sillitoe on behalf of the Wellcome Sanger Institute COVID-19 Surveillance Team |
| EPI_ISL_2718436 | Lighthouse Lab in Alderley Park | Wellcome Sanger Institute for the COVID-19 Genomics UK (COG-UK) Consortium | Jacquelyn Wynn, Mairead Hyland, The Lighthouse Lab in Alderley Park and Alex Alderton, Roberto Amato, Jeffrey Barrett, Sonia Goncalves, Ewan Harrison, David K. Jackson, Ian Johnston, Dominic Kwiatkowski, Cordelia Langford, John Sillitoe on behalf of the Wellcome Sanger Institute COVID-19 Surveillance Team |
| EPI_ISL_2718437 | Lighthouse Laboratory Plymouth | Wellcome Sanger Institute for the COVID-19 Genomics UK (COG-UK) Consortium | Lighthouse Laboratory Plymouth and Alex Alderton, Roberto Amato, Jeffrey Barrett, Sonia Goncalves, Ewan Harrison, David K. Jackson, Ian Johnston, Dominic Kwiatkowski, Cordelia Langford, John Sillitoe on behalf of the Wellcome Sanger Institute COVID-19 Surveillance Team |
| EPI_ISL_2718438, EPI_ISL_2718439, EPI_ISL_2718440 | Lighthouse Lab in Glasgow | Wellcome Sanger Institute for the COVID-19 Genomics UK (COG-UK) Consortium | Harper VanSteenhouse, Yumi Kasai, David Gray, Carol Clugston, Anna Dominiczak and Alex Alderton, Roberto Amato, Jeffrey Barrett, Sonia Goncalves, Ewan Harrison, David K. Jackson, Ian Johnston, Dominic Kwiatkowski, Cordelia Langford, John Sillitoe on behalf of the Wellcome Sanger Institute COVID-19 Surveillance Team |
| EPI_ISL_2718442, EPI_ISL_2718443 | Lighthouse Lab in Alderley Park | Wellcome Sanger Institute for the COVID-19 Genomics UK (COG-UK) Consortium | Jacquelyn Wynn, Mairead Hyland, The Lighthouse Lab in Alderley Park and Alex Alderton, Roberto Amato, Jeffrey Barrett, Sonia Goncalves, Ewan Harrison, David K. Jackson, Ian Johnston, Dominic Kwiatkowski, Cordelia Langford, John Sillitoe on behalf of the Wellcome Sanger Institute COVID-19 Surveillance Team |
| EPI_ISL_2718444 | Lighthouse Laboratory Plymouth | Wellcome Sanger Institute for the COVID-19 Genomics UK (COG-UK) Consortium | Lighthouse Laboratory Plymouth and Alex Alderton, Roberto Amato, Jeffrey Barrett, Sonia Goncalves, Ewan Harrison, David K. Jackson, Ian Johnston, Dominic Kwiatkowski, Cordelia Langford, John Sillitoe on behalf of the Wellcome Sanger Institute COVID-19 Surveillance Team |
| EPI_ISL_2718445, EPI_ISL_2718446 | Lighthouse Lab in Glasgow | Wellcome Sanger Institute for the COVID-19 Genomics UK (COG-UK) Consortium | Harper VanSteenhouse, Yumi Kasai, David Gray, Carol Clugston, Anna Dominiczak and Alex Alderton, Roberto Amato, Jeffrey Barrett, Sonia Goncalves, Ewan Harrison, David K. Jackson, Ian Johnston, Dominic Kwiatkowski, Cordelia Langford, John Sillitoe on behalf of the Wellcome Sanger Institute COVID-19 Surveillance Team |
| EPI_ISL_2718448, EPI_ISL_2718449 | Lighthouse Lab in Alderley Park | Wellcome Sanger Institute for the COVID-19 Genomics UK (COG-UK) Consortium | Jacquelyn Wynn, Mairead Hyland, The Lighthouse Lab in Alderley Park and Alex Alderton, Roberto Amato, Jeffrey Barrett, Sonia Goncalves, Ewan Harrison, David K. Jackson, Ian Johnston, Dominic Kwiatkowski, Cordelia Langford, John Sillitoe on behalf of the Wellcome Sanger Institute COVID-19 Surveillance Team |
| EPI_ISL_2718450 | Lighthouse Laboratory Plymouth | Wellcome Sanger Institute for the COVID-19 Genomics UK (COG-UK) Consortium | Lighthouse Laboratory Plymouth and Alex Alderton, Roberto Amato, Jeffrey Barrett, Sonia Goncalves, Ewan Harrison, David K. Jackson, Ian Johnston, Dominic Kwiatkowski, Cordelia Langford, John Sillitoe on behalf of the Wellcome Sanger Institute COVID-19 Surveillance Team |
| EPI_ISL_2718451 | Lighthouse Lab in Glasgow | Wellcome Sanger Institute for the COVID-19 Genomics UK (COG-UK) Consortium | Harper VanSteenhouse, Yumi Kasai, David Gray, Carol Clugston, Anna Dominiczak and Alex Alderton, Roberto Amato, Jeffrey Barrett, Sonia Goncalves, Ewan Harrison, David K. Jackson, Ian Johnston, Dominic Kwiatkowski, Cordelia Langford, John Sillitoe on behalf of the Wellcome Sanger Institute COVID-19 Surveillance Team |
| EPI_ISL_2718452 | Lighthouse Laboratory Plymouth | Wellcome Sanger Institute for the COVID-19 Genomics UK (COG-UK) Consortium | Lighthouse Laboratory Plymouth and Alex Alderton, Roberto Amato, Jeffrey Barrett, Sonia Goncalves, Ewan Harrison, David K. Jackson, Ian Johnston, Dominic Kwiatkowski, Cordelia Langford, John Sillitoe on behalf of the Wellcome Sanger Institute COVID-19 Surveillance Team |
| EPI_ISL_2718453 | Lighthouse Lab in Glasgow | Wellcome Sanger Institute for the COVID-19 Genomics UK (COG-UK) Consortium | Harper VanSteenhouse, Yumi Kasai, David Gray, Carol Clugston, Anna Dominiczak and Alex Alderton, Roberto Amato, Jeffrey Barrett, Sonia Goncalves, Ewan Harrison, David K. Jackson, Ian Johnston, Dominic Kwiatkowski, Cordelia Langford, John Sillitoe on behalf of the Wellcome Sanger Institute COVID-19 Surveillance Team |
| EPI_ISL_2718454, EPI_ISL_2718455, EPI_ISL_2718456 | Lighthouse Lab in Alderley Park | Wellcome Sanger Institute for the COVID-19 Genomics UK (COG-UK) Consortium | Jacquelyn Wynn, Mairead Hyland, The Lighthouse Lab in Alderley Park and Alex Alderton, Roberto Amato, Jeffrey Barrett, Sonia Goncalves, Ewan Harrison, David K. Jackson, Ian Johnston, Dominic Kwiatkowski, Cordelia Langford, John Sillitoe on behalf of the Wellcome Sanger Institute COVID-19 Surveillance Team |
| EPI_ISL_2718457 | Lighthouse Laboratory Plymouth | Wellcome Sanger Institute for the COVID-19 Genomics UK (COG-UK) Consortium | Lighthouse Laboratory Plymouth and Alex Alderton, Roberto Amato, Jeffrey Barrett, Sonia Goncalves, Ewan Harrison, David K. Jackson, Ian Johnston, Dominic Kwiatkowski, Cordelia Langford, John Sillitoe on behalf of the Wellcome Sanger Institute COVID-19 Surveillance Team |
| EPI_ISL_2718458 | Lighthouse Lab in Glasgow | Wellcome Sanger Institute for the COVID-19 Genomics UK (COG-UK) Consortium | Harper VanSteenhouse, Yumi Kasai, David Gray, Carol Clugston, Anna Dominiczak and Alex Alderton, Roberto Amato, Jeffrey Barrett, Sonia Goncalves, Ewan Harrison, David K. Jackson, Ian Johnston, Dominic Kwiatkowski, Cordelia Langford, John Sillitoe on behalf of the Wellcome Sanger Institute COVID-19 Surveillance Team |
| EPI_ISL_2718459, EPI_ISL_2718460 | Lighthouse Laboratory Plymouth | Wellcome Sanger Institute for the COVID-19 Genomics UK (COG-UK) Consortium | Lighthouse Laboratory Plymouth and Alex Alderton, Roberto Amato, Jeffrey Barrett, Sonia Goncalves, Ewan Harrison, David K. Jackson, Ian Johnston, Dominic Kwiatkowski, Cordelia Langford, John Sillitoe on behalf of the Wellcome Sanger Institute COVID-19 Surveillance Team |
| EPI_ISL_2718461 | Lighthouse Lab in Alderley Park | Wellcome Sanger Institute for the COVID-19 Genomics UK (COG-UK) Consortium | Jacquelyn Wynn, Mairead Hyland, The Lighthouse Lab in Alderley Park and Alex Alderton, Roberto Amato, Jeffrey Barrett, Sonia Goncalves, Ewan Harrison, David K. Jackson, Ian Johnston, Dominic Kwiatkowski, Cordelia Langford, John Sillitoe on behalf of the Wellcome Sanger Institute COVID-19 Surveillance Team |
| EPI_ISL_2718462 | Lighthouse Lab in Glasgow | Wellcome Sanger Institute for the COVID-19 Genomics UK (COG-UK) Consortium | Harper VanSteenhouse, Yumi Kasai, David Gray, Carol Clugston, Anna Dominiczak and Alex Alderton, Roberto Amato, Jeffrey Barrett, Sonia Goncalves, Ewan Harrison, David K. Jackson, Ian Johnston, Dominic Kwiatkowski, Cordelia Langford, John Sillitoe on behalf of the Wellcome Sanger Institute COVID-19 Surveillance Team |
| EPI_ISL_2718463 | Lighthouse Lab in Alderley Park | Wellcome Sanger Institute for the COVID-19 Genomics UK (COG-UK) Consortium | Jacquelyn Wynn, Mairead Hyland, The Lighthouse Lab in Alderley Park and Alex Alderton, Roberto Amato, Jeffrey Barrett, Sonia Goncalves, Ewan Harrison, David K. Jackson, Ian Johnston, Dominic Kwiatkowski, Cordelia Langford, John Sillitoe on behalf of the Wellcome Sanger Institute COVID-19 Surveillance Team |
| EPI_ISL_2718464 | Lighthouse Laboratory Plymouth | Wellcome Sanger Institute for the COVID-19 Genomics UK (COG-UK) Consortium | Lighthouse Laboratory Plymouth and Alex Alderton, Roberto Amato, Jeffrey Barrett, Sonia Goncalves, Ewan Harrison, David K. Jackson, Ian Johnston, Dominic Kwiatkowski, Cordelia Langford, John Sillitoe on behalf of the Wellcome Sanger Institute COVID-19 Surveillance Team |
| EPI_ISL_2718465 | Lighthouse Lab in Alderley Park | Wellcome Sanger Institute for the COVID-19 Genomics UK (COG-UK) Consortium | Jacquelyn Wynn, Mairead Hyland, The Lighthouse Lab in Alderley Park and Alex Alderton, Roberto Amato, Jeffrey Barrett, Sonia Goncalves, Ewan Harrison, David K. Jackson, Ian Johnston, Dominic Kwiatkowski, Cordelia Langford, John Sillitoe on behalf of the Wellcome Sanger Institute COVID-19 Surveillance Team |
| EPI_ISL_2718466, EPI_ISL_2718467 | Lighthouse Lab in Glasgow | Wellcome Sanger Institute for the COVID-19 Genomics UK (COG-UK) Consortium | Harper VanSteenhouse, Yumi Kasai, David Gray, Carol Clugston, Anna Dominiczak and Alex Alderton, Roberto Amato, Jeffrey Barrett, Sonia Goncalves, Ewan Harrison, David K. Jackson, Ian Johnston, Dominic Kwiatkowski, Cordelia Langford, John Sillitoe on behalf of the Wellcome Sanger Institute COVID-19 Surveillance Team |
| EPI_ISL_2718468 | Lighthouse Laboratory Plymouth | Wellcome Sanger Institute for the COVID-19 Genomics UK (COG-UK) Consortium | Lighthouse Laboratory Plymouth and Alex Alderton, Roberto Amato, Jeffrey Barrett, Sonia Goncalves, Ewan Harrison, David K. Jackson, Ian Johnston, Dominic Kwiatkowski, Cordelia Langford, John Sillitoe on behalf of the Wellcome Sanger Institute COVID-19 Surveillance Team |
| EPI_ISL_2718469 | Lighthouse Lab in Alderley Park | Wellcome Sanger Institute for the COVID-19 Genomics UK (COG-UK) Consortium | Jacquelyn Wynn, Mairead Hyland, The Lighthouse Lab in Alderley Park and Alex Alderton, Roberto Amato, Jeffrey Barrett, Sonia Goncalves, Ewan Harrison, David K. Jackson, Ian Johnston, Dominic Kwiatkowski, Cordelia Langford, John Sillitoe on behalf of the Wellcome Sanger Institute COVID-19 Surveillance Team |
| EPI_ISL_2718470 | Lighthouse Lab in Glasgow | Wellcome Sanger Institute for the COVID-19 Genomics UK (COG-UK) Consortium | Harper VanSteenhouse, Yumi Kasai, David Gray, Carol Clugston, Anna Dominiczak and Alex Alderton, Roberto Amato, Jeffrey Barrett, Sonia Goncalves, Ewan Harrison, David K. Jackson, Ian Johnston, Dominic Kwiatkowski, Cordelia Langford, John Sillitoe on behalf of the Wellcome Sanger Institute COVID-19 Surveillance Team |
| EPI_ISL_2718471, EPI_ISL_2718472 | Lighthouse Lab in Alderley Park | Wellcome Sanger Institute for the COVID-19 Genomics UK (COG-UK) Consortium | Jacquelyn Wynn, Mairead Hyland, The Lighthouse Lab in Alderley Park and Alex Alderton, Roberto Amato, Jeffrey Barrett, Sonia Goncalves, Ewan Harrison, David K. Jackson, Ian Johnston, Dominic Kwiatkowski, Cordelia Langford, John Sillitoe on behalf of the Wellcome Sanger Institute COVID-19 Surveillance Team |
| EPI_ISL_2718473 | Lighthouse Laboratory Plymouth | Wellcome Sanger Institute for the COVID-19 Genomics UK (COG-UK) Consortium | Lighthouse Laboratory Plymouth and Alex Alderton, Roberto Amato, Jeffrey Barrett, Sonia Goncalves, Ewan Harrison, David K. Jackson, Ian Johnston, Dominic Kwiatkowski, Cordelia Langford, John Sillitoe on behalf of the Wellcome Sanger Institute COVID-19 Surveillance Team |
| EPI_ISL_2718474, EPI_ISL_2718475, EPI_ISL_2718476 | Lighthouse Lab in Alderley Park | Wellcome Sanger Institute for the COVID-19 Genomics UK (COG-UK) Consortium | Jacquelyn Wynn, Mairead Hyland, The Lighthouse Lab in Alderley Park and Alex Alderton, Roberto Amato, Jeffrey Barrett, Sonia Goncalves, Ewan Harrison, David K. Jackson, Ian Johnston, Dominic Kwiatkowski, Cordelia Langford, John Sillitoe on behalf of the Wellcome Sanger Institute COVID-19 Surveillance Team |
| EPI_ISL_2718477, EPI_ISL_2718478 | Lighthouse Lab in Glasgow | Wellcome Sanger Institute for the COVID-19 Genomics UK (COG-UK) Consortium | Harper VanSteenhouse, Yumi Kasai, David Gray, Carol Clugston, Anna Dominiczak and Alex Alderton, Roberto Amato, Jeffrey Barrett, Sonia Goncalves, Ewan Harrison, David K. Jackson, Ian Johnston, Dominic Kwiatkowski, Cordelia Langford, John Sillitoe on behalf of the Wellcome Sanger Institute COVID-19 Surveillance Team |
| EPI_ISL_2718479 | Lighthouse Lab in Alderley Park | Wellcome Sanger Institute for the COVID-19 Genomics UK (COG-UK) Consortium | Jacquelyn Wynn, Mairead Hyland, The Lighthouse Lab in Alderley Park and Alex Alderton, Roberto Amato, Jeffrey Barrett, Sonia Goncalves, Ewan Harrison, David K. Jackson, Ian Johnston, Dominic Kwiatkowski, Cordelia Langford, John Sillitoe on behalf of the Wellcome Sanger Institute COVID-19 Surveillance Team |
| EPI_ISL_2718480 | Lighthouse Lab in Glasgow | Wellcome Sanger Institute for the COVID-19 Genomics UK (COG-UK) Consortium | Harper VanSteenhouse, Yumi Kasai, David Gray, Carol Clugston, Anna Dominiczak and Alex Alderton, Roberto Amato, Jeffrey Barrett, Sonia Goncalves, Ewan Harrison, David K. Jackson, Ian Johnston, Dominic Kwiatkowski, Cordelia Langford, John Sillitoe on behalf of the Wellcome Sanger Institute COVID-19 Surveillance Team |
| EPI_ISL_2718481, | Lighthouse Laboratory Plymouth | Wellcome Sanger Institute for the COVID-19 Genomics UK (COG-UK) | Lighthouse Laboratory Plymouth and Alex Alderton, Roberto Amato, Jeffrey Barrett, Sonia Goncalves, Ewan Harrison, David K. Jackson, Ian Johnston, Dominic Kwiatkowski, Cordelia Langford, John Sillitoe on behalf of the Wellcome Sanger |

| | | | |
|---|---|---|---|
| EPI_ISL_2718482, EPI_ISL_2718483 | | Consortium | Institute COVID-19 Surveillance Team |
| EPI_ISL_2718484 | Lighthouse Lab in Alderley Park | Wellcome Sanger Institute for the COVID-19 Genomics UK (COG-UK) Consortium | Jacquelyn Wynn, Mairead Hyland, The Lighthouse Lab in Alderley Park and Alex Alderton, Roberto Amato, Jeffrey Barrett, Sonia Goncalves, Ewan Harrison, David K. Jackson, Ian Johnston, Dominic Kwiatkowski, Cordelia Langford, John Sillitoe on behalf of the Wellcome Sanger Institute COVID-19 Surveillance Team |
| EPI_ISL_2718485 | Lighthouse Laboratory Plymouth | Wellcome Sanger Institute for the COVID-19 Genomics UK (COG-UK) Consortium | Lighthouse Laboratory Plymouth and Alex Alderton, Roberto Amato, Jeffrey Barrett, Sonia Goncalves, Ewan Harrison, David K. Jackson, Ian Johnston, Dominic Kwiatkowski, Cordelia Langford, John Sillitoe on behalf of the Wellcome Sanger Institute COVID-19 Surveillance Team |
| EPI_ISL_2718486 | Lighthouse Lab in Glasgow | Wellcome Sanger Institute for the COVID-19 Genomics UK (COG-UK) Consortium | Harper VanSteenhouse, Yumi Kasai, David Gray, Carol Clugston, Anna Dominiczak and Alex Alderton, Roberto Amato, Jeffrey Barrett, Sonia Goncalves, Ewan Harrison, David K. Jackson, Ian Johnston, Dominic Kwiatkowski, Cordelia Langford, John Sillitoe on behalf of the Wellcome Sanger Institute COVID-19 Surveillance Team |
| EPI_ISL_2718487 | Lighthouse Lab in Alderley Park | Wellcome Sanger Institute for the COVID-19 Genomics UK (COG-UK) Consortium | Jacquelyn Wynn, Mairead Hyland, The Lighthouse Lab in Alderley Park and Alex Alderton, Roberto Amato, Jeffrey Barrett, Sonia Goncalves, Ewan Harrison, David K. Jackson, Ian Johnston, Dominic Kwiatkowski, Cordelia Langford, John Sillitoe on behalf of the Wellcome Sanger Institute COVID-19 Surveillance Team |
| EPI_ISL_2718488 | Lighthouse Lab in Glasgow | Wellcome Sanger Institute for the COVID-19 Genomics UK (COG-UK) Consortium | Harper VanSteenhouse, Yumi Kasai, David Gray, Carol Clugston, Anna Dominiczak and Alex Alderton, Roberto Amato, Jeffrey Barrett, Sonia Goncalves, Ewan Harrison, David K. Jackson, Ian Johnston, Dominic Kwiatkowski, Cordelia Langford, John Sillitoe on behalf of the Wellcome Sanger Institute COVID-19 Surveillance Team |
| EPI_ISL_2718490 | Lighthouse Lab in Alderley Park | Wellcome Sanger Institute for the COVID-19 Genomics UK (COG-UK) Consortium | Jacquelyn Wynn, Mairead Hyland, The Lighthouse Lab in Alderley Park and Alex Alderton, Roberto Amato, Jeffrey Barrett, Sonia Goncalves, Ewan Harrison, David K. Jackson, Ian Johnston, Dominic Kwiatkowski, Cordelia Langford, John Sillitoe on behalf of the Wellcome Sanger Institute COVID-19 Surveillance Team |
| EPI_ISL_2718491, EPI_ISL_2718492 | Lighthouse Laboratory Plymouth | Wellcome Sanger Institute for the COVID-19 Genomics UK (COG-UK) Consortium | Lighthouse Laboratory Plymouth and Alex Alderton, Roberto Amato, Jeffrey Barrett, Sonia Goncalves, Ewan Harrison, David K. Jackson, Ian Johnston, Dominic Kwiatkowski, Cordelia Langford, John Sillitoe on behalf of the Wellcome Sanger Institute COVID-19 Surveillance Team |
| EPI_ISL_2718493, EPI_ISL_2718495 | Lighthouse Lab in Alderley Park | Wellcome Sanger Institute for the COVID-19 Genomics UK (COG-UK) Consortium | Jacquelyn Wynn, Mairead Hyland, The Lighthouse Lab in Alderley Park and Alex Alderton, Roberto Amato, Jeffrey Barrett, Sonia Goncalves, Ewan Harrison, David K. Jackson, Ian Johnston, Dominic Kwiatkowski, Cordelia Langford, John Sillitoe on behalf of the Wellcome Sanger Institute COVID-19 Surveillance Team |
| EPI_ISL_2718497, EPI_ISL_2718498 | Lighthouse Laboratory Plymouth | Wellcome Sanger Institute for the COVID-19 Genomics UK (COG-UK) Consortium | Lighthouse Laboratory Plymouth and Alex Alderton, Roberto Amato, Jeffrey Barrett, Sonia Goncalves, Ewan Harrison, David K. Jackson, Ian Johnston, Dominic Kwiatkowski, Cordelia Langford, John Sillitoe on behalf of the Wellcome Sanger Institute COVID-19 Surveillance Team |
| EPI_ISL_2718499 | Lighthouse Lab in Alderley Park | Wellcome Sanger Institute for the COVID-19 Genomics UK (COG-UK) Consortium | Jacquelyn Wynn, Mairead Hyland, The Lighthouse Lab in Alderley Park and Alex Alderton, Roberto Amato, Jeffrey Barrett, Sonia Goncalves, Ewan Harrison, David K. Jackson, Ian Johnston, Dominic Kwiatkowski, Cordelia Langford, John Sillitoe on behalf of the Wellcome Sanger Institute COVID-19 Surveillance Team |
| EPI_ISL_2718500, EPI_ISL_2718501 | Lighthouse Laboratory Plymouth | Wellcome Sanger Institute for the COVID-19 Genomics UK (COG-UK) Consortium | Lighthouse Laboratory Plymouth and Alex Alderton, Roberto Amato, Jeffrey Barrett, Sonia Goncalves, Ewan Harrison, David K. Jackson, Ian Johnston, Dominic Kwiatkowski, Cordelia Langford, John Sillitoe on behalf of the Wellcome Sanger Institute COVID-19 Surveillance Team |
| EPI_ISL_2718502 | Lighthouse Lab in Alderley Park | Wellcome Sanger Institute for the COVID-19 Genomics UK (COG-UK) Consortium | Jacquelyn Wynn, Mairead Hyland, The Lighthouse Lab in Alderley Park and Alex Alderton, Roberto Amato, Jeffrey Barrett, Sonia Goncalves, Ewan Harrison, David K. Jackson, Ian Johnston, Dominic Kwiatkowski, Cordelia Langford, John Sillitoe on behalf of the Wellcome Sanger Institute COVID-19 Surveillance Team |
| EPI_ISL_2718503 | Lighthouse Laboratory Plymouth | Wellcome Sanger Institute for the COVID-19 Genomics UK (COG-UK) Consortium | Lighthouse Laboratory Plymouth and Alex Alderton, Roberto Amato, Jeffrey Barrett, Sonia Goncalves, Ewan Harrison, David K. Jackson, Ian Johnston, Dominic Kwiatkowski, Cordelia Langford, John Sillitoe on behalf of the Wellcome Sanger Institute COVID-19 Surveillance Team |
| EPI_ISL_2718504 | Lighthouse Lab in Glasgow | Wellcome Sanger Institute for the COVID-19 Genomics UK (COG-UK) Consortium | Harper VanSteenhouse, Yumi Kasai, David Gray, Carol Clugston, Anna Dominiczak and Alex Alderton, Roberto Amato, Jeffrey Barrett, Sonia Goncalves, Ewan Harrison, David K. Jackson, Ian Johnston, Dominic Kwiatkowski, Cordelia Langford, John Sillitoe on behalf of the Wellcome Sanger Institute COVID-19 Surveillance Team |
| EPI_ISL_2718505 | Lighthouse Laboratory Plymouth | Wellcome Sanger Institute for the COVID-19 Genomics UK (COG-UK) Consortium | Lighthouse Laboratory Plymouth and Alex Alderton, Roberto Amato, Jeffrey Barrett, Sonia Goncalves, Ewan Harrison, David K. Jackson, Ian Johnston, Dominic Kwiatkowski, Cordelia Langford, John Sillitoe on behalf of the Wellcome Sanger Institute COVID-19 Surveillance Team |
| EPI_ISL_2718507 | Lighthouse Lab in Glasgow | Wellcome Sanger Institute for the COVID-19 Genomics UK (COG-UK) Consortium | Harper VanSteenhouse, Yumi Kasai, David Gray, Carol Clugston, Anna Dominiczak and Alex Alderton, Roberto Amato, Jeffrey Barrett, Sonia Goncalves, Ewan Harrison, David K. Jackson, Ian Johnston, Dominic Kwiatkowski, Cordelia Langford, John Sillitoe on behalf of the Wellcome Sanger Institute COVID-19 Surveillance Team |
| EPI_ISL_2718508 | Lighthouse Laboratory Plymouth | Wellcome Sanger Institute for the COVID-19 Genomics UK (COG-UK) Consortium | Lighthouse Laboratory Plymouth and Alex Alderton, Roberto Amato, Jeffrey Barrett, Sonia Goncalves, Ewan Harrison, David K. Jackson, Ian Johnston, Dominic Kwiatkowski, Cordelia Langford, John Sillitoe on behalf of the Wellcome Sanger Institute COVID-19 Surveillance Team |
| EPI_ISL_2718509 | Lighthouse Lab in Alderley Park | Wellcome Sanger Institute for the COVID-19 Genomics UK (COG-UK) Consortium | Jacquelyn Wynn, Mairead Hyland, The Lighthouse Lab in Alderley Park and Alex Alderton, Roberto Amato, Jeffrey Barrett, Sonia Goncalves, Ewan Harrison, David K. Jackson, Ian Johnston, Dominic Kwiatkowski, Cordelia Langford, John Sillitoe on behalf of the Wellcome Sanger Institute COVID-19 Surveillance Team |
| EPI_ISL_2718510 | Lighthouse Laboratory Plymouth | Wellcome Sanger Institute for the COVID-19 Genomics UK (COG-UK) Consortium | Lighthouse Laboratory Plymouth and Alex Alderton, Roberto Amato, Jeffrey Barrett, Sonia Goncalves, Ewan Harrison, David K. Jackson, Ian Johnston, Dominic Kwiatkowski, Cordelia Langford, John Sillitoe on behalf of the Wellcome Sanger Institute COVID-19 Surveillance Team |
| EPI_ISL_2718511 | Lighthouse Lab in Glasgow | Wellcome Sanger Institute for the COVID-19 Genomics UK (COG-UK) Consortium | Harper VanSteenhouse, Yumi Kasai, David Gray, Carol Clugston, Anna Dominiczak and Alex Alderton, Roberto Amato, Jeffrey Barrett, Sonia Goncalves, Ewan Harrison, David K. Jackson, Ian Johnston, Dominic Kwiatkowski, Cordelia Langford, John Sillitoe on behalf of the Wellcome Sanger Institute COVID-19 Surveillance Team |
| EPI_ISL_2718512 | Lighthouse Lab in Alderley Park | Wellcome Sanger Institute for the COVID-19 Genomics UK (COG-UK) Consortium | Jacquelyn Wynn, Mairead Hyland, The Lighthouse Lab in Alderley Park and Alex Alderton, Roberto Amato, Jeffrey Barrett, Sonia Goncalves, Ewan Harrison, David K. Jackson, Ian Johnston, Dominic Kwiatkowski, Cordelia Langford, John Sillitoe on behalf of the Wellcome Sanger Institute COVID-19 Surveillance Team |
| EPI_ISL_2718513 | Lighthouse Lab in Glasgow | Wellcome Sanger Institute for the COVID-19 Genomics UK (COG-UK) Consortium | Harper VanSteenhouse, Yumi Kasai, David Gray, Carol Clugston, Anna Dominiczak and Alex Alderton, Roberto Amato, Jeffrey Barrett, Sonia Goncalves, Ewan Harrison, David K. Jackson, Ian Johnston, Dominic Kwiatkowski, Cordelia Langford, John Sillitoe on behalf of the Wellcome Sanger Institute COVID-19 Surveillance Team |
| EPI_ISL_2718514 | Lighthouse Laboratory Plymouth | Wellcome Sanger Institute for the COVID-19 Genomics UK (COG-UK) Consortium | Lighthouse Laboratory Plymouth and Alex Alderton, Roberto Amato, Jeffrey Barrett, Sonia Goncalves, Ewan Harrison, David K. Jackson, Ian Johnston, Dominic Kwiatkowski, Cordelia Langford, John Sillitoe on behalf of the Wellcome Sanger Institute COVID-19 Surveillance Team |
| EPI_ISL_2718515 | Lighthouse Lab in Glasgow | Wellcome Sanger Institute for the COVID-19 Genomics UK (COG-UK) Consortium | Harper VanSteenhouse, Yumi Kasai, David Gray, Carol Clugston, Anna Dominiczak and Alex Alderton, Roberto Amato, Jeffrey Barrett, Sonia Goncalves, Ewan Harrison, David K. Jackson, Ian Johnston, Dominic Kwiatkowski, Cordelia Langford, John Sillitoe on behalf of the Wellcome Sanger Institute COVID-19 Surveillance Team |
| EPI_ISL_2718516, EPI_ISL_2718517 | Lighthouse Laboratory Plymouth | Wellcome Sanger Institute for the COVID-19 Genomics UK (COG-UK) Consortium | Lighthouse Laboratory Plymouth and Alex Alderton, Roberto Amato, Jeffrey Barrett, Sonia Goncalves, Ewan Harrison, David K. Jackson, Ian Johnston, Dominic Kwiatkowski, Cordelia Langford, John Sillitoe on behalf of the Wellcome Sanger Institute COVID-19 Surveillance Team |
| EPI_ISL_2718518, EPI_ISL_2718519, EPI_ISL_2718520, EPI_ISL_2718521, EPI_ISL_2718522 | Lighthouse Lab in Alderley Park | Wellcome Sanger Institute for the COVID-19 Genomics UK (COG-UK) Consortium | Jacquelyn Wynn, Mairead Hyland, The Lighthouse Lab in Alderley Park and Alex Alderton, Roberto Amato, Jeffrey Barrett, Sonia Goncalves, Ewan Harrison, David K. Jackson, Ian Johnston, Dominic Kwiatkowski, Cordelia Langford, John Sillitoe on behalf of the Wellcome Sanger Institute COVID-19 Surveillance Team |
| EPI_ISL_2718523, EPI_ISL_2718524, EPI_ISL_2718526 | Lighthouse Laboratory Plymouth | Wellcome Sanger Institute for the COVID-19 Genomics UK (COG-UK) Consortium | Lighthouse Laboratory Plymouth and Alex Alderton, Roberto Amato, Jeffrey Barrett, Sonia Goncalves, Ewan Harrison, David K. Jackson, Ian Johnston, Dominic Kwiatkowski, Cordelia Langford, John Sillitoe on behalf of the Wellcome Sanger Institute COVID-19 Surveillance Team |
| EPI_ISL_2718527, EPI_ISL_2718528 | Lighthouse Lab in Alderley Park | Wellcome Sanger Institute for the COVID-19 Genomics UK (COG-UK) Consortium | Jacquelyn Wynn, Mairead Hyland, The Lighthouse Lab in Alderley Park and Alex Alderton, Roberto Amato, Jeffrey Barrett, Sonia Goncalves, Ewan Harrison, David K. Jackson, Ian Johnston, Dominic Kwiatkowski, Cordelia Langford, John Sillitoe on behalf of the Wellcome Sanger Institute COVID-19 Surveillance Team |
| EPI_ISL_2718529 | Lighthouse Laboratory Plymouth | Wellcome Sanger Institute for the COVID-19 Genomics UK (COG-UK) Consortium | Lighthouse Laboratory Plymouth and Alex Alderton, Roberto Amato, Jeffrey Barrett, Sonia Goncalves, Ewan Harrison, David K. Jackson, Ian Johnston, Dominic Kwiatkowski, Cordelia Langford, John Sillitoe on behalf of the Wellcome Sanger Institute COVID-19 Surveillance Team |
| EPI_ISL_2718530 | Lighthouse Lab in Alderley Park | Wellcome Sanger Institute for the COVID-19 Genomics UK (COG-UK) Consortium | Jacquelyn Wynn, Mairead Hyland, The Lighthouse Lab in Alderley Park and Alex Alderton, Roberto Amato, Jeffrey Barrett, Sonia Goncalves, Ewan Harrison, David K. Jackson, Ian Johnston, Dominic Kwiatkowski, Cordelia Langford, John Sillitoe on behalf of the Wellcome Sanger Institute COVID-19 Surveillance Team |
| EPI_ISL_2718531 | Lighthouse Laboratory Plymouth | Wellcome Sanger Institute for the COVID-19 Genomics UK (COG-UK) Consortium | Lighthouse Laboratory Plymouth and Alex Alderton, Roberto Amato, Jeffrey Barrett, Sonia Goncalves, Ewan Harrison, David K. Jackson, Ian Johnston, Dominic Kwiatkowski, Cordelia Langford, John Sillitoe on behalf of the Wellcome Sanger Institute COVID-19 Surveillance Team |
| EPI_ISL_2718541, EPI_ISL_2718546 | Molecular Diagnostics Pathology Department Mater Dei Hospital Malta | Molecular Diagnostics Pathology Department Mater Dei Hospital Malta | G Zahra, R Borg, C Cilia, L Grech, M Briffa |
| EPI_ISL_2718550, EPI_ISL_2718554, EPI_ISL_2718557, EPI_ISL_2718560, EPI_ISL_2718562, EPI_ISL_2718563, EPI_ISL_2718564, EPI_ISL_2718569, EPI_ISL_2718570, EPI_ISL_2718572, EPI_ISL_2718573, EPI_ISL_2718575, EPI_ISL_2718577, EPI_ISL_2718578, EPI_ISL_2718579, EPI_ISL_2718582, EPI_ISL_2718585, EPI_ISL_2718586, EPI_ISL_2718588, EPI_ISL_2718589, EPI_ISL_2718593, EPI_ISL_2718594, EPI_ISL_2718596, EPI_ISL_2718597, EPI_ISL_2718600, EPI_ISL_2718601, EPI_ISL_2718605, EPI_ISL_2718608, EPI_ISL_2718609, EPI_ISL_2718611, EPI_ISL_2718613, EPI_ISL_2718615, EPI_ISL_2718618, EPI_ISL_2718620, EPI_ISL_2718621, EPI_ISL_2718622, EPI_ISL_2718623, EPI_ISL_2718625, EPI_ISL_2718626, EPI_ISL_2718631, EPI_ISL_2718633, EPI_ISL_2718634 | | | |
| see above | Microbiology Department, Laboratori Clínic Metropolitana Nord. Hospital Universitari Germans Trias i Pujol | Can Ruti SARS-CoV-2 Sequencing Hub (HUGTiP/IrsiCaixa/IGTP) | Marc Noguera-Julian, Pilar Armengol, Ignacio Blanco, Antoni E Bordoy, Francesc Catala-Moll, Pere-Joan Cardona, Maria Casadellà, Cristina Casañ, Gemma Clara, Bonaventura Clotet, Cristina Esteban, Montserrat Giménez, Mercedes Guerrero, Anna Not, Roger Paredes, Mariona Parera, Verónica Saludes, Alba Sánchez, and Elisa Martró on behalf of the Can Ruti SARS-CoV-2 Sequencing Hub |
| EPI_ISL_2719041, EPI_ISL_2719042, EPI_ISL_2719043, EPI_ISL_2719048, EPI_ISL_2719261, EPI_ISL_2719268, EPI_ISL_2719339, EPI_ISL_2719387, EPI_ISL_2719389, EPI_ISL_2719392, EPI_ISL_2719401, EPI_ISL_2719407, EPI_ISL_2719427, EPI_ISL_2719459, EPI_ISL_2719483, EPI_ISL_2719510, EPI_ISL_2719559, EPI_ISL_2719571, EPI_ISL_2719705, EPI_ISL_2719706, EPI_ISL_2719732, EPI_ISL_2719808, EPI_ISL_2719825, EPI_ISL_2719829, EPI_ISL_2719872, EPI_ISL_2719873, EPI_ISL_2719879, EPI_ISL_2719883, EPI_ISL_2719897, EPI_ISL_2719905, EPI_ISL_2719923, EPI_ISL_2719931, EPI_ISL_2719960, EPI_ISL_2719961, EPI_ISL_2719983, EPI_ISL_2719984, EPI_ISL_2719994, EPI_ISL_2720222, EPI_ISL_2720228, EPI_ISL_2720230, EPI_ISL_2720291, EPI_ISL_2720376, EPI_ISL_2720378, EPI_ISL_2720387, EPI_ISL_2720393, EPI_ISL_2720396, EPI_ISL_2720398, EPI_ISL_2720418, EPI_ISL_2720423, EPI_ISL_2720442, EPI_ISL_2720445, EPI_ISL_2720609, EPI_ISL_2720610, EPI_ISL_2720639, EPI_ISL_2720679, EPI_ISL_2720689, EPI_ISL_2720708, EPI_ISL_2720714, EPI_ISL_2720718, EPI_ISL_2720720, EPI_ISL_2720773, EPI_ISL_2721073, EPI_ISL_2721074, EPI_ISL_2721075, EPI_ISL_2721120, EPI_ISL_2721139, EPI_ISL_2721140 | | | |
| see above | BCCDC Public Health Laboratory | BCCDC Public Health Laboratory | Prystajecky Natalie, Linda Hoang, Dan Fornika, John Tyson, Shannon Russell, Kim Macdonald, Kimia Kamelian, Ana Pacagnella, Corrinne Ng, Loretta Janz, Robert Azana, Mel Krajden |
| EPI_ISL_2721143, EPI_ISL_2721145, EPI_ISL_2721146, EPI_ISL_2721148, EPI_ISL_2721150, EPI_ISL_2721152, EPI_ISL_2721155, EPI_ISL_2721157, EPI_ISL_2721159, EPI_ISL_2721162 | Quadram Institute Bioscience | COVID-19 Genomics UK (COG-UK) Consortium | Dave J. Baker, Gemma L. Kay, Alp Aydin, Thanh Le-Viet, Steven Rudder, Ana P. Tedim, Anastasia Kolyva, Maria Diaz, Leonardo de Oliveira Martins, Nabil-Fareed Alikhan, Lizzie Meadows, Rachael Stanley, Ngozi Elumogo, Muhammed Yasir, Nicholas M. Thomson, Alexander J Trotter, Rachel Gilroy, Samuel Bloomfield, Claire Stuart, Andrew Bell, Reenesh Prakash, Samir Dervisevic, Alison E. Mather, John Wain, Mark Webber, Andrew J. Page, Justin O'Grady |
| EPI_ISL_2721182 | A.S.L. TO5 | Fondazione del Piemonte per l'Oncologia IRCCS | Antonino Sottile, Silvia Brossa, Paola Marino, Giorgio Giardina |
| EPI_ISL_2721223, EPI_ISL_2721230 | A.S.L. CN1 | Fondazione del Piemonte per l'Oncologia IRCCS | Antonino Sottile, Silvia Brossa, Paola Marino, Giorgio Giardina |
| EPI_ISL_2721281, | A.S.L. CN2 | Fondazione del Piemonte per l'Oncologia IRCCS | Antonino Sottile, Silvia Brossa, Paola Marino, Giorgio Giardina |

| | | | |
|---|---|---|---|
| EPI_ISL_2721283 | | | |
| EPI_ISL_2721288 | A.S.L. TO4 | Fondazione del Piemonte per l'Oncologia IRCCS | Antonino Sottile, Silvia Brossa, Paola Marino, Giorgio Giardina |
| EPI_ISL_2721355 | BCCDC Public Health Laboratory | BCCDC Public Health Laboratory | Prystajecky Natalie, Linda Hoang, Dan Fornika, John Tyson, Shannon Russell, Kim Macdonald, Kimia Kamelian, Ana Pacagnella, Corrinne Ng, Loretta Janz, Robert Azana, Mel Krajden |
| EPI_ISL_2721376 | Pathology North - Hunter - NSW Health Pathology | NSW Health Pathology - Institute of Clinical Pathology and Medical Research; Westmead Hospital; University of Sydney | CIDM-PH et al. |
| EPI_ISL_2721378, EPI_ISL_2721398, EPI_ISL_2721424, EPI_ISL_2721428, EPI_ISL_2721429, EPI_ISL_2721488, EPI_ISL_2721494, EPI_ISL_2721514, EPI_ISL_2721532, EPI_ISL_2721548, EPI_ISL_2721551, EPI_ISL_2721566, EPI_ISL_2721583, EPI_ISL_2721636, EPI_ISL_2721653, EPI_ISL_2721679, EPI_ISL_2721716, EPI_ISL_2721717, EPI_ISL_2721731, EPI_ISL_2721739, EPI_ISL_2721753, EPI_ISL_2721759, EPI_ISL_2721762 | | | |
| see above | BCCDC Public Health Laboratory | BCCDC Public Health Laboratory | Prystajecky Natalie, Linda Hoang, Dan Fornika, John Tyson, Shannon Russell, Kim Macdonald, Kimia Kamelian, Ana Pacagnella, Corrinne Ng, Loretta Janz, Robert Azana, Mel Krajden |
| EPI_ISL_2721853, EPI_ISL_2721855, EPI_ISL_2721858, EPI_ISL_2721860, EPI_ISL_2721863, EPI_ISL_2721865, EPI_ISL_2721866, EPI_ISL_2721868, EPI_ISL_2721870, EPI_ISL_2721871, EPI_ISL_2721873, EPI_ISL_2721874, EPI_ISL_2721876, EPI_ISL_2721878, EPI_ISL_2721880, EPI_ISL_2721881, EPI_ISL_2721884, EPI_ISL_2721886, EPI_ISL_2721887, EPI_ISL_2721889, EPI_ISL_2721891, EPI_ISL_2721893, EPI_ISL_2721895, EPI_ISL_2721896, EPI_ISL_2721898, EPI_ISL_2721900, EPI_ISL_2721901, EPI_ISL_2721903, EPI_ISL_2721905, EPI_ISL_2721906, EPI_ISL_2721908, EPI_ISL_2721910, EPI_ISL_2721911, EPI_ISL_2721913, EPI_ISL_2721914, EPI_ISL_2721915, EPI_ISL_2721917, EPI_ISL_2721919, EPI_ISL_2721921, EPI_ISL_2721922, EPI_ISL_2721924, EPI_ISL_2721926, EPI_ISL_2721927, EPI_ISL_2721931, EPI_ISL_2721935, EPI_ISL_2721937, EPI_ISL_2721939, EPI_ISL_2721940, EPI_ISL_2721943, EPI_ISL_2721944, EPI_ISL_2721946, EPI_ISL_2721948, EPI_ISL_2721949, EPI_ISL_2721951, EPI_ISL_2721954, EPI_ISL_2721956, EPI_ISL_2721960, EPI_ISL_2721961, EPI_ISL_2721963, EPI_ISL_2721965, EPI_ISL_2721966, EPI_ISL_2721968, EPI_ISL_2721971, EPI_ISL_2721973, EPI_ISL_2721974, EPI_ISL_2721976, EPI_ISL_2721977, EPI_ISL_2721979, EPI_ISL_2721982, EPI_ISL_2721984, EPI_ISL_2721985, EPI_ISL_2721987, EPI_ISL_2721989, EPI_ISL_2721990, EPI_ISL_2721992, EPI_ISL_2721993, EPI_ISL_2721995, EPI_ISL_2721996, EPI_ISL_2721998, EPI_ISL_2721999, EPI_ISL_2722001, EPI_ISL_2722003, EPI_ISL_2722008, EPI_ISL_2722009, EPI_ISL_2722013, EPI_ISL_2722015, EPI_ISL_2722016, EPI_ISL_2722018, EPI_ISL_2722020, EPI_ISL_2722021, EPI_ISL_2722023, EPI_ISL_2722025, EPI_ISL_2722026, EPI_ISL_2722030, EPI_ISL_2722032, EPI_ISL_2722036, EPI_ISL_2722037, EPI_ISL_2722039, EPI_ISL_2722041, EPI_ISL_2722042, EPI_ISL_2722044, EPI_ISL_2722046, EPI_ISL_2722048, EPI_ISL_2722049, EPI_ISL_2722050, EPI_ISL_2722051, EPI_ISL_2722052, EPI_ISL_2722053, EPI_ISL_2722054, EPI_ISL_2722055, EPI_ISL_2722056, EPI_ISL_2722057, EPI_ISL_2722058, EPI_ISL_2722059, EPI_ISL_2722060, EPI_ISL_2722061, EPI_ISL_2722062, EPI_ISL_2722063, EPI_ISL_2722064, EPI_ISL_2722065, EPI_ISL_2722066, EPI_ISL_2722067, EPI_ISL_2722068, EPI_ISL_2722069, EPI_ISL_2722070, EPI_ISL_2722071, EPI_ISL_2722072, EPI_ISL_2722073, EPI_ISL_2722074, EPI_ISL_2722075, EPI_ISL_2722076, EPI_ISL_2722077, EPI_ISL_2722078, EPI_ISL_2722079, EPI_ISL_2722080, EPI_ISL_2722081, EPI_ISL_2722082, EPI_ISL_2722083, EPI_ISL_2722084, EPI_ISL_2722085, EPI_ISL_2722086, EPI_ISL_2722087, EPI_ISL_2722088, EPI_ISL_2722089, EPI_ISL_2722090, EPI_ISL_2722091, EPI_ISL_2722092, EPI_ISL_2722093, EPI_ISL_2722094, EPI_ISL_2722095, EPI_ISL_2722096, EPI_ISL_2722097, EPI_ISL_2722098, EPI_ISL_2722099, EPI_ISL_2722101, EPI_ISL_2722102, EPI_ISL_2722103, EPI_ISL_2722104 | | | |
| see above | University of Birmingham | COVID-19 Genomics UK (COG-UK) Consortium | Institute of Microbiology, University of Birmingham: Claire McMurray, Joanne Stockton, Samuel Nicholls, Radoslaw Poplawski, Will Rowe, Josh Quick, Nicholas Loman. University of Birmingham Testing Laboratory: Celina M Whalley, Andrew Bosworth, Charlotte Poxon, Kasun Wanigasooriya, Oliver Pickles, Mike Kidd, Alex Richter, Andrew D Beggs PHE Heartlands Lab: Husam Osman, Andrew Bosworth. Queen Elizabeth Hospital: Anna Casey |
| EPI_ISL_2722106, EPI_ISL_2722107, EPI_ISL_2722108, EPI_ISL_2722109, EPI_ISL_2722110, EPI_ISL_2722111, EPI_ISL_2722112, EPI_ISL_2722113, EPI_ISL_2722114, EPI_ISL_2722115, EPI_ISL_2722116, EPI_ISL_2722118, EPI_ISL_2722120, EPI_ISL_2722121, EPI_ISL_2722122, EPI_ISL_2722123, EPI_ISL_2722124, EPI_ISL_2722125, EPI_ISL_2722126, EPI_ISL_2722127, EPI_ISL_2722128, EPI_ISL_2722129, EPI_ISL_2722131, EPI_ISL_2722132, EPI_ISL_2722133, EPI_ISL_2722134, EPI_ISL_2722135, EPI_ISL_2722136, EPI_ISL_2722137, EPI_ISL_2722138, EPI_ISL_2722139, EPI_ISL_2722140, EPI_ISL_2722141, EPI_ISL_2722142, EPI_ISL_2722143, EPI_ISL_2722144, EPI_ISL_2722145, EPI_ISL_2722146, EPI_ISL_2722147, EPI_ISL_2722148, EPI_ISL_2722149, EPI_ISL_2722150, EPI_ISL_2722151, EPI_ISL_2722153, EPI_ISL_2722155, EPI_ISL_2722157, EPI_ISL_2722158, EPI_ISL_2722159, EPI_ISL_2722160, EPI_ISL_2722161, EPI_ISL_2722162, EPI_ISL_2722163, EPI_ISL_2722164, EPI_ISL_2722165, EPI_ISL_2722166, EPI_ISL_2722167, EPI_ISL_2722168, EPI_ISL_2722169, EPI_ISL_2722170, EPI_ISL_2722171, EPI_ISL_2722172, EPI_ISL_2722173, EPI_ISL_2722174, EPI_ISL_2722175, EPI_ISL_2722177, EPI_ISL_2722178, EPI_ISL_2722179, EPI_ISL_2722180, EPI_ISL_2722181, EPI_ISL_2722183, EPI_ISL_2722184, EPI_ISL_2722185, EPI_ISL_2722186, EPI_ISL_2722187, EPI_ISL_2722188, EPI_ISL_2722189, EPI_ISL_2722191, EPI_ISL_2722192, EPI_ISL_2722193, EPI_ISL_2722194, EPI_ISL_2722195, EPI_ISL_2722196, EPI_ISL_2722197, EPI_ISL_2722199, EPI_ISL_2722200, EPI_ISL_2722201, EPI_ISL_2722202, EPI_ISL_2722203, EPI_ISL_2722204, EPI_ISL_2722205, EPI_ISL_2722206, EPI_ISL_2722207, EPI_ISL_2722208, EPI_ISL_2722209, EPI_ISL_2722210, EPI_ISL_2722211, EPI_ISL_2722212, EPI_ISL_2722213, EPI_ISL_2722215, EPI_ISL_2722216, EPI_ISL_2722217, EPI_ISL_2722218, EPI_ISL_2722219, EPI_ISL_2722220, EPI_ISL_2722221, EPI_ISL_2722222, EPI_ISL_2722223, EPI_ISL_2722224, EPI_ISL_2722226, EPI_ISL_2722227, EPI_ISL_2722228, EPI_ISL_2722229, EPI_ISL_2722230, EPI_ISL_2722231, EPI_ISL_2722232, EPI_ISL_2722233, EPI_ISL_2722234, EPI_ISL_2722235, EPI_ISL_2722236, EPI_ISL_2722237, EPI_ISL_2722238, EPI_ISL_2722239, EPI_ISL_2722240, EPI_ISL_2722241, EPI_ISL_2722242, EPI_ISL_2722243, EPI_ISL_2722245, EPI_ISL_2722246, EPI_ISL_2722247, EPI_ISL_2722248, EPI_ISL_2722249, EPI_ISL_2722250, EPI_ISL_2722251, EPI_ISL_2722252, EPI_ISL_2722253, EPI_ISL_2722254, EPI_ISL_2722255, EPI_ISL_2722256, EPI_ISL_2722257, EPI_ISL_2722258, EPI_ISL_2722259, EPI_ISL_2722260, EPI_ISL_2722261, EPI_ISL_2722262, EPI_ISL_2722263, EPI_ISL_2722264, EPI_ISL_2722265, EPI_ISL_2722266, EPI_ISL_2722268, EPI_ISL_2722269, EPI_ISL_2722270, EPI_ISL_2722271, EPI_ISL_2722272, EPI_ISL_2722273, EPI_ISL_2722274, EPI_ISL_2722275, EPI_ISL_2722276, EPI_ISL_2722277, EPI_ISL_2722279, EPI_ISL_2722280, EPI_ISL_2722281, EPI_ISL_2722282, EPI_ISL_2722283, EPI_ISL_2722284, EPI_ISL_2722285, EPI_ISL_2722286, EPI_ISL_2722287, EPI_ISL_2722288, EPI_ISL_2722289, EPI_ISL_2722290, EPI_ISL_2722291, EPI_ISL_2722292, EPI_ISL_2722293, EPI_ISL_2722294, EPI_ISL_2722295, EPI_ISL_2722296, EPI_ISL_2722297, EPI_ISL_2722298, EPI_ISL_2722299, EPI_ISL_2722300, EPI_ISL_2722301, EPI_ISL_2722302, EPI_ISL_2722303, EPI_ISL_2722304, EPI_ISL_2722305, EPI_ISL_2722306, EPI_ISL_2722307, EPI_ISL_2722308, EPI_ISL_2722309, EPI_ISL_2722310, EPI_ISL_2722311, EPI_ISL_2722312, EPI_ISL_2722313, EPI_ISL_2722314, EPI_ISL_2722315, EPI_ISL_2722316, EPI_ISL_2722317 | | | |
| see above | Department of Pathology, University of Cambridge | COVID-19 Genomics UK (COG-UK) Consortium | Aminu S. Jahun, Yasmin Chaudhry, Iliana Georgana, Myra Hosmillo, Rhys Izuagbe, William L. Hamilton, Martin D. Curran, Surendra Parmar, Ian Goodfellow |
| EPI_ISL_2722318 | Intibios Karawang | Indonesian Institute of Sciences (LIPI)/National Agency of Research and Innovation (BRIN) and National Institute of Health Research and Development (NIHRD) | Indriawati, Idris, Masrukhin, Syam B. Iryanto, Anik. B. Dharmayanthi, Isa Nuryana, Syaiful Rizal, Ratih A. Ningrum, Wien Kusharyoto, Yayuk Sri Rahayu, Subangkit, Vivi Setiawaty, Sugiyono Saputra |
| EPI_ISL_2722319 | Department of Pathology, University of Cambridge | COVID-19 Genomics UK (COG-UK) Consortium | Aminu S. Jahun, Yasmin Chaudhry, Iliana Georgana, Myra Hosmillo, Rhys Izuagbe, William L. Hamilton, Martin D. Curran, Surendra Parmar, Ian Goodfellow |
| EPI_ISL_2722320 | Intibios Karawang | Indonesian Institute of Sciences (LIPI)/National Agency of Research and Innovation (BRIN) and National Institute of Health Research and Development (NIHRD) | Idris, Isa Nuryana, Erwin Fajar Hasrianda, Indriawati, Masrukhin, Gita Syahputra, Syam B. Iryanto, Anik. B. Dharmayanthi, Ratih A. Ningrum, Wien Kusharyoto, Yayuk Sri Rahayu, Subangkit, Vivi Setiawaty, Puspita Lisdiyanti |
| EPI_ISL_2722321 | Department of Pathology, University of Cambridge | COVID-19 Genomics UK (COG-UK) Consortium | Aminu S. Jahun, Yasmin Chaudhry, Iliana Georgana, Myra Hosmillo, Rhys Izuagbe, William L. Hamilton, Martin D. Curran, Surendra Parmar, Ian Goodfellow |
| EPI_ISL_2722322 | Intibios Karawang | Indonesian Institute of Sciences (LIPI)/National Agency of Research and Innovation (BRIN) and National Institute of Health Research and Development (NIHRD) | Idris, Isa Nuryana, Erwin Fajar Hasrianda, Indriawati, Masrukhin, Gita Syahputra, Syam B. Iryanto, Anik. B. Dharmayanthi, Ratih A. Ningrum, Wien Kusharyoto, Yayuk Sri Rahayu, Subangkit, Vivi Setiawaty, Sugiyono Saputra |
| EPI_ISL_2722323 | Department of Pathology, University of Cambridge | COVID-19 Genomics UK (COG-UK) Consortium | Aminu S. Jahun, Yasmin Chaudhry, Iliana Georgana, Myra Hosmillo, Rhys Izuagbe, William L. Hamilton, Martin D. Curran, Surendra Parmar, Ian Goodfellow |
| EPI_ISL_2722324 | Intibios Karawang | Indonesian Institute of Sciences (LIPI)/National Agency of Research and Innovation (BRIN) and National Institute of Health Research and Development (NIHRD) | Syaiful Rizal, Erwin Fajar Hasrianda, Indriawati, Idris, Gita Syahputra, Syam B. Iryanto, Anggia Prasetyoputri, Ratih A. Ningrum, Wien Kusharyoto, Yayuk Sri Rahayu, Subangkit, Vivi Setiawaty, Sugiyono Saputra |
| EPI_ISL_2722325 | Department of Pathology, University of Cambridge | COVID-19 Genomics UK (COG-UK) Consortium | Aminu S. Jahun, Yasmin Chaudhry, Iliana Georgana, Myra Hosmillo, Rhys Izuagbe, William L. Hamilton, Martin D. Curran, Surendra Parmar, Ian Goodfellow |
| EPI_ISL_2722326 | Intibios Karawang | Indonesian Institute of Sciences (LIPI)/National Agency of Research and Innovation (BRIN) and National Institute of Health Research and Development (NIHRD) | Masrukhin, Isa Nuryana, Erwin Fajar Hasrianda, Indriawati, Idris, Gita Syahputra, Syam B. Iryanto, Anik. B. Dharmayanthi, Ratih A. Ningrum, Wien Kusharyoto, Yayuk Sri Rahayu, Subangkit, Vivi Setiawaty, Sugiyono Saputra |
| EPI_ISL_2722327 | Department of Pathology, University of Cambridge | COVID-19 Genomics UK (COG-UK) Consortium | Aminu S. Jahun, Yasmin Chaudhry, Iliana Georgana, Myra Hosmillo, Rhys Izuagbe, William L. Hamilton, Martin D. Curran, Surendra Parmar, Ian Goodfellow |
| EPI_ISL_2722328 | Intibios Karawang | Indonesian Institute of Sciences (LIPI)/National Agency of Research and Innovation (BRIN) and National Institute of Health Research and Development (NIHRD) | Syaiful Rizal, Erwin Fajar Hasrianda, Indriawati, Idris, Gita Syahputra, Syam B. Iryanto, Anggia Prasetyoputri, Ratih A. Ningrum, Wien Kusharyoto, Yayuk Sri Rahayu, Subangkit, Vivi Setiawaty, Sugiyono Saputra |
| EPI_ISL_2722330 | Department of Pathology, University of Cambridge | COVID-19 Genomics UK (COG-UK) Consortium | Aminu S. Jahun, Yasmin Chaudhry, Iliana Georgana, Myra Hosmillo, Rhys Izuagbe, William L. Hamilton, Martin D. Curran, Surendra Parmar, Ian Goodfellow |
| EPI_ISL_2722331 | Intibios Karawang | Indonesian Institute of Sciences (LIPI)/National Agency of Research and Innovation (BRIN) and National Institute of Health Research and Development (NIHRD) | Isa Nuryana, Syaiful Rizal, Indriawati, Idris, Masrukhin, Syam B. Iryanto, Andri Wardiana, Ratih A. Ningrum, Wien Kusharyoto, Yayuk Sri Rahayu, Subangkit, Vivi Setiawaty, Sugiyono Saputra |
| EPI_ISL_2722332, EPI_ISL_2722334, EPI_ISL_2722336 | Department of Pathology, University of Cambridge | COVID-19 Genomics UK (COG-UK) Consortium | Aminu S. Jahun, Yasmin Chaudhry, Iliana Georgana, Myra Hosmillo, Rhys Izuagbe, William L. Hamilton, Martin D. Curran, Surendra Parmar, Ian Goodfellow |
| EPI_ISL_2722337 | Intibios Karawang | Indonesian Institute of Sciences (LIPI)/National Agency of Research and Innovation (BRIN) and National Institute of Health Research and Development (NIHRD) | Syaiful Rizal, Erwin Fajar Hasrianda, Indriawati, Idris, Gita Syahputra, Syam B. Iryanto, Anggia Prasetyoputri, Ratih A. Ningrum, Wien Kusharyoto, Yayuk Sri Rahayu, Subangkit, Vivi Setiawaty, Sugiyono Saputra |
| EPI_ISL_2722338, EPI_ISL_2722340 | Department of Pathology, University of Cambridge | COVID-19 Genomics UK (COG-UK) Consortium | Aminu S. Jahun, Yasmin Chaudhry, Iliana Georgana, Myra Hosmillo, Rhys Izuagbe, William L. Hamilton, Martin D. Curran, Surendra Parmar, Ian Goodfellow |
| EPI_ISL_2722341 | Intibios Karawang | Indonesian Institute of Sciences (LIPI)/National Agency of Research and Innovation (BRIN) and National Institute of Health Research and Development (NIHRD) | Masrukhin, Isa Nuryana, Erwin Fajar Hasrianda, Indriawati, Idris, Gita Syahputra, Syam B. Iryanto, Anik. B. Dharmayanthi, Ratih A. Ningrum, Wien Kusharyoto, Yayuk Sri Rahayu, Subangkit, Vivi Setiawaty, Sugiyono Saputra |
| EPI_ISL_2722342 | Intibios Karawang | Indonesian Institute of Sciences (LIPI)/National Agency of Research and Innovation (BRIN) and National Institute of Health Research and Development (NIHRD) | Indriawati, Idris, Masrukhin, Syam B. Iryanto, Anik. B. Dharmayanthi, Isa Nuryana, Syaiful Rizal, Ratih A. Ningrum, Wien Kusharyoto, Yayuk Sri Rahayu, Subangkit, Vivi Setiawaty, Sugiyono Saputra |
| EPI_ISL_2722343 | Department of Pathology, University of Cambridge | COVID-19 Genomics UK (COG-UK) Consortium | Aminu S. Jahun, Yasmin Chaudhry, Iliana Georgana, Myra Hosmillo, Rhys Izuagbe, William L. Hamilton, Martin D. Curran, Surendra Parmar, Ian Goodfellow |
| EPI_ISL_2722344 | Intibios Karawang | Indonesian Institute of Sciences (LIPI)/National Agency of Research and Innovation (BRIN) and National Institute of Health Research and Development (NIHRD) | Erwin Fajar Hasrianda, Indriawati, Masrukhin, Isa Nuryana, Gita Syahputra, Syam B. Iryanto, Andri Wardiana, Ratih A. Ningrum, Wien Kusharyoto, Yayuk Sri Rahayu, Subangkit, Vivi Setiawaty, Puspita Lisdiyanti |
| EPI_ISL_2722345 | Department of Pathology, University of Cambridge | COVID-19 Genomics UK (COG-UK) Consortium | Aminu S. Jahun, Yasmin Chaudhry, Iliana Georgana, Myra Hosmillo, Rhys Izuagbe, William L. Hamilton, Martin D. Curran, Surendra Parmar, Ian Goodfellow |
| EPI_ISL_2722346 | Intibios Karawang | Indonesian Institute of Sciences (LIPI)/National Agency of Research and Innovation (BRIN) and National Institute of Health Research and Development (NIHRD) | Erwin Fajar Hasrianda, Indriawati, Masrukhin, Isa Nuryana, Gita Syahputra, Syam B. Iryanto, Andri Wardiana, Ratih A. Ningrum, Wien Kusharyoto, Yayuk Sri Rahayu, Subangkit, Vivi Setiawaty, Puspita Lisdiyanti |
| EPI_ISL_2722347, EPI_ISL_2722348, EPI_ISL_2722349, EPI_ISL_2722350, EPI_ISL_2722351, EPI_ISL_2722352, EPI_ISL_2722353, EPI_ISL_2722354, EPI_ISL_2722355, EPI_ISL_2722356, EPI_ISL_2722357, EPI_ISL_2722358, EPI_ISL_2722359, EPI_ISL_2722360, EPI_ISL_2722361, EPI_ISL_2722362, EPI_ISL_2722363, EPI_ISL_2722364, EPI_ISL_2722365, EPI_ISL_2722366, EPI_ISL_2722367, EPI_ISL_2722368, EPI_ISL_2722369, EPI_ISL_2722370, EPI_ISL_2722371, EPI_ISL_2722372, EPI_ISL_2722373, EPI_ISL_2722374, EPI_ISL_2722375, EPI_ISL_2722376, EPI_ISL_2722377, EPI_ISL_2722378, EPI_ISL_2722379, EPI_ISL_2722380, EPI_ISL_2722381, EPI_ISL_2722382, EPI_ISL_2722383, EPI_ISL_2722384, EPI_ISL_2722385, EPI_ISL_2722386, EPI_ISL_2722387, EPI_ISL_2722388, EPI_ISL_2722389, EPI_ISL_2722390, EPI_ISL_2722391, EPI_ISL_2722392, EPI_ISL_2722393, EPI_ISL_2722394, EPI_ISL_2722395, EPI_ISL_2722396, EPI_ISL_2722397, EPI_ISL_2722398, EPI_ISL_2722399, EPI_ISL_2722400, EPI_ISL_2722401, EPI_ISL_2722402, EPI_ISL_2722403, EPI_ISL_2722404, EPI_ISL_2722405, EPI_ISL_2722406, EPI_ISL_2722407, EPI_ISL_2722408, EPI_ISL_2722409, EPI_ISL_2722410, EPI_ISL_2722411, EPI_ISL_2722412, EPI_ISL_2722413, EPI_ISL_2722414, EPI_ISL_2722415, EPI_ISL_2722416, EPI_ISL_2722417, EPI_ISL_2722418, EPI_ISL_2722419, EPI_ISL_2722420, EPI_ISL_2722421, EPI_ISL_2722422, EPI_ISL_2722423, EPI_ISL_2722424, EPI_ISL_2722425, EPI_ISL_2722426, EPI_ISL_2722427, EPI_ISL_2722428, EPI_ISL_2722429, EPI_ISL_2722430, EPI_ISL_2722431, EPI_ISL_2722432, EPI_ISL_2722433, EPI_ISL_2722434, EPI_ISL_2722435, EPI_ISL_2722436, EPI_ISL_2722437, EPI_ISL_2722438, EPI_ISL_2722440, EPI_ISL_2722441, EPI_ISL_2722442, EPI_ISL_2722443, EPI_ISL_2722444, EPI_ISL_2722445, EPI_ISL_2722446, EPI_ISL_2722447, EPI_ISL_2722448, EPI_ISL_2722449, EPI_ISL_2722450, EPI_ISL_2722451, EPI_ISL_2722452, EPI_ISL_2722453, EPI_ISL_2722454, EPI_ISL_2722455, EPI_ISL_2722456, EPI_ISL_2722457, EPI_ISL_2722458, EPI_ISL_2722459, EPI_ISL_2722460, EPI_ISL_2722461, EPI_ISL_2722462, EPI_ISL_2722463, EPI_ISL_2722464, EPI_ISL_2722465, EPI_ISL_2722466, EPI_ISL_2722467, EPI_ISL_2722468, EPI_ISL_2722469, EPI_ISL_2722470, EPI_ISL_2722471, EPI_ISL_2722472, EPI_ISL_2722473, EPI_ISL_2722474, EPI_ISL_2722475, EPI_ISL_2722476, EPI_ISL_2722477, EPI_ISL_2722478, EPI_ISL_2722479, EPI_ISL_2722480, EPI_ISL_2722481, EPI_ISL_2722482, EPI_ISL_2722483, EPI_ISL_2722484, EPI_ISL_2722485, EPI_ISL_2722486, EPI_ISL_2722487, EPI_ISL_2722488, EPI_ISL_2722489, EPI_ISL_2722490, EPI_ISL_2722491, EPI_ISL_2722492, EPI_ISL_2722493, EPI_ISL_2722494, EPI_ISL_2722495, EPI_ISL_2722496, EPI_ISL_2722497, EPI_ISL_2722498, EPI_ISL_2722499, EPI_ISL_2722500, EPI_ISL_2722501, EPI_ISL_2722502, EPI_ISL_2722503, EPI_ISL_2722504, EPI_ISL_2722505, EPI_ISL_2722506, EPI_ISL_2722507, EPI_ISL_2722508, EPI_ISL_2722509, EPI_ISL_2722510, EPI_ISL_2722511, EPI_ISL_2722512, EPI_ISL_2722513, EPI_ISL_2722514, EPI_ISL_2722515, EPI_ISL_2722516, EPI_ISL_2722517, EPI_ISL_2722518, EPI_ISL_2722519, EPI_ISL_2722520, EPI_ISL_2722521, EPI_ISL_2722522, EPI_ISL_2722523, EPI_ISL_2722524, EPI_ISL_2722525, EPI_ISL_2722526, EPI_ISL_2722527, EPI_ISL_2722528, EPI_ISL_2722529, EPI_ISL_2722530, EPI_ISL_2722531, EPI_ISL_2722532, EPI_ISL_2722533, EPI_ISL_2722534, EPI_ISL_2722535, EPI_ISL_2722536, EPI_ISL_2722537, EPI_ISL_2722538, EPI_ISL_2722539, EPI_ISL_2722540, EPI_ISL_2722541, EPI_ISL_2722542, EPI_ISL_2722543, EPI_ISL_2722544, EPI_ISL_2722545, EPI_ISL_2722546, EPI_ISL_2722547, EPI_ISL_2722548, EPI_ISL_2722549, EPI_ISL_2722550, EPI_ISL_2722551, EPI_ISL_2722552, EPI_ISL_2722553, EPI_ISL_2722554, EPI_ISL_2722555, EPI_ISL_2722556, EPI_ISL_2722557, EPI_ISL_2722558, EPI_ISL_2722559, EPI_ISL_2722560, EPI_ISL_2722561, EPI_ISL_2722562, EPI_ISL_2722563, EPI_ISL_2722564, EPI_ISL_2722565, EPI_ISL_2722566, EPI_ISL_2722567, EPI_ISL_2722568, EPI_ISL_2722569, EPI_ISL_2722570, EPI_ISL_2722571, EPI_ISL_2722572, EPI_ISL_2722573, EPI_ISL_2722574, EPI_ISL_2722575, EPI_ISL_2722576, EPI_ISL_2722577, EPI_ISL_2722578, EPI_ISL_2722579, EPI_ISL_2722580, EPI_ISL_2722581, EPI_ISL_2722582, EPI_ISL_2722583, EPI_ISL_2722584, EPI_ISL_2722585, EPI_ISL_2722586, EPI_ISL_2722587, EPI_ISL_2722588, EPI_ISL_2722589, EPI_ISL_2722590, EPI_ISL_2722591, EPI_ISL_2722592, EPI_ISL_2722593, EPI_ISL_2722594, EPI_ISL_2722595, EPI_ISL_2722596, EPI_ISL_2722597, EPI_ISL_2722598, EPI_ISL_2722599, EPI_ISL_2722600, EPI_ISL_2722601, EPI_ISL_2722602, EPI_ISL_2722603, EPI_ISL_2722604, EPI_ISL_2722605, EPI_ISL_2722606, EPI_ISL_2722607, EPI_ISL_2722608, EPI_ISL_2722609, EPI_ISL_2722610, EPI_ISL_2722611, EPI_ISL_2722612, EPI_ISL_2722613, EPI_ISL_2722614, EPI_ISL_2722615, EPI_ISL_2722616, EPI_ISL_2722617, EPI_ISL_2722618, EPI_ISL_2722619, EPI_ISL_2722620, EPI_ISL_2722621, EPI_ISL_2722622, EPI_ISL_2722623, EPI_ISL_2722624, EPI_ISL_2722625, EPI_ISL_2722626, EPI_ISL_2722627, EPI_ISL_2722628, EPI_ISL_2722629, EPI_ISL_2722630, EPI_ISL_2722631 | | | |
| see above | Department of Pathology, University of Cambridge | COVID-19 Genomics UK (COG-UK) Consortium | Aminu S. Jahun, Yasmin Chaudhry, Iliana Georgana, Myra Hosmillo, Rhys Izuagbe, William L. Hamilton, Martin D. Curran, Surendra Parmar, Ian Goodfellow |
| EPI_ISL_2722633, EPI_ISL_2722634, EPI_ISL_2722636, EPI_ISL_2722640, EPI_ISL_2722641, EPI_ISL_2722642, EPI_ISL_2722643, EPI_ISL_2722644, EPI_ISL_2722645, EPI_ISL_2722646, EPI_ISL_2722648, EPI_ISL_2722651, EPI_ISL_2722652, EPI_ISL_2722654, EPI_ISL_2722655, EPI_ISL_2722656, EPI_ISL_2722657, EPI_ISL_2722658, EPI_ISL_2722659, EPI_ISL_2722660, EPI_ISL_2722661, EPI_ISL_2722663, EPI_ISL_2722664, EPI_ISL_2722665, EPI_ISL_2722666, EPI_ISL_2722667, EPI_ISL_2722668, EPI_ISL_2722669, EPI_ISL_2722670, EPI_ISL_2722671, EPI_ISL_2722672, EPI_ISL_2722673, EPI_ISL_2722674, EPI_ISL_2722675, EPI_ISL_2722677, EPI_ISL_2722678 | | | |
| see above | West of Scotland Specialist Virology Centre, NHSGGC / MRC- | COVID-19 Genomics UK (COG-UK) Consortium | Ana da Silva Filipe, Natasha Johnson, Kathy Smollett, Daniel Mair, Stephen Carmichael, Alice Broos, Lily Tong, Jenna Nichols, Kyriaki Nomikou; Sarah McDonald; Ioulia Tsatsani, Richard Orton, Joseph Hughes, Sreenu Vattipally, David L |

| | | | |
|---|---|---|---|
| | University of Glasgow Centre for Virus Research | | Robertson; Alasdair MacLean, Rory Gunson; Sharif Shaaban, Matthew Holden; Rachel Blacow, Guy Mollett, Kathy Li, James Shepherd, Antonia Ho, Emma Thomson |
| EPI_ISL_2722999, EPI_ISL_2723000, EPI_ISL_2723001, EPI_ISL_2723002, EPI_ISL_2723003, EPI_ISL_2723004, EPI_ISL_2723005, EPI_ISL_2723006, EPI_ISL_2723007, EPI_ISL_2723008, EPI_ISL_2723009, EPI_ISL_2723010, EPI_ISL_2723011, EPI_ISL_2723012, EPI_ISL_2723013, EPI_ISL_2723014, EPI_ISL_2723015, EPI_ISL_2723016, EPI_ISL_2723017, EPI_ISL_2723018, EPI_ISL_2723019, EPI_ISL_2723020, EPI_ISL_2723021, EPI_ISL_2723022, EPI_ISL_2723023, EPI_ISL_2723024, EPI_ISL_2723025, EPI_ISL_2723026, EPI_ISL_2723027, EPI_ISL_2723028, EPI_ISL_2723029, EPI_ISL_2723030, EPI_ISL_2723031, EPI_ISL_2723032, EPI_ISL_2723033, EPI_ISL_2723034, EPI_ISL_2723035, EPI_ISL_2723036, EPI_ISL_2723037, EPI_ISL_2723039, EPI_ISL_2723040, EPI_ISL_2723041, EPI_ISL_2723042, EPI_ISL_2723043, EPI_ISL_2723044, EPI_ISL_2723045, EPI_ISL_2723046, EPI_ISL_2723047, EPI_ISL_2723048, EPI_ISL_2723049, EPI_ISL_2723050, EPI_ISL_2723051, EPI_ISL_2723052, EPI_ISL_2723053, EPI_ISL_2723054, EPI_ISL_2723055, EPI_ISL_2723056, EPI_ISL_2723057, EPI_ISL_2723058, EPI_ISL_2723059, EPI_ISL_2723060, EPI_ISL_2723061, EPI_ISL_2723062, EPI_ISL_2723063, EPI_ISL_2723064, EPI_ISL_2723065, EPI_ISL_2723066, EPI_ISL_2723067, EPI_ISL_2723068, EPI_ISL_2723069, EPI_ISL_2723070, EPI_ISL_2723071, EPI_ISL_2723072, EPI_ISL_2723073, EPI_ISL_2723074, EPI_ISL_2723075, EPI_ISL_2723076, EPI_ISL_2723077, EPI_ISL_2723078, EPI_ISL_2723079, EPI_ISL_2723080, EPI_ISL_2723081, EPI_ISL_2723082, EPI_ISL_2723083, EPI_ISL_2723084, EPI_ISL_2723085, EPI_ISL_2723086, EPI_ISL_2723087, EPI_ISL_2723088, EPI_ISL_2723089, EPI_ISL_2723090, EPI_ISL_2723091, EPI_ISL_2723092, EPI_ISL_2723093, EPI_ISL_2723094, EPI_ISL_2723095, EPI_ISL_2723096, EPI_ISL_2723097, EPI_ISL_2723098, EPI_ISL_2723099, EPI_ISL_2723100, EPI_ISL_2723101, EPI_ISL_2723102, EPI_ISL_2723103, EPI_ISL_2723104, EPI_ISL_2723105, EPI_ISL_2723106, EPI_ISL_2723107, EPI_ISL_2723108, EPI_ISL_2723109, EPI_ISL_2723110, EPI_ISL_2723111, EPI_ISL_2723112, EPI_ISL_2723113, EPI_ISL_2723114, EPI_ISL_2723115, EPI_ISL_2723116, EPI_ISL_2723117, EPI_ISL_2723118, EPI_ISL_2723119, EPI_ISL_2723120, EPI_ISL_2723121, EPI_ISL_2723122, EPI_ISL_2723123, EPI_ISL_2723124, EPI_ISL_2723125, EPI_ISL_2723126, EPI_ISL_2723127, EPI_ISL_2723128, EPI_ISL_2723129, EPI_ISL_2723131, EPI_ISL_2723132, EPI_ISL_2723134, EPI_ISL_2723135, EPI_ISL_2723136, EPI_ISL_2723137, EPI_ISL_2723138, EPI_ISL_2723139, EPI_ISL_2723140, EPI_ISL_2723141, EPI_ISL_2723142, EPI_ISL_2723143, EPI_ISL_2723144, EPI_ISL_2723145, EPI_ISL_2723146, EPI_ISL_2723147, EPI_ISL_2723148, EPI_ISL_2723149, EPI_ISL_2723150, EPI_ISL_2723151, EPI_ISL_2723152, EPI_ISL_2723153, EPI_ISL_2723154, EPI_ISL_2723155, EPI_ISL_2723156, EPI_ISL_2723158, EPI_ISL_2723159, EPI_ISL_2723160, EPI_ISL_2723161, EPI_ISL_2723163 | | | |
| see above | University College London, Great Ormond Street Hospital for Children NHS Foundation Trust, Imperial College Healthcare NHS Trust | COVID-19 Genomics UK (COG-UK) Consortium | Sergi Castellano, Rachel Williams, Mark Kristiansen, Paola Resende Silva, Sunando Roy, Tony Brooks, Helena Tutill, Paola Niola, Patricia Dyal, Charlotte Williams, Leysa Forrest, Yasmin Panchbhaya, Jacqueline Findlay, Samuel Weeks, Julianne Brown, Kathryn Harris, Paul Randell, James Price, Alison Holmes, Judith Breuer |
| EPI_ISL_2723164, EPI_ISL_2723165, EPI_ISL_2723166, EPI_ISL_2723167, EPI_ISL_2723168, EPI_ISL_2723169, EPI_ISL_2723170, EPI_ISL_2723171, EPI_ISL_2723172, EPI_ISL_2723173, EPI_ISL_2723174, EPI_ISL_2723175, EPI_ISL_2723176, EPI_ISL_2723177, EPI_ISL_2723178, EPI_ISL_2723179, EPI_ISL_2723180, EPI_ISL_2723181, EPI_ISL_2723182, EPI_ISL_2723184, EPI_ISL_2723185, EPI_ISL_2723186, EPI_ISL_2723187, EPI_ISL_2723188, EPI_ISL_2723189, EPI_ISL_2723190, EPI_ISL_2723191, EPI_ISL_2723192, EPI_ISL_2723193, EPI_ISL_2723194, EPI_ISL_2723195, EPI_ISL_2723196, EPI_ISL_2723197, EPI_ISL_2723198, EPI_ISL_2723199, EPI_ISL_2723200, EPI_ISL_2723201, EPI_ISL_2723202, EPI_ISL_2723203, EPI_ISL_2723204, EPI_ISL_2723205, EPI_ISL_2723206, EPI_ISL_2723207, EPI_ISL_2723208, EPI_ISL_2723209, EPI_ISL_2723210, EPI_ISL_2723211, EPI_ISL_2723212, EPI_ISL_2723213, EPI_ISL_2723214, EPI_ISL_2723215, EPI_ISL_2723216, EPI_ISL_2723217, EPI_ISL_2723218, EPI_ISL_2723219, EPI_ISL_2723221, EPI_ISL_2723222, EPI_ISL_2723223, EPI_ISL_2723224, EPI_ISL_2723225, EPI_ISL_2723226, EPI_ISL_2723227, EPI_ISL_2723228, EPI_ISL_2723229, EPI_ISL_2723230, EPI_ISL_2723231, EPI_ISL_2723232, EPI_ISL_2723233, EPI_ISL_2723234, EPI_ISL_2723235, EPI_ISL_2723236, EPI_ISL_2723237, EPI_ISL_2723238, EPI_ISL_2723239, EPI_ISL_2723240, EPI_ISL_2723241, EPI_ISL_2723242, EPI_ISL_2723243, EPI_ISL_2723244, EPI_ISL_2723245, EPI_ISL_2723246, EPI_ISL_2723247, EPI_ISL_2723248, EPI_ISL_2723249, EPI_ISL_2723250, EPI_ISL_2723251, EPI_ISL_2723252, EPI_ISL_2723253, EPI_ISL_2723254, EPI_ISL_2723255, EPI_ISL_2723256, EPI_ISL_2723257, EPI_ISL_2723258, EPI_ISL_2723259, EPI_ISL_2723260, EPI_ISL_2723261, EPI_ISL_2723262, EPI_ISL_2723263, EPI_ISL_2723264, EPI_ISL_2723265, EPI_ISL_2723266, EPI_ISL_2723267, EPI_ISL_2723268, EPI_ISL_2723269, EPI_ISL_2723270, EPI_ISL_2723271, EPI_ISL_2723272, EPI_ISL_2723273, EPI_ISL_2723274, EPI_ISL_2723275, EPI_ISL_2723276, EPI_ISL_2723277, EPI_ISL_2723278, EPI_ISL_2723279, EPI_ISL_2723280, EPI_ISL_2723281, EPI_ISL_2723282, EPI_ISL_2723283, EPI_ISL_2723284, EPI_ISL_2723285, EPI_ISL_2723286, EPI_ISL_2723287, EPI_ISL_2723288, EPI_ISL_2723289, EPI_ISL_2723290, EPI_ISL_2723291, EPI_ISL_2723292, EPI_ISL_2723293, EPI_ISL_2723294, EPI_ISL_2723295, EPI_ISL_2723296, EPI_ISL_2723297, EPI_ISL_2723298, EPI_ISL_2723299, EPI_ISL_2723300, EPI_ISL_2723301, EPI_ISL_2723302, EPI_ISL_2723303, EPI_ISL_2723304, EPI_ISL_2723305, EPI_ISL_2723306, EPI_ISL_2723307, EPI_ISL_2723308, EPI_ISL_2723309, EPI_ISL_2723310, EPI_ISL_2723311, EPI_ISL_2723312, EPI_ISL_2723313, EPI_ISL_2723314, EPI_ISL_2723315, EPI_ISL_2723316, EPI_ISL_2723317, EPI_ISL_2723318, EPI_ISL_2723319, EPI_ISL_2723320, EPI_ISL_2723321, EPI_ISL_2723322, EPI_ISL_2723323, EPI_ISL_2723324, EPI_ISL_2723325, EPI_ISL_2723326, EPI_ISL_2723327, EPI_ISL_2723328, EPI_ISL_2723329, EPI_ISL_2723330, EPI_ISL_2723331, EPI_ISL_2723332, EPI_ISL_2723333, EPI_ISL_2723334, EPI_ISL_2723335, EPI_ISL_2723336, EPI_ISL_2723337, EPI_ISL_2723338, EPI_ISL_2723339, EPI_ISL_2723340, EPI_ISL_2723341, EPI_ISL_2723342, EPI_ISL_2723343, EPI_ISL_2723344, EPI_ISL_2723345, EPI_ISL_2723346, EPI_ISL_2723347, EPI_ISL_2723349, EPI_ISL_2723350, EPI_ISL_2723351, EPI_ISL_2723352, EPI_ISL_2723353, EPI_ISL_2723354, EPI_ISL_2723355, EPI_ISL_2723356, EPI_ISL_2723357, EPI_ISL_2723358, EPI_ISL_2723359, EPI_ISL_2723360, EPI_ISL_2723361, EPI_ISL_2723362, EPI_ISL_2723363, EPI_ISL_2723364, EPI_ISL_2723365, EPI_ISL_2723366, EPI_ISL_2723367, EPI_ISL_2723368, EPI_ISL_2723369, EPI_ISL_2723370, EPI_ISL_2723371, EPI_ISL_2723372, EPI_ISL_2723373, EPI_ISL_2723374, EPI_ISL_2723375, EPI_ISL_2723376, EPI_ISL_2723377, EPI_ISL_2723378, EPI_ISL_2723379, EPI_ISL_2723380, EPI_ISL_2723381, EPI_ISL_2723382, EPI_ISL_2723383, EPI_ISL_2723384, EPI_ISL_2723385, EPI_ISL_2723386, EPI_ISL_2723387, EPI_ISL_2723388, EPI_ISL_2723389, EPI_ISL_2723390, EPI_ISL_2723391, EPI_ISL_2723392, EPI_ISL_2723393, EPI_ISL_2723394, EPI_ISL_2723395, EPI_ISL_2723396, EPI_ISL_2723397, EPI_ISL_2723398, EPI_ISL_2723399, EPI_ISL_2723400, EPI_ISL_2723401, EPI_ISL_2723402, EPI_ISL_2723403, EPI_ISL_2723404, EPI_ISL_2723405, EPI_ISL_2723406, EPI_ISL_2723408, EPI_ISL_2723409, EPI_ISL_2723410, EPI_ISL_2723411, EPI_ISL_2723412, EPI_ISL_2723413, EPI_ISL_2723414, EPI_ISL_2723415, EPI_ISL_2723416, EPI_ISL_2723417, EPI_ISL_2723418, EPI_ISL_2723419, EPI_ISL_2723420, EPI_ISL_2723421, EPI_ISL_2723422, EPI_ISL_2723423, EPI_ISL_2723424, EPI_ISL_2723425, EPI_ISL_2723426, EPI_ISL_2723427, EPI_ISL_2723428, EPI_ISL_2723429, EPI_ISL_2723430, EPI_ISL_2723431, EPI_ISL_2723432, EPI_ISL_2723433, EPI_ISL_2723434, EPI_ISL_2723435, EPI_ISL_2723436, EPI_ISL_2723437, EPI_ISL_2723438, EPI_ISL_2723439, EPI_ISL_2723440, EPI_ISL_2723441, EPI_ISL_2723442, EPI_ISL_2723443, EPI_ISL_2723444, EPI_ISL_2723445, EPI_ISL_2723446, EPI_ISL_2723447, EPI_ISL_2723448, EPI_ISL_2723449, EPI_ISL_2723450, EPI_ISL_2723451, EPI_ISL_2723452, EPI_ISL_2723453, EPI_ISL_2723454, EPI_ISL_2723455, EPI_ISL_2723456, EPI_ISL_2723457, EPI_ISL_2723458, EPI_ISL_2723459, EPI_ISL_2723460, EPI_ISL_2723461, EPI_ISL_2723462, EPI_ISL_2723463, EPI_ISL_2723464, EPI_ISL_2723465, EPI_ISL_2723466, EPI_ISL_2723467, EPI_ISL_2723468, EPI_ISL_2723469, EPI_ISL_2723470, EPI_ISL_2723471, EPI_ISL_2723472, EPI_ISL_2723473, EPI_ISL_2723474, EPI_ISL_2723475, EPI_ISL_2723476, EPI_ISL_2723477, EPI_ISL_2723478, EPI_ISL_2723479, EPI_ISL_2723480, EPI_ISL_2723481, EPI_ISL_2723482, EPI_ISL_2723483, EPI_ISL_2723484, EPI_ISL_2723485, EPI_ISL_2723486, EPI_ISL_2723487, EPI_ISL_2723488, EPI_ISL_2723489, EPI_ISL_2723490, EPI_ISL_2723491, EPI_ISL_2723492, EPI_ISL_2723493, EPI_ISL_2723494, EPI_ISL_2723495, EPI_ISL_2723496, EPI_ISL_2723497, EPI_ISL_2723498, EPI_ISL_2723499, EPI_ISL_2723500, EPI_ISL_2723501, EPI_ISL_2723502, EPI_ISL_2723503, EPI_ISL_2723504, EPI_ISL_2723505, EPI_ISL_2723506, EPI_ISL_2723507, EPI_ISL_2723508, EPI_ISL_2723509, EPI_ISL_2723510, EPI_ISL_2723511, EPI_ISL_2723512, EPI_ISL_2723513, EPI_ISL_2723514, EPI_ISL_2723515, EPI_ISL_2723516, EPI_ISL_2723517, EPI_ISL_2723518, EPI_ISL_2723519, EPI_ISL_2723520, EPI_ISL_2723521, EPI_ISL_2723522, EPI_ISL_2723523, EPI_ISL_2723524, EPI_ISL_2723525, EPI_ISL_2723526, EPI_ISL_2723527, EPI_ISL_2723528, EPI_ISL_2723529, EPI_ISL_2723530, EPI_ISL_2723531, EPI_ISL_2723532, EPI_ISL_2723533, EPI_ISL_2723534, EPI_ISL_2723535, EPI_ISL_2723536, EPI_ISL_2723537, EPI_ISL_2723538, EPI_ISL_2723539, EPI_ISL_2723540, EPI_ISL_2723541, EPI_ISL_2723542, EPI_ISL_2723543, EPI_ISL_2723544, EPI_ISL_2723545, EPI_ISL_2723546, EPI_ISL_2723547 | | | |
| see above | Northumbria University / South Tees Hospitals NHS Foundation Trust / North Cumbria Integrated Care NHS Foundation Trust / North Tees and Hartlepool NHS Foundation Trust / Newcastle Hospitals NHS Foundation Trust | COVID-19 Genomics UK (COG-UK) Consortium | Darren L Smith,Andrew Nelson,Matthew Bashton,Greg R Young,Joshua Loh,John Allan,Mohammad A Tariq,Giles S Holt,Gary Black,Wen C Yew,Lynn Dover,Paul Baker,Steve Liggett,Sarah Essex,Jane Greenaway,Debra Padgett,Clive Graham,Garren Scott,Edward Barton,Emma Swindells,Brendan Payne,Jennifer Collins,Yusri Taha,Gary Eltringham |
| EPI_ISL_2723562, EPI_ISL_2723563, EPI_ISL_2723564 | Department of Acute Infectious Diseases Control and Prevention, Yunnan Provincial Center for Disease Control and Prevention | Department of Acute Infectious Diseases Control and Prevention, Yunnan Provincial Center for Disease Control and Prevention | Meiling Zhang, Senquan Jia, Jienan Zhou, Zhaosheng Liu, Xiaoqing Fu |
| EPI_ISL_2723565, EPI_ISL_2723566, EPI_ISL_2723567, EPI_ISL_2723568, EPI_ISL_2723569 | Tokyo Metropolitan Institute of Public Health | Tokyo Metropolitan Institute of Public Health | Masaki Hayashi, Takako Yamazaki, Akane Negishi, Hirofumi Miyake, Kenshirou Kuroki, Maya Isogai, Takushi Fujiwara, Mami Nagashima, Jun Suzuki, Kenji Sadamasu |
| EPI_ISL_2723570, EPI_ISL_2723571, EPI_ISL_2723572, EPI_ISL_2723573, EPI_ISL_2723574, EPI_ISL_2723578, EPI_ISL_2723581, EPI_ISL_2723582 | SC (UCO) Igiene e Sanità Pubblica, ASUGI, Trieste | ARGO Laboratorio Genomica ed Epigenomica | Licastro D, Dal Monego S, Degasperi M, Marcello A, Segat L, Piscianz E, Busetti M, Fontana F, Braida C, Barbone F, D'Agaro P |
| EPI_ISL_2723583, EPI_ISL_2723584 | SC (UCO) Igiene e Sanità Pubblica, ASUGI, Trieste | ARGO Laboratorio Genomica ed Epigenomica | Licastro D, Dal Monego S, Degasperi M, Marcello A, Segat L, Piscianz E, Pipan C, Braida C, Barbone F, D'Agaro P |
| EPI_ISL_2723585, EPI_ISL_2723586, EPI_ISL_2723588, EPI_ISL_2723589, EPI_ISL_2723590, EPI_ISL_2723591 | SC (UCO) Igiene e Sanità Pubblica, ASUGI, Trieste | ARGO Laboratorio Genomica ed Epigenomica | Licastro D, Dal Monego S, Degasperi M, Marcello A, Segat L, Piscianz E, Busetti M, Fontana F, Braida C, Barbone F, D'Agaro P |
| EPI_ISL_2723600, EPI_ISL_2723604 | RS PELNI | Genomik Solidaritas Indonesia Laboratorium | Alfin Mohammad Abdillah, Dhahlia Agustina Cahyono, Normastuti Adhini Tantyo, Vania Gavrila Wikasa, Carissa Sintca Wijaya, Mohammad Anang Syarifudin, Annisa Muthiah Sukirman, Gracia Felias Enos Korompis, Widyastuti, Dwi Oktavia, Tiranti Vindha, Ngabila Salama, Anuraj Shankar, Meutia Ayuputeri Kumaheri, Ariel Pradipta |
| EPI_ISL_2723605, EPI_ISL_2723606, EPI_ISL_2723607, EPI_ISL_2723608, EPI_ISL_2723609, EPI_ISL_2723610, EPI_ISL_2723611, EPI_ISL_2723612, EPI_ISL_2723613, EPI_ISL_2723614, EPI_ISL_2723615, EPI_ISL_2723616, EPI_ISL_2723617, EPI_ISL_2723618, EPI_ISL_2723619 | | | |
| see above | SPEEDLAB | Genomik Solidaritas Indonesia Laboratorium | Alfin Mohammad Abdillah, Dhahlia Agustina Cahyono, Normastuti Adhini Tantyo, Vania Gavrila Wikasa, Carissa Sintca Wijaya, Mohammad Anang Syarifudin, Annisa Muthiah Sukirman, Gracia Felias Enos Korompis, Widyastuti, Dwi Oktavia, Tiranti Vindha, Ngabila Salama, Anuraj Shankar, Meutia Ayuputeri Kumaheri, Ariel Pradipta |
| EPI_ISL_2723620, EPI_ISL_2723621, EPI_ISL_2723622, EPI_ISL_2723623, EPI_ISL_2723624 | LABKESDA DKI JAKARTA | Genomik Solidaritas Indonesia Laboratorium | Alfin Mohammad Abdillah, Dhahlia Agustina Cahyono, Normastuti Adhini Tantyo, Vania Gavrila Wikasa, Carissa Sintca Wijaya, Mohammad Anang Syarifudin, Annisa Muthiah Sukirman, Gracia Felias Enos Korompis, Widyastuti, Dwi Oktavia, Tiranti Vindha, Ngabila Salama, Anuraj Shankar, Meutia Ayuputeri Kumaheri, Ariel Pradipta |
| EPI_ISL_2723625, EPI_ISL_2723626, EPI_ISL_2723627, EPI_ISL_2723628, EPI_ISL_2723629, EPI_ISL_2723630, EPI_ISL_2723631, EPI_ISL_2723632, EPI_ISL_2723633, EPI_ISL_2723634, EPI_ISL_2723635, EPI_ISL_2723636, EPI_ISL_2723637, EPI_ISL_2723638, EPI_ISL_2723639, EPI_ISL_2723640, EPI_ISL_2723641, EPI_ISL_2723642, EPI_ISL_2723643, EPI_ISL_2723644, EPI_ISL_2723645, EPI_ISL_2723646, EPI_ISL_2723647, EPI_ISL_2723648 | | | |
| see above | LABKESDA DKI JAKARTA | Genomik Solidaritas Indonesia Laboratorium | Dhahlia Agustina Cahyono, Alfin Mohammad Abdillah, Normastuti Adhini Tantyo, Vania Gavrila Wikasa, Carissa Sintca Wijaya, Mohammad Anang Syarifudin, Annisa Muthiah Sukirman, Gracia Felias Enos Korompis, Widyastuti, Dwi Oktavia, Tiranti Vindha, Ngabila Salama, Anuraj Shankar, Meutia Ayuputeri Kumaheri, Ariel Pradipta |
| EPI_ISL_2723769 | Department of Health Technology and Informatics, The Hong Kong Polytechnic University | Department of Health Technology and Informatics, The Hong Kong Polytechnic University | Gilman Kit-Hang Siu, Lam-Kwong Lee, Kenneth Siu-Sing Leung, Jake Siu-Lun Leung, Wing-Hei Lo, Timothy Ting-Leung Ng, Chloe Toi-Mei Chan, Hoi-Ching Jim, Ivan Tak-Fai Wong, Kingsley King-Gee Tam, Hiu-Yin Lao, Denise Sze-Hang Wong, Alan Ka-Lun Wu, Miranda Chong-Yee Yau, Yvette Wai-Man Lai, Kitty Sau-Chun Fung, Sandy Ka-Yee Chau, Barry Kin-Chung Wong, Wing-Kin To, Kristine Luk, Alex Yat-Man Ho, Tak-Lun Que, Kam-Tong Yip, Wing Cheong Yam, David Ho-Keung Shum, Shea Ping Yip |
| EPI_ISL_2723777, EPI_ISL_2723778, EPI_ISL_2723779, EPI_ISL_2723780, EPI_ISL_2723781, EPI_ISL_2723782, EPI_ISL_2723783, EPI_ISL_2723784, EPI_ISL_2723787, EPI_ISL_2723788, EPI_ISL_2723790, EPI_ISL_2723791, EPI_ISL_2723792, EPI_ISL_2723794, EPI_ISL_2723795, EPI_ISL_2723796, EPI_ISL_2723797, EPI_ISL_2723798, EPI_ISL_2723799, EPI_ISL_2723800, EPI_ISL_2723801, EPI_ISL_2723804, EPI_ISL_2723805, EPI_ISL_2723806, EPI_ISL_2723807, EPI_ISL_2723809, EPI_ISL_2723810, EPI_ISL_2723812, EPI_ISL_2723813 | | | |
| see above | Dr S raju, Director of public heath and preventive medicine | inStem NCBS – INSACOG | Uma Ramakrishnan Dasaradhi Palakodeti Aswin SaiNarain |
| EPI_ISL_2723814, EPI_ISL_2723815 | Dr. Madhusudhan, MBBD, MD, ICMR VRDL | inStem NCBS – INSACOG | Uma Ramakrishnan Dasaradhi Palakodeti Aswin SaiNarain |
| EPI_ISL_2723817, EPI_ISL_2723818, EPI_ISL_2723819, EPI_ISL_2723820, EPI_ISL_2723821, EPI_ISL_2723822 | JIPMER Asia / India / Puducherry / Puducherry | inStem NCBS – INSACOG | Uma Ramakrishnan Dasaradhi Palakodeti Aswin SaiNarain |
| EPI_ISL_2723823, EPI_ISL_2723824, EPI_ISL_2723825, EPI_ISL_2723826, EPI_ISL_2723828, EPI_ISL_2723829 | Dr. Madhusudhan, MBBD, MD, ICMR VRDL | inStem NCBS – INSACOG | Uma Ramakrishnan Dasaradhi Palakodeti Aswin SaiNarain |
| EPI_ISL_2723830, EPI_ISL_2723831, EPI_ISL_2723834, EPI_ISL_2723835, EPI_ISL_2723836, EPI_ISL_2723837, EPI_ISL_2723838, EPI_ISL_2723843, EPI_ISL_2723845, EPI_ISL_2723847, EPI_ISL_2723848, EPI_ISL_2723849, EPI_ISL_2723850, EPI_ISL_2723851, EPI_ISL_2723852, EPI_ISL_2723853, EPI_ISL_2723857, EPI_ISL_2723858, EPI_ISL_2723862, EPI_ISL_2723863, EPI_ISL_2723864, EPI_ISL_2723865, EPI_ISL_2723866, EPI_ISL_2723870, EPI_ISL_2723871, EPI_ISL_2723872, EPI_ISL_2723874, EPI_ISL_2723875, EPI_ISL_2723876, EPI_ISL_2723878, EPI_ISL_2723879, EPI_ISL_2723880, EPI_ISL_2723881, EPI_ISL_2723882, EPI_ISL_2723883, EPI_ISL_2723884, EPI_ISL_2723885, EPI_ISL_2723886, EPI_ISL_2723887, EPI_ISL_2723888, EPI_ISL_2723889, EPI_ISL_2723890, EPI_ISL_2723891, EPI_ISL_2723892, EPI_ISL_2723893, EPI_ISL_2723894, EPI_ISL_2723895, EPI_ISL_2723896, EPI_ISL_2723897, EPI_ISL_2723898, EPI_ISL_2723900, EPI_ISL_2723901, EPI_ISL_2723902, EPI_ISL_2723903, EPI_ISL_2723904, EPI_ISL_2723905, EPI_ISL_2723906, EPI_ISL_2723907, EPI_ISL_2723908, EPI_ISL_2723909, EPI_ISL_2723910, EPI_ISL_2723911, EPI_ISL_2723912, EPI_ISL_2723913, EPI_ISL_2723914, EPI_ISL_2723915, EPI_ISL_2723916, EPI_ISL_2723917, EPI_ISL_2723918, EPI_ISL_2723919, EPI_ISL_2723920, EPI_ISL_2723922, EPI_ISL_2723923, EPI_ISL_2723924, EPI_ISL_2723925, EPI_ISL_2723926, EPI_ISL_2723927, EPI_ISL_2723928, EPI_ISL_2723929, EPI_ISL_2723930, EPI_ISL_2723931, EPI_ISL_2723932, EPI_ISL_2723933, EPI_ISL_2723934, EPI_ISL_2723935, EPI_ISL_2723936, EPI_ISL_2723937, EPI_ISL_2723938, EPI_ISL_2723940, EPI_ISL_2723941, EPI_ISL_2723942, EPI_ISL_2723943, EPI_ISL_2723944, EPI_ISL_2723945, EPI_ISL_2723946, EPI_ISL_2723949, EPI_ISL_2723950, EPI_ISL_2723951, EPI_ISL_2723952, EPI_ISL_2723953, EPI_ISL_2723954, EPI_ISL_2723955, EPI_ISL_2723956, EPI_ISL_2723957, EPI_ISL_2723958, EPI_ISL_2723959, EPI_ISL_2723960, EPI_ISL_2723961, EPI_ISL_2723962, EPI_ISL_2723963, EPI_ISL_2723964, EPI_ISL_2723966, EPI_ISL_2723967, EPI_ISL_2723968, EPI_ISL_2723969, EPI_ISL_2723970, EPI_ISL_2723971, EPI_ISL_2723972, EPI_ISL_2723973, EPI_ISL_2723974, EPI_ISL_2723975, EPI_ISL_2723976, EPI_ISL_2723977, EPI_ISL_2723978, EPI_ISL_2723979 | | | |
| see above | Dr S raju, Director of public heath and preventive medicine | inStem NCBS – INSACOG | Uma Ramakrishnan Dasaradhi Palakodeti Aswin SaiNarain |
| EPI_ISL_2723980, EPI_ISL_2723981, EPI_ISL_2723982, EPI_ISL_2723983, EPI_ISL_2723984, EPI_ISL_2723985, EPI_ISL_2723986, EPI_ISL_2723987, EPI_ISL_2723988, EPI_ISL_2723989, EPI_ISL_2723990, EPI_ISL_2723992, EPI_ISL_2723994, EPI_ISL_2723995, EPI_ISL_2723998, EPI_ISL_2723999, EPI_ISL_2724000, EPI_ISL_2724001, EPI_ISL_2724002 | | | |

| | | | |
|---|---|---|---|
| see above | Dr. Madhusudhan, MBBD, MD, ICMR VRDL | inStem NCBS – INSACOG | Uma Ramakrishnan Dasaradhi Palakodeti Aswin SaiNarain |
| EPI_ISL_2724009 | Molecular Biology Laboratory, Infectious Diseases Department, Instituto Nacional de Ciencias Medicas y Nutricion Salvador Zubiran | Instituto Nacional de Ciencias Medicas y Nutricion Infectious Diseases | Pilar Ramos Cervantes, Fernando Ledesma Barrientos, Violeta Ibarra Gonzalez, Patricia Leal, Eric Ochoa, Arturo Galindo, Guillermo Ruiz-Palacios y Santos |
| EPI_ISL_2724010, EPI_ISL_2724011, EPI_ISL_2724012, EPI_ISL_2724013, EPI_ISL_2724014, EPI_ISL_2724015, EPI_ISL_2724016, EPI_ISL_2724017, EPI_ISL_2724018, EPI_ISL_2724019, EPI_ISL_2724020, EPI_ISL_2724021, EPI_ISL_2724023, EPI_ISL_2724025, EPI_ISL_2724031, EPI_ISL_2724032, EPI_ISL_2724033, EPI_ISL_2724034, EPI_ISL_2724035, EPI_ISL_2724036, EPI_ISL_2724037, EPI_ISL_2724038, EPI_ISL_2724039, EPI_ISL_2724040, EPI_ISL_2724041, EPI_ISL_2724042, EPI_ISL_2724043, EPI_ISL_2724044, EPI_ISL_2724045, EPI_ISL_2724046, EPI_ISL_2724047, EPI_ISL_2724048, EPI_ISL_2724049, EPI_ISL_2724050, EPI_ISL_2724051, EPI_ISL_2724052, EPI_ISL_2724054, EPI_ISL_2724055, EPI_ISL_2724057, EPI_ISL_2724059, EPI_ISL_2724060, EPI_ISL_2724061, EPI_ISL_2724063, EPI_ISL_2724064, EPI_ISL_2724065, EPI_ISL_2724066, EPI_ISL_2724067, EPI_ISL_2724068, EPI_ISL_2724070, EPI_ISL_2724071, EPI_ISL_2724072, EPI_ISL_2724073, EPI_ISL_2724074, EPI_ISL_2724075, EPI_ISL_2724076, EPI_ISL_2724077, EPI_ISL_2724078, EPI_ISL_2724079, EPI_ISL_2724080, EPI_ISL_2724081, EPI_ISL_2724082, EPI_ISL_2724083, EPI_ISL_2724084, EPI_ISL_2724085, EPI_ISL_2724087, EPI_ISL_2724088, EPI_ISL_2724089, EPI_ISL_2724090, EPI_ISL_2724091, EPI_ISL_2724092, EPI_ISL_2724093, EPI_ISL_2724094, EPI_ISL_2724095, EPI_ISL_2724096, EPI_ISL_2724097, EPI_ISL_2724098, EPI_ISL_2724099, EPI_ISL_2724100, EPI_ISL_2724101, EPI_ISL_2724102, EPI_ISL_2724103, EPI_ISL_2724104, EPI_ISL_2724105, EPI_ISL_2724106, EPI_ISL_2724107, EPI_ISL_2724108, EPI_ISL_2724109, EPI_ISL_2724110, EPI_ISL_2724112, EPI_ISL_2724113, EPI_ISL_2724114, EPI_ISL_2724115, EPI_ISL_2724116, EPI_ISL_2724117, EPI_ISL_2724118, EPI_ISL_2724119, EPI_ISL_2724120, EPI_ISL_2724121, EPI_ISL_2724122, EPI_ISL_2724123, EPI_ISL_2724124, EPI_ISL_2724125, EPI_ISL_2724127, EPI_ISL_2724128, EPI_ISL_2724129, EPI_ISL_2724130, EPI_ISL_2724132, EPI_ISL_2724133, EPI_ISL_2724134, EPI_ISL_2724135, EPI_ISL_2724136, EPI_ISL_2724137, EPI_ISL_2724139, EPI_ISL_2724140, EPI_ISL_2724141, EPI_ISL_2724142, EPI_ISL_2724143, EPI_ISL_2724144, EPI_ISL_2724145, EPI_ISL_2724146, EPI_ISL_2724147, EPI_ISL_2724148, EPI_ISL_2724149, EPI_ISL_2724150, EPI_ISL_2724151, EPI_ISL_2724152, EPI_ISL_2724153, EPI_ISL_2724154, EPI_ISL_2724155, EPI_ISL_2724157, EPI_ISL_2724158, EPI_ISL_2724159, EPI_ISL_2724160, EPI_ISL_2724161, EPI_ISL_2724162, EPI_ISL_2724163, EPI_ISL_2724164, EPI_ISL_2724165, EPI_ISL_2724166, EPI_ISL_2724167, EPI_ISL_2724169, EPI_ISL_2724170, EPI_ISL_2724171, EPI_ISL_2724172, EPI_ISL_2724173, EPI_ISL_2724174, EPI_ISL_2724175, EPI_ISL_2724176, EPI_ISL_2724177, EPI_ISL_2724178, EPI_ISL_2724179, EPI_ISL_2724180, EPI_ISL_2724181, EPI_ISL_2724182, EPI_ISL_2724183, EPI_ISL_2724184, EPI_ISL_2724185, EPI_ISL_2724186, EPI_ISL_2724187, EPI_ISL_2724188, EPI_ISL_2724189, EPI_ISL_2724190, EPI_ISL_2724191, EPI_ISL_2724192, EPI_ISL_2724193, EPI_ISL_2724195, EPI_ISL_2724196, EPI_ISL_2724197, EPI_ISL_2724198, EPI_ISL_2724199, EPI_ISL_2724200, EPI_ISL_2724201, EPI_ISL_2724203, EPI_ISL_2724204, EPI_ISL_2724205, EPI_ISL_2724206, EPI_ISL_2724207, EPI_ISL_2724208, EPI_ISL_2724209, EPI_ISL_2724210, EPI_ISL_2724211, EPI_ISL_2724212, EPI_ISL_2724213, EPI_ISL_2724214, EPI_ISL_2724215, EPI_ISL_2724216, EPI_ISL_2724217, EPI_ISL_2724218, EPI_ISL_2724221, EPI_ISL_2724222, EPI_ISL_2724223, EPI_ISL_2724225, EPI_ISL_2724227, EPI_ISL_2724228, EPI_ISL_2724229, EPI_ISL_2724231, EPI_ISL_2724232, EPI_ISL_2724233, EPI_ISL_2724234, EPI_ISL_2724235, EPI_ISL_2724236, EPI_ISL_2724237, EPI_ISL_2724238, EPI_ISL_2724239, EPI_ISL_2724240, EPI_ISL_2724243, EPI_ISL_2724244, EPI_ISL_2724247, EPI_ISL_2724249, EPI_ISL_2724251, EPI_ISL_2724252, EPI_ISL_2724253, EPI_ISL_2724254, EPI_ISL_2724255, EPI_ISL_2724256, EPI_ISL_2724257, EPI_ISL_2724258, EPI_ISL_2724259, EPI_ISL_2724260, EPI_ISL_2724261, EPI_ISL_2724262, EPI_ISL_2724263, EPI_ISL_2724264, EPI_ISL_2724267, EPI_ISL_2724268, EPI_ISL_2724269, EPI_ISL_2724270, EPI_ISL_2724271, EPI_ISL_2724272, EPI_ISL_2724273, EPI_ISL_2724275, EPI_ISL_2724276, EPI_ISL_2724277, EPI_ISL_2724278, EPI_ISL_2724279, EPI_ISL_2724280, EPI_ISL_2724281, EPI_ISL_2724283, EPI_ISL_2724284, EPI_ISL_2724285, EPI_ISL_2724286 | | | |
| see above | Dr S raju, Director of public heath and preventive medicine | inStem NCBS – INSACOG | Uma Ramakrishnan Dasaradhi Palakodeti Aswin SaiNarain |
| EPI_ISL_2724287, EPI_ISL_2724288, EPI_ISL_2724289, EPI_ISL_2724290, EPI_ISL_2724291, EPI_ISL_2724296, EPI_ISL_2724297, EPI_ISL_2724299, EPI_ISL_2724300, EPI_ISL_2724301, EPI_ISL_2724302, EPI_ISL_2724303, EPI_ISL_2724305, EPI_ISL_2724306, EPI_ISL_2724307, EPI_ISL_2724308, EPI_ISL_2724309, EPI_ISL_2724310, EPI_ISL_2724311, EPI_ISL_2724312, EPI_ISL_2724313, EPI_ISL_2724314, EPI_ISL_2724315, EPI_ISL_2724316, EPI_ISL_2724317 | | | |
| see above | Bangalore Baptist Hospital | inStem NCBS – INSACOG | Uma Ramakrishnan Dasaradhi Palakodeti Aswin SaiNarain |
| EPI_ISL_2724318, EPI_ISL_2724319, EPI_ISL_2724320, EPI_ISL_2724321, EPI_ISL_2724322, EPI_ISL_2724323, EPI_ISL_2724324, EPI_ISL_2724326, EPI_ISL_2724327, EPI_ISL_2724328, EPI_ISL_2724329, EPI_ISL_2724330, EPI_ISL_2724331, EPI_ISL_2724332, EPI_ISL_2724333, EPI_ISL_2724334, EPI_ISL_2724335, EPI_ISL_2724347, EPI_ISL_2724348, EPI_ISL_2724349 | | | |
| see above | Dr S raju, Director of public heath and preventive medicine | inStem NCBS – INSACOG | Uma Ramakrishnan Dasaradhi Palakodeti Aswin SaiNarain |
| EPI_ISL_2724352, EPI_ISL_2724353, EPI_ISL_2724354, EPI_ISL_2724355, EPI_ISL_2724356, EPI_ISL_2724357 | Dr. Madhusudhan, MBBD, MD, ICMR VRDL | inStem NCBS – INSACOG | Uma Ramakrishnan Dasaradhi Palakodeti Aswin SaiNarain |
| EPI_ISL_2724385, EPI_ISL_2724386, EPI_ISL_2724387, EPI_ISL_2724389, EPI_ISL_2724396, EPI_ISL_2724400, EPI_ISL_2724401, EPI_ISL_2724410, EPI_ISL_2724423, EPI_ISL_2724426, EPI_ISL_2724430, EPI_ISL_2724445, EPI_ISL_2724446, EPI_ISL_2724449, EPI_ISL_2724450, EPI_ISL_2724452, EPI_ISL_2724454, EPI_ISL_2724455, EPI_ISL_2724456, EPI_ISL_2724465, EPI_ISL_2724481, EPI_ISL_2724484, EPI_ISL_2724486, EPI_ISL_2724487, EPI_ISL_2724488, EPI_ISL_2724501, EPI_ISL_2724509, EPI_ISL_2724510 | | | |
| see above | Viollier AG | Department of Biosystems Science and Engineering, ETH Zürich | Christian Beisel, Sarah Nadeau, Chaoran Chen, Ivan Topolsky, Kim Philipp Jablonski, Lara Fuhrmann, Rebecca Denes, Mirjam Feldkamp, Ina Nissen, Natascha Santacroce, Elodie Burcklen, Christiane Beckmann, Maurice Redondo, Olivier Kobel, Christoph Noppen, Niko Beerenwinkel, Tanja Stadler |
| EPI_ISL_2724672, EPI_ISL_2724756, EPI_ISL_2724793, EPI_ISL_2724835, EPI_ISL_2724843, EPI_ISL_2724851, EPI_ISL_2724853, EPI_ISL_2724859, EPI_ISL_2724861, EPI_ISL_2724867, EPI_ISL_2724871, EPI_ISL_2724872, EPI_ISL_2724883, EPI_ISL_2724903 | | | |
| see above | State Laboratories Division, Hawaii State Department of Health | State Laboratories Division, Hawaii State Department of Health | Pamela O'Brien, Drew Kuwazaki, Ayana Garnet, Razvan Sultana, Edward Desmond |
| EPI_ISL_2724915, EPI_ISL_2724916, EPI_ISL_2724921, EPI_ISL_2724922, EPI_ISL_2724924, EPI_ISL_2724925, EPI_ISL_2724926, EPI_ISL_2724927, EPI_ISL_2724929, EPI_ISL_2724935, EPI_ISL_2724936, EPI_ISL_2724945 | | | |
| see above | Servicio Microbiología Hospital La Paz | Servicio Microbiología Hospital La Paz | Fernando Lázaro, Rubén Cáceres, Jesús Mingorance Cruz, Elie Dahdouh |
| EPI_ISL_2724975 | Department of Medical Microbiology - section Molde, Molde Hospital | Norwegian Institute of Public Health, Department of Virology | Kathrine Stene-Johansen, Kamilla Heddeland Instefjord, Hilde Elshaug, Garcia Llorente Ignacio, Jon Bråte, Engebretsen Serina Beate,Pedersen Benedikte Nevjen, Line Victoria Moen, Debech Nadia, Atiya R Ali,Marie Paulsen Madsen, Rasmus Riis Kopperud, Hilde Vollan, Karoline Bragstad, Olav Hungnes |
| EPI_ISL_2724999, EPI_ISL_2725002 | Ostfold Hospital Trust - Kalnes, Centre for Laboratory Medicine, Section for gene technology and infection serology | Norwegian Institute of Public Health, Department of Virology | Kathrine Stene-Johansen, Kamilla Heddeland Instefjord, Hilde Elshaug, Garcia Llorente Ignacio, Jon Bråte, Engebretsen Serina Beate,Pedersen Benedikte Nevjen, Line Victoria Moen, Debech Nadia, Atiya R Ali,Marie Paulsen Madsen, Rasmus Riis Kopperud, Hilde Vollan, Karoline Bragstad, Olav Hungnes |
| EPI_ISL_2725007 | Furst Medical Laboratory | Norwegian Institute of Public Health, Department of Virology | Kathrine Stene-Johansen, Kamilla Heddeland Instefjord, Hilde Elshaug, Garcia Llorente Ignacio, Jon Bråte, Engebretsen Serina Beate,Pedersen Benedikte Nevjen, Line Victoria Moen, Debech Nadia, Atiya R Ali,Marie Paulsen Madsen, Rasmus Riis Kopperud, Hilde Vollan, Karoline Bragstad, Olav Hungnes |
| EPI_ISL_2725043, EPI_ISL_2725045, EPI_ISL_2725048, EPI_ISL_2725049, EPI_ISL_2725050, EPI_ISL_2725051, EPI_ISL_2725053, EPI_ISL_2725054, EPI_ISL_2725055, EPI_ISL_2725056, EPI_ISL_2725062, EPI_ISL_2725067, EPI_ISL_2725068, EPI_ISL_2725069 | | | |
| see above | IN State Department of Health Laboratory Services | IN State Department of Health Laboratory Services | Cassandra Campion, Jamie Yeadon, Brian Pope, Lixia Liu, Kyle Brownlee, Melissa Hindenlang, Mark Glazier |
| EPI_ISL_2725072 | DNAGYN | LGBio - Laboratório de Genética & Biodiversidade | Mariana Pires de Campos Telles, Daniela de Melo e Silva, Elisangela de Paula Silveira Lacerda, Renata de Oliveira Dias, Rhewter Nunes, Cintia Pelegrineti Targueta de Azevedo Brito, Ramilla dos Santos Braga, Thais Guimarães Castro, Thays Millena Alves Pedroso, Amanda Alves de Melo, Aparecido Divino da Cruz, Luiz Augusto Pereira, Thais Cidália Vieira Gigonzac, Marc Alexandre Duarte Gigonzac, Alex Honda Bernardes, Francyelli Mello Andrade, Juliana Santana de Curcio |
| EPI_ISL_2725077 | Vestfold Hospital, Toensberg Department of Microbiology | Norwegian Institute of Public Health, Department of Virology | Kathrine Stene-Johansen, Kamilla Heddeland Instefjord, Hilde Elshaug, Garcia Llorente Ignacio, Jon Bråte, Engebretsen Serina Beate,Pedersen Benedikte Nevjen, Line Victoria Moen, Debech Nadia, Atiya R Ali,Marie Paulsen Madsen, Rasmus Riis Kopperud, Hilde Vollan, Karoline Bragstad, Olav Hungnes |
| EPI_ISL_2725080, EPI_ISL_2725083, EPI_ISL_2725084 | Akershus University Hospital, Department for Microbiology and Infectious Disease Control | Norwegian Institute of Public Health, Department of Virology | Kathrine Stene-Johansen, Kamilla Heddeland Instefjord, Hilde Elshaug, Garcia Llorente Ignacio, Jon Bråte, Engebretsen Serina Beate,Pedersen Benedikte Nevjen, Line Victoria Moen, Debech Nadia, Atiya R Ali,Marie Paulsen Madsen, Rasmus Riis Kopperud, Hilde Vollan, Karoline Bragstad, Olav Hungnes |
| EPI_ISL_2725088, EPI_ISL_2725089, EPI_ISL_2725090 | Furst Medical Laboratory | Norwegian Institute of Public Health, Department of Virology | Kathrine Stene-Johansen, Kamilla Heddeland Instefjord, Hilde Elshaug, Garcia Llorente Ignacio, Jon Bråte, Engebretsen Serina Beate,Pedersen Benedikte Nevjen, Line Victoria Moen, Debech Nadia, Atiya R Ali,Marie Paulsen Madsen, Rasmus Riis Kopperud, Hilde Vollan, Karoline Bragstad, Olav Hungnes |
| EPI_ISL_2725101 | Nordland Hospital - Bodo, Laboratory Department, Molecular Biology Unit | Norwegian Institute of Public Health, Department of Virology | Kathrine Stene-Johansen, Kamilla Heddeland Instefjord, Hilde Elshaug, Garcia Llorente Ignacio, Jon Bråte, Engebretsen Serina Beate,Pedersen Benedikte Nevjen, Line Victoria Moen, Debech Nadia, Atiya R Ali,Marie Paulsen Madsen, Rasmus Riis Kopperud, Hilde Vollan, Karoline Bragstad, Olav Hungnes |
| EPI_ISL_2725113, EPI_ISL_2725114, EPI_ISL_2725117, EPI_ISL_2725118, EPI_ISL_2725119, EPI_ISL_2725120, EPI_ISL_2725138, EPI_ISL_2725139, EPI_ISL_2725140, EPI_ISL_2725141, EPI_ISL_2725142, EPI_ISL_2725143, EPI_ISL_2725144, EPI_ISL_2725145 | | | |
| see above | Istituto Zooprofilattico Sperimentale del Mezzogiorno | Telethon Institute of Genetics and Medicine (TIGEM) | Antonio Grimaldi Patrizia Annunziata Francesco Panariello Biancamaria Pierri Claudia Tiberio Teresa Giuliano Valentina Bouche Chiara Colantuono Maria Concetta Cuomo Denise Di Concilio Lucio Di Filippo Anna Manfredi Marcello Salvi Antonio Limone Luigi Atripaldi Pellegrino Cerino Andrea Ballabio Davide Cacchiarelli |
| EPI_ISL_2725147, EPI_ISL_2725148, EPI_ISL_2725149, EPI_ISL_2725154, EPI_ISL_2725155, EPI_ISL_2725156, EPI_ISL_2725157, EPI_ISL_2725158, EPI_ISL_2725159, EPI_ISL_2725161, EPI_ISL_2725164, EPI_ISL_2725165, EPI_ISL_2725166, EPI_ISL_2725168, EPI_ISL_2725169, EPI_ISL_2725173, EPI_ISL_2725174, EPI_ISL_2725179, EPI_ISL_2725181, EPI_ISL_2725192, EPI_ISL_2725196, EPI_ISL_2725197, EPI_ISL_2725199, EPI_ISL_2725200, EPI_ISL_2725205, EPI_ISL_2725217, EPI_ISL_2725218, EPI_ISL_2725222, EPI_ISL_2725227, EPI_ISL_2725237, EPI_ISL_2725242, EPI_ISL_2725255, EPI_ISL_2725258, EPI_ISL_2725259 | | | |
| see above | Lab. Microbiologia e Virologia Cotugno A.O. dei Colli - Istituto Zooprofilattico Sperimentale del Mezzogiorno | Telethon Institute of Genetics and Medicine (TIGEM) | Antonio Grimaldi Patrizia Annunziata Francesco Panariello Biancamaria Pierri Claudia Tiberio Teresa Giuliano Valentina Bouche Chiara Colantuono Maria Concetta Cuomo Denise Di Concilio Lucio Di Filippo Anna Manfredi Marcello Salvi Antonio Limone Luigi Atripaldi Pellegrino Cerino Andrea Ballabio Davide Cacchiarelli |
| EPI_ISL_2725301, EPI_ISL_2725320, EPI_ISL_2725321, EPI_ISL_2725322, EPI_ISL_2725323, EPI_ISL_2725324, EPI_ISL_2725325, EPI_ISL_2725326, EPI_ISL_2725328, EPI_ISL_2725329, EPI_ISL_2725330, EPI_ISL_2725334, EPI_ISL_2725335 | | | |
| see above | Istituto Zooprofilattico Sperimentale del Mezzogiorno - Azienda Ospedaliera "Pugliese Ciaccio" di Catanzaro | Telethon Institute of Genetics and Medicine (TIGEM) | Antonio Grimaldi Patrizia Annunziata Francesco Panariello Biancamaria Pierri Claudia Tiberio Teresa Giuliano Valentina Bouche Chiara Colantuono Maria Concetta Cuomo Denise Di Concilio Lucio Di Filippo Anna Manfredi Pasquale Minchella Marcello Salvi Antonio Limone Luigi Atripaldi Pellegrino Cerino Andrea Ballabio Davide Cacchiarelli |
| EPI_ISL_2725359, EPI_ISL_2725399, EPI_ISL_2725400, EPI_ISL_2725401, EPI_ISL_2725402, EPI_ISL_2725403, EPI_ISL_2725405, EPI_ISL_2725408, EPI_ISL_2725409 | Istituto Zooprofilattico Sperimentale del Mezzogiorno | Telethon Institute of Genetics and Medicine (TIGEM) | Antonio Grimaldi Patrizia Annunziata Francesco Panariello Biancamaria Pierri Claudia Tiberio Teresa Giuliano Valentina Bouche Chiara Colantuono Maria Concetta Cuomo Denise Di Concilio Lucio Di Filippo Anna Manfredi Marcello Salvi Antonio Limone Luigi Atripaldi Pellegrino Cerino Andrea Ballabio Davide Cacchiarelli |
| EPI_ISL_2725440 | Lab. Microbiologia e Virologia Cotugno A.O. dei Colli - Istituto Zooprofilattico Sperimentale del Mezzogiorno | Telethon Institute of Genetics and Medicine (TIGEM) | Antonio Grimaldi Patrizia Annunziata Francesco Panariello Biancamaria Pierri Claudia Tiberio Teresa Giuliano Valentina Bouche Chiara Colantuono Maria Concetta Cuomo Denise Di Concilio Lucio Di Filippo Anna Manfredi Marcello Salvi Antonio Limone Luigi Atripaldi Pellegrino Cerino Andrea Ballabio Davide Cacchiarelli |
| EPI_ISL_2725488, EPI_ISL_2725489, EPI_ISL_2725491, EPI_ISL_2725501, EPI_ISL_2725506, EPI_ISL_2725507, EPI_ISL_2725510 | AZDelta | AZ Delta Medical Laboratories in Roeselare, Belgium | Geert Martens, Dieter De Smet, Merijn Vanhee, on behalf of AZ Delta COVID-19 Genomics core (member of Genomic surveillance of SARS-CoV-2 in Belgium network) |
| EPI_ISL_2725525, EPI_ISL_2725526, EPI_ISL_2725527, EPI_ISL_2725528 | BBMP Urban | INSACOG-KA, NIMHANS | Chitra Pattabiraman, Anson Kunjumon George,Darshan Sreenivas,Ellango Ramasamy, Sony Sharma, Mahesh Kumar.C.S, Ananthapadmanabha Kotambail, Chetan G K, Gautham Arunachal Udupi, Anita S Desai, V Ravi |
| EPI_ISL_2725529 | RAILWAY HOSPITAL | INSACOG-KA, NIMHANS | Chitra Pattabiraman, Anson Kunjumon George,Darshan Sreenivas,Ellango Ramasamy, Sony Sharma, Mahesh Kumar.C.S, Ananthapadmanabha Kotambail, Chetan G K, Gautham Arunachal Udupi, Anita S Desai, V Ravi |
| EPI_ISL_2725530, EPI_ISL_2725531, EPI_ISL_2725532, EPI_ISL_2725534, EPI_ISL_2725535, EPI_ISL_2725536 | BBMP Urban | INSACOG-KA, NIMHANS | Chitra Pattabiraman, Anson Kunjumon George,Darshan Sreenivas,Ellango Ramasamy, Sony Sharma, Mahesh Kumar.C.S, Ananthapadmanabha Kotambail, Chetan G K, Gautham Arunachal Udupi, Anita S Desai, V Ravi |

| | | | |
|---|---|---|---|
| EPI_ISL_2725537 | NIMHANS | INSACOG-KA, NIMHANS | Chitra Pattabiraman, Anson Kunjumon George,Darshan Sreenivas,Ellango Ramasamy, Sony Sharma, Mahesh Kumar.C.S, Ananthapadmanabha Kotambail, Chetan G K, Gautham Arunachal Udupi, Anita S Desai, V Ravi |
| EPI_ISL_2725538, EPI_ISL_2725539, EPI_ISL_2725540, EPI_ISL_2725543, EPI_ISL_2725544, EPI_ISL_2725550, EPI_ISL_2725552 | Azienda Sanitaria dell'Alto Adige - Laboratorio Aziendale di Microbiologia e Virologia | Azienda Sanitaria dell'Alto Adige | Irene Bianconi |
| EPI_ISL_2725597, EPI_ISL_2725611, EPI_ISL_2725612, EPI_ISL_2725613, EPI_ISL_2725614, EPI_ISL_2725615, EPI_ISL_2725616, EPI_ISL_2725620, EPI_ISL_2725621, EPI_ISL_2725622, EPI_ISL_2725638, EPI_ISL_2725653, EPI_ISL_2725659, EPI_ISL_2725661, EPI_ISL_2725662, EPI_ISL_2725663, EPI_ISL_2725664, EPI_ISL_2725665, EPI_ISL_2725700, EPI_ISL_2725709, EPI_ISL_2726311, EPI_ISL_2726312, EPI_ISL_2726502, EPI_ISL_2726503, EPI_ISL_2726504, EPI_ISL_2726516, EPI_ISL_2726517, EPI_ISL_2726518, EPI_ISL_2726519, EPI_ISL_2726520, EPI_ISL_2726521, EPI_ISL_2726522, EPI_ISL_2726523, EPI_ISL_2726524, EPI_ISL_2726525, EPI_ISL_2726526, EPI_ISL_2726527 | | | |
| see above | KU Leuven, Rega Institute, Clinical and Epidemiological Virology | KU Leuven, Rega Institute, Clinical and Epidemiological Virology | Tony Wawina-Bokalanga, Bert Vanmechelen, Joan Marti-Carerras, Piet Maes |
| EPI_ISL_2726672, EPI_ISL_2726673, EPI_ISL_2726674, EPI_ISL_2726676, EPI_ISL_2726679, EPI_ISL_2726680, EPI_ISL_2726681, EPI_ISL_2726682, EPI_ISL_2726683, EPI_ISL_2726685, EPI_ISL_2726686, EPI_ISL_2726687, EPI_ISL_2726688, EPI_ISL_2726690, EPI_ISL_2726691, EPI_ISL_2726692, EPI_ISL_2726693, EPI_ISL_2726694, EPI_ISL_2726695 | | | |
| see above | Kansas Health and Environmental Lab | Kansas Health and Environmental Lab | Mike Grose, Katherine Wiggins, Jonathan Barnell, Ben Olsen, and Phil Adam |
| EPI_ISL_2726705, EPI_ISL_2726706 | Eurofins-NMDL | Eurofins-NMDL | Thierry Janssens, Anne Vogel, Lisa Dreesens, Roy Masius, Katja Plathan, Anco Molijn, Maurine Leversteijn-van Hall, Thomas van der Toorn, Cornel Brouwer, Diviana Lakker, Hamza Saleem, Layonel Jacobus, Rosalie Van der Stap, Marisol Williams, Chris van Koppen, Senay Ozturk, Marvin Ruiter, Mark de Jong, Alexandra Hansen, Coen de Jong, Madhavi Kharagjitsing, Ramon Koers, Rens Driel, Thomas van der Toorn, Bram Weijers, Chantal Schenker, Humaira Yousaf, Jelmer Eerland, Melek Aras, Rousjan Amir, Enes Sekerci, Goyaert Roosen, Laura Weijenberg, Mathijs Wielen, Nienke Meijer, Tobias van der Heijde, Chinmoy Saha, Glenn Bohms, Iman El-Idrioui, Jorian Flik, Lida Cen, Simon Lansu |
| EPI_ISL_2726782 | GenXOne SA | 1. National Institute of Public Health - National Institute of Hygiene, Warsaw, Poland 2. Biobank Lab, University of Lodz 3. Laboratory of Respiratory Viruses, Teaching and Clinical Center of the Medical University of Lodz | Dominik Strapagiel, Marta Sobalska-Kwapis, Klaudyna Królikowska, Jakub Lach, Marcin Słomka, Tomasz Płoszaj, Magdalena Traczyk-Borszyńska, Izabela Dróżdż, Maciej Borowiec, Katarzyna Zacharczuk, Magdalena Nowakowska, Małgorzata Sadkowska-Todys, Tomasz Wołkowicz |
| EPI_ISL_2726789 | Laboratorium Diagnostyczno-Analityczne Ośrodek Profilaktyki i Epidemiologii Nowotworów im. A. Pieńkowskiej | 1. National Institute of Public Health - National Institute of Hygiene, Warsaw, Poland 2. Biobank Lab, University of Lodz 3. Laboratory of Respiratory Viruses, Teaching and Clinical Center of the Medical University of Lodz | Dominik Strapagiel, Marta Sobalska-Kwapis, Klaudyna Królikowska, Jakub Lach, Marcin Słomka, Tomasz Płoszaj, Magdalena Traczyk-Borszyńska, Izabela Dróżdż, Maciej Borowiec, Katarzyna Zacharczuk, Magdalena Nowakowska, Małgorzata Sadkowska-Todys, Tomasz Wołkowicz |
| EPI_ISL_2726793 | UM im. Karola Marcinkowskiego w Poznaniu Laboratorium UCA_Covid-19 Centrum Biologii Medycznej | 1. National Institute of Public Health - National Institute of Hygiene, Warsaw, Poland 2. Biobank Lab, University of Lodz 3. Laboratory of Respiratory Viruses, Teaching and Clinical Center of the Medical University of Lodz | Dominik Strapagiel, Marta Sobalska-Kwapis, Klaudyna Królikowska, Jakub Lach, Marcin Słomka, Tomasz Płoszaj, Magdalena Traczyk-Borszyńska, Izabela Dróżdż, Maciej Borowiec, Katarzyna Zacharczuk, Magdalena Nowakowska, Małgorzata Sadkowska-Todys, Tomasz Wołkowicz |
| EPI_ISL_2726813, EPI_ISL_2726814 | SPWSZ w Szczecinie | 1. National Institute of Public Health - National Institute of Hygiene, Warsaw, Poland 2. Biobank Lab, University of Lodz 3. Laboratory of Respiratory Viruses, Teaching and Clinical Center of the Medical University of Lodz | Dominik Strapagiel, Marta Sobalska-Kwapis, Klaudyna Królikowska, Jakub Lach, Marcin Słomka, Tomasz Płoszaj, Magdalena Traczyk-Borszyńska, Izabela Dróżdż, Maciej Borowiec, Katarzyna Zacharczuk, Magdalena Nowakowska, Małgorzata Sadkowska-Todys, Tomasz Wołkowicz |
| EPI_ISL_2726832, EPI_ISL_2726833, EPI_ISL_2726834, EPI_ISL_2726836, EPI_ISL_2726837, EPI_ISL_2726838, EPI_ISL_2726839, EPI_ISL_2726840, EPI_ISL_2726843, EPI_ISL_2726846, EPI_ISL_2726847, EPI_ISL_2726848, EPI_ISL_2726849 | | | |
| see above | Wyoming Public Health Laboratory | Wyoming Public Health Laboratory | Jim Mildenberger, Taylor Fearing, Channing Weber, Ashley Norberg, Chayse Rowley, Marley Goetz, Brian Dominguez, Elliot Thomasson, Sam Britz, Cari Sloma, Robert Petit, and Rob Christensen |
| EPI_ISL_2726858, EPI_ISL_2726859, EPI_ISL_2726861, EPI_ISL_2726863, EPI_ISL_2726864, EPI_ISL_2726870, EPI_ISL_2726878, EPI_ISL_2726881, EPI_ISL_2726883, EPI_ISL_2726890, EPI_ISL_2726902, EPI_ISL_2726908, EPI_ISL_2726910, EPI_ISL_2726915, EPI_ISL_2726917, EPI_ISL_2726922, EPI_ISL_2726923, EPI_ISL_2726925, EPI_ISL_2726927 | | | |
| see above | Lifebrain Covid Labor GmbH | Dept. of Laboratory Medicine | Harald Esterbauer, Anna Gschaider, Petra Jurkowitsch, Fabian Konig, Sabina Plumer, Robert Strassl, Claudia Weber, Oswald Wagner |
| EPI_ISL_2726936 | UW Virology Lab | UW Virology Lab | Pavitra Roychoudhury, Hong Xie, Lasata Shrestha, Shah Mohamed Bakhash, Tien V. Nguyen, Noah R. Baker, Sean Ellis, Meei-Li Huang, Keith R Jerome, Alexander Greninger |
| EPI_ISL_2726943 | ISDH | Quantigen Biosciences | Paul J. Childress |
| EPI_ISL_2726978, EPI_ISL_2726980, EPI_ISL_2726987, EPI_ISL_2726991, EPI_ISL_2726992, EPI_ISL_2726993, EPI_ISL_2727018 | National Institute of Public Health | National Institute of Public Health | Helena Jirincova, Jaromira Vecerova, Timotej Suri, Dusan Trnka, Alexander Nagy |
| EPI_ISL_2727055, EPI_ISL_2727058, EPI_ISL_2727059 | MD PHL | MD PHL | Maryland Department of Health Laboratories Administration |
| EPI_ISL_2727158, EPI_ISL_2727163, EPI_ISL_2727172 | AZDelta | AZ Delta Medical Laboratories in Roeselare, Belgium | Geert Martens, Dieter De Smet, Merijn Vanhee, on behalf of AZ Delta COVID-19 Genomics core (member of Genomic surveillance of SARS-CoV-2 in Belgium network) |
| EPI_ISL_2727200, EPI_ISL_2727202, EPI_ISL_2727209, EPI_ISL_2727210, EPI_ISL_2727213, EPI_ISL_2727217, EPI_ISL_2727218, EPI_ISL_2727219, EPI_ISL_2727223, EPI_ISL_2727225, EPI_ISL_2727227, EPI_ISL_2727232 | | | |
| see above | NHLS_VIRO Eastern Cape | KRISP, KZn Research Innovation and Sequencing Platform | Laguda-Akingba O, Giandhari J, Pillay S, Lessells R, Mdlalose K, York D, Khan S, Emmanuel SJ, Tegally H, Wilkinson E, de Oliveira T |
| EPI_ISL_2727233, EPI_ISL_2727234, EPI_ISL_2727235, EPI_ISL_2727236, EPI_ISL_2727237, EPI_ISL_2727238 | Ampath Greenacres Hospital | KRISP, KZn Research Innovation and Sequencing Platform | Giandhari Jennifer, Pillay Sureshnee, Yajna Ramphal, Naidoo Yeshnee, Tshabuila Derek,Tegally Houriiyah, San James, Wilkinson Eduan, de Oliveira Tulio |
| EPI_ISL_2727323 | AZDelta | AZ Delta Medical Laboratories in Roeselare, Belgium | Geert Martens, Dieter De Smet, Merijn Vanhee, on behalf of AZ Delta COVID-19 Genomics core (member of Genomic surveillance of SARS-CoV-2 in Belgium network) |
| EPI_ISL_2727329, EPI_ISL_2727335, EPI_ISL_2727347, EPI_ISL_2727351, EPI_ISL_2727352, EPI_ISL_2727353, EPI_ISL_2727354, EPI_ISL_2727357, EPI_ISL_2727358, EPI_ISL_2727359, EPI_ISL_2727360, EPI_ISL_2727361, EPI_ISL_2727362, EPI_ISL_2727363, EPI_ISL_2727364, EPI_ISL_2727365, EPI_ISL_2727366, EPI_ISL_2727369, EPI_ISL_2727370, EPI_ISL_2727371, EPI_ISL_2727373, EPI_ISL_2727374, EPI_ISL_2727375, EPI_ISL_2727376, EPI_ISL_2727377, EPI_ISL_2727378, EPI_ISL_2727379 | | | |
| see above | ZARV/NHLS, Department Medical Virology, University of Pretoria | KRISP, KZn Research Innovation and Sequencing Platform | Amy Strydom, Adriano Mendes, Micheala Davids, Sim Mayaphi and Marietjie Venter, Giandhari J, Pillay S, Naidoo Y, Ramphal U, Lessells R, Emmanuel SJ, Tegally H, Wilkinson E, de Oliveira T |
| EPI_ISL_2727380, EPI_ISL_2727381, EPI_ISL_2727383, EPI_ISL_2727384, EPI_ISL_2727386, EPI_ISL_2727387, EPI_ISL_2727388, EPI_ISL_2727389, EPI_ISL_2727391, EPI_ISL_2727392, EPI_ISL_2727393, EPI_ISL_2727394, EPI_ISL_2727395, EPI_ISL_2727396, EPI_ISL_2727397, EPI_ISL_2727398, EPI_ISL_2727399, EPI_ISL_2727400, EPI_ISL_2727401, EPI_ISL_2727402, EPI_ISL_2727403 | | | |
| see above | NHLS_VIRO | KRISP, KZn Research Innovation and Sequencing Platform | Giandhari Jennifer, Pillay Sureshnee, Yajna Ramphal, Naidoo Yeshnee, Tshabuila Derek,Tegally Houriiyah, San James, Wilkinson Eduan, de Oliveira Tulio |
| EPI_ISL_2727404 | Netcare Umhlanga Hospital | KRISP, KZn Research Innovation and Sequencing Platform | Giandhari Jennifer, Pillay Sureshnee, Yajna Ramphal, Naidoo Yeshnee, Tshabuila Derek,Tegally Houriiyah, San James, Wilkinson Eduan, de Oliveira Tulio |
| EPI_ISL_2727444 | Wyoming Public Health Laboratory | Wyoming Public Health Laboratory | Jim Mildenberger, Taylor Fearing, Channing Weber, Ashley Norberg, Chayse Rowley, Marley Goetz, Brian Dominguez, Elliot Thomasson, Sam Britz, Cari Sloma, Robert Petit, and Rob Christensen |
| EPI_ISL_2727474 | Oregon State Public Health Laboratory | Oregon State Public Health Laboratory | Rafia Razzaque, Eugene Yeboah, Vanda Makris, Laura Tsaknaridis, John Fontana and Shane Sevey |
| EPI_ISL_2727568 | Cantacuzino National Military-Medical Institute, Viral Respiratory Infections Laboratory | Cantacuzino Institute Virology | Carmen Cherciu, Luiza Ustea, Nicoleta Paraschiv, Sorin Dinu, Mihaela Oprea, Mihaela Lazar |
| EPI_ISL_2727569, EPI_ISL_2727570, EPI_ISL_2727571, EPI_ISL_2727574, EPI_ISL_2727575, EPI_ISL_2727576, EPI_ISL_2727581, EPI_ISL_2727592, EPI_ISL_2727599, EPI_ISL_2727601, EPI_ISL_2727602, EPI_ISL_2727603, EPI_ISL_2727604, EPI_ISL_2727605, EPI_ISL_2727606, EPI_ISL_2727607, EPI_ISL_2727608, EPI_ISL_2727609, EPI_ISL_2727611, EPI_ISL_2727614 | | | |
| see above | NJDOH, Public Health and Environmental Laboratories | NJ_PHEL | Lindsey Bodnar, Shiv K. Verma, Jacquelyn Deverell, Ryan Pachucki, Dana Woell, Allison Roder, Byeong Jeong |
| EPI_ISL_2727636, EPI_ISL_2727638, EPI_ISL_2727639, EPI_ISL_2727640, EPI_ISL_2727641, EPI_ISL_2727642 | Queensland Health Forensic and Scientific Services | Queensland Health Forensic and Scientific Services | Son Nguyen |
| EPI_ISL_2727654 | Genome Information Research Center, Research Institute for Microbial Diseases, Osaka University | Genome Information Research Center, Research Institute for Microbial Diseases, Osaka University | Daisuke Motooka, Kentaro Tanaka, Hiroya Oki, and Shota Nakamura |
| EPI_ISL_2727663 | Bundeswehrkrankenhaus Hamburg | Bundeswehr Institute of Microbiology | Markus Antwerpen, Alexandra Rehn, Mathias Walter, Malena Bestehorn-Willmann, Mike Pillukat, Sabine Zange, Enrico Georgi, Roman Wölfel |
| EPI_ISL_2727680, EPI_ISL_2727683, EPI_ISL_2727685, EPI_ISL_2727686, EPI_ISL_2727730, EPI_ISL_2727736, EPI_ISL_2727738, EPI_ISL_2727758, EPI_ISL_2727759, EPI_ISL_2727761, EPI_ISL_2727768, EPI_ISL_2727769, EPI_ISL_2727771, EPI_ISL_2727777, EPI_ISL_2727778, EPI_ISL_2727779, EPI_ISL_2727780, EPI_ISL_2727781, EPI_ISL_2727785, EPI_ISL_2727789, EPI_ISL_2727797, EPI_ISL_2727798, EPI_ISL_2727806, EPI_ISL_2727808, EPI_ISL_2727809, EPI_ISL_2727812, EPI_ISL_2727818, EPI_ISL_2727825, EPI_ISL_2727828, EPI_ISL_2727830, EPI_ISL_2727834 | | | |
| see above | Wisconsin State Laboratory of Hygiene Communicable Disease Division | Wisconsin State Laboratory of Hygiene Communicable Disease Division | Abigail C. Shockey, Alicia J. Mooney, Erika M. Hanson, Tonya Danz, Richard Griesser, Sara Wagner, Kelsey R. Florek |
| EPI_ISL_2727836 | Hospital Angkatan Tentera Kota Kinabalu | Institute for Medical Research, Infectious Disease Research Centre, National Institutes of Health, Ministry of Health Malaysia | Suppiah J, Kamel K, Mohd Zawawi Z, Azizan MA, Ramly N, Robert F, Thayan R |
| EPI_ISL_2727837 | Hospital Angkatan Tentera Wilayah Kota Kinabalu | Institute for Medical Research, Infectious Disease Research Centre, National Institutes of Health, Ministry of Health Malaysia | Suppiah J, Kamel K, Mohd Zawawi Z, Azizan MA, Ramly N, Robert F, Thayan R |
| EPI_ISL_2727840, EPI_ISL_2727843, | Minnesota Department of Health, Public Health Laboratory | Minnesota Department of Health, Public Health Laboratory | Alexandra Lorentz, Jacob Garfin, Matt Plumb, and Xiong Wang |

| | | | |
|---|---|---|---|
| EPI_ISL_2727853 | | | |
| EPI_ISL_2727862, EPI_ISL_2727863, EPI_ISL_2727864 | M Health Fairview | Minnesota Department of Health, Public Health Laboratory | Alexandra Lorentz, Jacob Garfin, Matt Plumb, and Xiong Wang |
| EPI_ISL_2727876 | ZOL | Jessa | Marijke Raymaekers et al. on behalf of the Jessa_cmdLab |
| EPI_ISL_2728130, EPI_ISL_2728133, EPI_ISL_2728137, EPI_ISL_2728138, EPI_ISL_2728140, EPI_ISL_2728141, EPI_ISL_2728142, EPI_ISL_2728143 | Wyoming Public Health Laboratory | Wyoming Public Health Laboratory | Jim Mildenberger, Taylor Fearing, Channing Weber, Ashley Norberg, Chayse Rowley, Marley Goetz, Brian Dominguez, Elliot Thomasson, Sam Britz, Cari Sloma, Robert Petit, and Rob Christensen |
| EPI_ISL_2728230, EPI_ISL_2728231 | BBMP Urban | INSACOG-KA, NIMHANS | Chitra Pattabiraman, Anson Kunjumon George,Darshan Sreenivas,Ellango Ramasamy, Sony Sharma, Mahesh Kumar.C.S, Ananthapadmanabha Kotambail, Chetan G K, Gautham Arunachal Udupi, Anita S Desai, V Ravi |
| EPI_ISL_2728232 | NIMHANS | INSACOG-KA, NIMHANS | Chitra Pattabiraman, Anson Kunjumon George,Darshan Sreenivas,Ellango Ramasamy, Sony Sharma, Mahesh Kumar.C.S, Ananthapadmanabha Kotambail, Chetan G K, Gautham Arunachal Udupi, Anita S Desai, V Ravi |
| EPI_ISL_2728233, EPI_ISL_2728234, EPI_ISL_2728235 | BBMP Urban | INSACOG-KA, NIMHANS | Chitra Pattabiraman, Anson Kunjumon George,Darshan Sreenivas,Ellango Ramasamy, Sony Sharma, Mahesh Kumar.C.S, Ananthapadmanabha Kotambail, Chetan G K, Gautham Arunachal Udupi, Anita S Desai, V Ravi |
| EPI_ISL_2728236 | SRI JAYADEVA INSTITUTE OF CARDIOLOGY | INSACOG-KA, NIMHANS | Chitra Pattabiraman, Anson Kunjumon George,Darshan Sreenivas,Ellango Ramasamy, Sony Sharma, Mahesh Kumar.C.S, Ananthapadmanabha Kotambail, Chetan G K, Gautham Arunachal Udupi, Anita S Desai, V Ravi |
| EPI_ISL_2728237, EPI_ISL_2728238, EPI_ISL_2728239, EPI_ISL_2728240, EPI_ISL_2728241, EPI_ISL_2728242, EPI_ISL_2728243, EPI_ISL_2728244, EPI_ISL_2728245, EPI_ISL_2728246, EPI_ISL_2728247, EPI_ISL_2728248 | | | |
| see above | BBMP Urban | INSACOG-KA, NIMHANS | Chitra Pattabiraman, Anson Kunjumon George,Darshan Sreenivas,Ellango Ramasamy, Sony Sharma, Mahesh Kumar.C.S, Ananthapadmanabha Kotambail, Chetan G K, Gautham Arunachal Udupi, Anita S Desai, V Ravi |
| EPI_ISL_2728249 | NIMHANS | INSACOG-KA, NIMHANS | Chitra Pattabiraman, Anson Kunjumon George,Darshan Sreenivas,Ellango Ramasamy, Sony Sharma, Mahesh Kumar.C.S, Ananthapadmanabha Kotambail, Chetan G K, Gautham Arunachal Udupi, Anita S Desai, V Ravi |
| EPI_ISL_2728250, EPI_ISL_2728251, EPI_ISL_2728252, EPI_ISL_2728253, EPI_ISL_2728254, EPI_ISL_2728255, EPI_ISL_2728256, EPI_ISL_2728257, EPI_ISL_2728258, EPI_ISL_2728259, EPI_ISL_2728260, EPI_ISL_2728261, EPI_ISL_2728262, EPI_ISL_2728263, EPI_ISL_2728264 | | | |
| see above | BBMP Urban | INSACOG-KA, NIMHANS | Chitra Pattabiraman, Anson Kunjumon George,Darshan Sreenivas,Ellango Ramasamy, Sony Sharma, Mahesh Kumar.C.S, Ananthapadmanabha Kotambail, Chetan G K, Gautham Arunachal Udupi, Anita S Desai, V Ravi |
| EPI_ISL_2728265 | RAILWAY HOSPITAL | INSACOG-KA, NIMHANS | Chitra Pattabiraman, Anson Kunjumon George,Darshan Sreenivas,Ellango Ramasamy, Sony Sharma, Mahesh Kumar.C.S, Ananthapadmanabha Kotambail, Chetan G K, Gautham Arunachal Udupi, Anita S Desai, V Ravi |
| EPI_ISL_2728266, EPI_ISL_2728267, EPI_ISL_2728268, EPI_ISL_2728269 | BBMP Urban | INSACOG-KA, NIMHANS | Chitra Pattabiraman, Anson Kunjumon George,Darshan Sreenivas,Ellango Ramasamy, Sony Sharma, Mahesh Kumar.C.S, Ananthapadmanabha Kotambail, Chetan G K, Gautham Arunachal Udupi, Anita S Desai, V Ravi |
| EPI_ISL_2728270, EPI_ISL_2728271 | SRI JAYADEVA INSTITUTE OF CARDIOLOGY | INSACOG-KA, NIMHANS | Chitra Pattabiraman, Anson Kunjumon George,Darshan Sreenivas,Ellango Ramasamy, Sony Sharma, Mahesh Kumar.C.S, Ananthapadmanabha Kotambail, Chetan G K, Gautham Arunachal Udupi, Anita S Desai, V Ravi |
| EPI_ISL_2728272, EPI_ISL_2728273, EPI_ISL_2728274, EPI_ISL_2728275, EPI_ISL_2728276, EPI_ISL_2728277, EPI_ISL_2728278, EPI_ISL_2728280, EPI_ISL_2728281, EPI_ISL_2728282, EPI_ISL_2728283, EPI_ISL_2728284, EPI_ISL_2728285, EPI_ISL_2728286, EPI_ISL_2728287, EPI_ISL_2728288, EPI_ISL_2728289, EPI_ISL_2728290 | | | |
| see above | BBMP Urban | INSACOG-KA, NIMHANS | Chitra Pattabiraman, Anson Kunjumon George,Darshan Sreenivas,Ellango Ramasamy, Sony Sharma, Mahesh Kumar.C.S, Ananthapadmanabha Kotambail, Chetan G K, Gautham Arunachal Udupi, Anita S Desai, V Ravi |
| EPI_ISL_2728291, EPI_ISL_2728292 | NIMHANS | INSACOG-KA, NIMHANS | Chitra Pattabiraman, Anson Kunjumon George,Darshan Sreenivas,Ellango Ramasamy, Sony Sharma, Mahesh Kumar.C.S, Ananthapadmanabha Kotambail, Chetan G K, Gautham Arunachal Udupi, Anita S Desai, V Ravi |
| EPI_ISL_2728293 | SANJAY GANDHI INSTITUTE OF TRAUMA AND ORTHOPAEDICS | INSACOG-KA, NIMHANS | Chitra Pattabiraman, Anson Kunjumon George,Darshan Sreenivas,Ellango Ramasamy, Sony Sharma, Mahesh Kumar.C.S, Ananthapadmanabha Kotambail, Chetan G K, Gautham Arunachal Udupi, Anita S Desai, V Ravi |
| EPI_ISL_2728294, EPI_ISL_2728295, EPI_ISL_2728296, EPI_ISL_2728297, EPI_ISL_2728300, EPI_ISL_2728302, EPI_ISL_2728303, EPI_ISL_2728304, EPI_ISL_2728305, EPI_ISL_2728306, EPI_ISL_2728307 | | | |
| see above | BBMP Urban | INSACOG-KA, NIMHANS | Chitra Pattabiraman, Anson Kunjumon George,Darshan Sreenivas,Ellango Ramasamy, Sony Sharma, Mahesh Kumar.C.S, Ananthapadmanabha Kotambail, Chetan G K, Gautham Arunachal Udupi, Anita S Desai, V Ravi |
| EPI_ISL_2728308 | DSU HASSAN | INSACOG-KA, NIMHANS | Chitra Pattabiraman, Anson Kunjumon George,Darshan Sreenivas,Ellango Ramasamy, Sony Sharma, Mahesh Kumar.C.S, Ananthapadmanabha Kotambail, Chetan G K, Gautham Arunachal Udupi, Anita S Desai, V Ravi |
| EPI_ISL_2728309, EPI_ISL_2728310, EPI_ISL_2728311 | BBMP Urban | INSACOG-KA, NIMHANS | Chitra Pattabiraman, Anson Kunjumon George,Darshan Sreenivas,Ellango Ramasamy, Sony Sharma, Mahesh Kumar.C.S, Ananthapadmanabha Kotambail, Chetan G K, Gautham Arunachal Udupi, Anita S Desai, V Ravi |
| EPI_ISL_2728312 | NIMHANS | INSACOG-KA, NIMHANS | Chitra Pattabiraman, Anson Kunjumon George,Darshan Sreenivas,Ellango Ramasamy, Sony Sharma, Mahesh Kumar.C.S, Ananthapadmanabha Kotambail, Chetan G K, Gautham Arunachal Udupi, Anita S Desai, V Ravi |
| EPI_ISL_2728313 | BBMP Urban | INSACOG-KA, NIMHANS | Chitra Pattabiraman, Anson Kunjumon George,Darshan Sreenivas,Ellango Ramasamy, Sony Sharma, Mahesh Kumar.C.S, Ananthapadmanabha Kotambail, Chetan G K, Gautham Arunachal Udupi, Anita S Desai, V Ravi |
| EPI_ISL_2728314 | SANJAY GANDHI INSTITUTE OF TRAUMA AND ORTHOPAEDICS | INSACOG-KA, NIMHANS | Chitra Pattabiraman, Anson Kunjumon George,Darshan Sreenivas,Ellango Ramasamy, Sony Sharma, Mahesh Kumar.C.S, Ananthapadmanabha Kotambail, Chetan G K, Gautham Arunachal Udupi, Anita S Desai, V Ravi |
| EPI_ISL_2728315 | BBMP Urban | INSACOG-KA, NIMHANS | Chitra Pattabiraman, Anson Kunjumon George,Darshan Sreenivas,Ellango Ramasamy, Sony Sharma, Mahesh Kumar.C.S, Ananthapadmanabha Kotambail, Chetan G K, Gautham Arunachal Udupi, Anita S Desai, V Ravi |
| EPI_ISL_2728316 | NIMHANS | INSACOG-KA, NIMHANS | Chitra Pattabiraman, Anson Kunjumon George,Darshan Sreenivas,Ellango Ramasamy, Sony Sharma, Mahesh Kumar.C.S, Ananthapadmanabha Kotambail, Chetan G K, Gautham Arunachal Udupi, Anita S Desai, V Ravi |
| EPI_ISL_2728317, EPI_ISL_2728318, EPI_ISL_2728319, EPI_ISL_2728320, EPI_ISL_2728321, EPI_ISL_2728322, EPI_ISL_2728323, EPI_ISL_2728324, EPI_ISL_2728325, EPI_ISL_2728326, EPI_ISL_2728327, EPI_ISL_2728328, EPI_ISL_2728329 | | | |
| see above | DSU HASSAN | INSACOG-KA, NIMHANS | Chitra Pattabiraman, Anson Kunjumon George,Darshan Sreenivas,Ellango Ramasamy, Sony Sharma, Mahesh Kumar.C.S, Ananthapadmanabha Kotambail, Chetan G K, Gautham Arunachal Udupi, Anita S Desai, V Ravi |
| EPI_ISL_2728330, EPI_ISL_2728331, EPI_ISL_2728332, EPI_ISL_2728333, EPI_ISL_2728334, EPI_ISL_2728335, EPI_ISL_2728336, EPI_ISL_2728337, EPI_ISL_2728338, EPI_ISL_2728339, EPI_ISL_2728340, EPI_ISL_2728341 | | | |
| see above | DSU KALABURAGI | INSACOG-KA, NIMHANS | Chitra Pattabiraman, Anson Kunjumon George,Darshan Sreenivas,Ellango Ramasamy, Sony Sharma, Mahesh Kumar.C.S, Ananthapadmanabha Kotambail, Chetan G K, Gautham Arunachal Udupi, Anita S Desai, V Ravi |
| EPI_ISL_2728342 | BBMP Urban | INSACOG-KA, NIMHANS | Chitra Pattabiraman, Anson Kunjumon George,Darshan Sreenivas,Ellango Ramasamy, Sony Sharma, Mahesh Kumar.C.S, Ananthapadmanabha Kotambail, Chetan G K, Gautham Arunachal Udupi, Anita S Desai, V Ravi |
| EPI_ISL_2728343, EPI_ISL_2728344, EPI_ISL_2728345, EPI_ISL_2728346, EPI_ISL_2728347 | DSU DAVANAGERE | INSACOG-KA, NIMHANS | Chitra Pattabiraman, Anson Kunjumon George,Darshan Sreenivas,Ellango Ramasamy, Sony Sharma, Mahesh Kumar.C.S, Ananthapadmanabha Kotambail, Chetan G K, Gautham Arunachal Udupi, Anita S Desai, V Ravi |
| EPI_ISL_2728348 | RAILWAY HOSPITAL | INSACOG-KA, NIMHANS | Chitra Pattabiraman, Anson Kunjumon George,Darshan Sreenivas,Ellango Ramasamy, Sony Sharma, Mahesh Kumar.C.S, Ananthapadmanabha Kotambail, Chetan G K, Gautham Arunachal Udupi, Anita S Desai, V Ravi |
| EPI_ISL_2728349, EPI_ISL_2728351, EPI_ISL_2728352, EPI_ISL_2728353, EPI_ISL_2728354, EPI_ISL_2728355, EPI_ISL_2728356, EPI_ISL_2728357, EPI_ISL_2728359, EPI_ISL_2728360, EPI_ISL_2728361, EPI_ISL_2728362, EPI_ISL_2728363, EPI_ISL_2728364, EPI_ISL_2728365, EPI_ISL_2728366, EPI_ISL_2728367, EPI_ISL_2728368, EPI_ISL_2728370, EPI_ISL_2728371, EPI_ISL_2728372, EPI_ISL_2728373, EPI_ISL_2728374, EPI_ISL_2728375, EPI_ISL_2728376, EPI_ISL_2728377, EPI_ISL_2728378, EPI_ISL_2728379, EPI_ISL_2728380, EPI_ISL_2728383, EPI_ISL_2728384, EPI_ISL_2728385, EPI_ISL_2728388, EPI_ISL_2728389, EPI_ISL_2728390, EPI_ISL_2728391, EPI_ISL_2728392, EPI_ISL_2728393, EPI_ISL_2728394, EPI_ISL_2728396, EPI_ISL_2728397, EPI_ISL_2728398, EPI_ISL_2728399, EPI_ISL_2728400, EPI_ISL_2728401, EPI_ISL_2728402, EPI_ISL_2728403, EPI_ISL_2728405, EPI_ISL_2728406, EPI_ISL_2728408, EPI_ISL_2728409, EPI_ISL_2728410, EPI_ISL_2728412, EPI_ISL_2728413, EPI_ISL_2728414, EPI_ISL_2728415, EPI_ISL_2728416, EPI_ISL_2728417, EPI_ISL_2728418, EPI_ISL_2728419, EPI_ISL_2728420, EPI_ISL_2728421, EPI_ISL_2728422, EPI_ISL_2728423, EPI_ISL_2728424, EPI_ISL_2728425, EPI_ISL_2728426, EPI_ISL_2728427 | | | |
| see above | DSU VIJAYAPURA | INSACOG-KA, NIMHANS | Chitra Pattabiraman, Anson Kunjumon George,Darshan Sreenivas,Ellango Ramasamy, Sony Sharma, Mahesh Kumar.C.S, Ananthapadmanabha Kotambail, Chetan G K, Gautham Arunachal Udupi, Anita S Desai, V Ravi |
| EPI_ISL_2728428 | NIMHANS | INSACOG-KA, NIMHANS | Chitra Pattabiraman, Anson Kunjumon George,Darshan Sreenivas,Ellango Ramasamy, Sony Sharma, Mahesh Kumar.C.S, Ananthapadmanabha Kotambail, Chetan G K, Gautham Arunachal Udupi, Anita S Desai, V Ravi |
| EPI_ISL_2728430, EPI_ISL_2728431, EPI_ISL_2728432, EPI_ISL_2728433, EPI_ISL_2728434, EPI_ISL_2728435, EPI_ISL_2728436, EPI_ISL_2728437, EPI_ISL_2728438 | BBMP Urban | INSACOG-KA, NIMHANS | Chitra Pattabiraman, Anson Kunjumon George,Darshan Sreenivas,Ellango Ramasamy, Sony Sharma, Mahesh Kumar.C.S, Ananthapadmanabha Kotambail, Chetan G K, Gautham Arunachal Udupi, Anita S Desai, V Ravi |
| EPI_ISL_2728444, EPI_ISL_2728455, EPI_ISL_2728456 | NIMHANS | INSACOG-KA, NIMHANS | Chitra Pattabiraman, Anson Kunjumon George,Darshan Sreenivas,Ellango Ramasamy, Sony Sharma, Mahesh Kumar.C.S, Ananthapadmanabha Kotambail, Chetan G K, Gautham Arunachal Udupi, Anita S Desai, V Ravi |
| EPI_ISL_2728459 | RAILWAY HOSPITAL | INSACOG-KA, NIMHANS | Chitra Pattabiraman, Anson Kunjumon George,Darshan Sreenivas,Ellango Ramasamy, Sony Sharma, Mahesh Kumar.C.S, Ananthapadmanabha Kotambail, Chetan G K, Gautham Arunachal Udupi, Anita S Desai, V Ravi |
| EPI_ISL_2728465, EPI_ISL_2728466, EPI_ISL_2728469 | BBMP Urban | INSACOG-KA, NIMHANS | Chitra Pattabiraman, Anson Kunjumon George,Darshan Sreenivas,Ellango Ramasamy, Sony Sharma, Mahesh Kumar.C.S, Ananthapadmanabha Kotambail, Chetan G K, Gautham Arunachal Udupi, Anita S Desai, V Ravi |
| EPI_ISL_2728471 | SRI JAYADEVA INSTITUTE OF CARDIOLOGY | INSACOG-KA, NIMHANS | Chitra Pattabiraman, Anson Kunjumon George,Darshan Sreenivas,Ellango Ramasamy, Sony Sharma, Mahesh Kumar.C.S, Ananthapadmanabha Kotambail, Chetan G K, Gautham Arunachal Udupi, Anita S Desai, V Ravi |
| EPI_ISL_2728472, EPI_ISL_2728473, EPI_ISL_2728475, EPI_ISL_2728476, EPI_ISL_2728477 | BBMP Urban | INSACOG-KA, NIMHANS | Chitra Pattabiraman, Anson Kunjumon George,Darshan Sreenivas,Ellango Ramasamy, Sony Sharma, Mahesh Kumar.C.S, Ananthapadmanabha Kotambail, Chetan G K, Gautham Arunachal Udupi, Anita S Desai, V Ravi |
| EPI_ISL_2728478 | RAILWAY HOSPITAL | INSACOG-KA, NIMHANS | Chitra Pattabiraman, Anson Kunjumon George,Darshan Sreenivas,Ellango Ramasamy, Sony Sharma, Mahesh Kumar.C.S, Ananthapadmanabha Kotambail, Chetan G K, Gautham Arunachal Udupi, Anita S Desai, V Ravi |
| EPI_ISL_2728479 | BBMP Urban | INSACOG-KA, NIMHANS | Chitra Pattabiraman, Anson Kunjumon George,Darshan Sreenivas,Ellango Ramasamy, Sony Sharma, Mahesh Kumar.C.S, Ananthapadmanabha Kotambail, Chetan G K, Gautham Arunachal Udupi, Anita S Desai, V Ravi |

| | | | |
|---|---|---|---|
| EPI_ISL_2728480 | NIMHANS | INSACOG-KA, NIMHANS | Chitra Pattabiraman, Anson Kunjumon George,Darshan Sreenivas,Ellango Ramasamy, Sony Sharma, Mahesh Kumar.C.S, Ananthapadmanabha Kotambail, Chetan G K, Gautham Arunachal Udupi, Anita S Desai, V Ravi |
| EPI_ISL_2728481, EPI_ISL_2728482 | RAILWAY HOSPITAL | INSACOG-KA, NIMHANS | Chitra Pattabiraman, Anson Kunjumon George,Darshan Sreenivas,Ellango Ramasamy, Sony Sharma, Mahesh Kumar.C.S, Ananthapadmanabha Kotambail, Chetan G K, Gautham Arunachal Udupi, Anita S Desai, V Ravi |
| EPI_ISL_2728484, EPI_ISL_2728485, EPI_ISL_2728486 | NIMHANS | INSACOG-KA, NIMHANS | Chitra Pattabiraman, Anson Kunjumon George,Darshan Sreenivas,Ellango Ramasamy, Sony Sharma, Mahesh Kumar.C.S, Ananthapadmanabha Kotambail, Chetan G K, Gautham Arunachal Udupi, Anita S Desai, V Ravi |
| EPI_ISL_2728487, EPI_ISL_2728488, EPI_ISL_2728489, EPI_ISL_2728492, EPI_ISL_2728493, EPI_ISL_2728494 | RAILWAY HOSPITAL | INSACOG-KA, NIMHANS | Chitra Pattabiraman, Anson Kunjumon George,Darshan Sreenivas,Ellango Ramasamy, Sony Sharma, Mahesh Kumar.C.S, Ananthapadmanabha Kotambail, Chetan G K, Gautham Arunachal Udupi, Anita S Desai, V Ravi |
| EPI_ISL_2728495, EPI_ISL_2728496 | NIMHANS | INSACOG-KA, NIMHANS | Chitra Pattabiraman, Anson Kunjumon George,Darshan Sreenivas,Ellango Ramasamy, Sony Sharma, Mahesh Kumar.C.S, Ananthapadmanabha Kotambail, Chetan G K, Gautham Arunachal Udupi, Anita S Desai, V Ravi |
| EPI_ISL_2728498, EPI_ISL_2728499 | RAILWAY HOSPITAL | INSACOG-KA, NIMHANS | Chitra Pattabiraman, Anson Kunjumon George,Darshan Sreenivas,Ellango Ramasamy, Sony Sharma, Mahesh Kumar.C.S, Ananthapadmanabha Kotambail, Chetan G K, Gautham Arunachal Udupi, Anita S Desai, V Ravi |
| EPI_ISL_2728500, EPI_ISL_2728501 | NIMHANS | INSACOG-KA, NIMHANS | Chitra Pattabiraman, Anson Kunjumon George,Darshan Sreenivas,Ellango Ramasamy, Sony Sharma, Mahesh Kumar.C.S, Ananthapadmanabha Kotambail, Chetan G K, Gautham Arunachal Udupi, Anita S Desai, V Ravi |
| EPI_ISL_2728503 | RAILWAY HOSPITAL | INSACOG-KA, NIMHANS | Chitra Pattabiraman, Anson Kunjumon George,Darshan Sreenivas,Ellango Ramasamy, Sony Sharma, Mahesh Kumar.C.S, Ananthapadmanabha Kotambail, Chetan G K, Gautham Arunachal Udupi, Anita S Desai, V Ravi |
| EPI_ISL_2728504, EPI_ISL_2728505, EPI_ISL_2728507, EPI_ISL_2728509, EPI_ISL_2728510, EPI_ISL_2728511, EPI_ISL_2728512, EPI_ISL_2728513, EPI_ISL_2728514, EPI_ISL_2728515, EPI_ISL_2728516, EPI_ISL_2728517, EPI_ISL_2728518, EPI_ISL_2728519, EPI_ISL_2728520, EPI_ISL_2728521, EPI_ISL_2728522 | | | |
| see above | NIMHANS | INSACOG-KA, NIMHANS | Chitra Pattabiraman, Anson Kunjumon George,Darshan Sreenivas,Ellango Ramasamy, Sony Sharma, Mahesh Kumar.C.S, Ananthapadmanabha Kotambail, Chetan G K, Gautham Arunachal Udupi, Anita S Desai, V Ravi |
| EPI_ISL_2728525, EPI_ISL_2728526, EPI_ISL_2728527, EPI_ISL_2728528 | RAILWAY HOSPITAL | INSACOG-KA, NIMHANS | Chitra Pattabiraman, Anson Kunjumon George,Darshan Sreenivas,Ellango Ramasamy, Sony Sharma, Mahesh Kumar.C.S, Ananthapadmanabha Kotambail, Chetan G K, Gautham Arunachal Udupi, Anita S Desai, V Ravi |
| EPI_ISL_2728529 | NIMHANS | INSACOG-KA, NIMHANS | Chitra Pattabiraman, Anson Kunjumon George,Darshan Sreenivas,Ellango Ramasamy, Sony Sharma, Mahesh Kumar.C.S, Ananthapadmanabha Kotambail, Chetan G K, Gautham Arunachal Udupi, Anita S Desai, V Ravi |
| EPI_ISL_2728608, EPI_ISL_2728610, EPI_ISL_2728614, EPI_ISL_2728618 | Fondazione IRCCS Ca' Granda Ospedale Maggiore Policlinico | Fondazione IRCCS Ca' Granda Ospedale Maggiore Policlinico | Ferruccio Ceriotti, Sara Uceda Renteria |
| EPI_ISL_2728623, EPI_ISL_2728627, EPI_ISL_2728630, EPI_ISL_2728632, EPI_ISL_2728633, EPI_ISL_2728638 | SYNLAB | GIGA Medical Genomics | Keith Durkin, Maria Artesi, Bouchra Boujemla, Nathalie Renotte, Nadine Cambisano, Sébastien Bontems, Cécile Meex, Claire Gourzonès, Olivier Ek, Laurent Gillet, Marie-Pierre Hayette, Vincent Bours |
| EPI_ISL_2728639 | Department of Clinical Microbiology | GIGA Medical Genomics | Keith Durkin, Maria Artesi, Bouchra Boujemla, Nathalie Renotte, Nadine Cambisano, Sébastien Bontems, Cécile Meex, Claire Gourzonès, Olivier Ek, Laurent Gillet, Marie-Pierre Hayette, Vincent Bours |
| EPI_ISL_2728642, EPI_ISL_2728643, EPI_ISL_2728645 | SYNLAB | GIGA Medical Genomics | Keith Durkin, Maria Artesi, Bouchra Boujemla, Nathalie Renotte, Nadine Cambisano, Sébastien Bontems, Cécile Meex, Claire Gourzonès, Olivier Ek, Laurent Gillet, Marie-Pierre Hayette, Vincent Bours |
| EPI_ISL_2728649, EPI_ISL_2728650, EPI_ISL_2728652, EPI_ISL_2728653, EPI_ISL_2728654, EPI_ISL_2728657, EPI_ISL_2728658, EPI_ISL_2728659, EPI_ISL_2728660, EPI_ISL_2728661, EPI_ISL_2728662, EPI_ISL_2728663, EPI_ISL_2728664, EPI_ISL_2728665, EPI_ISL_2728667, EPI_ISL_2728668, EPI_ISL_2728670, EPI_ISL_2728674, EPI_ISL_2728675, EPI_ISL_2728676, EPI_ISL_2728679, EPI_ISL_2728680 | | | |
| see above | University of Liège COVID-19 testing center | GIGA Medical Genomics | Keith Durkin, Maria Artesi, Bouchra Boujemla, Nathalie Renotte, Nadine Cambisano, Sébastien Bontems, Cécile Meex, Claire Gourzonès, Olivier Ek, Laurent Gillet, Marie-Pierre Hayette, Vincent Bours |
| EPI_ISL_2728683 | Lighthouse Laboratory Plymouth | Wellcome Sanger Institute for the COVID-19 Genomics UK (COG-UK) Consortium | Lighthouse Laboratory Plymouth and Alex Alderton, Roberto Amato, Jeffrey Barrett, Sonia Goncalves, Ewan Harrison, David K. Jackson, Ian Johnston, Dominic Kwiatkowski, Cordelia Langford, John Sillitoe on behalf of the Wellcome Sanger Institute COVID-19 Surveillance Team |
| EPI_ISL_2728684 | Lighthouse Lab in Glasgow | Wellcome Sanger Institute for the COVID-19 Genomics UK (COG-UK) Consortium | Harper VanSteenhouse, Yumi Kasai, David Gray, Carol Clugston, Anna Dominiczak and Alex Alderton, Roberto Amato, Jeffrey Barrett, Sonia Goncalves, Ewan Harrison, David K. Jackson, Ian Johnston, Dominic Kwiatkowski, Cordelia Langford, John Sillitoe on behalf of the Wellcome Sanger Institute COVID-19 Surveillance Team |
| EPI_ISL_2728685 | Lighthouse Laboratory Plymouth | Wellcome Sanger Institute for the COVID-19 Genomics UK (COG-UK) Consortium | Lighthouse Laboratory Plymouth and Alex Alderton, Roberto Amato, Jeffrey Barrett, Sonia Goncalves, Ewan Harrison, David K. Jackson, Ian Johnston, Dominic Kwiatkowski, Cordelia Langford, John Sillitoe on behalf of the Wellcome Sanger Institute COVID-19 Surveillance Team |
| EPI_ISL_2728686, EPI_ISL_2728687 | Lighthouse Lab in Milton Keynes | Wellcome Sanger Institute for the COVID-19 Genomics UK (COG-UK) Consortium | The Lighthouse Lab in Milton Keynes and Alex Alderton, Roberto Amato, Jeffrey Barrett, Sonia Goncalves, Ewan Harrison, David K. Jackson, Ian Johnston, Dominic Kwiatkowski, Cordelia Langford, John Sillitoe on behalf of the Wellcome Sanger Institute COVID-19 Surveillance Team |
| EPI_ISL_2728688 | Lighthouse Lab in Glasgow | Wellcome Sanger Institute for the COVID-19 Genomics UK (COG-UK) Consortium | Harper VanSteenhouse, Yumi Kasai, David Gray, Carol Clugston, Anna Dominiczak and Alex Alderton, Roberto Amato, Jeffrey Barrett, Sonia Goncalves, Ewan Harrison, David K. Jackson, Ian Johnston, Dominic Kwiatkowski, Cordelia Langford, John Sillitoe on behalf of the Wellcome Sanger Institute COVID-19 Surveillance Team |
| EPI_ISL_2728689 | Lighthouse Laboratory Plymouth | Wellcome Sanger Institute for the COVID-19 Genomics UK (COG-UK) Consortium | Lighthouse Laboratory Plymouth and Alex Alderton, Roberto Amato, Jeffrey Barrett, Sonia Goncalves, Ewan Harrison, David K. Jackson, Ian Johnston, Dominic Kwiatkowski, Cordelia Langford, John Sillitoe on behalf of the Wellcome Sanger Institute COVID-19 Surveillance Team |
| EPI_ISL_2728690, EPI_ISL_2728691, EPI_ISL_2728692, EPI_ISL_2728693, EPI_ISL_2728694, EPI_ISL_2728695 | Lighthouse Lab in Glasgow | Wellcome Sanger Institute for the COVID-19 Genomics UK (COG-UK) Consortium | Harper VanSteenhouse, Yumi Kasai, David Gray, Carol Clugston, Anna Dominiczak and Alex Alderton, Roberto Amato, Jeffrey Barrett, Sonia Goncalves, Ewan Harrison, David K. Jackson, Ian Johnston, Dominic Kwiatkowski, Cordelia Langford, John Sillitoe on behalf of the Wellcome Sanger Institute COVID-19 Surveillance Team |
| EPI_ISL_2728696 | Lighthouse Lab in Milton Keynes | Wellcome Sanger Institute for the COVID-19 Genomics UK (COG-UK) Consortium | The Lighthouse Lab in Milton Keynes and Alex Alderton, Roberto Amato, Jeffrey Barrett, Sonia Goncalves, Ewan Harrison, David K. Jackson, Ian Johnston, Dominic Kwiatkowski, Cordelia Langford, John Sillitoe on behalf of the Wellcome Sanger Institute COVID-19 Surveillance Team |
| EPI_ISL_2728697 | Lighthouse Lab in Glasgow | Wellcome Sanger Institute for the COVID-19 Genomics UK (COG-UK) Consortium | Harper VanSteenhouse, Yumi Kasai, David Gray, Carol Clugston, Anna Dominiczak and Alex Alderton, Roberto Amato, Jeffrey Barrett, Sonia Goncalves, Ewan Harrison, David K. Jackson, Ian Johnston, Dominic Kwiatkowski, Cordelia Langford, John Sillitoe on behalf of the Wellcome Sanger Institute COVID-19 Surveillance Team |
| EPI_ISL_2728698 | Lighthouse Lab in Milton Keynes | Wellcome Sanger Institute for the COVID-19 Genomics UK (COG-UK) Consortium | The Lighthouse Lab in Milton Keynes and Alex Alderton, Roberto Amato, Jeffrey Barrett, Sonia Goncalves, Ewan Harrison, David K. Jackson, Ian Johnston, Dominic Kwiatkowski, Cordelia Langford, John Sillitoe on behalf of the Wellcome Sanger Institute COVID-19 Surveillance Team |
| EPI_ISL_2728699 | Lighthouse Lab in Glasgow | Wellcome Sanger Institute for the COVID-19 Genomics UK (COG-UK) Consortium | Harper VanSteenhouse, Yumi Kasai, David Gray, Carol Clugston, Anna Dominiczak and Alex Alderton, Roberto Amato, Jeffrey Barrett, Sonia Goncalves, Ewan Harrison, David K. Jackson, Ian Johnston, Dominic Kwiatkowski, Cordelia Langford, John Sillitoe on behalf of the Wellcome Sanger Institute COVID-19 Surveillance Team |
| EPI_ISL_2728700 | Lighthouse Lab in Milton Keynes | Wellcome Sanger Institute for the COVID-19 Genomics UK (COG-UK) Consortium | The Lighthouse Lab in Milton Keynes and Alex Alderton, Roberto Amato, Jeffrey Barrett, Sonia Goncalves, Ewan Harrison, David K. Jackson, Ian Johnston, Dominic Kwiatkowski, Cordelia Langford, John Sillitoe on behalf of the Wellcome Sanger Institute COVID-19 Surveillance Team |
| EPI_ISL_2728701 | Lighthouse Lab in Glasgow | Wellcome Sanger Institute for the COVID-19 Genomics UK (COG-UK) Consortium | Harper VanSteenhouse, Yumi Kasai, David Gray, Carol Clugston, Anna Dominiczak and Alex Alderton, Roberto Amato, Jeffrey Barrett, Sonia Goncalves, Ewan Harrison, David K. Jackson, Ian Johnston, Dominic Kwiatkowski, Cordelia Langford, John Sillitoe on behalf of the Wellcome Sanger Institute COVID-19 Surveillance Team |
| EPI_ISL_2728702 | Lighthouse Lab in Milton Keynes | Wellcome Sanger Institute for the COVID-19 Genomics UK (COG-UK) Consortium | The Lighthouse Lab in Milton Keynes and Alex Alderton, Roberto Amato, Jeffrey Barrett, Sonia Goncalves, Ewan Harrison, David K. Jackson, Ian Johnston, Dominic Kwiatkowski, Cordelia Langford, John Sillitoe on behalf of the Wellcome Sanger Institute COVID-19 Surveillance Team |
| EPI_ISL_2728703 | Lighthouse Lab in Glasgow | Wellcome Sanger Institute for the COVID-19 Genomics UK (COG-UK) Consortium | Harper VanSteenhouse, Yumi Kasai, David Gray, Carol Clugston, Anna Dominiczak and Alex Alderton, Roberto Amato, Jeffrey Barrett, Sonia Goncalves, Ewan Harrison, David K. Jackson, Ian Johnston, Dominic Kwiatkowski, Cordelia Langford, John Sillitoe on behalf of the Wellcome Sanger Institute COVID-19 Surveillance Team |
| EPI_ISL_2728704 | Lighthouse Lab in Milton Keynes | Wellcome Sanger Institute for the COVID-19 Genomics UK (COG-UK) Consortium | The Lighthouse Lab in Milton Keynes and Alex Alderton, Roberto Amato, Jeffrey Barrett, Sonia Goncalves, Ewan Harrison, David K. Jackson, Ian Johnston, Dominic Kwiatkowski, Cordelia Langford, John Sillitoe on behalf of the Wellcome Sanger Institute COVID-19 Surveillance Team |
| EPI_ISL_2728705 | Lighthouse Laboratory Plymouth | Wellcome Sanger Institute for the COVID-19 Genomics UK (COG-UK) Consortium | Lighthouse Laboratory Plymouth and Alex Alderton, Roberto Amato, Jeffrey Barrett, Sonia Goncalves, Ewan Harrison, David K. Jackson, Ian Johnston, Dominic Kwiatkowski, Cordelia Langford, John Sillitoe on behalf of the Wellcome Sanger Institute COVID-19 Surveillance Team |
| EPI_ISL_2728706 | Lighthouse Lab in Milton Keynes | Wellcome Sanger Institute for the COVID-19 Genomics UK (COG-UK) Consortium | The Lighthouse Lab in Milton Keynes and Alex Alderton, Roberto Amato, Jeffrey Barrett, Sonia Goncalves, Ewan Harrison, David K. Jackson, Ian Johnston, Dominic Kwiatkowski, Cordelia Langford, John Sillitoe on behalf of the Wellcome Sanger Institute COVID-19 Surveillance Team |
| EPI_ISL_2728707, EPI_ISL_2728708, EPI_ISL_2728710 | Lighthouse Lab in Glasgow | Wellcome Sanger Institute for the COVID-19 Genomics UK (COG-UK) Consortium | Harper VanSteenhouse, Yumi Kasai, David Gray, Carol Clugston, Anna Dominiczak and Alex Alderton, Roberto Amato, Jeffrey Barrett, Sonia Goncalves, Ewan Harrison, David K. Jackson, Ian Johnston, Dominic Kwiatkowski, Cordelia Langford, John Sillitoe on behalf of the Wellcome Sanger Institute COVID-19 Surveillance Team |
| EPI_ISL_2728711, EPI_ISL_2728712 | Lighthouse Laboratory Plymouth | Wellcome Sanger Institute for the COVID-19 Genomics UK (COG-UK) Consortium | Lighthouse Laboratory Plymouth and Alex Alderton, Roberto Amato, Jeffrey Barrett, Sonia Goncalves, Ewan Harrison, David K. Jackson, Ian Johnston, Dominic Kwiatkowski, Cordelia Langford, John Sillitoe on behalf of the Wellcome Sanger Institute COVID-19 Surveillance Team |
| EPI_ISL_2728713 | Lighthouse Lab in Milton Keynes | Wellcome Sanger Institute for the COVID-19 Genomics UK (COG-UK) Consortium | The Lighthouse Lab in Milton Keynes and Alex Alderton, Roberto Amato, Jeffrey Barrett, Sonia Goncalves, Ewan Harrison, David K. Jackson, Ian Johnston, Dominic Kwiatkowski, Cordelia Langford, John Sillitoe on behalf of the Wellcome Sanger Institute COVID-19 Surveillance Team |
| EPI_ISL_2728714, EPI_ISL_2728715, EPI_ISL_2728716, | Lighthouse Laboratory Plymouth | Wellcome Sanger Institute for the COVID-19 Genomics UK (COG-UK) Consortium | Lighthouse Laboratory Plymouth and Alex Alderton, Roberto Amato, Jeffrey Barrett, Sonia Goncalves, Ewan Harrison, David K. Jackson, Ian Johnston, Dominic Kwiatkowski, Cordelia Langford, John Sillitoe on behalf of the Wellcome Sanger Institute COVID-19 Surveillance Team |

| | | | |
|---|---|---|---|
| EPI_ISL_2728718 | | | |
| EPI_ISL_2728719 | Lighthouse Lab in Glasgow | Wellcome Sanger Institute for the COVID-19 Genomics UK (COG-UK) Consortium | Harper VanSteenhouse, Yumi Kasai, David Gray, Carol Clugston, Anna Dominiczak and Alex Alderton, Roberto Amato, Jeffrey Barrett, Sonia Goncalves, Ewan Harrison, David K. Jackson, Ian Johnston, Dominic Kwiatkowski, Cordelia Langford, John Sillitoe on behalf of the Wellcome Sanger Institute COVID-19 Surveillance Team |
| EPI_ISL_2728720, EPI_ISL_2728721 | Lighthouse Lab in Milton Keynes | Wellcome Sanger Institute for the COVID-19 Genomics UK (COG-UK) Consortium | The Lighthouse Lab in Milton Keynes and Alex Alderton, Roberto Amato, Jeffrey Barrett, Sonia Goncalves, Ewan Harrison, David K. Jackson, Ian Johnston, Dominic Kwiatkowski, Cordelia Langford, John Sillitoe on behalf of the Wellcome Sanger Institute COVID-19 Surveillance Team |
| EPI_ISL_2728722, EPI_ISL_2728723 | Lighthouse Lab in Glasgow | Wellcome Sanger Institute for the COVID-19 Genomics UK (COG-UK) Consortium | Harper VanSteenhouse, Yumi Kasai, David Gray, Carol Clugston, Anna Dominiczak and Alex Alderton, Roberto Amato, Jeffrey Barrett, Sonia Goncalves, Ewan Harrison, David K. Jackson, Ian Johnston, Dominic Kwiatkowski, Cordelia Langford, John Sillitoe on behalf of the Wellcome Sanger Institute COVID-19 Surveillance Team |
| EPI_ISL_2728724, EPI_ISL_2728725 | Lighthouse Lab in Milton Keynes | Wellcome Sanger Institute for the COVID-19 Genomics UK (COG-UK) Consortium | The Lighthouse Lab in Milton Keynes and Alex Alderton, Roberto Amato, Jeffrey Barrett, Sonia Goncalves, Ewan Harrison, David K. Jackson, Ian Johnston, Dominic Kwiatkowski, Cordelia Langford, John Sillitoe on behalf of the Wellcome Sanger Institute COVID-19 Surveillance Team |
| EPI_ISL_2728726, EPI_ISL_2728727 | Lighthouse Laboratory Plymouth | Wellcome Sanger Institute for the COVID-19 Genomics UK (COG-UK) Consortium | Lighthouse Laboratory Plymouth and Alex Alderton, Roberto Amato, Jeffrey Barrett, Sonia Goncalves, Ewan Harrison, David K. Jackson, Ian Johnston, Dominic Kwiatkowski, Cordelia Langford, John Sillitoe on behalf of the Wellcome Sanger Institute COVID-19 Surveillance Team |
| EPI_ISL_2728728, EPI_ISL_2728729, EPI_ISL_2728730, EPI_ISL_2728731, EPI_ISL_2728732, EPI_ISL_2728733, EPI_ISL_2728734, EPI_ISL_2728735 | Lighthouse Lab in Glasgow | Wellcome Sanger Institute for the COVID-19 Genomics UK (COG-UK) Consortium | Harper VanSteenhouse, Yumi Kasai, David Gray, Carol Clugston, Anna Dominiczak and Alex Alderton, Roberto Amato, Jeffrey Barrett, Sonia Goncalves, Ewan Harrison, David K. Jackson, Ian Johnston, Dominic Kwiatkowski, Cordelia Langford, John Sillitoe on behalf of the Wellcome Sanger Institute COVID-19 Surveillance Team |
| EPI_ISL_2728736 | Lighthouse Laboratory Plymouth | Wellcome Sanger Institute for the COVID-19 Genomics UK (COG-UK) Consortium | Lighthouse Laboratory Plymouth and Alex Alderton, Roberto Amato, Jeffrey Barrett, Sonia Goncalves, Ewan Harrison, David K. Jackson, Ian Johnston, Dominic Kwiatkowski, Cordelia Langford, John Sillitoe on behalf of the Wellcome Sanger Institute COVID-19 Surveillance Team |
| EPI_ISL_2728737, EPI_ISL_2728738 | Lighthouse Lab in Glasgow | Wellcome Sanger Institute for the COVID-19 Genomics UK (COG-UK) Consortium | Harper VanSteenhouse, Yumi Kasai, David Gray, Carol Clugston, Anna Dominiczak and Alex Alderton, Roberto Amato, Jeffrey Barrett, Sonia Goncalves, Ewan Harrison, David K. Jackson, Ian Johnston, Dominic Kwiatkowski, Cordelia Langford, John Sillitoe on behalf of the Wellcome Sanger Institute COVID-19 Surveillance Team |
| EPI_ISL_2728739 | Lighthouse Lab in Milton Keynes | Wellcome Sanger Institute for the COVID-19 Genomics UK (COG-UK) Consortium | The Lighthouse Lab in Milton Keynes and Alex Alderton, Roberto Amato, Jeffrey Barrett, Sonia Goncalves, Ewan Harrison, David K. Jackson, Ian Johnston, Dominic Kwiatkowski, Cordelia Langford, John Sillitoe on behalf of the Wellcome Sanger Institute COVID-19 Surveillance Team |
| EPI_ISL_2728740, EPI_ISL_2728741 | Lighthouse Lab in Glasgow | Wellcome Sanger Institute for the COVID-19 Genomics UK (COG-UK) Consortium | Harper VanSteenhouse, Yumi Kasai, David Gray, Carol Clugston, Anna Dominiczak and Alex Alderton, Roberto Amato, Jeffrey Barrett, Sonia Goncalves, Ewan Harrison, David K. Jackson, Ian Johnston, Dominic Kwiatkowski, Cordelia Langford, John Sillitoe on behalf of the Wellcome Sanger Institute COVID-19 Surveillance Team |
| EPI_ISL_2728742 | Lighthouse Lab in Milton Keynes | Wellcome Sanger Institute for the COVID-19 Genomics UK (COG-UK) Consortium | The Lighthouse Lab in Milton Keynes and Alex Alderton, Roberto Amato, Jeffrey Barrett, Sonia Goncalves, Ewan Harrison, David K. Jackson, Ian Johnston, Dominic Kwiatkowski, Cordelia Langford, John Sillitoe on behalf of the Wellcome Sanger Institute COVID-19 Surveillance Team |
| EPI_ISL_2728743 | Lighthouse Laboratory Plymouth | Wellcome Sanger Institute for the COVID-19 Genomics UK (COG-UK) Consortium | Lighthouse Laboratory Plymouth and Alex Alderton, Roberto Amato, Jeffrey Barrett, Sonia Goncalves, Ewan Harrison, David K. Jackson, Ian Johnston, Dominic Kwiatkowski, Cordelia Langford, John Sillitoe on behalf of the Wellcome Sanger Institute COVID-19 Surveillance Team |
| EPI_ISL_2728744 | Lighthouse Lab in Glasgow | Wellcome Sanger Institute for the COVID-19 Genomics UK (COG-UK) Consortium | Harper VanSteenhouse, Yumi Kasai, David Gray, Carol Clugston, Anna Dominiczak and Alex Alderton, Roberto Amato, Jeffrey Barrett, Sonia Goncalves, Ewan Harrison, David K. Jackson, Ian Johnston, Dominic Kwiatkowski, Cordelia Langford, John Sillitoe on behalf of the Wellcome Sanger Institute COVID-19 Surveillance Team |
| EPI_ISL_2728745 | Lighthouse Lab in Milton Keynes | Wellcome Sanger Institute for the COVID-19 Genomics UK (COG-UK) Consortium | The Lighthouse Lab in Milton Keynes and Alex Alderton, Roberto Amato, Jeffrey Barrett, Sonia Goncalves, Ewan Harrison, David K. Jackson, Ian Johnston, Dominic Kwiatkowski, Cordelia Langford, John Sillitoe on behalf of the Wellcome Sanger Institute COVID-19 Surveillance Team |
| EPI_ISL_2728746 | Lighthouse Laboratory Plymouth | Wellcome Sanger Institute for the COVID-19 Genomics UK (COG-UK) Consortium | Lighthouse Laboratory Plymouth and Alex Alderton, Roberto Amato, Jeffrey Barrett, Sonia Goncalves, Ewan Harrison, David K. Jackson, Ian Johnston, Dominic Kwiatkowski, Cordelia Langford, John Sillitoe on behalf of the Wellcome Sanger Institute COVID-19 Surveillance Team |
| EPI_ISL_2728747, EPI_ISL_2728748 | Lighthouse Lab in Glasgow | Wellcome Sanger Institute for the COVID-19 Genomics UK (COG-UK) Consortium | Harper VanSteenhouse, Yumi Kasai, David Gray, Carol Clugston, Anna Dominiczak and Alex Alderton, Roberto Amato, Jeffrey Barrett, Sonia Goncalves, Ewan Harrison, David K. Jackson, Ian Johnston, Dominic Kwiatkowski, Cordelia Langford, John Sillitoe on behalf of the Wellcome Sanger Institute COVID-19 Surveillance Team |
| EPI_ISL_2728749 | Lighthouse Laboratory Plymouth | Wellcome Sanger Institute for the COVID-19 Genomics UK (COG-UK) Consortium | Lighthouse Laboratory Plymouth and Alex Alderton, Roberto Amato, Jeffrey Barrett, Sonia Goncalves, Ewan Harrison, David K. Jackson, Ian Johnston, Dominic Kwiatkowski, Cordelia Langford, John Sillitoe on behalf of the Wellcome Sanger Institute COVID-19 Surveillance Team |
| EPI_ISL_2728750 | Lighthouse Lab in Milton Keynes | Wellcome Sanger Institute for the COVID-19 Genomics UK (COG-UK) Consortium | The Lighthouse Lab in Milton Keynes and Alex Alderton, Roberto Amato, Jeffrey Barrett, Sonia Goncalves, Ewan Harrison, David K. Jackson, Ian Johnston, Dominic Kwiatkowski, Cordelia Langford, John Sillitoe on behalf of the Wellcome Sanger Institute COVID-19 Surveillance Team |
| EPI_ISL_2728751 | Lighthouse Laboratory Plymouth | Wellcome Sanger Institute for the COVID-19 Genomics UK (COG-UK) Consortium | Lighthouse Laboratory Plymouth and Alex Alderton, Roberto Amato, Jeffrey Barrett, Sonia Goncalves, Ewan Harrison, David K. Jackson, Ian Johnston, Dominic Kwiatkowski, Cordelia Langford, John Sillitoe on behalf of the Wellcome Sanger Institute COVID-19 Surveillance Team |
| EPI_ISL_2728752, EPI_ISL_2728753 | Lighthouse Lab in Glasgow | Wellcome Sanger Institute for the COVID-19 Genomics UK (COG-UK) Consortium | Harper VanSteenhouse, Yumi Kasai, David Gray, Carol Clugston, Anna Dominiczak and Alex Alderton, Roberto Amato, Jeffrey Barrett, Sonia Goncalves, Ewan Harrison, David K. Jackson, Ian Johnston, Dominic Kwiatkowski, Cordelia Langford, John Sillitoe on behalf of the Wellcome Sanger Institute COVID-19 Surveillance Team |
| EPI_ISL_2728755, EPI_ISL_2728756, EPI_ISL_2728757 | Lighthouse Laboratory Plymouth | Wellcome Sanger Institute for the COVID-19 Genomics UK (COG-UK) Consortium | Lighthouse Laboratory Plymouth and Alex Alderton, Roberto Amato, Jeffrey Barrett, Sonia Goncalves, Ewan Harrison, David K. Jackson, Ian Johnston, Dominic Kwiatkowski, Cordelia Langford, John Sillitoe on behalf of the Wellcome Sanger Institute COVID-19 Surveillance Team |
| EPI_ISL_2728758 | Lighthouse Lab in Milton Keynes | Wellcome Sanger Institute for the COVID-19 Genomics UK (COG-UK) Consortium | The Lighthouse Lab in Milton Keynes and Alex Alderton, Roberto Amato, Jeffrey Barrett, Sonia Goncalves, Ewan Harrison, David K. Jackson, Ian Johnston, Dominic Kwiatkowski, Cordelia Langford, John Sillitoe on behalf of the Wellcome Sanger Institute COVID-19 Surveillance Team |
| EPI_ISL_2728759, EPI_ISL_2728760 | Lighthouse Laboratory Plymouth | Wellcome Sanger Institute for the COVID-19 Genomics UK (COG-UK) Consortium | Lighthouse Laboratory Plymouth and Alex Alderton, Roberto Amato, Jeffrey Barrett, Sonia Goncalves, Ewan Harrison, David K. Jackson, Ian Johnston, Dominic Kwiatkowski, Cordelia Langford, John Sillitoe on behalf of the Wellcome Sanger Institute COVID-19 Surveillance Team |
| EPI_ISL_2728761, EPI_ISL_2728762 | Lighthouse Lab in Milton Keynes | Wellcome Sanger Institute for the COVID-19 Genomics UK (COG-UK) Consortium | The Lighthouse Lab in Milton Keynes and Alex Alderton, Roberto Amato, Jeffrey Barrett, Sonia Goncalves, Ewan Harrison, David K. Jackson, Ian Johnston, Dominic Kwiatkowski, Cordelia Langford, John Sillitoe on behalf of the Wellcome Sanger Institute COVID-19 Surveillance Team |
| EPI_ISL_2728763 | Lighthouse Lab in Glasgow | Wellcome Sanger Institute for the COVID-19 Genomics UK (COG-UK) Consortium | Harper VanSteenhouse, Yumi Kasai, David Gray, Carol Clugston, Anna Dominiczak and Alex Alderton, Roberto Amato, Jeffrey Barrett, Sonia Goncalves, Ewan Harrison, David K. Jackson, Ian Johnston, Dominic Kwiatkowski, Cordelia Langford, John Sillitoe on behalf of the Wellcome Sanger Institute COVID-19 Surveillance Team |
| EPI_ISL_2728764 | Lighthouse Lab in Milton Keynes | Wellcome Sanger Institute for the COVID-19 Genomics UK (COG-UK) Consortium | The Lighthouse Lab in Milton Keynes and Alex Alderton, Roberto Amato, Jeffrey Barrett, Sonia Goncalves, Ewan Harrison, David K. Jackson, Ian Johnston, Dominic Kwiatkowski, Cordelia Langford, John Sillitoe on behalf of the Wellcome Sanger Institute COVID-19 Surveillance Team |
| EPI_ISL_2728765, EPI_ISL_2728766 | Lighthouse Lab in Glasgow | Wellcome Sanger Institute for the COVID-19 Genomics UK (COG-UK) Consortium | Harper VanSteenhouse, Yumi Kasai, David Gray, Carol Clugston, Anna Dominiczak and Alex Alderton, Roberto Amato, Jeffrey Barrett, Sonia Goncalves, Ewan Harrison, David K. Jackson, Ian Johnston, Dominic Kwiatkowski, Cordelia Langford, John Sillitoe on behalf of the Wellcome Sanger Institute COVID-19 Surveillance Team |
| EPI_ISL_2728767 | Lighthouse Lab in Milton Keynes | Wellcome Sanger Institute for the COVID-19 Genomics UK (COG-UK) Consortium | The Lighthouse Lab in Milton Keynes and Alex Alderton, Roberto Amato, Jeffrey Barrett, Sonia Goncalves, Ewan Harrison, David K. Jackson, Ian Johnston, Dominic Kwiatkowski, Cordelia Langford, John Sillitoe on behalf of the Wellcome Sanger Institute COVID-19 Surveillance Team |
| EPI_ISL_2728768 | Lighthouse Lab in Glasgow | Wellcome Sanger Institute for the COVID-19 Genomics UK (COG-UK) Consortium | Harper VanSteenhouse, Yumi Kasai, David Gray, Carol Clugston, Anna Dominiczak and Alex Alderton, Roberto Amato, Jeffrey Barrett, Sonia Goncalves, Ewan Harrison, David K. Jackson, Ian Johnston, Dominic Kwiatkowski, Cordelia Langford, John Sillitoe on behalf of the Wellcome Sanger Institute COVID-19 Surveillance Team |
| EPI_ISL_2728769 | Lighthouse Laboratory Plymouth | Wellcome Sanger Institute for the COVID-19 Genomics UK (COG-UK) Consortium | Lighthouse Laboratory Plymouth and Alex Alderton, Roberto Amato, Jeffrey Barrett, Sonia Goncalves, Ewan Harrison, David K. Jackson, Ian Johnston, Dominic Kwiatkowski, Cordelia Langford, John Sillitoe on behalf of the Wellcome Sanger Institute COVID-19 Surveillance Team |
| EPI_ISL_2728770, EPI_ISL_2728771, EPI_ISL_2728772, EPI_ISL_2728773 | Lighthouse Lab in Glasgow | Wellcome Sanger Institute for the COVID-19 Genomics UK (COG-UK) Consortium | Harper VanSteenhouse, Yumi Kasai, David Gray, Carol Clugston, Anna Dominiczak and Alex Alderton, Roberto Amato, Jeffrey Barrett, Sonia Goncalves, Ewan Harrison, David K. Jackson, Ian Johnston, Dominic Kwiatkowski, Cordelia Langford, John Sillitoe on behalf of the Wellcome Sanger Institute COVID-19 Surveillance Team |
| EPI_ISL_2728774 | Lighthouse Laboratory Plymouth | Wellcome Sanger Institute for the COVID-19 Genomics UK (COG-UK) Consortium | Lighthouse Laboratory Plymouth and Alex Alderton, Roberto Amato, Jeffrey Barrett, Sonia Goncalves, Ewan Harrison, David K. Jackson, Ian Johnston, Dominic Kwiatkowski, Cordelia Langford, John Sillitoe on behalf of the Wellcome Sanger Institute COVID-19 Surveillance Team |
| EPI_ISL_2728775 | Lighthouse Lab in Glasgow | Wellcome Sanger Institute for the COVID-19 Genomics UK (COG-UK) Consortium | Harper VanSteenhouse, Yumi Kasai, David Gray, Carol Clugston, Anna Dominiczak and Alex Alderton, Roberto Amato, Jeffrey Barrett, Sonia Goncalves, Ewan Harrison, David K. Jackson, Ian Johnston, Dominic Kwiatkowski, Cordelia Langford, John Sillitoe on behalf of the Wellcome Sanger Institute COVID-19 Surveillance Team |
| EPI_ISL_2728776 | Lighthouse Laboratory Plymouth | Wellcome Sanger Institute for the COVID-19 Genomics UK (COG-UK) Consortium | Lighthouse Laboratory Plymouth and Alex Alderton, Roberto Amato, Jeffrey Barrett, Sonia Goncalves, Ewan Harrison, David K. Jackson, Ian Johnston, Dominic Kwiatkowski, Cordelia Langford, John Sillitoe on behalf of the Wellcome Sanger Institute COVID-19 Surveillance Team |
| EPI_ISL_2728777 | Lighthouse Lab in Glasgow | Wellcome Sanger Institute for the COVID-19 Genomics UK (COG-UK) Consortium | Harper VanSteenhouse, Yumi Kasai, David Gray, Carol Clugston, Anna Dominiczak and Alex Alderton, Roberto Amato, Jeffrey Barrett, Sonia Goncalves, Ewan Harrison, David K. Jackson, Ian Johnston, Dominic Kwiatkowski, Cordelia Langford, John Sillitoe on behalf of the Wellcome Sanger Institute COVID-19 Surveillance Team |
| EPI_ISL_2728778 | Lighthouse Lab in Milton Keynes | Wellcome Sanger Institute for the COVID-19 Genomics UK (COG-UK) Consortium | The Lighthouse Lab in Milton Keynes and Alex Alderton, Roberto Amato, Jeffrey Barrett, Sonia Goncalves, Ewan Harrison, David K. Jackson, Ian Johnston, Dominic Kwiatkowski, Cordelia Langford, John Sillitoe on behalf of the Wellcome Sanger Institute COVID-19 Surveillance Team |
| EPI_ISL_2728779 | Lighthouse Laboratory Plymouth | Wellcome Sanger Institute for the COVID-19 Genomics UK (COG-UK) Consortium | Lighthouse Laboratory Plymouth and Alex Alderton, Roberto Amato, Jeffrey Barrett, Sonia Goncalves, Ewan Harrison, David K. Jackson, Ian Johnston, Dominic Kwiatkowski, Cordelia Langford, John Sillitoe on behalf of the Wellcome Sanger Institute COVID-19 Surveillance Team |
| EPI_ISL_2728780 | Lighthouse Lab in Glasgow | Wellcome Sanger Institute for the COVID-19 Genomics UK (COG-UK) Consortium | Harper VanSteenhouse, Yumi Kasai, David Gray, Carol Clugston, Anna Dominiczak and Alex Alderton, Roberto Amato, Jeffrey Barrett, Sonia Goncalves, Ewan Harrison, David K. Jackson, Ian Johnston, Dominic Kwiatkowski, Cordelia Langford, John Sillitoe on behalf of the Wellcome Sanger Institute COVID-19 Surveillance Team |
| EPI_ISL_2728781 | Lighthouse Lab in Milton Keynes | Wellcome Sanger Institute for the COVID-19 Genomics UK (COG-UK) Consortium | The Lighthouse Lab in Milton Keynes and Alex Alderton, Roberto Amato, Jeffrey Barrett, Sonia Goncalves, Ewan Harrison, David K. Jackson, Ian Johnston, Dominic Kwiatkowski, Cordelia Langford, John Sillitoe on behalf of the Wellcome Sanger Institute COVID-19 Surveillance Team |
| EPI_ISL_2728782, EPI_ISL_2728783 | Lighthouse Lab in Glasgow | Wellcome Sanger Institute for the COVID-19 Genomics UK (COG-UK) Consortium | Harper VanSteenhouse, Yumi Kasai, David Gray, Carol Clugston, Anna Dominiczak and Alex Alderton, Roberto Amato, Jeffrey Barrett, Sonia Goncalves, Ewan Harrison, David K. Jackson, Ian Johnston, Dominic Kwiatkowski, Cordelia Langford, John Sillitoe on behalf of the Wellcome Sanger Institute COVID-19 Surveillance Team |
| EPI_ISL_2728784 | Lighthouse Lab in Milton Keynes | Wellcome Sanger Institute for the COVID-19 Genomics UK (COG-UK) Consortium | The Lighthouse Lab in Milton Keynes and Alex Alderton, Roberto Amato, Jeffrey Barrett, Sonia Goncalves, Ewan Harrison, David K. Jackson, Ian Johnston, Dominic Kwiatkowski, Cordelia Langford, John Sillitoe on behalf of the Wellcome Sanger Institute COVID-19 Surveillance Team |

| | | | |
|---|---|---|---|
| EPI_ISL_2728785, EPI_ISL_2728786 | Lighthouse Lab in Glasgow | Wellcome Sanger Institute for the COVID-19 Genomics UK (COG-UK) Consortium | Harper VanSteenhouse, Yumi Kasai, David Gray, Carol Clugston, Anna Dominiczak and Alex Alderton, Roberto Amato, Jeffrey Barrett, Sonia Goncalves, Ewan Harrison, David K. Jackson, Ian Johnston, Dominic Kwiatkowski, Cordelia Langford, John Sillitoe on behalf of the Wellcome Sanger Institute COVID-19 Surveillance Team |
| EPI_ISL_2728787 | Lighthouse Lab in Milton Keynes | Wellcome Sanger Institute for the COVID-19 Genomics UK (COG-UK) Consortium | The Lighthouse Lab in Milton Keynes and Alex Alderton, Roberto Amato, Jeffrey Barrett, Sonia Goncalves, Ewan Harrison, David K. Jackson, Ian Johnston, Dominic Kwiatkowski, Cordelia Langford, John Sillitoe on behalf of the Wellcome Sanger Institute COVID-19 Surveillance Team |
| EPI_ISL_2728788 | Lighthouse Lab in Glasgow | Wellcome Sanger Institute for the COVID-19 Genomics UK (COG-UK) Consortium | Harper VanSteenhouse, Yumi Kasai, David Gray, Carol Clugston, Anna Dominiczak and Alex Alderton, Roberto Amato, Jeffrey Barrett, Sonia Goncalves, Ewan Harrison, David K. Jackson, Ian Johnston, Dominic Kwiatkowski, Cordelia Langford, John Sillitoe on behalf of the Wellcome Sanger Institute COVID-19 Surveillance Team |
| EPI_ISL_2728789, EPI_ISL_2728790 | Lighthouse Laboratory Plymouth | Wellcome Sanger Institute for the COVID-19 Genomics UK (COG-UK) Consortium | Lighthouse Laboratory Plymouth and Alex Alderton, Roberto Amato, Jeffrey Barrett, Sonia Goncalves, Ewan Harrison, David K. Jackson, Ian Johnston, Dominic Kwiatkowski, Cordelia Langford, John Sillitoe on behalf of the Wellcome Sanger Institute COVID-19 Surveillance Team |
| EPI_ISL_2728791 | Lighthouse Lab in Glasgow | Wellcome Sanger Institute for the COVID-19 Genomics UK (COG-UK) Consortium | Harper VanSteenhouse, Yumi Kasai, David Gray, Carol Clugston, Anna Dominiczak and Alex Alderton, Roberto Amato, Jeffrey Barrett, Sonia Goncalves, Ewan Harrison, David K. Jackson, Ian Johnston, Dominic Kwiatkowski, Cordelia Langford, John Sillitoe on behalf of the Wellcome Sanger Institute COVID-19 Surveillance Team |
| EPI_ISL_2728792, EPI_ISL_2728793, EPI_ISL_2728794 | Lighthouse Lab in Milton Keynes | Wellcome Sanger Institute for the COVID-19 Genomics UK (COG-UK) Consortium | The Lighthouse Lab in Milton Keynes and Alex Alderton, Roberto Amato, Jeffrey Barrett, Sonia Goncalves, Ewan Harrison, David K. Jackson, Ian Johnston, Dominic Kwiatkowski, Cordelia Langford, John Sillitoe on behalf of the Wellcome Sanger Institute COVID-19 Surveillance Team |
| EPI_ISL_2728795 | Lighthouse Laboratory Plymouth | Wellcome Sanger Institute for the COVID-19 Genomics UK (COG-UK) Consortium | Lighthouse Laboratory Plymouth and Alex Alderton, Roberto Amato, Jeffrey Barrett, Sonia Goncalves, Ewan Harrison, David K. Jackson, Ian Johnston, Dominic Kwiatkowski, Cordelia Langford, John Sillitoe on behalf of the Wellcome Sanger Institute COVID-19 Surveillance Team |
| EPI_ISL_2728796 | Lighthouse Lab in Milton Keynes | Wellcome Sanger Institute for the COVID-19 Genomics UK (COG-UK) Consortium | The Lighthouse Lab in Milton Keynes and Alex Alderton, Roberto Amato, Jeffrey Barrett, Sonia Goncalves, Ewan Harrison, David K. Jackson, Ian Johnston, Dominic Kwiatkowski, Cordelia Langford, John Sillitoe on behalf of the Wellcome Sanger Institute COVID-19 Surveillance Team |
| EPI_ISL_2728797, EPI_ISL_2728798 | Lighthouse Laboratory Plymouth | Wellcome Sanger Institute for the COVID-19 Genomics UK (COG-UK) Consortium | Lighthouse Laboratory Plymouth and Alex Alderton, Roberto Amato, Jeffrey Barrett, Sonia Goncalves, Ewan Harrison, David K. Jackson, Ian Johnston, Dominic Kwiatkowski, Cordelia Langford, John Sillitoe on behalf of the Wellcome Sanger Institute COVID-19 Surveillance Team |
| EPI_ISL_2728799, EPI_ISL_2728800, EPI_ISL_2728801, EPI_ISL_2728802, EPI_ISL_2728803, EPI_ISL_2728804 | Lighthouse Lab in Glasgow | Wellcome Sanger Institute for the COVID-19 Genomics UK (COG-UK) Consortium | Harper VanSteenhouse, Yumi Kasai, David Gray, Carol Clugston, Anna Dominiczak and Alex Alderton, Roberto Amato, Jeffrey Barrett, Sonia Goncalves, Ewan Harrison, David K. Jackson, Ian Johnston, Dominic Kwiatkowski, Cordelia Langford, John Sillitoe on behalf of the Wellcome Sanger Institute COVID-19 Surveillance Team |
| EPI_ISL_2728805 | Lighthouse Laboratory Plymouth | Wellcome Sanger Institute for the COVID-19 Genomics UK (COG-UK) Consortium | Lighthouse Laboratory Plymouth and Alex Alderton, Roberto Amato, Jeffrey Barrett, Sonia Goncalves, Ewan Harrison, David K. Jackson, Ian Johnston, Dominic Kwiatkowski, Cordelia Langford, John Sillitoe on behalf of the Wellcome Sanger Institute COVID-19 Surveillance Team |
| EPI_ISL_2728806 | Lighthouse Lab in Glasgow | Wellcome Sanger Institute for the COVID-19 Genomics UK (COG-UK) Consortium | Harper VanSteenhouse, Yumi Kasai, David Gray, Carol Clugston, Anna Dominiczak and Alex Alderton, Roberto Amato, Jeffrey Barrett, Sonia Goncalves, Ewan Harrison, David K. Jackson, Ian Johnston, Dominic Kwiatkowski, Cordelia Langford, John Sillitoe on behalf of the Wellcome Sanger Institute COVID-19 Surveillance Team |
| EPI_ISL_2728807 | Lighthouse Laboratory Plymouth | Wellcome Sanger Institute for the COVID-19 Genomics UK (COG-UK) Consortium | Lighthouse Laboratory Plymouth and Alex Alderton, Roberto Amato, Jeffrey Barrett, Sonia Goncalves, Ewan Harrison, David K. Jackson, Ian Johnston, Dominic Kwiatkowski, Cordelia Langford, John Sillitoe on behalf of the Wellcome Sanger Institute COVID-19 Surveillance Team |
| EPI_ISL_2728808 | Lighthouse Lab in Glasgow | Wellcome Sanger Institute for the COVID-19 Genomics UK (COG-UK) Consortium | Harper VanSteenhouse, Yumi Kasai, David Gray, Carol Clugston, Anna Dominiczak and Alex Alderton, Roberto Amato, Jeffrey Barrett, Sonia Goncalves, Ewan Harrison, David K. Jackson, Ian Johnston, Dominic Kwiatkowski, Cordelia Langford, John Sillitoe on behalf of the Wellcome Sanger Institute COVID-19 Surveillance Team |
| EPI_ISL_2728809 | Lighthouse Lab in Milton Keynes | Wellcome Sanger Institute for the COVID-19 Genomics UK (COG-UK) Consortium | The Lighthouse Lab in Milton Keynes and Alex Alderton, Roberto Amato, Jeffrey Barrett, Sonia Goncalves, Ewan Harrison, David K. Jackson, Ian Johnston, Dominic Kwiatkowski, Cordelia Langford, John Sillitoe on behalf of the Wellcome Sanger Institute COVID-19 Surveillance Team |
| EPI_ISL_2728810 | Lighthouse Lab in Glasgow | Wellcome Sanger Institute for the COVID-19 Genomics UK (COG-UK) Consortium | Harper VanSteenhouse, Yumi Kasai, David Gray, Carol Clugston, Anna Dominiczak and Alex Alderton, Roberto Amato, Jeffrey Barrett, Sonia Goncalves, Ewan Harrison, David K. Jackson, Ian Johnston, Dominic Kwiatkowski, Cordelia Langford, John Sillitoe on behalf of the Wellcome Sanger Institute COVID-19 Surveillance Team |
| EPI_ISL_2728811 | Lighthouse Laboratory Plymouth | Wellcome Sanger Institute for the COVID-19 Genomics UK (COG-UK) Consortium | Lighthouse Laboratory Plymouth and Alex Alderton, Roberto Amato, Jeffrey Barrett, Sonia Goncalves, Ewan Harrison, David K. Jackson, Ian Johnston, Dominic Kwiatkowski, Cordelia Langford, John Sillitoe on behalf of the Wellcome Sanger Institute COVID-19 Surveillance Team |
| EPI_ISL_2728812 | Lighthouse Lab in Milton Keynes | Wellcome Sanger Institute for the COVID-19 Genomics UK (COG-UK) Consortium | The Lighthouse Lab in Milton Keynes and Alex Alderton, Roberto Amato, Jeffrey Barrett, Sonia Goncalves, Ewan Harrison, David K. Jackson, Ian Johnston, Dominic Kwiatkowski, Cordelia Langford, John Sillitoe on behalf of the Wellcome Sanger Institute COVID-19 Surveillance Team |
| EPI_ISL_2728814, EPI_ISL_2728815 | Lighthouse Lab in Glasgow | Wellcome Sanger Institute for the COVID-19 Genomics UK (COG-UK) Consortium | Harper VanSteenhouse, Yumi Kasai, David Gray, Carol Clugston, Anna Dominiczak and Alex Alderton, Roberto Amato, Jeffrey Barrett, Sonia Goncalves, Ewan Harrison, David K. Jackson, Ian Johnston, Dominic Kwiatkowski, Cordelia Langford, John Sillitoe on behalf of the Wellcome Sanger Institute COVID-19 Surveillance Team |
| EPI_ISL_2728816 | Lighthouse Lab in Milton Keynes | Wellcome Sanger Institute for the COVID-19 Genomics UK (COG-UK) Consortium | The Lighthouse Lab in Milton Keynes and Alex Alderton, Roberto Amato, Jeffrey Barrett, Sonia Goncalves, Ewan Harrison, David K. Jackson, Ian Johnston, Dominic Kwiatkowski, Cordelia Langford, John Sillitoe on behalf of the Wellcome Sanger Institute COVID-19 Surveillance Team |
| EPI_ISL_2728817 | Lighthouse Laboratory Plymouth | Wellcome Sanger Institute for the COVID-19 Genomics UK (COG-UK) Consortium | Lighthouse Laboratory Plymouth and Alex Alderton, Roberto Amato, Jeffrey Barrett, Sonia Goncalves, Ewan Harrison, David K. Jackson, Ian Johnston, Dominic Kwiatkowski, Cordelia Langford, John Sillitoe on behalf of the Wellcome Sanger Institute COVID-19 Surveillance Team |
| EPI_ISL_2728818 | Lighthouse Lab in Milton Keynes | Wellcome Sanger Institute for the COVID-19 Genomics UK (COG-UK) Consortium | The Lighthouse Lab in Milton Keynes and Alex Alderton, Roberto Amato, Jeffrey Barrett, Sonia Goncalves, Ewan Harrison, David K. Jackson, Ian Johnston, Dominic Kwiatkowski, Cordelia Langford, John Sillitoe on behalf of the Wellcome Sanger Institute COVID-19 Surveillance Team |
| EPI_ISL_2728819 | Lighthouse Laboratory Plymouth | Wellcome Sanger Institute for the COVID-19 Genomics UK (COG-UK) Consortium | Lighthouse Laboratory Plymouth and Alex Alderton, Roberto Amato, Jeffrey Barrett, Sonia Goncalves, Ewan Harrison, David K. Jackson, Ian Johnston, Dominic Kwiatkowski, Cordelia Langford, John Sillitoe on behalf of the Wellcome Sanger Institute COVID-19 Surveillance Team |
| EPI_ISL_2728820, EPI_ISL_2728821 | Lighthouse Lab in Glasgow | Wellcome Sanger Institute for the COVID-19 Genomics UK (COG-UK) Consortium | Harper VanSteenhouse, Yumi Kasai, David Gray, Carol Clugston, Anna Dominiczak and Alex Alderton, Roberto Amato, Jeffrey Barrett, Sonia Goncalves, Ewan Harrison, David K. Jackson, Ian Johnston, Dominic Kwiatkowski, Cordelia Langford, John Sillitoe on behalf of the Wellcome Sanger Institute COVID-19 Surveillance Team |
| EPI_ISL_2728822 | Lighthouse Lab in Milton Keynes | Wellcome Sanger Institute for the COVID-19 Genomics UK (COG-UK) Consortium | The Lighthouse Lab in Milton Keynes and Alex Alderton, Roberto Amato, Jeffrey Barrett, Sonia Goncalves, Ewan Harrison, David K. Jackson, Ian Johnston, Dominic Kwiatkowski, Cordelia Langford, John Sillitoe on behalf of the Wellcome Sanger Institute COVID-19 Surveillance Team |
| EPI_ISL_2728823 | Lighthouse Laboratory Plymouth | Wellcome Sanger Institute for the COVID-19 Genomics UK (COG-UK) Consortium | Lighthouse Laboratory Plymouth and Alex Alderton, Roberto Amato, Jeffrey Barrett, Sonia Goncalves, Ewan Harrison, David K. Jackson, Ian Johnston, Dominic Kwiatkowski, Cordelia Langford, John Sillitoe on behalf of the Wellcome Sanger Institute COVID-19 Surveillance Team |
| EPI_ISL_2728824, EPI_ISL_2728825, EPI_ISL_2728826, EPI_ISL_2728828 | Lighthouse Lab in Glasgow | Wellcome Sanger Institute for the COVID-19 Genomics UK (COG-UK) Consortium | Harper VanSteenhouse, Yumi Kasai, David Gray, Carol Clugston, Anna Dominiczak and Alex Alderton, Roberto Amato, Jeffrey Barrett, Sonia Goncalves, Ewan Harrison, David K. Jackson, Ian Johnston, Dominic Kwiatkowski, Cordelia Langford, John Sillitoe on behalf of the Wellcome Sanger Institute COVID-19 Surveillance Team |
| EPI_ISL_2728829 | Lighthouse Laboratory Plymouth | Wellcome Sanger Institute for the COVID-19 Genomics UK (COG-UK) Consortium | Lighthouse Laboratory Plymouth and Alex Alderton, Roberto Amato, Jeffrey Barrett, Sonia Goncalves, Ewan Harrison, David K. Jackson, Ian Johnston, Dominic Kwiatkowski, Cordelia Langford, John Sillitoe on behalf of the Wellcome Sanger Institute COVID-19 Surveillance Team |
| EPI_ISL_2728830, EPI_ISL_2728831 | Lighthouse Lab in Glasgow | Wellcome Sanger Institute for the COVID-19 Genomics UK (COG-UK) Consortium | Harper VanSteenhouse, Yumi Kasai, David Gray, Carol Clugston, Anna Dominiczak and Alex Alderton, Roberto Amato, Jeffrey Barrett, Sonia Goncalves, Ewan Harrison, David K. Jackson, Ian Johnston, Dominic Kwiatkowski, Cordelia Langford, John Sillitoe on behalf of the Wellcome Sanger Institute COVID-19 Surveillance Team |
| EPI_ISL_2728832, EPI_ISL_2728833 | Lighthouse Lab in Milton Keynes | Wellcome Sanger Institute for the COVID-19 Genomics UK (COG-UK) Consortium | The Lighthouse Lab in Milton Keynes and Alex Alderton, Roberto Amato, Jeffrey Barrett, Sonia Goncalves, Ewan Harrison, David K. Jackson, Ian Johnston, Dominic Kwiatkowski, Cordelia Langford, John Sillitoe on behalf of the Wellcome Sanger Institute COVID-19 Surveillance Team |
| EPI_ISL_2728834, EPI_ISL_2728835, EPI_ISL_2728836 | Lighthouse Laboratory Plymouth | Wellcome Sanger Institute for the COVID-19 Genomics UK (COG-UK) Consortium | Lighthouse Laboratory Plymouth and Alex Alderton, Roberto Amato, Jeffrey Barrett, Sonia Goncalves, Ewan Harrison, David K. Jackson, Ian Johnston, Dominic Kwiatkowski, Cordelia Langford, John Sillitoe on behalf of the Wellcome Sanger Institute COVID-19 Surveillance Team |
| EPI_ISL_2728837 | Lighthouse Lab in Glasgow | Wellcome Sanger Institute for the COVID-19 Genomics UK (COG-UK) Consortium | Harper VanSteenhouse, Yumi Kasai, David Gray, Carol Clugston, Anna Dominiczak and Alex Alderton, Roberto Amato, Jeffrey Barrett, Sonia Goncalves, Ewan Harrison, David K. Jackson, Ian Johnston, Dominic Kwiatkowski, Cordelia Langford, John Sillitoe on behalf of the Wellcome Sanger Institute COVID-19 Surveillance Team |
| EPI_ISL_2728838 | Lighthouse Lab in Milton Keynes | Wellcome Sanger Institute for the COVID-19 Genomics UK (COG-UK) Consortium | The Lighthouse Lab in Milton Keynes and Alex Alderton, Roberto Amato, Jeffrey Barrett, Sonia Goncalves, Ewan Harrison, David K. Jackson, Ian Johnston, Dominic Kwiatkowski, Cordelia Langford, John Sillitoe on behalf of the Wellcome Sanger Institute COVID-19 Surveillance Team |
| EPI_ISL_2728839, EPI_ISL_2728840, EPI_ISL_2728841 | Lighthouse Lab in Glasgow | Wellcome Sanger Institute for the COVID-19 Genomics UK (COG-UK) Consortium | Harper VanSteenhouse, Yumi Kasai, David Gray, Carol Clugston, Anna Dominiczak and Alex Alderton, Roberto Amato, Jeffrey Barrett, Sonia Goncalves, Ewan Harrison, David K. Jackson, Ian Johnston, Dominic Kwiatkowski, Cordelia Langford, John Sillitoe on behalf of the Wellcome Sanger Institute COVID-19 Surveillance Team |
| EPI_ISL_2728842 | Lighthouse Laboratory Plymouth | Wellcome Sanger Institute for the COVID-19 Genomics UK (COG-UK) Consortium | Lighthouse Laboratory Plymouth and Alex Alderton, Roberto Amato, Jeffrey Barrett, Sonia Goncalves, Ewan Harrison, David K. Jackson, Ian Johnston, Dominic Kwiatkowski, Cordelia Langford, John Sillitoe on behalf of the Wellcome Sanger Institute COVID-19 Surveillance Team |
| EPI_ISL_2728843 | Lighthouse Lab in Milton Keynes | Wellcome Sanger Institute for the COVID-19 Genomics UK (COG-UK) Consortium | The Lighthouse Lab in Milton Keynes and Alex Alderton, Roberto Amato, Jeffrey Barrett, Sonia Goncalves, Ewan Harrison, David K. Jackson, Ian Johnston, Dominic Kwiatkowski, Cordelia Langford, John Sillitoe on behalf of the Wellcome Sanger Institute COVID-19 Surveillance Team |
| EPI_ISL_2728844 | Lighthouse Laboratory Plymouth | Wellcome Sanger Institute for the COVID-19 Genomics UK (COG-UK) Consortium | Lighthouse Laboratory Plymouth and Alex Alderton, Roberto Amato, Jeffrey Barrett, Sonia Goncalves, Ewan Harrison, David K. Jackson, Ian Johnston, Dominic Kwiatkowski, Cordelia Langford, John Sillitoe on behalf of the Wellcome Sanger Institute COVID-19 Surveillance Team |
| EPI_ISL_2728845 | Lighthouse Lab in Milton Keynes | Wellcome Sanger Institute for the COVID-19 Genomics UK (COG-UK) Consortium | The Lighthouse Lab in Milton Keynes and Alex Alderton, Roberto Amato, Jeffrey Barrett, Sonia Goncalves, Ewan Harrison, David K. Jackson, Ian Johnston, Dominic Kwiatkowski, Cordelia Langford, John Sillitoe on behalf of the Wellcome Sanger Institute COVID-19 Surveillance Team |
| EPI_ISL_2728846, EPI_ISL_2728847 | Lighthouse Laboratory Plymouth | Wellcome Sanger Institute for the COVID-19 Genomics UK (COG-UK) Consortium | Lighthouse Laboratory Plymouth and Alex Alderton, Roberto Amato, Jeffrey Barrett, Sonia Goncalves, Ewan Harrison, David K. Jackson, Ian Johnston, Dominic Kwiatkowski, Cordelia Langford, John Sillitoe on behalf of the Wellcome Sanger Institute COVID-19 Surveillance Team |
| EPI_ISL_2728848, EPI_ISL_2728849, EPI_ISL_2728850 | Lighthouse Lab in Milton Keynes | Wellcome Sanger Institute for the COVID-19 Genomics UK (COG-UK) Consortium | The Lighthouse Lab in Milton Keynes and Alex Alderton, Roberto Amato, Jeffrey Barrett, Sonia Goncalves, Ewan Harrison, David K. Jackson, Ian Johnston, Dominic Kwiatkowski, Cordelia Langford, John Sillitoe on behalf of the Wellcome Sanger Institute COVID-19 Surveillance Team |
| EPI_ISL_2728851, EPI_ISL_2728852 | Lighthouse Lab in Glasgow | Wellcome Sanger Institute for the COVID-19 Genomics UK (COG-UK) Consortium | Harper VanSteenhouse, Yumi Kasai, David Gray, Carol Clugston, Anna Dominiczak and Alex Alderton, Roberto Amato, Jeffrey Barrett, Sonia Goncalves, Ewan Harrison, David K. Jackson, Ian Johnston, Dominic Kwiatkowski, Cordelia Langford, John Sillitoe on behalf of the Wellcome Sanger Institute COVID-19 Surveillance Team |

| | | | |
|---|---|---|---|
| EPI_ISL_2728853 | Lighthouse Laboratory Plymouth | Wellcome Sanger Institute for the COVID-19 Genomics UK (COG-UK) Consortium | Lighthouse Laboratory Plymouth and Alex Alderton, Roberto Amato, Jeffrey Barrett, Sonia Goncalves, Ewan Harrison, David K. Jackson, Ian Johnston, Dominic Kwiatkowski, Cordelia Langford, John Sillitoe on behalf of the Wellcome Sanger Institute COVID-19 Surveillance Team |
| EPI_ISL_2728854, EPI_ISL_2728855 | Lighthouse Lab in Glasgow | Wellcome Sanger Institute for the COVID-19 Genomics UK (COG-UK) Consortium | Harper VanSteenhouse, Yumi Kasai, David Gray, Carol Clugston, Anna Dominiczak and Alex Alderton, Roberto Amato, Jeffrey Barrett, Sonia Goncalves, Ewan Harrison, David K. Jackson, Ian Johnston, Dominic Kwiatkowski, Cordelia Langford, John Sillitoe on behalf of the Wellcome Sanger Institute COVID-19 Surveillance Team |
| EPI_ISL_2728856 | Lighthouse Laboratory Plymouth | Wellcome Sanger Institute for the COVID-19 Genomics UK (COG-UK) Consortium | Lighthouse Laboratory Plymouth and Alex Alderton, Roberto Amato, Jeffrey Barrett, Sonia Goncalves, Ewan Harrison, David K. Jackson, Ian Johnston, Dominic Kwiatkowski, Cordelia Langford, John Sillitoe on behalf of the Wellcome Sanger Institute COVID-19 Surveillance Team |
| EPI_ISL_2728857 | Lighthouse Lab in Milton Keynes | Wellcome Sanger Institute for the COVID-19 Genomics UK (COG-UK) Consortium | The Lighthouse Lab in Milton Keynes and Alex Alderton, Roberto Amato, Jeffrey Barrett, Sonia Goncalves, Ewan Harrison, David K. Jackson, Ian Johnston, Dominic Kwiatkowski, Cordelia Langford, John Sillitoe on behalf of the Wellcome Sanger Institute COVID-19 Surveillance Team |
| EPI_ISL_2728858 | Lighthouse Laboratory Plymouth | Wellcome Sanger Institute for the COVID-19 Genomics UK (COG-UK) Consortium | Lighthouse Laboratory Plymouth and Alex Alderton, Roberto Amato, Jeffrey Barrett, Sonia Goncalves, Ewan Harrison, David K. Jackson, Ian Johnston, Dominic Kwiatkowski, Cordelia Langford, John Sillitoe on behalf of the Wellcome Sanger Institute COVID-19 Surveillance Team |
| EPI_ISL_2728859 | Lighthouse Lab in Glasgow | Wellcome Sanger Institute for the COVID-19 Genomics UK (COG-UK) Consortium | Harper VanSteenhouse, Yumi Kasai, David Gray, Carol Clugston, Anna Dominiczak and Alex Alderton, Roberto Amato, Jeffrey Barrett, Sonia Goncalves, Ewan Harrison, David K. Jackson, Ian Johnston, Dominic Kwiatkowski, Cordelia Langford, John Sillitoe on behalf of the Wellcome Sanger Institute COVID-19 Surveillance Team |
| EPI_ISL_2728860, EPI_ISL_2728861 | Lighthouse Laboratory Plymouth | Wellcome Sanger Institute for the COVID-19 Genomics UK (COG-UK) Consortium | Lighthouse Laboratory Plymouth and Alex Alderton, Roberto Amato, Jeffrey Barrett, Sonia Goncalves, Ewan Harrison, David K. Jackson, Ian Johnston, Dominic Kwiatkowski, Cordelia Langford, John Sillitoe on behalf of the Wellcome Sanger Institute COVID-19 Surveillance Team |
| EPI_ISL_2728862 | Lighthouse Lab in Glasgow | Wellcome Sanger Institute for the COVID-19 Genomics UK (COG-UK) Consortium | Harper VanSteenhouse, Yumi Kasai, David Gray, Carol Clugston, Anna Dominiczak and Alex Alderton, Roberto Amato, Jeffrey Barrett, Sonia Goncalves, Ewan Harrison, David K. Jackson, Ian Johnston, Dominic Kwiatkowski, Cordelia Langford, John Sillitoe on behalf of the Wellcome Sanger Institute COVID-19 Surveillance Team |
| EPI_ISL_2728863 | Lighthouse Laboratory Plymouth | Wellcome Sanger Institute for the COVID-19 Genomics UK (COG-UK) Consortium | Lighthouse Laboratory Plymouth and Alex Alderton, Roberto Amato, Jeffrey Barrett, Sonia Goncalves, Ewan Harrison, David K. Jackson, Ian Johnston, Dominic Kwiatkowski, Cordelia Langford, John Sillitoe on behalf of the Wellcome Sanger Institute COVID-19 Surveillance Team |
| EPI_ISL_2728864, EPI_ISL_2728865 | Lighthouse Lab in Glasgow | Wellcome Sanger Institute for the COVID-19 Genomics UK (COG-UK) Consortium | Harper VanSteenhouse, Yumi Kasai, David Gray, Carol Clugston, Anna Dominiczak and Alex Alderton, Roberto Amato, Jeffrey Barrett, Sonia Goncalves, Ewan Harrison, David K. Jackson, Ian Johnston, Dominic Kwiatkowski, Cordelia Langford, John Sillitoe on behalf of the Wellcome Sanger Institute COVID-19 Surveillance Team |
| EPI_ISL_2728866 | Lighthouse Laboratory Plymouth | Wellcome Sanger Institute for the COVID-19 Genomics UK (COG-UK) Consortium | Lighthouse Laboratory Plymouth and Alex Alderton, Roberto Amato, Jeffrey Barrett, Sonia Goncalves, Ewan Harrison, David K. Jackson, Ian Johnston, Dominic Kwiatkowski, Cordelia Langford, John Sillitoe on behalf of the Wellcome Sanger Institute COVID-19 Surveillance Team |
| EPI_ISL_2728867 | Lighthouse Lab in Milton Keynes | Wellcome Sanger Institute for the COVID-19 Genomics UK (COG-UK) Consortium | The Lighthouse Lab in Milton Keynes and Alex Alderton, Roberto Amato, Jeffrey Barrett, Sonia Goncalves, Ewan Harrison, David K. Jackson, Ian Johnston, Dominic Kwiatkowski, Cordelia Langford, John Sillitoe on behalf of the Wellcome Sanger Institute COVID-19 Surveillance Team |
| EPI_ISL_2728868, EPI_ISL_2728869 | Lighthouse Lab in Glasgow | Wellcome Sanger Institute for the COVID-19 Genomics UK (COG-UK) Consortium | Harper VanSteenhouse, Yumi Kasai, David Gray, Carol Clugston, Anna Dominiczak and Alex Alderton, Roberto Amato, Jeffrey Barrett, Sonia Goncalves, Ewan Harrison, David K. Jackson, Ian Johnston, Dominic Kwiatkowski, Cordelia Langford, John Sillitoe on behalf of the Wellcome Sanger Institute COVID-19 Surveillance Team |
| EPI_ISL_2728870 | Lighthouse Lab in Milton Keynes | Wellcome Sanger Institute for the COVID-19 Genomics UK (COG-UK) Consortium | The Lighthouse Lab in Milton Keynes and Alex Alderton, Roberto Amato, Jeffrey Barrett, Sonia Goncalves, Ewan Harrison, David K. Jackson, Ian Johnston, Dominic Kwiatkowski, Cordelia Langford, John Sillitoe on behalf of the Wellcome Sanger Institute COVID-19 Surveillance Team |
| EPI_ISL_2728871 | Lighthouse Lab in Glasgow | Wellcome Sanger Institute for the COVID-19 Genomics UK (COG-UK) Consortium | Harper VanSteenhouse, Yumi Kasai, David Gray, Carol Clugston, Anna Dominiczak and Alex Alderton, Roberto Amato, Jeffrey Barrett, Sonia Goncalves, Ewan Harrison, David K. Jackson, Ian Johnston, Dominic Kwiatkowski, Cordelia Langford, John Sillitoe on behalf of the Wellcome Sanger Institute COVID-19 Surveillance Team |
| EPI_ISL_2728872, EPI_ISL_2728873 | Lighthouse Lab in Milton Keynes | Wellcome Sanger Institute for the COVID-19 Genomics UK (COG-UK) Consortium | The Lighthouse Lab in Milton Keynes and Alex Alderton, Roberto Amato, Jeffrey Barrett, Sonia Goncalves, Ewan Harrison, David K. Jackson, Ian Johnston, Dominic Kwiatkowski, Cordelia Langford, John Sillitoe on behalf of the Wellcome Sanger Institute COVID-19 Surveillance Team |
| EPI_ISL_2728874, EPI_ISL_2728875, EPI_ISL_2728876 | Lighthouse Lab in Glasgow | Wellcome Sanger Institute for the COVID-19 Genomics UK (COG-UK) Consortium | Harper VanSteenhouse, Yumi Kasai, David Gray, Carol Clugston, Anna Dominiczak and Alex Alderton, Roberto Amato, Jeffrey Barrett, Sonia Goncalves, Ewan Harrison, David K. Jackson, Ian Johnston, Dominic Kwiatkowski, Cordelia Langford, John Sillitoe on behalf of the Wellcome Sanger Institute COVID-19 Surveillance Team |
| EPI_ISL_2728877 | Lighthouse Laboratory Plymouth | Wellcome Sanger Institute for the COVID-19 Genomics UK (COG-UK) Consortium | Lighthouse Laboratory Plymouth and Alex Alderton, Roberto Amato, Jeffrey Barrett, Sonia Goncalves, Ewan Harrison, David K. Jackson, Ian Johnston, Dominic Kwiatkowski, Cordelia Langford, John Sillitoe on behalf of the Wellcome Sanger Institute COVID-19 Surveillance Team |
| EPI_ISL_2728878, EPI_ISL_2728879, EPI_ISL_2728880, EPI_ISL_2728881 | Lighthouse Lab in Glasgow | Wellcome Sanger Institute for the COVID-19 Genomics UK (COG-UK) Consortium | Harper VanSteenhouse, Yumi Kasai, David Gray, Carol Clugston, Anna Dominiczak and Alex Alderton, Roberto Amato, Jeffrey Barrett, Sonia Goncalves, Ewan Harrison, David K. Jackson, Ian Johnston, Dominic Kwiatkowski, Cordelia Langford, John Sillitoe on behalf of the Wellcome Sanger Institute COVID-19 Surveillance Team |
| EPI_ISL_2728882 | Lighthouse Lab in Milton Keynes | Wellcome Sanger Institute for the COVID-19 Genomics UK (COG-UK) Consortium | The Lighthouse Lab in Milton Keynes and Alex Alderton, Roberto Amato, Jeffrey Barrett, Sonia Goncalves, Ewan Harrison, David K. Jackson, Ian Johnston, Dominic Kwiatkowski, Cordelia Langford, John Sillitoe on behalf of the Wellcome Sanger Institute COVID-19 Surveillance Team |
| EPI_ISL_2728883 | Lighthouse Lab in Glasgow | Wellcome Sanger Institute for the COVID-19 Genomics UK (COG-UK) Consortium | Harper VanSteenhouse, Yumi Kasai, David Gray, Carol Clugston, Anna Dominiczak and Alex Alderton, Roberto Amato, Jeffrey Barrett, Sonia Goncalves, Ewan Harrison, David K. Jackson, Ian Johnston, Dominic Kwiatkowski, Cordelia Langford, John Sillitoe on behalf of the Wellcome Sanger Institute COVID-19 Surveillance Team |
| EPI_ISL_2728884 | Lighthouse Lab in Milton Keynes | Wellcome Sanger Institute for the COVID-19 Genomics UK (COG-UK) Consortium | The Lighthouse Lab in Milton Keynes and Alex Alderton, Roberto Amato, Jeffrey Barrett, Sonia Goncalves, Ewan Harrison, David K. Jackson, Ian Johnston, Dominic Kwiatkowski, Cordelia Langford, John Sillitoe on behalf of the Wellcome Sanger Institute COVID-19 Surveillance Team |
| EPI_ISL_2728885, EPI_ISL_2728886, EPI_ISL_2728887, EPI_ISL_2728888, EPI_ISL_2728889, EPI_ISL_2728890 | Lighthouse Lab in Glasgow | Wellcome Sanger Institute for the COVID-19 Genomics UK (COG-UK) Consortium | Harper VanSteenhouse, Yumi Kasai, David Gray, Carol Clugston, Anna Dominiczak and Alex Alderton, Roberto Amato, Jeffrey Barrett, Sonia Goncalves, Ewan Harrison, David K. Jackson, Ian Johnston, Dominic Kwiatkowski, Cordelia Langford, John Sillitoe on behalf of the Wellcome Sanger Institute COVID-19 Surveillance Team |
| EPI_ISL_2728891 | Lighthouse Laboratory Plymouth | Wellcome Sanger Institute for the COVID-19 Genomics UK (COG-UK) Consortium | Lighthouse Laboratory Plymouth and Alex Alderton, Roberto Amato, Jeffrey Barrett, Sonia Goncalves, Ewan Harrison, David K. Jackson, Ian Johnston, Dominic Kwiatkowski, Cordelia Langford, John Sillitoe on behalf of the Wellcome Sanger Institute COVID-19 Surveillance Team |
| EPI_ISL_2728892 | Lighthouse Lab in Milton Keynes | Wellcome Sanger Institute for the COVID-19 Genomics UK (COG-UK) Consortium | The Lighthouse Lab in Milton Keynes and Alex Alderton, Roberto Amato, Jeffrey Barrett, Sonia Goncalves, Ewan Harrison, David K. Jackson, Ian Johnston, Dominic Kwiatkowski, Cordelia Langford, John Sillitoe on behalf of the Wellcome Sanger Institute COVID-19 Surveillance Team |
| EPI_ISL_2728893 | Lighthouse Laboratory Plymouth | Wellcome Sanger Institute for the COVID-19 Genomics UK (COG-UK) Consortium | Lighthouse Laboratory Plymouth and Alex Alderton, Roberto Amato, Jeffrey Barrett, Sonia Goncalves, Ewan Harrison, David K. Jackson, Ian Johnston, Dominic Kwiatkowski, Cordelia Langford, John Sillitoe on behalf of the Wellcome Sanger Institute COVID-19 Surveillance Team |
| EPI_ISL_2728894 | Lighthouse Lab in Milton Keynes | Wellcome Sanger Institute for the COVID-19 Genomics UK (COG-UK) Consortium | The Lighthouse Lab in Milton Keynes and Alex Alderton, Roberto Amato, Jeffrey Barrett, Sonia Goncalves, Ewan Harrison, David K. Jackson, Ian Johnston, Dominic Kwiatkowski, Cordelia Langford, John Sillitoe on behalf of the Wellcome Sanger Institute COVID-19 Surveillance Team |
| EPI_ISL_2728895, EPI_ISL_2728896 | Lighthouse Laboratory Plymouth | Wellcome Sanger Institute for the COVID-19 Genomics UK (COG-UK) Consortium | Lighthouse Laboratory Plymouth and Alex Alderton, Roberto Amato, Jeffrey Barrett, Sonia Goncalves, Ewan Harrison, David K. Jackson, Ian Johnston, Dominic Kwiatkowski, Cordelia Langford, John Sillitoe on behalf of the Wellcome Sanger Institute COVID-19 Surveillance Team |
| EPI_ISL_2728897 | Lighthouse Lab in Milton Keynes | Wellcome Sanger Institute for the COVID-19 Genomics UK (COG-UK) Consortium | The Lighthouse Lab in Milton Keynes and Alex Alderton, Roberto Amato, Jeffrey Barrett, Sonia Goncalves, Ewan Harrison, David K. Jackson, Ian Johnston, Dominic Kwiatkowski, Cordelia Langford, John Sillitoe on behalf of the Wellcome Sanger Institute COVID-19 Surveillance Team |
| EPI_ISL_2728898, EPI_ISL_2728899 | Lighthouse Lab in Glasgow | Wellcome Sanger Institute for the COVID-19 Genomics UK (COG-UK) Consortium | Harper VanSteenhouse, Yumi Kasai, David Gray, Carol Clugston, Anna Dominiczak and Alex Alderton, Roberto Amato, Jeffrey Barrett, Sonia Goncalves, Ewan Harrison, David K. Jackson, Ian Johnston, Dominic Kwiatkowski, Cordelia Langford, John Sillitoe on behalf of the Wellcome Sanger Institute COVID-19 Surveillance Team |
| EPI_ISL_2728900, EPI_ISL_2728901, EPI_ISL_2728902 | Lighthouse Lab in Milton Keynes | Wellcome Sanger Institute for the COVID-19 Genomics UK (COG-UK) Consortium | The Lighthouse Lab in Milton Keynes and Alex Alderton, Roberto Amato, Jeffrey Barrett, Sonia Goncalves, Ewan Harrison, David K. Jackson, Ian Johnston, Dominic Kwiatkowski, Cordelia Langford, John Sillitoe on behalf of the Wellcome Sanger Institute COVID-19 Surveillance Team |
| EPI_ISL_2728903 | Lighthouse Lab in Glasgow | Wellcome Sanger Institute for the COVID-19 Genomics UK (COG-UK) Consortium | Harper VanSteenhouse, Yumi Kasai, David Gray, Carol Clugston, Anna Dominiczak and Alex Alderton, Roberto Amato, Jeffrey Barrett, Sonia Goncalves, Ewan Harrison, David K. Jackson, Ian Johnston, Dominic Kwiatkowski, Cordelia Langford, John Sillitoe on behalf of the Wellcome Sanger Institute COVID-19 Surveillance Team |
| EPI_ISL_2728904, EPI_ISL_2728905 | Lighthouse Laboratory Plymouth | Wellcome Sanger Institute for the COVID-19 Genomics UK (COG-UK) Consortium | Lighthouse Laboratory Plymouth and Alex Alderton, Roberto Amato, Jeffrey Barrett, Sonia Goncalves, Ewan Harrison, David K. Jackson, Ian Johnston, Dominic Kwiatkowski, Cordelia Langford, John Sillitoe on behalf of the Wellcome Sanger Institute COVID-19 Surveillance Team |
| EPI_ISL_2728906 | Lighthouse Lab in Glasgow | Wellcome Sanger Institute for the COVID-19 Genomics UK (COG-UK) Consortium | Harper VanSteenhouse, Yumi Kasai, David Gray, Carol Clugston, Anna Dominiczak and Alex Alderton, Roberto Amato, Jeffrey Barrett, Sonia Goncalves, Ewan Harrison, David K. Jackson, Ian Johnston, Dominic Kwiatkowski, Cordelia Langford, John Sillitoe on behalf of the Wellcome Sanger Institute COVID-19 Surveillance Team |
| EPI_ISL_2728907 | Lighthouse Lab in Milton Keynes | Wellcome Sanger Institute for the COVID-19 Genomics UK (COG-UK) Consortium | The Lighthouse Lab in Milton Keynes and Alex Alderton, Roberto Amato, Jeffrey Barrett, Sonia Goncalves, Ewan Harrison, David K. Jackson, Ian Johnston, Dominic Kwiatkowski, Cordelia Langford, John Sillitoe on behalf of the Wellcome Sanger Institute COVID-19 Surveillance Team |
| EPI_ISL_2728909 | Lighthouse Laboratory Plymouth | Wellcome Sanger Institute for the COVID-19 Genomics UK (COG-UK) Consortium | Lighthouse Laboratory Plymouth and Alex Alderton, Roberto Amato, Jeffrey Barrett, Sonia Goncalves, Ewan Harrison, David K. Jackson, Ian Johnston, Dominic Kwiatkowski, Cordelia Langford, John Sillitoe on behalf of the Wellcome Sanger Institute COVID-19 Surveillance Team |
| EPI_ISL_2728910 | Lighthouse Lab in Glasgow | Wellcome Sanger Institute for the COVID-19 Genomics UK (COG-UK) Consortium | Harper VanSteenhouse, Yumi Kasai, David Gray, Carol Clugston, Anna Dominiczak and Alex Alderton, Roberto Amato, Jeffrey Barrett, Sonia Goncalves, Ewan Harrison, David K. Jackson, Ian Johnston, Dominic Kwiatkowski, Cordelia Langford, John Sillitoe on behalf of the Wellcome Sanger Institute COVID-19 Surveillance Team |
| EPI_ISL_2728911 | Lighthouse Lab in Milton Keynes | Wellcome Sanger Institute for the COVID-19 Genomics UK (COG-UK) Consortium | The Lighthouse Lab in Milton Keynes and Alex Alderton, Roberto Amato, Jeffrey Barrett, Sonia Goncalves, Ewan Harrison, David K. Jackson, Ian Johnston, Dominic Kwiatkowski, Cordelia Langford, John Sillitoe on behalf of the Wellcome Sanger Institute COVID-19 Surveillance Team |
| EPI_ISL_2728912 | Lighthouse Laboratory Plymouth | Wellcome Sanger Institute for the COVID-19 Genomics UK (COG-UK) Consortium | Lighthouse Laboratory Plymouth and Alex Alderton, Roberto Amato, Jeffrey Barrett, Sonia Goncalves, Ewan Harrison, David K. Jackson, Ian Johnston, Dominic Kwiatkowski, Cordelia Langford, John Sillitoe on behalf of the Wellcome Sanger Institute COVID-19 Surveillance Team |
| EPI_ISL_2728913 | Lighthouse Lab in Glasgow | Wellcome Sanger Institute for the COVID-19 Genomics UK (COG-UK) Consortium | Harper VanSteenhouse, Yumi Kasai, David Gray, Carol Clugston, Anna Dominiczak and Alex Alderton, Roberto Amato, Jeffrey Barrett, Sonia Goncalves, Ewan Harrison, David K. Jackson, Ian Johnston, Dominic Kwiatkowski, Cordelia Langford, John Sillitoe on behalf of the Wellcome Sanger Institute COVID-19 Surveillance Team |
| EPI_ISL_2728914 | Lighthouse Laboratory Plymouth | Wellcome Sanger Institute for the COVID-19 Genomics UK (COG-UK) Consortium | Lighthouse Laboratory Plymouth and Alex Alderton, Roberto Amato, Jeffrey Barrett, Sonia Goncalves, Ewan Harrison, David K. Jackson, Ian Johnston, Dominic Kwiatkowski, Cordelia Langford, John Sillitoe on behalf of the Wellcome Sanger Institute COVID-19 Surveillance Team |
| EPI_ISL_2728915 | Lighthouse Lab in Glasgow | Wellcome Sanger Institute for the COVID-19 Genomics UK (COG-UK) Consortium | Harper VanSteenhouse, Yumi Kasai, David Gray, Carol Clugston, Anna Dominiczak and Alex Alderton, Roberto Amato, Jeffrey Barrett, Sonia Goncalves, Ewan Harrison, David K. Jackson, Ian Johnston, Dominic Kwiatkowski, Cordelia Langford, John Sillitoe on behalf of the Wellcome Sanger Institute COVID-19 Surveillance Team |

| | | | |
|---|---|---|---|
| EPI_ISL_2728916 | Lighthouse Lab in Milton Keynes | Wellcome Sanger Institute for the COVID-19 Genomics UK (COG-UK) Consortium | The Lighthouse Lab in Milton Keynes and Alex Alderton, Roberto Amato, Jeffrey Barrett, Sonia Goncalves, Ewan Harrison, David K. Jackson, Ian Johnston, Dominic Kwiatkowski, Cordelia Langford, John Sillitoe on behalf of the Wellcome Sanger Institute COVID-19 Surveillance Team |
| EPI_ISL_2728917 | Lighthouse Laboratory Plymouth | Wellcome Sanger Institute for the COVID-19 Genomics UK (COG-UK) Consortium | Lighthouse Laboratory Plymouth and Alex Alderton, Roberto Amato, Jeffrey Barrett, Sonia Goncalves, Ewan Harrison, David K. Jackson, Ian Johnston, Dominic Kwiatkowski, Cordelia Langford, John Sillitoe on behalf of the Wellcome Sanger Institute COVID-19 Surveillance Team |
| EPI_ISL_2728918 | Lighthouse Lab in Glasgow | Wellcome Sanger Institute for the COVID-19 Genomics UK (COG-UK) Consortium | Harper VanSteenhouse, Yumi Kasai, David Gray, Carol Clugston, Anna Dominiczak and Alex Alderton, Roberto Amato, Jeffrey Barrett, Sonia Goncalves, Ewan Harrison, David K. Jackson, Ian Johnston, Dominic Kwiatkowski, Cordelia Langford, John Sillitoe on behalf of the Wellcome Sanger Institute COVID-19 Surveillance Team |
| EPI_ISL_2728919 | Lighthouse Lab in Milton Keynes | Wellcome Sanger Institute for the COVID-19 Genomics UK (COG-UK) Consortium | The Lighthouse Lab in Milton Keynes and Alex Alderton, Roberto Amato, Jeffrey Barrett, Sonia Goncalves, Ewan Harrison, David K. Jackson, Ian Johnston, Dominic Kwiatkowski, Cordelia Langford, John Sillitoe on behalf of the Wellcome Sanger Institute COVID-19 Surveillance Team |
| EPI_ISL_2728920, EPI_ISL_2728921 | Lighthouse Lab in Glasgow | Wellcome Sanger Institute for the COVID-19 Genomics UK (COG-UK) Consortium | Harper VanSteenhouse, Yumi Kasai, David Gray, Carol Clugston, Anna Dominiczak and Alex Alderton, Roberto Amato, Jeffrey Barrett, Sonia Goncalves, Ewan Harrison, David K. Jackson, Ian Johnston, Dominic Kwiatkowski, Cordelia Langford, John Sillitoe on behalf of the Wellcome Sanger Institute COVID-19 Surveillance Team |
| EPI_ISL_2728922 | Lighthouse Laboratory Plymouth | Wellcome Sanger Institute for the COVID-19 Genomics UK (COG-UK) Consortium | Lighthouse Laboratory Plymouth and Alex Alderton, Roberto Amato, Jeffrey Barrett, Sonia Goncalves, Ewan Harrison, David K. Jackson, Ian Johnston, Dominic Kwiatkowski, Cordelia Langford, John Sillitoe on behalf of the Wellcome Sanger Institute COVID-19 Surveillance Team |
| EPI_ISL_2728923, EPI_ISL_2728924 | Lighthouse Lab in Milton Keynes | Wellcome Sanger Institute for the COVID-19 Genomics UK (COG-UK) Consortium | The Lighthouse Lab in Milton Keynes and Alex Alderton, Roberto Amato, Jeffrey Barrett, Sonia Goncalves, Ewan Harrison, David K. Jackson, Ian Johnston, Dominic Kwiatkowski, Cordelia Langford, John Sillitoe on behalf of the Wellcome Sanger Institute COVID-19 Surveillance Team |
| EPI_ISL_2728925, EPI_ISL_2728926 | Lighthouse Lab in Glasgow | Wellcome Sanger Institute for the COVID-19 Genomics UK (COG-UK) Consortium | Harper VanSteenhouse, Yumi Kasai, David Gray, Carol Clugston, Anna Dominiczak and Alex Alderton, Roberto Amato, Jeffrey Barrett, Sonia Goncalves, Ewan Harrison, David K. Jackson, Ian Johnston, Dominic Kwiatkowski, Cordelia Langford, John Sillitoe on behalf of the Wellcome Sanger Institute COVID-19 Surveillance Team |
| EPI_ISL_2728927 | Lighthouse Laboratory Plymouth | Wellcome Sanger Institute for the COVID-19 Genomics UK (COG-UK) Consortium | Lighthouse Laboratory Plymouth and Alex Alderton, Roberto Amato, Jeffrey Barrett, Sonia Goncalves, Ewan Harrison, David K. Jackson, Ian Johnston, Dominic Kwiatkowski, Cordelia Langford, John Sillitoe on behalf of the Wellcome Sanger Institute COVID-19 Surveillance Team |
| EPI_ISL_2728928 | Lighthouse Lab in Glasgow | Wellcome Sanger Institute for the COVID-19 Genomics UK (COG-UK) Consortium | Harper VanSteenhouse, Yumi Kasai, David Gray, Carol Clugston, Anna Dominiczak and Alex Alderton, Roberto Amato, Jeffrey Barrett, Sonia Goncalves, Ewan Harrison, David K. Jackson, Ian Johnston, Dominic Kwiatkowski, Cordelia Langford, John Sillitoe on behalf of the Wellcome Sanger Institute COVID-19 Surveillance Team |
| EPI_ISL_2728929, EPI_ISL_2728930 | Lighthouse Lab in Milton Keynes | Wellcome Sanger Institute for the COVID-19 Genomics UK (COG-UK) Consortium | The Lighthouse Lab in Milton Keynes and Alex Alderton, Roberto Amato, Jeffrey Barrett, Sonia Goncalves, Ewan Harrison, David K. Jackson, Ian Johnston, Dominic Kwiatkowski, Cordelia Langford, John Sillitoe on behalf of the Wellcome Sanger Institute COVID-19 Surveillance Team |
| EPI_ISL_2728931 | Lighthouse Lab in Glasgow | Wellcome Sanger Institute for the COVID-19 Genomics UK (COG-UK) Consortium | Harper VanSteenhouse, Yumi Kasai, David Gray, Carol Clugston, Anna Dominiczak and Alex Alderton, Roberto Amato, Jeffrey Barrett, Sonia Goncalves, Ewan Harrison, David K. Jackson, Ian Johnston, Dominic Kwiatkowski, Cordelia Langford, John Sillitoe on behalf of the Wellcome Sanger Institute COVID-19 Surveillance Team |
| EPI_ISL_2728932 | Lighthouse Lab in Milton Keynes | Wellcome Sanger Institute for the COVID-19 Genomics UK (COG-UK) Consortium | The Lighthouse Lab in Milton Keynes and Alex Alderton, Roberto Amato, Jeffrey Barrett, Sonia Goncalves, Ewan Harrison, David K. Jackson, Ian Johnston, Dominic Kwiatkowski, Cordelia Langford, John Sillitoe on behalf of the Wellcome Sanger Institute COVID-19 Surveillance Team |
| EPI_ISL_2728933 | Lighthouse Laboratory Plymouth | Wellcome Sanger Institute for the COVID-19 Genomics UK (COG-UK) Consortium | Lighthouse Laboratory Plymouth and Alex Alderton, Roberto Amato, Jeffrey Barrett, Sonia Goncalves, Ewan Harrison, David K. Jackson, Ian Johnston, Dominic Kwiatkowski, Cordelia Langford, John Sillitoe on behalf of the Wellcome Sanger Institute COVID-19 Surveillance Team |
| EPI_ISL_2728934, EPI_ISL_2728935, EPI_ISL_2728936 | Lighthouse Lab in Glasgow | Wellcome Sanger Institute for the COVID-19 Genomics UK (COG-UK) Consortium | Harper VanSteenhouse, Yumi Kasai, David Gray, Carol Clugston, Anna Dominiczak and Alex Alderton, Roberto Amato, Jeffrey Barrett, Sonia Goncalves, Ewan Harrison, David K. Jackson, Ian Johnston, Dominic Kwiatkowski, Cordelia Langford, John Sillitoe on behalf of the Wellcome Sanger Institute COVID-19 Surveillance Team |
| EPI_ISL_2728937, EPI_ISL_2728938 | Lighthouse Laboratory Plymouth | Wellcome Sanger Institute for the COVID-19 Genomics UK (COG-UK) Consortium | Lighthouse Laboratory Plymouth and Alex Alderton, Roberto Amato, Jeffrey Barrett, Sonia Goncalves, Ewan Harrison, David K. Jackson, Ian Johnston, Dominic Kwiatkowski, Cordelia Langford, John Sillitoe on behalf of the Wellcome Sanger Institute COVID-19 Surveillance Team |
| EPI_ISL_2728939, EPI_ISL_2728940, EPI_ISL_2728941 | Lighthouse Lab in Glasgow | Wellcome Sanger Institute for the COVID-19 Genomics UK (COG-UK) Consortium | Harper VanSteenhouse, Yumi Kasai, David Gray, Carol Clugston, Anna Dominiczak and Alex Alderton, Roberto Amato, Jeffrey Barrett, Sonia Goncalves, Ewan Harrison, David K. Jackson, Ian Johnston, Dominic Kwiatkowski, Cordelia Langford, John Sillitoe on behalf of the Wellcome Sanger Institute COVID-19 Surveillance Team |
| EPI_ISL_2728942 | Lighthouse Laboratory Plymouth | Wellcome Sanger Institute for the COVID-19 Genomics UK (COG-UK) Consortium | Lighthouse Laboratory Plymouth and Alex Alderton, Roberto Amato, Jeffrey Barrett, Sonia Goncalves, Ewan Harrison, David K. Jackson, Ian Johnston, Dominic Kwiatkowski, Cordelia Langford, John Sillitoe on behalf of the Wellcome Sanger Institute COVID-19 Surveillance Team |
| EPI_ISL_2728943, EPI_ISL_2728944, EPI_ISL_2728945 | Lighthouse Lab in Glasgow | Wellcome Sanger Institute for the COVID-19 Genomics UK (COG-UK) Consortium | Harper VanSteenhouse, Yumi Kasai, David Gray, Carol Clugston, Anna Dominiczak and Alex Alderton, Roberto Amato, Jeffrey Barrett, Sonia Goncalves, Ewan Harrison, David K. Jackson, Ian Johnston, Dominic Kwiatkowski, Cordelia Langford, John Sillitoe on behalf of the Wellcome Sanger Institute COVID-19 Surveillance Team |
| EPI_ISL_2728946 | Lighthouse Lab in Milton Keynes | Wellcome Sanger Institute for the COVID-19 Genomics UK (COG-UK) Consortium | The Lighthouse Lab in Milton Keynes and Alex Alderton, Roberto Amato, Jeffrey Barrett, Sonia Goncalves, Ewan Harrison, David K. Jackson, Ian Johnston, Dominic Kwiatkowski, Cordelia Langford, John Sillitoe on behalf of the Wellcome Sanger Institute COVID-19 Surveillance Team |
| EPI_ISL_2728947 | Lighthouse Laboratory Plymouth | Wellcome Sanger Institute for the COVID-19 Genomics UK (COG-UK) Consortium | Lighthouse Laboratory Plymouth and Alex Alderton, Roberto Amato, Jeffrey Barrett, Sonia Goncalves, Ewan Harrison, David K. Jackson, Ian Johnston, Dominic Kwiatkowski, Cordelia Langford, John Sillitoe on behalf of the Wellcome Sanger Institute COVID-19 Surveillance Team |
| EPI_ISL_2728948 | Lighthouse Lab in Glasgow | Wellcome Sanger Institute for the COVID-19 Genomics UK (COG-UK) Consortium | Harper VanSteenhouse, Yumi Kasai, David Gray, Carol Clugston, Anna Dominiczak and Alex Alderton, Roberto Amato, Jeffrey Barrett, Sonia Goncalves, Ewan Harrison, David K. Jackson, Ian Johnston, Dominic Kwiatkowski, Cordelia Langford, John Sillitoe on behalf of the Wellcome Sanger Institute COVID-19 Surveillance Team |
| EPI_ISL_2728949 | Lighthouse Laboratory Plymouth | Wellcome Sanger Institute for the COVID-19 Genomics UK (COG-UK) Consortium | Lighthouse Laboratory Plymouth and Alex Alderton, Roberto Amato, Jeffrey Barrett, Sonia Goncalves, Ewan Harrison, David K. Jackson, Ian Johnston, Dominic Kwiatkowski, Cordelia Langford, John Sillitoe on behalf of the Wellcome Sanger Institute COVID-19 Surveillance Team |
| EPI_ISL_2728950, EPI_ISL_2728951, EPI_ISL_2728952, EPI_ISL_2728953, EPI_ISL_2728954 | Lighthouse Lab in Glasgow | Wellcome Sanger Institute for the COVID-19 Genomics UK (COG-UK) Consortium | Harper VanSteenhouse, Yumi Kasai, David Gray, Carol Clugston, Anna Dominiczak and Alex Alderton, Roberto Amato, Jeffrey Barrett, Sonia Goncalves, Ewan Harrison, David K. Jackson, Ian Johnston, Dominic Kwiatkowski, Cordelia Langford, John Sillitoe on behalf of the Wellcome Sanger Institute COVID-19 Surveillance Team |
| EPI_ISL_2728955 | Lighthouse Laboratory Plymouth | Wellcome Sanger Institute for the COVID-19 Genomics UK (COG-UK) Consortium | Lighthouse Laboratory Plymouth and Alex Alderton, Roberto Amato, Jeffrey Barrett, Sonia Goncalves, Ewan Harrison, David K. Jackson, Ian Johnston, Dominic Kwiatkowski, Cordelia Langford, John Sillitoe on behalf of the Wellcome Sanger Institute COVID-19 Surveillance Team |
| EPI_ISL_2728956, EPI_ISL_2728957 | Lighthouse Lab in Milton Keynes | Wellcome Sanger Institute for the COVID-19 Genomics UK (COG-UK) Consortium | The Lighthouse Lab in Milton Keynes and Alex Alderton, Roberto Amato, Jeffrey Barrett, Sonia Goncalves, Ewan Harrison, David K. Jackson, Ian Johnston, Dominic Kwiatkowski, Cordelia Langford, John Sillitoe on behalf of the Wellcome Sanger Institute COVID-19 Surveillance Team |
| EPI_ISL_2728958 | Lighthouse Lab in Glasgow | Wellcome Sanger Institute for the COVID-19 Genomics UK (COG-UK) Consortium | Harper VanSteenhouse, Yumi Kasai, David Gray, Carol Clugston, Anna Dominiczak and Alex Alderton, Roberto Amato, Jeffrey Barrett, Sonia Goncalves, Ewan Harrison, David K. Jackson, Ian Johnston, Dominic Kwiatkowski, Cordelia Langford, John Sillitoe on behalf of the Wellcome Sanger Institute COVID-19 Surveillance Team |
| EPI_ISL_2728959 | Lighthouse Lab in Milton Keynes | Wellcome Sanger Institute for the COVID-19 Genomics UK (COG-UK) Consortium | The Lighthouse Lab in Milton Keynes and Alex Alderton, Roberto Amato, Jeffrey Barrett, Sonia Goncalves, Ewan Harrison, David K. Jackson, Ian Johnston, Dominic Kwiatkowski, Cordelia Langford, John Sillitoe on behalf of the Wellcome Sanger Institute COVID-19 Surveillance Team |
| EPI_ISL_2728960, EPI_ISL_2728961, EPI_ISL_2728962 | Lighthouse Lab in Glasgow | Wellcome Sanger Institute for the COVID-19 Genomics UK (COG-UK) Consortium | Harper VanSteenhouse, Yumi Kasai, David Gray, Carol Clugston, Anna Dominiczak and Alex Alderton, Roberto Amato, Jeffrey Barrett, Sonia Goncalves, Ewan Harrison, David K. Jackson, Ian Johnston, Dominic Kwiatkowski, Cordelia Langford, John Sillitoe on behalf of the Wellcome Sanger Institute COVID-19 Surveillance Team |
| EPI_ISL_2728963 | Lighthouse Laboratory Plymouth | Wellcome Sanger Institute for the COVID-19 Genomics UK (COG-UK) Consortium | Lighthouse Laboratory Plymouth and Alex Alderton, Roberto Amato, Jeffrey Barrett, Sonia Goncalves, Ewan Harrison, David K. Jackson, Ian Johnston, Dominic Kwiatkowski, Cordelia Langford, John Sillitoe on behalf of the Wellcome Sanger Institute COVID-19 Surveillance Team |
| EPI_ISL_2728964, EPI_ISL_2728965, EPI_ISL_2728966, EPI_ISL_2728967 | Lighthouse Lab in Glasgow | Wellcome Sanger Institute for the COVID-19 Genomics UK (COG-UK) Consortium | Harper VanSteenhouse, Yumi Kasai, David Gray, Carol Clugston, Anna Dominiczak and Alex Alderton, Roberto Amato, Jeffrey Barrett, Sonia Goncalves, Ewan Harrison, David K. Jackson, Ian Johnston, Dominic Kwiatkowski, Cordelia Langford, John Sillitoe on behalf of the Wellcome Sanger Institute COVID-19 Surveillance Team |
| EPI_ISL_2728968 | Lighthouse Laboratory Plymouth | Wellcome Sanger Institute for the COVID-19 Genomics UK (COG-UK) Consortium | Lighthouse Laboratory Plymouth and Alex Alderton, Roberto Amato, Jeffrey Barrett, Sonia Goncalves, Ewan Harrison, David K. Jackson, Ian Johnston, Dominic Kwiatkowski, Cordelia Langford, John Sillitoe on behalf of the Wellcome Sanger Institute COVID-19 Surveillance Team |
| EPI_ISL_2728969 | Lighthouse Lab in Glasgow | Wellcome Sanger Institute for the COVID-19 Genomics UK (COG-UK) Consortium | Harper VanSteenhouse, Yumi Kasai, David Gray, Carol Clugston, Anna Dominiczak and Alex Alderton, Roberto Amato, Jeffrey Barrett, Sonia Goncalves, Ewan Harrison, David K. Jackson, Ian Johnston, Dominic Kwiatkowski, Cordelia Langford, John Sillitoe on behalf of the Wellcome Sanger Institute COVID-19 Surveillance Team |
| EPI_ISL_2728970 | Lighthouse Lab in Milton Keynes | Wellcome Sanger Institute for the COVID-19 Genomics UK (COG-UK) Consortium | The Lighthouse Lab in Milton Keynes and Alex Alderton, Roberto Amato, Jeffrey Barrett, Sonia Goncalves, Ewan Harrison, David K. Jackson, Ian Johnston, Dominic Kwiatkowski, Cordelia Langford, John Sillitoe on behalf of the Wellcome Sanger Institute COVID-19 Surveillance Team |
| EPI_ISL_2728971, EPI_ISL_2728972, EPI_ISL_2728973, EPI_ISL_2728974 | Lighthouse Lab in Glasgow | Wellcome Sanger Institute for the COVID-19 Genomics UK (COG-UK) Consortium | Harper VanSteenhouse, Yumi Kasai, David Gray, Carol Clugston, Anna Dominiczak and Alex Alderton, Roberto Amato, Jeffrey Barrett, Sonia Goncalves, Ewan Harrison, David K. Jackson, Ian Johnston, Dominic Kwiatkowski, Cordelia Langford, John Sillitoe on behalf of the Wellcome Sanger Institute COVID-19 Surveillance Team |
| EPI_ISL_2728975 | Lighthouse Lab in Milton Keynes | Wellcome Sanger Institute for the COVID-19 Genomics UK (COG-UK) Consortium | The Lighthouse Lab in Milton Keynes and Alex Alderton, Roberto Amato, Jeffrey Barrett, Sonia Goncalves, Ewan Harrison, David K. Jackson, Ian Johnston, Dominic Kwiatkowski, Cordelia Langford, John Sillitoe on behalf of the Wellcome Sanger Institute COVID-19 Surveillance Team |
| EPI_ISL_2728976, EPI_ISL_2728977, EPI_ISL_2728978 | Lighthouse Lab in Glasgow | Wellcome Sanger Institute for the COVID-19 Genomics UK (COG-UK) Consortium | Harper VanSteenhouse, Yumi Kasai, David Gray, Carol Clugston, Anna Dominiczak and Alex Alderton, Roberto Amato, Jeffrey Barrett, Sonia Goncalves, Ewan Harrison, David K. Jackson, Ian Johnston, Dominic Kwiatkowski, Cordelia Langford, John Sillitoe on behalf of the Wellcome Sanger Institute COVID-19 Surveillance Team |
| EPI_ISL_2728979, EPI_ISL_2728980 | Lighthouse Lab in Milton Keynes | Wellcome Sanger Institute for the COVID-19 Genomics UK (COG-UK) Consortium | The Lighthouse Lab in Milton Keynes and Alex Alderton, Roberto Amato, Jeffrey Barrett, Sonia Goncalves, Ewan Harrison, David K. Jackson, Ian Johnston, Dominic Kwiatkowski, Cordelia Langford, John Sillitoe on behalf of the Wellcome Sanger Institute COVID-19 Surveillance Team |
| EPI_ISL_2728981, EPI_ISL_2728982, EPI_ISL_2728983, EPI_ISL_2728984 | Lighthouse Lab in Glasgow | Wellcome Sanger Institute for the COVID-19 Genomics UK (COG-UK) Consortium | Harper VanSteenhouse, Yumi Kasai, David Gray, Carol Clugston, Anna Dominiczak and Alex Alderton, Roberto Amato, Jeffrey Barrett, Sonia Goncalves, Ewan Harrison, David K. Jackson, Ian Johnston, Dominic Kwiatkowski, Cordelia Langford, John Sillitoe on behalf of the Wellcome Sanger Institute COVID-19 Surveillance Team |
| EPI_ISL_2728985 | Lighthouse Lab in Milton Keynes | Wellcome Sanger Institute for the COVID-19 Genomics UK (COG-UK) | The Lighthouse Lab in Milton Keynes and Alex Alderton, Roberto Amato, Jeffrey Barrett, Sonia Goncalves, Ewan Harrison, David K. Jackson, Ian Johnston, Dominic Kwiatkowski, Cordelia Langford, John Sillitoe on behalf of the Wellcome Sanger |

| | | | |
|---|---|---|---|
| | | Consortium | Institute COVID-19 Surveillance Team |
| EPI_ISL_2728986 | Lighthouse Lab in Glasgow | Wellcome Sanger Institute for the COVID-19 Genomics UK (COG-UK) Consortium | Harper VanSteenhouse, Yumi Kasai, David Gray, Carol Clugston, Anna Dominiczak and Alex Alderton, Roberto Amato, Jeffrey Barrett, Sonia Goncalves, Ewan Harrison, David K. Jackson, Ian Johnston, Dominic Kwiatkowski, Cordelia Langford, John Sillitoe on behalf of the Wellcome Sanger Institute COVID-19 Surveillance Team |
| EPI_ISL_2728987 | Lighthouse Lab in Milton Keynes | Wellcome Sanger Institute for the COVID-19 Genomics UK (COG-UK) Consortium | The Lighthouse Lab in Milton Keynes and Alex Alderton, Roberto Amato, Jeffrey Barrett, Sonia Goncalves, Ewan Harrison, David K. Jackson, Ian Johnston, Dominic Kwiatkowski, Cordelia Langford, John Sillitoe on behalf of the Wellcome Sanger Institute COVID-19 Surveillance Team |
| EPI_ISL_2728988, EPI_ISL_2728989 | Lighthouse Laboratory Plymouth | Wellcome Sanger Institute for the COVID-19 Genomics UK (COG-UK) Consortium | Lighthouse Laboratory Plymouth and Alex Alderton, Roberto Amato, Jeffrey Barrett, Sonia Goncalves, Ewan Harrison, David K. Jackson, Ian Johnston, Dominic Kwiatkowski, Cordelia Langford, John Sillitoe on behalf of the Wellcome Sanger Institute COVID-19 Surveillance Team |
| EPI_ISL_2728990 | Lighthouse Lab in Milton Keynes | Wellcome Sanger Institute for the COVID-19 Genomics UK (COG-UK) Consortium | The Lighthouse Lab in Milton Keynes and Alex Alderton, Roberto Amato, Jeffrey Barrett, Sonia Goncalves, Ewan Harrison, David K. Jackson, Ian Johnston, Dominic Kwiatkowski, Cordelia Langford, John Sillitoe on behalf of the Wellcome Sanger Institute COVID-19 Surveillance Team |
| EPI_ISL_2728991 | Lighthouse Laboratory Plymouth | Wellcome Sanger Institute for the COVID-19 Genomics UK (COG-UK) Consortium | Lighthouse Laboratory Plymouth and Alex Alderton, Roberto Amato, Jeffrey Barrett, Sonia Goncalves, Ewan Harrison, David K. Jackson, Ian Johnston, Dominic Kwiatkowski, Cordelia Langford, John Sillitoe on behalf of the Wellcome Sanger Institute COVID-19 Surveillance Team |
| EPI_ISL_2728992, EPI_ISL_2728993 | Lighthouse Lab in Glasgow | Wellcome Sanger Institute for the COVID-19 Genomics UK (COG-UK) Consortium | Harper VanSteenhouse, Yumi Kasai, David Gray, Carol Clugston, Anna Dominiczak and Alex Alderton, Roberto Amato, Jeffrey Barrett, Sonia Goncalves, Ewan Harrison, David K. Jackson, Ian Johnston, Dominic Kwiatkowski, Cordelia Langford, John Sillitoe on behalf of the Wellcome Sanger Institute COVID-19 Surveillance Team |
| EPI_ISL_2728995 | Lighthouse Laboratory Plymouth | Wellcome Sanger Institute for the COVID-19 Genomics UK (COG-UK) Consortium | Lighthouse Laboratory Plymouth and Alex Alderton, Roberto Amato, Jeffrey Barrett, Sonia Goncalves, Ewan Harrison, David K. Jackson, Ian Johnston, Dominic Kwiatkowski, Cordelia Langford, John Sillitoe on behalf of the Wellcome Sanger Institute COVID-19 Surveillance Team |
| EPI_ISL_2728996 | Lighthouse Lab in Milton Keynes | Wellcome Sanger Institute for the COVID-19 Genomics UK (COG-UK) Consortium | The Lighthouse Lab in Milton Keynes and Alex Alderton, Roberto Amato, Jeffrey Barrett, Sonia Goncalves, Ewan Harrison, David K. Jackson, Ian Johnston, Dominic Kwiatkowski, Cordelia Langford, John Sillitoe on behalf of the Wellcome Sanger Institute COVID-19 Surveillance Team |
| EPI_ISL_2728997, EPI_ISL_2728998 | Lighthouse Lab in Glasgow | Wellcome Sanger Institute for the COVID-19 Genomics UK (COG-UK) Consortium | Harper VanSteenhouse, Yumi Kasai, David Gray, Carol Clugston, Anna Dominiczak and Alex Alderton, Roberto Amato, Jeffrey Barrett, Sonia Goncalves, Ewan Harrison, David K. Jackson, Ian Johnston, Dominic Kwiatkowski, Cordelia Langford, John Sillitoe on behalf of the Wellcome Sanger Institute COVID-19 Surveillance Team |
| EPI_ISL_2728999 | Lighthouse Laboratory Plymouth | Wellcome Sanger Institute for the COVID-19 Genomics UK (COG-UK) Consortium | Lighthouse Laboratory Plymouth and Alex Alderton, Roberto Amato, Jeffrey Barrett, Sonia Goncalves, Ewan Harrison, David K. Jackson, Ian Johnston, Dominic Kwiatkowski, Cordelia Langford, John Sillitoe on behalf of the Wellcome Sanger Institute COVID-19 Surveillance Team |
| EPI_ISL_2729000 | Lighthouse Lab in Glasgow | Wellcome Sanger Institute for the COVID-19 Genomics UK (COG-UK) Consortium | Harper VanSteenhouse, Yumi Kasai, David Gray, Carol Clugston, Anna Dominiczak and Alex Alderton, Roberto Amato, Jeffrey Barrett, Sonia Goncalves, Ewan Harrison, David K. Jackson, Ian Johnston, Dominic Kwiatkowski, Cordelia Langford, John Sillitoe on behalf of the Wellcome Sanger Institute COVID-19 Surveillance Team |
| EPI_ISL_2729001 | Lighthouse Laboratory Plymouth | Wellcome Sanger Institute for the COVID-19 Genomics UK (COG-UK) Consortium | Lighthouse Laboratory Plymouth and Alex Alderton, Roberto Amato, Jeffrey Barrett, Sonia Goncalves, Ewan Harrison, David K. Jackson, Ian Johnston, Dominic Kwiatkowski, Cordelia Langford, John Sillitoe on behalf of the Wellcome Sanger Institute COVID-19 Surveillance Team |
| EPI_ISL_2729002, EPI_ISL_2729003 | Lighthouse Lab in Glasgow | Wellcome Sanger Institute for the COVID-19 Genomics UK (COG-UK) Consortium | Harper VanSteenhouse, Yumi Kasai, David Gray, Carol Clugston, Anna Dominiczak and Alex Alderton, Roberto Amato, Jeffrey Barrett, Sonia Goncalves, Ewan Harrison, David K. Jackson, Ian Johnston, Dominic Kwiatkowski, Cordelia Langford, John Sillitoe on behalf of the Wellcome Sanger Institute COVID-19 Surveillance Team |
| EPI_ISL_2729004 | Lighthouse Lab in Milton Keynes | Wellcome Sanger Institute for the COVID-19 Genomics UK (COG-UK) Consortium | The Lighthouse Lab in Milton Keynes and Alex Alderton, Roberto Amato, Jeffrey Barrett, Sonia Goncalves, Ewan Harrison, David K. Jackson, Ian Johnston, Dominic Kwiatkowski, Cordelia Langford, John Sillitoe on behalf of the Wellcome Sanger Institute COVID-19 Surveillance Team |
| EPI_ISL_2729006, EPI_ISL_2729007, EPI_ISL_2729008, EPI_ISL_2729009 | Lighthouse Lab in Glasgow | Wellcome Sanger Institute for the COVID-19 Genomics UK (COG-UK) Consortium | Harper VanSteenhouse, Yumi Kasai, David Gray, Carol Clugston, Anna Dominiczak and Alex Alderton, Roberto Amato, Jeffrey Barrett, Sonia Goncalves, Ewan Harrison, David K. Jackson, Ian Johnston, Dominic Kwiatkowski, Cordelia Langford, John Sillitoe on behalf of the Wellcome Sanger Institute COVID-19 Surveillance Team |
| EPI_ISL_2729010 | Lighthouse Laboratory Plymouth | Wellcome Sanger Institute for the COVID-19 Genomics UK (COG-UK) Consortium | Lighthouse Laboratory Plymouth and Alex Alderton, Roberto Amato, Jeffrey Barrett, Sonia Goncalves, Ewan Harrison, David K. Jackson, Ian Johnston, Dominic Kwiatkowski, Cordelia Langford, John Sillitoe on behalf of the Wellcome Sanger Institute COVID-19 Surveillance Team |
| EPI_ISL_2729011, EPI_ISL_2729012, EPI_ISL_2729013 | Lighthouse Lab in Glasgow | Wellcome Sanger Institute for the COVID-19 Genomics UK (COG-UK) Consortium | Harper VanSteenhouse, Yumi Kasai, David Gray, Carol Clugston, Anna Dominiczak and Alex Alderton, Roberto Amato, Jeffrey Barrett, Sonia Goncalves, Ewan Harrison, David K. Jackson, Ian Johnston, Dominic Kwiatkowski, Cordelia Langford, John Sillitoe on behalf of the Wellcome Sanger Institute COVID-19 Surveillance Team |
| EPI_ISL_2729014 | Lighthouse Laboratory Plymouth | Wellcome Sanger Institute for the COVID-19 Genomics UK (COG-UK) Consortium | Lighthouse Laboratory Plymouth and Alex Alderton, Roberto Amato, Jeffrey Barrett, Sonia Goncalves, Ewan Harrison, David K. Jackson, Ian Johnston, Dominic Kwiatkowski, Cordelia Langford, John Sillitoe on behalf of the Wellcome Sanger Institute COVID-19 Surveillance Team |
| EPI_ISL_2729015 | Lighthouse Lab in Glasgow | Wellcome Sanger Institute for the COVID-19 Genomics UK (COG-UK) Consortium | Harper VanSteenhouse, Yumi Kasai, David Gray, Carol Clugston, Anna Dominiczak and Alex Alderton, Roberto Amato, Jeffrey Barrett, Sonia Goncalves, Ewan Harrison, David K. Jackson, Ian Johnston, Dominic Kwiatkowski, Cordelia Langford, John Sillitoe on behalf of the Wellcome Sanger Institute COVID-19 Surveillance Team |
| EPI_ISL_2729016, EPI_ISL_2729017 | Lighthouse Laboratory Plymouth | Wellcome Sanger Institute for the COVID-19 Genomics UK (COG-UK) Consortium | Lighthouse Laboratory Plymouth and Alex Alderton, Roberto Amato, Jeffrey Barrett, Sonia Goncalves, Ewan Harrison, David K. Jackson, Ian Johnston, Dominic Kwiatkowski, Cordelia Langford, John Sillitoe on behalf of the Wellcome Sanger Institute COVID-19 Surveillance Team |
| EPI_ISL_2729018, EPI_ISL_2729019 | Lighthouse Lab in Glasgow | Wellcome Sanger Institute for the COVID-19 Genomics UK (COG-UK) Consortium | Harper VanSteenhouse, Yumi Kasai, David Gray, Carol Clugston, Anna Dominiczak and Alex Alderton, Roberto Amato, Jeffrey Barrett, Sonia Goncalves, Ewan Harrison, David K. Jackson, Ian Johnston, Dominic Kwiatkowski, Cordelia Langford, John Sillitoe on behalf of the Wellcome Sanger Institute COVID-19 Surveillance Team |
| EPI_ISL_2729020, EPI_ISL_2729021, EPI_ISL_2729022 | Lighthouse Lab in Milton Keynes | Wellcome Sanger Institute for the COVID-19 Genomics UK (COG-UK) Consortium | The Lighthouse Lab in Milton Keynes and Alex Alderton, Roberto Amato, Jeffrey Barrett, Sonia Goncalves, Ewan Harrison, David K. Jackson, Ian Johnston, Dominic Kwiatkowski, Cordelia Langford, John Sillitoe on behalf of the Wellcome Sanger Institute COVID-19 Surveillance Team |
| EPI_ISL_2729023, EPI_ISL_2729024, EPI_ISL_2729025, EPI_ISL_2729026, EPI_ISL_2729027, EPI_ISL_2729028 | Lighthouse Lab in Glasgow | Wellcome Sanger Institute for the COVID-19 Genomics UK (COG-UK) Consortium | Harper VanSteenhouse, Yumi Kasai, David Gray, Carol Clugston, Anna Dominiczak and Alex Alderton, Roberto Amato, Jeffrey Barrett, Sonia Goncalves, Ewan Harrison, David K. Jackson, Ian Johnston, Dominic Kwiatkowski, Cordelia Langford, John Sillitoe on behalf of the Wellcome Sanger Institute COVID-19 Surveillance Team |
| EPI_ISL_2729029 | Lighthouse Laboratory Plymouth | Wellcome Sanger Institute for the COVID-19 Genomics UK (COG-UK) Consortium | Lighthouse Laboratory Plymouth and Alex Alderton, Roberto Amato, Jeffrey Barrett, Sonia Goncalves, Ewan Harrison, David K. Jackson, Ian Johnston, Dominic Kwiatkowski, Cordelia Langford, John Sillitoe on behalf of the Wellcome Sanger Institute COVID-19 Surveillance Team |
| EPI_ISL_2729030 | Lighthouse Lab in Milton Keynes | Wellcome Sanger Institute for the COVID-19 Genomics UK (COG-UK) Consortium | The Lighthouse Lab in Milton Keynes and Alex Alderton, Roberto Amato, Jeffrey Barrett, Sonia Goncalves, Ewan Harrison, David K. Jackson, Ian Johnston, Dominic Kwiatkowski, Cordelia Langford, John Sillitoe on behalf of the Wellcome Sanger Institute COVID-19 Surveillance Team |
| EPI_ISL_2729031 | Lighthouse Laboratory Plymouth | Wellcome Sanger Institute for the COVID-19 Genomics UK (COG-UK) Consortium | Lighthouse Laboratory Plymouth and Alex Alderton, Roberto Amato, Jeffrey Barrett, Sonia Goncalves, Ewan Harrison, David K. Jackson, Ian Johnston, Dominic Kwiatkowski, Cordelia Langford, John Sillitoe on behalf of the Wellcome Sanger Institute COVID-19 Surveillance Team |
| EPI_ISL_2729032 | Lighthouse Lab in Milton Keynes | Wellcome Sanger Institute for the COVID-19 Genomics UK (COG-UK) Consortium | The Lighthouse Lab in Milton Keynes and Alex Alderton, Roberto Amato, Jeffrey Barrett, Sonia Goncalves, Ewan Harrison, David K. Jackson, Ian Johnston, Dominic Kwiatkowski, Cordelia Langford, John Sillitoe on behalf of the Wellcome Sanger Institute COVID-19 Surveillance Team |
| EPI_ISL_2729033, EPI_ISL_2729034, EPI_ISL_2729035 | Lighthouse Lab in Glasgow | Wellcome Sanger Institute for the COVID-19 Genomics UK (COG-UK) Consortium | Harper VanSteenhouse, Yumi Kasai, David Gray, Carol Clugston, Anna Dominiczak and Alex Alderton, Roberto Amato, Jeffrey Barrett, Sonia Goncalves, Ewan Harrison, David K. Jackson, Ian Johnston, Dominic Kwiatkowski, Cordelia Langford, John Sillitoe on behalf of the Wellcome Sanger Institute COVID-19 Surveillance Team |
| EPI_ISL_2729036 | Lighthouse Laboratory Plymouth | Wellcome Sanger Institute for the COVID-19 Genomics UK (COG-UK) Consortium | Lighthouse Laboratory Plymouth and Alex Alderton, Roberto Amato, Jeffrey Barrett, Sonia Goncalves, Ewan Harrison, David K. Jackson, Ian Johnston, Dominic Kwiatkowski, Cordelia Langford, John Sillitoe on behalf of the Wellcome Sanger Institute COVID-19 Surveillance Team |
| EPI_ISL_2729038 | Lighthouse Lab in Glasgow | Wellcome Sanger Institute for the COVID-19 Genomics UK (COG-UK) Consortium | Harper VanSteenhouse, Yumi Kasai, David Gray, Carol Clugston, Anna Dominiczak and Alex Alderton, Roberto Amato, Jeffrey Barrett, Sonia Goncalves, Ewan Harrison, David K. Jackson, Ian Johnston, Dominic Kwiatkowski, Cordelia Langford, John Sillitoe on behalf of the Wellcome Sanger Institute COVID-19 Surveillance Team |
| EPI_ISL_2729039 | Lighthouse Lab in Milton Keynes | Wellcome Sanger Institute for the COVID-19 Genomics UK (COG-UK) Consortium | The Lighthouse Lab in Milton Keynes and Alex Alderton, Roberto Amato, Jeffrey Barrett, Sonia Goncalves, Ewan Harrison, David K. Jackson, Ian Johnston, Dominic Kwiatkowski, Cordelia Langford, John Sillitoe on behalf of the Wellcome Sanger Institute COVID-19 Surveillance Team |
| EPI_ISL_2729040 | Lighthouse Lab in Glasgow | Wellcome Sanger Institute for the COVID-19 Genomics UK (COG-UK) Consortium | Harper VanSteenhouse, Yumi Kasai, David Gray, Carol Clugston, Anna Dominiczak and Alex Alderton, Roberto Amato, Jeffrey Barrett, Sonia Goncalves, Ewan Harrison, David K. Jackson, Ian Johnston, Dominic Kwiatkowski, Cordelia Langford, John Sillitoe on behalf of the Wellcome Sanger Institute COVID-19 Surveillance Team |
| EPI_ISL_2729041 | Lighthouse Laboratory Plymouth | Wellcome Sanger Institute for the COVID-19 Genomics UK (COG-UK) Consortium | Lighthouse Laboratory Plymouth and Alex Alderton, Roberto Amato, Jeffrey Barrett, Sonia Goncalves, Ewan Harrison, David K. Jackson, Ian Johnston, Dominic Kwiatkowski, Cordelia Langford, John Sillitoe on behalf of the Wellcome Sanger Institute COVID-19 Surveillance Team |
| EPI_ISL_2729042, EPI_ISL_2729043 | Lighthouse Lab in Glasgow | Wellcome Sanger Institute for the COVID-19 Genomics UK (COG-UK) Consortium | Harper VanSteenhouse, Yumi Kasai, David Gray, Carol Clugston, Anna Dominiczak and Alex Alderton, Roberto Amato, Jeffrey Barrett, Sonia Goncalves, Ewan Harrison, David K. Jackson, Ian Johnston, Dominic Kwiatkowski, Cordelia Langford, John Sillitoe on behalf of the Wellcome Sanger Institute COVID-19 Surveillance Team |
| EPI_ISL_2729044 | Lighthouse Lab in Milton Keynes | Wellcome Sanger Institute for the COVID-19 Genomics UK (COG-UK) Consortium | The Lighthouse Lab in Milton Keynes and Alex Alderton, Roberto Amato, Jeffrey Barrett, Sonia Goncalves, Ewan Harrison, David K. Jackson, Ian Johnston, Dominic Kwiatkowski, Cordelia Langford, John Sillitoe on behalf of the Wellcome Sanger Institute COVID-19 Surveillance Team |
| EPI_ISL_2729045 | Lighthouse Lab in Glasgow | Wellcome Sanger Institute for the COVID-19 Genomics UK (COG-UK) Consortium | Harper VanSteenhouse, Yumi Kasai, David Gray, Carol Clugston, Anna Dominiczak and Alex Alderton, Roberto Amato, Jeffrey Barrett, Sonia Goncalves, Ewan Harrison, David K. Jackson, Ian Johnston, Dominic Kwiatkowski, Cordelia Langford, John Sillitoe on behalf of the Wellcome Sanger Institute COVID-19 Surveillance Team |
| EPI_ISL_2729046 | Lighthouse Lab in Alderley Park | Wellcome Sanger Institute for the COVID-19 Genomics UK (COG-UK) Consortium | Jacquelyn Wynn, Mairead Hyland, The Lighthouse Lab in Alderley Park and Alex Alderton, Roberto Amato, Jeffrey Barrett, Sonia Goncalves, Ewan Harrison, David K. Jackson, Ian Johnston, Dominic Kwiatkowski, Cordelia Langford, John Sillitoe on behalf of the Wellcome Sanger Institute COVID-19 Surveillance Team |
| EPI_ISL_2729047 | Lighthouse Lab in Glasgow | Wellcome Sanger Institute for the COVID-19 Genomics UK (COG-UK) Consortium | Harper VanSteenhouse, Yumi Kasai, David Gray, Carol Clugston, Anna Dominiczak and Alex Alderton, Roberto Amato, Jeffrey Barrett, Sonia Goncalves, Ewan Harrison, David K. Jackson, Ian Johnston, Dominic Kwiatkowski, Cordelia Langford, John Sillitoe on behalf of the Wellcome Sanger Institute COVID-19 Surveillance Team |
| EPI_ISL_2729048, EPI_ISL_2729049 | Lighthouse Lab in Milton Keynes | Wellcome Sanger Institute for the COVID-19 Genomics UK (COG-UK) Consortium | The Lighthouse Lab in Milton Keynes and Alex Alderton, Roberto Amato, Jeffrey Barrett, Sonia Goncalves, Ewan Harrison, David K. Jackson, Ian Johnston, Dominic Kwiatkowski, Cordelia Langford, John Sillitoe on behalf of the Wellcome Sanger Institute COVID-19 Surveillance Team |
| EPI_ISL_2729050 | Lighthouse Lab in Glasgow | Wellcome Sanger Institute for the COVID-19 Genomics UK (COG-UK) Consortium | Harper VanSteenhouse, Yumi Kasai, David Gray, Carol Clugston, Anna Dominiczak and Alex Alderton, Roberto Amato, Jeffrey Barrett, Sonia Goncalves, Ewan Harrison, David K. Jackson, Ian Johnston, Dominic Kwiatkowski, Cordelia Langford, John Sillitoe on behalf of the Wellcome Sanger Institute COVID-19 Surveillance Team |
| EPI_ISL_2729051 | Lighthouse Lab in Milton Keynes | Wellcome Sanger Institute for the COVID-19 Genomics UK (COG-UK) Consortium | The Lighthouse Lab in Milton Keynes and Alex Alderton, Roberto Amato, Jeffrey Barrett, Sonia Goncalves, Ewan Harrison, David K. Jackson, Ian Johnston, Dominic Kwiatkowski, Cordelia Langford, John Sillitoe on behalf of the Wellcome Sanger Institute COVID-19 Surveillance Team |

| | | | |
|---|---|---|---|
| EPI_ISL_2729052, EPI_ISL_2729053 | Lighthouse Lab in Glasgow | Wellcome Sanger Institute for the COVID-19 Genomics UK (COG-UK) Consortium | Harper VanSteenhouse, Yumi Kasai, David Gray, Carol Clugston, Anna Dominiczak and Alex Alderton, Roberto Amato, Jeffrey Barrett, Sonia Goncalves, Ewan Harrison, David K. Jackson, Ian Johnston, Dominic Kwiatkowski, Cordelia Langford, John Sillitoe on behalf of the Wellcome Sanger Institute COVID-19 Surveillance Team |
| EPI_ISL_2729054, EPI_ISL_2729055, EPI_ISL_2729056 | Lighthouse Lab in Alderley Park | Wellcome Sanger Institute for the COVID-19 Genomics UK (COG-UK) Consortium | Jacquelyn Wynn, Mairead Hyland, The Lighthouse Lab in Alderley Park and Alex Alderton, Roberto Amato, Jeffrey Barrett, Sonia Goncalves, Ewan Harrison, David K. Jackson, Ian Johnston, Dominic Kwiatkowski, Cordelia Langford, John Sillitoe on behalf of the Wellcome Sanger Institute COVID-19 Surveillance Team |
| EPI_ISL_2729057 | Lighthouse Lab in Milton Keynes | Wellcome Sanger Institute for the COVID-19 Genomics UK (COG-UK) Consortium | The Lighthouse Lab in Milton Keynes and Alex Alderton, Roberto Amato, Jeffrey Barrett, Sonia Goncalves, Ewan Harrison, David K. Jackson, Ian Johnston, Dominic Kwiatkowski, Cordelia Langford, John Sillitoe on behalf of the Wellcome Sanger Institute COVID-19 Surveillance Team |
| EPI_ISL_2729058 | Lighthouse Lab in Alderley Park | Wellcome Sanger Institute for the COVID-19 Genomics UK (COG-UK) Consortium | Jacquelyn Wynn, Mairead Hyland, The Lighthouse Lab in Alderley Park and Alex Alderton, Roberto Amato, Jeffrey Barrett, Sonia Goncalves, Ewan Harrison, David K. Jackson, Ian Johnston, Dominic Kwiatkowski, Cordelia Langford, John Sillitoe on behalf of the Wellcome Sanger Institute COVID-19 Surveillance Team |
| EPI_ISL_2729059 | Lighthouse Lab in Glasgow | Wellcome Sanger Institute for the COVID-19 Genomics UK (COG-UK) Consortium | Harper VanSteenhouse, Yumi Kasai, David Gray, Carol Clugston, Anna Dominiczak and Alex Alderton, Roberto Amato, Jeffrey Barrett, Sonia Goncalves, Ewan Harrison, David K. Jackson, Ian Johnston, Dominic Kwiatkowski, Cordelia Langford, John Sillitoe on behalf of the Wellcome Sanger Institute COVID-19 Surveillance Team |
| EPI_ISL_2729060 | Lighthouse Lab in Milton Keynes | Wellcome Sanger Institute for the COVID-19 Genomics UK (COG-UK) Consortium | The Lighthouse Lab in Milton Keynes and Alex Alderton, Roberto Amato, Jeffrey Barrett, Sonia Goncalves, Ewan Harrison, David K. Jackson, Ian Johnston, Dominic Kwiatkowski, Cordelia Langford, John Sillitoe on behalf of the Wellcome Sanger Institute COVID-19 Surveillance Team |
| EPI_ISL_2729061 | Lighthouse Lab in Alderley Park | Wellcome Sanger Institute for the COVID-19 Genomics UK (COG-UK) Consortium | Jacquelyn Wynn, Mairead Hyland, The Lighthouse Lab in Alderley Park and Alex Alderton, Roberto Amato, Jeffrey Barrett, Sonia Goncalves, Ewan Harrison, David K. Jackson, Ian Johnston, Dominic Kwiatkowski, Cordelia Langford, John Sillitoe on behalf of the Wellcome Sanger Institute COVID-19 Surveillance Team |
| EPI_ISL_2729062 | Lighthouse Lab in Milton Keynes | Wellcome Sanger Institute for the COVID-19 Genomics UK (COG-UK) Consortium | The Lighthouse Lab in Milton Keynes and Alex Alderton, Roberto Amato, Jeffrey Barrett, Sonia Goncalves, Ewan Harrison, David K. Jackson, Ian Johnston, Dominic Kwiatkowski, Cordelia Langford, John Sillitoe on behalf of the Wellcome Sanger Institute COVID-19 Surveillance Team |
| EPI_ISL_2729063 | Lighthouse Lab in Alderley Park | Wellcome Sanger Institute for the COVID-19 Genomics UK (COG-UK) Consortium | Jacquelyn Wynn, Mairead Hyland, The Lighthouse Lab in Alderley Park and Alex Alderton, Roberto Amato, Jeffrey Barrett, Sonia Goncalves, Ewan Harrison, David K. Jackson, Ian Johnston, Dominic Kwiatkowski, Cordelia Langford, John Sillitoe on behalf of the Wellcome Sanger Institute COVID-19 Surveillance Team |
| EPI_ISL_2729064 | Lighthouse Lab in Glasgow | Wellcome Sanger Institute for the COVID-19 Genomics UK (COG-UK) Consortium | Harper VanSteenhouse, Yumi Kasai, David Gray, Carol Clugston, Anna Dominiczak and Alex Alderton, Roberto Amato, Jeffrey Barrett, Sonia Goncalves, Ewan Harrison, David K. Jackson, Ian Johnston, Dominic Kwiatkowski, Cordelia Langford, John Sillitoe on behalf of the Wellcome Sanger Institute COVID-19 Surveillance Team |
| EPI_ISL_2729065, EPI_ISL_2729066, EPI_ISL_2729067, EPI_ISL_2729068 | Lighthouse Lab in Alderley Park | Wellcome Sanger Institute for the COVID-19 Genomics UK (COG-UK) Consortium | Jacquelyn Wynn, Mairead Hyland, The Lighthouse Lab in Alderley Park and Alex Alderton, Roberto Amato, Jeffrey Barrett, Sonia Goncalves, Ewan Harrison, David K. Jackson, Ian Johnston, Dominic Kwiatkowski, Cordelia Langford, John Sillitoe on behalf of the Wellcome Sanger Institute COVID-19 Surveillance Team |
| EPI_ISL_2729069 | Lighthouse Lab in Milton Keynes | Wellcome Sanger Institute for the COVID-19 Genomics UK (COG-UK) Consortium | The Lighthouse Lab in Milton Keynes and Alex Alderton, Roberto Amato, Jeffrey Barrett, Sonia Goncalves, Ewan Harrison, David K. Jackson, Ian Johnston, Dominic Kwiatkowski, Cordelia Langford, John Sillitoe on behalf of the Wellcome Sanger Institute COVID-19 Surveillance Team |
| EPI_ISL_2729070, EPI_ISL_2729071 | Lighthouse Lab in Alderley Park | Wellcome Sanger Institute for the COVID-19 Genomics UK (COG-UK) Consortium | Jacquelyn Wynn, Mairead Hyland, The Lighthouse Lab in Alderley Park and Alex Alderton, Roberto Amato, Jeffrey Barrett, Sonia Goncalves, Ewan Harrison, David K. Jackson, Ian Johnston, Dominic Kwiatkowski, Cordelia Langford, John Sillitoe on behalf of the Wellcome Sanger Institute COVID-19 Surveillance Team |
| EPI_ISL_2729072, EPI_ISL_2729073 | Lighthouse Lab in Glasgow | Wellcome Sanger Institute for the COVID-19 Genomics UK (COG-UK) Consortium | Harper VanSteenhouse, Yumi Kasai, David Gray, Carol Clugston, Anna Dominiczak and Alex Alderton, Roberto Amato, Jeffrey Barrett, Sonia Goncalves, Ewan Harrison, David K. Jackson, Ian Johnston, Dominic Kwiatkowski, Cordelia Langford, John Sillitoe on behalf of the Wellcome Sanger Institute COVID-19 Surveillance Team |
| EPI_ISL_2729074, EPI_ISL_2729075 | Lighthouse Lab in Alderley Park | Wellcome Sanger Institute for the COVID-19 Genomics UK (COG-UK) Consortium | Jacquelyn Wynn, Mairead Hyland, The Lighthouse Lab in Alderley Park and Alex Alderton, Roberto Amato, Jeffrey Barrett, Sonia Goncalves, Ewan Harrison, David K. Jackson, Ian Johnston, Dominic Kwiatkowski, Cordelia Langford, John Sillitoe on behalf of the Wellcome Sanger Institute COVID-19 Surveillance Team |
| EPI_ISL_2729076 | Lighthouse Lab in Glasgow | Wellcome Sanger Institute for the COVID-19 Genomics UK (COG-UK) Consortium | Harper VanSteenhouse, Yumi Kasai, David Gray, Carol Clugston, Anna Dominiczak and Alex Alderton, Roberto Amato, Jeffrey Barrett, Sonia Goncalves, Ewan Harrison, David K. Jackson, Ian Johnston, Dominic Kwiatkowski, Cordelia Langford, John Sillitoe on behalf of the Wellcome Sanger Institute COVID-19 Surveillance Team |
| EPI_ISL_2729077, EPI_ISL_2729078, EPI_ISL_2729079, EPI_ISL_2729080, EPI_ISL_2729081 | Lighthouse Lab in Alderley Park | Wellcome Sanger Institute for the COVID-19 Genomics UK (COG-UK) Consortium | Jacquelyn Wynn, Mairead Hyland, The Lighthouse Lab in Alderley Park and Alex Alderton, Roberto Amato, Jeffrey Barrett, Sonia Goncalves, Ewan Harrison, David K. Jackson, Ian Johnston, Dominic Kwiatkowski, Cordelia Langford, John Sillitoe on behalf of the Wellcome Sanger Institute COVID-19 Surveillance Team |
| EPI_ISL_2729082 | Lighthouse Lab in Milton Keynes | Wellcome Sanger Institute for the COVID-19 Genomics UK (COG-UK) Consortium | The Lighthouse Lab in Milton Keynes and Alex Alderton, Roberto Amato, Jeffrey Barrett, Sonia Goncalves, Ewan Harrison, David K. Jackson, Ian Johnston, Dominic Kwiatkowski, Cordelia Langford, John Sillitoe on behalf of the Wellcome Sanger Institute COVID-19 Surveillance Team |
| EPI_ISL_2729083, EPI_ISL_2729084 | Lighthouse Lab in Alderley Park | Wellcome Sanger Institute for the COVID-19 Genomics UK (COG-UK) Consortium | Jacquelyn Wynn, Mairead Hyland, The Lighthouse Lab in Alderley Park and Alex Alderton, Roberto Amato, Jeffrey Barrett, Sonia Goncalves, Ewan Harrison, David K. Jackson, Ian Johnston, Dominic Kwiatkowski, Cordelia Langford, John Sillitoe on behalf of the Wellcome Sanger Institute COVID-19 Surveillance Team |
| EPI_ISL_2729085 | Lighthouse Lab in Milton Keynes | Wellcome Sanger Institute for the COVID-19 Genomics UK (COG-UK) Consortium | The Lighthouse Lab in Milton Keynes and Alex Alderton, Roberto Amato, Jeffrey Barrett, Sonia Goncalves, Ewan Harrison, David K. Jackson, Ian Johnston, Dominic Kwiatkowski, Cordelia Langford, John Sillitoe on behalf of the Wellcome Sanger Institute COVID-19 Surveillance Team |
| EPI_ISL_2729086 | Lighthouse Lab in Alderley Park | Wellcome Sanger Institute for the COVID-19 Genomics UK (COG-UK) Consortium | Jacquelyn Wynn, Mairead Hyland, The Lighthouse Lab in Alderley Park and Alex Alderton, Roberto Amato, Jeffrey Barrett, Sonia Goncalves, Ewan Harrison, David K. Jackson, Ian Johnston, Dominic Kwiatkowski, Cordelia Langford, John Sillitoe on behalf of the Wellcome Sanger Institute COVID-19 Surveillance Team |
| EPI_ISL_2729087 | Lighthouse Lab in Glasgow | Wellcome Sanger Institute for the COVID-19 Genomics UK (COG-UK) Consortium | Harper VanSteenhouse, Yumi Kasai, David Gray, Carol Clugston, Anna Dominiczak and Alex Alderton, Roberto Amato, Jeffrey Barrett, Sonia Goncalves, Ewan Harrison, David K. Jackson, Ian Johnston, Dominic Kwiatkowski, Cordelia Langford, John Sillitoe on behalf of the Wellcome Sanger Institute COVID-19 Surveillance Team |
| EPI_ISL_2729088 | Lighthouse Lab in Milton Keynes | Wellcome Sanger Institute for the COVID-19 Genomics UK (COG-UK) Consortium | The Lighthouse Lab in Milton Keynes and Alex Alderton, Roberto Amato, Jeffrey Barrett, Sonia Goncalves, Ewan Harrison, David K. Jackson, Ian Johnston, Dominic Kwiatkowski, Cordelia Langford, John Sillitoe on behalf of the Wellcome Sanger Institute COVID-19 Surveillance Team |
| EPI_ISL_2729089 | Lighthouse Lab in Alderley Park | Wellcome Sanger Institute for the COVID-19 Genomics UK (COG-UK) Consortium | Jacquelyn Wynn, Mairead Hyland, The Lighthouse Lab in Alderley Park and Alex Alderton, Roberto Amato, Jeffrey Barrett, Sonia Goncalves, Ewan Harrison, David K. Jackson, Ian Johnston, Dominic Kwiatkowski, Cordelia Langford, John Sillitoe on behalf of the Wellcome Sanger Institute COVID-19 Surveillance Team |
| EPI_ISL_2729090 | Lighthouse Lab in Glasgow | Wellcome Sanger Institute for the COVID-19 Genomics UK (COG-UK) Consortium | Harper VanSteenhouse, Yumi Kasai, David Gray, Carol Clugston, Anna Dominiczak and Alex Alderton, Roberto Amato, Jeffrey Barrett, Sonia Goncalves, Ewan Harrison, David K. Jackson, Ian Johnston, Dominic Kwiatkowski, Cordelia Langford, John Sillitoe on behalf of the Wellcome Sanger Institute COVID-19 Surveillance Team |
| EPI_ISL_2729091, EPI_ISL_2729092 | Lighthouse Lab in Milton Keynes | Wellcome Sanger Institute for the COVID-19 Genomics UK (COG-UK) Consortium | The Lighthouse Lab in Milton Keynes and Alex Alderton, Roberto Amato, Jeffrey Barrett, Sonia Goncalves, Ewan Harrison, David K. Jackson, Ian Johnston, Dominic Kwiatkowski, Cordelia Langford, John Sillitoe on behalf of the Wellcome Sanger Institute COVID-19 Surveillance Team |
| EPI_ISL_2729093, EPI_ISL_2729094 | Lighthouse Lab in Alderley Park | Wellcome Sanger Institute for the COVID-19 Genomics UK (COG-UK) Consortium | Jacquelyn Wynn, Mairead Hyland, The Lighthouse Lab in Alderley Park and Alex Alderton, Roberto Amato, Jeffrey Barrett, Sonia Goncalves, Ewan Harrison, David K. Jackson, Ian Johnston, Dominic Kwiatkowski, Cordelia Langford, John Sillitoe on behalf of the Wellcome Sanger Institute COVID-19 Surveillance Team |
| EPI_ISL_2729095 | Lighthouse Lab in Glasgow | Wellcome Sanger Institute for the COVID-19 Genomics UK (COG-UK) Consortium | Harper VanSteenhouse, Yumi Kasai, David Gray, Carol Clugston, Anna Dominiczak and Alex Alderton, Roberto Amato, Jeffrey Barrett, Sonia Goncalves, Ewan Harrison, David K. Jackson, Ian Johnston, Dominic Kwiatkowski, Cordelia Langford, John Sillitoe on behalf of the Wellcome Sanger Institute COVID-19 Surveillance Team |
| EPI_ISL_2729096, EPI_ISL_2729097, EPI_ISL_2729098, EPI_ISL_2729099, EPI_ISL_2729100 | Lighthouse Lab in Alderley Park | Wellcome Sanger Institute for the COVID-19 Genomics UK (COG-UK) Consortium | Jacquelyn Wynn, Mairead Hyland, The Lighthouse Lab in Alderley Park and Alex Alderton, Roberto Amato, Jeffrey Barrett, Sonia Goncalves, Ewan Harrison, David K. Jackson, Ian Johnston, Dominic Kwiatkowski, Cordelia Langford, John Sillitoe on behalf of the Wellcome Sanger Institute COVID-19 Surveillance Team |
| EPI_ISL_2729101 | Lighthouse Lab in Glasgow | Wellcome Sanger Institute for the COVID-19 Genomics UK (COG-UK) Consortium | Harper VanSteenhouse, Yumi Kasai, David Gray, Carol Clugston, Anna Dominiczak and Alex Alderton, Roberto Amato, Jeffrey Barrett, Sonia Goncalves, Ewan Harrison, David K. Jackson, Ian Johnston, Dominic Kwiatkowski, Cordelia Langford, John Sillitoe on behalf of the Wellcome Sanger Institute COVID-19 Surveillance Team |
| EPI_ISL_2729102 | Lighthouse Lab in Alderley Park | Wellcome Sanger Institute for the COVID-19 Genomics UK (COG-UK) Consortium | Jacquelyn Wynn, Mairead Hyland, The Lighthouse Lab in Alderley Park and Alex Alderton, Roberto Amato, Jeffrey Barrett, Sonia Goncalves, Ewan Harrison, David K. Jackson, Ian Johnston, Dominic Kwiatkowski, Cordelia Langford, John Sillitoe on behalf of the Wellcome Sanger Institute COVID-19 Surveillance Team |
| EPI_ISL_2729103 | Lighthouse Lab in Glasgow | Wellcome Sanger Institute for the COVID-19 Genomics UK (COG-UK) Consortium | Harper VanSteenhouse, Yumi Kasai, David Gray, Carol Clugston, Anna Dominiczak and Alex Alderton, Roberto Amato, Jeffrey Barrett, Sonia Goncalves, Ewan Harrison, David K. Jackson, Ian Johnston, Dominic Kwiatkowski, Cordelia Langford, John Sillitoe on behalf of the Wellcome Sanger Institute COVID-19 Surveillance Team |
| EPI_ISL_2729104 | Lighthouse Lab in Alderley Park | Wellcome Sanger Institute for the COVID-19 Genomics UK (COG-UK) Consortium | Jacquelyn Wynn, Mairead Hyland, The Lighthouse Lab in Alderley Park and Alex Alderton, Roberto Amato, Jeffrey Barrett, Sonia Goncalves, Ewan Harrison, David K. Jackson, Ian Johnston, Dominic Kwiatkowski, Cordelia Langford, John Sillitoe on behalf of the Wellcome Sanger Institute COVID-19 Surveillance Team |
| EPI_ISL_2729105, EPI_ISL_2729106 | Lighthouse Lab in Glasgow | Wellcome Sanger Institute for the COVID-19 Genomics UK (COG-UK) Consortium | Harper VanSteenhouse, Yumi Kasai, David Gray, Carol Clugston, Anna Dominiczak and Alex Alderton, Roberto Amato, Jeffrey Barrett, Sonia Goncalves, Ewan Harrison, David K. Jackson, Ian Johnston, Dominic Kwiatkowski, Cordelia Langford, John Sillitoe on behalf of the Wellcome Sanger Institute COVID-19 Surveillance Team |
| EPI_ISL_2729107, EPI_ISL_2729108 | Lighthouse Lab in Alderley Park | Wellcome Sanger Institute for the COVID-19 Genomics UK (COG-UK) Consortium | Jacquelyn Wynn, Mairead Hyland, The Lighthouse Lab in Alderley Park and Alex Alderton, Roberto Amato, Jeffrey Barrett, Sonia Goncalves, Ewan Harrison, David K. Jackson, Ian Johnston, Dominic Kwiatkowski, Cordelia Langford, John Sillitoe on behalf of the Wellcome Sanger Institute COVID-19 Surveillance Team |
| EPI_ISL_2729109 | Lighthouse Lab in Milton Keynes | Wellcome Sanger Institute for the COVID-19 Genomics UK (COG-UK) Consortium | The Lighthouse Lab in Milton Keynes and Alex Alderton, Roberto Amato, Jeffrey Barrett, Sonia Goncalves, Ewan Harrison, David K. Jackson, Ian Johnston, Dominic Kwiatkowski, Cordelia Langford, John Sillitoe on behalf of the Wellcome Sanger Institute COVID-19 Surveillance Team |
| EPI_ISL_2729110 | Lighthouse Lab in Alderley Park | Wellcome Sanger Institute for the COVID-19 Genomics UK (COG-UK) Consortium | Jacquelyn Wynn, Mairead Hyland, The Lighthouse Lab in Alderley Park and Alex Alderton, Roberto Amato, Jeffrey Barrett, Sonia Goncalves, Ewan Harrison, David K. Jackson, Ian Johnston, Dominic Kwiatkowski, Cordelia Langford, John Sillitoe on behalf of the Wellcome Sanger Institute COVID-19 Surveillance Team |
| EPI_ISL_2729111 | Lighthouse Lab in Glasgow | Wellcome Sanger Institute for the COVID-19 Genomics UK (COG-UK) Consortium | Harper VanSteenhouse, Yumi Kasai, David Gray, Carol Clugston, Anna Dominiczak and Alex Alderton, Roberto Amato, Jeffrey Barrett, Sonia Goncalves, Ewan Harrison, David K. Jackson, Ian Johnston, Dominic Kwiatkowski, Cordelia Langford, John Sillitoe on behalf of the Wellcome Sanger Institute COVID-19 Surveillance Team |
| EPI_ISL_2729112 | Lighthouse Lab in Alderley Park | Wellcome Sanger Institute for the COVID-19 Genomics UK (COG-UK) Consortium | Jacquelyn Wynn, Mairead Hyland, The Lighthouse Lab in Alderley Park and Alex Alderton, Roberto Amato, Jeffrey Barrett, Sonia Goncalves, Ewan Harrison, David K. Jackson, Ian Johnston, Dominic Kwiatkowski, Cordelia Langford, John Sillitoe on behalf of the Wellcome Sanger Institute COVID-19 Surveillance Team |
| EPI_ISL_2729113 | Lighthouse Lab in Glasgow | Wellcome Sanger Institute for the COVID-19 Genomics UK (COG-UK) Consortium | Harper VanSteenhouse, Yumi Kasai, David Gray, Carol Clugston, Anna Dominiczak and Alex Alderton, Roberto Amato, Jeffrey Barrett, Sonia Goncalves, Ewan Harrison, David K. Jackson, Ian Johnston, Dominic Kwiatkowski, Cordelia Langford, John Sillitoe on behalf of the Wellcome Sanger Institute COVID-19 Surveillance Team |
| EPI_ISL_2729114, EPI_ISL_2729115 | Lighthouse Lab in Alderley Park | Wellcome Sanger Institute for the COVID-19 Genomics UK (COG-UK) Consortium | Jacquelyn Wynn, Mairead Hyland, The Lighthouse Lab in Alderley Park and Alex Alderton, Roberto Amato, Jeffrey Barrett, Sonia Goncalves, Ewan Harrison, David K. Jackson, Ian Johnston, Dominic Kwiatkowski, Cordelia Langford, John Sillitoe on behalf of the Wellcome Sanger Institute COVID-19 Surveillance Team |

| | | | |
|---|---|---|---|
| EPI_ISL_2729116 | Lighthouse Lab in Glasgow | Wellcome Sanger Institute for the COVID-19 Genomics UK (COG-UK) Consortium | Harper VanSteenhouse, Yumi Kasai, David Gray, Carol Clugston, Anna Dominiczak and Alex Alderton, Roberto Amato, Jeffrey Barrett, Sonia Goncalves, Ewan Harrison, David K. Jackson, Ian Johnston, Dominic Kwiatkowski, Cordelia Langford, John Sillitoe on behalf of the Wellcome Sanger Institute COVID-19 Surveillance Team |
| EPI_ISL_2729117 | Lighthouse Lab in Milton Keynes | Wellcome Sanger Institute for the COVID-19 Genomics UK (COG-UK) Consortium | The Lighthouse Lab in Milton Keynes and Alex Alderton, Roberto Amato, Jeffrey Barrett, Sonia Goncalves, Ewan Harrison, David K. Jackson, Ian Johnston, Dominic Kwiatkowski, Cordelia Langford, John Sillitoe on behalf of the Wellcome Sanger Institute COVID-19 Surveillance Team |
| EPI_ISL_2729118 | Lighthouse Lab in Alderley Park | Wellcome Sanger Institute for the COVID-19 Genomics UK (COG-UK) Consortium | Jacquelyn Wynn, Mairead Hyland, The Lighthouse Lab in Alderley Park and Alex Alderton, Roberto Amato, Jeffrey Barrett, Sonia Goncalves, Ewan Harrison, David K. Jackson, Ian Johnston, Dominic Kwiatkowski, Cordelia Langford, John Sillitoe on behalf of the Wellcome Sanger Institute COVID-19 Surveillance Team |
| EPI_ISL_2729119 | Lighthouse Lab in Glasgow | Wellcome Sanger Institute for the COVID-19 Genomics UK (COG-UK) Consortium | Harper VanSteenhouse, Yumi Kasai, David Gray, Carol Clugston, Anna Dominiczak and Alex Alderton, Roberto Amato, Jeffrey Barrett, Sonia Goncalves, Ewan Harrison, David K. Jackson, Ian Johnston, Dominic Kwiatkowski, Cordelia Langford, John Sillitoe on behalf of the Wellcome Sanger Institute COVID-19 Surveillance Team |
| EPI_ISL_2729120, EPI_ISL_2729121 | Lighthouse Lab in Alderley Park | Wellcome Sanger Institute for the COVID-19 Genomics UK (COG-UK) Consortium | Jacquelyn Wynn, Mairead Hyland, The Lighthouse Lab in Alderley Park and Alex Alderton, Roberto Amato, Jeffrey Barrett, Sonia Goncalves, Ewan Harrison, David K. Jackson, Ian Johnston, Dominic Kwiatkowski, Cordelia Langford, John Sillitoe on behalf of the Wellcome Sanger Institute COVID-19 Surveillance Team |
| EPI_ISL_2729122 | Lighthouse Lab in Glasgow | Wellcome Sanger Institute for the COVID-19 Genomics UK (COG-UK) Consortium | Harper VanSteenhouse, Yumi Kasai, David Gray, Carol Clugston, Anna Dominiczak and Alex Alderton, Roberto Amato, Jeffrey Barrett, Sonia Goncalves, Ewan Harrison, David K. Jackson, Ian Johnston, Dominic Kwiatkowski, Cordelia Langford, John Sillitoe on behalf of the Wellcome Sanger Institute COVID-19 Surveillance Team |
| EPI_ISL_2729123 | Lighthouse Lab in Alderley Park | Wellcome Sanger Institute for the COVID-19 Genomics UK (COG-UK) Consortium | Jacquelyn Wynn, Mairead Hyland, The Lighthouse Lab in Alderley Park and Alex Alderton, Roberto Amato, Jeffrey Barrett, Sonia Goncalves, Ewan Harrison, David K. Jackson, Ian Johnston, Dominic Kwiatkowski, Cordelia Langford, John Sillitoe on behalf of the Wellcome Sanger Institute COVID-19 Surveillance Team |
| EPI_ISL_2729124 | Lighthouse Lab in Glasgow | Wellcome Sanger Institute for the COVID-19 Genomics UK (COG-UK) Consortium | Harper VanSteenhouse, Yumi Kasai, David Gray, Carol Clugston, Anna Dominiczak and Alex Alderton, Roberto Amato, Jeffrey Barrett, Sonia Goncalves, Ewan Harrison, David K. Jackson, Ian Johnston, Dominic Kwiatkowski, Cordelia Langford, John Sillitoe on behalf of the Wellcome Sanger Institute COVID-19 Surveillance Team |
| EPI_ISL_2729125 | Lighthouse Lab in Alderley Park | Wellcome Sanger Institute for the COVID-19 Genomics UK (COG-UK) Consortium | Jacquelyn Wynn, Mairead Hyland, The Lighthouse Lab in Alderley Park and Alex Alderton, Roberto Amato, Jeffrey Barrett, Sonia Goncalves, Ewan Harrison, David K. Jackson, Ian Johnston, Dominic Kwiatkowski, Cordelia Langford, John Sillitoe on behalf of the Wellcome Sanger Institute COVID-19 Surveillance Team |
| EPI_ISL_2729126, EPI_ISL_2729127, EPI_ISL_2729128, EPI_ISL_2729129 | Lighthouse Lab in Glasgow | Wellcome Sanger Institute for the COVID-19 Genomics UK (COG-UK) Consortium | Harper VanSteenhouse, Yumi Kasai, David Gray, Carol Clugston, Anna Dominiczak and Alex Alderton, Roberto Amato, Jeffrey Barrett, Sonia Goncalves, Ewan Harrison, David K. Jackson, Ian Johnston, Dominic Kwiatkowski, Cordelia Langford, John Sillitoe on behalf of the Wellcome Sanger Institute COVID-19 Surveillance Team |
| EPI_ISL_2729130, EPI_ISL_2729131, EPI_ISL_2729132 | Lighthouse Lab in Alderley Park | Wellcome Sanger Institute for the COVID-19 Genomics UK (COG-UK) Consortium | Jacquelyn Wynn, Mairead Hyland, The Lighthouse Lab in Alderley Park and Alex Alderton, Roberto Amato, Jeffrey Barrett, Sonia Goncalves, Ewan Harrison, David K. Jackson, Ian Johnston, Dominic Kwiatkowski, Cordelia Langford, John Sillitoe on behalf of the Wellcome Sanger Institute COVID-19 Surveillance Team |
| EPI_ISL_2729133 | Lighthouse Lab in Milton Keynes | Wellcome Sanger Institute for the COVID-19 Genomics UK (COG-UK) Consortium | The Lighthouse Lab in Milton Keynes and Alex Alderton, Roberto Amato, Jeffrey Barrett, Sonia Goncalves, Ewan Harrison, David K. Jackson, Ian Johnston, Dominic Kwiatkowski, Cordelia Langford, John Sillitoe on behalf of the Wellcome Sanger Institute COVID-19 Surveillance Team |
| EPI_ISL_2729134, EPI_ISL_2729135 | Lighthouse Lab in Alderley Park | Wellcome Sanger Institute for the COVID-19 Genomics UK (COG-UK) Consortium | Jacquelyn Wynn, Mairead Hyland, The Lighthouse Lab in Alderley Park and Alex Alderton, Roberto Amato, Jeffrey Barrett, Sonia Goncalves, Ewan Harrison, David K. Jackson, Ian Johnston, Dominic Kwiatkowski, Cordelia Langford, John Sillitoe on behalf of the Wellcome Sanger Institute COVID-19 Surveillance Team |
| EPI_ISL_2729136 | Lighthouse Lab in Milton Keynes | Wellcome Sanger Institute for the COVID-19 Genomics UK (COG-UK) Consortium | The Lighthouse Lab in Milton Keynes and Alex Alderton, Roberto Amato, Jeffrey Barrett, Sonia Goncalves, Ewan Harrison, David K. Jackson, Ian Johnston, Dominic Kwiatkowski, Cordelia Langford, John Sillitoe on behalf of the Wellcome Sanger Institute COVID-19 Surveillance Team |
| EPI_ISL_2729137 | Lighthouse Lab in Glasgow | Wellcome Sanger Institute for the COVID-19 Genomics UK (COG-UK) Consortium | Harper VanSteenhouse, Yumi Kasai, David Gray, Carol Clugston, Anna Dominiczak and Alex Alderton, Roberto Amato, Jeffrey Barrett, Sonia Goncalves, Ewan Harrison, David K. Jackson, Ian Johnston, Dominic Kwiatkowski, Cordelia Langford, John Sillitoe on behalf of the Wellcome Sanger Institute COVID-19 Surveillance Team |
| EPI_ISL_2729138 | Lighthouse Lab in Milton Keynes | Wellcome Sanger Institute for the COVID-19 Genomics UK (COG-UK) Consortium | The Lighthouse Lab in Milton Keynes and Alex Alderton, Roberto Amato, Jeffrey Barrett, Sonia Goncalves, Ewan Harrison, David K. Jackson, Ian Johnston, Dominic Kwiatkowski, Cordelia Langford, John Sillitoe on behalf of the Wellcome Sanger Institute COVID-19 Surveillance Team |
| EPI_ISL_2729139 | Lighthouse Lab in Glasgow | Wellcome Sanger Institute for the COVID-19 Genomics UK (COG-UK) Consortium | Harper VanSteenhouse, Yumi Kasai, David Gray, Carol Clugston, Anna Dominiczak and Alex Alderton, Roberto Amato, Jeffrey Barrett, Sonia Goncalves, Ewan Harrison, David K. Jackson, Ian Johnston, Dominic Kwiatkowski, Cordelia Langford, John Sillitoe on behalf of the Wellcome Sanger Institute COVID-19 Surveillance Team |
| EPI_ISL_2729140 | Lighthouse Lab in Milton Keynes | Wellcome Sanger Institute for the COVID-19 Genomics UK (COG-UK) Consortium | The Lighthouse Lab in Milton Keynes and Alex Alderton, Roberto Amato, Jeffrey Barrett, Sonia Goncalves, Ewan Harrison, David K. Jackson, Ian Johnston, Dominic Kwiatkowski, Cordelia Langford, John Sillitoe on behalf of the Wellcome Sanger Institute COVID-19 Surveillance Team |
| EPI_ISL_2729141, EPI_ISL_2729142, EPI_ISL_2729143 | Lighthouse Lab in Alderley Park | Wellcome Sanger Institute for the COVID-19 Genomics UK (COG-UK) Consortium | Jacquelyn Wynn, Mairead Hyland, The Lighthouse Lab in Alderley Park and Alex Alderton, Roberto Amato, Jeffrey Barrett, Sonia Goncalves, Ewan Harrison, David K. Jackson, Ian Johnston, Dominic Kwiatkowski, Cordelia Langford, John Sillitoe on behalf of the Wellcome Sanger Institute COVID-19 Surveillance Team |
| EPI_ISL_2729144 | Lighthouse Lab in Glasgow | Wellcome Sanger Institute for the COVID-19 Genomics UK (COG-UK) Consortium | Harper VanSteenhouse, Yumi Kasai, David Gray, Carol Clugston, Anna Dominiczak and Alex Alderton, Roberto Amato, Jeffrey Barrett, Sonia Goncalves, Ewan Harrison, David K. Jackson, Ian Johnston, Dominic Kwiatkowski, Cordelia Langford, John Sillitoe on behalf of the Wellcome Sanger Institute COVID-19 Surveillance Team |
| EPI_ISL_2729145 | Lighthouse Lab in Alderley Park | Wellcome Sanger Institute for the COVID-19 Genomics UK (COG-UK) Consortium | Jacquelyn Wynn, Mairead Hyland, The Lighthouse Lab in Alderley Park and Alex Alderton, Roberto Amato, Jeffrey Barrett, Sonia Goncalves, Ewan Harrison, David K. Jackson, Ian Johnston, Dominic Kwiatkowski, Cordelia Langford, John Sillitoe on behalf of the Wellcome Sanger Institute COVID-19 Surveillance Team |
| EPI_ISL_2729146 | Lighthouse Lab in Glasgow | Wellcome Sanger Institute for the COVID-19 Genomics UK (COG-UK) Consortium | Harper VanSteenhouse, Yumi Kasai, David Gray, Carol Clugston, Anna Dominiczak and Alex Alderton, Roberto Amato, Jeffrey Barrett, Sonia Goncalves, Ewan Harrison, David K. Jackson, Ian Johnston, Dominic Kwiatkowski, Cordelia Langford, John Sillitoe on behalf of the Wellcome Sanger Institute COVID-19 Surveillance Team |
| EPI_ISL_2729147 | Lighthouse Lab in Alderley Park | Wellcome Sanger Institute for the COVID-19 Genomics UK (COG-UK) Consortium | Jacquelyn Wynn, Mairead Hyland, The Lighthouse Lab in Alderley Park and Alex Alderton, Roberto Amato, Jeffrey Barrett, Sonia Goncalves, Ewan Harrison, David K. Jackson, Ian Johnston, Dominic Kwiatkowski, Cordelia Langford, John Sillitoe on behalf of the Wellcome Sanger Institute COVID-19 Surveillance Team |
| EPI_ISL_2729148 | Lighthouse Lab in Glasgow | Wellcome Sanger Institute for the COVID-19 Genomics UK (COG-UK) Consortium | Harper VanSteenhouse, Yumi Kasai, David Gray, Carol Clugston, Anna Dominiczak and Alex Alderton, Roberto Amato, Jeffrey Barrett, Sonia Goncalves, Ewan Harrison, David K. Jackson, Ian Johnston, Dominic Kwiatkowski, Cordelia Langford, John Sillitoe on behalf of the Wellcome Sanger Institute COVID-19 Surveillance Team |
| EPI_ISL_2729149 | Lighthouse Lab in Alderley Park | Wellcome Sanger Institute for the COVID-19 Genomics UK (COG-UK) Consortium | Jacquelyn Wynn, Mairead Hyland, The Lighthouse Lab in Alderley Park and Alex Alderton, Roberto Amato, Jeffrey Barrett, Sonia Goncalves, Ewan Harrison, David K. Jackson, Ian Johnston, Dominic Kwiatkowski, Cordelia Langford, John Sillitoe on behalf of the Wellcome Sanger Institute COVID-19 Surveillance Team |
| EPI_ISL_2729150 | Lighthouse Lab in Glasgow | Wellcome Sanger Institute for the COVID-19 Genomics UK (COG-UK) Consortium | Harper VanSteenhouse, Yumi Kasai, David Gray, Carol Clugston, Anna Dominiczak and Alex Alderton, Roberto Amato, Jeffrey Barrett, Sonia Goncalves, Ewan Harrison, David K. Jackson, Ian Johnston, Dominic Kwiatkowski, Cordelia Langford, John Sillitoe on behalf of the Wellcome Sanger Institute COVID-19 Surveillance Team |
| EPI_ISL_2729151, EPI_ISL_2729152, EPI_ISL_2729153 | Lighthouse Lab in Alderley Park | Wellcome Sanger Institute for the COVID-19 Genomics UK (COG-UK) Consortium | Jacquelyn Wynn, Mairead Hyland, The Lighthouse Lab in Alderley Park and Alex Alderton, Roberto Amato, Jeffrey Barrett, Sonia Goncalves, Ewan Harrison, David K. Jackson, Ian Johnston, Dominic Kwiatkowski, Cordelia Langford, John Sillitoe on behalf of the Wellcome Sanger Institute COVID-19 Surveillance Team |
| EPI_ISL_2729154 | Lighthouse Lab in Glasgow | Wellcome Sanger Institute for the COVID-19 Genomics UK (COG-UK) Consortium | Harper VanSteenhouse, Yumi Kasai, David Gray, Carol Clugston, Anna Dominiczak and Alex Alderton, Roberto Amato, Jeffrey Barrett, Sonia Goncalves, Ewan Harrison, David K. Jackson, Ian Johnston, Dominic Kwiatkowski, Cordelia Langford, John Sillitoe on behalf of the Wellcome Sanger Institute COVID-19 Surveillance Team |
| EPI_ISL_2729155 | Lighthouse Lab in Milton Keynes | Wellcome Sanger Institute for the COVID-19 Genomics UK (COG-UK) Consortium | The Lighthouse Lab in Milton Keynes and Alex Alderton, Roberto Amato, Jeffrey Barrett, Sonia Goncalves, Ewan Harrison, David K. Jackson, Ian Johnston, Dominic Kwiatkowski, Cordelia Langford, John Sillitoe on behalf of the Wellcome Sanger Institute COVID-19 Surveillance Team |
| EPI_ISL_2729156, EPI_ISL_2729157, EPI_ISL_2729158, EPI_ISL_2729159 | Lighthouse Lab in Alderley Park | Wellcome Sanger Institute for the COVID-19 Genomics UK (COG-UK) Consortium | Jacquelyn Wynn, Mairead Hyland, The Lighthouse Lab in Alderley Park and Alex Alderton, Roberto Amato, Jeffrey Barrett, Sonia Goncalves, Ewan Harrison, David K. Jackson, Ian Johnston, Dominic Kwiatkowski, Cordelia Langford, John Sillitoe on behalf of the Wellcome Sanger Institute COVID-19 Surveillance Team |
| EPI_ISL_2729160, EPI_ISL_2729161 | Lighthouse Lab in Glasgow | Wellcome Sanger Institute for the COVID-19 Genomics UK (COG-UK) Consortium | Harper VanSteenhouse, Yumi Kasai, David Gray, Carol Clugston, Anna Dominiczak and Alex Alderton, Roberto Amato, Jeffrey Barrett, Sonia Goncalves, Ewan Harrison, David K. Jackson, Ian Johnston, Dominic Kwiatkowski, Cordelia Langford, John Sillitoe on behalf of the Wellcome Sanger Institute COVID-19 Surveillance Team |
| EPI_ISL_2729162 | Lighthouse Lab in Alderley Park | Wellcome Sanger Institute for the COVID-19 Genomics UK (COG-UK) Consortium | Jacquelyn Wynn, Mairead Hyland, The Lighthouse Lab in Alderley Park and Alex Alderton, Roberto Amato, Jeffrey Barrett, Sonia Goncalves, Ewan Harrison, David K. Jackson, Ian Johnston, Dominic Kwiatkowski, Cordelia Langford, John Sillitoe on behalf of the Wellcome Sanger Institute COVID-19 Surveillance Team |
| EPI_ISL_2729163 | Lighthouse Lab in Milton Keynes | Wellcome Sanger Institute for the COVID-19 Genomics UK (COG-UK) Consortium | The Lighthouse Lab in Milton Keynes and Alex Alderton, Roberto Amato, Jeffrey Barrett, Sonia Goncalves, Ewan Harrison, David K. Jackson, Ian Johnston, Dominic Kwiatkowski, Cordelia Langford, John Sillitoe on behalf of the Wellcome Sanger Institute COVID-19 Surveillance Team |
| EPI_ISL_2729164, EPI_ISL_2729165, EPI_ISL_2729166 | Lighthouse Lab in Glasgow | Wellcome Sanger Institute for the COVID-19 Genomics UK (COG-UK) Consortium | Harper VanSteenhouse, Yumi Kasai, David Gray, Carol Clugston, Anna Dominiczak and Alex Alderton, Roberto Amato, Jeffrey Barrett, Sonia Goncalves, Ewan Harrison, David K. Jackson, Ian Johnston, Dominic Kwiatkowski, Cordelia Langford, John Sillitoe on behalf of the Wellcome Sanger Institute COVID-19 Surveillance Team |
| EPI_ISL_2729167, EPI_ISL_2729168, EPI_ISL_2729169 | Lighthouse Lab in Alderley Park | Wellcome Sanger Institute for the COVID-19 Genomics UK (COG-UK) Consortium | Jacquelyn Wynn, Mairead Hyland, The Lighthouse Lab in Alderley Park and Alex Alderton, Roberto Amato, Jeffrey Barrett, Sonia Goncalves, Ewan Harrison, David K. Jackson, Ian Johnston, Dominic Kwiatkowski, Cordelia Langford, John Sillitoe on behalf of the Wellcome Sanger Institute COVID-19 Surveillance Team |
| EPI_ISL_2729170, EPI_ISL_2729172 | Lighthouse Lab in Glasgow | Wellcome Sanger Institute for the COVID-19 Genomics UK (COG-UK) Consortium | Harper VanSteenhouse, Yumi Kasai, David Gray, Carol Clugston, Anna Dominiczak and Alex Alderton, Roberto Amato, Jeffrey Barrett, Sonia Goncalves, Ewan Harrison, David K. Jackson, Ian Johnston, Dominic Kwiatkowski, Cordelia Langford, John Sillitoe on behalf of the Wellcome Sanger Institute COVID-19 Surveillance Team |
| EPI_ISL_2729173 | Lighthouse Lab in Alderley Park | Wellcome Sanger Institute for the COVID-19 Genomics UK (COG-UK) Consortium | Jacquelyn Wynn, Mairead Hyland, The Lighthouse Lab in Alderley Park and Alex Alderton, Roberto Amato, Jeffrey Barrett, Sonia Goncalves, Ewan Harrison, David K. Jackson, Ian Johnston, Dominic Kwiatkowski, Cordelia Langford, John Sillitoe on behalf of the Wellcome Sanger Institute COVID-19 Surveillance Team |
| EPI_ISL_2729174 | Lighthouse Lab in Milton Keynes | Wellcome Sanger Institute for the COVID-19 Genomics UK (COG-UK) Consortium | The Lighthouse Lab in Milton Keynes and Alex Alderton, Roberto Amato, Jeffrey Barrett, Sonia Goncalves, Ewan Harrison, David K. Jackson, Ian Johnston, Dominic Kwiatkowski, Cordelia Langford, John Sillitoe on behalf of the Wellcome Sanger Institute COVID-19 Surveillance Team |
| EPI_ISL_2729175 | Lighthouse Lab in Alderley Park | Wellcome Sanger Institute for the COVID-19 Genomics UK (COG-UK) Consortium | Jacquelyn Wynn, Mairead Hyland, The Lighthouse Lab in Alderley Park and Alex Alderton, Roberto Amato, Jeffrey Barrett, Sonia Goncalves, Ewan Harrison, David K. Jackson, Ian Johnston, Dominic Kwiatkowski, Cordelia Langford, John Sillitoe on behalf of the Wellcome Sanger Institute COVID-19 Surveillance Team |
| EPI_ISL_2729176 | Lighthouse Lab in Glasgow | Wellcome Sanger Institute for the COVID-19 Genomics UK (COG-UK) Consortium | Harper VanSteenhouse, Yumi Kasai, David Gray, Carol Clugston, Anna Dominiczak and Alex Alderton, Roberto Amato, Jeffrey Barrett, Sonia Goncalves, Ewan Harrison, David K. Jackson, Ian Johnston, Dominic Kwiatkowski, Cordelia Langford, John Sillitoe on behalf of the Wellcome Sanger Institute COVID-19 Surveillance Team |
| EPI_ISL_2729177 | Lighthouse Lab in Milton Keynes | Wellcome Sanger Institute for the COVID-19 Genomics UK (COG-UK) Consortium | The Lighthouse Lab in Milton Keynes and Alex Alderton, Roberto Amato, Jeffrey Barrett, Sonia Goncalves, Ewan Harrison, David K. Jackson, Ian Johnston, Dominic Kwiatkowski, Cordelia Langford, John Sillitoe on behalf of the Wellcome Sanger Institute COVID-19 Surveillance Team |

| | | | |
|---|---|---|---|
| EPI_ISL_2729178, EPI_ISL_2729179, EPI_ISL_2729180, EPI_ISL_2729181, EPI_ISL_2729182, EPI_ISL_2729183, EPI_ISL_2729184 | Lighthouse Lab in Alderley Park | Wellcome Sanger Institute for the COVID-19 Genomics UK (COG-UK) Consortium | Jacquelyn Wynn, Mairead Hyland, The Lighthouse Lab in Alderley Park and Alex Alderton, Roberto Amato, Jeffrey Barrett, Sonia Goncalves, Ewan Harrison, David K. Jackson, Ian Johnston, Dominic Kwiatkowski, Cordelia Langford, John Sillitoe on behalf of the Wellcome Sanger Institute COVID-19 Surveillance Team |
| EPI_ISL_2729185 | Lighthouse Lab in Glasgow | Wellcome Sanger Institute for the COVID-19 Genomics UK (COG-UK) Consortium | Harper VanSteenhouse, Yumi Kasai, David Gray, Carol Clugston, Anna Dominiczak and Alex Alderton, Roberto Amato, Jeffrey Barrett, Sonia Goncalves, Ewan Harrison, David K. Jackson, Ian Johnston, Dominic Kwiatkowski, Cordelia Langford, John Sillitoe on behalf of the Wellcome Sanger Institute COVID-19 Surveillance Team |
| EPI_ISL_2729186 | Lighthouse Lab in Milton Keynes | Wellcome Sanger Institute for the COVID-19 Genomics UK (COG-UK) Consortium | The Lighthouse Lab in Milton Keynes and Alex Alderton, Roberto Amato, Jeffrey Barrett, Sonia Goncalves, Ewan Harrison, David K. Jackson, Ian Johnston, Dominic Kwiatkowski, Cordelia Langford, John Sillitoe on behalf of the Wellcome Sanger Institute COVID-19 Surveillance Team |
| EPI_ISL_2729187, EPI_ISL_2729188, EPI_ISL_2729189 | Lighthouse Lab in Alderley Park | Wellcome Sanger Institute for the COVID-19 Genomics UK (COG-UK) Consortium | Jacquelyn Wynn, Mairead Hyland, The Lighthouse Lab in Alderley Park and Alex Alderton, Roberto Amato, Jeffrey Barrett, Sonia Goncalves, Ewan Harrison, David K. Jackson, Ian Johnston, Dominic Kwiatkowski, Cordelia Langford, John Sillitoe on behalf of the Wellcome Sanger Institute COVID-19 Surveillance Team |
| EPI_ISL_2729190 | Lighthouse Lab in Milton Keynes | Wellcome Sanger Institute for the COVID-19 Genomics UK (COG-UK) Consortium | The Lighthouse Lab in Milton Keynes and Alex Alderton, Roberto Amato, Jeffrey Barrett, Sonia Goncalves, Ewan Harrison, David K. Jackson, Ian Johnston, Dominic Kwiatkowski, Cordelia Langford, John Sillitoe on behalf of the Wellcome Sanger Institute COVID-19 Surveillance Team |
| EPI_ISL_2729191 | Lighthouse Lab in Glasgow | Wellcome Sanger Institute for the COVID-19 Genomics UK (COG-UK) Consortium | Harper VanSteenhouse, Yumi Kasai, David Gray, Carol Clugston, Anna Dominiczak and Alex Alderton, Roberto Amato, Jeffrey Barrett, Sonia Goncalves, Ewan Harrison, David K. Jackson, Ian Johnston, Dominic Kwiatkowski, Cordelia Langford, John Sillitoe on behalf of the Wellcome Sanger Institute COVID-19 Surveillance Team |
| EPI_ISL_2729192, EPI_ISL_2729193 | Lighthouse Lab in Alderley Park | Wellcome Sanger Institute for the COVID-19 Genomics UK (COG-UK) Consortium | Jacquelyn Wynn, Mairead Hyland, The Lighthouse Lab in Alderley Park and Alex Alderton, Roberto Amato, Jeffrey Barrett, Sonia Goncalves, Ewan Harrison, David K. Jackson, Ian Johnston, Dominic Kwiatkowski, Cordelia Langford, John Sillitoe on behalf of the Wellcome Sanger Institute COVID-19 Surveillance Team |
| EPI_ISL_2729194, EPI_ISL_2729195 | Lighthouse Lab in Milton Keynes | Wellcome Sanger Institute for the COVID-19 Genomics UK (COG-UK) Consortium | The Lighthouse Lab in Milton Keynes and Alex Alderton, Roberto Amato, Jeffrey Barrett, Sonia Goncalves, Ewan Harrison, David K. Jackson, Ian Johnston, Dominic Kwiatkowski, Cordelia Langford, John Sillitoe on behalf of the Wellcome Sanger Institute COVID-19 Surveillance Team |
| EPI_ISL_2729196 | Lighthouse Lab in Alderley Park | Wellcome Sanger Institute for the COVID-19 Genomics UK (COG-UK) Consortium | Jacquelyn Wynn, Mairead Hyland, The Lighthouse Lab in Alderley Park and Alex Alderton, Roberto Amato, Jeffrey Barrett, Sonia Goncalves, Ewan Harrison, David K. Jackson, Ian Johnston, Dominic Kwiatkowski, Cordelia Langford, John Sillitoe on behalf of the Wellcome Sanger Institute COVID-19 Surveillance Team |
| EPI_ISL_2729197, EPI_ISL_2729198, EPI_ISL_2729199 | Lighthouse Lab in Glasgow | Wellcome Sanger Institute for the COVID-19 Genomics UK (COG-UK) Consortium | Harper VanSteenhouse, Yumi Kasai, David Gray, Carol Clugston, Anna Dominiczak and Alex Alderton, Roberto Amato, Jeffrey Barrett, Sonia Goncalves, Ewan Harrison, David K. Jackson, Ian Johnston, Dominic Kwiatkowski, Cordelia Langford, John Sillitoe on behalf of the Wellcome Sanger Institute COVID-19 Surveillance Team |
| EPI_ISL_2729200, EPI_ISL_2729201, EPI_ISL_2729202, EPI_ISL_2729203 | Lighthouse Lab in Alderley Park | Wellcome Sanger Institute for the COVID-19 Genomics UK (COG-UK) Consortium | Jacquelyn Wynn, Mairead Hyland, The Lighthouse Lab in Alderley Park and Alex Alderton, Roberto Amato, Jeffrey Barrett, Sonia Goncalves, Ewan Harrison, David K. Jackson, Ian Johnston, Dominic Kwiatkowski, Cordelia Langford, John Sillitoe on behalf of the Wellcome Sanger Institute COVID-19 Surveillance Team |
| EPI_ISL_2729204 | Lighthouse Lab in Glasgow | Wellcome Sanger Institute for the COVID-19 Genomics UK (COG-UK) Consortium | Harper VanSteenhouse, Yumi Kasai, David Gray, Carol Clugston, Anna Dominiczak and Alex Alderton, Roberto Amato, Jeffrey Barrett, Sonia Goncalves, Ewan Harrison, David K. Jackson, Ian Johnston, Dominic Kwiatkowski, Cordelia Langford, John Sillitoe on behalf of the Wellcome Sanger Institute COVID-19 Surveillance Team |
| EPI_ISL_2729205 | Lighthouse Lab in Alderley Park | Wellcome Sanger Institute for the COVID-19 Genomics UK (COG-UK) Consortium | Jacquelyn Wynn, Mairead Hyland, The Lighthouse Lab in Alderley Park and Alex Alderton, Roberto Amato, Jeffrey Barrett, Sonia Goncalves, Ewan Harrison, David K. Jackson, Ian Johnston, Dominic Kwiatkowski, Cordelia Langford, John Sillitoe on behalf of the Wellcome Sanger Institute COVID-19 Surveillance Team |
| EPI_ISL_2729206 | Lighthouse Lab in Milton Keynes | Wellcome Sanger Institute for the COVID-19 Genomics UK (COG-UK) Consortium | The Lighthouse Lab in Milton Keynes and Alex Alderton, Roberto Amato, Jeffrey Barrett, Sonia Goncalves, Ewan Harrison, David K. Jackson, Ian Johnston, Dominic Kwiatkowski, Cordelia Langford, John Sillitoe on behalf of the Wellcome Sanger Institute COVID-19 Surveillance Team |
| EPI_ISL_2729207, EPI_ISL_2729208 | Lighthouse Lab in Alderley Park | Wellcome Sanger Institute for the COVID-19 Genomics UK (COG-UK) Consortium | Jacquelyn Wynn, Mairead Hyland, The Lighthouse Lab in Alderley Park and Alex Alderton, Roberto Amato, Jeffrey Barrett, Sonia Goncalves, Ewan Harrison, David K. Jackson, Ian Johnston, Dominic Kwiatkowski, Cordelia Langford, John Sillitoe on behalf of the Wellcome Sanger Institute COVID-19 Surveillance Team |
| EPI_ISL_2729209 | Lighthouse Lab in Glasgow | Wellcome Sanger Institute for the COVID-19 Genomics UK (COG-UK) Consortium | Harper VanSteenhouse, Yumi Kasai, David Gray, Carol Clugston, Anna Dominiczak and Alex Alderton, Roberto Amato, Jeffrey Barrett, Sonia Goncalves, Ewan Harrison, David K. Jackson, Ian Johnston, Dominic Kwiatkowski, Cordelia Langford, John Sillitoe on behalf of the Wellcome Sanger Institute COVID-19 Surveillance Team |
| EPI_ISL_2729210 | Lighthouse Lab in Milton Keynes | Wellcome Sanger Institute for the COVID-19 Genomics UK (COG-UK) Consortium | The Lighthouse Lab in Milton Keynes and Alex Alderton, Roberto Amato, Jeffrey Barrett, Sonia Goncalves, Ewan Harrison, David K. Jackson, Ian Johnston, Dominic Kwiatkowski, Cordelia Langford, John Sillitoe on behalf of the Wellcome Sanger Institute COVID-19 Surveillance Team |
| EPI_ISL_2729211 | Lighthouse Lab in Glasgow | Wellcome Sanger Institute for the COVID-19 Genomics UK (COG-UK) Consortium | Harper VanSteenhouse, Yumi Kasai, David Gray, Carol Clugston, Anna Dominiczak and Alex Alderton, Roberto Amato, Jeffrey Barrett, Sonia Goncalves, Ewan Harrison, David K. Jackson, Ian Johnston, Dominic Kwiatkowski, Cordelia Langford, John Sillitoe on behalf of the Wellcome Sanger Institute COVID-19 Surveillance Team |
| EPI_ISL_2729212 | Lighthouse Lab in Alderley Park | Wellcome Sanger Institute for the COVID-19 Genomics UK (COG-UK) Consortium | Jacquelyn Wynn, Mairead Hyland, The Lighthouse Lab in Alderley Park and Alex Alderton, Roberto Amato, Jeffrey Barrett, Sonia Goncalves, Ewan Harrison, David K. Jackson, Ian Johnston, Dominic Kwiatkowski, Cordelia Langford, John Sillitoe on behalf of the Wellcome Sanger Institute COVID-19 Surveillance Team |
| EPI_ISL_2729213 | Lighthouse Lab in Glasgow | Wellcome Sanger Institute for the COVID-19 Genomics UK (COG-UK) Consortium | Harper VanSteenhouse, Yumi Kasai, David Gray, Carol Clugston, Anna Dominiczak and Alex Alderton, Roberto Amato, Jeffrey Barrett, Sonia Goncalves, Ewan Harrison, David K. Jackson, Ian Johnston, Dominic Kwiatkowski, Cordelia Langford, John Sillitoe on behalf of the Wellcome Sanger Institute COVID-19 Surveillance Team |
| EPI_ISL_2729214 | Lighthouse Lab in Milton Keynes | Wellcome Sanger Institute for the COVID-19 Genomics UK (COG-UK) Consortium | The Lighthouse Lab in Milton Keynes and Alex Alderton, Roberto Amato, Jeffrey Barrett, Sonia Goncalves, Ewan Harrison, David K. Jackson, Ian Johnston, Dominic Kwiatkowski, Cordelia Langford, John Sillitoe on behalf of the Wellcome Sanger Institute COVID-19 Surveillance Team |
| EPI_ISL_2729215 | Lighthouse Lab in Alderley Park | Wellcome Sanger Institute for the COVID-19 Genomics UK (COG-UK) Consortium | Jacquelyn Wynn, Mairead Hyland, The Lighthouse Lab in Alderley Park and Alex Alderton, Roberto Amato, Jeffrey Barrett, Sonia Goncalves, Ewan Harrison, David K. Jackson, Ian Johnston, Dominic Kwiatkowski, Cordelia Langford, John Sillitoe on behalf of the Wellcome Sanger Institute COVID-19 Surveillance Team |
| EPI_ISL_2729216 | Lighthouse Lab in Milton Keynes | Wellcome Sanger Institute for the COVID-19 Genomics UK (COG-UK) Consortium | The Lighthouse Lab in Milton Keynes and Alex Alderton, Roberto Amato, Jeffrey Barrett, Sonia Goncalves, Ewan Harrison, David K. Jackson, Ian Johnston, Dominic Kwiatkowski, Cordelia Langford, John Sillitoe on behalf of the Wellcome Sanger Institute COVID-19 Surveillance Team |
| EPI_ISL_2729217, EPI_ISL_2729218 | Lighthouse Lab in Alderley Park | Wellcome Sanger Institute for the COVID-19 Genomics UK (COG-UK) Consortium | Jacquelyn Wynn, Mairead Hyland, The Lighthouse Lab in Alderley Park and Alex Alderton, Roberto Amato, Jeffrey Barrett, Sonia Goncalves, Ewan Harrison, David K. Jackson, Ian Johnston, Dominic Kwiatkowski, Cordelia Langford, John Sillitoe on behalf of the Wellcome Sanger Institute COVID-19 Surveillance Team |
| EPI_ISL_2729219 | Lighthouse Lab in Milton Keynes | Wellcome Sanger Institute for the COVID-19 Genomics UK (COG-UK) Consortium | The Lighthouse Lab in Milton Keynes and Alex Alderton, Roberto Amato, Jeffrey Barrett, Sonia Goncalves, Ewan Harrison, David K. Jackson, Ian Johnston, Dominic Kwiatkowski, Cordelia Langford, John Sillitoe on behalf of the Wellcome Sanger Institute COVID-19 Surveillance Team |
| EPI_ISL_2729220 | Lighthouse Lab in Alderley Park | Wellcome Sanger Institute for the COVID-19 Genomics UK (COG-UK) Consortium | Jacquelyn Wynn, Mairead Hyland, The Lighthouse Lab in Alderley Park and Alex Alderton, Roberto Amato, Jeffrey Barrett, Sonia Goncalves, Ewan Harrison, David K. Jackson, Ian Johnston, Dominic Kwiatkowski, Cordelia Langford, John Sillitoe on behalf of the Wellcome Sanger Institute COVID-19 Surveillance Team |
| EPI_ISL_2729221 | Lighthouse Lab in Milton Keynes | Wellcome Sanger Institute for the COVID-19 Genomics UK (COG-UK) Consortium | The Lighthouse Lab in Milton Keynes and Alex Alderton, Roberto Amato, Jeffrey Barrett, Sonia Goncalves, Ewan Harrison, David K. Jackson, Ian Johnston, Dominic Kwiatkowski, Cordelia Langford, John Sillitoe on behalf of the Wellcome Sanger Institute COVID-19 Surveillance Team |
| EPI_ISL_2729222, EPI_ISL_2729223 | Lighthouse Lab in Glasgow | Wellcome Sanger Institute for the COVID-19 Genomics UK (COG-UK) Consortium | Harper VanSteenhouse, Yumi Kasai, David Gray, Carol Clugston, Anna Dominiczak and Alex Alderton, Roberto Amato, Jeffrey Barrett, Sonia Goncalves, Ewan Harrison, David K. Jackson, Ian Johnston, Dominic Kwiatkowski, Cordelia Langford, John Sillitoe on behalf of the Wellcome Sanger Institute COVID-19 Surveillance Team |
| EPI_ISL_2729224 | Lighthouse Lab in Milton Keynes | Wellcome Sanger Institute for the COVID-19 Genomics UK (COG-UK) Consortium | The Lighthouse Lab in Milton Keynes and Alex Alderton, Roberto Amato, Jeffrey Barrett, Sonia Goncalves, Ewan Harrison, David K. Jackson, Ian Johnston, Dominic Kwiatkowski, Cordelia Langford, John Sillitoe on behalf of the Wellcome Sanger Institute COVID-19 Surveillance Team |
| EPI_ISL_2729225 | Lighthouse Lab in Glasgow | Wellcome Sanger Institute for the COVID-19 Genomics UK (COG-UK) Consortium | Harper VanSteenhouse, Yumi Kasai, David Gray, Carol Clugston, Anna Dominiczak and Alex Alderton, Roberto Amato, Jeffrey Barrett, Sonia Goncalves, Ewan Harrison, David K. Jackson, Ian Johnston, Dominic Kwiatkowski, Cordelia Langford, John Sillitoe on behalf of the Wellcome Sanger Institute COVID-19 Surveillance Team |
| EPI_ISL_2729226 | Lighthouse Lab in Alderley Park | Wellcome Sanger Institute for the COVID-19 Genomics UK (COG-UK) Consortium | Jacquelyn Wynn, Mairead Hyland, The Lighthouse Lab in Alderley Park and Alex Alderton, Roberto Amato, Jeffrey Barrett, Sonia Goncalves, Ewan Harrison, David K. Jackson, Ian Johnston, Dominic Kwiatkowski, Cordelia Langford, John Sillitoe on behalf of the Wellcome Sanger Institute COVID-19 Surveillance Team |
| EPI_ISL_2729227 | Lighthouse Lab in Milton Keynes | Wellcome Sanger Institute for the COVID-19 Genomics UK (COG-UK) Consortium | The Lighthouse Lab in Milton Keynes and Alex Alderton, Roberto Amato, Jeffrey Barrett, Sonia Goncalves, Ewan Harrison, David K. Jackson, Ian Johnston, Dominic Kwiatkowski, Cordelia Langford, John Sillitoe on behalf of the Wellcome Sanger Institute COVID-19 Surveillance Team |
| EPI_ISL_2729228, EPI_ISL_2729229 | Lighthouse Lab in Alderley Park | Wellcome Sanger Institute for the COVID-19 Genomics UK (COG-UK) Consortium | Jacquelyn Wynn, Mairead Hyland, The Lighthouse Lab in Alderley Park and Alex Alderton, Roberto Amato, Jeffrey Barrett, Sonia Goncalves, Ewan Harrison, David K. Jackson, Ian Johnston, Dominic Kwiatkowski, Cordelia Langford, John Sillitoe on behalf of the Wellcome Sanger Institute COVID-19 Surveillance Team |
| EPI_ISL_2729230, EPI_ISL_2729231 | Lighthouse Lab in Milton Keynes | Wellcome Sanger Institute for the COVID-19 Genomics UK (COG-UK) Consortium | The Lighthouse Lab in Milton Keynes and Alex Alderton, Roberto Amato, Jeffrey Barrett, Sonia Goncalves, Ewan Harrison, David K. Jackson, Ian Johnston, Dominic Kwiatkowski, Cordelia Langford, John Sillitoe on behalf of the Wellcome Sanger Institute COVID-19 Surveillance Team |
| EPI_ISL_2729232, EPI_ISL_2729233, EPI_ISL_2729234 | Lighthouse Lab in Alderley Park | Wellcome Sanger Institute for the COVID-19 Genomics UK (COG-UK) Consortium | Jacquelyn Wynn, Mairead Hyland, The Lighthouse Lab in Alderley Park and Alex Alderton, Roberto Amato, Jeffrey Barrett, Sonia Goncalves, Ewan Harrison, David K. Jackson, Ian Johnston, Dominic Kwiatkowski, Cordelia Langford, John Sillitoe on behalf of the Wellcome Sanger Institute COVID-19 Surveillance Team |
| EPI_ISL_2729235, EPI_ISL_2729236 | Lighthouse Lab in Milton Keynes | Wellcome Sanger Institute for the COVID-19 Genomics UK (COG-UK) Consortium | The Lighthouse Lab in Milton Keynes and Alex Alderton, Roberto Amato, Jeffrey Barrett, Sonia Goncalves, Ewan Harrison, David K. Jackson, Ian Johnston, Dominic Kwiatkowski, Cordelia Langford, John Sillitoe on behalf of the Wellcome Sanger Institute COVID-19 Surveillance Team |
| EPI_ISL_2729237 | Lighthouse Lab in Alderley Park | Wellcome Sanger Institute for the COVID-19 Genomics UK (COG-UK) Consortium | Jacquelyn Wynn, Mairead Hyland, The Lighthouse Lab in Alderley Park and Alex Alderton, Roberto Amato, Jeffrey Barrett, Sonia Goncalves, Ewan Harrison, David K. Jackson, Ian Johnston, Dominic Kwiatkowski, Cordelia Langford, John Sillitoe on behalf of the Wellcome Sanger Institute COVID-19 Surveillance Team |
| EPI_ISL_2729238 | Lighthouse Lab in Milton Keynes | Wellcome Sanger Institute for the COVID-19 Genomics UK (COG-UK) Consortium | The Lighthouse Lab in Milton Keynes and Alex Alderton, Roberto Amato, Jeffrey Barrett, Sonia Goncalves, Ewan Harrison, David K. Jackson, Ian Johnston, Dominic Kwiatkowski, Cordelia Langford, John Sillitoe on behalf of the Wellcome Sanger Institute COVID-19 Surveillance Team |
| EPI_ISL_2729239, EPI_ISL_2729240 | Lighthouse Lab in Alderley Park | Wellcome Sanger Institute for the COVID-19 Genomics UK (COG-UK) Consortium | Jacquelyn Wynn, Mairead Hyland, The Lighthouse Lab in Alderley Park and Alex Alderton, Roberto Amato, Jeffrey Barrett, Sonia Goncalves, Ewan Harrison, David K. Jackson, Ian Johnston, Dominic Kwiatkowski, Cordelia Langford, John Sillitoe on behalf of the Wellcome Sanger Institute COVID-19 Surveillance Team |
| EPI_ISL_2729241 | Lighthouse Lab in Milton Keynes | Wellcome Sanger Institute for the COVID-19 Genomics UK (COG-UK) Consortium | The Lighthouse Lab in Milton Keynes and Alex Alderton, Roberto Amato, Jeffrey Barrett, Sonia Goncalves, Ewan Harrison, David K. Jackson, Ian Johnston, Dominic Kwiatkowski, Cordelia Langford, John Sillitoe on behalf of the Wellcome Sanger Institute COVID-19 Surveillance Team |
| EPI_ISL_2729242 | Lighthouse Lab in Glasgow | Wellcome Sanger Institute for the COVID-19 Genomics UK (COG-UK) | Harper VanSteenhouse, Yumi Kasai, David Gray, Carol Clugston, Anna Dominiczak and Alex Alderton, Roberto Amato, Jeffrey Barrett, Sonia Goncalves, Ewan Harrison, David K. Jackson, Ian Johnston, Dominic Kwiatkowski, Cordelia Langford, |

| | | | |
|---|---|---|---|
| | | Consortium | John Sillitoe on behalf of the Wellcome Sanger Institute COVID-19 Surveillance Team |
| EPI_ISL_2729243 | Lighthouse Lab in Milton Keynes | Wellcome Sanger Institute for the COVID-19 Genomics UK (COG-UK) Consortium | The Lighthouse Lab in Milton Keynes and Alex Alderton, Roberto Amato, Jeffrey Barrett, Sonia Goncalves, Ewan Harrison, David K. Jackson, Ian Johnston, Dominic Kwiatkowski, Cordelia Langford, John Sillitoe on behalf of the Wellcome Sanger Institute COVID-19 Surveillance Team |
| EPI_ISL_2729244, EPI_ISL_2729245, EPI_ISL_2729246, EPI_ISL_2729247 | Lighthouse Lab in Alderley Park | Wellcome Sanger Institute for the COVID-19 Genomics UK (COG-UK) Consortium | Jacquelyn Wynn, Mairead Hyland, The Lighthouse Lab in Alderley Park and Alex Alderton, Roberto Amato, Jeffrey Barrett, Sonia Goncalves, Ewan Harrison, David K. Jackson, Ian Johnston, Dominic Kwiatkowski, Cordelia Langford, John Sillitoe on behalf of the Wellcome Sanger Institute COVID-19 Surveillance Team |
| EPI_ISL_2729248 | Lighthouse Lab in Milton Keynes | Wellcome Sanger Institute for the COVID-19 Genomics UK (COG-UK) Consortium | The Lighthouse Lab in Milton Keynes and Alex Alderton, Roberto Amato, Jeffrey Barrett, Sonia Goncalves, Ewan Harrison, David K. Jackson, Ian Johnston, Dominic Kwiatkowski, Cordelia Langford, John Sillitoe on behalf of the Wellcome Sanger Institute COVID-19 Surveillance Team |
| EPI_ISL_2729249, EPI_ISL_2729250, EPI_ISL_2729251, EPI_ISL_2729252 | Lighthouse Lab in Alderley Park | Wellcome Sanger Institute for the COVID-19 Genomics UK (COG-UK) Consortium | Jacquelyn Wynn, Mairead Hyland, The Lighthouse Lab in Alderley Park and Alex Alderton, Roberto Amato, Jeffrey Barrett, Sonia Goncalves, Ewan Harrison, David K. Jackson, Ian Johnston, Dominic Kwiatkowski, Cordelia Langford, John Sillitoe on behalf of the Wellcome Sanger Institute COVID-19 Surveillance Team |
| EPI_ISL_2729253 | Lighthouse Lab in Glasgow | Wellcome Sanger Institute for the COVID-19 Genomics UK (COG-UK) Consortium | Harper VanSteenhouse, Yumi Kasai, David Gray, Carol Clugston, Anna Dominiczak and Alex Alderton, Roberto Amato, Jeffrey Barrett, Sonia Goncalves, Ewan Harrison, David K. Jackson, Ian Johnston, Dominic Kwiatkowski, Cordelia Langford, John Sillitoe on behalf of the Wellcome Sanger Institute COVID-19 Surveillance Team |
| EPI_ISL_2729254, EPI_ISL_2729255 | Lighthouse Lab in Milton Keynes | Wellcome Sanger Institute for the COVID-19 Genomics UK (COG-UK) Consortium | The Lighthouse Lab in Milton Keynes and Alex Alderton, Roberto Amato, Jeffrey Barrett, Sonia Goncalves, Ewan Harrison, David K. Jackson, Ian Johnston, Dominic Kwiatkowski, Cordelia Langford, John Sillitoe on behalf of the Wellcome Sanger Institute COVID-19 Surveillance Team |
| EPI_ISL_2729256 | Lighthouse Lab in Alderley Park | Wellcome Sanger Institute for the COVID-19 Genomics UK (COG-UK) Consortium | Jacquelyn Wynn, Mairead Hyland, The Lighthouse Lab in Alderley Park and Alex Alderton, Roberto Amato, Jeffrey Barrett, Sonia Goncalves, Ewan Harrison, David K. Jackson, Ian Johnston, Dominic Kwiatkowski, Cordelia Langford, John Sillitoe on behalf of the Wellcome Sanger Institute COVID-19 Surveillance Team |
| EPI_ISL_2729257, EPI_ISL_2729258 | Lighthouse Lab in Glasgow | Wellcome Sanger Institute for the COVID-19 Genomics UK (COG-UK) Consortium | Harper VanSteenhouse, Yumi Kasai, David Gray, Carol Clugston, Anna Dominiczak and Alex Alderton, Roberto Amato, Jeffrey Barrett, Sonia Goncalves, Ewan Harrison, David K. Jackson, Ian Johnston, Dominic Kwiatkowski, Cordelia Langford, John Sillitoe on behalf of the Wellcome Sanger Institute COVID-19 Surveillance Team |
| EPI_ISL_2729259, EPI_ISL_2729260 | Lighthouse Lab in Milton Keynes | Wellcome Sanger Institute for the COVID-19 Genomics UK (COG-UK) Consortium | The Lighthouse Lab in Milton Keynes and Alex Alderton, Roberto Amato, Jeffrey Barrett, Sonia Goncalves, Ewan Harrison, David K. Jackson, Ian Johnston, Dominic Kwiatkowski, Cordelia Langford, John Sillitoe on behalf of the Wellcome Sanger Institute COVID-19 Surveillance Team |
| EPI_ISL_2729261 | Lighthouse Lab in Glasgow | Wellcome Sanger Institute for the COVID-19 Genomics UK (COG-UK) Consortium | Harper VanSteenhouse, Yumi Kasai, David Gray, Carol Clugston, Anna Dominiczak and Alex Alderton, Roberto Amato, Jeffrey Barrett, Sonia Goncalves, Ewan Harrison, David K. Jackson, Ian Johnston, Dominic Kwiatkowski, Cordelia Langford, John Sillitoe on behalf of the Wellcome Sanger Institute COVID-19 Surveillance Team |
| EPI_ISL_2729262, EPI_ISL_2729263 | Lighthouse Lab in Milton Keynes | Wellcome Sanger Institute for the COVID-19 Genomics UK (COG-UK) Consortium | The Lighthouse Lab in Milton Keynes and Alex Alderton, Roberto Amato, Jeffrey Barrett, Sonia Goncalves, Ewan Harrison, David K. Jackson, Ian Johnston, Dominic Kwiatkowski, Cordelia Langford, John Sillitoe on behalf of the Wellcome Sanger Institute COVID-19 Surveillance Team |
| EPI_ISL_2729264 | Lighthouse Lab in Glasgow | Wellcome Sanger Institute for the COVID-19 Genomics UK (COG-UK) Consortium | Harper VanSteenhouse, Yumi Kasai, David Gray, Carol Clugston, Anna Dominiczak and Alex Alderton, Roberto Amato, Jeffrey Barrett, Sonia Goncalves, Ewan Harrison, David K. Jackson, Ian Johnston, Dominic Kwiatkowski, Cordelia Langford, John Sillitoe on behalf of the Wellcome Sanger Institute COVID-19 Surveillance Team |
| EPI_ISL_2729265 | Lighthouse Lab in Alderley Park | Wellcome Sanger Institute for the COVID-19 Genomics UK (COG-UK) Consortium | Jacquelyn Wynn, Mairead Hyland, The Lighthouse Lab in Alderley Park and Alex Alderton, Roberto Amato, Jeffrey Barrett, Sonia Goncalves, Ewan Harrison, David K. Jackson, Ian Johnston, Dominic Kwiatkowski, Cordelia Langford, John Sillitoe on behalf of the Wellcome Sanger Institute COVID-19 Surveillance Team |
| EPI_ISL_2729266 | Lighthouse Lab in Milton Keynes | Wellcome Sanger Institute for the COVID-19 Genomics UK (COG-UK) Consortium | The Lighthouse Lab in Milton Keynes and Alex Alderton, Roberto Amato, Jeffrey Barrett, Sonia Goncalves, Ewan Harrison, David K. Jackson, Ian Johnston, Dominic Kwiatkowski, Cordelia Langford, John Sillitoe on behalf of the Wellcome Sanger Institute COVID-19 Surveillance Team |
| EPI_ISL_2729267, EPI_ISL_2729268, EPI_ISL_2729269, EPI_ISL_2729270 | Lighthouse Lab in Alderley Park | Wellcome Sanger Institute for the COVID-19 Genomics UK (COG-UK) Consortium | Jacquelyn Wynn, Mairead Hyland, The Lighthouse Lab in Alderley Park and Alex Alderton, Roberto Amato, Jeffrey Barrett, Sonia Goncalves, Ewan Harrison, David K. Jackson, Ian Johnston, Dominic Kwiatkowski, Cordelia Langford, John Sillitoe on behalf of the Wellcome Sanger Institute COVID-19 Surveillance Team |
| EPI_ISL_2729271 | Lighthouse Lab in Milton Keynes | Wellcome Sanger Institute for the COVID-19 Genomics UK (COG-UK) Consortium | The Lighthouse Lab in Milton Keynes and Alex Alderton, Roberto Amato, Jeffrey Barrett, Sonia Goncalves, Ewan Harrison, David K. Jackson, Ian Johnston, Dominic Kwiatkowski, Cordelia Langford, John Sillitoe on behalf of the Wellcome Sanger Institute COVID-19 Surveillance Team |
| EPI_ISL_2729273, EPI_ISL_2729274 | Lighthouse Lab in Alderley Park | Wellcome Sanger Institute for the COVID-19 Genomics UK (COG-UK) Consortium | Jacquelyn Wynn, Mairead Hyland, The Lighthouse Lab in Alderley Park and Alex Alderton, Roberto Amato, Jeffrey Barrett, Sonia Goncalves, Ewan Harrison, David K. Jackson, Ian Johnston, Dominic Kwiatkowski, Cordelia Langford, John Sillitoe on behalf of the Wellcome Sanger Institute COVID-19 Surveillance Team |
| EPI_ISL_2729275 | Lighthouse Lab in Glasgow | Wellcome Sanger Institute for the COVID-19 Genomics UK (COG-UK) Consortium | Harper VanSteenhouse, Yumi Kasai, David Gray, Carol Clugston, Anna Dominiczak and Alex Alderton, Roberto Amato, Jeffrey Barrett, Sonia Goncalves, Ewan Harrison, David K. Jackson, Ian Johnston, Dominic Kwiatkowski, Cordelia Langford, John Sillitoe on behalf of the Wellcome Sanger Institute COVID-19 Surveillance Team |
| EPI_ISL_2729276 | Lighthouse Lab in Alderley Park | Wellcome Sanger Institute for the COVID-19 Genomics UK (COG-UK) Consortium | Jacquelyn Wynn, Mairead Hyland, The Lighthouse Lab in Alderley Park and Alex Alderton, Roberto Amato, Jeffrey Barrett, Sonia Goncalves, Ewan Harrison, David K. Jackson, Ian Johnston, Dominic Kwiatkowski, Cordelia Langford, John Sillitoe on behalf of the Wellcome Sanger Institute COVID-19 Surveillance Team |
| EPI_ISL_2729277 | Lighthouse Lab in Milton Keynes | Wellcome Sanger Institute for the COVID-19 Genomics UK (COG-UK) Consortium | The Lighthouse Lab in Milton Keynes and Alex Alderton, Roberto Amato, Jeffrey Barrett, Sonia Goncalves, Ewan Harrison, David K. Jackson, Ian Johnston, Dominic Kwiatkowski, Cordelia Langford, John Sillitoe on behalf of the Wellcome Sanger Institute COVID-19 Surveillance Team |
| EPI_ISL_2729278 | Lighthouse Lab in Alderley Park | Wellcome Sanger Institute for the COVID-19 Genomics UK (COG-UK) Consortium | Jacquelyn Wynn, Mairead Hyland, The Lighthouse Lab in Alderley Park and Alex Alderton, Roberto Amato, Jeffrey Barrett, Sonia Goncalves, Ewan Harrison, David K. Jackson, Ian Johnston, Dominic Kwiatkowski, Cordelia Langford, John Sillitoe on behalf of the Wellcome Sanger Institute COVID-19 Surveillance Team |
| EPI_ISL_2729279 | Lighthouse Lab in Milton Keynes | Wellcome Sanger Institute for the COVID-19 Genomics UK (COG-UK) Consortium | The Lighthouse Lab in Milton Keynes and Alex Alderton, Roberto Amato, Jeffrey Barrett, Sonia Goncalves, Ewan Harrison, David K. Jackson, Ian Johnston, Dominic Kwiatkowski, Cordelia Langford, John Sillitoe on behalf of the Wellcome Sanger Institute COVID-19 Surveillance Team |
| EPI_ISL_2729280 | Lighthouse Lab in Alderley Park | Wellcome Sanger Institute for the COVID-19 Genomics UK (COG-UK) Consortium | Jacquelyn Wynn, Mairead Hyland, The Lighthouse Lab in Alderley Park and Alex Alderton, Roberto Amato, Jeffrey Barrett, Sonia Goncalves, Ewan Harrison, David K. Jackson, Ian Johnston, Dominic Kwiatkowski, Cordelia Langford, John Sillitoe on behalf of the Wellcome Sanger Institute COVID-19 Surveillance Team |
| EPI_ISL_2729281 | Lighthouse Lab in Milton Keynes | Wellcome Sanger Institute for the COVID-19 Genomics UK (COG-UK) Consortium | The Lighthouse Lab in Milton Keynes and Alex Alderton, Roberto Amato, Jeffrey Barrett, Sonia Goncalves, Ewan Harrison, David K. Jackson, Ian Johnston, Dominic Kwiatkowski, Cordelia Langford, John Sillitoe on behalf of the Wellcome Sanger Institute COVID-19 Surveillance Team |
| EPI_ISL_2729282 | Lighthouse Lab in Glasgow | Wellcome Sanger Institute for the COVID-19 Genomics UK (COG-UK) Consortium | Harper VanSteenhouse, Yumi Kasai, David Gray, Carol Clugston, Anna Dominiczak and Alex Alderton, Roberto Amato, Jeffrey Barrett, Sonia Goncalves, Ewan Harrison, David K. Jackson, Ian Johnston, Dominic Kwiatkowski, Cordelia Langford, John Sillitoe on behalf of the Wellcome Sanger Institute COVID-19 Surveillance Team |
| EPI_ISL_2729284 | Lighthouse Lab in Alderley Park | Wellcome Sanger Institute for the COVID-19 Genomics UK (COG-UK) Consortium | Jacquelyn Wynn, Mairead Hyland, The Lighthouse Lab in Alderley Park and Alex Alderton, Roberto Amato, Jeffrey Barrett, Sonia Goncalves, Ewan Harrison, David K. Jackson, Ian Johnston, Dominic Kwiatkowski, Cordelia Langford, John Sillitoe on behalf of the Wellcome Sanger Institute COVID-19 Surveillance Team |
| EPI_ISL_2729285, EPI_ISL_2729286 | Lighthouse Lab in Milton Keynes | Wellcome Sanger Institute for the COVID-19 Genomics UK (COG-UK) Consortium | The Lighthouse Lab in Milton Keynes and Alex Alderton, Roberto Amato, Jeffrey Barrett, Sonia Goncalves, Ewan Harrison, David K. Jackson, Ian Johnston, Dominic Kwiatkowski, Cordelia Langford, John Sillitoe on behalf of the Wellcome Sanger Institute COVID-19 Surveillance Team |
| EPI_ISL_2729287, EPI_ISL_2729288 | Lighthouse Lab in Glasgow | Wellcome Sanger Institute for the COVID-19 Genomics UK (COG-UK) Consortium | Harper VanSteenhouse, Yumi Kasai, David Gray, Carol Clugston, Anna Dominiczak and Alex Alderton, Roberto Amato, Jeffrey Barrett, Sonia Goncalves, Ewan Harrison, David K. Jackson, Ian Johnston, Dominic Kwiatkowski, Cordelia Langford, John Sillitoe on behalf of the Wellcome Sanger Institute COVID-19 Surveillance Team |
| EPI_ISL_2729289 | Lighthouse Lab in Alderley Park | Wellcome Sanger Institute for the COVID-19 Genomics UK (COG-UK) Consortium | Jacquelyn Wynn, Mairead Hyland, The Lighthouse Lab in Alderley Park and Alex Alderton, Roberto Amato, Jeffrey Barrett, Sonia Goncalves, Ewan Harrison, David K. Jackson, Ian Johnston, Dominic Kwiatkowski, Cordelia Langford, John Sillitoe on behalf of the Wellcome Sanger Institute COVID-19 Surveillance Team |
| EPI_ISL_2729290 | Lighthouse Lab in Milton Keynes | Wellcome Sanger Institute for the COVID-19 Genomics UK (COG-UK) Consortium | The Lighthouse Lab in Milton Keynes and Alex Alderton, Roberto Amato, Jeffrey Barrett, Sonia Goncalves, Ewan Harrison, David K. Jackson, Ian Johnston, Dominic Kwiatkowski, Cordelia Langford, John Sillitoe on behalf of the Wellcome Sanger Institute COVID-19 Surveillance Team |
| EPI_ISL_2729291 | Lighthouse Lab in Alderley Park | Wellcome Sanger Institute for the COVID-19 Genomics UK (COG-UK) Consortium | Jacquelyn Wynn, Mairead Hyland, The Lighthouse Lab in Alderley Park and Alex Alderton, Roberto Amato, Jeffrey Barrett, Sonia Goncalves, Ewan Harrison, David K. Jackson, Ian Johnston, Dominic Kwiatkowski, Cordelia Langford, John Sillitoe on behalf of the Wellcome Sanger Institute COVID-19 Surveillance Team |
| EPI_ISL_2729292 | Lighthouse Lab in Glasgow | Wellcome Sanger Institute for the COVID-19 Genomics UK (COG-UK) Consortium | Harper VanSteenhouse, Yumi Kasai, David Gray, Carol Clugston, Anna Dominiczak and Alex Alderton, Roberto Amato, Jeffrey Barrett, Sonia Goncalves, Ewan Harrison, David K. Jackson, Ian Johnston, Dominic Kwiatkowski, Cordelia Langford, John Sillitoe on behalf of the Wellcome Sanger Institute COVID-19 Surveillance Team |
| EPI_ISL_2729293, EPI_ISL_2729294, EPI_ISL_2729295 | Lighthouse Lab in Alderley Park | Wellcome Sanger Institute for the COVID-19 Genomics UK (COG-UK) Consortium | Jacquelyn Wynn, Mairead Hyland, The Lighthouse Lab in Alderley Park and Alex Alderton, Roberto Amato, Jeffrey Barrett, Sonia Goncalves, Ewan Harrison, David K. Jackson, Ian Johnston, Dominic Kwiatkowski, Cordelia Langford, John Sillitoe on behalf of the Wellcome Sanger Institute COVID-19 Surveillance Team |
| EPI_ISL_2729296 | Lighthouse Lab in Glasgow | Wellcome Sanger Institute for the COVID-19 Genomics UK (COG-UK) Consortium | Harper VanSteenhouse, Yumi Kasai, David Gray, Carol Clugston, Anna Dominiczak and Alex Alderton, Roberto Amato, Jeffrey Barrett, Sonia Goncalves, Ewan Harrison, David K. Jackson, Ian Johnston, Dominic Kwiatkowski, Cordelia Langford, John Sillitoe on behalf of the Wellcome Sanger Institute COVID-19 Surveillance Team |
| EPI_ISL_2729297 | Lighthouse Lab in Alderley Park | Wellcome Sanger Institute for the COVID-19 Genomics UK (COG-UK) Consortium | Jacquelyn Wynn, Mairead Hyland, The Lighthouse Lab in Alderley Park and Alex Alderton, Roberto Amato, Jeffrey Barrett, Sonia Goncalves, Ewan Harrison, David K. Jackson, Ian Johnston, Dominic Kwiatkowski, Cordelia Langford, John Sillitoe on behalf of the Wellcome Sanger Institute COVID-19 Surveillance Team |
| EPI_ISL_2729298 | Lighthouse Lab in Milton Keynes | Wellcome Sanger Institute for the COVID-19 Genomics UK (COG-UK) Consortium | The Lighthouse Lab in Milton Keynes and Alex Alderton, Roberto Amato, Jeffrey Barrett, Sonia Goncalves, Ewan Harrison, David K. Jackson, Ian Johnston, Dominic Kwiatkowski, Cordelia Langford, John Sillitoe on behalf of the Wellcome Sanger Institute COVID-19 Surveillance Team |
| EPI_ISL_2729299 | Lighthouse Lab in Glasgow | Wellcome Sanger Institute for the COVID-19 Genomics UK (COG-UK) Consortium | Harper VanSteenhouse, Yumi Kasai, David Gray, Carol Clugston, Anna Dominiczak and Alex Alderton, Roberto Amato, Jeffrey Barrett, Sonia Goncalves, Ewan Harrison, David K. Jackson, Ian Johnston, Dominic Kwiatkowski, Cordelia Langford, John Sillitoe on behalf of the Wellcome Sanger Institute COVID-19 Surveillance Team |
| EPI_ISL_2729300, EPI_ISL_2729301, EPI_ISL_2729302 | Lighthouse Lab in Alderley Park | Wellcome Sanger Institute for the COVID-19 Genomics UK (COG-UK) Consortium | Jacquelyn Wynn, Mairead Hyland, The Lighthouse Lab in Alderley Park and Alex Alderton, Roberto Amato, Jeffrey Barrett, Sonia Goncalves, Ewan Harrison, David K. Jackson, Ian Johnston, Dominic Kwiatkowski, Cordelia Langford, John Sillitoe on behalf of the Wellcome Sanger Institute COVID-19 Surveillance Team |
| EPI_ISL_2729303, EPI_ISL_2729304 | Lighthouse Lab in Milton Keynes | Wellcome Sanger Institute for the COVID-19 Genomics UK (COG-UK) Consortium | The Lighthouse Lab in Milton Keynes and Alex Alderton, Roberto Amato, Jeffrey Barrett, Sonia Goncalves, Ewan Harrison, David K. Jackson, Ian Johnston, Dominic Kwiatkowski, Cordelia Langford, John Sillitoe on behalf of the Wellcome Sanger Institute COVID-19 Surveillance Team |
| EPI_ISL_2729305 | Lighthouse Lab in Alderley Park | Wellcome Sanger Institute for the COVID-19 Genomics UK (COG-UK) Consortium | Jacquelyn Wynn, Mairead Hyland, The Lighthouse Lab in Alderley Park and Alex Alderton, Roberto Amato, Jeffrey Barrett, Sonia Goncalves, Ewan Harrison, David K. Jackson, Ian Johnston, Dominic Kwiatkowski, Cordelia Langford, John Sillitoe on behalf of the Wellcome Sanger Institute COVID-19 Surveillance Team |
| EPI_ISL_2729306 | Lighthouse Lab in Glasgow | Wellcome Sanger Institute for the COVID-19 Genomics UK (COG-UK) Consortium | Harper VanSteenhouse, Yumi Kasai, David Gray, Carol Clugston, Anna Dominiczak and Alex Alderton, Roberto Amato, Jeffrey Barrett, Sonia Goncalves, Ewan Harrison, David K. Jackson, Ian Johnston, Dominic Kwiatkowski, Cordelia Langford, John Sillitoe on behalf of the Wellcome Sanger Institute COVID-19 Surveillance Team |
| EPI_ISL_2729307 | Lighthouse Lab in Milton Keynes | Wellcome Sanger Institute for the COVID-19 Genomics UK (COG-UK) | The Lighthouse Lab in Milton Keynes and Alex Alderton, Roberto Amato, Jeffrey Barrett, Sonia Goncalves, Ewan Harrison, David K. Jackson, Ian Johnston, Dominic Kwiatkowski, Cordelia Langford, John Sillitoe on behalf of the Wellcome Sanger |

| | | | |
|---|---|---|---|
| | | Consortium | Institute COVID-19 Surveillance Team |
| EPI_ISL_2729308, EPI_ISL_2729309 | Lighthouse Lab in Alderley Park | Wellcome Sanger Institute for the COVID-19 Genomics UK (COG-UK) Consortium | Jacquelyn Wynn, Mairead Hyland, The Lighthouse Lab in Alderley Park and Alex Alderton, Roberto Amato, Jeffrey Barrett, Sonia Goncalves, Ewan Harrison, David K. Jackson, Ian Johnston, Dominic Kwiatkowski, Cordelia Langford, John Sillitoe on behalf of the Wellcome Sanger Institute COVID-19 Surveillance Team |
| EPI_ISL_2729310 | Lighthouse Lab in Milton Keynes | Wellcome Sanger Institute for the COVID-19 Genomics UK (COG-UK) Consortium | The Lighthouse Lab in Milton Keynes and Alex Alderton, Roberto Amato, Jeffrey Barrett, Sonia Goncalves, Ewan Harrison, David K. Jackson, Ian Johnston, Dominic Kwiatkowski, Cordelia Langford, John Sillitoe on behalf of the Wellcome Sanger Institute COVID-19 Surveillance Team |
| EPI_ISL_2729311, EPI_ISL_2729312 | Lighthouse Lab in Alderley Park | Wellcome Sanger Institute for the COVID-19 Genomics UK (COG-UK) Consortium | Jacquelyn Wynn, Mairead Hyland, The Lighthouse Lab in Alderley Park and Alex Alderton, Roberto Amato, Jeffrey Barrett, Sonia Goncalves, Ewan Harrison, David K. Jackson, Ian Johnston, Dominic Kwiatkowski, Cordelia Langford, John Sillitoe on behalf of the Wellcome Sanger Institute COVID-19 Surveillance Team |
| EPI_ISL_2729313 | Lighthouse Lab in Glasgow | Wellcome Sanger Institute for the COVID-19 Genomics UK (COG-UK) Consortium | Harper VanSteenhouse, Yumi Kasai, David Gray, Carol Clugston, Anna Dominiczak and Alex Alderton, Roberto Amato, Jeffrey Barrett, Sonia Goncalves, Ewan Harrison, David K. Jackson, Ian Johnston, Dominic Kwiatkowski, Cordelia Langford, John Sillitoe on behalf of the Wellcome Sanger Institute COVID-19 Surveillance Team |
| EPI_ISL_2729314, EPI_ISL_2729315, EPI_ISL_2729316 | Lighthouse Lab in Alderley Park | Wellcome Sanger Institute for the COVID-19 Genomics UK (COG-UK) Consortium | Jacquelyn Wynn, Mairead Hyland, The Lighthouse Lab in Alderley Park and Alex Alderton, Roberto Amato, Jeffrey Barrett, Sonia Goncalves, Ewan Harrison, David K. Jackson, Ian Johnston, Dominic Kwiatkowski, Cordelia Langford, John Sillitoe on behalf of the Wellcome Sanger Institute COVID-19 Surveillance Team |
| EPI_ISL_2729317 | Lighthouse Lab in Glasgow | Wellcome Sanger Institute for the COVID-19 Genomics UK (COG-UK) Consortium | Harper VanSteenhouse, Yumi Kasai, David Gray, Carol Clugston, Anna Dominiczak and Alex Alderton, Roberto Amato, Jeffrey Barrett, Sonia Goncalves, Ewan Harrison, David K. Jackson, Ian Johnston, Dominic Kwiatkowski, Cordelia Langford, John Sillitoe on behalf of the Wellcome Sanger Institute COVID-19 Surveillance Team |
| EPI_ISL_2729318, EPI_ISL_2729319 | Lighthouse Lab in Milton Keynes | Wellcome Sanger Institute for the COVID-19 Genomics UK (COG-UK) Consortium | The Lighthouse Lab in Milton Keynes and Alex Alderton, Roberto Amato, Jeffrey Barrett, Sonia Goncalves, Ewan Harrison, David K. Jackson, Ian Johnston, Dominic Kwiatkowski, Cordelia Langford, John Sillitoe on behalf of the Wellcome Sanger Institute COVID-19 Surveillance Team |
| EPI_ISL_2729320, EPI_ISL_2729321, EPI_ISL_2729322 | Lighthouse Lab in Glasgow | Wellcome Sanger Institute for the COVID-19 Genomics UK (COG-UK) Consortium | Harper VanSteenhouse, Yumi Kasai, David Gray, Carol Clugston, Anna Dominiczak and Alex Alderton, Roberto Amato, Jeffrey Barrett, Sonia Goncalves, Ewan Harrison, David K. Jackson, Ian Johnston, Dominic Kwiatkowski, Cordelia Langford, John Sillitoe on behalf of the Wellcome Sanger Institute COVID-19 Surveillance Team |
| EPI_ISL_2729323 | Lighthouse Lab in Milton Keynes | Wellcome Sanger Institute for the COVID-19 Genomics UK (COG-UK) Consortium | The Lighthouse Lab in Milton Keynes and Alex Alderton, Roberto Amato, Jeffrey Barrett, Sonia Goncalves, Ewan Harrison, David K. Jackson, Ian Johnston, Dominic Kwiatkowski, Cordelia Langford, John Sillitoe on behalf of the Wellcome Sanger Institute COVID-19 Surveillance Team |
| EPI_ISL_2729324, EPI_ISL_2729325, EPI_ISL_2729326, EPI_ISL_2729327, EPI_ISL_2729328, EPI_ISL_2729329 | Lighthouse Lab in Alderley Park | Wellcome Sanger Institute for the COVID-19 Genomics UK (COG-UK) Consortium | Jacquelyn Wynn, Mairead Hyland, The Lighthouse Lab in Alderley Park and Alex Alderton, Roberto Amato, Jeffrey Barrett, Sonia Goncalves, Ewan Harrison, David K. Jackson, Ian Johnston, Dominic Kwiatkowski, Cordelia Langford, John Sillitoe on behalf of the Wellcome Sanger Institute COVID-19 Surveillance Team |
| EPI_ISL_2729330, EPI_ISL_2729331 | Lighthouse Lab in Milton Keynes | Wellcome Sanger Institute for the COVID-19 Genomics UK (COG-UK) Consortium | The Lighthouse Lab in Milton Keynes and Alex Alderton, Roberto Amato, Jeffrey Barrett, Sonia Goncalves, Ewan Harrison, David K. Jackson, Ian Johnston, Dominic Kwiatkowski, Cordelia Langford, John Sillitoe on behalf of the Wellcome Sanger Institute COVID-19 Surveillance Team |
| EPI_ISL_2729332 | Lighthouse Lab in Glasgow | Wellcome Sanger Institute for the COVID-19 Genomics UK (COG-UK) Consortium | Harper VanSteenhouse, Yumi Kasai, David Gray, Carol Clugston, Anna Dominiczak and Alex Alderton, Roberto Amato, Jeffrey Barrett, Sonia Goncalves, Ewan Harrison, David K. Jackson, Ian Johnston, Dominic Kwiatkowski, Cordelia Langford, John Sillitoe on behalf of the Wellcome Sanger Institute COVID-19 Surveillance Team |
| EPI_ISL_2729333, EPI_ISL_2729334, EPI_ISL_2729335, EPI_ISL_2729336, EPI_ISL_2729337, EPI_ISL_2729338, EPI_ISL_2729339 | Lighthouse Lab in Milton Keynes | Wellcome Sanger Institute for the COVID-19 Genomics UK (COG-UK) Consortium | The Lighthouse Lab in Milton Keynes and Alex Alderton, Roberto Amato, Jeffrey Barrett, Sonia Goncalves, Ewan Harrison, David K. Jackson, Ian Johnston, Dominic Kwiatkowski, Cordelia Langford, John Sillitoe on behalf of the Wellcome Sanger Institute COVID-19 Surveillance Team |
| EPI_ISL_2729340 | Lighthouse Lab in Glasgow | Wellcome Sanger Institute for the COVID-19 Genomics UK (COG-UK) Consortium | Harper VanSteenhouse, Yumi Kasai, David Gray, Carol Clugston, Anna Dominiczak and Alex Alderton, Roberto Amato, Jeffrey Barrett, Sonia Goncalves, Ewan Harrison, David K. Jackson, Ian Johnston, Dominic Kwiatkowski, Cordelia Langford, John Sillitoe on behalf of the Wellcome Sanger Institute COVID-19 Surveillance Team |
| EPI_ISL_2729341, EPI_ISL_2729342, EPI_ISL_2729343 | Lighthouse Lab in Alderley Park | Wellcome Sanger Institute for the COVID-19 Genomics UK (COG-UK) Consortium | Jacquelyn Wynn, Mairead Hyland, The Lighthouse Lab in Alderley Park and Alex Alderton, Roberto Amato, Jeffrey Barrett, Sonia Goncalves, Ewan Harrison, David K. Jackson, Ian Johnston, Dominic Kwiatkowski, Cordelia Langford, John Sillitoe on behalf of the Wellcome Sanger Institute COVID-19 Surveillance Team |
| EPI_ISL_2729344, EPI_ISL_2729345 | Lighthouse Lab in Milton Keynes | Wellcome Sanger Institute for the COVID-19 Genomics UK (COG-UK) Consortium | The Lighthouse Lab in Milton Keynes and Alex Alderton, Roberto Amato, Jeffrey Barrett, Sonia Goncalves, Ewan Harrison, David K. Jackson, Ian Johnston, Dominic Kwiatkowski, Cordelia Langford, John Sillitoe on behalf of the Wellcome Sanger Institute COVID-19 Surveillance Team |
| EPI_ISL_2729346, EPI_ISL_2729347 | Lighthouse Lab in Alderley Park | Wellcome Sanger Institute for the COVID-19 Genomics UK (COG-UK) Consortium | Jacquelyn Wynn, Mairead Hyland, The Lighthouse Lab in Alderley Park and Alex Alderton, Roberto Amato, Jeffrey Barrett, Sonia Goncalves, Ewan Harrison, David K. Jackson, Ian Johnston, Dominic Kwiatkowski, Cordelia Langford, John Sillitoe on behalf of the Wellcome Sanger Institute COVID-19 Surveillance Team |
| EPI_ISL_2729348 | Lighthouse Lab in Milton Keynes | Wellcome Sanger Institute for the COVID-19 Genomics UK (COG-UK) Consortium | The Lighthouse Lab in Milton Keynes and Alex Alderton, Roberto Amato, Jeffrey Barrett, Sonia Goncalves, Ewan Harrison, David K. Jackson, Ian Johnston, Dominic Kwiatkowski, Cordelia Langford, John Sillitoe on behalf of the Wellcome Sanger Institute COVID-19 Surveillance Team |
| EPI_ISL_2729349 | Lighthouse Lab in Alderley Park | Wellcome Sanger Institute for the COVID-19 Genomics UK (COG-UK) Consortium | Jacquelyn Wynn, Mairead Hyland, The Lighthouse Lab in Alderley Park and Alex Alderton, Roberto Amato, Jeffrey Barrett, Sonia Goncalves, Ewan Harrison, David K. Jackson, Ian Johnston, Dominic Kwiatkowski, Cordelia Langford, John Sillitoe on behalf of the Wellcome Sanger Institute COVID-19 Surveillance Team |
| EPI_ISL_2729350 | Lighthouse Lab in Glasgow | Wellcome Sanger Institute for the COVID-19 Genomics UK (COG-UK) Consortium | Harper VanSteenhouse, Yumi Kasai, David Gray, Carol Clugston, Anna Dominiczak and Alex Alderton, Roberto Amato, Jeffrey Barrett, Sonia Goncalves, Ewan Harrison, David K. Jackson, Ian Johnston, Dominic Kwiatkowski, Cordelia Langford, John Sillitoe on behalf of the Wellcome Sanger Institute COVID-19 Surveillance Team |
| EPI_ISL_2729351 | Lighthouse Lab in Alderley Park | Wellcome Sanger Institute for the COVID-19 Genomics UK (COG-UK) Consortium | Jacquelyn Wynn, Mairead Hyland, The Lighthouse Lab in Alderley Park and Alex Alderton, Roberto Amato, Jeffrey Barrett, Sonia Goncalves, Ewan Harrison, David K. Jackson, Ian Johnston, Dominic Kwiatkowski, Cordelia Langford, John Sillitoe on behalf of the Wellcome Sanger Institute COVID-19 Surveillance Team |
| EPI_ISL_2729352, EPI_ISL_2729353, EPI_ISL_2729354 | Lighthouse Lab in Glasgow | Wellcome Sanger Institute for the COVID-19 Genomics UK (COG-UK) Consortium | Harper VanSteenhouse, Yumi Kasai, David Gray, Carol Clugston, Anna Dominiczak and Alex Alderton, Roberto Amato, Jeffrey Barrett, Sonia Goncalves, Ewan Harrison, David K. Jackson, Ian Johnston, Dominic Kwiatkowski, Cordelia Langford, John Sillitoe on behalf of the Wellcome Sanger Institute COVID-19 Surveillance Team |
| EPI_ISL_2729355 | Lighthouse Lab in Alderley Park | Wellcome Sanger Institute for the COVID-19 Genomics UK (COG-UK) Consortium | Jacquelyn Wynn, Mairead Hyland, The Lighthouse Lab in Alderley Park and Alex Alderton, Roberto Amato, Jeffrey Barrett, Sonia Goncalves, Ewan Harrison, David K. Jackson, Ian Johnston, Dominic Kwiatkowski, Cordelia Langford, John Sillitoe on behalf of the Wellcome Sanger Institute COVID-19 Surveillance Team |
| EPI_ISL_2729356 | Lighthouse Lab in Glasgow | Wellcome Sanger Institute for the COVID-19 Genomics UK (COG-UK) Consortium | Harper VanSteenhouse, Yumi Kasai, David Gray, Carol Clugston, Anna Dominiczak and Alex Alderton, Roberto Amato, Jeffrey Barrett, Sonia Goncalves, Ewan Harrison, David K. Jackson, Ian Johnston, Dominic Kwiatkowski, Cordelia Langford, John Sillitoe on behalf of the Wellcome Sanger Institute COVID-19 Surveillance Team |
| EPI_ISL_2729357 | Lighthouse Lab in Milton Keynes | Wellcome Sanger Institute for the COVID-19 Genomics UK (COG-UK) Consortium | The Lighthouse Lab in Milton Keynes and Alex Alderton, Roberto Amato, Jeffrey Barrett, Sonia Goncalves, Ewan Harrison, David K. Jackson, Ian Johnston, Dominic Kwiatkowski, Cordelia Langford, John Sillitoe on behalf of the Wellcome Sanger Institute COVID-19 Surveillance Team |
| EPI_ISL_2729358, EPI_ISL_2729359, EPI_ISL_2729360, EPI_ISL_2729361 | Lighthouse Lab in Glasgow | Wellcome Sanger Institute for the COVID-19 Genomics UK (COG-UK) Consortium | Harper VanSteenhouse, Yumi Kasai, David Gray, Carol Clugston, Anna Dominiczak and Alex Alderton, Roberto Amato, Jeffrey Barrett, Sonia Goncalves, Ewan Harrison, David K. Jackson, Ian Johnston, Dominic Kwiatkowski, Cordelia Langford, John Sillitoe on behalf of the Wellcome Sanger Institute COVID-19 Surveillance Team |
| EPI_ISL_2729362 | Lighthouse Lab in Alderley Park | Wellcome Sanger Institute for the COVID-19 Genomics UK (COG-UK) Consortium | Jacquelyn Wynn, Mairead Hyland, The Lighthouse Lab in Alderley Park and Alex Alderton, Roberto Amato, Jeffrey Barrett, Sonia Goncalves, Ewan Harrison, David K. Jackson, Ian Johnston, Dominic Kwiatkowski, Cordelia Langford, John Sillitoe on behalf of the Wellcome Sanger Institute COVID-19 Surveillance Team |
| EPI_ISL_2729363, EPI_ISL_2729364 | Lighthouse Lab in Glasgow | Wellcome Sanger Institute for the COVID-19 Genomics UK (COG-UK) Consortium | Harper VanSteenhouse, Yumi Kasai, David Gray, Carol Clugston, Anna Dominiczak and Alex Alderton, Roberto Amato, Jeffrey Barrett, Sonia Goncalves, Ewan Harrison, David K. Jackson, Ian Johnston, Dominic Kwiatkowski, Cordelia Langford, John Sillitoe on behalf of the Wellcome Sanger Institute COVID-19 Surveillance Team |
| EPI_ISL_2729365 | Lighthouse Lab in Milton Keynes | Wellcome Sanger Institute for the COVID-19 Genomics UK (COG-UK) Consortium | The Lighthouse Lab in Milton Keynes and Alex Alderton, Roberto Amato, Jeffrey Barrett, Sonia Goncalves, Ewan Harrison, David K. Jackson, Ian Johnston, Dominic Kwiatkowski, Cordelia Langford, John Sillitoe on behalf of the Wellcome Sanger Institute COVID-19 Surveillance Team |
| EPI_ISL_2729366 | Lighthouse Lab in Alderley Park | Wellcome Sanger Institute for the COVID-19 Genomics UK (COG-UK) Consortium | Jacquelyn Wynn, Mairead Hyland, The Lighthouse Lab in Alderley Park and Alex Alderton, Roberto Amato, Jeffrey Barrett, Sonia Goncalves, Ewan Harrison, David K. Jackson, Ian Johnston, Dominic Kwiatkowski, Cordelia Langford, John Sillitoe on behalf of the Wellcome Sanger Institute COVID-19 Surveillance Team |
| EPI_ISL_2729367 | Lighthouse Lab in Glasgow | Wellcome Sanger Institute for the COVID-19 Genomics UK (COG-UK) Consortium | Harper VanSteenhouse, Yumi Kasai, David Gray, Carol Clugston, Anna Dominiczak and Alex Alderton, Roberto Amato, Jeffrey Barrett, Sonia Goncalves, Ewan Harrison, David K. Jackson, Ian Johnston, Dominic Kwiatkowski, Cordelia Langford, John Sillitoe on behalf of the Wellcome Sanger Institute COVID-19 Surveillance Team |
| EPI_ISL_2729368 | Lighthouse Lab in Alderley Park | Wellcome Sanger Institute for the COVID-19 Genomics UK (COG-UK) Consortium | Jacquelyn Wynn, Mairead Hyland, The Lighthouse Lab in Alderley Park and Alex Alderton, Roberto Amato, Jeffrey Barrett, Sonia Goncalves, Ewan Harrison, David K. Jackson, Ian Johnston, Dominic Kwiatkowski, Cordelia Langford, John Sillitoe on behalf of the Wellcome Sanger Institute COVID-19 Surveillance Team |
| EPI_ISL_2729369 | Lighthouse Lab in Glasgow | Wellcome Sanger Institute for the COVID-19 Genomics UK (COG-UK) Consortium | Harper VanSteenhouse, Yumi Kasai, David Gray, Carol Clugston, Anna Dominiczak and Alex Alderton, Roberto Amato, Jeffrey Barrett, Sonia Goncalves, Ewan Harrison, David K. Jackson, Ian Johnston, Dominic Kwiatkowski, Cordelia Langford, John Sillitoe on behalf of the Wellcome Sanger Institute COVID-19 Surveillance Team |
| EPI_ISL_2729371, EPI_ISL_2729372 | Lighthouse Lab in Alderley Park | Wellcome Sanger Institute for the COVID-19 Genomics UK (COG-UK) Consortium | Jacquelyn Wynn, Mairead Hyland, The Lighthouse Lab in Alderley Park and Alex Alderton, Roberto Amato, Jeffrey Barrett, Sonia Goncalves, Ewan Harrison, David K. Jackson, Ian Johnston, Dominic Kwiatkowski, Cordelia Langford, John Sillitoe on behalf of the Wellcome Sanger Institute COVID-19 Surveillance Team |
| EPI_ISL_2729373 | Lighthouse Lab in Glasgow | Wellcome Sanger Institute for the COVID-19 Genomics UK (COG-UK) Consortium | Harper VanSteenhouse, Yumi Kasai, David Gray, Carol Clugston, Anna Dominiczak and Alex Alderton, Roberto Amato, Jeffrey Barrett, Sonia Goncalves, Ewan Harrison, David K. Jackson, Ian Johnston, Dominic Kwiatkowski, Cordelia Langford, John Sillitoe on behalf of the Wellcome Sanger Institute COVID-19 Surveillance Team |
| EPI_ISL_2729374, EPI_ISL_2729375 | Lighthouse Lab in Alderley Park | Wellcome Sanger Institute for the COVID-19 Genomics UK (COG-UK) Consortium | Jacquelyn Wynn, Mairead Hyland, The Lighthouse Lab in Alderley Park and Alex Alderton, Roberto Amato, Jeffrey Barrett, Sonia Goncalves, Ewan Harrison, David K. Jackson, Ian Johnston, Dominic Kwiatkowski, Cordelia Langford, John Sillitoe on behalf of the Wellcome Sanger Institute COVID-19 Surveillance Team |
| EPI_ISL_2729376 | Lighthouse Lab in Milton Keynes | Wellcome Sanger Institute for the COVID-19 Genomics UK (COG-UK) Consortium | The Lighthouse Lab in Milton Keynes and Alex Alderton, Roberto Amato, Jeffrey Barrett, Sonia Goncalves, Ewan Harrison, David K. Jackson, Ian Johnston, Dominic Kwiatkowski, Cordelia Langford, John Sillitoe on behalf of the Wellcome Sanger Institute COVID-19 Surveillance Team |
| EPI_ISL_2729377 | Lighthouse Lab in Glasgow | Wellcome Sanger Institute for the COVID-19 Genomics UK (COG-UK) Consortium | Harper VanSteenhouse, Yumi Kasai, David Gray, Carol Clugston, Anna Dominiczak and Alex Alderton, Roberto Amato, Jeffrey Barrett, Sonia Goncalves, Ewan Harrison, David K. Jackson, Ian Johnston, Dominic Kwiatkowski, Cordelia Langford, John Sillitoe on behalf of the Wellcome Sanger Institute COVID-19 Surveillance Team |
| EPI_ISL_2729378 | Lighthouse Lab in Alderley Park | Wellcome Sanger Institute for the COVID-19 Genomics UK (COG-UK) | Jacquelyn Wynn, Mairead Hyland, The Lighthouse Lab in Alderley Park and Alex Alderton, Roberto Amato, Jeffrey Barrett, Sonia Goncalves, Ewan Harrison, David K. Jackson, Ian Johnston, Dominic Kwiatkowski, Cordelia Langford, John Sillitoe on |

| | | | |
|---|---|---|---|
| | | Consortium | behalf of the Wellcome Sanger Institute COVID-19 Surveillance Team |
| EPI_ISL_2729379, EPI_ISL_2729380 | Lighthouse Lab in Milton Keynes | Wellcome Sanger Institute for the COVID-19 Genomics UK (COG-UK) Consortium | The Lighthouse Lab in Milton Keynes and Alex Alderton, Roberto Amato, Jeffrey Barrett, Sonia Goncalves, Ewan Harrison, David K. Jackson, Ian Johnston, Dominic Kwiatkowski, Cordelia Langford, John Sillitoe on behalf of the Wellcome Sanger Institute COVID-19 Surveillance Team |
| EPI_ISL_2729381, EPI_ISL_2729382, EPI_ISL_2729383 | Lighthouse Lab in Alderley Park | Wellcome Sanger Institute for the COVID-19 Genomics UK (COG-UK) Consortium | Jacquelyn Wynn, Mairead Hyland, The Lighthouse Lab in Alderley Park and Alex Alderton, Roberto Amato, Jeffrey Barrett, Sonia Goncalves, Ewan Harrison, David K. Jackson, Ian Johnston, Dominic Kwiatkowski, Cordelia Langford, John Sillitoe on behalf of the Wellcome Sanger Institute COVID-19 Surveillance Team |
| EPI_ISL_2729384, EPI_ISL_2729385, EPI_ISL_2729386 | Lighthouse Lab in Milton Keynes | Wellcome Sanger Institute for the COVID-19 Genomics UK (COG-UK) Consortium | The Lighthouse Lab in Milton Keynes and Alex Alderton, Roberto Amato, Jeffrey Barrett, Sonia Goncalves, Ewan Harrison, David K. Jackson, Ian Johnston, Dominic Kwiatkowski, Cordelia Langford, John Sillitoe on behalf of the Wellcome Sanger Institute COVID-19 Surveillance Team |
| EPI_ISL_2729387 | Lighthouse Lab in Alderley Park | Wellcome Sanger Institute for the COVID-19 Genomics UK (COG-UK) Consortium | Jacquelyn Wynn, Mairead Hyland, The Lighthouse Lab in Alderley Park and Alex Alderton, Roberto Amato, Jeffrey Barrett, Sonia Goncalves, Ewan Harrison, David K. Jackson, Ian Johnston, Dominic Kwiatkowski, Cordelia Langford, John Sillitoe on behalf of the Wellcome Sanger Institute COVID-19 Surveillance Team |
| EPI_ISL_2729388, EPI_ISL_2729389 | Lighthouse Lab in Milton Keynes | Wellcome Sanger Institute for the COVID-19 Genomics UK (COG-UK) Consortium | The Lighthouse Lab in Milton Keynes and Alex Alderton, Roberto Amato, Jeffrey Barrett, Sonia Goncalves, Ewan Harrison, David K. Jackson, Ian Johnston, Dominic Kwiatkowski, Cordelia Langford, John Sillitoe on behalf of the Wellcome Sanger Institute COVID-19 Surveillance Team |
| EPI_ISL_2729390, EPI_ISL_2729391, EPI_ISL_2729392, EPI_ISL_2729393 | Lighthouse Lab in Alderley Park | Wellcome Sanger Institute for the COVID-19 Genomics UK (COG-UK) Consortium | Jacquelyn Wynn, Mairead Hyland, The Lighthouse Lab in Alderley Park and Alex Alderton, Roberto Amato, Jeffrey Barrett, Sonia Goncalves, Ewan Harrison, David K. Jackson, Ian Johnston, Dominic Kwiatkowski, Cordelia Langford, John Sillitoe on behalf of the Wellcome Sanger Institute COVID-19 Surveillance Team |
| EPI_ISL_2729394, EPI_ISL_2729395, EPI_ISL_2729396, EPI_ISL_2729397, EPI_ISL_2729398 | Lighthouse Lab in Glasgow | Wellcome Sanger Institute for the COVID-19 Genomics UK (COG-UK) Consortium | Harper VanSteenhouse, Yumi Kasai, David Gray, Carol Clugston, Anna Dominiczak and Alex Alderton, Roberto Amato, Jeffrey Barrett, Sonia Goncalves, Ewan Harrison, David K. Jackson, Ian Johnston, Dominic Kwiatkowski, Cordelia Langford, John Sillitoe on behalf of the Wellcome Sanger Institute COVID-19 Surveillance Team |
| EPI_ISL_2729399 | Lighthouse Lab in Alderley Park | Wellcome Sanger Institute for the COVID-19 Genomics UK (COG-UK) Consortium | Jacquelyn Wynn, Mairead Hyland, The Lighthouse Lab in Alderley Park and Alex Alderton, Roberto Amato, Jeffrey Barrett, Sonia Goncalves, Ewan Harrison, David K. Jackson, Ian Johnston, Dominic Kwiatkowski, Cordelia Langford, John Sillitoe on behalf of the Wellcome Sanger Institute COVID-19 Surveillance Team |
| EPI_ISL_2729400 | Lighthouse Lab in Glasgow | Wellcome Sanger Institute for the COVID-19 Genomics UK (COG-UK) Consortium | Harper VanSteenhouse, Yumi Kasai, David Gray, Carol Clugston, Anna Dominiczak and Alex Alderton, Roberto Amato, Jeffrey Barrett, Sonia Goncalves, Ewan Harrison, David K. Jackson, Ian Johnston, Dominic Kwiatkowski, Cordelia Langford, John Sillitoe on behalf of the Wellcome Sanger Institute COVID-19 Surveillance Team |
| EPI_ISL_2729401, EPI_ISL_2729402, EPI_ISL_2729403 | Lighthouse Lab in Milton Keynes | Wellcome Sanger Institute for the COVID-19 Genomics UK (COG-UK) Consortium | The Lighthouse Lab in Milton Keynes and Alex Alderton, Roberto Amato, Jeffrey Barrett, Sonia Goncalves, Ewan Harrison, David K. Jackson, Ian Johnston, Dominic Kwiatkowski, Cordelia Langford, John Sillitoe on behalf of the Wellcome Sanger Institute COVID-19 Surveillance Team |
| EPI_ISL_2729404, EPI_ISL_2729405 | Lighthouse Lab in Glasgow | Wellcome Sanger Institute for the COVID-19 Genomics UK (COG-UK) Consortium | Harper VanSteenhouse, Yumi Kasai, David Gray, Carol Clugston, Anna Dominiczak and Alex Alderton, Roberto Amato, Jeffrey Barrett, Sonia Goncalves, Ewan Harrison, David K. Jackson, Ian Johnston, Dominic Kwiatkowski, Cordelia Langford, John Sillitoe on behalf of the Wellcome Sanger Institute COVID-19 Surveillance Team |
| EPI_ISL_2729406 | Lighthouse Lab in Milton Keynes | Wellcome Sanger Institute for the COVID-19 Genomics UK (COG-UK) Consortium | The Lighthouse Lab in Milton Keynes and Alex Alderton, Roberto Amato, Jeffrey Barrett, Sonia Goncalves, Ewan Harrison, David K. Jackson, Ian Johnston, Dominic Kwiatkowski, Cordelia Langford, John Sillitoe on behalf of the Wellcome Sanger Institute COVID-19 Surveillance Team |
| EPI_ISL_2729407, EPI_ISL_2729408 | Lighthouse Lab in Alderley Park | Wellcome Sanger Institute for the COVID-19 Genomics UK (COG-UK) Consortium | Jacquelyn Wynn, Mairead Hyland, The Lighthouse Lab in Alderley Park and Alex Alderton, Roberto Amato, Jeffrey Barrett, Sonia Goncalves, Ewan Harrison, David K. Jackson, Ian Johnston, Dominic Kwiatkowski, Cordelia Langford, John Sillitoe on behalf of the Wellcome Sanger Institute COVID-19 Surveillance Team |
| EPI_ISL_2729409 | Lighthouse Lab in Glasgow | Wellcome Sanger Institute for the COVID-19 Genomics UK (COG-UK) Consortium | Harper VanSteenhouse, Yumi Kasai, David Gray, Carol Clugston, Anna Dominiczak and Alex Alderton, Roberto Amato, Jeffrey Barrett, Sonia Goncalves, Ewan Harrison, David K. Jackson, Ian Johnston, Dominic Kwiatkowski, Cordelia Langford, John Sillitoe on behalf of the Wellcome Sanger Institute COVID-19 Surveillance Team |
| EPI_ISL_2729410, EPI_ISL_2729412 | Lighthouse Lab in Milton Keynes | Wellcome Sanger Institute for the COVID-19 Genomics UK (COG-UK) Consortium | The Lighthouse Lab in Milton Keynes and Alex Alderton, Roberto Amato, Jeffrey Barrett, Sonia Goncalves, Ewan Harrison, David K. Jackson, Ian Johnston, Dominic Kwiatkowski, Cordelia Langford, John Sillitoe on behalf of the Wellcome Sanger Institute COVID-19 Surveillance Team |
| EPI_ISL_2729413 | Lighthouse Lab in Glasgow | Wellcome Sanger Institute for the COVID-19 Genomics UK (COG-UK) Consortium | Harper VanSteenhouse, Yumi Kasai, David Gray, Carol Clugston, Anna Dominiczak and Alex Alderton, Roberto Amato, Jeffrey Barrett, Sonia Goncalves, Ewan Harrison, David K. Jackson, Ian Johnston, Dominic Kwiatkowski, Cordelia Langford, John Sillitoe on behalf of the Wellcome Sanger Institute COVID-19 Surveillance Team |
| EPI_ISL_2729414 | Lighthouse Lab in Milton Keynes | Wellcome Sanger Institute for the COVID-19 Genomics UK (COG-UK) Consortium | The Lighthouse Lab in Milton Keynes and Alex Alderton, Roberto Amato, Jeffrey Barrett, Sonia Goncalves, Ewan Harrison, David K. Jackson, Ian Johnston, Dominic Kwiatkowski, Cordelia Langford, John Sillitoe on behalf of the Wellcome Sanger Institute COVID-19 Surveillance Team |
| EPI_ISL_2729415, EPI_ISL_2729416 | Lighthouse Lab in Glasgow | Wellcome Sanger Institute for the COVID-19 Genomics UK (COG-UK) Consortium | Harper VanSteenhouse, Yumi Kasai, David Gray, Carol Clugston, Anna Dominiczak and Alex Alderton, Roberto Amato, Jeffrey Barrett, Sonia Goncalves, Ewan Harrison, David K. Jackson, Ian Johnston, Dominic Kwiatkowski, Cordelia Langford, John Sillitoe on behalf of the Wellcome Sanger Institute COVID-19 Surveillance Team |
| EPI_ISL_2729417 | Lighthouse Lab in Alderley Park | Wellcome Sanger Institute for the COVID-19 Genomics UK (COG-UK) Consortium | Jacquelyn Wynn, Mairead Hyland, The Lighthouse Lab in Alderley Park and Alex Alderton, Roberto Amato, Jeffrey Barrett, Sonia Goncalves, Ewan Harrison, David K. Jackson, Ian Johnston, Dominic Kwiatkowski, Cordelia Langford, John Sillitoe on behalf of the Wellcome Sanger Institute COVID-19 Surveillance Team |
| EPI_ISL_2729418 | Lighthouse Lab in Glasgow | Wellcome Sanger Institute for the COVID-19 Genomics UK (COG-UK) Consortium | Harper VanSteenhouse, Yumi Kasai, David Gray, Carol Clugston, Anna Dominiczak and Alex Alderton, Roberto Amato, Jeffrey Barrett, Sonia Goncalves, Ewan Harrison, David K. Jackson, Ian Johnston, Dominic Kwiatkowski, Cordelia Langford, John Sillitoe on behalf of the Wellcome Sanger Institute COVID-19 Surveillance Team |
| EPI_ISL_2729419 | Lighthouse Lab in Alderley Park | Wellcome Sanger Institute for the COVID-19 Genomics UK (COG-UK) Consortium | Jacquelyn Wynn, Mairead Hyland, The Lighthouse Lab in Alderley Park and Alex Alderton, Roberto Amato, Jeffrey Barrett, Sonia Goncalves, Ewan Harrison, David K. Jackson, Ian Johnston, Dominic Kwiatkowski, Cordelia Langford, John Sillitoe on behalf of the Wellcome Sanger Institute COVID-19 Surveillance Team |
| EPI_ISL_2729420 | Lighthouse Lab in Milton Keynes | Wellcome Sanger Institute for the COVID-19 Genomics UK (COG-UK) Consortium | The Lighthouse Lab in Milton Keynes and Alex Alderton, Roberto Amato, Jeffrey Barrett, Sonia Goncalves, Ewan Harrison, David K. Jackson, Ian Johnston, Dominic Kwiatkowski, Cordelia Langford, John Sillitoe on behalf of the Wellcome Sanger Institute COVID-19 Surveillance Team |
| EPI_ISL_2729421 | Lighthouse Lab in Alderley Park | Wellcome Sanger Institute for the COVID-19 Genomics UK (COG-UK) Consortium | Jacquelyn Wynn, Mairead Hyland, The Lighthouse Lab in Alderley Park and Alex Alderton, Roberto Amato, Jeffrey Barrett, Sonia Goncalves, Ewan Harrison, David K. Jackson, Ian Johnston, Dominic Kwiatkowski, Cordelia Langford, John Sillitoe on behalf of the Wellcome Sanger Institute COVID-19 Surveillance Team |
| EPI_ISL_2729422 | Lighthouse Lab in Glasgow | Wellcome Sanger Institute for the COVID-19 Genomics UK (COG-UK) Consortium | Harper VanSteenhouse, Yumi Kasai, David Gray, Carol Clugston, Anna Dominiczak and Alex Alderton, Roberto Amato, Jeffrey Barrett, Sonia Goncalves, Ewan Harrison, David K. Jackson, Ian Johnston, Dominic Kwiatkowski, Cordelia Langford, John Sillitoe on behalf of the Wellcome Sanger Institute COVID-19 Surveillance Team |
| EPI_ISL_2729423 | Lighthouse Lab in Milton Keynes | Wellcome Sanger Institute for the COVID-19 Genomics UK (COG-UK) Consortium | The Lighthouse Lab in Milton Keynes and Alex Alderton, Roberto Amato, Jeffrey Barrett, Sonia Goncalves, Ewan Harrison, David K. Jackson, Ian Johnston, Dominic Kwiatkowski, Cordelia Langford, John Sillitoe on behalf of the Wellcome Sanger Institute COVID-19 Surveillance Team |
| EPI_ISL_2729424 | Lighthouse Lab in Glasgow | Wellcome Sanger Institute for the COVID-19 Genomics UK (COG-UK) Consortium | Harper VanSteenhouse, Yumi Kasai, David Gray, Carol Clugston, Anna Dominiczak and Alex Alderton, Roberto Amato, Jeffrey Barrett, Sonia Goncalves, Ewan Harrison, David K. Jackson, Ian Johnston, Dominic Kwiatkowski, Cordelia Langford, John Sillitoe on behalf of the Wellcome Sanger Institute COVID-19 Surveillance Team |
| EPI_ISL_2729425 | Lighthouse Lab in Milton Keynes | Wellcome Sanger Institute for the COVID-19 Genomics UK (COG-UK) Consortium | The Lighthouse Lab in Milton Keynes and Alex Alderton, Roberto Amato, Jeffrey Barrett, Sonia Goncalves, Ewan Harrison, David K. Jackson, Ian Johnston, Dominic Kwiatkowski, Cordelia Langford, John Sillitoe on behalf of the Wellcome Sanger Institute COVID-19 Surveillance Team |
| EPI_ISL_2729426 | Lighthouse Lab in Glasgow | Wellcome Sanger Institute for the COVID-19 Genomics UK (COG-UK) Consortium | Harper VanSteenhouse, Yumi Kasai, David Gray, Carol Clugston, Anna Dominiczak and Alex Alderton, Roberto Amato, Jeffrey Barrett, Sonia Goncalves, Ewan Harrison, David K. Jackson, Ian Johnston, Dominic Kwiatkowski, Cordelia Langford, John Sillitoe on behalf of the Wellcome Sanger Institute COVID-19 Surveillance Team |
| EPI_ISL_2729427 | Lighthouse Lab in Milton Keynes | Wellcome Sanger Institute for the COVID-19 Genomics UK (COG-UK) Consortium | The Lighthouse Lab in Milton Keynes and Alex Alderton, Roberto Amato, Jeffrey Barrett, Sonia Goncalves, Ewan Harrison, David K. Jackson, Ian Johnston, Dominic Kwiatkowski, Cordelia Langford, John Sillitoe on behalf of the Wellcome Sanger Institute COVID-19 Surveillance Team |
| EPI_ISL_2729428 | Lighthouse Lab in Glasgow | Wellcome Sanger Institute for the COVID-19 Genomics UK (COG-UK) Consortium | Harper VanSteenhouse, Yumi Kasai, David Gray, Carol Clugston, Anna Dominiczak and Alex Alderton, Roberto Amato, Jeffrey Barrett, Sonia Goncalves, Ewan Harrison, David K. Jackson, Ian Johnston, Dominic Kwiatkowski, Cordelia Langford, John Sillitoe on behalf of the Wellcome Sanger Institute COVID-19 Surveillance Team |
| EPI_ISL_2729430, EPI_ISL_2729431 | Lighthouse Lab in Alderley Park | Wellcome Sanger Institute for the COVID-19 Genomics UK (COG-UK) Consortium | Jacquelyn Wynn, Mairead Hyland, The Lighthouse Lab in Alderley Park and Alex Alderton, Roberto Amato, Jeffrey Barrett, Sonia Goncalves, Ewan Harrison, David K. Jackson, Ian Johnston, Dominic Kwiatkowski, Cordelia Langford, John Sillitoe on behalf of the Wellcome Sanger Institute COVID-19 Surveillance Team |
| EPI_ISL_2729432, EPI_ISL_2729433 | Lighthouse Lab in Milton Keynes | Wellcome Sanger Institute for the COVID-19 Genomics UK (COG-UK) Consortium | The Lighthouse Lab in Milton Keynes and Alex Alderton, Roberto Amato, Jeffrey Barrett, Sonia Goncalves, Ewan Harrison, David K. Jackson, Ian Johnston, Dominic Kwiatkowski, Cordelia Langford, John Sillitoe on behalf of the Wellcome Sanger Institute COVID-19 Surveillance Team |
| EPI_ISL_2729434 | Lighthouse Lab in Glasgow | Wellcome Sanger Institute for the COVID-19 Genomics UK (COG-UK) Consortium | Harper VanSteenhouse, Yumi Kasai, David Gray, Carol Clugston, Anna Dominiczak and Alex Alderton, Roberto Amato, Jeffrey Barrett, Sonia Goncalves, Ewan Harrison, David K. Jackson, Ian Johnston, Dominic Kwiatkowski, Cordelia Langford, John Sillitoe on behalf of the Wellcome Sanger Institute COVID-19 Surveillance Team |
| EPI_ISL_2729435 | Lighthouse Lab in Milton Keynes | Wellcome Sanger Institute for the COVID-19 Genomics UK (COG-UK) Consortium | The Lighthouse Lab in Milton Keynes and Alex Alderton, Roberto Amato, Jeffrey Barrett, Sonia Goncalves, Ewan Harrison, David K. Jackson, Ian Johnston, Dominic Kwiatkowski, Cordelia Langford, John Sillitoe on behalf of the Wellcome Sanger Institute COVID-19 Surveillance Team |
| EPI_ISL_2729436 | Lighthouse Lab in Alderley Park | Wellcome Sanger Institute for the COVID-19 Genomics UK (COG-UK) Consortium | Jacquelyn Wynn, Mairead Hyland, The Lighthouse Lab in Alderley Park and Alex Alderton, Roberto Amato, Jeffrey Barrett, Sonia Goncalves, Ewan Harrison, David K. Jackson, Ian Johnston, Dominic Kwiatkowski, Cordelia Langford, John Sillitoe on behalf of the Wellcome Sanger Institute COVID-19 Surveillance Team |
| EPI_ISL_2729437 | Lighthouse Lab in Milton Keynes | Wellcome Sanger Institute for the COVID-19 Genomics UK (COG-UK) Consortium | The Lighthouse Lab in Milton Keynes and Alex Alderton, Roberto Amato, Jeffrey Barrett, Sonia Goncalves, Ewan Harrison, David K. Jackson, Ian Johnston, Dominic Kwiatkowski, Cordelia Langford, John Sillitoe on behalf of the Wellcome Sanger Institute COVID-19 Surveillance Team |
| EPI_ISL_2729438 | Lighthouse Lab in Alderley Park | Wellcome Sanger Institute for the COVID-19 Genomics UK (COG-UK) Consortium | Jacquelyn Wynn, Mairead Hyland, The Lighthouse Lab in Alderley Park and Alex Alderton, Roberto Amato, Jeffrey Barrett, Sonia Goncalves, Ewan Harrison, David K. Jackson, Ian Johnston, Dominic Kwiatkowski, Cordelia Langford, John Sillitoe on behalf of the Wellcome Sanger Institute COVID-19 Surveillance Team |
| EPI_ISL_2729439 | Lighthouse Lab in Glasgow | Wellcome Sanger Institute for the COVID-19 Genomics UK (COG-UK) Consortium | Harper VanSteenhouse, Yumi Kasai, David Gray, Carol Clugston, Anna Dominiczak and Alex Alderton, Roberto Amato, Jeffrey Barrett, Sonia Goncalves, Ewan Harrison, David K. Jackson, Ian Johnston, Dominic Kwiatkowski, Cordelia Langford, John Sillitoe on behalf of the Wellcome Sanger Institute COVID-19 Surveillance Team |
| EPI_ISL_2729440, EPI_ISL_2729441 | Lighthouse Lab in Alderley Park | Wellcome Sanger Institute for the COVID-19 Genomics UK (COG-UK) Consortium | Jacquelyn Wynn, Mairead Hyland, The Lighthouse Lab in Alderley Park and Alex Alderton, Roberto Amato, Jeffrey Barrett, Sonia Goncalves, Ewan Harrison, David K. Jackson, Ian Johnston, Dominic Kwiatkowski, Cordelia Langford, John Sillitoe on behalf of the Wellcome Sanger Institute COVID-19 Surveillance Team |
| EPI_ISL_2729442 | Lighthouse Lab in Milton Keynes | Wellcome Sanger Institute for the COVID-19 Genomics UK (COG-UK) Consortium | The Lighthouse Lab in Milton Keynes and Alex Alderton, Roberto Amato, Jeffrey Barrett, Sonia Goncalves, Ewan Harrison, David K. Jackson, Ian Johnston, Dominic Kwiatkowski, Cordelia Langford, John Sillitoe on behalf of the Wellcome Sanger Institute COVID-19 Surveillance Team |
| EPI_ISL_2729443 | Lighthouse Lab in Glasgow | Wellcome Sanger Institute for the COVID-19 Genomics UK (COG-UK) Consortium | Harper VanSteenhouse, Yumi Kasai, David Gray, Carol Clugston, Anna Dominiczak and Alex Alderton, Roberto Amato, Jeffrey Barrett, Sonia Goncalves, Ewan Harrison, David K. Jackson, Ian Johnston, Dominic Kwiatkowski, Cordelia Langford, John Sillitoe on behalf of the Wellcome Sanger Institute COVID-19 Surveillance Team |

| | | | |
|---|---|---|---|
| EPI_ISL_2729445, EPI_ISL_2729446 | Lighthouse Lab in Alderley Park | Wellcome Sanger Institute for the COVID-19 Genomics UK (COG-UK) Consortium | Jacquelyn Wynn, Mairead Hyland, The Lighthouse Lab in Alderley Park and Alex Alderton, Roberto Amato, Jeffrey Barrett, Sonia Goncalves, Ewan Harrison, David K. Jackson, Ian Johnston, Dominic Kwiatkowski, Cordelia Langford, John Sillitoe on behalf of the Wellcome Sanger Institute COVID-19 Surveillance Team |
| EPI_ISL_2729447, EPI_ISL_2729448 | Lighthouse Lab in Glasgow | Wellcome Sanger Institute for the COVID-19 Genomics UK (COG-UK) Consortium | Harper VanSteenhouse, Yumi Kasai, David Gray, Carol Clugston, Anna Dominiczak and Alex Alderton, Roberto Amato, Jeffrey Barrett, Sonia Goncalves, Ewan Harrison, David K. Jackson, Ian Johnston, Dominic Kwiatkowski, Cordelia Langford, John Sillitoe on behalf of the Wellcome Sanger Institute COVID-19 Surveillance Team |
| EPI_ISL_2729449 | Lighthouse Lab in Alderley Park | Wellcome Sanger Institute for the COVID-19 Genomics UK (COG-UK) Consortium | Jacquelyn Wynn, Mairead Hyland, The Lighthouse Lab in Alderley Park and Alex Alderton, Roberto Amato, Jeffrey Barrett, Sonia Goncalves, Ewan Harrison, David K. Jackson, Ian Johnston, Dominic Kwiatkowski, Cordelia Langford, John Sillitoe on behalf of the Wellcome Sanger Institute COVID-19 Surveillance Team |
| EPI_ISL_2729450 | Lighthouse Lab in Glasgow | Wellcome Sanger Institute for the COVID-19 Genomics UK (COG-UK) Consortium | Harper VanSteenhouse, Yumi Kasai, David Gray, Carol Clugston, Anna Dominiczak and Alex Alderton, Roberto Amato, Jeffrey Barrett, Sonia Goncalves, Ewan Harrison, David K. Jackson, Ian Johnston, Dominic Kwiatkowski, Cordelia Langford, John Sillitoe on behalf of the Wellcome Sanger Institute COVID-19 Surveillance Team |
| EPI_ISL_2729451 | Lighthouse Lab in Milton Keynes | Wellcome Sanger Institute for the COVID-19 Genomics UK (COG-UK) Consortium | The Lighthouse Lab in Milton Keynes and Alex Alderton, Roberto Amato, Jeffrey Barrett, Sonia Goncalves, Ewan Harrison, David K. Jackson, Ian Johnston, Dominic Kwiatkowski, Cordelia Langford, John Sillitoe on behalf of the Wellcome Sanger Institute COVID-19 Surveillance Team |
| EPI_ISL_2729452 | Lighthouse Lab in Glasgow | Wellcome Sanger Institute for the COVID-19 Genomics UK (COG-UK) Consortium | Harper VanSteenhouse, Yumi Kasai, David Gray, Carol Clugston, Anna Dominiczak and Alex Alderton, Roberto Amato, Jeffrey Barrett, Sonia Goncalves, Ewan Harrison, David K. Jackson, Ian Johnston, Dominic Kwiatkowski, Cordelia Langford, John Sillitoe on behalf of the Wellcome Sanger Institute COVID-19 Surveillance Team |
| EPI_ISL_2729453, EPI_ISL_2729454 | Lighthouse Lab in Milton Keynes | Wellcome Sanger Institute for the COVID-19 Genomics UK (COG-UK) Consortium | The Lighthouse Lab in Milton Keynes and Alex Alderton, Roberto Amato, Jeffrey Barrett, Sonia Goncalves, Ewan Harrison, David K. Jackson, Ian Johnston, Dominic Kwiatkowski, Cordelia Langford, John Sillitoe on behalf of the Wellcome Sanger Institute COVID-19 Surveillance Team |
| EPI_ISL_2729455 | Lighthouse Lab in Glasgow | Wellcome Sanger Institute for the COVID-19 Genomics UK (COG-UK) Consortium | Harper VanSteenhouse, Yumi Kasai, David Gray, Carol Clugston, Anna Dominiczak and Alex Alderton, Roberto Amato, Jeffrey Barrett, Sonia Goncalves, Ewan Harrison, David K. Jackson, Ian Johnston, Dominic Kwiatkowski, Cordelia Langford, John Sillitoe on behalf of the Wellcome Sanger Institute COVID-19 Surveillance Team |
| EPI_ISL_2729456, EPI_ISL_2729457, EPI_ISL_2729458 | Lighthouse Lab in Alderley Park | Wellcome Sanger Institute for the COVID-19 Genomics UK (COG-UK) Consortium | Jacquelyn Wynn, Mairead Hyland, The Lighthouse Lab in Alderley Park and Alex Alderton, Roberto Amato, Jeffrey Barrett, Sonia Goncalves, Ewan Harrison, David K. Jackson, Ian Johnston, Dominic Kwiatkowski, Cordelia Langford, John Sillitoe on behalf of the Wellcome Sanger Institute COVID-19 Surveillance Team |
| EPI_ISL_2729459, EPI_ISL_2729460 | Lighthouse Lab in Milton Keynes | Wellcome Sanger Institute for the COVID-19 Genomics UK (COG-UK) Consortium | The Lighthouse Lab in Milton Keynes and Alex Alderton, Roberto Amato, Jeffrey Barrett, Sonia Goncalves, Ewan Harrison, David K. Jackson, Ian Johnston, Dominic Kwiatkowski, Cordelia Langford, John Sillitoe on behalf of the Wellcome Sanger Institute COVID-19 Surveillance Team |
| EPI_ISL_2729461 | Lighthouse Lab in Alderley Park | Wellcome Sanger Institute for the COVID-19 Genomics UK (COG-UK) Consortium | Jacquelyn Wynn, Mairead Hyland, The Lighthouse Lab in Alderley Park and Alex Alderton, Roberto Amato, Jeffrey Barrett, Sonia Goncalves, Ewan Harrison, David K. Jackson, Ian Johnston, Dominic Kwiatkowski, Cordelia Langford, John Sillitoe on behalf of the Wellcome Sanger Institute COVID-19 Surveillance Team |
| EPI_ISL_2729462 | Lighthouse Lab in Glasgow | Wellcome Sanger Institute for the COVID-19 Genomics UK (COG-UK) Consortium | Harper VanSteenhouse, Yumi Kasai, David Gray, Carol Clugston, Anna Dominiczak and Alex Alderton, Roberto Amato, Jeffrey Barrett, Sonia Goncalves, Ewan Harrison, David K. Jackson, Ian Johnston, Dominic Kwiatkowski, Cordelia Langford, John Sillitoe on behalf of the Wellcome Sanger Institute COVID-19 Surveillance Team |
| EPI_ISL_2729463, EPI_ISL_2729464, EPI_ISL_2729465 | Lighthouse Lab in Alderley Park | Wellcome Sanger Institute for the COVID-19 Genomics UK (COG-UK) Consortium | Jacquelyn Wynn, Mairead Hyland, The Lighthouse Lab in Alderley Park and Alex Alderton, Roberto Amato, Jeffrey Barrett, Sonia Goncalves, Ewan Harrison, David K. Jackson, Ian Johnston, Dominic Kwiatkowski, Cordelia Langford, John Sillitoe on behalf of the Wellcome Sanger Institute COVID-19 Surveillance Team |
| EPI_ISL_2729466, EPI_ISL_2729467 | Lighthouse Lab in Glasgow | Wellcome Sanger Institute for the COVID-19 Genomics UK (COG-UK) Consortium | Harper VanSteenhouse, Yumi Kasai, David Gray, Carol Clugston, Anna Dominiczak and Alex Alderton, Roberto Amato, Jeffrey Barrett, Sonia Goncalves, Ewan Harrison, David K. Jackson, Ian Johnston, Dominic Kwiatkowski, Cordelia Langford, John Sillitoe on behalf of the Wellcome Sanger Institute COVID-19 Surveillance Team |
| EPI_ISL_2729468 | Lighthouse Lab in Milton Keynes | Wellcome Sanger Institute for the COVID-19 Genomics UK (COG-UK) Consortium | The Lighthouse Lab in Milton Keynes and Alex Alderton, Roberto Amato, Jeffrey Barrett, Sonia Goncalves, Ewan Harrison, David K. Jackson, Ian Johnston, Dominic Kwiatkowski, Cordelia Langford, John Sillitoe on behalf of the Wellcome Sanger Institute COVID-19 Surveillance Team |
| EPI_ISL_2729469, EPI_ISL_2729470, EPI_ISL_2729471, EPI_ISL_2729472, EPI_ISL_2729473 | Lighthouse Lab in Glasgow | Wellcome Sanger Institute for the COVID-19 Genomics UK (COG-UK) Consortium | Harper VanSteenhouse, Yumi Kasai, David Gray, Carol Clugston, Anna Dominiczak and Alex Alderton, Roberto Amato, Jeffrey Barrett, Sonia Goncalves, Ewan Harrison, David K. Jackson, Ian Johnston, Dominic Kwiatkowski, Cordelia Langford, John Sillitoe on behalf of the Wellcome Sanger Institute COVID-19 Surveillance Team |
| EPI_ISL_2729474 | Lighthouse Lab in Milton Keynes | Wellcome Sanger Institute for the COVID-19 Genomics UK (COG-UK) Consortium | The Lighthouse Lab in Milton Keynes and Alex Alderton, Roberto Amato, Jeffrey Barrett, Sonia Goncalves, Ewan Harrison, David K. Jackson, Ian Johnston, Dominic Kwiatkowski, Cordelia Langford, John Sillitoe on behalf of the Wellcome Sanger Institute COVID-19 Surveillance Team |
| EPI_ISL_2729475 | Lighthouse Lab in Alderley Park | Wellcome Sanger Institute for the COVID-19 Genomics UK (COG-UK) Consortium | Jacquelyn Wynn, Mairead Hyland, The Lighthouse Lab in Alderley Park and Alex Alderton, Roberto Amato, Jeffrey Barrett, Sonia Goncalves, Ewan Harrison, David K. Jackson, Ian Johnston, Dominic Kwiatkowski, Cordelia Langford, John Sillitoe on behalf of the Wellcome Sanger Institute COVID-19 Surveillance Team |
| EPI_ISL_2729477 | Lighthouse Lab in Milton Keynes | Wellcome Sanger Institute for the COVID-19 Genomics UK (COG-UK) Consortium | The Lighthouse Lab in Milton Keynes and Alex Alderton, Roberto Amato, Jeffrey Barrett, Sonia Goncalves, Ewan Harrison, David K. Jackson, Ian Johnston, Dominic Kwiatkowski, Cordelia Langford, John Sillitoe on behalf of the Wellcome Sanger Institute COVID-19 Surveillance Team |
| EPI_ISL_2729478, EPI_ISL_2729479 | Lighthouse Lab in Glasgow | Wellcome Sanger Institute for the COVID-19 Genomics UK (COG-UK) Consortium | Harper VanSteenhouse, Yumi Kasai, David Gray, Carol Clugston, Anna Dominiczak and Alex Alderton, Roberto Amato, Jeffrey Barrett, Sonia Goncalves, Ewan Harrison, David K. Jackson, Ian Johnston, Dominic Kwiatkowski, Cordelia Langford, John Sillitoe on behalf of the Wellcome Sanger Institute COVID-19 Surveillance Team |
| EPI_ISL_2729480, EPI_ISL_2729481 | Lighthouse Lab in Alderley Park | Wellcome Sanger Institute for the COVID-19 Genomics UK (COG-UK) Consortium | Jacquelyn Wynn, Mairead Hyland, The Lighthouse Lab in Alderley Park and Alex Alderton, Roberto Amato, Jeffrey Barrett, Sonia Goncalves, Ewan Harrison, David K. Jackson, Ian Johnston, Dominic Kwiatkowski, Cordelia Langford, John Sillitoe on behalf of the Wellcome Sanger Institute COVID-19 Surveillance Team |
| EPI_ISL_2729482 | Lighthouse Lab in Milton Keynes | Wellcome Sanger Institute for the COVID-19 Genomics UK (COG-UK) Consortium | The Lighthouse Lab in Milton Keynes and Alex Alderton, Roberto Amato, Jeffrey Barrett, Sonia Goncalves, Ewan Harrison, David K. Jackson, Ian Johnston, Dominic Kwiatkowski, Cordelia Langford, John Sillitoe on behalf of the Wellcome Sanger Institute COVID-19 Surveillance Team |
| EPI_ISL_2729483 | Lighthouse Lab in Glasgow | Wellcome Sanger Institute for the COVID-19 Genomics UK (COG-UK) Consortium | Harper VanSteenhouse, Yumi Kasai, David Gray, Carol Clugston, Anna Dominiczak and Alex Alderton, Roberto Amato, Jeffrey Barrett, Sonia Goncalves, Ewan Harrison, David K. Jackson, Ian Johnston, Dominic Kwiatkowski, Cordelia Langford, John Sillitoe on behalf of the Wellcome Sanger Institute COVID-19 Surveillance Team |
| EPI_ISL_2729484, EPI_ISL_2729485, EPI_ISL_2729486, EPI_ISL_2729487 | Lighthouse Lab in Milton Keynes | Wellcome Sanger Institute for the COVID-19 Genomics UK (COG-UK) Consortium | The Lighthouse Lab in Milton Keynes and Alex Alderton, Roberto Amato, Jeffrey Barrett, Sonia Goncalves, Ewan Harrison, David K. Jackson, Ian Johnston, Dominic Kwiatkowski, Cordelia Langford, John Sillitoe on behalf of the Wellcome Sanger Institute COVID-19 Surveillance Team |
| EPI_ISL_2729488 | Lighthouse Lab in Glasgow | Wellcome Sanger Institute for the COVID-19 Genomics UK (COG-UK) Consortium | Harper VanSteenhouse, Yumi Kasai, David Gray, Carol Clugston, Anna Dominiczak and Alex Alderton, Roberto Amato, Jeffrey Barrett, Sonia Goncalves, Ewan Harrison, David K. Jackson, Ian Johnston, Dominic Kwiatkowski, Cordelia Langford, John Sillitoe on behalf of the Wellcome Sanger Institute COVID-19 Surveillance Team |
| EPI_ISL_2729489, EPI_ISL_2729490, EPI_ISL_2729491 | Lighthouse Lab in Alderley Park | Wellcome Sanger Institute for the COVID-19 Genomics UK (COG-UK) Consortium | Jacquelyn Wynn, Mairead Hyland, The Lighthouse Lab in Alderley Park and Alex Alderton, Roberto Amato, Jeffrey Barrett, Sonia Goncalves, Ewan Harrison, David K. Jackson, Ian Johnston, Dominic Kwiatkowski, Cordelia Langford, John Sillitoe on behalf of the Wellcome Sanger Institute COVID-19 Surveillance Team |
| EPI_ISL_2729492 | Lighthouse Lab in Glasgow | Wellcome Sanger Institute for the COVID-19 Genomics UK (COG-UK) Consortium | Harper VanSteenhouse, Yumi Kasai, David Gray, Carol Clugston, Anna Dominiczak and Alex Alderton, Roberto Amato, Jeffrey Barrett, Sonia Goncalves, Ewan Harrison, David K. Jackson, Ian Johnston, Dominic Kwiatkowski, Cordelia Langford, John Sillitoe on behalf of the Wellcome Sanger Institute COVID-19 Surveillance Team |
| EPI_ISL_2729493, EPI_ISL_2729494 | Lighthouse Lab in Milton Keynes | Wellcome Sanger Institute for the COVID-19 Genomics UK (COG-UK) Consortium | The Lighthouse Lab in Milton Keynes and Alex Alderton, Roberto Amato, Jeffrey Barrett, Sonia Goncalves, Ewan Harrison, David K. Jackson, Ian Johnston, Dominic Kwiatkowski, Cordelia Langford, John Sillitoe on behalf of the Wellcome Sanger Institute COVID-19 Surveillance Team |
| EPI_ISL_2729495 | Lighthouse Lab in Glasgow | Wellcome Sanger Institute for the COVID-19 Genomics UK (COG-UK) Consortium | Harper VanSteenhouse, Yumi Kasai, David Gray, Carol Clugston, Anna Dominiczak and Alex Alderton, Roberto Amato, Jeffrey Barrett, Sonia Goncalves, Ewan Harrison, David K. Jackson, Ian Johnston, Dominic Kwiatkowski, Cordelia Langford, John Sillitoe on behalf of the Wellcome Sanger Institute COVID-19 Surveillance Team |
| EPI_ISL_2729496 | Lighthouse Lab in Milton Keynes | Wellcome Sanger Institute for the COVID-19 Genomics UK (COG-UK) Consortium | The Lighthouse Lab in Milton Keynes and Alex Alderton, Roberto Amato, Jeffrey Barrett, Sonia Goncalves, Ewan Harrison, David K. Jackson, Ian Johnston, Dominic Kwiatkowski, Cordelia Langford, John Sillitoe on behalf of the Wellcome Sanger Institute COVID-19 Surveillance Team |
| EPI_ISL_2729497, EPI_ISL_2729498 | Lighthouse Lab in Alderley Park | Wellcome Sanger Institute for the COVID-19 Genomics UK (COG-UK) Consortium | Jacquelyn Wynn, Mairead Hyland, The Lighthouse Lab in Alderley Park and Alex Alderton, Roberto Amato, Jeffrey Barrett, Sonia Goncalves, Ewan Harrison, David K. Jackson, Ian Johnston, Dominic Kwiatkowski, Cordelia Langford, John Sillitoe on behalf of the Wellcome Sanger Institute COVID-19 Surveillance Team |
| EPI_ISL_2729499, EPI_ISL_2729500, EPI_ISL_2729501 | Lighthouse Lab in Glasgow | Wellcome Sanger Institute for the COVID-19 Genomics UK (COG-UK) Consortium | Harper VanSteenhouse, Yumi Kasai, David Gray, Carol Clugston, Anna Dominiczak and Alex Alderton, Roberto Amato, Jeffrey Barrett, Sonia Goncalves, Ewan Harrison, David K. Jackson, Ian Johnston, Dominic Kwiatkowski, Cordelia Langford, John Sillitoe on behalf of the Wellcome Sanger Institute COVID-19 Surveillance Team |
| EPI_ISL_2729502 | Lighthouse Lab in Alderley Park | Wellcome Sanger Institute for the COVID-19 Genomics UK (COG-UK) Consortium | Jacquelyn Wynn, Mairead Hyland, The Lighthouse Lab in Alderley Park and Alex Alderton, Roberto Amato, Jeffrey Barrett, Sonia Goncalves, Ewan Harrison, David K. Jackson, Ian Johnston, Dominic Kwiatkowski, Cordelia Langford, John Sillitoe on behalf of the Wellcome Sanger Institute COVID-19 Surveillance Team |
| EPI_ISL_2729503 | Lighthouse Lab in Glasgow | Wellcome Sanger Institute for the COVID-19 Genomics UK (COG-UK) Consortium | Harper VanSteenhouse, Yumi Kasai, David Gray, Carol Clugston, Anna Dominiczak and Alex Alderton, Roberto Amato, Jeffrey Barrett, Sonia Goncalves, Ewan Harrison, David K. Jackson, Ian Johnston, Dominic Kwiatkowski, Cordelia Langford, John Sillitoe on behalf of the Wellcome Sanger Institute COVID-19 Surveillance Team |
| EPI_ISL_2729504 | Lighthouse Lab in Alderley Park | Wellcome Sanger Institute for the COVID-19 Genomics UK (COG-UK) Consortium | Jacquelyn Wynn, Mairead Hyland, The Lighthouse Lab in Alderley Park and Alex Alderton, Roberto Amato, Jeffrey Barrett, Sonia Goncalves, Ewan Harrison, David K. Jackson, Ian Johnston, Dominic Kwiatkowski, Cordelia Langford, John Sillitoe on behalf of the Wellcome Sanger Institute COVID-19 Surveillance Team |
| EPI_ISL_2729505 | Lighthouse Lab in Milton Keynes | Wellcome Sanger Institute for the COVID-19 Genomics UK (COG-UK) Consortium | The Lighthouse Lab in Milton Keynes and Alex Alderton, Roberto Amato, Jeffrey Barrett, Sonia Goncalves, Ewan Harrison, David K. Jackson, Ian Johnston, Dominic Kwiatkowski, Cordelia Langford, John Sillitoe on behalf of the Wellcome Sanger Institute COVID-19 Surveillance Team |
| EPI_ISL_2729506 | Lighthouse Lab in Glasgow | Wellcome Sanger Institute for the COVID-19 Genomics UK (COG-UK) Consortium | Harper VanSteenhouse, Yumi Kasai, David Gray, Carol Clugston, Anna Dominiczak and Alex Alderton, Roberto Amato, Jeffrey Barrett, Sonia Goncalves, Ewan Harrison, David K. Jackson, Ian Johnston, Dominic Kwiatkowski, Cordelia Langford, John Sillitoe on behalf of the Wellcome Sanger Institute COVID-19 Surveillance Team |
| EPI_ISL_2729507 | Lighthouse Lab in Alderley Park | Wellcome Sanger Institute for the COVID-19 Genomics UK (COG-UK) Consortium | Jacquelyn Wynn, Mairead Hyland, The Lighthouse Lab in Alderley Park and Alex Alderton, Roberto Amato, Jeffrey Barrett, Sonia Goncalves, Ewan Harrison, David K. Jackson, Ian Johnston, Dominic Kwiatkowski, Cordelia Langford, John Sillitoe on behalf of the Wellcome Sanger Institute COVID-19 Surveillance Team |
| EPI_ISL_2729508 | Lighthouse Lab in Glasgow | Wellcome Sanger Institute for the COVID-19 Genomics UK (COG-UK) Consortium | Harper VanSteenhouse, Yumi Kasai, David Gray, Carol Clugston, Anna Dominiczak and Alex Alderton, Roberto Amato, Jeffrey Barrett, Sonia Goncalves, Ewan Harrison, David K. Jackson, Ian Johnston, Dominic Kwiatkowski, Cordelia Langford, John Sillitoe on behalf of the Wellcome Sanger Institute COVID-19 Surveillance Team |
| EPI_ISL_2729509 | Lighthouse Lab in Milton Keynes | Wellcome Sanger Institute for the COVID-19 Genomics UK (COG-UK) Consortium | The Lighthouse Lab in Milton Keynes and Alex Alderton, Roberto Amato, Jeffrey Barrett, Sonia Goncalves, Ewan Harrison, David K. Jackson, Ian Johnston, Dominic Kwiatkowski, Cordelia Langford, John Sillitoe on behalf of the Wellcome Sanger Institute COVID-19 Surveillance Team |
| EPI_ISL_2729510 | Lighthouse Lab in Glasgow | Wellcome Sanger Institute for the COVID-19 Genomics UK (COG-UK) Consortium | Harper VanSteenhouse, Yumi Kasai, David Gray, Carol Clugston, Anna Dominiczak and Alex Alderton, Roberto Amato, Jeffrey Barrett, Sonia Goncalves, Ewan Harrison, David K. Jackson, Ian Johnston, Dominic Kwiatkowski, Cordelia Langford, John Sillitoe on behalf of the Wellcome Sanger Institute COVID-19 Surveillance Team |
| EPI_ISL_2729511, | Lighthouse Lab in Alderley Park | Wellcome Sanger Institute for the COVID-19 Genomics UK (COG-UK) | Jacquelyn Wynn, Mairead Hyland, The Lighthouse Lab in Alderley Park and Alex Alderton, Roberto Amato, Jeffrey Barrett, Sonia Goncalves, Ewan Harrison, David K. Jackson, Ian Johnston, Dominic Kwiatkowski, Cordelia Langford, John Sillitoe on |

| | | | |
|---|---|---|---|
| EPI_ISL_2729512 | | Consortium | behalf of the Wellcome Sanger Institute COVID-19 Surveillance Team |
| EPI_ISL_2729513, EPI_ISL_2729514, EPI_ISL_2729515 | Lighthouse Lab in Glasgow | Wellcome Sanger Institute for the COVID-19 Genomics UK (COG-UK) Consortium | Harper VanSteenhouse, Yumi Kasai, David Gray, Carol Clugston, Anna Dominiczak and Alex Alderton, Roberto Amato, Jeffrey Barrett, Sonia Goncalves, Ewan Harrison, David K. Jackson, Ian Johnston, Dominic Kwiatkowski, Cordelia Langford, John Sillitoe on behalf of the Wellcome Sanger Institute COVID-19 Surveillance Team |
| EPI_ISL_2729516, EPI_ISL_2729517 | Lighthouse Lab in Alderley Park | Wellcome Sanger Institute for the COVID-19 Genomics UK (COG-UK) Consortium | Jacquelyn Wynn, Mairead Hyland, The Lighthouse Lab in Alderley Park and Alex Alderton, Roberto Amato, Jeffrey Barrett, Sonia Goncalves, Ewan Harrison, David K. Jackson, Ian Johnston, Dominic Kwiatkowski, Cordelia Langford, John Sillitoe on behalf of the Wellcome Sanger Institute COVID-19 Surveillance Team |
| EPI_ISL_2729518 | Lighthouse Lab in Milton Keynes | Wellcome Sanger Institute for the COVID-19 Genomics UK (COG-UK) Consortium | The Lighthouse Lab in Milton Keynes and Alex Alderton, Roberto Amato, Jeffrey Barrett, Sonia Goncalves, Ewan Harrison, David K. Jackson, Ian Johnston, Dominic Kwiatkowski, Cordelia Langford, John Sillitoe on behalf of the Wellcome Sanger Institute COVID-19 Surveillance Team |
| EPI_ISL_2729519 | Lighthouse Lab in Alderley Park | Wellcome Sanger Institute for the COVID-19 Genomics UK (COG-UK) Consortium | Jacquelyn Wynn, Mairead Hyland, The Lighthouse Lab in Alderley Park and Alex Alderton, Roberto Amato, Jeffrey Barrett, Sonia Goncalves, Ewan Harrison, David K. Jackson, Ian Johnston, Dominic Kwiatkowski, Cordelia Langford, John Sillitoe on behalf of the Wellcome Sanger Institute COVID-19 Surveillance Team |
| EPI_ISL_2729520 | Lighthouse Lab in Glasgow | Wellcome Sanger Institute for the COVID-19 Genomics UK (COG-UK) Consortium | Harper VanSteenhouse, Yumi Kasai, David Gray, Carol Clugston, Anna Dominiczak and Alex Alderton, Roberto Amato, Jeffrey Barrett, Sonia Goncalves, Ewan Harrison, David K. Jackson, Ian Johnston, Dominic Kwiatkowski, Cordelia Langford, John Sillitoe on behalf of the Wellcome Sanger Institute COVID-19 Surveillance Team |
| EPI_ISL_2729521, EPI_ISL_2729522 | Lighthouse Lab in Milton Keynes | Wellcome Sanger Institute for the COVID-19 Genomics UK (COG-UK) Consortium | The Lighthouse Lab in Milton Keynes and Alex Alderton, Roberto Amato, Jeffrey Barrett, Sonia Goncalves, Ewan Harrison, David K. Jackson, Ian Johnston, Dominic Kwiatkowski, Cordelia Langford, John Sillitoe on behalf of the Wellcome Sanger Institute COVID-19 Surveillance Team |
| EPI_ISL_2729523, EPI_ISL_2729525, EPI_ISL_2729527, EPI_ISL_2729528, EPI_ISL_2729529 | Lighthouse Lab in Alderley Park | Wellcome Sanger Institute for the COVID-19 Genomics UK (COG-UK) Consortium | Jacquelyn Wynn, Mairead Hyland, The Lighthouse Lab in Alderley Park and Alex Alderton, Roberto Amato, Jeffrey Barrett, Sonia Goncalves, Ewan Harrison, David K. Jackson, Ian Johnston, Dominic Kwiatkowski, Cordelia Langford, John Sillitoe on behalf of the Wellcome Sanger Institute COVID-19 Surveillance Team |
| EPI_ISL_2729530 | Lighthouse Lab in Glasgow | Wellcome Sanger Institute for the COVID-19 Genomics UK (COG-UK) Consortium | Harper VanSteenhouse, Yumi Kasai, David Gray, Carol Clugston, Anna Dominiczak and Alex Alderton, Roberto Amato, Jeffrey Barrett, Sonia Goncalves, Ewan Harrison, David K. Jackson, Ian Johnston, Dominic Kwiatkowski, Cordelia Langford, John Sillitoe on behalf of the Wellcome Sanger Institute COVID-19 Surveillance Team |
| EPI_ISL_2729531, EPI_ISL_2729532 | Lighthouse Lab in Alderley Park | Wellcome Sanger Institute for the COVID-19 Genomics UK (COG-UK) Consortium | Jacquelyn Wynn, Mairead Hyland, The Lighthouse Lab in Alderley Park and Alex Alderton, Roberto Amato, Jeffrey Barrett, Sonia Goncalves, Ewan Harrison, David K. Jackson, Ian Johnston, Dominic Kwiatkowski, Cordelia Langford, John Sillitoe on behalf of the Wellcome Sanger Institute COVID-19 Surveillance Team |
| EPI_ISL_2729533, EPI_ISL_2729534 | Lighthouse Lab in Milton Keynes | Wellcome Sanger Institute for the COVID-19 Genomics UK (COG-UK) Consortium | The Lighthouse Lab in Milton Keynes and Alex Alderton, Roberto Amato, Jeffrey Barrett, Sonia Goncalves, Ewan Harrison, David K. Jackson, Ian Johnston, Dominic Kwiatkowski, Cordelia Langford, John Sillitoe on behalf of the Wellcome Sanger Institute COVID-19 Surveillance Team |
| EPI_ISL_2729535 | Lighthouse Lab in Glasgow | Wellcome Sanger Institute for the COVID-19 Genomics UK (COG-UK) Consortium | Harper VanSteenhouse, Yumi Kasai, David Gray, Carol Clugston, Anna Dominiczak and Alex Alderton, Roberto Amato, Jeffrey Barrett, Sonia Goncalves, Ewan Harrison, David K. Jackson, Ian Johnston, Dominic Kwiatkowski, Cordelia Langford, John Sillitoe on behalf of the Wellcome Sanger Institute COVID-19 Surveillance Team |
| EPI_ISL_2729536 | Lighthouse Lab in Alderley Park | Wellcome Sanger Institute for the COVID-19 Genomics UK (COG-UK) Consortium | Jacquelyn Wynn, Mairead Hyland, The Lighthouse Lab in Alderley Park and Alex Alderton, Roberto Amato, Jeffrey Barrett, Sonia Goncalves, Ewan Harrison, David K. Jackson, Ian Johnston, Dominic Kwiatkowski, Cordelia Langford, John Sillitoe on behalf of the Wellcome Sanger Institute COVID-19 Surveillance Team |
| EPI_ISL_2729537 | Lighthouse Lab in Milton Keynes | Wellcome Sanger Institute for the COVID-19 Genomics UK (COG-UK) Consortium | The Lighthouse Lab in Milton Keynes and Alex Alderton, Roberto Amato, Jeffrey Barrett, Sonia Goncalves, Ewan Harrison, David K. Jackson, Ian Johnston, Dominic Kwiatkowski, Cordelia Langford, John Sillitoe on behalf of the Wellcome Sanger Institute COVID-19 Surveillance Team |
| EPI_ISL_2729538 | Lighthouse Lab in Glasgow | Wellcome Sanger Institute for the COVID-19 Genomics UK (COG-UK) Consortium | Harper VanSteenhouse, Yumi Kasai, David Gray, Carol Clugston, Anna Dominiczak and Alex Alderton, Roberto Amato, Jeffrey Barrett, Sonia Goncalves, Ewan Harrison, David K. Jackson, Ian Johnston, Dominic Kwiatkowski, Cordelia Langford, John Sillitoe on behalf of the Wellcome Sanger Institute COVID-19 Surveillance Team |
| EPI_ISL_2729539 | Lighthouse Lab in Milton Keynes | Wellcome Sanger Institute for the COVID-19 Genomics UK (COG-UK) Consortium | The Lighthouse Lab in Milton Keynes and Alex Alderton, Roberto Amato, Jeffrey Barrett, Sonia Goncalves, Ewan Harrison, David K. Jackson, Ian Johnston, Dominic Kwiatkowski, Cordelia Langford, John Sillitoe on behalf of the Wellcome Sanger Institute COVID-19 Surveillance Team |
| EPI_ISL_2729540 | Lighthouse Lab in Alderley Park | Wellcome Sanger Institute for the COVID-19 Genomics UK (COG-UK) Consortium | Jacquelyn Wynn, Mairead Hyland, The Lighthouse Lab in Alderley Park and Alex Alderton, Roberto Amato, Jeffrey Barrett, Sonia Goncalves, Ewan Harrison, David K. Jackson, Ian Johnston, Dominic Kwiatkowski, Cordelia Langford, John Sillitoe on behalf of the Wellcome Sanger Institute COVID-19 Surveillance Team |
| EPI_ISL_2729541 | Lighthouse Lab in Milton Keynes | Wellcome Sanger Institute for the COVID-19 Genomics UK (COG-UK) Consortium | The Lighthouse Lab in Milton Keynes and Alex Alderton, Roberto Amato, Jeffrey Barrett, Sonia Goncalves, Ewan Harrison, David K. Jackson, Ian Johnston, Dominic Kwiatkowski, Cordelia Langford, John Sillitoe on behalf of the Wellcome Sanger Institute COVID-19 Surveillance Team |
| EPI_ISL_2729542 | Lighthouse Lab in Alderley Park | Wellcome Sanger Institute for the COVID-19 Genomics UK (COG-UK) Consortium | Jacquelyn Wynn, Mairead Hyland, The Lighthouse Lab in Alderley Park and Alex Alderton, Roberto Amato, Jeffrey Barrett, Sonia Goncalves, Ewan Harrison, David K. Jackson, Ian Johnston, Dominic Kwiatkowski, Cordelia Langford, John Sillitoe on behalf of the Wellcome Sanger Institute COVID-19 Surveillance Team |
| EPI_ISL_2729543, EPI_ISL_2729544 | Lighthouse Lab in Milton Keynes | Wellcome Sanger Institute for the COVID-19 Genomics UK (COG-UK) Consortium | The Lighthouse Lab in Milton Keynes and Alex Alderton, Roberto Amato, Jeffrey Barrett, Sonia Goncalves, Ewan Harrison, David K. Jackson, Ian Johnston, Dominic Kwiatkowski, Cordelia Langford, John Sillitoe on behalf of the Wellcome Sanger Institute COVID-19 Surveillance Team |
| EPI_ISL_2729545 | Lighthouse Lab in Alderley Park | Wellcome Sanger Institute for the COVID-19 Genomics UK (COG-UK) Consortium | Jacquelyn Wynn, Mairead Hyland, The Lighthouse Lab in Alderley Park and Alex Alderton, Roberto Amato, Jeffrey Barrett, Sonia Goncalves, Ewan Harrison, David K. Jackson, Ian Johnston, Dominic Kwiatkowski, Cordelia Langford, John Sillitoe on behalf of the Wellcome Sanger Institute COVID-19 Surveillance Team |
| EPI_ISL_2729546 | Lighthouse Lab in Milton Keynes | Wellcome Sanger Institute for the COVID-19 Genomics UK (COG-UK) Consortium | The Lighthouse Lab in Milton Keynes and Alex Alderton, Roberto Amato, Jeffrey Barrett, Sonia Goncalves, Ewan Harrison, David K. Jackson, Ian Johnston, Dominic Kwiatkowski, Cordelia Langford, John Sillitoe on behalf of the Wellcome Sanger Institute COVID-19 Surveillance Team |
| EPI_ISL_2729547, EPI_ISL_2729548 | Lighthouse Lab in Alderley Park | Wellcome Sanger Institute for the COVID-19 Genomics UK (COG-UK) Consortium | Jacquelyn Wynn, Mairead Hyland, The Lighthouse Lab in Alderley Park and Alex Alderton, Roberto Amato, Jeffrey Barrett, Sonia Goncalves, Ewan Harrison, David K. Jackson, Ian Johnston, Dominic Kwiatkowski, Cordelia Langford, John Sillitoe on behalf of the Wellcome Sanger Institute COVID-19 Surveillance Team |
| EPI_ISL_2729549 | Lighthouse Lab in Glasgow | Wellcome Sanger Institute for the COVID-19 Genomics UK (COG-UK) Consortium | Harper VanSteenhouse, Yumi Kasai, David Gray, Carol Clugston, Anna Dominiczak and Alex Alderton, Roberto Amato, Jeffrey Barrett, Sonia Goncalves, Ewan Harrison, David K. Jackson, Ian Johnston, Dominic Kwiatkowski, Cordelia Langford, John Sillitoe on behalf of the Wellcome Sanger Institute COVID-19 Surveillance Team |
| EPI_ISL_2729550 | Lighthouse Lab in Milton Keynes | Wellcome Sanger Institute for the COVID-19 Genomics UK (COG-UK) Consortium | The Lighthouse Lab in Milton Keynes and Alex Alderton, Roberto Amato, Jeffrey Barrett, Sonia Goncalves, Ewan Harrison, David K. Jackson, Ian Johnston, Dominic Kwiatkowski, Cordelia Langford, John Sillitoe on behalf of the Wellcome Sanger Institute COVID-19 Surveillance Team |
| EPI_ISL_2729551, EPI_ISL_2729552 | Lighthouse Lab in Alderley Park | Wellcome Sanger Institute for the COVID-19 Genomics UK (COG-UK) Consortium | Jacquelyn Wynn, Mairead Hyland, The Lighthouse Lab in Alderley Park and Alex Alderton, Roberto Amato, Jeffrey Barrett, Sonia Goncalves, Ewan Harrison, David K. Jackson, Ian Johnston, Dominic Kwiatkowski, Cordelia Langford, John Sillitoe on behalf of the Wellcome Sanger Institute COVID-19 Surveillance Team |
| EPI_ISL_2729553 | Lighthouse Lab in Milton Keynes | Wellcome Sanger Institute for the COVID-19 Genomics UK (COG-UK) Consortium | The Lighthouse Lab in Milton Keynes and Alex Alderton, Roberto Amato, Jeffrey Barrett, Sonia Goncalves, Ewan Harrison, David K. Jackson, Ian Johnston, Dominic Kwiatkowski, Cordelia Langford, John Sillitoe on behalf of the Wellcome Sanger Institute COVID-19 Surveillance Team |
| EPI_ISL_2729554 | Lighthouse Lab in Glasgow | Wellcome Sanger Institute for the COVID-19 Genomics UK (COG-UK) Consortium | Harper VanSteenhouse, Yumi Kasai, David Gray, Carol Clugston, Anna Dominiczak and Alex Alderton, Roberto Amato, Jeffrey Barrett, Sonia Goncalves, Ewan Harrison, David K. Jackson, Ian Johnston, Dominic Kwiatkowski, Cordelia Langford, John Sillitoe on behalf of the Wellcome Sanger Institute COVID-19 Surveillance Team |
| EPI_ISL_2729555 | Lighthouse Lab in Milton Keynes | Wellcome Sanger Institute for the COVID-19 Genomics UK (COG-UK) Consortium | The Lighthouse Lab in Milton Keynes and Alex Alderton, Roberto Amato, Jeffrey Barrett, Sonia Goncalves, Ewan Harrison, David K. Jackson, Ian Johnston, Dominic Kwiatkowski, Cordelia Langford, John Sillitoe on behalf of the Wellcome Sanger Institute COVID-19 Surveillance Team |
| EPI_ISL_2729556 | Lighthouse Lab in Glasgow | Wellcome Sanger Institute for the COVID-19 Genomics UK (COG-UK) Consortium | Harper VanSteenhouse, Yumi Kasai, David Gray, Carol Clugston, Anna Dominiczak and Alex Alderton, Roberto Amato, Jeffrey Barrett, Sonia Goncalves, Ewan Harrison, David K. Jackson, Ian Johnston, Dominic Kwiatkowski, Cordelia Langford, John Sillitoe on behalf of the Wellcome Sanger Institute COVID-19 Surveillance Team |
| EPI_ISL_2729557 | Lighthouse Lab in Alderley Park | Wellcome Sanger Institute for the COVID-19 Genomics UK (COG-UK) Consortium | Jacquelyn Wynn, Mairead Hyland, The Lighthouse Lab in Alderley Park and Alex Alderton, Roberto Amato, Jeffrey Barrett, Sonia Goncalves, Ewan Harrison, David K. Jackson, Ian Johnston, Dominic Kwiatkowski, Cordelia Langford, John Sillitoe on behalf of the Wellcome Sanger Institute COVID-19 Surveillance Team |
| EPI_ISL_2729558 | Lighthouse Lab in Glasgow | Wellcome Sanger Institute for the COVID-19 Genomics UK (COG-UK) Consortium | Harper VanSteenhouse, Yumi Kasai, David Gray, Carol Clugston, Anna Dominiczak and Alex Alderton, Roberto Amato, Jeffrey Barrett, Sonia Goncalves, Ewan Harrison, David K. Jackson, Ian Johnston, Dominic Kwiatkowski, Cordelia Langford, John Sillitoe on behalf of the Wellcome Sanger Institute COVID-19 Surveillance Team |
| EPI_ISL_2729559, EPI_ISL_2729560 | Lighthouse Lab in Milton Keynes | Wellcome Sanger Institute for the COVID-19 Genomics UK (COG-UK) Consortium | The Lighthouse Lab in Milton Keynes and Alex Alderton, Roberto Amato, Jeffrey Barrett, Sonia Goncalves, Ewan Harrison, David K. Jackson, Ian Johnston, Dominic Kwiatkowski, Cordelia Langford, John Sillitoe on behalf of the Wellcome Sanger Institute COVID-19 Surveillance Team |
| EPI_ISL_2729561 | Lighthouse Lab in Alderley Park | Wellcome Sanger Institute for the COVID-19 Genomics UK (COG-UK) Consortium | Jacquelyn Wynn, Mairead Hyland, The Lighthouse Lab in Alderley Park and Alex Alderton, Roberto Amato, Jeffrey Barrett, Sonia Goncalves, Ewan Harrison, David K. Jackson, Ian Johnston, Dominic Kwiatkowski, Cordelia Langford, John Sillitoe on behalf of the Wellcome Sanger Institute COVID-19 Surveillance Team |
| EPI_ISL_2729562 | Lighthouse Lab in Glasgow | Wellcome Sanger Institute for the COVID-19 Genomics UK (COG-UK) Consortium | Harper VanSteenhouse, Yumi Kasai, David Gray, Carol Clugston, Anna Dominiczak and Alex Alderton, Roberto Amato, Jeffrey Barrett, Sonia Goncalves, Ewan Harrison, David K. Jackson, Ian Johnston, Dominic Kwiatkowski, Cordelia Langford, John Sillitoe on behalf of the Wellcome Sanger Institute COVID-19 Surveillance Team |
| EPI_ISL_2729563 | Lighthouse Lab in Alderley Park | Wellcome Sanger Institute for the COVID-19 Genomics UK (COG-UK) Consortium | Jacquelyn Wynn, Mairead Hyland, The Lighthouse Lab in Alderley Park and Alex Alderton, Roberto Amato, Jeffrey Barrett, Sonia Goncalves, Ewan Harrison, David K. Jackson, Ian Johnston, Dominic Kwiatkowski, Cordelia Langford, John Sillitoe on behalf of the Wellcome Sanger Institute COVID-19 Surveillance Team |
| EPI_ISL_2729564 | Lighthouse Lab in Milton Keynes | Wellcome Sanger Institute for the COVID-19 Genomics UK (COG-UK) Consortium | The Lighthouse Lab in Milton Keynes and Alex Alderton, Roberto Amato, Jeffrey Barrett, Sonia Goncalves, Ewan Harrison, David K. Jackson, Ian Johnston, Dominic Kwiatkowski, Cordelia Langford, John Sillitoe on behalf of the Wellcome Sanger Institute COVID-19 Surveillance Team |
| EPI_ISL_2729565 | Lighthouse Lab in Glasgow | Wellcome Sanger Institute for the COVID-19 Genomics UK (COG-UK) Consortium | Harper VanSteenhouse, Yumi Kasai, David Gray, Carol Clugston, Anna Dominiczak and Alex Alderton, Roberto Amato, Jeffrey Barrett, Sonia Goncalves, Ewan Harrison, David K. Jackson, Ian Johnston, Dominic Kwiatkowski, Cordelia Langford, John Sillitoe on behalf of the Wellcome Sanger Institute COVID-19 Surveillance Team |
| EPI_ISL_2729566, EPI_ISL_2729567 | Lighthouse Lab in Milton Keynes | Wellcome Sanger Institute for the COVID-19 Genomics UK (COG-UK) Consortium | The Lighthouse Lab in Milton Keynes and Alex Alderton, Roberto Amato, Jeffrey Barrett, Sonia Goncalves, Ewan Harrison, David K. Jackson, Ian Johnston, Dominic Kwiatkowski, Cordelia Langford, John Sillitoe on behalf of the Wellcome Sanger Institute COVID-19 Surveillance Team |
| EPI_ISL_2729568 | Lighthouse Lab in Glasgow | Wellcome Sanger Institute for the COVID-19 Genomics UK (COG-UK) Consortium | Harper VanSteenhouse, Yumi Kasai, David Gray, Carol Clugston, Anna Dominiczak and Alex Alderton, Roberto Amato, Jeffrey Barrett, Sonia Goncalves, Ewan Harrison, David K. Jackson, Ian Johnston, Dominic Kwiatkowski, Cordelia Langford, John Sillitoe on behalf of the Wellcome Sanger Institute COVID-19 Surveillance Team |
| EPI_ISL_2729569, EPI_ISL_2729570, EPI_ISL_2729572 | Lighthouse Lab in Alderley Park | Wellcome Sanger Institute for the COVID-19 Genomics UK (COG-UK) Consortium | Jacquelyn Wynn, Mairead Hyland, The Lighthouse Lab in Alderley Park and Alex Alderton, Roberto Amato, Jeffrey Barrett, Sonia Goncalves, Ewan Harrison, David K. Jackson, Ian Johnston, Dominic Kwiatkowski, Cordelia Langford, John Sillitoe on behalf of the Wellcome Sanger Institute COVID-19 Surveillance Team |
| EPI_ISL_2729573 | Lighthouse Lab in Milton Keynes | Wellcome Sanger Institute for the COVID-19 Genomics UK (COG-UK) Consortium | The Lighthouse Lab in Milton Keynes and Alex Alderton, Roberto Amato, Jeffrey Barrett, Sonia Goncalves, Ewan Harrison, David K. Jackson, Ian Johnston, Dominic Kwiatkowski, Cordelia Langford, John Sillitoe on behalf of the Wellcome Sanger Institute COVID-19 Surveillance Team |
| EPI_ISL_2729574 | Lighthouse Lab in Glasgow | Wellcome Sanger Institute for the COVID-19 Genomics UK (COG-UK) Consortium | Harper VanSteenhouse, Yumi Kasai, David Gray, Carol Clugston, Anna Dominiczak and Alex Alderton, Roberto Amato, Jeffrey Barrett, Sonia Goncalves, Ewan Harrison, David K. Jackson, Ian Johnston, Dominic Kwiatkowski, Cordelia Langford, John Sillitoe on behalf of the Wellcome Sanger Institute COVID-19 Surveillance Team |
| EPI_ISL_2729575 | Lighthouse Lab in Alderley Park | Wellcome Sanger Institute for the COVID-19 Genomics UK (COG-UK) | Jacquelyn Wynn, Mairead Hyland, The Lighthouse Lab in Alderley Park and Alex Alderton, Roberto Amato, Jeffrey Barrett, Sonia Goncalves, Ewan Harrison, David K. Jackson, Ian Johnston, Dominic Kwiatkowski, Cordelia Langford, John Sillitoe on |

| | | | |
|---|---|---|---|
| | | Consortium | behalf of the Wellcome Sanger Institute COVID-19 Surveillance Team |
| EPI_ISL_2729576 | Lighthouse Lab in Glasgow | Wellcome Sanger Institute for the COVID-19 Genomics UK (COG-UK) Consortium | Harper VanSteenhouse, Yumi Kasai, David Gray, Carol Clugston, Anna Dominiczak and Alex Alderton, Roberto Amato, Jeffrey Barrett, Sonia Goncalves, Ewan Harrison, David K. Jackson, Ian Johnston, Dominic Kwiatkowski, Cordelia Langford, John Sillitoe on behalf of the Wellcome Sanger Institute COVID-19 Surveillance Team |
| EPI_ISL_2729577 | Lighthouse Lab in Milton Keynes | Wellcome Sanger Institute for the COVID-19 Genomics UK (COG-UK) Consortium | The Lighthouse Lab in Milton Keynes and Alex Alderton, Roberto Amato, Jeffrey Barrett, Sonia Goncalves, Ewan Harrison, David K. Jackson, Ian Johnston, Dominic Kwiatkowski, Cordelia Langford, John Sillitoe on behalf of the Wellcome Sanger Institute COVID-19 Surveillance Team |
| EPI_ISL_2729578, EPI_ISL_2729579, EPI_ISL_2729580 | Lighthouse Lab in Glasgow | Wellcome Sanger Institute for the COVID-19 Genomics UK (COG-UK) Consortium | Harper VanSteenhouse, Yumi Kasai, David Gray, Carol Clugston, Anna Dominiczak and Alex Alderton, Roberto Amato, Jeffrey Barrett, Sonia Goncalves, Ewan Harrison, David K. Jackson, Ian Johnston, Dominic Kwiatkowski, Cordelia Langford, John Sillitoe on behalf of the Wellcome Sanger Institute COVID-19 Surveillance Team |
| EPI_ISL_2729581 | Lighthouse Lab in Milton Keynes | Wellcome Sanger Institute for the COVID-19 Genomics UK (COG-UK) Consortium | The Lighthouse Lab in Milton Keynes and Alex Alderton, Roberto Amato, Jeffrey Barrett, Sonia Goncalves, Ewan Harrison, David K. Jackson, Ian Johnston, Dominic Kwiatkowski, Cordelia Langford, John Sillitoe on behalf of the Wellcome Sanger Institute COVID-19 Surveillance Team |
| EPI_ISL_2729582, EPI_ISL_2729583 | Lighthouse Lab in Glasgow | Wellcome Sanger Institute for the COVID-19 Genomics UK (COG-UK) Consortium | Harper VanSteenhouse, Yumi Kasai, David Gray, Carol Clugston, Anna Dominiczak and Alex Alderton, Roberto Amato, Jeffrey Barrett, Sonia Goncalves, Ewan Harrison, David K. Jackson, Ian Johnston, Dominic Kwiatkowski, Cordelia Langford, John Sillitoe on behalf of the Wellcome Sanger Institute COVID-19 Surveillance Team |
| EPI_ISL_2729584 | Lighthouse Lab in Alderley Park | Wellcome Sanger Institute for the COVID-19 Genomics UK (COG-UK) Consortium | Jacquelyn Wynn, Mairead Hyland, The Lighthouse Lab in Alderley Park and Alex Alderton, Roberto Amato, Jeffrey Barrett, Sonia Goncalves, Ewan Harrison, David K. Jackson, Ian Johnston, Dominic Kwiatkowski, Cordelia Langford, John Sillitoe on behalf of the Wellcome Sanger Institute COVID-19 Surveillance Team |
| EPI_ISL_2729585, EPI_ISL_2729586 | Lighthouse Lab in Milton Keynes | Wellcome Sanger Institute for the COVID-19 Genomics UK (COG-UK) Consortium | The Lighthouse Lab in Milton Keynes and Alex Alderton, Roberto Amato, Jeffrey Barrett, Sonia Goncalves, Ewan Harrison, David K. Jackson, Ian Johnston, Dominic Kwiatkowski, Cordelia Langford, John Sillitoe on behalf of the Wellcome Sanger Institute COVID-19 Surveillance Team |
| EPI_ISL_2729587 | Lighthouse Lab in Alderley Park | Wellcome Sanger Institute for the COVID-19 Genomics UK (COG-UK) Consortium | Jacquelyn Wynn, Mairead Hyland, The Lighthouse Lab in Alderley Park and Alex Alderton, Roberto Amato, Jeffrey Barrett, Sonia Goncalves, Ewan Harrison, David K. Jackson, Ian Johnston, Dominic Kwiatkowski, Cordelia Langford, John Sillitoe on behalf of the Wellcome Sanger Institute COVID-19 Surveillance Team |
| EPI_ISL_2729588 | Lighthouse Lab in Glasgow | Wellcome Sanger Institute for the COVID-19 Genomics UK (COG-UK) Consortium | Harper VanSteenhouse, Yumi Kasai, David Gray, Carol Clugston, Anna Dominiczak and Alex Alderton, Roberto Amato, Jeffrey Barrett, Sonia Goncalves, Ewan Harrison, David K. Jackson, Ian Johnston, Dominic Kwiatkowski, Cordelia Langford, John Sillitoe on behalf of the Wellcome Sanger Institute COVID-19 Surveillance Team |
| EPI_ISL_2729589, EPI_ISL_2729590 | Lighthouse Lab in Milton Keynes | Wellcome Sanger Institute for the COVID-19 Genomics UK (COG-UK) Consortium | The Lighthouse Lab in Milton Keynes and Alex Alderton, Roberto Amato, Jeffrey Barrett, Sonia Goncalves, Ewan Harrison, David K. Jackson, Ian Johnston, Dominic Kwiatkowski, Cordelia Langford, John Sillitoe on behalf of the Wellcome Sanger Institute COVID-19 Surveillance Team |
| EPI_ISL_2729591 | Lighthouse Lab in Glasgow | Wellcome Sanger Institute for the COVID-19 Genomics UK (COG-UK) Consortium | Harper VanSteenhouse, Yumi Kasai, David Gray, Carol Clugston, Anna Dominiczak and Alex Alderton, Roberto Amato, Jeffrey Barrett, Sonia Goncalves, Ewan Harrison, David K. Jackson, Ian Johnston, Dominic Kwiatkowski, Cordelia Langford, John Sillitoe on behalf of the Wellcome Sanger Institute COVID-19 Surveillance Team |
| EPI_ISL_2729592 | Lighthouse Lab in Alderley Park | Wellcome Sanger Institute for the COVID-19 Genomics UK (COG-UK) Consortium | Jacquelyn Wynn, Mairead Hyland, The Lighthouse Lab in Alderley Park and Alex Alderton, Roberto Amato, Jeffrey Barrett, Sonia Goncalves, Ewan Harrison, David K. Jackson, Ian Johnston, Dominic Kwiatkowski, Cordelia Langford, John Sillitoe on behalf of the Wellcome Sanger Institute COVID-19 Surveillance Team |
| EPI_ISL_2729593 | Lighthouse Lab in Glasgow | Wellcome Sanger Institute for the COVID-19 Genomics UK (COG-UK) Consortium | Harper VanSteenhouse, Yumi Kasai, David Gray, Carol Clugston, Anna Dominiczak and Alex Alderton, Roberto Amato, Jeffrey Barrett, Sonia Goncalves, Ewan Harrison, David K. Jackson, Ian Johnston, Dominic Kwiatkowski, Cordelia Langford, John Sillitoe on behalf of the Wellcome Sanger Institute COVID-19 Surveillance Team |
| EPI_ISL_2729594 | Lighthouse Lab in Alderley Park | Wellcome Sanger Institute for the COVID-19 Genomics UK (COG-UK) Consortium | Jacquelyn Wynn, Mairead Hyland, The Lighthouse Lab in Alderley Park and Alex Alderton, Roberto Amato, Jeffrey Barrett, Sonia Goncalves, Ewan Harrison, David K. Jackson, Ian Johnston, Dominic Kwiatkowski, Cordelia Langford, John Sillitoe on behalf of the Wellcome Sanger Institute COVID-19 Surveillance Team |
| EPI_ISL_2729595, EPI_ISL_2729596 | Lighthouse Lab in Milton Keynes | Wellcome Sanger Institute for the COVID-19 Genomics UK (COG-UK) Consortium | The Lighthouse Lab in Milton Keynes and Alex Alderton, Roberto Amato, Jeffrey Barrett, Sonia Goncalves, Ewan Harrison, David K. Jackson, Ian Johnston, Dominic Kwiatkowski, Cordelia Langford, John Sillitoe on behalf of the Wellcome Sanger Institute COVID-19 Surveillance Team |
| EPI_ISL_2729597 | Lighthouse Lab in Alderley Park | Wellcome Sanger Institute for the COVID-19 Genomics UK (COG-UK) Consortium | Jacquelyn Wynn, Mairead Hyland, The Lighthouse Lab in Alderley Park and Alex Alderton, Roberto Amato, Jeffrey Barrett, Sonia Goncalves, Ewan Harrison, David K. Jackson, Ian Johnston, Dominic Kwiatkowski, Cordelia Langford, John Sillitoe on behalf of the Wellcome Sanger Institute COVID-19 Surveillance Team |
| EPI_ISL_2729598 | Lighthouse Lab in Milton Keynes | Wellcome Sanger Institute for the COVID-19 Genomics UK (COG-UK) Consortium | The Lighthouse Lab in Milton Keynes and Alex Alderton, Roberto Amato, Jeffrey Barrett, Sonia Goncalves, Ewan Harrison, David K. Jackson, Ian Johnston, Dominic Kwiatkowski, Cordelia Langford, John Sillitoe on behalf of the Wellcome Sanger Institute COVID-19 Surveillance Team |
| EPI_ISL_2729599 | Lighthouse Lab in Alderley Park | Wellcome Sanger Institute for the COVID-19 Genomics UK (COG-UK) Consortium | Jacquelyn Wynn, Mairead Hyland, The Lighthouse Lab in Alderley Park and Alex Alderton, Roberto Amato, Jeffrey Barrett, Sonia Goncalves, Ewan Harrison, David K. Jackson, Ian Johnston, Dominic Kwiatkowski, Cordelia Langford, John Sillitoe on behalf of the Wellcome Sanger Institute COVID-19 Surveillance Team |
| EPI_ISL_2729600 | Lighthouse Lab in Milton Keynes | Wellcome Sanger Institute for the COVID-19 Genomics UK (COG-UK) Consortium | The Lighthouse Lab in Milton Keynes and Alex Alderton, Roberto Amato, Jeffrey Barrett, Sonia Goncalves, Ewan Harrison, David K. Jackson, Ian Johnston, Dominic Kwiatkowski, Cordelia Langford, John Sillitoe on behalf of the Wellcome Sanger Institute COVID-19 Surveillance Team |
| EPI_ISL_2729601 | Lighthouse Lab in Alderley Park | Wellcome Sanger Institute for the COVID-19 Genomics UK (COG-UK) Consortium | Jacquelyn Wynn, Mairead Hyland, The Lighthouse Lab in Alderley Park and Alex Alderton, Roberto Amato, Jeffrey Barrett, Sonia Goncalves, Ewan Harrison, David K. Jackson, Ian Johnston, Dominic Kwiatkowski, Cordelia Langford, John Sillitoe on behalf of the Wellcome Sanger Institute COVID-19 Surveillance Team |
| EPI_ISL_2729602 | Lighthouse Lab in Milton Keynes | Wellcome Sanger Institute for the COVID-19 Genomics UK (COG-UK) Consortium | The Lighthouse Lab in Milton Keynes and Alex Alderton, Roberto Amato, Jeffrey Barrett, Sonia Goncalves, Ewan Harrison, David K. Jackson, Ian Johnston, Dominic Kwiatkowski, Cordelia Langford, John Sillitoe on behalf of the Wellcome Sanger Institute COVID-19 Surveillance Team |
| EPI_ISL_2729603 | Lighthouse Lab in Glasgow | Wellcome Sanger Institute for the COVID-19 Genomics UK (COG-UK) Consortium | Harper VanSteenhouse, Yumi Kasai, David Gray, Carol Clugston, Anna Dominiczak and Alex Alderton, Roberto Amato, Jeffrey Barrett, Sonia Goncalves, Ewan Harrison, David K. Jackson, Ian Johnston, Dominic Kwiatkowski, Cordelia Langford, John Sillitoe on behalf of the Wellcome Sanger Institute COVID-19 Surveillance Team |
| EPI_ISL_2729604 | Lighthouse Lab in Milton Keynes | Wellcome Sanger Institute for the COVID-19 Genomics UK (COG-UK) Consortium | The Lighthouse Lab in Milton Keynes and Alex Alderton, Roberto Amato, Jeffrey Barrett, Sonia Goncalves, Ewan Harrison, David K. Jackson, Ian Johnston, Dominic Kwiatkowski, Cordelia Langford, John Sillitoe on behalf of the Wellcome Sanger Institute COVID-19 Surveillance Team |
| EPI_ISL_2729605 | Lighthouse Lab in Glasgow | Wellcome Sanger Institute for the COVID-19 Genomics UK (COG-UK) Consortium | Harper VanSteenhouse, Yumi Kasai, David Gray, Carol Clugston, Anna Dominiczak and Alex Alderton, Roberto Amato, Jeffrey Barrett, Sonia Goncalves, Ewan Harrison, David K. Jackson, Ian Johnston, Dominic Kwiatkowski, Cordelia Langford, John Sillitoe on behalf of the Wellcome Sanger Institute COVID-19 Surveillance Team |
| EPI_ISL_2729606 | Lighthouse Lab in Milton Keynes | Wellcome Sanger Institute for the COVID-19 Genomics UK (COG-UK) Consortium | The Lighthouse Lab in Milton Keynes and Alex Alderton, Roberto Amato, Jeffrey Barrett, Sonia Goncalves, Ewan Harrison, David K. Jackson, Ian Johnston, Dominic Kwiatkowski, Cordelia Langford, John Sillitoe on behalf of the Wellcome Sanger Institute COVID-19 Surveillance Team |
| EPI_ISL_2729607, EPI_ISL_2729608 | Lighthouse Lab in Alderley Park | Wellcome Sanger Institute for the COVID-19 Genomics UK (COG-UK) Consortium | Jacquelyn Wynn, Mairead Hyland, The Lighthouse Lab in Alderley Park and Alex Alderton, Roberto Amato, Jeffrey Barrett, Sonia Goncalves, Ewan Harrison, David K. Jackson, Ian Johnston, Dominic Kwiatkowski, Cordelia Langford, John Sillitoe on behalf of the Wellcome Sanger Institute COVID-19 Surveillance Team |
| EPI_ISL_2729609, EPI_ISL_2729610 | Lighthouse Lab in Milton Keynes | Wellcome Sanger Institute for the COVID-19 Genomics UK (COG-UK) Consortium | The Lighthouse Lab in Milton Keynes and Alex Alderton, Roberto Amato, Jeffrey Barrett, Sonia Goncalves, Ewan Harrison, David K. Jackson, Ian Johnston, Dominic Kwiatkowski, Cordelia Langford, John Sillitoe on behalf of the Wellcome Sanger Institute COVID-19 Surveillance Team |
| EPI_ISL_2729611 | Lighthouse Lab in Glasgow | Wellcome Sanger Institute for the COVID-19 Genomics UK (COG-UK) Consortium | Harper VanSteenhouse, Yumi Kasai, David Gray, Carol Clugston, Anna Dominiczak and Alex Alderton, Roberto Amato, Jeffrey Barrett, Sonia Goncalves, Ewan Harrison, David K. Jackson, Ian Johnston, Dominic Kwiatkowski, Cordelia Langford, John Sillitoe on behalf of the Wellcome Sanger Institute COVID-19 Surveillance Team |
| EPI_ISL_2729612, EPI_ISL_2729613, EPI_ISL_2729614 | Lighthouse Lab in Alderley Park | Wellcome Sanger Institute for the COVID-19 Genomics UK (COG-UK) Consortium | Jacquelyn Wynn, Mairead Hyland, The Lighthouse Lab in Alderley Park and Alex Alderton, Roberto Amato, Jeffrey Barrett, Sonia Goncalves, Ewan Harrison, David K. Jackson, Ian Johnston, Dominic Kwiatkowski, Cordelia Langford, John Sillitoe on behalf of the Wellcome Sanger Institute COVID-19 Surveillance Team |
| EPI_ISL_2729615 | Lighthouse Lab in Milton Keynes | Wellcome Sanger Institute for the COVID-19 Genomics UK (COG-UK) Consortium | The Lighthouse Lab in Milton Keynes and Alex Alderton, Roberto Amato, Jeffrey Barrett, Sonia Goncalves, Ewan Harrison, David K. Jackson, Ian Johnston, Dominic Kwiatkowski, Cordelia Langford, John Sillitoe on behalf of the Wellcome Sanger Institute COVID-19 Surveillance Team |
| EPI_ISL_2729616, EPI_ISL_2729617, EPI_ISL_2729618, EPI_ISL_2729619 | Lighthouse Lab in Alderley Park | Wellcome Sanger Institute for the COVID-19 Genomics UK (COG-UK) Consortium | Jacquelyn Wynn, Mairead Hyland, The Lighthouse Lab in Alderley Park and Alex Alderton, Roberto Amato, Jeffrey Barrett, Sonia Goncalves, Ewan Harrison, David K. Jackson, Ian Johnston, Dominic Kwiatkowski, Cordelia Langford, John Sillitoe on behalf of the Wellcome Sanger Institute COVID-19 Surveillance Team |
| EPI_ISL_2729620, EPI_ISL_2729621 | Lighthouse Lab in Milton Keynes | Wellcome Sanger Institute for the COVID-19 Genomics UK (COG-UK) Consortium | The Lighthouse Lab in Milton Keynes and Alex Alderton, Roberto Amato, Jeffrey Barrett, Sonia Goncalves, Ewan Harrison, David K. Jackson, Ian Johnston, Dominic Kwiatkowski, Cordelia Langford, John Sillitoe on behalf of the Wellcome Sanger Institute COVID-19 Surveillance Team |
| EPI_ISL_2729622, EPI_ISL_2729623 | Lighthouse Lab in Alderley Park | Wellcome Sanger Institute for the COVID-19 Genomics UK (COG-UK) Consortium | Jacquelyn Wynn, Mairead Hyland, The Lighthouse Lab in Alderley Park and Alex Alderton, Roberto Amato, Jeffrey Barrett, Sonia Goncalves, Ewan Harrison, David K. Jackson, Ian Johnston, Dominic Kwiatkowski, Cordelia Langford, John Sillitoe on behalf of the Wellcome Sanger Institute COVID-19 Surveillance Team |
| EPI_ISL_2729624 | Lighthouse Lab in Glasgow | Wellcome Sanger Institute for the COVID-19 Genomics UK (COG-UK) Consortium | Harper VanSteenhouse, Yumi Kasai, David Gray, Carol Clugston, Anna Dominiczak and Alex Alderton, Roberto Amato, Jeffrey Barrett, Sonia Goncalves, Ewan Harrison, David K. Jackson, Ian Johnston, Dominic Kwiatkowski, Cordelia Langford, John Sillitoe on behalf of the Wellcome Sanger Institute COVID-19 Surveillance Team |
| EPI_ISL_2729625 | Lighthouse Lab in Milton Keynes | Wellcome Sanger Institute for the COVID-19 Genomics UK (COG-UK) Consortium | The Lighthouse Lab in Milton Keynes and Alex Alderton, Roberto Amato, Jeffrey Barrett, Sonia Goncalves, Ewan Harrison, David K. Jackson, Ian Johnston, Dominic Kwiatkowski, Cordelia Langford, John Sillitoe on behalf of the Wellcome Sanger Institute COVID-19 Surveillance Team |
| EPI_ISL_2729626 | Lighthouse Lab in Glasgow | Wellcome Sanger Institute for the COVID-19 Genomics UK (COG-UK) Consortium | Harper VanSteenhouse, Yumi Kasai, David Gray, Carol Clugston, Anna Dominiczak and Alex Alderton, Roberto Amato, Jeffrey Barrett, Sonia Goncalves, Ewan Harrison, David K. Jackson, Ian Johnston, Dominic Kwiatkowski, Cordelia Langford, John Sillitoe on behalf of the Wellcome Sanger Institute COVID-19 Surveillance Team |
| EPI_ISL_2729627 | Lighthouse Lab in Alderley Park | Wellcome Sanger Institute for the COVID-19 Genomics UK (COG-UK) Consortium | Jacquelyn Wynn, Mairead Hyland, The Lighthouse Lab in Alderley Park and Alex Alderton, Roberto Amato, Jeffrey Barrett, Sonia Goncalves, Ewan Harrison, David K. Jackson, Ian Johnston, Dominic Kwiatkowski, Cordelia Langford, John Sillitoe on behalf of the Wellcome Sanger Institute COVID-19 Surveillance Team |
| EPI_ISL_2729628 | Lighthouse Lab in Glasgow | Wellcome Sanger Institute for the COVID-19 Genomics UK (COG-UK) Consortium | Harper VanSteenhouse, Yumi Kasai, David Gray, Carol Clugston, Anna Dominiczak and Alex Alderton, Roberto Amato, Jeffrey Barrett, Sonia Goncalves, Ewan Harrison, David K. Jackson, Ian Johnston, Dominic Kwiatkowski, Cordelia Langford, John Sillitoe on behalf of the Wellcome Sanger Institute COVID-19 Surveillance Team |
| EPI_ISL_2729629, EPI_ISL_2729630 | Lighthouse Lab in Alderley Park | Wellcome Sanger Institute for the COVID-19 Genomics UK (COG-UK) Consortium | Jacquelyn Wynn, Mairead Hyland, The Lighthouse Lab in Alderley Park and Alex Alderton, Roberto Amato, Jeffrey Barrett, Sonia Goncalves, Ewan Harrison, David K. Jackson, Ian Johnston, Dominic Kwiatkowski, Cordelia Langford, John Sillitoe on behalf of the Wellcome Sanger Institute COVID-19 Surveillance Team |
| EPI_ISL_2729631, EPI_ISL_2729632 | Lighthouse Lab in Milton Keynes | Wellcome Sanger Institute for the COVID-19 Genomics UK (COG-UK) Consortium | The Lighthouse Lab in Milton Keynes and Alex Alderton, Roberto Amato, Jeffrey Barrett, Sonia Goncalves, Ewan Harrison, David K. Jackson, Ian Johnston, Dominic Kwiatkowski, Cordelia Langford, John Sillitoe on behalf of the Wellcome Sanger Institute COVID-19 Surveillance Team |
| EPI_ISL_2729633 | Lighthouse Lab in Glasgow | Wellcome Sanger Institute for the COVID-19 Genomics UK (COG-UK) Consortium | Harper VanSteenhouse, Yumi Kasai, David Gray, Carol Clugston, Anna Dominiczak and Alex Alderton, Roberto Amato, Jeffrey Barrett, Sonia Goncalves, Ewan Harrison, David K. Jackson, Ian Johnston, Dominic Kwiatkowski, Cordelia Langford, John Sillitoe on behalf of the Wellcome Sanger Institute COVID-19 Surveillance Team |
| EPI_ISL_2729634, EPI_ISL_2729635 | Lighthouse Lab in Alderley Park | Wellcome Sanger Institute for the COVID-19 Genomics UK (COG-UK) Consortium | Jacquelyn Wynn, Mairead Hyland, The Lighthouse Lab in Alderley Park and Alex Alderton, Roberto Amato, Jeffrey Barrett, Sonia Goncalves, Ewan Harrison, David K. Jackson, Ian Johnston, Dominic Kwiatkowski, Cordelia Langford, John Sillitoe on behalf of the Wellcome Sanger Institute COVID-19 Surveillance Team |
| EPI_ISL_2729636, EPI_ISL_2729637, | Lighthouse Lab in Glasgow | Wellcome Sanger Institute for the COVID-19 Genomics UK (COG-UK) Consortium | Harper VanSteenhouse, Yumi Kasai, David Gray, Carol Clugston, Anna Dominiczak and Alex Alderton, Roberto Amato, Jeffrey Barrett, Sonia Goncalves, Ewan Harrison, David K. Jackson, Ian Johnston, Dominic Kwiatkowski, Cordelia Langford, John Sillitoe on behalf of the Wellcome Sanger Institute COVID-19 Surveillance Team |

| | | | |
|---|---|---|---|
| EPI_ISL_2729638, EPI_ISL_2729639 | | | |
| EPI_ISL_2729640 | Lighthouse Lab in Milton Keynes | Wellcome Sanger Institute for the COVID-19 Genomics UK (COG-UK) Consortium | The Lighthouse Lab in Milton Keynes and Alex Alderton, Roberto Amato, Jeffrey Barrett, Sonia Goncalves, Ewan Harrison, David K. Jackson, Ian Johnston, Dominic Kwiatkowski, Cordelia Langford, John Sillitoe on behalf of the Wellcome Sanger Institute COVID-19 Surveillance Team |
| EPI_ISL_2729641 | Lighthouse Lab in Glasgow | Wellcome Sanger Institute for the COVID-19 Genomics UK (COG-UK) Consortium | Harper VanSteenhouse, Yumi Kasai, David Gray, Carol Clugston, Anna Dominiczak and Alex Alderton, Roberto Amato, Jeffrey Barrett, Sonia Goncalves, Ewan Harrison, David K. Jackson, Ian Johnston, Dominic Kwiatkowski, Cordelia Langford, John Sillitoe on behalf of the Wellcome Sanger Institute COVID-19 Surveillance Team |
| EPI_ISL_2729642 | Lighthouse Lab in Milton Keynes | Wellcome Sanger Institute for the COVID-19 Genomics UK (COG-UK) Consortium | The Lighthouse Lab in Milton Keynes and Alex Alderton, Roberto Amato, Jeffrey Barrett, Sonia Goncalves, Ewan Harrison, David K. Jackson, Ian Johnston, Dominic Kwiatkowski, Cordelia Langford, John Sillitoe on behalf of the Wellcome Sanger Institute COVID-19 Surveillance Team |
| EPI_ISL_2729643 | Lighthouse Lab in Alderley Park | Wellcome Sanger Institute for the COVID-19 Genomics UK (COG-UK) Consortium | Jacquelyn Wynn, Mairead Hyland, The Lighthouse Lab in Alderley Park and Alex Alderton, Roberto Amato, Jeffrey Barrett, Sonia Goncalves, Ewan Harrison, David K. Jackson, Ian Johnston, Dominic Kwiatkowski, Cordelia Langford, John Sillitoe on behalf of the Wellcome Sanger Institute COVID-19 Surveillance Team |
| EPI_ISL_2729644, EPI_ISL_2729645, EPI_ISL_2729646 | Lighthouse Lab in Milton Keynes | Wellcome Sanger Institute for the COVID-19 Genomics UK (COG-UK) Consortium | The Lighthouse Lab in Milton Keynes and Alex Alderton, Roberto Amato, Jeffrey Barrett, Sonia Goncalves, Ewan Harrison, David K. Jackson, Ian Johnston, Dominic Kwiatkowski, Cordelia Langford, John Sillitoe on behalf of the Wellcome Sanger Institute COVID-19 Surveillance Team |
| EPI_ISL_2729647, EPI_ISL_2729648 | Lighthouse Lab in Alderley Park | Wellcome Sanger Institute for the COVID-19 Genomics UK (COG-UK) Consortium | Jacquelyn Wynn, Mairead Hyland, The Lighthouse Lab in Alderley Park and Alex Alderton, Roberto Amato, Jeffrey Barrett, Sonia Goncalves, Ewan Harrison, David K. Jackson, Ian Johnston, Dominic Kwiatkowski, Cordelia Langford, John Sillitoe on behalf of the Wellcome Sanger Institute COVID-19 Surveillance Team |
| EPI_ISL_2729649 | Lighthouse Lab in Milton Keynes | Wellcome Sanger Institute for the COVID-19 Genomics UK (COG-UK) Consortium | The Lighthouse Lab in Milton Keynes and Alex Alderton, Roberto Amato, Jeffrey Barrett, Sonia Goncalves, Ewan Harrison, David K. Jackson, Ian Johnston, Dominic Kwiatkowski, Cordelia Langford, John Sillitoe on behalf of the Wellcome Sanger Institute COVID-19 Surveillance Team |
| EPI_ISL_2729650 | Lighthouse Lab in Glasgow | Wellcome Sanger Institute for the COVID-19 Genomics UK (COG-UK) Consortium | Harper VanSteenhouse, Yumi Kasai, David Gray, Carol Clugston, Anna Dominiczak and Alex Alderton, Roberto Amato, Jeffrey Barrett, Sonia Goncalves, Ewan Harrison, David K. Jackson, Ian Johnston, Dominic Kwiatkowski, Cordelia Langford, John Sillitoe on behalf of the Wellcome Sanger Institute COVID-19 Surveillance Team |
| EPI_ISL_2729651, EPI_ISL_2729652 | Lighthouse Lab in Alderley Park | Wellcome Sanger Institute for the COVID-19 Genomics UK (COG-UK) Consortium | Jacquelyn Wynn, Mairead Hyland, The Lighthouse Lab in Alderley Park and Alex Alderton, Roberto Amato, Jeffrey Barrett, Sonia Goncalves, Ewan Harrison, David K. Jackson, Ian Johnston, Dominic Kwiatkowski, Cordelia Langford, John Sillitoe on behalf of the Wellcome Sanger Institute COVID-19 Surveillance Team |
| EPI_ISL_2729653 | Lighthouse Lab in Milton Keynes | Wellcome Sanger Institute for the COVID-19 Genomics UK (COG-UK) Consortium | The Lighthouse Lab in Milton Keynes and Alex Alderton, Roberto Amato, Jeffrey Barrett, Sonia Goncalves, Ewan Harrison, David K. Jackson, Ian Johnston, Dominic Kwiatkowski, Cordelia Langford, John Sillitoe on behalf of the Wellcome Sanger Institute COVID-19 Surveillance Team |
| EPI_ISL_2729654 | Lighthouse Lab in Glasgow | Wellcome Sanger Institute for the COVID-19 Genomics UK (COG-UK) Consortium | Harper VanSteenhouse, Yumi Kasai, David Gray, Carol Clugston, Anna Dominiczak and Alex Alderton, Roberto Amato, Jeffrey Barrett, Sonia Goncalves, Ewan Harrison, David K. Jackson, Ian Johnston, Dominic Kwiatkowski, Cordelia Langford, John Sillitoe on behalf of the Wellcome Sanger Institute COVID-19 Surveillance Team |
| EPI_ISL_2729655 | Lighthouse Lab in Milton Keynes | Wellcome Sanger Institute for the COVID-19 Genomics UK (COG-UK) Consortium | The Lighthouse Lab in Milton Keynes and Alex Alderton, Roberto Amato, Jeffrey Barrett, Sonia Goncalves, Ewan Harrison, David K. Jackson, Ian Johnston, Dominic Kwiatkowski, Cordelia Langford, John Sillitoe on behalf of the Wellcome Sanger Institute COVID-19 Surveillance Team |
| EPI_ISL_2729656 | Lighthouse Laboratory Plymouth | Wellcome Sanger Institute for the COVID-19 Genomics UK (COG-UK) Consortium | Lighthouse Laboratory Plymouth and Alex Alderton, Roberto Amato, Jeffrey Barrett, Sonia Goncalves, Ewan Harrison, David K. Jackson, Ian Johnston, Dominic Kwiatkowski, Cordelia Langford, John Sillitoe on behalf of the Wellcome Sanger Institute COVID-19 Surveillance Team |
| EPI_ISL_2729657, EPI_ISL_2729658, EPI_ISL_2729659 | Lighthouse Lab in Alderley Park | Wellcome Sanger Institute for the COVID-19 Genomics UK (COG-UK) Consortium | Jacquelyn Wynn, Mairead Hyland, The Lighthouse Lab in Alderley Park and Alex Alderton, Roberto Amato, Jeffrey Barrett, Sonia Goncalves, Ewan Harrison, David K. Jackson, Ian Johnston, Dominic Kwiatkowski, Cordelia Langford, John Sillitoe on behalf of the Wellcome Sanger Institute COVID-19 Surveillance Team |
| EPI_ISL_2729660 | Lighthouse Laboratory Plymouth | Wellcome Sanger Institute for the COVID-19 Genomics UK (COG-UK) Consortium | Lighthouse Laboratory Plymouth and Alex Alderton, Roberto Amato, Jeffrey Barrett, Sonia Goncalves, Ewan Harrison, David K. Jackson, Ian Johnston, Dominic Kwiatkowski, Cordelia Langford, John Sillitoe on behalf of the Wellcome Sanger Institute COVID-19 Surveillance Team |
| EPI_ISL_2729661 | Lighthouse Lab in Milton Keynes | Wellcome Sanger Institute for the COVID-19 Genomics UK (COG-UK) Consortium | The Lighthouse Lab in Milton Keynes and Alex Alderton, Roberto Amato, Jeffrey Barrett, Sonia Goncalves, Ewan Harrison, David K. Jackson, Ian Johnston, Dominic Kwiatkowski, Cordelia Langford, John Sillitoe on behalf of the Wellcome Sanger Institute COVID-19 Surveillance Team |
| EPI_ISL_2729662 | Lighthouse Laboratory Plymouth | Wellcome Sanger Institute for the COVID-19 Genomics UK (COG-UK) Consortium | Lighthouse Laboratory Plymouth and Alex Alderton, Roberto Amato, Jeffrey Barrett, Sonia Goncalves, Ewan Harrison, David K. Jackson, Ian Johnston, Dominic Kwiatkowski, Cordelia Langford, John Sillitoe on behalf of the Wellcome Sanger Institute COVID-19 Surveillance Team |
| EPI_ISL_2729663, EPI_ISL_2729664 | Lighthouse Lab in Alderley Park | Wellcome Sanger Institute for the COVID-19 Genomics UK (COG-UK) Consortium | Jacquelyn Wynn, Mairead Hyland, The Lighthouse Lab in Alderley Park and Alex Alderton, Roberto Amato, Jeffrey Barrett, Sonia Goncalves, Ewan Harrison, David K. Jackson, Ian Johnston, Dominic Kwiatkowski, Cordelia Langford, John Sillitoe on behalf of the Wellcome Sanger Institute COVID-19 Surveillance Team |
| EPI_ISL_2729665, EPI_ISL_2729666 | Lighthouse Laboratory Plymouth | Wellcome Sanger Institute for the COVID-19 Genomics UK (COG-UK) Consortium | Lighthouse Laboratory Plymouth and Alex Alderton, Roberto Amato, Jeffrey Barrett, Sonia Goncalves, Ewan Harrison, David K. Jackson, Ian Johnston, Dominic Kwiatkowski, Cordelia Langford, John Sillitoe on behalf of the Wellcome Sanger Institute COVID-19 Surveillance Team |
| EPI_ISL_2729667 | Lighthouse Lab in Alderley Park | Wellcome Sanger Institute for the COVID-19 Genomics UK (COG-UK) Consortium | Jacquelyn Wynn, Mairead Hyland, The Lighthouse Lab in Alderley Park and Alex Alderton, Roberto Amato, Jeffrey Barrett, Sonia Goncalves, Ewan Harrison, David K. Jackson, Ian Johnston, Dominic Kwiatkowski, Cordelia Langford, John Sillitoe on behalf of the Wellcome Sanger Institute COVID-19 Surveillance Team |
| EPI_ISL_2729668 | Lighthouse Laboratory Plymouth | Wellcome Sanger Institute for the COVID-19 Genomics UK (COG-UK) Consortium | Lighthouse Laboratory Plymouth and Alex Alderton, Roberto Amato, Jeffrey Barrett, Sonia Goncalves, Ewan Harrison, David K. Jackson, Ian Johnston, Dominic Kwiatkowski, Cordelia Langford, John Sillitoe on behalf of the Wellcome Sanger Institute COVID-19 Surveillance Team |
| EPI_ISL_2729669, EPI_ISL_2729670, EPI_ISL_2729671 | Lighthouse Lab in Milton Keynes | Wellcome Sanger Institute for the COVID-19 Genomics UK (COG-UK) Consortium | The Lighthouse Lab in Milton Keynes and Alex Alderton, Roberto Amato, Jeffrey Barrett, Sonia Goncalves, Ewan Harrison, David K. Jackson, Ian Johnston, Dominic Kwiatkowski, Cordelia Langford, John Sillitoe on behalf of the Wellcome Sanger Institute COVID-19 Surveillance Team |
| EPI_ISL_2729672, EPI_ISL_2729673 | Lighthouse Lab in Alderley Park | Wellcome Sanger Institute for the COVID-19 Genomics UK (COG-UK) Consortium | Jacquelyn Wynn, Mairead Hyland, The Lighthouse Lab in Alderley Park and Alex Alderton, Roberto Amato, Jeffrey Barrett, Sonia Goncalves, Ewan Harrison, David K. Jackson, Ian Johnston, Dominic Kwiatkowski, Cordelia Langford, John Sillitoe on behalf of the Wellcome Sanger Institute COVID-19 Surveillance Team |
| EPI_ISL_2729674 | Lighthouse Lab in Glasgow | Wellcome Sanger Institute for the COVID-19 Genomics UK (COG-UK) Consortium | Harper VanSteenhouse, Yumi Kasai, David Gray, Carol Clugston, Anna Dominiczak and Alex Alderton, Roberto Amato, Jeffrey Barrett, Sonia Goncalves, Ewan Harrison, David K. Jackson, Ian Johnston, Dominic Kwiatkowski, Cordelia Langford, John Sillitoe on behalf of the Wellcome Sanger Institute COVID-19 Surveillance Team |
| EPI_ISL_2729675, EPI_ISL_2729676 | Lighthouse Laboratory Plymouth | Wellcome Sanger Institute for the COVID-19 Genomics UK (COG-UK) Consortium | Lighthouse Laboratory Plymouth and Alex Alderton, Roberto Amato, Jeffrey Barrett, Sonia Goncalves, Ewan Harrison, David K. Jackson, Ian Johnston, Dominic Kwiatkowski, Cordelia Langford, John Sillitoe on behalf of the Wellcome Sanger Institute COVID-19 Surveillance Team |
| EPI_ISL_2729677 | Lighthouse Lab in Alderley Park | Wellcome Sanger Institute for the COVID-19 Genomics UK (COG-UK) Consortium | Jacquelyn Wynn, Mairead Hyland, The Lighthouse Lab in Alderley Park and Alex Alderton, Roberto Amato, Jeffrey Barrett, Sonia Goncalves, Ewan Harrison, David K. Jackson, Ian Johnston, Dominic Kwiatkowski, Cordelia Langford, John Sillitoe on behalf of the Wellcome Sanger Institute COVID-19 Surveillance Team |
| EPI_ISL_2729678, EPI_ISL_2729679 | Lighthouse Lab in Milton Keynes | Wellcome Sanger Institute for the COVID-19 Genomics UK (COG-UK) Consortium | The Lighthouse Lab in Milton Keynes and Alex Alderton, Roberto Amato, Jeffrey Barrett, Sonia Goncalves, Ewan Harrison, David K. Jackson, Ian Johnston, Dominic Kwiatkowski, Cordelia Langford, John Sillitoe on behalf of the Wellcome Sanger Institute COVID-19 Surveillance Team |
| EPI_ISL_2729680 | Lighthouse Laboratory Plymouth | Wellcome Sanger Institute for the COVID-19 Genomics UK (COG-UK) Consortium | Lighthouse Laboratory Plymouth and Alex Alderton, Roberto Amato, Jeffrey Barrett, Sonia Goncalves, Ewan Harrison, David K. Jackson, Ian Johnston, Dominic Kwiatkowski, Cordelia Langford, John Sillitoe on behalf of the Wellcome Sanger Institute COVID-19 Surveillance Team |
| EPI_ISL_2729681, EPI_ISL_2729682 | Lighthouse Lab in Alderley Park | Wellcome Sanger Institute for the COVID-19 Genomics UK (COG-UK) Consortium | Jacquelyn Wynn, Mairead Hyland, The Lighthouse Lab in Alderley Park and Alex Alderton, Roberto Amato, Jeffrey Barrett, Sonia Goncalves, Ewan Harrison, David K. Jackson, Ian Johnston, Dominic Kwiatkowski, Cordelia Langford, John Sillitoe on behalf of the Wellcome Sanger Institute COVID-19 Surveillance Team |
| EPI_ISL_2729683 | Lighthouse Lab in Glasgow | Wellcome Sanger Institute for the COVID-19 Genomics UK (COG-UK) Consortium | Harper VanSteenhouse, Yumi Kasai, David Gray, Carol Clugston, Anna Dominiczak and Alex Alderton, Roberto Amato, Jeffrey Barrett, Sonia Goncalves, Ewan Harrison, David K. Jackson, Ian Johnston, Dominic Kwiatkowski, Cordelia Langford, John Sillitoe on behalf of the Wellcome Sanger Institute COVID-19 Surveillance Team |
| EPI_ISL_2729684 | Lighthouse Laboratory Plymouth | Wellcome Sanger Institute for the COVID-19 Genomics UK (COG-UK) Consortium | Lighthouse Laboratory Plymouth and Alex Alderton, Roberto Amato, Jeffrey Barrett, Sonia Goncalves, Ewan Harrison, David K. Jackson, Ian Johnston, Dominic Kwiatkowski, Cordelia Langford, John Sillitoe on behalf of the Wellcome Sanger Institute COVID-19 Surveillance Team |
| EPI_ISL_2729685 | Lighthouse Lab in Glasgow | Wellcome Sanger Institute for the COVID-19 Genomics UK (COG-UK) Consortium | Harper VanSteenhouse, Yumi Kasai, David Gray, Carol Clugston, Anna Dominiczak and Alex Alderton, Roberto Amato, Jeffrey Barrett, Sonia Goncalves, Ewan Harrison, David K. Jackson, Ian Johnston, Dominic Kwiatkowski, Cordelia Langford, John Sillitoe on behalf of the Wellcome Sanger Institute COVID-19 Surveillance Team |
| EPI_ISL_2729686 | Lighthouse Lab in Milton Keynes | Wellcome Sanger Institute for the COVID-19 Genomics UK (COG-UK) Consortium | The Lighthouse Lab in Milton Keynes and Alex Alderton, Roberto Amato, Jeffrey Barrett, Sonia Goncalves, Ewan Harrison, David K. Jackson, Ian Johnston, Dominic Kwiatkowski, Cordelia Langford, John Sillitoe on behalf of the Wellcome Sanger Institute COVID-19 Surveillance Team |
| EPI_ISL_2729687 | Lighthouse Lab in Alderley Park | Wellcome Sanger Institute for the COVID-19 Genomics UK (COG-UK) Consortium | Jacquelyn Wynn, Mairead Hyland, The Lighthouse Lab in Alderley Park and Alex Alderton, Roberto Amato, Jeffrey Barrett, Sonia Goncalves, Ewan Harrison, David K. Jackson, Ian Johnston, Dominic Kwiatkowski, Cordelia Langford, John Sillitoe on behalf of the Wellcome Sanger Institute COVID-19 Surveillance Team |
| EPI_ISL_2729688 | Lighthouse Lab in Milton Keynes | Wellcome Sanger Institute for the COVID-19 Genomics UK (COG-UK) Consortium | The Lighthouse Lab in Milton Keynes and Alex Alderton, Roberto Amato, Jeffrey Barrett, Sonia Goncalves, Ewan Harrison, David K. Jackson, Ian Johnston, Dominic Kwiatkowski, Cordelia Langford, John Sillitoe on behalf of the Wellcome Sanger Institute COVID-19 Surveillance Team |
| EPI_ISL_2729689 | Lighthouse Laboratory Plymouth | Wellcome Sanger Institute for the COVID-19 Genomics UK (COG-UK) Consortium | Lighthouse Laboratory Plymouth and Alex Alderton, Roberto Amato, Jeffrey Barrett, Sonia Goncalves, Ewan Harrison, David K. Jackson, Ian Johnston, Dominic Kwiatkowski, Cordelia Langford, John Sillitoe on behalf of the Wellcome Sanger Institute COVID-19 Surveillance Team |
| EPI_ISL_2729690, EPI_ISL_2729691 | Lighthouse Lab in Alderley Park | Wellcome Sanger Institute for the COVID-19 Genomics UK (COG-UK) Consortium | Jacquelyn Wynn, Mairead Hyland, The Lighthouse Lab in Alderley Park and Alex Alderton, Roberto Amato, Jeffrey Barrett, Sonia Goncalves, Ewan Harrison, David K. Jackson, Ian Johnston, Dominic Kwiatkowski, Cordelia Langford, John Sillitoe on behalf of the Wellcome Sanger Institute COVID-19 Surveillance Team |
| EPI_ISL_2729692 | Lighthouse Lab in Milton Keynes | Wellcome Sanger Institute for the COVID-19 Genomics UK (COG-UK) Consortium | The Lighthouse Lab in Milton Keynes and Alex Alderton, Roberto Amato, Jeffrey Barrett, Sonia Goncalves, Ewan Harrison, David K. Jackson, Ian Johnston, Dominic Kwiatkowski, Cordelia Langford, John Sillitoe on behalf of the Wellcome Sanger Institute COVID-19 Surveillance Team |
| EPI_ISL_2729693, EPI_ISL_2729694 | Lighthouse Lab in Alderley Park | Wellcome Sanger Institute for the COVID-19 Genomics UK (COG-UK) Consortium | Jacquelyn Wynn, Mairead Hyland, The Lighthouse Lab in Alderley Park and Alex Alderton, Roberto Amato, Jeffrey Barrett, Sonia Goncalves, Ewan Harrison, David K. Jackson, Ian Johnston, Dominic Kwiatkowski, Cordelia Langford, John Sillitoe on behalf of the Wellcome Sanger Institute COVID-19 Surveillance Team |
| EPI_ISL_2729695, EPI_ISL_2729696 | Lighthouse Lab in Milton Keynes | Wellcome Sanger Institute for the COVID-19 Genomics UK (COG-UK) Consortium | The Lighthouse Lab in Milton Keynes and Alex Alderton, Roberto Amato, Jeffrey Barrett, Sonia Goncalves, Ewan Harrison, David K. Jackson, Ian Johnston, Dominic Kwiatkowski, Cordelia Langford, John Sillitoe on behalf of the Wellcome Sanger Institute COVID-19 Surveillance Team |
| EPI_ISL_2729697, EPI_ISL_2729698, EPI_ISL_2729699 | Lighthouse Lab in Alderley Park | Wellcome Sanger Institute for the COVID-19 Genomics UK (COG-UK) Consortium | Jacquelyn Wynn, Mairead Hyland, The Lighthouse Lab in Alderley Park and Alex Alderton, Roberto Amato, Jeffrey Barrett, Sonia Goncalves, Ewan Harrison, David K. Jackson, Ian Johnston, Dominic Kwiatkowski, Cordelia Langford, John Sillitoe on behalf of the Wellcome Sanger Institute COVID-19 Surveillance Team |
| EPI_ISL_2729700 | Lighthouse Laboratory Plymouth | Wellcome Sanger Institute for the COVID-19 Genomics UK (COG-UK) Consortium | Lighthouse Laboratory Plymouth and Alex Alderton, Roberto Amato, Jeffrey Barrett, Sonia Goncalves, Ewan Harrison, David K. Jackson, Ian Johnston, Dominic Kwiatkowski, Cordelia Langford, John Sillitoe on behalf of the Wellcome Sanger Institute COVID-19 Surveillance Team |
| EPI_ISL_2729701, | Lighthouse Lab in Alderley Park | Wellcome Sanger Institute for the COVID-19 Genomics UK (COG-UK) | Jacquelyn Wynn, Mairead Hyland, The Lighthouse Lab in Alderley Park and Alex Alderton, Roberto Amato, Jeffrey Barrett, Sonia Goncalves, Ewan Harrison, David K. Jackson, Ian Johnston, Dominic Kwiatkowski, Cordelia Langford, John Sillitoe on |

| | | | |
|---|---|---|---|
| EPI_ISL_2729702 | | Consortium | behalf of the Wellcome Sanger Institute COVID-19 Surveillance Team |
| EPI_ISL_2729703 | Lighthouse Laboratory Plymouth | Wellcome Sanger Institute for the COVID-19 Genomics UK (COG-UK) Consortium | Lighthouse Laboratory Plymouth and Alex Alderton, Roberto Amato, Jeffrey Barrett, Sonia Goncalves, Ewan Harrison, David K. Jackson, Ian Johnston, Dominic Kwiatkowski, Cordelia Langford, John Sillitoe on behalf of the Wellcome Sanger Institute COVID-19 Surveillance Team |
| EPI_ISL_2729704 | Lighthouse Lab in Alderley Park | Wellcome Sanger Institute for the COVID-19 Genomics UK (COG-UK) Consortium | Jacquelyn Wynn, Mairead Hyland, The Lighthouse Lab in Alderley Park and Alex Alderton, Roberto Amato, Jeffrey Barrett, Sonia Goncalves, Ewan Harrison, David K. Jackson, Ian Johnston, Dominic Kwiatkowski, Cordelia Langford, John Sillitoe on behalf of the Wellcome Sanger Institute COVID-19 Surveillance Team |
| EPI_ISL_2729705 | Lighthouse Laboratory Plymouth | Wellcome Sanger Institute for the COVID-19 Genomics UK (COG-UK) Consortium | Lighthouse Laboratory Plymouth and Alex Alderton, Roberto Amato, Jeffrey Barrett, Sonia Goncalves, Ewan Harrison, David K. Jackson, Ian Johnston, Dominic Kwiatkowski, Cordelia Langford, John Sillitoe on behalf of the Wellcome Sanger Institute COVID-19 Surveillance Team |
| EPI_ISL_2729706 | Lighthouse Lab in Alderley Park | Wellcome Sanger Institute for the COVID-19 Genomics UK (COG-UK) Consortium | Jacquelyn Wynn, Mairead Hyland, The Lighthouse Lab in Alderley Park and Alex Alderton, Roberto Amato, Jeffrey Barrett, Sonia Goncalves, Ewan Harrison, David K. Jackson, Ian Johnston, Dominic Kwiatkowski, Cordelia Langford, John Sillitoe on behalf of the Wellcome Sanger Institute COVID-19 Surveillance Team |
| EPI_ISL_2729707, EPI_ISL_2729708 | Lighthouse Laboratory Plymouth | Wellcome Sanger Institute for the COVID-19 Genomics UK (COG-UK) Consortium | Lighthouse Laboratory Plymouth and Alex Alderton, Roberto Amato, Jeffrey Barrett, Sonia Goncalves, Ewan Harrison, David K. Jackson, Ian Johnston, Dominic Kwiatkowski, Cordelia Langford, John Sillitoe on behalf of the Wellcome Sanger Institute COVID-19 Surveillance Team |
| EPI_ISL_2729709 | Lighthouse Lab in Alderley Park | Wellcome Sanger Institute for the COVID-19 Genomics UK (COG-UK) Consortium | Jacquelyn Wynn, Mairead Hyland, The Lighthouse Lab in Alderley Park and Alex Alderton, Roberto Amato, Jeffrey Barrett, Sonia Goncalves, Ewan Harrison, David K. Jackson, Ian Johnston, Dominic Kwiatkowski, Cordelia Langford, John Sillitoe on behalf of the Wellcome Sanger Institute COVID-19 Surveillance Team |
| EPI_ISL_2729710 | Lighthouse Laboratory Plymouth | Wellcome Sanger Institute for the COVID-19 Genomics UK (COG-UK) Consortium | Lighthouse Laboratory Plymouth and Alex Alderton, Roberto Amato, Jeffrey Barrett, Sonia Goncalves, Ewan Harrison, David K. Jackson, Ian Johnston, Dominic Kwiatkowski, Cordelia Langford, John Sillitoe on behalf of the Wellcome Sanger Institute COVID-19 Surveillance Team |
| EPI_ISL_2729711 | Lighthouse Lab in Milton Keynes | Wellcome Sanger Institute for the COVID-19 Genomics UK (COG-UK) Consortium | The Lighthouse Lab in Milton Keynes and Alex Alderton, Roberto Amato, Jeffrey Barrett, Sonia Goncalves, Ewan Harrison, David K. Jackson, Ian Johnston, Dominic Kwiatkowski, Cordelia Langford, John Sillitoe on behalf of the Wellcome Sanger Institute COVID-19 Surveillance Team |
| EPI_ISL_2729712 | Lighthouse Lab in Alderley Park | Wellcome Sanger Institute for the COVID-19 Genomics UK (COG-UK) Consortium | Jacquelyn Wynn, Mairead Hyland, The Lighthouse Lab in Alderley Park and Alex Alderton, Roberto Amato, Jeffrey Barrett, Sonia Goncalves, Ewan Harrison, David K. Jackson, Ian Johnston, Dominic Kwiatkowski, Cordelia Langford, John Sillitoe on behalf of the Wellcome Sanger Institute COVID-19 Surveillance Team |
| EPI_ISL_2729713, EPI_ISL_2729714 | Lighthouse Lab in Milton Keynes | Wellcome Sanger Institute for the COVID-19 Genomics UK (COG-UK) Consortium | The Lighthouse Lab in Milton Keynes and Alex Alderton, Roberto Amato, Jeffrey Barrett, Sonia Goncalves, Ewan Harrison, David K. Jackson, Ian Johnston, Dominic Kwiatkowski, Cordelia Langford, John Sillitoe on behalf of the Wellcome Sanger Institute COVID-19 Surveillance Team |
| EPI_ISL_2729715, EPI_ISL_2729718, EPI_ISL_2729719 | Lighthouse Laboratory Plymouth | Wellcome Sanger Institute for the COVID-19 Genomics UK (COG-UK) Consortium | Lighthouse Laboratory Plymouth and Alex Alderton, Roberto Amato, Jeffrey Barrett, Sonia Goncalves, Ewan Harrison, David K. Jackson, Ian Johnston, Dominic Kwiatkowski, Cordelia Langford, John Sillitoe on behalf of the Wellcome Sanger Institute COVID-19 Surveillance Team |
| EPI_ISL_2729720 | Lighthouse Lab in Glasgow | Wellcome Sanger Institute for the COVID-19 Genomics UK (COG-UK) Consortium | Harper VanSteenhouse, Yumi Kasai, David Gray, Carol Clugston, Anna Dominiczak and Alex Alderton, Roberto Amato, Jeffrey Barrett, Sonia Goncalves, Ewan Harrison, David K. Jackson, Ian Johnston, Dominic Kwiatkowski, Cordelia Langford, John Sillitoe on behalf of the Wellcome Sanger Institute COVID-19 Surveillance Team |
| EPI_ISL_2729721 | Lighthouse Lab in Alderley Park | Wellcome Sanger Institute for the COVID-19 Genomics UK (COG-UK) Consortium | Jacquelyn Wynn, Mairead Hyland, The Lighthouse Lab in Alderley Park and Alex Alderton, Roberto Amato, Jeffrey Barrett, Sonia Goncalves, Ewan Harrison, David K. Jackson, Ian Johnston, Dominic Kwiatkowski, Cordelia Langford, John Sillitoe on behalf of the Wellcome Sanger Institute COVID-19 Surveillance Team |
| EPI_ISL_2729722 | Lighthouse Laboratory Plymouth | Wellcome Sanger Institute for the COVID-19 Genomics UK (COG-UK) Consortium | Lighthouse Laboratory Plymouth and Alex Alderton, Roberto Amato, Jeffrey Barrett, Sonia Goncalves, Ewan Harrison, David K. Jackson, Ian Johnston, Dominic Kwiatkowski, Cordelia Langford, John Sillitoe on behalf of the Wellcome Sanger Institute COVID-19 Surveillance Team |
| EPI_ISL_2729723, EPI_ISL_2729724 | Lighthouse Lab in Milton Keynes | Wellcome Sanger Institute for the COVID-19 Genomics UK (COG-UK) Consortium | The Lighthouse Lab in Milton Keynes and Alex Alderton, Roberto Amato, Jeffrey Barrett, Sonia Goncalves, Ewan Harrison, David K. Jackson, Ian Johnston, Dominic Kwiatkowski, Cordelia Langford, John Sillitoe on behalf of the Wellcome Sanger Institute COVID-19 Surveillance Team |
| EPI_ISL_2729725, EPI_ISL_2729726, EPI_ISL_2729727, EPI_ISL_2729728 | Lighthouse Lab in Alderley Park | Wellcome Sanger Institute for the COVID-19 Genomics UK (COG-UK) Consortium | Jacquelyn Wynn, Mairead Hyland, The Lighthouse Lab in Alderley Park and Alex Alderton, Roberto Amato, Jeffrey Barrett, Sonia Goncalves, Ewan Harrison, David K. Jackson, Ian Johnston, Dominic Kwiatkowski, Cordelia Langford, John Sillitoe on behalf of the Wellcome Sanger Institute COVID-19 Surveillance Team |
| EPI_ISL_2729729, EPI_ISL_2729730 | Lighthouse Lab in Glasgow | Wellcome Sanger Institute for the COVID-19 Genomics UK (COG-UK) Consortium | Harper VanSteenhouse, Yumi Kasai, David Gray, Carol Clugston, Anna Dominiczak and Alex Alderton, Roberto Amato, Jeffrey Barrett, Sonia Goncalves, Ewan Harrison, David K. Jackson, Ian Johnston, Dominic Kwiatkowski, Cordelia Langford, John Sillitoe on behalf of the Wellcome Sanger Institute COVID-19 Surveillance Team |
| EPI_ISL_2729731 | Lighthouse Lab in Milton Keynes | Wellcome Sanger Institute for the COVID-19 Genomics UK (COG-UK) Consortium | The Lighthouse Lab in Milton Keynes and Alex Alderton, Roberto Amato, Jeffrey Barrett, Sonia Goncalves, Ewan Harrison, David K. Jackson, Ian Johnston, Dominic Kwiatkowski, Cordelia Langford, John Sillitoe on behalf of the Wellcome Sanger Institute COVID-19 Surveillance Team |
| EPI_ISL_2729732 | Lighthouse Laboratory Plymouth | Wellcome Sanger Institute for the COVID-19 Genomics UK (COG-UK) Consortium | Lighthouse Laboratory Plymouth and Alex Alderton, Roberto Amato, Jeffrey Barrett, Sonia Goncalves, Ewan Harrison, David K. Jackson, Ian Johnston, Dominic Kwiatkowski, Cordelia Langford, John Sillitoe on behalf of the Wellcome Sanger Institute COVID-19 Surveillance Team |
| EPI_ISL_2729733 | Lighthouse Lab in Glasgow | Wellcome Sanger Institute for the COVID-19 Genomics UK (COG-UK) Consortium | Harper VanSteenhouse, Yumi Kasai, David Gray, Carol Clugston, Anna Dominiczak and Alex Alderton, Roberto Amato, Jeffrey Barrett, Sonia Goncalves, Ewan Harrison, David K. Jackson, Ian Johnston, Dominic Kwiatkowski, Cordelia Langford, John Sillitoe on behalf of the Wellcome Sanger Institute COVID-19 Surveillance Team |
| EPI_ISL_2729734 | Lighthouse Lab in Alderley Park | Wellcome Sanger Institute for the COVID-19 Genomics UK (COG-UK) Consortium | Jacquelyn Wynn, Mairead Hyland, The Lighthouse Lab in Alderley Park and Alex Alderton, Roberto Amato, Jeffrey Barrett, Sonia Goncalves, Ewan Harrison, David K. Jackson, Ian Johnston, Dominic Kwiatkowski, Cordelia Langford, John Sillitoe on behalf of the Wellcome Sanger Institute COVID-19 Surveillance Team |
| EPI_ISL_2729735 | Lighthouse Laboratory Plymouth | Wellcome Sanger Institute for the COVID-19 Genomics UK (COG-UK) Consortium | Lighthouse Laboratory Plymouth and Alex Alderton, Roberto Amato, Jeffrey Barrett, Sonia Goncalves, Ewan Harrison, David K. Jackson, Ian Johnston, Dominic Kwiatkowski, Cordelia Langford, John Sillitoe on behalf of the Wellcome Sanger Institute COVID-19 Surveillance Team |
| EPI_ISL_2729736 | Lighthouse Lab in Milton Keynes | Wellcome Sanger Institute for the COVID-19 Genomics UK (COG-UK) Consortium | The Lighthouse Lab in Milton Keynes and Alex Alderton, Roberto Amato, Jeffrey Barrett, Sonia Goncalves, Ewan Harrison, David K. Jackson, Ian Johnston, Dominic Kwiatkowski, Cordelia Langford, John Sillitoe on behalf of the Wellcome Sanger Institute COVID-19 Surveillance Team |
| EPI_ISL_2729737 | Lighthouse Lab in Alderley Park | Wellcome Sanger Institute for the COVID-19 Genomics UK (COG-UK) Consortium | Jacquelyn Wynn, Mairead Hyland, The Lighthouse Lab in Alderley Park and Alex Alderton, Roberto Amato, Jeffrey Barrett, Sonia Goncalves, Ewan Harrison, David K. Jackson, Ian Johnston, Dominic Kwiatkowski, Cordelia Langford, John Sillitoe on behalf of the Wellcome Sanger Institute COVID-19 Surveillance Team |
| EPI_ISL_2729738 | Lighthouse Laboratory Plymouth | Wellcome Sanger Institute for the COVID-19 Genomics UK (COG-UK) Consortium | Lighthouse Laboratory Plymouth and Alex Alderton, Roberto Amato, Jeffrey Barrett, Sonia Goncalves, Ewan Harrison, David K. Jackson, Ian Johnston, Dominic Kwiatkowski, Cordelia Langford, John Sillitoe on behalf of the Wellcome Sanger Institute COVID-19 Surveillance Team |
| EPI_ISL_2729739, EPI_ISL_2729740 | Lighthouse Lab in Glasgow | Wellcome Sanger Institute for the COVID-19 Genomics UK (COG-UK) Consortium | Harper VanSteenhouse, Yumi Kasai, David Gray, Carol Clugston, Anna Dominiczak and Alex Alderton, Roberto Amato, Jeffrey Barrett, Sonia Goncalves, Ewan Harrison, David K. Jackson, Ian Johnston, Dominic Kwiatkowski, Cordelia Langford, John Sillitoe on behalf of the Wellcome Sanger Institute COVID-19 Surveillance Team |
| EPI_ISL_2729741 | Lighthouse Laboratory Plymouth | Wellcome Sanger Institute for the COVID-19 Genomics UK (COG-UK) Consortium | Lighthouse Laboratory Plymouth and Alex Alderton, Roberto Amato, Jeffrey Barrett, Sonia Goncalves, Ewan Harrison, David K. Jackson, Ian Johnston, Dominic Kwiatkowski, Cordelia Langford, John Sillitoe on behalf of the Wellcome Sanger Institute COVID-19 Surveillance Team |
| EPI_ISL_2729742 | Lighthouse Lab in Glasgow | Wellcome Sanger Institute for the COVID-19 Genomics UK (COG-UK) Consortium | Harper VanSteenhouse, Yumi Kasai, David Gray, Carol Clugston, Anna Dominiczak and Alex Alderton, Roberto Amato, Jeffrey Barrett, Sonia Goncalves, Ewan Harrison, David K. Jackson, Ian Johnston, Dominic Kwiatkowski, Cordelia Langford, John Sillitoe on behalf of the Wellcome Sanger Institute COVID-19 Surveillance Team |
| EPI_ISL_2729743 | Lighthouse Lab in Milton Keynes | Wellcome Sanger Institute for the COVID-19 Genomics UK (COG-UK) Consortium | The Lighthouse Lab in Milton Keynes and Alex Alderton, Roberto Amato, Jeffrey Barrett, Sonia Goncalves, Ewan Harrison, David K. Jackson, Ian Johnston, Dominic Kwiatkowski, Cordelia Langford, John Sillitoe on behalf of the Wellcome Sanger Institute COVID-19 Surveillance Team |
| EPI_ISL_2729744 | Lighthouse Lab in Alderley Park | Wellcome Sanger Institute for the COVID-19 Genomics UK (COG-UK) Consortium | Jacquelyn Wynn, Mairead Hyland, The Lighthouse Lab in Alderley Park and Alex Alderton, Roberto Amato, Jeffrey Barrett, Sonia Goncalves, Ewan Harrison, David K. Jackson, Ian Johnston, Dominic Kwiatkowski, Cordelia Langford, John Sillitoe on behalf of the Wellcome Sanger Institute COVID-19 Surveillance Team |
| EPI_ISL_2729745, EPI_ISL_2729746, EPI_ISL_2729747, EPI_ISL_2729748, EPI_ISL_2729749 | Lighthouse Laboratory Plymouth | Wellcome Sanger Institute for the COVID-19 Genomics UK (COG-UK) Consortium | Lighthouse Laboratory Plymouth and Alex Alderton, Roberto Amato, Jeffrey Barrett, Sonia Goncalves, Ewan Harrison, David K. Jackson, Ian Johnston, Dominic Kwiatkowski, Cordelia Langford, John Sillitoe on behalf of the Wellcome Sanger Institute COVID-19 Surveillance Team |
| EPI_ISL_2729750, EPI_ISL_2729751 | Lighthouse Lab in Milton Keynes | Wellcome Sanger Institute for the COVID-19 Genomics UK (COG-UK) Consortium | The Lighthouse Lab in Milton Keynes and Alex Alderton, Roberto Amato, Jeffrey Barrett, Sonia Goncalves, Ewan Harrison, David K. Jackson, Ian Johnston, Dominic Kwiatkowski, Cordelia Langford, John Sillitoe on behalf of the Wellcome Sanger Institute COVID-19 Surveillance Team |
| EPI_ISL_2729752 | Lighthouse Laboratory Plymouth | Wellcome Sanger Institute for the COVID-19 Genomics UK (COG-UK) Consortium | Lighthouse Laboratory Plymouth and Alex Alderton, Roberto Amato, Jeffrey Barrett, Sonia Goncalves, Ewan Harrison, David K. Jackson, Ian Johnston, Dominic Kwiatkowski, Cordelia Langford, John Sillitoe on behalf of the Wellcome Sanger Institute COVID-19 Surveillance Team |
| EPI_ISL_2729753 | Lighthouse Lab in Alderley Park | Wellcome Sanger Institute for the COVID-19 Genomics UK (COG-UK) Consortium | Jacquelyn Wynn, Mairead Hyland, The Lighthouse Lab in Alderley Park and Alex Alderton, Roberto Amato, Jeffrey Barrett, Sonia Goncalves, Ewan Harrison, David K. Jackson, Ian Johnston, Dominic Kwiatkowski, Cordelia Langford, John Sillitoe on behalf of the Wellcome Sanger Institute COVID-19 Surveillance Team |
| EPI_ISL_2729754 | Lighthouse Lab in Glasgow | Wellcome Sanger Institute for the COVID-19 Genomics UK (COG-UK) Consortium | Harper VanSteenhouse, Yumi Kasai, David Gray, Carol Clugston, Anna Dominiczak and Alex Alderton, Roberto Amato, Jeffrey Barrett, Sonia Goncalves, Ewan Harrison, David K. Jackson, Ian Johnston, Dominic Kwiatkowski, Cordelia Langford, John Sillitoe on behalf of the Wellcome Sanger Institute COVID-19 Surveillance Team |
| EPI_ISL_2729755 | Lighthouse Lab in Alderley Park | Wellcome Sanger Institute for the COVID-19 Genomics UK (COG-UK) Consortium | Jacquelyn Wynn, Mairead Hyland, The Lighthouse Lab in Alderley Park and Alex Alderton, Roberto Amato, Jeffrey Barrett, Sonia Goncalves, Ewan Harrison, David K. Jackson, Ian Johnston, Dominic Kwiatkowski, Cordelia Langford, John Sillitoe on behalf of the Wellcome Sanger Institute COVID-19 Surveillance Team |
| EPI_ISL_2729756 | Lighthouse Lab in Milton Keynes | Wellcome Sanger Institute for the COVID-19 Genomics UK (COG-UK) Consortium | The Lighthouse Lab in Milton Keynes and Alex Alderton, Roberto Amato, Jeffrey Barrett, Sonia Goncalves, Ewan Harrison, David K. Jackson, Ian Johnston, Dominic Kwiatkowski, Cordelia Langford, John Sillitoe on behalf of the Wellcome Sanger Institute COVID-19 Surveillance Team |
| EPI_ISL_2729757, EPI_ISL_2729758 | Lighthouse Laboratory Plymouth | Wellcome Sanger Institute for the COVID-19 Genomics UK (COG-UK) Consortium | Lighthouse Laboratory Plymouth and Alex Alderton, Roberto Amato, Jeffrey Barrett, Sonia Goncalves, Ewan Harrison, David K. Jackson, Ian Johnston, Dominic Kwiatkowski, Cordelia Langford, John Sillitoe on behalf of the Wellcome Sanger Institute COVID-19 Surveillance Team |
| EPI_ISL_2729759 | Lighthouse Lab in Milton Keynes | Wellcome Sanger Institute for the COVID-19 Genomics UK (COG-UK) Consortium | The Lighthouse Lab in Milton Keynes and Alex Alderton, Roberto Amato, Jeffrey Barrett, Sonia Goncalves, Ewan Harrison, David K. Jackson, Ian Johnston, Dominic Kwiatkowski, Cordelia Langford, John Sillitoe on behalf of the Wellcome Sanger Institute COVID-19 Surveillance Team |
| EPI_ISL_2729760 | Lighthouse Lab in Glasgow | Wellcome Sanger Institute for the COVID-19 Genomics UK (COG-UK) Consortium | Harper VanSteenhouse, Yumi Kasai, David Gray, Carol Clugston, Anna Dominiczak and Alex Alderton, Roberto Amato, Jeffrey Barrett, Sonia Goncalves, Ewan Harrison, David K. Jackson, Ian Johnston, Dominic Kwiatkowski, Cordelia Langford, John Sillitoe on behalf of the Wellcome Sanger Institute COVID-19 Surveillance Team |
| EPI_ISL_2729761, EPI_ISL_2729762, EPI_ISL_2729763 | Lighthouse Laboratory Plymouth | Wellcome Sanger Institute for the COVID-19 Genomics UK (COG-UK) Consortium | Lighthouse Laboratory Plymouth and Alex Alderton, Roberto Amato, Jeffrey Barrett, Sonia Goncalves, Ewan Harrison, David K. Jackson, Ian Johnston, Dominic Kwiatkowski, Cordelia Langford, John Sillitoe on behalf of the Wellcome Sanger Institute COVID-19 Surveillance Team |
| EPI_ISL_2729764 | Lighthouse Lab in Milton Keynes | Wellcome Sanger Institute for the COVID-19 Genomics UK (COG-UK) Consortium | The Lighthouse Lab in Milton Keynes and Alex Alderton, Roberto Amato, Jeffrey Barrett, Sonia Goncalves, Ewan Harrison, David K. Jackson, Ian Johnston, Dominic Kwiatkowski, Cordelia Langford, John Sillitoe on behalf of the Wellcome Sanger Institute COVID-19 Surveillance Team |

| | | | |
|---|---|---|---|
| EPI_ISL_2729765, EPI_ISL_2729766 | Lighthouse Laboratory Plymouth | Wellcome Sanger Institute for the COVID-19 Genomics UK (COG-UK) Consortium | Lighthouse Laboratory Plymouth and Alex Alderton, Roberto Amato, Jeffrey Barrett, Sonia Goncalves, Ewan Harrison, David K. Jackson, Ian Johnston, Dominic Kwiatkowski, Cordelia Langford, John Sillitoe on behalf of the Wellcome Sanger Institute COVID-19 Surveillance Team |
| EPI_ISL_2729767 | Lighthouse Lab in Alderley Park | Wellcome Sanger Institute for the COVID-19 Genomics UK (COG-UK) Consortium | Jacquelyn Wynn, Mairead Hyland, The Lighthouse Lab in Alderley Park and Alex Alderton, Roberto Amato, Jeffrey Barrett, Sonia Goncalves, Ewan Harrison, David K. Jackson, Ian Johnston, Dominic Kwiatkowski, Cordelia Langford, John Sillitoe on behalf of the Wellcome Sanger Institute COVID-19 Surveillance Team |
| EPI_ISL_2729768, EPI_ISL_2729769, EPI_ISL_2729770 | Lighthouse Lab in Milton Keynes | Wellcome Sanger Institute for the COVID-19 Genomics UK (COG-UK) Consortium | The Lighthouse Lab in Milton Keynes and Alex Alderton, Roberto Amato, Jeffrey Barrett, Sonia Goncalves, Ewan Harrison, David K. Jackson, Ian Johnston, Dominic Kwiatkowski, Cordelia Langford, John Sillitoe on behalf of the Wellcome Sanger Institute COVID-19 Surveillance Team |
| EPI_ISL_2729771, EPI_ISL_2729772 | Lighthouse Lab in Alderley Park | Wellcome Sanger Institute for the COVID-19 Genomics UK (COG-UK) Consortium | Jacquelyn Wynn, Mairead Hyland, The Lighthouse Lab in Alderley Park and Alex Alderton, Roberto Amato, Jeffrey Barrett, Sonia Goncalves, Ewan Harrison, David K. Jackson, Ian Johnston, Dominic Kwiatkowski, Cordelia Langford, John Sillitoe on behalf of the Wellcome Sanger Institute COVID-19 Surveillance Team |
| EPI_ISL_2729773, EPI_ISL_2729774 | Lighthouse Lab in Milton Keynes | Wellcome Sanger Institute for the COVID-19 Genomics UK (COG-UK) Consortium | The Lighthouse Lab in Milton Keynes and Alex Alderton, Roberto Amato, Jeffrey Barrett, Sonia Goncalves, Ewan Harrison, David K. Jackson, Ian Johnston, Dominic Kwiatkowski, Cordelia Langford, John Sillitoe on behalf of the Wellcome Sanger Institute COVID-19 Surveillance Team |
| EPI_ISL_2729775 | Lighthouse Laboratory Plymouth | Wellcome Sanger Institute for the COVID-19 Genomics UK (COG-UK) Consortium | Lighthouse Laboratory Plymouth and Alex Alderton, Roberto Amato, Jeffrey Barrett, Sonia Goncalves, Ewan Harrison, David K. Jackson, Ian Johnston, Dominic Kwiatkowski, Cordelia Langford, John Sillitoe on behalf of the Wellcome Sanger Institute COVID-19 Surveillance Team |
| EPI_ISL_2729776, EPI_ISL_2729777, EPI_ISL_2729778 | Lighthouse Lab in Milton Keynes | Wellcome Sanger Institute for the COVID-19 Genomics UK (COG-UK) Consortium | The Lighthouse Lab in Milton Keynes and Alex Alderton, Roberto Amato, Jeffrey Barrett, Sonia Goncalves, Ewan Harrison, David K. Jackson, Ian Johnston, Dominic Kwiatkowski, Cordelia Langford, John Sillitoe on behalf of the Wellcome Sanger Institute COVID-19 Surveillance Team |
| EPI_ISL_2729779 | Lighthouse Laboratory Plymouth | Wellcome Sanger Institute for the COVID-19 Genomics UK (COG-UK) Consortium | Lighthouse Laboratory Plymouth and Alex Alderton, Roberto Amato, Jeffrey Barrett, Sonia Goncalves, Ewan Harrison, David K. Jackson, Ian Johnston, Dominic Kwiatkowski, Cordelia Langford, John Sillitoe on behalf of the Wellcome Sanger Institute COVID-19 Surveillance Team |
| EPI_ISL_2729780 | Lighthouse Lab in Glasgow | Wellcome Sanger Institute for the COVID-19 Genomics UK (COG-UK) Consortium | Harper VanSteenhouse, Yumi Kasai, David Gray, Carol Clugston, Anna Dominiczak and Alex Alderton, Roberto Amato, Jeffrey Barrett, Sonia Goncalves, Ewan Harrison, David K. Jackson, Ian Johnston, Dominic Kwiatkowski, Cordelia Langford, John Sillitoe on behalf of the Wellcome Sanger Institute COVID-19 Surveillance Team |
| EPI_ISL_2729781 | Lighthouse Laboratory Plymouth | Wellcome Sanger Institute for the COVID-19 Genomics UK (COG-UK) Consortium | Lighthouse Laboratory Plymouth and Alex Alderton, Roberto Amato, Jeffrey Barrett, Sonia Goncalves, Ewan Harrison, David K. Jackson, Ian Johnston, Dominic Kwiatkowski, Cordelia Langford, John Sillitoe on behalf of the Wellcome Sanger Institute COVID-19 Surveillance Team |
| EPI_ISL_2729782 | Lighthouse Lab in Glasgow | Wellcome Sanger Institute for the COVID-19 Genomics UK (COG-UK) Consortium | Harper VanSteenhouse, Yumi Kasai, David Gray, Carol Clugston, Anna Dominiczak and Alex Alderton, Roberto Amato, Jeffrey Barrett, Sonia Goncalves, Ewan Harrison, David K. Jackson, Ian Johnston, Dominic Kwiatkowski, Cordelia Langford, John Sillitoe on behalf of the Wellcome Sanger Institute COVID-19 Surveillance Team |
| EPI_ISL_2729783 | Lighthouse Lab in Milton Keynes | Wellcome Sanger Institute for the COVID-19 Genomics UK (COG-UK) Consortium | The Lighthouse Lab in Milton Keynes and Alex Alderton, Roberto Amato, Jeffrey Barrett, Sonia Goncalves, Ewan Harrison, David K. Jackson, Ian Johnston, Dominic Kwiatkowski, Cordelia Langford, John Sillitoe on behalf of the Wellcome Sanger Institute COVID-19 Surveillance Team |
| EPI_ISL_2729784, EPI_ISL_2729785 | Lighthouse Laboratory Plymouth | Wellcome Sanger Institute for the COVID-19 Genomics UK (COG-UK) Consortium | Lighthouse Laboratory Plymouth and Alex Alderton, Roberto Amato, Jeffrey Barrett, Sonia Goncalves, Ewan Harrison, David K. Jackson, Ian Johnston, Dominic Kwiatkowski, Cordelia Langford, John Sillitoe on behalf of the Wellcome Sanger Institute COVID-19 Surveillance Team |
| EPI_ISL_2729786 | Lighthouse Lab in Alderley Park | Wellcome Sanger Institute for the COVID-19 Genomics UK (COG-UK) Consortium | Jacquelyn Wynn, Mairead Hyland, The Lighthouse Lab in Alderley Park and Alex Alderton, Roberto Amato, Jeffrey Barrett, Sonia Goncalves, Ewan Harrison, David K. Jackson, Ian Johnston, Dominic Kwiatkowski, Cordelia Langford, John Sillitoe on behalf of the Wellcome Sanger Institute COVID-19 Surveillance Team |
| EPI_ISL_2729787 | Lighthouse Lab in Milton Keynes | Wellcome Sanger Institute for the COVID-19 Genomics UK (COG-UK) Consortium | The Lighthouse Lab in Milton Keynes and Alex Alderton, Roberto Amato, Jeffrey Barrett, Sonia Goncalves, Ewan Harrison, David K. Jackson, Ian Johnston, Dominic Kwiatkowski, Cordelia Langford, John Sillitoe on behalf of the Wellcome Sanger Institute COVID-19 Surveillance Team |
| EPI_ISL_2729788 | Lighthouse Laboratory Plymouth | Wellcome Sanger Institute for the COVID-19 Genomics UK (COG-UK) Consortium | Lighthouse Laboratory Plymouth and Alex Alderton, Roberto Amato, Jeffrey Barrett, Sonia Goncalves, Ewan Harrison, David K. Jackson, Ian Johnston, Dominic Kwiatkowski, Cordelia Langford, John Sillitoe on behalf of the Wellcome Sanger Institute COVID-19 Surveillance Team |
| EPI_ISL_2729789 | Lighthouse Lab in Alderley Park | Wellcome Sanger Institute for the COVID-19 Genomics UK (COG-UK) Consortium | Jacquelyn Wynn, Mairead Hyland, The Lighthouse Lab in Alderley Park and Alex Alderton, Roberto Amato, Jeffrey Barrett, Sonia Goncalves, Ewan Harrison, David K. Jackson, Ian Johnston, Dominic Kwiatkowski, Cordelia Langford, John Sillitoe on behalf of the Wellcome Sanger Institute COVID-19 Surveillance Team |
| EPI_ISL_2729790 | Lighthouse Laboratory Plymouth | Wellcome Sanger Institute for the COVID-19 Genomics UK (COG-UK) Consortium | Lighthouse Laboratory Plymouth and Alex Alderton, Roberto Amato, Jeffrey Barrett, Sonia Goncalves, Ewan Harrison, David K. Jackson, Ian Johnston, Dominic Kwiatkowski, Cordelia Langford, John Sillitoe on behalf of the Wellcome Sanger Institute COVID-19 Surveillance Team |
| EPI_ISL_2729791 | Lighthouse Lab in Milton Keynes | Wellcome Sanger Institute for the COVID-19 Genomics UK (COG-UK) Consortium | The Lighthouse Lab in Milton Keynes and Alex Alderton, Roberto Amato, Jeffrey Barrett, Sonia Goncalves, Ewan Harrison, David K. Jackson, Ian Johnston, Dominic Kwiatkowski, Cordelia Langford, John Sillitoe on behalf of the Wellcome Sanger Institute COVID-19 Surveillance Team |
| EPI_ISL_2729792, EPI_ISL_2729793 | Lighthouse Laboratory Plymouth | Wellcome Sanger Institute for the COVID-19 Genomics UK (COG-UK) Consortium | Lighthouse Laboratory Plymouth and Alex Alderton, Roberto Amato, Jeffrey Barrett, Sonia Goncalves, Ewan Harrison, David K. Jackson, Ian Johnston, Dominic Kwiatkowski, Cordelia Langford, John Sillitoe on behalf of the Wellcome Sanger Institute COVID-19 Surveillance Team |
| EPI_ISL_2729794 | Lighthouse Lab in Alderley Park | Wellcome Sanger Institute for the COVID-19 Genomics UK (COG-UK) Consortium | Jacquelyn Wynn, Mairead Hyland, The Lighthouse Lab in Alderley Park and Alex Alderton, Roberto Amato, Jeffrey Barrett, Sonia Goncalves, Ewan Harrison, David K. Jackson, Ian Johnston, Dominic Kwiatkowski, Cordelia Langford, John Sillitoe on behalf of the Wellcome Sanger Institute COVID-19 Surveillance Team |
| EPI_ISL_2729795 | Lighthouse Lab in Milton Keynes | Wellcome Sanger Institute for the COVID-19 Genomics UK (COG-UK) Consortium | The Lighthouse Lab in Milton Keynes and Alex Alderton, Roberto Amato, Jeffrey Barrett, Sonia Goncalves, Ewan Harrison, David K. Jackson, Ian Johnston, Dominic Kwiatkowski, Cordelia Langford, John Sillitoe on behalf of the Wellcome Sanger Institute COVID-19 Surveillance Team |
| EPI_ISL_2729796, EPI_ISL_2729797, EPI_ISL_2729798 | Lighthouse Laboratory Plymouth | Wellcome Sanger Institute for the COVID-19 Genomics UK (COG-UK) Consortium | Lighthouse Laboratory Plymouth and Alex Alderton, Roberto Amato, Jeffrey Barrett, Sonia Goncalves, Ewan Harrison, David K. Jackson, Ian Johnston, Dominic Kwiatkowski, Cordelia Langford, John Sillitoe on behalf of the Wellcome Sanger Institute COVID-19 Surveillance Team |
| EPI_ISL_2729799 | Lighthouse Lab in Milton Keynes | Wellcome Sanger Institute for the COVID-19 Genomics UK (COG-UK) Consortium | The Lighthouse Lab in Milton Keynes and Alex Alderton, Roberto Amato, Jeffrey Barrett, Sonia Goncalves, Ewan Harrison, David K. Jackson, Ian Johnston, Dominic Kwiatkowski, Cordelia Langford, John Sillitoe on behalf of the Wellcome Sanger Institute COVID-19 Surveillance Team |
| EPI_ISL_2729800, EPI_ISL_2729801, EPI_ISL_2729802 | Lighthouse Lab in Alderley Park | Wellcome Sanger Institute for the COVID-19 Genomics UK (COG-UK) Consortium | Jacquelyn Wynn, Mairead Hyland, The Lighthouse Lab in Alderley Park and Alex Alderton, Roberto Amato, Jeffrey Barrett, Sonia Goncalves, Ewan Harrison, David K. Jackson, Ian Johnston, Dominic Kwiatkowski, Cordelia Langford, John Sillitoe on behalf of the Wellcome Sanger Institute COVID-19 Surveillance Team |
| EPI_ISL_2729803, EPI_ISL_2729804 | Lighthouse Lab in Milton Keynes | Wellcome Sanger Institute for the COVID-19 Genomics UK (COG-UK) Consortium | The Lighthouse Lab in Milton Keynes and Alex Alderton, Roberto Amato, Jeffrey Barrett, Sonia Goncalves, Ewan Harrison, David K. Jackson, Ian Johnston, Dominic Kwiatkowski, Cordelia Langford, John Sillitoe on behalf of the Wellcome Sanger Institute COVID-19 Surveillance Team |
| EPI_ISL_2729805, EPI_ISL_2729806 | Lighthouse Lab in Alderley Park | Wellcome Sanger Institute for the COVID-19 Genomics UK (COG-UK) Consortium | Jacquelyn Wynn, Mairead Hyland, The Lighthouse Lab in Alderley Park and Alex Alderton, Roberto Amato, Jeffrey Barrett, Sonia Goncalves, Ewan Harrison, David K. Jackson, Ian Johnston, Dominic Kwiatkowski, Cordelia Langford, John Sillitoe on behalf of the Wellcome Sanger Institute COVID-19 Surveillance Team |
| EPI_ISL_2729807 | Lighthouse Lab in Milton Keynes | Wellcome Sanger Institute for the COVID-19 Genomics UK (COG-UK) Consortium | The Lighthouse Lab in Milton Keynes and Alex Alderton, Roberto Amato, Jeffrey Barrett, Sonia Goncalves, Ewan Harrison, David K. Jackson, Ian Johnston, Dominic Kwiatkowski, Cordelia Langford, John Sillitoe on behalf of the Wellcome Sanger Institute COVID-19 Surveillance Team |
| EPI_ISL_2729808, EPI_ISL_2729809 | Lighthouse Laboratory Plymouth | Wellcome Sanger Institute for the COVID-19 Genomics UK (COG-UK) Consortium | Lighthouse Laboratory Plymouth and Alex Alderton, Roberto Amato, Jeffrey Barrett, Sonia Goncalves, Ewan Harrison, David K. Jackson, Ian Johnston, Dominic Kwiatkowski, Cordelia Langford, John Sillitoe on behalf of the Wellcome Sanger Institute COVID-19 Surveillance Team |
| EPI_ISL_2729810, EPI_ISL_2729811, EPI_ISL_2729812, EPI_ISL_2729813, EPI_ISL_2729814 | Lighthouse Lab in Alderley Park | Wellcome Sanger Institute for the COVID-19 Genomics UK (COG-UK) Consortium | Jacquelyn Wynn, Mairead Hyland, The Lighthouse Lab in Alderley Park and Alex Alderton, Roberto Amato, Jeffrey Barrett, Sonia Goncalves, Ewan Harrison, David K. Jackson, Ian Johnston, Dominic Kwiatkowski, Cordelia Langford, John Sillitoe on behalf of the Wellcome Sanger Institute COVID-19 Surveillance Team |
| EPI_ISL_2729815 | Lighthouse Lab in Milton Keynes | Wellcome Sanger Institute for the COVID-19 Genomics UK (COG-UK) Consortium | The Lighthouse Lab in Milton Keynes and Alex Alderton, Roberto Amato, Jeffrey Barrett, Sonia Goncalves, Ewan Harrison, David K. Jackson, Ian Johnston, Dominic Kwiatkowski, Cordelia Langford, John Sillitoe on behalf of the Wellcome Sanger Institute COVID-19 Surveillance Team |
| EPI_ISL_2729816 | Lighthouse Laboratory Plymouth | Wellcome Sanger Institute for the COVID-19 Genomics UK (COG-UK) Consortium | Lighthouse Laboratory Plymouth and Alex Alderton, Roberto Amato, Jeffrey Barrett, Sonia Goncalves, Ewan Harrison, David K. Jackson, Ian Johnston, Dominic Kwiatkowski, Cordelia Langford, John Sillitoe on behalf of the Wellcome Sanger Institute COVID-19 Surveillance Team |
| EPI_ISL_2729817, EPI_ISL_2729818 | Lighthouse Lab in Milton Keynes | Wellcome Sanger Institute for the COVID-19 Genomics UK (COG-UK) Consortium | The Lighthouse Lab in Milton Keynes and Alex Alderton, Roberto Amato, Jeffrey Barrett, Sonia Goncalves, Ewan Harrison, David K. Jackson, Ian Johnston, Dominic Kwiatkowski, Cordelia Langford, John Sillitoe on behalf of the Wellcome Sanger Institute COVID-19 Surveillance Team |
| EPI_ISL_2729819, EPI_ISL_2729820 | Lighthouse Lab in Alderley Park | Wellcome Sanger Institute for the COVID-19 Genomics UK (COG-UK) Consortium | Jacquelyn Wynn, Mairead Hyland, The Lighthouse Lab in Alderley Park and Alex Alderton, Roberto Amato, Jeffrey Barrett, Sonia Goncalves, Ewan Harrison, David K. Jackson, Ian Johnston, Dominic Kwiatkowski, Cordelia Langford, John Sillitoe on behalf of the Wellcome Sanger Institute COVID-19 Surveillance Team |
| EPI_ISL_2729821 | Lighthouse Lab in Milton Keynes | Wellcome Sanger Institute for the COVID-19 Genomics UK (COG-UK) Consortium | The Lighthouse Lab in Milton Keynes and Alex Alderton, Roberto Amato, Jeffrey Barrett, Sonia Goncalves, Ewan Harrison, David K. Jackson, Ian Johnston, Dominic Kwiatkowski, Cordelia Langford, John Sillitoe on behalf of the Wellcome Sanger Institute COVID-19 Surveillance Team |
| EPI_ISL_2729822 | Lighthouse Lab in Alderley Park | Wellcome Sanger Institute for the COVID-19 Genomics UK (COG-UK) Consortium | Jacquelyn Wynn, Mairead Hyland, The Lighthouse Lab in Alderley Park and Alex Alderton, Roberto Amato, Jeffrey Barrett, Sonia Goncalves, Ewan Harrison, David K. Jackson, Ian Johnston, Dominic Kwiatkowski, Cordelia Langford, John Sillitoe on behalf of the Wellcome Sanger Institute COVID-19 Surveillance Team |
| EPI_ISL_2729823 | Lighthouse Lab in Glasgow | Wellcome Sanger Institute for the COVID-19 Genomics UK (COG-UK) Consortium | Harper VanSteenhouse, Yumi Kasai, David Gray, Carol Clugston, Anna Dominiczak and Alex Alderton, Roberto Amato, Jeffrey Barrett, Sonia Goncalves, Ewan Harrison, David K. Jackson, Ian Johnston, Dominic Kwiatkowski, Cordelia Langford, John Sillitoe on behalf of the Wellcome Sanger Institute COVID-19 Surveillance Team |
| EPI_ISL_2729824 | Lighthouse Lab in Alderley Park | Wellcome Sanger Institute for the COVID-19 Genomics UK (COG-UK) Consortium | Jacquelyn Wynn, Mairead Hyland, The Lighthouse Lab in Alderley Park and Alex Alderton, Roberto Amato, Jeffrey Barrett, Sonia Goncalves, Ewan Harrison, David K. Jackson, Ian Johnston, Dominic Kwiatkowski, Cordelia Langford, John Sillitoe on behalf of the Wellcome Sanger Institute COVID-19 Surveillance Team |
| EPI_ISL_2729825 | Lighthouse Lab in Milton Keynes | Wellcome Sanger Institute for the COVID-19 Genomics UK (COG-UK) Consortium | The Lighthouse Lab in Milton Keynes and Alex Alderton, Roberto Amato, Jeffrey Barrett, Sonia Goncalves, Ewan Harrison, David K. Jackson, Ian Johnston, Dominic Kwiatkowski, Cordelia Langford, John Sillitoe on behalf of the Wellcome Sanger Institute COVID-19 Surveillance Team |
| EPI_ISL_2729826 | Lighthouse Laboratory Plymouth | Wellcome Sanger Institute for the COVID-19 Genomics UK (COG-UK) Consortium | Lighthouse Laboratory Plymouth and Alex Alderton, Roberto Amato, Jeffrey Barrett, Sonia Goncalves, Ewan Harrison, David K. Jackson, Ian Johnston, Dominic Kwiatkowski, Cordelia Langford, John Sillitoe on behalf of the Wellcome Sanger Institute COVID-19 Surveillance Team |
| EPI_ISL_2729827 | Lighthouse Lab in Milton Keynes | Wellcome Sanger Institute for the COVID-19 Genomics UK (COG-UK) Consortium | The Lighthouse Lab in Milton Keynes and Alex Alderton, Roberto Amato, Jeffrey Barrett, Sonia Goncalves, Ewan Harrison, David K. Jackson, Ian Johnston, Dominic Kwiatkowski, Cordelia Langford, John Sillitoe on behalf of the Wellcome Sanger Institute COVID-19 Surveillance Team |
| EPI_ISL_2729828, EPI_ISL_2729829 | Lighthouse Laboratory Plymouth | Wellcome Sanger Institute for the COVID-19 Genomics UK (COG-UK) Consortium | Lighthouse Laboratory Plymouth and Alex Alderton, Roberto Amato, Jeffrey Barrett, Sonia Goncalves, Ewan Harrison, David K. Jackson, Ian Johnston, Dominic Kwiatkowski, Cordelia Langford, John Sillitoe on behalf of the Wellcome Sanger Institute COVID-19 Surveillance Team |

| | | | |
|---|---|---|---|
| EPI_ISL_2729830 | Lighthouse Lab in Milton Keynes | Wellcome Sanger Institute for the COVID-19 Genomics UK (COG-UK) Consortium | The Lighthouse Lab in Milton Keynes and Alex Alderton, Roberto Amato, Jeffrey Barrett, Sonia Goncalves, Ewan Harrison, David K. Jackson, Ian Johnston, Dominic Kwiatkowski, Cordelia Langford, John Sillitoe on behalf of the Wellcome Sanger Institute COVID-19 Surveillance Team |
| EPI_ISL_2729831 | Lighthouse Lab in Alderley Park | Wellcome Sanger Institute for the COVID-19 Genomics UK (COG-UK) Consortium | Jacquelyn Wynn, Mairead Hyland, The Lighthouse Lab in Alderley Park and Alex Alderton, Roberto Amato, Jeffrey Barrett, Sonia Goncalves, Ewan Harrison, David K. Jackson, Ian Johnston, Dominic Kwiatkowski, Cordelia Langford, John Sillitoe on behalf of the Wellcome Sanger Institute COVID-19 Surveillance Team |
| EPI_ISL_2729832 | Lighthouse Lab in Glasgow | Wellcome Sanger Institute for the COVID-19 Genomics UK (COG-UK) Consortium | Harper VanSteenhouse, Yumi Kasai, David Gray, Carol Clugston, Anna Dominiczak and Alex Alderton, Roberto Amato, Jeffrey Barrett, Sonia Goncalves, Ewan Harrison, David K. Jackson, Ian Johnston, Dominic Kwiatkowski, Cordelia Langford, John Sillitoe on behalf of the Wellcome Sanger Institute COVID-19 Surveillance Team |
| EPI_ISL_2729833, EPI_ISL_2729834 | Lighthouse Laboratory Plymouth | Wellcome Sanger Institute for the COVID-19 Genomics UK (COG-UK) Consortium | Lighthouse Laboratory Plymouth and Alex Alderton, Roberto Amato, Jeffrey Barrett, Sonia Goncalves, Ewan Harrison, David K. Jackson, Ian Johnston, Dominic Kwiatkowski, Cordelia Langford, John Sillitoe on behalf of the Wellcome Sanger Institute COVID-19 Surveillance Team |
| EPI_ISL_2729835 | Lighthouse Lab in Alderley Park | Wellcome Sanger Institute for the COVID-19 Genomics UK (COG-UK) Consortium | Jacquelyn Wynn, Mairead Hyland, The Lighthouse Lab in Alderley Park and Alex Alderton, Roberto Amato, Jeffrey Barrett, Sonia Goncalves, Ewan Harrison, David K. Jackson, Ian Johnston, Dominic Kwiatkowski, Cordelia Langford, John Sillitoe on behalf of the Wellcome Sanger Institute COVID-19 Surveillance Team |
| EPI_ISL_2729836 | Lighthouse Laboratory Plymouth | Wellcome Sanger Institute for the COVID-19 Genomics UK (COG-UK) Consortium | Lighthouse Laboratory Plymouth and Alex Alderton, Roberto Amato, Jeffrey Barrett, Sonia Goncalves, Ewan Harrison, David K. Jackson, Ian Johnston, Dominic Kwiatkowski, Cordelia Langford, John Sillitoe on behalf of the Wellcome Sanger Institute COVID-19 Surveillance Team |
| EPI_ISL_2729837 | Lighthouse Lab in Milton Keynes | Wellcome Sanger Institute for the COVID-19 Genomics UK (COG-UK) Consortium | The Lighthouse Lab in Milton Keynes and Alex Alderton, Roberto Amato, Jeffrey Barrett, Sonia Goncalves, Ewan Harrison, David K. Jackson, Ian Johnston, Dominic Kwiatkowski, Cordelia Langford, John Sillitoe on behalf of the Wellcome Sanger Institute COVID-19 Surveillance Team |
| EPI_ISL_2729838 | Lighthouse Lab in Alderley Park | Wellcome Sanger Institute for the COVID-19 Genomics UK (COG-UK) Consortium | Jacquelyn Wynn, Mairead Hyland, The Lighthouse Lab in Alderley Park and Alex Alderton, Roberto Amato, Jeffrey Barrett, Sonia Goncalves, Ewan Harrison, David K. Jackson, Ian Johnston, Dominic Kwiatkowski, Cordelia Langford, John Sillitoe on behalf of the Wellcome Sanger Institute COVID-19 Surveillance Team |
| EPI_ISL_2729839 | Lighthouse Laboratory Plymouth | Wellcome Sanger Institute for the COVID-19 Genomics UK (COG-UK) Consortium | Lighthouse Laboratory Plymouth and Alex Alderton, Roberto Amato, Jeffrey Barrett, Sonia Goncalves, Ewan Harrison, David K. Jackson, Ian Johnston, Dominic Kwiatkowski, Cordelia Langford, John Sillitoe on behalf of the Wellcome Sanger Institute COVID-19 Surveillance Team |
| EPI_ISL_2729840 | Lighthouse Lab in Milton Keynes | Wellcome Sanger Institute for the COVID-19 Genomics UK (COG-UK) Consortium | The Lighthouse Lab in Milton Keynes and Alex Alderton, Roberto Amato, Jeffrey Barrett, Sonia Goncalves, Ewan Harrison, David K. Jackson, Ian Johnston, Dominic Kwiatkowski, Cordelia Langford, John Sillitoe on behalf of the Wellcome Sanger Institute COVID-19 Surveillance Team |
| EPI_ISL_2729841 | Lighthouse Laboratory Plymouth | Wellcome Sanger Institute for the COVID-19 Genomics UK (COG-UK) Consortium | Lighthouse Laboratory Plymouth and Alex Alderton, Roberto Amato, Jeffrey Barrett, Sonia Goncalves, Ewan Harrison, David K. Jackson, Ian Johnston, Dominic Kwiatkowski, Cordelia Langford, John Sillitoe on behalf of the Wellcome Sanger Institute COVID-19 Surveillance Team |
| EPI_ISL_2729842 | Lighthouse Lab in Milton Keynes | Wellcome Sanger Institute for the COVID-19 Genomics UK (COG-UK) Consortium | The Lighthouse Lab in Milton Keynes and Alex Alderton, Roberto Amato, Jeffrey Barrett, Sonia Goncalves, Ewan Harrison, David K. Jackson, Ian Johnston, Dominic Kwiatkowski, Cordelia Langford, John Sillitoe on behalf of the Wellcome Sanger Institute COVID-19 Surveillance Team |
| EPI_ISL_2729843 | Lighthouse Lab in Alderley Park | Wellcome Sanger Institute for the COVID-19 Genomics UK (COG-UK) Consortium | Jacquelyn Wynn, Mairead Hyland, The Lighthouse Lab in Alderley Park and Alex Alderton, Roberto Amato, Jeffrey Barrett, Sonia Goncalves, Ewan Harrison, David K. Jackson, Ian Johnston, Dominic Kwiatkowski, Cordelia Langford, John Sillitoe on behalf of the Wellcome Sanger Institute COVID-19 Surveillance Team |
| EPI_ISL_2729844 | Lighthouse Laboratory Plymouth | Wellcome Sanger Institute for the COVID-19 Genomics UK (COG-UK) Consortium | Lighthouse Laboratory Plymouth and Alex Alderton, Roberto Amato, Jeffrey Barrett, Sonia Goncalves, Ewan Harrison, David K. Jackson, Ian Johnston, Dominic Kwiatkowski, Cordelia Langford, John Sillitoe on behalf of the Wellcome Sanger Institute COVID-19 Surveillance Team |
| EPI_ISL_2729845 | Lighthouse Lab in Milton Keynes | Wellcome Sanger Institute for the COVID-19 Genomics UK (COG-UK) Consortium | The Lighthouse Lab in Milton Keynes and Alex Alderton, Roberto Amato, Jeffrey Barrett, Sonia Goncalves, Ewan Harrison, David K. Jackson, Ian Johnston, Dominic Kwiatkowski, Cordelia Langford, John Sillitoe on behalf of the Wellcome Sanger Institute COVID-19 Surveillance Team |
| EPI_ISL_2729846 | Lighthouse Lab in Alderley Park | Wellcome Sanger Institute for the COVID-19 Genomics UK (COG-UK) Consortium | Jacquelyn Wynn, Mairead Hyland, The Lighthouse Lab in Alderley Park and Alex Alderton, Roberto Amato, Jeffrey Barrett, Sonia Goncalves, Ewan Harrison, David K. Jackson, Ian Johnston, Dominic Kwiatkowski, Cordelia Langford, John Sillitoe on behalf of the Wellcome Sanger Institute COVID-19 Surveillance Team |
| EPI_ISL_2729847 | Lighthouse Lab in Milton Keynes | Wellcome Sanger Institute for the COVID-19 Genomics UK (COG-UK) Consortium | The Lighthouse Lab in Milton Keynes and Alex Alderton, Roberto Amato, Jeffrey Barrett, Sonia Goncalves, Ewan Harrison, David K. Jackson, Ian Johnston, Dominic Kwiatkowski, Cordelia Langford, John Sillitoe on behalf of the Wellcome Sanger Institute COVID-19 Surveillance Team |
| EPI_ISL_2729848 | Lighthouse Lab in Alderley Park | Wellcome Sanger Institute for the COVID-19 Genomics UK (COG-UK) Consortium | Jacquelyn Wynn, Mairead Hyland, The Lighthouse Lab in Alderley Park and Alex Alderton, Roberto Amato, Jeffrey Barrett, Sonia Goncalves, Ewan Harrison, David K. Jackson, Ian Johnston, Dominic Kwiatkowski, Cordelia Langford, John Sillitoe on behalf of the Wellcome Sanger Institute COVID-19 Surveillance Team |
| EPI_ISL_2729849, EPI_ISL_2729850, EPI_ISL_2729851 | Lighthouse Lab in Milton Keynes | Wellcome Sanger Institute for the COVID-19 Genomics UK (COG-UK) Consortium | The Lighthouse Lab in Milton Keynes and Alex Alderton, Roberto Amato, Jeffrey Barrett, Sonia Goncalves, Ewan Harrison, David K. Jackson, Ian Johnston, Dominic Kwiatkowski, Cordelia Langford, John Sillitoe on behalf of the Wellcome Sanger Institute COVID-19 Surveillance Team |
| EPI_ISL_2729852, EPI_ISL_2729853 | Lighthouse Lab in Alderley Park | Wellcome Sanger Institute for the COVID-19 Genomics UK (COG-UK) Consortium | Jacquelyn Wynn, Mairead Hyland, The Lighthouse Lab in Alderley Park and Alex Alderton, Roberto Amato, Jeffrey Barrett, Sonia Goncalves, Ewan Harrison, David K. Jackson, Ian Johnston, Dominic Kwiatkowski, Cordelia Langford, John Sillitoe on behalf of the Wellcome Sanger Institute COVID-19 Surveillance Team |
| EPI_ISL_2729854 | Lighthouse Lab in Glasgow | Wellcome Sanger Institute for the COVID-19 Genomics UK (COG-UK) Consortium | Harper VanSteenhouse, Yumi Kasai, David Gray, Carol Clugston, Anna Dominiczak and Alex Alderton, Roberto Amato, Jeffrey Barrett, Sonia Goncalves, Ewan Harrison, David K. Jackson, Ian Johnston, Dominic Kwiatkowski, Cordelia Langford, John Sillitoe on behalf of the Wellcome Sanger Institute COVID-19 Surveillance Team |
| EPI_ISL_2729855 | Lighthouse Lab in Milton Keynes | Wellcome Sanger Institute for the COVID-19 Genomics UK (COG-UK) Consortium | The Lighthouse Lab in Milton Keynes and Alex Alderton, Roberto Amato, Jeffrey Barrett, Sonia Goncalves, Ewan Harrison, David K. Jackson, Ian Johnston, Dominic Kwiatkowski, Cordelia Langford, John Sillitoe on behalf of the Wellcome Sanger Institute COVID-19 Surveillance Team |
| EPI_ISL_2729856, EPI_ISL_2729857 | Lighthouse Lab in Alderley Park | Wellcome Sanger Institute for the COVID-19 Genomics UK (COG-UK) Consortium | Jacquelyn Wynn, Mairead Hyland, The Lighthouse Lab in Alderley Park and Alex Alderton, Roberto Amato, Jeffrey Barrett, Sonia Goncalves, Ewan Harrison, David K. Jackson, Ian Johnston, Dominic Kwiatkowski, Cordelia Langford, John Sillitoe on behalf of the Wellcome Sanger Institute COVID-19 Surveillance Team |
| EPI_ISL_2729858, EPI_ISL_2729859 | Lighthouse Laboratory Plymouth | Wellcome Sanger Institute for the COVID-19 Genomics UK (COG-UK) Consortium | Lighthouse Laboratory Plymouth and Alex Alderton, Roberto Amato, Jeffrey Barrett, Sonia Goncalves, Ewan Harrison, David K. Jackson, Ian Johnston, Dominic Kwiatkowski, Cordelia Langford, John Sillitoe on behalf of the Wellcome Sanger Institute COVID-19 Surveillance Team |
| EPI_ISL_2729860 | Lighthouse Lab in Milton Keynes | Wellcome Sanger Institute for the COVID-19 Genomics UK (COG-UK) Consortium | The Lighthouse Lab in Milton Keynes and Alex Alderton, Roberto Amato, Jeffrey Barrett, Sonia Goncalves, Ewan Harrison, David K. Jackson, Ian Johnston, Dominic Kwiatkowski, Cordelia Langford, John Sillitoe on behalf of the Wellcome Sanger Institute COVID-19 Surveillance Team |
| EPI_ISL_2729861, EPI_ISL_2729862 | Lighthouse Laboratory Plymouth | Wellcome Sanger Institute for the COVID-19 Genomics UK (COG-UK) Consortium | Lighthouse Laboratory Plymouth and Alex Alderton, Roberto Amato, Jeffrey Barrett, Sonia Goncalves, Ewan Harrison, David K. Jackson, Ian Johnston, Dominic Kwiatkowski, Cordelia Langford, John Sillitoe on behalf of the Wellcome Sanger Institute COVID-19 Surveillance Team |
| EPI_ISL_2729863 | Lighthouse Lab in Milton Keynes | Wellcome Sanger Institute for the COVID-19 Genomics UK (COG-UK) Consortium | The Lighthouse Lab in Milton Keynes and Alex Alderton, Roberto Amato, Jeffrey Barrett, Sonia Goncalves, Ewan Harrison, David K. Jackson, Ian Johnston, Dominic Kwiatkowski, Cordelia Langford, John Sillitoe on behalf of the Wellcome Sanger Institute COVID-19 Surveillance Team |
| EPI_ISL_2729865 | Lighthouse Lab in Glasgow | Wellcome Sanger Institute for the COVID-19 Genomics UK (COG-UK) Consortium | Harper VanSteenhouse, Yumi Kasai, David Gray, Carol Clugston, Anna Dominiczak and Alex Alderton, Roberto Amato, Jeffrey Barrett, Sonia Goncalves, Ewan Harrison, David K. Jackson, Ian Johnston, Dominic Kwiatkowski, Cordelia Langford, John Sillitoe on behalf of the Wellcome Sanger Institute COVID-19 Surveillance Team |
| EPI_ISL_2729866 | Lighthouse Lab in Milton Keynes | Wellcome Sanger Institute for the COVID-19 Genomics UK (COG-UK) Consortium | The Lighthouse Lab in Milton Keynes and Alex Alderton, Roberto Amato, Jeffrey Barrett, Sonia Goncalves, Ewan Harrison, David K. Jackson, Ian Johnston, Dominic Kwiatkowski, Cordelia Langford, John Sillitoe on behalf of the Wellcome Sanger Institute COVID-19 Surveillance Team |
| EPI_ISL_2729867 | Lighthouse Lab in Glasgow | Wellcome Sanger Institute for the COVID-19 Genomics UK (COG-UK) Consortium | Harper VanSteenhouse, Yumi Kasai, David Gray, Carol Clugston, Anna Dominiczak and Alex Alderton, Roberto Amato, Jeffrey Barrett, Sonia Goncalves, Ewan Harrison, David K. Jackson, Ian Johnston, Dominic Kwiatkowski, Cordelia Langford, John Sillitoe on behalf of the Wellcome Sanger Institute COVID-19 Surveillance Team |
| EPI_ISL_2729868, EPI_ISL_2729869 | Lighthouse Lab in Milton Keynes | Wellcome Sanger Institute for the COVID-19 Genomics UK (COG-UK) Consortium | The Lighthouse Lab in Milton Keynes and Alex Alderton, Roberto Amato, Jeffrey Barrett, Sonia Goncalves, Ewan Harrison, David K. Jackson, Ian Johnston, Dominic Kwiatkowski, Cordelia Langford, John Sillitoe on behalf of the Wellcome Sanger Institute COVID-19 Surveillance Team |
| EPI_ISL_2729870 | Lighthouse Lab in Alderley Park | Wellcome Sanger Institute for the COVID-19 Genomics UK (COG-UK) Consortium | Jacquelyn Wynn, Mairead Hyland, The Lighthouse Lab in Alderley Park and Alex Alderton, Roberto Amato, Jeffrey Barrett, Sonia Goncalves, Ewan Harrison, David K. Jackson, Ian Johnston, Dominic Kwiatkowski, Cordelia Langford, John Sillitoe on behalf of the Wellcome Sanger Institute COVID-19 Surveillance Team |
| EPI_ISL_2729871, EPI_ISL_2729872 | Lighthouse Laboratory Plymouth | Wellcome Sanger Institute for the COVID-19 Genomics UK (COG-UK) Consortium | Lighthouse Laboratory Plymouth and Alex Alderton, Roberto Amato, Jeffrey Barrett, Sonia Goncalves, Ewan Harrison, David K. Jackson, Ian Johnston, Dominic Kwiatkowski, Cordelia Langford, John Sillitoe on behalf of the Wellcome Sanger Institute COVID-19 Surveillance Team |
| EPI_ISL_2729873 | Lighthouse Lab in Milton Keynes | Wellcome Sanger Institute for the COVID-19 Genomics UK (COG-UK) Consortium | The Lighthouse Lab in Milton Keynes and Alex Alderton, Roberto Amato, Jeffrey Barrett, Sonia Goncalves, Ewan Harrison, David K. Jackson, Ian Johnston, Dominic Kwiatkowski, Cordelia Langford, John Sillitoe on behalf of the Wellcome Sanger Institute COVID-19 Surveillance Team |
| EPI_ISL_2729874 | Lighthouse Lab in Glasgow | Wellcome Sanger Institute for the COVID-19 Genomics UK (COG-UK) Consortium | Harper VanSteenhouse, Yumi Kasai, David Gray, Carol Clugston, Anna Dominiczak and Alex Alderton, Roberto Amato, Jeffrey Barrett, Sonia Goncalves, Ewan Harrison, David K. Jackson, Ian Johnston, Dominic Kwiatkowski, Cordelia Langford, John Sillitoe on behalf of the Wellcome Sanger Institute COVID-19 Surveillance Team |
| EPI_ISL_2729875 | Lighthouse Lab in Alderley Park | Wellcome Sanger Institute for the COVID-19 Genomics UK (COG-UK) Consortium | Jacquelyn Wynn, Mairead Hyland, The Lighthouse Lab in Alderley Park and Alex Alderton, Roberto Amato, Jeffrey Barrett, Sonia Goncalves, Ewan Harrison, David K. Jackson, Ian Johnston, Dominic Kwiatkowski, Cordelia Langford, John Sillitoe on behalf of the Wellcome Sanger Institute COVID-19 Surveillance Team |
| EPI_ISL_2729876, EPI_ISL_2729877 | Lighthouse Lab in Milton Keynes | Wellcome Sanger Institute for the COVID-19 Genomics UK (COG-UK) Consortium | The Lighthouse Lab in Milton Keynes and Alex Alderton, Roberto Amato, Jeffrey Barrett, Sonia Goncalves, Ewan Harrison, David K. Jackson, Ian Johnston, Dominic Kwiatkowski, Cordelia Langford, John Sillitoe on behalf of the Wellcome Sanger Institute COVID-19 Surveillance Team |
| EPI_ISL_2729878 | Lighthouse Laboratory Plymouth | Wellcome Sanger Institute for the COVID-19 Genomics UK (COG-UK) Consortium | Lighthouse Laboratory Plymouth and Alex Alderton, Roberto Amato, Jeffrey Barrett, Sonia Goncalves, Ewan Harrison, David K. Jackson, Ian Johnston, Dominic Kwiatkowski, Cordelia Langford, John Sillitoe on behalf of the Wellcome Sanger Institute COVID-19 Surveillance Team |
| EPI_ISL_2729879 | Lighthouse Lab in Milton Keynes | Wellcome Sanger Institute for the COVID-19 Genomics UK (COG-UK) Consortium | The Lighthouse Lab in Milton Keynes and Alex Alderton, Roberto Amato, Jeffrey Barrett, Sonia Goncalves, Ewan Harrison, David K. Jackson, Ian Johnston, Dominic Kwiatkowski, Cordelia Langford, John Sillitoe on behalf of the Wellcome Sanger Institute COVID-19 Surveillance Team |
| EPI_ISL_2729880, EPI_ISL_2729881 | Lighthouse Lab in Alderley Park | Wellcome Sanger Institute for the COVID-19 Genomics UK (COG-UK) Consortium | Jacquelyn Wynn, Mairead Hyland, The Lighthouse Lab in Alderley Park and Alex Alderton, Roberto Amato, Jeffrey Barrett, Sonia Goncalves, Ewan Harrison, David K. Jackson, Ian Johnston, Dominic Kwiatkowski, Cordelia Langford, John Sillitoe on behalf of the Wellcome Sanger Institute COVID-19 Surveillance Team |
| EPI_ISL_2729882 | Lighthouse Laboratory Plymouth | Wellcome Sanger Institute for the COVID-19 Genomics UK (COG-UK) Consortium | Lighthouse Laboratory Plymouth and Alex Alderton, Roberto Amato, Jeffrey Barrett, Sonia Goncalves, Ewan Harrison, David K. Jackson, Ian Johnston, Dominic Kwiatkowski, Cordelia Langford, John Sillitoe on behalf of the Wellcome Sanger Institute COVID-19 Surveillance Team |
| EPI_ISL_2729883 | Lighthouse Lab in Milton Keynes | Wellcome Sanger Institute for the COVID-19 Genomics UK (COG-UK) Consortium | The Lighthouse Lab in Milton Keynes and Alex Alderton, Roberto Amato, Jeffrey Barrett, Sonia Goncalves, Ewan Harrison, David K. Jackson, Ian Johnston, Dominic Kwiatkowski, Cordelia Langford, John Sillitoe on behalf of the Wellcome Sanger Institute COVID-19 Surveillance Team |
| EPI_ISL_2729884 | Lighthouse Lab in Alderley Park | Wellcome Sanger Institute for the COVID-19 Genomics UK (COG-UK) Consortium | Jacquelyn Wynn, Mairead Hyland, The Lighthouse Lab in Alderley Park and Alex Alderton, Roberto Amato, Jeffrey Barrett, Sonia Goncalves, Ewan Harrison, David K. Jackson, Ian Johnston, Dominic Kwiatkowski, Cordelia Langford, John Sillitoe on behalf of the Wellcome Sanger Institute COVID-19 Surveillance Team |
| EPI_ISL_2729885 | Lighthouse Lab in Glasgow | Wellcome Sanger Institute for the COVID-19 Genomics UK (COG-UK) Consortium | Harper VanSteenhouse, Yumi Kasai, David Gray, Carol Clugston, Anna Dominiczak and Alex Alderton, Roberto Amato, Jeffrey Barrett, Sonia Goncalves, Ewan Harrison, David K. Jackson, Ian Johnston, Dominic Kwiatkowski, Cordelia Langford, John Sillitoe on behalf of the Wellcome Sanger Institute COVID-19 Surveillance Team |

| | | | |
|---|---|---|---|
| EPI_ISL_2729886 | Lighthouse Lab in Alderley Park | Wellcome Sanger Institute for the COVID-19 Genomics UK (COG-UK) Consortium | Jacquelyn Wynn, Mairead Hyland, The Lighthouse Lab in Alderley Park and Alex Alderton, Roberto Amato, Jeffrey Barrett, Sonia Goncalves, Ewan Harrison, David K. Jackson, Ian Johnston, Dominic Kwiatkowski, Cordelia Langford, John Sillitoe on behalf of the Wellcome Sanger Institute COVID-19 Surveillance Team |
| EPI_ISL_2729887, EPI_ISL_2729888 | Lighthouse Laboratory Plymouth | Wellcome Sanger Institute for the COVID-19 Genomics UK (COG-UK) Consortium | Lighthouse Laboratory Plymouth and Alex Alderton, Roberto Amato, Jeffrey Barrett, Sonia Goncalves, Ewan Harrison, David K. Jackson, Ian Johnston, Dominic Kwiatkowski, Cordelia Langford, John Sillitoe on behalf of the Wellcome Sanger Institute COVID-19 Surveillance Team |
| EPI_ISL_2729889, EPI_ISL_2729890 | Lighthouse Lab in Milton Keynes | Wellcome Sanger Institute for the COVID-19 Genomics UK (COG-UK) Consortium | The Lighthouse Lab in Milton Keynes and Alex Alderton, Roberto Amato, Jeffrey Barrett, Sonia Goncalves, Ewan Harrison, David K. Jackson, Ian Johnston, Dominic Kwiatkowski, Cordelia Langford, John Sillitoe on behalf of the Wellcome Sanger Institute COVID-19 Surveillance Team |
| EPI_ISL_2729891, EPI_ISL_2729892 | Lighthouse Lab in Alderley Park | Wellcome Sanger Institute for the COVID-19 Genomics UK (COG-UK) Consortium | Jacquelyn Wynn, Mairead Hyland, The Lighthouse Lab in Alderley Park and Alex Alderton, Roberto Amato, Jeffrey Barrett, Sonia Goncalves, Ewan Harrison, David K. Jackson, Ian Johnston, Dominic Kwiatkowski, Cordelia Langford, John Sillitoe on behalf of the Wellcome Sanger Institute COVID-19 Surveillance Team |
| EPI_ISL_2729894 | Lighthouse Laboratory Plymouth | Wellcome Sanger Institute for the COVID-19 Genomics UK (COG-UK) Consortium | Lighthouse Laboratory Plymouth and Alex Alderton, Roberto Amato, Jeffrey Barrett, Sonia Goncalves, Ewan Harrison, David K. Jackson, Ian Johnston, Dominic Kwiatkowski, Cordelia Langford, John Sillitoe on behalf of the Wellcome Sanger Institute COVID-19 Surveillance Team |
| EPI_ISL_2729895 | Lighthouse Lab in Alderley Park | Wellcome Sanger Institute for the COVID-19 Genomics UK (COG-UK) Consortium | Jacquelyn Wynn, Mairead Hyland, The Lighthouse Lab in Alderley Park and Alex Alderton, Roberto Amato, Jeffrey Barrett, Sonia Goncalves, Ewan Harrison, David K. Jackson, Ian Johnston, Dominic Kwiatkowski, Cordelia Langford, John Sillitoe on behalf of the Wellcome Sanger Institute COVID-19 Surveillance Team |
| EPI_ISL_2729896, EPI_ISL_2729897, EPI_ISL_2729898 | Lighthouse Laboratory Plymouth | Wellcome Sanger Institute for the COVID-19 Genomics UK (COG-UK) Consortium | Lighthouse Laboratory Plymouth and Alex Alderton, Roberto Amato, Jeffrey Barrett, Sonia Goncalves, Ewan Harrison, David K. Jackson, Ian Johnston, Dominic Kwiatkowski, Cordelia Langford, John Sillitoe on behalf of the Wellcome Sanger Institute COVID-19 Surveillance Team |
| EPI_ISL_2729899, EPI_ISL_2729900 | Lighthouse Lab in Milton Keynes | Wellcome Sanger Institute for the COVID-19 Genomics UK (COG-UK) Consortium | The Lighthouse Lab in Milton Keynes and Alex Alderton, Roberto Amato, Jeffrey Barrett, Sonia Goncalves, Ewan Harrison, David K. Jackson, Ian Johnston, Dominic Kwiatkowski, Cordelia Langford, John Sillitoe on behalf of the Wellcome Sanger Institute COVID-19 Surveillance Team |
| EPI_ISL_2729901 | Lighthouse Laboratory Plymouth | Wellcome Sanger Institute for the COVID-19 Genomics UK (COG-UK) Consortium | Lighthouse Laboratory Plymouth and Alex Alderton, Roberto Amato, Jeffrey Barrett, Sonia Goncalves, Ewan Harrison, David K. Jackson, Ian Johnston, Dominic Kwiatkowski, Cordelia Langford, John Sillitoe on behalf of the Wellcome Sanger Institute COVID-19 Surveillance Team |
| EPI_ISL_2729902, EPI_ISL_2729903 | Lighthouse Lab in Milton Keynes | Wellcome Sanger Institute for the COVID-19 Genomics UK (COG-UK) Consortium | The Lighthouse Lab in Milton Keynes and Alex Alderton, Roberto Amato, Jeffrey Barrett, Sonia Goncalves, Ewan Harrison, David K. Jackson, Ian Johnston, Dominic Kwiatkowski, Cordelia Langford, John Sillitoe on behalf of the Wellcome Sanger Institute COVID-19 Surveillance Team |
| EPI_ISL_2729904 | Lighthouse Laboratory Plymouth | Wellcome Sanger Institute for the COVID-19 Genomics UK (COG-UK) Consortium | Lighthouse Laboratory Plymouth and Alex Alderton, Roberto Amato, Jeffrey Barrett, Sonia Goncalves, Ewan Harrison, David K. Jackson, Ian Johnston, Dominic Kwiatkowski, Cordelia Langford, John Sillitoe on behalf of the Wellcome Sanger Institute COVID-19 Surveillance Team |
| EPI_ISL_2729905 | Lighthouse Lab in Milton Keynes | Wellcome Sanger Institute for the COVID-19 Genomics UK (COG-UK) Consortium | The Lighthouse Lab in Milton Keynes and Alex Alderton, Roberto Amato, Jeffrey Barrett, Sonia Goncalves, Ewan Harrison, David K. Jackson, Ian Johnston, Dominic Kwiatkowski, Cordelia Langford, John Sillitoe on behalf of the Wellcome Sanger Institute COVID-19 Surveillance Team |
| EPI_ISL_2729906 | Lighthouse Lab in Alderley Park | Wellcome Sanger Institute for the COVID-19 Genomics UK (COG-UK) Consortium | Jacquelyn Wynn, Mairead Hyland, The Lighthouse Lab in Alderley Park and Alex Alderton, Roberto Amato, Jeffrey Barrett, Sonia Goncalves, Ewan Harrison, David K. Jackson, Ian Johnston, Dominic Kwiatkowski, Cordelia Langford, John Sillitoe on behalf of the Wellcome Sanger Institute COVID-19 Surveillance Team |
| EPI_ISL_2729907 | Lighthouse Laboratory Plymouth | Wellcome Sanger Institute for the COVID-19 Genomics UK (COG-UK) Consortium | Lighthouse Laboratory Plymouth and Alex Alderton, Roberto Amato, Jeffrey Barrett, Sonia Goncalves, Ewan Harrison, David K. Jackson, Ian Johnston, Dominic Kwiatkowski, Cordelia Langford, John Sillitoe on behalf of the Wellcome Sanger Institute COVID-19 Surveillance Team |
| EPI_ISL_2729908 | Lighthouse Lab in Alderley Park | Wellcome Sanger Institute for the COVID-19 Genomics UK (COG-UK) Consortium | Jacquelyn Wynn, Mairead Hyland, The Lighthouse Lab in Alderley Park and Alex Alderton, Roberto Amato, Jeffrey Barrett, Sonia Goncalves, Ewan Harrison, David K. Jackson, Ian Johnston, Dominic Kwiatkowski, Cordelia Langford, John Sillitoe on behalf of the Wellcome Sanger Institute COVID-19 Surveillance Team |
| EPI_ISL_2729909 | Lighthouse Lab in Milton Keynes | Wellcome Sanger Institute for the COVID-19 Genomics UK (COG-UK) Consortium | The Lighthouse Lab in Milton Keynes and Alex Alderton, Roberto Amato, Jeffrey Barrett, Sonia Goncalves, Ewan Harrison, David K. Jackson, Ian Johnston, Dominic Kwiatkowski, Cordelia Langford, John Sillitoe on behalf of the Wellcome Sanger Institute COVID-19 Surveillance Team |
| EPI_ISL_2729910 | Lighthouse Lab in Alderley Park | Wellcome Sanger Institute for the COVID-19 Genomics UK (COG-UK) Consortium | Jacquelyn Wynn, Mairead Hyland, The Lighthouse Lab in Alderley Park and Alex Alderton, Roberto Amato, Jeffrey Barrett, Sonia Goncalves, Ewan Harrison, David K. Jackson, Ian Johnston, Dominic Kwiatkowski, Cordelia Langford, John Sillitoe on behalf of the Wellcome Sanger Institute COVID-19 Surveillance Team |
| EPI_ISL_2729911, EPI_ISL_2729912 | Lighthouse Laboratory Plymouth | Wellcome Sanger Institute for the COVID-19 Genomics UK (COG-UK) Consortium | Lighthouse Laboratory Plymouth and Alex Alderton, Roberto Amato, Jeffrey Barrett, Sonia Goncalves, Ewan Harrison, David K. Jackson, Ian Johnston, Dominic Kwiatkowski, Cordelia Langford, John Sillitoe on behalf of the Wellcome Sanger Institute COVID-19 Surveillance Team |
| EPI_ISL_2729913 | Lighthouse Lab in Milton Keynes | Wellcome Sanger Institute for the COVID-19 Genomics UK (COG-UK) Consortium | The Lighthouse Lab in Milton Keynes and Alex Alderton, Roberto Amato, Jeffrey Barrett, Sonia Goncalves, Ewan Harrison, David K. Jackson, Ian Johnston, Dominic Kwiatkowski, Cordelia Langford, John Sillitoe on behalf of the Wellcome Sanger Institute COVID-19 Surveillance Team |
| EPI_ISL_2729914 | Lighthouse Lab in Alderley Park | Wellcome Sanger Institute for the COVID-19 Genomics UK (COG-UK) Consortium | Jacquelyn Wynn, Mairead Hyland, The Lighthouse Lab in Alderley Park and Alex Alderton, Roberto Amato, Jeffrey Barrett, Sonia Goncalves, Ewan Harrison, David K. Jackson, Ian Johnston, Dominic Kwiatkowski, Cordelia Langford, John Sillitoe on behalf of the Wellcome Sanger Institute COVID-19 Surveillance Team |
| EPI_ISL_2729915 | Lighthouse Laboratory Plymouth | Wellcome Sanger Institute for the COVID-19 Genomics UK (COG-UK) Consortium | Lighthouse Laboratory Plymouth and Alex Alderton, Roberto Amato, Jeffrey Barrett, Sonia Goncalves, Ewan Harrison, David K. Jackson, Ian Johnston, Dominic Kwiatkowski, Cordelia Langford, John Sillitoe on behalf of the Wellcome Sanger Institute COVID-19 Surveillance Team |
| EPI_ISL_2729916 | Lighthouse Lab in Milton Keynes | Wellcome Sanger Institute for the COVID-19 Genomics UK (COG-UK) Consortium | The Lighthouse Lab in Milton Keynes and Alex Alderton, Roberto Amato, Jeffrey Barrett, Sonia Goncalves, Ewan Harrison, David K. Jackson, Ian Johnston, Dominic Kwiatkowski, Cordelia Langford, John Sillitoe on behalf of the Wellcome Sanger Institute COVID-19 Surveillance Team |
| EPI_ISL_2729917, EPI_ISL_2729918 | Lighthouse Lab in Glasgow | Wellcome Sanger Institute for the COVID-19 Genomics UK (COG-UK) Consortium | Harper VanSteenhouse, Yumi Kasai, David Gray, Carol Clugston, Anna Dominiczak and Alex Alderton, Roberto Amato, Jeffrey Barrett, Sonia Goncalves, Ewan Harrison, David K. Jackson, Ian Johnston, Dominic Kwiatkowski, Cordelia Langford, John Sillitoe on behalf of the Wellcome Sanger Institute COVID-19 Surveillance Team |
| EPI_ISL_2729919 | Lighthouse Lab in Alderley Park | Wellcome Sanger Institute for the COVID-19 Genomics UK (COG-UK) Consortium | Jacquelyn Wynn, Mairead Hyland, The Lighthouse Lab in Alderley Park and Alex Alderton, Roberto Amato, Jeffrey Barrett, Sonia Goncalves, Ewan Harrison, David K. Jackson, Ian Johnston, Dominic Kwiatkowski, Cordelia Langford, John Sillitoe on behalf of the Wellcome Sanger Institute COVID-19 Surveillance Team |
| EPI_ISL_2729920 | Lighthouse Lab in Glasgow | Wellcome Sanger Institute for the COVID-19 Genomics UK (COG-UK) Consortium | Harper VanSteenhouse, Yumi Kasai, David Gray, Carol Clugston, Anna Dominiczak and Alex Alderton, Roberto Amato, Jeffrey Barrett, Sonia Goncalves, Ewan Harrison, David K. Jackson, Ian Johnston, Dominic Kwiatkowski, Cordelia Langford, John Sillitoe on behalf of the Wellcome Sanger Institute COVID-19 Surveillance Team |
| EPI_ISL_2729921 | Lighthouse Lab in Milton Keynes | Wellcome Sanger Institute for the COVID-19 Genomics UK (COG-UK) Consortium | The Lighthouse Lab in Milton Keynes and Alex Alderton, Roberto Amato, Jeffrey Barrett, Sonia Goncalves, Ewan Harrison, David K. Jackson, Ian Johnston, Dominic Kwiatkowski, Cordelia Langford, John Sillitoe on behalf of the Wellcome Sanger Institute COVID-19 Surveillance Team |
| EPI_ISL_2729922 | Lighthouse Lab in Alderley Park | Wellcome Sanger Institute for the COVID-19 Genomics UK (COG-UK) Consortium | Jacquelyn Wynn, Mairead Hyland, The Lighthouse Lab in Alderley Park and Alex Alderton, Roberto Amato, Jeffrey Barrett, Sonia Goncalves, Ewan Harrison, David K. Jackson, Ian Johnston, Dominic Kwiatkowski, Cordelia Langford, John Sillitoe on behalf of the Wellcome Sanger Institute COVID-19 Surveillance Team |
| EPI_ISL_2729923, EPI_ISL_2729924 | Lighthouse Laboratory Plymouth | Wellcome Sanger Institute for the COVID-19 Genomics UK (COG-UK) Consortium | Lighthouse Laboratory Plymouth and Alex Alderton, Roberto Amato, Jeffrey Barrett, Sonia Goncalves, Ewan Harrison, David K. Jackson, Ian Johnston, Dominic Kwiatkowski, Cordelia Langford, John Sillitoe on behalf of the Wellcome Sanger Institute COVID-19 Surveillance Team |
| EPI_ISL_2729925 | Lighthouse Lab in Milton Keynes | Wellcome Sanger Institute for the COVID-19 Genomics UK (COG-UK) Consortium | The Lighthouse Lab in Milton Keynes and Alex Alderton, Roberto Amato, Jeffrey Barrett, Sonia Goncalves, Ewan Harrison, David K. Jackson, Ian Johnston, Dominic Kwiatkowski, Cordelia Langford, John Sillitoe on behalf of the Wellcome Sanger Institute COVID-19 Surveillance Team |
| EPI_ISL_2729926, EPI_ISL_2729927 | Lighthouse Lab in Alderley Park | Wellcome Sanger Institute for the COVID-19 Genomics UK (COG-UK) Consortium | Jacquelyn Wynn, Mairead Hyland, The Lighthouse Lab in Alderley Park and Alex Alderton, Roberto Amato, Jeffrey Barrett, Sonia Goncalves, Ewan Harrison, David K. Jackson, Ian Johnston, Dominic Kwiatkowski, Cordelia Langford, John Sillitoe on behalf of the Wellcome Sanger Institute COVID-19 Surveillance Team |
| EPI_ISL_2729928, EPI_ISL_2729929, EPI_ISL_2729930 | Lighthouse Laboratory Plymouth | Wellcome Sanger Institute for the COVID-19 Genomics UK (COG-UK) Consortium | Lighthouse Laboratory Plymouth and Alex Alderton, Roberto Amato, Jeffrey Barrett, Sonia Goncalves, Ewan Harrison, David K. Jackson, Ian Johnston, Dominic Kwiatkowski, Cordelia Langford, John Sillitoe on behalf of the Wellcome Sanger Institute COVID-19 Surveillance Team |
| EPI_ISL_2729931 | Lighthouse Lab in Glasgow | Wellcome Sanger Institute for the COVID-19 Genomics UK (COG-UK) Consortium | Harper VanSteenhouse, Yumi Kasai, David Gray, Carol Clugston, Anna Dominiczak and Alex Alderton, Roberto Amato, Jeffrey Barrett, Sonia Goncalves, Ewan Harrison, David K. Jackson, Ian Johnston, Dominic Kwiatkowski, Cordelia Langford, John Sillitoe on behalf of the Wellcome Sanger Institute COVID-19 Surveillance Team |
| EPI_ISL_2729932, EPI_ISL_2729933 | Lighthouse Lab in Milton Keynes | Wellcome Sanger Institute for the COVID-19 Genomics UK (COG-UK) Consortium | The Lighthouse Lab in Milton Keynes and Alex Alderton, Roberto Amato, Jeffrey Barrett, Sonia Goncalves, Ewan Harrison, David K. Jackson, Ian Johnston, Dominic Kwiatkowski, Cordelia Langford, John Sillitoe on behalf of the Wellcome Sanger Institute COVID-19 Surveillance Team |
| EPI_ISL_2729934 | Lighthouse Laboratory Plymouth | Wellcome Sanger Institute for the COVID-19 Genomics UK (COG-UK) Consortium | Lighthouse Laboratory Plymouth and Alex Alderton, Roberto Amato, Jeffrey Barrett, Sonia Goncalves, Ewan Harrison, David K. Jackson, Ian Johnston, Dominic Kwiatkowski, Cordelia Langford, John Sillitoe on behalf of the Wellcome Sanger Institute COVID-19 Surveillance Team |
| EPI_ISL_2729935 | Lighthouse Lab in Alderley Park | Wellcome Sanger Institute for the COVID-19 Genomics UK (COG-UK) Consortium | Jacquelyn Wynn, Mairead Hyland, The Lighthouse Lab in Alderley Park and Alex Alderton, Roberto Amato, Jeffrey Barrett, Sonia Goncalves, Ewan Harrison, David K. Jackson, Ian Johnston, Dominic Kwiatkowski, Cordelia Langford, John Sillitoe on behalf of the Wellcome Sanger Institute COVID-19 Surveillance Team |
| EPI_ISL_2729936 | Lighthouse Lab in Milton Keynes | Wellcome Sanger Institute for the COVID-19 Genomics UK (COG-UK) Consortium | The Lighthouse Lab in Milton Keynes and Alex Alderton, Roberto Amato, Jeffrey Barrett, Sonia Goncalves, Ewan Harrison, David K. Jackson, Ian Johnston, Dominic Kwiatkowski, Cordelia Langford, John Sillitoe on behalf of the Wellcome Sanger Institute COVID-19 Surveillance Team |
| EPI_ISL_2729937, EPI_ISL_2729938 | Lighthouse Lab in Alderley Park | Wellcome Sanger Institute for the COVID-19 Genomics UK (COG-UK) Consortium | Jacquelyn Wynn, Mairead Hyland, The Lighthouse Lab in Alderley Park and Alex Alderton, Roberto Amato, Jeffrey Barrett, Sonia Goncalves, Ewan Harrison, David K. Jackson, Ian Johnston, Dominic Kwiatkowski, Cordelia Langford, John Sillitoe on behalf of the Wellcome Sanger Institute COVID-19 Surveillance Team |
| EPI_ISL_2729939 | Lighthouse Lab in Glasgow | Wellcome Sanger Institute for the COVID-19 Genomics UK (COG-UK) Consortium | Harper VanSteenhouse, Yumi Kasai, David Gray, Carol Clugston, Anna Dominiczak and Alex Alderton, Roberto Amato, Jeffrey Barrett, Sonia Goncalves, Ewan Harrison, David K. Jackson, Ian Johnston, Dominic Kwiatkowski, Cordelia Langford, John Sillitoe on behalf of the Wellcome Sanger Institute COVID-19 Surveillance Team |
| EPI_ISL_2729940, EPI_ISL_2729941 | Lighthouse Lab in Alderley Park | Wellcome Sanger Institute for the COVID-19 Genomics UK (COG-UK) Consortium | Jacquelyn Wynn, Mairead Hyland, The Lighthouse Lab in Alderley Park and Alex Alderton, Roberto Amato, Jeffrey Barrett, Sonia Goncalves, Ewan Harrison, David K. Jackson, Ian Johnston, Dominic Kwiatkowski, Cordelia Langford, John Sillitoe on behalf of the Wellcome Sanger Institute COVID-19 Surveillance Team |
| EPI_ISL_2729942 | Lighthouse Lab in Milton Keynes | Wellcome Sanger Institute for the COVID-19 Genomics UK (COG-UK) Consortium | The Lighthouse Lab in Milton Keynes and Alex Alderton, Roberto Amato, Jeffrey Barrett, Sonia Goncalves, Ewan Harrison, David K. Jackson, Ian Johnston, Dominic Kwiatkowski, Cordelia Langford, John Sillitoe on behalf of the Wellcome Sanger Institute COVID-19 Surveillance Team |
| EPI_ISL_2729943 | Lighthouse Lab in Alderley Park | Wellcome Sanger Institute for the COVID-19 Genomics UK (COG-UK) Consortium | Jacquelyn Wynn, Mairead Hyland, The Lighthouse Lab in Alderley Park and Alex Alderton, Roberto Amato, Jeffrey Barrett, Sonia Goncalves, Ewan Harrison, David K. Jackson, Ian Johnston, Dominic Kwiatkowski, Cordelia Langford, John Sillitoe on behalf of the Wellcome Sanger Institute COVID-19 Surveillance Team |
| EPI_ISL_2729944, EPI_ISL_2729945 | Lighthouse Lab in Glasgow | Wellcome Sanger Institute for the COVID-19 Genomics UK (COG-UK) Consortium | Harper VanSteenhouse, Yumi Kasai, David Gray, Carol Clugston, Anna Dominiczak and Alex Alderton, Roberto Amato, Jeffrey Barrett, Sonia Goncalves, Ewan Harrison, David K. Jackson, Ian Johnston, Dominic Kwiatkowski, Cordelia Langford, John Sillitoe on behalf of the Wellcome Sanger Institute COVID-19 Surveillance Team |
| EPI_ISL_2729946 | Lighthouse Lab in Alderley Park | Wellcome Sanger Institute for the COVID-19 Genomics UK (COG-UK) Consortium | Jacquelyn Wynn, Mairead Hyland, The Lighthouse Lab in Alderley Park and Alex Alderton, Roberto Amato, Jeffrey Barrett, Sonia Goncalves, Ewan Harrison, David K. Jackson, Ian Johnston, Dominic Kwiatkowski, Cordelia Langford, John Sillitoe on behalf of the Wellcome Sanger Institute COVID-19 Surveillance Team |
| EPI_ISL_2729947 | Lighthouse Laboratory Plymouth | Wellcome Sanger Institute for the COVID-19 Genomics UK (COG-UK) Consortium | Lighthouse Laboratory Plymouth and Alex Alderton, Roberto Amato, Jeffrey Barrett, Sonia Goncalves, Ewan Harrison, David K. Jackson, Ian Johnston, Dominic Kwiatkowski, Cordelia Langford, John Sillitoe on behalf of the Wellcome Sanger Institute COVID-19 Surveillance Team |

| | | | |
|---|---|---|---|
| EPI_ISL_2729948 | Lighthouse Lab in Milton Keynes | Wellcome Sanger Institute for the COVID-19 Genomics UK (COG-UK) Consortium | The Lighthouse Lab in Milton Keynes and Alex Alderton, Roberto Amato, Jeffrey Barrett, Sonia Goncalves, Ewan Harrison, David K. Jackson, Ian Johnston, Dominic Kwiatkowski, Cordelia Langford, John Sillitoe on behalf of the Wellcome Sanger Institute COVID-19 Surveillance Team |
| EPI_ISL_2729949, EPI_ISL_2729950 | Lighthouse Laboratory Plymouth | Wellcome Sanger Institute for the COVID-19 Genomics UK (COG-UK) Consortium | Lighthouse Laboratory Plymouth and Alex Alderton, Roberto Amato, Jeffrey Barrett, Sonia Goncalves, Ewan Harrison, David K. Jackson, Ian Johnston, Dominic Kwiatkowski, Cordelia Langford, John Sillitoe on behalf of the Wellcome Sanger Institute COVID-19 Surveillance Team |
| EPI_ISL_2729951 | Lighthouse Lab in Milton Keynes | Wellcome Sanger Institute for the COVID-19 Genomics UK (COG-UK) Consortium | The Lighthouse Lab in Milton Keynes and Alex Alderton, Roberto Amato, Jeffrey Barrett, Sonia Goncalves, Ewan Harrison, David K. Jackson, Ian Johnston, Dominic Kwiatkowski, Cordelia Langford, John Sillitoe on behalf of the Wellcome Sanger Institute COVID-19 Surveillance Team |
| EPI_ISL_2729952 | Lighthouse Lab in Alderley Park | Wellcome Sanger Institute for the COVID-19 Genomics UK (COG-UK) Consortium | Jacquelyn Wynn, Mairead Hyland, The Lighthouse Lab in Alderley Park and Alex Alderton, Roberto Amato, Jeffrey Barrett, Sonia Goncalves, Ewan Harrison, David K. Jackson, Ian Johnston, Dominic Kwiatkowski, Cordelia Langford, John Sillitoe on behalf of the Wellcome Sanger Institute COVID-19 Surveillance Team |
| EPI_ISL_2729953 | Lighthouse Lab in Glasgow | Wellcome Sanger Institute for the COVID-19 Genomics UK (COG-UK) Consortium | Harper VanSteenhouse, Yumi Kasai, David Gray, Carol Clugston, Anna Dominiczak and Alex Alderton, Roberto Amato, Jeffrey Barrett, Sonia Goncalves, Ewan Harrison, David K. Jackson, Ian Johnston, Dominic Kwiatkowski, Cordelia Langford, John Sillitoe on behalf of the Wellcome Sanger Institute COVID-19 Surveillance Team |
| EPI_ISL_2729954 | Lighthouse Laboratory Plymouth | Wellcome Sanger Institute for the COVID-19 Genomics UK (COG-UK) Consortium | Lighthouse Laboratory Plymouth and Alex Alderton, Roberto Amato, Jeffrey Barrett, Sonia Goncalves, Ewan Harrison, David K. Jackson, Ian Johnston, Dominic Kwiatkowski, Cordelia Langford, John Sillitoe on behalf of the Wellcome Sanger Institute COVID-19 Surveillance Team |
| EPI_ISL_2729955 | Lighthouse Lab in Glasgow | Wellcome Sanger Institute for the COVID-19 Genomics UK (COG-UK) Consortium | Harper VanSteenhouse, Yumi Kasai, David Gray, Carol Clugston, Anna Dominiczak and Alex Alderton, Roberto Amato, Jeffrey Barrett, Sonia Goncalves, Ewan Harrison, David K. Jackson, Ian Johnston, Dominic Kwiatkowski, Cordelia Langford, John Sillitoe on behalf of the Wellcome Sanger Institute COVID-19 Surveillance Team |
| EPI_ISL_2729956 | Lighthouse Lab in Milton Keynes | Wellcome Sanger Institute for the COVID-19 Genomics UK (COG-UK) Consortium | The Lighthouse Lab in Milton Keynes and Alex Alderton, Roberto Amato, Jeffrey Barrett, Sonia Goncalves, Ewan Harrison, David K. Jackson, Ian Johnston, Dominic Kwiatkowski, Cordelia Langford, John Sillitoe on behalf of the Wellcome Sanger Institute COVID-19 Surveillance Team |
| EPI_ISL_2729957 | Lighthouse Laboratory Plymouth | Wellcome Sanger Institute for the COVID-19 Genomics UK (COG-UK) Consortium | Lighthouse Laboratory Plymouth and Alex Alderton, Roberto Amato, Jeffrey Barrett, Sonia Goncalves, Ewan Harrison, David K. Jackson, Ian Johnston, Dominic Kwiatkowski, Cordelia Langford, John Sillitoe on behalf of the Wellcome Sanger Institute COVID-19 Surveillance Team |
| EPI_ISL_2729958 | Lighthouse Lab in Glasgow | Wellcome Sanger Institute for the COVID-19 Genomics UK (COG-UK) Consortium | Harper VanSteenhouse, Yumi Kasai, David Gray, Carol Clugston, Anna Dominiczak and Alex Alderton, Roberto Amato, Jeffrey Barrett, Sonia Goncalves, Ewan Harrison, David K. Jackson, Ian Johnston, Dominic Kwiatkowski, Cordelia Langford, John Sillitoe on behalf of the Wellcome Sanger Institute COVID-19 Surveillance Team |
| EPI_ISL_2729959, EPI_ISL_2729960 | Lighthouse Lab in Alderley Park | Wellcome Sanger Institute for the COVID-19 Genomics UK (COG-UK) Consortium | Jacquelyn Wynn, Mairead Hyland, The Lighthouse Lab in Alderley Park and Alex Alderton, Roberto Amato, Jeffrey Barrett, Sonia Goncalves, Ewan Harrison, David K. Jackson, Ian Johnston, Dominic Kwiatkowski, Cordelia Langford, John Sillitoe on behalf of the Wellcome Sanger Institute COVID-19 Surveillance Team |
| EPI_ISL_2729961 | Lighthouse Lab in Glasgow | Wellcome Sanger Institute for the COVID-19 Genomics UK (COG-UK) Consortium | Harper VanSteenhouse, Yumi Kasai, David Gray, Carol Clugston, Anna Dominiczak and Alex Alderton, Roberto Amato, Jeffrey Barrett, Sonia Goncalves, Ewan Harrison, David K. Jackson, Ian Johnston, Dominic Kwiatkowski, Cordelia Langford, John Sillitoe on behalf of the Wellcome Sanger Institute COVID-19 Surveillance Team |
| EPI_ISL_2729962 | Lighthouse Laboratory Plymouth | Wellcome Sanger Institute for the COVID-19 Genomics UK (COG-UK) Consortium | Lighthouse Laboratory Plymouth and Alex Alderton, Roberto Amato, Jeffrey Barrett, Sonia Goncalves, Ewan Harrison, David K. Jackson, Ian Johnston, Dominic Kwiatkowski, Cordelia Langford, John Sillitoe on behalf of the Wellcome Sanger Institute COVID-19 Surveillance Team |
| EPI_ISL_2729963, EPI_ISL_2729964 | Lighthouse Lab in Milton Keynes | Wellcome Sanger Institute for the COVID-19 Genomics UK (COG-UK) Consortium | The Lighthouse Lab in Milton Keynes and Alex Alderton, Roberto Amato, Jeffrey Barrett, Sonia Goncalves, Ewan Harrison, David K. Jackson, Ian Johnston, Dominic Kwiatkowski, Cordelia Langford, John Sillitoe on behalf of the Wellcome Sanger Institute COVID-19 Surveillance Team |
| EPI_ISL_2729965, EPI_ISL_2729966, EPI_ISL_2729967 | Lighthouse Lab in Glasgow | Wellcome Sanger Institute for the COVID-19 Genomics UK (COG-UK) Consortium | Harper VanSteenhouse, Yumi Kasai, David Gray, Carol Clugston, Anna Dominiczak and Alex Alderton, Roberto Amato, Jeffrey Barrett, Sonia Goncalves, Ewan Harrison, David K. Jackson, Ian Johnston, Dominic Kwiatkowski, Cordelia Langford, John Sillitoe on behalf of the Wellcome Sanger Institute COVID-19 Surveillance Team |
| EPI_ISL_2729968 | Lighthouse Lab in Milton Keynes | Wellcome Sanger Institute for the COVID-19 Genomics UK (COG-UK) Consortium | The Lighthouse Lab in Milton Keynes and Alex Alderton, Roberto Amato, Jeffrey Barrett, Sonia Goncalves, Ewan Harrison, David K. Jackson, Ian Johnston, Dominic Kwiatkowski, Cordelia Langford, John Sillitoe on behalf of the Wellcome Sanger Institute COVID-19 Surveillance Team |
| EPI_ISL_2729969 | Lighthouse Laboratory Plymouth | Wellcome Sanger Institute for the COVID-19 Genomics UK (COG-UK) Consortium | Lighthouse Laboratory Plymouth and Alex Alderton, Roberto Amato, Jeffrey Barrett, Sonia Goncalves, Ewan Harrison, David K. Jackson, Ian Johnston, Dominic Kwiatkowski, Cordelia Langford, John Sillitoe on behalf of the Wellcome Sanger Institute COVID-19 Surveillance Team |
| EPI_ISL_2729970 | Lighthouse Lab in Milton Keynes | Wellcome Sanger Institute for the COVID-19 Genomics UK (COG-UK) Consortium | The Lighthouse Lab in Milton Keynes and Alex Alderton, Roberto Amato, Jeffrey Barrett, Sonia Goncalves, Ewan Harrison, David K. Jackson, Ian Johnston, Dominic Kwiatkowski, Cordelia Langford, John Sillitoe on behalf of the Wellcome Sanger Institute COVID-19 Surveillance Team |
| EPI_ISL_2729971 | Lighthouse Laboratory Plymouth | Wellcome Sanger Institute for the COVID-19 Genomics UK (COG-UK) Consortium | Lighthouse Laboratory Plymouth and Alex Alderton, Roberto Amato, Jeffrey Barrett, Sonia Goncalves, Ewan Harrison, David K. Jackson, Ian Johnston, Dominic Kwiatkowski, Cordelia Langford, John Sillitoe on behalf of the Wellcome Sanger Institute COVID-19 Surveillance Team |
| EPI_ISL_2729972 | Lighthouse Lab in Milton Keynes | Wellcome Sanger Institute for the COVID-19 Genomics UK (COG-UK) Consortium | The Lighthouse Lab in Milton Keynes and Alex Alderton, Roberto Amato, Jeffrey Barrett, Sonia Goncalves, Ewan Harrison, David K. Jackson, Ian Johnston, Dominic Kwiatkowski, Cordelia Langford, John Sillitoe on behalf of the Wellcome Sanger Institute COVID-19 Surveillance Team |
| EPI_ISL_2729973 | Lighthouse Lab in Glasgow | Wellcome Sanger Institute for the COVID-19 Genomics UK (COG-UK) Consortium | Harper VanSteenhouse, Yumi Kasai, David Gray, Carol Clugston, Anna Dominiczak and Alex Alderton, Roberto Amato, Jeffrey Barrett, Sonia Goncalves, Ewan Harrison, David K. Jackson, Ian Johnston, Dominic Kwiatkowski, Cordelia Langford, John Sillitoe on behalf of the Wellcome Sanger Institute COVID-19 Surveillance Team |
| EPI_ISL_2729974 | Lighthouse Laboratory Plymouth | Wellcome Sanger Institute for the COVID-19 Genomics UK (COG-UK) Consortium | Lighthouse Laboratory Plymouth and Alex Alderton, Roberto Amato, Jeffrey Barrett, Sonia Goncalves, Ewan Harrison, David K. Jackson, Ian Johnston, Dominic Kwiatkowski, Cordelia Langford, John Sillitoe on behalf of the Wellcome Sanger Institute COVID-19 Surveillance Team |
| EPI_ISL_2729975 | Lighthouse Lab in Glasgow | Wellcome Sanger Institute for the COVID-19 Genomics UK (COG-UK) Consortium | Harper VanSteenhouse, Yumi Kasai, David Gray, Carol Clugston, Anna Dominiczak and Alex Alderton, Roberto Amato, Jeffrey Barrett, Sonia Goncalves, Ewan Harrison, David K. Jackson, Ian Johnston, Dominic Kwiatkowski, Cordelia Langford, John Sillitoe on behalf of the Wellcome Sanger Institute COVID-19 Surveillance Team |
| EPI_ISL_2729976 | Lighthouse Lab in Milton Keynes | Wellcome Sanger Institute for the COVID-19 Genomics UK (COG-UK) Consortium | The Lighthouse Lab in Milton Keynes and Alex Alderton, Roberto Amato, Jeffrey Barrett, Sonia Goncalves, Ewan Harrison, David K. Jackson, Ian Johnston, Dominic Kwiatkowski, Cordelia Langford, John Sillitoe on behalf of the Wellcome Sanger Institute COVID-19 Surveillance Team |
| EPI_ISL_2729977 | Lighthouse Lab in Glasgow | Wellcome Sanger Institute for the COVID-19 Genomics UK (COG-UK) Consortium | Harper VanSteenhouse, Yumi Kasai, David Gray, Carol Clugston, Anna Dominiczak and Alex Alderton, Roberto Amato, Jeffrey Barrett, Sonia Goncalves, Ewan Harrison, David K. Jackson, Ian Johnston, Dominic Kwiatkowski, Cordelia Langford, John Sillitoe on behalf of the Wellcome Sanger Institute COVID-19 Surveillance Team |
| EPI_ISL_2729978 | Lighthouse Laboratory Plymouth | Wellcome Sanger Institute for the COVID-19 Genomics UK (COG-UK) Consortium | Lighthouse Laboratory Plymouth and Alex Alderton, Roberto Amato, Jeffrey Barrett, Sonia Goncalves, Ewan Harrison, David K. Jackson, Ian Johnston, Dominic Kwiatkowski, Cordelia Langford, John Sillitoe on behalf of the Wellcome Sanger Institute COVID-19 Surveillance Team |
| EPI_ISL_2729979 | Lighthouse Lab in Glasgow | Wellcome Sanger Institute for the COVID-19 Genomics UK (COG-UK) Consortium | Harper VanSteenhouse, Yumi Kasai, David Gray, Carol Clugston, Anna Dominiczak and Alex Alderton, Roberto Amato, Jeffrey Barrett, Sonia Goncalves, Ewan Harrison, David K. Jackson, Ian Johnston, Dominic Kwiatkowski, Cordelia Langford, John Sillitoe on behalf of the Wellcome Sanger Institute COVID-19 Surveillance Team |
| EPI_ISL_2729980 | Lighthouse Laboratory Plymouth | Wellcome Sanger Institute for the COVID-19 Genomics UK (COG-UK) Consortium | Lighthouse Laboratory Plymouth and Alex Alderton, Roberto Amato, Jeffrey Barrett, Sonia Goncalves, Ewan Harrison, David K. Jackson, Ian Johnston, Dominic Kwiatkowski, Cordelia Langford, John Sillitoe on behalf of the Wellcome Sanger Institute COVID-19 Surveillance Team |
| EPI_ISL_2729981 | Lighthouse Lab in Glasgow | Wellcome Sanger Institute for the COVID-19 Genomics UK (COG-UK) Consortium | Harper VanSteenhouse, Yumi Kasai, David Gray, Carol Clugston, Anna Dominiczak and Alex Alderton, Roberto Amato, Jeffrey Barrett, Sonia Goncalves, Ewan Harrison, David K. Jackson, Ian Johnston, Dominic Kwiatkowski, Cordelia Langford, John Sillitoe on behalf of the Wellcome Sanger Institute COVID-19 Surveillance Team |
| EPI_ISL_2729982 | Lighthouse Lab in Alderley Park | Wellcome Sanger Institute for the COVID-19 Genomics UK (COG-UK) Consortium | Jacquelyn Wynn, Mairead Hyland, The Lighthouse Lab in Alderley Park and Alex Alderton, Roberto Amato, Jeffrey Barrett, Sonia Goncalves, Ewan Harrison, David K. Jackson, Ian Johnston, Dominic Kwiatkowski, Cordelia Langford, John Sillitoe on behalf of the Wellcome Sanger Institute COVID-19 Surveillance Team |
| EPI_ISL_2729983 | Lighthouse Lab in Glasgow | Wellcome Sanger Institute for the COVID-19 Genomics UK (COG-UK) Consortium | Harper VanSteenhouse, Yumi Kasai, David Gray, Carol Clugston, Anna Dominiczak and Alex Alderton, Roberto Amato, Jeffrey Barrett, Sonia Goncalves, Ewan Harrison, David K. Jackson, Ian Johnston, Dominic Kwiatkowski, Cordelia Langford, John Sillitoe on behalf of the Wellcome Sanger Institute COVID-19 Surveillance Team |
| EPI_ISL_2729984 | Lighthouse Lab in Alderley Park | Wellcome Sanger Institute for the COVID-19 Genomics UK (COG-UK) Consortium | Jacquelyn Wynn, Mairead Hyland, The Lighthouse Lab in Alderley Park and Alex Alderton, Roberto Amato, Jeffrey Barrett, Sonia Goncalves, Ewan Harrison, David K. Jackson, Ian Johnston, Dominic Kwiatkowski, Cordelia Langford, John Sillitoe on behalf of the Wellcome Sanger Institute COVID-19 Surveillance Team |
| EPI_ISL_2729985 | Lighthouse Lab in Milton Keynes | Wellcome Sanger Institute for the COVID-19 Genomics UK (COG-UK) Consortium | The Lighthouse Lab in Milton Keynes and Alex Alderton, Roberto Amato, Jeffrey Barrett, Sonia Goncalves, Ewan Harrison, David K. Jackson, Ian Johnston, Dominic Kwiatkowski, Cordelia Langford, John Sillitoe on behalf of the Wellcome Sanger Institute COVID-19 Surveillance Team |
| EPI_ISL_2729986, EPI_ISL_2729987 | Lighthouse Lab in Alderley Park | Wellcome Sanger Institute for the COVID-19 Genomics UK (COG-UK) Consortium | Jacquelyn Wynn, Mairead Hyland, The Lighthouse Lab in Alderley Park and Alex Alderton, Roberto Amato, Jeffrey Barrett, Sonia Goncalves, Ewan Harrison, David K. Jackson, Ian Johnston, Dominic Kwiatkowski, Cordelia Langford, John Sillitoe on behalf of the Wellcome Sanger Institute COVID-19 Surveillance Team |
| EPI_ISL_2729988, EPI_ISL_2729989 | Lighthouse Lab in Glasgow | Wellcome Sanger Institute for the COVID-19 Genomics UK (COG-UK) Consortium | Harper VanSteenhouse, Yumi Kasai, David Gray, Carol Clugston, Anna Dominiczak and Alex Alderton, Roberto Amato, Jeffrey Barrett, Sonia Goncalves, Ewan Harrison, David K. Jackson, Ian Johnston, Dominic Kwiatkowski, Cordelia Langford, John Sillitoe on behalf of the Wellcome Sanger Institute COVID-19 Surveillance Team |
| EPI_ISL_2729990 | Lighthouse Lab in Alderley Park | Wellcome Sanger Institute for the COVID-19 Genomics UK (COG-UK) Consortium | Jacquelyn Wynn, Mairead Hyland, The Lighthouse Lab in Alderley Park and Alex Alderton, Roberto Amato, Jeffrey Barrett, Sonia Goncalves, Ewan Harrison, David K. Jackson, Ian Johnston, Dominic Kwiatkowski, Cordelia Langford, John Sillitoe on behalf of the Wellcome Sanger Institute COVID-19 Surveillance Team |
| EPI_ISL_2729991 | Lighthouse Lab in Milton Keynes | Wellcome Sanger Institute for the COVID-19 Genomics UK (COG-UK) Consortium | The Lighthouse Lab in Milton Keynes and Alex Alderton, Roberto Amato, Jeffrey Barrett, Sonia Goncalves, Ewan Harrison, David K. Jackson, Ian Johnston, Dominic Kwiatkowski, Cordelia Langford, John Sillitoe on behalf of the Wellcome Sanger Institute COVID-19 Surveillance Team |
| EPI_ISL_2729992 | Lighthouse Lab in Alderley Park | Wellcome Sanger Institute for the COVID-19 Genomics UK (COG-UK) Consortium | Jacquelyn Wynn, Mairead Hyland, The Lighthouse Lab in Alderley Park and Alex Alderton, Roberto Amato, Jeffrey Barrett, Sonia Goncalves, Ewan Harrison, David K. Jackson, Ian Johnston, Dominic Kwiatkowski, Cordelia Langford, John Sillitoe on behalf of the Wellcome Sanger Institute COVID-19 Surveillance Team |
| EPI_ISL_2729993 | Lighthouse Lab in Glasgow | Wellcome Sanger Institute for the COVID-19 Genomics UK (COG-UK) Consortium | Harper VanSteenhouse, Yumi Kasai, David Gray, Carol Clugston, Anna Dominiczak and Alex Alderton, Roberto Amato, Jeffrey Barrett, Sonia Goncalves, Ewan Harrison, David K. Jackson, Ian Johnston, Dominic Kwiatkowski, Cordelia Langford, John Sillitoe on behalf of the Wellcome Sanger Institute COVID-19 Surveillance Team |
| EPI_ISL_2729994 | Lighthouse Lab in Alderley Park | Wellcome Sanger Institute for the COVID-19 Genomics UK (COG-UK) Consortium | Jacquelyn Wynn, Mairead Hyland, The Lighthouse Lab in Alderley Park and Alex Alderton, Roberto Amato, Jeffrey Barrett, Sonia Goncalves, Ewan Harrison, David K. Jackson, Ian Johnston, Dominic Kwiatkowski, Cordelia Langford, John Sillitoe on behalf of the Wellcome Sanger Institute COVID-19 Surveillance Team |
| EPI_ISL_2729995 | Lighthouse Laboratory Plymouth | Wellcome Sanger Institute for the COVID-19 Genomics UK (COG-UK) Consortium | Lighthouse Laboratory Plymouth and Alex Alderton, Roberto Amato, Jeffrey Barrett, Sonia Goncalves, Ewan Harrison, David K. Jackson, Ian Johnston, Dominic Kwiatkowski, Cordelia Langford, John Sillitoe on behalf of the Wellcome Sanger Institute COVID-19 Surveillance Team |
| EPI_ISL_2729996, EPI_ISL_2729997, EPI_ISL_2729998, EPI_ISL_2729999 | Lighthouse Lab in Milton Keynes | Wellcome Sanger Institute for the COVID-19 Genomics UK (COG-UK) Consortium | The Lighthouse Lab in Milton Keynes and Alex Alderton, Roberto Amato, Jeffrey Barrett, Sonia Goncalves, Ewan Harrison, David K. Jackson, Ian Johnston, Dominic Kwiatkowski, Cordelia Langford, John Sillitoe on behalf of the Wellcome Sanger Institute COVID-19 Surveillance Team |
| EPI_ISL_2730000 | Lighthouse Lab in Alderley Park | Wellcome Sanger Institute for the COVID-19 Genomics UK (COG-UK) Consortium | Jacquelyn Wynn, Mairead Hyland, The Lighthouse Lab in Alderley Park and Alex Alderton, Roberto Amato, Jeffrey Barrett, Sonia Goncalves, Ewan Harrison, David K. Jackson, Ian Johnston, Dominic Kwiatkowski, Cordelia Langford, John Sillitoe on behalf of the Wellcome Sanger Institute COVID-19 Surveillance Team |
| EPI_ISL_2730001 | Lighthouse Laboratory Plymouth | Wellcome Sanger Institute for the COVID-19 Genomics UK (COG-UK) Consortium | Lighthouse Laboratory Plymouth and Alex Alderton, Roberto Amato, Jeffrey Barrett, Sonia Goncalves, Ewan Harrison, David K. Jackson, Ian Johnston, Dominic Kwiatkowski, Cordelia Langford, John Sillitoe on behalf of the Wellcome Sanger Institute COVID-19 Surveillance Team |

| | | | |
|---|---|---|---|
| EPI_ISL_2730002 | Lighthouse Lab in Alderley Park | Wellcome Sanger Institute for the COVID-19 Genomics UK (COG-UK) Consortium | Jacquelyn Wynn, Mairead Hyland, The Lighthouse Lab in Alderley Park and Alex Alderton, Roberto Amato, Jeffrey Barrett, Sonia Goncalves, Ewan Harrison, David K. Jackson, Ian Johnston, Dominic Kwiatkowski, Cordelia Langford, John Sillitoe on behalf of the Wellcome Sanger Institute COVID-19 Surveillance Team |
| EPI_ISL_2730003 | Lighthouse Lab in Glasgow | Wellcome Sanger Institute for the COVID-19 Genomics UK (COG-UK) Consortium | Harper VanSteenhouse, Yumi Kasai, David Gray, Carol Clugston, Anna Dominiczak and Alex Alderton, Roberto Amato, Jeffrey Barrett, Sonia Goncalves, Ewan Harrison, David K. Jackson, Ian Johnston, Dominic Kwiatkowski, Cordelia Langford, John Sillitoe on behalf of the Wellcome Sanger Institute COVID-19 Surveillance Team |
| EPI_ISL_2730004 | Lighthouse Lab in Milton Keynes | Wellcome Sanger Institute for the COVID-19 Genomics UK (COG-UK) Consortium | The Lighthouse Lab in Milton Keynes and Alex Alderton, Roberto Amato, Jeffrey Barrett, Sonia Goncalves, Ewan Harrison, David K. Jackson, Ian Johnston, Dominic Kwiatkowski, Cordelia Langford, John Sillitoe on behalf of the Wellcome Sanger Institute COVID-19 Surveillance Team |
| EPI_ISL_2730005 | Lighthouse Lab in Glasgow | Wellcome Sanger Institute for the COVID-19 Genomics UK (COG-UK) Consortium | Harper VanSteenhouse, Yumi Kasai, David Gray, Carol Clugston, Anna Dominiczak and Alex Alderton, Roberto Amato, Jeffrey Barrett, Sonia Goncalves, Ewan Harrison, David K. Jackson, Ian Johnston, Dominic Kwiatkowski, Cordelia Langford, John Sillitoe on behalf of the Wellcome Sanger Institute COVID-19 Surveillance Team |
| EPI_ISL_2730006 | Lighthouse Lab in Milton Keynes | Wellcome Sanger Institute for the COVID-19 Genomics UK (COG-UK) Consortium | The Lighthouse Lab in Milton Keynes and Alex Alderton, Roberto Amato, Jeffrey Barrett, Sonia Goncalves, Ewan Harrison, David K. Jackson, Ian Johnston, Dominic Kwiatkowski, Cordelia Langford, John Sillitoe on behalf of the Wellcome Sanger Institute COVID-19 Surveillance Team |
| EPI_ISL_2730007 | Lighthouse Lab in Glasgow | Wellcome Sanger Institute for the COVID-19 Genomics UK (COG-UK) Consortium | Harper VanSteenhouse, Yumi Kasai, David Gray, Carol Clugston, Anna Dominiczak and Alex Alderton, Roberto Amato, Jeffrey Barrett, Sonia Goncalves, Ewan Harrison, David K. Jackson, Ian Johnston, Dominic Kwiatkowski, Cordelia Langford, John Sillitoe on behalf of the Wellcome Sanger Institute COVID-19 Surveillance Team |
| EPI_ISL_2730008 | Lighthouse Laboratory Plymouth | Wellcome Sanger Institute for the COVID-19 Genomics UK (COG-UK) Consortium | Lighthouse Laboratory Plymouth and Alex Alderton, Roberto Amato, Jeffrey Barrett, Sonia Goncalves, Ewan Harrison, David K. Jackson, Ian Johnston, Dominic Kwiatkowski, Cordelia Langford, John Sillitoe on behalf of the Wellcome Sanger Institute COVID-19 Surveillance Team |
| EPI_ISL_2730009 | Lighthouse Lab in Milton Keynes | Wellcome Sanger Institute for the COVID-19 Genomics UK (COG-UK) Consortium | The Lighthouse Lab in Milton Keynes and Alex Alderton, Roberto Amato, Jeffrey Barrett, Sonia Goncalves, Ewan Harrison, David K. Jackson, Ian Johnston, Dominic Kwiatkowski, Cordelia Langford, John Sillitoe on behalf of the Wellcome Sanger Institute COVID-19 Surveillance Team |
| EPI_ISL_2730010, EPI_ISL_2730011, EPI_ISL_2730012, EPI_ISL_2730013 | Lighthouse Lab in Glasgow | Wellcome Sanger Institute for the COVID-19 Genomics UK (COG-UK) Consortium | Harper VanSteenhouse, Yumi Kasai, David Gray, Carol Clugston, Anna Dominiczak and Alex Alderton, Roberto Amato, Jeffrey Barrett, Sonia Goncalves, Ewan Harrison, David K. Jackson, Ian Johnston, Dominic Kwiatkowski, Cordelia Langford, John Sillitoe on behalf of the Wellcome Sanger Institute COVID-19 Surveillance Team |
| EPI_ISL_2730014 | Lighthouse Lab in Alderley Park | Wellcome Sanger Institute for the COVID-19 Genomics UK (COG-UK) Consortium | Jacquelyn Wynn, Mairead Hyland, The Lighthouse Lab in Alderley Park and Alex Alderton, Roberto Amato, Jeffrey Barrett, Sonia Goncalves, Ewan Harrison, David K. Jackson, Ian Johnston, Dominic Kwiatkowski, Cordelia Langford, John Sillitoe on behalf of the Wellcome Sanger Institute COVID-19 Surveillance Team |
| EPI_ISL_2730015, EPI_ISL_2730016, EPI_ISL_2730017 | Lighthouse Lab in Glasgow | Wellcome Sanger Institute for the COVID-19 Genomics UK (COG-UK) Consortium | Harper VanSteenhouse, Yumi Kasai, David Gray, Carol Clugston, Anna Dominiczak and Alex Alderton, Roberto Amato, Jeffrey Barrett, Sonia Goncalves, Ewan Harrison, David K. Jackson, Ian Johnston, Dominic Kwiatkowski, Cordelia Langford, John Sillitoe on behalf of the Wellcome Sanger Institute COVID-19 Surveillance Team |
| EPI_ISL_2730018, EPI_ISL_2730019 | Lighthouse Lab in Milton Keynes | Wellcome Sanger Institute for the COVID-19 Genomics UK (COG-UK) Consortium | The Lighthouse Lab in Milton Keynes and Alex Alderton, Roberto Amato, Jeffrey Barrett, Sonia Goncalves, Ewan Harrison, David K. Jackson, Ian Johnston, Dominic Kwiatkowski, Cordelia Langford, John Sillitoe on behalf of the Wellcome Sanger Institute COVID-19 Surveillance Team |
| EPI_ISL_2730020 | Lighthouse Lab in Glasgow | Wellcome Sanger Institute for the COVID-19 Genomics UK (COG-UK) Consortium | Harper VanSteenhouse, Yumi Kasai, David Gray, Carol Clugston, Anna Dominiczak and Alex Alderton, Roberto Amato, Jeffrey Barrett, Sonia Goncalves, Ewan Harrison, David K. Jackson, Ian Johnston, Dominic Kwiatkowski, Cordelia Langford, John Sillitoe on behalf of the Wellcome Sanger Institute COVID-19 Surveillance Team |
| EPI_ISL_2730021, EPI_ISL_2730022 | Lighthouse Lab in Alderley Park | Wellcome Sanger Institute for the COVID-19 Genomics UK (COG-UK) Consortium | Jacquelyn Wynn, Mairead Hyland, The Lighthouse Lab in Alderley Park and Alex Alderton, Roberto Amato, Jeffrey Barrett, Sonia Goncalves, Ewan Harrison, David K. Jackson, Ian Johnston, Dominic Kwiatkowski, Cordelia Langford, John Sillitoe on behalf of the Wellcome Sanger Institute COVID-19 Surveillance Team |
| EPI_ISL_2730023 | Lighthouse Laboratory Plymouth | Wellcome Sanger Institute for the COVID-19 Genomics UK (COG-UK) Consortium | Lighthouse Laboratory Plymouth and Alex Alderton, Roberto Amato, Jeffrey Barrett, Sonia Goncalves, Ewan Harrison, David K. Jackson, Ian Johnston, Dominic Kwiatkowski, Cordelia Langford, John Sillitoe on behalf of the Wellcome Sanger Institute COVID-19 Surveillance Team |
| EPI_ISL_2730024 | Lighthouse Lab in Milton Keynes | Wellcome Sanger Institute for the COVID-19 Genomics UK (COG-UK) Consortium | The Lighthouse Lab in Milton Keynes and Alex Alderton, Roberto Amato, Jeffrey Barrett, Sonia Goncalves, Ewan Harrison, David K. Jackson, Ian Johnston, Dominic Kwiatkowski, Cordelia Langford, John Sillitoe on behalf of the Wellcome Sanger Institute COVID-19 Surveillance Team |
| EPI_ISL_2730025 | Lighthouse Lab in Alderley Park | Wellcome Sanger Institute for the COVID-19 Genomics UK (COG-UK) Consortium | Jacquelyn Wynn, Mairead Hyland, The Lighthouse Lab in Alderley Park and Alex Alderton, Roberto Amato, Jeffrey Barrett, Sonia Goncalves, Ewan Harrison, David K. Jackson, Ian Johnston, Dominic Kwiatkowski, Cordelia Langford, John Sillitoe on behalf of the Wellcome Sanger Institute COVID-19 Surveillance Team |
| EPI_ISL_2730026 | Lighthouse Lab in Glasgow | Wellcome Sanger Institute for the COVID-19 Genomics UK (COG-UK) Consortium | Harper VanSteenhouse, Yumi Kasai, David Gray, Carol Clugston, Anna Dominiczak and Alex Alderton, Roberto Amato, Jeffrey Barrett, Sonia Goncalves, Ewan Harrison, David K. Jackson, Ian Johnston, Dominic Kwiatkowski, Cordelia Langford, John Sillitoe on behalf of the Wellcome Sanger Institute COVID-19 Surveillance Team |
| EPI_ISL_2730027 | Lighthouse Laboratory Plymouth | Wellcome Sanger Institute for the COVID-19 Genomics UK (COG-UK) Consortium | Lighthouse Laboratory Plymouth and Alex Alderton, Roberto Amato, Jeffrey Barrett, Sonia Goncalves, Ewan Harrison, David K. Jackson, Ian Johnston, Dominic Kwiatkowski, Cordelia Langford, John Sillitoe on behalf of the Wellcome Sanger Institute COVID-19 Surveillance Team |
| EPI_ISL_2730028 | Lighthouse Lab in Milton Keynes | Wellcome Sanger Institute for the COVID-19 Genomics UK (COG-UK) Consortium | The Lighthouse Lab in Milton Keynes and Alex Alderton, Roberto Amato, Jeffrey Barrett, Sonia Goncalves, Ewan Harrison, David K. Jackson, Ian Johnston, Dominic Kwiatkowski, Cordelia Langford, John Sillitoe on behalf of the Wellcome Sanger Institute COVID-19 Surveillance Team |
| EPI_ISL_2730029 | Lighthouse Lab in Alderley Park | Wellcome Sanger Institute for the COVID-19 Genomics UK (COG-UK) Consortium | Jacquelyn Wynn, Mairead Hyland, The Lighthouse Lab in Alderley Park and Alex Alderton, Roberto Amato, Jeffrey Barrett, Sonia Goncalves, Ewan Harrison, David K. Jackson, Ian Johnston, Dominic Kwiatkowski, Cordelia Langford, John Sillitoe on behalf of the Wellcome Sanger Institute COVID-19 Surveillance Team |
| EPI_ISL_2730030 | Lighthouse Lab in Milton Keynes | Wellcome Sanger Institute for the COVID-19 Genomics UK (COG-UK) Consortium | The Lighthouse Lab in Milton Keynes and Alex Alderton, Roberto Amato, Jeffrey Barrett, Sonia Goncalves, Ewan Harrison, David K. Jackson, Ian Johnston, Dominic Kwiatkowski, Cordelia Langford, John Sillitoe on behalf of the Wellcome Sanger Institute COVID-19 Surveillance Team |
| EPI_ISL_2730031 | Lighthouse Lab in Alderley Park | Wellcome Sanger Institute for the COVID-19 Genomics UK (COG-UK) Consortium | Jacquelyn Wynn, Mairead Hyland, The Lighthouse Lab in Alderley Park and Alex Alderton, Roberto Amato, Jeffrey Barrett, Sonia Goncalves, Ewan Harrison, David K. Jackson, Ian Johnston, Dominic Kwiatkowski, Cordelia Langford, John Sillitoe on behalf of the Wellcome Sanger Institute COVID-19 Surveillance Team |
| EPI_ISL_2730032 | Lighthouse Lab in Glasgow | Wellcome Sanger Institute for the COVID-19 Genomics UK (COG-UK) Consortium | Harper VanSteenhouse, Yumi Kasai, David Gray, Carol Clugston, Anna Dominiczak and Alex Alderton, Roberto Amato, Jeffrey Barrett, Sonia Goncalves, Ewan Harrison, David K. Jackson, Ian Johnston, Dominic Kwiatkowski, Cordelia Langford, John Sillitoe on behalf of the Wellcome Sanger Institute COVID-19 Surveillance Team |
| EPI_ISL_2730033 | Lighthouse Lab in Milton Keynes | Wellcome Sanger Institute for the COVID-19 Genomics UK (COG-UK) Consortium | The Lighthouse Lab in Milton Keynes and Alex Alderton, Roberto Amato, Jeffrey Barrett, Sonia Goncalves, Ewan Harrison, David K. Jackson, Ian Johnston, Dominic Kwiatkowski, Cordelia Langford, John Sillitoe on behalf of the Wellcome Sanger Institute COVID-19 Surveillance Team |
| EPI_ISL_2730034 | Lighthouse Lab in Alderley Park | Wellcome Sanger Institute for the COVID-19 Genomics UK (COG-UK) Consortium | Jacquelyn Wynn, Mairead Hyland, The Lighthouse Lab in Alderley Park and Alex Alderton, Roberto Amato, Jeffrey Barrett, Sonia Goncalves, Ewan Harrison, David K. Jackson, Ian Johnston, Dominic Kwiatkowski, Cordelia Langford, John Sillitoe on behalf of the Wellcome Sanger Institute COVID-19 Surveillance Team |
| EPI_ISL_2730035 | Lighthouse Lab in Milton Keynes | Wellcome Sanger Institute for the COVID-19 Genomics UK (COG-UK) Consortium | The Lighthouse Lab in Milton Keynes and Alex Alderton, Roberto Amato, Jeffrey Barrett, Sonia Goncalves, Ewan Harrison, David K. Jackson, Ian Johnston, Dominic Kwiatkowski, Cordelia Langford, John Sillitoe on behalf of the Wellcome Sanger Institute COVID-19 Surveillance Team |
| EPI_ISL_2730036 | Lighthouse Laboratory Plymouth | Wellcome Sanger Institute for the COVID-19 Genomics UK (COG-UK) Consortium | Lighthouse Laboratory Plymouth and Alex Alderton, Roberto Amato, Jeffrey Barrett, Sonia Goncalves, Ewan Harrison, David K. Jackson, Ian Johnston, Dominic Kwiatkowski, Cordelia Langford, John Sillitoe on behalf of the Wellcome Sanger Institute COVID-19 Surveillance Team |
| EPI_ISL_2730037 | Lighthouse Lab in Glasgow | Wellcome Sanger Institute for the COVID-19 Genomics UK (COG-UK) Consortium | Harper VanSteenhouse, Yumi Kasai, David Gray, Carol Clugston, Anna Dominiczak and Alex Alderton, Roberto Amato, Jeffrey Barrett, Sonia Goncalves, Ewan Harrison, David K. Jackson, Ian Johnston, Dominic Kwiatkowski, Cordelia Langford, John Sillitoe on behalf of the Wellcome Sanger Institute COVID-19 Surveillance Team |
| EPI_ISL_2730038 | Lighthouse Lab in Milton Keynes | Wellcome Sanger Institute for the COVID-19 Genomics UK (COG-UK) Consortium | The Lighthouse Lab in Milton Keynes and Alex Alderton, Roberto Amato, Jeffrey Barrett, Sonia Goncalves, Ewan Harrison, David K. Jackson, Ian Johnston, Dominic Kwiatkowski, Cordelia Langford, John Sillitoe on behalf of the Wellcome Sanger Institute COVID-19 Surveillance Team |
| EPI_ISL_2730039 | Lighthouse Lab in Alderley Park | Wellcome Sanger Institute for the COVID-19 Genomics UK (COG-UK) Consortium | Jacquelyn Wynn, Mairead Hyland, The Lighthouse Lab in Alderley Park and Alex Alderton, Roberto Amato, Jeffrey Barrett, Sonia Goncalves, Ewan Harrison, David K. Jackson, Ian Johnston, Dominic Kwiatkowski, Cordelia Langford, John Sillitoe on behalf of the Wellcome Sanger Institute COVID-19 Surveillance Team |
| EPI_ISL_2730040, EPI_ISL_2730041 | Lighthouse Lab in Glasgow | Wellcome Sanger Institute for the COVID-19 Genomics UK (COG-UK) Consortium | Harper VanSteenhouse, Yumi Kasai, David Gray, Carol Clugston, Anna Dominiczak and Alex Alderton, Roberto Amato, Jeffrey Barrett, Sonia Goncalves, Ewan Harrison, David K. Jackson, Ian Johnston, Dominic Kwiatkowski, Cordelia Langford, John Sillitoe on behalf of the Wellcome Sanger Institute COVID-19 Surveillance Team |
| EPI_ISL_2730042 | Lighthouse Lab in Milton Keynes | Wellcome Sanger Institute for the COVID-19 Genomics UK (COG-UK) Consortium | The Lighthouse Lab in Milton Keynes and Alex Alderton, Roberto Amato, Jeffrey Barrett, Sonia Goncalves, Ewan Harrison, David K. Jackson, Ian Johnston, Dominic Kwiatkowski, Cordelia Langford, John Sillitoe on behalf of the Wellcome Sanger Institute COVID-19 Surveillance Team |
| EPI_ISL_2730043 | Lighthouse Laboratory Plymouth | Wellcome Sanger Institute for the COVID-19 Genomics UK (COG-UK) Consortium | Lighthouse Laboratory Plymouth and Alex Alderton, Roberto Amato, Jeffrey Barrett, Sonia Goncalves, Ewan Harrison, David K. Jackson, Ian Johnston, Dominic Kwiatkowski, Cordelia Langford, John Sillitoe on behalf of the Wellcome Sanger Institute COVID-19 Surveillance Team |
| EPI_ISL_2730044 | Lighthouse Lab in Glasgow | Wellcome Sanger Institute for the COVID-19 Genomics UK (COG-UK) Consortium | Harper VanSteenhouse, Yumi Kasai, David Gray, Carol Clugston, Anna Dominiczak and Alex Alderton, Roberto Amato, Jeffrey Barrett, Sonia Goncalves, Ewan Harrison, David K. Jackson, Ian Johnston, Dominic Kwiatkowski, Cordelia Langford, John Sillitoe on behalf of the Wellcome Sanger Institute COVID-19 Surveillance Team |
| EPI_ISL_2730045 | Lighthouse Lab in Milton Keynes | Wellcome Sanger Institute for the COVID-19 Genomics UK (COG-UK) Consortium | The Lighthouse Lab in Milton Keynes and Alex Alderton, Roberto Amato, Jeffrey Barrett, Sonia Goncalves, Ewan Harrison, David K. Jackson, Ian Johnston, Dominic Kwiatkowski, Cordelia Langford, John Sillitoe on behalf of the Wellcome Sanger Institute COVID-19 Surveillance Team |
| EPI_ISL_2730046 | Lighthouse Laboratory Plymouth | Wellcome Sanger Institute for the COVID-19 Genomics UK (COG-UK) Consortium | Lighthouse Laboratory Plymouth and Alex Alderton, Roberto Amato, Jeffrey Barrett, Sonia Goncalves, Ewan Harrison, David K. Jackson, Ian Johnston, Dominic Kwiatkowski, Cordelia Langford, John Sillitoe on behalf of the Wellcome Sanger Institute COVID-19 Surveillance Team |
| EPI_ISL_2730047 | Lighthouse Lab in Alderley Park | Wellcome Sanger Institute for the COVID-19 Genomics UK (COG-UK) Consortium | Jacquelyn Wynn, Mairead Hyland, The Lighthouse Lab in Alderley Park and Alex Alderton, Roberto Amato, Jeffrey Barrett, Sonia Goncalves, Ewan Harrison, David K. Jackson, Ian Johnston, Dominic Kwiatkowski, Cordelia Langford, John Sillitoe on behalf of the Wellcome Sanger Institute COVID-19 Surveillance Team |
| EPI_ISL_2730048 | Lighthouse Lab in Milton Keynes | Wellcome Sanger Institute for the COVID-19 Genomics UK (COG-UK) Consortium | The Lighthouse Lab in Milton Keynes and Alex Alderton, Roberto Amato, Jeffrey Barrett, Sonia Goncalves, Ewan Harrison, David K. Jackson, Ian Johnston, Dominic Kwiatkowski, Cordelia Langford, John Sillitoe on behalf of the Wellcome Sanger Institute COVID-19 Surveillance Team |
| EPI_ISL_2730049, EPI_ISL_2730050, EPI_ISL_2730051 | Lighthouse Laboratory Plymouth | Wellcome Sanger Institute for the COVID-19 Genomics UK (COG-UK) Consortium | Lighthouse Laboratory Plymouth and Alex Alderton, Roberto Amato, Jeffrey Barrett, Sonia Goncalves, Ewan Harrison, David K. Jackson, Ian Johnston, Dominic Kwiatkowski, Cordelia Langford, John Sillitoe on behalf of the Wellcome Sanger Institute COVID-19 Surveillance Team |
| EPI_ISL_2730052 | Lighthouse Lab in Glasgow | Wellcome Sanger Institute for the COVID-19 Genomics UK (COG-UK) Consortium | Harper VanSteenhouse, Yumi Kasai, David Gray, Carol Clugston, Anna Dominiczak and Alex Alderton, Roberto Amato, Jeffrey Barrett, Sonia Goncalves, Ewan Harrison, David K. Jackson, Ian Johnston, Dominic Kwiatkowski, Cordelia Langford, John Sillitoe on behalf of the Wellcome Sanger Institute COVID-19 Surveillance Team |
| EPI_ISL_2730053 | Lighthouse Lab in Alderley Park | Wellcome Sanger Institute for the COVID-19 Genomics UK (COG-UK) Consortium | Jacquelyn Wynn, Mairead Hyland, The Lighthouse Lab in Alderley Park and Alex Alderton, Roberto Amato, Jeffrey Barrett, Sonia Goncalves, Ewan Harrison, David K. Jackson, Ian Johnston, Dominic Kwiatkowski, Cordelia Langford, John Sillitoe on behalf of the Wellcome Sanger Institute COVID-19 Surveillance Team |
| EPI_ISL_2730054 | Lighthouse Laboratory Plymouth | Wellcome Sanger Institute for the COVID-19 Genomics UK (COG-UK) Consortium | Lighthouse Laboratory Plymouth and Alex Alderton, Roberto Amato, Jeffrey Barrett, Sonia Goncalves, Ewan Harrison, David K. Jackson, Ian Johnston, Dominic Kwiatkowski, Cordelia Langford, John Sillitoe on behalf of the Wellcome Sanger Institute COVID-19 Surveillance Team |
| EPI_ISL_2730055 | Lighthouse Lab in Alderley Park | Wellcome Sanger Institute for the COVID-19 Genomics UK (COG-UK) | Jacquelyn Wynn, Mairead Hyland, The Lighthouse Lab in Alderley Park and Alex Alderton, Roberto Amato, Jeffrey Barrett, Sonia Goncalves, Ewan Harrison, David K. Jackson, Ian Johnston, Dominic Kwiatkowski, Cordelia Langford, John Sillitoe on |

| | | | |
|---|---|---|---|
| | | Consortium | behalf of the Wellcome Sanger Institute COVID-19 Surveillance Team |
| EPI_ISL_2730056 | Lighthouse Lab in Glasgow | Wellcome Sanger Institute for the COVID-19 Genomics UK (COG-UK) Consortium | Harper VanSteenhouse, Yumi Kasai, David Gray, Carol Clugston, Anna Dominiczak and Alex Alderton, Roberto Amato, Jeffrey Barrett, Sonia Goncalves, Ewan Harrison, David K. Jackson, Ian Johnston, Dominic Kwiatkowski, Cordelia Langford, John Sillitoe on behalf of the Wellcome Sanger Institute COVID-19 Surveillance Team |
| EPI_ISL_2730057 | Lighthouse Laboratory Plymouth | Wellcome Sanger Institute for the COVID-19 Genomics UK (COG-UK) Consortium | Lighthouse Laboratory Plymouth and Alex Alderton, Roberto Amato, Jeffrey Barrett, Sonia Goncalves, Ewan Harrison, David K. Jackson, Ian Johnston, Dominic Kwiatkowski, Cordelia Langford, John Sillitoe on behalf of the Wellcome Sanger Institute COVID-19 Surveillance Team |
| EPI_ISL_2730058 | Lighthouse Lab in Glasgow | Wellcome Sanger Institute for the COVID-19 Genomics UK (COG-UK) Consortium | Harper VanSteenhouse, Yumi Kasai, David Gray, Carol Clugston, Anna Dominiczak and Alex Alderton, Roberto Amato, Jeffrey Barrett, Sonia Goncalves, Ewan Harrison, David K. Jackson, Ian Johnston, Dominic Kwiatkowski, Cordelia Langford, John Sillitoe on behalf of the Wellcome Sanger Institute COVID-19 Surveillance Team |
| EPI_ISL_2730060 | Lighthouse Laboratory Plymouth | Wellcome Sanger Institute for the COVID-19 Genomics UK (COG-UK) Consortium | Lighthouse Laboratory Plymouth and Alex Alderton, Roberto Amato, Jeffrey Barrett, Sonia Goncalves, Ewan Harrison, David K. Jackson, Ian Johnston, Dominic Kwiatkowski, Cordelia Langford, John Sillitoe on behalf of the Wellcome Sanger Institute COVID-19 Surveillance Team |
| EPI_ISL_2730061 | Lighthouse Lab in Alderley Park | Wellcome Sanger Institute for the COVID-19 Genomics UK (COG-UK) Consortium | Jacquelyn Wynn, Mairead Hyland, The Lighthouse Lab in Alderley Park and Alex Alderton, Roberto Amato, Jeffrey Barrett, Sonia Goncalves, Ewan Harrison, David K. Jackson, Ian Johnston, Dominic Kwiatkowski, Cordelia Langford, John Sillitoe on behalf of the Wellcome Sanger Institute COVID-19 Surveillance Team |
| EPI_ISL_2730062 | Lighthouse Lab in Milton Keynes | Wellcome Sanger Institute for the COVID-19 Genomics UK (COG-UK) Consortium | The Lighthouse Lab in Milton Keynes and Alex Alderton, Roberto Amato, Jeffrey Barrett, Sonia Goncalves, Ewan Harrison, David K. Jackson, Ian Johnston, Dominic Kwiatkowski, Cordelia Langford, John Sillitoe on behalf of the Wellcome Sanger Institute COVID-19 Surveillance Team |
| EPI_ISL_2730063 | Lighthouse Lab in Glasgow | Wellcome Sanger Institute for the COVID-19 Genomics UK (COG-UK) Consortium | Harper VanSteenhouse, Yumi Kasai, David Gray, Carol Clugston, Anna Dominiczak and Alex Alderton, Roberto Amato, Jeffrey Barrett, Sonia Goncalves, Ewan Harrison, David K. Jackson, Ian Johnston, Dominic Kwiatkowski, Cordelia Langford, John Sillitoe on behalf of the Wellcome Sanger Institute COVID-19 Surveillance Team |
| EPI_ISL_2730064, EPI_ISL_2730065, EPI_ISL_2730066, EPI_ISL_2730067, EPI_ISL_2730068 | Lighthouse Lab in Milton Keynes | Wellcome Sanger Institute for the COVID-19 Genomics UK (COG-UK) Consortium | The Lighthouse Lab in Milton Keynes and Alex Alderton, Roberto Amato, Jeffrey Barrett, Sonia Goncalves, Ewan Harrison, David K. Jackson, Ian Johnston, Dominic Kwiatkowski, Cordelia Langford, John Sillitoe on behalf of the Wellcome Sanger Institute COVID-19 Surveillance Team |
| EPI_ISL_2730069, EPI_ISL_2730070 | Lighthouse Lab in Alderley Park | Wellcome Sanger Institute for the COVID-19 Genomics UK (COG-UK) Consortium | Jacquelyn Wynn, Mairead Hyland, The Lighthouse Lab in Alderley Park and Alex Alderton, Roberto Amato, Jeffrey Barrett, Sonia Goncalves, Ewan Harrison, David K. Jackson, Ian Johnston, Dominic Kwiatkowski, Cordelia Langford, John Sillitoe on behalf of the Wellcome Sanger Institute COVID-19 Surveillance Team |
| EPI_ISL_2730071 | Lighthouse Laboratory Plymouth | Wellcome Sanger Institute for the COVID-19 Genomics UK (COG-UK) Consortium | Lighthouse Laboratory Plymouth and Alex Alderton, Roberto Amato, Jeffrey Barrett, Sonia Goncalves, Ewan Harrison, David K. Jackson, Ian Johnston, Dominic Kwiatkowski, Cordelia Langford, John Sillitoe on behalf of the Wellcome Sanger Institute COVID-19 Surveillance Team |
| EPI_ISL_2730072, EPI_ISL_2730073 | Lighthouse Lab in Glasgow | Wellcome Sanger Institute for the COVID-19 Genomics UK (COG-UK) Consortium | Harper VanSteenhouse, Yumi Kasai, David Gray, Carol Clugston, Anna Dominiczak and Alex Alderton, Roberto Amato, Jeffrey Barrett, Sonia Goncalves, Ewan Harrison, David K. Jackson, Ian Johnston, Dominic Kwiatkowski, Cordelia Langford, John Sillitoe on behalf of the Wellcome Sanger Institute COVID-19 Surveillance Team |
| EPI_ISL_2730074 | Lighthouse Lab in Alderley Park | Wellcome Sanger Institute for the COVID-19 Genomics UK (COG-UK) Consortium | Jacquelyn Wynn, Mairead Hyland, The Lighthouse Lab in Alderley Park and Alex Alderton, Roberto Amato, Jeffrey Barrett, Sonia Goncalves, Ewan Harrison, David K. Jackson, Ian Johnston, Dominic Kwiatkowski, Cordelia Langford, John Sillitoe on behalf of the Wellcome Sanger Institute COVID-19 Surveillance Team |
| EPI_ISL_2730075 | Lighthouse Lab in Milton Keynes | Wellcome Sanger Institute for the COVID-19 Genomics UK (COG-UK) Consortium | The Lighthouse Lab in Milton Keynes and Alex Alderton, Roberto Amato, Jeffrey Barrett, Sonia Goncalves, Ewan Harrison, David K. Jackson, Ian Johnston, Dominic Kwiatkowski, Cordelia Langford, John Sillitoe on behalf of the Wellcome Sanger Institute COVID-19 Surveillance Team |
| EPI_ISL_2730076 | Lighthouse Laboratory Plymouth | Wellcome Sanger Institute for the COVID-19 Genomics UK (COG-UK) Consortium | Lighthouse Laboratory Plymouth and Alex Alderton, Roberto Amato, Jeffrey Barrett, Sonia Goncalves, Ewan Harrison, David K. Jackson, Ian Johnston, Dominic Kwiatkowski, Cordelia Langford, John Sillitoe on behalf of the Wellcome Sanger Institute COVID-19 Surveillance Team |
| EPI_ISL_2730077 | Lighthouse Lab in Glasgow | Wellcome Sanger Institute for the COVID-19 Genomics UK (COG-UK) Consortium | Harper VanSteenhouse, Yumi Kasai, David Gray, Carol Clugston, Anna Dominiczak and Alex Alderton, Roberto Amato, Jeffrey Barrett, Sonia Goncalves, Ewan Harrison, David K. Jackson, Ian Johnston, Dominic Kwiatkowski, Cordelia Langford, John Sillitoe on behalf of the Wellcome Sanger Institute COVID-19 Surveillance Team |
| EPI_ISL_2730078 | Lighthouse Laboratory Plymouth | Wellcome Sanger Institute for the COVID-19 Genomics UK (COG-UK) Consortium | Lighthouse Laboratory Plymouth and Alex Alderton, Roberto Amato, Jeffrey Barrett, Sonia Goncalves, Ewan Harrison, David K. Jackson, Ian Johnston, Dominic Kwiatkowski, Cordelia Langford, John Sillitoe on behalf of the Wellcome Sanger Institute COVID-19 Surveillance Team |
| EPI_ISL_2730079, EPI_ISL_2730080 | Lighthouse Lab in Milton Keynes | Wellcome Sanger Institute for the COVID-19 Genomics UK (COG-UK) Consortium | The Lighthouse Lab in Milton Keynes and Alex Alderton, Roberto Amato, Jeffrey Barrett, Sonia Goncalves, Ewan Harrison, David K. Jackson, Ian Johnston, Dominic Kwiatkowski, Cordelia Langford, John Sillitoe on behalf of the Wellcome Sanger Institute COVID-19 Surveillance Team |
| EPI_ISL_2730081 | Lighthouse Laboratory Plymouth | Wellcome Sanger Institute for the COVID-19 Genomics UK (COG-UK) Consortium | Lighthouse Laboratory Plymouth and Alex Alderton, Roberto Amato, Jeffrey Barrett, Sonia Goncalves, Ewan Harrison, David K. Jackson, Ian Johnston, Dominic Kwiatkowski, Cordelia Langford, John Sillitoe on behalf of the Wellcome Sanger Institute COVID-19 Surveillance Team |
| EPI_ISL_2730082 | Lighthouse Lab in Alderley Park | Wellcome Sanger Institute for the COVID-19 Genomics UK (COG-UK) Consortium | Jacquelyn Wynn, Mairead Hyland, The Lighthouse Lab in Alderley Park and Alex Alderton, Roberto Amato, Jeffrey Barrett, Sonia Goncalves, Ewan Harrison, David K. Jackson, Ian Johnston, Dominic Kwiatkowski, Cordelia Langford, John Sillitoe on behalf of the Wellcome Sanger Institute COVID-19 Surveillance Team |
| EPI_ISL_2730083 | Lighthouse Laboratory Plymouth | Wellcome Sanger Institute for the COVID-19 Genomics UK (COG-UK) Consortium | Lighthouse Laboratory Plymouth and Alex Alderton, Roberto Amato, Jeffrey Barrett, Sonia Goncalves, Ewan Harrison, David K. Jackson, Ian Johnston, Dominic Kwiatkowski, Cordelia Langford, John Sillitoe on behalf of the Wellcome Sanger Institute COVID-19 Surveillance Team |
| EPI_ISL_2730084 | Lighthouse Lab in Glasgow | Wellcome Sanger Institute for the COVID-19 Genomics UK (COG-UK) Consortium | Harper VanSteenhouse, Yumi Kasai, David Gray, Carol Clugston, Anna Dominiczak and Alex Alderton, Roberto Amato, Jeffrey Barrett, Sonia Goncalves, Ewan Harrison, David K. Jackson, Ian Johnston, Dominic Kwiatkowski, Cordelia Langford, John Sillitoe on behalf of the Wellcome Sanger Institute COVID-19 Surveillance Team |
| EPI_ISL_2730085 | Lighthouse Laboratory Plymouth | Wellcome Sanger Institute for the COVID-19 Genomics UK (COG-UK) Consortium | Lighthouse Laboratory Plymouth and Alex Alderton, Roberto Amato, Jeffrey Barrett, Sonia Goncalves, Ewan Harrison, David K. Jackson, Ian Johnston, Dominic Kwiatkowski, Cordelia Langford, John Sillitoe on behalf of the Wellcome Sanger Institute COVID-19 Surveillance Team |
| EPI_ISL_2730086 | Lighthouse Lab in Glasgow | Wellcome Sanger Institute for the COVID-19 Genomics UK (COG-UK) Consortium | Harper VanSteenhouse, Yumi Kasai, David Gray, Carol Clugston, Anna Dominiczak and Alex Alderton, Roberto Amato, Jeffrey Barrett, Sonia Goncalves, Ewan Harrison, David K. Jackson, Ian Johnston, Dominic Kwiatkowski, Cordelia Langford, John Sillitoe on behalf of the Wellcome Sanger Institute COVID-19 Surveillance Team |
| EPI_ISL_2730087 | Lighthouse Laboratory Plymouth | Wellcome Sanger Institute for the COVID-19 Genomics UK (COG-UK) Consortium | Lighthouse Laboratory Plymouth and Alex Alderton, Roberto Amato, Jeffrey Barrett, Sonia Goncalves, Ewan Harrison, David K. Jackson, Ian Johnston, Dominic Kwiatkowski, Cordelia Langford, John Sillitoe on behalf of the Wellcome Sanger Institute COVID-19 Surveillance Team |
| EPI_ISL_2730088 | Lighthouse Lab in Alderley Park | Wellcome Sanger Institute for the COVID-19 Genomics UK (COG-UK) Consortium | Jacquelyn Wynn, Mairead Hyland, The Lighthouse Lab in Alderley Park and Alex Alderton, Roberto Amato, Jeffrey Barrett, Sonia Goncalves, Ewan Harrison, David K. Jackson, Ian Johnston, Dominic Kwiatkowski, Cordelia Langford, John Sillitoe on behalf of the Wellcome Sanger Institute COVID-19 Surveillance Team |
| EPI_ISL_2730089 | Lighthouse Laboratory Plymouth | Wellcome Sanger Institute for the COVID-19 Genomics UK (COG-UK) Consortium | Lighthouse Laboratory Plymouth and Alex Alderton, Roberto Amato, Jeffrey Barrett, Sonia Goncalves, Ewan Harrison, David K. Jackson, Ian Johnston, Dominic Kwiatkowski, Cordelia Langford, John Sillitoe on behalf of the Wellcome Sanger Institute COVID-19 Surveillance Team |
| EPI_ISL_2730090 | Lighthouse Lab in Glasgow | Wellcome Sanger Institute for the COVID-19 Genomics UK (COG-UK) Consortium | Harper VanSteenhouse, Yumi Kasai, David Gray, Carol Clugston, Anna Dominiczak and Alex Alderton, Roberto Amato, Jeffrey Barrett, Sonia Goncalves, Ewan Harrison, David K. Jackson, Ian Johnston, Dominic Kwiatkowski, Cordelia Langford, John Sillitoe on behalf of the Wellcome Sanger Institute COVID-19 Surveillance Team |
| EPI_ISL_2730091, EPI_ISL_2730092 | Lighthouse Lab in Milton Keynes | Wellcome Sanger Institute for the COVID-19 Genomics UK (COG-UK) Consortium | The Lighthouse Lab in Milton Keynes and Alex Alderton, Roberto Amato, Jeffrey Barrett, Sonia Goncalves, Ewan Harrison, David K. Jackson, Ian Johnston, Dominic Kwiatkowski, Cordelia Langford, John Sillitoe on behalf of the Wellcome Sanger Institute COVID-19 Surveillance Team |
| EPI_ISL_2730093, EPI_ISL_2730094 | Lighthouse Laboratory Plymouth | Wellcome Sanger Institute for the COVID-19 Genomics UK (COG-UK) Consortium | Lighthouse Laboratory Plymouth and Alex Alderton, Roberto Amato, Jeffrey Barrett, Sonia Goncalves, Ewan Harrison, David K. Jackson, Ian Johnston, Dominic Kwiatkowski, Cordelia Langford, John Sillitoe on behalf of the Wellcome Sanger Institute COVID-19 Surveillance Team |
| EPI_ISL_2730095 | Lighthouse Lab in Alderley Park | Wellcome Sanger Institute for the COVID-19 Genomics UK (COG-UK) Consortium | Jacquelyn Wynn, Mairead Hyland, The Lighthouse Lab in Alderley Park and Alex Alderton, Roberto Amato, Jeffrey Barrett, Sonia Goncalves, Ewan Harrison, David K. Jackson, Ian Johnston, Dominic Kwiatkowski, Cordelia Langford, John Sillitoe on behalf of the Wellcome Sanger Institute COVID-19 Surveillance Team |
| EPI_ISL_2730096, EPI_ISL_2730097 | Lighthouse Lab in Glasgow | Wellcome Sanger Institute for the COVID-19 Genomics UK (COG-UK) Consortium | Harper VanSteenhouse, Yumi Kasai, David Gray, Carol Clugston, Anna Dominiczak and Alex Alderton, Roberto Amato, Jeffrey Barrett, Sonia Goncalves, Ewan Harrison, David K. Jackson, Ian Johnston, Dominic Kwiatkowski, Cordelia Langford, John Sillitoe on behalf of the Wellcome Sanger Institute COVID-19 Surveillance Team |
| EPI_ISL_2730098, EPI_ISL_2730099 | Lighthouse Laboratory Plymouth | Wellcome Sanger Institute for the COVID-19 Genomics UK (COG-UK) Consortium | Lighthouse Laboratory Plymouth and Alex Alderton, Roberto Amato, Jeffrey Barrett, Sonia Goncalves, Ewan Harrison, David K. Jackson, Ian Johnston, Dominic Kwiatkowski, Cordelia Langford, John Sillitoe on behalf of the Wellcome Sanger Institute COVID-19 Surveillance Team |
| EPI_ISL_2730100 | Lighthouse Lab in Glasgow | Wellcome Sanger Institute for the COVID-19 Genomics UK (COG-UK) Consortium | Harper VanSteenhouse, Yumi Kasai, David Gray, Carol Clugston, Anna Dominiczak and Alex Alderton, Roberto Amato, Jeffrey Barrett, Sonia Goncalves, Ewan Harrison, David K. Jackson, Ian Johnston, Dominic Kwiatkowski, Cordelia Langford, John Sillitoe on behalf of the Wellcome Sanger Institute COVID-19 Surveillance Team |
| EPI_ISL_2730101 | Lighthouse Lab in Alderley Park | Wellcome Sanger Institute for the COVID-19 Genomics UK (COG-UK) Consortium | Jacquelyn Wynn, Mairead Hyland, The Lighthouse Lab in Alderley Park and Alex Alderton, Roberto Amato, Jeffrey Barrett, Sonia Goncalves, Ewan Harrison, David K. Jackson, Ian Johnston, Dominic Kwiatkowski, Cordelia Langford, John Sillitoe on behalf of the Wellcome Sanger Institute COVID-19 Surveillance Team |
| EPI_ISL_2730102, EPI_ISL_2730103, EPI_ISL_2730104, EPI_ISL_2730105 | Lighthouse Laboratory Plymouth | Wellcome Sanger Institute for the COVID-19 Genomics UK (COG-UK) Consortium | Lighthouse Laboratory Plymouth and Alex Alderton, Roberto Amato, Jeffrey Barrett, Sonia Goncalves, Ewan Harrison, David K. Jackson, Ian Johnston, Dominic Kwiatkowski, Cordelia Langford, John Sillitoe on behalf of the Wellcome Sanger Institute COVID-19 Surveillance Team |
| EPI_ISL_2730106 | Lighthouse Lab in Alderley Park | Wellcome Sanger Institute for the COVID-19 Genomics UK (COG-UK) Consortium | Jacquelyn Wynn, Mairead Hyland, The Lighthouse Lab in Alderley Park and Alex Alderton, Roberto Amato, Jeffrey Barrett, Sonia Goncalves, Ewan Harrison, David K. Jackson, Ian Johnston, Dominic Kwiatkowski, Cordelia Langford, John Sillitoe on behalf of the Wellcome Sanger Institute COVID-19 Surveillance Team |
| EPI_ISL_2730107, EPI_ISL_2730108 | Lighthouse Laboratory Plymouth | Wellcome Sanger Institute for the COVID-19 Genomics UK (COG-UK) Consortium | Lighthouse Laboratory Plymouth and Alex Alderton, Roberto Amato, Jeffrey Barrett, Sonia Goncalves, Ewan Harrison, David K. Jackson, Ian Johnston, Dominic Kwiatkowski, Cordelia Langford, John Sillitoe on behalf of the Wellcome Sanger Institute COVID-19 Surveillance Team |
| EPI_ISL_2730109, EPI_ISL_2730110, EPI_ISL_2730111 | Lighthouse Lab in Glasgow | Wellcome Sanger Institute for the COVID-19 Genomics UK (COG-UK) Consortium | Harper VanSteenhouse, Yumi Kasai, David Gray, Carol Clugston, Anna Dominiczak and Alex Alderton, Roberto Amato, Jeffrey Barrett, Sonia Goncalves, Ewan Harrison, David K. Jackson, Ian Johnston, Dominic Kwiatkowski, Cordelia Langford, John Sillitoe on behalf of the Wellcome Sanger Institute COVID-19 Surveillance Team |
| EPI_ISL_2730112 | Lighthouse Laboratory Plymouth | Wellcome Sanger Institute for the COVID-19 Genomics UK (COG-UK) Consortium | Lighthouse Laboratory Plymouth and Alex Alderton, Roberto Amato, Jeffrey Barrett, Sonia Goncalves, Ewan Harrison, David K. Jackson, Ian Johnston, Dominic Kwiatkowski, Cordelia Langford, John Sillitoe on behalf of the Wellcome Sanger Institute COVID-19 Surveillance Team |
| EPI_ISL_2730113, EPI_ISL_2730114, EPI_ISL_2730115 | Lighthouse Lab in Milton Keynes | Wellcome Sanger Institute for the COVID-19 Genomics UK (COG-UK) Consortium | The Lighthouse Lab in Milton Keynes and Alex Alderton, Roberto Amato, Jeffrey Barrett, Sonia Goncalves, Ewan Harrison, David K. Jackson, Ian Johnston, Dominic Kwiatkowski, Cordelia Langford, John Sillitoe on behalf of the Wellcome Sanger Institute COVID-19 Surveillance Team |
| EPI_ISL_2730116 | Lighthouse Lab in Alderley Park | Wellcome Sanger Institute for the COVID-19 Genomics UK (COG-UK) Consortium | Jacquelyn Wynn, Mairead Hyland, The Lighthouse Lab in Alderley Park and Alex Alderton, Roberto Amato, Jeffrey Barrett, Sonia Goncalves, Ewan Harrison, David K. Jackson, Ian Johnston, Dominic Kwiatkowski, Cordelia Langford, John Sillitoe on behalf of the Wellcome Sanger Institute COVID-19 Surveillance Team |
| EPI_ISL_2730117 | Lighthouse Lab in Glasgow | Wellcome Sanger Institute for the COVID-19 Genomics UK (COG-UK) | Harper VanSteenhouse, Yumi Kasai, David Gray, Carol Clugston, Anna Dominiczak and Alex Alderton, Roberto Amato, Jeffrey Barrett, Sonia Goncalves, Ewan Harrison, David K. Jackson, Ian Johnston, Dominic Kwiatkowski, Cordelia Langford, |

| | | | |
|---|---|---|---|
| | | Consortium | John Sillitoe on behalf of the Wellcome Sanger Institute COVID-19 Surveillance Team |
| EPI_ISL_2730118, EPI_ISL_2730119 | Lighthouse Lab in Milton Keynes | Wellcome Sanger Institute for the COVID-19 Genomics UK (COG-UK) Consortium | The Lighthouse Lab in Milton Keynes and Alex Alderton, Roberto Amato, Jeffrey Barrett, Sonia Goncalves, Ewan Harrison, David K. Jackson, Ian Johnston, Dominic Kwiatkowski, Cordelia Langford, John Sillitoe on behalf of the Wellcome Sanger Institute COVID-19 Surveillance Team |
| EPI_ISL_2730120 | Lighthouse Lab in Glasgow | Wellcome Sanger Institute for the COVID-19 Genomics UK (COG-UK) Consortium | Harper VanSteenhouse, Yumi Kasai, David Gray, Carol Clugston, Anna Dominiczak and Alex Alderton, Roberto Amato, Jeffrey Barrett, Sonia Goncalves, Ewan Harrison, David K. Jackson, Ian Johnston, Dominic Kwiatkowski, Cordelia Langford, John Sillitoe on behalf of the Wellcome Sanger Institute COVID-19 Surveillance Team |
| EPI_ISL_2730121 | Lighthouse Lab in Milton Keynes | Wellcome Sanger Institute for the COVID-19 Genomics UK (COG-UK) Consortium | The Lighthouse Lab in Milton Keynes and Alex Alderton, Roberto Amato, Jeffrey Barrett, Sonia Goncalves, Ewan Harrison, David K. Jackson, Ian Johnston, Dominic Kwiatkowski, Cordelia Langford, John Sillitoe on behalf of the Wellcome Sanger Institute COVID-19 Surveillance Team |
| EPI_ISL_2730122, EPI_ISL_2730123 | Lighthouse Lab in Alderley Park | Wellcome Sanger Institute for the COVID-19 Genomics UK (COG-UK) Consortium | Jacquelyn Wynn, Mairead Hyland, The Lighthouse Lab in Alderley Park and Alex Alderton, Roberto Amato, Jeffrey Barrett, Sonia Goncalves, Ewan Harrison, David K. Jackson, Ian Johnston, Dominic Kwiatkowski, Cordelia Langford, John Sillitoe on behalf of the Wellcome Sanger Institute COVID-19 Surveillance Team |
| EPI_ISL_2730124, EPI_ISL_2730125 | Lighthouse Lab in Milton Keynes | Wellcome Sanger Institute for the COVID-19 Genomics UK (COG-UK) Consortium | The Lighthouse Lab in Milton Keynes and Alex Alderton, Roberto Amato, Jeffrey Barrett, Sonia Goncalves, Ewan Harrison, David K. Jackson, Ian Johnston, Dominic Kwiatkowski, Cordelia Langford, John Sillitoe on behalf of the Wellcome Sanger Institute COVID-19 Surveillance Team |
| EPI_ISL_2730126 | Lighthouse Lab in Glasgow | Wellcome Sanger Institute for the COVID-19 Genomics UK (COG-UK) Consortium | Harper VanSteenhouse, Yumi Kasai, David Gray, Carol Clugston, Anna Dominiczak and Alex Alderton, Roberto Amato, Jeffrey Barrett, Sonia Goncalves, Ewan Harrison, David K. Jackson, Ian Johnston, Dominic Kwiatkowski, Cordelia Langford, John Sillitoe on behalf of the Wellcome Sanger Institute COVID-19 Surveillance Team |
| EPI_ISL_2730127 | Lighthouse Lab in Alderley Park | Wellcome Sanger Institute for the COVID-19 Genomics UK (COG-UK) Consortium | Jacquelyn Wynn, Mairead Hyland, The Lighthouse Lab in Alderley Park and Alex Alderton, Roberto Amato, Jeffrey Barrett, Sonia Goncalves, Ewan Harrison, David K. Jackson, Ian Johnston, Dominic Kwiatkowski, Cordelia Langford, John Sillitoe on behalf of the Wellcome Sanger Institute COVID-19 Surveillance Team |
| EPI_ISL_2730128, EPI_ISL_2730129 | Lighthouse Laboratory Plymouth | Wellcome Sanger Institute for the COVID-19 Genomics UK (COG-UK) Consortium | Lighthouse Laboratory Plymouth and Alex Alderton, Roberto Amato, Jeffrey Barrett, Sonia Goncalves, Ewan Harrison, David K. Jackson, Ian Johnston, Dominic Kwiatkowski, Cordelia Langford, John Sillitoe on behalf of the Wellcome Sanger Institute COVID-19 Surveillance Team |
| EPI_ISL_2730130 | Lighthouse Lab in Glasgow | Wellcome Sanger Institute for the COVID-19 Genomics UK (COG-UK) Consortium | Harper VanSteenhouse, Yumi Kasai, David Gray, Carol Clugston, Anna Dominiczak and Alex Alderton, Roberto Amato, Jeffrey Barrett, Sonia Goncalves, Ewan Harrison, David K. Jackson, Ian Johnston, Dominic Kwiatkowski, Cordelia Langford, John Sillitoe on behalf of the Wellcome Sanger Institute COVID-19 Surveillance Team |
| EPI_ISL_2730131 | Lighthouse Lab in Alderley Park | Wellcome Sanger Institute for the COVID-19 Genomics UK (COG-UK) Consortium | Jacquelyn Wynn, Mairead Hyland, The Lighthouse Lab in Alderley Park and Alex Alderton, Roberto Amato, Jeffrey Barrett, Sonia Goncalves, Ewan Harrison, David K. Jackson, Ian Johnston, Dominic Kwiatkowski, Cordelia Langford, John Sillitoe on behalf of the Wellcome Sanger Institute COVID-19 Surveillance Team |
| EPI_ISL_2730132 | Lighthouse Lab in Glasgow | Wellcome Sanger Institute for the COVID-19 Genomics UK (COG-UK) Consortium | Harper VanSteenhouse, Yumi Kasai, David Gray, Carol Clugston, Anna Dominiczak and Alex Alderton, Roberto Amato, Jeffrey Barrett, Sonia Goncalves, Ewan Harrison, David K. Jackson, Ian Johnston, Dominic Kwiatkowski, Cordelia Langford, John Sillitoe on behalf of the Wellcome Sanger Institute COVID-19 Surveillance Team |
| EPI_ISL_2730133 | Lighthouse Lab in Milton Keynes | Wellcome Sanger Institute for the COVID-19 Genomics UK (COG-UK) Consortium | The Lighthouse Lab in Milton Keynes and Alex Alderton, Roberto Amato, Jeffrey Barrett, Sonia Goncalves, Ewan Harrison, David K. Jackson, Ian Johnston, Dominic Kwiatkowski, Cordelia Langford, John Sillitoe on behalf of the Wellcome Sanger Institute COVID-19 Surveillance Team |
| EPI_ISL_2730134 | Lighthouse Lab in Glasgow | Wellcome Sanger Institute for the COVID-19 Genomics UK (COG-UK) Consortium | Harper VanSteenhouse, Yumi Kasai, David Gray, Carol Clugston, Anna Dominiczak and Alex Alderton, Roberto Amato, Jeffrey Barrett, Sonia Goncalves, Ewan Harrison, David K. Jackson, Ian Johnston, Dominic Kwiatkowski, Cordelia Langford, John Sillitoe on behalf of the Wellcome Sanger Institute COVID-19 Surveillance Team |
| EPI_ISL_2730135, EPI_ISL_2730136, EPI_ISL_2730137 | Lighthouse Lab in Milton Keynes | Wellcome Sanger Institute for the COVID-19 Genomics UK (COG-UK) Consortium | The Lighthouse Lab in Milton Keynes and Alex Alderton, Roberto Amato, Jeffrey Barrett, Sonia Goncalves, Ewan Harrison, David K. Jackson, Ian Johnston, Dominic Kwiatkowski, Cordelia Langford, John Sillitoe on behalf of the Wellcome Sanger Institute COVID-19 Surveillance Team |
| EPI_ISL_2730138 | Lighthouse Laboratory Plymouth | Wellcome Sanger Institute for the COVID-19 Genomics UK (COG-UK) Consortium | Lighthouse Laboratory Plymouth and Alex Alderton, Roberto Amato, Jeffrey Barrett, Sonia Goncalves, Ewan Harrison, David K. Jackson, Ian Johnston, Dominic Kwiatkowski, Cordelia Langford, John Sillitoe on behalf of the Wellcome Sanger Institute COVID-19 Surveillance Team |
| EPI_ISL_2730139, EPI_ISL_2730140 | Lighthouse Lab in Milton Keynes | Wellcome Sanger Institute for the COVID-19 Genomics UK (COG-UK) Consortium | The Lighthouse Lab in Milton Keynes and Alex Alderton, Roberto Amato, Jeffrey Barrett, Sonia Goncalves, Ewan Harrison, David K. Jackson, Ian Johnston, Dominic Kwiatkowski, Cordelia Langford, John Sillitoe on behalf of the Wellcome Sanger Institute COVID-19 Surveillance Team |
| EPI_ISL_2730141 | Lighthouse Laboratory Plymouth | Wellcome Sanger Institute for the COVID-19 Genomics UK (COG-UK) Consortium | Lighthouse Laboratory Plymouth and Alex Alderton, Roberto Amato, Jeffrey Barrett, Sonia Goncalves, Ewan Harrison, David K. Jackson, Ian Johnston, Dominic Kwiatkowski, Cordelia Langford, John Sillitoe on behalf of the Wellcome Sanger Institute COVID-19 Surveillance Team |
| EPI_ISL_2730142 | Lighthouse Lab in Alderley Park | Wellcome Sanger Institute for the COVID-19 Genomics UK (COG-UK) Consortium | Jacquelyn Wynn, Mairead Hyland, The Lighthouse Lab in Alderley Park and Alex Alderton, Roberto Amato, Jeffrey Barrett, Sonia Goncalves, Ewan Harrison, David K. Jackson, Ian Johnston, Dominic Kwiatkowski, Cordelia Langford, John Sillitoe on behalf of the Wellcome Sanger Institute COVID-19 Surveillance Team |
| EPI_ISL_2730143, EPI_ISL_2730144, EPI_ISL_2730145 | Lighthouse Lab in Glasgow | Wellcome Sanger Institute for the COVID-19 Genomics UK (COG-UK) Consortium | Harper VanSteenhouse, Yumi Kasai, David Gray, Carol Clugston, Anna Dominiczak and Alex Alderton, Roberto Amato, Jeffrey Barrett, Sonia Goncalves, Ewan Harrison, David K. Jackson, Ian Johnston, Dominic Kwiatkowski, Cordelia Langford, John Sillitoe on behalf of the Wellcome Sanger Institute COVID-19 Surveillance Team |
| EPI_ISL_2730146 | Lighthouse Laboratory Plymouth | Wellcome Sanger Institute for the COVID-19 Genomics UK (COG-UK) Consortium | Lighthouse Laboratory Plymouth and Alex Alderton, Roberto Amato, Jeffrey Barrett, Sonia Goncalves, Ewan Harrison, David K. Jackson, Ian Johnston, Dominic Kwiatkowski, Cordelia Langford, John Sillitoe on behalf of the Wellcome Sanger Institute COVID-19 Surveillance Team |
| EPI_ISL_2730147 | Lighthouse Lab in Glasgow | Wellcome Sanger Institute for the COVID-19 Genomics UK (COG-UK) Consortium | Harper VanSteenhouse, Yumi Kasai, David Gray, Carol Clugston, Anna Dominiczak and Alex Alderton, Roberto Amato, Jeffrey Barrett, Sonia Goncalves, Ewan Harrison, David K. Jackson, Ian Johnston, Dominic Kwiatkowski, Cordelia Langford, John Sillitoe on behalf of the Wellcome Sanger Institute COVID-19 Surveillance Team |
| EPI_ISL_2730148 | Lighthouse Lab in Alderley Park | Wellcome Sanger Institute for the COVID-19 Genomics UK (COG-UK) Consortium | Jacquelyn Wynn, Mairead Hyland, The Lighthouse Lab in Alderley Park and Alex Alderton, Roberto Amato, Jeffrey Barrett, Sonia Goncalves, Ewan Harrison, David K. Jackson, Ian Johnston, Dominic Kwiatkowski, Cordelia Langford, John Sillitoe on behalf of the Wellcome Sanger Institute COVID-19 Surveillance Team |
| EPI_ISL_2730149 | Lighthouse Lab in Glasgow | Wellcome Sanger Institute for the COVID-19 Genomics UK (COG-UK) Consortium | Harper VanSteenhouse, Yumi Kasai, David Gray, Carol Clugston, Anna Dominiczak and Alex Alderton, Roberto Amato, Jeffrey Barrett, Sonia Goncalves, Ewan Harrison, David K. Jackson, Ian Johnston, Dominic Kwiatkowski, Cordelia Langford, John Sillitoe on behalf of the Wellcome Sanger Institute COVID-19 Surveillance Team |
| EPI_ISL_2730150 | Lighthouse Lab in Alderley Park | Wellcome Sanger Institute for the COVID-19 Genomics UK (COG-UK) Consortium | Jacquelyn Wynn, Mairead Hyland, The Lighthouse Lab in Alderley Park and Alex Alderton, Roberto Amato, Jeffrey Barrett, Sonia Goncalves, Ewan Harrison, David K. Jackson, Ian Johnston, Dominic Kwiatkowski, Cordelia Langford, John Sillitoe on behalf of the Wellcome Sanger Institute COVID-19 Surveillance Team |
| EPI_ISL_2730151 | Lighthouse Laboratory Plymouth | Wellcome Sanger Institute for the COVID-19 Genomics UK (COG-UK) Consortium | Lighthouse Laboratory Plymouth and Alex Alderton, Roberto Amato, Jeffrey Barrett, Sonia Goncalves, Ewan Harrison, David K. Jackson, Ian Johnston, Dominic Kwiatkowski, Cordelia Langford, John Sillitoe on behalf of the Wellcome Sanger Institute COVID-19 Surveillance Team |
| EPI_ISL_2730152, EPI_ISL_2730153 | Lighthouse Lab in Milton Keynes | Wellcome Sanger Institute for the COVID-19 Genomics UK (COG-UK) Consortium | The Lighthouse Lab in Milton Keynes and Alex Alderton, Roberto Amato, Jeffrey Barrett, Sonia Goncalves, Ewan Harrison, David K. Jackson, Ian Johnston, Dominic Kwiatkowski, Cordelia Langford, John Sillitoe on behalf of the Wellcome Sanger Institute COVID-19 Surveillance Team |
| EPI_ISL_2730154 | Lighthouse Lab in Alderley Park | Wellcome Sanger Institute for the COVID-19 Genomics UK (COG-UK) Consortium | Jacquelyn Wynn, Mairead Hyland, The Lighthouse Lab in Alderley Park and Alex Alderton, Roberto Amato, Jeffrey Barrett, Sonia Goncalves, Ewan Harrison, David K. Jackson, Ian Johnston, Dominic Kwiatkowski, Cordelia Langford, John Sillitoe on behalf of the Wellcome Sanger Institute COVID-19 Surveillance Team |
| EPI_ISL_2730155 | Lighthouse Lab in Glasgow | Wellcome Sanger Institute for the COVID-19 Genomics UK (COG-UK) Consortium | Harper VanSteenhouse, Yumi Kasai, David Gray, Carol Clugston, Anna Dominiczak and Alex Alderton, Roberto Amato, Jeffrey Barrett, Sonia Goncalves, Ewan Harrison, David K. Jackson, Ian Johnston, Dominic Kwiatkowski, Cordelia Langford, John Sillitoe on behalf of the Wellcome Sanger Institute COVID-19 Surveillance Team |
| EPI_ISL_2730156 | Lighthouse Lab in Milton Keynes | Wellcome Sanger Institute for the COVID-19 Genomics UK (COG-UK) Consortium | The Lighthouse Lab in Milton Keynes and Alex Alderton, Roberto Amato, Jeffrey Barrett, Sonia Goncalves, Ewan Harrison, David K. Jackson, Ian Johnston, Dominic Kwiatkowski, Cordelia Langford, John Sillitoe on behalf of the Wellcome Sanger Institute COVID-19 Surveillance Team |
| EPI_ISL_2730157, EPI_ISL_2730158, EPI_ISL_2730159 | Lighthouse Lab in Alderley Park | Wellcome Sanger Institute for the COVID-19 Genomics UK (COG-UK) Consortium | Jacquelyn Wynn, Mairead Hyland, The Lighthouse Lab in Alderley Park and Alex Alderton, Roberto Amato, Jeffrey Barrett, Sonia Goncalves, Ewan Harrison, David K. Jackson, Ian Johnston, Dominic Kwiatkowski, Cordelia Langford, John Sillitoe on behalf of the Wellcome Sanger Institute COVID-19 Surveillance Team |
| EPI_ISL_2730160 | Lighthouse Laboratory Plymouth | Wellcome Sanger Institute for the COVID-19 Genomics UK (COG-UK) Consortium | Lighthouse Laboratory Plymouth and Alex Alderton, Roberto Amato, Jeffrey Barrett, Sonia Goncalves, Ewan Harrison, David K. Jackson, Ian Johnston, Dominic Kwiatkowski, Cordelia Langford, John Sillitoe on behalf of the Wellcome Sanger Institute COVID-19 Surveillance Team |
| EPI_ISL_2730161, EPI_ISL_2730162 | Lighthouse Lab in Glasgow | Wellcome Sanger Institute for the COVID-19 Genomics UK (COG-UK) Consortium | Harper VanSteenhouse, Yumi Kasai, David Gray, Carol Clugston, Anna Dominiczak and Alex Alderton, Roberto Amato, Jeffrey Barrett, Sonia Goncalves, Ewan Harrison, David K. Jackson, Ian Johnston, Dominic Kwiatkowski, Cordelia Langford, John Sillitoe on behalf of the Wellcome Sanger Institute COVID-19 Surveillance Team |
| EPI_ISL_2730163 | Lighthouse Lab in Milton Keynes | Wellcome Sanger Institute for the COVID-19 Genomics UK (COG-UK) Consortium | The Lighthouse Lab in Milton Keynes and Alex Alderton, Roberto Amato, Jeffrey Barrett, Sonia Goncalves, Ewan Harrison, David K. Jackson, Ian Johnston, Dominic Kwiatkowski, Cordelia Langford, John Sillitoe on behalf of the Wellcome Sanger Institute COVID-19 Surveillance Team |
| EPI_ISL_2730164 | Lighthouse Lab in Glasgow | Wellcome Sanger Institute for the COVID-19 Genomics UK (COG-UK) Consortium | Harper VanSteenhouse, Yumi Kasai, David Gray, Carol Clugston, Anna Dominiczak and Alex Alderton, Roberto Amato, Jeffrey Barrett, Sonia Goncalves, Ewan Harrison, David K. Jackson, Ian Johnston, Dominic Kwiatkowski, Cordelia Langford, John Sillitoe on behalf of the Wellcome Sanger Institute COVID-19 Surveillance Team |
| EPI_ISL_2730165, EPI_ISL_2730166, EPI_ISL_2730167, EPI_ISL_2730168 | Lighthouse Lab in Alderley Park | Wellcome Sanger Institute for the COVID-19 Genomics UK (COG-UK) Consortium | Jacquelyn Wynn, Mairead Hyland, The Lighthouse Lab in Alderley Park and Alex Alderton, Roberto Amato, Jeffrey Barrett, Sonia Goncalves, Ewan Harrison, David K. Jackson, Ian Johnston, Dominic Kwiatkowski, Cordelia Langford, John Sillitoe on behalf of the Wellcome Sanger Institute COVID-19 Surveillance Team |
| EPI_ISL_2730169, EPI_ISL_2730170, EPI_ISL_2730171, EPI_ISL_2730172 | Lighthouse Lab in Milton Keynes | Wellcome Sanger Institute for the COVID-19 Genomics UK (COG-UK) Consortium | The Lighthouse Lab in Milton Keynes and Alex Alderton, Roberto Amato, Jeffrey Barrett, Sonia Goncalves, Ewan Harrison, David K. Jackson, Ian Johnston, Dominic Kwiatkowski, Cordelia Langford, John Sillitoe on behalf of the Wellcome Sanger Institute COVID-19 Surveillance Team |
| EPI_ISL_2730173 | Lighthouse Lab in Glasgow | Wellcome Sanger Institute for the COVID-19 Genomics UK (COG-UK) Consortium | Harper VanSteenhouse, Yumi Kasai, David Gray, Carol Clugston, Anna Dominiczak and Alex Alderton, Roberto Amato, Jeffrey Barrett, Sonia Goncalves, Ewan Harrison, David K. Jackson, Ian Johnston, Dominic Kwiatkowski, Cordelia Langford, John Sillitoe on behalf of the Wellcome Sanger Institute COVID-19 Surveillance Team |
| EPI_ISL_2730174 | Lighthouse Lab in Alderley Park | Wellcome Sanger Institute for the COVID-19 Genomics UK (COG-UK) Consortium | Jacquelyn Wynn, Mairead Hyland, The Lighthouse Lab in Alderley Park and Alex Alderton, Roberto Amato, Jeffrey Barrett, Sonia Goncalves, Ewan Harrison, David K. Jackson, Ian Johnston, Dominic Kwiatkowski, Cordelia Langford, John Sillitoe on behalf of the Wellcome Sanger Institute COVID-19 Surveillance Team |
| EPI_ISL_2730175 | Lighthouse Laboratory Plymouth | Wellcome Sanger Institute for the COVID-19 Genomics UK (COG-UK) Consortium | Lighthouse Laboratory Plymouth and Alex Alderton, Roberto Amato, Jeffrey Barrett, Sonia Goncalves, Ewan Harrison, David K. Jackson, Ian Johnston, Dominic Kwiatkowski, Cordelia Langford, John Sillitoe on behalf of the Wellcome Sanger Institute COVID-19 Surveillance Team |
| EPI_ISL_2730176 | Lighthouse Lab in Milton Keynes | Wellcome Sanger Institute for the COVID-19 Genomics UK (COG-UK) Consortium | The Lighthouse Lab in Milton Keynes and Alex Alderton, Roberto Amato, Jeffrey Barrett, Sonia Goncalves, Ewan Harrison, David K. Jackson, Ian Johnston, Dominic Kwiatkowski, Cordelia Langford, John Sillitoe on behalf of the Wellcome Sanger Institute COVID-19 Surveillance Team |
| EPI_ISL_2730177 | Lighthouse Lab in Alderley Park | Wellcome Sanger Institute for the COVID-19 Genomics UK (COG-UK) Consortium | Jacquelyn Wynn, Mairead Hyland, The Lighthouse Lab in Alderley Park and Alex Alderton, Roberto Amato, Jeffrey Barrett, Sonia Goncalves, Ewan Harrison, David K. Jackson, Ian Johnston, Dominic Kwiatkowski, Cordelia Langford, John Sillitoe on behalf of the Wellcome Sanger Institute COVID-19 Surveillance Team |
| EPI_ISL_2730178 | Lighthouse Lab in Milton Keynes | Wellcome Sanger Institute for the COVID-19 Genomics UK (COG-UK) | The Lighthouse Lab in Milton Keynes and Alex Alderton, Roberto Amato, Jeffrey Barrett, Sonia Goncalves, Ewan Harrison, David K. Jackson, Ian Johnston, Dominic Kwiatkowski, Cordelia Langford, John Sillitoe on behalf of the Wellcome Sanger |

| | | | |
|---|---|---|---|
| | | Consortium | Institute COVID-19 Surveillance Team |
| EPI_ISL_2730179 | Lighthouse Lab in Glasgow | Wellcome Sanger Institute for the COVID-19 Genomics UK (COG-UK) Consortium | Harper VanSteenhouse, Yumi Kasai, David Gray, Carol Clugston, Anna Dominiczak and Alex Alderton, Roberto Amato, Jeffrey Barrett, Sonia Goncalves, Ewan Harrison, David K. Jackson, Ian Johnston, Dominic Kwiatkowski, Cordelia Langford, John Sillitoe on behalf of the Wellcome Sanger Institute COVID-19 Surveillance Team |
| EPI_ISL_2730180 | Lighthouse Lab in Milton Keynes | Wellcome Sanger Institute for the COVID-19 Genomics UK (COG-UK) Consortium | The Lighthouse Lab in Milton Keynes and Alex Alderton, Roberto Amato, Jeffrey Barrett, Sonia Goncalves, Ewan Harrison, David K. Jackson, Ian Johnston, Dominic Kwiatkowski, Cordelia Langford, John Sillitoe on behalf of the Wellcome Sanger Institute COVID-19 Surveillance Team |
| EPI_ISL_2730181 | Lighthouse Lab in Glasgow | Wellcome Sanger Institute for the COVID-19 Genomics UK (COG-UK) Consortium | Harper VanSteenhouse, Yumi Kasai, David Gray, Carol Clugston, Anna Dominiczak and Alex Alderton, Roberto Amato, Jeffrey Barrett, Sonia Goncalves, Ewan Harrison, David K. Jackson, Ian Johnston, Dominic Kwiatkowski, Cordelia Langford, John Sillitoe on behalf of the Wellcome Sanger Institute COVID-19 Surveillance Team |
| EPI_ISL_2730182 | Lighthouse Lab in Milton Keynes | Wellcome Sanger Institute for the COVID-19 Genomics UK (COG-UK) Consortium | The Lighthouse Lab in Milton Keynes and Alex Alderton, Roberto Amato, Jeffrey Barrett, Sonia Goncalves, Ewan Harrison, David K. Jackson, Ian Johnston, Dominic Kwiatkowski, Cordelia Langford, John Sillitoe on behalf of the Wellcome Sanger Institute COVID-19 Surveillance Team |
| EPI_ISL_2730183 | Lighthouse Lab in Alderley Park | Wellcome Sanger Institute for the COVID-19 Genomics UK (COG-UK) Consortium | Jacquelyn Wynn, Mairead Hyland, The Lighthouse Lab in Alderley Park and Alex Alderton, Roberto Amato, Jeffrey Barrett, Sonia Goncalves, Ewan Harrison, David K. Jackson, Ian Johnston, Dominic Kwiatkowski, Cordelia Langford, John Sillitoe on behalf of the Wellcome Sanger Institute COVID-19 Surveillance Team |
| EPI_ISL_2730184 | Lighthouse Lab in Milton Keynes | Wellcome Sanger Institute for the COVID-19 Genomics UK (COG-UK) Consortium | The Lighthouse Lab in Milton Keynes and Alex Alderton, Roberto Amato, Jeffrey Barrett, Sonia Goncalves, Ewan Harrison, David K. Jackson, Ian Johnston, Dominic Kwiatkowski, Cordelia Langford, John Sillitoe on behalf of the Wellcome Sanger Institute COVID-19 Surveillance Team |
| EPI_ISL_2730185 | Lighthouse Laboratory Plymouth | Wellcome Sanger Institute for the COVID-19 Genomics UK (COG-UK) Consortium | Lighthouse Laboratory Plymouth and Alex Alderton, Roberto Amato, Jeffrey Barrett, Sonia Goncalves, Ewan Harrison, David K. Jackson, Ian Johnston, Dominic Kwiatkowski, Cordelia Langford, John Sillitoe on behalf of the Wellcome Sanger Institute COVID-19 Surveillance Team |
| EPI_ISL_2730186 | Lighthouse Lab in Milton Keynes | Wellcome Sanger Institute for the COVID-19 Genomics UK (COG-UK) Consortium | The Lighthouse Lab in Milton Keynes and Alex Alderton, Roberto Amato, Jeffrey Barrett, Sonia Goncalves, Ewan Harrison, David K. Jackson, Ian Johnston, Dominic Kwiatkowski, Cordelia Langford, John Sillitoe on behalf of the Wellcome Sanger Institute COVID-19 Surveillance Team |
| EPI_ISL_2730187, EPI_ISL_2730188, EPI_ISL_2730189 | Lighthouse Lab in Glasgow | Wellcome Sanger Institute for the COVID-19 Genomics UK (COG-UK) Consortium | Harper VanSteenhouse, Yumi Kasai, David Gray, Carol Clugston, Anna Dominiczak and Alex Alderton, Roberto Amato, Jeffrey Barrett, Sonia Goncalves, Ewan Harrison, David K. Jackson, Ian Johnston, Dominic Kwiatkowski, Cordelia Langford, John Sillitoe on behalf of the Wellcome Sanger Institute COVID-19 Surveillance Team |
| EPI_ISL_2730190 | Lighthouse Laboratory Plymouth | Wellcome Sanger Institute for the COVID-19 Genomics UK (COG-UK) Consortium | Lighthouse Laboratory Plymouth and Alex Alderton, Roberto Amato, Jeffrey Barrett, Sonia Goncalves, Ewan Harrison, David K. Jackson, Ian Johnston, Dominic Kwiatkowski, Cordelia Langford, John Sillitoe on behalf of the Wellcome Sanger Institute COVID-19 Surveillance Team |
| EPI_ISL_2730191, EPI_ISL_2730192 | Lighthouse Lab in Milton Keynes | Wellcome Sanger Institute for the COVID-19 Genomics UK (COG-UK) Consortium | The Lighthouse Lab in Milton Keynes and Alex Alderton, Roberto Amato, Jeffrey Barrett, Sonia Goncalves, Ewan Harrison, David K. Jackson, Ian Johnston, Dominic Kwiatkowski, Cordelia Langford, John Sillitoe on behalf of the Wellcome Sanger Institute COVID-19 Surveillance Team |
| EPI_ISL_2730193, EPI_ISL_2730194 | Lighthouse Lab in Alderley Park | Wellcome Sanger Institute for the COVID-19 Genomics UK (COG-UK) Consortium | Jacquelyn Wynn, Mairead Hyland, The Lighthouse Lab in Alderley Park and Alex Alderton, Roberto Amato, Jeffrey Barrett, Sonia Goncalves, Ewan Harrison, David K. Jackson, Ian Johnston, Dominic Kwiatkowski, Cordelia Langford, John Sillitoe on behalf of the Wellcome Sanger Institute COVID-19 Surveillance Team |
| EPI_ISL_2730195 | Lighthouse Lab in Milton Keynes | Wellcome Sanger Institute for the COVID-19 Genomics UK (COG-UK) Consortium | The Lighthouse Lab in Milton Keynes and Alex Alderton, Roberto Amato, Jeffrey Barrett, Sonia Goncalves, Ewan Harrison, David K. Jackson, Ian Johnston, Dominic Kwiatkowski, Cordelia Langford, John Sillitoe on behalf of the Wellcome Sanger Institute COVID-19 Surveillance Team |
| EPI_ISL_2730196 | Lighthouse Lab in Glasgow | Wellcome Sanger Institute for the COVID-19 Genomics UK (COG-UK) Consortium | Harper VanSteenhouse, Yumi Kasai, David Gray, Carol Clugston, Anna Dominiczak and Alex Alderton, Roberto Amato, Jeffrey Barrett, Sonia Goncalves, Ewan Harrison, David K. Jackson, Ian Johnston, Dominic Kwiatkowski, Cordelia Langford, John Sillitoe on behalf of the Wellcome Sanger Institute COVID-19 Surveillance Team |
| EPI_ISL_2730197 | Lighthouse Lab in Milton Keynes | Wellcome Sanger Institute for the COVID-19 Genomics UK (COG-UK) Consortium | The Lighthouse Lab in Milton Keynes and Alex Alderton, Roberto Amato, Jeffrey Barrett, Sonia Goncalves, Ewan Harrison, David K. Jackson, Ian Johnston, Dominic Kwiatkowski, Cordelia Langford, John Sillitoe on behalf of the Wellcome Sanger Institute COVID-19 Surveillance Team |
| EPI_ISL_2730198 | Lighthouse Lab in Alderley Park | Wellcome Sanger Institute for the COVID-19 Genomics UK (COG-UK) Consortium | Jacquelyn Wynn, Mairead Hyland, The Lighthouse Lab in Alderley Park and Alex Alderton, Roberto Amato, Jeffrey Barrett, Sonia Goncalves, Ewan Harrison, David K. Jackson, Ian Johnston, Dominic Kwiatkowski, Cordelia Langford, John Sillitoe on behalf of the Wellcome Sanger Institute COVID-19 Surveillance Team |
| EPI_ISL_2730199 | Lighthouse Lab in Glasgow | Wellcome Sanger Institute for the COVID-19 Genomics UK (COG-UK) Consortium | Harper VanSteenhouse, Yumi Kasai, David Gray, Carol Clugston, Anna Dominiczak and Alex Alderton, Roberto Amato, Jeffrey Barrett, Sonia Goncalves, Ewan Harrison, David K. Jackson, Ian Johnston, Dominic Kwiatkowski, Cordelia Langford, John Sillitoe on behalf of the Wellcome Sanger Institute COVID-19 Surveillance Team |
| EPI_ISL_2730200 | Lighthouse Lab in Milton Keynes | Wellcome Sanger Institute for the COVID-19 Genomics UK (COG-UK) Consortium | The Lighthouse Lab in Milton Keynes and Alex Alderton, Roberto Amato, Jeffrey Barrett, Sonia Goncalves, Ewan Harrison, David K. Jackson, Ian Johnston, Dominic Kwiatkowski, Cordelia Langford, John Sillitoe on behalf of the Wellcome Sanger Institute COVID-19 Surveillance Team |
| EPI_ISL_2730201 | Lighthouse Lab in Glasgow | Wellcome Sanger Institute for the COVID-19 Genomics UK (COG-UK) Consortium | Harper VanSteenhouse, Yumi Kasai, David Gray, Carol Clugston, Anna Dominiczak and Alex Alderton, Roberto Amato, Jeffrey Barrett, Sonia Goncalves, Ewan Harrison, David K. Jackson, Ian Johnston, Dominic Kwiatkowski, Cordelia Langford, John Sillitoe on behalf of the Wellcome Sanger Institute COVID-19 Surveillance Team |
| EPI_ISL_2730202, EPI_ISL_2730203 | Lighthouse Laboratory Plymouth | Wellcome Sanger Institute for the COVID-19 Genomics UK (COG-UK) Consortium | Lighthouse Laboratory Plymouth and Alex Alderton, Roberto Amato, Jeffrey Barrett, Sonia Goncalves, Ewan Harrison, David K. Jackson, Ian Johnston, Dominic Kwiatkowski, Cordelia Langford, John Sillitoe on behalf of the Wellcome Sanger Institute COVID-19 Surveillance Team |
| EPI_ISL_2730204 | Lighthouse Lab in Alderley Park | Wellcome Sanger Institute for the COVID-19 Genomics UK (COG-UK) Consortium | Jacquelyn Wynn, Mairead Hyland, The Lighthouse Lab in Alderley Park and Alex Alderton, Roberto Amato, Jeffrey Barrett, Sonia Goncalves, Ewan Harrison, David K. Jackson, Ian Johnston, Dominic Kwiatkowski, Cordelia Langford, John Sillitoe on behalf of the Wellcome Sanger Institute COVID-19 Surveillance Team |
| EPI_ISL_2730205 | Lighthouse Lab in Milton Keynes | Wellcome Sanger Institute for the COVID-19 Genomics UK (COG-UK) Consortium | The Lighthouse Lab in Milton Keynes and Alex Alderton, Roberto Amato, Jeffrey Barrett, Sonia Goncalves, Ewan Harrison, David K. Jackson, Ian Johnston, Dominic Kwiatkowski, Cordelia Langford, John Sillitoe on behalf of the Wellcome Sanger Institute COVID-19 Surveillance Team |
| EPI_ISL_2730206 | Lighthouse Laboratory Plymouth | Wellcome Sanger Institute for the COVID-19 Genomics UK (COG-UK) Consortium | Lighthouse Laboratory Plymouth and Alex Alderton, Roberto Amato, Jeffrey Barrett, Sonia Goncalves, Ewan Harrison, David K. Jackson, Ian Johnston, Dominic Kwiatkowski, Cordelia Langford, John Sillitoe on behalf of the Wellcome Sanger Institute COVID-19 Surveillance Team |
| EPI_ISL_2730207 | Lighthouse Lab in Milton Keynes | Wellcome Sanger Institute for the COVID-19 Genomics UK (COG-UK) Consortium | The Lighthouse Lab in Milton Keynes and Alex Alderton, Roberto Amato, Jeffrey Barrett, Sonia Goncalves, Ewan Harrison, David K. Jackson, Ian Johnston, Dominic Kwiatkowski, Cordelia Langford, John Sillitoe on behalf of the Wellcome Sanger Institute COVID-19 Surveillance Team |
| EPI_ISL_2730208 | Lighthouse Lab in Glasgow | Wellcome Sanger Institute for the COVID-19 Genomics UK (COG-UK) Consortium | Harper VanSteenhouse, Yumi Kasai, David Gray, Carol Clugston, Anna Dominiczak and Alex Alderton, Roberto Amato, Jeffrey Barrett, Sonia Goncalves, Ewan Harrison, David K. Jackson, Ian Johnston, Dominic Kwiatkowski, Cordelia Langford, John Sillitoe on behalf of the Wellcome Sanger Institute COVID-19 Surveillance Team |
| EPI_ISL_2730209 | Lighthouse Laboratory Plymouth | Wellcome Sanger Institute for the COVID-19 Genomics UK (COG-UK) Consortium | Lighthouse Laboratory Plymouth and Alex Alderton, Roberto Amato, Jeffrey Barrett, Sonia Goncalves, Ewan Harrison, David K. Jackson, Ian Johnston, Dominic Kwiatkowski, Cordelia Langford, John Sillitoe on behalf of the Wellcome Sanger Institute COVID-19 Surveillance Team |
| EPI_ISL_2730210 | Lighthouse Lab in Milton Keynes | Wellcome Sanger Institute for the COVID-19 Genomics UK (COG-UK) Consortium | The Lighthouse Lab in Milton Keynes and Alex Alderton, Roberto Amato, Jeffrey Barrett, Sonia Goncalves, Ewan Harrison, David K. Jackson, Ian Johnston, Dominic Kwiatkowski, Cordelia Langford, John Sillitoe on behalf of the Wellcome Sanger Institute COVID-19 Surveillance Team |
| EPI_ISL_2730211, EPI_ISL_2730212 | Lighthouse Lab in Glasgow | Wellcome Sanger Institute for the COVID-19 Genomics UK (COG-UK) Consortium | Harper VanSteenhouse, Yumi Kasai, David Gray, Carol Clugston, Anna Dominiczak and Alex Alderton, Roberto Amato, Jeffrey Barrett, Sonia Goncalves, Ewan Harrison, David K. Jackson, Ian Johnston, Dominic Kwiatkowski, Cordelia Langford, John Sillitoe on behalf of the Wellcome Sanger Institute COVID-19 Surveillance Team |
| EPI_ISL_2730213 | Lighthouse Lab in Alderley Park | Wellcome Sanger Institute for the COVID-19 Genomics UK (COG-UK) Consortium | Jacquelyn Wynn, Mairead Hyland, The Lighthouse Lab in Alderley Park and Alex Alderton, Roberto Amato, Jeffrey Barrett, Sonia Goncalves, Ewan Harrison, David K. Jackson, Ian Johnston, Dominic Kwiatkowski, Cordelia Langford, John Sillitoe on behalf of the Wellcome Sanger Institute COVID-19 Surveillance Team |
| EPI_ISL_2730214 | Lighthouse Lab in Glasgow | Wellcome Sanger Institute for the COVID-19 Genomics UK (COG-UK) Consortium | Harper VanSteenhouse, Yumi Kasai, David Gray, Carol Clugston, Anna Dominiczak and Alex Alderton, Roberto Amato, Jeffrey Barrett, Sonia Goncalves, Ewan Harrison, David K. Jackson, Ian Johnston, Dominic Kwiatkowski, Cordelia Langford, John Sillitoe on behalf of the Wellcome Sanger Institute COVID-19 Surveillance Team |
| EPI_ISL_2730215, EPI_ISL_2730216 | Lighthouse Lab in Milton Keynes | Wellcome Sanger Institute for the COVID-19 Genomics UK (COG-UK) Consortium | The Lighthouse Lab in Milton Keynes and Alex Alderton, Roberto Amato, Jeffrey Barrett, Sonia Goncalves, Ewan Harrison, David K. Jackson, Ian Johnston, Dominic Kwiatkowski, Cordelia Langford, John Sillitoe on behalf of the Wellcome Sanger Institute COVID-19 Surveillance Team |
| EPI_ISL_2730217 | Lighthouse Lab in Alderley Park | Wellcome Sanger Institute for the COVID-19 Genomics UK (COG-UK) Consortium | Jacquelyn Wynn, Mairead Hyland, The Lighthouse Lab in Alderley Park and Alex Alderton, Roberto Amato, Jeffrey Barrett, Sonia Goncalves, Ewan Harrison, David K. Jackson, Ian Johnston, Dominic Kwiatkowski, Cordelia Langford, John Sillitoe on behalf of the Wellcome Sanger Institute COVID-19 Surveillance Team |
| EPI_ISL_2730218 | Lighthouse Lab in Glasgow | Wellcome Sanger Institute for the COVID-19 Genomics UK (COG-UK) Consortium | Harper VanSteenhouse, Yumi Kasai, David Gray, Carol Clugston, Anna Dominiczak and Alex Alderton, Roberto Amato, Jeffrey Barrett, Sonia Goncalves, Ewan Harrison, David K. Jackson, Ian Johnston, Dominic Kwiatkowski, Cordelia Langford, John Sillitoe on behalf of the Wellcome Sanger Institute COVID-19 Surveillance Team |
| EPI_ISL_2730219 | Lighthouse Lab in Alderley Park | Wellcome Sanger Institute for the COVID-19 Genomics UK (COG-UK) Consortium | Jacquelyn Wynn, Mairead Hyland, The Lighthouse Lab in Alderley Park and Alex Alderton, Roberto Amato, Jeffrey Barrett, Sonia Goncalves, Ewan Harrison, David K. Jackson, Ian Johnston, Dominic Kwiatkowski, Cordelia Langford, John Sillitoe on behalf of the Wellcome Sanger Institute COVID-19 Surveillance Team |
| EPI_ISL_2730220 | Lighthouse Lab in Glasgow | Wellcome Sanger Institute for the COVID-19 Genomics UK (COG-UK) Consortium | Harper VanSteenhouse, Yumi Kasai, David Gray, Carol Clugston, Anna Dominiczak and Alex Alderton, Roberto Amato, Jeffrey Barrett, Sonia Goncalves, Ewan Harrison, David K. Jackson, Ian Johnston, Dominic Kwiatkowski, Cordelia Langford, John Sillitoe on behalf of the Wellcome Sanger Institute COVID-19 Surveillance Team |
| EPI_ISL_2730221, EPI_ISL_2730222, EPI_ISL_2730223 | Lighthouse Lab in Alderley Park | Wellcome Sanger Institute for the COVID-19 Genomics UK (COG-UK) Consortium | Jacquelyn Wynn, Mairead Hyland, The Lighthouse Lab in Alderley Park and Alex Alderton, Roberto Amato, Jeffrey Barrett, Sonia Goncalves, Ewan Harrison, David K. Jackson, Ian Johnston, Dominic Kwiatkowski, Cordelia Langford, John Sillitoe on behalf of the Wellcome Sanger Institute COVID-19 Surveillance Team |
| EPI_ISL_2730224, EPI_ISL_2730225 | Lighthouse Lab in Milton Keynes | Wellcome Sanger Institute for the COVID-19 Genomics UK (COG-UK) Consortium | The Lighthouse Lab in Milton Keynes and Alex Alderton, Roberto Amato, Jeffrey Barrett, Sonia Goncalves, Ewan Harrison, David K. Jackson, Ian Johnston, Dominic Kwiatkowski, Cordelia Langford, John Sillitoe on behalf of the Wellcome Sanger Institute COVID-19 Surveillance Team |
| EPI_ISL_2730226, EPI_ISL_2730227, EPI_ISL_2730228 | Lighthouse Lab in Glasgow | Wellcome Sanger Institute for the COVID-19 Genomics UK (COG-UK) Consortium | Harper VanSteenhouse, Yumi Kasai, David Gray, Carol Clugston, Anna Dominiczak and Alex Alderton, Roberto Amato, Jeffrey Barrett, Sonia Goncalves, Ewan Harrison, David K. Jackson, Ian Johnston, Dominic Kwiatkowski, Cordelia Langford, John Sillitoe on behalf of the Wellcome Sanger Institute COVID-19 Surveillance Team |
| EPI_ISL_2730229 | Lighthouse Lab in Milton Keynes | Wellcome Sanger Institute for the COVID-19 Genomics UK (COG-UK) Consortium | The Lighthouse Lab in Milton Keynes and Alex Alderton, Roberto Amato, Jeffrey Barrett, Sonia Goncalves, Ewan Harrison, David K. Jackson, Ian Johnston, Dominic Kwiatkowski, Cordelia Langford, John Sillitoe on behalf of the Wellcome Sanger Institute COVID-19 Surveillance Team |
| EPI_ISL_2730230, EPI_ISL_2730231, EPI_ISL_2730232, EPI_ISL_2730233 | Lighthouse Laboratory Plymouth | Wellcome Sanger Institute for the COVID-19 Genomics UK (COG-UK) Consortium | Lighthouse Laboratory Plymouth and Alex Alderton, Roberto Amato, Jeffrey Barrett, Sonia Goncalves, Ewan Harrison, David K. Jackson, Ian Johnston, Dominic Kwiatkowski, Cordelia Langford, John Sillitoe on behalf of the Wellcome Sanger Institute COVID-19 Surveillance Team |
| EPI_ISL_2730234 | Lighthouse Lab in Alderley Park | Wellcome Sanger Institute for the COVID-19 Genomics UK (COG-UK) Consortium | Jacquelyn Wynn, Mairead Hyland, The Lighthouse Lab in Alderley Park and Alex Alderton, Roberto Amato, Jeffrey Barrett, Sonia Goncalves, Ewan Harrison, David K. Jackson, Ian Johnston, Dominic Kwiatkowski, Cordelia Langford, John Sillitoe on behalf of the Wellcome Sanger Institute COVID-19 Surveillance Team |
| EPI_ISL_2730235 | Lighthouse Laboratory Plymouth | Wellcome Sanger Institute for the COVID-19 Genomics UK (COG-UK) Consortium | Lighthouse Laboratory Plymouth and Alex Alderton, Roberto Amato, Jeffrey Barrett, Sonia Goncalves, Ewan Harrison, David K. Jackson, Ian Johnston, Dominic Kwiatkowski, Cordelia Langford, John Sillitoe on behalf of the Wellcome Sanger Institute COVID-19 Surveillance Team |
| EPI_ISL_2730236 | Lighthouse Lab in Glasgow | Wellcome Sanger Institute for the COVID-19 Genomics UK (COG-UK) | Harper VanSteenhouse, Yumi Kasai, David Gray, Carol Clugston, Anna Dominiczak and Alex Alderton, Roberto Amato, Jeffrey Barrett, Sonia Goncalves, Ewan Harrison, David K. Jackson, Ian Johnston, Dominic Kwiatkowski, Cordelia Langford, |

| | | | |
|---|---|---|---|
| | | Consortium | John Sillitoe on behalf of the Wellcome Sanger Institute COVID-19 Surveillance Team |
| EPI_ISL_2730237, EPI_ISL_2730238, EPI_ISL_2730239, EPI_ISL_2730240 | Lighthouse Laboratory Plymouth | Wellcome Sanger Institute for the COVID-19 Genomics UK (COG-UK) Consortium | Lighthouse Laboratory Plymouth and Alex Alderton, Roberto Amato, Jeffrey Barrett, Sonia Goncalves, Ewan Harrison, David K. Jackson, Ian Johnston, Dominic Kwiatkowski, Cordelia Langford, John Sillitoe on behalf of the Wellcome Sanger Institute COVID-19 Surveillance Team |
| EPI_ISL_2730241 | Lighthouse Lab in Glasgow | Wellcome Sanger Institute for the COVID-19 Genomics UK (COG-UK) Consortium | Harper VanSteenhouse, Yumi Kasai, David Gray, Carol Clugston, Anna Dominiczak and Alex Alderton, Roberto Amato, Jeffrey Barrett, Sonia Goncalves, Ewan Harrison, David K. Jackson, Ian Johnston, Dominic Kwiatkowski, Cordelia Langford, John Sillitoe on behalf of the Wellcome Sanger Institute COVID-19 Surveillance Team |
| EPI_ISL_2730242, EPI_ISL_2730243, EPI_ISL_2730244 | Lighthouse Lab in Milton Keynes | Wellcome Sanger Institute for the COVID-19 Genomics UK (COG-UK) Consortium | The Lighthouse Lab in Milton Keynes and Alex Alderton, Roberto Amato, Jeffrey Barrett, Sonia Goncalves, Ewan Harrison, David K. Jackson, Ian Johnston, Dominic Kwiatkowski, Cordelia Langford, John Sillitoe on behalf of the Wellcome Sanger Institute COVID-19 Surveillance Team |
| EPI_ISL_2730245 | Lighthouse Lab in Alderley Park | Wellcome Sanger Institute for the COVID-19 Genomics UK (COG-UK) Consortium | Jacquelyn Wynn, Mairead Hyland, The Lighthouse Lab in Alderley Park and Alex Alderton, Roberto Amato, Jeffrey Barrett, Sonia Goncalves, Ewan Harrison, David K. Jackson, Ian Johnston, Dominic Kwiatkowski, Cordelia Langford, John Sillitoe on behalf of the Wellcome Sanger Institute COVID-19 Surveillance Team |
| EPI_ISL_2730246 | Lighthouse Laboratory Plymouth | Wellcome Sanger Institute for the COVID-19 Genomics UK (COG-UK) Consortium | Lighthouse Laboratory Plymouth and Alex Alderton, Roberto Amato, Jeffrey Barrett, Sonia Goncalves, Ewan Harrison, David K. Jackson, Ian Johnston, Dominic Kwiatkowski, Cordelia Langford, John Sillitoe on behalf of the Wellcome Sanger Institute COVID-19 Surveillance Team |
| EPI_ISL_2730247 | Lighthouse Lab in Alderley Park | Wellcome Sanger Institute for the COVID-19 Genomics UK (COG-UK) Consortium | Jacquelyn Wynn, Mairead Hyland, The Lighthouse Lab in Alderley Park and Alex Alderton, Roberto Amato, Jeffrey Barrett, Sonia Goncalves, Ewan Harrison, David K. Jackson, Ian Johnston, Dominic Kwiatkowski, Cordelia Langford, John Sillitoe on behalf of the Wellcome Sanger Institute COVID-19 Surveillance Team |
| EPI_ISL_2730248 | Lighthouse Lab in Milton Keynes | Wellcome Sanger Institute for the COVID-19 Genomics UK (COG-UK) Consortium | The Lighthouse Lab in Milton Keynes and Alex Alderton, Roberto Amato, Jeffrey Barrett, Sonia Goncalves, Ewan Harrison, David K. Jackson, Ian Johnston, Dominic Kwiatkowski, Cordelia Langford, John Sillitoe on behalf of the Wellcome Sanger Institute COVID-19 Surveillance Team |
| EPI_ISL_2730249 | Lighthouse Lab in Alderley Park | Wellcome Sanger Institute for the COVID-19 Genomics UK (COG-UK) Consortium | Jacquelyn Wynn, Mairead Hyland, The Lighthouse Lab in Alderley Park and Alex Alderton, Roberto Amato, Jeffrey Barrett, Sonia Goncalves, Ewan Harrison, David K. Jackson, Ian Johnston, Dominic Kwiatkowski, Cordelia Langford, John Sillitoe on behalf of the Wellcome Sanger Institute COVID-19 Surveillance Team |
| EPI_ISL_2730250 | Lighthouse Lab in Milton Keynes | Wellcome Sanger Institute for the COVID-19 Genomics UK (COG-UK) Consortium | The Lighthouse Lab in Milton Keynes and Alex Alderton, Roberto Amato, Jeffrey Barrett, Sonia Goncalves, Ewan Harrison, David K. Jackson, Ian Johnston, Dominic Kwiatkowski, Cordelia Langford, John Sillitoe on behalf of the Wellcome Sanger Institute COVID-19 Surveillance Team |
| EPI_ISL_2730251 | Lighthouse Lab in Glasgow | Wellcome Sanger Institute for the COVID-19 Genomics UK (COG-UK) Consortium | Harper VanSteenhouse, Yumi Kasai, David Gray, Carol Clugston, Anna Dominiczak and Alex Alderton, Roberto Amato, Jeffrey Barrett, Sonia Goncalves, Ewan Harrison, David K. Jackson, Ian Johnston, Dominic Kwiatkowski, Cordelia Langford, John Sillitoe on behalf of the Wellcome Sanger Institute COVID-19 Surveillance Team |
| EPI_ISL_2730252 | Lighthouse Laboratory Plymouth | Wellcome Sanger Institute for the COVID-19 Genomics UK (COG-UK) Consortium | Lighthouse Laboratory Plymouth and Alex Alderton, Roberto Amato, Jeffrey Barrett, Sonia Goncalves, Ewan Harrison, David K. Jackson, Ian Johnston, Dominic Kwiatkowski, Cordelia Langford, John Sillitoe on behalf of the Wellcome Sanger Institute COVID-19 Surveillance Team |
| EPI_ISL_2730253 | Lighthouse Lab in Milton Keynes | Wellcome Sanger Institute for the COVID-19 Genomics UK (COG-UK) Consortium | The Lighthouse Lab in Milton Keynes and Alex Alderton, Roberto Amato, Jeffrey Barrett, Sonia Goncalves, Ewan Harrison, David K. Jackson, Ian Johnston, Dominic Kwiatkowski, Cordelia Langford, John Sillitoe on behalf of the Wellcome Sanger Institute COVID-19 Surveillance Team |
| EPI_ISL_2730254 | Lighthouse Lab in Alderley Park | Wellcome Sanger Institute for the COVID-19 Genomics UK (COG-UK) Consortium | Jacquelyn Wynn, Mairead Hyland, The Lighthouse Lab in Alderley Park and Alex Alderton, Roberto Amato, Jeffrey Barrett, Sonia Goncalves, Ewan Harrison, David K. Jackson, Ian Johnston, Dominic Kwiatkowski, Cordelia Langford, John Sillitoe on behalf of the Wellcome Sanger Institute COVID-19 Surveillance Team |
| EPI_ISL_2730255 | Lighthouse Laboratory Plymouth | Wellcome Sanger Institute for the COVID-19 Genomics UK (COG-UK) Consortium | Lighthouse Laboratory Plymouth and Alex Alderton, Roberto Amato, Jeffrey Barrett, Sonia Goncalves, Ewan Harrison, David K. Jackson, Ian Johnston, Dominic Kwiatkowski, Cordelia Langford, John Sillitoe on behalf of the Wellcome Sanger Institute COVID-19 Surveillance Team |
| EPI_ISL_2730256, EPI_ISL_2730257 | Lighthouse Lab in Alderley Park | Wellcome Sanger Institute for the COVID-19 Genomics UK (COG-UK) Consortium | Jacquelyn Wynn, Mairead Hyland, The Lighthouse Lab in Alderley Park and Alex Alderton, Roberto Amato, Jeffrey Barrett, Sonia Goncalves, Ewan Harrison, David K. Jackson, Ian Johnston, Dominic Kwiatkowski, Cordelia Langford, John Sillitoe on behalf of the Wellcome Sanger Institute COVID-19 Surveillance Team |
| EPI_ISL_2730258 | Lighthouse Lab in Glasgow | Wellcome Sanger Institute for the COVID-19 Genomics UK (COG-UK) Consortium | Harper VanSteenhouse, Yumi Kasai, David Gray, Carol Clugston, Anna Dominiczak and Alex Alderton, Roberto Amato, Jeffrey Barrett, Sonia Goncalves, Ewan Harrison, David K. Jackson, Ian Johnston, Dominic Kwiatkowski, Cordelia Langford, John Sillitoe on behalf of the Wellcome Sanger Institute COVID-19 Surveillance Team |
| EPI_ISL_2730259, EPI_ISL_2730260, EPI_ISL_2730261, EPI_ISL_2730262 | Lighthouse Laboratory Plymouth | Wellcome Sanger Institute for the COVID-19 Genomics UK (COG-UK) Consortium | Lighthouse Laboratory Plymouth and Alex Alderton, Roberto Amato, Jeffrey Barrett, Sonia Goncalves, Ewan Harrison, David K. Jackson, Ian Johnston, Dominic Kwiatkowski, Cordelia Langford, John Sillitoe on behalf of the Wellcome Sanger Institute COVID-19 Surveillance Team |
| EPI_ISL_2730263 | Lighthouse Lab in Alderley Park | Wellcome Sanger Institute for the COVID-19 Genomics UK (COG-UK) Consortium | Jacquelyn Wynn, Mairead Hyland, The Lighthouse Lab in Alderley Park and Alex Alderton, Roberto Amato, Jeffrey Barrett, Sonia Goncalves, Ewan Harrison, David K. Jackson, Ian Johnston, Dominic Kwiatkowski, Cordelia Langford, John Sillitoe on behalf of the Wellcome Sanger Institute COVID-19 Surveillance Team |
| EPI_ISL_2730264, EPI_ISL_2730265 | Lighthouse Laboratory Plymouth | Wellcome Sanger Institute for the COVID-19 Genomics UK (COG-UK) Consortium | Lighthouse Laboratory Plymouth and Alex Alderton, Roberto Amato, Jeffrey Barrett, Sonia Goncalves, Ewan Harrison, David K. Jackson, Ian Johnston, Dominic Kwiatkowski, Cordelia Langford, John Sillitoe on behalf of the Wellcome Sanger Institute COVID-19 Surveillance Team |
| EPI_ISL_2730266 | Lighthouse Lab in Milton Keynes | Wellcome Sanger Institute for the COVID-19 Genomics UK (COG-UK) Consortium | The Lighthouse Lab in Milton Keynes and Alex Alderton, Roberto Amato, Jeffrey Barrett, Sonia Goncalves, Ewan Harrison, David K. Jackson, Ian Johnston, Dominic Kwiatkowski, Cordelia Langford, John Sillitoe on behalf of the Wellcome Sanger Institute COVID-19 Surveillance Team |
| EPI_ISL_2730267 | Lighthouse Lab in Glasgow | Wellcome Sanger Institute for the COVID-19 Genomics UK (COG-UK) Consortium | Harper VanSteenhouse, Yumi Kasai, David Gray, Carol Clugston, Anna Dominiczak and Alex Alderton, Roberto Amato, Jeffrey Barrett, Sonia Goncalves, Ewan Harrison, David K. Jackson, Ian Johnston, Dominic Kwiatkowski, Cordelia Langford, John Sillitoe on behalf of the Wellcome Sanger Institute COVID-19 Surveillance Team |
| EPI_ISL_2730268, EPI_ISL_2730269 | Lighthouse Laboratory Plymouth | Wellcome Sanger Institute for the COVID-19 Genomics UK (COG-UK) Consortium | Lighthouse Laboratory Plymouth and Alex Alderton, Roberto Amato, Jeffrey Barrett, Sonia Goncalves, Ewan Harrison, David K. Jackson, Ian Johnston, Dominic Kwiatkowski, Cordelia Langford, John Sillitoe on behalf of the Wellcome Sanger Institute COVID-19 Surveillance Team |
| EPI_ISL_2730270 | Lighthouse Lab in Alderley Park | Wellcome Sanger Institute for the COVID-19 Genomics UK (COG-UK) Consortium | Jacquelyn Wynn, Mairead Hyland, The Lighthouse Lab in Alderley Park and Alex Alderton, Roberto Amato, Jeffrey Barrett, Sonia Goncalves, Ewan Harrison, David K. Jackson, Ian Johnston, Dominic Kwiatkowski, Cordelia Langford, John Sillitoe on behalf of the Wellcome Sanger Institute COVID-19 Surveillance Team |
| EPI_ISL_2730271 | Lighthouse Lab in Milton Keynes | Wellcome Sanger Institute for the COVID-19 Genomics UK (COG-UK) Consortium | The Lighthouse Lab in Milton Keynes and Alex Alderton, Roberto Amato, Jeffrey Barrett, Sonia Goncalves, Ewan Harrison, David K. Jackson, Ian Johnston, Dominic Kwiatkowski, Cordelia Langford, John Sillitoe on behalf of the Wellcome Sanger Institute COVID-19 Surveillance Team |
| EPI_ISL_2730272 | Lighthouse Lab in Alderley Park | Wellcome Sanger Institute for the COVID-19 Genomics UK (COG-UK) Consortium | Jacquelyn Wynn, Mairead Hyland, The Lighthouse Lab in Alderley Park and Alex Alderton, Roberto Amato, Jeffrey Barrett, Sonia Goncalves, Ewan Harrison, David K. Jackson, Ian Johnston, Dominic Kwiatkowski, Cordelia Langford, John Sillitoe on behalf of the Wellcome Sanger Institute COVID-19 Surveillance Team |
| EPI_ISL_2730273, EPI_ISL_2730274 | Lighthouse Laboratory Plymouth | Wellcome Sanger Institute for the COVID-19 Genomics UK (COG-UK) Consortium | Lighthouse Laboratory Plymouth and Alex Alderton, Roberto Amato, Jeffrey Barrett, Sonia Goncalves, Ewan Harrison, David K. Jackson, Ian Johnston, Dominic Kwiatkowski, Cordelia Langford, John Sillitoe on behalf of the Wellcome Sanger Institute COVID-19 Surveillance Team |
| EPI_ISL_2730275 | Lighthouse Lab in Alderley Park | Wellcome Sanger Institute for the COVID-19 Genomics UK (COG-UK) Consortium | Jacquelyn Wynn, Mairead Hyland, The Lighthouse Lab in Alderley Park and Alex Alderton, Roberto Amato, Jeffrey Barrett, Sonia Goncalves, Ewan Harrison, David K. Jackson, Ian Johnston, Dominic Kwiatkowski, Cordelia Langford, John Sillitoe on behalf of the Wellcome Sanger Institute COVID-19 Surveillance Team |
| EPI_ISL_2730276 | Lighthouse Laboratory Plymouth | Wellcome Sanger Institute for the COVID-19 Genomics UK (COG-UK) Consortium | Lighthouse Laboratory Plymouth and Alex Alderton, Roberto Amato, Jeffrey Barrett, Sonia Goncalves, Ewan Harrison, David K. Jackson, Ian Johnston, Dominic Kwiatkowski, Cordelia Langford, John Sillitoe on behalf of the Wellcome Sanger Institute COVID-19 Surveillance Team |
| EPI_ISL_2730277 | Lighthouse Lab in Milton Keynes | Wellcome Sanger Institute for the COVID-19 Genomics UK (COG-UK) Consortium | The Lighthouse Lab in Milton Keynes and Alex Alderton, Roberto Amato, Jeffrey Barrett, Sonia Goncalves, Ewan Harrison, David K. Jackson, Ian Johnston, Dominic Kwiatkowski, Cordelia Langford, John Sillitoe on behalf of the Wellcome Sanger Institute COVID-19 Surveillance Team |
| EPI_ISL_2730278 | Lighthouse Lab in Alderley Park | Wellcome Sanger Institute for the COVID-19 Genomics UK (COG-UK) Consortium | Jacquelyn Wynn, Mairead Hyland, The Lighthouse Lab in Alderley Park and Alex Alderton, Roberto Amato, Jeffrey Barrett, Sonia Goncalves, Ewan Harrison, David K. Jackson, Ian Johnston, Dominic Kwiatkowski, Cordelia Langford, John Sillitoe on behalf of the Wellcome Sanger Institute COVID-19 Surveillance Team |
| EPI_ISL_2730279 | Lighthouse Laboratory Plymouth | Wellcome Sanger Institute for the COVID-19 Genomics UK (COG-UK) Consortium | Lighthouse Laboratory Plymouth and Alex Alderton, Roberto Amato, Jeffrey Barrett, Sonia Goncalves, Ewan Harrison, David K. Jackson, Ian Johnston, Dominic Kwiatkowski, Cordelia Langford, John Sillitoe on behalf of the Wellcome Sanger Institute COVID-19 Surveillance Team |
| EPI_ISL_2730280 | Lighthouse Lab in Milton Keynes | Wellcome Sanger Institute for the COVID-19 Genomics UK (COG-UK) Consortium | The Lighthouse Lab in Milton Keynes and Alex Alderton, Roberto Amato, Jeffrey Barrett, Sonia Goncalves, Ewan Harrison, David K. Jackson, Ian Johnston, Dominic Kwiatkowski, Cordelia Langford, John Sillitoe on behalf of the Wellcome Sanger Institute COVID-19 Surveillance Team |
| EPI_ISL_2730281 | Lighthouse Laboratory Plymouth | Wellcome Sanger Institute for the COVID-19 Genomics UK (COG-UK) Consortium | Lighthouse Laboratory Plymouth and Alex Alderton, Roberto Amato, Jeffrey Barrett, Sonia Goncalves, Ewan Harrison, David K. Jackson, Ian Johnston, Dominic Kwiatkowski, Cordelia Langford, John Sillitoe on behalf of the Wellcome Sanger Institute COVID-19 Surveillance Team |
| EPI_ISL_2730282 | Lighthouse Lab in Glasgow | Wellcome Sanger Institute for the COVID-19 Genomics UK (COG-UK) Consortium | Harper VanSteenhouse, Yumi Kasai, David Gray, Carol Clugston, Anna Dominiczak and Alex Alderton, Roberto Amato, Jeffrey Barrett, Sonia Goncalves, Ewan Harrison, David K. Jackson, Ian Johnston, Dominic Kwiatkowski, Cordelia Langford, John Sillitoe on behalf of the Wellcome Sanger Institute COVID-19 Surveillance Team |
| EPI_ISL_2730283, EPI_ISL_2730284 | Lighthouse Laboratory Plymouth | Wellcome Sanger Institute for the COVID-19 Genomics UK (COG-UK) Consortium | Lighthouse Laboratory Plymouth and Alex Alderton, Roberto Amato, Jeffrey Barrett, Sonia Goncalves, Ewan Harrison, David K. Jackson, Ian Johnston, Dominic Kwiatkowski, Cordelia Langford, John Sillitoe on behalf of the Wellcome Sanger Institute COVID-19 Surveillance Team |
| EPI_ISL_2730285 | Lighthouse Lab in Milton Keynes | Wellcome Sanger Institute for the COVID-19 Genomics UK (COG-UK) Consortium | The Lighthouse Lab in Milton Keynes and Alex Alderton, Roberto Amato, Jeffrey Barrett, Sonia Goncalves, Ewan Harrison, David K. Jackson, Ian Johnston, Dominic Kwiatkowski, Cordelia Langford, John Sillitoe on behalf of the Wellcome Sanger Institute COVID-19 Surveillance Team |
| EPI_ISL_2730286, EPI_ISL_2730287, EPI_ISL_2730288 | Lighthouse Lab in Alderley Park | Wellcome Sanger Institute for the COVID-19 Genomics UK (COG-UK) Consortium | Jacquelyn Wynn, Mairead Hyland, The Lighthouse Lab in Alderley Park and Alex Alderton, Roberto Amato, Jeffrey Barrett, Sonia Goncalves, Ewan Harrison, David K. Jackson, Ian Johnston, Dominic Kwiatkowski, Cordelia Langford, John Sillitoe on behalf of the Wellcome Sanger Institute COVID-19 Surveillance Team |
| EPI_ISL_2730289, EPI_ISL_2730290, EPI_ISL_2730291 | Lighthouse Lab in Glasgow | Wellcome Sanger Institute for the COVID-19 Genomics UK (COG-UK) Consortium | Harper VanSteenhouse, Yumi Kasai, David Gray, Carol Clugston, Anna Dominiczak and Alex Alderton, Roberto Amato, Jeffrey Barrett, Sonia Goncalves, Ewan Harrison, David K. Jackson, Ian Johnston, Dominic Kwiatkowski, Cordelia Langford, John Sillitoe on behalf of the Wellcome Sanger Institute COVID-19 Surveillance Team |
| EPI_ISL_2730292 | Lighthouse Lab in Alderley Park | Wellcome Sanger Institute for the COVID-19 Genomics UK (COG-UK) Consortium | Jacquelyn Wynn, Mairead Hyland, The Lighthouse Lab in Alderley Park and Alex Alderton, Roberto Amato, Jeffrey Barrett, Sonia Goncalves, Ewan Harrison, David K. Jackson, Ian Johnston, Dominic Kwiatkowski, Cordelia Langford, John Sillitoe on behalf of the Wellcome Sanger Institute COVID-19 Surveillance Team |
| EPI_ISL_2730293 | Lighthouse Lab in Milton Keynes | Wellcome Sanger Institute for the COVID-19 Genomics UK (COG-UK) Consortium | The Lighthouse Lab in Milton Keynes and Alex Alderton, Roberto Amato, Jeffrey Barrett, Sonia Goncalves, Ewan Harrison, David K. Jackson, Ian Johnston, Dominic Kwiatkowski, Cordelia Langford, John Sillitoe on behalf of the Wellcome Sanger Institute COVID-19 Surveillance Team |
| EPI_ISL_2730294, EPI_ISL_2730295, EPI_ISL_2730296, | Lighthouse Lab in Glasgow | Wellcome Sanger Institute for the COVID-19 Genomics UK (COG-UK) Consortium | Harper VanSteenhouse, Yumi Kasai, David Gray, Carol Clugston, Anna Dominiczak and Alex Alderton, Roberto Amato, Jeffrey Barrett, Sonia Goncalves, Ewan Harrison, David K. Jackson, Ian Johnston, Dominic Kwiatkowski, Cordelia Langford, John Sillitoe on behalf of the Wellcome Sanger Institute COVID-19 Surveillance Team |

| | | | |
|---|---|---|---|
| EPI_ISL_2730297, EPI_ISL_2730298 | | | |
| EPI_ISL_2730299 | Lighthouse Lab in Milton Keynes | Wellcome Sanger Institute for the COVID-19 Genomics UK (COG-UK) Consortium | The Lighthouse Lab in Milton Keynes and Alex Alderton, Roberto Amato, Jeffrey Barrett, Sonia Goncalves, Ewan Harrison, David K. Jackson, Ian Johnston, Dominic Kwiatkowski, Cordelia Langford, John Sillitoe on behalf of the Wellcome Sanger Institute COVID-19 Surveillance Team |
| EPI_ISL_2730300 | Lighthouse Lab in Alderley Park | Wellcome Sanger Institute for the COVID-19 Genomics UK (COG-UK) Consortium | Jacquelyn Wynn, Mairead Hyland, The Lighthouse Lab in Alderley Park and Alex Alderton, Roberto Amato, Jeffrey Barrett, Sonia Goncalves, Ewan Harrison, David K. Jackson, Ian Johnston, Dominic Kwiatkowski, Cordelia Langford, John Sillitoe on behalf of the Wellcome Sanger Institute COVID-19 Surveillance Team |
| EPI_ISL_2730301, EPI_ISL_2730303, EPI_ISL_2730304, EPI_ISL_2730305, EPI_ISL_2730306 | Lighthouse Lab in Glasgow | Wellcome Sanger Institute for the COVID-19 Genomics UK (COG-UK) Consortium | Harper VanSteenhouse, Yumi Kasai, David Gray, Carol Clugston, Anna Dominiczak and Alex Alderton, Roberto Amato, Jeffrey Barrett, Sonia Goncalves, Ewan Harrison, David K. Jackson, Ian Johnston, Dominic Kwiatkowski, Cordelia Langford, John Sillitoe on behalf of the Wellcome Sanger Institute COVID-19 Surveillance Team |
| EPI_ISL_2730307, EPI_ISL_2730308 | Lighthouse Lab in Alderley Park | Wellcome Sanger Institute for the COVID-19 Genomics UK (COG-UK) Consortium | Jacquelyn Wynn, Mairead Hyland, The Lighthouse Lab in Alderley Park and Alex Alderton, Roberto Amato, Jeffrey Barrett, Sonia Goncalves, Ewan Harrison, David K. Jackson, Ian Johnston, Dominic Kwiatkowski, Cordelia Langford, John Sillitoe on behalf of the Wellcome Sanger Institute COVID-19 Surveillance Team |
| EPI_ISL_2730309, EPI_ISL_2730310 | Lighthouse Lab in Glasgow | Wellcome Sanger Institute for the COVID-19 Genomics UK (COG-UK) Consortium | Harper VanSteenhouse, Yumi Kasai, David Gray, Carol Clugston, Anna Dominiczak and Alex Alderton, Roberto Amato, Jeffrey Barrett, Sonia Goncalves, Ewan Harrison, David K. Jackson, Ian Johnston, Dominic Kwiatkowski, Cordelia Langford, John Sillitoe on behalf of the Wellcome Sanger Institute COVID-19 Surveillance Team |
| EPI_ISL_2730311 | Lighthouse Lab in Milton Keynes | Wellcome Sanger Institute for the COVID-19 Genomics UK (COG-UK) Consortium | The Lighthouse Lab in Milton Keynes and Alex Alderton, Roberto Amato, Jeffrey Barrett, Sonia Goncalves, Ewan Harrison, David K. Jackson, Ian Johnston, Dominic Kwiatkowski, Cordelia Langford, John Sillitoe on behalf of the Wellcome Sanger Institute COVID-19 Surveillance Team |
| EPI_ISL_2730312, EPI_ISL_2730313, EPI_ISL_2730314, EPI_ISL_2730315, EPI_ISL_2730316 | Lighthouse Lab in Glasgow | Wellcome Sanger Institute for the COVID-19 Genomics UK (COG-UK) Consortium | Harper VanSteenhouse, Yumi Kasai, David Gray, Carol Clugston, Anna Dominiczak and Alex Alderton, Roberto Amato, Jeffrey Barrett, Sonia Goncalves, Ewan Harrison, David K. Jackson, Ian Johnston, Dominic Kwiatkowski, Cordelia Langford, John Sillitoe on behalf of the Wellcome Sanger Institute COVID-19 Surveillance Team |
| EPI_ISL_2730317 | Lighthouse Lab in Alderley Park | Wellcome Sanger Institute for the COVID-19 Genomics UK (COG-UK) Consortium | Jacquelyn Wynn, Mairead Hyland, The Lighthouse Lab in Alderley Park and Alex Alderton, Roberto Amato, Jeffrey Barrett, Sonia Goncalves, Ewan Harrison, David K. Jackson, Ian Johnston, Dominic Kwiatkowski, Cordelia Langford, John Sillitoe on behalf of the Wellcome Sanger Institute COVID-19 Surveillance Team |
| EPI_ISL_2730318 | Lighthouse Lab in Milton Keynes | Wellcome Sanger Institute for the COVID-19 Genomics UK (COG-UK) Consortium | The Lighthouse Lab in Milton Keynes and Alex Alderton, Roberto Amato, Jeffrey Barrett, Sonia Goncalves, Ewan Harrison, David K. Jackson, Ian Johnston, Dominic Kwiatkowski, Cordelia Langford, John Sillitoe on behalf of the Wellcome Sanger Institute COVID-19 Surveillance Team |
| EPI_ISL_2730319 | Lighthouse Lab in Glasgow | Wellcome Sanger Institute for the COVID-19 Genomics UK (COG-UK) Consortium | Harper VanSteenhouse, Yumi Kasai, David Gray, Carol Clugston, Anna Dominiczak and Alex Alderton, Roberto Amato, Jeffrey Barrett, Sonia Goncalves, Ewan Harrison, David K. Jackson, Ian Johnston, Dominic Kwiatkowski, Cordelia Langford, John Sillitoe on behalf of the Wellcome Sanger Institute COVID-19 Surveillance Team |
| EPI_ISL_2730320 | Lighthouse Lab in Milton Keynes | Wellcome Sanger Institute for the COVID-19 Genomics UK (COG-UK) Consortium | The Lighthouse Lab in Milton Keynes and Alex Alderton, Roberto Amato, Jeffrey Barrett, Sonia Goncalves, Ewan Harrison, David K. Jackson, Ian Johnston, Dominic Kwiatkowski, Cordelia Langford, John Sillitoe on behalf of the Wellcome Sanger Institute COVID-19 Surveillance Team |
| EPI_ISL_2730321, EPI_ISL_2730322, EPI_ISL_2730323, EPI_ISL_2730324, EPI_ISL_2730325 | Lighthouse Lab in Glasgow | Wellcome Sanger Institute for the COVID-19 Genomics UK (COG-UK) Consortium | Harper VanSteenhouse, Yumi Kasai, David Gray, Carol Clugston, Anna Dominiczak and Alex Alderton, Roberto Amato, Jeffrey Barrett, Sonia Goncalves, Ewan Harrison, David K. Jackson, Ian Johnston, Dominic Kwiatkowski, Cordelia Langford, John Sillitoe on behalf of the Wellcome Sanger Institute COVID-19 Surveillance Team |
| EPI_ISL_2730326 | Lighthouse Lab in Alderley Park | Wellcome Sanger Institute for the COVID-19 Genomics UK (COG-UK) Consortium | Jacquelyn Wynn, Mairead Hyland, The Lighthouse Lab in Alderley Park and Alex Alderton, Roberto Amato, Jeffrey Barrett, Sonia Goncalves, Ewan Harrison, David K. Jackson, Ian Johnston, Dominic Kwiatkowski, Cordelia Langford, John Sillitoe on behalf of the Wellcome Sanger Institute COVID-19 Surveillance Team |
| EPI_ISL_2730327 | Lighthouse Lab in Glasgow | Wellcome Sanger Institute for the COVID-19 Genomics UK (COG-UK) Consortium | Harper VanSteenhouse, Yumi Kasai, David Gray, Carol Clugston, Anna Dominiczak and Alex Alderton, Roberto Amato, Jeffrey Barrett, Sonia Goncalves, Ewan Harrison, David K. Jackson, Ian Johnston, Dominic Kwiatkowski, Cordelia Langford, John Sillitoe on behalf of the Wellcome Sanger Institute COVID-19 Surveillance Team |
| EPI_ISL_2730328, EPI_ISL_2730329 | Lighthouse Lab in Alderley Park | Wellcome Sanger Institute for the COVID-19 Genomics UK (COG-UK) Consortium | Jacquelyn Wynn, Mairead Hyland, The Lighthouse Lab in Alderley Park and Alex Alderton, Roberto Amato, Jeffrey Barrett, Sonia Goncalves, Ewan Harrison, David K. Jackson, Ian Johnston, Dominic Kwiatkowski, Cordelia Langford, John Sillitoe on behalf of the Wellcome Sanger Institute COVID-19 Surveillance Team |
| EPI_ISL_2730330 | Lighthouse Lab in Milton Keynes | Wellcome Sanger Institute for the COVID-19 Genomics UK (COG-UK) Consortium | The Lighthouse Lab in Milton Keynes and Alex Alderton, Roberto Amato, Jeffrey Barrett, Sonia Goncalves, Ewan Harrison, David K. Jackson, Ian Johnston, Dominic Kwiatkowski, Cordelia Langford, John Sillitoe on behalf of the Wellcome Sanger Institute COVID-19 Surveillance Team |
| EPI_ISL_2730331 | Lighthouse Lab in Alderley Park | Wellcome Sanger Institute for the COVID-19 Genomics UK (COG-UK) Consortium | Jacquelyn Wynn, Mairead Hyland, The Lighthouse Lab in Alderley Park and Alex Alderton, Roberto Amato, Jeffrey Barrett, Sonia Goncalves, Ewan Harrison, David K. Jackson, Ian Johnston, Dominic Kwiatkowski, Cordelia Langford, John Sillitoe on behalf of the Wellcome Sanger Institute COVID-19 Surveillance Team |
| EPI_ISL_2730332 | Lighthouse Lab in Milton Keynes | Wellcome Sanger Institute for the COVID-19 Genomics UK (COG-UK) Consortium | The Lighthouse Lab in Milton Keynes and Alex Alderton, Roberto Amato, Jeffrey Barrett, Sonia Goncalves, Ewan Harrison, David K. Jackson, Ian Johnston, Dominic Kwiatkowski, Cordelia Langford, John Sillitoe on behalf of the Wellcome Sanger Institute COVID-19 Surveillance Team |
| EPI_ISL_2730333 | Lighthouse Lab in Glasgow | Wellcome Sanger Institute for the COVID-19 Genomics UK (COG-UK) Consortium | Harper VanSteenhouse, Yumi Kasai, David Gray, Carol Clugston, Anna Dominiczak and Alex Alderton, Roberto Amato, Jeffrey Barrett, Sonia Goncalves, Ewan Harrison, David K. Jackson, Ian Johnston, Dominic Kwiatkowski, Cordelia Langford, John Sillitoe on behalf of the Wellcome Sanger Institute COVID-19 Surveillance Team |
| EPI_ISL_2730334 | Lighthouse Lab in Milton Keynes | Wellcome Sanger Institute for the COVID-19 Genomics UK (COG-UK) Consortium | The Lighthouse Lab in Milton Keynes and Alex Alderton, Roberto Amato, Jeffrey Barrett, Sonia Goncalves, Ewan Harrison, David K. Jackson, Ian Johnston, Dominic Kwiatkowski, Cordelia Langford, John Sillitoe on behalf of the Wellcome Sanger Institute COVID-19 Surveillance Team |
| EPI_ISL_2730335, EPI_ISL_2730336 | Lighthouse Lab in Alderley Park | Wellcome Sanger Institute for the COVID-19 Genomics UK (COG-UK) Consortium | Jacquelyn Wynn, Mairead Hyland, The Lighthouse Lab in Alderley Park and Alex Alderton, Roberto Amato, Jeffrey Barrett, Sonia Goncalves, Ewan Harrison, David K. Jackson, Ian Johnston, Dominic Kwiatkowski, Cordelia Langford, John Sillitoe on behalf of the Wellcome Sanger Institute COVID-19 Surveillance Team |
| EPI_ISL_2730337 | Lighthouse Lab in Glasgow | Wellcome Sanger Institute for the COVID-19 Genomics UK (COG-UK) Consortium | Harper VanSteenhouse, Yumi Kasai, David Gray, Carol Clugston, Anna Dominiczak and Alex Alderton, Roberto Amato, Jeffrey Barrett, Sonia Goncalves, Ewan Harrison, David K. Jackson, Ian Johnston, Dominic Kwiatkowski, Cordelia Langford, John Sillitoe on behalf of the Wellcome Sanger Institute COVID-19 Surveillance Team |
| EPI_ISL_2730338, EPI_ISL_2730339, EPI_ISL_2730340, EPI_ISL_2730341 | Lighthouse Lab in Alderley Park | Wellcome Sanger Institute for the COVID-19 Genomics UK (COG-UK) Consortium | Jacquelyn Wynn, Mairead Hyland, The Lighthouse Lab in Alderley Park and Alex Alderton, Roberto Amato, Jeffrey Barrett, Sonia Goncalves, Ewan Harrison, David K. Jackson, Ian Johnston, Dominic Kwiatkowski, Cordelia Langford, John Sillitoe on behalf of the Wellcome Sanger Institute COVID-19 Surveillance Team |
| EPI_ISL_2730342 | Lighthouse Lab in Milton Keynes | Wellcome Sanger Institute for the COVID-19 Genomics UK (COG-UK) Consortium | The Lighthouse Lab in Milton Keynes and Alex Alderton, Roberto Amato, Jeffrey Barrett, Sonia Goncalves, Ewan Harrison, David K. Jackson, Ian Johnston, Dominic Kwiatkowski, Cordelia Langford, John Sillitoe on behalf of the Wellcome Sanger Institute COVID-19 Surveillance Team |
| EPI_ISL_2730343, EPI_ISL_2730344 | Lighthouse Lab in Glasgow | Wellcome Sanger Institute for the COVID-19 Genomics UK (COG-UK) Consortium | Harper VanSteenhouse, Yumi Kasai, David Gray, Carol Clugston, Anna Dominiczak and Alex Alderton, Roberto Amato, Jeffrey Barrett, Sonia Goncalves, Ewan Harrison, David K. Jackson, Ian Johnston, Dominic Kwiatkowski, Cordelia Langford, John Sillitoe on behalf of the Wellcome Sanger Institute COVID-19 Surveillance Team |
| EPI_ISL_2730345 | Lighthouse Lab in Alderley Park | Wellcome Sanger Institute for the COVID-19 Genomics UK (COG-UK) Consortium | Jacquelyn Wynn, Mairead Hyland, The Lighthouse Lab in Alderley Park and Alex Alderton, Roberto Amato, Jeffrey Barrett, Sonia Goncalves, Ewan Harrison, David K. Jackson, Ian Johnston, Dominic Kwiatkowski, Cordelia Langford, John Sillitoe on behalf of the Wellcome Sanger Institute COVID-19 Surveillance Team |
| EPI_ISL_2730346, EPI_ISL_2730347, EPI_ISL_2730348 | Lighthouse Lab in Glasgow | Wellcome Sanger Institute for the COVID-19 Genomics UK (COG-UK) Consortium | Harper VanSteenhouse, Yumi Kasai, David Gray, Carol Clugston, Anna Dominiczak and Alex Alderton, Roberto Amato, Jeffrey Barrett, Sonia Goncalves, Ewan Harrison, David K. Jackson, Ian Johnston, Dominic Kwiatkowski, Cordelia Langford, John Sillitoe on behalf of the Wellcome Sanger Institute COVID-19 Surveillance Team |
| EPI_ISL_2730349 | Lighthouse Lab in Milton Keynes | Wellcome Sanger Institute for the COVID-19 Genomics UK (COG-UK) Consortium | The Lighthouse Lab in Milton Keynes and Alex Alderton, Roberto Amato, Jeffrey Barrett, Sonia Goncalves, Ewan Harrison, David K. Jackson, Ian Johnston, Dominic Kwiatkowski, Cordelia Langford, John Sillitoe on behalf of the Wellcome Sanger Institute COVID-19 Surveillance Team |
| EPI_ISL_2730350 | Lighthouse Lab in Glasgow | Wellcome Sanger Institute for the COVID-19 Genomics UK (COG-UK) Consortium | Harper VanSteenhouse, Yumi Kasai, David Gray, Carol Clugston, Anna Dominiczak and Alex Alderton, Roberto Amato, Jeffrey Barrett, Sonia Goncalves, Ewan Harrison, David K. Jackson, Ian Johnston, Dominic Kwiatkowski, Cordelia Langford, John Sillitoe on behalf of the Wellcome Sanger Institute COVID-19 Surveillance Team |
| EPI_ISL_2730351, EPI_ISL_2730352 | Lighthouse Lab in Alderley Park | Wellcome Sanger Institute for the COVID-19 Genomics UK (COG-UK) Consortium | Jacquelyn Wynn, Mairead Hyland, The Lighthouse Lab in Alderley Park and Alex Alderton, Roberto Amato, Jeffrey Barrett, Sonia Goncalves, Ewan Harrison, David K. Jackson, Ian Johnston, Dominic Kwiatkowski, Cordelia Langford, John Sillitoe on behalf of the Wellcome Sanger Institute COVID-19 Surveillance Team |
| EPI_ISL_2730353, EPI_ISL_2730354, EPI_ISL_2730355, EPI_ISL_2730356 | Lighthouse Lab in Glasgow | Wellcome Sanger Institute for the COVID-19 Genomics UK (COG-UK) Consortium | Harper VanSteenhouse, Yumi Kasai, David Gray, Carol Clugston, Anna Dominiczak and Alex Alderton, Roberto Amato, Jeffrey Barrett, Sonia Goncalves, Ewan Harrison, David K. Jackson, Ian Johnston, Dominic Kwiatkowski, Cordelia Langford, John Sillitoe on behalf of the Wellcome Sanger Institute COVID-19 Surveillance Team |
| EPI_ISL_2730357 | Lighthouse Lab in Milton Keynes | Wellcome Sanger Institute for the COVID-19 Genomics UK (COG-UK) Consortium | The Lighthouse Lab in Milton Keynes and Alex Alderton, Roberto Amato, Jeffrey Barrett, Sonia Goncalves, Ewan Harrison, David K. Jackson, Ian Johnston, Dominic Kwiatkowski, Cordelia Langford, John Sillitoe on behalf of the Wellcome Sanger Institute COVID-19 Surveillance Team |
| EPI_ISL_2730358 | Lighthouse Lab in Glasgow | Wellcome Sanger Institute for the COVID-19 Genomics UK (COG-UK) Consortium | Harper VanSteenhouse, Yumi Kasai, David Gray, Carol Clugston, Anna Dominiczak and Alex Alderton, Roberto Amato, Jeffrey Barrett, Sonia Goncalves, Ewan Harrison, David K. Jackson, Ian Johnston, Dominic Kwiatkowski, Cordelia Langford, John Sillitoe on behalf of the Wellcome Sanger Institute COVID-19 Surveillance Team |
| EPI_ISL_2730359 | Lighthouse Lab in Milton Keynes | Wellcome Sanger Institute for the COVID-19 Genomics UK (COG-UK) Consortium | The Lighthouse Lab in Milton Keynes and Alex Alderton, Roberto Amato, Jeffrey Barrett, Sonia Goncalves, Ewan Harrison, David K. Jackson, Ian Johnston, Dominic Kwiatkowski, Cordelia Langford, John Sillitoe on behalf of the Wellcome Sanger Institute COVID-19 Surveillance Team |
| EPI_ISL_2730360, EPI_ISL_2730361, EPI_ISL_2730362, EPI_ISL_2730363, EPI_ISL_2730364, EPI_ISL_2730366 | Lighthouse Lab in Glasgow | Wellcome Sanger Institute for the COVID-19 Genomics UK (COG-UK) Consortium | Harper VanSteenhouse, Yumi Kasai, David Gray, Carol Clugston, Anna Dominiczak and Alex Alderton, Roberto Amato, Jeffrey Barrett, Sonia Goncalves, Ewan Harrison, David K. Jackson, Ian Johnston, Dominic Kwiatkowski, Cordelia Langford, John Sillitoe on behalf of the Wellcome Sanger Institute COVID-19 Surveillance Team |
| EPI_ISL_2730367 | Lighthouse Lab in Alderley Park | Wellcome Sanger Institute for the COVID-19 Genomics UK (COG-UK) Consortium | Jacquelyn Wynn, Mairead Hyland, The Lighthouse Lab in Alderley Park and Alex Alderton, Roberto Amato, Jeffrey Barrett, Sonia Goncalves, Ewan Harrison, David K. Jackson, Ian Johnston, Dominic Kwiatkowski, Cordelia Langford, John Sillitoe on behalf of the Wellcome Sanger Institute COVID-19 Surveillance Team |
| EPI_ISL_2730368 | Lighthouse Lab in Milton Keynes | Wellcome Sanger Institute for the COVID-19 Genomics UK (COG-UK) Consortium | The Lighthouse Lab in Milton Keynes and Alex Alderton, Roberto Amato, Jeffrey Barrett, Sonia Goncalves, Ewan Harrison, David K. Jackson, Ian Johnston, Dominic Kwiatkowski, Cordelia Langford, John Sillitoe on behalf of the Wellcome Sanger Institute COVID-19 Surveillance Team |

| | | | |
|---|---|---|---|
| EPI_ISL_2730369 | Lighthouse Lab in Glasgow | Wellcome Sanger Institute for the COVID-19 Genomics UK (COG-UK) Consortium | Harper VanSteenhouse, Yumi Kasai, David Gray, Carol Clugston, Anna Dominiczak and Alex Alderton, Roberto Amato, Jeffrey Barrett, Sonia Goncalves, Ewan Harrison, David K. Jackson, Ian Johnston, Dominic Kwiatkowski, Cordelia Langford, John Sillitoe on behalf of the Wellcome Sanger Institute COVID-19 Surveillance Team |
| EPI_ISL_2730370 | Lighthouse Lab in Alderley Park | Wellcome Sanger Institute for the COVID-19 Genomics UK (COG-UK) Consortium | Jacquelyn Wynn, Mairead Hyland, The Lighthouse Lab in Alderley Park and Alex Alderton, Roberto Amato, Jeffrey Barrett, Sonia Goncalves, Ewan Harrison, David K. Jackson, Ian Johnston, Dominic Kwiatkowski, Cordelia Langford, John Sillitoe on behalf of the Wellcome Sanger Institute COVID-19 Surveillance Team |
| EPI_ISL_2730371 | Lighthouse Lab in Glasgow | Wellcome Sanger Institute for the COVID-19 Genomics UK (COG-UK) Consortium | Harper VanSteenhouse, Yumi Kasai, David Gray, Carol Clugston, Anna Dominiczak and Alex Alderton, Roberto Amato, Jeffrey Barrett, Sonia Goncalves, Ewan Harrison, David K. Jackson, Ian Johnston, Dominic Kwiatkowski, Cordelia Langford, John Sillitoe on behalf of the Wellcome Sanger Institute COVID-19 Surveillance Team |
| EPI_ISL_2730372 | Lighthouse Lab in Milton Keynes | Wellcome Sanger Institute for the COVID-19 Genomics UK (COG-UK) Consortium | The Lighthouse Lab in Milton Keynes and Alex Alderton, Roberto Amato, Jeffrey Barrett, Sonia Goncalves, Ewan Harrison, David K. Jackson, Ian Johnston, Dominic Kwiatkowski, Cordelia Langford, John Sillitoe on behalf of the Wellcome Sanger Institute COVID-19 Surveillance Team |
| EPI_ISL_2730373 | Lighthouse Lab in Alderley Park | Wellcome Sanger Institute for the COVID-19 Genomics UK (COG-UK) Consortium | Jacquelyn Wynn, Mairead Hyland, The Lighthouse Lab in Alderley Park and Alex Alderton, Roberto Amato, Jeffrey Barrett, Sonia Goncalves, Ewan Harrison, David K. Jackson, Ian Johnston, Dominic Kwiatkowski, Cordelia Langford, John Sillitoe on behalf of the Wellcome Sanger Institute COVID-19 Surveillance Team |
| EPI_ISL_2730374, EPI_ISL_2730375 | Lighthouse Lab in Glasgow | Wellcome Sanger Institute for the COVID-19 Genomics UK (COG-UK) Consortium | Harper VanSteenhouse, Yumi Kasai, David Gray, Carol Clugston, Anna Dominiczak and Alex Alderton, Roberto Amato, Jeffrey Barrett, Sonia Goncalves, Ewan Harrison, David K. Jackson, Ian Johnston, Dominic Kwiatkowski, Cordelia Langford, John Sillitoe on behalf of the Wellcome Sanger Institute COVID-19 Surveillance Team |
| EPI_ISL_2730376 | Lighthouse Lab in Milton Keynes | Wellcome Sanger Institute for the COVID-19 Genomics UK (COG-UK) Consortium | The Lighthouse Lab in Milton Keynes and Alex Alderton, Roberto Amato, Jeffrey Barrett, Sonia Goncalves, Ewan Harrison, David K. Jackson, Ian Johnston, Dominic Kwiatkowski, Cordelia Langford, John Sillitoe on behalf of the Wellcome Sanger Institute COVID-19 Surveillance Team |
| EPI_ISL_2730377, EPI_ISL_2730378, EPI_ISL_2730379, EPI_ISL_2730380, EPI_ISL_2730381 | Lighthouse Lab in Glasgow | Wellcome Sanger Institute for the COVID-19 Genomics UK (COG-UK) Consortium | Harper VanSteenhouse, Yumi Kasai, David Gray, Carol Clugston, Anna Dominiczak and Alex Alderton, Roberto Amato, Jeffrey Barrett, Sonia Goncalves, Ewan Harrison, David K. Jackson, Ian Johnston, Dominic Kwiatkowski, Cordelia Langford, John Sillitoe on behalf of the Wellcome Sanger Institute COVID-19 Surveillance Team |
| EPI_ISL_2730382 | Lighthouse Lab in Alderley Park | Wellcome Sanger Institute for the COVID-19 Genomics UK (COG-UK) Consortium | Jacquelyn Wynn, Mairead Hyland, The Lighthouse Lab in Alderley Park and Alex Alderton, Roberto Amato, Jeffrey Barrett, Sonia Goncalves, Ewan Harrison, David K. Jackson, Ian Johnston, Dominic Kwiatkowski, Cordelia Langford, John Sillitoe on behalf of the Wellcome Sanger Institute COVID-19 Surveillance Team |
| EPI_ISL_2730383 | Lighthouse Lab in Glasgow | Wellcome Sanger Institute for the COVID-19 Genomics UK (COG-UK) Consortium | Harper VanSteenhouse, Yumi Kasai, David Gray, Carol Clugston, Anna Dominiczak and Alex Alderton, Roberto Amato, Jeffrey Barrett, Sonia Goncalves, Ewan Harrison, David K. Jackson, Ian Johnston, Dominic Kwiatkowski, Cordelia Langford, John Sillitoe on behalf of the Wellcome Sanger Institute COVID-19 Surveillance Team |
| EPI_ISL_2730384, EPI_ISL_2730385 | Lighthouse Lab in Alderley Park | Wellcome Sanger Institute for the COVID-19 Genomics UK (COG-UK) Consortium | Jacquelyn Wynn, Mairead Hyland, The Lighthouse Lab in Alderley Park and Alex Alderton, Roberto Amato, Jeffrey Barrett, Sonia Goncalves, Ewan Harrison, David K. Jackson, Ian Johnston, Dominic Kwiatkowski, Cordelia Langford, John Sillitoe on behalf of the Wellcome Sanger Institute COVID-19 Surveillance Team |
| EPI_ISL_2730386, EPI_ISL_2730387, EPI_ISL_2730388, EPI_ISL_2730390 | Lighthouse Lab in Glasgow | Wellcome Sanger Institute for the COVID-19 Genomics UK (COG-UK) Consortium | Harper VanSteenhouse, Yumi Kasai, David Gray, Carol Clugston, Anna Dominiczak and Alex Alderton, Roberto Amato, Jeffrey Barrett, Sonia Goncalves, Ewan Harrison, David K. Jackson, Ian Johnston, Dominic Kwiatkowski, Cordelia Langford, John Sillitoe on behalf of the Wellcome Sanger Institute COVID-19 Surveillance Team |
| EPI_ISL_2730391 | Lighthouse Lab in Alderley Park | Wellcome Sanger Institute for the COVID-19 Genomics UK (COG-UK) Consortium | Jacquelyn Wynn, Mairead Hyland, The Lighthouse Lab in Alderley Park and Alex Alderton, Roberto Amato, Jeffrey Barrett, Sonia Goncalves, Ewan Harrison, David K. Jackson, Ian Johnston, Dominic Kwiatkowski, Cordelia Langford, John Sillitoe on behalf of the Wellcome Sanger Institute COVID-19 Surveillance Team |
| EPI_ISL_2730392 | Lighthouse Lab in Glasgow | Wellcome Sanger Institute for the COVID-19 Genomics UK (COG-UK) Consortium | Harper VanSteenhouse, Yumi Kasai, David Gray, Carol Clugston, Anna Dominiczak and Alex Alderton, Roberto Amato, Jeffrey Barrett, Sonia Goncalves, Ewan Harrison, David K. Jackson, Ian Johnston, Dominic Kwiatkowski, Cordelia Langford, John Sillitoe on behalf of the Wellcome Sanger Institute COVID-19 Surveillance Team |
| EPI_ISL_2730393, EPI_ISL_2730394 | Lighthouse Lab in Alderley Park | Wellcome Sanger Institute for the COVID-19 Genomics UK (COG-UK) Consortium | Jacquelyn Wynn, Mairead Hyland, The Lighthouse Lab in Alderley Park and Alex Alderton, Roberto Amato, Jeffrey Barrett, Sonia Goncalves, Ewan Harrison, David K. Jackson, Ian Johnston, Dominic Kwiatkowski, Cordelia Langford, John Sillitoe on behalf of the Wellcome Sanger Institute COVID-19 Surveillance Team |
| EPI_ISL_2730395 | Lighthouse Lab in Milton Keynes | Wellcome Sanger Institute for the COVID-19 Genomics UK (COG-UK) Consortium | The Lighthouse Lab in Milton Keynes and Alex Alderton, Roberto Amato, Jeffrey Barrett, Sonia Goncalves, Ewan Harrison, David K. Jackson, Ian Johnston, Dominic Kwiatkowski, Cordelia Langford, John Sillitoe on behalf of the Wellcome Sanger Institute COVID-19 Surveillance Team |
| EPI_ISL_2730396 | Lighthouse Lab in Alderley Park | Wellcome Sanger Institute for the COVID-19 Genomics UK (COG-UK) Consortium | Jacquelyn Wynn, Mairead Hyland, The Lighthouse Lab in Alderley Park and Alex Alderton, Roberto Amato, Jeffrey Barrett, Sonia Goncalves, Ewan Harrison, David K. Jackson, Ian Johnston, Dominic Kwiatkowski, Cordelia Langford, John Sillitoe on behalf of the Wellcome Sanger Institute COVID-19 Surveillance Team |
| EPI_ISL_2730397, EPI_ISL_2730398, EPI_ISL_2730399, EPI_ISL_2730400 | Lighthouse Lab in Milton Keynes | Wellcome Sanger Institute for the COVID-19 Genomics UK (COG-UK) Consortium | The Lighthouse Lab in Milton Keynes and Alex Alderton, Roberto Amato, Jeffrey Barrett, Sonia Goncalves, Ewan Harrison, David K. Jackson, Ian Johnston, Dominic Kwiatkowski, Cordelia Langford, John Sillitoe on behalf of the Wellcome Sanger Institute COVID-19 Surveillance Team |
| EPI_ISL_2730401, EPI_ISL_2730402 | Lighthouse Lab in Alderley Park | Wellcome Sanger Institute for the COVID-19 Genomics UK (COG-UK) Consortium | Jacquelyn Wynn, Mairead Hyland, The Lighthouse Lab in Alderley Park and Alex Alderton, Roberto Amato, Jeffrey Barrett, Sonia Goncalves, Ewan Harrison, David K. Jackson, Ian Johnston, Dominic Kwiatkowski, Cordelia Langford, John Sillitoe on behalf of the Wellcome Sanger Institute COVID-19 Surveillance Team |
| EPI_ISL_2730403, EPI_ISL_2730404, EPI_ISL_2730405 | Lighthouse Lab in Glasgow | Wellcome Sanger Institute for the COVID-19 Genomics UK (COG-UK) Consortium | Harper VanSteenhouse, Yumi Kasai, David Gray, Carol Clugston, Anna Dominiczak and Alex Alderton, Roberto Amato, Jeffrey Barrett, Sonia Goncalves, Ewan Harrison, David K. Jackson, Ian Johnston, Dominic Kwiatkowski, Cordelia Langford, John Sillitoe on behalf of the Wellcome Sanger Institute COVID-19 Surveillance Team |
| EPI_ISL_2730406, EPI_ISL_2730407, EPI_ISL_2730408 | Lighthouse Lab in Milton Keynes | Wellcome Sanger Institute for the COVID-19 Genomics UK (COG-UK) Consortium | The Lighthouse Lab in Milton Keynes and Alex Alderton, Roberto Amato, Jeffrey Barrett, Sonia Goncalves, Ewan Harrison, David K. Jackson, Ian Johnston, Dominic Kwiatkowski, Cordelia Langford, John Sillitoe on behalf of the Wellcome Sanger Institute COVID-19 Surveillance Team |
| EPI_ISL_2730409 | Lighthouse Lab in Glasgow | Wellcome Sanger Institute for the COVID-19 Genomics UK (COG-UK) Consortium | Harper VanSteenhouse, Yumi Kasai, David Gray, Carol Clugston, Anna Dominiczak and Alex Alderton, Roberto Amato, Jeffrey Barrett, Sonia Goncalves, Ewan Harrison, David K. Jackson, Ian Johnston, Dominic Kwiatkowski, Cordelia Langford, John Sillitoe on behalf of the Wellcome Sanger Institute COVID-19 Surveillance Team |
| EPI_ISL_2730410 | Lighthouse Lab in Alderley Park | Wellcome Sanger Institute for the COVID-19 Genomics UK (COG-UK) Consortium | Jacquelyn Wynn, Mairead Hyland, The Lighthouse Lab in Alderley Park and Alex Alderton, Roberto Amato, Jeffrey Barrett, Sonia Goncalves, Ewan Harrison, David K. Jackson, Ian Johnston, Dominic Kwiatkowski, Cordelia Langford, John Sillitoe on behalf of the Wellcome Sanger Institute COVID-19 Surveillance Team |
| EPI_ISL_2730411 | Lighthouse Lab in Glasgow | Wellcome Sanger Institute for the COVID-19 Genomics UK (COG-UK) Consortium | Harper VanSteenhouse, Yumi Kasai, David Gray, Carol Clugston, Anna Dominiczak and Alex Alderton, Roberto Amato, Jeffrey Barrett, Sonia Goncalves, Ewan Harrison, David K. Jackson, Ian Johnston, Dominic Kwiatkowski, Cordelia Langford, John Sillitoe on behalf of the Wellcome Sanger Institute COVID-19 Surveillance Team |
| EPI_ISL_2730412 | Lighthouse Lab in Alderley Park | Wellcome Sanger Institute for the COVID-19 Genomics UK (COG-UK) Consortium | Jacquelyn Wynn, Mairead Hyland, The Lighthouse Lab in Alderley Park and Alex Alderton, Roberto Amato, Jeffrey Barrett, Sonia Goncalves, Ewan Harrison, David K. Jackson, Ian Johnston, Dominic Kwiatkowski, Cordelia Langford, John Sillitoe on behalf of the Wellcome Sanger Institute COVID-19 Surveillance Team |
| EPI_ISL_2730413 | Lighthouse Lab in Milton Keynes | Wellcome Sanger Institute for the COVID-19 Genomics UK (COG-UK) Consortium | The Lighthouse Lab in Milton Keynes and Alex Alderton, Roberto Amato, Jeffrey Barrett, Sonia Goncalves, Ewan Harrison, David K. Jackson, Ian Johnston, Dominic Kwiatkowski, Cordelia Langford, John Sillitoe on behalf of the Wellcome Sanger Institute COVID-19 Surveillance Team |
| EPI_ISL_2730414 | Lighthouse Lab in Alderley Park | Wellcome Sanger Institute for the COVID-19 Genomics UK (COG-UK) Consortium | Jacquelyn Wynn, Mairead Hyland, The Lighthouse Lab in Alderley Park and Alex Alderton, Roberto Amato, Jeffrey Barrett, Sonia Goncalves, Ewan Harrison, David K. Jackson, Ian Johnston, Dominic Kwiatkowski, Cordelia Langford, John Sillitoe on behalf of the Wellcome Sanger Institute COVID-19 Surveillance Team |
| EPI_ISL_2730415 | Lighthouse Lab in Glasgow | Wellcome Sanger Institute for the COVID-19 Genomics UK (COG-UK) Consortium | Harper VanSteenhouse, Yumi Kasai, David Gray, Carol Clugston, Anna Dominiczak and Alex Alderton, Roberto Amato, Jeffrey Barrett, Sonia Goncalves, Ewan Harrison, David K. Jackson, Ian Johnston, Dominic Kwiatkowski, Cordelia Langford, John Sillitoe on behalf of the Wellcome Sanger Institute COVID-19 Surveillance Team |
| EPI_ISL_2730416 | Lighthouse Lab in Alderley Park | Wellcome Sanger Institute for the COVID-19 Genomics UK (COG-UK) Consortium | Jacquelyn Wynn, Mairead Hyland, The Lighthouse Lab in Alderley Park and Alex Alderton, Roberto Amato, Jeffrey Barrett, Sonia Goncalves, Ewan Harrison, David K. Jackson, Ian Johnston, Dominic Kwiatkowski, Cordelia Langford, John Sillitoe on behalf of the Wellcome Sanger Institute COVID-19 Surveillance Team |
| EPI_ISL_2730417 | Lighthouse Lab in Milton Keynes | Wellcome Sanger Institute for the COVID-19 Genomics UK (COG-UK) Consortium | The Lighthouse Lab in Milton Keynes and Alex Alderton, Roberto Amato, Jeffrey Barrett, Sonia Goncalves, Ewan Harrison, David K. Jackson, Ian Johnston, Dominic Kwiatkowski, Cordelia Langford, John Sillitoe on behalf of the Wellcome Sanger Institute COVID-19 Surveillance Team |
| EPI_ISL_2730418, EPI_ISL_2730419 | Lighthouse Lab in Glasgow | Wellcome Sanger Institute for the COVID-19 Genomics UK (COG-UK) Consortium | Harper VanSteenhouse, Yumi Kasai, David Gray, Carol Clugston, Anna Dominiczak and Alex Alderton, Roberto Amato, Jeffrey Barrett, Sonia Goncalves, Ewan Harrison, David K. Jackson, Ian Johnston, Dominic Kwiatkowski, Cordelia Langford, John Sillitoe on behalf of the Wellcome Sanger Institute COVID-19 Surveillance Team |
| EPI_ISL_2730420 | Lighthouse Lab in Alderley Park | Wellcome Sanger Institute for the COVID-19 Genomics UK (COG-UK) Consortium | Jacquelyn Wynn, Mairead Hyland, The Lighthouse Lab in Alderley Park and Alex Alderton, Roberto Amato, Jeffrey Barrett, Sonia Goncalves, Ewan Harrison, David K. Jackson, Ian Johnston, Dominic Kwiatkowski, Cordelia Langford, John Sillitoe on behalf of the Wellcome Sanger Institute COVID-19 Surveillance Team |
| EPI_ISL_2730421, EPI_ISL_2730422 | Lighthouse Lab in Glasgow | Wellcome Sanger Institute for the COVID-19 Genomics UK (COG-UK) Consortium | Harper VanSteenhouse, Yumi Kasai, David Gray, Carol Clugston, Anna Dominiczak and Alex Alderton, Roberto Amato, Jeffrey Barrett, Sonia Goncalves, Ewan Harrison, David K. Jackson, Ian Johnston, Dominic Kwiatkowski, Cordelia Langford, John Sillitoe on behalf of the Wellcome Sanger Institute COVID-19 Surveillance Team |
| EPI_ISL_2730423, EPI_ISL_2730424 | Lighthouse Lab in Alderley Park | Wellcome Sanger Institute for the COVID-19 Genomics UK (COG-UK) Consortium | Jacquelyn Wynn, Mairead Hyland, The Lighthouse Lab in Alderley Park and Alex Alderton, Roberto Amato, Jeffrey Barrett, Sonia Goncalves, Ewan Harrison, David K. Jackson, Ian Johnston, Dominic Kwiatkowski, Cordelia Langford, John Sillitoe on behalf of the Wellcome Sanger Institute COVID-19 Surveillance Team |
| EPI_ISL_2730425 | Lighthouse Lab in Milton Keynes | Wellcome Sanger Institute for the COVID-19 Genomics UK (COG-UK) Consortium | The Lighthouse Lab in Milton Keynes and Alex Alderton, Roberto Amato, Jeffrey Barrett, Sonia Goncalves, Ewan Harrison, David K. Jackson, Ian Johnston, Dominic Kwiatkowski, Cordelia Langford, John Sillitoe on behalf of the Wellcome Sanger Institute COVID-19 Surveillance Team |
| EPI_ISL_2730426 | Lighthouse Lab in Glasgow | Wellcome Sanger Institute for the COVID-19 Genomics UK (COG-UK) Consortium | Harper VanSteenhouse, Yumi Kasai, David Gray, Carol Clugston, Anna Dominiczak and Alex Alderton, Roberto Amato, Jeffrey Barrett, Sonia Goncalves, Ewan Harrison, David K. Jackson, Ian Johnston, Dominic Kwiatkowski, Cordelia Langford, John Sillitoe on behalf of the Wellcome Sanger Institute COVID-19 Surveillance Team |
| EPI_ISL_2730427 | Lighthouse Lab in Milton Keynes | Wellcome Sanger Institute for the COVID-19 Genomics UK (COG-UK) Consortium | The Lighthouse Lab in Milton Keynes and Alex Alderton, Roberto Amato, Jeffrey Barrett, Sonia Goncalves, Ewan Harrison, David K. Jackson, Ian Johnston, Dominic Kwiatkowski, Cordelia Langford, John Sillitoe on behalf of the Wellcome Sanger Institute COVID-19 Surveillance Team |
| EPI_ISL_2730428, EPI_ISL_2730429 | Lighthouse Lab in Alderley Park | Wellcome Sanger Institute for the COVID-19 Genomics UK (COG-UK) Consortium | Jacquelyn Wynn, Mairead Hyland, The Lighthouse Lab in Alderley Park and Alex Alderton, Roberto Amato, Jeffrey Barrett, Sonia Goncalves, Ewan Harrison, David K. Jackson, Ian Johnston, Dominic Kwiatkowski, Cordelia Langford, John Sillitoe on behalf of the Wellcome Sanger Institute COVID-19 Surveillance Team |
| EPI_ISL_2730430 | Lighthouse Lab in Glasgow | Wellcome Sanger Institute for the COVID-19 Genomics UK (COG-UK) Consortium | Harper VanSteenhouse, Yumi Kasai, David Gray, Carol Clugston, Anna Dominiczak and Alex Alderton, Roberto Amato, Jeffrey Barrett, Sonia Goncalves, Ewan Harrison, David K. Jackson, Ian Johnston, Dominic Kwiatkowski, Cordelia Langford, John Sillitoe on behalf of the Wellcome Sanger Institute COVID-19 Surveillance Team |
| EPI_ISL_2730431 | Lighthouse Lab in Alderley Park | Wellcome Sanger Institute for the COVID-19 Genomics UK (COG-UK) Consortium | Jacquelyn Wynn, Mairead Hyland, The Lighthouse Lab in Alderley Park and Alex Alderton, Roberto Amato, Jeffrey Barrett, Sonia Goncalves, Ewan Harrison, David K. Jackson, Ian Johnston, Dominic Kwiatkowski, Cordelia Langford, John Sillitoe on behalf of the Wellcome Sanger Institute COVID-19 Surveillance Team |
| EPI_ISL_2730432, EPI_ISL_2730434, EPI_ISL_2730435, | Lighthouse Lab in Glasgow | Wellcome Sanger Institute for the COVID-19 Genomics UK (COG-UK) Consortium | Harper VanSteenhouse, Yumi Kasai, David Gray, Carol Clugston, Anna Dominiczak and Alex Alderton, Roberto Amato, Jeffrey Barrett, Sonia Goncalves, Ewan Harrison, David K. Jackson, Ian Johnston, Dominic Kwiatkowski, Cordelia Langford, John Sillitoe on behalf of the Wellcome Sanger Institute COVID-19 Surveillance Team |

| | | | |
|---|---|---|---|
| EPI_ISL_2730436 | | | |
| EPI_ISL_2730437 | Lighthouse Lab in Milton Keynes | Wellcome Sanger Institute for the COVID-19 Genomics UK (COG-UK) Consortium | The Lighthouse Lab in Milton Keynes and Alex Alderton, Roberto Amato, Jeffrey Barrett, Sonia Goncalves, Ewan Harrison, David K. Jackson, Ian Johnston, Dominic Kwiatkowski, Cordelia Langford, John Sillitoe on behalf of the Wellcome Sanger Institute COVID-19 Surveillance Team |
| EPI_ISL_2730438, EPI_ISL_2730439, EPI_ISL_2730440 | Lighthouse Lab in Glasgow | Wellcome Sanger Institute for the COVID-19 Genomics UK (COG-UK) Consortium | Harper VanSteenhouse, Yumi Kasai, David Gray, Carol Clugston, Anna Dominiczak and Alex Alderton, Roberto Amato, Jeffrey Barrett, Sonia Goncalves, Ewan Harrison, David K. Jackson, Ian Johnston, Dominic Kwiatkowski, Cordelia Langford, John Sillitoe on behalf of the Wellcome Sanger Institute COVID-19 Surveillance Team |
| EPI_ISL_2730441 | Lighthouse Lab in Milton Keynes | Wellcome Sanger Institute for the COVID-19 Genomics UK (COG-UK) Consortium | The Lighthouse Lab in Milton Keynes and Alex Alderton, Roberto Amato, Jeffrey Barrett, Sonia Goncalves, Ewan Harrison, David K. Jackson, Ian Johnston, Dominic Kwiatkowski, Cordelia Langford, John Sillitoe on behalf of the Wellcome Sanger Institute COVID-19 Surveillance Team |
| EPI_ISL_2730442, EPI_ISL_2730443, EPI_ISL_2730444 | Lighthouse Lab in Glasgow | Wellcome Sanger Institute for the COVID-19 Genomics UK (COG-UK) Consortium | Harper VanSteenhouse, Yumi Kasai, David Gray, Carol Clugston, Anna Dominiczak and Alex Alderton, Roberto Amato, Jeffrey Barrett, Sonia Goncalves, Ewan Harrison, David K. Jackson, Ian Johnston, Dominic Kwiatkowski, Cordelia Langford, John Sillitoe on behalf of the Wellcome Sanger Institute COVID-19 Surveillance Team |
| EPI_ISL_2730445 | Lighthouse Lab in Milton Keynes | Wellcome Sanger Institute for the COVID-19 Genomics UK (COG-UK) Consortium | The Lighthouse Lab in Milton Keynes and Alex Alderton, Roberto Amato, Jeffrey Barrett, Sonia Goncalves, Ewan Harrison, David K. Jackson, Ian Johnston, Dominic Kwiatkowski, Cordelia Langford, John Sillitoe on behalf of the Wellcome Sanger Institute COVID-19 Surveillance Team |
| EPI_ISL_2730446 | Lighthouse Lab in Glasgow | Wellcome Sanger Institute for the COVID-19 Genomics UK (COG-UK) Consortium | Harper VanSteenhouse, Yumi Kasai, David Gray, Carol Clugston, Anna Dominiczak and Alex Alderton, Roberto Amato, Jeffrey Barrett, Sonia Goncalves, Ewan Harrison, David K. Jackson, Ian Johnston, Dominic Kwiatkowski, Cordelia Langford, John Sillitoe on behalf of the Wellcome Sanger Institute COVID-19 Surveillance Team |
| EPI_ISL_2730447 | Lighthouse Lab in Alderley Park | Wellcome Sanger Institute for the COVID-19 Genomics UK (COG-UK) Consortium | Jacquelyn Wynn, Mairead Hyland, The Lighthouse Lab in Alderley Park and Alex Alderton, Roberto Amato, Jeffrey Barrett, Sonia Goncalves, Ewan Harrison, David K. Jackson, Ian Johnston, Dominic Kwiatkowski, Cordelia Langford, John Sillitoe on behalf of the Wellcome Sanger Institute COVID-19 Surveillance Team |
| EPI_ISL_2730449 | Lighthouse Lab in Glasgow | Wellcome Sanger Institute for the COVID-19 Genomics UK (COG-UK) Consortium | Harper VanSteenhouse, Yumi Kasai, David Gray, Carol Clugston, Anna Dominiczak and Alex Alderton, Roberto Amato, Jeffrey Barrett, Sonia Goncalves, Ewan Harrison, David K. Jackson, Ian Johnston, Dominic Kwiatkowski, Cordelia Langford, John Sillitoe on behalf of the Wellcome Sanger Institute COVID-19 Surveillance Team |
| EPI_ISL_2730450 | Lighthouse Lab in Alderley Park | Wellcome Sanger Institute for the COVID-19 Genomics UK (COG-UK) Consortium | Jacquelyn Wynn, Mairead Hyland, The Lighthouse Lab in Alderley Park and Alex Alderton, Roberto Amato, Jeffrey Barrett, Sonia Goncalves, Ewan Harrison, David K. Jackson, Ian Johnston, Dominic Kwiatkowski, Cordelia Langford, John Sillitoe on behalf of the Wellcome Sanger Institute COVID-19 Surveillance Team |
| EPI_ISL_2730451 | Lighthouse Lab in Milton Keynes | Wellcome Sanger Institute for the COVID-19 Genomics UK (COG-UK) Consortium | The Lighthouse Lab in Milton Keynes and Alex Alderton, Roberto Amato, Jeffrey Barrett, Sonia Goncalves, Ewan Harrison, David K. Jackson, Ian Johnston, Dominic Kwiatkowski, Cordelia Langford, John Sillitoe on behalf of the Wellcome Sanger Institute COVID-19 Surveillance Team |
| EPI_ISL_2730452, EPI_ISL_2730453 | Lighthouse Lab in Alderley Park | Wellcome Sanger Institute for the COVID-19 Genomics UK (COG-UK) Consortium | Jacquelyn Wynn, Mairead Hyland, The Lighthouse Lab in Alderley Park and Alex Alderton, Roberto Amato, Jeffrey Barrett, Sonia Goncalves, Ewan Harrison, David K. Jackson, Ian Johnston, Dominic Kwiatkowski, Cordelia Langford, John Sillitoe on behalf of the Wellcome Sanger Institute COVID-19 Surveillance Team |
| EPI_ISL_2730454 | Lighthouse Lab in Milton Keynes | Wellcome Sanger Institute for the COVID-19 Genomics UK (COG-UK) Consortium | The Lighthouse Lab in Milton Keynes and Alex Alderton, Roberto Amato, Jeffrey Barrett, Sonia Goncalves, Ewan Harrison, David K. Jackson, Ian Johnston, Dominic Kwiatkowski, Cordelia Langford, John Sillitoe on behalf of the Wellcome Sanger Institute COVID-19 Surveillance Team |
| EPI_ISL_2730455, EPI_ISL_2730456 | Lighthouse Lab in Glasgow | Wellcome Sanger Institute for the COVID-19 Genomics UK (COG-UK) Consortium | Harper VanSteenhouse, Yumi Kasai, David Gray, Carol Clugston, Anna Dominiczak and Alex Alderton, Roberto Amato, Jeffrey Barrett, Sonia Goncalves, Ewan Harrison, David K. Jackson, Ian Johnston, Dominic Kwiatkowski, Cordelia Langford, John Sillitoe on behalf of the Wellcome Sanger Institute COVID-19 Surveillance Team |
| EPI_ISL_2730457 | Lighthouse Lab in Alderley Park | Wellcome Sanger Institute for the COVID-19 Genomics UK (COG-UK) Consortium | Jacquelyn Wynn, Mairead Hyland, The Lighthouse Lab in Alderley Park and Alex Alderton, Roberto Amato, Jeffrey Barrett, Sonia Goncalves, Ewan Harrison, David K. Jackson, Ian Johnston, Dominic Kwiatkowski, Cordelia Langford, John Sillitoe on behalf of the Wellcome Sanger Institute COVID-19 Surveillance Team |
| EPI_ISL_2730458, EPI_ISL_2730459 | Lighthouse Lab in Glasgow | Wellcome Sanger Institute for the COVID-19 Genomics UK (COG-UK) Consortium | Harper VanSteenhouse, Yumi Kasai, David Gray, Carol Clugston, Anna Dominiczak and Alex Alderton, Roberto Amato, Jeffrey Barrett, Sonia Goncalves, Ewan Harrison, David K. Jackson, Ian Johnston, Dominic Kwiatkowski, Cordelia Langford, John Sillitoe on behalf of the Wellcome Sanger Institute COVID-19 Surveillance Team |
| EPI_ISL_2730460 | Lighthouse Lab in Milton Keynes | Wellcome Sanger Institute for the COVID-19 Genomics UK (COG-UK) Consortium | The Lighthouse Lab in Milton Keynes and Alex Alderton, Roberto Amato, Jeffrey Barrett, Sonia Goncalves, Ewan Harrison, David K. Jackson, Ian Johnston, Dominic Kwiatkowski, Cordelia Langford, John Sillitoe on behalf of the Wellcome Sanger Institute COVID-19 Surveillance Team |
| EPI_ISL_2730461 | Lighthouse Lab in Glasgow | Wellcome Sanger Institute for the COVID-19 Genomics UK (COG-UK) Consortium | Harper VanSteenhouse, Yumi Kasai, David Gray, Carol Clugston, Anna Dominiczak and Alex Alderton, Roberto Amato, Jeffrey Barrett, Sonia Goncalves, Ewan Harrison, David K. Jackson, Ian Johnston, Dominic Kwiatkowski, Cordelia Langford, John Sillitoe on behalf of the Wellcome Sanger Institute COVID-19 Surveillance Team |
| EPI_ISL_2730462 | Lighthouse Lab in Alderley Park | Wellcome Sanger Institute for the COVID-19 Genomics UK (COG-UK) Consortium | Jacquelyn Wynn, Mairead Hyland, The Lighthouse Lab in Alderley Park and Alex Alderton, Roberto Amato, Jeffrey Barrett, Sonia Goncalves, Ewan Harrison, David K. Jackson, Ian Johnston, Dominic Kwiatkowski, Cordelia Langford, John Sillitoe on behalf of the Wellcome Sanger Institute COVID-19 Surveillance Team |
| EPI_ISL_2730463, EPI_ISL_2730464 | Lighthouse Lab in Milton Keynes | Wellcome Sanger Institute for the COVID-19 Genomics UK (COG-UK) Consortium | The Lighthouse Lab in Milton Keynes and Alex Alderton, Roberto Amato, Jeffrey Barrett, Sonia Goncalves, Ewan Harrison, David K. Jackson, Ian Johnston, Dominic Kwiatkowski, Cordelia Langford, John Sillitoe on behalf of the Wellcome Sanger Institute COVID-19 Surveillance Team |
| EPI_ISL_2730465, EPI_ISL_2730466, EPI_ISL_2730467, EPI_ISL_2730468 | Lighthouse Lab in Glasgow | Wellcome Sanger Institute for the COVID-19 Genomics UK (COG-UK) Consortium | Harper VanSteenhouse, Yumi Kasai, David Gray, Carol Clugston, Anna Dominiczak and Alex Alderton, Roberto Amato, Jeffrey Barrett, Sonia Goncalves, Ewan Harrison, David K. Jackson, Ian Johnston, Dominic Kwiatkowski, Cordelia Langford, John Sillitoe on behalf of the Wellcome Sanger Institute COVID-19 Surveillance Team |
| EPI_ISL_2730469 | Lighthouse Lab in Alderley Park | Wellcome Sanger Institute for the COVID-19 Genomics UK (COG-UK) Consortium | Jacquelyn Wynn, Mairead Hyland, The Lighthouse Lab in Alderley Park and Alex Alderton, Roberto Amato, Jeffrey Barrett, Sonia Goncalves, Ewan Harrison, David K. Jackson, Ian Johnston, Dominic Kwiatkowski, Cordelia Langford, John Sillitoe on behalf of the Wellcome Sanger Institute COVID-19 Surveillance Team |
| EPI_ISL_2730470 | Lighthouse Lab in Milton Keynes | Wellcome Sanger Institute for the COVID-19 Genomics UK (COG-UK) Consortium | The Lighthouse Lab in Milton Keynes and Alex Alderton, Roberto Amato, Jeffrey Barrett, Sonia Goncalves, Ewan Harrison, David K. Jackson, Ian Johnston, Dominic Kwiatkowski, Cordelia Langford, John Sillitoe on behalf of the Wellcome Sanger Institute COVID-19 Surveillance Team |
| EPI_ISL_2730471 | Lighthouse Lab in Glasgow | Wellcome Sanger Institute for the COVID-19 Genomics UK (COG-UK) Consortium | Harper VanSteenhouse, Yumi Kasai, David Gray, Carol Clugston, Anna Dominiczak and Alex Alderton, Roberto Amato, Jeffrey Barrett, Sonia Goncalves, Ewan Harrison, David K. Jackson, Ian Johnston, Dominic Kwiatkowski, Cordelia Langford, John Sillitoe on behalf of the Wellcome Sanger Institute COVID-19 Surveillance Team |
| EPI_ISL_2730472 | Lighthouse Lab in Alderley Park | Wellcome Sanger Institute for the COVID-19 Genomics UK (COG-UK) Consortium | Jacquelyn Wynn, Mairead Hyland, The Lighthouse Lab in Alderley Park and Alex Alderton, Roberto Amato, Jeffrey Barrett, Sonia Goncalves, Ewan Harrison, David K. Jackson, Ian Johnston, Dominic Kwiatkowski, Cordelia Langford, John Sillitoe on behalf of the Wellcome Sanger Institute COVID-19 Surveillance Team |
| EPI_ISL_2730473 | Lighthouse Lab in Glasgow | Wellcome Sanger Institute for the COVID-19 Genomics UK (COG-UK) Consortium | Harper VanSteenhouse, Yumi Kasai, David Gray, Carol Clugston, Anna Dominiczak and Alex Alderton, Roberto Amato, Jeffrey Barrett, Sonia Goncalves, Ewan Harrison, David K. Jackson, Ian Johnston, Dominic Kwiatkowski, Cordelia Langford, John Sillitoe on behalf of the Wellcome Sanger Institute COVID-19 Surveillance Team |
| EPI_ISL_2730474 | Lighthouse Lab in Alderley Park | Wellcome Sanger Institute for the COVID-19 Genomics UK (COG-UK) Consortium | Jacquelyn Wynn, Mairead Hyland, The Lighthouse Lab in Alderley Park and Alex Alderton, Roberto Amato, Jeffrey Barrett, Sonia Goncalves, Ewan Harrison, David K. Jackson, Ian Johnston, Dominic Kwiatkowski, Cordelia Langford, John Sillitoe on behalf of the Wellcome Sanger Institute COVID-19 Surveillance Team |
| EPI_ISL_2730475, EPI_ISL_2730476 | Lighthouse Lab in Milton Keynes | Wellcome Sanger Institute for the COVID-19 Genomics UK (COG-UK) Consortium | The Lighthouse Lab in Milton Keynes and Alex Alderton, Roberto Amato, Jeffrey Barrett, Sonia Goncalves, Ewan Harrison, David K. Jackson, Ian Johnston, Dominic Kwiatkowski, Cordelia Langford, John Sillitoe on behalf of the Wellcome Sanger Institute COVID-19 Surveillance Team |
| EPI_ISL_2730477, EPI_ISL_2730478, EPI_ISL_2730479, EPI_ISL_2730480, EPI_ISL_2730481 | Lighthouse Lab in Glasgow | Wellcome Sanger Institute for the COVID-19 Genomics UK (COG-UK) Consortium | Harper VanSteenhouse, Yumi Kasai, David Gray, Carol Clugston, Anna Dominiczak and Alex Alderton, Roberto Amato, Jeffrey Barrett, Sonia Goncalves, Ewan Harrison, David K. Jackson, Ian Johnston, Dominic Kwiatkowski, Cordelia Langford, John Sillitoe on behalf of the Wellcome Sanger Institute COVID-19 Surveillance Team |
| EPI_ISL_2730482 | Lighthouse Lab in Milton Keynes | Wellcome Sanger Institute for the COVID-19 Genomics UK (COG-UK) Consortium | The Lighthouse Lab in Milton Keynes and Alex Alderton, Roberto Amato, Jeffrey Barrett, Sonia Goncalves, Ewan Harrison, David K. Jackson, Ian Johnston, Dominic Kwiatkowski, Cordelia Langford, John Sillitoe on behalf of the Wellcome Sanger Institute COVID-19 Surveillance Team |
| EPI_ISL_2730483 | Lighthouse Lab in Glasgow | Wellcome Sanger Institute for the COVID-19 Genomics UK (COG-UK) Consortium | Harper VanSteenhouse, Yumi Kasai, David Gray, Carol Clugston, Anna Dominiczak and Alex Alderton, Roberto Amato, Jeffrey Barrett, Sonia Goncalves, Ewan Harrison, David K. Jackson, Ian Johnston, Dominic Kwiatkowski, Cordelia Langford, John Sillitoe on behalf of the Wellcome Sanger Institute COVID-19 Surveillance Team |
| EPI_ISL_2730484, EPI_ISL_2730485 | Lighthouse Lab in Alderley Park | Wellcome Sanger Institute for the COVID-19 Genomics UK (COG-UK) Consortium | Jacquelyn Wynn, Mairead Hyland, The Lighthouse Lab in Alderley Park and Alex Alderton, Roberto Amato, Jeffrey Barrett, Sonia Goncalves, Ewan Harrison, David K. Jackson, Ian Johnston, Dominic Kwiatkowski, Cordelia Langford, John Sillitoe on behalf of the Wellcome Sanger Institute COVID-19 Surveillance Team |
| EPI_ISL_2730486, EPI_ISL_2730488, EPI_ISL_2730489 | Lighthouse Lab in Glasgow | Wellcome Sanger Institute for the COVID-19 Genomics UK (COG-UK) Consortium | Harper VanSteenhouse, Yumi Kasai, David Gray, Carol Clugston, Anna Dominiczak and Alex Alderton, Roberto Amato, Jeffrey Barrett, Sonia Goncalves, Ewan Harrison, David K. Jackson, Ian Johnston, Dominic Kwiatkowski, Cordelia Langford, John Sillitoe on behalf of the Wellcome Sanger Institute COVID-19 Surveillance Team |
| EPI_ISL_2730490 | Lighthouse Lab in Alderley Park | Wellcome Sanger Institute for the COVID-19 Genomics UK (COG-UK) Consortium | Jacquelyn Wynn, Mairead Hyland, The Lighthouse Lab in Alderley Park and Alex Alderton, Roberto Amato, Jeffrey Barrett, Sonia Goncalves, Ewan Harrison, David K. Jackson, Ian Johnston, Dominic Kwiatkowski, Cordelia Langford, John Sillitoe on behalf of the Wellcome Sanger Institute COVID-19 Surveillance Team |
| EPI_ISL_2730491, EPI_ISL_2730492 | Lighthouse Lab in Glasgow | Wellcome Sanger Institute for the COVID-19 Genomics UK (COG-UK) Consortium | Harper VanSteenhouse, Yumi Kasai, David Gray, Carol Clugston, Anna Dominiczak and Alex Alderton, Roberto Amato, Jeffrey Barrett, Sonia Goncalves, Ewan Harrison, David K. Jackson, Ian Johnston, Dominic Kwiatkowski, Cordelia Langford, John Sillitoe on behalf of the Wellcome Sanger Institute COVID-19 Surveillance Team |
| EPI_ISL_2730493 | Lighthouse Lab in Milton Keynes | Wellcome Sanger Institute for the COVID-19 Genomics UK (COG-UK) Consortium | The Lighthouse Lab in Milton Keynes and Alex Alderton, Roberto Amato, Jeffrey Barrett, Sonia Goncalves, Ewan Harrison, David K. Jackson, Ian Johnston, Dominic Kwiatkowski, Cordelia Langford, John Sillitoe on behalf of the Wellcome Sanger Institute COVID-19 Surveillance Team |
| EPI_ISL_2730494, EPI_ISL_2730495 | Lighthouse Lab in Glasgow | Wellcome Sanger Institute for the COVID-19 Genomics UK (COG-UK) Consortium | Harper VanSteenhouse, Yumi Kasai, David Gray, Carol Clugston, Anna Dominiczak and Alex Alderton, Roberto Amato, Jeffrey Barrett, Sonia Goncalves, Ewan Harrison, David K. Jackson, Ian Johnston, Dominic Kwiatkowski, Cordelia Langford, John Sillitoe on behalf of the Wellcome Sanger Institute COVID-19 Surveillance Team |
| EPI_ISL_2730496 | Lighthouse Lab in Milton Keynes | Wellcome Sanger Institute for the COVID-19 Genomics UK (COG-UK) Consortium | The Lighthouse Lab in Milton Keynes and Alex Alderton, Roberto Amato, Jeffrey Barrett, Sonia Goncalves, Ewan Harrison, David K. Jackson, Ian Johnston, Dominic Kwiatkowski, Cordelia Langford, John Sillitoe on behalf of the Wellcome Sanger Institute COVID-19 Surveillance Team |
| EPI_ISL_2730497 | Lighthouse Lab in Alderley Park | Wellcome Sanger Institute for the COVID-19 Genomics UK (COG-UK) Consortium | Jacquelyn Wynn, Mairead Hyland, The Lighthouse Lab in Alderley Park and Alex Alderton, Roberto Amato, Jeffrey Barrett, Sonia Goncalves, Ewan Harrison, David K. Jackson, Ian Johnston, Dominic Kwiatkowski, Cordelia Langford, John Sillitoe on behalf of the Wellcome Sanger Institute COVID-19 Surveillance Team |
| EPI_ISL_2730498 | Lighthouse Lab in Milton Keynes | Wellcome Sanger Institute for the COVID-19 Genomics UK (COG-UK) Consortium | The Lighthouse Lab in Milton Keynes and Alex Alderton, Roberto Amato, Jeffrey Barrett, Sonia Goncalves, Ewan Harrison, David K. Jackson, Ian Johnston, Dominic Kwiatkowski, Cordelia Langford, John Sillitoe on behalf of the Wellcome Sanger Institute COVID-19 Surveillance Team |
| EPI_ISL_2730499 | Lighthouse Lab in Alderley Park | Wellcome Sanger Institute for the COVID-19 Genomics UK (COG-UK) Consortium | Jacquelyn Wynn, Mairead Hyland, The Lighthouse Lab in Alderley Park and Alex Alderton, Roberto Amato, Jeffrey Barrett, Sonia Goncalves, Ewan Harrison, David K. Jackson, Ian Johnston, Dominic Kwiatkowski, Cordelia Langford, John Sillitoe on behalf of the Wellcome Sanger Institute COVID-19 Surveillance Team |
| EPI_ISL_2730500, EPI_ISL_2730501, EPI_ISL_2730502 | Lighthouse Lab in Glasgow | Wellcome Sanger Institute for the COVID-19 Genomics UK (COG-UK) Consortium | Harper VanSteenhouse, Yumi Kasai, David Gray, Carol Clugston, Anna Dominiczak and Alex Alderton, Roberto Amato, Jeffrey Barrett, Sonia Goncalves, Ewan Harrison, David K. Jackson, Ian Johnston, Dominic Kwiatkowski, Cordelia Langford, John Sillitoe on behalf of the Wellcome Sanger Institute COVID-19 Surveillance Team |

| | | | |
|---|---|---|---|
| EPI_ISL_2730503 | Lighthouse Lab in Milton Keynes | Wellcome Sanger Institute for the COVID-19 Genomics UK (COG-UK) Consortium | The Lighthouse Lab in Milton Keynes and Alex Alderton, Roberto Amato, Jeffrey Barrett, Sonia Goncalves, Ewan Harrison, David K. Jackson, Ian Johnston, Dominic Kwiatkowski, Cordelia Langford, John Sillitoe on behalf of the Wellcome Sanger Institute COVID-19 Surveillance Team |
| EPI_ISL_2730504 | Lighthouse Lab in Glasgow | Wellcome Sanger Institute for the COVID-19 Genomics UK (COG-UK) Consortium | Harper VanSteenhouse, Yumi Kasai, David Gray, Carol Clugston, Anna Dominiczak and Alex Alderton, Roberto Amato, Jeffrey Barrett, Sonia Goncalves, Ewan Harrison, David K. Jackson, Ian Johnston, Dominic Kwiatkowski, Cordelia Langford, John Sillitoe on behalf of the Wellcome Sanger Institute COVID-19 Surveillance Team |
| EPI_ISL_2730505 | Lighthouse Lab in Milton Keynes | Wellcome Sanger Institute for the COVID-19 Genomics UK (COG-UK) Consortium | The Lighthouse Lab in Milton Keynes and Alex Alderton, Roberto Amato, Jeffrey Barrett, Sonia Goncalves, Ewan Harrison, David K. Jackson, Ian Johnston, Dominic Kwiatkowski, Cordelia Langford, John Sillitoe on behalf of the Wellcome Sanger Institute COVID-19 Surveillance Team |
| EPI_ISL_2730506, EPI_ISL_2730507, EPI_ISL_2730508 | Lighthouse Lab in Glasgow | Wellcome Sanger Institute for the COVID-19 Genomics UK (COG-UK) Consortium | Harper VanSteenhouse, Yumi Kasai, David Gray, Carol Clugston, Anna Dominiczak and Alex Alderton, Roberto Amato, Jeffrey Barrett, Sonia Goncalves, Ewan Harrison, David K. Jackson, Ian Johnston, Dominic Kwiatkowski, Cordelia Langford, John Sillitoe on behalf of the Wellcome Sanger Institute COVID-19 Surveillance Team |
| EPI_ISL_2730509 | Lighthouse Lab in Alderley Park | Wellcome Sanger Institute for the COVID-19 Genomics UK (COG-UK) Consortium | Jacquelyn Wynn, Mairead Hyland, The Lighthouse Lab in Alderley Park and Alex Alderton, Roberto Amato, Jeffrey Barrett, Sonia Goncalves, Ewan Harrison, David K. Jackson, Ian Johnston, Dominic Kwiatkowski, Cordelia Langford, John Sillitoe on behalf of the Wellcome Sanger Institute COVID-19 Surveillance Team |
| EPI_ISL_2730510, EPI_ISL_2730511 | Lighthouse Lab in Milton Keynes | Wellcome Sanger Institute for the COVID-19 Genomics UK (COG-UK) Consortium | The Lighthouse Lab in Milton Keynes and Alex Alderton, Roberto Amato, Jeffrey Barrett, Sonia Goncalves, Ewan Harrison, David K. Jackson, Ian Johnston, Dominic Kwiatkowski, Cordelia Langford, John Sillitoe on behalf of the Wellcome Sanger Institute COVID-19 Surveillance Team |
| EPI_ISL_2730512, EPI_ISL_2730513, EPI_ISL_2730514, EPI_ISL_2730515, EPI_ISL_2730516 | Lighthouse Lab in Glasgow | Wellcome Sanger Institute for the COVID-19 Genomics UK (COG-UK) Consortium | Harper VanSteenhouse, Yumi Kasai, David Gray, Carol Clugston, Anna Dominiczak and Alex Alderton, Roberto Amato, Jeffrey Barrett, Sonia Goncalves, Ewan Harrison, David K. Jackson, Ian Johnston, Dominic Kwiatkowski, Cordelia Langford, John Sillitoe on behalf of the Wellcome Sanger Institute COVID-19 Surveillance Team |
| EPI_ISL_2730517 | Lighthouse Lab in Alderley Park | Wellcome Sanger Institute for the COVID-19 Genomics UK (COG-UK) Consortium | Jacquelyn Wynn, Mairead Hyland, The Lighthouse Lab in Alderley Park and Alex Alderton, Roberto Amato, Jeffrey Barrett, Sonia Goncalves, Ewan Harrison, David K. Jackson, Ian Johnston, Dominic Kwiatkowski, Cordelia Langford, John Sillitoe on behalf of the Wellcome Sanger Institute COVID-19 Surveillance Team |
| EPI_ISL_2730518 | Lighthouse Lab in Glasgow | Wellcome Sanger Institute for the COVID-19 Genomics UK (COG-UK) Consortium | Harper VanSteenhouse, Yumi Kasai, David Gray, Carol Clugston, Anna Dominiczak and Alex Alderton, Roberto Amato, Jeffrey Barrett, Sonia Goncalves, Ewan Harrison, David K. Jackson, Ian Johnston, Dominic Kwiatkowski, Cordelia Langford, John Sillitoe on behalf of the Wellcome Sanger Institute COVID-19 Surveillance Team |
| EPI_ISL_2730519, EPI_ISL_2730520 | Lighthouse Lab in Alderley Park | Wellcome Sanger Institute for the COVID-19 Genomics UK (COG-UK) Consortium | Jacquelyn Wynn, Mairead Hyland, The Lighthouse Lab in Alderley Park and Alex Alderton, Roberto Amato, Jeffrey Barrett, Sonia Goncalves, Ewan Harrison, David K. Jackson, Ian Johnston, Dominic Kwiatkowski, Cordelia Langford, John Sillitoe on behalf of the Wellcome Sanger Institute COVID-19 Surveillance Team |
| EPI_ISL_2730521 | Lighthouse Lab in Milton Keynes | Wellcome Sanger Institute for the COVID-19 Genomics UK (COG-UK) Consortium | The Lighthouse Lab in Milton Keynes and Alex Alderton, Roberto Amato, Jeffrey Barrett, Sonia Goncalves, Ewan Harrison, David K. Jackson, Ian Johnston, Dominic Kwiatkowski, Cordelia Langford, John Sillitoe on behalf of the Wellcome Sanger Institute COVID-19 Surveillance Team |
| EPI_ISL_2730522 | Lighthouse Lab in Alderley Park | Wellcome Sanger Institute for the COVID-19 Genomics UK (COG-UK) Consortium | Jacquelyn Wynn, Mairead Hyland, The Lighthouse Lab in Alderley Park and Alex Alderton, Roberto Amato, Jeffrey Barrett, Sonia Goncalves, Ewan Harrison, David K. Jackson, Ian Johnston, Dominic Kwiatkowski, Cordelia Langford, John Sillitoe on behalf of the Wellcome Sanger Institute COVID-19 Surveillance Team |
| EPI_ISL_2730523 | Lighthouse Lab in Glasgow | Wellcome Sanger Institute for the COVID-19 Genomics UK (COG-UK) Consortium | Harper VanSteenhouse, Yumi Kasai, David Gray, Carol Clugston, Anna Dominiczak and Alex Alderton, Roberto Amato, Jeffrey Barrett, Sonia Goncalves, Ewan Harrison, David K. Jackson, Ian Johnston, Dominic Kwiatkowski, Cordelia Langford, John Sillitoe on behalf of the Wellcome Sanger Institute COVID-19 Surveillance Team |
| EPI_ISL_2730524 | Lighthouse Lab in Milton Keynes | Wellcome Sanger Institute for the COVID-19 Genomics UK (COG-UK) Consortium | The Lighthouse Lab in Milton Keynes and Alex Alderton, Roberto Amato, Jeffrey Barrett, Sonia Goncalves, Ewan Harrison, David K. Jackson, Ian Johnston, Dominic Kwiatkowski, Cordelia Langford, John Sillitoe on behalf of the Wellcome Sanger Institute COVID-19 Surveillance Team |
| EPI_ISL_2730525, EPI_ISL_2730526 | Lighthouse Lab in Glasgow | Wellcome Sanger Institute for the COVID-19 Genomics UK (COG-UK) Consortium | Harper VanSteenhouse, Yumi Kasai, David Gray, Carol Clugston, Anna Dominiczak and Alex Alderton, Roberto Amato, Jeffrey Barrett, Sonia Goncalves, Ewan Harrison, David K. Jackson, Ian Johnston, Dominic Kwiatkowski, Cordelia Langford, John Sillitoe on behalf of the Wellcome Sanger Institute COVID-19 Surveillance Team |
| EPI_ISL_2730527 | Lighthouse Lab in Alderley Park | Wellcome Sanger Institute for the COVID-19 Genomics UK (COG-UK) Consortium | Jacquelyn Wynn, Mairead Hyland, The Lighthouse Lab in Alderley Park and Alex Alderton, Roberto Amato, Jeffrey Barrett, Sonia Goncalves, Ewan Harrison, David K. Jackson, Ian Johnston, Dominic Kwiatkowski, Cordelia Langford, John Sillitoe on behalf of the Wellcome Sanger Institute COVID-19 Surveillance Team |
| EPI_ISL_2730528 | Lighthouse Lab in Glasgow | Wellcome Sanger Institute for the COVID-19 Genomics UK (COG-UK) Consortium | Harper VanSteenhouse, Yumi Kasai, David Gray, Carol Clugston, Anna Dominiczak and Alex Alderton, Roberto Amato, Jeffrey Barrett, Sonia Goncalves, Ewan Harrison, David K. Jackson, Ian Johnston, Dominic Kwiatkowski, Cordelia Langford, John Sillitoe on behalf of the Wellcome Sanger Institute COVID-19 Surveillance Team |
| EPI_ISL_2730529 | Lighthouse Lab in Alderley Park | Wellcome Sanger Institute for the COVID-19 Genomics UK (COG-UK) Consortium | Jacquelyn Wynn, Mairead Hyland, The Lighthouse Lab in Alderley Park and Alex Alderton, Roberto Amato, Jeffrey Barrett, Sonia Goncalves, Ewan Harrison, David K. Jackson, Ian Johnston, Dominic Kwiatkowski, Cordelia Langford, John Sillitoe on behalf of the Wellcome Sanger Institute COVID-19 Surveillance Team |
| EPI_ISL_2730530, EPI_ISL_2730531 | Lighthouse Lab in Milton Keynes | Wellcome Sanger Institute for the COVID-19 Genomics UK (COG-UK) Consortium | The Lighthouse Lab in Milton Keynes and Alex Alderton, Roberto Amato, Jeffrey Barrett, Sonia Goncalves, Ewan Harrison, David K. Jackson, Ian Johnston, Dominic Kwiatkowski, Cordelia Langford, John Sillitoe on behalf of the Wellcome Sanger Institute COVID-19 Surveillance Team |
| EPI_ISL_2730532 | Lighthouse Lab in Alderley Park | Wellcome Sanger Institute for the COVID-19 Genomics UK (COG-UK) Consortium | Jacquelyn Wynn, Mairead Hyland, The Lighthouse Lab in Alderley Park and Alex Alderton, Roberto Amato, Jeffrey Barrett, Sonia Goncalves, Ewan Harrison, David K. Jackson, Ian Johnston, Dominic Kwiatkowski, Cordelia Langford, John Sillitoe on behalf of the Wellcome Sanger Institute COVID-19 Surveillance Team |
| EPI_ISL_2730533, EPI_ISL_2730534 | Lighthouse Lab in Glasgow | Wellcome Sanger Institute for the COVID-19 Genomics UK (COG-UK) Consortium | Harper VanSteenhouse, Yumi Kasai, David Gray, Carol Clugston, Anna Dominiczak and Alex Alderton, Roberto Amato, Jeffrey Barrett, Sonia Goncalves, Ewan Harrison, David K. Jackson, Ian Johnston, Dominic Kwiatkowski, Cordelia Langford, John Sillitoe on behalf of the Wellcome Sanger Institute COVID-19 Surveillance Team |
| EPI_ISL_2730535, EPI_ISL_2730536 | Lighthouse Lab in Milton Keynes | Wellcome Sanger Institute for the COVID-19 Genomics UK (COG-UK) Consortium | The Lighthouse Lab in Milton Keynes and Alex Alderton, Roberto Amato, Jeffrey Barrett, Sonia Goncalves, Ewan Harrison, David K. Jackson, Ian Johnston, Dominic Kwiatkowski, Cordelia Langford, John Sillitoe on behalf of the Wellcome Sanger Institute COVID-19 Surveillance Team |
| EPI_ISL_2730537, EPI_ISL_2730538, EPI_ISL_2730539, EPI_ISL_2730540 | Lighthouse Lab in Glasgow | Wellcome Sanger Institute for the COVID-19 Genomics UK (COG-UK) Consortium | Harper VanSteenhouse, Yumi Kasai, David Gray, Carol Clugston, Anna Dominiczak and Alex Alderton, Roberto Amato, Jeffrey Barrett, Sonia Goncalves, Ewan Harrison, David K. Jackson, Ian Johnston, Dominic Kwiatkowski, Cordelia Langford, John Sillitoe on behalf of the Wellcome Sanger Institute COVID-19 Surveillance Team |
| EPI_ISL_2730542, EPI_ISL_2730543 | Lighthouse Lab in Alderley Park | Wellcome Sanger Institute for the COVID-19 Genomics UK (COG-UK) Consortium | Jacquelyn Wynn, Mairead Hyland, The Lighthouse Lab in Alderley Park and Alex Alderton, Roberto Amato, Jeffrey Barrett, Sonia Goncalves, Ewan Harrison, David K. Jackson, Ian Johnston, Dominic Kwiatkowski, Cordelia Langford, John Sillitoe on behalf of the Wellcome Sanger Institute COVID-19 Surveillance Team |
| EPI_ISL_2730544, EPI_ISL_2730545, EPI_ISL_2730546 | Lighthouse Lab in Milton Keynes | Wellcome Sanger Institute for the COVID-19 Genomics UK (COG-UK) Consortium | The Lighthouse Lab in Milton Keynes and Alex Alderton, Roberto Amato, Jeffrey Barrett, Sonia Goncalves, Ewan Harrison, David K. Jackson, Ian Johnston, Dominic Kwiatkowski, Cordelia Langford, John Sillitoe on behalf of the Wellcome Sanger Institute COVID-19 Surveillance Team |
| EPI_ISL_2730547, EPI_ISL_2730548 | Lighthouse Lab in Alderley Park | Wellcome Sanger Institute for the COVID-19 Genomics UK (COG-UK) Consortium | Jacquelyn Wynn, Mairead Hyland, The Lighthouse Lab in Alderley Park and Alex Alderton, Roberto Amato, Jeffrey Barrett, Sonia Goncalves, Ewan Harrison, David K. Jackson, Ian Johnston, Dominic Kwiatkowski, Cordelia Langford, John Sillitoe on behalf of the Wellcome Sanger Institute COVID-19 Surveillance Team |
| EPI_ISL_2730549 | Lighthouse Lab in Glasgow | Wellcome Sanger Institute for the COVID-19 Genomics UK (COG-UK) Consortium | Harper VanSteenhouse, Yumi Kasai, David Gray, Carol Clugston, Anna Dominiczak and Alex Alderton, Roberto Amato, Jeffrey Barrett, Sonia Goncalves, Ewan Harrison, David K. Jackson, Ian Johnston, Dominic Kwiatkowski, Cordelia Langford, John Sillitoe on behalf of the Wellcome Sanger Institute COVID-19 Surveillance Team |
| EPI_ISL_2730550 | Lighthouse Lab in Alderley Park | Wellcome Sanger Institute for the COVID-19 Genomics UK (COG-UK) Consortium | Jacquelyn Wynn, Mairead Hyland, The Lighthouse Lab in Alderley Park and Alex Alderton, Roberto Amato, Jeffrey Barrett, Sonia Goncalves, Ewan Harrison, David K. Jackson, Ian Johnston, Dominic Kwiatkowski, Cordelia Langford, John Sillitoe on behalf of the Wellcome Sanger Institute COVID-19 Surveillance Team |
| EPI_ISL_2730551 | Lighthouse Lab in Milton Keynes | Wellcome Sanger Institute for the COVID-19 Genomics UK (COG-UK) Consortium | The Lighthouse Lab in Milton Keynes and Alex Alderton, Roberto Amato, Jeffrey Barrett, Sonia Goncalves, Ewan Harrison, David K. Jackson, Ian Johnston, Dominic Kwiatkowski, Cordelia Langford, John Sillitoe on behalf of the Wellcome Sanger Institute COVID-19 Surveillance Team |
| EPI_ISL_2730552 | Lighthouse Lab in Glasgow | Wellcome Sanger Institute for the COVID-19 Genomics UK (COG-UK) Consortium | Harper VanSteenhouse, Yumi Kasai, David Gray, Carol Clugston, Anna Dominiczak and Alex Alderton, Roberto Amato, Jeffrey Barrett, Sonia Goncalves, Ewan Harrison, David K. Jackson, Ian Johnston, Dominic Kwiatkowski, Cordelia Langford, John Sillitoe on behalf of the Wellcome Sanger Institute COVID-19 Surveillance Team |
| EPI_ISL_2730553 | Lighthouse Lab in Alderley Park | Wellcome Sanger Institute for the COVID-19 Genomics UK (COG-UK) Consortium | Jacquelyn Wynn, Mairead Hyland, The Lighthouse Lab in Alderley Park and Alex Alderton, Roberto Amato, Jeffrey Barrett, Sonia Goncalves, Ewan Harrison, David K. Jackson, Ian Johnston, Dominic Kwiatkowski, Cordelia Langford, John Sillitoe on behalf of the Wellcome Sanger Institute COVID-19 Surveillance Team |
| EPI_ISL_2730554 | Lighthouse Lab in Glasgow | Wellcome Sanger Institute for the COVID-19 Genomics UK (COG-UK) Consortium | Harper VanSteenhouse, Yumi Kasai, David Gray, Carol Clugston, Anna Dominiczak and Alex Alderton, Roberto Amato, Jeffrey Barrett, Sonia Goncalves, Ewan Harrison, David K. Jackson, Ian Johnston, Dominic Kwiatkowski, Cordelia Langford, John Sillitoe on behalf of the Wellcome Sanger Institute COVID-19 Surveillance Team |
| EPI_ISL_2730555 | Lighthouse Lab in Milton Keynes | Wellcome Sanger Institute for the COVID-19 Genomics UK (COG-UK) Consortium | The Lighthouse Lab in Milton Keynes and Alex Alderton, Roberto Amato, Jeffrey Barrett, Sonia Goncalves, Ewan Harrison, David K. Jackson, Ian Johnston, Dominic Kwiatkowski, Cordelia Langford, John Sillitoe on behalf of the Wellcome Sanger Institute COVID-19 Surveillance Team |
| EPI_ISL_2730556 | Lighthouse Lab in Alderley Park | Wellcome Sanger Institute for the COVID-19 Genomics UK (COG-UK) Consortium | Jacquelyn Wynn, Mairead Hyland, The Lighthouse Lab in Alderley Park and Alex Alderton, Roberto Amato, Jeffrey Barrett, Sonia Goncalves, Ewan Harrison, David K. Jackson, Ian Johnston, Dominic Kwiatkowski, Cordelia Langford, John Sillitoe on behalf of the Wellcome Sanger Institute COVID-19 Surveillance Team |
| EPI_ISL_2730557 | Lighthouse Lab in Glasgow | Wellcome Sanger Institute for the COVID-19 Genomics UK (COG-UK) Consortium | Harper VanSteenhouse, Yumi Kasai, David Gray, Carol Clugston, Anna Dominiczak and Alex Alderton, Roberto Amato, Jeffrey Barrett, Sonia Goncalves, Ewan Harrison, David K. Jackson, Ian Johnston, Dominic Kwiatkowski, Cordelia Langford, John Sillitoe on behalf of the Wellcome Sanger Institute COVID-19 Surveillance Team |
| EPI_ISL_2730558 | Lighthouse Lab in Milton Keynes | Wellcome Sanger Institute for the COVID-19 Genomics UK (COG-UK) Consortium | The Lighthouse Lab in Milton Keynes and Alex Alderton, Roberto Amato, Jeffrey Barrett, Sonia Goncalves, Ewan Harrison, David K. Jackson, Ian Johnston, Dominic Kwiatkowski, Cordelia Langford, John Sillitoe on behalf of the Wellcome Sanger Institute COVID-19 Surveillance Team |
| EPI_ISL_2730559 | Lighthouse Lab in Glasgow | Wellcome Sanger Institute for the COVID-19 Genomics UK (COG-UK) Consortium | Harper VanSteenhouse, Yumi Kasai, David Gray, Carol Clugston, Anna Dominiczak and Alex Alderton, Roberto Amato, Jeffrey Barrett, Sonia Goncalves, Ewan Harrison, David K. Jackson, Ian Johnston, Dominic Kwiatkowski, Cordelia Langford, John Sillitoe on behalf of the Wellcome Sanger Institute COVID-19 Surveillance Team |
| EPI_ISL_2730560 | Lighthouse Lab in Alderley Park | Wellcome Sanger Institute for the COVID-19 Genomics UK (COG-UK) Consortium | Jacquelyn Wynn, Mairead Hyland, The Lighthouse Lab in Alderley Park and Alex Alderton, Roberto Amato, Jeffrey Barrett, Sonia Goncalves, Ewan Harrison, David K. Jackson, Ian Johnston, Dominic Kwiatkowski, Cordelia Langford, John Sillitoe on behalf of the Wellcome Sanger Institute COVID-19 Surveillance Team |
| EPI_ISL_2730561 | Lighthouse Lab in Glasgow | Wellcome Sanger Institute for the COVID-19 Genomics UK (COG-UK) Consortium | Harper VanSteenhouse, Yumi Kasai, David Gray, Carol Clugston, Anna Dominiczak and Alex Alderton, Roberto Amato, Jeffrey Barrett, Sonia Goncalves, Ewan Harrison, David K. Jackson, Ian Johnston, Dominic Kwiatkowski, Cordelia Langford, John Sillitoe on behalf of the Wellcome Sanger Institute COVID-19 Surveillance Team |
| EPI_ISL_2730562 | Lighthouse Lab in Alderley Park | Wellcome Sanger Institute for the COVID-19 Genomics UK (COG-UK) Consortium | Jacquelyn Wynn, Mairead Hyland, The Lighthouse Lab in Alderley Park and Alex Alderton, Roberto Amato, Jeffrey Barrett, Sonia Goncalves, Ewan Harrison, David K. Jackson, Ian Johnston, Dominic Kwiatkowski, Cordelia Langford, John Sillitoe on behalf of the Wellcome Sanger Institute COVID-19 Surveillance Team |
| EPI_ISL_2730563 | Lighthouse Lab in Glasgow | Wellcome Sanger Institute for the COVID-19 Genomics UK (COG-UK) Consortium | Harper VanSteenhouse, Yumi Kasai, David Gray, Carol Clugston, Anna Dominiczak and Alex Alderton, Roberto Amato, Jeffrey Barrett, Sonia Goncalves, Ewan Harrison, David K. Jackson, Ian Johnston, Dominic Kwiatkowski, Cordelia Langford, John Sillitoe on behalf of the Wellcome Sanger Institute COVID-19 Surveillance Team |
| EPI_ISL_2730564, EPI_ISL_2730565 | Lighthouse Lab in Milton Keynes | Wellcome Sanger Institute for the COVID-19 Genomics UK (COG-UK) Consortium | The Lighthouse Lab in Milton Keynes and Alex Alderton, Roberto Amato, Jeffrey Barrett, Sonia Goncalves, Ewan Harrison, David K. Jackson, Ian Johnston, Dominic Kwiatkowski, Cordelia Langford, John Sillitoe on behalf of the Wellcome Sanger Institute COVID-19 Surveillance Team |

| | | | |
|---|---|---|---|
| EPI_ISL_2730566 | Lighthouse Lab in Glasgow | Wellcome Sanger Institute for the COVID-19 Genomics UK (COG-UK) Consortium | Harper VanSteenhouse, Yumi Kasai, David Gray, Carol Clugston, Anna Dominiczak and Alex Alderton, Roberto Amato, Jeffrey Barrett, Sonia Goncalves, Ewan Harrison, David K. Jackson, Ian Johnston, Dominic Kwiatkowski, Cordelia Langford, John Sillitoe on behalf of the Wellcome Sanger Institute COVID-19 Surveillance Team |
| EPI_ISL_2730567 | Lighthouse Lab in Alderley Park | Wellcome Sanger Institute for the COVID-19 Genomics UK (COG-UK) Consortium | Jacquelyn Wynn, Mairead Hyland, The Lighthouse Lab in Alderley Park and Alex Alderton, Roberto Amato, Jeffrey Barrett, Sonia Goncalves, Ewan Harrison, David K. Jackson, Ian Johnston, Dominic Kwiatkowski, Cordelia Langford, John Sillitoe on behalf of the Wellcome Sanger Institute COVID-19 Surveillance Team |
| EPI_ISL_2730568 | Lighthouse Lab in Milton Keynes | Wellcome Sanger Institute for the COVID-19 Genomics UK (COG-UK) Consortium | The Lighthouse Lab in Milton Keynes and Alex Alderton, Roberto Amato, Jeffrey Barrett, Sonia Goncalves, Ewan Harrison, David K. Jackson, Ian Johnston, Dominic Kwiatkowski, Cordelia Langford, John Sillitoe on behalf of the Wellcome Sanger Institute COVID-19 Surveillance Team |
| EPI_ISL_2730569 | Lighthouse Lab in Alderley Park | Wellcome Sanger Institute for the COVID-19 Genomics UK (COG-UK) Consortium | Jacquelyn Wynn, Mairead Hyland, The Lighthouse Lab in Alderley Park and Alex Alderton, Roberto Amato, Jeffrey Barrett, Sonia Goncalves, Ewan Harrison, David K. Jackson, Ian Johnston, Dominic Kwiatkowski, Cordelia Langford, John Sillitoe on behalf of the Wellcome Sanger Institute COVID-19 Surveillance Team |
| EPI_ISL_2730570, EPI_ISL_2730571 | Lighthouse Lab in Glasgow | Wellcome Sanger Institute for the COVID-19 Genomics UK (COG-UK) Consortium | Harper VanSteenhouse, Yumi Kasai, David Gray, Carol Clugston, Anna Dominiczak and Alex Alderton, Roberto Amato, Jeffrey Barrett, Sonia Goncalves, Ewan Harrison, David K. Jackson, Ian Johnston, Dominic Kwiatkowski, Cordelia Langford, John Sillitoe on behalf of the Wellcome Sanger Institute COVID-19 Surveillance Team |
| EPI_ISL_2730572 | Lighthouse Lab in Alderley Park | Wellcome Sanger Institute for the COVID-19 Genomics UK (COG-UK) Consortium | Jacquelyn Wynn, Mairead Hyland, The Lighthouse Lab in Alderley Park and Alex Alderton, Roberto Amato, Jeffrey Barrett, Sonia Goncalves, Ewan Harrison, David K. Jackson, Ian Johnston, Dominic Kwiatkowski, Cordelia Langford, John Sillitoe on behalf of the Wellcome Sanger Institute COVID-19 Surveillance Team |
| EPI_ISL_2730573 | Lighthouse Lab in Milton Keynes | Wellcome Sanger Institute for the COVID-19 Genomics UK (COG-UK) Consortium | The Lighthouse Lab in Milton Keynes and Alex Alderton, Roberto Amato, Jeffrey Barrett, Sonia Goncalves, Ewan Harrison, David K. Jackson, Ian Johnston, Dominic Kwiatkowski, Cordelia Langford, John Sillitoe on behalf of the Wellcome Sanger Institute COVID-19 Surveillance Team |
| EPI_ISL_2730574 | Lighthouse Lab in Glasgow | Wellcome Sanger Institute for the COVID-19 Genomics UK (COG-UK) Consortium | Harper VanSteenhouse, Yumi Kasai, David Gray, Carol Clugston, Anna Dominiczak and Alex Alderton, Roberto Amato, Jeffrey Barrett, Sonia Goncalves, Ewan Harrison, David K. Jackson, Ian Johnston, Dominic Kwiatkowski, Cordelia Langford, John Sillitoe on behalf of the Wellcome Sanger Institute COVID-19 Surveillance Team |
| EPI_ISL_2730575 | Lighthouse Lab in Milton Keynes | Wellcome Sanger Institute for the COVID-19 Genomics UK (COG-UK) Consortium | The Lighthouse Lab in Milton Keynes and Alex Alderton, Roberto Amato, Jeffrey Barrett, Sonia Goncalves, Ewan Harrison, David K. Jackson, Ian Johnston, Dominic Kwiatkowski, Cordelia Langford, John Sillitoe on behalf of the Wellcome Sanger Institute COVID-19 Surveillance Team |
| EPI_ISL_2730576 | Lighthouse Lab in Glasgow | Wellcome Sanger Institute for the COVID-19 Genomics UK (COG-UK) Consortium | Harper VanSteenhouse, Yumi Kasai, David Gray, Carol Clugston, Anna Dominiczak and Alex Alderton, Roberto Amato, Jeffrey Barrett, Sonia Goncalves, Ewan Harrison, David K. Jackson, Ian Johnston, Dominic Kwiatkowski, Cordelia Langford, John Sillitoe on behalf of the Wellcome Sanger Institute COVID-19 Surveillance Team |
| EPI_ISL_2730577 | Lighthouse Lab in Alderley Park | Wellcome Sanger Institute for the COVID-19 Genomics UK (COG-UK) Consortium | Jacquelyn Wynn, Mairead Hyland, The Lighthouse Lab in Alderley Park and Alex Alderton, Roberto Amato, Jeffrey Barrett, Sonia Goncalves, Ewan Harrison, David K. Jackson, Ian Johnston, Dominic Kwiatkowski, Cordelia Langford, John Sillitoe on behalf of the Wellcome Sanger Institute COVID-19 Surveillance Team |
| EPI_ISL_2730578 | Lighthouse Lab in Milton Keynes | Wellcome Sanger Institute for the COVID-19 Genomics UK (COG-UK) Consortium | The Lighthouse Lab in Milton Keynes and Alex Alderton, Roberto Amato, Jeffrey Barrett, Sonia Goncalves, Ewan Harrison, David K. Jackson, Ian Johnston, Dominic Kwiatkowski, Cordelia Langford, John Sillitoe on behalf of the Wellcome Sanger Institute COVID-19 Surveillance Team |
| EPI_ISL_2730579 | Lighthouse Lab in Glasgow | Wellcome Sanger Institute for the COVID-19 Genomics UK (COG-UK) Consortium | Harper VanSteenhouse, Yumi Kasai, David Gray, Carol Clugston, Anna Dominiczak and Alex Alderton, Roberto Amato, Jeffrey Barrett, Sonia Goncalves, Ewan Harrison, David K. Jackson, Ian Johnston, Dominic Kwiatkowski, Cordelia Langford, John Sillitoe on behalf of the Wellcome Sanger Institute COVID-19 Surveillance Team |
| EPI_ISL_2730580 | Lighthouse Lab in Alderley Park | Wellcome Sanger Institute for the COVID-19 Genomics UK (COG-UK) Consortium | Jacquelyn Wynn, Mairead Hyland, The Lighthouse Lab in Alderley Park and Alex Alderton, Roberto Amato, Jeffrey Barrett, Sonia Goncalves, Ewan Harrison, David K. Jackson, Ian Johnston, Dominic Kwiatkowski, Cordelia Langford, John Sillitoe on behalf of the Wellcome Sanger Institute COVID-19 Surveillance Team |
| EPI_ISL_2730581 | Lighthouse Lab in Milton Keynes | Wellcome Sanger Institute for the COVID-19 Genomics UK (COG-UK) Consortium | The Lighthouse Lab in Milton Keynes and Alex Alderton, Roberto Amato, Jeffrey Barrett, Sonia Goncalves, Ewan Harrison, David K. Jackson, Ian Johnston, Dominic Kwiatkowski, Cordelia Langford, John Sillitoe on behalf of the Wellcome Sanger Institute COVID-19 Surveillance Team |
| EPI_ISL_2730582 | Lighthouse Lab in Glasgow | Wellcome Sanger Institute for the COVID-19 Genomics UK (COG-UK) Consortium | Harper VanSteenhouse, Yumi Kasai, David Gray, Carol Clugston, Anna Dominiczak and Alex Alderton, Roberto Amato, Jeffrey Barrett, Sonia Goncalves, Ewan Harrison, David K. Jackson, Ian Johnston, Dominic Kwiatkowski, Cordelia Langford, John Sillitoe on behalf of the Wellcome Sanger Institute COVID-19 Surveillance Team |
| EPI_ISL_2730583, EPI_ISL_2730584 | Lighthouse Lab in Milton Keynes | Wellcome Sanger Institute for the COVID-19 Genomics UK (COG-UK) Consortium | The Lighthouse Lab in Milton Keynes and Alex Alderton, Roberto Amato, Jeffrey Barrett, Sonia Goncalves, Ewan Harrison, David K. Jackson, Ian Johnston, Dominic Kwiatkowski, Cordelia Langford, John Sillitoe on behalf of the Wellcome Sanger Institute COVID-19 Surveillance Team |
| EPI_ISL_2730585 | Lighthouse Lab in Glasgow | Wellcome Sanger Institute for the COVID-19 Genomics UK (COG-UK) Consortium | Harper VanSteenhouse, Yumi Kasai, David Gray, Carol Clugston, Anna Dominiczak and Alex Alderton, Roberto Amato, Jeffrey Barrett, Sonia Goncalves, Ewan Harrison, David K. Jackson, Ian Johnston, Dominic Kwiatkowski, Cordelia Langford, John Sillitoe on behalf of the Wellcome Sanger Institute COVID-19 Surveillance Team |
| EPI_ISL_2730586 | Lighthouse Lab in Milton Keynes | Wellcome Sanger Institute for the COVID-19 Genomics UK (COG-UK) Consortium | The Lighthouse Lab in Milton Keynes and Alex Alderton, Roberto Amato, Jeffrey Barrett, Sonia Goncalves, Ewan Harrison, David K. Jackson, Ian Johnston, Dominic Kwiatkowski, Cordelia Langford, John Sillitoe on behalf of the Wellcome Sanger Institute COVID-19 Surveillance Team |
| EPI_ISL_2730587, EPI_ISL_2730588 | Lighthouse Lab in Glasgow | Wellcome Sanger Institute for the COVID-19 Genomics UK (COG-UK) Consortium | Harper VanSteenhouse, Yumi Kasai, David Gray, Carol Clugston, Anna Dominiczak and Alex Alderton, Roberto Amato, Jeffrey Barrett, Sonia Goncalves, Ewan Harrison, David K. Jackson, Ian Johnston, Dominic Kwiatkowski, Cordelia Langford, John Sillitoe on behalf of the Wellcome Sanger Institute COVID-19 Surveillance Team |
| EPI_ISL_2730590 | Lighthouse Lab in Milton Keynes | Wellcome Sanger Institute for the COVID-19 Genomics UK (COG-UK) Consortium | The Lighthouse Lab in Milton Keynes and Alex Alderton, Roberto Amato, Jeffrey Barrett, Sonia Goncalves, Ewan Harrison, David K. Jackson, Ian Johnston, Dominic Kwiatkowski, Cordelia Langford, John Sillitoe on behalf of the Wellcome Sanger Institute COVID-19 Surveillance Team |
| EPI_ISL_2730591, EPI_ISL_2730592 | Lighthouse Lab in Glasgow | Wellcome Sanger Institute for the COVID-19 Genomics UK (COG-UK) Consortium | Harper VanSteenhouse, Yumi Kasai, David Gray, Carol Clugston, Anna Dominiczak and Alex Alderton, Roberto Amato, Jeffrey Barrett, Sonia Goncalves, Ewan Harrison, David K. Jackson, Ian Johnston, Dominic Kwiatkowski, Cordelia Langford, John Sillitoe on behalf of the Wellcome Sanger Institute COVID-19 Surveillance Team |
| EPI_ISL_2730593 | Lighthouse Lab in Milton Keynes | Wellcome Sanger Institute for the COVID-19 Genomics UK (COG-UK) Consortium | The Lighthouse Lab in Milton Keynes and Alex Alderton, Roberto Amato, Jeffrey Barrett, Sonia Goncalves, Ewan Harrison, David K. Jackson, Ian Johnston, Dominic Kwiatkowski, Cordelia Langford, John Sillitoe on behalf of the Wellcome Sanger Institute COVID-19 Surveillance Team |
| EPI_ISL_2730594, EPI_ISL_2730595, EPI_ISL_2730596, EPI_ISL_2730597, EPI_ISL_2730598, EPI_ISL_2730599, EPI_ISL_2730600 | Lighthouse Lab in Glasgow | Wellcome Sanger Institute for the COVID-19 Genomics UK (COG-UK) Consortium | Harper VanSteenhouse, Yumi Kasai, David Gray, Carol Clugston, Anna Dominiczak and Alex Alderton, Roberto Amato, Jeffrey Barrett, Sonia Goncalves, Ewan Harrison, David K. Jackson, Ian Johnston, Dominic Kwiatkowski, Cordelia Langford, John Sillitoe on behalf of the Wellcome Sanger Institute COVID-19 Surveillance Team |
| EPI_ISL_2730601 | Lighthouse Lab in Milton Keynes | Wellcome Sanger Institute for the COVID-19 Genomics UK (COG-UK) Consortium | The Lighthouse Lab in Milton Keynes and Alex Alderton, Roberto Amato, Jeffrey Barrett, Sonia Goncalves, Ewan Harrison, David K. Jackson, Ian Johnston, Dominic Kwiatkowski, Cordelia Langford, John Sillitoe on behalf of the Wellcome Sanger Institute COVID-19 Surveillance Team |
| EPI_ISL_2730602, EPI_ISL_2730603 | Lighthouse Lab in Alderley Park | Wellcome Sanger Institute for the COVID-19 Genomics UK (COG-UK) Consortium | Jacquelyn Wynn, Mairead Hyland, The Lighthouse Lab in Alderley Park and Alex Alderton, Roberto Amato, Jeffrey Barrett, Sonia Goncalves, Ewan Harrison, David K. Jackson, Ian Johnston, Dominic Kwiatkowski, Cordelia Langford, John Sillitoe on behalf of the Wellcome Sanger Institute COVID-19 Surveillance Team |
| EPI_ISL_2730604 | Lighthouse Lab in Milton Keynes | Wellcome Sanger Institute for the COVID-19 Genomics UK (COG-UK) Consortium | The Lighthouse Lab in Milton Keynes and Alex Alderton, Roberto Amato, Jeffrey Barrett, Sonia Goncalves, Ewan Harrison, David K. Jackson, Ian Johnston, Dominic Kwiatkowski, Cordelia Langford, John Sillitoe on behalf of the Wellcome Sanger Institute COVID-19 Surveillance Team |
| EPI_ISL_2730605, EPI_ISL_2730606 | Lighthouse Lab in Alderley Park | Wellcome Sanger Institute for the COVID-19 Genomics UK (COG-UK) Consortium | Jacquelyn Wynn, Mairead Hyland, The Lighthouse Lab in Alderley Park and Alex Alderton, Roberto Amato, Jeffrey Barrett, Sonia Goncalves, Ewan Harrison, David K. Jackson, Ian Johnston, Dominic Kwiatkowski, Cordelia Langford, John Sillitoe on behalf of the Wellcome Sanger Institute COVID-19 Surveillance Team |
| EPI_ISL_2730607, EPI_ISL_2730608 | Lighthouse Lab in Milton Keynes | Wellcome Sanger Institute for the COVID-19 Genomics UK (COG-UK) Consortium | The Lighthouse Lab in Milton Keynes and Alex Alderton, Roberto Amato, Jeffrey Barrett, Sonia Goncalves, Ewan Harrison, David K. Jackson, Ian Johnston, Dominic Kwiatkowski, Cordelia Langford, John Sillitoe on behalf of the Wellcome Sanger Institute COVID-19 Surveillance Team |
| EPI_ISL_2730609, EPI_ISL_2730610 | Lighthouse Lab in Glasgow | Wellcome Sanger Institute for the COVID-19 Genomics UK (COG-UK) Consortium | Harper VanSteenhouse, Yumi Kasai, David Gray, Carol Clugston, Anna Dominiczak and Alex Alderton, Roberto Amato, Jeffrey Barrett, Sonia Goncalves, Ewan Harrison, David K. Jackson, Ian Johnston, Dominic Kwiatkowski, Cordelia Langford, John Sillitoe on behalf of the Wellcome Sanger Institute COVID-19 Surveillance Team |
| EPI_ISL_2730611 | Lighthouse Lab in Milton Keynes | Wellcome Sanger Institute for the COVID-19 Genomics UK (COG-UK) Consortium | The Lighthouse Lab in Milton Keynes and Alex Alderton, Roberto Amato, Jeffrey Barrett, Sonia Goncalves, Ewan Harrison, David K. Jackson, Ian Johnston, Dominic Kwiatkowski, Cordelia Langford, John Sillitoe on behalf of the Wellcome Sanger Institute COVID-19 Surveillance Team |
| EPI_ISL_2730612, EPI_ISL_2730614, EPI_ISL_2730615, EPI_ISL_2730616 | Lighthouse Lab in Glasgow | Wellcome Sanger Institute for the COVID-19 Genomics UK (COG-UK) Consortium | Harper VanSteenhouse, Yumi Kasai, David Gray, Carol Clugston, Anna Dominiczak and Alex Alderton, Roberto Amato, Jeffrey Barrett, Sonia Goncalves, Ewan Harrison, David K. Jackson, Ian Johnston, Dominic Kwiatkowski, Cordelia Langford, John Sillitoe on behalf of the Wellcome Sanger Institute COVID-19 Surveillance Team |
| EPI_ISL_2730617 | Lighthouse Lab in Milton Keynes | Wellcome Sanger Institute for the COVID-19 Genomics UK (COG-UK) Consortium | The Lighthouse Lab in Milton Keynes and Alex Alderton, Roberto Amato, Jeffrey Barrett, Sonia Goncalves, Ewan Harrison, David K. Jackson, Ian Johnston, Dominic Kwiatkowski, Cordelia Langford, John Sillitoe on behalf of the Wellcome Sanger Institute COVID-19 Surveillance Team |
| EPI_ISL_2730618 | Lighthouse Lab in Glasgow | Wellcome Sanger Institute for the COVID-19 Genomics UK (COG-UK) Consortium | Harper VanSteenhouse, Yumi Kasai, David Gray, Carol Clugston, Anna Dominiczak and Alex Alderton, Roberto Amato, Jeffrey Barrett, Sonia Goncalves, Ewan Harrison, David K. Jackson, Ian Johnston, Dominic Kwiatkowski, Cordelia Langford, John Sillitoe on behalf of the Wellcome Sanger Institute COVID-19 Surveillance Team |
| EPI_ISL_2730619 | Lighthouse Lab in Alderley Park | Wellcome Sanger Institute for the COVID-19 Genomics UK (COG-UK) Consortium | Jacquelyn Wynn, Mairead Hyland, The Lighthouse Lab in Alderley Park and Alex Alderton, Roberto Amato, Jeffrey Barrett, Sonia Goncalves, Ewan Harrison, David K. Jackson, Ian Johnston, Dominic Kwiatkowski, Cordelia Langford, John Sillitoe on behalf of the Wellcome Sanger Institute COVID-19 Surveillance Team |
| EPI_ISL_2730620, EPI_ISL_2730621 | Lighthouse Lab in Glasgow | Wellcome Sanger Institute for the COVID-19 Genomics UK (COG-UK) Consortium | Harper VanSteenhouse, Yumi Kasai, David Gray, Carol Clugston, Anna Dominiczak and Alex Alderton, Roberto Amato, Jeffrey Barrett, Sonia Goncalves, Ewan Harrison, David K. Jackson, Ian Johnston, Dominic Kwiatkowski, Cordelia Langford, John Sillitoe on behalf of the Wellcome Sanger Institute COVID-19 Surveillance Team |
| EPI_ISL_2730622 | Lighthouse Lab in Milton Keynes | Wellcome Sanger Institute for the COVID-19 Genomics UK (COG-UK) Consortium | The Lighthouse Lab in Milton Keynes and Alex Alderton, Roberto Amato, Jeffrey Barrett, Sonia Goncalves, Ewan Harrison, David K. Jackson, Ian Johnston, Dominic Kwiatkowski, Cordelia Langford, John Sillitoe on behalf of the Wellcome Sanger Institute COVID-19 Surveillance Team |
| EPI_ISL_2730623 | Lighthouse Lab in Glasgow | Wellcome Sanger Institute for the COVID-19 Genomics UK (COG-UK) Consortium | Harper VanSteenhouse, Yumi Kasai, David Gray, Carol Clugston, Anna Dominiczak and Alex Alderton, Roberto Amato, Jeffrey Barrett, Sonia Goncalves, Ewan Harrison, David K. Jackson, Ian Johnston, Dominic Kwiatkowski, Cordelia Langford, John Sillitoe on behalf of the Wellcome Sanger Institute COVID-19 Surveillance Team |
| EPI_ISL_2730624 | Lighthouse Lab in Alderley Park | Wellcome Sanger Institute for the COVID-19 Genomics UK (COG-UK) Consortium | Jacquelyn Wynn, Mairead Hyland, The Lighthouse Lab in Alderley Park and Alex Alderton, Roberto Amato, Jeffrey Barrett, Sonia Goncalves, Ewan Harrison, David K. Jackson, Ian Johnston, Dominic Kwiatkowski, Cordelia Langford, John Sillitoe on behalf of the Wellcome Sanger Institute COVID-19 Surveillance Team |
| EPI_ISL_2730625 | Lighthouse Lab in Milton Keynes | Wellcome Sanger Institute for the COVID-19 Genomics UK (COG-UK) Consortium | The Lighthouse Lab in Milton Keynes and Alex Alderton, Roberto Amato, Jeffrey Barrett, Sonia Goncalves, Ewan Harrison, David K. Jackson, Ian Johnston, Dominic Kwiatkowski, Cordelia Langford, John Sillitoe on behalf of the Wellcome Sanger Institute COVID-19 Surveillance Team |
| EPI_ISL_2730626, EPI_ISL_2730627 | Lighthouse Lab in Glasgow | Wellcome Sanger Institute for the COVID-19 Genomics UK (COG-UK) Consortium | Harper VanSteenhouse, Yumi Kasai, David Gray, Carol Clugston, Anna Dominiczak and Alex Alderton, Roberto Amato, Jeffrey Barrett, Sonia Goncalves, Ewan Harrison, David K. Jackson, Ian Johnston, Dominic Kwiatkowski, Cordelia Langford, John Sillitoe on behalf of the Wellcome Sanger Institute COVID-19 Surveillance Team |
| EPI_ISL_2730628 | Lighthouse Lab in Milton Keynes | Wellcome Sanger Institute for the COVID-19 Genomics UK (COG-UK) Consortium | The Lighthouse Lab in Milton Keynes and Alex Alderton, Roberto Amato, Jeffrey Barrett, Sonia Goncalves, Ewan Harrison, David K. Jackson, Ian Johnston, Dominic Kwiatkowski, Cordelia Langford, John Sillitoe on behalf of the Wellcome Sanger Institute COVID-19 Surveillance Team |
| EPI_ISL_2730629 | Lighthouse Lab in Glasgow | Wellcome Sanger Institute for the COVID-19 Genomics UK (COG-UK) | Harper VanSteenhouse, Yumi Kasai, David Gray, Carol Clugston, Anna Dominiczak and Alex Alderton, Roberto Amato, Jeffrey Barrett, Sonia Goncalves, Ewan Harrison, David K. Jackson, Ian Johnston, Dominic Kwiatkowski, Cordelia Langford, |

| | | | |
|---|---|---|---|
| | | Consortium | John Sillitoe on behalf of the Wellcome Sanger Institute COVID-19 Surveillance Team |
| EPI_ISL_2730630 | Lighthouse Lab in Milton Keynes | Wellcome Sanger Institute for the COVID-19 Genomics UK (COG-UK) Consortium | The Lighthouse Lab in Milton Keynes and Alex Alderton, Roberto Amato, Jeffrey Barrett, Sonia Goncalves, Ewan Harrison, David K. Jackson, Ian Johnston, Dominic Kwiatkowski, Cordelia Langford, John Sillitoe on behalf of the Wellcome Sanger Institute COVID-19 Surveillance Team |
| EPI_ISL_2730631 | Lighthouse Lab in Glasgow | Wellcome Sanger Institute for the COVID-19 Genomics UK (COG-UK) Consortium | Harper VanSteenhouse, Yumi Kasai, David Gray, Carol Clugston, Anna Dominiczak and Alex Alderton, Roberto Amato, Jeffrey Barrett, Sonia Goncalves, Ewan Harrison, David K. Jackson, Ian Johnston, Dominic Kwiatkowski, Cordelia Langford, John Sillitoe on behalf of the Wellcome Sanger Institute COVID-19 Surveillance Team |
| EPI_ISL_2730632 | Lighthouse Lab in Milton Keynes | Wellcome Sanger Institute for the COVID-19 Genomics UK (COG-UK) Consortium | The Lighthouse Lab in Milton Keynes and Alex Alderton, Roberto Amato, Jeffrey Barrett, Sonia Goncalves, Ewan Harrison, David K. Jackson, Ian Johnston, Dominic Kwiatkowski, Cordelia Langford, John Sillitoe on behalf of the Wellcome Sanger Institute COVID-19 Surveillance Team |
| EPI_ISL_2730633 | Lighthouse Lab in Alderley Park | Wellcome Sanger Institute for the COVID-19 Genomics UK (COG-UK) Consortium | Jacquelyn Wynn, Mairead Hyland, The Lighthouse Lab in Alderley Park and Alex Alderton, Roberto Amato, Jeffrey Barrett, Sonia Goncalves, Ewan Harrison, David K. Jackson, Ian Johnston, Dominic Kwiatkowski, Cordelia Langford, John Sillitoe on behalf of the Wellcome Sanger Institute COVID-19 Surveillance Team |
| EPI_ISL_2730634 | Lighthouse Lab in Glasgow | Wellcome Sanger Institute for the COVID-19 Genomics UK (COG-UK) Consortium | Harper VanSteenhouse, Yumi Kasai, David Gray, Carol Clugston, Anna Dominiczak and Alex Alderton, Roberto Amato, Jeffrey Barrett, Sonia Goncalves, Ewan Harrison, David K. Jackson, Ian Johnston, Dominic Kwiatkowski, Cordelia Langford, John Sillitoe on behalf of the Wellcome Sanger Institute COVID-19 Surveillance Team |
| EPI_ISL_2730635 | Lighthouse Lab in Milton Keynes | Wellcome Sanger Institute for the COVID-19 Genomics UK (COG-UK) Consortium | The Lighthouse Lab in Milton Keynes and Alex Alderton, Roberto Amato, Jeffrey Barrett, Sonia Goncalves, Ewan Harrison, David K. Jackson, Ian Johnston, Dominic Kwiatkowski, Cordelia Langford, John Sillitoe on behalf of the Wellcome Sanger Institute COVID-19 Surveillance Team |
| EPI_ISL_2730636 | Lighthouse Lab in Alderley Park | Wellcome Sanger Institute for the COVID-19 Genomics UK (COG-UK) Consortium | Jacquelyn Wynn, Mairead Hyland, The Lighthouse Lab in Alderley Park and Alex Alderton, Roberto Amato, Jeffrey Barrett, Sonia Goncalves, Ewan Harrison, David K. Jackson, Ian Johnston, Dominic Kwiatkowski, Cordelia Langford, John Sillitoe on behalf of the Wellcome Sanger Institute COVID-19 Surveillance Team |
| EPI_ISL_2730637 | Lighthouse Lab in Glasgow | Wellcome Sanger Institute for the COVID-19 Genomics UK (COG-UK) Consortium | Harper VanSteenhouse, Yumi Kasai, David Gray, Carol Clugston, Anna Dominiczak and Alex Alderton, Roberto Amato, Jeffrey Barrett, Sonia Goncalves, Ewan Harrison, David K. Jackson, Ian Johnston, Dominic Kwiatkowski, Cordelia Langford, John Sillitoe on behalf of the Wellcome Sanger Institute COVID-19 Surveillance Team |
| EPI_ISL_2730638 | Lighthouse Lab in Milton Keynes | Wellcome Sanger Institute for the COVID-19 Genomics UK (COG-UK) Consortium | The Lighthouse Lab in Milton Keynes and Alex Alderton, Roberto Amato, Jeffrey Barrett, Sonia Goncalves, Ewan Harrison, David K. Jackson, Ian Johnston, Dominic Kwiatkowski, Cordelia Langford, John Sillitoe on behalf of the Wellcome Sanger Institute COVID-19 Surveillance Team |
| EPI_ISL_2730639, EPI_ISL_2730640, EPI_ISL_2730641, EPI_ISL_2730642 | Lighthouse Lab in Glasgow | Wellcome Sanger Institute for the COVID-19 Genomics UK (COG-UK) Consortium | Harper VanSteenhouse, Yumi Kasai, David Gray, Carol Clugston, Anna Dominiczak and Alex Alderton, Roberto Amato, Jeffrey Barrett, Sonia Goncalves, Ewan Harrison, David K. Jackson, Ian Johnston, Dominic Kwiatkowski, Cordelia Langford, John Sillitoe on behalf of the Wellcome Sanger Institute COVID-19 Surveillance Team |
| EPI_ISL_2730643 | Lighthouse Lab in Milton Keynes | Wellcome Sanger Institute for the COVID-19 Genomics UK (COG-UK) Consortium | The Lighthouse Lab in Milton Keynes and Alex Alderton, Roberto Amato, Jeffrey Barrett, Sonia Goncalves, Ewan Harrison, David K. Jackson, Ian Johnston, Dominic Kwiatkowski, Cordelia Langford, John Sillitoe on behalf of the Wellcome Sanger Institute COVID-19 Surveillance Team |
| EPI_ISL_2730644 | Lighthouse Lab in Glasgow | Wellcome Sanger Institute for the COVID-19 Genomics UK (COG-UK) Consortium | Harper VanSteenhouse, Yumi Kasai, David Gray, Carol Clugston, Anna Dominiczak and Alex Alderton, Roberto Amato, Jeffrey Barrett, Sonia Goncalves, Ewan Harrison, David K. Jackson, Ian Johnston, Dominic Kwiatkowski, Cordelia Langford, John Sillitoe on behalf of the Wellcome Sanger Institute COVID-19 Surveillance Team |
| EPI_ISL_2730645 | Lighthouse Lab in Alderley Park | Wellcome Sanger Institute for the COVID-19 Genomics UK (COG-UK) Consortium | Jacquelyn Wynn, Mairead Hyland, The Lighthouse Lab in Alderley Park and Alex Alderton, Roberto Amato, Jeffrey Barrett, Sonia Goncalves, Ewan Harrison, David K. Jackson, Ian Johnston, Dominic Kwiatkowski, Cordelia Langford, John Sillitoe on behalf of the Wellcome Sanger Institute COVID-19 Surveillance Team |
| EPI_ISL_2730646, EPI_ISL_2730647 | Lighthouse Lab in Milton Keynes | Wellcome Sanger Institute for the COVID-19 Genomics UK (COG-UK) Consortium | The Lighthouse Lab in Milton Keynes and Alex Alderton, Roberto Amato, Jeffrey Barrett, Sonia Goncalves, Ewan Harrison, David K. Jackson, Ian Johnston, Dominic Kwiatkowski, Cordelia Langford, John Sillitoe on behalf of the Wellcome Sanger Institute COVID-19 Surveillance Team |
| EPI_ISL_2730648, EPI_ISL_2730649, EPI_ISL_2730650, EPI_ISL_2730651, EPI_ISL_2730652, EPI_ISL_2730653, EPI_ISL_2730654, EPI_ISL_2730655, EPI_ISL_2730656, EPI_ISL_2730657, EPI_ISL_2730658, EPI_ISL_2730659, EPI_ISL_2730660, EPI_ISL_2730661, EPI_ISL_2730662, EPI_ISL_2730663, EPI_ISL_2730664, EPI_ISL_2730665, EPI_ISL_2730666, EPI_ISL_2730667, EPI_ISL_2730668, EPI_ISL_2730669, EPI_ISL_2730670, EPI_ISL_2730671, EPI_ISL_2730672, EPI_ISL_2730673, EPI_ISL_2730674, EPI_ISL_2730675, EPI_ISL_2730676, EPI_ISL_2730677, EPI_ISL_2730678, EPI_ISL_2730679, EPI_ISL_2730680, EPI_ISL_2730681, EPI_ISL_2730682, EPI_ISL_2730683, EPI_ISL_2730684, EPI_ISL_2730685, EPI_ISL_2730686, EPI_ISL_2730687, EPI_ISL_2730688, EPI_ISL_2730689, EPI_ISL_2730690, EPI_ISL_2730691, EPI_ISL_2730692, EPI_ISL_2730693, EPI_ISL_2730694, EPI_ISL_2730695, EPI_ISL_2730696, EPI_ISL_2730697, EPI_ISL_2730698, EPI_ISL_2730699, EPI_ISL_2730700, EPI_ISL_2730701, EPI_ISL_2730702, EPI_ISL_2730703, EPI_ISL_2730704, EPI_ISL_2730705, EPI_ISL_2730706, EPI_ISL_2730707, EPI_ISL_2730708, EPI_ISL_2730709, EPI_ISL_2730710, EPI_ISL_2730711, EPI_ISL_2730712, EPI_ISL_2730713, EPI_ISL_2730714, EPI_ISL_2730715, EPI_ISL_2730716, EPI_ISL_2730717, EPI_ISL_2730718, EPI_ISL_2730719, EPI_ISL_2730720, EPI_ISL_2730721, EPI_ISL_2730722, EPI_ISL_2730723, EPI_ISL_2730724, EPI_ISL_2730726, EPI_ISL_2730727, EPI_ISL_2730728, EPI_ISL_2730729, EPI_ISL_2730730, EPI_ISL_2730731, EPI_ISL_2730732, EPI_ISL_2730733, EPI_ISL_2730734, EPI_ISL_2730735, EPI_ISL_2730736, EPI_ISL_2730737, EPI_ISL_2730738, EPI_ISL_2730739, EPI_ISL_2730740, EPI_ISL_2730741, EPI_ISL_2730742, EPI_ISL_2730743, EPI_ISL_2730744, EPI_ISL_2730745, EPI_ISL_2730746, EPI_ISL_2730747, EPI_ISL_2730748, EPI_ISL_2730749, EPI_ISL_2730750, EPI_ISL_2730751, EPI_ISL_2730752, EPI_ISL_2730753, EPI_ISL_2730754, EPI_ISL_2730755, EPI_ISL_2730756, EPI_ISL_2730757, EPI_ISL_2730758, EPI_ISL_2730759, EPI_ISL_2730760, EPI_ISL_2730761, EPI_ISL_2730762, EPI_ISL_2730763, EPI_ISL_2730764, EPI_ISL_2730765, EPI_ISL_2730766, EPI_ISL_2730767, EPI_ISL_2730768, EPI_ISL_2730769, EPI_ISL_2730770, EPI_ISL_2730771, EPI_ISL_2730772, EPI_ISL_2730773, EPI_ISL_2730774, EPI_ISL_2730775, EPI_ISL_2730776, EPI_ISL_2730777, EPI_ISL_2730778, EPI_ISL_2730779, EPI_ISL_2730780, EPI_ISL_2730781, EPI_ISL_2730782, EPI_ISL_2730783, EPI_ISL_2730784, EPI_ISL_2730785, EPI_ISL_2730786, EPI_ISL_2730787, EPI_ISL_2730788, EPI_ISL_2730789, EPI_ISL_2730790, EPI_ISL_2730791, EPI_ISL_2730792, EPI_ISL_2730793, EPI_ISL_2730794, EPI_ISL_2730795, EPI_ISL_2730796, EPI_ISL_2730797, EPI_ISL_2730798, EPI_ISL_2730799, EPI_ISL_2730800, EPI_ISL_2730801, EPI_ISL_2730802, EPI_ISL_2730803, EPI_ISL_2730804, EPI_ISL_2730805, EPI_ISL_2730806, EPI_ISL_2730807, EPI_ISL_2730808, EPI_ISL_2730809, EPI_ISL_2730810, EPI_ISL_2730811, EPI_ISL_2730812, EPI_ISL_2730813, EPI_ISL_2730814, EPI_ISL_2730815, EPI_ISL_2730816, EPI_ISL_2730817, EPI_ISL_2730818, EPI_ISL_2730819, EPI_ISL_2730820, EPI_ISL_2730821, EPI_ISL_2730822, EPI_ISL_2730823, EPI_ISL_2730824, EPI_ISL_2730825, EPI_ISL_2730826, EPI_ISL_2730827, EPI_ISL_2730828, EPI_ISL_2730829, EPI_ISL_2730830, EPI_ISL_2730831, EPI_ISL_2730832, EPI_ISL_2730833, EPI_ISL_2730834, EPI_ISL_2730835, EPI_ISL_2730836, EPI_ISL_2730837, EPI_ISL_2730838, EPI_ISL_2730839, EPI_ISL_2730840, EPI_ISL_2730841, EPI_ISL_2730844, EPI_ISL_2730845, EPI_ISL_2730846, EPI_ISL_2730847, EPI_ISL_2730848, EPI_ISL_2730849, EPI_ISL_2730850, EPI_ISL_2730851, EPI_ISL_2730852, EPI_ISL_2730853, EPI_ISL_2730854, EPI_ISL_2730855, EPI_ISL_2730856, EPI_ISL_2730857, EPI_ISL_2730858, EPI_ISL_2730859, EPI_ISL_2730860, EPI_ISL_2730861, EPI_ISL_2730862, EPI_ISL_2730863, EPI_ISL_2730864, EPI_ISL_2730865, EPI_ISL_2730866, EPI_ISL_2730867, EPI_ISL_2730868, EPI_ISL_2730869, EPI_ISL_2730870, EPI_ISL_2730871, EPI_ISL_2730872, EPI_ISL_2730873, EPI_ISL_2730874, EPI_ISL_2730875, EPI_ISL_2730876, EPI_ISL_2730877, EPI_ISL_2730878, EPI_ISL_2730879, EPI_ISL_2730880, EPI_ISL_2730881, EPI_ISL_2730882, EPI_ISL_2730883, EPI_ISL_2730884, EPI_ISL_2730885, EPI_ISL_2730886, EPI_ISL_2730887, EPI_ISL_2730888, EPI_ISL_2730889, EPI_ISL_2730890, EPI_ISL_2730891, EPI_ISL_2730892, EPI_ISL_2730893, EPI_ISL_2730894, EPI_ISL_2730895, EPI_ISL_2730896, EPI_ISL_2730897, EPI_ISL_2730898, EPI_ISL_2730899, EPI_ISL_2730900, EPI_ISL_2730901, EPI_ISL_2730902, EPI_ISL_2730903, EPI_ISL_2730904, EPI_ISL_2730905, EPI_ISL_2730906, EPI_ISL_2730907, EPI_ISL_2730908, EPI_ISL_2730910, EPI_ISL_2730911, EPI_ISL_2730912, EPI_ISL_2730913, EPI_ISL_2730914, EPI_ISL_2730915, EPI_ISL_2730916, EPI_ISL_2730917, EPI_ISL_2730918, EPI_ISL_2730919, EPI_ISL_2730920, EPI_ISL_2730921, EPI_ISL_2730922, EPI_ISL_2730923, EPI_ISL_2730924, EPI_ISL_2730925 | | | |
| see above | Lighthouse Lab in Glasgow | Wellcome Sanger Institute for the COVID-19 Genomics UK (COG-UK) Consortium | Harper VanSteenhouse, Yumi Kasai, David Gray, Carol Clugston, Anna Dominiczak and Alex Alderton, Roberto Amato, Jeffrey Barrett, Sonia Goncalves, Ewan Harrison, David K. Jackson, Ian Johnston, Dominic Kwiatkowski, Cordelia Langford, John Sillitoe on behalf of the Wellcome Sanger Institute COVID-19 Surveillance Team |
| EPI_ISL_2730926 | Lighthouse Lab in Milton Keynes | Wellcome Sanger Institute for the COVID-19 Genomics UK (COG-UK) Consortium | The Lighthouse Lab in Milton Keynes and Alex Alderton, Roberto Amato, Jeffrey Barrett, Sonia Goncalves, Ewan Harrison, David K. Jackson, Ian Johnston, Dominic Kwiatkowski, Cordelia Langford, John Sillitoe on behalf of the Wellcome Sanger Institute COVID-19 Surveillance Team |
| EPI_ISL_2730928, EPI_ISL_2730929, EPI_ISL_2730930, EPI_ISL_2730931, EPI_ISL_2730932, EPI_ISL_2730933, EPI_ISL_2730934, EPI_ISL_2730935, EPI_ISL_2730936, EPI_ISL_2730937, EPI_ISL_2730938, EPI_ISL_2730939, EPI_ISL_2730940, EPI_ISL_2730941, EPI_ISL_2730942, EPI_ISL_2730944, EPI_ISL_2730945, EPI_ISL_2730946, EPI_ISL_2730947, EPI_ISL_2730948, EPI_ISL_2730949, EPI_ISL_2730950, EPI_ISL_2730951, EPI_ISL_2730952, EPI_ISL_2730953, EPI_ISL_2730954, EPI_ISL_2730955, EPI_ISL_2730956, EPI_ISL_2730957, EPI_ISL_2730958, EPI_ISL_2730959, EPI_ISL_2730960, EPI_ISL_2730961, EPI_ISL_2730962, EPI_ISL_2730963, EPI_ISL_2730964, EPI_ISL_2730965, EPI_ISL_2730966, EPI_ISL_2730967, EPI_ISL_2730968, EPI_ISL_2730969, EPI_ISL_2730970, EPI_ISL_2730971, EPI_ISL_2730972, EPI_ISL_2730973, EPI_ISL_2730974 | | | |
| see above | Lighthouse Lab in Glasgow | Wellcome Sanger Institute for the COVID-19 Genomics UK (COG-UK) Consortium | Harper VanSteenhouse, Yumi Kasai, David Gray, Carol Clugston, Anna Dominiczak and Alex Alderton, Roberto Amato, Jeffrey Barrett, Sonia Goncalves, Ewan Harrison, David K. Jackson, Ian Johnston, Dominic Kwiatkowski, Cordelia Langford, John Sillitoe on behalf of the Wellcome Sanger Institute COVID-19 Surveillance Team |
| EPI_ISL_2730975 | Lighthouse Lab in Milton Keynes | Wellcome Sanger Institute for the COVID-19 Genomics UK (COG-UK) Consortium | The Lighthouse Lab in Milton Keynes and Alex Alderton, Roberto Amato, Jeffrey Barrett, Sonia Goncalves, Ewan Harrison, David K. Jackson, Ian Johnston, Dominic Kwiatkowski, Cordelia Langford, John Sillitoe on behalf of the Wellcome Sanger Institute COVID-19 Surveillance Team |
| EPI_ISL_2730976, EPI_ISL_2730977, EPI_ISL_2730978, EPI_ISL_2730979, EPI_ISL_2730980, EPI_ISL_2730981, EPI_ISL_2730982, EPI_ISL_2730983, EPI_ISL_2730984, EPI_ISL_2730985, EPI_ISL_2730986, EPI_ISL_2730987, EPI_ISL_2730988, EPI_ISL_2730989, EPI_ISL_2730990, EPI_ISL_2730991, EPI_ISL_2730992, EPI_ISL_2730993, EPI_ISL_2730994, EPI_ISL_2730995, EPI_ISL_2730996, EPI_ISL_2730997, EPI_ISL_2730998, EPI_ISL_2730999, EPI_ISL_2731000, EPI_ISL_2731001, EPI_ISL_2731002, EPI_ISL_2731003, EPI_ISL_2731004, EPI_ISL_2731005, EPI_ISL_2731006, EPI_ISL_2731007, EPI_ISL_2731008, EPI_ISL_2731009, EPI_ISL_2731010, EPI_ISL_2731011, EPI_ISL_2731012, EPI_ISL_2731013, EPI_ISL_2731014, EPI_ISL_2731015, EPI_ISL_2731016, EPI_ISL_2731017, EPI_ISL_2731018, EPI_ISL_2731019, EPI_ISL_2731020, EPI_ISL_2731021, EPI_ISL_2731022, EPI_ISL_2731023, EPI_ISL_2731024, EPI_ISL_2731025, EPI_ISL_2731026, EPI_ISL_2731027, EPI_ISL_2731028, EPI_ISL_2731029, EPI_ISL_2731030, EPI_ISL_2731031, EPI_ISL_2731032, EPI_ISL_2731033, EPI_ISL_2731034, EPI_ISL_2731035, EPI_ISL_2731036, EPI_ISL_2731037, EPI_ISL_2731038, EPI_ISL_2731039, EPI_ISL_2731040, EPI_ISL_2731041, EPI_ISL_2731042, EPI_ISL_2731043, EPI_ISL_2731044, EPI_ISL_2731045, EPI_ISL_2731046, EPI_ISL_2731047, EPI_ISL_2731048, EPI_ISL_2731049, EPI_ISL_2731050, EPI_ISL_2731051, EPI_ISL_2731052, EPI_ISL_2731053, EPI_ISL_2731054, EPI_ISL_2731055, EPI_ISL_2731056, EPI_ISL_2731057, EPI_ISL_2731058, EPI_ISL_2731059, EPI_ISL_2731060, EPI_ISL_2731061, EPI_ISL_2731062, EPI_ISL_2731063, EPI_ISL_2731064, EPI_ISL_2731065, EPI_ISL_2731066, EPI_ISL_2731067, EPI_ISL_2731068, EPI_ISL_2731069, EPI_ISL_2731070, EPI_ISL_2731071, EPI_ISL_2731072, EPI_ISL_2731073, EPI_ISL_2731074, EPI_ISL_2731075, EPI_ISL_2731076, EPI_ISL_2731077, EPI_ISL_2731078, EPI_ISL_2731079, EPI_ISL_2731080, EPI_ISL_2731081, EPI_ISL_2731082, EPI_ISL_2731083, EPI_ISL_2731084, EPI_ISL_2731085, EPI_ISL_2731086, EPI_ISL_2731087, EPI_ISL_2731088, EPI_ISL_2731089, EPI_ISL_2731090, EPI_ISL_2731091, EPI_ISL_2731092, EPI_ISL_2731093, EPI_ISL_2731094, EPI_ISL_2731095, EPI_ISL_2731096, EPI_ISL_2731097, EPI_ISL_2731098, EPI_ISL_2731099, EPI_ISL_2731100, EPI_ISL_2731101, EPI_ISL_2731102, EPI_ISL_2731103, EPI_ISL_2731104, EPI_ISL_2731105, EPI_ISL_2731106, EPI_ISL_2731107, EPI_ISL_2731108, EPI_ISL_2731109, EPI_ISL_2731110, EPI_ISL_2731111, EPI_ISL_2731112, EPI_ISL_2731113, EPI_ISL_2731114, EPI_ISL_2731115, EPI_ISL_2731116, EPI_ISL_2731117, EPI_ISL_2731118, EPI_ISL_2731119, EPI_ISL_2731120, EPI_ISL_2731121, EPI_ISL_2731122, EPI_ISL_2731123, EPI_ISL_2731124, EPI_ISL_2731125, EPI_ISL_2731126, EPI_ISL_2731127, EPI_ISL_2731128, EPI_ISL_2731129, EPI_ISL_2731130, EPI_ISL_2731131, EPI_ISL_2731132, EPI_ISL_2731133, EPI_ISL_2731134, EPI_ISL_2731135, EPI_ISL_2731136, EPI_ISL_2731137, EPI_ISL_2731138, EPI_ISL_2731139, EPI_ISL_2731140, EPI_ISL_2731141, EPI_ISL_2731142, EPI_ISL_2731143, EPI_ISL_2731144, EPI_ISL_2731145, EPI_ISL_2731146, EPI_ISL_2731147, EPI_ISL_2731148, EPI_ISL_2731149, EPI_ISL_2731150, EPI_ISL_2731151, EPI_ISL_2731152, EPI_ISL_2731153, EPI_ISL_2731155, EPI_ISL_2731156, EPI_ISL_2731157, EPI_ISL_2731158, EPI_ISL_2731159, EPI_ISL_2731160, EPI_ISL_2731161, EPI_ISL_2731162, EPI_ISL_2731163, EPI_ISL_2731164, EPI_ISL_2731165, EPI_ISL_2731167, EPI_ISL_2731168, EPI_ISL_2731169, EPI_ISL_2731170, EPI_ISL_2731171, EPI_ISL_2731172, EPI_ISL_2731173, EPI_ISL_2731174, EPI_ISL_2731175, EPI_ISL_2731176, EPI_ISL_2731177, EPI_ISL_2731178, EPI_ISL_2731179, EPI_ISL_2731180, EPI_ISL_2731181, EPI_ISL_2731182, EPI_ISL_2731183, EPI_ISL_2731185, EPI_ISL_2731186 | | | |
| see above | Lighthouse Lab in Glasgow | Wellcome Sanger Institute for the COVID-19 Genomics UK (COG-UK) Consortium | Harper VanSteenhouse, Yumi Kasai, David Gray, Carol Clugston, Anna Dominiczak and Alex Alderton, Roberto Amato, Jeffrey Barrett, Sonia Goncalves, Ewan Harrison, David K. Jackson, Ian Johnston, Dominic Kwiatkowski, Cordelia Langford, John Sillitoe on behalf of the Wellcome Sanger Institute COVID-19 Surveillance Team |
| EPI_ISL_2731187 | Lighthouse Laboratory Plymouth | Wellcome Sanger Institute for the COVID-19 Genomics UK (COG-UK) Consortium | Lighthouse Laboratory Plymouth and Alex Alderton, Roberto Amato, Jeffrey Barrett, Sonia Goncalves, Ewan Harrison, David K. Jackson, Ian Johnston, Dominic Kwiatkowski, Cordelia Langford, John Sillitoe on behalf of the Wellcome Sanger Institute COVID-19 Surveillance Team |
| EPI_ISL_2731188, EPI_ISL_2731189, EPI_ISL_2731190, EPI_ISL_2731191, EPI_ISL_2731192, EPI_ISL_2731193, EPI_ISL_2731194, EPI_ISL_2731195, EPI_ISL_2731196, EPI_ISL_2731197, EPI_ISL_2731198, EPI_ISL_2731199, EPI_ISL_2731200, EPI_ISL_2731201, EPI_ISL_2731202, EPI_ISL_2731203, EPI_ISL_2731204, EPI_ISL_2731205, EPI_ISL_2731207, EPI_ISL_2731208, EPI_ISL_2731209, EPI_ISL_2731210, EPI_ISL_2731211, EPI_ISL_2731212, EPI_ISL_2731213, EPI_ISL_2731214, EPI_ISL_2731215, EPI_ISL_2731216, EPI_ISL_2731217, EPI_ISL_2731218, EPI_ISL_2731219, EPI_ISL_2731220, EPI_ISL_2731221, EPI_ISL_2731222, EPI_ISL_2731223, EPI_ISL_2731224, EPI_ISL_2731225, EPI_ISL_2731226, EPI_ISL_2731227, EPI_ISL_2731228, EPI_ISL_2731229, EPI_ISL_2731231, EPI_ISL_2731232, EPI_ISL_2731233, EPI_ISL_2731234, EPI_ISL_2731235, EPI_ISL_2731236, EPI_ISL_2731237, EPI_ISL_2731238, EPI_ISL_2731240, EPI_ISL_2731241, EPI_ISL_2731242, EPI_ISL_2731243, EPI_ISL_2731244, EPI_ISL_2731245, EPI_ISL_2731246, EPI_ISL_2731247, EPI_ISL_2731248, EPI_ISL_2731250, EPI_ISL_2731251, EPI_ISL_2731252, EPI_ISL_2731253, EPI_ISL_2731254, EPI_ISL_2731255, EPI_ISL_2731256, EPI_ISL_2731261, EPI_ISL_2731262, EPI_ISL_2731263, EPI_ISL_2731264, EPI_ISL_2731265, EPI_ISL_2731266, EPI_ISL_2731267, EPI_ISL_2731268, EPI_ISL_2731269 | | | |
| see above | Lighthouse Lab in Glasgow | Wellcome Sanger Institute for the COVID-19 Genomics UK (COG-UK) Consortium | Harper VanSteenhouse, Yumi Kasai, David Gray, Carol Clugston, Anna Dominiczak and Alex Alderton, Roberto Amato, Jeffrey Barrett, Sonia Goncalves, Ewan Harrison, David K. Jackson, Ian Johnston, Dominic Kwiatkowski, Cordelia Langford, John Sillitoe on behalf of the Wellcome Sanger Institute COVID-19 Surveillance Team |
| EPI_ISL_2731277, EPI_ISL_2731278, EPI_ISL_2731280, EPI_ISL_2731281, EPI_ISL_2731286, EPI_ISL_2731287, EPI_ISL_2731291 | Valais Hospital, Central Institute | Valais Hospital, Central Institute | Alexis Dumoulin, Lorenzo Cerutti, Henri Pegeot, Melyssa Elies, Deborah Penet, Keith Harshman, Ioannis Xenarios, Emmanouil Dermitzakis |
| EPI_ISL_2731326 | Microvida | Microvida | Suzan D. Pas, Jaco J. Verweij, Joep J. J. M. Stohr |
| EPI_ISL_2731419, EPI_ISL_2731421, EPI_ISL_2731427, EPI_ISL_2731435, EPI_ISL_2731436 | Montana Public Health Laboratory | Montana Public Health Laboratory | Joy Ritter, Michelle Mozer, Carrie Biskupiak, Deborah Gibson, Michael Dills |
| EPI_ISL_2731451, EPI_ISL_2731452 | HLAGYN-Laboratório de Imunologia de Transplantes de Goiás | Laboratory of Respiratory Viruses and Measles, Oswaldo Cruz Institute, FIOCRUZ | Paola Resende, Luciana Appolinario, Fernando Motta, Anna Carolina Paixao, Ana Carolina Mendonca, Alice Sampaio Rocha, Taina Venas, Elisa Cavalcante Pereira, Renata Serrano Lopes, Marilda Siqueira on behalf of the Fiocruz COVID-19 Genomic Surveillance Network |
| EPI_ISL_2733016, EPI_ISL_2733027 | Respiratory Virus Unit, Microbiology Services Colindale, Public Health England | COVID-19 Genomics UK (COG-UK) Consortium | PHE Covid Sequencing Team |
| EPI_ISL_2733071 | IRCCSSanta Lucia Foundation | National Institute for Infectious Diseases (INMI) L. Spallanzani I.R.C.C.S | CEM Gruber, F Messina, F Santini, M Rueca, G Bonfiglio, O Butera, B Bartolini, E Giombini, MR Capobianchi, A Di Caro |
| EPI_ISL_2733074 | Istituto figlie di S.Camillo Ospedale MADRE GIUSEPPINA VANNINI | National Institute for Infectious Diseases (INMI) L. Spallanzani I.R.C.C.S | CEM Gruber, F Messina, F Santini, M Rueca, G Bonfiglio, O Butera, B Bartolini, E Giombini, MR Capobianchi, A Di Caro |

| | | | |
|---|---|---|---|
| EPI_ISL_2733103 | Respiratory Virus Unit, Microbiology Services Colindale, Public Health England | COVID-19 Genomics UK (COG-UK) Consortium | PHE Covid Sequencing Team |
| EPI_ISL_2733919, EPI_ISL_2733920, EPI_ISL_2733921, EPI_ISL_2733922, EPI_ISL_2733923, EPI_ISL_2733924, EPI_ISL_2733925, EPI_ISL_2733926, EPI_ISL_2733927, EPI_ISL_2733928, EPI_ISL_2733929, EPI_ISL_2733930, EPI_ISL_2733931, EPI_ISL_2733932, EPI_ISL_2733933, EPI_ISL_2733934, EPI_ISL_2733935, EPI_ISL_2733936, EPI_ISL_2733937, EPI_ISL_2733938, EPI_ISL_2733939, EPI_ISL_2733940, EPI_ISL_2733941, EPI_ISL_2733942, EPI_ISL_2733943, EPI_ISL_2733944, EPI_ISL_2733945, EPI_ISL_2733946, EPI_ISL_2733947, EPI_ISL_2733948, EPI_ISL_2733949, EPI_ISL_2733950, EPI_ISL_2733951, EPI_ISL_2733952, EPI_ISL_2733953, EPI_ISL_2733954, EPI_ISL_2733955, EPI_ISL_2733956, EPI_ISL_2733957, EPI_ISL_2733958, EPI_ISL_2733959, EPI_ISL_2733960, EPI_ISL_2733961, EPI_ISL_2733962, EPI_ISL_2733963, EPI_ISL_2733964, EPI_ISL_2733965, EPI_ISL_2733966, EPI_ISL_2733967, EPI_ISL_2733968, EPI_ISL_2733969, EPI_ISL_2733970, EPI_ISL_2733971, EPI_ISL_2733972, EPI_ISL_2733973, EPI_ISL_2733974, EPI_ISL_2733975, EPI_ISL_2733976, EPI_ISL_2733977, EPI_ISL_2733978, EPI_ISL_2733979, EPI_ISL_2733980, EPI_ISL_2733981, EPI_ISL_2733982, EPI_ISL_2733983, EPI_ISL_2733984, EPI_ISL_2733985, EPI_ISL_2733986, EPI_ISL_2733987, EPI_ISL_2733988, EPI_ISL_2733989, EPI_ISL_2733990, EPI_ISL_2733991, EPI_ISL_2733992, EPI_ISL_2733993 | | | |
| see above | Department of Pathology, University of Cambridge | COVID-19 Genomics UK (COG-UK) Consortium | Aminu S. Jahun, Yasmin Chaudhry, Iliana Georgana, Myra Hosmillo, Rhys Izuagbe, William L. Hamilton, Martin D. Curran, Surendra Parmar, Ian Goodfellow |
| EPI_ISL_2734268, EPI_ISL_2734269, EPI_ISL_2734270, EPI_ISL_2734271, EPI_ISL_2734272, EPI_ISL_2734273, EPI_ISL_2734274, EPI_ISL_2734275, EPI_ISL_2734276, EPI_ISL_2734277, EPI_ISL_2734278, EPI_ISL_2734279, EPI_ISL_2734280, EPI_ISL_2734282, EPI_ISL_2734283, EPI_ISL_2734284, EPI_ISL_2734285, EPI_ISL_2734286, EPI_ISL_2734287, EPI_ISL_2734288, EPI_ISL_2734289, EPI_ISL_2734290, EPI_ISL_2734294, EPI_ISL_2734295, EPI_ISL_2734296, EPI_ISL_2734297, EPI_ISL_2734298, EPI_ISL_2734299, EPI_ISL_2734300, EPI_ISL_2734301, EPI_ISL_2734302, EPI_ISL_2734303, EPI_ISL_2734304, EPI_ISL_2734305, EPI_ISL_2734306, EPI_ISL_2734307, EPI_ISL_2734308, EPI_ISL_2734310, EPI_ISL_2734311, EPI_ISL_2734312, EPI_ISL_2734313, EPI_ISL_2734314, EPI_ISL_2734316, EPI_ISL_2734317, EPI_ISL_2734318 | | | |
| see above | West of Scotland Specialist Virology Centre, NHSGGC / MRC-University of Glasgow Centre for Virus Research | COVID-19 Genomics UK (COG-UK) Consortium | Ana da Silva Filipe, Natasha Johnson, Kathy Smollett, Daniel Mair, Stephen Carmichael, Alice Broos, Lily Tong, Jenna Nichols, Kyriaki Nomikou; Sarah McDonald; Ioulia Tsatsani, Richard Orton, Joseph Hughes, Sreenu Vattipally, David L Robertson; Alasdair MacLean, Rory Gunson; Sharif Shaaban, Matthew Holden; Rachel Blacow, Guy Mollett, Kathy Li, James Shepherd, Antonia Ho, Emma Thomson |
| EPI_ISL_2734319, EPI_ISL_2734320, EPI_ISL_2734321, EPI_ISL_2734322, EPI_ISL_2734323, EPI_ISL_2734324, EPI_ISL_2734325, EPI_ISL_2734326, EPI_ISL_2734327, EPI_ISL_2734328, EPI_ISL_2734329, EPI_ISL_2734331, EPI_ISL_2734332, EPI_ISL_2734333, EPI_ISL_2734334, EPI_ISL_2734335, EPI_ISL_2734336, EPI_ISL_2734338, EPI_ISL_2734339, EPI_ISL_2734340, EPI_ISL_2734341, EPI_ISL_2734342, EPI_ISL_2734343, EPI_ISL_2734344, EPI_ISL_2734345, EPI_ISL_2734346, EPI_ISL_2734347, EPI_ISL_2734348, EPI_ISL_2734349, EPI_ISL_2734350, EPI_ISL_2734351, EPI_ISL_2734352, EPI_ISL_2734353, EPI_ISL_2734354, EPI_ISL_2734355, EPI_ISL_2734356, EPI_ISL_2734357, EPI_ISL_2734359, EPI_ISL_2734360, EPI_ISL_2734361, EPI_ISL_2734362, EPI_ISL_2734363, EPI_ISL_2734364, EPI_ISL_2734365, EPI_ISL_2734366, EPI_ISL_2734367, EPI_ISL_2734368, EPI_ISL_2734369, EPI_ISL_2734370, EPI_ISL_2734371, EPI_ISL_2734372, EPI_ISL_2734373, EPI_ISL_2734374, EPI_ISL_2734375, EPI_ISL_2734376, EPI_ISL_2734377, EPI_ISL_2734378, EPI_ISL_2734379, EPI_ISL_2734380, EPI_ISL_2734381, EPI_ISL_2734383, EPI_ISL_2734384, EPI_ISL_2734385, EPI_ISL_2734386, EPI_ISL_2734387, EPI_ISL_2734388, EPI_ISL_2734389, EPI_ISL_2734390, EPI_ISL_2734391, EPI_ISL_2734392, EPI_ISL_2734393, EPI_ISL_2734394, EPI_ISL_2734396, EPI_ISL_2734397, EPI_ISL_2734398, EPI_ISL_2734399, EPI_ISL_2734400, EPI_ISL_2734401, EPI_ISL_2734402, EPI_ISL_2734403, EPI_ISL_2734404, EPI_ISL_2734405, EPI_ISL_2734406, EPI_ISL_2734407, EPI_ISL_2734408, EPI_ISL_2734409, EPI_ISL_2734410, EPI_ISL_2734411, EPI_ISL_2734412, EPI_ISL_2734413, EPI_ISL_2734414, EPI_ISL_2734415, EPI_ISL_2734416, EPI_ISL_2734417, EPI_ISL_2734419, EPI_ISL_2734420, EPI_ISL_2734421, EPI_ISL_2734422, EPI_ISL_2734423, EPI_ISL_2734424, EPI_ISL_2734425, EPI_ISL_2734426, EPI_ISL_2734427, EPI_ISL_2734428, EPI_ISL_2734430, EPI_ISL_2734431, EPI_ISL_2734432, EPI_ISL_2734434, EPI_ISL_2734435 | | | |
| see above | Virology Department, Royal Infirmary of Edinburgh, NHS Lothian / School of Biological Sciences, University of Edinburgh | COVID-19 Genomics UK (COG-UK) Consortium | McHugh M, Dewar R, Cotton S, Gallagher A, Maloney D, Rooke S, O'Toole Á, Scher E, Hill V, McCrone JT, Colquhoun R, Yu X, Jackson B, Rambaut A, Templeton K |
| EPI_ISL_2734939, EPI_ISL_2734942, EPI_ISL_2734943, EPI_ISL_2734944, EPI_ISL_2734945 | Respiratory Virus Unit, Microbiology Services Colindale, Public Health England | COVID-19 Genomics UK (COG-UK) Consortium | PHE Covid Sequencing Team |
| EPI_ISL_2734990, EPI_ISL_2734991, EPI_ISL_2734992, EPI_ISL_2734993, EPI_ISL_2734994, EPI_ISL_2734995, EPI_ISL_2734996, EPI_ISL_2734997, EPI_ISL_2734998, EPI_ISL_2734999, EPI_ISL_2735000, EPI_ISL_2735001, EPI_ISL_2735002, EPI_ISL_2735003, EPI_ISL_2735004, EPI_ISL_2735005, EPI_ISL_2735006, EPI_ISL_2735007, EPI_ISL_2735008, EPI_ISL_2735009, EPI_ISL_2735010, EPI_ISL_2735011, EPI_ISL_2735012, EPI_ISL_2735013, EPI_ISL_2735014, EPI_ISL_2735015, EPI_ISL_2735016, EPI_ISL_2735017, EPI_ISL_2735018, EPI_ISL_2735019, EPI_ISL_2735020, EPI_ISL_2735021, EPI_ISL_2735022, EPI_ISL_2735023, EPI_ISL_2735024, EPI_ISL_2735025, EPI_ISL_2735026, EPI_ISL_2735027, EPI_ISL_2735028, EPI_ISL_2735029, EPI_ISL_2735030, EPI_ISL_2735031, EPI_ISL_2735032, EPI_ISL_2735033, EPI_ISL_2735034, EPI_ISL_2735035, EPI_ISL_2735036, EPI_ISL_2735037, EPI_ISL_2735038, EPI_ISL_2735039, EPI_ISL_2735040, EPI_ISL_2735041, EPI_ISL_2735042, EPI_ISL_2735043, EPI_ISL_2735044, EPI_ISL_2735045, EPI_ISL_2735046, EPI_ISL_2735047, EPI_ISL_2735048, EPI_ISL_2735049, EPI_ISL_2735050, EPI_ISL_2735051, EPI_ISL_2735052, EPI_ISL_2735053, EPI_ISL_2735054, EPI_ISL_2735055, EPI_ISL_2735056, EPI_ISL_2735057, EPI_ISL_2735058, EPI_ISL_2735059, EPI_ISL_2735060, EPI_ISL_2735061 | | | |
| see above | University College London, Great Ormond Street Hospital for Children NHS Foundation Trust, Imperial College Healthcare NHS Trust | COVID-19 Genomics UK (COG-UK) Consortium | Sergi Castellano, Rachel Williams, Mark Kristiansen, Paola Resende Silva, Sunando Roy, Tony Brooks, Helena Tutill, Paola Niola, Patricia Dyal, Charlotte Williams, Leysa Forrest, Yasmin Panchbhaya, Jacqueline Findlay, Samuel Weeks, Julianne Brown, Kathryn Harris, Paul Randell, James Price, Alison Holmes, Judith Breuer |
| EPI_ISL_2735679, EPI_ISL_2735680, EPI_ISL_2735681, EPI_ISL_2735683, EPI_ISL_2735684, EPI_ISL_2735686, EPI_ISL_2735687, EPI_ISL_2735688, EPI_ISL_2735689, EPI_ISL_2735690, EPI_ISL_2735692, EPI_ISL_2735693, EPI_ISL_2735695, EPI_ISL_2735696, EPI_ISL_2735697, EPI_ISL_2735699, EPI_ISL_2735700, EPI_ISL_2735701, EPI_ISL_2735704, EPI_ISL_2735705, EPI_ISL_2735706, EPI_ISL_2735707 | | | |
| see above | Oxford Viromics, NDM, University of Oxford; Oxford University Hospitals; Basingstoke and North Hampshire Hospital | COVID-19 Genomics UK (COG-UK) Consortium | Tanya Golubchik, David Bonsall, George Macintyre, Amy Trebes, Mariateresa de Cesare, Catrin Moore, Alex Mobbs, Anita Justice, Robert Shaw, Monique Andersson, Timothy Peto, Emma Wise, Nathan Moore, Jessica Lynch, Nick Cortes, Matilde Mori, Stephen Kidd, David Buck, John Todd, Christophe Fraser |
| EPI_ISL_2735897, EPI_ISL_2735929, EPI_ISL_2735940, EPI_ISL_2735941, EPI_ISL_2735943, EPI_ISL_2736012 | Seattle Flu Study | Seattle Flu Study | Deborah A. Nickerson, Chris D. Frazar, Jover Lee, Benjamin Pelle, Erica Ryke, Matthew Richardson, Amanda Adler, Elisabeth Brandstetter, Peter D. Han, Kairsten Fay, Misja Ilcisin, Kirsten Lacombe, Thomas R. Sibley, Melissa Truong, Caitlin R. Wolf, Karen Cowgill, Stephanie Schrag, Jeff Duchin, Michael Boeckh, Janet A. Englund, Michael Famulare, Barry R. Lutz, Mark J. Rieder, Lea M. Starita, Matthew Thompson, Helen Y. Chu, Trevor Bedford, Jay Shendure |
| EPI_ISL_2736196, EPI_ISL_2736207 | Northwest Laboratory | Seattle Flu Study | Deborah A. Nickerson, Chris D. Frazar, Jover Lee, Benjamin Pelle, Erica Ryke, Matthew Richardson, Amanda Adler, Elisabeth Brandstetter, Peter D. Han, Kairsten Fay, Misja Ilcisin, Kirsten Lacombe, Thomas R. Sibley, Melissa Truong, Caitlin R. Wolf, Romesh Gautom, Geoff Melly, Brian Hiatt, Philip Dykema, Scott Lindquist, Michael Boeckh, Janet A. Englund, Michael Famulare, Barry R. Lutz, Mark J. Rieder, Lea M. Starita, Matthew Thompson, Helen Y. Chu, Jay Shendure, Trevor Bedford |
| EPI_ISL_2736253, EPI_ISL_2736259, EPI_ISL_2736285, EPI_ISL_2736287, EPI_ISL_2736301, EPI_ISL_2736304, EPI_ISL_2736310, EPI_ISL_2736320, EPI_ISL_2736330, EPI_ISL_2736336 | Atlas Genomics | Seattle Flu Study | Deborah A. Nickerson, Chris D. Frazar, Jover Lee, Benjamin Pelle, Erica Ryke, Matthew Richardson, Amanda Adler, Elisabeth Brandstetter, Peter D. Han, Kairsten Fay, Misja Ilcisin, Kirsten Lacombe, Thomas R. Sibley, Melissa Truong, Caitlin R. Wolf, Romesh Gautom, Geoff Melly, Brian Hiatt, Philip Dykema, Scott Lindquist, Michael Boeckh, Janet A. Englund, Michael Famulare, Barry R. Lutz, Mark J. Rieder, Lea M. Starita, Matthew Thompson, Helen Y. Chu, Jay Shendure, Trevor Bedford |
| EPI_ISL_2736461, EPI_ISL_2736470 | Seattle Flu Study | Seattle Flu Study | Deborah A. Nickerson, Chris D. Frazar, Jover Lee, Benjamin Pelle, Erica Ryke, Matthew Richardson, Amanda Adler, Elisabeth Brandstetter, Peter D. Han, Kairsten Fay, Misja Ilcisin, Kirsten Lacombe, Thomas R. Sibley, Melissa Truong, Caitlin R. Wolf, Michael Boeckh, Janet A. Englund, Michael Famulare, Barry R. Lutz, Mark J. Rieder, Lea M. Starita, Matthew Thompson, Jay Shendure, Trevor Bedford, Helen Y. Chu |
| EPI_ISL_2736780, EPI_ISL_2736781, EPI_ISL_2736783 | LESP Sinaloa | Instituto de Diagnostico y Referencia Epidemiologicos (INDRE) | Claudia Wong-Arambula, Abril Rodriguez-Maldonado, Vanessa Rivero-Arredondo, Ariadna Medina-Benitez, Joaquin Quiroz-Mercado, Sergio Rangel-Guerrero, Natividad Cruz-Ortiz, Tatiana Nunez-Garcia, Gisela Barrera-Badillo, Lucia Hernandez-Rivas, Irma Lopez-Martinez, Ernesto Ramirez-Gonzalez. |
| EPI_ISL_2736784 | LESP Michoacan | Instituto de Diagnostico y Referencia Epidemiologicos (INDRE) | Claudia Wong-Arambula, Abril Rodriguez-Maldonado, Vanessa Rivero-Arredondo, Ariadna Medina-Benitez, Joaquin Quiroz-Mercado, Sergio Rangel-Guerrero, Natividad Cruz-Ortiz, Tatiana Nunez-Garcia, Gisela Barrera-Badillo, Lucia Hernandez-Rivas, Irma Lopez-Martinez, Ernesto Ramirez-Gonzalez. |
| EPI_ISL_2736785 | LESP Puebla | Instituto de Diagnostico y Referencia Epidemiologicos (INDRE) | Claudia Wong-Arambula, Abril Rodriguez-Maldonado, Vanessa Rivero-Arredondo, Ariadna Medina-Benitez, Joaquin Quiroz-Mercado, Sergio Rangel-Guerrero, Natividad Cruz-Ortiz, Tatiana Nunez-Garcia, Gisela Barrera-Badillo, Lucia Hernandez-Rivas, Irma Lopez-Martinez, Ernesto Ramirez-Gonzalez. |
| EPI_ISL_2736787 | LESP Guanajuato | Instituto de Diagnostico y Referencia Epidemiologicos (INDRE) | Claudia Wong-Arambula, Abril Rodriguez-Maldonado, Vanessa Rivero-Arredondo, Ariadna Medina-Benitez, Joaquin Quiroz-Mercado, Sergio Rangel-Guerrero, Natividad Cruz-Ortiz, Tatiana Nunez-Garcia, Gisela Barrera-Badillo, Lucia Hernandez-Rivas, Irma Lopez-Martinez, Ernesto Ramirez-Gonzalez. |
| EPI_ISL_2738504, EPI_ISL_2738567 | Hackensack Medical Center | New York Genome Center | Michael Zody, Andre Corvelo, Dayna M. Oschwald, Samantha Fennessey, Tom Maniatis, Liang Chen, Jose Mediavilla, Marcus Cunningham, Kaelea Composto, Kar Chow, David Perlin, Barry Kreiswirth |
| EPI_ISL_2740797, EPI_ISL_2740891, EPI_ISL_2741099, EPI_ISL_2741124, EPI_ISL_2741166, EPI_ISL_2741169, EPI_ISL_2741268, EPI_ISL_2741307, EPI_ISL_2741327, EPI_ISL_2741350, EPI_ISL_2741434, EPI_ISL_2741444, EPI_ISL_2741482, EPI_ISL_2741531, EPI_ISL_2741538, EPI_ISL_2741628, EPI_ISL_2741630, EPI_ISL_2741632, EPI_ISL_2741634, EPI_ISL_2741636, EPI_ISL_2741638, EPI_ISL_2741640, EPI_ISL_2741642, EPI_ISL_2741643, EPI_ISL_2741645, EPI_ISL_2741647, EPI_ISL_2741649 | | | |
| see above | Respiratory Virus Unit, Microbiology Services Colindale, Public Health England | COVID-19 Genomics UK (COG-UK) Consortium | PHE Covid Sequencing Team |
| EPI_ISL_2741650 | A.S.L. CN2 | Fondazione del Piemonte per l'Oncologia IRCCS | Antonino Sottile, Silvia Brossa, Paola Marino, Giorgio Giardina |
| EPI_ISL_2741664 | A.S.L. VERCELLI | Fondazione del Piemonte per l'Oncologia IRCCS | Antonino Sottile, Silvia Brossa, Paola Marino, Giorgio Giardina |
| EPI_ISL_2741677 | A.S.L. BIELLA | Fondazione del Piemonte per l'Oncologia IRCCS | Antonino Sottile, Silvia Brossa, Paola Marino, Giorgio Giardina |
| EPI_ISL_2741700 | A.S.L CN1 | Fondazione del Piemonte per l'Oncologia IRCCS | Antonino Sottile, Silvia Brossa, Paola Marino, Giorgio Giardina |
| EPI_ISL_2741708 | A.S.L. VERCELLI | Fondazione del Piemonte per l'Oncologia IRCCS | Antonino Sottile, Silvia Brossa, Paola Marino, Giorgio Giardina |
| EPI_ISL_2741731, EPI_ISL_2741756, EPI_ISL_2741757, EPI_ISL_2741758, EPI_ISL_2741808, EPI_ISL_2741812, EPI_ISL_2741819, EPI_ISL_2741824, EPI_ISL_2741868, EPI_ISL_2741888, EPI_ISL_2741889, EPI_ISL_2741902, EPI_ISL_2741910, EPI_ISL_2741922, EPI_ISL_2741937, EPI_ISL_2741944, EPI_ISL_2741945, EPI_ISL_2741955, EPI_ISL_2741958, EPI_ISL_2741961, EPI_ISL_2741967, EPI_ISL_2741976, EPI_ISL_2741981, EPI_ISL_2741984, EPI_ISL_2741985, EPI_ISL_2741989, EPI_ISL_2742015, EPI_ISL_2742016, EPI_ISL_2742020, EPI_ISL_2742028, EPI_ISL_2742029, EPI_ISL_2742030, EPI_ISL_2742035, EPI_ISL_2742037, EPI_ISL_2742039, EPI_ISL_2742040, EPI_ISL_2742046, EPI_ISL_2742051, EPI_ISL_2742064, EPI_ISL_2742071, EPI_ISL_2742074, EPI_ISL_2742089, EPI_ISL_2742090, EPI_ISL_2742097, EPI_ISL_2742106, EPI_ISL_2742113, EPI_ISL_2742114, EPI_ISL_2742118, EPI_ISL_2742122, EPI_ISL_2742132, EPI_ISL_2742160, EPI_ISL_2742191, EPI_ISL_2742192, EPI_ISL_2742193, EPI_ISL_2742194, EPI_ISL_2742196, EPI_ISL_2742199, EPI_ISL_2742200, EPI_ISL_2742201, EPI_ISL_2742202, EPI_ISL_2742203, EPI_ISL_2742204, EPI_ISL_2742207, EPI_ISL_2742208, EPI_ISL_2742210, EPI_ISL_2742211, EPI_ISL_2742212, EPI_ISL_2742224, EPI_ISL_2742225, EPI_ISL_2742228, EPI_ISL_2742229, EPI_ISL_2742230, EPI_ISL_2742231, EPI_ISL_2742233, EPI_ISL_2742235, EPI_ISL_2742236, EPI_ISL_2742237, EPI_ISL_2742242, EPI_ISL_2742244, EPI_ISL_2742253, EPI_ISL_2742254, EPI_ISL_2742256, EPI_ISL_2742259, EPI_ISL_2742262, EPI_ISL_2742263, EPI_ISL_2742267, EPI_ISL_2742287, EPI_ISL_2742288, EPI_ISL_2742302, EPI_ISL_2742303, EPI_ISL_2742310, EPI_ISL_2742312, EPI_ISL_2742315, EPI_ISL_2742316, EPI_ISL_2742317, EPI_ISL_2742322, EPI_ISL_2742331, EPI_ISL_2742333, EPI_ISL_2742347, EPI_ISL_2742357, EPI_ISL_2742360, EPI_ISL_2742370, EPI_ISL_2742371, EPI_ISL_2742374, EPI_ISL_2742378, EPI_ISL_2742381, EPI_ISL_2742387, EPI_ISL_2742388, EPI_ISL_2742390, EPI_ISL_2742391, EPI_ISL_2742392, EPI_ISL_2742397, EPI_ISL_2742399, EPI_ISL_2742402, EPI_ISL_2742404, EPI_ISL_2742405, EPI_ISL_2742407, EPI_ISL_2742408, EPI_ISL_2742411, EPI_ISL_2742415, EPI_ISL_2742422, EPI_ISL_2742423, EPI_ISL_2742426, EPI_ISL_2742428, EPI_ISL_2742432, EPI_ISL_2742435, EPI_ISL_2742438, EPI_ISL_2742439, EPI_ISL_2742441, EPI_ISL_2742442, EPI_ISL_2742443, EPI_ISL_2742444, EPI_ISL_2742452, EPI_ISL_2742459, EPI_ISL_2742463, EPI_ISL_2742467, EPI_ISL_2742468, EPI_ISL_2742471, EPI_ISL_2742472, EPI_ISL_2742475, EPI_ISL_2742476, EPI_ISL_2742477, EPI_ISL_2742478, EPI_ISL_2742480, EPI_ISL_2742481, EPI_ISL_2742482, EPI_ISL_2742486, EPI_ISL_2742488, EPI_ISL_2742493, EPI_ISL_2742494, EPI_ISL_2742495, EPI_ISL_2742496, EPI_ISL_2742498, EPI_ISL_2742501, EPI_ISL_2742504, EPI_ISL_2742509, EPI_ISL_2742514, EPI_ISL_2742515, EPI_ISL_2742516, EPI_ISL_2742518, EPI_ISL_2742519, EPI_ISL_2742520, EPI_ISL_2742521, EPI_ISL_2742522, EPI_ISL_2742523, EPI_ISL_2742525, EPI_ISL_2742526, EPI_ISL_2742527, EPI_ISL_2742529, EPI_ISL_2742530, EPI_ISL_2742531, EPI_ISL_2742532, EPI_ISL_2742534, EPI_ISL_2742535, EPI_ISL_2742538, EPI_ISL_2742539, EPI_ISL_2742542, EPI_ISL_2742544, EPI_ISL_2742545, EPI_ISL_2742548, EPI_ISL_2742553, EPI_ISL_2742554, EPI_ISL_2742560, EPI_ISL_2742562, EPI_ISL_2742564, EPI_ISL_2742565, EPI_ISL_2742566, EPI_ISL_2742568, EPI_ISL_2742569, EPI_ISL_2742570, EPI_ISL_2742572, EPI_ISL_2742573, EPI_ISL_2742574, EPI_ISL_2742575, EPI_ISL_2742576, EPI_ISL_2742577, EPI_ISL_2742578, EPI_ISL_2742579, EPI_ISL_2742580, EPI_ISL_2742581, EPI_ISL_2742582, EPI_ISL_2742583, EPI_ISL_2742584, EPI_ISL_2742585, EPI_ISL_2742587, EPI_ISL_2742590, EPI_ISL_2742591, EPI_ISL_2742593, EPI_ISL_2742594, EPI_ISL_2742595, EPI_ISL_2742598, EPI_ISL_2742599, EPI_ISL_2742601, EPI_ISL_2742602, EPI_ISL_2742603, EPI_ISL_2742607, EPI_ISL_2742609, EPI_ISL_2742617, EPI_ISL_2742618, EPI_ISL_2742619, EPI_ISL_2742620, EPI_ISL_2742621, EPI_ISL_2742629, EPI_ISL_2742630, EPI_ISL_2742631, EPI_ISL_2742633, EPI_ISL_2742635, EPI_ISL_2742636, EPI_ISL_2742637, EPI_ISL_2742638, EPI_ISL_2742639, EPI_ISL_2742640, EPI_ISL_2742643, EPI_ISL_2742645, EPI_ISL_2742650, EPI_ISL_2742651, EPI_ISL_2742652, EPI_ISL_2742656, EPI_ISL_2742657, EPI_ISL_2742658, EPI_ISL_2742659, EPI_ISL_2742662, EPI_ISL_2742664, EPI_ISL_2742665, EPI_ISL_2742666, EPI_ISL_2742668, EPI_ISL_2742670, EPI_ISL_2742671, EPI_ISL_2742672, EPI_ISL_2742676, EPI_ISL_2742679, EPI_ISL_2742680, EPI_ISL_2742681, EPI_ISL_2742684, EPI_ISL_2742687, EPI_ISL_2742688, EPI_ISL_2742689, EPI_ISL_2742691, EPI_ISL_2742694, EPI_ISL_2742695, EPI_ISL_2742697, EPI_ISL_2742698, EPI_ISL_2742699, EPI_ISL_2742700, EPI_ISL_2742701, EPI_ISL_2742703, EPI_ISL_2742704, EPI_ISL_2742708, EPI_ISL_2742709, EPI_ISL_2742710, EPI_ISL_2742711, EPI_ISL_2742715, EPI_ISL_2742716, EPI_ISL_2742723, EPI_ISL_2742725, EPI_ISL_2742726, EPI_ISL_2742727, EPI_ISL_2742728, EPI_ISL_2742730, EPI_ISL_2742732, EPI_ISL_2742733, EPI_ISL_2742736, EPI_ISL_2742737, EPI_ISL_2742738, EPI_ISL_2742739, EPI_ISL_2742742, EPI_ISL_2742743, EPI_ISL_2742748, EPI_ISL_2742749, EPI_ISL_2742750, EPI_ISL_2742754, EPI_ISL_2742755, EPI_ISL_2742758, EPI_ISL_2742761, EPI_ISL_2742762, EPI_ISL_2742763, EPI_ISL_2742767, EPI_ISL_2742769, EPI_ISL_2742771, EPI_ISL_2742774, EPI_ISL_2742775, EPI_ISL_2742776, EPI_ISL_2742777, EPI_ISL_2742778, EPI_ISL_2742779, EPI_ISL_2742780, EPI_ISL_2742783, EPI_ISL_2742785, EPI_ISL_2742788, EPI_ISL_2742789, EPI_ISL_2742791, EPI_ISL_2742795, EPI_ISL_2742796, EPI_ISL_2742798, EPI_ISL_2742800, EPI_ISL_2742801, EPI_ISL_2742803, EPI_ISL_2742804, EPI_ISL_2742805, EPI_ISL_2742806, EPI_ISL_2742809, EPI_ISL_2742810, EPI_ISL_2742812, EPI_ISL_2742815, EPI_ISL_2742817, EPI_ISL_2742820, EPI_ISL_2742823, EPI_ISL_2742825, EPI_ISL_2742828, EPI_ISL_2742829, EPI_ISL_2742831, EPI_ISL_2742833, EPI_ISL_2742835, EPI_ISL_2742836, EPI_ISL_2742837, EPI_ISL_2742838, EPI_ISL_2742843, EPI_ISL_2742844, EPI_ISL_2742845, EPI_ISL_2742846, EPI_ISL_2742847, EPI_ISL_2742848, EPI_ISL_2742850, EPI_ISL_2742851, EPI_ISL_2742852, EPI_ISL_2742856, EPI_ISL_2742858, EPI_ISL_2742860, EPI_ISL_2742861, EPI_ISL_2742862, EPI_ISL_2742864, EPI_ISL_2742865, EPI_ISL_2742866, EPI_ISL_2742867, EPI_ISL_2742868, EPI_ISL_2742869, EPI_ISL_2742870, EPI_ISL_2742871, EPI_ISL_2742872, EPI_ISL_2742873, EPI_ISL_2742874, EPI_ISL_2742875, EPI_ISL_2742876, EPI_ISL_2742877, EPI_ISL_2742878, EPI_ISL_2742879, EPI_ISL_2742880, EPI_ISL_2742881, EPI_ISL_2742882, EPI_ISL_2742884, EPI_ISL_2742885, EPI_ISL_2742886, EPI_ISL_2742887, EPI_ISL_2742888, EPI_ISL_2742889, EPI_ISL_2742890, EPI_ISL_2742891, EPI_ISL_2742892, EPI_ISL_2742894, EPI_ISL_2742899, EPI_ISL_2742900, EPI_ISL_2742901, EPI_ISL_2742903, EPI_ISL_2742904, EPI_ISL_2742905, EPI_ISL_2742906, EPI_ISL_2742907, EPI_ISL_2742908, EPI_ISL_2742909, EPI_ISL_2742910, EPI_ISL_2742911, EPI_ISL_2742912, EPI_ISL_2742913, EPI_ISL_2742914, EPI_ISL_2742915, EPI_ISL_2742916, EPI_ISL_2742917, EPI_ISL_2742919, EPI_ISL_2742920, EPI_ISL_2742921, EPI_ISL_2742922, EPI_ISL_2742924, EPI_ISL_2742926, EPI_ISL_2742928, EPI_ISL_2742929, EPI_ISL_2742930, EPI_ISL_2742934, EPI_ISL_2742935, EPI_ISL_2742938, EPI_ISL_2742939, EPI_ISL_2742943, EPI_ISL_2742946, EPI_ISL_2742947, EPI_ISL_2742949, EPI_ISL_2742950, EPI_ISL_2742952, EPI_ISL_2742956, EPI_ISL_2742957, EPI_ISL_2742958, EPI_ISL_2742963, EPI_ISL_2742964, EPI_ISL_2742965, EPI_ISL_2742966, EPI_ISL_2742967, EPI_ISL_2742970, EPI_ISL_2742972, EPI_ISL_2742975, EPI_ISL_2742976, EPI_ISL_2742977, EPI_ISL_2742978, EPI_ISL_2742979, EPI_ISL_2742980, EPI_ISL_2742981, EPI_ISL_2742982, EPI_ISL_2742983, EPI_ISL_2742984, EPI_ISL_2742985, EPI_ISL_2742986, EPI_ISL_2742989, EPI_ISL_2742990, EPI_ISL_2742992, EPI_ISL_2742993, EPI_ISL_2742994, EPI_ISL_2742995, EPI_ISL_2742996, EPI_ISL_2742997, EPI_ISL_2742999, EPI_ISL_2743000, EPI_ISL_2743001, EPI_ISL_2743002, EPI_ISL_2743003, EPI_ISL_2743004, EPI_ISL_2743005, EPI_ISL_2743006, EPI_ISL_2743008, EPI_ISL_2743009, EPI_ISL_2743011, EPI_ISL_2743012, EPI_ISL_2743013, EPI_ISL_2743014, EPI_ISL_2743015, EPI_ISL_2743016, EPI_ISL_2743017, EPI_ISL_2743018, EPI_ISL_2743019, EPI_ISL_2743021, EPI_ISL_2743022, EPI_ISL_2743024, EPI_ISL_2743025, EPI_ISL_2743028, EPI_ISL_2743029, EPI_ISL_2743030, EPI_ISL_2743032, EPI_ISL_2743034, EPI_ISL_2743036, EPI_ISL_2743037, EPI_ISL_2743038, EPI_ISL_2743039, EPI_ISL_2743040, EPI_ISL_2743041, EPI_ISL_2743042, EPI_ISL_2743043, EPI_ISL_2743044, EPI_ISL_2743048, EPI_ISL_2743051, EPI_ISL_2743055, EPI_ISL_2743056, EPI_ISL_2743057, EPI_ISL_2743060, EPI_ISL_2743061, EPI_ISL_2743062, EPI_ISL_2743063, EPI_ISL_2743064, EPI_ISL_2743065, EPI_ISL_2743066, EPI_ISL_2743067, EPI_ISL_2743068, EPI_ISL_2743069, EPI_ISL_2743070, EPI_ISL_2743071, EPI_ISL_2743072, EPI_ISL_2743073, EPI_ISL_2743078, EPI_ISL_2743081, EPI_ISL_2743082, EPI_ISL_2743083, | | | |

EPI_ISL_2743085, EPI_ISL_2743086, EPI_ISL_2743089, EPI_ISL_2743090, EPI_ISL_2743091, EPI_ISL_2743092, EPI_ISL_2743095, EPI_ISL_2743096, EPI_ISL_2743097, EPI_ISL_2743099, EPI_ISL_2743103, EPI_ISL_2743104, EPI_ISL_2743110, EPI_ISL_2743111, EPI_ISL_2743113, EPI_ISL_2743114, EPI_ISL_2743115, EPI_ISL_2743117, EPI_ISL_2743118, EPI_ISL_2743119, EPI_ISL_2743120, EPI_ISL_2743123, EPI_ISL_2743124, EPI_ISL_2743125, EPI_ISL_2743126, EPI_ISL_2743127, EPI_ISL_2743128, EPI_ISL_2743130, EPI_ISL_2743132, EPI_ISL_2743133, EPI_ISL_2743134, EPI_ISL_2743135, EPI_ISL_2743136, EPI_ISL_2743140, EPI_ISL_2743142, EPI_ISL_2743143, EPI_ISL_2743146, EPI_ISL_2743148, EPI_ISL_2743149, EPI_ISL_2743151, EPI_ISL_2743153, EPI_ISL_2743154, EPI_ISL_2743155, EPI_ISL_2743156, EPI_ISL_2743157, EPI_ISL_2743158, EPI_ISL_2743160, EPI_ISL_2743163, EPI_ISL_2743164, EPI_ISL_2743165, EPI_ISL_2743166, EPI_ISL_2743167, EPI_ISL_2743168, EPI_ISL_2743169, EPI_ISL_2743172, EPI_ISL_2743173, EPI_ISL_2743174, EPI_ISL_2743176, EPI_ISL_2743177, EPI_ISL_2743179, EPI_ISL_2743180, EPI_ISL_2743183, EPI_ISL_2743185, EPI_ISL_2743186, EPI_ISL_2743187, EPI_ISL_2743188, EPI_ISL_2743189, EPI_ISL_2743190, EPI_ISL_2743191, EPI_ISL_2743192, EPI_ISL_2743193, EPI_ISL_2743194, EPI_ISL_2743195, EPI_ISL_2743196, EPI_ISL_2743197, EPI_ISL_2743200, EPI_ISL_2743201, EPI_ISL_2743202, EPI_ISL_2743203, EPI_ISL_2743204, EPI_ISL_2743205, EPI_ISL_2743206, EPI_ISL_2743207, EPI_ISL_2743211, EPI_ISL_2743212, EPI_ISL_2743213, EPI_ISL_2743214, EPI_ISL_2743217, EPI_ISL_2743218, EPI_ISL_2743220, EPI_ISL_2743221, EPI_ISL_2743222, EPI_ISL_2743223, EPI_ISL_2743230, EPI_ISL_2743232, EPI_ISL_2743233, EPI_ISL_2743234, EPI_ISL_2743235, EPI_ISL_2743236, EPI_ISL_2743237, EPI_ISL_2743238, EPI_ISL_2743239, EPI_ISL_2743240, EPI_ISL_2743241, EPI_ISL_2743242, EPI_ISL_2743243, EPI_ISL_2743244, EPI_ISL_2743245, EPI_ISL_2743246, EPI_ISL_2743247, EPI_ISL_2743248, EPI_ISL_2743249, EPI_ISL_2743250, EPI_ISL_2743251, EPI_ISL_2743252, EPI_ISL_2743256, EPI_ISL_2743257, EPI_ISL_2743258, EPI_ISL_2743259, EPI_ISL_2743261, EPI_ISL_2743262, EPI_ISL_2743263, EPI_ISL_2743264, EPI_ISL_2743266, EPI_ISL_2743267, EPI_ISL_2743270, EPI_ISL_2743271, EPI_ISL_2743272, EPI_ISL_2743273, EPI_ISL_2743276, EPI_ISL_2743277, EPI_ISL_2743279, EPI_ISL_2743280, EPI_ISL_2743281, EPI_ISL_2743282, EPI_ISL_2743283, EPI_ISL_2743287, EPI_ISL_2743288, EPI_ISL_2743289, EPI_ISL_2743293, EPI_ISL_2743294, EPI_ISL_2743295, EPI_ISL_2743296, EPI_ISL_2743297, EPI_ISL_2743298, EPI_ISL_2743299, EPI_ISL_2743300, EPI_ISL_2743302, EPI_ISL_2743303, EPI_ISL_2743304, EPI_ISL_2743305, EPI_ISL_2743307, EPI_ISL_2743308, EPI_ISL_2743309, EPI_ISL_2743310, EPI_ISL_2743311, EPI_ISL_2743312, EPI_ISL_2743313, EPI_ISL_2743315, EPI_ISL_2743316, EPI_ISL_2743317, EPI_ISL_2743318, EPI_ISL_2743319, EPI_ISL_2743320, EPI_ISL_2743321, EPI_ISL_2743322, EPI_ISL_2743324, EPI_ISL_2743325, EPI_ISL_2743326, EPI_ISL_2743328, EPI_ISL_2743329, EPI_ISL_2743330, EPI_ISL_2743332, EPI_ISL_2743333, EPI_ISL_2743334, EPI_ISL_2743336, EPI_ISL_2743337, EPI_ISL_2743338, EPI_ISL_2743339, EPI_ISL_2743340, EPI_ISL_2743341, EPI_ISL_2743342, EPI_ISL_2743343, EPI_ISL_2743344, EPI_ISL_2743345, EPI_ISL_2743346, EPI_ISL_2743348, EPI_ISL_2743349, EPI_ISL_2743350, EPI_ISL_2743351, EPI_ISL_2743352, EPI_ISL_2743353, EPI_ISL_2743354, EPI_ISL_2743355, EPI_ISL_2743356, EPI_ISL_2743357, EPI_ISL_2743358, EPI_ISL_2743359, EPI_ISL_2743360, EPI_ISL_2743361, EPI_ISL_2743362, EPI_ISL_2743363, EPI_ISL_2743364, EPI_ISL_2743368, EPI_ISL_2743369, EPI_ISL_2743370, EPI_ISL_2743371, EPI_ISL_2743372, EPI_ISL_2743374, EPI_ISL_2743375, EPI_ISL_2743376, EPI_ISL_2743377, EPI_ISL_2743378, EPI_ISL_2743380, EPI_ISL_2743381, EPI_ISL_2743382, EPI_ISL_2743387, EPI_ISL_2743388, EPI_ISL_2743389, EPI_ISL_2743390, EPI_ISL_2743391, EPI_ISL_2743392, EPI_ISL_2743393, EPI_ISL_2743394, EPI_ISL_2743395, EPI_ISL_2743396, EPI_ISL_2743397, EPI_ISL_2743398, EPI_ISL_2743399, EPI_ISL_2743401, EPI_ISL_2743402, EPI_ISL_2743403, EPI_ISL_2743404, EPI_ISL_2743405, EPI_ISL_2743406, EPI_ISL_2743407, EPI_ISL_2743408, EPI_ISL_2743409, EPI_ISL_2743410, EPI_ISL_2743411, EPI_ISL_2743412, EPI_ISL_2743413, EPI_ISL_2743416, EPI_ISL_2743417, EPI_ISL_2743418, EPI_ISL_2743419, EPI_ISL_2743420, EPI_ISL_2743421, EPI_ISL_2743422, EPI_ISL_2743423, EPI_ISL_2743424, EPI_ISL_2743427, EPI_ISL_2743428, EPI_ISL_2743436, EPI_ISL_2743437, EPI_ISL_2743438, EPI_ISL_2743439, EPI_ISL_2743440, EPI_ISL_2743441, EPI_ISL_2743442, EPI_ISL_2743443, EPI_ISL_2743444, EPI_ISL_2743445, EPI_ISL_2743446, EPI_ISL_2743447, EPI_ISL_2743448, EPI_ISL_2743449, EPI_ISL_2743450, EPI_ISL_2743451, EPI_ISL_2743452, EPI_ISL_2743453, EPI_ISL_2743454, EPI_ISL_2743455, EPI_ISL_2743456, EPI_ISL_2743457, EPI_ISL_2743459, EPI_ISL_2743460, EPI_ISL_2743461, EPI_ISL_2743462, EPI_ISL_2743463, EPI_ISL_2743464, EPI_ISL_2743465, EPI_ISL_2743466, EPI_ISL_2743467, EPI_ISL_2743468, EPI_ISL_2743470, EPI_ISL_2743473, EPI_ISL_2743474, EPI_ISL_2743475, EPI_ISL_2743476, EPI_ISL_2743477, EPI_ISL_2743478, EPI_ISL_2743479, EPI_ISL_2743480, EPI_ISL_2743481, EPI_ISL_2743483, EPI_ISL_2743484, EPI_ISL_2743485, EPI_ISL_2743486, EPI_ISL_2743487, EPI_ISL_2743488, EPI_ISL_2743489, EPI_ISL_2743490, EPI_ISL_2743491, EPI_ISL_2743492, EPI_ISL_2743493, EPI_ISL_2743494, EPI_ISL_2743495, EPI_ISL_2743496, EPI_ISL_2743497, EPI_ISL_2743498, EPI_ISL_2743499, EPI_ISL_2743500, EPI_ISL_2743501, EPI_ISL_2743502, EPI_ISL_2743504, EPI_ISL_2743505, EPI_ISL_2743506, EPI_ISL_2743507, EPI_ISL_2743508, EPI_ISL_2743509, EPI_ISL_2743510, EPI_ISL_2743511, EPI_ISL_2743512, EPI_ISL_2743513, EPI_ISL_2743514, EPI_ISL_2743515, EPI_ISL_2743516, EPI_ISL_2743517, EPI_ISL_2743518, EPI_ISL_2743519, EPI_ISL_2743520, EPI_ISL_2743521, EPI_ISL_2743522, EPI_ISL_2743523, EPI_ISL_2743524, EPI_ISL_2743525, EPI_ISL_2743527, EPI_ISL_2743529, EPI_ISL_2743530, EPI_ISL_2743531, EPI_ISL_2743533, EPI_ISL_2743534, EPI_ISL_2743535, EPI_ISL_2743536, EPI_ISL_2743537, EPI_ISL_2743539, EPI_ISL_2743540, EPI_ISL_2743541, EPI_ISL_2743542, EPI_ISL_2743543, EPI_ISL_2743545, EPI_ISL_2743546, EPI_ISL_2743547, EPI_ISL_2743548, EPI_ISL_2743550, EPI_ISL_2743551, EPI_ISL_2743555, EPI_ISL_2743556, EPI_ISL_2743557, EPI_ISL_2743558, EPI_ISL_2743559, EPI_ISL_2743560, EPI_ISL_2743562, EPI_ISL_2743563, EPI_ISL_2743564, EPI_ISL_2743565, EPI_ISL_2743566, EPI_ISL_2743567, EPI_ISL_2743568, EPI_ISL_2743569, EPI_ISL_2743570, EPI_ISL_2743573, EPI_ISL_2743574, EPI_ISL_2743575, EPI_ISL_2743576, EPI_ISL_2743577, EPI_ISL_2743578, EPI_ISL_2743580, EPI_ISL_2743581, EPI_ISL_2743582, EPI_ISL_2743587, EPI_ISL_2743588, EPI_ISL_2743589, EPI_ISL_2743590, EPI_ISL_2743591, EPI_ISL_2743592, EPI_ISL_2743594, EPI_ISL_2743596, EPI_ISL_2743600, EPI_ISL_2743601, EPI_ISL_2743602, EPI_ISL_2743603, EPI_ISL_2743604, EPI_ISL_2743605, EPI_ISL_2743606, EPI_ISL_2743607, EPI_ISL_2743608, EPI_ISL_2743609, EPI_ISL_2743610, EPI_ISL_2743611, EPI_ISL_2743613, EPI_ISL_2743614, EPI_ISL_2743615, EPI_ISL_2743616, EPI_ISL_2743617, EPI_ISL_2743618, EPI_ISL_2743619, EPI_ISL_2743620, EPI_ISL_2743621, EPI_ISL_2743622, EPI_ISL_2743623, EPI_ISL_2743624, EPI_ISL_2743625, EPI_ISL_2743626, EPI_ISL_2743627, EPI_ISL_2743628, EPI_ISL_2743629, EPI_ISL_2743630, EPI_ISL_2743633, EPI_ISL_2743634, EPI_ISL_2743635, EPI_ISL_2743636, EPI_ISL_2743637, EPI_ISL_2743638, EPI_ISL_2743639, EPI_ISL_2743641, EPI_ISL_2743642, EPI_ISL_2743643, EPI_ISL_2743648, EPI_ISL_2743649, EPI_ISL_2743650, EPI_ISL_2743651, EPI_ISL_2743652, EPI_ISL_2743653, EPI_ISL_2743655, EPI_ISL_2743656, EPI_ISL_2743657, EPI_ISL_2743658, EPI_ISL_2743660, EPI_ISL_2743661, EPI_ISL_2743662, EPI_ISL_2743664, EPI_ISL_2743665, EPI_ISL_2743666, EPI_ISL_2743667, EPI_ISL_2743668, EPI_ISL_2743670, EPI_ISL_2743671, EPI_ISL_2743672, EPI_ISL_2743676, EPI_ISL_2743677, EPI_ISL_2743678, EPI_ISL_2743679, EPI_ISL_2743680, EPI_ISL_2743681, EPI_ISL_2743682, EPI_ISL_2743683, EPI_ISL_2743684, EPI_ISL_2743686, EPI_ISL_2743687, EPI_ISL_2743689, EPI_ISL_2743690, EPI_ISL_2743691, EPI_ISL_2743692, EPI_ISL_2743693, EPI_ISL_2743694, EPI_ISL_2743695, EPI_ISL_2743696, EPI_ISL_2743697, EPI_ISL_2743698, EPI_ISL_2743699, EPI_ISL_2743700, EPI_ISL_2743701, EPI_ISL_2743702, EPI_ISL_2743703, EPI_ISL_2743704, EPI_ISL_2743705, EPI_ISL_2743706, EPI_ISL_2743707, EPI_ISL_2743708, EPI_ISL_2743709, EPI_ISL_2743710, EPI_ISL_2743714, EPI_ISL_2743715, EPI_ISL_2743716, EPI_ISL_2743718, EPI_ISL_2743720, EPI_ISL_2743724, EPI_ISL_2743727, EPI_ISL_2743728, EPI_ISL_2743729, EPI_ISL_2743730, EPI_ISL_2743731, EPI_ISL_2743732, EPI_ISL_2743734, EPI_ISL_2743735, EPI_ISL_2743736, EPI_ISL_2743739, EPI_ISL_2743740, EPI_ISL_2743741, EPI_ISL_2743742, EPI_ISL_2743743, EPI_ISL_2743745, EPI_ISL_2743747, EPI_ISL_2743748, EPI_ISL_2743749, EPI_ISL_2743750, EPI_ISL_2743751, EPI_ISL_2743752, EPI_ISL_2743754, EPI_ISL_2743755, EPI_ISL_2743758, EPI_ISL_2743760, EPI_ISL_2743761, EPI_ISL_2743762, EPI_ISL_2743763, EPI_ISL_2743764, EPI_ISL_2743765, EPI_ISL_2743766, EPI_ISL_2743767, EPI_ISL_2743768, EPI_ISL_2743769, EPI_ISL_2743770, EPI_ISL_2743771, EPI_ISL_2743772, EPI_ISL_2743773, EPI_ISL_2743774, EPI_ISL_2743775, EPI_ISL_2743776, EPI_ISL_2743777, EPI_ISL_2743778, EPI_ISL_2743779, EPI_ISL_2743780, EPI_ISL_2743781, EPI_ISL_2743782, EPI_ISL_2743783, EPI_ISL_2743785, EPI_ISL_2743786, EPI_ISL_2743787, EPI_ISL_2743788, EPI_ISL_2743789, EPI_ISL_2743790, EPI_ISL_2743792, EPI_ISL_2743793, EPI_ISL_2743794, EPI_ISL_2743795, EPI_ISL_2743796, EPI_ISL_2743797, EPI_ISL_2743798, EPI_ISL_2743799, EPI_ISL_2743800, EPI_ISL_2743801, EPI_ISL_2743802, EPI_ISL_2743803, EPI_ISL_2743804, EPI_ISL_2743805, EPI_ISL_2743807, EPI_ISL_2743810, EPI_ISL_2743811, EPI_ISL_2743812, EPI_ISL_2743814, EPI_ISL_2743815, EPI_ISL_2743816, EPI_ISL_2743817, EPI_ISL_2743818, EPI_ISL_2743819, EPI_ISL_2743822, EPI_ISL_2743823, EPI_ISL_2743824, EPI_ISL_2743825, EPI_ISL_2743826, EPI_ISL_2743829, EPI_ISL_2743830, EPI_ISL_2743831, EPI_ISL_2743832, EPI_ISL_2743833, EPI_ISL_2743834, EPI_ISL_2743835, EPI_ISL_2743836, EPI_ISL_2743837, EPI_ISL_2743838, EPI_ISL_2743840, EPI_ISL_2743841, EPI_ISL_2743842, EPI_ISL_2743843, EPI_ISL_2743844, EPI_ISL_2743845, EPI_ISL_2743847, EPI_ISL_2743850, EPI_ISL_2743851, EPI_ISL_2743852, EPI_ISL_2743855, EPI_ISL_2743856, EPI_ISL_2743857, EPI_ISL_2743858, EPI_ISL_2743859, EPI_ISL_2743860, EPI_ISL_2743861, EPI_ISL_2743862, EPI_ISL_2743863, EPI_ISL_2743864, EPI_ISL_2743865, EPI_ISL_2743866, EPI_ISL_2743867, EPI_ISL_2743868, EPI_ISL_2743869, EPI_ISL_2743870, EPI_ISL_2743871, EPI_ISL_2743872, EPI_ISL_2743875, EPI_ISL_2743876, EPI_ISL_2743878, EPI_ISL_2743879, EPI_ISL_2743880, EPI_ISL_2743882, EPI_ISL_2743883, EPI_ISL_2743884, EPI_ISL_2743885, EPI_ISL_2743886, EPI_ISL_2743889, EPI_ISL_2743890, EPI_ISL_2743892, EPI_ISL_2743893, EPI_ISL_2743894, EPI_ISL_2743895, EPI_ISL_2743896, EPI_ISL_2743897, EPI_ISL_2743898, EPI_ISL_2743899, EPI_ISL_2743900, EPI_ISL_2743901, EPI_ISL_2743904, EPI_ISL_2743905, EPI_ISL_2743906, EPI_ISL_2743907, EPI_ISL_2743908, EPI_ISL_2743909, EPI_ISL_2743910, EPI_ISL_2743911, EPI_ISL_2743912, EPI_ISL_2743913, EPI_ISL_2743914, EPI_ISL_2743915, EPI_ISL_2743916, EPI_ISL_2743917, EPI_ISL_2743918, EPI_ISL_2743919, EPI_ISL_2743920, EPI_ISL_2743921, EPI_ISL_2743922, EPI_ISL_2743923, EPI_ISL_2743925, EPI_ISL_2743927, EPI_ISL_2743928, EPI_ISL_2743929, EPI_ISL_2743930, EPI_ISL_2743931, EPI_ISL_2743932, EPI_ISL_2743933, EPI_ISL_2743934, EPI_ISL_2743935, EPI_ISL_2743936, EPI_ISL_2743938, EPI_ISL_2743939, EPI_ISL_2743941, EPI_ISL_2743942, EPI_ISL_2743943, EPI_ISL_2743944, EPI_ISL_2743945, EPI_ISL_2743946, EPI_ISL_2743947, EPI_ISL_2743948, EPI_ISL_2743951, EPI_ISL_2743952, EPI_ISL_2743953, EPI_ISL_2743954, EPI_ISL_2743956, EPI_ISL_2743957, EPI_ISL_2743958, EPI_ISL_2743959, EPI_ISL_2743960, EPI_ISL_2743961, EPI_ISL_2743962, EPI_ISL_2743963, EPI_ISL_2743964, EPI_ISL_2743965, EPI_ISL_2743966, EPI_ISL_2743967, EPI_ISL_2743968, EPI_ISL_2743969, EPI_ISL_2743971, EPI_ISL_2743973, EPI_ISL_2743974, EPI_ISL_2743975, EPI_ISL_2743976, EPI_ISL_2743977, EPI_ISL_2743978, EPI_ISL_2743979, EPI_ISL_2743980, EPI_ISL_2743981, EPI_ISL_2743982, EPI_ISL_2743983, EPI_ISL_2743984, EPI_ISL_2743985, EPI_ISL_2743986, EPI_ISL_2743987, EPI_ISL_2743989, EPI_ISL_2743990, EPI_ISL_2743991, EPI_ISL_2743993, EPI_ISL_2743994, EPI_ISL_2743995, EPI_ISL_2743996, EPI_ISL_2743997, EPI_ISL_2743999, EPI_ISL_2744001, EPI_ISL_2744003, EPI_ISL_2744004, EPI_ISL_2744005, EPI_ISL_2744006, EPI_ISL_2744007, EPI_ISL_2744008, EPI_ISL_2744009, EPI_ISL_2744010, EPI_ISL_2744011, EPI_ISL_2744012, EPI_ISL_2744013, EPI_ISL_2744014, EPI_ISL_2744015, EPI_ISL_2744016, EPI_ISL_2744017, EPI_ISL_2744018, EPI_ISL_2744019, EPI_ISL_2744020, EPI_ISL_2744023, EPI_ISL_2744024, EPI_ISL_2744025, EPI_ISL_2744026, EPI_ISL_2744027, EPI_ISL_2744028, EPI_ISL_2744029, EPI_ISL_2744030, EPI_ISL_2744032, EPI_ISL_2744034, EPI_ISL_2744035, EPI_ISL_2744036, EPI_ISL_2744037, EPI_ISL_2744038, EPI_ISL_2744039

| | | | |
|---|---|---|---|
| see above | Respiratory Virus Unit, Microbiology Services Colindale, Public Health England | COVID-19 Genomics UK (COG-UK) Consortium | PHE Covid Sequencing Team |
| EPI_ISL_2744040 | Department of Bacteria, Parasites and Fungi, Statens Serum Institut, Copenhagen, Denmark | Statens Serum Institut Bioinformatics and Microbial Genomics | Danish Covid-19 Genome Consortium |
| EPI_ISL_2744041, EPI_ISL_2744043 | Respiratory Virus Unit, Microbiology Services Colindale, Public Health England | COVID-19 Genomics UK (COG-UK) Consortium | PHE Covid Sequencing Team |
| EPI_ISL_2744044 | Department of Bacteria, Parasites and Fungi, Statens Serum Institut, Copenhagen, Denmark | Statens Serum Institut Bioinformatics and Microbial Genomics | Danish Covid-19 Genome Consortium |
| EPI_ISL_2744045 | Respiratory Virus Unit, Microbiology Services Colindale, Public Health England | COVID-19 Genomics UK (COG-UK) Consortium | PHE Covid Sequencing Team |
| EPI_ISL_2744046 | Department of Bacteria, Parasites and Fungi, Statens Serum Institut, Copenhagen, Denmark | Statens Serum Institut Bioinformatics and Microbial Genomics | Danish Covid-19 Genome Consortium |
| EPI_ISL_2744047 | Respiratory Virus Unit, Microbiology Services Colindale, Public Health England | COVID-19 Genomics UK (COG-UK) Consortium | PHE Covid Sequencing Team |
| EPI_ISL_2744048 | Department of Bacteria, Parasites and Fungi, Statens Serum Institut, Copenhagen, Denmark | Statens Serum Institut Bioinformatics and Microbial Genomics | Danish Covid-19 Genome Consortium |
| EPI_ISL_2744049 | Respiratory Virus Unit, Microbiology Services Colindale, Public Health England | COVID-19 Genomics UK (COG-UK) Consortium | PHE Covid Sequencing Team |
| EPI_ISL_2744050 | Department of Bacteria, Parasites and Fungi, Statens Serum Institut, Copenhagen, Denmark | Statens Serum Institut Bioinformatics and Microbial Genomics | Danish Covid-19 Genome Consortium |
| EPI_ISL_2744051 | Respiratory Virus Unit, Microbiology Services Colindale, Public Health England | COVID-19 Genomics UK (COG-UK) Consortium | PHE Covid Sequencing Team |
| EPI_ISL_2744052 | Department of Bacteria, Parasites and Fungi, Statens Serum Institut, Copenhagen, Denmark | Statens Serum Institut Bioinformatics and Microbial Genomics | Danish Covid-19 Genome Consortium |
| EPI_ISL_2744053 | Respiratory Virus Unit, Microbiology Services Colindale, Public Health England | COVID-19 Genomics UK (COG-UK) Consortium | PHE Covid Sequencing Team |
| EPI_ISL_2744054 | Department of Bacteria, Parasites and Fungi, Statens Serum Institut, Copenhagen, Denmark | Statens Serum Institut Bioinformatics and Microbial Genomics | Danish Covid-19 Genome Consortium |
| EPI_ISL_2744055 | Respiratory Virus Unit, Microbiology Services Colindale, Public Health England | COVID-19 Genomics UK (COG-UK) Consortium | PHE Covid Sequencing Team |
| EPI_ISL_2744056 | Department of Bacteria, Parasites and Fungi, Statens Serum Institut, Copenhagen, Denmark | Statens Serum Institut Bioinformatics and Microbial Genomics | Danish Covid-19 Genome Consortium |
| EPI_ISL_2744058, EPI_ISL_2744060, EPI_ISL_2744062, EPI_ISL_2744064, EPI_ISL_2744066, EPI_ISL_2744068 | Respiratory Virus Unit, Microbiology Services Colindale, Public Health England | COVID-19 Genomics UK (COG-UK) Consortium | PHE Covid Sequencing Team |
| EPI_ISL_2744069 | Department of Clinical Microbiology and Center for Genomic Medicine, Rigshospitalet, Copenhagen, Denmark | Statens Serum Institut Bioinformatics and Microbial Genomics | Danish Covid-19 Genome Consortium |
| EPI_ISL_2744070, EPI_ISL_2744072, EPI_ISL_2744074, EPI_ISL_2744076, EPI_ISL_2744078, EPI_ISL_2744080 | Respiratory Virus Unit, Microbiology Services Colindale, Public Health England | COVID-19 Genomics UK (COG-UK) Consortium | PHE Covid Sequencing Team |
| EPI_ISL_2744081 | Department of Bacteria, Parasites and Fungi, Statens Serum Institut, Copenhagen, Denmark | Statens Serum Institut Bioinformatics and Microbial Genomics | Danish Covid-19 Genome Consortium |
| EPI_ISL_2744082 | Respiratory Virus Unit, Microbiology Services Colindale, Public Health England | COVID-19 Genomics UK (COG-UK) Consortium | PHE Covid Sequencing Team |
| EPI_ISL_2744083 | Department of Bacteria, Parasites and Fungi, Statens Serum Institut, Copenhagen, Denmark | Statens Serum Institut Bioinformatics and Microbial Genomics | Danish Covid-19 Genome Consortium |
| EPI_ISL_2744084, EPI_ISL_2744086, EPI_ISL_2744088, EPI_ISL_2744090 | Respiratory Virus Unit, Microbiology Services Colindale, Public Health England | COVID-19 Genomics UK (COG-UK) Consortium | PHE Covid Sequencing Team |
| EPI_ISL_2744091 | Department of Bacteria, Parasites and Fungi, Statens Serum Institut, Copenhagen, Denmark | Statens Serum Institut Bioinformatics and Microbial Genomics | Danish Covid-19 Genome Consortium |

EPI_ISL_2744092, EPI_ISL_2744094, EPI_ISL_2744095, EPI_ISL_2744096, EPI_ISL_2744097, EPI_ISL_2744098, EPI_ISL_2744099, EPI_ISL_2744100, EPI_ISL_2744101, EPI_ISL_2744102, EPI_ISL_2744103, EPI_ISL_2744104, EPI_ISL_2744105, EPI_ISL_2744106, EPI_ISL_2744107, EPI_ISL_2744109, EPI_ISL_2744110, EPI_ISL_2744111, EPI_ISL_2744112, EPI_ISL_2744113, EPI_ISL_2744114, EPI_ISL_2744115, EPI_ISL_2744116, EPI_ISL_2744117, EPI_ISL_2744118, EPI_ISL_2744119, EPI_ISL_2744120, EPI_ISL_2744121, EPI_ISL_2744122, EPI_ISL_2744124, EPI_ISL_2744125, EPI_ISL_2744126, EPI_ISL_2744127, EPI_ISL_2744128, EPI_ISL_2744129, EPI_ISL_2744130, EPI_ISL_2744131, EPI_ISL_2744133, EPI_ISL_2744134, EPI_ISL_2744135, EPI_ISL_2744136, EPI_ISL_2744137, EPI_ISL_2744138, EPI_ISL_2744139, EPI_ISL_2744140, EPI_ISL_2744141, EPI_ISL_2744142, EPI_ISL_2744143, EPI_ISL_2744144, EPI_ISL_2744146, EPI_ISL_2744148, EPI_ISL_2744150, EPI_ISL_2744152, EPI_ISL_2744153, EPI_ISL_2744155, EPI_ISL_2744156, EPI_ISL_2744157, EPI_ISL_2744159, EPI_ISL_2744160, EPI_ISL_2744162, EPI_ISL_2744163, EPI_ISL_2744164, EPI_ISL_2744165, EPI_ISL_2744166, EPI_ISL_2744167, EPI_ISL_2744168, EPI_ISL_2744169, EPI_ISL_2744170, EPI_ISL_2744171, EPI_ISL_2744172, EPI_ISL_2744173, EPI_ISL_2744174, EPI_ISL_2744175, EPI_ISL_2744177, EPI_ISL_2744178, EPI_ISL_2744179, EPI_ISL_2744180, EPI_ISL_2744181, EPI_ISL_2744182, EPI_ISL_2744183, EPI_ISL_2744185, EPI_ISL_2744186, EPI_ISL_2744187, EPI_ISL_2744188, EPI_ISL_2744189, EPI_ISL_2744192, EPI_ISL_2744193, EPI_ISL_2744194, EPI_ISL_2744195, EPI_ISL_2744196, EPI_ISL_2744199, EPI_ISL_2744200, EPI_ISL_2744202, EPI_ISL_2744203, EPI_ISL_2744204, EPI_ISL_2744205, EPI_ISL_2744206, EPI_ISL_2744207, EPI_ISL_2744208, EPI_ISL_2744209, EPI_ISL_2744210, EPI_ISL_2744211, EPI_ISL_2744212, EPI_ISL_2744214, EPI_ISL_2744215, EPI_ISL_2744216, EPI_ISL_2744218, EPI_ISL_2744219, EPI_ISL_2744220, EPI_ISL_2744222, EPI_ISL_2744223, EPI_ISL_2744224, EPI_ISL_2744225, EPI_ISL_2744226, EPI_ISL_2744227, EPI_ISL_2744229, EPI_ISL_2744235, EPI_ISL_2744236, EPI_ISL_2744237, EPI_ISL_2744238, EPI_ISL_2744239, EPI_ISL_2744241, EPI_ISL_2744242, EPI_ISL_2744243, EPI_ISL_2744244, EPI_ISL_2744245, EPI_ISL_2744247, EPI_ISL_2744249, EPI_ISL_2744256, EPI_ISL_2744258, EPI_ISL_2744259, EPI_ISL_2744261,

EPI_ISL_2744265, EPI_ISL_2744268, EPI_ISL_2744269, EPI_ISL_2744270, EPI_ISL_2744271, EPI_ISL_2744272, EPI_ISL_2744273, EPI_ISL_2744274, EPI_ISL_2744275, EPI_ISL_2744276, EPI_ISL_2744277, EPI_ISL_2744279, EPI_ISL_2744292, EPI_ISL_2744293, EPI_ISL_2744295, EPI_ISL_2744296, EPI_ISL_2744297, EPI_ISL_2744298, EPI_ISL_2744299, EPI_ISL_2744300, EPI_ISL_2744306, EPI_ISL_2744307,
EPI_ISL_2744308, EPI_ISL_2744309, EPI_ISL_2744310, EPI_ISL_2744311, EPI_ISL_2744312, EPI_ISL_2744313, EPI_ISL_2744315, EPI_ISL_2744316, EPI_ISL_2744317, EPI_ISL_2744319, EPI_ISL_2744322, EPI_ISL_2744323, EPI_ISL_2744324, EPI_ISL_2744325, EPI_ISL_2744326, EPI_ISL_2744327, EPI_ISL_2744328, EPI_ISL_2744329, EPI_ISL_2744330, EPI_ISL_2744331, EPI_ISL_2744332, EPI_ISL_2744333,
EPI_ISL_2744334, EPI_ISL_2744335, EPI_ISL_2744336, EPI_ISL_2744337, EPI_ISL_2744338, EPI_ISL_2744339, EPI_ISL_2744340, EPI_ISL_2744341, EPI_ISL_2744342, EPI_ISL_2744344, EPI_ISL_2744345, EPI_ISL_2744347, EPI_ISL_2744348, EPI_ISL_2744349, EPI_ISL_2744350, EPI_ISL_2744351, EPI_ISL_2744352, EPI_ISL_2744353, EPI_ISL_2744354, EPI_ISL_2744355, EPI_ISL_2744356, EPI_ISL_2744357,
EPI_ISL_2744358, EPI_ISL_2744359, EPI_ISL_2744360, EPI_ISL_2744361, EPI_ISL_2744362, EPI_ISL_2744363, EPI_ISL_2744364, EPI_ISL_2744365, EPI_ISL_2744366, EPI_ISL_2744367, EPI_ISL_2744368, EPI_ISL_2744369, EPI_ISL_2744370, EPI_ISL_2744371, EPI_ISL_2744372, EPI_ISL_2744373, EPI_ISL_2744374, EPI_ISL_2744375, EPI_ISL_2744376, EPI_ISL_2744377, EPI_ISL_2744378, EPI_ISL_2744379,
EPI_ISL_2744380, EPI_ISL_2744381, EPI_ISL_2744382, EPI_ISL_2744383, EPI_ISL_2744384, EPI_ISL_2744385, EPI_ISL_2744386, EPI_ISL_2744387, EPI_ISL_2744388, EPI_ISL_2744389, EPI_ISL_2744390, EPI_ISL_2744391, EPI_ISL_2744392, EPI_ISL_2744394, EPI_ISL_2744395, EPI_ISL_2744396, EPI_ISL_2744397, EPI_ISL_2744398, EPI_ISL_2744399, EPI_ISL_2744400, EPI_ISL_2744401, EPI_ISL_2744402,
EPI_ISL_2744403, EPI_ISL_2744404, EPI_ISL_2744405, EPI_ISL_2744406, EPI_ISL_2744408, EPI_ISL_2744409, EPI_ISL_2744410, EPI_ISL_2744411, EPI_ISL_2744412, EPI_ISL_2744413, EPI_ISL_2744414, EPI_ISL_2744415, EPI_ISL_2744416, EPI_ISL_2744417, EPI_ISL_2744418, EPI_ISL_2744419, EPI_ISL_2744420, EPI_ISL_2744421, EPI_ISL_2744422, EPI_ISL_2744423, EPI_ISL_2744424, EPI_ISL_2744426,
EPI_ISL_2744427, EPI_ISL_2744428, EPI_ISL_2744432, EPI_ISL_2744433, EPI_ISL_2744434, EPI_ISL_2744435, EPI_ISL_2744436, EPI_ISL_2744437, EPI_ISL_2744438, EPI_ISL_2744439, EPI_ISL_2744440, EPI_ISL_2744443, EPI_ISL_2744444, EPI_ISL_2744445, EPI_ISL_2744446, EPI_ISL_2744447, EPI_ISL_2744448, EPI_ISL_2744449, EPI_ISL_2744450, EPI_ISL_2744451, EPI_ISL_2744452, EPI_ISL_2744453,
EPI_ISL_2744454, EPI_ISL_2744455, EPI_ISL_2744456, EPI_ISL_2744457, EPI_ISL_2744458, EPI_ISL_2744459, EPI_ISL_2744460, EPI_ISL_2744461, EPI_ISL_2744462, EPI_ISL_2744463, EPI_ISL_2744464, EPI_ISL_2744465, EPI_ISL_2744466, EPI_ISL_2744467, EPI_ISL_2744468, EPI_ISL_2744469, EPI_ISL_2744470, EPI_ISL_2744471, EPI_ISL_2744472, EPI_ISL_2744473, EPI_ISL_2744474, EPI_ISL_2744475,
EPI_ISL_2744476, EPI_ISL_2744477, EPI_ISL_2744478, EPI_ISL_2744479, EPI_ISL_2744480, EPI_ISL_2744481, EPI_ISL_2744482, EPI_ISL_2744483, EPI_ISL_2744484, EPI_ISL_2744485, EPI_ISL_2744486, EPI_ISL_2744487, EPI_ISL_2744488, EPI_ISL_2744489, EPI_ISL_2744490, EPI_ISL_2744491, EPI_ISL_2744492, EPI_ISL_2744493, EPI_ISL_2744494, EPI_ISL_2744495, EPI_ISL_2744496, EPI_ISL_2744497,
EPI_ISL_2744499, EPI_ISL_2744500, EPI_ISL_2744501, EPI_ISL_2744502, EPI_ISL_2744504, EPI_ISL_2744506, EPI_ISL_2744509, EPI_ISL_2744512, EPI_ISL_2744514, EPI_ISL_2744518, EPI_ISL_2744520, EPI_ISL_2744521, EPI_ISL_2744522, EPI_ISL_2744523, EPI_ISL_2744526, EPI_ISL_2744527, EPI_ISL_2744528, EPI_ISL_2744529, EPI_ISL_2744530, EPI_ISL_2744531, EPI_ISL_2744532, EPI_ISL_2744533,
EPI_ISL_2744534, EPI_ISL_2744535, EPI_ISL_2744536, EPI_ISL_2744537, EPI_ISL_2744538, EPI_ISL_2744539, EPI_ISL_2744540, EPI_ISL_2744541, EPI_ISL_2744542, EPI_ISL_2744544, EPI_ISL_2744545, EPI_ISL_2744546, EPI_ISL_2744551, EPI_ISL_2744553, EPI_ISL_2744554, EPI_ISL_2744555, EPI_ISL_2744556, EPI_ISL_2744557, EPI_ISL_2744558, EPI_ISL_2744559, EPI_ISL_2744560, EPI_ISL_2744561,
EPI_ISL_2744562, EPI_ISL_2744563, EPI_ISL_2744565, EPI_ISL_2744566, EPI_ISL_2744567, EPI_ISL_2744568, EPI_ISL_2744569, EPI_ISL_2744570, EPI_ISL_2744571, EPI_ISL_2744572, EPI_ISL_2744573, EPI_ISL_2744574, EPI_ISL_2744575, EPI_ISL_2744576, EPI_ISL_2744577, EPI_ISL_2744581, EPI_ISL_2744582, EPI_ISL_2744584, EPI_ISL_2744586, EPI_ISL_2744588, EPI_ISL_2744589, EPI_ISL_2744590,
EPI_ISL_2744592, EPI_ISL_2744593, EPI_ISL_2744594, EPI_ISL_2744595, EPI_ISL_2744596, EPI_ISL_2744597, EPI_ISL_2744598, EPI_ISL_2744599, EPI_ISL_2744600, EPI_ISL_2744601, EPI_ISL_2744602, EPI_ISL_2744603, EPI_ISL_2744604, EPI_ISL_2744605, EPI_ISL_2744606, EPI_ISL_2744607, EPI_ISL_2744608, EPI_ISL_2744609, EPI_ISL_2744610, EPI_ISL_2744612, EPI_ISL_2744613, EPI_ISL_2744614,
EPI_ISL_2744615, EPI_ISL_2744616, EPI_ISL_2744618, EPI_ISL_2744619, EPI_ISL_2744620, EPI_ISL_2744621, EPI_ISL_2744622, EPI_ISL_2744623, EPI_ISL_2744624, EPI_ISL_2744626, EPI_ISL_2744627, EPI_ISL_2744628, EPI_ISL_2744629, EPI_ISL_2744630, EPI_ISL_2744631, EPI_ISL_2744632, EPI_ISL_2744633, EPI_ISL_2744634, EPI_ISL_2744635, EPI_ISL_2744636, EPI_ISL_2744637, EPI_ISL_2744639,
EPI_ISL_2744640, EPI_ISL_2744641, EPI_ISL_2744642, EPI_ISL_2744643, EPI_ISL_2744644, EPI_ISL_2744645, EPI_ISL_2744646, EPI_ISL_2744647, EPI_ISL_2744648, EPI_ISL_2744649, EPI_ISL_2744650, EPI_ISL_2744651, EPI_ISL_2744652, EPI_ISL_2744653, EPI_ISL_2744654, EPI_ISL_2744655, EPI_ISL_2744656, EPI_ISL_2744657, EPI_ISL_2744658, EPI_ISL_2744659, EPI_ISL_2744660, EPI_ISL_2744661,
EPI_ISL_2744662, EPI_ISL_2744663, EPI_ISL_2744664, EPI_ISL_2744665, EPI_ISL_2744666, EPI_ISL_2744667, EPI_ISL_2744668, EPI_ISL_2744669, EPI_ISL_2744671, EPI_ISL_2744672, EPI_ISL_2744673, EPI_ISL_2744674, EPI_ISL_2744675, EPI_ISL_2744676, EPI_ISL_2744677, EPI_ISL_2744678, EPI_ISL_2744679, EPI_ISL_2744680, EPI_ISL_2744681, EPI_ISL_2744682, EPI_ISL_2744683, EPI_ISL_2744684,
EPI_ISL_2744685, EPI_ISL_2744686, EPI_ISL_2744687, EPI_ISL_2744688, EPI_ISL_2744689, EPI_ISL_2744690, EPI_ISL_2744692, EPI_ISL_2744693, EPI_ISL_2744694, EPI_ISL_2744695, EPI_ISL_2744696, EPI_ISL_2744697, EPI_ISL_2744698, EPI_ISL_2744699, EPI_ISL_2744700, EPI_ISL_2744701, EPI_ISL_2744702, EPI_ISL_2744703, EPI_ISL_2744704, EPI_ISL_2744706, EPI_ISL_2744707, EPI_ISL_2744708,
EPI_ISL_2744709, EPI_ISL_2744710, EPI_ISL_2744711, EPI_ISL_2744712, EPI_ISL_2744714, EPI_ISL_2744715, EPI_ISL_2744719, EPI_ISL_2744720, EPI_ISL_2744721, EPI_ISL_2744722, EPI_ISL_2744724, EPI_ISL_2744725, EPI_ISL_2744727, EPI_ISL_2744728, EPI_ISL_2744729, EPI_ISL_2744730, EPI_ISL_2744731, EPI_ISL_2744732, EPI_ISL_2744733, EPI_ISL_2744735, EPI_ISL_2744736, EPI_ISL_2744737,
EPI_ISL_2744738, EPI_ISL_2744739, EPI_ISL_2744740, EPI_ISL_2744741, EPI_ISL_2744742, EPI_ISL_2744743, EPI_ISL_2744744, EPI_ISL_2744745, EPI_ISL_2744746, EPI_ISL_2744747, EPI_ISL_2744748, EPI_ISL_2744749, EPI_ISL_2744750, EPI_ISL_2744753, EPI_ISL_2744754, EPI_ISL_2744755, EPI_ISL_2744756, EPI_ISL_2744758, EPI_ISL_2744759, EPI_ISL_2744761, EPI_ISL_2744762, EPI_ISL_2744763,
EPI_ISL_2744764, EPI_ISL_2744765, EPI_ISL_2744766, EPI_ISL_2744767, EPI_ISL_2744768, EPI_ISL_2744769, EPI_ISL_2744770, EPI_ISL_2744771, EPI_ISL_2744772, EPI_ISL_2744773, EPI_ISL_2744774, EPI_ISL_2744775, EPI_ISL_2744776, EPI_ISL_2744777, EPI_ISL_2744778, EPI_ISL_2744780, EPI_ISL_2744781, EPI_ISL_2744782, EPI_ISL_2744783, EPI_ISL_2744784, EPI_ISL_2744785, EPI_ISL_2744786,
EPI_ISL_2744787, EPI_ISL_2744788, EPI_ISL_2744789, EPI_ISL_2744790, EPI_ISL_2744791, EPI_ISL_2744792, EPI_ISL_2744793, EPI_ISL_2744794, EPI_ISL_2744795, EPI_ISL_2744796, EPI_ISL_2744798, EPI_ISL_2744799, EPI_ISL_2744800, EPI_ISL_2744801, EPI_ISL_2744802, EPI_ISL_2744803, EPI_ISL_2744804, EPI_ISL_2744805, EPI_ISL_2744807, EPI_ISL_2744808, EPI_ISL_2744809, EPI_ISL_2744810,
EPI_ISL_2744811, EPI_ISL_2744812, EPI_ISL_2744813, EPI_ISL_2744814, EPI_ISL_2744815, EPI_ISL_2744816, EPI_ISL_2744819, EPI_ISL_2744820, EPI_ISL_2744822, EPI_ISL_2744823, EPI_ISL_2744824, EPI_ISL_2744825, EPI_ISL_2744826, EPI_ISL_2744827, EPI_ISL_2744828, EPI_ISL_2744829, EPI_ISL_2744830, EPI_ISL_2744831, EPI_ISL_2744832, EPI_ISL_2744833, EPI_ISL_2744834, EPI_ISL_2744835,
EPI_ISL_2744836, EPI_ISL_2744837, EPI_ISL_2744838, EPI_ISL_2744839, EPI_ISL_2744840, EPI_ISL_2744841, EPI_ISL_2744842, EPI_ISL_2744843, EPI_ISL_2744845, EPI_ISL_2744846, EPI_ISL_2744847, EPI_ISL_2744848, EPI_ISL_2744849, EPI_ISL_2744851, EPI_ISL_2744852, EPI_ISL_2744853, EPI_ISL_2744854, EPI_ISL_2744855, EPI_ISL_2744856, EPI_ISL_2744857, EPI_ISL_2744858, EPI_ISL_2744859,
EPI_ISL_2744860, EPI_ISL_2744861, EPI_ISL_2744862, EPI_ISL_2744863, EPI_ISL_2744864, EPI_ISL_2744865, EPI_ISL_2744866, EPI_ISL_2744867, EPI_ISL_2744868, EPI_ISL_2744869, EPI_ISL_2744870, EPI_ISL_2744871, EPI_ISL_2744872, EPI_ISL_2744873, EPI_ISL_2744874, EPI_ISL_2744875, EPI_ISL_2744876, EPI_ISL_2744877, EPI_ISL_2744878, EPI_ISL_2744879, EPI_ISL_2744880, EPI_ISL_2744881,
EPI_ISL_2744882, EPI_ISL_2744883, EPI_ISL_2744884, EPI_ISL_2744885, EPI_ISL_2744886, EPI_ISL_2744887, EPI_ISL_2744888, EPI_ISL_2744889, EPI_ISL_2744890, EPI_ISL_2744891, EPI_ISL_2744892, EPI_ISL_2744893, EPI_ISL_2744894, EPI_ISL_2744895, EPI_ISL_2744896, EPI_ISL_2744897, EPI_ISL_2744898, EPI_ISL_2744899, EPI_ISL_2744900, EPI_ISL_2744901, EPI_ISL_2744902, EPI_ISL_2744903,
EPI_ISL_2744904, EPI_ISL_2744905, EPI_ISL_2744906, EPI_ISL_2744907, EPI_ISL_2744908, EPI_ISL_2744909, EPI_ISL_2744910, EPI_ISL_2744911, EPI_ISL_2744912, EPI_ISL_2744913, EPI_ISL_2744914, EPI_ISL_2744916, EPI_ISL_2744918, EPI_ISL_2744921, EPI_ISL_2744922, EPI_ISL_2744923, EPI_ISL_2744924, EPI_ISL_2744925, EPI_ISL_2744926, EPI_ISL_2744927, EPI_ISL_2744928, EPI_ISL_2744929,
EPI_ISL_2744930, EPI_ISL_2744931, EPI_ISL_2744932, EPI_ISL_2744933, EPI_ISL_2744934, EPI_ISL_2744935, EPI_ISL_2744936, EPI_ISL_2744937, EPI_ISL_2744938, EPI_ISL_2744940, EPI_ISL_2744941, EPI_ISL_2744942, EPI_ISL_2744943, EPI_ISL_2744944, EPI_ISL_2744947, EPI_ISL_2744948, EPI_ISL_2744949, EPI_ISL_2744950, EPI_ISL_2744951, EPI_ISL_2744952, EPI_ISL_2744953, EPI_ISL_2744955,
EPI_ISL_2744956, EPI_ISL_2744957, EPI_ISL_2744958, EPI_ISL_2744959, EPI_ISL_2744960, EPI_ISL_2744961, EPI_ISL_2744962, EPI_ISL_2744963, EPI_ISL_2744964, EPI_ISL_2744965, EPI_ISL_2744966, EPI_ISL_2744967, EPI_ISL_2744968, EPI_ISL_2744969, EPI_ISL_2744970, EPI_ISL_2744971, EPI_ISL_2744972, EPI_ISL_2744973, EPI_ISL_2744974, EPI_ISL_2744975, EPI_ISL_2744976, EPI_ISL_2744977,
EPI_ISL_2744978, EPI_ISL_2744979, EPI_ISL_2744980, EPI_ISL_2744982, EPI_ISL_2744984, EPI_ISL_2744985, EPI_ISL_2744986, EPI_ISL_2744987, EPI_ISL_2744988, EPI_ISL_2744989, EPI_ISL_2744990, EPI_ISL_2744991, EPI_ISL_2744992, EPI_ISL_2744993, EPI_ISL_2744994, EPI_ISL_2744995, EPI_ISL_2744996, EPI_ISL_2744997, EPI_ISL_2744998, EPI_ISL_2744999, EPI_ISL_2745000, EPI_ISL_2745001,
EPI_ISL_2745002, EPI_ISL_2745003, EPI_ISL_2745004, EPI_ISL_2745005, EPI_ISL_2745006, EPI_ISL_2745007, EPI_ISL_2745008, EPI_ISL_2745009, EPI_ISL_2745010, EPI_ISL_2745011, EPI_ISL_2745012, EPI_ISL_2745013, EPI_ISL_2745014, EPI_ISL_2745015, EPI_ISL_2745016, EPI_ISL_2745017, EPI_ISL_2745018, EPI_ISL_2745019, EPI_ISL_2745020, EPI_ISL_2745021, EPI_ISL_2745022, EPI_ISL_2745023,
EPI_ISL_2745024, EPI_ISL_2745026, EPI_ISL_2745027, EPI_ISL_2745028, EPI_ISL_2745029, EPI_ISL_2745030, EPI_ISL_2745031, EPI_ISL_2745032, EPI_ISL_2745033, EPI_ISL_2745034, EPI_ISL_2745035, EPI_ISL_2745036, EPI_ISL_2745037, EPI_ISL_2745039, EPI_ISL_2745040, EPI_ISL_2745041, EPI_ISL_2745042, EPI_ISL_2745043, EPI_ISL_2745044, EPI_ISL_2745045, EPI_ISL_2745046, EPI_ISL_2745047,
EPI_ISL_2745048, EPI_ISL_2745049, EPI_ISL_2745050, EPI_ISL_2745051, EPI_ISL_2745052, EPI_ISL_2745053, EPI_ISL_2745054, EPI_ISL_2745055, EPI_ISL_2745056, EPI_ISL_2745057, EPI_ISL_2745058, EPI_ISL_2745059, EPI_ISL_2745060, EPI_ISL_2745061, EPI_ISL_2745062, EPI_ISL_2745063, EPI_ISL_2745064, EPI_ISL_2745065, EPI_ISL_2745066, EPI_ISL_2745067, EPI_ISL_2745068, EPI_ISL_2745069,
EPI_ISL_2745070, EPI_ISL_2745071, EPI_ISL_2745072, EPI_ISL_2745073, EPI_ISL_2745074, EPI_ISL_2745075, EPI_ISL_2745076, EPI_ISL_2745077, EPI_ISL_2745078, EPI_ISL_2745079, EPI_ISL_2745080, EPI_ISL_2745081, EPI_ISL_2745082, EPI_ISL_2745083, EPI_ISL_2745084, EPI_ISL_2745085, EPI_ISL_2745086, EPI_ISL_2745087, EPI_ISL_2745088, EPI_ISL_2745090, EPI_ISL_2745091, EPI_ISL_2745092,
EPI_ISL_2745094, EPI_ISL_2745096, EPI_ISL_2745097, EPI_ISL_2745098, EPI_ISL_2745099, EPI_ISL_2745100, EPI_ISL_2745101, EPI_ISL_2745102, EPI_ISL_2745103, EPI_ISL_2745104, EPI_ISL_2745105, EPI_ISL_2745106, EPI_ISL_2745107, EPI_ISL_2745108, EPI_ISL_2745109, EPI_ISL_2745110, EPI_ISL_2745111, EPI_ISL_2745112, EPI_ISL_2745113, EPI_ISL_2745114, EPI_ISL_2745115, EPI_ISL_2745116,
EPI_ISL_2745117, EPI_ISL_2745118, EPI_ISL_2745119, EPI_ISL_2745120, EPI_ISL_2745121, EPI_ISL_2745122, EPI_ISL_2745123, EPI_ISL_2745124, EPI_ISL_2745125, EPI_ISL_2745126, EPI_ISL_2745127, EPI_ISL_2745128, EPI_ISL_2745129, EPI_ISL_2745131, EPI_ISL_2745132, EPI_ISL_2745133, EPI_ISL_2745134, EPI_ISL_2745135, EPI_ISL_2745136, EPI_ISL_2745137, EPI_ISL_2745138, EPI_ISL_2745139,
EPI_ISL_2745140, EPI_ISL_2745141, EPI_ISL_2745142, EPI_ISL_2745143, EPI_ISL_2745144, EPI_ISL_2745145, EPI_ISL_2745146, EPI_ISL_2745147, EPI_ISL_2745148, EPI_ISL_2745149, EPI_ISL_2745150, EPI_ISL_2745151, EPI_ISL_2745152, EPI_ISL_2745153, EPI_ISL_2745154, EPI_ISL_2745155, EPI_ISL_2745156, EPI_ISL_2745157, EPI_ISL_2745159, EPI_ISL_2745160, EPI_ISL_2745161, EPI_ISL_2745162,
EPI_ISL_2745163, EPI_ISL_2745164, EPI_ISL_2745165, EPI_ISL_2745166, EPI_ISL_2745167, EPI_ISL_2745168, EPI_ISL_2745169, EPI_ISL_2745170, EPI_ISL_2745171, EPI_ISL_2745173, EPI_ISL_2745174, EPI_ISL_2745176, EPI_ISL_2745177, EPI_ISL_2745178, EPI_ISL_2745179, EPI_ISL_2745180, EPI_ISL_2745182, EPI_ISL_2745183, EPI_ISL_2745184, EPI_ISL_2745185, EPI_ISL_2745186, EPI_ISL_2745188,
EPI_ISL_2745190, EPI_ISL_2745191, EPI_ISL_2745192, EPI_ISL_2745193, EPI_ISL_2745194, EPI_ISL_2745196, EPI_ISL_2745197, EPI_ISL_2745198, EPI_ISL_2745199, EPI_ISL_2745200, EPI_ISL_2745201, EPI_ISL_2745202, EPI_ISL_2745203, EPI_ISL_2745204, EPI_ISL_2745205, EPI_ISL_2745206, EPI_ISL_2745207, EPI_ISL_2745208, EPI_ISL_2745209, EPI_ISL_2745210, EPI_ISL_2745211, EPI_ISL_2745212,
EPI_ISL_2745213, EPI_ISL_2745214, EPI_ISL_2745215, EPI_ISL_2745217, EPI_ISL_2745219, EPI_ISL_2745222, EPI_ISL_2745224, EPI_ISL_2745227, EPI_ISL_2745228, EPI_ISL_2745230, EPI_ISL_2745231, EPI_ISL_2745232, EPI_ISL_2745233, EPI_ISL_2745235, EPI_ISL_2745236, EPI_ISL_2745237, EPI_ISL_2745238, EPI_ISL_2745239, EPI_ISL_2745240, EPI_ISL_2745241, EPI_ISL_2745242, EPI_ISL_2745243,
EPI_ISL_2745245, EPI_ISL_2745246, EPI_ISL_2745247, EPI_ISL_2745248, EPI_ISL_2745249, EPI_ISL_2745250, EPI_ISL_2745251, EPI_ISL_2745252, EPI_ISL_2745253, EPI_ISL_2745254, EPI_ISL_2745255, EPI_ISL_2745256, EPI_ISL_2745257, EPI_ISL_2745258, EPI_ISL_2745260, EPI_ISL_2745261, EPI_ISL_2745262, EPI_ISL_2745263, EPI_ISL_2745264, EPI_ISL_2745265, EPI_ISL_2745266, EPI_ISL_2745267,
EPI_ISL_2745268, EPI_ISL_2745269, EPI_ISL_2745270, EPI_ISL_2745271, EPI_ISL_2745272, EPI_ISL_2745273, EPI_ISL_2745274, EPI_ISL_2745275, EPI_ISL_2745276, EPI_ISL_2745277, EPI_ISL_2745278, EPI_ISL_2745279, EPI_ISL_2745280, EPI_ISL_2745281, EPI_ISL_2745282, EPI_ISL_2745284, EPI_ISL_2745286, EPI_ISL_2745287, EPI_ISL_2745288, EPI_ISL_2745289, EPI_ISL_2745290, EPI_ISL_2745291,
EPI_ISL_2745292, EPI_ISL_2745293, EPI_ISL_2745294, EPI_ISL_2745295, EPI_ISL_2745296, EPI_ISL_2745297, EPI_ISL_2745298, EPI_ISL_2745299, EPI_ISL_2745300, EPI_ISL_2745301, EPI_ISL_2745302, EPI_ISL_2745303, EPI_ISL_2745304, EPI_ISL_2745305, EPI_ISL_2745306, EPI_ISL_2745307, EPI_ISL_2745308, EPI_ISL_2745309, EPI_ISL_2745311, EPI_ISL_2745313, EPI_ISL_2745314, EPI_ISL_2745315,
EPI_ISL_2745316, EPI_ISL_2745317, EPI_ISL_2745318, EPI_ISL_2745319, EPI_ISL_2745320, EPI_ISL_2745322, EPI_ISL_2745323, EPI_ISL_2745324, EPI_ISL_2745325, EPI_ISL_2745326, EPI_ISL_2745327, EPI_ISL_2745328, EPI_ISL_2745329, EPI_ISL_2745330, EPI_ISL_2745331, EPI_ISL_2745332, EPI_ISL_2745333, EPI_ISL_2745334, EPI_ISL_2745335, EPI_ISL_2745336, EPI_ISL_2745337,
EPI_ISL_2745338, EPI_ISL_2745339, EPI_ISL_2745340, EPI_ISL_2745341, EPI_ISL_2745342, EPI_ISL_2745344, EPI_ISL_2745346, EPI_ISL_2745347, EPI_ISL_2745348, EPI_ISL_2745349, EPI_ISL_2745350, EPI_ISL_2745351, EPI_ISL_2745353, EPI_ISL_2745354, EPI_ISL_2745355, EPI_ISL_2745356, EPI_ISL_2745357, EPI_ISL_2745358, EPI_ISL_2745361, EPI_ISL_2745362, EPI_ISL_2745363, EPI_ISL_2745364,
EPI_ISL_2745365, EPI_ISL_2745366, EPI_ISL_2745367, EPI_ISL_2745368, EPI_ISL_2745369, EPI_ISL_2745370, EPI_ISL_2745371, EPI_ISL_2745372, EPI_ISL_2745373, EPI_ISL_2745374, EPI_ISL_2745375, EPI_ISL_2745376, EPI_ISL_2745377, EPI_ISL_2745378, EPI_ISL_2745379, EPI_ISL_2745381, EPI_ISL_2745382, EPI_ISL_2745383, EPI_ISL_2745384, EPI_ISL_2745385, EPI_ISL_2745386, EPI_ISL_2745387,
EPI_ISL_2745388, EPI_ISL_2745389, EPI_ISL_2745390, EPI_ISL_2745391, EPI_ISL_2745392, EPI_ISL_2745393, EPI_ISL_2745394, EPI_ISL_2745395, EPI_ISL_2745396, EPI_ISL_2745397, EPI_ISL_2745398, EPI_ISL_2745399, EPI_ISL_2745400, EPI_ISL_2745401, EPI_ISL_2745402, EPI_ISL_2745403, EPI_ISL_2745404, EPI_ISL_2745405, EPI_ISL_2745406, EPI_ISL_2745407, EPI_ISL_2745408, EPI_ISL_2745409,
EPI_ISL_2745410, EPI_ISL_2745411, EPI_ISL_2745412, EPI_ISL_2745413, EPI_ISL_2745414, EPI_ISL_2745416, EPI_ISL_2745419, EPI_ISL_2745421, EPI_ISL_2745422, EPI_ISL_2745423, EPI_ISL_2745424, EPI_ISL_2745426, EPI_ISL_2745427, EPI_ISL_2745429, EPI_ISL_2745430, EPI_ISL_2745432, EPI_ISL_2745434, EPI_ISL_2745435, EPI_ISL_2745436, EPI_ISL_2745437, EPI_ISL_2745438, EPI_ISL_2745439,
EPI_ISL_2745440, EPI_ISL_2745441, EPI_ISL_2745442, EPI_ISL_2745443, EPI_ISL_2745444, EPI_ISL_2745445, EPI_ISL_2745446, EPI_ISL_2745447, EPI_ISL_2745448, EPI_ISL_2745449, EPI_ISL_2745451, EPI_ISL_2745452, EPI_ISL_2745453, EPI_ISL_2745454, EPI_ISL_2745455, EPI_ISL_2745456, EPI_ISL_2745457, EPI_ISL_2745458, EPI_ISL_2745459, EPI_ISL_2745460, EPI_ISL_2745461, EPI_ISL_2745462,
EPI_ISL_2745463, EPI_ISL_2745464, EPI_ISL_2745465, EPI_ISL_2745466, EPI_ISL_2745467, EPI_ISL_2745468, EPI_ISL_2745469, EPI_ISL_2745470, EPI_ISL_2745471, EPI_ISL_2745472, EPI_ISL_2745473, EPI_ISL_2745474, EPI_ISL_2745475, EPI_ISL_2745476, EPI_ISL_2745477, EPI_ISL_2745478, EPI_ISL_2745480, EPI_ISL_2745481, EPI_ISL_2745482, EPI_ISL_2745483, EPI_ISL_2745484, EPI_ISL_2745485,
EPI_ISL_2745486, EPI_ISL_2745487, EPI_ISL_2745488, EPI_ISL_2745489, EPI_ISL_2745490, EPI_ISL_2745491, EPI_ISL_2745492, EPI_ISL_2745493, EPI_ISL_2745494, EPI_ISL_2745495, EPI_ISL_2745496, EPI_ISL_2745497, EPI_ISL_2745498, EPI_ISL_2745499, EPI_ISL_2745500, EPI_ISL_2745501, EPI_ISL_2745502, EPI_ISL_2745503, EPI_ISL_2745504, EPI_ISL_2745505, EPI_ISL_2745506, EPI_ISL_2745507,
EPI_ISL_2745508, EPI_ISL_2745509, EPI_ISL_2745510, EPI_ISL_2745511, EPI_ISL_2745512, EPI_ISL_2745514, EPI_ISL_2745515, EPI_ISL_2745516, EPI_ISL_2745517, EPI_ISL_2745518, EPI_ISL_2745519, EPI_ISL_2745520, EPI_ISL_2745521, EPI_ISL_2745522, EPI_ISL_2745523, EPI_ISL_2745525, EPI_ISL_2745526, EPI_ISL_2745527, EPI_ISL_2745528, EPI_ISL_2745530, EPI_ISL_2745531,
EPI_ISL_2745532, EPI_ISL_2745533, EPI_ISL_2745534, EPI_ISL_2745535, EPI_ISL_2745536, EPI_ISL_2745537, EPI_ISL_2745538, EPI_ISL_2745539, EPI_ISL_2745540, EPI_ISL_2745541, EPI_ISL_2745542, EPI_ISL_2745543, EPI_ISL_2745544, EPI_ISL_2745545, EPI_ISL_2745546, EPI_ISL_2745547, EPI_ISL_2745548, EPI_ISL_2745549, EPI_ISL_2745550, EPI_ISL_2745551, EPI_ISL_2745552, EPI_ISL_2745553,
EPI_ISL_2745554, EPI_ISL_2745555, EPI_ISL_2745556, EPI_ISL_2745557, EPI_ISL_2745558, EPI_ISL_2745559, EPI_ISL_2745560, EPI_ISL_2745561, EPI_ISL_2745562, EPI_ISL_2745563, EPI_ISL_2745564, EPI_ISL_2745565, EPI_ISL_2745566, EPI_ISL_2745567, EPI_ISL_2745568, EPI_ISL_2745569, EPI_ISL_2745570, EPI_ISL_2745571, EPI_ISL_2745572, EPI_ISL_2745573, EPI_ISL_2745574, EPI_ISL_2745575,
EPI_ISL_2745576, EPI_ISL_2745577, EPI_ISL_2745578, EPI_ISL_2745579, EPI_ISL_2745580, EPI_ISL_2745581, EPI_ISL_2745582, EPI_ISL_2745583, EPI_ISL_2745584, EPI_ISL_2745585, EPI_ISL_2745586, EPI_ISL_2745587, EPI_ISL_2745588, EPI_ISL_2745589, EPI_ISL_2745590, EPI_ISL_2745591, EPI_ISL_2745592, EPI_ISL_2745593, EPI_ISL_2745594, EPI_ISL_2745595, EPI_ISL_2745596, EPI_ISL_2745597,
EPI_ISL_2745598, EPI_ISL_2745599, EPI_ISL_2745600, EPI_ISL_2745601, EPI_ISL_2745602, EPI_ISL_2745603, EPI_ISL_2745604, EPI_ISL_2745605, EPI_ISL_2745606, EPI_ISL_2745607, EPI_ISL_2745608, EPI_ISL_2745609, EPI_ISL_2745610, EPI_ISL_2745611, EPI_ISL_2745612, EPI_ISL_2745613, EPI_ISL_2745614, EPI_ISL_2745615, EPI_ISL_2745616, EPI_ISL_2745617, EPI_ISL_2745618, EPI_ISL_2745619,
EPI_ISL_2745620, EPI_ISL_2745621, EPI_ISL_2745622, EPI_ISL_2745623, EPI_ISL_2745624, EPI_ISL_2745625, EPI_ISL_2745626, EPI_ISL_2745627, EPI_ISL_2745628, EPI_ISL_2745629, EPI_ISL_2745630, EPI_ISL_2745631, EPI_ISL_2745632, EPI_ISL_2745633, EPI_ISL_2745634, EPI_ISL_2745635, EPI_ISL_2745636, EPI_ISL_2745637, EPI_ISL_2745638, EPI_ISL_2745639, EPI_ISL_2745640, EPI_ISL_2745642,
EPI_ISL_2745643, EPI_ISL_2745644, EPI_ISL_2745646, EPI_ISL_2745647, EPI_ISL_2745648, EPI_ISL_2745649, EPI_ISL_2745650, EPI_ISL_2745651, EPI_ISL_2745652, EPI_ISL_2745653, EPI_ISL_2745654, EPI_ISL_2745655, EPI_ISL_2745656, EPI_ISL_2745657, EPI_ISL_2745658, EPI_ISL_2745659, EPI_ISL_2745660, EPI_ISL_2745661, EPI_ISL_2745662, EPI_ISL_2745663, EPI_ISL_2745664, EPI_ISL_2745665,
EPI_ISL_2745666, EPI_ISL_2745667, EPI_ISL_2745668, EPI_ISL_2745669, EPI_ISL_2745670, EPI_ISL_2745672, EPI_ISL_2745673, EPI_ISL_2745674, EPI_ISL_2745675, EPI_ISL_2745676, EPI_ISL_2745677, EPI_ISL_2745678, EPI_ISL_2745679, EPI_ISL_2745680, EPI_ISL_2745681, EPI_ISL_2745682, EPI_ISL_2745683, EPI_ISL_2745685, EPI_ISL_2745686, EPI_ISL_2745687, EPI_ISL_2745688,
EPI_ISL_2745689, EPI_ISL_2745690, EPI_ISL_2745691, EPI_ISL_2745692, EPI_ISL_2745693, EPI_ISL_2745695, EPI_ISL_2745696, EPI_ISL_2745697, EPI_ISL_2745698, EPI_ISL_2745699, EPI_ISL_2745700, EPI_ISL_2745701, EPI_ISL_2745702, EPI_ISL_2745703, EPI_ISL_2745704, EPI_ISL_2745705, EPI_ISL_2745706, EPI_ISL_2745707, EPI_ISL_2745708, EPI_ISL_2745709, EPI_ISL_2745710, EPI_ISL_2745711,
EPI_ISL_2745712, EPI_ISL_2745713, EPI_ISL_2745714, EPI_ISL_2745715, EPI_ISL_2745716, EPI_ISL_2745718, EPI_ISL_2745719, EPI_ISL_2745720, EPI_ISL_2745721, EPI_ISL_2745722, EPI_ISL_2745723, EPI_ISL_2745724, EPI_ISL_2745725, EPI_ISL_2745726, EPI_ISL_2745727, EPI_ISL_2745728, EPI_ISL_2745729, EPI_ISL_2745730, EPI_ISL_2745731, EPI_ISL_2745732, EPI_ISL_2745733, EPI_ISL_2745734,
EPI_ISL_2745735, EPI_ISL_2745736, EPI_ISL_2745737, EPI_ISL_2745738, EPI_ISL_2745739, EPI_ISL_2745740, EPI_ISL_2745741, EPI_ISL_2745742, EPI_ISL_2745743, EPI_ISL_2745744, EPI_ISL_2745745, EPI_ISL_2745746, EPI_ISL_2745747, EPI_ISL_2745748, EPI_ISL_2745749, EPI_ISL_2745750, EPI_ISL_2745751, EPI_ISL_2745752, EPI_ISL_2745753, EPI_ISL_2745754, EPI_ISL_2745755, EPI_ISL_2745756,
EPI_ISL_2745757, EPI_ISL_2745758, EPI_ISL_2745759, EPI_ISL_2745760, EPI_ISL_2745761, EPI_ISL_2745762, EPI_ISL_2745763, EPI_ISL_2745764, EPI_ISL_2745765, EPI_ISL_2745766, EPI_ISL_2745767, EPI_ISL_2745769, EPI_ISL_2745770, EPI_ISL_2745771, EPI_ISL_2745772, EPI_ISL_2745773, EPI_ISL_2745774, EPI_ISL_2745775, EPI_ISL_2745776, EPI_ISL_2745777, EPI_ISL_2745778, EPI_ISL_2745779,
EPI_ISL_2745780, EPI_ISL_2745781, EPI_ISL_2745783, EPI_ISL_2745784, EPI_ISL_2745786, EPI_ISL_2745787, EPI_ISL_2745788, EPI_ISL_2745789, EPI_ISL_2745790, EPI_ISL_2745791, EPI_ISL_2745792, EPI_ISL_2745793, EPI_ISL_2745794, EPI_ISL_2745795, EPI_ISL_2745796, EPI_ISL_2745797, EPI_ISL_2745798, EPI_ISL_2745799, EPI_ISL_2745800, EPI_ISL_2745801, EPI_ISL_2745802, EPI_ISL_2745803,
EPI_ISL_2745804, EPI_ISL_2745805, EPI_ISL_2745806, EPI_ISL_2745807, EPI_ISL_2745808, EPI_ISL_2745809, EPI_ISL_2745810, EPI_ISL_2745811, EPI_ISL_2745812, EPI_ISL_2745813, EPI_ISL_2745814, EPI_ISL_2745815, EPI_ISL_2745816, EPI_ISL_2745817, EPI_ISL_2745818, EPI_ISL_2745819, EPI_ISL_2745820, EPI_ISL_2745821, EPI_ISL_2745822, EPI_ISL_2745823, EPI_ISL_2745824, EPI_ISL_2745825,
EPI_ISL_2745826, EPI_ISL_2745827, EPI_ISL_2745828, EPI_ISL_2745829, EPI_ISL_2745830, EPI_ISL_2745831, EPI_ISL_2745832, EPI_ISL_2745833, EPI_ISL_2745834, EPI_ISL_2745836, EPI_ISL_2745837, EPI_ISL_2745838, EPI_ISL_2745839, EPI_ISL_2745840, EPI_ISL_2745841, EPI_ISL_2745842, EPI_ISL_2745843, EPI_ISL_2745844, EPI_ISL_2745845, EPI_ISL_2745846, EPI_ISL_2745847, EPI_ISL_2745848,
EPI_ISL_2745849, EPI_ISL_2745851, EPI_ISL_2745852, EPI_ISL_2745853, EPI_ISL_2745854, EPI_ISL_2745855, EPI_ISL_2745856, EPI_ISL_2745857, EPI_ISL_2745858, EPI_ISL_2745859, EPI_ISL_2745860, EPI_ISL_2745861, EPI_ISL_2745862, EPI_ISL_2745863, EPI_ISL_2745864, EPI_ISL_2745865, EPI_ISL_2745866, EPI_ISL_2745867, EPI_ISL_2745869, EPI_ISL_2745870, EPI_ISL_2745871, EPI_ISL_2745872,
EPI_ISL_2745873, EPI_ISL_2745874, EPI_ISL_2745875, EPI_ISL_2745876, EPI_ISL_2745877, EPI_ISL_2745878, EPI_ISL_2745879, EPI_ISL_2745880, EPI_ISL_2745881, EPI_ISL_2745882, EPI_ISL_2745883, EPI_ISL_2745884, EPI_ISL_2745885, EPI_ISL_2745886, EPI_ISL_2745887, EPI_ISL_2745889, EPI_ISL_2745890, EPI_ISL_2745891, EPI_ISL_2745892, EPI_ISL_2745894, EPI_ISL_2745895, EPI_ISL_2745896,
EPI_ISL_2745897, EPI_ISL_2745898, EPI_ISL_2745899, EPI_ISL_2745900, EPI_ISL_2745901, EPI_ISL_2745904, EPI_ISL_2745905, EPI_ISL_2745906, EPI_ISL_2745907, EPI_ISL_2745908, EPI_ISL_2745909, EPI_ISL_2745910, EPI_ISL_2745911, EPI_ISL_2745912, EPI_ISL_2745913, EPI_ISL_2745914, EPI_ISL_2745915, EPI_ISL_2745916, EPI_ISL_2745917, EPI_ISL_2745918, EPI_ISL_2745919,
EPI_ISL_2745920, EPI_ISL_2745921, EPI_ISL_2745922, EPI_ISL_2745923, EPI_ISL_2745924, EPI_ISL_2745925, EPI_ISL_2745926, EPI_ISL_2745927, EPI_ISL_2745928, EPI_ISL_2745929, EPI_ISL_2745930, EPI_ISL_2745931, EPI_ISL_2745932, EPI_ISL_2745933, EPI_ISL_2745934, EPI_ISL_2745935, EPI_ISL_2745936, EPI_ISL_2745937, EPI_ISL_2745938, EPI_ISL_2745939, EPI_ISL_2745940, EPI_ISL_2745941,
EPI_ISL_2745942, EPI_ISL_2745943, EPI_ISL_2745944, EPI_ISL_2745945, EPI_ISL_2745946, EPI_ISL_2745947, EPI_ISL_2745948, EPI_ISL_2745949, EPI_ISL_2745950, EPI_ISL_2745951, EPI_ISL_2745952, EPI_ISL_2745953, EPI_ISL_2745954, EPI_ISL_2745955, EPI_ISL_2745956, EPI_ISL_2745957, EPI_ISL_2745958, EPI_ISL_2745959, EPI_ISL_2745960, EPI_ISL_2745961, EPI_ISL_2745962, EPI_ISL_2745963,
EPI_ISL_2745964, EPI_ISL_2745965, EPI_ISL_2745966, EPI_ISL_2745967, EPI_ISL_2745968, EPI_ISL_2745969, EPI_ISL_2745970, EPI_ISL_2745971, EPI_ISL_2745972, EPI_ISL_2745973, EPI_ISL_2745974, EPI_ISL_2745975, EPI_ISL_2745976, EPI_ISL_2745977, EPI_ISL_2745979, EPI_ISL_2745980, EPI_ISL_2745981, EPI_ISL_2745982, EPI_ISL_2745983, EPI_ISL_2745984, EPI_ISL_2745985, EPI_ISL_2745986,
EPI_ISL_2745987, EPI_ISL_2745988, EPI_ISL_2745989, EPI_ISL_2745990, EPI_ISL_2745991, EPI_ISL_2745992, EPI_ISL_2745993, EPI_ISL_2745994, EPI_ISL_2745995, EPI_ISL_2745996, EPI_ISL_2745997, EPI_ISL_2745998, EPI_ISL_2745999, EPI_ISL_2746000, EPI_ISL_2746001, EPI_ISL_2746002, EPI_ISL_2746003, EPI_ISL_2746004, EPI_ISL_2746005, EPI_ISL_2746006, EPI_ISL_2746007, EPI_ISL_2746008,
EPI_ISL_2746009, EPI_ISL_2746010, EPI_ISL_2746011, EPI_ISL_2746012, EPI_ISL_2746013, EPI_ISL_2746014, EPI_ISL_2746015, EPI_ISL_2746016, EPI_ISL_2746017, EPI_ISL_2746018, EPI_ISL_2746019, EPI_ISL_2746020, EPI_ISL_2746021, EPI_ISL_2746022, EPI_ISL_2746023, EPI_ISL_2746024, EPI_ISL_2746025, EPI_ISL_2746027, EPI_ISL_2746028, EPI_ISL_2746029, EPI_ISL_2746030,
EPI_ISL_2746032, EPI_ISL_2746033, EPI_ISL_2746034, EPI_ISL_2746035, EPI_ISL_2746037, EPI_ISL_2746038, EPI_ISL_2746039, EPI_ISL_2746040, EPI_ISL_2746041, EPI_ISL_2746042, EPI_ISL_2746043, EPI_ISL_2746044, EPI_ISL_2746045, EPI_ISL_2746046, EPI_ISL_2746047, EPI_ISL_2746048, EPI_ISL_2746049, EPI_ISL_2746050, EPI_ISL_2746051, EPI_ISL_2746052, EPI_ISL_2746053, EPI_ISL_2746054,
EPI_ISL_2746055, EPI_ISL_2746056, EPI_ISL_2746057, EPI_ISL_2746058, EPI_ISL_2746059, EPI_ISL_2746060, EPI_ISL_2746061, EPI_ISL_2746063, EPI_ISL_2746065, EPI_ISL_2746066, EPI_ISL_2746067, EPI_ISL_2746068, EPI_ISL_2746069, EPI_ISL_2746070, EPI_ISL_2746071, EPI_ISL_2746072, EPI_ISL_2746073, EPI_ISL_2746074, EPI_ISL_2746075, EPI_ISL_2746076, EPI_ISL_2746077, EPI_ISL_2746078,
EPI_ISL_2746079, EPI_ISL_2746080, EPI_ISL_2746081, EPI_ISL_2746082, EPI_ISL_2746083, EPI_ISL_2746084, EPI_ISL_2746085, EPI_ISL_2746086, EPI_ISL_2746087, EPI_ISL_2746088, EPI_ISL_2746089, EPI_ISL_2746090, EPI_ISL_2746091, EPI_ISL_2746092, EPI_ISL_2746093, EPI_ISL_2746094, EPI_ISL_2746095, EPI_ISL_2746096, EPI_ISL_2746097, EPI_ISL_2746098, EPI_ISL_2746099, EPI_ISL_2746100,
EPI_ISL_2746101, EPI_ISL_2746102, EPI_ISL_2746103, EPI_ISL_2746104, EPI_ISL_2746105, EPI_ISL_2746106, EPI_ISL_2746107, EPI_ISL_2746108, EPI_ISL_2746109, EPI_ISL_2746110, EPI_ISL_2746111, EPI_ISL_2746112, EPI_ISL_2746113, EPI_ISL_2746114, EPI_ISL_2746115, EPI_ISL_2746116, EPI_ISL_2746117, EPI_ISL_2746118, EPI_ISL_2746119, EPI_ISL_2746120, EPI_ISL_2746121, EPI_ISL_2746122,
EPI_ISL_2746123, EPI_ISL_2746124, EPI_ISL_2746125, EPI_ISL_2746126, EPI_ISL_2746127, EPI_ISL_2746128, EPI_ISL_2746129, EPI_ISL_2746130, EPI_ISL_2746131, EPI_ISL_2746132, EPI_ISL_2746133, EPI_ISL_2746134, EPI_ISL_2746135, EPI_ISL_2746136, EPI_ISL_2746138, EPI_ISL_2746139, EPI_ISL_2746140, EPI_ISL_2746141, EPI_ISL_2746142, EPI_ISL_2746144, EPI_ISL_2746145,
EPI_ISL_2746146, EPI_ISL_2746147, EPI_ISL_2746148, EPI_ISL_2746149, EPI_ISL_2746151, EPI_ISL_2746152, EPI_ISL_2746153, EPI_ISL_2746154, EPI_ISL_2746156, EPI_ISL_2746157, EPI_ISL_2746158, EPI_ISL_2746159, EPI_ISL_2746160, EPI_ISL_2746161, EPI_ISL_2746162, EPI_ISL_2746164, EPI_ISL_2746165, EPI_ISL_2746166, EPI_ISL_2746167, EPI_ISL_2746168, EPI_ISL_2746169, EPI_ISL_2746170,
EPI_ISL_2746171, EPI_ISL_2746172, EPI_ISL_2746173, EPI_ISL_2746174, EPI_ISL_2746176, EPI_ISL_2746177, EPI_ISL_2746179, EPI_ISL_2746180, EPI_ISL_2746182, EPI_ISL_2746183, EPI_ISL_2746184, EPI_ISL_2746186, EPI_ISL_2746187, EPI_ISL_2746188, EPI_ISL_2746189, EPI_ISL_2746190, EPI_ISL_2746191, EPI_ISL_2746192, EPI_ISL_2746193, EPI_ISL_2746194, EPI_ISL_2746195, EPI_ISL_2746196,
EPI_ISL_2746197, EPI_ISL_2746198, EPI_ISL_2746199, EPI_ISL_2746200, EPI_ISL_2746201, EPI_ISL_2746202, EPI_ISL_2746203, EPI_ISL_2746204, EPI_ISL_2746205, EPI_ISL_2746206, EPI_ISL_2746207, EPI_ISL_2746208, EPI_ISL_2746209, EPI_ISL_2746210, EPI_ISL_2746211, EPI_ISL_2746212, EPI_ISL_2746213, EPI_ISL_2746214, EPI_ISL_2746215, EPI_ISL_2746216, EPI_ISL_2746217, EPI_ISL_2746218,
EPI_ISL_2746219, EPI_ISL_2746220, EPI_ISL_2746221, EPI_ISL_2746222, EPI_ISL_2746223, EPI_ISL_2746224, EPI_ISL_2746226, EPI_ISL_2746229, EPI_ISL_2746230, EPI_ISL_2746231, EPI_ISL_2746232, EPI_ISL_2746233, EPI_ISL_2746234, EPI_ISL_2746235, EPI_ISL_2746236, EPI_ISL_2746237, EPI_ISL_2746238, EPI_ISL_2746239, EPI_ISL_2746240, EPI_ISL_2746241, EPI_ISL_2746242, EPI_ISL_2746243,
EPI_ISL_2746245, EPI_ISL_2746246, EPI_ISL_2746247, EPI_ISL_2746248, EPI_ISL_2746249, EPI_ISL_2746250, EPI_ISL_2746251, EPI_ISL_2746252, EPI_ISL_2746253, EPI_ISL_2746254, EPI_ISL_2746255, EPI_ISL_2746256, EPI_ISL_2746257, EPI_ISL_2746258, EPI_ISL_2746259, EPI_ISL_2746261, EPI_ISL_2746262, EPI_ISL_2746263, EPI_ISL_2746264, EPI_ISL_2746265, EPI_ISL_2746266, EPI_ISL_2746267,
EPI_ISL_2746268, EPI_ISL_2746270, EPI_ISL_2746271, EPI_ISL_2746272, EPI_ISL_2746273, EPI_ISL_2746274, EPI_ISL_2746275, EPI_ISL_2746276, EPI_ISL_2746277, EPI_ISL_2746278, EPI_ISL_2746279, EPI_ISL_2746280, EPI_ISL_2746281, EPI_ISL_2746282, EPI_ISL_2746283, EPI_ISL_2746284, EPI_ISL_2746285, EPI_ISL_2746286, EPI_ISL_2746287, EPI_ISL_2746288, EPI_ISL_2746289, EPI_ISL_2746290,
EPI_ISL_2746291, EPI_ISL_2746292, EPI_ISL_2746293, EPI_ISL_2746294, EPI_ISL_2746295, EPI_ISL_2746296, EPI_ISL_2746297, EPI_ISL_2746298, EPI_ISL_2746299, EPI_ISL_2746300, EPI_ISL_2746301, EPI_ISL_2746302, EPI_ISL_2746303, EPI_ISL_2746304, EPI_ISL_2746305, EPI_ISL_2746306, EPI_ISL_2746307, EPI_ISL_2746308, EPI_ISL_2746309, EPI_ISL_2746310, EPI_ISL_2746311, EPI_ISL_2746312,
EPI_ISL_2746313, EPI_ISL_2746314, EPI_ISL_2746315, EPI_ISL_2746316, EPI_ISL_2746317, EPI_ISL_2746318, EPI_ISL_2746319, EPI_ISL_2746320, EPI_ISL_2746321, EPI_ISL_2746322, EPI_ISL_2746323, EPI_ISL_2746324, EPI_ISL_2746325, EPI_ISL_2746326, EPI_ISL_2746327, EPI_ISL_2746328, EPI_ISL_2746329, EPI_ISL_2746330, EPI_ISL_2746331, EPI_ISL_2746332, EPI_ISL_2746333, EPI_ISL_2746334,
EPI_ISL_2746335, EPI_ISL_2746336, EPI_ISL_2746337, EPI_ISL_2746338, EPI_ISL_2746339, EPI_ISL_2746340, EPI_ISL_2746341, EPI_ISL_2746342, EPI_ISL_2746343, EPI_ISL_2746344, EPI_ISL_2746345, EPI_ISL_2746346, EPI_ISL_2746347, EPI_ISL_2746348, EPI_ISL_2746349, EPI_ISL_2746350, EPI_ISL_2746351, EPI_ISL_2746352, EPI_ISL_2746353, EPI_ISL_2746354, EPI_ISL_2746355, EPI_ISL_2746357,
EPI_ISL_2746358, EPI_ISL_2746359, EPI_ISL_2746360, EPI_ISL_2746361, EPI_ISL_2746362, EPI_ISL_2746363, EPI_ISL_2746364, EPI_ISL_2746365, EPI_ISL_2746366, EPI_ISL_2746367, EPI_ISL_2746368, EPI_ISL_2746369, EPI_ISL_2746370, EPI_ISL_2746371, EPI_ISL_2746372, EPI_ISL_2746373, EPI_ISL_2746374, EPI_ISL_2746375, EPI_ISL_2746376, EPI_ISL_2746377, EPI_ISL_2746378, EPI_ISL_2746379,
EPI_ISL_2746380, EPI_ISL_2746381, EPI_ISL_2746383, EPI_ISL_2746384, EPI_ISL_2746385, EPI_ISL_2746386, EPI_ISL_2746387, EPI_ISL_2746388, EPI_ISL_2746389, EPI_ISL_2746390, EPI_ISL_2746391, EPI_ISL_2746392, EPI_ISL_2746393, EPI_ISL_2746394, EPI_ISL_2746395, EPI_ISL_2746396, EPI_ISL_2746397, EPI_ISL_2746398, EPI_ISL_2746399, EPI_ISL_2746400, EPI_ISL_2746401, EPI_ISL_2746402,
EPI_ISL_2746403, EPI_ISL_2746404, EPI_ISL_2746405, EPI_ISL_2746406, EPI_ISL_2746407, EPI_ISL_2746408, EPI_ISL_2746409, EPI_ISL_2746410, EPI_ISL_2746411, EPI_ISL_2746412, EPI_ISL_2746413, EPI_ISL_2746414, EPI_ISL_2746415, EPI_ISL_2746416, EPI_ISL_2746417, EPI_ISL_2746418, EPI_ISL_2746419, EPI_ISL_2746420, EPI_ISL_2746421, EPI_ISL_2746422, EPI_ISL_2746423, EPI_ISL_2746424,
EPI_ISL_2746425, EPI_ISL_2746426, EPI_ISL_2746427, EPI_ISL_2746428, EPI_ISL_2746429, EPI_ISL_2746430, EPI_ISL_2746431, EPI_ISL_2746432, EPI_ISL_2746433, EPI_ISL_2746434, EPI_ISL_2746435, EPI_ISL_2746436, EPI_ISL_2746437, EPI_ISL_2746438, EPI_ISL_2746439, EPI_ISL_2746440, EPI_ISL_2746441, EPI_ISL_2746442, EPI_ISL_2746444, EPI_ISL_2746445, EPI_ISL_2746446, EPI_ISL_2746447,
EPI_ISL_2746448, EPI_ISL_2746449, EPI_ISL_2746450, EPI_ISL_2746451, EPI_ISL_2746452, EPI_ISL_2746453, EPI_ISL_2746454, EPI_ISL_2746455, EPI_ISL_2746456, EPI_ISL_2746457, EPI_ISL_2746458, EPI_ISL_2746461, EPI_ISL_2746462, EPI_ISL_2746463, EPI_ISL_2746465, EPI_ISL_2746466, EPI_ISL_2746467, EPI_ISL_2746468, EPI_ISL_2746469, EPI_ISL_2746470, EPI_ISL_2746471, EPI_ISL_2746473,
EPI_ISL_2746474, EPI_ISL_2746475, EPI_ISL_2746476, EPI_ISL_2746477, EPI_ISL_2746478, EPI_ISL_2746479, EPI_ISL_2746480, EPI_ISL_2746481, EPI_ISL_2746482, EPI_ISL_2746483, EPI_ISL_2746484, EPI_ISL_2746485, EPI_ISL_2746486, EPI_ISL_2746487, EPI_ISL_2746488, EPI_ISL_2746489, EPI_ISL_2746490, EPI_ISL_2746491, EPI_ISL_2746492, EPI_ISL_2746493, EPI_ISL_2746494, EPI_ISL_2746495,
EPI_ISL_2746496, EPI_ISL_2746497, EPI_ISL_2746498, EPI_ISL_2746499, EPI_ISL_2746500, EPI_ISL_2746501, EPI_ISL_2746502, EPI_ISL_2746503, EPI_ISL_2746504, EPI_ISL_2746505, EPI_ISL_2746506, EPI_ISL_2746507, EPI_ISL_2746508, EPI_ISL_2746509, EPI_ISL_2746510, EPI_ISL_2746512, EPI_ISL_2746513, EPI_ISL_2746514, EPI_ISL_2746515, EPI_ISL_2746516, EPI_ISL_2746517, EPI_ISL_2746518,
EPI_ISL_2746519, EPI_ISL_2746520, EPI_ISL_2746521, EPI_ISL_2746522, EPI_ISL_2746524, EPI_ISL_2746525, EPI_ISL_2746526, EPI_ISL_2746527, EPI_ISL_2746528, EPI_ISL_2746529, EPI_ISL_2746530, EPI_ISL_2746531, EPI_ISL_2746532, EPI_ISL_2746533, EPI_ISL_2746534, EPI_ISL_2746535, EPI_ISL_2746536, EPI_ISL_2746537, EPI_ISL_2746538, EPI_ISL_2746540, EPI_ISL_2746541, EPI_ISL_2746542,
EPI_ISL_2746543, EPI_ISL_2746544, EPI_ISL_2746545, EPI_ISL_2746546, EPI_ISL_2746547, EPI_ISL_2746548, EPI_ISL_2746549, EPI_ISL_2746550, EPI_ISL_2746551, EPI_ISL_2746552, EPI_ISL_2746553, EPI_ISL_2746555, EPI_ISL_2746556, EPI_ISL_2746557, EPI_ISL_2746558, EPI_ISL_2746559, EPI_ISL_2746560, EPI_ISL_2746561, EPI_ISL_2746562, EPI_ISL_2746563, EPI_ISL_2746564, EPI_ISL_2746565,
EPI_ISL_2746566, EPI_ISL_2746567, EPI_ISL_2746568, EPI_ISL_2746569, EPI_ISL_2746570, EPI_ISL_2746571, EPI_ISL_2746572, EPI_ISL_2746573, EPI_ISL_2746574, EPI_ISL_2746575, EPI_ISL_2746576, EPI_ISL_2746577, EPI_ISL_2746578, EPI_ISL_2746579, EPI_ISL_2746580, EPI_ISL_2746581, EPI_ISL_2746582, EPI_ISL_2746583, EPI_ISL_2746584, EPI_ISL_2746585, EPI_ISL_2746586, EPI_ISL_2746587,
EPI_ISL_2746588, EPI_ISL_2746589, EPI_ISL_2746590, EPI_ISL_2746591, EPI_ISL_2746592, EPI_ISL_2746593, EPI_ISL_2746594, EPI_ISL_2746595, EPI_ISL_2746596, EPI_ISL_2746597, EPI_ISL_2746598, EPI_ISL_2746599, EPI_ISL_2746600, EPI_ISL_2746601, EPI_ISL_2746602, EPI_ISL_2746603, EPI_ISL_2746604, EPI_ISL_2746605, EPI_ISL_2746606, EPI_ISL_2746608, EPI_ISL_2746609, EPI_ISL_2746610,
EPI_ISL_2746611, EPI_ISL_2746612, EPI_ISL_2746613, EPI_ISL_2746615, EPI_ISL_2746616, EPI_ISL_2746617, EPI_ISL_2746618, EPI_ISL_2746619, EPI_ISL_2746620, EPI_ISL_2746621, EPI_ISL_2746622, EPI_ISL_2746623, EPI_ISL_2746624, EPI_ISL_2746625, EPI_ISL_2746626, EPI_ISL_2746627, EPI_ISL_2746628, EPI_ISL_2746629, EPI_ISL_2746630, EPI_ISL_2746631, EPI_ISL_2746632, EPI_ISL_2746633,
EPI_ISL_2746635, EPI_ISL_2746636, EPI_ISL_2746637, EPI_ISL_2746638, EPI_ISL_2746639, EPI_ISL_2746642, EPI_ISL_2746643, EPI_ISL_2746644, EPI_ISL_2746645, EPI_ISL_2746646, EPI_ISL_2746647, EPI_ISL_2746648, EPI_ISL_2746649, EPI_ISL_2746650, EPI_ISL_2746651, EPI_ISL_2746652, EPI_ISL_2746653, EPI_ISL_2746654, EPI_ISL_2746655, EPI_ISL_2746656, EPI_ISL_2746657, EPI_ISL_2746658,
EPI_ISL_2746659, EPI_ISL_2746660, EPI_ISL_2746661, EPI_ISL_2746662, EPI_ISL_2746663, EPI_ISL_2746664, EPI_ISL_2746665, EPI_ISL_2746666, EPI_ISL_2746667, EPI_ISL_2746668, EPI_ISL_2746669, EPI_ISL_2746670, EPI_ISL_2746671, EPI_ISL_2746672, EPI_ISL_2746673, EPI_ISL_2746674, EPI_ISL_2746675, EPI_ISL_2746676, EPI_ISL_2746677, EPI_ISL_2746678, EPI_ISL_2746679, EPI_ISL_2746680,
EPI_ISL_2746681, EPI_ISL_2746682, EPI_ISL_2746683, EPI_ISL_2746684, EPI_ISL_2746685, EPI_ISL_2746686, EPI_ISL_2746687, EPI_ISL_2746688, EPI_ISL_2746689, EPI_ISL_2746691, EPI_ISL_2746692, EPI_ISL_2746693, EPI_ISL_2746694, EPI_ISL_2746695, EPI_ISL_2746696, EPI_ISL_2746697, EPI_ISL_2746698, EPI_ISL_2746699, EPI_ISL_2746700, EPI_ISL_2746701, EPI_ISL_2746702, EPI_ISL_2746703,
EPI_ISL_2746704, EPI_ISL_2746705, EPI_ISL_2746706, EPI_ISL_2746707, EPI_ISL_2746709, EPI_ISL_2746710, EPI_ISL_2746711, EPI_ISL_2746712, EPI_ISL_2746713, EPI_ISL_2746714, EPI_ISL_2746715, EPI_ISL_2746716, EPI_ISL_2746717, EPI_ISL_2746718, EPI_ISL_2746719, EPI_ISL_2746720, EPI_ISL_2746721, EPI_ISL_2746722, EPI_ISL_2746723, EPI_ISL_2746724, EPI_ISL_2746725, EPI_ISL_2746726,
EPI_ISL_2746727, EPI_ISL_2746728, EPI_ISL_2746729, EPI_ISL_2746730, EPI_ISL_2746731, EPI_ISL_2746732, EPI_ISL_2746733, EPI_ISL_2746734, EPI_ISL_2746735, EPI_ISL_2746736, EPI_ISL_2746737, EPI_ISL_2746738, EPI_ISL_2746739, EPI_ISL_2746740, EPI_ISL_2746741, EPI_ISL_2746742, EPI_ISL_2746743, EPI_ISL_2746744, EPI_ISL_2746745, EPI_ISL_2746746, EPI_ISL_2746747, EPI_ISL_2746748,
EPI_ISL_2746749, EPI_ISL_2746750, EPI_ISL_2746751, EPI_ISL_2746752, EPI_ISL_2746753, EPI_ISL_2746754, EPI_ISL_2746755, EPI_ISL_2746756, EPI_ISL_2746757, EPI_ISL_2746758, EPI_ISL_2746759, EPI_ISL_2746760, EPI_ISL_2746761, EPI_ISL_2746762, EPI_ISL_2746763, EPI_ISL_2746764, EPI_ISL_2746765, EPI_ISL_2746766, EPI_ISL_2746767, EPI_ISL_2746769, EPI_ISL_2746770, EPI_ISL_2746771,
EPI_ISL_2746772, EPI_ISL_2746773, EPI_ISL_2746775, EPI_ISL_2746776, EPI_ISL_2746777, EPI_ISL_2746778, EPI_ISL_2746779, EPI_ISL_2746780, EPI_ISL_2746781, EPI_ISL_2746782, EPI_ISL_2746783, EPI_ISL_2746784, EPI_ISL_2746785, EPI_ISL_2746786, EPI_ISL_2746787, EPI_ISL_2746788, EPI_ISL_2746789, EPI_ISL_2746790, EPI_ISL_2746791, EPI_ISL_2746792, EPI_ISL_2746793, EPI_ISL_2746794,
EPI_ISL_2746795, EPI_ISL_2746796, EPI_ISL_2746797, EPI_ISL_2746798, EPI_ISL_2746799, EPI_ISL_2746800, EPI_ISL_2746801, EPI_ISL_2746802, EPI_ISL_2746803, EPI_ISL_2746804, EPI_ISL_2746805, EPI_ISL_2746806, EPI_ISL_2746807, EPI_ISL_2746808, EPI_ISL_2746809, EPI_ISL_2746810, EPI_ISL_2746811, EPI_ISL_2746812, EPI_ISL_2746813, EPI_ISL_2746814, EPI_ISL_2746815, EPI_ISL_2746816,
EPI_ISL_2746817, EPI_ISL_2746818, EPI_ISL_2746819, EPI_ISL_2746820, EPI_ISL_2746821, EPI_ISL_2746823, EPI_ISL_2746824, EPI_ISL_2746825, EPI_ISL_2746826, EPI_ISL_2746827, EPI_ISL_2746828, EPI_ISL_2746829, EPI_ISL_2746830, EPI_ISL_2746831, EPI_ISL_2746832, EPI_ISL_2746833, EPI_ISL_2746834, EPI_ISL_2746835, EPI_ISL_2746836, EPI_ISL_2746837, EPI_ISL_2746838,
EPI_ISL_2746839, EPI_ISL_2746840, EPI_ISL_2746841, EPI_ISL_2746842, EPI_ISL_2746843, EPI_ISL_2746844, EPI_ISL_2746845, EPI_ISL_2746846, EPI_ISL_2746847, EPI_ISL_2746848, EPI_ISL_2746849, EPI_ISL_2746850, EPI_ISL_2746851, EPI_ISL_2746852, EPI_ISL_2746853, EPI_ISL_2746854, EPI_ISL_2746855, EPI_ISL_2746856, EPI_ISL_2746857, EPI_ISL_2746858, EPI_ISL_2746859, EPI_ISL_2746860,
EPI_ISL_2746861, EPI_ISL_2746862, EPI_ISL_2746863, EPI_ISL_2746864, EPI_ISL_2746865, EPI_ISL_2746866, EPI_ISL_2746867, EPI_ISL_2746868, EPI_ISL_2746869, EPI_ISL_2746870, EPI_ISL_2746871, EPI_ISL_2746872, EPI_ISL_2746873, EPI_ISL_2746874, EPI_ISL_2746875, EPI_ISL_2746876, EPI_ISL_2746877, EPI_ISL_2746878, EPI_ISL_2746879, EPI_ISL_2746880, EPI_ISL_2746881, EPI_ISL_2746882,

EPI_ISL_2746883, EPI_ISL_2746884, EPI_ISL_2746885, EPI_ISL_2746886, EPI_ISL_2746887, EPI_ISL_2746888, EPI_ISL_2746889, EPI_ISL_2746890, EPI_ISL_2746891, EPI_ISL_2746892, EPI_ISL_2746893, EPI_ISL_2746894, EPI_ISL_2746895, EPI_ISL_2746896, EPI_ISL_2746897, EPI_ISL_2746898, EPI_ISL_2746899, EPI_ISL_2746900, EPI_ISL_2746901, EPI_ISL_2746902, EPI_ISL_2746903, EPI_ISL_2746904, EPI_ISL_2746905, EPI_ISL_2746906, EPI_ISL_2746907, EPI_ISL_2746908, EPI_ISL_2746909, EPI_ISL_2746910, EPI_ISL_2746911, EPI_ISL_2746912, EPI_ISL_2746913, EPI_ISL_2746914, EPI_ISL_2746915, EPI_ISL_2746916, EPI_ISL_2746917, EPI_ISL_2746918, EPI_ISL_2746919, EPI_ISL_2746920, EPI_ISL_2746921, EPI_ISL_2746922, EPI_ISL_2746923, EPI_ISL_2746924, EPI_ISL_2746925, EPI_ISL_2746926, EPI_ISL_2746927, EPI_ISL_2746928, EPI_ISL_2746929, EPI_ISL_2746931, EPI_ISL_2746932, EPI_ISL_2746933, EPI_ISL_2746934, EPI_ISL_2746935, EPI_ISL_2746936, EPI_ISL_2746937, EPI_ISL_2746938, EPI_ISL_2746939, EPI_ISL_2746940, EPI_ISL_2746941, EPI_ISL_2746942, EPI_ISL_2746943, EPI_ISL_2746944, EPI_ISL_2746945, EPI_ISL_2746946, EPI_ISL_2746947, EPI_ISL_2746948, EPI_ISL_2746949, EPI_ISL_2746950, EPI_ISL_2746951, EPI_ISL_2746952, EPI_ISL_2746953, EPI_ISL_2746954, EPI_ISL_2746955, EPI_ISL_2746956, EPI_ISL_2746957, EPI_ISL_2746958, EPI_ISL_2746959, EPI_ISL_2746960, EPI_ISL_2746961, EPI_ISL_2746962, EPI_ISL_2746963, EPI_ISL_2746964, EPI_ISL_2746965, EPI_ISL_2746966, EPI_ISL_2746967, EPI_ISL_2746968, EPI_ISL_2746969, EPI_ISL_2746970, EPI_ISL_2746971, EPI_ISL_2746972, EPI_ISL_2746973, EPI_ISL_2746974, EPI_ISL_2746975, EPI_ISL_2746976, EPI_ISL_2746977, EPI_ISL_2746978, EPI_ISL_2746979, EPI_ISL_2746980, EPI_ISL_2746981, EPI_ISL_2746982, EPI_ISL_2746983, EPI_ISL_2746984, EPI_ISL_2746985, EPI_ISL_2746986, EPI_ISL_2746987, EPI_ISL_2746988, EPI_ISL_2746989, EPI_ISL_2746990, EPI_ISL_2746991, EPI_ISL_2746992, EPI_ISL_2746993, EPI_ISL_2746994, EPI_ISL_2746995, EPI_ISL_2746996, EPI_ISL_2746998, EPI_ISL_2746999, EPI_ISL_2747000, EPI_ISL_2747001, EPI_ISL_2747002, EPI_ISL_2747003, EPI_ISL_2747004, EPI_ISL_2747005, EPI_ISL_2747006, EPI_ISL_2747007, EPI_ISL_2747008, EPI_ISL_2747009, EPI_ISL_2747010, EPI_ISL_2747011, EPI_ISL_2747012, EPI_ISL_2747013, EPI_ISL_2747014, EPI_ISL_2747015, EPI_ISL_2747016, EPI_ISL_2747017, EPI_ISL_2747018, EPI_ISL_2747019, EPI_ISL_2747020, EPI_ISL_2747021, EPI_ISL_2747022, EPI_ISL_2747023, EPI_ISL_2747024, EPI_ISL_2747025, EPI_ISL_2747026, EPI_ISL_2747027, EPI_ISL_2747028, EPI_ISL_2747029, EPI_ISL_2747030, EPI_ISL_2747031, EPI_ISL_2747032, EPI_ISL_2747033, EPI_ISL_2747034, EPI_ISL_2747035, EPI_ISL_2747036, EPI_ISL_2747037, EPI_ISL_2747038, EPI_ISL_2747039, EPI_ISL_2747040, EPI_ISL_2747041, EPI_ISL_2747042, EPI_ISL_2747043, EPI_ISL_2747044, EPI_ISL_2747045, EPI_ISL_2747046, EPI_ISL_2747047, EPI_ISL_2747048, EPI_ISL_2747049, EPI_ISL_2747050, EPI_ISL_2747051, EPI_ISL_2747052, EPI_ISL_2747053, EPI_ISL_2747054, EPI_ISL_2747055, EPI_ISL_2747057, EPI_ISL_2747058, EPI_ISL_2747059, EPI_ISL_2747060, EPI_ISL_2747061, EPI_ISL_2747062, EPI_ISL_2747063, EPI_ISL_2747064, EPI_ISL_2747065, EPI_ISL_2747066, EPI_ISL_2747067, EPI_ISL_2747068, EPI_ISL_2747069, EPI_ISL_2747070, EPI_ISL_2747071, EPI_ISL_2747072, EPI_ISL_2747073, EPI_ISL_2747074, EPI_ISL_2747075, EPI_ISL_2747076, EPI_ISL_2747077, EPI_ISL_2747079, EPI_ISL_2747080, EPI_ISL_2747081, EPI_ISL_2747082, EPI_ISL_2747083, EPI_ISL_2747084, EPI_ISL_2747085, EPI_ISL_2747086, EPI_ISL_2747087, EPI_ISL_2747088, EPI_ISL_2747089, EPI_ISL_2747090, EPI_ISL_2747091, EPI_ISL_2747092, EPI_ISL_2747093, EPI_ISL_2747094, EPI_ISL_2747095, EPI_ISL_2747096, EPI_ISL_2747097, EPI_ISL_2747098, EPI_ISL_2747099, EPI_ISL_2747100, EPI_ISL_2747101, EPI_ISL_2747102, EPI_ISL_2747103, EPI_ISL_2747104, EPI_ISL_2747105, EPI_ISL_2747106, EPI_ISL_2747107, EPI_ISL_2747108, EPI_ISL_2747109, EPI_ISL_2747110, EPI_ISL_2747111, EPI_ISL_2747113, EPI_ISL_2747114, EPI_ISL_2747115, EPI_ISL_2747116, EPI_ISL_2747117, EPI_ISL_2747118, EPI_ISL_2747119, EPI_ISL_2747120, EPI_ISL_2747121, EPI_ISL_2747122, EPI_ISL_2747123, EPI_ISL_2747124, EPI_ISL_2747125, EPI_ISL_2747126, EPI_ISL_2747127, EPI_ISL_2747128, EPI_ISL_2747129, EPI_ISL_2747130, EPI_ISL_2747131, EPI_ISL_2747132, EPI_ISL_2747133, EPI_ISL_2747134, EPI_ISL_2747135, EPI_ISL_2747136, EPI_ISL_2747137, EPI_ISL_2747138, EPI_ISL_2747139, EPI_ISL_2747140, EPI_ISL_2747141, EPI_ISL_2747142, EPI_ISL_2747143, EPI_ISL_2747144, EPI_ISL_2747145, EPI_ISL_2747146, EPI_ISL_2747147, EPI_ISL_2747148, EPI_ISL_2747149, EPI_ISL_2747151, EPI_ISL_2747152, EPI_ISL_2747153, EPI_ISL_2747160, EPI_ISL_2747161, EPI_ISL_2747162, EPI_ISL_2747163, EPI_ISL_2747164, EPI_ISL_2747165, EPI_ISL_2747166, EPI_ISL_2747167, EPI_ISL_2747168, EPI_ISL_2747169, EPI_ISL_2747170, EPI_ISL_2747171, EPI_ISL_2747172, EPI_ISL_2747173, EPI_ISL_2747174, EPI_ISL_2747176, EPI_ISL_2747177, EPI_ISL_2747178, EPI_ISL_2747180, EPI_ISL_2747181, EPI_ISL_2747182, EPI_ISL_2747183, EPI_ISL_2747184, EPI_ISL_2747186, EPI_ISL_2747187, EPI_ISL_2747188, EPI_ISL_2747189, EPI_ISL_2747190, EPI_ISL_2747191, EPI_ISL_2747192, EPI_ISL_2747193, EPI_ISL_2747194, EPI_ISL_2747195, EPI_ISL_2747196, EPI_ISL_2747197, EPI_ISL_2747198, EPI_ISL_2747199, EPI_ISL_2747200, EPI_ISL_2747201, EPI_ISL_2747202, EPI_ISL_2747203, EPI_ISL_2747205, EPI_ISL_2747206, EPI_ISL_2747207, EPI_ISL_2747208, EPI_ISL_2747209, EPI_ISL_2747211, EPI_ISL_2747212, EPI_ISL_2747213, EPI_ISL_2747214, EPI_ISL_2747215, EPI_ISL_2747216, EPI_ISL_2747217, EPI_ISL_2747218, EPI_ISL_2747219, EPI_ISL_2747220, EPI_ISL_2747221, EPI_ISL_2747222, EPI_ISL_2747223, EPI_ISL_2747224, EPI_ISL_2747225, EPI_ISL_2747226, EPI_ISL_2747227, EPI_ISL_2747228, EPI_ISL_2747229, EPI_ISL_2747230, EPI_ISL_2747231, EPI_ISL_2747232, EPI_ISL_2747233, EPI_ISL_2747234, EPI_ISL_2747235, EPI_ISL_2747236, EPI_ISL_2747237, EPI_ISL_2747238, EPI_ISL_2747239, EPI_ISL_2747240, EPI_ISL_2747241, EPI_ISL_2747242, EPI_ISL_2747243, EPI_ISL_2747244, EPI_ISL_2747245, EPI_ISL_2747246, EPI_ISL_2747247, EPI_ISL_2747248, EPI_ISL_2747249, EPI_ISL_2747250, EPI_ISL_2747251, EPI_ISL_2747252, EPI_ISL_2747253, EPI_ISL_2747254, EPI_ISL_2747255, EPI_ISL_2747256, EPI_ISL_2747257, EPI_ISL_2747258, EPI_ISL_2747259, EPI_ISL_2747260, EPI_ISL_2747261, EPI_ISL_2747262, EPI_ISL_2747263, EPI_ISL_2747264, EPI_ISL_2747265, EPI_ISL_2747266, EPI_ISL_2747267, EPI_ISL_2747268, EPI_ISL_2747269, EPI_ISL_2747270, EPI_ISL_2747271, EPI_ISL_2747272, EPI_ISL_2747273, EPI_ISL_2747274, EPI_ISL_2747275, EPI_ISL_2747276, EPI_ISL_2747277, EPI_ISL_2747278, EPI_ISL_2747279, EPI_ISL_2747280, EPI_ISL_2747281, EPI_ISL_2747282, EPI_ISL_2747283, EPI_ISL_2747284, EPI_ISL_2747285, EPI_ISL_2747286, EPI_ISL_2747287, EPI_ISL_2747288, EPI_ISL_2747289, EPI_ISL_2747290, EPI_ISL_2747291, EPI_ISL_2747292, EPI_ISL_2747293, EPI_ISL_2747294, EPI_ISL_2747295, EPI_ISL_2747296, EPI_ISL_2747297, EPI_ISL_2747298, EPI_ISL_2747299, EPI_ISL_2747300, EPI_ISL_2747301, EPI_ISL_2747302, EPI_ISL_2747303, EPI_ISL_2747304, EPI_ISL_2747305, EPI_ISL_2747306, EPI_ISL_2747308, EPI_ISL_2747309, EPI_ISL_2747310, EPI_ISL_2747311, EPI_ISL_2747312, EPI_ISL_2747313, EPI_ISL_2747314, EPI_ISL_2747315, EPI_ISL_2747316, EPI_ISL_2747317, EPI_ISL_2747318, EPI_ISL_2747319, EPI_ISL_2747320, EPI_ISL_2747321, EPI_ISL_2747322, EPI_ISL_2747323, EPI_ISL_2747324, EPI_ISL_2747325, EPI_ISL_2747326, EPI_ISL_2747328, EPI_ISL_2747329, EPI_ISL_2747330, EPI_ISL_2747331, EPI_ISL_2747332, EPI_ISL_2747333, EPI_ISL_2747334, EPI_ISL_2747335, EPI_ISL_2747336, EPI_ISL_2747337, EPI_ISL_2747338, EPI_ISL_2747339, EPI_ISL_2747340, EPI_ISL_2747341, EPI_ISL_2747342, EPI_ISL_2747343, EPI_ISL_2747344, EPI_ISL_2747345, EPI_ISL_2747347, EPI_ISL_2747348, EPI_ISL_2747349, EPI_ISL_2747350, EPI_ISL_2747351, EPI_ISL_2747352, EPI_ISL_2747353, EPI_ISL_2747354, EPI_ISL_2747355, EPI_ISL_2747356, EPI_ISL_2747357, EPI_ISL_2747358, EPI_ISL_2747359, EPI_ISL_2747360, EPI_ISL_2747361, EPI_ISL_2747362, EPI_ISL_2747363, EPI_ISL_2747364, EPI_ISL_2747365, EPI_ISL_2747366, EPI_ISL_2747367, EPI_ISL_2747368, EPI_ISL_2747369, EPI_ISL_2747370, EPI_ISL_2747371, EPI_ISL_2747372, EPI_ISL_2747373, EPI_ISL_2747374, EPI_ISL_2747375, EPI_ISL_2747376, EPI_ISL_2747377, EPI_ISL_2747378, EPI_ISL_2747379, EPI_ISL_2747380, EPI_ISL_2747381, EPI_ISL_2747382, EPI_ISL_2747383, EPI_ISL_2747384, EPI_ISL_2747385, EPI_ISL_2747386, EPI_ISL_2747387, EPI_ISL_2747388, EPI_ISL_2747389, EPI_ISL_2747390, EPI_ISL_2747392, EPI_ISL_2747393, EPI_ISL_2747394, EPI_ISL_2747395, EPI_ISL_2747396, EPI_ISL_2747397, EPI_ISL_2747398, EPI_ISL_2747399, EPI_ISL_2747400, EPI_ISL_2747401, EPI_ISL_2747402, EPI_ISL_2747403, EPI_ISL_2747404, EPI_ISL_2747405, EPI_ISL_2747406, EPI_ISL_2747407, EPI_ISL_2747408, EPI_ISL_2747409, EPI_ISL_2747410, EPI_ISL_2747411, EPI_ISL_2747412, EPI_ISL_2747413, EPI_ISL_2747414, EPI_ISL_2747415, EPI_ISL_2747416, EPI_ISL_2747417, EPI_ISL_2747418, EPI_ISL_2747419, EPI_ISL_2747420, EPI_ISL_2747421, EPI_ISL_2747422, EPI_ISL_2747423, EPI_ISL_2747425, EPI_ISL_2747426, EPI_ISL_2747427, EPI_ISL_2747428, EPI_ISL_2747429, EPI_ISL_2747430, EPI_ISL_2747431, EPI_ISL_2747432, EPI_ISL_2747433, EPI_ISL_2747434, EPI_ISL_2747435, EPI_ISL_2747436, EPI_ISL_2747437, EPI_ISL_2747438, EPI_ISL_2747439, EPI_ISL_2747440, EPI_ISL_2747441, EPI_ISL_2747442, EPI_ISL_2747443, EPI_ISL_2747444, EPI_ISL_2747445, EPI_ISL_2747447, EPI_ISL_2747448, EPI_ISL_2747449, EPI_ISL_2747450, EPI_ISL_2747451, EPI_ISL_2747452, EPI_ISL_2747453, EPI_ISL_2747454, EPI_ISL_2747455, EPI_ISL_2747456, EPI_ISL_2747457, EPI_ISL_2747459, EPI_ISL_2747461

| | | | |
|---|---|---|---|
| see above | Respiratory Virus Unit, Microbiology Services Colindale, Public Health England | COVID-19 Genomics UK (COG-UK) Consortium | PHE Covid Sequencing Team |
| EPI_ISL_2747462 | Department of Bacteria, Parasites and Fungi, Statens Serum Institut, Copenhagen, Denmark | Statens Serum Institut Bioinformatics and Microbial Genomics | Danish Covid-19 Genome Consortium |

EPI_ISL_2747464, EPI_ISL_2747465, EPI_ISL_2747466, EPI_ISL_2747467, EPI_ISL_2747469, EPI_ISL_2747471, EPI_ISL_2747473, EPI_ISL_2747475, EPI_ISL_2747477, EPI_ISL_2747479, EPI_ISL_2747481, EPI_ISL_2747483, EPI_ISL_2747485, EPI_ISL_2747487

| | | | |
|---|---|---|---|
| see above | Respiratory Virus Unit, Microbiology Services Colindale, Public Health England | COVID-19 Genomics UK (COG-UK) Consortium | PHE Covid Sequencing Team |
| EPI_ISL_2747488 | Department of Bacteria, Parasites and Fungi, Statens Serum Institut, Copenhagen, Denmark | Statens Serum Institut Bioinformatics and Microbial Genomics | Danish Covid-19 Genome Consortium |
| EPI_ISL_2747489, EPI_ISL_2747490, EPI_ISL_2747492, EPI_ISL_2747495, EPI_ISL_2747497, EPI_ISL_2747499, EPI_ISL_2747501 | Respiratory Virus Unit, Microbiology Services Colindale, Public Health England | COVID-19 Genomics UK (COG-UK) Consortium | PHE Covid Sequencing Team |
| EPI_ISL_2747502 | Department of Bacteria, Parasites and Fungi, Statens Serum Institut, Copenhagen, Denmark | Statens Serum Institut Bioinformatics and Microbial Genomics | Danish Covid-19 Genome Consortium |
| EPI_ISL_2747503, EPI_ISL_2747505 | Respiratory Virus Unit, Microbiology Services Colindale, Public Health England | COVID-19 Genomics UK (COG-UK) Consortium | PHE Covid Sequencing Team |
| EPI_ISL_2747506 | Department of Bacteria, Parasites and Fungi, Statens Serum Institut, Copenhagen, Denmark | Statens Serum Institut Bioinformatics and Microbial Genomics | Danish Covid-19 Genome Consortium |
| EPI_ISL_2747507, EPI_ISL_2747509, EPI_ISL_2747511, EPI_ISL_2747513, EPI_ISL_2747515, EPI_ISL_2747516, EPI_ISL_2747518, EPI_ISL_2747520 | Respiratory Virus Unit, Microbiology Services Colindale, Public Health England | COVID-19 Genomics UK (COG-UK) Consortium | PHE Covid Sequencing Team |
| EPI_ISL_2747522 | Department of Bacteria, Parasites and Fungi, Statens Serum Institut, Copenhagen, Denmark | Statens Serum Institut Bioinformatics and Microbial Genomics | Danish Covid-19 Genome Consortium |
| EPI_ISL_2747523, EPI_ISL_2747525, EPI_ISL_2747527 | Respiratory Virus Unit, Microbiology Services Colindale, Public Health England | COVID-19 Genomics UK (COG-UK) Consortium | PHE Covid Sequencing Team |
| EPI_ISL_2747528 | Department of Bacteria, Parasites and Fungi, Statens Serum Institut, Copenhagen, Denmark | Statens Serum Institut Bioinformatics and Microbial Genomics | Danish Covid-19 Genome Consortium |
| EPI_ISL_2747529, EPI_ISL_2747530, EPI_ISL_2747533, EPI_ISL_2747535, EPI_ISL_2747537, EPI_ISL_2747539, EPI_ISL_2747541 | Respiratory Virus Unit, Microbiology Services Colindale, Public Health England | COVID-19 Genomics UK (COG-UK) Consortium | PHE Covid Sequencing Team |
| EPI_ISL_2747542 | Department of Bacteria, Parasites and Fungi, Statens Serum Institut, Copenhagen, Denmark | Statens Serum Institut Bioinformatics and Microbial Genomics | Danish Covid-19 Genome Consortium |
| EPI_ISL_2747543, EPI_ISL_2747545 | Respiratory Virus Unit, Microbiology Services Colindale, Public Health England | COVID-19 Genomics UK (COG-UK) Consortium | PHE Covid Sequencing Team |
| EPI_ISL_2747546 | Department of Bacteria, Parasites and Fungi, Statens Serum Institut, Copenhagen, Denmark | Statens Serum Institut Bioinformatics and Microbial Genomics | Danish Covid-19 Genome Consortium |
| EPI_ISL_2747547 | Respiratory Virus Unit, Microbiology Services Colindale, Public Health England | COVID-19 Genomics UK (COG-UK) Consortium | PHE Covid Sequencing Team |
| EPI_ISL_2747548 | Department of Bacteria, Parasites and Fungi, Statens Serum Institut, Copenhagen, Denmark | Statens Serum Institut Bioinformatics and Microbial Genomics | Danish Covid-19 Genome Consortium |
| EPI_ISL_2747549 | Respiratory Virus Unit, Microbiology Services Colindale, Public Health England | COVID-19 Genomics UK (COG-UK) Consortium | PHE Covid Sequencing Team |
| EPI_ISL_2747550 | Department of Bacteria, Parasites and Fungi, Statens Serum Institut, Copenhagen, Denmark | Statens Serum Institut Bioinformatics and Microbial Genomics | Danish Covid-19 Genome Consortium |
| EPI_ISL_2747551 | Respiratory Virus Unit, Microbiology Services Colindale, Public Health England | COVID-19 Genomics UK (COG-UK) Consortium | PHE Covid Sequencing Team |
| EPI_ISL_2747556 | Department of Bacteria, Parasites and Fungi, Statens Serum Institut, Copenhagen, Denmark | Statens Serum Institut Bioinformatics and Microbial Genomics | Danish Covid-19 Genome Consortium |
| EPI_ISL_2747557 | Respiratory Virus Unit, Microbiology Services Colindale, Public Health England | COVID-19 Genomics UK (COG-UK) Consortium | PHE Covid Sequencing Team |
| EPI_ISL_2747559 | Department of Bacteria, Parasites and Fungi, Statens Serum Institut, Copenhagen, Denmark | Statens Serum Institut Bioinformatics and Microbial Genomics | Danish Covid-19 Genome Consortium |
| EPI_ISL_2747560 | Respiratory Virus Unit, Microbiology Services Colindale, Public Health England | COVID-19 Genomics UK (COG-UK) Consortium | PHE Covid Sequencing Team |
| EPI_ISL_2747561 | Department of Bacteria, Parasites and Fungi, Statens Serum Institut, Copenhagen, Denmark | Statens Serum Institut Bioinformatics and Microbial Genomics | Danish Covid-19 Genome Consortium |
| EPI_ISL_2747562 | Respiratory Virus Unit, Microbiology Services Colindale, Public Health England | COVID-19 Genomics UK (COG-UK) Consortium | PHE Covid Sequencing Team |

| | | | |
|---|---|---|---|
| EPI_ISL_2747563 | Department of Bacteria, Parasites and Fungi, Statens Serum Institut, Copenhagen, Denmark | Statens Serum Institut Bioinformatics and Microbial Genomics | Danish Covid-19 Genome Consortium |
| EPI_ISL_2747564 | Respiratory Virus Unit, Microbiology Services Colindale, Public Health England | COVID-19 Genomics UK (COG-UK) Consortium | PHE Covid Sequencing Team |
| EPI_ISL_2747565, EPI_ISL_2747566 | Department of Bacteria, Parasites and Fungi, Statens Serum Institut, Copenhagen, Denmark | Statens Serum Institut Bioinformatics and Microbial Genomics | Danish Covid-19 Genome Consortium |
| EPI_ISL_2747567, EPI_ISL_2747568 | Respiratory Virus Unit, Microbiology Services Colindale, Public Health England | COVID-19 Genomics UK (COG-UK) Consortium | PHE Covid Sequencing Team |
| EPI_ISL_2747569 | Department of Bacteria, Parasites and Fungi, Statens Serum Institut, Copenhagen, Denmark | Statens Serum Institut Bioinformatics and Microbial Genomics | Danish Covid-19 Genome Consortium |
| EPI_ISL_2747570 | Respiratory Virus Unit, Microbiology Services Colindale, Public Health England | COVID-19 Genomics UK (COG-UK) Consortium | PHE Covid Sequencing Team |
| EPI_ISL_2747571 | Department of Bacteria, Parasites and Fungi, Statens Serum Institut, Copenhagen, Denmark | Statens Serum Institut Bioinformatics and Microbial Genomics | Danish Covid-19 Genome Consortium |
| EPI_ISL_2747573, EPI_ISL_2747575, EPI_ISL_2747577, EPI_ISL_2747579 | Respiratory Virus Unit, Microbiology Services Colindale, Public Health England | COVID-19 Genomics UK (COG-UK) Consortium | PHE Covid Sequencing Team |
| EPI_ISL_2747580 | Department of Bacteria, Parasites and Fungi, Statens Serum Institut, Copenhagen, Denmark | Statens Serum Institut Bioinformatics and Microbial Genomics | Danish Covid-19 Genome Consortium |
| EPI_ISL_2747581 | Respiratory Virus Unit, Microbiology Services Colindale, Public Health England | COVID-19 Genomics UK (COG-UK) Consortium | PHE Covid Sequencing Team |
| EPI_ISL_2747582 | Department of Bacteria, Parasites and Fungi, Statens Serum Institut, Copenhagen, Denmark | Statens Serum Institut Bioinformatics and Microbial Genomics | Danish Covid-19 Genome Consortium |
| EPI_ISL_2747583, EPI_ISL_2747585 | Respiratory Virus Unit, Microbiology Services Colindale, Public Health England | COVID-19 Genomics UK (COG-UK) Consortium | PHE Covid Sequencing Team |
| EPI_ISL_2747586 | Department of Bacteria, Parasites and Fungi, Statens Serum Institut, Copenhagen, Denmark | Statens Serum Institut Bioinformatics and Microbial Genomics | Danish Covid-19 Genome Consortium |
| EPI_ISL_2747587, EPI_ISL_2747589, EPI_ISL_2747592, EPI_ISL_2747594 | Respiratory Virus Unit, Microbiology Services Colindale, Public Health England | COVID-19 Genomics UK (COG-UK) Consortium | PHE Covid Sequencing Team |
| EPI_ISL_2747595 | Department of Bacteria, Parasites and Fungi, Statens Serum Institut, Copenhagen, Denmark | Statens Serum Institut Bioinformatics and Microbial Genomics | Danish Covid-19 Genome Consortium |
| EPI_ISL_2747596, EPI_ISL_2747598, EPI_ISL_2747600, EPI_ISL_2747602, EPI_ISL_2747604 | Respiratory Virus Unit, Microbiology Services Colindale, Public Health England | COVID-19 Genomics UK (COG-UK) Consortium | PHE Covid Sequencing Team |
| EPI_ISL_2747605 | Department of Bacteria, Parasites and Fungi, Statens Serum Institut, Copenhagen, Denmark | Statens Serum Institut Bioinformatics and Microbial Genomics | Danish Covid-19 Genome Consortium |
| EPI_ISL_2747606, EPI_ISL_2747608, EPI_ISL_2747610, EPI_ISL_2747612, EPI_ISL_2747614, EPI_ISL_2747616, EPI_ISL_2747618, EPI_ISL_2747619 | Respiratory Virus Unit, Microbiology Services Colindale, Public Health England | COVID-19 Genomics UK (COG-UK) Consortium | PHE Covid Sequencing Team |
| EPI_ISL_2747621 | Department of Bacteria, Parasites and Fungi, Statens Serum Institut, Copenhagen, Denmark | Statens Serum Institut Bioinformatics and Microbial Genomics | Danish Covid-19 Genome Consortium |
| EPI_ISL_2747622, EPI_ISL_2747624, EPI_ISL_2747625 | Respiratory Virus Unit, Microbiology Services Colindale, Public Health England | COVID-19 Genomics UK (COG-UK) Consortium | PHE Covid Sequencing Team |
| EPI_ISL_2747626 | Department of Bacteria, Parasites and Fungi, Statens Serum Institut, Copenhagen, Denmark | Statens Serum Institut Bioinformatics and Microbial Genomics | Danish Covid-19 Genome Consortium |
| EPI_ISL_2747627 | Respiratory Virus Unit, Microbiology Services Colindale, Public Health England | COVID-19 Genomics UK (COG-UK) Consortium | PHE Covid Sequencing Team |
| EPI_ISL_2747628 | Department of Bacteria, Parasites and Fungi, Statens Serum Institut, Copenhagen, Denmark | Statens Serum Institut Bioinformatics and Microbial Genomics | Danish Covid-19 Genome Consortium |
| EPI_ISL_2747630 | Respiratory Virus Unit, Microbiology Services Colindale, Public Health England | COVID-19 Genomics UK (COG-UK) Consortium | PHE Covid Sequencing Team |
| EPI_ISL_2747631 | Department of Bacteria, Parasites and Fungi, Statens Serum Institut, Copenhagen, Denmark | Statens Serum Institut Bioinformatics and Microbial Genomics | Danish Covid-19 Genome Consortium |
| EPI_ISL_2747632, EPI_ISL_2747634 | Respiratory Virus Unit, Microbiology Services Colindale, Public Health England | COVID-19 Genomics UK (COG-UK) Consortium | PHE Covid Sequencing Team |
| EPI_ISL_2747635 | Department of Bacteria, Parasites and Fungi, Statens Serum Institut, Copenhagen, Denmark | Statens Serum Institut Bioinformatics and Microbial Genomics | Danish Covid-19 Genome Consortium |
| EPI_ISL_2747636, EPI_ISL_2747638 | Respiratory Virus Unit, Microbiology Services Colindale, Public Health England | COVID-19 Genomics UK (COG-UK) Consortium | PHE Covid Sequencing Team |
| EPI_ISL_2747639 | Department of Bacteria, Parasites and Fungi, Statens Serum Institut, Copenhagen, Denmark | Statens Serum Institut Bioinformatics and Microbial Genomics | Danish Covid-19 Genome Consortium |
| EPI_ISL_2747640, EPI_ISL_2747642, EPI_ISL_2747644 | Respiratory Virus Unit, Microbiology Services Colindale, Public Health England | COVID-19 Genomics UK (COG-UK) Consortium | PHE Covid Sequencing Team |
| EPI_ISL_2747646 | Department of Bacteria, Parasites and Fungi, Statens Serum Institut, Copenhagen, Denmark | Statens Serum Institut Bioinformatics and Microbial Genomics | Danish Covid-19 Genome Consortium |
| EPI_ISL_2747647, EPI_ISL_2747649 | Respiratory Virus Unit, Microbiology Services Colindale, Public Health England | COVID-19 Genomics UK (COG-UK) Consortium | PHE Covid Sequencing Team |
| EPI_ISL_2747650 | Department of Bacteria, Parasites and Fungi, Statens Serum Institut, Copenhagen, Denmark | Statens Serum Institut Bioinformatics and Microbial Genomics | Danish Covid-19 Genome Consortium |
| EPI_ISL_2747651 | Respiratory Virus Unit, Microbiology Services Colindale, Public Health England | COVID-19 Genomics UK (COG-UK) Consortium | PHE Covid Sequencing Team |
| EPI_ISL_2747652 | Department of Bacteria, Parasites and Fungi, Statens Serum Institut, Copenhagen, Denmark | Statens Serum Institut Bioinformatics and Microbial Genomics | Danish Covid-19 Genome Consortium |
| EPI_ISL_2747653 | Respiratory Virus Unit, Microbiology Services Colindale, Public Health England | COVID-19 Genomics UK (COG-UK) Consortium | PHE Covid Sequencing Team |
| EPI_ISL_2747654 | Department of Bacteria, Parasites and Fungi, Statens Serum Institut, Copenhagen, Denmark | Statens Serum Institut Bioinformatics and Microbial Genomics | Danish Covid-19 Genome Consortium |
| EPI_ISL_2747655, EPI_ISL_2747657 | Respiratory Virus Unit, Microbiology Services Colindale, Public Health England | COVID-19 Genomics UK (COG-UK) Consortium | PHE Covid Sequencing Team |
| EPI_ISL_2747658 | Department of Bacteria, Parasites and Fungi, Statens Serum Institut, Copenhagen, Denmark | Statens Serum Institut Bioinformatics and Microbial Genomics | Danish Covid-19 Genome Consortium |

| | | | |
|---|---|---|---|
| EPI_ISL_2747659, EPI_ISL_2747660 | Respiratory Virus Unit, Microbiology Services Colindale, Public Health England | COVID-19 Genomics UK (COG-UK) Consortium | PHE Covid Sequencing Team |
| EPI_ISL_2747661 | Department of Bacteria, Parasites and Fungi, Statens Serum Institut, Copenhagen, Denmark | Statens Serum Institut Bioinformatics and Microbial Genomics | Danish Covid-19 Genome Consortium |
| EPI_ISL_2747662, EPI_ISL_2747665, EPI_ISL_2747667, EPI_ISL_2747669, EPI_ISL_2747673 | Respiratory Virus Unit, Microbiology Services Colindale, Public Health England | COVID-19 Genomics UK (COG-UK) Consortium | PHE Covid Sequencing Team |
| EPI_ISL_2747674 | Department of Bacteria, Parasites and Fungi, Statens Serum Institut, Copenhagen, Denmark | Statens Serum Institut Bioinformatics and Microbial Genomics | Danish Covid-19 Genome Consortium |
| EPI_ISL_2747675, EPI_ISL_2747677, EPI_ISL_2747679, EPI_ISL_2747681, EPI_ISL_2747683, EPI_ISL_2747685, EPI_ISL_2747687 | Respiratory Virus Unit, Microbiology Services Colindale, Public Health England | COVID-19 Genomics UK (COG-UK) Consortium | PHE Covid Sequencing Team |
| EPI_ISL_2747688 | Department of Clinical Microbiology and Center for Genomic Medicine, Rigshospitalet, Copenhagen, Denmark | Statens Serum Institut Bioinformatics and Microbial Genomics | Danish Covid-19 Genome Consortium |
| EPI_ISL_2747689, EPI_ISL_2747691 | Respiratory Virus Unit, Microbiology Services Colindale, Public Health England | COVID-19 Genomics UK (COG-UK) Consortium | PHE Covid Sequencing Team |
| EPI_ISL_2747692 | Department of Bacteria, Parasites and Fungi, Statens Serum Institut, Copenhagen, Denmark | Statens Serum Institut Bioinformatics and Microbial Genomics | Danish Covid-19 Genome Consortium |
| EPI_ISL_2747693 | Respiratory Virus Unit, Microbiology Services Colindale, Public Health England | COVID-19 Genomics UK (COG-UK) Consortium | PHE Covid Sequencing Team |
| EPI_ISL_2747694 | Department of Bacteria, Parasites and Fungi, Statens Serum Institut, Copenhagen, Denmark | Statens Serum Institut Bioinformatics and Microbial Genomics | Danish Covid-19 Genome Consortium |
| EPI_ISL_2747695, EPI_ISL_2747697, EPI_ISL_2747699 | Respiratory Virus Unit, Microbiology Services Colindale, Public Health England | COVID-19 Genomics UK (COG-UK) Consortium | PHE Covid Sequencing Team |
| EPI_ISL_2747700 | Department of Bacteria, Parasites and Fungi, Statens Serum Institut, Copenhagen, Denmark | Statens Serum Institut Bioinformatics and Microbial Genomics | Danish Covid-19 Genome Consortium |
| EPI_ISL_2747701 | Respiratory Virus Unit, Microbiology Services Colindale, Public Health England | COVID-19 Genomics UK (COG-UK) Consortium | PHE Covid Sequencing Team |
| EPI_ISL_2747702 | Department of Bacteria, Parasites and Fungi, Statens Serum Institut, Copenhagen, Denmark | Statens Serum Institut Bioinformatics and Microbial Genomics | Danish Covid-19 Genome Consortium |
| EPI_ISL_2747703, EPI_ISL_2747705 | Respiratory Virus Unit, Microbiology Services Colindale, Public Health England | COVID-19 Genomics UK (COG-UK) Consortium | PHE Covid Sequencing Team |
| EPI_ISL_2747706 | Department of Bacteria, Parasites and Fungi, Statens Serum Institut, Copenhagen, Denmark | Statens Serum Institut Bioinformatics and Microbial Genomics | Danish Covid-19 Genome Consortium |
| EPI_ISL_2747707, EPI_ISL_2747709, EPI_ISL_2747711 | Respiratory Virus Unit, Microbiology Services Colindale, Public Health England | COVID-19 Genomics UK (COG-UK) Consortium | PHE Covid Sequencing Team |
| EPI_ISL_2747712 | Department of Bacteria, Parasites and Fungi, Statens Serum Institut, Copenhagen, Denmark | Statens Serum Institut Bioinformatics and Microbial Genomics | Danish Covid-19 Genome Consortium |
| EPI_ISL_2747713, EPI_ISL_2747715, EPI_ISL_2747717, EPI_ISL_2747719, EPI_ISL_2747721, EPI_ISL_2747723, EPI_ISL_2747725, EPI_ISL_2747727 | Respiratory Virus Unit, Microbiology Services Colindale, Public Health England | COVID-19 Genomics UK (COG-UK) Consortium | PHE Covid Sequencing Team |
| EPI_ISL_2747728 | Department of Bacteria, Parasites and Fungi, Statens Serum Institut, Copenhagen, Denmark | Statens Serum Institut Bioinformatics and Microbial Genomics | Danish Covid-19 Genome Consortium |
| EPI_ISL_2747729, EPI_ISL_2747731 | Respiratory Virus Unit, Microbiology Services Colindale, Public Health England | COVID-19 Genomics UK (COG-UK) Consortium | PHE Covid Sequencing Team |
| EPI_ISL_2747732 | Department of Bacteria, Parasites and Fungi, Statens Serum Institut, Copenhagen, Denmark | Statens Serum Institut Bioinformatics and Microbial Genomics | Danish Covid-19 Genome Consortium |
| EPI_ISL_2747733, EPI_ISL_2747735 | Respiratory Virus Unit, Microbiology Services Colindale, Public Health England | COVID-19 Genomics UK (COG-UK) Consortium | PHE Covid Sequencing Team |
| EPI_ISL_2747736 | Department of Bacteria, Parasites and Fungi, Statens Serum Institut, Copenhagen, Denmark | Statens Serum Institut Bioinformatics and Microbial Genomics | Danish Covid-19 Genome Consortium |
| EPI_ISL_2747737, EPI_ISL_2747739 | Respiratory Virus Unit, Microbiology Services Colindale, Public Health England | COVID-19 Genomics UK (COG-UK) Consortium | PHE Covid Sequencing Team |
| EPI_ISL_2747740 | Department of Bacteria, Parasites and Fungi, Statens Serum Institut, Copenhagen, Denmark | Statens Serum Institut Bioinformatics and Microbial Genomics | Danish Covid-19 Genome Consortium |
| EPI_ISL_2747741 | Respiratory Virus Unit, Microbiology Services Colindale, Public Health England | COVID-19 Genomics UK (COG-UK) Consortium | PHE Covid Sequencing Team |
| EPI_ISL_2747742 | Department of Bacteria, Parasites and Fungi, Statens Serum Institut, Copenhagen, Denmark | Statens Serum Institut Bioinformatics and Microbial Genomics | Danish Covid-19 Genome Consortium |
| EPI_ISL_2747743, EPI_ISL_2747745 | Respiratory Virus Unit, Microbiology Services Colindale, Public Health England | COVID-19 Genomics UK (COG-UK) Consortium | PHE Covid Sequencing Team |
| EPI_ISL_2747746 | Department of Bacteria, Parasites and Fungi, Statens Serum Institut, Copenhagen, Denmark | Statens Serum Institut Bioinformatics and Microbial Genomics | Danish Covid-19 Genome Consortium |
| EPI_ISL_2747747 | Respiratory Virus Unit, Microbiology Services Colindale, Public Health England | COVID-19 Genomics UK (COG-UK) Consortium | PHE Covid Sequencing Team |
| EPI_ISL_2747748 | Department of Bacteria, Parasites and Fungi, Statens Serum Institut, Copenhagen, Denmark | Statens Serum Institut Bioinformatics and Microbial Genomics | Danish Covid-19 Genome Consortium |
| EPI_ISL_2747749, EPI_ISL_2747750, EPI_ISL_2747752 | Respiratory Virus Unit, Microbiology Services Colindale, Public Health England | COVID-19 Genomics UK (COG-UK) Consortium | PHE Covid Sequencing Team |
| EPI_ISL_2747754 | Department of Bacteria, Parasites and Fungi, Statens Serum Institut, Copenhagen, Denmark | Statens Serum Institut Bioinformatics and Microbial Genomics | Danish Covid-19 Genome Consortium |
| EPI_ISL_2747755, EPI_ISL_2747756 | Respiratory Virus Unit, Microbiology Services Colindale, Public Health England | COVID-19 Genomics UK (COG-UK) Consortium | PHE Covid Sequencing Team |
| EPI_ISL_2747757 | Department of Bacteria, Parasites and Fungi, Statens Serum Institut, Copenhagen, Denmark | Statens Serum Institut Bioinformatics and Microbial Genomics | Danish Covid-19 Genome Consortium |
| EPI_ISL_2747758, EPI_ISL_2747760, EPI_ISL_2747762, EPI_ISL_2747764 | Respiratory Virus Unit, Microbiology Services Colindale, Public Health England | COVID-19 Genomics UK (COG-UK) Consortium | PHE Covid Sequencing Team |

| | | | |
|---|---|---|---|
| EPI_ISL_2747765 | Department of Bacteria, Parasites and Fungi, Statens Serum Institut, Copenhagen, Denmark | Statens Serum Institut Bioinformatics and Microbial Genomics | Danish Covid-19 Genome Consortium |
| EPI_ISL_2747766 | Respiratory Virus Unit, Microbiology Services Colindale, Public Health England | COVID-19 Genomics UK (COG-UK) Consortium | PHE Covid Sequencing Team |
| EPI_ISL_2747767 | Department of Bacteria, Parasites and Fungi, Statens Serum Institut, Copenhagen, Denmark | Statens Serum Institut Bioinformatics and Microbial Genomics | Danish Covid-19 Genome Consortium |
| EPI_ISL_2747768 | Respiratory Virus Unit, Microbiology Services Colindale, Public Health England | COVID-19 Genomics UK (COG-UK) Consortium | PHE Covid Sequencing Team |
| EPI_ISL_2747769 | Department of Bacteria, Parasites and Fungi, Statens Serum Institut, Copenhagen, Denmark | Statens Serum Institut Bioinformatics and Microbial Genomics | Danish Covid-19 Genome Consortium |
| EPI_ISL_2747770 | Respiratory Virus Unit, Microbiology Services Colindale, Public Health England | COVID-19 Genomics UK (COG-UK) Consortium | PHE Covid Sequencing Team |
| EPI_ISL_2747771 | Department of Bacteria, Parasites and Fungi, Statens Serum Institut, Copenhagen, Denmark | Statens Serum Institut Bioinformatics and Microbial Genomics | Danish Covid-19 Genome Consortium |
| EPI_ISL_2747772, EPI_ISL_2747774, EPI_ISL_2747776, EPI_ISL_2747778 | Respiratory Virus Unit, Microbiology Services Colindale, Public Health England | COVID-19 Genomics UK (COG-UK) Consortium | PHE Covid Sequencing Team |
| EPI_ISL_2747779 | Department of Bacteria, Parasites and Fungi, Statens Serum Institut, Copenhagen, Denmark | Statens Serum Institut Bioinformatics and Microbial Genomics | Danish Covid-19 Genome Consortium |
| EPI_ISL_2747780, EPI_ISL_2747782 | Respiratory Virus Unit, Microbiology Services Colindale, Public Health England | COVID-19 Genomics UK (COG-UK) Consortium | PHE Covid Sequencing Team |
| EPI_ISL_2747783 | Department of Bacteria, Parasites and Fungi, Statens Serum Institut, Copenhagen, Denmark | Statens Serum Institut Bioinformatics and Microbial Genomics | Danish Covid-19 Genome Consortium |
| EPI_ISL_2747784, EPI_ISL_2747786, EPI_ISL_2747788, EPI_ISL_2747790, EPI_ISL_2747792, EPI_ISL_2747794, EPI_ISL_2747796, EPI_ISL_2747798 | Respiratory Virus Unit, Microbiology Services Colindale, Public Health England | COVID-19 Genomics UK (COG-UK) Consortium | PHE Covid Sequencing Team |
| EPI_ISL_2747799 | Department of Bacteria, Parasites and Fungi, Statens Serum Institut, Copenhagen, Denmark | Statens Serum Institut Bioinformatics and Microbial Genomics | Danish Covid-19 Genome Consortium |
| EPI_ISL_2747800, EPI_ISL_2747802 | Respiratory Virus Unit, Microbiology Services Colindale, Public Health England | COVID-19 Genomics UK (COG-UK) Consortium | PHE Covid Sequencing Team |
| EPI_ISL_2747803 | Department of Bacteria, Parasites and Fungi, Statens Serum Institut, Copenhagen, Denmark | Statens Serum Institut Bioinformatics and Microbial Genomics | Danish Covid-19 Genome Consortium |
| EPI_ISL_2747804, EPI_ISL_2747806 | Respiratory Virus Unit, Microbiology Services Colindale, Public Health England | COVID-19 Genomics UK (COG-UK) Consortium | PHE Covid Sequencing Team |
| EPI_ISL_2747807 | Department of Bacteria, Parasites and Fungi, Statens Serum Institut, Copenhagen, Denmark | Statens Serum Institut Bioinformatics and Microbial Genomics | Danish Covid-19 Genome Consortium |
| EPI_ISL_2747808 | Respiratory Virus Unit, Microbiology Services Colindale, Public Health England | COVID-19 Genomics UK (COG-UK) Consortium | PHE Covid Sequencing Team |
| EPI_ISL_2747809 | Department of Bacteria, Parasites and Fungi, Statens Serum Institut, Copenhagen, Denmark | Statens Serum Institut Bioinformatics and Microbial Genomics | Danish Covid-19 Genome Consortium |
| EPI_ISL_2747810 | Respiratory Virus Unit, Microbiology Services Colindale, Public Health England | COVID-19 Genomics UK (COG-UK) Consortium | PHE Covid Sequencing Team |
| EPI_ISL_2747811 | Department of Bacteria, Parasites and Fungi, Statens Serum Institut, Copenhagen, Denmark | Statens Serum Institut Bioinformatics and Microbial Genomics | Danish Covid-19 Genome Consortium |
| EPI_ISL_2747812 | Respiratory Virus Unit, Microbiology Services Colindale, Public Health England | COVID-19 Genomics UK (COG-UK) Consortium | PHE Covid Sequencing Team |
| EPI_ISL_2747813 | Department of Bacteria, Parasites and Fungi, Statens Serum Institut, Copenhagen, Denmark | Statens Serum Institut Bioinformatics and Microbial Genomics | Danish Covid-19 Genome Consortium |
| EPI_ISL_2747814 | Respiratory Virus Unit, Microbiology Services Colindale, Public Health England | COVID-19 Genomics UK (COG-UK) Consortium | PHE Covid Sequencing Team |
| EPI_ISL_2747815 | Department of Bacteria, Parasites and Fungi, Statens Serum Institut, Copenhagen, Denmark | Statens Serum Institut Bioinformatics and Microbial Genomics | Danish Covid-19 Genome Consortium |
| EPI_ISL_2747816, EPI_ISL_2747818, EPI_ISL_2747820 | Respiratory Virus Unit, Microbiology Services Colindale, Public Health England | COVID-19 Genomics UK (COG-UK) Consortium | PHE Covid Sequencing Team |
| EPI_ISL_2747821 | Department of Bacteria, Parasites and Fungi, Statens Serum Institut, Copenhagen, Denmark | Statens Serum Institut Bioinformatics and Microbial Genomics | Danish Covid-19 Genome Consortium |
| EPI_ISL_2747822 | Respiratory Virus Unit, Microbiology Services Colindale, Public Health England | COVID-19 Genomics UK (COG-UK) Consortium | PHE Covid Sequencing Team |
| EPI_ISL_2747824 | Department of Bacteria, Parasites and Fungi, Statens Serum Institut, Copenhagen, Denmark | Statens Serum Institut Bioinformatics and Microbial Genomics | Danish Covid-19 Genome Consortium |
| EPI_ISL_2747825 | Respiratory Virus Unit, Microbiology Services Colindale, Public Health England | COVID-19 Genomics UK (COG-UK) Consortium | PHE Covid Sequencing Team |
| EPI_ISL_2747826, EPI_ISL_2747828 | Department of Bacteria, Parasites and Fungi, Statens Serum Institut, Copenhagen, Denmark | Statens Serum Institut Bioinformatics and Microbial Genomics | Danish Covid-19 Genome Consortium |
| EPI_ISL_2747829 | Respiratory Virus Unit, Microbiology Services Colindale, Public Health England | COVID-19 Genomics UK (COG-UK) Consortium | PHE Covid Sequencing Team |
| EPI_ISL_2747830 | Department of Bacteria, Parasites and Fungi, Statens Serum Institut, Copenhagen, Denmark | Statens Serum Institut Bioinformatics and Microbial Genomics | Danish Covid-19 Genome Consortium |
| EPI_ISL_2747831, EPI_ISL_2747832, EPI_ISL_2747834 | Respiratory Virus Unit, Microbiology Services Colindale, Public Health England | COVID-19 Genomics UK (COG-UK) Consortium | PHE Covid Sequencing Team |
| EPI_ISL_2747835 | Department of Bacteria, Parasites and Fungi, Statens Serum Institut, Copenhagen, Denmark | Statens Serum Institut Bioinformatics and Microbial Genomics | Danish Covid-19 Genome Consortium |
| EPI_ISL_2747837 | Respiratory Virus Unit, Microbiology Services Colindale, Public Health England | COVID-19 Genomics UK (COG-UK) Consortium | PHE Covid Sequencing Team |
| EPI_ISL_2747838 | Department of Bacteria, Parasites and Fungi, Statens Serum Institut, Copenhagen, Denmark | Statens Serum Institut Bioinformatics and Microbial Genomics | Danish Covid-19 Genome Consortium |
| EPI_ISL_2747839 | Respiratory Virus Unit, Microbiology Services Colindale, Public Health England | COVID-19 Genomics UK (COG-UK) Consortium | PHE Covid Sequencing Team |
| EPI_ISL_2747840 | Department of Bacteria, Parasites and Fungi, Statens Serum Institut, Copenhagen, Denmark | Statens Serum Institut Bioinformatics and Microbial Genomics | Danish Covid-19 Genome Consortium |
| EPI_ISL_2747841, EPI_ISL_2747843, EPI_ISL_2747845, EPI_ISL_2747847 | Respiratory Virus Unit, Microbiology Services Colindale, Public Health England | COVID-19 Genomics UK (COG-UK) Consortium | PHE Covid Sequencing Team |
| EPI_ISL_2747848 | Department of Bacteria, Parasites and Fungi, Statens Serum | Statens Serum Institut Bioinformatics and Microbial Genomics | Danish Covid-19 Genome Consortium |

| | | | |
|---|---|---|---|
| | Institut, Copenhagen, Denmark | | |
| EPI_ISL_2747849, EPI_ISL_2747851, EPI_ISL_2747853 | Respiratory Virus Unit, Microbiology Services Colindale, Public Health England | COVID-19 Genomics UK (COG-UK) Consortium | PHE Covid Sequencing Team |
| EPI_ISL_2747854 | Department of Bacteria, Parasites and Fungi, Statens Serum Institut, Copenhagen, Denmark | Statens Serum Institut Bioinformatics and Microbial Genomics | Danish Covid-19 Genome Consortium |
| EPI_ISL_2747855, EPI_ISL_2747857, EPI_ISL_2747859, EPI_ISL_2747861, EPI_ISL_2747863, EPI_ISL_2747865, EPI_ISL_2747867, EPI_ISL_2747869, EPI_ISL_2747871, EPI_ISL_2747873, EPI_ISL_2747875, EPI_ISL_2747877, EPI_ISL_2747878 | | | |
| see above | Respiratory Virus Unit, Microbiology Services Colindale, Public Health England | COVID-19 Genomics UK (COG-UK) Consortium | PHE Covid Sequencing Team |
| EPI_ISL_2747879 | Department of Bacteria, Parasites and Fungi, Statens Serum Institut, Copenhagen, Denmark | Statens Serum Institut Bioinformatics and Microbial Genomics | Danish Covid-19 Genome Consortium |
| EPI_ISL_2747880 | Respiratory Virus Unit, Microbiology Services Colindale, Public Health England | COVID-19 Genomics UK (COG-UK) Consortium | PHE Covid Sequencing Team |
| EPI_ISL_2747881 | Department of Bacteria, Parasites and Fungi, Statens Serum Institut, Copenhagen, Denmark | Statens Serum Institut Bioinformatics and Microbial Genomics | Danish Covid-19 Genome Consortium |
| EPI_ISL_2747882, EPI_ISL_2747884 | Respiratory Virus Unit, Microbiology Services Colindale, Public Health England | COVID-19 Genomics UK (COG-UK) Consortium | PHE Covid Sequencing Team |
| EPI_ISL_2747885 | Department of Bacteria, Parasites and Fungi, Statens Serum Institut, Copenhagen, Denmark | Statens Serum Institut Bioinformatics and Microbial Genomics | Danish Covid-19 Genome Consortium |
| EPI_ISL_2747886, EPI_ISL_2747888, EPI_ISL_2747890 | Respiratory Virus Unit, Microbiology Services Colindale, Public Health England | COVID-19 Genomics UK (COG-UK) Consortium | PHE Covid Sequencing Team |
| EPI_ISL_2747891 | Department of Bacteria, Parasites and Fungi, Statens Serum Institut, Copenhagen, Denmark | Statens Serum Institut Bioinformatics and Microbial Genomics | Danish Covid-19 Genome Consortium |
| EPI_ISL_2747892, EPI_ISL_2747894, EPI_ISL_2747896, EPI_ISL_2747898, EPI_ISL_2747900, EPI_ISL_2747902 | Respiratory Virus Unit, Microbiology Services Colindale, Public Health England | COVID-19 Genomics UK (COG-UK) Consortium | PHE Covid Sequencing Team |
| EPI_ISL_2747903 | Department of Bacteria, Parasites and Fungi, Statens Serum Institut, Copenhagen, Denmark | Statens Serum Institut Bioinformatics and Microbial Genomics | Danish Covid-19 Genome Consortium |
| EPI_ISL_2747904 | Respiratory Virus Unit, Microbiology Services Colindale, Public Health England | COVID-19 Genomics UK (COG-UK) Consortium | PHE Covid Sequencing Team |
| EPI_ISL_2747905 | Department of Bacteria, Parasites and Fungi, Statens Serum Institut, Copenhagen, Denmark | Statens Serum Institut Bioinformatics and Microbial Genomics | Danish Covid-19 Genome Consortium |
| EPI_ISL_2747906 | Respiratory Virus Unit, Microbiology Services Colindale, Public Health England | COVID-19 Genomics UK (COG-UK) Consortium | PHE Covid Sequencing Team |
| EPI_ISL_2747907 | Department of Bacteria, Parasites and Fungi, Statens Serum Institut, Copenhagen, Denmark | Statens Serum Institut Bioinformatics and Microbial Genomics | Danish Covid-19 Genome Consortium |
| EPI_ISL_2747908, EPI_ISL_2747910, EPI_ISL_2747912 | Respiratory Virus Unit, Microbiology Services Colindale, Public Health England | COVID-19 Genomics UK (COG-UK) Consortium | PHE Covid Sequencing Team |
| EPI_ISL_2747913 | Department of Bacteria, Parasites and Fungi, Statens Serum Institut, Copenhagen, Denmark | Statens Serum Institut Bioinformatics and Microbial Genomics | Danish Covid-19 Genome Consortium |
| EPI_ISL_2747914, EPI_ISL_2747916, EPI_ISL_2747918, EPI_ISL_2747920, EPI_ISL_2747922, EPI_ISL_2747924, EPI_ISL_2747926, EPI_ISL_2747928, EPI_ISL_2747930, EPI_ISL_2747932, EPI_ISL_2747934, EPI_ISL_2747936, EPI_ISL_2747938, EPI_ISL_2747940, EPI_ISL_2747942, EPI_ISL_2747944, EPI_ISL_2747946 | | | |
| see above | Respiratory Virus Unit, Microbiology Services Colindale, Public Health England | COVID-19 Genomics UK (COG-UK) Consortium | PHE Covid Sequencing Team |
| EPI_ISL_2747947 | Department of Bacteria, Parasites and Fungi, Statens Serum Institut, Copenhagen, Denmark | Statens Serum Institut Bioinformatics and Microbial Genomics | Danish Covid-19 Genome Consortium |
| EPI_ISL_2747948, EPI_ISL_2747951, EPI_ISL_2747953, EPI_ISL_2747955 | Respiratory Virus Unit, Microbiology Services Colindale, Public Health England | COVID-19 Genomics UK (COG-UK) Consortium | PHE Covid Sequencing Team |
| EPI_ISL_2747956 | Department of Bacteria, Parasites and Fungi, Statens Serum Institut, Copenhagen, Denmark | Statens Serum Institut Bioinformatics and Microbial Genomics | Danish Covid-19 Genome Consortium |
| EPI_ISL_2747957, EPI_ISL_2747959 | Respiratory Virus Unit, Microbiology Services Colindale, Public Health England | COVID-19 Genomics UK (COG-UK) Consortium | PHE Covid Sequencing Team |
| EPI_ISL_2747960 | Department of Bacteria, Parasites and Fungi, Statens Serum Institut, Copenhagen, Denmark | Statens Serum Institut Bioinformatics and Microbial Genomics | Danish Covid-19 Genome Consortium |
| EPI_ISL_2747961 | Respiratory Virus Unit, Microbiology Services Colindale, Public Health England | COVID-19 Genomics UK (COG-UK) Consortium | PHE Covid Sequencing Team |
| EPI_ISL_2747962 | Department of Bacteria, Parasites and Fungi, Statens Serum Institut, Copenhagen, Denmark | Statens Serum Institut Bioinformatics and Microbial Genomics | Danish Covid-19 Genome Consortium |
| EPI_ISL_2747963 | Respiratory Virus Unit, Microbiology Services Colindale, Public Health England | COVID-19 Genomics UK (COG-UK) Consortium | PHE Covid Sequencing Team |
| EPI_ISL_2747964 | Department of Bacteria, Parasites and Fungi, Statens Serum Institut, Copenhagen, Denmark | Statens Serum Institut Bioinformatics and Microbial Genomics | Danish Covid-19 Genome Consortium |
| EPI_ISL_2747965, EPI_ISL_2747967 | Respiratory Virus Unit, Microbiology Services Colindale, Public Health England | COVID-19 Genomics UK (COG-UK) Consortium | PHE Covid Sequencing Team |
| EPI_ISL_2747968 | Department of Bacteria, Parasites and Fungi, Statens Serum Institut, Copenhagen, Denmark | Statens Serum Institut Bioinformatics and Microbial Genomics | Danish Covid-19 Genome Consortium |
| EPI_ISL_2747969 | Respiratory Virus Unit, Microbiology Services Colindale, Public Health England | COVID-19 Genomics UK (COG-UK) Consortium | PHE Covid Sequencing Team |
| EPI_ISL_2747970 | Department of Bacteria, Parasites and Fungi, Statens Serum Institut, Copenhagen, Denmark | Statens Serum Institut Bioinformatics and Microbial Genomics | Danish Covid-19 Genome Consortium |
| EPI_ISL_2747971 | Respiratory Virus Unit, Microbiology Services Colindale, Public Health England | COVID-19 Genomics UK (COG-UK) Consortium | PHE Covid Sequencing Team |
| EPI_ISL_2747972 | Department of Bacteria, Parasites and Fungi, Statens Serum Institut, Copenhagen, Denmark | Statens Serum Institut Bioinformatics and Microbial Genomics | Danish Covid-19 Genome Consortium |
| EPI_ISL_2747973 | Respiratory Virus Unit, Microbiology Services Colindale, Public Health England | COVID-19 Genomics UK (COG-UK) Consortium | PHE Covid Sequencing Team |
| EPI_ISL_2747974 | Department of Bacteria, Parasites and Fungi, Statens Serum Institut, Copenhagen, Denmark | Statens Serum Institut Bioinformatics and Microbial Genomics | Danish Covid-19 Genome Consortium |
| EPI_ISL_2747975, EPI_ISL_2747977, EPI_ISL_2747979 | Respiratory Virus Unit, Microbiology Services Colindale, Public Health England | COVID-19 Genomics UK (COG-UK) Consortium | PHE Covid Sequencing Team |
| EPI_ISL_2747981, EPI_ISL_2747983, EPI_ISL_2747985 | Department of Bacteria, Parasites and Fungi, Statens Serum Institut, Copenhagen, Denmark | Statens Serum Institut Bioinformatics and Microbial Genomics | Danish Covid-19 Genome Consortium |
| EPI_ISL_2747987, EPI_ISL_2747990, | Respiratory Virus Unit, Microbiology Services Colindale, Public Health England | COVID-19 Genomics UK (COG-UK) Consortium | PHE Covid Sequencing Team |

| | | | |
|---|---|---|---|
| EPI_ISL_2747992, EPI_ISL_2747994, EPI_ISL_2747996, EPI_ISL_2747998, EPI_ISL_2748000, EPI_ISL_2748002, EPI_ISL_2748004 | | | |
| EPI_ISL_2748005 | Department of Bacteria, Parasites and Fungi, Statens Serum Institut, Copenhagen, Denmark | Statens Serum Institut Bioinformatics and Microbial Genomics | Danish Covid-19 Genome Consortium |
| EPI_ISL_2748006 | Respiratory Virus Unit, Microbiology Services Colindale, Public Health England | COVID-19 Genomics UK (COG-UK) Consortium | PHE Covid Sequencing Team |
| EPI_ISL_2748007 | Department of Bacteria, Parasites and Fungi, Statens Serum Institut, Copenhagen, Denmark | Statens Serum Institut Bioinformatics and Microbial Genomics | Danish Covid-19 Genome Consortium |
| EPI_ISL_2748008, EPI_ISL_2748009 | Respiratory Virus Unit, Microbiology Services Colindale, Public Health England | COVID-19 Genomics UK (COG-UK) Consortium | PHE Covid Sequencing Team |
| EPI_ISL_2748010 | Department of Bacteria, Parasites and Fungi, Statens Serum Institut, Copenhagen, Denmark | Statens Serum Institut Bioinformatics and Microbial Genomics | Danish Covid-19 Genome Consortium |
| EPI_ISL_2748011, EPI_ISL_2748013, EPI_ISL_2748015, EPI_ISL_2748017, EPI_ISL_2748019 | Respiratory Virus Unit, Microbiology Services Colindale, Public Health England | COVID-19 Genomics UK (COG-UK) Consortium | PHE Covid Sequencing Team |
| EPI_ISL_2748020 | Department of Bacteria, Parasites and Fungi, Statens Serum Institut, Copenhagen, Denmark | Statens Serum Institut Bioinformatics and Microbial Genomics | Danish Covid-19 Genome Consortium |
| EPI_ISL_2748021 | Respiratory Virus Unit, Microbiology Services Colindale, Public Health England | COVID-19 Genomics UK (COG-UK) Consortium | PHE Covid Sequencing Team |
| EPI_ISL_2748022 | Department of Bacteria, Parasites and Fungi, Statens Serum Institut, Copenhagen, Denmark | Statens Serum Institut Bioinformatics and Microbial Genomics | Danish Covid-19 Genome Consortium |
| EPI_ISL_2748023 | Respiratory Virus Unit, Microbiology Services Colindale, Public Health England | COVID-19 Genomics UK (COG-UK) Consortium | PHE Covid Sequencing Team |
| EPI_ISL_2748024 | Department of Bacteria, Parasites and Fungi, Statens Serum Institut, Copenhagen, Denmark | Statens Serum Institut Bioinformatics and Microbial Genomics | Danish Covid-19 Genome Consortium |
| EPI_ISL_2748026, EPI_ISL_2748027 | Respiratory Virus Unit, Microbiology Services Colindale, Public Health England | COVID-19 Genomics UK (COG-UK) Consortium | PHE Covid Sequencing Team |
| EPI_ISL_2748028 | Department of Bacteria, Parasites and Fungi, Statens Serum Institut, Copenhagen, Denmark | Statens Serum Institut Bioinformatics and Microbial Genomics | Danish Covid-19 Genome Consortium |
| EPI_ISL_2748029, EPI_ISL_2748031 | Respiratory Virus Unit, Microbiology Services Colindale, Public Health England | COVID-19 Genomics UK (COG-UK) Consortium | PHE Covid Sequencing Team |
| EPI_ISL_2748032, EPI_ISL_2748033 | Department of Bacteria, Parasites and Fungi, Statens Serum Institut, Copenhagen, Denmark | Statens Serum Institut Bioinformatics and Microbial Genomics | Danish Covid-19 Genome Consortium |
| EPI_ISL_2748034, EPI_ISL_2748036, EPI_ISL_2748038 | Respiratory Virus Unit, Microbiology Services Colindale, Public Health England | COVID-19 Genomics UK (COG-UK) Consortium | PHE Covid Sequencing Team |
| EPI_ISL_2748039 | Department of Bacteria, Parasites and Fungi, Statens Serum Institut, Copenhagen, Denmark | Statens Serum Institut Bioinformatics and Microbial Genomics | Danish Covid-19 Genome Consortium |
| EPI_ISL_2748040 | Respiratory Virus Unit, Microbiology Services Colindale, Public Health England | COVID-19 Genomics UK (COG-UK) Consortium | PHE Covid Sequencing Team |
| EPI_ISL_2748041 | Department of Bacteria, Parasites and Fungi, Statens Serum Institut, Copenhagen, Denmark | Statens Serum Institut Bioinformatics and Microbial Genomics | Danish Covid-19 Genome Consortium |
| EPI_ISL_2748043, EPI_ISL_2748045 | Respiratory Virus Unit, Microbiology Services Colindale, Public Health England | COVID-19 Genomics UK (COG-UK) Consortium | PHE Covid Sequencing Team |
| EPI_ISL_2748047 | Department of Bacteria, Parasites and Fungi, Statens Serum Institut, Copenhagen, Denmark | Statens Serum Institut Bioinformatics and Microbial Genomics | Danish Covid-19 Genome Consortium |
| EPI_ISL_2748048, EPI_ISL_2748050, EPI_ISL_2748052, EPI_ISL_2748054, EPI_ISL_2748056, EPI_ISL_2748058, EPI_ISL_2748062, EPI_ISL_2748064 | Respiratory Virus Unit, Microbiology Services Colindale, Public Health England | COVID-19 Genomics UK (COG-UK) Consortium | PHE Covid Sequencing Team |
| EPI_ISL_2748065 | Department of Bacteria, Parasites and Fungi, Statens Serum Institut, Copenhagen, Denmark | Statens Serum Institut Bioinformatics and Microbial Genomics | Danish Covid-19 Genome Consortium |
| EPI_ISL_2748066, EPI_ISL_2748067, EPI_ISL_2748069, EPI_ISL_2748071, EPI_ISL_2748073, EPI_ISL_2748075, EPI_ISL_2748078, EPI_ISL_2748080, EPI_ISL_2748082, EPI_ISL_2748084, EPI_ISL_2748085 | | | |
| see above | Respiratory Virus Unit, Microbiology Services Colindale, Public Health England | COVID-19 Genomics UK (COG-UK) Consortium | PHE Covid Sequencing Team |
| EPI_ISL_2748087 | Department of Bacteria, Parasites and Fungi, Statens Serum Institut, Copenhagen, Denmark | Statens Serum Institut Bioinformatics and Microbial Genomics | Danish Covid-19 Genome Consortium |
| EPI_ISL_2748088, EPI_ISL_2748089, EPI_ISL_2748092, EPI_ISL_2748093 | Respiratory Virus Unit, Microbiology Services Colindale, Public Health England | COVID-19 Genomics UK (COG-UK) Consortium | PHE Covid Sequencing Team |
| EPI_ISL_2748094, EPI_ISL_2748095 | Department of Bacteria, Parasites and Fungi, Statens Serum Institut, Copenhagen, Denmark | Statens Serum Institut Bioinformatics and Microbial Genomics | Danish Covid-19 Genome Consortium |
| EPI_ISL_2748096, EPI_ISL_2748097, EPI_ISL_2748099 | Respiratory Virus Unit, Microbiology Services Colindale, Public Health England | COVID-19 Genomics UK (COG-UK) Consortium | PHE Covid Sequencing Team |
| EPI_ISL_2748100 | Department of Bacteria, Parasites and Fungi, Statens Serum Institut, Copenhagen, Denmark | Statens Serum Institut Bioinformatics and Microbial Genomics | Danish Covid-19 Genome Consortium |
| EPI_ISL_2748101 | Respiratory Virus Unit, Microbiology Services Colindale, Public Health England | COVID-19 Genomics UK (COG-UK) Consortium | PHE Covid Sequencing Team |
| EPI_ISL_2748103 | Department of Bacteria, Parasites and Fungi, Statens Serum Institut, Copenhagen, Denmark | Statens Serum Institut Bioinformatics and Microbial Genomics | Danish Covid-19 Genome Consortium |
| EPI_ISL_2748104, EPI_ISL_2748106, EPI_ISL_2748108 | Respiratory Virus Unit, Microbiology Services Colindale, Public Health England | COVID-19 Genomics UK (COG-UK) Consortium | PHE Covid Sequencing Team |
| EPI_ISL_2748109 | Department of Bacteria, Parasites and Fungi, Statens Serum Institut, Copenhagen, Denmark | Statens Serum Institut Bioinformatics and Microbial Genomics | Danish Covid-19 Genome Consortium |
| EPI_ISL_2748110 | Respiratory Virus Unit, Microbiology Services Colindale, Public Health England | COVID-19 Genomics UK (COG-UK) Consortium | PHE Covid Sequencing Team |
| EPI_ISL_2748111 | Department of Bacteria, Parasites and Fungi, Statens Serum Institut, Copenhagen, Denmark | Statens Serum Institut Bioinformatics and Microbial Genomics | Danish Covid-19 Genome Consortium |
| EPI_ISL_2748112, | Respiratory Virus Unit, Microbiology Services Colindale, Public | COVID-19 Genomics UK (COG-UK) Consortium | PHE Covid Sequencing Team |

| | | | |
|---|---|---|---|
| EPI_ISL_2748114, EPI_ISL_2748116, EPI_ISL_2748118 | Health England | | |
| EPI_ISL_2748119 | Department of Bacteria, Parasites and Fungi, Statens Serum Institut, Copenhagen, Denmark | Statens Serum Institut Bioinformatics and Microbial Genomics | Danish Covid-19 Genome Consortium |
| EPI_ISL_2748120 | Respiratory Virus Unit, Microbiology Services Colindale, Public Health England | COVID-19 Genomics UK (COG-UK) Consortium | PHE Covid Sequencing Team |
| EPI_ISL_2748121 | Department of Bacteria, Parasites and Fungi, Statens Serum Institut, Copenhagen, Denmark | Statens Serum Institut Bioinformatics and Microbial Genomics | Danish Covid-19 Genome Consortium |
| EPI_ISL_2748122 | Respiratory Virus Unit, Microbiology Services Colindale, Public Health England | COVID-19 Genomics UK (COG-UK) Consortium | PHE Covid Sequencing Team |
| EPI_ISL_2748123 | Department of Bacteria, Parasites and Fungi, Statens Serum Institut, Copenhagen, Denmark | Statens Serum Institut Bioinformatics and Microbial Genomics | Danish Covid-19 Genome Consortium |
| EPI_ISL_2748124 | Respiratory Virus Unit, Microbiology Services Colindale, Public Health England | COVID-19 Genomics UK (COG-UK) Consortium | PHE Covid Sequencing Team |
| EPI_ISL_2748125 | Department of Bacteria, Parasites and Fungi, Statens Serum Institut, Copenhagen, Denmark | Statens Serum Institut Bioinformatics and Microbial Genomics | Danish Covid-19 Genome Consortium |
| EPI_ISL_2748126, EPI_ISL_2748128, EPI_ISL_2748130, EPI_ISL_2748132, EPI_ISL_2748134, EPI_ISL_2748136, EPI_ISL_2748138, EPI_ISL_2748139, EPI_ISL_2748142, EPI_ISL_2748144, EPI_ISL_2748146, EPI_ISL_2748148, EPI_ISL_2748150, EPI_ISL_2748152, EPI_ISL_2748154, EPI_ISL_2748156, EPI_ISL_2748158, EPI_ISL_2748160, EPI_ISL_2748162, EPI_ISL_2748163, EPI_ISL_2748165 | | | |
| see above | Respiratory Virus Unit, Microbiology Services Colindale, Public Health England | COVID-19 Genomics UK (COG-UK) Consortium | PHE Covid Sequencing Team |
| EPI_ISL_2748166 | Department of Bacteria, Parasites and Fungi, Statens Serum Institut, Copenhagen, Denmark | Statens Serum Institut Bioinformatics and Microbial Genomics | Danish Covid-19 Genome Consortium |
| EPI_ISL_2748167 | Respiratory Virus Unit, Microbiology Services Colindale, Public Health England | COVID-19 Genomics UK (COG-UK) Consortium | PHE Covid Sequencing Team |
| EPI_ISL_2748168 | Department of Bacteria, Parasites and Fungi, Statens Serum Institut, Copenhagen, Denmark | Statens Serum Institut Bioinformatics and Microbial Genomics | Danish Covid-19 Genome Consortium |
| EPI_ISL_2748169 | Respiratory Virus Unit, Microbiology Services Colindale, Public Health England | COVID-19 Genomics UK (COG-UK) Consortium | PHE Covid Sequencing Team |
| EPI_ISL_2748170 | Department of Bacteria, Parasites and Fungi, Statens Serum Institut, Copenhagen, Denmark | Statens Serum Institut Bioinformatics and Microbial Genomics | Danish Covid-19 Genome Consortium |
| EPI_ISL_2748171 | Respiratory Virus Unit, Microbiology Services Colindale, Public Health England | COVID-19 Genomics UK (COG-UK) Consortium | PHE Covid Sequencing Team |
| EPI_ISL_2748172 | Department of Bacteria, Parasites and Fungi, Statens Serum Institut, Copenhagen, Denmark | Statens Serum Institut Bioinformatics and Microbial Genomics | Danish Covid-19 Genome Consortium |
| EPI_ISL_2748173 | Respiratory Virus Unit, Microbiology Services Colindale, Public Health England | COVID-19 Genomics UK (COG-UK) Consortium | PHE Covid Sequencing Team |
| EPI_ISL_2748174 | Department of Bacteria, Parasites and Fungi, Statens Serum Institut, Copenhagen, Denmark | Statens Serum Institut Bioinformatics and Microbial Genomics | Danish Covid-19 Genome Consortium |
| EPI_ISL_2748175, EPI_ISL_2748177 | Respiratory Virus Unit, Microbiology Services Colindale, Public Health England | COVID-19 Genomics UK (COG-UK) Consortium | PHE Covid Sequencing Team |
| EPI_ISL_2748178 | Department of Bacteria, Parasites and Fungi, Statens Serum Institut, Copenhagen, Denmark | Statens Serum Institut Bioinformatics and Microbial Genomics | Danish Covid-19 Genome Consortium |
| EPI_ISL_2748179, EPI_ISL_2748180 | Respiratory Virus Unit, Microbiology Services Colindale, Public Health England | COVID-19 Genomics UK (COG-UK) Consortium | PHE Covid Sequencing Team |
| EPI_ISL_2748194, EPI_ISL_2748196, EPI_ISL_2748198, EPI_ISL_2748200 | Department of Bacteria, Parasites and Fungi, Statens Serum Institut, Copenhagen, Denmark | Statens Serum Institut Bioinformatics and Microbial Genomics | Danish Covid-19 Genome Consortium |
| EPI_ISL_2748219 | Respiratory Virus Unit, Microbiology Services Colindale, Public Health England | COVID-19 Genomics UK (COG-UK) Consortium | PHE Covid Sequencing Team |
| EPI_ISL_2748220, EPI_ISL_2748224, EPI_ISL_2748234, EPI_ISL_2748236 | Department of Bacteria, Parasites and Fungi, Statens Serum Institut, Copenhagen, Denmark | Statens Serum Institut Bioinformatics and Microbial Genomics | Danish Covid-19 Genome Consortium |
| EPI_ISL_2748237, EPI_ISL_2748245 | Respiratory Virus Unit, Microbiology Services Colindale, Public Health England | COVID-19 Genomics UK (COG-UK) Consortium | PHE Covid Sequencing Team |
| EPI_ISL_2748256, EPI_ISL_2748259, EPI_ISL_2748263 | Department of Bacteria, Parasites and Fungi, Statens Serum Institut, Copenhagen, Denmark | Statens Serum Institut Bioinformatics and Microbial Genomics | Danish Covid-19 Genome Consortium |
| EPI_ISL_2748278 | Respiratory Virus Unit, Microbiology Services Colindale, Public Health England | COVID-19 Genomics UK (COG-UK) Consortium | PHE Covid Sequencing Team |
| EPI_ISL_2748297, EPI_ISL_2748307, EPI_ISL_2748311, EPI_ISL_2748313 | Department of Bacteria, Parasites and Fungi, Statens Serum Institut, Copenhagen, Denmark | Statens Serum Institut Bioinformatics and Microbial Genomics | Danish Covid-19 Genome Consortium |
| EPI_ISL_2748314 | Respiratory Virus Unit, Microbiology Services Colindale, Public Health England | COVID-19 Genomics UK (COG-UK) Consortium | PHE Covid Sequencing Team |
| EPI_ISL_2748315, EPI_ISL_2748321, EPI_ISL_2748329 | Department of Bacteria, Parasites and Fungi, Statens Serum Institut, Copenhagen, Denmark | Statens Serum Institut Bioinformatics and Microbial Genomics | Danish Covid-19 Genome Consortium |
| EPI_ISL_2748332 | Respiratory Virus Unit, Microbiology Services Colindale, Public Health England | COVID-19 Genomics UK (COG-UK) Consortium | PHE Covid Sequencing Team |
| EPI_ISL_2748341, EPI_ISL_2748345, EPI_ISL_2748348, EPI_ISL_2748354 | Department of Bacteria, Parasites and Fungi, Statens Serum Institut, Copenhagen, Denmark | Statens Serum Institut Bioinformatics and Microbial Genomics | Danish Covid-19 Genome Consortium |
| EPI_ISL_2748364 | Respiratory Virus Unit, Microbiology Services Colindale, Public Health England | COVID-19 Genomics UK (COG-UK) Consortium | PHE Covid Sequencing Team |
| EPI_ISL_2748366 | Department of Bacteria, Parasites and Fungi, Statens Serum Institut, Copenhagen, Denmark | Statens Serum Institut Bioinformatics and Microbial Genomics | Danish Covid-19 Genome Consortium |
| EPI_ISL_2748367 | Respiratory Virus Unit, Microbiology Services Colindale, Public Health England | COVID-19 Genomics UK (COG-UK) Consortium | PHE Covid Sequencing Team |
| EPI_ISL_2748370 | Department of Bacteria, Parasites and Fungi, Statens Serum Institut, Copenhagen, Denmark | Statens Serum Institut Bioinformatics and Microbial Genomics | Danish Covid-19 Genome Consortium |
| EPI_ISL_2748374 | Department of Clinical Microbiology and Center for Genomic Medicine, Rigshospitalet, Copenhagen, Denmark | Statens Serum Institut Bioinformatics and Microbial Genomics | Danish Covid-19 Genome Consortium |
| EPI_ISL_2748378 | Respiratory Virus Unit, Microbiology Services Colindale, Public Health England | COVID-19 Genomics UK (COG-UK) Consortium | PHE Covid Sequencing Team |
| EPI_ISL_2748382, EPI_ISL_2748388, EPI_ISL_2748394, EPI_ISL_2748401, | Department of Bacteria, Parasites and Fungi, Statens Serum Institut, Copenhagen, Denmark | Statens Serum Institut Bioinformatics and Microbial Genomics | Danish Covid-19 Genome Consortium |

| | | | |
|---|---|---|---|
| EPI_ISL_2748403, EPI_ISL_2748411, EPI_ISL_2748418 | | | |
| EPI_ISL_2748419 | Respiratory Virus Unit, Microbiology Services Colindale, Public Health England | COVID-19 Genomics UK (COG-UK) Consortium | PHE Covid Sequencing Team |
| EPI_ISL_2748420 | Department of Bacteria, Parasites and Fungi, Statens Serum Institut, Copenhagen, Denmark | Statens Serum Institut Bioinformatics and Microbial Genomics | Danish Covid-19 Genome Consortium |
| EPI_ISL_2748421 | Respiratory Virus Unit, Microbiology Services Colindale, Public Health England | COVID-19 Genomics UK (COG-UK) Consortium | PHE Covid Sequencing Team |
| EPI_ISL_2748422 | Department of Bacteria, Parasites and Fungi, Statens Serum Institut, Copenhagen, Denmark | Statens Serum Institut Bioinformatics and Microbial Genomics | Danish Covid-19 Genome Consortium |
| EPI_ISL_2748425 | Respiratory Virus Unit, Microbiology Services Colindale, Public Health England | COVID-19 Genomics UK (COG-UK) Consortium | PHE Covid Sequencing Team |
| EPI_ISL_2748428, EPI_ISL_2748429 | Department of Bacteria, Parasites and Fungi, Statens Serum Institut, Copenhagen, Denmark | Statens Serum Institut Bioinformatics and Microbial Genomics | Danish Covid-19 Genome Consortium |
| EPI_ISL_2748430 | Respiratory Virus Unit, Microbiology Services Colindale, Public Health England | COVID-19 Genomics UK (COG-UK) Consortium | PHE Covid Sequencing Team |
| EPI_ISL_2748433, EPI_ISL_2748439, EPI_ISL_2748441, EPI_ISL_2748443, EPI_ISL_2748447 | Department of Bacteria, Parasites and Fungi, Statens Serum Institut, Copenhagen, Denmark | Statens Serum Institut Bioinformatics and Microbial Genomics | Danish Covid-19 Genome Consortium |
| EPI_ISL_2748451, EPI_ISL_2748455, EPI_ISL_2748457, EPI_ISL_2748462, EPI_ISL_2748463, EPI_ISL_2748468, EPI_ISL_2748480, EPI_ISL_2748485, EPI_ISL_2748491, EPI_ISL_2748495, EPI_ISL_2748501, EPI_ISL_2748521, EPI_ISL_2748523, EPI_ISL_2748527, EPI_ISL_2748529, EPI_ISL_2748530, EPI_ISL_2748540, EPI_ISL_2748542, EPI_ISL_2748544, EPI_ISL_2748551, EPI_ISL_2748552, EPI_ISL_2748560, EPI_ISL_2748572, EPI_ISL_2748579, EPI_ISL_2748580, EPI_ISL_2748586, EPI_ISL_2748588, EPI_ISL_2748591, EPI_ISL_2748595, EPI_ISL_2748596, EPI_ISL_2748597, EPI_ISL_2748599, EPI_ISL_2748601, EPI_ISL_2748602, EPI_ISL_2748605, EPI_ISL_2748606, EPI_ISL_2748607, EPI_ISL_2748612, EPI_ISL_2748613, EPI_ISL_2748614, EPI_ISL_2748617, EPI_ISL_2748619, EPI_ISL_2748621, EPI_ISL_2748631, EPI_ISL_2748638, EPI_ISL_2748646, EPI_ISL_2748648, EPI_ISL_2748658, EPI_ISL_2748659, EPI_ISL_2748663, EPI_ISL_2748666, EPI_ISL_2748671, EPI_ISL_2748674, EPI_ISL_2748675, EPI_ISL_2748680, EPI_ISL_2748682, EPI_ISL_2748684, EPI_ISL_2748691, EPI_ISL_2748696, EPI_ISL_2748697, EPI_ISL_2748698, EPI_ISL_2748699, EPI_ISL_2748700, EPI_ISL_2748701, EPI_ISL_2748702, EPI_ISL_2748703, EPI_ISL_2748704, EPI_ISL_2748712, EPI_ISL_2748714, EPI_ISL_2748716, EPI_ISL_2748722, EPI_ISL_2748737, EPI_ISL_2748751, EPI_ISL_2748774, EPI_ISL_2748788, EPI_ISL_2748790, EPI_ISL_2748791, EPI_ISL_2748794, EPI_ISL_2748799, EPI_ISL_2748809, EPI_ISL_2748811, EPI_ISL_2748812, EPI_ISL_2748813, EPI_ISL_2748818, EPI_ISL_2748820, EPI_ISL_2748821, EPI_ISL_2748823, EPI_ISL_2748835, EPI_ISL_2748840, EPI_ISL_2748843, EPI_ISL_2748844, EPI_ISL_2748845, EPI_ISL_2748855, EPI_ISL_2748886, EPI_ISL_2748897, EPI_ISL_2748914, EPI_ISL_2748922, EPI_ISL_2748924, EPI_ISL_2748933, EPI_ISL_2748934, EPI_ISL_2748935, EPI_ISL_2748936, EPI_ISL_2748938, EPI_ISL_2748939, EPI_ISL_2748942, EPI_ISL_2748947, EPI_ISL_2748948, EPI_ISL_2748960, EPI_ISL_2748962, EPI_ISL_2748966, EPI_ISL_2748972, EPI_ISL_2748985, EPI_ISL_2748994, EPI_ISL_2748997, EPI_ISL_2748999, EPI_ISL_2749003, EPI_ISL_2749012, EPI_ISL_2749014, EPI_ISL_2749017, EPI_ISL_2749022, EPI_ISL_2749029, EPI_ISL_2749030, EPI_ISL_2749031, EPI_ISL_2749098, EPI_ISL_2749106, EPI_ISL_2749114, EPI_ISL_2749128, EPI_ISL_2749131, EPI_ISL_2749140, EPI_ISL_2749144, EPI_ISL_2749145, EPI_ISL_2749151, EPI_ISL_2749153, EPI_ISL_2749154, EPI_ISL_2749155, EPI_ISL_2749157, EPI_ISL_2749160, EPI_ISL_2749162, EPI_ISL_2749165, EPI_ISL_2749175, EPI_ISL_2749179, EPI_ISL_2749180, EPI_ISL_2749188, EPI_ISL_2749189, EPI_ISL_2749190, EPI_ISL_2749191, EPI_ISL_2749192, EPI_ISL_2749193, EPI_ISL_2749194, EPI_ISL_2749196, EPI_ISL_2749198, EPI_ISL_2749199, EPI_ISL_2749200, EPI_ISL_2749203, EPI_ISL_2749204, EPI_ISL_2749206, EPI_ISL_2749207, EPI_ISL_2749211, EPI_ISL_2749213, EPI_ISL_2749214, EPI_ISL_2749215, EPI_ISL_2749218, EPI_ISL_2749219, EPI_ISL_2749222, EPI_ISL_2749228, EPI_ISL_2749229, EPI_ISL_2749232, EPI_ISL_2749233, EPI_ISL_2749234, EPI_ISL_2749235, EPI_ISL_2749236, EPI_ISL_2749237, EPI_ISL_2749239, EPI_ISL_2749240, EPI_ISL_2749241, EPI_ISL_2749242, EPI_ISL_2749243, EPI_ISL_2749244, EPI_ISL_2749245, EPI_ISL_2749248, EPI_ISL_2749250, EPI_ISL_2749253, EPI_ISL_2749255, EPI_ISL_2749256, EPI_ISL_2749257, EPI_ISL_2749258, EPI_ISL_2749260, EPI_ISL_2749261, EPI_ISL_2749263, EPI_ISL_2749264, EPI_ISL_2749265, EPI_ISL_2749266, EPI_ISL_2749267, EPI_ISL_2749268, EPI_ISL_2749269, EPI_ISL_2749270, EPI_ISL_2749271, EPI_ISL_2749275, EPI_ISL_2749276, EPI_ISL_2749277, EPI_ISL_2749278, EPI_ISL_2749279, EPI_ISL_2749280, EPI_ISL_2749281, EPI_ISL_2749282, EPI_ISL_2749283, EPI_ISL_2749284, EPI_ISL_2749285, EPI_ISL_2749286, EPI_ISL_2749288, EPI_ISL_2749289, EPI_ISL_2749290, EPI_ISL_2749291, EPI_ISL_2749292, EPI_ISL_2749295, EPI_ISL_2749297, EPI_ISL_2749298, EPI_ISL_2749299, EPI_ISL_2749302, EPI_ISL_2749303, EPI_ISL_2749305, EPI_ISL_2749306, EPI_ISL_2749308, EPI_ISL_2749309, EPI_ISL_2749310, EPI_ISL_2749312, EPI_ISL_2749314, EPI_ISL_2749315, EPI_ISL_2749325, EPI_ISL_2749326, EPI_ISL_2749327, EPI_ISL_2749330, EPI_ISL_2749331, EPI_ISL_2749333, EPI_ISL_2749334, EPI_ISL_2749336, EPI_ISL_2749337, EPI_ISL_2749339, EPI_ISL_2749340, EPI_ISL_2749341, EPI_ISL_2749342, EPI_ISL_2749343, EPI_ISL_2749344, EPI_ISL_2749345, EPI_ISL_2749346, EPI_ISL_2749347, EPI_ISL_2749348, EPI_ISL_2749349, EPI_ISL_2749350, EPI_ISL_2749351, EPI_ISL_2749352, EPI_ISL_2749353, EPI_ISL_2749354, EPI_ISL_2749355, EPI_ISL_2749356, EPI_ISL_2749357, EPI_ISL_2749358, EPI_ISL_2749360, EPI_ISL_2749361, EPI_ISL_2749362, EPI_ISL_2749363, EPI_ISL_2749364, EPI_ISL_2749365, EPI_ISL_2749367, EPI_ISL_2749368, EPI_ISL_2749369, EPI_ISL_2749370, EPI_ISL_2749371, EPI_ISL_2749372, EPI_ISL_2749373, EPI_ISL_2749374, EPI_ISL_2749375, EPI_ISL_2749376, EPI_ISL_2749377, EPI_ISL_2749378, EPI_ISL_2749380, EPI_ISL_2749381, EPI_ISL_2749382, EPI_ISL_2749383, EPI_ISL_2749384, EPI_ISL_2749385, EPI_ISL_2749386, EPI_ISL_2749387, EPI_ISL_2749388, EPI_ISL_2749389, EPI_ISL_2749390, EPI_ISL_2749391, EPI_ISL_2749392, EPI_ISL_2749393, EPI_ISL_2749454, EPI_ISL_2749456, EPI_ISL_2749589, EPI_ISL_2749590, EPI_ISL_2749610, EPI_ISL_2749622, EPI_ISL_2749647, EPI_ISL_2749686, EPI_ISL_2749694 | | | |
| see above | Respiratory Virus Unit, Microbiology Services Colindale, Public Health England | COVID-19 Genomics UK (COG-UK) Consortium | PHE Covid Sequencing Team |
| EPI_ISL_2749952, EPI_ISL_2749953, EPI_ISL_2749954, EPI_ISL_2749955, EPI_ISL_2749956, EPI_ISL_2749957, EPI_ISL_2749958, EPI_ISL_2749959, EPI_ISL_2749960, EPI_ISL_2749961, EPI_ISL_2749962, EPI_ISL_2749963, EPI_ISL_2749964, EPI_ISL_2749965, EPI_ISL_2749966, EPI_ISL_2749967, EPI_ISL_2749968, EPI_ISL_2749969, EPI_ISL_2749970, EPI_ISL_2749971, EPI_ISL_2749972, EPI_ISL_2749973, EPI_ISL_2749974, EPI_ISL_2749975, EPI_ISL_2749976, EPI_ISL_2749977, EPI_ISL_2749978, EPI_ISL_2749979, EPI_ISL_2749980, EPI_ISL_2749981, EPI_ISL_2749982, EPI_ISL_2749983, EPI_ISL_2749984, EPI_ISL_2749985, EPI_ISL_2749986, EPI_ISL_2749987, EPI_ISL_2749988, EPI_ISL_2749989, EPI_ISL_2749990, EPI_ISL_2749991, EPI_ISL_2749992, EPI_ISL_2749993, EPI_ISL_2749994, EPI_ISL_2749995, EPI_ISL_2749996, EPI_ISL_2749997, EPI_ISL_2749998, EPI_ISL_2749999, EPI_ISL_2750000, EPI_ISL_2750002, EPI_ISL_2750003, EPI_ISL_2750004, EPI_ISL_2750005, EPI_ISL_2750006, EPI_ISL_2750007, EPI_ISL_2750008, EPI_ISL_2750009, EPI_ISL_2750010, EPI_ISL_2750011, EPI_ISL_2750012, EPI_ISL_2750013, EPI_ISL_2750014, EPI_ISL_2750015, EPI_ISL_2750016, EPI_ISL_2750017, EPI_ISL_2750018, EPI_ISL_2750019, EPI_ISL_2750020, EPI_ISL_2750021, EPI_ISL_2750022, EPI_ISL_2750023, EPI_ISL_2750024, EPI_ISL_2750025, EPI_ISL_2750026, EPI_ISL_2750027, EPI_ISL_2750028, EPI_ISL_2750029, EPI_ISL_2750030, EPI_ISL_2750031, EPI_ISL_2750032, EPI_ISL_2750033, EPI_ISL_2750034, EPI_ISL_2750035, EPI_ISL_2750036, EPI_ISL_2750038, EPI_ISL_2750039, EPI_ISL_2750040, EPI_ISL_2750041, EPI_ISL_2750042, EPI_ISL_2750043, EPI_ISL_2750044, EPI_ISL_2750045, EPI_ISL_2750046, EPI_ISL_2750047, EPI_ISL_2750048, EPI_ISL_2750049, EPI_ISL_2750051, EPI_ISL_2750052, EPI_ISL_2750054, EPI_ISL_2750055, EPI_ISL_2750056, EPI_ISL_2750057, EPI_ISL_2750058, EPI_ISL_2750059, EPI_ISL_2750060, EPI_ISL_2750061, EPI_ISL_2750062, EPI_ISL_2750063, EPI_ISL_2750064, EPI_ISL_2750065, EPI_ISL_2750066, EPI_ISL_2750067, EPI_ISL_2750069, EPI_ISL_2750070, EPI_ISL_2750071, EPI_ISL_2750072, EPI_ISL_2750073, EPI_ISL_2750074, EPI_ISL_2750075, EPI_ISL_2750076, EPI_ISL_2750077, EPI_ISL_2750078, EPI_ISL_2750080, EPI_ISL_2750081, EPI_ISL_2750082, EPI_ISL_2750083, EPI_ISL_2750084, EPI_ISL_2750085, EPI_ISL_2750086, EPI_ISL_2750087, EPI_ISL_2750088, EPI_ISL_2750089, EPI_ISL_2750090, EPI_ISL_2750091, EPI_ISL_2750092, EPI_ISL_2750093, EPI_ISL_2750094, EPI_ISL_2750095, EPI_ISL_2750096, EPI_ISL_2750097, EPI_ISL_2750098, EPI_ISL_2750099, EPI_ISL_2750100, EPI_ISL_2750101, EPI_ISL_2750102, EPI_ISL_2750103, EPI_ISL_2750105, EPI_ISL_2750106, EPI_ISL_2750107, EPI_ISL_2750108, EPI_ISL_2750109, EPI_ISL_2750110, EPI_ISL_2750111, EPI_ISL_2750112, EPI_ISL_2750113, EPI_ISL_2750114, EPI_ISL_2750115, EPI_ISL_2750116, EPI_ISL_2750117, EPI_ISL_2750118, EPI_ISL_2750120, EPI_ISL_2750125, EPI_ISL_2750126, EPI_ISL_2750127, EPI_ISL_2750128, EPI_ISL_2750130, EPI_ISL_2750131, EPI_ISL_2750133, EPI_ISL_2750134, EPI_ISL_2750135, EPI_ISL_2750136, EPI_ISL_2750137, EPI_ISL_2750138, EPI_ISL_2750139, EPI_ISL_2750140, EPI_ISL_2750141, EPI_ISL_2750142, EPI_ISL_2750143, EPI_ISL_2750144, EPI_ISL_2750145, EPI_ISL_2750146, EPI_ISL_2750147, EPI_ISL_2750148, EPI_ISL_2750149, EPI_ISL_2750150, EPI_ISL_2750151, EPI_ISL_2750152, EPI_ISL_2750153, EPI_ISL_2750154, EPI_ISL_2750155, EPI_ISL_2750156, EPI_ISL_2750157, EPI_ISL_2750158, EPI_ISL_2750159, EPI_ISL_2750160, EPI_ISL_2750161, EPI_ISL_2750162, EPI_ISL_2750163, EPI_ISL_2750164, EPI_ISL_2750165, EPI_ISL_2750166, EPI_ISL_2750167, EPI_ISL_2750168, EPI_ISL_2750169, EPI_ISL_2750170, EPI_ISL_2750171, EPI_ISL_2750172, EPI_ISL_2750173, EPI_ISL_2750174, EPI_ISL_2750175, EPI_ISL_2750176, EPI_ISL_2750177, EPI_ISL_2750178, EPI_ISL_2750179, EPI_ISL_2750180, EPI_ISL_2750181, EPI_ISL_2750182, EPI_ISL_2750183, EPI_ISL_2750184, EPI_ISL_2750185, EPI_ISL_2750186, EPI_ISL_2750187, EPI_ISL_2750188, EPI_ISL_2750189, EPI_ISL_2750190, EPI_ISL_2750191, EPI_ISL_2750192, EPI_ISL_2750193, EPI_ISL_2750194, EPI_ISL_2750195, EPI_ISL_2750196, EPI_ISL_2750197, EPI_ISL_2750198, EPI_ISL_2750199, EPI_ISL_2750200, EPI_ISL_2750201, EPI_ISL_2750202, EPI_ISL_2750203, EPI_ISL_2750204, EPI_ISL_2750205, EPI_ISL_2750206, EPI_ISL_2750207, EPI_ISL_2750208, EPI_ISL_2750209, EPI_ISL_2750210, EPI_ISL_2750212, EPI_ISL_2750215, EPI_ISL_2750217, EPI_ISL_2750219, EPI_ISL_2750220, EPI_ISL_2750221, EPI_ISL_2750223, EPI_ISL_2750227, EPI_ISL_2750229, EPI_ISL_2750230, EPI_ISL_2750231, EPI_ISL_2750232, EPI_ISL_2750233, EPI_ISL_2750235, EPI_ISL_2750236, EPI_ISL_2750237, EPI_ISL_2750238, EPI_ISL_2750239, EPI_ISL_2750240, EPI_ISL_2750241, EPI_ISL_2750242, EPI_ISL_2750243, EPI_ISL_2750244, EPI_ISL_2750245, EPI_ISL_2750246, EPI_ISL_2750247, EPI_ISL_2750248, EPI_ISL_2750249, EPI_ISL_2750250, EPI_ISL_2750251, EPI_ISL_2750252, EPI_ISL_2750253, EPI_ISL_2750254, EPI_ISL_2750255, EPI_ISL_2750256, EPI_ISL_2750257, EPI_ISL_2750258, EPI_ISL_2750259, EPI_ISL_2750260, EPI_ISL_2750261, EPI_ISL_2750262, EPI_ISL_2750263, EPI_ISL_2750264, EPI_ISL_2750265, EPI_ISL_2750266, EPI_ISL_2750267, EPI_ISL_2750268, EPI_ISL_2750269, EPI_ISL_2750270, EPI_ISL_2750271, EPI_ISL_2750272, EPI_ISL_2750273, EPI_ISL_2750274, EPI_ISL_2750275, EPI_ISL_2750276, EPI_ISL_2750277, EPI_ISL_2750278, EPI_ISL_2750279, EPI_ISL_2750280, EPI_ISL_2750281, EPI_ISL_2750282, EPI_ISL_2750283, EPI_ISL_2750284, EPI_ISL_2750285, EPI_ISL_2750286, EPI_ISL_2750287, EPI_ISL_2750288, EPI_ISL_2750289, EPI_ISL_2750290, EPI_ISL_2750291, EPI_ISL_2750292, EPI_ISL_2750293, EPI_ISL_2750294, EPI_ISL_2750295, EPI_ISL_2750296, EPI_ISL_2750297, EPI_ISL_2750298, EPI_ISL_2750299, EPI_ISL_2750301, EPI_ISL_2750302, EPI_ISL_2750303, EPI_ISL_2750305, EPI_ISL_2750306, EPI_ISL_2750307, EPI_ISL_2750308, EPI_ISL_2750309, EPI_ISL_2750310, EPI_ISL_2750311, EPI_ISL_2750312, EPI_ISL_2750313, EPI_ISL_2750314, EPI_ISL_2750315, EPI_ISL_2750316, EPI_ISL_2750317, EPI_ISL_2750318, EPI_ISL_2750319, EPI_ISL_2750320, EPI_ISL_2750321, EPI_ISL_2750322, EPI_ISL_2750323, EPI_ISL_2750324, EPI_ISL_2750325, EPI_ISL_2750326, EPI_ISL_2750327, EPI_ISL_2750328, EPI_ISL_2750329, EPI_ISL_2750330, EPI_ISL_2750331, EPI_ISL_2750332, EPI_ISL_2750333, EPI_ISL_2750334, EPI_ISL_2750335, EPI_ISL_2750336, EPI_ISL_2750337, EPI_ISL_2750338, EPI_ISL_2750339, EPI_ISL_2750340, EPI_ISL_2750341, EPI_ISL_2750342, EPI_ISL_2750343, EPI_ISL_2750344, EPI_ISL_2750345, EPI_ISL_2750346, EPI_ISL_2750347, EPI_ISL_2750348, EPI_ISL_2750349, EPI_ISL_2750350, EPI_ISL_2750351, EPI_ISL_2750352, EPI_ISL_2750353, EPI_ISL_2750354, EPI_ISL_2750355, EPI_ISL_2750356, EPI_ISL_2750357, EPI_ISL_2750358, EPI_ISL_2750359, EPI_ISL_2750360, EPI_ISL_2750361, EPI_ISL_2750362, EPI_ISL_2750363, EPI_ISL_2750364, EPI_ISL_2750365, EPI_ISL_2750366, EPI_ISL_2750367, EPI_ISL_2750368, EPI_ISL_2750369, EPI_ISL_2750370, EPI_ISL_2750371, EPI_ISL_2750372, EPI_ISL_2750373, EPI_ISL_2750374, EPI_ISL_2750375, EPI_ISL_2750376, EPI_ISL_2750377, EPI_ISL_2750378, EPI_ISL_2750379, EPI_ISL_2750380, EPI_ISL_2750381, EPI_ISL_2750382, EPI_ISL_2750383, EPI_ISL_2750384, EPI_ISL_2750385, EPI_ISL_2750386, EPI_ISL_2750387, EPI_ISL_2750388, EPI_ISL_2750389, EPI_ISL_2750390, EPI_ISL_2750391, EPI_ISL_2750392, EPI_ISL_2750393, EPI_ISL_2750394, EPI_ISL_2750395, EPI_ISL_2750396, EPI_ISL_2750397, EPI_ISL_2750398, EPI_ISL_2750399, EPI_ISL_2750400, EPI_ISL_2750401, EPI_ISL_2750402, EPI_ISL_2750403, EPI_ISL_2750404, EPI_ISL_2750405, EPI_ISL_2750406, EPI_ISL_2750407, EPI_ISL_2750408, EPI_ISL_2750409, EPI_ISL_2750410, EPI_ISL_2750411, EPI_ISL_2750412, EPI_ISL_2750413, EPI_ISL_2750414, EPI_ISL_2750415, EPI_ISL_2750416, EPI_ISL_2750417, EPI_ISL_2750418, EPI_ISL_2750419, EPI_ISL_2750420, EPI_ISL_2750421, EPI_ISL_2750422, EPI_ISL_2750423, EPI_ISL_2750424, EPI_ISL_2750425, EPI_ISL_2750426, EPI_ISL_2750427, EPI_ISL_2750428, EPI_ISL_2750429, EPI_ISL_2750430, EPI_ISL_2750431, EPI_ISL_2750432, EPI_ISL_2750433, EPI_ISL_2750434, EPI_ISL_2750435, EPI_ISL_2750436, EPI_ISL_2750437, EPI_ISL_2750438, EPI_ISL_2750439, EPI_ISL_2750440, EPI_ISL_2750441, EPI_ISL_2750442, EPI_ISL_2750443, EPI_ISL_2750444, EPI_ISL_2750445, EPI_ISL_2750446, EPI_ISL_2750447, EPI_ISL_2750448, EPI_ISL_2750449, EPI_ISL_2750450, EPI_ISL_2750451, EPI_ISL_2750452, EPI_ISL_2750453, EPI_ISL_2750454, EPI_ISL_2750455, EPI_ISL_2750456, EPI_ISL_2750457, EPI_ISL_2750458, EPI_ISL_2750459, EPI_ISL_2750460, EPI_ISL_2750461, EPI_ISL_2750462, EPI_ISL_2750463, EPI_ISL_2750464, EPI_ISL_2750465, EPI_ISL_2750466, EPI_ISL_2750467, EPI_ISL_2750468, EPI_ISL_2750469, EPI_ISL_2750470, EPI_ISL_2750471, EPI_ISL_2750472, EPI_ISL_2750473, EPI_ISL_2750474, EPI_ISL_2750475, EPI_ISL_2750476, EPI_ISL_2750477, EPI_ISL_2750478, EPI_ISL_2750479, EPI_ISL_2750480, EPI_ISL_2750481, EPI_ISL_2750482, EPI_ISL_2750483, EPI_ISL_2750484, EPI_ISL_2750485, EPI_ISL_2750486, EPI_ISL_2750487, EPI_ISL_2750488, EPI_ISL_2750489, EPI_ISL_2750490, EPI_ISL_2750491, EPI_ISL_2750493, EPI_ISL_2750494, EPI_ISL_2750495, EPI_ISL_2750496, EPI_ISL_2750497, EPI_ISL_2750498, EPI_ISL_2750499, EPI_ISL_2750500, EPI_ISL_2750501, EPI_ISL_2750502, EPI_ISL_2750503, EPI_ISL_2750504, EPI_ISL_2750505, EPI_ISL_2750506, EPI_ISL_2750507, EPI_ISL_2750508, EPI_ISL_2750509, EPI_ISL_2750510, EPI_ISL_2750511, EPI_ISL_2750512, EPI_ISL_2750513, EPI_ISL_2750514, EPI_ISL_2750515, EPI_ISL_2750516, EPI_ISL_2750517, EPI_ISL_2750518, EPI_ISL_2750519, EPI_ISL_2750520, EPI_ISL_2750521, EPI_ISL_2750522, EPI_ISL_2750523, EPI_ISL_2750524, EPI_ISL_2750525, EPI_ISL_2750526, EPI_ISL_2750527, EPI_ISL_2750528, EPI_ISL_2750529, EPI_ISL_2750531, EPI_ISL_2750532, EPI_ISL_2750533, EPI_ISL_2750534, EPI_ISL_2750535, EPI_ISL_2750536, EPI_ISL_2750537, EPI_ISL_2750539, EPI_ISL_2750540, EPI_ISL_2750541, EPI_ISL_2750542, EPI_ISL_2750543, EPI_ISL_2750544, EPI_ISL_2750546, EPI_ISL_2750547, EPI_ISL_2750548, EPI_ISL_2750549, EPI_ISL_2750550, EPI_ISL_2750551, EPI_ISL_2750552, EPI_ISL_2750553, EPI_ISL_2750555, EPI_ISL_2750556, EPI_ISL_2750557, EPI_ISL_2750558, EPI_ISL_2750559, EPI_ISL_2750560, EPI_ISL_2750561, EPI_ISL_2750562, EPI_ISL_2750563, EPI_ISL_2750564, EPI_ISL_2750565, EPI_ISL_2750566, EPI_ISL_2750567, EPI_ISL_2750568, EPI_ISL_2750570, EPI_ISL_2750571, EPI_ISL_2750572, EPI_ISL_2750573, EPI_ISL_2750574, EPI_ISL_2750575, EPI_ISL_2750576, EPI_ISL_2750578, EPI_ISL_2750579, EPI_ISL_2750580, EPI_ISL_2750581, EPI_ISL_2750582, EPI_ISL_2750583, EPI_ISL_2750584, EPI_ISL_2750585, EPI_ISL_2750586, EPI_ISL_2750587, EPI_ISL_2750588, EPI_ISL_2750589, EPI_ISL_2750590, EPI_ISL_2750591, EPI_ISL_2750592, EPI_ISL_2750593, EPI_ISL_2750594, EPI_ISL_2750595, EPI_ISL_2750596, EPI_ISL_2750597, EPI_ISL_2750598, EPI_ISL_2750599, EPI_ISL_2750600, EPI_ISL_2750601, EPI_ISL_2750602, EPI_ISL_2750603, EPI_ISL_2750604, EPI_ISL_2750605, EPI_ISL_2750606, EPI_ISL_2750607, EPI_ISL_2750608, EPI_ISL_2750609, EPI_ISL_2750610, EPI_ISL_2750611, EPI_ISL_2750612, EPI_ISL_2750613, EPI_ISL_2750614, EPI_ISL_2750615, EPI_ISL_2750616, EPI_ISL_2750617, EPI_ISL_2750618, EPI_ISL_2750619, EPI_ISL_2750620, EPI_ISL_2750621, EPI_ISL_2750622, EPI_ISL_2750623, EPI_ISL_2750624, EPI_ISL_2750625, EPI_ISL_2750626, EPI_ISL_2750627, EPI_ISL_2750628, EPI_ISL_2750629, EPI_ISL_2750630, EPI_ISL_2750631, EPI_ISL_2750632, EPI_ISL_2750633, EPI_ISL_2750634, EPI_ISL_2750635, EPI_ISL_2750636, EPI_ISL_2750637, EPI_ISL_2750638, EPI_ISL_2750639, EPI_ISL_2750640, EPI_ISL_2750641, EPI_ISL_2750642, EPI_ISL_2750643, EPI_ISL_2750644, EPI_ISL_2750645, EPI_ISL_2750646, EPI_ISL_2750647, EPI_ISL_2750648, EPI_ISL_2750649, EPI_ISL_2750650, EPI_ISL_2750651, EPI_ISL_2750652, EPI_ISL_2750653, EPI_ISL_2750654, EPI_ISL_2750655, EPI_ISL_2750656, EPI_ISL_2750657, EPI_ISL_2750658, EPI_ISL_2750659, EPI_ISL_2750660, EPI_ISL_2750661, EPI_ISL_2750662, EPI_ISL_2750663, EPI_ISL_2750664, EPI_ISL_2750665, EPI_ISL_2750666, EPI_ISL_2750667, EPI_ISL_2750668, EPI_ISL_2750669, EPI_ISL_2750670, EPI_ISL_2750671, EPI_ISL_2750672, EPI_ISL_2750673, EPI_ISL_2750674, EPI_ISL_2750675, EPI_ISL_2750676, EPI_ISL_2750677, EPI_ISL_2750678, EPI_ISL_2750679, EPI_ISL_2750680, EPI_ISL_2750681, EPI_ISL_2750682, EPI_ISL_2750683, EPI_ISL_2750684, EPI_ISL_2750685, EPI_ISL_2750686, EPI_ISL_2750687, EPI_ISL_2750688, EPI_ISL_2750689 | | | |
| see above | Originating lab: Wales Specialist Virology Centre Sequencing lab: Pathogen Genomics Unit | Public Health Wales Microbiology Cardiff Wales Specialist Virology Centre | Catherine Moore, Johnathan Evans, Laura Gifford, Malorie Perry, Simon Cottrell, Angela Marchbank, Alec Birchley, Alexander Adams, Amy Gaskin, Bree Gatica-Wilcox, Jason Coombes, Joel Southgate, Lauren Gilbert, Lee Graham, Nicole Pacchiarini, Sara Kumziene-Summerhayes, Sarah Taylor, Sophie Jones, Sara Rey, Matthew Bull, Joanne Watkins, Sally Corden, Tom Connor |
| EPI_ISL_2750845 | Center for Laboratory Medicine | Center for Laboratory Medicine | Yannick Gerth |
| EPI_ISL_2752169 | Ohio Department of Health Laboratory | Ohio Department of Health Laboratory | Holmes, Jennifer; Eric Brandt, Keoni Omura, Glen McGillivary, Caitlin McDonnell, Jade Mowery, Stephanie Mccracken, Tyler Payne, Kirtana Ramadugu, Erica Leasure, Brent Lee, Kelsey Florek, Heather Blankenship, Allison Black, Quanta Brown, and Tammy Bannerman |
| EPI_ISL_2753615, EPI_ISL_2753616, EPI_ISL_2753617, EPI_ISL_2753619, EPI_ISL_2753623, EPI_ISL_2753624, EPI_ISL_2753627, EPI_ISL_2753630, EPI_ISL_2753631, EPI_ISL_2753632, EPI_ISL_2753633, EPI_ISL_2753634, EPI_ISL_2753635, EPI_ISL_2753637, EPI_ISL_2753638, EPI_ISL_2753639, EPI_ISL_2753640, EPI_ISL_2753641, EPI_ISL_2753642, EPI_ISL_2753643, EPI_ISL_2753644, EPI_ISL_2753649, EPI_ISL_2753650, EPI_ISL_2753651, EPI_ISL_2753652, EPI_ISL_2753653, EPI_ISL_2753654, EPI_ISL_2753655, EPI_ISL_2753656, EPI_ISL_2753657, EPI_ISL_2753658, EPI_ISL_2753659, EPI_ISL_2753660, EPI_ISL_2753661, EPI_ISL_2753662, EPI_ISL_2753663, EPI_ISL_2753664, EPI_ISL_2753665, EPI_ISL_2753666, EPI_ISL_2753667, EPI_ISL_2753668, EPI_ISL_2753669, EPI_ISL_2753670, EPI_ISL_2753672, EPI_ISL_2753673, EPI_ISL_2753674, EPI_ISL_2753675, EPI_ISL_2753676, EPI_ISL_2753677, EPI_ISL_2753678, EPI_ISL_2753679, EPI_ISL_2753680, EPI_ISL_2753681, EPI_ISL_2753682, EPI_ISL_2753683, EPI_ISL_2753684, EPI_ISL_2753685, EPI_ISL_2753686, EPI_ISL_2753687, EPI_ISL_2753690, EPI_ISL_2753691, EPI_ISL_2753692, EPI_ISL_2753693, EPI_ISL_2753718, EPI_ISL_2753719, EPI_ISL_2753720, EPI_ISL_2753721, EPI_ISL_2753722, EPI_ISL_2753723, EPI_ISL_2753724, EPI_ISL_2753725, EPI_ISL_2753726, EPI_ISL_2753727, EPI_ISL_2753729, EPI_ISL_2753730, EPI_ISL_2753731, EPI_ISL_2753732, EPI_ISL_2753733, EPI_ISL_2753734, EPI_ISL_2753735, EPI_ISL_2753736, EPI_ISL_2753737, EPI_ISL_2753738, EPI_ISL_2753739, EPI_ISL_2753740, EPI_ISL_2753741, EPI_ISL_2753742, EPI_ISL_2753743, EPI_ISL_2753744, EPI_ISL_2753745, EPI_ISL_2753746, EPI_ISL_2753747, EPI_ISL_2753748, EPI_ISL_2753749, EPI_ISL_2753750, EPI_ISL_2753751, EPI_ISL_2753752, EPI_ISL_2753753, EPI_ISL_2753754, EPI_ISL_2753755, EPI_ISL_2753757, EPI_ISL_2753758, EPI_ISL_2753759, EPI_ISL_2753760, EPI_ISL_2753762, EPI_ISL_2753763, EPI_ISL_2753765, EPI_ISL_2753766, EPI_ISL_2753767, EPI_ISL_2753772, EPI_ISL_2753773, EPI_ISL_2753775, EPI_ISL_2753780, EPI_ISL_2753783, EPI_ISL_2753784, EPI_ISL_2753785, EPI_ISL_2753786, EPI_ISL_2753787, EPI_ISL_2753788, EPI_ISL_2753789, EPI_ISL_2753790, EPI_ISL_2753791, EPI_ISL_2753792, EPI_ISL_2753793, EPI_ISL_2753794, EPI_ISL_2753795, EPI_ISL_2753796, EPI_ISL_2753797, EPI_ISL_2753798, EPI_ISL_2753799, EPI_ISL_2753800, EPI_ISL_2753801, EPI_ISL_2753802, EPI_ISL_2753804, EPI_ISL_2753805, EPI_ISL_2753806, EPI_ISL_2753807, EPI_ISL_2753808, EPI_ISL_2753809, EPI_ISL_2753810, EPI_ISL_2753811, EPI_ISL_2753812, EPI_ISL_2753813, EPI_ISL_2753814, EPI_ISL_2753815, EPI_ISL_2753816, EPI_ISL_2753817, EPI_ISL_2753818, EPI_ISL_2753819, EPI_ISL_2753820, EPI_ISL_2753821, EPI_ISL_2753822, EPI_ISL_2753823, EPI_ISL_2753824, EPI_ISL_2753825, EPI_ISL_2753826, EPI_ISL_2753827, EPI_ISL_2753828, EPI_ISL_2753829, EPI_ISL_2753830, EPI_ISL_2753831, EPI_ISL_2753832, EPI_ISL_2753833, EPI_ISL_2753834, EPI_ISL_2753835, EPI_ISL_2753836, EPI_ISL_2753837, EPI_ISL_2753838, EPI_ISL_2753839, EPI_ISL_2753840, EPI_ISL_2753841, EPI_ISL_2753842, EPI_ISL_2753843, EPI_ISL_2753844, EPI_ISL_2753845, EPI_ISL_2753846 | | | |
| see above | Missouri State Public Health Laboratory | Missouri State Public Health Laboratory | Matthew Sinn, Joshua Barry, Ashley New |
| EPI_ISL_2753940, EPI_ISL_2753942, EPI_ISL_2753943, EPI_ISL_2753944, EPI_ISL_2753951 | NYU Langone Health | Departments of Pathology and Medicine, New York University School of Medicine | Adriana Heguy, Dacia Dimartino, Emily Guzman, Christian Marier, Peter Meyn, Sitharam Ramaswami, Gael Westby, Paul Zappile, Yutong Zhang, Paolo Cotzia, Guiqing Wang |
| EPI_ISL_2753984, EPI_ISL_2753985, EPI_ISL_2753986, EPI_ISL_2753987, EPI_ISL_2753988 | PAMF Palo Alto | Santa Clara County Public Health Laboratory | Santa Clara County Public Health Department |
| EPI_ISL_2754006, EPI_ISL_2754009, EPI_ISL_2754012, EPI_ISL_2754013, EPI_ISL_2754014, EPI_ISL_2754015, EPI_ISL_2754019 | Kaiser Regional | Santa Clara County Public Health Laboratory | Santa Clara County Public Health Department |

| | | | |
|---|---|---|---|
| EPI_ISL_2754045, EPI_ISL_2754046, EPI_ISL_2754047, EPI_ISL_2754050, EPI_ISL_2754051, EPI_ISL_2754052, EPI_ISL_2754053, EPI_ISL_2754054, EPI_ISL_2754055, EPI_ISL_2754056, EPI_ISL_2754058, EPI_ISL_2754060, EPI_ISL_2754061, EPI_ISL_2754071 | | | |
| see above | Wyoming Public Health Laboratory | Wyoming Public Health Laboratory | Jim Mildenberger, Taylor Fearing, Channing Weber, Ashley Norberg, Chayse Rowley, Marley Goetz, Brian Dominguez, Elliot Thomasson, Sam Britz, Cari Sloma, Robert Petit, and Rob Christensen |
| EPI_ISL_2754086, EPI_ISL_2754088, EPI_ISL_2754089, EPI_ISL_2754091, EPI_ISL_2754092, EPI_ISL_2754093, EPI_ISL_2754094 | Nebraska Public Health Laboratory | NPHL COVID-19 Response Team | NPHL COVID-19 Response Team |
| EPI_ISL_2754097 | Kansas Health and Environmental Lab | Wichita State University - Molecular Diagnostics Lab | Katherine Wiggins, Mike Grose, Jonathan Barnell, Ben Olsen, and Phil Adam |
| EPI_ISL_2754100, EPI_ISL_2754103, EPI_ISL_2754105, EPI_ISL_2754106, EPI_ISL_2754107, EPI_ISL_2754109, EPI_ISL_2754110, EPI_ISL_2754111, EPI_ISL_2754112, EPI_ISL_2754113 | Kansas Health and Environmental Lab | Clinical Reference Laboratory | Katherine Wiggins, Mike Grose, Jonathan Barnell, Ben Olsen, and Phil Adam |
| EPI_ISL_2754116 | Laverty Pathology | NSW Health Pathology - Institute of Clinical Pathology and Medical Research; Westmead Hospital; University of Sydney | CIDM-PH et al. |
| EPI_ISL_2754117, EPI_ISL_2754118, EPI_ISL_2754119 | Douglass Hanly Moir Pathology | NSW Health Pathology - Institute of Clinical Pathology and Medical Research; Westmead Hospital; University of Sydney | CIDM-PH et al. |
| EPI_ISL_2754120 | Laverty Pathology | NSW Health Pathology - Institute of Clinical Pathology and Medical Research; Westmead Hospital; University of Sydney | CIDM-PH et al. |
| EPI_ISL_2754121 | St Vincent's Pathology (SydPath) | NSW Health Pathology - Institute of Clinical Pathology and Medical Research; Westmead Hospital; University of Sydney | CIDM-PH et al. |
| EPI_ISL_2754122 | Histopath | NSW Health Pathology - Institute of Clinical Pathology and Medical Research; Westmead Hospital; University of Sydney | CIDM-PH et al. |
| EPI_ISL_2754123, EPI_ISL_2754124 | Laverty Pathology | NSW Health Pathology - Institute of Clinical Pathology and Medical Research; Westmead Hospital; University of Sydney | CIDM-PH et al. |
| EPI_ISL_2754125 | 4Cyte Pathology | NSW Health Pathology - Institute of Clinical Pathology and Medical Research; Westmead Hospital; University of Sydney | CIDM-PH et al. |
| EPI_ISL_2754126, EPI_ISL_2754127 | Histopath | NSW Health Pathology - Institute of Clinical Pathology and Medical Research; Westmead Hospital; University of Sydney | CIDM-PH et al. |
| EPI_ISL_2754128 | Medlab Pathology | NSW Health Pathology - Institute of Clinical Pathology and Medical Research; Westmead Hospital; University of Sydney | CIDM-PH et al. |
| EPI_ISL_2754196 | S.C Microbiologia e virologia - Speciale Centro influenza -AOU di Sassari | Laboratorio Biologia Molecolare Sars Cov2 - UOC Laboratorio Analisi - Servizio Medicina di Laboratorio, Ospedale "San Francesco" - ATS-ASSL Nuoro and Laboratorio specialistico UOC Ematologia - Ospedale "San Francesco" - ATS-ASSL Nuoro | Asproni Rosanna, Piras Giovanna, Malune Paolo, Pira Giovanna, Fancello Patrizia, Rosu Valentina, Carta Franco, Fiamma Maura, Mameli Giuseppe, Garau Maria Cristina , Palmas Angelo Domenico,Rimini Elena ,Serra Caterina, Salvatore Rubino |
| EPI_ISL_2754197 | S.C Microbiologia e virologia - Speciale Centro influenza -AOU di Sassari | Laboratorio Biologia Molecolare Sars Cov2 - UOC Laboratorio Analisi - Servizio Medicina di Laboratorio, Ospedale "San Francesco" - ATS-ASSL Nuoro and Laboratorio specialistico UOC Ematologia - Ospedale "San Francesco" - ATS-ASSL Nuoro | Asproni Rosanna, Piras Giovanna, Malune Paolo, Pira Giovanna, Fancello Patrizia, Rosu Valentina, Carta Franco, Fiamma Maura, Mameli Giuseppe, Garau Maria Cristina , Palmas Angelo Domenico,Rimini Elena ,Serra Caterina, Rubino Salvatore |
| EPI_ISL_2754198, EPI_ISL_2754199, EPI_ISL_2754200, EPI_ISL_2754202 | S.C Microbiologia e virologia - Speciale Centro influenza -AOU di Sassari | Laboratorio Biologia Molecolare Sars Cov2 - UOC Laboratorio Analisi - Servizio Medicina di Laboratorio, Ospedale "San Francesco" - ATS-ASSL Nuoro and Laboratorio specialistico UOC Ematologia - Ospedale "San Francesco" - ATS-ASSL Nuoro | Asproni Rosanna, Piras Giovanna, Malune Paolo, Pira Giovanna, Fancello Patrizia, Rosu Valentina, Carta Franco, Fiamma Maura, Mameli Giuseppe, Garau Maria Cristina , Palmas Angelo Domenico,Rimini Elena ,Serra Caterina, Salvatore Rubino |
| EPI_ISL_2754296, EPI_ISL_2754298, EPI_ISL_2754299, EPI_ISL_2754302, EPI_ISL_2754303, EPI_ISL_2754304, EPI_ISL_2754308, EPI_ISL_2754309 | Arizona State Public Health Laboratory | Arizona State Public Health Laboratory | Trung Huynh, Jessica Escobar, Katherine Fullerton, Nobuko Fukushima, Matthew Contursi, Stacy White, Linda Getsinger, Victor Waddell |
| EPI_ISL_2754858, EPI_ISL_2754870, EPI_ISL_2754871, EPI_ISL_2754872, EPI_ISL_2754876, EPI_ISL_2754877, EPI_ISL_2754878, EPI_ISL_2754887, EPI_ISL_2754895, EPI_ISL_2754897, EPI_ISL_2754898, EPI_ISL_2754899, EPI_ISL_2754900, EPI_ISL_2754902 | | | |
| see above | Nebraska Public Health Laboratory | NPHL COVID-19 Response Team | NPHL COVID-19 Response Team |
| EPI_ISL_2754911 | GA Department of Public Health Laboratory | Genomics and Discovery, Respiratory Viruses Branch, Division of Viral Diseases, Centers for Disease Control and Prevention | Anna Kelleher, Ying Tao, Yan Li, Jing Zhang, Brian Lynch, Krista Queen, Anna Uehara, Peter Cook, Han Jia Justin Ng, Rachel Marine, Clinton R. Paden, Dhwani Batra, Haibin Wang, Tara Coalter, Jasmine Padilla, Morgan Davis, Mili Sheth, Sarah Nobles, Mark Burroughs, Justin Lee, Adam Retchless, Suxiang Tong |
| EPI_ISL_2754913 | AR Dept. of Health-PHL, Molecular Diagnostics | Genomics and Discovery, Respiratory Viruses Branch, Division of Viral Diseases, Centers for Disease Control and Prevention | Anna Kelleher, Ying Tao, Yan Li, Jing Zhang, Brian Lynch, Krista Queen, Anna Uehara, Peter Cook, Han Jia Justin Ng, Rachel Marine, Clinton R. Paden, Dhwani Batra, Haibin Wang, Tara Coalter, Jasmine Padilla, Morgan Davis, Mili Sheth, Sarah Nobles, Mark Burroughs, Justin Lee, Adam Retchless, Suxiang Tong |
| EPI_ISL_2754914 | NJ Public Health and Environmental Laboratories | Genomics and Discovery, Respiratory Viruses Branch, Division of Viral Diseases, Centers for Disease Control and Prevention | Anna Kelleher, Ying Tao, Yan Li, Jing Zhang, Brian Lynch, Krista Queen, Anna Uehara, Peter Cook, Han Jia Justin Ng, Rachel Marine, Clinton R. Paden, Dhwani Batra, Haibin Wang, Tara Coalter, Jasmine Padilla, Morgan Davis, Mili Sheth, Sarah Nobles, Mark Burroughs, Justin Lee, Adam Retchless, Suxiang Tong |
| EPI_ISL_2755034, EPI_ISL_2755117, EPI_ISL_2755119, EPI_ISL_2755122, EPI_ISL_2755159, EPI_ISL_2755176, EPI_ISL_2755181, EPI_ISL_2755207, EPI_ISL_2755213, EPI_ISL_2755223, EPI_ISL_2755267, EPI_ISL_2755293, EPI_ISL_2755317, EPI_ISL_2755319, EPI_ISL_2755321, EPI_ISL_2755334, EPI_ISL_2755335, EPI_ISL_2755354, EPI_ISL_2755356, EPI_ISL_2755364, EPI_ISL_2755367, EPI_ISL_2755394, EPI_ISL_2755409, EPI_ISL_2755411, EPI_ISL_2755439, EPI_ISL_2755447, EPI_ISL_2755448, EPI_ISL_2755449, EPI_ISL_2755450, EPI_ISL_2755459, EPI_ISL_2755464, EPI_ISL_2755467, EPI_ISL_2755468, EPI_ISL_2755472, EPI_ISL_2755788, EPI_ISL_2755837, EPI_ISL_2755905, EPI_ISL_2755935 | | | |
| see above | Curative Labs | Curative Labs | Mikhail Hanewich-Hollatz, J.Cesar Ignacio-Espinoza, Elias L. Salfati, Nabjot Sandhu, George Way, Zhiyi Xie, Eugenia Khorosheva, Sophia Quasem, Janet Chen, Vladimir Slepnev |
| EPI_ISL_2756159 | Genetica Molecular and Subdepartamento de Virologia ISP Chile | Instituto de Salud Publica de Chile | Karen Orostica, Constanza Campano, Barbara Parra, Loredana Arata, Gisselle Barra, Patricia Bustos, Rodrigo Fasce, Javier Tognarelli, Andres Castillo, Soledad Ulloa, Jorge Fernandez |
| EPI_ISL_2756206, EPI_ISL_2756208, EPI_ISL_2756210, EPI_ISL_2756211, EPI_ISL_2756215, EPI_ISL_2756216, EPI_ISL_2756217, EPI_ISL_2756218 | National Platform bis UMONS/Jolimont | National Platform bis UMONS/Jolimont | François Dufrasne, Guillaume Bayon-Vicente, Florian Juszczak, Eric Tarantino, Gautier Detry, Ruddy Wattiez |
| EPI_ISL_2756496, EPI_ISL_2756497, EPI_ISL_2756514, EPI_ISL_2756519 | Montana Public Health Laboratory | Montana Public Health Laboratory | Joy Ritter, Michelle Mozer, Carrie Biskupiak, Deborah Gibson, Michael Dills |
| EPI_ISL_2756523, EPI_ISL_2756524, EPI_ISL_2756525, EPI_ISL_2756527, EPI_ISL_2756528, EPI_ISL_2756529, EPI_ISL_2756530, EPI_ISL_2756531, EPI_ISL_2756533, EPI_ISL_2756535, EPI_ISL_2756536, EPI_ISL_2756537, EPI_ISL_2756538, EPI_ISL_2756539, EPI_ISL_2756540 | | | |
| see above | Wichita State University - Molecular Diagnostics Lab | Kansas Health and Environmental Lab | Katherine Wiggins, Mike Grose, Jonathan Barnell, Ben Olsen, and Phil Adam |
| EPI_ISL_2757080 | MEPHI, Aix Marseille University | MEPHI, Aix Marseille University | Anthony LEVASSEUR |
| EPI_ISL_2757655, EPI_ISL_2757656, EPI_ISL_2757657 | PathWest Laboratory Medicine WA | PathWest Laboratory Medicine WA Microbial Surveillance Unit | PathWest Laboratory Medicine WA Microbial Surveillance Unit |
| EPI_ISL_2757658 | Hospital | National Reference Center for Viruses of Respiratory Infections, Institut Pasteur, Paris | Marion Barbet, Sylvie Behillil, Méline Bizard, Angela Brisebarre, Camille Capel, Vincent Enouf, Louise Lefrançois, Frédéric Lemoine, Christophe Malabat, Corinne Maufrais, Etienne Simon-Lorière, Maud Vanpeene, Sylvie Van der Werf ,JéRome Guinard |
| EPI_ISL_2757659 | Labo Analyses Med | National Reference Center for Viruses of Respiratory Infections, Institut Pasteur, Paris | Marion Barbet, Sylvie Behillil, Méline Bizard, Angela Brisebarre, Camille Capel, Vincent Enouf, Louise Lefrançois, Frédéric Lemoine, Christophe Malabat, Corinne Maufrais, Emmanuelle Permal, Etienne Simon-Lorière, Maud Vanpeene, Sylvie Van der Werf ,GréGoire Potiron |
| EPI_ISL_2757660 | Hospital | National Reference Center for Viruses of Respiratory Infections, Institut Pasteur, Paris | Marion Barbet, Sylvie Behillil, Méline Bizard, Angela Brisebarre, Camille Capel, Vincent Enouf, Louise Lefrançois, Frédéric Lemoine, Christophe Malabat, Corinne Maufrais, Etienne Simon-Lorière, Maud Vanpeene, Sylvie Van der Werf ,Eric Farfour |

| | | | |
|---|---|---|---|
| EPI_ISL_2757662 | Hospital | National Reference Center for Viruses of Respiratory Infections, Institut Pasteur, Paris | Marion Barbet, Sylvie Behillil, Méline Bizard, Angela Brisebarre, Camille Capel, Vincent Enouf, Louise Lefrançois, Frédéric Lemoine, Christophe Malabat, Corinne Maufrais, Adrien Pain, Etienne Simon-Lorière, Maud Vanpeene, Sylvie Van der Werf ,Laurence Courdavault |
| EPI_ISL_2757663, EPI_ISL_2757664, EPI_ISL_2757665 | Hospital | National Reference Center for Viruses of Respiratory Infections, Institut Pasteur, Paris | Marion Barbet, Sylvie Behillil, Méline Bizard, Angela Brisebarre, Camille Capel, Vincent Enouf, Louise Lefrançois, Frédéric Lemoine, Christophe Malabat, Corinne Maufrais, Etienne Simon-Lorière, Maud Vanpeene, Sylvie Van der Werf ,JéRôMe Durivault |
| EPI_ISL_2757666 | Hospital | National Reference Center for Viruses of Respiratory Infections, Institut Pasteur, Paris | Marion Barbet, Sylvie Behillil, Méline Bizard, Angela Brisebarre, Camille Capel, Vincent Enouf, Louise Lefrançois, Frédéric Lemoine, Christophe Malabat, Corinne Maufrais, Adrien Pain, Etienne Simon-Lorière, Maud Vanpeene, Sylvie Van der Werf ,JéRôMe Durivault |
| EPI_ISL_2757667, EPI_ISL_2757668, EPI_ISL_2757669, EPI_ISL_2757670, EPI_ISL_2757671, EPI_ISL_2757672, EPI_ISL_2757673 | Hospital | National Reference Center for Viruses of Respiratory Infections, Institut Pasteur, Paris | Marion Barbet, Sylvie Behillil, Méline Bizard, Angela Brisebarre, Camille Capel, Vincent Enouf, Louise Lefrançois, Frédéric Lemoine, Christophe Malabat, Corinne Maufrais, Etienne Simon-Lorière, Maud Vanpeene, Sylvie Van der Werf ,JéRôMe Durivault |
| EPI_ISL_2757674, EPI_ISL_2757675 | Hospital | National Reference Center for Viruses of Respiratory Infections, Institut Pasteur, Paris | Marion Barbet, Sylvie Behillil, Méline Bizard, Angela Brisebarre, Camille Capel, Vincent Enouf, Louise Lefrançois, Frédéric Lemoine, Christophe Malabat, Corinne Maufrais, Amaury Vaysse, Etienne Simon-Lorière, Maud Vanpeene, Sylvie Van der Werf ,JéRôMe Durivault |
| EPI_ISL_2757676, EPI_ISL_2757677, EPI_ISL_2757678 | Hospital | National Reference Center for Viruses of Respiratory Infections, Institut Pasteur, Paris | Marion Barbet, Sylvie Behillil, Méline Bizard, Angela Brisebarre, Camille Capel, Vincent Enouf, Louise Lefrançois, Frédéric Lemoine, Christophe Malabat, Corinne Maufrais, Etienne Simon-Lorière, Maud Vanpeene, Sylvie Van der Werf ,JéRôMe Durivault |
| EPI_ISL_2757680, EPI_ISL_2757681 | Labo Analyses Med | National Reference Center for Viruses of Respiratory Infections, Institut Pasteur, Paris | Marion Barbet, Sylvie Behillil, Méline Bizard, Angela Brisebarre, Camille Capel, Vincent Enouf, Louise Lefrançois, Frédéric Lemoine, Christophe Malabat, Corinne Maufrais, Etienne Simon-Lorière, Maud Vanpeene, Sylvie Van der Werf ,Nader Kherbek |
| EPI_ISL_2757682 | Labo Analyses Med | National Reference Center for Viruses of Respiratory Infections, Institut Pasteur, Paris | Marion Barbet, Sylvie Behillil, Méline Bizard, Angela Brisebarre, Camille Capel, Vincent Enouf, Louise Lefrançois, Frédéric Lemoine, Christophe Malabat, Corinne Maufrais, Amaury Vaysse, Etienne Simon-Lorière, Maud Vanpeene, Sylvie Van der Werf ,GréGoire Potiron |
| EPI_ISL_2757685, EPI_ISL_2757686, EPI_ISL_2757687, EPI_ISL_2757688 | Presidio di Brindisi Di Summa - Perrino | Istituto Zooprofilattico Sperimentale della Puglia e della Basilicata | Parisi A., Bianco A., Capozzi L., Del Sambro L., Simone D., Difato L., Santoro A., Ridolfi D., Giannico A. |
| EPI_ISL_2757693, EPI_ISL_2757694 | Ospedale Santa Caterina Novella | Istituto Zooprofilattico Sperimentale della Puglia e della Basilicata | Parisi A., Bianco A., Capozzi L., Del Sambro L., Simone D., Difato L., Bruno A. R. |
| EPI_ISL_2757696 | Ospedale Cardinale G. Panico | Istituto Zooprofilattico Sperimentale della Puglia e della Basilicata | Parisi A., Bianco A., Capozzi L., Del Sambro L., Simone D., Difato L., Bruno A. R. |
| EPI_ISL_2757698, EPI_ISL_2757701 | Ospedale Vito Fazzi | Istituto Zooprofilattico Sperimentale della Puglia e della Basilicata | Parisi A., Bianco A., Capozzi L., Del Sambro L., Simone D., Difato L., Lobreglio G. |
| EPI_ISL_2757702, EPI_ISL_2757703, EPI_ISL_2757705 | Ospedale Santa Caterina Novella | Istituto Zooprofilattico Sperimentale della Puglia e della Basilicata | Parisi A., Bianco A., Capozzi L., Del Sambro L., Simone D., Difato L., Bruno A. R. |
| EPI_ISL_2757706, EPI_ISL_2757707, EPI_ISL_2757712 | Presidio di Brindisi Di Summa - Perrino | Istituto Zooprofilattico Sperimentale della Puglia e della Basilicata | Parisi A., Bianco A., Capozzi L., Del Sambro L., Simone D., Difato L., Santoro A., Ridolfi D., Giannico A. |
| EPI_ISL_2757713 | Ospedale Santissima Annunziata | Istituto Zooprofilattico Sperimentale della Puglia e della Basilicata | Parisi A., Bianco A., Capozzi L., Del Sambro L., Simone D., Difato L., Ridolfi D., Giannico A, Tampoia M. |
| EPI_ISL_2757720 | Ospedale Santa Caterina Novella | Istituto Zooprofilattico Sperimentale della Puglia e della Basilicata | Parisi A., Bianco A., Capozzi L., Del Sambro L., Simone D., Difato L., Bruno A. R. |
| EPI_ISL_2757722 | Dipartimento di prevenzione Bari | Istituto Zooprofilattico Sperimentale della Puglia e della Basilicata | Parisi A., Bianco A., Capozzi L., Del Sambro L., Simone D., Difato L., Ridolfi D., Giannico A., Lagravinese D. |
| EPI_ISL_2757734, EPI_ISL_2757736, EPI_ISL_2757737, EPI_ISL_2757738, EPI_ISL_2757739, EPI_ISL_2757741, EPI_ISL_2757743, EPI_ISL_2757744, EPI_ISL_2757756, EPI_ISL_2757757, EPI_ISL_2757758, EPI_ISL_2757759, EPI_ISL_2757760, EPI_ISL_2757761, EPI_ISL_2757762 | | | |
| see above | Department of Virology | Department of Virology | Massab Umair, Aamer Ikram, Muhammad Salman, Nazish Badar, Zaira Rehman, Muhammad Ammar, Syed Adnan Haider |
| EPI_ISL_2757811, EPI_ISL_2757812, EPI_ISL_2757813, EPI_ISL_2757814, EPI_ISL_2757815, EPI_ISL_2757816, EPI_ISL_2757817 | OSPEDALE CIVILE TERAMO - CENTRO TRASFUSIONALE | Istituto Zooprofilattico Sperimentale dell'Abruzzo e Molise "G. Caporale" | Lorusso A, Marcacci M, Di Domenico M, Ancora M, Curini V, Di Lollo Valeria, Mangone I, Rinaldi A, Delli Compagni E, Scialabba S, Caporale M, Di Pasquale A, Cammà C, Puglia I, Calistri P, Savini G |
| EPI_ISL_2757819, EPI_ISL_2757820, EPI_ISL_2757821 | SIESP CHIETI - DRIVE IN LANCIANO | Istituto Zooprofilattico Sperimentale dell'Abruzzo e Molise "G. Caporale" | Lorusso A, Marcacci M, Di Domenico M, Ancora M, Curini V, Di Lollo Valeria, Mangone I, Rinaldi A, Delli Compagni E, Scialabba S, Caporale M, Di Pasquale A, Cammà C, Puglia I, Calistri P, Savini G |
| EPI_ISL_2757822, EPI_ISL_2757823 | OSPEDALE CIVILE TERAMO - CENTRO TRASFUSIONALE | Istituto Zooprofilattico Sperimentale dell'Abruzzo e Molise "G. Caporale" | Lorusso A, Marcacci M, Di Domenico M, Ancora M, Curini V, Di Lollo Valeria, Mangone I, Rinaldi A, Delli Compagni E, Scialabba S, Caporale M, Di Pasquale A, Cammà C, Puglia I, Calistri P, Savini G |
| EPI_ISL_2757829, EPI_ISL_2757830, EPI_ISL_2757831 | SIESP CHIETI - DRIVE IN LANCIANO | Istituto Zooprofilattico Sperimentale dell'Abruzzo e Molise "G. Caporale" | Lorusso A, Marcacci M, Di Domenico M, Ancora M, Curini V, Di Lollo Valeria, Mangone I, Rinaldi A, Delli Compagni E, Scialabba S, Caporale M, Di Pasquale A, Cammà C, Puglia I, Calistri P, Savini G |
| EPI_ISL_2757832 | OSPEDALE CIVILE TERAMO - CENTRO TRASFUSIONALE | Istituto Zooprofilattico Sperimentale dell'Abruzzo e Molise "G. Caporale" | Lorusso A, Marcacci M, Di Domenico M, Ancora M, Curini V, Di Lollo Valeria, Mangone I, Rinaldi A, Delli Compagni E, Scialabba S, Caporale M, Di Pasquale A, Cammà C, Puglia I, Calistri P, Savini G |
| EPI_ISL_2757862, EPI_ISL_2757863, EPI_ISL_2757865, EPI_ISL_2757873, EPI_ISL_2757874, EPI_ISL_2757876, EPI_ISL_2757878, EPI_ISL_2757879, EPI_ISL_2757880, EPI_ISL_2757882, EPI_ISL_2757885, EPI_ISL_2757886, EPI_ISL_2757887, EPI_ISL_2757892, EPI_ISL_2757895, EPI_ISL_2757896, EPI_ISL_2757897, EPI_ISL_2757898, EPI_ISL_2757902, EPI_ISL_2757903, EPI_ISL_2757907, EPI_ISL_2757909, EPI_ISL_2757911, EPI_ISL_2757912, EPI_ISL_2757913, EPI_ISL_2757915, EPI_ISL_2757916, EPI_ISL_2757917, EPI_ISL_2757918, EPI_ISL_2757921, EPI_ISL_2757922, EPI_ISL_2757923, EPI_ISL_2757927, EPI_ISL_2757929 | | | |
| see above | Lifebrain Covid Labor GmbH | Dept. of Laboratory Medicine | Harald Esterbauer, Anna Gschaider, Petra Jurkowitsch, Fabian Konig, Sabina Plumer, Robert Strassl, Claudia Weber, Oswald Wagner |
| EPI_ISL_2757931 | Labuan Hospital | Institute for Medical Research, Infectious Disease Research Centre, National Institutes of Health, Ministry of Health Malaysia | Suppiah J, Kamel K, Mohd Zawawi Z, Azizan MA, Ramly N, Robert F, Thayan R |
| EPI_ISL_2757940 | ISDH | Quantigen Biosciences | Paul J. Childress |
| EPI_ISL_2757953, EPI_ISL_2757954, EPI_ISL_2757955, EPI_ISL_2757956, EPI_ISL_2757957, EPI_ISL_2757958, EPI_ISL_2757959, EPI_ISL_2757966, EPI_ISL_2757970, EPI_ISL_2757972, EPI_ISL_2757977, EPI_ISL_2757978, EPI_ISL_2757979, EPI_ISL_2757980, EPI_ISL_2757981, EPI_ISL_2757982, EPI_ISL_2757983, EPI_ISL_2757984, EPI_ISL_2757990, EPI_ISL_2757992, EPI_ISL_2758002, EPI_ISL_2758003, EPI_ISL_2758005, EPI_ISL_2758006, EPI_ISL_2758024, EPI_ISL_2758025, EPI_ISL_2758026, EPI_ISL_2758029, EPI_ISL_2758030, EPI_ISL_2758031, EPI_ISL_2758032, EPI_ISL_2758033, EPI_ISL_2758034, EPI_ISL_2758035, EPI_ISL_2758036, EPI_ISL_2758037, EPI_ISL_2758038, EPI_ISL_2758039, EPI_ISL_2758040, EPI_ISL_2758041, EPI_ISL_2758043, EPI_ISL_2758044, EPI_ISL_2758045, EPI_ISL_2758046, EPI_ISL_2758047, EPI_ISL_2758048, EPI_ISL_2758049, EPI_ISL_2758050, EPI_ISL_2758051, EPI_ISL_2758052, EPI_ISL_2758053, EPI_ISL_2758054, EPI_ISL_2758056, EPI_ISL_2758057, EPI_ISL_2758058, EPI_ISL_2758059, EPI_ISL_2758060, EPI_ISL_2758061, EPI_ISL_2758063, EPI_ISL_2758064, EPI_ISL_2758065, EPI_ISL_2758066, EPI_ISL_2758067, EPI_ISL_2758068, EPI_ISL_2758069, EPI_ISL_2758070, EPI_ISL_2758071, EPI_ISL_2758072, EPI_ISL_2758073, EPI_ISL_2758074, EPI_ISL_2758075, EPI_ISL_2758076, EPI_ISL_2758077, EPI_ISL_2758078, EPI_ISL_2758079, EPI_ISL_2758080, EPI_ISL_2758082, EPI_ISL_2758083, EPI_ISL_2758084, EPI_ISL_2758085, EPI_ISL_2758086, EPI_ISL_2758087, EPI_ISL_2758088, EPI_ISL_2758089, EPI_ISL_2758090, EPI_ISL_2758091, EPI_ISL_2758092, EPI_ISL_2758093, EPI_ISL_2758094, EPI_ISL_2758095, EPI_ISL_2758096, EPI_ISL_2758097, EPI_ISL_2758098, EPI_ISL_2758099, EPI_ISL_2758100, EPI_ISL_2758101, EPI_ISL_2758102, EPI_ISL_2758103, EPI_ISL_2758104, EPI_ISL_2758105, EPI_ISL_2758106, EPI_ISL_2758107, EPI_ISL_2758108, EPI_ISL_2758109, EPI_ISL_2758110, EPI_ISL_2758111, EPI_ISL_2758112, EPI_ISL_2758113, EPI_ISL_2758114, EPI_ISL_2758115, EPI_ISL_2758116, EPI_ISL_2758117, EPI_ISL_2758118, EPI_ISL_2758119, EPI_ISL_2758120, EPI_ISL_2758121, EPI_ISL_2758122, EPI_ISL_2758123, EPI_ISL_2758124, EPI_ISL_2758125, EPI_ISL_2758126, EPI_ISL_2758127, EPI_ISL_2758128, EPI_ISL_2758130, EPI_ISL_2758133, EPI_ISL_2758137, EPI_ISL_2758138, EPI_ISL_2758139, EPI_ISL_2758140, EPI_ISL_2758141, EPI_ISL_2758142, EPI_ISL_2758143, EPI_ISL_2758144, EPI_ISL_2758146, EPI_ISL_2758147, EPI_ISL_2758148, EPI_ISL_2758151, EPI_ISL_2758153, EPI_ISL_2758154, EPI_ISL_2758156, EPI_ISL_2758158, EPI_ISL_2758163, EPI_ISL_2758164, EPI_ISL_2758165, EPI_ISL_2758167, EPI_ISL_2758170, EPI_ISL_2758171, EPI_ISL_2758173, EPI_ISL_2758174 | | | |
| see above | Austrian Agency for Health and Food Safety (AGES) | Bergthaler laboratory, CeMM Research Center for Molecular Medicine of the Austrian Academy of Sciences | Lukas Endler, Anna Schedl, Fabian Amman, Petr Triska, Matthew Thornton, Thomas Penz, Benedikt Agerer, Maelle Le Moing, Michael Schuster, Bekir Erguner, Jan Laine, Martin Senekowitsch, Christoph Bock, Andreas Bergthaler |
| EPI_ISL_2758180 | ILV Kärnten | Bergthaler laboratory, CeMM Research Center for Molecular Medicine of the Austrian Academy of Sciences | Lukas Endler, Anna Schedl, Fabian Amman, Petr Triska, Matthew Thornton, Thomas Penz, Benedikt Agerer, Maelle Le Moing, Michael Schuster, Bekir Erguner, Jan Laine, Martin Senekowitsch, Christoph Bock, Andreas Bergthaler |
| EPI_ISL_2758186, EPI_ISL_2758188 | Institute for Laboratory Diagnostics and Microbiology, Klinikum Klagenfurt am Wörthersee | Bergthaler laboratory, CeMM Research Center for Molecular Medicine of the Austrian Academy of Sciences | Lukas Endler, Anna Schedl, Fabian Amman, Petr Triska, Matthew Thornton, Thomas Penz, Benedikt Agerer, Maelle Le Moing, Michael Schuster, Bekir Erguner, Jan Laine, Martin Senekowitsch, Christoph Bock, Andreas Bergthaler |
| EPI_ISL_2758190 | Klinikum Wels-Grieskirchen | Bergthaler laboratory, CeMM Research Center for Molecular Medicine of the Austrian Academy of Sciences | Lukas Endler, Anna Schedl, Fabian Amman, Petr Triska, Matthew Thornton, Thomas Penz, Benedikt Agerer, Maelle Le Moing, Michael Schuster, Bekir Erguner, Jan Laine, Martin Senekowitsch, Christoph Bock, Andreas Bergthaler |
| EPI_ISL_2758229, EPI_ISL_2758230, EPI_ISL_2758234 | Weill Cornell Medicine | New York Genome Center | Michael Zody, Andre Corvelo, Dayna M. Oschwald, Samantha Fennessey, Tom Maniatis, Melissa Cushing, Olivier Elemento, Margaret Elizabeth Ross, Chris Mason, Priya Velu, Hanna Rennert, Arryn Craney, Lars F Westblade |
| EPI_ISL_2758272 | J.W. Ruby Memorial Hospital | WVU and Marshall University Combined Genomics Core Facilities | James Denvir, Peter Stoilov, Peter Perrotta, Wesley Kimble, Ryan Percifield |
| EPI_ISL_2758374 | Hampshire Memorial Hospital | WVU and Marshall University Combined Genomics Core Facilities | James Denvir, Peter Stoilov, Peter Perrotta, Wesley Kimble, Ryan Percifield |
| EPI_ISL_2758377 | Berkeley Medical Center | WVU and Marshall University Combined Genomics Core Facilities | James Denvir, Peter Stoilov, Peter Perrotta, Wesley Kimble, Ryan Percifield |
| EPI_ISL_2758396, EPI_ISL_2758397, EPI_ISL_2758398, EPI_ISL_2758399 | QLabs | WVU and Marshall University Combined Genomics Core Facilities | James Denvir, Peter Stoilov, Peter Perrotta, Wesley Kimble, Ryan Percifield |
| EPI_ISL_2758551, EPI_ISL_2758555, | Servicio de Microbiologia Hospital Ramon y Cajal | Servicio de Microbiologia Hospital Ramon y Cajal | Ponce M, Galan JC, Martinez L. Abreu M, y Gonzalez-Alba JM |

| | | | |
|---|---|---|---|
| EPI_ISL_2758557, EPI_ISL_2758576, EPI_ISL_2758582 | | | |
| EPI_ISL_2758635 | DOHMH Morrisania | New York City Public Health Laboratory | Jade Wang, et al. |
| EPI_ISL_2758636 | DOHMH Fort Greene | New York City Public Health Laboratory | Jade Wang, et al. |
| EPI_ISL_2758637 | DOHMH Riverside | New York City Public Health Laboratory | Jade Wang, et al. |
| EPI_ISL_2758638 | DOHMH Chelsea | New York City Public Health Laboratory | Jade Wang, et al. |
| EPI_ISL_2758639 | DOHMH Riverside | New York City Public Health Laboratory | Jade Wang, et al. |
| EPI_ISL_2758640, EPI_ISL_2758641, EPI_ISL_2758642, EPI_ISL_2758643, EPI_ISL_2758644 | Quest Diagnostics | New York City Public Health Laboratory | Jade Wang, et al. |
| EPI_ISL_2758645 | OCME Office Of Chief Medical Examiner | New York City Public Health Laboratory | Jade Wang, et al. |
| EPI_ISL_2758809 | PathWest Laboratory Medicine WA | PathWest Laboratory Medicine WA | PathWest Laboratory Medicine WA Microbial Surveillance Unit |
| EPI_ISL_2758922 | The Ohio State University Applied Microbiology Services Laboratory | The Ohio State University Applied Microbiology Services Laboratory | Seth A. Faith PhD |
| EPI_ISL_2759093 | MOH - Jaber Al-Ahmad Hospital (Innovation Research Laboratory) | MOH - Jaber Al-Ahmad Hospital (Innovation Research Laboratory) | Salman Al-Sabah , Mohammad Alghounaim |
| EPI_ISL_2759188 | Population Medicine and Diagnostic Sciences, Cornell University | Population Medicine and Diagnostic Sciences, Cornell University | Mitchell,P.K., Caserta,L.C., Cronk,B.D., Anderson,R.R., Laverack,M., Venugopalan,R., Plocharczyk,E., Diel,D.G. |
| EPI_ISL_2759660, EPI_ISL_2759668, EPI_ISL_2759671 | SYNLAB MVZ Leverkusen | Robert Koch Institute | unknown |
| EPI_ISL_2759672 | SYNLAB MVZ Weiden | Robert Koch Institute | unknown |
| EPI_ISL_2759675, EPI_ISL_2759677 | SYNLAB MVZ Leinfelden-Echterdingen | Robert Koch Institute | unknown |
| EPI_ISL_2759679 | SYNLAB MVZ Leverkusen | Robert Koch Institute | unknown |
| EPI_ISL_2759680, EPI_ISL_2759681, EPI_ISL_2759682, EPI_ISL_2759684, EPI_ISL_2759685 | SYNLAB MVZ Leinfelden-Echterdingen | Robert Koch Institute | unknown |
| EPI_ISL_2759689 | SYNLAB Labor München Zentrum LMZ | Robert Koch Institute | unknown |
| EPI_ISL_2759693, EPI_ISL_2759697, EPI_ISL_2759699 | SYNLAB MVZ Leinfelden-Echterdingen | Robert Koch Institute | unknown |
| EPI_ISL_2759721 | SYNLAB MVZ Weiden | Robert Koch Institute | unknown |
| EPI_ISL_2759724, EPI_ISL_2759731 | SYNLAB MVZ Leinfelden-Echterdingen | Robert Koch Institute | unknown |
| EPI_ISL_2759735 | SYNLAB Labor München Zentrum LMZ | Robert Koch Institute | unknown |
| EPI_ISL_2759737 | SYNLAB MVZ Leverkusen | Robert Koch Institute | unknown |
| EPI_ISL_2759738 | SYNLAB Labor München Zentrum LMZ | Robert Koch Institute | unknown |
| EPI_ISL_2759740 | SYNLAB MVZ Trier | Robert Koch Institute | unknown |
| EPI_ISL_2759741 | SYNLAB MVZ Neustadt a. d. Weinstraße | Robert Koch Institute | unknown |
| EPI_ISL_2759743, EPI_ISL_2759745 | SYNLAB MVZ Leverkusen | Robert Koch Institute | unknown |
| EPI_ISL_2759757 | SYNLAB MVZ Weiden | Robert Koch Institute | unknown |
| EPI_ISL_2759759, EPI_ISL_2759764, EPI_ISL_2759771 | SYNLAB MVZ Leverkusen | Robert Koch Institute | unknown |
| EPI_ISL_2759772 | SYNLAB MVZ Leinfelden-Echterdingen | Robert Koch Institute | unknown |
| EPI_ISL_2759778 | SYNLAB MVZ Leverkusen | Robert Koch Institute | unknown |
| EPI_ISL_2759817 | Universitätsklinikum Bonn - Institut für Virologie | Robert Koch Institute | unknown |
| EPI_ISL_2759818 | Uniklinikum Regensburg; Institut für klinische Mikrobiologie und Hygiene | Robert Koch Institute | unknown |
| EPI_ISL_2759829, EPI_ISL_2759842 | Bioscientia Labor Wermsdorf | Robert Koch Institute | unknown |
| EPI_ISL_2759844 | Sonic - Labor Dr. von Froreich GmbH | Robert Koch Institute | unknown |
| EPI_ISL_2759847 | Sonic - MVZ Medizinisches Labor Bremen GmbH | Robert Koch Institute | unknown |
| EPI_ISL_2759850 | Sonic - Labor Staber Bayreuth | Robert Koch Institute | unknown |
| EPI_ISL_2759877, EPI_ISL_2759878, EPI_ISL_2759879, EPI_ISL_2759882, EPI_ISL_2759883, EPI_ISL_2759884 | Washington State Department of Health Public Health Laboratories | Washington State Department of Health Public Health Laboratories | Drew MacKellar, Philip Dykema, Denny Russell, Joenice Gonzalez, Hannah Gray, Geoff Melly, Vanessa De Los Santos, Darren Lucas, JohnAric Peterson, Avi Singh, Rebecca Cao |
| EPI_ISL_2759889, EPI_ISL_2759904 | SYNLAB MVZ Neustadt a. d. Weinstraße | Robert Koch Institute | unknown |
| EPI_ISL_2759909 | SYNLAB MVZ Weiden | Robert Koch Institute | unknown |
| EPI_ISL_2759913 | SYNLAB MVZ Heidelberg | Robert Koch Institute | unknown |
| EPI_ISL_2759915 | SYNLAB MVZ Leinfelden-Echterdingen | Robert Koch Institute | unknown |
| EPI_ISL_2759916 | SYNLAB MVZ Leverkusen | Robert Koch Institute | unknown |
| EPI_ISL_2759922 | SYNLAB MVZ Leinfelden-Echterdingen | Robert Koch Institute | unknown |
| EPI_ISL_2759929 | SYNLAB MVZ Leverkusen | Robert Koch Institute | unknown |
| EPI_ISL_2759931 | SYNLAB MVZ Berlin | Robert Koch Institute | unknown |
| EPI_ISL_2759937 | SYNLAB MVZ Leinfelden-Echterdingen | Robert Koch Institute | unknown |
| EPI_ISL_2759940 | SYNLAB MVZ Trier | Robert Koch Institute | unknown |
| EPI_ISL_2759941 | SYNLAB MVZ Leinfelden-Echterdingen | Robert Koch Institute | unknown |
| EPI_ISL_2759944 | SYNLAB MVZ Trier | Robert Koch Institute | unknown |
| EPI_ISL_2759952, EPI_ISL_2759954, EPI_ISL_2759963, EPI_ISL_2759966 | SYNLAB MVZ Leinfelden-Echterdingen | Robert Koch Institute | unknown |
| EPI_ISL_2759972, EPI_ISL_2759983 | SYNLAB MVZ Leverkusen | Robert Koch Institute | unknown |
| EPI_ISL_2759987 | SYNLAB Labor München Zentrum LMZ | Robert Koch Institute | unknown |

| | | | |
|---|---|---|---|
| EPI_ISL_2759988 | SYNLAB MVZ Leinfelden-Echterdingen | Robert Koch Institute | unknown |
| EPI_ISL_2759989 | SYNLAB Labor München Zentrum LMZ | Robert Koch Institute | unknown |
| EPI_ISL_2759995 | SYNLAB MVZ Leinfelden-Echterdingen | Robert Koch Institute | unknown |
| EPI_ISL_2759999, EPI_ISL_2760000, EPI_ISL_2760001 | SYNLAB MVZ Weiden | Robert Koch Institute | unknown |
| EPI_ISL_2760005, EPI_ISL_2760008 | SYNLAB MVZ Leinfelden-Echterdingen | Robert Koch Institute | unknown |
| EPI_ISL_2760009, EPI_ISL_2760012, EPI_ISL_2760019 | SYNLAB MVZ Heidelberg | Robert Koch Institute | unknown |
| EPI_ISL_2760023, EPI_ISL_2760025 | Bayerisches Landesamt für Gesundheit und Lebensmittelsicherheit (LGL) | Robert Koch Institute | unknown |
| EPI_ISL_2760083, EPI_ISL_2760085 | Pathogen Genomics Center, National Institute of Infectious Diseases | Pathogen Genomics Center, National Institute of Infectious Diseases | Tsuyoshi Sekizuka, Kentaro Itokawa, Rina Tanaka, Masanori Hashino, Makoto Kuroda |
| EPI_ISL_2760100 | SARS-CoV-2 testing team, National Institute of Infectious Diseases | Pathogen Genomics Center, National Institute of Infectious Diseases | Tsuyoshi Sekizuka, Kentaro Itokawa, Rina Tanaka, Masanori Hashino, Nozomu Hanaoka, Masumichi Saito, Naomi Nojiri, Hazuka Y Furihata, Hiromizu Takahashi, Tsuguto Fujimoto, Makoto Kuroda |
| EPI_ISL_2760117, EPI_ISL_2760131 | Bayerisches Landesamt für Gesundheit und Lebensmittelsicherheit (LGL) | Robert Koch Institute | unknown |
| EPI_ISL_2760153, EPI_ISL_2760155, EPI_ISL_2760157, EPI_ISL_2760159, EPI_ISL_2760161, EPI_ISL_2760164, EPI_ISL_2760166, EPI_ISL_2760168, EPI_ISL_2760171, EPI_ISL_2760174, EPI_ISL_2760176, EPI_ISL_2760178, EPI_ISL_2760181, EPI_ISL_2760184, EPI_ISL_2760186, EPI_ISL_2760189 | | | |
| see above | Landesgesundheitsamt Baden-Württemberg | Robert Koch Institute | unknown |
| EPI_ISL_2760207 | Pathogen Genomics Center, National Institute of Infectious Diseases | Pathogen Genomics Center, National Institute of Infectious Diseases | Tsuyoshi Sekizuka, Kentaro Itokawa, Rina Tanaka, Masanori Hashino, Makoto Kuroda |
| EPI_ISL_2760212, EPI_ISL_2760213, EPI_ISL_2760215, EPI_ISL_2760216 | Osaka Institute of Public Health, Morinomiya Center | Pathogen Genomics Center, National Institute of Infectious Diseases | Tsuyoshi Sekizuka, Kentaro Itokawa, Rina Tanaka, Masanori Hashino, Makoto Kuroda |
| EPI_ISL_2760315, EPI_ISL_2760320 | MVZ Labor Krone GbR | Robert Koch Institute | unknown |
| EPI_ISL_2760374 | Pathogen Genomics Center, National Institute of Infectious Diseases | Pathogen Genomics Center, National Institute of Infectious Diseases | Tsuyoshi Sekizuka, Kentaro Itokawa, Rina Tanaka, Masanori Hashino, Makoto Kuroda |
| EPI_ISL_2760394, EPI_ISL_2760396 | Labormedizin Darmstadt | Robert Koch Institute | unknown |
| EPI_ISL_2760399 | MVZ Dr. Eberhard & Partner Dortmund | Robert Koch Institute | unknown |
| EPI_ISL_2760405 | Pathogen Genomics Center, National Institute of Infectious Diseases | Pathogen Genomics Center, National Institute of Infectious Diseases | Tsuyoshi Sekizuka, Kentaro Itokawa, Rina Tanaka, Masanori Hashino, Makoto Kuroda |
| EPI_ISL_2760409 | MVZ für Laboratoriumsmedizin und Mikrobiologie Würzburg | Robert Koch Institute | unknown |
| EPI_ISL_2760412 | MVZ Dr. Eberhard & Partner Dortmund | Robert Koch Institute | unknown |
| EPI_ISL_2760414, EPI_ISL_2760417 | MVZ für Laboratoriumsmedizin und Mikrobiologie Würzburg | Robert Koch Institute | unknown |
| EPI_ISL_2760418 | Pathogen Genomics Center, National Institute of Infectious Diseases | Pathogen Genomics Center, National Institute of Infectious Diseases | Tsuyoshi Sekizuka, Kentaro Itokawa, Rina Tanaka, Masanori Hashino, Makoto Kuroda |
| EPI_ISL_2760419 | Labor Lübeck bzw. Laborärztliche Gemeinschaftspraxis Lübeck | Robert Koch Institute | unknown |
| EPI_ISL_2760420, EPI_ISL_2760423 | Pathogen Genomics Center, National Institute of Infectious Diseases | Pathogen Genomics Center, National Institute of Infectious Diseases | Tsuyoshi Sekizuka, Kentaro Itokawa, Rina Tanaka, Masanori Hashino, Makoto Kuroda |
| EPI_ISL_2760426, EPI_ISL_2760439 | MVZ für Laboratoriumsmedizin und Mikrobiologie Würzburg | Robert Koch Institute | unknown |
| EPI_ISL_2760440 | Pathogen Genomics Center, National Institute of Infectious Diseases | Pathogen Genomics Center, National Institute of Infectious Diseases | Tsuyoshi Sekizuka, Kentaro Itokawa, Rina Tanaka, Masanori Hashino, Makoto Kuroda |
| EPI_ISL_2760447 | MVZ Dr. Eberhard & Partner Dortmund | Robert Koch Institute | unknown |
| EPI_ISL_2760451, EPI_ISL_2760454 | Pathogen Genomics Center, National Institute of Infectious Diseases | Pathogen Genomics Center, National Institute of Infectious Diseases | Tsuyoshi Sekizuka, Kentaro Itokawa, Rina Tanaka, Masanori Hashino, Makoto Kuroda |
| EPI_ISL_2760455 | amedes MVZ Hannover | Robert Koch Institute | unknown |
| EPI_ISL_2760458 | Pathogen Genomics Center, National Institute of Infectious Diseases | Pathogen Genomics Center, National Institute of Infectious Diseases | Tsuyoshi Sekizuka, Kentaro Itokawa, Rina Tanaka, Masanori Hashino, Makoto Kuroda |
| EPI_ISL_2760463 | amedes MVZ für Laboratoriumsdiagnostik Raubling GmbH | Robert Koch Institute | unknown |
| EPI_ISL_2760467 | SARS-CoV-2 testing team, National Institute of Infectious Diseases | Pathogen Genomics Center, National Institute of Infectious Diseases | Tsuyoshi Sekizuka, Kentaro Itokawa, Rina Tanaka, Masanori Hashino, Nozomu Hanaoka, Masumichi Saito, Naomi Nojiri, Hazuka Y Furihata, Hiromizu Takahashi, Tsuguto Fujimoto, Makoto Kuroda |
| EPI_ISL_2760471, EPI_ISL_2760478 | nordlab - Partnerschaftspraxis für Laboratoriumsmedizin | Robert Koch Institute | unknown |
| EPI_ISL_2760480 | amedes MVZ für Laboratoriumsmedizin Rhein/Ruhr | Robert Koch Institute | unknown |
| EPI_ISL_2760485, EPI_ISL_2760488 | nordlab - Partnerschaftspraxis für Laboratoriumsmedizin | Robert Koch Institute | unknown |
| EPI_ISL_2760493 | amedes MVZ DIAMEDIS Sennestadt | Robert Koch Institute | unknown |
| EPI_ISL_2760496, EPI_ISL_2760501 | nordlab - Partnerschaftspraxis für Laboratoriumsmedizin | Robert Koch Institute | unknown |
| EPI_ISL_2760504 | amedes MVZ DIAMEDIS Sennestadt | Robert Koch Institute | unknown |
| EPI_ISL_2760506 | amedes MVZ für Laboratoriumsmedizin Fürstenfeldbruck | Robert Koch Institute | unknown |
| EPI_ISL_2760513 | nordlab - Partnerschaftspraxis für Laboratoriumsmedizin | Robert Koch Institute | unknown |
| EPI_ISL_2760525 | amedes MVZ für Labordiagnostik Rhein-Main | Robert Koch Institute | unknown |
| EPI_ISL_2760528 | nordlab - Partnerschaftspraxis für Laboratoriumsmedizin | Robert Koch Institute | unknown |
| EPI_ISL_2760530 | amedes MVZ für Labordiagnostik Rhein-Main | Robert Koch Institute | unknown |
| EPI_ISL_2760533 | nordlab - Partnerschaftspraxis für Laboratoriumsmedizin | Robert Koch Institute | unknown |
| EPI_ISL_2760535 | amedes MVZ Hannover | Robert Koch Institute | unknown |
| EPI_ISL_2760538 | amedes MVZ für Labordiagnostik Rhein-Main | Robert Koch Institute | unknown |
| EPI_ISL_2760543 | amedes MVZ für Laboratoriumsmedizin München | Robert Koch Institute | unknown |
| EPI_ISL_2760545, EPI_ISL_2760547, EPI_ISL_2760549, EPI_ISL_2760553, EPI_ISL_2760555 | Niedersächsisches Landesgesundheitsamt (NLGA) | Robert Koch Institute | unknown |
| EPI_ISL_2760558, EPI_ISL_2760560, EPI_ISL_2760576, EPI_ISL_2760594, EPI_ISL_2760611 | MVZ Labor Dr. Limbach & Kollegen GbR | Robert Koch Institute | unknown |
| EPI_ISL_2760623, EPI_ISL_2760625, | Medizinische Laboratorien Düsseldorf | Robert Koch Institute | unknown |

| | | | |
|---|---|---|---|
| EPI_ISL_2760630, EPI_ISL_2760642 | | | |
| EPI_ISL_2760666 | SARS-CoV-2 testing team, National Institute of Infectious Diseases | Pathogen Genomics Center, National Institute of Infectious Diseases | Tsuyoshi Sekizuka, Kentaro Itokawa, Rina Tanaka, Masanori Hashino, Nozomu Hanaoka, Masumichi Saito, Naomi Nojiri, Hazuka Y Furihata, Hiromizu Takahashi, Tsuguto Fujimoto, Makoto Kuroda |
| EPI_ISL_2760667 | SYNLAB MVZ Leinfelden-Echterdingen | Robert Koch Institute | unknown |
| EPI_ISL_2760668 | SARS-CoV-2 testing team, National Institute of Infectious Diseases | Pathogen Genomics Center, National Institute of Infectious Diseases | Tsuyoshi Sekizuka, Kentaro Itokawa, Rina Tanaka, Masanori Hashino, Nozomu Hanaoka, Masumichi Saito, Naomi Nojiri, Hazuka Y Furihata, Hiromizu Takahashi, Tsuguto Fujimoto, Makoto Kuroda |
| EPI_ISL_2760674 | SYNLAB MVZ Leinfelden-Echterdingen | Robert Koch Institute | unknown |
| EPI_ISL_2760676 | SYNLAB MVZ Leverkusen | Robert Koch Institute | unknown |
| EPI_ISL_2760693 | SYNLAB MVZ Leinfelden-Echterdingen | Robert Koch Institute | unknown |
| EPI_ISL_2760700 | SYNLAB Labor München Zentrum LMZ | Robert Koch Institute | unknown |
| EPI_ISL_2760720 | SYNLAB MVZ Leinfelden-Echterdingen | Robert Koch Institute | unknown |
| EPI_ISL_2760722 | SYNLAB Labor München Zentrum LMZ | Robert Koch Institute | unknown |
| EPI_ISL_2760724, EPI_ISL_2760743 | SYNLAB MVZ Leinfelden-Echterdingen | Robert Koch Institute | unknown |
| EPI_ISL_2760751 | SYNLAB MVZ Weiden | Robert Koch Institute | unknown |
| EPI_ISL_2760763 | SYNLAB MVZ Trier | Robert Koch Institute | unknown |
| EPI_ISL_2760791 | SYNLAB MVZ Leinfelden-Echterdingen | Robert Koch Institute | unknown |
| EPI_ISL_2760811 | SYNLAB MVZ Weiden | Robert Koch Institute | unknown |
| EPI_ISL_2760816 | SYNLAB MVZ Leinfelden-Echterdingen | Robert Koch Institute | unknown |
| EPI_ISL_2760830, EPI_ISL_2760835, EPI_ISL_2760839 | Chiba Prefectural Institute of Public Health | Pathogen Genomics Center, National Institute of Infectious Diseases | Tsuyoshi Sekizuka, Kentaro Itokawa, Rina Tanaka, Masanori Hashino, Makoto Kuroda |
| EPI_ISL_2760840, EPI_ISL_2760842, EPI_ISL_2760844 | Osaka Institute of Public Health, Morinomiya Center | Pathogen Genomics Center, National Institute of Infectious Diseases | Tsuyoshi Sekizuka, Kentaro Itokawa, Rina Tanaka, Masanori Hashino, Makoto Kuroda |
| EPI_ISL_2760870 | SYNLAB MVZ Leverkusen | Robert Koch Institute | unknown |
| EPI_ISL_2760886, EPI_ISL_2760887, EPI_ISL_2760888, EPI_ISL_2760889, EPI_ISL_2760890 | SARS-CoV-2 testing team, National Institute of Infectious Diseases | Pathogen Genomics Center, National Institute of Infectious Diseases | Tsuyoshi Sekizuka, Kentaro Itokawa, Rina Tanaka, Masanori Hashino, Nozomu Hanaoka, Masumichi Saito, Naomi Nojiri, Hazuka Y Furihata, Hiromizu Takahashi, Tsuguto Fujimoto, Makoto Kuroda |
| EPI_ISL_2761070 | Bioscientia Labor Wermsdorf | Robert Koch Institute | unknown |
| EPI_ISL_2761072, EPI_ISL_2761074, EPI_ISL_2761075, EPI_ISL_2761077, EPI_ISL_2761079 | SARS-CoV-2 testing team, National Institute of Infectious Diseases | Pathogen Genomics Center, National Institute of Infectious Diseases | Tsuyoshi Sekizuka, Kentaro Itokawa, Rina Tanaka, Masanori Hashino, Nozomu Hanaoka, Masumichi Saito, Naomi Nojiri, Hazuka Y Furihata, Hiromizu Takahashi, Tsuguto Fujimoto, Makoto Kuroda |
| EPI_ISL_2761080 | LabKom - Labor Augsburg MVZ GmbH | Robert Koch Institute | unknown |
| EPI_ISL_2761081 | SARS-CoV-2 testing team, National Institute of Infectious Diseases | Pathogen Genomics Center, National Institute of Infectious Diseases | Tsuyoshi Sekizuka, Kentaro Itokawa, Rina Tanaka, Masanori Hashino, Nozomu Hanaoka, Masumichi Saito, Naomi Nojiri, Hazuka Y Furihata, Hiromizu Takahashi, Tsuguto Fujimoto, Makoto Kuroda |
| EPI_ISL_2761086, EPI_ISL_2761091, EPI_ISL_2761093 | Bioscientia Labor Wermsdorf | Robert Koch Institute | unknown |
| EPI_ISL_2761096 | Sonic - Labor Dr. von Froreich GmbH | Robert Koch Institute | unknown |
| EPI_ISL_2761100 | Sonic - MVZ Medizinisches Labor Bremen GmbH | Robert Koch Institute | unknown |
| EPI_ISL_2761108, EPI_ISL_2761122, EPI_ISL_2761127, EPI_ISL_2761132 | Bioscientia Labor Wermsdorf | Robert Koch Institute | unknown |
| EPI_ISL_2761138 | Sonic - Bioscientia - MVZ Labor Saar GmbH | Robert Koch Institute | unknown |
| EPI_ISL_2761159, EPI_ISL_2761178 | Bioscientia Labor Wermsdorf | Robert Koch Institute | unknown |
| EPI_ISL_2761180 | Bioscientia MVZ Labor Karlsruhe GmbH | Robert Koch Institute | unknown |
| EPI_ISL_2761182 | Bioscientia Labor Wermsdorf | Robert Koch Institute | unknown |
| EPI_ISL_2761213 | Sonic - Labor Staber Bayreuth | Robert Koch Institute | unknown |
| EPI_ISL_2761219 | Sonic - MVZ Medizinisches Labor Bremen GmbH | Robert Koch Institute | unknown |
| EPI_ISL_2761232 | Bioscientia Labor Wermsdorf | Robert Koch Institute | unknown |
| EPI_ISL_2761242 | LabKom - Labor Augsburg MVZ GmbH | Robert Koch Institute | unknown |
| EPI_ISL_2761250, EPI_ISL_2761259, EPI_ISL_2761269 | Bioscientia Labor Wermsdorf | Robert Koch Institute | unknown |
| EPI_ISL_2761285 | Bioscientia MVZ Labor Karlsruhe GmbH | Robert Koch Institute | unknown |
| EPI_ISL_2761292, EPI_ISL_2761297, EPI_ISL_2761299 | Bioscientia Labor Wermsdorf | Robert Koch Institute | unknown |
| EPI_ISL_2761302 | Labor 28 MVZ GmbH | Robert Koch Institute | unknown |
| EPI_ISL_2761312 | Robert-Bosch-Krankenhaus | Robert Koch Institute | unknown |
| EPI_ISL_2761318, EPI_ISL_2761321, EPI_ISL_2761328, EPI_ISL_2761349, EPI_ISL_2761360, EPI_ISL_2761367, EPI_ISL_2761374, EPI_ISL_2761379, EPI_ISL_2761389, EPI_ISL_2761395, EPI_ISL_2761400 | | | |
| see above | Bioscientia Labor Wermsdorf | Robert Koch Institute | unknown |
| EPI_ISL_2761405 | Bioscientia MVZ Labor Karlsruhe GmbH | Robert Koch Institute | unknown |
| EPI_ISL_2761408 | Sonic - Dr. Staber & Kollegen GmbH München | Robert Koch Institute | unknown |
| EPI_ISL_2761413 | Bioscientia Labor Wermsdorf | Robert Koch Institute | unknown |
| EPI_ISL_2761415 | Sonic - Labor Dr. von Froreich GmbH | Robert Koch Institute | unknown |
| EPI_ISL_2761417 | Bioscientia Labor Wermsdorf | Robert Koch Institute | unknown |
| EPI_ISL_2761457 | Sonic - Labor Dr. von Froreich GmbH | Robert Koch Institute | unknown |
| EPI_ISL_2761460, EPI_ISL_2761477, EPI_ISL_2761493 | Bioscientia Labor Wermsdorf | Robert Koch Institute | unknown |
| EPI_ISL_2761508 | Bioscientia MVZ Labor Karlsruhe GmbH | Robert Koch Institute | unknown |
| EPI_ISL_2761510 | Bioscientia Labor Wermsdorf | Robert Koch Institute | unknown |
| EPI_ISL_2761513 | Klinikum Fulda gAG | Robert Koch Institute | unknown |
| EPI_ISL_2761528, EPI_ISL_2761536 | Bioscientia Labor Wermsdorf | Robert Koch Institute | unknown |
| EPI_ISL_2761554 | LabKom - Labor Augsburg MVZ GmbH | Robert Koch Institute | unknown |
| EPI_ISL_2761563 | Sonic - MVZ Medizinisches Labor Bremen GmbH | Robert Koch Institute | unknown |

| | | | |
|---|---|---|---|
| EPI_ISL_2761568 | Bioscientia Labor Wermsdorf | Robert Koch Institute | unknown |
| EPI_ISL_2761576 | LabKom - Labor Augsburg MVZ GmbH | Robert Koch Institute | unknown |
| EPI_ISL_2761578 | Bioscientia Labor Wermsdorf | Robert Koch Institute | unknown |
| EPI_ISL_2761583 | Labor 28 MVZ GmbH | Robert Koch Institute | unknown |
| EPI_ISL_2761586 | Sonic - Bioscientia - MVZ Labor Saar GmbH | Robert Koch Institute | unknown |
| EPI_ISL_2761598, EPI_ISL_2761600 | Bioscientia Labor Wermsdorf | Robert Koch Institute | unknown |
| EPI_ISL_2761602 | Bioscientia MVZ Labor Karlsruhe GmbH | Robert Koch Institute | unknown |
| EPI_ISL_2761603 | Bioscientia Labor Wermsdorf | Robert Koch Institute | unknown |
| EPI_ISL_2761607 | Labor Mönchengladbach MVZ Dr. Stein + Kollegen GbR | Robert Koch Institute | unknown |
| EPI_ISL_2761608 | MVZ Labor Passau GbR | Robert Koch Institute | unknown |
| EPI_ISL_2761611, EPI_ISL_2761613, EPI_ISL_2761614 | Labor Mönchengladbach MVZ Dr. Stein + Kollegen GbR | Robert Koch Institute | unknown |
| EPI_ISL_2761615 | MVZ Labor Dr. Limbach & Kollegen GbR | Robert Koch Institute | unknown |
| EPI_ISL_2761625 | MVZ Labor Dr. Reising-Ackermann und Kollegen | Robert Koch Institute | unknown |
| EPI_ISL_2761627 | MDI Limbach Berlin GmbH; MVZ Labor Berlin | Robert Koch Institute | unknown |
| EPI_ISL_2761629 | Labor Mönchengladbach MVZ Dr. Stein + Kollegen GbR | Robert Koch Institute | unknown |
| EPI_ISL_2761631 | MVZ Labor Dr. Limbach & Kollegen GbR | Robert Koch Institute | unknown |
| EPI_ISL_2761636 | MVZ Labor Passau GbR | Robert Koch Institute | unknown |
| EPI_ISL_2761642, EPI_ISL_2761643, EPI_ISL_2761644, EPI_ISL_2761645, EPI_ISL_2761646, EPI_ISL_2761647, EPI_ISL_2761648, EPI_ISL_2761649, EPI_ISL_2761650, EPI_ISL_2761651, EPI_ISL_2761652, EPI_ISL_2761653, EPI_ISL_2761654, EPI_ISL_2761655, EPI_ISL_2761656, EPI_ISL_2761657, EPI_ISL_2761660, EPI_ISL_2761661, EPI_ISL_2761662, EPI_ISL_2761663 | | | |
| see above | Landesgesundheitsamt Baden-Württemberg | Robert Koch Institute | unknown |
| EPI_ISL_2761664, EPI_ISL_2761665, EPI_ISL_2761668, EPI_ISL_2761673, EPI_ISL_2761679, EPI_ISL_2761682 | MVZ Labor Dr. Fenner und Kollegen (Standort Hamburg) | Robert Koch Institute | unknown |
| EPI_ISL_2761684, EPI_ISL_2761687, EPI_ISL_2761689, EPI_ISL_2761692, EPI_ISL_2761695, EPI_ISL_2761696, EPI_ISL_2761698, EPI_ISL_2761699, EPI_ISL_2761700, EPI_ISL_2761701, EPI_ISL_2761702, EPI_ISL_2761704, EPI_ISL_2761705, EPI_ISL_2761706, EPI_ISL_2761707 | | | |
| see above | Labor Dr. Heidrich & Kollegen MVZ GmbH Hamburg | Robert Koch Institute | unknown |
| EPI_ISL_2761708, EPI_ISL_2761712, EPI_ISL_2761713, EPI_ISL_2761714, EPI_ISL_2761716, EPI_ISL_2761718, EPI_ISL_2761719, EPI_ISL_2761720, EPI_ISL_2761721, EPI_ISL_2761722, EPI_ISL_2761723, EPI_ISL_2761726, EPI_ISL_2761727 | | | |
| see above | Labor Mönchengladbach MVZ Dr. Stein + Kollegen GbR | Robert Koch Institute | unknown |
| EPI_ISL_2761730, EPI_ISL_2761731, EPI_ISL_2761733, EPI_ISL_2761734 | Institut für Labormedizin Mikrobiologie und Hygiene | Robert Koch Institute | unknown |
| EPI_ISL_2761745, EPI_ISL_2761746, EPI_ISL_2761757, EPI_ISL_2761759, EPI_ISL_2761760, EPI_ISL_2761762, EPI_ISL_2761763, EPI_ISL_2761767, EPI_ISL_2761768, EPI_ISL_2761770, EPI_ISL_2761786, EPI_ISL_2761788, EPI_ISL_2761789, EPI_ISL_2761794, EPI_ISL_2761796, EPI_ISL_2761797, EPI_ISL_2761805, EPI_ISL_2761813, EPI_ISL_2761815, EPI_ISL_2761816, EPI_ISL_2761819, EPI_ISL_2761820, EPI_ISL_2761825, EPI_ISL_2761826, EPI_ISL_2761830, EPI_ISL_2761831, EPI_ISL_2761833, EPI_ISL_2761839, EPI_ISL_2761843, EPI_ISL_2761846, EPI_ISL_2761847, EPI_ISL_2761849, EPI_ISL_2761852, EPI_ISL_2761853, EPI_ISL_2761856, EPI_ISL_2761858 | | | |
| see above | CENTOGENE Frankfurt Laboratory: Niederlassung Industriepark Höchst | Robert Koch Institute | unknown |
| EPI_ISL_2761868 | MVZ für Laboratoriumsmedizin und Mikrobiologie Koblenz-Mittelrhein (Labor Koblenz) | Robert Koch Institute | unknown |
| EPI_ISL_2761870 | MDI Limbach Berlin GmbH; MVZ Labor Berlin | Robert Koch Institute | unknown |
| EPI_ISL_2761874 | MVZ Medizinisches Labor Hannover GmbH | Robert Koch Institute | unknown |
| EPI_ISL_2761875 | MVZ Labor Dr. Limbach & Kollegen GbR | Robert Koch Institute | unknown |
| EPI_ISL_2761876 | MVZ Labor Passau GbR | Robert Koch Institute | unknown |
| EPI_ISL_2761878, EPI_ISL_2761879 | MVZ für Laboratoriumsmedizin und Mikrobiologie Koblenz-Mittelrhein (Labor Koblenz) | Robert Koch Institute | unknown |
| EPI_ISL_2761882, EPI_ISL_2761883 | MDI Limbach Berlin GmbH; MVZ Labor Berlin | Robert Koch Institute | unknown |
| EPI_ISL_2761884, EPI_ISL_2761885, EPI_ISL_2761887 | MVZ für Laboratoriumsmedizin und Mikrobiologie Koblenz-Mittelrhein (Labor Koblenz) | Robert Koch Institute | unknown |
| EPI_ISL_2761888 | MDI Limbach Berlin GmbH; MVZ Labor Berlin | Robert Koch Institute | unknown |
| EPI_ISL_2761889, EPI_ISL_2761890, EPI_ISL_2761891 | MVZ für Laboratoriumsmedizin und Mikrobiologie Koblenz-Mittelrhein (Labor Koblenz) | Robert Koch Institute | unknown |
| EPI_ISL_2761892 | MDI Limbach Berlin GmbH; MVZ Labor Berlin | Robert Koch Institute | unknown |
| EPI_ISL_2761893 | MVZ Labor Dr. Limbach & Kollegen GbR | Robert Koch Institute | unknown |
| EPI_ISL_2761894 | MDI Limbach Berlin GmbH; MVZ Labor Berlin | Robert Koch Institute | unknown |
| EPI_ISL_2761895 | MVZ für Laboratoriumsmedizin und Mikrobiologie Koblenz-Mittelrhein (Labor Koblenz) | Robert Koch Institute | unknown |
| EPI_ISL_2761896 | MVZ Labor Passau GbR | Robert Koch Institute | unknown |
| EPI_ISL_2761899, EPI_ISL_2761901, EPI_ISL_2761902, EPI_ISL_2761904, EPI_ISL_2761906 | MVZ für Laboratoriumsmedizin und Mikrobiologie Koblenz-Mittelrhein (Labor Koblenz) | Robert Koch Institute | unknown |
| EPI_ISL_2761907, EPI_ISL_2761909, EPI_ISL_2761912, EPI_ISL_2761922 | Medizinisch-Diagnostisches Labor Kempten allgäulab | Robert Koch Institute | unknown |
| EPI_ISL_2761923 | MVZ Dr. Eberhard & Partner Dortmund | Robert Koch Institute | unknown |
| EPI_ISL_2761924 | Medizinisches Versorgungszentrum für Labormedizin und Mikrobiologie Ruhr GmbH - mvzlm RUHR GmbH | Robert Koch Institute | unknown |
| EPI_ISL_2761925 | Laborarztpraxis Osnabrück | Robert Koch Institute | unknown |
| EPI_ISL_2761926 | Procomcure Biotech Germany GmbH | Robert Koch Institute | unknown |
| EPI_ISL_2761927 | MVZ Dr. Eberhard & Partner Dortmund | Robert Koch Institute | unknown |
| EPI_ISL_2761931 | Laborarztpraxis Osnabrück | Robert Koch Institute | unknown |
| EPI_ISL_2761933 | MVZ für Laboratoriumsmedizin und Mikrobiologie Würzburg | Robert Koch Institute | unknown |
| EPI_ISL_2761935 | Laborarztpraxis Osnabrück | Robert Koch Institute | unknown |
| EPI_ISL_2761937 | Procomcure Biotech Germany GmbH | Robert Koch Institute | unknown |
| EPI_ISL_2761940 | Institut für Klinische Chemie Transfusions- und Laboratoriumsmedizin Berufsgenossenschaftliches Universitätsklinikum Bergmannsheil | Robert Koch Institute | unknown |
| EPI_ISL_2761941 | MVZ Dr. Eberhard & Partner Dortmund | Robert Koch Institute | unknown |

| | | | |
|---|---|---|---|
| EPI_ISL_2761942 | Laborarztpraxis Osnabrück | Robert Koch Institute | unknown |
| EPI_ISL_2761944, EPI_ISL_2761948, EPI_ISL_2761955, EPI_ISL_2761956, EPI_ISL_2761957 | labopart - Medizinische Laboratorien Dresden | Robert Koch Institute | unknown |
| EPI_ISL_2761960, EPI_ISL_2761961, EPI_ISL_2761962 | Universitätsklinikum Frankfurt - Institut für Medizinische Virologie | Robert Koch Institute | unknown |
| EPI_ISL_2761965 | Klinikum Stuttgart | Robert Koch Institute | unknown |
| EPI_ISL_2761971 | LabKom - Labor Mainz MVZ GmbH | Robert Koch Institute | unknown |
| EPI_ISL_2761973 | LabKom - Labor an der Salzbrücke MVZ GmbH | Robert Koch Institute | unknown |
| EPI_ISL_2761976, EPI_ISL_2761979 | Bioscientia Labor Wermsdorf | Robert Koch Institute | unknown |
| EPI_ISL_2761980 | LabKom - Labor an der Salzbrücke MVZ GmbH | Robert Koch Institute | unknown |
| EPI_ISL_2761981 | Sonic - MVZ Medizinisches Labor Bremen GmbH | Robert Koch Institute | unknown |
| EPI_ISL_2761983 | Bioscientia Labor Wermsdorf | Robert Koch Institute | unknown |
| EPI_ISL_2761984 | Sonic - MVZ Medizinisches Labor Bremen GmbH | Robert Koch Institute | unknown |
| EPI_ISL_2761985 | Bioscientia MVZ Labor Karlsruhe GmbH | Robert Koch Institute | unknown |
| EPI_ISL_2761986 | Sonic - Dr. Staber & Kollegen GmbH München | Robert Koch Institute | unknown |
| EPI_ISL_2761990, EPI_ISL_2761992 | Bioscientia Labor Wermsdorf | Robert Koch Institute | unknown |
| EPI_ISL_2761999 | Sonic - MVZ Medizinisches Labor Bremen GmbH | Robert Koch Institute | unknown |
| EPI_ISL_2762000 | Bioscientia MVZ Labor Karlsruhe GmbH | Robert Koch Institute | unknown |
| EPI_ISL_2762001 | LabKom - Labor Hamburg-Lübeck | Robert Koch Institute | unknown |
| EPI_ISL_2762002 | Sonic - MVZ Medizinisches Labor Bremen GmbH | Robert Koch Institute | unknown |
| EPI_ISL_2762003 | Bioscientia Labor Wermsdorf | Robert Koch Institute | unknown |
| EPI_ISL_2762007 | Bioscientia MVZ Labor Karlsruhe GmbH | Robert Koch Institute | unknown |
| EPI_ISL_2762008 | Sonic - MVZ Medizinisches Labor Bremen GmbH | Robert Koch Institute | unknown |
| EPI_ISL_2762009 | Bioscientia Labor Wermsdorf | Robert Koch Institute | unknown |
| EPI_ISL_2762011, EPI_ISL_2762014, EPI_ISL_2762015 | Sonic - MVZ Medizinisches Labor Bremen GmbH | Robert Koch Institute | unknown |
| EPI_ISL_2762017, EPI_ISL_2762019, EPI_ISL_2762021, EPI_ISL_2762023, EPI_ISL_2762026 | Bioscientia Labor Wermsdorf | Robert Koch Institute | unknown |
| EPI_ISL_2762028 | Sonic - MVZ Medizinisches Labor Bremen GmbH | Robert Koch Institute | unknown |
| EPI_ISL_2762029 | LabKom - Labor Hannover MVZ GmbH | Robert Koch Institute | unknown |
| EPI_ISL_2762030 | Bioscientia MVZ Labor Karlsruhe GmbH | Robert Koch Institute | unknown |
| EPI_ISL_2762032 | Robert-Bosch-Krankenhaus | Robert Koch Institute | unknown |
| EPI_ISL_2762034 | Bioscientia Labor Wermsdorf | Robert Koch Institute | unknown |
| EPI_ISL_2762039 | LabKom - MVZ Labor Bochum MLB GmbH | Robert Koch Institute | unknown |
| EPI_ISL_2762040 | Sonic - MVZ Medizinisches Labor Bremen GmbH | Robert Koch Institute | unknown |
| EPI_ISL_2762041 | Bioscientia MVZ Labor Karlsruhe GmbH | Robert Koch Institute | unknown |
| EPI_ISL_2762042 | LabKom - Labor an der Salzbrücke MVZ GmbH | Robert Koch Institute | unknown |
| EPI_ISL_2762049 | Bioscientia Labor Wermsdorf | Robert Koch Institute | unknown |
| EPI_ISL_2762050 | Sonic - MVZ Medizinisches Labor Bremen GmbH | Robert Koch Institute | unknown |
| EPI_ISL_2762054, EPI_ISL_2762055 | Bioscientia Labor Wermsdorf | Robert Koch Institute | unknown |
| EPI_ISL_2762057 | Sonic - MVZ Medizinisches Labor Bremen GmbH | Robert Koch Institute | unknown |
| EPI_ISL_2762058 | Sonic - Labor Staber Dresden (Klipphausen) | Robert Koch Institute | unknown |
| EPI_ISL_2762059 | Sonic - Labor Staber Nürnberg | Robert Koch Institute | unknown |
| EPI_ISL_2762062 | Bioscientia Labor Wermsdorf | Robert Koch Institute | unknown |
| EPI_ISL_2762063 | Sonic - MVZ Medizinisches Labor Bremen GmbH | Robert Koch Institute | unknown |
| EPI_ISL_2762064, EPI_ISL_2762065 | Bioscientia Labor Wermsdorf | Robert Koch Institute | unknown |
| EPI_ISL_2762066 | Sonic - MVZ Medizinisches Labor Bremen GmbH | Robert Koch Institute | unknown |
| EPI_ISL_2762070, EPI_ISL_2762071, EPI_ISL_2762075, EPI_ISL_2762076, EPI_ISL_2762078, EPI_ISL_2762079, EPI_ISL_2762080, EPI_ISL_2762081, EPI_ISL_2762082, EPI_ISL_2762084, EPI_ISL_2762089 | | | |
| see above | Bioscientia Labor Wermsdorf | Robert Koch Institute | unknown |
| EPI_ISL_2762094 | Bioscientia MVZ Labor Karlsruhe GmbH | Robert Koch Institute | unknown |
| EPI_ISL_2762095 | Sonic - MVZ Medizinisches Labor Oldenburg | Robert Koch Institute | unknown |
| EPI_ISL_2762098, EPI_ISL_2762102 | Bioscientia Labor Wermsdorf | Robert Koch Institute | unknown |
| EPI_ISL_2762103 | Sonic - Dr. Staber & Kollegen GmbH München | Robert Koch Institute | unknown |
| EPI_ISL_2762106 | LabKom - MVZ Labor Bochum MLB GmbH | Robert Koch Institute | unknown |
| EPI_ISL_2762109, EPI_ISL_2762110, EPI_ISL_2762111, EPI_ISL_2762112, EPI_ISL_2762113, EPI_ISL_2762115, EPI_ISL_2762116, EPI_ISL_2762117, EPI_ISL_2762118, EPI_ISL_2762119, EPI_ISL_2762120, EPI_ISL_2762122, EPI_ISL_2762123, EPI_ISL_2762125, EPI_ISL_2762126, EPI_ISL_2762127, EPI_ISL_2762128, EPI_ISL_2762129 | | | |
| see above | MVZ Labor Krone GbR | Robert Koch Institute | unknown |
| EPI_ISL_2762135 | Eurofins MVZ Labor Gelsenkirchen | Robert Koch Institute | unknown |
| EPI_ISL_2762138 | IMD - MVZ Labor Greifswald | Robert Koch Institute | unknown |
| EPI_ISL_2762139 | IMD - MVZ Labor Martinsried | Robert Koch Institute | unknown |
| EPI_ISL_2762141, EPI_ISL_2762142, EPI_ISL_2762144, EPI_ISL_2762146, EPI_ISL_2762148, EPI_ISL_2762149, EPI_ISL_2762150, EPI_ISL_2762151, EPI_ISL_2762152, EPI_ISL_2762155, EPI_ISL_2762156, EPI_ISL_2762159, EPI_ISL_2762160, EPI_ISL_2762161, EPI_ISL_2762163, EPI_ISL_2762165 | | | |
| see above | Eurofins LifeCodexx GmbH | Robert Koch Institute | unknown |
| EPI_ISL_2762166 | Fraunhofer-Institut für Zelltherapie und Immunologie IZI AG Next-Generation Diagnostics | Robert Koch Institute | unknown |
| EPI_ISL_2762170, EPI_ISL_2762171, EPI_ISL_2762172, EPI_ISL_2762173 | Institut für Hygiene und Umwelt Hamburg | Robert Koch Institute | unknown |
| EPI_ISL_2762174, EPI_ISL_2762175 | alphaomega Labor GbR (Medizinische Kooperationsgemeinschaft Dres. Ackermann/Tennert/Zaiß) | Robert Koch Institute | unknown |
| EPI_ISL_2762176 | MVZ Labor Dr. Fenner und Kollegen (Standort Hamburg) | Robert Koch Institute | unknown |

| | | | |
|---|---|---|---|
| EPI_ISL_2762180, EPI_ISL_2762187 | DKMS Life Science Lab GmbH Dresden | Robert Koch Institute | unknown |
| EPI_ISL_2762190 | Städtisches Klinikum Dresden Institut für Klinische Chemie und Laboratoriumsmedizin | Robert Koch Institute | unknown |
| EPI_ISL_2762192 | Uniklinikum Carl Gustav Carus an der TU Dresden; Institut für Virologie | Robert Koch Institute | unknown |
| EPI_ISL_2762197 | Medizinisches Labor Ostsachsen MVZ GbR Nebenbetriebsstätte Dresden | Robert Koch Institute | unknown |
| EPI_ISL_2762200 | Uniklinikum Carl Gustav Carus an der TU Dresden; Institut für Virologie | Robert Koch Institute | unknown |
| EPI_ISL_2762209, EPI_ISL_2762218, EPI_ISL_2762223, EPI_ISL_2762229, EPI_ISL_2762237, EPI_ISL_2762241, EPI_ISL_2762246, EPI_ISL_2762247, EPI_ISL_2762251, EPI_ISL_2762252, EPI_ISL_2762253, EPI_ISL_2762256, EPI_ISL_2762264, EPI_ISL_2762267, EPI_ISL_2762268, EPI_ISL_2762274 | | | |
| see above | Bayerisches Landesamt für Gesundheit und Lebensmittelsicherheit (LGL) | Robert Koch Institute | unknown |
| EPI_ISL_2762276 | MVZ Labor Krone GbR | Robert Koch Institute | unknown |
| EPI_ISL_2762279 | Laborarztpraxis Dres. med. Walther Weindel & Kollegen | Robert Koch Institute | unknown |
| EPI_ISL_2762284, EPI_ISL_2762286, EPI_ISL_2762288, EPI_ISL_2762289, EPI_ISL_2762291, EPI_ISL_2762292, EPI_ISL_2762293, EPI_ISL_2762295, EPI_ISL_2762298, EPI_ISL_2762299, EPI_ISL_2762301 | | | |
| see above | MVZ Labor Dr. Fenner und Kollegen (Standort Hamburg) | Robert Koch Institute | unknown |
| EPI_ISL_2762307, EPI_ISL_2762326, EPI_ISL_2762329, EPI_ISL_2762342, EPI_ISL_2762343, EPI_ISL_2762351 | Diagnosticum - Labor Neukirchen | Robert Koch Institute | unknown |
| EPI_ISL_2762357, EPI_ISL_2762358, EPI_ISL_2762360, EPI_ISL_2762362, EPI_ISL_2762364, EPI_ISL_2762366, EPI_ISL_2762367 | SARS-CoV-2 testing team, National Institute of Infectious Diseases | Pathogen Genomics Center, National Institute of Infectious Diseases | Tsuyoshi Sekizuka, Kentaro Itokawa, Rina Tanaka, Masanori Hashino, Nozomu Hanaoka, Masumichi Saito, Naomi Nojiri, Hazuka Y Furihata, Hiromizu Takahashi, Tsuguto Fujimoto, Makoto Kuroda |
| EPI_ISL_2762377, EPI_ISL_2762385, EPI_ISL_2762388 | Diagnosticum - Labor Neukirchen | Robert Koch Institute | unknown |
| EPI_ISL_2762452, EPI_ISL_2762456 | Labor für medizinische Mikrobiologie Bakteriologie Mykologie Virologie & Infektionsserologie - Labor Mölbis | Robert Koch Institute | unknown |
| EPI_ISL_2762465, EPI_ISL_2762477, EPI_ISL_2762479, EPI_ISL_2762481, EPI_ISL_2762483, EPI_ISL_2762485, EPI_ISL_2762488, EPI_ISL_2762492, EPI_ISL_2762494, EPI_ISL_2762497, EPI_ISL_2762508, EPI_ISL_2762513, EPI_ISL_2762515 | | | |
| see above | MVZ Labor Dr. Limbach & Kollegen GbR | Robert Koch Institute | unknown |
| EPI_ISL_2762517 | MVZ Medizinisches Labor Hannover GmbH | Robert Koch Institute | unknown |
| EPI_ISL_2762520 | MVZ Labor Dr. Limbach & Kollegen GbR | Robert Koch Institute | unknown |
| EPI_ISL_2762534 | MVZ für Laboratoriumsmedizin und Mikrobiologie Koblenz-Mittelrhein (Labor Koblenz) | Robert Koch Institute | unknown |
| EPI_ISL_2762539 | Labor Prof. Dr. G. Enders MVZ GbR | Robert Koch Institute | unknown |
| EPI_ISL_2762541 | MVZ für Laboratoriumsmedizin und Mikrobiologie Koblenz-Mittelrhein (Labor Koblenz) | Robert Koch Institute | unknown |
| EPI_ISL_2762548 | Labor Prof. Dr. G. Enders MVZ GbR | Robert Koch Institute | unknown |
| EPI_ISL_2762555 | MVZ für Laboratoriumsmedizin und Mikrobiologie Koblenz-Mittelrhein (Labor Koblenz) | Robert Koch Institute | unknown |
| EPI_ISL_2762561, EPI_ISL_2762564 | Labor Prof. Dr. G. Enders MVZ GbR | Robert Koch Institute | unknown |
| EPI_ISL_2762566 | MVZ Labor Dr. Limbach & Kollegen GbR | Robert Koch Institute | unknown |
| EPI_ISL_2762571 | MVZ Medizinisches Labor Hannover GmbH | Robert Koch Institute | unknown |
| EPI_ISL_2762573, EPI_ISL_2762575, EPI_ISL_2762577, EPI_ISL_2762580 | MVZ für Laboratoriumsmedizin und Mikrobiologie Koblenz-Mittelrhein (Labor Koblenz) | Robert Koch Institute | unknown |
| EPI_ISL_2762584 | MDI Limbach Berlin GmbH; MVZ Labor Berlin | Robert Koch Institute | unknown |
| EPI_ISL_2762587 | MVZ Medizinisches Labor Hannover GmbH | Robert Koch Institute | unknown |
| EPI_ISL_2762589 | MVZ für Laboratoriumsmedizin und Mikrobiologie Koblenz-Mittelrhein (Labor Koblenz) | Robert Koch Institute | unknown |
| EPI_ISL_2762591 | MVZ Medizinisches Labor Hannover GmbH | Robert Koch Institute | unknown |
| EPI_ISL_2762594 | MVZ für Laboratoriumsmedizin und Mikrobiologie Koblenz-Mittelrhein (Labor Koblenz) | Robert Koch Institute | unknown |
| EPI_ISL_2762596 | MVZ Medizinisches Labor Hannover GmbH | Robert Koch Institute | unknown |
| EPI_ISL_2762601 | MDI Limbach Berlin GmbH; MVZ Labor Berlin | Robert Koch Institute | unknown |
| EPI_ISL_2762603 | MVZ Medizinisches Labor Hannover GmbH | Robert Koch Institute | unknown |
| EPI_ISL_2762605, EPI_ISL_2762609, EPI_ISL_2762611, EPI_ISL_2762613, EPI_ISL_2762616 | MDI Limbach Berlin GmbH; MVZ Labor Berlin | Robert Koch Institute | unknown |
| EPI_ISL_2762618 | MVZ Labor Dr. Limbach & Kollegen GbR | Robert Koch Institute | unknown |
| EPI_ISL_2762623 | MDI Limbach Berlin GmbH; MVZ Labor Berlin | Robert Koch Institute | unknown |
| EPI_ISL_2762625, EPI_ISL_2762630 | Limbach - MVZ Labor Ludwigsburg | Robert Koch Institute | unknown |
| EPI_ISL_2762633 | MDI Limbach Berlin GmbH; MVZ Labor Berlin | Robert Koch Institute | unknown |
| EPI_ISL_2762636 | Limbach - MVZ Labor Ludwigsburg | Robert Koch Institute | unknown |
| EPI_ISL_2762638 | Klinikum Darmstadt GmbH | Robert Koch Institute | unknown |
| EPI_ISL_2762641 | Bioscientia Labor Wermsdorf | Robert Koch Institute | unknown |
| EPI_ISL_2762649 | LabKom - MVZ Labor Bochum MLB GmbH | Robert Koch Institute | unknown |
| EPI_ISL_2762654, EPI_ISL_2762656 | Bioscientia Labor Wermsdorf | Robert Koch Institute | unknown |
| EPI_ISL_2762659 | Labor Dr. Schumacher MVZ | Robert Koch Institute | unknown |
| EPI_ISL_2762664, EPI_ISL_2762666, EPI_ISL_2762669, EPI_ISL_2762671, EPI_ISL_2762673, EPI_ISL_2762684, EPI_ISL_2762686, | Bioscientia Labor Wermsdorf | Robert Koch Institute | unknown |

| | | | |
|---|---|---|---|
| EPI_ISL_2762693, EPI_ISL_2762695 | | | |
| EPI_ISL_2762698 | Labor 28 MVZ GmbH | Robert Koch Institute | unknown |
| EPI_ISL_2762703, EPI_ISL_2762705, EPI_ISL_2762710, EPI_ISL_2762712, EPI_ISL_2762718, EPI_ISL_2762720 | Bioscientia Labor Wermsdorf | Robert Koch Institute | unknown |
| EPI_ISL_2762725 | Bioscientia MVZ Labor Karlsruhe GmbH | Robert Koch Institute | unknown |
| EPI_ISL_2762727, EPI_ISL_2762729, EPI_ISL_2762735, EPI_ISL_2762737, EPI_ISL_2762739, EPI_ISL_2762742, EPI_ISL_2762744, EPI_ISL_2762747, EPI_ISL_2762751, EPI_ISL_2762753, EPI_ISL_2762755, EPI_ISL_2762758, EPI_ISL_2762760, EPI_ISL_2762762, EPI_ISL_2762766, EPI_ISL_2762768, EPI_ISL_2762770 | | | |
| see above | Bioscientia Labor Wermsdorf | Robert Koch Institute | unknown |
| EPI_ISL_2762773 | Sonic - Labor Staber Nürnberg | Robert Koch Institute | unknown |
| EPI_ISL_2762775, EPI_ISL_2762778 | Bioscientia Labor Wermsdorf | Robert Koch Institute | unknown |
| EPI_ISL_2762780 | Sonic - Labor Staber Nürnberg | Robert Koch Institute | unknown |
| EPI_ISL_2762785, EPI_ISL_2762788, EPI_ISL_2762791, EPI_ISL_2762793 | Bioscientia Labor Wermsdorf | Robert Koch Institute | unknown |
| EPI_ISL_2762796 | Bioscientia MVZ Labor Karlsruhe GmbH | Robert Koch Institute | unknown |
| EPI_ISL_2762798 | Bioscientia Labor Wermsdorf | Robert Koch Institute | unknown |
| EPI_ISL_2762808 | Sonic - Bioscientia - MVZ Labor Saar GmbH | Robert Koch Institute | unknown |
| EPI_ISL_2762812, EPI_ISL_2762815, EPI_ISL_2762817, EPI_ISL_2762820, EPI_ISL_2762822 | Bioscientia Labor Wermsdorf | Robert Koch Institute | unknown |
| EPI_ISL_2762825 | Medizinisch-Diagnostisches Labor Kempten allgäulab | Robert Koch Institute | unknown |
| EPI_ISL_2762961 | Pathogen Genomics Center, National Institute of Infectious Diseases | Pathogen Genomics Center, National Institute of Infectious Diseases | Tsuyoshi Sekizuka, Kentaro Itokawa, Rina Tanaka, Masanori Hashino, Makoto Kuroda |
| EPI_ISL_2763624 | MOH - Jaber Al-Ahmad Hospital (Innovation Research Laboratory) | MOH - Jaber Al-Ahmad Hospital (Innovation Research Laboratory) | Salman Al-Sabah , Mohammad Alghounaim |
| EPI_ISL_2763970, EPI_ISL_2763987, EPI_ISL_2763989, EPI_ISL_2764011, EPI_ISL_2764017, EPI_ISL_2764018, EPI_ISL_2764101, EPI_ISL_2764102, EPI_ISL_2764103 | Pathogen Genomics Center, National Institute of Infectious Diseases | Pathogen Genomics Center, National Institute of Infectious Diseases | Tsuyoshi Sekizuka, Kentaro Itokawa, Rina Tanaka, Masanori Hashino, Makoto Kuroda |
| EPI_ISL_2764904 | SARS-CoV-2 testing team, National Institute of Infectious Diseases | Pathogen Genomics Center, National Institute of Infectious Diseases | Tsuyoshi Sekizuka, Kentaro Itokawa, Rina Tanaka, Masanori Hashino, Nozomu Hanaoka, Masumichi Saito, Naomi Nojiri, Hazuka Y Furihata, Hiromizu Takahashi, Tsuguto Fujimoto, Makoto Kuroda |
| EPI_ISL_2764950 | Pathogen Genomics Center, National Institute of Infectious Diseases | Pathogen Genomics Center, National Institute of Infectious Diseases | Tsuyoshi Sekizuka, Kentaro Itokawa, Rina Tanaka, Masanori Hashino, Makoto Kuroda |
| EPI_ISL_2765167, EPI_ISL_2765502, EPI_ISL_2765503, EPI_ISL_2765504 | SARS-CoV-2 testing team, National Institute of Infectious Diseases | Pathogen Genomics Center, National Institute of Infectious Diseases | Tsuyoshi Sekizuka, Kentaro Itokawa, Rina Tanaka, Masanori Hashino, Nozomu Hanaoka, Masumichi Saito, Naomi Nojiri, Hazuka Y Furihata, Hiromizu Takahashi, Tsuguto Fujimoto, Makoto Kuroda |
| EPI_ISL_2765548 | Laboratorio di Riferimento Regionale della Sicilia Occidentale per l'Emergenza COVID-19 | Laboratorio di Riferimento Regionale della Sicilia Occidentale per l'Emergenza COVID-19 | Fabio Tramuto, Carmelo Massimo Maida, Daniela Di Naro, Giulia Randazzo, Walter Mazzucco, Giorgio Graziano, Vincenzo Restivo, Claudio Costantino, Francesco Vitale |
| EPI_ISL_2765570, EPI_ISL_2765571, EPI_ISL_2765572, EPI_ISL_2765573, EPI_ISL_2765574 | Lighthouse Lab in Alderley Park | Wellcome Sanger Institute for the COVID-19 Genomics UK (COG-UK) Consortium | Jacquelyn Wynn, Mairead Hyland, The Lighthouse Lab in Alderley Park and Alex Alderton, Roberto Amato, Jeffrey Barrett, Sonia Goncalves, Ewan Harrison, David K. Jackson, Ian Johnston, Dominic Kwiatkowski, Cordelia Langford, John Sillitoe on behalf of the Wellcome Sanger Institute COVID-19 Surveillance Team |
| EPI_ISL_2765575 | Lighthouse Lab in Milton Keynes | Wellcome Sanger Institute for the COVID-19 Genomics UK (COG-UK) Consortium | The Lighthouse Lab in Milton Keynes and Alex Alderton, Roberto Amato, Jeffrey Barrett, Sonia Goncalves, Ewan Harrison, David K. Jackson, Ian Johnston, Dominic Kwiatkowski, Cordelia Langford, John Sillitoe on behalf of the Wellcome Sanger Institute COVID-19 Surveillance Team |
| EPI_ISL_2765576, EPI_ISL_2765577, EPI_ISL_2765578, EPI_ISL_2765579 | Lighthouse Lab in Alderley Park | Wellcome Sanger Institute for the COVID-19 Genomics UK (COG-UK) Consortium | Jacquelyn Wynn, Mairead Hyland, The Lighthouse Lab in Alderley Park and Alex Alderton, Roberto Amato, Jeffrey Barrett, Sonia Goncalves, Ewan Harrison, David K. Jackson, Ian Johnston, Dominic Kwiatkowski, Cordelia Langford, John Sillitoe on behalf of the Wellcome Sanger Institute COVID-19 Surveillance Team |
| EPI_ISL_2765580 | Lighthouse Lab in Milton Keynes | Wellcome Sanger Institute for the COVID-19 Genomics UK (COG-UK) Consortium | The Lighthouse Lab in Milton Keynes and Alex Alderton, Roberto Amato, Jeffrey Barrett, Sonia Goncalves, Ewan Harrison, David K. Jackson, Ian Johnston, Dominic Kwiatkowski, Cordelia Langford, John Sillitoe on behalf of the Wellcome Sanger Institute COVID-19 Surveillance Team |
| EPI_ISL_2765581 | Lighthouse Lab in Alderley Park | Wellcome Sanger Institute for the COVID-19 Genomics UK (COG-UK) Consortium | Jacquelyn Wynn, Mairead Hyland, The Lighthouse Lab in Alderley Park and Alex Alderton, Roberto Amato, Jeffrey Barrett, Sonia Goncalves, Ewan Harrison, David K. Jackson, Ian Johnston, Dominic Kwiatkowski, Cordelia Langford, John Sillitoe on behalf of the Wellcome Sanger Institute COVID-19 Surveillance Team |
| EPI_ISL_2765582, EPI_ISL_2765583 | Lighthouse Laboratory Plymouth | Wellcome Sanger Institute for the COVID-19 Genomics UK (COG-UK) Consortium | Lighthouse Laboratory Plymouth and Alex Alderton, Roberto Amato, Jeffrey Barrett, Sonia Goncalves, Ewan Harrison, David K. Jackson, Ian Johnston, Dominic Kwiatkowski, Cordelia Langford, John Sillitoe on behalf of the Wellcome Sanger Institute COVID-19 Surveillance Team |
| EPI_ISL_2765584 | Lighthouse Lab in Milton Keynes | Wellcome Sanger Institute for the COVID-19 Genomics UK (COG-UK) Consortium | The Lighthouse Lab in Milton Keynes and Alex Alderton, Roberto Amato, Jeffrey Barrett, Sonia Goncalves, Ewan Harrison, David K. Jackson, Ian Johnston, Dominic Kwiatkowski, Cordelia Langford, John Sillitoe on behalf of the Wellcome Sanger Institute COVID-19 Surveillance Team |
| EPI_ISL_2765585, EPI_ISL_2765586 | Lighthouse Lab in Alderley Park | Wellcome Sanger Institute for the COVID-19 Genomics UK (COG-UK) Consortium | Jacquelyn Wynn, Mairead Hyland, The Lighthouse Lab in Alderley Park and Alex Alderton, Roberto Amato, Jeffrey Barrett, Sonia Goncalves, Ewan Harrison, David K. Jackson, Ian Johnston, Dominic Kwiatkowski, Cordelia Langford, John Sillitoe on behalf of the Wellcome Sanger Institute COVID-19 Surveillance Team |
| EPI_ISL_2765587 | Lighthouse Lab in Glasgow | Wellcome Sanger Institute for the COVID-19 Genomics UK (COG-UK) Consortium | Harper VanSteenhouse, Yumi Kasai, David Gray, Carol Clugston, Anna Dominiczak and Alex Alderton, Roberto Amato, Jeffrey Barrett, Sonia Goncalves, Ewan Harrison, David K. Jackson, Ian Johnston, Dominic Kwiatkowski, Cordelia Langford, John Sillitoe on behalf of the Wellcome Sanger Institute COVID-19 Surveillance Team |
| EPI_ISL_2765588 | Lighthouse Lab in Milton Keynes | Wellcome Sanger Institute for the COVID-19 Genomics UK (COG-UK) Consortium | The Lighthouse Lab in Milton Keynes and Alex Alderton, Roberto Amato, Jeffrey Barrett, Sonia Goncalves, Ewan Harrison, David K. Jackson, Ian Johnston, Dominic Kwiatkowski, Cordelia Langford, John Sillitoe on behalf of the Wellcome Sanger Institute COVID-19 Surveillance Team |
| EPI_ISL_2765589 | Lighthouse Lab in Alderley Park | Wellcome Sanger Institute for the COVID-19 Genomics UK (COG-UK) Consortium | Jacquelyn Wynn, Mairead Hyland, The Lighthouse Lab in Alderley Park and Alex Alderton, Roberto Amato, Jeffrey Barrett, Sonia Goncalves, Ewan Harrison, David K. Jackson, Ian Johnston, Dominic Kwiatkowski, Cordelia Langford, John Sillitoe on behalf of the Wellcome Sanger Institute COVID-19 Surveillance Team |
| EPI_ISL_2765590 | Lighthouse Lab in Milton Keynes | Wellcome Sanger Institute for the COVID-19 Genomics UK (COG-UK) Consortium | The Lighthouse Lab in Milton Keynes and Alex Alderton, Roberto Amato, Jeffrey Barrett, Sonia Goncalves, Ewan Harrison, David K. Jackson, Ian Johnston, Dominic Kwiatkowski, Cordelia Langford, John Sillitoe on behalf of the Wellcome Sanger Institute COVID-19 Surveillance Team |
| EPI_ISL_2765591 | Lighthouse Laboratory Plymouth | Wellcome Sanger Institute for the COVID-19 Genomics UK (COG-UK) Consortium | Lighthouse Laboratory Plymouth and Alex Alderton, Roberto Amato, Jeffrey Barrett, Sonia Goncalves, Ewan Harrison, David K. Jackson, Ian Johnston, Dominic Kwiatkowski, Cordelia Langford, John Sillitoe on behalf of the Wellcome Sanger Institute COVID-19 Surveillance Team |
| EPI_ISL_2765592, EPI_ISL_2765593, EPI_ISL_2765594 | Lighthouse Lab in Alderley Park | Wellcome Sanger Institute for the COVID-19 Genomics UK (COG-UK) Consortium | Jacquelyn Wynn, Mairead Hyland, The Lighthouse Lab in Alderley Park and Alex Alderton, Roberto Amato, Jeffrey Barrett, Sonia Goncalves, Ewan Harrison, David K. Jackson, Ian Johnston, Dominic Kwiatkowski, Cordelia Langford, John Sillitoe on behalf of the Wellcome Sanger Institute COVID-19 Surveillance Team |
| EPI_ISL_2765595 | Lighthouse Laboratory Plymouth | Wellcome Sanger Institute for the COVID-19 Genomics UK (COG-UK) Consortium | Lighthouse Laboratory Plymouth and Alex Alderton, Roberto Amato, Jeffrey Barrett, Sonia Goncalves, Ewan Harrison, David K. Jackson, Ian Johnston, Dominic Kwiatkowski, Cordelia Langford, John Sillitoe on behalf of the Wellcome Sanger Institute COVID-19 Surveillance Team |
| EPI_ISL_2765596, EPI_ISL_2765597 | Lighthouse Lab in Alderley Park | Wellcome Sanger Institute for the COVID-19 Genomics UK (COG-UK) Consortium | Jacquelyn Wynn, Mairead Hyland, The Lighthouse Lab in Alderley Park and Alex Alderton, Roberto Amato, Jeffrey Barrett, Sonia Goncalves, Ewan Harrison, David K. Jackson, Ian Johnston, Dominic Kwiatkowski, Cordelia Langford, John Sillitoe on behalf of the Wellcome Sanger Institute COVID-19 Surveillance Team |
| EPI_ISL_2765598 | Lighthouse Lab in Milton Keynes | Wellcome Sanger Institute for the COVID-19 Genomics UK (COG-UK) Consortium | The Lighthouse Lab in Milton Keynes and Alex Alderton, Roberto Amato, Jeffrey Barrett, Sonia Goncalves, Ewan Harrison, David K. Jackson, Ian Johnston, Dominic Kwiatkowski, Cordelia Langford, John Sillitoe on behalf of the Wellcome Sanger Institute COVID-19 Surveillance Team |
| EPI_ISL_2765599, | Lighthouse Lab in Alderley Park | Wellcome Sanger Institute for the COVID-19 Genomics UK (COG-UK) | Jacquelyn Wynn, Mairead Hyland, The Lighthouse Lab in Alderley Park and Alex Alderton, Roberto Amato, Jeffrey Barrett, Sonia Goncalves, Ewan Harrison, David K. Jackson, Ian Johnston, Dominic Kwiatkowski, Cordelia Langford, John Sillitoe on |

| | | | |
|---|---|---|---|
| EPI_ISL_2765600, EPI_ISL_2765601 | | Consortium | behalf of the Wellcome Sanger Institute COVID-19 Surveillance Team |
| EPI_ISL_2765602 | Lighthouse Laboratory Plymouth | Wellcome Sanger Institute for the COVID-19 Genomics UK (COG-UK) Consortium | Lighthouse Laboratory Plymouth and Alex Alderton, Roberto Amato, Jeffrey Barrett, Sonia Goncalves, Ewan Harrison, David K. Jackson, Ian Johnston, Dominic Kwiatkowski, Cordelia Langford, John Sillitoe on behalf of the Wellcome Sanger Institute COVID-19 Surveillance Team |
| EPI_ISL_2765603 | Lighthouse Lab in Alderley Park | Wellcome Sanger Institute for the COVID-19 Genomics UK (COG-UK) Consortium | Jacquelyn Wynn, Mairead Hyland, The Lighthouse Lab in Alderley Park and Alex Alderton, Roberto Amato, Jeffrey Barrett, Sonia Goncalves, Ewan Harrison, David K. Jackson, Ian Johnston, Dominic Kwiatkowski, Cordelia Langford, John Sillitoe on behalf of the Wellcome Sanger Institute COVID-19 Surveillance Team |
| EPI_ISL_2765604 | Lighthouse Lab in Milton Keynes | Wellcome Sanger Institute for the COVID-19 Genomics UK (COG-UK) Consortium | The Lighthouse Lab in Milton Keynes and Alex Alderton, Roberto Amato, Jeffrey Barrett, Sonia Goncalves, Ewan Harrison, David K. Jackson, Ian Johnston, Dominic Kwiatkowski, Cordelia Langford, John Sillitoe on behalf of the Wellcome Sanger Institute COVID-19 Surveillance Team |
| EPI_ISL_2765605 | Lighthouse Lab in Alderley Park | Wellcome Sanger Institute for the COVID-19 Genomics UK (COG-UK) Consortium | Jacquelyn Wynn, Mairead Hyland, The Lighthouse Lab in Alderley Park and Alex Alderton, Roberto Amato, Jeffrey Barrett, Sonia Goncalves, Ewan Harrison, David K. Jackson, Ian Johnston, Dominic Kwiatkowski, Cordelia Langford, John Sillitoe on behalf of the Wellcome Sanger Institute COVID-19 Surveillance Team |
| EPI_ISL_2765606 | Lighthouse Lab in Milton Keynes | Wellcome Sanger Institute for the COVID-19 Genomics UK (COG-UK) Consortium | The Lighthouse Lab in Milton Keynes and Alex Alderton, Roberto Amato, Jeffrey Barrett, Sonia Goncalves, Ewan Harrison, David K. Jackson, Ian Johnston, Dominic Kwiatkowski, Cordelia Langford, John Sillitoe on behalf of the Wellcome Sanger Institute COVID-19 Surveillance Team |
| EPI_ISL_2765607, EPI_ISL_2765608, EPI_ISL_2765609, EPI_ISL_2765610 | Lighthouse Lab in Alderley Park | Wellcome Sanger Institute for the COVID-19 Genomics UK (COG-UK) Consortium | Jacquelyn Wynn, Mairead Hyland, The Lighthouse Lab in Alderley Park and Alex Alderton, Roberto Amato, Jeffrey Barrett, Sonia Goncalves, Ewan Harrison, David K. Jackson, Ian Johnston, Dominic Kwiatkowski, Cordelia Langford, John Sillitoe on behalf of the Wellcome Sanger Institute COVID-19 Surveillance Team |
| EPI_ISL_2765611 | Lighthouse Lab in Glasgow | Wellcome Sanger Institute for the COVID-19 Genomics UK (COG-UK) Consortium | Harper VanSteenhouse, Yumi Kasai, David Gray, Carol Clugston, Anna Dominiczak and Alex Alderton, Roberto Amato, Jeffrey Barrett, Sonia Goncalves, Ewan Harrison, David K. Jackson, Ian Johnston, Dominic Kwiatkowski, Cordelia Langford, John Sillitoe on behalf of the Wellcome Sanger Institute COVID-19 Surveillance Team |
| EPI_ISL_2765612, EPI_ISL_2765613 | Lighthouse Lab in Milton Keynes | Wellcome Sanger Institute for the COVID-19 Genomics UK (COG-UK) Consortium | The Lighthouse Lab in Milton Keynes and Alex Alderton, Roberto Amato, Jeffrey Barrett, Sonia Goncalves, Ewan Harrison, David K. Jackson, Ian Johnston, Dominic Kwiatkowski, Cordelia Langford, John Sillitoe on behalf of the Wellcome Sanger Institute COVID-19 Surveillance Team |
| EPI_ISL_2765614, EPI_ISL_2765615, EPI_ISL_2765616, EPI_ISL_2765617 | Lighthouse Lab in Alderley Park | Wellcome Sanger Institute for the COVID-19 Genomics UK (COG-UK) Consortium | Jacquelyn Wynn, Mairead Hyland, The Lighthouse Lab in Alderley Park and Alex Alderton, Roberto Amato, Jeffrey Barrett, Sonia Goncalves, Ewan Harrison, David K. Jackson, Ian Johnston, Dominic Kwiatkowski, Cordelia Langford, John Sillitoe on behalf of the Wellcome Sanger Institute COVID-19 Surveillance Team |
| EPI_ISL_2765618 | Lighthouse Laboratory Plymouth | Wellcome Sanger Institute for the COVID-19 Genomics UK (COG-UK) Consortium | Lighthouse Laboratory Plymouth and Alex Alderton, Roberto Amato, Jeffrey Barrett, Sonia Goncalves, Ewan Harrison, David K. Jackson, Ian Johnston, Dominic Kwiatkowski, Cordelia Langford, John Sillitoe on behalf of the Wellcome Sanger Institute COVID-19 Surveillance Team |
| EPI_ISL_2765619, EPI_ISL_2765620 | Lighthouse Lab in Alderley Park | Wellcome Sanger Institute for the COVID-19 Genomics UK (COG-UK) Consortium | Jacquelyn Wynn, Mairead Hyland, The Lighthouse Lab in Alderley Park and Alex Alderton, Roberto Amato, Jeffrey Barrett, Sonia Goncalves, Ewan Harrison, David K. Jackson, Ian Johnston, Dominic Kwiatkowski, Cordelia Langford, John Sillitoe on behalf of the Wellcome Sanger Institute COVID-19 Surveillance Team |
| EPI_ISL_2765621, EPI_ISL_2765622 | Lighthouse Laboratory Plymouth | Wellcome Sanger Institute for the COVID-19 Genomics UK (COG-UK) Consortium | Lighthouse Laboratory Plymouth and Alex Alderton, Roberto Amato, Jeffrey Barrett, Sonia Goncalves, Ewan Harrison, David K. Jackson, Ian Johnston, Dominic Kwiatkowski, Cordelia Langford, John Sillitoe on behalf of the Wellcome Sanger Institute COVID-19 Surveillance Team |
| EPI_ISL_2765623, EPI_ISL_2765624 | Lighthouse Lab in Alderley Park | Wellcome Sanger Institute for the COVID-19 Genomics UK (COG-UK) Consortium | Jacquelyn Wynn, Mairead Hyland, The Lighthouse Lab in Alderley Park and Alex Alderton, Roberto Amato, Jeffrey Barrett, Sonia Goncalves, Ewan Harrison, David K. Jackson, Ian Johnston, Dominic Kwiatkowski, Cordelia Langford, John Sillitoe on behalf of the Wellcome Sanger Institute COVID-19 Surveillance Team |
| EPI_ISL_2765625 | Lighthouse Laboratory Plymouth | Wellcome Sanger Institute for the COVID-19 Genomics UK (COG-UK) Consortium | Lighthouse Laboratory Plymouth and Alex Alderton, Roberto Amato, Jeffrey Barrett, Sonia Goncalves, Ewan Harrison, David K. Jackson, Ian Johnston, Dominic Kwiatkowski, Cordelia Langford, John Sillitoe on behalf of the Wellcome Sanger Institute COVID-19 Surveillance Team |
| EPI_ISL_2765626 | Lighthouse Lab in Alderley Park | Wellcome Sanger Institute for the COVID-19 Genomics UK (COG-UK) Consortium | Jacquelyn Wynn, Mairead Hyland, The Lighthouse Lab in Alderley Park and Alex Alderton, Roberto Amato, Jeffrey Barrett, Sonia Goncalves, Ewan Harrison, David K. Jackson, Ian Johnston, Dominic Kwiatkowski, Cordelia Langford, John Sillitoe on behalf of the Wellcome Sanger Institute COVID-19 Surveillance Team |
| EPI_ISL_2765627 | Lighthouse Laboratory Plymouth | Wellcome Sanger Institute for the COVID-19 Genomics UK (COG-UK) Consortium | Lighthouse Laboratory Plymouth and Alex Alderton, Roberto Amato, Jeffrey Barrett, Sonia Goncalves, Ewan Harrison, David K. Jackson, Ian Johnston, Dominic Kwiatkowski, Cordelia Langford, John Sillitoe on behalf of the Wellcome Sanger Institute COVID-19 Surveillance Team |
| EPI_ISL_2765628 | Lighthouse Lab in Alderley Park | Wellcome Sanger Institute for the COVID-19 Genomics UK (COG-UK) Consortium | Jacquelyn Wynn, Mairead Hyland, The Lighthouse Lab in Alderley Park and Alex Alderton, Roberto Amato, Jeffrey Barrett, Sonia Goncalves, Ewan Harrison, David K. Jackson, Ian Johnston, Dominic Kwiatkowski, Cordelia Langford, John Sillitoe on behalf of the Wellcome Sanger Institute COVID-19 Surveillance Team |
| EPI_ISL_2765629 | Lighthouse Lab in Milton Keynes | Wellcome Sanger Institute for the COVID-19 Genomics UK (COG-UK) Consortium | The Lighthouse Lab in Milton Keynes and Alex Alderton, Roberto Amato, Jeffrey Barrett, Sonia Goncalves, Ewan Harrison, David K. Jackson, Ian Johnston, Dominic Kwiatkowski, Cordelia Langford, John Sillitoe on behalf of the Wellcome Sanger Institute COVID-19 Surveillance Team |
| EPI_ISL_2765630 | Lighthouse Lab in Alderley Park | Wellcome Sanger Institute for the COVID-19 Genomics UK (COG-UK) Consortium | Jacquelyn Wynn, Mairead Hyland, The Lighthouse Lab in Alderley Park and Alex Alderton, Roberto Amato, Jeffrey Barrett, Sonia Goncalves, Ewan Harrison, David K. Jackson, Ian Johnston, Dominic Kwiatkowski, Cordelia Langford, John Sillitoe on behalf of the Wellcome Sanger Institute COVID-19 Surveillance Team |
| EPI_ISL_2765631 | Lighthouse Lab in Milton Keynes | Wellcome Sanger Institute for the COVID-19 Genomics UK (COG-UK) Consortium | The Lighthouse Lab in Milton Keynes and Alex Alderton, Roberto Amato, Jeffrey Barrett, Sonia Goncalves, Ewan Harrison, David K. Jackson, Ian Johnston, Dominic Kwiatkowski, Cordelia Langford, John Sillitoe on behalf of the Wellcome Sanger Institute COVID-19 Surveillance Team |
| EPI_ISL_2765632 | Lighthouse Lab in Glasgow | Wellcome Sanger Institute for the COVID-19 Genomics UK (COG-UK) Consortium | Harper VanSteenhouse, Yumi Kasai, David Gray, Carol Clugston, Anna Dominiczak and Alex Alderton, Roberto Amato, Jeffrey Barrett, Sonia Goncalves, Ewan Harrison, David K. Jackson, Ian Johnston, Dominic Kwiatkowski, Cordelia Langford, John Sillitoe on behalf of the Wellcome Sanger Institute COVID-19 Surveillance Team |
| EPI_ISL_2765633 | Lighthouse Lab in Milton Keynes | Wellcome Sanger Institute for the COVID-19 Genomics UK (COG-UK) Consortium | The Lighthouse Lab in Milton Keynes and Alex Alderton, Roberto Amato, Jeffrey Barrett, Sonia Goncalves, Ewan Harrison, David K. Jackson, Ian Johnston, Dominic Kwiatkowski, Cordelia Langford, John Sillitoe on behalf of the Wellcome Sanger Institute COVID-19 Surveillance Team |
| EPI_ISL_2765634, EPI_ISL_2765635, EPI_ISL_2765636 | Lighthouse Lab in Alderley Park | Wellcome Sanger Institute for the COVID-19 Genomics UK (COG-UK) Consortium | Jacquelyn Wynn, Mairead Hyland, The Lighthouse Lab in Alderley Park and Alex Alderton, Roberto Amato, Jeffrey Barrett, Sonia Goncalves, Ewan Harrison, David K. Jackson, Ian Johnston, Dominic Kwiatkowski, Cordelia Langford, John Sillitoe on behalf of the Wellcome Sanger Institute COVID-19 Surveillance Team |
| EPI_ISL_2765637 | Lighthouse Lab in Milton Keynes | Wellcome Sanger Institute for the COVID-19 Genomics UK (COG-UK) Consortium | The Lighthouse Lab in Milton Keynes and Alex Alderton, Roberto Amato, Jeffrey Barrett, Sonia Goncalves, Ewan Harrison, David K. Jackson, Ian Johnston, Dominic Kwiatkowski, Cordelia Langford, John Sillitoe on behalf of the Wellcome Sanger Institute COVID-19 Surveillance Team |
| EPI_ISL_2765638, EPI_ISL_2765639 | Lighthouse Lab in Alderley Park | Wellcome Sanger Institute for the COVID-19 Genomics UK (COG-UK) Consortium | Jacquelyn Wynn, Mairead Hyland, The Lighthouse Lab in Alderley Park and Alex Alderton, Roberto Amato, Jeffrey Barrett, Sonia Goncalves, Ewan Harrison, David K. Jackson, Ian Johnston, Dominic Kwiatkowski, Cordelia Langford, John Sillitoe on behalf of the Wellcome Sanger Institute COVID-19 Surveillance Team |
| EPI_ISL_2765640 | Lighthouse Laboratory Plymouth | Wellcome Sanger Institute for the COVID-19 Genomics UK (COG-UK) Consortium | Lighthouse Laboratory Plymouth and Alex Alderton, Roberto Amato, Jeffrey Barrett, Sonia Goncalves, Ewan Harrison, David K. Jackson, Ian Johnston, Dominic Kwiatkowski, Cordelia Langford, John Sillitoe on behalf of the Wellcome Sanger Institute COVID-19 Surveillance Team |
| EPI_ISL_2765641 | Lighthouse Lab in Milton Keynes | Wellcome Sanger Institute for the COVID-19 Genomics UK (COG-UK) Consortium | The Lighthouse Lab in Milton Keynes and Alex Alderton, Roberto Amato, Jeffrey Barrett, Sonia Goncalves, Ewan Harrison, David K. Jackson, Ian Johnston, Dominic Kwiatkowski, Cordelia Langford, John Sillitoe on behalf of the Wellcome Sanger Institute COVID-19 Surveillance Team |
| EPI_ISL_2765642, EPI_ISL_2765643 | Lighthouse Lab in Alderley Park | Wellcome Sanger Institute for the COVID-19 Genomics UK (COG-UK) Consortium | Jacquelyn Wynn, Mairead Hyland, The Lighthouse Lab in Alderley Park and Alex Alderton, Roberto Amato, Jeffrey Barrett, Sonia Goncalves, Ewan Harrison, David K. Jackson, Ian Johnston, Dominic Kwiatkowski, Cordelia Langford, John Sillitoe on behalf of the Wellcome Sanger Institute COVID-19 Surveillance Team |
| EPI_ISL_2765644, EPI_ISL_2765645 | Lighthouse Lab in Milton Keynes | Wellcome Sanger Institute for the COVID-19 Genomics UK (COG-UK) Consortium | The Lighthouse Lab in Milton Keynes and Alex Alderton, Roberto Amato, Jeffrey Barrett, Sonia Goncalves, Ewan Harrison, David K. Jackson, Ian Johnston, Dominic Kwiatkowski, Cordelia Langford, John Sillitoe on behalf of the Wellcome Sanger Institute COVID-19 Surveillance Team |
| EPI_ISL_2765646, EPI_ISL_2765647 | Lighthouse Lab in Alderley Park | Wellcome Sanger Institute for the COVID-19 Genomics UK (COG-UK) Consortium | Jacquelyn Wynn, Mairead Hyland, The Lighthouse Lab in Alderley Park and Alex Alderton, Roberto Amato, Jeffrey Barrett, Sonia Goncalves, Ewan Harrison, David K. Jackson, Ian Johnston, Dominic Kwiatkowski, Cordelia Langford, John Sillitoe on behalf of the Wellcome Sanger Institute COVID-19 Surveillance Team |
| EPI_ISL_2765648, EPI_ISL_2765649 | Lighthouse Lab in Milton Keynes | Wellcome Sanger Institute for the COVID-19 Genomics UK (COG-UK) Consortium | The Lighthouse Lab in Milton Keynes and Alex Alderton, Roberto Amato, Jeffrey Barrett, Sonia Goncalves, Ewan Harrison, David K. Jackson, Ian Johnston, Dominic Kwiatkowski, Cordelia Langford, John Sillitoe on behalf of the Wellcome Sanger Institute COVID-19 Surveillance Team |
| EPI_ISL_2765650, EPI_ISL_2765651 | Lighthouse Lab in Alderley Park | Wellcome Sanger Institute for the COVID-19 Genomics UK (COG-UK) Consortium | Jacquelyn Wynn, Mairead Hyland, The Lighthouse Lab in Alderley Park and Alex Alderton, Roberto Amato, Jeffrey Barrett, Sonia Goncalves, Ewan Harrison, David K. Jackson, Ian Johnston, Dominic Kwiatkowski, Cordelia Langford, John Sillitoe on behalf of the Wellcome Sanger Institute COVID-19 Surveillance Team |
| EPI_ISL_2765652 | Lighthouse Laboratory Plymouth | Wellcome Sanger Institute for the COVID-19 Genomics UK (COG-UK) Consortium | Lighthouse Laboratory Plymouth and Alex Alderton, Roberto Amato, Jeffrey Barrett, Sonia Goncalves, Ewan Harrison, David K. Jackson, Ian Johnston, Dominic Kwiatkowski, Cordelia Langford, John Sillitoe on behalf of the Wellcome Sanger Institute COVID-19 Surveillance Team |
| EPI_ISL_2765653 | Lighthouse Lab in Milton Keynes | Wellcome Sanger Institute for the COVID-19 Genomics UK (COG-UK) Consortium | The Lighthouse Lab in Milton Keynes and Alex Alderton, Roberto Amato, Jeffrey Barrett, Sonia Goncalves, Ewan Harrison, David K. Jackson, Ian Johnston, Dominic Kwiatkowski, Cordelia Langford, John Sillitoe on behalf of the Wellcome Sanger Institute COVID-19 Surveillance Team |
| EPI_ISL_2765654, EPI_ISL_2765655 | Lighthouse Lab in Alderley Park | Wellcome Sanger Institute for the COVID-19 Genomics UK (COG-UK) Consortium | Jacquelyn Wynn, Mairead Hyland, The Lighthouse Lab in Alderley Park and Alex Alderton, Roberto Amato, Jeffrey Barrett, Sonia Goncalves, Ewan Harrison, David K. Jackson, Ian Johnston, Dominic Kwiatkowski, Cordelia Langford, John Sillitoe on behalf of the Wellcome Sanger Institute COVID-19 Surveillance Team |
| EPI_ISL_2765656, EPI_ISL_2765657, EPI_ISL_2765658 | Lighthouse Lab in Milton Keynes | Wellcome Sanger Institute for the COVID-19 Genomics UK (COG-UK) Consortium | The Lighthouse Lab in Milton Keynes and Alex Alderton, Roberto Amato, Jeffrey Barrett, Sonia Goncalves, Ewan Harrison, David K. Jackson, Ian Johnston, Dominic Kwiatkowski, Cordelia Langford, John Sillitoe on behalf of the Wellcome Sanger Institute COVID-19 Surveillance Team |
| EPI_ISL_2765659, EPI_ISL_2765660, EPI_ISL_2765661, EPI_ISL_2765662 | Lighthouse Lab in Alderley Park | Wellcome Sanger Institute for the COVID-19 Genomics UK (COG-UK) Consortium | Jacquelyn Wynn, Mairead Hyland, The Lighthouse Lab in Alderley Park and Alex Alderton, Roberto Amato, Jeffrey Barrett, Sonia Goncalves, Ewan Harrison, David K. Jackson, Ian Johnston, Dominic Kwiatkowski, Cordelia Langford, John Sillitoe on behalf of the Wellcome Sanger Institute COVID-19 Surveillance Team |
| EPI_ISL_2765663 | Lighthouse Lab in Milton Keynes | Wellcome Sanger Institute for the COVID-19 Genomics UK (COG-UK) Consortium | The Lighthouse Lab in Milton Keynes and Alex Alderton, Roberto Amato, Jeffrey Barrett, Sonia Goncalves, Ewan Harrison, David K. Jackson, Ian Johnston, Dominic Kwiatkowski, Cordelia Langford, John Sillitoe on behalf of the Wellcome Sanger Institute COVID-19 Surveillance Team |
| EPI_ISL_2765664, EPI_ISL_2765665 | Lighthouse Laboratory Plymouth | Wellcome Sanger Institute for the COVID-19 Genomics UK (COG-UK) Consortium | Lighthouse Laboratory Plymouth and Alex Alderton, Roberto Amato, Jeffrey Barrett, Sonia Goncalves, Ewan Harrison, David K. Jackson, Ian Johnston, Dominic Kwiatkowski, Cordelia Langford, John Sillitoe on behalf of the Wellcome Sanger Institute COVID-19 Surveillance Team |
| EPI_ISL_2765666 | Lighthouse Lab in Milton Keynes | Wellcome Sanger Institute for the COVID-19 Genomics UK (COG-UK) Consortium | The Lighthouse Lab in Milton Keynes and Alex Alderton, Roberto Amato, Jeffrey Barrett, Sonia Goncalves, Ewan Harrison, David K. Jackson, Ian Johnston, Dominic Kwiatkowski, Cordelia Langford, John Sillitoe on behalf of the Wellcome Sanger Institute COVID-19 Surveillance Team |
| EPI_ISL_2765667, EPI_ISL_2765668, | Lighthouse Lab in Alderley Park | Wellcome Sanger Institute for the COVID-19 Genomics UK (COG-UK) Consortium | Jacquelyn Wynn, Mairead Hyland, The Lighthouse Lab in Alderley Park and Alex Alderton, Roberto Amato, Jeffrey Barrett, Sonia Goncalves, Ewan Harrison, David K. Jackson, Ian Johnston, Dominic Kwiatkowski, Cordelia Langford, John Sillitoe on behalf of the Wellcome Sanger Institute COVID-19 Surveillance Team |

| | | | |
|---|---|---|---|
| EPI_ISL_2765669 | | | |
| EPI_ISL_2765670 | Lighthouse Lab in Milton Keynes | Wellcome Sanger Institute for the COVID-19 Genomics UK (COG-UK) Consortium | The Lighthouse Lab in Milton Keynes and Alex Alderton, Roberto Amato, Jeffrey Barrett, Sonia Goncalves, Ewan Harrison, David K. Jackson, Ian Johnston, Dominic Kwiatkowski, Cordelia Langford, John Sillitoe on behalf of the Wellcome Sanger Institute COVID-19 Surveillance Team |
| EPI_ISL_2765671, EPI_ISL_2765672 | Lighthouse Lab in Alderley Park | Wellcome Sanger Institute for the COVID-19 Genomics UK (COG-UK) Consortium | Jacquelyn Wynn, Mairead Hyland, The Lighthouse Lab in Alderley Park and Alex Alderton, Roberto Amato, Jeffrey Barrett, Sonia Goncalves, Ewan Harrison, David K. Jackson, Ian Johnston, Dominic Kwiatkowski, Cordelia Langford, John Sillitoe on behalf of the Wellcome Sanger Institute COVID-19 Surveillance Team |
| EPI_ISL_2765673, EPI_ISL_2765674 | Lighthouse Lab in Milton Keynes | Wellcome Sanger Institute for the COVID-19 Genomics UK (COG-UK) Consortium | The Lighthouse Lab in Milton Keynes and Alex Alderton, Roberto Amato, Jeffrey Barrett, Sonia Goncalves, Ewan Harrison, David K. Jackson, Ian Johnston, Dominic Kwiatkowski, Cordelia Langford, John Sillitoe on behalf of the Wellcome Sanger Institute COVID-19 Surveillance Team |
| EPI_ISL_2765675 | Lighthouse Laboratory Plymouth | Wellcome Sanger Institute for the COVID-19 Genomics UK (COG-UK) Consortium | Lighthouse Laboratory Plymouth and Alex Alderton, Roberto Amato, Jeffrey Barrett, Sonia Goncalves, Ewan Harrison, David K. Jackson, Ian Johnston, Dominic Kwiatkowski, Cordelia Langford, John Sillitoe on behalf of the Wellcome Sanger Institute COVID-19 Surveillance Team |
| EPI_ISL_2765676 | Lighthouse Lab in Milton Keynes | Wellcome Sanger Institute for the COVID-19 Genomics UK (COG-UK) Consortium | The Lighthouse Lab in Milton Keynes and Alex Alderton, Roberto Amato, Jeffrey Barrett, Sonia Goncalves, Ewan Harrison, David K. Jackson, Ian Johnston, Dominic Kwiatkowski, Cordelia Langford, John Sillitoe on behalf of the Wellcome Sanger Institute COVID-19 Surveillance Team |
| EPI_ISL_2765677, EPI_ISL_2765679, EPI_ISL_2765680, EPI_ISL_2765681 | Lighthouse Lab in Alderley Park | Wellcome Sanger Institute for the COVID-19 Genomics UK (COG-UK) Consortium | Jacquelyn Wynn, Mairead Hyland, The Lighthouse Lab in Alderley Park and Alex Alderton, Roberto Amato, Jeffrey Barrett, Sonia Goncalves, Ewan Harrison, David K. Jackson, Ian Johnston, Dominic Kwiatkowski, Cordelia Langford, John Sillitoe on behalf of the Wellcome Sanger Institute COVID-19 Surveillance Team |
| EPI_ISL_2765682 | Lighthouse Lab in Milton Keynes | Wellcome Sanger Institute for the COVID-19 Genomics UK (COG-UK) Consortium | The Lighthouse Lab in Milton Keynes and Alex Alderton, Roberto Amato, Jeffrey Barrett, Sonia Goncalves, Ewan Harrison, David K. Jackson, Ian Johnston, Dominic Kwiatkowski, Cordelia Langford, John Sillitoe on behalf of the Wellcome Sanger Institute COVID-19 Surveillance Team |
| EPI_ISL_2765683, EPI_ISL_2765684 | Lighthouse Lab in Alderley Park | Wellcome Sanger Institute for the COVID-19 Genomics UK (COG-UK) Consortium | Jacquelyn Wynn, Mairead Hyland, The Lighthouse Lab in Alderley Park and Alex Alderton, Roberto Amato, Jeffrey Barrett, Sonia Goncalves, Ewan Harrison, David K. Jackson, Ian Johnston, Dominic Kwiatkowski, Cordelia Langford, John Sillitoe on behalf of the Wellcome Sanger Institute COVID-19 Surveillance Team |
| EPI_ISL_2765685 | Lighthouse Lab in Milton Keynes | Wellcome Sanger Institute for the COVID-19 Genomics UK (COG-UK) Consortium | The Lighthouse Lab in Milton Keynes and Alex Alderton, Roberto Amato, Jeffrey Barrett, Sonia Goncalves, Ewan Harrison, David K. Jackson, Ian Johnston, Dominic Kwiatkowski, Cordelia Langford, John Sillitoe on behalf of the Wellcome Sanger Institute COVID-19 Surveillance Team |
| EPI_ISL_2765686 | Lighthouse Lab in Alderley Park | Wellcome Sanger Institute for the COVID-19 Genomics UK (COG-UK) Consortium | Jacquelyn Wynn, Mairead Hyland, The Lighthouse Lab in Alderley Park and Alex Alderton, Roberto Amato, Jeffrey Barrett, Sonia Goncalves, Ewan Harrison, David K. Jackson, Ian Johnston, Dominic Kwiatkowski, Cordelia Langford, John Sillitoe on behalf of the Wellcome Sanger Institute COVID-19 Surveillance Team |
| EPI_ISL_2765687 | Lighthouse Lab in Milton Keynes | Wellcome Sanger Institute for the COVID-19 Genomics UK (COG-UK) Consortium | The Lighthouse Lab in Milton Keynes and Alex Alderton, Roberto Amato, Jeffrey Barrett, Sonia Goncalves, Ewan Harrison, David K. Jackson, Ian Johnston, Dominic Kwiatkowski, Cordelia Langford, John Sillitoe on behalf of the Wellcome Sanger Institute COVID-19 Surveillance Team |
| EPI_ISL_2765688, EPI_ISL_2765689, EPI_ISL_2765690 | Lighthouse Lab in Alderley Park | Wellcome Sanger Institute for the COVID-19 Genomics UK (COG-UK) Consortium | Jacquelyn Wynn, Mairead Hyland, The Lighthouse Lab in Alderley Park and Alex Alderton, Roberto Amato, Jeffrey Barrett, Sonia Goncalves, Ewan Harrison, David K. Jackson, Ian Johnston, Dominic Kwiatkowski, Cordelia Langford, John Sillitoe on behalf of the Wellcome Sanger Institute COVID-19 Surveillance Team |
| EPI_ISL_2765691 | Lighthouse Lab in Milton Keynes | Wellcome Sanger Institute for the COVID-19 Genomics UK (COG-UK) Consortium | The Lighthouse Lab in Milton Keynes and Alex Alderton, Roberto Amato, Jeffrey Barrett, Sonia Goncalves, Ewan Harrison, David K. Jackson, Ian Johnston, Dominic Kwiatkowski, Cordelia Langford, John Sillitoe on behalf of the Wellcome Sanger Institute COVID-19 Surveillance Team |
| EPI_ISL_2765692 | Lighthouse Lab in Alderley Park | Wellcome Sanger Institute for the COVID-19 Genomics UK (COG-UK) Consortium | Jacquelyn Wynn, Mairead Hyland, The Lighthouse Lab in Alderley Park and Alex Alderton, Roberto Amato, Jeffrey Barrett, Sonia Goncalves, Ewan Harrison, David K. Jackson, Ian Johnston, Dominic Kwiatkowski, Cordelia Langford, John Sillitoe on behalf of the Wellcome Sanger Institute COVID-19 Surveillance Team |
| EPI_ISL_2765693, EPI_ISL_2765694, EPI_ISL_2765695, EPI_ISL_2765696, EPI_ISL_2765697 | Lighthouse Lab in Milton Keynes | Wellcome Sanger Institute for the COVID-19 Genomics UK (COG-UK) Consortium | The Lighthouse Lab in Milton Keynes and Alex Alderton, Roberto Amato, Jeffrey Barrett, Sonia Goncalves, Ewan Harrison, David K. Jackson, Ian Johnston, Dominic Kwiatkowski, Cordelia Langford, John Sillitoe on behalf of the Wellcome Sanger Institute COVID-19 Surveillance Team |
| EPI_ISL_2765698, EPI_ISL_2765699 | Lighthouse Lab in Alderley Park | Wellcome Sanger Institute for the COVID-19 Genomics UK (COG-UK) Consortium | Jacquelyn Wynn, Mairead Hyland, The Lighthouse Lab in Alderley Park and Alex Alderton, Roberto Amato, Jeffrey Barrett, Sonia Goncalves, Ewan Harrison, David K. Jackson, Ian Johnston, Dominic Kwiatkowski, Cordelia Langford, John Sillitoe on behalf of the Wellcome Sanger Institute COVID-19 Surveillance Team |
| EPI_ISL_2765700 | Lighthouse Laboratory Plymouth | Wellcome Sanger Institute for the COVID-19 Genomics UK (COG-UK) Consortium | Lighthouse Laboratory Plymouth and Alex Alderton, Roberto Amato, Jeffrey Barrett, Sonia Goncalves, Ewan Harrison, David K. Jackson, Ian Johnston, Dominic Kwiatkowski, Cordelia Langford, John Sillitoe on behalf of the Wellcome Sanger Institute COVID-19 Surveillance Team |
| EPI_ISL_2765701 | Lighthouse Lab in Milton Keynes | Wellcome Sanger Institute for the COVID-19 Genomics UK (COG-UK) Consortium | The Lighthouse Lab in Milton Keynes and Alex Alderton, Roberto Amato, Jeffrey Barrett, Sonia Goncalves, Ewan Harrison, David K. Jackson, Ian Johnston, Dominic Kwiatkowski, Cordelia Langford, John Sillitoe on behalf of the Wellcome Sanger Institute COVID-19 Surveillance Team |
| EPI_ISL_2765702 | Lighthouse Lab in Alderley Park | Wellcome Sanger Institute for the COVID-19 Genomics UK (COG-UK) Consortium | Jacquelyn Wynn, Mairead Hyland, The Lighthouse Lab in Alderley Park and Alex Alderton, Roberto Amato, Jeffrey Barrett, Sonia Goncalves, Ewan Harrison, David K. Jackson, Ian Johnston, Dominic Kwiatkowski, Cordelia Langford, John Sillitoe on behalf of the Wellcome Sanger Institute COVID-19 Surveillance Team |
| EPI_ISL_2765703 | Lighthouse Lab in Milton Keynes | Wellcome Sanger Institute for the COVID-19 Genomics UK (COG-UK) Consortium | The Lighthouse Lab in Milton Keynes and Alex Alderton, Roberto Amato, Jeffrey Barrett, Sonia Goncalves, Ewan Harrison, David K. Jackson, Ian Johnston, Dominic Kwiatkowski, Cordelia Langford, John Sillitoe on behalf of the Wellcome Sanger Institute COVID-19 Surveillance Team |
| EPI_ISL_2765704, EPI_ISL_2765705, EPI_ISL_2765706 | Lighthouse Lab in Alderley Park | Wellcome Sanger Institute for the COVID-19 Genomics UK (COG-UK) Consortium | Jacquelyn Wynn, Mairead Hyland, The Lighthouse Lab in Alderley Park and Alex Alderton, Roberto Amato, Jeffrey Barrett, Sonia Goncalves, Ewan Harrison, David K. Jackson, Ian Johnston, Dominic Kwiatkowski, Cordelia Langford, John Sillitoe on behalf of the Wellcome Sanger Institute COVID-19 Surveillance Team |
| EPI_ISL_2765707, EPI_ISL_2765709 | Lighthouse Lab in Milton Keynes | Wellcome Sanger Institute for the COVID-19 Genomics UK (COG-UK) Consortium | The Lighthouse Lab in Milton Keynes and Alex Alderton, Roberto Amato, Jeffrey Barrett, Sonia Goncalves, Ewan Harrison, David K. Jackson, Ian Johnston, Dominic Kwiatkowski, Cordelia Langford, John Sillitoe on behalf of the Wellcome Sanger Institute COVID-19 Surveillance Team |
| EPI_ISL_2765710 | Lighthouse Lab in Alderley Park | Wellcome Sanger Institute for the COVID-19 Genomics UK (COG-UK) Consortium | Jacquelyn Wynn, Mairead Hyland, The Lighthouse Lab in Alderley Park and Alex Alderton, Roberto Amato, Jeffrey Barrett, Sonia Goncalves, Ewan Harrison, David K. Jackson, Ian Johnston, Dominic Kwiatkowski, Cordelia Langford, John Sillitoe on behalf of the Wellcome Sanger Institute COVID-19 Surveillance Team |
| EPI_ISL_2765711, EPI_ISL_2765712 | Lighthouse Lab in Milton Keynes | Wellcome Sanger Institute for the COVID-19 Genomics UK (COG-UK) Consortium | The Lighthouse Lab in Milton Keynes and Alex Alderton, Roberto Amato, Jeffrey Barrett, Sonia Goncalves, Ewan Harrison, David K. Jackson, Ian Johnston, Dominic Kwiatkowski, Cordelia Langford, John Sillitoe on behalf of the Wellcome Sanger Institute COVID-19 Surveillance Team |
| EPI_ISL_2765713, EPI_ISL_2765714 | Lighthouse Laboratory Plymouth | Wellcome Sanger Institute for the COVID-19 Genomics UK (COG-UK) Consortium | Lighthouse Laboratory Plymouth and Alex Alderton, Roberto Amato, Jeffrey Barrett, Sonia Goncalves, Ewan Harrison, David K. Jackson, Ian Johnston, Dominic Kwiatkowski, Cordelia Langford, John Sillitoe on behalf of the Wellcome Sanger Institute COVID-19 Surveillance Team |
| EPI_ISL_2765715, EPI_ISL_2765716, EPI_ISL_2765717 | Lighthouse Lab in Alderley Park | Wellcome Sanger Institute for the COVID-19 Genomics UK (COG-UK) Consortium | Jacquelyn Wynn, Mairead Hyland, The Lighthouse Lab in Alderley Park and Alex Alderton, Roberto Amato, Jeffrey Barrett, Sonia Goncalves, Ewan Harrison, David K. Jackson, Ian Johnston, Dominic Kwiatkowski, Cordelia Langford, John Sillitoe on behalf of the Wellcome Sanger Institute COVID-19 Surveillance Team |
| EPI_ISL_2765718, EPI_ISL_2765719 | Lighthouse Lab in Milton Keynes | Wellcome Sanger Institute for the COVID-19 Genomics UK (COG-UK) Consortium | The Lighthouse Lab in Milton Keynes and Alex Alderton, Roberto Amato, Jeffrey Barrett, Sonia Goncalves, Ewan Harrison, David K. Jackson, Ian Johnston, Dominic Kwiatkowski, Cordelia Langford, John Sillitoe on behalf of the Wellcome Sanger Institute COVID-19 Surveillance Team |
| EPI_ISL_2765720 | Lighthouse Laboratory Plymouth | Wellcome Sanger Institute for the COVID-19 Genomics UK (COG-UK) Consortium | Lighthouse Laboratory Plymouth and Alex Alderton, Roberto Amato, Jeffrey Barrett, Sonia Goncalves, Ewan Harrison, David K. Jackson, Ian Johnston, Dominic Kwiatkowski, Cordelia Langford, John Sillitoe on behalf of the Wellcome Sanger Institute COVID-19 Surveillance Team |
| EPI_ISL_2765721 | Lighthouse Lab in Alderley Park | Wellcome Sanger Institute for the COVID-19 Genomics UK (COG-UK) Consortium | Jacquelyn Wynn, Mairead Hyland, The Lighthouse Lab in Alderley Park and Alex Alderton, Roberto Amato, Jeffrey Barrett, Sonia Goncalves, Ewan Harrison, David K. Jackson, Ian Johnston, Dominic Kwiatkowski, Cordelia Langford, John Sillitoe on behalf of the Wellcome Sanger Institute COVID-19 Surveillance Team |
| EPI_ISL_2765722 | Lighthouse Lab in Milton Keynes | Wellcome Sanger Institute for the COVID-19 Genomics UK (COG-UK) Consortium | The Lighthouse Lab in Milton Keynes and Alex Alderton, Roberto Amato, Jeffrey Barrett, Sonia Goncalves, Ewan Harrison, David K. Jackson, Ian Johnston, Dominic Kwiatkowski, Cordelia Langford, John Sillitoe on behalf of the Wellcome Sanger Institute COVID-19 Surveillance Team |
| EPI_ISL_2765723, EPI_ISL_2765724 | Lighthouse Lab in Alderley Park | Wellcome Sanger Institute for the COVID-19 Genomics UK (COG-UK) Consortium | Jacquelyn Wynn, Mairead Hyland, The Lighthouse Lab in Alderley Park and Alex Alderton, Roberto Amato, Jeffrey Barrett, Sonia Goncalves, Ewan Harrison, David K. Jackson, Ian Johnston, Dominic Kwiatkowski, Cordelia Langford, John Sillitoe on behalf of the Wellcome Sanger Institute COVID-19 Surveillance Team |
| EPI_ISL_2765725, EPI_ISL_2765726 | Lighthouse Lab in Milton Keynes | Wellcome Sanger Institute for the COVID-19 Genomics UK (COG-UK) Consortium | The Lighthouse Lab in Milton Keynes and Alex Alderton, Roberto Amato, Jeffrey Barrett, Sonia Goncalves, Ewan Harrison, David K. Jackson, Ian Johnston, Dominic Kwiatkowski, Cordelia Langford, John Sillitoe on behalf of the Wellcome Sanger Institute COVID-19 Surveillance Team |
| EPI_ISL_2765727, EPI_ISL_2765729 | Lighthouse Lab in Alderley Park | Wellcome Sanger Institute for the COVID-19 Genomics UK (COG-UK) Consortium | Jacquelyn Wynn, Mairead Hyland, The Lighthouse Lab in Alderley Park and Alex Alderton, Roberto Amato, Jeffrey Barrett, Sonia Goncalves, Ewan Harrison, David K. Jackson, Ian Johnston, Dominic Kwiatkowski, Cordelia Langford, John Sillitoe on behalf of the Wellcome Sanger Institute COVID-19 Surveillance Team |
| EPI_ISL_2765730 | Lighthouse Laboratory Plymouth | Wellcome Sanger Institute for the COVID-19 Genomics UK (COG-UK) Consortium | Lighthouse Laboratory Plymouth and Alex Alderton, Roberto Amato, Jeffrey Barrett, Sonia Goncalves, Ewan Harrison, David K. Jackson, Ian Johnston, Dominic Kwiatkowski, Cordelia Langford, John Sillitoe on behalf of the Wellcome Sanger Institute COVID-19 Surveillance Team |
| EPI_ISL_2765731, EPI_ISL_2765732, EPI_ISL_2765733, EPI_ISL_2765734, EPI_ISL_2765735 | Lighthouse Lab in Alderley Park | Wellcome Sanger Institute for the COVID-19 Genomics UK (COG-UK) Consortium | Jacquelyn Wynn, Mairead Hyland, The Lighthouse Lab in Alderley Park and Alex Alderton, Roberto Amato, Jeffrey Barrett, Sonia Goncalves, Ewan Harrison, David K. Jackson, Ian Johnston, Dominic Kwiatkowski, Cordelia Langford, John Sillitoe on behalf of the Wellcome Sanger Institute COVID-19 Surveillance Team |
| EPI_ISL_2765736 | Lighthouse Lab in Milton Keynes | Wellcome Sanger Institute for the COVID-19 Genomics UK (COG-UK) Consortium | The Lighthouse Lab in Milton Keynes and Alex Alderton, Roberto Amato, Jeffrey Barrett, Sonia Goncalves, Ewan Harrison, David K. Jackson, Ian Johnston, Dominic Kwiatkowski, Cordelia Langford, John Sillitoe on behalf of the Wellcome Sanger Institute COVID-19 Surveillance Team |
| EPI_ISL_2765737 | Lighthouse Lab in Alderley Park | Wellcome Sanger Institute for the COVID-19 Genomics UK (COG-UK) Consortium | Jacquelyn Wynn, Mairead Hyland, The Lighthouse Lab in Alderley Park and Alex Alderton, Roberto Amato, Jeffrey Barrett, Sonia Goncalves, Ewan Harrison, David K. Jackson, Ian Johnston, Dominic Kwiatkowski, Cordelia Langford, John Sillitoe on behalf of the Wellcome Sanger Institute COVID-19 Surveillance Team |
| EPI_ISL_2765738 | Lighthouse Lab in Milton Keynes | Wellcome Sanger Institute for the COVID-19 Genomics UK (COG-UK) Consortium | The Lighthouse Lab in Milton Keynes and Alex Alderton, Roberto Amato, Jeffrey Barrett, Sonia Goncalves, Ewan Harrison, David K. Jackson, Ian Johnston, Dominic Kwiatkowski, Cordelia Langford, John Sillitoe on behalf of the Wellcome Sanger Institute COVID-19 Surveillance Team |
| EPI_ISL_2765739, EPI_ISL_2765740 | Lighthouse Lab in Alderley Park | Wellcome Sanger Institute for the COVID-19 Genomics UK (COG-UK) Consortium | Jacquelyn Wynn, Mairead Hyland, The Lighthouse Lab in Alderley Park and Alex Alderton, Roberto Amato, Jeffrey Barrett, Sonia Goncalves, Ewan Harrison, David K. Jackson, Ian Johnston, Dominic Kwiatkowski, Cordelia Langford, John Sillitoe on behalf of the Wellcome Sanger Institute COVID-19 Surveillance Team |
| EPI_ISL_2765741 | Lighthouse Lab in Milton Keynes | Wellcome Sanger Institute for the COVID-19 Genomics UK (COG-UK) Consortium | The Lighthouse Lab in Milton Keynes and Alex Alderton, Roberto Amato, Jeffrey Barrett, Sonia Goncalves, Ewan Harrison, David K. Jackson, Ian Johnston, Dominic Kwiatkowski, Cordelia Langford, John Sillitoe on behalf of the Wellcome Sanger Institute COVID-19 Surveillance Team |

| | | | |
|---|---|---|---|
| EPI_ISL_2765742, EPI_ISL_2765743 | Lighthouse Laboratory Plymouth | Wellcome Sanger Institute for the COVID-19 Genomics UK (COG-UK) Consortium | Lighthouse Laboratory Plymouth and Alex Alderton, Roberto Amato, Jeffrey Barrett, Sonia Goncalves, Ewan Harrison, David K. Jackson, Ian Johnston, Dominic Kwiatkowski, Cordelia Langford, John Sillitoe on behalf of the Wellcome Sanger Institute COVID-19 Surveillance Team |
| EPI_ISL_2765744 | Lighthouse Lab in Milton Keynes | Wellcome Sanger Institute for the COVID-19 Genomics UK (COG-UK) Consortium | The Lighthouse Lab in Milton Keynes and Alex Alderton, Roberto Amato, Jeffrey Barrett, Sonia Goncalves, Ewan Harrison, David K. Jackson, Ian Johnston, Dominic Kwiatkowski, Cordelia Langford, John Sillitoe on behalf of the Wellcome Sanger Institute COVID-19 Surveillance Team |
| EPI_ISL_2765745 | Lighthouse Laboratory Plymouth | Wellcome Sanger Institute for the COVID-19 Genomics UK (COG-UK) Consortium | Lighthouse Laboratory Plymouth and Alex Alderton, Roberto Amato, Jeffrey Barrett, Sonia Goncalves, Ewan Harrison, David K. Jackson, Ian Johnston, Dominic Kwiatkowski, Cordelia Langford, John Sillitoe on behalf of the Wellcome Sanger Institute COVID-19 Surveillance Team |
| EPI_ISL_2765746, EPI_ISL_2765747, EPI_ISL_2765748, EPI_ISL_2765749 | Lighthouse Lab in Alderley Park | Wellcome Sanger Institute for the COVID-19 Genomics UK (COG-UK) Consortium | Jacquelyn Wynn, Mairead Hyland, The Lighthouse Lab in Alderley Park and Alex Alderton, Roberto Amato, Jeffrey Barrett, Sonia Goncalves, Ewan Harrison, David K. Jackson, Ian Johnston, Dominic Kwiatkowski, Cordelia Langford, John Sillitoe on behalf of the Wellcome Sanger Institute COVID-19 Surveillance Team |
| EPI_ISL_2765750, EPI_ISL_2765751, EPI_ISL_2765752 | Lighthouse Lab in Milton Keynes | Wellcome Sanger Institute for the COVID-19 Genomics UK (COG-UK) Consortium | The Lighthouse Lab in Milton Keynes and Alex Alderton, Roberto Amato, Jeffrey Barrett, Sonia Goncalves, Ewan Harrison, David K. Jackson, Ian Johnston, Dominic Kwiatkowski, Cordelia Langford, John Sillitoe on behalf of the Wellcome Sanger Institute COVID-19 Surveillance Team |
| EPI_ISL_2765753 | Lighthouse Laboratory Plymouth | Wellcome Sanger Institute for the COVID-19 Genomics UK (COG-UK) Consortium | Lighthouse Laboratory Plymouth and Alex Alderton, Roberto Amato, Jeffrey Barrett, Sonia Goncalves, Ewan Harrison, David K. Jackson, Ian Johnston, Dominic Kwiatkowski, Cordelia Langford, John Sillitoe on behalf of the Wellcome Sanger Institute COVID-19 Surveillance Team |
| EPI_ISL_2765754, EPI_ISL_2765755, EPI_ISL_2765756 | Lighthouse Lab in Alderley Park | Wellcome Sanger Institute for the COVID-19 Genomics UK (COG-UK) Consortium | Jacquelyn Wynn, Mairead Hyland, The Lighthouse Lab in Alderley Park and Alex Alderton, Roberto Amato, Jeffrey Barrett, Sonia Goncalves, Ewan Harrison, David K. Jackson, Ian Johnston, Dominic Kwiatkowski, Cordelia Langford, John Sillitoe on behalf of the Wellcome Sanger Institute COVID-19 Surveillance Team |
| EPI_ISL_2765757 | Lighthouse Lab in Milton Keynes | Wellcome Sanger Institute for the COVID-19 Genomics UK (COG-UK) Consortium | The Lighthouse Lab in Milton Keynes and Alex Alderton, Roberto Amato, Jeffrey Barrett, Sonia Goncalves, Ewan Harrison, David K. Jackson, Ian Johnston, Dominic Kwiatkowski, Cordelia Langford, John Sillitoe on behalf of the Wellcome Sanger Institute COVID-19 Surveillance Team |
| EPI_ISL_2765758 | Lighthouse Lab in Alderley Park | Wellcome Sanger Institute for the COVID-19 Genomics UK (COG-UK) Consortium | Jacquelyn Wynn, Mairead Hyland, The Lighthouse Lab in Alderley Park and Alex Alderton, Roberto Amato, Jeffrey Barrett, Sonia Goncalves, Ewan Harrison, David K. Jackson, Ian Johnston, Dominic Kwiatkowski, Cordelia Langford, John Sillitoe on behalf of the Wellcome Sanger Institute COVID-19 Surveillance Team |
| EPI_ISL_2765759 | Lighthouse Laboratory Plymouth | Wellcome Sanger Institute for the COVID-19 Genomics UK (COG-UK) Consortium | Lighthouse Laboratory Plymouth and Alex Alderton, Roberto Amato, Jeffrey Barrett, Sonia Goncalves, Ewan Harrison, David K. Jackson, Ian Johnston, Dominic Kwiatkowski, Cordelia Langford, John Sillitoe on behalf of the Wellcome Sanger Institute COVID-19 Surveillance Team |
| EPI_ISL_2765760, EPI_ISL_2765761, EPI_ISL_2765762 | Lighthouse Lab in Alderley Park | Wellcome Sanger Institute for the COVID-19 Genomics UK (COG-UK) Consortium | Jacquelyn Wynn, Mairead Hyland, The Lighthouse Lab in Alderley Park and Alex Alderton, Roberto Amato, Jeffrey Barrett, Sonia Goncalves, Ewan Harrison, David K. Jackson, Ian Johnston, Dominic Kwiatkowski, Cordelia Langford, John Sillitoe on behalf of the Wellcome Sanger Institute COVID-19 Surveillance Team |
| EPI_ISL_2765763 | Lighthouse Lab in Milton Keynes | Wellcome Sanger Institute for the COVID-19 Genomics UK (COG-UK) Consortium | The Lighthouse Lab in Milton Keynes and Alex Alderton, Roberto Amato, Jeffrey Barrett, Sonia Goncalves, Ewan Harrison, David K. Jackson, Ian Johnston, Dominic Kwiatkowski, Cordelia Langford, John Sillitoe on behalf of the Wellcome Sanger Institute COVID-19 Surveillance Team |
| EPI_ISL_2765764, EPI_ISL_2765765, EPI_ISL_2765766 | Lighthouse Lab in Alderley Park | Wellcome Sanger Institute for the COVID-19 Genomics UK (COG-UK) Consortium | Jacquelyn Wynn, Mairead Hyland, The Lighthouse Lab in Alderley Park and Alex Alderton, Roberto Amato, Jeffrey Barrett, Sonia Goncalves, Ewan Harrison, David K. Jackson, Ian Johnston, Dominic Kwiatkowski, Cordelia Langford, John Sillitoe on behalf of the Wellcome Sanger Institute COVID-19 Surveillance Team |
| EPI_ISL_2765767 | Lighthouse Laboratory Plymouth | Wellcome Sanger Institute for the COVID-19 Genomics UK (COG-UK) Consortium | Lighthouse Laboratory Plymouth and Alex Alderton, Roberto Amato, Jeffrey Barrett, Sonia Goncalves, Ewan Harrison, David K. Jackson, Ian Johnston, Dominic Kwiatkowski, Cordelia Langford, John Sillitoe on behalf of the Wellcome Sanger Institute COVID-19 Surveillance Team |
| EPI_ISL_2765768 | Lighthouse Lab in Milton Keynes | Wellcome Sanger Institute for the COVID-19 Genomics UK (COG-UK) Consortium | The Lighthouse Lab in Milton Keynes and Alex Alderton, Roberto Amato, Jeffrey Barrett, Sonia Goncalves, Ewan Harrison, David K. Jackson, Ian Johnston, Dominic Kwiatkowski, Cordelia Langford, John Sillitoe on behalf of the Wellcome Sanger Institute COVID-19 Surveillance Team |
| EPI_ISL_2765769 | Lighthouse Lab in Alderley Park | Wellcome Sanger Institute for the COVID-19 Genomics UK (COG-UK) Consortium | Jacquelyn Wynn, Mairead Hyland, The Lighthouse Lab in Alderley Park and Alex Alderton, Roberto Amato, Jeffrey Barrett, Sonia Goncalves, Ewan Harrison, David K. Jackson, Ian Johnston, Dominic Kwiatkowski, Cordelia Langford, John Sillitoe on behalf of the Wellcome Sanger Institute COVID-19 Surveillance Team |
| EPI_ISL_2765771 | Lighthouse Lab in Glasgow | Wellcome Sanger Institute for the COVID-19 Genomics UK (COG-UK) Consortium | Harper VanSteenhouse, Yumi Kasai, David Gray, Carol Clugston, Anna Dominiczak and Alex Alderton, Roberto Amato, Jeffrey Barrett, Sonia Goncalves, Ewan Harrison, David K. Jackson, Ian Johnston, Dominic Kwiatkowski, Cordelia Langford, John Sillitoe on behalf of the Wellcome Sanger Institute COVID-19 Surveillance Team |
| EPI_ISL_2765772, EPI_ISL_2765773, EPI_ISL_2765774, EPI_ISL_2765775, EPI_ISL_2765776 | Lighthouse Lab in Alderley Park | Wellcome Sanger Institute for the COVID-19 Genomics UK (COG-UK) Consortium | Jacquelyn Wynn, Mairead Hyland, The Lighthouse Lab in Alderley Park and Alex Alderton, Roberto Amato, Jeffrey Barrett, Sonia Goncalves, Ewan Harrison, David K. Jackson, Ian Johnston, Dominic Kwiatkowski, Cordelia Langford, John Sillitoe on behalf of the Wellcome Sanger Institute COVID-19 Surveillance Team |
| EPI_ISL_2765777 | Lighthouse Lab in Milton Keynes | Wellcome Sanger Institute for the COVID-19 Genomics UK (COG-UK) Consortium | The Lighthouse Lab in Milton Keynes and Alex Alderton, Roberto Amato, Jeffrey Barrett, Sonia Goncalves, Ewan Harrison, David K. Jackson, Ian Johnston, Dominic Kwiatkowski, Cordelia Langford, John Sillitoe on behalf of the Wellcome Sanger Institute COVID-19 Surveillance Team |
| EPI_ISL_2765778, EPI_ISL_2765779, EPI_ISL_2765780 | Lighthouse Lab in Alderley Park | Wellcome Sanger Institute for the COVID-19 Genomics UK (COG-UK) Consortium | Jacquelyn Wynn, Mairead Hyland, The Lighthouse Lab in Alderley Park and Alex Alderton, Roberto Amato, Jeffrey Barrett, Sonia Goncalves, Ewan Harrison, David K. Jackson, Ian Johnston, Dominic Kwiatkowski, Cordelia Langford, John Sillitoe on behalf of the Wellcome Sanger Institute COVID-19 Surveillance Team |
| EPI_ISL_2765781 | Lighthouse Laboratory Plymouth | Wellcome Sanger Institute for the COVID-19 Genomics UK (COG-UK) Consortium | Lighthouse Laboratory Plymouth and Alex Alderton, Roberto Amato, Jeffrey Barrett, Sonia Goncalves, Ewan Harrison, David K. Jackson, Ian Johnston, Dominic Kwiatkowski, Cordelia Langford, John Sillitoe on behalf of the Wellcome Sanger Institute COVID-19 Surveillance Team |
| EPI_ISL_2765782, EPI_ISL_2765783 | Lighthouse Lab in Alderley Park | Wellcome Sanger Institute for the COVID-19 Genomics UK (COG-UK) Consortium | Jacquelyn Wynn, Mairead Hyland, The Lighthouse Lab in Alderley Park and Alex Alderton, Roberto Amato, Jeffrey Barrett, Sonia Goncalves, Ewan Harrison, David K. Jackson, Ian Johnston, Dominic Kwiatkowski, Cordelia Langford, John Sillitoe on behalf of the Wellcome Sanger Institute COVID-19 Surveillance Team |
| EPI_ISL_2765784 | Lighthouse Lab in Milton Keynes | Wellcome Sanger Institute for the COVID-19 Genomics UK (COG-UK) Consortium | The Lighthouse Lab in Milton Keynes and Alex Alderton, Roberto Amato, Jeffrey Barrett, Sonia Goncalves, Ewan Harrison, David K. Jackson, Ian Johnston, Dominic Kwiatkowski, Cordelia Langford, John Sillitoe on behalf of the Wellcome Sanger Institute COVID-19 Surveillance Team |
| EPI_ISL_2765785, EPI_ISL_2765786, EPI_ISL_2765787 | Lighthouse Lab in Alderley Park | Wellcome Sanger Institute for the COVID-19 Genomics UK (COG-UK) Consortium | Jacquelyn Wynn, Mairead Hyland, The Lighthouse Lab in Alderley Park and Alex Alderton, Roberto Amato, Jeffrey Barrett, Sonia Goncalves, Ewan Harrison, David K. Jackson, Ian Johnston, Dominic Kwiatkowski, Cordelia Langford, John Sillitoe on behalf of the Wellcome Sanger Institute COVID-19 Surveillance Team |
| EPI_ISL_2765788 | Lighthouse Laboratory Plymouth | Wellcome Sanger Institute for the COVID-19 Genomics UK (COG-UK) Consortium | Lighthouse Laboratory Plymouth and Alex Alderton, Roberto Amato, Jeffrey Barrett, Sonia Goncalves, Ewan Harrison, David K. Jackson, Ian Johnston, Dominic Kwiatkowski, Cordelia Langford, John Sillitoe on behalf of the Wellcome Sanger Institute COVID-19 Surveillance Team |
| EPI_ISL_2765789, EPI_ISL_2765790 | Lighthouse Lab in Milton Keynes | Wellcome Sanger Institute for the COVID-19 Genomics UK (COG-UK) Consortium | The Lighthouse Lab in Milton Keynes and Alex Alderton, Roberto Amato, Jeffrey Barrett, Sonia Goncalves, Ewan Harrison, David K. Jackson, Ian Johnston, Dominic Kwiatkowski, Cordelia Langford, John Sillitoe on behalf of the Wellcome Sanger Institute COVID-19 Surveillance Team |
| EPI_ISL_2765791 | Lighthouse Lab in Alderley Park | Wellcome Sanger Institute for the COVID-19 Genomics UK (COG-UK) Consortium | Jacquelyn Wynn, Mairead Hyland, The Lighthouse Lab in Alderley Park and Alex Alderton, Roberto Amato, Jeffrey Barrett, Sonia Goncalves, Ewan Harrison, David K. Jackson, Ian Johnston, Dominic Kwiatkowski, Cordelia Langford, John Sillitoe on behalf of the Wellcome Sanger Institute COVID-19 Surveillance Team |
| EPI_ISL_2765792 | Lighthouse Lab in Milton Keynes | Wellcome Sanger Institute for the COVID-19 Genomics UK (COG-UK) Consortium | The Lighthouse Lab in Milton Keynes and Alex Alderton, Roberto Amato, Jeffrey Barrett, Sonia Goncalves, Ewan Harrison, David K. Jackson, Ian Johnston, Dominic Kwiatkowski, Cordelia Langford, John Sillitoe on behalf of the Wellcome Sanger Institute COVID-19 Surveillance Team |
| EPI_ISL_2765793, EPI_ISL_2765794 | Lighthouse Lab in Alderley Park | Wellcome Sanger Institute for the COVID-19 Genomics UK (COG-UK) Consortium | Jacquelyn Wynn, Mairead Hyland, The Lighthouse Lab in Alderley Park and Alex Alderton, Roberto Amato, Jeffrey Barrett, Sonia Goncalves, Ewan Harrison, David K. Jackson, Ian Johnston, Dominic Kwiatkowski, Cordelia Langford, John Sillitoe on behalf of the Wellcome Sanger Institute COVID-19 Surveillance Team |
| EPI_ISL_2765795 | Lighthouse Laboratory Plymouth | Wellcome Sanger Institute for the COVID-19 Genomics UK (COG-UK) Consortium | Lighthouse Laboratory Plymouth and Alex Alderton, Roberto Amato, Jeffrey Barrett, Sonia Goncalves, Ewan Harrison, David K. Jackson, Ian Johnston, Dominic Kwiatkowski, Cordelia Langford, John Sillitoe on behalf of the Wellcome Sanger Institute COVID-19 Surveillance Team |
| EPI_ISL_2765796 | Lighthouse Lab in Alderley Park | Wellcome Sanger Institute for the COVID-19 Genomics UK (COG-UK) Consortium | Jacquelyn Wynn, Mairead Hyland, The Lighthouse Lab in Alderley Park and Alex Alderton, Roberto Amato, Jeffrey Barrett, Sonia Goncalves, Ewan Harrison, David K. Jackson, Ian Johnston, Dominic Kwiatkowski, Cordelia Langford, John Sillitoe on behalf of the Wellcome Sanger Institute COVID-19 Surveillance Team |
| EPI_ISL_2765797 | Lighthouse Lab in Milton Keynes | Wellcome Sanger Institute for the COVID-19 Genomics UK (COG-UK) Consortium | The Lighthouse Lab in Milton Keynes and Alex Alderton, Roberto Amato, Jeffrey Barrett, Sonia Goncalves, Ewan Harrison, David K. Jackson, Ian Johnston, Dominic Kwiatkowski, Cordelia Langford, John Sillitoe on behalf of the Wellcome Sanger Institute COVID-19 Surveillance Team |
| EPI_ISL_2765798, EPI_ISL_2765799 | Lighthouse Laboratory Plymouth | Wellcome Sanger Institute for the COVID-19 Genomics UK (COG-UK) Consortium | Lighthouse Laboratory Plymouth and Alex Alderton, Roberto Amato, Jeffrey Barrett, Sonia Goncalves, Ewan Harrison, David K. Jackson, Ian Johnston, Dominic Kwiatkowski, Cordelia Langford, John Sillitoe on behalf of the Wellcome Sanger Institute COVID-19 Surveillance Team |
| EPI_ISL_2765800, EPI_ISL_2765801, EPI_ISL_2765802 | Lighthouse Lab in Alderley Park | Wellcome Sanger Institute for the COVID-19 Genomics UK (COG-UK) Consortium | Jacquelyn Wynn, Mairead Hyland, The Lighthouse Lab in Alderley Park and Alex Alderton, Roberto Amato, Jeffrey Barrett, Sonia Goncalves, Ewan Harrison, David K. Jackson, Ian Johnston, Dominic Kwiatkowski, Cordelia Langford, John Sillitoe on behalf of the Wellcome Sanger Institute COVID-19 Surveillance Team |
| EPI_ISL_2765803 | Lighthouse Lab in Milton Keynes | Wellcome Sanger Institute for the COVID-19 Genomics UK (COG-UK) Consortium | The Lighthouse Lab in Milton Keynes and Alex Alderton, Roberto Amato, Jeffrey Barrett, Sonia Goncalves, Ewan Harrison, David K. Jackson, Ian Johnston, Dominic Kwiatkowski, Cordelia Langford, John Sillitoe on behalf of the Wellcome Sanger Institute COVID-19 Surveillance Team |
| EPI_ISL_2765804, EPI_ISL_2765805, EPI_ISL_2765806, EPI_ISL_2765807, EPI_ISL_2765808, EPI_ISL_2765809, EPI_ISL_2765810, EPI_ISL_2765811, EPI_ISL_2765812 | Lighthouse Lab in Alderley Park | Wellcome Sanger Institute for the COVID-19 Genomics UK (COG-UK) Consortium | Jacquelyn Wynn, Mairead Hyland, The Lighthouse Lab in Alderley Park and Alex Alderton, Roberto Amato, Jeffrey Barrett, Sonia Goncalves, Ewan Harrison, David K. Jackson, Ian Johnston, Dominic Kwiatkowski, Cordelia Langford, John Sillitoe on behalf of the Wellcome Sanger Institute COVID-19 Surveillance Team |
| EPI_ISL_2765813 | Lighthouse Laboratory Plymouth | Wellcome Sanger Institute for the COVID-19 Genomics UK (COG-UK) Consortium | Lighthouse Laboratory Plymouth and Alex Alderton, Roberto Amato, Jeffrey Barrett, Sonia Goncalves, Ewan Harrison, David K. Jackson, Ian Johnston, Dominic Kwiatkowski, Cordelia Langford, John Sillitoe on behalf of the Wellcome Sanger Institute COVID-19 Surveillance Team |
| EPI_ISL_2765814 | Lighthouse Lab in Alderley Park | Wellcome Sanger Institute for the COVID-19 Genomics UK (COG-UK) | Jacquelyn Wynn, Mairead Hyland, The Lighthouse Lab in Alderley Park and Alex Alderton, Roberto Amato, Jeffrey Barrett, Sonia Goncalves, Ewan Harrison, David K. Jackson, Ian Johnston, Dominic Kwiatkowski, Cordelia Langford, John Sillitoe on |

| | | | |
|---|---|---|---|
| | | Consortium | behalf of the Wellcome Sanger Institute COVID-19 Surveillance Team |
| EPI_ISL_2765815 | Lighthouse Lab in Glasgow | Wellcome Sanger Institute for the COVID-19 Genomics UK (COG-UK) Consortium | Harper VanSteenhouse, Yumi Kasai, David Gray, Carol Clugston, Anna Dominiczak and Alex Alderton, Roberto Amato, Jeffrey Barrett, Sonia Goncalves, Ewan Harrison, David K. Jackson, Ian Johnston, Dominic Kwiatkowski, Cordelia Langford, John Sillitoe on behalf of the Wellcome Sanger Institute COVID-19 Surveillance Team |
| EPI_ISL_2765816 | Lighthouse Lab in Alderley Park | Wellcome Sanger Institute for the COVID-19 Genomics UK (COG-UK) Consortium | Jacquelyn Wynn, Mairead Hyland, The Lighthouse Lab in Alderley Park and Alex Alderton, Roberto Amato, Jeffrey Barrett, Sonia Goncalves, Ewan Harrison, David K. Jackson, Ian Johnston, Dominic Kwiatkowski, Cordelia Langford, John Sillitoe on behalf of the Wellcome Sanger Institute COVID-19 Surveillance Team |
| EPI_ISL_2765817 | Lighthouse Lab in Milton Keynes | Wellcome Sanger Institute for the COVID-19 Genomics UK (COG-UK) Consortium | The Lighthouse Lab in Milton Keynes and Alex Alderton, Roberto Amato, Jeffrey Barrett, Sonia Goncalves, Ewan Harrison, David K. Jackson, Ian Johnston, Dominic Kwiatkowski, Cordelia Langford, John Sillitoe on behalf of the Wellcome Sanger Institute COVID-19 Surveillance Team |
| EPI_ISL_2765818 | Lighthouse Lab in Alderley Park | Wellcome Sanger Institute for the COVID-19 Genomics UK (COG-UK) Consortium | Jacquelyn Wynn, Mairead Hyland, The Lighthouse Lab in Alderley Park and Alex Alderton, Roberto Amato, Jeffrey Barrett, Sonia Goncalves, Ewan Harrison, David K. Jackson, Ian Johnston, Dominic Kwiatkowski, Cordelia Langford, John Sillitoe on behalf of the Wellcome Sanger Institute COVID-19 Surveillance Team |
| EPI_ISL_2765819 | Lighthouse Lab in Milton Keynes | Wellcome Sanger Institute for the COVID-19 Genomics UK (COG-UK) Consortium | The Lighthouse Lab in Milton Keynes and Alex Alderton, Roberto Amato, Jeffrey Barrett, Sonia Goncalves, Ewan Harrison, David K. Jackson, Ian Johnston, Dominic Kwiatkowski, Cordelia Langford, John Sillitoe on behalf of the Wellcome Sanger Institute COVID-19 Surveillance Team |
| EPI_ISL_2765820 | Lighthouse Lab in Alderley Park | Wellcome Sanger Institute for the COVID-19 Genomics UK (COG-UK) Consortium | Jacquelyn Wynn, Mairead Hyland, The Lighthouse Lab in Alderley Park and Alex Alderton, Roberto Amato, Jeffrey Barrett, Sonia Goncalves, Ewan Harrison, David K. Jackson, Ian Johnston, Dominic Kwiatkowski, Cordelia Langford, John Sillitoe on behalf of the Wellcome Sanger Institute COVID-19 Surveillance Team |
| EPI_ISL_2765821 | Lighthouse Laboratory Plymouth | Wellcome Sanger Institute for the COVID-19 Genomics UK (COG-UK) Consortium | Lighthouse Laboratory Plymouth and Alex Alderton, Roberto Amato, Jeffrey Barrett, Sonia Goncalves, Ewan Harrison, David K. Jackson, Ian Johnston, Dominic Kwiatkowski, Cordelia Langford, John Sillitoe on behalf of the Wellcome Sanger Institute COVID-19 Surveillance Team |
| EPI_ISL_2765822, EPI_ISL_2765823, EPI_ISL_2765824, EPI_ISL_2765825 | Lighthouse Lab in Alderley Park | Wellcome Sanger Institute for the COVID-19 Genomics UK (COG-UK) Consortium | Jacquelyn Wynn, Mairead Hyland, The Lighthouse Lab in Alderley Park and Alex Alderton, Roberto Amato, Jeffrey Barrett, Sonia Goncalves, Ewan Harrison, David K. Jackson, Ian Johnston, Dominic Kwiatkowski, Cordelia Langford, John Sillitoe on behalf of the Wellcome Sanger Institute COVID-19 Surveillance Team |
| EPI_ISL_2765826 | Lighthouse Lab in Milton Keynes | Wellcome Sanger Institute for the COVID-19 Genomics UK (COG-UK) Consortium | The Lighthouse Lab in Milton Keynes and Alex Alderton, Roberto Amato, Jeffrey Barrett, Sonia Goncalves, Ewan Harrison, David K. Jackson, Ian Johnston, Dominic Kwiatkowski, Cordelia Langford, John Sillitoe on behalf of the Wellcome Sanger Institute COVID-19 Surveillance Team |
| EPI_ISL_2765827 | Lighthouse Laboratory Plymouth | Wellcome Sanger Institute for the COVID-19 Genomics UK (COG-UK) Consortium | Lighthouse Laboratory Plymouth and Alex Alderton, Roberto Amato, Jeffrey Barrett, Sonia Goncalves, Ewan Harrison, David K. Jackson, Ian Johnston, Dominic Kwiatkowski, Cordelia Langford, John Sillitoe on behalf of the Wellcome Sanger Institute COVID-19 Surveillance Team |
| EPI_ISL_2765828, EPI_ISL_2765829 | Lighthouse Lab in Alderley Park | Wellcome Sanger Institute for the COVID-19 Genomics UK (COG-UK) Consortium | Jacquelyn Wynn, Mairead Hyland, The Lighthouse Lab in Alderley Park and Alex Alderton, Roberto Amato, Jeffrey Barrett, Sonia Goncalves, Ewan Harrison, David K. Jackson, Ian Johnston, Dominic Kwiatkowski, Cordelia Langford, John Sillitoe on behalf of the Wellcome Sanger Institute COVID-19 Surveillance Team |
| EPI_ISL_2765830 | Lighthouse Lab in Milton Keynes | Wellcome Sanger Institute for the COVID-19 Genomics UK (COG-UK) Consortium | The Lighthouse Lab in Milton Keynes and Alex Alderton, Roberto Amato, Jeffrey Barrett, Sonia Goncalves, Ewan Harrison, David K. Jackson, Ian Johnston, Dominic Kwiatkowski, Cordelia Langford, John Sillitoe on behalf of the Wellcome Sanger Institute COVID-19 Surveillance Team |
| EPI_ISL_2765831 | Lighthouse Lab in Alderley Park | Wellcome Sanger Institute for the COVID-19 Genomics UK (COG-UK) Consortium | Jacquelyn Wynn, Mairead Hyland, The Lighthouse Lab in Alderley Park and Alex Alderton, Roberto Amato, Jeffrey Barrett, Sonia Goncalves, Ewan Harrison, David K. Jackson, Ian Johnston, Dominic Kwiatkowski, Cordelia Langford, John Sillitoe on behalf of the Wellcome Sanger Institute COVID-19 Surveillance Team |
| EPI_ISL_2765832, EPI_ISL_2765833, EPI_ISL_2765834 | Lighthouse Laboratory Plymouth | Wellcome Sanger Institute for the COVID-19 Genomics UK (COG-UK) Consortium | Lighthouse Laboratory Plymouth and Alex Alderton, Roberto Amato, Jeffrey Barrett, Sonia Goncalves, Ewan Harrison, David K. Jackson, Ian Johnston, Dominic Kwiatkowski, Cordelia Langford, John Sillitoe on behalf of the Wellcome Sanger Institute COVID-19 Surveillance Team |
| EPI_ISL_2765835, EPI_ISL_2765836 | Lighthouse Lab in Alderley Park | Wellcome Sanger Institute for the COVID-19 Genomics UK (COG-UK) Consortium | Jacquelyn Wynn, Mairead Hyland, The Lighthouse Lab in Alderley Park and Alex Alderton, Roberto Amato, Jeffrey Barrett, Sonia Goncalves, Ewan Harrison, David K. Jackson, Ian Johnston, Dominic Kwiatkowski, Cordelia Langford, John Sillitoe on behalf of the Wellcome Sanger Institute COVID-19 Surveillance Team |
| EPI_ISL_2765837 | Lighthouse Laboratory Plymouth | Wellcome Sanger Institute for the COVID-19 Genomics UK (COG-UK) Consortium | Lighthouse Laboratory Plymouth and Alex Alderton, Roberto Amato, Jeffrey Barrett, Sonia Goncalves, Ewan Harrison, David K. Jackson, Ian Johnston, Dominic Kwiatkowski, Cordelia Langford, John Sillitoe on behalf of the Wellcome Sanger Institute COVID-19 Surveillance Team |
| EPI_ISL_2765838 | Lighthouse Lab in Alderley Park | Wellcome Sanger Institute for the COVID-19 Genomics UK (COG-UK) Consortium | Jacquelyn Wynn, Mairead Hyland, The Lighthouse Lab in Alderley Park and Alex Alderton, Roberto Amato, Jeffrey Barrett, Sonia Goncalves, Ewan Harrison, David K. Jackson, Ian Johnston, Dominic Kwiatkowski, Cordelia Langford, John Sillitoe on behalf of the Wellcome Sanger Institute COVID-19 Surveillance Team |
| EPI_ISL_2765839 | Lighthouse Lab in Milton Keynes | Wellcome Sanger Institute for the COVID-19 Genomics UK (COG-UK) Consortium | The Lighthouse Lab in Milton Keynes and Alex Alderton, Roberto Amato, Jeffrey Barrett, Sonia Goncalves, Ewan Harrison, David K. Jackson, Ian Johnston, Dominic Kwiatkowski, Cordelia Langford, John Sillitoe on behalf of the Wellcome Sanger Institute COVID-19 Surveillance Team |
| EPI_ISL_2765840, EPI_ISL_2765841 | Lighthouse Lab in Alderley Park | Wellcome Sanger Institute for the COVID-19 Genomics UK (COG-UK) Consortium | Jacquelyn Wynn, Mairead Hyland, The Lighthouse Lab in Alderley Park and Alex Alderton, Roberto Amato, Jeffrey Barrett, Sonia Goncalves, Ewan Harrison, David K. Jackson, Ian Johnston, Dominic Kwiatkowski, Cordelia Langford, John Sillitoe on behalf of the Wellcome Sanger Institute COVID-19 Surveillance Team |
| EPI_ISL_2765842 | Lighthouse Lab in Milton Keynes | Wellcome Sanger Institute for the COVID-19 Genomics UK (COG-UK) Consortium | The Lighthouse Lab in Milton Keynes and Alex Alderton, Roberto Amato, Jeffrey Barrett, Sonia Goncalves, Ewan Harrison, David K. Jackson, Ian Johnston, Dominic Kwiatkowski, Cordelia Langford, John Sillitoe on behalf of the Wellcome Sanger Institute COVID-19 Surveillance Team |
| EPI_ISL_2765843 | Lighthouse Lab in Alderley Park | Wellcome Sanger Institute for the COVID-19 Genomics UK (COG-UK) Consortium | Jacquelyn Wynn, Mairead Hyland, The Lighthouse Lab in Alderley Park and Alex Alderton, Roberto Amato, Jeffrey Barrett, Sonia Goncalves, Ewan Harrison, David K. Jackson, Ian Johnston, Dominic Kwiatkowski, Cordelia Langford, John Sillitoe on behalf of the Wellcome Sanger Institute COVID-19 Surveillance Team |
| EPI_ISL_2765844 | Lighthouse Lab in Milton Keynes | Wellcome Sanger Institute for the COVID-19 Genomics UK (COG-UK) Consortium | The Lighthouse Lab in Milton Keynes and Alex Alderton, Roberto Amato, Jeffrey Barrett, Sonia Goncalves, Ewan Harrison, David K. Jackson, Ian Johnston, Dominic Kwiatkowski, Cordelia Langford, John Sillitoe on behalf of the Wellcome Sanger Institute COVID-19 Surveillance Team |
| EPI_ISL_2765845, EPI_ISL_2765846, EPI_ISL_2765847, EPI_ISL_2765848 | Lighthouse Lab in Alderley Park | Wellcome Sanger Institute for the COVID-19 Genomics UK (COG-UK) Consortium | Jacquelyn Wynn, Mairead Hyland, The Lighthouse Lab in Alderley Park and Alex Alderton, Roberto Amato, Jeffrey Barrett, Sonia Goncalves, Ewan Harrison, David K. Jackson, Ian Johnston, Dominic Kwiatkowski, Cordelia Langford, John Sillitoe on behalf of the Wellcome Sanger Institute COVID-19 Surveillance Team |
| EPI_ISL_2765849 | Lighthouse Laboratory Plymouth | Wellcome Sanger Institute for the COVID-19 Genomics UK (COG-UK) Consortium | Lighthouse Laboratory Plymouth and Alex Alderton, Roberto Amato, Jeffrey Barrett, Sonia Goncalves, Ewan Harrison, David K. Jackson, Ian Johnston, Dominic Kwiatkowski, Cordelia Langford, John Sillitoe on behalf of the Wellcome Sanger Institute COVID-19 Surveillance Team |
| EPI_ISL_2765850, EPI_ISL_2765851, EPI_ISL_2765852 | Lighthouse Lab in Milton Keynes | Wellcome Sanger Institute for the COVID-19 Genomics UK (COG-UK) Consortium | The Lighthouse Lab in Milton Keynes and Alex Alderton, Roberto Amato, Jeffrey Barrett, Sonia Goncalves, Ewan Harrison, David K. Jackson, Ian Johnston, Dominic Kwiatkowski, Cordelia Langford, John Sillitoe on behalf of the Wellcome Sanger Institute COVID-19 Surveillance Team |
| EPI_ISL_2765853, EPI_ISL_2765854, EPI_ISL_2765855, EPI_ISL_2765856, EPI_ISL_2765857, EPI_ISL_2765858 | Lighthouse Lab in Alderley Park | Wellcome Sanger Institute for the COVID-19 Genomics UK (COG-UK) Consortium | Jacquelyn Wynn, Mairead Hyland, The Lighthouse Lab in Alderley Park and Alex Alderton, Roberto Amato, Jeffrey Barrett, Sonia Goncalves, Ewan Harrison, David K. Jackson, Ian Johnston, Dominic Kwiatkowski, Cordelia Langford, John Sillitoe on behalf of the Wellcome Sanger Institute COVID-19 Surveillance Team |
| EPI_ISL_2765860 | Lighthouse Lab in Milton Keynes | Wellcome Sanger Institute for the COVID-19 Genomics UK (COG-UK) Consortium | The Lighthouse Lab in Milton Keynes and Alex Alderton, Roberto Amato, Jeffrey Barrett, Sonia Goncalves, Ewan Harrison, David K. Jackson, Ian Johnston, Dominic Kwiatkowski, Cordelia Langford, John Sillitoe on behalf of the Wellcome Sanger Institute COVID-19 Surveillance Team |
| EPI_ISL_2765861 | Lighthouse Lab in Alderley Park | Wellcome Sanger Institute for the COVID-19 Genomics UK (COG-UK) Consortium | Jacquelyn Wynn, Mairead Hyland, The Lighthouse Lab in Alderley Park and Alex Alderton, Roberto Amato, Jeffrey Barrett, Sonia Goncalves, Ewan Harrison, David K. Jackson, Ian Johnston, Dominic Kwiatkowski, Cordelia Langford, John Sillitoe on behalf of the Wellcome Sanger Institute COVID-19 Surveillance Team |
| EPI_ISL_2765862 | Lighthouse Lab in Milton Keynes | Wellcome Sanger Institute for the COVID-19 Genomics UK (COG-UK) Consortium | The Lighthouse Lab in Milton Keynes and Alex Alderton, Roberto Amato, Jeffrey Barrett, Sonia Goncalves, Ewan Harrison, David K. Jackson, Ian Johnston, Dominic Kwiatkowski, Cordelia Langford, John Sillitoe on behalf of the Wellcome Sanger Institute COVID-19 Surveillance Team |
| EPI_ISL_2765863, EPI_ISL_2765865, EPI_ISL_2765866, EPI_ISL_2765867, EPI_ISL_2765868, EPI_ISL_2765869, EPI_ISL_2765870 | Lighthouse Lab in Alderley Park | Wellcome Sanger Institute for the COVID-19 Genomics UK (COG-UK) Consortium | Jacquelyn Wynn, Mairead Hyland, The Lighthouse Lab in Alderley Park and Alex Alderton, Roberto Amato, Jeffrey Barrett, Sonia Goncalves, Ewan Harrison, David K. Jackson, Ian Johnston, Dominic Kwiatkowski, Cordelia Langford, John Sillitoe on behalf of the Wellcome Sanger Institute COVID-19 Surveillance Team |
| EPI_ISL_2765871 | Health Services Laboratories | Wellcome Sanger Institute for the COVID-19 Genomics UK (COG-UK) Consortium | Health Services Laboratories and Alex Alderton, Roberto Amato, Jeffrey Barrett, Sonia Goncalves, Ewan Harrison, David K. Jackson, Ian Johnston, Dominic Kwiatkowski, Cordelia Langford, John Sillitoe on behalf of the Wellcome Sanger Institute COVID-19 Surveillance Team |
| EPI_ISL_2765872, EPI_ISL_2765873, EPI_ISL_2765874, EPI_ISL_2765875, EPI_ISL_2765876, EPI_ISL_2765877, EPI_ISL_2765878 | Lighthouse Lab in Alderley Park | Wellcome Sanger Institute for the COVID-19 Genomics UK (COG-UK) Consortium | Jacquelyn Wynn, Mairead Hyland, The Lighthouse Lab in Alderley Park and Alex Alderton, Roberto Amato, Jeffrey Barrett, Sonia Goncalves, Ewan Harrison, David K. Jackson, Ian Johnston, Dominic Kwiatkowski, Cordelia Langford, John Sillitoe on behalf of the Wellcome Sanger Institute COVID-19 Surveillance Team |
| EPI_ISL_2765879 | Health Services Laboratories | Wellcome Sanger Institute for the COVID-19 Genomics UK (COG-UK) Consortium | Health Services Laboratories and Alex Alderton, Roberto Amato, Jeffrey Barrett, Sonia Goncalves, Ewan Harrison, David K. Jackson, Ian Johnston, Dominic Kwiatkowski, Cordelia Langford, John Sillitoe on behalf of the Wellcome Sanger Institute COVID-19 Surveillance Team |
| EPI_ISL_2765880, EPI_ISL_2765881 | Lighthouse Lab in Alderley Park | Wellcome Sanger Institute for the COVID-19 Genomics UK (COG-UK) Consortium | Jacquelyn Wynn, Mairead Hyland, The Lighthouse Lab in Alderley Park and Alex Alderton, Roberto Amato, Jeffrey Barrett, Sonia Goncalves, Ewan Harrison, David K. Jackson, Ian Johnston, Dominic Kwiatkowski, Cordelia Langford, John Sillitoe on behalf of the Wellcome Sanger Institute COVID-19 Surveillance Team |
| EPI_ISL_2765882 | Health Services Laboratories | Wellcome Sanger Institute for the COVID-19 Genomics UK (COG-UK) Consortium | Health Services Laboratories and Alex Alderton, Roberto Amato, Jeffrey Barrett, Sonia Goncalves, Ewan Harrison, David K. Jackson, Ian Johnston, Dominic Kwiatkowski, Cordelia Langford, John Sillitoe on behalf of the Wellcome Sanger Institute COVID-19 Surveillance Team |
| EPI_ISL_2765883, EPI_ISL_2765884, EPI_ISL_2765885, EPI_ISL_2765886, EPI_ISL_2765887, EPI_ISL_2765888, EPI_ISL_2765889, EPI_ISL_2765890, EPI_ISL_2765891, EPI_ISL_2765892, EPI_ISL_2765893, EPI_ISL_2765894, EPI_ISL_2765895, EPI_ISL_2765896, EPI_ISL_2765897, EPI_ISL_2765898 | | | |
| see above | Lighthouse Lab in Alderley Park | Wellcome Sanger Institute for the COVID-19 Genomics UK (COG-UK) Consortium | Jacquelyn Wynn, Mairead Hyland, The Lighthouse Lab in Alderley Park and Alex Alderton, Roberto Amato, Jeffrey Barrett, Sonia Goncalves, Ewan Harrison, David K. Jackson, Ian Johnston, Dominic Kwiatkowski, Cordelia Langford, John Sillitoe on behalf of the Wellcome Sanger Institute COVID-19 Surveillance Team |

| | | | |
|---|---|---|---|
| EPI_ISL_2765899 | Health Services Laboratories | Wellcome Sanger Institute for the COVID-19 Genomics UK (COG-UK) Consortium | Health Services Laboratories and Alex Alderton, Roberto Amato, Jeffrey Barrett, Sonia Goncalves, Ewan Harrison, David K. Jackson, Ian Johnston, Dominic Kwiatkowski, Cordelia Langford, John Sillitoe on behalf of the Wellcome Sanger Institute COVID-19 Surveillance Team |
| EPI_ISL_2765900, EPI_ISL_2765901, EPI_ISL_2765902, EPI_ISL_2765903 | Lighthouse Lab in Alderley Park | Wellcome Sanger Institute for the COVID-19 Genomics UK (COG-UK) Consortium | Jacquelyn Wynn, Mairead Hyland, The Lighthouse Lab in Alderley Park and Alex Alderton, Roberto Amato, Jeffrey Barrett, Sonia Goncalves, Ewan Harrison, David K. Jackson, Ian Johnston, Dominic Kwiatkowski, Cordelia Langford, John Sillitoe on behalf of the Wellcome Sanger Institute COVID-19 Surveillance Team |
| EPI_ISL_2765904 | Health Services Laboratories | Wellcome Sanger Institute for the COVID-19 Genomics UK (COG-UK) Consortium | Health Services Laboratories and Alex Alderton, Roberto Amato, Jeffrey Barrett, Sonia Goncalves, Ewan Harrison, David K. Jackson, Ian Johnston, Dominic Kwiatkowski, Cordelia Langford, John Sillitoe on behalf of the Wellcome Sanger Institute COVID-19 Surveillance Team |
| EPI_ISL_2765905, EPI_ISL_2765906, EPI_ISL_2765907 | Lighthouse Lab in Alderley Park | Wellcome Sanger Institute for the COVID-19 Genomics UK (COG-UK) Consortium | Jacquelyn Wynn, Mairead Hyland, The Lighthouse Lab in Alderley Park and Alex Alderton, Roberto Amato, Jeffrey Barrett, Sonia Goncalves, Ewan Harrison, David K. Jackson, Ian Johnston, Dominic Kwiatkowski, Cordelia Langford, John Sillitoe on behalf of the Wellcome Sanger Institute COVID-19 Surveillance Team |
| EPI_ISL_2765908 | Health Services Laboratories | Wellcome Sanger Institute for the COVID-19 Genomics UK (COG-UK) Consortium | Health Services Laboratories and Alex Alderton, Roberto Amato, Jeffrey Barrett, Sonia Goncalves, Ewan Harrison, David K. Jackson, Ian Johnston, Dominic Kwiatkowski, Cordelia Langford, John Sillitoe on behalf of the Wellcome Sanger Institute COVID-19 Surveillance Team |
| EPI_ISL_2765909, EPI_ISL_2765910, EPI_ISL_2765911, EPI_ISL_2765912, EPI_ISL_2765913, EPI_ISL_2765914, EPI_ISL_2765915 | Lighthouse Lab in Alderley Park | Wellcome Sanger Institute for the COVID-19 Genomics UK (COG-UK) Consortium | Jacquelyn Wynn, Mairead Hyland, The Lighthouse Lab in Alderley Park and Alex Alderton, Roberto Amato, Jeffrey Barrett, Sonia Goncalves, Ewan Harrison, David K. Jackson, Ian Johnston, Dominic Kwiatkowski, Cordelia Langford, John Sillitoe on behalf of the Wellcome Sanger Institute COVID-19 Surveillance Team |
| EPI_ISL_2765916 | Health Services Laboratories | Wellcome Sanger Institute for the COVID-19 Genomics UK (COG-UK) Consortium | Health Services Laboratories and Alex Alderton, Roberto Amato, Jeffrey Barrett, Sonia Goncalves, Ewan Harrison, David K. Jackson, Ian Johnston, Dominic Kwiatkowski, Cordelia Langford, John Sillitoe on behalf of the Wellcome Sanger Institute COVID-19 Surveillance Team |
| EPI_ISL_2765917, EPI_ISL_2765918, EPI_ISL_2765919, EPI_ISL_2765920, EPI_ISL_2765921, EPI_ISL_2765922, EPI_ISL_2765923 | Lighthouse Lab in Alderley Park | Wellcome Sanger Institute for the COVID-19 Genomics UK (COG-UK) Consortium | Jacquelyn Wynn, Mairead Hyland, The Lighthouse Lab in Alderley Park and Alex Alderton, Roberto Amato, Jeffrey Barrett, Sonia Goncalves, Ewan Harrison, David K. Jackson, Ian Johnston, Dominic Kwiatkowski, Cordelia Langford, John Sillitoe on behalf of the Wellcome Sanger Institute COVID-19 Surveillance Team |
| EPI_ISL_2765924, EPI_ISL_2765925, EPI_ISL_2765926, EPI_ISL_2765927 | Health Services Laboratories | Wellcome Sanger Institute for the COVID-19 Genomics UK (COG-UK) Consortium | Health Services Laboratories and Alex Alderton, Roberto Amato, Jeffrey Barrett, Sonia Goncalves, Ewan Harrison, David K. Jackson, Ian Johnston, Dominic Kwiatkowski, Cordelia Langford, John Sillitoe on behalf of the Wellcome Sanger Institute COVID-19 Surveillance Team |
| EPI_ISL_2765928 | Lighthouse Lab in Alderley Park | Wellcome Sanger Institute for the COVID-19 Genomics UK (COG-UK) Consortium | Jacquelyn Wynn, Mairead Hyland, The Lighthouse Lab in Alderley Park and Alex Alderton, Roberto Amato, Jeffrey Barrett, Sonia Goncalves, Ewan Harrison, David K. Jackson, Ian Johnston, Dominic Kwiatkowski, Cordelia Langford, John Sillitoe on behalf of the Wellcome Sanger Institute COVID-19 Surveillance Team |
| EPI_ISL_2765929 | Health Services Laboratories | Wellcome Sanger Institute for the COVID-19 Genomics UK (COG-UK) Consortium | Health Services Laboratories and Alex Alderton, Roberto Amato, Jeffrey Barrett, Sonia Goncalves, Ewan Harrison, David K. Jackson, Ian Johnston, Dominic Kwiatkowski, Cordelia Langford, John Sillitoe on behalf of the Wellcome Sanger Institute COVID-19 Surveillance Team |
| EPI_ISL_2765930, EPI_ISL_2765931 | Lighthouse Lab in Alderley Park | Wellcome Sanger Institute for the COVID-19 Genomics UK (COG-UK) Consortium | Jacquelyn Wynn, Mairead Hyland, The Lighthouse Lab in Alderley Park and Alex Alderton, Roberto Amato, Jeffrey Barrett, Sonia Goncalves, Ewan Harrison, David K. Jackson, Ian Johnston, Dominic Kwiatkowski, Cordelia Langford, John Sillitoe on behalf of the Wellcome Sanger Institute COVID-19 Surveillance Team |
| EPI_ISL_2765932 | Health Services Laboratories | Wellcome Sanger Institute for the COVID-19 Genomics UK (COG-UK) Consortium | Health Services Laboratories and Alex Alderton, Roberto Amato, Jeffrey Barrett, Sonia Goncalves, Ewan Harrison, David K. Jackson, Ian Johnston, Dominic Kwiatkowski, Cordelia Langford, John Sillitoe on behalf of the Wellcome Sanger Institute COVID-19 Surveillance Team |
| EPI_ISL_2765933 | Lighthouse Lab in Alderley Park | Wellcome Sanger Institute for the COVID-19 Genomics UK (COG-UK) Consortium | Jacquelyn Wynn, Mairead Hyland, The Lighthouse Lab in Alderley Park and Alex Alderton, Roberto Amato, Jeffrey Barrett, Sonia Goncalves, Ewan Harrison, David K. Jackson, Ian Johnston, Dominic Kwiatkowski, Cordelia Langford, John Sillitoe on behalf of the Wellcome Sanger Institute COVID-19 Surveillance Team |
| EPI_ISL_2765934, EPI_ISL_2765935 | Health Services Laboratories | Wellcome Sanger Institute for the COVID-19 Genomics UK (COG-UK) Consortium | Health Services Laboratories and Alex Alderton, Roberto Amato, Jeffrey Barrett, Sonia Goncalves, Ewan Harrison, David K. Jackson, Ian Johnston, Dominic Kwiatkowski, Cordelia Langford, John Sillitoe on behalf of the Wellcome Sanger Institute COVID-19 Surveillance Team |
| EPI_ISL_2765936 | Lighthouse Lab in Alderley Park | Wellcome Sanger Institute for the COVID-19 Genomics UK (COG-UK) Consortium | Jacquelyn Wynn, Mairead Hyland, The Lighthouse Lab in Alderley Park and Alex Alderton, Roberto Amato, Jeffrey Barrett, Sonia Goncalves, Ewan Harrison, David K. Jackson, Ian Johnston, Dominic Kwiatkowski, Cordelia Langford, John Sillitoe on behalf of the Wellcome Sanger Institute COVID-19 Surveillance Team |
| EPI_ISL_2765937 | Health Services Laboratories | Wellcome Sanger Institute for the COVID-19 Genomics UK (COG-UK) Consortium | Health Services Laboratories and Alex Alderton, Roberto Amato, Jeffrey Barrett, Sonia Goncalves, Ewan Harrison, David K. Jackson, Ian Johnston, Dominic Kwiatkowski, Cordelia Langford, John Sillitoe on behalf of the Wellcome Sanger Institute COVID-19 Surveillance Team |
| EPI_ISL_2765938, EPI_ISL_2765939, EPI_ISL_2765940, EPI_ISL_2765941, EPI_ISL_2765942, EPI_ISL_2765943, EPI_ISL_2765944 | Lighthouse Lab in Alderley Park | Wellcome Sanger Institute for the COVID-19 Genomics UK (COG-UK) Consortium | Jacquelyn Wynn, Mairead Hyland, The Lighthouse Lab in Alderley Park and Alex Alderton, Roberto Amato, Jeffrey Barrett, Sonia Goncalves, Ewan Harrison, David K. Jackson, Ian Johnston, Dominic Kwiatkowski, Cordelia Langford, John Sillitoe on behalf of the Wellcome Sanger Institute COVID-19 Surveillance Team |
| EPI_ISL_2765945 | Health Services Laboratories | Wellcome Sanger Institute for the COVID-19 Genomics UK (COG-UK) Consortium | Health Services Laboratories and Alex Alderton, Roberto Amato, Jeffrey Barrett, Sonia Goncalves, Ewan Harrison, David K. Jackson, Ian Johnston, Dominic Kwiatkowski, Cordelia Langford, John Sillitoe on behalf of the Wellcome Sanger Institute COVID-19 Surveillance Team |
| EPI_ISL_2765946, EPI_ISL_2765947, EPI_ISL_2765948 | Lighthouse Lab in Alderley Park | Wellcome Sanger Institute for the COVID-19 Genomics UK (COG-UK) Consortium | Jacquelyn Wynn, Mairead Hyland, The Lighthouse Lab in Alderley Park and Alex Alderton, Roberto Amato, Jeffrey Barrett, Sonia Goncalves, Ewan Harrison, David K. Jackson, Ian Johnston, Dominic Kwiatkowski, Cordelia Langford, John Sillitoe on behalf of the Wellcome Sanger Institute COVID-19 Surveillance Team |
| EPI_ISL_2765949, EPI_ISL_2765950 | Health Services Laboratories | Wellcome Sanger Institute for the COVID-19 Genomics UK (COG-UK) Consortium | Health Services Laboratories and Alex Alderton, Roberto Amato, Jeffrey Barrett, Sonia Goncalves, Ewan Harrison, David K. Jackson, Ian Johnston, Dominic Kwiatkowski, Cordelia Langford, John Sillitoe on behalf of the Wellcome Sanger Institute COVID-19 Surveillance Team |
| EPI_ISL_2765951, EPI_ISL_2765952, EPI_ISL_2765953 | Lighthouse Lab in Alderley Park | Wellcome Sanger Institute for the COVID-19 Genomics UK (COG-UK) Consortium | Jacquelyn Wynn, Mairead Hyland, The Lighthouse Lab in Alderley Park and Alex Alderton, Roberto Amato, Jeffrey Barrett, Sonia Goncalves, Ewan Harrison, David K. Jackson, Ian Johnston, Dominic Kwiatkowski, Cordelia Langford, John Sillitoe on behalf of the Wellcome Sanger Institute COVID-19 Surveillance Team |
| EPI_ISL_2765954, EPI_ISL_2765955 | Health Services Laboratories | Wellcome Sanger Institute for the COVID-19 Genomics UK (COG-UK) Consortium | Health Services Laboratories and Alex Alderton, Roberto Amato, Jeffrey Barrett, Sonia Goncalves, Ewan Harrison, David K. Jackson, Ian Johnston, Dominic Kwiatkowski, Cordelia Langford, John Sillitoe on behalf of the Wellcome Sanger Institute COVID-19 Surveillance Team |
| EPI_ISL_2765956, EPI_ISL_2765957, EPI_ISL_2765958, EPI_ISL_2765959 | Lighthouse Lab in Alderley Park | Wellcome Sanger Institute for the COVID-19 Genomics UK (COG-UK) Consortium | Jacquelyn Wynn, Mairead Hyland, The Lighthouse Lab in Alderley Park and Alex Alderton, Roberto Amato, Jeffrey Barrett, Sonia Goncalves, Ewan Harrison, David K. Jackson, Ian Johnston, Dominic Kwiatkowski, Cordelia Langford, John Sillitoe on behalf of the Wellcome Sanger Institute COVID-19 Surveillance Team |
| EPI_ISL_2765960, EPI_ISL_2765961 | Health Services Laboratories | Wellcome Sanger Institute for the COVID-19 Genomics UK (COG-UK) Consortium | Health Services Laboratories and Alex Alderton, Roberto Amato, Jeffrey Barrett, Sonia Goncalves, Ewan Harrison, David K. Jackson, Ian Johnston, Dominic Kwiatkowski, Cordelia Langford, John Sillitoe on behalf of the Wellcome Sanger Institute COVID-19 Surveillance Team |
| EPI_ISL_2765962, EPI_ISL_2765963 | Lighthouse Lab in Alderley Park | Wellcome Sanger Institute for the COVID-19 Genomics UK (COG-UK) Consortium | Jacquelyn Wynn, Mairead Hyland, The Lighthouse Lab in Alderley Park and Alex Alderton, Roberto Amato, Jeffrey Barrett, Sonia Goncalves, Ewan Harrison, David K. Jackson, Ian Johnston, Dominic Kwiatkowski, Cordelia Langford, John Sillitoe on behalf of the Wellcome Sanger Institute COVID-19 Surveillance Team |
| EPI_ISL_2765964, EPI_ISL_2765965, EPI_ISL_2765966, EPI_ISL_2765967 | Health Services Laboratories | Wellcome Sanger Institute for the COVID-19 Genomics UK (COG-UK) Consortium | Health Services Laboratories and Alex Alderton, Roberto Amato, Jeffrey Barrett, Sonia Goncalves, Ewan Harrison, David K. Jackson, Ian Johnston, Dominic Kwiatkowski, Cordelia Langford, John Sillitoe on behalf of the Wellcome Sanger Institute COVID-19 Surveillance Team |
| EPI_ISL_2765968 | Lighthouse Lab in Alderley Park | Wellcome Sanger Institute for the COVID-19 Genomics UK (COG-UK) Consortium | Jacquelyn Wynn, Mairead Hyland, The Lighthouse Lab in Alderley Park and Alex Alderton, Roberto Amato, Jeffrey Barrett, Sonia Goncalves, Ewan Harrison, David K. Jackson, Ian Johnston, Dominic Kwiatkowski, Cordelia Langford, John Sillitoe on behalf of the Wellcome Sanger Institute COVID-19 Surveillance Team |
| EPI_ISL_2765969 | Health Services Laboratories | Wellcome Sanger Institute for the COVID-19 Genomics UK (COG-UK) Consortium | Health Services Laboratories and Alex Alderton, Roberto Amato, Jeffrey Barrett, Sonia Goncalves, Ewan Harrison, David K. Jackson, Ian Johnston, Dominic Kwiatkowski, Cordelia Langford, John Sillitoe on behalf of the Wellcome Sanger Institute COVID-19 Surveillance Team |
| EPI_ISL_2765970 | Lighthouse Lab in Alderley Park | Wellcome Sanger Institute for the COVID-19 Genomics UK (COG-UK) Consortium | Jacquelyn Wynn, Mairead Hyland, The Lighthouse Lab in Alderley Park and Alex Alderton, Roberto Amato, Jeffrey Barrett, Sonia Goncalves, Ewan Harrison, David K. Jackson, Ian Johnston, Dominic Kwiatkowski, Cordelia Langford, John Sillitoe on behalf of the Wellcome Sanger Institute COVID-19 Surveillance Team |
| EPI_ISL_2765971 | Health Services Laboratories | Wellcome Sanger Institute for the COVID-19 Genomics UK (COG-UK) Consortium | Health Services Laboratories and Alex Alderton, Roberto Amato, Jeffrey Barrett, Sonia Goncalves, Ewan Harrison, David K. Jackson, Ian Johnston, Dominic Kwiatkowski, Cordelia Langford, John Sillitoe on behalf of the Wellcome Sanger Institute COVID-19 Surveillance Team |
| EPI_ISL_2765972, EPI_ISL_2765973, EPI_ISL_2765974, EPI_ISL_2765975, EPI_ISL_2765976 | Lighthouse Lab in Alderley Park | Wellcome Sanger Institute for the COVID-19 Genomics UK (COG-UK) Consortium | Jacquelyn Wynn, Mairead Hyland, The Lighthouse Lab in Alderley Park and Alex Alderton, Roberto Amato, Jeffrey Barrett, Sonia Goncalves, Ewan Harrison, David K. Jackson, Ian Johnston, Dominic Kwiatkowski, Cordelia Langford, John Sillitoe on behalf of the Wellcome Sanger Institute COVID-19 Surveillance Team |
| EPI_ISL_2765977 | Health Services Laboratories | Wellcome Sanger Institute for the COVID-19 Genomics UK (COG-UK) Consortium | Health Services Laboratories and Alex Alderton, Roberto Amato, Jeffrey Barrett, Sonia Goncalves, Ewan Harrison, David K. Jackson, Ian Johnston, Dominic Kwiatkowski, Cordelia Langford, John Sillitoe on behalf of the Wellcome Sanger Institute COVID-19 Surveillance Team |

| | | | |
|---|---|---|---|
| EPI_ISL_2765978, EPI_ISL_2765979, EPI_ISL_2765980, EPI_ISL_2765981, EPI_ISL_2765982 | Lighthouse Lab in Alderley Park | Wellcome Sanger Institute for the COVID-19 Genomics UK (COG-UK) Consortium | Jacquelyn Wynn, Mairead Hyland, The Lighthouse Lab in Alderley Park and Alex Alderton, Roberto Amato, Jeffrey Barrett, Sonia Goncalves, Ewan Harrison, David K. Jackson, Ian Johnston, Dominic Kwiatkowski, Cordelia Langford, John Sillitoe on behalf of the Wellcome Sanger Institute COVID-19 Surveillance Team |
| EPI_ISL_2765983, EPI_ISL_2765984 | Health Services Laboratories | Wellcome Sanger Institute for the COVID-19 Genomics UK (COG-UK) Consortium | Health Services Laboratories and Alex Alderton, Roberto Amato, Jeffrey Barrett, Sonia Goncalves, Ewan Harrison, David K. Jackson, Ian Johnston, Dominic Kwiatkowski, Cordelia Langford, John Sillitoe on behalf of the Wellcome Sanger Institute COVID-19 Surveillance Team |
| EPI_ISL_2765985, EPI_ISL_2765986 | Lighthouse Lab in Alderley Park | Wellcome Sanger Institute for the COVID-19 Genomics UK (COG-UK) Consortium | Jacquelyn Wynn, Mairead Hyland, The Lighthouse Lab in Alderley Park and Alex Alderton, Roberto Amato, Jeffrey Barrett, Sonia Goncalves, Ewan Harrison, David K. Jackson, Ian Johnston, Dominic Kwiatkowski, Cordelia Langford, John Sillitoe on behalf of the Wellcome Sanger Institute COVID-19 Surveillance Team |
| EPI_ISL_2765987 | Health Services Laboratories | Wellcome Sanger Institute for the COVID-19 Genomics UK (COG-UK) Consortium | Health Services Laboratories and Alex Alderton, Roberto Amato, Jeffrey Barrett, Sonia Goncalves, Ewan Harrison, David K. Jackson, Ian Johnston, Dominic Kwiatkowski, Cordelia Langford, John Sillitoe on behalf of the Wellcome Sanger Institute COVID-19 Surveillance Team |
| EPI_ISL_2765988 | Lighthouse Lab in Alderley Park | Wellcome Sanger Institute for the COVID-19 Genomics UK (COG-UK) Consortium | Jacquelyn Wynn, Mairead Hyland, The Lighthouse Lab in Alderley Park and Alex Alderton, Roberto Amato, Jeffrey Barrett, Sonia Goncalves, Ewan Harrison, David K. Jackson, Ian Johnston, Dominic Kwiatkowski, Cordelia Langford, John Sillitoe on behalf of the Wellcome Sanger Institute COVID-19 Surveillance Team |
| EPI_ISL_2765989 | Health Services Laboratories | Wellcome Sanger Institute for the COVID-19 Genomics UK (COG-UK) Consortium | Health Services Laboratories and Alex Alderton, Roberto Amato, Jeffrey Barrett, Sonia Goncalves, Ewan Harrison, David K. Jackson, Ian Johnston, Dominic Kwiatkowski, Cordelia Langford, John Sillitoe on behalf of the Wellcome Sanger Institute COVID-19 Surveillance Team |
| EPI_ISL_2765990 | Lighthouse Lab in Alderley Park | Wellcome Sanger Institute for the COVID-19 Genomics UK (COG-UK) Consortium | Jacquelyn Wynn, Mairead Hyland, The Lighthouse Lab in Alderley Park and Alex Alderton, Roberto Amato, Jeffrey Barrett, Sonia Goncalves, Ewan Harrison, David K. Jackson, Ian Johnston, Dominic Kwiatkowski, Cordelia Langford, John Sillitoe on behalf of the Wellcome Sanger Institute COVID-19 Surveillance Team |
| EPI_ISL_2765991, EPI_ISL_2765992 | Health Services Laboratories | Wellcome Sanger Institute for the COVID-19 Genomics UK (COG-UK) Consortium | Health Services Laboratories and Alex Alderton, Roberto Amato, Jeffrey Barrett, Sonia Goncalves, Ewan Harrison, David K. Jackson, Ian Johnston, Dominic Kwiatkowski, Cordelia Langford, John Sillitoe on behalf of the Wellcome Sanger Institute COVID-19 Surveillance Team |
| EPI_ISL_2765993 | Lighthouse Lab in Alderley Park | Wellcome Sanger Institute for the COVID-19 Genomics UK (COG-UK) Consortium | Jacquelyn Wynn, Mairead Hyland, The Lighthouse Lab in Alderley Park and Alex Alderton, Roberto Amato, Jeffrey Barrett, Sonia Goncalves, Ewan Harrison, David K. Jackson, Ian Johnston, Dominic Kwiatkowski, Cordelia Langford, John Sillitoe on behalf of the Wellcome Sanger Institute COVID-19 Surveillance Team |
| EPI_ISL_2765994, EPI_ISL_2765995 | Health Services Laboratories | Wellcome Sanger Institute for the COVID-19 Genomics UK (COG-UK) Consortium | Health Services Laboratories and Alex Alderton, Roberto Amato, Jeffrey Barrett, Sonia Goncalves, Ewan Harrison, David K. Jackson, Ian Johnston, Dominic Kwiatkowski, Cordelia Langford, John Sillitoe on behalf of the Wellcome Sanger Institute COVID-19 Surveillance Team |
| EPI_ISL_2765996 | Lighthouse Lab in Alderley Park | Wellcome Sanger Institute for the COVID-19 Genomics UK (COG-UK) Consortium | Jacquelyn Wynn, Mairead Hyland, The Lighthouse Lab in Alderley Park and Alex Alderton, Roberto Amato, Jeffrey Barrett, Sonia Goncalves, Ewan Harrison, David K. Jackson, Ian Johnston, Dominic Kwiatkowski, Cordelia Langford, John Sillitoe on behalf of the Wellcome Sanger Institute COVID-19 Surveillance Team |
| EPI_ISL_2765997, EPI_ISL_2765998 | Health Services Laboratories | Wellcome Sanger Institute for the COVID-19 Genomics UK (COG-UK) Consortium | Health Services Laboratories and Alex Alderton, Roberto Amato, Jeffrey Barrett, Sonia Goncalves, Ewan Harrison, David K. Jackson, Ian Johnston, Dominic Kwiatkowski, Cordelia Langford, John Sillitoe on behalf of the Wellcome Sanger Institute COVID-19 Surveillance Team |
| EPI_ISL_2765999, EPI_ISL_2766000 | Lighthouse Lab in Alderley Park | Wellcome Sanger Institute for the COVID-19 Genomics UK (COG-UK) Consortium | Jacquelyn Wynn, Mairead Hyland, The Lighthouse Lab in Alderley Park and Alex Alderton, Roberto Amato, Jeffrey Barrett, Sonia Goncalves, Ewan Harrison, David K. Jackson, Ian Johnston, Dominic Kwiatkowski, Cordelia Langford, John Sillitoe on behalf of the Wellcome Sanger Institute COVID-19 Surveillance Team |
| EPI_ISL_2766001 | Health Services Laboratories | Wellcome Sanger Institute for the COVID-19 Genomics UK (COG-UK) Consortium | Health Services Laboratories and Alex Alderton, Roberto Amato, Jeffrey Barrett, Sonia Goncalves, Ewan Harrison, David K. Jackson, Ian Johnston, Dominic Kwiatkowski, Cordelia Langford, John Sillitoe on behalf of the Wellcome Sanger Institute COVID-19 Surveillance Team |
| EPI_ISL_2766002 | Lighthouse Lab in Alderley Park | Wellcome Sanger Institute for the COVID-19 Genomics UK (COG-UK) Consortium | Jacquelyn Wynn, Mairead Hyland, The Lighthouse Lab in Alderley Park and Alex Alderton, Roberto Amato, Jeffrey Barrett, Sonia Goncalves, Ewan Harrison, David K. Jackson, Ian Johnston, Dominic Kwiatkowski, Cordelia Langford, John Sillitoe on behalf of the Wellcome Sanger Institute COVID-19 Surveillance Team |
| EPI_ISL_2766003 | Health Services Laboratories | Wellcome Sanger Institute for the COVID-19 Genomics UK (COG-UK) Consortium | Health Services Laboratories and Alex Alderton, Roberto Amato, Jeffrey Barrett, Sonia Goncalves, Ewan Harrison, David K. Jackson, Ian Johnston, Dominic Kwiatkowski, Cordelia Langford, John Sillitoe on behalf of the Wellcome Sanger Institute COVID-19 Surveillance Team |
| EPI_ISL_2766004, EPI_ISL_2766005, EPI_ISL_2766006, EPI_ISL_2766007, EPI_ISL_2766008 | Lighthouse Lab in Alderley Park | Wellcome Sanger Institute for the COVID-19 Genomics UK (COG-UK) Consortium | Jacquelyn Wynn, Mairead Hyland, The Lighthouse Lab in Alderley Park and Alex Alderton, Roberto Amato, Jeffrey Barrett, Sonia Goncalves, Ewan Harrison, David K. Jackson, Ian Johnston, Dominic Kwiatkowski, Cordelia Langford, John Sillitoe on behalf of the Wellcome Sanger Institute COVID-19 Surveillance Team |
| EPI_ISL_2766009 | Health Services Laboratories | Wellcome Sanger Institute for the COVID-19 Genomics UK (COG-UK) Consortium | Health Services Laboratories and Alex Alderton, Roberto Amato, Jeffrey Barrett, Sonia Goncalves, Ewan Harrison, David K. Jackson, Ian Johnston, Dominic Kwiatkowski, Cordelia Langford, John Sillitoe on behalf of the Wellcome Sanger Institute COVID-19 Surveillance Team |
| EPI_ISL_2766010, EPI_ISL_2766011, EPI_ISL_2766012, EPI_ISL_2766013, EPI_ISL_2766014, EPI_ISL_2766015, EPI_ISL_2766016, EPI_ISL_2766017, EPI_ISL_2766018, EPI_ISL_2766019 | Lighthouse Lab in Alderley Park | Wellcome Sanger Institute for the COVID-19 Genomics UK (COG-UK) Consortium | Jacquelyn Wynn, Mairead Hyland, The Lighthouse Lab in Alderley Park and Alex Alderton, Roberto Amato, Jeffrey Barrett, Sonia Goncalves, Ewan Harrison, David K. Jackson, Ian Johnston, Dominic Kwiatkowski, Cordelia Langford, John Sillitoe on behalf of the Wellcome Sanger Institute COVID-19 Surveillance Team |
| EPI_ISL_2766020 | Health Services Laboratories | Wellcome Sanger Institute for the COVID-19 Genomics UK (COG-UK) Consortium | Health Services Laboratories and Alex Alderton, Roberto Amato, Jeffrey Barrett, Sonia Goncalves, Ewan Harrison, David K. Jackson, Ian Johnston, Dominic Kwiatkowski, Cordelia Langford, John Sillitoe on behalf of the Wellcome Sanger Institute COVID-19 Surveillance Team |
| EPI_ISL_2766021, EPI_ISL_2766022, EPI_ISL_2766023, EPI_ISL_2766024, EPI_ISL_2766025, EPI_ISL_2766026, EPI_ISL_2766027 | Lighthouse Lab in Alderley Park | Wellcome Sanger Institute for the COVID-19 Genomics UK (COG-UK) Consortium | Jacquelyn Wynn, Mairead Hyland, The Lighthouse Lab in Alderley Park and Alex Alderton, Roberto Amato, Jeffrey Barrett, Sonia Goncalves, Ewan Harrison, David K. Jackson, Ian Johnston, Dominic Kwiatkowski, Cordelia Langford, John Sillitoe on behalf of the Wellcome Sanger Institute COVID-19 Surveillance Team |
| EPI_ISL_2766028 | Health Services Laboratories | Wellcome Sanger Institute for the COVID-19 Genomics UK (COG-UK) Consortium | Health Services Laboratories and Alex Alderton, Roberto Amato, Jeffrey Barrett, Sonia Goncalves, Ewan Harrison, David K. Jackson, Ian Johnston, Dominic Kwiatkowski, Cordelia Langford, John Sillitoe on behalf of the Wellcome Sanger Institute COVID-19 Surveillance Team |
| EPI_ISL_2766029, EPI_ISL_2766030 | Lighthouse Lab in Alderley Park | Wellcome Sanger Institute for the COVID-19 Genomics UK (COG-UK) Consortium | Jacquelyn Wynn, Mairead Hyland, The Lighthouse Lab in Alderley Park and Alex Alderton, Roberto Amato, Jeffrey Barrett, Sonia Goncalves, Ewan Harrison, David K. Jackson, Ian Johnston, Dominic Kwiatkowski, Cordelia Langford, John Sillitoe on behalf of the Wellcome Sanger Institute COVID-19 Surveillance Team |
| EPI_ISL_2766031, EPI_ISL_2766032, EPI_ISL_2766033 | Health Services Laboratories | Wellcome Sanger Institute for the COVID-19 Genomics UK (COG-UK) Consortium | Health Services Laboratories and Alex Alderton, Roberto Amato, Jeffrey Barrett, Sonia Goncalves, Ewan Harrison, David K. Jackson, Ian Johnston, Dominic Kwiatkowski, Cordelia Langford, John Sillitoe on behalf of the Wellcome Sanger Institute COVID-19 Surveillance Team |
| EPI_ISL_2766034, EPI_ISL_2766035, EPI_ISL_2766036, EPI_ISL_2766037 | Lighthouse Lab in Alderley Park | Wellcome Sanger Institute for the COVID-19 Genomics UK (COG-UK) Consortium | Jacquelyn Wynn, Mairead Hyland, The Lighthouse Lab in Alderley Park and Alex Alderton, Roberto Amato, Jeffrey Barrett, Sonia Goncalves, Ewan Harrison, David K. Jackson, Ian Johnston, Dominic Kwiatkowski, Cordelia Langford, John Sillitoe on behalf of the Wellcome Sanger Institute COVID-19 Surveillance Team |
| EPI_ISL_2766038, EPI_ISL_2766039, EPI_ISL_2766040 | Health Services Laboratories | Wellcome Sanger Institute for the COVID-19 Genomics UK (COG-UK) Consortium | Health Services Laboratories and Alex Alderton, Roberto Amato, Jeffrey Barrett, Sonia Goncalves, Ewan Harrison, David K. Jackson, Ian Johnston, Dominic Kwiatkowski, Cordelia Langford, John Sillitoe on behalf of the Wellcome Sanger Institute COVID-19 Surveillance Team |
| EPI_ISL_2766041 | Lighthouse Lab in Alderley Park | Wellcome Sanger Institute for the COVID-19 Genomics UK (COG-UK) Consortium | Jacquelyn Wynn, Mairead Hyland, The Lighthouse Lab in Alderley Park and Alex Alderton, Roberto Amato, Jeffrey Barrett, Sonia Goncalves, Ewan Harrison, David K. Jackson, Ian Johnston, Dominic Kwiatkowski, Cordelia Langford, John Sillitoe on behalf of the Wellcome Sanger Institute COVID-19 Surveillance Team |
| EPI_ISL_2766042 | Health Services Laboratories | Wellcome Sanger Institute for the COVID-19 Genomics UK (COG-UK) Consortium | Health Services Laboratories and Alex Alderton, Roberto Amato, Jeffrey Barrett, Sonia Goncalves, Ewan Harrison, David K. Jackson, Ian Johnston, Dominic Kwiatkowski, Cordelia Langford, John Sillitoe on behalf of the Wellcome Sanger Institute COVID-19 Surveillance Team |
| EPI_ISL_2766043, EPI_ISL_2766044, EPI_ISL_2766045, EPI_ISL_2766046, EPI_ISL_2766047, EPI_ISL_2766048, EPI_ISL_2766049, EPI_ISL_2766050, EPI_ISL_2766051, EPI_ISL_2766052, EPI_ISL_2766053, EPI_ISL_2766054, EPI_ISL_2766055, EPI_ISL_2766056, EPI_ISL_2766057, EPI_ISL_2766058, EPI_ISL_2766059, EPI_ISL_2766060 | | | |
| see above | Lighthouse Lab in Alderley Park | Wellcome Sanger Institute for the COVID-19 Genomics UK (COG-UK) Consortium | Jacquelyn Wynn, Mairead Hyland, The Lighthouse Lab in Alderley Park and Alex Alderton, Roberto Amato, Jeffrey Barrett, Sonia Goncalves, Ewan Harrison, David K. Jackson, Ian Johnston, Dominic Kwiatkowski, Cordelia Langford, John Sillitoe on behalf of the Wellcome Sanger Institute COVID-19 Surveillance Team |
| EPI_ISL_2766061 | Health Services Laboratories | Wellcome Sanger Institute for the COVID-19 Genomics UK (COG-UK) Consortium | Health Services Laboratories and Alex Alderton, Roberto Amato, Jeffrey Barrett, Sonia Goncalves, Ewan Harrison, David K. Jackson, Ian Johnston, Dominic Kwiatkowski, Cordelia Langford, John Sillitoe on behalf of the Wellcome Sanger Institute COVID-19 Surveillance Team |
| EPI_ISL_2766062, EPI_ISL_2766063, EPI_ISL_2766064, EPI_ISL_2766065, EPI_ISL_2766066, EPI_ISL_2766067, EPI_ISL_2766068 | Lighthouse Lab in Alderley Park | Wellcome Sanger Institute for the COVID-19 Genomics UK (COG-UK) Consortium | Jacquelyn Wynn, Mairead Hyland, The Lighthouse Lab in Alderley Park and Alex Alderton, Roberto Amato, Jeffrey Barrett, Sonia Goncalves, Ewan Harrison, David K. Jackson, Ian Johnston, Dominic Kwiatkowski, Cordelia Langford, John Sillitoe on behalf of the Wellcome Sanger Institute COVID-19 Surveillance Team |
| EPI_ISL_2766069 | Health Services Laboratories | Wellcome Sanger Institute for the COVID-19 Genomics UK (COG-UK) Consortium | Health Services Laboratories and Alex Alderton, Roberto Amato, Jeffrey Barrett, Sonia Goncalves, Ewan Harrison, David K. Jackson, Ian Johnston, Dominic Kwiatkowski, Cordelia Langford, John Sillitoe on behalf of the Wellcome Sanger Institute COVID-19 Surveillance Team |
| EPI_ISL_2766070, EPI_ISL_2766071, EPI_ISL_2766072, | Lighthouse Lab in Alderley Park | Wellcome Sanger Institute for the COVID-19 Genomics UK (COG-UK) Consortium | Jacquelyn Wynn, Mairead Hyland, The Lighthouse Lab in Alderley Park and Alex Alderton, Roberto Amato, Jeffrey Barrett, Sonia Goncalves, Ewan Harrison, David K. Jackson, Ian Johnston, Dominic Kwiatkowski, Cordelia Langford, John Sillitoe on behalf of the Wellcome Sanger Institute COVID-19 Surveillance Team |

| | | | |
|---|---|---|---|
| EPI_ISL_2766073, EPI_ISL_2766074 | | | |
| EPI_ISL_2766075 | Health Services Laboratories | Wellcome Sanger Institute for the COVID-19 Genomics UK (COG-UK) Consortium | Health Services Laboratories and Alex Alderton, Roberto Amato, Jeffrey Barrett, Sonia Goncalves, Ewan Harrison, David K. Jackson, Ian Johnston, Dominic Kwiatkowski, Cordelia Langford, John Sillitoe on behalf of the Wellcome Sanger Institute COVID-19 Surveillance Team |
| EPI_ISL_2766076, EPI_ISL_2766078, EPI_ISL_2766079, EPI_ISL_2766080 | Lighthouse Lab in Alderley Park | Wellcome Sanger Institute for the COVID-19 Genomics UK (COG-UK) Consortium | Jacquelyn Wynn, Mairead Hyland, The Lighthouse Lab in Alderley Park and Alex Alderton, Roberto Amato, Jeffrey Barrett, Sonia Goncalves, Ewan Harrison, David K. Jackson, Ian Johnston, Dominic Kwiatkowski, Cordelia Langford, John Sillitoe on behalf of the Wellcome Sanger Institute COVID-19 Surveillance Team |
| EPI_ISL_2766081 | Health Services Laboratories | Wellcome Sanger Institute for the COVID-19 Genomics UK (COG-UK) Consortium | Health Services Laboratories and Alex Alderton, Roberto Amato, Jeffrey Barrett, Sonia Goncalves, Ewan Harrison, David K. Jackson, Ian Johnston, Dominic Kwiatkowski, Cordelia Langford, John Sillitoe on behalf of the Wellcome Sanger Institute COVID-19 Surveillance Team |
| EPI_ISL_2766082, EPI_ISL_2766083, EPI_ISL_2766084 | Lighthouse Lab in Alderley Park | Wellcome Sanger Institute for the COVID-19 Genomics UK (COG-UK) Consortium | Jacquelyn Wynn, Mairead Hyland, The Lighthouse Lab in Alderley Park and Alex Alderton, Roberto Amato, Jeffrey Barrett, Sonia Goncalves, Ewan Harrison, David K. Jackson, Ian Johnston, Dominic Kwiatkowski, Cordelia Langford, John Sillitoe on behalf of the Wellcome Sanger Institute COVID-19 Surveillance Team |
| EPI_ISL_2766085, EPI_ISL_2766086 | Health Services Laboratories | Wellcome Sanger Institute for the COVID-19 Genomics UK (COG-UK) Consortium | Health Services Laboratories and Alex Alderton, Roberto Amato, Jeffrey Barrett, Sonia Goncalves, Ewan Harrison, David K. Jackson, Ian Johnston, Dominic Kwiatkowski, Cordelia Langford, John Sillitoe on behalf of the Wellcome Sanger Institute COVID-19 Surveillance Team |
| EPI_ISL_2766087, EPI_ISL_2766088, EPI_ISL_2766089, EPI_ISL_2766090 | Lighthouse Lab in Alderley Park | Wellcome Sanger Institute for the COVID-19 Genomics UK (COG-UK) Consortium | Jacquelyn Wynn, Mairead Hyland, The Lighthouse Lab in Alderley Park and Alex Alderton, Roberto Amato, Jeffrey Barrett, Sonia Goncalves, Ewan Harrison, David K. Jackson, Ian Johnston, Dominic Kwiatkowski, Cordelia Langford, John Sillitoe on behalf of the Wellcome Sanger Institute COVID-19 Surveillance Team |
| EPI_ISL_2766091 | Health Services Laboratories | Wellcome Sanger Institute for the COVID-19 Genomics UK (COG-UK) Consortium | Health Services Laboratories and Alex Alderton, Roberto Amato, Jeffrey Barrett, Sonia Goncalves, Ewan Harrison, David K. Jackson, Ian Johnston, Dominic Kwiatkowski, Cordelia Langford, John Sillitoe on behalf of the Wellcome Sanger Institute COVID-19 Surveillance Team |
| EPI_ISL_2766092, EPI_ISL_2766093, EPI_ISL_2766094, EPI_ISL_2766095, EPI_ISL_2766096 | Lighthouse Lab in Alderley Park | Wellcome Sanger Institute for the COVID-19 Genomics UK (COG-UK) Consortium | Jacquelyn Wynn, Mairead Hyland, The Lighthouse Lab in Alderley Park and Alex Alderton, Roberto Amato, Jeffrey Barrett, Sonia Goncalves, Ewan Harrison, David K. Jackson, Ian Johnston, Dominic Kwiatkowski, Cordelia Langford, John Sillitoe on behalf of the Wellcome Sanger Institute COVID-19 Surveillance Team |
| EPI_ISL_2766097, EPI_ISL_2766098 | Health Services Laboratories | Wellcome Sanger Institute for the COVID-19 Genomics UK (COG-UK) Consortium | Health Services Laboratories and Alex Alderton, Roberto Amato, Jeffrey Barrett, Sonia Goncalves, Ewan Harrison, David K. Jackson, Ian Johnston, Dominic Kwiatkowski, Cordelia Langford, John Sillitoe on behalf of the Wellcome Sanger Institute COVID-19 Surveillance Team |
| EPI_ISL_2766099, EPI_ISL_2766100, EPI_ISL_2766101, EPI_ISL_2766102, EPI_ISL_2766103, EPI_ISL_2766104, EPI_ISL_2766105 | Lighthouse Lab in Alderley Park | Wellcome Sanger Institute for the COVID-19 Genomics UK (COG-UK) Consortium | Jacquelyn Wynn, Mairead Hyland, The Lighthouse Lab in Alderley Park and Alex Alderton, Roberto Amato, Jeffrey Barrett, Sonia Goncalves, Ewan Harrison, David K. Jackson, Ian Johnston, Dominic Kwiatkowski, Cordelia Langford, John Sillitoe on behalf of the Wellcome Sanger Institute COVID-19 Surveillance Team |
| EPI_ISL_2766106 | Health Services Laboratories | Wellcome Sanger Institute for the COVID-19 Genomics UK (COG-UK) Consortium | Health Services Laboratories and Alex Alderton, Roberto Amato, Jeffrey Barrett, Sonia Goncalves, Ewan Harrison, David K. Jackson, Ian Johnston, Dominic Kwiatkowski, Cordelia Langford, John Sillitoe on behalf of the Wellcome Sanger Institute COVID-19 Surveillance Team |
| EPI_ISL_2766107, EPI_ISL_2766108, EPI_ISL_2766109 | Lighthouse Lab in Alderley Park | Wellcome Sanger Institute for the COVID-19 Genomics UK (COG-UK) Consortium | Jacquelyn Wynn, Mairead Hyland, The Lighthouse Lab in Alderley Park and Alex Alderton, Roberto Amato, Jeffrey Barrett, Sonia Goncalves, Ewan Harrison, David K. Jackson, Ian Johnston, Dominic Kwiatkowski, Cordelia Langford, John Sillitoe on behalf of the Wellcome Sanger Institute COVID-19 Surveillance Team |
| EPI_ISL_2766110, EPI_ISL_2766111, EPI_ISL_2766112, EPI_ISL_2766113 | Health Services Laboratories | Wellcome Sanger Institute for the COVID-19 Genomics UK (COG-UK) Consortium | Health Services Laboratories and Alex Alderton, Roberto Amato, Jeffrey Barrett, Sonia Goncalves, Ewan Harrison, David K. Jackson, Ian Johnston, Dominic Kwiatkowski, Cordelia Langford, John Sillitoe on behalf of the Wellcome Sanger Institute COVID-19 Surveillance Team |
| EPI_ISL_2766114, EPI_ISL_2766115, EPI_ISL_2766116, EPI_ISL_2766117 | Lighthouse Lab in Alderley Park | Wellcome Sanger Institute for the COVID-19 Genomics UK (COG-UK) Consortium | Jacquelyn Wynn, Mairead Hyland, The Lighthouse Lab in Alderley Park and Alex Alderton, Roberto Amato, Jeffrey Barrett, Sonia Goncalves, Ewan Harrison, David K. Jackson, Ian Johnston, Dominic Kwiatkowski, Cordelia Langford, John Sillitoe on behalf of the Wellcome Sanger Institute COVID-19 Surveillance Team |
| EPI_ISL_2766118, EPI_ISL_2766119 | Health Services Laboratories | Wellcome Sanger Institute for the COVID-19 Genomics UK (COG-UK) Consortium | Health Services Laboratories and Alex Alderton, Roberto Amato, Jeffrey Barrett, Sonia Goncalves, Ewan Harrison, David K. Jackson, Ian Johnston, Dominic Kwiatkowski, Cordelia Langford, John Sillitoe on behalf of the Wellcome Sanger Institute COVID-19 Surveillance Team |
| EPI_ISL_2766120, EPI_ISL_2766121, EPI_ISL_2766122, EPI_ISL_2766123, EPI_ISL_2766124, EPI_ISL_2766125, EPI_ISL_2766126, EPI_ISL_2766127, EPI_ISL_2766128 | Lighthouse Lab in Alderley Park | Wellcome Sanger Institute for the COVID-19 Genomics UK (COG-UK) Consortium | Jacquelyn Wynn, Mairead Hyland, The Lighthouse Lab in Alderley Park and Alex Alderton, Roberto Amato, Jeffrey Barrett, Sonia Goncalves, Ewan Harrison, David K. Jackson, Ian Johnston, Dominic Kwiatkowski, Cordelia Langford, John Sillitoe on behalf of the Wellcome Sanger Institute COVID-19 Surveillance Team |
| EPI_ISL_2766129 | Health Services Laboratories | Wellcome Sanger Institute for the COVID-19 Genomics UK (COG-UK) Consortium | Health Services Laboratories and Alex Alderton, Roberto Amato, Jeffrey Barrett, Sonia Goncalves, Ewan Harrison, David K. Jackson, Ian Johnston, Dominic Kwiatkowski, Cordelia Langford, John Sillitoe on behalf of the Wellcome Sanger Institute COVID-19 Surveillance Team |
| EPI_ISL_2766130, EPI_ISL_2766131, EPI_ISL_2766132, EPI_ISL_2766133, EPI_ISL_2766134 | Lighthouse Lab in Alderley Park | Wellcome Sanger Institute for the COVID-19 Genomics UK (COG-UK) Consortium | Jacquelyn Wynn, Mairead Hyland, The Lighthouse Lab in Alderley Park and Alex Alderton, Roberto Amato, Jeffrey Barrett, Sonia Goncalves, Ewan Harrison, David K. Jackson, Ian Johnston, Dominic Kwiatkowski, Cordelia Langford, John Sillitoe on behalf of the Wellcome Sanger Institute COVID-19 Surveillance Team |
| EPI_ISL_2766135 | Health Services Laboratories | Wellcome Sanger Institute for the COVID-19 Genomics UK (COG-UK) Consortium | Health Services Laboratories and Alex Alderton, Roberto Amato, Jeffrey Barrett, Sonia Goncalves, Ewan Harrison, David K. Jackson, Ian Johnston, Dominic Kwiatkowski, Cordelia Langford, John Sillitoe on behalf of the Wellcome Sanger Institute COVID-19 Surveillance Team |
| EPI_ISL_2766136 | Lighthouse Lab in Alderley Park | Wellcome Sanger Institute for the COVID-19 Genomics UK (COG-UK) Consortium | Jacquelyn Wynn, Mairead Hyland, The Lighthouse Lab in Alderley Park and Alex Alderton, Roberto Amato, Jeffrey Barrett, Sonia Goncalves, Ewan Harrison, David K. Jackson, Ian Johnston, Dominic Kwiatkowski, Cordelia Langford, John Sillitoe on behalf of the Wellcome Sanger Institute COVID-19 Surveillance Team |
| EPI_ISL_2766137 | Health Services Laboratories | Wellcome Sanger Institute for the COVID-19 Genomics UK (COG-UK) Consortium | Health Services Laboratories and Alex Alderton, Roberto Amato, Jeffrey Barrett, Sonia Goncalves, Ewan Harrison, David K. Jackson, Ian Johnston, Dominic Kwiatkowski, Cordelia Langford, John Sillitoe on behalf of the Wellcome Sanger Institute COVID-19 Surveillance Team |
| EPI_ISL_2766138, EPI_ISL_2766139, EPI_ISL_2766140, EPI_ISL_2766141 | Lighthouse Lab in Alderley Park | Wellcome Sanger Institute for the COVID-19 Genomics UK (COG-UK) Consortium | Jacquelyn Wynn, Mairead Hyland, The Lighthouse Lab in Alderley Park and Alex Alderton, Roberto Amato, Jeffrey Barrett, Sonia Goncalves, Ewan Harrison, David K. Jackson, Ian Johnston, Dominic Kwiatkowski, Cordelia Langford, John Sillitoe on behalf of the Wellcome Sanger Institute COVID-19 Surveillance Team |
| EPI_ISL_2766142, EPI_ISL_2766143, EPI_ISL_2766144, EPI_ISL_2766145 | Health Services Laboratories | Wellcome Sanger Institute for the COVID-19 Genomics UK (COG-UK) Consortium | Health Services Laboratories and Alex Alderton, Roberto Amato, Jeffrey Barrett, Sonia Goncalves, Ewan Harrison, David K. Jackson, Ian Johnston, Dominic Kwiatkowski, Cordelia Langford, John Sillitoe on behalf of the Wellcome Sanger Institute COVID-19 Surveillance Team |
| EPI_ISL_2766146, EPI_ISL_2766147 | Lighthouse Lab in Alderley Park | Wellcome Sanger Institute for the COVID-19 Genomics UK (COG-UK) Consortium | Jacquelyn Wynn, Mairead Hyland, The Lighthouse Lab in Alderley Park and Alex Alderton, Roberto Amato, Jeffrey Barrett, Sonia Goncalves, Ewan Harrison, David K. Jackson, Ian Johnston, Dominic Kwiatkowski, Cordelia Langford, John Sillitoe on behalf of the Wellcome Sanger Institute COVID-19 Surveillance Team |
| EPI_ISL_2766148, EPI_ISL_2766149 | Health Services Laboratories | Wellcome Sanger Institute for the COVID-19 Genomics UK (COG-UK) Consortium | Health Services Laboratories and Alex Alderton, Roberto Amato, Jeffrey Barrett, Sonia Goncalves, Ewan Harrison, David K. Jackson, Ian Johnston, Dominic Kwiatkowski, Cordelia Langford, John Sillitoe on behalf of the Wellcome Sanger Institute COVID-19 Surveillance Team |
| EPI_ISL_2766150, EPI_ISL_2766151, EPI_ISL_2766152, EPI_ISL_2766153, EPI_ISL_2766154, EPI_ISL_2766155 | Lighthouse Lab in Alderley Park | Wellcome Sanger Institute for the COVID-19 Genomics UK (COG-UK) Consortium | Jacquelyn Wynn, Mairead Hyland, The Lighthouse Lab in Alderley Park and Alex Alderton, Roberto Amato, Jeffrey Barrett, Sonia Goncalves, Ewan Harrison, David K. Jackson, Ian Johnston, Dominic Kwiatkowski, Cordelia Langford, John Sillitoe on behalf of the Wellcome Sanger Institute COVID-19 Surveillance Team |
| EPI_ISL_2766156 | Health Services Laboratories | Wellcome Sanger Institute for the COVID-19 Genomics UK (COG-UK) Consortium | Health Services Laboratories and Alex Alderton, Roberto Amato, Jeffrey Barrett, Sonia Goncalves, Ewan Harrison, David K. Jackson, Ian Johnston, Dominic Kwiatkowski, Cordelia Langford, John Sillitoe on behalf of the Wellcome Sanger Institute COVID-19 Surveillance Team |
| EPI_ISL_2766157, EPI_ISL_2766158 | Lighthouse Lab in Alderley Park | Wellcome Sanger Institute for the COVID-19 Genomics UK (COG-UK) Consortium | Jacquelyn Wynn, Mairead Hyland, The Lighthouse Lab in Alderley Park and Alex Alderton, Roberto Amato, Jeffrey Barrett, Sonia Goncalves, Ewan Harrison, David K. Jackson, Ian Johnston, Dominic Kwiatkowski, Cordelia Langford, John Sillitoe on behalf of the Wellcome Sanger Institute COVID-19 Surveillance Team |
| EPI_ISL_2766159, EPI_ISL_2766160 | Health Services Laboratories | Wellcome Sanger Institute for the COVID-19 Genomics UK (COG-UK) Consortium | Health Services Laboratories and Alex Alderton, Roberto Amato, Jeffrey Barrett, Sonia Goncalves, Ewan Harrison, David K. Jackson, Ian Johnston, Dominic Kwiatkowski, Cordelia Langford, John Sillitoe on behalf of the Wellcome Sanger Institute COVID-19 Surveillance Team |
| EPI_ISL_2766161 | Lighthouse Lab in Alderley Park | Wellcome Sanger Institute for the COVID-19 Genomics UK (COG-UK) | Jacquelyn Wynn, Mairead Hyland, The Lighthouse Lab in Alderley Park and Alex Alderton, Roberto Amato, Jeffrey Barrett, Sonia Goncalves, Ewan Harrison, David K. Jackson, Ian Johnston, Dominic Kwiatkowski, Cordelia Langford, John Sillitoe on |

| | | | |
|---|---|---|---|
| | | Consortium | behalf of the Wellcome Sanger Institute COVID-19 Surveillance Team |
| EPI_ISL_2766162 | Health Services Laboratories | Wellcome Sanger Institute for the COVID-19 Genomics UK (COG-UK) Consortium | Health Services Laboratories and Alex Alderton, Roberto Amato, Jeffrey Barrett, Sonia Goncalves, Ewan Harrison, David K. Jackson, Ian Johnston, Dominic Kwiatkowski, Cordelia Langford, John Sillitoe on behalf of the Wellcome Sanger Institute COVID-19 Surveillance Team |
| EPI_ISL_2766163, EPI_ISL_2766164, EPI_ISL_2766165, EPI_ISL_2766166, EPI_ISL_2766167, EPI_ISL_2766168 | Lighthouse Lab in Alderley Park | Wellcome Sanger Institute for the COVID-19 Genomics UK (COG-UK) Consortium | Jacquelyn Wynn, Mairead Hyland, The Lighthouse Lab in Alderley Park and Alex Alderton, Roberto Amato, Jeffrey Barrett, Sonia Goncalves, Ewan Harrison, David K. Jackson, Ian Johnston, Dominic Kwiatkowski, Cordelia Langford, John Sillitoe on behalf of the Wellcome Sanger Institute COVID-19 Surveillance Team |
| EPI_ISL_2766169, EPI_ISL_2766170 | Health Services Laboratories | Wellcome Sanger Institute for the COVID-19 Genomics UK (COG-UK) Consortium | Health Services Laboratories and Alex Alderton, Roberto Amato, Jeffrey Barrett, Sonia Goncalves, Ewan Harrison, David K. Jackson, Ian Johnston, Dominic Kwiatkowski, Cordelia Langford, John Sillitoe on behalf of the Wellcome Sanger Institute COVID-19 Surveillance Team |
| EPI_ISL_2766171, EPI_ISL_2766172, EPI_ISL_2766173 | Lighthouse Lab in Alderley Park | Wellcome Sanger Institute for the COVID-19 Genomics UK (COG-UK) Consortium | Jacquelyn Wynn, Mairead Hyland, The Lighthouse Lab in Alderley Park and Alex Alderton, Roberto Amato, Jeffrey Barrett, Sonia Goncalves, Ewan Harrison, David K. Jackson, Ian Johnston, Dominic Kwiatkowski, Cordelia Langford, John Sillitoe on behalf of the Wellcome Sanger Institute COVID-19 Surveillance Team |
| EPI_ISL_2766174 | Health Services Laboratories | Wellcome Sanger Institute for the COVID-19 Genomics UK (COG-UK) Consortium | Health Services Laboratories and Alex Alderton, Roberto Amato, Jeffrey Barrett, Sonia Goncalves, Ewan Harrison, David K. Jackson, Ian Johnston, Dominic Kwiatkowski, Cordelia Langford, John Sillitoe on behalf of the Wellcome Sanger Institute COVID-19 Surveillance Team |
| EPI_ISL_2766175, EPI_ISL_2766176, EPI_ISL_2766177 | Lighthouse Lab in Alderley Park | Wellcome Sanger Institute for the COVID-19 Genomics UK (COG-UK) Consortium | Jacquelyn Wynn, Mairead Hyland, The Lighthouse Lab in Alderley Park and Alex Alderton, Roberto Amato, Jeffrey Barrett, Sonia Goncalves, Ewan Harrison, David K. Jackson, Ian Johnston, Dominic Kwiatkowski, Cordelia Langford, John Sillitoe on behalf of the Wellcome Sanger Institute COVID-19 Surveillance Team |
| EPI_ISL_2766178 | Health Services Laboratories | Wellcome Sanger Institute for the COVID-19 Genomics UK (COG-UK) Consortium | Health Services Laboratories and Alex Alderton, Roberto Amato, Jeffrey Barrett, Sonia Goncalves, Ewan Harrison, David K. Jackson, Ian Johnston, Dominic Kwiatkowski, Cordelia Langford, John Sillitoe on behalf of the Wellcome Sanger Institute COVID-19 Surveillance Team |
| EPI_ISL_2766179, EPI_ISL_2766180, EPI_ISL_2766181, EPI_ISL_2766182, EPI_ISL_2766183 | Lighthouse Lab in Alderley Park | Wellcome Sanger Institute for the COVID-19 Genomics UK (COG-UK) Consortium | Jacquelyn Wynn, Mairead Hyland, The Lighthouse Lab in Alderley Park and Alex Alderton, Roberto Amato, Jeffrey Barrett, Sonia Goncalves, Ewan Harrison, David K. Jackson, Ian Johnston, Dominic Kwiatkowski, Cordelia Langford, John Sillitoe on behalf of the Wellcome Sanger Institute COVID-19 Surveillance Team |
| EPI_ISL_2766184 | Health Services Laboratories | Wellcome Sanger Institute for the COVID-19 Genomics UK (COG-UK) Consortium | Health Services Laboratories and Alex Alderton, Roberto Amato, Jeffrey Barrett, Sonia Goncalves, Ewan Harrison, David K. Jackson, Ian Johnston, Dominic Kwiatkowski, Cordelia Langford, John Sillitoe on behalf of the Wellcome Sanger Institute COVID-19 Surveillance Team |
| EPI_ISL_2766185, EPI_ISL_2766186 | Lighthouse Lab in Alderley Park | Wellcome Sanger Institute for the COVID-19 Genomics UK (COG-UK) Consortium | Jacquelyn Wynn, Mairead Hyland, The Lighthouse Lab in Alderley Park and Alex Alderton, Roberto Amato, Jeffrey Barrett, Sonia Goncalves, Ewan Harrison, David K. Jackson, Ian Johnston, Dominic Kwiatkowski, Cordelia Langford, John Sillitoe on behalf of the Wellcome Sanger Institute COVID-19 Surveillance Team |
| EPI_ISL_2766187 | Health Services Laboratories | Wellcome Sanger Institute for the COVID-19 Genomics UK (COG-UK) Consortium | Health Services Laboratories and Alex Alderton, Roberto Amato, Jeffrey Barrett, Sonia Goncalves, Ewan Harrison, David K. Jackson, Ian Johnston, Dominic Kwiatkowski, Cordelia Langford, John Sillitoe on behalf of the Wellcome Sanger Institute COVID-19 Surveillance Team |
| EPI_ISL_2766188, EPI_ISL_2766189, EPI_ISL_2766190, EPI_ISL_2766191, EPI_ISL_2766192, EPI_ISL_2766193, EPI_ISL_2766194 | Lighthouse Lab in Alderley Park | Wellcome Sanger Institute for the COVID-19 Genomics UK (COG-UK) Consortium | Jacquelyn Wynn, Mairead Hyland, The Lighthouse Lab in Alderley Park and Alex Alderton, Roberto Amato, Jeffrey Barrett, Sonia Goncalves, Ewan Harrison, David K. Jackson, Ian Johnston, Dominic Kwiatkowski, Cordelia Langford, John Sillitoe on behalf of the Wellcome Sanger Institute COVID-19 Surveillance Team |
| EPI_ISL_2766195, EPI_ISL_2766196 | Lighthouse Laboratory Plymouth | Wellcome Sanger Institute for the COVID-19 Genomics UK (COG-UK) Consortium | Lighthouse Laboratory Plymouth and Alex Alderton, Roberto Amato, Jeffrey Barrett, Sonia Goncalves, Ewan Harrison, David K. Jackson, Ian Johnston, Dominic Kwiatkowski, Cordelia Langford, John Sillitoe on behalf of the Wellcome Sanger Institute COVID-19 Surveillance Team |
| EPI_ISL_2766197 | Health Services Laboratories | Wellcome Sanger Institute for the COVID-19 Genomics UK (COG-UK) Consortium | Health Services Laboratories and Alex Alderton, Roberto Amato, Jeffrey Barrett, Sonia Goncalves, Ewan Harrison, David K. Jackson, Ian Johnston, Dominic Kwiatkowski, Cordelia Langford, John Sillitoe on behalf of the Wellcome Sanger Institute COVID-19 Surveillance Team |
| EPI_ISL_2766198 | Lighthouse Laboratory Plymouth | Wellcome Sanger Institute for the COVID-19 Genomics UK (COG-UK) Consortium | Lighthouse Laboratory Plymouth and Alex Alderton, Roberto Amato, Jeffrey Barrett, Sonia Goncalves, Ewan Harrison, David K. Jackson, Ian Johnston, Dominic Kwiatkowski, Cordelia Langford, John Sillitoe on behalf of the Wellcome Sanger Institute COVID-19 Surveillance Team |
| EPI_ISL_2766199 | Health Services Laboratories | Wellcome Sanger Institute for the COVID-19 Genomics UK (COG-UK) Consortium | Health Services Laboratories and Alex Alderton, Roberto Amato, Jeffrey Barrett, Sonia Goncalves, Ewan Harrison, David K. Jackson, Ian Johnston, Dominic Kwiatkowski, Cordelia Langford, John Sillitoe on behalf of the Wellcome Sanger Institute COVID-19 Surveillance Team |
| EPI_ISL_2766200 | Lighthouse Laboratory Plymouth | Wellcome Sanger Institute for the COVID-19 Genomics UK (COG-UK) Consortium | Lighthouse Laboratory Plymouth and Alex Alderton, Roberto Amato, Jeffrey Barrett, Sonia Goncalves, Ewan Harrison, David K. Jackson, Ian Johnston, Dominic Kwiatkowski, Cordelia Langford, John Sillitoe on behalf of the Wellcome Sanger Institute COVID-19 Surveillance Team |
| EPI_ISL_2766201, EPI_ISL_2766202 | Health Services Laboratories | Wellcome Sanger Institute for the COVID-19 Genomics UK (COG-UK) Consortium | Health Services Laboratories and Alex Alderton, Roberto Amato, Jeffrey Barrett, Sonia Goncalves, Ewan Harrison, David K. Jackson, Ian Johnston, Dominic Kwiatkowski, Cordelia Langford, John Sillitoe on behalf of the Wellcome Sanger Institute COVID-19 Surveillance Team |
| EPI_ISL_2766203, EPI_ISL_2766204, EPI_ISL_2766205, EPI_ISL_2766206, EPI_ISL_2766207, EPI_ISL_2766208, EPI_ISL_2766209, EPI_ISL_2766210, EPI_ISL_2766211 | Lighthouse Laboratory Plymouth | Wellcome Sanger Institute for the COVID-19 Genomics UK (COG-UK) Consortium | Lighthouse Laboratory Plymouth and Alex Alderton, Roberto Amato, Jeffrey Barrett, Sonia Goncalves, Ewan Harrison, David K. Jackson, Ian Johnston, Dominic Kwiatkowski, Cordelia Langford, John Sillitoe on behalf of the Wellcome Sanger Institute COVID-19 Surveillance Team |
| EPI_ISL_2766212 | Health Services Laboratories | Wellcome Sanger Institute for the COVID-19 Genomics UK (COG-UK) Consortium | Health Services Laboratories and Alex Alderton, Roberto Amato, Jeffrey Barrett, Sonia Goncalves, Ewan Harrison, David K. Jackson, Ian Johnston, Dominic Kwiatkowski, Cordelia Langford, John Sillitoe on behalf of the Wellcome Sanger Institute COVID-19 Surveillance Team |
| EPI_ISL_2766213 | Lighthouse Laboratory Plymouth | Wellcome Sanger Institute for the COVID-19 Genomics UK (COG-UK) Consortium | Lighthouse Laboratory Plymouth and Alex Alderton, Roberto Amato, Jeffrey Barrett, Sonia Goncalves, Ewan Harrison, David K. Jackson, Ian Johnston, Dominic Kwiatkowski, Cordelia Langford, John Sillitoe on behalf of the Wellcome Sanger Institute COVID-19 Surveillance Team |
| EPI_ISL_2766214 | Health Services Laboratories | Wellcome Sanger Institute for the COVID-19 Genomics UK (COG-UK) Consortium | Health Services Laboratories and Alex Alderton, Roberto Amato, Jeffrey Barrett, Sonia Goncalves, Ewan Harrison, David K. Jackson, Ian Johnston, Dominic Kwiatkowski, Cordelia Langford, John Sillitoe on behalf of the Wellcome Sanger Institute COVID-19 Surveillance Team |
| EPI_ISL_2766215, EPI_ISL_2766216, EPI_ISL_2766217, EPI_ISL_2766218, EPI_ISL_2766219 | Lighthouse Laboratory Plymouth | Wellcome Sanger Institute for the COVID-19 Genomics UK (COG-UK) Consortium | Lighthouse Laboratory Plymouth and Alex Alderton, Roberto Amato, Jeffrey Barrett, Sonia Goncalves, Ewan Harrison, David K. Jackson, Ian Johnston, Dominic Kwiatkowski, Cordelia Langford, John Sillitoe on behalf of the Wellcome Sanger Institute COVID-19 Surveillance Team |
| EPI_ISL_2766220, EPI_ISL_2766221, EPI_ISL_2766222 | Health Services Laboratories | Wellcome Sanger Institute for the COVID-19 Genomics UK (COG-UK) Consortium | Health Services Laboratories and Alex Alderton, Roberto Amato, Jeffrey Barrett, Sonia Goncalves, Ewan Harrison, David K. Jackson, Ian Johnston, Dominic Kwiatkowski, Cordelia Langford, John Sillitoe on behalf of the Wellcome Sanger Institute COVID-19 Surveillance Team |
| EPI_ISL_2766223, EPI_ISL_2766224 | Lighthouse Laboratory Plymouth | Wellcome Sanger Institute for the COVID-19 Genomics UK (COG-UK) Consortium | Lighthouse Laboratory Plymouth and Alex Alderton, Roberto Amato, Jeffrey Barrett, Sonia Goncalves, Ewan Harrison, David K. Jackson, Ian Johnston, Dominic Kwiatkowski, Cordelia Langford, John Sillitoe on behalf of the Wellcome Sanger Institute COVID-19 Surveillance Team |
| EPI_ISL_2766225 | Health Services Laboratories | Wellcome Sanger Institute for the COVID-19 Genomics UK (COG-UK) Consortium | Health Services Laboratories and Alex Alderton, Roberto Amato, Jeffrey Barrett, Sonia Goncalves, Ewan Harrison, David K. Jackson, Ian Johnston, Dominic Kwiatkowski, Cordelia Langford, John Sillitoe on behalf of the Wellcome Sanger Institute COVID-19 Surveillance Team |
| EPI_ISL_2766226, EPI_ISL_2766227 | Lighthouse Laboratory Plymouth | Wellcome Sanger Institute for the COVID-19 Genomics UK (COG-UK) Consortium | Lighthouse Laboratory Plymouth and Alex Alderton, Roberto Amato, Jeffrey Barrett, Sonia Goncalves, Ewan Harrison, David K. Jackson, Ian Johnston, Dominic Kwiatkowski, Cordelia Langford, John Sillitoe on behalf of the Wellcome Sanger Institute COVID-19 Surveillance Team |
| EPI_ISL_2766228 | Health Services Laboratories | Wellcome Sanger Institute for the COVID-19 Genomics UK (COG-UK) Consortium | Health Services Laboratories and Alex Alderton, Roberto Amato, Jeffrey Barrett, Sonia Goncalves, Ewan Harrison, David K. Jackson, Ian Johnston, Dominic Kwiatkowski, Cordelia Langford, John Sillitoe on behalf of the Wellcome Sanger Institute COVID-19 Surveillance Team |
| EPI_ISL_2766229, EPI_ISL_2766230, EPI_ISL_2766231, EPI_ISL_2766232, EPI_ISL_2766233, EPI_ISL_2766234, EPI_ISL_2766235, EPI_ISL_2766236, EPI_ISL_2766237, EPI_ISL_2766238, EPI_ISL_2766239, EPI_ISL_2766240, EPI_ISL_2766241, EPI_ISL_2766242 | | | |
| see above | Lighthouse Laboratory Plymouth | Wellcome Sanger Institute for the COVID-19 Genomics UK (COG-UK) Consortium | Lighthouse Laboratory Plymouth and Alex Alderton, Roberto Amato, Jeffrey Barrett, Sonia Goncalves, Ewan Harrison, David K. Jackson, Ian Johnston, Dominic Kwiatkowski, Cordelia Langford, John Sillitoe on behalf of the Wellcome Sanger Institute COVID-19 Surveillance Team |
| EPI_ISL_2766243, EPI_ISL_2766244 | Health Services Laboratories | Wellcome Sanger Institute for the COVID-19 Genomics UK (COG-UK) Consortium | Health Services Laboratories and Alex Alderton, Roberto Amato, Jeffrey Barrett, Sonia Goncalves, Ewan Harrison, David K. Jackson, Ian Johnston, Dominic Kwiatkowski, Cordelia Langford, John Sillitoe on behalf of the Wellcome Sanger Institute COVID-19 Surveillance Team |
| EPI_ISL_2766245, EPI_ISL_2766246, EPI_ISL_2766247, EPI_ISL_2766248, EPI_ISL_2766249 | Lighthouse Laboratory Plymouth | Wellcome Sanger Institute for the COVID-19 Genomics UK (COG-UK) Consortium | Lighthouse Laboratory Plymouth and Alex Alderton, Roberto Amato, Jeffrey Barrett, Sonia Goncalves, Ewan Harrison, David K. Jackson, Ian Johnston, Dominic Kwiatkowski, Cordelia Langford, John Sillitoe on behalf of the Wellcome Sanger Institute COVID-19 Surveillance Team |
| EPI_ISL_2766250 | Health Services Laboratories | Wellcome Sanger Institute for the COVID-19 Genomics UK (COG-UK) Consortium | Health Services Laboratories and Alex Alderton, Roberto Amato, Jeffrey Barrett, Sonia Goncalves, Ewan Harrison, David K. Jackson, Ian Johnston, Dominic Kwiatkowski, Cordelia Langford, John Sillitoe on behalf of the Wellcome Sanger Institute COVID-19 Surveillance Team |
| EPI_ISL_2766251, | Lighthouse Laboratory Plymouth | Wellcome Sanger Institute for the COVID-19 Genomics UK (COG-UK) | Lighthouse Laboratory Plymouth and Alex Alderton, Roberto Amato, Jeffrey Barrett, Sonia Goncalves, Ewan Harrison, David K. Jackson, Ian Johnston, Dominic Kwiatkowski, Cordelia Langford, John Sillitoe on behalf of the Wellcome Sanger |

| | | | |
|---|---|---|---|
| EPI_ISL_2766252, EPI_ISL_2766253, EPI_ISL_2766254 | | Consortium | Institute COVID-19 Surveillance Team |
| EPI_ISL_2766255 | Health Services Laboratories | Wellcome Sanger Institute for the COVID-19 Genomics UK (COG-UK) Consortium | Health Services Laboratories and Alex Alderton, Roberto Amato, Jeffrey Barrett, Sonia Goncalves, Ewan Harrison, David K. Jackson, Ian Johnston, Dominic Kwiatkowski, Cordelia Langford, John Sillitoe on behalf of the Wellcome Sanger Institute COVID-19 Surveillance Team |
| EPI_ISL_2766257, EPI_ISL_2766258, EPI_ISL_2766259, EPI_ISL_2766260 | Lighthouse Laboratory Plymouth | Wellcome Sanger Institute for the COVID-19 Genomics UK (COG-UK) Consortium | Lighthouse Laboratory Plymouth and Alex Alderton, Roberto Amato, Jeffrey Barrett, Sonia Goncalves, Ewan Harrison, David K. Jackson, Ian Johnston, Dominic Kwiatkowski, Cordelia Langford, John Sillitoe on behalf of the Wellcome Sanger Institute COVID-19 Surveillance Team |
| EPI_ISL_2766261 | Health Services Laboratories | Wellcome Sanger Institute for the COVID-19 Genomics UK (COG-UK) Consortium | Health Services Laboratories and Alex Alderton, Roberto Amato, Jeffrey Barrett, Sonia Goncalves, Ewan Harrison, David K. Jackson, Ian Johnston, Dominic Kwiatkowski, Cordelia Langford, John Sillitoe on behalf of the Wellcome Sanger Institute COVID-19 Surveillance Team |
| EPI_ISL_2766262, EPI_ISL_2766263, EPI_ISL_2766264 | Lighthouse Laboratory Plymouth | Wellcome Sanger Institute for the COVID-19 Genomics UK (COG-UK) Consortium | Lighthouse Laboratory Plymouth and Alex Alderton, Roberto Amato, Jeffrey Barrett, Sonia Goncalves, Ewan Harrison, David K. Jackson, Ian Johnston, Dominic Kwiatkowski, Cordelia Langford, John Sillitoe on behalf of the Wellcome Sanger Institute COVID-19 Surveillance Team |
| EPI_ISL_2766265 | Health Services Laboratories | Wellcome Sanger Institute for the COVID-19 Genomics UK (COG-UK) Consortium | Health Services Laboratories and Alex Alderton, Roberto Amato, Jeffrey Barrett, Sonia Goncalves, Ewan Harrison, David K. Jackson, Ian Johnston, Dominic Kwiatkowski, Cordelia Langford, John Sillitoe on behalf of the Wellcome Sanger Institute COVID-19 Surveillance Team |
| EPI_ISL_2766266, EPI_ISL_2766267, EPI_ISL_2766268, EPI_ISL_2766269, EPI_ISL_2766270, EPI_ISL_2766271, EPI_ISL_2766272 | Lighthouse Laboratory Plymouth | Wellcome Sanger Institute for the COVID-19 Genomics UK (COG-UK) Consortium | Lighthouse Laboratory Plymouth and Alex Alderton, Roberto Amato, Jeffrey Barrett, Sonia Goncalves, Ewan Harrison, David K. Jackson, Ian Johnston, Dominic Kwiatkowski, Cordelia Langford, John Sillitoe on behalf of the Wellcome Sanger Institute COVID-19 Surveillance Team |
| EPI_ISL_2766273, EPI_ISL_2766274 | Health Services Laboratories | Wellcome Sanger Institute for the COVID-19 Genomics UK (COG-UK) Consortium | Health Services Laboratories and Alex Alderton, Roberto Amato, Jeffrey Barrett, Sonia Goncalves, Ewan Harrison, David K. Jackson, Ian Johnston, Dominic Kwiatkowski, Cordelia Langford, John Sillitoe on behalf of the Wellcome Sanger Institute COVID-19 Surveillance Team |
| EPI_ISL_2766275, EPI_ISL_2766276, EPI_ISL_2766277, EPI_ISL_2766278, EPI_ISL_2766279 | Lighthouse Laboratory Plymouth | Wellcome Sanger Institute for the COVID-19 Genomics UK (COG-UK) Consortium | Lighthouse Laboratory Plymouth and Alex Alderton, Roberto Amato, Jeffrey Barrett, Sonia Goncalves, Ewan Harrison, David K. Jackson, Ian Johnston, Dominic Kwiatkowski, Cordelia Langford, John Sillitoe on behalf of the Wellcome Sanger Institute COVID-19 Surveillance Team |
| EPI_ISL_2766280 | Health Services Laboratories | Wellcome Sanger Institute for the COVID-19 Genomics UK (COG-UK) Consortium | Health Services Laboratories and Alex Alderton, Roberto Amato, Jeffrey Barrett, Sonia Goncalves, Ewan Harrison, David K. Jackson, Ian Johnston, Dominic Kwiatkowski, Cordelia Langford, John Sillitoe on behalf of the Wellcome Sanger Institute COVID-19 Surveillance Team |
| EPI_ISL_2766281 | Lighthouse Laboratory Plymouth | Wellcome Sanger Institute for the COVID-19 Genomics UK (COG-UK) Consortium | Lighthouse Laboratory Plymouth and Alex Alderton, Roberto Amato, Jeffrey Barrett, Sonia Goncalves, Ewan Harrison, David K. Jackson, Ian Johnston, Dominic Kwiatkowski, Cordelia Langford, John Sillitoe on behalf of the Wellcome Sanger Institute COVID-19 Surveillance Team |
| EPI_ISL_2766282, EPI_ISL_2766283, EPI_ISL_2766284, EPI_ISL_2766285 | Health Services Laboratories | Wellcome Sanger Institute for the COVID-19 Genomics UK (COG-UK) Consortium | Health Services Laboratories and Alex Alderton, Roberto Amato, Jeffrey Barrett, Sonia Goncalves, Ewan Harrison, David K. Jackson, Ian Johnston, Dominic Kwiatkowski, Cordelia Langford, John Sillitoe on behalf of the Wellcome Sanger Institute COVID-19 Surveillance Team |
| EPI_ISL_2766286, EPI_ISL_2766287 | Lighthouse Laboratory Plymouth | Wellcome Sanger Institute for the COVID-19 Genomics UK (COG-UK) Consortium | Lighthouse Laboratory Plymouth and Alex Alderton, Roberto Amato, Jeffrey Barrett, Sonia Goncalves, Ewan Harrison, David K. Jackson, Ian Johnston, Dominic Kwiatkowski, Cordelia Langford, John Sillitoe on behalf of the Wellcome Sanger Institute COVID-19 Surveillance Team |
| EPI_ISL_2766288 | Health Services Laboratories | Wellcome Sanger Institute for the COVID-19 Genomics UK (COG-UK) Consortium | Health Services Laboratories and Alex Alderton, Roberto Amato, Jeffrey Barrett, Sonia Goncalves, Ewan Harrison, David K. Jackson, Ian Johnston, Dominic Kwiatkowski, Cordelia Langford, John Sillitoe on behalf of the Wellcome Sanger Institute COVID-19 Surveillance Team |
| EPI_ISL_2766289, EPI_ISL_2766290, EPI_ISL_2766291, EPI_ISL_2766292, EPI_ISL_2766293, EPI_ISL_2766294, EPI_ISL_2766295, EPI_ISL_2766296 | Lighthouse Laboratory Plymouth | Wellcome Sanger Institute for the COVID-19 Genomics UK (COG-UK) Consortium | Lighthouse Laboratory Plymouth and Alex Alderton, Roberto Amato, Jeffrey Barrett, Sonia Goncalves, Ewan Harrison, David K. Jackson, Ian Johnston, Dominic Kwiatkowski, Cordelia Langford, John Sillitoe on behalf of the Wellcome Sanger Institute COVID-19 Surveillance Team |
| EPI_ISL_2766297 | Health Services Laboratories | Wellcome Sanger Institute for the COVID-19 Genomics UK (COG-UK) Consortium | Health Services Laboratories and Alex Alderton, Roberto Amato, Jeffrey Barrett, Sonia Goncalves, Ewan Harrison, David K. Jackson, Ian Johnston, Dominic Kwiatkowski, Cordelia Langford, John Sillitoe on behalf of the Wellcome Sanger Institute COVID-19 Surveillance Team |
| EPI_ISL_2766298, EPI_ISL_2766299 | Lighthouse Laboratory Plymouth | Wellcome Sanger Institute for the COVID-19 Genomics UK (COG-UK) Consortium | Lighthouse Laboratory Plymouth and Alex Alderton, Roberto Amato, Jeffrey Barrett, Sonia Goncalves, Ewan Harrison, David K. Jackson, Ian Johnston, Dominic Kwiatkowski, Cordelia Langford, John Sillitoe on behalf of the Wellcome Sanger Institute COVID-19 Surveillance Team |
| EPI_ISL_2766300 | Health Services Laboratories | Wellcome Sanger Institute for the COVID-19 Genomics UK (COG-UK) Consortium | Health Services Laboratories and Alex Alderton, Roberto Amato, Jeffrey Barrett, Sonia Goncalves, Ewan Harrison, David K. Jackson, Ian Johnston, Dominic Kwiatkowski, Cordelia Langford, John Sillitoe on behalf of the Wellcome Sanger Institute COVID-19 Surveillance Team |
| EPI_ISL_2766301, EPI_ISL_2766302, EPI_ISL_2766303, EPI_ISL_2766304, EPI_ISL_2766305, EPI_ISL_2766306, EPI_ISL_2766307, EPI_ISL_2766308, EPI_ISL_2766309, EPI_ISL_2766310 | Lighthouse Laboratory Plymouth | Wellcome Sanger Institute for the COVID-19 Genomics UK (COG-UK) Consortium | Lighthouse Laboratory Plymouth and Alex Alderton, Roberto Amato, Jeffrey Barrett, Sonia Goncalves, Ewan Harrison, David K. Jackson, Ian Johnston, Dominic Kwiatkowski, Cordelia Langford, John Sillitoe on behalf of the Wellcome Sanger Institute COVID-19 Surveillance Team |
| EPI_ISL_2766311 | Health Services Laboratories | Wellcome Sanger Institute for the COVID-19 Genomics UK (COG-UK) Consortium | Health Services Laboratories and Alex Alderton, Roberto Amato, Jeffrey Barrett, Sonia Goncalves, Ewan Harrison, David K. Jackson, Ian Johnston, Dominic Kwiatkowski, Cordelia Langford, John Sillitoe on behalf of the Wellcome Sanger Institute COVID-19 Surveillance Team |
| EPI_ISL_2766312 | Lighthouse Laboratory Plymouth | Wellcome Sanger Institute for the COVID-19 Genomics UK (COG-UK) Consortium | Lighthouse Laboratory Plymouth and Alex Alderton, Roberto Amato, Jeffrey Barrett, Sonia Goncalves, Ewan Harrison, David K. Jackson, Ian Johnston, Dominic Kwiatkowski, Cordelia Langford, John Sillitoe on behalf of the Wellcome Sanger Institute COVID-19 Surveillance Team |
| EPI_ISL_2766313 | Health Services Laboratories | Wellcome Sanger Institute for the COVID-19 Genomics UK (COG-UK) Consortium | Health Services Laboratories and Alex Alderton, Roberto Amato, Jeffrey Barrett, Sonia Goncalves, Ewan Harrison, David K. Jackson, Ian Johnston, Dominic Kwiatkowski, Cordelia Langford, John Sillitoe on behalf of the Wellcome Sanger Institute COVID-19 Surveillance Team |
| EPI_ISL_2766314 | Lighthouse Laboratory Plymouth | Wellcome Sanger Institute for the COVID-19 Genomics UK (COG-UK) Consortium | Lighthouse Laboratory Plymouth and Alex Alderton, Roberto Amato, Jeffrey Barrett, Sonia Goncalves, Ewan Harrison, David K. Jackson, Ian Johnston, Dominic Kwiatkowski, Cordelia Langford, John Sillitoe on behalf of the Wellcome Sanger Institute COVID-19 Surveillance Team |
| EPI_ISL_2766315 | Health Services Laboratories | Wellcome Sanger Institute for the COVID-19 Genomics UK (COG-UK) Consortium | Health Services Laboratories and Alex Alderton, Roberto Amato, Jeffrey Barrett, Sonia Goncalves, Ewan Harrison, David K. Jackson, Ian Johnston, Dominic Kwiatkowski, Cordelia Langford, John Sillitoe on behalf of the Wellcome Sanger Institute COVID-19 Surveillance Team |
| EPI_ISL_2766316 | Lighthouse Laboratory Plymouth | Wellcome Sanger Institute for the COVID-19 Genomics UK (COG-UK) Consortium | Lighthouse Laboratory Plymouth and Alex Alderton, Roberto Amato, Jeffrey Barrett, Sonia Goncalves, Ewan Harrison, David K. Jackson, Ian Johnston, Dominic Kwiatkowski, Cordelia Langford, John Sillitoe on behalf of the Wellcome Sanger Institute COVID-19 Surveillance Team |
| EPI_ISL_2766317, EPI_ISL_2766318, EPI_ISL_2766319, EPI_ISL_2766320, EPI_ISL_2766321, EPI_ISL_2766322 | Health Services Laboratories | Wellcome Sanger Institute for the COVID-19 Genomics UK (COG-UK) Consortium | Health Services Laboratories and Alex Alderton, Roberto Amato, Jeffrey Barrett, Sonia Goncalves, Ewan Harrison, David K. Jackson, Ian Johnston, Dominic Kwiatkowski, Cordelia Langford, John Sillitoe on behalf of the Wellcome Sanger Institute COVID-19 Surveillance Team |
| EPI_ISL_2766323 | Lighthouse Laboratory Plymouth | Wellcome Sanger Institute for the COVID-19 Genomics UK (COG-UK) Consortium | Lighthouse Laboratory Plymouth and Alex Alderton, Roberto Amato, Jeffrey Barrett, Sonia Goncalves, Ewan Harrison, David K. Jackson, Ian Johnston, Dominic Kwiatkowski, Cordelia Langford, John Sillitoe on behalf of the Wellcome Sanger Institute COVID-19 Surveillance Team |
| EPI_ISL_2766324 | Health Services Laboratories | Wellcome Sanger Institute for the COVID-19 Genomics UK (COG-UK) Consortium | Health Services Laboratories and Alex Alderton, Roberto Amato, Jeffrey Barrett, Sonia Goncalves, Ewan Harrison, David K. Jackson, Ian Johnston, Dominic Kwiatkowski, Cordelia Langford, John Sillitoe on behalf of the Wellcome Sanger Institute COVID-19 Surveillance Team |
| EPI_ISL_2766325, EPI_ISL_2766326 | Lighthouse Laboratory Plymouth | Wellcome Sanger Institute for the COVID-19 Genomics UK (COG-UK) Consortium | Lighthouse Laboratory Plymouth and Alex Alderton, Roberto Amato, Jeffrey Barrett, Sonia Goncalves, Ewan Harrison, David K. Jackson, Ian Johnston, Dominic Kwiatkowski, Cordelia Langford, John Sillitoe on behalf of the Wellcome Sanger Institute COVID-19 Surveillance Team |
| EPI_ISL_2766327 | Health Services Laboratories | Wellcome Sanger Institute for the COVID-19 Genomics UK (COG-UK) Consortium | Health Services Laboratories and Alex Alderton, Roberto Amato, Jeffrey Barrett, Sonia Goncalves, Ewan Harrison, David K. Jackson, Ian Johnston, Dominic Kwiatkowski, Cordelia Langford, John Sillitoe on behalf of the Wellcome Sanger Institute COVID-19 Surveillance Team |
| EPI_ISL_2766328, EPI_ISL_2766329, EPI_ISL_2766330, EPI_ISL_2766331, EPI_ISL_2766332, EPI_ISL_2766333, EPI_ISL_2766334, EPI_ISL_2766335, EPI_ISL_2766336, EPI_ISL_2766337, EPI_ISL_2766338, EPI_ISL_2766339, EPI_ISL_2766340 | | | |
| see above | Lighthouse Laboratory Plymouth | Wellcome Sanger Institute for the COVID-19 Genomics UK (COG-UK) Consortium | Lighthouse Laboratory Plymouth and Alex Alderton, Roberto Amato, Jeffrey Barrett, Sonia Goncalves, Ewan Harrison, David K. Jackson, Ian Johnston, Dominic Kwiatkowski, Cordelia Langford, John Sillitoe on behalf of the Wellcome Sanger Institute COVID-19 Surveillance Team |
| EPI_ISL_2766341 | Health Services Laboratories | Wellcome Sanger Institute for the COVID-19 Genomics UK (COG-UK) | Health Services Laboratories and Alex Alderton, Roberto Amato, Jeffrey Barrett, Sonia Goncalves, Ewan Harrison, David K. Jackson, Ian Johnston, Dominic Kwiatkowski, Cordelia Langford, John Sillitoe on behalf of the Wellcome Sanger Institute |

| | | | |
|---|---|---|---|
| | | Consortium | COVID-19 Surveillance Team |
| EPI_ISL_2766342, EPI_ISL_2766343 | Lighthouse Laboratory Plymouth | Wellcome Sanger Institute for the COVID-19 Genomics UK (COG-UK) Consortium | Lighthouse Laboratory Plymouth and Alex Alderton, Roberto Amato, Jeffrey Barrett, Sonia Goncalves, Ewan Harrison, David K. Jackson, Ian Johnston, Dominic Kwiatkowski, Cordelia Langford, John Sillitoe on behalf of the Wellcome Sanger Institute COVID-19 Surveillance Team |
| EPI_ISL_2766344 | Health Services Laboratories | Wellcome Sanger Institute for the COVID-19 Genomics UK (COG-UK) Consortium | Health Services Laboratories and Alex Alderton, Roberto Amato, Jeffrey Barrett, Sonia Goncalves, Ewan Harrison, David K. Jackson, Ian Johnston, Dominic Kwiatkowski, Cordelia Langford, John Sillitoe on behalf of the Wellcome Sanger Institute COVID-19 Surveillance Team |
| EPI_ISL_2766345, EPI_ISL_2766346, EPI_ISL_2766347, EPI_ISL_2766348 | Lighthouse Laboratory Plymouth | Wellcome Sanger Institute for the COVID-19 Genomics UK (COG-UK) Consortium | Lighthouse Laboratory Plymouth and Alex Alderton, Roberto Amato, Jeffrey Barrett, Sonia Goncalves, Ewan Harrison, David K. Jackson, Ian Johnston, Dominic Kwiatkowski, Cordelia Langford, John Sillitoe on behalf of the Wellcome Sanger Institute COVID-19 Surveillance Team |
| EPI_ISL_2766349 | Health Services Laboratories | Wellcome Sanger Institute for the COVID-19 Genomics UK (COG-UK) Consortium | Health Services Laboratories and Alex Alderton, Roberto Amato, Jeffrey Barrett, Sonia Goncalves, Ewan Harrison, David K. Jackson, Ian Johnston, Dominic Kwiatkowski, Cordelia Langford, John Sillitoe on behalf of the Wellcome Sanger Institute COVID-19 Surveillance Team |
| EPI_ISL_2766350, EPI_ISL_2766351, EPI_ISL_2766352, EPI_ISL_2766353, EPI_ISL_2766354 | Lighthouse Laboratory Plymouth | Wellcome Sanger Institute for the COVID-19 Genomics UK (COG-UK) Consortium | Lighthouse Laboratory Plymouth and Alex Alderton, Roberto Amato, Jeffrey Barrett, Sonia Goncalves, Ewan Harrison, David K. Jackson, Ian Johnston, Dominic Kwiatkowski, Cordelia Langford, John Sillitoe on behalf of the Wellcome Sanger Institute COVID-19 Surveillance Team |
| EPI_ISL_2766355 | Health Services Laboratories | Wellcome Sanger Institute for the COVID-19 Genomics UK (COG-UK) Consortium | Health Services Laboratories and Alex Alderton, Roberto Amato, Jeffrey Barrett, Sonia Goncalves, Ewan Harrison, David K. Jackson, Ian Johnston, Dominic Kwiatkowski, Cordelia Langford, John Sillitoe on behalf of the Wellcome Sanger Institute COVID-19 Surveillance Team |
| EPI_ISL_2766356 | Lighthouse Laboratory Plymouth | Wellcome Sanger Institute for the COVID-19 Genomics UK (COG-UK) Consortium | Lighthouse Laboratory Plymouth and Alex Alderton, Roberto Amato, Jeffrey Barrett, Sonia Goncalves, Ewan Harrison, David K. Jackson, Ian Johnston, Dominic Kwiatkowski, Cordelia Langford, John Sillitoe on behalf of the Wellcome Sanger Institute COVID-19 Surveillance Team |
| EPI_ISL_2766357, EPI_ISL_2766358 | Health Services Laboratories | Wellcome Sanger Institute for the COVID-19 Genomics UK (COG-UK) Consortium | Health Services Laboratories and Alex Alderton, Roberto Amato, Jeffrey Barrett, Sonia Goncalves, Ewan Harrison, David K. Jackson, Ian Johnston, Dominic Kwiatkowski, Cordelia Langford, John Sillitoe on behalf of the Wellcome Sanger Institute COVID-19 Surveillance Team |
| EPI_ISL_2766359 | Lighthouse Laboratory Plymouth | Wellcome Sanger Institute for the COVID-19 Genomics UK (COG-UK) Consortium | Lighthouse Laboratory Plymouth and Alex Alderton, Roberto Amato, Jeffrey Barrett, Sonia Goncalves, Ewan Harrison, David K. Jackson, Ian Johnston, Dominic Kwiatkowski, Cordelia Langford, John Sillitoe on behalf of the Wellcome Sanger Institute COVID-19 Surveillance Team |
| EPI_ISL_2766360, EPI_ISL_2766361 | Health Services Laboratories | Wellcome Sanger Institute for the COVID-19 Genomics UK (COG-UK) Consortium | Health Services Laboratories and Alex Alderton, Roberto Amato, Jeffrey Barrett, Sonia Goncalves, Ewan Harrison, David K. Jackson, Ian Johnston, Dominic Kwiatkowski, Cordelia Langford, John Sillitoe on behalf of the Wellcome Sanger Institute COVID-19 Surveillance Team |
| EPI_ISL_2766362, EPI_ISL_2766363, EPI_ISL_2766364, EPI_ISL_2766365, EPI_ISL_2766366 | Lighthouse Laboratory Plymouth | Wellcome Sanger Institute for the COVID-19 Genomics UK (COG-UK) Consortium | Lighthouse Laboratory Plymouth and Alex Alderton, Roberto Amato, Jeffrey Barrett, Sonia Goncalves, Ewan Harrison, David K. Jackson, Ian Johnston, Dominic Kwiatkowski, Cordelia Langford, John Sillitoe on behalf of the Wellcome Sanger Institute COVID-19 Surveillance Team |
| EPI_ISL_2766367, EPI_ISL_2766368 | Health Services Laboratories | Wellcome Sanger Institute for the COVID-19 Genomics UK (COG-UK) Consortium | Health Services Laboratories and Alex Alderton, Roberto Amato, Jeffrey Barrett, Sonia Goncalves, Ewan Harrison, David K. Jackson, Ian Johnston, Dominic Kwiatkowski, Cordelia Langford, John Sillitoe on behalf of the Wellcome Sanger Institute COVID-19 Surveillance Team |
| EPI_ISL_2766369, EPI_ISL_2766370 | Lighthouse Laboratory Plymouth | Wellcome Sanger Institute for the COVID-19 Genomics UK (COG-UK) Consortium | Lighthouse Laboratory Plymouth and Alex Alderton, Roberto Amato, Jeffrey Barrett, Sonia Goncalves, Ewan Harrison, David K. Jackson, Ian Johnston, Dominic Kwiatkowski, Cordelia Langford, John Sillitoe on behalf of the Wellcome Sanger Institute COVID-19 Surveillance Team |
| EPI_ISL_2766371 | Health Services Laboratories | Wellcome Sanger Institute for the COVID-19 Genomics UK (COG-UK) Consortium | Health Services Laboratories and Alex Alderton, Roberto Amato, Jeffrey Barrett, Sonia Goncalves, Ewan Harrison, David K. Jackson, Ian Johnston, Dominic Kwiatkowski, Cordelia Langford, John Sillitoe on behalf of the Wellcome Sanger Institute COVID-19 Surveillance Team |
| EPI_ISL_2766372, EPI_ISL_2766373, EPI_ISL_2766374, EPI_ISL_2766375, EPI_ISL_2766376, EPI_ISL_2766377, EPI_ISL_2766378, EPI_ISL_2766379, EPI_ISL_2766380 | Lighthouse Laboratory Plymouth | Wellcome Sanger Institute for the COVID-19 Genomics UK (COG-UK) Consortium | Lighthouse Laboratory Plymouth and Alex Alderton, Roberto Amato, Jeffrey Barrett, Sonia Goncalves, Ewan Harrison, David K. Jackson, Ian Johnston, Dominic Kwiatkowski, Cordelia Langford, John Sillitoe on behalf of the Wellcome Sanger Institute COVID-19 Surveillance Team |
| EPI_ISL_2766381 | Health Services Laboratories | Wellcome Sanger Institute for the COVID-19 Genomics UK (COG-UK) Consortium | Health Services Laboratories and Alex Alderton, Roberto Amato, Jeffrey Barrett, Sonia Goncalves, Ewan Harrison, David K. Jackson, Ian Johnston, Dominic Kwiatkowski, Cordelia Langford, John Sillitoe on behalf of the Wellcome Sanger Institute COVID-19 Surveillance Team |
| EPI_ISL_2766382, EPI_ISL_2766383, EPI_ISL_2766384, EPI_ISL_2766385, EPI_ISL_2766386, EPI_ISL_2766387 | Lighthouse Laboratory Plymouth | Wellcome Sanger Institute for the COVID-19 Genomics UK (COG-UK) Consortium | Lighthouse Laboratory Plymouth and Alex Alderton, Roberto Amato, Jeffrey Barrett, Sonia Goncalves, Ewan Harrison, David K. Jackson, Ian Johnston, Dominic Kwiatkowski, Cordelia Langford, John Sillitoe on behalf of the Wellcome Sanger Institute COVID-19 Surveillance Team |
| EPI_ISL_2766388 | Health Services Laboratories | Wellcome Sanger Institute for the COVID-19 Genomics UK (COG-UK) Consortium | Health Services Laboratories and Alex Alderton, Roberto Amato, Jeffrey Barrett, Sonia Goncalves, Ewan Harrison, David K. Jackson, Ian Johnston, Dominic Kwiatkowski, Cordelia Langford, John Sillitoe on behalf of the Wellcome Sanger Institute COVID-19 Surveillance Team |
| EPI_ISL_2766389, EPI_ISL_2766390, EPI_ISL_2766391, EPI_ISL_2766392, EPI_ISL_2766393, EPI_ISL_2766394, EPI_ISL_2766395 | Lighthouse Laboratory Plymouth | Wellcome Sanger Institute for the COVID-19 Genomics UK (COG-UK) Consortium | Lighthouse Laboratory Plymouth and Alex Alderton, Roberto Amato, Jeffrey Barrett, Sonia Goncalves, Ewan Harrison, David K. Jackson, Ian Johnston, Dominic Kwiatkowski, Cordelia Langford, John Sillitoe on behalf of the Wellcome Sanger Institute COVID-19 Surveillance Team |
| EPI_ISL_2766396 | Health Services Laboratories | Wellcome Sanger Institute for the COVID-19 Genomics UK (COG-UK) Consortium | Health Services Laboratories and Alex Alderton, Roberto Amato, Jeffrey Barrett, Sonia Goncalves, Ewan Harrison, David K. Jackson, Ian Johnston, Dominic Kwiatkowski, Cordelia Langford, John Sillitoe on behalf of the Wellcome Sanger Institute COVID-19 Surveillance Team |
| EPI_ISL_2766397, EPI_ISL_2766398, EPI_ISL_2766399, EPI_ISL_2766400, EPI_ISL_2766401 | Lighthouse Laboratory Plymouth | Wellcome Sanger Institute for the COVID-19 Genomics UK (COG-UK) Consortium | Lighthouse Laboratory Plymouth and Alex Alderton, Roberto Amato, Jeffrey Barrett, Sonia Goncalves, Ewan Harrison, David K. Jackson, Ian Johnston, Dominic Kwiatkowski, Cordelia Langford, John Sillitoe on behalf of the Wellcome Sanger Institute COVID-19 Surveillance Team |
| EPI_ISL_2766402, EPI_ISL_2766403 | Health Services Laboratories | Wellcome Sanger Institute for the COVID-19 Genomics UK (COG-UK) Consortium | Health Services Laboratories and Alex Alderton, Roberto Amato, Jeffrey Barrett, Sonia Goncalves, Ewan Harrison, David K. Jackson, Ian Johnston, Dominic Kwiatkowski, Cordelia Langford, John Sillitoe on behalf of the Wellcome Sanger Institute COVID-19 Surveillance Team |
| EPI_ISL_2766404, EPI_ISL_2766405, EPI_ISL_2766406, EPI_ISL_2766407, EPI_ISL_2766408, EPI_ISL_2766409 | Lighthouse Laboratory Plymouth | Wellcome Sanger Institute for the COVID-19 Genomics UK (COG-UK) Consortium | Lighthouse Laboratory Plymouth and Alex Alderton, Roberto Amato, Jeffrey Barrett, Sonia Goncalves, Ewan Harrison, David K. Jackson, Ian Johnston, Dominic Kwiatkowski, Cordelia Langford, John Sillitoe on behalf of the Wellcome Sanger Institute COVID-19 Surveillance Team |
| EPI_ISL_2766410 | Health Services Laboratories | Wellcome Sanger Institute for the COVID-19 Genomics UK (COG-UK) Consortium | Health Services Laboratories and Alex Alderton, Roberto Amato, Jeffrey Barrett, Sonia Goncalves, Ewan Harrison, David K. Jackson, Ian Johnston, Dominic Kwiatkowski, Cordelia Langford, John Sillitoe on behalf of the Wellcome Sanger Institute COVID-19 Surveillance Team |
| EPI_ISL_2766411 | Lighthouse Laboratory Plymouth | Wellcome Sanger Institute for the COVID-19 Genomics UK (COG-UK) Consortium | Lighthouse Laboratory Plymouth and Alex Alderton, Roberto Amato, Jeffrey Barrett, Sonia Goncalves, Ewan Harrison, David K. Jackson, Ian Johnston, Dominic Kwiatkowski, Cordelia Langford, John Sillitoe on behalf of the Wellcome Sanger Institute COVID-19 Surveillance Team |
| EPI_ISL_2766412 | Health Services Laboratories | Wellcome Sanger Institute for the COVID-19 Genomics UK (COG-UK) Consortium | Health Services Laboratories and Alex Alderton, Roberto Amato, Jeffrey Barrett, Sonia Goncalves, Ewan Harrison, David K. Jackson, Ian Johnston, Dominic Kwiatkowski, Cordelia Langford, John Sillitoe on behalf of the Wellcome Sanger Institute COVID-19 Surveillance Team |
| EPI_ISL_2766413, EPI_ISL_2766414 | Lighthouse Laboratory Plymouth | Wellcome Sanger Institute for the COVID-19 Genomics UK (COG-UK) Consortium | Lighthouse Laboratory Plymouth and Alex Alderton, Roberto Amato, Jeffrey Barrett, Sonia Goncalves, Ewan Harrison, David K. Jackson, Ian Johnston, Dominic Kwiatkowski, Cordelia Langford, John Sillitoe on behalf of the Wellcome Sanger Institute COVID-19 Surveillance Team |
| EPI_ISL_2766415 | Health Services Laboratories | Wellcome Sanger Institute for the COVID-19 Genomics UK (COG-UK) Consortium | Health Services Laboratories and Alex Alderton, Roberto Amato, Jeffrey Barrett, Sonia Goncalves, Ewan Harrison, David K. Jackson, Ian Johnston, Dominic Kwiatkowski, Cordelia Langford, John Sillitoe on behalf of the Wellcome Sanger Institute COVID-19 Surveillance Team |
| EPI_ISL_2766416 | Lighthouse Laboratory Plymouth | Wellcome Sanger Institute for the COVID-19 Genomics UK (COG-UK) Consortium | Lighthouse Laboratory Plymouth and Alex Alderton, Roberto Amato, Jeffrey Barrett, Sonia Goncalves, Ewan Harrison, David K. Jackson, Ian Johnston, Dominic Kwiatkowski, Cordelia Langford, John Sillitoe on behalf of the Wellcome Sanger Institute COVID-19 Surveillance Team |
| EPI_ISL_2766417 | Health Services Laboratories | Wellcome Sanger Institute for the COVID-19 Genomics UK (COG-UK) Consortium | Health Services Laboratories and Alex Alderton, Roberto Amato, Jeffrey Barrett, Sonia Goncalves, Ewan Harrison, David K. Jackson, Ian Johnston, Dominic Kwiatkowski, Cordelia Langford, John Sillitoe on behalf of the Wellcome Sanger Institute COVID-19 Surveillance Team |
| EPI_ISL_2766418, EPI_ISL_2766419 | Lighthouse Laboratory Plymouth | Wellcome Sanger Institute for the COVID-19 Genomics UK (COG-UK) Consortium | Lighthouse Laboratory Plymouth and Alex Alderton, Roberto Amato, Jeffrey Barrett, Sonia Goncalves, Ewan Harrison, David K. Jackson, Ian Johnston, Dominic Kwiatkowski, Cordelia Langford, John Sillitoe on behalf of the Wellcome Sanger Institute COVID-19 Surveillance Team |
| EPI_ISL_2766420 | Health Services Laboratories | Wellcome Sanger Institute for the COVID-19 Genomics UK (COG-UK) | Health Services Laboratories and Alex Alderton, Roberto Amato, Jeffrey Barrett, Sonia Goncalves, Ewan Harrison, David K. Jackson, Ian Johnston, Dominic Kwiatkowski, Cordelia Langford, John Sillitoe on behalf of the Wellcome Sanger Institute |

| | | | |
|---|---|---|---|
| | | Consortium | COVID-19 Surveillance Team |
| EPI_ISL_2766421 | Lighthouse Laboratory Plymouth | Wellcome Sanger Institute for the COVID-19 Genomics UK (COG-UK) Consortium | Lighthouse Laboratory Plymouth and Alex Alderton, Roberto Amato, Jeffrey Barrett, Sonia Goncalves, Ewan Harrison, David K. Jackson, Ian Johnston, Dominic Kwiatkowski, Cordelia Langford, John Sillitoe on behalf of the Wellcome Sanger Institute COVID-19 Surveillance Team |
| EPI_ISL_2766422 | Health Services Laboratories | Wellcome Sanger Institute for the COVID-19 Genomics UK (COG-UK) Consortium | Health Services Laboratories and Alex Alderton, Roberto Amato, Jeffrey Barrett, Sonia Goncalves, Ewan Harrison, David K. Jackson, Ian Johnston, Dominic Kwiatkowski, Cordelia Langford, John Sillitoe on behalf of the Wellcome Sanger Institute COVID-19 Surveillance Team |
| EPI_ISL_2766423 | Lighthouse Laboratory Plymouth | Wellcome Sanger Institute for the COVID-19 Genomics UK (COG-UK) Consortium | Lighthouse Laboratory Plymouth and Alex Alderton, Roberto Amato, Jeffrey Barrett, Sonia Goncalves, Ewan Harrison, David K. Jackson, Ian Johnston, Dominic Kwiatkowski, Cordelia Langford, John Sillitoe on behalf of the Wellcome Sanger Institute COVID-19 Surveillance Team |
| EPI_ISL_2766424 | Health Services Laboratories | Wellcome Sanger Institute for the COVID-19 Genomics UK (COG-UK) Consortium | Health Services Laboratories and Alex Alderton, Roberto Amato, Jeffrey Barrett, Sonia Goncalves, Ewan Harrison, David K. Jackson, Ian Johnston, Dominic Kwiatkowski, Cordelia Langford, John Sillitoe on behalf of the Wellcome Sanger Institute COVID-19 Surveillance Team |
| EPI_ISL_2766425, EPI_ISL_2766426, EPI_ISL_2766427 | Lighthouse Laboratory Plymouth | Wellcome Sanger Institute for the COVID-19 Genomics UK (COG-UK) Consortium | Lighthouse Laboratory Plymouth and Alex Alderton, Roberto Amato, Jeffrey Barrett, Sonia Goncalves, Ewan Harrison, David K. Jackson, Ian Johnston, Dominic Kwiatkowski, Cordelia Langford, John Sillitoe on behalf of the Wellcome Sanger Institute COVID-19 Surveillance Team |
| EPI_ISL_2766428 | Health Services Laboratories | Wellcome Sanger Institute for the COVID-19 Genomics UK (COG-UK) Consortium | Health Services Laboratories and Alex Alderton, Roberto Amato, Jeffrey Barrett, Sonia Goncalves, Ewan Harrison, David K. Jackson, Ian Johnston, Dominic Kwiatkowski, Cordelia Langford, John Sillitoe on behalf of the Wellcome Sanger Institute COVID-19 Surveillance Team |
| EPI_ISL_2766429, EPI_ISL_2766430, EPI_ISL_2766431, EPI_ISL_2766432, EPI_ISL_2766433, EPI_ISL_2766434 | Lighthouse Laboratory Plymouth | Wellcome Sanger Institute for the COVID-19 Genomics UK (COG-UK) Consortium | Lighthouse Laboratory Plymouth and Alex Alderton, Roberto Amato, Jeffrey Barrett, Sonia Goncalves, Ewan Harrison, David K. Jackson, Ian Johnston, Dominic Kwiatkowski, Cordelia Langford, John Sillitoe on behalf of the Wellcome Sanger Institute COVID-19 Surveillance Team |
| EPI_ISL_2766435 | Health Services Laboratories | Wellcome Sanger Institute for the COVID-19 Genomics UK (COG-UK) Consortium | Health Services Laboratories and Alex Alderton, Roberto Amato, Jeffrey Barrett, Sonia Goncalves, Ewan Harrison, David K. Jackson, Ian Johnston, Dominic Kwiatkowski, Cordelia Langford, John Sillitoe on behalf of the Wellcome Sanger Institute COVID-19 Surveillance Team |
| EPI_ISL_2766436, EPI_ISL_2766437 | Lighthouse Laboratory Plymouth | Wellcome Sanger Institute for the COVID-19 Genomics UK (COG-UK) Consortium | Lighthouse Laboratory Plymouth and Alex Alderton, Roberto Amato, Jeffrey Barrett, Sonia Goncalves, Ewan Harrison, David K. Jackson, Ian Johnston, Dominic Kwiatkowski, Cordelia Langford, John Sillitoe on behalf of the Wellcome Sanger Institute COVID-19 Surveillance Team |
| EPI_ISL_2766438 | Health Services Laboratories | Wellcome Sanger Institute for the COVID-19 Genomics UK (COG-UK) Consortium | Health Services Laboratories and Alex Alderton, Roberto Amato, Jeffrey Barrett, Sonia Goncalves, Ewan Harrison, David K. Jackson, Ian Johnston, Dominic Kwiatkowski, Cordelia Langford, John Sillitoe on behalf of the Wellcome Sanger Institute COVID-19 Surveillance Team |
| EPI_ISL_2766439, EPI_ISL_2766440, EPI_ISL_2766441, EPI_ISL_2766442, EPI_ISL_2766443, EPI_ISL_2766444, EPI_ISL_2766445, EPI_ISL_2766446, EPI_ISL_2766447, EPI_ISL_2766448 | Lighthouse Laboratory Plymouth | Wellcome Sanger Institute for the COVID-19 Genomics UK (COG-UK) Consortium | Lighthouse Laboratory Plymouth and Alex Alderton, Roberto Amato, Jeffrey Barrett, Sonia Goncalves, Ewan Harrison, David K. Jackson, Ian Johnston, Dominic Kwiatkowski, Cordelia Langford, John Sillitoe on behalf of the Wellcome Sanger Institute COVID-19 Surveillance Team |
| EPI_ISL_2766449, EPI_ISL_2766450 | Health Services Laboratories | Wellcome Sanger Institute for the COVID-19 Genomics UK (COG-UK) Consortium | Health Services Laboratories and Alex Alderton, Roberto Amato, Jeffrey Barrett, Sonia Goncalves, Ewan Harrison, David K. Jackson, Ian Johnston, Dominic Kwiatkowski, Cordelia Langford, John Sillitoe on behalf of the Wellcome Sanger Institute COVID-19 Surveillance Team |
| EPI_ISL_2766451 | Lighthouse Laboratory Plymouth | Wellcome Sanger Institute for the COVID-19 Genomics UK (COG-UK) Consortium | Lighthouse Laboratory Plymouth and Alex Alderton, Roberto Amato, Jeffrey Barrett, Sonia Goncalves, Ewan Harrison, David K. Jackson, Ian Johnston, Dominic Kwiatkowski, Cordelia Langford, John Sillitoe on behalf of the Wellcome Sanger Institute COVID-19 Surveillance Team |
| EPI_ISL_2766452 | Health Services Laboratories | Wellcome Sanger Institute for the COVID-19 Genomics UK (COG-UK) Consortium | Health Services Laboratories and Alex Alderton, Roberto Amato, Jeffrey Barrett, Sonia Goncalves, Ewan Harrison, David K. Jackson, Ian Johnston, Dominic Kwiatkowski, Cordelia Langford, John Sillitoe on behalf of the Wellcome Sanger Institute COVID-19 Surveillance Team |
| EPI_ISL_2766454, EPI_ISL_2766455, EPI_ISL_2766456, EPI_ISL_2766457, EPI_ISL_2766458, EPI_ISL_2766459, EPI_ISL_2766460, EPI_ISL_2766461, EPI_ISL_2766462, EPI_ISL_2766463, EPI_ISL_2766464, EPI_ISL_2766465 | | | |
| see above | Lighthouse Laboratory Plymouth | Wellcome Sanger Institute for the COVID-19 Genomics UK (COG-UK) Consortium | Lighthouse Laboratory Plymouth and Alex Alderton, Roberto Amato, Jeffrey Barrett, Sonia Goncalves, Ewan Harrison, David K. Jackson, Ian Johnston, Dominic Kwiatkowski, Cordelia Langford, John Sillitoe on behalf of the Wellcome Sanger Institute COVID-19 Surveillance Team |
| EPI_ISL_2766466 | Health Services Laboratories | Wellcome Sanger Institute for the COVID-19 Genomics UK (COG-UK) Consortium | Health Services Laboratories and Alex Alderton, Roberto Amato, Jeffrey Barrett, Sonia Goncalves, Ewan Harrison, David K. Jackson, Ian Johnston, Dominic Kwiatkowski, Cordelia Langford, John Sillitoe on behalf of the Wellcome Sanger Institute COVID-19 Surveillance Team |
| EPI_ISL_2766467, EPI_ISL_2766468, EPI_ISL_2766469, EPI_ISL_2766470 | Lighthouse Laboratory Plymouth | Wellcome Sanger Institute for the COVID-19 Genomics UK (COG-UK) Consortium | Lighthouse Laboratory Plymouth and Alex Alderton, Roberto Amato, Jeffrey Barrett, Sonia Goncalves, Ewan Harrison, David K. Jackson, Ian Johnston, Dominic Kwiatkowski, Cordelia Langford, John Sillitoe on behalf of the Wellcome Sanger Institute COVID-19 Surveillance Team |
| EPI_ISL_2766471 | Health Services Laboratories | Wellcome Sanger Institute for the COVID-19 Genomics UK (COG-UK) Consortium | Health Services Laboratories and Alex Alderton, Roberto Amato, Jeffrey Barrett, Sonia Goncalves, Ewan Harrison, David K. Jackson, Ian Johnston, Dominic Kwiatkowski, Cordelia Langford, John Sillitoe on behalf of the Wellcome Sanger Institute COVID-19 Surveillance Team |
| EPI_ISL_2766472, EPI_ISL_2766473, EPI_ISL_2766474 | Lighthouse Laboratory Plymouth | Wellcome Sanger Institute for the COVID-19 Genomics UK (COG-UK) Consortium | Lighthouse Laboratory Plymouth and Alex Alderton, Roberto Amato, Jeffrey Barrett, Sonia Goncalves, Ewan Harrison, David K. Jackson, Ian Johnston, Dominic Kwiatkowski, Cordelia Langford, John Sillitoe on behalf of the Wellcome Sanger Institute COVID-19 Surveillance Team |
| EPI_ISL_2766475, EPI_ISL_2766476 | Health Services Laboratories | Wellcome Sanger Institute for the COVID-19 Genomics UK (COG-UK) Consortium | Health Services Laboratories and Alex Alderton, Roberto Amato, Jeffrey Barrett, Sonia Goncalves, Ewan Harrison, David K. Jackson, Ian Johnston, Dominic Kwiatkowski, Cordelia Langford, John Sillitoe on behalf of the Wellcome Sanger Institute COVID-19 Surveillance Team |
| EPI_ISL_2766477 | Lighthouse Laboratory Plymouth | Wellcome Sanger Institute for the COVID-19 Genomics UK (COG-UK) Consortium | Lighthouse Laboratory Plymouth and Alex Alderton, Roberto Amato, Jeffrey Barrett, Sonia Goncalves, Ewan Harrison, David K. Jackson, Ian Johnston, Dominic Kwiatkowski, Cordelia Langford, John Sillitoe on behalf of the Wellcome Sanger Institute COVID-19 Surveillance Team |
| EPI_ISL_2766478 | Health Services Laboratories | Wellcome Sanger Institute for the COVID-19 Genomics UK (COG-UK) Consortium | Health Services Laboratories and Alex Alderton, Roberto Amato, Jeffrey Barrett, Sonia Goncalves, Ewan Harrison, David K. Jackson, Ian Johnston, Dominic Kwiatkowski, Cordelia Langford, John Sillitoe on behalf of the Wellcome Sanger Institute COVID-19 Surveillance Team |
| EPI_ISL_2766479, EPI_ISL_2766480 | Lighthouse Laboratory Plymouth | Wellcome Sanger Institute for the COVID-19 Genomics UK (COG-UK) Consortium | Lighthouse Laboratory Plymouth and Alex Alderton, Roberto Amato, Jeffrey Barrett, Sonia Goncalves, Ewan Harrison, David K. Jackson, Ian Johnston, Dominic Kwiatkowski, Cordelia Langford, John Sillitoe on behalf of the Wellcome Sanger Institute COVID-19 Surveillance Team |
| EPI_ISL_2766481 | Health Services Laboratories | Wellcome Sanger Institute for the COVID-19 Genomics UK (COG-UK) Consortium | Health Services Laboratories and Alex Alderton, Roberto Amato, Jeffrey Barrett, Sonia Goncalves, Ewan Harrison, David K. Jackson, Ian Johnston, Dominic Kwiatkowski, Cordelia Langford, John Sillitoe on behalf of the Wellcome Sanger Institute COVID-19 Surveillance Team |
| EPI_ISL_2766482, EPI_ISL_2766483 | Lighthouse Laboratory Plymouth | Wellcome Sanger Institute for the COVID-19 Genomics UK (COG-UK) Consortium | Lighthouse Laboratory Plymouth and Alex Alderton, Roberto Amato, Jeffrey Barrett, Sonia Goncalves, Ewan Harrison, David K. Jackson, Ian Johnston, Dominic Kwiatkowski, Cordelia Langford, John Sillitoe on behalf of the Wellcome Sanger Institute COVID-19 Surveillance Team |
| EPI_ISL_2766484 | Health Services Laboratories | Wellcome Sanger Institute for the COVID-19 Genomics UK (COG-UK) Consortium | Health Services Laboratories and Alex Alderton, Roberto Amato, Jeffrey Barrett, Sonia Goncalves, Ewan Harrison, David K. Jackson, Ian Johnston, Dominic Kwiatkowski, Cordelia Langford, John Sillitoe on behalf of the Wellcome Sanger Institute COVID-19 Surveillance Team |
| EPI_ISL_2766485, EPI_ISL_2766486, EPI_ISL_2766487, EPI_ISL_2766488, EPI_ISL_2766489, EPI_ISL_2766490, EPI_ISL_2766491, EPI_ISL_2766492, EPI_ISL_2766493, EPI_ISL_2766494, EPI_ISL_2766495 | | | |
| see above | Lighthouse Laboratory Plymouth | Wellcome Sanger Institute for the COVID-19 Genomics UK (COG-UK) Consortium | Lighthouse Laboratory Plymouth and Alex Alderton, Roberto Amato, Jeffrey Barrett, Sonia Goncalves, Ewan Harrison, David K. Jackson, Ian Johnston, Dominic Kwiatkowski, Cordelia Langford, John Sillitoe on behalf of the Wellcome Sanger Institute COVID-19 Surveillance Team |
| EPI_ISL_2766496 | Health Services Laboratories | Wellcome Sanger Institute for the COVID-19 Genomics UK (COG-UK) Consortium | Health Services Laboratories and Alex Alderton, Roberto Amato, Jeffrey Barrett, Sonia Goncalves, Ewan Harrison, David K. Jackson, Ian Johnston, Dominic Kwiatkowski, Cordelia Langford, John Sillitoe on behalf of the Wellcome Sanger Institute COVID-19 Surveillance Team |
| EPI_ISL_2766497, EPI_ISL_2766498 | Lighthouse Laboratory Plymouth | Wellcome Sanger Institute for the COVID-19 Genomics UK (COG-UK) Consortium | Lighthouse Laboratory Plymouth and Alex Alderton, Roberto Amato, Jeffrey Barrett, Sonia Goncalves, Ewan Harrison, David K. Jackson, Ian Johnston, Dominic Kwiatkowski, Cordelia Langford, John Sillitoe on behalf of the Wellcome Sanger Institute COVID-19 Surveillance Team |
| EPI_ISL_2766499 | Health Services Laboratories | Wellcome Sanger Institute for the COVID-19 Genomics UK (COG-UK) Consortium | Health Services Laboratories and Alex Alderton, Roberto Amato, Jeffrey Barrett, Sonia Goncalves, Ewan Harrison, David K. Jackson, Ian Johnston, Dominic Kwiatkowski, Cordelia Langford, John Sillitoe on behalf of the Wellcome Sanger Institute COVID-19 Surveillance Team |
| EPI_ISL_2766500, EPI_ISL_2766501, EPI_ISL_2766502, EPI_ISL_2766503, EPI_ISL_2766504, EPI_ISL_2766505, EPI_ISL_2766506, EPI_ISL_2766507 | Lighthouse Laboratory Plymouth | Wellcome Sanger Institute for the COVID-19 Genomics UK (COG-UK) Consortium | Lighthouse Laboratory Plymouth and Alex Alderton, Roberto Amato, Jeffrey Barrett, Sonia Goncalves, Ewan Harrison, David K. Jackson, Ian Johnston, Dominic Kwiatkowski, Cordelia Langford, John Sillitoe on behalf of the Wellcome Sanger Institute COVID-19 Surveillance Team |
| EPI_ISL_2766508 | Health Services Laboratories | Wellcome Sanger Institute for the COVID-19 Genomics UK (COG-UK) Consortium | Health Services Laboratories and Alex Alderton, Roberto Amato, Jeffrey Barrett, Sonia Goncalves, Ewan Harrison, David K. Jackson, Ian Johnston, Dominic Kwiatkowski, Cordelia Langford, John Sillitoe on behalf of the Wellcome Sanger Institute COVID-19 Surveillance Team |
| EPI_ISL_2766509, EPI_ISL_2766510 | Lighthouse Laboratory Plymouth | Wellcome Sanger Institute for the COVID-19 Genomics UK (COG-UK) Consortium | Lighthouse Laboratory Plymouth and Alex Alderton, Roberto Amato, Jeffrey Barrett, Sonia Goncalves, Ewan Harrison, David K. Jackson, Ian Johnston, Dominic Kwiatkowski, Cordelia Langford, John Sillitoe on behalf of the Wellcome Sanger Institute COVID-19 Surveillance Team |
| EPI_ISL_2766511, EPI_ISL_2766512, EPI_ISL_2766513 | Health Services Laboratories | Wellcome Sanger Institute for the COVID-19 Genomics UK (COG-UK) Consortium | Health Services Laboratories and Alex Alderton, Roberto Amato, Jeffrey Barrett, Sonia Goncalves, Ewan Harrison, David K. Jackson, Ian Johnston, Dominic Kwiatkowski, Cordelia Langford, John Sillitoe on behalf of the Wellcome Sanger Institute COVID-19 Surveillance Team |

| | | | |
|---|---|---|---|
| EPI_ISL_2766514, EPI_ISL_2766515, EPI_ISL_2766516, EPI_ISL_2766517, EPI_ISL_2766518 | Lighthouse Laboratory Plymouth | Wellcome Sanger Institute for the COVID-19 Genomics UK (COG-UK) Consortium | Lighthouse Laboratory Plymouth and Alex Alderton, Roberto Amato, Jeffrey Barrett, Sonia Goncalves, Ewan Harrison, David K. Jackson, Ian Johnston, Dominic Kwiatkowski, Cordelia Langford, John Sillitoe on behalf of the Wellcome Sanger Institute COVID-19 Surveillance Team |
| EPI_ISL_2766519 | Health Services Laboratories | Wellcome Sanger Institute for the COVID-19 Genomics UK (COG-UK) Consortium | Health Services Laboratories and Alex Alderton, Roberto Amato, Jeffrey Barrett, Sonia Goncalves, Ewan Harrison, David K. Jackson, Ian Johnston, Dominic Kwiatkowski, Cordelia Langford, John Sillitoe on behalf of the Wellcome Sanger Institute COVID-19 Surveillance Team |
| EPI_ISL_2766520 | Lighthouse Laboratory Plymouth | Wellcome Sanger Institute for the COVID-19 Genomics UK (COG-UK) Consortium | Lighthouse Laboratory Plymouth and Alex Alderton, Roberto Amato, Jeffrey Barrett, Sonia Goncalves, Ewan Harrison, David K. Jackson, Ian Johnston, Dominic Kwiatkowski, Cordelia Langford, John Sillitoe on behalf of the Wellcome Sanger Institute COVID-19 Surveillance Team |
| EPI_ISL_2766521, EPI_ISL_2766522 | Health Services Laboratories | Wellcome Sanger Institute for the COVID-19 Genomics UK (COG-UK) Consortium | Health Services Laboratories and Alex Alderton, Roberto Amato, Jeffrey Barrett, Sonia Goncalves, Ewan Harrison, David K. Jackson, Ian Johnston, Dominic Kwiatkowski, Cordelia Langford, John Sillitoe on behalf of the Wellcome Sanger Institute COVID-19 Surveillance Team |
| EPI_ISL_2766523, EPI_ISL_2766524, EPI_ISL_2766525 | Lighthouse Laboratory Plymouth | Wellcome Sanger Institute for the COVID-19 Genomics UK (COG-UK) Consortium | Lighthouse Laboratory Plymouth and Alex Alderton, Roberto Amato, Jeffrey Barrett, Sonia Goncalves, Ewan Harrison, David K. Jackson, Ian Johnston, Dominic Kwiatkowski, Cordelia Langford, John Sillitoe on behalf of the Wellcome Sanger Institute COVID-19 Surveillance Team |
| EPI_ISL_2766526 | Health Services Laboratories | Wellcome Sanger Institute for the COVID-19 Genomics UK (COG-UK) Consortium | Health Services Laboratories and Alex Alderton, Roberto Amato, Jeffrey Barrett, Sonia Goncalves, Ewan Harrison, David K. Jackson, Ian Johnston, Dominic Kwiatkowski, Cordelia Langford, John Sillitoe on behalf of the Wellcome Sanger Institute COVID-19 Surveillance Team |
| EPI_ISL_2766527, EPI_ISL_2766528, EPI_ISL_2766529, EPI_ISL_2766530 | Lighthouse Laboratory Plymouth | Wellcome Sanger Institute for the COVID-19 Genomics UK (COG-UK) Consortium | Lighthouse Laboratory Plymouth and Alex Alderton, Roberto Amato, Jeffrey Barrett, Sonia Goncalves, Ewan Harrison, David K. Jackson, Ian Johnston, Dominic Kwiatkowski, Cordelia Langford, John Sillitoe on behalf of the Wellcome Sanger Institute COVID-19 Surveillance Team |
| EPI_ISL_2766531, EPI_ISL_2766532 | Health Services Laboratories | Wellcome Sanger Institute for the COVID-19 Genomics UK (COG-UK) Consortium | Health Services Laboratories and Alex Alderton, Roberto Amato, Jeffrey Barrett, Sonia Goncalves, Ewan Harrison, David K. Jackson, Ian Johnston, Dominic Kwiatkowski, Cordelia Langford, John Sillitoe on behalf of the Wellcome Sanger Institute COVID-19 Surveillance Team |
| EPI_ISL_2766533, EPI_ISL_2766534, EPI_ISL_2766535 | Lighthouse Laboratory Plymouth | Wellcome Sanger Institute for the COVID-19 Genomics UK (COG-UK) Consortium | Lighthouse Laboratory Plymouth and Alex Alderton, Roberto Amato, Jeffrey Barrett, Sonia Goncalves, Ewan Harrison, David K. Jackson, Ian Johnston, Dominic Kwiatkowski, Cordelia Langford, John Sillitoe on behalf of the Wellcome Sanger Institute COVID-19 Surveillance Team |
| EPI_ISL_2766536 | Health Services Laboratories | Wellcome Sanger Institute for the COVID-19 Genomics UK (COG-UK) Consortium | Health Services Laboratories and Alex Alderton, Roberto Amato, Jeffrey Barrett, Sonia Goncalves, Ewan Harrison, David K. Jackson, Ian Johnston, Dominic Kwiatkowski, Cordelia Langford, John Sillitoe on behalf of the Wellcome Sanger Institute COVID-19 Surveillance Team |
| EPI_ISL_2766537, EPI_ISL_2766538, EPI_ISL_2766539 | Lighthouse Laboratory Plymouth | Wellcome Sanger Institute for the COVID-19 Genomics UK (COG-UK) Consortium | Lighthouse Laboratory Plymouth and Alex Alderton, Roberto Amato, Jeffrey Barrett, Sonia Goncalves, Ewan Harrison, David K. Jackson, Ian Johnston, Dominic Kwiatkowski, Cordelia Langford, John Sillitoe on behalf of the Wellcome Sanger Institute COVID-19 Surveillance Team |
| EPI_ISL_2766540 | Health Services Laboratories | Wellcome Sanger Institute for the COVID-19 Genomics UK (COG-UK) Consortium | Health Services Laboratories and Alex Alderton, Roberto Amato, Jeffrey Barrett, Sonia Goncalves, Ewan Harrison, David K. Jackson, Ian Johnston, Dominic Kwiatkowski, Cordelia Langford, John Sillitoe on behalf of the Wellcome Sanger Institute COVID-19 Surveillance Team |
| EPI_ISL_2766541, EPI_ISL_2766542, EPI_ISL_2766543 | Lighthouse Laboratory Plymouth | Wellcome Sanger Institute for the COVID-19 Genomics UK (COG-UK) Consortium | Lighthouse Laboratory Plymouth and Alex Alderton, Roberto Amato, Jeffrey Barrett, Sonia Goncalves, Ewan Harrison, David K. Jackson, Ian Johnston, Dominic Kwiatkowski, Cordelia Langford, John Sillitoe on behalf of the Wellcome Sanger Institute COVID-19 Surveillance Team |
| EPI_ISL_2766544 | Health Services Laboratories | Wellcome Sanger Institute for the COVID-19 Genomics UK (COG-UK) Consortium | Health Services Laboratories and Alex Alderton, Roberto Amato, Jeffrey Barrett, Sonia Goncalves, Ewan Harrison, David K. Jackson, Ian Johnston, Dominic Kwiatkowski, Cordelia Langford, John Sillitoe on behalf of the Wellcome Sanger Institute COVID-19 Surveillance Team |
| EPI_ISL_2766545, EPI_ISL_2766546 | Lighthouse Laboratory Plymouth | Wellcome Sanger Institute for the COVID-19 Genomics UK (COG-UK) Consortium | Lighthouse Laboratory Plymouth and Alex Alderton, Roberto Amato, Jeffrey Barrett, Sonia Goncalves, Ewan Harrison, David K. Jackson, Ian Johnston, Dominic Kwiatkowski, Cordelia Langford, John Sillitoe on behalf of the Wellcome Sanger Institute COVID-19 Surveillance Team |
| EPI_ISL_2766547 | Lighthouse Lab in Alderley Park | Wellcome Sanger Institute for the COVID-19 Genomics UK (COG-UK) Consortium | Jacquelyn Wynn, Mairead Hyland, The Lighthouse Lab in Alderley Park and Alex Alderton, Roberto Amato, Jeffrey Barrett, Sonia Goncalves, Ewan Harrison, David K. Jackson, Ian Johnston, Dominic Kwiatkowski, Cordelia Langford, John Sillitoe on behalf of the Wellcome Sanger Institute COVID-19 Surveillance Team |
| EPI_ISL_2766548, EPI_ISL_2766549 | Lighthouse Laboratory Plymouth | Wellcome Sanger Institute for the COVID-19 Genomics UK (COG-UK) Consortium | Lighthouse Laboratory Plymouth and Alex Alderton, Roberto Amato, Jeffrey Barrett, Sonia Goncalves, Ewan Harrison, David K. Jackson, Ian Johnston, Dominic Kwiatkowski, Cordelia Langford, John Sillitoe on behalf of the Wellcome Sanger Institute COVID-19 Surveillance Team |
| EPI_ISL_2766550, EPI_ISL_2766551, EPI_ISL_2766552, EPI_ISL_2766553 | Lighthouse Lab in Alderley Park | Wellcome Sanger Institute for the COVID-19 Genomics UK (COG-UK) Consortium | Jacquelyn Wynn, Mairead Hyland, The Lighthouse Lab in Alderley Park and Alex Alderton, Roberto Amato, Jeffrey Barrett, Sonia Goncalves, Ewan Harrison, David K. Jackson, Ian Johnston, Dominic Kwiatkowski, Cordelia Langford, John Sillitoe on behalf of the Wellcome Sanger Institute COVID-19 Surveillance Team |
| EPI_ISL_2766554 | Lighthouse Laboratory Plymouth | Wellcome Sanger Institute for the COVID-19 Genomics UK (COG-UK) Consortium | Lighthouse Laboratory Plymouth and Alex Alderton, Roberto Amato, Jeffrey Barrett, Sonia Goncalves, Ewan Harrison, David K. Jackson, Ian Johnston, Dominic Kwiatkowski, Cordelia Langford, John Sillitoe on behalf of the Wellcome Sanger Institute COVID-19 Surveillance Team |
| EPI_ISL_2766555 | Lighthouse Lab in Milton Keynes | Wellcome Sanger Institute for the COVID-19 Genomics UK (COG-UK) Consortium | The Lighthouse Lab in Milton Keynes and Alex Alderton, Roberto Amato, Jeffrey Barrett, Sonia Goncalves, Ewan Harrison, David K. Jackson, Ian Johnston, Dominic Kwiatkowski, Cordelia Langford, John Sillitoe on behalf of the Wellcome Sanger Institute COVID-19 Surveillance Team |
| EPI_ISL_2766556, EPI_ISL_2766557 | Lighthouse Lab in Alderley Park | Wellcome Sanger Institute for the COVID-19 Genomics UK (COG-UK) Consortium | Jacquelyn Wynn, Mairead Hyland, The Lighthouse Lab in Alderley Park and Alex Alderton, Roberto Amato, Jeffrey Barrett, Sonia Goncalves, Ewan Harrison, David K. Jackson, Ian Johnston, Dominic Kwiatkowski, Cordelia Langford, John Sillitoe on behalf of the Wellcome Sanger Institute COVID-19 Surveillance Team |
| EPI_ISL_2766558 | Lighthouse Laboratory Plymouth | Wellcome Sanger Institute for the COVID-19 Genomics UK (COG-UK) Consortium | Lighthouse Laboratory Plymouth and Alex Alderton, Roberto Amato, Jeffrey Barrett, Sonia Goncalves, Ewan Harrison, David K. Jackson, Ian Johnston, Dominic Kwiatkowski, Cordelia Langford, John Sillitoe on behalf of the Wellcome Sanger Institute COVID-19 Surveillance Team |
| EPI_ISL_2766559, EPI_ISL_2766560, EPI_ISL_2766561 | Lighthouse Lab in Milton Keynes | Wellcome Sanger Institute for the COVID-19 Genomics UK (COG-UK) Consortium | The Lighthouse Lab in Milton Keynes and Alex Alderton, Roberto Amato, Jeffrey Barrett, Sonia Goncalves, Ewan Harrison, David K. Jackson, Ian Johnston, Dominic Kwiatkowski, Cordelia Langford, John Sillitoe on behalf of the Wellcome Sanger Institute COVID-19 Surveillance Team |
| EPI_ISL_2766562, EPI_ISL_2766563, EPI_ISL_2766564, EPI_ISL_2766565 | Lighthouse Lab in Alderley Park | Wellcome Sanger Institute for the COVID-19 Genomics UK (COG-UK) Consortium | Jacquelyn Wynn, Mairead Hyland, The Lighthouse Lab in Alderley Park and Alex Alderton, Roberto Amato, Jeffrey Barrett, Sonia Goncalves, Ewan Harrison, David K. Jackson, Ian Johnston, Dominic Kwiatkowski, Cordelia Langford, John Sillitoe on behalf of the Wellcome Sanger Institute COVID-19 Surveillance Team |
| EPI_ISL_2766566 | Lighthouse Lab in Milton Keynes | Wellcome Sanger Institute for the COVID-19 Genomics UK (COG-UK) Consortium | The Lighthouse Lab in Milton Keynes and Alex Alderton, Roberto Amato, Jeffrey Barrett, Sonia Goncalves, Ewan Harrison, David K. Jackson, Ian Johnston, Dominic Kwiatkowski, Cordelia Langford, John Sillitoe on behalf of the Wellcome Sanger Institute COVID-19 Surveillance Team |
| EPI_ISL_2766567 | Lighthouse Lab in Alderley Park | Wellcome Sanger Institute for the COVID-19 Genomics UK (COG-UK) Consortium | Jacquelyn Wynn, Mairead Hyland, The Lighthouse Lab in Alderley Park and Alex Alderton, Roberto Amato, Jeffrey Barrett, Sonia Goncalves, Ewan Harrison, David K. Jackson, Ian Johnston, Dominic Kwiatkowski, Cordelia Langford, John Sillitoe on behalf of the Wellcome Sanger Institute COVID-19 Surveillance Team |
| EPI_ISL_2766568 | Lighthouse Laboratory Plymouth | Wellcome Sanger Institute for the COVID-19 Genomics UK (COG-UK) Consortium | Lighthouse Laboratory Plymouth and Alex Alderton, Roberto Amato, Jeffrey Barrett, Sonia Goncalves, Ewan Harrison, David K. Jackson, Ian Johnston, Dominic Kwiatkowski, Cordelia Langford, John Sillitoe on behalf of the Wellcome Sanger Institute COVID-19 Surveillance Team |
| EPI_ISL_2766569 | Lighthouse Lab in Milton Keynes | Wellcome Sanger Institute for the COVID-19 Genomics UK (COG-UK) Consortium | The Lighthouse Lab in Milton Keynes and Alex Alderton, Roberto Amato, Jeffrey Barrett, Sonia Goncalves, Ewan Harrison, David K. Jackson, Ian Johnston, Dominic Kwiatkowski, Cordelia Langford, John Sillitoe on behalf of the Wellcome Sanger Institute COVID-19 Surveillance Team |
| EPI_ISL_2766570 | Lighthouse Lab in Alderley Park | Wellcome Sanger Institute for the COVID-19 Genomics UK (COG-UK) Consortium | Jacquelyn Wynn, Mairead Hyland, The Lighthouse Lab in Alderley Park and Alex Alderton, Roberto Amato, Jeffrey Barrett, Sonia Goncalves, Ewan Harrison, David K. Jackson, Ian Johnston, Dominic Kwiatkowski, Cordelia Langford, John Sillitoe on behalf of the Wellcome Sanger Institute COVID-19 Surveillance Team |
| EPI_ISL_2766571 | Lighthouse Lab in Milton Keynes | Wellcome Sanger Institute for the COVID-19 Genomics UK (COG-UK) Consortium | The Lighthouse Lab in Milton Keynes and Alex Alderton, Roberto Amato, Jeffrey Barrett, Sonia Goncalves, Ewan Harrison, David K. Jackson, Ian Johnston, Dominic Kwiatkowski, Cordelia Langford, John Sillitoe on behalf of the Wellcome Sanger Institute COVID-19 Surveillance Team |
| EPI_ISL_2766572, EPI_ISL_2766573 | Lighthouse Lab in Alderley Park | Wellcome Sanger Institute for the COVID-19 Genomics UK (COG-UK) Consortium | Jacquelyn Wynn, Mairead Hyland, The Lighthouse Lab in Alderley Park and Alex Alderton, Roberto Amato, Jeffrey Barrett, Sonia Goncalves, Ewan Harrison, David K. Jackson, Ian Johnston, Dominic Kwiatkowski, Cordelia Langford, John Sillitoe on behalf of the Wellcome Sanger Institute COVID-19 Surveillance Team |
| EPI_ISL_2766574 | Lighthouse Lab in Milton Keynes | Wellcome Sanger Institute for the COVID-19 Genomics UK (COG-UK) Consortium | The Lighthouse Lab in Milton Keynes and Alex Alderton, Roberto Amato, Jeffrey Barrett, Sonia Goncalves, Ewan Harrison, David K. Jackson, Ian Johnston, Dominic Kwiatkowski, Cordelia Langford, John Sillitoe on behalf of the Wellcome Sanger Institute COVID-19 Surveillance Team |
| EPI_ISL_2766575, EPI_ISL_2766576 | Lighthouse Lab in Alderley Park | Wellcome Sanger Institute for the COVID-19 Genomics UK (COG-UK) Consortium | Jacquelyn Wynn, Mairead Hyland, The Lighthouse Lab in Alderley Park and Alex Alderton, Roberto Amato, Jeffrey Barrett, Sonia Goncalves, Ewan Harrison, David K. Jackson, Ian Johnston, Dominic Kwiatkowski, Cordelia Langford, John Sillitoe on behalf of the Wellcome Sanger Institute COVID-19 Surveillance Team |
| EPI_ISL_2766577, EPI_ISL_2766578 | Lighthouse Lab in Milton Keynes | Wellcome Sanger Institute for the COVID-19 Genomics UK (COG-UK) Consortium | The Lighthouse Lab in Milton Keynes and Alex Alderton, Roberto Amato, Jeffrey Barrett, Sonia Goncalves, Ewan Harrison, David K. Jackson, Ian Johnston, Dominic Kwiatkowski, Cordelia Langford, John Sillitoe on behalf of the Wellcome Sanger Institute COVID-19 Surveillance Team |
| EPI_ISL_2766579, EPI_ISL_2766580 | Lighthouse Lab in Alderley Park | Wellcome Sanger Institute for the COVID-19 Genomics UK (COG-UK) Consortium | Jacquelyn Wynn, Mairead Hyland, The Lighthouse Lab in Alderley Park and Alex Alderton, Roberto Amato, Jeffrey Barrett, Sonia Goncalves, Ewan Harrison, David K. Jackson, Ian Johnston, Dominic Kwiatkowski, Cordelia Langford, John Sillitoe on behalf of the Wellcome Sanger Institute COVID-19 Surveillance Team |
| EPI_ISL_2766581 | Lighthouse Laboratory Plymouth | Wellcome Sanger Institute for the COVID-19 Genomics UK (COG-UK) Consortium | Lighthouse Laboratory Plymouth and Alex Alderton, Roberto Amato, Jeffrey Barrett, Sonia Goncalves, Ewan Harrison, David K. Jackson, Ian Johnston, Dominic Kwiatkowski, Cordelia Langford, John Sillitoe on behalf of the Wellcome Sanger Institute COVID-19 Surveillance Team |
| EPI_ISL_2766582, EPI_ISL_2766583 | Lighthouse Lab in Alderley Park | Wellcome Sanger Institute for the COVID-19 Genomics UK (COG-UK) Consortium | Jacquelyn Wynn, Mairead Hyland, The Lighthouse Lab in Alderley Park and Alex Alderton, Roberto Amato, Jeffrey Barrett, Sonia Goncalves, Ewan Harrison, David K. Jackson, Ian Johnston, Dominic Kwiatkowski, Cordelia Langford, John Sillitoe on behalf of the Wellcome Sanger Institute COVID-19 Surveillance Team |
| EPI_ISL_2766584, EPI_ISL_2766585 | Lighthouse Laboratory Plymouth | Wellcome Sanger Institute for the COVID-19 Genomics UK (COG-UK) Consortium | Lighthouse Laboratory Plymouth and Alex Alderton, Roberto Amato, Jeffrey Barrett, Sonia Goncalves, Ewan Harrison, David K. Jackson, Ian Johnston, Dominic Kwiatkowski, Cordelia Langford, John Sillitoe on behalf of the Wellcome Sanger Institute COVID-19 Surveillance Team |
| EPI_ISL_2766586 | Lighthouse Lab in Milton Keynes | Wellcome Sanger Institute for the COVID-19 Genomics UK (COG-UK) Consortium | The Lighthouse Lab in Milton Keynes and Alex Alderton, Roberto Amato, Jeffrey Barrett, Sonia Goncalves, Ewan Harrison, David K. Jackson, Ian Johnston, Dominic Kwiatkowski, Cordelia Langford, John Sillitoe on behalf of the Wellcome Sanger Institute COVID-19 Surveillance Team |

| | | | |
|---|---|---|---|
| EPI_ISL_2766587, EPI_ISL_2766588 | Lighthouse Lab in Alderley Park | Wellcome Sanger Institute for the COVID-19 Genomics UK (COG-UK) Consortium | Jacquelyn Wynn, Mairead Hyland, The Lighthouse Lab in Alderley Park and Alex Alderton, Roberto Amato, Jeffrey Barrett, Sonia Goncalves, Ewan Harrison, David K. Jackson, Ian Johnston, Dominic Kwiatkowski, Cordelia Langford, John Sillitoe on behalf of the Wellcome Sanger Institute COVID-19 Surveillance Team |
| EPI_ISL_2766589 | Lighthouse Lab in Milton Keynes | Wellcome Sanger Institute for the COVID-19 Genomics UK (COG-UK) Consortium | The Lighthouse Lab in Milton Keynes and Alex Alderton, Roberto Amato, Jeffrey Barrett, Sonia Goncalves, Ewan Harrison, David K. Jackson, Ian Johnston, Dominic Kwiatkowski, Cordelia Langford, John Sillitoe on behalf of the Wellcome Sanger Institute COVID-19 Surveillance Team |
| EPI_ISL_2766590 | Lighthouse Laboratory Plymouth | Wellcome Sanger Institute for the COVID-19 Genomics UK (COG-UK) Consortium | Lighthouse Laboratory Plymouth and Alex Alderton, Roberto Amato, Jeffrey Barrett, Sonia Goncalves, Ewan Harrison, David K. Jackson, Ian Johnston, Dominic Kwiatkowski, Cordelia Langford, John Sillitoe on behalf of the Wellcome Sanger Institute COVID-19 Surveillance Team |
| EPI_ISL_2766591 | Lighthouse Lab in Alderley Park | Wellcome Sanger Institute for the COVID-19 Genomics UK (COG-UK) Consortium | Jacquelyn Wynn, Mairead Hyland, The Lighthouse Lab in Alderley Park and Alex Alderton, Roberto Amato, Jeffrey Barrett, Sonia Goncalves, Ewan Harrison, David K. Jackson, Ian Johnston, Dominic Kwiatkowski, Cordelia Langford, John Sillitoe on behalf of the Wellcome Sanger Institute COVID-19 Surveillance Team |
| EPI_ISL_2766592, EPI_ISL_2766593 | Lighthouse Laboratory Plymouth | Wellcome Sanger Institute for the COVID-19 Genomics UK (COG-UK) Consortium | Lighthouse Laboratory Plymouth and Alex Alderton, Roberto Amato, Jeffrey Barrett, Sonia Goncalves, Ewan Harrison, David K. Jackson, Ian Johnston, Dominic Kwiatkowski, Cordelia Langford, John Sillitoe on behalf of the Wellcome Sanger Institute COVID-19 Surveillance Team |
| EPI_ISL_2766594 | Lighthouse Lab in Alderley Park | Wellcome Sanger Institute for the COVID-19 Genomics UK (COG-UK) Consortium | Jacquelyn Wynn, Mairead Hyland, The Lighthouse Lab in Alderley Park and Alex Alderton, Roberto Amato, Jeffrey Barrett, Sonia Goncalves, Ewan Harrison, David K. Jackson, Ian Johnston, Dominic Kwiatkowski, Cordelia Langford, John Sillitoe on behalf of the Wellcome Sanger Institute COVID-19 Surveillance Team |
| EPI_ISL_2766595 | Lighthouse Lab in Milton Keynes | Wellcome Sanger Institute for the COVID-19 Genomics UK (COG-UK) Consortium | The Lighthouse Lab in Milton Keynes and Alex Alderton, Roberto Amato, Jeffrey Barrett, Sonia Goncalves, Ewan Harrison, David K. Jackson, Ian Johnston, Dominic Kwiatkowski, Cordelia Langford, John Sillitoe on behalf of the Wellcome Sanger Institute COVID-19 Surveillance Team |
| EPI_ISL_2766596 | Lighthouse Lab in Alderley Park | Wellcome Sanger Institute for the COVID-19 Genomics UK (COG-UK) Consortium | Jacquelyn Wynn, Mairead Hyland, The Lighthouse Lab in Alderley Park and Alex Alderton, Roberto Amato, Jeffrey Barrett, Sonia Goncalves, Ewan Harrison, David K. Jackson, Ian Johnston, Dominic Kwiatkowski, Cordelia Langford, John Sillitoe on behalf of the Wellcome Sanger Institute COVID-19 Surveillance Team |
| EPI_ISL_2766597 | Lighthouse Lab in Milton Keynes | Wellcome Sanger Institute for the COVID-19 Genomics UK (COG-UK) Consortium | The Lighthouse Lab in Milton Keynes and Alex Alderton, Roberto Amato, Jeffrey Barrett, Sonia Goncalves, Ewan Harrison, David K. Jackson, Ian Johnston, Dominic Kwiatkowski, Cordelia Langford, John Sillitoe on behalf of the Wellcome Sanger Institute COVID-19 Surveillance Team |
| EPI_ISL_2766598 | Lighthouse Lab in Alderley Park | Wellcome Sanger Institute for the COVID-19 Genomics UK (COG-UK) Consortium | Jacquelyn Wynn, Mairead Hyland, The Lighthouse Lab in Alderley Park and Alex Alderton, Roberto Amato, Jeffrey Barrett, Sonia Goncalves, Ewan Harrison, David K. Jackson, Ian Johnston, Dominic Kwiatkowski, Cordelia Langford, John Sillitoe on behalf of the Wellcome Sanger Institute COVID-19 Surveillance Team |
| EPI_ISL_2766599 | Lighthouse Lab in Milton Keynes | Wellcome Sanger Institute for the COVID-19 Genomics UK (COG-UK) Consortium | The Lighthouse Lab in Milton Keynes and Alex Alderton, Roberto Amato, Jeffrey Barrett, Sonia Goncalves, Ewan Harrison, David K. Jackson, Ian Johnston, Dominic Kwiatkowski, Cordelia Langford, John Sillitoe on behalf of the Wellcome Sanger Institute COVID-19 Surveillance Team |
| EPI_ISL_2766600 | Lighthouse Lab in Alderley Park | Wellcome Sanger Institute for the COVID-19 Genomics UK (COG-UK) Consortium | Jacquelyn Wynn, Mairead Hyland, The Lighthouse Lab in Alderley Park and Alex Alderton, Roberto Amato, Jeffrey Barrett, Sonia Goncalves, Ewan Harrison, David K. Jackson, Ian Johnston, Dominic Kwiatkowski, Cordelia Langford, John Sillitoe on behalf of the Wellcome Sanger Institute COVID-19 Surveillance Team |
| EPI_ISL_2766601 | Lighthouse Lab in Milton Keynes | Wellcome Sanger Institute for the COVID-19 Genomics UK (COG-UK) Consortium | The Lighthouse Lab in Milton Keynes and Alex Alderton, Roberto Amato, Jeffrey Barrett, Sonia Goncalves, Ewan Harrison, David K. Jackson, Ian Johnston, Dominic Kwiatkowski, Cordelia Langford, John Sillitoe on behalf of the Wellcome Sanger Institute COVID-19 Surveillance Team |
| EPI_ISL_2766602 | Lighthouse Laboratory Plymouth | Wellcome Sanger Institute for the COVID-19 Genomics UK (COG-UK) Consortium | Lighthouse Laboratory Plymouth and Alex Alderton, Roberto Amato, Jeffrey Barrett, Sonia Goncalves, Ewan Harrison, David K. Jackson, Ian Johnston, Dominic Kwiatkowski, Cordelia Langford, John Sillitoe on behalf of the Wellcome Sanger Institute COVID-19 Surveillance Team |
| EPI_ISL_2766603, EPI_ISL_2766604, EPI_ISL_2766605 | Lighthouse Lab in Alderley Park | Wellcome Sanger Institute for the COVID-19 Genomics UK (COG-UK) Consortium | Jacquelyn Wynn, Mairead Hyland, The Lighthouse Lab in Alderley Park and Alex Alderton, Roberto Amato, Jeffrey Barrett, Sonia Goncalves, Ewan Harrison, David K. Jackson, Ian Johnston, Dominic Kwiatkowski, Cordelia Langford, John Sillitoe on behalf of the Wellcome Sanger Institute COVID-19 Surveillance Team |
| EPI_ISL_2766606 | Lighthouse Laboratory Plymouth | Wellcome Sanger Institute for the COVID-19 Genomics UK (COG-UK) Consortium | Lighthouse Laboratory Plymouth and Alex Alderton, Roberto Amato, Jeffrey Barrett, Sonia Goncalves, Ewan Harrison, David K. Jackson, Ian Johnston, Dominic Kwiatkowski, Cordelia Langford, John Sillitoe on behalf of the Wellcome Sanger Institute COVID-19 Surveillance Team |
| EPI_ISL_2766607, EPI_ISL_2766608 | Lighthouse Lab in Alderley Park | Wellcome Sanger Institute for the COVID-19 Genomics UK (COG-UK) Consortium | Jacquelyn Wynn, Mairead Hyland, The Lighthouse Lab in Alderley Park and Alex Alderton, Roberto Amato, Jeffrey Barrett, Sonia Goncalves, Ewan Harrison, David K. Jackson, Ian Johnston, Dominic Kwiatkowski, Cordelia Langford, John Sillitoe on behalf of the Wellcome Sanger Institute COVID-19 Surveillance Team |
| EPI_ISL_2766609 | Lighthouse Laboratory Plymouth | Wellcome Sanger Institute for the COVID-19 Genomics UK (COG-UK) Consortium | Lighthouse Laboratory Plymouth and Alex Alderton, Roberto Amato, Jeffrey Barrett, Sonia Goncalves, Ewan Harrison, David K. Jackson, Ian Johnston, Dominic Kwiatkowski, Cordelia Langford, John Sillitoe on behalf of the Wellcome Sanger Institute COVID-19 Surveillance Team |
| EPI_ISL_2766610 | Lighthouse Lab in Alderley Park | Wellcome Sanger Institute for the COVID-19 Genomics UK (COG-UK) Consortium | Jacquelyn Wynn, Mairead Hyland, The Lighthouse Lab in Alderley Park and Alex Alderton, Roberto Amato, Jeffrey Barrett, Sonia Goncalves, Ewan Harrison, David K. Jackson, Ian Johnston, Dominic Kwiatkowski, Cordelia Langford, John Sillitoe on behalf of the Wellcome Sanger Institute COVID-19 Surveillance Team |
| EPI_ISL_2766611 | Lighthouse Lab in Milton Keynes | Wellcome Sanger Institute for the COVID-19 Genomics UK (COG-UK) Consortium | The Lighthouse Lab in Milton Keynes and Alex Alderton, Roberto Amato, Jeffrey Barrett, Sonia Goncalves, Ewan Harrison, David K. Jackson, Ian Johnston, Dominic Kwiatkowski, Cordelia Langford, John Sillitoe on behalf of the Wellcome Sanger Institute COVID-19 Surveillance Team |
| EPI_ISL_2766612, EPI_ISL_2766613, EPI_ISL_2766614 | Lighthouse Lab in Alderley Park | Wellcome Sanger Institute for the COVID-19 Genomics UK (COG-UK) Consortium | Jacquelyn Wynn, Mairead Hyland, The Lighthouse Lab in Alderley Park and Alex Alderton, Roberto Amato, Jeffrey Barrett, Sonia Goncalves, Ewan Harrison, David K. Jackson, Ian Johnston, Dominic Kwiatkowski, Cordelia Langford, John Sillitoe on behalf of the Wellcome Sanger Institute COVID-19 Surveillance Team |
| EPI_ISL_2766615 | Lighthouse Lab in Milton Keynes | Wellcome Sanger Institute for the COVID-19 Genomics UK (COG-UK) Consortium | The Lighthouse Lab in Milton Keynes and Alex Alderton, Roberto Amato, Jeffrey Barrett, Sonia Goncalves, Ewan Harrison, David K. Jackson, Ian Johnston, Dominic Kwiatkowski, Cordelia Langford, John Sillitoe on behalf of the Wellcome Sanger Institute COVID-19 Surveillance Team |
| EPI_ISL_2766616 | Lighthouse Laboratory Plymouth | Wellcome Sanger Institute for the COVID-19 Genomics UK (COG-UK) Consortium | Lighthouse Laboratory Plymouth and Alex Alderton, Roberto Amato, Jeffrey Barrett, Sonia Goncalves, Ewan Harrison, David K. Jackson, Ian Johnston, Dominic Kwiatkowski, Cordelia Langford, John Sillitoe on behalf of the Wellcome Sanger Institute COVID-19 Surveillance Team |
| EPI_ISL_2766617 | Lighthouse Lab in Alderley Park | Wellcome Sanger Institute for the COVID-19 Genomics UK (COG-UK) Consortium | Jacquelyn Wynn, Mairead Hyland, The Lighthouse Lab in Alderley Park and Alex Alderton, Roberto Amato, Jeffrey Barrett, Sonia Goncalves, Ewan Harrison, David K. Jackson, Ian Johnston, Dominic Kwiatkowski, Cordelia Langford, John Sillitoe on behalf of the Wellcome Sanger Institute COVID-19 Surveillance Team |
| EPI_ISL_2766618 | Lighthouse Lab in Milton Keynes | Wellcome Sanger Institute for the COVID-19 Genomics UK (COG-UK) Consortium | The Lighthouse Lab in Milton Keynes and Alex Alderton, Roberto Amato, Jeffrey Barrett, Sonia Goncalves, Ewan Harrison, David K. Jackson, Ian Johnston, Dominic Kwiatkowski, Cordelia Langford, John Sillitoe on behalf of the Wellcome Sanger Institute COVID-19 Surveillance Team |
| EPI_ISL_2766619 | Lighthouse Lab in Alderley Park | Wellcome Sanger Institute for the COVID-19 Genomics UK (COG-UK) Consortium | Jacquelyn Wynn, Mairead Hyland, The Lighthouse Lab in Alderley Park and Alex Alderton, Roberto Amato, Jeffrey Barrett, Sonia Goncalves, Ewan Harrison, David K. Jackson, Ian Johnston, Dominic Kwiatkowski, Cordelia Langford, John Sillitoe on behalf of the Wellcome Sanger Institute COVID-19 Surveillance Team |
| EPI_ISL_2766620 | Lighthouse Lab in Milton Keynes | Wellcome Sanger Institute for the COVID-19 Genomics UK (COG-UK) Consortium | The Lighthouse Lab in Milton Keynes and Alex Alderton, Roberto Amato, Jeffrey Barrett, Sonia Goncalves, Ewan Harrison, David K. Jackson, Ian Johnston, Dominic Kwiatkowski, Cordelia Langford, John Sillitoe on behalf of the Wellcome Sanger Institute COVID-19 Surveillance Team |
| EPI_ISL_2766621, EPI_ISL_2766622 | Lighthouse Lab in Alderley Park | Wellcome Sanger Institute for the COVID-19 Genomics UK (COG-UK) Consortium | Jacquelyn Wynn, Mairead Hyland, The Lighthouse Lab in Alderley Park and Alex Alderton, Roberto Amato, Jeffrey Barrett, Sonia Goncalves, Ewan Harrison, David K. Jackson, Ian Johnston, Dominic Kwiatkowski, Cordelia Langford, John Sillitoe on behalf of the Wellcome Sanger Institute COVID-19 Surveillance Team |
| EPI_ISL_2766623 | Lighthouse Laboratory Plymouth | Wellcome Sanger Institute for the COVID-19 Genomics UK (COG-UK) Consortium | Lighthouse Laboratory Plymouth and Alex Alderton, Roberto Amato, Jeffrey Barrett, Sonia Goncalves, Ewan Harrison, David K. Jackson, Ian Johnston, Dominic Kwiatkowski, Cordelia Langford, John Sillitoe on behalf of the Wellcome Sanger Institute COVID-19 Surveillance Team |
| EPI_ISL_2766624 | Lighthouse Lab in Alderley Park | Wellcome Sanger Institute for the COVID-19 Genomics UK (COG-UK) Consortium | Jacquelyn Wynn, Mairead Hyland, The Lighthouse Lab in Alderley Park and Alex Alderton, Roberto Amato, Jeffrey Barrett, Sonia Goncalves, Ewan Harrison, David K. Jackson, Ian Johnston, Dominic Kwiatkowski, Cordelia Langford, John Sillitoe on behalf of the Wellcome Sanger Institute COVID-19 Surveillance Team |
| EPI_ISL_2766625 | Lighthouse Lab in Milton Keynes | Wellcome Sanger Institute for the COVID-19 Genomics UK (COG-UK) Consortium | The Lighthouse Lab in Milton Keynes and Alex Alderton, Roberto Amato, Jeffrey Barrett, Sonia Goncalves, Ewan Harrison, David K. Jackson, Ian Johnston, Dominic Kwiatkowski, Cordelia Langford, John Sillitoe on behalf of the Wellcome Sanger Institute COVID-19 Surveillance Team |
| EPI_ISL_2766626 | Lighthouse Lab in Alderley Park | Wellcome Sanger Institute for the COVID-19 Genomics UK (COG-UK) Consortium | Jacquelyn Wynn, Mairead Hyland, The Lighthouse Lab in Alderley Park and Alex Alderton, Roberto Amato, Jeffrey Barrett, Sonia Goncalves, Ewan Harrison, David K. Jackson, Ian Johnston, Dominic Kwiatkowski, Cordelia Langford, John Sillitoe on behalf of the Wellcome Sanger Institute COVID-19 Surveillance Team |
| EPI_ISL_2766627 | Lighthouse Lab in Milton Keynes | Wellcome Sanger Institute for the COVID-19 Genomics UK (COG-UK) Consortium | The Lighthouse Lab in Milton Keynes and Alex Alderton, Roberto Amato, Jeffrey Barrett, Sonia Goncalves, Ewan Harrison, David K. Jackson, Ian Johnston, Dominic Kwiatkowski, Cordelia Langford, John Sillitoe on behalf of the Wellcome Sanger Institute COVID-19 Surveillance Team |
| EPI_ISL_2766628 | Lighthouse Lab in Alderley Park | Wellcome Sanger Institute for the COVID-19 Genomics UK (COG-UK) Consortium | Jacquelyn Wynn, Mairead Hyland, The Lighthouse Lab in Alderley Park and Alex Alderton, Roberto Amato, Jeffrey Barrett, Sonia Goncalves, Ewan Harrison, David K. Jackson, Ian Johnston, Dominic Kwiatkowski, Cordelia Langford, John Sillitoe on behalf of the Wellcome Sanger Institute COVID-19 Surveillance Team |
| EPI_ISL_2766629 | Lighthouse Lab in Milton Keynes | Wellcome Sanger Institute for the COVID-19 Genomics UK (COG-UK) Consortium | The Lighthouse Lab in Milton Keynes and Alex Alderton, Roberto Amato, Jeffrey Barrett, Sonia Goncalves, Ewan Harrison, David K. Jackson, Ian Johnston, Dominic Kwiatkowski, Cordelia Langford, John Sillitoe on behalf of the Wellcome Sanger Institute COVID-19 Surveillance Team |
| EPI_ISL_2766630 | Lighthouse Lab in Alderley Park | Wellcome Sanger Institute for the COVID-19 Genomics UK (COG-UK) Consortium | Jacquelyn Wynn, Mairead Hyland, The Lighthouse Lab in Alderley Park and Alex Alderton, Roberto Amato, Jeffrey Barrett, Sonia Goncalves, Ewan Harrison, David K. Jackson, Ian Johnston, Dominic Kwiatkowski, Cordelia Langford, John Sillitoe on behalf of the Wellcome Sanger Institute COVID-19 Surveillance Team |
| EPI_ISL_2766631 | Lighthouse Lab in Milton Keynes | Wellcome Sanger Institute for the COVID-19 Genomics UK (COG-UK) Consortium | The Lighthouse Lab in Milton Keynes and Alex Alderton, Roberto Amato, Jeffrey Barrett, Sonia Goncalves, Ewan Harrison, David K. Jackson, Ian Johnston, Dominic Kwiatkowski, Cordelia Langford, John Sillitoe on behalf of the Wellcome Sanger Institute COVID-19 Surveillance Team |
| EPI_ISL_2766632, EPI_ISL_2766633 | Lighthouse Lab in Alderley Park | Wellcome Sanger Institute for the COVID-19 Genomics UK (COG-UK) Consortium | Jacquelyn Wynn, Mairead Hyland, The Lighthouse Lab in Alderley Park and Alex Alderton, Roberto Amato, Jeffrey Barrett, Sonia Goncalves, Ewan Harrison, David K. Jackson, Ian Johnston, Dominic Kwiatkowski, Cordelia Langford, John Sillitoe on behalf of the Wellcome Sanger Institute COVID-19 Surveillance Team |
| EPI_ISL_2766634, EPI_ISL_2766635 | Lighthouse Lab in Milton Keynes | Wellcome Sanger Institute for the COVID-19 Genomics UK (COG-UK) Consortium | The Lighthouse Lab in Milton Keynes and Alex Alderton, Roberto Amato, Jeffrey Barrett, Sonia Goncalves, Ewan Harrison, David K. Jackson, Ian Johnston, Dominic Kwiatkowski, Cordelia Langford, John Sillitoe on behalf of the Wellcome Sanger Institute COVID-19 Surveillance Team |
| EPI_ISL_2766636, EPI_ISL_2766637 | Lighthouse Lab in Alderley Park | Wellcome Sanger Institute for the COVID-19 Genomics UK (COG-UK) Consortium | Jacquelyn Wynn, Mairead Hyland, The Lighthouse Lab in Alderley Park and Alex Alderton, Roberto Amato, Jeffrey Barrett, Sonia Goncalves, Ewan Harrison, David K. Jackson, Ian Johnston, Dominic Kwiatkowski, Cordelia Langford, John Sillitoe on behalf of the Wellcome Sanger Institute COVID-19 Surveillance Team |
| EPI_ISL_2766638, EPI_ISL_2766639 | Lighthouse Laboratory Plymouth | Wellcome Sanger Institute for the COVID-19 Genomics UK (COG-UK) Consortium | Lighthouse Laboratory Plymouth and Alex Alderton, Roberto Amato, Jeffrey Barrett, Sonia Goncalves, Ewan Harrison, David K. Jackson, Ian Johnston, Dominic Kwiatkowski, Cordelia Langford, John Sillitoe on behalf of the Wellcome Sanger Institute COVID-19 Surveillance Team |
| EPI_ISL_2766640, EPI_ISL_2766641 | Lighthouse Lab in Alderley Park | Wellcome Sanger Institute for the COVID-19 Genomics UK (COG-UK) Consortium | Jacquelyn Wynn, Mairead Hyland, The Lighthouse Lab in Alderley Park and Alex Alderton, Roberto Amato, Jeffrey Barrett, Sonia Goncalves, Ewan Harrison, David K. Jackson, Ian Johnston, Dominic Kwiatkowski, Cordelia Langford, John Sillitoe on behalf of the Wellcome Sanger Institute COVID-19 Surveillance Team |
| EPI_ISL_2766642 | Lighthouse Lab in Milton Keynes | Wellcome Sanger Institute for the COVID-19 Genomics UK (COG-UK) Consortium | The Lighthouse Lab in Milton Keynes and Alex Alderton, Roberto Amato, Jeffrey Barrett, Sonia Goncalves, Ewan Harrison, David K. Jackson, Ian Johnston, Dominic Kwiatkowski, Cordelia Langford, John Sillitoe on behalf of the Wellcome Sanger Institute COVID-19 Surveillance Team |
| EPI_ISL_2766643 | Lighthouse Lab in Alderley Park | Wellcome Sanger Institute for the COVID-19 Genomics UK (COG-UK) Consortium | Jacquelyn Wynn, Mairead Hyland, The Lighthouse Lab in Alderley Park and Alex Alderton, Roberto Amato, Jeffrey Barrett, Sonia Goncalves, Ewan Harrison, David K. Jackson, Ian Johnston, Dominic Kwiatkowski, Cordelia Langford, John Sillitoe on behalf of the Wellcome Sanger Institute COVID-19 Surveillance Team |
| EPI_ISL_2766644 | Lighthouse Lab in Milton Keynes | Wellcome Sanger Institute for the COVID-19 Genomics UK (COG-UK) | The Lighthouse Lab in Milton Keynes and Alex Alderton, Roberto Amato, Jeffrey Barrett, Sonia Goncalves, Ewan Harrison, David K. Jackson, Ian Johnston, Dominic Kwiatkowski, Cordelia Langford, John Sillitoe on behalf of the Wellcome Sanger |

| | | | |
|---|---|---|---|
| | | Consortium | Institute COVID-19 Surveillance Team |
| EPI_ISL_2766645 | Lighthouse Lab in Alderley Park | Wellcome Sanger Institute for the COVID-19 Genomics UK (COG-UK) Consortium | Jacquelyn Wynn, Mairead Hyland, The Lighthouse Lab in Alderley Park and Alex Alderton, Roberto Amato, Jeffrey Barrett, Sonia Goncalves, Ewan Harrison, David K. Jackson, Ian Johnston, Dominic Kwiatkowski, Cordelia Langford, John Sillitoe on behalf of the Wellcome Sanger Institute COVID-19 Surveillance Team |
| EPI_ISL_2766646 | Lighthouse Lab in Milton Keynes | Wellcome Sanger Institute for the COVID-19 Genomics UK (COG-UK) Consortium | The Lighthouse Lab in Milton Keynes and Alex Alderton, Roberto Amato, Jeffrey Barrett, Sonia Goncalves, Ewan Harrison, David K. Jackson, Ian Johnston, Dominic Kwiatkowski, Cordelia Langford, John Sillitoe on behalf of the Wellcome Sanger Institute COVID-19 Surveillance Team |
| EPI_ISL_2766647 | Lighthouse Lab in Alderley Park | Wellcome Sanger Institute for the COVID-19 Genomics UK (COG-UK) Consortium | Jacquelyn Wynn, Mairead Hyland, The Lighthouse Lab in Alderley Park and Alex Alderton, Roberto Amato, Jeffrey Barrett, Sonia Goncalves, Ewan Harrison, David K. Jackson, Ian Johnston, Dominic Kwiatkowski, Cordelia Langford, John Sillitoe on behalf of the Wellcome Sanger Institute COVID-19 Surveillance Team |
| EPI_ISL_2766648, EPI_ISL_2766649 | Lighthouse Lab in Milton Keynes | Wellcome Sanger Institute for the COVID-19 Genomics UK (COG-UK) Consortium | The Lighthouse Lab in Milton Keynes and Alex Alderton, Roberto Amato, Jeffrey Barrett, Sonia Goncalves, Ewan Harrison, David K. Jackson, Ian Johnston, Dominic Kwiatkowski, Cordelia Langford, John Sillitoe on behalf of the Wellcome Sanger Institute COVID-19 Surveillance Team |
| EPI_ISL_2766650 | National Institute of Laboratory Medicine and Referral Center | Genomic Research Lab, BCSIR | Md. Murshed Hasan Sarkar, Mohammad Samir Uzzaman, Eshrar Osman, Md. Ahasan Habib, Shahina Akter, Tanjina Akhter Banu, Barna Goswami, Iffat Jahan, Mohammad Mohi Uddin, Md. Kamrul Islam, Tasnim Nafisa, Md. Maruf Ahmed Molla, Mahmuda Yeasmin, Asish Kumar Ghosh, Arifa Akram, Md. Salim Khan |
| EPI_ISL_2766651, EPI_ISL_2766652, EPI_ISL_2766653 | Lighthouse Lab in Alderley Park | Wellcome Sanger Institute for the COVID-19 Genomics UK (COG-UK) Consortium | Jacquelyn Wynn, Mairead Hyland, The Lighthouse Lab in Alderley Park and Alex Alderton, Roberto Amato, Jeffrey Barrett, Sonia Goncalves, Ewan Harrison, David K. Jackson, Ian Johnston, Dominic Kwiatkowski, Cordelia Langford, John Sillitoe on behalf of the Wellcome Sanger Institute COVID-19 Surveillance Team |
| EPI_ISL_2766654 | Lighthouse Lab in Milton Keynes | Wellcome Sanger Institute for the COVID-19 Genomics UK (COG-UK) Consortium | The Lighthouse Lab in Milton Keynes and Alex Alderton, Roberto Amato, Jeffrey Barrett, Sonia Goncalves, Ewan Harrison, David K. Jackson, Ian Johnston, Dominic Kwiatkowski, Cordelia Langford, John Sillitoe on behalf of the Wellcome Sanger Institute COVID-19 Surveillance Team |
| EPI_ISL_2766655 | Lighthouse Lab in Alderley Park | Wellcome Sanger Institute for the COVID-19 Genomics UK (COG-UK) Consortium | Jacquelyn Wynn, Mairead Hyland, The Lighthouse Lab in Alderley Park and Alex Alderton, Roberto Amato, Jeffrey Barrett, Sonia Goncalves, Ewan Harrison, David K. Jackson, Ian Johnston, Dominic Kwiatkowski, Cordelia Langford, John Sillitoe on behalf of the Wellcome Sanger Institute COVID-19 Surveillance Team |
| EPI_ISL_2766656 | Lighthouse Lab in Milton Keynes | Wellcome Sanger Institute for the COVID-19 Genomics UK (COG-UK) Consortium | The Lighthouse Lab in Milton Keynes and Alex Alderton, Roberto Amato, Jeffrey Barrett, Sonia Goncalves, Ewan Harrison, David K. Jackson, Ian Johnston, Dominic Kwiatkowski, Cordelia Langford, John Sillitoe on behalf of the Wellcome Sanger Institute COVID-19 Surveillance Team |
| EPI_ISL_2766657, EPI_ISL_2766658, EPI_ISL_2766659 | Lighthouse Lab in Alderley Park | Wellcome Sanger Institute for the COVID-19 Genomics UK (COG-UK) Consortium | Jacquelyn Wynn, Mairead Hyland, The Lighthouse Lab in Alderley Park and Alex Alderton, Roberto Amato, Jeffrey Barrett, Sonia Goncalves, Ewan Harrison, David K. Jackson, Ian Johnston, Dominic Kwiatkowski, Cordelia Langford, John Sillitoe on behalf of the Wellcome Sanger Institute COVID-19 Surveillance Team |
| EPI_ISL_2766660 | Lighthouse Laboratory Plymouth | Wellcome Sanger Institute for the COVID-19 Genomics UK (COG-UK) Consortium | Lighthouse Laboratory Plymouth and Alex Alderton, Roberto Amato, Jeffrey Barrett, Sonia Goncalves, Ewan Harrison, David K. Jackson, Ian Johnston, Dominic Kwiatkowski, Cordelia Langford, John Sillitoe on behalf of the Wellcome Sanger Institute COVID-19 Surveillance Team |
| EPI_ISL_2766661, EPI_ISL_2766662 | Lighthouse Lab in Alderley Park | Wellcome Sanger Institute for the COVID-19 Genomics UK (COG-UK) Consortium | Jacquelyn Wynn, Mairead Hyland, The Lighthouse Lab in Alderley Park and Alex Alderton, Roberto Amato, Jeffrey Barrett, Sonia Goncalves, Ewan Harrison, David K. Jackson, Ian Johnston, Dominic Kwiatkowski, Cordelia Langford, John Sillitoe on behalf of the Wellcome Sanger Institute COVID-19 Surveillance Team |
| EPI_ISL_2766663 | Lighthouse Laboratory Plymouth | Wellcome Sanger Institute for the COVID-19 Genomics UK (COG-UK) Consortium | Lighthouse Laboratory Plymouth and Alex Alderton, Roberto Amato, Jeffrey Barrett, Sonia Goncalves, Ewan Harrison, David K. Jackson, Ian Johnston, Dominic Kwiatkowski, Cordelia Langford, John Sillitoe on behalf of the Wellcome Sanger Institute COVID-19 Surveillance Team |
| EPI_ISL_2766664 | Lighthouse Lab in Alderley Park | Wellcome Sanger Institute for the COVID-19 Genomics UK (COG-UK) Consortium | Jacquelyn Wynn, Mairead Hyland, The Lighthouse Lab in Alderley Park and Alex Alderton, Roberto Amato, Jeffrey Barrett, Sonia Goncalves, Ewan Harrison, David K. Jackson, Ian Johnston, Dominic Kwiatkowski, Cordelia Langford, John Sillitoe on behalf of the Wellcome Sanger Institute COVID-19 Surveillance Team |
| EPI_ISL_2766665, EPI_ISL_2766666, EPI_ISL_2766667 | Lighthouse Lab in Milton Keynes | Wellcome Sanger Institute for the COVID-19 Genomics UK (COG-UK) Consortium | The Lighthouse Lab in Milton Keynes and Alex Alderton, Roberto Amato, Jeffrey Barrett, Sonia Goncalves, Ewan Harrison, David K. Jackson, Ian Johnston, Dominic Kwiatkowski, Cordelia Langford, John Sillitoe on behalf of the Wellcome Sanger Institute COVID-19 Surveillance Team |
| EPI_ISL_2766668 | Lighthouse Lab in Alderley Park | Wellcome Sanger Institute for the COVID-19 Genomics UK (COG-UK) Consortium | Jacquelyn Wynn, Mairead Hyland, The Lighthouse Lab in Alderley Park and Alex Alderton, Roberto Amato, Jeffrey Barrett, Sonia Goncalves, Ewan Harrison, David K. Jackson, Ian Johnston, Dominic Kwiatkowski, Cordelia Langford, John Sillitoe on behalf of the Wellcome Sanger Institute COVID-19 Surveillance Team |
| EPI_ISL_2766669, EPI_ISL_2766670 | Lighthouse Laboratory Plymouth | Wellcome Sanger Institute for the COVID-19 Genomics UK (COG-UK) Consortium | Lighthouse Laboratory Plymouth and Alex Alderton, Roberto Amato, Jeffrey Barrett, Sonia Goncalves, Ewan Harrison, David K. Jackson, Ian Johnston, Dominic Kwiatkowski, Cordelia Langford, John Sillitoe on behalf of the Wellcome Sanger Institute COVID-19 Surveillance Team |
| EPI_ISL_2766671 | Lighthouse Lab in Milton Keynes | Wellcome Sanger Institute for the COVID-19 Genomics UK (COG-UK) Consortium | The Lighthouse Lab in Milton Keynes and Alex Alderton, Roberto Amato, Jeffrey Barrett, Sonia Goncalves, Ewan Harrison, David K. Jackson, Ian Johnston, Dominic Kwiatkowski, Cordelia Langford, John Sillitoe on behalf of the Wellcome Sanger Institute COVID-19 Surveillance Team |
| EPI_ISL_2766672, EPI_ISL_2766673, EPI_ISL_2766674 | Lighthouse Lab in Alderley Park | Wellcome Sanger Institute for the COVID-19 Genomics UK (COG-UK) Consortium | Jacquelyn Wynn, Mairead Hyland, The Lighthouse Lab in Alderley Park and Alex Alderton, Roberto Amato, Jeffrey Barrett, Sonia Goncalves, Ewan Harrison, David K. Jackson, Ian Johnston, Dominic Kwiatkowski, Cordelia Langford, John Sillitoe on behalf of the Wellcome Sanger Institute COVID-19 Surveillance Team |
| EPI_ISL_2766675 | Lighthouse Lab in Milton Keynes | Wellcome Sanger Institute for the COVID-19 Genomics UK (COG-UK) Consortium | The Lighthouse Lab in Milton Keynes and Alex Alderton, Roberto Amato, Jeffrey Barrett, Sonia Goncalves, Ewan Harrison, David K. Jackson, Ian Johnston, Dominic Kwiatkowski, Cordelia Langford, John Sillitoe on behalf of the Wellcome Sanger Institute COVID-19 Surveillance Team |
| EPI_ISL_2766676, EPI_ISL_2766677 | Lighthouse Lab in Alderley Park | Wellcome Sanger Institute for the COVID-19 Genomics UK (COG-UK) Consortium | Jacquelyn Wynn, Mairead Hyland, The Lighthouse Lab in Alderley Park and Alex Alderton, Roberto Amato, Jeffrey Barrett, Sonia Goncalves, Ewan Harrison, David K. Jackson, Ian Johnston, Dominic Kwiatkowski, Cordelia Langford, John Sillitoe on behalf of the Wellcome Sanger Institute COVID-19 Surveillance Team |
| EPI_ISL_2766678 | Lighthouse Lab in Milton Keynes | Wellcome Sanger Institute for the COVID-19 Genomics UK (COG-UK) Consortium | The Lighthouse Lab in Milton Keynes and Alex Alderton, Roberto Amato, Jeffrey Barrett, Sonia Goncalves, Ewan Harrison, David K. Jackson, Ian Johnston, Dominic Kwiatkowski, Cordelia Langford, John Sillitoe on behalf of the Wellcome Sanger Institute COVID-19 Surveillance Team |
| EPI_ISL_2766679, EPI_ISL_2766680 | Lighthouse Laboratory Plymouth | Wellcome Sanger Institute for the COVID-19 Genomics UK (COG-UK) Consortium | Lighthouse Laboratory Plymouth and Alex Alderton, Roberto Amato, Jeffrey Barrett, Sonia Goncalves, Ewan Harrison, David K. Jackson, Ian Johnston, Dominic Kwiatkowski, Cordelia Langford, John Sillitoe on behalf of the Wellcome Sanger Institute COVID-19 Surveillance Team |
| EPI_ISL_2766681 | Lighthouse Lab in Milton Keynes | Wellcome Sanger Institute for the COVID-19 Genomics UK (COG-UK) Consortium | The Lighthouse Lab in Milton Keynes and Alex Alderton, Roberto Amato, Jeffrey Barrett, Sonia Goncalves, Ewan Harrison, David K. Jackson, Ian Johnston, Dominic Kwiatkowski, Cordelia Langford, John Sillitoe on behalf of the Wellcome Sanger Institute COVID-19 Surveillance Team |
| EPI_ISL_2766682, EPI_ISL_2766683, EPI_ISL_2766684, EPI_ISL_2766685, EPI_ISL_2766686 | Lighthouse Lab in Alderley Park | Wellcome Sanger Institute for the COVID-19 Genomics UK (COG-UK) Consortium | Jacquelyn Wynn, Mairead Hyland, The Lighthouse Lab in Alderley Park and Alex Alderton, Roberto Amato, Jeffrey Barrett, Sonia Goncalves, Ewan Harrison, David K. Jackson, Ian Johnston, Dominic Kwiatkowski, Cordelia Langford, John Sillitoe on behalf of the Wellcome Sanger Institute COVID-19 Surveillance Team |
| EPI_ISL_2766687 | Lighthouse Lab in Milton Keynes | Wellcome Sanger Institute for the COVID-19 Genomics UK (COG-UK) Consortium | The Lighthouse Lab in Milton Keynes and Alex Alderton, Roberto Amato, Jeffrey Barrett, Sonia Goncalves, Ewan Harrison, David K. Jackson, Ian Johnston, Dominic Kwiatkowski, Cordelia Langford, John Sillitoe on behalf of the Wellcome Sanger Institute COVID-19 Surveillance Team |
| EPI_ISL_2766688 | Lighthouse Lab in Alderley Park | Wellcome Sanger Institute for the COVID-19 Genomics UK (COG-UK) Consortium | Jacquelyn Wynn, Mairead Hyland, The Lighthouse Lab in Alderley Park and Alex Alderton, Roberto Amato, Jeffrey Barrett, Sonia Goncalves, Ewan Harrison, David K. Jackson, Ian Johnston, Dominic Kwiatkowski, Cordelia Langford, John Sillitoe on behalf of the Wellcome Sanger Institute COVID-19 Surveillance Team |
| EPI_ISL_2766689 | Lighthouse Laboratory Plymouth | Wellcome Sanger Institute for the COVID-19 Genomics UK (COG-UK) Consortium | Lighthouse Laboratory Plymouth and Alex Alderton, Roberto Amato, Jeffrey Barrett, Sonia Goncalves, Ewan Harrison, David K. Jackson, Ian Johnston, Dominic Kwiatkowski, Cordelia Langford, John Sillitoe on behalf of the Wellcome Sanger Institute COVID-19 Surveillance Team |
| EPI_ISL_2766690 | Lighthouse Lab in Alderley Park | Wellcome Sanger Institute for the COVID-19 Genomics UK (COG-UK) Consortium | Jacquelyn Wynn, Mairead Hyland, The Lighthouse Lab in Alderley Park and Alex Alderton, Roberto Amato, Jeffrey Barrett, Sonia Goncalves, Ewan Harrison, David K. Jackson, Ian Johnston, Dominic Kwiatkowski, Cordelia Langford, John Sillitoe on behalf of the Wellcome Sanger Institute COVID-19 Surveillance Team |
| EPI_ISL_2766691, EPI_ISL_2766692 | Lighthouse Lab in Milton Keynes | Wellcome Sanger Institute for the COVID-19 Genomics UK (COG-UK) Consortium | The Lighthouse Lab in Milton Keynes and Alex Alderton, Roberto Amato, Jeffrey Barrett, Sonia Goncalves, Ewan Harrison, David K. Jackson, Ian Johnston, Dominic Kwiatkowski, Cordelia Langford, John Sillitoe on behalf of the Wellcome Sanger Institute COVID-19 Surveillance Team |
| EPI_ISL_2766693 | Lighthouse Lab in Alderley Park | Wellcome Sanger Institute for the COVID-19 Genomics UK (COG-UK) Consortium | Jacquelyn Wynn, Mairead Hyland, The Lighthouse Lab in Alderley Park and Alex Alderton, Roberto Amato, Jeffrey Barrett, Sonia Goncalves, Ewan Harrison, David K. Jackson, Ian Johnston, Dominic Kwiatkowski, Cordelia Langford, John Sillitoe on behalf of the Wellcome Sanger Institute COVID-19 Surveillance Team |
| EPI_ISL_2766694 | Lighthouse Laboratory Plymouth | Wellcome Sanger Institute for the COVID-19 Genomics UK (COG-UK) Consortium | Lighthouse Laboratory Plymouth and Alex Alderton, Roberto Amato, Jeffrey Barrett, Sonia Goncalves, Ewan Harrison, David K. Jackson, Ian Johnston, Dominic Kwiatkowski, Cordelia Langford, John Sillitoe on behalf of the Wellcome Sanger Institute COVID-19 Surveillance Team |
| EPI_ISL_2766695, EPI_ISL_2766696, EPI_ISL_2766697 | Lighthouse Lab in Alderley Park | Wellcome Sanger Institute for the COVID-19 Genomics UK (COG-UK) Consortium | Jacquelyn Wynn, Mairead Hyland, The Lighthouse Lab in Alderley Park and Alex Alderton, Roberto Amato, Jeffrey Barrett, Sonia Goncalves, Ewan Harrison, David K. Jackson, Ian Johnston, Dominic Kwiatkowski, Cordelia Langford, John Sillitoe on behalf of the Wellcome Sanger Institute COVID-19 Surveillance Team |
| EPI_ISL_2766698 | Lighthouse Lab in Milton Keynes | Wellcome Sanger Institute for the COVID-19 Genomics UK (COG-UK) Consortium | The Lighthouse Lab in Milton Keynes and Alex Alderton, Roberto Amato, Jeffrey Barrett, Sonia Goncalves, Ewan Harrison, David K. Jackson, Ian Johnston, Dominic Kwiatkowski, Cordelia Langford, John Sillitoe on behalf of the Wellcome Sanger Institute COVID-19 Surveillance Team |
| EPI_ISL_2766699 | Lighthouse Lab in Alderley Park | Wellcome Sanger Institute for the COVID-19 Genomics UK (COG-UK) Consortium | Jacquelyn Wynn, Mairead Hyland, The Lighthouse Lab in Alderley Park and Alex Alderton, Roberto Amato, Jeffrey Barrett, Sonia Goncalves, Ewan Harrison, David K. Jackson, Ian Johnston, Dominic Kwiatkowski, Cordelia Langford, John Sillitoe on behalf of the Wellcome Sanger Institute COVID-19 Surveillance Team |
| EPI_ISL_2766700, EPI_ISL_2766701 | Lighthouse Lab in Milton Keynes | Wellcome Sanger Institute for the COVID-19 Genomics UK (COG-UK) Consortium | The Lighthouse Lab in Milton Keynes and Alex Alderton, Roberto Amato, Jeffrey Barrett, Sonia Goncalves, Ewan Harrison, David K. Jackson, Ian Johnston, Dominic Kwiatkowski, Cordelia Langford, John Sillitoe on behalf of the Wellcome Sanger Institute COVID-19 Surveillance Team |
| EPI_ISL_2766702 | Lighthouse Lab in Alderley Park | Wellcome Sanger Institute for the COVID-19 Genomics UK (COG-UK) Consortium | Jacquelyn Wynn, Mairead Hyland, The Lighthouse Lab in Alderley Park and Alex Alderton, Roberto Amato, Jeffrey Barrett, Sonia Goncalves, Ewan Harrison, David K. Jackson, Ian Johnston, Dominic Kwiatkowski, Cordelia Langford, John Sillitoe on behalf of the Wellcome Sanger Institute COVID-19 Surveillance Team |
| EPI_ISL_2766703, EPI_ISL_2766704 | Lighthouse Laboratory Plymouth | Wellcome Sanger Institute for the COVID-19 Genomics UK (COG-UK) Consortium | Lighthouse Laboratory Plymouth and Alex Alderton, Roberto Amato, Jeffrey Barrett, Sonia Goncalves, Ewan Harrison, David K. Jackson, Ian Johnston, Dominic Kwiatkowski, Cordelia Langford, John Sillitoe on behalf of the Wellcome Sanger Institute COVID-19 Surveillance Team |
| EPI_ISL_2766705, EPI_ISL_2766706, EPI_ISL_2766707 | Lighthouse Lab in Alderley Park | Wellcome Sanger Institute for the COVID-19 Genomics UK (COG-UK) Consortium | Jacquelyn Wynn, Mairead Hyland, The Lighthouse Lab in Alderley Park and Alex Alderton, Roberto Amato, Jeffrey Barrett, Sonia Goncalves, Ewan Harrison, David K. Jackson, Ian Johnston, Dominic Kwiatkowski, Cordelia Langford, John Sillitoe on behalf of the Wellcome Sanger Institute COVID-19 Surveillance Team |
| EPI_ISL_2766708 | Lighthouse Laboratory Plymouth | Wellcome Sanger Institute for the COVID-19 Genomics UK (COG-UK) Consortium | Lighthouse Laboratory Plymouth and Alex Alderton, Roberto Amato, Jeffrey Barrett, Sonia Goncalves, Ewan Harrison, David K. Jackson, Ian Johnston, Dominic Kwiatkowski, Cordelia Langford, John Sillitoe on behalf of the Wellcome Sanger Institute COVID-19 Surveillance Team |
| EPI_ISL_2766709, EPI_ISL_2766710 | Lighthouse Lab in Milton Keynes | Wellcome Sanger Institute for the COVID-19 Genomics UK (COG-UK) Consortium | The Lighthouse Lab in Milton Keynes and Alex Alderton, Roberto Amato, Jeffrey Barrett, Sonia Goncalves, Ewan Harrison, David K. Jackson, Ian Johnston, Dominic Kwiatkowski, Cordelia Langford, John Sillitoe on behalf of the Wellcome Sanger Institute COVID-19 Surveillance Team |

| | | | |
|---|---|---|---|
| EPI_ISL_2766711 | Lighthouse Lab in Alderley Park | Wellcome Sanger Institute for the COVID-19 Genomics UK (COG-UK) Consortium | Jacquelyn Wynn, Mairead Hyland, The Lighthouse Lab in Alderley Park and Alex Alderton, Roberto Amato, Jeffrey Barrett, Sonia Goncalves, Ewan Harrison, David K. Jackson, Ian Johnston, Dominic Kwiatkowski, Cordelia Langford, John Sillitoe on behalf of the Wellcome Sanger Institute COVID-19 Surveillance Team |
| EPI_ISL_2766712, EPI_ISL_2766713 | Lighthouse Lab in Milton Keynes | Wellcome Sanger Institute for the COVID-19 Genomics UK (COG-UK) Consortium | The Lighthouse Lab in Milton Keynes and Alex Alderton, Roberto Amato, Jeffrey Barrett, Sonia Goncalves, Ewan Harrison, David K. Jackson, Ian Johnston, Dominic Kwiatkowski, Cordelia Langford, John Sillitoe on behalf of the Wellcome Sanger Institute COVID-19 Surveillance Team |
| EPI_ISL_2766714, EPI_ISL_2766715, EPI_ISL_2766716, EPI_ISL_2766717, EPI_ISL_2766718, EPI_ISL_2766719 | Lighthouse Lab in Alderley Park | Wellcome Sanger Institute for the COVID-19 Genomics UK (COG-UK) Consortium | Jacquelyn Wynn, Mairead Hyland, The Lighthouse Lab in Alderley Park and Alex Alderton, Roberto Amato, Jeffrey Barrett, Sonia Goncalves, Ewan Harrison, David K. Jackson, Ian Johnston, Dominic Kwiatkowski, Cordelia Langford, John Sillitoe on behalf of the Wellcome Sanger Institute COVID-19 Surveillance Team |
| EPI_ISL_2766720 | Lighthouse Lab in Milton Keynes | Wellcome Sanger Institute for the COVID-19 Genomics UK (COG-UK) Consortium | The Lighthouse Lab in Milton Keynes and Alex Alderton, Roberto Amato, Jeffrey Barrett, Sonia Goncalves, Ewan Harrison, David K. Jackson, Ian Johnston, Dominic Kwiatkowski, Cordelia Langford, John Sillitoe on behalf of the Wellcome Sanger Institute COVID-19 Surveillance Team |
| EPI_ISL_2766721 | Lighthouse Lab in Alderley Park | Wellcome Sanger Institute for the COVID-19 Genomics UK (COG-UK) Consortium | Jacquelyn Wynn, Mairead Hyland, The Lighthouse Lab in Alderley Park and Alex Alderton, Roberto Amato, Jeffrey Barrett, Sonia Goncalves, Ewan Harrison, David K. Jackson, Ian Johnston, Dominic Kwiatkowski, Cordelia Langford, John Sillitoe on behalf of the Wellcome Sanger Institute COVID-19 Surveillance Team |
| EPI_ISL_2766722, EPI_ISL_2766723 | Lighthouse Lab in Milton Keynes | Wellcome Sanger Institute for the COVID-19 Genomics UK (COG-UK) Consortium | The Lighthouse Lab in Milton Keynes and Alex Alderton, Roberto Amato, Jeffrey Barrett, Sonia Goncalves, Ewan Harrison, David K. Jackson, Ian Johnston, Dominic Kwiatkowski, Cordelia Langford, John Sillitoe on behalf of the Wellcome Sanger Institute COVID-19 Surveillance Team |
| EPI_ISL_2766724 | Lighthouse Laboratory Plymouth | Wellcome Sanger Institute for the COVID-19 Genomics UK (COG-UK) Consortium | Lighthouse Laboratory Plymouth and Alex Alderton, Roberto Amato, Jeffrey Barrett, Sonia Goncalves, Ewan Harrison, David K. Jackson, Ian Johnston, Dominic Kwiatkowski, Cordelia Langford, John Sillitoe on behalf of the Wellcome Sanger Institute COVID-19 Surveillance Team |
| EPI_ISL_2766725, EPI_ISL_2766726, EPI_ISL_2766727 | Lighthouse Lab in Milton Keynes | Wellcome Sanger Institute for the COVID-19 Genomics UK (COG-UK) Consortium | The Lighthouse Lab in Milton Keynes and Alex Alderton, Roberto Amato, Jeffrey Barrett, Sonia Goncalves, Ewan Harrison, David K. Jackson, Ian Johnston, Dominic Kwiatkowski, Cordelia Langford, John Sillitoe on behalf of the Wellcome Sanger Institute COVID-19 Surveillance Team |
| EPI_ISL_2766728 | Lighthouse Lab in Alderley Park | Wellcome Sanger Institute for the COVID-19 Genomics UK (COG-UK) Consortium | Jacquelyn Wynn, Mairead Hyland, The Lighthouse Lab in Alderley Park and Alex Alderton, Roberto Amato, Jeffrey Barrett, Sonia Goncalves, Ewan Harrison, David K. Jackson, Ian Johnston, Dominic Kwiatkowski, Cordelia Langford, John Sillitoe on behalf of the Wellcome Sanger Institute COVID-19 Surveillance Team |
| EPI_ISL_2766729 | Lighthouse Lab in Milton Keynes | Wellcome Sanger Institute for the COVID-19 Genomics UK (COG-UK) Consortium | The Lighthouse Lab in Milton Keynes and Alex Alderton, Roberto Amato, Jeffrey Barrett, Sonia Goncalves, Ewan Harrison, David K. Jackson, Ian Johnston, Dominic Kwiatkowski, Cordelia Langford, John Sillitoe on behalf of the Wellcome Sanger Institute COVID-19 Surveillance Team |
| EPI_ISL_2766730 | Lighthouse Lab in Alderley Park | Wellcome Sanger Institute for the COVID-19 Genomics UK (COG-UK) Consortium | Jacquelyn Wynn, Mairead Hyland, The Lighthouse Lab in Alderley Park and Alex Alderton, Roberto Amato, Jeffrey Barrett, Sonia Goncalves, Ewan Harrison, David K. Jackson, Ian Johnston, Dominic Kwiatkowski, Cordelia Langford, John Sillitoe on behalf of the Wellcome Sanger Institute COVID-19 Surveillance Team |
| EPI_ISL_2766731, EPI_ISL_2766732, EPI_ISL_2766733 | Lighthouse Lab in Milton Keynes | Wellcome Sanger Institute for the COVID-19 Genomics UK (COG-UK) Consortium | The Lighthouse Lab in Milton Keynes and Alex Alderton, Roberto Amato, Jeffrey Barrett, Sonia Goncalves, Ewan Harrison, David K. Jackson, Ian Johnston, Dominic Kwiatkowski, Cordelia Langford, John Sillitoe on behalf of the Wellcome Sanger Institute COVID-19 Surveillance Team |
| EPI_ISL_2766734 | Lighthouse Laboratory Plymouth | Wellcome Sanger Institute for the COVID-19 Genomics UK (COG-UK) Consortium | Lighthouse Laboratory Plymouth and Alex Alderton, Roberto Amato, Jeffrey Barrett, Sonia Goncalves, Ewan Harrison, David K. Jackson, Ian Johnston, Dominic Kwiatkowski, Cordelia Langford, John Sillitoe on behalf of the Wellcome Sanger Institute COVID-19 Surveillance Team |
| EPI_ISL_2766735 | Lighthouse Lab in Alderley Park | Wellcome Sanger Institute for the COVID-19 Genomics UK (COG-UK) Consortium | Jacquelyn Wynn, Mairead Hyland, The Lighthouse Lab in Alderley Park and Alex Alderton, Roberto Amato, Jeffrey Barrett, Sonia Goncalves, Ewan Harrison, David K. Jackson, Ian Johnston, Dominic Kwiatkowski, Cordelia Langford, John Sillitoe on behalf of the Wellcome Sanger Institute COVID-19 Surveillance Team |
| EPI_ISL_2766736 | Lighthouse Laboratory Plymouth | Wellcome Sanger Institute for the COVID-19 Genomics UK (COG-UK) Consortium | Lighthouse Laboratory Plymouth and Alex Alderton, Roberto Amato, Jeffrey Barrett, Sonia Goncalves, Ewan Harrison, David K. Jackson, Ian Johnston, Dominic Kwiatkowski, Cordelia Langford, John Sillitoe on behalf of the Wellcome Sanger Institute COVID-19 Surveillance Team |
| EPI_ISL_2766737 | Lighthouse Lab in Alderley Park | Wellcome Sanger Institute for the COVID-19 Genomics UK (COG-UK) Consortium | Jacquelyn Wynn, Mairead Hyland, The Lighthouse Lab in Alderley Park and Alex Alderton, Roberto Amato, Jeffrey Barrett, Sonia Goncalves, Ewan Harrison, David K. Jackson, Ian Johnston, Dominic Kwiatkowski, Cordelia Langford, John Sillitoe on behalf of the Wellcome Sanger Institute COVID-19 Surveillance Team |
| EPI_ISL_2766738 | Lighthouse Laboratory Plymouth | Wellcome Sanger Institute for the COVID-19 Genomics UK (COG-UK) Consortium | Lighthouse Laboratory Plymouth and Alex Alderton, Roberto Amato, Jeffrey Barrett, Sonia Goncalves, Ewan Harrison, David K. Jackson, Ian Johnston, Dominic Kwiatkowski, Cordelia Langford, John Sillitoe on behalf of the Wellcome Sanger Institute COVID-19 Surveillance Team |
| EPI_ISL_2766739, EPI_ISL_2766740 | Lighthouse Lab in Milton Keynes | Wellcome Sanger Institute for the COVID-19 Genomics UK (COG-UK) Consortium | The Lighthouse Lab in Milton Keynes and Alex Alderton, Roberto Amato, Jeffrey Barrett, Sonia Goncalves, Ewan Harrison, David K. Jackson, Ian Johnston, Dominic Kwiatkowski, Cordelia Langford, John Sillitoe on behalf of the Wellcome Sanger Institute COVID-19 Surveillance Team |
| EPI_ISL_2766741 | Lighthouse Lab in Alderley Park | Wellcome Sanger Institute for the COVID-19 Genomics UK (COG-UK) Consortium | Jacquelyn Wynn, Mairead Hyland, The Lighthouse Lab in Alderley Park and Alex Alderton, Roberto Amato, Jeffrey Barrett, Sonia Goncalves, Ewan Harrison, David K. Jackson, Ian Johnston, Dominic Kwiatkowski, Cordelia Langford, John Sillitoe on behalf of the Wellcome Sanger Institute COVID-19 Surveillance Team |
| EPI_ISL_2766742 | Lighthouse Laboratory Plymouth | Wellcome Sanger Institute for the COVID-19 Genomics UK (COG-UK) Consortium | Lighthouse Laboratory Plymouth and Alex Alderton, Roberto Amato, Jeffrey Barrett, Sonia Goncalves, Ewan Harrison, David K. Jackson, Ian Johnston, Dominic Kwiatkowski, Cordelia Langford, John Sillitoe on behalf of the Wellcome Sanger Institute COVID-19 Surveillance Team |
| EPI_ISL_2766743 | Lighthouse Lab in Alderley Park | Wellcome Sanger Institute for the COVID-19 Genomics UK (COG-UK) Consortium | Jacquelyn Wynn, Mairead Hyland, The Lighthouse Lab in Alderley Park and Alex Alderton, Roberto Amato, Jeffrey Barrett, Sonia Goncalves, Ewan Harrison, David K. Jackson, Ian Johnston, Dominic Kwiatkowski, Cordelia Langford, John Sillitoe on behalf of the Wellcome Sanger Institute COVID-19 Surveillance Team |
| EPI_ISL_2766744 | Lighthouse Lab in Milton Keynes | Wellcome Sanger Institute for the COVID-19 Genomics UK (COG-UK) Consortium | The Lighthouse Lab in Milton Keynes and Alex Alderton, Roberto Amato, Jeffrey Barrett, Sonia Goncalves, Ewan Harrison, David K. Jackson, Ian Johnston, Dominic Kwiatkowski, Cordelia Langford, John Sillitoe on behalf of the Wellcome Sanger Institute COVID-19 Surveillance Team |
| EPI_ISL_2766745 | Lighthouse Lab in Alderley Park | Wellcome Sanger Institute for the COVID-19 Genomics UK (COG-UK) Consortium | Jacquelyn Wynn, Mairead Hyland, The Lighthouse Lab in Alderley Park and Alex Alderton, Roberto Amato, Jeffrey Barrett, Sonia Goncalves, Ewan Harrison, David K. Jackson, Ian Johnston, Dominic Kwiatkowski, Cordelia Langford, John Sillitoe on behalf of the Wellcome Sanger Institute COVID-19 Surveillance Team |
| EPI_ISL_2766746, EPI_ISL_2766747 | Lighthouse Lab in Milton Keynes | Wellcome Sanger Institute for the COVID-19 Genomics UK (COG-UK) Consortium | The Lighthouse Lab in Milton Keynes and Alex Alderton, Roberto Amato, Jeffrey Barrett, Sonia Goncalves, Ewan Harrison, David K. Jackson, Ian Johnston, Dominic Kwiatkowski, Cordelia Langford, John Sillitoe on behalf of the Wellcome Sanger Institute COVID-19 Surveillance Team |
| EPI_ISL_2766748 | Lighthouse Laboratory Plymouth | Wellcome Sanger Institute for the COVID-19 Genomics UK (COG-UK) Consortium | Lighthouse Laboratory Plymouth and Alex Alderton, Roberto Amato, Jeffrey Barrett, Sonia Goncalves, Ewan Harrison, David K. Jackson, Ian Johnston, Dominic Kwiatkowski, Cordelia Langford, John Sillitoe on behalf of the Wellcome Sanger Institute COVID-19 Surveillance Team |
| EPI_ISL_2766749 | Lighthouse Lab in Milton Keynes | Wellcome Sanger Institute for the COVID-19 Genomics UK (COG-UK) Consortium | The Lighthouse Lab in Milton Keynes and Alex Alderton, Roberto Amato, Jeffrey Barrett, Sonia Goncalves, Ewan Harrison, David K. Jackson, Ian Johnston, Dominic Kwiatkowski, Cordelia Langford, John Sillitoe on behalf of the Wellcome Sanger Institute COVID-19 Surveillance Team |
| EPI_ISL_2766750, EPI_ISL_2766751 | Lighthouse Lab in Alderley Park | Wellcome Sanger Institute for the COVID-19 Genomics UK (COG-UK) Consortium | Jacquelyn Wynn, Mairead Hyland, The Lighthouse Lab in Alderley Park and Alex Alderton, Roberto Amato, Jeffrey Barrett, Sonia Goncalves, Ewan Harrison, David K. Jackson, Ian Johnston, Dominic Kwiatkowski, Cordelia Langford, John Sillitoe on behalf of the Wellcome Sanger Institute COVID-19 Surveillance Team |
| EPI_ISL_2766752, EPI_ISL_2766753 | Lighthouse Lab in Milton Keynes | Wellcome Sanger Institute for the COVID-19 Genomics UK (COG-UK) Consortium | The Lighthouse Lab in Milton Keynes and Alex Alderton, Roberto Amato, Jeffrey Barrett, Sonia Goncalves, Ewan Harrison, David K. Jackson, Ian Johnston, Dominic Kwiatkowski, Cordelia Langford, John Sillitoe on behalf of the Wellcome Sanger Institute COVID-19 Surveillance Team |
| EPI_ISL_2766754 | Lighthouse Laboratory Plymouth | Wellcome Sanger Institute for the COVID-19 Genomics UK (COG-UK) Consortium | Lighthouse Laboratory Plymouth and Alex Alderton, Roberto Amato, Jeffrey Barrett, Sonia Goncalves, Ewan Harrison, David K. Jackson, Ian Johnston, Dominic Kwiatkowski, Cordelia Langford, John Sillitoe on behalf of the Wellcome Sanger Institute COVID-19 Surveillance Team |
| EPI_ISL_2766755, EPI_ISL_2766756, EPI_ISL_2766757, EPI_ISL_2766758, EPI_ISL_2766759, EPI_ISL_2766760, EPI_ISL_2766761, EPI_ISL_2766762, EPI_ISL_2766763 | Lighthouse Lab in Alderley Park | Wellcome Sanger Institute for the COVID-19 Genomics UK (COG-UK) Consortium | Jacquelyn Wynn, Mairead Hyland, The Lighthouse Lab in Alderley Park and Alex Alderton, Roberto Amato, Jeffrey Barrett, Sonia Goncalves, Ewan Harrison, David K. Jackson, Ian Johnston, Dominic Kwiatkowski, Cordelia Langford, John Sillitoe on behalf of the Wellcome Sanger Institute COVID-19 Surveillance Team |
| EPI_ISL_2766764, EPI_ISL_2766765, EPI_ISL_2766766 | Lighthouse Lab in Milton Keynes | Wellcome Sanger Institute for the COVID-19 Genomics UK (COG-UK) Consortium | The Lighthouse Lab in Milton Keynes and Alex Alderton, Roberto Amato, Jeffrey Barrett, Sonia Goncalves, Ewan Harrison, David K. Jackson, Ian Johnston, Dominic Kwiatkowski, Cordelia Langford, John Sillitoe on behalf of the Wellcome Sanger Institute COVID-19 Surveillance Team |
| EPI_ISL_2766767, EPI_ISL_2766768 | Lighthouse Lab in Alderley Park | Wellcome Sanger Institute for the COVID-19 Genomics UK (COG-UK) Consortium | Jacquelyn Wynn, Mairead Hyland, The Lighthouse Lab in Alderley Park and Alex Alderton, Roberto Amato, Jeffrey Barrett, Sonia Goncalves, Ewan Harrison, David K. Jackson, Ian Johnston, Dominic Kwiatkowski, Cordelia Langford, John Sillitoe on behalf of the Wellcome Sanger Institute COVID-19 Surveillance Team |
| EPI_ISL_2766769 | Lighthouse Lab in Milton Keynes | Wellcome Sanger Institute for the COVID-19 Genomics UK (COG-UK) Consortium | The Lighthouse Lab in Milton Keynes and Alex Alderton, Roberto Amato, Jeffrey Barrett, Sonia Goncalves, Ewan Harrison, David K. Jackson, Ian Johnston, Dominic Kwiatkowski, Cordelia Langford, John Sillitoe on behalf of the Wellcome Sanger Institute COVID-19 Surveillance Team |
| EPI_ISL_2766770 | Lighthouse Laboratory Plymouth | Wellcome Sanger Institute for the COVID-19 Genomics UK (COG-UK) Consortium | Lighthouse Laboratory Plymouth and Alex Alderton, Roberto Amato, Jeffrey Barrett, Sonia Goncalves, Ewan Harrison, David K. Jackson, Ian Johnston, Dominic Kwiatkowski, Cordelia Langford, John Sillitoe on behalf of the Wellcome Sanger Institute COVID-19 Surveillance Team |
| EPI_ISL_2766771 | Lighthouse Lab in Milton Keynes | Wellcome Sanger Institute for the COVID-19 Genomics UK (COG-UK) Consortium | The Lighthouse Lab in Milton Keynes and Alex Alderton, Roberto Amato, Jeffrey Barrett, Sonia Goncalves, Ewan Harrison, David K. Jackson, Ian Johnston, Dominic Kwiatkowski, Cordelia Langford, John Sillitoe on behalf of the Wellcome Sanger Institute COVID-19 Surveillance Team |
| EPI_ISL_2766772, EPI_ISL_2766773 | Lighthouse Lab in Alderley Park | Wellcome Sanger Institute for the COVID-19 Genomics UK (COG-UK) Consortium | Jacquelyn Wynn, Mairead Hyland, The Lighthouse Lab in Alderley Park and Alex Alderton, Roberto Amato, Jeffrey Barrett, Sonia Goncalves, Ewan Harrison, David K. Jackson, Ian Johnston, Dominic Kwiatkowski, Cordelia Langford, John Sillitoe on behalf of the Wellcome Sanger Institute COVID-19 Surveillance Team |
| EPI_ISL_2766774 | Lighthouse Lab in Milton Keynes | Wellcome Sanger Institute for the COVID-19 Genomics UK (COG-UK) Consortium | The Lighthouse Lab in Milton Keynes and Alex Alderton, Roberto Amato, Jeffrey Barrett, Sonia Goncalves, Ewan Harrison, David K. Jackson, Ian Johnston, Dominic Kwiatkowski, Cordelia Langford, John Sillitoe on behalf of the Wellcome Sanger Institute COVID-19 Surveillance Team |
| EPI_ISL_2766775 | Lighthouse Laboratory Plymouth | Wellcome Sanger Institute for the COVID-19 Genomics UK (COG-UK) Consortium | Lighthouse Laboratory Plymouth and Alex Alderton, Roberto Amato, Jeffrey Barrett, Sonia Goncalves, Ewan Harrison, David K. Jackson, Ian Johnston, Dominic Kwiatkowski, Cordelia Langford, John Sillitoe on behalf of the Wellcome Sanger Institute COVID-19 Surveillance Team |
| EPI_ISL_2766776 | Lighthouse Lab in Milton Keynes | Wellcome Sanger Institute for the COVID-19 Genomics UK (COG-UK) Consortium | The Lighthouse Lab in Milton Keynes and Alex Alderton, Roberto Amato, Jeffrey Barrett, Sonia Goncalves, Ewan Harrison, David K. Jackson, Ian Johnston, Dominic Kwiatkowski, Cordelia Langford, John Sillitoe on behalf of the Wellcome Sanger Institute COVID-19 Surveillance Team |

| | | | |
|---|---|---|---|
| EPI_ISL_2766777 | Lighthouse Lab in Alderley Park | Wellcome Sanger Institute for the COVID-19 Genomics UK (COG-UK) Consortium | Jacquelyn Wynn, Mairead Hyland, The Lighthouse Lab in Alderley Park and Alex Alderton, Roberto Amato, Jeffrey Barrett, Sonia Goncalves, Ewan Harrison, David K. Jackson, Ian Johnston, Dominic Kwiatkowski, Cordelia Langford, John Sillitoe on behalf of the Wellcome Sanger Institute COVID-19 Surveillance Team |
| EPI_ISL_2766778 | Lighthouse Lab in Milton Keynes | Wellcome Sanger Institute for the COVID-19 Genomics UK (COG-UK) Consortium | The Lighthouse Lab in Milton Keynes and Alex Alderton, Roberto Amato, Jeffrey Barrett, Sonia Goncalves, Ewan Harrison, David K. Jackson, Ian Johnston, Dominic Kwiatkowski, Cordelia Langford, John Sillitoe on behalf of the Wellcome Sanger Institute COVID-19 Surveillance Team |
| EPI_ISL_2766779, EPI_ISL_2766780 | Lighthouse Lab in Alderley Park | Wellcome Sanger Institute for the COVID-19 Genomics UK (COG-UK) Consortium | Jacquelyn Wynn, Mairead Hyland, The Lighthouse Lab in Alderley Park and Alex Alderton, Roberto Amato, Jeffrey Barrett, Sonia Goncalves, Ewan Harrison, David K. Jackson, Ian Johnston, Dominic Kwiatkowski, Cordelia Langford, John Sillitoe on behalf of the Wellcome Sanger Institute COVID-19 Surveillance Team |
| EPI_ISL_2766781, EPI_ISL_2766782 | Lighthouse Laboratory Plymouth | Wellcome Sanger Institute for the COVID-19 Genomics UK (COG-UK) Consortium | Lighthouse Laboratory Plymouth and Alex Alderton, Roberto Amato, Jeffrey Barrett, Sonia Goncalves, Ewan Harrison, David K. Jackson, Ian Johnston, Dominic Kwiatkowski, Cordelia Langford, John Sillitoe on behalf of the Wellcome Sanger Institute COVID-19 Surveillance Team |
| EPI_ISL_2766783, EPI_ISL_2766784, EPI_ISL_2766785, EPI_ISL_2766786, EPI_ISL_2766787, EPI_ISL_2766788 | Lighthouse Lab in Alderley Park | Wellcome Sanger Institute for the COVID-19 Genomics UK (COG-UK) Consortium | Jacquelyn Wynn, Mairead Hyland, The Lighthouse Lab in Alderley Park and Alex Alderton, Roberto Amato, Jeffrey Barrett, Sonia Goncalves, Ewan Harrison, David K. Jackson, Ian Johnston, Dominic Kwiatkowski, Cordelia Langford, John Sillitoe on behalf of the Wellcome Sanger Institute COVID-19 Surveillance Team |
| EPI_ISL_2766789 | Lighthouse Lab in Milton Keynes | Wellcome Sanger Institute for the COVID-19 Genomics UK (COG-UK) Consortium | The Lighthouse Lab in Milton Keynes and Alex Alderton, Roberto Amato, Jeffrey Barrett, Sonia Goncalves, Ewan Harrison, David K. Jackson, Ian Johnston, Dominic Kwiatkowski, Cordelia Langford, John Sillitoe on behalf of the Wellcome Sanger Institute COVID-19 Surveillance Team |
| EPI_ISL_2766790 | Lighthouse Laboratory Plymouth | Wellcome Sanger Institute for the COVID-19 Genomics UK (COG-UK) Consortium | Lighthouse Laboratory Plymouth and Alex Alderton, Roberto Amato, Jeffrey Barrett, Sonia Goncalves, Ewan Harrison, David K. Jackson, Ian Johnston, Dominic Kwiatkowski, Cordelia Langford, John Sillitoe on behalf of the Wellcome Sanger Institute COVID-19 Surveillance Team |
| EPI_ISL_2766791, EPI_ISL_2766792 | Lighthouse Lab in Milton Keynes | Wellcome Sanger Institute for the COVID-19 Genomics UK (COG-UK) Consortium | The Lighthouse Lab in Milton Keynes and Alex Alderton, Roberto Amato, Jeffrey Barrett, Sonia Goncalves, Ewan Harrison, David K. Jackson, Ian Johnston, Dominic Kwiatkowski, Cordelia Langford, John Sillitoe on behalf of the Wellcome Sanger Institute COVID-19 Surveillance Team |
| EPI_ISL_2766793 | Lighthouse Lab in Alderley Park | Wellcome Sanger Institute for the COVID-19 Genomics UK (COG-UK) Consortium | Jacquelyn Wynn, Mairead Hyland, The Lighthouse Lab in Alderley Park and Alex Alderton, Roberto Amato, Jeffrey Barrett, Sonia Goncalves, Ewan Harrison, David K. Jackson, Ian Johnston, Dominic Kwiatkowski, Cordelia Langford, John Sillitoe on behalf of the Wellcome Sanger Institute COVID-19 Surveillance Team |
| EPI_ISL_2766794 | Lighthouse Laboratory Plymouth | Wellcome Sanger Institute for the COVID-19 Genomics UK (COG-UK) Consortium | Lighthouse Laboratory Plymouth and Alex Alderton, Roberto Amato, Jeffrey Barrett, Sonia Goncalves, Ewan Harrison, David K. Jackson, Ian Johnston, Dominic Kwiatkowski, Cordelia Langford, John Sillitoe on behalf of the Wellcome Sanger Institute COVID-19 Surveillance Team |
| EPI_ISL_2766795 | Lighthouse Lab in Milton Keynes | Wellcome Sanger Institute for the COVID-19 Genomics UK (COG-UK) Consortium | The Lighthouse Lab in Milton Keynes and Alex Alderton, Roberto Amato, Jeffrey Barrett, Sonia Goncalves, Ewan Harrison, David K. Jackson, Ian Johnston, Dominic Kwiatkowski, Cordelia Langford, John Sillitoe on behalf of the Wellcome Sanger Institute COVID-19 Surveillance Team |
| EPI_ISL_2766796 | Lighthouse Laboratory Plymouth | Wellcome Sanger Institute for the COVID-19 Genomics UK (COG-UK) Consortium | Lighthouse Laboratory Plymouth and Alex Alderton, Roberto Amato, Jeffrey Barrett, Sonia Goncalves, Ewan Harrison, David K. Jackson, Ian Johnston, Dominic Kwiatkowski, Cordelia Langford, John Sillitoe on behalf of the Wellcome Sanger Institute COVID-19 Surveillance Team |
| EPI_ISL_2766797 | Lighthouse Lab in Milton Keynes | Wellcome Sanger Institute for the COVID-19 Genomics UK (COG-UK) Consortium | The Lighthouse Lab in Milton Keynes and Alex Alderton, Roberto Amato, Jeffrey Barrett, Sonia Goncalves, Ewan Harrison, David K. Jackson, Ian Johnston, Dominic Kwiatkowski, Cordelia Langford, John Sillitoe on behalf of the Wellcome Sanger Institute COVID-19 Surveillance Team |
| EPI_ISL_2766798 | Lighthouse Lab in Alderley Park | Wellcome Sanger Institute for the COVID-19 Genomics UK (COG-UK) Consortium | Jacquelyn Wynn, Mairead Hyland, The Lighthouse Lab in Alderley Park and Alex Alderton, Roberto Amato, Jeffrey Barrett, Sonia Goncalves, Ewan Harrison, David K. Jackson, Ian Johnston, Dominic Kwiatkowski, Cordelia Langford, John Sillitoe on behalf of the Wellcome Sanger Institute COVID-19 Surveillance Team |
| EPI_ISL_2766799, EPI_ISL_2766800 | Lighthouse Lab in Milton Keynes | Wellcome Sanger Institute for the COVID-19 Genomics UK (COG-UK) Consortium | The Lighthouse Lab in Milton Keynes and Alex Alderton, Roberto Amato, Jeffrey Barrett, Sonia Goncalves, Ewan Harrison, David K. Jackson, Ian Johnston, Dominic Kwiatkowski, Cordelia Langford, John Sillitoe on behalf of the Wellcome Sanger Institute COVID-19 Surveillance Team |
| EPI_ISL_2766801, EPI_ISL_2766802 | Lighthouse Lab in Alderley Park | Wellcome Sanger Institute for the COVID-19 Genomics UK (COG-UK) Consortium | Jacquelyn Wynn, Mairead Hyland, The Lighthouse Lab in Alderley Park and Alex Alderton, Roberto Amato, Jeffrey Barrett, Sonia Goncalves, Ewan Harrison, David K. Jackson, Ian Johnston, Dominic Kwiatkowski, Cordelia Langford, John Sillitoe on behalf of the Wellcome Sanger Institute COVID-19 Surveillance Team |
| EPI_ISL_2766803 | Lighthouse Lab in Milton Keynes | Wellcome Sanger Institute for the COVID-19 Genomics UK (COG-UK) Consortium | The Lighthouse Lab in Milton Keynes and Alex Alderton, Roberto Amato, Jeffrey Barrett, Sonia Goncalves, Ewan Harrison, David K. Jackson, Ian Johnston, Dominic Kwiatkowski, Cordelia Langford, John Sillitoe on behalf of the Wellcome Sanger Institute COVID-19 Surveillance Team |
| EPI_ISL_2766804 | Lighthouse Lab in Alderley Park | Wellcome Sanger Institute for the COVID-19 Genomics UK (COG-UK) Consortium | Jacquelyn Wynn, Mairead Hyland, The Lighthouse Lab in Alderley Park and Alex Alderton, Roberto Amato, Jeffrey Barrett, Sonia Goncalves, Ewan Harrison, David K. Jackson, Ian Johnston, Dominic Kwiatkowski, Cordelia Langford, John Sillitoe on behalf of the Wellcome Sanger Institute COVID-19 Surveillance Team |
| EPI_ISL_2766805 | Lighthouse Lab in Milton Keynes | Wellcome Sanger Institute for the COVID-19 Genomics UK (COG-UK) Consortium | The Lighthouse Lab in Milton Keynes and Alex Alderton, Roberto Amato, Jeffrey Barrett, Sonia Goncalves, Ewan Harrison, David K. Jackson, Ian Johnston, Dominic Kwiatkowski, Cordelia Langford, John Sillitoe on behalf of the Wellcome Sanger Institute COVID-19 Surveillance Team |
| EPI_ISL_2766806, EPI_ISL_2766807 | Lighthouse Lab in Alderley Park | Wellcome Sanger Institute for the COVID-19 Genomics UK (COG-UK) Consortium | Jacquelyn Wynn, Mairead Hyland, The Lighthouse Lab in Alderley Park and Alex Alderton, Roberto Amato, Jeffrey Barrett, Sonia Goncalves, Ewan Harrison, David K. Jackson, Ian Johnston, Dominic Kwiatkowski, Cordelia Langford, John Sillitoe on behalf of the Wellcome Sanger Institute COVID-19 Surveillance Team |
| EPI_ISL_2766808 | Lighthouse Lab in Milton Keynes | Wellcome Sanger Institute for the COVID-19 Genomics UK (COG-UK) Consortium | The Lighthouse Lab in Milton Keynes and Alex Alderton, Roberto Amato, Jeffrey Barrett, Sonia Goncalves, Ewan Harrison, David K. Jackson, Ian Johnston, Dominic Kwiatkowski, Cordelia Langford, John Sillitoe on behalf of the Wellcome Sanger Institute COVID-19 Surveillance Team |
| EPI_ISL_2766809, EPI_ISL_2766810 | Lighthouse Laboratory Plymouth | Wellcome Sanger Institute for the COVID-19 Genomics UK (COG-UK) Consortium | Lighthouse Laboratory Plymouth and Alex Alderton, Roberto Amato, Jeffrey Barrett, Sonia Goncalves, Ewan Harrison, David K. Jackson, Ian Johnston, Dominic Kwiatkowski, Cordelia Langford, John Sillitoe on behalf of the Wellcome Sanger Institute COVID-19 Surveillance Team |
| EPI_ISL_2766811, EPI_ISL_2766812 | Lighthouse Lab in Alderley Park | Wellcome Sanger Institute for the COVID-19 Genomics UK (COG-UK) Consortium | Jacquelyn Wynn, Mairead Hyland, The Lighthouse Lab in Alderley Park and Alex Alderton, Roberto Amato, Jeffrey Barrett, Sonia Goncalves, Ewan Harrison, David K. Jackson, Ian Johnston, Dominic Kwiatkowski, Cordelia Langford, John Sillitoe on behalf of the Wellcome Sanger Institute COVID-19 Surveillance Team |
| EPI_ISL_2766813 | Lighthouse Lab in Milton Keynes | Wellcome Sanger Institute for the COVID-19 Genomics UK (COG-UK) Consortium | The Lighthouse Lab in Milton Keynes and Alex Alderton, Roberto Amato, Jeffrey Barrett, Sonia Goncalves, Ewan Harrison, David K. Jackson, Ian Johnston, Dominic Kwiatkowski, Cordelia Langford, John Sillitoe on behalf of the Wellcome Sanger Institute COVID-19 Surveillance Team |
| EPI_ISL_2766814, EPI_ISL_2766815 | Lighthouse Laboratory Plymouth | Wellcome Sanger Institute for the COVID-19 Genomics UK (COG-UK) Consortium | Lighthouse Laboratory Plymouth and Alex Alderton, Roberto Amato, Jeffrey Barrett, Sonia Goncalves, Ewan Harrison, David K. Jackson, Ian Johnston, Dominic Kwiatkowski, Cordelia Langford, John Sillitoe on behalf of the Wellcome Sanger Institute COVID-19 Surveillance Team |
| EPI_ISL_2766816 | Lighthouse Lab in Alderley Park | Wellcome Sanger Institute for the COVID-19 Genomics UK (COG-UK) Consortium | Jacquelyn Wynn, Mairead Hyland, The Lighthouse Lab in Alderley Park and Alex Alderton, Roberto Amato, Jeffrey Barrett, Sonia Goncalves, Ewan Harrison, David K. Jackson, Ian Johnston, Dominic Kwiatkowski, Cordelia Langford, John Sillitoe on behalf of the Wellcome Sanger Institute COVID-19 Surveillance Team |
| EPI_ISL_2766817 | Lighthouse Lab in Milton Keynes | Wellcome Sanger Institute for the COVID-19 Genomics UK (COG-UK) Consortium | The Lighthouse Lab in Milton Keynes and Alex Alderton, Roberto Amato, Jeffrey Barrett, Sonia Goncalves, Ewan Harrison, David K. Jackson, Ian Johnston, Dominic Kwiatkowski, Cordelia Langford, John Sillitoe on behalf of the Wellcome Sanger Institute COVID-19 Surveillance Team |
| EPI_ISL_2766818 | Lighthouse Lab in Alderley Park | Wellcome Sanger Institute for the COVID-19 Genomics UK (COG-UK) Consortium | Jacquelyn Wynn, Mairead Hyland, The Lighthouse Lab in Alderley Park and Alex Alderton, Roberto Amato, Jeffrey Barrett, Sonia Goncalves, Ewan Harrison, David K. Jackson, Ian Johnston, Dominic Kwiatkowski, Cordelia Langford, John Sillitoe on behalf of the Wellcome Sanger Institute COVID-19 Surveillance Team |
| EPI_ISL_2766819 | Lighthouse Lab in Milton Keynes | Wellcome Sanger Institute for the COVID-19 Genomics UK (COG-UK) Consortium | The Lighthouse Lab in Milton Keynes and Alex Alderton, Roberto Amato, Jeffrey Barrett, Sonia Goncalves, Ewan Harrison, David K. Jackson, Ian Johnston, Dominic Kwiatkowski, Cordelia Langford, John Sillitoe on behalf of the Wellcome Sanger Institute COVID-19 Surveillance Team |
| EPI_ISL_2766820, EPI_ISL_2766821, EPI_ISL_2766822 | Lighthouse Lab in Alderley Park | Wellcome Sanger Institute for the COVID-19 Genomics UK (COG-UK) Consortium | Jacquelyn Wynn, Mairead Hyland, The Lighthouse Lab in Alderley Park and Alex Alderton, Roberto Amato, Jeffrey Barrett, Sonia Goncalves, Ewan Harrison, David K. Jackson, Ian Johnston, Dominic Kwiatkowski, Cordelia Langford, John Sillitoe on behalf of the Wellcome Sanger Institute COVID-19 Surveillance Team |
| EPI_ISL_2766823, EPI_ISL_2766824 | Lighthouse Lab in Milton Keynes | Wellcome Sanger Institute for the COVID-19 Genomics UK (COG-UK) Consortium | The Lighthouse Lab in Milton Keynes and Alex Alderton, Roberto Amato, Jeffrey Barrett, Sonia Goncalves, Ewan Harrison, David K. Jackson, Ian Johnston, Dominic Kwiatkowski, Cordelia Langford, John Sillitoe on behalf of the Wellcome Sanger Institute COVID-19 Surveillance Team |
| EPI_ISL_2766825, EPI_ISL_2766826, EPI_ISL_2766827, EPI_ISL_2766828 | Lighthouse Lab in Alderley Park | Wellcome Sanger Institute for the COVID-19 Genomics UK (COG-UK) Consortium | Jacquelyn Wynn, Mairead Hyland, The Lighthouse Lab in Alderley Park and Alex Alderton, Roberto Amato, Jeffrey Barrett, Sonia Goncalves, Ewan Harrison, David K. Jackson, Ian Johnston, Dominic Kwiatkowski, Cordelia Langford, John Sillitoe on behalf of the Wellcome Sanger Institute COVID-19 Surveillance Team |
| EPI_ISL_2766829 | Lighthouse Laboratory Plymouth | Wellcome Sanger Institute for the COVID-19 Genomics UK (COG-UK) Consortium | Lighthouse Laboratory Plymouth and Alex Alderton, Roberto Amato, Jeffrey Barrett, Sonia Goncalves, Ewan Harrison, David K. Jackson, Ian Johnston, Dominic Kwiatkowski, Cordelia Langford, John Sillitoe on behalf of the Wellcome Sanger Institute COVID-19 Surveillance Team |
| EPI_ISL_2766830 | Lighthouse Lab in Alderley Park | Wellcome Sanger Institute for the COVID-19 Genomics UK (COG-UK) Consortium | Jacquelyn Wynn, Mairead Hyland, The Lighthouse Lab in Alderley Park and Alex Alderton, Roberto Amato, Jeffrey Barrett, Sonia Goncalves, Ewan Harrison, David K. Jackson, Ian Johnston, Dominic Kwiatkowski, Cordelia Langford, John Sillitoe on behalf of the Wellcome Sanger Institute COVID-19 Surveillance Team |
| EPI_ISL_2766831 | Lighthouse Lab in Milton Keynes | Wellcome Sanger Institute for the COVID-19 Genomics UK (COG-UK) Consortium | The Lighthouse Lab in Milton Keynes and Alex Alderton, Roberto Amato, Jeffrey Barrett, Sonia Goncalves, Ewan Harrison, David K. Jackson, Ian Johnston, Dominic Kwiatkowski, Cordelia Langford, John Sillitoe on behalf of the Wellcome Sanger Institute COVID-19 Surveillance Team |
| EPI_ISL_2766832 | Lighthouse Lab in Alderley Park | Wellcome Sanger Institute for the COVID-19 Genomics UK (COG-UK) Consortium | Jacquelyn Wynn, Mairead Hyland, The Lighthouse Lab in Alderley Park and Alex Alderton, Roberto Amato, Jeffrey Barrett, Sonia Goncalves, Ewan Harrison, David K. Jackson, Ian Johnston, Dominic Kwiatkowski, Cordelia Langford, John Sillitoe on behalf of the Wellcome Sanger Institute COVID-19 Surveillance Team |
| EPI_ISL_2766833 | Lighthouse Lab in Milton Keynes | Wellcome Sanger Institute for the COVID-19 Genomics UK (COG-UK) Consortium | The Lighthouse Lab in Milton Keynes and Alex Alderton, Roberto Amato, Jeffrey Barrett, Sonia Goncalves, Ewan Harrison, David K. Jackson, Ian Johnston, Dominic Kwiatkowski, Cordelia Langford, John Sillitoe on behalf of the Wellcome Sanger Institute COVID-19 Surveillance Team |
| EPI_ISL_2766834, EPI_ISL_2766835 | Lighthouse Lab in Alderley Park | Wellcome Sanger Institute for the COVID-19 Genomics UK (COG-UK) Consortium | Jacquelyn Wynn, Mairead Hyland, The Lighthouse Lab in Alderley Park and Alex Alderton, Roberto Amato, Jeffrey Barrett, Sonia Goncalves, Ewan Harrison, David K. Jackson, Ian Johnston, Dominic Kwiatkowski, Cordelia Langford, John Sillitoe on behalf of the Wellcome Sanger Institute COVID-19 Surveillance Team |
| EPI_ISL_2766836 | Lighthouse Lab in Milton Keynes | Wellcome Sanger Institute for the COVID-19 Genomics UK (COG-UK) Consortium | The Lighthouse Lab in Milton Keynes and Alex Alderton, Roberto Amato, Jeffrey Barrett, Sonia Goncalves, Ewan Harrison, David K. Jackson, Ian Johnston, Dominic Kwiatkowski, Cordelia Langford, John Sillitoe on behalf of the Wellcome Sanger Institute COVID-19 Surveillance Team |
| EPI_ISL_2766837, EPI_ISL_2766838 | Lighthouse Lab in Alderley Park | Wellcome Sanger Institute for the COVID-19 Genomics UK (COG-UK) Consortium | Jacquelyn Wynn, Mairead Hyland, The Lighthouse Lab in Alderley Park and Alex Alderton, Roberto Amato, Jeffrey Barrett, Sonia Goncalves, Ewan Harrison, David K. Jackson, Ian Johnston, Dominic Kwiatkowski, Cordelia Langford, John Sillitoe on behalf of the Wellcome Sanger Institute COVID-19 Surveillance Team |
| EPI_ISL_2766839 | Lighthouse Lab in Milton Keynes | Wellcome Sanger Institute for the COVID-19 Genomics UK (COG-UK) Consortium | The Lighthouse Lab in Milton Keynes and Alex Alderton, Roberto Amato, Jeffrey Barrett, Sonia Goncalves, Ewan Harrison, David K. Jackson, Ian Johnston, Dominic Kwiatkowski, Cordelia Langford, John Sillitoe on behalf of the Wellcome Sanger Institute COVID-19 Surveillance Team |
| EPI_ISL_2766840 | Lighthouse Laboratory Plymouth | Wellcome Sanger Institute for the COVID-19 Genomics UK (COG-UK) Consortium | Lighthouse Laboratory Plymouth and Alex Alderton, Roberto Amato, Jeffrey Barrett, Sonia Goncalves, Ewan Harrison, David K. Jackson, Ian Johnston, Dominic Kwiatkowski, Cordelia Langford, John Sillitoe on behalf of the Wellcome Sanger Institute COVID-19 Surveillance Team |
| EPI_ISL_2766841 | Lighthouse Lab in Milton Keynes | Wellcome Sanger Institute for the COVID-19 Genomics UK (COG-UK) | The Lighthouse Lab in Milton Keynes and Alex Alderton, Roberto Amato, Jeffrey Barrett, Sonia Goncalves, Ewan Harrison, David K. Jackson, Ian Johnston, Dominic Kwiatkowski, Cordelia Langford, John Sillitoe on behalf of the Wellcome Sanger |

| | | | |
|---|---|---|---|
| | | Consortium | Institute COVID-19 Surveillance Team |
| EPI_ISL_2766842 | Lighthouse Laboratory Plymouth | Wellcome Sanger Institute for the COVID-19 Genomics UK (COG-UK) Consortium | Lighthouse Laboratory Plymouth and Alex Alderton, Roberto Amato, Jeffrey Barrett, Sonia Goncalves, Ewan Harrison, David K. Jackson, Ian Johnston, Dominic Kwiatkowski, Cordelia Langford, John Sillitoe on behalf of the Wellcome Sanger Institute COVID-19 Surveillance Team |
| EPI_ISL_2766843 | Lighthouse Lab in Milton Keynes | Wellcome Sanger Institute for the COVID-19 Genomics UK (COG-UK) Consortium | The Lighthouse Lab in Milton Keynes and Alex Alderton, Roberto Amato, Jeffrey Barrett, Sonia Goncalves, Ewan Harrison, David K. Jackson, Ian Johnston, Dominic Kwiatkowski, Cordelia Langford, John Sillitoe on behalf of the Wellcome Sanger Institute COVID-19 Surveillance Team |
| EPI_ISL_2766844, EPI_ISL_2766845 | Lighthouse Lab in Alderley Park | Wellcome Sanger Institute for the COVID-19 Genomics UK (COG-UK) Consortium | Jacquelyn Wynn, Mairead Hyland, The Lighthouse Lab in Alderley Park and Alex Alderton, Roberto Amato, Jeffrey Barrett, Sonia Goncalves, Ewan Harrison, David K. Jackson, Ian Johnston, Dominic Kwiatkowski, Cordelia Langford, John Sillitoe on behalf of the Wellcome Sanger Institute COVID-19 Surveillance Team |
| EPI_ISL_2766846 | Lighthouse Laboratory Plymouth | Wellcome Sanger Institute for the COVID-19 Genomics UK (COG-UK) Consortium | Lighthouse Laboratory Plymouth and Alex Alderton, Roberto Amato, Jeffrey Barrett, Sonia Goncalves, Ewan Harrison, David K. Jackson, Ian Johnston, Dominic Kwiatkowski, Cordelia Langford, John Sillitoe on behalf of the Wellcome Sanger Institute COVID-19 Surveillance Team |
| EPI_ISL_2766847, EPI_ISL_2766848 | Lighthouse Lab in Alderley Park | Wellcome Sanger Institute for the COVID-19 Genomics UK (COG-UK) Consortium | Jacquelyn Wynn, Mairead Hyland, The Lighthouse Lab in Alderley Park and Alex Alderton, Roberto Amato, Jeffrey Barrett, Sonia Goncalves, Ewan Harrison, David K. Jackson, Ian Johnston, Dominic Kwiatkowski, Cordelia Langford, John Sillitoe on behalf of the Wellcome Sanger Institute COVID-19 Surveillance Team |
| EPI_ISL_2766849 | Lighthouse Laboratory Plymouth | Wellcome Sanger Institute for the COVID-19 Genomics UK (COG-UK) Consortium | Lighthouse Laboratory Plymouth and Alex Alderton, Roberto Amato, Jeffrey Barrett, Sonia Goncalves, Ewan Harrison, David K. Jackson, Ian Johnston, Dominic Kwiatkowski, Cordelia Langford, John Sillitoe on behalf of the Wellcome Sanger Institute COVID-19 Surveillance Team |
| EPI_ISL_2766850 | Lighthouse Lab in Milton Keynes | Wellcome Sanger Institute for the COVID-19 Genomics UK (COG-UK) Consortium | The Lighthouse Lab in Milton Keynes and Alex Alderton, Roberto Amato, Jeffrey Barrett, Sonia Goncalves, Ewan Harrison, David K. Jackson, Ian Johnston, Dominic Kwiatkowski, Cordelia Langford, John Sillitoe on behalf of the Wellcome Sanger Institute COVID-19 Surveillance Team |
| EPI_ISL_2766851, EPI_ISL_2766852 | Lighthouse Lab in Alderley Park | Wellcome Sanger Institute for the COVID-19 Genomics UK (COG-UK) Consortium | Jacquelyn Wynn, Mairead Hyland, The Lighthouse Lab in Alderley Park and Alex Alderton, Roberto Amato, Jeffrey Barrett, Sonia Goncalves, Ewan Harrison, David K. Jackson, Ian Johnston, Dominic Kwiatkowski, Cordelia Langford, John Sillitoe on behalf of the Wellcome Sanger Institute COVID-19 Surveillance Team |
| EPI_ISL_2766853, EPI_ISL_2766854, EPI_ISL_2766855 | Lighthouse Lab in Milton Keynes | Wellcome Sanger Institute for the COVID-19 Genomics UK (COG-UK) Consortium | The Lighthouse Lab in Milton Keynes and Alex Alderton, Roberto Amato, Jeffrey Barrett, Sonia Goncalves, Ewan Harrison, David K. Jackson, Ian Johnston, Dominic Kwiatkowski, Cordelia Langford, John Sillitoe on behalf of the Wellcome Sanger Institute COVID-19 Surveillance Team |
| EPI_ISL_2766856, EPI_ISL_2766857 | Lighthouse Lab in Alderley Park | Wellcome Sanger Institute for the COVID-19 Genomics UK (COG-UK) Consortium | Jacquelyn Wynn, Mairead Hyland, The Lighthouse Lab in Alderley Park and Alex Alderton, Roberto Amato, Jeffrey Barrett, Sonia Goncalves, Ewan Harrison, David K. Jackson, Ian Johnston, Dominic Kwiatkowski, Cordelia Langford, John Sillitoe on behalf of the Wellcome Sanger Institute COVID-19 Surveillance Team |
| EPI_ISL_2766858, EPI_ISL_2766859, EPI_ISL_2766860 | Lighthouse Lab in Milton Keynes | Wellcome Sanger Institute for the COVID-19 Genomics UK (COG-UK) Consortium | The Lighthouse Lab in Milton Keynes and Alex Alderton, Roberto Amato, Jeffrey Barrett, Sonia Goncalves, Ewan Harrison, David K. Jackson, Ian Johnston, Dominic Kwiatkowski, Cordelia Langford, John Sillitoe on behalf of the Wellcome Sanger Institute COVID-19 Surveillance Team |
| EPI_ISL_2766861 | Lighthouse Lab in Alderley Park | Wellcome Sanger Institute for the COVID-19 Genomics UK (COG-UK) Consortium | Jacquelyn Wynn, Mairead Hyland, The Lighthouse Lab in Alderley Park and Alex Alderton, Roberto Amato, Jeffrey Barrett, Sonia Goncalves, Ewan Harrison, David K. Jackson, Ian Johnston, Dominic Kwiatkowski, Cordelia Langford, John Sillitoe on behalf of the Wellcome Sanger Institute COVID-19 Surveillance Team |
| EPI_ISL_2766862 | Lighthouse Laboratory Plymouth | Wellcome Sanger Institute for the COVID-19 Genomics UK (COG-UK) Consortium | Lighthouse Laboratory Plymouth and Alex Alderton, Roberto Amato, Jeffrey Barrett, Sonia Goncalves, Ewan Harrison, David K. Jackson, Ian Johnston, Dominic Kwiatkowski, Cordelia Langford, John Sillitoe on behalf of the Wellcome Sanger Institute COVID-19 Surveillance Team |
| EPI_ISL_2766863, EPI_ISL_2766864, EPI_ISL_2766865 | Lighthouse Lab in Alderley Park | Wellcome Sanger Institute for the COVID-19 Genomics UK (COG-UK) Consortium | Jacquelyn Wynn, Mairead Hyland, The Lighthouse Lab in Alderley Park and Alex Alderton, Roberto Amato, Jeffrey Barrett, Sonia Goncalves, Ewan Harrison, David K. Jackson, Ian Johnston, Dominic Kwiatkowski, Cordelia Langford, John Sillitoe on behalf of the Wellcome Sanger Institute COVID-19 Surveillance Team |
| EPI_ISL_2766866, EPI_ISL_2766867 | Lighthouse Lab in Milton Keynes | Wellcome Sanger Institute for the COVID-19 Genomics UK (COG-UK) Consortium | The Lighthouse Lab in Milton Keynes and Alex Alderton, Roberto Amato, Jeffrey Barrett, Sonia Goncalves, Ewan Harrison, David K. Jackson, Ian Johnston, Dominic Kwiatkowski, Cordelia Langford, John Sillitoe on behalf of the Wellcome Sanger Institute COVID-19 Surveillance Team |
| EPI_ISL_2766868 | Lighthouse Lab in Alderley Park | Wellcome Sanger Institute for the COVID-19 Genomics UK (COG-UK) Consortium | Jacquelyn Wynn, Mairead Hyland, The Lighthouse Lab in Alderley Park and Alex Alderton, Roberto Amato, Jeffrey Barrett, Sonia Goncalves, Ewan Harrison, David K. Jackson, Ian Johnston, Dominic Kwiatkowski, Cordelia Langford, John Sillitoe on behalf of the Wellcome Sanger Institute COVID-19 Surveillance Team |
| EPI_ISL_2766869 | Lighthouse Lab in Milton Keynes | Wellcome Sanger Institute for the COVID-19 Genomics UK (COG-UK) Consortium | The Lighthouse Lab in Milton Keynes and Alex Alderton, Roberto Amato, Jeffrey Barrett, Sonia Goncalves, Ewan Harrison, David K. Jackson, Ian Johnston, Dominic Kwiatkowski, Cordelia Langford, John Sillitoe on behalf of the Wellcome Sanger Institute COVID-19 Surveillance Team |
| EPI_ISL_2766870 | Lighthouse Lab in Alderley Park | Wellcome Sanger Institute for the COVID-19 Genomics UK (COG-UK) Consortium | Jacquelyn Wynn, Mairead Hyland, The Lighthouse Lab in Alderley Park and Alex Alderton, Roberto Amato, Jeffrey Barrett, Sonia Goncalves, Ewan Harrison, David K. Jackson, Ian Johnston, Dominic Kwiatkowski, Cordelia Langford, John Sillitoe on behalf of the Wellcome Sanger Institute COVID-19 Surveillance Team |
| EPI_ISL_2766871 | Lighthouse Laboratory Plymouth | Wellcome Sanger Institute for the COVID-19 Genomics UK (COG-UK) Consortium | Lighthouse Laboratory Plymouth and Alex Alderton, Roberto Amato, Jeffrey Barrett, Sonia Goncalves, Ewan Harrison, David K. Jackson, Ian Johnston, Dominic Kwiatkowski, Cordelia Langford, John Sillitoe on behalf of the Wellcome Sanger Institute COVID-19 Surveillance Team |
| EPI_ISL_2766872 | Lighthouse Lab in Alderley Park | Wellcome Sanger Institute for the COVID-19 Genomics UK (COG-UK) Consortium | Jacquelyn Wynn, Mairead Hyland, The Lighthouse Lab in Alderley Park and Alex Alderton, Roberto Amato, Jeffrey Barrett, Sonia Goncalves, Ewan Harrison, David K. Jackson, Ian Johnston, Dominic Kwiatkowski, Cordelia Langford, John Sillitoe on behalf of the Wellcome Sanger Institute COVID-19 Surveillance Team |
| EPI_ISL_2766873 | Lighthouse Laboratory Plymouth | Wellcome Sanger Institute for the COVID-19 Genomics UK (COG-UK) Consortium | Lighthouse Laboratory Plymouth and Alex Alderton, Roberto Amato, Jeffrey Barrett, Sonia Goncalves, Ewan Harrison, David K. Jackson, Ian Johnston, Dominic Kwiatkowski, Cordelia Langford, John Sillitoe on behalf of the Wellcome Sanger Institute COVID-19 Surveillance Team |
| EPI_ISL_2766874 | Lighthouse Lab in Milton Keynes | Wellcome Sanger Institute for the COVID-19 Genomics UK (COG-UK) Consortium | The Lighthouse Lab in Milton Keynes and Alex Alderton, Roberto Amato, Jeffrey Barrett, Sonia Goncalves, Ewan Harrison, David K. Jackson, Ian Johnston, Dominic Kwiatkowski, Cordelia Langford, John Sillitoe on behalf of the Wellcome Sanger Institute COVID-19 Surveillance Team |
| EPI_ISL_2766875 | Lighthouse Lab in Alderley Park | Wellcome Sanger Institute for the COVID-19 Genomics UK (COG-UK) Consortium | Jacquelyn Wynn, Mairead Hyland, The Lighthouse Lab in Alderley Park and Alex Alderton, Roberto Amato, Jeffrey Barrett, Sonia Goncalves, Ewan Harrison, David K. Jackson, Ian Johnston, Dominic Kwiatkowski, Cordelia Langford, John Sillitoe on behalf of the Wellcome Sanger Institute COVID-19 Surveillance Team |
| EPI_ISL_2766876 | Lighthouse Lab in Milton Keynes | Wellcome Sanger Institute for the COVID-19 Genomics UK (COG-UK) Consortium | The Lighthouse Lab in Milton Keynes and Alex Alderton, Roberto Amato, Jeffrey Barrett, Sonia Goncalves, Ewan Harrison, David K. Jackson, Ian Johnston, Dominic Kwiatkowski, Cordelia Langford, John Sillitoe on behalf of the Wellcome Sanger Institute COVID-19 Surveillance Team |
| EPI_ISL_2766877 | Lighthouse Lab in Alderley Park | Wellcome Sanger Institute for the COVID-19 Genomics UK (COG-UK) Consortium | Jacquelyn Wynn, Mairead Hyland, The Lighthouse Lab in Alderley Park and Alex Alderton, Roberto Amato, Jeffrey Barrett, Sonia Goncalves, Ewan Harrison, David K. Jackson, Ian Johnston, Dominic Kwiatkowski, Cordelia Langford, John Sillitoe on behalf of the Wellcome Sanger Institute COVID-19 Surveillance Team |
| EPI_ISL_2766878 | Lighthouse Lab in Milton Keynes | Wellcome Sanger Institute for the COVID-19 Genomics UK (COG-UK) Consortium | The Lighthouse Lab in Milton Keynes and Alex Alderton, Roberto Amato, Jeffrey Barrett, Sonia Goncalves, Ewan Harrison, David K. Jackson, Ian Johnston, Dominic Kwiatkowski, Cordelia Langford, John Sillitoe on behalf of the Wellcome Sanger Institute COVID-19 Surveillance Team |
| EPI_ISL_2766879, EPI_ISL_2766880, EPI_ISL_2766881, EPI_ISL_2766882 | Lighthouse Lab in Alderley Park | Wellcome Sanger Institute for the COVID-19 Genomics UK (COG-UK) Consortium | Jacquelyn Wynn, Mairead Hyland, The Lighthouse Lab in Alderley Park and Alex Alderton, Roberto Amato, Jeffrey Barrett, Sonia Goncalves, Ewan Harrison, David K. Jackson, Ian Johnston, Dominic Kwiatkowski, Cordelia Langford, John Sillitoe on behalf of the Wellcome Sanger Institute COVID-19 Surveillance Team |
| EPI_ISL_2766883 | Lighthouse Lab in Milton Keynes | Wellcome Sanger Institute for the COVID-19 Genomics UK (COG-UK) Consortium | The Lighthouse Lab in Milton Keynes and Alex Alderton, Roberto Amato, Jeffrey Barrett, Sonia Goncalves, Ewan Harrison, David K. Jackson, Ian Johnston, Dominic Kwiatkowski, Cordelia Langford, John Sillitoe on behalf of the Wellcome Sanger Institute COVID-19 Surveillance Team |
| EPI_ISL_2766884, EPI_ISL_2766885 | Lighthouse Lab in Alderley Park | Wellcome Sanger Institute for the COVID-19 Genomics UK (COG-UK) Consortium | Jacquelyn Wynn, Mairead Hyland, The Lighthouse Lab in Alderley Park and Alex Alderton, Roberto Amato, Jeffrey Barrett, Sonia Goncalves, Ewan Harrison, David K. Jackson, Ian Johnston, Dominic Kwiatkowski, Cordelia Langford, John Sillitoe on behalf of the Wellcome Sanger Institute COVID-19 Surveillance Team |
| EPI_ISL_2766886 | Lighthouse Lab in Milton Keynes | Wellcome Sanger Institute for the COVID-19 Genomics UK (COG-UK) Consortium | The Lighthouse Lab in Milton Keynes and Alex Alderton, Roberto Amato, Jeffrey Barrett, Sonia Goncalves, Ewan Harrison, David K. Jackson, Ian Johnston, Dominic Kwiatkowski, Cordelia Langford, John Sillitoe on behalf of the Wellcome Sanger Institute COVID-19 Surveillance Team |
| EPI_ISL_2766887 | Lighthouse Lab in Alderley Park | Wellcome Sanger Institute for the COVID-19 Genomics UK (COG-UK) Consortium | Jacquelyn Wynn, Mairead Hyland, The Lighthouse Lab in Alderley Park and Alex Alderton, Roberto Amato, Jeffrey Barrett, Sonia Goncalves, Ewan Harrison, David K. Jackson, Ian Johnston, Dominic Kwiatkowski, Cordelia Langford, John Sillitoe on behalf of the Wellcome Sanger Institute COVID-19 Surveillance Team |
| EPI_ISL_2766888, EPI_ISL_2766889, EPI_ISL_2766890 | Lighthouse Lab in Milton Keynes | Wellcome Sanger Institute for the COVID-19 Genomics UK (COG-UK) Consortium | The Lighthouse Lab in Milton Keynes and Alex Alderton, Roberto Amato, Jeffrey Barrett, Sonia Goncalves, Ewan Harrison, David K. Jackson, Ian Johnston, Dominic Kwiatkowski, Cordelia Langford, John Sillitoe on behalf of the Wellcome Sanger Institute COVID-19 Surveillance Team |
| EPI_ISL_2766891, EPI_ISL_2766892 | Lighthouse Lab in Alderley Park | Wellcome Sanger Institute for the COVID-19 Genomics UK (COG-UK) Consortium | Jacquelyn Wynn, Mairead Hyland, The Lighthouse Lab in Alderley Park and Alex Alderton, Roberto Amato, Jeffrey Barrett, Sonia Goncalves, Ewan Harrison, David K. Jackson, Ian Johnston, Dominic Kwiatkowski, Cordelia Langford, John Sillitoe on behalf of the Wellcome Sanger Institute COVID-19 Surveillance Team |
| EPI_ISL_2766893 | Lighthouse Lab in Milton Keynes | Wellcome Sanger Institute for the COVID-19 Genomics UK (COG-UK) Consortium | The Lighthouse Lab in Milton Keynes and Alex Alderton, Roberto Amato, Jeffrey Barrett, Sonia Goncalves, Ewan Harrison, David K. Jackson, Ian Johnston, Dominic Kwiatkowski, Cordelia Langford, John Sillitoe on behalf of the Wellcome Sanger Institute COVID-19 Surveillance Team |
| EPI_ISL_2766894 | Lighthouse Lab in Alderley Park | Wellcome Sanger Institute for the COVID-19 Genomics UK (COG-UK) Consortium | Jacquelyn Wynn, Mairead Hyland, The Lighthouse Lab in Alderley Park and Alex Alderton, Roberto Amato, Jeffrey Barrett, Sonia Goncalves, Ewan Harrison, David K. Jackson, Ian Johnston, Dominic Kwiatkowski, Cordelia Langford, John Sillitoe on behalf of the Wellcome Sanger Institute COVID-19 Surveillance Team |
| EPI_ISL_2766895 | Lighthouse Lab in Milton Keynes | Wellcome Sanger Institute for the COVID-19 Genomics UK (COG-UK) Consortium | The Lighthouse Lab in Milton Keynes and Alex Alderton, Roberto Amato, Jeffrey Barrett, Sonia Goncalves, Ewan Harrison, David K. Jackson, Ian Johnston, Dominic Kwiatkowski, Cordelia Langford, John Sillitoe on behalf of the Wellcome Sanger Institute COVID-19 Surveillance Team |
| EPI_ISL_2766896 | Lighthouse Lab in Alderley Park | Wellcome Sanger Institute for the COVID-19 Genomics UK (COG-UK) Consortium | Jacquelyn Wynn, Mairead Hyland, The Lighthouse Lab in Alderley Park and Alex Alderton, Roberto Amato, Jeffrey Barrett, Sonia Goncalves, Ewan Harrison, David K. Jackson, Ian Johnston, Dominic Kwiatkowski, Cordelia Langford, John Sillitoe on behalf of the Wellcome Sanger Institute COVID-19 Surveillance Team |
| EPI_ISL_2766897 | Lighthouse Lab in Milton Keynes | Wellcome Sanger Institute for the COVID-19 Genomics UK (COG-UK) Consortium | The Lighthouse Lab in Milton Keynes and Alex Alderton, Roberto Amato, Jeffrey Barrett, Sonia Goncalves, Ewan Harrison, David K. Jackson, Ian Johnston, Dominic Kwiatkowski, Cordelia Langford, John Sillitoe on behalf of the Wellcome Sanger Institute COVID-19 Surveillance Team |
| EPI_ISL_2766898, EPI_ISL_2766899, EPI_ISL_2766900, EPI_ISL_2766901, EPI_ISL_2766902, EPI_ISL_2766903, EPI_ISL_2766904 | Lighthouse Lab in Alderley Park | Wellcome Sanger Institute for the COVID-19 Genomics UK (COG-UK) Consortium | Jacquelyn Wynn, Mairead Hyland, The Lighthouse Lab in Alderley Park and Alex Alderton, Roberto Amato, Jeffrey Barrett, Sonia Goncalves, Ewan Harrison, David K. Jackson, Ian Johnston, Dominic Kwiatkowski, Cordelia Langford, John Sillitoe on behalf of the Wellcome Sanger Institute COVID-19 Surveillance Team |
| EPI_ISL_2766905 | Lighthouse Lab in Milton Keynes | Wellcome Sanger Institute for the COVID-19 Genomics UK (COG-UK) Consortium | The Lighthouse Lab in Milton Keynes and Alex Alderton, Roberto Amato, Jeffrey Barrett, Sonia Goncalves, Ewan Harrison, David K. Jackson, Ian Johnston, Dominic Kwiatkowski, Cordelia Langford, John Sillitoe on behalf of the Wellcome Sanger Institute COVID-19 Surveillance Team |

| | | | |
|---|---|---|---|
| EPI_ISL_2766906, EPI_ISL_2766907 | Lighthouse Lab in Alderley Park | Wellcome Sanger Institute for the COVID-19 Genomics UK (COG-UK) Consortium | Jacquelyn Wynn, Mairead Hyland, The Lighthouse Lab in Alderley Park and Alex Alderton, Roberto Amato, Jeffrey Barrett, Sonia Goncalves, Ewan Harrison, David K. Jackson, Ian Johnston, Dominic Kwiatkowski, Cordelia Langford, John Sillitoe on behalf of the Wellcome Sanger Institute COVID-19 Surveillance Team |
| EPI_ISL_2766908 | Lighthouse Lab in Milton Keynes | Wellcome Sanger Institute for the COVID-19 Genomics UK (COG-UK) Consortium | The Lighthouse Lab in Milton Keynes and Alex Alderton, Roberto Amato, Jeffrey Barrett, Sonia Goncalves, Ewan Harrison, David K. Jackson, Ian Johnston, Dominic Kwiatkowski, Cordelia Langford, John Sillitoe on behalf of the Wellcome Sanger Institute COVID-19 Surveillance Team |
| EPI_ISL_2766909, EPI_ISL_2766910, EPI_ISL_2766911, EPI_ISL_2766912, EPI_ISL_2766913, EPI_ISL_2766914, EPI_ISL_2766915, EPI_ISL_2766916, EPI_ISL_2766917 | Lighthouse Lab in Alderley Park | Wellcome Sanger Institute for the COVID-19 Genomics UK (COG-UK) Consortium | Jacquelyn Wynn, Mairead Hyland, The Lighthouse Lab in Alderley Park and Alex Alderton, Roberto Amato, Jeffrey Barrett, Sonia Goncalves, Ewan Harrison, David K. Jackson, Ian Johnston, Dominic Kwiatkowski, Cordelia Langford, John Sillitoe on behalf of the Wellcome Sanger Institute COVID-19 Surveillance Team |
| EPI_ISL_2766918 | Lighthouse Lab in Milton Keynes | Wellcome Sanger Institute for the COVID-19 Genomics UK (COG-UK) Consortium | The Lighthouse Lab in Milton Keynes and Alex Alderton, Roberto Amato, Jeffrey Barrett, Sonia Goncalves, Ewan Harrison, David K. Jackson, Ian Johnston, Dominic Kwiatkowski, Cordelia Langford, John Sillitoe on behalf of the Wellcome Sanger Institute COVID-19 Surveillance Team |
| EPI_ISL_2766919, EPI_ISL_2766920, EPI_ISL_2766921, EPI_ISL_2766922 | Lighthouse Lab in Alderley Park | Wellcome Sanger Institute for the COVID-19 Genomics UK (COG-UK) Consortium | Jacquelyn Wynn, Mairead Hyland, The Lighthouse Lab in Alderley Park and Alex Alderton, Roberto Amato, Jeffrey Barrett, Sonia Goncalves, Ewan Harrison, David K. Jackson, Ian Johnston, Dominic Kwiatkowski, Cordelia Langford, John Sillitoe on behalf of the Wellcome Sanger Institute COVID-19 Surveillance Team |
| EPI_ISL_2766923, EPI_ISL_2766924 | Lighthouse Lab in Milton Keynes | Wellcome Sanger Institute for the COVID-19 Genomics UK (COG-UK) Consortium | The Lighthouse Lab in Milton Keynes and Alex Alderton, Roberto Amato, Jeffrey Barrett, Sonia Goncalves, Ewan Harrison, David K. Jackson, Ian Johnston, Dominic Kwiatkowski, Cordelia Langford, John Sillitoe on behalf of the Wellcome Sanger Institute COVID-19 Surveillance Team |
| EPI_ISL_2766925 | Lighthouse Lab in Alderley Park | Wellcome Sanger Institute for the COVID-19 Genomics UK (COG-UK) Consortium | Jacquelyn Wynn, Mairead Hyland, The Lighthouse Lab in Alderley Park and Alex Alderton, Roberto Amato, Jeffrey Barrett, Sonia Goncalves, Ewan Harrison, David K. Jackson, Ian Johnston, Dominic Kwiatkowski, Cordelia Langford, John Sillitoe on behalf of the Wellcome Sanger Institute COVID-19 Surveillance Team |
| EPI_ISL_2766926, EPI_ISL_2766927, EPI_ISL_2766928 | Lighthouse Lab in Milton Keynes | Wellcome Sanger Institute for the COVID-19 Genomics UK (COG-UK) Consortium | The Lighthouse Lab in Milton Keynes and Alex Alderton, Roberto Amato, Jeffrey Barrett, Sonia Goncalves, Ewan Harrison, David K. Jackson, Ian Johnston, Dominic Kwiatkowski, Cordelia Langford, John Sillitoe on behalf of the Wellcome Sanger Institute COVID-19 Surveillance Team |
| EPI_ISL_2766929 | Lighthouse Lab in Alderley Park | Wellcome Sanger Institute for the COVID-19 Genomics UK (COG-UK) Consortium | Jacquelyn Wynn, Mairead Hyland, The Lighthouse Lab in Alderley Park and Alex Alderton, Roberto Amato, Jeffrey Barrett, Sonia Goncalves, Ewan Harrison, David K. Jackson, Ian Johnston, Dominic Kwiatkowski, Cordelia Langford, John Sillitoe on behalf of the Wellcome Sanger Institute COVID-19 Surveillance Team |
| EPI_ISL_2766930, EPI_ISL_2766931 | Lighthouse Lab in Milton Keynes | Wellcome Sanger Institute for the COVID-19 Genomics UK (COG-UK) Consortium | The Lighthouse Lab in Milton Keynes and Alex Alderton, Roberto Amato, Jeffrey Barrett, Sonia Goncalves, Ewan Harrison, David K. Jackson, Ian Johnston, Dominic Kwiatkowski, Cordelia Langford, John Sillitoe on behalf of the Wellcome Sanger Institute COVID-19 Surveillance Team |
| EPI_ISL_2766932, EPI_ISL_2766933, EPI_ISL_2766934, EPI_ISL_2766935, EPI_ISL_2766936, EPI_ISL_2766937, EPI_ISL_2766938, EPI_ISL_2766939, EPI_ISL_2766940 | Lighthouse Lab in Alderley Park | Wellcome Sanger Institute for the COVID-19 Genomics UK (COG-UK) Consortium | Jacquelyn Wynn, Mairead Hyland, The Lighthouse Lab in Alderley Park and Alex Alderton, Roberto Amato, Jeffrey Barrett, Sonia Goncalves, Ewan Harrison, David K. Jackson, Ian Johnston, Dominic Kwiatkowski, Cordelia Langford, John Sillitoe on behalf of the Wellcome Sanger Institute COVID-19 Surveillance Team |
| EPI_ISL_2766941, EPI_ISL_2766942 | Lighthouse Lab in Milton Keynes | Wellcome Sanger Institute for the COVID-19 Genomics UK (COG-UK) Consortium | The Lighthouse Lab in Milton Keynes and Alex Alderton, Roberto Amato, Jeffrey Barrett, Sonia Goncalves, Ewan Harrison, David K. Jackson, Ian Johnston, Dominic Kwiatkowski, Cordelia Langford, John Sillitoe on behalf of the Wellcome Sanger Institute COVID-19 Surveillance Team |
| EPI_ISL_2766943, EPI_ISL_2766944, EPI_ISL_2766945, EPI_ISL_2766946, EPI_ISL_2766947 | Lighthouse Lab in Alderley Park | Wellcome Sanger Institute for the COVID-19 Genomics UK (COG-UK) Consortium | Jacquelyn Wynn, Mairead Hyland, The Lighthouse Lab in Alderley Park and Alex Alderton, Roberto Amato, Jeffrey Barrett, Sonia Goncalves, Ewan Harrison, David K. Jackson, Ian Johnston, Dominic Kwiatkowski, Cordelia Langford, John Sillitoe on behalf of the Wellcome Sanger Institute COVID-19 Surveillance Team |
| EPI_ISL_2766948, EPI_ISL_2766949 | Lighthouse Lab in Milton Keynes | Wellcome Sanger Institute for the COVID-19 Genomics UK (COG-UK) Consortium | The Lighthouse Lab in Milton Keynes and Alex Alderton, Roberto Amato, Jeffrey Barrett, Sonia Goncalves, Ewan Harrison, David K. Jackson, Ian Johnston, Dominic Kwiatkowski, Cordelia Langford, John Sillitoe on behalf of the Wellcome Sanger Institute COVID-19 Surveillance Team |
| EPI_ISL_2766950, EPI_ISL_2766951, EPI_ISL_2766952, EPI_ISL_2766953, EPI_ISL_2766954, EPI_ISL_2766955, EPI_ISL_2766956, EPI_ISL_2766958 | Lighthouse Lab in Alderley Park | Wellcome Sanger Institute for the COVID-19 Genomics UK (COG-UK) Consortium | Jacquelyn Wynn, Mairead Hyland, The Lighthouse Lab in Alderley Park and Alex Alderton, Roberto Amato, Jeffrey Barrett, Sonia Goncalves, Ewan Harrison, David K. Jackson, Ian Johnston, Dominic Kwiatkowski, Cordelia Langford, John Sillitoe on behalf of the Wellcome Sanger Institute COVID-19 Surveillance Team |
| EPI_ISL_2766959 | Lighthouse Lab in Milton Keynes | Wellcome Sanger Institute for the COVID-19 Genomics UK (COG-UK) Consortium | The Lighthouse Lab in Milton Keynes and Alex Alderton, Roberto Amato, Jeffrey Barrett, Sonia Goncalves, Ewan Harrison, David K. Jackson, Ian Johnston, Dominic Kwiatkowski, Cordelia Langford, John Sillitoe on behalf of the Wellcome Sanger Institute COVID-19 Surveillance Team |
| EPI_ISL_2766960, EPI_ISL_2766961 | Lighthouse Lab in Alderley Park | Wellcome Sanger Institute for the COVID-19 Genomics UK (COG-UK) Consortium | Jacquelyn Wynn, Mairead Hyland, The Lighthouse Lab in Alderley Park and Alex Alderton, Roberto Amato, Jeffrey Barrett, Sonia Goncalves, Ewan Harrison, David K. Jackson, Ian Johnston, Dominic Kwiatkowski, Cordelia Langford, John Sillitoe on behalf of the Wellcome Sanger Institute COVID-19 Surveillance Team |
| EPI_ISL_2766962 | Lighthouse Lab in Milton Keynes | Wellcome Sanger Institute for the COVID-19 Genomics UK (COG-UK) Consortium | The Lighthouse Lab in Milton Keynes and Alex Alderton, Roberto Amato, Jeffrey Barrett, Sonia Goncalves, Ewan Harrison, David K. Jackson, Ian Johnston, Dominic Kwiatkowski, Cordelia Langford, John Sillitoe on behalf of the Wellcome Sanger Institute COVID-19 Surveillance Team |
| EPI_ISL_2766964 | Lighthouse Lab in Alderley Park | Wellcome Sanger Institute for the COVID-19 Genomics UK (COG-UK) Consortium | Jacquelyn Wynn, Mairead Hyland, The Lighthouse Lab in Alderley Park and Alex Alderton, Roberto Amato, Jeffrey Barrett, Sonia Goncalves, Ewan Harrison, David K. Jackson, Ian Johnston, Dominic Kwiatkowski, Cordelia Langford, John Sillitoe on behalf of the Wellcome Sanger Institute COVID-19 Surveillance Team |
| EPI_ISL_2766965, EPI_ISL_2766966 | Lighthouse Lab in Milton Keynes | Wellcome Sanger Institute for the COVID-19 Genomics UK (COG-UK) Consortium | The Lighthouse Lab in Milton Keynes and Alex Alderton, Roberto Amato, Jeffrey Barrett, Sonia Goncalves, Ewan Harrison, David K. Jackson, Ian Johnston, Dominic Kwiatkowski, Cordelia Langford, John Sillitoe on behalf of the Wellcome Sanger Institute COVID-19 Surveillance Team |
| EPI_ISL_2766967, EPI_ISL_2766968, EPI_ISL_2766969, EPI_ISL_2766970 | Lighthouse Lab in Alderley Park | Wellcome Sanger Institute for the COVID-19 Genomics UK (COG-UK) Consortium | Jacquelyn Wynn, Mairead Hyland, The Lighthouse Lab in Alderley Park and Alex Alderton, Roberto Amato, Jeffrey Barrett, Sonia Goncalves, Ewan Harrison, David K. Jackson, Ian Johnston, Dominic Kwiatkowski, Cordelia Langford, John Sillitoe on behalf of the Wellcome Sanger Institute COVID-19 Surveillance Team |
| EPI_ISL_2766971 | Lighthouse Lab in Milton Keynes | Wellcome Sanger Institute for the COVID-19 Genomics UK (COG-UK) Consortium | The Lighthouse Lab in Milton Keynes and Alex Alderton, Roberto Amato, Jeffrey Barrett, Sonia Goncalves, Ewan Harrison, David K. Jackson, Ian Johnston, Dominic Kwiatkowski, Cordelia Langford, John Sillitoe on behalf of the Wellcome Sanger Institute COVID-19 Surveillance Team |
| EPI_ISL_2766972, EPI_ISL_2766973, EPI_ISL_2766974, EPI_ISL_2766975, EPI_ISL_2766976 | Lighthouse Lab in Alderley Park | Wellcome Sanger Institute for the COVID-19 Genomics UK (COG-UK) Consortium | Jacquelyn Wynn, Mairead Hyland, The Lighthouse Lab in Alderley Park and Alex Alderton, Roberto Amato, Jeffrey Barrett, Sonia Goncalves, Ewan Harrison, David K. Jackson, Ian Johnston, Dominic Kwiatkowski, Cordelia Langford, John Sillitoe on behalf of the Wellcome Sanger Institute COVID-19 Surveillance Team |
| EPI_ISL_2766977 | Lighthouse Lab in Milton Keynes | Wellcome Sanger Institute for the COVID-19 Genomics UK (COG-UK) Consortium | The Lighthouse Lab in Milton Keynes and Alex Alderton, Roberto Amato, Jeffrey Barrett, Sonia Goncalves, Ewan Harrison, David K. Jackson, Ian Johnston, Dominic Kwiatkowski, Cordelia Langford, John Sillitoe on behalf of the Wellcome Sanger Institute COVID-19 Surveillance Team |
| EPI_ISL_2766978, EPI_ISL_2766979, EPI_ISL_2766980, EPI_ISL_2766981, EPI_ISL_2766982, EPI_ISL_2766983, EPI_ISL_2766984 | Lighthouse Lab in Alderley Park | Wellcome Sanger Institute for the COVID-19 Genomics UK (COG-UK) Consortium | Jacquelyn Wynn, Mairead Hyland, The Lighthouse Lab in Alderley Park and Alex Alderton, Roberto Amato, Jeffrey Barrett, Sonia Goncalves, Ewan Harrison, David K. Jackson, Ian Johnston, Dominic Kwiatkowski, Cordelia Langford, John Sillitoe on behalf of the Wellcome Sanger Institute COVID-19 Surveillance Team |
| EPI_ISL_2766985 | Lighthouse Lab in Milton Keynes | Wellcome Sanger Institute for the COVID-19 Genomics UK (COG-UK) Consortium | The Lighthouse Lab in Milton Keynes and Alex Alderton, Roberto Amato, Jeffrey Barrett, Sonia Goncalves, Ewan Harrison, David K. Jackson, Ian Johnston, Dominic Kwiatkowski, Cordelia Langford, John Sillitoe on behalf of the Wellcome Sanger Institute COVID-19 Surveillance Team |
| EPI_ISL_2766986, EPI_ISL_2766987, EPI_ISL_2766988, EPI_ISL_2766989, EPI_ISL_2766990, EPI_ISL_2766991, EPI_ISL_2766992, EPI_ISL_2766993, EPI_ISL_2766994, EPI_ISL_2766995 | Lighthouse Lab in Alderley Park | Wellcome Sanger Institute for the COVID-19 Genomics UK (COG-UK) Consortium | Jacquelyn Wynn, Mairead Hyland, The Lighthouse Lab in Alderley Park and Alex Alderton, Roberto Amato, Jeffrey Barrett, Sonia Goncalves, Ewan Harrison, David K. Jackson, Ian Johnston, Dominic Kwiatkowski, Cordelia Langford, John Sillitoe on behalf of the Wellcome Sanger Institute COVID-19 Surveillance Team |

| | | | |
|---|---|---|---|
| EPI_ISL_2766996 | Lighthouse Lab in Milton Keynes | Wellcome Sanger Institute for the COVID-19 Genomics UK (COG-UK) Consortium | The Lighthouse Lab in Milton Keynes and Alex Alderton, Roberto Amato, Jeffrey Barrett, Sonia Goncalves, Ewan Harrison, David K. Jackson, Ian Johnston, Dominic Kwiatkowski, Cordelia Langford, John Sillitoe on behalf of the Wellcome Sanger Institute COVID-19 Surveillance Team |
| EPI_ISL_2766997, EPI_ISL_2766998, EPI_ISL_2766999, EPI_ISL_2767000, EPI_ISL_2767001, EPI_ISL_2767002, EPI_ISL_2767003 | Lighthouse Lab in Alderley Park | Wellcome Sanger Institute for the COVID-19 Genomics UK (COG-UK) Consortium | Jacquelyn Wynn, Mairead Hyland, The Lighthouse Lab in Alderley Park and Alex Alderton, Roberto Amato, Jeffrey Barrett, Sonia Goncalves, Ewan Harrison, David K. Jackson, Ian Johnston, Dominic Kwiatkowski, Cordelia Langford, John Sillitoe on behalf of the Wellcome Sanger Institute COVID-19 Surveillance Team |
| EPI_ISL_2767004 | Lighthouse Lab in Milton Keynes | Wellcome Sanger Institute for the COVID-19 Genomics UK (COG-UK) Consortium | The Lighthouse Lab in Milton Keynes and Alex Alderton, Roberto Amato, Jeffrey Barrett, Sonia Goncalves, Ewan Harrison, David K. Jackson, Ian Johnston, Dominic Kwiatkowski, Cordelia Langford, John Sillitoe on behalf of the Wellcome Sanger Institute COVID-19 Surveillance Team |
| EPI_ISL_2767005, EPI_ISL_2767006, EPI_ISL_2767007, EPI_ISL_2767008 | Lighthouse Lab in Alderley Park | Wellcome Sanger Institute for the COVID-19 Genomics UK (COG-UK) Consortium | Jacquelyn Wynn, Mairead Hyland, The Lighthouse Lab in Alderley Park and Alex Alderton, Roberto Amato, Jeffrey Barrett, Sonia Goncalves, Ewan Harrison, David K. Jackson, Ian Johnston, Dominic Kwiatkowski, Cordelia Langford, John Sillitoe on behalf of the Wellcome Sanger Institute COVID-19 Surveillance Team |
| EPI_ISL_2767009, EPI_ISL_2767010 | Lighthouse Lab in Milton Keynes | Wellcome Sanger Institute for the COVID-19 Genomics UK (COG-UK) Consortium | The Lighthouse Lab in Milton Keynes and Alex Alderton, Roberto Amato, Jeffrey Barrett, Sonia Goncalves, Ewan Harrison, David K. Jackson, Ian Johnston, Dominic Kwiatkowski, Cordelia Langford, John Sillitoe on behalf of the Wellcome Sanger Institute COVID-19 Surveillance Team |
| EPI_ISL_2767011, EPI_ISL_2767012 | Lighthouse Lab in Alderley Park | Wellcome Sanger Institute for the COVID-19 Genomics UK (COG-UK) Consortium | Jacquelyn Wynn, Mairead Hyland, The Lighthouse Lab in Alderley Park and Alex Alderton, Roberto Amato, Jeffrey Barrett, Sonia Goncalves, Ewan Harrison, David K. Jackson, Ian Johnston, Dominic Kwiatkowski, Cordelia Langford, John Sillitoe on behalf of the Wellcome Sanger Institute COVID-19 Surveillance Team |
| EPI_ISL_2767013 | Lighthouse Lab in Milton Keynes | Wellcome Sanger Institute for the COVID-19 Genomics UK (COG-UK) Consortium | The Lighthouse Lab in Milton Keynes and Alex Alderton, Roberto Amato, Jeffrey Barrett, Sonia Goncalves, Ewan Harrison, David K. Jackson, Ian Johnston, Dominic Kwiatkowski, Cordelia Langford, John Sillitoe on behalf of the Wellcome Sanger Institute COVID-19 Surveillance Team |
| EPI_ISL_2767014, EPI_ISL_2767015, EPI_ISL_2767016, EPI_ISL_2767017 | Lighthouse Lab in Alderley Park | Wellcome Sanger Institute for the COVID-19 Genomics UK (COG-UK) Consortium | Jacquelyn Wynn, Mairead Hyland, The Lighthouse Lab in Alderley Park and Alex Alderton, Roberto Amato, Jeffrey Barrett, Sonia Goncalves, Ewan Harrison, David K. Jackson, Ian Johnston, Dominic Kwiatkowski, Cordelia Langford, John Sillitoe on behalf of the Wellcome Sanger Institute COVID-19 Surveillance Team |
| EPI_ISL_2767018 | Lighthouse Lab in Milton Keynes | Wellcome Sanger Institute for the COVID-19 Genomics UK (COG-UK) Consortium | The Lighthouse Lab in Milton Keynes and Alex Alderton, Roberto Amato, Jeffrey Barrett, Sonia Goncalves, Ewan Harrison, David K. Jackson, Ian Johnston, Dominic Kwiatkowski, Cordelia Langford, John Sillitoe on behalf of the Wellcome Sanger Institute COVID-19 Surveillance Team |
| EPI_ISL_2767019, EPI_ISL_2767020 | Lighthouse Lab in Alderley Park | Wellcome Sanger Institute for the COVID-19 Genomics UK (COG-UK) Consortium | Jacquelyn Wynn, Mairead Hyland, The Lighthouse Lab in Alderley Park and Alex Alderton, Roberto Amato, Jeffrey Barrett, Sonia Goncalves, Ewan Harrison, David K. Jackson, Ian Johnston, Dominic Kwiatkowski, Cordelia Langford, John Sillitoe on behalf of the Wellcome Sanger Institute COVID-19 Surveillance Team |
| EPI_ISL_2767021 | Lighthouse Lab in Milton Keynes | Wellcome Sanger Institute for the COVID-19 Genomics UK (COG-UK) Consortium | The Lighthouse Lab in Milton Keynes and Alex Alderton, Roberto Amato, Jeffrey Barrett, Sonia Goncalves, Ewan Harrison, David K. Jackson, Ian Johnston, Dominic Kwiatkowski, Cordelia Langford, John Sillitoe on behalf of the Wellcome Sanger Institute COVID-19 Surveillance Team |
| EPI_ISL_2767022 | Lighthouse Lab in Alderley Park | Wellcome Sanger Institute for the COVID-19 Genomics UK (COG-UK) Consortium | Jacquelyn Wynn, Mairead Hyland, The Lighthouse Lab in Alderley Park and Alex Alderton, Roberto Amato, Jeffrey Barrett, Sonia Goncalves, Ewan Harrison, David K. Jackson, Ian Johnston, Dominic Kwiatkowski, Cordelia Langford, John Sillitoe on behalf of the Wellcome Sanger Institute COVID-19 Surveillance Team |
| EPI_ISL_2767023 | Lighthouse Lab in Milton Keynes | Wellcome Sanger Institute for the COVID-19 Genomics UK (COG-UK) Consortium | The Lighthouse Lab in Milton Keynes and Alex Alderton, Roberto Amato, Jeffrey Barrett, Sonia Goncalves, Ewan Harrison, David K. Jackson, Ian Johnston, Dominic Kwiatkowski, Cordelia Langford, John Sillitoe on behalf of the Wellcome Sanger Institute COVID-19 Surveillance Team |
| EPI_ISL_2767024, EPI_ISL_2767025, EPI_ISL_2767026, EPI_ISL_2767027 | Lighthouse Lab in Alderley Park | Wellcome Sanger Institute for the COVID-19 Genomics UK (COG-UK) Consortium | Jacquelyn Wynn, Mairead Hyland, The Lighthouse Lab in Alderley Park and Alex Alderton, Roberto Amato, Jeffrey Barrett, Sonia Goncalves, Ewan Harrison, David K. Jackson, Ian Johnston, Dominic Kwiatkowski, Cordelia Langford, John Sillitoe on behalf of the Wellcome Sanger Institute COVID-19 Surveillance Team |
| EPI_ISL_2767028 | Lighthouse Lab in Milton Keynes | Wellcome Sanger Institute for the COVID-19 Genomics UK (COG-UK) Consortium | The Lighthouse Lab in Milton Keynes and Alex Alderton, Roberto Amato, Jeffrey Barrett, Sonia Goncalves, Ewan Harrison, David K. Jackson, Ian Johnston, Dominic Kwiatkowski, Cordelia Langford, John Sillitoe on behalf of the Wellcome Sanger Institute COVID-19 Surveillance Team |
| EPI_ISL_2767029, EPI_ISL_2767030 | Lighthouse Lab in Alderley Park | Wellcome Sanger Institute for the COVID-19 Genomics UK (COG-UK) Consortium | Jacquelyn Wynn, Mairead Hyland, The Lighthouse Lab in Alderley Park and Alex Alderton, Roberto Amato, Jeffrey Barrett, Sonia Goncalves, Ewan Harrison, David K. Jackson, Ian Johnston, Dominic Kwiatkowski, Cordelia Langford, John Sillitoe on behalf of the Wellcome Sanger Institute COVID-19 Surveillance Team |
| EPI_ISL_2767031, EPI_ISL_2767032 | Lighthouse Lab in Milton Keynes | Wellcome Sanger Institute for the COVID-19 Genomics UK (COG-UK) Consortium | The Lighthouse Lab in Milton Keynes and Alex Alderton, Roberto Amato, Jeffrey Barrett, Sonia Goncalves, Ewan Harrison, David K. Jackson, Ian Johnston, Dominic Kwiatkowski, Cordelia Langford, John Sillitoe on behalf of the Wellcome Sanger Institute COVID-19 Surveillance Team |
| EPI_ISL_2767033, EPI_ISL_2767034, EPI_ISL_2767035, EPI_ISL_2767036, EPI_ISL_2767037, EPI_ISL_2767038 | Lighthouse Lab in Alderley Park | Wellcome Sanger Institute for the COVID-19 Genomics UK (COG-UK) Consortium | Jacquelyn Wynn, Mairead Hyland, The Lighthouse Lab in Alderley Park and Alex Alderton, Roberto Amato, Jeffrey Barrett, Sonia Goncalves, Ewan Harrison, David K. Jackson, Ian Johnston, Dominic Kwiatkowski, Cordelia Langford, John Sillitoe on behalf of the Wellcome Sanger Institute COVID-19 Surveillance Team |
| EPI_ISL_2767039, EPI_ISL_2767040 | Lighthouse Lab in Milton Keynes | Wellcome Sanger Institute for the COVID-19 Genomics UK (COG-UK) Consortium | The Lighthouse Lab in Milton Keynes and Alex Alderton, Roberto Amato, Jeffrey Barrett, Sonia Goncalves, Ewan Harrison, David K. Jackson, Ian Johnston, Dominic Kwiatkowski, Cordelia Langford, John Sillitoe on behalf of the Wellcome Sanger Institute COVID-19 Surveillance Team |
| EPI_ISL_2767041, EPI_ISL_2767042, EPI_ISL_2767043, EPI_ISL_2767044 | Lighthouse Lab in Alderley Park | Wellcome Sanger Institute for the COVID-19 Genomics UK (COG-UK) Consortium | Jacquelyn Wynn, Mairead Hyland, The Lighthouse Lab in Alderley Park and Alex Alderton, Roberto Amato, Jeffrey Barrett, Sonia Goncalves, Ewan Harrison, David K. Jackson, Ian Johnston, Dominic Kwiatkowski, Cordelia Langford, John Sillitoe on behalf of the Wellcome Sanger Institute COVID-19 Surveillance Team |
| EPI_ISL_2767046, EPI_ISL_2767047 | Lighthouse Lab in Milton Keynes | Wellcome Sanger Institute for the COVID-19 Genomics UK (COG-UK) Consortium | The Lighthouse Lab in Milton Keynes and Alex Alderton, Roberto Amato, Jeffrey Barrett, Sonia Goncalves, Ewan Harrison, David K. Jackson, Ian Johnston, Dominic Kwiatkowski, Cordelia Langford, John Sillitoe on behalf of the Wellcome Sanger Institute COVID-19 Surveillance Team |
| EPI_ISL_2767048, EPI_ISL_2767049, EPI_ISL_2767050 | Lighthouse Lab in Alderley Park | Wellcome Sanger Institute for the COVID-19 Genomics UK (COG-UK) Consortium | Jacquelyn Wynn, Mairead Hyland, The Lighthouse Lab in Alderley Park and Alex Alderton, Roberto Amato, Jeffrey Barrett, Sonia Goncalves, Ewan Harrison, David K. Jackson, Ian Johnston, Dominic Kwiatkowski, Cordelia Langford, John Sillitoe on behalf of the Wellcome Sanger Institute COVID-19 Surveillance Team |
| EPI_ISL_2767051 | Lighthouse Lab in Milton Keynes | Wellcome Sanger Institute for the COVID-19 Genomics UK (COG-UK) Consortium | The Lighthouse Lab in Milton Keynes and Alex Alderton, Roberto Amato, Jeffrey Barrett, Sonia Goncalves, Ewan Harrison, David K. Jackson, Ian Johnston, Dominic Kwiatkowski, Cordelia Langford, John Sillitoe on behalf of the Wellcome Sanger Institute COVID-19 Surveillance Team |
| EPI_ISL_2767052, EPI_ISL_2767053, EPI_ISL_2767054, EPI_ISL_2767055, EPI_ISL_2767056, EPI_ISL_2767057 | Lighthouse Lab in Alderley Park | Wellcome Sanger Institute for the COVID-19 Genomics UK (COG-UK) Consortium | Jacquelyn Wynn, Mairead Hyland, The Lighthouse Lab in Alderley Park and Alex Alderton, Roberto Amato, Jeffrey Barrett, Sonia Goncalves, Ewan Harrison, David K. Jackson, Ian Johnston, Dominic Kwiatkowski, Cordelia Langford, John Sillitoe on behalf of the Wellcome Sanger Institute COVID-19 Surveillance Team |
| EPI_ISL_2767058 | Lighthouse Lab in Milton Keynes | Wellcome Sanger Institute for the COVID-19 Genomics UK (COG-UK) Consortium | The Lighthouse Lab in Milton Keynes and Alex Alderton, Roberto Amato, Jeffrey Barrett, Sonia Goncalves, Ewan Harrison, David K. Jackson, Ian Johnston, Dominic Kwiatkowski, Cordelia Langford, John Sillitoe on behalf of the Wellcome Sanger Institute COVID-19 Surveillance Team |
| EPI_ISL_2767059, EPI_ISL_2767060, EPI_ISL_2767061, EPI_ISL_2767062, EPI_ISL_2767063, EPI_ISL_2767064, EPI_ISL_2767065 | Lighthouse Lab in Alderley Park | Wellcome Sanger Institute for the COVID-19 Genomics UK (COG-UK) Consortium | Jacquelyn Wynn, Mairead Hyland, The Lighthouse Lab in Alderley Park and Alex Alderton, Roberto Amato, Jeffrey Barrett, Sonia Goncalves, Ewan Harrison, David K. Jackson, Ian Johnston, Dominic Kwiatkowski, Cordelia Langford, John Sillitoe on behalf of the Wellcome Sanger Institute COVID-19 Surveillance Team |
| EPI_ISL_2767066 | Lighthouse Lab in Milton Keynes | Wellcome Sanger Institute for the COVID-19 Genomics UK (COG-UK) Consortium | The Lighthouse Lab in Milton Keynes and Alex Alderton, Roberto Amato, Jeffrey Barrett, Sonia Goncalves, Ewan Harrison, David K. Jackson, Ian Johnston, Dominic Kwiatkowski, Cordelia Langford, John Sillitoe on behalf of the Wellcome Sanger Institute COVID-19 Surveillance Team |
| EPI_ISL_2767067, EPI_ISL_2767068, EPI_ISL_2767069, EPI_ISL_2767070, EPI_ISL_2767071 | Lighthouse Lab in Alderley Park | Wellcome Sanger Institute for the COVID-19 Genomics UK (COG-UK) Consortium | Jacquelyn Wynn, Mairead Hyland, The Lighthouse Lab in Alderley Park and Alex Alderton, Roberto Amato, Jeffrey Barrett, Sonia Goncalves, Ewan Harrison, David K. Jackson, Ian Johnston, Dominic Kwiatkowski, Cordelia Langford, John Sillitoe on behalf of the Wellcome Sanger Institute COVID-19 Surveillance Team |
| EPI_ISL_2767072 | Lighthouse Lab in Milton Keynes | Wellcome Sanger Institute for the COVID-19 Genomics UK (COG-UK) Consortium | The Lighthouse Lab in Milton Keynes and Alex Alderton, Roberto Amato, Jeffrey Barrett, Sonia Goncalves, Ewan Harrison, David K. Jackson, Ian Johnston, Dominic Kwiatkowski, Cordelia Langford, John Sillitoe on behalf of the Wellcome Sanger Institute COVID-19 Surveillance Team |
| EPI_ISL_2767073, EPI_ISL_2767074, EPI_ISL_2767075, EPI_ISL_2767076, EPI_ISL_2767077, EPI_ISL_2767078, EPI_ISL_2767079, EPI_ISL_2767080 | Lighthouse Lab in Alderley Park | Wellcome Sanger Institute for the COVID-19 Genomics UK (COG-UK) Consortium | Jacquelyn Wynn, Mairead Hyland, The Lighthouse Lab in Alderley Park and Alex Alderton, Roberto Amato, Jeffrey Barrett, Sonia Goncalves, Ewan Harrison, David K. Jackson, Ian Johnston, Dominic Kwiatkowski, Cordelia Langford, John Sillitoe on behalf of the Wellcome Sanger Institute COVID-19 Surveillance Team |

| | | | |
|---|---|---|---|
| EPI_ISL_2767081 | Lighthouse Lab in Milton Keynes | Wellcome Sanger Institute for the COVID-19 Genomics UK (COG-UK) Consortium | The Lighthouse Lab in Milton Keynes and Alex Alderton, Roberto Amato, Jeffrey Barrett, Sonia Goncalves, Ewan Harrison, David K. Jackson, Ian Johnston, Dominic Kwiatkowski, Cordelia Langford, John Sillitoe on behalf of the Wellcome Sanger Institute COVID-19 Surveillance Team |
| EPI_ISL_2767082 | Lighthouse Lab in Alderley Park | Wellcome Sanger Institute for the COVID-19 Genomics UK (COG-UK) Consortium | Jacquelyn Wynn, Mairead Hyland, The Lighthouse Lab in Alderley Park and Alex Alderton, Roberto Amato, Jeffrey Barrett, Sonia Goncalves, Ewan Harrison, David K. Jackson, Ian Johnston, Dominic Kwiatkowski, Cordelia Langford, John Sillitoe on behalf of the Wellcome Sanger Institute COVID-19 Surveillance Team |
| EPI_ISL_2767083, EPI_ISL_2767084 | Lighthouse Lab in Milton Keynes | Wellcome Sanger Institute for the COVID-19 Genomics UK (COG-UK) Consortium | The Lighthouse Lab in Milton Keynes and Alex Alderton, Roberto Amato, Jeffrey Barrett, Sonia Goncalves, Ewan Harrison, David K. Jackson, Ian Johnston, Dominic Kwiatkowski, Cordelia Langford, John Sillitoe on behalf of the Wellcome Sanger Institute COVID-19 Surveillance Team |
| EPI_ISL_2767085, EPI_ISL_2767086, EPI_ISL_2767087 | Lighthouse Lab in Alderley Park | Wellcome Sanger Institute for the COVID-19 Genomics UK (COG-UK) Consortium | Jacquelyn Wynn, Mairead Hyland, The Lighthouse Lab in Alderley Park and Alex Alderton, Roberto Amato, Jeffrey Barrett, Sonia Goncalves, Ewan Harrison, David K. Jackson, Ian Johnston, Dominic Kwiatkowski, Cordelia Langford, John Sillitoe on behalf of the Wellcome Sanger Institute COVID-19 Surveillance Team |
| EPI_ISL_2767088, EPI_ISL_2767089 | Lighthouse Lab in Milton Keynes | Wellcome Sanger Institute for the COVID-19 Genomics UK (COG-UK) Consortium | The Lighthouse Lab in Milton Keynes and Alex Alderton, Roberto Amato, Jeffrey Barrett, Sonia Goncalves, Ewan Harrison, David K. Jackson, Ian Johnston, Dominic Kwiatkowski, Cordelia Langford, John Sillitoe on behalf of the Wellcome Sanger Institute COVID-19 Surveillance Team |
| EPI_ISL_2767090 | Lighthouse Lab in Alderley Park | Wellcome Sanger Institute for the COVID-19 Genomics UK (COG-UK) Consortium | Jacquelyn Wynn, Mairead Hyland, The Lighthouse Lab in Alderley Park and Alex Alderton, Roberto Amato, Jeffrey Barrett, Sonia Goncalves, Ewan Harrison, David K. Jackson, Ian Johnston, Dominic Kwiatkowski, Cordelia Langford, John Sillitoe on behalf of the Wellcome Sanger Institute COVID-19 Surveillance Team |
| EPI_ISL_2767091, EPI_ISL_2767092 | Lighthouse Lab in Milton Keynes | Wellcome Sanger Institute for the COVID-19 Genomics UK (COG-UK) Consortium | The Lighthouse Lab in Milton Keynes and Alex Alderton, Roberto Amato, Jeffrey Barrett, Sonia Goncalves, Ewan Harrison, David K. Jackson, Ian Johnston, Dominic Kwiatkowski, Cordelia Langford, John Sillitoe on behalf of the Wellcome Sanger Institute COVID-19 Surveillance Team |
| EPI_ISL_2767093, EPI_ISL_2767094, EPI_ISL_2767095, EPI_ISL_2767096, EPI_ISL_2767097, EPI_ISL_2767098 | Lighthouse Lab in Alderley Park | Wellcome Sanger Institute for the COVID-19 Genomics UK (COG-UK) Consortium | Jacquelyn Wynn, Mairead Hyland, The Lighthouse Lab in Alderley Park and Alex Alderton, Roberto Amato, Jeffrey Barrett, Sonia Goncalves, Ewan Harrison, David K. Jackson, Ian Johnston, Dominic Kwiatkowski, Cordelia Langford, John Sillitoe on behalf of the Wellcome Sanger Institute COVID-19 Surveillance Team |
| EPI_ISL_2767099 | Lighthouse Lab in Milton Keynes | Wellcome Sanger Institute for the COVID-19 Genomics UK (COG-UK) Consortium | The Lighthouse Lab in Milton Keynes and Alex Alderton, Roberto Amato, Jeffrey Barrett, Sonia Goncalves, Ewan Harrison, David K. Jackson, Ian Johnston, Dominic Kwiatkowski, Cordelia Langford, John Sillitoe on behalf of the Wellcome Sanger Institute COVID-19 Surveillance Team |
| EPI_ISL_2767100 | Lighthouse Lab in Alderley Park | Wellcome Sanger Institute for the COVID-19 Genomics UK (COG-UK) Consortium | Jacquelyn Wynn, Mairead Hyland, The Lighthouse Lab in Alderley Park and Alex Alderton, Roberto Amato, Jeffrey Barrett, Sonia Goncalves, Ewan Harrison, David K. Jackson, Ian Johnston, Dominic Kwiatkowski, Cordelia Langford, John Sillitoe on behalf of the Wellcome Sanger Institute COVID-19 Surveillance Team |
| EPI_ISL_2767101 | Lighthouse Lab in Milton Keynes | Wellcome Sanger Institute for the COVID-19 Genomics UK (COG-UK) Consortium | The Lighthouse Lab in Milton Keynes and Alex Alderton, Roberto Amato, Jeffrey Barrett, Sonia Goncalves, Ewan Harrison, David K. Jackson, Ian Johnston, Dominic Kwiatkowski, Cordelia Langford, John Sillitoe on behalf of the Wellcome Sanger Institute COVID-19 Surveillance Team |
| EPI_ISL_2767102, EPI_ISL_2767103, EPI_ISL_2767104, EPI_ISL_2767105, EPI_ISL_2767106, EPI_ISL_2767107, EPI_ISL_2767108 | Lighthouse Lab in Alderley Park | Wellcome Sanger Institute for the COVID-19 Genomics UK (COG-UK) Consortium | Jacquelyn Wynn, Mairead Hyland, The Lighthouse Lab in Alderley Park and Alex Alderton, Roberto Amato, Jeffrey Barrett, Sonia Goncalves, Ewan Harrison, David K. Jackson, Ian Johnston, Dominic Kwiatkowski, Cordelia Langford, John Sillitoe on behalf of the Wellcome Sanger Institute COVID-19 Surveillance Team |
| EPI_ISL_2767109 | Lighthouse Lab in Milton Keynes | Wellcome Sanger Institute for the COVID-19 Genomics UK (COG-UK) Consortium | The Lighthouse Lab in Milton Keynes and Alex Alderton, Roberto Amato, Jeffrey Barrett, Sonia Goncalves, Ewan Harrison, David K. Jackson, Ian Johnston, Dominic Kwiatkowski, Cordelia Langford, John Sillitoe on behalf of the Wellcome Sanger Institute COVID-19 Surveillance Team |
| EPI_ISL_2767110, EPI_ISL_2767111, EPI_ISL_2767112 | Lighthouse Lab in Alderley Park | Wellcome Sanger Institute for the COVID-19 Genomics UK (COG-UK) Consortium | Jacquelyn Wynn, Mairead Hyland, The Lighthouse Lab in Alderley Park and Alex Alderton, Roberto Amato, Jeffrey Barrett, Sonia Goncalves, Ewan Harrison, David K. Jackson, Ian Johnston, Dominic Kwiatkowski, Cordelia Langford, John Sillitoe on behalf of the Wellcome Sanger Institute COVID-19 Surveillance Team |
| EPI_ISL_2767113 | Lighthouse Lab in Milton Keynes | Wellcome Sanger Institute for the COVID-19 Genomics UK (COG-UK) Consortium | The Lighthouse Lab in Milton Keynes and Alex Alderton, Roberto Amato, Jeffrey Barrett, Sonia Goncalves, Ewan Harrison, David K. Jackson, Ian Johnston, Dominic Kwiatkowski, Cordelia Langford, John Sillitoe on behalf of the Wellcome Sanger Institute COVID-19 Surveillance Team |
| EPI_ISL_2767114, EPI_ISL_2767115 | Lighthouse Lab in Alderley Park | Wellcome Sanger Institute for the COVID-19 Genomics UK (COG-UK) Consortium | Jacquelyn Wynn, Mairead Hyland, The Lighthouse Lab in Alderley Park and Alex Alderton, Roberto Amato, Jeffrey Barrett, Sonia Goncalves, Ewan Harrison, David K. Jackson, Ian Johnston, Dominic Kwiatkowski, Cordelia Langford, John Sillitoe on behalf of the Wellcome Sanger Institute COVID-19 Surveillance Team |
| EPI_ISL_2767116 | Lighthouse Lab in Milton Keynes | Wellcome Sanger Institute for the COVID-19 Genomics UK (COG-UK) Consortium | The Lighthouse Lab in Milton Keynes and Alex Alderton, Roberto Amato, Jeffrey Barrett, Sonia Goncalves, Ewan Harrison, David K. Jackson, Ian Johnston, Dominic Kwiatkowski, Cordelia Langford, John Sillitoe on behalf of the Wellcome Sanger Institute COVID-19 Surveillance Team |
| EPI_ISL_2767117 | Lighthouse Lab in Alderley Park | Wellcome Sanger Institute for the COVID-19 Genomics UK (COG-UK) Consortium | Jacquelyn Wynn, Mairead Hyland, The Lighthouse Lab in Alderley Park and Alex Alderton, Roberto Amato, Jeffrey Barrett, Sonia Goncalves, Ewan Harrison, David K. Jackson, Ian Johnston, Dominic Kwiatkowski, Cordelia Langford, John Sillitoe on behalf of the Wellcome Sanger Institute COVID-19 Surveillance Team |
| EPI_ISL_2767118 | Lighthouse Lab in Milton Keynes | Wellcome Sanger Institute for the COVID-19 Genomics UK (COG-UK) Consortium | The Lighthouse Lab in Milton Keynes and Alex Alderton, Roberto Amato, Jeffrey Barrett, Sonia Goncalves, Ewan Harrison, David K. Jackson, Ian Johnston, Dominic Kwiatkowski, Cordelia Langford, John Sillitoe on behalf of the Wellcome Sanger Institute COVID-19 Surveillance Team |
| EPI_ISL_2767119, EPI_ISL_2767120, EPI_ISL_2767121, EPI_ISL_2767122, EPI_ISL_2767123 | Lighthouse Lab in Alderley Park | Wellcome Sanger Institute for the COVID-19 Genomics UK (COG-UK) Consortium | Jacquelyn Wynn, Mairead Hyland, The Lighthouse Lab in Alderley Park and Alex Alderton, Roberto Amato, Jeffrey Barrett, Sonia Goncalves, Ewan Harrison, David K. Jackson, Ian Johnston, Dominic Kwiatkowski, Cordelia Langford, John Sillitoe on behalf of the Wellcome Sanger Institute COVID-19 Surveillance Team |
| EPI_ISL_2767124 | Lighthouse Lab in Milton Keynes | Wellcome Sanger Institute for the COVID-19 Genomics UK (COG-UK) Consortium | The Lighthouse Lab in Milton Keynes and Alex Alderton, Roberto Amato, Jeffrey Barrett, Sonia Goncalves, Ewan Harrison, David K. Jackson, Ian Johnston, Dominic Kwiatkowski, Cordelia Langford, John Sillitoe on behalf of the Wellcome Sanger Institute COVID-19 Surveillance Team |
| EPI_ISL_2767125, EPI_ISL_2767126, EPI_ISL_2767127, EPI_ISL_2767128, EPI_ISL_2767129, EPI_ISL_2767130 | Lighthouse Lab in Alderley Park | Wellcome Sanger Institute for the COVID-19 Genomics UK (COG-UK) Consortium | Jacquelyn Wynn, Mairead Hyland, The Lighthouse Lab in Alderley Park and Alex Alderton, Roberto Amato, Jeffrey Barrett, Sonia Goncalves, Ewan Harrison, David K. Jackson, Ian Johnston, Dominic Kwiatkowski, Cordelia Langford, John Sillitoe on behalf of the Wellcome Sanger Institute COVID-19 Surveillance Team |
| EPI_ISL_2767131 | Lighthouse Lab in Milton Keynes | Wellcome Sanger Institute for the COVID-19 Genomics UK (COG-UK) Consortium | The Lighthouse Lab in Milton Keynes and Alex Alderton, Roberto Amato, Jeffrey Barrett, Sonia Goncalves, Ewan Harrison, David K. Jackson, Ian Johnston, Dominic Kwiatkowski, Cordelia Langford, John Sillitoe on behalf of the Wellcome Sanger Institute COVID-19 Surveillance Team |
| EPI_ISL_2767132, EPI_ISL_2767133, EPI_ISL_2767134 | Lighthouse Lab in Alderley Park | Wellcome Sanger Institute for the COVID-19 Genomics UK (COG-UK) Consortium | Jacquelyn Wynn, Mairead Hyland, The Lighthouse Lab in Alderley Park and Alex Alderton, Roberto Amato, Jeffrey Barrett, Sonia Goncalves, Ewan Harrison, David K. Jackson, Ian Johnston, Dominic Kwiatkowski, Cordelia Langford, John Sillitoe on behalf of the Wellcome Sanger Institute COVID-19 Surveillance Team |
| EPI_ISL_2767135, EPI_ISL_2767136 | Lighthouse Lab in Milton Keynes | Wellcome Sanger Institute for the COVID-19 Genomics UK (COG-UK) Consortium | The Lighthouse Lab in Milton Keynes and Alex Alderton, Roberto Amato, Jeffrey Barrett, Sonia Goncalves, Ewan Harrison, David K. Jackson, Ian Johnston, Dominic Kwiatkowski, Cordelia Langford, John Sillitoe on behalf of the Wellcome Sanger Institute COVID-19 Surveillance Team |
| EPI_ISL_2767137 | Lighthouse Lab in Alderley Park | Wellcome Sanger Institute for the COVID-19 Genomics UK (COG-UK) Consortium | Jacquelyn Wynn, Mairead Hyland, The Lighthouse Lab in Alderley Park and Alex Alderton, Roberto Amato, Jeffrey Barrett, Sonia Goncalves, Ewan Harrison, David K. Jackson, Ian Johnston, Dominic Kwiatkowski, Cordelia Langford, John Sillitoe on behalf of the Wellcome Sanger Institute COVID-19 Surveillance Team |
| EPI_ISL_2767138, EPI_ISL_2767139 | Lighthouse Lab in Milton Keynes | Wellcome Sanger Institute for the COVID-19 Genomics UK (COG-UK) Consortium | The Lighthouse Lab in Milton Keynes and Alex Alderton, Roberto Amato, Jeffrey Barrett, Sonia Goncalves, Ewan Harrison, David K. Jackson, Ian Johnston, Dominic Kwiatkowski, Cordelia Langford, John Sillitoe on behalf of the Wellcome Sanger Institute COVID-19 Surveillance Team |
| EPI_ISL_2767140, EPI_ISL_2767141, EPI_ISL_2767142 | Lighthouse Lab in Alderley Park | Wellcome Sanger Institute for the COVID-19 Genomics UK (COG-UK) Consortium | Jacquelyn Wynn, Mairead Hyland, The Lighthouse Lab in Alderley Park and Alex Alderton, Roberto Amato, Jeffrey Barrett, Sonia Goncalves, Ewan Harrison, David K. Jackson, Ian Johnston, Dominic Kwiatkowski, Cordelia Langford, John Sillitoe on behalf of the Wellcome Sanger Institute COVID-19 Surveillance Team |
| EPI_ISL_2767143 | Lighthouse Lab in Milton Keynes | Wellcome Sanger Institute for the COVID-19 Genomics UK (COG-UK) Consortium | The Lighthouse Lab in Milton Keynes and Alex Alderton, Roberto Amato, Jeffrey Barrett, Sonia Goncalves, Ewan Harrison, David K. Jackson, Ian Johnston, Dominic Kwiatkowski, Cordelia Langford, John Sillitoe on behalf of the Wellcome Sanger Institute COVID-19 Surveillance Team |
| EPI_ISL_2767144, EPI_ISL_2767145, EPI_ISL_2767146, EPI_ISL_2767147, EPI_ISL_2767148, EPI_ISL_2767149, EPI_ISL_2767150, EPI_ISL_2767151 | Lighthouse Lab in Alderley Park | Wellcome Sanger Institute for the COVID-19 Genomics UK (COG-UK) Consortium | Jacquelyn Wynn, Mairead Hyland, The Lighthouse Lab in Alderley Park and Alex Alderton, Roberto Amato, Jeffrey Barrett, Sonia Goncalves, Ewan Harrison, David K. Jackson, Ian Johnston, Dominic Kwiatkowski, Cordelia Langford, John Sillitoe on behalf of the Wellcome Sanger Institute COVID-19 Surveillance Team |
| EPI_ISL_2767152, EPI_ISL_2767153 | Lighthouse Lab in Milton Keynes | Wellcome Sanger Institute for the COVID-19 Genomics UK (COG-UK) Consortium | The Lighthouse Lab in Milton Keynes and Alex Alderton, Roberto Amato, Jeffrey Barrett, Sonia Goncalves, Ewan Harrison, David K. Jackson, Ian Johnston, Dominic Kwiatkowski, Cordelia Langford, John Sillitoe on behalf of the Wellcome Sanger Institute COVID-19 Surveillance Team |
| EPI_ISL_2767154, EPI_ISL_2767155, EPI_ISL_2767156, EPI_ISL_2767157, EPI_ISL_2767158 | Lighthouse Lab in Alderley Park | Wellcome Sanger Institute for the COVID-19 Genomics UK (COG-UK) Consortium | Jacquelyn Wynn, Mairead Hyland, The Lighthouse Lab in Alderley Park and Alex Alderton, Roberto Amato, Jeffrey Barrett, Sonia Goncalves, Ewan Harrison, David K. Jackson, Ian Johnston, Dominic Kwiatkowski, Cordelia Langford, John Sillitoe on behalf of the Wellcome Sanger Institute COVID-19 Surveillance Team |
| EPI_ISL_2767159 | Lighthouse Lab in Milton Keynes | Wellcome Sanger Institute for the COVID-19 Genomics UK (COG-UK) Consortium | The Lighthouse Lab in Milton Keynes and Alex Alderton, Roberto Amato, Jeffrey Barrett, Sonia Goncalves, Ewan Harrison, David K. Jackson, Ian Johnston, Dominic Kwiatkowski, Cordelia Langford, John Sillitoe on behalf of the Wellcome Sanger Institute COVID-19 Surveillance Team |
| EPI_ISL_2767160, | Lighthouse Lab in Alderley Park | Wellcome Sanger Institute for the COVID-19 Genomics UK (COG-UK) Consortium | Jacquelyn Wynn, Mairead Hyland, The Lighthouse Lab in Alderley Park and Alex Alderton, Roberto Amato, Jeffrey Barrett, Sonia Goncalves, Ewan Harrison, David K. Jackson, Ian Johnston, Dominic Kwiatkowski, Cordelia Langford, John Sillitoe on |

| | | | |
|---|---|---|---|
| EPI_ISL_2767161, EPI_ISL_2767162, EPI_ISL_2767163 | | Consortium | behalf of the Wellcome Sanger Institute COVID-19 Surveillance Team |
| EPI_ISL_2767164 | Lighthouse Lab in Milton Keynes | Wellcome Sanger Institute for the COVID-19 Genomics UK (COG-UK) Consortium | The Lighthouse Lab in Milton Keynes and Alex Alderton, Roberto Amato, Jeffrey Barrett, Sonia Goncalves, Ewan Harrison, David K. Jackson, Ian Johnston, Dominic Kwiatkowski, Cordelia Langford, John Sillitoe on behalf of the Wellcome Sanger Institute COVID-19 Surveillance Team |
| EPI_ISL_2767165, EPI_ISL_2767166 | Lighthouse Lab in Alderley Park | Wellcome Sanger Institute for the COVID-19 Genomics UK (COG-UK) Consortium | Jacquelyn Wynn, Mairead Hyland, The Lighthouse Lab in Alderley Park and Alex Alderton, Roberto Amato, Jeffrey Barrett, Sonia Goncalves, Ewan Harrison, David K. Jackson, Ian Johnston, Dominic Kwiatkowski, Cordelia Langford, John Sillitoe on behalf of the Wellcome Sanger Institute COVID-19 Surveillance Team |
| EPI_ISL_2767167 | Lighthouse Lab in Milton Keynes | Wellcome Sanger Institute for the COVID-19 Genomics UK (COG-UK) Consortium | The Lighthouse Lab in Milton Keynes and Alex Alderton, Roberto Amato, Jeffrey Barrett, Sonia Goncalves, Ewan Harrison, David K. Jackson, Ian Johnston, Dominic Kwiatkowski, Cordelia Langford, John Sillitoe on behalf of the Wellcome Sanger Institute COVID-19 Surveillance Team |
| EPI_ISL_2767168, EPI_ISL_2767169 | Lighthouse Lab in Alderley Park | Wellcome Sanger Institute for the COVID-19 Genomics UK (COG-UK) Consortium | Jacquelyn Wynn, Mairead Hyland, The Lighthouse Lab in Alderley Park and Alex Alderton, Roberto Amato, Jeffrey Barrett, Sonia Goncalves, Ewan Harrison, David K. Jackson, Ian Johnston, Dominic Kwiatkowski, Cordelia Langford, John Sillitoe on behalf of the Wellcome Sanger Institute COVID-19 Surveillance Team |
| EPI_ISL_2767170, EPI_ISL_2767171 | Lighthouse Lab in Milton Keynes | Wellcome Sanger Institute for the COVID-19 Genomics UK (COG-UK) Consortium | The Lighthouse Lab in Milton Keynes and Alex Alderton, Roberto Amato, Jeffrey Barrett, Sonia Goncalves, Ewan Harrison, David K. Jackson, Ian Johnston, Dominic Kwiatkowski, Cordelia Langford, John Sillitoe on behalf of the Wellcome Sanger Institute COVID-19 Surveillance Team |
| EPI_ISL_2767172, EPI_ISL_2767173, EPI_ISL_2767174, EPI_ISL_2767176 | Lighthouse Lab in Alderley Park | Wellcome Sanger Institute for the COVID-19 Genomics UK (COG-UK) Consortium | Jacquelyn Wynn, Mairead Hyland, The Lighthouse Lab in Alderley Park and Alex Alderton, Roberto Amato, Jeffrey Barrett, Sonia Goncalves, Ewan Harrison, David K. Jackson, Ian Johnston, Dominic Kwiatkowski, Cordelia Langford, John Sillitoe on behalf of the Wellcome Sanger Institute COVID-19 Surveillance Team |
| EPI_ISL_2767177 | Lighthouse Lab in Milton Keynes | Wellcome Sanger Institute for the COVID-19 Genomics UK (COG-UK) Consortium | The Lighthouse Lab in Milton Keynes and Alex Alderton, Roberto Amato, Jeffrey Barrett, Sonia Goncalves, Ewan Harrison, David K. Jackson, Ian Johnston, Dominic Kwiatkowski, Cordelia Langford, John Sillitoe on behalf of the Wellcome Sanger Institute COVID-19 Surveillance Team |
| EPI_ISL_2767178 | Lighthouse Lab in Alderley Park | Wellcome Sanger Institute for the COVID-19 Genomics UK (COG-UK) Consortium | Jacquelyn Wynn, Mairead Hyland, The Lighthouse Lab in Alderley Park and Alex Alderton, Roberto Amato, Jeffrey Barrett, Sonia Goncalves, Ewan Harrison, David K. Jackson, Ian Johnston, Dominic Kwiatkowski, Cordelia Langford, John Sillitoe on behalf of the Wellcome Sanger Institute COVID-19 Surveillance Team |
| EPI_ISL_2767179, EPI_ISL_2767180 | Lighthouse Lab in Milton Keynes | Wellcome Sanger Institute for the COVID-19 Genomics UK (COG-UK) Consortium | The Lighthouse Lab in Milton Keynes and Alex Alderton, Roberto Amato, Jeffrey Barrett, Sonia Goncalves, Ewan Harrison, David K. Jackson, Ian Johnston, Dominic Kwiatkowski, Cordelia Langford, John Sillitoe on behalf of the Wellcome Sanger Institute COVID-19 Surveillance Team |
| EPI_ISL_2767181, EPI_ISL_2767182, EPI_ISL_2767183, EPI_ISL_2767184, EPI_ISL_2767185, EPI_ISL_2767186, EPI_ISL_2767187, EPI_ISL_2767188, EPI_ISL_2767189, EPI_ISL_2767190 | Lighthouse Lab in Alderley Park | Wellcome Sanger Institute for the COVID-19 Genomics UK (COG-UK) Consortium | Jacquelyn Wynn, Mairead Hyland, The Lighthouse Lab in Alderley Park and Alex Alderton, Roberto Amato, Jeffrey Barrett, Sonia Goncalves, Ewan Harrison, David K. Jackson, Ian Johnston, Dominic Kwiatkowski, Cordelia Langford, John Sillitoe on behalf of the Wellcome Sanger Institute COVID-19 Surveillance Team |
| EPI_ISL_2767191 | Lighthouse Lab in Milton Keynes | Wellcome Sanger Institute for the COVID-19 Genomics UK (COG-UK) Consortium | The Lighthouse Lab in Milton Keynes and Alex Alderton, Roberto Amato, Jeffrey Barrett, Sonia Goncalves, Ewan Harrison, David K. Jackson, Ian Johnston, Dominic Kwiatkowski, Cordelia Langford, John Sillitoe on behalf of the Wellcome Sanger Institute COVID-19 Surveillance Team |
| EPI_ISL_2767192 | Lighthouse Lab in Alderley Park | Wellcome Sanger Institute for the COVID-19 Genomics UK (COG-UK) Consortium | Jacquelyn Wynn, Mairead Hyland, The Lighthouse Lab in Alderley Park and Alex Alderton, Roberto Amato, Jeffrey Barrett, Sonia Goncalves, Ewan Harrison, David K. Jackson, Ian Johnston, Dominic Kwiatkowski, Cordelia Langford, John Sillitoe on behalf of the Wellcome Sanger Institute COVID-19 Surveillance Team |
| EPI_ISL_2767193 | Lighthouse Lab in Milton Keynes | Wellcome Sanger Institute for the COVID-19 Genomics UK (COG-UK) Consortium | The Lighthouse Lab in Milton Keynes and Alex Alderton, Roberto Amato, Jeffrey Barrett, Sonia Goncalves, Ewan Harrison, David K. Jackson, Ian Johnston, Dominic Kwiatkowski, Cordelia Langford, John Sillitoe on behalf of the Wellcome Sanger Institute COVID-19 Surveillance Team |
| EPI_ISL_2767194, EPI_ISL_2767195 | Lighthouse Lab in Alderley Park | Wellcome Sanger Institute for the COVID-19 Genomics UK (COG-UK) Consortium | Jacquelyn Wynn, Mairead Hyland, The Lighthouse Lab in Alderley Park and Alex Alderton, Roberto Amato, Jeffrey Barrett, Sonia Goncalves, Ewan Harrison, David K. Jackson, Ian Johnston, Dominic Kwiatkowski, Cordelia Langford, John Sillitoe on behalf of the Wellcome Sanger Institute COVID-19 Surveillance Team |
| EPI_ISL_2767196 | Lighthouse Lab in Milton Keynes | Wellcome Sanger Institute for the COVID-19 Genomics UK (COG-UK) Consortium | The Lighthouse Lab in Milton Keynes and Alex Alderton, Roberto Amato, Jeffrey Barrett, Sonia Goncalves, Ewan Harrison, David K. Jackson, Ian Johnston, Dominic Kwiatkowski, Cordelia Langford, John Sillitoe on behalf of the Wellcome Sanger Institute COVID-19 Surveillance Team |
| EPI_ISL_2767197, EPI_ISL_2767198, EPI_ISL_2767199, EPI_ISL_2767200, EPI_ISL_2767201 | Lighthouse Lab in Alderley Park | Wellcome Sanger Institute for the COVID-19 Genomics UK (COG-UK) Consortium | Jacquelyn Wynn, Mairead Hyland, The Lighthouse Lab in Alderley Park and Alex Alderton, Roberto Amato, Jeffrey Barrett, Sonia Goncalves, Ewan Harrison, David K. Jackson, Ian Johnston, Dominic Kwiatkowski, Cordelia Langford, John Sillitoe on behalf of the Wellcome Sanger Institute COVID-19 Surveillance Team |
| EPI_ISL_2767202 | Lighthouse Lab in Milton Keynes | Wellcome Sanger Institute for the COVID-19 Genomics UK (COG-UK) Consortium | The Lighthouse Lab in Milton Keynes and Alex Alderton, Roberto Amato, Jeffrey Barrett, Sonia Goncalves, Ewan Harrison, David K. Jackson, Ian Johnston, Dominic Kwiatkowski, Cordelia Langford, John Sillitoe on behalf of the Wellcome Sanger Institute COVID-19 Surveillance Team |
| EPI_ISL_2767203 | Lighthouse Lab in Alderley Park | Wellcome Sanger Institute for the COVID-19 Genomics UK (COG-UK) Consortium | Jacquelyn Wynn, Mairead Hyland, The Lighthouse Lab in Alderley Park and Alex Alderton, Roberto Amato, Jeffrey Barrett, Sonia Goncalves, Ewan Harrison, David K. Jackson, Ian Johnston, Dominic Kwiatkowski, Cordelia Langford, John Sillitoe on behalf of the Wellcome Sanger Institute COVID-19 Surveillance Team |
| EPI_ISL_2767204 | Lighthouse Lab in Milton Keynes | Wellcome Sanger Institute for the COVID-19 Genomics UK (COG-UK) Consortium | The Lighthouse Lab in Milton Keynes and Alex Alderton, Roberto Amato, Jeffrey Barrett, Sonia Goncalves, Ewan Harrison, David K. Jackson, Ian Johnston, Dominic Kwiatkowski, Cordelia Langford, John Sillitoe on behalf of the Wellcome Sanger Institute COVID-19 Surveillance Team |
| EPI_ISL_2767206, EPI_ISL_2767207 | Lighthouse Lab in Alderley Park | Wellcome Sanger Institute for the COVID-19 Genomics UK (COG-UK) Consortium | Jacquelyn Wynn, Mairead Hyland, The Lighthouse Lab in Alderley Park and Alex Alderton, Roberto Amato, Jeffrey Barrett, Sonia Goncalves, Ewan Harrison, David K. Jackson, Ian Johnston, Dominic Kwiatkowski, Cordelia Langford, John Sillitoe on behalf of the Wellcome Sanger Institute COVID-19 Surveillance Team |
| EPI_ISL_2767208, EPI_ISL_2767209, EPI_ISL_2767210, EPI_ISL_2767211, EPI_ISL_2767212 | Lighthouse Lab in Milton Keynes | Wellcome Sanger Institute for the COVID-19 Genomics UK (COG-UK) Consortium | The Lighthouse Lab in Milton Keynes and Alex Alderton, Roberto Amato, Jeffrey Barrett, Sonia Goncalves, Ewan Harrison, David K. Jackson, Ian Johnston, Dominic Kwiatkowski, Cordelia Langford, John Sillitoe on behalf of the Wellcome Sanger Institute COVID-19 Surveillance Team |
| EPI_ISL_2767213, EPI_ISL_2767214, EPI_ISL_2767215 | Lighthouse Lab in Alderley Park | Wellcome Sanger Institute for the COVID-19 Genomics UK (COG-UK) Consortium | Jacquelyn Wynn, Mairead Hyland, The Lighthouse Lab in Alderley Park and Alex Alderton, Roberto Amato, Jeffrey Barrett, Sonia Goncalves, Ewan Harrison, David K. Jackson, Ian Johnston, Dominic Kwiatkowski, Cordelia Langford, John Sillitoe on behalf of the Wellcome Sanger Institute COVID-19 Surveillance Team |
| EPI_ISL_2767216 | Lighthouse Lab in Milton Keynes | Wellcome Sanger Institute for the COVID-19 Genomics UK (COG-UK) Consortium | The Lighthouse Lab in Milton Keynes and Alex Alderton, Roberto Amato, Jeffrey Barrett, Sonia Goncalves, Ewan Harrison, David K. Jackson, Ian Johnston, Dominic Kwiatkowski, Cordelia Langford, John Sillitoe on behalf of the Wellcome Sanger Institute COVID-19 Surveillance Team |
| EPI_ISL_2767217, EPI_ISL_2767218 | Lighthouse Lab in Alderley Park | Wellcome Sanger Institute for the COVID-19 Genomics UK (COG-UK) Consortium | Jacquelyn Wynn, Mairead Hyland, The Lighthouse Lab in Alderley Park and Alex Alderton, Roberto Amato, Jeffrey Barrett, Sonia Goncalves, Ewan Harrison, David K. Jackson, Ian Johnston, Dominic Kwiatkowski, Cordelia Langford, John Sillitoe on behalf of the Wellcome Sanger Institute COVID-19 Surveillance Team |
| EPI_ISL_2767219 | Lighthouse Lab in Milton Keynes | Wellcome Sanger Institute for the COVID-19 Genomics UK (COG-UK) Consortium | The Lighthouse Lab in Milton Keynes and Alex Alderton, Roberto Amato, Jeffrey Barrett, Sonia Goncalves, Ewan Harrison, David K. Jackson, Ian Johnston, Dominic Kwiatkowski, Cordelia Langford, John Sillitoe on behalf of the Wellcome Sanger Institute COVID-19 Surveillance Team |
| EPI_ISL_2767220, EPI_ISL_2767221, EPI_ISL_2767222, EPI_ISL_2767223, EPI_ISL_2767224, EPI_ISL_2767226 | Lighthouse Lab in Alderley Park | Wellcome Sanger Institute for the COVID-19 Genomics UK (COG-UK) Consortium | Jacquelyn Wynn, Mairead Hyland, The Lighthouse Lab in Alderley Park and Alex Alderton, Roberto Amato, Jeffrey Barrett, Sonia Goncalves, Ewan Harrison, David K. Jackson, Ian Johnston, Dominic Kwiatkowski, Cordelia Langford, John Sillitoe on behalf of the Wellcome Sanger Institute COVID-19 Surveillance Team |
| EPI_ISL_2767227 | Lighthouse Lab in Milton Keynes | Wellcome Sanger Institute for the COVID-19 Genomics UK (COG-UK) Consortium | The Lighthouse Lab in Milton Keynes and Alex Alderton, Roberto Amato, Jeffrey Barrett, Sonia Goncalves, Ewan Harrison, David K. Jackson, Ian Johnston, Dominic Kwiatkowski, Cordelia Langford, John Sillitoe on behalf of the Wellcome Sanger Institute COVID-19 Surveillance Team |
| EPI_ISL_2767228, EPI_ISL_2767229, EPI_ISL_2767230, EPI_ISL_2767231, EPI_ISL_2767232 | Lighthouse Lab in Alderley Park | Wellcome Sanger Institute for the COVID-19 Genomics UK (COG-UK) Consortium | Jacquelyn Wynn, Mairead Hyland, The Lighthouse Lab in Alderley Park and Alex Alderton, Roberto Amato, Jeffrey Barrett, Sonia Goncalves, Ewan Harrison, David K. Jackson, Ian Johnston, Dominic Kwiatkowski, Cordelia Langford, John Sillitoe on behalf of the Wellcome Sanger Institute COVID-19 Surveillance Team |
| EPI_ISL_2767233 | Lighthouse Lab in Milton Keynes | Wellcome Sanger Institute for the COVID-19 Genomics UK (COG-UK) Consortium | The Lighthouse Lab in Milton Keynes and Alex Alderton, Roberto Amato, Jeffrey Barrett, Sonia Goncalves, Ewan Harrison, David K. Jackson, Ian Johnston, Dominic Kwiatkowski, Cordelia Langford, John Sillitoe on behalf of the Wellcome Sanger Institute COVID-19 Surveillance Team |
| EPI_ISL_2767234, EPI_ISL_2767235, EPI_ISL_2767236, EPI_ISL_2767237, EPI_ISL_2767238 | Lighthouse Lab in Alderley Park | Wellcome Sanger Institute for the COVID-19 Genomics UK (COG-UK) Consortium | Jacquelyn Wynn, Mairead Hyland, The Lighthouse Lab in Alderley Park and Alex Alderton, Roberto Amato, Jeffrey Barrett, Sonia Goncalves, Ewan Harrison, David K. Jackson, Ian Johnston, Dominic Kwiatkowski, Cordelia Langford, John Sillitoe on behalf of the Wellcome Sanger Institute COVID-19 Surveillance Team |
| EPI_ISL_2767239 | Lighthouse Lab in Glasgow | Wellcome Sanger Institute for the COVID-19 Genomics UK (COG-UK) Consortium | Harper VanSteenhouse, Yumi Kasai, David Gray, Carol Clugston, Anna Dominiczak and Alex Alderton, Roberto Amato, Jeffrey Barrett, Sonia Goncalves, Ewan Harrison, David K. Jackson, Ian Johnston, Dominic Kwiatkowski, Cordelia Langford, John Sillitoe on behalf of the Wellcome Sanger Institute COVID-19 Surveillance Team |
| EPI_ISL_2767240, EPI_ISL_2767241, EPI_ISL_2767242, EPI_ISL_2767243, | Lighthouse Lab in Alderley Park | Wellcome Sanger Institute for the COVID-19 Genomics UK (COG-UK) Consortium | Jacquelyn Wynn, Mairead Hyland, The Lighthouse Lab in Alderley Park and Alex Alderton, Roberto Amato, Jeffrey Barrett, Sonia Goncalves, Ewan Harrison, David K. Jackson, Ian Johnston, Dominic Kwiatkowski, Cordelia Langford, John Sillitoe on behalf of the Wellcome Sanger Institute COVID-19 Surveillance Team |

| | | | |
|---|---|---|---|
| EPI_ISL_2767244 | | | |
| EPI_ISL_2767245, EPI_ISL_2767246, EPI_ISL_2767247 | Lighthouse Lab in Glasgow | Wellcome Sanger Institute for the COVID-19 Genomics UK (COG-UK) Consortium | Harper VanSteenhouse, Yumi Kasai, David Gray, Carol Clugston, Anna Dominiczak and Alex Alderton, Roberto Amato, Jeffrey Barrett, Sonia Goncalves, Ewan Harrison, David K. Jackson, Ian Johnston, Dominic Kwiatkowski, Cordelia Langford, John Sillitoe on behalf of the Wellcome Sanger Institute COVID-19 Surveillance Team |
| EPI_ISL_2767248, EPI_ISL_2767249 | Lighthouse Lab in Alderley Park | Wellcome Sanger Institute for the COVID-19 Genomics UK (COG-UK) Consortium | Jacquelyn Wynn, Mairead Hyland, The Lighthouse Lab in Alderley Park and Alex Alderton, Roberto Amato, Jeffrey Barrett, Sonia Goncalves, Ewan Harrison, David K. Jackson, Ian Johnston, Dominic Kwiatkowski, Cordelia Langford, John Sillitoe on behalf of the Wellcome Sanger Institute COVID-19 Surveillance Team |
| EPI_ISL_2767250, EPI_ISL_2767251 | Lighthouse Lab in Glasgow | Wellcome Sanger Institute for the COVID-19 Genomics UK (COG-UK) Consortium | Harper VanSteenhouse, Yumi Kasai, David Gray, Carol Clugston, Anna Dominiczak and Alex Alderton, Roberto Amato, Jeffrey Barrett, Sonia Goncalves, Ewan Harrison, David K. Jackson, Ian Johnston, Dominic Kwiatkowski, Cordelia Langford, John Sillitoe on behalf of the Wellcome Sanger Institute COVID-19 Surveillance Team |
| EPI_ISL_2767252, EPI_ISL_2767253, EPI_ISL_2767254 | Lighthouse Lab in Alderley Park | Wellcome Sanger Institute for the COVID-19 Genomics UK (COG-UK) Consortium | Jacquelyn Wynn, Mairead Hyland, The Lighthouse Lab in Alderley Park and Alex Alderton, Roberto Amato, Jeffrey Barrett, Sonia Goncalves, Ewan Harrison, David K. Jackson, Ian Johnston, Dominic Kwiatkowski, Cordelia Langford, John Sillitoe on behalf of the Wellcome Sanger Institute COVID-19 Surveillance Team |
| EPI_ISL_2767255 | Lighthouse Lab in Glasgow | Wellcome Sanger Institute for the COVID-19 Genomics UK (COG-UK) Consortium | Harper VanSteenhouse, Yumi Kasai, David Gray, Carol Clugston, Anna Dominiczak and Alex Alderton, Roberto Amato, Jeffrey Barrett, Sonia Goncalves, Ewan Harrison, David K. Jackson, Ian Johnston, Dominic Kwiatkowski, Cordelia Langford, John Sillitoe on behalf of the Wellcome Sanger Institute COVID-19 Surveillance Team |
| EPI_ISL_2767256, EPI_ISL_2767257 | Lighthouse Lab in Alderley Park | Wellcome Sanger Institute for the COVID-19 Genomics UK (COG-UK) Consortium | Jacquelyn Wynn, Mairead Hyland, The Lighthouse Lab in Alderley Park and Alex Alderton, Roberto Amato, Jeffrey Barrett, Sonia Goncalves, Ewan Harrison, David K. Jackson, Ian Johnston, Dominic Kwiatkowski, Cordelia Langford, John Sillitoe on behalf of the Wellcome Sanger Institute COVID-19 Surveillance Team |
| EPI_ISL_2767258 | Lighthouse Lab in Glasgow | Wellcome Sanger Institute for the COVID-19 Genomics UK (COG-UK) Consortium | Harper VanSteenhouse, Yumi Kasai, David Gray, Carol Clugston, Anna Dominiczak and Alex Alderton, Roberto Amato, Jeffrey Barrett, Sonia Goncalves, Ewan Harrison, David K. Jackson, Ian Johnston, Dominic Kwiatkowski, Cordelia Langford, John Sillitoe on behalf of the Wellcome Sanger Institute COVID-19 Surveillance Team |
| EPI_ISL_2767259, EPI_ISL_2767260 | Lighthouse Lab in Alderley Park | Wellcome Sanger Institute for the COVID-19 Genomics UK (COG-UK) Consortium | Jacquelyn Wynn, Mairead Hyland, The Lighthouse Lab in Alderley Park and Alex Alderton, Roberto Amato, Jeffrey Barrett, Sonia Goncalves, Ewan Harrison, David K. Jackson, Ian Johnston, Dominic Kwiatkowski, Cordelia Langford, John Sillitoe on behalf of the Wellcome Sanger Institute COVID-19 Surveillance Team |
| EPI_ISL_2767261 | Lighthouse Lab in Glasgow | Wellcome Sanger Institute for the COVID-19 Genomics UK (COG-UK) Consortium | Harper VanSteenhouse, Yumi Kasai, David Gray, Carol Clugston, Anna Dominiczak and Alex Alderton, Roberto Amato, Jeffrey Barrett, Sonia Goncalves, Ewan Harrison, David K. Jackson, Ian Johnston, Dominic Kwiatkowski, Cordelia Langford, John Sillitoe on behalf of the Wellcome Sanger Institute COVID-19 Surveillance Team |
| EPI_ISL_2767262, EPI_ISL_2767263, EPI_ISL_2767264, EPI_ISL_2767265 | Lighthouse Lab in Alderley Park | Wellcome Sanger Institute for the COVID-19 Genomics UK (COG-UK) Consortium | Jacquelyn Wynn, Mairead Hyland, The Lighthouse Lab in Alderley Park and Alex Alderton, Roberto Amato, Jeffrey Barrett, Sonia Goncalves, Ewan Harrison, David K. Jackson, Ian Johnston, Dominic Kwiatkowski, Cordelia Langford, John Sillitoe on behalf of the Wellcome Sanger Institute COVID-19 Surveillance Team |
| EPI_ISL_2767266 | Lighthouse Lab in Glasgow | Wellcome Sanger Institute for the COVID-19 Genomics UK (COG-UK) Consortium | Harper VanSteenhouse, Yumi Kasai, David Gray, Carol Clugston, Anna Dominiczak and Alex Alderton, Roberto Amato, Jeffrey Barrett, Sonia Goncalves, Ewan Harrison, David K. Jackson, Ian Johnston, Dominic Kwiatkowski, Cordelia Langford, John Sillitoe on behalf of the Wellcome Sanger Institute COVID-19 Surveillance Team |
| EPI_ISL_2767267, EPI_ISL_2767268, EPI_ISL_2767269, EPI_ISL_2767270 | Lighthouse Lab in Alderley Park | Wellcome Sanger Institute for the COVID-19 Genomics UK (COG-UK) Consortium | Jacquelyn Wynn, Mairead Hyland, The Lighthouse Lab in Alderley Park and Alex Alderton, Roberto Amato, Jeffrey Barrett, Sonia Goncalves, Ewan Harrison, David K. Jackson, Ian Johnston, Dominic Kwiatkowski, Cordelia Langford, John Sillitoe on behalf of the Wellcome Sanger Institute COVID-19 Surveillance Team |
| EPI_ISL_2767271, EPI_ISL_2767272, EPI_ISL_2767273 | Lighthouse Lab in Glasgow | Wellcome Sanger Institute for the COVID-19 Genomics UK (COG-UK) Consortium | Harper VanSteenhouse, Yumi Kasai, David Gray, Carol Clugston, Anna Dominiczak and Alex Alderton, Roberto Amato, Jeffrey Barrett, Sonia Goncalves, Ewan Harrison, David K. Jackson, Ian Johnston, Dominic Kwiatkowski, Cordelia Langford, John Sillitoe on behalf of the Wellcome Sanger Institute COVID-19 Surveillance Team |
| EPI_ISL_2767274, EPI_ISL_2767275, EPI_ISL_2767276 | Lighthouse Lab in Alderley Park | Wellcome Sanger Institute for the COVID-19 Genomics UK (COG-UK) Consortium | Jacquelyn Wynn, Mairead Hyland, The Lighthouse Lab in Alderley Park and Alex Alderton, Roberto Amato, Jeffrey Barrett, Sonia Goncalves, Ewan Harrison, David K. Jackson, Ian Johnston, Dominic Kwiatkowski, Cordelia Langford, John Sillitoe on behalf of the Wellcome Sanger Institute COVID-19 Surveillance Team |
| EPI_ISL_2767277, EPI_ISL_2767278 | Lighthouse Lab in Glasgow | Wellcome Sanger Institute for the COVID-19 Genomics UK (COG-UK) Consortium | Harper VanSteenhouse, Yumi Kasai, David Gray, Carol Clugston, Anna Dominiczak and Alex Alderton, Roberto Amato, Jeffrey Barrett, Sonia Goncalves, Ewan Harrison, David K. Jackson, Ian Johnston, Dominic Kwiatkowski, Cordelia Langford, John Sillitoe on behalf of the Wellcome Sanger Institute COVID-19 Surveillance Team |
| EPI_ISL_2767279, EPI_ISL_2767280, EPI_ISL_2767281, EPI_ISL_2767282, EPI_ISL_2767283 | Lighthouse Lab in Alderley Park | Wellcome Sanger Institute for the COVID-19 Genomics UK (COG-UK) Consortium | Jacquelyn Wynn, Mairead Hyland, The Lighthouse Lab in Alderley Park and Alex Alderton, Roberto Amato, Jeffrey Barrett, Sonia Goncalves, Ewan Harrison, David K. Jackson, Ian Johnston, Dominic Kwiatkowski, Cordelia Langford, John Sillitoe on behalf of the Wellcome Sanger Institute COVID-19 Surveillance Team |
| EPI_ISL_2767284 | Lighthouse Lab in Glasgow | Wellcome Sanger Institute for the COVID-19 Genomics UK (COG-UK) Consortium | Harper VanSteenhouse, Yumi Kasai, David Gray, Carol Clugston, Anna Dominiczak and Alex Alderton, Roberto Amato, Jeffrey Barrett, Sonia Goncalves, Ewan Harrison, David K. Jackson, Ian Johnston, Dominic Kwiatkowski, Cordelia Langford, John Sillitoe on behalf of the Wellcome Sanger Institute COVID-19 Surveillance Team |
| EPI_ISL_2767285 | Lighthouse Lab in Alderley Park | Wellcome Sanger Institute for the COVID-19 Genomics UK (COG-UK) Consortium | Jacquelyn Wynn, Mairead Hyland, The Lighthouse Lab in Alderley Park and Alex Alderton, Roberto Amato, Jeffrey Barrett, Sonia Goncalves, Ewan Harrison, David K. Jackson, Ian Johnston, Dominic Kwiatkowski, Cordelia Langford, John Sillitoe on behalf of the Wellcome Sanger Institute COVID-19 Surveillance Team |
| EPI_ISL_2767286 | Lighthouse Lab in Glasgow | Wellcome Sanger Institute for the COVID-19 Genomics UK (COG-UK) Consortium | Harper VanSteenhouse, Yumi Kasai, David Gray, Carol Clugston, Anna Dominiczak and Alex Alderton, Roberto Amato, Jeffrey Barrett, Sonia Goncalves, Ewan Harrison, David K. Jackson, Ian Johnston, Dominic Kwiatkowski, Cordelia Langford, John Sillitoe on behalf of the Wellcome Sanger Institute COVID-19 Surveillance Team |
| EPI_ISL_2767287, EPI_ISL_2767288, EPI_ISL_2767289, EPI_ISL_2767290 | Lighthouse Lab in Alderley Park | Wellcome Sanger Institute for the COVID-19 Genomics UK (COG-UK) Consortium | Jacquelyn Wynn, Mairead Hyland, The Lighthouse Lab in Alderley Park and Alex Alderton, Roberto Amato, Jeffrey Barrett, Sonia Goncalves, Ewan Harrison, David K. Jackson, Ian Johnston, Dominic Kwiatkowski, Cordelia Langford, John Sillitoe on behalf of the Wellcome Sanger Institute COVID-19 Surveillance Team |
| EPI_ISL_2767291 | Lighthouse Lab in Glasgow | Wellcome Sanger Institute for the COVID-19 Genomics UK (COG-UK) Consortium | Harper VanSteenhouse, Yumi Kasai, David Gray, Carol Clugston, Anna Dominiczak and Alex Alderton, Roberto Amato, Jeffrey Barrett, Sonia Goncalves, Ewan Harrison, David K. Jackson, Ian Johnston, Dominic Kwiatkowski, Cordelia Langford, John Sillitoe on behalf of the Wellcome Sanger Institute COVID-19 Surveillance Team |
| EPI_ISL_2767292 | Lighthouse Lab in Alderley Park | Wellcome Sanger Institute for the COVID-19 Genomics UK (COG-UK) Consortium | Jacquelyn Wynn, Mairead Hyland, The Lighthouse Lab in Alderley Park and Alex Alderton, Roberto Amato, Jeffrey Barrett, Sonia Goncalves, Ewan Harrison, David K. Jackson, Ian Johnston, Dominic Kwiatkowski, Cordelia Langford, John Sillitoe on behalf of the Wellcome Sanger Institute COVID-19 Surveillance Team |
| EPI_ISL_2767293, EPI_ISL_2767294 | Lighthouse Lab in Glasgow | Wellcome Sanger Institute for the COVID-19 Genomics UK (COG-UK) Consortium | Harper VanSteenhouse, Yumi Kasai, David Gray, Carol Clugston, Anna Dominiczak and Alex Alderton, Roberto Amato, Jeffrey Barrett, Sonia Goncalves, Ewan Harrison, David K. Jackson, Ian Johnston, Dominic Kwiatkowski, Cordelia Langford, John Sillitoe on behalf of the Wellcome Sanger Institute COVID-19 Surveillance Team |
| EPI_ISL_2767295, EPI_ISL_2767296, EPI_ISL_2767298, EPI_ISL_2767299, EPI_ISL_2767300, EPI_ISL_2767301, EPI_ISL_2767302 | Lighthouse Lab in Alderley Park | Wellcome Sanger Institute for the COVID-19 Genomics UK (COG-UK) Consortium | Jacquelyn Wynn, Mairead Hyland, The Lighthouse Lab in Alderley Park and Alex Alderton, Roberto Amato, Jeffrey Barrett, Sonia Goncalves, Ewan Harrison, David K. Jackson, Ian Johnston, Dominic Kwiatkowski, Cordelia Langford, John Sillitoe on behalf of the Wellcome Sanger Institute COVID-19 Surveillance Team |
| EPI_ISL_2767303, EPI_ISL_2767304, EPI_ISL_2767305 | Lighthouse Lab in Glasgow | Wellcome Sanger Institute for the COVID-19 Genomics UK (COG-UK) Consortium | Harper VanSteenhouse, Yumi Kasai, David Gray, Carol Clugston, Anna Dominiczak and Alex Alderton, Roberto Amato, Jeffrey Barrett, Sonia Goncalves, Ewan Harrison, David K. Jackson, Ian Johnston, Dominic Kwiatkowski, Cordelia Langford, John Sillitoe on behalf of the Wellcome Sanger Institute COVID-19 Surveillance Team |
| EPI_ISL_2767306, EPI_ISL_2767307, EPI_ISL_2767308, EPI_ISL_2767309, EPI_ISL_2767310, EPI_ISL_2767311, EPI_ISL_2767312, EPI_ISL_2767313, EPI_ISL_2767314, EPI_ISL_2767315 | Lighthouse Lab in Alderley Park | Wellcome Sanger Institute for the COVID-19 Genomics UK (COG-UK) Consortium | Jacquelyn Wynn, Mairead Hyland, The Lighthouse Lab in Alderley Park and Alex Alderton, Roberto Amato, Jeffrey Barrett, Sonia Goncalves, Ewan Harrison, David K. Jackson, Ian Johnston, Dominic Kwiatkowski, Cordelia Langford, John Sillitoe on behalf of the Wellcome Sanger Institute COVID-19 Surveillance Team |
| EPI_ISL_2767316 | Lighthouse Lab in Glasgow | Wellcome Sanger Institute for the COVID-19 Genomics UK (COG-UK) Consortium | Harper VanSteenhouse, Yumi Kasai, David Gray, Carol Clugston, Anna Dominiczak and Alex Alderton, Roberto Amato, Jeffrey Barrett, Sonia Goncalves, Ewan Harrison, David K. Jackson, Ian Johnston, Dominic Kwiatkowski, Cordelia Langford, John Sillitoe on behalf of the Wellcome Sanger Institute COVID-19 Surveillance Team |
| EPI_ISL_2767317 | Lighthouse Lab in Alderley Park | Wellcome Sanger Institute for the COVID-19 Genomics UK (COG-UK) Consortium | Jacquelyn Wynn, Mairead Hyland, The Lighthouse Lab in Alderley Park and Alex Alderton, Roberto Amato, Jeffrey Barrett, Sonia Goncalves, Ewan Harrison, David K. Jackson, Ian Johnston, Dominic Kwiatkowski, Cordelia Langford, John Sillitoe on behalf of the Wellcome Sanger Institute COVID-19 Surveillance Team |
| EPI_ISL_2767318, EPI_ISL_2767319 | Lighthouse Lab in Glasgow | Wellcome Sanger Institute for the COVID-19 Genomics UK (COG-UK) Consortium | Harper VanSteenhouse, Yumi Kasai, David Gray, Carol Clugston, Anna Dominiczak and Alex Alderton, Roberto Amato, Jeffrey Barrett, Sonia Goncalves, Ewan Harrison, David K. Jackson, Ian Johnston, Dominic Kwiatkowski, Cordelia Langford, John Sillitoe on behalf of the Wellcome Sanger Institute COVID-19 Surveillance Team |
| EPI_ISL_2767320, EPI_ISL_2767321, EPI_ISL_2767322, EPI_ISL_2767323, EPI_ISL_2767324, EPI_ISL_2767325, EPI_ISL_2767326, EPI_ISL_2767327, EPI_ISL_2767328 | Lighthouse Lab in Alderley Park | Wellcome Sanger Institute for the COVID-19 Genomics UK (COG-UK) Consortium | Jacquelyn Wynn, Mairead Hyland, The Lighthouse Lab in Alderley Park and Alex Alderton, Roberto Amato, Jeffrey Barrett, Sonia Goncalves, Ewan Harrison, David K. Jackson, Ian Johnston, Dominic Kwiatkowski, Cordelia Langford, John Sillitoe on behalf of the Wellcome Sanger Institute COVID-19 Surveillance Team |
| EPI_ISL_2767329 | Lighthouse Lab in Glasgow | Wellcome Sanger Institute for the COVID-19 Genomics UK (COG-UK) Consortium | Harper VanSteenhouse, Yumi Kasai, David Gray, Carol Clugston, Anna Dominiczak and Alex Alderton, Roberto Amato, Jeffrey Barrett, Sonia Goncalves, Ewan Harrison, David K. Jackson, Ian Johnston, Dominic Kwiatkowski, Cordelia Langford, John Sillitoe on behalf of the Wellcome Sanger Institute COVID-19 Surveillance Team |

| | | | |
|---|---|---|---|
| EPI_ISL_2767330, EPI_ISL_2767331, EPI_ISL_2767332 | Lighthouse Lab in Alderley Park | Wellcome Sanger Institute for the COVID-19 Genomics UK (COG-UK) Consortium | Jacquelyn Wynn, Mairead Hyland, The Lighthouse Lab in Alderley Park and Alex Alderton, Roberto Amato, Jeffrey Barrett, Sonia Goncalves, Ewan Harrison, David K. Jackson, Ian Johnston, Dominic Kwiatkowski, Cordelia Langford, John Sillitoe on behalf of the Wellcome Sanger Institute COVID-19 Surveillance Team |
| EPI_ISL_2767333 | Lighthouse Lab in Glasgow | Wellcome Sanger Institute for the COVID-19 Genomics UK (COG-UK) Consortium | Harper VanSteenhouse, Yumi Kasai, David Gray, Carol Clugston, Anna Dominiczak and Alex Alderton, Roberto Amato, Jeffrey Barrett, Sonia Goncalves, Ewan Harrison, David K. Jackson, Ian Johnston, Dominic Kwiatkowski, Cordelia Langford, John Sillitoe on behalf of the Wellcome Sanger Institute COVID-19 Surveillance Team |
| EPI_ISL_2767334, EPI_ISL_2767335, EPI_ISL_2767336, EPI_ISL_2767337 | Lighthouse Lab in Alderley Park | Wellcome Sanger Institute for the COVID-19 Genomics UK (COG-UK) Consortium | Jacquelyn Wynn, Mairead Hyland, The Lighthouse Lab in Alderley Park and Alex Alderton, Roberto Amato, Jeffrey Barrett, Sonia Goncalves, Ewan Harrison, David K. Jackson, Ian Johnston, Dominic Kwiatkowski, Cordelia Langford, John Sillitoe on behalf of the Wellcome Sanger Institute COVID-19 Surveillance Team |
| EPI_ISL_2767338 | Lighthouse Lab in Glasgow | Wellcome Sanger Institute for the COVID-19 Genomics UK (COG-UK) Consortium | Harper VanSteenhouse, Yumi Kasai, David Gray, Carol Clugston, Anna Dominiczak and Alex Alderton, Roberto Amato, Jeffrey Barrett, Sonia Goncalves, Ewan Harrison, David K. Jackson, Ian Johnston, Dominic Kwiatkowski, Cordelia Langford, John Sillitoe on behalf of the Wellcome Sanger Institute COVID-19 Surveillance Team |
| EPI_ISL_2767339, EPI_ISL_2767340, EPI_ISL_2767341 | Lighthouse Lab in Alderley Park | Wellcome Sanger Institute for the COVID-19 Genomics UK (COG-UK) Consortium | Jacquelyn Wynn, Mairead Hyland, The Lighthouse Lab in Alderley Park and Alex Alderton, Roberto Amato, Jeffrey Barrett, Sonia Goncalves, Ewan Harrison, David K. Jackson, Ian Johnston, Dominic Kwiatkowski, Cordelia Langford, John Sillitoe on behalf of the Wellcome Sanger Institute COVID-19 Surveillance Team |
| EPI_ISL_2767342 | Lighthouse Lab in Glasgow | Wellcome Sanger Institute for the COVID-19 Genomics UK (COG-UK) Consortium | Harper VanSteenhouse, Yumi Kasai, David Gray, Carol Clugston, Anna Dominiczak and Alex Alderton, Roberto Amato, Jeffrey Barrett, Sonia Goncalves, Ewan Harrison, David K. Jackson, Ian Johnston, Dominic Kwiatkowski, Cordelia Langford, John Sillitoe on behalf of the Wellcome Sanger Institute COVID-19 Surveillance Team |
| EPI_ISL_2767343 | Lighthouse Lab in Alderley Park | Wellcome Sanger Institute for the COVID-19 Genomics UK (COG-UK) Consortium | Jacquelyn Wynn, Mairead Hyland, The Lighthouse Lab in Alderley Park and Alex Alderton, Roberto Amato, Jeffrey Barrett, Sonia Goncalves, Ewan Harrison, David K. Jackson, Ian Johnston, Dominic Kwiatkowski, Cordelia Langford, John Sillitoe on behalf of the Wellcome Sanger Institute COVID-19 Surveillance Team |
| EPI_ISL_2767344 | Lighthouse Lab in Glasgow | Wellcome Sanger Institute for the COVID-19 Genomics UK (COG-UK) Consortium | Harper VanSteenhouse, Yumi Kasai, David Gray, Carol Clugston, Anna Dominiczak and Alex Alderton, Roberto Amato, Jeffrey Barrett, Sonia Goncalves, Ewan Harrison, David K. Jackson, Ian Johnston, Dominic Kwiatkowski, Cordelia Langford, John Sillitoe on behalf of the Wellcome Sanger Institute COVID-19 Surveillance Team |
| EPI_ISL_2767345, EPI_ISL_2767346 | Lighthouse Lab in Alderley Park | Wellcome Sanger Institute for the COVID-19 Genomics UK (COG-UK) Consortium | Jacquelyn Wynn, Mairead Hyland, The Lighthouse Lab in Alderley Park and Alex Alderton, Roberto Amato, Jeffrey Barrett, Sonia Goncalves, Ewan Harrison, David K. Jackson, Ian Johnston, Dominic Kwiatkowski, Cordelia Langford, John Sillitoe on behalf of the Wellcome Sanger Institute COVID-19 Surveillance Team |
| EPI_ISL_2767347 | Lighthouse Lab in Glasgow | Wellcome Sanger Institute for the COVID-19 Genomics UK (COG-UK) Consortium | Harper VanSteenhouse, Yumi Kasai, David Gray, Carol Clugston, Anna Dominiczak and Alex Alderton, Roberto Amato, Jeffrey Barrett, Sonia Goncalves, Ewan Harrison, David K. Jackson, Ian Johnston, Dominic Kwiatkowski, Cordelia Langford, John Sillitoe on behalf of the Wellcome Sanger Institute COVID-19 Surveillance Team |
| EPI_ISL_2767348, EPI_ISL_2767349, EPI_ISL_2767350 | Lighthouse Lab in Alderley Park | Wellcome Sanger Institute for the COVID-19 Genomics UK (COG-UK) Consortium | Jacquelyn Wynn, Mairead Hyland, The Lighthouse Lab in Alderley Park and Alex Alderton, Roberto Amato, Jeffrey Barrett, Sonia Goncalves, Ewan Harrison, David K. Jackson, Ian Johnston, Dominic Kwiatkowski, Cordelia Langford, John Sillitoe on behalf of the Wellcome Sanger Institute COVID-19 Surveillance Team |
| EPI_ISL_2767351 | Lighthouse Lab in Milton Keynes | Wellcome Sanger Institute for the COVID-19 Genomics UK (COG-UK) Consortium | The Lighthouse Lab in Milton Keynes and Alex Alderton, Roberto Amato, Jeffrey Barrett, Sonia Goncalves, Ewan Harrison, David K. Jackson, Ian Johnston, Dominic Kwiatkowski, Cordelia Langford, John Sillitoe on behalf of the Wellcome Sanger Institute COVID-19 Surveillance Team |
| EPI_ISL_2767352 | Lighthouse Laboratory Plymouth | Wellcome Sanger Institute for the COVID-19 Genomics UK (COG-UK) Consortium | Lighthouse Laboratory Plymouth and Alex Alderton, Roberto Amato, Jeffrey Barrett, Sonia Goncalves, Ewan Harrison, David K. Jackson, Ian Johnston, Dominic Kwiatkowski, Cordelia Langford, John Sillitoe on behalf of the Wellcome Sanger Institute COVID-19 Surveillance Team |
| EPI_ISL_2767353 | Lighthouse Lab in Milton Keynes | Wellcome Sanger Institute for the COVID-19 Genomics UK (COG-UK) Consortium | The Lighthouse Lab in Milton Keynes and Alex Alderton, Roberto Amato, Jeffrey Barrett, Sonia Goncalves, Ewan Harrison, David K. Jackson, Ian Johnston, Dominic Kwiatkowski, Cordelia Langford, John Sillitoe on behalf of the Wellcome Sanger Institute COVID-19 Surveillance Team |
| EPI_ISL_2767354 | Lighthouse Lab in Alderley Park | Wellcome Sanger Institute for the COVID-19 Genomics UK (COG-UK) Consortium | Jacquelyn Wynn, Mairead Hyland, The Lighthouse Lab in Alderley Park and Alex Alderton, Roberto Amato, Jeffrey Barrett, Sonia Goncalves, Ewan Harrison, David K. Jackson, Ian Johnston, Dominic Kwiatkowski, Cordelia Langford, John Sillitoe on behalf of the Wellcome Sanger Institute COVID-19 Surveillance Team |
| EPI_ISL_2767355 | Lighthouse Lab in Milton Keynes | Wellcome Sanger Institute for the COVID-19 Genomics UK (COG-UK) Consortium | The Lighthouse Lab in Milton Keynes and Alex Alderton, Roberto Amato, Jeffrey Barrett, Sonia Goncalves, Ewan Harrison, David K. Jackson, Ian Johnston, Dominic Kwiatkowski, Cordelia Langford, John Sillitoe on behalf of the Wellcome Sanger Institute COVID-19 Surveillance Team |
| EPI_ISL_2767356 | Lighthouse Laboratory Plymouth | Wellcome Sanger Institute for the COVID-19 Genomics UK (COG-UK) Consortium | Lighthouse Laboratory Plymouth and Alex Alderton, Roberto Amato, Jeffrey Barrett, Sonia Goncalves, Ewan Harrison, David K. Jackson, Ian Johnston, Dominic Kwiatkowski, Cordelia Langford, John Sillitoe on behalf of the Wellcome Sanger Institute COVID-19 Surveillance Team |
| EPI_ISL_2767357, EPI_ISL_2767358 | Lighthouse Lab in Milton Keynes | Wellcome Sanger Institute for the COVID-19 Genomics UK (COG-UK) Consortium | The Lighthouse Lab in Milton Keynes and Alex Alderton, Roberto Amato, Jeffrey Barrett, Sonia Goncalves, Ewan Harrison, David K. Jackson, Ian Johnston, Dominic Kwiatkowski, Cordelia Langford, John Sillitoe on behalf of the Wellcome Sanger Institute COVID-19 Surveillance Team |
| EPI_ISL_2767359 | Lighthouse Laboratory Plymouth | Wellcome Sanger Institute for the COVID-19 Genomics UK (COG-UK) Consortium | Lighthouse Laboratory Plymouth and Alex Alderton, Roberto Amato, Jeffrey Barrett, Sonia Goncalves, Ewan Harrison, David K. Jackson, Ian Johnston, Dominic Kwiatkowski, Cordelia Langford, John Sillitoe on behalf of the Wellcome Sanger Institute COVID-19 Surveillance Team |
| EPI_ISL_2767360 | Lighthouse Lab in Alderley Park | Wellcome Sanger Institute for the COVID-19 Genomics UK (COG-UK) Consortium | Jacquelyn Wynn, Mairead Hyland, The Lighthouse Lab in Alderley Park and Alex Alderton, Roberto Amato, Jeffrey Barrett, Sonia Goncalves, Ewan Harrison, David K. Jackson, Ian Johnston, Dominic Kwiatkowski, Cordelia Langford, John Sillitoe on behalf of the Wellcome Sanger Institute COVID-19 Surveillance Team |
| EPI_ISL_2767361, EPI_ISL_2767362 | Lighthouse Laboratory Plymouth | Wellcome Sanger Institute for the COVID-19 Genomics UK (COG-UK) Consortium | Lighthouse Laboratory Plymouth and Alex Alderton, Roberto Amato, Jeffrey Barrett, Sonia Goncalves, Ewan Harrison, David K. Jackson, Ian Johnston, Dominic Kwiatkowski, Cordelia Langford, John Sillitoe on behalf of the Wellcome Sanger Institute COVID-19 Surveillance Team |
| EPI_ISL_2767363 | Lighthouse Lab in Milton Keynes | Wellcome Sanger Institute for the COVID-19 Genomics UK (COG-UK) Consortium | The Lighthouse Lab in Milton Keynes and Alex Alderton, Roberto Amato, Jeffrey Barrett, Sonia Goncalves, Ewan Harrison, David K. Jackson, Ian Johnston, Dominic Kwiatkowski, Cordelia Langford, John Sillitoe on behalf of the Wellcome Sanger Institute COVID-19 Surveillance Team |
| EPI_ISL_2767364 | Lighthouse Laboratory Plymouth | Wellcome Sanger Institute for the COVID-19 Genomics UK (COG-UK) Consortium | Lighthouse Laboratory Plymouth and Alex Alderton, Roberto Amato, Jeffrey Barrett, Sonia Goncalves, Ewan Harrison, David K. Jackson, Ian Johnston, Dominic Kwiatkowski, Cordelia Langford, John Sillitoe on behalf of the Wellcome Sanger Institute COVID-19 Surveillance Team |
| EPI_ISL_2767365 | Lighthouse Lab in Milton Keynes | Wellcome Sanger Institute for the COVID-19 Genomics UK (COG-UK) Consortium | The Lighthouse Lab in Milton Keynes and Alex Alderton, Roberto Amato, Jeffrey Barrett, Sonia Goncalves, Ewan Harrison, David K. Jackson, Ian Johnston, Dominic Kwiatkowski, Cordelia Langford, John Sillitoe on behalf of the Wellcome Sanger Institute COVID-19 Surveillance Team |
| EPI_ISL_2767366, EPI_ISL_2767367 | Lighthouse Lab in Alderley Park | Wellcome Sanger Institute for the COVID-19 Genomics UK (COG-UK) Consortium | Jacquelyn Wynn, Mairead Hyland, The Lighthouse Lab in Alderley Park and Alex Alderton, Roberto Amato, Jeffrey Barrett, Sonia Goncalves, Ewan Harrison, David K. Jackson, Ian Johnston, Dominic Kwiatkowski, Cordelia Langford, John Sillitoe on behalf of the Wellcome Sanger Institute COVID-19 Surveillance Team |
| EPI_ISL_2767368 | Lighthouse Lab in Milton Keynes | Wellcome Sanger Institute for the COVID-19 Genomics UK (COG-UK) Consortium | The Lighthouse Lab in Milton Keynes and Alex Alderton, Roberto Amato, Jeffrey Barrett, Sonia Goncalves, Ewan Harrison, David K. Jackson, Ian Johnston, Dominic Kwiatkowski, Cordelia Langford, John Sillitoe on behalf of the Wellcome Sanger Institute COVID-19 Surveillance Team |
| EPI_ISL_2767369 | Lighthouse Laboratory Plymouth | Wellcome Sanger Institute for the COVID-19 Genomics UK (COG-UK) Consortium | Lighthouse Laboratory Plymouth and Alex Alderton, Roberto Amato, Jeffrey Barrett, Sonia Goncalves, Ewan Harrison, David K. Jackson, Ian Johnston, Dominic Kwiatkowski, Cordelia Langford, John Sillitoe on behalf of the Wellcome Sanger Institute COVID-19 Surveillance Team |
| EPI_ISL_2767370 | Lighthouse Lab in Milton Keynes | Wellcome Sanger Institute for the COVID-19 Genomics UK (COG-UK) Consortium | The Lighthouse Lab in Milton Keynes and Alex Alderton, Roberto Amato, Jeffrey Barrett, Sonia Goncalves, Ewan Harrison, David K. Jackson, Ian Johnston, Dominic Kwiatkowski, Cordelia Langford, John Sillitoe on behalf of the Wellcome Sanger Institute COVID-19 Surveillance Team |
| EPI_ISL_2767371, EPI_ISL_2767372 | Lighthouse Lab in Alderley Park | Wellcome Sanger Institute for the COVID-19 Genomics UK (COG-UK) Consortium | Jacquelyn Wynn, Mairead Hyland, The Lighthouse Lab in Alderley Park and Alex Alderton, Roberto Amato, Jeffrey Barrett, Sonia Goncalves, Ewan Harrison, David K. Jackson, Ian Johnston, Dominic Kwiatkowski, Cordelia Langford, John Sillitoe on behalf of the Wellcome Sanger Institute COVID-19 Surveillance Team |
| EPI_ISL_2767373 | Lighthouse Laboratory Plymouth | Wellcome Sanger Institute for the COVID-19 Genomics UK (COG-UK) Consortium | Lighthouse Laboratory Plymouth and Alex Alderton, Roberto Amato, Jeffrey Barrett, Sonia Goncalves, Ewan Harrison, David K. Jackson, Ian Johnston, Dominic Kwiatkowski, Cordelia Langford, John Sillitoe on behalf of the Wellcome Sanger Institute COVID-19 Surveillance Team |
| EPI_ISL_2767374, EPI_ISL_2767375, EPI_ISL_2767376 | Lighthouse Lab in Alderley Park | Wellcome Sanger Institute for the COVID-19 Genomics UK (COG-UK) Consortium | Jacquelyn Wynn, Mairead Hyland, The Lighthouse Lab in Alderley Park and Alex Alderton, Roberto Amato, Jeffrey Barrett, Sonia Goncalves, Ewan Harrison, David K. Jackson, Ian Johnston, Dominic Kwiatkowski, Cordelia Langford, John Sillitoe on behalf of the Wellcome Sanger Institute COVID-19 Surveillance Team |
| EPI_ISL_2767377 | Lighthouse Lab in Milton Keynes | Wellcome Sanger Institute for the COVID-19 Genomics UK (COG-UK) Consortium | The Lighthouse Lab in Milton Keynes and Alex Alderton, Roberto Amato, Jeffrey Barrett, Sonia Goncalves, Ewan Harrison, David K. Jackson, Ian Johnston, Dominic Kwiatkowski, Cordelia Langford, John Sillitoe on behalf of the Wellcome Sanger Institute COVID-19 Surveillance Team |
| EPI_ISL_2767378 | Lighthouse Laboratory Plymouth | Wellcome Sanger Institute for the COVID-19 Genomics UK (COG-UK) Consortium | Lighthouse Laboratory Plymouth and Alex Alderton, Roberto Amato, Jeffrey Barrett, Sonia Goncalves, Ewan Harrison, David K. Jackson, Ian Johnston, Dominic Kwiatkowski, Cordelia Langford, John Sillitoe on behalf of the Wellcome Sanger Institute COVID-19 Surveillance Team |
| EPI_ISL_2767379 | Lighthouse Lab in Milton Keynes | Wellcome Sanger Institute for the COVID-19 Genomics UK (COG-UK) Consortium | The Lighthouse Lab in Milton Keynes and Alex Alderton, Roberto Amato, Jeffrey Barrett, Sonia Goncalves, Ewan Harrison, David K. Jackson, Ian Johnston, Dominic Kwiatkowski, Cordelia Langford, John Sillitoe on behalf of the Wellcome Sanger Institute COVID-19 Surveillance Team |
| EPI_ISL_2767380, EPI_ISL_2767381 | Lighthouse Lab in Alderley Park | Wellcome Sanger Institute for the COVID-19 Genomics UK (COG-UK) Consortium | Jacquelyn Wynn, Mairead Hyland, The Lighthouse Lab in Alderley Park and Alex Alderton, Roberto Amato, Jeffrey Barrett, Sonia Goncalves, Ewan Harrison, David K. Jackson, Ian Johnston, Dominic Kwiatkowski, Cordelia Langford, John Sillitoe on behalf of the Wellcome Sanger Institute COVID-19 Surveillance Team |
| EPI_ISL_2767382 | Lighthouse Lab in Milton Keynes | Wellcome Sanger Institute for the COVID-19 Genomics UK (COG-UK) Consortium | The Lighthouse Lab in Milton Keynes and Alex Alderton, Roberto Amato, Jeffrey Barrett, Sonia Goncalves, Ewan Harrison, David K. Jackson, Ian Johnston, Dominic Kwiatkowski, Cordelia Langford, John Sillitoe on behalf of the Wellcome Sanger Institute COVID-19 Surveillance Team |
| EPI_ISL_2767383 | Lighthouse Laboratory Plymouth | Wellcome Sanger Institute for the COVID-19 Genomics UK (COG-UK) Consortium | Lighthouse Laboratory Plymouth and Alex Alderton, Roberto Amato, Jeffrey Barrett, Sonia Goncalves, Ewan Harrison, David K. Jackson, Ian Johnston, Dominic Kwiatkowski, Cordelia Langford, John Sillitoe on behalf of the Wellcome Sanger Institute COVID-19 Surveillance Team |
| EPI_ISL_2767384 | Lighthouse Lab in Glasgow | Wellcome Sanger Institute for the COVID-19 Genomics UK (COG-UK) Consortium | Harper VanSteenhouse, Yumi Kasai, David Gray, Carol Clugston, Anna Dominiczak and Alex Alderton, Roberto Amato, Jeffrey Barrett, Sonia Goncalves, Ewan Harrison, David K. Jackson, Ian Johnston, Dominic Kwiatkowski, Cordelia Langford, John Sillitoe on behalf of the Wellcome Sanger Institute COVID-19 Surveillance Team |
| EPI_ISL_2767385, EPI_ISL_2767386, EPI_ISL_2767387 | Lighthouse Lab in Milton Keynes | Wellcome Sanger Institute for the COVID-19 Genomics UK (COG-UK) Consortium | The Lighthouse Lab in Milton Keynes and Alex Alderton, Roberto Amato, Jeffrey Barrett, Sonia Goncalves, Ewan Harrison, David K. Jackson, Ian Johnston, Dominic Kwiatkowski, Cordelia Langford, John Sillitoe on behalf of the Wellcome Sanger Institute COVID-19 Surveillance Team |
| EPI_ISL_2767388 | Lighthouse Lab in Glasgow | Wellcome Sanger Institute for the COVID-19 Genomics UK (COG-UK) Consortium | Harper VanSteenhouse, Yumi Kasai, David Gray, Carol Clugston, Anna Dominiczak and Alex Alderton, Roberto Amato, Jeffrey Barrett, Sonia Goncalves, Ewan Harrison, David K. Jackson, Ian Johnston, Dominic Kwiatkowski, Cordelia Langford, John Sillitoe on behalf of the Wellcome Sanger Institute COVID-19 Surveillance Team |
| EPI_ISL_2767389 | Lighthouse Lab in Milton Keynes | Wellcome Sanger Institute for the COVID-19 Genomics UK (COG-UK) Consortium | The Lighthouse Lab in Milton Keynes and Alex Alderton, Roberto Amato, Jeffrey Barrett, Sonia Goncalves, Ewan Harrison, David K. Jackson, Ian Johnston, Dominic Kwiatkowski, Cordelia Langford, John Sillitoe on behalf of the Wellcome Sanger Institute COVID-19 Surveillance Team |
| EPI_ISL_2767390 | Lighthouse Laboratory Plymouth | Wellcome Sanger Institute for the COVID-19 Genomics UK (COG-UK) Consortium | Lighthouse Laboratory Plymouth and Alex Alderton, Roberto Amato, Jeffrey Barrett, Sonia Goncalves, Ewan Harrison, David K. Jackson, Ian Johnston, Dominic Kwiatkowski, Cordelia Langford, John Sillitoe on behalf of the Wellcome Sanger Institute COVID-19 Surveillance Team |

| | | | |
|---|---|---|---|
| EPI_ISL_2767391 | Lighthouse Lab in Glasgow | Wellcome Sanger Institute for the COVID-19 Genomics UK (COG-UK) Consortium | Harper VanSteenhouse, Yumi Kasai, David Gray, Carol Clugston, Anna Dominiczak and Alex Alderton, Roberto Amato, Jeffrey Barrett, Sonia Goncalves, Ewan Harrison, David K. Jackson, Ian Johnston, Dominic Kwiatkowski, Cordelia Langford, John Sillitoe on behalf of the Wellcome Sanger Institute COVID-19 Surveillance Team |
| EPI_ISL_2767392 | Lighthouse Lab in Alderley Park | Wellcome Sanger Institute for the COVID-19 Genomics UK (COG-UK) Consortium | Jacquelyn Wynn, Mairead Hyland, The Lighthouse Lab in Alderley Park and Alex Alderton, Roberto Amato, Jeffrey Barrett, Sonia Goncalves, Ewan Harrison, David K. Jackson, Ian Johnston, Dominic Kwiatkowski, Cordelia Langford, John Sillitoe on behalf of the Wellcome Sanger Institute COVID-19 Surveillance Team |
| EPI_ISL_2767393 | Lighthouse Lab in Milton Keynes | Wellcome Sanger Institute for the COVID-19 Genomics UK (COG-UK) Consortium | The Lighthouse Lab in Milton Keynes and Alex Alderton, Roberto Amato, Jeffrey Barrett, Sonia Goncalves, Ewan Harrison, David K. Jackson, Ian Johnston, Dominic Kwiatkowski, Cordelia Langford, John Sillitoe on behalf of the Wellcome Sanger Institute COVID-19 Surveillance Team |
| EPI_ISL_2767394 | Lighthouse Laboratory Plymouth | Wellcome Sanger Institute for the COVID-19 Genomics UK (COG-UK) Consortium | Lighthouse Laboratory Plymouth and Alex Alderton, Roberto Amato, Jeffrey Barrett, Sonia Goncalves, Ewan Harrison, David K. Jackson, Ian Johnston, Dominic Kwiatkowski, Cordelia Langford, John Sillitoe on behalf of the Wellcome Sanger Institute COVID-19 Surveillance Team |
| EPI_ISL_2767395, EPI_ISL_2767396 | Lighthouse Lab in Milton Keynes | Wellcome Sanger Institute for the COVID-19 Genomics UK (COG-UK) Consortium | The Lighthouse Lab in Milton Keynes and Alex Alderton, Roberto Amato, Jeffrey Barrett, Sonia Goncalves, Ewan Harrison, David K. Jackson, Ian Johnston, Dominic Kwiatkowski, Cordelia Langford, John Sillitoe on behalf of the Wellcome Sanger Institute COVID-19 Surveillance Team |
| EPI_ISL_2767397 | Lighthouse Lab in Alderley Park | Wellcome Sanger Institute for the COVID-19 Genomics UK (COG-UK) Consortium | Jacquelyn Wynn, Mairead Hyland, The Lighthouse Lab in Alderley Park and Alex Alderton, Roberto Amato, Jeffrey Barrett, Sonia Goncalves, Ewan Harrison, David K. Jackson, Ian Johnston, Dominic Kwiatkowski, Cordelia Langford, John Sillitoe on behalf of the Wellcome Sanger Institute COVID-19 Surveillance Team |
| EPI_ISL_2767398 | Lighthouse Lab in Glasgow | Wellcome Sanger Institute for the COVID-19 Genomics UK (COG-UK) Consortium | Harper VanSteenhouse, Yumi Kasai, David Gray, Carol Clugston, Anna Dominiczak and Alex Alderton, Roberto Amato, Jeffrey Barrett, Sonia Goncalves, Ewan Harrison, David K. Jackson, Ian Johnston, Dominic Kwiatkowski, Cordelia Langford, John Sillitoe on behalf of the Wellcome Sanger Institute COVID-19 Surveillance Team |
| EPI_ISL_2767399 | Lighthouse Lab in Milton Keynes | Wellcome Sanger Institute for the COVID-19 Genomics UK (COG-UK) Consortium | The Lighthouse Lab in Milton Keynes and Alex Alderton, Roberto Amato, Jeffrey Barrett, Sonia Goncalves, Ewan Harrison, David K. Jackson, Ian Johnston, Dominic Kwiatkowski, Cordelia Langford, John Sillitoe on behalf of the Wellcome Sanger Institute COVID-19 Surveillance Team |
| EPI_ISL_2767400 | Lighthouse Laboratory Plymouth | Wellcome Sanger Institute for the COVID-19 Genomics UK (COG-UK) Consortium | Lighthouse Laboratory Plymouth and Alex Alderton, Roberto Amato, Jeffrey Barrett, Sonia Goncalves, Ewan Harrison, David K. Jackson, Ian Johnston, Dominic Kwiatkowski, Cordelia Langford, John Sillitoe on behalf of the Wellcome Sanger Institute COVID-19 Surveillance Team |
| EPI_ISL_2767401 | Lighthouse Lab in Glasgow | Wellcome Sanger Institute for the COVID-19 Genomics UK (COG-UK) Consortium | Harper VanSteenhouse, Yumi Kasai, David Gray, Carol Clugston, Anna Dominiczak and Alex Alderton, Roberto Amato, Jeffrey Barrett, Sonia Goncalves, Ewan Harrison, David K. Jackson, Ian Johnston, Dominic Kwiatkowski, Cordelia Langford, John Sillitoe on behalf of the Wellcome Sanger Institute COVID-19 Surveillance Team |
| EPI_ISL_2767402 | Lighthouse Laboratory Plymouth | Wellcome Sanger Institute for the COVID-19 Genomics UK (COG-UK) Consortium | Lighthouse Laboratory Plymouth and Alex Alderton, Roberto Amato, Jeffrey Barrett, Sonia Goncalves, Ewan Harrison, David K. Jackson, Ian Johnston, Dominic Kwiatkowski, Cordelia Langford, John Sillitoe on behalf of the Wellcome Sanger Institute COVID-19 Surveillance Team |
| EPI_ISL_2767403, EPI_ISL_2767404, EPI_ISL_2767405 | Lighthouse Lab in Milton Keynes | Wellcome Sanger Institute for the COVID-19 Genomics UK (COG-UK) Consortium | The Lighthouse Lab in Milton Keynes and Alex Alderton, Roberto Amato, Jeffrey Barrett, Sonia Goncalves, Ewan Harrison, David K. Jackson, Ian Johnston, Dominic Kwiatkowski, Cordelia Langford, John Sillitoe on behalf of the Wellcome Sanger Institute COVID-19 Surveillance Team |
| EPI_ISL_2767406, EPI_ISL_2767407 | Lighthouse Lab in Alderley Park | Wellcome Sanger Institute for the COVID-19 Genomics UK (COG-UK) Consortium | Jacquelyn Wynn, Mairead Hyland, The Lighthouse Lab in Alderley Park and Alex Alderton, Roberto Amato, Jeffrey Barrett, Sonia Goncalves, Ewan Harrison, David K. Jackson, Ian Johnston, Dominic Kwiatkowski, Cordelia Langford, John Sillitoe on behalf of the Wellcome Sanger Institute COVID-19 Surveillance Team |
| EPI_ISL_2767408 | Lighthouse Lab in Milton Keynes | Wellcome Sanger Institute for the COVID-19 Genomics UK (COG-UK) Consortium | The Lighthouse Lab in Milton Keynes and Alex Alderton, Roberto Amato, Jeffrey Barrett, Sonia Goncalves, Ewan Harrison, David K. Jackson, Ian Johnston, Dominic Kwiatkowski, Cordelia Langford, John Sillitoe on behalf of the Wellcome Sanger Institute COVID-19 Surveillance Team |
| EPI_ISL_2767409 | Lighthouse Lab in Alderley Park | Wellcome Sanger Institute for the COVID-19 Genomics UK (COG-UK) Consortium | Jacquelyn Wynn, Mairead Hyland, The Lighthouse Lab in Alderley Park and Alex Alderton, Roberto Amato, Jeffrey Barrett, Sonia Goncalves, Ewan Harrison, David K. Jackson, Ian Johnston, Dominic Kwiatkowski, Cordelia Langford, John Sillitoe on behalf of the Wellcome Sanger Institute COVID-19 Surveillance Team |
| EPI_ISL_2767410 | Lighthouse Lab in Milton Keynes | Wellcome Sanger Institute for the COVID-19 Genomics UK (COG-UK) Consortium | The Lighthouse Lab in Milton Keynes and Alex Alderton, Roberto Amato, Jeffrey Barrett, Sonia Goncalves, Ewan Harrison, David K. Jackson, Ian Johnston, Dominic Kwiatkowski, Cordelia Langford, John Sillitoe on behalf of the Wellcome Sanger Institute COVID-19 Surveillance Team |
| EPI_ISL_2767411 | Lighthouse Lab in Alderley Park | Wellcome Sanger Institute for the COVID-19 Genomics UK (COG-UK) Consortium | Jacquelyn Wynn, Mairead Hyland, The Lighthouse Lab in Alderley Park and Alex Alderton, Roberto Amato, Jeffrey Barrett, Sonia Goncalves, Ewan Harrison, David K. Jackson, Ian Johnston, Dominic Kwiatkowski, Cordelia Langford, John Sillitoe on behalf of the Wellcome Sanger Institute COVID-19 Surveillance Team |
| EPI_ISL_2767412, EPI_ISL_2767413, EPI_ISL_2767414, EPI_ISL_2767415 | Lighthouse Laboratory Plymouth | Wellcome Sanger Institute for the COVID-19 Genomics UK (COG-UK) Consortium | Lighthouse Laboratory Plymouth and Alex Alderton, Roberto Amato, Jeffrey Barrett, Sonia Goncalves, Ewan Harrison, David K. Jackson, Ian Johnston, Dominic Kwiatkowski, Cordelia Langford, John Sillitoe on behalf of the Wellcome Sanger Institute COVID-19 Surveillance Team |
| EPI_ISL_2767416 | Lighthouse Lab in Glasgow | Wellcome Sanger Institute for the COVID-19 Genomics UK (COG-UK) Consortium | Harper VanSteenhouse, Yumi Kasai, David Gray, Carol Clugston, Anna Dominiczak and Alex Alderton, Roberto Amato, Jeffrey Barrett, Sonia Goncalves, Ewan Harrison, David K. Jackson, Ian Johnston, Dominic Kwiatkowski, Cordelia Langford, John Sillitoe on behalf of the Wellcome Sanger Institute COVID-19 Surveillance Team |
| EPI_ISL_2767417 | Lighthouse Laboratory Plymouth | Wellcome Sanger Institute for the COVID-19 Genomics UK (COG-UK) Consortium | Lighthouse Laboratory Plymouth and Alex Alderton, Roberto Amato, Jeffrey Barrett, Sonia Goncalves, Ewan Harrison, David K. Jackson, Ian Johnston, Dominic Kwiatkowski, Cordelia Langford, John Sillitoe on behalf of the Wellcome Sanger Institute COVID-19 Surveillance Team |
| EPI_ISL_2767418, EPI_ISL_2767419 | Lighthouse Lab in Milton Keynes | Wellcome Sanger Institute for the COVID-19 Genomics UK (COG-UK) Consortium | The Lighthouse Lab in Milton Keynes and Alex Alderton, Roberto Amato, Jeffrey Barrett, Sonia Goncalves, Ewan Harrison, David K. Jackson, Ian Johnston, Dominic Kwiatkowski, Cordelia Langford, John Sillitoe on behalf of the Wellcome Sanger Institute COVID-19 Surveillance Team |
| EPI_ISL_2767420 | Lighthouse Laboratory Plymouth | Wellcome Sanger Institute for the COVID-19 Genomics UK (COG-UK) Consortium | Lighthouse Laboratory Plymouth and Alex Alderton, Roberto Amato, Jeffrey Barrett, Sonia Goncalves, Ewan Harrison, David K. Jackson, Ian Johnston, Dominic Kwiatkowski, Cordelia Langford, John Sillitoe on behalf of the Wellcome Sanger Institute COVID-19 Surveillance Team |
| EPI_ISL_2767421 | Lighthouse Lab in Alderley Park | Wellcome Sanger Institute for the COVID-19 Genomics UK (COG-UK) Consortium | Jacquelyn Wynn, Mairead Hyland, The Lighthouse Lab in Alderley Park and Alex Alderton, Roberto Amato, Jeffrey Barrett, Sonia Goncalves, Ewan Harrison, David K. Jackson, Ian Johnston, Dominic Kwiatkowski, Cordelia Langford, John Sillitoe on behalf of the Wellcome Sanger Institute COVID-19 Surveillance Team |
| EPI_ISL_2767422 | Lighthouse Lab in Milton Keynes | Wellcome Sanger Institute for the COVID-19 Genomics UK (COG-UK) Consortium | The Lighthouse Lab in Milton Keynes and Alex Alderton, Roberto Amato, Jeffrey Barrett, Sonia Goncalves, Ewan Harrison, David K. Jackson, Ian Johnston, Dominic Kwiatkowski, Cordelia Langford, John Sillitoe on behalf of the Wellcome Sanger Institute COVID-19 Surveillance Team |
| EPI_ISL_2767423 | Lighthouse Laboratory Plymouth | Wellcome Sanger Institute for the COVID-19 Genomics UK (COG-UK) Consortium | Lighthouse Laboratory Plymouth and Alex Alderton, Roberto Amato, Jeffrey Barrett, Sonia Goncalves, Ewan Harrison, David K. Jackson, Ian Johnston, Dominic Kwiatkowski, Cordelia Langford, John Sillitoe on behalf of the Wellcome Sanger Institute COVID-19 Surveillance Team |
| EPI_ISL_2767424, EPI_ISL_2767425 | Lighthouse Lab in Alderley Park | Wellcome Sanger Institute for the COVID-19 Genomics UK (COG-UK) Consortium | Jacquelyn Wynn, Mairead Hyland, The Lighthouse Lab in Alderley Park and Alex Alderton, Roberto Amato, Jeffrey Barrett, Sonia Goncalves, Ewan Harrison, David K. Jackson, Ian Johnston, Dominic Kwiatkowski, Cordelia Langford, John Sillitoe on behalf of the Wellcome Sanger Institute COVID-19 Surveillance Team |
| EPI_ISL_2767426 | Lighthouse Laboratory Plymouth | Wellcome Sanger Institute for the COVID-19 Genomics UK (COG-UK) Consortium | Lighthouse Laboratory Plymouth and Alex Alderton, Roberto Amato, Jeffrey Barrett, Sonia Goncalves, Ewan Harrison, David K. Jackson, Ian Johnston, Dominic Kwiatkowski, Cordelia Langford, John Sillitoe on behalf of the Wellcome Sanger Institute COVID-19 Surveillance Team |
| EPI_ISL_2767427 | Lighthouse Lab in Glasgow | Wellcome Sanger Institute for the COVID-19 Genomics UK (COG-UK) Consortium | Harper VanSteenhouse, Yumi Kasai, David Gray, Carol Clugston, Anna Dominiczak and Alex Alderton, Roberto Amato, Jeffrey Barrett, Sonia Goncalves, Ewan Harrison, David K. Jackson, Ian Johnston, Dominic Kwiatkowski, Cordelia Langford, John Sillitoe on behalf of the Wellcome Sanger Institute COVID-19 Surveillance Team |
| EPI_ISL_2767428 | Lighthouse Lab in Milton Keynes | Wellcome Sanger Institute for the COVID-19 Genomics UK (COG-UK) Consortium | The Lighthouse Lab in Milton Keynes and Alex Alderton, Roberto Amato, Jeffrey Barrett, Sonia Goncalves, Ewan Harrison, David K. Jackson, Ian Johnston, Dominic Kwiatkowski, Cordelia Langford, John Sillitoe on behalf of the Wellcome Sanger Institute COVID-19 Surveillance Team |
| EPI_ISL_2767429 | Lighthouse Lab in Glasgow | Wellcome Sanger Institute for the COVID-19 Genomics UK (COG-UK) Consortium | Harper VanSteenhouse, Yumi Kasai, David Gray, Carol Clugston, Anna Dominiczak and Alex Alderton, Roberto Amato, Jeffrey Barrett, Sonia Goncalves, Ewan Harrison, David K. Jackson, Ian Johnston, Dominic Kwiatkowski, Cordelia Langford, John Sillitoe on behalf of the Wellcome Sanger Institute COVID-19 Surveillance Team |
| EPI_ISL_2767430 | Lighthouse Lab in Milton Keynes | Wellcome Sanger Institute for the COVID-19 Genomics UK (COG-UK) Consortium | The Lighthouse Lab in Milton Keynes and Alex Alderton, Roberto Amato, Jeffrey Barrett, Sonia Goncalves, Ewan Harrison, David K. Jackson, Ian Johnston, Dominic Kwiatkowski, Cordelia Langford, John Sillitoe on behalf of the Wellcome Sanger Institute COVID-19 Surveillance Team |
| EPI_ISL_2767431 | Lighthouse Laboratory Plymouth | Wellcome Sanger Institute for the COVID-19 Genomics UK (COG-UK) Consortium | Lighthouse Laboratory Plymouth and Alex Alderton, Roberto Amato, Jeffrey Barrett, Sonia Goncalves, Ewan Harrison, David K. Jackson, Ian Johnston, Dominic Kwiatkowski, Cordelia Langford, John Sillitoe on behalf of the Wellcome Sanger Institute COVID-19 Surveillance Team |
| EPI_ISL_2767432 | Lighthouse Lab in Alderley Park | Wellcome Sanger Institute for the COVID-19 Genomics UK (COG-UK) Consortium | Jacquelyn Wynn, Mairead Hyland, The Lighthouse Lab in Alderley Park and Alex Alderton, Roberto Amato, Jeffrey Barrett, Sonia Goncalves, Ewan Harrison, David K. Jackson, Ian Johnston, Dominic Kwiatkowski, Cordelia Langford, John Sillitoe on behalf of the Wellcome Sanger Institute COVID-19 Surveillance Team |
| EPI_ISL_2767433 | Lighthouse Lab in Milton Keynes | Wellcome Sanger Institute for the COVID-19 Genomics UK (COG-UK) Consortium | The Lighthouse Lab in Milton Keynes and Alex Alderton, Roberto Amato, Jeffrey Barrett, Sonia Goncalves, Ewan Harrison, David K. Jackson, Ian Johnston, Dominic Kwiatkowski, Cordelia Langford, John Sillitoe on behalf of the Wellcome Sanger Institute COVID-19 Surveillance Team |
| EPI_ISL_2767434 | Lighthouse Laboratory Plymouth | Wellcome Sanger Institute for the COVID-19 Genomics UK (COG-UK) Consortium | Lighthouse Laboratory Plymouth and Alex Alderton, Roberto Amato, Jeffrey Barrett, Sonia Goncalves, Ewan Harrison, David K. Jackson, Ian Johnston, Dominic Kwiatkowski, Cordelia Langford, John Sillitoe on behalf of the Wellcome Sanger Institute COVID-19 Surveillance Team |
| EPI_ISL_2767435 | Lighthouse Lab in Milton Keynes | Wellcome Sanger Institute for the COVID-19 Genomics UK (COG-UK) Consortium | The Lighthouse Lab in Milton Keynes and Alex Alderton, Roberto Amato, Jeffrey Barrett, Sonia Goncalves, Ewan Harrison, David K. Jackson, Ian Johnston, Dominic Kwiatkowski, Cordelia Langford, John Sillitoe on behalf of the Wellcome Sanger Institute COVID-19 Surveillance Team |
| EPI_ISL_2767436, EPI_ISL_2767437, EPI_ISL_2767438 | Lighthouse Lab in Alderley Park | Wellcome Sanger Institute for the COVID-19 Genomics UK (COG-UK) Consortium | Jacquelyn Wynn, Mairead Hyland, The Lighthouse Lab in Alderley Park and Alex Alderton, Roberto Amato, Jeffrey Barrett, Sonia Goncalves, Ewan Harrison, David K. Jackson, Ian Johnston, Dominic Kwiatkowski, Cordelia Langford, John Sillitoe on behalf of the Wellcome Sanger Institute COVID-19 Surveillance Team |
| EPI_ISL_2767439 | Lighthouse Laboratory Plymouth | Wellcome Sanger Institute for the COVID-19 Genomics UK (COG-UK) Consortium | Lighthouse Laboratory Plymouth and Alex Alderton, Roberto Amato, Jeffrey Barrett, Sonia Goncalves, Ewan Harrison, David K. Jackson, Ian Johnston, Dominic Kwiatkowski, Cordelia Langford, John Sillitoe on behalf of the Wellcome Sanger Institute COVID-19 Surveillance Team |
| EPI_ISL_2767440 | Lighthouse Lab in Alderley Park | Wellcome Sanger Institute for the COVID-19 Genomics UK (COG-UK) Consortium | Jacquelyn Wynn, Mairead Hyland, The Lighthouse Lab in Alderley Park and Alex Alderton, Roberto Amato, Jeffrey Barrett, Sonia Goncalves, Ewan Harrison, David K. Jackson, Ian Johnston, Dominic Kwiatkowski, Cordelia Langford, John Sillitoe on behalf of the Wellcome Sanger Institute COVID-19 Surveillance Team |
| EPI_ISL_2767441 | Lighthouse Laboratory Plymouth | Wellcome Sanger Institute for the COVID-19 Genomics UK (COG-UK) Consortium | Lighthouse Laboratory Plymouth and Alex Alderton, Roberto Amato, Jeffrey Barrett, Sonia Goncalves, Ewan Harrison, David K. Jackson, Ian Johnston, Dominic Kwiatkowski, Cordelia Langford, John Sillitoe on behalf of the Wellcome Sanger Institute COVID-19 Surveillance Team |
| EPI_ISL_2767442 | Lighthouse Lab in Alderley Park | Wellcome Sanger Institute for the COVID-19 Genomics UK (COG-UK) Consortium | Jacquelyn Wynn, Mairead Hyland, The Lighthouse Lab in Alderley Park and Alex Alderton, Roberto Amato, Jeffrey Barrett, Sonia Goncalves, Ewan Harrison, David K. Jackson, Ian Johnston, Dominic Kwiatkowski, Cordelia Langford, John Sillitoe on behalf of the Wellcome Sanger Institute COVID-19 Surveillance Team |
| EPI_ISL_2767443, EPI_ISL_2767444, EPI_ISL_2767445, EPI_ISL_2767446 | Lighthouse Laboratory Plymouth | Wellcome Sanger Institute for the COVID-19 Genomics UK (COG-UK) Consortium | Lighthouse Laboratory Plymouth and Alex Alderton, Roberto Amato, Jeffrey Barrett, Sonia Goncalves, Ewan Harrison, David K. Jackson, Ian Johnston, Dominic Kwiatkowski, Cordelia Langford, John Sillitoe on behalf of the Wellcome Sanger Institute COVID-19 Surveillance Team |
| EPI_ISL_2767447 | Lighthouse Lab in Milton Keynes | Wellcome Sanger Institute for the COVID-19 Genomics UK (COG-UK) | The Lighthouse Lab in Milton Keynes and Alex Alderton, Roberto Amato, Jeffrey Barrett, Sonia Goncalves, Ewan Harrison, David K. Jackson, Ian Johnston, Dominic Kwiatkowski, Cordelia Langford, John Sillitoe on behalf of the Wellcome Sanger |

| | | | |
|---|---|---|---|
| | | Consortium | Institute COVID-19 Surveillance Team |
| EPI_ISL_2767448, EPI_ISL_2767449 | Lighthouse Lab in Alderley Park | Wellcome Sanger Institute for the COVID-19 Genomics UK (COG-UK) Consortium | Jacquelyn Wynn, Mairead Hyland, The Lighthouse Lab in Alderley Park and Alex Alderton, Roberto Amato, Jeffrey Barrett, Sonia Goncalves, Ewan Harrison, David K. Jackson, Ian Johnston, Dominic Kwiatkowski, Cordelia Langford, John Sillitoe on behalf of the Wellcome Sanger Institute COVID-19 Surveillance Team |
| EPI_ISL_2767450 | Lighthouse Laboratory Plymouth | Wellcome Sanger Institute for the COVID-19 Genomics UK (COG-UK) Consortium | Lighthouse Laboratory Plymouth and Alex Alderton, Roberto Amato, Jeffrey Barrett, Sonia Goncalves, Ewan Harrison, David K. Jackson, Ian Johnston, Dominic Kwiatkowski, Cordelia Langford, John Sillitoe on behalf of the Wellcome Sanger Institute COVID-19 Surveillance Team |
| EPI_ISL_2767451 | Lighthouse Lab in Milton Keynes | Wellcome Sanger Institute for the COVID-19 Genomics UK (COG-UK) Consortium | The Lighthouse Lab in Milton Keynes and Alex Alderton, Roberto Amato, Jeffrey Barrett, Sonia Goncalves, Ewan Harrison, David K. Jackson, Ian Johnston, Dominic Kwiatkowski, Cordelia Langford, John Sillitoe on behalf of the Wellcome Sanger Institute COVID-19 Surveillance Team |
| EPI_ISL_2767452, EPI_ISL_2767453 | Lighthouse Lab in Alderley Park | Wellcome Sanger Institute for the COVID-19 Genomics UK (COG-UK) Consortium | Jacquelyn Wynn, Mairead Hyland, The Lighthouse Lab in Alderley Park and Alex Alderton, Roberto Amato, Jeffrey Barrett, Sonia Goncalves, Ewan Harrison, David K. Jackson, Ian Johnston, Dominic Kwiatkowski, Cordelia Langford, John Sillitoe on behalf of the Wellcome Sanger Institute COVID-19 Surveillance Team |
| EPI_ISL_2767454 | Lighthouse Laboratory Plymouth | Wellcome Sanger Institute for the COVID-19 Genomics UK (COG-UK) Consortium | Lighthouse Laboratory Plymouth and Alex Alderton, Roberto Amato, Jeffrey Barrett, Sonia Goncalves, Ewan Harrison, David K. Jackson, Ian Johnston, Dominic Kwiatkowski, Cordelia Langford, John Sillitoe on behalf of the Wellcome Sanger Institute COVID-19 Surveillance Team |
| EPI_ISL_2767455 | Lighthouse Lab in Alderley Park | Wellcome Sanger Institute for the COVID-19 Genomics UK (COG-UK) Consortium | Jacquelyn Wynn, Mairead Hyland, The Lighthouse Lab in Alderley Park and Alex Alderton, Roberto Amato, Jeffrey Barrett, Sonia Goncalves, Ewan Harrison, David K. Jackson, Ian Johnston, Dominic Kwiatkowski, Cordelia Langford, John Sillitoe on behalf of the Wellcome Sanger Institute COVID-19 Surveillance Team |
| EPI_ISL_2767456 | Lighthouse Lab in Milton Keynes | Wellcome Sanger Institute for the COVID-19 Genomics UK (COG-UK) Consortium | The Lighthouse Lab in Milton Keynes and Alex Alderton, Roberto Amato, Jeffrey Barrett, Sonia Goncalves, Ewan Harrison, David K. Jackson, Ian Johnston, Dominic Kwiatkowski, Cordelia Langford, John Sillitoe on behalf of the Wellcome Sanger Institute COVID-19 Surveillance Team |
| EPI_ISL_2767457 | Lighthouse Laboratory Plymouth | Wellcome Sanger Institute for the COVID-19 Genomics UK (COG-UK) Consortium | Lighthouse Laboratory Plymouth and Alex Alderton, Roberto Amato, Jeffrey Barrett, Sonia Goncalves, Ewan Harrison, David K. Jackson, Ian Johnston, Dominic Kwiatkowski, Cordelia Langford, John Sillitoe on behalf of the Wellcome Sanger Institute COVID-19 Surveillance Team |
| EPI_ISL_2767458 | Lighthouse Lab in Milton Keynes | Wellcome Sanger Institute for the COVID-19 Genomics UK (COG-UK) Consortium | The Lighthouse Lab in Milton Keynes and Alex Alderton, Roberto Amato, Jeffrey Barrett, Sonia Goncalves, Ewan Harrison, David K. Jackson, Ian Johnston, Dominic Kwiatkowski, Cordelia Langford, John Sillitoe on behalf of the Wellcome Sanger Institute COVID-19 Surveillance Team |
| EPI_ISL_2767459 | Lighthouse Lab in Glasgow | Wellcome Sanger Institute for the COVID-19 Genomics UK (COG-UK) Consortium | Harper VanSteenhouse, Yumi Kasai, David Gray, Carol Clugston, Anna Dominiczak and Alex Alderton, Roberto Amato, Jeffrey Barrett, Sonia Goncalves, Ewan Harrison, David K. Jackson, Ian Johnston, Dominic Kwiatkowski, Cordelia Langford, John Sillitoe on behalf of the Wellcome Sanger Institute COVID-19 Surveillance Team |
| EPI_ISL_2767460 | Lighthouse Laboratory Plymouth | Wellcome Sanger Institute for the COVID-19 Genomics UK (COG-UK) Consortium | Lighthouse Laboratory Plymouth and Alex Alderton, Roberto Amato, Jeffrey Barrett, Sonia Goncalves, Ewan Harrison, David K. Jackson, Ian Johnston, Dominic Kwiatkowski, Cordelia Langford, John Sillitoe on behalf of the Wellcome Sanger Institute COVID-19 Surveillance Team |
| EPI_ISL_2767461, EPI_ISL_2767462 | Lighthouse Lab in Alderley Park | Wellcome Sanger Institute for the COVID-19 Genomics UK (COG-UK) Consortium | Jacquelyn Wynn, Mairead Hyland, The Lighthouse Lab in Alderley Park and Alex Alderton, Roberto Amato, Jeffrey Barrett, Sonia Goncalves, Ewan Harrison, David K. Jackson, Ian Johnston, Dominic Kwiatkowski, Cordelia Langford, John Sillitoe on behalf of the Wellcome Sanger Institute COVID-19 Surveillance Team |
| EPI_ISL_2767463 | Lighthouse Laboratory Plymouth | Wellcome Sanger Institute for the COVID-19 Genomics UK (COG-UK) Consortium | Lighthouse Laboratory Plymouth and Alex Alderton, Roberto Amato, Jeffrey Barrett, Sonia Goncalves, Ewan Harrison, David K. Jackson, Ian Johnston, Dominic Kwiatkowski, Cordelia Langford, John Sillitoe on behalf of the Wellcome Sanger Institute COVID-19 Surveillance Team |
| EPI_ISL_2767464 | Lighthouse Lab in Glasgow | Wellcome Sanger Institute for the COVID-19 Genomics UK (COG-UK) Consortium | Harper VanSteenhouse, Yumi Kasai, David Gray, Carol Clugston, Anna Dominiczak and Alex Alderton, Roberto Amato, Jeffrey Barrett, Sonia Goncalves, Ewan Harrison, David K. Jackson, Ian Johnston, Dominic Kwiatkowski, Cordelia Langford, John Sillitoe on behalf of the Wellcome Sanger Institute COVID-19 Surveillance Team |
| EPI_ISL_2767465, EPI_ISL_2767466 | Lighthouse Lab in Alderley Park | Wellcome Sanger Institute for the COVID-19 Genomics UK (COG-UK) Consortium | Jacquelyn Wynn, Mairead Hyland, The Lighthouse Lab in Alderley Park and Alex Alderton, Roberto Amato, Jeffrey Barrett, Sonia Goncalves, Ewan Harrison, David K. Jackson, Ian Johnston, Dominic Kwiatkowski, Cordelia Langford, John Sillitoe on behalf of the Wellcome Sanger Institute COVID-19 Surveillance Team |
| EPI_ISL_2767467 | Lighthouse Laboratory Plymouth | Wellcome Sanger Institute for the COVID-19 Genomics UK (COG-UK) Consortium | Lighthouse Laboratory Plymouth and Alex Alderton, Roberto Amato, Jeffrey Barrett, Sonia Goncalves, Ewan Harrison, David K. Jackson, Ian Johnston, Dominic Kwiatkowski, Cordelia Langford, John Sillitoe on behalf of the Wellcome Sanger Institute COVID-19 Surveillance Team |
| EPI_ISL_2767468 | Lighthouse Lab in Alderley Park | Wellcome Sanger Institute for the COVID-19 Genomics UK (COG-UK) Consortium | Jacquelyn Wynn, Mairead Hyland, The Lighthouse Lab in Alderley Park and Alex Alderton, Roberto Amato, Jeffrey Barrett, Sonia Goncalves, Ewan Harrison, David K. Jackson, Ian Johnston, Dominic Kwiatkowski, Cordelia Langford, John Sillitoe on behalf of the Wellcome Sanger Institute COVID-19 Surveillance Team |
| EPI_ISL_2767469 | Lighthouse Lab in Glasgow | Wellcome Sanger Institute for the COVID-19 Genomics UK (COG-UK) Consortium | Harper VanSteenhouse, Yumi Kasai, David Gray, Carol Clugston, Anna Dominiczak and Alex Alderton, Roberto Amato, Jeffrey Barrett, Sonia Goncalves, Ewan Harrison, David K. Jackson, Ian Johnston, Dominic Kwiatkowski, Cordelia Langford, John Sillitoe on behalf of the Wellcome Sanger Institute COVID-19 Surveillance Team |
| EPI_ISL_2767470, EPI_ISL_2767471 | Lighthouse Laboratory Plymouth | Wellcome Sanger Institute for the COVID-19 Genomics UK (COG-UK) Consortium | Lighthouse Laboratory Plymouth and Alex Alderton, Roberto Amato, Jeffrey Barrett, Sonia Goncalves, Ewan Harrison, David K. Jackson, Ian Johnston, Dominic Kwiatkowski, Cordelia Langford, John Sillitoe on behalf of the Wellcome Sanger Institute COVID-19 Surveillance Team |
| EPI_ISL_2767472 | Lighthouse Lab in Milton Keynes | Wellcome Sanger Institute for the COVID-19 Genomics UK (COG-UK) Consortium | The Lighthouse Lab in Milton Keynes and Alex Alderton, Roberto Amato, Jeffrey Barrett, Sonia Goncalves, Ewan Harrison, David K. Jackson, Ian Johnston, Dominic Kwiatkowski, Cordelia Langford, John Sillitoe on behalf of the Wellcome Sanger Institute COVID-19 Surveillance Team |
| EPI_ISL_2767473 | Lighthouse Laboratory Plymouth | Wellcome Sanger Institute for the COVID-19 Genomics UK (COG-UK) Consortium | Lighthouse Laboratory Plymouth and Alex Alderton, Roberto Amato, Jeffrey Barrett, Sonia Goncalves, Ewan Harrison, David K. Jackson, Ian Johnston, Dominic Kwiatkowski, Cordelia Langford, John Sillitoe on behalf of the Wellcome Sanger Institute COVID-19 Surveillance Team |
| EPI_ISL_2767474 | Lighthouse Lab in Milton Keynes | Wellcome Sanger Institute for the COVID-19 Genomics UK (COG-UK) Consortium | The Lighthouse Lab in Milton Keynes and Alex Alderton, Roberto Amato, Jeffrey Barrett, Sonia Goncalves, Ewan Harrison, David K. Jackson, Ian Johnston, Dominic Kwiatkowski, Cordelia Langford, John Sillitoe on behalf of the Wellcome Sanger Institute COVID-19 Surveillance Team |
| EPI_ISL_2767475 | Lighthouse Lab in Alderley Park | Wellcome Sanger Institute for the COVID-19 Genomics UK (COG-UK) Consortium | Jacquelyn Wynn, Mairead Hyland, The Lighthouse Lab in Alderley Park and Alex Alderton, Roberto Amato, Jeffrey Barrett, Sonia Goncalves, Ewan Harrison, David K. Jackson, Ian Johnston, Dominic Kwiatkowski, Cordelia Langford, John Sillitoe on behalf of the Wellcome Sanger Institute COVID-19 Surveillance Team |
| EPI_ISL_2767476 | Lighthouse Laboratory Plymouth | Wellcome Sanger Institute for the COVID-19 Genomics UK (COG-UK) Consortium | Lighthouse Laboratory Plymouth and Alex Alderton, Roberto Amato, Jeffrey Barrett, Sonia Goncalves, Ewan Harrison, David K. Jackson, Ian Johnston, Dominic Kwiatkowski, Cordelia Langford, John Sillitoe on behalf of the Wellcome Sanger Institute COVID-19 Surveillance Team |
| EPI_ISL_2767477 | Lighthouse Lab in Alderley Park | Wellcome Sanger Institute for the COVID-19 Genomics UK (COG-UK) Consortium | Jacquelyn Wynn, Mairead Hyland, The Lighthouse Lab in Alderley Park and Alex Alderton, Roberto Amato, Jeffrey Barrett, Sonia Goncalves, Ewan Harrison, David K. Jackson, Ian Johnston, Dominic Kwiatkowski, Cordelia Langford, John Sillitoe on behalf of the Wellcome Sanger Institute COVID-19 Surveillance Team |
| EPI_ISL_2767478 | Lighthouse Laboratory Plymouth | Wellcome Sanger Institute for the COVID-19 Genomics UK (COG-UK) Consortium | Lighthouse Laboratory Plymouth and Alex Alderton, Roberto Amato, Jeffrey Barrett, Sonia Goncalves, Ewan Harrison, David K. Jackson, Ian Johnston, Dominic Kwiatkowski, Cordelia Langford, John Sillitoe on behalf of the Wellcome Sanger Institute COVID-19 Surveillance Team |
| EPI_ISL_2767479 | Lighthouse Lab in Glasgow | Wellcome Sanger Institute for the COVID-19 Genomics UK (COG-UK) Consortium | Harper VanSteenhouse, Yumi Kasai, David Gray, Carol Clugston, Anna Dominiczak and Alex Alderton, Roberto Amato, Jeffrey Barrett, Sonia Goncalves, Ewan Harrison, David K. Jackson, Ian Johnston, Dominic Kwiatkowski, Cordelia Langford, John Sillitoe on behalf of the Wellcome Sanger Institute COVID-19 Surveillance Team |
| EPI_ISL_2767480 | Lighthouse Lab in Alderley Park | Wellcome Sanger Institute for the COVID-19 Genomics UK (COG-UK) Consortium | Jacquelyn Wynn, Mairead Hyland, The Lighthouse Lab in Alderley Park and Alex Alderton, Roberto Amato, Jeffrey Barrett, Sonia Goncalves, Ewan Harrison, David K. Jackson, Ian Johnston, Dominic Kwiatkowski, Cordelia Langford, John Sillitoe on behalf of the Wellcome Sanger Institute COVID-19 Surveillance Team |
| EPI_ISL_2767481 | Lighthouse Laboratory Plymouth | Wellcome Sanger Institute for the COVID-19 Genomics UK (COG-UK) Consortium | Lighthouse Laboratory Plymouth and Alex Alderton, Roberto Amato, Jeffrey Barrett, Sonia Goncalves, Ewan Harrison, David K. Jackson, Ian Johnston, Dominic Kwiatkowski, Cordelia Langford, John Sillitoe on behalf of the Wellcome Sanger Institute COVID-19 Surveillance Team |
| EPI_ISL_2767482 | Lighthouse Lab in Alderley Park | Wellcome Sanger Institute for the COVID-19 Genomics UK (COG-UK) Consortium | Jacquelyn Wynn, Mairead Hyland, The Lighthouse Lab in Alderley Park and Alex Alderton, Roberto Amato, Jeffrey Barrett, Sonia Goncalves, Ewan Harrison, David K. Jackson, Ian Johnston, Dominic Kwiatkowski, Cordelia Langford, John Sillitoe on behalf of the Wellcome Sanger Institute COVID-19 Surveillance Team |
| EPI_ISL_2767483 | Lighthouse Lab in Glasgow | Wellcome Sanger Institute for the COVID-19 Genomics UK (COG-UK) Consortium | Harper VanSteenhouse, Yumi Kasai, David Gray, Carol Clugston, Anna Dominiczak and Alex Alderton, Roberto Amato, Jeffrey Barrett, Sonia Goncalves, Ewan Harrison, David K. Jackson, Ian Johnston, Dominic Kwiatkowski, Cordelia Langford, John Sillitoe on behalf of the Wellcome Sanger Institute COVID-19 Surveillance Team |
| EPI_ISL_2767484 | Lighthouse Lab in Alderley Park | Wellcome Sanger Institute for the COVID-19 Genomics UK (COG-UK) Consortium | Jacquelyn Wynn, Mairead Hyland, The Lighthouse Lab in Alderley Park and Alex Alderton, Roberto Amato, Jeffrey Barrett, Sonia Goncalves, Ewan Harrison, David K. Jackson, Ian Johnston, Dominic Kwiatkowski, Cordelia Langford, John Sillitoe on behalf of the Wellcome Sanger Institute COVID-19 Surveillance Team |
| EPI_ISL_2767485 | Lighthouse Laboratory Plymouth | Wellcome Sanger Institute for the COVID-19 Genomics UK (COG-UK) Consortium | Lighthouse Laboratory Plymouth and Alex Alderton, Roberto Amato, Jeffrey Barrett, Sonia Goncalves, Ewan Harrison, David K. Jackson, Ian Johnston, Dominic Kwiatkowski, Cordelia Langford, John Sillitoe on behalf of the Wellcome Sanger Institute COVID-19 Surveillance Team |
| EPI_ISL_2767486 | Lighthouse Lab in Alderley Park | Wellcome Sanger Institute for the COVID-19 Genomics UK (COG-UK) Consortium | Jacquelyn Wynn, Mairead Hyland, The Lighthouse Lab in Alderley Park and Alex Alderton, Roberto Amato, Jeffrey Barrett, Sonia Goncalves, Ewan Harrison, David K. Jackson, Ian Johnston, Dominic Kwiatkowski, Cordelia Langford, John Sillitoe on behalf of the Wellcome Sanger Institute COVID-19 Surveillance Team |
| EPI_ISL_2767487 | Lighthouse Lab in Milton Keynes | Wellcome Sanger Institute for the COVID-19 Genomics UK (COG-UK) Consortium | The Lighthouse Lab in Milton Keynes and Alex Alderton, Roberto Amato, Jeffrey Barrett, Sonia Goncalves, Ewan Harrison, David K. Jackson, Ian Johnston, Dominic Kwiatkowski, Cordelia Langford, John Sillitoe on behalf of the Wellcome Sanger Institute COVID-19 Surveillance Team |
| EPI_ISL_2767488 | Lighthouse Lab in Glasgow | Wellcome Sanger Institute for the COVID-19 Genomics UK (COG-UK) Consortium | Harper VanSteenhouse, Yumi Kasai, David Gray, Carol Clugston, Anna Dominiczak and Alex Alderton, Roberto Amato, Jeffrey Barrett, Sonia Goncalves, Ewan Harrison, David K. Jackson, Ian Johnston, Dominic Kwiatkowski, Cordelia Langford, John Sillitoe on behalf of the Wellcome Sanger Institute COVID-19 Surveillance Team |
| EPI_ISL_2767489, EPI_ISL_2767490 | Lighthouse Lab in Alderley Park | Wellcome Sanger Institute for the COVID-19 Genomics UK (COG-UK) Consortium | Jacquelyn Wynn, Mairead Hyland, The Lighthouse Lab in Alderley Park and Alex Alderton, Roberto Amato, Jeffrey Barrett, Sonia Goncalves, Ewan Harrison, David K. Jackson, Ian Johnston, Dominic Kwiatkowski, Cordelia Langford, John Sillitoe on behalf of the Wellcome Sanger Institute COVID-19 Surveillance Team |
| EPI_ISL_2767491, EPI_ISL_2767492 | Lighthouse Lab in Milton Keynes | Wellcome Sanger Institute for the COVID-19 Genomics UK (COG-UK) Consortium | The Lighthouse Lab in Milton Keynes and Alex Alderton, Roberto Amato, Jeffrey Barrett, Sonia Goncalves, Ewan Harrison, David K. Jackson, Ian Johnston, Dominic Kwiatkowski, Cordelia Langford, John Sillitoe on behalf of the Wellcome Sanger Institute COVID-19 Surveillance Team |
| EPI_ISL_2767493, EPI_ISL_2767494, EPI_ISL_2767495 | Lighthouse Lab in Alderley Park | Wellcome Sanger Institute for the COVID-19 Genomics UK (COG-UK) Consortium | Jacquelyn Wynn, Mairead Hyland, The Lighthouse Lab in Alderley Park and Alex Alderton, Roberto Amato, Jeffrey Barrett, Sonia Goncalves, Ewan Harrison, David K. Jackson, Ian Johnston, Dominic Kwiatkowski, Cordelia Langford, John Sillitoe on behalf of the Wellcome Sanger Institute COVID-19 Surveillance Team |
| EPI_ISL_2767496 | Lighthouse Lab in Milton Keynes | Wellcome Sanger Institute for the COVID-19 Genomics UK (COG-UK) Consortium | The Lighthouse Lab in Milton Keynes and Alex Alderton, Roberto Amato, Jeffrey Barrett, Sonia Goncalves, Ewan Harrison, David K. Jackson, Ian Johnston, Dominic Kwiatkowski, Cordelia Langford, John Sillitoe on behalf of the Wellcome Sanger Institute COVID-19 Surveillance Team |
| EPI_ISL_2767497 | Lighthouse Laboratory Plymouth | Wellcome Sanger Institute for the COVID-19 Genomics UK (COG-UK) Consortium | Lighthouse Laboratory Plymouth and Alex Alderton, Roberto Amato, Jeffrey Barrett, Sonia Goncalves, Ewan Harrison, David K. Jackson, Ian Johnston, Dominic Kwiatkowski, Cordelia Langford, John Sillitoe on behalf of the Wellcome Sanger Institute COVID-19 Surveillance Team |
| EPI_ISL_2767498 | Lighthouse Lab in Milton Keynes | Wellcome Sanger Institute for the COVID-19 Genomics UK (COG-UK) Consortium | The Lighthouse Lab in Milton Keynes and Alex Alderton, Roberto Amato, Jeffrey Barrett, Sonia Goncalves, Ewan Harrison, David K. Jackson, Ian Johnston, Dominic Kwiatkowski, Cordelia Langford, John Sillitoe on behalf of the Wellcome Sanger Institute COVID-19 Surveillance Team |
| EPI_ISL_2767499, EPI_ISL_2767500 | Lighthouse Lab in Alderley Park | Wellcome Sanger Institute for the COVID-19 Genomics UK (COG-UK) Consortium | Jacquelyn Wynn, Mairead Hyland, The Lighthouse Lab in Alderley Park and Alex Alderton, Roberto Amato, Jeffrey Barrett, Sonia Goncalves, Ewan Harrison, David K. Jackson, Ian Johnston, Dominic Kwiatkowski, Cordelia Langford, John Sillitoe on behalf of the Wellcome Sanger Institute COVID-19 Surveillance Team |
| EPI_ISL_2767501, EPI_ISL_2767502 | Lighthouse Laboratory Plymouth | Wellcome Sanger Institute for the COVID-19 Genomics UK (COG-UK) Consortium | Lighthouse Laboratory Plymouth and Alex Alderton, Roberto Amato, Jeffrey Barrett, Sonia Goncalves, Ewan Harrison, David K. Jackson, Ian Johnston, Dominic Kwiatkowski, Cordelia Langford, John Sillitoe on behalf of the Wellcome Sanger Institute COVID-19 Surveillance Team |

| | | | |
|---|---|---|---|
| EPI_ISL_2767503, EPI_ISL_2767504 | Lighthouse Lab in Alderley Park | Wellcome Sanger Institute for the COVID-19 Genomics UK (COG-UK) Consortium | Jacquelyn Wynn, Mairead Hyland, The Lighthouse Lab in Alderley Park and Alex Alderton, Roberto Amato, Jeffrey Barrett, Sonia Goncalves, Ewan Harrison, David K. Jackson, Ian Johnston, Dominic Kwiatkowski, Cordelia Langford, John Sillitoe on behalf of the Wellcome Sanger Institute COVID-19 Surveillance Team |
| EPI_ISL_2767505 | Lighthouse Lab in Milton Keynes | Wellcome Sanger Institute for the COVID-19 Genomics UK (COG-UK) Consortium | The Lighthouse Lab in Milton Keynes and Alex Alderton, Roberto Amato, Jeffrey Barrett, Sonia Goncalves, Ewan Harrison, David K. Jackson, Ian Johnston, Dominic Kwiatkowski, Cordelia Langford, John Sillitoe on behalf of the Wellcome Sanger Institute COVID-19 Surveillance Team |
| EPI_ISL_2767506 | Lighthouse Laboratory Plymouth | Wellcome Sanger Institute for the COVID-19 Genomics UK (COG-UK) Consortium | Lighthouse Laboratory Plymouth and Alex Alderton, Roberto Amato, Jeffrey Barrett, Sonia Goncalves, Ewan Harrison, David K. Jackson, Ian Johnston, Dominic Kwiatkowski, Cordelia Langford, John Sillitoe on behalf of the Wellcome Sanger Institute COVID-19 Surveillance Team |
| EPI_ISL_2767508 | Lighthouse Lab in Glasgow | Wellcome Sanger Institute for the COVID-19 Genomics UK (COG-UK) Consortium | Harper VanSteenhouse, Yumi Kasai, David Gray, Carol Clugston, Anna Dominiczak and Alex Alderton, Roberto Amato, Jeffrey Barrett, Sonia Goncalves, Ewan Harrison, David K. Jackson, Ian Johnston, Dominic Kwiatkowski, Cordelia Langford, John Sillitoe on behalf of the Wellcome Sanger Institute COVID-19 Surveillance Team |
| EPI_ISL_2767509 | Lighthouse Laboratory Plymouth | Wellcome Sanger Institute for the COVID-19 Genomics UK (COG-UK) Consortium | Lighthouse Laboratory Plymouth and Alex Alderton, Roberto Amato, Jeffrey Barrett, Sonia Goncalves, Ewan Harrison, David K. Jackson, Ian Johnston, Dominic Kwiatkowski, Cordelia Langford, John Sillitoe on behalf of the Wellcome Sanger Institute COVID-19 Surveillance Team |
| EPI_ISL_2767510, EPI_ISL_2767511, EPI_ISL_2767512 | Lighthouse Lab in Milton Keynes | Wellcome Sanger Institute for the COVID-19 Genomics UK (COG-UK) Consortium | The Lighthouse Lab in Milton Keynes and Alex Alderton, Roberto Amato, Jeffrey Barrett, Sonia Goncalves, Ewan Harrison, David K. Jackson, Ian Johnston, Dominic Kwiatkowski, Cordelia Langford, John Sillitoe on behalf of the Wellcome Sanger Institute COVID-19 Surveillance Team |
| EPI_ISL_2767513, EPI_ISL_2767514 | Lighthouse Lab in Alderley Park | Wellcome Sanger Institute for the COVID-19 Genomics UK (COG-UK) Consortium | Jacquelyn Wynn, Mairead Hyland, The Lighthouse Lab in Alderley Park and Alex Alderton, Roberto Amato, Jeffrey Barrett, Sonia Goncalves, Ewan Harrison, David K. Jackson, Ian Johnston, Dominic Kwiatkowski, Cordelia Langford, John Sillitoe on behalf of the Wellcome Sanger Institute COVID-19 Surveillance Team |
| EPI_ISL_2767515 | Lighthouse Lab in Milton Keynes | Wellcome Sanger Institute for the COVID-19 Genomics UK (COG-UK) Consortium | The Lighthouse Lab in Milton Keynes and Alex Alderton, Roberto Amato, Jeffrey Barrett, Sonia Goncalves, Ewan Harrison, David K. Jackson, Ian Johnston, Dominic Kwiatkowski, Cordelia Langford, John Sillitoe on behalf of the Wellcome Sanger Institute COVID-19 Surveillance Team |
| EPI_ISL_2767516, EPI_ISL_2767517 | Lighthouse Lab in Alderley Park | Wellcome Sanger Institute for the COVID-19 Genomics UK (COG-UK) Consortium | Jacquelyn Wynn, Mairead Hyland, The Lighthouse Lab in Alderley Park and Alex Alderton, Roberto Amato, Jeffrey Barrett, Sonia Goncalves, Ewan Harrison, David K. Jackson, Ian Johnston, Dominic Kwiatkowski, Cordelia Langford, John Sillitoe on behalf of the Wellcome Sanger Institute COVID-19 Surveillance Team |
| EPI_ISL_2767518 | Lighthouse Lab in Glasgow | Wellcome Sanger Institute for the COVID-19 Genomics UK (COG-UK) Consortium | Harper VanSteenhouse, Yumi Kasai, David Gray, Carol Clugston, Anna Dominiczak and Alex Alderton, Roberto Amato, Jeffrey Barrett, Sonia Goncalves, Ewan Harrison, David K. Jackson, Ian Johnston, Dominic Kwiatkowski, Cordelia Langford, John Sillitoe on behalf of the Wellcome Sanger Institute COVID-19 Surveillance Team |
| EPI_ISL_2767519 | Lighthouse Lab in Milton Keynes | Wellcome Sanger Institute for the COVID-19 Genomics UK (COG-UK) Consortium | The Lighthouse Lab in Milton Keynes and Alex Alderton, Roberto Amato, Jeffrey Barrett, Sonia Goncalves, Ewan Harrison, David K. Jackson, Ian Johnston, Dominic Kwiatkowski, Cordelia Langford, John Sillitoe on behalf of the Wellcome Sanger Institute COVID-19 Surveillance Team |
| EPI_ISL_2767520 | Lighthouse Laboratory Plymouth | Wellcome Sanger Institute for the COVID-19 Genomics UK (COG-UK) Consortium | Lighthouse Laboratory Plymouth and Alex Alderton, Roberto Amato, Jeffrey Barrett, Sonia Goncalves, Ewan Harrison, David K. Jackson, Ian Johnston, Dominic Kwiatkowski, Cordelia Langford, John Sillitoe on behalf of the Wellcome Sanger Institute COVID-19 Surveillance Team |
| EPI_ISL_2767521 | Lighthouse Lab in Alderley Park | Wellcome Sanger Institute for the COVID-19 Genomics UK (COG-UK) Consortium | Jacquelyn Wynn, Mairead Hyland, The Lighthouse Lab in Alderley Park and Alex Alderton, Roberto Amato, Jeffrey Barrett, Sonia Goncalves, Ewan Harrison, David K. Jackson, Ian Johnston, Dominic Kwiatkowski, Cordelia Langford, John Sillitoe on behalf of the Wellcome Sanger Institute COVID-19 Surveillance Team |
| EPI_ISL_2767522, EPI_ISL_2767523 | Lighthouse Lab in Milton Keynes | Wellcome Sanger Institute for the COVID-19 Genomics UK (COG-UK) Consortium | The Lighthouse Lab in Milton Keynes and Alex Alderton, Roberto Amato, Jeffrey Barrett, Sonia Goncalves, Ewan Harrison, David K. Jackson, Ian Johnston, Dominic Kwiatkowski, Cordelia Langford, John Sillitoe on behalf of the Wellcome Sanger Institute COVID-19 Surveillance Team |
| EPI_ISL_2767524 | Lighthouse Lab in Alderley Park | Wellcome Sanger Institute for the COVID-19 Genomics UK (COG-UK) Consortium | Jacquelyn Wynn, Mairead Hyland, The Lighthouse Lab in Alderley Park and Alex Alderton, Roberto Amato, Jeffrey Barrett, Sonia Goncalves, Ewan Harrison, David K. Jackson, Ian Johnston, Dominic Kwiatkowski, Cordelia Langford, John Sillitoe on behalf of the Wellcome Sanger Institute COVID-19 Surveillance Team |
| EPI_ISL_2767525 | Lighthouse Lab in Milton Keynes | Wellcome Sanger Institute for the COVID-19 Genomics UK (COG-UK) Consortium | The Lighthouse Lab in Milton Keynes and Alex Alderton, Roberto Amato, Jeffrey Barrett, Sonia Goncalves, Ewan Harrison, David K. Jackson, Ian Johnston, Dominic Kwiatkowski, Cordelia Langford, John Sillitoe on behalf of the Wellcome Sanger Institute COVID-19 Surveillance Team |
| EPI_ISL_2767526 | Lighthouse Laboratory Plymouth | Wellcome Sanger Institute for the COVID-19 Genomics UK (COG-UK) Consortium | Lighthouse Laboratory Plymouth and Alex Alderton, Roberto Amato, Jeffrey Barrett, Sonia Goncalves, Ewan Harrison, David K. Jackson, Ian Johnston, Dominic Kwiatkowski, Cordelia Langford, John Sillitoe on behalf of the Wellcome Sanger Institute COVID-19 Surveillance Team |
| EPI_ISL_2767527, EPI_ISL_2767528 | Lighthouse Lab in Glasgow | Wellcome Sanger Institute for the COVID-19 Genomics UK (COG-UK) Consortium | Harper VanSteenhouse, Yumi Kasai, David Gray, Carol Clugston, Anna Dominiczak and Alex Alderton, Roberto Amato, Jeffrey Barrett, Sonia Goncalves, Ewan Harrison, David K. Jackson, Ian Johnston, Dominic Kwiatkowski, Cordelia Langford, John Sillitoe on behalf of the Wellcome Sanger Institute COVID-19 Surveillance Team |
| EPI_ISL_2767529, EPI_ISL_2767530 | Lighthouse Lab in Alderley Park | Wellcome Sanger Institute for the COVID-19 Genomics UK (COG-UK) Consortium | Jacquelyn Wynn, Mairead Hyland, The Lighthouse Lab in Alderley Park and Alex Alderton, Roberto Amato, Jeffrey Barrett, Sonia Goncalves, Ewan Harrison, David K. Jackson, Ian Johnston, Dominic Kwiatkowski, Cordelia Langford, John Sillitoe on behalf of the Wellcome Sanger Institute COVID-19 Surveillance Team |
| EPI_ISL_2767531 | Lighthouse Laboratory Plymouth | Wellcome Sanger Institute for the COVID-19 Genomics UK (COG-UK) Consortium | Lighthouse Laboratory Plymouth and Alex Alderton, Roberto Amato, Jeffrey Barrett, Sonia Goncalves, Ewan Harrison, David K. Jackson, Ian Johnston, Dominic Kwiatkowski, Cordelia Langford, John Sillitoe on behalf of the Wellcome Sanger Institute COVID-19 Surveillance Team |
| EPI_ISL_2767532 | Lighthouse Lab in Milton Keynes | Wellcome Sanger Institute for the COVID-19 Genomics UK (COG-UK) Consortium | The Lighthouse Lab in Milton Keynes and Alex Alderton, Roberto Amato, Jeffrey Barrett, Sonia Goncalves, Ewan Harrison, David K. Jackson, Ian Johnston, Dominic Kwiatkowski, Cordelia Langford, John Sillitoe on behalf of the Wellcome Sanger Institute COVID-19 Surveillance Team |
| EPI_ISL_2767533 | Lighthouse Laboratory Plymouth | Wellcome Sanger Institute for the COVID-19 Genomics UK (COG-UK) Consortium | Lighthouse Laboratory Plymouth and Alex Alderton, Roberto Amato, Jeffrey Barrett, Sonia Goncalves, Ewan Harrison, David K. Jackson, Ian Johnston, Dominic Kwiatkowski, Cordelia Langford, John Sillitoe on behalf of the Wellcome Sanger Institute COVID-19 Surveillance Team |
| EPI_ISL_2767534 | Lighthouse Lab in Milton Keynes | Wellcome Sanger Institute for the COVID-19 Genomics UK (COG-UK) Consortium | The Lighthouse Lab in Milton Keynes and Alex Alderton, Roberto Amato, Jeffrey Barrett, Sonia Goncalves, Ewan Harrison, David K. Jackson, Ian Johnston, Dominic Kwiatkowski, Cordelia Langford, John Sillitoe on behalf of the Wellcome Sanger Institute COVID-19 Surveillance Team |
| EPI_ISL_2767535 | Lighthouse Laboratory Plymouth | Wellcome Sanger Institute for the COVID-19 Genomics UK (COG-UK) Consortium | Lighthouse Laboratory Plymouth and Alex Alderton, Roberto Amato, Jeffrey Barrett, Sonia Goncalves, Ewan Harrison, David K. Jackson, Ian Johnston, Dominic Kwiatkowski, Cordelia Langford, John Sillitoe on behalf of the Wellcome Sanger Institute COVID-19 Surveillance Team |
| EPI_ISL_2767536 | Lighthouse Lab in Glasgow | Wellcome Sanger Institute for the COVID-19 Genomics UK (COG-UK) Consortium | Harper VanSteenhouse, Yumi Kasai, David Gray, Carol Clugston, Anna Dominiczak and Alex Alderton, Roberto Amato, Jeffrey Barrett, Sonia Goncalves, Ewan Harrison, David K. Jackson, Ian Johnston, Dominic Kwiatkowski, Cordelia Langford, John Sillitoe on behalf of the Wellcome Sanger Institute COVID-19 Surveillance Team |
| EPI_ISL_2767537 | Lighthouse Lab in Alderley Park | Wellcome Sanger Institute for the COVID-19 Genomics UK (COG-UK) Consortium | Jacquelyn Wynn, Mairead Hyland, The Lighthouse Lab in Alderley Park and Alex Alderton, Roberto Amato, Jeffrey Barrett, Sonia Goncalves, Ewan Harrison, David K. Jackson, Ian Johnston, Dominic Kwiatkowski, Cordelia Langford, John Sillitoe on behalf of the Wellcome Sanger Institute COVID-19 Surveillance Team |
| EPI_ISL_2767538, EPI_ISL_2767539 | Lighthouse Lab in Milton Keynes | Wellcome Sanger Institute for the COVID-19 Genomics UK (COG-UK) Consortium | The Lighthouse Lab in Milton Keynes and Alex Alderton, Roberto Amato, Jeffrey Barrett, Sonia Goncalves, Ewan Harrison, David K. Jackson, Ian Johnston, Dominic Kwiatkowski, Cordelia Langford, John Sillitoe on behalf of the Wellcome Sanger Institute COVID-19 Surveillance Team |
| EPI_ISL_2767540 | Lighthouse Lab in Glasgow | Wellcome Sanger Institute for the COVID-19 Genomics UK (COG-UK) Consortium | Harper VanSteenhouse, Yumi Kasai, David Gray, Carol Clugston, Anna Dominiczak and Alex Alderton, Roberto Amato, Jeffrey Barrett, Sonia Goncalves, Ewan Harrison, David K. Jackson, Ian Johnston, Dominic Kwiatkowski, Cordelia Langford, John Sillitoe on behalf of the Wellcome Sanger Institute COVID-19 Surveillance Team |
| EPI_ISL_2767541 | Lighthouse Lab in Milton Keynes | Wellcome Sanger Institute for the COVID-19 Genomics UK (COG-UK) Consortium | The Lighthouse Lab in Milton Keynes and Alex Alderton, Roberto Amato, Jeffrey Barrett, Sonia Goncalves, Ewan Harrison, David K. Jackson, Ian Johnston, Dominic Kwiatkowski, Cordelia Langford, John Sillitoe on behalf of the Wellcome Sanger Institute COVID-19 Surveillance Team |
| EPI_ISL_2767542, EPI_ISL_2767543, EPI_ISL_2767544 | Lighthouse Lab in Alderley Park | Wellcome Sanger Institute for the COVID-19 Genomics UK (COG-UK) Consortium | Jacquelyn Wynn, Mairead Hyland, The Lighthouse Lab in Alderley Park and Alex Alderton, Roberto Amato, Jeffrey Barrett, Sonia Goncalves, Ewan Harrison, David K. Jackson, Ian Johnston, Dominic Kwiatkowski, Cordelia Langford, John Sillitoe on behalf of the Wellcome Sanger Institute COVID-19 Surveillance Team |
| EPI_ISL_2767545 | Lighthouse Laboratory Plymouth | Wellcome Sanger Institute for the COVID-19 Genomics UK (COG-UK) Consortium | Lighthouse Laboratory Plymouth and Alex Alderton, Roberto Amato, Jeffrey Barrett, Sonia Goncalves, Ewan Harrison, David K. Jackson, Ian Johnston, Dominic Kwiatkowski, Cordelia Langford, John Sillitoe on behalf of the Wellcome Sanger Institute COVID-19 Surveillance Team |
| EPI_ISL_2767546, EPI_ISL_2767547 | Lighthouse Lab in Milton Keynes | Wellcome Sanger Institute for the COVID-19 Genomics UK (COG-UK) Consortium | The Lighthouse Lab in Milton Keynes and Alex Alderton, Roberto Amato, Jeffrey Barrett, Sonia Goncalves, Ewan Harrison, David K. Jackson, Ian Johnston, Dominic Kwiatkowski, Cordelia Langford, John Sillitoe on behalf of the Wellcome Sanger Institute COVID-19 Surveillance Team |
| EPI_ISL_2767548 | Lighthouse Laboratory Plymouth | Wellcome Sanger Institute for the COVID-19 Genomics UK (COG-UK) Consortium | Lighthouse Laboratory Plymouth and Alex Alderton, Roberto Amato, Jeffrey Barrett, Sonia Goncalves, Ewan Harrison, David K. Jackson, Ian Johnston, Dominic Kwiatkowski, Cordelia Langford, John Sillitoe on behalf of the Wellcome Sanger Institute COVID-19 Surveillance Team |
| EPI_ISL_2767549 | Lighthouse Lab in Milton Keynes | Wellcome Sanger Institute for the COVID-19 Genomics UK (COG-UK) Consortium | The Lighthouse Lab in Milton Keynes and Alex Alderton, Roberto Amato, Jeffrey Barrett, Sonia Goncalves, Ewan Harrison, David K. Jackson, Ian Johnston, Dominic Kwiatkowski, Cordelia Langford, John Sillitoe on behalf of the Wellcome Sanger Institute COVID-19 Surveillance Team |
| EPI_ISL_2767550 | Lighthouse Laboratory Plymouth | Wellcome Sanger Institute for the COVID-19 Genomics UK (COG-UK) Consortium | Lighthouse Laboratory Plymouth and Alex Alderton, Roberto Amato, Jeffrey Barrett, Sonia Goncalves, Ewan Harrison, David K. Jackson, Ian Johnston, Dominic Kwiatkowski, Cordelia Langford, John Sillitoe on behalf of the Wellcome Sanger Institute COVID-19 Surveillance Team |
| EPI_ISL_2767551 | Lighthouse Lab in Alderley Park | Wellcome Sanger Institute for the COVID-19 Genomics UK (COG-UK) Consortium | Jacquelyn Wynn, Mairead Hyland, The Lighthouse Lab in Alderley Park and Alex Alderton, Roberto Amato, Jeffrey Barrett, Sonia Goncalves, Ewan Harrison, David K. Jackson, Ian Johnston, Dominic Kwiatkowski, Cordelia Langford, John Sillitoe on behalf of the Wellcome Sanger Institute COVID-19 Surveillance Team |
| EPI_ISL_2767552 | Lighthouse Lab in Glasgow | Wellcome Sanger Institute for the COVID-19 Genomics UK (COG-UK) Consortium | Harper VanSteenhouse, Yumi Kasai, David Gray, Carol Clugston, Anna Dominiczak and Alex Alderton, Roberto Amato, Jeffrey Barrett, Sonia Goncalves, Ewan Harrison, David K. Jackson, Ian Johnston, Dominic Kwiatkowski, Cordelia Langford, John Sillitoe on behalf of the Wellcome Sanger Institute COVID-19 Surveillance Team |
| EPI_ISL_2767553 | Lighthouse Laboratory Plymouth | Wellcome Sanger Institute for the COVID-19 Genomics UK (COG-UK) Consortium | Lighthouse Laboratory Plymouth and Alex Alderton, Roberto Amato, Jeffrey Barrett, Sonia Goncalves, Ewan Harrison, David K. Jackson, Ian Johnston, Dominic Kwiatkowski, Cordelia Langford, John Sillitoe on behalf of the Wellcome Sanger Institute COVID-19 Surveillance Team |
| EPI_ISL_2767554, EPI_ISL_2767555, EPI_ISL_2767556 | Lighthouse Lab in Milton Keynes | Wellcome Sanger Institute for the COVID-19 Genomics UK (COG-UK) Consortium | The Lighthouse Lab in Milton Keynes and Alex Alderton, Roberto Amato, Jeffrey Barrett, Sonia Goncalves, Ewan Harrison, David K. Jackson, Ian Johnston, Dominic Kwiatkowski, Cordelia Langford, John Sillitoe on behalf of the Wellcome Sanger Institute COVID-19 Surveillance Team |
| EPI_ISL_2767557 | Lighthouse Laboratory Plymouth | Wellcome Sanger Institute for the COVID-19 Genomics UK (COG-UK) Consortium | Lighthouse Laboratory Plymouth and Alex Alderton, Roberto Amato, Jeffrey Barrett, Sonia Goncalves, Ewan Harrison, David K. Jackson, Ian Johnston, Dominic Kwiatkowski, Cordelia Langford, John Sillitoe on behalf of the Wellcome Sanger Institute COVID-19 Surveillance Team |
| EPI_ISL_2767558, EPI_ISL_2767559, EPI_ISL_2767560 | Lighthouse Lab in Alderley Park | Wellcome Sanger Institute for the COVID-19 Genomics UK (COG-UK) Consortium | Jacquelyn Wynn, Mairead Hyland, The Lighthouse Lab in Alderley Park and Alex Alderton, Roberto Amato, Jeffrey Barrett, Sonia Goncalves, Ewan Harrison, David K. Jackson, Ian Johnston, Dominic Kwiatkowski, Cordelia Langford, John Sillitoe on behalf of the Wellcome Sanger Institute COVID-19 Surveillance Team |
| EPI_ISL_2767561 | Lighthouse Laboratory Plymouth | Wellcome Sanger Institute for the COVID-19 Genomics UK (COG-UK) Consortium | Lighthouse Laboratory Plymouth and Alex Alderton, Roberto Amato, Jeffrey Barrett, Sonia Goncalves, Ewan Harrison, David K. Jackson, Ian Johnston, Dominic Kwiatkowski, Cordelia Langford, John Sillitoe on behalf of the Wellcome Sanger Institute COVID-19 Surveillance Team |
| EPI_ISL_2767562, EPI_ISL_2767563, EPI_ISL_2767564 | Lighthouse Lab in Milton Keynes | Wellcome Sanger Institute for the COVID-19 Genomics UK (COG-UK) Consortium | The Lighthouse Lab in Milton Keynes and Alex Alderton, Roberto Amato, Jeffrey Barrett, Sonia Goncalves, Ewan Harrison, David K. Jackson, Ian Johnston, Dominic Kwiatkowski, Cordelia Langford, John Sillitoe on behalf of the Wellcome Sanger Institute COVID-19 Surveillance Team |

| | | | |
|---|---|---|---|
| EPI_ISL_2767565 | Lighthouse Lab in Glasgow | Wellcome Sanger Institute for the COVID-19 Genomics UK (COG-UK) Consortium | Harper VanSteenhouse, Yumi Kasai, David Gray, Carol Clugston, Anna Dominiczak and Alex Alderton, Roberto Amato, Jeffrey Barrett, Sonia Goncalves, Ewan Harrison, David K. Jackson, Ian Johnston, Dominic Kwiatkowski, Cordelia Langford, John Sillitoe on behalf of the Wellcome Sanger Institute COVID-19 Surveillance Team |
| EPI_ISL_2767566 | Lighthouse Lab in Milton Keynes | Wellcome Sanger Institute for the COVID-19 Genomics UK (COG-UK) Consortium | The Lighthouse Lab in Milton Keynes and Alex Alderton, Roberto Amato, Jeffrey Barrett, Sonia Goncalves, Ewan Harrison, David K. Jackson, Ian Johnston, Dominic Kwiatkowski, Cordelia Langford, John Sillitoe on behalf of the Wellcome Sanger Institute COVID-19 Surveillance Team |
| EPI_ISL_2767567 | Lighthouse Lab in Alderley Park | Wellcome Sanger Institute for the COVID-19 Genomics UK (COG-UK) Consortium | Jacquelyn Wynn, Mairead Hyland, The Lighthouse Lab in Alderley Park and Alex Alderton, Roberto Amato, Jeffrey Barrett, Sonia Goncalves, Ewan Harrison, David K. Jackson, Ian Johnston, Dominic Kwiatkowski, Cordelia Langford, John Sillitoe on behalf of the Wellcome Sanger Institute COVID-19 Surveillance Team |
| EPI_ISL_2767568 | Lighthouse Laboratory Plymouth | Wellcome Sanger Institute for the COVID-19 Genomics UK (COG-UK) Consortium | Lighthouse Laboratory Plymouth and Alex Alderton, Roberto Amato, Jeffrey Barrett, Sonia Goncalves, Ewan Harrison, David K. Jackson, Ian Johnston, Dominic Kwiatkowski, Cordelia Langford, John Sillitoe on behalf of the Wellcome Sanger Institute COVID-19 Surveillance Team |
| EPI_ISL_2767569, EPI_ISL_2767570 | Lighthouse Lab in Milton Keynes | Wellcome Sanger Institute for the COVID-19 Genomics UK (COG-UK) Consortium | The Lighthouse Lab in Milton Keynes and Alex Alderton, Roberto Amato, Jeffrey Barrett, Sonia Goncalves, Ewan Harrison, David K. Jackson, Ian Johnston, Dominic Kwiatkowski, Cordelia Langford, John Sillitoe on behalf of the Wellcome Sanger Institute COVID-19 Surveillance Team |
| EPI_ISL_2767571 | Lighthouse Laboratory Plymouth | Wellcome Sanger Institute for the COVID-19 Genomics UK (COG-UK) Consortium | Lighthouse Laboratory Plymouth and Alex Alderton, Roberto Amato, Jeffrey Barrett, Sonia Goncalves, Ewan Harrison, David K. Jackson, Ian Johnston, Dominic Kwiatkowski, Cordelia Langford, John Sillitoe on behalf of the Wellcome Sanger Institute COVID-19 Surveillance Team |
| EPI_ISL_2767572, EPI_ISL_2767573, EPI_ISL_2767574 | Lighthouse Lab in Alderley Park | Wellcome Sanger Institute for the COVID-19 Genomics UK (COG-UK) Consortium | Jacquelyn Wynn, Mairead Hyland, The Lighthouse Lab in Alderley Park and Alex Alderton, Roberto Amato, Jeffrey Barrett, Sonia Goncalves, Ewan Harrison, David K. Jackson, Ian Johnston, Dominic Kwiatkowski, Cordelia Langford, John Sillitoe on behalf of the Wellcome Sanger Institute COVID-19 Surveillance Team |
| EPI_ISL_2767575 | Lighthouse Lab in Milton Keynes | Wellcome Sanger Institute for the COVID-19 Genomics UK (COG-UK) Consortium | The Lighthouse Lab in Milton Keynes and Alex Alderton, Roberto Amato, Jeffrey Barrett, Sonia Goncalves, Ewan Harrison, David K. Jackson, Ian Johnston, Dominic Kwiatkowski, Cordelia Langford, John Sillitoe on behalf of the Wellcome Sanger Institute COVID-19 Surveillance Team |
| EPI_ISL_2767576 | Lighthouse Laboratory Plymouth | Wellcome Sanger Institute for the COVID-19 Genomics UK (COG-UK) Consortium | Lighthouse Laboratory Plymouth and Alex Alderton, Roberto Amato, Jeffrey Barrett, Sonia Goncalves, Ewan Harrison, David K. Jackson, Ian Johnston, Dominic Kwiatkowski, Cordelia Langford, John Sillitoe on behalf of the Wellcome Sanger Institute COVID-19 Surveillance Team |
| EPI_ISL_2767577 | Lighthouse Lab in Alderley Park | Wellcome Sanger Institute for the COVID-19 Genomics UK (COG-UK) Consortium | Jacquelyn Wynn, Mairead Hyland, The Lighthouse Lab in Alderley Park and Alex Alderton, Roberto Amato, Jeffrey Barrett, Sonia Goncalves, Ewan Harrison, David K. Jackson, Ian Johnston, Dominic Kwiatkowski, Cordelia Langford, John Sillitoe on behalf of the Wellcome Sanger Institute COVID-19 Surveillance Team |
| EPI_ISL_2767578 | Lighthouse Laboratory Plymouth | Wellcome Sanger Institute for the COVID-19 Genomics UK (COG-UK) Consortium | Lighthouse Laboratory Plymouth and Alex Alderton, Roberto Amato, Jeffrey Barrett, Sonia Goncalves, Ewan Harrison, David K. Jackson, Ian Johnston, Dominic Kwiatkowski, Cordelia Langford, John Sillitoe on behalf of the Wellcome Sanger Institute COVID-19 Surveillance Team |
| EPI_ISL_2767579 | Lighthouse Lab in Alderley Park | Wellcome Sanger Institute for the COVID-19 Genomics UK (COG-UK) Consortium | Jacquelyn Wynn, Mairead Hyland, The Lighthouse Lab in Alderley Park and Alex Alderton, Roberto Amato, Jeffrey Barrett, Sonia Goncalves, Ewan Harrison, David K. Jackson, Ian Johnston, Dominic Kwiatkowski, Cordelia Langford, John Sillitoe on behalf of the Wellcome Sanger Institute COVID-19 Surveillance Team |
| EPI_ISL_2767580 | Lighthouse Laboratory Plymouth | Wellcome Sanger Institute for the COVID-19 Genomics UK (COG-UK) Consortium | Lighthouse Laboratory Plymouth and Alex Alderton, Roberto Amato, Jeffrey Barrett, Sonia Goncalves, Ewan Harrison, David K. Jackson, Ian Johnston, Dominic Kwiatkowski, Cordelia Langford, John Sillitoe on behalf of the Wellcome Sanger Institute COVID-19 Surveillance Team |
| EPI_ISL_2767581, EPI_ISL_2767582 | Lighthouse Lab in Alderley Park | Wellcome Sanger Institute for the COVID-19 Genomics UK (COG-UK) Consortium | Jacquelyn Wynn, Mairead Hyland, The Lighthouse Lab in Alderley Park and Alex Alderton, Roberto Amato, Jeffrey Barrett, Sonia Goncalves, Ewan Harrison, David K. Jackson, Ian Johnston, Dominic Kwiatkowski, Cordelia Langford, John Sillitoe on behalf of the Wellcome Sanger Institute COVID-19 Surveillance Team |
| EPI_ISL_2767583, EPI_ISL_2767584 | Lighthouse Lab in Milton Keynes | Wellcome Sanger Institute for the COVID-19 Genomics UK (COG-UK) Consortium | The Lighthouse Lab in Milton Keynes and Alex Alderton, Roberto Amato, Jeffrey Barrett, Sonia Goncalves, Ewan Harrison, David K. Jackson, Ian Johnston, Dominic Kwiatkowski, Cordelia Langford, John Sillitoe on behalf of the Wellcome Sanger Institute COVID-19 Surveillance Team |
| EPI_ISL_2767585 | Lighthouse Lab in Alderley Park | Wellcome Sanger Institute for the COVID-19 Genomics UK (COG-UK) Consortium | Jacquelyn Wynn, Mairead Hyland, The Lighthouse Lab in Alderley Park and Alex Alderton, Roberto Amato, Jeffrey Barrett, Sonia Goncalves, Ewan Harrison, David K. Jackson, Ian Johnston, Dominic Kwiatkowski, Cordelia Langford, John Sillitoe on behalf of the Wellcome Sanger Institute COVID-19 Surveillance Team |
| EPI_ISL_2767586 | Lighthouse Lab in Milton Keynes | Wellcome Sanger Institute for the COVID-19 Genomics UK (COG-UK) Consortium | The Lighthouse Lab in Milton Keynes and Alex Alderton, Roberto Amato, Jeffrey Barrett, Sonia Goncalves, Ewan Harrison, David K. Jackson, Ian Johnston, Dominic Kwiatkowski, Cordelia Langford, John Sillitoe on behalf of the Wellcome Sanger Institute COVID-19 Surveillance Team |
| EPI_ISL_2767587 | Lighthouse Laboratory Plymouth | Wellcome Sanger Institute for the COVID-19 Genomics UK (COG-UK) Consortium | Lighthouse Laboratory Plymouth and Alex Alderton, Roberto Amato, Jeffrey Barrett, Sonia Goncalves, Ewan Harrison, David K. Jackson, Ian Johnston, Dominic Kwiatkowski, Cordelia Langford, John Sillitoe on behalf of the Wellcome Sanger Institute COVID-19 Surveillance Team |
| EPI_ISL_2767588, EPI_ISL_2767589, EPI_ISL_2767590 | Lighthouse Lab in Milton Keynes | Wellcome Sanger Institute for the COVID-19 Genomics UK (COG-UK) Consortium | The Lighthouse Lab in Milton Keynes and Alex Alderton, Roberto Amato, Jeffrey Barrett, Sonia Goncalves, Ewan Harrison, David K. Jackson, Ian Johnston, Dominic Kwiatkowski, Cordelia Langford, John Sillitoe on behalf of the Wellcome Sanger Institute COVID-19 Surveillance Team |
| EPI_ISL_2767591 | Lighthouse Laboratory Plymouth | Wellcome Sanger Institute for the COVID-19 Genomics UK (COG-UK) Consortium | Lighthouse Laboratory Plymouth and Alex Alderton, Roberto Amato, Jeffrey Barrett, Sonia Goncalves, Ewan Harrison, David K. Jackson, Ian Johnston, Dominic Kwiatkowski, Cordelia Langford, John Sillitoe on behalf of the Wellcome Sanger Institute COVID-19 Surveillance Team |
| EPI_ISL_2767592 | Lighthouse Lab in Glasgow | Wellcome Sanger Institute for the COVID-19 Genomics UK (COG-UK) Consortium | Harper VanSteenhouse, Yumi Kasai, David Gray, Carol Clugston, Anna Dominiczak and Alex Alderton, Roberto Amato, Jeffrey Barrett, Sonia Goncalves, Ewan Harrison, David K. Jackson, Ian Johnston, Dominic Kwiatkowski, Cordelia Langford, John Sillitoe on behalf of the Wellcome Sanger Institute COVID-19 Surveillance Team |
| EPI_ISL_2767593 | Lighthouse Laboratory Plymouth | Wellcome Sanger Institute for the COVID-19 Genomics UK (COG-UK) Consortium | Lighthouse Laboratory Plymouth and Alex Alderton, Roberto Amato, Jeffrey Barrett, Sonia Goncalves, Ewan Harrison, David K. Jackson, Ian Johnston, Dominic Kwiatkowski, Cordelia Langford, John Sillitoe on behalf of the Wellcome Sanger Institute COVID-19 Surveillance Team |
| EPI_ISL_2767594 | Lighthouse Lab in Milton Keynes | Wellcome Sanger Institute for the COVID-19 Genomics UK (COG-UK) Consortium | The Lighthouse Lab in Milton Keynes and Alex Alderton, Roberto Amato, Jeffrey Barrett, Sonia Goncalves, Ewan Harrison, David K. Jackson, Ian Johnston, Dominic Kwiatkowski, Cordelia Langford, John Sillitoe on behalf of the Wellcome Sanger Institute COVID-19 Surveillance Team |
| EPI_ISL_2767595, EPI_ISL_2767596, EPI_ISL_2767597 | Lighthouse Lab in Alderley Park | Wellcome Sanger Institute for the COVID-19 Genomics UK (COG-UK) Consortium | Jacquelyn Wynn, Mairead Hyland, The Lighthouse Lab in Alderley Park and Alex Alderton, Roberto Amato, Jeffrey Barrett, Sonia Goncalves, Ewan Harrison, David K. Jackson, Ian Johnston, Dominic Kwiatkowski, Cordelia Langford, John Sillitoe on behalf of the Wellcome Sanger Institute COVID-19 Surveillance Team |
| EPI_ISL_2767598 | Lighthouse Lab in Glasgow | Wellcome Sanger Institute for the COVID-19 Genomics UK (COG-UK) Consortium | Harper VanSteenhouse, Yumi Kasai, David Gray, Carol Clugston, Anna Dominiczak and Alex Alderton, Roberto Amato, Jeffrey Barrett, Sonia Goncalves, Ewan Harrison, David K. Jackson, Ian Johnston, Dominic Kwiatkowski, Cordelia Langford, John Sillitoe on behalf of the Wellcome Sanger Institute COVID-19 Surveillance Team |
| EPI_ISL_2767599, EPI_ISL_2767600 | Lighthouse Lab in Alderley Park | Wellcome Sanger Institute for the COVID-19 Genomics UK (COG-UK) Consortium | Jacquelyn Wynn, Mairead Hyland, The Lighthouse Lab in Alderley Park and Alex Alderton, Roberto Amato, Jeffrey Barrett, Sonia Goncalves, Ewan Harrison, David K. Jackson, Ian Johnston, Dominic Kwiatkowski, Cordelia Langford, John Sillitoe on behalf of the Wellcome Sanger Institute COVID-19 Surveillance Team |
| EPI_ISL_2767601 | Lighthouse Lab in Milton Keynes | Wellcome Sanger Institute for the COVID-19 Genomics UK (COG-UK) Consortium | The Lighthouse Lab in Milton Keynes and Alex Alderton, Roberto Amato, Jeffrey Barrett, Sonia Goncalves, Ewan Harrison, David K. Jackson, Ian Johnston, Dominic Kwiatkowski, Cordelia Langford, John Sillitoe on behalf of the Wellcome Sanger Institute COVID-19 Surveillance Team |
| EPI_ISL_2767602, EPI_ISL_2767603, EPI_ISL_2767604 | Lighthouse Lab in Glasgow | Wellcome Sanger Institute for the COVID-19 Genomics UK (COG-UK) Consortium | Harper VanSteenhouse, Yumi Kasai, David Gray, Carol Clugston, Anna Dominiczak and Alex Alderton, Roberto Amato, Jeffrey Barrett, Sonia Goncalves, Ewan Harrison, David K. Jackson, Ian Johnston, Dominic Kwiatkowski, Cordelia Langford, John Sillitoe on behalf of the Wellcome Sanger Institute COVID-19 Surveillance Team |
| EPI_ISL_2767605, EPI_ISL_2767606 | Lighthouse Lab in Alderley Park | Wellcome Sanger Institute for the COVID-19 Genomics UK (COG-UK) Consortium | Jacquelyn Wynn, Mairead Hyland, The Lighthouse Lab in Alderley Park and Alex Alderton, Roberto Amato, Jeffrey Barrett, Sonia Goncalves, Ewan Harrison, David K. Jackson, Ian Johnston, Dominic Kwiatkowski, Cordelia Langford, John Sillitoe on behalf of the Wellcome Sanger Institute COVID-19 Surveillance Team |
| EPI_ISL_2767607 | Lighthouse Lab in Milton Keynes | Wellcome Sanger Institute for the COVID-19 Genomics UK (COG-UK) Consortium | The Lighthouse Lab in Milton Keynes and Alex Alderton, Roberto Amato, Jeffrey Barrett, Sonia Goncalves, Ewan Harrison, David K. Jackson, Ian Johnston, Dominic Kwiatkowski, Cordelia Langford, John Sillitoe on behalf of the Wellcome Sanger Institute COVID-19 Surveillance Team |
| EPI_ISL_2767608 | Lighthouse Laboratory Plymouth | Wellcome Sanger Institute for the COVID-19 Genomics UK (COG-UK) Consortium | Lighthouse Laboratory Plymouth and Alex Alderton, Roberto Amato, Jeffrey Barrett, Sonia Goncalves, Ewan Harrison, David K. Jackson, Ian Johnston, Dominic Kwiatkowski, Cordelia Langford, John Sillitoe on behalf of the Wellcome Sanger Institute COVID-19 Surveillance Team |
| EPI_ISL_2767609, EPI_ISL_2767610 | Lighthouse Lab in Alderley Park | Wellcome Sanger Institute for the COVID-19 Genomics UK (COG-UK) Consortium | Jacquelyn Wynn, Mairead Hyland, The Lighthouse Lab in Alderley Park and Alex Alderton, Roberto Amato, Jeffrey Barrett, Sonia Goncalves, Ewan Harrison, David K. Jackson, Ian Johnston, Dominic Kwiatkowski, Cordelia Langford, John Sillitoe on behalf of the Wellcome Sanger Institute COVID-19 Surveillance Team |
| EPI_ISL_2767611 | Lighthouse Lab in Milton Keynes | Wellcome Sanger Institute for the COVID-19 Genomics UK (COG-UK) Consortium | The Lighthouse Lab in Milton Keynes and Alex Alderton, Roberto Amato, Jeffrey Barrett, Sonia Goncalves, Ewan Harrison, David K. Jackson, Ian Johnston, Dominic Kwiatkowski, Cordelia Langford, John Sillitoe on behalf of the Wellcome Sanger Institute COVID-19 Surveillance Team |
| EPI_ISL_2767612 | Lighthouse Laboratory Plymouth | Wellcome Sanger Institute for the COVID-19 Genomics UK (COG-UK) Consortium | Lighthouse Laboratory Plymouth and Alex Alderton, Roberto Amato, Jeffrey Barrett, Sonia Goncalves, Ewan Harrison, David K. Jackson, Ian Johnston, Dominic Kwiatkowski, Cordelia Langford, John Sillitoe on behalf of the Wellcome Sanger Institute COVID-19 Surveillance Team |
| EPI_ISL_2767613 | Lighthouse Lab in Alderley Park | Wellcome Sanger Institute for the COVID-19 Genomics UK (COG-UK) Consortium | Jacquelyn Wynn, Mairead Hyland, The Lighthouse Lab in Alderley Park and Alex Alderton, Roberto Amato, Jeffrey Barrett, Sonia Goncalves, Ewan Harrison, David K. Jackson, Ian Johnston, Dominic Kwiatkowski, Cordelia Langford, John Sillitoe on behalf of the Wellcome Sanger Institute COVID-19 Surveillance Team |
| EPI_ISL_2767614 | Lighthouse Lab in Milton Keynes | Wellcome Sanger Institute for the COVID-19 Genomics UK (COG-UK) Consortium | The Lighthouse Lab in Milton Keynes and Alex Alderton, Roberto Amato, Jeffrey Barrett, Sonia Goncalves, Ewan Harrison, David K. Jackson, Ian Johnston, Dominic Kwiatkowski, Cordelia Langford, John Sillitoe on behalf of the Wellcome Sanger Institute COVID-19 Surveillance Team |
| EPI_ISL_2767615 | Lighthouse Lab in Alderley Park | Wellcome Sanger Institute for the COVID-19 Genomics UK (COG-UK) Consortium | Jacquelyn Wynn, Mairead Hyland, The Lighthouse Lab in Alderley Park and Alex Alderton, Roberto Amato, Jeffrey Barrett, Sonia Goncalves, Ewan Harrison, David K. Jackson, Ian Johnston, Dominic Kwiatkowski, Cordelia Langford, John Sillitoe on behalf of the Wellcome Sanger Institute COVID-19 Surveillance Team |
| EPI_ISL_2767616 | Lighthouse Laboratory Plymouth | Wellcome Sanger Institute for the COVID-19 Genomics UK (COG-UK) Consortium | Lighthouse Laboratory Plymouth and Alex Alderton, Roberto Amato, Jeffrey Barrett, Sonia Goncalves, Ewan Harrison, David K. Jackson, Ian Johnston, Dominic Kwiatkowski, Cordelia Langford, John Sillitoe on behalf of the Wellcome Sanger Institute COVID-19 Surveillance Team |
| EPI_ISL_2767617, EPI_ISL_2767618, EPI_ISL_2767619 | Lighthouse Lab in Milton Keynes | Wellcome Sanger Institute for the COVID-19 Genomics UK (COG-UK) Consortium | The Lighthouse Lab in Milton Keynes and Alex Alderton, Roberto Amato, Jeffrey Barrett, Sonia Goncalves, Ewan Harrison, David K. Jackson, Ian Johnston, Dominic Kwiatkowski, Cordelia Langford, John Sillitoe on behalf of the Wellcome Sanger Institute COVID-19 Surveillance Team |
| EPI_ISL_2767620 | Lighthouse Lab in Glasgow | Wellcome Sanger Institute for the COVID-19 Genomics UK (COG-UK) Consortium | Harper VanSteenhouse, Yumi Kasai, David Gray, Carol Clugston, Anna Dominiczak and Alex Alderton, Roberto Amato, Jeffrey Barrett, Sonia Goncalves, Ewan Harrison, David K. Jackson, Ian Johnston, Dominic Kwiatkowski, Cordelia Langford, John Sillitoe on behalf of the Wellcome Sanger Institute COVID-19 Surveillance Team |
| EPI_ISL_2767621 | Lighthouse Laboratory Plymouth | Wellcome Sanger Institute for the COVID-19 Genomics UK (COG-UK) Consortium | Lighthouse Laboratory Plymouth and Alex Alderton, Roberto Amato, Jeffrey Barrett, Sonia Goncalves, Ewan Harrison, David K. Jackson, Ian Johnston, Dominic Kwiatkowski, Cordelia Langford, John Sillitoe on behalf of the Wellcome Sanger Institute COVID-19 Surveillance Team |
| EPI_ISL_2767622 | Lighthouse Lab in Milton Keynes | Wellcome Sanger Institute for the COVID-19 Genomics UK (COG-UK) Consortium | The Lighthouse Lab in Milton Keynes and Alex Alderton, Roberto Amato, Jeffrey Barrett, Sonia Goncalves, Ewan Harrison, David K. Jackson, Ian Johnston, Dominic Kwiatkowski, Cordelia Langford, John Sillitoe on behalf of the Wellcome Sanger Institute COVID-19 Surveillance Team |
| EPI_ISL_2767623 | Lighthouse Lab in Alderley Park | Wellcome Sanger Institute for the COVID-19 Genomics UK (COG-UK) Consortium | Jacquelyn Wynn, Mairead Hyland, The Lighthouse Lab in Alderley Park and Alex Alderton, Roberto Amato, Jeffrey Barrett, Sonia Goncalves, Ewan Harrison, David K. Jackson, Ian Johnston, Dominic Kwiatkowski, Cordelia Langford, John Sillitoe on behalf of the Wellcome Sanger Institute COVID-19 Surveillance Team |

| | | | |
|---|---|---|---|
| EPI_ISL_2767624 | Lighthouse Lab in Milton Keynes | Wellcome Sanger Institute for the COVID-19 Genomics UK (COG-UK) Consortium | The Lighthouse Lab in Milton Keynes and Alex Alderton, Roberto Amato, Jeffrey Barrett, Sonia Goncalves, Ewan Harrison, David K. Jackson, Ian Johnston, Dominic Kwiatkowski, Cordelia Langford, John Sillitoe on behalf of the Wellcome Sanger Institute COVID-19 Surveillance Team |
| EPI_ISL_2767625 | Lighthouse Lab in Glasgow | Wellcome Sanger Institute for the COVID-19 Genomics UK (COG-UK) Consortium | Harper VanSteenhouse, Yumi Kasai, David Gray, Carol Clugston, Anna Dominiczak and Alex Alderton, Roberto Amato, Jeffrey Barrett, Sonia Goncalves, Ewan Harrison, David K. Jackson, Ian Johnston, Dominic Kwiatkowski, Cordelia Langford, John Sillitoe on behalf of the Wellcome Sanger Institute COVID-19 Surveillance Team |
| EPI_ISL_2767626 | Lighthouse Lab in Alderley Park | Wellcome Sanger Institute for the COVID-19 Genomics UK (COG-UK) Consortium | Jacquelyn Wynn, Mairead Hyland, The Lighthouse Lab in Alderley Park and Alex Alderton, Roberto Amato, Jeffrey Barrett, Sonia Goncalves, Ewan Harrison, David K. Jackson, Ian Johnston, Dominic Kwiatkowski, Cordelia Langford, John Sillitoe on behalf of the Wellcome Sanger Institute COVID-19 Surveillance Team |
| EPI_ISL_2767627 | Lighthouse Lab in Glasgow | Wellcome Sanger Institute for the COVID-19 Genomics UK (COG-UK) Consortium | Harper VanSteenhouse, Yumi Kasai, David Gray, Carol Clugston, Anna Dominiczak and Alex Alderton, Roberto Amato, Jeffrey Barrett, Sonia Goncalves, Ewan Harrison, David K. Jackson, Ian Johnston, Dominic Kwiatkowski, Cordelia Langford, John Sillitoe on behalf of the Wellcome Sanger Institute COVID-19 Surveillance Team |
| EPI_ISL_2767628, EPI_ISL_2767629 | Lighthouse Lab in Alderley Park | Wellcome Sanger Institute for the COVID-19 Genomics UK (COG-UK) Consortium | Jacquelyn Wynn, Mairead Hyland, The Lighthouse Lab in Alderley Park and Alex Alderton, Roberto Amato, Jeffrey Barrett, Sonia Goncalves, Ewan Harrison, David K. Jackson, Ian Johnston, Dominic Kwiatkowski, Cordelia Langford, John Sillitoe on behalf of the Wellcome Sanger Institute COVID-19 Surveillance Team |
| EPI_ISL_2767630 | Lighthouse Lab in Milton Keynes | Wellcome Sanger Institute for the COVID-19 Genomics UK (COG-UK) Consortium | The Lighthouse Lab in Milton Keynes and Alex Alderton, Roberto Amato, Jeffrey Barrett, Sonia Goncalves, Ewan Harrison, David K. Jackson, Ian Johnston, Dominic Kwiatkowski, Cordelia Langford, John Sillitoe on behalf of the Wellcome Sanger Institute COVID-19 Surveillance Team |
| EPI_ISL_2767631, EPI_ISL_2767632 | Lighthouse Laboratory Plymouth | Wellcome Sanger Institute for the COVID-19 Genomics UK (COG-UK) Consortium | Lighthouse Laboratory Plymouth and Alex Alderton, Roberto Amato, Jeffrey Barrett, Sonia Goncalves, Ewan Harrison, David K. Jackson, Ian Johnston, Dominic Kwiatkowski, Cordelia Langford, John Sillitoe on behalf of the Wellcome Sanger Institute COVID-19 Surveillance Team |
| EPI_ISL_2767633, EPI_ISL_2767634 | Lighthouse Lab in Alderley Park | Wellcome Sanger Institute for the COVID-19 Genomics UK (COG-UK) Consortium | Jacquelyn Wynn, Mairead Hyland, The Lighthouse Lab in Alderley Park and Alex Alderton, Roberto Amato, Jeffrey Barrett, Sonia Goncalves, Ewan Harrison, David K. Jackson, Ian Johnston, Dominic Kwiatkowski, Cordelia Langford, John Sillitoe on behalf of the Wellcome Sanger Institute COVID-19 Surveillance Team |
| EPI_ISL_2767635 | Lighthouse Lab in Glasgow | Wellcome Sanger Institute for the COVID-19 Genomics UK (COG-UK) Consortium | Harper VanSteenhouse, Yumi Kasai, David Gray, Carol Clugston, Anna Dominiczak and Alex Alderton, Roberto Amato, Jeffrey Barrett, Sonia Goncalves, Ewan Harrison, David K. Jackson, Ian Johnston, Dominic Kwiatkowski, Cordelia Langford, John Sillitoe on behalf of the Wellcome Sanger Institute COVID-19 Surveillance Team |
| EPI_ISL_2767636 | Lighthouse Laboratory Plymouth | Wellcome Sanger Institute for the COVID-19 Genomics UK (COG-UK) Consortium | Lighthouse Laboratory Plymouth and Alex Alderton, Roberto Amato, Jeffrey Barrett, Sonia Goncalves, Ewan Harrison, David K. Jackson, Ian Johnston, Dominic Kwiatkowski, Cordelia Langford, John Sillitoe on behalf of the Wellcome Sanger Institute COVID-19 Surveillance Team |
| EPI_ISL_2767637, EPI_ISL_2767638 | Lighthouse Lab in Milton Keynes | Wellcome Sanger Institute for the COVID-19 Genomics UK (COG-UK) Consortium | The Lighthouse Lab in Milton Keynes and Alex Alderton, Roberto Amato, Jeffrey Barrett, Sonia Goncalves, Ewan Harrison, David K. Jackson, Ian Johnston, Dominic Kwiatkowski, Cordelia Langford, John Sillitoe on behalf of the Wellcome Sanger Institute COVID-19 Surveillance Team |
| EPI_ISL_2767639 | Lighthouse Lab in Glasgow | Wellcome Sanger Institute for the COVID-19 Genomics UK (COG-UK) Consortium | Harper VanSteenhouse, Yumi Kasai, David Gray, Carol Clugston, Anna Dominiczak and Alex Alderton, Roberto Amato, Jeffrey Barrett, Sonia Goncalves, Ewan Harrison, David K. Jackson, Ian Johnston, Dominic Kwiatkowski, Cordelia Langford, John Sillitoe on behalf of the Wellcome Sanger Institute COVID-19 Surveillance Team |
| EPI_ISL_2767640 | Lighthouse Laboratory Plymouth | Wellcome Sanger Institute for the COVID-19 Genomics UK (COG-UK) Consortium | Lighthouse Laboratory Plymouth and Alex Alderton, Roberto Amato, Jeffrey Barrett, Sonia Goncalves, Ewan Harrison, David K. Jackson, Ian Johnston, Dominic Kwiatkowski, Cordelia Langford, John Sillitoe on behalf of the Wellcome Sanger Institute COVID-19 Surveillance Team |
| EPI_ISL_2767641 | Lighthouse Lab in Alderley Park | Wellcome Sanger Institute for the COVID-19 Genomics UK (COG-UK) Consortium | Jacquelyn Wynn, Mairead Hyland, The Lighthouse Lab in Alderley Park and Alex Alderton, Roberto Amato, Jeffrey Barrett, Sonia Goncalves, Ewan Harrison, David K. Jackson, Ian Johnston, Dominic Kwiatkowski, Cordelia Langford, John Sillitoe on behalf of the Wellcome Sanger Institute COVID-19 Surveillance Team |
| EPI_ISL_2767642 | Lighthouse Laboratory Plymouth | Wellcome Sanger Institute for the COVID-19 Genomics UK (COG-UK) Consortium | Lighthouse Laboratory Plymouth and Alex Alderton, Roberto Amato, Jeffrey Barrett, Sonia Goncalves, Ewan Harrison, David K. Jackson, Ian Johnston, Dominic Kwiatkowski, Cordelia Langford, John Sillitoe on behalf of the Wellcome Sanger Institute COVID-19 Surveillance Team |
| EPI_ISL_2767643 | Lighthouse Lab in Milton Keynes | Wellcome Sanger Institute for the COVID-19 Genomics UK (COG-UK) Consortium | The Lighthouse Lab in Milton Keynes and Alex Alderton, Roberto Amato, Jeffrey Barrett, Sonia Goncalves, Ewan Harrison, David K. Jackson, Ian Johnston, Dominic Kwiatkowski, Cordelia Langford, John Sillitoe on behalf of the Wellcome Sanger Institute COVID-19 Surveillance Team |
| EPI_ISL_2767644 | Lighthouse Lab in Alderley Park | Wellcome Sanger Institute for the COVID-19 Genomics UK (COG-UK) Consortium | Jacquelyn Wynn, Mairead Hyland, The Lighthouse Lab in Alderley Park and Alex Alderton, Roberto Amato, Jeffrey Barrett, Sonia Goncalves, Ewan Harrison, David K. Jackson, Ian Johnston, Dominic Kwiatkowski, Cordelia Langford, John Sillitoe on behalf of the Wellcome Sanger Institute COVID-19 Surveillance Team |
| EPI_ISL_2767645, EPI_ISL_2767646, EPI_ISL_2767647, EPI_ISL_2767648 | Lighthouse Lab in Milton Keynes | Wellcome Sanger Institute for the COVID-19 Genomics UK (COG-UK) Consortium | The Lighthouse Lab in Milton Keynes and Alex Alderton, Roberto Amato, Jeffrey Barrett, Sonia Goncalves, Ewan Harrison, David K. Jackson, Ian Johnston, Dominic Kwiatkowski, Cordelia Langford, John Sillitoe on behalf of the Wellcome Sanger Institute COVID-19 Surveillance Team |
| EPI_ISL_2767649, EPI_ISL_2767650 | Lighthouse Lab in Alderley Park | Wellcome Sanger Institute for the COVID-19 Genomics UK (COG-UK) Consortium | Jacquelyn Wynn, Mairead Hyland, The Lighthouse Lab in Alderley Park and Alex Alderton, Roberto Amato, Jeffrey Barrett, Sonia Goncalves, Ewan Harrison, David K. Jackson, Ian Johnston, Dominic Kwiatkowski, Cordelia Langford, John Sillitoe on behalf of the Wellcome Sanger Institute COVID-19 Surveillance Team |
| EPI_ISL_2767651 | Lighthouse Lab in Milton Keynes | Wellcome Sanger Institute for the COVID-19 Genomics UK (COG-UK) Consortium | The Lighthouse Lab in Milton Keynes and Alex Alderton, Roberto Amato, Jeffrey Barrett, Sonia Goncalves, Ewan Harrison, David K. Jackson, Ian Johnston, Dominic Kwiatkowski, Cordelia Langford, John Sillitoe on behalf of the Wellcome Sanger Institute COVID-19 Surveillance Team |
| EPI_ISL_2767652, EPI_ISL_2767653 | Lighthouse Lab in Alderley Park | Wellcome Sanger Institute for the COVID-19 Genomics UK (COG-UK) Consortium | Jacquelyn Wynn, Mairead Hyland, The Lighthouse Lab in Alderley Park and Alex Alderton, Roberto Amato, Jeffrey Barrett, Sonia Goncalves, Ewan Harrison, David K. Jackson, Ian Johnston, Dominic Kwiatkowski, Cordelia Langford, John Sillitoe on behalf of the Wellcome Sanger Institute COVID-19 Surveillance Team |
| EPI_ISL_2767654 | Lighthouse Lab in Milton Keynes | Wellcome Sanger Institute for the COVID-19 Genomics UK (COG-UK) Consortium | The Lighthouse Lab in Milton Keynes and Alex Alderton, Roberto Amato, Jeffrey Barrett, Sonia Goncalves, Ewan Harrison, David K. Jackson, Ian Johnston, Dominic Kwiatkowski, Cordelia Langford, John Sillitoe on behalf of the Wellcome Sanger Institute COVID-19 Surveillance Team |
| EPI_ISL_2767655, EPI_ISL_2767656, EPI_ISL_2767657, EPI_ISL_2767658 | Lighthouse Lab in Alderley Park | Wellcome Sanger Institute for the COVID-19 Genomics UK (COG-UK) Consortium | Jacquelyn Wynn, Mairead Hyland, The Lighthouse Lab in Alderley Park and Alex Alderton, Roberto Amato, Jeffrey Barrett, Sonia Goncalves, Ewan Harrison, David K. Jackson, Ian Johnston, Dominic Kwiatkowski, Cordelia Langford, John Sillitoe on behalf of the Wellcome Sanger Institute COVID-19 Surveillance Team |
| EPI_ISL_2767659 | Lighthouse Lab in Milton Keynes | Wellcome Sanger Institute for the COVID-19 Genomics UK (COG-UK) Consortium | The Lighthouse Lab in Milton Keynes and Alex Alderton, Roberto Amato, Jeffrey Barrett, Sonia Goncalves, Ewan Harrison, David K. Jackson, Ian Johnston, Dominic Kwiatkowski, Cordelia Langford, John Sillitoe on behalf of the Wellcome Sanger Institute COVID-19 Surveillance Team |
| EPI_ISL_2767660, EPI_ISL_2767661 | Lighthouse Lab in Alderley Park | Wellcome Sanger Institute for the COVID-19 Genomics UK (COG-UK) Consortium | Jacquelyn Wynn, Mairead Hyland, The Lighthouse Lab in Alderley Park and Alex Alderton, Roberto Amato, Jeffrey Barrett, Sonia Goncalves, Ewan Harrison, David K. Jackson, Ian Johnston, Dominic Kwiatkowski, Cordelia Langford, John Sillitoe on behalf of the Wellcome Sanger Institute COVID-19 Surveillance Team |
| EPI_ISL_2767662 | Lighthouse Lab in Milton Keynes | Wellcome Sanger Institute for the COVID-19 Genomics UK (COG-UK) Consortium | The Lighthouse Lab in Milton Keynes and Alex Alderton, Roberto Amato, Jeffrey Barrett, Sonia Goncalves, Ewan Harrison, David K. Jackson, Ian Johnston, Dominic Kwiatkowski, Cordelia Langford, John Sillitoe on behalf of the Wellcome Sanger Institute COVID-19 Surveillance Team |
| EPI_ISL_2767663, EPI_ISL_2767664 | Lighthouse Lab in Alderley Park | Wellcome Sanger Institute for the COVID-19 Genomics UK (COG-UK) Consortium | Jacquelyn Wynn, Mairead Hyland, The Lighthouse Lab in Alderley Park and Alex Alderton, Roberto Amato, Jeffrey Barrett, Sonia Goncalves, Ewan Harrison, David K. Jackson, Ian Johnston, Dominic Kwiatkowski, Cordelia Langford, John Sillitoe on behalf of the Wellcome Sanger Institute COVID-19 Surveillance Team |
| EPI_ISL_2767665 | Lighthouse Lab in Milton Keynes | Wellcome Sanger Institute for the COVID-19 Genomics UK (COG-UK) Consortium | The Lighthouse Lab in Milton Keynes and Alex Alderton, Roberto Amato, Jeffrey Barrett, Sonia Goncalves, Ewan Harrison, David K. Jackson, Ian Johnston, Dominic Kwiatkowski, Cordelia Langford, John Sillitoe on behalf of the Wellcome Sanger Institute COVID-19 Surveillance Team |
| EPI_ISL_2767666, EPI_ISL_2767667 | Lighthouse Lab in Alderley Park | Wellcome Sanger Institute for the COVID-19 Genomics UK (COG-UK) Consortium | Jacquelyn Wynn, Mairead Hyland, The Lighthouse Lab in Alderley Park and Alex Alderton, Roberto Amato, Jeffrey Barrett, Sonia Goncalves, Ewan Harrison, David K. Jackson, Ian Johnston, Dominic Kwiatkowski, Cordelia Langford, John Sillitoe on behalf of the Wellcome Sanger Institute COVID-19 Surveillance Team |
| EPI_ISL_2767668, EPI_ISL_2767670, EPI_ISL_2767671, EPI_ISL_2767672, EPI_ISL_2767673 | Lighthouse Lab in Milton Keynes | Wellcome Sanger Institute for the COVID-19 Genomics UK (COG-UK) Consortium | The Lighthouse Lab in Milton Keynes and Alex Alderton, Roberto Amato, Jeffrey Barrett, Sonia Goncalves, Ewan Harrison, David K. Jackson, Ian Johnston, Dominic Kwiatkowski, Cordelia Langford, John Sillitoe on behalf of the Wellcome Sanger Institute COVID-19 Surveillance Team |
| EPI_ISL_2767674, EPI_ISL_2767675, EPI_ISL_2767676, EPI_ISL_2767677 | Lighthouse Lab in Alderley Park | Wellcome Sanger Institute for the COVID-19 Genomics UK (COG-UK) Consortium | Jacquelyn Wynn, Mairead Hyland, The Lighthouse Lab in Alderley Park and Alex Alderton, Roberto Amato, Jeffrey Barrett, Sonia Goncalves, Ewan Harrison, David K. Jackson, Ian Johnston, Dominic Kwiatkowski, Cordelia Langford, John Sillitoe on behalf of the Wellcome Sanger Institute COVID-19 Surveillance Team |
| EPI_ISL_2767678 | Lighthouse Lab in Milton Keynes | Wellcome Sanger Institute for the COVID-19 Genomics UK (COG-UK) Consortium | The Lighthouse Lab in Milton Keynes and Alex Alderton, Roberto Amato, Jeffrey Barrett, Sonia Goncalves, Ewan Harrison, David K. Jackson, Ian Johnston, Dominic Kwiatkowski, Cordelia Langford, John Sillitoe on behalf of the Wellcome Sanger Institute COVID-19 Surveillance Team |
| EPI_ISL_2767679 | Lighthouse Lab in Alderley Park | Wellcome Sanger Institute for the COVID-19 Genomics UK (COG-UK) Consortium | Jacquelyn Wynn, Mairead Hyland, The Lighthouse Lab in Alderley Park and Alex Alderton, Roberto Amato, Jeffrey Barrett, Sonia Goncalves, Ewan Harrison, David K. Jackson, Ian Johnston, Dominic Kwiatkowski, Cordelia Langford, John Sillitoe on behalf of the Wellcome Sanger Institute COVID-19 Surveillance Team |
| EPI_ISL_2767680, EPI_ISL_2767681 | Lighthouse Lab in Milton Keynes | Wellcome Sanger Institute for the COVID-19 Genomics UK (COG-UK) Consortium | The Lighthouse Lab in Milton Keynes and Alex Alderton, Roberto Amato, Jeffrey Barrett, Sonia Goncalves, Ewan Harrison, David K. Jackson, Ian Johnston, Dominic Kwiatkowski, Cordelia Langford, John Sillitoe on behalf of the Wellcome Sanger Institute COVID-19 Surveillance Team |
| EPI_ISL_2767682 | Lighthouse Lab in Alderley Park | Wellcome Sanger Institute for the COVID-19 Genomics UK (COG-UK) Consortium | Jacquelyn Wynn, Mairead Hyland, The Lighthouse Lab in Alderley Park and Alex Alderton, Roberto Amato, Jeffrey Barrett, Sonia Goncalves, Ewan Harrison, David K. Jackson, Ian Johnston, Dominic Kwiatkowski, Cordelia Langford, John Sillitoe on behalf of the Wellcome Sanger Institute COVID-19 Surveillance Team |
| EPI_ISL_2767683 | Lighthouse Lab in Milton Keynes | Wellcome Sanger Institute for the COVID-19 Genomics UK (COG-UK) Consortium | The Lighthouse Lab in Milton Keynes and Alex Alderton, Roberto Amato, Jeffrey Barrett, Sonia Goncalves, Ewan Harrison, David K. Jackson, Ian Johnston, Dominic Kwiatkowski, Cordelia Langford, John Sillitoe on behalf of the Wellcome Sanger Institute COVID-19 Surveillance Team |
| EPI_ISL_2767684 | Lighthouse Lab in Alderley Park | Wellcome Sanger Institute for the COVID-19 Genomics UK (COG-UK) Consortium | Jacquelyn Wynn, Mairead Hyland, The Lighthouse Lab in Alderley Park and Alex Alderton, Roberto Amato, Jeffrey Barrett, Sonia Goncalves, Ewan Harrison, David K. Jackson, Ian Johnston, Dominic Kwiatkowski, Cordelia Langford, John Sillitoe on behalf of the Wellcome Sanger Institute COVID-19 Surveillance Team |
| EPI_ISL_2767685, EPI_ISL_2767686 | Lighthouse Lab in Milton Keynes | Wellcome Sanger Institute for the COVID-19 Genomics UK (COG-UK) Consortium | The Lighthouse Lab in Milton Keynes and Alex Alderton, Roberto Amato, Jeffrey Barrett, Sonia Goncalves, Ewan Harrison, David K. Jackson, Ian Johnston, Dominic Kwiatkowski, Cordelia Langford, John Sillitoe on behalf of the Wellcome Sanger Institute COVID-19 Surveillance Team |
| EPI_ISL_2767687 | Lighthouse Lab in Alderley Park | Wellcome Sanger Institute for the COVID-19 Genomics UK (COG-UK) Consortium | Jacquelyn Wynn, Mairead Hyland, The Lighthouse Lab in Alderley Park and Alex Alderton, Roberto Amato, Jeffrey Barrett, Sonia Goncalves, Ewan Harrison, David K. Jackson, Ian Johnston, Dominic Kwiatkowski, Cordelia Langford, John Sillitoe on behalf of the Wellcome Sanger Institute COVID-19 Surveillance Team |
| EPI_ISL_2767688 | Lighthouse Lab in Milton Keynes | Wellcome Sanger Institute for the COVID-19 Genomics UK (COG-UK) Consortium | The Lighthouse Lab in Milton Keynes and Alex Alderton, Roberto Amato, Jeffrey Barrett, Sonia Goncalves, Ewan Harrison, David K. Jackson, Ian Johnston, Dominic Kwiatkowski, Cordelia Langford, John Sillitoe on behalf of the Wellcome Sanger Institute COVID-19 Surveillance Team |
| EPI_ISL_2767689, EPI_ISL_2767690, | Lighthouse Lab in Alderley Park | Wellcome Sanger Institute for the COVID-19 Genomics UK (COG-UK) Consortium | Jacquelyn Wynn, Mairead Hyland, The Lighthouse Lab in Alderley Park and Alex Alderton, Roberto Amato, Jeffrey Barrett, Sonia Goncalves, Ewan Harrison, David K. Jackson, Ian Johnston, Dominic Kwiatkowski, Cordelia Langford, John Sillitoe on behalf of the Wellcome Sanger Institute COVID-19 Surveillance Team |

| | | | |
|---|---|---|---|
| EPI_ISL_2767691, EPI_ISL_2767692, EPI_ISL_2767693 | | | |
| EPI_ISL_2767694 | Lighthouse Lab in Milton Keynes | Wellcome Sanger Institute for the COVID-19 Genomics UK (COG-UK) Consortium | The Lighthouse Lab in Milton Keynes and Alex Alderton, Roberto Amato, Jeffrey Barrett, Sonia Goncalves, Ewan Harrison, David K. Jackson, Ian Johnston, Dominic Kwiatkowski, Cordelia Langford, John Sillitoe on behalf of the Wellcome Sanger Institute COVID-19 Surveillance Team |
| EPI_ISL_2767695 | Lighthouse Lab in Alderley Park | Wellcome Sanger Institute for the COVID-19 Genomics UK (COG-UK) Consortium | Jacquelyn Wynn, Mairead Hyland, The Lighthouse Lab in Alderley Park and Alex Alderton, Roberto Amato, Jeffrey Barrett, Sonia Goncalves, Ewan Harrison, David K. Jackson, Ian Johnston, Dominic Kwiatkowski, Cordelia Langford, John Sillitoe on behalf of the Wellcome Sanger Institute COVID-19 Surveillance Team |
| EPI_ISL_2767696, EPI_ISL_2767697 | Lighthouse Lab in Milton Keynes | Wellcome Sanger Institute for the COVID-19 Genomics UK (COG-UK) Consortium | The Lighthouse Lab in Milton Keynes and Alex Alderton, Roberto Amato, Jeffrey Barrett, Sonia Goncalves, Ewan Harrison, David K. Jackson, Ian Johnston, Dominic Kwiatkowski, Cordelia Langford, John Sillitoe on behalf of the Wellcome Sanger Institute COVID-19 Surveillance Team |
| EPI_ISL_2767698, EPI_ISL_2767699, EPI_ISL_2767700, EPI_ISL_2767701, EPI_ISL_2767702, EPI_ISL_2767703 | Lighthouse Lab in Alderley Park | Wellcome Sanger Institute for the COVID-19 Genomics UK (COG-UK) Consortium | Jacquelyn Wynn, Mairead Hyland, The Lighthouse Lab in Alderley Park and Alex Alderton, Roberto Amato, Jeffrey Barrett, Sonia Goncalves, Ewan Harrison, David K. Jackson, Ian Johnston, Dominic Kwiatkowski, Cordelia Langford, John Sillitoe on behalf of the Wellcome Sanger Institute COVID-19 Surveillance Team |
| EPI_ISL_2767704, EPI_ISL_2767705, EPI_ISL_2767706, EPI_ISL_2767707, EPI_ISL_2767708 | Lighthouse Lab in Milton Keynes | Wellcome Sanger Institute for the COVID-19 Genomics UK (COG-UK) Consortium | The Lighthouse Lab in Milton Keynes and Alex Alderton, Roberto Amato, Jeffrey Barrett, Sonia Goncalves, Ewan Harrison, David K. Jackson, Ian Johnston, Dominic Kwiatkowski, Cordelia Langford, John Sillitoe on behalf of the Wellcome Sanger Institute COVID-19 Surveillance Team |
| EPI_ISL_2767709 | Lighthouse Lab in Alderley Park | Wellcome Sanger Institute for the COVID-19 Genomics UK (COG-UK) Consortium | Jacquelyn Wynn, Mairead Hyland, The Lighthouse Lab in Alderley Park and Alex Alderton, Roberto Amato, Jeffrey Barrett, Sonia Goncalves, Ewan Harrison, David K. Jackson, Ian Johnston, Dominic Kwiatkowski, Cordelia Langford, John Sillitoe on behalf of the Wellcome Sanger Institute COVID-19 Surveillance Team |
| EPI_ISL_2767710 | Lighthouse Lab in Milton Keynes | Wellcome Sanger Institute for the COVID-19 Genomics UK (COG-UK) Consortium | The Lighthouse Lab in Milton Keynes and Alex Alderton, Roberto Amato, Jeffrey Barrett, Sonia Goncalves, Ewan Harrison, David K. Jackson, Ian Johnston, Dominic Kwiatkowski, Cordelia Langford, John Sillitoe on behalf of the Wellcome Sanger Institute COVID-19 Surveillance Team |
| EPI_ISL_2767711, EPI_ISL_2767712, EPI_ISL_2767713 | Lighthouse Lab in Alderley Park | Wellcome Sanger Institute for the COVID-19 Genomics UK (COG-UK) Consortium | Jacquelyn Wynn, Mairead Hyland, The Lighthouse Lab in Alderley Park and Alex Alderton, Roberto Amato, Jeffrey Barrett, Sonia Goncalves, Ewan Harrison, David K. Jackson, Ian Johnston, Dominic Kwiatkowski, Cordelia Langford, John Sillitoe on behalf of the Wellcome Sanger Institute COVID-19 Surveillance Team |
| EPI_ISL_2767714, EPI_ISL_2767715 | Lighthouse Lab in Milton Keynes | Wellcome Sanger Institute for the COVID-19 Genomics UK (COG-UK) Consortium | The Lighthouse Lab in Milton Keynes and Alex Alderton, Roberto Amato, Jeffrey Barrett, Sonia Goncalves, Ewan Harrison, David K. Jackson, Ian Johnston, Dominic Kwiatkowski, Cordelia Langford, John Sillitoe on behalf of the Wellcome Sanger Institute COVID-19 Surveillance Team |
| EPI_ISL_2767716 | Lighthouse Lab in Alderley Park | Wellcome Sanger Institute for the COVID-19 Genomics UK (COG-UK) Consortium | Jacquelyn Wynn, Mairead Hyland, The Lighthouse Lab in Alderley Park and Alex Alderton, Roberto Amato, Jeffrey Barrett, Sonia Goncalves, Ewan Harrison, David K. Jackson, Ian Johnston, Dominic Kwiatkowski, Cordelia Langford, John Sillitoe on behalf of the Wellcome Sanger Institute COVID-19 Surveillance Team |
| EPI_ISL_2767717, EPI_ISL_2767718 | Lighthouse Lab in Milton Keynes | Wellcome Sanger Institute for the COVID-19 Genomics UK (COG-UK) Consortium | The Lighthouse Lab in Milton Keynes and Alex Alderton, Roberto Amato, Jeffrey Barrett, Sonia Goncalves, Ewan Harrison, David K. Jackson, Ian Johnston, Dominic Kwiatkowski, Cordelia Langford, John Sillitoe on behalf of the Wellcome Sanger Institute COVID-19 Surveillance Team |
| EPI_ISL_2767719, EPI_ISL_2767720, EPI_ISL_2767721 | Lighthouse Lab in Alderley Park | Wellcome Sanger Institute for the COVID-19 Genomics UK (COG-UK) Consortium | Jacquelyn Wynn, Mairead Hyland, The Lighthouse Lab in Alderley Park and Alex Alderton, Roberto Amato, Jeffrey Barrett, Sonia Goncalves, Ewan Harrison, David K. Jackson, Ian Johnston, Dominic Kwiatkowski, Cordelia Langford, John Sillitoe on behalf of the Wellcome Sanger Institute COVID-19 Surveillance Team |
| EPI_ISL_2767722 | Lighthouse Lab in Milton Keynes | Wellcome Sanger Institute for the COVID-19 Genomics UK (COG-UK) Consortium | The Lighthouse Lab in Milton Keynes and Alex Alderton, Roberto Amato, Jeffrey Barrett, Sonia Goncalves, Ewan Harrison, David K. Jackson, Ian Johnston, Dominic Kwiatkowski, Cordelia Langford, John Sillitoe on behalf of the Wellcome Sanger Institute COVID-19 Surveillance Team |
| EPI_ISL_2767723, EPI_ISL_2767724 | Lighthouse Lab in Alderley Park | Wellcome Sanger Institute for the COVID-19 Genomics UK (COG-UK) Consortium | Jacquelyn Wynn, Mairead Hyland, The Lighthouse Lab in Alderley Park and Alex Alderton, Roberto Amato, Jeffrey Barrett, Sonia Goncalves, Ewan Harrison, David K. Jackson, Ian Johnston, Dominic Kwiatkowski, Cordelia Langford, John Sillitoe on behalf of the Wellcome Sanger Institute COVID-19 Surveillance Team |
| EPI_ISL_2767725, EPI_ISL_2767726, EPI_ISL_2767727, EPI_ISL_2767728 | Lighthouse Lab in Milton Keynes | Wellcome Sanger Institute for the COVID-19 Genomics UK (COG-UK) Consortium | The Lighthouse Lab in Milton Keynes and Alex Alderton, Roberto Amato, Jeffrey Barrett, Sonia Goncalves, Ewan Harrison, David K. Jackson, Ian Johnston, Dominic Kwiatkowski, Cordelia Langford, John Sillitoe on behalf of the Wellcome Sanger Institute COVID-19 Surveillance Team |
| EPI_ISL_2767729, EPI_ISL_2767730, EPI_ISL_2767731 | Lighthouse Lab in Alderley Park | Wellcome Sanger Institute for the COVID-19 Genomics UK (COG-UK) Consortium | Jacquelyn Wynn, Mairead Hyland, The Lighthouse Lab in Alderley Park and Alex Alderton, Roberto Amato, Jeffrey Barrett, Sonia Goncalves, Ewan Harrison, David K. Jackson, Ian Johnston, Dominic Kwiatkowski, Cordelia Langford, John Sillitoe on behalf of the Wellcome Sanger Institute COVID-19 Surveillance Team |
| EPI_ISL_2767732, EPI_ISL_2767733, EPI_ISL_2767734 | Lighthouse Lab in Milton Keynes | Wellcome Sanger Institute for the COVID-19 Genomics UK (COG-UK) Consortium | The Lighthouse Lab in Milton Keynes and Alex Alderton, Roberto Amato, Jeffrey Barrett, Sonia Goncalves, Ewan Harrison, David K. Jackson, Ian Johnston, Dominic Kwiatkowski, Cordelia Langford, John Sillitoe on behalf of the Wellcome Sanger Institute COVID-19 Surveillance Team |
| EPI_ISL_2767735, EPI_ISL_2767736 | Lighthouse Lab in Alderley Park | Wellcome Sanger Institute for the COVID-19 Genomics UK (COG-UK) Consortium | Jacquelyn Wynn, Mairead Hyland, The Lighthouse Lab in Alderley Park and Alex Alderton, Roberto Amato, Jeffrey Barrett, Sonia Goncalves, Ewan Harrison, David K. Jackson, Ian Johnston, Dominic Kwiatkowski, Cordelia Langford, John Sillitoe on behalf of the Wellcome Sanger Institute COVID-19 Surveillance Team |
| EPI_ISL_2767737 | Lighthouse Lab in Milton Keynes | Wellcome Sanger Institute for the COVID-19 Genomics UK (COG-UK) Consortium | The Lighthouse Lab in Milton Keynes and Alex Alderton, Roberto Amato, Jeffrey Barrett, Sonia Goncalves, Ewan Harrison, David K. Jackson, Ian Johnston, Dominic Kwiatkowski, Cordelia Langford, John Sillitoe on behalf of the Wellcome Sanger Institute COVID-19 Surveillance Team |
| EPI_ISL_2767738, EPI_ISL_2767739, EPI_ISL_2767740 | Lighthouse Lab in Alderley Park | Wellcome Sanger Institute for the COVID-19 Genomics UK (COG-UK) Consortium | Jacquelyn Wynn, Mairead Hyland, The Lighthouse Lab in Alderley Park and Alex Alderton, Roberto Amato, Jeffrey Barrett, Sonia Goncalves, Ewan Harrison, David K. Jackson, Ian Johnston, Dominic Kwiatkowski, Cordelia Langford, John Sillitoe on behalf of the Wellcome Sanger Institute COVID-19 Surveillance Team |
| EPI_ISL_2767741, EPI_ISL_2767742, EPI_ISL_2767743 | Lighthouse Lab in Milton Keynes | Wellcome Sanger Institute for the COVID-19 Genomics UK (COG-UK) Consortium | The Lighthouse Lab in Milton Keynes and Alex Alderton, Roberto Amato, Jeffrey Barrett, Sonia Goncalves, Ewan Harrison, David K. Jackson, Ian Johnston, Dominic Kwiatkowski, Cordelia Langford, John Sillitoe on behalf of the Wellcome Sanger Institute COVID-19 Surveillance Team |
| EPI_ISL_2767744 | Lighthouse Lab in Alderley Park | Wellcome Sanger Institute for the COVID-19 Genomics UK (COG-UK) Consortium | Jacquelyn Wynn, Mairead Hyland, The Lighthouse Lab in Alderley Park and Alex Alderton, Roberto Amato, Jeffrey Barrett, Sonia Goncalves, Ewan Harrison, David K. Jackson, Ian Johnston, Dominic Kwiatkowski, Cordelia Langford, John Sillitoe on behalf of the Wellcome Sanger Institute COVID-19 Surveillance Team |
| EPI_ISL_2767745, EPI_ISL_2767746 | Lighthouse Lab in Milton Keynes | Wellcome Sanger Institute for the COVID-19 Genomics UK (COG-UK) Consortium | The Lighthouse Lab in Milton Keynes and Alex Alderton, Roberto Amato, Jeffrey Barrett, Sonia Goncalves, Ewan Harrison, David K. Jackson, Ian Johnston, Dominic Kwiatkowski, Cordelia Langford, John Sillitoe on behalf of the Wellcome Sanger Institute COVID-19 Surveillance Team |
| EPI_ISL_2767747, EPI_ISL_2767748, EPI_ISL_2767749 | Lighthouse Lab in Alderley Park | Wellcome Sanger Institute for the COVID-19 Genomics UK (COG-UK) Consortium | Jacquelyn Wynn, Mairead Hyland, The Lighthouse Lab in Alderley Park and Alex Alderton, Roberto Amato, Jeffrey Barrett, Sonia Goncalves, Ewan Harrison, David K. Jackson, Ian Johnston, Dominic Kwiatkowski, Cordelia Langford, John Sillitoe on behalf of the Wellcome Sanger Institute COVID-19 Surveillance Team |
| EPI_ISL_2767750, EPI_ISL_2767751 | Lighthouse Lab in Milton Keynes | Wellcome Sanger Institute for the COVID-19 Genomics UK (COG-UK) Consortium | The Lighthouse Lab in Milton Keynes and Alex Alderton, Roberto Amato, Jeffrey Barrett, Sonia Goncalves, Ewan Harrison, David K. Jackson, Ian Johnston, Dominic Kwiatkowski, Cordelia Langford, John Sillitoe on behalf of the Wellcome Sanger Institute COVID-19 Surveillance Team |
| EPI_ISL_2767752 | Lighthouse Lab in Alderley Park | Wellcome Sanger Institute for the COVID-19 Genomics UK (COG-UK) Consortium | Jacquelyn Wynn, Mairead Hyland, The Lighthouse Lab in Alderley Park and Alex Alderton, Roberto Amato, Jeffrey Barrett, Sonia Goncalves, Ewan Harrison, David K. Jackson, Ian Johnston, Dominic Kwiatkowski, Cordelia Langford, John Sillitoe on behalf of the Wellcome Sanger Institute COVID-19 Surveillance Team |
| EPI_ISL_2767753, EPI_ISL_2767754, EPI_ISL_2767755 | Lighthouse Lab in Milton Keynes | Wellcome Sanger Institute for the COVID-19 Genomics UK (COG-UK) Consortium | The Lighthouse Lab in Milton Keynes and Alex Alderton, Roberto Amato, Jeffrey Barrett, Sonia Goncalves, Ewan Harrison, David K. Jackson, Ian Johnston, Dominic Kwiatkowski, Cordelia Langford, John Sillitoe on behalf of the Wellcome Sanger Institute COVID-19 Surveillance Team |
| EPI_ISL_2767756 | Lighthouse Lab in Alderley Park | Wellcome Sanger Institute for the COVID-19 Genomics UK (COG-UK) Consortium | Jacquelyn Wynn, Mairead Hyland, The Lighthouse Lab in Alderley Park and Alex Alderton, Roberto Amato, Jeffrey Barrett, Sonia Goncalves, Ewan Harrison, David K. Jackson, Ian Johnston, Dominic Kwiatkowski, Cordelia Langford, John Sillitoe on behalf of the Wellcome Sanger Institute COVID-19 Surveillance Team |
| EPI_ISL_2767757 | Lighthouse Lab in Milton Keynes | Wellcome Sanger Institute for the COVID-19 Genomics UK (COG-UK) Consortium | The Lighthouse Lab in Milton Keynes and Alex Alderton, Roberto Amato, Jeffrey Barrett, Sonia Goncalves, Ewan Harrison, David K. Jackson, Ian Johnston, Dominic Kwiatkowski, Cordelia Langford, John Sillitoe on behalf of the Wellcome Sanger Institute COVID-19 Surveillance Team |
| EPI_ISL_2767758, EPI_ISL_2767759, EPI_ISL_2767760 | Lighthouse Lab in Alderley Park | Wellcome Sanger Institute for the COVID-19 Genomics UK (COG-UK) Consortium | Jacquelyn Wynn, Mairead Hyland, The Lighthouse Lab in Alderley Park and Alex Alderton, Roberto Amato, Jeffrey Barrett, Sonia Goncalves, Ewan Harrison, David K. Jackson, Ian Johnston, Dominic Kwiatkowski, Cordelia Langford, John Sillitoe on behalf of the Wellcome Sanger Institute COVID-19 Surveillance Team |
| EPI_ISL_2767762, EPI_ISL_2767763, EPI_ISL_2767764 | Lighthouse Lab in Milton Keynes | Wellcome Sanger Institute for the COVID-19 Genomics UK (COG-UK) Consortium | The Lighthouse Lab in Milton Keynes and Alex Alderton, Roberto Amato, Jeffrey Barrett, Sonia Goncalves, Ewan Harrison, David K. Jackson, Ian Johnston, Dominic Kwiatkowski, Cordelia Langford, John Sillitoe on behalf of the Wellcome Sanger Institute COVID-19 Surveillance Team |
| EPI_ISL_2767765 | Lighthouse Lab in Alderley Park | Wellcome Sanger Institute for the COVID-19 Genomics UK (COG-UK) Consortium | Jacquelyn Wynn, Mairead Hyland, The Lighthouse Lab in Alderley Park and Alex Alderton, Roberto Amato, Jeffrey Barrett, Sonia Goncalves, Ewan Harrison, David K. Jackson, Ian Johnston, Dominic Kwiatkowski, Cordelia Langford, John Sillitoe on behalf of the Wellcome Sanger Institute COVID-19 Surveillance Team |
| EPI_ISL_2767766, EPI_ISL_2767767 | Lighthouse Lab in Milton Keynes | Wellcome Sanger Institute for the COVID-19 Genomics UK (COG-UK) Consortium | The Lighthouse Lab in Milton Keynes and Alex Alderton, Roberto Amato, Jeffrey Barrett, Sonia Goncalves, Ewan Harrison, David K. Jackson, Ian Johnston, Dominic Kwiatkowski, Cordelia Langford, John Sillitoe on behalf of the Wellcome Sanger Institute COVID-19 Surveillance Team |
| EPI_ISL_2767768, EPI_ISL_2767769 | Lighthouse Lab in Alderley Park | Wellcome Sanger Institute for the COVID-19 Genomics UK (COG-UK) Consortium | Jacquelyn Wynn, Mairead Hyland, The Lighthouse Lab in Alderley Park and Alex Alderton, Roberto Amato, Jeffrey Barrett, Sonia Goncalves, Ewan Harrison, David K. Jackson, Ian Johnston, Dominic Kwiatkowski, Cordelia Langford, John Sillitoe on behalf of the Wellcome Sanger Institute COVID-19 Surveillance Team |
| EPI_ISL_2767770 | Lighthouse Lab in Milton Keynes | Wellcome Sanger Institute for the COVID-19 Genomics UK (COG-UK) Consortium | The Lighthouse Lab in Milton Keynes and Alex Alderton, Roberto Amato, Jeffrey Barrett, Sonia Goncalves, Ewan Harrison, David K. Jackson, Ian Johnston, Dominic Kwiatkowski, Cordelia Langford, John Sillitoe on behalf of the Wellcome Sanger Institute COVID-19 Surveillance Team |
| EPI_ISL_2767771, EPI_ISL_2767772, | Lighthouse Lab in Alderley Park | Wellcome Sanger Institute for the COVID-19 Genomics UK (COG-UK) Consortium | Jacquelyn Wynn, Mairead Hyland, The Lighthouse Lab in Alderley Park and Alex Alderton, Roberto Amato, Jeffrey Barrett, Sonia Goncalves, Ewan Harrison, David K. Jackson, Ian Johnston, Dominic Kwiatkowski, Cordelia Langford, John Sillitoe on behalf of the Wellcome Sanger Institute COVID-19 Surveillance Team |

| | | | |
|---|---|---|---|
| EPI_ISL_2767773 | | | |
| EPI_ISL_2767774 | Lighthouse Lab in Milton Keynes | Wellcome Sanger Institute for the COVID-19 Genomics UK (COG-UK) Consortium | The Lighthouse Lab in Milton Keynes and Alex Alderton, Roberto Amato, Jeffrey Barrett, Sonia Goncalves, Ewan Harrison, David K. Jackson, Ian Johnston, Dominic Kwiatkowski, Cordelia Langford, John Sillitoe on behalf of the Wellcome Sanger Institute COVID-19 Surveillance Team |
| EPI_ISL_2767775, EPI_ISL_2767776 | Lighthouse Lab in Alderley Park | Wellcome Sanger Institute for the COVID-19 Genomics UK (COG-UK) Consortium | Jacquelyn Wynn, Mairead Hyland, The Lighthouse Lab in Alderley Park and Alex Alderton, Roberto Amato, Jeffrey Barrett, Sonia Goncalves, Ewan Harrison, David K. Jackson, Ian Johnston, Dominic Kwiatkowski, Cordelia Langford, John Sillitoe on behalf of the Wellcome Sanger Institute COVID-19 Surveillance Team |
| EPI_ISL_2767777, EPI_ISL_2767778 | Lighthouse Lab in Milton Keynes | Wellcome Sanger Institute for the COVID-19 Genomics UK (COG-UK) Consortium | The Lighthouse Lab in Milton Keynes and Alex Alderton, Roberto Amato, Jeffrey Barrett, Sonia Goncalves, Ewan Harrison, David K. Jackson, Ian Johnston, Dominic Kwiatkowski, Cordelia Langford, John Sillitoe on behalf of the Wellcome Sanger Institute COVID-19 Surveillance Team |
| EPI_ISL_2767779 | Lighthouse Lab in Alderley Park | Wellcome Sanger Institute for the COVID-19 Genomics UK (COG-UK) Consortium | Jacquelyn Wynn, Mairead Hyland, The Lighthouse Lab in Alderley Park and Alex Alderton, Roberto Amato, Jeffrey Barrett, Sonia Goncalves, Ewan Harrison, David K. Jackson, Ian Johnston, Dominic Kwiatkowski, Cordelia Langford, John Sillitoe on behalf of the Wellcome Sanger Institute COVID-19 Surveillance Team |
| EPI_ISL_2767780 | Lighthouse Lab in Milton Keynes | Wellcome Sanger Institute for the COVID-19 Genomics UK (COG-UK) Consortium | The Lighthouse Lab in Milton Keynes and Alex Alderton, Roberto Amato, Jeffrey Barrett, Sonia Goncalves, Ewan Harrison, David K. Jackson, Ian Johnston, Dominic Kwiatkowski, Cordelia Langford, John Sillitoe on behalf of the Wellcome Sanger Institute COVID-19 Surveillance Team |
| EPI_ISL_2767781, EPI_ISL_2767782 | Lighthouse Lab in Alderley Park | Wellcome Sanger Institute for the COVID-19 Genomics UK (COG-UK) Consortium | Jacquelyn Wynn, Mairead Hyland, The Lighthouse Lab in Alderley Park and Alex Alderton, Roberto Amato, Jeffrey Barrett, Sonia Goncalves, Ewan Harrison, David K. Jackson, Ian Johnston, Dominic Kwiatkowski, Cordelia Langford, John Sillitoe on behalf of the Wellcome Sanger Institute COVID-19 Surveillance Team |
| EPI_ISL_2767783 | Lighthouse Lab in Milton Keynes | Wellcome Sanger Institute for the COVID-19 Genomics UK (COG-UK) Consortium | The Lighthouse Lab in Milton Keynes and Alex Alderton, Roberto Amato, Jeffrey Barrett, Sonia Goncalves, Ewan Harrison, David K. Jackson, Ian Johnston, Dominic Kwiatkowski, Cordelia Langford, John Sillitoe on behalf of the Wellcome Sanger Institute COVID-19 Surveillance Team |
| EPI_ISL_2767785 | Lighthouse Lab in Alderley Park | Wellcome Sanger Institute for the COVID-19 Genomics UK (COG-UK) Consortium | Jacquelyn Wynn, Mairead Hyland, The Lighthouse Lab in Alderley Park and Alex Alderton, Roberto Amato, Jeffrey Barrett, Sonia Goncalves, Ewan Harrison, David K. Jackson, Ian Johnston, Dominic Kwiatkowski, Cordelia Langford, John Sillitoe on behalf of the Wellcome Sanger Institute COVID-19 Surveillance Team |
| EPI_ISL_2767786 | Lighthouse Lab in Milton Keynes | Wellcome Sanger Institute for the COVID-19 Genomics UK (COG-UK) Consortium | The Lighthouse Lab in Milton Keynes and Alex Alderton, Roberto Amato, Jeffrey Barrett, Sonia Goncalves, Ewan Harrison, David K. Jackson, Ian Johnston, Dominic Kwiatkowski, Cordelia Langford, John Sillitoe on behalf of the Wellcome Sanger Institute COVID-19 Surveillance Team |
| EPI_ISL_2767787, EPI_ISL_2767788 | Lighthouse Lab in Alderley Park | Wellcome Sanger Institute for the COVID-19 Genomics UK (COG-UK) Consortium | Jacquelyn Wynn, Mairead Hyland, The Lighthouse Lab in Alderley Park and Alex Alderton, Roberto Amato, Jeffrey Barrett, Sonia Goncalves, Ewan Harrison, David K. Jackson, Ian Johnston, Dominic Kwiatkowski, Cordelia Langford, John Sillitoe on behalf of the Wellcome Sanger Institute COVID-19 Surveillance Team |
| EPI_ISL_2767789, EPI_ISL_2767790, EPI_ISL_2767791, EPI_ISL_2767792 | Lighthouse Lab in Milton Keynes | Wellcome Sanger Institute for the COVID-19 Genomics UK (COG-UK) Consortium | The Lighthouse Lab in Milton Keynes and Alex Alderton, Roberto Amato, Jeffrey Barrett, Sonia Goncalves, Ewan Harrison, David K. Jackson, Ian Johnston, Dominic Kwiatkowski, Cordelia Langford, John Sillitoe on behalf of the Wellcome Sanger Institute COVID-19 Surveillance Team |
| EPI_ISL_2767793 | Lighthouse Lab in Alderley Park | Wellcome Sanger Institute for the COVID-19 Genomics UK (COG-UK) Consortium | Jacquelyn Wynn, Mairead Hyland, The Lighthouse Lab in Alderley Park and Alex Alderton, Roberto Amato, Jeffrey Barrett, Sonia Goncalves, Ewan Harrison, David K. Jackson, Ian Johnston, Dominic Kwiatkowski, Cordelia Langford, John Sillitoe on behalf of the Wellcome Sanger Institute COVID-19 Surveillance Team |
| EPI_ISL_2767794, EPI_ISL_2767795, EPI_ISL_2767796 | Lighthouse Lab in Milton Keynes | Wellcome Sanger Institute for the COVID-19 Genomics UK (COG-UK) Consortium | The Lighthouse Lab in Milton Keynes and Alex Alderton, Roberto Amato, Jeffrey Barrett, Sonia Goncalves, Ewan Harrison, David K. Jackson, Ian Johnston, Dominic Kwiatkowski, Cordelia Langford, John Sillitoe on behalf of the Wellcome Sanger Institute COVID-19 Surveillance Team |
| EPI_ISL_2767797 | Lighthouse Lab in Alderley Park | Wellcome Sanger Institute for the COVID-19 Genomics UK (COG-UK) Consortium | Jacquelyn Wynn, Mairead Hyland, The Lighthouse Lab in Alderley Park and Alex Alderton, Roberto Amato, Jeffrey Barrett, Sonia Goncalves, Ewan Harrison, David K. Jackson, Ian Johnston, Dominic Kwiatkowski, Cordelia Langford, John Sillitoe on behalf of the Wellcome Sanger Institute COVID-19 Surveillance Team |
| EPI_ISL_2767798, EPI_ISL_2767799 | Lighthouse Lab in Milton Keynes | Wellcome Sanger Institute for the COVID-19 Genomics UK (COG-UK) Consortium | The Lighthouse Lab in Milton Keynes and Alex Alderton, Roberto Amato, Jeffrey Barrett, Sonia Goncalves, Ewan Harrison, David K. Jackson, Ian Johnston, Dominic Kwiatkowski, Cordelia Langford, John Sillitoe on behalf of the Wellcome Sanger Institute COVID-19 Surveillance Team |
| EPI_ISL_2767800 | Lighthouse Lab in Alderley Park | Wellcome Sanger Institute for the COVID-19 Genomics UK (COG-UK) Consortium | Jacquelyn Wynn, Mairead Hyland, The Lighthouse Lab in Alderley Park and Alex Alderton, Roberto Amato, Jeffrey Barrett, Sonia Goncalves, Ewan Harrison, David K. Jackson, Ian Johnston, Dominic Kwiatkowski, Cordelia Langford, John Sillitoe on behalf of the Wellcome Sanger Institute COVID-19 Surveillance Team |
| EPI_ISL_2767801, EPI_ISL_2767802, EPI_ISL_2767803 | Lighthouse Lab in Milton Keynes | Wellcome Sanger Institute for the COVID-19 Genomics UK (COG-UK) Consortium | The Lighthouse Lab in Milton Keynes and Alex Alderton, Roberto Amato, Jeffrey Barrett, Sonia Goncalves, Ewan Harrison, David K. Jackson, Ian Johnston, Dominic Kwiatkowski, Cordelia Langford, John Sillitoe on behalf of the Wellcome Sanger Institute COVID-19 Surveillance Team |
| EPI_ISL_2767804 | Lighthouse Lab in Alderley Park | Wellcome Sanger Institute for the COVID-19 Genomics UK (COG-UK) Consortium | Jacquelyn Wynn, Mairead Hyland, The Lighthouse Lab in Alderley Park and Alex Alderton, Roberto Amato, Jeffrey Barrett, Sonia Goncalves, Ewan Harrison, David K. Jackson, Ian Johnston, Dominic Kwiatkowski, Cordelia Langford, John Sillitoe on behalf of the Wellcome Sanger Institute COVID-19 Surveillance Team |
| EPI_ISL_2767805 | Lighthouse Lab in Milton Keynes | Wellcome Sanger Institute for the COVID-19 Genomics UK (COG-UK) Consortium | The Lighthouse Lab in Milton Keynes and Alex Alderton, Roberto Amato, Jeffrey Barrett, Sonia Goncalves, Ewan Harrison, David K. Jackson, Ian Johnston, Dominic Kwiatkowski, Cordelia Langford, John Sillitoe on behalf of the Wellcome Sanger Institute COVID-19 Surveillance Team |
| EPI_ISL_2767806 | Lighthouse Lab in Alderley Park | Wellcome Sanger Institute for the COVID-19 Genomics UK (COG-UK) Consortium | Jacquelyn Wynn, Mairead Hyland, The Lighthouse Lab in Alderley Park and Alex Alderton, Roberto Amato, Jeffrey Barrett, Sonia Goncalves, Ewan Harrison, David K. Jackson, Ian Johnston, Dominic Kwiatkowski, Cordelia Langford, John Sillitoe on behalf of the Wellcome Sanger Institute COVID-19 Surveillance Team |
| EPI_ISL_2767807 | Lighthouse Lab in Milton Keynes | Wellcome Sanger Institute for the COVID-19 Genomics UK (COG-UK) Consortium | The Lighthouse Lab in Milton Keynes and Alex Alderton, Roberto Amato, Jeffrey Barrett, Sonia Goncalves, Ewan Harrison, David K. Jackson, Ian Johnston, Dominic Kwiatkowski, Cordelia Langford, John Sillitoe on behalf of the Wellcome Sanger Institute COVID-19 Surveillance Team |
| EPI_ISL_2767808, EPI_ISL_2767809 | Lighthouse Lab in Alderley Park | Wellcome Sanger Institute for the COVID-19 Genomics UK (COG-UK) Consortium | Jacquelyn Wynn, Mairead Hyland, The Lighthouse Lab in Alderley Park and Alex Alderton, Roberto Amato, Jeffrey Barrett, Sonia Goncalves, Ewan Harrison, David K. Jackson, Ian Johnston, Dominic Kwiatkowski, Cordelia Langford, John Sillitoe on behalf of the Wellcome Sanger Institute COVID-19 Surveillance Team |
| EPI_ISL_2767810 | Lighthouse Lab in Milton Keynes | Wellcome Sanger Institute for the COVID-19 Genomics UK (COG-UK) Consortium | The Lighthouse Lab in Milton Keynes and Alex Alderton, Roberto Amato, Jeffrey Barrett, Sonia Goncalves, Ewan Harrison, David K. Jackson, Ian Johnston, Dominic Kwiatkowski, Cordelia Langford, John Sillitoe on behalf of the Wellcome Sanger Institute COVID-19 Surveillance Team |
| EPI_ISL_2767811, EPI_ISL_2767812, EPI_ISL_2767813 | Lighthouse Lab in Alderley Park | Wellcome Sanger Institute for the COVID-19 Genomics UK (COG-UK) Consortium | Jacquelyn Wynn, Mairead Hyland, The Lighthouse Lab in Alderley Park and Alex Alderton, Roberto Amato, Jeffrey Barrett, Sonia Goncalves, Ewan Harrison, David K. Jackson, Ian Johnston, Dominic Kwiatkowski, Cordelia Langford, John Sillitoe on behalf of the Wellcome Sanger Institute COVID-19 Surveillance Team |
| EPI_ISL_2767814 | Lighthouse Lab in Milton Keynes | Wellcome Sanger Institute for the COVID-19 Genomics UK (COG-UK) Consortium | The Lighthouse Lab in Milton Keynes and Alex Alderton, Roberto Amato, Jeffrey Barrett, Sonia Goncalves, Ewan Harrison, David K. Jackson, Ian Johnston, Dominic Kwiatkowski, Cordelia Langford, John Sillitoe on behalf of the Wellcome Sanger Institute COVID-19 Surveillance Team |
| EPI_ISL_2767815, EPI_ISL_2767816, EPI_ISL_2767817, EPI_ISL_2767818 | Lighthouse Lab in Alderley Park | Wellcome Sanger Institute for the COVID-19 Genomics UK (COG-UK) Consortium | Jacquelyn Wynn, Mairead Hyland, The Lighthouse Lab in Alderley Park and Alex Alderton, Roberto Amato, Jeffrey Barrett, Sonia Goncalves, Ewan Harrison, David K. Jackson, Ian Johnston, Dominic Kwiatkowski, Cordelia Langford, John Sillitoe on behalf of the Wellcome Sanger Institute COVID-19 Surveillance Team |
| EPI_ISL_2767819, EPI_ISL_2767820, EPI_ISL_2767821 | Lighthouse Lab in Milton Keynes | Wellcome Sanger Institute for the COVID-19 Genomics UK (COG-UK) Consortium | The Lighthouse Lab in Milton Keynes and Alex Alderton, Roberto Amato, Jeffrey Barrett, Sonia Goncalves, Ewan Harrison, David K. Jackson, Ian Johnston, Dominic Kwiatkowski, Cordelia Langford, John Sillitoe on behalf of the Wellcome Sanger Institute COVID-19 Surveillance Team |
| EPI_ISL_2767822 | Lighthouse Lab in Alderley Park | Wellcome Sanger Institute for the COVID-19 Genomics UK (COG-UK) Consortium | Jacquelyn Wynn, Mairead Hyland, The Lighthouse Lab in Alderley Park and Alex Alderton, Roberto Amato, Jeffrey Barrett, Sonia Goncalves, Ewan Harrison, David K. Jackson, Ian Johnston, Dominic Kwiatkowski, Cordelia Langford, John Sillitoe on behalf of the Wellcome Sanger Institute COVID-19 Surveillance Team |
| EPI_ISL_2767823 | Lighthouse Lab in Milton Keynes | Wellcome Sanger Institute for the COVID-19 Genomics UK (COG-UK) Consortium | The Lighthouse Lab in Milton Keynes and Alex Alderton, Roberto Amato, Jeffrey Barrett, Sonia Goncalves, Ewan Harrison, David K. Jackson, Ian Johnston, Dominic Kwiatkowski, Cordelia Langford, John Sillitoe on behalf of the Wellcome Sanger Institute COVID-19 Surveillance Team |
| EPI_ISL_2767824, EPI_ISL_2767825, EPI_ISL_2767826 | Lighthouse Lab in Alderley Park | Wellcome Sanger Institute for the COVID-19 Genomics UK (COG-UK) Consortium | Jacquelyn Wynn, Mairead Hyland, The Lighthouse Lab in Alderley Park and Alex Alderton, Roberto Amato, Jeffrey Barrett, Sonia Goncalves, Ewan Harrison, David K. Jackson, Ian Johnston, Dominic Kwiatkowski, Cordelia Langford, John Sillitoe on behalf of the Wellcome Sanger Institute COVID-19 Surveillance Team |
| EPI_ISL_2767827, EPI_ISL_2767828 | Lighthouse Lab in Milton Keynes | Wellcome Sanger Institute for the COVID-19 Genomics UK (COG-UK) Consortium | The Lighthouse Lab in Milton Keynes and Alex Alderton, Roberto Amato, Jeffrey Barrett, Sonia Goncalves, Ewan Harrison, David K. Jackson, Ian Johnston, Dominic Kwiatkowski, Cordelia Langford, John Sillitoe on behalf of the Wellcome Sanger Institute COVID-19 Surveillance Team |
| EPI_ISL_2767829, EPI_ISL_2767830, EPI_ISL_2767831 | Lighthouse Lab in Alderley Park | Wellcome Sanger Institute for the COVID-19 Genomics UK (COG-UK) Consortium | Jacquelyn Wynn, Mairead Hyland, The Lighthouse Lab in Alderley Park and Alex Alderton, Roberto Amato, Jeffrey Barrett, Sonia Goncalves, Ewan Harrison, David K. Jackson, Ian Johnston, Dominic Kwiatkowski, Cordelia Langford, John Sillitoe on behalf of the Wellcome Sanger Institute COVID-19 Surveillance Team |
| EPI_ISL_2767832 | Lighthouse Lab in Milton Keynes | Wellcome Sanger Institute for the COVID-19 Genomics UK (COG-UK) Consortium | The Lighthouse Lab in Milton Keynes and Alex Alderton, Roberto Amato, Jeffrey Barrett, Sonia Goncalves, Ewan Harrison, David K. Jackson, Ian Johnston, Dominic Kwiatkowski, Cordelia Langford, John Sillitoe on behalf of the Wellcome Sanger Institute COVID-19 Surveillance Team |
| EPI_ISL_2767833 | Lighthouse Lab in Alderley Park | Wellcome Sanger Institute for the COVID-19 Genomics UK (COG-UK) Consortium | Jacquelyn Wynn, Mairead Hyland, The Lighthouse Lab in Alderley Park and Alex Alderton, Roberto Amato, Jeffrey Barrett, Sonia Goncalves, Ewan Harrison, David K. Jackson, Ian Johnston, Dominic Kwiatkowski, Cordelia Langford, John Sillitoe on behalf of the Wellcome Sanger Institute COVID-19 Surveillance Team |
| EPI_ISL_2767834, EPI_ISL_2767835 | Lighthouse Lab in Milton Keynes | Wellcome Sanger Institute for the COVID-19 Genomics UK (COG-UK) Consortium | The Lighthouse Lab in Milton Keynes and Alex Alderton, Roberto Amato, Jeffrey Barrett, Sonia Goncalves, Ewan Harrison, David K. Jackson, Ian Johnston, Dominic Kwiatkowski, Cordelia Langford, John Sillitoe on behalf of the Wellcome Sanger Institute COVID-19 Surveillance Team |
| EPI_ISL_2767836 | Lighthouse Lab in Alderley Park | Wellcome Sanger Institute for the COVID-19 Genomics UK (COG-UK) Consortium | Jacquelyn Wynn, Mairead Hyland, The Lighthouse Lab in Alderley Park and Alex Alderton, Roberto Amato, Jeffrey Barrett, Sonia Goncalves, Ewan Harrison, David K. Jackson, Ian Johnston, Dominic Kwiatkowski, Cordelia Langford, John Sillitoe on behalf of the Wellcome Sanger Institute COVID-19 Surveillance Team |
| EPI_ISL_2767837 | Lighthouse Lab in Milton Keynes | Wellcome Sanger Institute for the COVID-19 Genomics UK (COG-UK) Consortium | The Lighthouse Lab in Milton Keynes and Alex Alderton, Roberto Amato, Jeffrey Barrett, Sonia Goncalves, Ewan Harrison, David K. Jackson, Ian Johnston, Dominic Kwiatkowski, Cordelia Langford, John Sillitoe on behalf of the Wellcome Sanger Institute COVID-19 Surveillance Team |
| EPI_ISL_2767838 | Lighthouse Lab in Alderley Park | Wellcome Sanger Institute for the COVID-19 Genomics UK (COG-UK) Consortium | Jacquelyn Wynn, Mairead Hyland, The Lighthouse Lab in Alderley Park and Alex Alderton, Roberto Amato, Jeffrey Barrett, Sonia Goncalves, Ewan Harrison, David K. Jackson, Ian Johnston, Dominic Kwiatkowski, Cordelia Langford, John Sillitoe on behalf of the Wellcome Sanger Institute COVID-19 Surveillance Team |
| EPI_ISL_2767839, EPI_ISL_2767840, EPI_ISL_2767841, EPI_ISL_2767842, EPI_ISL_2767843 | Lighthouse Lab in Milton Keynes | Wellcome Sanger Institute for the COVID-19 Genomics UK (COG-UK) Consortium | The Lighthouse Lab in Milton Keynes and Alex Alderton, Roberto Amato, Jeffrey Barrett, Sonia Goncalves, Ewan Harrison, David K. Jackson, Ian Johnston, Dominic Kwiatkowski, Cordelia Langford, John Sillitoe on behalf of the Wellcome Sanger Institute COVID-19 Surveillance Team |

| | | | |
|---|---|---|---|
| EPI_ISL_2767844 | Lighthouse Lab in Alderley Park | Wellcome Sanger Institute for the COVID-19 Genomics UK (COG-UK) Consortium | Jacquelyn Wynn, Mairead Hyland, The Lighthouse Lab in Alderley Park and Alex Alderton, Roberto Amato, Jeffrey Barrett, Sonia Goncalves, Ewan Harrison, David K. Jackson, Ian Johnston, Dominic Kwiatkowski, Cordelia Langford, John Sillitoe on behalf of the Wellcome Sanger Institute COVID-19 Surveillance Team |
| EPI_ISL_2767845, EPI_ISL_2767846, EPI_ISL_2767847 | Lighthouse Lab in Milton Keynes | Wellcome Sanger Institute for the COVID-19 Genomics UK (COG-UK) Consortium | The Lighthouse Lab in Milton Keynes and Alex Alderton, Roberto Amato, Jeffrey Barrett, Sonia Goncalves, Ewan Harrison, David K. Jackson, Ian Johnston, Dominic Kwiatkowski, Cordelia Langford, John Sillitoe on behalf of the Wellcome Sanger Institute COVID-19 Surveillance Team |
| EPI_ISL_2767848 | Lighthouse Lab in Alderley Park | Wellcome Sanger Institute for the COVID-19 Genomics UK (COG-UK) Consortium | Jacquelyn Wynn, Mairead Hyland, The Lighthouse Lab in Alderley Park and Alex Alderton, Roberto Amato, Jeffrey Barrett, Sonia Goncalves, Ewan Harrison, David K. Jackson, Ian Johnston, Dominic Kwiatkowski, Cordelia Langford, John Sillitoe on behalf of the Wellcome Sanger Institute COVID-19 Surveillance Team |
| EPI_ISL_2767849 | Lighthouse Lab in Milton Keynes | Wellcome Sanger Institute for the COVID-19 Genomics UK (COG-UK) Consortium | The Lighthouse Lab in Milton Keynes and Alex Alderton, Roberto Amato, Jeffrey Barrett, Sonia Goncalves, Ewan Harrison, David K. Jackson, Ian Johnston, Dominic Kwiatkowski, Cordelia Langford, John Sillitoe on behalf of the Wellcome Sanger Institute COVID-19 Surveillance Team |
| EPI_ISL_2767850, EPI_ISL_2767851, EPI_ISL_2767852, EPI_ISL_2767853 | Lighthouse Lab in Alderley Park | Wellcome Sanger Institute for the COVID-19 Genomics UK (COG-UK) Consortium | Jacquelyn Wynn, Mairead Hyland, The Lighthouse Lab in Alderley Park and Alex Alderton, Roberto Amato, Jeffrey Barrett, Sonia Goncalves, Ewan Harrison, David K. Jackson, Ian Johnston, Dominic Kwiatkowski, Cordelia Langford, John Sillitoe on behalf of the Wellcome Sanger Institute COVID-19 Surveillance Team |
| EPI_ISL_2767854 | Lighthouse Lab in Milton Keynes | Wellcome Sanger Institute for the COVID-19 Genomics UK (COG-UK) Consortium | The Lighthouse Lab in Milton Keynes and Alex Alderton, Roberto Amato, Jeffrey Barrett, Sonia Goncalves, Ewan Harrison, David K. Jackson, Ian Johnston, Dominic Kwiatkowski, Cordelia Langford, John Sillitoe on behalf of the Wellcome Sanger Institute COVID-19 Surveillance Team |
| EPI_ISL_2767855, EPI_ISL_2767856, EPI_ISL_2767857, EPI_ISL_2767858 | Lighthouse Lab in Alderley Park | Wellcome Sanger Institute for the COVID-19 Genomics UK (COG-UK) Consortium | Jacquelyn Wynn, Mairead Hyland, The Lighthouse Lab in Alderley Park and Alex Alderton, Roberto Amato, Jeffrey Barrett, Sonia Goncalves, Ewan Harrison, David K. Jackson, Ian Johnston, Dominic Kwiatkowski, Cordelia Langford, John Sillitoe on behalf of the Wellcome Sanger Institute COVID-19 Surveillance Team |
| EPI_ISL_2767859 | Lighthouse Lab in Milton Keynes | Wellcome Sanger Institute for the COVID-19 Genomics UK (COG-UK) Consortium | The Lighthouse Lab in Milton Keynes and Alex Alderton, Roberto Amato, Jeffrey Barrett, Sonia Goncalves, Ewan Harrison, David K. Jackson, Ian Johnston, Dominic Kwiatkowski, Cordelia Langford, John Sillitoe on behalf of the Wellcome Sanger Institute COVID-19 Surveillance Team |
| EPI_ISL_2767860 | Lighthouse Lab in Alderley Park | Wellcome Sanger Institute for the COVID-19 Genomics UK (COG-UK) Consortium | Jacquelyn Wynn, Mairead Hyland, The Lighthouse Lab in Alderley Park and Alex Alderton, Roberto Amato, Jeffrey Barrett, Sonia Goncalves, Ewan Harrison, David K. Jackson, Ian Johnston, Dominic Kwiatkowski, Cordelia Langford, John Sillitoe on behalf of the Wellcome Sanger Institute COVID-19 Surveillance Team |
| EPI_ISL_2767861 | Lighthouse Lab in Milton Keynes | Wellcome Sanger Institute for the COVID-19 Genomics UK (COG-UK) Consortium | The Lighthouse Lab in Milton Keynes and Alex Alderton, Roberto Amato, Jeffrey Barrett, Sonia Goncalves, Ewan Harrison, David K. Jackson, Ian Johnston, Dominic Kwiatkowski, Cordelia Langford, John Sillitoe on behalf of the Wellcome Sanger Institute COVID-19 Surveillance Team |
| EPI_ISL_2767862 | Lighthouse Lab in Alderley Park | Wellcome Sanger Institute for the COVID-19 Genomics UK (COG-UK) Consortium | Jacquelyn Wynn, Mairead Hyland, The Lighthouse Lab in Alderley Park and Alex Alderton, Roberto Amato, Jeffrey Barrett, Sonia Goncalves, Ewan Harrison, David K. Jackson, Ian Johnston, Dominic Kwiatkowski, Cordelia Langford, John Sillitoe on behalf of the Wellcome Sanger Institute COVID-19 Surveillance Team |
| EPI_ISL_2767863, EPI_ISL_2767864, EPI_ISL_2767865 | Lighthouse Lab in Milton Keynes | Wellcome Sanger Institute for the COVID-19 Genomics UK (COG-UK) Consortium | The Lighthouse Lab in Milton Keynes and Alex Alderton, Roberto Amato, Jeffrey Barrett, Sonia Goncalves, Ewan Harrison, David K. Jackson, Ian Johnston, Dominic Kwiatkowski, Cordelia Langford, John Sillitoe on behalf of the Wellcome Sanger Institute COVID-19 Surveillance Team |
| EPI_ISL_2767866, EPI_ISL_2767867 | Lighthouse Lab in Alderley Park | Wellcome Sanger Institute for the COVID-19 Genomics UK (COG-UK) Consortium | Jacquelyn Wynn, Mairead Hyland, The Lighthouse Lab in Alderley Park and Alex Alderton, Roberto Amato, Jeffrey Barrett, Sonia Goncalves, Ewan Harrison, David K. Jackson, Ian Johnston, Dominic Kwiatkowski, Cordelia Langford, John Sillitoe on behalf of the Wellcome Sanger Institute COVID-19 Surveillance Team |
| EPI_ISL_2767869, EPI_ISL_2767870 | Lighthouse Lab in Milton Keynes | Wellcome Sanger Institute for the COVID-19 Genomics UK (COG-UK) Consortium | The Lighthouse Lab in Milton Keynes and Alex Alderton, Roberto Amato, Jeffrey Barrett, Sonia Goncalves, Ewan Harrison, David K. Jackson, Ian Johnston, Dominic Kwiatkowski, Cordelia Langford, John Sillitoe on behalf of the Wellcome Sanger Institute COVID-19 Surveillance Team |
| EPI_ISL_2767871, EPI_ISL_2767872 | Lighthouse Lab in Alderley Park | Wellcome Sanger Institute for the COVID-19 Genomics UK (COG-UK) Consortium | Jacquelyn Wynn, Mairead Hyland, The Lighthouse Lab in Alderley Park and Alex Alderton, Roberto Amato, Jeffrey Barrett, Sonia Goncalves, Ewan Harrison, David K. Jackson, Ian Johnston, Dominic Kwiatkowski, Cordelia Langford, John Sillitoe on behalf of the Wellcome Sanger Institute COVID-19 Surveillance Team |
| EPI_ISL_2767873, EPI_ISL_2767874 | Lighthouse Lab in Milton Keynes | Wellcome Sanger Institute for the COVID-19 Genomics UK (COG-UK) Consortium | The Lighthouse Lab in Milton Keynes and Alex Alderton, Roberto Amato, Jeffrey Barrett, Sonia Goncalves, Ewan Harrison, David K. Jackson, Ian Johnston, Dominic Kwiatkowski, Cordelia Langford, John Sillitoe on behalf of the Wellcome Sanger Institute COVID-19 Surveillance Team |
| EPI_ISL_2767875 | Lighthouse Lab in Alderley Park | Wellcome Sanger Institute for the COVID-19 Genomics UK (COG-UK) Consortium | Jacquelyn Wynn, Mairead Hyland, The Lighthouse Lab in Alderley Park and Alex Alderton, Roberto Amato, Jeffrey Barrett, Sonia Goncalves, Ewan Harrison, David K. Jackson, Ian Johnston, Dominic Kwiatkowski, Cordelia Langford, John Sillitoe on behalf of the Wellcome Sanger Institute COVID-19 Surveillance Team |
| EPI_ISL_2767876 | Lighthouse Lab in Milton Keynes | Wellcome Sanger Institute for the COVID-19 Genomics UK (COG-UK) Consortium | The Lighthouse Lab in Milton Keynes and Alex Alderton, Roberto Amato, Jeffrey Barrett, Sonia Goncalves, Ewan Harrison, David K. Jackson, Ian Johnston, Dominic Kwiatkowski, Cordelia Langford, John Sillitoe on behalf of the Wellcome Sanger Institute COVID-19 Surveillance Team |
| EPI_ISL_2767877, EPI_ISL_2767878 | Lighthouse Lab in Alderley Park | Wellcome Sanger Institute for the COVID-19 Genomics UK (COG-UK) Consortium | Jacquelyn Wynn, Mairead Hyland, The Lighthouse Lab in Alderley Park and Alex Alderton, Roberto Amato, Jeffrey Barrett, Sonia Goncalves, Ewan Harrison, David K. Jackson, Ian Johnston, Dominic Kwiatkowski, Cordelia Langford, John Sillitoe on behalf of the Wellcome Sanger Institute COVID-19 Surveillance Team |
| EPI_ISL_2767879 | Lighthouse Lab in Milton Keynes | Wellcome Sanger Institute for the COVID-19 Genomics UK (COG-UK) Consortium | The Lighthouse Lab in Milton Keynes and Alex Alderton, Roberto Amato, Jeffrey Barrett, Sonia Goncalves, Ewan Harrison, David K. Jackson, Ian Johnston, Dominic Kwiatkowski, Cordelia Langford, John Sillitoe on behalf of the Wellcome Sanger Institute COVID-19 Surveillance Team |
| EPI_ISL_2767880 | Lighthouse Lab in Alderley Park | Wellcome Sanger Institute for the COVID-19 Genomics UK (COG-UK) Consortium | Jacquelyn Wynn, Mairead Hyland, The Lighthouse Lab in Alderley Park and Alex Alderton, Roberto Amato, Jeffrey Barrett, Sonia Goncalves, Ewan Harrison, David K. Jackson, Ian Johnston, Dominic Kwiatkowski, Cordelia Langford, John Sillitoe on behalf of the Wellcome Sanger Institute COVID-19 Surveillance Team |
| EPI_ISL_2767881 | Lighthouse Lab in Milton Keynes | Wellcome Sanger Institute for the COVID-19 Genomics UK (COG-UK) Consortium | The Lighthouse Lab in Milton Keynes and Alex Alderton, Roberto Amato, Jeffrey Barrett, Sonia Goncalves, Ewan Harrison, David K. Jackson, Ian Johnston, Dominic Kwiatkowski, Cordelia Langford, John Sillitoe on behalf of the Wellcome Sanger Institute COVID-19 Surveillance Team |
| EPI_ISL_2767882 | Lighthouse Lab in Alderley Park | Wellcome Sanger Institute for the COVID-19 Genomics UK (COG-UK) Consortium | Jacquelyn Wynn, Mairead Hyland, The Lighthouse Lab in Alderley Park and Alex Alderton, Roberto Amato, Jeffrey Barrett, Sonia Goncalves, Ewan Harrison, David K. Jackson, Ian Johnston, Dominic Kwiatkowski, Cordelia Langford, John Sillitoe on behalf of the Wellcome Sanger Institute COVID-19 Surveillance Team |
| EPI_ISL_2767883, EPI_ISL_2767884 | Lighthouse Lab in Milton Keynes | Wellcome Sanger Institute for the COVID-19 Genomics UK (COG-UK) Consortium | The Lighthouse Lab in Milton Keynes and Alex Alderton, Roberto Amato, Jeffrey Barrett, Sonia Goncalves, Ewan Harrison, David K. Jackson, Ian Johnston, Dominic Kwiatkowski, Cordelia Langford, John Sillitoe on behalf of the Wellcome Sanger Institute COVID-19 Surveillance Team |
| EPI_ISL_2767885 | Lighthouse Lab in Alderley Park | Wellcome Sanger Institute for the COVID-19 Genomics UK (COG-UK) Consortium | Jacquelyn Wynn, Mairead Hyland, The Lighthouse Lab in Alderley Park and Alex Alderton, Roberto Amato, Jeffrey Barrett, Sonia Goncalves, Ewan Harrison, David K. Jackson, Ian Johnston, Dominic Kwiatkowski, Cordelia Langford, John Sillitoe on behalf of the Wellcome Sanger Institute COVID-19 Surveillance Team |
| EPI_ISL_2767886, EPI_ISL_2767887, EPI_ISL_2767888, EPI_ISL_2767889 | Lighthouse Lab in Milton Keynes | Wellcome Sanger Institute for the COVID-19 Genomics UK (COG-UK) Consortium | The Lighthouse Lab in Milton Keynes and Alex Alderton, Roberto Amato, Jeffrey Barrett, Sonia Goncalves, Ewan Harrison, David K. Jackson, Ian Johnston, Dominic Kwiatkowski, Cordelia Langford, John Sillitoe on behalf of the Wellcome Sanger Institute COVID-19 Surveillance Team |
| EPI_ISL_2767890, EPI_ISL_2767891, EPI_ISL_2767892 | Lighthouse Lab in Alderley Park | Wellcome Sanger Institute for the COVID-19 Genomics UK (COG-UK) Consortium | Jacquelyn Wynn, Mairead Hyland, The Lighthouse Lab in Alderley Park and Alex Alderton, Roberto Amato, Jeffrey Barrett, Sonia Goncalves, Ewan Harrison, David K. Jackson, Ian Johnston, Dominic Kwiatkowski, Cordelia Langford, John Sillitoe on behalf of the Wellcome Sanger Institute COVID-19 Surveillance Team |
| EPI_ISL_2767893, EPI_ISL_2767894 | Lighthouse Lab in Milton Keynes | Wellcome Sanger Institute for the COVID-19 Genomics UK (COG-UK) Consortium | The Lighthouse Lab in Milton Keynes and Alex Alderton, Roberto Amato, Jeffrey Barrett, Sonia Goncalves, Ewan Harrison, David K. Jackson, Ian Johnston, Dominic Kwiatkowski, Cordelia Langford, John Sillitoe on behalf of the Wellcome Sanger Institute COVID-19 Surveillance Team |
| EPI_ISL_2767895 | Lighthouse Lab in Alderley Park | Wellcome Sanger Institute for the COVID-19 Genomics UK (COG-UK) Consortium | Jacquelyn Wynn, Mairead Hyland, The Lighthouse Lab in Alderley Park and Alex Alderton, Roberto Amato, Jeffrey Barrett, Sonia Goncalves, Ewan Harrison, David K. Jackson, Ian Johnston, Dominic Kwiatkowski, Cordelia Langford, John Sillitoe on behalf of the Wellcome Sanger Institute COVID-19 Surveillance Team |
| EPI_ISL_2767896 | Lighthouse Lab in Milton Keynes | Wellcome Sanger Institute for the COVID-19 Genomics UK (COG-UK) Consortium | The Lighthouse Lab in Milton Keynes and Alex Alderton, Roberto Amato, Jeffrey Barrett, Sonia Goncalves, Ewan Harrison, David K. Jackson, Ian Johnston, Dominic Kwiatkowski, Cordelia Langford, John Sillitoe on behalf of the Wellcome Sanger Institute COVID-19 Surveillance Team |
| EPI_ISL_2767898 | Lighthouse Lab in Alderley Park | Wellcome Sanger Institute for the COVID-19 Genomics UK (COG-UK) Consortium | Jacquelyn Wynn, Mairead Hyland, The Lighthouse Lab in Alderley Park and Alex Alderton, Roberto Amato, Jeffrey Barrett, Sonia Goncalves, Ewan Harrison, David K. Jackson, Ian Johnston, Dominic Kwiatkowski, Cordelia Langford, John Sillitoe on behalf of the Wellcome Sanger Institute COVID-19 Surveillance Team |
| EPI_ISL_2767899, EPI_ISL_2767900 | Lighthouse Lab in Milton Keynes | Wellcome Sanger Institute for the COVID-19 Genomics UK (COG-UK) Consortium | The Lighthouse Lab in Milton Keynes and Alex Alderton, Roberto Amato, Jeffrey Barrett, Sonia Goncalves, Ewan Harrison, David K. Jackson, Ian Johnston, Dominic Kwiatkowski, Cordelia Langford, John Sillitoe on behalf of the Wellcome Sanger Institute COVID-19 Surveillance Team |
| EPI_ISL_2767901 | Lighthouse Lab in Alderley Park | Wellcome Sanger Institute for the COVID-19 Genomics UK (COG-UK) Consortium | Jacquelyn Wynn, Mairead Hyland, The Lighthouse Lab in Alderley Park and Alex Alderton, Roberto Amato, Jeffrey Barrett, Sonia Goncalves, Ewan Harrison, David K. Jackson, Ian Johnston, Dominic Kwiatkowski, Cordelia Langford, John Sillitoe on behalf of the Wellcome Sanger Institute COVID-19 Surveillance Team |
| EPI_ISL_2767902, EPI_ISL_2767903 | Lighthouse Lab in Milton Keynes | Wellcome Sanger Institute for the COVID-19 Genomics UK (COG-UK) Consortium | The Lighthouse Lab in Milton Keynes and Alex Alderton, Roberto Amato, Jeffrey Barrett, Sonia Goncalves, Ewan Harrison, David K. Jackson, Ian Johnston, Dominic Kwiatkowski, Cordelia Langford, John Sillitoe on behalf of the Wellcome Sanger Institute COVID-19 Surveillance Team |
| EPI_ISL_2767904, EPI_ISL_2767905 | Lighthouse Lab in Alderley Park | Wellcome Sanger Institute for the COVID-19 Genomics UK (COG-UK) Consortium | Jacquelyn Wynn, Mairead Hyland, The Lighthouse Lab in Alderley Park and Alex Alderton, Roberto Amato, Jeffrey Barrett, Sonia Goncalves, Ewan Harrison, David K. Jackson, Ian Johnston, Dominic Kwiatkowski, Cordelia Langford, John Sillitoe on behalf of the Wellcome Sanger Institute COVID-19 Surveillance Team |
| EPI_ISL_2767906, EPI_ISL_2767907 | Lighthouse Lab in Milton Keynes | Wellcome Sanger Institute for the COVID-19 Genomics UK (COG-UK) Consortium | The Lighthouse Lab in Milton Keynes and Alex Alderton, Roberto Amato, Jeffrey Barrett, Sonia Goncalves, Ewan Harrison, David K. Jackson, Ian Johnston, Dominic Kwiatkowski, Cordelia Langford, John Sillitoe on behalf of the Wellcome Sanger Institute COVID-19 Surveillance Team |
| EPI_ISL_2767908, EPI_ISL_2767909 | Lighthouse Lab in Alderley Park | Wellcome Sanger Institute for the COVID-19 Genomics UK (COG-UK) Consortium | Jacquelyn Wynn, Mairead Hyland, The Lighthouse Lab in Alderley Park and Alex Alderton, Roberto Amato, Jeffrey Barrett, Sonia Goncalves, Ewan Harrison, David K. Jackson, Ian Johnston, Dominic Kwiatkowski, Cordelia Langford, John Sillitoe on behalf of the Wellcome Sanger Institute COVID-19 Surveillance Team |
| EPI_ISL_2767910 | Lighthouse Lab in Milton Keynes | Wellcome Sanger Institute for the COVID-19 Genomics UK (COG-UK) Consortium | The Lighthouse Lab in Milton Keynes and Alex Alderton, Roberto Amato, Jeffrey Barrett, Sonia Goncalves, Ewan Harrison, David K. Jackson, Ian Johnston, Dominic Kwiatkowski, Cordelia Langford, John Sillitoe on behalf of the Wellcome Sanger Institute COVID-19 Surveillance Team |
| EPI_ISL_2767911, EPI_ISL_2767912 | Lighthouse Lab in Alderley Park | Wellcome Sanger Institute for the COVID-19 Genomics UK (COG-UK) Consortium | Jacquelyn Wynn, Mairead Hyland, The Lighthouse Lab in Alderley Park and Alex Alderton, Roberto Amato, Jeffrey Barrett, Sonia Goncalves, Ewan Harrison, David K. Jackson, Ian Johnston, Dominic Kwiatkowski, Cordelia Langford, John Sillitoe on behalf of the Wellcome Sanger Institute COVID-19 Surveillance Team |
| EPI_ISL_2767913, EPI_ISL_2767914, EPI_ISL_2767915 | Lighthouse Lab in Milton Keynes | Wellcome Sanger Institute for the COVID-19 Genomics UK (COG-UK) Consortium | The Lighthouse Lab in Milton Keynes and Alex Alderton, Roberto Amato, Jeffrey Barrett, Sonia Goncalves, Ewan Harrison, David K. Jackson, Ian Johnston, Dominic Kwiatkowski, Cordelia Langford, John Sillitoe on behalf of the Wellcome Sanger Institute COVID-19 Surveillance Team |
| EPI_ISL_2767916 | Lighthouse Lab in Alderley Park | Wellcome Sanger Institute for the COVID-19 Genomics UK (COG-UK) Consortium | Jacquelyn Wynn, Mairead Hyland, The Lighthouse Lab in Alderley Park and Alex Alderton, Roberto Amato, Jeffrey Barrett, Sonia Goncalves, Ewan Harrison, David K. Jackson, Ian Johnston, Dominic Kwiatkowski, Cordelia Langford, John Sillitoe on behalf of the Wellcome Sanger Institute COVID-19 Surveillance Team |

| | | | |
|---|---|---|---|
| EPI_ISL_2767917 | Lighthouse Lab in Milton Keynes | Wellcome Sanger Institute for the COVID-19 Genomics UK (COG-UK) Consortium | The Lighthouse Lab in Milton Keynes and Alex Alderton, Roberto Amato, Jeffrey Barrett, Sonia Goncalves, Ewan Harrison, David K. Jackson, Ian Johnston, Dominic Kwiatkowski, Cordelia Langford, John Sillitoe on behalf of the Wellcome Sanger Institute COVID-19 Surveillance Team |
| EPI_ISL_2767918, EPI_ISL_2767919, EPI_ISL_2767920 | Lighthouse Lab in Alderley Park | Wellcome Sanger Institute for the COVID-19 Genomics UK (COG-UK) Consortium | Jacquelyn Wynn, Mairead Hyland, The Lighthouse Lab in Alderley Park and Alex Alderton, Roberto Amato, Jeffrey Barrett, Sonia Goncalves, Ewan Harrison, David K. Jackson, Ian Johnston, Dominic Kwiatkowski, Cordelia Langford, John Sillitoe on behalf of the Wellcome Sanger Institute COVID-19 Surveillance Team |
| EPI_ISL_2767921 | Lighthouse Lab in Milton Keynes | Wellcome Sanger Institute for the COVID-19 Genomics UK (COG-UK) Consortium | The Lighthouse Lab in Milton Keynes and Alex Alderton, Roberto Amato, Jeffrey Barrett, Sonia Goncalves, Ewan Harrison, David K. Jackson, Ian Johnston, Dominic Kwiatkowski, Cordelia Langford, John Sillitoe on behalf of the Wellcome Sanger Institute COVID-19 Surveillance Team |
| EPI_ISL_2767922 | Lighthouse Lab in Alderley Park | Wellcome Sanger Institute for the COVID-19 Genomics UK (COG-UK) Consortium | Jacquelyn Wynn, Mairead Hyland, The Lighthouse Lab in Alderley Park and Alex Alderton, Roberto Amato, Jeffrey Barrett, Sonia Goncalves, Ewan Harrison, David K. Jackson, Ian Johnston, Dominic Kwiatkowski, Cordelia Langford, John Sillitoe on behalf of the Wellcome Sanger Institute COVID-19 Surveillance Team |
| EPI_ISL_2767923 | Lighthouse Lab in Milton Keynes | Wellcome Sanger Institute for the COVID-19 Genomics UK (COG-UK) Consortium | The Lighthouse Lab in Milton Keynes and Alex Alderton, Roberto Amato, Jeffrey Barrett, Sonia Goncalves, Ewan Harrison, David K. Jackson, Ian Johnston, Dominic Kwiatkowski, Cordelia Langford, John Sillitoe on behalf of the Wellcome Sanger Institute COVID-19 Surveillance Team |
| EPI_ISL_2767924 | Lighthouse Lab in Alderley Park | Wellcome Sanger Institute for the COVID-19 Genomics UK (COG-UK) Consortium | Jacquelyn Wynn, Mairead Hyland, The Lighthouse Lab in Alderley Park and Alex Alderton, Roberto Amato, Jeffrey Barrett, Sonia Goncalves, Ewan Harrison, David K. Jackson, Ian Johnston, Dominic Kwiatkowski, Cordelia Langford, John Sillitoe on behalf of the Wellcome Sanger Institute COVID-19 Surveillance Team |
| EPI_ISL_2767925, EPI_ISL_2767926, EPI_ISL_2767927 | Lighthouse Lab in Milton Keynes | Wellcome Sanger Institute for the COVID-19 Genomics UK (COG-UK) Consortium | The Lighthouse Lab in Milton Keynes and Alex Alderton, Roberto Amato, Jeffrey Barrett, Sonia Goncalves, Ewan Harrison, David K. Jackson, Ian Johnston, Dominic Kwiatkowski, Cordelia Langford, John Sillitoe on behalf of the Wellcome Sanger Institute COVID-19 Surveillance Team |
| EPI_ISL_2767928, EPI_ISL_2767929, EPI_ISL_2767930, EPI_ISL_2767931, EPI_ISL_2767932, EPI_ISL_2767933, EPI_ISL_2767934, EPI_ISL_2767935, EPI_ISL_2767936 | Lighthouse Lab in Alderley Park | Wellcome Sanger Institute for the COVID-19 Genomics UK (COG-UK) Consortium | Jacquelyn Wynn, Mairead Hyland, The Lighthouse Lab in Alderley Park and Alex Alderton, Roberto Amato, Jeffrey Barrett, Sonia Goncalves, Ewan Harrison, David K. Jackson, Ian Johnston, Dominic Kwiatkowski, Cordelia Langford, John Sillitoe on behalf of the Wellcome Sanger Institute COVID-19 Surveillance Team |
| EPI_ISL_2767937, EPI_ISL_2767938, EPI_ISL_2767939 | Lighthouse Lab in Milton Keynes | Wellcome Sanger Institute for the COVID-19 Genomics UK (COG-UK) Consortium | The Lighthouse Lab in Milton Keynes and Alex Alderton, Roberto Amato, Jeffrey Barrett, Sonia Goncalves, Ewan Harrison, David K. Jackson, Ian Johnston, Dominic Kwiatkowski, Cordelia Langford, John Sillitoe on behalf of the Wellcome Sanger Institute COVID-19 Surveillance Team |
| EPI_ISL_2767940, EPI_ISL_2767941, EPI_ISL_2767942 | Lighthouse Lab in Alderley Park | Wellcome Sanger Institute for the COVID-19 Genomics UK (COG-UK) Consortium | Jacquelyn Wynn, Mairead Hyland, The Lighthouse Lab in Alderley Park and Alex Alderton, Roberto Amato, Jeffrey Barrett, Sonia Goncalves, Ewan Harrison, David K. Jackson, Ian Johnston, Dominic Kwiatkowski, Cordelia Langford, John Sillitoe on behalf of the Wellcome Sanger Institute COVID-19 Surveillance Team |
| EPI_ISL_2767943 | Lighthouse Lab in Milton Keynes | Wellcome Sanger Institute for the COVID-19 Genomics UK (COG-UK) Consortium | The Lighthouse Lab in Milton Keynes and Alex Alderton, Roberto Amato, Jeffrey Barrett, Sonia Goncalves, Ewan Harrison, David K. Jackson, Ian Johnston, Dominic Kwiatkowski, Cordelia Langford, John Sillitoe on behalf of the Wellcome Sanger Institute COVID-19 Surveillance Team |
| EPI_ISL_2767944 | Lighthouse Lab in Alderley Park | Wellcome Sanger Institute for the COVID-19 Genomics UK (COG-UK) Consortium | Jacquelyn Wynn, Mairead Hyland, The Lighthouse Lab in Alderley Park and Alex Alderton, Roberto Amato, Jeffrey Barrett, Sonia Goncalves, Ewan Harrison, David K. Jackson, Ian Johnston, Dominic Kwiatkowski, Cordelia Langford, John Sillitoe on behalf of the Wellcome Sanger Institute COVID-19 Surveillance Team |
| EPI_ISL_2767945 | Lighthouse Lab in Milton Keynes | Wellcome Sanger Institute for the COVID-19 Genomics UK (COG-UK) Consortium | The Lighthouse Lab in Milton Keynes and Alex Alderton, Roberto Amato, Jeffrey Barrett, Sonia Goncalves, Ewan Harrison, David K. Jackson, Ian Johnston, Dominic Kwiatkowski, Cordelia Langford, John Sillitoe on behalf of the Wellcome Sanger Institute COVID-19 Surveillance Team |
| EPI_ISL_2767946 | Lighthouse Lab in Alderley Park | Wellcome Sanger Institute for the COVID-19 Genomics UK (COG-UK) Consortium | Jacquelyn Wynn, Mairead Hyland, The Lighthouse Lab in Alderley Park and Alex Alderton, Roberto Amato, Jeffrey Barrett, Sonia Goncalves, Ewan Harrison, David K. Jackson, Ian Johnston, Dominic Kwiatkowski, Cordelia Langford, John Sillitoe on behalf of the Wellcome Sanger Institute COVID-19 Surveillance Team |
| EPI_ISL_2767947 | Lighthouse Lab in Milton Keynes | Wellcome Sanger Institute for the COVID-19 Genomics UK (COG-UK) Consortium | The Lighthouse Lab in Milton Keynes and Alex Alderton, Roberto Amato, Jeffrey Barrett, Sonia Goncalves, Ewan Harrison, David K. Jackson, Ian Johnston, Dominic Kwiatkowski, Cordelia Langford, John Sillitoe on behalf of the Wellcome Sanger Institute COVID-19 Surveillance Team |
| EPI_ISL_2767948, EPI_ISL_2767949, EPI_ISL_2767950, EPI_ISL_2767951, EPI_ISL_2767952, EPI_ISL_2767953 | Lighthouse Lab in Alderley Park | Wellcome Sanger Institute for the COVID-19 Genomics UK (COG-UK) Consortium | Jacquelyn Wynn, Mairead Hyland, The Lighthouse Lab in Alderley Park and Alex Alderton, Roberto Amato, Jeffrey Barrett, Sonia Goncalves, Ewan Harrison, David K. Jackson, Ian Johnston, Dominic Kwiatkowski, Cordelia Langford, John Sillitoe on behalf of the Wellcome Sanger Institute COVID-19 Surveillance Team |
| EPI_ISL_2767954 | Lighthouse Lab in Milton Keynes | Wellcome Sanger Institute for the COVID-19 Genomics UK (COG-UK) Consortium | The Lighthouse Lab in Milton Keynes and Alex Alderton, Roberto Amato, Jeffrey Barrett, Sonia Goncalves, Ewan Harrison, David K. Jackson, Ian Johnston, Dominic Kwiatkowski, Cordelia Langford, John Sillitoe on behalf of the Wellcome Sanger Institute COVID-19 Surveillance Team |
| EPI_ISL_2767955, EPI_ISL_2767956 | Lighthouse Lab in Alderley Park | Wellcome Sanger Institute for the COVID-19 Genomics UK (COG-UK) Consortium | Jacquelyn Wynn, Mairead Hyland, The Lighthouse Lab in Alderley Park and Alex Alderton, Roberto Amato, Jeffrey Barrett, Sonia Goncalves, Ewan Harrison, David K. Jackson, Ian Johnston, Dominic Kwiatkowski, Cordelia Langford, John Sillitoe on behalf of the Wellcome Sanger Institute COVID-19 Surveillance Team |
| EPI_ISL_2767957, EPI_ISL_2767958, EPI_ISL_2767959 | Lighthouse Lab in Milton Keynes | Wellcome Sanger Institute for the COVID-19 Genomics UK (COG-UK) Consortium | The Lighthouse Lab in Milton Keynes and Alex Alderton, Roberto Amato, Jeffrey Barrett, Sonia Goncalves, Ewan Harrison, David K. Jackson, Ian Johnston, Dominic Kwiatkowski, Cordelia Langford, John Sillitoe on behalf of the Wellcome Sanger Institute COVID-19 Surveillance Team |
| EPI_ISL_2767960 | Lighthouse Lab in Alderley Park | Wellcome Sanger Institute for the COVID-19 Genomics UK (COG-UK) Consortium | Jacquelyn Wynn, Mairead Hyland, The Lighthouse Lab in Alderley Park and Alex Alderton, Roberto Amato, Jeffrey Barrett, Sonia Goncalves, Ewan Harrison, David K. Jackson, Ian Johnston, Dominic Kwiatkowski, Cordelia Langford, John Sillitoe on behalf of the Wellcome Sanger Institute COVID-19 Surveillance Team |
| EPI_ISL_2767961, EPI_ISL_2767962 | Lighthouse Lab in Milton Keynes | Wellcome Sanger Institute for the COVID-19 Genomics UK (COG-UK) Consortium | The Lighthouse Lab in Milton Keynes and Alex Alderton, Roberto Amato, Jeffrey Barrett, Sonia Goncalves, Ewan Harrison, David K. Jackson, Ian Johnston, Dominic Kwiatkowski, Cordelia Langford, John Sillitoe on behalf of the Wellcome Sanger Institute COVID-19 Surveillance Team |
| EPI_ISL_2767963 | Lighthouse Lab in Alderley Park | Wellcome Sanger Institute for the COVID-19 Genomics UK (COG-UK) Consortium | Jacquelyn Wynn, Mairead Hyland, The Lighthouse Lab in Alderley Park and Alex Alderton, Roberto Amato, Jeffrey Barrett, Sonia Goncalves, Ewan Harrison, David K. Jackson, Ian Johnston, Dominic Kwiatkowski, Cordelia Langford, John Sillitoe on behalf of the Wellcome Sanger Institute COVID-19 Surveillance Team |
| EPI_ISL_2767964, EPI_ISL_2767965, EPI_ISL_2767966, EPI_ISL_2767967 | Lighthouse Lab in Milton Keynes | Wellcome Sanger Institute for the COVID-19 Genomics UK (COG-UK) Consortium | The Lighthouse Lab in Milton Keynes and Alex Alderton, Roberto Amato, Jeffrey Barrett, Sonia Goncalves, Ewan Harrison, David K. Jackson, Ian Johnston, Dominic Kwiatkowski, Cordelia Langford, John Sillitoe on behalf of the Wellcome Sanger Institute COVID-19 Surveillance Team |
| EPI_ISL_2767968 | Lighthouse Lab in Alderley Park | Wellcome Sanger Institute for the COVID-19 Genomics UK (COG-UK) Consortium | Jacquelyn Wynn, Mairead Hyland, The Lighthouse Lab in Alderley Park and Alex Alderton, Roberto Amato, Jeffrey Barrett, Sonia Goncalves, Ewan Harrison, David K. Jackson, Ian Johnston, Dominic Kwiatkowski, Cordelia Langford, John Sillitoe on behalf of the Wellcome Sanger Institute COVID-19 Surveillance Team |
| EPI_ISL_2767969, EPI_ISL_2767970 | Lighthouse Lab in Milton Keynes | Wellcome Sanger Institute for the COVID-19 Genomics UK (COG-UK) Consortium | The Lighthouse Lab in Milton Keynes and Alex Alderton, Roberto Amato, Jeffrey Barrett, Sonia Goncalves, Ewan Harrison, David K. Jackson, Ian Johnston, Dominic Kwiatkowski, Cordelia Langford, John Sillitoe on behalf of the Wellcome Sanger Institute COVID-19 Surveillance Team |
| EPI_ISL_2767971, EPI_ISL_2767972 | Lighthouse Lab in Alderley Park | Wellcome Sanger Institute for the COVID-19 Genomics UK (COG-UK) Consortium | Jacquelyn Wynn, Mairead Hyland, The Lighthouse Lab in Alderley Park and Alex Alderton, Roberto Amato, Jeffrey Barrett, Sonia Goncalves, Ewan Harrison, David K. Jackson, Ian Johnston, Dominic Kwiatkowski, Cordelia Langford, John Sillitoe on behalf of the Wellcome Sanger Institute COVID-19 Surveillance Team |
| EPI_ISL_2767973, EPI_ISL_2767974, EPI_ISL_2767975, EPI_ISL_2767976, EPI_ISL_2767977, EPI_ISL_2767978, EPI_ISL_2767979 | Lighthouse Lab in Milton Keynes | Wellcome Sanger Institute for the COVID-19 Genomics UK (COG-UK) Consortium | The Lighthouse Lab in Milton Keynes and Alex Alderton, Roberto Amato, Jeffrey Barrett, Sonia Goncalves, Ewan Harrison, David K. Jackson, Ian Johnston, Dominic Kwiatkowski, Cordelia Langford, John Sillitoe on behalf of the Wellcome Sanger Institute COVID-19 Surveillance Team |
| EPI_ISL_2767980 | Lighthouse Lab in Alderley Park | Wellcome Sanger Institute for the COVID-19 Genomics UK (COG-UK) Consortium | Jacquelyn Wynn, Mairead Hyland, The Lighthouse Lab in Alderley Park and Alex Alderton, Roberto Amato, Jeffrey Barrett, Sonia Goncalves, Ewan Harrison, David K. Jackson, Ian Johnston, Dominic Kwiatkowski, Cordelia Langford, John Sillitoe on behalf of the Wellcome Sanger Institute COVID-19 Surveillance Team |
| EPI_ISL_2767981, EPI_ISL_2767982, EPI_ISL_2767983 | Lighthouse Lab in Milton Keynes | Wellcome Sanger Institute for the COVID-19 Genomics UK (COG-UK) Consortium | The Lighthouse Lab in Milton Keynes and Alex Alderton, Roberto Amato, Jeffrey Barrett, Sonia Goncalves, Ewan Harrison, David K. Jackson, Ian Johnston, Dominic Kwiatkowski, Cordelia Langford, John Sillitoe on behalf of the Wellcome Sanger Institute COVID-19 Surveillance Team |
| EPI_ISL_2767984 | Lighthouse Lab in Alderley Park | Wellcome Sanger Institute for the COVID-19 Genomics UK (COG-UK) Consortium | Jacquelyn Wynn, Mairead Hyland, The Lighthouse Lab in Alderley Park and Alex Alderton, Roberto Amato, Jeffrey Barrett, Sonia Goncalves, Ewan Harrison, David K. Jackson, Ian Johnston, Dominic Kwiatkowski, Cordelia Langford, John Sillitoe on behalf of the Wellcome Sanger Institute COVID-19 Surveillance Team |
| EPI_ISL_2767985 | Lighthouse Lab in Milton Keynes | Wellcome Sanger Institute for the COVID-19 Genomics UK (COG-UK) Consortium | The Lighthouse Lab in Milton Keynes and Alex Alderton, Roberto Amato, Jeffrey Barrett, Sonia Goncalves, Ewan Harrison, David K. Jackson, Ian Johnston, Dominic Kwiatkowski, Cordelia Langford, John Sillitoe on behalf of the Wellcome Sanger Institute COVID-19 Surveillance Team |
| EPI_ISL_2767986, EPI_ISL_2767987, EPI_ISL_2767988, EPI_ISL_2767989, EPI_ISL_2767990 | Lighthouse Lab in Alderley Park | Wellcome Sanger Institute for the COVID-19 Genomics UK (COG-UK) Consortium | Jacquelyn Wynn, Mairead Hyland, The Lighthouse Lab in Alderley Park and Alex Alderton, Roberto Amato, Jeffrey Barrett, Sonia Goncalves, Ewan Harrison, David K. Jackson, Ian Johnston, Dominic Kwiatkowski, Cordelia Langford, John Sillitoe on behalf of the Wellcome Sanger Institute COVID-19 Surveillance Team |
| EPI_ISL_2767991, EPI_ISL_2767992, EPI_ISL_2767993 | Lighthouse Lab in Milton Keynes | Wellcome Sanger Institute for the COVID-19 Genomics UK (COG-UK) Consortium | The Lighthouse Lab in Milton Keynes and Alex Alderton, Roberto Amato, Jeffrey Barrett, Sonia Goncalves, Ewan Harrison, David K. Jackson, Ian Johnston, Dominic Kwiatkowski, Cordelia Langford, John Sillitoe on behalf of the Wellcome Sanger Institute COVID-19 Surveillance Team |
| EPI_ISL_2767994 | Lighthouse Lab in Alderley Park | Wellcome Sanger Institute for the COVID-19 Genomics UK (COG-UK) | Jacquelyn Wynn, Mairead Hyland, The Lighthouse Lab in Alderley Park and Alex Alderton, Roberto Amato, Jeffrey Barrett, Sonia Goncalves, Ewan Harrison, David K. Jackson, Ian Johnston, Dominic Kwiatkowski, Cordelia Langford, John Sillitoe on |

| | | | |
|---|---|---|---|
| | | Consortium | behalf of the Wellcome Sanger Institute COVID-19 Surveillance Team |
| EPI_ISL_2767995 | Lighthouse Lab in Milton Keynes | Wellcome Sanger Institute for the COVID-19 Genomics UK (COG-UK) Consortium | The Lighthouse Lab in Milton Keynes and Alex Alderton, Roberto Amato, Jeffrey Barrett, Sonia Goncalves, Ewan Harrison, David K. Jackson, Ian Johnston, Dominic Kwiatkowski, Cordelia Langford, John Sillitoe on behalf of the Wellcome Sanger Institute COVID-19 Surveillance Team |
| EPI_ISL_2767996 | Lighthouse Lab in Alderley Park | Wellcome Sanger Institute for the COVID-19 Genomics UK (COG-UK) Consortium | Jacquelyn Wynn, Mairead Hyland, The Lighthouse Lab in Alderley Park and Alex Alderton, Roberto Amato, Jeffrey Barrett, Sonia Goncalves, Ewan Harrison, David K. Jackson, Ian Johnston, Dominic Kwiatkowski, Cordelia Langford, John Sillitoe on behalf of the Wellcome Sanger Institute COVID-19 Surveillance Team |
| EPI_ISL_2767997, EPI_ISL_2767998 | Lighthouse Lab in Milton Keynes | Wellcome Sanger Institute for the COVID-19 Genomics UK (COG-UK) Consortium | The Lighthouse Lab in Milton Keynes and Alex Alderton, Roberto Amato, Jeffrey Barrett, Sonia Goncalves, Ewan Harrison, David K. Jackson, Ian Johnston, Dominic Kwiatkowski, Cordelia Langford, John Sillitoe on behalf of the Wellcome Sanger Institute COVID-19 Surveillance Team |
| EPI_ISL_2767999, EPI_ISL_2768000, EPI_ISL_2768001, EPI_ISL_2768002, EPI_ISL_2768003, EPI_ISL_2768004, EPI_ISL_2768005, EPI_ISL_2768006, EPI_ISL_2768007, EPI_ISL_2768008 | Lighthouse Lab in Alderley Park | Wellcome Sanger Institute for the COVID-19 Genomics UK (COG-UK) Consortium | Jacquelyn Wynn, Mairead Hyland, The Lighthouse Lab in Alderley Park and Alex Alderton, Roberto Amato, Jeffrey Barrett, Sonia Goncalves, Ewan Harrison, David K. Jackson, Ian Johnston, Dominic Kwiatkowski, Cordelia Langford, John Sillitoe on behalf of the Wellcome Sanger Institute COVID-19 Surveillance Team |
| EPI_ISL_2768009, EPI_ISL_2768010, EPI_ISL_2768011, EPI_ISL_2768012, EPI_ISL_2768013 | Lighthouse Laboratory Plymouth | Wellcome Sanger Institute for the COVID-19 Genomics UK (COG-UK) Consortium | Lighthouse Laboratory Plymouth and Alex Alderton, Roberto Amato, Jeffrey Barrett, Sonia Goncalves, Ewan Harrison, David K. Jackson, Ian Johnston, Dominic Kwiatkowski, Cordelia Langford, John Sillitoe on behalf of the Wellcome Sanger Institute COVID-19 Surveillance Team |
| EPI_ISL_2768037, EPI_ISL_2768041 | Clinical Virology | Clinical Bacteriology | Tim Roloff, Fanny Wegner, Helena MB Seth-Smith, Alfredo Mari, Karoline Leuzinger, Julia Bielicki, Manuel Battegay, Hans Hirsch, Adrian Egli |
| EPI_ISL_2768052, EPI_ISL_2768056, EPI_ISL_2768064, EPI_ISL_2768068, EPI_ISL_2768069, EPI_ISL_2768078 | IN State Department of Health Laboratory Services | IN State Department of Health Laboratory Services | Cassandra Campion, Jamie Yeadon, Brian Pope, Lixia Liu, Kyle Brownlee, Melissa Hindenlang, Mark Glazier |
| EPI_ISL_2768083 | Clinical Virology | Clinical Bacteriology | Tim Roloff, Fanny Wegner, Helena MB Seth-Smith, Alfredo Mari, Karoline Leuzinger, Julia Bielicki, Manuel Battegay, Hans Hirsch, Adrian Egli |
| EPI_ISL_2768092 | Synlab Suisse SA | Clinical Bacteriology | Tim Roloff, Fanny Wegner, Helena MB Seth-Smith, Alfredo Mari, Karoline Leuzinger, Julia Bielicki, Manuel Battegay, Hans Hirsch, Adrian Egli |
| EPI_ISL_2768093, EPI_ISL_2768094 | Rothen Medizinische Laboratorien AG | Clinical Bacteriology | Tim Roloff, Fanny Wegner, Helena MB Seth-Smith, Alfredo Mari, Karoline Leuzinger, Julia Bielicki, Ingrid Steffen, Manuel Battegay, Hans Hirsch, Adrian Egli |
| EPI_ISL_2768236, EPI_ISL_2768313, EPI_ISL_2768314, EPI_ISL_2768318, EPI_ISL_2768322, EPI_ISL_2768323, EPI_ISL_2768324, EPI_ISL_2768326, EPI_ISL_2768336, EPI_ISL_2768338, EPI_ISL_2768341, EPI_ISL_2768345, EPI_ISL_2768355, EPI_ISL_2768357, EPI_ISL_2768359, EPI_ISL_2768363, EPI_ISL_2768364, EPI_ISL_2768366, EPI_ISL_2768368, EPI_ISL_2768370, EPI_ISL_2768371, EPI_ISL_2768372, EPI_ISL_2768375 | | | |
| see above | Pathogen Genomics Center, National Institute of Infectious Diseases | Pathogen Genomics Center, National Institute of Infectious Diseases | Tsuyoshi Sekizuka, Kentaro Itokawa, Rina Tanaka, Masanori Hashino, Makoto Kuroda |
| EPI_ISL_2768526, EPI_ISL_2768527 | SARS-CoV-2 testing team, National Institute of Infectious Diseases | Pathogen Genomics Center, National Institute of Infectious Diseases | Tsuyoshi Sekizuka, Kentaro Itokawa, Rina Tanaka, Masanori Hashino, Nozomu Hanaoka, Masumichi Saito, Naomi Nojiri, Hazuka Y Furihata, Hiromizu Takahashi, Tsuguto Fujimoto, Makoto Kuroda |
| EPI_ISL_2768839, EPI_ISL_2768840, EPI_ISL_2768841, EPI_ISL_2768842, EPI_ISL_2768843 | Osaka Institute of Public Health, Morinomiya Center | Pathogen Genomics Center, National Institute of Infectious Diseases | Tsuyoshi Sekizuka, Kentaro Itokawa, Rina Tanaka, Masanori Hashino, Makoto Kuroda |
| EPI_ISL_2768849, EPI_ISL_2768850 | Pathogen Genomics Center, National Institute of Infectious Diseases | Pathogen Genomics Center, National Institute of Infectious Diseases | Tsuyoshi Sekizuka, Kentaro Itokawa, Rina Tanaka, Masanori Hashino, Makoto Kuroda |
| EPI_ISL_2769109 | Unilabs | Institute of Medical Virology | Verena Kufner, Gabriela Ziltener, Maryam Zaheri, Stefan Schmutz, Annette Audigé, Maria Grünberg, Kevin Steiner, Jon Huder, Cyril Shah, Riccarda Capaul, Guido Bloemberg, Jürg Böni, Michael Huber, Alexandra Trkola |
| EPI_ISL_2769117 | UniversitätsSpital Zürich | Institute of Medical Virology | Verena Kufner, Gabriela Ziltener, Maryam Zaheri, Stefan Schmutz, Annette Audigé, Maria Grünberg, Kevin Steiner, Jon Huder, Cyril Shah, Riccarda Capaul, Guido Bloemberg, Jürg Böni, Michael Huber, Alexandra Trkola |
| EPI_ISL_2769149 | Spital Männedorf AG | Institute of Medical Virology | Verena Kufner, Gabriela Ziltener, Maryam Zaheri, Stefan Schmutz, Annette Audigé, Maria Grünberg, Kevin Steiner, Jon Huder, Cyril Shah, Riccarda Capaul, Guido Bloemberg, Jürg Böni, Michael Huber, Alexandra Trkola |
| EPI_ISL_2769163, EPI_ISL_2769195, EPI_ISL_2769230, EPI_ISL_2769235, EPI_ISL_2769251, EPI_ISL_2769264, EPI_ISL_2769286, EPI_ISL_2769287, EPI_ISL_2769299 | TGen North | TGen North | "Jolene Bowers, Heather Centner, Brett Van Tassel, Chris French, Hayley Yaglom, Darrin Lemmer, Dave Engelthaler" |
| EPI_ISL_2769624, EPI_ISL_2769825, EPI_ISL_2769826, EPI_ISL_2769918, EPI_ISL_2769919, EPI_ISL_2769961, EPI_ISL_2770144, EPI_ISL_2770145, EPI_ISL_2770197, EPI_ISL_2770198, EPI_ISL_2770202, EPI_ISL_2770203, EPI_ISL_2770204, EPI_ISL_2770205, EPI_ISL_2770206, EPI_ISL_2770207, EPI_ISL_2770208, EPI_ISL_2770209, EPI_ISL_2770210, EPI_ISL_2770211, EPI_ISL_2770212, EPI_ISL_2770213, EPI_ISL_2770214, EPI_ISL_2770215 | | | |
| see above | SARS-CoV-2 testing team, National Institute of Infectious Diseases | Pathogen Genomics Center, National Institute of Infectious Diseases | Tsuyoshi Sekizuka, Kentaro Itokawa, Rina Tanaka, Masanori Hashino, Nozomu Hanaoka, Masumichi Saito, Naomi Nojiri, Hazuka Y Furihata, Hiromizu Takahashi, Tsuguto Fujimoto, Makoto Kuroda |
| EPI_ISL_2770316, EPI_ISL_2770333 | Pathogen Genomics Center, National Institute of Infectious Diseases | Pathogen Genomics Center, National Institute of Infectious Diseases | Tsuyoshi Sekizuka, Kentaro Itokawa, Rina Tanaka, Masanori Hashino, Makoto Kuroda |
| EPI_ISL_2770439, EPI_ISL_2770440, EPI_ISL_2770444, EPI_ISL_2770445, EPI_ISL_2770447, EPI_ISL_2770453, EPI_ISL_2770457, EPI_ISL_2770458, EPI_ISL_2770459, EPI_ISL_2770460, EPI_ISL_2770461, EPI_ISL_2770463, EPI_ISL_2770464, EPI_ISL_2770465, EPI_ISL_2770466, EPI_ISL_2770467, EPI_ISL_2770469, EPI_ISL_2770473, EPI_ISL_2770479, EPI_ISL_2770480, EPI_ISL_2770486, EPI_ISL_2770488 | | | |
| see above | NHLS Charlotte Maxeke Johannesburg Academic Hospital and the University of the Witwatersrand | KRISP, KZN Research Innovation and Sequencing Platform | Florette Treurnicht, Bulelani Manene, Kathleen Subramoney, Giandhari Jennifer, Pillay Sureshnee, Yajna Ramphal, Naidoo Yeshnee, Tshabuila Derek,Tegally Houriiyah, San James, Wilkinson Eduan, de Oliveira Tulio |
| EPI_ISL_2770683, EPI_ISL_2770685, EPI_ISL_2770687, EPI_ISL_2770688, EPI_ISL_2770689, EPI_ISL_2770690, EPI_ISL_2770696, EPI_ISL_2770698, EPI_ISL_2770700, EPI_ISL_2770702, EPI_ISL_2770703, EPI_ISL_2770704, EPI_ISL_2770705, EPI_ISL_2770742 | | | |
| see above | Pathogen Genomics Center, National Institute of Infectious Diseases | Pathogen Genomics Center, National Institute of Infectious Diseases | Tsuyoshi Sekizuka, Kentaro Itokawa, Rina Tanaka, Masanori Hashino, Makoto Kuroda |
| EPI_ISL_2770819, EPI_ISL_2771183, EPI_ISL_2771184, EPI_ISL_2771592, EPI_ISL_2771593, EPI_ISL_2772002, EPI_ISL_2772003, EPI_ISL_2772017, EPI_ISL_2772018, EPI_ISL_2772019, EPI_ISL_2772020, EPI_ISL_2772021, EPI_ISL_2772022, EPI_ISL_2772023 | | | |
| see above | SARS-CoV-2 testing team, National Institute of Infectious Diseases | Pathogen Genomics Center, National Institute of Infectious Diseases | Tsuyoshi Sekizuka, Kentaro Itokawa, Rina Tanaka, Masanori Hashino, Nozomu Hanaoka, Masumichi Saito, Naomi Nojiri, Hazuka Y Furihata, Hiromizu Takahashi, Tsuguto Fujimoto, Makoto Kuroda |
| EPI_ISL_2772109, EPI_ISL_2772110, EPI_ISL_2772111, EPI_ISL_2772112, EPI_ISL_2772113, EPI_ISL_2772114, EPI_ISL_2772116, EPI_ISL_2772117, EPI_ISL_2772118, EPI_ISL_2772119, EPI_ISL_2772121, EPI_ISL_2772123, EPI_ISL_2772124, EPI_ISL_2772127, EPI_ISL_2772128 | | | |
| see above | Pathogen Genomics Center, National Institute of Infectious Diseases | Pathogen Genomics Center, National Institute of Infectious Diseases | Tsuyoshi Sekizuka, Kentaro Itokawa, Rina Tanaka, Masanori Hashino, Makoto Kuroda |
| EPI_ISL_2772503, EPI_ISL_2772504 | California Hospital Medical Center | Los Angeles County Public Health Laboratories | P. Hemarajata et al. |
| EPI_ISL_2772515, EPI_ISL_2772517 | UCLA Clinical Micro Lab | Los Angeles County Public Health Laboratories | P. Hemarajata et al. |
| EPI_ISL_2772525 | Palmdale Regional Medical Center | Los Angeles County Public Health Laboratories | P. Hemarajata et al. |
| EPI_ISL_2772526, EPI_ISL_2772527, EPI_ISL_2772529, EPI_ISL_2772530, EPI_ISL_2772531 | National Center of Infectious and Parasitic Diseases | National Center of Infectious and Parasitic Diseases | Alexiev et al |
| EPI_ISL_2772532, EPI_ISL_2772534, EPI_ISL_2772535, EPI_ISL_2772536, EPI_ISL_2772537, EPI_ISL_2772538, EPI_ISL_2772539, EPI_ISL_2772540, EPI_ISL_2772541, EPI_ISL_2772542, EPI_ISL_2772543, EPI_ISL_2772544, EPI_ISL_2772545, EPI_ISL_2772546, EPI_ISL_2772547, EPI_ISL_2772548, EPI_ISL_2772549, EPI_ISL_2772550, EPI_ISL_2772551, EPI_ISL_2772552, EPI_ISL_2772553, EPI_ISL_2772554, EPI_ISL_2772555, EPI_ISL_2772556, EPI_ISL_2772557, EPI_ISL_2772558, EPI_ISL_2772559, EPI_ISL_2772560, EPI_ISL_2772561, EPI_ISL_2772562, EPI_ISL_2772563, EPI_ISL_2772564, EPI_ISL_2772565, EPI_ISL_2772566, EPI_ISL_2772567, EPI_ISL_2772568, EPI_ISL_2772569, EPI_ISL_2772570, EPI_ISL_2772571, EPI_ISL_2772572, EPI_ISL_2772573, EPI_ISL_2772574, EPI_ISL_2772575, EPI_ISL_2772576, EPI_ISL_2772577, EPI_ISL_2772578, EPI_ISL_2772579, EPI_ISL_2772580, EPI_ISL_2772581, EPI_ISL_2772582 | | | |
| see above | Kansas Health and Environmental Lab | Kansas Health and Environmental Lab | Katherine Wiggins, Mike Grose, Jonathan Barnell, Ben Olsen, and Phil Adam |
| EPI_ISL_2772587, EPI_ISL_2772594, EPI_ISL_2772610, EPI_ISL_2772613, EPI_ISL_2772620, EPI_ISL_2772622, EPI_ISL_2772624, EPI_ISL_2772625, EPI_ISL_2772626, EPI_ISL_2772628, EPI_ISL_2772635, EPI_ISL_2772637, EPI_ISL_2772642, EPI_ISL_2772645, EPI_ISL_2772649 | | | |
| see above | Dept. of Microbiology and Infection Control, Akershus University | Dept. of Microbiology and Infection Control, Akershus University | Hege Vangstein Aamot, Alexander Hesselberg Løvestad |

| | | | |
|---|---|---|---|
| | Hospital HF | Hospital HF | |
| EPI_ISL_2772665, EPI_ISL_2772666, EPI_ISL_2772667, EPI_ISL_2772692 | Max von Pettenkofer Institute, Virology, National Reference Center for Retroviruses, LMU Munich | Laboratory for Functional Genome Analysis; Dept. Genomics; Gene Center of the LMU Munich | Max Muenchhoff; Stefan Krebs; Alexander Graf; Oliver Keppler; Helmut Blum |
| EPI_ISL_2772699, EPI_ISL_2772700 | Max von Pettenkofer Institute, Virology, National Reference Center for Retroviruses, LMU Munich | Laboratory for Functional Genome Analysis; Dept. Genomics; Gene Center of the LMU Munich | Marcel Stern; Stefan Krebs; Alexander Graf; Oliver Keppler; Helmut Blum |
| EPI_ISL_2772702, EPI_ISL_2772703, EPI_ISL_2772705 | Max von Pettenkofer Institute, Virology, National Reference Center for Retroviruses, LMU Munich | Laboratory for Functional Genome Analysis; Dept. Genomics; Gene Center of the LMU Munich | Max Muenchhoff; Stefan Krebs; Alexander Graf; Oliver Keppler; Helmut Blum |
| EPI_ISL_2773859 | University Hospital Brno, OKMI | University Hospital Brno, CMBG | Matej Bezdicek, Jan Svaton, Martina Lengerova, Monika Dolejska |
| EPI_ISL_2773860, EPI_ISL_2773861 | St. Anne´s University Hospital Brno | University Hospital Brno, CMBG | Matej Bezdicek, Jan Svaton, Martina Lengerova, Monika Dolejska |
| EPI_ISL_2773862 | University Hospital Brno, OKMI | University Hospital Brno, CMBG | Matej Bezdicek, Jan Svaton, Martina Lengerova, Monika Dolejska |
| EPI_ISL_2773867, EPI_ISL_2773868, EPI_ISL_2773869, EPI_ISL_2773945, EPI_ISL_2773946 | Genome Information Research Center, Research Institute for Microbial Diseases, Osaka University | Genome Information Research Center, Research Institute for Microbial Diseases, Osaka University | Daisuke Motooka, Kentaro Tanaka, Hiroya Oki, and Shota Nakamura |
| EPI_ISL_2773986, EPI_ISL_2773988, EPI_ISL_2773989, EPI_ISL_2773990, EPI_ISL_2773991, EPI_ISL_2773993, EPI_ISL_2773994, EPI_ISL_2773995, EPI_ISL_2773996, EPI_ISL_2773997, EPI_ISL_2773998, EPI_ISL_2773999, EPI_ISL_2774001, EPI_ISL_2774002 | | | |
| see above | OLVZ Aalst | OLVZ Aalst | Astrid Holderbeke |
| EPI_ISL_2774008, EPI_ISL_2774010 | Cantacuzino National Military-Medical Institute, Viral Respiratory Infections Laboratory | Cantacuzino Institute Virology | Luiza Ustea, Nicoleta Paraschiv, Sorin Dinu, Mihaela Oprea, Mihaela Lazar |
| EPI_ISL_2774036 | Clinical Emergency County Hospital, Craiova | Cantacuzino Institute Virology | Luiza Ustea, Nicoleta Paraschiv, Sorin Dinu, Mihaela Oprea, Mihaela Lazar |
| EPI_ISL_2774038, EPI_ISL_2774039, EPI_ISL_2774040, EPI_ISL_2774042, EPI_ISL_2774043, EPI_ISL_2774044, EPI_ISL_2774045, EPI_ISL_2774046, EPI_ISL_2774048, EPI_ISL_2774049, EPI_ISL_2774050, EPI_ISL_2774051, EPI_ISL_2774052, EPI_ISL_2774053, EPI_ISL_2774054, EPI_ISL_2774055, EPI_ISL_2774056, EPI_ISL_2774057, EPI_ISL_2774058, EPI_ISL_2774059, EPI_ISL_2774060, EPI_ISL_2774061, EPI_ISL_2774062, EPI_ISL_2774063, EPI_ISL_2774064, EPI_ISL_2774065, EPI_ISL_2774066, EPI_ISL_2774067, EPI_ISL_2774068, EPI_ISL_2774069, EPI_ISL_2774070, EPI_ISL_2774071, EPI_ISL_2774072, EPI_ISL_2774073, EPI_ISL_2774074, EPI_ISL_2774075 | | | |
| see above | Institute of Epidemiology, Disease Control and Research (IEDCR) | IEDCR-ideSHi-icddr,b | Hassan Afrad, Sadia Rahman, Manjur Hossain Khan, Firdausi Qadri, Tahmina Shirin |
| EPI_ISL_2774076 | Institute for Developing Science and Health Initiatives (ideSHi) | IEDCR-ideSHi-icddr,b | Hassan Afrad, Sadia Rahman, Manjur Hossain Khan, Firdausi Qadri, Tahmina Shirin |
| EPI_ISL_2774088 | Multiplex DX | Multiplex DX | Diana Drobna, Veronika Mancikova, Silvia Rybecka, Jakub Kovac, Pavol Cekan |
| EPI_ISL_2774122 | Microbiology Department, University Hospital Donostia | Microbiology Department, University Hospital Donostia | Marimon JM, Montes M, Piñeiro L, Martin-Peñaranda T, Gomez M, Sorarrain A, Cilla G. |
| EPI_ISL_2774133 | ZOL | Jessa | Marijke Raymaekers et al. on behalf of the Jessa_cmdLab |
| EPI_ISL_2774142, EPI_ISL_2774143 | Jessa | Jessa | Marijke Raymaekers et al. on behalf of the Jessa_cmdLab |
| EPI_ISL_2774149 | IRCCS San Gallicano Dermatological Institute | IRCCS Regina Elena National Cancer Institute | Francesca De Nicola, Giulia Orlandi, Alice Massacci, Martina Pontone, Fabrizio Ensoli, Fulvia Pimpinelli, Maurizio Fanciulli, Giovanni Blandino, Aldo Morrone, Gennaro Ciliberto |
| EPI_ISL_2774164 | IRCCS San Gallicano Dermatological Institute | IRCCS Regina Elena National Cancer Institute | Francesca De Nicola, Giulia Orlandi, Eleonora Sperandio, Giovanna D'agosto, Fabrizio Ensoli, Fulvia Pimpinelli, Maurizio Fanciulli, Giovanni Blandino, Aldo Morrone, Gennaro Ciliberto |
| EPI_ISL_2774165 | IRCCS San Gallicano Dermatological Institute | IRCCS Regina Elena National Cancer Institute | Ludovica Ciuffreda, Sara Donzelli, Alice Massacci, Carla Mottini, Fabrizio Ensoli, Fulvia Pimpinelli, Maurizio Fanciulli, Giovanni Blandino, Aldo Morrone, Gennaro Ciliberto |
| EPI_ISL_2774173 | IRCCS San Gallicano Dermatological Institute | IRCCS Regina Elena National Cancer Institute | Sabrina Strano, Sara Donzelli, Matteo Pallocca, Lorenzo Furzi, Fabrizio Ensoli, Fulvia Pimpinelli, Maurizio Fanciulli, Giovanni Blandino, Aldo Morrone, Gennaro Ciliberto |
| EPI_ISL_2774204 | IRCCS San Gallicano Dermatological Institute | IRCCS Regina Elena National Cancer Institute | Sabrina Strano, Sara Donzelli, Alice Massacci, Arianna Mastrofrancesco, Fabrizio Ensoli, Fulvia Pimpinelli, Maurizio Fanciulli, Giovanni Blandino, Aldo Morrone, Gennaro Ciliberto |
| EPI_ISL_2774212 | IRCCS San Gallicano Dermatological Institute | IRCCS Regina Elena National Cancer Institute | Ludovica Ciuffreda, Sara Donzelli, Matteo Pallocca, Antonio Federico, Fabrizio Ensoli, Fulvia Pimpinelli, Maurizio Fanciulli, Giovanni Blandino, Aldo Morrone, Gennaro Ciliberto |
| EPI_ISL_2774215 | IRCCS San Gallicano Dermatological Institute | IRCCS Regina Elena National Cancer Institute | Ludovica Ciuffreda, Sara Donzelli, Eleonora Sperandio, Francesca Maione, Fabrizio Ensoli, Fulvia Pimpinelli, Maurizio Fanciulli, Giovanni Blandino, Aldo Morrone, Gennaro Ciliberto |
| EPI_ISL_2774223 | IRCCS San Gallicano Dermatological Institute | IRCCS Regina Elena National Cancer Institute | Frauke Goeman, Giulia Orlandi, Eleonora Sperandio, Giovanna D'agosto, Fabrizio Ensoli, Fulvia Pimpinelli, Maurizio Fanciulli, Giovanni Blandino, Aldo Morrone, Gennaro Ciliberto |
| EPI_ISL_2774228 | IRCCS San Gallicano Dermatological Institute | IRCCS Regina Elena National Cancer Institute | Sabrina Strano, Sara Donzelli, Matteo Pallocca, Francesca Sivori, Fabrizio Ensoli, Fulvia Pimpinelli, Maurizio Fanciulli, Giovanni Blandino, Aldo Morrone, Gennaro Ciliberto |
| EPI_ISL_2774230 | IRCCS San Gallicano Dermatological Institute | IRCCS Regina Elena National Cancer Institute | Ludovica Ciuffreda, Sara Donzelli, Alice Massacci, Ilaria Celesti, Fabrizio Ensoli, Fulvia Pimpinelli, Maurizio Fanciulli, Giovanni Blandino, Aldo Morrone, Gennaro Ciliberto |
| EPI_ISL_2774232 | IRCCS San Gallicano Dermatological Institute | IRCCS Regina Elena National Cancer Institute | Sabrina Strano, Sara Donzelli, Eleonora Sperandio, Antonio Federico, Fabrizio Ensoli, Fulvia Pimpinelli, Maurizio Fanciulli, Giovanni Blandino, Aldo Morrone, Gennaro Ciliberto |
| EPI_ISL_2774233 | IRCCS San Gallicano Dermatological Institute | IRCCS Regina Elena National Cancer Institute | Francesca De Nicola, Giulia Orlandi, Alice Massacci, Fulvia Fraticelli, Fabrizio Ensoli, Fulvia Pimpinelli, Maurizio Fanciulli, Giovanni Blandino, Aldo Morrone, Gennaro Ciliberto |
| EPI_ISL_2774234 | IRCCS San Gallicano Dermatological Institute | IRCCS Regina Elena National Cancer Institute | Ludovica Ciuffreda, Sara Donzelli, Matteo Pallocca, Lorenzo Furzi, Fabrizio Ensoli, Fulvia Pimpinelli, Maurizio Fanciulli, Giovanni Blandino, Aldo Morrone, Gennaro Ciliberto |
| EPI_ISL_2774238 | IRCCS San Gallicano Dermatological Institute | IRCCS Regina Elena National Cancer Institute | Ludovica Ciuffreda, Sara Donzelli, Eleonora Sperandio, Silvia Paluzzi, Fabrizio Ensoli, Fulvia Pimpinelli, Maurizio Fanciulli, Giovanni Blandino, Aldo Morrone, Gennaro Ciliberto |
| EPI_ISL_2774239 | IRCCS San Gallicano Dermatological Institute | IRCCS Regina Elena National Cancer Institute | Frauke Goeman, Giulia Orlandi, Alice Massacci, Sara Petrolo, Fabrizio Ensoli, Fulvia Pimpinelli, Maurizio Fanciulli, Giovanni Blandino, Aldo Morrone, Gennaro Ciliberto |
| EPI_ISL_2774240 | IRCCS San Gallicano Dermatological Institute | IRCCS Regina Elena National Cancer Institute | Sabrina Strano, Sara Donzelli, Matteo Pallocca, Valentina Ricca, Fabrizio Ensoli, Fulvia Pimpinelli, Maurizio Fanciulli, Giovanni Blandino, Aldo Morrone, Gennaro Ciliberto |
| EPI_ISL_2774241, EPI_ISL_2774242, EPI_ISL_2774245, EPI_ISL_2774249, EPI_ISL_2774254, EPI_ISL_2774258, EPI_ISL_2774260, EPI_ISL_2774261, EPI_ISL_2774262, EPI_ISL_2774264, EPI_ISL_2774266, EPI_ISL_2774267, EPI_ISL_2774270, EPI_ISL_2774277, EPI_ISL_2774278, EPI_ISL_2774279, EPI_ISL_2774290, EPI_ISL_2774291, EPI_ISL_2774292, EPI_ISL_2774293, EPI_ISL_2774295, EPI_ISL_2774300, EPI_ISL_2774302, EPI_ISL_2774304 | | | |
| see above | Baylor Scott & White-Temple | Baylor Scott & White-Temple | Ari Rao, Linden Morales, Kimberly Walker, Marcus Volz, Shelby Johnson |
| EPI_ISL_2774307, EPI_ISL_2774308, EPI_ISL_2774309, EPI_ISL_2774310, EPI_ISL_2774311 | Baylor Scott & White-Irving | Baylor Scott & White-Temple | Ari Rao, Linden Morales, Kimberly Walker, Marcus Volz, Shelby Johnson |
| EPI_ISL_2774329, EPI_ISL_2774337, EPI_ISL_2774341, EPI_ISL_2774356, EPI_ISL_2774374, EPI_ISL_2774376, EPI_ISL_2774386, EPI_ISL_2774388, EPI_ISL_2774390, EPI_ISL_2774393, EPI_ISL_2774394, EPI_ISL_2774395 | | | |
| see above | Illinois Department of Public Health - Springfield Lab | Illinois Department of Public Health - Springfield Lab | Bryan Sim, Gordon McCall |
| EPI_ISL_2774515 | ZOL | Jessa | Marijke Raymaekers et al. on behalf of the Jessa_cmdLab |
| EPI_ISL_2774535, EPI_ISL_2774543, EPI_ISL_2774544, EPI_ISL_2774545, EPI_ISL_2774547, EPI_ISL_2774548, EPI_ISL_2774549, EPI_ISL_2774553, EPI_ISL_2774554 | Nebraska Public Health Laboratory | NPHL COVID-19 Response Team | NPHL COVID-19 Response Team |
| EPI_ISL_2774577, EPI_ISL_2774595, EPI_ISL_2774603, EPI_ISL_2774606, EPI_ISL_2774607, EPI_ISL_2774609, EPI_ISL_2774616 | Santa Clara Valley Medical Center | Santa Clara County Public Health Laboratory | Santa Clara County Public Health Department |
| EPI_ISL_2774622, EPI_ISL_2774623 | PAMF Palo Alto | Santa Clara County Public Health Laboratory | Santa Clara County Public Health Department |
| EPI_ISL_2774627, EPI_ISL_2774628, EPI_ISL_2774629, EPI_ISL_2774631, EPI_ISL_2774634 | Santa Clara Valley Medical Center | Santa Clara County Public Health Laboratory | Santa Clara County Public Health Department |
| EPI_ISL_2774658 | Mapmygenome | CSIR-Centre for Cellular and Molecular Biology-INSACOG | Lamuk Zaveri,Tulasi Nagabandi,Payel Mukherjee,Onkar Kulkarni, Sofia Banu, Jandhyala Sai Krishna,Ara Sreenivas,Shreekant Verma, Amareshwar Vodapalli ,B Himasri,Valli Nagalakshmi Undamatla, Sumedha Avadhanula, Priya Nurkurthy, Vidhyadhari Methuku, Archana Bharadwaj Siva, Karthik Bharadwaj Tallapaka,Rakesh K Mishra,Divya Tej Sowpati |
| EPI_ISL_2774666, EPI_ISL_2774678, EPI_ISL_2774679 | Mapmygenome | CSIR-Centre for Cellular and Molecular Biology-INSACOG | Tulasi Nagabandi,Lamuk Zaveri,Payel Mukherjee,Onkar Kulkarni,Jandhyala Sai Krishna, Sofia Banu, Ara Sreenivas,Shreekant Verma, Amareshwar Vodapalli ,B Himasri,Valli Nagalakshmi Undamatla, Priya Nurkurthy,Sumedha Avadhanula, Vidhyadhari Methuku, Archana Bharadwaj Siva, Karthik Bharadwaj Tallapaka,Rakesh K Mishra,Divya Tej Sowpati |
| EPI_ISL_2774681, | Mapmygenome | CSIR-Centre for Cellular and Molecular Biology-INSACOG | Ara Sreenivas,Shreekant Verma, Amareshwar Vodapalli ,B Himasri,Valli Nagalakshmi Undamatla,Payel Mukherjee, Lamuk Zaveri, Onkar Kulkarni,Jandhyala Sai Krishna,Tulasi Nagabandi, Sofia Banu, ,Priya Nurkurthy, Vidhyadhari Methuku, |

| | | | |
|---|---|---|---|
| EPI_ISL_2774682, EPI_ISL_2774683, EPI_ISL_2774696, EPI_ISL_2774698 | | | Sumedha Avadhanula, Archana Bharadwaj Siva, Karthik Bharadwaj Tallapaka,Rakesh K Mishra,Divya Tej Sowpati |
| EPI_ISL_2774700, EPI_ISL_2774701, EPI_ISL_2774707, EPI_ISL_2774708, EPI_ISL_2774709, EPI_ISL_2774710, EPI_ISL_2774713, EPI_ISL_2774714 | Mapmygenome | CSIR-Centre for Cellular and Molecular Biology-INSACOG | Shreekant Verma, Amareshwar Vodapalli ,Ara Sreenivas,B Himasri,Valli Nagalakshmi Undamatla,Payel Mukherjee, Lamuk Zaveri, Onkar Kulkarni,Tulasi Nagabandi, Sofia Banu, Jandhyala Sai Krishna,Sumedha Avadhanula, Priya Nurkurthy, Vidhyadhari Methuku, Archana Bharadwaj Siva, Karthik Bharadwaj Tallapaka,Rakesh K Mishra,Divya Tej Sowpati |
| EPI_ISL_2774724, EPI_ISL_2774725, EPI_ISL_2774730, EPI_ISL_2774731, EPI_ISL_2774732, EPI_ISL_2774733, EPI_ISL_2774734, EPI_ISL_2774735, EPI_ISL_2774736, EPI_ISL_2774737, EPI_ISL_2774738, EPI_ISL_2774739, EPI_ISL_2774740, EPI_ISL_2774741 | | | |
| see above | AP SSO | CSIR-Centre for Cellular and Molecular Biology-INSACOG | Amareshwar Vodapalli,Ara Sreenivas,Shreekant Verma, B Himasri,Valli Nagalakshmi Undamatla,Payel Mukherjee, Lamuk Zaveri, Onkar Kulkarni,Tulasi Nagabandi, Sofia Banu, Jandhyala Sai Krishna,Priya Nurkurthy, Vidhyadhari Methuku, Sumedha Avadhanula, Archana Bharadwaj Siva, Karthik Bharadwaj Tallapaka,Rakesh K Mishra,Divya Tej Sowpati |
| EPI_ISL_2774742, EPI_ISL_2774743, EPI_ISL_2774744, EPI_ISL_2774745, EPI_ISL_2774747, EPI_ISL_2774748, EPI_ISL_2774749, EPI_ISL_2774750, EPI_ISL_2774751, EPI_ISL_2774752, EPI_ISL_2774753, EPI_ISL_2774754, EPI_ISL_2774755, EPI_ISL_2774756, EPI_ISL_2774757, EPI_ISL_2774758 | | | |
| see above | AP SSO | CSIR-Centre for Cellular and Molecular Biology-INSACOG | B Himasri,Amareshwar Vodapalli,Ara Sreenivas,Shreekant Verma,Valli Nagalakshmi Undamatla,Payel Mukherjee, Lamuk Zaveri, Onkar Kulkarni,Tulasi Nagabandi, Sofia Banu, Vidhyadhari Methuku,Priya Nurkurthy, Jandhyala Sai Krishna,Sumedha Avadhanula, Archana Bharadwaj Siva, Karthik Bharadwaj Tallapaka,Rakesh K Mishra,Divya Tej Sowpati |
| EPI_ISL_2774759, EPI_ISL_2774762, EPI_ISL_2774763, EPI_ISL_2774765, EPI_ISL_2774766, EPI_ISL_2774767, EPI_ISL_2774768, EPI_ISL_2774769, EPI_ISL_2774770, EPI_ISL_2774771, EPI_ISL_2774772, EPI_ISL_2774773, EPI_ISL_2774774, EPI_ISL_2774775, EPI_ISL_2774776, EPI_ISL_2774777, EPI_ISL_2774778 | | | |
| see above | AP SSO | CSIR-Centre for Cellular and Molecular Biology-INSACOG | Sofia Banu,Payel Mukherjee, Lamuk Zaveri, Onkar Kulkarni,Tulasi Nagabandi, Jandhyala Sai Krishna, Ara Sreenivas,Shreekant Verma, Amareshwar Vodapalli ,B Himasri,Valli Nagalakshmi Undamatla, Sumedha Avadhanula, Vidhyadhari Methuku, Priya Nurkurthy, Archana Bharadwaj Siva, Karthik Bharadwaj Tallapaka,Rakesh K Mishra,Divya Tej Sowpati |
| EPI_ISL_2774779, EPI_ISL_2774780, EPI_ISL_2774781, EPI_ISL_2774782, EPI_ISL_2774783, EPI_ISL_2774784, EPI_ISL_2774785, EPI_ISL_2774786, EPI_ISL_2774787, EPI_ISL_2774788, EPI_ISL_2774789, EPI_ISL_2774790, EPI_ISL_2774791, EPI_ISL_2774792, EPI_ISL_2774793, EPI_ISL_2774794, EPI_ISL_2774795, EPI_ISL_2774796 | | | |
| see above | AP SSO | CSIR-Centre for Cellular and Molecular Biology-INSACOG | Sumedha Avadhanula, Lamuk Zaveri, Tulasi Nagabandi, Ara Sreenivas,Shreekant Verma, Amareshwar Vodapalli ,B Himasri,Valli Nagalakshmi Undamatla, Vidhyadhari Methuku, Priya Nurkurthy, Payel Mukherjee,Jandhyala Sai Krishna,Onkar Kulkarni, Sofia Banu, Archana Bharadwaj Siva, Karthik Bharadwaj Tallapaka,Rakesh K Mishra,Divya Tej Sowpati |
| EPI_ISL_2774797, EPI_ISL_2774798, EPI_ISL_2774799, EPI_ISL_2774800, EPI_ISL_2774801, EPI_ISL_2774802, EPI_ISL_2774803, EPI_ISL_2774804, EPI_ISL_2774805, EPI_ISL_2774806, EPI_ISL_2774807, EPI_ISL_2774808, EPI_ISL_2774809, EPI_ISL_2774810, EPI_ISL_2774811, EPI_ISL_2774812, EPI_ISL_2774813, EPI_ISL_2774814, EPI_ISL_2774815, EPI_ISL_2774816, EPI_ISL_2774817 | | | |
| see above | AP SSO | CSIR-Centre for Cellular and Molecular Biology-INSACOG | Priya Nurkurthy,Ara Sreenivas,Shreekant Verma, Amareshwar Vodapalli ,Lamuk Zaveri,Tulasi Nagabandi,B Himasri,Valli Nagalakshmi Undamatla, Sumedha Avadhanula, Vidhyadhari Methuku,Payel Mukherjee,Onkar Kulkarni, Jandhyala Sai Krishna,Sofia Banu, Archana Bharadwaj Siva, Karthik Bharadwaj Tallapaka,Rakesh K Mishra,Divya Tej Sowpati |
| EPI_ISL_2774818, EPI_ISL_2774819, EPI_ISL_2774820, EPI_ISL_2774821, EPI_ISL_2774822, EPI_ISL_2774823, EPI_ISL_2774824, EPI_ISL_2774825, EPI_ISL_2774826, EPI_ISL_2774827, EPI_ISL_2774828, EPI_ISL_2774829, EPI_ISL_2774830, EPI_ISL_2774831, EPI_ISL_2774832, EPI_ISL_2774833 | | | |
| see above | AP SSO | CSIR-Centre for Cellular and Molecular Biology-INSACOG | Vidhyadhari Methuku,Tulasi Nagabandi,Lamuk Zaveri, Ara Sreenivas,Shreekant Verma, Amareshwar Vodapalli ,B Himasri,Valli Nagalakshmi Undamatla, Priya Nurkurthy,Sumedha Avadhanula, Archana Bharadwaj Siva,Payel Mukherjee,Onkar Kulkarni, Jandhyala Sai Krishna,Sofia Banu, Karthik Bharadwaj Tallapaka,Rakesh K Mishra,Divya Tej Sowpati |
| EPI_ISL_2774835, EPI_ISL_2774836 | CSIR-Centre for Cellular and Molecular Biology | CSIR-Centre for Cellular and Molecular Biology-INSACOG | Vidhyadhari Methuku,Tulasi Nagabandi,Lamuk Zaveri, Ara Sreenivas,Shreekant Verma, Amareshwar Vodapalli ,B Himasri,Valli Nagalakshmi Undamatla, Priya Nurkurthy,Sumedha Avadhanula, Archana Bharadwaj Siva,Payel Mukherjee,Onkar Kulkarni, Jandhyala Sai Krishna,Sofia Banu, Karthik Bharadwaj Tallapaka,Rakesh K Mishra,Divya Tej Sowpati |
| EPI_ISL_2774839 | CSIR-Centre for Cellular and Molecular Biology | CSIR-Centre for Cellular and Molecular Biology-INSACOG | Sumedha Avadhanula, Lamuk Zaveri, Tulasi Nagabandi, Ara Sreenivas,Shreekant Verma, Amareshwar Vodapalli ,B Himasri,Valli Nagalakshmi Undamatla, Vidhyadhari Methuku, Priya Nurkurthy, Payel Mukherjee,Jandhyala Sai Krishna,Onkar Kulkarni, Sofia Banu, Archana Bharadwaj Siva, Karthik Bharadwaj Tallapaka,Rakesh K Mishra,Divya Tej Sowpati |
| EPI_ISL_2774844, EPI_ISL_2774846 | CSIR-Centre for Cellular and Molecular Biology | CSIR-Centre for Cellular and Molecular Biology-INSACOG | Payel Mukherjee, Lamuk Zaveri, Onkar Kulkarni,Jandhyala Sai Krishna,Tulasi Nagabandi, Sofia Banu, Ara Sreenivas,Shreekant Verma, Amareshwar Vodapalli ,B Himasri,Valli Nagalakshmi Undamatla, Sumedha Avadhanula, Vidhyadhari Methuku, Priya Nurkurthy, Archana Bharadwaj Siva, Karthik Bharadwaj Tallapaka,Rakesh K Mishra,Divya Tej Sowpati |
| EPI_ISL_2774858, EPI_ISL_2774863, EPI_ISL_2774868 | CSIR-Centre for Cellular and Molecular Biology | CSIR-Centre for Cellular and Molecular Biology-INSACOG | Lamuk Zaveri,Tulasi Nagabandi,Payel Mukherjee,Onkar Kulkarni, Sofia Banu, Jandhyala Sai Krishna,Ara Sreenivas,Shreekant Verma, Amareshwar Vodapalli ,B Himasri,Valli Nagalakshmi Undamatla, Sumedha Avadhanula, Priya Nurkurthy, Vidhyadhari Methuku, Archana Bharadwaj Siva, Karthik Bharadwaj Tallapaka,Rakesh K Mishra,Divya Tej Sowpati |
| EPI_ISL_2774878, EPI_ISL_2774881, EPI_ISL_2774882 | CSIR-Centre for Cellular and Molecular Biology | CSIR-Centre for Cellular and Molecular Biology-INSACOG | Payel Mukherjee, Lamuk Zaveri, Onkar Kulkarni,Jandhyala Sai Krishna,Tulasi Nagabandi, Sofia Banu, Ara Sreenivas,Shreekant Verma, Amareshwar Vodapalli ,B Himasri,Valli Nagalakshmi Undamatla, Sumedha Avadhanula, Vidhyadhari Methuku, Priya Nurkurthy, Archana Bharadwaj Siva, Karthik Bharadwaj Tallapaka,Rakesh K Mishra,Divya Tej Sowpati |
| EPI_ISL_2774893, EPI_ISL_2774897, EPI_ISL_2774898, EPI_ISL_2774899, EPI_ISL_2774901, EPI_ISL_2774902, EPI_ISL_2774903, EPI_ISL_2774905 | CSIR-Centre for Cellular and Molecular Biology | CSIR-Centre for Cellular and Molecular Biology-INSACOG | Lamuk Zaveri,Tulasi Nagabandi,Payel Mukherjee,Onkar Kulkarni, Sofia Banu, Jandhyala Sai Krishna,Ara Sreenivas,Shreekant Verma, Amareshwar Vodapalli ,B Himasri,Valli Nagalakshmi Undamatla, Sumedha Avadhanula, Priya Nurkurthy, Vidhyadhari Methuku, Archana Bharadwaj Siva, Karthik Bharadwaj Tallapaka,Rakesh K Mishra,Divya Tej Sowpati |
| EPI_ISL_2774909, EPI_ISL_2774913, EPI_ISL_2774915, EPI_ISL_2774916, EPI_ISL_2774917, EPI_ISL_2774918, EPI_ISL_2774919, EPI_ISL_2774920, EPI_ISL_2774921, EPI_ISL_2774922, EPI_ISL_2774923 | | | |
| see above | CSIR-Centre for Cellular and Molecular Biology | CSIR-Centre for Cellular and Molecular Biology-INSACOG | Tulasi Nagabandi,Lamuk Zaveri,Payel Mukherjee,Onkar Kulkarni,Jandhyala Sai Krishna, Sofia Banu, Ara Sreenivas,Shreekant Verma, Amareshwar Vodapalli ,B Himasri,Valli Nagalakshmi Undamatla, Priya Nurkurthy,Sumedha Avadhanula, Vidhyadhari Methuku, Archana Bharadwaj Siva, Karthik Bharadwaj Tallapaka,Rakesh K Mishra,Divya Tej Sowpati |
| EPI_ISL_2774924, EPI_ISL_2774925, EPI_ISL_2774926, EPI_ISL_2774927, EPI_ISL_2774928, EPI_ISL_2774929, EPI_ISL_2774930, EPI_ISL_2774931, EPI_ISL_2774932, EPI_ISL_2774933, EPI_ISL_2774934, EPI_ISL_2774935, EPI_ISL_2774936, EPI_ISL_2774938, EPI_ISL_2774939, EPI_ISL_2774940 | | | |
| see above | CSIR-Centre for Cellular and Molecular Biology | CSIR-Centre for Cellular and Molecular Biology-INSACOG | Ara Sreenivas,Shreekant Verma, Amareshwar Vodapalli ,B Himasri,Valli Nagalakshmi Undamatla,Payel Mukherjee, Lamuk Zaveri, Onkar Kulkarni,Jandhyala Sai Krishna,Tulasi Nagabandi, Sofia Banu, ,Priya Nurkurthy, Vidhyadhari Methuku, Sumedha Avadhanula, Archana Bharadwaj Siva, Karthik Bharadwaj Tallapaka,Rakesh K Mishra,Divya Tej Sowpati |
| EPI_ISL_2774941, EPI_ISL_2774942, EPI_ISL_2774943, EPI_ISL_2774944, EPI_ISL_2774945, EPI_ISL_2774946, EPI_ISL_2774947, EPI_ISL_2774948, EPI_ISL_2774949, EPI_ISL_2774950, EPI_ISL_2774953, EPI_ISL_2774954, EPI_ISL_2774955, EPI_ISL_2774956, EPI_ISL_2774957, EPI_ISL_2774958 | | | |
| see above | CSIR-Centre for Cellular and Molecular Biology | CSIR-Centre for Cellular and Molecular Biology-INSACOG | Shreekant Verma, Amareshwar Vodapalli ,Ara Sreenivas,B Himasri,Valli Nagalakshmi Undamatla,Payel Mukherjee, Lamuk Zaveri, Onkar Kulkarni,Tulasi Nagabandi, Sofia Banu, Jandhyala Sai Krishna,Sumedha Avadhanula, Priya Nurkurthy, Vidhyadhari Methuku, Archana Bharadwaj Siva, Karthik Bharadwaj Tallapaka,Rakesh K Mishra,Divya Tej Sowpati |
| EPI_ISL_2774959, EPI_ISL_2774960, EPI_ISL_2774961, EPI_ISL_2774962, EPI_ISL_2774963, EPI_ISL_2774964, EPI_ISL_2774965, EPI_ISL_2774966, EPI_ISL_2774967, EPI_ISL_2774968, EPI_ISL_2774969, EPI_ISL_2774970, EPI_ISL_2774971, EPI_ISL_2774973, EPI_ISL_2774974, EPI_ISL_2774975, EPI_ISL_2774976, EPI_ISL_2774977, EPI_ISL_2774978, EPI_ISL_2774979, EPI_ISL_2774980, EPI_ISL_2774981, EPI_ISL_2774982, EPI_ISL_2774983 | | | |
| see above | CSIR-Centre for Cellular and Molecular Biology | CSIR-Centre for Cellular and Molecular Biology-INSACOG | Amareshwar Vodapalli,Ara Sreenivas,Shreekant Verma, B Himasri,Valli Nagalakshmi Undamatla,Payel Mukherjee, Lamuk Zaveri, Onkar Kulkarni,Tulasi Nagabandi, Sofia Banu, Jandhyala Sai Krishna,Priya Nurkurthy, Vidhyadhari Methuku, Sumedha Avadhanula, Archana Bharadwaj Siva, Karthik Bharadwaj Tallapaka,Rakesh K Mishra,Divya Tej Sowpati |
| EPI_ISL_2774984, EPI_ISL_2774985, EPI_ISL_2774986, EPI_ISL_2774987, EPI_ISL_2774988, EPI_ISL_2774989, EPI_ISL_2774990, EPI_ISL_2774991, EPI_ISL_2774993, EPI_ISL_2774994, EPI_ISL_2774995, EPI_ISL_2774996, EPI_ISL_2774997, EPI_ISL_2774998, EPI_ISL_2774999, EPI_ISL_2775000, EPI_ISL_2775001 | | | |
| see above | CSIR-Centre for Cellular and Molecular Biology | CSIR-Centre for Cellular and Molecular Biology-INSACOG | B Himasri,Amareshwar Vodapalli,Ara Sreenivas,Shreekant Verma,Valli Nagalakshmi Undamatla,Payel Mukherjee, Lamuk Zaveri, Onkar Kulkarni,Tulasi Nagabandi, Sofia Banu, Vidhyadhari Methuku,Priya Nurkurthy, Jandhyala Sai Krishna,Sumedha Avadhanula, Archana Bharadwaj Siva, Karthik Bharadwaj Tallapaka,Rakesh K Mishra,Divya Tej Sowpati |
| EPI_ISL_2775002, EPI_ISL_2775003, EPI_ISL_2775004, EPI_ISL_2775005, EPI_ISL_2775006, EPI_ISL_2775007, EPI_ISL_2775008, EPI_ISL_2775009, EPI_ISL_2775010, EPI_ISL_2775011, EPI_ISL_2775012, EPI_ISL_2775013, EPI_ISL_2775014, EPI_ISL_2775015, EPI_ISL_2775016, EPI_ISL_2775017, EPI_ISL_2775018, EPI_ISL_2775019, EPI_ISL_2775020 | | | |
| see above | CSIR-Centre for Cellular and Molecular Biology | CSIR-Centre for Cellular and Molecular Biology-INSACOG | Sofia Banu,Payel Mukherjee, Lamuk Zaveri, Onkar Kulkarni,Tulasi Nagabandi, Jandhyala Sai Krishna, Ara Sreenivas,Shreekant Verma, Amareshwar Vodapalli ,B Himasri,Valli Nagalakshmi Undamatla, Sumedha Avadhanula, Vidhyadhari Methuku, Priya Nurkurthy, Archana Bharadwaj Siva, Karthik Bharadwaj Tallapaka,Rakesh K Mishra,Divya Tej Sowpati |
| EPI_ISL_2775021, EPI_ISL_2775022, EPI_ISL_2775023, EPI_ISL_2775024, EPI_ISL_2775025, EPI_ISL_2775026, EPI_ISL_2775027, EPI_ISL_2775028, EPI_ISL_2775029, EPI_ISL_2775030, EPI_ISL_2775031, EPI_ISL_2775032, EPI_ISL_2775034, EPI_ISL_2775035, EPI_ISL_2775036, EPI_ISL_2775037, EPI_ISL_2775038 | | | |
| see above | CSIR-Centre for Cellular and Molecular Biology | CSIR-Centre for Cellular and Molecular Biology-INSACOG | Sumedha Avadhanula, Lamuk Zaveri, Tulasi Nagabandi, Ara Sreenivas,Shreekant Verma, Amareshwar Vodapalli ,B Himasri,Valli Nagalakshmi Undamatla, Vidhyadhari Methuku, Priya Nurkurthy, Payel Mukherjee,Jandhyala Sai Krishna,Onkar Kulkarni, Sofia Banu, Archana Bharadwaj Siva, Karthik Bharadwaj Tallapaka,Rakesh K Mishra,Divya Tej Sowpati |
| EPI_ISL_2775039, EPI_ISL_2775040, EPI_ISL_2775041, EPI_ISL_2775042, EPI_ISL_2775043, EPI_ISL_2775044, EPI_ISL_2775045, EPI_ISL_2775046, EPI_ISL_2775047, EPI_ISL_2775048, EPI_ISL_2775050, EPI_ISL_2775051, EPI_ISL_2775052, EPI_ISL_2775053, EPI_ISL_2775054, EPI_ISL_2775055, EPI_ISL_2775056, EPI_ISL_2775057, EPI_ISL_2775058, EPI_ISL_2775059 | | | |
| see above | CSIR-Centre for Cellular and Molecular Biology | CSIR-Centre for Cellular and Molecular Biology-INSACOG | Priya Nurkurthy,Ara Sreenivas,Shreekant Verma, Amareshwar Vodapalli ,Lamuk Zaveri,Tulasi Nagabandi,B Himasri,Valli Nagalakshmi Undamatla, Sumedha Avadhanula, Vidhyadhari Methuku,Payel Mukherjee,Onkar Kulkarni, Jandhyala Sai Krishna,Sofia Banu, Archana Bharadwaj Siva, Karthik Bharadwaj Tallapaka,Rakesh K Mishra,Divya Tej Sowpati |
| EPI_ISL_2775060, EPI_ISL_2775061, EPI_ISL_2775062, EPI_ISL_2775063, EPI_ISL_2775064, EPI_ISL_2775065, EPI_ISL_2775066, EPI_ISL_2775067, EPI_ISL_2775068, EPI_ISL_2775069, EPI_ISL_2775070, EPI_ISL_2775071, EPI_ISL_2775072, EPI_ISL_2775073, EPI_ISL_2775074, EPI_ISL_2775075, EPI_ISL_2775076, EPI_ISL_2775077 | | | |
| see above | CSIR-Centre for Cellular and Molecular Biology | CSIR-Centre for Cellular and Molecular Biology-INSACOG | Vidhyadhari Methuku,Tulasi Nagabandi,Lamuk Zaveri, Ara Sreenivas,Shreekant Verma, Amareshwar Vodapalli ,B Himasri,Valli Nagalakshmi Undamatla, Priya Nurkurthy,Sumedha Avadhanula, Archana Bharadwaj Siva,Payel Mukherjee,Onkar Kulkarni, Jandhyala Sai Krishna,Sofia Banu, Karthik Bharadwaj Tallapaka,Rakesh K Mishra,Divya Tej Sowpati |
| EPI_ISL_2775079, EPI_ISL_2775080, EPI_ISL_2775081 | CSIR-Centre for Cellular and Molecular Biology | CSIR-Centre for Cellular and Molecular Biology-INSACOG | Sumedha Avadhanula, Lamuk Zaveri, Tulasi Nagabandi, Ara Sreenivas,Shreekant Verma, Amareshwar Vodapalli ,B Himasri,Valli Nagalakshmi Undamatla, Vidhyadhari Methuku, Priya Nurkurthy, Payel Mukherjee,Jandhyala Sai Krishna,Onkar Kulkarni, Sofia Banu, Archana Bharadwaj Siva, Karthik Bharadwaj Tallapaka,Rakesh K Mishra,Divya Tej Sowpati |
| EPI_ISL_2775082, EPI_ISL_2775083, EPI_ISL_2775084 | CSIR-Centre for Cellular and Molecular Biology | CSIR-Centre for Cellular and Molecular Biology-INSACOG | Priya Nurkurthy,Ara Sreenivas,Shreekant Verma, Amareshwar Vodapalli ,Lamuk Zaveri,Tulasi Nagabandi,B Himasri,Valli Nagalakshmi Undamatla, Sumedha Avadhanula, Vidhyadhari Methuku,Payel Mukherjee,Onkar Kulkarni, Jandhyala Sai Krishna,Sofia Banu, Archana Bharadwaj Siva, Karthik Bharadwaj Tallapaka,Rakesh K Mishra,Divya Tej Sowpati |
| EPI_ISL_2775085, EPI_ISL_2775086, EPI_ISL_2775087, EPI_ISL_2775088, EPI_ISL_2775089, EPI_ISL_2775090, EPI_ISL_2775091, EPI_ISL_2775092, EPI_ISL_2775093, EPI_ISL_2775094, EPI_ISL_2775095 | | | |
| see above | CSIR-Centre for Cellular and Molecular Biology | CSIR-Centre for Cellular and Molecular Biology-INSACOG | Payel Mukherjee, Lamuk Zaveri, Onkar Kulkarni,Jandhyala Sai Krishna,Tulasi Nagabandi, Sofia Banu, Ara Sreenivas,Shreekant Verma, Amareshwar Vodapalli ,B Himasri,Valli Nagalakshmi Undamatla, Sumedha Avadhanula, Vidhyadhari Methuku, Priya Nurkurthy, Archana Bharadwaj Siva, Karthik Bharadwaj Tallapaka,Rakesh K Mishra,Divya Tej Sowpati |
| EPI_ISL_2775096, EPI_ISL_2775097, EPI_ISL_2775098, EPI_ISL_2775099, EPI_ISL_2775100, EPI_ISL_2775101, EPI_ISL_2775102, EPI_ISL_2775103, EPI_ISL_2775104, EPI_ISL_2775105, EPI_ISL_2775106, EPI_ISL_2775108, EPI_ISL_2775109 | | | |
| see above | CSIR-Centre for Cellular and Molecular Biology | CSIR-Centre for Cellular and Molecular Biology-INSACOG | Lamuk Zaveri,Tulasi Nagabandi,Payel Mukherjee,Onkar Kulkarni, Sofia Banu, Jandhyala Sai Krishna,Ara Sreenivas,Shreekant Verma, Amareshwar Vodapalli ,B Himasri,Valli Nagalakshmi Undamatla, Sumedha Avadhanula, Priya Nurkurthy, Vidhyadhari Methuku, Archana Bharadwaj Siva, Karthik Bharadwaj Tallapaka,Rakesh K Mishra,Divya Tej Sowpati |
| EPI_ISL_2775112, | CSIR-Centre for Cellular and Molecular Biology | CSIR-Centre for Cellular and Molecular Biology-INSACOG | Tulasi Nagabandi,Lamuk Zaveri,Payel Mukherjee,Onkar Kulkarni,Jandhyala Sai Krishna, Sofia Banu, Ara Sreenivas,Shreekant Verma, Amareshwar Vodapalli ,B Himasri,Valli Nagalakshmi Undamatla, Priya Nurkurthy,Sumedha Avadhanula, |

| | | | |
|---|---|---|---|
| EPI_ISL_2775114, EPI_ISL_2775116, EPI_ISL_2775117 | | | Vidhyadhari Methuku, Archana Bharadwaj Siva, Karthik Bharadwaj Tallapaka,Rakesh K Mishra,Divya Tej Sowpati |
| EPI_ISL_2775119 | CSIR-Centre for Cellular and Molecular Biology | CSIR-Centre for Cellular and Molecular Biology-INSACOG | Amareshwar Vodapalli,Ara Sreenivas,Shreekant Verma, B Himasri,Valli Nagalakshmi Undamatla,Payel Mukherjee, Lamuk Zaveri, Onkar Kulkarni,Tulasi Nagabandi, Sofia Banu, Jandhyala Sai Krishna,Priya Nurkurthy, Vidhyadhari Methuku, Sumedha Avadhanula, Archana Bharadwaj Siva, Karthik Bharadwaj Tallapaka,Rakesh K Mishra,Divya Tej Sowpati |
| EPI_ISL_2775128 | Mapmygenome | CSIR-Centre for Cellular and Molecular Biology-INSACOG | Lamuk Zaveri,Tulasi Nagabandi,Payel Mukherjee,Onkar Kulkarni, Sofia Banu, Jandhyala Sai Krishna,Ara Sreenivas,Shreekant Verma, Amareshwar Vodapalli ,B Himasri,Valli Nagalakshmi Undamatla, Sumedha Avadhanula, Priya Nurkurthy, Vidhyadhari Methuku, Archana Bharadwaj Siva, Karthik Bharadwaj Tallapaka,Rakesh K Mishra,Divya Tej Sowpati |
| EPI_ISL_2775130 | Mapmygenome | CSIR-Centre for Cellular and Molecular Biology-INSACOG | Tulasi Nagabandi,Lamuk Zaveri,Payel Mukherjee,Onkar Kulkarni,Jandhyala Sai Krishna, Sofia Banu, Ara Sreenivas,Shreekant Verma, Amareshwar Vodapalli ,B Himasri,Valli Nagalakshmi Undamatla, Priya Nurkurthy,Sumedha Avadhanula, Vidhyadhari Methuku, Archana Bharadwaj Siva, Karthik Bharadwaj Tallapaka,Rakesh K Mishra,Divya Tej Sowpati |
| EPI_ISL_2775136 | Mapmygenome | CSIR-Centre for Cellular and Molecular Biology-INSACOG | Amareshwar Vodapalli,Ara Sreenivas,Shreekant Verma, B Himasri,Valli Nagalakshmi Undamatla,Payel Mukherjee, Lamuk Zaveri, Onkar Kulkarni,Tulasi Nagabandi, Sofia Banu, Jandhyala Sai Krishna,Priya Nurkurthy, Vidhyadhari Methuku, Sumedha Avadhanula, Archana Bharadwaj Siva, Karthik Bharadwaj Tallapaka,Rakesh K Mishra,Divya Tej Sowpati |
| EPI_ISL_2775149, EPI_ISL_2775150, EPI_ISL_2775151, EPI_ISL_2775152, EPI_ISL_2775154, EPI_ISL_2775155, EPI_ISL_2775157, EPI_ISL_2775159, EPI_ISL_2775160, EPI_ISL_2775161, EPI_ISL_2775162 | | | |
| see above | CSIR-Centre for Cellular and Molecular Biology | CSIR-Centre for Cellular and Molecular Biology-INSACOG | Payel Mukherjee, Lamuk Zaveri, Onkar Kulkarni,Jandhyala Sai Krishna,Tulasi Nagabandi, Sofia Banu, Ara Sreenivas,Shreekant Verma, Amareshwar Vodapalli ,B Himasri,Valli Nagalakshmi Undamatla, Sumedha Avadhanula, Vidhyadhari Methuku, Priya Nurkurthy, Archana Bharadwaj Siva, Karthik Bharadwaj Tallapaka,Rakesh K Mishra,Divya Tej Sowpati |
| EPI_ISL_2775163, EPI_ISL_2775164, EPI_ISL_2775165, EPI_ISL_2775166, EPI_ISL_2775167, EPI_ISL_2775168, EPI_ISL_2775169, EPI_ISL_2775170, EPI_ISL_2775171, EPI_ISL_2775172, EPI_ISL_2775173, EPI_ISL_2775174, EPI_ISL_2775175, EPI_ISL_2775176, EPI_ISL_2775177, EPI_ISL_2775179 | | | |
| see above | CSIR-Centre for Cellular and Molecular Biology | CSIR-Centre for Cellular and Molecular Biology-INSACOG | Lamuk Zaveri,Tulasi Nagabandi,Payel Mukherjee,Onkar Kulkarni, Sofia Banu, Jandhyala Sai Krishna,Ara Sreenivas,Shreekant Verma, Amareshwar Vodapalli ,B Himasri,Valli Nagalakshmi Undamatla, Sumedha Avadhanula, Priya Nurkurthy, Vidhyadhari Methuku, Archana Bharadwaj Siva, Karthik Bharadwaj Tallapaka,Rakesh K Mishra,Divya Tej Sowpati |
| EPI_ISL_2775180, EPI_ISL_2775181, EPI_ISL_2775182, EPI_ISL_2775183, EPI_ISL_2775184, EPI_ISL_2775185, EPI_ISL_2775186, EPI_ISL_2775187, EPI_ISL_2775188, EPI_ISL_2775190, EPI_ISL_2775191, EPI_ISL_2775192, EPI_ISL_2775193, EPI_ISL_2775194, EPI_ISL_2775195 | | | |
| see above | CSIR-Centre for Cellular and Molecular Biology | CSIR-Centre for Cellular and Molecular Biology-INSACOG | Tulasi Nagabandi,Lamuk Zaveri,Payel Mukherjee,Onkar Kulkarni,Jandhyala Sai Krishna, Sofia Banu, Ara Sreenivas,Shreekant Verma, Amareshwar Vodapalli ,B Himasri,Valli Nagalakshmi Undamatla, Priya Nurkurthy,Sumedha Avadhanula, Vidhyadhari Methuku, Archana Bharadwaj Siva, Karthik Bharadwaj Tallapaka,Rakesh K Mishra,Divya Tej Sowpati |
| EPI_ISL_2775196, EPI_ISL_2775198, EPI_ISL_2775199, EPI_ISL_2775201, EPI_ISL_2775202, EPI_ISL_2775203, EPI_ISL_2775204, EPI_ISL_2775205, EPI_ISL_2775206, EPI_ISL_2775207, EPI_ISL_2775208, EPI_ISL_2775209, EPI_ISL_2775210, EPI_ISL_2775211, EPI_ISL_2775212, EPI_ISL_2775213 | | | |
| see above | CSIR-Centre for Cellular and Molecular Biology | CSIR-Centre for Cellular and Molecular Biology-INSACOG | Ara Sreenivas,Shreekant Verma, Amareshwar Vodapalli ,B Himasri,Valli Nagalakshmi Undamatla,Payel Mukherjee, Lamuk Zaveri, Onkar Kulkarni,Jandhyala Sai Krishna,Tulasi Nagabandi, Sofia Banu, ,Priya Nurkurthy, Vidhyadhari Methuku, Sumedha Avadhanula, Archana Bharadwaj Siva, Karthik Bharadwaj Tallapaka,Rakesh K Mishra,Divya Tej Sowpati |
| EPI_ISL_2775214, EPI_ISL_2775215, EPI_ISL_2775216, EPI_ISL_2775217, EPI_ISL_2775218, EPI_ISL_2775219, EPI_ISL_2775220, EPI_ISL_2775221, EPI_ISL_2775222, EPI_ISL_2775223, EPI_ISL_2775224, EPI_ISL_2775225, EPI_ISL_2775226, EPI_ISL_2775227, EPI_ISL_2775228, EPI_ISL_2775229, EPI_ISL_2775230, EPI_ISL_2775231 | | | |
| see above | CSIR-Centre for Cellular and Molecular Biology | CSIR-Centre for Cellular and Molecular Biology-INSACOG | Shreekant Verma, Amareshwar Vodapalli ,Ara Sreenivas,B Himasri,Valli Nagalakshmi Undamatla,Payel Mukherjee, Lamuk Zaveri, Onkar Kulkarni,Tulasi Nagabandi, Sofia Banu, Jandhyala Sai Krishna,Sumedha Avadhanula, Priya Nurkurthy, Vidhyadhari Methuku, Archana Bharadwaj Siva, Karthik Bharadwaj Tallapaka,Rakesh K Mishra,Divya Tej Sowpati |
| EPI_ISL_2775232, EPI_ISL_2775233, EPI_ISL_2775234, EPI_ISL_2775235, EPI_ISL_2775236, EPI_ISL_2775237, EPI_ISL_2775238, EPI_ISL_2775239, EPI_ISL_2775240, EPI_ISL_2775241, EPI_ISL_2775242, EPI_ISL_2775243, EPI_ISL_2775244, EPI_ISL_2775245, EPI_ISL_2775246, EPI_ISL_2775247, EPI_ISL_2775248, EPI_ISL_2775249, EPI_ISL_2775250, EPI_ISL_2775251, EPI_ISL_2775252, EPI_ISL_2775253, EPI_ISL_2775254, EPI_ISL_2775255, EPI_ISL_2775256 | | | |
| see above | CSIR-Centre for Cellular and Molecular Biology | CSIR-Centre for Cellular and Molecular Biology-INSACOG | Amareshwar Vodapalli,Ara Sreenivas,Shreekant Verma, B Himasri,Valli Nagalakshmi Undamatla,Payel Mukherjee, Lamuk Zaveri, Onkar Kulkarni,Tulasi Nagabandi, Sofia Banu, Jandhyala Sai Krishna,Priya Nurkurthy, Vidhyadhari Methuku, Sumedha Avadhanula, Archana Bharadwaj Siva, Karthik Bharadwaj Tallapaka,Rakesh K Mishra,Divya Tej Sowpati |
| EPI_ISL_2775257, EPI_ISL_2775258, EPI_ISL_2775259, EPI_ISL_2775260, EPI_ISL_2775261, EPI_ISL_2775262, EPI_ISL_2775263, EPI_ISL_2775264, EPI_ISL_2775265, EPI_ISL_2775266, EPI_ISL_2775269, EPI_ISL_2775272 | | | |
| see above | CSIR-Centre for Cellular and Molecular Biology | CSIR-Centre for Cellular and Molecular Biology-INSACOG | B Himasri,Amareshwar Vodapalli,Ara Sreenivas,Shreekant Verma,Valli Nagalakshmi Undamatla,Payel Mukherjee, Lamuk Zaveri, Onkar Kulkarni,Tulasi Nagabandi, Sofia Banu, Vidhyadhari Methuku,Priya Nurkurthy, Jandhyala Sai Krishna,Sumedha Avadhanula, Archana Bharadwaj Siva, Karthik Bharadwaj Tallapaka,Rakesh K Mishra,Divya Tej Sowpati |
| EPI_ISL_2775275, EPI_ISL_2775278, EPI_ISL_2775279, EPI_ISL_2775280, EPI_ISL_2775281, EPI_ISL_2775282, EPI_ISL_2775283, EPI_ISL_2775284, EPI_ISL_2775285, EPI_ISL_2775286, EPI_ISL_2775287 | | | |
| see above | CSIR-Centre for Cellular and Molecular Biology | CSIR-Centre for Cellular and Molecular Biology-INSACOG | Sofia Banu,Payel Mukherjee, Lamuk Zaveri, Onkar Kulkarni,Tulasi Nagabandi, Jandhyala Sai Krishna, Ara Sreenivas,Shreekant Verma, Amareshwar Vodapalli ,B Himasri,Valli Nagalakshmi Undamatla, Sumedha Avadhanula, Vidhyadhari Methuku, Priya Nurkurthy, Archana Bharadwaj Siva, Karthik Bharadwaj Tallapaka,Rakesh K Mishra,Divya Tej Sowpati |
| EPI_ISL_2775288, EPI_ISL_2775289, EPI_ISL_2775290, EPI_ISL_2775291, EPI_ISL_2775292, EPI_ISL_2775293, EPI_ISL_2775294 | AP SSO | CSIR-Centre for Cellular and Molecular Biology-INSACOG | Sofia Banu,Payel Mukherjee, Lamuk Zaveri, Onkar Kulkarni,Tulasi Nagabandi, Jandhyala Sai Krishna, Ara Sreenivas,Shreekant Verma, Amareshwar Vodapalli ,B Himasri,Valli Nagalakshmi Undamatla, Sumedha Avadhanula, Vidhyadhari Methuku, Priya Nurkurthy, Archana Bharadwaj Siva, Karthik Bharadwaj Tallapaka,Rakesh K Mishra,Divya Tej Sowpati |
| EPI_ISL_2775295, EPI_ISL_2775296, EPI_ISL_2775297, EPI_ISL_2775299, EPI_ISL_2775300, EPI_ISL_2775301, EPI_ISL_2775302, EPI_ISL_2775303, EPI_ISL_2775304, EPI_ISL_2775305 | AP SSO | CSIR-Centre for Cellular and Molecular Biology-INSACOG | Sumedha Avadhanula, Lamuk Zaveri, Tulasi Nagabandi, Ara Sreenivas,Shreekant Verma, Amareshwar Vodapalli ,B Himasri,Valli Nagalakshmi Undamatla, Vidhyadhari Methuku, Priya Nurkurthy, Payel Mukherjee,Jandhyala Sai Krishna,Onkar Kulkarni, Sofia Banu, Archana Bharadwaj Siva, Karthik Bharadwaj Tallapaka,Rakesh K Mishra,Divya Tej Sowpati |
| EPI_ISL_2775314, EPI_ISL_2775315, EPI_ISL_2775321, EPI_ISL_2775322, EPI_ISL_2775325, EPI_ISL_2775326, EPI_ISL_2775327, EPI_ISL_2775328, EPI_ISL_2775329, EPI_ISL_2775330, EPI_ISL_2775331 | | | |
| see above | AP SSO | CSIR-Centre for Cellular and Molecular Biology-INSACOG | Priya Nurkurthy,Ara Sreenivas,Shreekant Verma, Amareshwar Vodapalli ,Lamuk Zaveri,Tulasi Nagabandi,B Himasri,Valli Nagalakshmi Undamatla, Sumedha Avadhanula, Vidhyadhari Methuku,Payel Mukherjee,Onkar Kulkarni, Jandhyala Sai Krishna,Sofia Banu, Archana Bharadwaj Siva, Karthik Bharadwaj Tallapaka,Rakesh K Mishra,Divya Tej Sowpati |
| EPI_ISL_2775332, EPI_ISL_2775333, EPI_ISL_2775334, EPI_ISL_2775335, EPI_ISL_2775336, EPI_ISL_2775337, EPI_ISL_2775338, EPI_ISL_2775339, EPI_ISL_2775340, EPI_ISL_2775341, EPI_ISL_2775342, EPI_ISL_2775343, EPI_ISL_2775344, EPI_ISL_2775345, EPI_ISL_2775346, EPI_ISL_2775347, EPI_ISL_2775348, EPI_ISL_2775349, EPI_ISL_2775350 | | | |
| see above | AP SSO | CSIR-Centre for Cellular and Molecular Biology-INSACOG | Vidhyadhari Methuku,Tulasi Nagabandi,Lamuk Zaveri, Ara Sreenivas,Shreekant Verma, Amareshwar Vodapalli ,B Himasri,Valli Nagalakshmi Undamatla, Priya Nurkurthy,Sumedha Avadhanula, Archana Bharadwaj Siva,Payel Mukherjee,Onkar Kulkarni, Jandhyala Sai Krishna,Sofia Banu, Karthik Bharadwaj Tallapaka,Rakesh K Mishra,Divya Tej Sowpati |
| EPI_ISL_2775351, EPI_ISL_2775352, EPI_ISL_2775353 | AP SSO | CSIR-Centre for Cellular and Molecular Biology-INSACOG | Sumedha Avadhanula, Lamuk Zaveri, Tulasi Nagabandi, Ara Sreenivas,Shreekant Verma, Amareshwar Vodapalli ,B Himasri,Valli Nagalakshmi Undamatla, Vidhyadhari Methuku, Priya Nurkurthy, Payel Mukherjee,Jandhyala Sai Krishna,Onkar Kulkarni, Sofia Banu, Archana Bharadwaj Siva, Karthik Bharadwaj Tallapaka,Rakesh K Mishra,Divya Tej Sowpati |
| EPI_ISL_2775354, EPI_ISL_2775355, EPI_ISL_2775356 | AP SSO | CSIR-Centre for Cellular and Molecular Biology-INSACOG | Priya Nurkurthy,Ara Sreenivas,Shreekant Verma, Amareshwar Vodapalli ,Lamuk Zaveri,Tulasi Nagabandi,B Himasri,Valli Nagalakshmi Undamatla, Sumedha Avadhanula, Vidhyadhari Methuku,Payel Mukherjee,Onkar Kulkarni, Jandhyala Sai Krishna,Sofia Banu, Archana Bharadwaj Siva, Karthik Bharadwaj Tallapaka,Rakesh K Mishra,Divya Tej Sowpati |
| EPI_ISL_2775357, EPI_ISL_2775358, EPI_ISL_2775359, EPI_ISL_2775360, EPI_ISL_2775361, EPI_ISL_2775362, EPI_ISL_2775363, EPI_ISL_2775364, EPI_ISL_2775365 | AP SSO | CSIR-Centre for Cellular and Molecular Biology-INSACOG | Payel Mukherjee, Lamuk Zaveri, Onkar Kulkarni,Jandhyala Sai Krishna,Tulasi Nagabandi, Sofia Banu, Ara Sreenivas,Shreekant Verma, Amareshwar Vodapalli ,B Himasri,Valli Nagalakshmi Undamatla, Sumedha Avadhanula, Vidhyadhari Methuku, Priya Nurkurthy, Archana Bharadwaj Siva, Karthik Bharadwaj Tallapaka,Rakesh K Mishra,Divya Tej Sowpati |
| EPI_ISL_2775368, EPI_ISL_2775369, EPI_ISL_2775370, EPI_ISL_2775371, EPI_ISL_2775372, EPI_ISL_2775373, EPI_ISL_2775374, EPI_ISL_2775375, EPI_ISL_2775376, EPI_ISL_2775379, EPI_ISL_2775381, EPI_ISL_2775382, EPI_ISL_2775383 | | | |
| see above | AP SSO | CSIR-Centre for Cellular and Molecular Biology-INSACOG | Lamuk Zaveri,Tulasi Nagabandi,Payel Mukherjee,Onkar Kulkarni, Sofia Banu, Jandhyala Sai Krishna,Ara Sreenivas,Shreekant Verma, Amareshwar Vodapalli ,B Himasri,Valli Nagalakshmi Undamatla, Sumedha Avadhanula, Priya Nurkurthy, Vidhyadhari Methuku, Archana Bharadwaj Siva, Karthik Bharadwaj Tallapaka,Rakesh K Mishra,Divya Tej Sowpati |
| EPI_ISL_2775385, EPI_ISL_2775386, EPI_ISL_2775387, EPI_ISL_2775389 | AP SSO | CSIR-Centre for Cellular and Molecular Biology-INSACOG | Tulasi Nagabandi,Lamuk Zaveri,Payel Mukherjee,Onkar Kulkarni,Jandhyala Sai Krishna, Sofia Banu, Ara Sreenivas,Shreekant Verma, Amareshwar Vodapalli ,B Himasri,Valli Nagalakshmi Undamatla, Priya Nurkurthy,Sumedha Avadhanula, Vidhyadhari Methuku, Archana Bharadwaj Siva, Karthik Bharadwaj Tallapaka,Rakesh K Mishra,Divya Tej Sowpati |
| EPI_ISL_2775579, EPI_ISL_2775580, EPI_ISL_2775581, EPI_ISL_2775583, EPI_ISL_2775597, EPI_ISL_2775600, EPI_ISL_2775601, EPI_ISL_2775602, EPI_ISL_2775606, EPI_ISL_2775607, EPI_ISL_2775647, EPI_ISL_2775648, EPI_ISL_2775649, EPI_ISL_2775650, EPI_ISL_2775651, EPI_ISL_2775652, EPI_ISL_2775653, EPI_ISL_2775654, EPI_ISL_2775655, EPI_ISL_2775656, EPI_ISL_2775657, EPI_ISL_2775658, EPI_ISL_2775659, EPI_ISL_2775661, EPI_ISL_2775662, EPI_ISL_2775663, EPI_ISL_2775668, EPI_ISL_2775669, EPI_ISL_2775671 | | | |
| see above | Wyoming Public Health Laboratory | Wyoming Public Health Laboratory | Jim Mildenberger, Taylor Fearing, Channing Weber, Ashley Norberg, Chayse Rowley, Marley Goetz, Brian Dominguez, Elliot Thomasson, Sam Britz, Cari Sloma, Robert Petit, and Rob Christensen |
| EPI_ISL_2775676 | Pathogen Genomics Center, National Institute of Infectious Diseases | Pathogen Genomics Center, National Institute of Infectious Diseases | Tsuyoshi Sekizuka, Kentaro Itokawa, Rina Tanaka, Masanori Hashino, Makoto Kuroda |
| EPI_ISL_2775926 | Laboratorio de Biología Molecular. Servicio de Análisis Clínicos y Microbiología. Hospital de Sagunto | SeqCOVID-SPAIN consortium/IBV(CSIC) | Dra. Victoria Simón García, Gonzalo Llop Furquet, Agustín Irazo Tatay, Sandra Moreno Marro, Noelia Lozano Rodríguez and SeqCOVID-SPAIN consortium |
| EPI_ISL_2775929, EPI_ISL_2775935, EPI_ISL_2775936, EPI_ISL_2775937, EPI_ISL_2775942, EPI_ISL_2775949, EPI_ISL_2775951, EPI_ISL_2775953, EPI_ISL_2775954, EPI_ISL_2775957, EPI_ISL_2775964, EPI_ISL_2775967, EPI_ISL_2775969, EPI_ISL_2775975, EPI_ISL_2775977, EPI_ISL_2775979, EPI_ISL_2775981, EPI_ISL_2775982, EPI_ISL_2775983, EPI_ISL_2775984, EPI_ISL_2775985 | | | |
| see above | Hospital Universitari i Politècnic La Fe de València | SeqCOVID-SPAIN consortium/IBV(CSIC) | María Dolores Gómez Ruiz, Eva González Barberá, Ana Gil Brusola, Salvador Giner Almaraz, José Luis López Hontangas and SeqCOVID-SPAIN consortium |
| EPI_ISL_2775999, EPI_ISL_2776003, EPI_ISL_2776005, | Servicio de Microbiología. Hospital General Universitario de Castellón | SeqCOVID-SPAIN consortium/IBV(CSIC) | Rosario Moreno Muñoz, María Dolores Tirado Balaguer and SeqCOVID-SPAIN consortium |

| | | | |
|---|---|---|---|
| EPI_ISL_2776006, EPI_ISL_2776008, EPI_ISL_2776010 | | | |
| EPI_ISL_2776014, EPI_ISL_2776033, EPI_ISL_2776035, EPI_ISL_2776037, EPI_ISL_2776055, EPI_ISL_2776063, EPI_ISL_2776064, EPI_ISL_2776065, EPI_ISL_2776066 | Virginia Division of Consolidated Laboratory Services | Virginia Division of Consolidated Laboratory Services | Virginia Division of Consolidated Laboratory Services |
| EPI_ISL_2776076, EPI_ISL_2776078, EPI_ISL_2776081 | Texas Children's Hospital | Texas Children's Microbiome Center | Ruth Ann Luna, Jennifer K. Spinler, James Dunn, James Versalovic, Ila Singh |
| EPI_ISL_2776137, EPI_ISL_2776138, EPI_ISL_2776139, EPI_ISL_2776140, EPI_ISL_2776142, EPI_ISL_2776143, EPI_ISL_2776144, EPI_ISL_2776145, EPI_ISL_2776147 | Wyoming Public Health Laboratory | Wyoming Public Health Laboratory | Jim Mildenberger, Taylor Fearing, Channing Weber, Ashley Norberg, Chayse Rowley, Marley Goetz, Brian Dominguez, Elliot Thomasson, Sam Britz, Cari Sloma, Robert Petit, and Rob Christensen |
| EPI_ISL_2776149, EPI_ISL_2776150 | Wojewodzka Stacja Sanitarno-Epidemiologiczna w Olsztynie, Laboratorium Badan Epidemiologiczno-Klinicznych | Wojewodzka Stacja Sanitarno-Epidemiologiczna w Olsztynie, Laboratorium Badan Epidemiologiczno-Klinicznych | Sylwia Krzetowska, Monika Czerminska, Ewa Liszewska, Tomasz Jakubczak, Marta Lukian, Patryk Bielecki, Aleksandra Kobiatko, Emilia Tarabasz, Paulina Rozycka, Barbara Dolinska |
| EPI_ISL_2776369 | Ministry of Health, Jaber Al-Ahmad Hospital | Virology Unit, Department of Microbiology, Faulty of Medicine, Kuwait University | Nada Madi, Hussain Safar, Ebaa Al-Awadhi, Anfal Al-Adwani |
| EPI_ISL_2776430 | Ministry of Health, Jaber Al-Ahmad Hospital | Virology Unit, Department of Microbiology, Faculty of Medicine, Kuwait University | Nada Madi, Hussain Safar, Ebaa Al-Awadhi, Anfal Al-Adwani |
| EPI_ISL_2776663, EPI_ISL_2776664, EPI_ISL_2776899, EPI_ISL_2776922 | SARS-CoV-2 testing team, National Institute of Infectious Diseases | Pathogen Genomics Center, National Institute of Infectious Diseases | Tsuyoshi Sekizuka, Kentaro Itokawa, Rina Tanaka, Masanori Hashino, Nozomu Hanaoka, Masumichi Saito, Naomi Nojiri, Hazuka Y Furihata, Hiromizu Takahashi, Tsuguto Fujimoto, Makoto Kuroda |
| EPI_ISL_2777214, EPI_ISL_2777216, EPI_ISL_2777219 | Instituto Nacional de Enfermedades Respiratorias (INER) | Centro de Investigación en Enfermedades Infecciosas (CIENI), Instituto Nacional de Enfermedades Respiratorias (INER) | Consorcio Mexicano de Vigilancia Genómica (CoViGen-Mex). Authors (in alphabetical order): Julio Elias Alvarado-Yaah, Carlos F. Arias, Santiago Ávila-Ríos, Eduardo Becerril-Vargas, Víctor Hugo Borja-Aburto, Celia Boukadida, Cristóbal Cháidez-Quiróz, Juan Bautista Chale-Dzul, Ricardo Ciria Merce, Andreu Comas-García, Célida Duque Molina, Julissa Enciso-Ibarra, José Antonio Enciso-Moreno, Gloria Elena Espinosa-Ayala, Fernando Fontove-Herrera, Daniel Fregoso-Rueda, Victor Eduardo Garcia-Arias, Alejandra Garcia-Gasca, Bruno Gomez-Gil, Jean Pierre González, Irvin González-López, Concepción Grajales-Muñiz, Ricardo Grande, Rosa María Gutierrez Rios, Alejandra Hernández-Terán, Alfredo Herrera-Estrella, Carla Ivón Herrera-Najera, Pavel Isa, Daniel Lira Morales, Susana Lopez, Antonio Loza Román, Brenda Irasema Maldonado-Meza, Bernardo Martínez-Miguel, José Arturo Martínez-Orozco, Célida Martínez- Rodríguez, Margarita Matías-Florentino, Fidencio Mejía-Nepomuceno, María Guadalupe de Jesús Mireles-Rivera, Gloria María Molina-Salinas, Hector Montoya-Fuentes, Mario Mújica-Sánchez, José Esteban Muñoz-Medina, José de Jesús Nuñez-Contreras, Alicia Ocaña-Mondragón, Luis Alberto Ochoa-Carrera, Hector Esteban Paz-Juárez, Marissa Pérez-García, Francisco Pulido, Helen Haydee Fernanda Ramírez-Plascencia, Jorge Salas-Hernández, Angel Gustavo Salas-Lais, Alejandro Sanchez-Flores, Clara Esperanza Santacruz-Tinoco, María Guadalupe Santiago-Mauricio, Selene Zárate, Nelly Sélem-Mojica, Blanca Taboada, Kathia Elizabeth Tapia-Diaz, Gloria Vazquez, Joel Armando Vázquez-Pérez. |
| EPI_ISL_2777222, EPI_ISL_2777224, EPI_ISL_2777225, EPI_ISL_2777226, EPI_ISL_2777229, EPI_ISL_2777230, EPI_ISL_2777231, EPI_ISL_2777232, EPI_ISL_2777233 | Instituto Nacional de Cancerología | Centro de Investigación en Enfermedades Infecciosas (CIENI), Instituto Nacional de Enfermedades Respiratorias (INER) | Consorcio Mexicano de Vigilancia Genómica (CoViGen-Mex). Authors (in alphabetical order): Julio Elias Alvarado-Yaah, Carlos F. Arias, Santiago Ávila-Ríos, Eduardo Becerril-Vargas, Víctor Hugo Borja-Aburto, Celia Boukadida, Cristóbal Cháidez-Quiróz, Juan Bautista Chale-Dzul, Ricardo Ciria Merce, Andreu Comas-García, Célida Duque Molina, Julissa Enciso-Ibarra, José Antonio Enciso-Moreno, Gloria Elena Espinosa-Ayala, Fernando Fontove-Herrera, Daniel Fregoso-Rueda, Victor Eduardo Garcia-Arias, Alejandra Garcia-Gasca, Bruno Gomez-Gil, Jean Pierre González, Irvin González-López, Concepción Grajales-Muñiz, Ricardo Grande, Rosa María Gutierrez Rios, Alejandra Hernández-Terán, Alfredo Herrera-Estrella, Carla Ivón Herrera-Najera, Pavel Isa, Daniel Lira Morales, Susana Lopez, Antonio Loza Román, Brenda Irasema Maldonado-Meza, Bernardo Martínez-Miguel, José Arturo Martínez-Orozco, Célida Martínez- Rodríguez, Margarita Matías-Florentino, Fidencio Mejía-Nepomuceno, María Guadalupe de Jesús Mireles-Rivera, Gloria María Molina-Salinas, Hector Montoya-Fuentes, Mario Mújica-Sánchez, José Esteban Muñoz-Medina, José de Jesús Nuñez-Contreras, Alicia Ocaña-Mondragón, Luis Alberto Ochoa-Carrera, Hector Esteban Paz-Juárez, Marissa Pérez-García, Francisco Pulido, Helen Haydee Fernanda Ramírez-Plascencia, Jorge Salas-Hernández, Angel Gustavo Salas-Lais, Alejandro Sanchez-Flores, Clara Esperanza Santacruz-Tinoco, María Guadalupe Santiago-Mauricio, Selene Zárate, Nelly Sélem-Mojica, Blanca Taboada, Kathia Elizabeth Tapia-Diaz, Gloria Vazquez, Joel Armando Vázquez-Pérez. |
| EPI_ISL_2777291 | A.S.L. NOVARA-BORGOMANERO | Fondazione del Piemonte per l'Oncologia IRCCS | Antonino Sottile, Silvia Brossa, Paola Marino, Giorgio Giardina |
| EPI_ISL_2778017 | Instituto Nacional de Medicina Genomica | Centro de Investigación en Enfermedades Infecciosas (CIENI), Instituto Nacional de Enfermedades Respiratorias (INER) | Matias-Florentino M, Perez-Garcia M, Hidalgo-Miranda A, Mendoza-Vargas A, Reyes-Grajeda JP, Cedro-Tanda A, Herrera-Montalvo LA, Reyes-Teran G, Avila-Rios S, Boukadida C |
| EPI_ISL_2778018, EPI_ISL_2778019, EPI_ISL_2778020, EPI_ISL_2778022, EPI_ISL_2778024, EPI_ISL_2778025, EPI_ISL_2778026, EPI_ISL_2778028, EPI_ISL_2778029, EPI_ISL_2778032, EPI_ISL_2778037, EPI_ISL_2778040, EPI_ISL_2778041, EPI_ISL_2778047, EPI_ISL_2778048, EPI_ISL_2778051, EPI_ISL_2778053, EPI_ISL_2778055, EPI_ISL_2778057, EPI_ISL_2778063, EPI_ISL_2778064, EPI_ISL_2778066, EPI_ISL_2778069, EPI_ISL_2778071, EPI_ISL_2778072, EPI_ISL_2778074, EPI_ISL_2778079, EPI_ISL_2778081, EPI_ISL_2778082, EPI_ISL_2778090 | | | |
| see above | Cliniques universitaires Saint-Luc | UCLouvain/IREC/MBLG | Jean Ruelle, Bertrand Bearzatto, Benoit Kabamba Mukadi |
| EPI_ISL_2778098 | Clinique Saint-Pierre Ottignies | UCLouvain/IREC/MBLG | Jean Ruelle, Bertrand Bearzatto, Benoit Kabamba Mukadi |
| EPI_ISL_2778814, EPI_ISL_2778817, EPI_ISL_2778831, EPI_ISL_2778833, EPI_ISL_2778873, EPI_ISL_2778886, EPI_ISL_2778903, EPI_ISL_2778937, EPI_ISL_2778944, EPI_ISL_2778951 | California Department of Public Health Valencia Branch Laboratory (CDPH VBL) | California Department of Public Health | CDPH-COVIDNet,UCLA Technology Center for Genomics & Bioinformatics |
| EPI_ISL_2778967 | Instituto Nacional de Medicina Genomica | Instituto Nacional de Medicina Genomica | Cedro-Tanda A, Hidalgo-Miranda A, Cisneros-Villanueva M, Orjuela-Rodríguez M, Mendoza-Vargas A, Reyes-Grajeda JP, Abraham Campos-Romero, Moreno-Camacho José Luis, Rodriguez-Gallegos Jorge, Luna-Ruiz Marco, Gonzalez-Barrera D, Rangel-DeLeon D, Munguia-Garza P, Ramirez-Vega O, Escobar-Arrazola, M, Herrera-Montalvo LA. |
| EPI_ISL_2778969, EPI_ISL_2778971, EPI_ISL_2778976, EPI_ISL_2778977 | Humboldt County Public Health Laboratory | Humboldt County Public Health Laboratory | Jeremy Corrigan |
| EPI_ISL_2779640 | Lighthouse Lab in Glasgow | Wellcome Sanger Institute for the COVID-19 Genomics UK (COG-UK) Consortium | Harper VanSteenhouse, Yumi Kasai, David Gray, Carol Clugston, Anna Dominiczak and Alex Alderton, Roberto Amato, Jeffrey Barrett, Sonia Goncalves, Ewan Harrison, David K. Jackson, Ian Johnston, Dominic Kwiatkowski, Cordelia Langford, John Sillitoe on behalf of the Wellcome Sanger Institute COVID-19 Surveillance Team |
| EPI_ISL_2779641, EPI_ISL_2779642, EPI_ISL_2779643 | Health Services Laboratories | Wellcome Sanger Institute for the COVID-19 Genomics UK (COG-UK) Consortium | Health Services Laboratories and Alex Alderton, Roberto Amato, Jeffrey Barrett, Sonia Goncalves, Ewan Harrison, David K. Jackson, Ian Johnston, Dominic Kwiatkowski, Cordelia Langford, John Sillitoe on behalf of the Wellcome Sanger Institute COVID-19 Surveillance Team |
| EPI_ISL_2779644 | Lighthouse Lab in Milton Keynes | Wellcome Sanger Institute for the COVID-19 Genomics UK (COG-UK) Consortium | The Lighthouse Lab in Milton Keynes and Alex Alderton, Roberto Amato, Jeffrey Barrett, Sonia Goncalves, Ewan Harrison, David K. Jackson, Ian Johnston, Dominic Kwiatkowski, Cordelia Langford, John Sillitoe on behalf of the Wellcome Sanger Institute COVID-19 Surveillance Team |
| EPI_ISL_2779645, EPI_ISL_2779646, EPI_ISL_2779647, EPI_ISL_2779648, EPI_ISL_2779649 | Health Services Laboratories | Wellcome Sanger Institute for the COVID-19 Genomics UK (COG-UK) Consortium | Health Services Laboratories and Alex Alderton, Roberto Amato, Jeffrey Barrett, Sonia Goncalves, Ewan Harrison, David K. Jackson, Ian Johnston, Dominic Kwiatkowski, Cordelia Langford, John Sillitoe on behalf of the Wellcome Sanger Institute COVID-19 Surveillance Team |
| EPI_ISL_2779650, EPI_ISL_2779651, EPI_ISL_2779652 | Lighthouse Lab in Glasgow | Wellcome Sanger Institute for the COVID-19 Genomics UK (COG-UK) Consortium | Harper VanSteenhouse, Yumi Kasai, David Gray, Carol Clugston, Anna Dominiczak and Alex Alderton, Roberto Amato, Jeffrey Barrett, Sonia Goncalves, Ewan Harrison, David K. Jackson, Ian Johnston, Dominic Kwiatkowski, Cordelia Langford, John Sillitoe on behalf of the Wellcome Sanger Institute COVID-19 Surveillance Team |
| EPI_ISL_2779653 | Health Services Laboratories | Wellcome Sanger Institute for the COVID-19 Genomics UK (COG-UK) Consortium | Health Services Laboratories and Alex Alderton, Roberto Amato, Jeffrey Barrett, Sonia Goncalves, Ewan Harrison, David K. Jackson, Ian Johnston, Dominic Kwiatkowski, Cordelia Langford, John Sillitoe on behalf of the Wellcome Sanger Institute COVID-19 Surveillance Team |
| EPI_ISL_2779654 | Lighthouse Lab in Milton Keynes | Wellcome Sanger Institute for the COVID-19 Genomics UK (COG-UK) Consortium | The Lighthouse Lab in Milton Keynes and Alex Alderton, Roberto Amato, Jeffrey Barrett, Sonia Goncalves, Ewan Harrison, David K. Jackson, Ian Johnston, Dominic Kwiatkowski, Cordelia Langford, John Sillitoe on behalf of the Wellcome Sanger Institute COVID-19 Surveillance Team |
| EPI_ISL_2779655 | Health Services Laboratories | Wellcome Sanger Institute for the COVID-19 Genomics UK (COG-UK) Consortium | Health Services Laboratories and Alex Alderton, Roberto Amato, Jeffrey Barrett, Sonia Goncalves, Ewan Harrison, David K. Jackson, Ian Johnston, Dominic Kwiatkowski, Cordelia Langford, John Sillitoe on behalf of the Wellcome Sanger Institute COVID-19 Surveillance Team |
| EPI_ISL_2779656, EPI_ISL_2779657, EPI_ISL_2779658 | Lighthouse Lab in Milton Keynes | Wellcome Sanger Institute for the COVID-19 Genomics UK (COG-UK) Consortium | The Lighthouse Lab in Milton Keynes and Alex Alderton, Roberto Amato, Jeffrey Barrett, Sonia Goncalves, Ewan Harrison, David K. Jackson, Ian Johnston, Dominic Kwiatkowski, Cordelia Langford, John Sillitoe on behalf of the Wellcome Sanger Institute COVID-19 Surveillance Team |

| | | | |
|---|---|---|---|
| EPI_ISL_2779659 | Lighthouse Lab in Glasgow | Wellcome Sanger Institute for the COVID-19 Genomics UK (COG-UK) Consortium | Harper VanSteenhouse, Yumi Kasai, David Gray, Carol Clugston, Anna Dominiczak and Alex Alderton, Roberto Amato, Jeffrey Barrett, Sonia Goncalves, Ewan Harrison, David K. Jackson, Ian Johnston, Dominic Kwiatkowski, Cordelia Langford, John Sillitoe on behalf of the Wellcome Sanger Institute COVID-19 Surveillance Team |
| EPI_ISL_2779661 | Lighthouse Lab in Milton Keynes | Wellcome Sanger Institute for the COVID-19 Genomics UK (COG-UK) Consortium | The Lighthouse Lab in Milton Keynes and Alex Alderton, Roberto Amato, Jeffrey Barrett, Sonia Goncalves, Ewan Harrison, David K. Jackson, Ian Johnston, Dominic Kwiatkowski, Cordelia Langford, John Sillitoe on behalf of the Wellcome Sanger Institute COVID-19 Surveillance Team |
| EPI_ISL_2779662, EPI_ISL_2779663, EPI_ISL_2779664 | Health Services Laboratories | Wellcome Sanger Institute for the COVID-19 Genomics UK (COG-UK) Consortium | Health Services Laboratories and Alex Alderton, Roberto Amato, Jeffrey Barrett, Sonia Goncalves, Ewan Harrison, David K. Jackson, Ian Johnston, Dominic Kwiatkowski, Cordelia Langford, John Sillitoe on behalf of the Wellcome Sanger Institute COVID-19 Surveillance Team |
| EPI_ISL_2779665 | Lighthouse Lab in Milton Keynes | Wellcome Sanger Institute for the COVID-19 Genomics UK (COG-UK) Consortium | The Lighthouse Lab in Milton Keynes and Alex Alderton, Roberto Amato, Jeffrey Barrett, Sonia Goncalves, Ewan Harrison, David K. Jackson, Ian Johnston, Dominic Kwiatkowski, Cordelia Langford, John Sillitoe on behalf of the Wellcome Sanger Institute COVID-19 Surveillance Team |
| EPI_ISL_2779666 | Health Services Laboratories | Wellcome Sanger Institute for the COVID-19 Genomics UK (COG-UK) Consortium | Health Services Laboratories and Alex Alderton, Roberto Amato, Jeffrey Barrett, Sonia Goncalves, Ewan Harrison, David K. Jackson, Ian Johnston, Dominic Kwiatkowski, Cordelia Langford, John Sillitoe on behalf of the Wellcome Sanger Institute COVID-19 Surveillance Team |
| EPI_ISL_2779667, EPI_ISL_2779668, EPI_ISL_2779669 | Lighthouse Lab in Milton Keynes | Wellcome Sanger Institute for the COVID-19 Genomics UK (COG-UK) Consortium | The Lighthouse Lab in Milton Keynes and Alex Alderton, Roberto Amato, Jeffrey Barrett, Sonia Goncalves, Ewan Harrison, David K. Jackson, Ian Johnston, Dominic Kwiatkowski, Cordelia Langford, John Sillitoe on behalf of the Wellcome Sanger Institute COVID-19 Surveillance Team |
| EPI_ISL_2779670, EPI_ISL_2779671 | Health Services Laboratories | Wellcome Sanger Institute for the COVID-19 Genomics UK (COG-UK) Consortium | Health Services Laboratories and Alex Alderton, Roberto Amato, Jeffrey Barrett, Sonia Goncalves, Ewan Harrison, David K. Jackson, Ian Johnston, Dominic Kwiatkowski, Cordelia Langford, John Sillitoe on behalf of the Wellcome Sanger Institute COVID-19 Surveillance Team |
| EPI_ISL_2779672 | Lighthouse Lab in Glasgow | Wellcome Sanger Institute for the COVID-19 Genomics UK (COG-UK) Consortium | Harper VanSteenhouse, Yumi Kasai, David Gray, Carol Clugston, Anna Dominiczak and Alex Alderton, Roberto Amato, Jeffrey Barrett, Sonia Goncalves, Ewan Harrison, David K. Jackson, Ian Johnston, Dominic Kwiatkowski, Cordelia Langford, John Sillitoe on behalf of the Wellcome Sanger Institute COVID-19 Surveillance Team |
| EPI_ISL_2779673, EPI_ISL_2779674, EPI_ISL_2779675 | Health Services Laboratories | Wellcome Sanger Institute for the COVID-19 Genomics UK (COG-UK) Consortium | Health Services Laboratories and Alex Alderton, Roberto Amato, Jeffrey Barrett, Sonia Goncalves, Ewan Harrison, David K. Jackson, Ian Johnston, Dominic Kwiatkowski, Cordelia Langford, John Sillitoe on behalf of the Wellcome Sanger Institute COVID-19 Surveillance Team |
| EPI_ISL_2779676 | Lighthouse Lab in Milton Keynes | Wellcome Sanger Institute for the COVID-19 Genomics UK (COG-UK) Consortium | The Lighthouse Lab in Milton Keynes and Alex Alderton, Roberto Amato, Jeffrey Barrett, Sonia Goncalves, Ewan Harrison, David K. Jackson, Ian Johnston, Dominic Kwiatkowski, Cordelia Langford, John Sillitoe on behalf of the Wellcome Sanger Institute COVID-19 Surveillance Team |
| EPI_ISL_2779677 | Lighthouse Lab in Glasgow | Wellcome Sanger Institute for the COVID-19 Genomics UK (COG-UK) Consortium | Harper VanSteenhouse, Yumi Kasai, David Gray, Carol Clugston, Anna Dominiczak and Alex Alderton, Roberto Amato, Jeffrey Barrett, Sonia Goncalves, Ewan Harrison, David K. Jackson, Ian Johnston, Dominic Kwiatkowski, Cordelia Langford, John Sillitoe on behalf of the Wellcome Sanger Institute COVID-19 Surveillance Team |
| EPI_ISL_2779678, EPI_ISL_2779679 | Lighthouse Lab in Milton Keynes | Wellcome Sanger Institute for the COVID-19 Genomics UK (COG-UK) Consortium | The Lighthouse Lab in Milton Keynes and Alex Alderton, Roberto Amato, Jeffrey Barrett, Sonia Goncalves, Ewan Harrison, David K. Jackson, Ian Johnston, Dominic Kwiatkowski, Cordelia Langford, John Sillitoe on behalf of the Wellcome Sanger Institute COVID-19 Surveillance Team |
| EPI_ISL_2779680, EPI_ISL_2779681 | Lighthouse Lab in Glasgow | Wellcome Sanger Institute for the COVID-19 Genomics UK (COG-UK) Consortium | Harper VanSteenhouse, Yumi Kasai, David Gray, Carol Clugston, Anna Dominiczak and Alex Alderton, Roberto Amato, Jeffrey Barrett, Sonia Goncalves, Ewan Harrison, David K. Jackson, Ian Johnston, Dominic Kwiatkowski, Cordelia Langford, John Sillitoe on behalf of the Wellcome Sanger Institute COVID-19 Surveillance Team |
| EPI_ISL_2779682, EPI_ISL_2779683, EPI_ISL_2779684, EPI_ISL_2779685 | Health Services Laboratories | Wellcome Sanger Institute for the COVID-19 Genomics UK (COG-UK) Consortium | Health Services Laboratories and Alex Alderton, Roberto Amato, Jeffrey Barrett, Sonia Goncalves, Ewan Harrison, David K. Jackson, Ian Johnston, Dominic Kwiatkowski, Cordelia Langford, John Sillitoe on behalf of the Wellcome Sanger Institute COVID-19 Surveillance Team |
| EPI_ISL_2779686 | Lighthouse Lab in Glasgow | Wellcome Sanger Institute for the COVID-19 Genomics UK (COG-UK) Consortium | Harper VanSteenhouse, Yumi Kasai, David Gray, Carol Clugston, Anna Dominiczak and Alex Alderton, Roberto Amato, Jeffrey Barrett, Sonia Goncalves, Ewan Harrison, David K. Jackson, Ian Johnston, Dominic Kwiatkowski, Cordelia Langford, John Sillitoe on behalf of the Wellcome Sanger Institute COVID-19 Surveillance Team |
| EPI_ISL_2779687 | Lighthouse Lab in Milton Keynes | Wellcome Sanger Institute for the COVID-19 Genomics UK (COG-UK) Consortium | The Lighthouse Lab in Milton Keynes and Alex Alderton, Roberto Amato, Jeffrey Barrett, Sonia Goncalves, Ewan Harrison, David K. Jackson, Ian Johnston, Dominic Kwiatkowski, Cordelia Langford, John Sillitoe on behalf of the Wellcome Sanger Institute COVID-19 Surveillance Team |
| EPI_ISL_2779688 | Lighthouse Lab in Glasgow | Wellcome Sanger Institute for the COVID-19 Genomics UK (COG-UK) Consortium | Harper VanSteenhouse, Yumi Kasai, David Gray, Carol Clugston, Anna Dominiczak and Alex Alderton, Roberto Amato, Jeffrey Barrett, Sonia Goncalves, Ewan Harrison, David K. Jackson, Ian Johnston, Dominic Kwiatkowski, Cordelia Langford, John Sillitoe on behalf of the Wellcome Sanger Institute COVID-19 Surveillance Team |
| EPI_ISL_2779689, EPI_ISL_2779690 | Lighthouse Lab in Milton Keynes | Wellcome Sanger Institute for the COVID-19 Genomics UK (COG-UK) Consortium | The Lighthouse Lab in Milton Keynes and Alex Alderton, Roberto Amato, Jeffrey Barrett, Sonia Goncalves, Ewan Harrison, David K. Jackson, Ian Johnston, Dominic Kwiatkowski, Cordelia Langford, John Sillitoe on behalf of the Wellcome Sanger Institute COVID-19 Surveillance Team |
| EPI_ISL_2779691 | Lighthouse Lab in Glasgow | Wellcome Sanger Institute for the COVID-19 Genomics UK (COG-UK) Consortium | Harper VanSteenhouse, Yumi Kasai, David Gray, Carol Clugston, Anna Dominiczak and Alex Alderton, Roberto Amato, Jeffrey Barrett, Sonia Goncalves, Ewan Harrison, David K. Jackson, Ian Johnston, Dominic Kwiatkowski, Cordelia Langford, John Sillitoe on behalf of the Wellcome Sanger Institute COVID-19 Surveillance Team |
| EPI_ISL_2779692 | Health Services Laboratories | Wellcome Sanger Institute for the COVID-19 Genomics UK (COG-UK) Consortium | Health Services Laboratories and Alex Alderton, Roberto Amato, Jeffrey Barrett, Sonia Goncalves, Ewan Harrison, David K. Jackson, Ian Johnston, Dominic Kwiatkowski, Cordelia Langford, John Sillitoe on behalf of the Wellcome Sanger Institute COVID-19 Surveillance Team |
| EPI_ISL_2779693, EPI_ISL_2779694 | Lighthouse Lab in Glasgow | Wellcome Sanger Institute for the COVID-19 Genomics UK (COG-UK) Consortium | Harper VanSteenhouse, Yumi Kasai, David Gray, Carol Clugston, Anna Dominiczak and Alex Alderton, Roberto Amato, Jeffrey Barrett, Sonia Goncalves, Ewan Harrison, David K. Jackson, Ian Johnston, Dominic Kwiatkowski, Cordelia Langford, John Sillitoe on behalf of the Wellcome Sanger Institute COVID-19 Surveillance Team |
| EPI_ISL_2779695 | Lighthouse Lab in Milton Keynes | Wellcome Sanger Institute for the COVID-19 Genomics UK (COG-UK) Consortium | The Lighthouse Lab in Milton Keynes and Alex Alderton, Roberto Amato, Jeffrey Barrett, Sonia Goncalves, Ewan Harrison, David K. Jackson, Ian Johnston, Dominic Kwiatkowski, Cordelia Langford, John Sillitoe on behalf of the Wellcome Sanger Institute COVID-19 Surveillance Team |
| EPI_ISL_2779696 | Lighthouse Lab in Glasgow | Wellcome Sanger Institute for the COVID-19 Genomics UK (COG-UK) Consortium | Harper VanSteenhouse, Yumi Kasai, David Gray, Carol Clugston, Anna Dominiczak and Alex Alderton, Roberto Amato, Jeffrey Barrett, Sonia Goncalves, Ewan Harrison, David K. Jackson, Ian Johnston, Dominic Kwiatkowski, Cordelia Langford, John Sillitoe on behalf of the Wellcome Sanger Institute COVID-19 Surveillance Team |
| EPI_ISL_2779697, EPI_ISL_2779698 | Lighthouse Lab in Milton Keynes | Wellcome Sanger Institute for the COVID-19 Genomics UK (COG-UK) Consortium | The Lighthouse Lab in Milton Keynes and Alex Alderton, Roberto Amato, Jeffrey Barrett, Sonia Goncalves, Ewan Harrison, David K. Jackson, Ian Johnston, Dominic Kwiatkowski, Cordelia Langford, John Sillitoe on behalf of the Wellcome Sanger Institute COVID-19 Surveillance Team |
| EPI_ISL_2779699 | Health Services Laboratories | Wellcome Sanger Institute for the COVID-19 Genomics UK (COG-UK) Consortium | Health Services Laboratories and Alex Alderton, Roberto Amato, Jeffrey Barrett, Sonia Goncalves, Ewan Harrison, David K. Jackson, Ian Johnston, Dominic Kwiatkowski, Cordelia Langford, John Sillitoe on behalf of the Wellcome Sanger Institute COVID-19 Surveillance Team |
| EPI_ISL_2779700 | Lighthouse Lab in Milton Keynes | Wellcome Sanger Institute for the COVID-19 Genomics UK (COG-UK) Consortium | The Lighthouse Lab in Milton Keynes and Alex Alderton, Roberto Amato, Jeffrey Barrett, Sonia Goncalves, Ewan Harrison, David K. Jackson, Ian Johnston, Dominic Kwiatkowski, Cordelia Langford, John Sillitoe on behalf of the Wellcome Sanger Institute COVID-19 Surveillance Team |
| EPI_ISL_2779701 | Lighthouse Lab in Glasgow | Wellcome Sanger Institute for the COVID-19 Genomics UK (COG-UK) Consortium | Harper VanSteenhouse, Yumi Kasai, David Gray, Carol Clugston, Anna Dominiczak and Alex Alderton, Roberto Amato, Jeffrey Barrett, Sonia Goncalves, Ewan Harrison, David K. Jackson, Ian Johnston, Dominic Kwiatkowski, Cordelia Langford, John Sillitoe on behalf of the Wellcome Sanger Institute COVID-19 Surveillance Team |
| EPI_ISL_2779702, EPI_ISL_2779703 | Lighthouse Lab in Milton Keynes | Wellcome Sanger Institute for the COVID-19 Genomics UK (COG-UK) Consortium | The Lighthouse Lab in Milton Keynes and Alex Alderton, Roberto Amato, Jeffrey Barrett, Sonia Goncalves, Ewan Harrison, David K. Jackson, Ian Johnston, Dominic Kwiatkowski, Cordelia Langford, John Sillitoe on behalf of the Wellcome Sanger Institute COVID-19 Surveillance Team |
| EPI_ISL_2779704 | Health Services Laboratories | Wellcome Sanger Institute for the COVID-19 Genomics UK (COG-UK) Consortium | Health Services Laboratories and Alex Alderton, Roberto Amato, Jeffrey Barrett, Sonia Goncalves, Ewan Harrison, David K. Jackson, Ian Johnston, Dominic Kwiatkowski, Cordelia Langford, John Sillitoe on behalf of the Wellcome Sanger Institute COVID-19 Surveillance Team |
| EPI_ISL_2779705 | Lighthouse Lab in Glasgow | Wellcome Sanger Institute for the COVID-19 Genomics UK (COG-UK) Consortium | Harper VanSteenhouse, Yumi Kasai, David Gray, Carol Clugston, Anna Dominiczak and Alex Alderton, Roberto Amato, Jeffrey Barrett, Sonia Goncalves, Ewan Harrison, David K. Jackson, Ian Johnston, Dominic Kwiatkowski, Cordelia Langford, John Sillitoe on behalf of the Wellcome Sanger Institute COVID-19 Surveillance Team |
| EPI_ISL_2779706, EPI_ISL_2779707, EPI_ISL_2779708 | Lighthouse Lab in Milton Keynes | Wellcome Sanger Institute for the COVID-19 Genomics UK (COG-UK) Consortium | The Lighthouse Lab in Milton Keynes and Alex Alderton, Roberto Amato, Jeffrey Barrett, Sonia Goncalves, Ewan Harrison, David K. Jackson, Ian Johnston, Dominic Kwiatkowski, Cordelia Langford, John Sillitoe on behalf of the Wellcome Sanger Institute COVID-19 Surveillance Team |
| EPI_ISL_2779709, EPI_ISL_2779710 | Health Services Laboratories | Wellcome Sanger Institute for the COVID-19 Genomics UK (COG-UK) Consortium | Health Services Laboratories and Alex Alderton, Roberto Amato, Jeffrey Barrett, Sonia Goncalves, Ewan Harrison, David K. Jackson, Ian Johnston, Dominic Kwiatkowski, Cordelia Langford, John Sillitoe on behalf of the Wellcome Sanger Institute COVID-19 Surveillance Team |
| EPI_ISL_2779711 | Lighthouse Lab in Glasgow | Wellcome Sanger Institute for the COVID-19 Genomics UK (COG-UK) Consortium | Harper VanSteenhouse, Yumi Kasai, David Gray, Carol Clugston, Anna Dominiczak and Alex Alderton, Roberto Amato, Jeffrey Barrett, Sonia Goncalves, Ewan Harrison, David K. Jackson, Ian Johnston, Dominic Kwiatkowski, Cordelia Langford, John Sillitoe on behalf of the Wellcome Sanger Institute COVID-19 Surveillance Team |
| EPI_ISL_2779712 | Lighthouse Lab in Milton Keynes | Wellcome Sanger Institute for the COVID-19 Genomics UK (COG-UK) Consortium | The Lighthouse Lab in Milton Keynes and Alex Alderton, Roberto Amato, Jeffrey Barrett, Sonia Goncalves, Ewan Harrison, David K. Jackson, Ian Johnston, Dominic Kwiatkowski, Cordelia Langford, John Sillitoe on behalf of the Wellcome Sanger Institute COVID-19 Surveillance Team |
| EPI_ISL_2779713 | Lighthouse Lab in Glasgow | Wellcome Sanger Institute for the COVID-19 Genomics UK (COG-UK) Consortium | Harper VanSteenhouse, Yumi Kasai, David Gray, Carol Clugston, Anna Dominiczak and Alex Alderton, Roberto Amato, Jeffrey Barrett, Sonia Goncalves, Ewan Harrison, David K. Jackson, Ian Johnston, Dominic Kwiatkowski, Cordelia Langford, John Sillitoe on behalf of the Wellcome Sanger Institute COVID-19 Surveillance Team |
| EPI_ISL_2779714 | Lighthouse Lab in Milton Keynes | Wellcome Sanger Institute for the COVID-19 Genomics UK (COG-UK) Consortium | The Lighthouse Lab in Milton Keynes and Alex Alderton, Roberto Amato, Jeffrey Barrett, Sonia Goncalves, Ewan Harrison, David K. Jackson, Ian Johnston, Dominic Kwiatkowski, Cordelia Langford, John Sillitoe on behalf of the Wellcome Sanger Institute COVID-19 Surveillance Team |
| EPI_ISL_2779715, EPI_ISL_2779716 | Lighthouse Lab in Glasgow | Wellcome Sanger Institute for the COVID-19 Genomics UK (COG-UK) Consortium | Harper VanSteenhouse, Yumi Kasai, David Gray, Carol Clugston, Anna Dominiczak and Alex Alderton, Roberto Amato, Jeffrey Barrett, Sonia Goncalves, Ewan Harrison, David K. Jackson, Ian Johnston, Dominic Kwiatkowski, Cordelia Langford, John Sillitoe on behalf of the Wellcome Sanger Institute COVID-19 Surveillance Team |
| EPI_ISL_2779717 | Lighthouse Lab in Milton Keynes | Wellcome Sanger Institute for the COVID-19 Genomics UK (COG-UK) Consortium | The Lighthouse Lab in Milton Keynes and Alex Alderton, Roberto Amato, Jeffrey Barrett, Sonia Goncalves, Ewan Harrison, David K. Jackson, Ian Johnston, Dominic Kwiatkowski, Cordelia Langford, John Sillitoe on behalf of the Wellcome Sanger Institute COVID-19 Surveillance Team |
| EPI_ISL_2779718 | Lighthouse Lab in Glasgow | Wellcome Sanger Institute for the COVID-19 Genomics UK (COG-UK) Consortium | Harper VanSteenhouse, Yumi Kasai, David Gray, Carol Clugston, Anna Dominiczak and Alex Alderton, Roberto Amato, Jeffrey Barrett, Sonia Goncalves, Ewan Harrison, David K. Jackson, Ian Johnston, Dominic Kwiatkowski, Cordelia Langford, John Sillitoe on behalf of the Wellcome Sanger Institute COVID-19 Surveillance Team |
| EPI_ISL_2779719, EPI_ISL_2779720, EPI_ISL_2779721 | Health Services Laboratories | Wellcome Sanger Institute for the COVID-19 Genomics UK (COG-UK) Consortium | Health Services Laboratories and Alex Alderton, Roberto Amato, Jeffrey Barrett, Sonia Goncalves, Ewan Harrison, David K. Jackson, Ian Johnston, Dominic Kwiatkowski, Cordelia Langford, John Sillitoe on behalf of the Wellcome Sanger Institute COVID-19 Surveillance Team |
| EPI_ISL_2779722 | Lighthouse Lab in Milton Keynes | Wellcome Sanger Institute for the COVID-19 Genomics UK (COG-UK) Consortium | The Lighthouse Lab in Milton Keynes and Alex Alderton, Roberto Amato, Jeffrey Barrett, Sonia Goncalves, Ewan Harrison, David K. Jackson, Ian Johnston, Dominic Kwiatkowski, Cordelia Langford, John Sillitoe on behalf of the Wellcome Sanger Institute COVID-19 Surveillance Team |
| EPI_ISL_2779723 | Health Services Laboratories | Wellcome Sanger Institute for the COVID-19 Genomics UK (COG-UK) Consortium | Health Services Laboratories and Alex Alderton, Roberto Amato, Jeffrey Barrett, Sonia Goncalves, Ewan Harrison, David K. Jackson, Ian Johnston, Dominic Kwiatkowski, Cordelia Langford, John Sillitoe on behalf of the Wellcome Sanger Institute COVID-19 Surveillance Team |
| EPI_ISL_2779724 | Lighthouse Lab in Glasgow | Wellcome Sanger Institute for the COVID-19 Genomics UK (COG-UK) | Harper VanSteenhouse, Yumi Kasai, David Gray, Carol Clugston, Anna Dominiczak and Alex Alderton, Roberto Amato, Jeffrey Barrett, Sonia Goncalves, Ewan Harrison, David K. Jackson, Ian Johnston, Dominic Kwiatkowski, Cordelia Langford, |

| | | | |
|---|---|---|---|
| | | Consortium | John Sillitoe on behalf of the Wellcome Sanger Institute COVID-19 Surveillance Team |
| EPI_ISL_2779725, EPI_ISL_2779726 | Health Services Laboratories | Wellcome Sanger Institute for the COVID-19 Genomics UK (COG-UK) Consortium | Health Services Laboratories and Alex Alderton, Roberto Amato, Jeffrey Barrett, Sonia Goncalves, Ewan Harrison, David K. Jackson, Ian Johnston, Dominic Kwiatkowski, Cordelia Langford, John Sillitoe on behalf of the Wellcome Sanger Institute COVID-19 Surveillance Team |
| EPI_ISL_2779727, EPI_ISL_2779728 | Lighthouse Lab in Glasgow | Wellcome Sanger Institute for the COVID-19 Genomics UK (COG-UK) Consortium | Harper VanSteenhouse, Yumi Kasai, David Gray, Carol Clugston, Anna Dominiczak and Alex Alderton, Roberto Amato, Jeffrey Barrett, Sonia Goncalves, Ewan Harrison, David K. Jackson, Ian Johnston, Dominic Kwiatkowski, Cordelia Langford, John Sillitoe on behalf of the Wellcome Sanger Institute COVID-19 Surveillance Team |
| EPI_ISL_2779729 | Lighthouse Lab in Milton Keynes | Wellcome Sanger Institute for the COVID-19 Genomics UK (COG-UK) Consortium | The Lighthouse Lab in Milton Keynes and Alex Alderton, Roberto Amato, Jeffrey Barrett, Sonia Goncalves, Ewan Harrison, David K. Jackson, Ian Johnston, Dominic Kwiatkowski, Cordelia Langford, John Sillitoe on behalf of the Wellcome Sanger Institute COVID-19 Surveillance Team |
| EPI_ISL_2779730 | Lighthouse Lab in Glasgow | Wellcome Sanger Institute for the COVID-19 Genomics UK (COG-UK) Consortium | Harper VanSteenhouse, Yumi Kasai, David Gray, Carol Clugston, Anna Dominiczak and Alex Alderton, Roberto Amato, Jeffrey Barrett, Sonia Goncalves, Ewan Harrison, David K. Jackson, Ian Johnston, Dominic Kwiatkowski, Cordelia Langford, John Sillitoe on behalf of the Wellcome Sanger Institute COVID-19 Surveillance Team |
| EPI_ISL_2779731 | Lighthouse Lab in Milton Keynes | Wellcome Sanger Institute for the COVID-19 Genomics UK (COG-UK) Consortium | The Lighthouse Lab in Milton Keynes and Alex Alderton, Roberto Amato, Jeffrey Barrett, Sonia Goncalves, Ewan Harrison, David K. Jackson, Ian Johnston, Dominic Kwiatkowski, Cordelia Langford, John Sillitoe on behalf of the Wellcome Sanger Institute COVID-19 Surveillance Team |
| EPI_ISL_2779732, EPI_ISL_2779733, EPI_ISL_2779734 | Health Services Laboratories | Wellcome Sanger Institute for the COVID-19 Genomics UK (COG-UK) Consortium | Health Services Laboratories and Alex Alderton, Roberto Amato, Jeffrey Barrett, Sonia Goncalves, Ewan Harrison, David K. Jackson, Ian Johnston, Dominic Kwiatkowski, Cordelia Langford, John Sillitoe on behalf of the Wellcome Sanger Institute COVID-19 Surveillance Team |
| EPI_ISL_2779735 | Lighthouse Lab in Glasgow | Wellcome Sanger Institute for the COVID-19 Genomics UK (COG-UK) Consortium | Harper VanSteenhouse, Yumi Kasai, David Gray, Carol Clugston, Anna Dominiczak and Alex Alderton, Roberto Amato, Jeffrey Barrett, Sonia Goncalves, Ewan Harrison, David K. Jackson, Ian Johnston, Dominic Kwiatkowski, Cordelia Langford, John Sillitoe on behalf of the Wellcome Sanger Institute COVID-19 Surveillance Team |
| EPI_ISL_2779736 | Lighthouse Lab in Milton Keynes | Wellcome Sanger Institute for the COVID-19 Genomics UK (COG-UK) Consortium | The Lighthouse Lab in Milton Keynes and Alex Alderton, Roberto Amato, Jeffrey Barrett, Sonia Goncalves, Ewan Harrison, David K. Jackson, Ian Johnston, Dominic Kwiatkowski, Cordelia Langford, John Sillitoe on behalf of the Wellcome Sanger Institute COVID-19 Surveillance Team |
| EPI_ISL_2779737, EPI_ISL_2779738 | Health Services Laboratories | Wellcome Sanger Institute for the COVID-19 Genomics UK (COG-UK) Consortium | Health Services Laboratories and Alex Alderton, Roberto Amato, Jeffrey Barrett, Sonia Goncalves, Ewan Harrison, David K. Jackson, Ian Johnston, Dominic Kwiatkowski, Cordelia Langford, John Sillitoe on behalf of the Wellcome Sanger Institute COVID-19 Surveillance Team |
| EPI_ISL_2779739 | Lighthouse Lab in Milton Keynes | Wellcome Sanger Institute for the COVID-19 Genomics UK (COG-UK) Consortium | The Lighthouse Lab in Milton Keynes and Alex Alderton, Roberto Amato, Jeffrey Barrett, Sonia Goncalves, Ewan Harrison, David K. Jackson, Ian Johnston, Dominic Kwiatkowski, Cordelia Langford, John Sillitoe on behalf of the Wellcome Sanger Institute COVID-19 Surveillance Team |
| EPI_ISL_2779740 | Lighthouse Lab in Glasgow | Wellcome Sanger Institute for the COVID-19 Genomics UK (COG-UK) Consortium | Harper VanSteenhouse, Yumi Kasai, David Gray, Carol Clugston, Anna Dominiczak and Alex Alderton, Roberto Amato, Jeffrey Barrett, Sonia Goncalves, Ewan Harrison, David K. Jackson, Ian Johnston, Dominic Kwiatkowski, Cordelia Langford, John Sillitoe on behalf of the Wellcome Sanger Institute COVID-19 Surveillance Team |
| EPI_ISL_2779741, EPI_ISL_2779742 | Health Services Laboratories | Wellcome Sanger Institute for the COVID-19 Genomics UK (COG-UK) Consortium | Health Services Laboratories and Alex Alderton, Roberto Amato, Jeffrey Barrett, Sonia Goncalves, Ewan Harrison, David K. Jackson, Ian Johnston, Dominic Kwiatkowski, Cordelia Langford, John Sillitoe on behalf of the Wellcome Sanger Institute COVID-19 Surveillance Team |
| EPI_ISL_2779743 | Lighthouse Lab in Glasgow | Wellcome Sanger Institute for the COVID-19 Genomics UK (COG-UK) Consortium | Harper VanSteenhouse, Yumi Kasai, David Gray, Carol Clugston, Anna Dominiczak and Alex Alderton, Roberto Amato, Jeffrey Barrett, Sonia Goncalves, Ewan Harrison, David K. Jackson, Ian Johnston, Dominic Kwiatkowski, Cordelia Langford, John Sillitoe on behalf of the Wellcome Sanger Institute COVID-19 Surveillance Team |
| EPI_ISL_2779744 | Lighthouse Lab in Milton Keynes | Wellcome Sanger Institute for the COVID-19 Genomics UK (COG-UK) Consortium | The Lighthouse Lab in Milton Keynes and Alex Alderton, Roberto Amato, Jeffrey Barrett, Sonia Goncalves, Ewan Harrison, David K. Jackson, Ian Johnston, Dominic Kwiatkowski, Cordelia Langford, John Sillitoe on behalf of the Wellcome Sanger Institute COVID-19 Surveillance Team |
| EPI_ISL_2779745, EPI_ISL_2779746 | Health Services Laboratories | Wellcome Sanger Institute for the COVID-19 Genomics UK (COG-UK) Consortium | Health Services Laboratories and Alex Alderton, Roberto Amato, Jeffrey Barrett, Sonia Goncalves, Ewan Harrison, David K. Jackson, Ian Johnston, Dominic Kwiatkowski, Cordelia Langford, John Sillitoe on behalf of the Wellcome Sanger Institute COVID-19 Surveillance Team |
| EPI_ISL_2779747 | Lighthouse Lab in Milton Keynes | Wellcome Sanger Institute for the COVID-19 Genomics UK (COG-UK) Consortium | The Lighthouse Lab in Milton Keynes and Alex Alderton, Roberto Amato, Jeffrey Barrett, Sonia Goncalves, Ewan Harrison, David K. Jackson, Ian Johnston, Dominic Kwiatkowski, Cordelia Langford, John Sillitoe on behalf of the Wellcome Sanger Institute COVID-19 Surveillance Team |
| EPI_ISL_2779748, EPI_ISL_2779749, EPI_ISL_2779750 | Health Services Laboratories | Wellcome Sanger Institute for the COVID-19 Genomics UK (COG-UK) Consortium | Health Services Laboratories and Alex Alderton, Roberto Amato, Jeffrey Barrett, Sonia Goncalves, Ewan Harrison, David K. Jackson, Ian Johnston, Dominic Kwiatkowski, Cordelia Langford, John Sillitoe on behalf of the Wellcome Sanger Institute COVID-19 Surveillance Team |
| EPI_ISL_2779751 | Lighthouse Lab in Milton Keynes | Wellcome Sanger Institute for the COVID-19 Genomics UK (COG-UK) Consortium | The Lighthouse Lab in Milton Keynes and Alex Alderton, Roberto Amato, Jeffrey Barrett, Sonia Goncalves, Ewan Harrison, David K. Jackson, Ian Johnston, Dominic Kwiatkowski, Cordelia Langford, John Sillitoe on behalf of the Wellcome Sanger Institute COVID-19 Surveillance Team |
| EPI_ISL_2779752 | Health Services Laboratories | Wellcome Sanger Institute for the COVID-19 Genomics UK (COG-UK) Consortium | Health Services Laboratories and Alex Alderton, Roberto Amato, Jeffrey Barrett, Sonia Goncalves, Ewan Harrison, David K. Jackson, Ian Johnston, Dominic Kwiatkowski, Cordelia Langford, John Sillitoe on behalf of the Wellcome Sanger Institute COVID-19 Surveillance Team |
| EPI_ISL_2779753 | Lighthouse Lab in Milton Keynes | Wellcome Sanger Institute for the COVID-19 Genomics UK (COG-UK) Consortium | The Lighthouse Lab in Milton Keynes and Alex Alderton, Roberto Amato, Jeffrey Barrett, Sonia Goncalves, Ewan Harrison, David K. Jackson, Ian Johnston, Dominic Kwiatkowski, Cordelia Langford, John Sillitoe on behalf of the Wellcome Sanger Institute COVID-19 Surveillance Team |
| EPI_ISL_2779754, EPI_ISL_2779755 | Lighthouse Lab in Glasgow | Wellcome Sanger Institute for the COVID-19 Genomics UK (COG-UK) Consortium | Harper VanSteenhouse, Yumi Kasai, David Gray, Carol Clugston, Anna Dominiczak and Alex Alderton, Roberto Amato, Jeffrey Barrett, Sonia Goncalves, Ewan Harrison, David K. Jackson, Ian Johnston, Dominic Kwiatkowski, Cordelia Langford, John Sillitoe on behalf of the Wellcome Sanger Institute COVID-19 Surveillance Team |
| EPI_ISL_2779756 | Health Services Laboratories | Wellcome Sanger Institute for the COVID-19 Genomics UK (COG-UK) Consortium | Health Services Laboratories and Alex Alderton, Roberto Amato, Jeffrey Barrett, Sonia Goncalves, Ewan Harrison, David K. Jackson, Ian Johnston, Dominic Kwiatkowski, Cordelia Langford, John Sillitoe on behalf of the Wellcome Sanger Institute COVID-19 Surveillance Team |
| EPI_ISL_2779757 | Lighthouse Lab in Glasgow | Wellcome Sanger Institute for the COVID-19 Genomics UK (COG-UK) Consortium | Harper VanSteenhouse, Yumi Kasai, David Gray, Carol Clugston, Anna Dominiczak and Alex Alderton, Roberto Amato, Jeffrey Barrett, Sonia Goncalves, Ewan Harrison, David K. Jackson, Ian Johnston, Dominic Kwiatkowski, Cordelia Langford, John Sillitoe on behalf of the Wellcome Sanger Institute COVID-19 Surveillance Team |
| EPI_ISL_2779758 | Health Services Laboratories | Wellcome Sanger Institute for the COVID-19 Genomics UK (COG-UK) Consortium | Health Services Laboratories and Alex Alderton, Roberto Amato, Jeffrey Barrett, Sonia Goncalves, Ewan Harrison, David K. Jackson, Ian Johnston, Dominic Kwiatkowski, Cordelia Langford, John Sillitoe on behalf of the Wellcome Sanger Institute COVID-19 Surveillance Team |
| EPI_ISL_2779759 | Lighthouse Lab in Glasgow | Wellcome Sanger Institute for the COVID-19 Genomics UK (COG-UK) Consortium | Harper VanSteenhouse, Yumi Kasai, David Gray, Carol Clugston, Anna Dominiczak and Alex Alderton, Roberto Amato, Jeffrey Barrett, Sonia Goncalves, Ewan Harrison, David K. Jackson, Ian Johnston, Dominic Kwiatkowski, Cordelia Langford, John Sillitoe on behalf of the Wellcome Sanger Institute COVID-19 Surveillance Team |
| EPI_ISL_2779760 | Lighthouse Lab in Milton Keynes | Wellcome Sanger Institute for the COVID-19 Genomics UK (COG-UK) Consortium | The Lighthouse Lab in Milton Keynes and Alex Alderton, Roberto Amato, Jeffrey Barrett, Sonia Goncalves, Ewan Harrison, David K. Jackson, Ian Johnston, Dominic Kwiatkowski, Cordelia Langford, John Sillitoe on behalf of the Wellcome Sanger Institute COVID-19 Surveillance Team |
| EPI_ISL_2779761, EPI_ISL_2779762, EPI_ISL_2779763, EPI_ISL_2779764 | Health Services Laboratories | Wellcome Sanger Institute for the COVID-19 Genomics UK (COG-UK) Consortium | Health Services Laboratories and Alex Alderton, Roberto Amato, Jeffrey Barrett, Sonia Goncalves, Ewan Harrison, David K. Jackson, Ian Johnston, Dominic Kwiatkowski, Cordelia Langford, John Sillitoe on behalf of the Wellcome Sanger Institute COVID-19 Surveillance Team |
| EPI_ISL_2779765 | Lighthouse Lab in Milton Keynes | Wellcome Sanger Institute for the COVID-19 Genomics UK (COG-UK) Consortium | The Lighthouse Lab in Milton Keynes and Alex Alderton, Roberto Amato, Jeffrey Barrett, Sonia Goncalves, Ewan Harrison, David K. Jackson, Ian Johnston, Dominic Kwiatkowski, Cordelia Langford, John Sillitoe on behalf of the Wellcome Sanger Institute COVID-19 Surveillance Team |
| EPI_ISL_2779766 | Health Services Laboratories | Wellcome Sanger Institute for the COVID-19 Genomics UK (COG-UK) Consortium | Health Services Laboratories and Alex Alderton, Roberto Amato, Jeffrey Barrett, Sonia Goncalves, Ewan Harrison, David K. Jackson, Ian Johnston, Dominic Kwiatkowski, Cordelia Langford, John Sillitoe on behalf of the Wellcome Sanger Institute COVID-19 Surveillance Team |
| EPI_ISL_2779767, EPI_ISL_2779768 | Lighthouse Lab in Glasgow | Wellcome Sanger Institute for the COVID-19 Genomics UK (COG-UK) Consortium | Harper VanSteenhouse, Yumi Kasai, David Gray, Carol Clugston, Anna Dominiczak and Alex Alderton, Roberto Amato, Jeffrey Barrett, Sonia Goncalves, Ewan Harrison, David K. Jackson, Ian Johnston, Dominic Kwiatkowski, Cordelia Langford, John Sillitoe on behalf of the Wellcome Sanger Institute COVID-19 Surveillance Team |
| EPI_ISL_2779769 | Lighthouse Lab in Milton Keynes | Wellcome Sanger Institute for the COVID-19 Genomics UK (COG-UK) Consortium | The Lighthouse Lab in Milton Keynes and Alex Alderton, Roberto Amato, Jeffrey Barrett, Sonia Goncalves, Ewan Harrison, David K. Jackson, Ian Johnston, Dominic Kwiatkowski, Cordelia Langford, John Sillitoe on behalf of the Wellcome Sanger Institute COVID-19 Surveillance Team |
| EPI_ISL_2779770 | Health Services Laboratories | Wellcome Sanger Institute for the COVID-19 Genomics UK (COG-UK) Consortium | Health Services Laboratories and Alex Alderton, Roberto Amato, Jeffrey Barrett, Sonia Goncalves, Ewan Harrison, David K. Jackson, Ian Johnston, Dominic Kwiatkowski, Cordelia Langford, John Sillitoe on behalf of the Wellcome Sanger Institute COVID-19 Surveillance Team |
| EPI_ISL_2779771 | Lighthouse Lab in Milton Keynes | Wellcome Sanger Institute for the COVID-19 Genomics UK (COG-UK) Consortium | The Lighthouse Lab in Milton Keynes and Alex Alderton, Roberto Amato, Jeffrey Barrett, Sonia Goncalves, Ewan Harrison, David K. Jackson, Ian Johnston, Dominic Kwiatkowski, Cordelia Langford, John Sillitoe on behalf of the Wellcome Sanger Institute COVID-19 Surveillance Team |
| EPI_ISL_2779772 | Health Services Laboratories | Wellcome Sanger Institute for the COVID-19 Genomics UK (COG-UK) Consortium | Health Services Laboratories and Alex Alderton, Roberto Amato, Jeffrey Barrett, Sonia Goncalves, Ewan Harrison, David K. Jackson, Ian Johnston, Dominic Kwiatkowski, Cordelia Langford, John Sillitoe on behalf of the Wellcome Sanger Institute COVID-19 Surveillance Team |
| EPI_ISL_2779773 | Lighthouse Lab in Glasgow | Wellcome Sanger Institute for the COVID-19 Genomics UK (COG-UK) Consortium | Harper VanSteenhouse, Yumi Kasai, David Gray, Carol Clugston, Anna Dominiczak and Alex Alderton, Roberto Amato, Jeffrey Barrett, Sonia Goncalves, Ewan Harrison, David K. Jackson, Ian Johnston, Dominic Kwiatkowski, Cordelia Langford, John Sillitoe on behalf of the Wellcome Sanger Institute COVID-19 Surveillance Team |
| EPI_ISL_2779774 | Health Services Laboratories | Wellcome Sanger Institute for the COVID-19 Genomics UK (COG-UK) Consortium | Health Services Laboratories and Alex Alderton, Roberto Amato, Jeffrey Barrett, Sonia Goncalves, Ewan Harrison, David K. Jackson, Ian Johnston, Dominic Kwiatkowski, Cordelia Langford, John Sillitoe on behalf of the Wellcome Sanger Institute COVID-19 Surveillance Team |
| EPI_ISL_2779775, EPI_ISL_2779776 | Lighthouse Lab in Milton Keynes | Wellcome Sanger Institute for the COVID-19 Genomics UK (COG-UK) Consortium | The Lighthouse Lab in Milton Keynes and Alex Alderton, Roberto Amato, Jeffrey Barrett, Sonia Goncalves, Ewan Harrison, David K. Jackson, Ian Johnston, Dominic Kwiatkowski, Cordelia Langford, John Sillitoe on behalf of the Wellcome Sanger Institute COVID-19 Surveillance Team |
| EPI_ISL_2779777 | Health Services Laboratories | Wellcome Sanger Institute for the COVID-19 Genomics UK (COG-UK) Consortium | Health Services Laboratories and Alex Alderton, Roberto Amato, Jeffrey Barrett, Sonia Goncalves, Ewan Harrison, David K. Jackson, Ian Johnston, Dominic Kwiatkowski, Cordelia Langford, John Sillitoe on behalf of the Wellcome Sanger Institute COVID-19 Surveillance Team |
| EPI_ISL_2779778, EPI_ISL_2779779 | Lighthouse Lab in Milton Keynes | Wellcome Sanger Institute for the COVID-19 Genomics UK (COG-UK) Consortium | The Lighthouse Lab in Milton Keynes and Alex Alderton, Roberto Amato, Jeffrey Barrett, Sonia Goncalves, Ewan Harrison, David K. Jackson, Ian Johnston, Dominic Kwiatkowski, Cordelia Langford, John Sillitoe on behalf of the Wellcome Sanger Institute COVID-19 Surveillance Team |
| EPI_ISL_2779780 | Health Services Laboratories | Wellcome Sanger Institute for the COVID-19 Genomics UK (COG-UK) Consortium | Health Services Laboratories and Alex Alderton, Roberto Amato, Jeffrey Barrett, Sonia Goncalves, Ewan Harrison, David K. Jackson, Ian Johnston, Dominic Kwiatkowski, Cordelia Langford, John Sillitoe on behalf of the Wellcome Sanger Institute COVID-19 Surveillance Team |
| EPI_ISL_2779781, EPI_ISL_2779782 | Lighthouse Lab in Glasgow | Wellcome Sanger Institute for the COVID-19 Genomics UK (COG-UK) Consortium | Harper VanSteenhouse, Yumi Kasai, David Gray, Carol Clugston, Anna Dominiczak and Alex Alderton, Roberto Amato, Jeffrey Barrett, Sonia Goncalves, Ewan Harrison, David K. Jackson, Ian Johnston, Dominic Kwiatkowski, Cordelia Langford, John Sillitoe on behalf of the Wellcome Sanger Institute COVID-19 Surveillance Team |
| EPI_ISL_2779783 | Lighthouse Lab in Milton Keynes | Wellcome Sanger Institute for the COVID-19 Genomics UK (COG-UK) Consortium | The Lighthouse Lab in Milton Keynes and Alex Alderton, Roberto Amato, Jeffrey Barrett, Sonia Goncalves, Ewan Harrison, David K. Jackson, Ian Johnston, Dominic Kwiatkowski, Cordelia Langford, John Sillitoe on behalf of the Wellcome Sanger Institute COVID-19 Surveillance Team |
| EPI_ISL_2779784, EPI_ISL_2779785 | Lighthouse Lab in Glasgow | Wellcome Sanger Institute for the COVID-19 Genomics UK (COG-UK) Consortium | Harper VanSteenhouse, Yumi Kasai, David Gray, Carol Clugston, Anna Dominiczak and Alex Alderton, Roberto Amato, Jeffrey Barrett, Sonia Goncalves, Ewan Harrison, David K. Jackson, Ian Johnston, Dominic Kwiatkowski, Cordelia Langford, John Sillitoe on behalf of the Wellcome Sanger Institute COVID-19 Surveillance Team |

| | | | |
|---|---|---|---|
| EPI_ISL_2779786 | Lighthouse Lab in Milton Keynes | Wellcome Sanger Institute for the COVID-19 Genomics UK (COG-UK) Consortium | The Lighthouse Lab in Milton Keynes and Alex Alderton, Roberto Amato, Jeffrey Barrett, Sonia Goncalves, Ewan Harrison, David K. Jackson, Ian Johnston, Dominic Kwiatkowski, Cordelia Langford, John Sillitoe on behalf of the Wellcome Sanger Institute COVID-19 Surveillance Team |
| EPI_ISL_2779787 | Health Services Laboratories | Wellcome Sanger Institute for the COVID-19 Genomics UK (COG-UK) Consortium | Health Services Laboratories and Alex Alderton, Roberto Amato, Jeffrey Barrett, Sonia Goncalves, Ewan Harrison, David K. Jackson, Ian Johnston, Dominic Kwiatkowski, Cordelia Langford, John Sillitoe on behalf of the Wellcome Sanger Institute COVID-19 Surveillance Team |
| EPI_ISL_2779788, EPI_ISL_2779789, EPI_ISL_2779790, EPI_ISL_2779791, EPI_ISL_2779792 | Lighthouse Lab in Milton Keynes | Wellcome Sanger Institute for the COVID-19 Genomics UK (COG-UK) Consortium | The Lighthouse Lab in Milton Keynes and Alex Alderton, Roberto Amato, Jeffrey Barrett, Sonia Goncalves, Ewan Harrison, David K. Jackson, Ian Johnston, Dominic Kwiatkowski, Cordelia Langford, John Sillitoe on behalf of the Wellcome Sanger Institute COVID-19 Surveillance Team |
| EPI_ISL_2779793, EPI_ISL_2779794 | Health Services Laboratories | Wellcome Sanger Institute for the COVID-19 Genomics UK (COG-UK) Consortium | Health Services Laboratories and Alex Alderton, Roberto Amato, Jeffrey Barrett, Sonia Goncalves, Ewan Harrison, David K. Jackson, Ian Johnston, Dominic Kwiatkowski, Cordelia Langford, John Sillitoe on behalf of the Wellcome Sanger Institute COVID-19 Surveillance Team |
| EPI_ISL_2779795 | Lighthouse Lab in Milton Keynes | Wellcome Sanger Institute for the COVID-19 Genomics UK (COG-UK) Consortium | The Lighthouse Lab in Milton Keynes and Alex Alderton, Roberto Amato, Jeffrey Barrett, Sonia Goncalves, Ewan Harrison, David K. Jackson, Ian Johnston, Dominic Kwiatkowski, Cordelia Langford, John Sillitoe on behalf of the Wellcome Sanger Institute COVID-19 Surveillance Team |
| EPI_ISL_2779796 | Health Services Laboratories | Wellcome Sanger Institute for the COVID-19 Genomics UK (COG-UK) Consortium | Health Services Laboratories and Alex Alderton, Roberto Amato, Jeffrey Barrett, Sonia Goncalves, Ewan Harrison, David K. Jackson, Ian Johnston, Dominic Kwiatkowski, Cordelia Langford, John Sillitoe on behalf of the Wellcome Sanger Institute COVID-19 Surveillance Team |
| EPI_ISL_2779797 | Lighthouse Lab in Milton Keynes | Wellcome Sanger Institute for the COVID-19 Genomics UK (COG-UK) Consortium | The Lighthouse Lab in Milton Keynes and Alex Alderton, Roberto Amato, Jeffrey Barrett, Sonia Goncalves, Ewan Harrison, David K. Jackson, Ian Johnston, Dominic Kwiatkowski, Cordelia Langford, John Sillitoe on behalf of the Wellcome Sanger Institute COVID-19 Surveillance Team |
| EPI_ISL_2779798 | Health Services Laboratories | Wellcome Sanger Institute for the COVID-19 Genomics UK (COG-UK) Consortium | Health Services Laboratories and Alex Alderton, Roberto Amato, Jeffrey Barrett, Sonia Goncalves, Ewan Harrison, David K. Jackson, Ian Johnston, Dominic Kwiatkowski, Cordelia Langford, John Sillitoe on behalf of the Wellcome Sanger Institute COVID-19 Surveillance Team |
| EPI_ISL_2779799, EPI_ISL_2779800, EPI_ISL_2779801 | Lighthouse Lab in Milton Keynes | Wellcome Sanger Institute for the COVID-19 Genomics UK (COG-UK) Consortium | The Lighthouse Lab in Milton Keynes and Alex Alderton, Roberto Amato, Jeffrey Barrett, Sonia Goncalves, Ewan Harrison, David K. Jackson, Ian Johnston, Dominic Kwiatkowski, Cordelia Langford, John Sillitoe on behalf of the Wellcome Sanger Institute COVID-19 Surveillance Team |
| EPI_ISL_2779802, EPI_ISL_2779803, EPI_ISL_2779804, EPI_ISL_2779805, EPI_ISL_2779806 | Health Services Laboratories | Wellcome Sanger Institute for the COVID-19 Genomics UK (COG-UK) Consortium | Health Services Laboratories and Alex Alderton, Roberto Amato, Jeffrey Barrett, Sonia Goncalves, Ewan Harrison, David K. Jackson, Ian Johnston, Dominic Kwiatkowski, Cordelia Langford, John Sillitoe on behalf of the Wellcome Sanger Institute COVID-19 Surveillance Team |
| EPI_ISL_2779807 | Lighthouse Lab in Glasgow | Wellcome Sanger Institute for the COVID-19 Genomics UK (COG-UK) Consortium | Harper VanSteenhouse, Yumi Kasai, David Gray, Carol Clugston, Anna Dominiczak and Alex Alderton, Roberto Amato, Jeffrey Barrett, Sonia Goncalves, Ewan Harrison, David K. Jackson, Ian Johnston, Dominic Kwiatkowski, Cordelia Langford, John Sillitoe on behalf of the Wellcome Sanger Institute COVID-19 Surveillance Team |
| EPI_ISL_2779808 | Lighthouse Lab in Milton Keynes | Wellcome Sanger Institute for the COVID-19 Genomics UK (COG-UK) Consortium | The Lighthouse Lab in Milton Keynes and Alex Alderton, Roberto Amato, Jeffrey Barrett, Sonia Goncalves, Ewan Harrison, David K. Jackson, Ian Johnston, Dominic Kwiatkowski, Cordelia Langford, John Sillitoe on behalf of the Wellcome Sanger Institute COVID-19 Surveillance Team |
| EPI_ISL_2779809, EPI_ISL_2779810 | Health Services Laboratories | Wellcome Sanger Institute for the COVID-19 Genomics UK (COG-UK) Consortium | Health Services Laboratories and Alex Alderton, Roberto Amato, Jeffrey Barrett, Sonia Goncalves, Ewan Harrison, David K. Jackson, Ian Johnston, Dominic Kwiatkowski, Cordelia Langford, John Sillitoe on behalf of the Wellcome Sanger Institute COVID-19 Surveillance Team |
| EPI_ISL_2779811 | Lighthouse Lab in Milton Keynes | Wellcome Sanger Institute for the COVID-19 Genomics UK (COG-UK) Consortium | The Lighthouse Lab in Milton Keynes and Alex Alderton, Roberto Amato, Jeffrey Barrett, Sonia Goncalves, Ewan Harrison, David K. Jackson, Ian Johnston, Dominic Kwiatkowski, Cordelia Langford, John Sillitoe on behalf of the Wellcome Sanger Institute COVID-19 Surveillance Team |
| EPI_ISL_2779812, EPI_ISL_2779813, EPI_ISL_2779814 | Health Services Laboratories | Wellcome Sanger Institute for the COVID-19 Genomics UK (COG-UK) Consortium | Health Services Laboratories and Alex Alderton, Roberto Amato, Jeffrey Barrett, Sonia Goncalves, Ewan Harrison, David K. Jackson, Ian Johnston, Dominic Kwiatkowski, Cordelia Langford, John Sillitoe on behalf of the Wellcome Sanger Institute COVID-19 Surveillance Team |
| EPI_ISL_2779815 | Lighthouse Lab in Milton Keynes | Wellcome Sanger Institute for the COVID-19 Genomics UK (COG-UK) Consortium | The Lighthouse Lab in Milton Keynes and Alex Alderton, Roberto Amato, Jeffrey Barrett, Sonia Goncalves, Ewan Harrison, David K. Jackson, Ian Johnston, Dominic Kwiatkowski, Cordelia Langford, John Sillitoe on behalf of the Wellcome Sanger Institute COVID-19 Surveillance Team |
| EPI_ISL_2779816 | Health Services Laboratories | Wellcome Sanger Institute for the COVID-19 Genomics UK (COG-UK) Consortium | Health Services Laboratories and Alex Alderton, Roberto Amato, Jeffrey Barrett, Sonia Goncalves, Ewan Harrison, David K. Jackson, Ian Johnston, Dominic Kwiatkowski, Cordelia Langford, John Sillitoe on behalf of the Wellcome Sanger Institute COVID-19 Surveillance Team |
| EPI_ISL_2779817 | Lighthouse Lab in Milton Keynes | Wellcome Sanger Institute for the COVID-19 Genomics UK (COG-UK) Consortium | The Lighthouse Lab in Milton Keynes and Alex Alderton, Roberto Amato, Jeffrey Barrett, Sonia Goncalves, Ewan Harrison, David K. Jackson, Ian Johnston, Dominic Kwiatkowski, Cordelia Langford, John Sillitoe on behalf of the Wellcome Sanger Institute COVID-19 Surveillance Team |
| EPI_ISL_2779818, EPI_ISL_2779819 | Lighthouse Lab in Glasgow | Wellcome Sanger Institute for the COVID-19 Genomics UK (COG-UK) Consortium | Harper VanSteenhouse, Yumi Kasai, David Gray, Carol Clugston, Anna Dominiczak and Alex Alderton, Roberto Amato, Jeffrey Barrett, Sonia Goncalves, Ewan Harrison, David K. Jackson, Ian Johnston, Dominic Kwiatkowski, Cordelia Langford, John Sillitoe on behalf of the Wellcome Sanger Institute COVID-19 Surveillance Team |
| EPI_ISL_2779820 | Health Services Laboratories | Wellcome Sanger Institute for the COVID-19 Genomics UK (COG-UK) Consortium | Health Services Laboratories and Alex Alderton, Roberto Amato, Jeffrey Barrett, Sonia Goncalves, Ewan Harrison, David K. Jackson, Ian Johnston, Dominic Kwiatkowski, Cordelia Langford, John Sillitoe on behalf of the Wellcome Sanger Institute COVID-19 Surveillance Team |
| EPI_ISL_2779821, EPI_ISL_2779822 | Lighthouse Lab in Glasgow | Wellcome Sanger Institute for the COVID-19 Genomics UK (COG-UK) Consortium | Harper VanSteenhouse, Yumi Kasai, David Gray, Carol Clugston, Anna Dominiczak and Alex Alderton, Roberto Amato, Jeffrey Barrett, Sonia Goncalves, Ewan Harrison, David K. Jackson, Ian Johnston, Dominic Kwiatkowski, Cordelia Langford, John Sillitoe on behalf of the Wellcome Sanger Institute COVID-19 Surveillance Team |
| EPI_ISL_2779823 | Lighthouse Lab in Milton Keynes | Wellcome Sanger Institute for the COVID-19 Genomics UK (COG-UK) Consortium | The Lighthouse Lab in Milton Keynes and Alex Alderton, Roberto Amato, Jeffrey Barrett, Sonia Goncalves, Ewan Harrison, David K. Jackson, Ian Johnston, Dominic Kwiatkowski, Cordelia Langford, John Sillitoe on behalf of the Wellcome Sanger Institute COVID-19 Surveillance Team |
| EPI_ISL_2779824 | Lighthouse Lab in Glasgow | Wellcome Sanger Institute for the COVID-19 Genomics UK (COG-UK) Consortium | Harper VanSteenhouse, Yumi Kasai, David Gray, Carol Clugston, Anna Dominiczak and Alex Alderton, Roberto Amato, Jeffrey Barrett, Sonia Goncalves, Ewan Harrison, David K. Jackson, Ian Johnston, Dominic Kwiatkowski, Cordelia Langford, John Sillitoe on behalf of the Wellcome Sanger Institute COVID-19 Surveillance Team |
| EPI_ISL_2779825, EPI_ISL_2779826, EPI_ISL_2779827 | Lighthouse Lab in Milton Keynes | Wellcome Sanger Institute for the COVID-19 Genomics UK (COG-UK) Consortium | The Lighthouse Lab in Milton Keynes and Alex Alderton, Roberto Amato, Jeffrey Barrett, Sonia Goncalves, Ewan Harrison, David K. Jackson, Ian Johnston, Dominic Kwiatkowski, Cordelia Langford, John Sillitoe on behalf of the Wellcome Sanger Institute COVID-19 Surveillance Team |
| EPI_ISL_2779828, EPI_ISL_2779829 | Health Services Laboratories | Wellcome Sanger Institute for the COVID-19 Genomics UK (COG-UK) Consortium | Health Services Laboratories and Alex Alderton, Roberto Amato, Jeffrey Barrett, Sonia Goncalves, Ewan Harrison, David K. Jackson, Ian Johnston, Dominic Kwiatkowski, Cordelia Langford, John Sillitoe on behalf of the Wellcome Sanger Institute COVID-19 Surveillance Team |
| EPI_ISL_2779830 | Lighthouse Lab in Milton Keynes | Wellcome Sanger Institute for the COVID-19 Genomics UK (COG-UK) Consortium | The Lighthouse Lab in Milton Keynes and Alex Alderton, Roberto Amato, Jeffrey Barrett, Sonia Goncalves, Ewan Harrison, David K. Jackson, Ian Johnston, Dominic Kwiatkowski, Cordelia Langford, John Sillitoe on behalf of the Wellcome Sanger Institute COVID-19 Surveillance Team |
| EPI_ISL_2779831 | Lighthouse Lab in Glasgow | Wellcome Sanger Institute for the COVID-19 Genomics UK (COG-UK) Consortium | Harper VanSteenhouse, Yumi Kasai, David Gray, Carol Clugston, Anna Dominiczak and Alex Alderton, Roberto Amato, Jeffrey Barrett, Sonia Goncalves, Ewan Harrison, David K. Jackson, Ian Johnston, Dominic Kwiatkowski, Cordelia Langford, John Sillitoe on behalf of the Wellcome Sanger Institute COVID-19 Surveillance Team |
| EPI_ISL_2779832, EPI_ISL_2779833 | Health Services Laboratories | Wellcome Sanger Institute for the COVID-19 Genomics UK (COG-UK) Consortium | Health Services Laboratories and Alex Alderton, Roberto Amato, Jeffrey Barrett, Sonia Goncalves, Ewan Harrison, David K. Jackson, Ian Johnston, Dominic Kwiatkowski, Cordelia Langford, John Sillitoe on behalf of the Wellcome Sanger Institute COVID-19 Surveillance Team |
| EPI_ISL_2779834 | Lighthouse Lab in Milton Keynes | Wellcome Sanger Institute for the COVID-19 Genomics UK (COG-UK) Consortium | The Lighthouse Lab in Milton Keynes and Alex Alderton, Roberto Amato, Jeffrey Barrett, Sonia Goncalves, Ewan Harrison, David K. Jackson, Ian Johnston, Dominic Kwiatkowski, Cordelia Langford, John Sillitoe on behalf of the Wellcome Sanger Institute COVID-19 Surveillance Team |
| EPI_ISL_2779835, EPI_ISL_2779836 | Lighthouse Lab in Glasgow | Wellcome Sanger Institute for the COVID-19 Genomics UK (COG-UK) Consortium | Harper VanSteenhouse, Yumi Kasai, David Gray, Carol Clugston, Anna Dominiczak and Alex Alderton, Roberto Amato, Jeffrey Barrett, Sonia Goncalves, Ewan Harrison, David K. Jackson, Ian Johnston, Dominic Kwiatkowski, Cordelia Langford, John Sillitoe on behalf of the Wellcome Sanger Institute COVID-19 Surveillance Team |
| EPI_ISL_2779837 | Health Services Laboratories | Wellcome Sanger Institute for the COVID-19 Genomics UK (COG-UK) Consortium | Health Services Laboratories and Alex Alderton, Roberto Amato, Jeffrey Barrett, Sonia Goncalves, Ewan Harrison, David K. Jackson, Ian Johnston, Dominic Kwiatkowski, Cordelia Langford, John Sillitoe on behalf of the Wellcome Sanger Institute COVID-19 Surveillance Team |
| EPI_ISL_2779838 | Lighthouse Lab in Glasgow | Wellcome Sanger Institute for the COVID-19 Genomics UK (COG-UK) Consortium | Harper VanSteenhouse, Yumi Kasai, David Gray, Carol Clugston, Anna Dominiczak and Alex Alderton, Roberto Amato, Jeffrey Barrett, Sonia Goncalves, Ewan Harrison, David K. Jackson, Ian Johnston, Dominic Kwiatkowski, Cordelia Langford, John Sillitoe on behalf of the Wellcome Sanger Institute COVID-19 Surveillance Team |
| EPI_ISL_2779839 | Lighthouse Lab in Milton Keynes | Wellcome Sanger Institute for the COVID-19 Genomics UK (COG-UK) Consortium | The Lighthouse Lab in Milton Keynes and Alex Alderton, Roberto Amato, Jeffrey Barrett, Sonia Goncalves, Ewan Harrison, David K. Jackson, Ian Johnston, Dominic Kwiatkowski, Cordelia Langford, John Sillitoe on behalf of the Wellcome Sanger Institute COVID-19 Surveillance Team |
| EPI_ISL_2779840 | Health Services Laboratories | Wellcome Sanger Institute for the COVID-19 Genomics UK (COG-UK) Consortium | Health Services Laboratories and Alex Alderton, Roberto Amato, Jeffrey Barrett, Sonia Goncalves, Ewan Harrison, David K. Jackson, Ian Johnston, Dominic Kwiatkowski, Cordelia Langford, John Sillitoe on behalf of the Wellcome Sanger Institute COVID-19 Surveillance Team |
| EPI_ISL_2779841 | Lighthouse Lab in Glasgow | Wellcome Sanger Institute for the COVID-19 Genomics UK (COG-UK) Consortium | Harper VanSteenhouse, Yumi Kasai, David Gray, Carol Clugston, Anna Dominiczak and Alex Alderton, Roberto Amato, Jeffrey Barrett, Sonia Goncalves, Ewan Harrison, David K. Jackson, Ian Johnston, Dominic Kwiatkowski, Cordelia Langford, John Sillitoe on behalf of the Wellcome Sanger Institute COVID-19 Surveillance Team |
| EPI_ISL_2779842 | Health Services Laboratories | Wellcome Sanger Institute for the COVID-19 Genomics UK (COG-UK) Consortium | Health Services Laboratories and Alex Alderton, Roberto Amato, Jeffrey Barrett, Sonia Goncalves, Ewan Harrison, David K. Jackson, Ian Johnston, Dominic Kwiatkowski, Cordelia Langford, John Sillitoe on behalf of the Wellcome Sanger Institute COVID-19 Surveillance Team |
| EPI_ISL_2779843, EPI_ISL_2779844 | Lighthouse Lab in Milton Keynes | Wellcome Sanger Institute for the COVID-19 Genomics UK (COG-UK) Consortium | The Lighthouse Lab in Milton Keynes and Alex Alderton, Roberto Amato, Jeffrey Barrett, Sonia Goncalves, Ewan Harrison, David K. Jackson, Ian Johnston, Dominic Kwiatkowski, Cordelia Langford, John Sillitoe on behalf of the Wellcome Sanger Institute COVID-19 Surveillance Team |
| EPI_ISL_2779845 | Health Services Laboratories | Wellcome Sanger Institute for the COVID-19 Genomics UK (COG-UK) Consortium | Health Services Laboratories and Alex Alderton, Roberto Amato, Jeffrey Barrett, Sonia Goncalves, Ewan Harrison, David K. Jackson, Ian Johnston, Dominic Kwiatkowski, Cordelia Langford, John Sillitoe on behalf of the Wellcome Sanger Institute COVID-19 Surveillance Team |
| EPI_ISL_2779846, EPI_ISL_2779847 | Lighthouse Lab in Milton Keynes | Wellcome Sanger Institute for the COVID-19 Genomics UK (COG-UK) Consortium | The Lighthouse Lab in Milton Keynes and Alex Alderton, Roberto Amato, Jeffrey Barrett, Sonia Goncalves, Ewan Harrison, David K. Jackson, Ian Johnston, Dominic Kwiatkowski, Cordelia Langford, John Sillitoe on behalf of the Wellcome Sanger Institute COVID-19 Surveillance Team |
| EPI_ISL_2779848 | Health Services Laboratories | Wellcome Sanger Institute for the COVID-19 Genomics UK (COG-UK) Consortium | Health Services Laboratories and Alex Alderton, Roberto Amato, Jeffrey Barrett, Sonia Goncalves, Ewan Harrison, David K. Jackson, Ian Johnston, Dominic Kwiatkowski, Cordelia Langford, John Sillitoe on behalf of the Wellcome Sanger Institute COVID-19 Surveillance Team |
| EPI_ISL_2779849 | Lighthouse Lab in Milton Keynes | Wellcome Sanger Institute for the COVID-19 Genomics UK (COG-UK) Consortium | The Lighthouse Lab in Milton Keynes and Alex Alderton, Roberto Amato, Jeffrey Barrett, Sonia Goncalves, Ewan Harrison, David K. Jackson, Ian Johnston, Dominic Kwiatkowski, Cordelia Langford, John Sillitoe on behalf of the Wellcome Sanger Institute COVID-19 Surveillance Team |
| EPI_ISL_2779850, | Health Services Laboratories | Wellcome Sanger Institute for the COVID-19 Genomics UK (COG-UK) | Health Services Laboratories and Alex Alderton, Roberto Amato, Jeffrey Barrett, Sonia Goncalves, Ewan Harrison, David K. Jackson, Ian Johnston, Dominic Kwiatkowski, Cordelia Langford, John Sillitoe on behalf of the Wellcome Sanger Institute |

| | | | |
|---|---|---|---|
| EPI_ISL_2779851 | | Consortium | COVID-19 Surveillance Team |
| EPI_ISL_2779852 | Lighthouse Lab in Milton Keynes | Wellcome Sanger Institute for the COVID-19 Genomics UK (COG-UK) Consortium | The Lighthouse Lab in Milton Keynes and Alex Alderton, Roberto Amato, Jeffrey Barrett, Sonia Goncalves, Ewan Harrison, David K. Jackson, Ian Johnston, Dominic Kwiatkowski, Cordelia Langford, John Sillitoe on behalf of the Wellcome Sanger Institute COVID-19 Surveillance Team |
| EPI_ISL_2779853 | Health Services Laboratories | Wellcome Sanger Institute for the COVID-19 Genomics UK (COG-UK) Consortium | Health Services Laboratories and Alex Alderton, Roberto Amato, Jeffrey Barrett, Sonia Goncalves, Ewan Harrison, David K. Jackson, Ian Johnston, Dominic Kwiatkowski, Cordelia Langford, John Sillitoe on behalf of the Wellcome Sanger Institute COVID-19 Surveillance Team |
| EPI_ISL_2779854 | Lighthouse Lab in Glasgow | Wellcome Sanger Institute for the COVID-19 Genomics UK (COG-UK) Consortium | Harper VanSteenhouse, Yumi Kasai, David Gray, Carol Clugston, Anna Dominiczak and Alex Alderton, Roberto Amato, Jeffrey Barrett, Sonia Goncalves, Ewan Harrison, David K. Jackson, Ian Johnston, Dominic Kwiatkowski, Cordelia Langford, John Sillitoe on behalf of the Wellcome Sanger Institute COVID-19 Surveillance Team |
| EPI_ISL_2779855 | Health Services Laboratories | Wellcome Sanger Institute for the COVID-19 Genomics UK (COG-UK) Consortium | Health Services Laboratories and Alex Alderton, Roberto Amato, Jeffrey Barrett, Sonia Goncalves, Ewan Harrison, David K. Jackson, Ian Johnston, Dominic Kwiatkowski, Cordelia Langford, John Sillitoe on behalf of the Wellcome Sanger Institute COVID-19 Surveillance Team |
| EPI_ISL_2779856 | Lighthouse Lab in Glasgow | Wellcome Sanger Institute for the COVID-19 Genomics UK (COG-UK) Consortium | Harper VanSteenhouse, Yumi Kasai, David Gray, Carol Clugston, Anna Dominiczak and Alex Alderton, Roberto Amato, Jeffrey Barrett, Sonia Goncalves, Ewan Harrison, David K. Jackson, Ian Johnston, Dominic Kwiatkowski, Cordelia Langford, John Sillitoe on behalf of the Wellcome Sanger Institute COVID-19 Surveillance Team |
| EPI_ISL_2779857 | Lighthouse Lab in Milton Keynes | Wellcome Sanger Institute for the COVID-19 Genomics UK (COG-UK) Consortium | The Lighthouse Lab in Milton Keynes and Alex Alderton, Roberto Amato, Jeffrey Barrett, Sonia Goncalves, Ewan Harrison, David K. Jackson, Ian Johnston, Dominic Kwiatkowski, Cordelia Langford, John Sillitoe on behalf of the Wellcome Sanger Institute COVID-19 Surveillance Team |
| EPI_ISL_2779858 | Lighthouse Lab in Glasgow | Wellcome Sanger Institute for the COVID-19 Genomics UK (COG-UK) Consortium | Harper VanSteenhouse, Yumi Kasai, David Gray, Carol Clugston, Anna Dominiczak and Alex Alderton, Roberto Amato, Jeffrey Barrett, Sonia Goncalves, Ewan Harrison, David K. Jackson, Ian Johnston, Dominic Kwiatkowski, Cordelia Langford, John Sillitoe on behalf of the Wellcome Sanger Institute COVID-19 Surveillance Team |
| EPI_ISL_2779859, EPI_ISL_2779860 | Health Services Laboratories | Wellcome Sanger Institute for the COVID-19 Genomics UK (COG-UK) Consortium | Health Services Laboratories and Alex Alderton, Roberto Amato, Jeffrey Barrett, Sonia Goncalves, Ewan Harrison, David K. Jackson, Ian Johnston, Dominic Kwiatkowski, Cordelia Langford, John Sillitoe on behalf of the Wellcome Sanger Institute COVID-19 Surveillance Team |
| EPI_ISL_2779861 | Lighthouse Lab in Milton Keynes | Wellcome Sanger Institute for the COVID-19 Genomics UK (COG-UK) Consortium | The Lighthouse Lab in Milton Keynes and Alex Alderton, Roberto Amato, Jeffrey Barrett, Sonia Goncalves, Ewan Harrison, David K. Jackson, Ian Johnston, Dominic Kwiatkowski, Cordelia Langford, John Sillitoe on behalf of the Wellcome Sanger Institute COVID-19 Surveillance Team |
| EPI_ISL_2779862 | Health Services Laboratories | Wellcome Sanger Institute for the COVID-19 Genomics UK (COG-UK) Consortium | Health Services Laboratories and Alex Alderton, Roberto Amato, Jeffrey Barrett, Sonia Goncalves, Ewan Harrison, David K. Jackson, Ian Johnston, Dominic Kwiatkowski, Cordelia Langford, John Sillitoe on behalf of the Wellcome Sanger Institute COVID-19 Surveillance Team |
| EPI_ISL_2779863 | Lighthouse Lab in Milton Keynes | Wellcome Sanger Institute for the COVID-19 Genomics UK (COG-UK) Consortium | The Lighthouse Lab in Milton Keynes and Alex Alderton, Roberto Amato, Jeffrey Barrett, Sonia Goncalves, Ewan Harrison, David K. Jackson, Ian Johnston, Dominic Kwiatkowski, Cordelia Langford, John Sillitoe on behalf of the Wellcome Sanger Institute COVID-19 Surveillance Team |
| EPI_ISL_2779864 | Health Services Laboratories | Wellcome Sanger Institute for the COVID-19 Genomics UK (COG-UK) Consortium | Health Services Laboratories and Alex Alderton, Roberto Amato, Jeffrey Barrett, Sonia Goncalves, Ewan Harrison, David K. Jackson, Ian Johnston, Dominic Kwiatkowski, Cordelia Langford, John Sillitoe on behalf of the Wellcome Sanger Institute COVID-19 Surveillance Team |
| EPI_ISL_2779865, EPI_ISL_2779866 | Lighthouse Lab in Milton Keynes | Wellcome Sanger Institute for the COVID-19 Genomics UK (COG-UK) Consortium | The Lighthouse Lab in Milton Keynes and Alex Alderton, Roberto Amato, Jeffrey Barrett, Sonia Goncalves, Ewan Harrison, David K. Jackson, Ian Johnston, Dominic Kwiatkowski, Cordelia Langford, John Sillitoe on behalf of the Wellcome Sanger Institute COVID-19 Surveillance Team |
| EPI_ISL_2779867, EPI_ISL_2779868 | Lighthouse Lab in Glasgow | Wellcome Sanger Institute for the COVID-19 Genomics UK (COG-UK) Consortium | Harper VanSteenhouse, Yumi Kasai, David Gray, Carol Clugston, Anna Dominiczak and Alex Alderton, Roberto Amato, Jeffrey Barrett, Sonia Goncalves, Ewan Harrison, David K. Jackson, Ian Johnston, Dominic Kwiatkowski, Cordelia Langford, John Sillitoe on behalf of the Wellcome Sanger Institute COVID-19 Surveillance Team |
| EPI_ISL_2779869, EPI_ISL_2779870 | Lighthouse Laboratory Plymouth | Wellcome Sanger Institute for the COVID-19 Genomics UK (COG-UK) Consortium | Lighthouse Laboratory Plymouth and Alex Alderton, Roberto Amato, Jeffrey Barrett, Sonia Goncalves, Ewan Harrison, David K. Jackson, Ian Johnston, Dominic Kwiatkowski, Cordelia Langford, John Sillitoe on behalf of the Wellcome Sanger Institute COVID-19 Surveillance Team |
| EPI_ISL_2779871 | Lighthouse Lab in Milton Keynes | Wellcome Sanger Institute for the COVID-19 Genomics UK (COG-UK) Consortium | The Lighthouse Lab in Milton Keynes and Alex Alderton, Roberto Amato, Jeffrey Barrett, Sonia Goncalves, Ewan Harrison, David K. Jackson, Ian Johnston, Dominic Kwiatkowski, Cordelia Langford, John Sillitoe on behalf of the Wellcome Sanger Institute COVID-19 Surveillance Team |
| EPI_ISL_2779872, EPI_ISL_2779873 | Lighthouse Laboratory Plymouth | Wellcome Sanger Institute for the COVID-19 Genomics UK (COG-UK) Consortium | Lighthouse Laboratory Plymouth and Alex Alderton, Roberto Amato, Jeffrey Barrett, Sonia Goncalves, Ewan Harrison, David K. Jackson, Ian Johnston, Dominic Kwiatkowski, Cordelia Langford, John Sillitoe on behalf of the Wellcome Sanger Institute COVID-19 Surveillance Team |
| EPI_ISL_2779874 | Lighthouse Lab in Glasgow | Wellcome Sanger Institute for the COVID-19 Genomics UK (COG-UK) Consortium | Harper VanSteenhouse, Yumi Kasai, David Gray, Carol Clugston, Anna Dominiczak and Alex Alderton, Roberto Amato, Jeffrey Barrett, Sonia Goncalves, Ewan Harrison, David K. Jackson, Ian Johnston, Dominic Kwiatkowski, Cordelia Langford, John Sillitoe on behalf of the Wellcome Sanger Institute COVID-19 Surveillance Team |
| EPI_ISL_2779875, EPI_ISL_2779876, EPI_ISL_2779877 | Lighthouse Laboratory Plymouth | Wellcome Sanger Institute for the COVID-19 Genomics UK (COG-UK) Consortium | Lighthouse Laboratory Plymouth and Alex Alderton, Roberto Amato, Jeffrey Barrett, Sonia Goncalves, Ewan Harrison, David K. Jackson, Ian Johnston, Dominic Kwiatkowski, Cordelia Langford, John Sillitoe on behalf of the Wellcome Sanger Institute COVID-19 Surveillance Team |
| EPI_ISL_2779878 | Lighthouse Lab in Glasgow | Wellcome Sanger Institute for the COVID-19 Genomics UK (COG-UK) Consortium | Harper VanSteenhouse, Yumi Kasai, David Gray, Carol Clugston, Anna Dominiczak and Alex Alderton, Roberto Amato, Jeffrey Barrett, Sonia Goncalves, Ewan Harrison, David K. Jackson, Ian Johnston, Dominic Kwiatkowski, Cordelia Langford, John Sillitoe on behalf of the Wellcome Sanger Institute COVID-19 Surveillance Team |
| EPI_ISL_2779879 | Lighthouse Laboratory Plymouth | Wellcome Sanger Institute for the COVID-19 Genomics UK (COG-UK) Consortium | Lighthouse Laboratory Plymouth and Alex Alderton, Roberto Amato, Jeffrey Barrett, Sonia Goncalves, Ewan Harrison, David K. Jackson, Ian Johnston, Dominic Kwiatkowski, Cordelia Langford, John Sillitoe on behalf of the Wellcome Sanger Institute COVID-19 Surveillance Team |
| EPI_ISL_2779880 | Lighthouse Lab in Milton Keynes | Wellcome Sanger Institute for the COVID-19 Genomics UK (COG-UK) Consortium | The Lighthouse Lab in Milton Keynes and Alex Alderton, Roberto Amato, Jeffrey Barrett, Sonia Goncalves, Ewan Harrison, David K. Jackson, Ian Johnston, Dominic Kwiatkowski, Cordelia Langford, John Sillitoe on behalf of the Wellcome Sanger Institute COVID-19 Surveillance Team |
| EPI_ISL_2779882 | Lighthouse Lab in Glasgow | Wellcome Sanger Institute for the COVID-19 Genomics UK (COG-UK) Consortium | Harper VanSteenhouse, Yumi Kasai, David Gray, Carol Clugston, Anna Dominiczak and Alex Alderton, Roberto Amato, Jeffrey Barrett, Sonia Goncalves, Ewan Harrison, David K. Jackson, Ian Johnston, Dominic Kwiatkowski, Cordelia Langford, John Sillitoe on behalf of the Wellcome Sanger Institute COVID-19 Surveillance Team |
| EPI_ISL_2779883 | Lighthouse Lab in Milton Keynes | Wellcome Sanger Institute for the COVID-19 Genomics UK (COG-UK) Consortium | The Lighthouse Lab in Milton Keynes and Alex Alderton, Roberto Amato, Jeffrey Barrett, Sonia Goncalves, Ewan Harrison, David K. Jackson, Ian Johnston, Dominic Kwiatkowski, Cordelia Langford, John Sillitoe on behalf of the Wellcome Sanger Institute COVID-19 Surveillance Team |
| EPI_ISL_2779884 | Lighthouse Laboratory Plymouth | Wellcome Sanger Institute for the COVID-19 Genomics UK (COG-UK) Consortium | Lighthouse Laboratory Plymouth and Alex Alderton, Roberto Amato, Jeffrey Barrett, Sonia Goncalves, Ewan Harrison, David K. Jackson, Ian Johnston, Dominic Kwiatkowski, Cordelia Langford, John Sillitoe on behalf of the Wellcome Sanger Institute COVID-19 Surveillance Team |
| EPI_ISL_2779885, EPI_ISL_2779886, EPI_ISL_2779887, EPI_ISL_2779888 | Lighthouse Lab in Glasgow | Wellcome Sanger Institute for the COVID-19 Genomics UK (COG-UK) Consortium | Harper VanSteenhouse, Yumi Kasai, David Gray, Carol Clugston, Anna Dominiczak and Alex Alderton, Roberto Amato, Jeffrey Barrett, Sonia Goncalves, Ewan Harrison, David K. Jackson, Ian Johnston, Dominic Kwiatkowski, Cordelia Langford, John Sillitoe on behalf of the Wellcome Sanger Institute COVID-19 Surveillance Team |
| EPI_ISL_2779889 | Lighthouse Laboratory Plymouth | Wellcome Sanger Institute for the COVID-19 Genomics UK (COG-UK) Consortium | Lighthouse Laboratory Plymouth and Alex Alderton, Roberto Amato, Jeffrey Barrett, Sonia Goncalves, Ewan Harrison, David K. Jackson, Ian Johnston, Dominic Kwiatkowski, Cordelia Langford, John Sillitoe on behalf of the Wellcome Sanger Institute COVID-19 Surveillance Team |
| EPI_ISL_2779890 | Lighthouse Lab in Milton Keynes | Wellcome Sanger Institute for the COVID-19 Genomics UK (COG-UK) Consortium | The Lighthouse Lab in Milton Keynes and Alex Alderton, Roberto Amato, Jeffrey Barrett, Sonia Goncalves, Ewan Harrison, David K. Jackson, Ian Johnston, Dominic Kwiatkowski, Cordelia Langford, John Sillitoe on behalf of the Wellcome Sanger Institute COVID-19 Surveillance Team |
| EPI_ISL_2779891 | Lighthouse Laboratory Plymouth | Wellcome Sanger Institute for the COVID-19 Genomics UK (COG-UK) Consortium | Lighthouse Laboratory Plymouth and Alex Alderton, Roberto Amato, Jeffrey Barrett, Sonia Goncalves, Ewan Harrison, David K. Jackson, Ian Johnston, Dominic Kwiatkowski, Cordelia Langford, John Sillitoe on behalf of the Wellcome Sanger Institute COVID-19 Surveillance Team |
| EPI_ISL_2779892 | Lighthouse Lab in Glasgow | Wellcome Sanger Institute for the COVID-19 Genomics UK (COG-UK) Consortium | Harper VanSteenhouse, Yumi Kasai, David Gray, Carol Clugston, Anna Dominiczak and Alex Alderton, Roberto Amato, Jeffrey Barrett, Sonia Goncalves, Ewan Harrison, David K. Jackson, Ian Johnston, Dominic Kwiatkowski, Cordelia Langford, John Sillitoe on behalf of the Wellcome Sanger Institute COVID-19 Surveillance Team |
| EPI_ISL_2779893 | Lighthouse Laboratory Plymouth | Wellcome Sanger Institute for the COVID-19 Genomics UK (COG-UK) Consortium | Lighthouse Laboratory Plymouth and Alex Alderton, Roberto Amato, Jeffrey Barrett, Sonia Goncalves, Ewan Harrison, David K. Jackson, Ian Johnston, Dominic Kwiatkowski, Cordelia Langford, John Sillitoe on behalf of the Wellcome Sanger Institute COVID-19 Surveillance Team |
| EPI_ISL_2779894, EPI_ISL_2779895 | Lighthouse Lab in Glasgow | Wellcome Sanger Institute for the COVID-19 Genomics UK (COG-UK) Consortium | Harper VanSteenhouse, Yumi Kasai, David Gray, Carol Clugston, Anna Dominiczak and Alex Alderton, Roberto Amato, Jeffrey Barrett, Sonia Goncalves, Ewan Harrison, David K. Jackson, Ian Johnston, Dominic Kwiatkowski, Cordelia Langford, John Sillitoe on behalf of the Wellcome Sanger Institute COVID-19 Surveillance Team |
| EPI_ISL_2779896, EPI_ISL_2779897, EPI_ISL_2779898 | Lighthouse Laboratory Plymouth | Wellcome Sanger Institute for the COVID-19 Genomics UK (COG-UK) Consortium | Lighthouse Laboratory Plymouth and Alex Alderton, Roberto Amato, Jeffrey Barrett, Sonia Goncalves, Ewan Harrison, David K. Jackson, Ian Johnston, Dominic Kwiatkowski, Cordelia Langford, John Sillitoe on behalf of the Wellcome Sanger Institute COVID-19 Surveillance Team |
| EPI_ISL_2779899, EPI_ISL_2779900 | Lighthouse Lab in Milton Keynes | Wellcome Sanger Institute for the COVID-19 Genomics UK (COG-UK) Consortium | The Lighthouse Lab in Milton Keynes and Alex Alderton, Roberto Amato, Jeffrey Barrett, Sonia Goncalves, Ewan Harrison, David K. Jackson, Ian Johnston, Dominic Kwiatkowski, Cordelia Langford, John Sillitoe on behalf of the Wellcome Sanger Institute COVID-19 Surveillance Team |
| EPI_ISL_2779901 | Lighthouse Laboratory Plymouth | Wellcome Sanger Institute for the COVID-19 Genomics UK (COG-UK) Consortium | Lighthouse Laboratory Plymouth and Alex Alderton, Roberto Amato, Jeffrey Barrett, Sonia Goncalves, Ewan Harrison, David K. Jackson, Ian Johnston, Dominic Kwiatkowski, Cordelia Langford, John Sillitoe on behalf of the Wellcome Sanger Institute COVID-19 Surveillance Team |
| EPI_ISL_2779902 | Lighthouse Lab in Milton Keynes | Wellcome Sanger Institute for the COVID-19 Genomics UK (COG-UK) Consortium | The Lighthouse Lab in Milton Keynes and Alex Alderton, Roberto Amato, Jeffrey Barrett, Sonia Goncalves, Ewan Harrison, David K. Jackson, Ian Johnston, Dominic Kwiatkowski, Cordelia Langford, John Sillitoe on behalf of the Wellcome Sanger Institute COVID-19 Surveillance Team |
| EPI_ISL_2779903 | Lighthouse Laboratory Plymouth | Wellcome Sanger Institute for the COVID-19 Genomics UK (COG-UK) Consortium | Lighthouse Laboratory Plymouth and Alex Alderton, Roberto Amato, Jeffrey Barrett, Sonia Goncalves, Ewan Harrison, David K. Jackson, Ian Johnston, Dominic Kwiatkowski, Cordelia Langford, John Sillitoe on behalf of the Wellcome Sanger Institute COVID-19 Surveillance Team |
| EPI_ISL_2779904, EPI_ISL_2779905 | Lighthouse Lab in Milton Keynes | Wellcome Sanger Institute for the COVID-19 Genomics UK (COG-UK) Consortium | The Lighthouse Lab in Milton Keynes and Alex Alderton, Roberto Amato, Jeffrey Barrett, Sonia Goncalves, Ewan Harrison, David K. Jackson, Ian Johnston, Dominic Kwiatkowski, Cordelia Langford, John Sillitoe on behalf of the Wellcome Sanger Institute COVID-19 Surveillance Team |
| EPI_ISL_2779906 | Lighthouse Laboratory Plymouth | Wellcome Sanger Institute for the COVID-19 Genomics UK (COG-UK) Consortium | Lighthouse Laboratory Plymouth and Alex Alderton, Roberto Amato, Jeffrey Barrett, Sonia Goncalves, Ewan Harrison, David K. Jackson, Ian Johnston, Dominic Kwiatkowski, Cordelia Langford, John Sillitoe on behalf of the Wellcome Sanger Institute COVID-19 Surveillance Team |
| EPI_ISL_2779907 | Lighthouse Lab in Glasgow | Wellcome Sanger Institute for the COVID-19 Genomics UK (COG-UK) Consortium | Harper VanSteenhouse, Yumi Kasai, David Gray, Carol Clugston, Anna Dominiczak and Alex Alderton, Roberto Amato, Jeffrey Barrett, Sonia Goncalves, Ewan Harrison, David K. Jackson, Ian Johnston, Dominic Kwiatkowski, Cordelia Langford, John Sillitoe on behalf of the Wellcome Sanger Institute COVID-19 Surveillance Team |
| EPI_ISL_2779908 | Lighthouse Lab in Milton Keynes | Wellcome Sanger Institute for the COVID-19 Genomics UK (COG-UK) Consortium | The Lighthouse Lab in Milton Keynes and Alex Alderton, Roberto Amato, Jeffrey Barrett, Sonia Goncalves, Ewan Harrison, David K. Jackson, Ian Johnston, Dominic Kwiatkowski, Cordelia Langford, John Sillitoe on behalf of the Wellcome Sanger Institute COVID-19 Surveillance Team |
| EPI_ISL_2779909, EPI_ISL_2779910, EPI_ISL_2779911 | Lighthouse Laboratory Plymouth | Wellcome Sanger Institute for the COVID-19 Genomics UK (COG-UK) Consortium | Lighthouse Laboratory Plymouth and Alex Alderton, Roberto Amato, Jeffrey Barrett, Sonia Goncalves, Ewan Harrison, David K. Jackson, Ian Johnston, Dominic Kwiatkowski, Cordelia Langford, John Sillitoe on behalf of the Wellcome Sanger Institute COVID-19 Surveillance Team |

| | | | |
|---|---|---|---|
| EPI_ISL_2779912 | Lighthouse Lab in Milton Keynes | Wellcome Sanger Institute for the COVID-19 Genomics UK (COG-UK) Consortium | The Lighthouse Lab in Milton Keynes and Alex Alderton, Roberto Amato, Jeffrey Barrett, Sonia Goncalves, Ewan Harrison, David K. Jackson, Ian Johnston, Dominic Kwiatkowski, Cordelia Langford, John Sillitoe on behalf of the Wellcome Sanger Institute COVID-19 Surveillance Team |
| EPI_ISL_2779913 | Lighthouse Lab in Glasgow | Wellcome Sanger Institute for the COVID-19 Genomics UK (COG-UK) Consortium | Harper VanSteenhouse, Yumi Kasai, David Gray, Carol Clugston, Anna Dominiczak and Alex Alderton, Roberto Amato, Jeffrey Barrett, Sonia Goncalves, Ewan Harrison, David K. Jackson, Ian Johnston, Dominic Kwiatkowski, Cordelia Langford, John Sillitoe on behalf of the Wellcome Sanger Institute COVID-19 Surveillance Team |
| EPI_ISL_2779914, EPI_ISL_2779915 | Lighthouse Laboratory Plymouth | Wellcome Sanger Institute for the COVID-19 Genomics UK (COG-UK) Consortium | Lighthouse Laboratory Plymouth and Alex Alderton, Roberto Amato, Jeffrey Barrett, Sonia Goncalves, Ewan Harrison, David K. Jackson, Ian Johnston, Dominic Kwiatkowski, Cordelia Langford, John Sillitoe on behalf of the Wellcome Sanger Institute COVID-19 Surveillance Team |
| EPI_ISL_2779916 | Lighthouse Lab in Glasgow | Wellcome Sanger Institute for the COVID-19 Genomics UK (COG-UK) Consortium | Harper VanSteenhouse, Yumi Kasai, David Gray, Carol Clugston, Anna Dominiczak and Alex Alderton, Roberto Amato, Jeffrey Barrett, Sonia Goncalves, Ewan Harrison, David K. Jackson, Ian Johnston, Dominic Kwiatkowski, Cordelia Langford, John Sillitoe on behalf of the Wellcome Sanger Institute COVID-19 Surveillance Team |
| EPI_ISL_2779917 | Lighthouse Laboratory Plymouth | Wellcome Sanger Institute for the COVID-19 Genomics UK (COG-UK) Consortium | Lighthouse Laboratory Plymouth and Alex Alderton, Roberto Amato, Jeffrey Barrett, Sonia Goncalves, Ewan Harrison, David K. Jackson, Ian Johnston, Dominic Kwiatkowski, Cordelia Langford, John Sillitoe on behalf of the Wellcome Sanger Institute COVID-19 Surveillance Team |
| EPI_ISL_2779918, EPI_ISL_2779919 | Lighthouse Lab in Glasgow | Wellcome Sanger Institute for the COVID-19 Genomics UK (COG-UK) Consortium | Harper VanSteenhouse, Yumi Kasai, David Gray, Carol Clugston, Anna Dominiczak and Alex Alderton, Roberto Amato, Jeffrey Barrett, Sonia Goncalves, Ewan Harrison, David K. Jackson, Ian Johnston, Dominic Kwiatkowski, Cordelia Langford, John Sillitoe on behalf of the Wellcome Sanger Institute COVID-19 Surveillance Team |
| EPI_ISL_2779920 | Lighthouse Laboratory Plymouth | Wellcome Sanger Institute for the COVID-19 Genomics UK (COG-UK) Consortium | Lighthouse Laboratory Plymouth and Alex Alderton, Roberto Amato, Jeffrey Barrett, Sonia Goncalves, Ewan Harrison, David K. Jackson, Ian Johnston, Dominic Kwiatkowski, Cordelia Langford, John Sillitoe on behalf of the Wellcome Sanger Institute COVID-19 Surveillance Team |
| EPI_ISL_2779921, EPI_ISL_2779922 | Lighthouse Lab in Glasgow | Wellcome Sanger Institute for the COVID-19 Genomics UK (COG-UK) Consortium | Harper VanSteenhouse, Yumi Kasai, David Gray, Carol Clugston, Anna Dominiczak and Alex Alderton, Roberto Amato, Jeffrey Barrett, Sonia Goncalves, Ewan Harrison, David K. Jackson, Ian Johnston, Dominic Kwiatkowski, Cordelia Langford, John Sillitoe on behalf of the Wellcome Sanger Institute COVID-19 Surveillance Team |
| EPI_ISL_2779923, EPI_ISL_2779924, EPI_ISL_2779925, EPI_ISL_2779926, EPI_ISL_2779927, EPI_ISL_2779928 | Lighthouse Laboratory Plymouth | Wellcome Sanger Institute for the COVID-19 Genomics UK (COG-UK) Consortium | Lighthouse Laboratory Plymouth and Alex Alderton, Roberto Amato, Jeffrey Barrett, Sonia Goncalves, Ewan Harrison, David K. Jackson, Ian Johnston, Dominic Kwiatkowski, Cordelia Langford, John Sillitoe on behalf of the Wellcome Sanger Institute COVID-19 Surveillance Team |
| EPI_ISL_2779929 | Lighthouse Lab in Milton Keynes | Wellcome Sanger Institute for the COVID-19 Genomics UK (COG-UK) Consortium | The Lighthouse Lab in Milton Keynes and Alex Alderton, Roberto Amato, Jeffrey Barrett, Sonia Goncalves, Ewan Harrison, David K. Jackson, Ian Johnston, Dominic Kwiatkowski, Cordelia Langford, John Sillitoe on behalf of the Wellcome Sanger Institute COVID-19 Surveillance Team |
| EPI_ISL_2779930 | Lighthouse Lab in Glasgow | Wellcome Sanger Institute for the COVID-19 Genomics UK (COG-UK) Consortium | Harper VanSteenhouse, Yumi Kasai, David Gray, Carol Clugston, Anna Dominiczak and Alex Alderton, Roberto Amato, Jeffrey Barrett, Sonia Goncalves, Ewan Harrison, David K. Jackson, Ian Johnston, Dominic Kwiatkowski, Cordelia Langford, John Sillitoe on behalf of the Wellcome Sanger Institute COVID-19 Surveillance Team |
| EPI_ISL_2779931 | Lighthouse Laboratory Plymouth | Wellcome Sanger Institute for the COVID-19 Genomics UK (COG-UK) Consortium | Lighthouse Laboratory Plymouth and Alex Alderton, Roberto Amato, Jeffrey Barrett, Sonia Goncalves, Ewan Harrison, David K. Jackson, Ian Johnston, Dominic Kwiatkowski, Cordelia Langford, John Sillitoe on behalf of the Wellcome Sanger Institute COVID-19 Surveillance Team |
| EPI_ISL_2779932 | Lighthouse Lab in Milton Keynes | Wellcome Sanger Institute for the COVID-19 Genomics UK (COG-UK) Consortium | The Lighthouse Lab in Milton Keynes and Alex Alderton, Roberto Amato, Jeffrey Barrett, Sonia Goncalves, Ewan Harrison, David K. Jackson, Ian Johnston, Dominic Kwiatkowski, Cordelia Langford, John Sillitoe on behalf of the Wellcome Sanger Institute COVID-19 Surveillance Team |
| EPI_ISL_2779933 | Lighthouse Lab in Glasgow | Wellcome Sanger Institute for the COVID-19 Genomics UK (COG-UK) Consortium | Harper VanSteenhouse, Yumi Kasai, David Gray, Carol Clugston, Anna Dominiczak and Alex Alderton, Roberto Amato, Jeffrey Barrett, Sonia Goncalves, Ewan Harrison, David K. Jackson, Ian Johnston, Dominic Kwiatkowski, Cordelia Langford, John Sillitoe on behalf of the Wellcome Sanger Institute COVID-19 Surveillance Team |
| EPI_ISL_2779934, EPI_ISL_2779935, EPI_ISL_2779936 | Lighthouse Lab in Milton Keynes | Wellcome Sanger Institute for the COVID-19 Genomics UK (COG-UK) Consortium | The Lighthouse Lab in Milton Keynes and Alex Alderton, Roberto Amato, Jeffrey Barrett, Sonia Goncalves, Ewan Harrison, David K. Jackson, Ian Johnston, Dominic Kwiatkowski, Cordelia Langford, John Sillitoe on behalf of the Wellcome Sanger Institute COVID-19 Surveillance Team |
| EPI_ISL_2779937, EPI_ISL_2779938 | Lighthouse Lab in Glasgow | Wellcome Sanger Institute for the COVID-19 Genomics UK (COG-UK) Consortium | Harper VanSteenhouse, Yumi Kasai, David Gray, Carol Clugston, Anna Dominiczak and Alex Alderton, Roberto Amato, Jeffrey Barrett, Sonia Goncalves, Ewan Harrison, David K. Jackson, Ian Johnston, Dominic Kwiatkowski, Cordelia Langford, John Sillitoe on behalf of the Wellcome Sanger Institute COVID-19 Surveillance Team |
| EPI_ISL_2779939 | Lighthouse Laboratory Plymouth | Wellcome Sanger Institute for the COVID-19 Genomics UK (COG-UK) Consortium | Lighthouse Laboratory Plymouth and Alex Alderton, Roberto Amato, Jeffrey Barrett, Sonia Goncalves, Ewan Harrison, David K. Jackson, Ian Johnston, Dominic Kwiatkowski, Cordelia Langford, John Sillitoe on behalf of the Wellcome Sanger Institute COVID-19 Surveillance Team |
| EPI_ISL_2779940, EPI_ISL_2779941 | Lighthouse Lab in Milton Keynes | Wellcome Sanger Institute for the COVID-19 Genomics UK (COG-UK) Consortium | The Lighthouse Lab in Milton Keynes and Alex Alderton, Roberto Amato, Jeffrey Barrett, Sonia Goncalves, Ewan Harrison, David K. Jackson, Ian Johnston, Dominic Kwiatkowski, Cordelia Langford, John Sillitoe on behalf of the Wellcome Sanger Institute COVID-19 Surveillance Team |
| EPI_ISL_2779942, EPI_ISL_2779943, EPI_ISL_2779944 | Lighthouse Laboratory Plymouth | Wellcome Sanger Institute for the COVID-19 Genomics UK (COG-UK) Consortium | Lighthouse Laboratory Plymouth and Alex Alderton, Roberto Amato, Jeffrey Barrett, Sonia Goncalves, Ewan Harrison, David K. Jackson, Ian Johnston, Dominic Kwiatkowski, Cordelia Langford, John Sillitoe on behalf of the Wellcome Sanger Institute COVID-19 Surveillance Team |
| EPI_ISL_2779945 | Lighthouse Lab in Glasgow | Wellcome Sanger Institute for the COVID-19 Genomics UK (COG-UK) Consortium | Harper VanSteenhouse, Yumi Kasai, David Gray, Carol Clugston, Anna Dominiczak and Alex Alderton, Roberto Amato, Jeffrey Barrett, Sonia Goncalves, Ewan Harrison, David K. Jackson, Ian Johnston, Dominic Kwiatkowski, Cordelia Langford, John Sillitoe on behalf of the Wellcome Sanger Institute COVID-19 Surveillance Team |
| EPI_ISL_2779946 | Lighthouse Laboratory Plymouth | Wellcome Sanger Institute for the COVID-19 Genomics UK (COG-UK) Consortium | Lighthouse Laboratory Plymouth and Alex Alderton, Roberto Amato, Jeffrey Barrett, Sonia Goncalves, Ewan Harrison, David K. Jackson, Ian Johnston, Dominic Kwiatkowski, Cordelia Langford, John Sillitoe on behalf of the Wellcome Sanger Institute COVID-19 Surveillance Team |
| EPI_ISL_2779947 | Lighthouse Lab in Glasgow | Wellcome Sanger Institute for the COVID-19 Genomics UK (COG-UK) Consortium | Harper VanSteenhouse, Yumi Kasai, David Gray, Carol Clugston, Anna Dominiczak and Alex Alderton, Roberto Amato, Jeffrey Barrett, Sonia Goncalves, Ewan Harrison, David K. Jackson, Ian Johnston, Dominic Kwiatkowski, Cordelia Langford, John Sillitoe on behalf of the Wellcome Sanger Institute COVID-19 Surveillance Team |
| EPI_ISL_2779948, EPI_ISL_2779949 | Lighthouse Laboratory Plymouth | Wellcome Sanger Institute for the COVID-19 Genomics UK (COG-UK) Consortium | Lighthouse Laboratory Plymouth and Alex Alderton, Roberto Amato, Jeffrey Barrett, Sonia Goncalves, Ewan Harrison, David K. Jackson, Ian Johnston, Dominic Kwiatkowski, Cordelia Langford, John Sillitoe on behalf of the Wellcome Sanger Institute COVID-19 Surveillance Team |
| EPI_ISL_2779950 | Lighthouse Lab in Milton Keynes | Wellcome Sanger Institute for the COVID-19 Genomics UK (COG-UK) Consortium | The Lighthouse Lab in Milton Keynes and Alex Alderton, Roberto Amato, Jeffrey Barrett, Sonia Goncalves, Ewan Harrison, David K. Jackson, Ian Johnston, Dominic Kwiatkowski, Cordelia Langford, John Sillitoe on behalf of the Wellcome Sanger Institute COVID-19 Surveillance Team |
| EPI_ISL_2779951, EPI_ISL_2779952 | Lighthouse Laboratory Plymouth | Wellcome Sanger Institute for the COVID-19 Genomics UK (COG-UK) Consortium | Lighthouse Laboratory Plymouth and Alex Alderton, Roberto Amato, Jeffrey Barrett, Sonia Goncalves, Ewan Harrison, David K. Jackson, Ian Johnston, Dominic Kwiatkowski, Cordelia Langford, John Sillitoe on behalf of the Wellcome Sanger Institute COVID-19 Surveillance Team |
| EPI_ISL_2779953 | Lighthouse Lab in Glasgow | Wellcome Sanger Institute for the COVID-19 Genomics UK (COG-UK) Consortium | Harper VanSteenhouse, Yumi Kasai, David Gray, Carol Clugston, Anna Dominiczak and Alex Alderton, Roberto Amato, Jeffrey Barrett, Sonia Goncalves, Ewan Harrison, David K. Jackson, Ian Johnston, Dominic Kwiatkowski, Cordelia Langford, John Sillitoe on behalf of the Wellcome Sanger Institute COVID-19 Surveillance Team |
| EPI_ISL_2779954, EPI_ISL_2779955 | Lighthouse Laboratory Plymouth | Wellcome Sanger Institute for the COVID-19 Genomics UK (COG-UK) Consortium | Lighthouse Laboratory Plymouth and Alex Alderton, Roberto Amato, Jeffrey Barrett, Sonia Goncalves, Ewan Harrison, David K. Jackson, Ian Johnston, Dominic Kwiatkowski, Cordelia Langford, John Sillitoe on behalf of the Wellcome Sanger Institute COVID-19 Surveillance Team |
| EPI_ISL_2779956 | Lighthouse Lab in Glasgow | Wellcome Sanger Institute for the COVID-19 Genomics UK (COG-UK) Consortium | Harper VanSteenhouse, Yumi Kasai, David Gray, Carol Clugston, Anna Dominiczak and Alex Alderton, Roberto Amato, Jeffrey Barrett, Sonia Goncalves, Ewan Harrison, David K. Jackson, Ian Johnston, Dominic Kwiatkowski, Cordelia Langford, John Sillitoe on behalf of the Wellcome Sanger Institute COVID-19 Surveillance Team |
| EPI_ISL_2779957 | Lighthouse Laboratory Plymouth | Wellcome Sanger Institute for the COVID-19 Genomics UK (COG-UK) Consortium | Lighthouse Laboratory Plymouth and Alex Alderton, Roberto Amato, Jeffrey Barrett, Sonia Goncalves, Ewan Harrison, David K. Jackson, Ian Johnston, Dominic Kwiatkowski, Cordelia Langford, John Sillitoe on behalf of the Wellcome Sanger Institute COVID-19 Surveillance Team |
| EPI_ISL_2779958 | Lighthouse Lab in Milton Keynes | Wellcome Sanger Institute for the COVID-19 Genomics UK (COG-UK) Consortium | The Lighthouse Lab in Milton Keynes and Alex Alderton, Roberto Amato, Jeffrey Barrett, Sonia Goncalves, Ewan Harrison, David K. Jackson, Ian Johnston, Dominic Kwiatkowski, Cordelia Langford, John Sillitoe on behalf of the Wellcome Sanger Institute COVID-19 Surveillance Team |
| EPI_ISL_2779959 | Lighthouse Lab in Glasgow | Wellcome Sanger Institute for the COVID-19 Genomics UK (COG-UK) Consortium | Harper VanSteenhouse, Yumi Kasai, David Gray, Carol Clugston, Anna Dominiczak and Alex Alderton, Roberto Amato, Jeffrey Barrett, Sonia Goncalves, Ewan Harrison, David K. Jackson, Ian Johnston, Dominic Kwiatkowski, Cordelia Langford, John Sillitoe on behalf of the Wellcome Sanger Institute COVID-19 Surveillance Team |
| EPI_ISL_2779960, EPI_ISL_2779961 | Lighthouse Laboratory Plymouth | Wellcome Sanger Institute for the COVID-19 Genomics UK (COG-UK) Consortium | Lighthouse Laboratory Plymouth and Alex Alderton, Roberto Amato, Jeffrey Barrett, Sonia Goncalves, Ewan Harrison, David K. Jackson, Ian Johnston, Dominic Kwiatkowski, Cordelia Langford, John Sillitoe on behalf of the Wellcome Sanger Institute COVID-19 Surveillance Team |
| EPI_ISL_2779962 | Lighthouse Lab in Glasgow | Wellcome Sanger Institute for the COVID-19 Genomics UK (COG-UK) Consortium | Harper VanSteenhouse, Yumi Kasai, David Gray, Carol Clugston, Anna Dominiczak and Alex Alderton, Roberto Amato, Jeffrey Barrett, Sonia Goncalves, Ewan Harrison, David K. Jackson, Ian Johnston, Dominic Kwiatkowski, Cordelia Langford, John Sillitoe on behalf of the Wellcome Sanger Institute COVID-19 Surveillance Team |
| EPI_ISL_2779963, EPI_ISL_2779964 | Lighthouse Laboratory Plymouth | Wellcome Sanger Institute for the COVID-19 Genomics UK (COG-UK) Consortium | Lighthouse Laboratory Plymouth and Alex Alderton, Roberto Amato, Jeffrey Barrett, Sonia Goncalves, Ewan Harrison, David K. Jackson, Ian Johnston, Dominic Kwiatkowski, Cordelia Langford, John Sillitoe on behalf of the Wellcome Sanger Institute COVID-19 Surveillance Team |
| EPI_ISL_2779965 | Lighthouse Lab in Glasgow | Wellcome Sanger Institute for the COVID-19 Genomics UK (COG-UK) Consortium | Harper VanSteenhouse, Yumi Kasai, David Gray, Carol Clugston, Anna Dominiczak and Alex Alderton, Roberto Amato, Jeffrey Barrett, Sonia Goncalves, Ewan Harrison, David K. Jackson, Ian Johnston, Dominic Kwiatkowski, Cordelia Langford, John Sillitoe on behalf of the Wellcome Sanger Institute COVID-19 Surveillance Team |
| EPI_ISL_2779966, EPI_ISL_2779967 | Lighthouse Lab in Milton Keynes | Wellcome Sanger Institute for the COVID-19 Genomics UK (COG-UK) Consortium | The Lighthouse Lab in Milton Keynes and Alex Alderton, Roberto Amato, Jeffrey Barrett, Sonia Goncalves, Ewan Harrison, David K. Jackson, Ian Johnston, Dominic Kwiatkowski, Cordelia Langford, John Sillitoe on behalf of the Wellcome Sanger Institute COVID-19 Surveillance Team |
| EPI_ISL_2779968, EPI_ISL_2779969 | Lighthouse Lab in Glasgow | Wellcome Sanger Institute for the COVID-19 Genomics UK (COG-UK) Consortium | Harper VanSteenhouse, Yumi Kasai, David Gray, Carol Clugston, Anna Dominiczak and Alex Alderton, Roberto Amato, Jeffrey Barrett, Sonia Goncalves, Ewan Harrison, David K. Jackson, Ian Johnston, Dominic Kwiatkowski, Cordelia Langford, John Sillitoe on behalf of the Wellcome Sanger Institute COVID-19 Surveillance Team |
| EPI_ISL_2779970, EPI_ISL_2779971, EPI_ISL_2779972 | Lighthouse Laboratory Plymouth | Wellcome Sanger Institute for the COVID-19 Genomics UK (COG-UK) Consortium | Lighthouse Laboratory Plymouth and Alex Alderton, Roberto Amato, Jeffrey Barrett, Sonia Goncalves, Ewan Harrison, David K. Jackson, Ian Johnston, Dominic Kwiatkowski, Cordelia Langford, John Sillitoe on behalf of the Wellcome Sanger Institute COVID-19 Surveillance Team |
| EPI_ISL_2779973 | Lighthouse Lab in Milton Keynes | Wellcome Sanger Institute for the COVID-19 Genomics UK (COG-UK) Consortium | The Lighthouse Lab in Milton Keynes and Alex Alderton, Roberto Amato, Jeffrey Barrett, Sonia Goncalves, Ewan Harrison, David K. Jackson, Ian Johnston, Dominic Kwiatkowski, Cordelia Langford, John Sillitoe on behalf of the Wellcome Sanger Institute COVID-19 Surveillance Team |
| EPI_ISL_2779974, EPI_ISL_2779975 | Lighthouse Lab in Glasgow | Wellcome Sanger Institute for the COVID-19 Genomics UK (COG-UK) Consortium | Harper VanSteenhouse, Yumi Kasai, David Gray, Carol Clugston, Anna Dominiczak and Alex Alderton, Roberto Amato, Jeffrey Barrett, Sonia Goncalves, Ewan Harrison, David K. Jackson, Ian Johnston, Dominic Kwiatkowski, Cordelia Langford, John Sillitoe on behalf of the Wellcome Sanger Institute COVID-19 Surveillance Team |
| EPI_ISL_2779976 | Lighthouse Laboratory Plymouth | Wellcome Sanger Institute for the COVID-19 Genomics UK (COG-UK) Consortium | Lighthouse Laboratory Plymouth and Alex Alderton, Roberto Amato, Jeffrey Barrett, Sonia Goncalves, Ewan Harrison, David K. Jackson, Ian Johnston, Dominic Kwiatkowski, Cordelia Langford, John Sillitoe on behalf of the Wellcome Sanger Institute COVID-19 Surveillance Team |
| EPI_ISL_2779977 | Lighthouse Lab in Milton Keynes | Wellcome Sanger Institute for the COVID-19 Genomics UK (COG-UK) Consortium | The Lighthouse Lab in Milton Keynes and Alex Alderton, Roberto Amato, Jeffrey Barrett, Sonia Goncalves, Ewan Harrison, David K. Jackson, Ian Johnston, Dominic Kwiatkowski, Cordelia Langford, John Sillitoe on behalf of the Wellcome Sanger Institute COVID-19 Surveillance Team |

| | | | |
|---|---|---|---|
| EPI_ISL_2779978 | Lighthouse Laboratory Plymouth | Wellcome Sanger Institute for the COVID-19 Genomics UK (COG-UK) Consortium | Lighthouse Laboratory Plymouth and Alex Alderton, Roberto Amato, Jeffrey Barrett, Sonia Goncalves, Ewan Harrison, David K. Jackson, Ian Johnston, Dominic Kwiatkowski, Cordelia Langford, John Sillitoe on behalf of the Wellcome Sanger Institute COVID-19 Surveillance Team |
| EPI_ISL_2779979 | Lighthouse Lab in Milton Keynes | Wellcome Sanger Institute for the COVID-19 Genomics UK (COG-UK) Consortium | The Lighthouse Lab in Milton Keynes and Alex Alderton, Roberto Amato, Jeffrey Barrett, Sonia Goncalves, Ewan Harrison, David K. Jackson, Ian Johnston, Dominic Kwiatkowski, Cordelia Langford, John Sillitoe on behalf of the Wellcome Sanger Institute COVID-19 Surveillance Team |
| EPI_ISL_2779980, EPI_ISL_2779981 | Lighthouse Lab in Glasgow | Wellcome Sanger Institute for the COVID-19 Genomics UK (COG-UK) Consortium | Harper VanSteenhouse, Yumi Kasai, David Gray, Carol Clugston, Anna Dominiczak and Alex Alderton, Roberto Amato, Jeffrey Barrett, Sonia Goncalves, Ewan Harrison, David K. Jackson, Ian Johnston, Dominic Kwiatkowski, Cordelia Langford, John Sillitoe on behalf of the Wellcome Sanger Institute COVID-19 Surveillance Team |
| EPI_ISL_2779982, EPI_ISL_2779983 | Lighthouse Laboratory Plymouth | Wellcome Sanger Institute for the COVID-19 Genomics UK (COG-UK) Consortium | Lighthouse Laboratory Plymouth and Alex Alderton, Roberto Amato, Jeffrey Barrett, Sonia Goncalves, Ewan Harrison, David K. Jackson, Ian Johnston, Dominic Kwiatkowski, Cordelia Langford, John Sillitoe on behalf of the Wellcome Sanger Institute COVID-19 Surveillance Team |
| EPI_ISL_2779984, EPI_ISL_2779985 | Lighthouse Lab in Milton Keynes | Wellcome Sanger Institute for the COVID-19 Genomics UK (COG-UK) Consortium | The Lighthouse Lab in Milton Keynes and Alex Alderton, Roberto Amato, Jeffrey Barrett, Sonia Goncalves, Ewan Harrison, David K. Jackson, Ian Johnston, Dominic Kwiatkowski, Cordelia Langford, John Sillitoe on behalf of the Wellcome Sanger Institute COVID-19 Surveillance Team |
| EPI_ISL_2779986 | Lighthouse Laboratory Plymouth | Wellcome Sanger Institute for the COVID-19 Genomics UK (COG-UK) Consortium | Lighthouse Laboratory Plymouth and Alex Alderton, Roberto Amato, Jeffrey Barrett, Sonia Goncalves, Ewan Harrison, David K. Jackson, Ian Johnston, Dominic Kwiatkowski, Cordelia Langford, John Sillitoe on behalf of the Wellcome Sanger Institute COVID-19 Surveillance Team |
| EPI_ISL_2779987, EPI_ISL_2779988 | Lighthouse Lab in Milton Keynes | Wellcome Sanger Institute for the COVID-19 Genomics UK (COG-UK) Consortium | The Lighthouse Lab in Milton Keynes and Alex Alderton, Roberto Amato, Jeffrey Barrett, Sonia Goncalves, Ewan Harrison, David K. Jackson, Ian Johnston, Dominic Kwiatkowski, Cordelia Langford, John Sillitoe on behalf of the Wellcome Sanger Institute COVID-19 Surveillance Team |
| EPI_ISL_2779989 | Lighthouse Laboratory Plymouth | Wellcome Sanger Institute for the COVID-19 Genomics UK (COG-UK) Consortium | Lighthouse Laboratory Plymouth and Alex Alderton, Roberto Amato, Jeffrey Barrett, Sonia Goncalves, Ewan Harrison, David K. Jackson, Ian Johnston, Dominic Kwiatkowski, Cordelia Langford, John Sillitoe on behalf of the Wellcome Sanger Institute COVID-19 Surveillance Team |
| EPI_ISL_2779990 | Lighthouse Lab in Milton Keynes | Wellcome Sanger Institute for the COVID-19 Genomics UK (COG-UK) Consortium | The Lighthouse Lab in Milton Keynes and Alex Alderton, Roberto Amato, Jeffrey Barrett, Sonia Goncalves, Ewan Harrison, David K. Jackson, Ian Johnston, Dominic Kwiatkowski, Cordelia Langford, John Sillitoe on behalf of the Wellcome Sanger Institute COVID-19 Surveillance Team |
| EPI_ISL_2779991 | Lighthouse Laboratory Plymouth | Wellcome Sanger Institute for the COVID-19 Genomics UK (COG-UK) Consortium | Lighthouse Laboratory Plymouth and Alex Alderton, Roberto Amato, Jeffrey Barrett, Sonia Goncalves, Ewan Harrison, David K. Jackson, Ian Johnston, Dominic Kwiatkowski, Cordelia Langford, John Sillitoe on behalf of the Wellcome Sanger Institute COVID-19 Surveillance Team |
| EPI_ISL_2779992 | Lighthouse Lab in Glasgow | Wellcome Sanger Institute for the COVID-19 Genomics UK (COG-UK) Consortium | Harper VanSteenhouse, Yumi Kasai, David Gray, Carol Clugston, Anna Dominiczak and Alex Alderton, Roberto Amato, Jeffrey Barrett, Sonia Goncalves, Ewan Harrison, David K. Jackson, Ian Johnston, Dominic Kwiatkowski, Cordelia Langford, John Sillitoe on behalf of the Wellcome Sanger Institute COVID-19 Surveillance Team |
| EPI_ISL_2779993 | Lighthouse Lab in Milton Keynes | Wellcome Sanger Institute for the COVID-19 Genomics UK (COG-UK) Consortium | The Lighthouse Lab in Milton Keynes and Alex Alderton, Roberto Amato, Jeffrey Barrett, Sonia Goncalves, Ewan Harrison, David K. Jackson, Ian Johnston, Dominic Kwiatkowski, Cordelia Langford, John Sillitoe on behalf of the Wellcome Sanger Institute COVID-19 Surveillance Team |
| EPI_ISL_2779994 | Lighthouse Laboratory Plymouth | Wellcome Sanger Institute for the COVID-19 Genomics UK (COG-UK) Consortium | Lighthouse Laboratory Plymouth and Alex Alderton, Roberto Amato, Jeffrey Barrett, Sonia Goncalves, Ewan Harrison, David K. Jackson, Ian Johnston, Dominic Kwiatkowski, Cordelia Langford, John Sillitoe on behalf of the Wellcome Sanger Institute COVID-19 Surveillance Team |
| EPI_ISL_2779995 | Lighthouse Lab in Milton Keynes | Wellcome Sanger Institute for the COVID-19 Genomics UK (COG-UK) Consortium | The Lighthouse Lab in Milton Keynes and Alex Alderton, Roberto Amato, Jeffrey Barrett, Sonia Goncalves, Ewan Harrison, David K. Jackson, Ian Johnston, Dominic Kwiatkowski, Cordelia Langford, John Sillitoe on behalf of the Wellcome Sanger Institute COVID-19 Surveillance Team |
| EPI_ISL_2779996 | Lighthouse Laboratory Plymouth | Wellcome Sanger Institute for the COVID-19 Genomics UK (COG-UK) Consortium | Lighthouse Laboratory Plymouth and Alex Alderton, Roberto Amato, Jeffrey Barrett, Sonia Goncalves, Ewan Harrison, David K. Jackson, Ian Johnston, Dominic Kwiatkowski, Cordelia Langford, John Sillitoe on behalf of the Wellcome Sanger Institute COVID-19 Surveillance Team |
| EPI_ISL_2779997 | Lighthouse Lab in Milton Keynes | Wellcome Sanger Institute for the COVID-19 Genomics UK (COG-UK) Consortium | The Lighthouse Lab in Milton Keynes and Alex Alderton, Roberto Amato, Jeffrey Barrett, Sonia Goncalves, Ewan Harrison, David K. Jackson, Ian Johnston, Dominic Kwiatkowski, Cordelia Langford, John Sillitoe on behalf of the Wellcome Sanger Institute COVID-19 Surveillance Team |
| EPI_ISL_2779998, EPI_ISL_2779999 | Lighthouse Laboratory Plymouth | Wellcome Sanger Institute for the COVID-19 Genomics UK (COG-UK) Consortium | Lighthouse Laboratory Plymouth and Alex Alderton, Roberto Amato, Jeffrey Barrett, Sonia Goncalves, Ewan Harrison, David K. Jackson, Ian Johnston, Dominic Kwiatkowski, Cordelia Langford, John Sillitoe on behalf of the Wellcome Sanger Institute COVID-19 Surveillance Team |
| EPI_ISL_2780000, EPI_ISL_2780001 | Lighthouse Lab in Glasgow | Wellcome Sanger Institute for the COVID-19 Genomics UK (COG-UK) Consortium | Harper VanSteenhouse, Yumi Kasai, David Gray, Carol Clugston, Anna Dominiczak and Alex Alderton, Roberto Amato, Jeffrey Barrett, Sonia Goncalves, Ewan Harrison, David K. Jackson, Ian Johnston, Dominic Kwiatkowski, Cordelia Langford, John Sillitoe on behalf of the Wellcome Sanger Institute COVID-19 Surveillance Team |
| EPI_ISL_2780002, EPI_ISL_2780003, EPI_ISL_2780004, EPI_ISL_2780005 | Lighthouse Lab in Milton Keynes | Wellcome Sanger Institute for the COVID-19 Genomics UK (COG-UK) Consortium | The Lighthouse Lab in Milton Keynes and Alex Alderton, Roberto Amato, Jeffrey Barrett, Sonia Goncalves, Ewan Harrison, David K. Jackson, Ian Johnston, Dominic Kwiatkowski, Cordelia Langford, John Sillitoe on behalf of the Wellcome Sanger Institute COVID-19 Surveillance Team |
| EPI_ISL_2780006 | Lighthouse Laboratory Plymouth | Wellcome Sanger Institute for the COVID-19 Genomics UK (COG-UK) Consortium | Lighthouse Laboratory Plymouth and Alex Alderton, Roberto Amato, Jeffrey Barrett, Sonia Goncalves, Ewan Harrison, David K. Jackson, Ian Johnston, Dominic Kwiatkowski, Cordelia Langford, John Sillitoe on behalf of the Wellcome Sanger Institute COVID-19 Surveillance Team |
| EPI_ISL_2780007 | Lighthouse Lab in Milton Keynes | Wellcome Sanger Institute for the COVID-19 Genomics UK (COG-UK) Consortium | The Lighthouse Lab in Milton Keynes and Alex Alderton, Roberto Amato, Jeffrey Barrett, Sonia Goncalves, Ewan Harrison, David K. Jackson, Ian Johnston, Dominic Kwiatkowski, Cordelia Langford, John Sillitoe on behalf of the Wellcome Sanger Institute COVID-19 Surveillance Team |
| EPI_ISL_2780008 | Lighthouse Laboratory Plymouth | Wellcome Sanger Institute for the COVID-19 Genomics UK (COG-UK) Consortium | Lighthouse Laboratory Plymouth and Alex Alderton, Roberto Amato, Jeffrey Barrett, Sonia Goncalves, Ewan Harrison, David K. Jackson, Ian Johnston, Dominic Kwiatkowski, Cordelia Langford, John Sillitoe on behalf of the Wellcome Sanger Institute COVID-19 Surveillance Team |
| EPI_ISL_2780009 | Lighthouse Lab in Glasgow | Wellcome Sanger Institute for the COVID-19 Genomics UK (COG-UK) Consortium | Harper VanSteenhouse, Yumi Kasai, David Gray, Carol Clugston, Anna Dominiczak and Alex Alderton, Roberto Amato, Jeffrey Barrett, Sonia Goncalves, Ewan Harrison, David K. Jackson, Ian Johnston, Dominic Kwiatkowski, Cordelia Langford, John Sillitoe on behalf of the Wellcome Sanger Institute COVID-19 Surveillance Team |
| EPI_ISL_2780010 | Lighthouse Lab in Milton Keynes | Wellcome Sanger Institute for the COVID-19 Genomics UK (COG-UK) Consortium | The Lighthouse Lab in Milton Keynes and Alex Alderton, Roberto Amato, Jeffrey Barrett, Sonia Goncalves, Ewan Harrison, David K. Jackson, Ian Johnston, Dominic Kwiatkowski, Cordelia Langford, John Sillitoe on behalf of the Wellcome Sanger Institute COVID-19 Surveillance Team |
| EPI_ISL_2780011 | Lighthouse Laboratory Plymouth | Wellcome Sanger Institute for the COVID-19 Genomics UK (COG-UK) Consortium | Lighthouse Laboratory Plymouth and Alex Alderton, Roberto Amato, Jeffrey Barrett, Sonia Goncalves, Ewan Harrison, David K. Jackson, Ian Johnston, Dominic Kwiatkowski, Cordelia Langford, John Sillitoe on behalf of the Wellcome Sanger Institute COVID-19 Surveillance Team |
| EPI_ISL_2780012 | Lighthouse Lab in Glasgow | Wellcome Sanger Institute for the COVID-19 Genomics UK (COG-UK) Consortium | Harper VanSteenhouse, Yumi Kasai, David Gray, Carol Clugston, Anna Dominiczak and Alex Alderton, Roberto Amato, Jeffrey Barrett, Sonia Goncalves, Ewan Harrison, David K. Jackson, Ian Johnston, Dominic Kwiatkowski, Cordelia Langford, John Sillitoe on behalf of the Wellcome Sanger Institute COVID-19 Surveillance Team |
| EPI_ISL_2780013 | Lighthouse Laboratory Plymouth | Wellcome Sanger Institute for the COVID-19 Genomics UK (COG-UK) Consortium | Lighthouse Laboratory Plymouth and Alex Alderton, Roberto Amato, Jeffrey Barrett, Sonia Goncalves, Ewan Harrison, David K. Jackson, Ian Johnston, Dominic Kwiatkowski, Cordelia Langford, John Sillitoe on behalf of the Wellcome Sanger Institute COVID-19 Surveillance Team |
| EPI_ISL_2780014 | Lighthouse Lab in Glasgow | Wellcome Sanger Institute for the COVID-19 Genomics UK (COG-UK) Consortium | Harper VanSteenhouse, Yumi Kasai, David Gray, Carol Clugston, Anna Dominiczak and Alex Alderton, Roberto Amato, Jeffrey Barrett, Sonia Goncalves, Ewan Harrison, David K. Jackson, Ian Johnston, Dominic Kwiatkowski, Cordelia Langford, John Sillitoe on behalf of the Wellcome Sanger Institute COVID-19 Surveillance Team |
| EPI_ISL_2780015 | Lighthouse Laboratory Plymouth | Wellcome Sanger Institute for the COVID-19 Genomics UK (COG-UK) Consortium | Lighthouse Laboratory Plymouth and Alex Alderton, Roberto Amato, Jeffrey Barrett, Sonia Goncalves, Ewan Harrison, David K. Jackson, Ian Johnston, Dominic Kwiatkowski, Cordelia Langford, John Sillitoe on behalf of the Wellcome Sanger Institute COVID-19 Surveillance Team |
| EPI_ISL_2780016 | Lighthouse Lab in Glasgow | Wellcome Sanger Institute for the COVID-19 Genomics UK (COG-UK) Consortium | Harper VanSteenhouse, Yumi Kasai, David Gray, Carol Clugston, Anna Dominiczak and Alex Alderton, Roberto Amato, Jeffrey Barrett, Sonia Goncalves, Ewan Harrison, David K. Jackson, Ian Johnston, Dominic Kwiatkowski, Cordelia Langford, John Sillitoe on behalf of the Wellcome Sanger Institute COVID-19 Surveillance Team |
| EPI_ISL_2780017, EPI_ISL_2780018 | Lighthouse Lab in Milton Keynes | Wellcome Sanger Institute for the COVID-19 Genomics UK (COG-UK) Consortium | The Lighthouse Lab in Milton Keynes and Alex Alderton, Roberto Amato, Jeffrey Barrett, Sonia Goncalves, Ewan Harrison, David K. Jackson, Ian Johnston, Dominic Kwiatkowski, Cordelia Langford, John Sillitoe on behalf of the Wellcome Sanger Institute COVID-19 Surveillance Team |
| EPI_ISL_2780019 | Lighthouse Laboratory Plymouth | Wellcome Sanger Institute for the COVID-19 Genomics UK (COG-UK) Consortium | Lighthouse Laboratory Plymouth and Alex Alderton, Roberto Amato, Jeffrey Barrett, Sonia Goncalves, Ewan Harrison, David K. Jackson, Ian Johnston, Dominic Kwiatkowski, Cordelia Langford, John Sillitoe on behalf of the Wellcome Sanger Institute COVID-19 Surveillance Team |
| EPI_ISL_2780020, EPI_ISL_2780021, EPI_ISL_2780022 | Lighthouse Lab in Milton Keynes | Wellcome Sanger Institute for the COVID-19 Genomics UK (COG-UK) Consortium | The Lighthouse Lab in Milton Keynes and Alex Alderton, Roberto Amato, Jeffrey Barrett, Sonia Goncalves, Ewan Harrison, David K. Jackson, Ian Johnston, Dominic Kwiatkowski, Cordelia Langford, John Sillitoe on behalf of the Wellcome Sanger Institute COVID-19 Surveillance Team |
| EPI_ISL_2780023 | Lighthouse Laboratory Plymouth | Wellcome Sanger Institute for the COVID-19 Genomics UK (COG-UK) Consortium | Lighthouse Laboratory Plymouth and Alex Alderton, Roberto Amato, Jeffrey Barrett, Sonia Goncalves, Ewan Harrison, David K. Jackson, Ian Johnston, Dominic Kwiatkowski, Cordelia Langford, John Sillitoe on behalf of the Wellcome Sanger Institute COVID-19 Surveillance Team |
| EPI_ISL_2780024 | Lighthouse Lab in Glasgow | Wellcome Sanger Institute for the COVID-19 Genomics UK (COG-UK) Consortium | Harper VanSteenhouse, Yumi Kasai, David Gray, Carol Clugston, Anna Dominiczak and Alex Alderton, Roberto Amato, Jeffrey Barrett, Sonia Goncalves, Ewan Harrison, David K. Jackson, Ian Johnston, Dominic Kwiatkowski, Cordelia Langford, John Sillitoe on behalf of the Wellcome Sanger Institute COVID-19 Surveillance Team |
| EPI_ISL_2780025 | Lighthouse Lab in Milton Keynes | Wellcome Sanger Institute for the COVID-19 Genomics UK (COG-UK) Consortium | The Lighthouse Lab in Milton Keynes and Alex Alderton, Roberto Amato, Jeffrey Barrett, Sonia Goncalves, Ewan Harrison, David K. Jackson, Ian Johnston, Dominic Kwiatkowski, Cordelia Langford, John Sillitoe on behalf of the Wellcome Sanger Institute COVID-19 Surveillance Team |
| EPI_ISL_2780026 | Lighthouse Lab in Glasgow | Wellcome Sanger Institute for the COVID-19 Genomics UK (COG-UK) Consortium | Harper VanSteenhouse, Yumi Kasai, David Gray, Carol Clugston, Anna Dominiczak and Alex Alderton, Roberto Amato, Jeffrey Barrett, Sonia Goncalves, Ewan Harrison, David K. Jackson, Ian Johnston, Dominic Kwiatkowski, Cordelia Langford, John Sillitoe on behalf of the Wellcome Sanger Institute COVID-19 Surveillance Team |
| EPI_ISL_2780027, EPI_ISL_2780028 | Lighthouse Laboratory Plymouth | Wellcome Sanger Institute for the COVID-19 Genomics UK (COG-UK) Consortium | Lighthouse Laboratory Plymouth and Alex Alderton, Roberto Amato, Jeffrey Barrett, Sonia Goncalves, Ewan Harrison, David K. Jackson, Ian Johnston, Dominic Kwiatkowski, Cordelia Langford, John Sillitoe on behalf of the Wellcome Sanger Institute COVID-19 Surveillance Team |
| EPI_ISL_2780029, EPI_ISL_2780030 | Lighthouse Lab in Milton Keynes | Wellcome Sanger Institute for the COVID-19 Genomics UK (COG-UK) Consortium | The Lighthouse Lab in Milton Keynes and Alex Alderton, Roberto Amato, Jeffrey Barrett, Sonia Goncalves, Ewan Harrison, David K. Jackson, Ian Johnston, Dominic Kwiatkowski, Cordelia Langford, John Sillitoe on behalf of the Wellcome Sanger Institute COVID-19 Surveillance Team |
| EPI_ISL_2780031 | Lighthouse Laboratory Plymouth | Wellcome Sanger Institute for the COVID-19 Genomics UK (COG-UK) Consortium | Lighthouse Laboratory Plymouth and Alex Alderton, Roberto Amato, Jeffrey Barrett, Sonia Goncalves, Ewan Harrison, David K. Jackson, Ian Johnston, Dominic Kwiatkowski, Cordelia Langford, John Sillitoe on behalf of the Wellcome Sanger Institute COVID-19 Surveillance Team |
| EPI_ISL_2780032 | Lighthouse Lab in Glasgow | Wellcome Sanger Institute for the COVID-19 Genomics UK (COG-UK) Consortium | Harper VanSteenhouse, Yumi Kasai, David Gray, Carol Clugston, Anna Dominiczak and Alex Alderton, Roberto Amato, Jeffrey Barrett, Sonia Goncalves, Ewan Harrison, David K. Jackson, Ian Johnston, Dominic Kwiatkowski, Cordelia Langford, John Sillitoe on behalf of the Wellcome Sanger Institute COVID-19 Surveillance Team |
| EPI_ISL_2780033 | Lighthouse Lab in Milton Keynes | Wellcome Sanger Institute for the COVID-19 Genomics UK (COG-UK) Consortium | The Lighthouse Lab in Milton Keynes and Alex Alderton, Roberto Amato, Jeffrey Barrett, Sonia Goncalves, Ewan Harrison, David K. Jackson, Ian Johnston, Dominic Kwiatkowski, Cordelia Langford, John Sillitoe on behalf of the Wellcome Sanger Institute COVID-19 Surveillance Team |
| EPI_ISL_2780034, EPI_ISL_2780035 | Lighthouse Laboratory Plymouth | Wellcome Sanger Institute for the COVID-19 Genomics UK (COG-UK) Consortium | Lighthouse Laboratory Plymouth and Alex Alderton, Roberto Amato, Jeffrey Barrett, Sonia Goncalves, Ewan Harrison, David K. Jackson, Ian Johnston, Dominic Kwiatkowski, Cordelia Langford, John Sillitoe on behalf of the Wellcome Sanger Institute COVID-19 Surveillance Team |
| EPI_ISL_2780036 | Lighthouse Lab in Glasgow | Wellcome Sanger Institute for the COVID-19 Genomics UK (COG-UK) Consortium | Harper VanSteenhouse, Yumi Kasai, David Gray, Carol Clugston, Anna Dominiczak and Alex Alderton, Roberto Amato, Jeffrey Barrett, Sonia Goncalves, Ewan Harrison, David K. Jackson, Ian Johnston, Dominic Kwiatkowski, Cordelia Langford, John Sillitoe on behalf of the Wellcome Sanger Institute COVID-19 Surveillance Team |

| | | | |
|---|---|---|---|
| EPI_ISL_2780037, EPI_ISL_2780038 | Lighthouse Lab in Milton Keynes | Wellcome Sanger Institute for the COVID-19 Genomics UK (COG-UK) Consortium | The Lighthouse Lab in Milton Keynes and Alex Alderton, Roberto Amato, Jeffrey Barrett, Sonia Goncalves, Ewan Harrison, David K. Jackson, Ian Johnston, Dominic Kwiatkowski, Cordelia Langford, John Sillitoe on behalf of the Wellcome Sanger Institute COVID-19 Surveillance Team |
| EPI_ISL_2780039, EPI_ISL_2780040 | Lighthouse Laboratory Plymouth | Wellcome Sanger Institute for the COVID-19 Genomics UK (COG-UK) Consortium | Lighthouse Laboratory Plymouth and Alex Alderton, Roberto Amato, Jeffrey Barrett, Sonia Goncalves, Ewan Harrison, David K. Jackson, Ian Johnston, Dominic Kwiatkowski, Cordelia Langford, John Sillitoe on behalf of the Wellcome Sanger Institute COVID-19 Surveillance Team |
| EPI_ISL_2780041, EPI_ISL_2780042 | Lighthouse Lab in Glasgow | Wellcome Sanger Institute for the COVID-19 Genomics UK (COG-UK) Consortium | Harper VanSteenhouse, Yumi Kasai, David Gray, Carol Clugston, Anna Dominiczak and Alex Alderton, Roberto Amato, Jeffrey Barrett, Sonia Goncalves, Ewan Harrison, David K. Jackson, Ian Johnston, Dominic Kwiatkowski, Cordelia Langford, John Sillitoe on behalf of the Wellcome Sanger Institute COVID-19 Surveillance Team |
| EPI_ISL_2780043 | Lighthouse Lab in Milton Keynes | Wellcome Sanger Institute for the COVID-19 Genomics UK (COG-UK) Consortium | The Lighthouse Lab in Milton Keynes and Alex Alderton, Roberto Amato, Jeffrey Barrett, Sonia Goncalves, Ewan Harrison, David K. Jackson, Ian Johnston, Dominic Kwiatkowski, Cordelia Langford, John Sillitoe on behalf of the Wellcome Sanger Institute COVID-19 Surveillance Team |
| EPI_ISL_2780044, EPI_ISL_2780045 | Lighthouse Lab in Glasgow | Wellcome Sanger Institute for the COVID-19 Genomics UK (COG-UK) Consortium | Harper VanSteenhouse, Yumi Kasai, David Gray, Carol Clugston, Anna Dominiczak and Alex Alderton, Roberto Amato, Jeffrey Barrett, Sonia Goncalves, Ewan Harrison, David K. Jackson, Ian Johnston, Dominic Kwiatkowski, Cordelia Langford, John Sillitoe on behalf of the Wellcome Sanger Institute COVID-19 Surveillance Team |
| EPI_ISL_2780046 | Lighthouse Lab in Milton Keynes | Wellcome Sanger Institute for the COVID-19 Genomics UK (COG-UK) Consortium | The Lighthouse Lab in Milton Keynes and Alex Alderton, Roberto Amato, Jeffrey Barrett, Sonia Goncalves, Ewan Harrison, David K. Jackson, Ian Johnston, Dominic Kwiatkowski, Cordelia Langford, John Sillitoe on behalf of the Wellcome Sanger Institute COVID-19 Surveillance Team |
| EPI_ISL_2780047 | Lighthouse Lab in Glasgow | Wellcome Sanger Institute for the COVID-19 Genomics UK (COG-UK) Consortium | Harper VanSteenhouse, Yumi Kasai, David Gray, Carol Clugston, Anna Dominiczak and Alex Alderton, Roberto Amato, Jeffrey Barrett, Sonia Goncalves, Ewan Harrison, David K. Jackson, Ian Johnston, Dominic Kwiatkowski, Cordelia Langford, John Sillitoe on behalf of the Wellcome Sanger Institute COVID-19 Surveillance Team |
| EPI_ISL_2780048 | Lighthouse Laboratory Plymouth | Wellcome Sanger Institute for the COVID-19 Genomics UK (COG-UK) Consortium | Lighthouse Laboratory Plymouth and Alex Alderton, Roberto Amato, Jeffrey Barrett, Sonia Goncalves, Ewan Harrison, David K. Jackson, Ian Johnston, Dominic Kwiatkowski, Cordelia Langford, John Sillitoe on behalf of the Wellcome Sanger Institute COVID-19 Surveillance Team |
| EPI_ISL_2780049 | Lighthouse Lab in Milton Keynes | Wellcome Sanger Institute for the COVID-19 Genomics UK (COG-UK) Consortium | The Lighthouse Lab in Milton Keynes and Alex Alderton, Roberto Amato, Jeffrey Barrett, Sonia Goncalves, Ewan Harrison, David K. Jackson, Ian Johnston, Dominic Kwiatkowski, Cordelia Langford, John Sillitoe on behalf of the Wellcome Sanger Institute COVID-19 Surveillance Team |
| EPI_ISL_2780050 | Lighthouse Laboratory Plymouth | Wellcome Sanger Institute for the COVID-19 Genomics UK (COG-UK) Consortium | Lighthouse Laboratory Plymouth and Alex Alderton, Roberto Amato, Jeffrey Barrett, Sonia Goncalves, Ewan Harrison, David K. Jackson, Ian Johnston, Dominic Kwiatkowski, Cordelia Langford, John Sillitoe on behalf of the Wellcome Sanger Institute COVID-19 Surveillance Team |
| EPI_ISL_2780051 | Lighthouse Lab in Glasgow | Wellcome Sanger Institute for the COVID-19 Genomics UK (COG-UK) Consortium | Harper VanSteenhouse, Yumi Kasai, David Gray, Carol Clugston, Anna Dominiczak and Alex Alderton, Roberto Amato, Jeffrey Barrett, Sonia Goncalves, Ewan Harrison, David K. Jackson, Ian Johnston, Dominic Kwiatkowski, Cordelia Langford, John Sillitoe on behalf of the Wellcome Sanger Institute COVID-19 Surveillance Team |
| EPI_ISL_2780052 | Lighthouse Laboratory Plymouth | Wellcome Sanger Institute for the COVID-19 Genomics UK (COG-UK) Consortium | Lighthouse Laboratory Plymouth and Alex Alderton, Roberto Amato, Jeffrey Barrett, Sonia Goncalves, Ewan Harrison, David K. Jackson, Ian Johnston, Dominic Kwiatkowski, Cordelia Langford, John Sillitoe on behalf of the Wellcome Sanger Institute COVID-19 Surveillance Team |
| EPI_ISL_2780053 | Lighthouse Lab in Milton Keynes | Wellcome Sanger Institute for the COVID-19 Genomics UK (COG-UK) Consortium | The Lighthouse Lab in Milton Keynes and Alex Alderton, Roberto Amato, Jeffrey Barrett, Sonia Goncalves, Ewan Harrison, David K. Jackson, Ian Johnston, Dominic Kwiatkowski, Cordelia Langford, John Sillitoe on behalf of the Wellcome Sanger Institute COVID-19 Surveillance Team |
| EPI_ISL_2780054 | Lighthouse Laboratory Plymouth | Wellcome Sanger Institute for the COVID-19 Genomics UK (COG-UK) Consortium | Lighthouse Laboratory Plymouth and Alex Alderton, Roberto Amato, Jeffrey Barrett, Sonia Goncalves, Ewan Harrison, David K. Jackson, Ian Johnston, Dominic Kwiatkowski, Cordelia Langford, John Sillitoe on behalf of the Wellcome Sanger Institute COVID-19 Surveillance Team |
| EPI_ISL_2780055, EPI_ISL_2780056, EPI_ISL_2780057 | Lighthouse Lab in Glasgow | Wellcome Sanger Institute for the COVID-19 Genomics UK (COG-UK) Consortium | Harper VanSteenhouse, Yumi Kasai, David Gray, Carol Clugston, Anna Dominiczak and Alex Alderton, Roberto Amato, Jeffrey Barrett, Sonia Goncalves, Ewan Harrison, David K. Jackson, Ian Johnston, Dominic Kwiatkowski, Cordelia Langford, John Sillitoe on behalf of the Wellcome Sanger Institute COVID-19 Surveillance Team |
| EPI_ISL_2780058 | Lighthouse Laboratory Plymouth | Wellcome Sanger Institute for the COVID-19 Genomics UK (COG-UK) Consortium | Lighthouse Laboratory Plymouth and Alex Alderton, Roberto Amato, Jeffrey Barrett, Sonia Goncalves, Ewan Harrison, David K. Jackson, Ian Johnston, Dominic Kwiatkowski, Cordelia Langford, John Sillitoe on behalf of the Wellcome Sanger Institute COVID-19 Surveillance Team |
| EPI_ISL_2780059 | Lighthouse Lab in Glasgow | Wellcome Sanger Institute for the COVID-19 Genomics UK (COG-UK) Consortium | Harper VanSteenhouse, Yumi Kasai, David Gray, Carol Clugston, Anna Dominiczak and Alex Alderton, Roberto Amato, Jeffrey Barrett, Sonia Goncalves, Ewan Harrison, David K. Jackson, Ian Johnston, Dominic Kwiatkowski, Cordelia Langford, John Sillitoe on behalf of the Wellcome Sanger Institute COVID-19 Surveillance Team |
| EPI_ISL_2780060 | Lighthouse Lab in Milton Keynes | Wellcome Sanger Institute for the COVID-19 Genomics UK (COG-UK) Consortium | The Lighthouse Lab in Milton Keynes and Alex Alderton, Roberto Amato, Jeffrey Barrett, Sonia Goncalves, Ewan Harrison, David K. Jackson, Ian Johnston, Dominic Kwiatkowski, Cordelia Langford, John Sillitoe on behalf of the Wellcome Sanger Institute COVID-19 Surveillance Team |
| EPI_ISL_2780061, EPI_ISL_2780062, EPI_ISL_2780063, EPI_ISL_2780064 | Lighthouse Laboratory Plymouth | Wellcome Sanger Institute for the COVID-19 Genomics UK (COG-UK) Consortium | Lighthouse Laboratory Plymouth and Alex Alderton, Roberto Amato, Jeffrey Barrett, Sonia Goncalves, Ewan Harrison, David K. Jackson, Ian Johnston, Dominic Kwiatkowski, Cordelia Langford, John Sillitoe on behalf of the Wellcome Sanger Institute COVID-19 Surveillance Team |
| EPI_ISL_2780065 | Lighthouse Lab in Glasgow | Wellcome Sanger Institute for the COVID-19 Genomics UK (COG-UK) Consortium | Harper VanSteenhouse, Yumi Kasai, David Gray, Carol Clugston, Anna Dominiczak and Alex Alderton, Roberto Amato, Jeffrey Barrett, Sonia Goncalves, Ewan Harrison, David K. Jackson, Ian Johnston, Dominic Kwiatkowski, Cordelia Langford, John Sillitoe on behalf of the Wellcome Sanger Institute COVID-19 Surveillance Team |
| EPI_ISL_2780066, EPI_ISL_2780067, EPI_ISL_2780068, EPI_ISL_2780069 | Lighthouse Lab in Milton Keynes | Wellcome Sanger Institute for the COVID-19 Genomics UK (COG-UK) Consortium | The Lighthouse Lab in Milton Keynes and Alex Alderton, Roberto Amato, Jeffrey Barrett, Sonia Goncalves, Ewan Harrison, David K. Jackson, Ian Johnston, Dominic Kwiatkowski, Cordelia Langford, John Sillitoe on behalf of the Wellcome Sanger Institute COVID-19 Surveillance Team |
| EPI_ISL_2780070 | Lighthouse Lab in Glasgow | Wellcome Sanger Institute for the COVID-19 Genomics UK (COG-UK) Consortium | Harper VanSteenhouse, Yumi Kasai, David Gray, Carol Clugston, Anna Dominiczak and Alex Alderton, Roberto Amato, Jeffrey Barrett, Sonia Goncalves, Ewan Harrison, David K. Jackson, Ian Johnston, Dominic Kwiatkowski, Cordelia Langford, John Sillitoe on behalf of the Wellcome Sanger Institute COVID-19 Surveillance Team |
| EPI_ISL_2780071 | Lighthouse Lab in Milton Keynes | Wellcome Sanger Institute for the COVID-19 Genomics UK (COG-UK) Consortium | The Lighthouse Lab in Milton Keynes and Alex Alderton, Roberto Amato, Jeffrey Barrett, Sonia Goncalves, Ewan Harrison, David K. Jackson, Ian Johnston, Dominic Kwiatkowski, Cordelia Langford, John Sillitoe on behalf of the Wellcome Sanger Institute COVID-19 Surveillance Team |
| EPI_ISL_2780072 | Lighthouse Lab in Glasgow | Wellcome Sanger Institute for the COVID-19 Genomics UK (COG-UK) Consortium | Harper VanSteenhouse, Yumi Kasai, David Gray, Carol Clugston, Anna Dominiczak and Alex Alderton, Roberto Amato, Jeffrey Barrett, Sonia Goncalves, Ewan Harrison, David K. Jackson, Ian Johnston, Dominic Kwiatkowski, Cordelia Langford, John Sillitoe on behalf of the Wellcome Sanger Institute COVID-19 Surveillance Team |
| EPI_ISL_2780073 | Lighthouse Lab in Milton Keynes | Wellcome Sanger Institute for the COVID-19 Genomics UK (COG-UK) Consortium | The Lighthouse Lab in Milton Keynes and Alex Alderton, Roberto Amato, Jeffrey Barrett, Sonia Goncalves, Ewan Harrison, David K. Jackson, Ian Johnston, Dominic Kwiatkowski, Cordelia Langford, John Sillitoe on behalf of the Wellcome Sanger Institute COVID-19 Surveillance Team |
| EPI_ISL_2780074 | Lighthouse Lab in Glasgow | Wellcome Sanger Institute for the COVID-19 Genomics UK (COG-UK) Consortium | Harper VanSteenhouse, Yumi Kasai, David Gray, Carol Clugston, Anna Dominiczak and Alex Alderton, Roberto Amato, Jeffrey Barrett, Sonia Goncalves, Ewan Harrison, David K. Jackson, Ian Johnston, Dominic Kwiatkowski, Cordelia Langford, John Sillitoe on behalf of the Wellcome Sanger Institute COVID-19 Surveillance Team |
| EPI_ISL_2780075, EPI_ISL_2780076 | Lighthouse Lab in Milton Keynes | Wellcome Sanger Institute for the COVID-19 Genomics UK (COG-UK) Consortium | The Lighthouse Lab in Milton Keynes and Alex Alderton, Roberto Amato, Jeffrey Barrett, Sonia Goncalves, Ewan Harrison, David K. Jackson, Ian Johnston, Dominic Kwiatkowski, Cordelia Langford, John Sillitoe on behalf of the Wellcome Sanger Institute COVID-19 Surveillance Team |
| EPI_ISL_2780077 | Lighthouse Lab in Glasgow | Wellcome Sanger Institute for the COVID-19 Genomics UK (COG-UK) Consortium | Harper VanSteenhouse, Yumi Kasai, David Gray, Carol Clugston, Anna Dominiczak and Alex Alderton, Roberto Amato, Jeffrey Barrett, Sonia Goncalves, Ewan Harrison, David K. Jackson, Ian Johnston, Dominic Kwiatkowski, Cordelia Langford, John Sillitoe on behalf of the Wellcome Sanger Institute COVID-19 Surveillance Team |
| EPI_ISL_2780078 | Lighthouse Laboratory Plymouth | Wellcome Sanger Institute for the COVID-19 Genomics UK (COG-UK) Consortium | Lighthouse Laboratory Plymouth and Alex Alderton, Roberto Amato, Jeffrey Barrett, Sonia Goncalves, Ewan Harrison, David K. Jackson, Ian Johnston, Dominic Kwiatkowski, Cordelia Langford, John Sillitoe on behalf of the Wellcome Sanger Institute COVID-19 Surveillance Team |
| EPI_ISL_2780079 | Lighthouse Lab in Milton Keynes | Wellcome Sanger Institute for the COVID-19 Genomics UK (COG-UK) Consortium | The Lighthouse Lab in Milton Keynes and Alex Alderton, Roberto Amato, Jeffrey Barrett, Sonia Goncalves, Ewan Harrison, David K. Jackson, Ian Johnston, Dominic Kwiatkowski, Cordelia Langford, John Sillitoe on behalf of the Wellcome Sanger Institute COVID-19 Surveillance Team |
| EPI_ISL_2780080 | Lighthouse Lab in Glasgow | Wellcome Sanger Institute for the COVID-19 Genomics UK (COG-UK) Consortium | Harper VanSteenhouse, Yumi Kasai, David Gray, Carol Clugston, Anna Dominiczak and Alex Alderton, Roberto Amato, Jeffrey Barrett, Sonia Goncalves, Ewan Harrison, David K. Jackson, Ian Johnston, Dominic Kwiatkowski, Cordelia Langford, John Sillitoe on behalf of the Wellcome Sanger Institute COVID-19 Surveillance Team |
| EPI_ISL_2780081 | Lighthouse Lab in Milton Keynes | Wellcome Sanger Institute for the COVID-19 Genomics UK (COG-UK) Consortium | The Lighthouse Lab in Milton Keynes and Alex Alderton, Roberto Amato, Jeffrey Barrett, Sonia Goncalves, Ewan Harrison, David K. Jackson, Ian Johnston, Dominic Kwiatkowski, Cordelia Langford, John Sillitoe on behalf of the Wellcome Sanger Institute COVID-19 Surveillance Team |
| EPI_ISL_2780082 | Lighthouse Lab in Glasgow | Wellcome Sanger Institute for the COVID-19 Genomics UK (COG-UK) Consortium | Harper VanSteenhouse, Yumi Kasai, David Gray, Carol Clugston, Anna Dominiczak and Alex Alderton, Roberto Amato, Jeffrey Barrett, Sonia Goncalves, Ewan Harrison, David K. Jackson, Ian Johnston, Dominic Kwiatkowski, Cordelia Langford, John Sillitoe on behalf of the Wellcome Sanger Institute COVID-19 Surveillance Team |
| EPI_ISL_2780083, EPI_ISL_2780084, EPI_ISL_2780085, EPI_ISL_2780086, EPI_ISL_2780087 | Lighthouse Laboratory Plymouth | Wellcome Sanger Institute for the COVID-19 Genomics UK (COG-UK) Consortium | Lighthouse Laboratory Plymouth and Alex Alderton, Roberto Amato, Jeffrey Barrett, Sonia Goncalves, Ewan Harrison, David K. Jackson, Ian Johnston, Dominic Kwiatkowski, Cordelia Langford, John Sillitoe on behalf of the Wellcome Sanger Institute COVID-19 Surveillance Team |
| EPI_ISL_2780088, EPI_ISL_2780089 | Lighthouse Lab in Milton Keynes | Wellcome Sanger Institute for the COVID-19 Genomics UK (COG-UK) Consortium | The Lighthouse Lab in Milton Keynes and Alex Alderton, Roberto Amato, Jeffrey Barrett, Sonia Goncalves, Ewan Harrison, David K. Jackson, Ian Johnston, Dominic Kwiatkowski, Cordelia Langford, John Sillitoe on behalf of the Wellcome Sanger Institute COVID-19 Surveillance Team |
| EPI_ISL_2780090 | Lighthouse Lab in Glasgow | Wellcome Sanger Institute for the COVID-19 Genomics UK (COG-UK) Consortium | Harper VanSteenhouse, Yumi Kasai, David Gray, Carol Clugston, Anna Dominiczak and Alex Alderton, Roberto Amato, Jeffrey Barrett, Sonia Goncalves, Ewan Harrison, David K. Jackson, Ian Johnston, Dominic Kwiatkowski, Cordelia Langford, John Sillitoe on behalf of the Wellcome Sanger Institute COVID-19 Surveillance Team |
| EPI_ISL_2780091 | Lighthouse Laboratory Plymouth | Wellcome Sanger Institute for the COVID-19 Genomics UK (COG-UK) Consortium | Lighthouse Laboratory Plymouth and Alex Alderton, Roberto Amato, Jeffrey Barrett, Sonia Goncalves, Ewan Harrison, David K. Jackson, Ian Johnston, Dominic Kwiatkowski, Cordelia Langford, John Sillitoe on behalf of the Wellcome Sanger Institute COVID-19 Surveillance Team |
| EPI_ISL_2780092, EPI_ISL_2780093 | Lighthouse Lab in Glasgow | Wellcome Sanger Institute for the COVID-19 Genomics UK (COG-UK) Consortium | Harper VanSteenhouse, Yumi Kasai, David Gray, Carol Clugston, Anna Dominiczak and Alex Alderton, Roberto Amato, Jeffrey Barrett, Sonia Goncalves, Ewan Harrison, David K. Jackson, Ian Johnston, Dominic Kwiatkowski, Cordelia Langford, John Sillitoe on behalf of the Wellcome Sanger Institute COVID-19 Surveillance Team |
| EPI_ISL_2780094 | Lighthouse Laboratory Plymouth | Wellcome Sanger Institute for the COVID-19 Genomics UK (COG-UK) Consortium | Lighthouse Laboratory Plymouth and Alex Alderton, Roberto Amato, Jeffrey Barrett, Sonia Goncalves, Ewan Harrison, David K. Jackson, Ian Johnston, Dominic Kwiatkowski, Cordelia Langford, John Sillitoe on behalf of the Wellcome Sanger Institute COVID-19 Surveillance Team |
| EPI_ISL_2780095 | Lighthouse Lab in Milton Keynes | Wellcome Sanger Institute for the COVID-19 Genomics UK (COG-UK) Consortium | The Lighthouse Lab in Milton Keynes and Alex Alderton, Roberto Amato, Jeffrey Barrett, Sonia Goncalves, Ewan Harrison, David K. Jackson, Ian Johnston, Dominic Kwiatkowski, Cordelia Langford, John Sillitoe on behalf of the Wellcome Sanger Institute COVID-19 Surveillance Team |
| EPI_ISL_2780096, EPI_ISL_2780097, EPI_ISL_2780098 | Lighthouse Laboratory Plymouth | Wellcome Sanger Institute for the COVID-19 Genomics UK (COG-UK) Consortium | Lighthouse Laboratory Plymouth and Alex Alderton, Roberto Amato, Jeffrey Barrett, Sonia Goncalves, Ewan Harrison, David K. Jackson, Ian Johnston, Dominic Kwiatkowski, Cordelia Langford, John Sillitoe on behalf of the Wellcome Sanger Institute COVID-19 Surveillance Team |
| EPI_ISL_2780099, | Lighthouse Lab in Milton Keynes | Wellcome Sanger Institute for the COVID-19 Genomics UK (COG-UK) | The Lighthouse Lab in Milton Keynes and Alex Alderton, Roberto Amato, Jeffrey Barrett, Sonia Goncalves, Ewan Harrison, David K. Jackson, Ian Johnston, Dominic Kwiatkowski, Cordelia Langford, John Sillitoe on behalf of the Wellcome Sanger |

| | | | |
|---|---|---|---|
| EPI_ISL_2780100 | | Consortium | Institute COVID-19 Surveillance Team |
| EPI_ISL_2780101 | Lighthouse Lab in Glasgow | Wellcome Sanger Institute for the COVID-19 Genomics UK (COG-UK) Consortium | Harper VanSteenhouse, Yumi Kasai, David Gray, Carol Clugston, Anna Dominiczak and Alex Alderton, Roberto Amato, Jeffrey Barrett, Sonia Goncalves, Ewan Harrison, David K. Jackson, Ian Johnston, Dominic Kwiatkowski, Cordelia Langford, John Sillitoe on behalf of the Wellcome Sanger Institute COVID-19 Surveillance Team |
| EPI_ISL_2780102 | Lighthouse Laboratory Plymouth | Wellcome Sanger Institute for the COVID-19 Genomics UK (COG-UK) Consortium | Lighthouse Laboratory Plymouth and Alex Alderton, Roberto Amato, Jeffrey Barrett, Sonia Goncalves, Ewan Harrison, David K. Jackson, Ian Johnston, Dominic Kwiatkowski, Cordelia Langford, John Sillitoe on behalf of the Wellcome Sanger Institute COVID-19 Surveillance Team |
| EPI_ISL_2780103, EPI_ISL_2780104 | Lighthouse Lab in Milton Keynes | Wellcome Sanger Institute for the COVID-19 Genomics UK (COG-UK) Consortium | The Lighthouse Lab in Milton Keynes and Alex Alderton, Roberto Amato, Jeffrey Barrett, Sonia Goncalves, Ewan Harrison, David K. Jackson, Ian Johnston, Dominic Kwiatkowski, Cordelia Langford, John Sillitoe on behalf of the Wellcome Sanger Institute COVID-19 Surveillance Team |
| EPI_ISL_2780105, EPI_ISL_2780106 | Lighthouse Lab in Glasgow | Wellcome Sanger Institute for the COVID-19 Genomics UK (COG-UK) Consortium | Harper VanSteenhouse, Yumi Kasai, David Gray, Carol Clugston, Anna Dominiczak and Alex Alderton, Roberto Amato, Jeffrey Barrett, Sonia Goncalves, Ewan Harrison, David K. Jackson, Ian Johnston, Dominic Kwiatkowski, Cordelia Langford, John Sillitoe on behalf of the Wellcome Sanger Institute COVID-19 Surveillance Team |
| EPI_ISL_2780107, EPI_ISL_2780108 | Lighthouse Laboratory Plymouth | Wellcome Sanger Institute for the COVID-19 Genomics UK (COG-UK) Consortium | Lighthouse Laboratory Plymouth and Alex Alderton, Roberto Amato, Jeffrey Barrett, Sonia Goncalves, Ewan Harrison, David K. Jackson, Ian Johnston, Dominic Kwiatkowski, Cordelia Langford, John Sillitoe on behalf of the Wellcome Sanger Institute COVID-19 Surveillance Team |
| EPI_ISL_2780109 | Lighthouse Lab in Glasgow | Wellcome Sanger Institute for the COVID-19 Genomics UK (COG-UK) Consortium | Harper VanSteenhouse, Yumi Kasai, David Gray, Carol Clugston, Anna Dominiczak and Alex Alderton, Roberto Amato, Jeffrey Barrett, Sonia Goncalves, Ewan Harrison, David K. Jackson, Ian Johnston, Dominic Kwiatkowski, Cordelia Langford, John Sillitoe on behalf of the Wellcome Sanger Institute COVID-19 Surveillance Team |
| EPI_ISL_2780110, EPI_ISL_2780111 | Lighthouse Lab in Milton Keynes | Wellcome Sanger Institute for the COVID-19 Genomics UK (COG-UK) Consortium | The Lighthouse Lab in Milton Keynes and Alex Alderton, Roberto Amato, Jeffrey Barrett, Sonia Goncalves, Ewan Harrison, David K. Jackson, Ian Johnston, Dominic Kwiatkowski, Cordelia Langford, John Sillitoe on behalf of the Wellcome Sanger Institute COVID-19 Surveillance Team |
| EPI_ISL_2780112, EPI_ISL_2780113 | Lighthouse Lab in Glasgow | Wellcome Sanger Institute for the COVID-19 Genomics UK (COG-UK) Consortium | Harper VanSteenhouse, Yumi Kasai, David Gray, Carol Clugston, Anna Dominiczak and Alex Alderton, Roberto Amato, Jeffrey Barrett, Sonia Goncalves, Ewan Harrison, David K. Jackson, Ian Johnston, Dominic Kwiatkowski, Cordelia Langford, John Sillitoe on behalf of the Wellcome Sanger Institute COVID-19 Surveillance Team |
| EPI_ISL_2780114 | Lighthouse Lab in Milton Keynes | Wellcome Sanger Institute for the COVID-19 Genomics UK (COG-UK) Consortium | The Lighthouse Lab in Milton Keynes and Alex Alderton, Roberto Amato, Jeffrey Barrett, Sonia Goncalves, Ewan Harrison, David K. Jackson, Ian Johnston, Dominic Kwiatkowski, Cordelia Langford, John Sillitoe on behalf of the Wellcome Sanger Institute COVID-19 Surveillance Team |
| EPI_ISL_2780115 | Lighthouse Laboratory Plymouth | Wellcome Sanger Institute for the COVID-19 Genomics UK (COG-UK) Consortium | Lighthouse Laboratory Plymouth and Alex Alderton, Roberto Amato, Jeffrey Barrett, Sonia Goncalves, Ewan Harrison, David K. Jackson, Ian Johnston, Dominic Kwiatkowski, Cordelia Langford, John Sillitoe on behalf of the Wellcome Sanger Institute COVID-19 Surveillance Team |
| EPI_ISL_2780116, EPI_ISL_2780117, EPI_ISL_2780118, EPI_ISL_2780119 | Lighthouse Lab in Milton Keynes | Wellcome Sanger Institute for the COVID-19 Genomics UK (COG-UK) Consortium | The Lighthouse Lab in Milton Keynes and Alex Alderton, Roberto Amato, Jeffrey Barrett, Sonia Goncalves, Ewan Harrison, David K. Jackson, Ian Johnston, Dominic Kwiatkowski, Cordelia Langford, John Sillitoe on behalf of the Wellcome Sanger Institute COVID-19 Surveillance Team |
| EPI_ISL_2780120, EPI_ISL_2780121 | Lighthouse Lab in Glasgow | Wellcome Sanger Institute for the COVID-19 Genomics UK (COG-UK) Consortium | Harper VanSteenhouse, Yumi Kasai, David Gray, Carol Clugston, Anna Dominiczak and Alex Alderton, Roberto Amato, Jeffrey Barrett, Sonia Goncalves, Ewan Harrison, David K. Jackson, Ian Johnston, Dominic Kwiatkowski, Cordelia Langford, John Sillitoe on behalf of the Wellcome Sanger Institute COVID-19 Surveillance Team |
| EPI_ISL_2780122, EPI_ISL_2780123 | Lighthouse Laboratory Plymouth | Wellcome Sanger Institute for the COVID-19 Genomics UK (COG-UK) Consortium | Lighthouse Laboratory Plymouth and Alex Alderton, Roberto Amato, Jeffrey Barrett, Sonia Goncalves, Ewan Harrison, David K. Jackson, Ian Johnston, Dominic Kwiatkowski, Cordelia Langford, John Sillitoe on behalf of the Wellcome Sanger Institute COVID-19 Surveillance Team |
| EPI_ISL_2780124 | Lighthouse Lab in Alderley Park | Wellcome Sanger Institute for the COVID-19 Genomics UK (COG-UK) Consortium | Jacquelyn Wynn, Mairead Hyland, The Lighthouse Lab in Alderley Park and Alex Alderton, Roberto Amato, Jeffrey Barrett, Sonia Goncalves, Ewan Harrison, David K. Jackson, Ian Johnston, Dominic Kwiatkowski, Cordelia Langford, John Sillitoe on behalf of the Wellcome Sanger Institute COVID-19 Surveillance Team |
| EPI_ISL_2780125 | Lighthouse Lab in Milton Keynes | Wellcome Sanger Institute for the COVID-19 Genomics UK (COG-UK) Consortium | The Lighthouse Lab in Milton Keynes and Alex Alderton, Roberto Amato, Jeffrey Barrett, Sonia Goncalves, Ewan Harrison, David K. Jackson, Ian Johnston, Dominic Kwiatkowski, Cordelia Langford, John Sillitoe on behalf of the Wellcome Sanger Institute COVID-19 Surveillance Team |
| EPI_ISL_2780126, EPI_ISL_2780127 | Lighthouse Lab in Alderley Park | Wellcome Sanger Institute for the COVID-19 Genomics UK (COG-UK) Consortium | Jacquelyn Wynn, Mairead Hyland, The Lighthouse Lab in Alderley Park and Alex Alderton, Roberto Amato, Jeffrey Barrett, Sonia Goncalves, Ewan Harrison, David K. Jackson, Ian Johnston, Dominic Kwiatkowski, Cordelia Langford, John Sillitoe on behalf of the Wellcome Sanger Institute COVID-19 Surveillance Team |
| EPI_ISL_2780128 | Lighthouse Lab in Milton Keynes | Wellcome Sanger Institute for the COVID-19 Genomics UK (COG-UK) Consortium | The Lighthouse Lab in Milton Keynes and Alex Alderton, Roberto Amato, Jeffrey Barrett, Sonia Goncalves, Ewan Harrison, David K. Jackson, Ian Johnston, Dominic Kwiatkowski, Cordelia Langford, John Sillitoe on behalf of the Wellcome Sanger Institute COVID-19 Surveillance Team |
| EPI_ISL_2780129 | Lighthouse Lab in Alderley Park | Wellcome Sanger Institute for the COVID-19 Genomics UK (COG-UK) Consortium | Jacquelyn Wynn, Mairead Hyland, The Lighthouse Lab in Alderley Park and Alex Alderton, Roberto Amato, Jeffrey Barrett, Sonia Goncalves, Ewan Harrison, David K. Jackson, Ian Johnston, Dominic Kwiatkowski, Cordelia Langford, John Sillitoe on behalf of the Wellcome Sanger Institute COVID-19 Surveillance Team |
| EPI_ISL_2780130 | Lighthouse Lab in Milton Keynes | Wellcome Sanger Institute for the COVID-19 Genomics UK (COG-UK) Consortium | The Lighthouse Lab in Milton Keynes and Alex Alderton, Roberto Amato, Jeffrey Barrett, Sonia Goncalves, Ewan Harrison, David K. Jackson, Ian Johnston, Dominic Kwiatkowski, Cordelia Langford, John Sillitoe on behalf of the Wellcome Sanger Institute COVID-19 Surveillance Team |
| EPI_ISL_2780131 | Lighthouse Lab in Alderley Park | Wellcome Sanger Institute for the COVID-19 Genomics UK (COG-UK) Consortium | Jacquelyn Wynn, Mairead Hyland, The Lighthouse Lab in Alderley Park and Alex Alderton, Roberto Amato, Jeffrey Barrett, Sonia Goncalves, Ewan Harrison, David K. Jackson, Ian Johnston, Dominic Kwiatkowski, Cordelia Langford, John Sillitoe on behalf of the Wellcome Sanger Institute COVID-19 Surveillance Team |
| EPI_ISL_2780132 | Lighthouse Lab in Milton Keynes | Wellcome Sanger Institute for the COVID-19 Genomics UK (COG-UK) Consortium | The Lighthouse Lab in Milton Keynes and Alex Alderton, Roberto Amato, Jeffrey Barrett, Sonia Goncalves, Ewan Harrison, David K. Jackson, Ian Johnston, Dominic Kwiatkowski, Cordelia Langford, John Sillitoe on behalf of the Wellcome Sanger Institute COVID-19 Surveillance Team |
| EPI_ISL_2780133, EPI_ISL_2780134, EPI_ISL_2780135, EPI_ISL_2780136 | Lighthouse Lab in Alderley Park | Wellcome Sanger Institute for the COVID-19 Genomics UK (COG-UK) Consortium | Jacquelyn Wynn, Mairead Hyland, The Lighthouse Lab in Alderley Park and Alex Alderton, Roberto Amato, Jeffrey Barrett, Sonia Goncalves, Ewan Harrison, David K. Jackson, Ian Johnston, Dominic Kwiatkowski, Cordelia Langford, John Sillitoe on behalf of the Wellcome Sanger Institute COVID-19 Surveillance Team |
| EPI_ISL_2780137 | Lighthouse Lab in Milton Keynes | Wellcome Sanger Institute for the COVID-19 Genomics UK (COG-UK) Consortium | The Lighthouse Lab in Milton Keynes and Alex Alderton, Roberto Amato, Jeffrey Barrett, Sonia Goncalves, Ewan Harrison, David K. Jackson, Ian Johnston, Dominic Kwiatkowski, Cordelia Langford, John Sillitoe on behalf of the Wellcome Sanger Institute COVID-19 Surveillance Team |
| EPI_ISL_2780138 | Lighthouse Laboratory Plymouth | Wellcome Sanger Institute for the COVID-19 Genomics UK (COG-UK) Consortium | Lighthouse Laboratory Plymouth and Alex Alderton, Roberto Amato, Jeffrey Barrett, Sonia Goncalves, Ewan Harrison, David K. Jackson, Ian Johnston, Dominic Kwiatkowski, Cordelia Langford, John Sillitoe on behalf of the Wellcome Sanger Institute COVID-19 Surveillance Team |
| EPI_ISL_2780139 | Lighthouse Lab in Alderley Park | Wellcome Sanger Institute for the COVID-19 Genomics UK (COG-UK) Consortium | Jacquelyn Wynn, Mairead Hyland, The Lighthouse Lab in Alderley Park and Alex Alderton, Roberto Amato, Jeffrey Barrett, Sonia Goncalves, Ewan Harrison, David K. Jackson, Ian Johnston, Dominic Kwiatkowski, Cordelia Langford, John Sillitoe on behalf of the Wellcome Sanger Institute COVID-19 Surveillance Team |
| EPI_ISL_2780140, EPI_ISL_2780141 | Lighthouse Laboratory Plymouth | Wellcome Sanger Institute for the COVID-19 Genomics UK (COG-UK) Consortium | Lighthouse Laboratory Plymouth and Alex Alderton, Roberto Amato, Jeffrey Barrett, Sonia Goncalves, Ewan Harrison, David K. Jackson, Ian Johnston, Dominic Kwiatkowski, Cordelia Langford, John Sillitoe on behalf of the Wellcome Sanger Institute COVID-19 Surveillance Team |
| EPI_ISL_2780142 | Lighthouse Lab in Alderley Park | Wellcome Sanger Institute for the COVID-19 Genomics UK (COG-UK) Consortium | Jacquelyn Wynn, Mairead Hyland, The Lighthouse Lab in Alderley Park and Alex Alderton, Roberto Amato, Jeffrey Barrett, Sonia Goncalves, Ewan Harrison, David K. Jackson, Ian Johnston, Dominic Kwiatkowski, Cordelia Langford, John Sillitoe on behalf of the Wellcome Sanger Institute COVID-19 Surveillance Team |
| EPI_ISL_2780143 | Lighthouse Laboratory Plymouth | Wellcome Sanger Institute for the COVID-19 Genomics UK (COG-UK) Consortium | Lighthouse Laboratory Plymouth and Alex Alderton, Roberto Amato, Jeffrey Barrett, Sonia Goncalves, Ewan Harrison, David K. Jackson, Ian Johnston, Dominic Kwiatkowski, Cordelia Langford, John Sillitoe on behalf of the Wellcome Sanger Institute COVID-19 Surveillance Team |
| EPI_ISL_2780144 | Lighthouse Lab in Milton Keynes | Wellcome Sanger Institute for the COVID-19 Genomics UK (COG-UK) Consortium | The Lighthouse Lab in Milton Keynes and Alex Alderton, Roberto Amato, Jeffrey Barrett, Sonia Goncalves, Ewan Harrison, David K. Jackson, Ian Johnston, Dominic Kwiatkowski, Cordelia Langford, John Sillitoe on behalf of the Wellcome Sanger Institute COVID-19 Surveillance Team |
| EPI_ISL_2780145 | Lighthouse Lab in Alderley Park | Wellcome Sanger Institute for the COVID-19 Genomics UK (COG-UK) Consortium | Jacquelyn Wynn, Mairead Hyland, The Lighthouse Lab in Alderley Park and Alex Alderton, Roberto Amato, Jeffrey Barrett, Sonia Goncalves, Ewan Harrison, David K. Jackson, Ian Johnston, Dominic Kwiatkowski, Cordelia Langford, John Sillitoe on behalf of the Wellcome Sanger Institute COVID-19 Surveillance Team |
| EPI_ISL_2780146 | Lighthouse Laboratory Plymouth | Wellcome Sanger Institute for the COVID-19 Genomics UK (COG-UK) Consortium | Lighthouse Laboratory Plymouth and Alex Alderton, Roberto Amato, Jeffrey Barrett, Sonia Goncalves, Ewan Harrison, David K. Jackson, Ian Johnston, Dominic Kwiatkowski, Cordelia Langford, John Sillitoe on behalf of the Wellcome Sanger Institute COVID-19 Surveillance Team |
| EPI_ISL_2780147, EPI_ISL_2780148, EPI_ISL_2780149, EPI_ISL_2780150 | Lighthouse Lab in Alderley Park | Wellcome Sanger Institute for the COVID-19 Genomics UK (COG-UK) Consortium | Jacquelyn Wynn, Mairead Hyland, The Lighthouse Lab in Alderley Park and Alex Alderton, Roberto Amato, Jeffrey Barrett, Sonia Goncalves, Ewan Harrison, David K. Jackson, Ian Johnston, Dominic Kwiatkowski, Cordelia Langford, John Sillitoe on behalf of the Wellcome Sanger Institute COVID-19 Surveillance Team |
| EPI_ISL_2780151 | Lighthouse Lab in Milton Keynes | Wellcome Sanger Institute for the COVID-19 Genomics UK (COG-UK) Consortium | The Lighthouse Lab in Milton Keynes and Alex Alderton, Roberto Amato, Jeffrey Barrett, Sonia Goncalves, Ewan Harrison, David K. Jackson, Ian Johnston, Dominic Kwiatkowski, Cordelia Langford, John Sillitoe on behalf of the Wellcome Sanger Institute COVID-19 Surveillance Team |
| EPI_ISL_2780152, EPI_ISL_2780153, EPI_ISL_2780154, EPI_ISL_2780155 | Lighthouse Lab in Alderley Park | Wellcome Sanger Institute for the COVID-19 Genomics UK (COG-UK) Consortium | Jacquelyn Wynn, Mairead Hyland, The Lighthouse Lab in Alderley Park and Alex Alderton, Roberto Amato, Jeffrey Barrett, Sonia Goncalves, Ewan Harrison, David K. Jackson, Ian Johnston, Dominic Kwiatkowski, Cordelia Langford, John Sillitoe on behalf of the Wellcome Sanger Institute COVID-19 Surveillance Team |
| EPI_ISL_2780156 | Lighthouse Laboratory Plymouth | Wellcome Sanger Institute for the COVID-19 Genomics UK (COG-UK) Consortium | Lighthouse Laboratory Plymouth and Alex Alderton, Roberto Amato, Jeffrey Barrett, Sonia Goncalves, Ewan Harrison, David K. Jackson, Ian Johnston, Dominic Kwiatkowski, Cordelia Langford, John Sillitoe on behalf of the Wellcome Sanger Institute COVID-19 Surveillance Team |
| EPI_ISL_2780157, EPI_ISL_2780158 | Lighthouse Lab in Alderley Park | Wellcome Sanger Institute for the COVID-19 Genomics UK (COG-UK) Consortium | Jacquelyn Wynn, Mairead Hyland, The Lighthouse Lab in Alderley Park and Alex Alderton, Roberto Amato, Jeffrey Barrett, Sonia Goncalves, Ewan Harrison, David K. Jackson, Ian Johnston, Dominic Kwiatkowski, Cordelia Langford, John Sillitoe on behalf of the Wellcome Sanger Institute COVID-19 Surveillance Team |
| EPI_ISL_2780159 | Lighthouse Lab in Milton Keynes | Wellcome Sanger Institute for the COVID-19 Genomics UK (COG-UK) Consortium | The Lighthouse Lab in Milton Keynes and Alex Alderton, Roberto Amato, Jeffrey Barrett, Sonia Goncalves, Ewan Harrison, David K. Jackson, Ian Johnston, Dominic Kwiatkowski, Cordelia Langford, John Sillitoe on behalf of the Wellcome Sanger Institute COVID-19 Surveillance Team |
| EPI_ISL_2780160 | Lighthouse Lab in Alderley Park | Wellcome Sanger Institute for the COVID-19 Genomics UK (COG-UK) Consortium | Jacquelyn Wynn, Mairead Hyland, The Lighthouse Lab in Alderley Park and Alex Alderton, Roberto Amato, Jeffrey Barrett, Sonia Goncalves, Ewan Harrison, David K. Jackson, Ian Johnston, Dominic Kwiatkowski, Cordelia Langford, John Sillitoe on behalf of the Wellcome Sanger Institute COVID-19 Surveillance Team |
| EPI_ISL_2780161, EPI_ISL_2780162 | Lighthouse Lab in Milton Keynes | Wellcome Sanger Institute for the COVID-19 Genomics UK (COG-UK) Consortium | The Lighthouse Lab in Milton Keynes and Alex Alderton, Roberto Amato, Jeffrey Barrett, Sonia Goncalves, Ewan Harrison, David K. Jackson, Ian Johnston, Dominic Kwiatkowski, Cordelia Langford, John Sillitoe on behalf of the Wellcome Sanger Institute COVID-19 Surveillance Team |
| EPI_ISL_2780163, EPI_ISL_2780164 | Lighthouse Lab in Alderley Park | Wellcome Sanger Institute for the COVID-19 Genomics UK (COG-UK) Consortium | Jacquelyn Wynn, Mairead Hyland, The Lighthouse Lab in Alderley Park and Alex Alderton, Roberto Amato, Jeffrey Barrett, Sonia Goncalves, Ewan Harrison, David K. Jackson, Ian Johnston, Dominic Kwiatkowski, Cordelia Langford, John Sillitoe on behalf of the Wellcome Sanger Institute COVID-19 Surveillance Team |
| EPI_ISL_2780165 | Lighthouse Lab in Milton Keynes | Wellcome Sanger Institute for the COVID-19 Genomics UK (COG-UK) Consortium | The Lighthouse Lab in Milton Keynes and Alex Alderton, Roberto Amato, Jeffrey Barrett, Sonia Goncalves, Ewan Harrison, David K. Jackson, Ian Johnston, Dominic Kwiatkowski, Cordelia Langford, John Sillitoe on behalf of the Wellcome Sanger Institute COVID-19 Surveillance Team |
| EPI_ISL_2780166 | Lighthouse Lab in Alderley Park | Wellcome Sanger Institute for the COVID-19 Genomics UK (COG-UK) | Jacquelyn Wynn, Mairead Hyland, The Lighthouse Lab in Alderley Park and Alex Alderton, Roberto Amato, Jeffrey Barrett, Sonia Goncalves, Ewan Harrison, David K. Jackson, Ian Johnston, Dominic Kwiatkowski, Cordelia Langford, John Sillitoe on |

| | | | |
|---|---|---|---|
| | | Consortium | behalf of the Wellcome Sanger Institute COVID-19 Surveillance Team |
| EPI_ISL_2780167 | Lighthouse Lab in Milton Keynes | Wellcome Sanger Institute for the COVID-19 Genomics UK (COG-UK) Consortium | The Lighthouse Lab in Milton Keynes and Alex Alderton, Roberto Amato, Jeffrey Barrett, Sonia Goncalves, Ewan Harrison, David K. Jackson, Ian Johnston, Dominic Kwiatkowski, Cordelia Langford, John Sillitoe on behalf of the Wellcome Sanger Institute COVID-19 Surveillance Team |
| EPI_ISL_2780168 | Lighthouse Laboratory Plymouth | Wellcome Sanger Institute for the COVID-19 Genomics UK (COG-UK) Consortium | Lighthouse Laboratory Plymouth and Alex Alderton, Roberto Amato, Jeffrey Barrett, Sonia Goncalves, Ewan Harrison, David K. Jackson, Ian Johnston, Dominic Kwiatkowski, Cordelia Langford, John Sillitoe on behalf of the Wellcome Sanger Institute COVID-19 Surveillance Team |
| EPI_ISL_2780169, EPI_ISL_2780170 | Lighthouse Lab in Alderley Park | Wellcome Sanger Institute for the COVID-19 Genomics UK (COG-UK) Consortium | Jacquelyn Wynn, Mairead Hyland, The Lighthouse Lab in Alderley Park and Alex Alderton, Roberto Amato, Jeffrey Barrett, Sonia Goncalves, Ewan Harrison, David K. Jackson, Ian Johnston, Dominic Kwiatkowski, Cordelia Langford, John Sillitoe on behalf of the Wellcome Sanger Institute COVID-19 Surveillance Team |
| EPI_ISL_2780171 | Lighthouse Lab in Milton Keynes | Wellcome Sanger Institute for the COVID-19 Genomics UK (COG-UK) Consortium | The Lighthouse Lab in Milton Keynes and Alex Alderton, Roberto Amato, Jeffrey Barrett, Sonia Goncalves, Ewan Harrison, David K. Jackson, Ian Johnston, Dominic Kwiatkowski, Cordelia Langford, John Sillitoe on behalf of the Wellcome Sanger Institute COVID-19 Surveillance Team |
| EPI_ISL_2780172, EPI_ISL_2780173, EPI_ISL_2780174, EPI_ISL_2780175, EPI_ISL_2780176 | Lighthouse Lab in Alderley Park | Wellcome Sanger Institute for the COVID-19 Genomics UK (COG-UK) Consortium | Jacquelyn Wynn, Mairead Hyland, The Lighthouse Lab in Alderley Park and Alex Alderton, Roberto Amato, Jeffrey Barrett, Sonia Goncalves, Ewan Harrison, David K. Jackson, Ian Johnston, Dominic Kwiatkowski, Cordelia Langford, John Sillitoe on behalf of the Wellcome Sanger Institute COVID-19 Surveillance Team |
| EPI_ISL_2780177, EPI_ISL_2780178 | Lighthouse Lab in Milton Keynes | Wellcome Sanger Institute for the COVID-19 Genomics UK (COG-UK) Consortium | The Lighthouse Lab in Milton Keynes and Alex Alderton, Roberto Amato, Jeffrey Barrett, Sonia Goncalves, Ewan Harrison, David K. Jackson, Ian Johnston, Dominic Kwiatkowski, Cordelia Langford, John Sillitoe on behalf of the Wellcome Sanger Institute COVID-19 Surveillance Team |
| EPI_ISL_2780179 | Lighthouse Laboratory Plymouth | Wellcome Sanger Institute for the COVID-19 Genomics UK (COG-UK) Consortium | Lighthouse Laboratory Plymouth and Alex Alderton, Roberto Amato, Jeffrey Barrett, Sonia Goncalves, Ewan Harrison, David K. Jackson, Ian Johnston, Dominic Kwiatkowski, Cordelia Langford, John Sillitoe on behalf of the Wellcome Sanger Institute COVID-19 Surveillance Team |
| EPI_ISL_2780180, EPI_ISL_2780181, EPI_ISL_2780182 | Lighthouse Lab in Alderley Park | Wellcome Sanger Institute for the COVID-19 Genomics UK (COG-UK) Consortium | Jacquelyn Wynn, Mairead Hyland, The Lighthouse Lab in Alderley Park and Alex Alderton, Roberto Amato, Jeffrey Barrett, Sonia Goncalves, Ewan Harrison, David K. Jackson, Ian Johnston, Dominic Kwiatkowski, Cordelia Langford, John Sillitoe on behalf of the Wellcome Sanger Institute COVID-19 Surveillance Team |
| EPI_ISL_2780183 | Lighthouse Laboratory Plymouth | Wellcome Sanger Institute for the COVID-19 Genomics UK (COG-UK) Consortium | Lighthouse Laboratory Plymouth and Alex Alderton, Roberto Amato, Jeffrey Barrett, Sonia Goncalves, Ewan Harrison, David K. Jackson, Ian Johnston, Dominic Kwiatkowski, Cordelia Langford, John Sillitoe on behalf of the Wellcome Sanger Institute COVID-19 Surveillance Team |
| EPI_ISL_2780184 | Lighthouse Lab in Alderley Park | Wellcome Sanger Institute for the COVID-19 Genomics UK (COG-UK) Consortium | Jacquelyn Wynn, Mairead Hyland, The Lighthouse Lab in Alderley Park and Alex Alderton, Roberto Amato, Jeffrey Barrett, Sonia Goncalves, Ewan Harrison, David K. Jackson, Ian Johnston, Dominic Kwiatkowski, Cordelia Langford, John Sillitoe on behalf of the Wellcome Sanger Institute COVID-19 Surveillance Team |
| EPI_ISL_2780185, EPI_ISL_2780186, EPI_ISL_2780187 | Lighthouse Lab in Milton Keynes | Wellcome Sanger Institute for the COVID-19 Genomics UK (COG-UK) Consortium | The Lighthouse Lab in Milton Keynes and Alex Alderton, Roberto Amato, Jeffrey Barrett, Sonia Goncalves, Ewan Harrison, David K. Jackson, Ian Johnston, Dominic Kwiatkowski, Cordelia Langford, John Sillitoe on behalf of the Wellcome Sanger Institute COVID-19 Surveillance Team |
| EPI_ISL_2780188 | Lighthouse Lab in Alderley Park | Wellcome Sanger Institute for the COVID-19 Genomics UK (COG-UK) Consortium | Jacquelyn Wynn, Mairead Hyland, The Lighthouse Lab in Alderley Park and Alex Alderton, Roberto Amato, Jeffrey Barrett, Sonia Goncalves, Ewan Harrison, David K. Jackson, Ian Johnston, Dominic Kwiatkowski, Cordelia Langford, John Sillitoe on behalf of the Wellcome Sanger Institute COVID-19 Surveillance Team |
| EPI_ISL_2780189 | Lighthouse Laboratory Plymouth | Wellcome Sanger Institute for the COVID-19 Genomics UK (COG-UK) Consortium | Lighthouse Laboratory Plymouth and Alex Alderton, Roberto Amato, Jeffrey Barrett, Sonia Goncalves, Ewan Harrison, David K. Jackson, Ian Johnston, Dominic Kwiatkowski, Cordelia Langford, John Sillitoe on behalf of the Wellcome Sanger Institute COVID-19 Surveillance Team |
| EPI_ISL_2780190, EPI_ISL_2780191, EPI_ISL_2780192 | Lighthouse Lab in Alderley Park | Wellcome Sanger Institute for the COVID-19 Genomics UK (COG-UK) Consortium | Jacquelyn Wynn, Mairead Hyland, The Lighthouse Lab in Alderley Park and Alex Alderton, Roberto Amato, Jeffrey Barrett, Sonia Goncalves, Ewan Harrison, David K. Jackson, Ian Johnston, Dominic Kwiatkowski, Cordelia Langford, John Sillitoe on behalf of the Wellcome Sanger Institute COVID-19 Surveillance Team |
| EPI_ISL_2780193 | Lighthouse Laboratory Plymouth | Wellcome Sanger Institute for the COVID-19 Genomics UK (COG-UK) Consortium | Lighthouse Laboratory Plymouth and Alex Alderton, Roberto Amato, Jeffrey Barrett, Sonia Goncalves, Ewan Harrison, David K. Jackson, Ian Johnston, Dominic Kwiatkowski, Cordelia Langford, John Sillitoe on behalf of the Wellcome Sanger Institute COVID-19 Surveillance Team |
| EPI_ISL_2780194, EPI_ISL_2780195, EPI_ISL_2780196 | Lighthouse Lab in Alderley Park | Wellcome Sanger Institute for the COVID-19 Genomics UK (COG-UK) Consortium | Jacquelyn Wynn, Mairead Hyland, The Lighthouse Lab in Alderley Park and Alex Alderton, Roberto Amato, Jeffrey Barrett, Sonia Goncalves, Ewan Harrison, David K. Jackson, Ian Johnston, Dominic Kwiatkowski, Cordelia Langford, John Sillitoe on behalf of the Wellcome Sanger Institute COVID-19 Surveillance Team |
| EPI_ISL_2780197, EPI_ISL_2780198, EPI_ISL_2780199 | Lighthouse Laboratory Plymouth | Wellcome Sanger Institute for the COVID-19 Genomics UK (COG-UK) Consortium | Lighthouse Laboratory Plymouth and Alex Alderton, Roberto Amato, Jeffrey Barrett, Sonia Goncalves, Ewan Harrison, David K. Jackson, Ian Johnston, Dominic Kwiatkowski, Cordelia Langford, John Sillitoe on behalf of the Wellcome Sanger Institute COVID-19 Surveillance Team |
| EPI_ISL_2780200, EPI_ISL_2780201 | Lighthouse Lab in Alderley Park | Wellcome Sanger Institute for the COVID-19 Genomics UK (COG-UK) Consortium | Jacquelyn Wynn, Mairead Hyland, The Lighthouse Lab in Alderley Park and Alex Alderton, Roberto Amato, Jeffrey Barrett, Sonia Goncalves, Ewan Harrison, David K. Jackson, Ian Johnston, Dominic Kwiatkowski, Cordelia Langford, John Sillitoe on behalf of the Wellcome Sanger Institute COVID-19 Surveillance Team |
| EPI_ISL_2780202, EPI_ISL_2780203 | Lighthouse Laboratory Plymouth | Wellcome Sanger Institute for the COVID-19 Genomics UK (COG-UK) Consortium | Lighthouse Laboratory Plymouth and Alex Alderton, Roberto Amato, Jeffrey Barrett, Sonia Goncalves, Ewan Harrison, David K. Jackson, Ian Johnston, Dominic Kwiatkowski, Cordelia Langford, John Sillitoe on behalf of the Wellcome Sanger Institute COVID-19 Surveillance Team |
| EPI_ISL_2780204, EPI_ISL_2780205 | Lighthouse Lab in Alderley Park | Wellcome Sanger Institute for the COVID-19 Genomics UK (COG-UK) Consortium | Jacquelyn Wynn, Mairead Hyland, The Lighthouse Lab in Alderley Park and Alex Alderton, Roberto Amato, Jeffrey Barrett, Sonia Goncalves, Ewan Harrison, David K. Jackson, Ian Johnston, Dominic Kwiatkowski, Cordelia Langford, John Sillitoe on behalf of the Wellcome Sanger Institute COVID-19 Surveillance Team |
| EPI_ISL_2780206 | Lighthouse Lab in Milton Keynes | Wellcome Sanger Institute for the COVID-19 Genomics UK (COG-UK) Consortium | The Lighthouse Lab in Milton Keynes and Alex Alderton, Roberto Amato, Jeffrey Barrett, Sonia Goncalves, Ewan Harrison, David K. Jackson, Ian Johnston, Dominic Kwiatkowski, Cordelia Langford, John Sillitoe on behalf of the Wellcome Sanger Institute COVID-19 Surveillance Team |
| EPI_ISL_2780207 | Lighthouse Laboratory Plymouth | Wellcome Sanger Institute for the COVID-19 Genomics UK (COG-UK) Consortium | Lighthouse Laboratory Plymouth and Alex Alderton, Roberto Amato, Jeffrey Barrett, Sonia Goncalves, Ewan Harrison, David K. Jackson, Ian Johnston, Dominic Kwiatkowski, Cordelia Langford, John Sillitoe on behalf of the Wellcome Sanger Institute COVID-19 Surveillance Team |
| EPI_ISL_2780208, EPI_ISL_2780209 | Lighthouse Lab in Alderley Park | Wellcome Sanger Institute for the COVID-19 Genomics UK (COG-UK) Consortium | Jacquelyn Wynn, Mairead Hyland, The Lighthouse Lab in Alderley Park and Alex Alderton, Roberto Amato, Jeffrey Barrett, Sonia Goncalves, Ewan Harrison, David K. Jackson, Ian Johnston, Dominic Kwiatkowski, Cordelia Langford, John Sillitoe on behalf of the Wellcome Sanger Institute COVID-19 Surveillance Team |
| EPI_ISL_2780210, EPI_ISL_2780211 | Lighthouse Lab in Milton Keynes | Wellcome Sanger Institute for the COVID-19 Genomics UK (COG-UK) Consortium | The Lighthouse Lab in Milton Keynes and Alex Alderton, Roberto Amato, Jeffrey Barrett, Sonia Goncalves, Ewan Harrison, David K. Jackson, Ian Johnston, Dominic Kwiatkowski, Cordelia Langford, John Sillitoe on behalf of the Wellcome Sanger Institute COVID-19 Surveillance Team |
| EPI_ISL_2780212 | Lighthouse Lab in Alderley Park | Wellcome Sanger Institute for the COVID-19 Genomics UK (COG-UK) Consortium | Jacquelyn Wynn, Mairead Hyland, The Lighthouse Lab in Alderley Park and Alex Alderton, Roberto Amato, Jeffrey Barrett, Sonia Goncalves, Ewan Harrison, David K. Jackson, Ian Johnston, Dominic Kwiatkowski, Cordelia Langford, John Sillitoe on behalf of the Wellcome Sanger Institute COVID-19 Surveillance Team |
| EPI_ISL_2780213 | Lighthouse Lab in Milton Keynes | Wellcome Sanger Institute for the COVID-19 Genomics UK (COG-UK) Consortium | The Lighthouse Lab in Milton Keynes and Alex Alderton, Roberto Amato, Jeffrey Barrett, Sonia Goncalves, Ewan Harrison, David K. Jackson, Ian Johnston, Dominic Kwiatkowski, Cordelia Langford, John Sillitoe on behalf of the Wellcome Sanger Institute COVID-19 Surveillance Team |
| EPI_ISL_2780214, EPI_ISL_2780215, EPI_ISL_2780216 | Lighthouse Lab in Alderley Park | Wellcome Sanger Institute for the COVID-19 Genomics UK (COG-UK) Consortium | Jacquelyn Wynn, Mairead Hyland, The Lighthouse Lab in Alderley Park and Alex Alderton, Roberto Amato, Jeffrey Barrett, Sonia Goncalves, Ewan Harrison, David K. Jackson, Ian Johnston, Dominic Kwiatkowski, Cordelia Langford, John Sillitoe on behalf of the Wellcome Sanger Institute COVID-19 Surveillance Team |
| EPI_ISL_2780217 | Lighthouse Laboratory Plymouth | Wellcome Sanger Institute for the COVID-19 Genomics UK (COG-UK) Consortium | Lighthouse Laboratory Plymouth and Alex Alderton, Roberto Amato, Jeffrey Barrett, Sonia Goncalves, Ewan Harrison, David K. Jackson, Ian Johnston, Dominic Kwiatkowski, Cordelia Langford, John Sillitoe on behalf of the Wellcome Sanger Institute COVID-19 Surveillance Team |
| EPI_ISL_2780218 | Lighthouse Lab in Alderley Park | Wellcome Sanger Institute for the COVID-19 Genomics UK (COG-UK) Consortium | Jacquelyn Wynn, Mairead Hyland, The Lighthouse Lab in Alderley Park and Alex Alderton, Roberto Amato, Jeffrey Barrett, Sonia Goncalves, Ewan Harrison, David K. Jackson, Ian Johnston, Dominic Kwiatkowski, Cordelia Langford, John Sillitoe on behalf of the Wellcome Sanger Institute COVID-19 Surveillance Team |
| EPI_ISL_2780219 | Lighthouse Lab in Milton Keynes | Wellcome Sanger Institute for the COVID-19 Genomics UK (COG-UK) Consortium | The Lighthouse Lab in Milton Keynes and Alex Alderton, Roberto Amato, Jeffrey Barrett, Sonia Goncalves, Ewan Harrison, David K. Jackson, Ian Johnston, Dominic Kwiatkowski, Cordelia Langford, John Sillitoe on behalf of the Wellcome Sanger Institute COVID-19 Surveillance Team |
| EPI_ISL_2780220, EPI_ISL_2780221 | Lighthouse Laboratory Plymouth | Wellcome Sanger Institute for the COVID-19 Genomics UK (COG-UK) Consortium | Lighthouse Laboratory Plymouth and Alex Alderton, Roberto Amato, Jeffrey Barrett, Sonia Goncalves, Ewan Harrison, David K. Jackson, Ian Johnston, Dominic Kwiatkowski, Cordelia Langford, John Sillitoe on behalf of the Wellcome Sanger Institute COVID-19 Surveillance Team |
| EPI_ISL_2780222 | Lighthouse Lab in Alderley Park | Wellcome Sanger Institute for the COVID-19 Genomics UK (COG-UK) Consortium | Jacquelyn Wynn, Mairead Hyland, The Lighthouse Lab in Alderley Park and Alex Alderton, Roberto Amato, Jeffrey Barrett, Sonia Goncalves, Ewan Harrison, David K. Jackson, Ian Johnston, Dominic Kwiatkowski, Cordelia Langford, John Sillitoe on behalf of the Wellcome Sanger Institute COVID-19 Surveillance Team |
| EPI_ISL_2780223 | Lighthouse Lab in Milton Keynes | Wellcome Sanger Institute for the COVID-19 Genomics UK (COG-UK) Consortium | The Lighthouse Lab in Milton Keynes and Alex Alderton, Roberto Amato, Jeffrey Barrett, Sonia Goncalves, Ewan Harrison, David K. Jackson, Ian Johnston, Dominic Kwiatkowski, Cordelia Langford, John Sillitoe on behalf of the Wellcome Sanger Institute COVID-19 Surveillance Team |
| EPI_ISL_2780225 | Lighthouse Lab in Alderley Park | Wellcome Sanger Institute for the COVID-19 Genomics UK (COG-UK) Consortium | Jacquelyn Wynn, Mairead Hyland, The Lighthouse Lab in Alderley Park and Alex Alderton, Roberto Amato, Jeffrey Barrett, Sonia Goncalves, Ewan Harrison, David K. Jackson, Ian Johnston, Dominic Kwiatkowski, Cordelia Langford, John Sillitoe on behalf of the Wellcome Sanger Institute COVID-19 Surveillance Team |
| EPI_ISL_2780226 | Lighthouse Lab in Milton Keynes | Wellcome Sanger Institute for the COVID-19 Genomics UK (COG-UK) Consortium | The Lighthouse Lab in Milton Keynes and Alex Alderton, Roberto Amato, Jeffrey Barrett, Sonia Goncalves, Ewan Harrison, David K. Jackson, Ian Johnston, Dominic Kwiatkowski, Cordelia Langford, John Sillitoe on behalf of the Wellcome Sanger Institute COVID-19 Surveillance Team |
| EPI_ISL_2780227, EPI_ISL_2780228 | Lighthouse Laboratory Plymouth | Wellcome Sanger Institute for the COVID-19 Genomics UK (COG-UK) Consortium | Lighthouse Laboratory Plymouth and Alex Alderton, Roberto Amato, Jeffrey Barrett, Sonia Goncalves, Ewan Harrison, David K. Jackson, Ian Johnston, Dominic Kwiatkowski, Cordelia Langford, John Sillitoe on behalf of the Wellcome Sanger Institute COVID-19 Surveillance Team |
| EPI_ISL_2780229, EPI_ISL_2780230 | Lighthouse Lab in Alderley Park | Wellcome Sanger Institute for the COVID-19 Genomics UK (COG-UK) Consortium | Jacquelyn Wynn, Mairead Hyland, The Lighthouse Lab in Alderley Park and Alex Alderton, Roberto Amato, Jeffrey Barrett, Sonia Goncalves, Ewan Harrison, David K. Jackson, Ian Johnston, Dominic Kwiatkowski, Cordelia Langford, John Sillitoe on behalf of the Wellcome Sanger Institute COVID-19 Surveillance Team |
| EPI_ISL_2780231, EPI_ISL_2780232 | Lighthouse Laboratory Plymouth | Wellcome Sanger Institute for the COVID-19 Genomics UK (COG-UK) Consortium | Lighthouse Laboratory Plymouth and Alex Alderton, Roberto Amato, Jeffrey Barrett, Sonia Goncalves, Ewan Harrison, David K. Jackson, Ian Johnston, Dominic Kwiatkowski, Cordelia Langford, John Sillitoe on behalf of the Wellcome Sanger Institute COVID-19 Surveillance Team |
| EPI_ISL_2780233, EPI_ISL_2780234 | Lighthouse Lab in Alderley Park | Wellcome Sanger Institute for the COVID-19 Genomics UK (COG-UK) Consortium | Jacquelyn Wynn, Mairead Hyland, The Lighthouse Lab in Alderley Park and Alex Alderton, Roberto Amato, Jeffrey Barrett, Sonia Goncalves, Ewan Harrison, David K. Jackson, Ian Johnston, Dominic Kwiatkowski, Cordelia Langford, John Sillitoe on behalf of the Wellcome Sanger Institute COVID-19 Surveillance Team |
| EPI_ISL_2780235 | Lighthouse Lab in Milton Keynes | Wellcome Sanger Institute for the COVID-19 Genomics UK (COG-UK) Consortium | The Lighthouse Lab in Milton Keynes and Alex Alderton, Roberto Amato, Jeffrey Barrett, Sonia Goncalves, Ewan Harrison, David K. Jackson, Ian Johnston, Dominic Kwiatkowski, Cordelia Langford, John Sillitoe on behalf of the Wellcome Sanger Institute COVID-19 Surveillance Team |
| EPI_ISL_2780236 | Lighthouse Laboratory Plymouth | Wellcome Sanger Institute for the COVID-19 Genomics UK (COG-UK) Consortium | Lighthouse Laboratory Plymouth and Alex Alderton, Roberto Amato, Jeffrey Barrett, Sonia Goncalves, Ewan Harrison, David K. Jackson, Ian Johnston, Dominic Kwiatkowski, Cordelia Langford, John Sillitoe on behalf of the Wellcome Sanger Institute COVID-19 Surveillance Team |

| | | | |
|---|---|---|---|
| EPI_ISL_2780237, EPI_ISL_2780238, EPI_ISL_2780239, EPI_ISL_2780240, EPI_ISL_2780241 | Lighthouse Lab in Alderley Park | Wellcome Sanger Institute for the COVID-19 Genomics UK (COG-UK) Consortium | Jacquelyn Wynn, Mairead Hyland, The Lighthouse Lab in Alderley Park and Alex Alderton, Roberto Amato, Jeffrey Barrett, Sonia Goncalves, Ewan Harrison, David K. Jackson, Ian Johnston, Dominic Kwiatkowski, Cordelia Langford, John Sillitoe on behalf of the Wellcome Sanger Institute COVID-19 Surveillance Team |
| EPI_ISL_2780242 | Lighthouse Lab in Milton Keynes | Wellcome Sanger Institute for the COVID-19 Genomics UK (COG-UK) Consortium | The Lighthouse Lab in Milton Keynes and Alex Alderton, Roberto Amato, Jeffrey Barrett, Sonia Goncalves, Ewan Harrison, David K. Jackson, Ian Johnston, Dominic Kwiatkowski, Cordelia Langford, John Sillitoe on behalf of the Wellcome Sanger Institute COVID-19 Surveillance Team |
| EPI_ISL_2780243, EPI_ISL_2780244, EPI_ISL_2780245 | Lighthouse Lab in Alderley Park | Wellcome Sanger Institute for the COVID-19 Genomics UK (COG-UK) Consortium | Jacquelyn Wynn, Mairead Hyland, The Lighthouse Lab in Alderley Park and Alex Alderton, Roberto Amato, Jeffrey Barrett, Sonia Goncalves, Ewan Harrison, David K. Jackson, Ian Johnston, Dominic Kwiatkowski, Cordelia Langford, John Sillitoe on behalf of the Wellcome Sanger Institute COVID-19 Surveillance Team |
| EPI_ISL_2780246, EPI_ISL_2780247 | Lighthouse Lab in Milton Keynes | Wellcome Sanger Institute for the COVID-19 Genomics UK (COG-UK) Consortium | The Lighthouse Lab in Milton Keynes and Alex Alderton, Roberto Amato, Jeffrey Barrett, Sonia Goncalves, Ewan Harrison, David K. Jackson, Ian Johnston, Dominic Kwiatkowski, Cordelia Langford, John Sillitoe on behalf of the Wellcome Sanger Institute COVID-19 Surveillance Team |
| EPI_ISL_2780248 | Lighthouse Lab in Alderley Park | Wellcome Sanger Institute for the COVID-19 Genomics UK (COG-UK) Consortium | Jacquelyn Wynn, Mairead Hyland, The Lighthouse Lab in Alderley Park and Alex Alderton, Roberto Amato, Jeffrey Barrett, Sonia Goncalves, Ewan Harrison, David K. Jackson, Ian Johnston, Dominic Kwiatkowski, Cordelia Langford, John Sillitoe on behalf of the Wellcome Sanger Institute COVID-19 Surveillance Team |
| EPI_ISL_2780249, EPI_ISL_2780251, EPI_ISL_2780252 | Lighthouse Lab in Milton Keynes | Wellcome Sanger Institute for the COVID-19 Genomics UK (COG-UK) Consortium | The Lighthouse Lab in Milton Keynes and Alex Alderton, Roberto Amato, Jeffrey Barrett, Sonia Goncalves, Ewan Harrison, David K. Jackson, Ian Johnston, Dominic Kwiatkowski, Cordelia Langford, John Sillitoe on behalf of the Wellcome Sanger Institute COVID-19 Surveillance Team |
| EPI_ISL_2780253, EPI_ISL_2780254, EPI_ISL_2780255 | Lighthouse Lab in Alderley Park | Wellcome Sanger Institute for the COVID-19 Genomics UK (COG-UK) Consortium | Jacquelyn Wynn, Mairead Hyland, The Lighthouse Lab in Alderley Park and Alex Alderton, Roberto Amato, Jeffrey Barrett, Sonia Goncalves, Ewan Harrison, David K. Jackson, Ian Johnston, Dominic Kwiatkowski, Cordelia Langford, John Sillitoe on behalf of the Wellcome Sanger Institute COVID-19 Surveillance Team |
| EPI_ISL_2780256 | Lighthouse Lab in Milton Keynes | Wellcome Sanger Institute for the COVID-19 Genomics UK (COG-UK) Consortium | The Lighthouse Lab in Milton Keynes and Alex Alderton, Roberto Amato, Jeffrey Barrett, Sonia Goncalves, Ewan Harrison, David K. Jackson, Ian Johnston, Dominic Kwiatkowski, Cordelia Langford, John Sillitoe on behalf of the Wellcome Sanger Institute COVID-19 Surveillance Team |
| EPI_ISL_2780257 | Lighthouse Lab in Alderley Park | Wellcome Sanger Institute for the COVID-19 Genomics UK (COG-UK) Consortium | Jacquelyn Wynn, Mairead Hyland, The Lighthouse Lab in Alderley Park and Alex Alderton, Roberto Amato, Jeffrey Barrett, Sonia Goncalves, Ewan Harrison, David K. Jackson, Ian Johnston, Dominic Kwiatkowski, Cordelia Langford, John Sillitoe on behalf of the Wellcome Sanger Institute COVID-19 Surveillance Team |
| EPI_ISL_2780258, EPI_ISL_2780259, EPI_ISL_2780260, EPI_ISL_2780261, EPI_ISL_2780262 | Lighthouse Laboratory Plymouth | Wellcome Sanger Institute for the COVID-19 Genomics UK (COG-UK) Consortium | Lighthouse Laboratory Plymouth and Alex Alderton, Roberto Amato, Jeffrey Barrett, Sonia Goncalves, Ewan Harrison, David K. Jackson, Ian Johnston, Dominic Kwiatkowski, Cordelia Langford, John Sillitoe on behalf of the Wellcome Sanger Institute COVID-19 Surveillance Team |
| EPI_ISL_2780263, EPI_ISL_2780264, EPI_ISL_2780265, EPI_ISL_2780266 | Lighthouse Lab in Alderley Park | Wellcome Sanger Institute for the COVID-19 Genomics UK (COG-UK) Consortium | Jacquelyn Wynn, Mairead Hyland, The Lighthouse Lab in Alderley Park and Alex Alderton, Roberto Amato, Jeffrey Barrett, Sonia Goncalves, Ewan Harrison, David K. Jackson, Ian Johnston, Dominic Kwiatkowski, Cordelia Langford, John Sillitoe on behalf of the Wellcome Sanger Institute COVID-19 Surveillance Team |
| EPI_ISL_2780267 | Lighthouse Laboratory Plymouth | Wellcome Sanger Institute for the COVID-19 Genomics UK (COG-UK) Consortium | Lighthouse Laboratory Plymouth and Alex Alderton, Roberto Amato, Jeffrey Barrett, Sonia Goncalves, Ewan Harrison, David K. Jackson, Ian Johnston, Dominic Kwiatkowski, Cordelia Langford, John Sillitoe on behalf of the Wellcome Sanger Institute COVID-19 Surveillance Team |
| EPI_ISL_2780268 | Lighthouse Lab in Alderley Park | Wellcome Sanger Institute for the COVID-19 Genomics UK (COG-UK) Consortium | Jacquelyn Wynn, Mairead Hyland, The Lighthouse Lab in Alderley Park and Alex Alderton, Roberto Amato, Jeffrey Barrett, Sonia Goncalves, Ewan Harrison, David K. Jackson, Ian Johnston, Dominic Kwiatkowski, Cordelia Langford, John Sillitoe on behalf of the Wellcome Sanger Institute COVID-19 Surveillance Team |
| EPI_ISL_2780269 | Lighthouse Laboratory Plymouth | Wellcome Sanger Institute for the COVID-19 Genomics UK (COG-UK) Consortium | Lighthouse Laboratory Plymouth and Alex Alderton, Roberto Amato, Jeffrey Barrett, Sonia Goncalves, Ewan Harrison, David K. Jackson, Ian Johnston, Dominic Kwiatkowski, Cordelia Langford, John Sillitoe on behalf of the Wellcome Sanger Institute COVID-19 Surveillance Team |
| EPI_ISL_2780270 | Lighthouse Lab in Alderley Park | Wellcome Sanger Institute for the COVID-19 Genomics UK (COG-UK) Consortium | Jacquelyn Wynn, Mairead Hyland, The Lighthouse Lab in Alderley Park and Alex Alderton, Roberto Amato, Jeffrey Barrett, Sonia Goncalves, Ewan Harrison, David K. Jackson, Ian Johnston, Dominic Kwiatkowski, Cordelia Langford, John Sillitoe on behalf of the Wellcome Sanger Institute COVID-19 Surveillance Team |
| EPI_ISL_2780271 | Lighthouse Laboratory Plymouth | Wellcome Sanger Institute for the COVID-19 Genomics UK (COG-UK) Consortium | Lighthouse Laboratory Plymouth and Alex Alderton, Roberto Amato, Jeffrey Barrett, Sonia Goncalves, Ewan Harrison, David K. Jackson, Ian Johnston, Dominic Kwiatkowski, Cordelia Langford, John Sillitoe on behalf of the Wellcome Sanger Institute COVID-19 Surveillance Team |
| EPI_ISL_2780272 | Lighthouse Lab in Milton Keynes | Wellcome Sanger Institute for the COVID-19 Genomics UK (COG-UK) Consortium | The Lighthouse Lab in Milton Keynes and Alex Alderton, Roberto Amato, Jeffrey Barrett, Sonia Goncalves, Ewan Harrison, David K. Jackson, Ian Johnston, Dominic Kwiatkowski, Cordelia Langford, John Sillitoe on behalf of the Wellcome Sanger Institute COVID-19 Surveillance Team |
| EPI_ISL_2780273 | Lighthouse Laboratory Plymouth | Wellcome Sanger Institute for the COVID-19 Genomics UK (COG-UK) Consortium | Lighthouse Laboratory Plymouth and Alex Alderton, Roberto Amato, Jeffrey Barrett, Sonia Goncalves, Ewan Harrison, David K. Jackson, Ian Johnston, Dominic Kwiatkowski, Cordelia Langford, John Sillitoe on behalf of the Wellcome Sanger Institute COVID-19 Surveillance Team |
| EPI_ISL_2780274, EPI_ISL_2780275 | Lighthouse Lab in Alderley Park | Wellcome Sanger Institute for the COVID-19 Genomics UK (COG-UK) Consortium | Jacquelyn Wynn, Mairead Hyland, The Lighthouse Lab in Alderley Park and Alex Alderton, Roberto Amato, Jeffrey Barrett, Sonia Goncalves, Ewan Harrison, David K. Jackson, Ian Johnston, Dominic Kwiatkowski, Cordelia Langford, John Sillitoe on behalf of the Wellcome Sanger Institute COVID-19 Surveillance Team |
| EPI_ISL_2780276, EPI_ISL_2780277 | Lighthouse Laboratory Plymouth | Wellcome Sanger Institute for the COVID-19 Genomics UK (COG-UK) Consortium | Lighthouse Laboratory Plymouth and Alex Alderton, Roberto Amato, Jeffrey Barrett, Sonia Goncalves, Ewan Harrison, David K. Jackson, Ian Johnston, Dominic Kwiatkowski, Cordelia Langford, John Sillitoe on behalf of the Wellcome Sanger Institute COVID-19 Surveillance Team |
| EPI_ISL_2780278 | Lighthouse Lab in Alderley Park | Wellcome Sanger Institute for the COVID-19 Genomics UK (COG-UK) Consortium | Jacquelyn Wynn, Mairead Hyland, The Lighthouse Lab in Alderley Park and Alex Alderton, Roberto Amato, Jeffrey Barrett, Sonia Goncalves, Ewan Harrison, David K. Jackson, Ian Johnston, Dominic Kwiatkowski, Cordelia Langford, John Sillitoe on behalf of the Wellcome Sanger Institute COVID-19 Surveillance Team |
| EPI_ISL_2780279 | Lighthouse Lab in Milton Keynes | Wellcome Sanger Institute for the COVID-19 Genomics UK (COG-UK) Consortium | The Lighthouse Lab in Milton Keynes and Alex Alderton, Roberto Amato, Jeffrey Barrett, Sonia Goncalves, Ewan Harrison, David K. Jackson, Ian Johnston, Dominic Kwiatkowski, Cordelia Langford, John Sillitoe on behalf of the Wellcome Sanger Institute COVID-19 Surveillance Team |
| EPI_ISL_2780280 | Lighthouse Lab in Alderley Park | Wellcome Sanger Institute for the COVID-19 Genomics UK (COG-UK) Consortium | Jacquelyn Wynn, Mairead Hyland, The Lighthouse Lab in Alderley Park and Alex Alderton, Roberto Amato, Jeffrey Barrett, Sonia Goncalves, Ewan Harrison, David K. Jackson, Ian Johnston, Dominic Kwiatkowski, Cordelia Langford, John Sillitoe on behalf of the Wellcome Sanger Institute COVID-19 Surveillance Team |
| EPI_ISL_2780281 | Lighthouse Laboratory Plymouth | Wellcome Sanger Institute for the COVID-19 Genomics UK (COG-UK) Consortium | Lighthouse Laboratory Plymouth and Alex Alderton, Roberto Amato, Jeffrey Barrett, Sonia Goncalves, Ewan Harrison, David K. Jackson, Ian Johnston, Dominic Kwiatkowski, Cordelia Langford, John Sillitoe on behalf of the Wellcome Sanger Institute COVID-19 Surveillance Team |
| EPI_ISL_2780282 | Lighthouse Lab in Alderley Park | Wellcome Sanger Institute for the COVID-19 Genomics UK (COG-UK) Consortium | Jacquelyn Wynn, Mairead Hyland, The Lighthouse Lab in Alderley Park and Alex Alderton, Roberto Amato, Jeffrey Barrett, Sonia Goncalves, Ewan Harrison, David K. Jackson, Ian Johnston, Dominic Kwiatkowski, Cordelia Langford, John Sillitoe on behalf of the Wellcome Sanger Institute COVID-19 Surveillance Team |
| EPI_ISL_2780283, EPI_ISL_2780284 | Lighthouse Laboratory Plymouth | Wellcome Sanger Institute for the COVID-19 Genomics UK (COG-UK) Consortium | Lighthouse Laboratory Plymouth and Alex Alderton, Roberto Amato, Jeffrey Barrett, Sonia Goncalves, Ewan Harrison, David K. Jackson, Ian Johnston, Dominic Kwiatkowski, Cordelia Langford, John Sillitoe on behalf of the Wellcome Sanger Institute COVID-19 Surveillance Team |
| EPI_ISL_2780285 | Lighthouse Lab in Alderley Park | Wellcome Sanger Institute for the COVID-19 Genomics UK (COG-UK) Consortium | Jacquelyn Wynn, Mairead Hyland, The Lighthouse Lab in Alderley Park and Alex Alderton, Roberto Amato, Jeffrey Barrett, Sonia Goncalves, Ewan Harrison, David K. Jackson, Ian Johnston, Dominic Kwiatkowski, Cordelia Langford, John Sillitoe on behalf of the Wellcome Sanger Institute COVID-19 Surveillance Team |
| EPI_ISL_2780286 | Lighthouse Lab in Milton Keynes | Wellcome Sanger Institute for the COVID-19 Genomics UK (COG-UK) Consortium | The Lighthouse Lab in Milton Keynes and Alex Alderton, Roberto Amato, Jeffrey Barrett, Sonia Goncalves, Ewan Harrison, David K. Jackson, Ian Johnston, Dominic Kwiatkowski, Cordelia Langford, John Sillitoe on behalf of the Wellcome Sanger Institute COVID-19 Surveillance Team |
| EPI_ISL_2780287 | Lighthouse Lab in Alderley Park | Wellcome Sanger Institute for the COVID-19 Genomics UK (COG-UK) Consortium | Jacquelyn Wynn, Mairead Hyland, The Lighthouse Lab in Alderley Park and Alex Alderton, Roberto Amato, Jeffrey Barrett, Sonia Goncalves, Ewan Harrison, David K. Jackson, Ian Johnston, Dominic Kwiatkowski, Cordelia Langford, John Sillitoe on behalf of the Wellcome Sanger Institute COVID-19 Surveillance Team |
| EPI_ISL_2780288, EPI_ISL_2780289 | Lighthouse Lab in Milton Keynes | Wellcome Sanger Institute for the COVID-19 Genomics UK (COG-UK) Consortium | The Lighthouse Lab in Milton Keynes and Alex Alderton, Roberto Amato, Jeffrey Barrett, Sonia Goncalves, Ewan Harrison, David K. Jackson, Ian Johnston, Dominic Kwiatkowski, Cordelia Langford, John Sillitoe on behalf of the Wellcome Sanger Institute COVID-19 Surveillance Team |
| EPI_ISL_2780290 | Lighthouse Lab in Alderley Park | Wellcome Sanger Institute for the COVID-19 Genomics UK (COG-UK) Consortium | Jacquelyn Wynn, Mairead Hyland, The Lighthouse Lab in Alderley Park and Alex Alderton, Roberto Amato, Jeffrey Barrett, Sonia Goncalves, Ewan Harrison, David K. Jackson, Ian Johnston, Dominic Kwiatkowski, Cordelia Langford, John Sillitoe on behalf of the Wellcome Sanger Institute COVID-19 Surveillance Team |
| EPI_ISL_2780291 | Lighthouse Lab in Milton Keynes | Wellcome Sanger Institute for the COVID-19 Genomics UK (COG-UK) Consortium | The Lighthouse Lab in Milton Keynes and Alex Alderton, Roberto Amato, Jeffrey Barrett, Sonia Goncalves, Ewan Harrison, David K. Jackson, Ian Johnston, Dominic Kwiatkowski, Cordelia Langford, John Sillitoe on behalf of the Wellcome Sanger Institute COVID-19 Surveillance Team |
| EPI_ISL_2780292, EPI_ISL_2780293 | Lighthouse Laboratory Plymouth | Wellcome Sanger Institute for the COVID-19 Genomics UK (COG-UK) Consortium | Lighthouse Laboratory Plymouth and Alex Alderton, Roberto Amato, Jeffrey Barrett, Sonia Goncalves, Ewan Harrison, David K. Jackson, Ian Johnston, Dominic Kwiatkowski, Cordelia Langford, John Sillitoe on behalf of the Wellcome Sanger Institute COVID-19 Surveillance Team |
| EPI_ISL_2780294, EPI_ISL_2780295, EPI_ISL_2780296 | Lighthouse Lab in Alderley Park | Wellcome Sanger Institute for the COVID-19 Genomics UK (COG-UK) Consortium | Jacquelyn Wynn, Mairead Hyland, The Lighthouse Lab in Alderley Park and Alex Alderton, Roberto Amato, Jeffrey Barrett, Sonia Goncalves, Ewan Harrison, David K. Jackson, Ian Johnston, Dominic Kwiatkowski, Cordelia Langford, John Sillitoe on behalf of the Wellcome Sanger Institute COVID-19 Surveillance Team |
| EPI_ISL_2780297 | Lighthouse Lab in Milton Keynes | Wellcome Sanger Institute for the COVID-19 Genomics UK (COG-UK) Consortium | The Lighthouse Lab in Milton Keynes and Alex Alderton, Roberto Amato, Jeffrey Barrett, Sonia Goncalves, Ewan Harrison, David K. Jackson, Ian Johnston, Dominic Kwiatkowski, Cordelia Langford, John Sillitoe on behalf of the Wellcome Sanger Institute COVID-19 Surveillance Team |
| EPI_ISL_2780298 | Lighthouse Laboratory Plymouth | Wellcome Sanger Institute for the COVID-19 Genomics UK (COG-UK) Consortium | Lighthouse Laboratory Plymouth and Alex Alderton, Roberto Amato, Jeffrey Barrett, Sonia Goncalves, Ewan Harrison, David K. Jackson, Ian Johnston, Dominic Kwiatkowski, Cordelia Langford, John Sillitoe on behalf of the Wellcome Sanger Institute COVID-19 Surveillance Team |
| EPI_ISL_2780299 | Lighthouse Lab in Milton Keynes | Wellcome Sanger Institute for the COVID-19 Genomics UK (COG-UK) Consortium | The Lighthouse Lab in Milton Keynes and Alex Alderton, Roberto Amato, Jeffrey Barrett, Sonia Goncalves, Ewan Harrison, David K. Jackson, Ian Johnston, Dominic Kwiatkowski, Cordelia Langford, John Sillitoe on behalf of the Wellcome Sanger Institute COVID-19 Surveillance Team |
| EPI_ISL_2780300, EPI_ISL_2780301, EPI_ISL_2780302, EPI_ISL_2780303, EPI_ISL_2780304, EPI_ISL_2780305, EPI_ISL_2780306, | Lighthouse Lab in Alderley Park | Wellcome Sanger Institute for the COVID-19 Genomics UK (COG-UK) Consortium | Jacquelyn Wynn, Mairead Hyland, The Lighthouse Lab in Alderley Park and Alex Alderton, Roberto Amato, Jeffrey Barrett, Sonia Goncalves, Ewan Harrison, David K. Jackson, Ian Johnston, Dominic Kwiatkowski, Cordelia Langford, John Sillitoe on behalf of the Wellcome Sanger Institute COVID-19 Surveillance Team |

| | | | |
|---|---|---|---|
| EPI_ISL_2780307, EPI_ISL_2780308, EPI_ISL_2780309 | | | |
| EPI_ISL_2780310, EPI_ISL_2780311, EPI_ISL_2780312 | Lighthouse Lab in Milton Keynes | Wellcome Sanger Institute for the COVID-19 Genomics UK (COG-UK) Consortium | The Lighthouse Lab in Milton Keynes and Alex Alderton, Roberto Amato, Jeffrey Barrett, Sonia Goncalves, Ewan Harrison, David K. Jackson, Ian Johnston, Dominic Kwiatkowski, Cordelia Langford, John Sillitoe on behalf of the Wellcome Sanger Institute COVID-19 Surveillance Team |
| EPI_ISL_2780313 | Lighthouse Laboratory Plymouth | Wellcome Sanger Institute for the COVID-19 Genomics UK (COG-UK) Consortium | Lighthouse Laboratory Plymouth and Alex Alderton, Roberto Amato, Jeffrey Barrett, Sonia Goncalves, Ewan Harrison, David K. Jackson, Ian Johnston, Dominic Kwiatkowski, Cordelia Langford, John Sillitoe on behalf of the Wellcome Sanger Institute COVID-19 Surveillance Team |
| EPI_ISL_2780314 | Lighthouse Lab in Milton Keynes | Wellcome Sanger Institute for the COVID-19 Genomics UK (COG-UK) Consortium | The Lighthouse Lab in Milton Keynes and Alex Alderton, Roberto Amato, Jeffrey Barrett, Sonia Goncalves, Ewan Harrison, David K. Jackson, Ian Johnston, Dominic Kwiatkowski, Cordelia Langford, John Sillitoe on behalf of the Wellcome Sanger Institute COVID-19 Surveillance Team |
| EPI_ISL_2780315, EPI_ISL_2780316, EPI_ISL_2780317 | Lighthouse Laboratory Plymouth | Wellcome Sanger Institute for the COVID-19 Genomics UK (COG-UK) Consortium | Lighthouse Laboratory Plymouth and Alex Alderton, Roberto Amato, Jeffrey Barrett, Sonia Goncalves, Ewan Harrison, David K. Jackson, Ian Johnston, Dominic Kwiatkowski, Cordelia Langford, John Sillitoe on behalf of the Wellcome Sanger Institute COVID-19 Surveillance Team |
| EPI_ISL_2780318 | Lighthouse Lab in Alderley Park | Wellcome Sanger Institute for the COVID-19 Genomics UK (COG-UK) Consortium | Jacquelyn Wynn, Mairead Hyland, The Lighthouse Lab in Alderley Park and Alex Alderton, Roberto Amato, Jeffrey Barrett, Sonia Goncalves, Ewan Harrison, David K. Jackson, Ian Johnston, Dominic Kwiatkowski, Cordelia Langford, John Sillitoe on behalf of the Wellcome Sanger Institute COVID-19 Surveillance Team |
| EPI_ISL_2780319, EPI_ISL_2780320 | Lighthouse Laboratory Plymouth | Wellcome Sanger Institute for the COVID-19 Genomics UK (COG-UK) Consortium | Lighthouse Laboratory Plymouth and Alex Alderton, Roberto Amato, Jeffrey Barrett, Sonia Goncalves, Ewan Harrison, David K. Jackson, Ian Johnston, Dominic Kwiatkowski, Cordelia Langford, John Sillitoe on behalf of the Wellcome Sanger Institute COVID-19 Surveillance Team |
| EPI_ISL_2780321 | Lighthouse Lab in Alderley Park | Wellcome Sanger Institute for the COVID-19 Genomics UK (COG-UK) Consortium | Jacquelyn Wynn, Mairead Hyland, The Lighthouse Lab in Alderley Park and Alex Alderton, Roberto Amato, Jeffrey Barrett, Sonia Goncalves, Ewan Harrison, David K. Jackson, Ian Johnston, Dominic Kwiatkowski, Cordelia Langford, John Sillitoe on behalf of the Wellcome Sanger Institute COVID-19 Surveillance Team |
| EPI_ISL_2780322 | Lighthouse Laboratory Plymouth | Wellcome Sanger Institute for the COVID-19 Genomics UK (COG-UK) Consortium | Lighthouse Laboratory Plymouth and Alex Alderton, Roberto Amato, Jeffrey Barrett, Sonia Goncalves, Ewan Harrison, David K. Jackson, Ian Johnston, Dominic Kwiatkowski, Cordelia Langford, John Sillitoe on behalf of the Wellcome Sanger Institute COVID-19 Surveillance Team |
| EPI_ISL_2780323 | Lighthouse Lab in Alderley Park | Wellcome Sanger Institute for the COVID-19 Genomics UK (COG-UK) Consortium | Jacquelyn Wynn, Mairead Hyland, The Lighthouse Lab in Alderley Park and Alex Alderton, Roberto Amato, Jeffrey Barrett, Sonia Goncalves, Ewan Harrison, David K. Jackson, Ian Johnston, Dominic Kwiatkowski, Cordelia Langford, John Sillitoe on behalf of the Wellcome Sanger Institute COVID-19 Surveillance Team |
| EPI_ISL_2780324 | Lighthouse Laboratory Plymouth | Wellcome Sanger Institute for the COVID-19 Genomics UK (COG-UK) Consortium | Lighthouse Laboratory Plymouth and Alex Alderton, Roberto Amato, Jeffrey Barrett, Sonia Goncalves, Ewan Harrison, David K. Jackson, Ian Johnston, Dominic Kwiatkowski, Cordelia Langford, John Sillitoe on behalf of the Wellcome Sanger Institute COVID-19 Surveillance Team |
| EPI_ISL_2780325 | Lighthouse Lab in Alderley Park | Wellcome Sanger Institute for the COVID-19 Genomics UK (COG-UK) Consortium | Jacquelyn Wynn, Mairead Hyland, The Lighthouse Lab in Alderley Park and Alex Alderton, Roberto Amato, Jeffrey Barrett, Sonia Goncalves, Ewan Harrison, David K. Jackson, Ian Johnston, Dominic Kwiatkowski, Cordelia Langford, John Sillitoe on behalf of the Wellcome Sanger Institute COVID-19 Surveillance Team |
| EPI_ISL_2780327, EPI_ISL_2780328 | Lighthouse Lab in Milton Keynes | Wellcome Sanger Institute for the COVID-19 Genomics UK (COG-UK) Consortium | The Lighthouse Lab in Milton Keynes and Alex Alderton, Roberto Amato, Jeffrey Barrett, Sonia Goncalves, Ewan Harrison, David K. Jackson, Ian Johnston, Dominic Kwiatkowski, Cordelia Langford, John Sillitoe on behalf of the Wellcome Sanger Institute COVID-19 Surveillance Team |
| EPI_ISL_2780329, EPI_ISL_2780330 | Lighthouse Laboratory Plymouth | Wellcome Sanger Institute for the COVID-19 Genomics UK (COG-UK) Consortium | Lighthouse Laboratory Plymouth and Alex Alderton, Roberto Amato, Jeffrey Barrett, Sonia Goncalves, Ewan Harrison, David K. Jackson, Ian Johnston, Dominic Kwiatkowski, Cordelia Langford, John Sillitoe on behalf of the Wellcome Sanger Institute COVID-19 Surveillance Team |
| EPI_ISL_2780331 | Lighthouse Lab in Alderley Park | Wellcome Sanger Institute for the COVID-19 Genomics UK (COG-UK) Consortium | Jacquelyn Wynn, Mairead Hyland, The Lighthouse Lab in Alderley Park and Alex Alderton, Roberto Amato, Jeffrey Barrett, Sonia Goncalves, Ewan Harrison, David K. Jackson, Ian Johnston, Dominic Kwiatkowski, Cordelia Langford, John Sillitoe on behalf of the Wellcome Sanger Institute COVID-19 Surveillance Team |
| EPI_ISL_2780332 | Lighthouse Lab in Milton Keynes | Wellcome Sanger Institute for the COVID-19 Genomics UK (COG-UK) Consortium | The Lighthouse Lab in Milton Keynes and Alex Alderton, Roberto Amato, Jeffrey Barrett, Sonia Goncalves, Ewan Harrison, David K. Jackson, Ian Johnston, Dominic Kwiatkowski, Cordelia Langford, John Sillitoe on behalf of the Wellcome Sanger Institute COVID-19 Surveillance Team |
| EPI_ISL_2780333 | Lighthouse Lab in Alderley Park | Wellcome Sanger Institute for the COVID-19 Genomics UK (COG-UK) Consortium | Jacquelyn Wynn, Mairead Hyland, The Lighthouse Lab in Alderley Park and Alex Alderton, Roberto Amato, Jeffrey Barrett, Sonia Goncalves, Ewan Harrison, David K. Jackson, Ian Johnston, Dominic Kwiatkowski, Cordelia Langford, John Sillitoe on behalf of the Wellcome Sanger Institute COVID-19 Surveillance Team |
| EPI_ISL_2780334 | Lighthouse Lab in Milton Keynes | Wellcome Sanger Institute for the COVID-19 Genomics UK (COG-UK) Consortium | The Lighthouse Lab in Milton Keynes and Alex Alderton, Roberto Amato, Jeffrey Barrett, Sonia Goncalves, Ewan Harrison, David K. Jackson, Ian Johnston, Dominic Kwiatkowski, Cordelia Langford, John Sillitoe on behalf of the Wellcome Sanger Institute COVID-19 Surveillance Team |
| EPI_ISL_2780335 | Lighthouse Laboratory Plymouth | Wellcome Sanger Institute for the COVID-19 Genomics UK (COG-UK) Consortium | Lighthouse Laboratory Plymouth and Alex Alderton, Roberto Amato, Jeffrey Barrett, Sonia Goncalves, Ewan Harrison, David K. Jackson, Ian Johnston, Dominic Kwiatkowski, Cordelia Langford, John Sillitoe on behalf of the Wellcome Sanger Institute COVID-19 Surveillance Team |
| EPI_ISL_2780336, EPI_ISL_2780337, EPI_ISL_2780338 | Lighthouse Lab in Alderley Park | Wellcome Sanger Institute for the COVID-19 Genomics UK (COG-UK) Consortium | Jacquelyn Wynn, Mairead Hyland, The Lighthouse Lab in Alderley Park and Alex Alderton, Roberto Amato, Jeffrey Barrett, Sonia Goncalves, Ewan Harrison, David K. Jackson, Ian Johnston, Dominic Kwiatkowski, Cordelia Langford, John Sillitoe on behalf of the Wellcome Sanger Institute COVID-19 Surveillance Team |
| EPI_ISL_2780339 | Lighthouse Laboratory Plymouth | Wellcome Sanger Institute for the COVID-19 Genomics UK (COG-UK) Consortium | Lighthouse Laboratory Plymouth and Alex Alderton, Roberto Amato, Jeffrey Barrett, Sonia Goncalves, Ewan Harrison, David K. Jackson, Ian Johnston, Dominic Kwiatkowski, Cordelia Langford, John Sillitoe on behalf of the Wellcome Sanger Institute COVID-19 Surveillance Team |
| EPI_ISL_2780340, EPI_ISL_2780341, EPI_ISL_2780342 | Lighthouse Lab in Alderley Park | Wellcome Sanger Institute for the COVID-19 Genomics UK (COG-UK) Consortium | Jacquelyn Wynn, Mairead Hyland, The Lighthouse Lab in Alderley Park and Alex Alderton, Roberto Amato, Jeffrey Barrett, Sonia Goncalves, Ewan Harrison, David K. Jackson, Ian Johnston, Dominic Kwiatkowski, Cordelia Langford, John Sillitoe on behalf of the Wellcome Sanger Institute COVID-19 Surveillance Team |
| EPI_ISL_2780343 | Lighthouse Lab in Milton Keynes | Wellcome Sanger Institute for the COVID-19 Genomics UK (COG-UK) Consortium | The Lighthouse Lab in Milton Keynes and Alex Alderton, Roberto Amato, Jeffrey Barrett, Sonia Goncalves, Ewan Harrison, David K. Jackson, Ian Johnston, Dominic Kwiatkowski, Cordelia Langford, John Sillitoe on behalf of the Wellcome Sanger Institute COVID-19 Surveillance Team |
| EPI_ISL_2780344, EPI_ISL_2780345, EPI_ISL_2780346 | Lighthouse Lab in Alderley Park | Wellcome Sanger Institute for the COVID-19 Genomics UK (COG-UK) Consortium | Jacquelyn Wynn, Mairead Hyland, The Lighthouse Lab in Alderley Park and Alex Alderton, Roberto Amato, Jeffrey Barrett, Sonia Goncalves, Ewan Harrison, David K. Jackson, Ian Johnston, Dominic Kwiatkowski, Cordelia Langford, John Sillitoe on behalf of the Wellcome Sanger Institute COVID-19 Surveillance Team |
| EPI_ISL_2780347 | Lighthouse Lab in Milton Keynes | Wellcome Sanger Institute for the COVID-19 Genomics UK (COG-UK) Consortium | The Lighthouse Lab in Milton Keynes and Alex Alderton, Roberto Amato, Jeffrey Barrett, Sonia Goncalves, Ewan Harrison, David K. Jackson, Ian Johnston, Dominic Kwiatkowski, Cordelia Langford, John Sillitoe on behalf of the Wellcome Sanger Institute COVID-19 Surveillance Team |
| EPI_ISL_2780348 | Lighthouse Lab in Alderley Park | Wellcome Sanger Institute for the COVID-19 Genomics UK (COG-UK) Consortium | Jacquelyn Wynn, Mairead Hyland, The Lighthouse Lab in Alderley Park and Alex Alderton, Roberto Amato, Jeffrey Barrett, Sonia Goncalves, Ewan Harrison, David K. Jackson, Ian Johnston, Dominic Kwiatkowski, Cordelia Langford, John Sillitoe on behalf of the Wellcome Sanger Institute COVID-19 Surveillance Team |
| EPI_ISL_2780349 | Lighthouse Laboratory Plymouth | Wellcome Sanger Institute for the COVID-19 Genomics UK (COG-UK) Consortium | Lighthouse Laboratory Plymouth and Alex Alderton, Roberto Amato, Jeffrey Barrett, Sonia Goncalves, Ewan Harrison, David K. Jackson, Ian Johnston, Dominic Kwiatkowski, Cordelia Langford, John Sillitoe on behalf of the Wellcome Sanger Institute COVID-19 Surveillance Team |
| EPI_ISL_2780350 | Lighthouse Lab in Milton Keynes | Wellcome Sanger Institute for the COVID-19 Genomics UK (COG-UK) Consortium | The Lighthouse Lab in Milton Keynes and Alex Alderton, Roberto Amato, Jeffrey Barrett, Sonia Goncalves, Ewan Harrison, David K. Jackson, Ian Johnston, Dominic Kwiatkowski, Cordelia Langford, John Sillitoe on behalf of the Wellcome Sanger Institute COVID-19 Surveillance Team |
| EPI_ISL_2780351, EPI_ISL_2780352 | Lighthouse Lab in Alderley Park | Wellcome Sanger Institute for the COVID-19 Genomics UK (COG-UK) Consortium | Jacquelyn Wynn, Mairead Hyland, The Lighthouse Lab in Alderley Park and Alex Alderton, Roberto Amato, Jeffrey Barrett, Sonia Goncalves, Ewan Harrison, David K. Jackson, Ian Johnston, Dominic Kwiatkowski, Cordelia Langford, John Sillitoe on behalf of the Wellcome Sanger Institute COVID-19 Surveillance Team |
| EPI_ISL_2780353 | Lighthouse Lab in Milton Keynes | Wellcome Sanger Institute for the COVID-19 Genomics UK (COG-UK) Consortium | The Lighthouse Lab in Milton Keynes and Alex Alderton, Roberto Amato, Jeffrey Barrett, Sonia Goncalves, Ewan Harrison, David K. Jackson, Ian Johnston, Dominic Kwiatkowski, Cordelia Langford, John Sillitoe on behalf of the Wellcome Sanger Institute COVID-19 Surveillance Team |
| EPI_ISL_2780354 | Lighthouse Laboratory Plymouth | Wellcome Sanger Institute for the COVID-19 Genomics UK (COG-UK) Consortium | Lighthouse Laboratory Plymouth and Alex Alderton, Roberto Amato, Jeffrey Barrett, Sonia Goncalves, Ewan Harrison, David K. Jackson, Ian Johnston, Dominic Kwiatkowski, Cordelia Langford, John Sillitoe on behalf of the Wellcome Sanger Institute COVID-19 Surveillance Team |
| EPI_ISL_2780355 | Lighthouse Lab in Milton Keynes | Wellcome Sanger Institute for the COVID-19 Genomics UK (COG-UK) Consortium | The Lighthouse Lab in Milton Keynes and Alex Alderton, Roberto Amato, Jeffrey Barrett, Sonia Goncalves, Ewan Harrison, David K. Jackson, Ian Johnston, Dominic Kwiatkowski, Cordelia Langford, John Sillitoe on behalf of the Wellcome Sanger Institute COVID-19 Surveillance Team |
| EPI_ISL_2780356 | Lighthouse Lab in Alderley Park | Wellcome Sanger Institute for the COVID-19 Genomics UK (COG-UK) Consortium | Jacquelyn Wynn, Mairead Hyland, The Lighthouse Lab in Alderley Park and Alex Alderton, Roberto Amato, Jeffrey Barrett, Sonia Goncalves, Ewan Harrison, David K. Jackson, Ian Johnston, Dominic Kwiatkowski, Cordelia Langford, John Sillitoe on behalf of the Wellcome Sanger Institute COVID-19 Surveillance Team |
| EPI_ISL_2780357, EPI_ISL_2780358 | Lighthouse Lab in Milton Keynes | Wellcome Sanger Institute for the COVID-19 Genomics UK (COG-UK) Consortium | The Lighthouse Lab in Milton Keynes and Alex Alderton, Roberto Amato, Jeffrey Barrett, Sonia Goncalves, Ewan Harrison, David K. Jackson, Ian Johnston, Dominic Kwiatkowski, Cordelia Langford, John Sillitoe on behalf of the Wellcome Sanger Institute COVID-19 Surveillance Team |
| EPI_ISL_2780359 | Lighthouse Laboratory Plymouth | Wellcome Sanger Institute for the COVID-19 Genomics UK (COG-UK) Consortium | Lighthouse Laboratory Plymouth and Alex Alderton, Roberto Amato, Jeffrey Barrett, Sonia Goncalves, Ewan Harrison, David K. Jackson, Ian Johnston, Dominic Kwiatkowski, Cordelia Langford, John Sillitoe on behalf of the Wellcome Sanger Institute COVID-19 Surveillance Team |
| EPI_ISL_2780360, EPI_ISL_2780361 | Lighthouse Lab in Alderley Park | Wellcome Sanger Institute for the COVID-19 Genomics UK (COG-UK) Consortium | Jacquelyn Wynn, Mairead Hyland, The Lighthouse Lab in Alderley Park and Alex Alderton, Roberto Amato, Jeffrey Barrett, Sonia Goncalves, Ewan Harrison, David K. Jackson, Ian Johnston, Dominic Kwiatkowski, Cordelia Langford, John Sillitoe on behalf of the Wellcome Sanger Institute COVID-19 Surveillance Team |
| EPI_ISL_2780362, EPI_ISL_2780363 | Lighthouse Laboratory Plymouth | Wellcome Sanger Institute for the COVID-19 Genomics UK (COG-UK) Consortium | Lighthouse Laboratory Plymouth and Alex Alderton, Roberto Amato, Jeffrey Barrett, Sonia Goncalves, Ewan Harrison, David K. Jackson, Ian Johnston, Dominic Kwiatkowski, Cordelia Langford, John Sillitoe on behalf of the Wellcome Sanger Institute COVID-19 Surveillance Team |
| EPI_ISL_2780364 | Lighthouse Lab in Alderley Park | Wellcome Sanger Institute for the COVID-19 Genomics UK (COG-UK) Consortium | Jacquelyn Wynn, Mairead Hyland, The Lighthouse Lab in Alderley Park and Alex Alderton, Roberto Amato, Jeffrey Barrett, Sonia Goncalves, Ewan Harrison, David K. Jackson, Ian Johnston, Dominic Kwiatkowski, Cordelia Langford, John Sillitoe on behalf of the Wellcome Sanger Institute COVID-19 Surveillance Team |
| EPI_ISL_2780365 | Lighthouse Laboratory Plymouth | Wellcome Sanger Institute for the COVID-19 Genomics UK (COG-UK) Consortium | Lighthouse Laboratory Plymouth and Alex Alderton, Roberto Amato, Jeffrey Barrett, Sonia Goncalves, Ewan Harrison, David K. Jackson, Ian Johnston, Dominic Kwiatkowski, Cordelia Langford, John Sillitoe on behalf of the Wellcome Sanger Institute COVID-19 Surveillance Team |
| EPI_ISL_2780366, EPI_ISL_2780367 | Lighthouse Lab in Alderley Park | Wellcome Sanger Institute for the COVID-19 Genomics UK (COG-UK) Consortium | Jacquelyn Wynn, Mairead Hyland, The Lighthouse Lab in Alderley Park and Alex Alderton, Roberto Amato, Jeffrey Barrett, Sonia Goncalves, Ewan Harrison, David K. Jackson, Ian Johnston, Dominic Kwiatkowski, Cordelia Langford, John Sillitoe on behalf of the Wellcome Sanger Institute COVID-19 Surveillance Team |
| EPI_ISL_2780368 | Lighthouse Lab in Milton Keynes | Wellcome Sanger Institute for the COVID-19 Genomics UK (COG-UK) Consortium | The Lighthouse Lab in Milton Keynes and Alex Alderton, Roberto Amato, Jeffrey Barrett, Sonia Goncalves, Ewan Harrison, David K. Jackson, Ian Johnston, Dominic Kwiatkowski, Cordelia Langford, John Sillitoe on behalf of the Wellcome Sanger Institute COVID-19 Surveillance Team |
| EPI_ISL_2780369 | Lighthouse Laboratory Plymouth | Wellcome Sanger Institute for the COVID-19 Genomics UK (COG-UK) Consortium | Lighthouse Laboratory Plymouth and Alex Alderton, Roberto Amato, Jeffrey Barrett, Sonia Goncalves, Ewan Harrison, David K. Jackson, Ian Johnston, Dominic Kwiatkowski, Cordelia Langford, John Sillitoe on behalf of the Wellcome Sanger Institute COVID-19 Surveillance Team |
| EPI_ISL_2780370 | Lighthouse Lab in Milton Keynes | Wellcome Sanger Institute for the COVID-19 Genomics UK (COG-UK) Consortium | The Lighthouse Lab in Milton Keynes and Alex Alderton, Roberto Amato, Jeffrey Barrett, Sonia Goncalves, Ewan Harrison, David K. Jackson, Ian Johnston, Dominic Kwiatkowski, Cordelia Langford, John Sillitoe on behalf of the Wellcome Sanger Institute COVID-19 Surveillance Team |

| | | | |
|---|---|---|---|
| EPI_ISL_2780371 | Lighthouse Laboratory Plymouth | Wellcome Sanger Institute for the COVID-19 Genomics UK (COG-UK) Consortium | Lighthouse Laboratory Plymouth and Alex Alderton, Roberto Amato, Jeffrey Barrett, Sonia Goncalves, Ewan Harrison, David K. Jackson, Ian Johnston, Dominic Kwiatkowski, Cordelia Langford, John Sillitoe on behalf of the Wellcome Sanger Institute COVID-19 Surveillance Team |
| EPI_ISL_2780372, EPI_ISL_2780373 | Lighthouse Lab in Alderley Park | Wellcome Sanger Institute for the COVID-19 Genomics UK (COG-UK) Consortium | Jacquelyn Wynn, Mairead Hyland, The Lighthouse Lab in Alderley Park and Alex Alderton, Roberto Amato, Jeffrey Barrett, Sonia Goncalves, Ewan Harrison, David K. Jackson, Ian Johnston, Dominic Kwiatkowski, Cordelia Langford, John Sillitoe on behalf of the Wellcome Sanger Institute COVID-19 Surveillance Team |
| EPI_ISL_2780374, EPI_ISL_2780375 | Lighthouse Laboratory Plymouth | Wellcome Sanger Institute for the COVID-19 Genomics UK (COG-UK) Consortium | Lighthouse Laboratory Plymouth and Alex Alderton, Roberto Amato, Jeffrey Barrett, Sonia Goncalves, Ewan Harrison, David K. Jackson, Ian Johnston, Dominic Kwiatkowski, Cordelia Langford, John Sillitoe on behalf of the Wellcome Sanger Institute COVID-19 Surveillance Team |
| EPI_ISL_2780376, EPI_ISL_2780377 | Lighthouse Lab in Milton Keynes | Wellcome Sanger Institute for the COVID-19 Genomics UK (COG-UK) Consortium | The Lighthouse Lab in Milton Keynes and Alex Alderton, Roberto Amato, Jeffrey Barrett, Sonia Goncalves, Ewan Harrison, David K. Jackson, Ian Johnston, Dominic Kwiatkowski, Cordelia Langford, John Sillitoe on behalf of the Wellcome Sanger Institute COVID-19 Surveillance Team |
| EPI_ISL_2780378 | Lighthouse Lab in Alderley Park | Wellcome Sanger Institute for the COVID-19 Genomics UK (COG-UK) Consortium | Jacquelyn Wynn, Mairead Hyland, The Lighthouse Lab in Alderley Park and Alex Alderton, Roberto Amato, Jeffrey Barrett, Sonia Goncalves, Ewan Harrison, David K. Jackson, Ian Johnston, Dominic Kwiatkowski, Cordelia Langford, John Sillitoe on behalf of the Wellcome Sanger Institute COVID-19 Surveillance Team |
| EPI_ISL_2780379 | Lighthouse Lab in Milton Keynes | Wellcome Sanger Institute for the COVID-19 Genomics UK (COG-UK) Consortium | The Lighthouse Lab in Milton Keynes and Alex Alderton, Roberto Amato, Jeffrey Barrett, Sonia Goncalves, Ewan Harrison, David K. Jackson, Ian Johnston, Dominic Kwiatkowski, Cordelia Langford, John Sillitoe on behalf of the Wellcome Sanger Institute COVID-19 Surveillance Team |
| EPI_ISL_2780380 | Lighthouse Lab in Alderley Park | Wellcome Sanger Institute for the COVID-19 Genomics UK (COG-UK) Consortium | Jacquelyn Wynn, Mairead Hyland, The Lighthouse Lab in Alderley Park and Alex Alderton, Roberto Amato, Jeffrey Barrett, Sonia Goncalves, Ewan Harrison, David K. Jackson, Ian Johnston, Dominic Kwiatkowski, Cordelia Langford, John Sillitoe on behalf of the Wellcome Sanger Institute COVID-19 Surveillance Team |
| EPI_ISL_2780381 | Lighthouse Laboratory Plymouth | Wellcome Sanger Institute for the COVID-19 Genomics UK (COG-UK) Consortium | Lighthouse Laboratory Plymouth and Alex Alderton, Roberto Amato, Jeffrey Barrett, Sonia Goncalves, Ewan Harrison, David K. Jackson, Ian Johnston, Dominic Kwiatkowski, Cordelia Langford, John Sillitoe on behalf of the Wellcome Sanger Institute COVID-19 Surveillance Team |
| EPI_ISL_2780382 | Lighthouse Lab in Alderley Park | Wellcome Sanger Institute for the COVID-19 Genomics UK (COG-UK) Consortium | Jacquelyn Wynn, Mairead Hyland, The Lighthouse Lab in Alderley Park and Alex Alderton, Roberto Amato, Jeffrey Barrett, Sonia Goncalves, Ewan Harrison, David K. Jackson, Ian Johnston, Dominic Kwiatkowski, Cordelia Langford, John Sillitoe on behalf of the Wellcome Sanger Institute COVID-19 Surveillance Team |
| EPI_ISL_2780383 | Lighthouse Laboratory Plymouth | Wellcome Sanger Institute for the COVID-19 Genomics UK (COG-UK) Consortium | Lighthouse Laboratory Plymouth and Alex Alderton, Roberto Amato, Jeffrey Barrett, Sonia Goncalves, Ewan Harrison, David K. Jackson, Ian Johnston, Dominic Kwiatkowski, Cordelia Langford, John Sillitoe on behalf of the Wellcome Sanger Institute COVID-19 Surveillance Team |
| EPI_ISL_2780384 | Lighthouse Lab in Alderley Park | Wellcome Sanger Institute for the COVID-19 Genomics UK (COG-UK) Consortium | Jacquelyn Wynn, Mairead Hyland, The Lighthouse Lab in Alderley Park and Alex Alderton, Roberto Amato, Jeffrey Barrett, Sonia Goncalves, Ewan Harrison, David K. Jackson, Ian Johnston, Dominic Kwiatkowski, Cordelia Langford, John Sillitoe on behalf of the Wellcome Sanger Institute COVID-19 Surveillance Team |
| EPI_ISL_2780385 | Lighthouse Laboratory Plymouth | Wellcome Sanger Institute for the COVID-19 Genomics UK (COG-UK) Consortium | Lighthouse Laboratory Plymouth and Alex Alderton, Roberto Amato, Jeffrey Barrett, Sonia Goncalves, Ewan Harrison, David K. Jackson, Ian Johnston, Dominic Kwiatkowski, Cordelia Langford, John Sillitoe on behalf of the Wellcome Sanger Institute COVID-19 Surveillance Team |
| EPI_ISL_2780386 | Lighthouse Lab in Milton Keynes | Wellcome Sanger Institute for the COVID-19 Genomics UK (COG-UK) Consortium | The Lighthouse Lab in Milton Keynes and Alex Alderton, Roberto Amato, Jeffrey Barrett, Sonia Goncalves, Ewan Harrison, David K. Jackson, Ian Johnston, Dominic Kwiatkowski, Cordelia Langford, John Sillitoe on behalf of the Wellcome Sanger Institute COVID-19 Surveillance Team |
| EPI_ISL_2780387 | Lighthouse Lab in Alderley Park | Wellcome Sanger Institute for the COVID-19 Genomics UK (COG-UK) Consortium | Jacquelyn Wynn, Mairead Hyland, The Lighthouse Lab in Alderley Park and Alex Alderton, Roberto Amato, Jeffrey Barrett, Sonia Goncalves, Ewan Harrison, David K. Jackson, Ian Johnston, Dominic Kwiatkowski, Cordelia Langford, John Sillitoe on behalf of the Wellcome Sanger Institute COVID-19 Surveillance Team |
| EPI_ISL_2780388 | Lighthouse Lab in Milton Keynes | Wellcome Sanger Institute for the COVID-19 Genomics UK (COG-UK) Consortium | The Lighthouse Lab in Milton Keynes and Alex Alderton, Roberto Amato, Jeffrey Barrett, Sonia Goncalves, Ewan Harrison, David K. Jackson, Ian Johnston, Dominic Kwiatkowski, Cordelia Langford, John Sillitoe on behalf of the Wellcome Sanger Institute COVID-19 Surveillance Team |
| EPI_ISL_2780389 | Lighthouse Laboratory Plymouth | Wellcome Sanger Institute for the COVID-19 Genomics UK (COG-UK) Consortium | Lighthouse Laboratory Plymouth and Alex Alderton, Roberto Amato, Jeffrey Barrett, Sonia Goncalves, Ewan Harrison, David K. Jackson, Ian Johnston, Dominic Kwiatkowski, Cordelia Langford, John Sillitoe on behalf of the Wellcome Sanger Institute COVID-19 Surveillance Team |
| EPI_ISL_2780390 | Lighthouse Lab in Alderley Park | Wellcome Sanger Institute for the COVID-19 Genomics UK (COG-UK) Consortium | Jacquelyn Wynn, Mairead Hyland, The Lighthouse Lab in Alderley Park and Alex Alderton, Roberto Amato, Jeffrey Barrett, Sonia Goncalves, Ewan Harrison, David K. Jackson, Ian Johnston, Dominic Kwiatkowski, Cordelia Langford, John Sillitoe on behalf of the Wellcome Sanger Institute COVID-19 Surveillance Team |
| EPI_ISL_2780391, EPI_ISL_2780392 | Lighthouse Laboratory Plymouth | Wellcome Sanger Institute for the COVID-19 Genomics UK (COG-UK) Consortium | Lighthouse Laboratory Plymouth and Alex Alderton, Roberto Amato, Jeffrey Barrett, Sonia Goncalves, Ewan Harrison, David K. Jackson, Ian Johnston, Dominic Kwiatkowski, Cordelia Langford, John Sillitoe on behalf of the Wellcome Sanger Institute COVID-19 Surveillance Team |
| EPI_ISL_2780393, EPI_ISL_2780394 | Lighthouse Lab in Alderley Park | Wellcome Sanger Institute for the COVID-19 Genomics UK (COG-UK) Consortium | Jacquelyn Wynn, Mairead Hyland, The Lighthouse Lab in Alderley Park and Alex Alderton, Roberto Amato, Jeffrey Barrett, Sonia Goncalves, Ewan Harrison, David K. Jackson, Ian Johnston, Dominic Kwiatkowski, Cordelia Langford, John Sillitoe on behalf of the Wellcome Sanger Institute COVID-19 Surveillance Team |
| EPI_ISL_2780395 | Lighthouse Laboratory Plymouth | Wellcome Sanger Institute for the COVID-19 Genomics UK (COG-UK) Consortium | Lighthouse Laboratory Plymouth and Alex Alderton, Roberto Amato, Jeffrey Barrett, Sonia Goncalves, Ewan Harrison, David K. Jackson, Ian Johnston, Dominic Kwiatkowski, Cordelia Langford, John Sillitoe on behalf of the Wellcome Sanger Institute COVID-19 Surveillance Team |
| EPI_ISL_2780396 | Lighthouse Lab in Alderley Park | Wellcome Sanger Institute for the COVID-19 Genomics UK (COG-UK) Consortium | Jacquelyn Wynn, Mairead Hyland, The Lighthouse Lab in Alderley Park and Alex Alderton, Roberto Amato, Jeffrey Barrett, Sonia Goncalves, Ewan Harrison, David K. Jackson, Ian Johnston, Dominic Kwiatkowski, Cordelia Langford, John Sillitoe on behalf of the Wellcome Sanger Institute COVID-19 Surveillance Team |
| EPI_ISL_2780397 | Lighthouse Laboratory Plymouth | Wellcome Sanger Institute for the COVID-19 Genomics UK (COG-UK) Consortium | Lighthouse Laboratory Plymouth and Alex Alderton, Roberto Amato, Jeffrey Barrett, Sonia Goncalves, Ewan Harrison, David K. Jackson, Ian Johnston, Dominic Kwiatkowski, Cordelia Langford, John Sillitoe on behalf of the Wellcome Sanger Institute COVID-19 Surveillance Team |
| EPI_ISL_2780398 | Lighthouse Lab in Milton Keynes | Wellcome Sanger Institute for the COVID-19 Genomics UK (COG-UK) Consortium | The Lighthouse Lab in Milton Keynes and Alex Alderton, Roberto Amato, Jeffrey Barrett, Sonia Goncalves, Ewan Harrison, David K. Jackson, Ian Johnston, Dominic Kwiatkowski, Cordelia Langford, John Sillitoe on behalf of the Wellcome Sanger Institute COVID-19 Surveillance Team |
| EPI_ISL_2780399, EPI_ISL_2780400 | Lighthouse Lab in Alderley Park | Wellcome Sanger Institute for the COVID-19 Genomics UK (COG-UK) Consortium | Jacquelyn Wynn, Mairead Hyland, The Lighthouse Lab in Alderley Park and Alex Alderton, Roberto Amato, Jeffrey Barrett, Sonia Goncalves, Ewan Harrison, David K. Jackson, Ian Johnston, Dominic Kwiatkowski, Cordelia Langford, John Sillitoe on behalf of the Wellcome Sanger Institute COVID-19 Surveillance Team |
| EPI_ISL_2780401, EPI_ISL_2780402 | Lighthouse Lab in Milton Keynes | Wellcome Sanger Institute for the COVID-19 Genomics UK (COG-UK) Consortium | The Lighthouse Lab in Milton Keynes and Alex Alderton, Roberto Amato, Jeffrey Barrett, Sonia Goncalves, Ewan Harrison, David K. Jackson, Ian Johnston, Dominic Kwiatkowski, Cordelia Langford, John Sillitoe on behalf of the Wellcome Sanger Institute COVID-19 Surveillance Team |
| EPI_ISL_2780403, EPI_ISL_2780404 | Lighthouse Laboratory Plymouth | Wellcome Sanger Institute for the COVID-19 Genomics UK (COG-UK) Consortium | Lighthouse Laboratory Plymouth and Alex Alderton, Roberto Amato, Jeffrey Barrett, Sonia Goncalves, Ewan Harrison, David K. Jackson, Ian Johnston, Dominic Kwiatkowski, Cordelia Langford, John Sillitoe on behalf of the Wellcome Sanger Institute COVID-19 Surveillance Team |
| EPI_ISL_2780405 | Lighthouse Lab in Alderley Park | Wellcome Sanger Institute for the COVID-19 Genomics UK (COG-UK) Consortium | Jacquelyn Wynn, Mairead Hyland, The Lighthouse Lab in Alderley Park and Alex Alderton, Roberto Amato, Jeffrey Barrett, Sonia Goncalves, Ewan Harrison, David K. Jackson, Ian Johnston, Dominic Kwiatkowski, Cordelia Langford, John Sillitoe on behalf of the Wellcome Sanger Institute COVID-19 Surveillance Team |
| EPI_ISL_2780406 | Lighthouse Lab in Milton Keynes | Wellcome Sanger Institute for the COVID-19 Genomics UK (COG-UK) Consortium | The Lighthouse Lab in Milton Keynes and Alex Alderton, Roberto Amato, Jeffrey Barrett, Sonia Goncalves, Ewan Harrison, David K. Jackson, Ian Johnston, Dominic Kwiatkowski, Cordelia Langford, John Sillitoe on behalf of the Wellcome Sanger Institute COVID-19 Surveillance Team |
| EPI_ISL_2780407 | Lighthouse Laboratory Plymouth | Wellcome Sanger Institute for the COVID-19 Genomics UK (COG-UK) Consortium | Lighthouse Laboratory Plymouth and Alex Alderton, Roberto Amato, Jeffrey Barrett, Sonia Goncalves, Ewan Harrison, David K. Jackson, Ian Johnston, Dominic Kwiatkowski, Cordelia Langford, John Sillitoe on behalf of the Wellcome Sanger Institute COVID-19 Surveillance Team |
| EPI_ISL_2780408, EPI_ISL_2780409, EPI_ISL_2780410 | Lighthouse Lab in Alderley Park | Wellcome Sanger Institute for the COVID-19 Genomics UK (COG-UK) Consortium | Jacquelyn Wynn, Mairead Hyland, The Lighthouse Lab in Alderley Park and Alex Alderton, Roberto Amato, Jeffrey Barrett, Sonia Goncalves, Ewan Harrison, David K. Jackson, Ian Johnston, Dominic Kwiatkowski, Cordelia Langford, John Sillitoe on behalf of the Wellcome Sanger Institute COVID-19 Surveillance Team |
| EPI_ISL_2780411 | Lighthouse Laboratory Plymouth | Wellcome Sanger Institute for the COVID-19 Genomics UK (COG-UK) Consortium | Lighthouse Laboratory Plymouth and Alex Alderton, Roberto Amato, Jeffrey Barrett, Sonia Goncalves, Ewan Harrison, David K. Jackson, Ian Johnston, Dominic Kwiatkowski, Cordelia Langford, John Sillitoe on behalf of the Wellcome Sanger Institute COVID-19 Surveillance Team |
| EPI_ISL_2780412, EPI_ISL_2780413 | Lighthouse Lab in Alderley Park | Wellcome Sanger Institute for the COVID-19 Genomics UK (COG-UK) Consortium | Jacquelyn Wynn, Mairead Hyland, The Lighthouse Lab in Alderley Park and Alex Alderton, Roberto Amato, Jeffrey Barrett, Sonia Goncalves, Ewan Harrison, David K. Jackson, Ian Johnston, Dominic Kwiatkowski, Cordelia Langford, John Sillitoe on behalf of the Wellcome Sanger Institute COVID-19 Surveillance Team |
| EPI_ISL_2780414 | Lighthouse Laboratory Plymouth | Wellcome Sanger Institute for the COVID-19 Genomics UK (COG-UK) Consortium | Lighthouse Laboratory Plymouth and Alex Alderton, Roberto Amato, Jeffrey Barrett, Sonia Goncalves, Ewan Harrison, David K. Jackson, Ian Johnston, Dominic Kwiatkowski, Cordelia Langford, John Sillitoe on behalf of the Wellcome Sanger Institute COVID-19 Surveillance Team |
| EPI_ISL_2780415, EPI_ISL_2780416, EPI_ISL_2780417 | Lighthouse Lab in Alderley Park | Wellcome Sanger Institute for the COVID-19 Genomics UK (COG-UK) Consortium | Jacquelyn Wynn, Mairead Hyland, The Lighthouse Lab in Alderley Park and Alex Alderton, Roberto Amato, Jeffrey Barrett, Sonia Goncalves, Ewan Harrison, David K. Jackson, Ian Johnston, Dominic Kwiatkowski, Cordelia Langford, John Sillitoe on behalf of the Wellcome Sanger Institute COVID-19 Surveillance Team |
| EPI_ISL_2780418, EPI_ISL_2780419, EPI_ISL_2780420, EPI_ISL_2780421 | Lighthouse Laboratory Plymouth | Wellcome Sanger Institute for the COVID-19 Genomics UK (COG-UK) Consortium | Lighthouse Laboratory Plymouth and Alex Alderton, Roberto Amato, Jeffrey Barrett, Sonia Goncalves, Ewan Harrison, David K. Jackson, Ian Johnston, Dominic Kwiatkowski, Cordelia Langford, John Sillitoe on behalf of the Wellcome Sanger Institute COVID-19 Surveillance Team |
| EPI_ISL_2780422 | Lighthouse Lab in Alderley Park | Wellcome Sanger Institute for the COVID-19 Genomics UK (COG-UK) Consortium | Jacquelyn Wynn, Mairead Hyland, The Lighthouse Lab in Alderley Park and Alex Alderton, Roberto Amato, Jeffrey Barrett, Sonia Goncalves, Ewan Harrison, David K. Jackson, Ian Johnston, Dominic Kwiatkowski, Cordelia Langford, John Sillitoe on behalf of the Wellcome Sanger Institute COVID-19 Surveillance Team |
| EPI_ISL_2780423 | Lighthouse Lab in Milton Keynes | Wellcome Sanger Institute for the COVID-19 Genomics UK (COG-UK) Consortium | The Lighthouse Lab in Milton Keynes and Alex Alderton, Roberto Amato, Jeffrey Barrett, Sonia Goncalves, Ewan Harrison, David K. Jackson, Ian Johnston, Dominic Kwiatkowski, Cordelia Langford, John Sillitoe on behalf of the Wellcome Sanger Institute COVID-19 Surveillance Team |
| EPI_ISL_2780424, EPI_ISL_2780425 | Lighthouse Lab in Alderley Park | Wellcome Sanger Institute for the COVID-19 Genomics UK (COG-UK) Consortium | Jacquelyn Wynn, Mairead Hyland, The Lighthouse Lab in Alderley Park and Alex Alderton, Roberto Amato, Jeffrey Barrett, Sonia Goncalves, Ewan Harrison, David K. Jackson, Ian Johnston, Dominic Kwiatkowski, Cordelia Langford, John Sillitoe on behalf of the Wellcome Sanger Institute COVID-19 Surveillance Team |
| EPI_ISL_2780426 | Lighthouse Lab in Milton Keynes | Wellcome Sanger Institute for the COVID-19 Genomics UK (COG-UK) Consortium | The Lighthouse Lab in Milton Keynes and Alex Alderton, Roberto Amato, Jeffrey Barrett, Sonia Goncalves, Ewan Harrison, David K. Jackson, Ian Johnston, Dominic Kwiatkowski, Cordelia Langford, John Sillitoe on behalf of the Wellcome Sanger Institute COVID-19 Surveillance Team |
| EPI_ISL_2780427 | Lighthouse Lab in Alderley Park | Wellcome Sanger Institute for the COVID-19 Genomics UK (COG-UK) Consortium | Jacquelyn Wynn, Mairead Hyland, The Lighthouse Lab in Alderley Park and Alex Alderton, Roberto Amato, Jeffrey Barrett, Sonia Goncalves, Ewan Harrison, David K. Jackson, Ian Johnston, Dominic Kwiatkowski, Cordelia Langford, John Sillitoe on behalf of the Wellcome Sanger Institute COVID-19 Surveillance Team |
| EPI_ISL_2780428 | Lighthouse Lab in Milton Keynes | Wellcome Sanger Institute for the COVID-19 Genomics UK (COG-UK) Consortium | The Lighthouse Lab in Milton Keynes and Alex Alderton, Roberto Amato, Jeffrey Barrett, Sonia Goncalves, Ewan Harrison, David K. Jackson, Ian Johnston, Dominic Kwiatkowski, Cordelia Langford, John Sillitoe on behalf of the Wellcome Sanger Institute COVID-19 Surveillance Team |
| EPI_ISL_2780429, EPI_ISL_2780430, EPI_ISL_2780431, EPI_ISL_2780432 | Lighthouse Lab in Alderley Park | Wellcome Sanger Institute for the COVID-19 Genomics UK (COG-UK) Consortium | Jacquelyn Wynn, Mairead Hyland, The Lighthouse Lab in Alderley Park and Alex Alderton, Roberto Amato, Jeffrey Barrett, Sonia Goncalves, Ewan Harrison, David K. Jackson, Ian Johnston, Dominic Kwiatkowski, Cordelia Langford, John Sillitoe on behalf of the Wellcome Sanger Institute COVID-19 Surveillance Team |
| EPI_ISL_2780433, | Lighthouse Lab in Milton Keynes | Wellcome Sanger Institute for the COVID-19 Genomics UK (COG-UK) | The Lighthouse Lab in Milton Keynes and Alex Alderton, Roberto Amato, Jeffrey Barrett, Sonia Goncalves, Ewan Harrison, David K. Jackson, Ian Johnston, Dominic Kwiatkowski, Cordelia Langford, John Sillitoe on behalf of the Wellcome Sanger |

| | | | |
|---|---|---|---|
| EPI_ISL_2780434 | | Consortium | Institute COVID-19 Surveillance Team |
| EPI_ISL_2780435, EPI_ISL_2780436 | Lighthouse Lab in Alderley Park | Wellcome Sanger Institute for the COVID-19 Genomics UK (COG-UK) Consortium | Jacquelyn Wynn, Mairead Hyland, The Lighthouse Lab in Alderley Park and Alex Alderton, Roberto Amato, Jeffrey Barrett, Sonia Goncalves, Ewan Harrison, David K. Jackson, Ian Johnston, Dominic Kwiatkowski, Cordelia Langford, John Sillitoe on behalf of the Wellcome Sanger Institute COVID-19 Surveillance Team |
| EPI_ISL_2780437 | Lighthouse Lab in Milton Keynes | Wellcome Sanger Institute for the COVID-19 Genomics UK (COG-UK) Consortium | The Lighthouse Lab in Milton Keynes and Alex Alderton, Roberto Amato, Jeffrey Barrett, Sonia Goncalves, Ewan Harrison, David K. Jackson, Ian Johnston, Dominic Kwiatkowski, Cordelia Langford, John Sillitoe on behalf of the Wellcome Sanger Institute COVID-19 Surveillance Team |
| EPI_ISL_2780438 | Lighthouse Lab in Alderley Park | Wellcome Sanger Institute for the COVID-19 Genomics UK (COG-UK) Consortium | Jacquelyn Wynn, Mairead Hyland, The Lighthouse Lab in Alderley Park and Alex Alderton, Roberto Amato, Jeffrey Barrett, Sonia Goncalves, Ewan Harrison, David K. Jackson, Ian Johnston, Dominic Kwiatkowski, Cordelia Langford, John Sillitoe on behalf of the Wellcome Sanger Institute COVID-19 Surveillance Team |
| EPI_ISL_2780439, EPI_ISL_2780440, EPI_ISL_2780441 | Lighthouse Lab in Milton Keynes | Wellcome Sanger Institute for the COVID-19 Genomics UK (COG-UK) Consortium | The Lighthouse Lab in Milton Keynes and Alex Alderton, Roberto Amato, Jeffrey Barrett, Sonia Goncalves, Ewan Harrison, David K. Jackson, Ian Johnston, Dominic Kwiatkowski, Cordelia Langford, John Sillitoe on behalf of the Wellcome Sanger Institute COVID-19 Surveillance Team |
| EPI_ISL_2780442, EPI_ISL_2780443 | Lighthouse Lab in Alderley Park | Wellcome Sanger Institute for the COVID-19 Genomics UK (COG-UK) Consortium | Jacquelyn Wynn, Mairead Hyland, The Lighthouse Lab in Alderley Park and Alex Alderton, Roberto Amato, Jeffrey Barrett, Sonia Goncalves, Ewan Harrison, David K. Jackson, Ian Johnston, Dominic Kwiatkowski, Cordelia Langford, John Sillitoe on behalf of the Wellcome Sanger Institute COVID-19 Surveillance Team |
| EPI_ISL_2780444 | Lighthouse Lab in Milton Keynes | Wellcome Sanger Institute for the COVID-19 Genomics UK (COG-UK) Consortium | The Lighthouse Lab in Milton Keynes and Alex Alderton, Roberto Amato, Jeffrey Barrett, Sonia Goncalves, Ewan Harrison, David K. Jackson, Ian Johnston, Dominic Kwiatkowski, Cordelia Langford, John Sillitoe on behalf of the Wellcome Sanger Institute COVID-19 Surveillance Team |
| EPI_ISL_2780445 | Lighthouse Lab in Alderley Park | Wellcome Sanger Institute for the COVID-19 Genomics UK (COG-UK) Consortium | Jacquelyn Wynn, Mairead Hyland, The Lighthouse Lab in Alderley Park and Alex Alderton, Roberto Amato, Jeffrey Barrett, Sonia Goncalves, Ewan Harrison, David K. Jackson, Ian Johnston, Dominic Kwiatkowski, Cordelia Langford, John Sillitoe on behalf of the Wellcome Sanger Institute COVID-19 Surveillance Team |
| EPI_ISL_2780446 | Lighthouse Lab in Milton Keynes | Wellcome Sanger Institute for the COVID-19 Genomics UK (COG-UK) Consortium | The Lighthouse Lab in Milton Keynes and Alex Alderton, Roberto Amato, Jeffrey Barrett, Sonia Goncalves, Ewan Harrison, David K. Jackson, Ian Johnston, Dominic Kwiatkowski, Cordelia Langford, John Sillitoe on behalf of the Wellcome Sanger Institute COVID-19 Surveillance Team |
| EPI_ISL_2780447 | Lighthouse Laboratory Plymouth | Wellcome Sanger Institute for the COVID-19 Genomics UK (COG-UK) Consortium | Lighthouse Laboratory Plymouth and Alex Alderton, Roberto Amato, Jeffrey Barrett, Sonia Goncalves, Ewan Harrison, David K. Jackson, Ian Johnston, Dominic Kwiatkowski, Cordelia Langford, John Sillitoe on behalf of the Wellcome Sanger Institute COVID-19 Surveillance Team |
| EPI_ISL_2780448 | Lighthouse Lab in Alderley Park | Wellcome Sanger Institute for the COVID-19 Genomics UK (COG-UK) Consortium | Jacquelyn Wynn, Mairead Hyland, The Lighthouse Lab in Alderley Park and Alex Alderton, Roberto Amato, Jeffrey Barrett, Sonia Goncalves, Ewan Harrison, David K. Jackson, Ian Johnston, Dominic Kwiatkowski, Cordelia Langford, John Sillitoe on behalf of the Wellcome Sanger Institute COVID-19 Surveillance Team |
| EPI_ISL_2780449, EPI_ISL_2780450, EPI_ISL_2780451 | Lighthouse Lab in Milton Keynes | Wellcome Sanger Institute for the COVID-19 Genomics UK (COG-UK) Consortium | The Lighthouse Lab in Milton Keynes and Alex Alderton, Roberto Amato, Jeffrey Barrett, Sonia Goncalves, Ewan Harrison, David K. Jackson, Ian Johnston, Dominic Kwiatkowski, Cordelia Langford, John Sillitoe on behalf of the Wellcome Sanger Institute COVID-19 Surveillance Team |
| EPI_ISL_2780452, EPI_ISL_2780453 | Lighthouse Lab in Alderley Park | Wellcome Sanger Institute for the COVID-19 Genomics UK (COG-UK) Consortium | Jacquelyn Wynn, Mairead Hyland, The Lighthouse Lab in Alderley Park and Alex Alderton, Roberto Amato, Jeffrey Barrett, Sonia Goncalves, Ewan Harrison, David K. Jackson, Ian Johnston, Dominic Kwiatkowski, Cordelia Langford, John Sillitoe on behalf of the Wellcome Sanger Institute COVID-19 Surveillance Team |
| EPI_ISL_2780454 | Lighthouse Laboratory Plymouth | Wellcome Sanger Institute for the COVID-19 Genomics UK (COG-UK) Consortium | Lighthouse Laboratory Plymouth and Alex Alderton, Roberto Amato, Jeffrey Barrett, Sonia Goncalves, Ewan Harrison, David K. Jackson, Ian Johnston, Dominic Kwiatkowski, Cordelia Langford, John Sillitoe on behalf of the Wellcome Sanger Institute COVID-19 Surveillance Team |
| EPI_ISL_2780455, EPI_ISL_2780456, EPI_ISL_2780457 | Lighthouse Lab in Alderley Park | Wellcome Sanger Institute for the COVID-19 Genomics UK (COG-UK) Consortium | Jacquelyn Wynn, Mairead Hyland, The Lighthouse Lab in Alderley Park and Alex Alderton, Roberto Amato, Jeffrey Barrett, Sonia Goncalves, Ewan Harrison, David K. Jackson, Ian Johnston, Dominic Kwiatkowski, Cordelia Langford, John Sillitoe on behalf of the Wellcome Sanger Institute COVID-19 Surveillance Team |
| EPI_ISL_2780458, EPI_ISL_2780459 | Lighthouse Lab in Milton Keynes | Wellcome Sanger Institute for the COVID-19 Genomics UK (COG-UK) Consortium | The Lighthouse Lab in Milton Keynes and Alex Alderton, Roberto Amato, Jeffrey Barrett, Sonia Goncalves, Ewan Harrison, David K. Jackson, Ian Johnston, Dominic Kwiatkowski, Cordelia Langford, John Sillitoe on behalf of the Wellcome Sanger Institute COVID-19 Surveillance Team |
| EPI_ISL_2780460, EPI_ISL_2780461 | Lighthouse Lab in Alderley Park | Wellcome Sanger Institute for the COVID-19 Genomics UK (COG-UK) Consortium | Jacquelyn Wynn, Mairead Hyland, The Lighthouse Lab in Alderley Park and Alex Alderton, Roberto Amato, Jeffrey Barrett, Sonia Goncalves, Ewan Harrison, David K. Jackson, Ian Johnston, Dominic Kwiatkowski, Cordelia Langford, John Sillitoe on behalf of the Wellcome Sanger Institute COVID-19 Surveillance Team |
| EPI_ISL_2780462, EPI_ISL_2780463 | Lighthouse Laboratory Plymouth | Wellcome Sanger Institute for the COVID-19 Genomics UK (COG-UK) Consortium | Lighthouse Laboratory Plymouth and Alex Alderton, Roberto Amato, Jeffrey Barrett, Sonia Goncalves, Ewan Harrison, David K. Jackson, Ian Johnston, Dominic Kwiatkowski, Cordelia Langford, John Sillitoe on behalf of the Wellcome Sanger Institute COVID-19 Surveillance Team |
| EPI_ISL_2780464 | Lighthouse Lab in Alderley Park | Wellcome Sanger Institute for the COVID-19 Genomics UK (COG-UK) Consortium | Jacquelyn Wynn, Mairead Hyland, The Lighthouse Lab in Alderley Park and Alex Alderton, Roberto Amato, Jeffrey Barrett, Sonia Goncalves, Ewan Harrison, David K. Jackson, Ian Johnston, Dominic Kwiatkowski, Cordelia Langford, John Sillitoe on behalf of the Wellcome Sanger Institute COVID-19 Surveillance Team |
| EPI_ISL_2780465 | Lighthouse Laboratory Plymouth | Wellcome Sanger Institute for the COVID-19 Genomics UK (COG-UK) Consortium | Lighthouse Laboratory Plymouth and Alex Alderton, Roberto Amato, Jeffrey Barrett, Sonia Goncalves, Ewan Harrison, David K. Jackson, Ian Johnston, Dominic Kwiatkowski, Cordelia Langford, John Sillitoe on behalf of the Wellcome Sanger Institute COVID-19 Surveillance Team |
| EPI_ISL_2780466 | Lighthouse Lab in Alderley Park | Wellcome Sanger Institute for the COVID-19 Genomics UK (COG-UK) Consortium | Jacquelyn Wynn, Mairead Hyland, The Lighthouse Lab in Alderley Park and Alex Alderton, Roberto Amato, Jeffrey Barrett, Sonia Goncalves, Ewan Harrison, David K. Jackson, Ian Johnston, Dominic Kwiatkowski, Cordelia Langford, John Sillitoe on behalf of the Wellcome Sanger Institute COVID-19 Surveillance Team |
| EPI_ISL_2780467, EPI_ISL_2780468 | Lighthouse Lab in Milton Keynes | Wellcome Sanger Institute for the COVID-19 Genomics UK (COG-UK) Consortium | The Lighthouse Lab in Milton Keynes and Alex Alderton, Roberto Amato, Jeffrey Barrett, Sonia Goncalves, Ewan Harrison, David K. Jackson, Ian Johnston, Dominic Kwiatkowski, Cordelia Langford, John Sillitoe on behalf of the Wellcome Sanger Institute COVID-19 Surveillance Team |
| EPI_ISL_2780469, EPI_ISL_2780470 | Lighthouse Laboratory Plymouth | Wellcome Sanger Institute for the COVID-19 Genomics UK (COG-UK) Consortium | Lighthouse Laboratory Plymouth and Alex Alderton, Roberto Amato, Jeffrey Barrett, Sonia Goncalves, Ewan Harrison, David K. Jackson, Ian Johnston, Dominic Kwiatkowski, Cordelia Langford, John Sillitoe on behalf of the Wellcome Sanger Institute COVID-19 Surveillance Team |
| EPI_ISL_2780471, EPI_ISL_2780472, EPI_ISL_2780473, EPI_ISL_2780474, EPI_ISL_2780475, EPI_ISL_2780476 | Lighthouse Lab in Milton Keynes | Wellcome Sanger Institute for the COVID-19 Genomics UK (COG-UK) Consortium | The Lighthouse Lab in Milton Keynes and Alex Alderton, Roberto Amato, Jeffrey Barrett, Sonia Goncalves, Ewan Harrison, David K. Jackson, Ian Johnston, Dominic Kwiatkowski, Cordelia Langford, John Sillitoe on behalf of the Wellcome Sanger Institute COVID-19 Surveillance Team |
| EPI_ISL_2780477 | Lighthouse Lab in Glasgow | Wellcome Sanger Institute for the COVID-19 Genomics UK (COG-UK) Consortium | Harper VanSteenhouse, Yumi Kasai, David Gray, Carol Clugston, Anna Dominiczak and Alex Alderton, Roberto Amato, Jeffrey Barrett, Sonia Goncalves, Ewan Harrison, David K. Jackson, Ian Johnston, Dominic Kwiatkowski, Cordelia Langford, John Sillitoe on behalf of the Wellcome Sanger Institute COVID-19 Surveillance Team |
| EPI_ISL_2780478, EPI_ISL_2780479 | Lighthouse Lab in Milton Keynes | Wellcome Sanger Institute for the COVID-19 Genomics UK (COG-UK) Consortium | The Lighthouse Lab in Milton Keynes and Alex Alderton, Roberto Amato, Jeffrey Barrett, Sonia Goncalves, Ewan Harrison, David K. Jackson, Ian Johnston, Dominic Kwiatkowski, Cordelia Langford, John Sillitoe on behalf of the Wellcome Sanger Institute COVID-19 Surveillance Team |
| EPI_ISL_2780480, EPI_ISL_2780481 | Lighthouse Lab in Glasgow | Wellcome Sanger Institute for the COVID-19 Genomics UK (COG-UK) Consortium | Harper VanSteenhouse, Yumi Kasai, David Gray, Carol Clugston, Anna Dominiczak and Alex Alderton, Roberto Amato, Jeffrey Barrett, Sonia Goncalves, Ewan Harrison, David K. Jackson, Ian Johnston, Dominic Kwiatkowski, Cordelia Langford, John Sillitoe on behalf of the Wellcome Sanger Institute COVID-19 Surveillance Team |
| EPI_ISL_2780482, EPI_ISL_2780483 | Lighthouse Lab in Milton Keynes | Wellcome Sanger Institute for the COVID-19 Genomics UK (COG-UK) Consortium | The Lighthouse Lab in Milton Keynes and Alex Alderton, Roberto Amato, Jeffrey Barrett, Sonia Goncalves, Ewan Harrison, David K. Jackson, Ian Johnston, Dominic Kwiatkowski, Cordelia Langford, John Sillitoe on behalf of the Wellcome Sanger Institute COVID-19 Surveillance Team |
| EPI_ISL_2780484 | Lighthouse Lab in Glasgow | Wellcome Sanger Institute for the COVID-19 Genomics UK (COG-UK) Consortium | Harper VanSteenhouse, Yumi Kasai, David Gray, Carol Clugston, Anna Dominiczak and Alex Alderton, Roberto Amato, Jeffrey Barrett, Sonia Goncalves, Ewan Harrison, David K. Jackson, Ian Johnston, Dominic Kwiatkowski, Cordelia Langford, John Sillitoe on behalf of the Wellcome Sanger Institute COVID-19 Surveillance Team |
| EPI_ISL_2780485, EPI_ISL_2780486, EPI_ISL_2780487, EPI_ISL_2780488 | Lighthouse Lab in Milton Keynes | Wellcome Sanger Institute for the COVID-19 Genomics UK (COG-UK) Consortium | The Lighthouse Lab in Milton Keynes and Alex Alderton, Roberto Amato, Jeffrey Barrett, Sonia Goncalves, Ewan Harrison, David K. Jackson, Ian Johnston, Dominic Kwiatkowski, Cordelia Langford, John Sillitoe on behalf of the Wellcome Sanger Institute COVID-19 Surveillance Team |
| EPI_ISL_2780489 | Lighthouse Lab in Glasgow | Wellcome Sanger Institute for the COVID-19 Genomics UK (COG-UK) Consortium | Harper VanSteenhouse, Yumi Kasai, David Gray, Carol Clugston, Anna Dominiczak and Alex Alderton, Roberto Amato, Jeffrey Barrett, Sonia Goncalves, Ewan Harrison, David K. Jackson, Ian Johnston, Dominic Kwiatkowski, Cordelia Langford, John Sillitoe on behalf of the Wellcome Sanger Institute COVID-19 Surveillance Team |
| EPI_ISL_2780490 | Lighthouse Lab in Milton Keynes | Wellcome Sanger Institute for the COVID-19 Genomics UK (COG-UK) Consortium | The Lighthouse Lab in Milton Keynes and Alex Alderton, Roberto Amato, Jeffrey Barrett, Sonia Goncalves, Ewan Harrison, David K. Jackson, Ian Johnston, Dominic Kwiatkowski, Cordelia Langford, John Sillitoe on behalf of the Wellcome Sanger Institute COVID-19 Surveillance Team |
| EPI_ISL_2780491 | Lighthouse Lab in Glasgow | Wellcome Sanger Institute for the COVID-19 Genomics UK (COG-UK) Consortium | Harper VanSteenhouse, Yumi Kasai, David Gray, Carol Clugston, Anna Dominiczak and Alex Alderton, Roberto Amato, Jeffrey Barrett, Sonia Goncalves, Ewan Harrison, David K. Jackson, Ian Johnston, Dominic Kwiatkowski, Cordelia Langford, John Sillitoe on behalf of the Wellcome Sanger Institute COVID-19 Surveillance Team |
| EPI_ISL_2780492, EPI_ISL_2780493, EPI_ISL_2780494, EPI_ISL_2780495, EPI_ISL_2780496, EPI_ISL_2780497, EPI_ISL_2780498, EPI_ISL_2780499, EPI_ISL_2780500, EPI_ISL_2780501, EPI_ISL_2780502, EPI_ISL_2780503, EPI_ISL_2780504, EPI_ISL_2780505 | | | |
| see above | Lighthouse Lab in Milton Keynes | Wellcome Sanger Institute for the COVID-19 Genomics UK (COG-UK) Consortium | The Lighthouse Lab in Milton Keynes and Alex Alderton, Roberto Amato, Jeffrey Barrett, Sonia Goncalves, Ewan Harrison, David K. Jackson, Ian Johnston, Dominic Kwiatkowski, Cordelia Langford, John Sillitoe on behalf of the Wellcome Sanger Institute COVID-19 Surveillance Team |
| EPI_ISL_2780506 | Lighthouse Lab in Glasgow | Wellcome Sanger Institute for the COVID-19 Genomics UK (COG-UK) Consortium | Harper VanSteenhouse, Yumi Kasai, David Gray, Carol Clugston, Anna Dominiczak and Alex Alderton, Roberto Amato, Jeffrey Barrett, Sonia Goncalves, Ewan Harrison, David K. Jackson, Ian Johnston, Dominic Kwiatkowski, Cordelia Langford, John Sillitoe on behalf of the Wellcome Sanger Institute COVID-19 Surveillance Team |
| EPI_ISL_2780507 | Lighthouse Lab in Milton Keynes | Wellcome Sanger Institute for the COVID-19 Genomics UK (COG-UK) Consortium | The Lighthouse Lab in Milton Keynes and Alex Alderton, Roberto Amato, Jeffrey Barrett, Sonia Goncalves, Ewan Harrison, David K. Jackson, Ian Johnston, Dominic Kwiatkowski, Cordelia Langford, John Sillitoe on behalf of the Wellcome Sanger Institute COVID-19 Surveillance Team |
| EPI_ISL_2780508 | Lighthouse Lab in Glasgow | Wellcome Sanger Institute for the COVID-19 Genomics UK (COG-UK) Consortium | Harper VanSteenhouse, Yumi Kasai, David Gray, Carol Clugston, Anna Dominiczak and Alex Alderton, Roberto Amato, Jeffrey Barrett, Sonia Goncalves, Ewan Harrison, David K. Jackson, Ian Johnston, Dominic Kwiatkowski, Cordelia Langford, John Sillitoe on behalf of the Wellcome Sanger Institute COVID-19 Surveillance Team |
| EPI_ISL_2780509, EPI_ISL_2780510, EPI_ISL_2780511, EPI_ISL_2780513, EPI_ISL_2780514 | Lighthouse Lab in Milton Keynes | Wellcome Sanger Institute for the COVID-19 Genomics UK (COG-UK) Consortium | The Lighthouse Lab in Milton Keynes and Alex Alderton, Roberto Amato, Jeffrey Barrett, Sonia Goncalves, Ewan Harrison, David K. Jackson, Ian Johnston, Dominic Kwiatkowski, Cordelia Langford, John Sillitoe on behalf of the Wellcome Sanger Institute COVID-19 Surveillance Team |
| EPI_ISL_2780515 | Lighthouse Lab in Glasgow | Wellcome Sanger Institute for the COVID-19 Genomics UK (COG-UK) Consortium | Harper VanSteenhouse, Yumi Kasai, David Gray, Carol Clugston, Anna Dominiczak and Alex Alderton, Roberto Amato, Jeffrey Barrett, Sonia Goncalves, Ewan Harrison, David K. Jackson, Ian Johnston, Dominic Kwiatkowski, Cordelia Langford, John Sillitoe on behalf of the Wellcome Sanger Institute COVID-19 Surveillance Team |
| EPI_ISL_2780516, EPI_ISL_2780517, EPI_ISL_2780518, EPI_ISL_2780519, EPI_ISL_2780520, EPI_ISL_2780521, EPI_ISL_2780522, EPI_ISL_2780523, EPI_ISL_2780524, EPI_ISL_2780525, EPI_ISL_2780526, EPI_ISL_2780527, EPI_ISL_2780529, EPI_ISL_2780530, EPI_ISL_2780531, EPI_ISL_2780532, EPI_ISL_2780533, EPI_ISL_2780534, EPI_ISL_2780535 | | | |

| | | | |
|---|---|---|---|
| see above | Lighthouse Lab in Milton Keynes | Wellcome Sanger Institute for the COVID-19 Genomics UK (COG-UK) Consortium | The Lighthouse Lab in Milton Keynes and Alex Alderton, Roberto Amato, Jeffrey Barrett, Sonia Goncalves, Ewan Harrison, David K. Jackson, Ian Johnston, Dominic Kwiatkowski, Cordelia Langford, John Sillitoe on behalf of the Wellcome Sanger Institute COVID-19 Surveillance Team |
| EPI_ISL_2780536 | Lighthouse Lab in Glasgow | Wellcome Sanger Institute for the COVID-19 Genomics UK (COG-UK) Consortium | Harper VanSteenhouse, Yumi Kasai, David Gray, Carol Clugston, Anna Dominiczak and Alex Alderton, Roberto Amato, Jeffrey Barrett, Sonia Goncalves, Ewan Harrison, David K. Jackson, Ian Johnston, Dominic Kwiatkowski, Cordelia Langford, John Sillitoe on behalf of the Wellcome Sanger Institute COVID-19 Surveillance Team |
| EPI_ISL_2780537 | Lighthouse Lab in Milton Keynes | Wellcome Sanger Institute for the COVID-19 Genomics UK (COG-UK) Consortium | The Lighthouse Lab in Milton Keynes and Alex Alderton, Roberto Amato, Jeffrey Barrett, Sonia Goncalves, Ewan Harrison, David K. Jackson, Ian Johnston, Dominic Kwiatkowski, Cordelia Langford, John Sillitoe on behalf of the Wellcome Sanger Institute COVID-19 Surveillance Team |
| EPI_ISL_2780538, EPI_ISL_2780539 | Lighthouse Lab in Glasgow | Wellcome Sanger Institute for the COVID-19 Genomics UK (COG-UK) Consortium | Harper VanSteenhouse, Yumi Kasai, David Gray, Carol Clugston, Anna Dominiczak and Alex Alderton, Roberto Amato, Jeffrey Barrett, Sonia Goncalves, Ewan Harrison, David K. Jackson, Ian Johnston, Dominic Kwiatkowski, Cordelia Langford, John Sillitoe on behalf of the Wellcome Sanger Institute COVID-19 Surveillance Team |
| EPI_ISL_2780540, EPI_ISL_2780541, EPI_ISL_2780542 | Lighthouse Lab in Milton Keynes | Wellcome Sanger Institute for the COVID-19 Genomics UK (COG-UK) Consortium | The Lighthouse Lab in Milton Keynes and Alex Alderton, Roberto Amato, Jeffrey Barrett, Sonia Goncalves, Ewan Harrison, David K. Jackson, Ian Johnston, Dominic Kwiatkowski, Cordelia Langford, John Sillitoe on behalf of the Wellcome Sanger Institute COVID-19 Surveillance Team |
| EPI_ISL_2780543 | Lighthouse Lab in Glasgow | Wellcome Sanger Institute for the COVID-19 Genomics UK (COG-UK) Consortium | Harper VanSteenhouse, Yumi Kasai, David Gray, Carol Clugston, Anna Dominiczak and Alex Alderton, Roberto Amato, Jeffrey Barrett, Sonia Goncalves, Ewan Harrison, David K. Jackson, Ian Johnston, Dominic Kwiatkowski, Cordelia Langford, John Sillitoe on behalf of the Wellcome Sanger Institute COVID-19 Surveillance Team |
| EPI_ISL_2780544 | Lighthouse Lab in Milton Keynes | Wellcome Sanger Institute for the COVID-19 Genomics UK (COG-UK) Consortium | The Lighthouse Lab in Milton Keynes and Alex Alderton, Roberto Amato, Jeffrey Barrett, Sonia Goncalves, Ewan Harrison, David K. Jackson, Ian Johnston, Dominic Kwiatkowski, Cordelia Langford, John Sillitoe on behalf of the Wellcome Sanger Institute COVID-19 Surveillance Team |
| EPI_ISL_2780545 | Lighthouse Lab in Glasgow | Wellcome Sanger Institute for the COVID-19 Genomics UK (COG-UK) Consortium | Harper VanSteenhouse, Yumi Kasai, David Gray, Carol Clugston, Anna Dominiczak and Alex Alderton, Roberto Amato, Jeffrey Barrett, Sonia Goncalves, Ewan Harrison, David K. Jackson, Ian Johnston, Dominic Kwiatkowski, Cordelia Langford, John Sillitoe on behalf of the Wellcome Sanger Institute COVID-19 Surveillance Team |
| EPI_ISL_2780546, EPI_ISL_2780547, EPI_ISL_2780549, EPI_ISL_2780550, EPI_ISL_2780551, EPI_ISL_2780552, EPI_ISL_2780553, EPI_ISL_2780554 | Lighthouse Lab in Milton Keynes | Wellcome Sanger Institute for the COVID-19 Genomics UK (COG-UK) Consortium | The Lighthouse Lab in Milton Keynes and Alex Alderton, Roberto Amato, Jeffrey Barrett, Sonia Goncalves, Ewan Harrison, David K. Jackson, Ian Johnston, Dominic Kwiatkowski, Cordelia Langford, John Sillitoe on behalf of the Wellcome Sanger Institute COVID-19 Surveillance Team |
| EPI_ISL_2780555, EPI_ISL_2780556 | Lighthouse Lab in Glasgow | Wellcome Sanger Institute for the COVID-19 Genomics UK (COG-UK) Consortium | Harper VanSteenhouse, Yumi Kasai, David Gray, Carol Clugston, Anna Dominiczak and Alex Alderton, Roberto Amato, Jeffrey Barrett, Sonia Goncalves, Ewan Harrison, David K. Jackson, Ian Johnston, Dominic Kwiatkowski, Cordelia Langford, John Sillitoe on behalf of the Wellcome Sanger Institute COVID-19 Surveillance Team |
| EPI_ISL_2780557, EPI_ISL_2780558, EPI_ISL_2780559 | Lighthouse Lab in Milton Keynes | Wellcome Sanger Institute for the COVID-19 Genomics UK (COG-UK) Consortium | The Lighthouse Lab in Milton Keynes and Alex Alderton, Roberto Amato, Jeffrey Barrett, Sonia Goncalves, Ewan Harrison, David K. Jackson, Ian Johnston, Dominic Kwiatkowski, Cordelia Langford, John Sillitoe on behalf of the Wellcome Sanger Institute COVID-19 Surveillance Team |
| EPI_ISL_2780560 | Lighthouse Lab in Glasgow | Wellcome Sanger Institute for the COVID-19 Genomics UK (COG-UK) Consortium | Harper VanSteenhouse, Yumi Kasai, David Gray, Carol Clugston, Anna Dominiczak and Alex Alderton, Roberto Amato, Jeffrey Barrett, Sonia Goncalves, Ewan Harrison, David K. Jackson, Ian Johnston, Dominic Kwiatkowski, Cordelia Langford, John Sillitoe on behalf of the Wellcome Sanger Institute COVID-19 Surveillance Team |
| EPI_ISL_2780561, EPI_ISL_2780562, EPI_ISL_2780563, EPI_ISL_2780564, EPI_ISL_2780565, EPI_ISL_2780566, EPI_ISL_2780567, EPI_ISL_2780568, EPI_ISL_2780569, EPI_ISL_2780570, EPI_ISL_2780571, EPI_ISL_2780572, EPI_ISL_2780573, EPI_ISL_2780574, EPI_ISL_2780575, EPI_ISL_2780577, EPI_ISL_2780578, EPI_ISL_2780579, EPI_ISL_2780580, EPI_ISL_2780581, EPI_ISL_2780582, EPI_ISL_2780583, EPI_ISL_2780584 | | | |
| see above | Lighthouse Lab in Milton Keynes | Wellcome Sanger Institute for the COVID-19 Genomics UK (COG-UK) Consortium | The Lighthouse Lab in Milton Keynes and Alex Alderton, Roberto Amato, Jeffrey Barrett, Sonia Goncalves, Ewan Harrison, David K. Jackson, Ian Johnston, Dominic Kwiatkowski, Cordelia Langford, John Sillitoe on behalf of the Wellcome Sanger Institute COVID-19 Surveillance Team |
| EPI_ISL_2780585 | Lighthouse Lab in Glasgow | Wellcome Sanger Institute for the COVID-19 Genomics UK (COG-UK) Consortium | Harper VanSteenhouse, Yumi Kasai, David Gray, Carol Clugston, Anna Dominiczak and Alex Alderton, Roberto Amato, Jeffrey Barrett, Sonia Goncalves, Ewan Harrison, David K. Jackson, Ian Johnston, Dominic Kwiatkowski, Cordelia Langford, John Sillitoe on behalf of the Wellcome Sanger Institute COVID-19 Surveillance Team |
| EPI_ISL_2780586 | Lighthouse Lab in Milton Keynes | Wellcome Sanger Institute for the COVID-19 Genomics UK (COG-UK) Consortium | The Lighthouse Lab in Milton Keynes and Alex Alderton, Roberto Amato, Jeffrey Barrett, Sonia Goncalves, Ewan Harrison, David K. Jackson, Ian Johnston, Dominic Kwiatkowski, Cordelia Langford, John Sillitoe on behalf of the Wellcome Sanger Institute COVID-19 Surveillance Team |
| EPI_ISL_2780587 | Lighthouse Lab in Glasgow | Wellcome Sanger Institute for the COVID-19 Genomics UK (COG-UK) Consortium | Harper VanSteenhouse, Yumi Kasai, David Gray, Carol Clugston, Anna Dominiczak and Alex Alderton, Roberto Amato, Jeffrey Barrett, Sonia Goncalves, Ewan Harrison, David K. Jackson, Ian Johnston, Dominic Kwiatkowski, Cordelia Langford, John Sillitoe on behalf of the Wellcome Sanger Institute COVID-19 Surveillance Team |
| EPI_ISL_2780588, EPI_ISL_2780589, EPI_ISL_2780590, EPI_ISL_2780591, EPI_ISL_2780592, EPI_ISL_2780593 | Lighthouse Lab in Milton Keynes | Wellcome Sanger Institute for the COVID-19 Genomics UK (COG-UK) Consortium | The Lighthouse Lab in Milton Keynes and Alex Alderton, Roberto Amato, Jeffrey Barrett, Sonia Goncalves, Ewan Harrison, David K. Jackson, Ian Johnston, Dominic Kwiatkowski, Cordelia Langford, John Sillitoe on behalf of the Wellcome Sanger Institute COVID-19 Surveillance Team |
| EPI_ISL_2780594 | Lighthouse Lab in Glasgow | Wellcome Sanger Institute for the COVID-19 Genomics UK (COG-UK) Consortium | Harper VanSteenhouse, Yumi Kasai, David Gray, Carol Clugston, Anna Dominiczak and Alex Alderton, Roberto Amato, Jeffrey Barrett, Sonia Goncalves, Ewan Harrison, David K. Jackson, Ian Johnston, Dominic Kwiatkowski, Cordelia Langford, John Sillitoe on behalf of the Wellcome Sanger Institute COVID-19 Surveillance Team |
| EPI_ISL_2780595, EPI_ISL_2780596 | Lighthouse Lab in Milton Keynes | Wellcome Sanger Institute for the COVID-19 Genomics UK (COG-UK) Consortium | The Lighthouse Lab in Milton Keynes and Alex Alderton, Roberto Amato, Jeffrey Barrett, Sonia Goncalves, Ewan Harrison, David K. Jackson, Ian Johnston, Dominic Kwiatkowski, Cordelia Langford, John Sillitoe on behalf of the Wellcome Sanger Institute COVID-19 Surveillance Team |
| EPI_ISL_2780597 | Lighthouse Lab in Glasgow | Wellcome Sanger Institute for the COVID-19 Genomics UK (COG-UK) Consortium | Harper VanSteenhouse, Yumi Kasai, David Gray, Carol Clugston, Anna Dominiczak and Alex Alderton, Roberto Amato, Jeffrey Barrett, Sonia Goncalves, Ewan Harrison, David K. Jackson, Ian Johnston, Dominic Kwiatkowski, Cordelia Langford, John Sillitoe on behalf of the Wellcome Sanger Institute COVID-19 Surveillance Team |
| EPI_ISL_2780598, EPI_ISL_2780599, EPI_ISL_2780600, EPI_ISL_2780601, EPI_ISL_2780602, EPI_ISL_2780603, EPI_ISL_2780604, EPI_ISL_2780605, EPI_ISL_2780606, EPI_ISL_2780607, EPI_ISL_2780608 | | | |
| see above | Lighthouse Lab in Milton Keynes | Wellcome Sanger Institute for the COVID-19 Genomics UK (COG-UK) Consortium | The Lighthouse Lab in Milton Keynes and Alex Alderton, Roberto Amato, Jeffrey Barrett, Sonia Goncalves, Ewan Harrison, David K. Jackson, Ian Johnston, Dominic Kwiatkowski, Cordelia Langford, John Sillitoe on behalf of the Wellcome Sanger Institute COVID-19 Surveillance Team |
| EPI_ISL_2780609 | Lighthouse Lab in Glasgow | Wellcome Sanger Institute for the COVID-19 Genomics UK (COG-UK) Consortium | Harper VanSteenhouse, Yumi Kasai, David Gray, Carol Clugston, Anna Dominiczak and Alex Alderton, Roberto Amato, Jeffrey Barrett, Sonia Goncalves, Ewan Harrison, David K. Jackson, Ian Johnston, Dominic Kwiatkowski, Cordelia Langford, John Sillitoe on behalf of the Wellcome Sanger Institute COVID-19 Surveillance Team |
| EPI_ISL_2780610, EPI_ISL_2780611, EPI_ISL_2780612, EPI_ISL_2780613, EPI_ISL_2780614, EPI_ISL_2780615, EPI_ISL_2780616, EPI_ISL_2780617 | Lighthouse Lab in Milton Keynes | Wellcome Sanger Institute for the COVID-19 Genomics UK (COG-UK) Consortium | The Lighthouse Lab in Milton Keynes and Alex Alderton, Roberto Amato, Jeffrey Barrett, Sonia Goncalves, Ewan Harrison, David K. Jackson, Ian Johnston, Dominic Kwiatkowski, Cordelia Langford, John Sillitoe on behalf of the Wellcome Sanger Institute COVID-19 Surveillance Team |
| EPI_ISL_2780618 | Lighthouse Lab in Glasgow | Wellcome Sanger Institute for the COVID-19 Genomics UK (COG-UK) Consortium | Harper VanSteenhouse, Yumi Kasai, David Gray, Carol Clugston, Anna Dominiczak and Alex Alderton, Roberto Amato, Jeffrey Barrett, Sonia Goncalves, Ewan Harrison, David K. Jackson, Ian Johnston, Dominic Kwiatkowski, Cordelia Langford, John Sillitoe on behalf of the Wellcome Sanger Institute COVID-19 Surveillance Team |
| EPI_ISL_2780619 | Lighthouse Lab in Milton Keynes | Wellcome Sanger Institute for the COVID-19 Genomics UK (COG-UK) Consortium | The Lighthouse Lab in Milton Keynes and Alex Alderton, Roberto Amato, Jeffrey Barrett, Sonia Goncalves, Ewan Harrison, David K. Jackson, Ian Johnston, Dominic Kwiatkowski, Cordelia Langford, John Sillitoe on behalf of the Wellcome Sanger Institute COVID-19 Surveillance Team |
| EPI_ISL_2780620 | Lighthouse Lab in Glasgow | Wellcome Sanger Institute for the COVID-19 Genomics UK (COG-UK) Consortium | Harper VanSteenhouse, Yumi Kasai, David Gray, Carol Clugston, Anna Dominiczak and Alex Alderton, Roberto Amato, Jeffrey Barrett, Sonia Goncalves, Ewan Harrison, David K. Jackson, Ian Johnston, Dominic Kwiatkowski, Cordelia Langford, John Sillitoe on behalf of the Wellcome Sanger Institute COVID-19 Surveillance Team |
| EPI_ISL_2780621, EPI_ISL_2780622 | Lighthouse Lab in Milton Keynes | Wellcome Sanger Institute for the COVID-19 Genomics UK (COG-UK) Consortium | The Lighthouse Lab in Milton Keynes and Alex Alderton, Roberto Amato, Jeffrey Barrett, Sonia Goncalves, Ewan Harrison, David K. Jackson, Ian Johnston, Dominic Kwiatkowski, Cordelia Langford, John Sillitoe on behalf of the Wellcome Sanger Institute COVID-19 Surveillance Team |
| EPI_ISL_2780623, EPI_ISL_2780624 | Lighthouse Lab in Glasgow | Wellcome Sanger Institute for the COVID-19 Genomics UK (COG-UK) Consortium | Harper VanSteenhouse, Yumi Kasai, David Gray, Carol Clugston, Anna Dominiczak and Alex Alderton, Roberto Amato, Jeffrey Barrett, Sonia Goncalves, Ewan Harrison, David K. Jackson, Ian Johnston, Dominic Kwiatkowski, Cordelia Langford, John Sillitoe on behalf of the Wellcome Sanger Institute COVID-19 Surveillance Team |
| EPI_ISL_2780625, EPI_ISL_2780626, EPI_ISL_2780627, EPI_ISL_2780629, EPI_ISL_2780630, EPI_ISL_2780631, EPI_ISL_2780632, EPI_ISL_2780633 | Lighthouse Lab in Milton Keynes | Wellcome Sanger Institute for the COVID-19 Genomics UK (COG-UK) Consortium | The Lighthouse Lab in Milton Keynes and Alex Alderton, Roberto Amato, Jeffrey Barrett, Sonia Goncalves, Ewan Harrison, David K. Jackson, Ian Johnston, Dominic Kwiatkowski, Cordelia Langford, John Sillitoe on behalf of the Wellcome Sanger Institute COVID-19 Surveillance Team |
| EPI_ISL_2780634 | Lighthouse Lab in Glasgow | Wellcome Sanger Institute for the COVID-19 Genomics UK (COG-UK) Consortium | Harper VanSteenhouse, Yumi Kasai, David Gray, Carol Clugston, Anna Dominiczak and Alex Alderton, Roberto Amato, Jeffrey Barrett, Sonia Goncalves, Ewan Harrison, David K. Jackson, Ian Johnston, Dominic Kwiatkowski, Cordelia Langford, John Sillitoe on behalf of the Wellcome Sanger Institute COVID-19 Surveillance Team |
| EPI_ISL_2780635, EPI_ISL_2780636 | Lighthouse Lab in Milton Keynes | Wellcome Sanger Institute for the COVID-19 Genomics UK (COG-UK) Consortium | The Lighthouse Lab in Milton Keynes and Alex Alderton, Roberto Amato, Jeffrey Barrett, Sonia Goncalves, Ewan Harrison, David K. Jackson, Ian Johnston, Dominic Kwiatkowski, Cordelia Langford, John Sillitoe on behalf of the Wellcome Sanger Institute COVID-19 Surveillance Team |
| EPI_ISL_2780637 | Lighthouse Lab in Glasgow | Wellcome Sanger Institute for the COVID-19 Genomics UK (COG-UK) Consortium | Harper VanSteenhouse, Yumi Kasai, David Gray, Carol Clugston, Anna Dominiczak and Alex Alderton, Roberto Amato, Jeffrey Barrett, Sonia Goncalves, Ewan Harrison, David K. Jackson, Ian Johnston, Dominic Kwiatkowski, Cordelia Langford, John Sillitoe on behalf of the Wellcome Sanger Institute COVID-19 Surveillance Team |
| EPI_ISL_2780638, EPI_ISL_2780639, EPI_ISL_2780640, EPI_ISL_2780641, EPI_ISL_2780642, EPI_ISL_2780643, EPI_ISL_2780644, EPI_ISL_2780645, EPI_ISL_2780646, EPI_ISL_2780647, EPI_ISL_2780648, EPI_ISL_2780649, EPI_ISL_2780650, EPI_ISL_2780651, EPI_ISL_2780652, EPI_ISL_2780653, EPI_ISL_2780654, EPI_ISL_2780655 | | | |
| see above | Lighthouse Lab in Milton Keynes | Wellcome Sanger Institute for the COVID-19 Genomics UK (COG-UK) Consortium | The Lighthouse Lab in Milton Keynes and Alex Alderton, Roberto Amato, Jeffrey Barrett, Sonia Goncalves, Ewan Harrison, David K. Jackson, Ian Johnston, Dominic Kwiatkowski, Cordelia Langford, John Sillitoe on behalf of the Wellcome Sanger Institute COVID-19 Surveillance Team |

| | | | |
|---|---|---|---|
| EPI_ISL_2780656 | Lighthouse Lab in Glasgow | Wellcome Sanger Institute for the COVID-19 Genomics UK (COG-UK) Consortium | Harper VanSteenhouse, Yumi Kasai, David Gray, Carol Clugston, Anna Dominiczak and Alex Alderton, Roberto Amato, Jeffrey Barrett, Sonia Goncalves, Ewan Harrison, David K. Jackson, Ian Johnston, Dominic Kwiatkowski, Cordelia Langford, John Sillitoe on behalf of the Wellcome Sanger Institute COVID-19 Surveillance Team |
| EPI_ISL_2780657, EPI_ISL_2780658, EPI_ISL_2780659, EPI_ISL_2780660, EPI_ISL_2780661, EPI_ISL_2780662, EPI_ISL_2780663, EPI_ISL_2780664, EPI_ISL_2780665, EPI_ISL_2780666, EPI_ISL_2780667, EPI_ISL_2780668, EPI_ISL_2780669, EPI_ISL_2780670, EPI_ISL_2780671, EPI_ISL_2780672, EPI_ISL_2780673 | | | |
| see above | Lighthouse Lab in Milton Keynes | Wellcome Sanger Institute for the COVID-19 Genomics UK (COG-UK) Consortium | The Lighthouse Lab in Milton Keynes and Alex Alderton, Roberto Amato, Jeffrey Barrett, Sonia Goncalves, Ewan Harrison, David K. Jackson, Ian Johnston, Dominic Kwiatkowski, Cordelia Langford, John Sillitoe on behalf of the Wellcome Sanger Institute COVID-19 Surveillance Team |
| EPI_ISL_2780674 | Lighthouse Lab in Glasgow | Wellcome Sanger Institute for the COVID-19 Genomics UK (COG-UK) Consortium | Harper VanSteenhouse, Yumi Kasai, David Gray, Carol Clugston, Anna Dominiczak and Alex Alderton, Roberto Amato, Jeffrey Barrett, Sonia Goncalves, Ewan Harrison, David K. Jackson, Ian Johnston, Dominic Kwiatkowski, Cordelia Langford, John Sillitoe on behalf of the Wellcome Sanger Institute COVID-19 Surveillance Team |
| EPI_ISL_2780675, EPI_ISL_2780676, EPI_ISL_2780677, EPI_ISL_2780678, EPI_ISL_2780679 | Lighthouse Lab in Milton Keynes | Wellcome Sanger Institute for the COVID-19 Genomics UK (COG-UK) Consortium | The Lighthouse Lab in Milton Keynes and Alex Alderton, Roberto Amato, Jeffrey Barrett, Sonia Goncalves, Ewan Harrison, David K. Jackson, Ian Johnston, Dominic Kwiatkowski, Cordelia Langford, John Sillitoe on behalf of the Wellcome Sanger Institute COVID-19 Surveillance Team |
| EPI_ISL_2780680 | Lighthouse Lab in Glasgow | Wellcome Sanger Institute for the COVID-19 Genomics UK (COG-UK) Consortium | Harper VanSteenhouse, Yumi Kasai, David Gray, Carol Clugston, Anna Dominiczak and Alex Alderton, Roberto Amato, Jeffrey Barrett, Sonia Goncalves, Ewan Harrison, David K. Jackson, Ian Johnston, Dominic Kwiatkowski, Cordelia Langford, John Sillitoe on behalf of the Wellcome Sanger Institute COVID-19 Surveillance Team |
| EPI_ISL_2780681, EPI_ISL_2780682, EPI_ISL_2780683, EPI_ISL_2780684, EPI_ISL_2780685, EPI_ISL_2780686, EPI_ISL_2780687, EPI_ISL_2780688, EPI_ISL_2780689, EPI_ISL_2780690, EPI_ISL_2780691, EPI_ISL_2780692, EPI_ISL_2780693, EPI_ISL_2780694, EPI_ISL_2780695, EPI_ISL_2780696, EPI_ISL_2780697, EPI_ISL_2780698 | | | |
| see above | Lighthouse Lab in Milton Keynes | Wellcome Sanger Institute for the COVID-19 Genomics UK (COG-UK) Consortium | The Lighthouse Lab in Milton Keynes and Alex Alderton, Roberto Amato, Jeffrey Barrett, Sonia Goncalves, Ewan Harrison, David K. Jackson, Ian Johnston, Dominic Kwiatkowski, Cordelia Langford, John Sillitoe on behalf of the Wellcome Sanger Institute COVID-19 Surveillance Team |
| EPI_ISL_2780699 | Lighthouse Lab in Glasgow | Wellcome Sanger Institute for the COVID-19 Genomics UK (COG-UK) Consortium | Harper VanSteenhouse, Yumi Kasai, David Gray, Carol Clugston, Anna Dominiczak and Alex Alderton, Roberto Amato, Jeffrey Barrett, Sonia Goncalves, Ewan Harrison, David K. Jackson, Ian Johnston, Dominic Kwiatkowski, Cordelia Langford, John Sillitoe on behalf of the Wellcome Sanger Institute COVID-19 Surveillance Team |
| EPI_ISL_2780702, EPI_ISL_2780703, EPI_ISL_2780704, EPI_ISL_2780705, EPI_ISL_2780706, EPI_ISL_2780707, EPI_ISL_2780708, EPI_ISL_2780709, EPI_ISL_2780710, EPI_ISL_2780711, EPI_ISL_2780712, EPI_ISL_2780713, EPI_ISL_2780714, EPI_ISL_2780715, EPI_ISL_2780716, EPI_ISL_2780717, EPI_ISL_2780718, EPI_ISL_2780719, EPI_ISL_2780720, EPI_ISL_2780721, EPI_ISL_2780722, EPI_ISL_2780723, EPI_ISL_2780724, EPI_ISL_2780725, EPI_ISL_2780726, EPI_ISL_2780727, EPI_ISL_2780728 | | | |
| see above | Lighthouse Lab in Milton Keynes | Wellcome Sanger Institute for the COVID-19 Genomics UK (COG-UK) Consortium | The Lighthouse Lab in Milton Keynes and Alex Alderton, Roberto Amato, Jeffrey Barrett, Sonia Goncalves, Ewan Harrison, David K. Jackson, Ian Johnston, Dominic Kwiatkowski, Cordelia Langford, John Sillitoe on behalf of the Wellcome Sanger Institute COVID-19 Surveillance Team |
| EPI_ISL_2780729 | Lighthouse Lab in Glasgow | Wellcome Sanger Institute for the COVID-19 Genomics UK (COG-UK) Consortium | Harper VanSteenhouse, Yumi Kasai, David Gray, Carol Clugston, Anna Dominiczak and Alex Alderton, Roberto Amato, Jeffrey Barrett, Sonia Goncalves, Ewan Harrison, David K. Jackson, Ian Johnston, Dominic Kwiatkowski, Cordelia Langford, John Sillitoe on behalf of the Wellcome Sanger Institute COVID-19 Surveillance Team |
| EPI_ISL_2780730, EPI_ISL_2780731, EPI_ISL_2780732, EPI_ISL_2780733, EPI_ISL_2780734, EPI_ISL_2780735, EPI_ISL_2780736, EPI_ISL_2780737, EPI_ISL_2780738 | Lighthouse Lab in Milton Keynes | Wellcome Sanger Institute for the COVID-19 Genomics UK (COG-UK) Consortium | The Lighthouse Lab in Milton Keynes and Alex Alderton, Roberto Amato, Jeffrey Barrett, Sonia Goncalves, Ewan Harrison, David K. Jackson, Ian Johnston, Dominic Kwiatkowski, Cordelia Langford, John Sillitoe on behalf of the Wellcome Sanger Institute COVID-19 Surveillance Team |
| EPI_ISL_2780739 | Lighthouse Lab in Glasgow | Wellcome Sanger Institute for the COVID-19 Genomics UK (COG-UK) Consortium | Harper VanSteenhouse, Yumi Kasai, David Gray, Carol Clugston, Anna Dominiczak and Alex Alderton, Roberto Amato, Jeffrey Barrett, Sonia Goncalves, Ewan Harrison, David K. Jackson, Ian Johnston, Dominic Kwiatkowski, Cordelia Langford, John Sillitoe on behalf of the Wellcome Sanger Institute COVID-19 Surveillance Team |
| EPI_ISL_2780740, EPI_ISL_2780741, EPI_ISL_2780742, EPI_ISL_2780743, EPI_ISL_2780744, EPI_ISL_2780745, EPI_ISL_2780746, EPI_ISL_2780747, EPI_ISL_2780748, EPI_ISL_2780749, EPI_ISL_2780750 | | | |
| see above | Lighthouse Lab in Milton Keynes | Wellcome Sanger Institute for the COVID-19 Genomics UK (COG-UK) Consortium | The Lighthouse Lab in Milton Keynes and Alex Alderton, Roberto Amato, Jeffrey Barrett, Sonia Goncalves, Ewan Harrison, David K. Jackson, Ian Johnston, Dominic Kwiatkowski, Cordelia Langford, John Sillitoe on behalf of the Wellcome Sanger Institute COVID-19 Surveillance Team |
| EPI_ISL_2780751 | Lighthouse Lab in Glasgow | Wellcome Sanger Institute for the COVID-19 Genomics UK (COG-UK) Consortium | Harper VanSteenhouse, Yumi Kasai, David Gray, Carol Clugston, Anna Dominiczak and Alex Alderton, Roberto Amato, Jeffrey Barrett, Sonia Goncalves, Ewan Harrison, David K. Jackson, Ian Johnston, Dominic Kwiatkowski, Cordelia Langford, John Sillitoe on behalf of the Wellcome Sanger Institute COVID-19 Surveillance Team |
| EPI_ISL_2780752, EPI_ISL_2780753, EPI_ISL_2780754, EPI_ISL_2780755, EPI_ISL_2780756, EPI_ISL_2780757, EPI_ISL_2780758, EPI_ISL_2780759, EPI_ISL_2780760, EPI_ISL_2780761, EPI_ISL_2780762, EPI_ISL_2780763, EPI_ISL_2780764, EPI_ISL_2780765, EPI_ISL_2780766, EPI_ISL_2780767, EPI_ISL_2780768, EPI_ISL_2780769, EPI_ISL_2780770, EPI_ISL_2780771, EPI_ISL_2780772, EPI_ISL_2780773 | | | |
| see above | Lighthouse Lab in Milton Keynes | Wellcome Sanger Institute for the COVID-19 Genomics UK (COG-UK) Consortium | The Lighthouse Lab in Milton Keynes and Alex Alderton, Roberto Amato, Jeffrey Barrett, Sonia Goncalves, Ewan Harrison, David K. Jackson, Ian Johnston, Dominic Kwiatkowski, Cordelia Langford, John Sillitoe on behalf of the Wellcome Sanger Institute COVID-19 Surveillance Team |
| EPI_ISL_2780774, EPI_ISL_2780775 | Lighthouse Lab in Alderley Park | Wellcome Sanger Institute for the COVID-19 Genomics UK (COG-UK) Consortium | Jacquelyn Wynn, Mairead Hyland, The Lighthouse Lab in Alderley Park and Alex Alderton, Roberto Amato, Jeffrey Barrett, Sonia Goncalves, Ewan Harrison, David K. Jackson, Ian Johnston, Dominic Kwiatkowski, Cordelia Langford, John Sillitoe on behalf of the Wellcome Sanger Institute COVID-19 Surveillance Team |
| EPI_ISL_2780776, EPI_ISL_2780777, EPI_ISL_2780778, EPI_ISL_2780779, EPI_ISL_2780780 | Lighthouse Lab in Milton Keynes | Wellcome Sanger Institute for the COVID-19 Genomics UK (COG-UK) Consortium | The Lighthouse Lab in Milton Keynes and Alex Alderton, Roberto Amato, Jeffrey Barrett, Sonia Goncalves, Ewan Harrison, David K. Jackson, Ian Johnston, Dominic Kwiatkowski, Cordelia Langford, John Sillitoe on behalf of the Wellcome Sanger Institute COVID-19 Surveillance Team |
| EPI_ISL_2780781 | Lighthouse Lab in Alderley Park | Wellcome Sanger Institute for the COVID-19 Genomics UK (COG-UK) Consortium | Jacquelyn Wynn, Mairead Hyland, The Lighthouse Lab in Alderley Park and Alex Alderton, Roberto Amato, Jeffrey Barrett, Sonia Goncalves, Ewan Harrison, David K. Jackson, Ian Johnston, Dominic Kwiatkowski, Cordelia Langford, John Sillitoe on behalf of the Wellcome Sanger Institute COVID-19 Surveillance Team |
| EPI_ISL_2780782, EPI_ISL_2780783, EPI_ISL_2780784, EPI_ISL_2780785, EPI_ISL_2780786, EPI_ISL_2780787, EPI_ISL_2780788, EPI_ISL_2780789 | Lighthouse Lab in Milton Keynes | Wellcome Sanger Institute for the COVID-19 Genomics UK (COG-UK) Consortium | The Lighthouse Lab in Milton Keynes and Alex Alderton, Roberto Amato, Jeffrey Barrett, Sonia Goncalves, Ewan Harrison, David K. Jackson, Ian Johnston, Dominic Kwiatkowski, Cordelia Langford, John Sillitoe on behalf of the Wellcome Sanger Institute COVID-19 Surveillance Team |
| EPI_ISL_2780790 | Lighthouse Lab in Alderley Park | Wellcome Sanger Institute for the COVID-19 Genomics UK (COG-UK) Consortium | Jacquelyn Wynn, Mairead Hyland, The Lighthouse Lab in Alderley Park and Alex Alderton, Roberto Amato, Jeffrey Barrett, Sonia Goncalves, Ewan Harrison, David K. Jackson, Ian Johnston, Dominic Kwiatkowski, Cordelia Langford, John Sillitoe on behalf of the Wellcome Sanger Institute COVID-19 Surveillance Team |
| EPI_ISL_2780791 | Lighthouse Lab in Milton Keynes | Wellcome Sanger Institute for the COVID-19 Genomics UK (COG-UK) Consortium | The Lighthouse Lab in Milton Keynes and Alex Alderton, Roberto Amato, Jeffrey Barrett, Sonia Goncalves, Ewan Harrison, David K. Jackson, Ian Johnston, Dominic Kwiatkowski, Cordelia Langford, John Sillitoe on behalf of the Wellcome Sanger Institute COVID-19 Surveillance Team |
| EPI_ISL_2780792, EPI_ISL_2780793 | Lighthouse Lab in Alderley Park | Wellcome Sanger Institute for the COVID-19 Genomics UK (COG-UK) Consortium | Jacquelyn Wynn, Mairead Hyland, The Lighthouse Lab in Alderley Park and Alex Alderton, Roberto Amato, Jeffrey Barrett, Sonia Goncalves, Ewan Harrison, David K. Jackson, Ian Johnston, Dominic Kwiatkowski, Cordelia Langford, John Sillitoe on behalf of the Wellcome Sanger Institute COVID-19 Surveillance Team |
| EPI_ISL_2780794, EPI_ISL_2780795, EPI_ISL_2780796 | Lighthouse Lab in Milton Keynes | Wellcome Sanger Institute for the COVID-19 Genomics UK (COG-UK) Consortium | The Lighthouse Lab in Milton Keynes and Alex Alderton, Roberto Amato, Jeffrey Barrett, Sonia Goncalves, Ewan Harrison, David K. Jackson, Ian Johnston, Dominic Kwiatkowski, Cordelia Langford, John Sillitoe on behalf of the Wellcome Sanger Institute COVID-19 Surveillance Team |
| EPI_ISL_2780797, EPI_ISL_2780798 | Lighthouse Lab in Alderley Park | Wellcome Sanger Institute for the COVID-19 Genomics UK (COG-UK) Consortium | Jacquelyn Wynn, Mairead Hyland, The Lighthouse Lab in Alderley Park and Alex Alderton, Roberto Amato, Jeffrey Barrett, Sonia Goncalves, Ewan Harrison, David K. Jackson, Ian Johnston, Dominic Kwiatkowski, Cordelia Langford, John Sillitoe on behalf of the Wellcome Sanger Institute COVID-19 Surveillance Team |
| EPI_ISL_2780799 | Lighthouse Lab in Milton Keynes | Wellcome Sanger Institute for the COVID-19 Genomics UK (COG-UK) Consortium | The Lighthouse Lab in Milton Keynes and Alex Alderton, Roberto Amato, Jeffrey Barrett, Sonia Goncalves, Ewan Harrison, David K. Jackson, Ian Johnston, Dominic Kwiatkowski, Cordelia Langford, John Sillitoe on behalf of the Wellcome Sanger Institute COVID-19 Surveillance Team |
| EPI_ISL_2780800 | Lighthouse Lab in Alderley Park | Wellcome Sanger Institute for the COVID-19 Genomics UK (COG-UK) Consortium | Jacquelyn Wynn, Mairead Hyland, The Lighthouse Lab in Alderley Park and Alex Alderton, Roberto Amato, Jeffrey Barrett, Sonia Goncalves, Ewan Harrison, David K. Jackson, Ian Johnston, Dominic Kwiatkowski, Cordelia Langford, John Sillitoe on behalf of the Wellcome Sanger Institute COVID-19 Surveillance Team |
| EPI_ISL_2780801, EPI_ISL_2780802, EPI_ISL_2780803 | Lighthouse Lab in Milton Keynes | Wellcome Sanger Institute for the COVID-19 Genomics UK (COG-UK) Consortium | The Lighthouse Lab in Milton Keynes and Alex Alderton, Roberto Amato, Jeffrey Barrett, Sonia Goncalves, Ewan Harrison, David K. Jackson, Ian Johnston, Dominic Kwiatkowski, Cordelia Langford, John Sillitoe on behalf of the Wellcome Sanger Institute COVID-19 Surveillance Team |
| EPI_ISL_2780804 | Lighthouse Lab in Alderley Park | Wellcome Sanger Institute for the COVID-19 Genomics UK (COG-UK) Consortium | Jacquelyn Wynn, Mairead Hyland, The Lighthouse Lab in Alderley Park and Alex Alderton, Roberto Amato, Jeffrey Barrett, Sonia Goncalves, Ewan Harrison, David K. Jackson, Ian Johnston, Dominic Kwiatkowski, Cordelia Langford, John Sillitoe on behalf of the Wellcome Sanger Institute COVID-19 Surveillance Team |
| EPI_ISL_2780805, EPI_ISL_2780806, EPI_ISL_2780807, EPI_ISL_2780808, EPI_ISL_2780809, EPI_ISL_2780810, EPI_ISL_2780811, EPI_ISL_2780812, EPI_ISL_2780813, EPI_ISL_2780814 | Lighthouse Lab in Milton Keynes | Wellcome Sanger Institute for the COVID-19 Genomics UK (COG-UK) Consortium | The Lighthouse Lab in Milton Keynes and Alex Alderton, Roberto Amato, Jeffrey Barrett, Sonia Goncalves, Ewan Harrison, David K. Jackson, Ian Johnston, Dominic Kwiatkowski, Cordelia Langford, John Sillitoe on behalf of the Wellcome Sanger Institute COVID-19 Surveillance Team |
| EPI_ISL_2780815 | Lighthouse Lab in Alderley Park | Wellcome Sanger Institute for the COVID-19 Genomics UK (COG-UK) Consortium | Jacquelyn Wynn, Mairead Hyland, The Lighthouse Lab in Alderley Park and Alex Alderton, Roberto Amato, Jeffrey Barrett, Sonia Goncalves, Ewan Harrison, David K. Jackson, Ian Johnston, Dominic Kwiatkowski, Cordelia Langford, John Sillitoe on behalf of the Wellcome Sanger Institute COVID-19 Surveillance Team |
| EPI_ISL_2780816 | Lighthouse Lab in Milton Keynes | Wellcome Sanger Institute for the COVID-19 Genomics UK (COG-UK) Consortium | The Lighthouse Lab in Milton Keynes and Alex Alderton, Roberto Amato, Jeffrey Barrett, Sonia Goncalves, Ewan Harrison, David K. Jackson, Ian Johnston, Dominic Kwiatkowski, Cordelia Langford, John Sillitoe on behalf of the Wellcome Sanger Institute COVID-19 Surveillance Team |

| | | | |
|---|---|---|---|
| EPI_ISL_2780817 | Lighthouse Lab in Alderley Park | Wellcome Sanger Institute for the COVID-19 Genomics UK (COG-UK) Consortium | Jacquelyn Wynn, Mairead Hyland, The Lighthouse Lab in Alderley Park and Alex Alderton, Roberto Amato, Jeffrey Barrett, Sonia Goncalves, Ewan Harrison, David K. Jackson, Ian Johnston, Dominic Kwiatkowski, Cordelia Langford, John Sillitoe on behalf of the Wellcome Sanger Institute COVID-19 Surveillance Team |
| EPI_ISL_2780818, EPI_ISL_2780819, EPI_ISL_2780820, EPI_ISL_2780821, EPI_ISL_2780822 | Lighthouse Lab in Milton Keynes | Wellcome Sanger Institute for the COVID-19 Genomics UK (COG-UK) Consortium | The Lighthouse Lab in Milton Keynes and Alex Alderton, Roberto Amato, Jeffrey Barrett, Sonia Goncalves, Ewan Harrison, David K. Jackson, Ian Johnston, Dominic Kwiatkowski, Cordelia Langford, John Sillitoe on behalf of the Wellcome Sanger Institute COVID-19 Surveillance Team |
| EPI_ISL_2780823 | Lighthouse Lab in Alderley Park | Wellcome Sanger Institute for the COVID-19 Genomics UK (COG-UK) Consortium | Jacquelyn Wynn, Mairead Hyland, The Lighthouse Lab in Alderley Park and Alex Alderton, Roberto Amato, Jeffrey Barrett, Sonia Goncalves, Ewan Harrison, David K. Jackson, Ian Johnston, Dominic Kwiatkowski, Cordelia Langford, John Sillitoe on behalf of the Wellcome Sanger Institute COVID-19 Surveillance Team |
| EPI_ISL_2780824, EPI_ISL_2780825, EPI_ISL_2780826, EPI_ISL_2780827, EPI_ISL_2780828, EPI_ISL_2780829, EPI_ISL_2780830, EPI_ISL_2780831, EPI_ISL_2780832, EPI_ISL_2780833, EPI_ISL_2780834, EPI_ISL_2780835, EPI_ISL_2780836, EPI_ISL_2780837, EPI_ISL_2780838 | | | |
| see above | Lighthouse Lab in Milton Keynes | Wellcome Sanger Institute for the COVID-19 Genomics UK (COG-UK) Consortium | The Lighthouse Lab in Milton Keynes and Alex Alderton, Roberto Amato, Jeffrey Barrett, Sonia Goncalves, Ewan Harrison, David K. Jackson, Ian Johnston, Dominic Kwiatkowski, Cordelia Langford, John Sillitoe on behalf of the Wellcome Sanger Institute COVID-19 Surveillance Team |
| EPI_ISL_2780839 | Lighthouse Lab in Alderley Park | Wellcome Sanger Institute for the COVID-19 Genomics UK (COG-UK) Consortium | Jacquelyn Wynn, Mairead Hyland, The Lighthouse Lab in Alderley Park and Alex Alderton, Roberto Amato, Jeffrey Barrett, Sonia Goncalves, Ewan Harrison, David K. Jackson, Ian Johnston, Dominic Kwiatkowski, Cordelia Langford, John Sillitoe on behalf of the Wellcome Sanger Institute COVID-19 Surveillance Team |
| EPI_ISL_2780840 | Lighthouse Lab in Milton Keynes | Wellcome Sanger Institute for the COVID-19 Genomics UK (COG-UK) Consortium | The Lighthouse Lab in Milton Keynes and Alex Alderton, Roberto Amato, Jeffrey Barrett, Sonia Goncalves, Ewan Harrison, David K. Jackson, Ian Johnston, Dominic Kwiatkowski, Cordelia Langford, John Sillitoe on behalf of the Wellcome Sanger Institute COVID-19 Surveillance Team |
| EPI_ISL_2780841 | Lighthouse Lab in Alderley Park | Wellcome Sanger Institute for the COVID-19 Genomics UK (COG-UK) Consortium | Jacquelyn Wynn, Mairead Hyland, The Lighthouse Lab in Alderley Park and Alex Alderton, Roberto Amato, Jeffrey Barrett, Sonia Goncalves, Ewan Harrison, David K. Jackson, Ian Johnston, Dominic Kwiatkowski, Cordelia Langford, John Sillitoe on behalf of the Wellcome Sanger Institute COVID-19 Surveillance Team |
| EPI_ISL_2780842, EPI_ISL_2780843 | Lighthouse Lab in Milton Keynes | Wellcome Sanger Institute for the COVID-19 Genomics UK (COG-UK) Consortium | The Lighthouse Lab in Milton Keynes and Alex Alderton, Roberto Amato, Jeffrey Barrett, Sonia Goncalves, Ewan Harrison, David K. Jackson, Ian Johnston, Dominic Kwiatkowski, Cordelia Langford, John Sillitoe on behalf of the Wellcome Sanger Institute COVID-19 Surveillance Team |
| EPI_ISL_2780844 | Lighthouse Lab in Alderley Park | Wellcome Sanger Institute for the COVID-19 Genomics UK (COG-UK) Consortium | Jacquelyn Wynn, Mairead Hyland, The Lighthouse Lab in Alderley Park and Alex Alderton, Roberto Amato, Jeffrey Barrett, Sonia Goncalves, Ewan Harrison, David K. Jackson, Ian Johnston, Dominic Kwiatkowski, Cordelia Langford, John Sillitoe on behalf of the Wellcome Sanger Institute COVID-19 Surveillance Team |
| EPI_ISL_2780845 | Lighthouse Lab in Milton Keynes | Wellcome Sanger Institute for the COVID-19 Genomics UK (COG-UK) Consortium | The Lighthouse Lab in Milton Keynes and Alex Alderton, Roberto Amato, Jeffrey Barrett, Sonia Goncalves, Ewan Harrison, David K. Jackson, Ian Johnston, Dominic Kwiatkowski, Cordelia Langford, John Sillitoe on behalf of the Wellcome Sanger Institute COVID-19 Surveillance Team |
| EPI_ISL_2780846 | Lighthouse Lab in Alderley Park | Wellcome Sanger Institute for the COVID-19 Genomics UK (COG-UK) Consortium | Jacquelyn Wynn, Mairead Hyland, The Lighthouse Lab in Alderley Park and Alex Alderton, Roberto Amato, Jeffrey Barrett, Sonia Goncalves, Ewan Harrison, David K. Jackson, Ian Johnston, Dominic Kwiatkowski, Cordelia Langford, John Sillitoe on behalf of the Wellcome Sanger Institute COVID-19 Surveillance Team |
| EPI_ISL_2780847 | Lighthouse Lab in Milton Keynes | Wellcome Sanger Institute for the COVID-19 Genomics UK (COG-UK) Consortium | The Lighthouse Lab in Milton Keynes and Alex Alderton, Roberto Amato, Jeffrey Barrett, Sonia Goncalves, Ewan Harrison, David K. Jackson, Ian Johnston, Dominic Kwiatkowski, Cordelia Langford, John Sillitoe on behalf of the Wellcome Sanger Institute COVID-19 Surveillance Team |
| EPI_ISL_2780848 | Lighthouse Lab in Alderley Park | Wellcome Sanger Institute for the COVID-19 Genomics UK (COG-UK) Consortium | Jacquelyn Wynn, Mairead Hyland, The Lighthouse Lab in Alderley Park and Alex Alderton, Roberto Amato, Jeffrey Barrett, Sonia Goncalves, Ewan Harrison, David K. Jackson, Ian Johnston, Dominic Kwiatkowski, Cordelia Langford, John Sillitoe on behalf of the Wellcome Sanger Institute COVID-19 Surveillance Team |
| EPI_ISL_2780849, EPI_ISL_2780850, EPI_ISL_2780851 | Lighthouse Lab in Milton Keynes | Wellcome Sanger Institute for the COVID-19 Genomics UK (COG-UK) Consortium | The Lighthouse Lab in Milton Keynes and Alex Alderton, Roberto Amato, Jeffrey Barrett, Sonia Goncalves, Ewan Harrison, David K. Jackson, Ian Johnston, Dominic Kwiatkowski, Cordelia Langford, John Sillitoe on behalf of the Wellcome Sanger Institute COVID-19 Surveillance Team |
| EPI_ISL_2780852, EPI_ISL_2780853, EPI_ISL_2780854 | Lighthouse Lab in Alderley Park | Wellcome Sanger Institute for the COVID-19 Genomics UK (COG-UK) Consortium | Jacquelyn Wynn, Mairead Hyland, The Lighthouse Lab in Alderley Park and Alex Alderton, Roberto Amato, Jeffrey Barrett, Sonia Goncalves, Ewan Harrison, David K. Jackson, Ian Johnston, Dominic Kwiatkowski, Cordelia Langford, John Sillitoe on behalf of the Wellcome Sanger Institute COVID-19 Surveillance Team |
| EPI_ISL_2780855 | Lighthouse Lab in Milton Keynes | Wellcome Sanger Institute for the COVID-19 Genomics UK (COG-UK) Consortium | The Lighthouse Lab in Milton Keynes and Alex Alderton, Roberto Amato, Jeffrey Barrett, Sonia Goncalves, Ewan Harrison, David K. Jackson, Ian Johnston, Dominic Kwiatkowski, Cordelia Langford, John Sillitoe on behalf of the Wellcome Sanger Institute COVID-19 Surveillance Team |
| EPI_ISL_2780856, EPI_ISL_2780857 | Lighthouse Lab in Alderley Park | Wellcome Sanger Institute for the COVID-19 Genomics UK (COG-UK) Consortium | Jacquelyn Wynn, Mairead Hyland, The Lighthouse Lab in Alderley Park and Alex Alderton, Roberto Amato, Jeffrey Barrett, Sonia Goncalves, Ewan Harrison, David K. Jackson, Ian Johnston, Dominic Kwiatkowski, Cordelia Langford, John Sillitoe on behalf of the Wellcome Sanger Institute COVID-19 Surveillance Team |
| EPI_ISL_2780858, EPI_ISL_2780859, EPI_ISL_2780860, EPI_ISL_2780861, EPI_ISL_2780862 | Lighthouse Lab in Milton Keynes | Wellcome Sanger Institute for the COVID-19 Genomics UK (COG-UK) Consortium | The Lighthouse Lab in Milton Keynes and Alex Alderton, Roberto Amato, Jeffrey Barrett, Sonia Goncalves, Ewan Harrison, David K. Jackson, Ian Johnston, Dominic Kwiatkowski, Cordelia Langford, John Sillitoe on behalf of the Wellcome Sanger Institute COVID-19 Surveillance Team |
| EPI_ISL_2780863 | Lighthouse Lab in Alderley Park | Wellcome Sanger Institute for the COVID-19 Genomics UK (COG-UK) Consortium | Jacquelyn Wynn, Mairead Hyland, The Lighthouse Lab in Alderley Park and Alex Alderton, Roberto Amato, Jeffrey Barrett, Sonia Goncalves, Ewan Harrison, David K. Jackson, Ian Johnston, Dominic Kwiatkowski, Cordelia Langford, John Sillitoe on behalf of the Wellcome Sanger Institute COVID-19 Surveillance Team |
| EPI_ISL_2780864, EPI_ISL_2780865, EPI_ISL_2780866 | Lighthouse Lab in Milton Keynes | Wellcome Sanger Institute for the COVID-19 Genomics UK (COG-UK) Consortium | The Lighthouse Lab in Milton Keynes and Alex Alderton, Roberto Amato, Jeffrey Barrett, Sonia Goncalves, Ewan Harrison, David K. Jackson, Ian Johnston, Dominic Kwiatkowski, Cordelia Langford, John Sillitoe on behalf of the Wellcome Sanger Institute COVID-19 Surveillance Team |
| EPI_ISL_2780867, EPI_ISL_2780868 | Lighthouse Lab in Alderley Park | Wellcome Sanger Institute for the COVID-19 Genomics UK (COG-UK) Consortium | Jacquelyn Wynn, Mairead Hyland, The Lighthouse Lab in Alderley Park and Alex Alderton, Roberto Amato, Jeffrey Barrett, Sonia Goncalves, Ewan Harrison, David K. Jackson, Ian Johnston, Dominic Kwiatkowski, Cordelia Langford, John Sillitoe on behalf of the Wellcome Sanger Institute COVID-19 Surveillance Team |
| EPI_ISL_2780869, EPI_ISL_2780870 | Lighthouse Lab in Milton Keynes | Wellcome Sanger Institute for the COVID-19 Genomics UK (COG-UK) Consortium | The Lighthouse Lab in Milton Keynes and Alex Alderton, Roberto Amato, Jeffrey Barrett, Sonia Goncalves, Ewan Harrison, David K. Jackson, Ian Johnston, Dominic Kwiatkowski, Cordelia Langford, John Sillitoe on behalf of the Wellcome Sanger Institute COVID-19 Surveillance Team |
| EPI_ISL_2780871 | Lighthouse Lab in Alderley Park | Wellcome Sanger Institute for the COVID-19 Genomics UK (COG-UK) Consortium | Jacquelyn Wynn, Mairead Hyland, The Lighthouse Lab in Alderley Park and Alex Alderton, Roberto Amato, Jeffrey Barrett, Sonia Goncalves, Ewan Harrison, David K. Jackson, Ian Johnston, Dominic Kwiatkowski, Cordelia Langford, John Sillitoe on behalf of the Wellcome Sanger Institute COVID-19 Surveillance Team |
| EPI_ISL_2780872, EPI_ISL_2780873, EPI_ISL_2780874, EPI_ISL_2780875, EPI_ISL_2780876, EPI_ISL_2780877, EPI_ISL_2780878, EPI_ISL_2780879 | Lighthouse Lab in Milton Keynes | Wellcome Sanger Institute for the COVID-19 Genomics UK (COG-UK) Consortium | The Lighthouse Lab in Milton Keynes and Alex Alderton, Roberto Amato, Jeffrey Barrett, Sonia Goncalves, Ewan Harrison, David K. Jackson, Ian Johnston, Dominic Kwiatkowski, Cordelia Langford, John Sillitoe on behalf of the Wellcome Sanger Institute COVID-19 Surveillance Team |
| EPI_ISL_2780881 | Lighthouse Lab in Alderley Park | Wellcome Sanger Institute for the COVID-19 Genomics UK (COG-UK) Consortium | Jacquelyn Wynn, Mairead Hyland, The Lighthouse Lab in Alderley Park and Alex Alderton, Roberto Amato, Jeffrey Barrett, Sonia Goncalves, Ewan Harrison, David K. Jackson, Ian Johnston, Dominic Kwiatkowski, Cordelia Langford, John Sillitoe on behalf of the Wellcome Sanger Institute COVID-19 Surveillance Team |
| EPI_ISL_2780882, EPI_ISL_2780883, EPI_ISL_2780884, EPI_ISL_2780885, EPI_ISL_2780886, EPI_ISL_2780887 | Lighthouse Lab in Milton Keynes | Wellcome Sanger Institute for the COVID-19 Genomics UK (COG-UK) Consortium | The Lighthouse Lab in Milton Keynes and Alex Alderton, Roberto Amato, Jeffrey Barrett, Sonia Goncalves, Ewan Harrison, David K. Jackson, Ian Johnston, Dominic Kwiatkowski, Cordelia Langford, John Sillitoe on behalf of the Wellcome Sanger Institute COVID-19 Surveillance Team |
| EPI_ISL_2780888 | Lighthouse Lab in Alderley Park | Wellcome Sanger Institute for the COVID-19 Genomics UK (COG-UK) Consortium | Jacquelyn Wynn, Mairead Hyland, The Lighthouse Lab in Alderley Park and Alex Alderton, Roberto Amato, Jeffrey Barrett, Sonia Goncalves, Ewan Harrison, David K. Jackson, Ian Johnston, Dominic Kwiatkowski, Cordelia Langford, John Sillitoe on behalf of the Wellcome Sanger Institute COVID-19 Surveillance Team |
| EPI_ISL_2780889, EPI_ISL_2780890, EPI_ISL_2780891, EPI_ISL_2780892, EPI_ISL_2780893, EPI_ISL_2780894, EPI_ISL_2780895, EPI_ISL_2780896 | Lighthouse Lab in Milton Keynes | Wellcome Sanger Institute for the COVID-19 Genomics UK (COG-UK) Consortium | The Lighthouse Lab in Milton Keynes and Alex Alderton, Roberto Amato, Jeffrey Barrett, Sonia Goncalves, Ewan Harrison, David K. Jackson, Ian Johnston, Dominic Kwiatkowski, Cordelia Langford, John Sillitoe on behalf of the Wellcome Sanger Institute COVID-19 Surveillance Team |
| EPI_ISL_2780897 | Lighthouse Lab in Alderley Park | Wellcome Sanger Institute for the COVID-19 Genomics UK (COG-UK) Consortium | Jacquelyn Wynn, Mairead Hyland, The Lighthouse Lab in Alderley Park and Alex Alderton, Roberto Amato, Jeffrey Barrett, Sonia Goncalves, Ewan Harrison, David K. Jackson, Ian Johnston, Dominic Kwiatkowski, Cordelia Langford, John Sillitoe on behalf of the Wellcome Sanger Institute COVID-19 Surveillance Team |
| EPI_ISL_2780898, EPI_ISL_2780899, EPI_ISL_2780900, EPI_ISL_2780901, EPI_ISL_2780902, EPI_ISL_2780903, EPI_ISL_2780904 | Lighthouse Lab in Milton Keynes | Wellcome Sanger Institute for the COVID-19 Genomics UK (COG-UK) Consortium | The Lighthouse Lab in Milton Keynes and Alex Alderton, Roberto Amato, Jeffrey Barrett, Sonia Goncalves, Ewan Harrison, David K. Jackson, Ian Johnston, Dominic Kwiatkowski, Cordelia Langford, John Sillitoe on behalf of the Wellcome Sanger Institute COVID-19 Surveillance Team |
| EPI_ISL_2780905 | Lighthouse Lab in Alderley Park | Wellcome Sanger Institute for the COVID-19 Genomics UK (COG-UK) Consortium | Jacquelyn Wynn, Mairead Hyland, The Lighthouse Lab in Alderley Park and Alex Alderton, Roberto Amato, Jeffrey Barrett, Sonia Goncalves, Ewan Harrison, David K. Jackson, Ian Johnston, Dominic Kwiatkowski, Cordelia Langford, John Sillitoe on behalf of the Wellcome Sanger Institute COVID-19 Surveillance Team |
| EPI_ISL_2780906, EPI_ISL_2780907, | Lighthouse Lab in Milton Keynes | Wellcome Sanger Institute for the COVID-19 Genomics UK (COG-UK) Consortium | The Lighthouse Lab in Milton Keynes and Alex Alderton, Roberto Amato, Jeffrey Barrett, Sonia Goncalves, Ewan Harrison, David K. Jackson, Ian Johnston, Dominic Kwiatkowski, Cordelia Langford, John Sillitoe on behalf of the Wellcome Sanger Institute COVID-19 Surveillance Team |

| | | | |
|---|---|---|---|
| EPI_ISL_2780908, EPI_ISL_2780909 | | | |
| EPI_ISL_2780910 | Lighthouse Lab in Alderley Park | Wellcome Sanger Institute for the COVID-19 Genomics UK (COG-UK) Consortium | Jacquelyn Wynn, Mairead Hyland, The Lighthouse Lab in Alderley Park and Alex Alderton, Roberto Amato, Jeffrey Barrett, Sonia Goncalves, Ewan Harrison, David K. Jackson, Ian Johnston, Dominic Kwiatkowski, Cordelia Langford, John Sillitoe on behalf of the Wellcome Sanger Institute COVID-19 Surveillance Team |
| EPI_ISL_2780911, EPI_ISL_2780912 | Lighthouse Lab in Milton Keynes | Wellcome Sanger Institute for the COVID-19 Genomics UK (COG-UK) Consortium | The Lighthouse Lab in Milton Keynes and Alex Alderton, Roberto Amato, Jeffrey Barrett, Sonia Goncalves, Ewan Harrison, David K. Jackson, Ian Johnston, Dominic Kwiatkowski, Cordelia Langford, John Sillitoe on behalf of the Wellcome Sanger Institute COVID-19 Surveillance Team |
| EPI_ISL_2780913 | Lighthouse Lab in Alderley Park | Wellcome Sanger Institute for the COVID-19 Genomics UK (COG-UK) Consortium | Jacquelyn Wynn, Mairead Hyland, The Lighthouse Lab in Alderley Park and Alex Alderton, Roberto Amato, Jeffrey Barrett, Sonia Goncalves, Ewan Harrison, David K. Jackson, Ian Johnston, Dominic Kwiatkowski, Cordelia Langford, John Sillitoe on behalf of the Wellcome Sanger Institute COVID-19 Surveillance Team |
| EPI_ISL_2780914 | Lighthouse Lab in Milton Keynes | Wellcome Sanger Institute for the COVID-19 Genomics UK (COG-UK) Consortium | The Lighthouse Lab in Milton Keynes and Alex Alderton, Roberto Amato, Jeffrey Barrett, Sonia Goncalves, Ewan Harrison, David K. Jackson, Ian Johnston, Dominic Kwiatkowski, Cordelia Langford, John Sillitoe on behalf of the Wellcome Sanger Institute COVID-19 Surveillance Team |
| EPI_ISL_2780915 | Lighthouse Lab in Alderley Park | Wellcome Sanger Institute for the COVID-19 Genomics UK (COG-UK) Consortium | Jacquelyn Wynn, Mairead Hyland, The Lighthouse Lab in Alderley Park and Alex Alderton, Roberto Amato, Jeffrey Barrett, Sonia Goncalves, Ewan Harrison, David K. Jackson, Ian Johnston, Dominic Kwiatkowski, Cordelia Langford, John Sillitoe on behalf of the Wellcome Sanger Institute COVID-19 Surveillance Team |
| EPI_ISL_2780916, EPI_ISL_2780917 | Lighthouse Lab in Milton Keynes | Wellcome Sanger Institute for the COVID-19 Genomics UK (COG-UK) Consortium | The Lighthouse Lab in Milton Keynes and Alex Alderton, Roberto Amato, Jeffrey Barrett, Sonia Goncalves, Ewan Harrison, David K. Jackson, Ian Johnston, Dominic Kwiatkowski, Cordelia Langford, John Sillitoe on behalf of the Wellcome Sanger Institute COVID-19 Surveillance Team |
| EPI_ISL_2780918, EPI_ISL_2780919 | Lighthouse Lab in Alderley Park | Wellcome Sanger Institute for the COVID-19 Genomics UK (COG-UK) Consortium | Jacquelyn Wynn, Mairead Hyland, The Lighthouse Lab in Alderley Park and Alex Alderton, Roberto Amato, Jeffrey Barrett, Sonia Goncalves, Ewan Harrison, David K. Jackson, Ian Johnston, Dominic Kwiatkowski, Cordelia Langford, John Sillitoe on behalf of the Wellcome Sanger Institute COVID-19 Surveillance Team |
| EPI_ISL_2780920, EPI_ISL_2780921, EPI_ISL_2780922 | Lighthouse Lab in Milton Keynes | Wellcome Sanger Institute for the COVID-19 Genomics UK (COG-UK) Consortium | The Lighthouse Lab in Milton Keynes and Alex Alderton, Roberto Amato, Jeffrey Barrett, Sonia Goncalves, Ewan Harrison, David K. Jackson, Ian Johnston, Dominic Kwiatkowski, Cordelia Langford, John Sillitoe on behalf of the Wellcome Sanger Institute COVID-19 Surveillance Team |
| EPI_ISL_2780923 | Lighthouse Lab in Alderley Park | Wellcome Sanger Institute for the COVID-19 Genomics UK (COG-UK) Consortium | Jacquelyn Wynn, Mairead Hyland, The Lighthouse Lab in Alderley Park and Alex Alderton, Roberto Amato, Jeffrey Barrett, Sonia Goncalves, Ewan Harrison, David K. Jackson, Ian Johnston, Dominic Kwiatkowski, Cordelia Langford, John Sillitoe on behalf of the Wellcome Sanger Institute COVID-19 Surveillance Team |
| EPI_ISL_2780924, EPI_ISL_2780925 | Lighthouse Lab in Milton Keynes | Wellcome Sanger Institute for the COVID-19 Genomics UK (COG-UK) Consortium | The Lighthouse Lab in Milton Keynes and Alex Alderton, Roberto Amato, Jeffrey Barrett, Sonia Goncalves, Ewan Harrison, David K. Jackson, Ian Johnston, Dominic Kwiatkowski, Cordelia Langford, John Sillitoe on behalf of the Wellcome Sanger Institute COVID-19 Surveillance Team |
| EPI_ISL_2780926, EPI_ISL_2780927 | Lighthouse Lab in Alderley Park | Wellcome Sanger Institute for the COVID-19 Genomics UK (COG-UK) Consortium | Jacquelyn Wynn, Mairead Hyland, The Lighthouse Lab in Alderley Park and Alex Alderton, Roberto Amato, Jeffrey Barrett, Sonia Goncalves, Ewan Harrison, David K. Jackson, Ian Johnston, Dominic Kwiatkowski, Cordelia Langford, John Sillitoe on behalf of the Wellcome Sanger Institute COVID-19 Surveillance Team |
| EPI_ISL_2780928 | Lighthouse Lab in Milton Keynes | Wellcome Sanger Institute for the COVID-19 Genomics UK (COG-UK) Consortium | The Lighthouse Lab in Milton Keynes and Alex Alderton, Roberto Amato, Jeffrey Barrett, Sonia Goncalves, Ewan Harrison, David K. Jackson, Ian Johnston, Dominic Kwiatkowski, Cordelia Langford, John Sillitoe on behalf of the Wellcome Sanger Institute COVID-19 Surveillance Team |
| EPI_ISL_2780929 | Lighthouse Lab in Alderley Park | Wellcome Sanger Institute for the COVID-19 Genomics UK (COG-UK) Consortium | Jacquelyn Wynn, Mairead Hyland, The Lighthouse Lab in Alderley Park and Alex Alderton, Roberto Amato, Jeffrey Barrett, Sonia Goncalves, Ewan Harrison, David K. Jackson, Ian Johnston, Dominic Kwiatkowski, Cordelia Langford, John Sillitoe on behalf of the Wellcome Sanger Institute COVID-19 Surveillance Team |
| EPI_ISL_2780930, EPI_ISL_2780931, EPI_ISL_2780932, EPI_ISL_2780933, EPI_ISL_2780934, EPI_ISL_2780935, EPI_ISL_2780936, EPI_ISL_2780937, EPI_ISL_2780938, EPI_ISL_2780939 | Lighthouse Lab in Milton Keynes | Wellcome Sanger Institute for the COVID-19 Genomics UK (COG-UK) Consortium | The Lighthouse Lab in Milton Keynes and Alex Alderton, Roberto Amato, Jeffrey Barrett, Sonia Goncalves, Ewan Harrison, David K. Jackson, Ian Johnston, Dominic Kwiatkowski, Cordelia Langford, John Sillitoe on behalf of the Wellcome Sanger Institute COVID-19 Surveillance Team |
| EPI_ISL_2780940, EPI_ISL_2780941, EPI_ISL_2780942 | Lighthouse Lab in Alderley Park | Wellcome Sanger Institute for the COVID-19 Genomics UK (COG-UK) Consortium | Jacquelyn Wynn, Mairead Hyland, The Lighthouse Lab in Alderley Park and Alex Alderton, Roberto Amato, Jeffrey Barrett, Sonia Goncalves, Ewan Harrison, David K. Jackson, Ian Johnston, Dominic Kwiatkowski, Cordelia Langford, John Sillitoe on behalf of the Wellcome Sanger Institute COVID-19 Surveillance Team |
| EPI_ISL_2780943 | Lighthouse Lab in Milton Keynes | Wellcome Sanger Institute for the COVID-19 Genomics UK (COG-UK) Consortium | The Lighthouse Lab in Milton Keynes and Alex Alderton, Roberto Amato, Jeffrey Barrett, Sonia Goncalves, Ewan Harrison, David K. Jackson, Ian Johnston, Dominic Kwiatkowski, Cordelia Langford, John Sillitoe on behalf of the Wellcome Sanger Institute COVID-19 Surveillance Team |
| EPI_ISL_2780944, EPI_ISL_2780945 | Lighthouse Lab in Alderley Park | Wellcome Sanger Institute for the COVID-19 Genomics UK (COG-UK) Consortium | Jacquelyn Wynn, Mairead Hyland, The Lighthouse Lab in Alderley Park and Alex Alderton, Roberto Amato, Jeffrey Barrett, Sonia Goncalves, Ewan Harrison, David K. Jackson, Ian Johnston, Dominic Kwiatkowski, Cordelia Langford, John Sillitoe on behalf of the Wellcome Sanger Institute COVID-19 Surveillance Team |
| EPI_ISL_2780946, EPI_ISL_2780947 | Lighthouse Lab in Milton Keynes | Wellcome Sanger Institute for the COVID-19 Genomics UK (COG-UK) Consortium | The Lighthouse Lab in Milton Keynes and Alex Alderton, Roberto Amato, Jeffrey Barrett, Sonia Goncalves, Ewan Harrison, David K. Jackson, Ian Johnston, Dominic Kwiatkowski, Cordelia Langford, John Sillitoe on behalf of the Wellcome Sanger Institute COVID-19 Surveillance Team |
| EPI_ISL_2780948, EPI_ISL_2780949, EPI_ISL_2780950 | Lighthouse Lab in Alderley Park | Wellcome Sanger Institute for the COVID-19 Genomics UK (COG-UK) Consortium | Jacquelyn Wynn, Mairead Hyland, The Lighthouse Lab in Alderley Park and Alex Alderton, Roberto Amato, Jeffrey Barrett, Sonia Goncalves, Ewan Harrison, David K. Jackson, Ian Johnston, Dominic Kwiatkowski, Cordelia Langford, John Sillitoe on behalf of the Wellcome Sanger Institute COVID-19 Surveillance Team |
| EPI_ISL_2780951 | Lighthouse Laboratory Plymouth | Wellcome Sanger Institute for the COVID-19 Genomics UK (COG-UK) Consortium | Lighthouse Laboratory Plymouth and Alex Alderton, Roberto Amato, Jeffrey Barrett, Sonia Goncalves, Ewan Harrison, David K. Jackson, Ian Johnston, Dominic Kwiatkowski, Cordelia Langford, John Sillitoe on behalf of the Wellcome Sanger Institute COVID-19 Surveillance Team |
| EPI_ISL_2780953 | Lighthouse Lab in Alderley Park | Wellcome Sanger Institute for the COVID-19 Genomics UK (COG-UK) Consortium | Jacquelyn Wynn, Mairead Hyland, The Lighthouse Lab in Alderley Park and Alex Alderton, Roberto Amato, Jeffrey Barrett, Sonia Goncalves, Ewan Harrison, David K. Jackson, Ian Johnston, Dominic Kwiatkowski, Cordelia Langford, John Sillitoe on behalf of the Wellcome Sanger Institute COVID-19 Surveillance Team |
| EPI_ISL_2780954 | Lighthouse Lab in Milton Keynes | Wellcome Sanger Institute for the COVID-19 Genomics UK (COG-UK) Consortium | The Lighthouse Lab in Milton Keynes and Alex Alderton, Roberto Amato, Jeffrey Barrett, Sonia Goncalves, Ewan Harrison, David K. Jackson, Ian Johnston, Dominic Kwiatkowski, Cordelia Langford, John Sillitoe on behalf of the Wellcome Sanger Institute COVID-19 Surveillance Team |
| EPI_ISL_2780955, EPI_ISL_2780956 | Lighthouse Lab in Alderley Park | Wellcome Sanger Institute for the COVID-19 Genomics UK (COG-UK) Consortium | Jacquelyn Wynn, Mairead Hyland, The Lighthouse Lab in Alderley Park and Alex Alderton, Roberto Amato, Jeffrey Barrett, Sonia Goncalves, Ewan Harrison, David K. Jackson, Ian Johnston, Dominic Kwiatkowski, Cordelia Langford, John Sillitoe on behalf of the Wellcome Sanger Institute COVID-19 Surveillance Team |
| EPI_ISL_2780957, EPI_ISL_2780958 | Lighthouse Lab in Milton Keynes | Wellcome Sanger Institute for the COVID-19 Genomics UK (COG-UK) Consortium | The Lighthouse Lab in Milton Keynes and Alex Alderton, Roberto Amato, Jeffrey Barrett, Sonia Goncalves, Ewan Harrison, David K. Jackson, Ian Johnston, Dominic Kwiatkowski, Cordelia Langford, John Sillitoe on behalf of the Wellcome Sanger Institute COVID-19 Surveillance Team |
| EPI_ISL_2780959 | Lighthouse Lab in Alderley Park | Wellcome Sanger Institute for the COVID-19 Genomics UK (COG-UK) Consortium | Jacquelyn Wynn, Mairead Hyland, The Lighthouse Lab in Alderley Park and Alex Alderton, Roberto Amato, Jeffrey Barrett, Sonia Goncalves, Ewan Harrison, David K. Jackson, Ian Johnston, Dominic Kwiatkowski, Cordelia Langford, John Sillitoe on behalf of the Wellcome Sanger Institute COVID-19 Surveillance Team |
| EPI_ISL_2780960, EPI_ISL_2780961, EPI_ISL_2780962, EPI_ISL_2780963, EPI_ISL_2780964 | Lighthouse Lab in Milton Keynes | Wellcome Sanger Institute for the COVID-19 Genomics UK (COG-UK) Consortium | The Lighthouse Lab in Milton Keynes and Alex Alderton, Roberto Amato, Jeffrey Barrett, Sonia Goncalves, Ewan Harrison, David K. Jackson, Ian Johnston, Dominic Kwiatkowski, Cordelia Langford, John Sillitoe on behalf of the Wellcome Sanger Institute COVID-19 Surveillance Team |
| EPI_ISL_2780965 | Lighthouse Lab in Alderley Park | Wellcome Sanger Institute for the COVID-19 Genomics UK (COG-UK) Consortium | Jacquelyn Wynn, Mairead Hyland, The Lighthouse Lab in Alderley Park and Alex Alderton, Roberto Amato, Jeffrey Barrett, Sonia Goncalves, Ewan Harrison, David K. Jackson, Ian Johnston, Dominic Kwiatkowski, Cordelia Langford, John Sillitoe on behalf of the Wellcome Sanger Institute COVID-19 Surveillance Team |
| EPI_ISL_2780966, EPI_ISL_2780967, EPI_ISL_2780968, EPI_ISL_2780969 | Lighthouse Lab in Milton Keynes | Wellcome Sanger Institute for the COVID-19 Genomics UK (COG-UK) Consortium | The Lighthouse Lab in Milton Keynes and Alex Alderton, Roberto Amato, Jeffrey Barrett, Sonia Goncalves, Ewan Harrison, David K. Jackson, Ian Johnston, Dominic Kwiatkowski, Cordelia Langford, John Sillitoe on behalf of the Wellcome Sanger Institute COVID-19 Surveillance Team |
| EPI_ISL_2780970, EPI_ISL_2780971 | Lighthouse Lab in Alderley Park | Wellcome Sanger Institute for the COVID-19 Genomics UK (COG-UK) Consortium | Jacquelyn Wynn, Mairead Hyland, The Lighthouse Lab in Alderley Park and Alex Alderton, Roberto Amato, Jeffrey Barrett, Sonia Goncalves, Ewan Harrison, David K. Jackson, Ian Johnston, Dominic Kwiatkowski, Cordelia Langford, John Sillitoe on behalf of the Wellcome Sanger Institute COVID-19 Surveillance Team |
| EPI_ISL_2780972, EPI_ISL_2780973, EPI_ISL_2780974, EPI_ISL_2780975 | Lighthouse Lab in Milton Keynes | Wellcome Sanger Institute for the COVID-19 Genomics UK (COG-UK) Consortium | The Lighthouse Lab in Milton Keynes and Alex Alderton, Roberto Amato, Jeffrey Barrett, Sonia Goncalves, Ewan Harrison, David K. Jackson, Ian Johnston, Dominic Kwiatkowski, Cordelia Langford, John Sillitoe on behalf of the Wellcome Sanger Institute COVID-19 Surveillance Team |
| EPI_ISL_2780976, EPI_ISL_2780977 | Lighthouse Lab in Alderley Park | Wellcome Sanger Institute for the COVID-19 Genomics UK (COG-UK) Consortium | Jacquelyn Wynn, Mairead Hyland, The Lighthouse Lab in Alderley Park and Alex Alderton, Roberto Amato, Jeffrey Barrett, Sonia Goncalves, Ewan Harrison, David K. Jackson, Ian Johnston, Dominic Kwiatkowski, Cordelia Langford, John Sillitoe on behalf of the Wellcome Sanger Institute COVID-19 Surveillance Team |
| EPI_ISL_2780978, EPI_ISL_2780979, EPI_ISL_2780980, EPI_ISL_2780981 | Lighthouse Lab in Milton Keynes | Wellcome Sanger Institute for the COVID-19 Genomics UK (COG-UK) Consortium | The Lighthouse Lab in Milton Keynes and Alex Alderton, Roberto Amato, Jeffrey Barrett, Sonia Goncalves, Ewan Harrison, David K. Jackson, Ian Johnston, Dominic Kwiatkowski, Cordelia Langford, John Sillitoe on behalf of the Wellcome Sanger Institute COVID-19 Surveillance Team |
| EPI_ISL_2780982 | Lighthouse Laboratory Plymouth | Wellcome Sanger Institute for the COVID-19 Genomics UK (COG-UK) Consortium | Lighthouse Laboratory Plymouth and Alex Alderton, Roberto Amato, Jeffrey Barrett, Sonia Goncalves, Ewan Harrison, David K. Jackson, Ian Johnston, Dominic Kwiatkowski, Cordelia Langford, John Sillitoe on behalf of the Wellcome Sanger Institute COVID-19 Surveillance Team |
| EPI_ISL_2780983 | Lighthouse Lab in Milton Keynes | Wellcome Sanger Institute for the COVID-19 Genomics UK (COG-UK) Consortium | The Lighthouse Lab in Milton Keynes and Alex Alderton, Roberto Amato, Jeffrey Barrett, Sonia Goncalves, Ewan Harrison, David K. Jackson, Ian Johnston, Dominic Kwiatkowski, Cordelia Langford, John Sillitoe on behalf of the Wellcome Sanger Institute COVID-19 Surveillance Team |
| EPI_ISL_2780984 | Lighthouse Lab in Alderley Park | Wellcome Sanger Institute for the COVID-19 Genomics UK (COG-UK) Consortium | Jacquelyn Wynn, Mairead Hyland, The Lighthouse Lab in Alderley Park and Alex Alderton, Roberto Amato, Jeffrey Barrett, Sonia Goncalves, Ewan Harrison, David K. Jackson, Ian Johnston, Dominic Kwiatkowski, Cordelia Langford, John Sillitoe on behalf of the Wellcome Sanger Institute COVID-19 Surveillance Team |
| EPI_ISL_2780985, EPI_ISL_2780986, | Lighthouse Lab in Milton Keynes | Wellcome Sanger Institute for the COVID-19 Genomics UK (COG-UK) Consortium | The Lighthouse Lab in Milton Keynes and Alex Alderton, Roberto Amato, Jeffrey Barrett, Sonia Goncalves, Ewan Harrison, David K. Jackson, Ian Johnston, Dominic Kwiatkowski, Cordelia Langford, John Sillitoe on behalf of the Wellcome Sanger Institute COVID-19 Surveillance Team |

| | | | |
|---|---|---|---|
| EPI_ISL_2780987 | | | |
| EPI_ISL_2780988, EPI_ISL_2780989, EPI_ISL_2780990 | Lighthouse Lab in Alderley Park | Wellcome Sanger Institute for the COVID-19 Genomics UK (COG-UK) Consortium | Jacquelyn Wynn, Mairead Hyland, The Lighthouse Lab in Alderley Park and Alex Alderton, Roberto Amato, Jeffrey Barrett, Sonia Goncalves, Ewan Harrison, David K. Jackson, Ian Johnston, Dominic Kwiatkowski, Cordelia Langford, John Sillitoe on behalf of the Wellcome Sanger Institute COVID-19 Surveillance Team |
| EPI_ISL_2780991, EPI_ISL_2780992, EPI_ISL_2780993, EPI_ISL_2780994, EPI_ISL_2780995, EPI_ISL_2780996 | Lighthouse Lab in Milton Keynes | Wellcome Sanger Institute for the COVID-19 Genomics UK (COG-UK) Consortium | The Lighthouse Lab in Milton Keynes and Alex Alderton, Roberto Amato, Jeffrey Barrett, Sonia Goncalves, Ewan Harrison, David K. Jackson, Ian Johnston, Dominic Kwiatkowski, Cordelia Langford, John Sillitoe on behalf of the Wellcome Sanger Institute COVID-19 Surveillance Team |
| EPI_ISL_2780997 | Lighthouse Lab in Alderley Park | Wellcome Sanger Institute for the COVID-19 Genomics UK (COG-UK) Consortium | Jacquelyn Wynn, Mairead Hyland, The Lighthouse Lab in Alderley Park and Alex Alderton, Roberto Amato, Jeffrey Barrett, Sonia Goncalves, Ewan Harrison, David K. Jackson, Ian Johnston, Dominic Kwiatkowski, Cordelia Langford, John Sillitoe on behalf of the Wellcome Sanger Institute COVID-19 Surveillance Team |
| EPI_ISL_2780998, EPI_ISL_2780999, EPI_ISL_2781000, EPI_ISL_2781001 | Lighthouse Lab in Milton Keynes | Wellcome Sanger Institute for the COVID-19 Genomics UK (COG-UK) Consortium | The Lighthouse Lab in Milton Keynes and Alex Alderton, Roberto Amato, Jeffrey Barrett, Sonia Goncalves, Ewan Harrison, David K. Jackson, Ian Johnston, Dominic Kwiatkowski, Cordelia Langford, John Sillitoe on behalf of the Wellcome Sanger Institute COVID-19 Surveillance Team |
| EPI_ISL_2781002 | Lighthouse Lab in Alderley Park | Wellcome Sanger Institute for the COVID-19 Genomics UK (COG-UK) Consortium | Jacquelyn Wynn, Mairead Hyland, The Lighthouse Lab in Alderley Park and Alex Alderton, Roberto Amato, Jeffrey Barrett, Sonia Goncalves, Ewan Harrison, David K. Jackson, Ian Johnston, Dominic Kwiatkowski, Cordelia Langford, John Sillitoe on behalf of the Wellcome Sanger Institute COVID-19 Surveillance Team |
| EPI_ISL_2781003, EPI_ISL_2781004, EPI_ISL_2781005, EPI_ISL_2781006, EPI_ISL_2781007, EPI_ISL_2781008 | Lighthouse Lab in Milton Keynes | Wellcome Sanger Institute for the COVID-19 Genomics UK (COG-UK) Consortium | The Lighthouse Lab in Milton Keynes and Alex Alderton, Roberto Amato, Jeffrey Barrett, Sonia Goncalves, Ewan Harrison, David K. Jackson, Ian Johnston, Dominic Kwiatkowski, Cordelia Langford, John Sillitoe on behalf of the Wellcome Sanger Institute COVID-19 Surveillance Team |
| EPI_ISL_2781009 | Lighthouse Lab in Alderley Park | Wellcome Sanger Institute for the COVID-19 Genomics UK (COG-UK) Consortium | Jacquelyn Wynn, Mairead Hyland, The Lighthouse Lab in Alderley Park and Alex Alderton, Roberto Amato, Jeffrey Barrett, Sonia Goncalves, Ewan Harrison, David K. Jackson, Ian Johnston, Dominic Kwiatkowski, Cordelia Langford, John Sillitoe on behalf of the Wellcome Sanger Institute COVID-19 Surveillance Team |
| EPI_ISL_2781010 | Lighthouse Lab in Milton Keynes | Wellcome Sanger Institute for the COVID-19 Genomics UK (COG-UK) Consortium | The Lighthouse Lab in Milton Keynes and Alex Alderton, Roberto Amato, Jeffrey Barrett, Sonia Goncalves, Ewan Harrison, David K. Jackson, Ian Johnston, Dominic Kwiatkowski, Cordelia Langford, John Sillitoe on behalf of the Wellcome Sanger Institute COVID-19 Surveillance Team |
| EPI_ISL_2781011 | Lighthouse Lab in Alderley Park | Wellcome Sanger Institute for the COVID-19 Genomics UK (COG-UK) Consortium | Jacquelyn Wynn, Mairead Hyland, The Lighthouse Lab in Alderley Park and Alex Alderton, Roberto Amato, Jeffrey Barrett, Sonia Goncalves, Ewan Harrison, David K. Jackson, Ian Johnston, Dominic Kwiatkowski, Cordelia Langford, John Sillitoe on behalf of the Wellcome Sanger Institute COVID-19 Surveillance Team |
| EPI_ISL_2781012, EPI_ISL_2781013, EPI_ISL_2781014, EPI_ISL_2781015, EPI_ISL_2781016, EPI_ISL_2781017, EPI_ISL_2781018, EPI_ISL_2781019, EPI_ISL_2781020, EPI_ISL_2781021, EPI_ISL_2781022, EPI_ISL_2781023, EPI_ISL_2781024, EPI_ISL_2781025, EPI_ISL_2781026, EPI_ISL_2781027, EPI_ISL_2781028 | | | |
| see above | Lighthouse Lab in Milton Keynes | Wellcome Sanger Institute for the COVID-19 Genomics UK (COG-UK) Consortium | The Lighthouse Lab in Milton Keynes and Alex Alderton, Roberto Amato, Jeffrey Barrett, Sonia Goncalves, Ewan Harrison, David K. Jackson, Ian Johnston, Dominic Kwiatkowski, Cordelia Langford, John Sillitoe on behalf of the Wellcome Sanger Institute COVID-19 Surveillance Team |
| EPI_ISL_2781029 | Lighthouse Lab in Alderley Park | Wellcome Sanger Institute for the COVID-19 Genomics UK (COG-UK) Consortium | Jacquelyn Wynn, Mairead Hyland, The Lighthouse Lab in Alderley Park and Alex Alderton, Roberto Amato, Jeffrey Barrett, Sonia Goncalves, Ewan Harrison, David K. Jackson, Ian Johnston, Dominic Kwiatkowski, Cordelia Langford, John Sillitoe on behalf of the Wellcome Sanger Institute COVID-19 Surveillance Team |
| EPI_ISL_2781030, EPI_ISL_2781031, EPI_ISL_2781032 | Lighthouse Lab in Milton Keynes | Wellcome Sanger Institute for the COVID-19 Genomics UK (COG-UK) Consortium | The Lighthouse Lab in Milton Keynes and Alex Alderton, Roberto Amato, Jeffrey Barrett, Sonia Goncalves, Ewan Harrison, David K. Jackson, Ian Johnston, Dominic Kwiatkowski, Cordelia Langford, John Sillitoe on behalf of the Wellcome Sanger Institute COVID-19 Surveillance Team |
| EPI_ISL_2781033 | Lighthouse Lab in Alderley Park | Wellcome Sanger Institute for the COVID-19 Genomics UK (COG-UK) Consortium | Jacquelyn Wynn, Mairead Hyland, The Lighthouse Lab in Alderley Park and Alex Alderton, Roberto Amato, Jeffrey Barrett, Sonia Goncalves, Ewan Harrison, David K. Jackson, Ian Johnston, Dominic Kwiatkowski, Cordelia Langford, John Sillitoe on behalf of the Wellcome Sanger Institute COVID-19 Surveillance Team |
| EPI_ISL_2781034 | Lighthouse Lab in Milton Keynes | Wellcome Sanger Institute for the COVID-19 Genomics UK (COG-UK) Consortium | The Lighthouse Lab in Milton Keynes and Alex Alderton, Roberto Amato, Jeffrey Barrett, Sonia Goncalves, Ewan Harrison, David K. Jackson, Ian Johnston, Dominic Kwiatkowski, Cordelia Langford, John Sillitoe on behalf of the Wellcome Sanger Institute COVID-19 Surveillance Team |
| EPI_ISL_2781035 | Lighthouse Lab in Alderley Park | Wellcome Sanger Institute for the COVID-19 Genomics UK (COG-UK) Consortium | Jacquelyn Wynn, Mairead Hyland, The Lighthouse Lab in Alderley Park and Alex Alderton, Roberto Amato, Jeffrey Barrett, Sonia Goncalves, Ewan Harrison, David K. Jackson, Ian Johnston, Dominic Kwiatkowski, Cordelia Langford, John Sillitoe on behalf of the Wellcome Sanger Institute COVID-19 Surveillance Team |
| EPI_ISL_2781036, EPI_ISL_2781037, EPI_ISL_2781038, EPI_ISL_2781039, EPI_ISL_2781040 | Lighthouse Lab in Milton Keynes | Wellcome Sanger Institute for the COVID-19 Genomics UK (COG-UK) Consortium | The Lighthouse Lab in Milton Keynes and Alex Alderton, Roberto Amato, Jeffrey Barrett, Sonia Goncalves, Ewan Harrison, David K. Jackson, Ian Johnston, Dominic Kwiatkowski, Cordelia Langford, John Sillitoe on behalf of the Wellcome Sanger Institute COVID-19 Surveillance Team |
| EPI_ISL_2781041 | Lighthouse Laboratory Plymouth | Wellcome Sanger Institute for the COVID-19 Genomics UK (COG-UK) Consortium | Lighthouse Laboratory Plymouth and Alex Alderton, Roberto Amato, Jeffrey Barrett, Sonia Goncalves, Ewan Harrison, David K. Jackson, Ian Johnston, Dominic Kwiatkowski, Cordelia Langford, John Sillitoe on behalf of the Wellcome Sanger Institute COVID-19 Surveillance Team |
| EPI_ISL_2781042, EPI_ISL_2781043 | Lighthouse Lab in Alderley Park | Wellcome Sanger Institute for the COVID-19 Genomics UK (COG-UK) Consortium | Jacquelyn Wynn, Mairead Hyland, The Lighthouse Lab in Alderley Park and Alex Alderton, Roberto Amato, Jeffrey Barrett, Sonia Goncalves, Ewan Harrison, David K. Jackson, Ian Johnston, Dominic Kwiatkowski, Cordelia Langford, John Sillitoe on behalf of the Wellcome Sanger Institute COVID-19 Surveillance Team |
| EPI_ISL_2781044, EPI_ISL_2781045, EPI_ISL_2781046, EPI_ISL_2781047, EPI_ISL_2781048, EPI_ISL_2781049, EPI_ISL_2781050, EPI_ISL_2781051, EPI_ISL_2781052, EPI_ISL_2781053, EPI_ISL_2781054 | | | |
| see above | Lighthouse Lab in Milton Keynes | Wellcome Sanger Institute for the COVID-19 Genomics UK (COG-UK) Consortium | The Lighthouse Lab in Milton Keynes and Alex Alderton, Roberto Amato, Jeffrey Barrett, Sonia Goncalves, Ewan Harrison, David K. Jackson, Ian Johnston, Dominic Kwiatkowski, Cordelia Langford, John Sillitoe on behalf of the Wellcome Sanger Institute COVID-19 Surveillance Team |
| EPI_ISL_2781055 | Lighthouse Lab in Alderley Park | Wellcome Sanger Institute for the COVID-19 Genomics UK (COG-UK) Consortium | Jacquelyn Wynn, Mairead Hyland, The Lighthouse Lab in Alderley Park and Alex Alderton, Roberto Amato, Jeffrey Barrett, Sonia Goncalves, Ewan Harrison, David K. Jackson, Ian Johnston, Dominic Kwiatkowski, Cordelia Langford, John Sillitoe on behalf of the Wellcome Sanger Institute COVID-19 Surveillance Team |
| EPI_ISL_2781056 | Lighthouse Lab in Milton Keynes | Wellcome Sanger Institute for the COVID-19 Genomics UK (COG-UK) Consortium | The Lighthouse Lab in Milton Keynes and Alex Alderton, Roberto Amato, Jeffrey Barrett, Sonia Goncalves, Ewan Harrison, David K. Jackson, Ian Johnston, Dominic Kwiatkowski, Cordelia Langford, John Sillitoe on behalf of the Wellcome Sanger Institute COVID-19 Surveillance Team |
| EPI_ISL_2781057 | Lighthouse Lab in Alderley Park | Wellcome Sanger Institute for the COVID-19 Genomics UK (COG-UK) Consortium | Jacquelyn Wynn, Mairead Hyland, The Lighthouse Lab in Alderley Park and Alex Alderton, Roberto Amato, Jeffrey Barrett, Sonia Goncalves, Ewan Harrison, David K. Jackson, Ian Johnston, Dominic Kwiatkowski, Cordelia Langford, John Sillitoe on behalf of the Wellcome Sanger Institute COVID-19 Surveillance Team |
| EPI_ISL_2781058 | Lighthouse Lab in Milton Keynes | Wellcome Sanger Institute for the COVID-19 Genomics UK (COG-UK) Consortium | The Lighthouse Lab in Milton Keynes and Alex Alderton, Roberto Amato, Jeffrey Barrett, Sonia Goncalves, Ewan Harrison, David K. Jackson, Ian Johnston, Dominic Kwiatkowski, Cordelia Langford, John Sillitoe on behalf of the Wellcome Sanger Institute COVID-19 Surveillance Team |
| EPI_ISL_2781059 | Lighthouse Lab in Alderley Park | Wellcome Sanger Institute for the COVID-19 Genomics UK (COG-UK) Consortium | Jacquelyn Wynn, Mairead Hyland, The Lighthouse Lab in Alderley Park and Alex Alderton, Roberto Amato, Jeffrey Barrett, Sonia Goncalves, Ewan Harrison, David K. Jackson, Ian Johnston, Dominic Kwiatkowski, Cordelia Langford, John Sillitoe on behalf of the Wellcome Sanger Institute COVID-19 Surveillance Team |
| EPI_ISL_2781060, EPI_ISL_2781061, EPI_ISL_2781062, EPI_ISL_2781063, EPI_ISL_2781064 | Lighthouse Lab in Milton Keynes | Wellcome Sanger Institute for the COVID-19 Genomics UK (COG-UK) Consortium | The Lighthouse Lab in Milton Keynes and Alex Alderton, Roberto Amato, Jeffrey Barrett, Sonia Goncalves, Ewan Harrison, David K. Jackson, Ian Johnston, Dominic Kwiatkowski, Cordelia Langford, John Sillitoe on behalf of the Wellcome Sanger Institute COVID-19 Surveillance Team |
| EPI_ISL_2781065 | Lighthouse Laboratory Plymouth | Wellcome Sanger Institute for the COVID-19 Genomics UK (COG-UK) Consortium | Lighthouse Laboratory Plymouth and Alex Alderton, Roberto Amato, Jeffrey Barrett, Sonia Goncalves, Ewan Harrison, David K. Jackson, Ian Johnston, Dominic Kwiatkowski, Cordelia Langford, John Sillitoe on behalf of the Wellcome Sanger Institute COVID-19 Surveillance Team |
| EPI_ISL_2781066 | Lighthouse Lab in Milton Keynes | Wellcome Sanger Institute for the COVID-19 Genomics UK (COG-UK) Consortium | The Lighthouse Lab in Milton Keynes and Alex Alderton, Roberto Amato, Jeffrey Barrett, Sonia Goncalves, Ewan Harrison, David K. Jackson, Ian Johnston, Dominic Kwiatkowski, Cordelia Langford, John Sillitoe on behalf of the Wellcome Sanger Institute COVID-19 Surveillance Team |
| EPI_ISL_2781067 | Lighthouse Lab in Alderley Park | Wellcome Sanger Institute for the COVID-19 Genomics UK (COG-UK) Consortium | Jacquelyn Wynn, Mairead Hyland, The Lighthouse Lab in Alderley Park and Alex Alderton, Roberto Amato, Jeffrey Barrett, Sonia Goncalves, Ewan Harrison, David K. Jackson, Ian Johnston, Dominic Kwiatkowski, Cordelia Langford, John Sillitoe on behalf of the Wellcome Sanger Institute COVID-19 Surveillance Team |
| EPI_ISL_2781068, EPI_ISL_2781069, EPI_ISL_2781070, EPI_ISL_2781071, EPI_ISL_2781072, EPI_ISL_2781073, EPI_ISL_2781074, EPI_ISL_2781075, EPI_ISL_2781076, EPI_ISL_2781077, EPI_ISL_2781078, EPI_ISL_2781079 | | | |
| see above | Lighthouse Lab in Milton Keynes | Wellcome Sanger Institute for the COVID-19 Genomics UK (COG-UK) Consortium | The Lighthouse Lab in Milton Keynes and Alex Alderton, Roberto Amato, Jeffrey Barrett, Sonia Goncalves, Ewan Harrison, David K. Jackson, Ian Johnston, Dominic Kwiatkowski, Cordelia Langford, John Sillitoe on behalf of the Wellcome Sanger Institute COVID-19 Surveillance Team |
| EPI_ISL_2781080 | Lighthouse Lab in Alderley Park | Wellcome Sanger Institute for the COVID-19 Genomics UK (COG-UK) Consortium | Jacquelyn Wynn, Mairead Hyland, The Lighthouse Lab in Alderley Park and Alex Alderton, Roberto Amato, Jeffrey Barrett, Sonia Goncalves, Ewan Harrison, David K. Jackson, Ian Johnston, Dominic Kwiatkowski, Cordelia Langford, John Sillitoe on behalf of the Wellcome Sanger Institute COVID-19 Surveillance Team |
| EPI_ISL_2781081 | Lighthouse Lab in Milton Keynes | Wellcome Sanger Institute for the COVID-19 Genomics UK (COG-UK) Consortium | The Lighthouse Lab in Milton Keynes and Alex Alderton, Roberto Amato, Jeffrey Barrett, Sonia Goncalves, Ewan Harrison, David K. Jackson, Ian Johnston, Dominic Kwiatkowski, Cordelia Langford, John Sillitoe on behalf of the Wellcome Sanger Institute COVID-19 Surveillance Team |
| EPI_ISL_2781082, EPI_ISL_2781083 | Lighthouse Lab in Alderley Park | Wellcome Sanger Institute for the COVID-19 Genomics UK (COG-UK) Consortium | Jacquelyn Wynn, Mairead Hyland, The Lighthouse Lab in Alderley Park and Alex Alderton, Roberto Amato, Jeffrey Barrett, Sonia Goncalves, Ewan Harrison, David K. Jackson, Ian Johnston, Dominic Kwiatkowski, Cordelia Langford, John Sillitoe on behalf of the Wellcome Sanger Institute COVID-19 Surveillance Team |
| EPI_ISL_2781084, EPI_ISL_2781085, EPI_ISL_2781086, EPI_ISL_2781087, EPI_ISL_2781088, EPI_ISL_2781089, EPI_ISL_2781090 | Lighthouse Lab in Milton Keynes | Wellcome Sanger Institute for the COVID-19 Genomics UK (COG-UK) Consortium | The Lighthouse Lab in Milton Keynes and Alex Alderton, Roberto Amato, Jeffrey Barrett, Sonia Goncalves, Ewan Harrison, David K. Jackson, Ian Johnston, Dominic Kwiatkowski, Cordelia Langford, John Sillitoe on behalf of the Wellcome Sanger Institute COVID-19 Surveillance Team |

| | | | |
|---|---|---|---|
| EPI_ISL_2781091, EPI_ISL_2781092 | Lighthouse Lab in Alderley Park | Wellcome Sanger Institute for the COVID-19 Genomics UK (COG-UK) Consortium | Jacquelyn Wynn, Mairead Hyland, The Lighthouse Lab in Alderley Park and Alex Alderton, Roberto Amato, Jeffrey Barrett, Sonia Goncalves, Ewan Harrison, David K. Jackson, Ian Johnston, Dominic Kwiatkowski, Cordelia Langford, John Sillitoe on behalf of the Wellcome Sanger Institute COVID-19 Surveillance Team |
| EPI_ISL_2781093, EPI_ISL_2781094, EPI_ISL_2781095 | Lighthouse Lab in Milton Keynes | Wellcome Sanger Institute for the COVID-19 Genomics UK (COG-UK) Consortium | The Lighthouse Lab in Milton Keynes and Alex Alderton, Roberto Amato, Jeffrey Barrett, Sonia Goncalves, Ewan Harrison, David K. Jackson, Ian Johnston, Dominic Kwiatkowski, Cordelia Langford, John Sillitoe on behalf of the Wellcome Sanger Institute COVID-19 Surveillance Team |
| EPI_ISL_2781096 | Lighthouse Lab in Alderley Park | Wellcome Sanger Institute for the COVID-19 Genomics UK (COG-UK) Consortium | Jacquelyn Wynn, Mairead Hyland, The Lighthouse Lab in Alderley Park and Alex Alderton, Roberto Amato, Jeffrey Barrett, Sonia Goncalves, Ewan Harrison, David K. Jackson, Ian Johnston, Dominic Kwiatkowski, Cordelia Langford, John Sillitoe on behalf of the Wellcome Sanger Institute COVID-19 Surveillance Team |
| EPI_ISL_2781097, EPI_ISL_2781098, EPI_ISL_2781099, EPI_ISL_2781100, EPI_ISL_2781101 | Lighthouse Lab in Milton Keynes | Wellcome Sanger Institute for the COVID-19 Genomics UK (COG-UK) Consortium | The Lighthouse Lab in Milton Keynes and Alex Alderton, Roberto Amato, Jeffrey Barrett, Sonia Goncalves, Ewan Harrison, David K. Jackson, Ian Johnston, Dominic Kwiatkowski, Cordelia Langford, John Sillitoe on behalf of the Wellcome Sanger Institute COVID-19 Surveillance Team |
| EPI_ISL_2781102, EPI_ISL_2781103 | Lighthouse Lab in Alderley Park | Wellcome Sanger Institute for the COVID-19 Genomics UK (COG-UK) Consortium | Jacquelyn Wynn, Mairead Hyland, The Lighthouse Lab in Alderley Park and Alex Alderton, Roberto Amato, Jeffrey Barrett, Sonia Goncalves, Ewan Harrison, David K. Jackson, Ian Johnston, Dominic Kwiatkowski, Cordelia Langford, John Sillitoe on behalf of the Wellcome Sanger Institute COVID-19 Surveillance Team |
| EPI_ISL_2781104, EPI_ISL_2781105, EPI_ISL_2781106, EPI_ISL_2781107, EPI_ISL_2781108, EPI_ISL_2781109, EPI_ISL_2781110, EPI_ISL_2781111, EPI_ISL_2781112, EPI_ISL_2781113, EPI_ISL_2781114, EPI_ISL_2781115 | | | |
| see above | Lighthouse Lab in Milton Keynes | Wellcome Sanger Institute for the COVID-19 Genomics UK (COG-UK) Consortium | The Lighthouse Lab in Milton Keynes and Alex Alderton, Roberto Amato, Jeffrey Barrett, Sonia Goncalves, Ewan Harrison, David K. Jackson, Ian Johnston, Dominic Kwiatkowski, Cordelia Langford, John Sillitoe on behalf of the Wellcome Sanger Institute COVID-19 Surveillance Team |
| EPI_ISL_2781116, EPI_ISL_2781117 | Lighthouse Lab in Alderley Park | Wellcome Sanger Institute for the COVID-19 Genomics UK (COG-UK) Consortium | Jacquelyn Wynn, Mairead Hyland, The Lighthouse Lab in Alderley Park and Alex Alderton, Roberto Amato, Jeffrey Barrett, Sonia Goncalves, Ewan Harrison, David K. Jackson, Ian Johnston, Dominic Kwiatkowski, Cordelia Langford, John Sillitoe on behalf of the Wellcome Sanger Institute COVID-19 Surveillance Team |
| EPI_ISL_2781118 | Lighthouse Lab in Milton Keynes | Wellcome Sanger Institute for the COVID-19 Genomics UK (COG-UK) Consortium | The Lighthouse Lab in Milton Keynes and Alex Alderton, Roberto Amato, Jeffrey Barrett, Sonia Goncalves, Ewan Harrison, David K. Jackson, Ian Johnston, Dominic Kwiatkowski, Cordelia Langford, John Sillitoe on behalf of the Wellcome Sanger Institute COVID-19 Surveillance Team |
| EPI_ISL_2781119 | Lighthouse Lab in Alderley Park | Wellcome Sanger Institute for the COVID-19 Genomics UK (COG-UK) Consortium | Jacquelyn Wynn, Mairead Hyland, The Lighthouse Lab in Alderley Park and Alex Alderton, Roberto Amato, Jeffrey Barrett, Sonia Goncalves, Ewan Harrison, David K. Jackson, Ian Johnston, Dominic Kwiatkowski, Cordelia Langford, John Sillitoe on behalf of the Wellcome Sanger Institute COVID-19 Surveillance Team |
| EPI_ISL_2781120, EPI_ISL_2781121, EPI_ISL_2781122, EPI_ISL_2781123, EPI_ISL_2781124, EPI_ISL_2781125 | Lighthouse Lab in Milton Keynes | Wellcome Sanger Institute for the COVID-19 Genomics UK (COG-UK) Consortium | The Lighthouse Lab in Milton Keynes and Alex Alderton, Roberto Amato, Jeffrey Barrett, Sonia Goncalves, Ewan Harrison, David K. Jackson, Ian Johnston, Dominic Kwiatkowski, Cordelia Langford, John Sillitoe on behalf of the Wellcome Sanger Institute COVID-19 Surveillance Team |
| EPI_ISL_2781126 | Lighthouse Lab in Alderley Park | Wellcome Sanger Institute for the COVID-19 Genomics UK (COG-UK) Consortium | Jacquelyn Wynn, Mairead Hyland, The Lighthouse Lab in Alderley Park and Alex Alderton, Roberto Amato, Jeffrey Barrett, Sonia Goncalves, Ewan Harrison, David K. Jackson, Ian Johnston, Dominic Kwiatkowski, Cordelia Langford, John Sillitoe on behalf of the Wellcome Sanger Institute COVID-19 Surveillance Team |
| EPI_ISL_2781127, EPI_ISL_2781128, EPI_ISL_2781129 | Lighthouse Lab in Milton Keynes | Wellcome Sanger Institute for the COVID-19 Genomics UK (COG-UK) Consortium | The Lighthouse Lab in Milton Keynes and Alex Alderton, Roberto Amato, Jeffrey Barrett, Sonia Goncalves, Ewan Harrison, David K. Jackson, Ian Johnston, Dominic Kwiatkowski, Cordelia Langford, John Sillitoe on behalf of the Wellcome Sanger Institute COVID-19 Surveillance Team |
| EPI_ISL_2781130 | Lighthouse Lab in Alderley Park | Wellcome Sanger Institute for the COVID-19 Genomics UK (COG-UK) Consortium | Jacquelyn Wynn, Mairead Hyland, The Lighthouse Lab in Alderley Park and Alex Alderton, Roberto Amato, Jeffrey Barrett, Sonia Goncalves, Ewan Harrison, David K. Jackson, Ian Johnston, Dominic Kwiatkowski, Cordelia Langford, John Sillitoe on behalf of the Wellcome Sanger Institute COVID-19 Surveillance Team |
| EPI_ISL_2781131, EPI_ISL_2781132, EPI_ISL_2781133, EPI_ISL_2781134 | Lighthouse Lab in Milton Keynes | Wellcome Sanger Institute for the COVID-19 Genomics UK (COG-UK) Consortium | The Lighthouse Lab in Milton Keynes and Alex Alderton, Roberto Amato, Jeffrey Barrett, Sonia Goncalves, Ewan Harrison, David K. Jackson, Ian Johnston, Dominic Kwiatkowski, Cordelia Langford, John Sillitoe on behalf of the Wellcome Sanger Institute COVID-19 Surveillance Team |
| EPI_ISL_2781135 | Lighthouse Lab in Alderley Park | Wellcome Sanger Institute for the COVID-19 Genomics UK (COG-UK) Consortium | Jacquelyn Wynn, Mairead Hyland, The Lighthouse Lab in Alderley Park and Alex Alderton, Roberto Amato, Jeffrey Barrett, Sonia Goncalves, Ewan Harrison, David K. Jackson, Ian Johnston, Dominic Kwiatkowski, Cordelia Langford, John Sillitoe on behalf of the Wellcome Sanger Institute COVID-19 Surveillance Team |
| EPI_ISL_2781136, EPI_ISL_2781137, EPI_ISL_2781138, EPI_ISL_2781139, EPI_ISL_2781140, EPI_ISL_2781141, EPI_ISL_2781142, EPI_ISL_2781143, EPI_ISL_2781144, EPI_ISL_2781145, EPI_ISL_2781146, EPI_ISL_2781147, EPI_ISL_2781148, EPI_ISL_2781149, EPI_ISL_2781150, EPI_ISL_2781151, EPI_ISL_2781152, EPI_ISL_2781153 | | | |
| see above | Lighthouse Lab in Milton Keynes | Wellcome Sanger Institute for the COVID-19 Genomics UK (COG-UK) Consortium | The Lighthouse Lab in Milton Keynes and Alex Alderton, Roberto Amato, Jeffrey Barrett, Sonia Goncalves, Ewan Harrison, David K. Jackson, Ian Johnston, Dominic Kwiatkowski, Cordelia Langford, John Sillitoe on behalf of the Wellcome Sanger Institute COVID-19 Surveillance Team |
| EPI_ISL_2781154, EPI_ISL_2781155 | Lighthouse Lab in Alderley Park | Wellcome Sanger Institute for the COVID-19 Genomics UK (COG-UK) Consortium | Jacquelyn Wynn, Mairead Hyland, The Lighthouse Lab in Alderley Park and Alex Alderton, Roberto Amato, Jeffrey Barrett, Sonia Goncalves, Ewan Harrison, David K. Jackson, Ian Johnston, Dominic Kwiatkowski, Cordelia Langford, John Sillitoe on behalf of the Wellcome Sanger Institute COVID-19 Surveillance Team |
| EPI_ISL_2781156, EPI_ISL_2781157, EPI_ISL_2781158, EPI_ISL_2781159, EPI_ISL_2781160 | Lighthouse Lab in Milton Keynes | Wellcome Sanger Institute for the COVID-19 Genomics UK (COG-UK) Consortium | The Lighthouse Lab in Milton Keynes and Alex Alderton, Roberto Amato, Jeffrey Barrett, Sonia Goncalves, Ewan Harrison, David K. Jackson, Ian Johnston, Dominic Kwiatkowski, Cordelia Langford, John Sillitoe on behalf of the Wellcome Sanger Institute COVID-19 Surveillance Team |
| EPI_ISL_2781161, EPI_ISL_2781162 | Lighthouse Lab in Alderley Park | Wellcome Sanger Institute for the COVID-19 Genomics UK (COG-UK) Consortium | Jacquelyn Wynn, Mairead Hyland, The Lighthouse Lab in Alderley Park and Alex Alderton, Roberto Amato, Jeffrey Barrett, Sonia Goncalves, Ewan Harrison, David K. Jackson, Ian Johnston, Dominic Kwiatkowski, Cordelia Langford, John Sillitoe on behalf of the Wellcome Sanger Institute COVID-19 Surveillance Team |
| EPI_ISL_2781163 | Lighthouse Lab in Milton Keynes | Wellcome Sanger Institute for the COVID-19 Genomics UK (COG-UK) Consortium | The Lighthouse Lab in Milton Keynes and Alex Alderton, Roberto Amato, Jeffrey Barrett, Sonia Goncalves, Ewan Harrison, David K. Jackson, Ian Johnston, Dominic Kwiatkowski, Cordelia Langford, John Sillitoe on behalf of the Wellcome Sanger Institute COVID-19 Surveillance Team |
| EPI_ISL_2781164 | Lighthouse Lab in Alderley Park | Wellcome Sanger Institute for the COVID-19 Genomics UK (COG-UK) Consortium | Jacquelyn Wynn, Mairead Hyland, The Lighthouse Lab in Alderley Park and Alex Alderton, Roberto Amato, Jeffrey Barrett, Sonia Goncalves, Ewan Harrison, David K. Jackson, Ian Johnston, Dominic Kwiatkowski, Cordelia Langford, John Sillitoe on behalf of the Wellcome Sanger Institute COVID-19 Surveillance Team |
| EPI_ISL_2781165, EPI_ISL_2781166, EPI_ISL_2781167, EPI_ISL_2781168, EPI_ISL_2781169, EPI_ISL_2781170, EPI_ISL_2781171 | Lighthouse Lab in Milton Keynes | Wellcome Sanger Institute for the COVID-19 Genomics UK (COG-UK) Consortium | The Lighthouse Lab in Milton Keynes and Alex Alderton, Roberto Amato, Jeffrey Barrett, Sonia Goncalves, Ewan Harrison, David K. Jackson, Ian Johnston, Dominic Kwiatkowski, Cordelia Langford, John Sillitoe on behalf of the Wellcome Sanger Institute COVID-19 Surveillance Team |
| EPI_ISL_2781172 | Lighthouse Lab in Alderley Park | Wellcome Sanger Institute for the COVID-19 Genomics UK (COG-UK) Consortium | Jacquelyn Wynn, Mairead Hyland, The Lighthouse Lab in Alderley Park and Alex Alderton, Roberto Amato, Jeffrey Barrett, Sonia Goncalves, Ewan Harrison, David K. Jackson, Ian Johnston, Dominic Kwiatkowski, Cordelia Langford, John Sillitoe on behalf of the Wellcome Sanger Institute COVID-19 Surveillance Team |
| EPI_ISL_2781173, EPI_ISL_2781174, EPI_ISL_2781175, EPI_ISL_2781176, EPI_ISL_2781177, EPI_ISL_2781178 | Lighthouse Lab in Milton Keynes | Wellcome Sanger Institute for the COVID-19 Genomics UK (COG-UK) Consortium | The Lighthouse Lab in Milton Keynes and Alex Alderton, Roberto Amato, Jeffrey Barrett, Sonia Goncalves, Ewan Harrison, David K. Jackson, Ian Johnston, Dominic Kwiatkowski, Cordelia Langford, John Sillitoe on behalf of the Wellcome Sanger Institute COVID-19 Surveillance Team |
| EPI_ISL_2781179 | Lighthouse Lab in Alderley Park | Wellcome Sanger Institute for the COVID-19 Genomics UK (COG-UK) Consortium | Jacquelyn Wynn, Mairead Hyland, The Lighthouse Lab in Alderley Park and Alex Alderton, Roberto Amato, Jeffrey Barrett, Sonia Goncalves, Ewan Harrison, David K. Jackson, Ian Johnston, Dominic Kwiatkowski, Cordelia Langford, John Sillitoe on behalf of the Wellcome Sanger Institute COVID-19 Surveillance Team |
| EPI_ISL_2781180 | Lighthouse Lab in Milton Keynes | Wellcome Sanger Institute for the COVID-19 Genomics UK (COG-UK) Consortium | The Lighthouse Lab in Milton Keynes and Alex Alderton, Roberto Amato, Jeffrey Barrett, Sonia Goncalves, Ewan Harrison, David K. Jackson, Ian Johnston, Dominic Kwiatkowski, Cordelia Langford, John Sillitoe on behalf of the Wellcome Sanger Institute COVID-19 Surveillance Team |
| EPI_ISL_2781181 | Lighthouse Lab in Alderley Park | Wellcome Sanger Institute for the COVID-19 Genomics UK (COG-UK) Consortium | Jacquelyn Wynn, Mairead Hyland, The Lighthouse Lab in Alderley Park and Alex Alderton, Roberto Amato, Jeffrey Barrett, Sonia Goncalves, Ewan Harrison, David K. Jackson, Ian Johnston, Dominic Kwiatkowski, Cordelia Langford, John Sillitoe on behalf of the Wellcome Sanger Institute COVID-19 Surveillance Team |
| EPI_ISL_2781182, EPI_ISL_2781183 | Lighthouse Lab in Milton Keynes | Wellcome Sanger Institute for the COVID-19 Genomics UK (COG-UK) Consortium | The Lighthouse Lab in Milton Keynes and Alex Alderton, Roberto Amato, Jeffrey Barrett, Sonia Goncalves, Ewan Harrison, David K. Jackson, Ian Johnston, Dominic Kwiatkowski, Cordelia Langford, John Sillitoe on behalf of the Wellcome Sanger Institute COVID-19 Surveillance Team |
| EPI_ISL_2781184 | Lighthouse Lab in Alderley Park | Wellcome Sanger Institute for the COVID-19 Genomics UK (COG-UK) Consortium | Jacquelyn Wynn, Mairead Hyland, The Lighthouse Lab in Alderley Park and Alex Alderton, Roberto Amato, Jeffrey Barrett, Sonia Goncalves, Ewan Harrison, David K. Jackson, Ian Johnston, Dominic Kwiatkowski, Cordelia Langford, John Sillitoe on behalf of the Wellcome Sanger Institute COVID-19 Surveillance Team |
| EPI_ISL_2781185 | Lighthouse Lab in Milton Keynes | Wellcome Sanger Institute for the COVID-19 Genomics UK (COG-UK) Consortium | The Lighthouse Lab in Milton Keynes and Alex Alderton, Roberto Amato, Jeffrey Barrett, Sonia Goncalves, Ewan Harrison, David K. Jackson, Ian Johnston, Dominic Kwiatkowski, Cordelia Langford, John Sillitoe on behalf of the Wellcome Sanger Institute COVID-19 Surveillance Team |
| EPI_ISL_2781186 | Lighthouse Lab in Alderley Park | Wellcome Sanger Institute for the COVID-19 Genomics UK (COG-UK) Consortium | Jacquelyn Wynn, Mairead Hyland, The Lighthouse Lab in Alderley Park and Alex Alderton, Roberto Amato, Jeffrey Barrett, Sonia Goncalves, Ewan Harrison, David K. Jackson, Ian Johnston, Dominic Kwiatkowski, Cordelia Langford, John Sillitoe on behalf of the Wellcome Sanger Institute COVID-19 Surveillance Team |
| EPI_ISL_2781187, EPI_ISL_2781188, EPI_ISL_2781189, EPI_ISL_2781190 | Lighthouse Lab in Milton Keynes | Wellcome Sanger Institute for the COVID-19 Genomics UK (COG-UK) Consortium | The Lighthouse Lab in Milton Keynes and Alex Alderton, Roberto Amato, Jeffrey Barrett, Sonia Goncalves, Ewan Harrison, David K. Jackson, Ian Johnston, Dominic Kwiatkowski, Cordelia Langford, John Sillitoe on behalf of the Wellcome Sanger Institute COVID-19 Surveillance Team |
| EPI_ISL_2781191 | Lighthouse Lab in Alderley Park | Wellcome Sanger Institute for the COVID-19 Genomics UK (COG-UK) Consortium | Jacquelyn Wynn, Mairead Hyland, The Lighthouse Lab in Alderley Park and Alex Alderton, Roberto Amato, Jeffrey Barrett, Sonia Goncalves, Ewan Harrison, David K. Jackson, Ian Johnston, Dominic Kwiatkowski, Cordelia Langford, John Sillitoe on behalf of the Wellcome Sanger Institute COVID-19 Surveillance Team |
| EPI_ISL_2781192 | Lighthouse Lab in Milton Keynes | Wellcome Sanger Institute for the COVID-19 Genomics UK (COG-UK) Consortium | The Lighthouse Lab in Milton Keynes and Alex Alderton, Roberto Amato, Jeffrey Barrett, Sonia Goncalves, Ewan Harrison, David K. Jackson, Ian Johnston, Dominic Kwiatkowski, Cordelia Langford, John Sillitoe on behalf of the Wellcome Sanger Institute COVID-19 Surveillance Team |

| | | | |
|---|---|---|---|
| EPI_ISL_2781193 | Lighthouse Lab in Alderley Park | Wellcome Sanger Institute for the COVID-19 Genomics UK (COG-UK) Consortium | Jacquelyn Wynn, Mairead Hyland, The Lighthouse Lab in Alderley Park and Alex Alderton, Roberto Amato, Jeffrey Barrett, Sonia Goncalves, Ewan Harrison, David K. Jackson, Ian Johnston, Dominic Kwiatkowski, Cordelia Langford, John Sillitoe on behalf of the Wellcome Sanger Institute COVID-19 Surveillance Team |
| EPI_ISL_2781194, EPI_ISL_2781195, EPI_ISL_2781196, EPI_ISL_2781197, EPI_ISL_2781198, EPI_ISL_2781199, EPI_ISL_2781200 | Lighthouse Lab in Milton Keynes | Wellcome Sanger Institute for the COVID-19 Genomics UK (COG-UK) Consortium | The Lighthouse Lab in Milton Keynes and Alex Alderton, Roberto Amato, Jeffrey Barrett, Sonia Goncalves, Ewan Harrison, David K. Jackson, Ian Johnston, Dominic Kwiatkowski, Cordelia Langford, John Sillitoe on behalf of the Wellcome Sanger Institute COVID-19 Surveillance Team |
| EPI_ISL_2781201 | Lighthouse Lab in Alderley Park | Wellcome Sanger Institute for the COVID-19 Genomics UK (COG-UK) Consortium | Jacquelyn Wynn, Mairead Hyland, The Lighthouse Lab in Alderley Park and Alex Alderton, Roberto Amato, Jeffrey Barrett, Sonia Goncalves, Ewan Harrison, David K. Jackson, Ian Johnston, Dominic Kwiatkowski, Cordelia Langford, John Sillitoe on behalf of the Wellcome Sanger Institute COVID-19 Surveillance Team |
| EPI_ISL_2781202, EPI_ISL_2781203, EPI_ISL_2781204, EPI_ISL_2781205, EPI_ISL_2781206, EPI_ISL_2781207, EPI_ISL_2781208, EPI_ISL_2781209 | Lighthouse Lab in Milton Keynes | Wellcome Sanger Institute for the COVID-19 Genomics UK (COG-UK) Consortium | The Lighthouse Lab in Milton Keynes and Alex Alderton, Roberto Amato, Jeffrey Barrett, Sonia Goncalves, Ewan Harrison, David K. Jackson, Ian Johnston, Dominic Kwiatkowski, Cordelia Langford, John Sillitoe on behalf of the Wellcome Sanger Institute COVID-19 Surveillance Team |
| EPI_ISL_2781210, EPI_ISL_2781211 | Lighthouse Lab in Alderley Park | Wellcome Sanger Institute for the COVID-19 Genomics UK (COG-UK) Consortium | Jacquelyn Wynn, Mairead Hyland, The Lighthouse Lab in Alderley Park and Alex Alderton, Roberto Amato, Jeffrey Barrett, Sonia Goncalves, Ewan Harrison, David K. Jackson, Ian Johnston, Dominic Kwiatkowski, Cordelia Langford, John Sillitoe on behalf of the Wellcome Sanger Institute COVID-19 Surveillance Team |
| EPI_ISL_2781212, EPI_ISL_2781213, EPI_ISL_2781214, EPI_ISL_2781215, EPI_ISL_2781216, EPI_ISL_2781217 | Lighthouse Lab in Milton Keynes | Wellcome Sanger Institute for the COVID-19 Genomics UK (COG-UK) Consortium | The Lighthouse Lab in Milton Keynes and Alex Alderton, Roberto Amato, Jeffrey Barrett, Sonia Goncalves, Ewan Harrison, David K. Jackson, Ian Johnston, Dominic Kwiatkowski, Cordelia Langford, John Sillitoe on behalf of the Wellcome Sanger Institute COVID-19 Surveillance Team |
| EPI_ISL_2781218 | Lighthouse Lab in Alderley Park | Wellcome Sanger Institute for the COVID-19 Genomics UK (COG-UK) Consortium | Jacquelyn Wynn, Mairead Hyland, The Lighthouse Lab in Alderley Park and Alex Alderton, Roberto Amato, Jeffrey Barrett, Sonia Goncalves, Ewan Harrison, David K. Jackson, Ian Johnston, Dominic Kwiatkowski, Cordelia Langford, John Sillitoe on behalf of the Wellcome Sanger Institute COVID-19 Surveillance Team |
| EPI_ISL_2781219, EPI_ISL_2781220, EPI_ISL_2781221 | Lighthouse Lab in Milton Keynes | Wellcome Sanger Institute for the COVID-19 Genomics UK (COG-UK) Consortium | The Lighthouse Lab in Milton Keynes and Alex Alderton, Roberto Amato, Jeffrey Barrett, Sonia Goncalves, Ewan Harrison, David K. Jackson, Ian Johnston, Dominic Kwiatkowski, Cordelia Langford, John Sillitoe on behalf of the Wellcome Sanger Institute COVID-19 Surveillance Team |
| EPI_ISL_2781222 | Lighthouse Lab in Alderley Park | Wellcome Sanger Institute for the COVID-19 Genomics UK (COG-UK) Consortium | Jacquelyn Wynn, Mairead Hyland, The Lighthouse Lab in Alderley Park and Alex Alderton, Roberto Amato, Jeffrey Barrett, Sonia Goncalves, Ewan Harrison, David K. Jackson, Ian Johnston, Dominic Kwiatkowski, Cordelia Langford, John Sillitoe on behalf of the Wellcome Sanger Institute COVID-19 Surveillance Team |
| EPI_ISL_2781223, EPI_ISL_2781224, EPI_ISL_2781225, EPI_ISL_2781226 | Lighthouse Lab in Milton Keynes | Wellcome Sanger Institute for the COVID-19 Genomics UK (COG-UK) Consortium | The Lighthouse Lab in Milton Keynes and Alex Alderton, Roberto Amato, Jeffrey Barrett, Sonia Goncalves, Ewan Harrison, David K. Jackson, Ian Johnston, Dominic Kwiatkowski, Cordelia Langford, John Sillitoe on behalf of the Wellcome Sanger Institute COVID-19 Surveillance Team |
| EPI_ISL_2781227 | Lighthouse Lab in Alderley Park | Wellcome Sanger Institute for the COVID-19 Genomics UK (COG-UK) Consortium | Jacquelyn Wynn, Mairead Hyland, The Lighthouse Lab in Alderley Park and Alex Alderton, Roberto Amato, Jeffrey Barrett, Sonia Goncalves, Ewan Harrison, David K. Jackson, Ian Johnston, Dominic Kwiatkowski, Cordelia Langford, John Sillitoe on behalf of the Wellcome Sanger Institute COVID-19 Surveillance Team |
| EPI_ISL_2781228 | Lighthouse Lab in Milton Keynes | Wellcome Sanger Institute for the COVID-19 Genomics UK (COG-UK) Consortium | The Lighthouse Lab in Milton Keynes and Alex Alderton, Roberto Amato, Jeffrey Barrett, Sonia Goncalves, Ewan Harrison, David K. Jackson, Ian Johnston, Dominic Kwiatkowski, Cordelia Langford, John Sillitoe on behalf of the Wellcome Sanger Institute COVID-19 Surveillance Team |
| EPI_ISL_2781229 | Lighthouse Lab in Alderley Park | Wellcome Sanger Institute for the COVID-19 Genomics UK (COG-UK) Consortium | Jacquelyn Wynn, Mairead Hyland, The Lighthouse Lab in Alderley Park and Alex Alderton, Roberto Amato, Jeffrey Barrett, Sonia Goncalves, Ewan Harrison, David K. Jackson, Ian Johnston, Dominic Kwiatkowski, Cordelia Langford, John Sillitoe on behalf of the Wellcome Sanger Institute COVID-19 Surveillance Team |
| EPI_ISL_2781230 | Lighthouse Lab in Milton Keynes | Wellcome Sanger Institute for the COVID-19 Genomics UK (COG-UK) Consortium | The Lighthouse Lab in Milton Keynes and Alex Alderton, Roberto Amato, Jeffrey Barrett, Sonia Goncalves, Ewan Harrison, David K. Jackson, Ian Johnston, Dominic Kwiatkowski, Cordelia Langford, John Sillitoe on behalf of the Wellcome Sanger Institute COVID-19 Surveillance Team |
| EPI_ISL_2781231, EPI_ISL_2781232, EPI_ISL_2781233 | Lighthouse Lab in Alderley Park | Wellcome Sanger Institute for the COVID-19 Genomics UK (COG-UK) Consortium | Jacquelyn Wynn, Mairead Hyland, The Lighthouse Lab in Alderley Park and Alex Alderton, Roberto Amato, Jeffrey Barrett, Sonia Goncalves, Ewan Harrison, David K. Jackson, Ian Johnston, Dominic Kwiatkowski, Cordelia Langford, John Sillitoe on behalf of the Wellcome Sanger Institute COVID-19 Surveillance Team |
| EPI_ISL_2781234, EPI_ISL_2781235 | Lighthouse Lab in Milton Keynes | Wellcome Sanger Institute for the COVID-19 Genomics UK (COG-UK) Consortium | The Lighthouse Lab in Milton Keynes and Alex Alderton, Roberto Amato, Jeffrey Barrett, Sonia Goncalves, Ewan Harrison, David K. Jackson, Ian Johnston, Dominic Kwiatkowski, Cordelia Langford, John Sillitoe on behalf of the Wellcome Sanger Institute COVID-19 Surveillance Team |
| EPI_ISL_2781236 | Lighthouse Lab in Alderley Park | Wellcome Sanger Institute for the COVID-19 Genomics UK (COG-UK) Consortium | Jacquelyn Wynn, Mairead Hyland, The Lighthouse Lab in Alderley Park and Alex Alderton, Roberto Amato, Jeffrey Barrett, Sonia Goncalves, Ewan Harrison, David K. Jackson, Ian Johnston, Dominic Kwiatkowski, Cordelia Langford, John Sillitoe on behalf of the Wellcome Sanger Institute COVID-19 Surveillance Team |
| EPI_ISL_2781237 | Lighthouse Lab in Milton Keynes | Wellcome Sanger Institute for the COVID-19 Genomics UK (COG-UK) Consortium | The Lighthouse Lab in Milton Keynes and Alex Alderton, Roberto Amato, Jeffrey Barrett, Sonia Goncalves, Ewan Harrison, David K. Jackson, Ian Johnston, Dominic Kwiatkowski, Cordelia Langford, John Sillitoe on behalf of the Wellcome Sanger Institute COVID-19 Surveillance Team |
| EPI_ISL_2781238 | Lighthouse Lab in Alderley Park | Wellcome Sanger Institute for the COVID-19 Genomics UK (COG-UK) Consortium | Jacquelyn Wynn, Mairead Hyland, The Lighthouse Lab in Alderley Park and Alex Alderton, Roberto Amato, Jeffrey Barrett, Sonia Goncalves, Ewan Harrison, David K. Jackson, Ian Johnston, Dominic Kwiatkowski, Cordelia Langford, John Sillitoe on behalf of the Wellcome Sanger Institute COVID-19 Surveillance Team |
| EPI_ISL_2781239, EPI_ISL_2781240, EPI_ISL_2781241, EPI_ISL_2781242, EPI_ISL_2781243, EPI_ISL_2781244, EPI_ISL_2781245, EPI_ISL_2781246, EPI_ISL_2781248 | Lighthouse Lab in Milton Keynes | Wellcome Sanger Institute for the COVID-19 Genomics UK (COG-UK) Consortium | The Lighthouse Lab in Milton Keynes and Alex Alderton, Roberto Amato, Jeffrey Barrett, Sonia Goncalves, Ewan Harrison, David K. Jackson, Ian Johnston, Dominic Kwiatkowski, Cordelia Langford, John Sillitoe on behalf of the Wellcome Sanger Institute COVID-19 Surveillance Team |
| EPI_ISL_2781249, EPI_ISL_2781250 | Lighthouse Lab in Alderley Park | Wellcome Sanger Institute for the COVID-19 Genomics UK (COG-UK) Consortium | Jacquelyn Wynn, Mairead Hyland, The Lighthouse Lab in Alderley Park and Alex Alderton, Roberto Amato, Jeffrey Barrett, Sonia Goncalves, Ewan Harrison, David K. Jackson, Ian Johnston, Dominic Kwiatkowski, Cordelia Langford, John Sillitoe on behalf of the Wellcome Sanger Institute COVID-19 Surveillance Team |
| EPI_ISL_2781251, EPI_ISL_2781252 | Lighthouse Lab in Milton Keynes | Wellcome Sanger Institute for the COVID-19 Genomics UK (COG-UK) Consortium | The Lighthouse Lab in Milton Keynes and Alex Alderton, Roberto Amato, Jeffrey Barrett, Sonia Goncalves, Ewan Harrison, David K. Jackson, Ian Johnston, Dominic Kwiatkowski, Cordelia Langford, John Sillitoe on behalf of the Wellcome Sanger Institute COVID-19 Surveillance Team |
| EPI_ISL_2781253 | Lighthouse Lab in Alderley Park | Wellcome Sanger Institute for the COVID-19 Genomics UK (COG-UK) Consortium | Jacquelyn Wynn, Mairead Hyland, The Lighthouse Lab in Alderley Park and Alex Alderton, Roberto Amato, Jeffrey Barrett, Sonia Goncalves, Ewan Harrison, David K. Jackson, Ian Johnston, Dominic Kwiatkowski, Cordelia Langford, John Sillitoe on behalf of the Wellcome Sanger Institute COVID-19 Surveillance Team |
| EPI_ISL_2781254, EPI_ISL_2781255, EPI_ISL_2781256, EPI_ISL_2781257, EPI_ISL_2781258 | Lighthouse Lab in Milton Keynes | Wellcome Sanger Institute for the COVID-19 Genomics UK (COG-UK) Consortium | The Lighthouse Lab in Milton Keynes and Alex Alderton, Roberto Amato, Jeffrey Barrett, Sonia Goncalves, Ewan Harrison, David K. Jackson, Ian Johnston, Dominic Kwiatkowski, Cordelia Langford, John Sillitoe on behalf of the Wellcome Sanger Institute COVID-19 Surveillance Team |
| EPI_ISL_2781259 | Lighthouse Lab in Alderley Park | Wellcome Sanger Institute for the COVID-19 Genomics UK (COG-UK) Consortium | Jacquelyn Wynn, Mairead Hyland, The Lighthouse Lab in Alderley Park and Alex Alderton, Roberto Amato, Jeffrey Barrett, Sonia Goncalves, Ewan Harrison, David K. Jackson, Ian Johnston, Dominic Kwiatkowski, Cordelia Langford, John Sillitoe on behalf of the Wellcome Sanger Institute COVID-19 Surveillance Team |
| EPI_ISL_2781260, EPI_ISL_2781261, EPI_ISL_2781262 | Lighthouse Lab in Milton Keynes | Wellcome Sanger Institute for the COVID-19 Genomics UK (COG-UK) Consortium | The Lighthouse Lab in Milton Keynes and Alex Alderton, Roberto Amato, Jeffrey Barrett, Sonia Goncalves, Ewan Harrison, David K. Jackson, Ian Johnston, Dominic Kwiatkowski, Cordelia Langford, John Sillitoe on behalf of the Wellcome Sanger Institute COVID-19 Surveillance Team |
| EPI_ISL_2781263 | Lighthouse Lab in Alderley Park | Wellcome Sanger Institute for the COVID-19 Genomics UK (COG-UK) Consortium | Jacquelyn Wynn, Mairead Hyland, The Lighthouse Lab in Alderley Park and Alex Alderton, Roberto Amato, Jeffrey Barrett, Sonia Goncalves, Ewan Harrison, David K. Jackson, Ian Johnston, Dominic Kwiatkowski, Cordelia Langford, John Sillitoe on behalf of the Wellcome Sanger Institute COVID-19 Surveillance Team |
| EPI_ISL_2781264, EPI_ISL_2781265, EPI_ISL_2781266, EPI_ISL_2781267 | Lighthouse Lab in Milton Keynes | Wellcome Sanger Institute for the COVID-19 Genomics UK (COG-UK) Consortium | The Lighthouse Lab in Milton Keynes and Alex Alderton, Roberto Amato, Jeffrey Barrett, Sonia Goncalves, Ewan Harrison, David K. Jackson, Ian Johnston, Dominic Kwiatkowski, Cordelia Langford, John Sillitoe on behalf of the Wellcome Sanger Institute COVID-19 Surveillance Team |
| EPI_ISL_2781268, EPI_ISL_2781269 | Lighthouse Lab in Alderley Park | Wellcome Sanger Institute for the COVID-19 Genomics UK (COG-UK) Consortium | Jacquelyn Wynn, Mairead Hyland, The Lighthouse Lab in Alderley Park and Alex Alderton, Roberto Amato, Jeffrey Barrett, Sonia Goncalves, Ewan Harrison, David K. Jackson, Ian Johnston, Dominic Kwiatkowski, Cordelia Langford, John Sillitoe on behalf of the Wellcome Sanger Institute COVID-19 Surveillance Team |
| EPI_ISL_2781270 | Lighthouse Lab in Milton Keynes | Wellcome Sanger Institute for the COVID-19 Genomics UK (COG-UK) Consortium | The Lighthouse Lab in Milton Keynes and Alex Alderton, Roberto Amato, Jeffrey Barrett, Sonia Goncalves, Ewan Harrison, David K. Jackson, Ian Johnston, Dominic Kwiatkowski, Cordelia Langford, John Sillitoe on behalf of the Wellcome Sanger Institute COVID-19 Surveillance Team |
| EPI_ISL_2781271 | Lighthouse Lab in Alderley Park | Wellcome Sanger Institute for the COVID-19 Genomics UK (COG-UK) Consortium | Jacquelyn Wynn, Mairead Hyland, The Lighthouse Lab in Alderley Park and Alex Alderton, Roberto Amato, Jeffrey Barrett, Sonia Goncalves, Ewan Harrison, David K. Jackson, Ian Johnston, Dominic Kwiatkowski, Cordelia Langford, John Sillitoe on behalf of the Wellcome Sanger Institute COVID-19 Surveillance Team |
| EPI_ISL_2781272, EPI_ISL_2781273, | Lighthouse Lab in Milton Keynes | Wellcome Sanger Institute for the COVID-19 Genomics UK (COG-UK) Consortium | The Lighthouse Lab in Milton Keynes and Alex Alderton, Roberto Amato, Jeffrey Barrett, Sonia Goncalves, Ewan Harrison, David K. Jackson, Ian Johnston, Dominic Kwiatkowski, Cordelia Langford, John Sillitoe on behalf of the Wellcome Sanger Institute COVID-19 Surveillance Team |

| | | | |
|---|---|---|---|
| EPI_ISL_2781274, EPI_ISL_2781276, EPI_ISL_2781277 | | | |
| EPI_ISL_2781278, EPI_ISL_2781279 | Lighthouse Lab in Alderley Park | Wellcome Sanger Institute for the COVID-19 Genomics UK (COG-UK) Consortium | Jacquelyn Wynn, Mairead Hyland, The Lighthouse Lab in Alderley Park and Alex Alderton, Roberto Amato, Jeffrey Barrett, Sonia Goncalves, Ewan Harrison, David K. Jackson, Ian Johnston, Dominic Kwiatkowski, Cordelia Langford, John Sillitoe on behalf of the Wellcome Sanger Institute COVID-19 Surveillance Team |
| EPI_ISL_2781280 | Lighthouse Lab in Milton Keynes | Wellcome Sanger Institute for the COVID-19 Genomics UK (COG-UK) Consortium | The Lighthouse Lab in Milton Keynes and Alex Alderton, Roberto Amato, Jeffrey Barrett, Sonia Goncalves, Ewan Harrison, David K. Jackson, Ian Johnston, Dominic Kwiatkowski, Cordelia Langford, John Sillitoe on behalf of the Wellcome Sanger Institute COVID-19 Surveillance Team |
| EPI_ISL_2781281 | Lighthouse Lab in Alderley Park | Wellcome Sanger Institute for the COVID-19 Genomics UK (COG-UK) Consortium | Jacquelyn Wynn, Mairead Hyland, The Lighthouse Lab in Alderley Park and Alex Alderton, Roberto Amato, Jeffrey Barrett, Sonia Goncalves, Ewan Harrison, David K. Jackson, Ian Johnston, Dominic Kwiatkowski, Cordelia Langford, John Sillitoe on behalf of the Wellcome Sanger Institute COVID-19 Surveillance Team |
| EPI_ISL_2781282, EPI_ISL_2781283 | Lighthouse Lab in Milton Keynes | Wellcome Sanger Institute for the COVID-19 Genomics UK (COG-UK) Consortium | The Lighthouse Lab in Milton Keynes and Alex Alderton, Roberto Amato, Jeffrey Barrett, Sonia Goncalves, Ewan Harrison, David K. Jackson, Ian Johnston, Dominic Kwiatkowski, Cordelia Langford, John Sillitoe on behalf of the Wellcome Sanger Institute COVID-19 Surveillance Team |
| EPI_ISL_2781284 | Lighthouse Lab in Alderley Park | Wellcome Sanger Institute for the COVID-19 Genomics UK (COG-UK) Consortium | Jacquelyn Wynn, Mairead Hyland, The Lighthouse Lab in Alderley Park and Alex Alderton, Roberto Amato, Jeffrey Barrett, Sonia Goncalves, Ewan Harrison, David K. Jackson, Ian Johnston, Dominic Kwiatkowski, Cordelia Langford, John Sillitoe on behalf of the Wellcome Sanger Institute COVID-19 Surveillance Team |
| EPI_ISL_2781285, EPI_ISL_2781286, EPI_ISL_2781287, EPI_ISL_2781288 | Lighthouse Lab in Milton Keynes | Wellcome Sanger Institute for the COVID-19 Genomics UK (COG-UK) Consortium | The Lighthouse Lab in Milton Keynes and Alex Alderton, Roberto Amato, Jeffrey Barrett, Sonia Goncalves, Ewan Harrison, David K. Jackson, Ian Johnston, Dominic Kwiatkowski, Cordelia Langford, John Sillitoe on behalf of the Wellcome Sanger Institute COVID-19 Surveillance Team |
| EPI_ISL_2781289 | Lighthouse Lab in Alderley Park | Wellcome Sanger Institute for the COVID-19 Genomics UK (COG-UK) Consortium | Jacquelyn Wynn, Mairead Hyland, The Lighthouse Lab in Alderley Park and Alex Alderton, Roberto Amato, Jeffrey Barrett, Sonia Goncalves, Ewan Harrison, David K. Jackson, Ian Johnston, Dominic Kwiatkowski, Cordelia Langford, John Sillitoe on behalf of the Wellcome Sanger Institute COVID-19 Surveillance Team |
| EPI_ISL_2781290, EPI_ISL_2781291, EPI_ISL_2781292 | Lighthouse Lab in Milton Keynes | Wellcome Sanger Institute for the COVID-19 Genomics UK (COG-UK) Consortium | The Lighthouse Lab in Milton Keynes and Alex Alderton, Roberto Amato, Jeffrey Barrett, Sonia Goncalves, Ewan Harrison, David K. Jackson, Ian Johnston, Dominic Kwiatkowski, Cordelia Langford, John Sillitoe on behalf of the Wellcome Sanger Institute COVID-19 Surveillance Team |
| EPI_ISL_2781293 | Lighthouse Lab in Alderley Park | Wellcome Sanger Institute for the COVID-19 Genomics UK (COG-UK) Consortium | Jacquelyn Wynn, Mairead Hyland, The Lighthouse Lab in Alderley Park and Alex Alderton, Roberto Amato, Jeffrey Barrett, Sonia Goncalves, Ewan Harrison, David K. Jackson, Ian Johnston, Dominic Kwiatkowski, Cordelia Langford, John Sillitoe on behalf of the Wellcome Sanger Institute COVID-19 Surveillance Team |
| EPI_ISL_2781294 | Lighthouse Lab in Milton Keynes | Wellcome Sanger Institute for the COVID-19 Genomics UK (COG-UK) Consortium | The Lighthouse Lab in Milton Keynes and Alex Alderton, Roberto Amato, Jeffrey Barrett, Sonia Goncalves, Ewan Harrison, David K. Jackson, Ian Johnston, Dominic Kwiatkowski, Cordelia Langford, John Sillitoe on behalf of the Wellcome Sanger Institute COVID-19 Surveillance Team |
| EPI_ISL_2781295 | Lighthouse Lab in Alderley Park | Wellcome Sanger Institute for the COVID-19 Genomics UK (COG-UK) Consortium | Jacquelyn Wynn, Mairead Hyland, The Lighthouse Lab in Alderley Park and Alex Alderton, Roberto Amato, Jeffrey Barrett, Sonia Goncalves, Ewan Harrison, David K. Jackson, Ian Johnston, Dominic Kwiatkowski, Cordelia Langford, John Sillitoe on behalf of the Wellcome Sanger Institute COVID-19 Surveillance Team |
| EPI_ISL_2781296, EPI_ISL_2781297, EPI_ISL_2781298 | Lighthouse Lab in Milton Keynes | Wellcome Sanger Institute for the COVID-19 Genomics UK (COG-UK) Consortium | The Lighthouse Lab in Milton Keynes and Alex Alderton, Roberto Amato, Jeffrey Barrett, Sonia Goncalves, Ewan Harrison, David K. Jackson, Ian Johnston, Dominic Kwiatkowski, Cordelia Langford, John Sillitoe on behalf of the Wellcome Sanger Institute COVID-19 Surveillance Team |
| EPI_ISL_2781299 | Lighthouse Lab in Alderley Park | Wellcome Sanger Institute for the COVID-19 Genomics UK (COG-UK) Consortium | Jacquelyn Wynn, Mairead Hyland, The Lighthouse Lab in Alderley Park and Alex Alderton, Roberto Amato, Jeffrey Barrett, Sonia Goncalves, Ewan Harrison, David K. Jackson, Ian Johnston, Dominic Kwiatkowski, Cordelia Langford, John Sillitoe on behalf of the Wellcome Sanger Institute COVID-19 Surveillance Team |
| EPI_ISL_2781300, EPI_ISL_2781301, EPI_ISL_2781302, EPI_ISL_2781303, EPI_ISL_2781304, EPI_ISL_2781305 | Lighthouse Lab in Milton Keynes | Wellcome Sanger Institute for the COVID-19 Genomics UK (COG-UK) Consortium | The Lighthouse Lab in Milton Keynes and Alex Alderton, Roberto Amato, Jeffrey Barrett, Sonia Goncalves, Ewan Harrison, David K. Jackson, Ian Johnston, Dominic Kwiatkowski, Cordelia Langford, John Sillitoe on behalf of the Wellcome Sanger Institute COVID-19 Surveillance Team |
| EPI_ISL_2781306 | Lighthouse Lab in Alderley Park | Wellcome Sanger Institute for the COVID-19 Genomics UK (COG-UK) Consortium | Jacquelyn Wynn, Mairead Hyland, The Lighthouse Lab in Alderley Park and Alex Alderton, Roberto Amato, Jeffrey Barrett, Sonia Goncalves, Ewan Harrison, David K. Jackson, Ian Johnston, Dominic Kwiatkowski, Cordelia Langford, John Sillitoe on behalf of the Wellcome Sanger Institute COVID-19 Surveillance Team |
| EPI_ISL_2781307, EPI_ISL_2781308, EPI_ISL_2781309, EPI_ISL_2781310 | Lighthouse Lab in Milton Keynes | Wellcome Sanger Institute for the COVID-19 Genomics UK (COG-UK) Consortium | The Lighthouse Lab in Milton Keynes and Alex Alderton, Roberto Amato, Jeffrey Barrett, Sonia Goncalves, Ewan Harrison, David K. Jackson, Ian Johnston, Dominic Kwiatkowski, Cordelia Langford, John Sillitoe on behalf of the Wellcome Sanger Institute COVID-19 Surveillance Team |
| EPI_ISL_2781311 | Lighthouse Lab in Alderley Park | Wellcome Sanger Institute for the COVID-19 Genomics UK (COG-UK) Consortium | Jacquelyn Wynn, Mairead Hyland, The Lighthouse Lab in Alderley Park and Alex Alderton, Roberto Amato, Jeffrey Barrett, Sonia Goncalves, Ewan Harrison, David K. Jackson, Ian Johnston, Dominic Kwiatkowski, Cordelia Langford, John Sillitoe on behalf of the Wellcome Sanger Institute COVID-19 Surveillance Team |
| EPI_ISL_2781312, EPI_ISL_2781313, EPI_ISL_2781314 | Lighthouse Lab in Milton Keynes | Wellcome Sanger Institute for the COVID-19 Genomics UK (COG-UK) Consortium | The Lighthouse Lab in Milton Keynes and Alex Alderton, Roberto Amato, Jeffrey Barrett, Sonia Goncalves, Ewan Harrison, David K. Jackson, Ian Johnston, Dominic Kwiatkowski, Cordelia Langford, John Sillitoe on behalf of the Wellcome Sanger Institute COVID-19 Surveillance Team |
| EPI_ISL_2781315 | Lighthouse Lab in Alderley Park | Wellcome Sanger Institute for the COVID-19 Genomics UK (COG-UK) Consortium | Jacquelyn Wynn, Mairead Hyland, The Lighthouse Lab in Alderley Park and Alex Alderton, Roberto Amato, Jeffrey Barrett, Sonia Goncalves, Ewan Harrison, David K. Jackson, Ian Johnston, Dominic Kwiatkowski, Cordelia Langford, John Sillitoe on behalf of the Wellcome Sanger Institute COVID-19 Surveillance Team |
| EPI_ISL_2781316, EPI_ISL_2781317, EPI_ISL_2781318, EPI_ISL_2781319, EPI_ISL_2781320, EPI_ISL_2781321 | Lighthouse Lab in Milton Keynes | Wellcome Sanger Institute for the COVID-19 Genomics UK (COG-UK) Consortium | The Lighthouse Lab in Milton Keynes and Alex Alderton, Roberto Amato, Jeffrey Barrett, Sonia Goncalves, Ewan Harrison, David K. Jackson, Ian Johnston, Dominic Kwiatkowski, Cordelia Langford, John Sillitoe on behalf of the Wellcome Sanger Institute COVID-19 Surveillance Team |
| EPI_ISL_2781322, EPI_ISL_2781323 | Lighthouse Lab in Alderley Park | Wellcome Sanger Institute for the COVID-19 Genomics UK (COG-UK) Consortium | Jacquelyn Wynn, Mairead Hyland, The Lighthouse Lab in Alderley Park and Alex Alderton, Roberto Amato, Jeffrey Barrett, Sonia Goncalves, Ewan Harrison, David K. Jackson, Ian Johnston, Dominic Kwiatkowski, Cordelia Langford, John Sillitoe on behalf of the Wellcome Sanger Institute COVID-19 Surveillance Team |
| EPI_ISL_2781324 | Lighthouse Lab in Milton Keynes | Wellcome Sanger Institute for the COVID-19 Genomics UK (COG-UK) Consortium | The Lighthouse Lab in Milton Keynes and Alex Alderton, Roberto Amato, Jeffrey Barrett, Sonia Goncalves, Ewan Harrison, David K. Jackson, Ian Johnston, Dominic Kwiatkowski, Cordelia Langford, John Sillitoe on behalf of the Wellcome Sanger Institute COVID-19 Surveillance Team |
| EPI_ISL_2781325, EPI_ISL_2781326, EPI_ISL_2781327 | Lighthouse Lab in Alderley Park | Wellcome Sanger Institute for the COVID-19 Genomics UK (COG-UK) Consortium | Jacquelyn Wynn, Mairead Hyland, The Lighthouse Lab in Alderley Park and Alex Alderton, Roberto Amato, Jeffrey Barrett, Sonia Goncalves, Ewan Harrison, David K. Jackson, Ian Johnston, Dominic Kwiatkowski, Cordelia Langford, John Sillitoe on behalf of the Wellcome Sanger Institute COVID-19 Surveillance Team |
| EPI_ISL_2781328, EPI_ISL_2781329 | Lighthouse Lab in Milton Keynes | Wellcome Sanger Institute for the COVID-19 Genomics UK (COG-UK) Consortium | The Lighthouse Lab in Milton Keynes and Alex Alderton, Roberto Amato, Jeffrey Barrett, Sonia Goncalves, Ewan Harrison, David K. Jackson, Ian Johnston, Dominic Kwiatkowski, Cordelia Langford, John Sillitoe on behalf of the Wellcome Sanger Institute COVID-19 Surveillance Team |
| EPI_ISL_2781330 | Lighthouse Lab in Alderley Park | Wellcome Sanger Institute for the COVID-19 Genomics UK (COG-UK) Consortium | Jacquelyn Wynn, Mairead Hyland, The Lighthouse Lab in Alderley Park and Alex Alderton, Roberto Amato, Jeffrey Barrett, Sonia Goncalves, Ewan Harrison, David K. Jackson, Ian Johnston, Dominic Kwiatkowski, Cordelia Langford, John Sillitoe on behalf of the Wellcome Sanger Institute COVID-19 Surveillance Team |
| EPI_ISL_2781331, EPI_ISL_2781332, EPI_ISL_2781333, EPI_ISL_2781334, EPI_ISL_2781335 | Lighthouse Lab in Milton Keynes | Wellcome Sanger Institute for the COVID-19 Genomics UK (COG-UK) Consortium | The Lighthouse Lab in Milton Keynes and Alex Alderton, Roberto Amato, Jeffrey Barrett, Sonia Goncalves, Ewan Harrison, David K. Jackson, Ian Johnston, Dominic Kwiatkowski, Cordelia Langford, John Sillitoe on behalf of the Wellcome Sanger Institute COVID-19 Surveillance Team |
| EPI_ISL_2781336 | Lighthouse Lab in Alderley Park | Wellcome Sanger Institute for the COVID-19 Genomics UK (COG-UK) Consortium | Jacquelyn Wynn, Mairead Hyland, The Lighthouse Lab in Alderley Park and Alex Alderton, Roberto Amato, Jeffrey Barrett, Sonia Goncalves, Ewan Harrison, David K. Jackson, Ian Johnston, Dominic Kwiatkowski, Cordelia Langford, John Sillitoe on behalf of the Wellcome Sanger Institute COVID-19 Surveillance Team |
| EPI_ISL_2781337 | Lighthouse Lab in Milton Keynes | Wellcome Sanger Institute for the COVID-19 Genomics UK (COG-UK) Consortium | The Lighthouse Lab in Milton Keynes and Alex Alderton, Roberto Amato, Jeffrey Barrett, Sonia Goncalves, Ewan Harrison, David K. Jackson, Ian Johnston, Dominic Kwiatkowski, Cordelia Langford, John Sillitoe on behalf of the Wellcome Sanger Institute COVID-19 Surveillance Team |
| EPI_ISL_2781338 | Lighthouse Lab in Alderley Park | Wellcome Sanger Institute for the COVID-19 Genomics UK (COG-UK) Consortium | Jacquelyn Wynn, Mairead Hyland, The Lighthouse Lab in Alderley Park and Alex Alderton, Roberto Amato, Jeffrey Barrett, Sonia Goncalves, Ewan Harrison, David K. Jackson, Ian Johnston, Dominic Kwiatkowski, Cordelia Langford, John Sillitoe on behalf of the Wellcome Sanger Institute COVID-19 Surveillance Team |
| EPI_ISL_2781339, EPI_ISL_2781340, EPI_ISL_2781341, EPI_ISL_2781342, EPI_ISL_2781343, EPI_ISL_2781344, EPI_ISL_2781345, EPI_ISL_2781346, EPI_ISL_2781347, EPI_ISL_2781349, EPI_ISL_2781350, EPI_ISL_2781351, EPI_ISL_2781352, EPI_ISL_2781353 | | | |
| see above | Lighthouse Lab in Milton Keynes | Wellcome Sanger Institute for the COVID-19 Genomics UK (COG-UK) Consortium | The Lighthouse Lab in Milton Keynes and Alex Alderton, Roberto Amato, Jeffrey Barrett, Sonia Goncalves, Ewan Harrison, David K. Jackson, Ian Johnston, Dominic Kwiatkowski, Cordelia Langford, John Sillitoe on behalf of the Wellcome Sanger Institute COVID-19 Surveillance Team |
| EPI_ISL_2781354 | Lighthouse Lab in Alderley Park | Wellcome Sanger Institute for the COVID-19 Genomics UK (COG-UK) Consortium | Jacquelyn Wynn, Mairead Hyland, The Lighthouse Lab in Alderley Park and Alex Alderton, Roberto Amato, Jeffrey Barrett, Sonia Goncalves, Ewan Harrison, David K. Jackson, Ian Johnston, Dominic Kwiatkowski, Cordelia Langford, John Sillitoe on behalf of the Wellcome Sanger Institute COVID-19 Surveillance Team |
| EPI_ISL_2781355, EPI_ISL_2781356, EPI_ISL_2781357, EPI_ISL_2781358, EPI_ISL_2781359, EPI_ISL_2781360, EPI_ISL_2781361, EPI_ISL_2781362, EPI_ISL_2781363, EPI_ISL_2781364, EPI_ISL_2781365, EPI_ISL_2781366, EPI_ISL_2781367, EPI_ISL_2781368, EPI_ISL_2781369, EPI_ISL_2781370, EPI_ISL_2781371, EPI_ISL_2781372, EPI_ISL_2781373 | | | |
| see above | Lighthouse Lab in Milton Keynes | Wellcome Sanger Institute for the COVID-19 Genomics UK (COG-UK) Consortium | The Lighthouse Lab in Milton Keynes and Alex Alderton, Roberto Amato, Jeffrey Barrett, Sonia Goncalves, Ewan Harrison, David K. Jackson, Ian Johnston, Dominic Kwiatkowski, Cordelia Langford, John Sillitoe on behalf of the Wellcome Sanger Institute COVID-19 Surveillance Team |
| EPI_ISL_2781374 | Lighthouse Lab in Alderley Park | Wellcome Sanger Institute for the COVID-19 Genomics UK (COG-UK) Consortium | Jacquelyn Wynn, Mairead Hyland, The Lighthouse Lab in Alderley Park and Alex Alderton, Roberto Amato, Jeffrey Barrett, Sonia Goncalves, Ewan Harrison, David K. Jackson, Ian Johnston, Dominic Kwiatkowski, Cordelia Langford, John Sillitoe on behalf of the Wellcome Sanger Institute COVID-19 Surveillance Team |
| EPI_ISL_2781375, EPI_ISL_2781376, EPI_ISL_2781377 | Lighthouse Lab in Milton Keynes | Wellcome Sanger Institute for the COVID-19 Genomics UK (COG-UK) Consortium | The Lighthouse Lab in Milton Keynes and Alex Alderton, Roberto Amato, Jeffrey Barrett, Sonia Goncalves, Ewan Harrison, David K. Jackson, Ian Johnston, Dominic Kwiatkowski, Cordelia Langford, John Sillitoe on behalf of the Wellcome Sanger Institute COVID-19 Surveillance Team |

| | | | |
|---|---|---|---|
| EPI_ISL_2781378 | Lighthouse Lab in Alderley Park | Wellcome Sanger Institute for the COVID-19 Genomics UK (COG-UK) Consortium | Jacquelyn Wynn, Mairead Hyland, The Lighthouse Lab in Alderley Park and Alex Alderton, Roberto Amato, Jeffrey Barrett, Sonia Goncalves, Ewan Harrison, David K. Jackson, Ian Johnston, Dominic Kwiatkowski, Cordelia Langford, John Sillitoe on behalf of the Wellcome Sanger Institute COVID-19 Surveillance Team |
| EPI_ISL_2781379 | Lighthouse Lab in Milton Keynes | Wellcome Sanger Institute for the COVID-19 Genomics UK (COG-UK) Consortium | The Lighthouse Lab in Milton Keynes and Alex Alderton, Roberto Amato, Jeffrey Barrett, Sonia Goncalves, Ewan Harrison, David K. Jackson, Ian Johnston, Dominic Kwiatkowski, Cordelia Langford, John Sillitoe on behalf of the Wellcome Sanger Institute COVID-19 Surveillance Team |
| EPI_ISL_2781380 | Lighthouse Lab in Alderley Park | Wellcome Sanger Institute for the COVID-19 Genomics UK (COG-UK) Consortium | Jacquelyn Wynn, Mairead Hyland, The Lighthouse Lab in Alderley Park and Alex Alderton, Roberto Amato, Jeffrey Barrett, Sonia Goncalves, Ewan Harrison, David K. Jackson, Ian Johnston, Dominic Kwiatkowski, Cordelia Langford, John Sillitoe on behalf of the Wellcome Sanger Institute COVID-19 Surveillance Team |
| EPI_ISL_2781381, EPI_ISL_2781382 | Lighthouse Lab in Milton Keynes | Wellcome Sanger Institute for the COVID-19 Genomics UK (COG-UK) Consortium | The Lighthouse Lab in Milton Keynes and Alex Alderton, Roberto Amato, Jeffrey Barrett, Sonia Goncalves, Ewan Harrison, David K. Jackson, Ian Johnston, Dominic Kwiatkowski, Cordelia Langford, John Sillitoe on behalf of the Wellcome Sanger Institute COVID-19 Surveillance Team |
| EPI_ISL_2781383, EPI_ISL_2781384 | Lighthouse Lab in Alderley Park | Wellcome Sanger Institute for the COVID-19 Genomics UK (COG-UK) Consortium | Jacquelyn Wynn, Mairead Hyland, The Lighthouse Lab in Alderley Park and Alex Alderton, Roberto Amato, Jeffrey Barrett, Sonia Goncalves, Ewan Harrison, David K. Jackson, Ian Johnston, Dominic Kwiatkowski, Cordelia Langford, John Sillitoe on behalf of the Wellcome Sanger Institute COVID-19 Surveillance Team |
| EPI_ISL_2781385 | Lighthouse Lab in Milton Keynes | Wellcome Sanger Institute for the COVID-19 Genomics UK (COG-UK) Consortium | The Lighthouse Lab in Milton Keynes and Alex Alderton, Roberto Amato, Jeffrey Barrett, Sonia Goncalves, Ewan Harrison, David K. Jackson, Ian Johnston, Dominic Kwiatkowski, Cordelia Langford, John Sillitoe on behalf of the Wellcome Sanger Institute COVID-19 Surveillance Team |
| EPI_ISL_2781386 | Lighthouse Lab in Alderley Park | Wellcome Sanger Institute for the COVID-19 Genomics UK (COG-UK) Consortium | Jacquelyn Wynn, Mairead Hyland, The Lighthouse Lab in Alderley Park and Alex Alderton, Roberto Amato, Jeffrey Barrett, Sonia Goncalves, Ewan Harrison, David K. Jackson, Ian Johnston, Dominic Kwiatkowski, Cordelia Langford, John Sillitoe on behalf of the Wellcome Sanger Institute COVID-19 Surveillance Team |
| EPI_ISL_2781387, EPI_ISL_2781388 | Lighthouse Lab in Milton Keynes | Wellcome Sanger Institute for the COVID-19 Genomics UK (COG-UK) Consortium | The Lighthouse Lab in Milton Keynes and Alex Alderton, Roberto Amato, Jeffrey Barrett, Sonia Goncalves, Ewan Harrison, David K. Jackson, Ian Johnston, Dominic Kwiatkowski, Cordelia Langford, John Sillitoe on behalf of the Wellcome Sanger Institute COVID-19 Surveillance Team |
| EPI_ISL_2781389, EPI_ISL_2781390 | Lighthouse Lab in Alderley Park | Wellcome Sanger Institute for the COVID-19 Genomics UK (COG-UK) Consortium | Jacquelyn Wynn, Mairead Hyland, The Lighthouse Lab in Alderley Park and Alex Alderton, Roberto Amato, Jeffrey Barrett, Sonia Goncalves, Ewan Harrison, David K. Jackson, Ian Johnston, Dominic Kwiatkowski, Cordelia Langford, John Sillitoe on behalf of the Wellcome Sanger Institute COVID-19 Surveillance Team |
| EPI_ISL_2781391, EPI_ISL_2781392, EPI_ISL_2781393, EPI_ISL_2781394, EPI_ISL_2781395, EPI_ISL_2781396, EPI_ISL_2781397 | Lighthouse Lab in Milton Keynes | Wellcome Sanger Institute for the COVID-19 Genomics UK (COG-UK) Consortium | The Lighthouse Lab in Milton Keynes and Alex Alderton, Roberto Amato, Jeffrey Barrett, Sonia Goncalves, Ewan Harrison, David K. Jackson, Ian Johnston, Dominic Kwiatkowski, Cordelia Langford, John Sillitoe on behalf of the Wellcome Sanger Institute COVID-19 Surveillance Team |
| EPI_ISL_2781398, EPI_ISL_2781399 | Lighthouse Lab in Alderley Park | Wellcome Sanger Institute for the COVID-19 Genomics UK (COG-UK) Consortium | Jacquelyn Wynn, Mairead Hyland, The Lighthouse Lab in Alderley Park and Alex Alderton, Roberto Amato, Jeffrey Barrett, Sonia Goncalves, Ewan Harrison, David K. Jackson, Ian Johnston, Dominic Kwiatkowski, Cordelia Langford, John Sillitoe on behalf of the Wellcome Sanger Institute COVID-19 Surveillance Team |
| EPI_ISL_2781400, EPI_ISL_2781401 | Lighthouse Lab in Milton Keynes | Wellcome Sanger Institute for the COVID-19 Genomics UK (COG-UK) Consortium | The Lighthouse Lab in Milton Keynes and Alex Alderton, Roberto Amato, Jeffrey Barrett, Sonia Goncalves, Ewan Harrison, David K. Jackson, Ian Johnston, Dominic Kwiatkowski, Cordelia Langford, John Sillitoe on behalf of the Wellcome Sanger Institute COVID-19 Surveillance Team |
| EPI_ISL_2781402, EPI_ISL_2781403 | Lighthouse Lab in Alderley Park | Wellcome Sanger Institute for the COVID-19 Genomics UK (COG-UK) Consortium | Jacquelyn Wynn, Mairead Hyland, The Lighthouse Lab in Alderley Park and Alex Alderton, Roberto Amato, Jeffrey Barrett, Sonia Goncalves, Ewan Harrison, David K. Jackson, Ian Johnston, Dominic Kwiatkowski, Cordelia Langford, John Sillitoe on behalf of the Wellcome Sanger Institute COVID-19 Surveillance Team |
| EPI_ISL_2781404, EPI_ISL_2781405, EPI_ISL_2781406 | Lighthouse Lab in Milton Keynes | Wellcome Sanger Institute for the COVID-19 Genomics UK (COG-UK) Consortium | The Lighthouse Lab in Milton Keynes and Alex Alderton, Roberto Amato, Jeffrey Barrett, Sonia Goncalves, Ewan Harrison, David K. Jackson, Ian Johnston, Dominic Kwiatkowski, Cordelia Langford, John Sillitoe on behalf of the Wellcome Sanger Institute COVID-19 Surveillance Team |
| EPI_ISL_2781407, EPI_ISL_2781408, EPI_ISL_2781409, EPI_ISL_2781410 | Lighthouse Lab in Alderley Park | Wellcome Sanger Institute for the COVID-19 Genomics UK (COG-UK) Consortium | Jacquelyn Wynn, Mairead Hyland, The Lighthouse Lab in Alderley Park and Alex Alderton, Roberto Amato, Jeffrey Barrett, Sonia Goncalves, Ewan Harrison, David K. Jackson, Ian Johnston, Dominic Kwiatkowski, Cordelia Langford, John Sillitoe on behalf of the Wellcome Sanger Institute COVID-19 Surveillance Team |
| EPI_ISL_2781411, EPI_ISL_2781412, EPI_ISL_2781413 | Lighthouse Lab in Milton Keynes | Wellcome Sanger Institute for the COVID-19 Genomics UK (COG-UK) Consortium | The Lighthouse Lab in Milton Keynes and Alex Alderton, Roberto Amato, Jeffrey Barrett, Sonia Goncalves, Ewan Harrison, David K. Jackson, Ian Johnston, Dominic Kwiatkowski, Cordelia Langford, John Sillitoe on behalf of the Wellcome Sanger Institute COVID-19 Surveillance Team |
| EPI_ISL_2781414 | Lighthouse Lab in Alderley Park | Wellcome Sanger Institute for the COVID-19 Genomics UK (COG-UK) Consortium | Jacquelyn Wynn, Mairead Hyland, The Lighthouse Lab in Alderley Park and Alex Alderton, Roberto Amato, Jeffrey Barrett, Sonia Goncalves, Ewan Harrison, David K. Jackson, Ian Johnston, Dominic Kwiatkowski, Cordelia Langford, John Sillitoe on behalf of the Wellcome Sanger Institute COVID-19 Surveillance Team |
| EPI_ISL_2781415, EPI_ISL_2781416 | Lighthouse Lab in Milton Keynes | Wellcome Sanger Institute for the COVID-19 Genomics UK (COG-UK) Consortium | The Lighthouse Lab in Milton Keynes and Alex Alderton, Roberto Amato, Jeffrey Barrett, Sonia Goncalves, Ewan Harrison, David K. Jackson, Ian Johnston, Dominic Kwiatkowski, Cordelia Langford, John Sillitoe on behalf of the Wellcome Sanger Institute COVID-19 Surveillance Team |
| EPI_ISL_2781417 | Lighthouse Lab in Alderley Park | Wellcome Sanger Institute for the COVID-19 Genomics UK (COG-UK) Consortium | Jacquelyn Wynn, Mairead Hyland, The Lighthouse Lab in Alderley Park and Alex Alderton, Roberto Amato, Jeffrey Barrett, Sonia Goncalves, Ewan Harrison, David K. Jackson, Ian Johnston, Dominic Kwiatkowski, Cordelia Langford, John Sillitoe on behalf of the Wellcome Sanger Institute COVID-19 Surveillance Team |
| EPI_ISL_2781418, EPI_ISL_2781419, EPI_ISL_2781420, EPI_ISL_2781421, EPI_ISL_2781422, EPI_ISL_2781423, EPI_ISL_2781424, EPI_ISL_2781425, EPI_ISL_2781426, EPI_ISL_2781427 | Lighthouse Lab in Milton Keynes | Wellcome Sanger Institute for the COVID-19 Genomics UK (COG-UK) Consortium | The Lighthouse Lab in Milton Keynes and Alex Alderton, Roberto Amato, Jeffrey Barrett, Sonia Goncalves, Ewan Harrison, David K. Jackson, Ian Johnston, Dominic Kwiatkowski, Cordelia Langford, John Sillitoe on behalf of the Wellcome Sanger Institute COVID-19 Surveillance Team |
| EPI_ISL_2781428 | Lighthouse Lab in Alderley Park | Wellcome Sanger Institute for the COVID-19 Genomics UK (COG-UK) Consortium | Jacquelyn Wynn, Mairead Hyland, The Lighthouse Lab in Alderley Park and Alex Alderton, Roberto Amato, Jeffrey Barrett, Sonia Goncalves, Ewan Harrison, David K. Jackson, Ian Johnston, Dominic Kwiatkowski, Cordelia Langford, John Sillitoe on behalf of the Wellcome Sanger Institute COVID-19 Surveillance Team |
| EPI_ISL_2781429, EPI_ISL_2781430, EPI_ISL_2781431, EPI_ISL_2781432, EPI_ISL_2781433 | Lighthouse Lab in Milton Keynes | Wellcome Sanger Institute for the COVID-19 Genomics UK (COG-UK) Consortium | The Lighthouse Lab in Milton Keynes and Alex Alderton, Roberto Amato, Jeffrey Barrett, Sonia Goncalves, Ewan Harrison, David K. Jackson, Ian Johnston, Dominic Kwiatkowski, Cordelia Langford, John Sillitoe on behalf of the Wellcome Sanger Institute COVID-19 Surveillance Team |
| EPI_ISL_2781434 | Lighthouse Lab in Alderley Park | Wellcome Sanger Institute for the COVID-19 Genomics UK (COG-UK) Consortium | Jacquelyn Wynn, Mairead Hyland, The Lighthouse Lab in Alderley Park and Alex Alderton, Roberto Amato, Jeffrey Barrett, Sonia Goncalves, Ewan Harrison, David K. Jackson, Ian Johnston, Dominic Kwiatkowski, Cordelia Langford, John Sillitoe on behalf of the Wellcome Sanger Institute COVID-19 Surveillance Team |
| EPI_ISL_2781435, EPI_ISL_2781436, EPI_ISL_2781437, EPI_ISL_2781438, EPI_ISL_2781439, EPI_ISL_2781440, EPI_ISL_2781441, EPI_ISL_2781442, EPI_ISL_2781443, EPI_ISL_2781444, EPI_ISL_2781445, EPI_ISL_2781446, EPI_ISL_2781447 | | | |
| see above | Lighthouse Lab in Milton Keynes | Wellcome Sanger Institute for the COVID-19 Genomics UK (COG-UK) Consortium | The Lighthouse Lab in Milton Keynes and Alex Alderton, Roberto Amato, Jeffrey Barrett, Sonia Goncalves, Ewan Harrison, David K. Jackson, Ian Johnston, Dominic Kwiatkowski, Cordelia Langford, John Sillitoe on behalf of the Wellcome Sanger Institute COVID-19 Surveillance Team |
| EPI_ISL_2781448 | Lighthouse Lab in Alderley Park | Wellcome Sanger Institute for the COVID-19 Genomics UK (COG-UK) Consortium | Jacquelyn Wynn, Mairead Hyland, The Lighthouse Lab in Alderley Park and Alex Alderton, Roberto Amato, Jeffrey Barrett, Sonia Goncalves, Ewan Harrison, David K. Jackson, Ian Johnston, Dominic Kwiatkowski, Cordelia Langford, John Sillitoe on behalf of the Wellcome Sanger Institute COVID-19 Surveillance Team |
| EPI_ISL_2781449 | Lighthouse Lab in Milton Keynes | Wellcome Sanger Institute for the COVID-19 Genomics UK (COG-UK) Consortium | The Lighthouse Lab in Milton Keynes and Alex Alderton, Roberto Amato, Jeffrey Barrett, Sonia Goncalves, Ewan Harrison, David K. Jackson, Ian Johnston, Dominic Kwiatkowski, Cordelia Langford, John Sillitoe on behalf of the Wellcome Sanger Institute COVID-19 Surveillance Team |
| EPI_ISL_2781450 | Lighthouse Lab in Alderley Park | Wellcome Sanger Institute for the COVID-19 Genomics UK (COG-UK) Consortium | Jacquelyn Wynn, Mairead Hyland, The Lighthouse Lab in Alderley Park and Alex Alderton, Roberto Amato, Jeffrey Barrett, Sonia Goncalves, Ewan Harrison, David K. Jackson, Ian Johnston, Dominic Kwiatkowski, Cordelia Langford, John Sillitoe on behalf of the Wellcome Sanger Institute COVID-19 Surveillance Team |
| EPI_ISL_2781451 | Lighthouse Lab in Milton Keynes | Wellcome Sanger Institute for the COVID-19 Genomics UK (COG-UK) Consortium | The Lighthouse Lab in Milton Keynes and Alex Alderton, Roberto Amato, Jeffrey Barrett, Sonia Goncalves, Ewan Harrison, David K. Jackson, Ian Johnston, Dominic Kwiatkowski, Cordelia Langford, John Sillitoe on behalf of the Wellcome Sanger Institute COVID-19 Surveillance Team |
| EPI_ISL_2781452 | Lighthouse Lab in Alderley Park | Wellcome Sanger Institute for the COVID-19 Genomics UK (COG-UK) Consortium | Jacquelyn Wynn, Mairead Hyland, The Lighthouse Lab in Alderley Park and Alex Alderton, Roberto Amato, Jeffrey Barrett, Sonia Goncalves, Ewan Harrison, David K. Jackson, Ian Johnston, Dominic Kwiatkowski, Cordelia Langford, John Sillitoe on behalf of the Wellcome Sanger Institute COVID-19 Surveillance Team |
| EPI_ISL_2781453, EPI_ISL_2781454 | Lighthouse Lab in Milton Keynes | Wellcome Sanger Institute for the COVID-19 Genomics UK (COG-UK) Consortium | The Lighthouse Lab in Milton Keynes and Alex Alderton, Roberto Amato, Jeffrey Barrett, Sonia Goncalves, Ewan Harrison, David K. Jackson, Ian Johnston, Dominic Kwiatkowski, Cordelia Langford, John Sillitoe on behalf of the Wellcome Sanger Institute COVID-19 Surveillance Team |
| EPI_ISL_2781455, EPI_ISL_2781456, EPI_ISL_2781457 | Lighthouse Lab in Alderley Park | Wellcome Sanger Institute for the COVID-19 Genomics UK (COG-UK) Consortium | Jacquelyn Wynn, Mairead Hyland, The Lighthouse Lab in Alderley Park and Alex Alderton, Roberto Amato, Jeffrey Barrett, Sonia Goncalves, Ewan Harrison, David K. Jackson, Ian Johnston, Dominic Kwiatkowski, Cordelia Langford, John Sillitoe on behalf of the Wellcome Sanger Institute COVID-19 Surveillance Team |
| EPI_ISL_2781458, EPI_ISL_2781459, EPI_ISL_2781460 | Lighthouse Lab in Milton Keynes | Wellcome Sanger Institute for the COVID-19 Genomics UK (COG-UK) Consortium | The Lighthouse Lab in Milton Keynes and Alex Alderton, Roberto Amato, Jeffrey Barrett, Sonia Goncalves, Ewan Harrison, David K. Jackson, Ian Johnston, Dominic Kwiatkowski, Cordelia Langford, John Sillitoe on behalf of the Wellcome Sanger Institute COVID-19 Surveillance Team |
| EPI_ISL_2781461 | Lighthouse Lab in Alderley Park | Wellcome Sanger Institute for the COVID-19 Genomics UK (COG-UK) Consortium | Jacquelyn Wynn, Mairead Hyland, The Lighthouse Lab in Alderley Park and Alex Alderton, Roberto Amato, Jeffrey Barrett, Sonia Goncalves, Ewan Harrison, David K. Jackson, Ian Johnston, Dominic Kwiatkowski, Cordelia Langford, John Sillitoe on behalf of the Wellcome Sanger Institute COVID-19 Surveillance Team |
| EPI_ISL_2781462, EPI_ISL_2781463, EPI_ISL_2781464, EPI_ISL_2781465 | Lighthouse Lab in Milton Keynes | Wellcome Sanger Institute for the COVID-19 Genomics UK (COG-UK) Consortium | The Lighthouse Lab in Milton Keynes and Alex Alderton, Roberto Amato, Jeffrey Barrett, Sonia Goncalves, Ewan Harrison, David K. Jackson, Ian Johnston, Dominic Kwiatkowski, Cordelia Langford, John Sillitoe on behalf of the Wellcome Sanger Institute COVID-19 Surveillance Team |
| EPI_ISL_2781466 | Lighthouse Lab in Alderley Park | Wellcome Sanger Institute for the COVID-19 Genomics UK (COG-UK) | Jacquelyn Wynn, Mairead Hyland, The Lighthouse Lab in Alderley Park and Alex Alderton, Roberto Amato, Jeffrey Barrett, Sonia Goncalves, Ewan Harrison, David K. Jackson, Ian Johnston, Dominic Kwiatkowski, Cordelia Langford, John Sillitoe on |

| | | | |
|---|---|---|---|
| | | Consortium | behalf of the Wellcome Sanger Institute COVID-19 Surveillance Team |
| EPI_ISL_2781467, EPI_ISL_2781468, EPI_ISL_2781469, EPI_ISL_2781470, EPI_ISL_2781471, EPI_ISL_2781472, EPI_ISL_2781473 | Lighthouse Lab in Milton Keynes | Wellcome Sanger Institute for the COVID-19 Genomics UK (COG-UK) Consortium | The Lighthouse Lab in Milton Keynes and Alex Alderton, Roberto Amato, Jeffrey Barrett, Sonia Goncalves, Ewan Harrison, David K. Jackson, Ian Johnston, Dominic Kwiatkowski, Cordelia Langford, John Sillitoe on behalf of the Wellcome Sanger Institute COVID-19 Surveillance Team |
| EPI_ISL_2781474 | Lighthouse Lab in Alderley Park | Wellcome Sanger Institute for the COVID-19 Genomics UK (COG-UK) Consortium | Jacquelyn Wynn, Mairead Hyland, The Lighthouse Lab in Alderley Park and Alex Alderton, Roberto Amato, Jeffrey Barrett, Sonia Goncalves, Ewan Harrison, David K. Jackson, Ian Johnston, Dominic Kwiatkowski, Cordelia Langford, John Sillitoe on behalf of the Wellcome Sanger Institute COVID-19 Surveillance Team |
| EPI_ISL_2781475 | Lighthouse Lab in Milton Keynes | Wellcome Sanger Institute for the COVID-19 Genomics UK (COG-UK) Consortium | The Lighthouse Lab in Milton Keynes and Alex Alderton, Roberto Amato, Jeffrey Barrett, Sonia Goncalves, Ewan Harrison, David K. Jackson, Ian Johnston, Dominic Kwiatkowski, Cordelia Langford, John Sillitoe on behalf of the Wellcome Sanger Institute COVID-19 Surveillance Team |
| EPI_ISL_2781476 | Lighthouse Lab in Alderley Park | Wellcome Sanger Institute for the COVID-19 Genomics UK (COG-UK) Consortium | Jacquelyn Wynn, Mairead Hyland, The Lighthouse Lab in Alderley Park and Alex Alderton, Roberto Amato, Jeffrey Barrett, Sonia Goncalves, Ewan Harrison, David K. Jackson, Ian Johnston, Dominic Kwiatkowski, Cordelia Langford, John Sillitoe on behalf of the Wellcome Sanger Institute COVID-19 Surveillance Team |
| EPI_ISL_2781477 | Lighthouse Lab in Milton Keynes | Wellcome Sanger Institute for the COVID-19 Genomics UK (COG-UK) Consortium | The Lighthouse Lab in Milton Keynes and Alex Alderton, Roberto Amato, Jeffrey Barrett, Sonia Goncalves, Ewan Harrison, David K. Jackson, Ian Johnston, Dominic Kwiatkowski, Cordelia Langford, John Sillitoe on behalf of the Wellcome Sanger Institute COVID-19 Surveillance Team |
| EPI_ISL_2781478 | Lighthouse Lab in Alderley Park | Wellcome Sanger Institute for the COVID-19 Genomics UK (COG-UK) Consortium | Jacquelyn Wynn, Mairead Hyland, The Lighthouse Lab in Alderley Park and Alex Alderton, Roberto Amato, Jeffrey Barrett, Sonia Goncalves, Ewan Harrison, David K. Jackson, Ian Johnston, Dominic Kwiatkowski, Cordelia Langford, John Sillitoe on behalf of the Wellcome Sanger Institute COVID-19 Surveillance Team |
| EPI_ISL_2781480, EPI_ISL_2781481, EPI_ISL_2781482, EPI_ISL_2781483, EPI_ISL_2781484 | Lighthouse Lab in Milton Keynes | Wellcome Sanger Institute for the COVID-19 Genomics UK (COG-UK) Consortium | The Lighthouse Lab in Milton Keynes and Alex Alderton, Roberto Amato, Jeffrey Barrett, Sonia Goncalves, Ewan Harrison, David K. Jackson, Ian Johnston, Dominic Kwiatkowski, Cordelia Langford, John Sillitoe on behalf of the Wellcome Sanger Institute COVID-19 Surveillance Team |
| EPI_ISL_2781485 | Lighthouse Lab in Alderley Park | Wellcome Sanger Institute for the COVID-19 Genomics UK (COG-UK) Consortium | Jacquelyn Wynn, Mairead Hyland, The Lighthouse Lab in Alderley Park and Alex Alderton, Roberto Amato, Jeffrey Barrett, Sonia Goncalves, Ewan Harrison, David K. Jackson, Ian Johnston, Dominic Kwiatkowski, Cordelia Langford, John Sillitoe on behalf of the Wellcome Sanger Institute COVID-19 Surveillance Team |
| EPI_ISL_2781486 | Lighthouse Lab in Milton Keynes | Wellcome Sanger Institute for the COVID-19 Genomics UK (COG-UK) Consortium | The Lighthouse Lab in Milton Keynes and Alex Alderton, Roberto Amato, Jeffrey Barrett, Sonia Goncalves, Ewan Harrison, David K. Jackson, Ian Johnston, Dominic Kwiatkowski, Cordelia Langford, John Sillitoe on behalf of the Wellcome Sanger Institute COVID-19 Surveillance Team |
| EPI_ISL_2781487, EPI_ISL_2781488, EPI_ISL_2781489 | Health Services Laboratories | Wellcome Sanger Institute for the COVID-19 Genomics UK (COG-UK) Consortium | Health Services Laboratories and Alex Alderton, Roberto Amato, Jeffrey Barrett, Sonia Goncalves, Ewan Harrison, David K. Jackson, Ian Johnston, Dominic Kwiatkowski, Cordelia Langford, John Sillitoe on behalf of the Wellcome Sanger Institute COVID-19 Surveillance Team |
| EPI_ISL_2781490 | Lighthouse Lab in Milton Keynes | Wellcome Sanger Institute for the COVID-19 Genomics UK (COG-UK) Consortium | The Lighthouse Lab in Milton Keynes and Alex Alderton, Roberto Amato, Jeffrey Barrett, Sonia Goncalves, Ewan Harrison, David K. Jackson, Ian Johnston, Dominic Kwiatkowski, Cordelia Langford, John Sillitoe on behalf of the Wellcome Sanger Institute COVID-19 Surveillance Team |
| EPI_ISL_2781491 | Lighthouse Lab in Alderley Park | Wellcome Sanger Institute for the COVID-19 Genomics UK (COG-UK) Consortium | Jacquelyn Wynn, Mairead Hyland, The Lighthouse Lab in Alderley Park and Alex Alderton, Roberto Amato, Jeffrey Barrett, Sonia Goncalves, Ewan Harrison, David K. Jackson, Ian Johnston, Dominic Kwiatkowski, Cordelia Langford, John Sillitoe on behalf of the Wellcome Sanger Institute COVID-19 Surveillance Team |
| EPI_ISL_2781492 | Health Services Laboratories | Wellcome Sanger Institute for the COVID-19 Genomics UK (COG-UK) Consortium | Health Services Laboratories and Alex Alderton, Roberto Amato, Jeffrey Barrett, Sonia Goncalves, Ewan Harrison, David K. Jackson, Ian Johnston, Dominic Kwiatkowski, Cordelia Langford, John Sillitoe on behalf of the Wellcome Sanger Institute COVID-19 Surveillance Team |
| EPI_ISL_2781493, EPI_ISL_2781494 | Lighthouse Lab in Alderley Park | Wellcome Sanger Institute for the COVID-19 Genomics UK (COG-UK) Consortium | Jacquelyn Wynn, Mairead Hyland, The Lighthouse Lab in Alderley Park and Alex Alderton, Roberto Amato, Jeffrey Barrett, Sonia Goncalves, Ewan Harrison, David K. Jackson, Ian Johnston, Dominic Kwiatkowski, Cordelia Langford, John Sillitoe on behalf of the Wellcome Sanger Institute COVID-19 Surveillance Team |
| EPI_ISL_2781495 | Lighthouse Laboratory Plymouth | Wellcome Sanger Institute for the COVID-19 Genomics UK (COG-UK) Consortium | Lighthouse Laboratory Plymouth and Alex Alderton, Roberto Amato, Jeffrey Barrett, Sonia Goncalves, Ewan Harrison, David K. Jackson, Ian Johnston, Dominic Kwiatkowski, Cordelia Langford, John Sillitoe on behalf of the Wellcome Sanger Institute COVID-19 Surveillance Team |
| EPI_ISL_2781496 | Health Services Laboratories | Wellcome Sanger Institute for the COVID-19 Genomics UK (COG-UK) Consortium | Health Services Laboratories and Alex Alderton, Roberto Amato, Jeffrey Barrett, Sonia Goncalves, Ewan Harrison, David K. Jackson, Ian Johnston, Dominic Kwiatkowski, Cordelia Langford, John Sillitoe on behalf of the Wellcome Sanger Institute COVID-19 Surveillance Team |
| EPI_ISL_2781497 | Lighthouse Lab in Alderley Park | Wellcome Sanger Institute for the COVID-19 Genomics UK (COG-UK) Consortium | Jacquelyn Wynn, Mairead Hyland, The Lighthouse Lab in Alderley Park and Alex Alderton, Roberto Amato, Jeffrey Barrett, Sonia Goncalves, Ewan Harrison, David K. Jackson, Ian Johnston, Dominic Kwiatkowski, Cordelia Langford, John Sillitoe on behalf of the Wellcome Sanger Institute COVID-19 Surveillance Team |
| EPI_ISL_2781498, EPI_ISL_2781499 | Health Services Laboratories | Wellcome Sanger Institute for the COVID-19 Genomics UK (COG-UK) Consortium | Health Services Laboratories and Alex Alderton, Roberto Amato, Jeffrey Barrett, Sonia Goncalves, Ewan Harrison, David K. Jackson, Ian Johnston, Dominic Kwiatkowski, Cordelia Langford, John Sillitoe on behalf of the Wellcome Sanger Institute COVID-19 Surveillance Team |
| EPI_ISL_2781500 | Lighthouse Lab in Milton Keynes | Wellcome Sanger Institute for the COVID-19 Genomics UK (COG-UK) Consortium | The Lighthouse Lab in Milton Keynes and Alex Alderton, Roberto Amato, Jeffrey Barrett, Sonia Goncalves, Ewan Harrison, David K. Jackson, Ian Johnston, Dominic Kwiatkowski, Cordelia Langford, John Sillitoe on behalf of the Wellcome Sanger Institute COVID-19 Surveillance Team |
| EPI_ISL_2781501 | Lighthouse Laboratory Plymouth | Wellcome Sanger Institute for the COVID-19 Genomics UK (COG-UK) Consortium | Lighthouse Laboratory Plymouth and Alex Alderton, Roberto Amato, Jeffrey Barrett, Sonia Goncalves, Ewan Harrison, David K. Jackson, Ian Johnston, Dominic Kwiatkowski, Cordelia Langford, John Sillitoe on behalf of the Wellcome Sanger Institute COVID-19 Surveillance Team |
| EPI_ISL_2781502, EPI_ISL_2781503, EPI_ISL_2781504, EPI_ISL_2781505 | Lighthouse Lab in Milton Keynes | Wellcome Sanger Institute for the COVID-19 Genomics UK (COG-UK) Consortium | The Lighthouse Lab in Milton Keynes and Alex Alderton, Roberto Amato, Jeffrey Barrett, Sonia Goncalves, Ewan Harrison, David K. Jackson, Ian Johnston, Dominic Kwiatkowski, Cordelia Langford, John Sillitoe on behalf of the Wellcome Sanger Institute COVID-19 Surveillance Team |
| EPI_ISL_2781506 | Health Services Laboratories | Wellcome Sanger Institute for the COVID-19 Genomics UK (COG-UK) Consortium | Health Services Laboratories and Alex Alderton, Roberto Amato, Jeffrey Barrett, Sonia Goncalves, Ewan Harrison, David K. Jackson, Ian Johnston, Dominic Kwiatkowski, Cordelia Langford, John Sillitoe on behalf of the Wellcome Sanger Institute COVID-19 Surveillance Team |
| EPI_ISL_2781507, EPI_ISL_2781508 | Lighthouse Lab in Milton Keynes | Wellcome Sanger Institute for the COVID-19 Genomics UK (COG-UK) Consortium | The Lighthouse Lab in Milton Keynes and Alex Alderton, Roberto Amato, Jeffrey Barrett, Sonia Goncalves, Ewan Harrison, David K. Jackson, Ian Johnston, Dominic Kwiatkowski, Cordelia Langford, John Sillitoe on behalf of the Wellcome Sanger Institute COVID-19 Surveillance Team |
| EPI_ISL_2781509, EPI_ISL_2781510, EPI_ISL_2781511 | Lighthouse Lab in Glasgow | Wellcome Sanger Institute for the COVID-19 Genomics UK (COG-UK) Consortium | Harper VanSteenhouse, Yumi Kasai, David Gray, Carol Clugston, Anna Dominiczak and Alex Alderton, Roberto Amato, Jeffrey Barrett, Sonia Goncalves, Ewan Harrison, David K. Jackson, Ian Johnston, Dominic Kwiatkowski, Cordelia Langford, John Sillitoe on behalf of the Wellcome Sanger Institute COVID-19 Surveillance Team |
| EPI_ISL_2781512, EPI_ISL_2781513 | Lighthouse Lab in Milton Keynes | Wellcome Sanger Institute for the COVID-19 Genomics UK (COG-UK) Consortium | The Lighthouse Lab in Milton Keynes and Alex Alderton, Roberto Amato, Jeffrey Barrett, Sonia Goncalves, Ewan Harrison, David K. Jackson, Ian Johnston, Dominic Kwiatkowski, Cordelia Langford, John Sillitoe on behalf of the Wellcome Sanger Institute COVID-19 Surveillance Team |
| EPI_ISL_2781514 | Lighthouse Lab in Alderley Park | Wellcome Sanger Institute for the COVID-19 Genomics UK (COG-UK) Consortium | Jacquelyn Wynn, Mairead Hyland, The Lighthouse Lab in Alderley Park and Alex Alderton, Roberto Amato, Jeffrey Barrett, Sonia Goncalves, Ewan Harrison, David K. Jackson, Ian Johnston, Dominic Kwiatkowski, Cordelia Langford, John Sillitoe on behalf of the Wellcome Sanger Institute COVID-19 Surveillance Team |
| EPI_ISL_2781515 | Health Services Laboratories | Wellcome Sanger Institute for the COVID-19 Genomics UK (COG-UK) Consortium | Health Services Laboratories and Alex Alderton, Roberto Amato, Jeffrey Barrett, Sonia Goncalves, Ewan Harrison, David K. Jackson, Ian Johnston, Dominic Kwiatkowski, Cordelia Langford, John Sillitoe on behalf of the Wellcome Sanger Institute COVID-19 Surveillance Team |
| EPI_ISL_2781516 | Lighthouse Lab in Milton Keynes | Wellcome Sanger Institute for the COVID-19 Genomics UK (COG-UK) Consortium | The Lighthouse Lab in Milton Keynes and Alex Alderton, Roberto Amato, Jeffrey Barrett, Sonia Goncalves, Ewan Harrison, David K. Jackson, Ian Johnston, Dominic Kwiatkowski, Cordelia Langford, John Sillitoe on behalf of the Wellcome Sanger Institute COVID-19 Surveillance Team |
| EPI_ISL_2781517 | Health Services Laboratories | Wellcome Sanger Institute for the COVID-19 Genomics UK (COG-UK) Consortium | Health Services Laboratories and Alex Alderton, Roberto Amato, Jeffrey Barrett, Sonia Goncalves, Ewan Harrison, David K. Jackson, Ian Johnston, Dominic Kwiatkowski, Cordelia Langford, John Sillitoe on behalf of the Wellcome Sanger Institute COVID-19 Surveillance Team |
| EPI_ISL_2781518 | Lighthouse Lab in Milton Keynes | Wellcome Sanger Institute for the COVID-19 Genomics UK (COG-UK) Consortium | The Lighthouse Lab in Milton Keynes and Alex Alderton, Roberto Amato, Jeffrey Barrett, Sonia Goncalves, Ewan Harrison, David K. Jackson, Ian Johnston, Dominic Kwiatkowski, Cordelia Langford, John Sillitoe on behalf of the Wellcome Sanger Institute COVID-19 Surveillance Team |
| EPI_ISL_2781519, EPI_ISL_2781520, EPI_ISL_2781521 | Lighthouse Lab in Alderley Park | Wellcome Sanger Institute for the COVID-19 Genomics UK (COG-UK) Consortium | Jacquelyn Wynn, Mairead Hyland, The Lighthouse Lab in Alderley Park and Alex Alderton, Roberto Amato, Jeffrey Barrett, Sonia Goncalves, Ewan Harrison, David K. Jackson, Ian Johnston, Dominic Kwiatkowski, Cordelia Langford, John Sillitoe on behalf of the Wellcome Sanger Institute COVID-19 Surveillance Team |
| EPI_ISL_2781522 | Lighthouse Laboratory Plymouth | Wellcome Sanger Institute for the COVID-19 Genomics UK (COG-UK) Consortium | Lighthouse Laboratory Plymouth and Alex Alderton, Roberto Amato, Jeffrey Barrett, Sonia Goncalves, Ewan Harrison, David K. Jackson, Ian Johnston, Dominic Kwiatkowski, Cordelia Langford, John Sillitoe on behalf of the Wellcome Sanger Institute COVID-19 Surveillance Team |
| EPI_ISL_2781523 | Lighthouse Lab in Milton Keynes | Wellcome Sanger Institute for the COVID-19 Genomics UK (COG-UK) Consortium | The Lighthouse Lab in Milton Keynes and Alex Alderton, Roberto Amato, Jeffrey Barrett, Sonia Goncalves, Ewan Harrison, David K. Jackson, Ian Johnston, Dominic Kwiatkowski, Cordelia Langford, John Sillitoe on behalf of the Wellcome Sanger Institute COVID-19 Surveillance Team |
| EPI_ISL_2781524 | Lighthouse Lab in Alderley Park | Wellcome Sanger Institute for the COVID-19 Genomics UK (COG-UK) Consortium | Jacquelyn Wynn, Mairead Hyland, The Lighthouse Lab in Alderley Park and Alex Alderton, Roberto Amato, Jeffrey Barrett, Sonia Goncalves, Ewan Harrison, David K. Jackson, Ian Johnston, Dominic Kwiatkowski, Cordelia Langford, John Sillitoe on behalf of the Wellcome Sanger Institute COVID-19 Surveillance Team |
| EPI_ISL_2781525, EPI_ISL_2781526, EPI_ISL_2781527 | Lighthouse Lab in Milton Keynes | Wellcome Sanger Institute for the COVID-19 Genomics UK (COG-UK) Consortium | The Lighthouse Lab in Milton Keynes and Alex Alderton, Roberto Amato, Jeffrey Barrett, Sonia Goncalves, Ewan Harrison, David K. Jackson, Ian Johnston, Dominic Kwiatkowski, Cordelia Langford, John Sillitoe on behalf of the Wellcome Sanger Institute COVID-19 Surveillance Team |
| EPI_ISL_2781528 | Lighthouse Lab in Glasgow | Wellcome Sanger Institute for the COVID-19 Genomics UK (COG-UK) Consortium | Harper VanSteenhouse, Yumi Kasai, David Gray, Carol Clugston, Anna Dominiczak and Alex Alderton, Roberto Amato, Jeffrey Barrett, Sonia Goncalves, Ewan Harrison, David K. Jackson, Ian Johnston, Dominic Kwiatkowski, Cordelia Langford, John Sillitoe on behalf of the Wellcome Sanger Institute COVID-19 Surveillance Team |
| EPI_ISL_2781529 | Lighthouse Lab in Alderley Park | Wellcome Sanger Institute for the COVID-19 Genomics UK (COG-UK) Consortium | Jacquelyn Wynn, Mairead Hyland, The Lighthouse Lab in Alderley Park and Alex Alderton, Roberto Amato, Jeffrey Barrett, Sonia Goncalves, Ewan Harrison, David K. Jackson, Ian Johnston, Dominic Kwiatkowski, Cordelia Langford, John Sillitoe on behalf of the Wellcome Sanger Institute COVID-19 Surveillance Team |
| EPI_ISL_2781530, EPI_ISL_2781531 | Health Services Laboratories | Wellcome Sanger Institute for the COVID-19 Genomics UK (COG-UK) Consortium | Health Services Laboratories and Alex Alderton, Roberto Amato, Jeffrey Barrett, Sonia Goncalves, Ewan Harrison, David K. Jackson, Ian Johnston, Dominic Kwiatkowski, Cordelia Langford, John Sillitoe on behalf of the Wellcome Sanger Institute COVID-19 Surveillance Team |
| EPI_ISL_2781532 | Lighthouse Lab in Glasgow | Wellcome Sanger Institute for the COVID-19 Genomics UK (COG-UK) Consortium | Harper VanSteenhouse, Yumi Kasai, David Gray, Carol Clugston, Anna Dominiczak and Alex Alderton, Roberto Amato, Jeffrey Barrett, Sonia Goncalves, Ewan Harrison, David K. Jackson, Ian Johnston, Dominic Kwiatkowski, Cordelia Langford, John Sillitoe on behalf of the Wellcome Sanger Institute COVID-19 Surveillance Team |

| | | | |
|---|---|---|---|
| EPI_ISL_2781533 | Health Services Laboratories | Wellcome Sanger Institute for the COVID-19 Genomics UK (COG-UK) Consortium | Health Services Laboratories and Alex Alderton, Roberto Amato, Jeffrey Barrett, Sonia Goncalves, Ewan Harrison, David K. Jackson, Ian Johnston, Dominic Kwiatkowski, Cordelia Langford, John Sillitoe on behalf of the Wellcome Sanger Institute COVID-19 Surveillance Team |
| EPI_ISL_2781534 | Lighthouse Lab in Glasgow | Wellcome Sanger Institute for the COVID-19 Genomics UK (COG-UK) Consortium | Harper VanSteenhouse, Yumi Kasai, David Gray, Carol Clugston, Anna Dominiczak and Alex Alderton, Roberto Amato, Jeffrey Barrett, Sonia Goncalves, Ewan Harrison, David K. Jackson, Ian Johnston, Dominic Kwiatkowski, Cordelia Langford, John Sillitoe on behalf of the Wellcome Sanger Institute COVID-19 Surveillance Team |
| EPI_ISL_2781535 | Health Services Laboratories | Wellcome Sanger Institute for the COVID-19 Genomics UK (COG-UK) Consortium | Health Services Laboratories and Alex Alderton, Roberto Amato, Jeffrey Barrett, Sonia Goncalves, Ewan Harrison, David K. Jackson, Ian Johnston, Dominic Kwiatkowski, Cordelia Langford, John Sillitoe on behalf of the Wellcome Sanger Institute COVID-19 Surveillance Team |
| EPI_ISL_2781536 | Lighthouse Lab in Milton Keynes | Wellcome Sanger Institute for the COVID-19 Genomics UK (COG-UK) Consortium | The Lighthouse Lab in Milton Keynes and Alex Alderton, Roberto Amato, Jeffrey Barrett, Sonia Goncalves, Ewan Harrison, David K. Jackson, Ian Johnston, Dominic Kwiatkowski, Cordelia Langford, John Sillitoe on behalf of the Wellcome Sanger Institute COVID-19 Surveillance Team |
| EPI_ISL_2781537 | Lighthouse Lab in Alderley Park | Wellcome Sanger Institute for the COVID-19 Genomics UK (COG-UK) Consortium | Jacquelyn Wynn, Mairead Hyland, The Lighthouse Lab in Alderley Park and Alex Alderton, Roberto Amato, Jeffrey Barrett, Sonia Goncalves, Ewan Harrison, David K. Jackson, Ian Johnston, Dominic Kwiatkowski, Cordelia Langford, John Sillitoe on behalf of the Wellcome Sanger Institute COVID-19 Surveillance Team |
| EPI_ISL_2781538 | Health Services Laboratories | Wellcome Sanger Institute for the COVID-19 Genomics UK (COG-UK) Consortium | Health Services Laboratories and Alex Alderton, Roberto Amato, Jeffrey Barrett, Sonia Goncalves, Ewan Harrison, David K. Jackson, Ian Johnston, Dominic Kwiatkowski, Cordelia Langford, John Sillitoe on behalf of the Wellcome Sanger Institute COVID-19 Surveillance Team |
| EPI_ISL_2781540 | Lighthouse Lab in Milton Keynes | Wellcome Sanger Institute for the COVID-19 Genomics UK (COG-UK) Consortium | The Lighthouse Lab in Milton Keynes and Alex Alderton, Roberto Amato, Jeffrey Barrett, Sonia Goncalves, Ewan Harrison, David K. Jackson, Ian Johnston, Dominic Kwiatkowski, Cordelia Langford, John Sillitoe on behalf of the Wellcome Sanger Institute COVID-19 Surveillance Team |
| EPI_ISL_2781541 | Health Services Laboratories | Wellcome Sanger Institute for the COVID-19 Genomics UK (COG-UK) Consortium | Health Services Laboratories and Alex Alderton, Roberto Amato, Jeffrey Barrett, Sonia Goncalves, Ewan Harrison, David K. Jackson, Ian Johnston, Dominic Kwiatkowski, Cordelia Langford, John Sillitoe on behalf of the Wellcome Sanger Institute COVID-19 Surveillance Team |
| EPI_ISL_2781542 | Lighthouse Lab in Milton Keynes | Wellcome Sanger Institute for the COVID-19 Genomics UK (COG-UK) Consortium | The Lighthouse Lab in Milton Keynes and Alex Alderton, Roberto Amato, Jeffrey Barrett, Sonia Goncalves, Ewan Harrison, David K. Jackson, Ian Johnston, Dominic Kwiatkowski, Cordelia Langford, John Sillitoe on behalf of the Wellcome Sanger Institute COVID-19 Surveillance Team |
| EPI_ISL_2781543 | Health Services Laboratories | Wellcome Sanger Institute for the COVID-19 Genomics UK (COG-UK) Consortium | Health Services Laboratories and Alex Alderton, Roberto Amato, Jeffrey Barrett, Sonia Goncalves, Ewan Harrison, David K. Jackson, Ian Johnston, Dominic Kwiatkowski, Cordelia Langford, John Sillitoe on behalf of the Wellcome Sanger Institute COVID-19 Surveillance Team |
| EPI_ISL_2781544, EPI_ISL_2781545, EPI_ISL_2781546 | Lighthouse Lab in Milton Keynes | Wellcome Sanger Institute for the COVID-19 Genomics UK (COG-UK) Consortium | The Lighthouse Lab in Milton Keynes and Alex Alderton, Roberto Amato, Jeffrey Barrett, Sonia Goncalves, Ewan Harrison, David K. Jackson, Ian Johnston, Dominic Kwiatkowski, Cordelia Langford, John Sillitoe on behalf of the Wellcome Sanger Institute COVID-19 Surveillance Team |
| EPI_ISL_2781547 | Health Services Laboratories | Wellcome Sanger Institute for the COVID-19 Genomics UK (COG-UK) Consortium | Health Services Laboratories and Alex Alderton, Roberto Amato, Jeffrey Barrett, Sonia Goncalves, Ewan Harrison, David K. Jackson, Ian Johnston, Dominic Kwiatkowski, Cordelia Langford, John Sillitoe on behalf of the Wellcome Sanger Institute COVID-19 Surveillance Team |
| EPI_ISL_2781548 | Lighthouse Lab in Milton Keynes | Wellcome Sanger Institute for the COVID-19 Genomics UK (COG-UK) Consortium | The Lighthouse Lab in Milton Keynes and Alex Alderton, Roberto Amato, Jeffrey Barrett, Sonia Goncalves, Ewan Harrison, David K. Jackson, Ian Johnston, Dominic Kwiatkowski, Cordelia Langford, John Sillitoe on behalf of the Wellcome Sanger Institute COVID-19 Surveillance Team |
| EPI_ISL_2781549, EPI_ISL_2781550, EPI_ISL_2781551, EPI_ISL_2781552, EPI_ISL_2781553 | Lighthouse Lab in Alderley Park | Wellcome Sanger Institute for the COVID-19 Genomics UK (COG-UK) Consortium | Jacquelyn Wynn, Mairead Hyland, The Lighthouse Lab in Alderley Park and Alex Alderton, Roberto Amato, Jeffrey Barrett, Sonia Goncalves, Ewan Harrison, David K. Jackson, Ian Johnston, Dominic Kwiatkowski, Cordelia Langford, John Sillitoe on behalf of the Wellcome Sanger Institute COVID-19 Surveillance Team |
| EPI_ISL_2781554 | Lighthouse Laboratory Plymouth | Wellcome Sanger Institute for the COVID-19 Genomics UK (COG-UK) Consortium | Lighthouse Laboratory Plymouth and Alex Alderton, Roberto Amato, Jeffrey Barrett, Sonia Goncalves, Ewan Harrison, David K. Jackson, Ian Johnston, Dominic Kwiatkowski, Cordelia Langford, John Sillitoe on behalf of the Wellcome Sanger Institute COVID-19 Surveillance Team |
| EPI_ISL_2781555, EPI_ISL_2781556 | Lighthouse Lab in Alderley Park | Wellcome Sanger Institute for the COVID-19 Genomics UK (COG-UK) Consortium | Jacquelyn Wynn, Mairead Hyland, The Lighthouse Lab in Alderley Park and Alex Alderton, Roberto Amato, Jeffrey Barrett, Sonia Goncalves, Ewan Harrison, David K. Jackson, Ian Johnston, Dominic Kwiatkowski, Cordelia Langford, John Sillitoe on behalf of the Wellcome Sanger Institute COVID-19 Surveillance Team |
| EPI_ISL_2781557 | Lighthouse Lab in Milton Keynes | Wellcome Sanger Institute for the COVID-19 Genomics UK (COG-UK) Consortium | The Lighthouse Lab in Milton Keynes and Alex Alderton, Roberto Amato, Jeffrey Barrett, Sonia Goncalves, Ewan Harrison, David K. Jackson, Ian Johnston, Dominic Kwiatkowski, Cordelia Langford, John Sillitoe on behalf of the Wellcome Sanger Institute COVID-19 Surveillance Team |
| EPI_ISL_2781558 | Health Services Laboratories | Wellcome Sanger Institute for the COVID-19 Genomics UK (COG-UK) Consortium | Health Services Laboratories and Alex Alderton, Roberto Amato, Jeffrey Barrett, Sonia Goncalves, Ewan Harrison, David K. Jackson, Ian Johnston, Dominic Kwiatkowski, Cordelia Langford, John Sillitoe on behalf of the Wellcome Sanger Institute COVID-19 Surveillance Team |
| EPI_ISL_2781559, EPI_ISL_2781560 | Lighthouse Lab in Alderley Park | Wellcome Sanger Institute for the COVID-19 Genomics UK (COG-UK) Consortium | Jacquelyn Wynn, Mairead Hyland, The Lighthouse Lab in Alderley Park and Alex Alderton, Roberto Amato, Jeffrey Barrett, Sonia Goncalves, Ewan Harrison, David K. Jackson, Ian Johnston, Dominic Kwiatkowski, Cordelia Langford, John Sillitoe on behalf of the Wellcome Sanger Institute COVID-19 Surveillance Team |
| EPI_ISL_2781561, EPI_ISL_2781562 | Health Services Laboratories | Wellcome Sanger Institute for the COVID-19 Genomics UK (COG-UK) Consortium | Health Services Laboratories and Alex Alderton, Roberto Amato, Jeffrey Barrett, Sonia Goncalves, Ewan Harrison, David K. Jackson, Ian Johnston, Dominic Kwiatkowski, Cordelia Langford, John Sillitoe on behalf of the Wellcome Sanger Institute COVID-19 Surveillance Team |
| EPI_ISL_2781563, EPI_ISL_2781564 | Lighthouse Lab in Alderley Park | Wellcome Sanger Institute for the COVID-19 Genomics UK (COG-UK) Consortium | Jacquelyn Wynn, Mairead Hyland, The Lighthouse Lab in Alderley Park and Alex Alderton, Roberto Amato, Jeffrey Barrett, Sonia Goncalves, Ewan Harrison, David K. Jackson, Ian Johnston, Dominic Kwiatkowski, Cordelia Langford, John Sillitoe on behalf of the Wellcome Sanger Institute COVID-19 Surveillance Team |
| EPI_ISL_2781565 | Health Services Laboratories | Wellcome Sanger Institute for the COVID-19 Genomics UK (COG-UK) Consortium | Health Services Laboratories and Alex Alderton, Roberto Amato, Jeffrey Barrett, Sonia Goncalves, Ewan Harrison, David K. Jackson, Ian Johnston, Dominic Kwiatkowski, Cordelia Langford, John Sillitoe on behalf of the Wellcome Sanger Institute COVID-19 Surveillance Team |
| EPI_ISL_2781566 | Lighthouse Lab in Alderley Park | Wellcome Sanger Institute for the COVID-19 Genomics UK (COG-UK) Consortium | Jacquelyn Wynn, Mairead Hyland, The Lighthouse Lab in Alderley Park and Alex Alderton, Roberto Amato, Jeffrey Barrett, Sonia Goncalves, Ewan Harrison, David K. Jackson, Ian Johnston, Dominic Kwiatkowski, Cordelia Langford, John Sillitoe on behalf of the Wellcome Sanger Institute COVID-19 Surveillance Team |
| EPI_ISL_2781567 | Health Services Laboratories | Wellcome Sanger Institute for the COVID-19 Genomics UK (COG-UK) Consortium | Health Services Laboratories and Alex Alderton, Roberto Amato, Jeffrey Barrett, Sonia Goncalves, Ewan Harrison, David K. Jackson, Ian Johnston, Dominic Kwiatkowski, Cordelia Langford, John Sillitoe on behalf of the Wellcome Sanger Institute COVID-19 Surveillance Team |
| EPI_ISL_2781568 | Lighthouse Lab in Milton Keynes | Wellcome Sanger Institute for the COVID-19 Genomics UK (COG-UK) Consortium | The Lighthouse Lab in Milton Keynes and Alex Alderton, Roberto Amato, Jeffrey Barrett, Sonia Goncalves, Ewan Harrison, David K. Jackson, Ian Johnston, Dominic Kwiatkowski, Cordelia Langford, John Sillitoe on behalf of the Wellcome Sanger Institute COVID-19 Surveillance Team |
| EPI_ISL_2781569 | Lighthouse Lab in Alderley Park | Wellcome Sanger Institute for the COVID-19 Genomics UK (COG-UK) Consortium | Jacquelyn Wynn, Mairead Hyland, The Lighthouse Lab in Alderley Park and Alex Alderton, Roberto Amato, Jeffrey Barrett, Sonia Goncalves, Ewan Harrison, David K. Jackson, Ian Johnston, Dominic Kwiatkowski, Cordelia Langford, John Sillitoe on behalf of the Wellcome Sanger Institute COVID-19 Surveillance Team |
| EPI_ISL_2781570 | Lighthouse Lab in Milton Keynes | Wellcome Sanger Institute for the COVID-19 Genomics UK (COG-UK) Consortium | The Lighthouse Lab in Milton Keynes and Alex Alderton, Roberto Amato, Jeffrey Barrett, Sonia Goncalves, Ewan Harrison, David K. Jackson, Ian Johnston, Dominic Kwiatkowski, Cordelia Langford, John Sillitoe on behalf of the Wellcome Sanger Institute COVID-19 Surveillance Team |
| EPI_ISL_2781571 | Lighthouse Lab in Alderley Park | Wellcome Sanger Institute for the COVID-19 Genomics UK (COG-UK) Consortium | Jacquelyn Wynn, Mairead Hyland, The Lighthouse Lab in Alderley Park and Alex Alderton, Roberto Amato, Jeffrey Barrett, Sonia Goncalves, Ewan Harrison, David K. Jackson, Ian Johnston, Dominic Kwiatkowski, Cordelia Langford, John Sillitoe on behalf of the Wellcome Sanger Institute COVID-19 Surveillance Team |
| EPI_ISL_2781572 | Lighthouse Laboratory Plymouth | Wellcome Sanger Institute for the COVID-19 Genomics UK (COG-UK) Consortium | Lighthouse Laboratory Plymouth and Alex Alderton, Roberto Amato, Jeffrey Barrett, Sonia Goncalves, Ewan Harrison, David K. Jackson, Ian Johnston, Dominic Kwiatkowski, Cordelia Langford, John Sillitoe on behalf of the Wellcome Sanger Institute COVID-19 Surveillance Team |
| EPI_ISL_2781573 | Lighthouse Lab in Glasgow | Wellcome Sanger Institute for the COVID-19 Genomics UK (COG-UK) Consortium | Harper VanSteenhouse, Yumi Kasai, David Gray, Carol Clugston, Anna Dominiczak and Alex Alderton, Roberto Amato, Jeffrey Barrett, Sonia Goncalves, Ewan Harrison, David K. Jackson, Ian Johnston, Dominic Kwiatkowski, Cordelia Langford, John Sillitoe on behalf of the Wellcome Sanger Institute COVID-19 Surveillance Team |
| EPI_ISL_2781574 | Health Services Laboratories | Wellcome Sanger Institute for the COVID-19 Genomics UK (COG-UK) Consortium | Health Services Laboratories and Alex Alderton, Roberto Amato, Jeffrey Barrett, Sonia Goncalves, Ewan Harrison, David K. Jackson, Ian Johnston, Dominic Kwiatkowski, Cordelia Langford, John Sillitoe on behalf of the Wellcome Sanger Institute COVID-19 Surveillance Team |
| EPI_ISL_2781575, EPI_ISL_2781576 | Lighthouse Lab in Milton Keynes | Wellcome Sanger Institute for the COVID-19 Genomics UK (COG-UK) Consortium | The Lighthouse Lab in Milton Keynes and Alex Alderton, Roberto Amato, Jeffrey Barrett, Sonia Goncalves, Ewan Harrison, David K. Jackson, Ian Johnston, Dominic Kwiatkowski, Cordelia Langford, John Sillitoe on behalf of the Wellcome Sanger Institute COVID-19 Surveillance Team |
| EPI_ISL_2781577, EPI_ISL_2781578 | Lighthouse Lab in Alderley Park | Wellcome Sanger Institute for the COVID-19 Genomics UK (COG-UK) Consortium | Jacquelyn Wynn, Mairead Hyland, The Lighthouse Lab in Alderley Park and Alex Alderton, Roberto Amato, Jeffrey Barrett, Sonia Goncalves, Ewan Harrison, David K. Jackson, Ian Johnston, Dominic Kwiatkowski, Cordelia Langford, John Sillitoe on behalf of the Wellcome Sanger Institute COVID-19 Surveillance Team |
| EPI_ISL_2781579 | Lighthouse Lab in Glasgow | Wellcome Sanger Institute for the COVID-19 Genomics UK (COG-UK) Consortium | Harper VanSteenhouse, Yumi Kasai, David Gray, Carol Clugston, Anna Dominiczak and Alex Alderton, Roberto Amato, Jeffrey Barrett, Sonia Goncalves, Ewan Harrison, David K. Jackson, Ian Johnston, Dominic Kwiatkowski, Cordelia Langford, John Sillitoe on behalf of the Wellcome Sanger Institute COVID-19 Surveillance Team |
| EPI_ISL_2781580, EPI_ISL_2781581 | Lighthouse Lab in Milton Keynes | Wellcome Sanger Institute for the COVID-19 Genomics UK (COG-UK) Consortium | The Lighthouse Lab in Milton Keynes and Alex Alderton, Roberto Amato, Jeffrey Barrett, Sonia Goncalves, Ewan Harrison, David K. Jackson, Ian Johnston, Dominic Kwiatkowski, Cordelia Langford, John Sillitoe on behalf of the Wellcome Sanger Institute COVID-19 Surveillance Team |
| EPI_ISL_2781582 | Health Services Laboratories | Wellcome Sanger Institute for the COVID-19 Genomics UK (COG-UK) Consortium | Health Services Laboratories and Alex Alderton, Roberto Amato, Jeffrey Barrett, Sonia Goncalves, Ewan Harrison, David K. Jackson, Ian Johnston, Dominic Kwiatkowski, Cordelia Langford, John Sillitoe on behalf of the Wellcome Sanger Institute COVID-19 Surveillance Team |
| EPI_ISL_2781583 | Lighthouse Laboratory Plymouth | Wellcome Sanger Institute for the COVID-19 Genomics UK (COG-UK) Consortium | Lighthouse Laboratory Plymouth and Alex Alderton, Roberto Amato, Jeffrey Barrett, Sonia Goncalves, Ewan Harrison, David K. Jackson, Ian Johnston, Dominic Kwiatkowski, Cordelia Langford, John Sillitoe on behalf of the Wellcome Sanger Institute COVID-19 Surveillance Team |
| EPI_ISL_2781584 | Lighthouse Lab in Alderley Park | Wellcome Sanger Institute for the COVID-19 Genomics UK (COG-UK) Consortium | Jacquelyn Wynn, Mairead Hyland, The Lighthouse Lab in Alderley Park and Alex Alderton, Roberto Amato, Jeffrey Barrett, Sonia Goncalves, Ewan Harrison, David K. Jackson, Ian Johnston, Dominic Kwiatkowski, Cordelia Langford, John Sillitoe on behalf of the Wellcome Sanger Institute COVID-19 Surveillance Team |
| EPI_ISL_2781585 | Lighthouse Lab in Glasgow | Wellcome Sanger Institute for the COVID-19 Genomics UK (COG-UK) Consortium | Harper VanSteenhouse, Yumi Kasai, David Gray, Carol Clugston, Anna Dominiczak and Alex Alderton, Roberto Amato, Jeffrey Barrett, Sonia Goncalves, Ewan Harrison, David K. Jackson, Ian Johnston, Dominic Kwiatkowski, Cordelia Langford, John Sillitoe on behalf of the Wellcome Sanger Institute COVID-19 Surveillance Team |
| EPI_ISL_2781586 | Lighthouse Lab in Alderley Park | Wellcome Sanger Institute for the COVID-19 Genomics UK (COG-UK) Consortium | Jacquelyn Wynn, Mairead Hyland, The Lighthouse Lab in Alderley Park and Alex Alderton, Roberto Amato, Jeffrey Barrett, Sonia Goncalves, Ewan Harrison, David K. Jackson, Ian Johnston, Dominic Kwiatkowski, Cordelia Langford, John Sillitoe on behalf of the Wellcome Sanger Institute COVID-19 Surveillance Team |
| EPI_ISL_2781587 | Lighthouse Lab in Milton Keynes | Wellcome Sanger Institute for the COVID-19 Genomics UK (COG-UK) Consortium | The Lighthouse Lab in Milton Keynes and Alex Alderton, Roberto Amato, Jeffrey Barrett, Sonia Goncalves, Ewan Harrison, David K. Jackson, Ian Johnston, Dominic Kwiatkowski, Cordelia Langford, John Sillitoe on behalf of the Wellcome Sanger Institute COVID-19 Surveillance Team |
| EPI_ISL_2781588 | Lighthouse Lab in Glasgow | Wellcome Sanger Institute for the COVID-19 Genomics UK (COG-UK) Consortium | Harper VanSteenhouse, Yumi Kasai, David Gray, Carol Clugston, Anna Dominiczak and Alex Alderton, Roberto Amato, Jeffrey Barrett, Sonia Goncalves, Ewan Harrison, David K. Jackson, Ian Johnston, Dominic Kwiatkowski, Cordelia Langford, John Sillitoe on behalf of the Wellcome Sanger Institute COVID-19 Surveillance Team |
| EPI_ISL_2781589 | Lighthouse Lab in Alderley Park | Wellcome Sanger Institute for the COVID-19 Genomics UK (COG-UK) Consortium | Jacquelyn Wynn, Mairead Hyland, The Lighthouse Lab in Alderley Park and Alex Alderton, Roberto Amato, Jeffrey Barrett, Sonia Goncalves, Ewan Harrison, David K. Jackson, Ian Johnston, Dominic Kwiatkowski, Cordelia Langford, John Sillitoe on behalf of the Wellcome Sanger Institute COVID-19 Surveillance Team |
| EPI_ISL_2781590 | Lighthouse Lab in Milton Keynes | Wellcome Sanger Institute for the COVID-19 Genomics UK (COG-UK) | The Lighthouse Lab in Milton Keynes and Alex Alderton, Roberto Amato, Jeffrey Barrett, Sonia Goncalves, Ewan Harrison, David K. Jackson, Ian Johnston, Dominic Kwiatkowski, Cordelia Langford, John Sillitoe on behalf of the Wellcome Sanger |

| | | | |
|---|---|---|---|
| | | Consortium | Institute COVID-19 Surveillance Team |
| EPI_ISL_2781591 | Lighthouse Laboratory Plymouth | Wellcome Sanger Institute for the COVID-19 Genomics UK (COG-UK) Consortium | Lighthouse Laboratory Plymouth and Alex Alderton, Roberto Amato, Jeffrey Barrett, Sonia Goncalves, Ewan Harrison, David K. Jackson, Ian Johnston, Dominic Kwiatkowski, Cordelia Langford, John Sillitoe on behalf of the Wellcome Sanger Institute COVID-19 Surveillance Team |
| EPI_ISL_2781592 | Lighthouse Lab in Glasgow | Wellcome Sanger Institute for the COVID-19 Genomics UK (COG-UK) Consortium | Harper VanSteenhouse, Yumi Kasai, David Gray, Carol Clugston, Anna Dominiczak and Alex Alderton, Roberto Amato, Jeffrey Barrett, Sonia Goncalves, Ewan Harrison, David K. Jackson, Ian Johnston, Dominic Kwiatkowski, Cordelia Langford, John Sillitoe on behalf of the Wellcome Sanger Institute COVID-19 Surveillance Team |
| EPI_ISL_2781593 | Lighthouse Lab in Milton Keynes | Wellcome Sanger Institute for the COVID-19 Genomics UK (COG-UK) Consortium | The Lighthouse Lab in Milton Keynes and Alex Alderton, Roberto Amato, Jeffrey Barrett, Sonia Goncalves, Ewan Harrison, David K. Jackson, Ian Johnston, Dominic Kwiatkowski, Cordelia Langford, John Sillitoe on behalf of the Wellcome Sanger Institute COVID-19 Surveillance Team |
| EPI_ISL_2781594 | Lighthouse Lab in Alderley Park | Wellcome Sanger Institute for the COVID-19 Genomics UK (COG-UK) Consortium | Jacquelyn Wynn, Mairead Hyland, The Lighthouse Lab in Alderley Park and Alex Alderton, Roberto Amato, Jeffrey Barrett, Sonia Goncalves, Ewan Harrison, David K. Jackson, Ian Johnston, Dominic Kwiatkowski, Cordelia Langford, John Sillitoe on behalf of the Wellcome Sanger Institute COVID-19 Surveillance Team |
| EPI_ISL_2781595 | Health Services Laboratories | Wellcome Sanger Institute for the COVID-19 Genomics UK (COG-UK) Consortium | Health Services Laboratories and Alex Alderton, Roberto Amato, Jeffrey Barrett, Sonia Goncalves, Ewan Harrison, David K. Jackson, Ian Johnston, Dominic Kwiatkowski, Cordelia Langford, John Sillitoe on behalf of the Wellcome Sanger Institute COVID-19 Surveillance Team |
| EPI_ISL_2781596, EPI_ISL_2781597 | Lighthouse Lab in Milton Keynes | Wellcome Sanger Institute for the COVID-19 Genomics UK (COG-UK) Consortium | The Lighthouse Lab in Milton Keynes and Alex Alderton, Roberto Amato, Jeffrey Barrett, Sonia Goncalves, Ewan Harrison, David K. Jackson, Ian Johnston, Dominic Kwiatkowski, Cordelia Langford, John Sillitoe on behalf of the Wellcome Sanger Institute COVID-19 Surveillance Team |
| EPI_ISL_2781598 | Lighthouse Lab in Alderley Park | Wellcome Sanger Institute for the COVID-19 Genomics UK (COG-UK) Consortium | Jacquelyn Wynn, Mairead Hyland, The Lighthouse Lab in Alderley Park and Alex Alderton, Roberto Amato, Jeffrey Barrett, Sonia Goncalves, Ewan Harrison, David K. Jackson, Ian Johnston, Dominic Kwiatkowski, Cordelia Langford, John Sillitoe on behalf of the Wellcome Sanger Institute COVID-19 Surveillance Team |
| EPI_ISL_2781599, EPI_ISL_2781600 | Lighthouse Laboratory Plymouth | Wellcome Sanger Institute for the COVID-19 Genomics UK (COG-UK) Consortium | Lighthouse Laboratory Plymouth and Alex Alderton, Roberto Amato, Jeffrey Barrett, Sonia Goncalves, Ewan Harrison, David K. Jackson, Ian Johnston, Dominic Kwiatkowski, Cordelia Langford, John Sillitoe on behalf of the Wellcome Sanger Institute COVID-19 Surveillance Team |
| EPI_ISL_2781601 | Lighthouse Lab in Alderley Park | Wellcome Sanger Institute for the COVID-19 Genomics UK (COG-UK) Consortium | Jacquelyn Wynn, Mairead Hyland, The Lighthouse Lab in Alderley Park and Alex Alderton, Roberto Amato, Jeffrey Barrett, Sonia Goncalves, Ewan Harrison, David K. Jackson, Ian Johnston, Dominic Kwiatkowski, Cordelia Langford, John Sillitoe on behalf of the Wellcome Sanger Institute COVID-19 Surveillance Team |
| EPI_ISL_2781602 | Health Services Laboratories | Wellcome Sanger Institute for the COVID-19 Genomics UK (COG-UK) Consortium | Health Services Laboratories and Alex Alderton, Roberto Amato, Jeffrey Barrett, Sonia Goncalves, Ewan Harrison, David K. Jackson, Ian Johnston, Dominic Kwiatkowski, Cordelia Langford, John Sillitoe on behalf of the Wellcome Sanger Institute COVID-19 Surveillance Team |
| EPI_ISL_2781603 | Lighthouse Lab in Milton Keynes | Wellcome Sanger Institute for the COVID-19 Genomics UK (COG-UK) Consortium | The Lighthouse Lab in Milton Keynes and Alex Alderton, Roberto Amato, Jeffrey Barrett, Sonia Goncalves, Ewan Harrison, David K. Jackson, Ian Johnston, Dominic Kwiatkowski, Cordelia Langford, John Sillitoe on behalf of the Wellcome Sanger Institute COVID-19 Surveillance Team |
| EPI_ISL_2781604 | Lighthouse Lab in Alderley Park | Wellcome Sanger Institute for the COVID-19 Genomics UK (COG-UK) Consortium | Jacquelyn Wynn, Mairead Hyland, The Lighthouse Lab in Alderley Park and Alex Alderton, Roberto Amato, Jeffrey Barrett, Sonia Goncalves, Ewan Harrison, David K. Jackson, Ian Johnston, Dominic Kwiatkowski, Cordelia Langford, John Sillitoe on behalf of the Wellcome Sanger Institute COVID-19 Surveillance Team |
| EPI_ISL_2781605 | Lighthouse Lab in Milton Keynes | Wellcome Sanger Institute for the COVID-19 Genomics UK (COG-UK) Consortium | The Lighthouse Lab in Milton Keynes and Alex Alderton, Roberto Amato, Jeffrey Barrett, Sonia Goncalves, Ewan Harrison, David K. Jackson, Ian Johnston, Dominic Kwiatkowski, Cordelia Langford, John Sillitoe on behalf of the Wellcome Sanger Institute COVID-19 Surveillance Team |
| EPI_ISL_2781606 | Lighthouse Lab in Alderley Park | Wellcome Sanger Institute for the COVID-19 Genomics UK (COG-UK) Consortium | Jacquelyn Wynn, Mairead Hyland, The Lighthouse Lab in Alderley Park and Alex Alderton, Roberto Amato, Jeffrey Barrett, Sonia Goncalves, Ewan Harrison, David K. Jackson, Ian Johnston, Dominic Kwiatkowski, Cordelia Langford, John Sillitoe on behalf of the Wellcome Sanger Institute COVID-19 Surveillance Team |
| EPI_ISL_2781607 | Lighthouse Lab in Milton Keynes | Wellcome Sanger Institute for the COVID-19 Genomics UK (COG-UK) Consortium | The Lighthouse Lab in Milton Keynes and Alex Alderton, Roberto Amato, Jeffrey Barrett, Sonia Goncalves, Ewan Harrison, David K. Jackson, Ian Johnston, Dominic Kwiatkowski, Cordelia Langford, John Sillitoe on behalf of the Wellcome Sanger Institute COVID-19 Surveillance Team |
| EPI_ISL_2781608 | Lighthouse Lab in Alderley Park | Wellcome Sanger Institute for the COVID-19 Genomics UK (COG-UK) Consortium | Jacquelyn Wynn, Mairead Hyland, The Lighthouse Lab in Alderley Park and Alex Alderton, Roberto Amato, Jeffrey Barrett, Sonia Goncalves, Ewan Harrison, David K. Jackson, Ian Johnston, Dominic Kwiatkowski, Cordelia Langford, John Sillitoe on behalf of the Wellcome Sanger Institute COVID-19 Surveillance Team |
| EPI_ISL_2781609 | Lighthouse Lab in Milton Keynes | Wellcome Sanger Institute for the COVID-19 Genomics UK (COG-UK) Consortium | The Lighthouse Lab in Milton Keynes and Alex Alderton, Roberto Amato, Jeffrey Barrett, Sonia Goncalves, Ewan Harrison, David K. Jackson, Ian Johnston, Dominic Kwiatkowski, Cordelia Langford, John Sillitoe on behalf of the Wellcome Sanger Institute COVID-19 Surveillance Team |
| EPI_ISL_2781610 | Health Services Laboratories | Wellcome Sanger Institute for the COVID-19 Genomics UK (COG-UK) Consortium | Health Services Laboratories and Alex Alderton, Roberto Amato, Jeffrey Barrett, Sonia Goncalves, Ewan Harrison, David K. Jackson, Ian Johnston, Dominic Kwiatkowski, Cordelia Langford, John Sillitoe on behalf of the Wellcome Sanger Institute COVID-19 Surveillance Team |
| EPI_ISL_2781611 | Lighthouse Laboratory Plymouth | Wellcome Sanger Institute for the COVID-19 Genomics UK (COG-UK) Consortium | Lighthouse Laboratory Plymouth and Alex Alderton, Roberto Amato, Jeffrey Barrett, Sonia Goncalves, Ewan Harrison, David K. Jackson, Ian Johnston, Dominic Kwiatkowski, Cordelia Langford, John Sillitoe on behalf of the Wellcome Sanger Institute COVID-19 Surveillance Team |
| EPI_ISL_2781612 | Lighthouse Lab in Alderley Park | Wellcome Sanger Institute for the COVID-19 Genomics UK (COG-UK) Consortium | Jacquelyn Wynn, Mairead Hyland, The Lighthouse Lab in Alderley Park and Alex Alderton, Roberto Amato, Jeffrey Barrett, Sonia Goncalves, Ewan Harrison, David K. Jackson, Ian Johnston, Dominic Kwiatkowski, Cordelia Langford, John Sillitoe on behalf of the Wellcome Sanger Institute COVID-19 Surveillance Team |
| EPI_ISL_2781613 | Health Services Laboratories | Wellcome Sanger Institute for the COVID-19 Genomics UK (COG-UK) Consortium | Health Services Laboratories and Alex Alderton, Roberto Amato, Jeffrey Barrett, Sonia Goncalves, Ewan Harrison, David K. Jackson, Ian Johnston, Dominic Kwiatkowski, Cordelia Langford, John Sillitoe on behalf of the Wellcome Sanger Institute COVID-19 Surveillance Team |
| EPI_ISL_2781614, EPI_ISL_2781615, EPI_ISL_2781616 | Lighthouse Lab in Alderley Park | Wellcome Sanger Institute for the COVID-19 Genomics UK (COG-UK) Consortium | Jacquelyn Wynn, Mairead Hyland, The Lighthouse Lab in Alderley Park and Alex Alderton, Roberto Amato, Jeffrey Barrett, Sonia Goncalves, Ewan Harrison, David K. Jackson, Ian Johnston, Dominic Kwiatkowski, Cordelia Langford, John Sillitoe on behalf of the Wellcome Sanger Institute COVID-19 Surveillance Team |
| EPI_ISL_2781617 | Health Services Laboratories | Wellcome Sanger Institute for the COVID-19 Genomics UK (COG-UK) Consortium | Health Services Laboratories and Alex Alderton, Roberto Amato, Jeffrey Barrett, Sonia Goncalves, Ewan Harrison, David K. Jackson, Ian Johnston, Dominic Kwiatkowski, Cordelia Langford, John Sillitoe on behalf of the Wellcome Sanger Institute COVID-19 Surveillance Team |
| EPI_ISL_2781618, EPI_ISL_2781619 | Lighthouse Lab in Milton Keynes | Wellcome Sanger Institute for the COVID-19 Genomics UK (COG-UK) Consortium | The Lighthouse Lab in Milton Keynes and Alex Alderton, Roberto Amato, Jeffrey Barrett, Sonia Goncalves, Ewan Harrison, David K. Jackson, Ian Johnston, Dominic Kwiatkowski, Cordelia Langford, John Sillitoe on behalf of the Wellcome Sanger Institute COVID-19 Surveillance Team |
| EPI_ISL_2781620, EPI_ISL_2781621 | Health Services Laboratories | Wellcome Sanger Institute for the COVID-19 Genomics UK (COG-UK) Consortium | Health Services Laboratories and Alex Alderton, Roberto Amato, Jeffrey Barrett, Sonia Goncalves, Ewan Harrison, David K. Jackson, Ian Johnston, Dominic Kwiatkowski, Cordelia Langford, John Sillitoe on behalf of the Wellcome Sanger Institute COVID-19 Surveillance Team |
| EPI_ISL_2781622 | Lighthouse Lab in Milton Keynes | Wellcome Sanger Institute for the COVID-19 Genomics UK (COG-UK) Consortium | The Lighthouse Lab in Milton Keynes and Alex Alderton, Roberto Amato, Jeffrey Barrett, Sonia Goncalves, Ewan Harrison, David K. Jackson, Ian Johnston, Dominic Kwiatkowski, Cordelia Langford, John Sillitoe on behalf of the Wellcome Sanger Institute COVID-19 Surveillance Team |
| EPI_ISL_2781623 | Health Services Laboratories | Wellcome Sanger Institute for the COVID-19 Genomics UK (COG-UK) Consortium | Health Services Laboratories and Alex Alderton, Roberto Amato, Jeffrey Barrett, Sonia Goncalves, Ewan Harrison, David K. Jackson, Ian Johnston, Dominic Kwiatkowski, Cordelia Langford, John Sillitoe on behalf of the Wellcome Sanger Institute COVID-19 Surveillance Team |
| EPI_ISL_2781624 | Lighthouse Lab in Alderley Park | Wellcome Sanger Institute for the COVID-19 Genomics UK (COG-UK) Consortium | Jacquelyn Wynn, Mairead Hyland, The Lighthouse Lab in Alderley Park and Alex Alderton, Roberto Amato, Jeffrey Barrett, Sonia Goncalves, Ewan Harrison, David K. Jackson, Ian Johnston, Dominic Kwiatkowski, Cordelia Langford, John Sillitoe on behalf of the Wellcome Sanger Institute COVID-19 Surveillance Team |
| EPI_ISL_2781625 | Health Services Laboratories | Wellcome Sanger Institute for the COVID-19 Genomics UK (COG-UK) Consortium | Health Services Laboratories and Alex Alderton, Roberto Amato, Jeffrey Barrett, Sonia Goncalves, Ewan Harrison, David K. Jackson, Ian Johnston, Dominic Kwiatkowski, Cordelia Langford, John Sillitoe on behalf of the Wellcome Sanger Institute COVID-19 Surveillance Team |
| EPI_ISL_2781626 | Lighthouse Lab in Milton Keynes | Wellcome Sanger Institute for the COVID-19 Genomics UK (COG-UK) Consortium | The Lighthouse Lab in Milton Keynes and Alex Alderton, Roberto Amato, Jeffrey Barrett, Sonia Goncalves, Ewan Harrison, David K. Jackson, Ian Johnston, Dominic Kwiatkowski, Cordelia Langford, John Sillitoe on behalf of the Wellcome Sanger Institute COVID-19 Surveillance Team |
| EPI_ISL_2781627 | Lighthouse Lab in Alderley Park | Wellcome Sanger Institute for the COVID-19 Genomics UK (COG-UK) Consortium | Jacquelyn Wynn, Mairead Hyland, The Lighthouse Lab in Alderley Park and Alex Alderton, Roberto Amato, Jeffrey Barrett, Sonia Goncalves, Ewan Harrison, David K. Jackson, Ian Johnston, Dominic Kwiatkowski, Cordelia Langford, John Sillitoe on behalf of the Wellcome Sanger Institute COVID-19 Surveillance Team |
| EPI_ISL_2781628 | Lighthouse Lab in Milton Keynes | Wellcome Sanger Institute for the COVID-19 Genomics UK (COG-UK) Consortium | The Lighthouse Lab in Milton Keynes and Alex Alderton, Roberto Amato, Jeffrey Barrett, Sonia Goncalves, Ewan Harrison, David K. Jackson, Ian Johnston, Dominic Kwiatkowski, Cordelia Langford, John Sillitoe on behalf of the Wellcome Sanger Institute COVID-19 Surveillance Team |
| EPI_ISL_2781629, EPI_ISL_2781630, EPI_ISL_2781631 | Health Services Laboratories | Wellcome Sanger Institute for the COVID-19 Genomics UK (COG-UK) Consortium | Health Services Laboratories and Alex Alderton, Roberto Amato, Jeffrey Barrett, Sonia Goncalves, Ewan Harrison, David K. Jackson, Ian Johnston, Dominic Kwiatkowski, Cordelia Langford, John Sillitoe on behalf of the Wellcome Sanger Institute COVID-19 Surveillance Team |
| EPI_ISL_2781632 | Lighthouse Lab in Milton Keynes | Wellcome Sanger Institute for the COVID-19 Genomics UK (COG-UK) Consortium | The Lighthouse Lab in Milton Keynes and Alex Alderton, Roberto Amato, Jeffrey Barrett, Sonia Goncalves, Ewan Harrison, David K. Jackson, Ian Johnston, Dominic Kwiatkowski, Cordelia Langford, John Sillitoe on behalf of the Wellcome Sanger Institute COVID-19 Surveillance Team |
| EPI_ISL_2781633, EPI_ISL_2781634 | Lighthouse Lab in Alderley Park | Wellcome Sanger Institute for the COVID-19 Genomics UK (COG-UK) Consortium | Jacquelyn Wynn, Mairead Hyland, The Lighthouse Lab in Alderley Park and Alex Alderton, Roberto Amato, Jeffrey Barrett, Sonia Goncalves, Ewan Harrison, David K. Jackson, Ian Johnston, Dominic Kwiatkowski, Cordelia Langford, John Sillitoe on behalf of the Wellcome Sanger Institute COVID-19 Surveillance Team |
| EPI_ISL_2781635 | Lighthouse Laboratory Plymouth | Wellcome Sanger Institute for the COVID-19 Genomics UK (COG-UK) Consortium | Lighthouse Laboratory Plymouth and Alex Alderton, Roberto Amato, Jeffrey Barrett, Sonia Goncalves, Ewan Harrison, David K. Jackson, Ian Johnston, Dominic Kwiatkowski, Cordelia Langford, John Sillitoe on behalf of the Wellcome Sanger Institute COVID-19 Surveillance Team |
| EPI_ISL_2781636, EPI_ISL_2781637 | Lighthouse Lab in Milton Keynes | Wellcome Sanger Institute for the COVID-19 Genomics UK (COG-UK) Consortium | The Lighthouse Lab in Milton Keynes and Alex Alderton, Roberto Amato, Jeffrey Barrett, Sonia Goncalves, Ewan Harrison, David K. Jackson, Ian Johnston, Dominic Kwiatkowski, Cordelia Langford, John Sillitoe on behalf of the Wellcome Sanger Institute COVID-19 Surveillance Team |
| EPI_ISL_2781638 | Health Services Laboratories | Wellcome Sanger Institute for the COVID-19 Genomics UK (COG-UK) Consortium | Health Services Laboratories and Alex Alderton, Roberto Amato, Jeffrey Barrett, Sonia Goncalves, Ewan Harrison, David K. Jackson, Ian Johnston, Dominic Kwiatkowski, Cordelia Langford, John Sillitoe on behalf of the Wellcome Sanger Institute COVID-19 Surveillance Team |
| EPI_ISL_2781639 | Lighthouse Lab in Alderley Park | Wellcome Sanger Institute for the COVID-19 Genomics UK (COG-UK) Consortium | Jacquelyn Wynn, Mairead Hyland, The Lighthouse Lab in Alderley Park and Alex Alderton, Roberto Amato, Jeffrey Barrett, Sonia Goncalves, Ewan Harrison, David K. Jackson, Ian Johnston, Dominic Kwiatkowski, Cordelia Langford, John Sillitoe on behalf of the Wellcome Sanger Institute COVID-19 Surveillance Team |
| EPI_ISL_2781640 | Health Services Laboratories | Wellcome Sanger Institute for the COVID-19 Genomics UK (COG-UK) Consortium | Health Services Laboratories and Alex Alderton, Roberto Amato, Jeffrey Barrett, Sonia Goncalves, Ewan Harrison, David K. Jackson, Ian Johnston, Dominic Kwiatkowski, Cordelia Langford, John Sillitoe on behalf of the Wellcome Sanger Institute COVID-19 Surveillance Team |
| EPI_ISL_2781641 | Lighthouse Lab in Alderley Park | Wellcome Sanger Institute for the COVID-19 Genomics UK (COG-UK) Consortium | Jacquelyn Wynn, Mairead Hyland, The Lighthouse Lab in Alderley Park and Alex Alderton, Roberto Amato, Jeffrey Barrett, Sonia Goncalves, Ewan Harrison, David K. Jackson, Ian Johnston, Dominic Kwiatkowski, Cordelia Langford, John Sillitoe on behalf of the Wellcome Sanger Institute COVID-19 Surveillance Team |
| EPI_ISL_2781642 | Lighthouse Laboratory Plymouth | Wellcome Sanger Institute for the COVID-19 Genomics UK (COG-UK) Consortium | Lighthouse Laboratory Plymouth and Alex Alderton, Roberto Amato, Jeffrey Barrett, Sonia Goncalves, Ewan Harrison, David K. Jackson, Ian Johnston, Dominic Kwiatkowski, Cordelia Langford, John Sillitoe on behalf of the Wellcome Sanger Institute COVID-19 Surveillance Team |
| EPI_ISL_2781643 | Lighthouse Lab in Alderley Park | Wellcome Sanger Institute for the COVID-19 Genomics UK (COG-UK) Consortium | Jacquelyn Wynn, Mairead Hyland, The Lighthouse Lab in Alderley Park and Alex Alderton, Roberto Amato, Jeffrey Barrett, Sonia Goncalves, Ewan Harrison, David K. Jackson, Ian Johnston, Dominic Kwiatkowski, Cordelia Langford, John Sillitoe on behalf of the Wellcome Sanger Institute COVID-19 Surveillance Team |
| EPI_ISL_2781644 | Lighthouse Lab in Glasgow | Wellcome Sanger Institute for the COVID-19 Genomics UK (COG-UK) | Harper VanSteenhouse, Yumi Kasai, David Gray, Carol Clugston, Anna Dominiczak and Alex Alderton, Roberto Amato, Jeffrey Barrett, Sonia Goncalves, Ewan Harrison, David K. Jackson, Ian Johnston, Dominic Kwiatkowski, Cordelia Langford, |

| | | | |
|---|---|---|---|
| | | Consortium | John Sillitoe on behalf of the Wellcome Sanger Institute COVID-19 Surveillance Team |
| EPI_ISL_2781645 | Health Services Laboratories | Wellcome Sanger Institute for the COVID-19 Genomics UK (COG-UK) Consortium | Health Services Laboratories and Alex Alderton, Roberto Amato, Jeffrey Barrett, Sonia Goncalves, Ewan Harrison, David K. Jackson, Ian Johnston, Dominic Kwiatkowski, Cordelia Langford, John Sillitoe on behalf of the Wellcome Sanger Institute COVID-19 Surveillance Team |
| EPI_ISL_2781646 | Lighthouse Lab in Milton Keynes | Wellcome Sanger Institute for the COVID-19 Genomics UK (COG-UK) Consortium | The Lighthouse Lab in Milton Keynes and Alex Alderton, Roberto Amato, Jeffrey Barrett, Sonia Goncalves, Ewan Harrison, David K. Jackson, Ian Johnston, Dominic Kwiatkowski, Cordelia Langford, John Sillitoe on behalf of the Wellcome Sanger Institute COVID-19 Surveillance Team |
| EPI_ISL_2781647 | Lighthouse Laboratory Plymouth | Wellcome Sanger Institute for the COVID-19 Genomics UK (COG-UK) Consortium | Lighthouse Laboratory Plymouth and Alex Alderton, Roberto Amato, Jeffrey Barrett, Sonia Goncalves, Ewan Harrison, David K. Jackson, Ian Johnston, Dominic Kwiatkowski, Cordelia Langford, John Sillitoe on behalf of the Wellcome Sanger Institute COVID-19 Surveillance Team |
| EPI_ISL_2781648 | Health Services Laboratories | Wellcome Sanger Institute for the COVID-19 Genomics UK (COG-UK) Consortium | Health Services Laboratories and Alex Alderton, Roberto Amato, Jeffrey Barrett, Sonia Goncalves, Ewan Harrison, David K. Jackson, Ian Johnston, Dominic Kwiatkowski, Cordelia Langford, John Sillitoe on behalf of the Wellcome Sanger Institute COVID-19 Surveillance Team |
| EPI_ISL_2781649 | Lighthouse Lab in Alderley Park | Wellcome Sanger Institute for the COVID-19 Genomics UK (COG-UK) Consortium | Jacquelyn Wynn, Mairead Hyland, The Lighthouse Lab in Alderley Park and Alex Alderton, Roberto Amato, Jeffrey Barrett, Sonia Goncalves, Ewan Harrison, David K. Jackson, Ian Johnston, Dominic Kwiatkowski, Cordelia Langford, John Sillitoe on behalf of the Wellcome Sanger Institute COVID-19 Surveillance Team |
| EPI_ISL_2781650 | Lighthouse Lab in Milton Keynes | Wellcome Sanger Institute for the COVID-19 Genomics UK (COG-UK) Consortium | The Lighthouse Lab in Milton Keynes and Alex Alderton, Roberto Amato, Jeffrey Barrett, Sonia Goncalves, Ewan Harrison, David K. Jackson, Ian Johnston, Dominic Kwiatkowski, Cordelia Langford, John Sillitoe on behalf of the Wellcome Sanger Institute COVID-19 Surveillance Team |
| EPI_ISL_2781651 | Lighthouse Lab in Glasgow | Wellcome Sanger Institute for the COVID-19 Genomics UK (COG-UK) Consortium | Harper VanSteenhouse, Yumi Kasai, David Gray, Carol Clugston, Anna Dominiczak and Alex Alderton, Roberto Amato, Jeffrey Barrett, Sonia Goncalves, Ewan Harrison, David K. Jackson, Ian Johnston, Dominic Kwiatkowski, Cordelia Langford, John Sillitoe on behalf of the Wellcome Sanger Institute COVID-19 Surveillance Team |
| EPI_ISL_2781652, EPI_ISL_2781653 | Lighthouse Lab in Alderley Park | Wellcome Sanger Institute for the COVID-19 Genomics UK (COG-UK) Consortium | Jacquelyn Wynn, Mairead Hyland, The Lighthouse Lab in Alderley Park and Alex Alderton, Roberto Amato, Jeffrey Barrett, Sonia Goncalves, Ewan Harrison, David K. Jackson, Ian Johnston, Dominic Kwiatkowski, Cordelia Langford, John Sillitoe on behalf of the Wellcome Sanger Institute COVID-19 Surveillance Team |
| EPI_ISL_2781654, EPI_ISL_2781655 | Lighthouse Lab in Milton Keynes | Wellcome Sanger Institute for the COVID-19 Genomics UK (COG-UK) Consortium | The Lighthouse Lab in Milton Keynes and Alex Alderton, Roberto Amato, Jeffrey Barrett, Sonia Goncalves, Ewan Harrison, David K. Jackson, Ian Johnston, Dominic Kwiatkowski, Cordelia Langford, John Sillitoe on behalf of the Wellcome Sanger Institute COVID-19 Surveillance Team |
| EPI_ISL_2781656 | Lighthouse Lab in Alderley Park | Wellcome Sanger Institute for the COVID-19 Genomics UK (COG-UK) Consortium | Jacquelyn Wynn, Mairead Hyland, The Lighthouse Lab in Alderley Park and Alex Alderton, Roberto Amato, Jeffrey Barrett, Sonia Goncalves, Ewan Harrison, David K. Jackson, Ian Johnston, Dominic Kwiatkowski, Cordelia Langford, John Sillitoe on behalf of the Wellcome Sanger Institute COVID-19 Surveillance Team |
| EPI_ISL_2781657, EPI_ISL_2781658 | Health Services Laboratories | Wellcome Sanger Institute for the COVID-19 Genomics UK (COG-UK) Consortium | Health Services Laboratories and Alex Alderton, Roberto Amato, Jeffrey Barrett, Sonia Goncalves, Ewan Harrison, David K. Jackson, Ian Johnston, Dominic Kwiatkowski, Cordelia Langford, John Sillitoe on behalf of the Wellcome Sanger Institute COVID-19 Surveillance Team |
| EPI_ISL_2781659 | Lighthouse Lab in Alderley Park | Wellcome Sanger Institute for the COVID-19 Genomics UK (COG-UK) Consortium | Jacquelyn Wynn, Mairead Hyland, The Lighthouse Lab in Alderley Park and Alex Alderton, Roberto Amato, Jeffrey Barrett, Sonia Goncalves, Ewan Harrison, David K. Jackson, Ian Johnston, Dominic Kwiatkowski, Cordelia Langford, John Sillitoe on behalf of the Wellcome Sanger Institute COVID-19 Surveillance Team |
| EPI_ISL_2781660 | Health Services Laboratories | Wellcome Sanger Institute for the COVID-19 Genomics UK (COG-UK) Consortium | Health Services Laboratories and Alex Alderton, Roberto Amato, Jeffrey Barrett, Sonia Goncalves, Ewan Harrison, David K. Jackson, Ian Johnston, Dominic Kwiatkowski, Cordelia Langford, John Sillitoe on behalf of the Wellcome Sanger Institute COVID-19 Surveillance Team |
| EPI_ISL_2781661 | Lighthouse Lab in Milton Keynes | Wellcome Sanger Institute for the COVID-19 Genomics UK (COG-UK) Consortium | The Lighthouse Lab in Milton Keynes and Alex Alderton, Roberto Amato, Jeffrey Barrett, Sonia Goncalves, Ewan Harrison, David K. Jackson, Ian Johnston, Dominic Kwiatkowski, Cordelia Langford, John Sillitoe on behalf of the Wellcome Sanger Institute COVID-19 Surveillance Team |
| EPI_ISL_2781662 | Health Services Laboratories | Wellcome Sanger Institute for the COVID-19 Genomics UK (COG-UK) Consortium | Health Services Laboratories and Alex Alderton, Roberto Amato, Jeffrey Barrett, Sonia Goncalves, Ewan Harrison, David K. Jackson, Ian Johnston, Dominic Kwiatkowski, Cordelia Langford, John Sillitoe on behalf of the Wellcome Sanger Institute COVID-19 Surveillance Team |
| EPI_ISL_2781663 | Lighthouse Lab in Alderley Park | Wellcome Sanger Institute for the COVID-19 Genomics UK (COG-UK) Consortium | Jacquelyn Wynn, Mairead Hyland, The Lighthouse Lab in Alderley Park and Alex Alderton, Roberto Amato, Jeffrey Barrett, Sonia Goncalves, Ewan Harrison, David K. Jackson, Ian Johnston, Dominic Kwiatkowski, Cordelia Langford, John Sillitoe on behalf of the Wellcome Sanger Institute COVID-19 Surveillance Team |
| EPI_ISL_2781664 | Lighthouse Lab in Milton Keynes | Wellcome Sanger Institute for the COVID-19 Genomics UK (COG-UK) Consortium | The Lighthouse Lab in Milton Keynes and Alex Alderton, Roberto Amato, Jeffrey Barrett, Sonia Goncalves, Ewan Harrison, David K. Jackson, Ian Johnston, Dominic Kwiatkowski, Cordelia Langford, John Sillitoe on behalf of the Wellcome Sanger Institute COVID-19 Surveillance Team |
| EPI_ISL_2781665 | Lighthouse Laboratory Plymouth | Wellcome Sanger Institute for the COVID-19 Genomics UK (COG-UK) Consortium | Lighthouse Laboratory Plymouth and Alex Alderton, Roberto Amato, Jeffrey Barrett, Sonia Goncalves, Ewan Harrison, David K. Jackson, Ian Johnston, Dominic Kwiatkowski, Cordelia Langford, John Sillitoe on behalf of the Wellcome Sanger Institute COVID-19 Surveillance Team |
| EPI_ISL_2781666 | Health Services Laboratories | Wellcome Sanger Institute for the COVID-19 Genomics UK (COG-UK) Consortium | Health Services Laboratories and Alex Alderton, Roberto Amato, Jeffrey Barrett, Sonia Goncalves, Ewan Harrison, David K. Jackson, Ian Johnston, Dominic Kwiatkowski, Cordelia Langford, John Sillitoe on behalf of the Wellcome Sanger Institute COVID-19 Surveillance Team |
| EPI_ISL_2781667 | Lighthouse Lab in Milton Keynes | Wellcome Sanger Institute for the COVID-19 Genomics UK (COG-UK) Consortium | The Lighthouse Lab in Milton Keynes and Alex Alderton, Roberto Amato, Jeffrey Barrett, Sonia Goncalves, Ewan Harrison, David K. Jackson, Ian Johnston, Dominic Kwiatkowski, Cordelia Langford, John Sillitoe on behalf of the Wellcome Sanger Institute COVID-19 Surveillance Team |
| EPI_ISL_2781668 | Lighthouse Lab in Alderley Park | Wellcome Sanger Institute for the COVID-19 Genomics UK (COG-UK) Consortium | Jacquelyn Wynn, Mairead Hyland, The Lighthouse Lab in Alderley Park and Alex Alderton, Roberto Amato, Jeffrey Barrett, Sonia Goncalves, Ewan Harrison, David K. Jackson, Ian Johnston, Dominic Kwiatkowski, Cordelia Langford, John Sillitoe on behalf of the Wellcome Sanger Institute COVID-19 Surveillance Team |
| EPI_ISL_2781669, EPI_ISL_2781670, EPI_ISL_2781671, EPI_ISL_2781672 | Health Services Laboratories | Wellcome Sanger Institute for the COVID-19 Genomics UK (COG-UK) Consortium | Health Services Laboratories and Alex Alderton, Roberto Amato, Jeffrey Barrett, Sonia Goncalves, Ewan Harrison, David K. Jackson, Ian Johnston, Dominic Kwiatkowski, Cordelia Langford, John Sillitoe on behalf of the Wellcome Sanger Institute COVID-19 Surveillance Team |
| EPI_ISL_2781673 | Lighthouse Lab in Alderley Park | Wellcome Sanger Institute for the COVID-19 Genomics UK (COG-UK) Consortium | Jacquelyn Wynn, Mairead Hyland, The Lighthouse Lab in Alderley Park and Alex Alderton, Roberto Amato, Jeffrey Barrett, Sonia Goncalves, Ewan Harrison, David K. Jackson, Ian Johnston, Dominic Kwiatkowski, Cordelia Langford, John Sillitoe on behalf of the Wellcome Sanger Institute COVID-19 Surveillance Team |
| EPI_ISL_2781674, EPI_ISL_2781675, EPI_ISL_2781676 | Lighthouse Lab in Milton Keynes | Wellcome Sanger Institute for the COVID-19 Genomics UK (COG-UK) Consortium | The Lighthouse Lab in Milton Keynes and Alex Alderton, Roberto Amato, Jeffrey Barrett, Sonia Goncalves, Ewan Harrison, David K. Jackson, Ian Johnston, Dominic Kwiatkowski, Cordelia Langford, John Sillitoe on behalf of the Wellcome Sanger Institute COVID-19 Surveillance Team |
| EPI_ISL_2781677 | Lighthouse Lab in Alderley Park | Wellcome Sanger Institute for the COVID-19 Genomics UK (COG-UK) Consortium | Jacquelyn Wynn, Mairead Hyland, The Lighthouse Lab in Alderley Park and Alex Alderton, Roberto Amato, Jeffrey Barrett, Sonia Goncalves, Ewan Harrison, David K. Jackson, Ian Johnston, Dominic Kwiatkowski, Cordelia Langford, John Sillitoe on behalf of the Wellcome Sanger Institute COVID-19 Surveillance Team |
| EPI_ISL_2781678 | Lighthouse Lab in Milton Keynes | Wellcome Sanger Institute for the COVID-19 Genomics UK (COG-UK) Consortium | The Lighthouse Lab in Milton Keynes and Alex Alderton, Roberto Amato, Jeffrey Barrett, Sonia Goncalves, Ewan Harrison, David K. Jackson, Ian Johnston, Dominic Kwiatkowski, Cordelia Langford, John Sillitoe on behalf of the Wellcome Sanger Institute COVID-19 Surveillance Team |
| EPI_ISL_2781679 | Lighthouse Lab in Alderley Park | Wellcome Sanger Institute for the COVID-19 Genomics UK (COG-UK) Consortium | Jacquelyn Wynn, Mairead Hyland, The Lighthouse Lab in Alderley Park and Alex Alderton, Roberto Amato, Jeffrey Barrett, Sonia Goncalves, Ewan Harrison, David K. Jackson, Ian Johnston, Dominic Kwiatkowski, Cordelia Langford, John Sillitoe on behalf of the Wellcome Sanger Institute COVID-19 Surveillance Team |
| EPI_ISL_2781680 | Lighthouse Laboratory Plymouth | Wellcome Sanger Institute for the COVID-19 Genomics UK (COG-UK) Consortium | Lighthouse Laboratory Plymouth and Alex Alderton, Roberto Amato, Jeffrey Barrett, Sonia Goncalves, Ewan Harrison, David K. Jackson, Ian Johnston, Dominic Kwiatkowski, Cordelia Langford, John Sillitoe on behalf of the Wellcome Sanger Institute COVID-19 Surveillance Team |
| EPI_ISL_2781681, EPI_ISL_2781682 | Health Services Laboratories | Wellcome Sanger Institute for the COVID-19 Genomics UK (COG-UK) Consortium | Health Services Laboratories and Alex Alderton, Roberto Amato, Jeffrey Barrett, Sonia Goncalves, Ewan Harrison, David K. Jackson, Ian Johnston, Dominic Kwiatkowski, Cordelia Langford, John Sillitoe on behalf of the Wellcome Sanger Institute COVID-19 Surveillance Team |
| EPI_ISL_2781683 | Lighthouse Lab in Alderley Park | Wellcome Sanger Institute for the COVID-19 Genomics UK (COG-UK) Consortium | Jacquelyn Wynn, Mairead Hyland, The Lighthouse Lab in Alderley Park and Alex Alderton, Roberto Amato, Jeffrey Barrett, Sonia Goncalves, Ewan Harrison, David K. Jackson, Ian Johnston, Dominic Kwiatkowski, Cordelia Langford, John Sillitoe on behalf of the Wellcome Sanger Institute COVID-19 Surveillance Team |
| EPI_ISL_2781684 | Lighthouse Laboratory Plymouth | Wellcome Sanger Institute for the COVID-19 Genomics UK (COG-UK) Consortium | Lighthouse Laboratory Plymouth and Alex Alderton, Roberto Amato, Jeffrey Barrett, Sonia Goncalves, Ewan Harrison, David K. Jackson, Ian Johnston, Dominic Kwiatkowski, Cordelia Langford, John Sillitoe on behalf of the Wellcome Sanger Institute COVID-19 Surveillance Team |
| EPI_ISL_2781685 | Lighthouse Lab in Milton Keynes | Wellcome Sanger Institute for the COVID-19 Genomics UK (COG-UK) Consortium | The Lighthouse Lab in Milton Keynes and Alex Alderton, Roberto Amato, Jeffrey Barrett, Sonia Goncalves, Ewan Harrison, David K. Jackson, Ian Johnston, Dominic Kwiatkowski, Cordelia Langford, John Sillitoe on behalf of the Wellcome Sanger Institute COVID-19 Surveillance Team |
| EPI_ISL_2781686 | Lighthouse Lab in Glasgow | Wellcome Sanger Institute for the COVID-19 Genomics UK (COG-UK) Consortium | Harper VanSteenhouse, Yumi Kasai, David Gray, Carol Clugston, Anna Dominiczak and Alex Alderton, Roberto Amato, Jeffrey Barrett, Sonia Goncalves, Ewan Harrison, David K. Jackson, Ian Johnston, Dominic Kwiatkowski, Cordelia Langford, John Sillitoe on behalf of the Wellcome Sanger Institute COVID-19 Surveillance Team |
| EPI_ISL_2781687 | Lighthouse Lab in Alderley Park | Wellcome Sanger Institute for the COVID-19 Genomics UK (COG-UK) Consortium | Jacquelyn Wynn, Mairead Hyland, The Lighthouse Lab in Alderley Park and Alex Alderton, Roberto Amato, Jeffrey Barrett, Sonia Goncalves, Ewan Harrison, David K. Jackson, Ian Johnston, Dominic Kwiatkowski, Cordelia Langford, John Sillitoe on behalf of the Wellcome Sanger Institute COVID-19 Surveillance Team |
| EPI_ISL_2781688 | Lighthouse Lab in Milton Keynes | Wellcome Sanger Institute for the COVID-19 Genomics UK (COG-UK) Consortium | The Lighthouse Lab in Milton Keynes and Alex Alderton, Roberto Amato, Jeffrey Barrett, Sonia Goncalves, Ewan Harrison, David K. Jackson, Ian Johnston, Dominic Kwiatkowski, Cordelia Langford, John Sillitoe on behalf of the Wellcome Sanger Institute COVID-19 Surveillance Team |
| EPI_ISL_2781689, EPI_ISL_2781690, EPI_ISL_2781691 | Lighthouse Laboratory Plymouth | Wellcome Sanger Institute for the COVID-19 Genomics UK (COG-UK) Consortium | Lighthouse Laboratory Plymouth and Alex Alderton, Roberto Amato, Jeffrey Barrett, Sonia Goncalves, Ewan Harrison, David K. Jackson, Ian Johnston, Dominic Kwiatkowski, Cordelia Langford, John Sillitoe on behalf of the Wellcome Sanger Institute COVID-19 Surveillance Team |
| EPI_ISL_2781692 | Lighthouse Lab in Milton Keynes | Wellcome Sanger Institute for the COVID-19 Genomics UK (COG-UK) Consortium | The Lighthouse Lab in Milton Keynes and Alex Alderton, Roberto Amato, Jeffrey Barrett, Sonia Goncalves, Ewan Harrison, David K. Jackson, Ian Johnston, Dominic Kwiatkowski, Cordelia Langford, John Sillitoe on behalf of the Wellcome Sanger Institute COVID-19 Surveillance Team |
| EPI_ISL_2781693 | Lighthouse Lab in Glasgow | Wellcome Sanger Institute for the COVID-19 Genomics UK (COG-UK) Consortium | Harper VanSteenhouse, Yumi Kasai, David Gray, Carol Clugston, Anna Dominiczak and Alex Alderton, Roberto Amato, Jeffrey Barrett, Sonia Goncalves, Ewan Harrison, David K. Jackson, Ian Johnston, Dominic Kwiatkowski, Cordelia Langford, John Sillitoe on behalf of the Wellcome Sanger Institute COVID-19 Surveillance Team |
| EPI_ISL_2781694 | Lighthouse Laboratory Plymouth | Wellcome Sanger Institute for the COVID-19 Genomics UK (COG-UK) Consortium | Lighthouse Laboratory Plymouth and Alex Alderton, Roberto Amato, Jeffrey Barrett, Sonia Goncalves, Ewan Harrison, David K. Jackson, Ian Johnston, Dominic Kwiatkowski, Cordelia Langford, John Sillitoe on behalf of the Wellcome Sanger Institute COVID-19 Surveillance Team |
| EPI_ISL_2781695 | Health Services Laboratories | Wellcome Sanger Institute for the COVID-19 Genomics UK (COG-UK) Consortium | Health Services Laboratories and Alex Alderton, Roberto Amato, Jeffrey Barrett, Sonia Goncalves, Ewan Harrison, David K. Jackson, Ian Johnston, Dominic Kwiatkowski, Cordelia Langford, John Sillitoe on behalf of the Wellcome Sanger Institute COVID-19 Surveillance Team |
| EPI_ISL_2781696 | Lighthouse Lab in Milton Keynes | Wellcome Sanger Institute for the COVID-19 Genomics UK (COG-UK) Consortium | The Lighthouse Lab in Milton Keynes and Alex Alderton, Roberto Amato, Jeffrey Barrett, Sonia Goncalves, Ewan Harrison, David K. Jackson, Ian Johnston, Dominic Kwiatkowski, Cordelia Langford, John Sillitoe on behalf of the Wellcome Sanger Institute COVID-19 Surveillance Team |
| EPI_ISL_2781697 | Lighthouse Lab in Alderley Park | Wellcome Sanger Institute for the COVID-19 Genomics UK (COG-UK) Consortium | Jacquelyn Wynn, Mairead Hyland, The Lighthouse Lab in Alderley Park and Alex Alderton, Roberto Amato, Jeffrey Barrett, Sonia Goncalves, Ewan Harrison, David K. Jackson, Ian Johnston, Dominic Kwiatkowski, Cordelia Langford, John Sillitoe on behalf of the Wellcome Sanger Institute COVID-19 Surveillance Team |
| EPI_ISL_2781698 | Health Services Laboratories | Wellcome Sanger Institute for the COVID-19 Genomics UK (COG-UK) Consortium | Health Services Laboratories and Alex Alderton, Roberto Amato, Jeffrey Barrett, Sonia Goncalves, Ewan Harrison, David K. Jackson, Ian Johnston, Dominic Kwiatkowski, Cordelia Langford, John Sillitoe on behalf of the Wellcome Sanger Institute COVID-19 Surveillance Team |

| | | | |
|---|---|---|---|
| EPI_ISL_2781699, EPI_ISL_2781700 | Lighthouse Lab in Alderley Park | Wellcome Sanger Institute for the COVID-19 Genomics UK (COG-UK) Consortium | Jacquelyn Wynn, Mairead Hyland, The Lighthouse Lab in Alderley Park and Alex Alderton, Roberto Amato, Jeffrey Barrett, Sonia Goncalves, Ewan Harrison, David K. Jackson, Ian Johnston, Dominic Kwiatkowski, Cordelia Langford, John Sillitoe on behalf of the Wellcome Sanger Institute COVID-19 Surveillance Team |
| EPI_ISL_2781701 | Health Services Laboratories | Wellcome Sanger Institute for the COVID-19 Genomics UK (COG-UK) Consortium | Health Services Laboratories and Alex Alderton, Roberto Amato, Jeffrey Barrett, Sonia Goncalves, Ewan Harrison, David K. Jackson, Ian Johnston, Dominic Kwiatkowski, Cordelia Langford, John Sillitoe on behalf of the Wellcome Sanger Institute COVID-19 Surveillance Team |
| EPI_ISL_2781702, EPI_ISL_2781703, EPI_ISL_2781704 | Lighthouse Lab in Milton Keynes | Wellcome Sanger Institute for the COVID-19 Genomics UK (COG-UK) Consortium | The Lighthouse Lab in Milton Keynes and Alex Alderton, Roberto Amato, Jeffrey Barrett, Sonia Goncalves, Ewan Harrison, David K. Jackson, Ian Johnston, Dominic Kwiatkowski, Cordelia Langford, John Sillitoe on behalf of the Wellcome Sanger Institute COVID-19 Surveillance Team |
| EPI_ISL_2781705 | Health Services Laboratories | Wellcome Sanger Institute for the COVID-19 Genomics UK (COG-UK) Consortium | Health Services Laboratories and Alex Alderton, Roberto Amato, Jeffrey Barrett, Sonia Goncalves, Ewan Harrison, David K. Jackson, Ian Johnston, Dominic Kwiatkowski, Cordelia Langford, John Sillitoe on behalf of the Wellcome Sanger Institute COVID-19 Surveillance Team |
| EPI_ISL_2781706, EPI_ISL_2781707, EPI_ISL_2781708 | Lighthouse Lab in Alderley Park | Wellcome Sanger Institute for the COVID-19 Genomics UK (COG-UK) Consortium | Jacquelyn Wynn, Mairead Hyland, The Lighthouse Lab in Alderley Park and Alex Alderton, Roberto Amato, Jeffrey Barrett, Sonia Goncalves, Ewan Harrison, David K. Jackson, Ian Johnston, Dominic Kwiatkowski, Cordelia Langford, John Sillitoe on behalf of the Wellcome Sanger Institute COVID-19 Surveillance Team |
| EPI_ISL_2781709 | Lighthouse Lab in Milton Keynes | Wellcome Sanger Institute for the COVID-19 Genomics UK (COG-UK) Consortium | The Lighthouse Lab in Milton Keynes and Alex Alderton, Roberto Amato, Jeffrey Barrett, Sonia Goncalves, Ewan Harrison, David K. Jackson, Ian Johnston, Dominic Kwiatkowski, Cordelia Langford, John Sillitoe on behalf of the Wellcome Sanger Institute COVID-19 Surveillance Team |
| EPI_ISL_2781710 | Lighthouse Lab in Glasgow | Wellcome Sanger Institute for the COVID-19 Genomics UK (COG-UK) Consortium | Harper VanSteenhouse, Yumi Kasai, David Gray, Carol Clugston, Anna Dominiczak and Alex Alderton, Roberto Amato, Jeffrey Barrett, Sonia Goncalves, Ewan Harrison, David K. Jackson, Ian Johnston, Dominic Kwiatkowski, Cordelia Langford, John Sillitoe on behalf of the Wellcome Sanger Institute COVID-19 Surveillance Team |
| EPI_ISL_2781711 | Lighthouse Lab in Alderley Park | Wellcome Sanger Institute for the COVID-19 Genomics UK (COG-UK) Consortium | Jacquelyn Wynn, Mairead Hyland, The Lighthouse Lab in Alderley Park and Alex Alderton, Roberto Amato, Jeffrey Barrett, Sonia Goncalves, Ewan Harrison, David K. Jackson, Ian Johnston, Dominic Kwiatkowski, Cordelia Langford, John Sillitoe on behalf of the Wellcome Sanger Institute COVID-19 Surveillance Team |
| EPI_ISL_2781712, EPI_ISL_2781713, EPI_ISL_2781714, EPI_ISL_2781715 | Lighthouse Lab in Milton Keynes | Wellcome Sanger Institute for the COVID-19 Genomics UK (COG-UK) Consortium | The Lighthouse Lab in Milton Keynes and Alex Alderton, Roberto Amato, Jeffrey Barrett, Sonia Goncalves, Ewan Harrison, David K. Jackson, Ian Johnston, Dominic Kwiatkowski, Cordelia Langford, John Sillitoe on behalf of the Wellcome Sanger Institute COVID-19 Surveillance Team |
| EPI_ISL_2781716 | Lighthouse Lab in Alderley Park | Wellcome Sanger Institute for the COVID-19 Genomics UK (COG-UK) Consortium | Jacquelyn Wynn, Mairead Hyland, The Lighthouse Lab in Alderley Park and Alex Alderton, Roberto Amato, Jeffrey Barrett, Sonia Goncalves, Ewan Harrison, David K. Jackson, Ian Johnston, Dominic Kwiatkowski, Cordelia Langford, John Sillitoe on behalf of the Wellcome Sanger Institute COVID-19 Surveillance Team |
| EPI_ISL_2781717 | Lighthouse Lab in Milton Keynes | Wellcome Sanger Institute for the COVID-19 Genomics UK (COG-UK) Consortium | The Lighthouse Lab in Milton Keynes and Alex Alderton, Roberto Amato, Jeffrey Barrett, Sonia Goncalves, Ewan Harrison, David K. Jackson, Ian Johnston, Dominic Kwiatkowski, Cordelia Langford, John Sillitoe on behalf of the Wellcome Sanger Institute COVID-19 Surveillance Team |
| EPI_ISL_2781718 | Lighthouse Lab in Alderley Park | Wellcome Sanger Institute for the COVID-19 Genomics UK (COG-UK) Consortium | Jacquelyn Wynn, Mairead Hyland, The Lighthouse Lab in Alderley Park and Alex Alderton, Roberto Amato, Jeffrey Barrett, Sonia Goncalves, Ewan Harrison, David K. Jackson, Ian Johnston, Dominic Kwiatkowski, Cordelia Langford, John Sillitoe on behalf of the Wellcome Sanger Institute COVID-19 Surveillance Team |
| EPI_ISL_2781719 | Lighthouse Laboratory Plymouth | Wellcome Sanger Institute for the COVID-19 Genomics UK (COG-UK) Consortium | Lighthouse Laboratory Plymouth and Alex Alderton, Roberto Amato, Jeffrey Barrett, Sonia Goncalves, Ewan Harrison, David K. Jackson, Ian Johnston, Dominic Kwiatkowski, Cordelia Langford, John Sillitoe on behalf of the Wellcome Sanger Institute COVID-19 Surveillance Team |
| EPI_ISL_2781720 | Health Services Laboratories | Wellcome Sanger Institute for the COVID-19 Genomics UK (COG-UK) Consortium | Health Services Laboratories and Alex Alderton, Roberto Amato, Jeffrey Barrett, Sonia Goncalves, Ewan Harrison, David K. Jackson, Ian Johnston, Dominic Kwiatkowski, Cordelia Langford, John Sillitoe on behalf of the Wellcome Sanger Institute COVID-19 Surveillance Team |
| EPI_ISL_2781721 | Lighthouse Lab in Alderley Park | Wellcome Sanger Institute for the COVID-19 Genomics UK (COG-UK) Consortium | Jacquelyn Wynn, Mairead Hyland, The Lighthouse Lab in Alderley Park and Alex Alderton, Roberto Amato, Jeffrey Barrett, Sonia Goncalves, Ewan Harrison, David K. Jackson, Ian Johnston, Dominic Kwiatkowski, Cordelia Langford, John Sillitoe on behalf of the Wellcome Sanger Institute COVID-19 Surveillance Team |
| EPI_ISL_2781722, EPI_ISL_2781723, EPI_ISL_2781724 | Health Services Laboratories | Wellcome Sanger Institute for the COVID-19 Genomics UK (COG-UK) Consortium | Health Services Laboratories and Alex Alderton, Roberto Amato, Jeffrey Barrett, Sonia Goncalves, Ewan Harrison, David K. Jackson, Ian Johnston, Dominic Kwiatkowski, Cordelia Langford, John Sillitoe on behalf of the Wellcome Sanger Institute COVID-19 Surveillance Team |
| EPI_ISL_2781725, EPI_ISL_2781726 | Lighthouse Lab in Milton Keynes | Wellcome Sanger Institute for the COVID-19 Genomics UK (COG-UK) Consortium | The Lighthouse Lab in Milton Keynes and Alex Alderton, Roberto Amato, Jeffrey Barrett, Sonia Goncalves, Ewan Harrison, David K. Jackson, Ian Johnston, Dominic Kwiatkowski, Cordelia Langford, John Sillitoe on behalf of the Wellcome Sanger Institute COVID-19 Surveillance Team |
| EPI_ISL_2781727 | Health Services Laboratories | Wellcome Sanger Institute for the COVID-19 Genomics UK (COG-UK) Consortium | Health Services Laboratories and Alex Alderton, Roberto Amato, Jeffrey Barrett, Sonia Goncalves, Ewan Harrison, David K. Jackson, Ian Johnston, Dominic Kwiatkowski, Cordelia Langford, John Sillitoe on behalf of the Wellcome Sanger Institute COVID-19 Surveillance Team |
| EPI_ISL_2781728 | Lighthouse Lab in Milton Keynes | Wellcome Sanger Institute for the COVID-19 Genomics UK (COG-UK) Consortium | The Lighthouse Lab in Milton Keynes and Alex Alderton, Roberto Amato, Jeffrey Barrett, Sonia Goncalves, Ewan Harrison, David K. Jackson, Ian Johnston, Dominic Kwiatkowski, Cordelia Langford, John Sillitoe on behalf of the Wellcome Sanger Institute COVID-19 Surveillance Team |
| EPI_ISL_2781729 | Lighthouse Lab in Alderley Park | Wellcome Sanger Institute for the COVID-19 Genomics UK (COG-UK) Consortium | Jacquelyn Wynn, Mairead Hyland, The Lighthouse Lab in Alderley Park and Alex Alderton, Roberto Amato, Jeffrey Barrett, Sonia Goncalves, Ewan Harrison, David K. Jackson, Ian Johnston, Dominic Kwiatkowski, Cordelia Langford, John Sillitoe on behalf of the Wellcome Sanger Institute COVID-19 Surveillance Team |
| EPI_ISL_2781730 | Health Services Laboratories | Wellcome Sanger Institute for the COVID-19 Genomics UK (COG-UK) Consortium | Health Services Laboratories and Alex Alderton, Roberto Amato, Jeffrey Barrett, Sonia Goncalves, Ewan Harrison, David K. Jackson, Ian Johnston, Dominic Kwiatkowski, Cordelia Langford, John Sillitoe on behalf of the Wellcome Sanger Institute COVID-19 Surveillance Team |
| EPI_ISL_2781731 | Lighthouse Lab in Alderley Park | Wellcome Sanger Institute for the COVID-19 Genomics UK (COG-UK) Consortium | Jacquelyn Wynn, Mairead Hyland, The Lighthouse Lab in Alderley Park and Alex Alderton, Roberto Amato, Jeffrey Barrett, Sonia Goncalves, Ewan Harrison, David K. Jackson, Ian Johnston, Dominic Kwiatkowski, Cordelia Langford, John Sillitoe on behalf of the Wellcome Sanger Institute COVID-19 Surveillance Team |
| EPI_ISL_2781732 | Lighthouse Lab in Milton Keynes | Wellcome Sanger Institute for the COVID-19 Genomics UK (COG-UK) Consortium | The Lighthouse Lab in Milton Keynes and Alex Alderton, Roberto Amato, Jeffrey Barrett, Sonia Goncalves, Ewan Harrison, David K. Jackson, Ian Johnston, Dominic Kwiatkowski, Cordelia Langford, John Sillitoe on behalf of the Wellcome Sanger Institute COVID-19 Surveillance Team |
| EPI_ISL_2781733 | Lighthouse Lab in Glasgow | Wellcome Sanger Institute for the COVID-19 Genomics UK (COG-UK) Consortium | Harper VanSteenhouse, Yumi Kasai, David Gray, Carol Clugston, Anna Dominiczak and Alex Alderton, Roberto Amato, Jeffrey Barrett, Sonia Goncalves, Ewan Harrison, David K. Jackson, Ian Johnston, Dominic Kwiatkowski, Cordelia Langford, John Sillitoe on behalf of the Wellcome Sanger Institute COVID-19 Surveillance Team |
| EPI_ISL_2781734 | Lighthouse Lab in Milton Keynes | Wellcome Sanger Institute for the COVID-19 Genomics UK (COG-UK) Consortium | The Lighthouse Lab in Milton Keynes and Alex Alderton, Roberto Amato, Jeffrey Barrett, Sonia Goncalves, Ewan Harrison, David K. Jackson, Ian Johnston, Dominic Kwiatkowski, Cordelia Langford, John Sillitoe on behalf of the Wellcome Sanger Institute COVID-19 Surveillance Team |
| EPI_ISL_2781735, EPI_ISL_2781736 | Lighthouse Lab in Alderley Park | Wellcome Sanger Institute for the COVID-19 Genomics UK (COG-UK) Consortium | Jacquelyn Wynn, Mairead Hyland, The Lighthouse Lab in Alderley Park and Alex Alderton, Roberto Amato, Jeffrey Barrett, Sonia Goncalves, Ewan Harrison, David K. Jackson, Ian Johnston, Dominic Kwiatkowski, Cordelia Langford, John Sillitoe on behalf of the Wellcome Sanger Institute COVID-19 Surveillance Team |
| EPI_ISL_2781737, EPI_ISL_2781738, EPI_ISL_2781739, EPI_ISL_2781740 | Lighthouse Lab in Milton Keynes | Wellcome Sanger Institute for the COVID-19 Genomics UK (COG-UK) Consortium | The Lighthouse Lab in Milton Keynes and Alex Alderton, Roberto Amato, Jeffrey Barrett, Sonia Goncalves, Ewan Harrison, David K. Jackson, Ian Johnston, Dominic Kwiatkowski, Cordelia Langford, John Sillitoe on behalf of the Wellcome Sanger Institute COVID-19 Surveillance Team |
| EPI_ISL_2781741 | Lighthouse Lab in Alderley Park | Wellcome Sanger Institute for the COVID-19 Genomics UK (COG-UK) Consortium | Jacquelyn Wynn, Mairead Hyland, The Lighthouse Lab in Alderley Park and Alex Alderton, Roberto Amato, Jeffrey Barrett, Sonia Goncalves, Ewan Harrison, David K. Jackson, Ian Johnston, Dominic Kwiatkowski, Cordelia Langford, John Sillitoe on behalf of the Wellcome Sanger Institute COVID-19 Surveillance Team |
| EPI_ISL_2781742 | Lighthouse Lab in Milton Keynes | Wellcome Sanger Institute for the COVID-19 Genomics UK (COG-UK) Consortium | The Lighthouse Lab in Milton Keynes and Alex Alderton, Roberto Amato, Jeffrey Barrett, Sonia Goncalves, Ewan Harrison, David K. Jackson, Ian Johnston, Dominic Kwiatkowski, Cordelia Langford, John Sillitoe on behalf of the Wellcome Sanger Institute COVID-19 Surveillance Team |
| EPI_ISL_2781743 | Lighthouse Lab in Alderley Park | Wellcome Sanger Institute for the COVID-19 Genomics UK (COG-UK) Consortium | Jacquelyn Wynn, Mairead Hyland, The Lighthouse Lab in Alderley Park and Alex Alderton, Roberto Amato, Jeffrey Barrett, Sonia Goncalves, Ewan Harrison, David K. Jackson, Ian Johnston, Dominic Kwiatkowski, Cordelia Langford, John Sillitoe on behalf of the Wellcome Sanger Institute COVID-19 Surveillance Team |
| EPI_ISL_2781744 | Health Services Laboratories | Wellcome Sanger Institute for the COVID-19 Genomics UK (COG-UK) Consortium | Health Services Laboratories and Alex Alderton, Roberto Amato, Jeffrey Barrett, Sonia Goncalves, Ewan Harrison, David K. Jackson, Ian Johnston, Dominic Kwiatkowski, Cordelia Langford, John Sillitoe on behalf of the Wellcome Sanger Institute COVID-19 Surveillance Team |
| EPI_ISL_2781745, EPI_ISL_2781746 | Lighthouse Lab in Alderley Park | Wellcome Sanger Institute for the COVID-19 Genomics UK (COG-UK) Consortium | Jacquelyn Wynn, Mairead Hyland, The Lighthouse Lab in Alderley Park and Alex Alderton, Roberto Amato, Jeffrey Barrett, Sonia Goncalves, Ewan Harrison, David K. Jackson, Ian Johnston, Dominic Kwiatkowski, Cordelia Langford, John Sillitoe on behalf of the Wellcome Sanger Institute COVID-19 Surveillance Team |
| EPI_ISL_2781747, EPI_ISL_2781748, EPI_ISL_2781749 | Lighthouse Lab in Milton Keynes | Wellcome Sanger Institute for the COVID-19 Genomics UK (COG-UK) Consortium | The Lighthouse Lab in Milton Keynes and Alex Alderton, Roberto Amato, Jeffrey Barrett, Sonia Goncalves, Ewan Harrison, David K. Jackson, Ian Johnston, Dominic Kwiatkowski, Cordelia Langford, John Sillitoe on behalf of the Wellcome Sanger Institute COVID-19 Surveillance Team |
| EPI_ISL_2781750, EPI_ISL_2781751 | Lighthouse Lab in Alderley Park | Wellcome Sanger Institute for the COVID-19 Genomics UK (COG-UK) Consortium | Jacquelyn Wynn, Mairead Hyland, The Lighthouse Lab in Alderley Park and Alex Alderton, Roberto Amato, Jeffrey Barrett, Sonia Goncalves, Ewan Harrison, David K. Jackson, Ian Johnston, Dominic Kwiatkowski, Cordelia Langford, John Sillitoe on behalf of the Wellcome Sanger Institute COVID-19 Surveillance Team |
| EPI_ISL_2781752 | Health Services Laboratories | Wellcome Sanger Institute for the COVID-19 Genomics UK (COG-UK) Consortium | Health Services Laboratories and Alex Alderton, Roberto Amato, Jeffrey Barrett, Sonia Goncalves, Ewan Harrison, David K. Jackson, Ian Johnston, Dominic Kwiatkowski, Cordelia Langford, John Sillitoe on behalf of the Wellcome Sanger Institute COVID-19 Surveillance Team |
| EPI_ISL_2781753 | Lighthouse Lab in Alderley Park | Wellcome Sanger Institute for the COVID-19 Genomics UK (COG-UK) Consortium | Jacquelyn Wynn, Mairead Hyland, The Lighthouse Lab in Alderley Park and Alex Alderton, Roberto Amato, Jeffrey Barrett, Sonia Goncalves, Ewan Harrison, David K. Jackson, Ian Johnston, Dominic Kwiatkowski, Cordelia Langford, John Sillitoe on behalf of the Wellcome Sanger Institute COVID-19 Surveillance Team |
| EPI_ISL_2781754 | Lighthouse Lab in Milton Keynes | Wellcome Sanger Institute for the COVID-19 Genomics UK (COG-UK) Consortium | The Lighthouse Lab in Milton Keynes and Alex Alderton, Roberto Amato, Jeffrey Barrett, Sonia Goncalves, Ewan Harrison, David K. Jackson, Ian Johnston, Dominic Kwiatkowski, Cordelia Langford, John Sillitoe on behalf of the Wellcome Sanger Institute COVID-19 Surveillance Team |
| EPI_ISL_2781755 | Lighthouse Lab in Alderley Park | Wellcome Sanger Institute for the COVID-19 Genomics UK (COG-UK) Consortium | Jacquelyn Wynn, Mairead Hyland, The Lighthouse Lab in Alderley Park and Alex Alderton, Roberto Amato, Jeffrey Barrett, Sonia Goncalves, Ewan Harrison, David K. Jackson, Ian Johnston, Dominic Kwiatkowski, Cordelia Langford, John Sillitoe on behalf of the Wellcome Sanger Institute COVID-19 Surveillance Team |
| EPI_ISL_2781756 | Lighthouse Laboratory Plymouth | Wellcome Sanger Institute for the COVID-19 Genomics UK (COG-UK) Consortium | Lighthouse Laboratory Plymouth and Alex Alderton, Roberto Amato, Jeffrey Barrett, Sonia Goncalves, Ewan Harrison, David K. Jackson, Ian Johnston, Dominic Kwiatkowski, Cordelia Langford, John Sillitoe on behalf of the Wellcome Sanger Institute COVID-19 Surveillance Team |
| EPI_ISL_2781757 | Health Services Laboratories | Wellcome Sanger Institute for the COVID-19 Genomics UK (COG-UK) Consortium | Health Services Laboratories and Alex Alderton, Roberto Amato, Jeffrey Barrett, Sonia Goncalves, Ewan Harrison, David K. Jackson, Ian Johnston, Dominic Kwiatkowski, Cordelia Langford, John Sillitoe on behalf of the Wellcome Sanger Institute COVID-19 Surveillance Team |
| EPI_ISL_2781758 | Lighthouse Lab in Milton Keynes | Wellcome Sanger Institute for the COVID-19 Genomics UK (COG-UK) Consortium | The Lighthouse Lab in Milton Keynes and Alex Alderton, Roberto Amato, Jeffrey Barrett, Sonia Goncalves, Ewan Harrison, David K. Jackson, Ian Johnston, Dominic Kwiatkowski, Cordelia Langford, John Sillitoe on behalf of the Wellcome Sanger Institute COVID-19 Surveillance Team |
| EPI_ISL_2781759 | Lighthouse Lab in Alderley Park | Wellcome Sanger Institute for the COVID-19 Genomics UK (COG-UK) | Jacquelyn Wynn, Mairead Hyland, The Lighthouse Lab in Alderley Park and Alex Alderton, Roberto Amato, Jeffrey Barrett, Sonia Goncalves, Ewan Harrison, David K. Jackson, Ian Johnston, Dominic Kwiatkowski, Cordelia Langford, John Sillitoe on |

| | | | |
|---|---|---|---|
| | | Consortium | behalf of the Wellcome Sanger Institute COVID-19 Surveillance Team |
| EPI_ISL_2781760 | Health Services Laboratories | Wellcome Sanger Institute for the COVID-19 Genomics UK (COG-UK) Consortium | Health Services Laboratories and Alex Alderton, Roberto Amato, Jeffrey Barrett, Sonia Goncalves, Ewan Harrison, David K. Jackson, Ian Johnston, Dominic Kwiatkowski, Cordelia Langford, John Sillitoe on behalf of the Wellcome Sanger Institute COVID-19 Surveillance Team |
| EPI_ISL_2781761, EPI_ISL_2781762 | Lighthouse Lab in Alderley Park | Wellcome Sanger Institute for the COVID-19 Genomics UK (COG-UK) Consortium | Jacquelyn Wynn, Mairead Hyland, The Lighthouse Lab in Alderley Park and Alex Alderton, Roberto Amato, Jeffrey Barrett, Sonia Goncalves, Ewan Harrison, David K. Jackson, Ian Johnston, Dominic Kwiatkowski, Cordelia Langford, John Sillitoe on behalf of the Wellcome Sanger Institute COVID-19 Surveillance Team |
| EPI_ISL_2781763 | Lighthouse Lab in Milton Keynes | Wellcome Sanger Institute for the COVID-19 Genomics UK (COG-UK) Consortium | The Lighthouse Lab in Milton Keynes and Alex Alderton, Roberto Amato, Jeffrey Barrett, Sonia Goncalves, Ewan Harrison, David K. Jackson, Ian Johnston, Dominic Kwiatkowski, Cordelia Langford, John Sillitoe on behalf of the Wellcome Sanger Institute COVID-19 Surveillance Team |
| EPI_ISL_2781764 | Lighthouse Laboratory Plymouth | Wellcome Sanger Institute for the COVID-19 Genomics UK (COG-UK) Consortium | Lighthouse Laboratory Plymouth and Alex Alderton, Roberto Amato, Jeffrey Barrett, Sonia Goncalves, Ewan Harrison, David K. Jackson, Ian Johnston, Dominic Kwiatkowski, Cordelia Langford, John Sillitoe on behalf of the Wellcome Sanger Institute COVID-19 Surveillance Team |
| EPI_ISL_2781765, EPI_ISL_2781766, EPI_ISL_2781767 | Lighthouse Lab in Alderley Park | Wellcome Sanger Institute for the COVID-19 Genomics UK (COG-UK) Consortium | Jacquelyn Wynn, Mairead Hyland, The Lighthouse Lab in Alderley Park and Alex Alderton, Roberto Amato, Jeffrey Barrett, Sonia Goncalves, Ewan Harrison, David K. Jackson, Ian Johnston, Dominic Kwiatkowski, Cordelia Langford, John Sillitoe on behalf of the Wellcome Sanger Institute COVID-19 Surveillance Team |
| EPI_ISL_2781768, EPI_ISL_2781769 | Health Services Laboratories | Wellcome Sanger Institute for the COVID-19 Genomics UK (COG-UK) Consortium | Health Services Laboratories and Alex Alderton, Roberto Amato, Jeffrey Barrett, Sonia Goncalves, Ewan Harrison, David K. Jackson, Ian Johnston, Dominic Kwiatkowski, Cordelia Langford, John Sillitoe on behalf of the Wellcome Sanger Institute COVID-19 Surveillance Team |
| EPI_ISL_2781770, EPI_ISL_2781771, EPI_ISL_2781772 | Lighthouse Lab in Alderley Park | Wellcome Sanger Institute for the COVID-19 Genomics UK (COG-UK) Consortium | Jacquelyn Wynn, Mairead Hyland, The Lighthouse Lab in Alderley Park and Alex Alderton, Roberto Amato, Jeffrey Barrett, Sonia Goncalves, Ewan Harrison, David K. Jackson, Ian Johnston, Dominic Kwiatkowski, Cordelia Langford, John Sillitoe on behalf of the Wellcome Sanger Institute COVID-19 Surveillance Team |
| EPI_ISL_2781773 | Lighthouse Laboratory Plymouth | Wellcome Sanger Institute for the COVID-19 Genomics UK (COG-UK) Consortium | Lighthouse Laboratory Plymouth and Alex Alderton, Roberto Amato, Jeffrey Barrett, Sonia Goncalves, Ewan Harrison, David K. Jackson, Ian Johnston, Dominic Kwiatkowski, Cordelia Langford, John Sillitoe on behalf of the Wellcome Sanger Institute COVID-19 Surveillance Team |
| EPI_ISL_2781774, EPI_ISL_2781775, EPI_ISL_2781776, EPI_ISL_2781777, EPI_ISL_2781778, EPI_ISL_2781780, EPI_ISL_2781781 | Lighthouse Lab in Alderley Park | Wellcome Sanger Institute for the COVID-19 Genomics UK (COG-UK) Consortium | Jacquelyn Wynn, Mairead Hyland, The Lighthouse Lab in Alderley Park and Alex Alderton, Roberto Amato, Jeffrey Barrett, Sonia Goncalves, Ewan Harrison, David K. Jackson, Ian Johnston, Dominic Kwiatkowski, Cordelia Langford, John Sillitoe on behalf of the Wellcome Sanger Institute COVID-19 Surveillance Team |
| EPI_ISL_2781782 | Lighthouse Lab in Milton Keynes | Wellcome Sanger Institute for the COVID-19 Genomics UK (COG-UK) Consortium | The Lighthouse Lab in Milton Keynes and Alex Alderton, Roberto Amato, Jeffrey Barrett, Sonia Goncalves, Ewan Harrison, David K. Jackson, Ian Johnston, Dominic Kwiatkowski, Cordelia Langford, John Sillitoe on behalf of the Wellcome Sanger Institute COVID-19 Surveillance Team |
| EPI_ISL_2781783 | Lighthouse Lab in Alderley Park | Wellcome Sanger Institute for the COVID-19 Genomics UK (COG-UK) Consortium | Jacquelyn Wynn, Mairead Hyland, The Lighthouse Lab in Alderley Park and Alex Alderton, Roberto Amato, Jeffrey Barrett, Sonia Goncalves, Ewan Harrison, David K. Jackson, Ian Johnston, Dominic Kwiatkowski, Cordelia Langford, John Sillitoe on behalf of the Wellcome Sanger Institute COVID-19 Surveillance Team |
| EPI_ISL_2781784 | Lighthouse Lab in Milton Keynes | Wellcome Sanger Institute for the COVID-19 Genomics UK (COG-UK) Consortium | The Lighthouse Lab in Milton Keynes and Alex Alderton, Roberto Amato, Jeffrey Barrett, Sonia Goncalves, Ewan Harrison, David K. Jackson, Ian Johnston, Dominic Kwiatkowski, Cordelia Langford, John Sillitoe on behalf of the Wellcome Sanger Institute COVID-19 Surveillance Team |
| EPI_ISL_2781785 | Lighthouse Lab in Alderley Park | Wellcome Sanger Institute for the COVID-19 Genomics UK (COG-UK) Consortium | Jacquelyn Wynn, Mairead Hyland, The Lighthouse Lab in Alderley Park and Alex Alderton, Roberto Amato, Jeffrey Barrett, Sonia Goncalves, Ewan Harrison, David K. Jackson, Ian Johnston, Dominic Kwiatkowski, Cordelia Langford, John Sillitoe on behalf of the Wellcome Sanger Institute COVID-19 Surveillance Team |
| EPI_ISL_2781786 | Lighthouse Lab in Milton Keynes | Wellcome Sanger Institute for the COVID-19 Genomics UK (COG-UK) Consortium | The Lighthouse Lab in Milton Keynes and Alex Alderton, Roberto Amato, Jeffrey Barrett, Sonia Goncalves, Ewan Harrison, David K. Jackson, Ian Johnston, Dominic Kwiatkowski, Cordelia Langford, John Sillitoe on behalf of the Wellcome Sanger Institute COVID-19 Surveillance Team |
| EPI_ISL_2781787, EPI_ISL_2781788 | Lighthouse Lab in Alderley Park | Wellcome Sanger Institute for the COVID-19 Genomics UK (COG-UK) Consortium | Jacquelyn Wynn, Mairead Hyland, The Lighthouse Lab in Alderley Park and Alex Alderton, Roberto Amato, Jeffrey Barrett, Sonia Goncalves, Ewan Harrison, David K. Jackson, Ian Johnston, Dominic Kwiatkowski, Cordelia Langford, John Sillitoe on behalf of the Wellcome Sanger Institute COVID-19 Surveillance Team |
| EPI_ISL_2781789, EPI_ISL_2781790 | Lighthouse Lab in Milton Keynes | Wellcome Sanger Institute for the COVID-19 Genomics UK (COG-UK) Consortium | The Lighthouse Lab in Milton Keynes and Alex Alderton, Roberto Amato, Jeffrey Barrett, Sonia Goncalves, Ewan Harrison, David K. Jackson, Ian Johnston, Dominic Kwiatkowski, Cordelia Langford, John Sillitoe on behalf of the Wellcome Sanger Institute COVID-19 Surveillance Team |
| EPI_ISL_2781791, EPI_ISL_2781792 | Lighthouse Lab in Alderley Park | Wellcome Sanger Institute for the COVID-19 Genomics UK (COG-UK) Consortium | Jacquelyn Wynn, Mairead Hyland, The Lighthouse Lab in Alderley Park and Alex Alderton, Roberto Amato, Jeffrey Barrett, Sonia Goncalves, Ewan Harrison, David K. Jackson, Ian Johnston, Dominic Kwiatkowski, Cordelia Langford, John Sillitoe on behalf of the Wellcome Sanger Institute COVID-19 Surveillance Team |
| EPI_ISL_2781793 | Lighthouse Lab in Milton Keynes | Wellcome Sanger Institute for the COVID-19 Genomics UK (COG-UK) Consortium | The Lighthouse Lab in Milton Keynes and Alex Alderton, Roberto Amato, Jeffrey Barrett, Sonia Goncalves, Ewan Harrison, David K. Jackson, Ian Johnston, Dominic Kwiatkowski, Cordelia Langford, John Sillitoe on behalf of the Wellcome Sanger Institute COVID-19 Surveillance Team |
| EPI_ISL_2781794 | Lighthouse Lab in Alderley Park | Wellcome Sanger Institute for the COVID-19 Genomics UK (COG-UK) Consortium | Jacquelyn Wynn, Mairead Hyland, The Lighthouse Lab in Alderley Park and Alex Alderton, Roberto Amato, Jeffrey Barrett, Sonia Goncalves, Ewan Harrison, David K. Jackson, Ian Johnston, Dominic Kwiatkowski, Cordelia Langford, John Sillitoe on behalf of the Wellcome Sanger Institute COVID-19 Surveillance Team |
| EPI_ISL_2781795 | Lighthouse Lab in Milton Keynes | Wellcome Sanger Institute for the COVID-19 Genomics UK (COG-UK) Consortium | The Lighthouse Lab in Milton Keynes and Alex Alderton, Roberto Amato, Jeffrey Barrett, Sonia Goncalves, Ewan Harrison, David K. Jackson, Ian Johnston, Dominic Kwiatkowski, Cordelia Langford, John Sillitoe on behalf of the Wellcome Sanger Institute COVID-19 Surveillance Team |
| EPI_ISL_2781796, EPI_ISL_2781797 | Lighthouse Lab in Alderley Park | Wellcome Sanger Institute for the COVID-19 Genomics UK (COG-UK) Consortium | Jacquelyn Wynn, Mairead Hyland, The Lighthouse Lab in Alderley Park and Alex Alderton, Roberto Amato, Jeffrey Barrett, Sonia Goncalves, Ewan Harrison, David K. Jackson, Ian Johnston, Dominic Kwiatkowski, Cordelia Langford, John Sillitoe on behalf of the Wellcome Sanger Institute COVID-19 Surveillance Team |
| EPI_ISL_2781798, EPI_ISL_2781799 | Lighthouse Lab in Milton Keynes | Wellcome Sanger Institute for the COVID-19 Genomics UK (COG-UK) Consortium | The Lighthouse Lab in Milton Keynes and Alex Alderton, Roberto Amato, Jeffrey Barrett, Sonia Goncalves, Ewan Harrison, David K. Jackson, Ian Johnston, Dominic Kwiatkowski, Cordelia Langford, John Sillitoe on behalf of the Wellcome Sanger Institute COVID-19 Surveillance Team |
| EPI_ISL_2781800, EPI_ISL_2781801 | Lighthouse Lab in Alderley Park | Wellcome Sanger Institute for the COVID-19 Genomics UK (COG-UK) Consortium | Jacquelyn Wynn, Mairead Hyland, The Lighthouse Lab in Alderley Park and Alex Alderton, Roberto Amato, Jeffrey Barrett, Sonia Goncalves, Ewan Harrison, David K. Jackson, Ian Johnston, Dominic Kwiatkowski, Cordelia Langford, John Sillitoe on behalf of the Wellcome Sanger Institute COVID-19 Surveillance Team |
| EPI_ISL_2781802 | Lighthouse Lab in Milton Keynes | Wellcome Sanger Institute for the COVID-19 Genomics UK (COG-UK) Consortium | The Lighthouse Lab in Milton Keynes and Alex Alderton, Roberto Amato, Jeffrey Barrett, Sonia Goncalves, Ewan Harrison, David K. Jackson, Ian Johnston, Dominic Kwiatkowski, Cordelia Langford, John Sillitoe on behalf of the Wellcome Sanger Institute COVID-19 Surveillance Team |
| EPI_ISL_2781803, EPI_ISL_2781804, EPI_ISL_2781805, EPI_ISL_2781806 | Lighthouse Lab in Alderley Park | Wellcome Sanger Institute for the COVID-19 Genomics UK (COG-UK) Consortium | Jacquelyn Wynn, Mairead Hyland, The Lighthouse Lab in Alderley Park and Alex Alderton, Roberto Amato, Jeffrey Barrett, Sonia Goncalves, Ewan Harrison, David K. Jackson, Ian Johnston, Dominic Kwiatkowski, Cordelia Langford, John Sillitoe on behalf of the Wellcome Sanger Institute COVID-19 Surveillance Team |
| EPI_ISL_2781807 | Lighthouse Lab in Milton Keynes | Wellcome Sanger Institute for the COVID-19 Genomics UK (COG-UK) Consortium | The Lighthouse Lab in Milton Keynes and Alex Alderton, Roberto Amato, Jeffrey Barrett, Sonia Goncalves, Ewan Harrison, David K. Jackson, Ian Johnston, Dominic Kwiatkowski, Cordelia Langford, John Sillitoe on behalf of the Wellcome Sanger Institute COVID-19 Surveillance Team |
| EPI_ISL_2781808 | Lighthouse Lab in Alderley Park | Wellcome Sanger Institute for the COVID-19 Genomics UK (COG-UK) Consortium | Jacquelyn Wynn, Mairead Hyland, The Lighthouse Lab in Alderley Park and Alex Alderton, Roberto Amato, Jeffrey Barrett, Sonia Goncalves, Ewan Harrison, David K. Jackson, Ian Johnston, Dominic Kwiatkowski, Cordelia Langford, John Sillitoe on behalf of the Wellcome Sanger Institute COVID-19 Surveillance Team |
| EPI_ISL_2781809 | Lighthouse Lab in Glasgow | Wellcome Sanger Institute for the COVID-19 Genomics UK (COG-UK) Consortium | Harper VanSteenhouse, Yumi Kasai, David Gray, Carol Clugston, Anna Dominiczak and Alex Alderton, Roberto Amato, Jeffrey Barrett, Sonia Goncalves, Ewan Harrison, David K. Jackson, Ian Johnston, Dominic Kwiatkowski, Cordelia Langford, John Sillitoe on behalf of the Wellcome Sanger Institute COVID-19 Surveillance Team |
| EPI_ISL_2781810, EPI_ISL_2781811, EPI_ISL_2781812 | Lighthouse Lab in Alderley Park | Wellcome Sanger Institute for the COVID-19 Genomics UK (COG-UK) Consortium | Jacquelyn Wynn, Mairead Hyland, The Lighthouse Lab in Alderley Park and Alex Alderton, Roberto Amato, Jeffrey Barrett, Sonia Goncalves, Ewan Harrison, David K. Jackson, Ian Johnston, Dominic Kwiatkowski, Cordelia Langford, John Sillitoe on behalf of the Wellcome Sanger Institute COVID-19 Surveillance Team |
| EPI_ISL_2781813 | Lighthouse Lab in Milton Keynes | Wellcome Sanger Institute for the COVID-19 Genomics UK (COG-UK) Consortium | The Lighthouse Lab in Milton Keynes and Alex Alderton, Roberto Amato, Jeffrey Barrett, Sonia Goncalves, Ewan Harrison, David K. Jackson, Ian Johnston, Dominic Kwiatkowski, Cordelia Langford, John Sillitoe on behalf of the Wellcome Sanger Institute COVID-19 Surveillance Team |
| EPI_ISL_2781814 | Lighthouse Lab in Alderley Park | Wellcome Sanger Institute for the COVID-19 Genomics UK (COG-UK) Consortium | Jacquelyn Wynn, Mairead Hyland, The Lighthouse Lab in Alderley Park and Alex Alderton, Roberto Amato, Jeffrey Barrett, Sonia Goncalves, Ewan Harrison, David K. Jackson, Ian Johnston, Dominic Kwiatkowski, Cordelia Langford, John Sillitoe on behalf of the Wellcome Sanger Institute COVID-19 Surveillance Team |
| EPI_ISL_2781815, EPI_ISL_2781816 | Lighthouse Lab in Glasgow | Wellcome Sanger Institute for the COVID-19 Genomics UK (COG-UK) Consortium | Harper VanSteenhouse, Yumi Kasai, David Gray, Carol Clugston, Anna Dominiczak and Alex Alderton, Roberto Amato, Jeffrey Barrett, Sonia Goncalves, Ewan Harrison, David K. Jackson, Ian Johnston, Dominic Kwiatkowski, Cordelia Langford, John Sillitoe on behalf of the Wellcome Sanger Institute COVID-19 Surveillance Team |
| EPI_ISL_2781817, EPI_ISL_2781818, EPI_ISL_2781819 | Lighthouse Lab in Alderley Park | Wellcome Sanger Institute for the COVID-19 Genomics UK (COG-UK) Consortium | Jacquelyn Wynn, Mairead Hyland, The Lighthouse Lab in Alderley Park and Alex Alderton, Roberto Amato, Jeffrey Barrett, Sonia Goncalves, Ewan Harrison, David K. Jackson, Ian Johnston, Dominic Kwiatkowski, Cordelia Langford, John Sillitoe on behalf of the Wellcome Sanger Institute COVID-19 Surveillance Team |
| EPI_ISL_2781820 | Lighthouse Lab in Milton Keynes | Wellcome Sanger Institute for the COVID-19 Genomics UK (COG-UK) Consortium | The Lighthouse Lab in Milton Keynes and Alex Alderton, Roberto Amato, Jeffrey Barrett, Sonia Goncalves, Ewan Harrison, David K. Jackson, Ian Johnston, Dominic Kwiatkowski, Cordelia Langford, John Sillitoe on behalf of the Wellcome Sanger Institute COVID-19 Surveillance Team |
| EPI_ISL_2781821, EPI_ISL_2781822, EPI_ISL_2781823 | Lighthouse Lab in Alderley Park | Wellcome Sanger Institute for the COVID-19 Genomics UK (COG-UK) Consortium | Jacquelyn Wynn, Mairead Hyland, The Lighthouse Lab in Alderley Park and Alex Alderton, Roberto Amato, Jeffrey Barrett, Sonia Goncalves, Ewan Harrison, David K. Jackson, Ian Johnston, Dominic Kwiatkowski, Cordelia Langford, John Sillitoe on behalf of the Wellcome Sanger Institute COVID-19 Surveillance Team |
| EPI_ISL_2781824, EPI_ISL_2781825 | Lighthouse Lab in Milton Keynes | Wellcome Sanger Institute for the COVID-19 Genomics UK (COG-UK) Consortium | The Lighthouse Lab in Milton Keynes and Alex Alderton, Roberto Amato, Jeffrey Barrett, Sonia Goncalves, Ewan Harrison, David K. Jackson, Ian Johnston, Dominic Kwiatkowski, Cordelia Langford, John Sillitoe on behalf of the Wellcome Sanger Institute COVID-19 Surveillance Team |
| EPI_ISL_2781826 | Lighthouse Lab in Alderley Park | Wellcome Sanger Institute for the COVID-19 Genomics UK (COG-UK) Consortium | Jacquelyn Wynn, Mairead Hyland, The Lighthouse Lab in Alderley Park and Alex Alderton, Roberto Amato, Jeffrey Barrett, Sonia Goncalves, Ewan Harrison, David K. Jackson, Ian Johnston, Dominic Kwiatkowski, Cordelia Langford, John Sillitoe on behalf of the Wellcome Sanger Institute COVID-19 Surveillance Team |
| EPI_ISL_2781827 | Lighthouse Lab in Milton Keynes | Wellcome Sanger Institute for the COVID-19 Genomics UK (COG-UK) Consortium | The Lighthouse Lab in Milton Keynes and Alex Alderton, Roberto Amato, Jeffrey Barrett, Sonia Goncalves, Ewan Harrison, David K. Jackson, Ian Johnston, Dominic Kwiatkowski, Cordelia Langford, John Sillitoe on behalf of the Wellcome Sanger Institute COVID-19 Surveillance Team |
| EPI_ISL_2781828, EPI_ISL_2781829, EPI_ISL_2781830 | Lighthouse Lab in Alderley Park | Wellcome Sanger Institute for the COVID-19 Genomics UK (COG-UK) Consortium | Jacquelyn Wynn, Mairead Hyland, The Lighthouse Lab in Alderley Park and Alex Alderton, Roberto Amato, Jeffrey Barrett, Sonia Goncalves, Ewan Harrison, David K. Jackson, Ian Johnston, Dominic Kwiatkowski, Cordelia Langford, John Sillitoe on behalf of the Wellcome Sanger Institute COVID-19 Surveillance Team |

| | | | |
|---|---|---|---|
| EPI_ISL_2781831, EPI_ISL_2781832, EPI_ISL_2781833 | Lighthouse Lab in Milton Keynes | Wellcome Sanger Institute for the COVID-19 Genomics UK (COG-UK) Consortium | The Lighthouse Lab in Milton Keynes and Alex Alderton, Roberto Amato, Jeffrey Barrett, Sonia Goncalves, Ewan Harrison, David K. Jackson, Ian Johnston, Dominic Kwiatkowski, Cordelia Langford, John Sillitoe on behalf of the Wellcome Sanger Institute COVID-19 Surveillance Team |
| EPI_ISL_2781834 | Lighthouse Lab in Glasgow | Wellcome Sanger Institute for the COVID-19 Genomics UK (COG-UK) Consortium | Harper VanSteenhouse, Yumi Kasai, David Gray, Carol Clugston, Anna Dominiczak and Alex Alderton, Roberto Amato, Jeffrey Barrett, Sonia Goncalves, Ewan Harrison, David K. Jackson, Ian Johnston, Dominic Kwiatkowski, Cordelia Langford, John Sillitoe on behalf of the Wellcome Sanger Institute COVID-19 Surveillance Team |
| EPI_ISL_2781835, EPI_ISL_2781836, EPI_ISL_2781837, EPI_ISL_2781838 | Lighthouse Lab in Alderley Park | Wellcome Sanger Institute for the COVID-19 Genomics UK (COG-UK) Consortium | Jacquelyn Wynn, Mairead Hyland, The Lighthouse Lab in Alderley Park and Alex Alderton, Roberto Amato, Jeffrey Barrett, Sonia Goncalves, Ewan Harrison, David K. Jackson, Ian Johnston, Dominic Kwiatkowski, Cordelia Langford, John Sillitoe on behalf of the Wellcome Sanger Institute COVID-19 Surveillance Team |
| EPI_ISL_2781839 | Lighthouse Lab in Milton Keynes | Wellcome Sanger Institute for the COVID-19 Genomics UK (COG-UK) Consortium | The Lighthouse Lab in Milton Keynes and Alex Alderton, Roberto Amato, Jeffrey Barrett, Sonia Goncalves, Ewan Harrison, David K. Jackson, Ian Johnston, Dominic Kwiatkowski, Cordelia Langford, John Sillitoe on behalf of the Wellcome Sanger Institute COVID-19 Surveillance Team |
| EPI_ISL_2781840, EPI_ISL_2781841, EPI_ISL_2781842, EPI_ISL_2781843, EPI_ISL_2781844, EPI_ISL_2781845 | Lighthouse Lab in Alderley Park | Wellcome Sanger Institute for the COVID-19 Genomics UK (COG-UK) Consortium | Jacquelyn Wynn, Mairead Hyland, The Lighthouse Lab in Alderley Park and Alex Alderton, Roberto Amato, Jeffrey Barrett, Sonia Goncalves, Ewan Harrison, David K. Jackson, Ian Johnston, Dominic Kwiatkowski, Cordelia Langford, John Sillitoe on behalf of the Wellcome Sanger Institute COVID-19 Surveillance Team |
| EPI_ISL_2781846, EPI_ISL_2781847 | Lighthouse Lab in Milton Keynes | Wellcome Sanger Institute for the COVID-19 Genomics UK (COG-UK) Consortium | The Lighthouse Lab in Milton Keynes and Alex Alderton, Roberto Amato, Jeffrey Barrett, Sonia Goncalves, Ewan Harrison, David K. Jackson, Ian Johnston, Dominic Kwiatkowski, Cordelia Langford, John Sillitoe on behalf of the Wellcome Sanger Institute COVID-19 Surveillance Team |
| EPI_ISL_2781848 | Lighthouse Lab in Alderley Park | Wellcome Sanger Institute for the COVID-19 Genomics UK (COG-UK) Consortium | Jacquelyn Wynn, Mairead Hyland, The Lighthouse Lab in Alderley Park and Alex Alderton, Roberto Amato, Jeffrey Barrett, Sonia Goncalves, Ewan Harrison, David K. Jackson, Ian Johnston, Dominic Kwiatkowski, Cordelia Langford, John Sillitoe on behalf of the Wellcome Sanger Institute COVID-19 Surveillance Team |
| EPI_ISL_2781849, EPI_ISL_2781850 | Lighthouse Lab in Milton Keynes | Wellcome Sanger Institute for the COVID-19 Genomics UK (COG-UK) Consortium | The Lighthouse Lab in Milton Keynes and Alex Alderton, Roberto Amato, Jeffrey Barrett, Sonia Goncalves, Ewan Harrison, David K. Jackson, Ian Johnston, Dominic Kwiatkowski, Cordelia Langford, John Sillitoe on behalf of the Wellcome Sanger Institute COVID-19 Surveillance Team |
| EPI_ISL_2781851, EPI_ISL_2781852, EPI_ISL_2781853 | Lighthouse Lab in Alderley Park | Wellcome Sanger Institute for the COVID-19 Genomics UK (COG-UK) Consortium | Jacquelyn Wynn, Mairead Hyland, The Lighthouse Lab in Alderley Park and Alex Alderton, Roberto Amato, Jeffrey Barrett, Sonia Goncalves, Ewan Harrison, David K. Jackson, Ian Johnston, Dominic Kwiatkowski, Cordelia Langford, John Sillitoe on behalf of the Wellcome Sanger Institute COVID-19 Surveillance Team |
| EPI_ISL_2781854 | Lighthouse Lab in Milton Keynes | Wellcome Sanger Institute for the COVID-19 Genomics UK (COG-UK) Consortium | The Lighthouse Lab in Milton Keynes and Alex Alderton, Roberto Amato, Jeffrey Barrett, Sonia Goncalves, Ewan Harrison, David K. Jackson, Ian Johnston, Dominic Kwiatkowski, Cordelia Langford, John Sillitoe on behalf of the Wellcome Sanger Institute COVID-19 Surveillance Team |
| EPI_ISL_2781855, EPI_ISL_2781856, EPI_ISL_2781857 | Lighthouse Lab in Alderley Park | Wellcome Sanger Institute for the COVID-19 Genomics UK (COG-UK) Consortium | Jacquelyn Wynn, Mairead Hyland, The Lighthouse Lab in Alderley Park and Alex Alderton, Roberto Amato, Jeffrey Barrett, Sonia Goncalves, Ewan Harrison, David K. Jackson, Ian Johnston, Dominic Kwiatkowski, Cordelia Langford, John Sillitoe on behalf of the Wellcome Sanger Institute COVID-19 Surveillance Team |
| EPI_ISL_2781858, EPI_ISL_2781859, EPI_ISL_2781860 | Lighthouse Lab in Milton Keynes | Wellcome Sanger Institute for the COVID-19 Genomics UK (COG-UK) Consortium | The Lighthouse Lab in Milton Keynes and Alex Alderton, Roberto Amato, Jeffrey Barrett, Sonia Goncalves, Ewan Harrison, David K. Jackson, Ian Johnston, Dominic Kwiatkowski, Cordelia Langford, John Sillitoe on behalf of the Wellcome Sanger Institute COVID-19 Surveillance Team |
| EPI_ISL_2781861, EPI_ISL_2781862 | Lighthouse Lab in Alderley Park | Wellcome Sanger Institute for the COVID-19 Genomics UK (COG-UK) Consortium | Jacquelyn Wynn, Mairead Hyland, The Lighthouse Lab in Alderley Park and Alex Alderton, Roberto Amato, Jeffrey Barrett, Sonia Goncalves, Ewan Harrison, David K. Jackson, Ian Johnston, Dominic Kwiatkowski, Cordelia Langford, John Sillitoe on behalf of the Wellcome Sanger Institute COVID-19 Surveillance Team |
| EPI_ISL_2781863 | Lighthouse Lab in Milton Keynes | Wellcome Sanger Institute for the COVID-19 Genomics UK (COG-UK) Consortium | The Lighthouse Lab in Milton Keynes and Alex Alderton, Roberto Amato, Jeffrey Barrett, Sonia Goncalves, Ewan Harrison, David K. Jackson, Ian Johnston, Dominic Kwiatkowski, Cordelia Langford, John Sillitoe on behalf of the Wellcome Sanger Institute COVID-19 Surveillance Team |
| EPI_ISL_2781864 | Lighthouse Lab in Alderley Park | Wellcome Sanger Institute for the COVID-19 Genomics UK (COG-UK) Consortium | Jacquelyn Wynn, Mairead Hyland, The Lighthouse Lab in Alderley Park and Alex Alderton, Roberto Amato, Jeffrey Barrett, Sonia Goncalves, Ewan Harrison, David K. Jackson, Ian Johnston, Dominic Kwiatkowski, Cordelia Langford, John Sillitoe on behalf of the Wellcome Sanger Institute COVID-19 Surveillance Team |
| EPI_ISL_2781865 | Lighthouse Lab in Glasgow | Wellcome Sanger Institute for the COVID-19 Genomics UK (COG-UK) Consortium | Harper VanSteenhouse, Yumi Kasai, David Gray, Carol Clugston, Anna Dominiczak and Alex Alderton, Roberto Amato, Jeffrey Barrett, Sonia Goncalves, Ewan Harrison, David K. Jackson, Ian Johnston, Dominic Kwiatkowski, Cordelia Langford, John Sillitoe on behalf of the Wellcome Sanger Institute COVID-19 Surveillance Team |
| EPI_ISL_2781866, EPI_ISL_2781867, EPI_ISL_2781868 | Lighthouse Lab in Alderley Park | Wellcome Sanger Institute for the COVID-19 Genomics UK (COG-UK) Consortium | Jacquelyn Wynn, Mairead Hyland, The Lighthouse Lab in Alderley Park and Alex Alderton, Roberto Amato, Jeffrey Barrett, Sonia Goncalves, Ewan Harrison, David K. Jackson, Ian Johnston, Dominic Kwiatkowski, Cordelia Langford, John Sillitoe on behalf of the Wellcome Sanger Institute COVID-19 Surveillance Team |
| EPI_ISL_2781869 | Lighthouse Lab in Milton Keynes | Wellcome Sanger Institute for the COVID-19 Genomics UK (COG-UK) Consortium | The Lighthouse Lab in Milton Keynes and Alex Alderton, Roberto Amato, Jeffrey Barrett, Sonia Goncalves, Ewan Harrison, David K. Jackson, Ian Johnston, Dominic Kwiatkowski, Cordelia Langford, John Sillitoe on behalf of the Wellcome Sanger Institute COVID-19 Surveillance Team |
| EPI_ISL_2781870, EPI_ISL_2781871, EPI_ISL_2781872, EPI_ISL_2781873, EPI_ISL_2781874, EPI_ISL_2781875 | Lighthouse Lab in Alderley Park | Wellcome Sanger Institute for the COVID-19 Genomics UK (COG-UK) Consortium | Jacquelyn Wynn, Mairead Hyland, The Lighthouse Lab in Alderley Park and Alex Alderton, Roberto Amato, Jeffrey Barrett, Sonia Goncalves, Ewan Harrison, David K. Jackson, Ian Johnston, Dominic Kwiatkowski, Cordelia Langford, John Sillitoe on behalf of the Wellcome Sanger Institute COVID-19 Surveillance Team |
| EPI_ISL_2781876 | Lighthouse Lab in Milton Keynes | Wellcome Sanger Institute for the COVID-19 Genomics UK (COG-UK) Consortium | The Lighthouse Lab in Milton Keynes and Alex Alderton, Roberto Amato, Jeffrey Barrett, Sonia Goncalves, Ewan Harrison, David K. Jackson, Ian Johnston, Dominic Kwiatkowski, Cordelia Langford, John Sillitoe on behalf of the Wellcome Sanger Institute COVID-19 Surveillance Team |
| EPI_ISL_2781877 | Lighthouse Lab in Alderley Park | Wellcome Sanger Institute for the COVID-19 Genomics UK (COG-UK) Consortium | Jacquelyn Wynn, Mairead Hyland, The Lighthouse Lab in Alderley Park and Alex Alderton, Roberto Amato, Jeffrey Barrett, Sonia Goncalves, Ewan Harrison, David K. Jackson, Ian Johnston, Dominic Kwiatkowski, Cordelia Langford, John Sillitoe on behalf of the Wellcome Sanger Institute COVID-19 Surveillance Team |
| EPI_ISL_2781878, EPI_ISL_2781879 | Lighthouse Lab in Milton Keynes | Wellcome Sanger Institute for the COVID-19 Genomics UK (COG-UK) Consortium | The Lighthouse Lab in Milton Keynes and Alex Alderton, Roberto Amato, Jeffrey Barrett, Sonia Goncalves, Ewan Harrison, David K. Jackson, Ian Johnston, Dominic Kwiatkowski, Cordelia Langford, John Sillitoe on behalf of the Wellcome Sanger Institute COVID-19 Surveillance Team |
| EPI_ISL_2781880, EPI_ISL_2781881, EPI_ISL_2781882 | Lighthouse Lab in Alderley Park | Wellcome Sanger Institute for the COVID-19 Genomics UK (COG-UK) Consortium | Jacquelyn Wynn, Mairead Hyland, The Lighthouse Lab in Alderley Park and Alex Alderton, Roberto Amato, Jeffrey Barrett, Sonia Goncalves, Ewan Harrison, David K. Jackson, Ian Johnston, Dominic Kwiatkowski, Cordelia Langford, John Sillitoe on behalf of the Wellcome Sanger Institute COVID-19 Surveillance Team |
| EPI_ISL_2781883, EPI_ISL_2781884 | Lighthouse Lab in Milton Keynes | Wellcome Sanger Institute for the COVID-19 Genomics UK (COG-UK) Consortium | The Lighthouse Lab in Milton Keynes and Alex Alderton, Roberto Amato, Jeffrey Barrett, Sonia Goncalves, Ewan Harrison, David K. Jackson, Ian Johnston, Dominic Kwiatkowski, Cordelia Langford, John Sillitoe on behalf of the Wellcome Sanger Institute COVID-19 Surveillance Team |
| EPI_ISL_2781885 | Lighthouse Lab in Alderley Park | Wellcome Sanger Institute for the COVID-19 Genomics UK (COG-UK) Consortium | Jacquelyn Wynn, Mairead Hyland, The Lighthouse Lab in Alderley Park and Alex Alderton, Roberto Amato, Jeffrey Barrett, Sonia Goncalves, Ewan Harrison, David K. Jackson, Ian Johnston, Dominic Kwiatkowski, Cordelia Langford, John Sillitoe on behalf of the Wellcome Sanger Institute COVID-19 Surveillance Team |
| EPI_ISL_2781886, EPI_ISL_2781887 | Lighthouse Lab in Milton Keynes | Wellcome Sanger Institute for the COVID-19 Genomics UK (COG-UK) Consortium | The Lighthouse Lab in Milton Keynes and Alex Alderton, Roberto Amato, Jeffrey Barrett, Sonia Goncalves, Ewan Harrison, David K. Jackson, Ian Johnston, Dominic Kwiatkowski, Cordelia Langford, John Sillitoe on behalf of the Wellcome Sanger Institute COVID-19 Surveillance Team |
| EPI_ISL_2781888, EPI_ISL_2781889 | Lighthouse Lab in Alderley Park | Wellcome Sanger Institute for the COVID-19 Genomics UK (COG-UK) Consortium | Jacquelyn Wynn, Mairead Hyland, The Lighthouse Lab in Alderley Park and Alex Alderton, Roberto Amato, Jeffrey Barrett, Sonia Goncalves, Ewan Harrison, David K. Jackson, Ian Johnston, Dominic Kwiatkowski, Cordelia Langford, John Sillitoe on behalf of the Wellcome Sanger Institute COVID-19 Surveillance Team |
| EPI_ISL_2781890, EPI_ISL_2781891 | Lighthouse Lab in Milton Keynes | Wellcome Sanger Institute for the COVID-19 Genomics UK (COG-UK) Consortium | The Lighthouse Lab in Milton Keynes and Alex Alderton, Roberto Amato, Jeffrey Barrett, Sonia Goncalves, Ewan Harrison, David K. Jackson, Ian Johnston, Dominic Kwiatkowski, Cordelia Langford, John Sillitoe on behalf of the Wellcome Sanger Institute COVID-19 Surveillance Team |
| EPI_ISL_2781892 | Lighthouse Lab in Alderley Park | Wellcome Sanger Institute for the COVID-19 Genomics UK (COG-UK) Consortium | Jacquelyn Wynn, Mairead Hyland, The Lighthouse Lab in Alderley Park and Alex Alderton, Roberto Amato, Jeffrey Barrett, Sonia Goncalves, Ewan Harrison, David K. Jackson, Ian Johnston, Dominic Kwiatkowski, Cordelia Langford, John Sillitoe on behalf of the Wellcome Sanger Institute COVID-19 Surveillance Team |
| EPI_ISL_2781893 | Lighthouse Lab in Milton Keynes | Wellcome Sanger Institute for the COVID-19 Genomics UK (COG-UK) Consortium | The Lighthouse Lab in Milton Keynes and Alex Alderton, Roberto Amato, Jeffrey Barrett, Sonia Goncalves, Ewan Harrison, David K. Jackson, Ian Johnston, Dominic Kwiatkowski, Cordelia Langford, John Sillitoe on behalf of the Wellcome Sanger Institute COVID-19 Surveillance Team |
| EPI_ISL_2781894, EPI_ISL_2781895, EPI_ISL_2781896, EPI_ISL_2781897 | Lighthouse Lab in Alderley Park | Wellcome Sanger Institute for the COVID-19 Genomics UK (COG-UK) Consortium | Jacquelyn Wynn, Mairead Hyland, The Lighthouse Lab in Alderley Park and Alex Alderton, Roberto Amato, Jeffrey Barrett, Sonia Goncalves, Ewan Harrison, David K. Jackson, Ian Johnston, Dominic Kwiatkowski, Cordelia Langford, John Sillitoe on behalf of the Wellcome Sanger Institute COVID-19 Surveillance Team |
| EPI_ISL_2781898 | Lighthouse Lab in Milton Keynes | Wellcome Sanger Institute for the COVID-19 Genomics UK (COG-UK) Consortium | The Lighthouse Lab in Milton Keynes and Alex Alderton, Roberto Amato, Jeffrey Barrett, Sonia Goncalves, Ewan Harrison, David K. Jackson, Ian Johnston, Dominic Kwiatkowski, Cordelia Langford, John Sillitoe on behalf of the Wellcome Sanger Institute COVID-19 Surveillance Team |
| EPI_ISL_2781899 | Lighthouse Lab in Glasgow | Wellcome Sanger Institute for the COVID-19 Genomics UK (COG-UK) Consortium | Harper VanSteenhouse, Yumi Kasai, David Gray, Carol Clugston, Anna Dominiczak and Alex Alderton, Roberto Amato, Jeffrey Barrett, Sonia Goncalves, Ewan Harrison, David K. Jackson, Ian Johnston, Dominic Kwiatkowski, Cordelia Langford, John Sillitoe on behalf of the Wellcome Sanger Institute COVID-19 Surveillance Team |
| EPI_ISL_2781900, EPI_ISL_2781901, EPI_ISL_2781902 | Lighthouse Lab in Alderley Park | Wellcome Sanger Institute for the COVID-19 Genomics UK (COG-UK) Consortium | Jacquelyn Wynn, Mairead Hyland, The Lighthouse Lab in Alderley Park and Alex Alderton, Roberto Amato, Jeffrey Barrett, Sonia Goncalves, Ewan Harrison, David K. Jackson, Ian Johnston, Dominic Kwiatkowski, Cordelia Langford, John Sillitoe on behalf of the Wellcome Sanger Institute COVID-19 Surveillance Team |
| EPI_ISL_2781903 | Lighthouse Lab in Milton Keynes | Wellcome Sanger Institute for the COVID-19 Genomics UK (COG-UK) Consortium | The Lighthouse Lab in Milton Keynes and Alex Alderton, Roberto Amato, Jeffrey Barrett, Sonia Goncalves, Ewan Harrison, David K. Jackson, Ian Johnston, Dominic Kwiatkowski, Cordelia Langford, John Sillitoe on behalf of the Wellcome Sanger Institute COVID-19 Surveillance Team |
| EPI_ISL_2781904, EPI_ISL_2781905 | Lighthouse Lab in Alderley Park | Wellcome Sanger Institute for the COVID-19 Genomics UK (COG-UK) Consortium | Jacquelyn Wynn, Mairead Hyland, The Lighthouse Lab in Alderley Park and Alex Alderton, Roberto Amato, Jeffrey Barrett, Sonia Goncalves, Ewan Harrison, David K. Jackson, Ian Johnston, Dominic Kwiatkowski, Cordelia Langford, John Sillitoe on behalf of the Wellcome Sanger Institute COVID-19 Surveillance Team |
| EPI_ISL_2781906 | Lighthouse Lab in Milton Keynes | Wellcome Sanger Institute for the COVID-19 Genomics UK (COG-UK) Consortium | The Lighthouse Lab in Milton Keynes and Alex Alderton, Roberto Amato, Jeffrey Barrett, Sonia Goncalves, Ewan Harrison, David K. Jackson, Ian Johnston, Dominic Kwiatkowski, Cordelia Langford, John Sillitoe on behalf of the Wellcome Sanger Institute COVID-19 Surveillance Team |
| EPI_ISL_2781907, | Lighthouse Lab in Alderley Park | Wellcome Sanger Institute for the COVID-19 Genomics UK (COG-UK) | Jacquelyn Wynn, Mairead Hyland, The Lighthouse Lab in Alderley Park and Alex Alderton, Roberto Amato, Jeffrey Barrett, Sonia Goncalves, Ewan Harrison, David K. Jackson, Ian Johnston, Dominic Kwiatkowski, Cordelia Langford, John Sillitoe on |

| | | | |
|---|---|---|---|
| EPI_ISL_2781908, EPI_ISL_2781909, EPI_ISL_2781910, EPI_ISL_2781911, EPI_ISL_2781912, EPI_ISL_2781913, EPI_ISL_2781914, EPI_ISL_2781915 | | Consortium | behalf of the Wellcome Sanger Institute COVID-19 Surveillance Team |
| EPI_ISL_2781916, EPI_ISL_2781917 | Lighthouse Lab in Milton Keynes | Wellcome Sanger Institute for the COVID-19 Genomics UK (COG-UK) Consortium | The Lighthouse Lab in Milton Keynes and Alex Alderton, Roberto Amato, Jeffrey Barrett, Sonia Goncalves, Ewan Harrison, David K. Jackson, Ian Johnston, Dominic Kwiatkowski, Cordelia Langford, John Sillitoe on behalf of the Wellcome Sanger Institute COVID-19 Surveillance Team |
| EPI_ISL_2781918 | Lighthouse Lab in Alderley Park | Wellcome Sanger Institute for the COVID-19 Genomics UK (COG-UK) Consortium | Jacquelyn Wynn, Mairead Hyland, The Lighthouse Lab in Alderley Park and Alex Alderton, Roberto Amato, Jeffrey Barrett, Sonia Goncalves, Ewan Harrison, David K. Jackson, Ian Johnston, Dominic Kwiatkowski, Cordelia Langford, John Sillitoe on behalf of the Wellcome Sanger Institute COVID-19 Surveillance Team |
| EPI_ISL_2781919 | Lighthouse Lab in Milton Keynes | Wellcome Sanger Institute for the COVID-19 Genomics UK (COG-UK) Consortium | The Lighthouse Lab in Milton Keynes and Alex Alderton, Roberto Amato, Jeffrey Barrett, Sonia Goncalves, Ewan Harrison, David K. Jackson, Ian Johnston, Dominic Kwiatkowski, Cordelia Langford, John Sillitoe on behalf of the Wellcome Sanger Institute COVID-19 Surveillance Team |
| EPI_ISL_2781920 | Lighthouse Lab in Alderley Park | Wellcome Sanger Institute for the COVID-19 Genomics UK (COG-UK) Consortium | Jacquelyn Wynn, Mairead Hyland, The Lighthouse Lab in Alderley Park and Alex Alderton, Roberto Amato, Jeffrey Barrett, Sonia Goncalves, Ewan Harrison, David K. Jackson, Ian Johnston, Dominic Kwiatkowski, Cordelia Langford, John Sillitoe on behalf of the Wellcome Sanger Institute COVID-19 Surveillance Team |
| EPI_ISL_2781921 | Lighthouse Lab in Glasgow | Wellcome Sanger Institute for the COVID-19 Genomics UK (COG-UK) Consortium | Harper VanSteenhouse, Yumi Kasai, David Gray, Carol Clugston, Anna Dominiczak and Alex Alderton, Roberto Amato, Jeffrey Barrett, Sonia Goncalves, Ewan Harrison, David K. Jackson, Ian Johnston, Dominic Kwiatkowski, Cordelia Langford, John Sillitoe on behalf of the Wellcome Sanger Institute COVID-19 Surveillance Team |
| EPI_ISL_2781922, EPI_ISL_2781923 | Lighthouse Lab in Milton Keynes | Wellcome Sanger Institute for the COVID-19 Genomics UK (COG-UK) Consortium | The Lighthouse Lab in Milton Keynes and Alex Alderton, Roberto Amato, Jeffrey Barrett, Sonia Goncalves, Ewan Harrison, David K. Jackson, Ian Johnston, Dominic Kwiatkowski, Cordelia Langford, John Sillitoe on behalf of the Wellcome Sanger Institute COVID-19 Surveillance Team |
| EPI_ISL_2781924 | Lighthouse Lab in Alderley Park | Wellcome Sanger Institute for the COVID-19 Genomics UK (COG-UK) Consortium | Jacquelyn Wynn, Mairead Hyland, The Lighthouse Lab in Alderley Park and Alex Alderton, Roberto Amato, Jeffrey Barrett, Sonia Goncalves, Ewan Harrison, David K. Jackson, Ian Johnston, Dominic Kwiatkowski, Cordelia Langford, John Sillitoe on behalf of the Wellcome Sanger Institute COVID-19 Surveillance Team |
| EPI_ISL_2781925 | Lighthouse Lab in Milton Keynes | Wellcome Sanger Institute for the COVID-19 Genomics UK (COG-UK) Consortium | The Lighthouse Lab in Milton Keynes and Alex Alderton, Roberto Amato, Jeffrey Barrett, Sonia Goncalves, Ewan Harrison, David K. Jackson, Ian Johnston, Dominic Kwiatkowski, Cordelia Langford, John Sillitoe on behalf of the Wellcome Sanger Institute COVID-19 Surveillance Team |
| EPI_ISL_2781926, EPI_ISL_2781927 | Lighthouse Lab in Alderley Park | Wellcome Sanger Institute for the COVID-19 Genomics UK (COG-UK) Consortium | Jacquelyn Wynn, Mairead Hyland, The Lighthouse Lab in Alderley Park and Alex Alderton, Roberto Amato, Jeffrey Barrett, Sonia Goncalves, Ewan Harrison, David K. Jackson, Ian Johnston, Dominic Kwiatkowski, Cordelia Langford, John Sillitoe on behalf of the Wellcome Sanger Institute COVID-19 Surveillance Team |
| EPI_ISL_2781928, EPI_ISL_2781929, EPI_ISL_2781930 | Lighthouse Lab in Milton Keynes | Wellcome Sanger Institute for the COVID-19 Genomics UK (COG-UK) Consortium | The Lighthouse Lab in Milton Keynes and Alex Alderton, Roberto Amato, Jeffrey Barrett, Sonia Goncalves, Ewan Harrison, David K. Jackson, Ian Johnston, Dominic Kwiatkowski, Cordelia Langford, John Sillitoe on behalf of the Wellcome Sanger Institute COVID-19 Surveillance Team |
| EPI_ISL_2781931, EPI_ISL_2781932, EPI_ISL_2781933, EPI_ISL_2781934 | Lighthouse Lab in Alderley Park | Wellcome Sanger Institute for the COVID-19 Genomics UK (COG-UK) Consortium | Jacquelyn Wynn, Mairead Hyland, The Lighthouse Lab in Alderley Park and Alex Alderton, Roberto Amato, Jeffrey Barrett, Sonia Goncalves, Ewan Harrison, David K. Jackson, Ian Johnston, Dominic Kwiatkowski, Cordelia Langford, John Sillitoe on behalf of the Wellcome Sanger Institute COVID-19 Surveillance Team |
| EPI_ISL_2781935 | Lighthouse Lab in Milton Keynes | Wellcome Sanger Institute for the COVID-19 Genomics UK (COG-UK) Consortium | The Lighthouse Lab in Milton Keynes and Alex Alderton, Roberto Amato, Jeffrey Barrett, Sonia Goncalves, Ewan Harrison, David K. Jackson, Ian Johnston, Dominic Kwiatkowski, Cordelia Langford, John Sillitoe on behalf of the Wellcome Sanger Institute COVID-19 Surveillance Team |
| EPI_ISL_2781936 | Lighthouse Lab in Alderley Park | Wellcome Sanger Institute for the COVID-19 Genomics UK (COG-UK) Consortium | Jacquelyn Wynn, Mairead Hyland, The Lighthouse Lab in Alderley Park and Alex Alderton, Roberto Amato, Jeffrey Barrett, Sonia Goncalves, Ewan Harrison, David K. Jackson, Ian Johnston, Dominic Kwiatkowski, Cordelia Langford, John Sillitoe on behalf of the Wellcome Sanger Institute COVID-19 Surveillance Team |
| EPI_ISL_2781937 | Lighthouse Lab in Glasgow | Wellcome Sanger Institute for the COVID-19 Genomics UK (COG-UK) Consortium | Harper VanSteenhouse, Yumi Kasai, David Gray, Carol Clugston, Anna Dominiczak and Alex Alderton, Roberto Amato, Jeffrey Barrett, Sonia Goncalves, Ewan Harrison, David K. Jackson, Ian Johnston, Dominic Kwiatkowski, Cordelia Langford, John Sillitoe on behalf of the Wellcome Sanger Institute COVID-19 Surveillance Team |
| EPI_ISL_2781938, EPI_ISL_2781939, EPI_ISL_2781940 | Lighthouse Lab in Alderley Park | Wellcome Sanger Institute for the COVID-19 Genomics UK (COG-UK) Consortium | Jacquelyn Wynn, Mairead Hyland, The Lighthouse Lab in Alderley Park and Alex Alderton, Roberto Amato, Jeffrey Barrett, Sonia Goncalves, Ewan Harrison, David K. Jackson, Ian Johnston, Dominic Kwiatkowski, Cordelia Langford, John Sillitoe on behalf of the Wellcome Sanger Institute COVID-19 Surveillance Team |
| EPI_ISL_2781941 | Lighthouse Lab in Milton Keynes | Wellcome Sanger Institute for the COVID-19 Genomics UK (COG-UK) Consortium | The Lighthouse Lab in Milton Keynes and Alex Alderton, Roberto Amato, Jeffrey Barrett, Sonia Goncalves, Ewan Harrison, David K. Jackson, Ian Johnston, Dominic Kwiatkowski, Cordelia Langford, John Sillitoe on behalf of the Wellcome Sanger Institute COVID-19 Surveillance Team |
| EPI_ISL_2781942, EPI_ISL_2781943, EPI_ISL_2781944 | Lighthouse Lab in Alderley Park | Wellcome Sanger Institute for the COVID-19 Genomics UK (COG-UK) Consortium | Jacquelyn Wynn, Mairead Hyland, The Lighthouse Lab in Alderley Park and Alex Alderton, Roberto Amato, Jeffrey Barrett, Sonia Goncalves, Ewan Harrison, David K. Jackson, Ian Johnston, Dominic Kwiatkowski, Cordelia Langford, John Sillitoe on behalf of the Wellcome Sanger Institute COVID-19 Surveillance Team |
| EPI_ISL_2781945, EPI_ISL_2781946, EPI_ISL_2781947 | Lighthouse Lab in Milton Keynes | Wellcome Sanger Institute for the COVID-19 Genomics UK (COG-UK) Consortium | The Lighthouse Lab in Milton Keynes and Alex Alderton, Roberto Amato, Jeffrey Barrett, Sonia Goncalves, Ewan Harrison, David K. Jackson, Ian Johnston, Dominic Kwiatkowski, Cordelia Langford, John Sillitoe on behalf of the Wellcome Sanger Institute COVID-19 Surveillance Team |
| EPI_ISL_2781948 | Lighthouse Lab in Alderley Park | Wellcome Sanger Institute for the COVID-19 Genomics UK (COG-UK) Consortium | Jacquelyn Wynn, Mairead Hyland, The Lighthouse Lab in Alderley Park and Alex Alderton, Roberto Amato, Jeffrey Barrett, Sonia Goncalves, Ewan Harrison, David K. Jackson, Ian Johnston, Dominic Kwiatkowski, Cordelia Langford, John Sillitoe on behalf of the Wellcome Sanger Institute COVID-19 Surveillance Team |
| EPI_ISL_2781949 | Lighthouse Lab in Milton Keynes | Wellcome Sanger Institute for the COVID-19 Genomics UK (COG-UK) Consortium | The Lighthouse Lab in Milton Keynes and Alex Alderton, Roberto Amato, Jeffrey Barrett, Sonia Goncalves, Ewan Harrison, David K. Jackson, Ian Johnston, Dominic Kwiatkowski, Cordelia Langford, John Sillitoe on behalf of the Wellcome Sanger Institute COVID-19 Surveillance Team |
| EPI_ISL_2781950, EPI_ISL_2781951, EPI_ISL_2781952 | Lighthouse Lab in Alderley Park | Wellcome Sanger Institute for the COVID-19 Genomics UK (COG-UK) Consortium | Jacquelyn Wynn, Mairead Hyland, The Lighthouse Lab in Alderley Park and Alex Alderton, Roberto Amato, Jeffrey Barrett, Sonia Goncalves, Ewan Harrison, David K. Jackson, Ian Johnston, Dominic Kwiatkowski, Cordelia Langford, John Sillitoe on behalf of the Wellcome Sanger Institute COVID-19 Surveillance Team |
| EPI_ISL_2781953 | Lighthouse Lab in Milton Keynes | Wellcome Sanger Institute for the COVID-19 Genomics UK (COG-UK) Consortium | The Lighthouse Lab in Milton Keynes and Alex Alderton, Roberto Amato, Jeffrey Barrett, Sonia Goncalves, Ewan Harrison, David K. Jackson, Ian Johnston, Dominic Kwiatkowski, Cordelia Langford, John Sillitoe on behalf of the Wellcome Sanger Institute COVID-19 Surveillance Team |
| EPI_ISL_2781954, EPI_ISL_2781955, EPI_ISL_2781956, EPI_ISL_2781957, EPI_ISL_2781958 | Lighthouse Lab in Alderley Park | Wellcome Sanger Institute for the COVID-19 Genomics UK (COG-UK) Consortium | Jacquelyn Wynn, Mairead Hyland, The Lighthouse Lab in Alderley Park and Alex Alderton, Roberto Amato, Jeffrey Barrett, Sonia Goncalves, Ewan Harrison, David K. Jackson, Ian Johnston, Dominic Kwiatkowski, Cordelia Langford, John Sillitoe on behalf of the Wellcome Sanger Institute COVID-19 Surveillance Team |
| EPI_ISL_2781959, EPI_ISL_2781960 | Lighthouse Lab in Milton Keynes | Wellcome Sanger Institute for the COVID-19 Genomics UK (COG-UK) Consortium | The Lighthouse Lab in Milton Keynes and Alex Alderton, Roberto Amato, Jeffrey Barrett, Sonia Goncalves, Ewan Harrison, David K. Jackson, Ian Johnston, Dominic Kwiatkowski, Cordelia Langford, John Sillitoe on behalf of the Wellcome Sanger Institute COVID-19 Surveillance Team |
| EPI_ISL_2781961 | Lighthouse Lab in Alderley Park | Wellcome Sanger Institute for the COVID-19 Genomics UK (COG-UK) Consortium | Jacquelyn Wynn, Mairead Hyland, The Lighthouse Lab in Alderley Park and Alex Alderton, Roberto Amato, Jeffrey Barrett, Sonia Goncalves, Ewan Harrison, David K. Jackson, Ian Johnston, Dominic Kwiatkowski, Cordelia Langford, John Sillitoe on behalf of the Wellcome Sanger Institute COVID-19 Surveillance Team |
| EPI_ISL_2781962, EPI_ISL_2781963 | Lighthouse Lab in Milton Keynes | Wellcome Sanger Institute for the COVID-19 Genomics UK (COG-UK) Consortium | The Lighthouse Lab in Milton Keynes and Alex Alderton, Roberto Amato, Jeffrey Barrett, Sonia Goncalves, Ewan Harrison, David K. Jackson, Ian Johnston, Dominic Kwiatkowski, Cordelia Langford, John Sillitoe on behalf of the Wellcome Sanger Institute COVID-19 Surveillance Team |
| EPI_ISL_2781964, EPI_ISL_2781965 | Lighthouse Lab in Alderley Park | Wellcome Sanger Institute for the COVID-19 Genomics UK (COG-UK) Consortium | Jacquelyn Wynn, Mairead Hyland, The Lighthouse Lab in Alderley Park and Alex Alderton, Roberto Amato, Jeffrey Barrett, Sonia Goncalves, Ewan Harrison, David K. Jackson, Ian Johnston, Dominic Kwiatkowski, Cordelia Langford, John Sillitoe on behalf of the Wellcome Sanger Institute COVID-19 Surveillance Team |
| EPI_ISL_2781966 | Lighthouse Lab in Glasgow | Wellcome Sanger Institute for the COVID-19 Genomics UK (COG-UK) Consortium | Harper VanSteenhouse, Yumi Kasai, David Gray, Carol Clugston, Anna Dominiczak and Alex Alderton, Roberto Amato, Jeffrey Barrett, Sonia Goncalves, Ewan Harrison, David K. Jackson, Ian Johnston, Dominic Kwiatkowski, Cordelia Langford, John Sillitoe on behalf of the Wellcome Sanger Institute COVID-19 Surveillance Team |
| EPI_ISL_2781967, EPI_ISL_2781968, EPI_ISL_2781969, EPI_ISL_2781970 | Lighthouse Lab in Alderley Park | Wellcome Sanger Institute for the COVID-19 Genomics UK (COG-UK) Consortium | Jacquelyn Wynn, Mairead Hyland, The Lighthouse Lab in Alderley Park and Alex Alderton, Roberto Amato, Jeffrey Barrett, Sonia Goncalves, Ewan Harrison, David K. Jackson, Ian Johnston, Dominic Kwiatkowski, Cordelia Langford, John Sillitoe on behalf of the Wellcome Sanger Institute COVID-19 Surveillance Team |
| EPI_ISL_2781971 | Lighthouse Lab in Glasgow | Wellcome Sanger Institute for the COVID-19 Genomics UK (COG-UK) Consortium | Harper VanSteenhouse, Yumi Kasai, David Gray, Carol Clugston, Anna Dominiczak and Alex Alderton, Roberto Amato, Jeffrey Barrett, Sonia Goncalves, Ewan Harrison, David K. Jackson, Ian Johnston, Dominic Kwiatkowski, Cordelia Langford, John Sillitoe on behalf of the Wellcome Sanger Institute COVID-19 Surveillance Team |
| EPI_ISL_2781972 | Lighthouse Lab in Alderley Park | Wellcome Sanger Institute for the COVID-19 Genomics UK (COG-UK) Consortium | Jacquelyn Wynn, Mairead Hyland, The Lighthouse Lab in Alderley Park and Alex Alderton, Roberto Amato, Jeffrey Barrett, Sonia Goncalves, Ewan Harrison, David K. Jackson, Ian Johnston, Dominic Kwiatkowski, Cordelia Langford, John Sillitoe on behalf of the Wellcome Sanger Institute COVID-19 Surveillance Team |
| EPI_ISL_2781973 | Lighthouse Lab in Milton Keynes | Wellcome Sanger Institute for the COVID-19 Genomics UK (COG-UK) Consortium | The Lighthouse Lab in Milton Keynes and Alex Alderton, Roberto Amato, Jeffrey Barrett, Sonia Goncalves, Ewan Harrison, David K. Jackson, Ian Johnston, Dominic Kwiatkowski, Cordelia Langford, John Sillitoe on behalf of the Wellcome Sanger Institute COVID-19 Surveillance Team |
| EPI_ISL_2781974, EPI_ISL_2781975, EPI_ISL_2781976, EPI_ISL_2781977, EPI_ISL_2781978 | Lighthouse Lab in Alderley Park | Wellcome Sanger Institute for the COVID-19 Genomics UK (COG-UK) Consortium | Jacquelyn Wynn, Mairead Hyland, The Lighthouse Lab in Alderley Park and Alex Alderton, Roberto Amato, Jeffrey Barrett, Sonia Goncalves, Ewan Harrison, David K. Jackson, Ian Johnston, Dominic Kwiatkowski, Cordelia Langford, John Sillitoe on behalf of the Wellcome Sanger Institute COVID-19 Surveillance Team |
| EPI_ISL_2781979, EPI_ISL_2781980, EPI_ISL_2781981 | Lighthouse Lab in Milton Keynes | Wellcome Sanger Institute for the COVID-19 Genomics UK (COG-UK) Consortium | The Lighthouse Lab in Milton Keynes and Alex Alderton, Roberto Amato, Jeffrey Barrett, Sonia Goncalves, Ewan Harrison, David K. Jackson, Ian Johnston, Dominic Kwiatkowski, Cordelia Langford, John Sillitoe on behalf of the Wellcome Sanger Institute COVID-19 Surveillance Team |
| EPI_ISL_2781982, EPI_ISL_2781983, EPI_ISL_2781984 | Lighthouse Lab in Alderley Park | Wellcome Sanger Institute for the COVID-19 Genomics UK (COG-UK) Consortium | Jacquelyn Wynn, Mairead Hyland, The Lighthouse Lab in Alderley Park and Alex Alderton, Roberto Amato, Jeffrey Barrett, Sonia Goncalves, Ewan Harrison, David K. Jackson, Ian Johnston, Dominic Kwiatkowski, Cordelia Langford, John Sillitoe on behalf of the Wellcome Sanger Institute COVID-19 Surveillance Team |

| | | | |
|---|---|---|---|
| EPI_ISL_2781985 | Lighthouse Lab in Glasgow | Wellcome Sanger Institute for the COVID-19 Genomics UK (COG-UK) Consortium | Harper VanSteenhouse, Yumi Kasai, David Gray, Carol Clugston, Anna Dominiczak and Alex Alderton, Roberto Amato, Jeffrey Barrett, Sonia Goncalves, Ewan Harrison, David K. Jackson, Ian Johnston, Dominic Kwiatkowski, Cordelia Langford, John Sillitoe on behalf of the Wellcome Sanger Institute COVID-19 Surveillance Team |
| EPI_ISL_2781986 | Lighthouse Lab in Alderley Park | Wellcome Sanger Institute for the COVID-19 Genomics UK (COG-UK) Consortium | Jacquelyn Wynn, Mairead Hyland, The Lighthouse Lab in Alderley Park and Alex Alderton, Roberto Amato, Jeffrey Barrett, Sonia Goncalves, Ewan Harrison, David K. Jackson, Ian Johnston, Dominic Kwiatkowski, Cordelia Langford, John Sillitoe on behalf of the Wellcome Sanger Institute COVID-19 Surveillance Team |
| EPI_ISL_2781987, EPI_ISL_2781988 | Lighthouse Lab in Milton Keynes | Wellcome Sanger Institute for the COVID-19 Genomics UK (COG-UK) Consortium | The Lighthouse Lab in Milton Keynes and Alex Alderton, Roberto Amato, Jeffrey Barrett, Sonia Goncalves, Ewan Harrison, David K. Jackson, Ian Johnston, Dominic Kwiatkowski, Cordelia Langford, John Sillitoe on behalf of the Wellcome Sanger Institute COVID-19 Surveillance Team |
| EPI_ISL_2781989 | Lighthouse Lab in Alderley Park | Wellcome Sanger Institute for the COVID-19 Genomics UK (COG-UK) Consortium | Jacquelyn Wynn, Mairead Hyland, The Lighthouse Lab in Alderley Park and Alex Alderton, Roberto Amato, Jeffrey Barrett, Sonia Goncalves, Ewan Harrison, David K. Jackson, Ian Johnston, Dominic Kwiatkowski, Cordelia Langford, John Sillitoe on behalf of the Wellcome Sanger Institute COVID-19 Surveillance Team |
| EPI_ISL_2781990 | Lighthouse Lab in Milton Keynes | Wellcome Sanger Institute for the COVID-19 Genomics UK (COG-UK) Consortium | The Lighthouse Lab in Milton Keynes and Alex Alderton, Roberto Amato, Jeffrey Barrett, Sonia Goncalves, Ewan Harrison, David K. Jackson, Ian Johnston, Dominic Kwiatkowski, Cordelia Langford, John Sillitoe on behalf of the Wellcome Sanger Institute COVID-19 Surveillance Team |
| EPI_ISL_2781991 | Lighthouse Lab in Alderley Park | Wellcome Sanger Institute for the COVID-19 Genomics UK (COG-UK) Consortium | Jacquelyn Wynn, Mairead Hyland, The Lighthouse Lab in Alderley Park and Alex Alderton, Roberto Amato, Jeffrey Barrett, Sonia Goncalves, Ewan Harrison, David K. Jackson, Ian Johnston, Dominic Kwiatkowski, Cordelia Langford, John Sillitoe on behalf of the Wellcome Sanger Institute COVID-19 Surveillance Team |
| EPI_ISL_2781992, EPI_ISL_2781993, EPI_ISL_2781994 | Lighthouse Lab in Milton Keynes | Wellcome Sanger Institute for the COVID-19 Genomics UK (COG-UK) Consortium | The Lighthouse Lab in Milton Keynes and Alex Alderton, Roberto Amato, Jeffrey Barrett, Sonia Goncalves, Ewan Harrison, David K. Jackson, Ian Johnston, Dominic Kwiatkowski, Cordelia Langford, John Sillitoe on behalf of the Wellcome Sanger Institute COVID-19 Surveillance Team |
| EPI_ISL_2781995 | Lighthouse Lab in Glasgow | Wellcome Sanger Institute for the COVID-19 Genomics UK (COG-UK) Consortium | Harper VanSteenhouse, Yumi Kasai, David Gray, Carol Clugston, Anna Dominiczak and Alex Alderton, Roberto Amato, Jeffrey Barrett, Sonia Goncalves, Ewan Harrison, David K. Jackson, Ian Johnston, Dominic Kwiatkowski, Cordelia Langford, John Sillitoe on behalf of the Wellcome Sanger Institute COVID-19 Surveillance Team |
| EPI_ISL_2781996 | Lighthouse Lab in Milton Keynes | Wellcome Sanger Institute for the COVID-19 Genomics UK (COG-UK) Consortium | The Lighthouse Lab in Milton Keynes and Alex Alderton, Roberto Amato, Jeffrey Barrett, Sonia Goncalves, Ewan Harrison, David K. Jackson, Ian Johnston, Dominic Kwiatkowski, Cordelia Langford, John Sillitoe on behalf of the Wellcome Sanger Institute COVID-19 Surveillance Team |
| EPI_ISL_2781997, EPI_ISL_2781998 | Lighthouse Lab in Alderley Park | Wellcome Sanger Institute for the COVID-19 Genomics UK (COG-UK) Consortium | Jacquelyn Wynn, Mairead Hyland, The Lighthouse Lab in Alderley Park and Alex Alderton, Roberto Amato, Jeffrey Barrett, Sonia Goncalves, Ewan Harrison, David K. Jackson, Ian Johnston, Dominic Kwiatkowski, Cordelia Langford, John Sillitoe on behalf of the Wellcome Sanger Institute COVID-19 Surveillance Team |
| EPI_ISL_2781999 | Lighthouse Lab in Milton Keynes | Wellcome Sanger Institute for the COVID-19 Genomics UK (COG-UK) Consortium | The Lighthouse Lab in Milton Keynes and Alex Alderton, Roberto Amato, Jeffrey Barrett, Sonia Goncalves, Ewan Harrison, David K. Jackson, Ian Johnston, Dominic Kwiatkowski, Cordelia Langford, John Sillitoe on behalf of the Wellcome Sanger Institute COVID-19 Surveillance Team |
| EPI_ISL_2782000, EPI_ISL_2782001, EPI_ISL_2782002, EPI_ISL_2782003 | Lighthouse Lab in Alderley Park | Wellcome Sanger Institute for the COVID-19 Genomics UK (COG-UK) Consortium | Jacquelyn Wynn, Mairead Hyland, The Lighthouse Lab in Alderley Park and Alex Alderton, Roberto Amato, Jeffrey Barrett, Sonia Goncalves, Ewan Harrison, David K. Jackson, Ian Johnston, Dominic Kwiatkowski, Cordelia Langford, John Sillitoe on behalf of the Wellcome Sanger Institute COVID-19 Surveillance Team |
| EPI_ISL_2782004, EPI_ISL_2782005 | Lighthouse Lab in Milton Keynes | Wellcome Sanger Institute for the COVID-19 Genomics UK (COG-UK) Consortium | The Lighthouse Lab in Milton Keynes and Alex Alderton, Roberto Amato, Jeffrey Barrett, Sonia Goncalves, Ewan Harrison, David K. Jackson, Ian Johnston, Dominic Kwiatkowski, Cordelia Langford, John Sillitoe on behalf of the Wellcome Sanger Institute COVID-19 Surveillance Team |
| EPI_ISL_2782006, EPI_ISL_2782007, EPI_ISL_2782008 | Lighthouse Lab in Alderley Park | Wellcome Sanger Institute for the COVID-19 Genomics UK (COG-UK) Consortium | Jacquelyn Wynn, Mairead Hyland, The Lighthouse Lab in Alderley Park and Alex Alderton, Roberto Amato, Jeffrey Barrett, Sonia Goncalves, Ewan Harrison, David K. Jackson, Ian Johnston, Dominic Kwiatkowski, Cordelia Langford, John Sillitoe on behalf of the Wellcome Sanger Institute COVID-19 Surveillance Team |
| EPI_ISL_2782009 | Lighthouse Lab in Glasgow | Wellcome Sanger Institute for the COVID-19 Genomics UK (COG-UK) Consortium | Harper VanSteenhouse, Yumi Kasai, David Gray, Carol Clugston, Anna Dominiczak and Alex Alderton, Roberto Amato, Jeffrey Barrett, Sonia Goncalves, Ewan Harrison, David K. Jackson, Ian Johnston, Dominic Kwiatkowski, Cordelia Langford, John Sillitoe on behalf of the Wellcome Sanger Institute COVID-19 Surveillance Team |
| EPI_ISL_2782010, EPI_ISL_2782011 | Lighthouse Lab in Alderley Park | Wellcome Sanger Institute for the COVID-19 Genomics UK (COG-UK) Consortium | Jacquelyn Wynn, Mairead Hyland, The Lighthouse Lab in Alderley Park and Alex Alderton, Roberto Amato, Jeffrey Barrett, Sonia Goncalves, Ewan Harrison, David K. Jackson, Ian Johnston, Dominic Kwiatkowski, Cordelia Langford, John Sillitoe on behalf of the Wellcome Sanger Institute COVID-19 Surveillance Team |
| EPI_ISL_2782012 | Lighthouse Lab in Glasgow | Wellcome Sanger Institute for the COVID-19 Genomics UK (COG-UK) Consortium | Harper VanSteenhouse, Yumi Kasai, David Gray, Carol Clugston, Anna Dominiczak and Alex Alderton, Roberto Amato, Jeffrey Barrett, Sonia Goncalves, Ewan Harrison, David K. Jackson, Ian Johnston, Dominic Kwiatkowski, Cordelia Langford, John Sillitoe on behalf of the Wellcome Sanger Institute COVID-19 Surveillance Team |
| EPI_ISL_2782013, EPI_ISL_2782014, EPI_ISL_2782015 | Lighthouse Lab in Alderley Park | Wellcome Sanger Institute for the COVID-19 Genomics UK (COG-UK) Consortium | Jacquelyn Wynn, Mairead Hyland, The Lighthouse Lab in Alderley Park and Alex Alderton, Roberto Amato, Jeffrey Barrett, Sonia Goncalves, Ewan Harrison, David K. Jackson, Ian Johnston, Dominic Kwiatkowski, Cordelia Langford, John Sillitoe on behalf of the Wellcome Sanger Institute COVID-19 Surveillance Team |
| EPI_ISL_2782016, EPI_ISL_2782017 | Lighthouse Lab in Milton Keynes | Wellcome Sanger Institute for the COVID-19 Genomics UK (COG-UK) Consortium | The Lighthouse Lab in Milton Keynes and Alex Alderton, Roberto Amato, Jeffrey Barrett, Sonia Goncalves, Ewan Harrison, David K. Jackson, Ian Johnston, Dominic Kwiatkowski, Cordelia Langford, John Sillitoe on behalf of the Wellcome Sanger Institute COVID-19 Surveillance Team |
| EPI_ISL_2782018, EPI_ISL_2782019, EPI_ISL_2782020 | Lighthouse Lab in Alderley Park | Wellcome Sanger Institute for the COVID-19 Genomics UK (COG-UK) Consortium | Jacquelyn Wynn, Mairead Hyland, The Lighthouse Lab in Alderley Park and Alex Alderton, Roberto Amato, Jeffrey Barrett, Sonia Goncalves, Ewan Harrison, David K. Jackson, Ian Johnston, Dominic Kwiatkowski, Cordelia Langford, John Sillitoe on behalf of the Wellcome Sanger Institute COVID-19 Surveillance Team |
| EPI_ISL_2782021 | Lighthouse Lab in Milton Keynes | Wellcome Sanger Institute for the COVID-19 Genomics UK (COG-UK) Consortium | The Lighthouse Lab in Milton Keynes and Alex Alderton, Roberto Amato, Jeffrey Barrett, Sonia Goncalves, Ewan Harrison, David K. Jackson, Ian Johnston, Dominic Kwiatkowski, Cordelia Langford, John Sillitoe on behalf of the Wellcome Sanger Institute COVID-19 Surveillance Team |
| EPI_ISL_2782022, EPI_ISL_2782023 | Lighthouse Lab in Alderley Park | Wellcome Sanger Institute for the COVID-19 Genomics UK (COG-UK) Consortium | Jacquelyn Wynn, Mairead Hyland, The Lighthouse Lab in Alderley Park and Alex Alderton, Roberto Amato, Jeffrey Barrett, Sonia Goncalves, Ewan Harrison, David K. Jackson, Ian Johnston, Dominic Kwiatkowski, Cordelia Langford, John Sillitoe on behalf of the Wellcome Sanger Institute COVID-19 Surveillance Team |
| EPI_ISL_2782024 | Lighthouse Lab in Milton Keynes | Wellcome Sanger Institute for the COVID-19 Genomics UK (COG-UK) Consortium | The Lighthouse Lab in Milton Keynes and Alex Alderton, Roberto Amato, Jeffrey Barrett, Sonia Goncalves, Ewan Harrison, David K. Jackson, Ian Johnston, Dominic Kwiatkowski, Cordelia Langford, John Sillitoe on behalf of the Wellcome Sanger Institute COVID-19 Surveillance Team |
| EPI_ISL_2782025, EPI_ISL_2782026 | Lighthouse Lab in Alderley Park | Wellcome Sanger Institute for the COVID-19 Genomics UK (COG-UK) Consortium | Jacquelyn Wynn, Mairead Hyland, The Lighthouse Lab in Alderley Park and Alex Alderton, Roberto Amato, Jeffrey Barrett, Sonia Goncalves, Ewan Harrison, David K. Jackson, Ian Johnston, Dominic Kwiatkowski, Cordelia Langford, John Sillitoe on behalf of the Wellcome Sanger Institute COVID-19 Surveillance Team |
| EPI_ISL_2782027, EPI_ISL_2782028 | Lighthouse Lab in Milton Keynes | Wellcome Sanger Institute for the COVID-19 Genomics UK (COG-UK) Consortium | The Lighthouse Lab in Milton Keynes and Alex Alderton, Roberto Amato, Jeffrey Barrett, Sonia Goncalves, Ewan Harrison, David K. Jackson, Ian Johnston, Dominic Kwiatkowski, Cordelia Langford, John Sillitoe on behalf of the Wellcome Sanger Institute COVID-19 Surveillance Team |
| EPI_ISL_2782029, EPI_ISL_2782030, EPI_ISL_2782031 | Lighthouse Lab in Alderley Park | Wellcome Sanger Institute for the COVID-19 Genomics UK (COG-UK) Consortium | Jacquelyn Wynn, Mairead Hyland, The Lighthouse Lab in Alderley Park and Alex Alderton, Roberto Amato, Jeffrey Barrett, Sonia Goncalves, Ewan Harrison, David K. Jackson, Ian Johnston, Dominic Kwiatkowski, Cordelia Langford, John Sillitoe on behalf of the Wellcome Sanger Institute COVID-19 Surveillance Team |
| EPI_ISL_2782032 | Lighthouse Lab in Glasgow | Wellcome Sanger Institute for the COVID-19 Genomics UK (COG-UK) Consortium | Harper VanSteenhouse, Yumi Kasai, David Gray, Carol Clugston, Anna Dominiczak and Alex Alderton, Roberto Amato, Jeffrey Barrett, Sonia Goncalves, Ewan Harrison, David K. Jackson, Ian Johnston, Dominic Kwiatkowski, Cordelia Langford, John Sillitoe on behalf of the Wellcome Sanger Institute COVID-19 Surveillance Team |
| EPI_ISL_2782033, EPI_ISL_2782034 | Lighthouse Lab in Alderley Park | Wellcome Sanger Institute for the COVID-19 Genomics UK (COG-UK) Consortium | Jacquelyn Wynn, Mairead Hyland, The Lighthouse Lab in Alderley Park and Alex Alderton, Roberto Amato, Jeffrey Barrett, Sonia Goncalves, Ewan Harrison, David K. Jackson, Ian Johnston, Dominic Kwiatkowski, Cordelia Langford, John Sillitoe on behalf of the Wellcome Sanger Institute COVID-19 Surveillance Team |
| EPI_ISL_2782035 | Lighthouse Lab in Milton Keynes | Wellcome Sanger Institute for the COVID-19 Genomics UK (COG-UK) Consortium | The Lighthouse Lab in Milton Keynes and Alex Alderton, Roberto Amato, Jeffrey Barrett, Sonia Goncalves, Ewan Harrison, David K. Jackson, Ian Johnston, Dominic Kwiatkowski, Cordelia Langford, John Sillitoe on behalf of the Wellcome Sanger Institute COVID-19 Surveillance Team |
| EPI_ISL_2782036 | Lighthouse Lab in Alderley Park | Wellcome Sanger Institute for the COVID-19 Genomics UK (COG-UK) Consortium | Jacquelyn Wynn, Mairead Hyland, The Lighthouse Lab in Alderley Park and Alex Alderton, Roberto Amato, Jeffrey Barrett, Sonia Goncalves, Ewan Harrison, David K. Jackson, Ian Johnston, Dominic Kwiatkowski, Cordelia Langford, John Sillitoe on behalf of the Wellcome Sanger Institute COVID-19 Surveillance Team |
| EPI_ISL_2782037 | Lighthouse Laboratory Plymouth | Wellcome Sanger Institute for the COVID-19 Genomics UK (COG-UK) Consortium | Lighthouse Laboratory Plymouth and Alex Alderton, Roberto Amato, Jeffrey Barrett, Sonia Goncalves, Ewan Harrison, David K. Jackson, Ian Johnston, Dominic Kwiatkowski, Cordelia Langford, John Sillitoe on behalf of the Wellcome Sanger Institute COVID-19 Surveillance Team |
| EPI_ISL_2782038 | Lighthouse Lab in Alderley Park | Wellcome Sanger Institute for the COVID-19 Genomics UK (COG-UK) Consortium | Jacquelyn Wynn, Mairead Hyland, The Lighthouse Lab in Alderley Park and Alex Alderton, Roberto Amato, Jeffrey Barrett, Sonia Goncalves, Ewan Harrison, David K. Jackson, Ian Johnston, Dominic Kwiatkowski, Cordelia Langford, John Sillitoe on behalf of the Wellcome Sanger Institute COVID-19 Surveillance Team |
| EPI_ISL_2782039, EPI_ISL_2782040, EPI_ISL_2782041 | Lighthouse Lab in Milton Keynes | Wellcome Sanger Institute for the COVID-19 Genomics UK (COG-UK) Consortium | The Lighthouse Lab in Milton Keynes and Alex Alderton, Roberto Amato, Jeffrey Barrett, Sonia Goncalves, Ewan Harrison, David K. Jackson, Ian Johnston, Dominic Kwiatkowski, Cordelia Langford, John Sillitoe on behalf of the Wellcome Sanger Institute COVID-19 Surveillance Team |
| EPI_ISL_2782042 | Lighthouse Lab in Alderley Park | Wellcome Sanger Institute for the COVID-19 Genomics UK (COG-UK) Consortium | Jacquelyn Wynn, Mairead Hyland, The Lighthouse Lab in Alderley Park and Alex Alderton, Roberto Amato, Jeffrey Barrett, Sonia Goncalves, Ewan Harrison, David K. Jackson, Ian Johnston, Dominic Kwiatkowski, Cordelia Langford, John Sillitoe on behalf of the Wellcome Sanger Institute COVID-19 Surveillance Team |
| EPI_ISL_2782043 | Lighthouse Lab in Milton Keynes | Wellcome Sanger Institute for the COVID-19 Genomics UK (COG-UK) Consortium | The Lighthouse Lab in Milton Keynes and Alex Alderton, Roberto Amato, Jeffrey Barrett, Sonia Goncalves, Ewan Harrison, David K. Jackson, Ian Johnston, Dominic Kwiatkowski, Cordelia Langford, John Sillitoe on behalf of the Wellcome Sanger Institute COVID-19 Surveillance Team |
| EPI_ISL_2782044 | Lighthouse Lab in Alderley Park | Wellcome Sanger Institute for the COVID-19 Genomics UK (COG-UK) Consortium | Jacquelyn Wynn, Mairead Hyland, The Lighthouse Lab in Alderley Park and Alex Alderton, Roberto Amato, Jeffrey Barrett, Sonia Goncalves, Ewan Harrison, David K. Jackson, Ian Johnston, Dominic Kwiatkowski, Cordelia Langford, John Sillitoe on behalf of the Wellcome Sanger Institute COVID-19 Surveillance Team |
| EPI_ISL_2782045 | Lighthouse Lab in Milton Keynes | Wellcome Sanger Institute for the COVID-19 Genomics UK (COG-UK) Consortium | The Lighthouse Lab in Milton Keynes and Alex Alderton, Roberto Amato, Jeffrey Barrett, Sonia Goncalves, Ewan Harrison, David K. Jackson, Ian Johnston, Dominic Kwiatkowski, Cordelia Langford, John Sillitoe on behalf of the Wellcome Sanger Institute COVID-19 Surveillance Team |
| EPI_ISL_2782046, EPI_ISL_2782047, EPI_ISL_2782048 | Lighthouse Laboratory Plymouth | Wellcome Sanger Institute for the COVID-19 Genomics UK (COG-UK) Consortium | Lighthouse Laboratory Plymouth and Alex Alderton, Roberto Amato, Jeffrey Barrett, Sonia Goncalves, Ewan Harrison, David K. Jackson, Ian Johnston, Dominic Kwiatkowski, Cordelia Langford, John Sillitoe on behalf of the Wellcome Sanger Institute COVID-19 Surveillance Team |
| EPI_ISL_2782049 | Lighthouse Lab in Glasgow | Wellcome Sanger Institute for the COVID-19 Genomics UK (COG-UK) Consortium | Harper VanSteenhouse, Yumi Kasai, David Gray, Carol Clugston, Anna Dominiczak and Alex Alderton, Roberto Amato, Jeffrey Barrett, Sonia Goncalves, Ewan Harrison, David K. Jackson, Ian Johnston, Dominic Kwiatkowski, Cordelia Langford, John Sillitoe on behalf of the Wellcome Sanger Institute COVID-19 Surveillance Team |
| EPI_ISL_2782050 | Lighthouse Laboratory Plymouth | Wellcome Sanger Institute for the COVID-19 Genomics UK (COG-UK) Consortium | Lighthouse Laboratory Plymouth and Alex Alderton, Roberto Amato, Jeffrey Barrett, Sonia Goncalves, Ewan Harrison, David K. Jackson, Ian Johnston, Dominic Kwiatkowski, Cordelia Langford, John Sillitoe on behalf of the Wellcome Sanger Institute COVID-19 Surveillance Team |
| EPI_ISL_2782051 | Lighthouse Lab in Milton Keynes | Wellcome Sanger Institute for the COVID-19 Genomics UK (COG-UK) Consortium | The Lighthouse Lab in Milton Keynes and Alex Alderton, Roberto Amato, Jeffrey Barrett, Sonia Goncalves, Ewan Harrison, David K. Jackson, Ian Johnston, Dominic Kwiatkowski, Cordelia Langford, John Sillitoe on behalf of the Wellcome Sanger Institute COVID-19 Surveillance Team |
| EPI_ISL_2782052 | Lighthouse Laboratory Plymouth | Wellcome Sanger Institute for the COVID-19 Genomics UK (COG-UK) | Lighthouse Laboratory Plymouth and Alex Alderton, Roberto Amato, Jeffrey Barrett, Sonia Goncalves, Ewan Harrison, David K. Jackson, Ian Johnston, Dominic Kwiatkowski, Cordelia Langford, John Sillitoe on behalf of the Wellcome Sanger |

| | | | |
|---|---|---|---|
| | | Consortium | Institute COVID-19 Surveillance Team |
| EPI_ISL_2782053 | Lighthouse Lab in Milton Keynes | Wellcome Sanger Institute for the COVID-19 Genomics UK (COG-UK) Consortium | The Lighthouse Lab in Milton Keynes and Alex Alderton, Roberto Amato, Jeffrey Barrett, Sonia Goncalves, Ewan Harrison, David K. Jackson, Ian Johnston, Dominic Kwiatkowski, Cordelia Langford, John Sillitoe on behalf of the Wellcome Sanger Institute COVID-19 Surveillance Team |
| EPI_ISL_2782054, EPI_ISL_2782055 | Lighthouse Lab in Alderley Park | Wellcome Sanger Institute for the COVID-19 Genomics UK (COG-UK) Consortium | Jacquelyn Wynn, Mairead Hyland, The Lighthouse Lab in Alderley Park and Alex Alderton, Roberto Amato, Jeffrey Barrett, Sonia Goncalves, Ewan Harrison, David K. Jackson, Ian Johnston, Dominic Kwiatkowski, Cordelia Langford, John Sillitoe on behalf of the Wellcome Sanger Institute COVID-19 Surveillance Team |
| EPI_ISL_2782056, EPI_ISL_2782057 | Lighthouse Laboratory Plymouth | Wellcome Sanger Institute for the COVID-19 Genomics UK (COG-UK) Consortium | Lighthouse Laboratory Plymouth and Alex Alderton, Roberto Amato, Jeffrey Barrett, Sonia Goncalves, Ewan Harrison, David K. Jackson, Ian Johnston, Dominic Kwiatkowski, Cordelia Langford, John Sillitoe on behalf of the Wellcome Sanger Institute COVID-19 Surveillance Team |
| EPI_ISL_2782058 | Lighthouse Lab in Alderley Park | Wellcome Sanger Institute for the COVID-19 Genomics UK (COG-UK) Consortium | Jacquelyn Wynn, Mairead Hyland, The Lighthouse Lab in Alderley Park and Alex Alderton, Roberto Amato, Jeffrey Barrett, Sonia Goncalves, Ewan Harrison, David K. Jackson, Ian Johnston, Dominic Kwiatkowski, Cordelia Langford, John Sillitoe on behalf of the Wellcome Sanger Institute COVID-19 Surveillance Team |
| EPI_ISL_2782059 | Lighthouse Lab in Milton Keynes | Wellcome Sanger Institute for the COVID-19 Genomics UK (COG-UK) Consortium | The Lighthouse Lab in Milton Keynes and Alex Alderton, Roberto Amato, Jeffrey Barrett, Sonia Goncalves, Ewan Harrison, David K. Jackson, Ian Johnston, Dominic Kwiatkowski, Cordelia Langford, John Sillitoe on behalf of the Wellcome Sanger Institute COVID-19 Surveillance Team |
| EPI_ISL_2782060 | Lighthouse Laboratory Plymouth | Wellcome Sanger Institute for the COVID-19 Genomics UK (COG-UK) Consortium | Lighthouse Laboratory Plymouth and Alex Alderton, Roberto Amato, Jeffrey Barrett, Sonia Goncalves, Ewan Harrison, David K. Jackson, Ian Johnston, Dominic Kwiatkowski, Cordelia Langford, John Sillitoe on behalf of the Wellcome Sanger Institute COVID-19 Surveillance Team |
| EPI_ISL_2782061, EPI_ISL_2782062 | Lighthouse Lab in Milton Keynes | Wellcome Sanger Institute for the COVID-19 Genomics UK (COG-UK) Consortium | The Lighthouse Lab in Milton Keynes and Alex Alderton, Roberto Amato, Jeffrey Barrett, Sonia Goncalves, Ewan Harrison, David K. Jackson, Ian Johnston, Dominic Kwiatkowski, Cordelia Langford, John Sillitoe on behalf of the Wellcome Sanger Institute COVID-19 Surveillance Team |
| EPI_ISL_2782063 | Lighthouse Lab in Alderley Park | Wellcome Sanger Institute for the COVID-19 Genomics UK (COG-UK) Consortium | Jacquelyn Wynn, Mairead Hyland, The Lighthouse Lab in Alderley Park and Alex Alderton, Roberto Amato, Jeffrey Barrett, Sonia Goncalves, Ewan Harrison, David K. Jackson, Ian Johnston, Dominic Kwiatkowski, Cordelia Langford, John Sillitoe on behalf of the Wellcome Sanger Institute COVID-19 Surveillance Team |
| EPI_ISL_2782064 | Lighthouse Laboratory Plymouth | Wellcome Sanger Institute for the COVID-19 Genomics UK (COG-UK) Consortium | Lighthouse Laboratory Plymouth and Alex Alderton, Roberto Amato, Jeffrey Barrett, Sonia Goncalves, Ewan Harrison, David K. Jackson, Ian Johnston, Dominic Kwiatkowski, Cordelia Langford, John Sillitoe on behalf of the Wellcome Sanger Institute COVID-19 Surveillance Team |
| EPI_ISL_2782065 | Lighthouse Lab in Milton Keynes | Wellcome Sanger Institute for the COVID-19 Genomics UK (COG-UK) Consortium | The Lighthouse Lab in Milton Keynes and Alex Alderton, Roberto Amato, Jeffrey Barrett, Sonia Goncalves, Ewan Harrison, David K. Jackson, Ian Johnston, Dominic Kwiatkowski, Cordelia Langford, John Sillitoe on behalf of the Wellcome Sanger Institute COVID-19 Surveillance Team |
| EPI_ISL_2782066 | Lighthouse Lab in Alderley Park | Wellcome Sanger Institute for the COVID-19 Genomics UK (COG-UK) Consortium | Jacquelyn Wynn, Mairead Hyland, The Lighthouse Lab in Alderley Park and Alex Alderton, Roberto Amato, Jeffrey Barrett, Sonia Goncalves, Ewan Harrison, David K. Jackson, Ian Johnston, Dominic Kwiatkowski, Cordelia Langford, John Sillitoe on behalf of the Wellcome Sanger Institute COVID-19 Surveillance Team |
| EPI_ISL_2782067 | Lighthouse Lab in Milton Keynes | Wellcome Sanger Institute for the COVID-19 Genomics UK (COG-UK) Consortium | The Lighthouse Lab in Milton Keynes and Alex Alderton, Roberto Amato, Jeffrey Barrett, Sonia Goncalves, Ewan Harrison, David K. Jackson, Ian Johnston, Dominic Kwiatkowski, Cordelia Langford, John Sillitoe on behalf of the Wellcome Sanger Institute COVID-19 Surveillance Team |
| EPI_ISL_2782068 | Lighthouse Lab in Alderley Park | Wellcome Sanger Institute for the COVID-19 Genomics UK (COG-UK) Consortium | Jacquelyn Wynn, Mairead Hyland, The Lighthouse Lab in Alderley Park and Alex Alderton, Roberto Amato, Jeffrey Barrett, Sonia Goncalves, Ewan Harrison, David K. Jackson, Ian Johnston, Dominic Kwiatkowski, Cordelia Langford, John Sillitoe on behalf of the Wellcome Sanger Institute COVID-19 Surveillance Team |
| EPI_ISL_2782069 | Lighthouse Lab in Milton Keynes | Wellcome Sanger Institute for the COVID-19 Genomics UK (COG-UK) Consortium | The Lighthouse Lab in Milton Keynes and Alex Alderton, Roberto Amato, Jeffrey Barrett, Sonia Goncalves, Ewan Harrison, David K. Jackson, Ian Johnston, Dominic Kwiatkowski, Cordelia Langford, John Sillitoe on behalf of the Wellcome Sanger Institute COVID-19 Surveillance Team |
| EPI_ISL_2782070, EPI_ISL_2782071 | Lighthouse Lab in Alderley Park | Wellcome Sanger Institute for the COVID-19 Genomics UK (COG-UK) Consortium | Jacquelyn Wynn, Mairead Hyland, The Lighthouse Lab in Alderley Park and Alex Alderton, Roberto Amato, Jeffrey Barrett, Sonia Goncalves, Ewan Harrison, David K. Jackson, Ian Johnston, Dominic Kwiatkowski, Cordelia Langford, John Sillitoe on behalf of the Wellcome Sanger Institute COVID-19 Surveillance Team |
| EPI_ISL_2782072 | Lighthouse Lab in Milton Keynes | Wellcome Sanger Institute for the COVID-19 Genomics UK (COG-UK) Consortium | The Lighthouse Lab in Milton Keynes and Alex Alderton, Roberto Amato, Jeffrey Barrett, Sonia Goncalves, Ewan Harrison, David K. Jackson, Ian Johnston, Dominic Kwiatkowski, Cordelia Langford, John Sillitoe on behalf of the Wellcome Sanger Institute COVID-19 Surveillance Team |
| EPI_ISL_2782073, EPI_ISL_2782074, EPI_ISL_2782075 | Lighthouse Laboratory Plymouth | Wellcome Sanger Institute for the COVID-19 Genomics UK (COG-UK) Consortium | Lighthouse Laboratory Plymouth and Alex Alderton, Roberto Amato, Jeffrey Barrett, Sonia Goncalves, Ewan Harrison, David K. Jackson, Ian Johnston, Dominic Kwiatkowski, Cordelia Langford, John Sillitoe on behalf of the Wellcome Sanger Institute COVID-19 Surveillance Team |
| EPI_ISL_2782076, EPI_ISL_2782077 | Lighthouse Lab in Alderley Park | Wellcome Sanger Institute for the COVID-19 Genomics UK (COG-UK) Consortium | Jacquelyn Wynn, Mairead Hyland, The Lighthouse Lab in Alderley Park and Alex Alderton, Roberto Amato, Jeffrey Barrett, Sonia Goncalves, Ewan Harrison, David K. Jackson, Ian Johnston, Dominic Kwiatkowski, Cordelia Langford, John Sillitoe on behalf of the Wellcome Sanger Institute COVID-19 Surveillance Team |
| EPI_ISL_2782078 | Lighthouse Laboratory Plymouth | Wellcome Sanger Institute for the COVID-19 Genomics UK (COG-UK) Consortium | Lighthouse Laboratory Plymouth and Alex Alderton, Roberto Amato, Jeffrey Barrett, Sonia Goncalves, Ewan Harrison, David K. Jackson, Ian Johnston, Dominic Kwiatkowski, Cordelia Langford, John Sillitoe on behalf of the Wellcome Sanger Institute COVID-19 Surveillance Team |
| EPI_ISL_2782079, EPI_ISL_2782080 | Lighthouse Lab in Alderley Park | Wellcome Sanger Institute for the COVID-19 Genomics UK (COG-UK) Consortium | Jacquelyn Wynn, Mairead Hyland, The Lighthouse Lab in Alderley Park and Alex Alderton, Roberto Amato, Jeffrey Barrett, Sonia Goncalves, Ewan Harrison, David K. Jackson, Ian Johnston, Dominic Kwiatkowski, Cordelia Langford, John Sillitoe on behalf of the Wellcome Sanger Institute COVID-19 Surveillance Team |
| EPI_ISL_2782081, EPI_ISL_2782082 | Lighthouse Lab in Milton Keynes | Wellcome Sanger Institute for the COVID-19 Genomics UK (COG-UK) Consortium | The Lighthouse Lab in Milton Keynes and Alex Alderton, Roberto Amato, Jeffrey Barrett, Sonia Goncalves, Ewan Harrison, David K. Jackson, Ian Johnston, Dominic Kwiatkowski, Cordelia Langford, John Sillitoe on behalf of the Wellcome Sanger Institute COVID-19 Surveillance Team |
| EPI_ISL_2782083, EPI_ISL_2782084, EPI_ISL_2782085 | Lighthouse Laboratory Plymouth | Wellcome Sanger Institute for the COVID-19 Genomics UK (COG-UK) Consortium | Lighthouse Laboratory Plymouth and Alex Alderton, Roberto Amato, Jeffrey Barrett, Sonia Goncalves, Ewan Harrison, David K. Jackson, Ian Johnston, Dominic Kwiatkowski, Cordelia Langford, John Sillitoe on behalf of the Wellcome Sanger Institute COVID-19 Surveillance Team |
| EPI_ISL_2782086 | Lighthouse Lab in Alderley Park | Wellcome Sanger Institute for the COVID-19 Genomics UK (COG-UK) Consortium | Jacquelyn Wynn, Mairead Hyland, The Lighthouse Lab in Alderley Park and Alex Alderton, Roberto Amato, Jeffrey Barrett, Sonia Goncalves, Ewan Harrison, David K. Jackson, Ian Johnston, Dominic Kwiatkowski, Cordelia Langford, John Sillitoe on behalf of the Wellcome Sanger Institute COVID-19 Surveillance Team |
| EPI_ISL_2782087 | Lighthouse Lab in Glasgow | Wellcome Sanger Institute for the COVID-19 Genomics UK (COG-UK) Consortium | Harper VanSteenhouse, Yumi Kasai, David Gray, Carol Clugston, Anna Dominiczak and Alex Alderton, Roberto Amato, Jeffrey Barrett, Sonia Goncalves, Ewan Harrison, David K. Jackson, Ian Johnston, Dominic Kwiatkowski, Cordelia Langford, John Sillitoe on behalf of the Wellcome Sanger Institute COVID-19 Surveillance Team |
| EPI_ISL_2782088, EPI_ISL_2782089 | Lighthouse Lab in Milton Keynes | Wellcome Sanger Institute for the COVID-19 Genomics UK (COG-UK) Consortium | The Lighthouse Lab in Milton Keynes and Alex Alderton, Roberto Amato, Jeffrey Barrett, Sonia Goncalves, Ewan Harrison, David K. Jackson, Ian Johnston, Dominic Kwiatkowski, Cordelia Langford, John Sillitoe on behalf of the Wellcome Sanger Institute COVID-19 Surveillance Team |
| EPI_ISL_2782090 | Lighthouse Lab in Alderley Park | Wellcome Sanger Institute for the COVID-19 Genomics UK (COG-UK) Consortium | Jacquelyn Wynn, Mairead Hyland, The Lighthouse Lab in Alderley Park and Alex Alderton, Roberto Amato, Jeffrey Barrett, Sonia Goncalves, Ewan Harrison, David K. Jackson, Ian Johnston, Dominic Kwiatkowski, Cordelia Langford, John Sillitoe on behalf of the Wellcome Sanger Institute COVID-19 Surveillance Team |
| EPI_ISL_2782091 | Lighthouse Lab in Milton Keynes | Wellcome Sanger Institute for the COVID-19 Genomics UK (COG-UK) Consortium | The Lighthouse Lab in Milton Keynes and Alex Alderton, Roberto Amato, Jeffrey Barrett, Sonia Goncalves, Ewan Harrison, David K. Jackson, Ian Johnston, Dominic Kwiatkowski, Cordelia Langford, John Sillitoe on behalf of the Wellcome Sanger Institute COVID-19 Surveillance Team |
| EPI_ISL_2782092 | Lighthouse Lab in Alderley Park | Wellcome Sanger Institute for the COVID-19 Genomics UK (COG-UK) Consortium | Jacquelyn Wynn, Mairead Hyland, The Lighthouse Lab in Alderley Park and Alex Alderton, Roberto Amato, Jeffrey Barrett, Sonia Goncalves, Ewan Harrison, David K. Jackson, Ian Johnston, Dominic Kwiatkowski, Cordelia Langford, John Sillitoe on behalf of the Wellcome Sanger Institute COVID-19 Surveillance Team |
| EPI_ISL_2782093 | Lighthouse Lab in Milton Keynes | Wellcome Sanger Institute for the COVID-19 Genomics UK (COG-UK) Consortium | The Lighthouse Lab in Milton Keynes and Alex Alderton, Roberto Amato, Jeffrey Barrett, Sonia Goncalves, Ewan Harrison, David K. Jackson, Ian Johnston, Dominic Kwiatkowski, Cordelia Langford, John Sillitoe on behalf of the Wellcome Sanger Institute COVID-19 Surveillance Team |
| EPI_ISL_2782094 | Lighthouse Lab in Glasgow | Wellcome Sanger Institute for the COVID-19 Genomics UK (COG-UK) Consortium | Harper VanSteenhouse, Yumi Kasai, David Gray, Carol Clugston, Anna Dominiczak and Alex Alderton, Roberto Amato, Jeffrey Barrett, Sonia Goncalves, Ewan Harrison, David K. Jackson, Ian Johnston, Dominic Kwiatkowski, Cordelia Langford, John Sillitoe on behalf of the Wellcome Sanger Institute COVID-19 Surveillance Team |
| EPI_ISL_2782095, EPI_ISL_2782096 | Lighthouse Lab in Alderley Park | Wellcome Sanger Institute for the COVID-19 Genomics UK (COG-UK) Consortium | Jacquelyn Wynn, Mairead Hyland, The Lighthouse Lab in Alderley Park and Alex Alderton, Roberto Amato, Jeffrey Barrett, Sonia Goncalves, Ewan Harrison, David K. Jackson, Ian Johnston, Dominic Kwiatkowski, Cordelia Langford, John Sillitoe on behalf of the Wellcome Sanger Institute COVID-19 Surveillance Team |
| EPI_ISL_2782097 | Lighthouse Lab in Glasgow | Wellcome Sanger Institute for the COVID-19 Genomics UK (COG-UK) Consortium | Harper VanSteenhouse, Yumi Kasai, David Gray, Carol Clugston, Anna Dominiczak and Alex Alderton, Roberto Amato, Jeffrey Barrett, Sonia Goncalves, Ewan Harrison, David K. Jackson, Ian Johnston, Dominic Kwiatkowski, Cordelia Langford, John Sillitoe on behalf of the Wellcome Sanger Institute COVID-19 Surveillance Team |
| EPI_ISL_2782098, EPI_ISL_2782099 | Lighthouse Laboratory Plymouth | Wellcome Sanger Institute for the COVID-19 Genomics UK (COG-UK) Consortium | Lighthouse Laboratory Plymouth and Alex Alderton, Roberto Amato, Jeffrey Barrett, Sonia Goncalves, Ewan Harrison, David K. Jackson, Ian Johnston, Dominic Kwiatkowski, Cordelia Langford, John Sillitoe on behalf of the Wellcome Sanger Institute COVID-19 Surveillance Team |
| EPI_ISL_2782100 | Lighthouse Lab in Alderley Park | Wellcome Sanger Institute for the COVID-19 Genomics UK (COG-UK) Consortium | Jacquelyn Wynn, Mairead Hyland, The Lighthouse Lab in Alderley Park and Alex Alderton, Roberto Amato, Jeffrey Barrett, Sonia Goncalves, Ewan Harrison, David K. Jackson, Ian Johnston, Dominic Kwiatkowski, Cordelia Langford, John Sillitoe on behalf of the Wellcome Sanger Institute COVID-19 Surveillance Team |
| EPI_ISL_2782101 | Lighthouse Lab in Glasgow | Wellcome Sanger Institute for the COVID-19 Genomics UK (COG-UK) Consortium | Harper VanSteenhouse, Yumi Kasai, David Gray, Carol Clugston, Anna Dominiczak and Alex Alderton, Roberto Amato, Jeffrey Barrett, Sonia Goncalves, Ewan Harrison, David K. Jackson, Ian Johnston, Dominic Kwiatkowski, Cordelia Langford, John Sillitoe on behalf of the Wellcome Sanger Institute COVID-19 Surveillance Team |
| EPI_ISL_2782102 | Lighthouse Laboratory Plymouth | Wellcome Sanger Institute for the COVID-19 Genomics UK (COG-UK) Consortium | Lighthouse Laboratory Plymouth and Alex Alderton, Roberto Amato, Jeffrey Barrett, Sonia Goncalves, Ewan Harrison, David K. Jackson, Ian Johnston, Dominic Kwiatkowski, Cordelia Langford, John Sillitoe on behalf of the Wellcome Sanger Institute COVID-19 Surveillance Team |
| EPI_ISL_2782103 | Lighthouse Lab in Milton Keynes | Wellcome Sanger Institute for the COVID-19 Genomics UK (COG-UK) Consortium | The Lighthouse Lab in Milton Keynes and Alex Alderton, Roberto Amato, Jeffrey Barrett, Sonia Goncalves, Ewan Harrison, David K. Jackson, Ian Johnston, Dominic Kwiatkowski, Cordelia Langford, John Sillitoe on behalf of the Wellcome Sanger Institute COVID-19 Surveillance Team |
| EPI_ISL_2782104, EPI_ISL_2782105 | Lighthouse Lab in Alderley Park | Wellcome Sanger Institute for the COVID-19 Genomics UK (COG-UK) Consortium | Jacquelyn Wynn, Mairead Hyland, The Lighthouse Lab in Alderley Park and Alex Alderton, Roberto Amato, Jeffrey Barrett, Sonia Goncalves, Ewan Harrison, David K. Jackson, Ian Johnston, Dominic Kwiatkowski, Cordelia Langford, John Sillitoe on behalf of the Wellcome Sanger Institute COVID-19 Surveillance Team |
| EPI_ISL_2782106, EPI_ISL_2782107 | Lighthouse Lab in Milton Keynes | Wellcome Sanger Institute for the COVID-19 Genomics UK (COG-UK) Consortium | The Lighthouse Lab in Milton Keynes and Alex Alderton, Roberto Amato, Jeffrey Barrett, Sonia Goncalves, Ewan Harrison, David K. Jackson, Ian Johnston, Dominic Kwiatkowski, Cordelia Langford, John Sillitoe on behalf of the Wellcome Sanger Institute COVID-19 Surveillance Team |
| EPI_ISL_2782108, EPI_ISL_2782109, EPI_ISL_2782110 | Lighthouse Laboratory Plymouth | Wellcome Sanger Institute for the COVID-19 Genomics UK (COG-UK) Consortium | Lighthouse Laboratory Plymouth and Alex Alderton, Roberto Amato, Jeffrey Barrett, Sonia Goncalves, Ewan Harrison, David K. Jackson, Ian Johnston, Dominic Kwiatkowski, Cordelia Langford, John Sillitoe on behalf of the Wellcome Sanger Institute COVID-19 Surveillance Team |
| EPI_ISL_2782111 | Lighthouse Lab in Milton Keynes | Wellcome Sanger Institute for the COVID-19 Genomics UK (COG-UK) Consortium | The Lighthouse Lab in Milton Keynes and Alex Alderton, Roberto Amato, Jeffrey Barrett, Sonia Goncalves, Ewan Harrison, David K. Jackson, Ian Johnston, Dominic Kwiatkowski, Cordelia Langford, John Sillitoe on behalf of the Wellcome Sanger Institute COVID-19 Surveillance Team |
| EPI_ISL_2782112 | Lighthouse Laboratory Plymouth | Wellcome Sanger Institute for the COVID-19 Genomics UK (COG-UK) Consortium | Lighthouse Laboratory Plymouth and Alex Alderton, Roberto Amato, Jeffrey Barrett, Sonia Goncalves, Ewan Harrison, David K. Jackson, Ian Johnston, Dominic Kwiatkowski, Cordelia Langford, John Sillitoe on behalf of the Wellcome Sanger Institute COVID-19 Surveillance Team |
| EPI_ISL_2782113 | Lighthouse Lab in Alderley Park | Wellcome Sanger Institute for the COVID-19 Genomics UK (COG-UK) Consortium | Jacquelyn Wynn, Mairead Hyland, The Lighthouse Lab in Alderley Park and Alex Alderton, Roberto Amato, Jeffrey Barrett, Sonia Goncalves, Ewan Harrison, David K. Jackson, Ian Johnston, Dominic Kwiatkowski, Cordelia Langford, John Sillitoe on behalf of the Wellcome Sanger Institute COVID-19 Surveillance Team |

| | | | |
|---|---|---|---|
| EPI_ISL_2782114, EPI_ISL_2782115, EPI_ISL_2782116 | Lighthouse Lab in Milton Keynes | Wellcome Sanger Institute for the COVID-19 Genomics UK (COG-UK) Consortium | The Lighthouse Lab in Milton Keynes and Alex Alderton, Roberto Amato, Jeffrey Barrett, Sonia Goncalves, Ewan Harrison, David K. Jackson, Ian Johnston, Dominic Kwiatkowski, Cordelia Langford, John Sillitoe on behalf of the Wellcome Sanger Institute COVID-19 Surveillance Team |
| EPI_ISL_2782117 | Lighthouse Lab in Alderley Park | Wellcome Sanger Institute for the COVID-19 Genomics UK (COG-UK) Consortium | Jacquelyn Wynn, Mairead Hyland, The Lighthouse Lab in Alderley Park and Alex Alderton, Roberto Amato, Jeffrey Barrett, Sonia Goncalves, Ewan Harrison, David K. Jackson, Ian Johnston, Dominic Kwiatkowski, Cordelia Langford, John Sillitoe on behalf of the Wellcome Sanger Institute COVID-19 Surveillance Team |
| EPI_ISL_2782118 | Lighthouse Lab in Glasgow | Wellcome Sanger Institute for the COVID-19 Genomics UK (COG-UK) Consortium | Harper VanSteenhouse, Yumi Kasai, David Gray, Carol Clugston, Anna Dominiczak and Alex Alderton, Roberto Amato, Jeffrey Barrett, Sonia Goncalves, Ewan Harrison, David K. Jackson, Ian Johnston, Dominic Kwiatkowski, Cordelia Langford, John Sillitoe on behalf of the Wellcome Sanger Institute COVID-19 Surveillance Team |
| EPI_ISL_2782119, EPI_ISL_2782120 | Lighthouse Lab in Milton Keynes | Wellcome Sanger Institute for the COVID-19 Genomics UK (COG-UK) Consortium | The Lighthouse Lab in Milton Keynes and Alex Alderton, Roberto Amato, Jeffrey Barrett, Sonia Goncalves, Ewan Harrison, David K. Jackson, Ian Johnston, Dominic Kwiatkowski, Cordelia Langford, John Sillitoe on behalf of the Wellcome Sanger Institute COVID-19 Surveillance Team |
| EPI_ISL_2782121 | Lighthouse Laboratory Plymouth | Wellcome Sanger Institute for the COVID-19 Genomics UK (COG-UK) Consortium | Lighthouse Laboratory Plymouth and Alex Alderton, Roberto Amato, Jeffrey Barrett, Sonia Goncalves, Ewan Harrison, David K. Jackson, Ian Johnston, Dominic Kwiatkowski, Cordelia Langford, John Sillitoe on behalf of the Wellcome Sanger Institute COVID-19 Surveillance Team |
| EPI_ISL_2782122 | Lighthouse Lab in Milton Keynes | Wellcome Sanger Institute for the COVID-19 Genomics UK (COG-UK) Consortium | The Lighthouse Lab in Milton Keynes and Alex Alderton, Roberto Amato, Jeffrey Barrett, Sonia Goncalves, Ewan Harrison, David K. Jackson, Ian Johnston, Dominic Kwiatkowski, Cordelia Langford, John Sillitoe on behalf of the Wellcome Sanger Institute COVID-19 Surveillance Team |
| EPI_ISL_2782123 | Lighthouse Laboratory Plymouth | Wellcome Sanger Institute for the COVID-19 Genomics UK (COG-UK) Consortium | Lighthouse Laboratory Plymouth and Alex Alderton, Roberto Amato, Jeffrey Barrett, Sonia Goncalves, Ewan Harrison, David K. Jackson, Ian Johnston, Dominic Kwiatkowski, Cordelia Langford, John Sillitoe on behalf of the Wellcome Sanger Institute COVID-19 Surveillance Team |
| EPI_ISL_2782124 | Lighthouse Lab in Alderley Park | Wellcome Sanger Institute for the COVID-19 Genomics UK (COG-UK) Consortium | Jacquelyn Wynn, Mairead Hyland, The Lighthouse Lab in Alderley Park and Alex Alderton, Roberto Amato, Jeffrey Barrett, Sonia Goncalves, Ewan Harrison, David K. Jackson, Ian Johnston, Dominic Kwiatkowski, Cordelia Langford, John Sillitoe on behalf of the Wellcome Sanger Institute COVID-19 Surveillance Team |
| EPI_ISL_2782125 | Lighthouse Laboratory Plymouth | Wellcome Sanger Institute for the COVID-19 Genomics UK (COG-UK) Consortium | Lighthouse Laboratory Plymouth and Alex Alderton, Roberto Amato, Jeffrey Barrett, Sonia Goncalves, Ewan Harrison, David K. Jackson, Ian Johnston, Dominic Kwiatkowski, Cordelia Langford, John Sillitoe on behalf of the Wellcome Sanger Institute COVID-19 Surveillance Team |
| EPI_ISL_2782126, EPI_ISL_2782127 | Lighthouse Lab in Alderley Park | Wellcome Sanger Institute for the COVID-19 Genomics UK (COG-UK) Consortium | Jacquelyn Wynn, Mairead Hyland, The Lighthouse Lab in Alderley Park and Alex Alderton, Roberto Amato, Jeffrey Barrett, Sonia Goncalves, Ewan Harrison, David K. Jackson, Ian Johnston, Dominic Kwiatkowski, Cordelia Langford, John Sillitoe on behalf of the Wellcome Sanger Institute COVID-19 Surveillance Team |
| EPI_ISL_2782128 | Lighthouse Lab in Milton Keynes | Wellcome Sanger Institute for the COVID-19 Genomics UK (COG-UK) Consortium | The Lighthouse Lab in Milton Keynes and Alex Alderton, Roberto Amato, Jeffrey Barrett, Sonia Goncalves, Ewan Harrison, David K. Jackson, Ian Johnston, Dominic Kwiatkowski, Cordelia Langford, John Sillitoe on behalf of the Wellcome Sanger Institute COVID-19 Surveillance Team |
| EPI_ISL_2782129 | Lighthouse Laboratory Plymouth | Wellcome Sanger Institute for the COVID-19 Genomics UK (COG-UK) Consortium | Lighthouse Laboratory Plymouth and Alex Alderton, Roberto Amato, Jeffrey Barrett, Sonia Goncalves, Ewan Harrison, David K. Jackson, Ian Johnston, Dominic Kwiatkowski, Cordelia Langford, John Sillitoe on behalf of the Wellcome Sanger Institute COVID-19 Surveillance Team |
| EPI_ISL_2782130 | Lighthouse Lab in Milton Keynes | Wellcome Sanger Institute for the COVID-19 Genomics UK (COG-UK) Consortium | The Lighthouse Lab in Milton Keynes and Alex Alderton, Roberto Amato, Jeffrey Barrett, Sonia Goncalves, Ewan Harrison, David K. Jackson, Ian Johnston, Dominic Kwiatkowski, Cordelia Langford, John Sillitoe on behalf of the Wellcome Sanger Institute COVID-19 Surveillance Team |
| EPI_ISL_2782131 | Lighthouse Lab in Glasgow | Wellcome Sanger Institute for the COVID-19 Genomics UK (COG-UK) Consortium | Harper VanSteenhouse, Yumi Kasai, David Gray, Carol Clugston, Anna Dominiczak and Alex Alderton, Roberto Amato, Jeffrey Barrett, Sonia Goncalves, Ewan Harrison, David K. Jackson, Ian Johnston, Dominic Kwiatkowski, Cordelia Langford, John Sillitoe on behalf of the Wellcome Sanger Institute COVID-19 Surveillance Team |
| EPI_ISL_2782132, EPI_ISL_2782133, EPI_ISL_2782134 | Lighthouse Laboratory Plymouth | Wellcome Sanger Institute for the COVID-19 Genomics UK (COG-UK) Consortium | Lighthouse Laboratory Plymouth and Alex Alderton, Roberto Amato, Jeffrey Barrett, Sonia Goncalves, Ewan Harrison, David K. Jackson, Ian Johnston, Dominic Kwiatkowski, Cordelia Langford, John Sillitoe on behalf of the Wellcome Sanger Institute COVID-19 Surveillance Team |
| EPI_ISL_2782135 | Lighthouse Lab in Milton Keynes | Wellcome Sanger Institute for the COVID-19 Genomics UK (COG-UK) Consortium | The Lighthouse Lab in Milton Keynes and Alex Alderton, Roberto Amato, Jeffrey Barrett, Sonia Goncalves, Ewan Harrison, David K. Jackson, Ian Johnston, Dominic Kwiatkowski, Cordelia Langford, John Sillitoe on behalf of the Wellcome Sanger Institute COVID-19 Surveillance Team |
| EPI_ISL_2782136 | Lighthouse Lab in Alderley Park | Wellcome Sanger Institute for the COVID-19 Genomics UK (COG-UK) Consortium | Jacquelyn Wynn, Mairead Hyland, The Lighthouse Lab in Alderley Park and Alex Alderton, Roberto Amato, Jeffrey Barrett, Sonia Goncalves, Ewan Harrison, David K. Jackson, Ian Johnston, Dominic Kwiatkowski, Cordelia Langford, John Sillitoe on behalf of the Wellcome Sanger Institute COVID-19 Surveillance Team |
| EPI_ISL_2782137 | Lighthouse Lab in Milton Keynes | Wellcome Sanger Institute for the COVID-19 Genomics UK (COG-UK) Consortium | The Lighthouse Lab in Milton Keynes and Alex Alderton, Roberto Amato, Jeffrey Barrett, Sonia Goncalves, Ewan Harrison, David K. Jackson, Ian Johnston, Dominic Kwiatkowski, Cordelia Langford, John Sillitoe on behalf of the Wellcome Sanger Institute COVID-19 Surveillance Team |
| EPI_ISL_2782138 | Lighthouse Laboratory Plymouth | Wellcome Sanger Institute for the COVID-19 Genomics UK (COG-UK) Consortium | Lighthouse Laboratory Plymouth and Alex Alderton, Roberto Amato, Jeffrey Barrett, Sonia Goncalves, Ewan Harrison, David K. Jackson, Ian Johnston, Dominic Kwiatkowski, Cordelia Langford, John Sillitoe on behalf of the Wellcome Sanger Institute COVID-19 Surveillance Team |
| EPI_ISL_2782139 | Lighthouse Lab in Milton Keynes | Wellcome Sanger Institute for the COVID-19 Genomics UK (COG-UK) Consortium | The Lighthouse Lab in Milton Keynes and Alex Alderton, Roberto Amato, Jeffrey Barrett, Sonia Goncalves, Ewan Harrison, David K. Jackson, Ian Johnston, Dominic Kwiatkowski, Cordelia Langford, John Sillitoe on behalf of the Wellcome Sanger Institute COVID-19 Surveillance Team |
| EPI_ISL_2782140 | Lighthouse Lab in Alderley Park | Wellcome Sanger Institute for the COVID-19 Genomics UK (COG-UK) Consortium | Jacquelyn Wynn, Mairead Hyland, The Lighthouse Lab in Alderley Park and Alex Alderton, Roberto Amato, Jeffrey Barrett, Sonia Goncalves, Ewan Harrison, David K. Jackson, Ian Johnston, Dominic Kwiatkowski, Cordelia Langford, John Sillitoe on behalf of the Wellcome Sanger Institute COVID-19 Surveillance Team |
| EPI_ISL_2782141, EPI_ISL_2782142 | Lighthouse Lab in Milton Keynes | Wellcome Sanger Institute for the COVID-19 Genomics UK (COG-UK) Consortium | The Lighthouse Lab in Milton Keynes and Alex Alderton, Roberto Amato, Jeffrey Barrett, Sonia Goncalves, Ewan Harrison, David K. Jackson, Ian Johnston, Dominic Kwiatkowski, Cordelia Langford, John Sillitoe on behalf of the Wellcome Sanger Institute COVID-19 Surveillance Team |
| EPI_ISL_2782143 | Lighthouse Laboratory Plymouth | Wellcome Sanger Institute for the COVID-19 Genomics UK (COG-UK) Consortium | Lighthouse Laboratory Plymouth and Alex Alderton, Roberto Amato, Jeffrey Barrett, Sonia Goncalves, Ewan Harrison, David K. Jackson, Ian Johnston, Dominic Kwiatkowski, Cordelia Langford, John Sillitoe on behalf of the Wellcome Sanger Institute COVID-19 Surveillance Team |
| EPI_ISL_2782144 | Lighthouse Lab in Milton Keynes | Wellcome Sanger Institute for the COVID-19 Genomics UK (COG-UK) Consortium | The Lighthouse Lab in Milton Keynes and Alex Alderton, Roberto Amato, Jeffrey Barrett, Sonia Goncalves, Ewan Harrison, David K. Jackson, Ian Johnston, Dominic Kwiatkowski, Cordelia Langford, John Sillitoe on behalf of the Wellcome Sanger Institute COVID-19 Surveillance Team |
| EPI_ISL_2782145 | Lighthouse Lab in Alderley Park | Wellcome Sanger Institute for the COVID-19 Genomics UK (COG-UK) Consortium | Jacquelyn Wynn, Mairead Hyland, The Lighthouse Lab in Alderley Park and Alex Alderton, Roberto Amato, Jeffrey Barrett, Sonia Goncalves, Ewan Harrison, David K. Jackson, Ian Johnston, Dominic Kwiatkowski, Cordelia Langford, John Sillitoe on behalf of the Wellcome Sanger Institute COVID-19 Surveillance Team |
| EPI_ISL_2782146 | Lighthouse Laboratory Plymouth | Wellcome Sanger Institute for the COVID-19 Genomics UK (COG-UK) Consortium | Lighthouse Laboratory Plymouth and Alex Alderton, Roberto Amato, Jeffrey Barrett, Sonia Goncalves, Ewan Harrison, David K. Jackson, Ian Johnston, Dominic Kwiatkowski, Cordelia Langford, John Sillitoe on behalf of the Wellcome Sanger Institute COVID-19 Surveillance Team |
| EPI_ISL_2782147 | Lighthouse Lab in Alderley Park | Wellcome Sanger Institute for the COVID-19 Genomics UK (COG-UK) Consortium | Jacquelyn Wynn, Mairead Hyland, The Lighthouse Lab in Alderley Park and Alex Alderton, Roberto Amato, Jeffrey Barrett, Sonia Goncalves, Ewan Harrison, David K. Jackson, Ian Johnston, Dominic Kwiatkowski, Cordelia Langford, John Sillitoe on behalf of the Wellcome Sanger Institute COVID-19 Surveillance Team |
| EPI_ISL_2782148 | Lighthouse Laboratory Plymouth | Wellcome Sanger Institute for the COVID-19 Genomics UK (COG-UK) Consortium | Lighthouse Laboratory Plymouth and Alex Alderton, Roberto Amato, Jeffrey Barrett, Sonia Goncalves, Ewan Harrison, David K. Jackson, Ian Johnston, Dominic Kwiatkowski, Cordelia Langford, John Sillitoe on behalf of the Wellcome Sanger Institute COVID-19 Surveillance Team |
| EPI_ISL_2782149, EPI_ISL_2782150, EPI_ISL_2782151, EPI_ISL_2782152, EPI_ISL_2782153 | Lighthouse Lab in Alderley Park | Wellcome Sanger Institute for the COVID-19 Genomics UK (COG-UK) Consortium | Jacquelyn Wynn, Mairead Hyland, The Lighthouse Lab in Alderley Park and Alex Alderton, Roberto Amato, Jeffrey Barrett, Sonia Goncalves, Ewan Harrison, David K. Jackson, Ian Johnston, Dominic Kwiatkowski, Cordelia Langford, John Sillitoe on behalf of the Wellcome Sanger Institute COVID-19 Surveillance Team |
| EPI_ISL_2782154 | Lighthouse Laboratory Plymouth | Wellcome Sanger Institute for the COVID-19 Genomics UK (COG-UK) Consortium | Lighthouse Laboratory Plymouth and Alex Alderton, Roberto Amato, Jeffrey Barrett, Sonia Goncalves, Ewan Harrison, David K. Jackson, Ian Johnston, Dominic Kwiatkowski, Cordelia Langford, John Sillitoe on behalf of the Wellcome Sanger Institute COVID-19 Surveillance Team |
| EPI_ISL_2782155 | Lighthouse Lab in Glasgow | Wellcome Sanger Institute for the COVID-19 Genomics UK (COG-UK) Consortium | Harper VanSteenhouse, Yumi Kasai, David Gray, Carol Clugston, Anna Dominiczak and Alex Alderton, Roberto Amato, Jeffrey Barrett, Sonia Goncalves, Ewan Harrison, David K. Jackson, Ian Johnston, Dominic Kwiatkowski, Cordelia Langford, John Sillitoe on behalf of the Wellcome Sanger Institute COVID-19 Surveillance Team |
| EPI_ISL_2782156 | Lighthouse Laboratory Plymouth | Wellcome Sanger Institute for the COVID-19 Genomics UK (COG-UK) Consortium | Lighthouse Laboratory Plymouth and Alex Alderton, Roberto Amato, Jeffrey Barrett, Sonia Goncalves, Ewan Harrison, David K. Jackson, Ian Johnston, Dominic Kwiatkowski, Cordelia Langford, John Sillitoe on behalf of the Wellcome Sanger Institute COVID-19 Surveillance Team |
| EPI_ISL_2782157 | Lighthouse Lab in Alderley Park | Wellcome Sanger Institute for the COVID-19 Genomics UK (COG-UK) Consortium | Jacquelyn Wynn, Mairead Hyland, The Lighthouse Lab in Alderley Park and Alex Alderton, Roberto Amato, Jeffrey Barrett, Sonia Goncalves, Ewan Harrison, David K. Jackson, Ian Johnston, Dominic Kwiatkowski, Cordelia Langford, John Sillitoe on behalf of the Wellcome Sanger Institute COVID-19 Surveillance Team |
| EPI_ISL_2782158 | Lighthouse Lab in Glasgow | Wellcome Sanger Institute for the COVID-19 Genomics UK (COG-UK) Consortium | Harper VanSteenhouse, Yumi Kasai, David Gray, Carol Clugston, Anna Dominiczak and Alex Alderton, Roberto Amato, Jeffrey Barrett, Sonia Goncalves, Ewan Harrison, David K. Jackson, Ian Johnston, Dominic Kwiatkowski, Cordelia Langford, John Sillitoe on behalf of the Wellcome Sanger Institute COVID-19 Surveillance Team |
| EPI_ISL_2782159 | Lighthouse Laboratory Plymouth | Wellcome Sanger Institute for the COVID-19 Genomics UK (COG-UK) Consortium | Lighthouse Laboratory Plymouth and Alex Alderton, Roberto Amato, Jeffrey Barrett, Sonia Goncalves, Ewan Harrison, David K. Jackson, Ian Johnston, Dominic Kwiatkowski, Cordelia Langford, John Sillitoe on behalf of the Wellcome Sanger Institute COVID-19 Surveillance Team |
| EPI_ISL_2782160 | Lighthouse Lab in Alderley Park | Wellcome Sanger Institute for the COVID-19 Genomics UK (COG-UK) Consortium | Jacquelyn Wynn, Mairead Hyland, The Lighthouse Lab in Alderley Park and Alex Alderton, Roberto Amato, Jeffrey Barrett, Sonia Goncalves, Ewan Harrison, David K. Jackson, Ian Johnston, Dominic Kwiatkowski, Cordelia Langford, John Sillitoe on behalf of the Wellcome Sanger Institute COVID-19 Surveillance Team |
| EPI_ISL_2782161 | Lighthouse Lab in Glasgow | Wellcome Sanger Institute for the COVID-19 Genomics UK (COG-UK) Consortium | Harper VanSteenhouse, Yumi Kasai, David Gray, Carol Clugston, Anna Dominiczak and Alex Alderton, Roberto Amato, Jeffrey Barrett, Sonia Goncalves, Ewan Harrison, David K. Jackson, Ian Johnston, Dominic Kwiatkowski, Cordelia Langford, John Sillitoe on behalf of the Wellcome Sanger Institute COVID-19 Surveillance Team |
| EPI_ISL_2782162 | Lighthouse Laboratory Plymouth | Wellcome Sanger Institute for the COVID-19 Genomics UK (COG-UK) Consortium | Lighthouse Laboratory Plymouth and Alex Alderton, Roberto Amato, Jeffrey Barrett, Sonia Goncalves, Ewan Harrison, David K. Jackson, Ian Johnston, Dominic Kwiatkowski, Cordelia Langford, John Sillitoe on behalf of the Wellcome Sanger Institute COVID-19 Surveillance Team |
| EPI_ISL_2782163 | Lighthouse Lab in Glasgow | Wellcome Sanger Institute for the COVID-19 Genomics UK (COG-UK) Consortium | Harper VanSteenhouse, Yumi Kasai, David Gray, Carol Clugston, Anna Dominiczak and Alex Alderton, Roberto Amato, Jeffrey Barrett, Sonia Goncalves, Ewan Harrison, David K. Jackson, Ian Johnston, Dominic Kwiatkowski, Cordelia Langford, John Sillitoe on behalf of the Wellcome Sanger Institute COVID-19 Surveillance Team |
| EPI_ISL_2782164 | Lighthouse Lab in Milton Keynes | Wellcome Sanger Institute for the COVID-19 Genomics UK (COG-UK) Consortium | The Lighthouse Lab in Milton Keynes and Alex Alderton, Roberto Amato, Jeffrey Barrett, Sonia Goncalves, Ewan Harrison, David K. Jackson, Ian Johnston, Dominic Kwiatkowski, Cordelia Langford, John Sillitoe on behalf of the Wellcome Sanger Institute COVID-19 Surveillance Team |
| EPI_ISL_2782166, EPI_ISL_2782167, EPI_ISL_2782168, EPI_ISL_2782169 | Lighthouse Laboratory Plymouth | Wellcome Sanger Institute for the COVID-19 Genomics UK (COG-UK) Consortium | Lighthouse Laboratory Plymouth and Alex Alderton, Roberto Amato, Jeffrey Barrett, Sonia Goncalves, Ewan Harrison, David K. Jackson, Ian Johnston, Dominic Kwiatkowski, Cordelia Langford, John Sillitoe on behalf of the Wellcome Sanger Institute COVID-19 Surveillance Team |
| EPI_ISL_2782170, EPI_ISL_2782171 | Lighthouse Lab in Alderley Park | Wellcome Sanger Institute for the COVID-19 Genomics UK (COG-UK) Consortium | Jacquelyn Wynn, Mairead Hyland, The Lighthouse Lab in Alderley Park and Alex Alderton, Roberto Amato, Jeffrey Barrett, Sonia Goncalves, Ewan Harrison, David K. Jackson, Ian Johnston, Dominic Kwiatkowski, Cordelia Langford, John Sillitoe on behalf of the Wellcome Sanger Institute COVID-19 Surveillance Team |
| EPI_ISL_2782172 | Lighthouse Lab in Milton Keynes | Wellcome Sanger Institute for the COVID-19 Genomics UK (COG-UK) | The Lighthouse Lab in Milton Keynes and Alex Alderton, Roberto Amato, Jeffrey Barrett, Sonia Goncalves, Ewan Harrison, David K. Jackson, Ian Johnston, Dominic Kwiatkowski, Cordelia Langford, John Sillitoe on behalf of the Wellcome Sanger |

| | | | |
|---|---|---|---|
| | | Consortium | Institute COVID-19 Surveillance Team |
| EPI_ISL_2782173, EPI_ISL_2782174, EPI_ISL_2782175 | Lighthouse Laboratory Plymouth | Wellcome Sanger Institute for the COVID-19 Genomics UK (COG-UK) Consortium | Lighthouse Laboratory Plymouth and Alex Alderton, Roberto Amato, Jeffrey Barrett, Sonia Goncalves, Ewan Harrison, David K. Jackson, Ian Johnston, Dominic Kwiatkowski, Cordelia Langford, John Sillitoe on behalf of the Wellcome Sanger Institute COVID-19 Surveillance Team |
| EPI_ISL_2782176, EPI_ISL_2782177 | Lighthouse Lab in Alderley Park | Wellcome Sanger Institute for the COVID-19 Genomics UK (COG-UK) Consortium | Jacquelyn Wynn, Mairead Hyland, The Lighthouse Lab in Alderley Park and Alex Alderton, Roberto Amato, Jeffrey Barrett, Sonia Goncalves, Ewan Harrison, David K. Jackson, Ian Johnston, Dominic Kwiatkowski, Cordelia Langford, John Sillitoe on behalf of the Wellcome Sanger Institute COVID-19 Surveillance Team |
| EPI_ISL_2782178, EPI_ISL_2782179 | Lighthouse Lab in Milton Keynes | Wellcome Sanger Institute for the COVID-19 Genomics UK (COG-UK) Consortium | The Lighthouse Lab in Milton Keynes and Alex Alderton, Roberto Amato, Jeffrey Barrett, Sonia Goncalves, Ewan Harrison, David K. Jackson, Ian Johnston, Dominic Kwiatkowski, Cordelia Langford, John Sillitoe on behalf of the Wellcome Sanger Institute COVID-19 Surveillance Team |
| EPI_ISL_2782180, EPI_ISL_2782181 | Lighthouse Lab in Alderley Park | Wellcome Sanger Institute for the COVID-19 Genomics UK (COG-UK) Consortium | Jacquelyn Wynn, Mairead Hyland, The Lighthouse Lab in Alderley Park and Alex Alderton, Roberto Amato, Jeffrey Barrett, Sonia Goncalves, Ewan Harrison, David K. Jackson, Ian Johnston, Dominic Kwiatkowski, Cordelia Langford, John Sillitoe on behalf of the Wellcome Sanger Institute COVID-19 Surveillance Team |
| EPI_ISL_2782182 | Lighthouse Lab in Glasgow | Wellcome Sanger Institute for the COVID-19 Genomics UK (COG-UK) Consortium | Harper VanSteenhouse, Yumi Kasai, David Gray, Carol Clugston, Anna Dominiczak and Alex Alderton, Roberto Amato, Jeffrey Barrett, Sonia Goncalves, Ewan Harrison, David K. Jackson, Ian Johnston, Dominic Kwiatkowski, Cordelia Langford, John Sillitoe on behalf of the Wellcome Sanger Institute COVID-19 Surveillance Team |
| EPI_ISL_2782183, EPI_ISL_2782184 | Lighthouse Lab in Milton Keynes | Wellcome Sanger Institute for the COVID-19 Genomics UK (COG-UK) Consortium | The Lighthouse Lab in Milton Keynes and Alex Alderton, Roberto Amato, Jeffrey Barrett, Sonia Goncalves, Ewan Harrison, David K. Jackson, Ian Johnston, Dominic Kwiatkowski, Cordelia Langford, John Sillitoe on behalf of the Wellcome Sanger Institute COVID-19 Surveillance Team |
| EPI_ISL_2782185 | Lighthouse Laboratory Plymouth | Wellcome Sanger Institute for the COVID-19 Genomics UK (COG-UK) Consortium | Lighthouse Laboratory Plymouth and Alex Alderton, Roberto Amato, Jeffrey Barrett, Sonia Goncalves, Ewan Harrison, David K. Jackson, Ian Johnston, Dominic Kwiatkowski, Cordelia Langford, John Sillitoe on behalf of the Wellcome Sanger Institute COVID-19 Surveillance Team |
| EPI_ISL_2782186, EPI_ISL_2782187 | Lighthouse Lab in Alderley Park | Wellcome Sanger Institute for the COVID-19 Genomics UK (COG-UK) Consortium | Jacquelyn Wynn, Mairead Hyland, The Lighthouse Lab in Alderley Park and Alex Alderton, Roberto Amato, Jeffrey Barrett, Sonia Goncalves, Ewan Harrison, David K. Jackson, Ian Johnston, Dominic Kwiatkowski, Cordelia Langford, John Sillitoe on behalf of the Wellcome Sanger Institute COVID-19 Surveillance Team |
| EPI_ISL_2782188 | Lighthouse Laboratory Plymouth | Wellcome Sanger Institute for the COVID-19 Genomics UK (COG-UK) Consortium | Lighthouse Laboratory Plymouth and Alex Alderton, Roberto Amato, Jeffrey Barrett, Sonia Goncalves, Ewan Harrison, David K. Jackson, Ian Johnston, Dominic Kwiatkowski, Cordelia Langford, John Sillitoe on behalf of the Wellcome Sanger Institute COVID-19 Surveillance Team |
| EPI_ISL_2782189, EPI_ISL_2782190 | Lighthouse Lab in Milton Keynes | Wellcome Sanger Institute for the COVID-19 Genomics UK (COG-UK) Consortium | The Lighthouse Lab in Milton Keynes and Alex Alderton, Roberto Amato, Jeffrey Barrett, Sonia Goncalves, Ewan Harrison, David K. Jackson, Ian Johnston, Dominic Kwiatkowski, Cordelia Langford, John Sillitoe on behalf of the Wellcome Sanger Institute COVID-19 Surveillance Team |
| EPI_ISL_2782191 | Lighthouse Lab in Glasgow | Wellcome Sanger Institute for the COVID-19 Genomics UK (COG-UK) Consortium | Harper VanSteenhouse, Yumi Kasai, David Gray, Carol Clugston, Anna Dominiczak and Alex Alderton, Roberto Amato, Jeffrey Barrett, Sonia Goncalves, Ewan Harrison, David K. Jackson, Ian Johnston, Dominic Kwiatkowski, Cordelia Langford, John Sillitoe on behalf of the Wellcome Sanger Institute COVID-19 Surveillance Team |
| EPI_ISL_2782192 | Lighthouse Lab in Milton Keynes | Wellcome Sanger Institute for the COVID-19 Genomics UK (COG-UK) Consortium | The Lighthouse Lab in Milton Keynes and Alex Alderton, Roberto Amato, Jeffrey Barrett, Sonia Goncalves, Ewan Harrison, David K. Jackson, Ian Johnston, Dominic Kwiatkowski, Cordelia Langford, John Sillitoe on behalf of the Wellcome Sanger Institute COVID-19 Surveillance Team |
| EPI_ISL_2782193 | Lighthouse Lab in Alderley Park | Wellcome Sanger Institute for the COVID-19 Genomics UK (COG-UK) Consortium | Jacquelyn Wynn, Mairead Hyland, The Lighthouse Lab in Alderley Park and Alex Alderton, Roberto Amato, Jeffrey Barrett, Sonia Goncalves, Ewan Harrison, David K. Jackson, Ian Johnston, Dominic Kwiatkowski, Cordelia Langford, John Sillitoe on behalf of the Wellcome Sanger Institute COVID-19 Surveillance Team |
| EPI_ISL_2782194, EPI_ISL_2782195 | Lighthouse Laboratory Plymouth | Wellcome Sanger Institute for the COVID-19 Genomics UK (COG-UK) Consortium | Lighthouse Laboratory Plymouth and Alex Alderton, Roberto Amato, Jeffrey Barrett, Sonia Goncalves, Ewan Harrison, David K. Jackson, Ian Johnston, Dominic Kwiatkowski, Cordelia Langford, John Sillitoe on behalf of the Wellcome Sanger Institute COVID-19 Surveillance Team |
| EPI_ISL_2782196 | Lighthouse Lab in Alderley Park | Wellcome Sanger Institute for the COVID-19 Genomics UK (COG-UK) Consortium | Jacquelyn Wynn, Mairead Hyland, The Lighthouse Lab in Alderley Park and Alex Alderton, Roberto Amato, Jeffrey Barrett, Sonia Goncalves, Ewan Harrison, David K. Jackson, Ian Johnston, Dominic Kwiatkowski, Cordelia Langford, John Sillitoe on behalf of the Wellcome Sanger Institute COVID-19 Surveillance Team |
| EPI_ISL_2782197, EPI_ISL_2782199, EPI_ISL_2782200, EPI_ISL_2782201 | Lighthouse Lab in Milton Keynes | Wellcome Sanger Institute for the COVID-19 Genomics UK (COG-UK) Consortium | The Lighthouse Lab in Milton Keynes and Alex Alderton, Roberto Amato, Jeffrey Barrett, Sonia Goncalves, Ewan Harrison, David K. Jackson, Ian Johnston, Dominic Kwiatkowski, Cordelia Langford, John Sillitoe on behalf of the Wellcome Sanger Institute COVID-19 Surveillance Team |
| EPI_ISL_2782202, EPI_ISL_2782203 | Lighthouse Lab in Alderley Park | Wellcome Sanger Institute for the COVID-19 Genomics UK (COG-UK) Consortium | Jacquelyn Wynn, Mairead Hyland, The Lighthouse Lab in Alderley Park and Alex Alderton, Roberto Amato, Jeffrey Barrett, Sonia Goncalves, Ewan Harrison, David K. Jackson, Ian Johnston, Dominic Kwiatkowski, Cordelia Langford, John Sillitoe on behalf of the Wellcome Sanger Institute COVID-19 Surveillance Team |
| EPI_ISL_2782204 | Lighthouse Lab in Milton Keynes | Wellcome Sanger Institute for the COVID-19 Genomics UK (COG-UK) Consortium | The Lighthouse Lab in Milton Keynes and Alex Alderton, Roberto Amato, Jeffrey Barrett, Sonia Goncalves, Ewan Harrison, David K. Jackson, Ian Johnston, Dominic Kwiatkowski, Cordelia Langford, John Sillitoe on behalf of the Wellcome Sanger Institute COVID-19 Surveillance Team |
| EPI_ISL_2782205 | Lighthouse Lab in Alderley Park | Wellcome Sanger Institute for the COVID-19 Genomics UK (COG-UK) Consortium | Jacquelyn Wynn, Mairead Hyland, The Lighthouse Lab in Alderley Park and Alex Alderton, Roberto Amato, Jeffrey Barrett, Sonia Goncalves, Ewan Harrison, David K. Jackson, Ian Johnston, Dominic Kwiatkowski, Cordelia Langford, John Sillitoe on behalf of the Wellcome Sanger Institute COVID-19 Surveillance Team |
| EPI_ISL_2782206, EPI_ISL_2782207 | Lighthouse Lab in Milton Keynes | Wellcome Sanger Institute for the COVID-19 Genomics UK (COG-UK) Consortium | The Lighthouse Lab in Milton Keynes and Alex Alderton, Roberto Amato, Jeffrey Barrett, Sonia Goncalves, Ewan Harrison, David K. Jackson, Ian Johnston, Dominic Kwiatkowski, Cordelia Langford, John Sillitoe on behalf of the Wellcome Sanger Institute COVID-19 Surveillance Team |
| EPI_ISL_2782208 | Lighthouse Lab in Glasgow | Wellcome Sanger Institute for the COVID-19 Genomics UK (COG-UK) Consortium | Harper VanSteenhouse, Yumi Kasai, David Gray, Carol Clugston, Anna Dominiczak and Alex Alderton, Roberto Amato, Jeffrey Barrett, Sonia Goncalves, Ewan Harrison, David K. Jackson, Ian Johnston, Dominic Kwiatkowski, Cordelia Langford, John Sillitoe on behalf of the Wellcome Sanger Institute COVID-19 Surveillance Team |
| EPI_ISL_2782209 | Lighthouse Lab in Alderley Park | Wellcome Sanger Institute for the COVID-19 Genomics UK (COG-UK) Consortium | Jacquelyn Wynn, Mairead Hyland, The Lighthouse Lab in Alderley Park and Alex Alderton, Roberto Amato, Jeffrey Barrett, Sonia Goncalves, Ewan Harrison, David K. Jackson, Ian Johnston, Dominic Kwiatkowski, Cordelia Langford, John Sillitoe on behalf of the Wellcome Sanger Institute COVID-19 Surveillance Team |
| EPI_ISL_2782210 | Lighthouse Lab in Milton Keynes | Wellcome Sanger Institute for the COVID-19 Genomics UK (COG-UK) Consortium | The Lighthouse Lab in Milton Keynes and Alex Alderton, Roberto Amato, Jeffrey Barrett, Sonia Goncalves, Ewan Harrison, David K. Jackson, Ian Johnston, Dominic Kwiatkowski, Cordelia Langford, John Sillitoe on behalf of the Wellcome Sanger Institute COVID-19 Surveillance Team |
| EPI_ISL_2782211, EPI_ISL_2782212 | Lighthouse Laboratory Plymouth | Wellcome Sanger Institute for the COVID-19 Genomics UK (COG-UK) Consortium | Lighthouse Laboratory Plymouth and Alex Alderton, Roberto Amato, Jeffrey Barrett, Sonia Goncalves, Ewan Harrison, David K. Jackson, Ian Johnston, Dominic Kwiatkowski, Cordelia Langford, John Sillitoe on behalf of the Wellcome Sanger Institute COVID-19 Surveillance Team |
| EPI_ISL_2782213, EPI_ISL_2782214 | Lighthouse Lab in Glasgow | Wellcome Sanger Institute for the COVID-19 Genomics UK (COG-UK) Consortium | Harper VanSteenhouse, Yumi Kasai, David Gray, Carol Clugston, Anna Dominiczak and Alex Alderton, Roberto Amato, Jeffrey Barrett, Sonia Goncalves, Ewan Harrison, David K. Jackson, Ian Johnston, Dominic Kwiatkowski, Cordelia Langford, John Sillitoe on behalf of the Wellcome Sanger Institute COVID-19 Surveillance Team |
| EPI_ISL_2782215, EPI_ISL_2782216 | Lighthouse Lab in Alderley Park | Wellcome Sanger Institute for the COVID-19 Genomics UK (COG-UK) Consortium | Jacquelyn Wynn, Mairead Hyland, The Lighthouse Lab in Alderley Park and Alex Alderton, Roberto Amato, Jeffrey Barrett, Sonia Goncalves, Ewan Harrison, David K. Jackson, Ian Johnston, Dominic Kwiatkowski, Cordelia Langford, John Sillitoe on behalf of the Wellcome Sanger Institute COVID-19 Surveillance Team |
| EPI_ISL_2782217 | Lighthouse Lab in Milton Keynes | Wellcome Sanger Institute for the COVID-19 Genomics UK (COG-UK) Consortium | The Lighthouse Lab in Milton Keynes and Alex Alderton, Roberto Amato, Jeffrey Barrett, Sonia Goncalves, Ewan Harrison, David K. Jackson, Ian Johnston, Dominic Kwiatkowski, Cordelia Langford, John Sillitoe on behalf of the Wellcome Sanger Institute COVID-19 Surveillance Team |
| EPI_ISL_2782218 | Lighthouse Lab in Alderley Park | Wellcome Sanger Institute for the COVID-19 Genomics UK (COG-UK) Consortium | Jacquelyn Wynn, Mairead Hyland, The Lighthouse Lab in Alderley Park and Alex Alderton, Roberto Amato, Jeffrey Barrett, Sonia Goncalves, Ewan Harrison, David K. Jackson, Ian Johnston, Dominic Kwiatkowski, Cordelia Langford, John Sillitoe on behalf of the Wellcome Sanger Institute COVID-19 Surveillance Team |
| EPI_ISL_2782219 | Lighthouse Lab in Milton Keynes | Wellcome Sanger Institute for the COVID-19 Genomics UK (COG-UK) Consortium | The Lighthouse Lab in Milton Keynes and Alex Alderton, Roberto Amato, Jeffrey Barrett, Sonia Goncalves, Ewan Harrison, David K. Jackson, Ian Johnston, Dominic Kwiatkowski, Cordelia Langford, John Sillitoe on behalf of the Wellcome Sanger Institute COVID-19 Surveillance Team |
| EPI_ISL_2782220 | Lighthouse Laboratory Plymouth | Wellcome Sanger Institute for the COVID-19 Genomics UK (COG-UK) Consortium | Lighthouse Laboratory Plymouth and Alex Alderton, Roberto Amato, Jeffrey Barrett, Sonia Goncalves, Ewan Harrison, David K. Jackson, Ian Johnston, Dominic Kwiatkowski, Cordelia Langford, John Sillitoe on behalf of the Wellcome Sanger Institute COVID-19 Surveillance Team |
| EPI_ISL_2782221 | Lighthouse Lab in Alderley Park | Wellcome Sanger Institute for the COVID-19 Genomics UK (COG-UK) Consortium | Jacquelyn Wynn, Mairead Hyland, The Lighthouse Lab in Alderley Park and Alex Alderton, Roberto Amato, Jeffrey Barrett, Sonia Goncalves, Ewan Harrison, David K. Jackson, Ian Johnston, Dominic Kwiatkowski, Cordelia Langford, John Sillitoe on behalf of the Wellcome Sanger Institute COVID-19 Surveillance Team |
| EPI_ISL_2782222 | Lighthouse Laboratory Plymouth | Wellcome Sanger Institute for the COVID-19 Genomics UK (COG-UK) Consortium | Lighthouse Laboratory Plymouth and Alex Alderton, Roberto Amato, Jeffrey Barrett, Sonia Goncalves, Ewan Harrison, David K. Jackson, Ian Johnston, Dominic Kwiatkowski, Cordelia Langford, John Sillitoe on behalf of the Wellcome Sanger Institute COVID-19 Surveillance Team |
| EPI_ISL_2782223, EPI_ISL_2782224, EPI_ISL_2782225, EPI_ISL_2782226 | Lighthouse Lab in Alderley Park | Wellcome Sanger Institute for the COVID-19 Genomics UK (COG-UK) Consortium | Jacquelyn Wynn, Mairead Hyland, The Lighthouse Lab in Alderley Park and Alex Alderton, Roberto Amato, Jeffrey Barrett, Sonia Goncalves, Ewan Harrison, David K. Jackson, Ian Johnston, Dominic Kwiatkowski, Cordelia Langford, John Sillitoe on behalf of the Wellcome Sanger Institute COVID-19 Surveillance Team |
| EPI_ISL_2782227, EPI_ISL_2782228, EPI_ISL_2782229 | Lighthouse Laboratory Plymouth | Wellcome Sanger Institute for the COVID-19 Genomics UK (COG-UK) Consortium | Lighthouse Laboratory Plymouth and Alex Alderton, Roberto Amato, Jeffrey Barrett, Sonia Goncalves, Ewan Harrison, David K. Jackson, Ian Johnston, Dominic Kwiatkowski, Cordelia Langford, John Sillitoe on behalf of the Wellcome Sanger Institute COVID-19 Surveillance Team |
| EPI_ISL_2782230 | Lighthouse Lab in Glasgow | Wellcome Sanger Institute for the COVID-19 Genomics UK (COG-UK) Consortium | Harper VanSteenhouse, Yumi Kasai, David Gray, Carol Clugston, Anna Dominiczak and Alex Alderton, Roberto Amato, Jeffrey Barrett, Sonia Goncalves, Ewan Harrison, David K. Jackson, Ian Johnston, Dominic Kwiatkowski, Cordelia Langford, John Sillitoe on behalf of the Wellcome Sanger Institute COVID-19 Surveillance Team |
| EPI_ISL_2782231, EPI_ISL_2782232 | Lighthouse Lab in Alderley Park | Wellcome Sanger Institute for the COVID-19 Genomics UK (COG-UK) Consortium | Jacquelyn Wynn, Mairead Hyland, The Lighthouse Lab in Alderley Park and Alex Alderton, Roberto Amato, Jeffrey Barrett, Sonia Goncalves, Ewan Harrison, David K. Jackson, Ian Johnston, Dominic Kwiatkowski, Cordelia Langford, John Sillitoe on behalf of the Wellcome Sanger Institute COVID-19 Surveillance Team |
| EPI_ISL_2782233 | Lighthouse Laboratory Plymouth | Wellcome Sanger Institute for the COVID-19 Genomics UK (COG-UK) Consortium | Lighthouse Laboratory Plymouth and Alex Alderton, Roberto Amato, Jeffrey Barrett, Sonia Goncalves, Ewan Harrison, David K. Jackson, Ian Johnston, Dominic Kwiatkowski, Cordelia Langford, John Sillitoe on behalf of the Wellcome Sanger Institute COVID-19 Surveillance Team |
| EPI_ISL_2782234 | Lighthouse Lab in Milton Keynes | Wellcome Sanger Institute for the COVID-19 Genomics UK (COG-UK) Consortium | The Lighthouse Lab in Milton Keynes and Alex Alderton, Roberto Amato, Jeffrey Barrett, Sonia Goncalves, Ewan Harrison, David K. Jackson, Ian Johnston, Dominic Kwiatkowski, Cordelia Langford, John Sillitoe on behalf of the Wellcome Sanger Institute COVID-19 Surveillance Team |
| EPI_ISL_2782235, EPI_ISL_2782236 | Lighthouse Laboratory Plymouth | Wellcome Sanger Institute for the COVID-19 Genomics UK (COG-UK) Consortium | Lighthouse Laboratory Plymouth and Alex Alderton, Roberto Amato, Jeffrey Barrett, Sonia Goncalves, Ewan Harrison, David K. Jackson, Ian Johnston, Dominic Kwiatkowski, Cordelia Langford, John Sillitoe on behalf of the Wellcome Sanger Institute COVID-19 Surveillance Team |
| EPI_ISL_2782237 | Lighthouse Lab in Alderley Park | Wellcome Sanger Institute for the COVID-19 Genomics UK (COG-UK) Consortium | Jacquelyn Wynn, Mairead Hyland, The Lighthouse Lab in Alderley Park and Alex Alderton, Roberto Amato, Jeffrey Barrett, Sonia Goncalves, Ewan Harrison, David K. Jackson, Ian Johnston, Dominic Kwiatkowski, Cordelia Langford, John Sillitoe on behalf of the Wellcome Sanger Institute COVID-19 Surveillance Team |
| EPI_ISL_2782238 | Lighthouse Lab in Milton Keynes | Wellcome Sanger Institute for the COVID-19 Genomics UK (COG-UK) Consortium | The Lighthouse Lab in Milton Keynes and Alex Alderton, Roberto Amato, Jeffrey Barrett, Sonia Goncalves, Ewan Harrison, David K. Jackson, Ian Johnston, Dominic Kwiatkowski, Cordelia Langford, John Sillitoe on behalf of the Wellcome Sanger Institute COVID-19 Surveillance Team |
| EPI_ISL_2782239 | Lighthouse Lab in Alderley Park | Wellcome Sanger Institute for the COVID-19 Genomics UK (COG-UK) Consortium | Jacquelyn Wynn, Mairead Hyland, The Lighthouse Lab in Alderley Park and Alex Alderton, Roberto Amato, Jeffrey Barrett, Sonia Goncalves, Ewan Harrison, David K. Jackson, Ian Johnston, Dominic Kwiatkowski, Cordelia Langford, John Sillitoe on behalf of the Wellcome Sanger Institute COVID-19 Surveillance Team |

| | | | |
|---|---|---|---|
| EPI_ISL_2782240 | Lighthouse Lab in Milton Keynes | Wellcome Sanger Institute for the COVID-19 Genomics UK (COG-UK) Consortium | The Lighthouse Lab in Milton Keynes and Alex Alderton, Roberto Amato, Jeffrey Barrett, Sonia Goncalves, Ewan Harrison, David K. Jackson, Ian Johnston, Dominic Kwiatkowski, Cordelia Langford, John Sillitoe on behalf of the Wellcome Sanger Institute COVID-19 Surveillance Team |
| EPI_ISL_2782241, EPI_ISL_2782242 | Lighthouse Lab in Alderley Park | Wellcome Sanger Institute for the COVID-19 Genomics UK (COG-UK) Consortium | Jacquelyn Wynn, Mairead Hyland, The Lighthouse Lab in Alderley Park and Alex Alderton, Roberto Amato, Jeffrey Barrett, Sonia Goncalves, Ewan Harrison, David K. Jackson, Ian Johnston, Dominic Kwiatkowski, Cordelia Langford, John Sillitoe on behalf of the Wellcome Sanger Institute COVID-19 Surveillance Team |
| EPI_ISL_2782243 | Lighthouse Laboratory Plymouth | Wellcome Sanger Institute for the COVID-19 Genomics UK (COG-UK) Consortium | Lighthouse Laboratory Plymouth and Alex Alderton, Roberto Amato, Jeffrey Barrett, Sonia Goncalves, Ewan Harrison, David K. Jackson, Ian Johnston, Dominic Kwiatkowski, Cordelia Langford, John Sillitoe on behalf of the Wellcome Sanger Institute COVID-19 Surveillance Team |
| EPI_ISL_2782244 | Lighthouse Lab in Milton Keynes | Wellcome Sanger Institute for the COVID-19 Genomics UK (COG-UK) Consortium | The Lighthouse Lab in Milton Keynes and Alex Alderton, Roberto Amato, Jeffrey Barrett, Sonia Goncalves, Ewan Harrison, David K. Jackson, Ian Johnston, Dominic Kwiatkowski, Cordelia Langford, John Sillitoe on behalf of the Wellcome Sanger Institute COVID-19 Surveillance Team |
| EPI_ISL_2782245 | Lighthouse Lab in Alderley Park | Wellcome Sanger Institute for the COVID-19 Genomics UK (COG-UK) Consortium | Jacquelyn Wynn, Mairead Hyland, The Lighthouse Lab in Alderley Park and Alex Alderton, Roberto Amato, Jeffrey Barrett, Sonia Goncalves, Ewan Harrison, David K. Jackson, Ian Johnston, Dominic Kwiatkowski, Cordelia Langford, John Sillitoe on behalf of the Wellcome Sanger Institute COVID-19 Surveillance Team |
| EPI_ISL_2782246 | Lighthouse Lab in Milton Keynes | Wellcome Sanger Institute for the COVID-19 Genomics UK (COG-UK) Consortium | The Lighthouse Lab in Milton Keynes and Alex Alderton, Roberto Amato, Jeffrey Barrett, Sonia Goncalves, Ewan Harrison, David K. Jackson, Ian Johnston, Dominic Kwiatkowski, Cordelia Langford, John Sillitoe on behalf of the Wellcome Sanger Institute COVID-19 Surveillance Team |
| EPI_ISL_2782247 | Lighthouse Lab in Alderley Park | Wellcome Sanger Institute for the COVID-19 Genomics UK (COG-UK) Consortium | Jacquelyn Wynn, Mairead Hyland, The Lighthouse Lab in Alderley Park and Alex Alderton, Roberto Amato, Jeffrey Barrett, Sonia Goncalves, Ewan Harrison, David K. Jackson, Ian Johnston, Dominic Kwiatkowski, Cordelia Langford, John Sillitoe on behalf of the Wellcome Sanger Institute COVID-19 Surveillance Team |
| EPI_ISL_2782248, EPI_ISL_2782249, EPI_ISL_2782250 | Lighthouse Lab in Milton Keynes | Wellcome Sanger Institute for the COVID-19 Genomics UK (COG-UK) Consortium | The Lighthouse Lab in Milton Keynes and Alex Alderton, Roberto Amato, Jeffrey Barrett, Sonia Goncalves, Ewan Harrison, David K. Jackson, Ian Johnston, Dominic Kwiatkowski, Cordelia Langford, John Sillitoe on behalf of the Wellcome Sanger Institute COVID-19 Surveillance Team |
| EPI_ISL_2782251 | Lighthouse Lab in Glasgow | Wellcome Sanger Institute for the COVID-19 Genomics UK (COG-UK) Consortium | Harper VanSteenhouse, Yumi Kasai, David Gray, Carol Clugston, Anna Dominiczak and Alex Alderton, Roberto Amato, Jeffrey Barrett, Sonia Goncalves, Ewan Harrison, David K. Jackson, Ian Johnston, Dominic Kwiatkowski, Cordelia Langford, John Sillitoe on behalf of the Wellcome Sanger Institute COVID-19 Surveillance Team |
| EPI_ISL_2782252 | Lighthouse Lab in Alderley Park | Wellcome Sanger Institute for the COVID-19 Genomics UK (COG-UK) Consortium | Jacquelyn Wynn, Mairead Hyland, The Lighthouse Lab in Alderley Park and Alex Alderton, Roberto Amato, Jeffrey Barrett, Sonia Goncalves, Ewan Harrison, David K. Jackson, Ian Johnston, Dominic Kwiatkowski, Cordelia Langford, John Sillitoe on behalf of the Wellcome Sanger Institute COVID-19 Surveillance Team |
| EPI_ISL_2782253 | Lighthouse Lab in Glasgow | Wellcome Sanger Institute for the COVID-19 Genomics UK (COG-UK) Consortium | Harper VanSteenhouse, Yumi Kasai, David Gray, Carol Clugston, Anna Dominiczak and Alex Alderton, Roberto Amato, Jeffrey Barrett, Sonia Goncalves, Ewan Harrison, David K. Jackson, Ian Johnston, Dominic Kwiatkowski, Cordelia Langford, John Sillitoe on behalf of the Wellcome Sanger Institute COVID-19 Surveillance Team |
| EPI_ISL_2782254 | Lighthouse Lab in Milton Keynes | Wellcome Sanger Institute for the COVID-19 Genomics UK (COG-UK) Consortium | The Lighthouse Lab in Milton Keynes and Alex Alderton, Roberto Amato, Jeffrey Barrett, Sonia Goncalves, Ewan Harrison, David K. Jackson, Ian Johnston, Dominic Kwiatkowski, Cordelia Langford, John Sillitoe on behalf of the Wellcome Sanger Institute COVID-19 Surveillance Team |
| EPI_ISL_2782255 | Lighthouse Laboratory Plymouth | Wellcome Sanger Institute for the COVID-19 Genomics UK (COG-UK) Consortium | Lighthouse Laboratory Plymouth and Alex Alderton, Roberto Amato, Jeffrey Barrett, Sonia Goncalves, Ewan Harrison, David K. Jackson, Ian Johnston, Dominic Kwiatkowski, Cordelia Langford, John Sillitoe on behalf of the Wellcome Sanger Institute COVID-19 Surveillance Team |
| EPI_ISL_2782256, EPI_ISL_2782257, EPI_ISL_2782258, EPI_ISL_2782259 | Lighthouse Lab in Alderley Park | Wellcome Sanger Institute for the COVID-19 Genomics UK (COG-UK) Consortium | Jacquelyn Wynn, Mairead Hyland, The Lighthouse Lab in Alderley Park and Alex Alderton, Roberto Amato, Jeffrey Barrett, Sonia Goncalves, Ewan Harrison, David K. Jackson, Ian Johnston, Dominic Kwiatkowski, Cordelia Langford, John Sillitoe on behalf of the Wellcome Sanger Institute COVID-19 Surveillance Team |
| EPI_ISL_2782260, EPI_ISL_2782261, EPI_ISL_2782262, EPI_ISL_2782263 | Lighthouse Lab in Milton Keynes | Wellcome Sanger Institute for the COVID-19 Genomics UK (COG-UK) Consortium | The Lighthouse Lab in Milton Keynes and Alex Alderton, Roberto Amato, Jeffrey Barrett, Sonia Goncalves, Ewan Harrison, David K. Jackson, Ian Johnston, Dominic Kwiatkowski, Cordelia Langford, John Sillitoe on behalf of the Wellcome Sanger Institute COVID-19 Surveillance Team |
| EPI_ISL_2782264 | Lighthouse Lab in Alderley Park | Wellcome Sanger Institute for the COVID-19 Genomics UK (COG-UK) Consortium | Jacquelyn Wynn, Mairead Hyland, The Lighthouse Lab in Alderley Park and Alex Alderton, Roberto Amato, Jeffrey Barrett, Sonia Goncalves, Ewan Harrison, David K. Jackson, Ian Johnston, Dominic Kwiatkowski, Cordelia Langford, John Sillitoe on behalf of the Wellcome Sanger Institute COVID-19 Surveillance Team |
| EPI_ISL_2782265 | Lighthouse Laboratory Plymouth | Wellcome Sanger Institute for the COVID-19 Genomics UK (COG-UK) Consortium | Lighthouse Laboratory Plymouth and Alex Alderton, Roberto Amato, Jeffrey Barrett, Sonia Goncalves, Ewan Harrison, David K. Jackson, Ian Johnston, Dominic Kwiatkowski, Cordelia Langford, John Sillitoe on behalf of the Wellcome Sanger Institute COVID-19 Surveillance Team |
| EPI_ISL_2782266 | Lighthouse Lab in Milton Keynes | Wellcome Sanger Institute for the COVID-19 Genomics UK (COG-UK) Consortium | The Lighthouse Lab in Milton Keynes and Alex Alderton, Roberto Amato, Jeffrey Barrett, Sonia Goncalves, Ewan Harrison, David K. Jackson, Ian Johnston, Dominic Kwiatkowski, Cordelia Langford, John Sillitoe on behalf of the Wellcome Sanger Institute COVID-19 Surveillance Team |
| EPI_ISL_2782267 | Lighthouse Lab in Alderley Park | Wellcome Sanger Institute for the COVID-19 Genomics UK (COG-UK) Consortium | Jacquelyn Wynn, Mairead Hyland, The Lighthouse Lab in Alderley Park and Alex Alderton, Roberto Amato, Jeffrey Barrett, Sonia Goncalves, Ewan Harrison, David K. Jackson, Ian Johnston, Dominic Kwiatkowski, Cordelia Langford, John Sillitoe on behalf of the Wellcome Sanger Institute COVID-19 Surveillance Team |
| EPI_ISL_2782268 | Lighthouse Lab in Milton Keynes | Wellcome Sanger Institute for the COVID-19 Genomics UK (COG-UK) Consortium | The Lighthouse Lab in Milton Keynes and Alex Alderton, Roberto Amato, Jeffrey Barrett, Sonia Goncalves, Ewan Harrison, David K. Jackson, Ian Johnston, Dominic Kwiatkowski, Cordelia Langford, John Sillitoe on behalf of the Wellcome Sanger Institute COVID-19 Surveillance Team |
| EPI_ISL_2782269, EPI_ISL_2782270 | Lighthouse Laboratory Plymouth | Wellcome Sanger Institute for the COVID-19 Genomics UK (COG-UK) Consortium | Lighthouse Laboratory Plymouth and Alex Alderton, Roberto Amato, Jeffrey Barrett, Sonia Goncalves, Ewan Harrison, David K. Jackson, Ian Johnston, Dominic Kwiatkowski, Cordelia Langford, John Sillitoe on behalf of the Wellcome Sanger Institute COVID-19 Surveillance Team |
| EPI_ISL_2782271 | Lighthouse Lab in Alderley Park | Wellcome Sanger Institute for the COVID-19 Genomics UK (COG-UK) Consortium | Jacquelyn Wynn, Mairead Hyland, The Lighthouse Lab in Alderley Park and Alex Alderton, Roberto Amato, Jeffrey Barrett, Sonia Goncalves, Ewan Harrison, David K. Jackson, Ian Johnston, Dominic Kwiatkowski, Cordelia Langford, John Sillitoe on behalf of the Wellcome Sanger Institute COVID-19 Surveillance Team |
| EPI_ISL_2782272, EPI_ISL_2782273 | Lighthouse Laboratory Plymouth | Wellcome Sanger Institute for the COVID-19 Genomics UK (COG-UK) Consortium | Lighthouse Laboratory Plymouth and Alex Alderton, Roberto Amato, Jeffrey Barrett, Sonia Goncalves, Ewan Harrison, David K. Jackson, Ian Johnston, Dominic Kwiatkowski, Cordelia Langford, John Sillitoe on behalf of the Wellcome Sanger Institute COVID-19 Surveillance Team |
| EPI_ISL_2782274 | Lighthouse Lab in Milton Keynes | Wellcome Sanger Institute for the COVID-19 Genomics UK (COG-UK) Consortium | The Lighthouse Lab in Milton Keynes and Alex Alderton, Roberto Amato, Jeffrey Barrett, Sonia Goncalves, Ewan Harrison, David K. Jackson, Ian Johnston, Dominic Kwiatkowski, Cordelia Langford, John Sillitoe on behalf of the Wellcome Sanger Institute COVID-19 Surveillance Team |
| EPI_ISL_2782275 | Lighthouse Lab in Glasgow | Wellcome Sanger Institute for the COVID-19 Genomics UK (COG-UK) Consortium | Harper VanSteenhouse, Yumi Kasai, David Gray, Carol Clugston, Anna Dominiczak and Alex Alderton, Roberto Amato, Jeffrey Barrett, Sonia Goncalves, Ewan Harrison, David K. Jackson, Ian Johnston, Dominic Kwiatkowski, Cordelia Langford, John Sillitoe on behalf of the Wellcome Sanger Institute COVID-19 Surveillance Team |
| EPI_ISL_2782276, EPI_ISL_2782277, EPI_ISL_2782278 | Lighthouse Laboratory Plymouth | Wellcome Sanger Institute for the COVID-19 Genomics UK (COG-UK) Consortium | Lighthouse Laboratory Plymouth and Alex Alderton, Roberto Amato, Jeffrey Barrett, Sonia Goncalves, Ewan Harrison, David K. Jackson, Ian Johnston, Dominic Kwiatkowski, Cordelia Langford, John Sillitoe on behalf of the Wellcome Sanger Institute COVID-19 Surveillance Team |
| EPI_ISL_2782279, EPI_ISL_2782280 | Lighthouse Lab in Milton Keynes | Wellcome Sanger Institute for the COVID-19 Genomics UK (COG-UK) Consortium | The Lighthouse Lab in Milton Keynes and Alex Alderton, Roberto Amato, Jeffrey Barrett, Sonia Goncalves, Ewan Harrison, David K. Jackson, Ian Johnston, Dominic Kwiatkowski, Cordelia Langford, John Sillitoe on behalf of the Wellcome Sanger Institute COVID-19 Surveillance Team |
| EPI_ISL_2782281 | Lighthouse Laboratory Plymouth | Wellcome Sanger Institute for the COVID-19 Genomics UK (COG-UK) Consortium | Lighthouse Laboratory Plymouth and Alex Alderton, Roberto Amato, Jeffrey Barrett, Sonia Goncalves, Ewan Harrison, David K. Jackson, Ian Johnston, Dominic Kwiatkowski, Cordelia Langford, John Sillitoe on behalf of the Wellcome Sanger Institute COVID-19 Surveillance Team |
| EPI_ISL_2782282 | Lighthouse Lab in Milton Keynes | Wellcome Sanger Institute for the COVID-19 Genomics UK (COG-UK) Consortium | The Lighthouse Lab in Milton Keynes and Alex Alderton, Roberto Amato, Jeffrey Barrett, Sonia Goncalves, Ewan Harrison, David K. Jackson, Ian Johnston, Dominic Kwiatkowski, Cordelia Langford, John Sillitoe on behalf of the Wellcome Sanger Institute COVID-19 Surveillance Team |
| EPI_ISL_2782283 | Lighthouse Laboratory Plymouth | Wellcome Sanger Institute for the COVID-19 Genomics UK (COG-UK) Consortium | Lighthouse Laboratory Plymouth and Alex Alderton, Roberto Amato, Jeffrey Barrett, Sonia Goncalves, Ewan Harrison, David K. Jackson, Ian Johnston, Dominic Kwiatkowski, Cordelia Langford, John Sillitoe on behalf of the Wellcome Sanger Institute COVID-19 Surveillance Team |
| EPI_ISL_2782284 | Lighthouse Lab in Glasgow | Wellcome Sanger Institute for the COVID-19 Genomics UK (COG-UK) Consortium | Harper VanSteenhouse, Yumi Kasai, David Gray, Carol Clugston, Anna Dominiczak and Alex Alderton, Roberto Amato, Jeffrey Barrett, Sonia Goncalves, Ewan Harrison, David K. Jackson, Ian Johnston, Dominic Kwiatkowski, Cordelia Langford, John Sillitoe on behalf of the Wellcome Sanger Institute COVID-19 Surveillance Team |
| EPI_ISL_2782285, EPI_ISL_2782286 | Lighthouse Laboratory Plymouth | Wellcome Sanger Institute for the COVID-19 Genomics UK (COG-UK) Consortium | Lighthouse Laboratory Plymouth and Alex Alderton, Roberto Amato, Jeffrey Barrett, Sonia Goncalves, Ewan Harrison, David K. Jackson, Ian Johnston, Dominic Kwiatkowski, Cordelia Langford, John Sillitoe on behalf of the Wellcome Sanger Institute COVID-19 Surveillance Team |
| EPI_ISL_2782287 | Lighthouse Lab in Alderley Park | Wellcome Sanger Institute for the COVID-19 Genomics UK (COG-UK) Consortium | Jacquelyn Wynn, Mairead Hyland, The Lighthouse Lab in Alderley Park and Alex Alderton, Roberto Amato, Jeffrey Barrett, Sonia Goncalves, Ewan Harrison, David K. Jackson, Ian Johnston, Dominic Kwiatkowski, Cordelia Langford, John Sillitoe on behalf of the Wellcome Sanger Institute COVID-19 Surveillance Team |
| EPI_ISL_2782288 | Lighthouse Laboratory Plymouth | Wellcome Sanger Institute for the COVID-19 Genomics UK (COG-UK) Consortium | Lighthouse Laboratory Plymouth and Alex Alderton, Roberto Amato, Jeffrey Barrett, Sonia Goncalves, Ewan Harrison, David K. Jackson, Ian Johnston, Dominic Kwiatkowski, Cordelia Langford, John Sillitoe on behalf of the Wellcome Sanger Institute COVID-19 Surveillance Team |
| EPI_ISL_2782289 | Lighthouse Lab in Alderley Park | Wellcome Sanger Institute for the COVID-19 Genomics UK (COG-UK) Consortium | Jacquelyn Wynn, Mairead Hyland, The Lighthouse Lab in Alderley Park and Alex Alderton, Roberto Amato, Jeffrey Barrett, Sonia Goncalves, Ewan Harrison, David K. Jackson, Ian Johnston, Dominic Kwiatkowski, Cordelia Langford, John Sillitoe on behalf of the Wellcome Sanger Institute COVID-19 Surveillance Team |
| EPI_ISL_2782290 | Lighthouse Lab in Milton Keynes | Wellcome Sanger Institute for the COVID-19 Genomics UK (COG-UK) Consortium | The Lighthouse Lab in Milton Keynes and Alex Alderton, Roberto Amato, Jeffrey Barrett, Sonia Goncalves, Ewan Harrison, David K. Jackson, Ian Johnston, Dominic Kwiatkowski, Cordelia Langford, John Sillitoe on behalf of the Wellcome Sanger Institute COVID-19 Surveillance Team |
| EPI_ISL_2782291 | Lighthouse Lab in Alderley Park | Wellcome Sanger Institute for the COVID-19 Genomics UK (COG-UK) Consortium | Jacquelyn Wynn, Mairead Hyland, The Lighthouse Lab in Alderley Park and Alex Alderton, Roberto Amato, Jeffrey Barrett, Sonia Goncalves, Ewan Harrison, David K. Jackson, Ian Johnston, Dominic Kwiatkowski, Cordelia Langford, John Sillitoe on behalf of the Wellcome Sanger Institute COVID-19 Surveillance Team |
| EPI_ISL_2782292 | Lighthouse Laboratory Plymouth | Wellcome Sanger Institute for the COVID-19 Genomics UK (COG-UK) Consortium | Lighthouse Laboratory Plymouth and Alex Alderton, Roberto Amato, Jeffrey Barrett, Sonia Goncalves, Ewan Harrison, David K. Jackson, Ian Johnston, Dominic Kwiatkowski, Cordelia Langford, John Sillitoe on behalf of the Wellcome Sanger Institute COVID-19 Surveillance Team |
| EPI_ISL_2782293 | Lighthouse Lab in Glasgow | Wellcome Sanger Institute for the COVID-19 Genomics UK (COG-UK) Consortium | Harper VanSteenhouse, Yumi Kasai, David Gray, Carol Clugston, Anna Dominiczak and Alex Alderton, Roberto Amato, Jeffrey Barrett, Sonia Goncalves, Ewan Harrison, David K. Jackson, Ian Johnston, Dominic Kwiatkowski, Cordelia Langford, John Sillitoe on behalf of the Wellcome Sanger Institute COVID-19 Surveillance Team |
| EPI_ISL_2782294 | Lighthouse Lab in Milton Keynes | Wellcome Sanger Institute for the COVID-19 Genomics UK (COG-UK) Consortium | The Lighthouse Lab in Milton Keynes and Alex Alderton, Roberto Amato, Jeffrey Barrett, Sonia Goncalves, Ewan Harrison, David K. Jackson, Ian Johnston, Dominic Kwiatkowski, Cordelia Langford, John Sillitoe on behalf of the Wellcome Sanger Institute COVID-19 Surveillance Team |
| EPI_ISL_2782295 | Lighthouse Lab in Alderley Park | Wellcome Sanger Institute for the COVID-19 Genomics UK (COG-UK) Consortium | Jacquelyn Wynn, Mairead Hyland, The Lighthouse Lab in Alderley Park and Alex Alderton, Roberto Amato, Jeffrey Barrett, Sonia Goncalves, Ewan Harrison, David K. Jackson, Ian Johnston, Dominic Kwiatkowski, Cordelia Langford, John Sillitoe on behalf of the Wellcome Sanger Institute COVID-19 Surveillance Team |
| EPI_ISL_2782296 | Lighthouse Laboratory Plymouth | Wellcome Sanger Institute for the COVID-19 Genomics UK (COG-UK) Consortium | Lighthouse Laboratory Plymouth and Alex Alderton, Roberto Amato, Jeffrey Barrett, Sonia Goncalves, Ewan Harrison, David K. Jackson, Ian Johnston, Dominic Kwiatkowski, Cordelia Langford, John Sillitoe on behalf of the Wellcome Sanger Institute COVID-19 Surveillance Team |
| EPI_ISL_2782297, EPI_ISL_2782298 | Lighthouse Lab in Milton Keynes | Wellcome Sanger Institute for the COVID-19 Genomics UK (COG-UK) Consortium | The Lighthouse Lab in Milton Keynes and Alex Alderton, Roberto Amato, Jeffrey Barrett, Sonia Goncalves, Ewan Harrison, David K. Jackson, Ian Johnston, Dominic Kwiatkowski, Cordelia Langford, John Sillitoe on behalf of the Wellcome Sanger Institute COVID-19 Surveillance Team |

| | | | |
|---|---|---|---|
| EPI_ISL_2782299 | Lighthouse Laboratory Plymouth | Wellcome Sanger Institute for the COVID-19 Genomics UK (COG-UK) Consortium | Lighthouse Laboratory Plymouth and Alex Alderton, Roberto Amato, Jeffrey Barrett, Sonia Goncalves, Ewan Harrison, David K. Jackson, Ian Johnston, Dominic Kwiatkowski, Cordelia Langford, John Sillitoe on behalf of the Wellcome Sanger Institute COVID-19 Surveillance Team |
| EPI_ISL_2782300 | Lighthouse Lab in Alderley Park | Wellcome Sanger Institute for the COVID-19 Genomics UK (COG-UK) Consortium | Jacquelyn Wynn, Mairead Hyland, The Lighthouse Lab in Alderley Park and Alex Alderton, Roberto Amato, Jeffrey Barrett, Sonia Goncalves, Ewan Harrison, David K. Jackson, Ian Johnston, Dominic Kwiatkowski, Cordelia Langford, John Sillitoe on behalf of the Wellcome Sanger Institute COVID-19 Surveillance Team |
| EPI_ISL_2782301 | Lighthouse Lab in Milton Keynes | Wellcome Sanger Institute for the COVID-19 Genomics UK (COG-UK) Consortium | The Lighthouse Lab in Milton Keynes and Alex Alderton, Roberto Amato, Jeffrey Barrett, Sonia Goncalves, Ewan Harrison, David K. Jackson, Ian Johnston, Dominic Kwiatkowski, Cordelia Langford, John Sillitoe on behalf of the Wellcome Sanger Institute COVID-19 Surveillance Team |
| EPI_ISL_2782302, EPI_ISL_2782303 | Lighthouse Lab in Alderley Park | Wellcome Sanger Institute for the COVID-19 Genomics UK (COG-UK) Consortium | Jacquelyn Wynn, Mairead Hyland, The Lighthouse Lab in Alderley Park and Alex Alderton, Roberto Amato, Jeffrey Barrett, Sonia Goncalves, Ewan Harrison, David K. Jackson, Ian Johnston, Dominic Kwiatkowski, Cordelia Langford, John Sillitoe on behalf of the Wellcome Sanger Institute COVID-19 Surveillance Team |
| EPI_ISL_2782304, EPI_ISL_2782305 | Lighthouse Laboratory Plymouth | Wellcome Sanger Institute for the COVID-19 Genomics UK (COG-UK) Consortium | Lighthouse Laboratory Plymouth and Alex Alderton, Roberto Amato, Jeffrey Barrett, Sonia Goncalves, Ewan Harrison, David K. Jackson, Ian Johnston, Dominic Kwiatkowski, Cordelia Langford, John Sillitoe on behalf of the Wellcome Sanger Institute COVID-19 Surveillance Team |
| EPI_ISL_2782306, EPI_ISL_2782307, EPI_ISL_2782308 | Lighthouse Lab in Milton Keynes | Wellcome Sanger Institute for the COVID-19 Genomics UK (COG-UK) Consortium | The Lighthouse Lab in Milton Keynes and Alex Alderton, Roberto Amato, Jeffrey Barrett, Sonia Goncalves, Ewan Harrison, David K. Jackson, Ian Johnston, Dominic Kwiatkowski, Cordelia Langford, John Sillitoe on behalf of the Wellcome Sanger Institute COVID-19 Surveillance Team |
| EPI_ISL_2782309, EPI_ISL_2782310, EPI_ISL_2782311 | Lighthouse Lab in Alderley Park | Wellcome Sanger Institute for the COVID-19 Genomics UK (COG-UK) Consortium | Jacquelyn Wynn, Mairead Hyland, The Lighthouse Lab in Alderley Park and Alex Alderton, Roberto Amato, Jeffrey Barrett, Sonia Goncalves, Ewan Harrison, David K. Jackson, Ian Johnston, Dominic Kwiatkowski, Cordelia Langford, John Sillitoe on behalf of the Wellcome Sanger Institute COVID-19 Surveillance Team |
| EPI_ISL_2782312 | Lighthouse Laboratory Plymouth | Wellcome Sanger Institute for the COVID-19 Genomics UK (COG-UK) Consortium | Lighthouse Laboratory Plymouth and Alex Alderton, Roberto Amato, Jeffrey Barrett, Sonia Goncalves, Ewan Harrison, David K. Jackson, Ian Johnston, Dominic Kwiatkowski, Cordelia Langford, John Sillitoe on behalf of the Wellcome Sanger Institute COVID-19 Surveillance Team |
| EPI_ISL_2782313 | Lighthouse Lab in Milton Keynes | Wellcome Sanger Institute for the COVID-19 Genomics UK (COG-UK) Consortium | The Lighthouse Lab in Milton Keynes and Alex Alderton, Roberto Amato, Jeffrey Barrett, Sonia Goncalves, Ewan Harrison, David K. Jackson, Ian Johnston, Dominic Kwiatkowski, Cordelia Langford, John Sillitoe on behalf of the Wellcome Sanger Institute COVID-19 Surveillance Team |
| EPI_ISL_2782318, EPI_ISL_2782319, EPI_ISL_2782320, EPI_ISL_2782322, EPI_ISL_2782323, EPI_ISL_2782326, EPI_ISL_2782329, EPI_ISL_2782330, EPI_ISL_2782333, EPI_ISL_2782336, EPI_ISL_2782339, EPI_ISL_2782341, EPI_ISL_2782344, EPI_ISL_2782346, EPI_ISL_2782358, EPI_ISL_2782359, EPI_ISL_2782361, EPI_ISL_2782364, EPI_ISL_2782366, EPI_ISL_2782370, EPI_ISL_2782376, EPI_ISL_2782378, EPI_ISL_2782380 | | | |
| see above | Hospital General Universitario Gregorio Marañón | Hospital General Universitario Gregorio Marañón | Sergio Buenestado Serrano, Pedro Sola Campoy, Laura Pérez-Lago, Cristina Rodriguez-Grande, Marta Herranz Martin, Victor Manuel de la Cueva, Julia Suárez, Pilar Catalán, Patricia Muñoz, Darío García de Viedma |
| EPI_ISL_2783636, EPI_ISL_2783656, EPI_ISL_2783657, EPI_ISL_2783662, EPI_ISL_2783736, EPI_ISL_2783751, EPI_ISL_2783792, EPI_ISL_2783795, EPI_ISL_2783859, EPI_ISL_2783873, EPI_ISL_2783903, EPI_ISL_2783926, EPI_ISL_2783976, EPI_ISL_2783979, EPI_ISL_2783981, EPI_ISL_2783982, EPI_ISL_2783985, EPI_ISL_2784026, EPI_ISL_2784109, EPI_ISL_2784123, EPI_ISL_2784131, EPI_ISL_2784163, EPI_ISL_2784169, EPI_ISL_2784173, EPI_ISL_2784188, EPI_ISL_2784193, EPI_ISL_2784199, EPI_ISL_2784203, EPI_ISL_2784207, EPI_ISL_2784210, EPI_ISL_2784211, EPI_ISL_2784216, EPI_ISL_2784217, EPI_ISL_2784225, EPI_ISL_2784228, EPI_ISL_2784232, EPI_ISL_2784234, EPI_ISL_2784243, EPI_ISL_2784246, EPI_ISL_2784247, EPI_ISL_2784249, EPI_ISL_2784250, EPI_ISL_2784252, EPI_ISL_2784254, EPI_ISL_2784265, EPI_ISL_2784267, EPI_ISL_2784268, EPI_ISL_2784270, EPI_ISL_2784274, EPI_ISL_2784277, EPI_ISL_2784278, EPI_ISL_2784280, EPI_ISL_2784282, EPI_ISL_2784288, EPI_ISL_2784290, EPI_ISL_2784291, EPI_ISL_2784301, EPI_ISL_2784302, EPI_ISL_2784304, EPI_ISL_2784316, EPI_ISL_2784319, EPI_ISL_2784326 | | | |
| see above | Curative Labs | Curative Labs | Mikhail Hanewich-Hollatz, J.Cesar Ignacio-Espinoza, Elias L. Salfati, Nabjot Sandhu, George Way, Zhiyi Xie, Eugenia Khorosheva, Sophia Quasem, Janet Chen, Vladimir Slepnev |
| EPI_ISL_2784330, EPI_ISL_2784331, EPI_ISL_2784332, EPI_ISL_2784333, EPI_ISL_2784334, EPI_ISL_2784335, EPI_ISL_2784336, EPI_ISL_2784337, EPI_ISL_2784338, EPI_ISL_2784339, EPI_ISL_2784340, EPI_ISL_2784341, EPI_ISL_2784342, EPI_ISL_2784343 | | | |
| see above | Tokyo Metropolitan Institute of Public Health | Tokyo Metropolitan Institute of Public Health | Masaki Hayashi, Takako Yamazaki, Akane Negishi,Hirofumi Miyake, Kenshirou Kuroki, Maya Isogai, Takushi Fujiwara, Mami Nagashima, Jun Suzuki, Kenji Sadamasu |
| EPI_ISL_2784344 | PathWest Laboratory Medicine WA | PathWest Laboratory Medicine WA Microbial Surveillance Unit | PathWest Laboratory Medicine WA Microbial Surveillance Unit |
| EPI_ISL_2784356, EPI_ISL_2784357, EPI_ISL_2784358, EPI_ISL_2784360, EPI_ISL_2784361, EPI_ISL_2784362, EPI_ISL_2784363, EPI_ISL_2784364, EPI_ISL_2784367, EPI_ISL_2784369, EPI_ISL_2784370, EPI_ISL_2784372, EPI_ISL_2784374, EPI_ISL_2784375, EPI_ISL_2784378, EPI_ISL_2784380, EPI_ISL_2784382, EPI_ISL_2784383, EPI_ISL_2784384, EPI_ISL_2784388, EPI_ISL_2784389, EPI_ISL_2784390, EPI_ISL_2784392, EPI_ISL_2784393 | | | |
| see above | Department of Bacteria, Parasites and Fungi, Statens Serum Institut, Copenhagen, Denmark | Statens Serum Institut Bioinformatics and Microbial Genomics | Danish Covid-19 Genome Consortium |
| EPI_ISL_2784394 | Department of Clinical Microbiology, Odense University Hospital, Odense, Denmark | Statens Serum Institut Bioinformatics and Microbial Genomics | Danish Covid-19 Genome Consortium |
| EPI_ISL_2784395, EPI_ISL_2784396, EPI_ISL_2784399, EPI_ISL_2784403, EPI_ISL_2784404, EPI_ISL_2784406, EPI_ISL_2784407, EPI_ISL_2784408, EPI_ISL_2784409, EPI_ISL_2784410, EPI_ISL_2784412, EPI_ISL_2784415, EPI_ISL_2784418, EPI_ISL_2784421, EPI_ISL_2784422, EPI_ISL_2784425, EPI_ISL_2784427, EPI_ISL_2784428, EPI_ISL_2784430, EPI_ISL_2784432, EPI_ISL_2784434, EPI_ISL_2784435, EPI_ISL_2784437, EPI_ISL_2784438, EPI_ISL_2784439, EPI_ISL_2784440, EPI_ISL_2784441, EPI_ISL_2784444, EPI_ISL_2784447, EPI_ISL_2784448, EPI_ISL_2784449, EPI_ISL_2784450, EPI_ISL_2784451, EPI_ISL_2784455, EPI_ISL_2784456, EPI_ISL_2784457, EPI_ISL_2784460, EPI_ISL_2784463, EPI_ISL_2784464, EPI_ISL_2784465, EPI_ISL_2784468, EPI_ISL_2784469, EPI_ISL_2784470, EPI_ISL_2784471, EPI_ISL_2784472, EPI_ISL_2784474, EPI_ISL_2784475, EPI_ISL_2784479, EPI_ISL_2784483, EPI_ISL_2784485, EPI_ISL_2784487 | | | |
| see above | Department of Bacteria, Parasites and Fungi, Statens Serum Institut, Copenhagen, Denmark | Statens Serum Institut Bioinformatics and Microbial Genomics | Danish Covid-19 Genome Consortium |
| EPI_ISL_2784491 | Molekylær Medicinsk Afdeling, Aarhus University Hospital, Aarhus, Denmark | Statens Serum Institut Bioinformatics and Microbial Genomics | Danish Covid-19 Genome Consortium |
| EPI_ISL_2784493, EPI_ISL_2784495, EPI_ISL_2784497, EPI_ISL_2784498, EPI_ISL_2784499, EPI_ISL_2784501, EPI_ISL_2784505, EPI_ISL_2784507, EPI_ISL_2784508, EPI_ISL_2784509, EPI_ISL_2784510, EPI_ISL_2784514, EPI_ISL_2784516, EPI_ISL_2784519, EPI_ISL_2784521, EPI_ISL_2784522, EPI_ISL_2784523, EPI_ISL_2784525, EPI_ISL_2784526, EPI_ISL_2784527, EPI_ISL_2784529, EPI_ISL_2784530, EPI_ISL_2784534, EPI_ISL_2784535, EPI_ISL_2784536, EPI_ISL_2784537, EPI_ISL_2784541, EPI_ISL_2784542, EPI_ISL_2784548, EPI_ISL_2784549, EPI_ISL_2784550, EPI_ISL_2784553, EPI_ISL_2784557, EPI_ISL_2784558, EPI_ISL_2784561, EPI_ISL_2784565, EPI_ISL_2784566, EPI_ISL_2784569, EPI_ISL_2784570, EPI_ISL_2784571, EPI_ISL_2784572, EPI_ISL_2784573, EPI_ISL_2784574, EPI_ISL_2784576, EPI_ISL_2784577, EPI_ISL_2784578, EPI_ISL_2784580, EPI_ISL_2784582, EPI_ISL_2784583, EPI_ISL_2784584, EPI_ISL_2784587, EPI_ISL_2784589, EPI_ISL_2784590, EPI_ISL_2784591, EPI_ISL_2784594, EPI_ISL_2784595, EPI_ISL_2784596, EPI_ISL_2784597, EPI_ISL_2784598, EPI_ISL_2784599, EPI_ISL_2784600, EPI_ISL_2784602, EPI_ISL_2784607, EPI_ISL_2784609, EPI_ISL_2784611, EPI_ISL_2784612, EPI_ISL_2784615, EPI_ISL_2784620, EPI_ISL_2784621, EPI_ISL_2784626, EPI_ISL_2784629, EPI_ISL_2784632, EPI_ISL_2784634, EPI_ISL_2784635, EPI_ISL_2784636, EPI_ISL_2784637, EPI_ISL_2784638, EPI_ISL_2784641, EPI_ISL_2784645, EPI_ISL_2784647, EPI_ISL_2784648, EPI_ISL_2784649, EPI_ISL_2784650, EPI_ISL_2784653, EPI_ISL_2784655, EPI_ISL_2784658, EPI_ISL_2784661, EPI_ISL_2784662, EPI_ISL_2784666, EPI_ISL_2784667, EPI_ISL_2784668, EPI_ISL_2784670, EPI_ISL_2784672, EPI_ISL_2784674, EPI_ISL_2784678, EPI_ISL_2784681, EPI_ISL_2784682, EPI_ISL_2784683 | | | |
| see above | Department of Bacteria, Parasites and Fungi, Statens Serum Institut, Copenhagen, Denmark | Statens Serum Institut Bioinformatics and Microbial Genomics | Danish Covid-19 Genome Consortium |
| EPI_ISL_2784684 | S Camillo | National Institute for Infectious Diseases (INMI) L. Spallanzani I.R.C.C.S | E Giombini, B Bartolini, M Rueca, G Bonfiglio, O Butera, CEM Gruber, F Santini, F Messina, A Di Caro, MR Capobianchi |
| EPI_ISL_2784685 | RM3 | National Institute for Infectious Diseases (INMI) L. Spallanzani I.R.C.C.S | E Giombini, B Bartolini, M Rueca, G Bonfiglio, O Butera, CEM Gruber, F Santini, F Messina, A Di Caro, MR Capobianchi |
| EPI_ISL_2784686 | Ifo Regina Elena | National Institute for Infectious Diseases (INMI) L. Spallanzani I.R.C.C.S | E Giombini, B Bartolini, M Rueca, G Bonfiglio, O Butera, CEM Gruber, F Santini, F Messina, A Di Caro, MR Capobianchi |
| EPI_ISL_2784687 | S Camillo | National Institute for Infectious Diseases (INMI) L. Spallanzani I.R.C.C.S | E Giombini, B Bartolini, M Rueca, G Bonfiglio, O Butera, CEM Gruber, F Santini, F Messina, A Di Caro, MR Capobianchi |
| EPI_ISL_2784688, EPI_ISL_2784691, EPI_ISL_2784693 | S Filippo Neri | National Institute for Infectious Diseases (INMI) L. Spallanzani I.R.C.C.S | E Giombini, B Bartolini, M Rueca, G Bonfiglio, O Butera, CEM Gruber, F Santini, F Messina, A Di Caro, MR Capobianchi |
| EPI_ISL_2784722 | National Platform bis UMONS/Jolimont | National Platform bis UMONS/Jolimont | François Dufrasne, Guillaume Bayon-Vicente, Florian Juszczak, Eric Tarantino, Gautier Detry, Ruddy Wattiez |
| EPI_ISL_2784728, EPI_ISL_2784733, EPI_ISL_2784734, EPI_ISL_2784735, EPI_ISL_2784736, EPI_ISL_2784737, EPI_ISL_2784739, EPI_ISL_2784740, EPI_ISL_2784743, EPI_ISL_2784746, EPI_ISL_2784752, EPI_ISL_2784753, EPI_ISL_2784757, EPI_ISL_2784758, EPI_ISL_2784760, EPI_ISL_2784762, EPI_ISL_2784768, EPI_ISL_2784769, EPI_ISL_2784770, EPI_ISL_2784772, EPI_ISL_2784774, EPI_ISL_2784775, EPI_ISL_2784778, EPI_ISL_2784779, EPI_ISL_2784780, EPI_ISL_2784781, EPI_ISL_2784783, EPI_ISL_2784786, EPI_ISL_2784787, EPI_ISL_2784788, EPI_ISL_2784789, EPI_ISL_2784790, EPI_ISL_2784797, EPI_ISL_2784799, EPI_ISL_2784801, EPI_ISL_2784802, EPI_ISL_2784804, EPI_ISL_2784807, EPI_ISL_2784809, EPI_ISL_2784810, EPI_ISL_2784811, EPI_ISL_2784814, EPI_ISL_2784816, EPI_ISL_2784818, EPI_ISL_2784819, EPI_ISL_2784821, EPI_ISL_2784822, EPI_ISL_2784823, EPI_ISL_2784826, EPI_ISL_2784829, EPI_ISL_2784832, EPI_ISL_2784836, EPI_ISL_2784840, EPI_ISL_2784841, EPI_ISL_2784842, EPI_ISL_2784844, EPI_ISL_2784847, EPI_ISL_2784857, EPI_ISL_2784863, EPI_ISL_2784864, EPI_ISL_2784866, EPI_ISL_2784871, EPI_ISL_2784872, EPI_ISL_2784874, EPI_ISL_2784875, EPI_ISL_2784878, EPI_ISL_2784879, EPI_ISL_2784883, EPI_ISL_2784885, EPI_ISL_2784886, EPI_ISL_2784890, EPI_ISL_2784891, EPI_ISL_2784893, EPI_ISL_2784894, EPI_ISL_2784898, EPI_ISL_2784899, EPI_ISL_2784903, EPI_ISL_2784905, EPI_ISL_2784906, EPI_ISL_2784907, EPI_ISL_2784908, EPI_ISL_2784912, EPI_ISL_2784913, EPI_ISL_2784915, EPI_ISL_2784920, EPI_ISL_2784922, EPI_ISL_2784924, EPI_ISL_2784926, EPI_ISL_2784930, EPI_ISL_2784932, EPI_ISL_2784934, EPI_ISL_2784938, EPI_ISL_2784941, EPI_ISL_2784945, EPI_ISL_2784947, EPI_ISL_2784948, EPI_ISL_2784949, EPI_ISL_2784950, EPI_ISL_2784955, EPI_ISL_2784958, EPI_ISL_2784962, EPI_ISL_2784963, EPI_ISL_2784966, EPI_ISL_2784968, EPI_ISL_2784969, EPI_ISL_2784970, EPI_ISL_2784971, EPI_ISL_2784972, EPI_ISL_2784976, EPI_ISL_2784978, EPI_ISL_2784986, EPI_ISL_2784987, EPI_ISL_2784992, EPI_ISL_2784998, EPI_ISL_2784999, EPI_ISL_2785000, EPI_ISL_2785004, EPI_ISL_2785006, EPI_ISL_2785007, EPI_ISL_2785009, EPI_ISL_2785014, EPI_ISL_2785019, EPI_ISL_2785020, EPI_ISL_2785022, EPI_ISL_2785024, EPI_ISL_2785026, EPI_ISL_2785029, EPI_ISL_2785030, EPI_ISL_2785033, EPI_ISL_2785037, EPI_ISL_2785053, EPI_ISL_2785054, EPI_ISL_2785056, EPI_ISL_2785057, EPI_ISL_2785059, EPI_ISL_2785060, EPI_ISL_2785062, EPI_ISL_2785063, EPI_ISL_2785064, EPI_ISL_2785068, EPI_ISL_2785071, EPI_ISL_2785073, EPI_ISL_2785076, EPI_ISL_2785078, EPI_ISL_2785080, EPI_ISL_2785081, EPI_ISL_2785086, EPI_ISL_2785087, EPI_ISL_2785093, EPI_ISL_2785094, EPI_ISL_2785095, EPI_ISL_2785096, EPI_ISL_2785104, EPI_ISL_2785105, EPI_ISL_2785106, EPI_ISL_2785107, EPI_ISL_2785110, EPI_ISL_2785113, EPI_ISL_2785114, EPI_ISL_2785118, EPI_ISL_2785121, EPI_ISL_2785124, EPI_ISL_2785128, EPI_ISL_2785132, EPI_ISL_2785133, EPI_ISL_2785134, EPI_ISL_2785140, EPI_ISL_2785145, EPI_ISL_2785147, EPI_ISL_2785148, EPI_ISL_2785149, EPI_ISL_2785150, EPI_ISL_2785151, EPI_ISL_2785153, EPI_ISL_2785154, EPI_ISL_2785155, EPI_ISL_2785156, EPI_ISL_2785158, EPI_ISL_2785161, EPI_ISL_2785163, EPI_ISL_2785165, EPI_ISL_2785169, EPI_ISL_2785171, EPI_ISL_2785172, EPI_ISL_2785173, EPI_ISL_2785177, EPI_ISL_2785178, EPI_ISL_2785179, EPI_ISL_2785182, EPI_ISL_2785184, EPI_ISL_2785187, EPI_ISL_2785188, EPI_ISL_2785189, EPI_ISL_2785190, EPI_ISL_2785191, EPI_ISL_2785192, EPI_ISL_2785194, EPI_ISL_2785203, EPI_ISL_2785205, EPI_ISL_2785207, EPI_ISL_2785208, EPI_ISL_2785209, EPI_ISL_2785211, EPI_ISL_2785213, EPI_ISL_2785217, EPI_ISL_2785219, EPI_ISL_2785222, EPI_ISL_2785223, EPI_ISL_2785224, EPI_ISL_2785225, EPI_ISL_2785232, EPI_ISL_2785235, EPI_ISL_2785239, EPI_ISL_2785243, EPI_ISL_2785244, EPI_ISL_2785246, EPI_ISL_2785250, EPI_ISL_2785255, EPI_ISL_2785256, EPI_ISL_2785259, EPI_ISL_2785261, EPI_ISL_2785262, EPI_ISL_2785263, EPI_ISL_2785266, EPI_ISL_2785270, EPI_ISL_2785277, EPI_ISL_2785279, EPI_ISL_2785281, EPI_ISL_2785282, EPI_ISL_2785284, EPI_ISL_2785288, EPI_ISL_2785289, EPI_ISL_2785292, EPI_ISL_2785293, EPI_ISL_2785304, EPI_ISL_2785307, EPI_ISL_2785308, EPI_ISL_2785312, EPI_ISL_2785314, EPI_ISL_2785321, EPI_ISL_2785328, EPI_ISL_2785336, EPI_ISL_2785337, EPI_ISL_2785340, EPI_ISL_2785345, EPI_ISL_2785346, EPI_ISL_2785347, EPI_ISL_2785348, EPI_ISL_2785351, EPI_ISL_2785352, EPI_ISL_2785357, EPI_ISL_2785378, EPI_ISL_2785380, EPI_ISL_2785386, EPI_ISL_2785387, EPI_ISL_2785388, EPI_ISL_2785389, EPI_ISL_2785390, EPI_ISL_2785391, EPI_ISL_2785395, EPI_ISL_2785401, EPI_ISL_2785406, EPI_ISL_2785407, EPI_ISL_2785409, EPI_ISL_2785410, EPI_ISL_2785412, EPI_ISL_2785416, EPI_ISL_2785417, EPI_ISL_2785426, EPI_ISL_2785431, EPI_ISL_2785432, EPI_ISL_2785446, EPI_ISL_2785447, EPI_ISL_2785448, EPI_ISL_2785449, EPI_ISL_2785450, EPI_ISL_2785455, EPI_ISL_2785460, EPI_ISL_2785462, EPI_ISL_2785467, EPI_ISL_2785480, EPI_ISL_2785483, EPI_ISL_2785501, EPI_ISL_2785503, EPI_ISL_2785505, EPI_ISL_2785506, EPI_ISL_2785517, EPI_ISL_2785518 | | | |
| see above | Aegis Sciences Corporation | Centers for Disease Control and Prevention Division of Viral Diseases, Pathogen Discovery | Dakota Howard, Dhwani Batra, Peter W. Cook, Kara Moser, Adrian Paskey, Jason Caravas, Benjamin Rambo-Martin, Shatavia Morrison, Christopher Gulvick, Scott Sammons, Yvette Unoarumhi, Darlene Wagner, Matthew Schmerer, Cyndi Clark, Patrick Campbell, Rob Case, Vikramsinha Ghorpade, Holly Houdeshell, Ola Kvalvaag, Dillon Nall, Ethan Sanders, Alec Vest, Shaun Westlund, Matthew Hardison, Clinton R. Paden, Duncan MacCannell |
| EPI_ISL_2785526, EPI_ISL_2785531, EPI_ISL_2785532, EPI_ISL_2785533, EPI_ISL_2785535, EPI_ISL_2785536, EPI_ISL_2785538, EPI_ISL_2785539, EPI_ISL_2785540, EPI_ISL_2785542, EPI_ISL_2785543, EPI_ISL_2785545, EPI_ISL_2785546, EPI_ISL_2785547, EPI_ISL_2785549, EPI_ISL_2785550, EPI_ISL_2785552, EPI_ISL_2785553, EPI_ISL_2785554, EPI_ISL_2785557, EPI_ISL_2785558, EPI_ISL_2785561, EPI_ISL_2785564, EPI_ISL_2785565, EPI_ISL_2785566, EPI_ISL_2785567, EPI_ISL_2785568, EPI_ISL_2785569, EPI_ISL_2785570, EPI_ISL_2785574, EPI_ISL_2785575, EPI_ISL_2785576, EPI_ISL_2785577, EPI_ISL_2785579, EPI_ISL_2785580, EPI_ISL_2785581, EPI_ISL_2785582, EPI_ISL_2785585, EPI_ISL_2785587, EPI_ISL_2785588, EPI_ISL_2785589, EPI_ISL_2785590, EPI_ISL_2785591, EPI_ISL_2785593, EPI_ISL_2785594, EPI_ISL_2785596, EPI_ISL_2785599, EPI_ISL_2785602, EPI_ISL_2785603, EPI_ISL_2785605, EPI_ISL_2785606, EPI_ISL_2785607, EPI_ISL_2785608, EPI_ISL_2785610, EPI_ISL_2785611, EPI_ISL_2785613, EPI_ISL_2785616, EPI_ISL_2785618, EPI_ISL_2785621, EPI_ISL_2785622, EPI_ISL_2785623, EPI_ISL_2785625, EPI_ISL_2785627, EPI_ISL_2785628, EPI_ISL_2785629, EPI_ISL_2785630, EPI_ISL_2785637, EPI_ISL_2785639, EPI_ISL_2785640, EPI_ISL_2785645, EPI_ISL_2785649, EPI_ISL_2785650, EPI_ISL_2785652, EPI_ISL_2785654, EPI_ISL_2785655, EPI_ISL_2785659, EPI_ISL_2785660, EPI_ISL_2785664, EPI_ISL_2785665, EPI_ISL_2785667, EPI_ISL_2785668, EPI_ISL_2785669, EPI_ISL_2785672, EPI_ISL_2785674, EPI_ISL_2785675, EPI_ISL_2785676, EPI_ISL_2785683, EPI_ISL_2785685, EPI_ISL_2785686, EPI_ISL_2785688, EPI_ISL_2785690, EPI_ISL_2785693, EPI_ISL_2785699, EPI_ISL_2785701, EPI_ISL_2785707, EPI_ISL_2785708, EPI_ISL_2785709, EPI_ISL_2785711, EPI_ISL_2785713, EPI_ISL_2785714, EPI_ISL_2785715, EPI_ISL_2785717, EPI_ISL_2785718, EPI_ISL_2785719, EPI_ISL_2785720, EPI_ISL_2785721, EPI_ISL_2785722, EPI_ISL_2785723, EPI_ISL_2785724, EPI_ISL_2785725, EPI_ISL_2785727, EPI_ISL_2785731, EPI_ISL_2785732, EPI_ISL_2785733, EPI_ISL_2785734, EPI_ISL_2785739, EPI_ISL_2785740, EPI_ISL_2785742, EPI_ISL_2785745, EPI_ISL_2785746, EPI_ISL_2785748, EPI_ISL_2785751, EPI_ISL_2785753, EPI_ISL_2785754, EPI_ISL_2785756, EPI_ISL_2785758, EPI_ISL_2785759, EPI_ISL_2785760, EPI_ISL_2785762, EPI_ISL_2785763, EPI_ISL_2785765, EPI_ISL_2785766, EPI_ISL_2785767, EPI_ISL_2785768, EPI_ISL_2785769, EPI_ISL_2785771, EPI_ISL_2785772, EPI_ISL_2785775, EPI_ISL_2785776, EPI_ISL_2785778, EPI_ISL_2785782, EPI_ISL_2785784, EPI_ISL_2785785, EPI_ISL_2785786, EPI_ISL_2785787, EPI_ISL_2785788, EPI_ISL_2785790, EPI_ISL_2785791, EPI_ISL_2785792, EPI_ISL_2785793, EPI_ISL_2785794, EPI_ISL_2785795, EPI_ISL_2785797, EPI_ISL_2785798, EPI_ISL_2785799, EPI_ISL_2785800, EPI_ISL_2785801, EPI_ISL_2785805, EPI_ISL_2785806, EPI_ISL_2785807, EPI_ISL_2785809, EPI_ISL_2785810, EPI_ISL_2785813, EPI_ISL_2785814, EPI_ISL_2785815, EPI_ISL_2785816, EPI_ISL_2785817, EPI_ISL_2785821, EPI_ISL_2785823, EPI_ISL_2785826, EPI_ISL_2785827, EPI_ISL_2785828, EPI_ISL_2785829, EPI_ISL_2785830, EPI_ISL_2785834, EPI_ISL_2785835, EPI_ISL_2785837, EPI_ISL_2785839, EPI_ISL_2785840, EPI_ISL_2785841, EPI_ISL_2785842, EPI_ISL_2785843, EPI_ISL_2785845, EPI_ISL_2785848, EPI_ISL_2785849, EPI_ISL_2785850, EPI_ISL_2785851, EPI_ISL_2785852, EPI_ISL_2785854, EPI_ISL_2785855, EPI_ISL_2785866, EPI_ISL_2785867, EPI_ISL_2785868, EPI_ISL_2785869, EPI_ISL_2785870, EPI_ISL_2785871, EPI_ISL_2785874, EPI_ISL_2785877, EPI_ISL_2785878, EPI_ISL_2785879, EPI_ISL_2785880, EPI_ISL_2785881, EPI_ISL_2785882, EPI_ISL_2785883, EPI_ISL_2785884, EPI_ISL_2785885, EPI_ISL_2785886, EPI_ISL_2785887, EPI_ISL_2785888, EPI_ISL_2785890, EPI_ISL_2785892, EPI_ISL_2785893, EPI_ISL_2785894, EPI_ISL_2785897, EPI_ISL_2785899, EPI_ISL_2785900, EPI_ISL_2785901, EPI_ISL_2785902, EPI_ISL_2785903, EPI_ISL_2785905, EPI_ISL_2785906, EPI_ISL_2785910, EPI_ISL_2785913, EPI_ISL_2785914, EPI_ISL_2785918, EPI_ISL_2785921, EPI_ISL_2785922, EPI_ISL_2785924, EPI_ISL_2785926, EPI_ISL_2785927, EPI_ISL_2785928, EPI_ISL_2785929, EPI_ISL_2785931, EPI_ISL_2785935, EPI_ISL_2785937, EPI_ISL_2785939, EPI_ISL_2785941, EPI_ISL_2785943, EPI_ISL_2785944, EPI_ISL_2785945, EPI_ISL_2785946, EPI_ISL_2785948, EPI_ISL_2785949, EPI_ISL_2785951, EPI_ISL_2785952, EPI_ISL_2785963 | | | |
| see above | Fulgent Genetics | Centers for Disease Control and Prevention Division of Viral Diseases, Pathogen Discovery | Dakota Howard, Dhwani Batra, Peter W. Cook, Kara Moser, Adrian Paskey, Jason Caravas, Benjamin Rambo-Martin, Shatavia Morrison, Christopher Gulvick, Scott Sammons, Yvette Unoarumhi, Darlene Wagner, Matthew Schmerer, Harry Gao, Mickey Li, John Gao, Joseph Fierro, Benafsh Sapra, Becky Tsai, Yan Meng, Doreen Ng, James Xie, Clinton R. Paden, Duncan MacCannell |
| EPI_ISL_2785972, EPI_ISL_2785973, EPI_ISL_2785978, EPI_ISL_2785979, EPI_ISL_2785980, EPI_ISL_2785990, EPI_ISL_2785992, EPI_ISL_2785994, EPI_ISL_2785995, EPI_ISL_2785999, EPI_ISL_2786000, EPI_ISL_2786001, EPI_ISL_2786005, EPI_ISL_2786011, EPI_ISL_2786013, EPI_ISL_2786019, EPI_ISL_2786023, EPI_ISL_2786025, EPI_ISL_2786028, EPI_ISL_2786029, EPI_ISL_2786031, EPI_ISL_2786037, EPI_ISL_2786039, EPI_ISL_2786041, EPI_ISL_2786047, EPI_ISL_2786052, EPI_ISL_2786053, EPI_ISL_2786055, EPI_ISL_2786061, EPI_ISL_2786065, EPI_ISL_2786071, EPI_ISL_2786072, EPI_ISL_2786074, EPI_ISL_2786078, EPI_ISL_2786079, EPI_ISL_2786080, EPI_ISL_2786081, EPI_ISL_2786082, EPI_ISL_2786083, EPI_ISL_2786084, EPI_ISL_2786087, EPI_ISL_2786088, EPI_ISL_2786090, EPI_ISL_2786092, EPI_ISL_2786096, EPI_ISL_2786097, EPI_ISL_2786098, EPI_ISL_2786104, EPI_ISL_2786105, EPI_ISL_2786108, EPI_ISL_2786118, EPI_ISL_2786122, EPI_ISL_2786126, EPI_ISL_2786130, EPI_ISL_2786132, EPI_ISL_2786134, EPI_ISL_2786149, EPI_ISL_2786152, EPI_ISL_2786154, EPI_ISL_2786157, EPI_ISL_2786166, EPI_ISL_2786167, EPI_ISL_2786176, EPI_ISL_2786180, EPI_ISL_2786188, EPI_ISL_2786189, | | | |

| | | | |
|---|---|---|---|
| EPI_ISL_2786193, EPI_ISL_2786199, EPI_ISL_2786201, EPI_ISL_2786204, EPI_ISL_2786206, EPI_ISL_2786218, EPI_ISL_2786219, EPI_ISL_2786221, EPI_ISL_2786223, EPI_ISL_2786237, EPI_ISL_2786239, EPI_ISL_2786240, EPI_ISL_2786241, EPI_ISL_2786242, EPI_ISL_2786243, EPI_ISL_2786244, EPI_ISL_2786246, EPI_ISL_2786247, EPI_ISL_2786248, EPI_ISL_2786252, EPI_ISL_2786253, EPI_ISL_2786255, EPI_ISL_2786256, EPI_ISL_2786258, EPI_ISL_2786259, EPI_ISL_2786260, EPI_ISL_2786261, EPI_ISL_2786263, EPI_ISL_2786264, EPI_ISL_2786265, EPI_ISL_2786269, EPI_ISL_2786272, EPI_ISL_2786273, EPI_ISL_2786277, EPI_ISL_2786278, EPI_ISL_2786279, EPI_ISL_2786280, EPI_ISL_2786281, EPI_ISL_2786282, EPI_ISL_2786284, EPI_ISL_2786285, EPI_ISL_2786288, EPI_ISL_2786293, EPI_ISL_2786295, EPI_ISL_2786298, EPI_ISL_2786301, EPI_ISL_2786302, EPI_ISL_2786307, EPI_ISL_2786310, EPI_ISL_2786317, EPI_ISL_2786319, EPI_ISL_2786320, EPI_ISL_2786321, EPI_ISL_2786323, EPI_ISL_2786324, EPI_ISL_2786327, EPI_ISL_2786328, EPI_ISL_2786330, EPI_ISL_2786332, EPI_ISL_2786333, EPI_ISL_2786337, EPI_ISL_2786338, EPI_ISL_2786340, EPI_ISL_2786343, EPI_ISL_2786346, EPI_ISL_2786349, EPI_ISL_2786352, EPI_ISL_2786355, EPI_ISL_2786357, EPI_ISL_2786362, EPI_ISL_2786363, EPI_ISL_2786365, EPI_ISL_2786366, EPI_ISL_2786367, EPI_ISL_2786371, EPI_ISL_2786372, EPI_ISL_2786373, EPI_ISL_2786376, EPI_ISL_2786377, EPI_ISL_2786378, EPI_ISL_2786380, EPI_ISL_2786381, EPI_ISL_2786384, EPI_ISL_2786386, EPI_ISL_2786388, EPI_ISL_2786389, EPI_ISL_2786392, EPI_ISL_2786394, EPI_ISL_2786395, EPI_ISL_2786397, EPI_ISL_2786398, EPI_ISL_2786399, EPI_ISL_2786403, EPI_ISL_2786405, EPI_ISL_2786407, EPI_ISL_2786409, EPI_ISL_2786410, EPI_ISL_2786412, EPI_ISL_2786415, EPI_ISL_2786416, EPI_ISL_2786418, EPI_ISL_2786419, EPI_ISL_2786431, EPI_ISL_2786435, EPI_ISL_2786436, EPI_ISL_2786438, EPI_ISL_2786439, EPI_ISL_2786444, EPI_ISL_2786445, EPI_ISL_2786450, EPI_ISL_2786451, EPI_ISL_2786452, EPI_ISL_2786453, EPI_ISL_2786454, EPI_ISL_2786455, EPI_ISL_2786456, EPI_ISL_2786457, EPI_ISL_2786459, EPI_ISL_2786461, EPI_ISL_2786462, EPI_ISL_2786467, EPI_ISL_2786469, EPI_ISL_2786471, EPI_ISL_2786472, EPI_ISL_2786475, EPI_ISL_2786477, EPI_ISL_2786478, EPI_ISL_2786481, EPI_ISL_2786483, EPI_ISL_2786484, EPI_ISL_2786487, EPI_ISL_2786488, EPI_ISL_2786489, EPI_ISL_2786490, EPI_ISL_2786491, EPI_ISL_2786492, EPI_ISL_2786494, EPI_ISL_2786496, EPI_ISL_2786497, EPI_ISL_2786500, EPI_ISL_2786501, EPI_ISL_2786504, EPI_ISL_2786505, EPI_ISL_2786506, EPI_ISL_2786509, EPI_ISL_2786512, EPI_ISL_2786513, EPI_ISL_2786514, EPI_ISL_2786515, EPI_ISL_2786517, EPI_ISL_2786518, EPI_ISL_2786522, EPI_ISL_2786525, EPI_ISL_2786528, EPI_ISL_2786529, EPI_ISL_2786530, EPI_ISL_2786532, EPI_ISL_2786537, EPI_ISL_2786546, EPI_ISL_2786555, EPI_ISL_2786557, EPI_ISL_2786562, EPI_ISL_2786563, EPI_ISL_2786567, EPI_ISL_2786568, EPI_ISL_2786569, EPI_ISL_2786572, EPI_ISL_2786573, EPI_ISL_2786578, EPI_ISL_2786579, EPI_ISL_2786583, EPI_ISL_2786587, EPI_ISL_2786590, EPI_ISL_2786591, EPI_ISL_2786597, EPI_ISL_2786599, EPI_ISL_2786600, EPI_ISL_2786601, EPI_ISL_2786603, EPI_ISL_2786605, EPI_ISL_2786613, EPI_ISL_2786619, EPI_ISL_2786620, EPI_ISL_2786622, EPI_ISL_2786623, EPI_ISL_2786624, EPI_ISL_2786626, EPI_ISL_2786627 | | | |
| see above | Mako Medical | Centers for Disease Control and Prevention Division of Viral Diseases, Pathogen Discovery | Dakota Howard, Dhwani Batra, Peter W. Cook, Kara Moser, Adrian Paskey, Jason Caravas, Benjamin Rambo-Martin, Shatavia Morrison, Christopher Gulvick, Scott Sammons, Yvette Unoarumhi, Darlene Wagner, Matthew Schmerer,Matthew Tugwell, Lauren Moon, Clinton R. Paden, Duncan MacCannell |
| EPI_ISL_2786634, EPI_ISL_2786648, EPI_ISL_2786655, EPI_ISL_2786657, EPI_ISL_2786666, EPI_ISL_2786667, EPI_ISL_2786671, EPI_ISL_2786676, EPI_ISL_2786677, EPI_ISL_2786682, EPI_ISL_2786685, EPI_ISL_2786688, EPI_ISL_2786693, EPI_ISL_2786694, EPI_ISL_2786697, EPI_ISL_2786698 | | | |
| see above | Infinity Biologix | Centers for Disease Control and Prevention Division of Viral Diseases, Pathogen Discovery | Dakota Howard, Dhwani Batra, Peter W. Cook, Kara Moser, Adrian Paskey, Jason Caravas, Benjamin Rambo-Martin, Shatavia Morrison, Christopher Gulvick, Scott Sammons, Yvette Unoarumhi, Darlene Wagner, Matthew Schmerer, Christian Bixby, Yihe Wang, Jonathan Schultz, Chirayu Goswami, Russ Hager, Robin Grimwood, Clinton R. Paden, Duncan MacCannell |
| EPI_ISL_2786930 | Platform BIS UZA/UAntwerpen | Labo Klinische Biologie, UZA | Marie Le Mercier, Jasmine Coppens, Basil Britto Xavier, Christine Lammens, Veerle Matheeussen, Herman Goossens |
| EPI_ISL_2786932 | Laverty Pathology | NSW Health Pathology - Institute of Clinical Pathology and Medical Research; Westmead Hospital; University of Sydney | CIDM-PH et al. |
| EPI_ISL_2786933, EPI_ISL_2786934 | St Vincent's Pathology (SydPath) | NSW Health Pathology - Institute of Clinical Pathology and Medical Research; Westmead Hospital; University of Sydney | CIDM-PH et al. |
| EPI_ISL_2786935 | Douglass Hanly Moir Pathology | NSW Health Pathology - Institute of Clinical Pathology and Medical Research; Westmead Hospital; University of Sydney | CIDM-PH et al. |
| EPI_ISL_2786991, EPI_ISL_2786992 | PORT HEALTH (HARBOUR) | Hong Kong Department of Health | Alan K.L. Tsang, Peter C.W. Yip, Ken H.L. Ng, Edman T.K. Lam, Rickjason C.W. Chan |
| EPI_ISL_2786993 | PRECIOUS BLOOD HOSPITAL | Hong Kong Department of Health | Alan K.L. Tsang, Peter C.W. Yip, Ken H.L. Ng, Edman T.K. Lam, Rickjason C.W. Chan |
| EPI_ISL_2786994 | private medical practitioners | Hong Kong Department of Health | Alan K.L. Tsang, Peter C.W. Yip, Ken H.L. Ng, Edman T.K. Lam, Rickjason C.W. Chan |
| EPI_ISL_2786995 | VICTORIA PUBLIC MORTUARY | Hong Kong Department of Health | Alan K.L. Tsang, Peter C.W. Yip, Ken H.L. Ng, Edman T.K. Lam, Rickjason C.W. Chan |
| EPI_ISL_2786996 | Queen Elizabeth Hospital | Hong Kong Department of Health | Alan K.L. Tsang, Peter C.W. Yip, Ken H.L. Ng, Edman T.K. Lam, Rickjason C.W. Chan |
| EPI_ISL_2786997, EPI_ISL_2786998, EPI_ISL_2786999, EPI_ISL_2787000, EPI_ISL_2787001 | North Lantau Hospital | Hong Kong Department of Health | Alan K.L. Tsang, Peter C.W. Yip, Ken H.L. Ng, Edman T.K. Lam, Rickjason C.W. Chan |
| EPI_ISL_2787002 | Queen Elizabeth Hospital | Hong Kong Department of Health | Alan K.L. Tsang, Peter C.W. Yip, Ken H.L. Ng, Edman T.K. Lam, Rickjason C.W. Chan |
| EPI_ISL_2787003, EPI_ISL_2787004, EPI_ISL_2787005, EPI_ISL_2787006, EPI_ISL_2787007, EPI_ISL_2787008, EPI_ISL_2787009 | Temporary Specimen Collection Centre at the AsiaWorld-Expo | Hong Kong Department of Health | Alan K.L. Tsang, Peter C.W. Yip, Ken H.L. Ng, Edman T.K. Lam, Rickjason C.W. Chan |
| EPI_ISL_2787010 | HOME QUARANTINE TASKFORCE | Hong Kong Department of Health | Alan K.L. Tsang, Peter C.W. Yip, Ken H.L. Ng, Edman T.K. Lam, Rickjason C.W. Chan |
| EPI_ISL_2787011, EPI_ISL_2787012 | QUARANTINE CAMP | Hong Kong Department of Health | Alan K.L. Tsang, Peter C.W. Yip, Ken H.L. Ng, Edman T.K. Lam, Rickjason C.W. Chan |
| EPI_ISL_2787013, EPI_ISL_2787014 | HOME QUARANTINE TASKFORCE | Hong Kong Department of Health | Alan K.L. Tsang, Peter C.W. Yip, Ken H.L. Ng, Edman T.K. Lam, Rickjason C.W. Chan |
| EPI_ISL_2787015 | Queen Elizabeth Hospital | Hong Kong Department of Health | Alan K.L. Tsang, Peter C.W. Yip, Ken H.L. Ng, Edman T.K. Lam, Rickjason C.W. Chan |
| EPI_ISL_2787079 | Ospedale San Camillo De Lellis Rieti | INMI Lazzaro Spallanzani IRCCS | CEM Gruber, F Santini, G Bonfiglio, O Butera, GF Inger, E Sperandio, D Di Fusco, D Cerini, B Bartolini |
| EPI_ISL_2787080 | Ospedale San Camillo De Lellis Rieti | INMI Lazzaro Spallanzani IRCCS | O Butera, M Rueca, F Santini, F Messina, L Ciuffreda, M Pallocca, D Di Fusco, D Cerini, E Giombini |
| EPI_ISL_2787081 | Ospedale San Filippo Neri | INMI Lazzaro Spallanzani IRCCS | F Santini, F Messina, O Butera, M Rueca, F De Nicola, M Pallocca, D Di Fusco, D Cerini, E Giombini |
| EPI_ISL_2787082 | Policlinico Universitario Campus Bio-Medico | INMI Lazzaro Spallanzani IRCCS | CEM Gruber, F Santini, G Bonfiglio, O Butera, G Orlandi, E Sperandio, D Di Fusco, D Cerini, E Giombini |
| EPI_ISL_2787083 | Fondazione Policlinico Tor Vergata | INMI Lazzaro Spallanzani IRCCS | M Rueca, G Bonfiglio, F Messina, CEM Gruber, S Donzelli, A Massacci, D Di Fusco, D Cerini, B Bartolini |
| EPI_ISL_2787084 | Fondazione Policlinico Tor Vergata | INMI Lazzaro Spallanzani IRCCS | G Bonfiglio, O Butera, CEM Gruber, F Santini, G Orlandi, E Sperandio, F Natoni, G Pessina, B Bartolini |
| EPI_ISL_2787085 | Fondazione Policlinico Tor Vergata | INMI Lazzaro Spallanzani IRCCS | F Santini, F Messina, O Butera, M Rueca, S Donzelli, M Pallocca, F Natoni, G Pessina, E Giombini |
| EPI_ISL_2787086 | Fondazione Policlinico Tor Vergata | INMI Lazzaro Spallanzani IRCCS | M Rueca, G Bonfiglio, F Messina, CEM Gruber, GF Inger, A Massacci, F Natoni, G Pessina, B Bartolini |
| EPI_ISL_2787087 | Fondazione Policlinico Tor Vergata | INMI Lazzaro Spallanzani IRCCS | CEM Gruber, F Santini, G Bonfiglio, O Butera, L Ciuffreda, E Sperandio, F Natoni, G Pessina, E Giombini |
| EPI_ISL_2787088 | Laboratorio Genzano - ASL RM 6 | INMI Lazzaro Spallanzani IRCCS | G Bonfiglio, O Butera, F Santini, F Messina, L Ciuffreda, E Sperandio, G Tramini, E Conti, E Giombini |
| EPI_ISL_2787089 | Azienda Sanitaria Locale Roma 5 | INMI Lazzaro Spallanzani IRCCS | G Bonfiglio, O Butera, CEM Gruber, F Santini, S Donzelli, E Sperandio, A Tamburro, M Meledandri, B Bartolini |
| EPI_ISL_2787090 | Azienda Sanitaria Locale Roma 5 | INMI Lazzaro Spallanzani IRCCS | F Santini, F Messina, O Butera, M Rueca, GF Inger, M Pallocca, M Meledandri, ML Schiavone, E Giombini |
| EPI_ISL_2787091 | Azienda Sanitaria Locale Roma 5 | INMI Lazzaro Spallanzani IRCCS | M Rueca, G Bonfiglio, F Messina, CEM Gruber, L Ciuffreda, A Massacci, ML Schiavone, A Tamburro, B Bartolini |
| EPI_ISL_2787092 | Azienda Sanitaria Locale Roma 5 | INMI Lazzaro Spallanzani IRCCS | O Butera, M Rueca, F Santini, F Messina, S Donzelli, M Pallocca, D Di Fusco, D Cerini, B Bartolini |
| EPI_ISL_2787093 | Fondazione Policlinico Tor Vergata | INMI Lazzaro Spallanzani IRCCS | F Messina, CEM Gruber, M Rueca, G Bonfiglio, F De Nicola, A Massacci, F Natoni, G Pessina, E Giombini |
| EPI_ISL_2787094 | Azienda Ospedaliera San Camillo Forlanini | INMI Lazzaro Spallanzani IRCCS | F Messina, CEM Gruber, M Rueca, G Bonfiglio, GF Inger, A Massacci, D Di Fusco, D Cerini, E Giombini |
| EPI_ISL_2787095 | Fondazione Policlinico Universitario “A. Gemelli” IRCCS | INMI Lazzaro Spallanzani IRCCS | G Bonfiglio, O Butera, CEM Gruber, F Santini, F De Nicola, E Sperandio, D Di Fusco, D Cerini, B Bartolini |
| EPI_ISL_2787096 | Ospedale Sandro Pertini ASL Roma2 | INMI Lazzaro Spallanzani IRCCS | M Rueca, G Bonfiglio, F Messina, CEM Gruber, GF Inger, A Massacci, F Natoni, G Pessina, B Bartolini |
| EPI_ISL_2787097 | Ospedale Sandro Pertini ASL Roma2 | INMI Lazzaro Spallanzani IRCCS | CEM Gruber, F Santini, G Bonfiglio, O Butera, L Ciuffreda, E Sperandio, F Natoni, G Pessina, E Giombini |
| EPI_ISL_2787098 | Ospedale Sandro Pertini ASL Roma2 | INMI Lazzaro Spallanzani IRCCS | F Messina, CEM Gruber, O Butera, M Rueca, F De Nicola, M Pallocca, G Tramini, E Conti, B Bartolini |
| EPI_ISL_2787132 | Dutch COVID-19 response team | National Institute for Public Health and the Environment (RIVM) | Adam Meijer, Harry Vennema, Dirk Eggink, Jeroen Cremer, Sharon van den Brink, Bas van der Veer, AnneMarie van den Brandt, Lisa Wijsman, Kim Freriks, Ryanne Jaarsma, Euníce Then, Lynn Aarts, Sanne Bos, Melissa van Tuil, Florian Zwagemaker, Dennis Schmitz, Annelies Kroneman, Karim Hajji, Chantal Reusken, on behalf of the national COVID-19 response team |
| EPI_ISL_2787325 | CA-Los Angeles County Public Health Laboratory | Centers for Disease Control and Prevention Division of Viral Diseases, Pathogen Discovery | Mili Sheth, Sarah Nobles, Jasmine Padilla, Mark Burroughs, Shoshona Le, Katie Dillon, Peter Cook, Clinton R. Paden, Dhwani Batra, Krista Queen, Kristen Knipe, Dakota Howard, Yvette Unoarumhi, Darlene Wagner, Matthew Schmerer, Ben L. Rambo-Martin, Kristine Lacek, Sam Shepard, Alison Laufer Halpin, Dave Wentworth, Vivien Dugan, Suxiang Tong, Justin Lee |
| EPI_ISL_2787331, EPI_ISL_2787336, EPI_ISL_2787339, EPI_ISL_2787342, EPI_ISL_2787348, EPI_ISL_2787351, EPI_ISL_2787354 | SD Public Health Laboratory | Centers for Disease Control and Prevention Division of Viral Diseases, Pathogen Discovery | Mili Sheth, Sarah Nobles, Jasmine Padilla, Mark Burroughs, Shoshona Le, Katie Dillon, Peter Cook, Clinton R. Paden, Dhwani Batra, Krista Queen, Kristen Knipe, Dakota Howard, Yvette Unoarumhi, Darlene Wagner, Matthew Schmerer, Ben L. Rambo-Martin, Kristine Lacek, Sam Shepard, Alison Laufer Halpin, Dave Wentworth, Vivien Dugan, Suxiang Tong, Justin Lee |
| EPI_ISL_2787359, EPI_ISL_2787372, EPI_ISL_2787375, EPI_ISL_2787395 | IA State Hygienic Laboratory | Centers for Disease Control and Prevention Division of Viral Diseases, Pathogen Discovery | Mili Sheth, Sarah Nobles, Jasmine Padilla, Mark Burroughs, Shoshona Le, Katie Dillon, Peter Cook, Clinton R. Paden, Dhwani Batra, Krista Queen, Kristen Knipe, Dakota Howard, Yvette Unoarumhi, Darlene Wagner, Matthew Schmerer, Ben L. Rambo-Martin, Kristine Lacek, Sam Shepard, Alison Laufer Halpin, Dave Wentworth, Vivien Dugan, Suxiang Tong, Justin Lee |
| EPI_ISL_2787400, EPI_ISL_2787403, EPI_ISL_2787414 | WY Public Health Laboratory | Centers for Disease Control and Prevention Division of Viral Diseases, Pathogen Discovery | Mili Sheth, Sarah Nobles, Jasmine Padilla, Mark Burroughs, Shoshona Le, Katie Dillon, Peter Cook, Clinton R. Paden, Dhwani Batra, Krista Queen, Kristen Knipe, Dakota Howard, Yvette Unoarumhi, Darlene Wagner, Matthew Schmerer, Ben L. Rambo-Martin, Kristine Lacek, Sam Shepard, Alison Laufer Halpin, Dave Wentworth, Vivien Dugan, Suxiang Tong, Justin Lee |
| EPI_ISL_2787477, EPI_ISL_2787508 | "NM Dept. Health, Scientific Laboratory Division " | Centers for Disease Control and Prevention Division of Viral Diseases, Pathogen Discovery | Mili Sheth, Sarah Nobles, Jasmine Padilla, Mark Burroughs, Shoshona Le, Katie Dillon, Peter Cook, Clinton R. Paden, Dhwani Batra, Krista Queen, Kristen Knipe, Dakota Howard, Yvette Unoarumhi, Darlene Wagner, Matthew Schmerer, Ben L. Rambo-Martin, Kristine Lacek, Sam Shepard, Alison Laufer Halpin, Dave Wentworth, Vivien Dugan, Suxiang Tong, Justin Lee |
| EPI_ISL_2787511 | "AK State Public Health Lab, State Health Department" | Centers for Disease Control and Prevention Division of Viral Diseases, Pathogen Discovery | Mili Sheth, Sarah Nobles, Jasmine Padilla, Mark Burroughs, Shoshona Le, Katie Dillon, Peter Cook, Clinton R. Paden, Dhwani Batra, Krista Queen, Kristen Knipe, Dakota Howard, Yvette Unoarumhi, Darlene Wagner, Matthew Schmerer, Ben L. Rambo-Martin, Kristine Lacek, Sam Shepard, Alison Laufer Halpin, Dave Wentworth, Vivien Dugan, Suxiang Tong, Justin Lee |
| EPI_ISL_2787562 | MS Public Health Laboratory | Centers for Disease Control and Prevention Division of Viral | Mili Sheth, Sarah Nobles, Jasmine Padilla, Mark Burroughs, Shoshona Le, Katie Dillon, Peter Cook, Clinton R. Paden, Dhwani Batra, Krista Queen, Kristen Knipe, Dakota Howard, Yvette Unoarumhi, Darlene Wagner, Matthew Schmerer, Ben L. |

| | | | |
|---|---|---|---|
| | | Diseases, Pathogen Discovery | Rambo-Martin, Kristine Lacek, Sam Shepard, Alison Laufer Halpin, Dave Wentworth, Vivien Dugan, Suxiang Tong, Justin Lee |
| EPI_ISL_2787738 | IN State Department of Health Laboratory Services | Centers for Disease Control and Prevention Division of Viral Diseases, Pathogen Discovery | Mili Sheth, Sarah Nobles, Jasmine Padilla, Mark Burroughs, Shoshona Le, Katie Dillon, Peter Cook, Clinton R. Paden, Dhwani Batra, Krista Queen, Kristen Knipe, Dakota Howard, Yvette Unoarumhi, Darlene Wagner, Matthew Schmerer, Ben L. Rambo-Martin, Kristine Lacek, Sam Shepard, Alison Laufer Halpin, Dave Wentworth, Vivien Dugan, Suxiang Tong, Justin Lee |
| EPI_ISL_2787741, EPI_ISL_2787744 | MO State Public Health Laboratory | Centers for Disease Control and Prevention Division of Viral Diseases, Pathogen Discovery | Mili Sheth, Sarah Nobles, Jasmine Padilla, Mark Burroughs, Shoshona Le, Katie Dillon, Peter Cook, Clinton R. Paden, Dhwani Batra, Krista Queen, Kristen Knipe, Dakota Howard, Yvette Unoarumhi, Darlene Wagner, Matthew Schmerer, Ben L. Rambo-Martin, Kristine Lacek, Sam Shepard, Alison Laufer Halpin, Dave Wentworth, Vivien Dugan, Suxiang Tong, Justin Lee |
| EPI_ISL_2787764 | AL Dept. of Public Health Bureau of Clinical Laboratories | Centers for Disease Control and Prevention Division of Viral Diseases, Pathogen Discovery | Mili Sheth, Sarah Nobles, Jasmine Padilla, Mark Burroughs, Shoshona Le, Katie Dillon, Peter Cook, Clinton R. Paden, Dhwani Batra, Krista Queen, Kristen Knipe, Dakota Howard, Yvette Unoarumhi, Darlene Wagner, Matthew Schmerer, Ben L. Rambo-Martin, Kristine Lacek, Sam Shepard, Alison Laufer Halpin, Dave Wentworth, Vivien Dugan, Suxiang Tong, Justin Lee |
| EPI_ISL_2787782, EPI_ISL_2787785, EPI_ISL_2787788, EPI_ISL_2787794, EPI_ISL_2787797 | KS Health and Environmental Laboratories | Centers for Disease Control and Prevention Division of Viral Diseases, Pathogen Discovery | Mili Sheth, Sarah Nobles, Jasmine Padilla, Mark Burroughs, Shoshona Le, Katie Dillon, Peter Cook, Clinton R. Paden, Dhwani Batra, Krista Queen, Kristen Knipe, Dakota Howard, Yvette Unoarumhi, Darlene Wagner, Matthew Schmerer, Ben L. Rambo-Martin, Kristine Lacek, Sam Shepard, Alison Laufer Halpin, Dave Wentworth, Vivien Dugan, Suxiang Tong, Justin Lee |
| EPI_ISL_2787805, EPI_ISL_2787811, EPI_ISL_2787820, EPI_ISL_2787822 | MT Public Health Laboratory | Centers for Disease Control and Prevention Division of Viral Diseases, Pathogen Discovery | Mili Sheth, Sarah Nobles, Jasmine Padilla, Mark Burroughs, Shoshona Le, Katie Dillon, Peter Cook, Clinton R. Paden, Dhwani Batra, Krista Queen, Kristen Knipe, Dakota Howard, Yvette Unoarumhi, Darlene Wagner, Matthew Schmerer, Ben L. Rambo-Martin, Kristine Lacek, Sam Shepard, Alison Laufer Halpin, Dave Wentworth, Vivien Dugan, Suxiang Tong, Justin Lee |
| EPI_ISL_2787831, EPI_ISL_2787834, EPI_ISL_2787840, EPI_ISL_2787846 | NV-Southern Nevada Public Health Laboratory | Centers for Disease Control and Prevention Division of Viral Diseases, Pathogen Discovery | Mili Sheth, Sarah Nobles, Jasmine Padilla, Mark Burroughs, Shoshona Le, Katie Dillon, Peter Cook, Clinton R. Paden, Dhwani Batra, Krista Queen, Kristen Knipe, Dakota Howard, Yvette Unoarumhi, Darlene Wagner, Matthew Schmerer, Ben L. Rambo-Martin, Kristine Lacek, Sam Shepard, Alison Laufer Halpin, Dave Wentworth, Vivien Dugan, Suxiang Tong, Justin Lee |
| EPI_ISL_2787860, EPI_ISL_2787879, EPI_ISL_2787892, EPI_ISL_2787904, EPI_ISL_2787923 | NJ Public Health and Environmental Laboratories | Centers for Disease Control and Prevention Division of Viral Diseases, Pathogen Discovery | Mili Sheth, Sarah Nobles, Jasmine Padilla, Mark Burroughs, Shoshona Le, Katie Dillon, Peter Cook, Clinton R. Paden, Dhwani Batra, Krista Queen, Kristen Knipe, Dakota Howard, Yvette Unoarumhi, Darlene Wagner, Matthew Schmerer, Ben L. Rambo-Martin, Kristine Lacek, Sam Shepard, Alison Laufer Halpin, Dave Wentworth, Vivien Dugan, Suxiang Tong, Justin Lee |
| EPI_ISL_2787970 | GA Department of Public Health Laboratory | Centers for Disease Control and Prevention Division of Viral Diseases, Pathogen Discovery | Mili Sheth, Sarah Nobles, Jasmine Padilla, Mark Burroughs, Shoshona Le, Katie Dillon, Peter Cook, Clinton R. Paden, Dhwani Batra, Krista Queen, Kristen Knipe, Dakota Howard, Yvette Unoarumhi, Darlene Wagner, Matthew Schmerer, Ben L. Rambo-Martin, Kristine Lacek, Sam Shepard, Alison Laufer Halpin, Dave Wentworth, Vivien Dugan, Suxiang Tong, Justin Lee |
| EPI_ISL_2788024, EPI_ISL_2788033 | WA State Department of Health | Centers for Disease Control and Prevention Division of Viral Diseases, Pathogen Discovery | Mili Sheth, Sarah Nobles, Jasmine Padilla, Mark Burroughs, Shoshona Le, Katie Dillon, Peter Cook, Clinton R. Paden, Dhwani Batra, Krista Queen, Kristen Knipe, Dakota Howard, Yvette Unoarumhi, Darlene Wagner, Matthew Schmerer, Ben L. Rambo-Martin, Kristine Lacek, Sam Shepard, Alison Laufer Halpin, Dave Wentworth, Vivien Dugan, Suxiang Tong, Justin Lee |
| EPI_ISL_2788051, EPI_ISL_2788056, EPI_ISL_2788061, EPI_ISL_2788067 | "AR Dept. of Health-PHL, Molecular Diagnostics" | Centers for Disease Control and Prevention Division of Viral Diseases, Pathogen Discovery | Mili Sheth, Sarah Nobles, Jasmine Padilla, Mark Burroughs, Shoshona Le, Katie Dillon, Peter Cook, Clinton R. Paden, Dhwani Batra, Krista Queen, Kristen Knipe, Dakota Howard, Yvette Unoarumhi, Darlene Wagner, Matthew Schmerer, Ben L. Rambo-Martin, Kristine Lacek, Sam Shepard, Alison Laufer Halpin, Dave Wentworth, Vivien Dugan, Suxiang Tong, Justin Lee |
| EPI_ISL_2788100, EPI_ISL_2788106, EPI_ISL_2788109 | "CDPH, Viral and Rickettsial Disease Laboratory" | Centers for Disease Control and Prevention Division of Viral Diseases, Pathogen Discovery | Mili Sheth, Sarah Nobles, Jasmine Padilla, Mark Burroughs, Shoshona Le, Katie Dillon, Peter Cook, Clinton R. Paden, Dhwani Batra, Krista Queen, Kristen Knipe, Dakota Howard, Yvette Unoarumhi, Darlene Wagner, Matthew Schmerer, Ben L. Rambo-Martin, Kristine Lacek, Sam Shepard, Alison Laufer Halpin, Dave Wentworth, Vivien Dugan, Suxiang Tong, Justin Lee |

EPI_ISL_2788156, EPI_ISL_2788157, EPI_ISL_2788158, EPI_ISL_2788159, EPI_ISL_2788160, EPI_ISL_2788161, EPI_ISL_2788162, EPI_ISL_2788163, EPI_ISL_2788164, EPI_ISL_2788165, EPI_ISL_2788166, EPI_ISL_2788167, EPI_ISL_2788168, EPI_ISL_2788169, EPI_ISL_2788170, EPI_ISL_2788174, EPI_ISL_2788176, EPI_ISL_2788190, EPI_ISL_2788191, EPI_ISL_2788192, EPI_ISL_2788193, EPI_ISL_2788194, EPI_ISL_2788195, EPI_ISL_2788196, EPI_ISL_2788197, EPI_ISL_2788198, EPI_ISL_2788199, EPI_ISL_2788200, EPI_ISL_2788201, EPI_ISL_2788202, EPI_ISL_2788203, EPI_ISL_2788204, EPI_ISL_2788205, EPI_ISL_2788206, EPI_ISL_2788207, EPI_ISL_2788208, EPI_ISL_2788209, EPI_ISL_2788210, EPI_ISL_2788211, EPI_ISL_2788212, EPI_ISL_2788213, EPI_ISL_2788214, EPI_ISL_2788215, EPI_ISL_2788216, EPI_ISL_2788217, EPI_ISL_2788218, EPI_ISL_2788219, EPI_ISL_2788220, EPI_ISL_2788221, EPI_ISL_2788238, EPI_ISL_2788239, EPI_ISL_2788240, EPI_ISL_2788241, EPI_ISL_2788242, EPI_ISL_2788243, EPI_ISL_2788244, EPI_ISL_2788245, EPI_ISL_2788246, EPI_ISL_2788247, EPI_ISL_2788248, EPI_ISL_2788249, EPI_ISL_2788250, EPI_ISL_2788251, EPI_ISL_2788252, EPI_ISL_2788253, EPI_ISL_2788254, EPI_ISL_2788255, EPI_ISL_2788256, EPI_ISL_2788257, EPI_ISL_2788258, EPI_ISL_2788259, EPI_ISL_2788260, EPI_ISL_2788261, EPI_ISL_2788262, EPI_ISL_2788263, EPI_ISL_2788264, EPI_ISL_2788265, EPI_ISL_2788266, EPI_ISL_2788267, EPI_ISL_2788268, EPI_ISL_2788269, EPI_ISL_2788270, EPI_ISL_2788271, EPI_ISL_2788272, EPI_ISL_2788273, EPI_ISL_2788274, EPI_ISL_2788275, EPI_ISL_2788276, EPI_ISL_2788277, EPI_ISL_2788278, EPI_ISL_2788279, EPI_ISL_2788280, EPI_ISL_2788281, EPI_ISL_2788282, EPI_ISL_2788283, EPI_ISL_2788284, EPI_ISL_2788285, EPI_ISL_2788286, EPI_ISL_2788287, EPI_ISL_2788288, EPI_ISL_2788289, EPI_ISL_2788290, EPI_ISL_2788291, EPI_ISL_2788292, EPI_ISL_2788293, EPI_ISL_2788294, EPI_ISL_2788295, EPI_ISL_2788296, EPI_ISL_2788297, EPI_ISL_2788298, EPI_ISL_2788299, EPI_ISL_2788300, EPI_ISL_2788301, EPI_ISL_2788302, EPI_ISL_2788303, EPI_ISL_2788304, EPI_ISL_2788305, EPI_ISL_2788306, EPI_ISL_2788307, EPI_ISL_2788308, EPI_ISL_2788309, EPI_ISL_2788310, EPI_ISL_2788311, EPI_ISL_2788312, EPI_ISL_2788313, EPI_ISL_2788314, EPI_ISL_2788315, EPI_ISL_2788316, EPI_ISL_2788317, EPI_ISL_2788318, EPI_ISL_2788319, EPI_ISL_2788320, EPI_ISL_2788321, EPI_ISL_2788322, EPI_ISL_2788323, EPI_ISL_2788324, EPI_ISL_2788325, EPI_ISL_2788326, EPI_ISL_2788327, EPI_ISL_2788328, EPI_ISL_2788329, EPI_ISL_2788330, EPI_ISL_2788331, EPI_ISL_2788332, EPI_ISL_2788333, EPI_ISL_2788334, EPI_ISL_2788335, EPI_ISL_2788336, EPI_ISL_2788337, EPI_ISL_2788338, EPI_ISL_2788339, EPI_ISL_2788340, EPI_ISL_2788341, EPI_ISL_2788342, EPI_ISL_2788343, EPI_ISL_2788344, EPI_ISL_2788345, EPI_ISL_2788346, EPI_ISL_2788347, EPI_ISL_2788348, EPI_ISL_2788349, EPI_ISL_2788350, EPI_ISL_2788351, EPI_ISL_2788352, EPI_ISL_2788353, EPI_ISL_2788354, EPI_ISL_2788355, EPI_ISL_2788356, EPI_ISL_2788357, EPI_ISL_2788358, EPI_ISL_2788359, EPI_ISL_2788360, EPI_ISL_2788361, EPI_ISL_2788362, EPI_ISL_2788363, EPI_ISL_2788364, EPI_ISL_2788365, EPI_ISL_2788366

| | | | |
|---|---|---|---|
| see above | Dutch COVID-19 response team | National Institute for Public Health and the Environment (RIVM) | Adam Meijer, Harry Vennema, Dirk Eggink, Jeroen Cremer, Sharon van den Brink, Bas van der Veer, AnneMarie van den Brandt, Lisa Wijsman, Kim Freriks, Ryanne Jaarsma, Euníce Then, Lynn Aarts, Sanne Bos, Melissa van Tuil, Florian Zwagemaker, Dennis Schmitz, Annelies Kroneman, Karim Hajji, Chantal Reusken, on behalf of the national COVID-19 response team |
| EPI_ISL_2788584, EPI_ISL_2788585, EPI_ISL_2788586, EPI_ISL_2788587, EPI_ISL_2788588 | Furst Medical Laboratory | Norwegian Institute of Public Health, Department of Virology | Kathrine Stene-Johansen, Kamilla Heddeland Instefjord, Hilde Elshaug, Garcia Llorente Ignacio, Jon Bråte, Engebretsen Serina Beate,Pedersen Benedikte Nevjen, Line Victoria Moen, Debech Nadia, Atiya R Ali,Marie Paulsen Madsen, Rasmus Riis Kopperud, Hilde Vollan, Karoline Bragstad, Olav Hungnes |
| EPI_ISL_2788589, EPI_ISL_2788590, EPI_ISL_2788591, EPI_ISL_2788592 | Akershus University Hospital, Department for Microbiology and Infectious Disease Control | Norwegian Institute of Public Health, Department of Virology | Kathrine Stene-Johansen, Kamilla Heddeland Instefjord, Hilde Elshaug, Garcia Llorente Ignacio, Jon Bråte, Engebretsen Serina Beate,Pedersen Benedikte Nevjen, Line Victoria Moen, Debech Nadia, Atiya R Ali,Marie Paulsen Madsen, Rasmus Riis Kopperud, Hilde Vollan, Karoline Bragstad, Olav Hungnes |
| EPI_ISL_2788593, EPI_ISL_2788594 | Furst Medical Laboratory | Norwegian Institute of Public Health, Department of Virology | Kathrine Stene-Johansen, Kamilla Heddeland Instefjord, Hilde Elshaug, Garcia Llorente Ignacio, Jon Bråte, Engebretsen Serina Beate,Pedersen Benedikte Nevjen, Line Victoria Moen, Debech Nadia, Atiya R Ali,Marie Paulsen Madsen, Rasmus Riis Kopperud, Hilde Vollan, Karoline Bragstad, Olav Hungnes |
| EPI_ISL_2788595 | Hospital of Southern Norway - Kristiansand, Department of Medical Microbiology | Norwegian Institute of Public Health, Department of Virology | Kathrine Stene-Johansen, Kamilla Heddeland Instefjord, Hilde Elshaug, Garcia Llorente Ignacio, Jon Bråte, Engebretsen Serina Beate,Pedersen Benedikte Nevjen, Line Victoria Moen, Debech Nadia, Atiya R Ali,Marie Paulsen Madsen, Rasmus Riis Kopperud, Hilde Vollan, Karoline Bragstad, Olav Hungnes |
| EPI_ISL_2788596 | Norwegian Institute of Public Health, Department of Virology | Oslo Helse | Kathrine Stene-Johansen, Kamilla Heddeland Instefjord, Hilde Elshaug, Garcia Llorente Ignacio, Jon Bråte, Engebretsen Serina Beate,Pedersen Benedikte Nevjen, Line Victoria Moen, Debech Nadia, Atiya R Ali,Marie Paulsen Madsen, Rasmus Riis Kopperud, Hilde Vollan, Karoline Bragstad, Olav Hungnes |
| EPI_ISL_2788597 | Medical Microbiology Unit, Department for Laboratory Medicine, Drammen Hospital, Vestre Viken Health Trust, | Norwegian Institute of Public Health, Department of Virology | Kathrine Stene-Johansen, Kamilla Heddeland Instefjord, Hilde Elshaug, Garcia Llorente Ignacio, Jon Bråte, Engebretsen Serina Beate,Pedersen Benedikte Nevjen, Line Victoria Moen, Debech Nadia, Atiya R Ali,Marie Paulsen Madsen, Rasmus Riis Kopperud, Hilde Vollan, Karoline Bragstad, Olav Hungnes |
| EPI_ISL_2788654, EPI_ISL_2788659, EPI_ISL_2788676, EPI_ISL_2788689, EPI_ISL_2788711, EPI_ISL_2788716, EPI_ISL_2788717, EPI_ISL_2788718, EPI_ISL_2788719 | Gunma Prefectural Institute of Public Health and Environmental Sciences | Gunma Prefectural Institute of Public Health and Environmental Sciences | Tsuyoshi Sekizuka, Kentaro Itokawa, Rina Tanaka, Masanori Hashino, Makoto Kuroda, Hiroyuki Tsukagoshi, Daisuke Shinoda, Mariko Saito |
| EPI_ISL_2788940 | National Laboratory for Health, Environment and Food, OMM, Maribor | NLZOH (National Laboratory for Health, Environment and Food) / CISLD (Clinical Institute of Special Laboratory Diagnostics), University Children's Hospital, University Medical Center Ljubljana | Sandra Janezic, Aleksander Mahnic, Maja Rupnik, Tjaša Žohar Čretnik, Alenka Štorman, Nika Gobec, Aleksander Kocuvan, Kaja Tominc, Maša Jarčič, Mojca Cimerman, Nika Volmajer, Andrej Golle / Jernej Kovač, Barbara Jenko Bizjan, Tine Tesovnik, Robert Šket, Katarina Kozmos, Ana Grom, Maruša Debeljak, Marko Pokorn, Tadej Battelino |
| EPI_ISL_2789321 | National Laboratory for Health, Environment and Food, OMM, Koper | NLZOH (National Laboratory for Health, Environment and Food) / CISLD (Clinical Institute of Special Laboratory Diagnostics), University Children's Hospital, University Medical Center Ljubljana | Sandra Janezic, Aleksander Mahnic, Maja Rupnik, Tjaša Žohar Čretnik, Alenka Štorman, Nika Gobec, Aleksander Kocuvan, Kaja Tominc, Maša Jarčič, Gašper Strugar, Tina Cvetković, Mitja Rak / Jernej Kovač, Barbara Jenko Bizjan, Tine Tesovnik, Robert Šket, Katarina Kozmos, Ana Grom, Maruša Debeljak, Marko Pokorn, Tadej Battelino |
| EPI_ISL_2789322 | National Laboratory for Health, Environment and Food, OMM, Celje | NLZOH (National Laboratory for Health, Environment and Food) / CISLD (Clinical Institute of Special Laboratory Diagnostics), University Children's Hospital, University Medical Center Ljubljana | Sandra Janezic, Aleksander Mahnic, Maja Rupnik, Tjaša Žohar Čretnik, Alenka Štorman, Nika Gobec, Aleksander Kocuvan, Kaja Tominc, Maša Jarčič, Mateja Borinc, Daša Kavka / Jernej Kovač, Barbara Jenko Bizjan, Tine Tesovnik, Robert Šket, Katarina Kozmos, Ana Grom, Maruša Debeljak, Marko Pokorn, Tadej Battelino |
| EPI_ISL_2789324 | National Laboratory for Health, Environment and Food, OMM, Maribor | NLZOH (National Laboratory for Health, Environment and Food) / CISLD (Clinical Institute of Special Laboratory Diagnostics), University Children's Hospital, University Medical Center Ljubljana | Sandra Janezic, Aleksander Mahnic, Maja Rupnik, Tjaša Žohar Čretnik, Alenka Štorman, Nika Gobec, Aleksander Kocuvan, Kaja Tominc, Maša Jarčič, Mojca Cimerman, Nika Volmajer, Andrej Golle / Jernej Kovač, Barbara Jenko Bizjan, Tine Tesovnik, Robert Šket, Katarina Kozmos, Ana Grom, Maruša Debeljak, Marko Pokorn, Tadej Battelino |
| EPI_ISL_2789325, EPI_ISL_2789326 | National Laboratory for Health, Environment and Food, OMM, Kranj | NLZOH (National Laboratory for Health, Environment and Food) / CISLD (Clinical Institute of Special Laboratory Diagnostics), University Children's Hospital, University Medical Center Ljubljana | Sandra Janezic, Aleksander Mahnic, Maja Rupnik, Tjaša Žohar Čretnik, Alenka Štorman, Nika Gobec, Aleksander Kocuvan, Kaja Tominc, Maša Jarčič, Mateja Ravnik, Monika Korošec, Marjana Petrevčič / Jernej Kovač, Barbara Jenko Bizjan, Tine Tesovnik, Robert Šket, Katarina Kozmos, Ana Grom, Maruša Debeljak, Marko Pokorn, Tadej Battelino |
| EPI_ISL_2789336 | OKMI | CMBG | Matej Bezdicek, Jan Svaton, Martina Lengerova, Monika Dolejska |
| EPI_ISL_2789347 | CSIR-Centre for Cellular and Molecular Biology | CSIR-Centre for Cellular and Molecular Biology-INSACOG | Sofia Banu,Payel Mukherjee, Lamuk Zaveri, Onkar Kulkarni,Tulasi Nagabandi, Jandhyala Sai Krishna, Ara Sreenivas,Shreekant Verma, Amareshwar Vodapalli ,B Himasri,Valli Nagalakshmi Undamatla, Sumedha Avadhanula, Vidhyadhari Methuku, Priya Nurkurthy, Archana Bharadwaj Siva, Karthik Bharadwaj Tallapaka,Rakesh K Mishra,Divya Tej Sowpati |
| EPI_ISL_2789349 | AP SSO | CSIR-Centre for Cellular and Molecular Biology-INSACOG | Priya Nurkurthy,Ara Sreenivas,Shreekant Verma, Amareshwar Vodapalli ,Lamuk Zaveri,Tulasi Nagabandi,B Himasri,Valli Nagalakshmi Undamatla, Sumedha Avadhanula, Vidhyadhari Methuku,Payel Mukherjee,Onkar Kulkarni, Jandhyala Sai Krishna,Sofia Banu, Archana Bharadwaj Siva, Karthik Bharadwaj Tallapaka,Rakesh K Mishra,Divya Tej Sowpati |
| EPI_ISL_2789350 | AP SSO | CSIR-Centre for Cellular and Molecular Biology-INSACOG | B Himasri,Amareshwar Vodapalli,Ara Sreenivas,Shreekant Verma,Valli Nagalakshmi Undamatla,Payel Mukherjee, Lamuk Zaveri, Onkar Kulkarni,Tulasi Nagabandi, Sofia Banu, Vidhyadhari Methuku,Priya Nurkurthy, Jandhyala Sai Krishna,Sumedha Avadhanula, Archana Bharadwaj Siva, Karthik Bharadwaj Tallapaka,Rakesh K Mishra,Divya Tej Sowpati |
| EPI_ISL_2789352 | Inovie AS MEDILAB 66 C0-1-2 (IASCM) | Inovie AS GenBio | Pauline LAURENT, Florian VERCRUYSSEN, Gwladys SEVRIN |
| EPI_ISL_2789353 | Inovie AS GenBio | Inovie AS GenBio | Pauline LAURENT, Florian VERCRUYSSEN, Gwladys SEVRIN |

| | | | |
|---|---|---|---|
| EPI_ISL_2789354 | Inovie AS MEDILAB 66 C0-1-2 (IASCM) | Inovie AS GenBio | Pauline LAURENT, Florian VERCRUYSSEN, Gwladys SEVRIN |
| EPI_ISL_2789355, EPI_ISL_2789356 | Inovie AS BIOMEDIVAL | Inovie AS GenBio | Pauline LAURENT, Florian VERCRUYSSEN, Gwladys SEVRIN |
| EPI_ISL_2789357 | Inovie AS MEDILAB 66 C0-1-2 (IASCM) | Inovie AS GenBio | Pauline LAURENT, Florian VERCRUYSSEN, Gwladys SEVRIN |
| EPI_ISL_2789358, EPI_ISL_2789359, EPI_ISL_2789360, EPI_ISL_2789361 | Inovie AS BIOFUSION | Inovie AS GenBio | Pauline LAURENT, Florian VERCRUYSSEN, Gwladys SEVRIN |
| EPI_ISL_2789362 | Inovie AS MEDILAB 66 C0-1-2 (IASCM) | Inovie AS GenBio | Pauline LAURENT, Florian VERCRUYSSEN, Gwladys SEVRIN |
| EPI_ISL_2789363 | Inovie AS BIOFUSION | Inovie AS GenBio | Pauline LAURENT, Florian VERCRUYSSEN, Gwladys SEVRIN |
| EPI_ISL_2789364 | Inovie AS BIOMEDIVAL | Inovie AS GenBio | Pauline LAURENT, Florian VERCRUYSSEN, Gwladys SEVRIN |
| EPI_ISL_2789365, EPI_ISL_2789366 | Inovie AS MEDILAB 66 C0-1-2 (IASCM) | Inovie AS GenBio | Pauline LAURENT, Florian VERCRUYSSEN, Gwladys SEVRIN |
| EPI_ISL_2789367 | Inovie AS BIOMEDIVAL | Inovie AS GenBio | Pauline LAURENT, Florian VERCRUYSSEN, Gwladys SEVRIN |
| EPI_ISL_2789368 | Inovie AS GenBio | Inovie AS GenBio | Pauline LAURENT, Florian VERCRUYSSEN, Gwladys SEVRIN |
| EPI_ISL_2789369 | Inovie AS BIOFUSION | Inovie AS GenBio | Pauline LAURENT, Florian VERCRUYSSEN, Gwladys SEVRIN |
| EPI_ISL_2789435, EPI_ISL_2789440, EPI_ISL_2789442 | National Virus Reference Laboratory | National Virus Reference Laboratory | Fiona Crispie, Calum Walsh, Matthew McCabe, Zoe Yandle, Charlene Bennet, Gabriel Gonzalez, Michael Carr, Jonathan Dean, Paul Cotter, Cillian F De Gascun |
| EPI_ISL_2789504, EPI_ISL_2789505, EPI_ISL_2789506, EPI_ISL_2789507, EPI_ISL_2789508, EPI_ISL_2789509, EPI_ISL_2789510, EPI_ISL_2789511, EPI_ISL_2789512, EPI_ISL_2789513, EPI_ISL_2789514, EPI_ISL_2789515, EPI_ISL_2789516, EPI_ISL_2789518, EPI_ISL_2789519, EPI_ISL_2789521, EPI_ISL_2789522, EPI_ISL_2789523, EPI_ISL_2789524, EPI_ISL_2789525, EPI_ISL_2789529, EPI_ISL_2789530, EPI_ISL_2789533, EPI_ISL_2789535, EPI_ISL_2789536, EPI_ISL_2789538, EPI_ISL_2789539, EPI_ISL_2789540, EPI_ISL_2789541, EPI_ISL_2789543, EPI_ISL_2789544, EPI_ISL_2789545, EPI_ISL_2789547, EPI_ISL_2789548, EPI_ISL_2789549, EPI_ISL_2789550, EPI_ISL_2789551, EPI_ISL_2789552, EPI_ISL_2789553, EPI_ISL_2789556, EPI_ISL_2789557, EPI_ISL_2789558, EPI_ISL_2789559, EPI_ISL_2789560, EPI_ISL_2789561, EPI_ISL_2789563, EPI_ISL_2789567, EPI_ISL_2789568, EPI_ISL_2789571, EPI_ISL_2789583, EPI_ISL_2789584, EPI_ISL_2789587, EPI_ISL_2789591, EPI_ISL_2789592, EPI_ISL_2789593, EPI_ISL_2789594, EPI_ISL_2789595, EPI_ISL_2789596, EPI_ISL_2789597, EPI_ISL_2789598, EPI_ISL_2789599, EPI_ISL_2789600, EPI_ISL_2789601, EPI_ISL_2789602, EPI_ISL_2789603, EPI_ISL_2789604, EPI_ISL_2789606, EPI_ISL_2789607, EPI_ISL_2789612, EPI_ISL_2789613, EPI_ISL_2789614, EPI_ISL_2789615, EPI_ISL_2789616, EPI_ISL_2789617, EPI_ISL_2789618, EPI_ISL_2789619, EPI_ISL_2789620, EPI_ISL_2789621, EPI_ISL_2789622, EPI_ISL_2789623, EPI_ISL_2789625, EPI_ISL_2789628, EPI_ISL_2789629, EPI_ISL_2789630, EPI_ISL_2789631, EPI_ISL_2789634, EPI_ISL_2789646, EPI_ISL_2789665, EPI_ISL_2789668, EPI_ISL_2789671, EPI_ISL_2789707, EPI_ISL_2789710, EPI_ISL_2789717 | | | |
| see above | National Virus Reference Laboratory | National Virus Reference Laboratory | Zoe Yandle, Charlene Bennett, Gabriel Gonzalez, Michael Carr, Jonathan Dean, Cillian F De Gascun |
| EPI_ISL_2789836 | Centre Hospitalier Universitaire Clermont-Ferrand | CHU Clermont-Ferrand, service de virologie | Bisseux Maxime, Mirand Audrey, Combes Patricia, Henquell Cécile |
| EPI_ISL_2789879, EPI_ISL_2789881, EPI_ISL_2789882, EPI_ISL_2789884, EPI_ISL_2789885, EPI_ISL_2789887, EPI_ISL_2789889, EPI_ISL_2789900, EPI_ISL_2789901, EPI_ISL_2789902, EPI_ISL_2789903, EPI_ISL_2789904, EPI_ISL_2789906, EPI_ISL_2789907, EPI_ISL_2789908 | | | |
| see above | IN State Department of Health Laboratory Services | IN State Department of Health Laboratory Services | Cassandra Campion, Jamie Yeadon, Brian Pope, Lixia Liu, Kyle Brownlee, Melissa Hindenlang, Mark Glazier |
| EPI_ISL_2789931, EPI_ISL_2789933, EPI_ISL_2789934, EPI_ISL_2789935 | Kansas Health and Environmental Lab | Kansas Health and Environmental Lab | Katherine Wiggins, Mike Grose, Jonathan Barnell, Ben Olsen, and Phil Adam |
| EPI_ISL_2789960 | IVY3 Central Lab, Vanderbilt University Medical Center | Lauring Lab, University of Michigan, Department of Microbiology and Immunology | Valesano |
| EPI_ISL_2790011, EPI_ISL_2790012, EPI_ISL_2790014 | Akershus University Hospital, Department for Microbiology and Infectious Disease Control | Norwegian Institute of Public Health, Department of Virology | Kathrine Stene-Johansen, Kamilla Heddeland Instefjord, Hilde Elshaug, Garcia Llorente Ignacio, Jon Bråte, Engebretsen Serina Beate,Pedersen Benedikte Nevjen, Line Victoria Moen, Debech Nadia, Atiya R Ali,Marie Paulsen Madsen, Rasmus Riis Kopperud, Hilde Vollan, Karoline Bragstad, Olav Hungnes |
| EPI_ISL_2790016, EPI_ISL_2790017 | Furst Medical Laboratory | Norwegian Institute of Public Health, Department of Virology | Kathrine Stene-Johansen, Kamilla Heddeland Instefjord, Hilde Elshaug, Garcia Llorente Ignacio, Jon Bråte, Engebretsen Serina Beate,Pedersen Benedikte Nevjen, Line Victoria Moen, Debech Nadia, Atiya R Ali,Marie Paulsen Madsen, Rasmus Riis Kopperud, Hilde Vollan, Karoline Bragstad, Olav Hungnes |
| EPI_ISL_2790020, EPI_ISL_2790022 | Vestfold Hospital, Toensberg Department of Microbiology | Norwegian Institute of Public Health, Department of Virology | Kathrine Stene-Johansen, Kamilla Heddeland Instefjord, Hilde Elshaug, Garcia Llorente Ignacio, Jon Bråte, Engebretsen Serina Beate,Pedersen Benedikte Nevjen, Line Victoria Moen, Debech Nadia, Atiya R Ali,Marie Paulsen Madsen, Rasmus Riis Kopperud, Hilde Vollan, Karoline Bragstad, Olav Hungnes |
| EPI_ISL_2790058, EPI_ISL_2790059, EPI_ISL_2790060 | MD PHL | MD PHL | Maryland Department of Health Laboratories Administration |
| EPI_ISL_2790082, EPI_ISL_2790084, EPI_ISL_2790088, EPI_ISL_2790091, EPI_ISL_2790092, EPI_ISL_2790093, EPI_ISL_2790094 | University Hospitals of Geneva, Laboratory of Virology | HUG, Laboratory of Virology and the Health2030 Genome Center | Samuel Cordey, Ana Rita Goncalves, Laurent Kaiser, Lorenzo Cerutti, Henri Pegeot, Melyssa Elies, Deborah Penet, Keith Harshman, Ioannis Xenarios, Emmanouil Dermitzakis |
| EPI_ISL_2790095, EPI_ISL_2790097, EPI_ISL_2790098, EPI_ISL_2790099, EPI_ISL_2790100, EPI_ISL_2790101, EPI_ISL_2790103, EPI_ISL_2790104, EPI_ISL_2790105, EPI_ISL_2790109, EPI_ISL_2790117, EPI_ISL_2790121, EPI_ISL_2790126, EPI_ISL_2790127, EPI_ISL_2790131 | | | |
| see above | AZDelta | AZ Delta Medical Laboratories in Roeselare, Belgium | Geert Martens, Dieter De Smet, Merijn Vanhee, on behalf of AZ Delta COVID-19 Genomics core (member of Genomic surveillance of SARS-CoV-2 in Belgium network) |
| EPI_ISL_2790186, EPI_ISL_2790204 | Centracare Laboratory Services | Minnesota Department of Health, Public Health Laboratory | Alexandra Lorentz, Jacob Garfin, Matt Plumb, and Xiong Wang |
| EPI_ISL_2790246 | Cabinet Priv | Institut de Pathologie et Genetique (IPG) | Pascale Hilbert, Jérémie Gras |
| EPI_ISL_2790247, EPI_ISL_2790248 | CHWAPI - SITE NOTRE DAME | Institut de Pathologie et Genetique (IPG) | Pascale Hilbert, Jérémie Gras |
| EPI_ISL_2790295, EPI_ISL_2790296, EPI_ISL_2790297, EPI_ISL_2790298, EPI_ISL_2790299, EPI_ISL_2790300, EPI_ISL_2790301, EPI_ISL_2790302, EPI_ISL_2790303, EPI_ISL_2790304, EPI_ISL_2790305 | | | |
| see above | National Public Health Laboratory, National Centre for Infectious Diseases | National Public Health Laboratory, National Centre for Infectious Diseases | Tze Minn Mak, Zhenyang Zhou, Royce Ang, Lin Cui, Raymond Tzer Pin Lin |
| EPI_ISL_2790329 | National Institute of Public Health | State Veterinary Institute Prague | Nagy,A;Cernikova,L;Stara,M;Suri,T;Vecerova,J;Jirincova,H |
| EPI_ISL_2790333, EPI_ISL_2790334, EPI_ISL_2790335, EPI_ISL_2790336, EPI_ISL_2790337, EPI_ISL_2790338, EPI_ISL_2790339, EPI_ISL_2790340, EPI_ISL_2790341, EPI_ISL_2790342, EPI_ISL_2790343, EPI_ISL_2790344, EPI_ISL_2790345, EPI_ISL_2790346, EPI_ISL_2790347, EPI_ISL_2790348, EPI_ISL_2790349, EPI_ISL_2790350, EPI_ISL_2790351, EPI_ISL_2790352, EPI_ISL_2790353, EPI_ISL_2790354, EPI_ISL_2790355, EPI_ISL_2790356, EPI_ISL_2790357, EPI_ISL_2790358, EPI_ISL_2790359, EPI_ISL_2790360, EPI_ISL_2790361, EPI_ISL_2790362, EPI_ISL_2790363, EPI_ISL_2790364, EPI_ISL_2790365, EPI_ISL_2790366, EPI_ISL_2790367, EPI_ISL_2790368, EPI_ISL_2790369, EPI_ISL_2790370, EPI_ISL_2790371, EPI_ISL_2790372, EPI_ISL_2790373, EPI_ISL_2790374, EPI_ISL_2790375, EPI_ISL_2790376, EPI_ISL_2790377, EPI_ISL_2790378, EPI_ISL_2790379, EPI_ISL_2790380, EPI_ISL_2790381, EPI_ISL_2790382, EPI_ISL_2790383, EPI_ISL_2790384, EPI_ISL_2790385, EPI_ISL_2790386, EPI_ISL_2790387 | | | |
| see above | National Public Health Laboratory, National Centre for Infectious Diseases | National Public Health Laboratory, National Centre for Infectious Diseases | Tze Minn Mak, Katherine Ching, Zhenyang Zhou, Royce Ang, Lin Cui, Raymond Tzer Pin Lin |
| EPI_ISL_2790551 | AZ Jan Portaels | Imelda Ziekenhuis | Johan Frans, Hanne Valgaeren, Dagmar Obbels |
| EPI_ISL_2790553 | Imelda Ziekenhuis | Imelda Ziekenhuis | Johan Frans, Hanne Valgaeren, Dagmar Obbels |
| EPI_ISL_2790626, EPI_ISL_2790631, EPI_ISL_2790646, EPI_ISL_2790649, EPI_ISL_2790651, EPI_ISL_2790652, EPI_ISL_2790678, EPI_ISL_2790684, EPI_ISL_2790685, EPI_ISL_2790689 | Colorado Department of Public Health and Environment | Colorado Department of Public Health and Environment | Laura Bankers, Molly C. Hetherington-Rauth, Diana Ir, Alexandria Rossheim, Michael Martin, Mandy Waters, Shannon R. Matzinger, Sarah Elizabeth Totten, Emily A. Travanty |
| EPI_ISL_2790716 | LAB MIKROBIOLOGI FKUI / DINAS KESEHATAN DKI JAKARTA | Genomik Solidaritas Indonesia Laboratorium | Reinhart Gabriel, Anna Christina Brazia, Vania Gavrila Wikasa, Carissa Sintca Wijaya, Dhahlia Agustina Cahyono, Alfin Mohammad Abdillah, Normastuti Adhini Tantyo, Annisa Muthiah Sukirman, Gracia Felias Enos Korompis, Yulia Rosa, Fera Ibrahim, Pratiwi Sudarmono, Widyastuti, Dwi Oktavia, Tiranti Vindha, Ngabila Salama, Anuraj Shankar, Meutia Ayuputeri Kumaheri, Ariel Pradipta |
| EPI_ISL_2790717, EPI_ISL_2790718 | LAB MIKROBIOLOGI FKUI / DINAS KESEHATAN DKI JAKARTA | Genomik Solidaritas Indonesia Laboratorium | Anna Christina Brazia, Reinhart Gabriel, Vania Gavrila Wikasa, Carissa Sintca Wijaya, Dhahlia Agustina Cahyono, Alfin Mohammad Abdillah, Normastuti Adhini Tantyo, Annisa Muthiah Sukirman, Gracia Felias Enos Korompis, Yulia Rosa, Fera Ibrahim, Pratiwi Sudarmono, Widyastuti, Dwi Oktavia, Tiranti Vindha, Ngabila Salama, Anuraj Shankar, Meutia Ayuputeri Kumaheri, Ariel Pradipta |
| EPI_ISL_2790719, EPI_ISL_2790720 | LAB MIKROBIOLOGI FKUI / DINAS KESEHATAN DKI JAKARTA | Genomik Solidaritas Indonesia Laboratorium | Vania Gavrila Wikasa, Anna Christina Brazia, Reinhart Gabriel, Carissa Sintca Wijaya, Dhahlia Agustina Cahyono, Alfin Mohammad Abdillah, Normastuti Adhini Tantyo, Annisa Muthiah Sukirman, Gracia Felias Enos Korompis, Yulia Rosa, Fera Ibrahim, Pratiwi Sudarmono, Widyastuti, Dwi Oktavia, Tiranti Vindha, Ngabila Salama, Anuraj Shankar, Meutia Ayuputeri Kumaheri, Ariel Pradipta |
| EPI_ISL_2790721, EPI_ISL_2790722 | RSUD KEBAYORAN BARU / DINAS KESEHATAN DKI JAKARTA | Genomik Solidaritas Indonesia Laboratorium | Carissa Sintca Wijaya, Vania Gavrila Wikasa, Anna Christina Brazia, Reinhart Gabriel, Dhahlia Agustina Cahyono, Alfin Mohammad Abdillah, Normastuti Adhini Tantyo, Annisa Muthiah Sukirman, Gracia Felias Enos Korompis, Widyastuti, Dwi Oktavia, Tiranti Vindha, Ngabila Salama, Anuraj Shankar, Meutia Ayuputeri Kumaheri, Ariel Pradipta |
| EPI_ISL_2790723 | RSUD KEBAYORAN BARU / DINAS KESEHATAN DKI JAKARTA | Genomik Solidaritas Indonesia Laboratorium | Reinhart Gabriel, Anna Christina Brazia, Vania Gavrila Wikasa, Carissa Sintca Wijaya, Dhahlia Agustina Cahyono, Alfin Mohammad Abdillah, Normastuti Adhini Tantyo, Annisa Muthiah Sukirman, Gracia Felias Enos Korompis, Widyastuti, Dwi Oktavia, Tiranti Vindha, Ngabila Salama, Anuraj Shankar, Meutia Ayuputeri Kumaheri, Ariel Pradipta |
| EPI_ISL_2790724, EPI_ISL_2790725 | RS MRCCC SILOAM SEMANGGI / DINAS KESEHATAN DKI JAKARTA | Genomik Solidaritas Indonesia Laboratorium | Anna Christina Brazia, Reinhart Gabriel, Vania Gavrila Wikasa, Carissa Sintca Wijaya, Dhahlia Agustina Cahyono, Alfin Mohammad Abdillah, Normastuti Adhini Tantyo, Annisa Muthiah Sukirman, Gracia Felias Enos Korompis, Widyastuti, Dwi Oktavia, Tiranti Vindha, Ngabila Salama, Anuraj Shankar, Meutia Ayuputeri Kumaheri, Ariel Pradipta |

| | | | |
|---|---|---|---|
| EPI_ISL_2790726, EPI_ISL_2790727 | RS MRCCC SILOAM SEMANGGI / DINAS KESEHATAN DKI JAKARTA | Genomik Solidaritas Indonesia Laboratorium | Vania Gavrila Wikasa, Anna Christina Brazia, Reinhart Gabriel, Carissa Sintca Wijaya, Dhahlia Agustina Cahyono, Alfin Mohammad Abdillah, Normastuti Adhini Tantyo, Annisa Muthiah Sukirman, Gracia Felias Enos Korompis, Widyastuti, Dwi Oktavia, Tiranti Vindha, Ngabila Salama, Anuraj Shankar, Meutia Ayuputeri Kumaheri, Ariel Pradipta |
| EPI_ISL_2790728, EPI_ISL_2790729 | RS MRCCC SILOAM SEMANGGI / DINAS KESEHATAN DKI JAKARTA | Genomik Solidaritas Indonesia Laboratorium | Carissa Sintca Wijaya, Vania Gavrila Wikasa, Anna Christina Brazia, Reinhart Gabriel, Dhahlia Agustina Cahyono, Alfin Mohammad Abdillah, Normastuti Adhini Tantyo, Annisa Muthiah Sukirman, Gracia Felias Enos Korompis, Widyastuti, Dwi Oktavia, Tiranti Vindha, Ngabila Salama, Anuraj Shankar, Meutia Ayuputeri Kumaheri, Ariel Pradipta |
| EPI_ISL_2790730 | RSUD KEBAYORAN BARU / DINAS KESEHATAN DKI JAKARTA | Genomik Solidaritas Indonesia Laboratorium | Vania Gavrila Wikasa, Anna Christina Brazia, Reinhart Gabriel, Carissa Sintca Wijaya, Dhahlia Agustina Cahyono, Alfin Mohammad Abdillah, Normastuti Adhini Tantyo, Annisa Muthiah Sukirman, Gracia Felias Enos Korompis, Widyastuti, Dwi Oktavia, Tiranti Vindha, Ngabila Salama, Anuraj Shankar, Meutia Ayuputeri Kumaheri, Ariel Pradipta |
| EPI_ISL_2790731 | RSUD PASAR MINGGU / DINAS KESEHATAN DKI JAKARTA | Genomik Solidaritas Indonesia Laboratorium | Carissa Sintca Wijaya, Vania Gavrila Wikasa, Anna Christina Brazia, Reinhart Gabriel, Dhahlia Agustina Cahyono, Alfin Mohammad Abdillah, Normastuti Adhini Tantyo, Annisa Muthiah Sukirman, Gracia Felias Enos Korompis, Widyastuti, Dwi Oktavia, Tiranti Vindha, Ngabila Salama, Anuraj Shankar, Meutia Ayuputeri Kumaheri, Ariel Pradipta |
| EPI_ISL_2790732, EPI_ISL_2790733 | Monterey County Public Health Laboratory | Monterey County Public Health Laboratory | Monterey County Public Health Laboratory |
| EPI_ISL_2790763, EPI_ISL_2790767, EPI_ISL_2790771, EPI_ISL_2790773, EPI_ISL_2790774, EPI_ISL_2790775, EPI_ISL_2790778, EPI_ISL_2790783, EPI_ISL_2790790, EPI_ISL_2790795, EPI_ISL_2790798, EPI_ISL_2790799 | | | |
| see above | Alaska State Virology Laboratory | Alaska State Virology Laboratory | Stephanie DeRonde, Elva House, Jacob Zidek, Lisa Smith, Ph.D., Jack Chen, Ph.D. |
| EPI_ISL_2790862, EPI_ISL_2790863 | Labo Analyses Med | National Reference Center for Viruses of Respiratory Infections, Institut Pasteur, Paris | Marion Barbet, Sylvie Behillil, Méline Bizard, Angela Brisebarre, Camille Capel, Vincent Enouf, Louise Lefrançois, Frédéric Lemoine, Christophe Malabat, Corinne Maufrais, Emmanuelle Permal, Etienne Simon-Lorière, Maud Vanpeene, Sylvie Van der Werf ,GréGoire Potiron |
| EPI_ISL_2790864 | Labo Analyses Med | National Reference Center for Viruses of Respiratory Infections, Institut Pasteur, Paris | Marion Barbet, Sylvie Behillil, Méline Bizard, Angela Brisebarre, Camille Capel, Vincent Enouf, Louise Lefrançois, Frédéric Lemoine, Christophe Malabat, Corinne Maufrais, Emmanuelle Permal, Etienne Simon-Lorière, Maud Vanpeene, Sylvie Van der Werf ,Sabine Trombert |
| EPI_ISL_2790865, EPI_ISL_2790866, EPI_ISL_2790867, EPI_ISL_2790868, EPI_ISL_2790869 | Labo Analyses Med | National Reference Center for Viruses of Respiratory Infections, Institut Pasteur, Paris | Marion Barbet, Sylvie Behillil, Méline Bizard, Angela Brisebarre, Camille Capel, Vincent Enouf, Louise Lefrançois, Frédéric Lemoine, Christophe Malabat, Corinne Maufrais, Emmanuelle Permal, Etienne Simon-Lorière, Maud Vanpeene, Sylvie Van der Werf ,BéNéDicte Roquebert |
| EPI_ISL_2790871 | Labo Analyses Med | National Reference Center for Viruses of Respiratory Infections, Institut Pasteur, Paris | Marion Barbet, Sylvie Behillil, Méline Bizard, Angela Brisebarre, Camille Capel, Vincent Enouf, Louise Lefrançois, Frédéric Lemoine, Christophe Malabat, Corinne Maufrais, Emmanuelle Permal, Etienne Simon-Lorière, Maud Vanpeene, Sylvie Van der Werf ,Sabine Trombert |
| EPI_ISL_2790910, EPI_ISL_2790911 | Hôpital | National Reference Center for Viruses of Respiratory Infections, Institut Pasteur, Paris | Marion Barbet, Sylvie Behillil, Méline Bizard, Angela Brisebarre, Camille Capel, Vincent Enouf, Louise Lefrançois, Frédéric Lemoine, Christophe Malabat, Corinne Maufrais, Emmanuelle Permal, Etienne Simon-Lorière, Maud Vanpeene, Sylvie Van der Werf ,Nabil Gastli |
| EPI_ISL_2790912, EPI_ISL_2790913, EPI_ISL_2790914, EPI_ISL_2790915 | Labo Analyses Med | National Reference Center for Viruses of Respiratory Infections, Institut Pasteur, Paris | Marion Barbet, Sylvie Behillil, Méline Bizard, Angela Brisebarre, Camille Capel, Vincent Enouf, Louise Lefrançois, Frédéric Lemoine, Christophe Malabat, Corinne Maufrais, Emmanuelle Permal, Etienne Simon-Lorière, Maud Vanpeene, Sylvie Van der Werf ,Nicolas Capron |
| EPI_ISL_2790924 | Labo Analyses Med | National Reference Center for Viruses of Respiratory Infections, Institut Pasteur, Paris | Marion Barbet, Sylvie Behillil, Méline Bizard, Angela Brisebarre, Camille Capel, Vincent Enouf, Louise Lefrançois, Frédéric Lemoine, Christophe Malabat, Corinne Maufrais, Emmanuelle Permal, Etienne Simon-Lorière, Maud Vanpeene, Sylvie Van der Werf ,Brieuc Gestin |
| EPI_ISL_2790927 | Hospital | National Reference Center for Viruses of Respiratory Infections, Institut Pasteur, Paris | Marion Barbet, Sylvie Behillil, Méline Bizard, Angela Brisebarre, Camille Capel, Vincent Enouf, Louise Lefrançois, Frédéric Lemoine, Christophe Malabat, Corinne Maufrais, Emmanuelle Permal, Etienne Simon-Lorière, Maud Vanpeene, Sylvie Van der Werf ,Laurence Louvet |
| EPI_ISL_2790929 | Labo Analyses Med | National Reference Center for Viruses of Respiratory Infections, Institut Pasteur, Paris | Marion Barbet, Sylvie Behillil, Méline Bizard, Angela Brisebarre, Camille Capel, Vincent Enouf, Louise Lefrançois, Frédéric Lemoine, Christophe Malabat, Corinne Maufrais, Emmanuelle Permal, Etienne Simon-Lorière, Maud Vanpeene, Sylvie Van der Werf ,BéNéDicte Roquebert |
| EPI_ISL_2790930, EPI_ISL_2790931, EPI_ISL_2790932, EPI_ISL_2790933, EPI_ISL_2790934, EPI_ISL_2790935, EPI_ISL_2790936, EPI_ISL_2790937, EPI_ISL_2790938, EPI_ISL_2790939, EPI_ISL_2790940, EPI_ISL_2790941, EPI_ISL_2790942, EPI_ISL_2790943, EPI_ISL_2790944, EPI_ISL_2790945, EPI_ISL_2790946, EPI_ISL_2790947, EPI_ISL_2790948, EPI_ISL_2790949, EPI_ISL_2790950, EPI_ISL_2790951, EPI_ISL_2790952, EPI_ISL_2790953, EPI_ISL_2790954, EPI_ISL_2790955, EPI_ISL_2790957, EPI_ISL_2790958, EPI_ISL_2790959, EPI_ISL_2790960, EPI_ISL_2790961, EPI_ISL_2790962, EPI_ISL_2790963, EPI_ISL_2790964, EPI_ISL_2790965, EPI_ISL_2790966 | | | |
| see above | Hôpital | National Reference Center for Viruses of Respiratory Infections, Institut Pasteur, Paris | Marion Barbet, Sylvie Behillil, Méline Bizard, Angela Brisebarre, Camille Capel, Vincent Enouf, Louise Lefrançois, Frédéric Lemoine, Christophe Malabat, Corinne Maufrais, Emmanuelle Permal, Etienne Simon-Lorière, Maud Vanpeene, Sylvie Van der Werf ,BéNéDicte Roquebert |
| EPI_ISL_2790969, EPI_ISL_2790972, EPI_ISL_2790973 | Labo Analyses Med | National Reference Center for Viruses of Respiratory Infections, Institut Pasteur, Paris | Marion Barbet, Sylvie Behillil, Méline Bizard, Angela Brisebarre, Camille Capel, Vincent Enouf, Louise Lefrançois, Frédéric Lemoine, Christophe Malabat, Corinne Maufrais, Emmanuelle Permal, Etienne Simon-Lorière, Maud Vanpeene, Sylvie Van der Werf ,Sabine Trombert |
| EPI_ISL_2790974 | Labo Analyses Med | National Reference Center for Viruses of Respiratory Infections, Institut Pasteur, Paris | Marion Barbet, Sylvie Behillil, Méline Bizard, Angela Brisebarre, Camille Capel, Vincent Enouf, Louise Lefrançois, Frédéric Lemoine, Christophe Malabat, Corinne Maufrais, Emmanuelle Permal, Etienne Simon-Lorière, Maud Vanpeene, Sylvie Van der Werf ,GréGoire Potiron |
| EPI_ISL_2790975, EPI_ISL_2790976, EPI_ISL_2790977 | Labo Analyses Med | National Reference Center for Viruses of Respiratory Infections, Institut Pasteur, Paris | Marion Barbet, Sylvie Behillil, Méline Bizard, Angela Brisebarre, Camille Capel, Vincent Enouf, Louise Lefrançois, Frédéric Lemoine, Christophe Malabat, Corinne Maufrais, Emmanuelle Permal, Etienne Simon-Lorière, Maud Vanpeene, Sylvie Van der Werf ,Eve Haguenoer |
| EPI_ISL_2790979, EPI_ISL_2790980 | Labo Analyses Med | National Reference Center for Viruses of Respiratory Infections, Institut Pasteur, Paris | Marion Barbet, Sylvie Behillil, Méline Bizard, Angela Brisebarre, Camille Capel, Vincent Enouf, Louise Lefrançois, Frédéric Lemoine, Christophe Malabat, Corinne Maufrais, Emmanuelle Permal, Etienne Simon-Lorière, Maud Vanpeene, Sylvie Van der Werf ,Nicolas Capron |
| EPI_ISL_2790981, EPI_ISL_2790982, EPI_ISL_2790983, EPI_ISL_2790984 | Hospital | National Reference Center for Viruses of Respiratory Infections, Institut Pasteur, Paris | Marion Barbet, Sylvie Behillil, Méline Bizard, Angela Brisebarre, Camille Capel, Vincent Enouf, Louise Lefrançois, Frédéric Lemoine, Christophe Malabat, Corinne Maufrais, Emmanuelle Permal, Etienne Simon-Lorière, Maud Vanpeene, Sylvie Van der Werf ,JéRôMe Durivault |
| EPI_ISL_2790989, EPI_ISL_2790990 | Labo Analyses Med | National Reference Center for Viruses of Respiratory Infections, Institut Pasteur, Paris | Marion Barbet, Sylvie Behillil, Méline Bizard, Angela Brisebarre, Camille Capel, Vincent Enouf, Louise Lefrançois, Frédéric Lemoine, Christophe Malabat, Corinne Maufrais, Emmanuelle Permal, Etienne Simon-Lorière, Maud Vanpeene, Sylvie Van der Werf ,BéNéDicte Roquebert |
| EPI_ISL_2791224, EPI_ISL_2791225, EPI_ISL_2791226 | Pathology and Laboratory Medicine Institute, Cleveland Clinic, Ohio, USA | Pathology and Laboratory Medicine Institute, Cleveland Clinic, Ohio, USA | Concetta Peck, Jennifer Starbuck, Joy Nakitandwe, Kristen McDonnell, David Plunkett, Zheng Jin Tu, Jay Brock, Yu-Wei Cheng, Gary Procop, Daniel Rhoads, Daniel H. Farkas, David Bosler |
| EPI_ISL_2791264 | BROUSSAIS | Department of Virology, Henri Mondor University Hospital, Assistance Publique Hôpitaux de Paris, Université Paris-Est Créteil, INSERM U955 | Christophe Rodriguez, Slim Fourati, Vanessa Demontant, Guillaume Gricourt, Melissa N'Debi, Alexandre Soulier, Elisabeth Trawinski, Jean-Michel Pawlotsky |
| EPI_ISL_2791268 | GH A.CHENEVIER-H.MONDOR | Department of Virology, Henri Mondor University Hospital, Assistance Publique Hôpitaux de Paris, Université Paris-Est Créteil, INSERM U955 | Christophe Rodriguez, Slim Fourati, Vanessa Demontant, Guillaume Gricourt, Melissa N'Debi, Alexandre Soulier, Elisabeth Trawinski, Jean-Michel Pawlotsky |
| EPI_ISL_2791271, EPI_ISL_2791273, EPI_ISL_2791290 | BROUSSAIS | Department of Virology, Henri Mondor University Hospital, Assistance Publique Hôpitaux de Paris, Université Paris-Est Créteil, INSERM U955 | Christophe Rodriguez, Slim Fourati, Vanessa Demontant, Guillaume Gricourt, Melissa N'Debi, Alexandre Soulier, Elisabeth Trawinski, Jean-Michel Pawlotsky |
| EPI_ISL_2791298, EPI_ISL_2791304 | GH A.CHENEVIER-H.MONDOR | Department of Virology, Henri Mondor University Hospital, Assistance Publique Hôpitaux de Paris, Université Paris-Est Créteil, INSERM U955 | Christophe Rodriguez, Slim Fourati, Vanessa Demontant, Guillaume Gricourt, Melissa N'Debi, Alexandre Soulier, Elisabeth Trawinski, Jean-Michel Pawlotsky |
| EPI_ISL_2791309 | CHU DE DIJON | Department of Virology, Henri Mondor University Hospital, Assistance Publique Hôpitaux de Paris, Université Paris-Est Créteil, INSERM U955 | Christophe Rodriguez, Slim Fourati, Vanessa Demontant, Guillaume Gricourt, Melissa N'Debi, Alexandre Soulier, Elisabeth Trawinski, Jean-Michel Pawlotsky |
| EPI_ISL_2791312 | BIOMNIS EUROFINS IVRY | Department of Virology, Henri Mondor University Hospital, Assistance Publique Hôpitaux de Paris, Université Paris-Est Créteil, INSERM U955 | Christophe Rodriguez, Slim Fourati, Vanessa Demontant, Guillaume Gricourt, Melissa N'Debi, Alexandre Soulier, Elisabeth Trawinski, Jean-Michel Pawlotsky |
| EPI_ISL_2791315, EPI_ISL_2791317 | BROUSSAIS | Department of Virology, Henri Mondor University Hospital, Assistance Publique Hôpitaux de Paris, Université Paris-Est Créteil, INSERM U955 | Christophe Rodriguez, Slim Fourati, Vanessa Demontant, Guillaume Gricourt, Melissa N'Debi, Alexandre Soulier, Elisabeth Trawinski, Jean-Michel Pawlotsky |
| EPI_ISL_2791320 | Hôpital Avicenne | Department of Virology, Henri Mondor University Hospital, Assistance Publique Hôpitaux de Paris, Université Paris-Est Créteil, INSERM U955 | Christophe Rodriguez, Slim Fourati, Vanessa Demontant, Guillaume Gricourt, Melissa N'Debi, Alexandre Soulier, Elisabeth Trawinski, Jean-Michel Pawlotsky |
| EPI_ISL_2791330 | BIOMNIS EUROFINS IVRY | Department of Virology, Henri Mondor University Hospital, Assistance Publique Hôpitaux de Paris, Université Paris-Est Créteil, INSERM U955 | Christophe Rodriguez, Slim Fourati, Vanessa Demontant, Guillaume Gricourt, Melissa N'Debi, Alexandre Soulier, Elisabeth Trawinski, Jean-Michel Pawlotsky |
| EPI_ISL_2791349, EPI_ISL_2791367 | BROUSSAIS | Department of Virology, Henri Mondor University Hospital, Assistance Publique Hôpitaux de Paris, Université Paris-Est Créteil, INSERM U955 | Christophe Rodriguez, Slim Fourati, Vanessa Demontant, Guillaume Gricourt, Melissa N'Debi, Alexandre Soulier, Elisabeth Trawinski, Jean-Michel Pawlotsky |
| EPI_ISL_2791463, EPI_ISL_2791465, EPI_ISL_2791466, EPI_ISL_2791467, EPI_ISL_2791469, EPI_ISL_2791470, EPI_ISL_2791471, | Alaska State Virology Laboratory | Alaska State Virology Laboratory | Stephanie DeRonde, Elva House, Jacob Zidek, Lisa Smith, Ph.D., Jack Chen, Ph.D. |

| | | | |
|---|---|---|---|
| EPI_ISL_2791477, EPI_ISL_2791479, EPI_ISL_2791486 | | | |
| EPI_ISL_2792366 | SARS-CoV-2 Sequencing Castilla y Leon-Spain Consortium | SARS-CoV-2 Sequencing Castilla y Leon-Spain Consortium | Antonio Orduña-Domingo, Marta Hernandez, David Abad, Marta Dominguez-Gil, Silvia Rojo, Laura Sánchez de Prada, Sonsoles Garcinuño Pérez, Carmen Aldea-Mansilla, Mª Fe Brezmes-Valdivieso, Gregoria Megías Lobón, María Antonia García Castro, Carmen Gimeno Crespo, Noelia Arenal Andrés, Carlos Fuster Foz, M. Isabel Fernandez-Natal, Jose María Eiros Bouza |
| EPI_ISL_2792567, EPI_ISL_2792572, EPI_ISL_2792577, EPI_ISL_2792627, EPI_ISL_2792631, EPI_ISL_2792650, EPI_ISL_2792651, EPI_ISL_2792654, EPI_ISL_2792658, EPI_ISL_2792665, EPI_ISL_2792676, EPI_ISL_2792687, EPI_ISL_2792688, EPI_ISL_2792690, EPI_ISL_2792704, EPI_ISL_2792706, EPI_ISL_2792710, EPI_ISL_2792711, EPI_ISL_2792713, EPI_ISL_2792715, EPI_ISL_2792717, EPI_ISL_2792718, EPI_ISL_2792720, EPI_ISL_2792730, EPI_ISL_2792737, EPI_ISL_2792744, EPI_ISL_2792749, EPI_ISL_2792751, EPI_ISL_2792755, EPI_ISL_2792772, EPI_ISL_2792773, EPI_ISL_2792774, EPI_ISL_2792776, EPI_ISL_2792777, EPI_ISL_2792790, EPI_ISL_2792791, EPI_ISL_2792792, EPI_ISL_2792794, EPI_ISL_2792798, EPI_ISL_2792800, EPI_ISL_2792803, EPI_ISL_2792809, EPI_ISL_2792811, EPI_ISL_2792819, EPI_ISL_2792820, EPI_ISL_2792825, EPI_ISL_2792829, EPI_ISL_2792831, EPI_ISL_2792832, EPI_ISL_2792836, EPI_ISL_2792861, EPI_ISL_2792863, EPI_ISL_2792868, EPI_ISL_2792869, EPI_ISL_2792872, EPI_ISL_2792878, EPI_ISL_2792879, EPI_ISL_2792881, EPI_ISL_2792886, EPI_ISL_2792888, EPI_ISL_2792889, EPI_ISL_2792891, EPI_ISL_2792894, EPI_ISL_2792901, EPI_ISL_2792903, EPI_ISL_2792920, EPI_ISL_2792924, EPI_ISL_2792925, EPI_ISL_2792926, EPI_ISL_2792928, EPI_ISL_2792930, EPI_ISL_2792934, EPI_ISL_2792936, EPI_ISL_2792937, EPI_ISL_2792941, EPI_ISL_2792943, EPI_ISL_2792949, EPI_ISL_2792950, EPI_ISL_2792953, EPI_ISL_2792957, EPI_ISL_2792958, EPI_ISL_2792962, EPI_ISL_2792968, EPI_ISL_2793059, EPI_ISL_2793064, EPI_ISL_2793066, EPI_ISL_2793067, EPI_ISL_2793069, EPI_ISL_2793073, EPI_ISL_2793079, EPI_ISL_2793080, EPI_ISL_2793081, EPI_ISL_2793085, EPI_ISL_2793086, EPI_ISL_2793089, EPI_ISL_2793092, EPI_ISL_2793093, EPI_ISL_2793101, EPI_ISL_2793110, EPI_ISL_2793111, EPI_ISL_2793116, EPI_ISL_2793117, EPI_ISL_2793118, EPI_ISL_2793122, EPI_ISL_2793123, EPI_ISL_2793125, EPI_ISL_2793127, EPI_ISL_2793131, EPI_ISL_2793132, EPI_ISL_2793133, EPI_ISL_2793135, EPI_ISL_2793136, EPI_ISL_2793138, EPI_ISL_2793140, EPI_ISL_2793141, EPI_ISL_2793143, EPI_ISL_2793149, EPI_ISL_2793153, EPI_ISL_2793154, EPI_ISL_2793156, EPI_ISL_2793157, EPI_ISL_2793160, EPI_ISL_2793161, EPI_ISL_2793162, EPI_ISL_2793167, EPI_ISL_2793169, EPI_ISL_2793181, EPI_ISL_2793184, EPI_ISL_2793191, EPI_ISL_2793196, EPI_ISL_2793203, EPI_ISL_2793210, EPI_ISL_2793212, EPI_ISL_2793214, EPI_ISL_2793216, EPI_ISL_2793218, EPI_ISL_2793221, EPI_ISL_2793222, EPI_ISL_2793224, EPI_ISL_2793230, EPI_ISL_2793233, EPI_ISL_2793234, EPI_ISL_2793238, EPI_ISL_2793241, EPI_ISL_2793266, EPI_ISL_2793267, EPI_ISL_2793268, EPI_ISL_2793269, EPI_ISL_2793270, EPI_ISL_2793271, EPI_ISL_2793272, EPI_ISL_2793273, EPI_ISL_2793274, EPI_ISL_2793275, EPI_ISL_2793276, EPI_ISL_2793277, EPI_ISL_2793278, EPI_ISL_2793279, EPI_ISL_2793280, EPI_ISL_2793281, EPI_ISL_2793316, EPI_ISL_2793321, EPI_ISL_2793322, EPI_ISL_2793330, EPI_ISL_2793351, EPI_ISL_2793361, EPI_ISL_2793429, EPI_ISL_2793430, EPI_ISL_2793431, EPI_ISL_2793432, EPI_ISL_2793433, EPI_ISL_2793434, EPI_ISL_2793435, EPI_ISL_2793436, EPI_ISL_2793437, EPI_ISL_2793438, EPI_ISL_2793439, EPI_ISL_2793440, EPI_ISL_2793441, EPI_ISL_2793442, EPI_ISL_2793443, EPI_ISL_2793444, EPI_ISL_2793445, EPI_ISL_2793446, EPI_ISL_2793447, EPI_ISL_2793448, EPI_ISL_2793449, EPI_ISL_2793451, EPI_ISL_2793452, EPI_ISL_2793453, EPI_ISL_2793454, EPI_ISL_2793455, EPI_ISL_2793456, EPI_ISL_2793457, EPI_ISL_2793458, EPI_ISL_2793459, EPI_ISL_2793460, EPI_ISL_2793461, EPI_ISL_2793462, EPI_ISL_2793463, EPI_ISL_2793464, EPI_ISL_2793465, EPI_ISL_2793466, EPI_ISL_2793467, EPI_ISL_2793468, EPI_ISL_2793469, EPI_ISL_2793470, EPI_ISL_2793474, EPI_ISL_2793475, EPI_ISL_2793476, EPI_ISL_2793477, EPI_ISL_2793478, EPI_ISL_2793479, EPI_ISL_2793480, EPI_ISL_2793481, EPI_ISL_2793482, EPI_ISL_2793483, EPI_ISL_2793484, EPI_ISL_2793485, EPI_ISL_2793486, EPI_ISL_2793487, EPI_ISL_2793488, EPI_ISL_2793489, EPI_ISL_2793490, EPI_ISL_2793491, EPI_ISL_2793492, EPI_ISL_2793493, EPI_ISL_2793496, EPI_ISL_2793497, EPI_ISL_2793499, EPI_ISL_2793500, EPI_ISL_2793501, EPI_ISL_2793502, EPI_ISL_2793503, EPI_ISL_2793504, EPI_ISL_2793505, EPI_ISL_2793506, EPI_ISL_2793507, EPI_ISL_2793508, EPI_ISL_2793509, EPI_ISL_2793510, EPI_ISL_2793511, EPI_ISL_2793512, EPI_ISL_2793513, EPI_ISL_2793514, EPI_ISL_2793515, EPI_ISL_2793516, EPI_ISL_2793517, EPI_ISL_2793518, EPI_ISL_2793519, EPI_ISL_2793520, EPI_ISL_2793521, EPI_ISL_2793523, EPI_ISL_2793525, EPI_ISL_2793526, EPI_ISL_2793529, EPI_ISL_2793530, EPI_ISL_2793531, EPI_ISL_2793532, EPI_ISL_2793533, EPI_ISL_2793535, EPI_ISL_2793537, EPI_ISL_2793538, EPI_ISL_2793539, EPI_ISL_2793540, EPI_ISL_2793541, EPI_ISL_2793542, EPI_ISL_2793543, EPI_ISL_2793544, EPI_ISL_2793545, EPI_ISL_2793546, EPI_ISL_2793547, EPI_ISL_2793548, EPI_ISL_2793549, EPI_ISL_2793550, EPI_ISL_2793551, EPI_ISL_2793552, EPI_ISL_2793553, EPI_ISL_2793554, EPI_ISL_2793555, EPI_ISL_2793557, EPI_ISL_2793558, EPI_ISL_2793559, EPI_ISL_2793560, EPI_ISL_2793561, EPI_ISL_2793562, EPI_ISL_2793563, EPI_ISL_2793564, EPI_ISL_2793565, EPI_ISL_2793566, EPI_ISL_2793567, EPI_ISL_2793568, EPI_ISL_2793570, EPI_ISL_2793571, EPI_ISL_2793572, EPI_ISL_2793573, EPI_ISL_2793574, EPI_ISL_2793575, EPI_ISL_2793576, EPI_ISL_2793577, EPI_ISL_2793578, EPI_ISL_2793579, EPI_ISL_2793580, EPI_ISL_2793581, EPI_ISL_2793582, EPI_ISL_2793583, EPI_ISL_2793584, EPI_ISL_2793585, EPI_ISL_2793586, EPI_ISL_2793587, EPI_ISL_2793588, EPI_ISL_2793589, EPI_ISL_2793590, EPI_ISL_2793591, EPI_ISL_2793592, EPI_ISL_2793593, EPI_ISL_2793594, EPI_ISL_2793595, EPI_ISL_2793597, EPI_ISL_2793598, EPI_ISL_2793599, EPI_ISL_2793600, EPI_ISL_2793601, EPI_ISL_2793602, EPI_ISL_2793603, EPI_ISL_2793604, EPI_ISL_2793605, EPI_ISL_2793606, EPI_ISL_2793607, EPI_ISL_2793608, EPI_ISL_2793609, EPI_ISL_2793611, EPI_ISL_2793612, EPI_ISL_2793613, EPI_ISL_2793614, EPI_ISL_2793615, EPI_ISL_2793616, EPI_ISL_2793617, EPI_ISL_2793618, EPI_ISL_2793619, EPI_ISL_2793621, EPI_ISL_2793623, EPI_ISL_2793624, EPI_ISL_2793625, EPI_ISL_2793626, EPI_ISL_2793627, EPI_ISL_2793628, EPI_ISL_2793629, EPI_ISL_2793631, EPI_ISL_2793632, EPI_ISL_2793633, EPI_ISL_2793634, EPI_ISL_2793635, EPI_ISL_2793636, EPI_ISL_2793637, EPI_ISL_2793639, EPI_ISL_2793641, EPI_ISL_2793643, EPI_ISL_2793644, EPI_ISL_2793645, EPI_ISL_2793646, EPI_ISL_2793647, EPI_ISL_2793648, EPI_ISL_2793649, EPI_ISL_2793650, EPI_ISL_2793651, EPI_ISL_2793652, EPI_ISL_2793653, EPI_ISL_2793654, EPI_ISL_2793655, EPI_ISL_2793656, EPI_ISL_2793657, EPI_ISL_2793658, EPI_ISL_2793659, EPI_ISL_2793662, EPI_ISL_2793663, EPI_ISL_2793664, EPI_ISL_2793665, EPI_ISL_2793666, EPI_ISL_2793667, EPI_ISL_2793668, EPI_ISL_2793669, EPI_ISL_2793670, EPI_ISL_2793671, EPI_ISL_2793672, EPI_ISL_2793673, EPI_ISL_2793674, EPI_ISL_2793675, EPI_ISL_2793676, EPI_ISL_2793678, EPI_ISL_2793679, EPI_ISL_2793680, EPI_ISL_2793681, EPI_ISL_2793682, EPI_ISL_2793683, EPI_ISL_2793684, EPI_ISL_2793685, EPI_ISL_2793686, EPI_ISL_2793687, EPI_ISL_2793688, EPI_ISL_2793689, EPI_ISL_2793690, EPI_ISL_2793691, EPI_ISL_2793692, EPI_ISL_2793693, EPI_ISL_2793694, EPI_ISL_2793695, EPI_ISL_2793696, EPI_ISL_2793697, EPI_ISL_2793698, EPI_ISL_2793699, EPI_ISL_2793700, EPI_ISL_2793701, EPI_ISL_2793703, EPI_ISL_2793704, EPI_ISL_2793705, EPI_ISL_2793706, EPI_ISL_2793707, EPI_ISL_2793708, EPI_ISL_2793709, EPI_ISL_2793710, EPI_ISL_2793711, EPI_ISL_2793712, EPI_ISL_2793713, EPI_ISL_2793716, EPI_ISL_2793717, EPI_ISL_2793718, EPI_ISL_2793719, EPI_ISL_2793720, EPI_ISL_2793721, EPI_ISL_2793722, EPI_ISL_2793723, EPI_ISL_2793724, EPI_ISL_2793725, EPI_ISL_2793726, EPI_ISL_2793727, EPI_ISL_2793728, EPI_ISL_2793729, EPI_ISL_2793730, EPI_ISL_2793731, EPI_ISL_2793732, EPI_ISL_2793733, EPI_ISL_2793734, EPI_ISL_2793735, EPI_ISL_2793736, EPI_ISL_2793737, EPI_ISL_2793738, EPI_ISL_2793739, EPI_ISL_2793740, EPI_ISL_2793742, EPI_ISL_2793743, EPI_ISL_2793744, EPI_ISL_2793745, EPI_ISL_2793746, EPI_ISL_2793747, EPI_ISL_2793748, EPI_ISL_2793749, EPI_ISL_2793750, EPI_ISL_2793751, EPI_ISL_2793752, EPI_ISL_2793753, EPI_ISL_2793754, EPI_ISL_2793755, EPI_ISL_2793756, EPI_ISL_2793757, EPI_ISL_2793763, EPI_ISL_2793764, EPI_ISL_2793765, EPI_ISL_2793766, EPI_ISL_2793767, EPI_ISL_2793768, EPI_ISL_2793769, EPI_ISL_2793770, EPI_ISL_2793771, EPI_ISL_2793772, EPI_ISL_2793773, EPI_ISL_2793774, EPI_ISL_2793775, EPI_ISL_2793776, EPI_ISL_2793777, EPI_ISL_2793778, EPI_ISL_2793779, EPI_ISL_2793780, EPI_ISL_2793781, EPI_ISL_2793782, EPI_ISL_2793783, EPI_ISL_2793784, EPI_ISL_2793785, EPI_ISL_2793786, EPI_ISL_2793787, EPI_ISL_2793788, EPI_ISL_2793789, EPI_ISL_2793790, EPI_ISL_2793791, EPI_ISL_2793792, EPI_ISL_2793793, EPI_ISL_2793794, EPI_ISL_2793795, EPI_ISL_2793796, EPI_ISL_2793797, EPI_ISL_2793798, EPI_ISL_2793799, EPI_ISL_2793800, EPI_ISL_2793801, EPI_ISL_2793802, EPI_ISL_2793803, EPI_ISL_2793804, EPI_ISL_2793805, EPI_ISL_2793806, EPI_ISL_2793807, EPI_ISL_2793808, EPI_ISL_2793809, EPI_ISL_2793810, EPI_ISL_2793811, EPI_ISL_2793812, EPI_ISL_2793813, EPI_ISL_2793814, EPI_ISL_2793815, EPI_ISL_2793816, EPI_ISL_2793817, EPI_ISL_2793818, EPI_ISL_2793819, EPI_ISL_2793820, EPI_ISL_2793821, EPI_ISL_2793822, EPI_ISL_2793823, EPI_ISL_2793824, EPI_ISL_2793825, EPI_ISL_2793826, EPI_ISL_2793827, EPI_ISL_2793828, EPI_ISL_2793829, EPI_ISL_2793830, EPI_ISL_2793831, EPI_ISL_2793832, EPI_ISL_2793834, EPI_ISL_2793835, EPI_ISL_2793836, EPI_ISL_2793837, EPI_ISL_2793838, EPI_ISL_2793839, EPI_ISL_2793840, EPI_ISL_2793841, EPI_ISL_2793842, EPI_ISL_2793843, EPI_ISL_2793844, EPI_ISL_2793845, EPI_ISL_2793846, EPI_ISL_2793847, EPI_ISL_2793848, EPI_ISL_2793849, EPI_ISL_2793850, EPI_ISL_2793851, EPI_ISL_2793852, EPI_ISL_2793853, EPI_ISL_2793854, EPI_ISL_2793855, EPI_ISL_2793856, EPI_ISL_2793857, EPI_ISL_2793858, EPI_ISL_2793859, EPI_ISL_2793860, EPI_ISL_2793861, EPI_ISL_2793862, EPI_ISL_2793863, EPI_ISL_2793864, EPI_ISL_2793865, EPI_ISL_2793866, EPI_ISL_2793868, EPI_ISL_2793869, EPI_ISL_2793870, EPI_ISL_2793871, EPI_ISL_2793872, EPI_ISL_2793873, EPI_ISL_2793874, EPI_ISL_2793875, EPI_ISL_2793876, EPI_ISL_2793877, EPI_ISL_2793878, EPI_ISL_2793879, EPI_ISL_2793880, EPI_ISL_2793881, EPI_ISL_2793882, EPI_ISL_2793883, EPI_ISL_2793884, EPI_ISL_2793885, EPI_ISL_2793886, EPI_ISL_2793887, EPI_ISL_2793888, EPI_ISL_2793889, EPI_ISL_2793890, EPI_ISL_2793891, EPI_ISL_2793892, EPI_ISL_2793893, EPI_ISL_2793894, EPI_ISL_2793895, EPI_ISL_2793896, EPI_ISL_2793897, EPI_ISL_2793898, EPI_ISL_2793899, EPI_ISL_2793900, EPI_ISL_2793901, EPI_ISL_2793902, EPI_ISL_2793903, EPI_ISL_2793904, EPI_ISL_2793905, EPI_ISL_2793906, EPI_ISL_2793907, EPI_ISL_2793908, EPI_ISL_2793909, EPI_ISL_2793910, EPI_ISL_2793911, EPI_ISL_2793912, EPI_ISL_2793913, EPI_ISL_2793914, EPI_ISL_2793915, EPI_ISL_2793916, EPI_ISL_2793917, EPI_ISL_2793918, EPI_ISL_2793919, EPI_ISL_2793920, EPI_ISL_2793921, EPI_ISL_2793922, EPI_ISL_2793923, EPI_ISL_2793924, EPI_ISL_2793925, EPI_ISL_2793926, EPI_ISL_2793927, EPI_ISL_2793928, EPI_ISL_2793929, EPI_ISL_2793930, EPI_ISL_2793931, EPI_ISL_2793932, EPI_ISL_2793933, EPI_ISL_2793934, EPI_ISL_2793935, EPI_ISL_2793936, EPI_ISL_2793937, EPI_ISL_2793938, EPI_ISL_2793939, EPI_ISL_2793940, EPI_ISL_2793941, EPI_ISL_2793942, EPI_ISL_2793943, EPI_ISL_2793944, EPI_ISL_2793945, EPI_ISL_2793946, EPI_ISL_2793948, EPI_ISL_2793949, EPI_ISL_2793950, EPI_ISL_2793951, EPI_ISL_2793952, EPI_ISL_2793953, EPI_ISL_2793954, EPI_ISL_2793955, EPI_ISL_2793956, EPI_ISL_2793957, EPI_ISL_2793983, EPI_ISL_2793986, EPI_ISL_2793991, EPI_ISL_2793992, EPI_ISL_2793995, EPI_ISL_2793996, EPI_ISL_2794006, EPI_ISL_2794008, EPI_ISL_2794009, EPI_ISL_2794011, EPI_ISL_2794070, EPI_ISL_2794072, EPI_ISL_2794077, EPI_ISL_2794078, EPI_ISL_2794079, EPI_ISL_2794083, EPI_ISL_2794087, EPI_ISL_2794218, EPI_ISL_2794219, EPI_ISL_2794220, EPI_ISL_2794223, EPI_ISL_2794224, EPI_ISL_2794225, EPI_ISL_2794226, EPI_ISL_2794227, EPI_ISL_2794267, EPI_ISL_2794330, EPI_ISL_2794354, EPI_ISL_2794398, EPI_ISL_2794440, EPI_ISL_2794477, EPI_ISL_2794860, EPI_ISL_2794944, EPI_ISL_2794949, EPI_ISL_2794959, EPI_ISL_2794960, EPI_ISL_2794962, EPI_ISL_2794968, EPI_ISL_2794995, EPI_ISL_2795205, EPI_ISL_2795210, EPI_ISL_2795212, EPI_ISL_2795215, EPI_ISL_2795226, EPI_ISL_2795239, EPI_ISL_2795249, EPI_ISL_2795250, EPI_ISL_2795251, EPI_ISL_2795255, EPI_ISL_2795261, EPI_ISL_2795265, EPI_ISL_2795308, EPI_ISL_2795309, EPI_ISL_2795310, EPI_ISL_2795311, EPI_ISL_2795312, EPI_ISL_2795313, EPI_ISL_2795324, EPI_ISL_2795325, EPI_ISL_2795326, EPI_ISL_2795327, EPI_ISL_2795328, EPI_ISL_2795329, EPI_ISL_2795332, EPI_ISL_2795333, EPI_ISL_2795334, EPI_ISL_2795336, EPI_ISL_2795337, EPI_ISL_2795338, EPI_ISL_2795340, EPI_ISL_2795341, EPI_ISL_2795342, EPI_ISL_2795343, EPI_ISL_2795344, EPI_ISL_2795346, EPI_ISL_2795347, EPI_ISL_2795348, EPI_ISL_2795349, EPI_ISL_2795350, EPI_ISL_2795351, EPI_ISL_2795352, EPI_ISL_2795357, EPI_ISL_2795359, EPI_ISL_2795361, EPI_ISL_2795364, EPI_ISL_2795365, EPI_ISL_2795369, EPI_ISL_2795370, EPI_ISL_2795371, EPI_ISL_2795372, EPI_ISL_2795374, EPI_ISL_2795375, EPI_ISL_2795378, EPI_ISL_2795384, EPI_ISL_2795389, EPI_ISL_2795392, EPI_ISL_2795393, EPI_ISL_2795394, EPI_ISL_2795397, EPI_ISL_2795399, EPI_ISL_2795400, EPI_ISL_2795402, EPI_ISL_2795404, EPI_ISL_2795405, EPI_ISL_2795406, EPI_ISL_2795407, EPI_ISL_2795408, EPI_ISL_2795409, EPI_ISL_2795410, EPI_ISL_2795411, EPI_ISL_2795413, EPI_ISL_2795414, EPI_ISL_2795415, EPI_ISL_2795416, EPI_ISL_2795417, EPI_ISL_2795418, EPI_ISL_2795419, EPI_ISL_2795420, EPI_ISL_2795421, EPI_ISL_2795422, EPI_ISL_2795423, EPI_ISL_2795424, EPI_ISL_2795426, EPI_ISL_2795427, EPI_ISL_2795431, EPI_ISL_2795433, EPI_ISL_2795434, EPI_ISL_2795435, EPI_ISL_2795436, EPI_ISL_2795437, EPI_ISL_2795438, EPI_ISL_2795439, EPI_ISL_2795440, EPI_ISL_2795441, EPI_ISL_2795443, EPI_ISL_2795444, EPI_ISL_2795445, EPI_ISL_2795446, EPI_ISL_2795447, EPI_ISL_2795448, EPI_ISL_2795449, EPI_ISL_2795450, EPI_ISL_2795451, EPI_ISL_2795452, EPI_ISL_2795453, EPI_ISL_2795456, EPI_ISL_2795459, EPI_ISL_2795460, EPI_ISL_2795464, EPI_ISL_2795465, EPI_ISL_2795466, EPI_ISL_2795467, EPI_ISL_2795468, EPI_ISL_2795469, EPI_ISL_2795502, EPI_ISL_2795503, EPI_ISL_2795504, EPI_ISL_2795505, EPI_ISL_2795506, EPI_ISL_2795507, EPI_ISL_2795508, EPI_ISL_2795509, EPI_ISL_2795510, EPI_ISL_2795511, EPI_ISL_2795512, EPI_ISL_2795513, EPI_ISL_2795514, EPI_ISL_2795515, EPI_ISL_2795516, EPI_ISL_2795517, EPI_ISL_2795518, EPI_ISL_2795519, EPI_ISL_2795520, EPI_ISL_2795521, EPI_ISL_2795522, EPI_ISL_2795523, EPI_ISL_2795524, EPI_ISL_2795525, EPI_ISL_2795526, EPI_ISL_2795527, EPI_ISL_2795528, EPI_ISL_2795529, EPI_ISL_2795530, EPI_ISL_2795555, EPI_ISL_2795580, EPI_ISL_2795581, EPI_ISL_2795596, EPI_ISL_2795600, EPI_ISL_2795601, EPI_ISL_2795611, EPI_ISL_2795614, EPI_ISL_2795629, EPI_ISL_2795630, EPI_ISL_2795640, EPI_ISL_2795648, EPI_ISL_2795652 | | | |
| see above | Swedish national genomic surveillance program of SARS-CoV-2 | The Public Health Agency of Sweden | Maximilian Riess, Maria Lind Karlberg, Alma Brolund, Swedish national genomic surveillance program of SARS-CoV-2 |
| EPI_ISL_2795663 | RM4 | National Institute for Infectious Diseases (INMI) L. Spallanzani I.R.C.C.S | E Giombini, B Bartolini, M Rueca, G Bonfiglio, O Butera, CEM Gruber, F Santini, F Messina, A Di Caro, MR Capobianchi |
| EPI_ISL_2795682 | Biogroup Bio Lam-LCD Saint-Denis | Department of Virology, Henri Mondor University Hospital, Assistance Publique Hôpitaux de Paris, Université Paris-Est Créteil, INSERM U955 | Christophe Rodriguez, Slim Fourati, Vanessa Demontant, Guillaume Gricourt, Melissa N'Debi, Alexandre Soulier, Elisabeth Trawinski, Jean-Michel Pawlotsky |
| EPI_ISL_2795683, EPI_ISL_2795684 | SELAS MEDILYS | Department of Virology, Henri Mondor University Hospital, Assistance Publique Hôpitaux de Paris, Université Paris-Est Créteil, INSERM U955 | Christophe Rodriguez, Slim Fourati, Vanessa Demontant, Guillaume Gricourt, Melissa N'Debi, Alexandre Soulier, Elisabeth Trawinski, Jean-Michel Pawlotsky |
| EPI_ISL_2795685, EPI_ISL_2795686 | Biogroup Bio Lam-LCD Saint-Denis | Department of Virology, Henri Mondor University Hospital, Assistance Publique Hôpitaux de Paris, Université Paris-Est Créteil, INSERM U955 | Christophe Rodriguez, Slim Fourati, Vanessa Demontant, Guillaume Gricourt, Melissa N'Debi, Alexandre Soulier, Elisabeth Trawinski, Jean-Michel Pawlotsky |
| EPI_ISL_2795689 | Plateau technique Biomer Metz Verriers | Department of Virology, Henri Mondor University Hospital, Assistance Publique Hôpitaux de Paris, Université Paris-Est Créteil, INSERM U955 | Christophe Rodriguez, Slim Fourati, Vanessa Demontant, Guillaume Gricourt, Melissa N'Debi, Alexandre Soulier, Elisabeth Trawinski, Jean-Michel Pawlotsky |
| EPI_ISL_2795690 | SELAS MEDILYS | Department of Virology, Henri Mondor University Hospital, Assistance Publique Hôpitaux de Paris, Université Paris-Est Créteil, INSERM U955 | Christophe Rodriguez, Slim Fourati, Vanessa Demontant, Guillaume Gricourt, Melissa N'Debi, Alexandre Soulier, Elisabeth Trawinski, Jean-Michel Pawlotsky |
| EPI_ISL_2795691 | Cerballiance Paris IDF est Saint-Denis | Department of Virology, Henri Mondor University Hospital, Assistance Publique Hôpitaux de Paris, Université Paris-Est Créteil, INSERM U955 | Christophe Rodriguez, Slim Fourati, Vanessa Demontant, Guillaume Gricourt, Melissa N'Debi, Alexandre Soulier, Elisabeth Trawinski, Jean-Michel Pawlotsky |
| EPI_ISL_2795696 | Biogroup Bio Lam-LCD Saint-Denis | Department of Virology, Henri Mondor University Hospital, Assistance Publique Hôpitaux de Paris, Université Paris-Est Créteil, INSERM U955 | Christophe Rodriguez, Slim Fourati, Vanessa Demontant, Guillaume Gricourt, Melissa N'Debi, Alexandre Soulier, Elisabeth Trawinski, Jean-Michel Pawlotsky |
| EPI_ISL_2795697 | CHU Rouen Normandie | Department of Virology, Henri Mondor University Hospital, Assistance Publique Hôpitaux de Paris, Université Paris-Est Créteil, INSERM U955 | Christophe Rodriguez, Slim Fourati, Vanessa Demontant, Guillaume Gricourt, Melissa N'Debi, Alexandre Soulier, Elisabeth Trawinski, Jean-Michel Pawlotsky |
| EPI_ISL_2795703 | Cerballiance Paris IDF est Saint-Denis | Department of Virology, Henri Mondor University Hospital, Assistance Publique Hôpitaux de Paris, Université Paris-Est Créteil, INSERM U955 | Christophe Rodriguez, Slim Fourati, Vanessa Demontant, Guillaume Gricourt, Melissa N'Debi, Alexandre Soulier, Elisabeth Trawinski, Jean-Michel Pawlotsky |
| EPI_ISL_2795704 | Laboratoire LENYS | Department of Virology, Henri Mondor University Hospital, Assistance Publique Hôpitaux de Paris, Université Paris-Est Créteil, INSERM U955 | Christophe Rodriguez, Slim Fourati, Vanessa Demontant, Guillaume Gricourt, Melissa N'Debi, Alexandre Soulier, Elisabeth Trawinski, Jean-Michel Pawlotsky |
| EPI_ISL_2795707 | Biogroup Bio Lam-LCD Saint-Denis | Department of Virology, Henri Mondor University Hospital, Assistance Publique Hôpitaux de Paris, Université Paris-Est Créteil, INSERM U955 | Christophe Rodriguez, Slim Fourati, Vanessa Demontant, Guillaume Gricourt, Melissa N'Debi, Alexandre Soulier, Elisabeth Trawinski, Jean-Michel Pawlotsky |
| EPI_ISL_2795708, EPI_ISL_2795709, EPI_ISL_2795710, EPI_ISL_2795711 | SELAS MEDILYS | Department of Virology, Henri Mondor University Hospital, Assistance Publique Hôpitaux de Paris, Université Paris-Est Créteil, INSERM U955 | Christophe Rodriguez, Slim Fourati, Vanessa Demontant, Guillaume Gricourt, Melissa N'Debi, Alexandre Soulier, Elisabeth Trawinski, Jean-Michel Pawlotsky |
| EPI_ISL_2795713 | BIOMNIS EUROFINS IVRY | Department of Virology, Henri Mondor University Hospital, Assistance Publique Hôpitaux de Paris, Université Paris-Est Créteil, INSERM U955 | Christophe Rodriguez, Slim Fourati, Vanessa Demontant, Guillaume Gricourt, Melissa N'Debi, Alexandre Soulier, Elisabeth Trawinski, Jean-Michel Pawlotsky |
| EPI_ISL_2795714 | Biogroup Bio Lam-LCD Saint-Denis | Department of Virology, Henri Mondor University Hospital, Assistance Publique Hôpitaux de Paris, Université Paris-Est Créteil, INSERM U955 | Christophe Rodriguez, Slim Fourati, Vanessa Demontant, Guillaume Gricourt, Melissa N'Debi, Alexandre Soulier, Elisabeth Trawinski, Jean-Michel Pawlotsky |
| EPI_ISL_2795715 | BIO67-BIOSPHERE | Department of Virology, Henri Mondor University Hospital, Assistance Publique Hôpitaux de Paris, Université Paris-Est Créteil, INSERM U955 | Christophe Rodriguez, Slim Fourati, Vanessa Demontant, Guillaume Gricourt, Melissa N'Debi, Alexandre Soulier, Elisabeth Trawinski, Jean-Michel Pawlotsky |
| EPI_ISL_2795716 | Biogroup Bio Lam-LCD Saint-Denis | Department of Virology, Henri Mondor University Hospital, Assistance Publique Hôpitaux de Paris, Université Paris-Est Créteil, INSERM U955 | Christophe Rodriguez, Slim Fourati, Vanessa Demontant, Guillaume Gricourt, Melissa N'Debi, Alexandre Soulier, Elisabeth Trawinski, Jean-Michel Pawlotsky |
| EPI_ISL_2795723 | CH SUD FRANCILIEN | Department of Virology, Henri Mondor University Hospital, Assistance Publique Hôpitaux de Paris, Université Paris-Est Créteil, INSERM U955 | Christophe Rodriguez, Slim Fourati, Vanessa Demontant, Guillaume Gricourt, Melissa N'Debi, Alexandre Soulier, Elisabeth Trawinski, Jean-Michel Pawlotsky |
| EPI_ISL_2795726 | laboratoire Belle Epine | Department of Virology, Henri Mondor University Hospital, | Christophe Rodriguez, Slim Fourati, Vanessa Demontant, Guillaume Gricourt, Melissa N'Debi, Alexandre Soulier, Elisabeth Trawinski, Jean-Michel Pawlotsky |

| | | | |
|---|---|---|---|
| | | Assistance Publique Hôpitaux de Paris, Université Paris-Est Créteil, INSERM U955 | |
| EPI_ISL_2795730 | BIOMNIS EUROFINS IVRY | Department of Virology, Henri Mondor University Hospital, Assistance Publique Hôpitaux de Paris, Université Paris-Est Créteil, INSERM U955 | Christophe Rodriguez, Slim Fourati, Vanessa Demontant, Guillaume Gricourt, Melissa N'Debi, Alexandre Soulier, Elisabeth Trawinski, Jean-Michel Pawlotsky |
| EPI_ISL_2795735, EPI_ISL_2795736 | Biogroup Bio Lam-LCD Saint-Denis | Department of Virology, Henri Mondor University Hospital, Assistance Publique Hôpitaux de Paris, Université Paris-Est Créteil, INSERM U955 | Christophe Rodriguez, Slim Fourati, Vanessa Demontant, Guillaume Gricourt, Melissa N'Debi, Alexandre Soulier, Elisabeth Trawinski, Jean-Michel Pawlotsky |
| EPI_ISL_2795740, EPI_ISL_2795742 | SELAS MEDILYS | Department of Virology, Henri Mondor University Hospital, Assistance Publique Hôpitaux de Paris, Université Paris-Est Créteil, INSERM U955 | Christophe Rodriguez, Slim Fourati, Vanessa Demontant, Guillaume Gricourt, Melissa N'Debi, Alexandre Soulier, Elisabeth Trawinski, Jean-Michel Pawlotsky |
| EPI_ISL_2795745 | Biogroup Bio Lam-LCD Saint-Denis | Department of Virology, Henri Mondor University Hospital, Assistance Publique Hôpitaux de Paris, Université Paris-Est Créteil, INSERM U955 | Christophe Rodriguez, Slim Fourati, Vanessa Demontant, Guillaume Gricourt, Melissa N'Debi, Alexandre Soulier, Elisabeth Trawinski, Jean-Michel Pawlotsky |
| EPI_ISL_2795746 | SELAS MEDILYS | Department of Virology, Henri Mondor University Hospital, Assistance Publique Hôpitaux de Paris, Université Paris-Est Créteil, INSERM U955 | Christophe Rodriguez, Slim Fourati, Vanessa Demontant, Guillaume Gricourt, Melissa N'Debi, Alexandre Soulier, Elisabeth Trawinski, Jean-Michel Pawlotsky |
| EPI_ISL_2795751, EPI_ISL_2795752 | Biogroup Bio Lam-LCD Saint-Denis | Department of Virology, Henri Mondor University Hospital, Assistance Publique Hôpitaux de Paris, Université Paris-Est Créteil, INSERM U955 | Christophe Rodriguez, Slim Fourati, Vanessa Demontant, Guillaume Gricourt, Melissa N'Debi, Alexandre Soulier, Elisabeth Trawinski, Jean-Michel Pawlotsky |
| EPI_ISL_2795753 | laboratoire Belle Epine | Department of Virology, Henri Mondor University Hospital, Assistance Publique Hôpitaux de Paris, Université Paris-Est Créteil, INSERM U955 | Christophe Rodriguez, Slim Fourati, Vanessa Demontant, Guillaume Gricourt, Melissa N'Debi, Alexandre Soulier, Elisabeth Trawinski, Jean-Michel Pawlotsky |
| EPI_ISL_2795755 | Biogroup Bio Lam-LCD Saint-Denis | Department of Virology, Henri Mondor University Hospital, Assistance Publique Hôpitaux de Paris, Université Paris-Est Créteil, INSERM U955 | Christophe Rodriguez, Slim Fourati, Vanessa Demontant, Guillaume Gricourt, Melissa N'Debi, Alexandre Soulier, Elisabeth Trawinski, Jean-Michel Pawlotsky |
| EPI_ISL_2795757 | SELAS MEDILYS | Department of Virology, Henri Mondor University Hospital, Assistance Publique Hôpitaux de Paris, Université Paris-Est Créteil, INSERM U955 | Christophe Rodriguez, Slim Fourati, Vanessa Demontant, Guillaume Gricourt, Melissa N'Debi, Alexandre Soulier, Elisabeth Trawinski, Jean-Michel Pawlotsky |
| EPI_ISL_2795764 | Biogroup Bio Lam-LCD Saint-Denis | Department of Virology, Henri Mondor University Hospital, Assistance Publique Hôpitaux de Paris, Université Paris-Est Créteil, INSERM U955 | Christophe Rodriguez, Slim Fourati, Vanessa Demontant, Guillaume Gricourt, Melissa N'Debi, Alexandre Soulier, Elisabeth Trawinski, Jean-Michel Pawlotsky |
| EPI_ISL_2795765 | BIO67-BIOSPHERE | Department of Virology, Henri Mondor University Hospital, Assistance Publique Hôpitaux de Paris, Université Paris-Est Créteil, INSERM U955 | Christophe Rodriguez, Slim Fourati, Vanessa Demontant, Guillaume Gricourt, Melissa N'Debi, Alexandre Soulier, Elisabeth Trawinski, Jean-Michel Pawlotsky |
| EPI_ISL_2795771 | Biogroup Bio Lam-LCD Saint-Denis | Department of Virology, Henri Mondor University Hospital, Assistance Publique Hôpitaux de Paris, Université Paris-Est Créteil, INSERM U955 | Christophe Rodriguez, Slim Fourati, Vanessa Demontant, Guillaume Gricourt, Melissa N'Debi, Alexandre Soulier, Elisabeth Trawinski, Jean-Michel Pawlotsky |
| EPI_ISL_2795772 | Cerballiance Paris IDF est Saint-Denis | Department of Virology, Henri Mondor University Hospital, Assistance Publique Hôpitaux de Paris, Université Paris-Est Créteil, INSERM U955 | Christophe Rodriguez, Slim Fourati, Vanessa Demontant, Guillaume Gricourt, Melissa N'Debi, Alexandre Soulier, Elisabeth Trawinski, Jean-Michel Pawlotsky |
| EPI_ISL_2795774 | SELAS MEDILYS | Department of Virology, Henri Mondor University Hospital, Assistance Publique Hôpitaux de Paris, Université Paris-Est Créteil, INSERM U955 | Christophe Rodriguez, Slim Fourati, Vanessa Demontant, Guillaume Gricourt, Melissa N'Debi, Alexandre Soulier, Elisabeth Trawinski, Jean-Michel Pawlotsky |
| EPI_ISL_2795776 | Biogroup Bio Lam-LCD Saint-Denis | Department of Virology, Henri Mondor University Hospital, Assistance Publique Hôpitaux de Paris, Université Paris-Est Créteil, INSERM U955 | Christophe Rodriguez, Slim Fourati, Vanessa Demontant, Guillaume Gricourt, Melissa N'Debi, Alexandre Soulier, Elisabeth Trawinski, Jean-Michel Pawlotsky |
| EPI_ISL_2795777 | CH SUD FRANCILIEN | Department of Virology, Henri Mondor University Hospital, Assistance Publique Hôpitaux de Paris, Université Paris-Est Créteil, INSERM U955 | Christophe Rodriguez, Slim Fourati, Vanessa Demontant, Guillaume Gricourt, Melissa N'Debi, Alexandre Soulier, Elisabeth Trawinski, Jean-Michel Pawlotsky |
| EPI_ISL_2795780 | Biogroup Bio Lam-LCD Saint-Denis | Department of Virology, Henri Mondor University Hospital, Assistance Publique Hôpitaux de Paris, Université Paris-Est Créteil, INSERM U955 | Christophe Rodriguez, Slim Fourati, Vanessa Demontant, Guillaume Gricourt, Melissa N'Debi, Alexandre Soulier, Elisabeth Trawinski, Jean-Michel Pawlotsky |
| EPI_ISL_2795781 | BIOLLAN Brognard | Department of Virology, Henri Mondor University Hospital, Assistance Publique Hôpitaux de Paris, Université Paris-Est Créteil, INSERM U955 | Christophe Rodriguez, Slim Fourati, Vanessa Demontant, Guillaume Gricourt, Melissa N'Debi, Alexandre Soulier, Elisabeth Trawinski, Jean-Michel Pawlotsky |
| EPI_ISL_2795782 | Biogroup Bio Lam-LCD Saint-Denis | Department of Virology, Henri Mondor University Hospital, Assistance Publique Hôpitaux de Paris, Université Paris-Est Créteil, INSERM U955 | Christophe Rodriguez, Slim Fourati, Vanessa Demontant, Guillaume Gricourt, Melissa N'Debi, Alexandre Soulier, Elisabeth Trawinski, Jean-Michel Pawlotsky |
| EPI_ISL_2795786 | Cerballiance Paris IDF est Saint-Denis | Department of Virology, Henri Mondor University Hospital, Assistance Publique Hôpitaux de Paris, Université Paris-Est Créteil, INSERM U955 | Christophe Rodriguez, Slim Fourati, Vanessa Demontant, Guillaume Gricourt, Melissa N'Debi, Alexandre Soulier, Elisabeth Trawinski, Jean-Michel Pawlotsky |
| EPI_ISL_2795787 | SELAS MEDILYS | Department of Virology, Henri Mondor University Hospital, Assistance Publique Hôpitaux de Paris, Université Paris-Est Créteil, INSERM U955 | Christophe Rodriguez, Slim Fourati, Vanessa Demontant, Guillaume Gricourt, Melissa N'Debi, Alexandre Soulier, Elisabeth Trawinski, Jean-Michel Pawlotsky |
| EPI_ISL_2795789 | Biogroup Bio Lam-LCD Saint-Denis | Department of Virology, Henri Mondor University Hospital, Assistance Publique Hôpitaux de Paris, Université Paris-Est Créteil, INSERM U955 | Christophe Rodriguez, Slim Fourati, Vanessa Demontant, Guillaume Gricourt, Melissa N'Debi, Alexandre Soulier, Elisabeth Trawinski, Jean-Michel Pawlotsky |
| EPI_ISL_2795791 | BIO67-BIOSPHERE | Department of Virology, Henri Mondor University Hospital, Assistance Publique Hôpitaux de Paris, Université Paris-Est Créteil, INSERM U955 | Christophe Rodriguez, Slim Fourati, Vanessa Demontant, Guillaume Gricourt, Melissa N'Debi, Alexandre Soulier, Elisabeth Trawinski, Jean-Michel Pawlotsky |
| EPI_ISL_2795793 | SELAS MEDILYS | Department of Virology, Henri Mondor University Hospital, Assistance Publique Hôpitaux de Paris, Université Paris-Est Créteil, INSERM U955 | Christophe Rodriguez, Slim Fourati, Vanessa Demontant, Guillaume Gricourt, Melissa N'Debi, Alexandre Soulier, Elisabeth Trawinski, Jean-Michel Pawlotsky |
| EPI_ISL_2795797 | laboratoire Belle Epine | Department of Virology, Henri Mondor University Hospital, Assistance Publique Hôpitaux de Paris, Université Paris-Est Créteil, INSERM U955 | Christophe Rodriguez, Slim Fourati, Vanessa Demontant, Guillaume Gricourt, Melissa N'Debi, Alexandre Soulier, Elisabeth Trawinski, Jean-Michel Pawlotsky |
| EPI_ISL_2795798 | Biogroup Bio Lam-LCD Saint-Denis | Department of Virology, Henri Mondor University Hospital, Assistance Publique Hôpitaux de Paris, Université Paris-Est Créteil, INSERM U955 | Christophe Rodriguez, Slim Fourati, Vanessa Demontant, Guillaume Gricourt, Melissa N'Debi, Alexandre Soulier, Elisabeth Trawinski, Jean-Michel Pawlotsky |
| EPI_ISL_2795799 | CH. E. MULLER | Department of Virology, Henri Mondor University Hospital, Assistance Publique Hôpitaux de Paris, Université Paris-Est Créteil, INSERM U955 | Christophe Rodriguez, Slim Fourati, Vanessa Demontant, Guillaume Gricourt, Melissa N'Debi, Alexandre Soulier, Elisabeth Trawinski, Jean-Michel Pawlotsky |
| EPI_ISL_2795801 | Laboratoire BioCarmes | Department of Virology, Henri Mondor University Hospital, Assistance Publique Hôpitaux de Paris, Université Paris-Est Créteil, INSERM U955 | Christophe Rodriguez, Slim Fourati, Vanessa Demontant, Guillaume Gricourt, Melissa N'Debi, Alexandre Soulier, Elisabeth Trawinski, Jean-Michel Pawlotsky |
| EPI_ISL_2795804 | SELAS MEDILYS | Department of Virology, Henri Mondor University Hospital, Assistance Publique Hôpitaux de Paris, Université Paris-Est Créteil, INSERM U955 | Christophe Rodriguez, Slim Fourati, Vanessa Demontant, Guillaume Gricourt, Melissa N'Debi, Alexandre Soulier, Elisabeth Trawinski, Jean-Michel Pawlotsky |
| EPI_ISL_2795805 | Biogroup Bio Lam-LCD Saint-Denis | Department of Virology, Henri Mondor University Hospital, Assistance Publique Hôpitaux de Paris, Université Paris-Est Créteil, INSERM U955 | Christophe Rodriguez, Slim Fourati, Vanessa Demontant, Guillaume Gricourt, Melissa N'Debi, Alexandre Soulier, Elisabeth Trawinski, Jean-Michel Pawlotsky |
| EPI_ISL_2795810 | BIO67-BIOSPHERE | Department of Virology, Henri Mondor University Hospital, Assistance Publique Hôpitaux de Paris, Université Paris-Est Créteil, INSERM U955 | Christophe Rodriguez, Slim Fourati, Vanessa Demontant, Guillaume Gricourt, Melissa N'Debi, Alexandre Soulier, Elisabeth Trawinski, Jean-Michel Pawlotsky |
| EPI_ISL_2795811 | Cerballiance Paris IDF est Saint-Denis | Department of Virology, Henri Mondor University Hospital, | Christophe Rodriguez, Slim Fourati, Vanessa Demontant, Guillaume Gricourt, Melissa N'Debi, Alexandre Soulier, Elisabeth Trawinski, Jean-Michel Pawlotsky |

| | | | |
|---|---|---|---|
| | | Assistance Publique Hôpitaux de Paris, Université Paris-Est Créteil, INSERM U955 | |
| EPI_ISL_2795812 | Biogroup Bio Lam-LCD Saint-Denis | Department of Virology, Henri Mondor University Hospital, Assistance Publique Hôpitaux de Paris, Université Paris-Est Créteil, INSERM U955 | Christophe Rodriguez, Slim Fourati, Vanessa Demontant, Guillaume Gricourt, Melissa N'Debi, Alexandre Soulier, Elisabeth Trawinski, Jean-Michel Pawlotsky |
| EPI_ISL_2795814 | Laboratoire LENYS | Department of Virology, Henri Mondor University Hospital, Assistance Publique Hôpitaux de Paris, Université Paris-Est Créteil, INSERM U955 | Christophe Rodriguez, Slim Fourati, Vanessa Demontant, Guillaume Gricourt, Melissa N'Debi, Alexandre Soulier, Elisabeth Trawinski, Jean-Michel Pawlotsky |
| EPI_ISL_2795816 | SELAS MEDILYS | Department of Virology, Henri Mondor University Hospital, Assistance Publique Hôpitaux de Paris, Université Paris-Est Créteil, INSERM U955 | Christophe Rodriguez, Slim Fourati, Vanessa Demontant, Guillaume Gricourt, Melissa N'Debi, Alexandre Soulier, Elisabeth Trawinski, Jean-Michel Pawlotsky |
| EPI_ISL_2795817, EPI_ISL_2795819, EPI_ISL_2795821 | Biogroup Bio Lam-LCD Saint-Denis | Department of Virology, Henri Mondor University Hospital, Assistance Publique Hôpitaux de Paris, Université Paris-Est Créteil, INSERM U955 | Christophe Rodriguez, Slim Fourati, Vanessa Demontant, Guillaume Gricourt, Melissa N'Debi, Alexandre Soulier, Elisabeth Trawinski, Jean-Michel Pawlotsky |
| EPI_ISL_2795824 | CH JURA SUD | Department of Virology, Henri Mondor University Hospital, Assistance Publique Hôpitaux de Paris, Université Paris-Est Créteil, INSERM U955 | Christophe Rodriguez, Slim Fourati, Vanessa Demontant, Guillaume Gricourt, Melissa N'Debi, Alexandre Soulier, Elisabeth Trawinski, Jean-Michel Pawlotsky |
| EPI_ISL_2795828 | Biogroup Bio Lam-LCD Saint-Denis | Department of Virology, Henri Mondor University Hospital, Assistance Publique Hôpitaux de Paris, Université Paris-Est Créteil, INSERM U955 | Christophe Rodriguez, Slim Fourati, Vanessa Demontant, Guillaume Gricourt, Melissa N'Debi, Alexandre Soulier, Elisabeth Trawinski, Jean-Michel Pawlotsky |
| EPI_ISL_2795832 | SELAS MEDILYS | Department of Virology, Henri Mondor University Hospital, Assistance Publique Hôpitaux de Paris, Université Paris-Est Créteil, INSERM U955 | Christophe Rodriguez, Slim Fourati, Vanessa Demontant, Guillaume Gricourt, Melissa N'Debi, Alexandre Soulier, Elisabeth Trawinski, Jean-Michel Pawlotsky |
| EPI_ISL_2795833 | Biogroup Bio Lam-LCD Saint-Denis | Department of Virology, Henri Mondor University Hospital, Assistance Publique Hôpitaux de Paris, Université Paris-Est Créteil, INSERM U955 | Christophe Rodriguez, Slim Fourati, Vanessa Demontant, Guillaume Gricourt, Melissa N'Debi, Alexandre Soulier, Elisabeth Trawinski, Jean-Michel Pawlotsky |
| EPI_ISL_2795834 | Cerballiance Paris IDF est Saint-Denis | Department of Virology, Henri Mondor University Hospital, Assistance Publique Hôpitaux de Paris, Université Paris-Est Créteil, INSERM U955 | Christophe Rodriguez, Slim Fourati, Vanessa Demontant, Guillaume Gricourt, Melissa N'Debi, Alexandre Soulier, Elisabeth Trawinski, Jean-Michel Pawlotsky |
| EPI_ISL_2795835 | Biogroup Bio Lam-LCD Saint-Denis | Department of Virology, Henri Mondor University Hospital, Assistance Publique Hôpitaux de Paris, Université Paris-Est Créteil, INSERM U955 | Christophe Rodriguez, Slim Fourati, Vanessa Demontant, Guillaume Gricourt, Melissa N'Debi, Alexandre Soulier, Elisabeth Trawinski, Jean-Michel Pawlotsky |
| EPI_ISL_2795836 | BIO67-BIOSPHERE | Department of Virology, Henri Mondor University Hospital, Assistance Publique Hôpitaux de Paris, Université Paris-Est Créteil, INSERM U955 | Christophe Rodriguez, Slim Fourati, Vanessa Demontant, Guillaume Gricourt, Melissa N'Debi, Alexandre Soulier, Elisabeth Trawinski, Jean-Michel Pawlotsky |
| EPI_ISL_2795838 | Biogroup Bio Lam-LCD Saint-Denis | Department of Virology, Henri Mondor University Hospital, Assistance Publique Hôpitaux de Paris, Université Paris-Est Créteil, INSERM U955 | Christophe Rodriguez, Slim Fourati, Vanessa Demontant, Guillaume Gricourt, Melissa N'Debi, Alexandre Soulier, Elisabeth Trawinski, Jean-Michel Pawlotsky |
| EPI_ISL_2795839 | SELAS MEDILYS | Department of Virology, Henri Mondor University Hospital, Assistance Publique Hôpitaux de Paris, Université Paris-Est Créteil, INSERM U955 | Christophe Rodriguez, Slim Fourati, Vanessa Demontant, Guillaume Gricourt, Melissa N'Debi, Alexandre Soulier, Elisabeth Trawinski, Jean-Michel Pawlotsky |
| EPI_ISL_2795841 | CH SUD FRANCILIEN | Department of Virology, Henri Mondor University Hospital, Assistance Publique Hôpitaux de Paris, Université Paris-Est Créteil, INSERM U955 | Christophe Rodriguez, Slim Fourati, Vanessa Demontant, Guillaume Gricourt, Melissa N'Debi, Alexandre Soulier, Elisabeth Trawinski, Jean-Michel Pawlotsky |
| EPI_ISL_2795842, EPI_ISL_2795843, EPI_ISL_2795844, EPI_ISL_2795845, EPI_ISL_2795848 | Biogroup Bio Lam-LCD Saint-Denis | Department of Virology, Henri Mondor University Hospital, Assistance Publique Hôpitaux de Paris, Université Paris-Est Créteil, INSERM U955 | Christophe Rodriguez, Slim Fourati, Vanessa Demontant, Guillaume Gricourt, Melissa N'Debi, Alexandre Soulier, Elisabeth Trawinski, Jean-Michel Pawlotsky |
| EPI_ISL_2795851 | SELAS CERBALLIANCE Normandie ouest | Department of Virology, Henri Mondor University Hospital, Assistance Publique Hôpitaux de Paris, Université Paris-Est Créteil, INSERM U955 | Christophe Rodriguez, Slim Fourati, Vanessa Demontant, Guillaume Gricourt, Melissa N'Debi, Alexandre Soulier, Elisabeth Trawinski, Jean-Michel Pawlotsky |
| EPI_ISL_2795855 | Biogroup Bio Lam-LCD Saint-Denis | Department of Virology, Henri Mondor University Hospital, Assistance Publique Hôpitaux de Paris, Université Paris-Est Créteil, INSERM U955 | Christophe Rodriguez, Slim Fourati, Vanessa Demontant, Guillaume Gricourt, Melissa N'Debi, Alexandre Soulier, Elisabeth Trawinski, Jean-Michel Pawlotsky |
| EPI_ISL_2795856 | Laboratoire BioCarmes | Department of Virology, Henri Mondor University Hospital, Assistance Publique Hôpitaux de Paris, Université Paris-Est Créteil, INSERM U955 | Christophe Rodriguez, Slim Fourati, Vanessa Demontant, Guillaume Gricourt, Melissa N'Debi, Alexandre Soulier, Elisabeth Trawinski, Jean-Michel Pawlotsky |
| EPI_ISL_2795857 | Biogroup Bio Lam-LCD Saint-Denis | Department of Virology, Henri Mondor University Hospital, Assistance Publique Hôpitaux de Paris, Université Paris-Est Créteil, INSERM U955 | Christophe Rodriguez, Slim Fourati, Vanessa Demontant, Guillaume Gricourt, Melissa N'Debi, Alexandre Soulier, Elisabeth Trawinski, Jean-Michel Pawlotsky |
| EPI_ISL_2795858 | BIO67-BIOSPHERE | Department of Virology, Henri Mondor University Hospital, Assistance Publique Hôpitaux de Paris, Université Paris-Est Créteil, INSERM U955 | Christophe Rodriguez, Slim Fourati, Vanessa Demontant, Guillaume Gricourt, Melissa N'Debi, Alexandre Soulier, Elisabeth Trawinski, Jean-Michel Pawlotsky |
| EPI_ISL_2795859 | Cerballiance Paris IDF est Saint-Denis | Department of Virology, Henri Mondor University Hospital, Assistance Publique Hôpitaux de Paris, Université Paris-Est Créteil, INSERM U955 | Christophe Rodriguez, Slim Fourati, Vanessa Demontant, Guillaume Gricourt, Melissa N'Debi, Alexandre Soulier, Elisabeth Trawinski, Jean-Michel Pawlotsky |
| EPI_ISL_2795860 | Biogroup Bio Lam-LCD Saint-Denis | Department of Virology, Henri Mondor University Hospital, Assistance Publique Hôpitaux de Paris, Université Paris-Est Créteil, INSERM U955 | Christophe Rodriguez, Slim Fourati, Vanessa Demontant, Guillaume Gricourt, Melissa N'Debi, Alexandre Soulier, Elisabeth Trawinski, Jean-Michel Pawlotsky |
| EPI_ISL_2795862 | SELAS MEDILYS | Department of Virology, Henri Mondor University Hospital, Assistance Publique Hôpitaux de Paris, Université Paris-Est Créteil, INSERM U955 | Christophe Rodriguez, Slim Fourati, Vanessa Demontant, Guillaume Gricourt, Melissa N'Debi, Alexandre Soulier, Elisabeth Trawinski, Jean-Michel Pawlotsky |
| EPI_ISL_2795864 | Biogroup Bio Lam-LCD Saint-Denis | Department of Virology, Henri Mondor University Hospital, Assistance Publique Hôpitaux de Paris, Université Paris-Est Créteil, INSERM U955 | Christophe Rodriguez, Slim Fourati, Vanessa Demontant, Guillaume Gricourt, Melissa N'Debi, Alexandre Soulier, Elisabeth Trawinski, Jean-Michel Pawlotsky |
| EPI_ISL_2795866 | Laboratoire LENYS | Department of Virology, Henri Mondor University Hospital, Assistance Publique Hôpitaux de Paris, Université Paris-Est Créteil, INSERM U955 | Christophe Rodriguez, Slim Fourati, Vanessa Demontant, Guillaume Gricourt, Melissa N'Debi, Alexandre Soulier, Elisabeth Trawinski, Jean-Michel Pawlotsky |
| EPI_ISL_2795867 | SELAS MEDILYS | Department of Virology, Henri Mondor University Hospital, Assistance Publique Hôpitaux de Paris, Université Paris-Est Créteil, INSERM U955 | Christophe Rodriguez, Slim Fourati, Vanessa Demontant, Guillaume Gricourt, Melissa N'Debi, Alexandre Soulier, Elisabeth Trawinski, Jean-Michel Pawlotsky |
| EPI_ISL_2795874 | Cerballiance Paris IDF est Saint-Denis | Department of Virology, Henri Mondor University Hospital, Assistance Publique Hôpitaux de Paris, Université Paris-Est Créteil, INSERM U955 | Christophe Rodriguez, Slim Fourati, Vanessa Demontant, Guillaume Gricourt, Melissa N'Debi, Alexandre Soulier, Elisabeth Trawinski, Jean-Michel Pawlotsky |
| EPI_ISL_2795876 | Biogroup Bio Lam-LCD Saint-Denis | Department of Virology, Henri Mondor University Hospital, Assistance Publique Hôpitaux de Paris, Université Paris-Est Créteil, INSERM U955 | Christophe Rodriguez, Slim Fourati, Vanessa Demontant, Guillaume Gricourt, Melissa N'Debi, Alexandre Soulier, Elisabeth Trawinski, Jean-Michel Pawlotsky |
| EPI_ISL_2795878 | SELAS MEDILYS | Department of Virology, Henri Mondor University Hospital, Assistance Publique Hôpitaux de Paris, Université Paris-Est Créteil, INSERM U955 | Christophe Rodriguez, Slim Fourati, Vanessa Demontant, Guillaume Gricourt, Melissa N'Debi, Alexandre Soulier, Elisabeth Trawinski, Jean-Michel Pawlotsky |
| EPI_ISL_2795879 | Laboratoire LENYS | Department of Virology, Henri Mondor University Hospital, Assistance Publique Hôpitaux de Paris, Université Paris-Est Créteil, INSERM U955 | Christophe Rodriguez, Slim Fourati, Vanessa Demontant, Guillaume Gricourt, Melissa N'Debi, Alexandre Soulier, Elisabeth Trawinski, Jean-Michel Pawlotsky |
| EPI_ISL_2795880 | Biogroup Bio Lam-LCD Saint-Denis | Department of Virology, Henri Mondor University Hospital, Assistance Publique Hôpitaux de Paris, Université Paris-Est Créteil, | Christophe Rodriguez, Slim Fourati, Vanessa Demontant, Guillaume Gricourt, Melissa N'Debi, Alexandre Soulier, Elisabeth Trawinski, Jean-Michel Pawlotsky |

| | | | |
|---|---|---|---|
| | | INSERM U955 | |
| EPI_ISL_2795886 | SELAS MEDILYS | Department of Virology, Henri Mondor University Hospital, Assistance Publique Hôpitaux de Paris, Université Paris-Est Créteil, INSERM U955 | Christophe Rodriguez, Slim Fourati, Vanessa Demontant, Guillaume Gricourt, Melissa N'Debi, Alexandre Soulier, Elisabeth Trawinski, Jean-Michel Pawlotsky |
| EPI_ISL_2795887, EPI_ISL_2795888 | Biogroup Bio Lam-LCD Saint-Denis | Department of Virology, Henri Mondor University Hospital, Assistance Publique Hôpitaux de Paris, Université Paris-Est Créteil, INSERM U955 | Christophe Rodriguez, Slim Fourati, Vanessa Demontant, Guillaume Gricourt, Melissa N'Debi, Alexandre Soulier, Elisabeth Trawinski, Jean-Michel Pawlotsky |
| EPI_ISL_2795889, EPI_ISL_2795890 | Cerballiance Paris IDF est Saint-Denis | Department of Virology, Henri Mondor University Hospital, Assistance Publique Hôpitaux de Paris, Université Paris-Est Créteil, INSERM U955 | Christophe Rodriguez, Slim Fourati, Vanessa Demontant, Guillaume Gricourt, Melissa N'Debi, Alexandre Soulier, Elisabeth Trawinski, Jean-Michel Pawlotsky |
| EPI_ISL_2795892 | Biogroup Bio Lam-LCD Saint-Denis | Department of Virology, Henri Mondor University Hospital, Assistance Publique Hôpitaux de Paris, Université Paris-Est Créteil, INSERM U955 | Christophe Rodriguez, Slim Fourati, Vanessa Demontant, Guillaume Gricourt, Melissa N'Debi, Alexandre Soulier, Elisabeth Trawinski, Jean-Michel Pawlotsky |
| EPI_ISL_2795893 | BIOMNIS EUROFINS IVRY | Department of Virology, Henri Mondor University Hospital, Assistance Publique Hôpitaux de Paris, Université Paris-Est Créteil, INSERM U955 | Christophe Rodriguez, Slim Fourati, Vanessa Demontant, Guillaume Gricourt, Melissa N'Debi, Alexandre Soulier, Elisabeth Trawinski, Jean-Michel Pawlotsky |
| EPI_ISL_2795895 | BIO67-BIOSPHERE | Department of Virology, Henri Mondor University Hospital, Assistance Publique Hôpitaux de Paris, Université Paris-Est Créteil, INSERM U955 | Christophe Rodriguez, Slim Fourati, Vanessa Demontant, Guillaume Gricourt, Melissa N'Debi, Alexandre Soulier, Elisabeth Trawinski, Jean-Michel Pawlotsky |
| EPI_ISL_2795896 | Plateau technique Biomer Metz Verriers | Department of Virology, Henri Mondor University Hospital, Assistance Publique Hôpitaux de Paris, Université Paris-Est Créteil, INSERM U955 | Christophe Rodriguez, Slim Fourati, Vanessa Demontant, Guillaume Gricourt, Melissa N'Debi, Alexandre Soulier, Elisabeth Trawinski, Jean-Michel Pawlotsky |
| EPI_ISL_2795897 | SELAS MEDILYS | Department of Virology, Henri Mondor University Hospital, Assistance Publique Hôpitaux de Paris, Université Paris-Est Créteil, INSERM U955 | Christophe Rodriguez, Slim Fourati, Vanessa Demontant, Guillaume Gricourt, Melissa N'Debi, Alexandre Soulier, Elisabeth Trawinski, Jean-Michel Pawlotsky |
| EPI_ISL_2795898 | Laboratoire LENYS | Department of Virology, Henri Mondor University Hospital, Assistance Publique Hôpitaux de Paris, Université Paris-Est Créteil, INSERM U955 | Christophe Rodriguez, Slim Fourati, Vanessa Demontant, Guillaume Gricourt, Melissa N'Debi, Alexandre Soulier, Elisabeth Trawinski, Jean-Michel Pawlotsky |
| EPI_ISL_2795900 | BIOMNIS EUROFINS IVRY | Department of Virology, Henri Mondor University Hospital, Assistance Publique Hôpitaux de Paris, Université Paris-Est Créteil, INSERM U955 | Christophe Rodriguez, Slim Fourati, Vanessa Demontant, Guillaume Gricourt, Melissa N'Debi, Alexandre Soulier, Elisabeth Trawinski, Jean-Michel Pawlotsky |
| EPI_ISL_2795903 | BIO67-BIOSPHERE | Department of Virology, Henri Mondor University Hospital, Assistance Publique Hôpitaux de Paris, Université Paris-Est Créteil, INSERM U955 | Christophe Rodriguez, Slim Fourati, Vanessa Demontant, Guillaume Gricourt, Melissa N'Debi, Alexandre Soulier, Elisabeth Trawinski, Jean-Michel Pawlotsky |
| EPI_ISL_2795904, EPI_ISL_2795906 | Biogroup Bio Lam-LCD Saint-Denis | Department of Virology, Henri Mondor University Hospital, Assistance Publique Hôpitaux de Paris, Université Paris-Est Créteil, INSERM U955 | Christophe Rodriguez, Slim Fourati, Vanessa Demontant, Guillaume Gricourt, Melissa N'Debi, Alexandre Soulier, Elisabeth Trawinski, Jean-Michel Pawlotsky |
| EPI_ISL_2795909 | CH SUD FRANCILIEN | Department of Virology, Henri Mondor University Hospital, Assistance Publique Hôpitaux de Paris, Université Paris-Est Créteil, INSERM U955 | Christophe Rodriguez, Slim Fourati, Vanessa Demontant, Guillaume Gricourt, Melissa N'Debi, Alexandre Soulier, Elisabeth Trawinski, Jean-Michel Pawlotsky |
| EPI_ISL_2795910 | BIO67-BIOSPHERE | Department of Virology, Henri Mondor University Hospital, Assistance Publique Hôpitaux de Paris, Université Paris-Est Créteil, INSERM U955 | Christophe Rodriguez, Slim Fourati, Vanessa Demontant, Guillaume Gricourt, Melissa N'Debi, Alexandre Soulier, Elisabeth Trawinski, Jean-Michel Pawlotsky |
| EPI_ISL_2795915, EPI_ISL_2795916, EPI_ISL_2795917 | Biogroup Bio Lam-LCD Saint-Denis | Department of Virology, Henri Mondor University Hospital, Assistance Publique Hôpitaux de Paris, Université Paris-Est Créteil, INSERM U955 | Christophe Rodriguez, Slim Fourati, Vanessa Demontant, Guillaume Gricourt, Melissa N'Debi, Alexandre Soulier, Elisabeth Trawinski, Jean-Michel Pawlotsky |
| EPI_ISL_2795918 | Laboratoire LENYS | Department of Virology, Henri Mondor University Hospital, Assistance Publique Hôpitaux de Paris, Université Paris-Est Créteil, INSERM U955 | Christophe Rodriguez, Slim Fourati, Vanessa Demontant, Guillaume Gricourt, Melissa N'Debi, Alexandre Soulier, Elisabeth Trawinski, Jean-Michel Pawlotsky |
| EPI_ISL_2795921, EPI_ISL_2795922, EPI_ISL_2795923, EPI_ISL_2795924, EPI_ISL_2795925, EPI_ISL_2795926, EPI_ISL_2795927, EPI_ISL_2795928, EPI_ISL_2795929, EPI_ISL_2795930, EPI_ISL_2795931, EPI_ISL_2795932, EPI_ISL_2795933, EPI_ISL_2795934, EPI_ISL_2795935, EPI_ISL_2795936, EPI_ISL_2795937, EPI_ISL_2795938, EPI_ISL_2795939, EPI_ISL_2795940, EPI_ISL_2795941, EPI_ISL_2795942, EPI_ISL_2795943, EPI_ISL_2795944, EPI_ISL_2795945, EPI_ISL_2795946, EPI_ISL_2795947, EPI_ISL_2795948, EPI_ISL_2795949, EPI_ISL_2795950, EPI_ISL_2795951, EPI_ISL_2795952, EPI_ISL_2795953, EPI_ISL_2795954, EPI_ISL_2795955, EPI_ISL_2795956, EPI_ISL_2795957, EPI_ISL_2795958, EPI_ISL_2795959, EPI_ISL_2795960, EPI_ISL_2795961, EPI_ISL_2795962, EPI_ISL_2795963, EPI_ISL_2795965 | | | |
| see above | Amazon Dx, AUV2 Laboratory | Amazon Dx, AUV2 Laboratory | Luke Meredith |
| EPI_ISL_2795968, EPI_ISL_2795969, EPI_ISL_2795970, EPI_ISL_2795971, EPI_ISL_2795972, EPI_ISL_2795973, EPI_ISL_2795974, EPI_ISL_2795975, EPI_ISL_2795976, EPI_ISL_2795978, EPI_ISL_2795979 | | | |
| see above | ABC Algarve | Instituto Nacional de Saude (INSA) and Faculdade de Medicina da Universidade de Coimbra | Borges et al |
| EPI_ISL_2795989 | LAC UCoimbra | Instituto Nacional de Saude (INSA) and Faculdade de Medicina da Universidade de Coimbra | Borges et al |
| EPI_ISL_2795990, EPI_ISL_2795991, EPI_ISL_2795992, EPI_ISL_2795993, EPI_ISL_2795994, EPI_ISL_2795995, EPI_ISL_2795996, EPI_ISL_2795997, EPI_ISL_2795998, EPI_ISL_2795999, EPI_ISL_2796000, EPI_ISL_2796001, EPI_ISL_2796002, EPI_ISL_2796003, EPI_ISL_2796004 | | | |
| see above | ULS Norte Alentejano | Instituto Nacional de Saude (INSA) and Faculdade de Medicina da Universidade de Coimbra | Borges et al |
| EPI_ISL_2796008, EPI_ISL_2796011, EPI_ISL_2796012, EPI_ISL_2796013, EPI_ISL_2796020, EPI_ISL_2796021, EPI_ISL_2796022, EPI_ISL_2796024, EPI_ISL_2796025, EPI_ISL_2796028, EPI_ISL_2796031, EPI_ISL_2796032, EPI_ISL_2796033, EPI_ISL_2796034, EPI_ISL_2796035, EPI_ISL_2796036, EPI_ISL_2796037, EPI_ISL_2796038, EPI_ISL_2796040, EPI_ISL_2796041, EPI_ISL_2796043, EPI_ISL_2796049, EPI_ISL_2796052, EPI_ISL_2796053, EPI_ISL_2796054, EPI_ISL_2796055, EPI_ISL_2796058, EPI_ISL_2796069, EPI_ISL_2796070, EPI_ISL_2796071, EPI_ISL_2796072, EPI_ISL_2796073, EPI_ISL_2796074, EPI_ISL_2796075, EPI_ISL_2796076, EPI_ISL_2796079, EPI_ISL_2796085, EPI_ISL_2796086, EPI_ISL_2796087, EPI_ISL_2796088, EPI_ISL_2796089, EPI_ISL_2796090, EPI_ISL_2796093, EPI_ISL_2796097, EPI_ISL_2796099, EPI_ISL_2796100, EPI_ISL_2796102, EPI_ISL_2796105, EPI_ISL_2796106, EPI_ISL_2796113, EPI_ISL_2796114, EPI_ISL_2796116, EPI_ISL_2796117, EPI_ISL_2796119, EPI_ISL_2796121, EPI_ISL_2796130, EPI_ISL_2796132, EPI_ISL_2796133, EPI_ISL_2796134, EPI_ISL_2796137, EPI_ISL_2796146, EPI_ISL_2796149, EPI_ISL_2796150, EPI_ISL_2796152, EPI_ISL_2796153, EPI_ISL_2796154, EPI_ISL_2796156, EPI_ISL_2796158, EPI_ISL_2796159, EPI_ISL_2796163, EPI_ISL_2796164, EPI_ISL_2796165, EPI_ISL_2796166, EPI_ISL_2796167, EPI_ISL_2796168, EPI_ISL_2796169, EPI_ISL_2796170, EPI_ISL_2796171, EPI_ISL_2796173, EPI_ISL_2796174, EPI_ISL_2796175, EPI_ISL_2796176, EPI_ISL_2796177, EPI_ISL_2796178, EPI_ISL_2796179, EPI_ISL_2796181, EPI_ISL_2796182, EPI_ISL_2796190 | | | |
| see above | UNILABS | Instituto Nacional de Saude (INSA) and Instituto Gulbenkian de Ciencia (IGC) | Borges et al |
| EPI_ISL_2796194, EPI_ISL_2796195 | ARS Algarve - Laboratorio Laura Ayres | Instituto Nacional de Saude (INSA) | Borges et al |
| EPI_ISL_2796199, EPI_ISL_2796200, EPI_ISL_2796201, EPI_ISL_2796202, EPI_ISL_2796203, EPI_ISL_2796204 | H Evora | Instituto Nacional de Saude (INSA) | Borges et al |
| EPI_ISL_2796205, EPI_ISL_2796206 | H Divino Espirito Santo - Ponta Delgada | Instituto Nacional de Saude (INSA) | Borges et al |
| EPI_ISL_2796207, EPI_ISL_2796208, EPI_ISL_2796209, EPI_ISL_2796210, EPI_ISL_2796211, EPI_ISL_2796212, EPI_ISL_2796213, EPI_ISL_2796214, EPI_ISL_2796215, EPI_ISL_2796216, EPI_ISL_2796217, EPI_ISL_2796218, EPI_ISL_2796219, EPI_ISL_2796220, EPI_ISL_2796221, EPI_ISL_2796222, EPI_ISL_2796223, EPI_ISL_2796224, EPI_ISL_2796225, EPI_ISL_2796226 | | | |
| see above | INSA | Instituto Nacional de Saude (INSA) | Borges et al |
| EPI_ISL_2796227, EPI_ISL_2796228 | CHLC | Instituto Nacional de Saude (INSA) | Borges et al |
| EPI_ISL_2796230, EPI_ISL_2796231, EPI_ISL_2796232 | Affidea | Instituto Nacional de Saude (INSA) | Borges et al |
| EPI_ISL_2796233, EPI_ISL_2796234 | Labeto - CAB - Leiria | Instituto Nacional de Saude (INSA) | Borges et al |
| EPI_ISL_2796235, EPI_ISL_2796236, EPI_ISL_2796237, EPI_ISL_2796238, EPI_ISL_2796239, EPI_ISL_2796240, EPI_ISL_2796241, EPI_ISL_2796242, EPI_ISL_2796243, EPI_ISL_2796244, EPI_ISL_2796246, EPI_ISL_2796247, EPI_ISL_2796248, EPI_ISL_2796249, EPI_ISL_2796250, EPI_ISL_2796252, EPI_ISL_2796253, EPI_ISL_2796254, EPI_ISL_2796255, EPI_ISL_2796256, EPI_ISL_2796257, EPI_ISL_2796258, EPI_ISL_2796259, EPI_ISL_2796260, EPI_ISL_2796261, EPI_ISL_2796262, EPI_ISL_2796263, EPI_ISL_2796264, EPI_ISL_2796265, EPI_ISL_2796266, EPI_ISL_2796267, EPI_ISL_2796268, EPI_ISL_2796269, EPI_ISL_2796270, EPI_ISL_2796271, EPI_ISL_2796272, EPI_ISL_2796273, EPI_ISL_2796274, EPI_ISL_2796275, EPI_ISL_2796276, EPI_ISL_2796277, EPI_ISL_2796278, EPI_ISL_2796279, EPI_ISL_2796280, EPI_ISL_2796281, EPI_ISL_2796282, EPI_ISL_2796283, EPI_ISL_2796284, EPI_ISL_2796285, EPI_ISL_2796286, EPI_ISL_2796288, EPI_ISL_2796290, EPI_ISL_2796291, EPI_ISL_2796292, EPI_ISL_2796293, EPI_ISL_2796294 | | | |
| see above | ARS Algarve - Laboratorio Laura Ayres | Instituto Nacional de Saude (INSA) | Borges et al |
| EPI_ISL_2796295, EPI_ISL_2796296, EPI_ISL_2796297, EPI_ISL_2796298, EPI_ISL_2796299, EPI_ISL_2796301, EPI_ISL_2796306, EPI_ISL_2796307, EPI_ISL_2796308, EPI_ISL_2796311, EPI_ISL_2796323, EPI_ISL_2796327, EPI_ISL_2796329, EPI_ISL_2796330, EPI_ISL_2796334, EPI_ISL_2796335, EPI_ISL_2796336, EPI_ISL_2796337, EPI_ISL_2796338, EPI_ISL_2796339, EPI_ISL_2796340, EPI_ISL_2796341, EPI_ISL_2796342, EPI_ISL_2796343, EPI_ISL_2796344, EPI_ISL_2796345, EPI_ISL_2796346, EPI_ISL_2796347, EPI_ISL_2796348, EPI_ISL_2796349, EPI_ISL_2796350, EPI_ISL_2796351, EPI_ISL_2796353, EPI_ISL_2796354, EPI_ISL_2796355, EPI_ISL_2796358, EPI_ISL_2796359, EPI_ISL_2796360, EPI_ISL_2796361, EPI_ISL_2796362, EPI_ISL_2796363, EPI_ISL_2796365, EPI_ISL_2796366, EPI_ISL_2796367, EPI_ISL_2796368, EPI_ISL_2796369, EPI_ISL_2796370, EPI_ISL_2796371, EPI_ISL_2796372, EPI_ISL_2796373, EPI_ISL_2796374, EPI_ISL_2796375, EPI_ISL_2796376, EPI_ISL_2796377, EPI_ISL_2796379, EPI_ISL_2796383, EPI_ISL_2796386, EPI_ISL_2796393, EPI_ISL_2796395, EPI_ISL_2796398, EPI_ISL_2796399, EPI_ISL_2796400, EPI_ISL_2796402, EPI_ISL_2796403, EPI_ISL_2796404, EPI_ISL_2796405, EPI_ISL_2796408, EPI_ISL_2796412, EPI_ISL_2796413, EPI_ISL_2796414, EPI_ISL_2796415, EPI_ISL_2796416, EPI_ISL_2796418, EPI_ISL_2796420, EPI_ISL_2796421, EPI_ISL_2796422, EPI_ISL_2796423, EPI_ISL_2796425, EPI_ISL_2796428, EPI_ISL_2796430, EPI_ISL_2796431, EPI_ISL_2796432, EPI_ISL_2796433, EPI_ISL_2796434, EPI_ISL_2796436, EPI_ISL_2796437, EPI_ISL_2796438, EPI_ISL_2796439, EPI_ISL_2796440, EPI_ISL_2796441, EPI_ISL_2796442, EPI_ISL_2796443, EPI_ISL_2796444, EPI_ISL_2796445, EPI_ISL_2796446, EPI_ISL_2796447, EPI_ISL_2796451, EPI_ISL_2796452, EPI_ISL_2796453, EPI_ISL_2796454, EPI_ISL_2796455, EPI_ISL_2796456, EPI_ISL_2796458, EPI_ISL_2796459, EPI_ISL_2796460, EPI_ISL_2796461, EPI_ISL_2796462, EPI_ISL_2796463, EPI_ISL_2796464, EPI_ISL_2796465 | | | |

| | | | |
|---|---|---|---|
| see above | UNILABS | Instituto Nacional de Saude (INSA) | Borges et al |
| EPI_ISL_2796466, EPI_ISL_2796467, EPI_ISL_2796469, EPI_ISL_2796470, EPI_ISL_2796471, EPI_ISL_2796472, EPI_ISL_2796473, EPI_ISL_2796474, EPI_ISL_2796475, EPI_ISL_2796476, EPI_ISL_2796477, EPI_ISL_2796479, EPI_ISL_2796480, EPI_ISL_2796482, EPI_ISL_2796483, EPI_ISL_2796484, EPI_ISL_2796485, EPI_ISL_2796486, EPI_ISL_2796487, EPI_ISL_2796488, EPI_ISL_2796489, EPI_ISL_2796490, EPI_ISL_2796492, EPI_ISL_2796493, EPI_ISL_2796495, EPI_ISL_2796496, EPI_ISL_2796497, EPI_ISL_2796498, EPI_ISL_2796499, EPI_ISL_2796500, EPI_ISL_2796501, EPI_ISL_2796502, EPI_ISL_2796503 | | | |
| see above | UNILABS | Instituto Nacional de Saude (INSA) and BioSystems & Integrative Sciences Institute (BioISI) Genomics Unit, FCUL | Borges et al |
| EPI_ISL_2796520, EPI_ISL_2796522 | National Institute of Laboratory Medicine and Referral Center | Genomic Research Lab, BCSIR | Md. Murshed Hasan Sarkar, Mohammad Samir Uzzaman, Eshrar Osman, Md. Ahasan Habib, Shahina Akter, Tanjina Akhter Banu, Barna Goswami, Iffat Jahan, Mohammad Mohi Uddin, Md. Kamrul Islam, Tasnim Nafisa, Md. Maruf Ahmed Molla, Mahmuda Yeasmin, Asish Kumar Ghosh, Arifa Akram, Md. Salim Khan |
| EPI_ISL_2796523, EPI_ISL_2796525 | National Institute of Laboratory Medicine and Referral Center | Genomic Research Lab, BCSIR | Tanjina Akhter Banu, Md. Murshed Hasan Sarkar, Mohammad Samir Uzzaman, Eshrar Osman, Md. Ahasan Habib, Shahina Akter, Barna Goswami, Iffat Jahan, Mohammad Mohi Uddin, Md. Kamrul Islam, Tasnim Nafisa, Md. Maruf Ahmed Molla, Mahmuda Yeasmin, Asish Kumar Ghosh, Arifa Akram, Md. Salim Khan |
| EPI_ISL_2796526 | National Institute of Laboratory Medicine and Referral Center | Genomic Research Lab, BCSIR | Shahina Akter, Md. Murshed Hasan Sarkar, Mohammad Samir Uzzaman, Eshrar Osman, Md. Ahasan Habib, Barna Goswami, Iffat Jahan, Mohammad Mohi Uddin, Md. Kamrul Islam, Tasnim Nafisa, Md. Maruf Ahmed Molla, Mahmuda Yeasmin, Asish Kumar Ghosh, Arifa Akram, Md. Salim Khan |
| EPI_ISL_2796527 | National Institute of Laboratory Medicine and Referral Center | Genomic Research Lab, BCSIR | Barna Goswami, Md. Murshed Hasan Sarkar, Mohammad Samir Uzzaman, Eshrar Osman, Md. Ahasan Habib, Iffat Jahan, Mohammad Mohi Uddin, Md. Kamrul Islam, Tasnim Nafisa, Md. Maruf Ahmed Molla, Mahmuda Yeasmin, Asish Kumar Ghosh, Arifa Akram, Md. Salim Khan |
| EPI_ISL_2796528 | National Institute of Laboratory Medicine and Referral Center | Genomic Research Lab, BCSIR | Iffat Jahan, Md. Murshed Hasan Sarkar, Mohammad Samir Uzzaman, Eshrar Osman, Md. Ahasan Habib, Mohammad Mohi Uddin, Md. Kamrul Islam, Tasnim Nafisa, Md. Maruf Ahmed Molla, Mahmuda Yeasmin, Asish Kumar Ghosh, Arifa Akram, Md. Salim Khan |
| EPI_ISL_2796530 | National Institute of Laboratory Medicine and Referral Center | Genomic Research Lab, BCSIR | Mohammad Mohi Uddin, Md. Murshed Hasan Sarkar, Mohammad Samir Uzzaman, Eshrar Osman, Md. Ahasan Habib, Shahina Akter, Tanjina Akhter Banu, Barna Goswami, Iffat Jahan, Md. Kamrul Islam, Tasnim Nafisa, Md. Maruf Ahmed Molla, Mahmuda Yeasmin, Asish Kumar Ghosh, Arifa Akram, Md. Salim Khan |
| EPI_ISL_2796531, EPI_ISL_2796710, EPI_ISL_2797382 | National Institute of Laboratory Medicine and Referral Center | Genomic Research Lab, BCSIR | Tasnim Nafisa, Md. Murshed Hasan Sarkar, Mohammad Samir Uzzaman, Eshrar Osman, Md. Ahasan Habib, Shahina Akter, Tanjina Akhter Banu, Barna Goswami, Iffat Jahan, Mohammad Mohi Uddin, Md. Kamrul Islam, Md. Maruf Ahmed Molla, Mahmuda Yeasmin, Asish Kumar Ghosh, Arifa Akram, Md. Salim Khan |
| EPI_ISL_2798352 | Riga East University Hospital, National Microbiology Reference Laboratory | Riga East University Hospital, National Microbiology Reference Laboratory; Eurofins Genomics Europe Sequencing GmbH | Ģirts Šķenders, Reinis Vangravs, Arzu Algulieva, Reinis Zeltmatis, Dārta Pūpola, Jevgenijs Bodrenko, Jūlija Čevere, Ilva Pole, Diāna Dušacka, Sergejs Ņikišins |
| EPI_ISL_2798753, EPI_ISL_2798754, EPI_ISL_2798755 | Centrālā Laboratorija, SIA | Riga East University Hospital, National Microbiology Reference Laboratory; Eurofins Genomics Europe Sequencing GmbH | Stella Lapiņa, Jana Osīte, Ģirts Šķenders, Reinis Vangravs, Arzu Algulieva, Reinis Zeltmatis, Dārta Pūpola, Jevgenijs Bodrenko, Jūlija Čevere, Ilva Pole, Diāna Dušacka, Sergejs Ņikišins |
| EPI_ISL_2798960 | Nacionālais medicīnas serviss - laboratorija, SIA | Riga East University Hospital, National Microbiology Reference Laboratory; Eurofins Genomics Europe Sequencing GmbH | Natalja Mikena, Inita Balta, Ģirts Šķenders, Reinis Vangravs, Arzu Algulieva, Reinis Zeltmatis, Dārta Pūpola, Jevgenijs Bodrenko, Jūlija Čevere, Ilva Pole, Diāna Dušacka, Sergejs Ņikišins |
| EPI_ISL_2798988 | Riga East University Hospital, National Microbiology Reference Laboratory | Riga East University Hospital, National Microbiology Reference Laboratory; Eurofins Genomics Europe Sequencing GmbH | Ģirts Šķenders, Reinis Vangravs, Arzu Algulieva, Reinis Zeltmatis, Dārta Pūpola, Jevgenijs Bodrenko, Jūlija Čevere, Ilva Pole, Diāna Dušacka, Sergejs Ņikišins |
| EPI_ISL_2798996, EPI_ISL_2799000, EPI_ISL_2799099, EPI_ISL_2799114, EPI_ISL_2799137, EPI_ISL_2799152, EPI_ISL_2799171, EPI_ISL_2799172, EPI_ISL_2799198, EPI_ISL_2799200, EPI_ISL_2799206, EPI_ISL_2799215, EPI_ISL_2799230, EPI_ISL_2799236, EPI_ISL_2799240, EPI_ISL_2799256 | | | |
| see above | Centrālā Laboratorija, SIA | Riga East University Hospital, National Microbiology Reference Laboratory; Eurofins Genomics Europe Sequencing GmbH | Stella Lapiņa, Jana Osīte, Ģirts Šķenders, Reinis Vangravs, Arzu Algulieva, Reinis Zeltmatis, Dārta Pūpola, Jevgenijs Bodrenko, Jūlija Čevere, Ilva Pole, Diāna Dušacka, Sergejs Ņikišins |
| EPI_ISL_2800346 | Vitalis Mostar | Alea Genetic Centre | Lana Salihefendic, Dino Pecar, Ivana Ceko, Rijad Konjhodzic |
| EPI_ISL_2800347, EPI_ISL_2800348, EPI_ISL_2800349, EPI_ISL_2800351, EPI_ISL_2800353, EPI_ISL_2800355, EPI_ISL_2800537, EPI_ISL_2800538, EPI_ISL_2800606, EPI_ISL_2800608, EPI_ISL_2800609, EPI_ISL_2800610, EPI_ISL_2800612, EPI_ISL_2800613, EPI_ISL_2800614, EPI_ISL_2800615, EPI_ISL_2800616, EPI_ISL_2800618, EPI_ISL_2800619, EPI_ISL_2800620, EPI_ISL_2800622, EPI_ISL_2800623, EPI_ISL_2800624, EPI_ISL_2800625, EPI_ISL_2800626, EPI_ISL_2800628, EPI_ISL_2800629, EPI_ISL_2800630, EPI_ISL_2800632, EPI_ISL_2800633, EPI_ISL_2800634, EPI_ISL_2800635, EPI_ISL_2800636, EPI_ISL_2800637, EPI_ISL_2800638, EPI_ISL_2800639, EPI_ISL_2800640, EPI_ISL_2800641, EPI_ISL_2800642, EPI_ISL_2800643, EPI_ISL_2800644, EPI_ISL_2800645, EPI_ISL_2800646, EPI_ISL_2800647, EPI_ISL_2800648, EPI_ISL_2800649, EPI_ISL_2800650, EPI_ISL_2800651, EPI_ISL_2800652, EPI_ISL_2800653, EPI_ISL_2800654, EPI_ISL_2800655, EPI_ISL_2800656, EPI_ISL_2800657, EPI_ISL_2800658, EPI_ISL_2800659, EPI_ISL_2800660, EPI_ISL_2800661, EPI_ISL_2800662, EPI_ISL_2800663, EPI_ISL_2800664, EPI_ISL_2800665, EPI_ISL_2800666, EPI_ISL_2800667, EPI_ISL_2800668, EPI_ISL_2800669, EPI_ISL_2800670, EPI_ISL_2800671, EPI_ISL_2800672, EPI_ISL_2800673, EPI_ISL_2800674, EPI_ISL_2800675, EPI_ISL_2800676, EPI_ISL_2800677, EPI_ISL_2800678, EPI_ISL_2800679, EPI_ISL_2800680 | | | |
| see above | State Testing Facility | Altius Institute for Biomedical Research | Daniel Bates, Rebecca Bruders, Michael Buckley, Mark Frerker, Amanda Gale, Clem Green, Muhammad Halimun, Kneshay Harper, Matt Hartman, Alex Isner, Audra Johnson, Jessica Kunder, Lauren Mitchell, Jemma Nelson, Alex Nguyen, Sofia Olsson, Sadie Patraw, Tobias Ragoczy, Joshua Richards, Jean Robinston, Jacob Rodriguez, John Stamatoyannopoulos, Eric Thorland, Julia Wald |
| EPI_ISL_2800728, EPI_ISL_2800766, EPI_ISL_2800824, EPI_ISL_2800868, EPI_ISL_2800870, EPI_ISL_2800871 | Illinois Department of Public Health | Illinois Department of Public Health - Chicago Lab | Vineet K. Dhiman, Ira Heimler, Joel Price |
| EPI_ISL_2800875, EPI_ISL_2800876, EPI_ISL_2800908, EPI_ISL_2800910, EPI_ISL_2800911, EPI_ISL_2800912 | Tampa General Hospital Esoteric Lab | Tampa General Hospital Esoteric Research & Development Lab | Jon Faughn, Deanna Becker, Dominic Uy, Jorge Mecias-Frias, Elaine Vendrone, Vicki Healer, Amorce Lima, Suzane Silbert |
| EPI_ISL_2800970, EPI_ISL_2800974, EPI_ISL_2800975, EPI_ISL_2800977, EPI_ISL_2800989, EPI_ISL_2800990, EPI_ISL_2800991 | Illinois Department of Public Health | Gagnon Lab, Southern Illinois University | Keith Gagnon |
| EPI_ISL_2800992, EPI_ISL_2800993 | Simple Laboratories | Gagnon Lab, Southern Illinois University | Keith Gagnon |
| EPI_ISL_2801110 | Illinois Department of Public Health | Gagnon Lab, Southern Illinois University | Keith Gagnon |
| EPI_ISL_2801162 | Simple Laboratories | Gagnon Lab, Southern Illinois University | Keith Gagnon |
| EPI_ISL_2801427 | LABORATORIUM BADAŃ KLINICZNYCH WSSE w OPOLU | National Institute of Public Health - National Institute of Hygiene | Wołkowicz Tomasz, Sadkowska-Todys Małgorzata, Gierczyńki Rafał |
| EPI_ISL_2801429 | Laboratorium Diagnostyczne Centrum Medycznego Luxmed | National Institute of Public Health - National Institute of Hygiene | Wołkowicz Tomasz, Sadkowska-Todys Małgorzata, Gierczyńki Rafał |
| EPI_ISL_2801430, EPI_ISL_2801431, EPI_ISL_2801432 | IHiT Pracownia Wirusologii Klinicznej | National Institute of Public Health - National Institute of Hygiene | Wołkowicz Tomasz, Sadkowska-Todys Małgorzata, Gierczyńki Rafał |
| EPI_ISL_2801438, EPI_ISL_2801441 | IVY3 Central Lab, Vanderbilt University Medical Center | Lauring Lab, University of Michigan, Department of Microbiology and Immunology | Valesano |
| EPI_ISL_2801478, EPI_ISL_2801479, EPI_ISL_2801480, EPI_ISL_2801481, EPI_ISL_2801482, EPI_ISL_2801483, EPI_ISL_2801484, EPI_ISL_2801485, EPI_ISL_2801486, EPI_ISL_2801487, EPI_ISL_2801488, EPI_ISL_2801489, EPI_ISL_2801490, EPI_ISL_2801491, EPI_ISL_2801492, EPI_ISL_2801493, EPI_ISL_2801494, EPI_ISL_2801499, EPI_ISL_2801500, EPI_ISL_2801501, EPI_ISL_2801502, EPI_ISL_2801503, EPI_ISL_2801504, EPI_ISL_2801505, EPI_ISL_2801506, EPI_ISL_2801509, EPI_ISL_2801511, EPI_ISL_2801512, EPI_ISL_2801513, EPI_ISL_2801514, EPI_ISL_2801515, EPI_ISL_2801516, EPI_ISL_2801517, EPI_ISL_2801518, EPI_ISL_2801519, EPI_ISL_2801520, EPI_ISL_2801521 | | | |
| see above | Nevada State Public Health Laboratory | Nevada State Public Health Laboratory | Andrew Gorzalski, Mark Pandori |
| EPI_ISL_2801552 | Wyoming Public Health Laboratory | Wyoming Public Health Laboratory | Jim Mildenberger, Taylor Fearing, Channing Weber, Ashley Norberg, Chayse Rowley, Marley Goetz, Brian Dominguez, Elliot Thomasson, Sam Britz, Cari Sloma, Robert Petit, and Rob Christensen |
| EPI_ISL_2801557, EPI_ISL_2801558, EPI_ISL_2801559, EPI_ISL_2801560, EPI_ISL_2801561, EPI_ISL_2801562, EPI_ISL_2801563 | DeRisi Lab, University of California, San Francisco | DeRisi Lab, University of California, San Francisco | Sabrina Mann, Anthea Mitchell, Matthias Hapte-Selassie, Jamin Liu, Sara Sunshine, Matthew Laurie, Genay Pilarowski, Amy Kistler, Manu Vanaerschot, Patrick Ayscue, Lucy Li, Eric Chow, IDseq Team, James Peng, Carina Marquez, Luis Rubio, Gabriel Chamie, Diane Jones, Jon Jacobo, Susana Rojas, Susy Rojas, Valerie Tulier-Laiwa, Douglas Black, Jackie Martinez, Maya Petersen, Diane Havlir, and Joseph DeRisi |
| EPI_ISL_2801578, EPI_ISL_2801593, EPI_ISL_2801595, EPI_ISL_2801630 | Unidad de Investigación Médica de Yucatán (UIMY) | Instituto de Biotecnología de la UNAM | Consorcio Mexicano de Vigilancia Genómica (CoViGen-Mex). Authors (in alphabetical order): Julio Elias Alvarado-Yaah, Carlos F. Arias, Santiago Ávila-Ríos, Eduardo Becerril-Vargas, Víctor Hugo Borja-Aburto, Celia Boukadida, Cristóbal Cháidez-Quiróz, Juan Bautista Chale-Dzul, Ricardo Ciria Merce, Andreu Comas-García, Célida Duque Molina, Julissa Enciso-Ibarra, José Antonio Enciso-Moreno, Gloria Elena Espinosa-Ayala, Fernando Fontove-Herrera, Daniel Fregoso-Rueda, Victor Eduardo García-Arias, Alejandra García-Gasca, Bruno Gómez-Gil, Jean Pierre González, Irvin González-López, Concepción Grajales-Muñiz, Ricardo Grande, Rosa María Gutiérrez Rios, Jesús Hernández, Alejandra Hernández-Terán, Alfredo Herrera-Estrella, Carla Ivón Herrera-Najera, Pavel Isa, Verónica Mata-Haro, Daniel Lira Morales, Susana Lopez, Antonio Loza Román, Brenda Irasema Maldonado-Meza, Bernardo Martínez-Miguel, José Arturo Martínez-Orozco, Célida Martínez-Rodríguez, Margarita Matías-Florentino, Fidencio Mejía-Nepomuceno, María Guadalupe de Jesús Mireles-Rivera, Gloria María Molina-Salinas, Hector Montoya-Fuentes, , Mario Mújica-Sánchez, José Esteban Muñoz-Medina, José de Jesús Nuñez-Contreras, Alicia Ocaña-Mondragón, Luis Alberto Ochoa-Carrera, Hector Esteban Paz-Juárez, Marissa Perez-Garcia, Francisco Pulido, Helen Haydee Fernanda Ramírez-Plascencia, Jorge Salas-Hernández, Angel Gustavo Salas-Lais, Alejandro Sánchez-Flores, Clara Esperanza Santacruz-Tinoco, María Guadalupe Santiago-Mauricio, Selene Zárate, Nelly Sélem-Mojica, Blanca Taboada, Gloria Vazquez, Joel Armando Vázquez-Pérez. |
| EPI_ISL_2801757, EPI_ISL_2801772, EPI_ISL_2801773, EPI_ISL_2801780, EPI_ISL_2801785, EPI_ISL_2801789, EPI_ISL_2801804, EPI_ISL_2801809, EPI_ISL_2801810, EPI_ISL_2801811, EPI_ISL_2801813, EPI_ISL_2801814, EPI_ISL_2801815, EPI_ISL_2801816, EPI_ISL_2801818, EPI_ISL_2801819, EPI_ISL_2801820, EPI_ISL_2801821, EPI_ISL_2801822, EPI_ISL_2801823, EPI_ISL_2801826, EPI_ISL_2801829, EPI_ISL_2801830, EPI_ISL_2801833, EPI_ISL_2801881 | | | |
| see above | Laboratorio Central de Epidemiología (LCE) | Instituto de Biotecnología de la UNAM | Consorcio Mexicano de Vigilancia Genómica (CoViGen-Mex). Authors (in alphabetical order): Julio Elias Alvarado-Yaah, Carlos F. Arias, Santiago Ávila-Ríos, Eduardo Becerril-Vargas, Víctor Hugo Borja-Aburto, Celia Boukadida, Cristóbal Cháidez-Quiróz, Juan Bautista Chale-Dzul, Ricardo Ciria Merce, Andreu Comas-García, Célida Duque Molina, Julissa Enciso-Ibarra, José Antonio Enciso-Moreno, Gloria Elena Espinosa-Ayala, Fernando Fontove-Herrera, Daniel Fregoso-Rueda, Victor Eduardo García-Arias, Alejandra García-Gasca, Bruno Gómez-Gil, Jean Pierre González, Irvin González-López, Concepción Grajales-Muñiz, Ricardo Grande, Rosa María Gutiérrez Rios, Jesús Hernández, Alejandra Hernández-Terán, Alfredo Herrera-Estrella, Carla Ivón Herrera-Najera, Pavel Isa, Verónica Mata-Haro, Daniel Lira Morales, Susana Lopez, Antonio Loza Román, Brenda Irasema Maldonado-Meza, Bernardo Martínez-Miguel, José Arturo Martínez-Orozco, Célida Martínez-Rodríguez, Margarita Matías-Florentino, Fidencio Mejía-Nepomuceno, María Guadalupe de Jesús Mireles-Rivera, Gloria María Molina-Salinas, Hector Montoya-Fuentes, , Mario Mújica-Sánchez, José Esteban Muñoz-Medina, José de Jesús Nuñez- |

| | | | |
|---|---|---|---|
| | | | Contreras, Alicia Ocaña-Mondragón, Luis Alberto Ochoa-Carrera, Hector Esteban Paz-Juárez, Marissa Perez-Garcia, Francisco Pulido, Helen Haydee Fernanda Ramírez-Plascencia, Jorge Salas-Hernández, Angel Gustavo Salas-Lais, Alejandro Sánchez-Flores, Clara Esperanza Santacruz-Tinoco, María Guadalupe Santiago-Mauricio, Selene Zárate, Nelly Sélem-Mojica, Blanca Taboada, Gloria Vazquez, Joel Armando Vázquez-Pérez. |
| EPI_ISL_2801892, EPI_ISL_2801893, EPI_ISL_2801894 | Banteay Meanchey Regional Laboratory | Virology Unit, Institut Pasteur du Cambodge | Cecile Troupin, Leakhena Pum, Jurre Y Siegers, Ly Sovann, Kraing Sidonn, Yi Sengdoeurn, Chin Savuth, Chau Darapheak, Veasna Duong, Erik A Karlsson |
| EPI_ISL_2801895 | Battambang Rapid Response Team | Virology Unit, Institut Pasteur du Cambodge | Cecile Troupin, Leakhena Pum, Jurre Y Siegers, Ly Sovann, Kraing Sidonn, Yi Sengdoeurn, Chin Savuth, Chau Darapheak, Veasna Duong, Erik A Karlsson |
| EPI_ISL_2801898 | Pursat Rapid Response Team | Virology Unit, Institut Pasteur du Cambodge | Cecile Troupin, Leakhena Pum, Jurre Y Siegers, Ly Sovann, Kraing Sidonn, Yi Sengdoeurn, Chin Savuth, Chau Darapheak, Veasna Duong, Erik A Karlsson |
| EPI_ISL_2801899, EPI_ISL_2801903 | Oddar Meanchey Rapid Response Team | Virology Unit, Institut Pasteur du Cambodge | Cecile Troupin, Leakhena Pum, Jurre Y Siegers, Ly Sovann, Kraing Sidonn, Yi Sengdoeurn, Chin Savuth, Chau Darapheak, Veasna Duong, Erik A Karlsson |
| EPI_ISL_2801904, EPI_ISL_2801905, EPI_ISL_2801906 | Tboung Khmum Rapid Response Team | Virology Unit, Institut Pasteur du Cambodge | Cecile Troupin, Leakhena Pum, Jurre Y Siegers, Ly Sovann, Kraing Sidonn, Yi Sengdoeurn, Chin Savuth, Chau Darapheak, Veasna Duong, Erik A Karlsson |
| EPI_ISL_2801922, EPI_ISL_2801923 | Virology Unit, Institut Pasteur du Cambodge | Virology Unit, Institut Pasteur du Cambodge | Cecile Troupin, Leakhena Pum, Jurre Y Siegers, Ly Sovann, Kraing Sidonn, Yi Sengdoeurn, Chin Savuth, Chau Darapheak, Veasna Duong, Erik A Karlsson |
| EPI_ISL_2801933 | FNUSA | CMBG | Matej Bezdicek, Jan Svaton, Martina Lengerova, Monika Dolejska |
| EPI_ISL_2801941, EPI_ISL_2801942, EPI_ISL_2801943, EPI_ISL_2801944, EPI_ISL_2801945, EPI_ISL_2801946, EPI_ISL_2801947, EPI_ISL_2801948, EPI_ISL_2801949, EPI_ISL_2801950, EPI_ISL_2801953, EPI_ISL_2801955, EPI_ISL_2801956, EPI_ISL_2801957, EPI_ISL_2801958, EPI_ISL_2801959, EPI_ISL_2801961, EPI_ISL_2801962, EPI_ISL_2801963, EPI_ISL_2801964, EPI_ISL_2801965, EPI_ISL_2801967, EPI_ISL_2801969, EPI_ISL_2801970, EPI_ISL_2801971, EPI_ISL_2801972, EPI_ISL_2801973, EPI_ISL_2801974, EPI_ISL_2801975, EPI_ISL_2801979, EPI_ISL_2801981, EPI_ISL_2801982, EPI_ISL_2801984 | | | |
| see above | Servicio Microbiología Hospital La Paz | Servicio Microbiología Hospital La Paz | Fernando Lázaro, Rubén Cáceres, Jesús Mingorance Cruz, Elie Dahdouh |
| EPI_ISL_2802008 | Institute of Virology, Biomedical Research Center of the Slovak Academy of Sciences, Bratislava | Faculty of Natural Sciences, Comenius University, Bratislava | Viktoria Cabanova, Kristina Borsova, Brona Brejova, Viktoria Hodorova, Sabina Fumacova Havlikova, Juraj Kopacek, Martina Lickova, Lubomira Lukacikova, Martina Nebohacova, Monika Slavikova, Tomas Vinar, Jozef Nosek, Boris Klempa |
| EPI_ISL_2802010 | Institute of Virology, Biomedical Research Center of the Slovak Academy of Sciences, Bratislava | Faculty of Natural Sciences, Comenius University, Bratislava | Brona Brejova, Viktoria Cabanova, Kristina Borsova, Viktoria Hodorova, Sabina Fumacova Havlikova, Juraj Kopacek, Martina Lickova, Lubomira Lukacikova, Martina Nebohacova, Monika Slavikova, Tomas Vinar, Jozef Nosek, Boris Klempa |
| EPI_ISL_2802112 | Tygerdal | NHLS/UCT | Arash Iranzadeh, Deelan Doolabh, Lynn Tyers, Bruna Galvao, Innocent Mudau, Marvin Hsiao, Gert Marais, Diana Hardie, Stephen Korsman, Carolyn Williamson |
| EPI_ISL_2802114 | Kraaifontein | NHLS/UCT | Arash Iranzadeh, Deelan Doolabh, Lynn Tyers, Bruna Galvao, Innocent Mudau, Marvin Hsiao, Gert Marais, Diana Hardie, Stephen Korsman, Carolyn Williamson |
| EPI_ISL_2802125, EPI_ISL_2802126, EPI_ISL_2802127 | Villiersdorp Clinic wc VDP | NHLS/UCT | Arash Iranzadeh, Deelan Doolabh, Lynn Tyers, Bruna Galvao, Innocent Mudau, Marvin Hsiao, Gert Marais, Diana Hardie, Stephen Korsman, Carolyn Williamson |
| EPI_ISL_2802180 | Sedgefield Clinic wc SGE | NHLS/UCT | Arash Iranzadeh, Deelan Doolabh, Lynn Tyers, Bruna Galvao, Innocent Mudau, Marvin Hsiao, Gert Marais, Diana Hardie, Stephen Korsman, Carolyn Williamson |
| EPI_ISL_2802181 | Thembalethu CDC wc THC | NHLS/UCT | Arash Iranzadeh, Deelan Doolabh, Lynn Tyers, Bruna Galvao, Innocent Mudau, Marvin Hsiao, Gert Marais, Diana Hardie, Stephen Korsman, Carolyn Williamson |
| EPI_ISL_2802190 | Swellendam PHC Clinic wc SHC | NHLS/UCT | Arash Iranzadeh, Deelan Doolabh, Lynn Tyers, Bruna Galvao, Innocent Mudau, Marvin Hsiao, Gert Marais, Diana Hardie, Stephen Korsman, Carolyn Williamson |
| EPI_ISL_2802194 | Groote Schuur Hospital wc GSH | NHLS/UCT | Arash Iranzadeh, Deelan Doolabh, Lynn Tyers, Bruna Galvao, Innocent Mudau, Marvin Hsiao, Gert Marais, Diana Hardie, Stephen Korsman, Carolyn Williamson |
| EPI_ISL_2802195 | Great Brak River Clinic wc GBC | NHLS/UCT | Arash Iranzadeh, Deelan Doolabh, Lynn Tyers, Bruna Galvao, Innocent Mudau, Marvin Hsiao, Gert Marais, Diana Hardie, Stephen Korsman, Carolyn Williamson |
| EPI_ISL_2802196 | NHLS/UCT | NHLS/UCT | Arash Iranzadeh, Deelan Doolabh, Lynn Tyers, Bruna Galvao, Innocent Mudau, Marvin Hsiao, Gert Marais, Diana Hardie, Stephen Korsman, Carolyn Williamson |
| EPI_ISL_2802199 | Groote Schuur Hospital wc GSH | NHLS/UCT | Arash Iranzadeh, Deelan Doolabh, Lynn Tyers, Bruna Galvao, Innocent Mudau, Marvin Hsiao, Gert Marais, Diana Hardie, Stephen Korsman, Carolyn Williamson |
| EPI_ISL_2802200 | Poliklinika Vitalis | Alea Genetic Centre | Rijad Konjhodzic, Dino Pecar, Lana Salihefendic, Ivana Ceko |
| EPI_ISL_2802622 | Maine Health and Environmental Testing Laboratory | Tewhey Lab, The Jackson Laboratory | Matluk,N., Dewey,H., Iosue,F., Barter,M., Lynch,R., Munger,H. and Tewhey,R. |
| EPI_ISL_2802831, EPI_ISL_2802832, EPI_ISL_2802834, EPI_ISL_2802835, EPI_ISL_2802841, EPI_ISL_2802844, EPI_ISL_2802846, EPI_ISL_2802848, EPI_ISL_2802850, EPI_ISL_2802851, EPI_ISL_2802853, EPI_ISL_2802854, EPI_ISL_2802855 | | | |
| see above | Ramathibodi Hospital | COVID-19 Network Investigations (CONI) Alliance | Elizabeth Batty, Wasun Chantratita, Thanat Chookajorn, Stefan Fernandez, Angkana Huang, Anthony R. Jones, Khajohn Joonlasak, Chonticha Klungtong, Theerarat Kochakarn, Namfon Kotanan, Krittikorn Kumpornsin, Duangkamon Loesbanluechai, Wudtichai Manasatienkij, Bhakbhoom Panthan, Ekawat Pasomsub, Kingkan Rakmanee, Insee Sensorn, Janjira Thaipadungpanit, Arporn Wangwiwatsin, Treewat Watthanachockchai |
| EPI_ISL_2802866, EPI_ISL_2802867, EPI_ISL_2802868, EPI_ISL_2802869, EPI_ISL_2802870, EPI_ISL_2802871, EPI_ISL_2802872, EPI_ISL_2802874, EPI_ISL_2802876, EPI_ISL_2802879, EPI_ISL_2802880, EPI_ISL_2802881, EPI_ISL_2802882, EPI_ISL_2802884, EPI_ISL_2802886, EPI_ISL_2802891, EPI_ISL_2802892, EPI_ISL_2802895, EPI_ISL_2802897, EPI_ISL_2802898, EPI_ISL_2802902, EPI_ISL_2802904, EPI_ISL_2802905, EPI_ISL_2802906, EPI_ISL_2802907, EPI_ISL_2802909, EPI_ISL_2802910, EPI_ISL_2802911, EPI_ISL_2802915, EPI_ISL_2802916, EPI_ISL_2802917, EPI_ISL_2802919, EPI_ISL_2802921, EPI_ISL_2802922, EPI_ISL_2802924, EPI_ISL_2802925, EPI_ISL_2802926, EPI_ISL_2802927, EPI_ISL_2802928, EPI_ISL_2802929, EPI_ISL_2802931, EPI_ISL_2802933, EPI_ISL_2802938, EPI_ISL_2802939, EPI_ISL_2802941, EPI_ISL_2802942, EPI_ISL_2802943, EPI_ISL_2802946, EPI_ISL_2802948, EPI_ISL_2802952, EPI_ISL_2802954, EPI_ISL_2802957, EPI_ISL_2802958, EPI_ISL_2802960, EPI_ISL_2802962, EPI_ISL_2802963, EPI_ISL_2802964, EPI_ISL_2802966, EPI_ISL_2802967, EPI_ISL_2802968, EPI_ISL_2802970, EPI_ISL_2802971, EPI_ISL_2802973, EPI_ISL_2802974, EPI_ISL_2802977, EPI_ISL_2802978, EPI_ISL_2802979, EPI_ISL_2802980, EPI_ISL_2802981, EPI_ISL_2802982, EPI_ISL_2802984, EPI_ISL_2802986, EPI_ISL_2802988, EPI_ISL_2802989, EPI_ISL_2802992, EPI_ISL_2802993, EPI_ISL_2802996, EPI_ISL_2802997, EPI_ISL_2803000, EPI_ISL_2803001, EPI_ISL_2803006, EPI_ISL_2803010, EPI_ISL_2803011, EPI_ISL_2803012, EPI_ISL_2803015, EPI_ISL_2803017, EPI_ISL_2803019, EPI_ISL_2803020, EPI_ISL_2803022, EPI_ISL_2803023, EPI_ISL_2803028, EPI_ISL_2803031, EPI_ISL_2803033, EPI_ISL_2803035, EPI_ISL_2803037, EPI_ISL_2803038, EPI_ISL_2803039, EPI_ISL_2803042, EPI_ISL_2803043, EPI_ISL_2803046, EPI_ISL_2803053, EPI_ISL_2803054, EPI_ISL_2803055, EPI_ISL_2803056, EPI_ISL_2803057, EPI_ISL_2803060, EPI_ISL_2803061, EPI_ISL_2803062, EPI_ISL_2803063, EPI_ISL_2803064, EPI_ISL_2803067, EPI_ISL_2803068, EPI_ISL_2803069, EPI_ISL_2803072, EPI_ISL_2803073, EPI_ISL_2803074, EPI_ISL_2803075, EPI_ISL_2803076, EPI_ISL_2803077, EPI_ISL_2803078, EPI_ISL_2803079, EPI_ISL_2803080, EPI_ISL_2803081, EPI_ISL_2803082, EPI_ISL_2803084, EPI_ISL_2803085, EPI_ISL_2803090, EPI_ISL_2803093, EPI_ISL_2803094, EPI_ISL_2803096, EPI_ISL_2803102, EPI_ISL_2803103, EPI_ISL_2803105, EPI_ISL_2803107, EPI_ISL_2803108, EPI_ISL_2803109, EPI_ISL_2803110, EPI_ISL_2803111, EPI_ISL_2803114, EPI_ISL_2803116, EPI_ISL_2803117, EPI_ISL_2803118, EPI_ISL_2803121, EPI_ISL_2803122, EPI_ISL_2803125, EPI_ISL_2803126, EPI_ISL_2803127, EPI_ISL_2803128, EPI_ISL_2803129, EPI_ISL_2803131, EPI_ISL_2803132, EPI_ISL_2803134, EPI_ISL_2803135, EPI_ISL_2803136, EPI_ISL_2803138, EPI_ISL_2803140, EPI_ISL_2803142, EPI_ISL_2803143, EPI_ISL_2803144, EPI_ISL_2803146, EPI_ISL_2803147, EPI_ISL_2803149, EPI_ISL_2803150, EPI_ISL_2803153, EPI_ISL_2803157, EPI_ISL_2803158, EPI_ISL_2803159, EPI_ISL_2803162, EPI_ISL_2803164, EPI_ISL_2803166, EPI_ISL_2803167 | | | |
| see above | UW Virology Lab | UW Virology Lab | Pavitra Roychoudhury, Hong Xie, Lasata Shrestha, Tien V. Nguyen, Shah Mohamed Bakhash, Michelle Lin, Noah R. Baker, Ricardo Perez, Sean Ellis, Nathan Breit, Robert J. Livingston, Meei-Li Huang, Keith R Jerome, Patrick Mathias, Alexander Greninger |
| EPI_ISL_2803176, EPI_ISL_2803187, EPI_ISL_2803191, EPI_ISL_2803196, EPI_ISL_2803227, EPI_ISL_2803228, EPI_ISL_2803243, EPI_ISL_2803244, EPI_ISL_2803248, EPI_ISL_2803270, EPI_ISL_2803271, EPI_ISL_2803275, EPI_ISL_2803285, EPI_ISL_2803292 | | | |
| see above | Centre for Dengue Research and AICBU, Department of Immunology and Molecular Medicine | Centre for Dengue Research and AICBU, Department of Immunology and Molecular Medicine | Chandima Jeewandara, Deshni Jayathilaka, Deshan Madhusanka, Diyanath Ranasinghe, Gathsaurie Neelika Malavige |
| EPI_ISL_2803306, EPI_ISL_2803310, EPI_ISL_2803311 | UW Virology Lab | UW Virology Lab | Pavitra Roychoudhury, Hong Xie, Lasata Shrestha, Shah Mohamed Bakhash, Tien V. Nguyen, Noah R. Baker, Sean Ellis, Meei-Li Huang, Keith R Jerome, Alexander Greninger |
| EPI_ISL_2803661, EPI_ISL_2803662, EPI_ISL_2803663, EPI_ISL_2803664, EPI_ISL_2803665, EPI_ISL_2803666, EPI_ISL_2803668, EPI_ISL_2803669, EPI_ISL_2803670, EPI_ISL_2803671, EPI_ISL_2803672, EPI_ISL_2803673, EPI_ISL_2803674, EPI_ISL_2803675, EPI_ISL_2803676, EPI_ISL_2803677, EPI_ISL_2803678, EPI_ISL_2803679, EPI_ISL_2803680, EPI_ISL_2803681, EPI_ISL_2803682, EPI_ISL_2803683, EPI_ISL_2803684, EPI_ISL_2803685, EPI_ISL_2803686, EPI_ISL_2803687, EPI_ISL_2803688, EPI_ISL_2803689, EPI_ISL_2803690, EPI_ISL_2803691, EPI_ISL_2803692, EPI_ISL_2803693, EPI_ISL_2803694, EPI_ISL_2803695, EPI_ISL_2803696, EPI_ISL_2803697, EPI_ISL_2803698, EPI_ISL_2803699, EPI_ISL_2803700, EPI_ISL_2803701, EPI_ISL_2803702, EPI_ISL_2803703, EPI_ISL_2803704, EPI_ISL_2803705, EPI_ISL_2803706, EPI_ISL_2803707, EPI_ISL_2803708, EPI_ISL_2803709, EPI_ISL_2803710, EPI_ISL_2803711, EPI_ISL_2803712, EPI_ISL_2803713, EPI_ISL_2803714, EPI_ISL_2803715, EPI_ISL_2803716, EPI_ISL_2803717, EPI_ISL_2803718, EPI_ISL_2803719, EPI_ISL_2803720, EPI_ISL_2803721, EPI_ISL_2803722, EPI_ISL_2803723, EPI_ISL_2803724, EPI_ISL_2803725, EPI_ISL_2803726, EPI_ISL_2803727, EPI_ISL_2803728, EPI_ISL_2803729, EPI_ISL_2803730, EPI_ISL_2803731, EPI_ISL_2803732, EPI_ISL_2803733, EPI_ISL_2803734, EPI_ISL_2803735, EPI_ISL_2803736, EPI_ISL_2803738, EPI_ISL_2803739, EPI_ISL_2803740, EPI_ISL_2803741, EPI_ISL_2803742, EPI_ISL_2803743, EPI_ISL_2803744 | | | |
| see above | University of Zambia, School of Veterinary Medicine | UNZAVET and PATH | Mulenga Mwenda-Chimfwembe, Ngonda Saasa, Daniel Bridges, ZNPHI and ZGSC |
| EPI_ISL_2803747 | Everight Diagnostic Laboratories | National Reference Laboratory, Nigeria Centre for Disease Control | Dr Ndodo Nnaemeka, Olusola Anuoluwapo Akanbi, Chimaobi Chukwu, Dr Omoare Adesuyi, Grace Esebanmen, Anthony Ahumibe, Catherine Okoi, Nwando Mba, Dr Chikwe Ihekweazu |
| EPI_ISL_2803757 | The Ohio State University Applied Microbiology Services Laboratory | The Ohio State University Applied Microbiology Services Laboratory | Seth A. Faith PhD |
| EPI_ISL_2803758, EPI_ISL_2803759 | National Institute of Laboratory Medicine and Referral Center | Genomic Research Lab, BCSIR | Md. Maruf Ahmed Molla, Md. Murshed Hasan Sarkar, Mohammad Samir Uzzaman, Eshrar Osman, Md. Ahasan Habib, Shahina Akter, Tanjina Akhter Banu, Barna Goswami, Iffat Jahan, Mohammad Mohi Uddin, Md. Kamrul Islam, Tasnim Nafisa, Mahmuda Yeasmin, Asish Kumar Ghosh, Arifa Akram, Md. Salim Khan |
| EPI_ISL_2803760, EPI_ISL_2803761, EPI_ISL_2803762, EPI_ISL_2803763 | National Institute of Laboratory Medicine and Referral Center | Genomic Research Lab, BCSIR | Md. Murshed Hasan Sarkar, Mohammad Samir Uzzaman, Eshrar Osman, Md. Ahasan Habib, Shahina Akter, Tanjina Akhter Banu, Barna Goswami, Iffat Jahan, Mohammad Mohi Uddin, Md. Kamrul Islam, Tasnim Nafisa, Mahmuda Yeasmin, Asish Kumar Ghosh, Arifa Akram, Md. Salim Khan |
| EPI_ISL_2803766, EPI_ISL_2803767, EPI_ISL_2803768 | Lighthouse Lab in Glasgow | Wellcome Sanger Institute for the COVID-19 Genomics UK (COG-UK) Consortium | Harper VanSteenhouse, Yumi Kasai, David Gray, Carol Clugston, Anna Dominiczak and Alex Alderton, Roberto Amato, Jeffrey Barrett, Sonia Goncalves, Ewan Harrison, David K. Jackson, Ian Johnston, Dominic Kwiatkowski, Cordelia Langford, John Sillitoe on behalf of the Wellcome Sanger Institute COVID-19 Surveillance Team |
| EPI_ISL_2803769, EPI_ISL_2803770, EPI_ISL_2803771, EPI_ISL_2803772 | Lighthouse Lab in Milton Keynes | Wellcome Sanger Institute for the COVID-19 Genomics UK (COG-UK) Consortium | The Lighthouse Lab in Milton Keynes and Alex Alderton, Roberto Amato, Jeffrey Barrett, Sonia Goncalves, Ewan Harrison, David K. Jackson, Ian Johnston, Dominic Kwiatkowski, Cordelia Langford, John Sillitoe on behalf of the Wellcome Sanger Institute COVID-19 Surveillance Team |
| EPI_ISL_2803773 | Lighthouse Lab in Glasgow | Wellcome Sanger Institute for the COVID-19 Genomics UK (COG-UK) Consortium | Harper VanSteenhouse, Yumi Kasai, David Gray, Carol Clugston, Anna Dominiczak and Alex Alderton, Roberto Amato, Jeffrey Barrett, Sonia Goncalves, Ewan Harrison, David K. Jackson, Ian Johnston, Dominic Kwiatkowski, Cordelia Langford, John Sillitoe on behalf of the Wellcome Sanger Institute COVID-19 Surveillance Team |
| EPI_ISL_2803774, EPI_ISL_2803775, EPI_ISL_2803776, EPI_ISL_2803777 | Lighthouse Lab in Milton Keynes | Wellcome Sanger Institute for the COVID-19 Genomics UK (COG-UK) Consortium | The Lighthouse Lab in Milton Keynes and Alex Alderton, Roberto Amato, Jeffrey Barrett, Sonia Goncalves, Ewan Harrison, David K. Jackson, Ian Johnston, Dominic Kwiatkowski, Cordelia Langford, John Sillitoe on behalf of the Wellcome Sanger Institute COVID-19 Surveillance Team |
| EPI_ISL_2803778 | Lighthouse Lab in Glasgow | Wellcome Sanger Institute for the COVID-19 Genomics UK (COG-UK) Consortium | Harper VanSteenhouse, Yumi Kasai, David Gray, Carol Clugston, Anna Dominiczak and Alex Alderton, Roberto Amato, Jeffrey Barrett, Sonia Goncalves, Ewan Harrison, David K. Jackson, Ian Johnston, Dominic Kwiatkowski, Cordelia Langford, John Sillitoe on behalf of the Wellcome Sanger Institute COVID-19 Surveillance Team |
| EPI_ISL_2803780, EPI_ISL_2803781, EPI_ISL_2803782, EPI_ISL_2803783, EPI_ISL_2803784 | Lighthouse Lab in Milton Keynes | Wellcome Sanger Institute for the COVID-19 Genomics UK (COG-UK) Consortium | The Lighthouse Lab in Milton Keynes and Alex Alderton, Roberto Amato, Jeffrey Barrett, Sonia Goncalves, Ewan Harrison, David K. Jackson, Ian Johnston, Dominic Kwiatkowski, Cordelia Langford, John Sillitoe on behalf of the Wellcome Sanger Institute COVID-19 Surveillance Team |
| EPI_ISL_2803785 | Lighthouse Lab in Glasgow | Wellcome Sanger Institute for the COVID-19 Genomics UK (COG-UK) Consortium | Harper VanSteenhouse, Yumi Kasai, David Gray, Carol Clugston, Anna Dominiczak and Alex Alderton, Roberto Amato, Jeffrey Barrett, Sonia Goncalves, Ewan Harrison, David K. Jackson, Ian Johnston, Dominic Kwiatkowski, Cordelia Langford, John Sillitoe on behalf of the Wellcome Sanger Institute COVID-19 Surveillance Team |

| | | | |
|---|---|---|---|
| EPI_ISL_2803786, EPI_ISL_2803787, EPI_ISL_2803788 | Lighthouse Lab in Milton Keynes | Wellcome Sanger Institute for the COVID-19 Genomics UK (COG-UK) Consortium | The Lighthouse Lab in Milton Keynes and Alex Alderton, Roberto Amato, Jeffrey Barrett, Sonia Goncalves, Ewan Harrison, David K. Jackson, Ian Johnston, Dominic Kwiatkowski, Cordelia Langford, John Sillitoe on behalf of the Wellcome Sanger Institute COVID-19 Surveillance Team |
| EPI_ISL_2803789, EPI_ISL_2803790 | Lighthouse Lab in Glasgow | Wellcome Sanger Institute for the COVID-19 Genomics UK (COG-UK) Consortium | Harper VanSteenhouse, Yumi Kasai, David Gray, Carol Clugston, Anna Dominiczak and Alex Alderton, Roberto Amato, Jeffrey Barrett, Sonia Goncalves, Ewan Harrison, David K. Jackson, Ian Johnston, Dominic Kwiatkowski, Cordelia Langford, John Sillitoe on behalf of the Wellcome Sanger Institute COVID-19 Surveillance Team |
| EPI_ISL_2803791, EPI_ISL_2803792, EPI_ISL_2803793 | Lighthouse Lab in Milton Keynes | Wellcome Sanger Institute for the COVID-19 Genomics UK (COG-UK) Consortium | The Lighthouse Lab in Milton Keynes and Alex Alderton, Roberto Amato, Jeffrey Barrett, Sonia Goncalves, Ewan Harrison, David K. Jackson, Ian Johnston, Dominic Kwiatkowski, Cordelia Langford, John Sillitoe on behalf of the Wellcome Sanger Institute COVID-19 Surveillance Team |
| EPI_ISL_2803794, EPI_ISL_2803795 | Lighthouse Lab in Glasgow | Wellcome Sanger Institute for the COVID-19 Genomics UK (COG-UK) Consortium | Harper VanSteenhouse, Yumi Kasai, David Gray, Carol Clugston, Anna Dominiczak and Alex Alderton, Roberto Amato, Jeffrey Barrett, Sonia Goncalves, Ewan Harrison, David K. Jackson, Ian Johnston, Dominic Kwiatkowski, Cordelia Langford, John Sillitoe on behalf of the Wellcome Sanger Institute COVID-19 Surveillance Team |
| EPI_ISL_2803796, EPI_ISL_2803797 | Lighthouse Lab in Milton Keynes | Wellcome Sanger Institute for the COVID-19 Genomics UK (COG-UK) Consortium | The Lighthouse Lab in Milton Keynes and Alex Alderton, Roberto Amato, Jeffrey Barrett, Sonia Goncalves, Ewan Harrison, David K. Jackson, Ian Johnston, Dominic Kwiatkowski, Cordelia Langford, John Sillitoe on behalf of the Wellcome Sanger Institute COVID-19 Surveillance Team |
| EPI_ISL_2803798, EPI_ISL_2803799 | Lighthouse Lab in Glasgow | Wellcome Sanger Institute for the COVID-19 Genomics UK (COG-UK) Consortium | Harper VanSteenhouse, Yumi Kasai, David Gray, Carol Clugston, Anna Dominiczak and Alex Alderton, Roberto Amato, Jeffrey Barrett, Sonia Goncalves, Ewan Harrison, David K. Jackson, Ian Johnston, Dominic Kwiatkowski, Cordelia Langford, John Sillitoe on behalf of the Wellcome Sanger Institute COVID-19 Surveillance Team |
| EPI_ISL_2803800 | Lighthouse Lab in Milton Keynes | Wellcome Sanger Institute for the COVID-19 Genomics UK (COG-UK) Consortium | The Lighthouse Lab in Milton Keynes and Alex Alderton, Roberto Amato, Jeffrey Barrett, Sonia Goncalves, Ewan Harrison, David K. Jackson, Ian Johnston, Dominic Kwiatkowski, Cordelia Langford, John Sillitoe on behalf of the Wellcome Sanger Institute COVID-19 Surveillance Team |
| EPI_ISL_2803801 | Lighthouse Lab in Glasgow | Wellcome Sanger Institute for the COVID-19 Genomics UK (COG-UK) Consortium | Harper VanSteenhouse, Yumi Kasai, David Gray, Carol Clugston, Anna Dominiczak and Alex Alderton, Roberto Amato, Jeffrey Barrett, Sonia Goncalves, Ewan Harrison, David K. Jackson, Ian Johnston, Dominic Kwiatkowski, Cordelia Langford, John Sillitoe on behalf of the Wellcome Sanger Institute COVID-19 Surveillance Team |
| EPI_ISL_2803802 | Lighthouse Lab in Milton Keynes | Wellcome Sanger Institute for the COVID-19 Genomics UK (COG-UK) Consortium | The Lighthouse Lab in Milton Keynes and Alex Alderton, Roberto Amato, Jeffrey Barrett, Sonia Goncalves, Ewan Harrison, David K. Jackson, Ian Johnston, Dominic Kwiatkowski, Cordelia Langford, John Sillitoe on behalf of the Wellcome Sanger Institute COVID-19 Surveillance Team |
| EPI_ISL_2803803 | Lighthouse Lab in Glasgow | Wellcome Sanger Institute for the COVID-19 Genomics UK (COG-UK) Consortium | Harper VanSteenhouse, Yumi Kasai, David Gray, Carol Clugston, Anna Dominiczak and Alex Alderton, Roberto Amato, Jeffrey Barrett, Sonia Goncalves, Ewan Harrison, David K. Jackson, Ian Johnston, Dominic Kwiatkowski, Cordelia Langford, John Sillitoe on behalf of the Wellcome Sanger Institute COVID-19 Surveillance Team |
| EPI_ISL_2803804, EPI_ISL_2803805, EPI_ISL_2803806, EPI_ISL_2803807 | Lighthouse Lab in Milton Keynes | Wellcome Sanger Institute for the COVID-19 Genomics UK (COG-UK) Consortium | The Lighthouse Lab in Milton Keynes and Alex Alderton, Roberto Amato, Jeffrey Barrett, Sonia Goncalves, Ewan Harrison, David K. Jackson, Ian Johnston, Dominic Kwiatkowski, Cordelia Langford, John Sillitoe on behalf of the Wellcome Sanger Institute COVID-19 Surveillance Team |
| EPI_ISL_2803808, EPI_ISL_2803809, EPI_ISL_2803810, EPI_ISL_2803811, EPI_ISL_2803812, EPI_ISL_2803813, EPI_ISL_2803814 | Lighthouse Lab in Glasgow | Wellcome Sanger Institute for the COVID-19 Genomics UK (COG-UK) Consortium | Harper VanSteenhouse, Yumi Kasai, David Gray, Carol Clugston, Anna Dominiczak and Alex Alderton, Roberto Amato, Jeffrey Barrett, Sonia Goncalves, Ewan Harrison, David K. Jackson, Ian Johnston, Dominic Kwiatkowski, Cordelia Langford, John Sillitoe on behalf of the Wellcome Sanger Institute COVID-19 Surveillance Team |
| EPI_ISL_2803816 | Lighthouse Lab in Milton Keynes | Wellcome Sanger Institute for the COVID-19 Genomics UK (COG-UK) Consortium | The Lighthouse Lab in Milton Keynes and Alex Alderton, Roberto Amato, Jeffrey Barrett, Sonia Goncalves, Ewan Harrison, David K. Jackson, Ian Johnston, Dominic Kwiatkowski, Cordelia Langford, John Sillitoe on behalf of the Wellcome Sanger Institute COVID-19 Surveillance Team |
| EPI_ISL_2803817, EPI_ISL_2803818, EPI_ISL_2803819 | Lighthouse Lab in Glasgow | Wellcome Sanger Institute for the COVID-19 Genomics UK (COG-UK) Consortium | Harper VanSteenhouse, Yumi Kasai, David Gray, Carol Clugston, Anna Dominiczak and Alex Alderton, Roberto Amato, Jeffrey Barrett, Sonia Goncalves, Ewan Harrison, David K. Jackson, Ian Johnston, Dominic Kwiatkowski, Cordelia Langford, John Sillitoe on behalf of the Wellcome Sanger Institute COVID-19 Surveillance Team |
| EPI_ISL_2803820 | Lighthouse Lab in Milton Keynes | Wellcome Sanger Institute for the COVID-19 Genomics UK (COG-UK) Consortium | The Lighthouse Lab in Milton Keynes and Alex Alderton, Roberto Amato, Jeffrey Barrett, Sonia Goncalves, Ewan Harrison, David K. Jackson, Ian Johnston, Dominic Kwiatkowski, Cordelia Langford, John Sillitoe on behalf of the Wellcome Sanger Institute COVID-19 Surveillance Team |
| EPI_ISL_2803821, EPI_ISL_2803822 | Lighthouse Lab in Glasgow | Wellcome Sanger Institute for the COVID-19 Genomics UK (COG-UK) Consortium | Harper VanSteenhouse, Yumi Kasai, David Gray, Carol Clugston, Anna Dominiczak and Alex Alderton, Roberto Amato, Jeffrey Barrett, Sonia Goncalves, Ewan Harrison, David K. Jackson, Ian Johnston, Dominic Kwiatkowski, Cordelia Langford, John Sillitoe on behalf of the Wellcome Sanger Institute COVID-19 Surveillance Team |
| EPI_ISL_2803823, EPI_ISL_2803824 | Lighthouse Lab in Milton Keynes | Wellcome Sanger Institute for the COVID-19 Genomics UK (COG-UK) Consortium | The Lighthouse Lab in Milton Keynes and Alex Alderton, Roberto Amato, Jeffrey Barrett, Sonia Goncalves, Ewan Harrison, David K. Jackson, Ian Johnston, Dominic Kwiatkowski, Cordelia Langford, John Sillitoe on behalf of the Wellcome Sanger Institute COVID-19 Surveillance Team |
| EPI_ISL_2803825, EPI_ISL_2803826, EPI_ISL_2803827 | Lighthouse Lab in Glasgow | Wellcome Sanger Institute for the COVID-19 Genomics UK (COG-UK) Consortium | Harper VanSteenhouse, Yumi Kasai, David Gray, Carol Clugston, Anna Dominiczak and Alex Alderton, Roberto Amato, Jeffrey Barrett, Sonia Goncalves, Ewan Harrison, David K. Jackson, Ian Johnston, Dominic Kwiatkowski, Cordelia Langford, John Sillitoe on behalf of the Wellcome Sanger Institute COVID-19 Surveillance Team |
| EPI_ISL_2803828, EPI_ISL_2803829 | Lighthouse Lab in Milton Keynes | Wellcome Sanger Institute for the COVID-19 Genomics UK (COG-UK) Consortium | The Lighthouse Lab in Milton Keynes and Alex Alderton, Roberto Amato, Jeffrey Barrett, Sonia Goncalves, Ewan Harrison, David K. Jackson, Ian Johnston, Dominic Kwiatkowski, Cordelia Langford, John Sillitoe on behalf of the Wellcome Sanger Institute COVID-19 Surveillance Team |
| EPI_ISL_2803830, EPI_ISL_2803831, EPI_ISL_2803832, EPI_ISL_2803833, EPI_ISL_2803834 | Lighthouse Lab in Glasgow | Wellcome Sanger Institute for the COVID-19 Genomics UK (COG-UK) Consortium | Harper VanSteenhouse, Yumi Kasai, David Gray, Carol Clugston, Anna Dominiczak and Alex Alderton, Roberto Amato, Jeffrey Barrett, Sonia Goncalves, Ewan Harrison, David K. Jackson, Ian Johnston, Dominic Kwiatkowski, Cordelia Langford, John Sillitoe on behalf of the Wellcome Sanger Institute COVID-19 Surveillance Team |
| EPI_ISL_2803835, EPI_ISL_2803836, EPI_ISL_2803837, EPI_ISL_2803838 | Lighthouse Lab in Milton Keynes | Wellcome Sanger Institute for the COVID-19 Genomics UK (COG-UK) Consortium | The Lighthouse Lab in Milton Keynes and Alex Alderton, Roberto Amato, Jeffrey Barrett, Sonia Goncalves, Ewan Harrison, David K. Jackson, Ian Johnston, Dominic Kwiatkowski, Cordelia Langford, John Sillitoe on behalf of the Wellcome Sanger Institute COVID-19 Surveillance Team |
| EPI_ISL_2803839 | Lighthouse Lab in Glasgow | Wellcome Sanger Institute for the COVID-19 Genomics UK (COG-UK) Consortium | Harper VanSteenhouse, Yumi Kasai, David Gray, Carol Clugston, Anna Dominiczak and Alex Alderton, Roberto Amato, Jeffrey Barrett, Sonia Goncalves, Ewan Harrison, David K. Jackson, Ian Johnston, Dominic Kwiatkowski, Cordelia Langford, John Sillitoe on behalf of the Wellcome Sanger Institute COVID-19 Surveillance Team |
| EPI_ISL_2803840 | Lighthouse Lab in Milton Keynes | Wellcome Sanger Institute for the COVID-19 Genomics UK (COG-UK) Consortium | The Lighthouse Lab in Milton Keynes and Alex Alderton, Roberto Amato, Jeffrey Barrett, Sonia Goncalves, Ewan Harrison, David K. Jackson, Ian Johnston, Dominic Kwiatkowski, Cordelia Langford, John Sillitoe on behalf of the Wellcome Sanger Institute COVID-19 Surveillance Team |
| EPI_ISL_2803842 | Lighthouse Lab in Glasgow | Wellcome Sanger Institute for the COVID-19 Genomics UK (COG-UK) Consortium | Harper VanSteenhouse, Yumi Kasai, David Gray, Carol Clugston, Anna Dominiczak and Alex Alderton, Roberto Amato, Jeffrey Barrett, Sonia Goncalves, Ewan Harrison, David K. Jackson, Ian Johnston, Dominic Kwiatkowski, Cordelia Langford, John Sillitoe on behalf of the Wellcome Sanger Institute COVID-19 Surveillance Team |
| EPI_ISL_2803843 | Lighthouse Lab in Milton Keynes | Wellcome Sanger Institute for the COVID-19 Genomics UK (COG-UK) Consortium | The Lighthouse Lab in Milton Keynes and Alex Alderton, Roberto Amato, Jeffrey Barrett, Sonia Goncalves, Ewan Harrison, David K. Jackson, Ian Johnston, Dominic Kwiatkowski, Cordelia Langford, John Sillitoe on behalf of the Wellcome Sanger Institute COVID-19 Surveillance Team |
| EPI_ISL_2803844, EPI_ISL_2803845 | Lighthouse Lab in Glasgow | Wellcome Sanger Institute for the COVID-19 Genomics UK (COG-UK) Consortium | Harper VanSteenhouse, Yumi Kasai, David Gray, Carol Clugston, Anna Dominiczak and Alex Alderton, Roberto Amato, Jeffrey Barrett, Sonia Goncalves, Ewan Harrison, David K. Jackson, Ian Johnston, Dominic Kwiatkowski, Cordelia Langford, John Sillitoe on behalf of the Wellcome Sanger Institute COVID-19 Surveillance Team |
| EPI_ISL_2803846 | Lighthouse Lab in Milton Keynes | Wellcome Sanger Institute for the COVID-19 Genomics UK (COG-UK) Consortium | The Lighthouse Lab in Milton Keynes and Alex Alderton, Roberto Amato, Jeffrey Barrett, Sonia Goncalves, Ewan Harrison, David K. Jackson, Ian Johnston, Dominic Kwiatkowski, Cordelia Langford, John Sillitoe on behalf of the Wellcome Sanger Institute COVID-19 Surveillance Team |
| EPI_ISL_2803847 | Lighthouse Lab in Glasgow | Wellcome Sanger Institute for the COVID-19 Genomics UK (COG-UK) Consortium | Harper VanSteenhouse, Yumi Kasai, David Gray, Carol Clugston, Anna Dominiczak and Alex Alderton, Roberto Amato, Jeffrey Barrett, Sonia Goncalves, Ewan Harrison, David K. Jackson, Ian Johnston, Dominic Kwiatkowski, Cordelia Langford, John Sillitoe on behalf of the Wellcome Sanger Institute COVID-19 Surveillance Team |
| EPI_ISL_2803848 | Lighthouse Lab in Milton Keynes | Wellcome Sanger Institute for the COVID-19 Genomics UK (COG-UK) Consortium | The Lighthouse Lab in Milton Keynes and Alex Alderton, Roberto Amato, Jeffrey Barrett, Sonia Goncalves, Ewan Harrison, David K. Jackson, Ian Johnston, Dominic Kwiatkowski, Cordelia Langford, John Sillitoe on behalf of the Wellcome Sanger Institute COVID-19 Surveillance Team |
| EPI_ISL_2803849 | Lighthouse Lab in Glasgow | Wellcome Sanger Institute for the COVID-19 Genomics UK (COG-UK) Consortium | Harper VanSteenhouse, Yumi Kasai, David Gray, Carol Clugston, Anna Dominiczak and Alex Alderton, Roberto Amato, Jeffrey Barrett, Sonia Goncalves, Ewan Harrison, David K. Jackson, Ian Johnston, Dominic Kwiatkowski, Cordelia Langford, John Sillitoe on behalf of the Wellcome Sanger Institute COVID-19 Surveillance Team |
| EPI_ISL_2803850 | Lighthouse Lab in Milton Keynes | Wellcome Sanger Institute for the COVID-19 Genomics UK (COG-UK) Consortium | The Lighthouse Lab in Milton Keynes and Alex Alderton, Roberto Amato, Jeffrey Barrett, Sonia Goncalves, Ewan Harrison, David K. Jackson, Ian Johnston, Dominic Kwiatkowski, Cordelia Langford, John Sillitoe on behalf of the Wellcome Sanger Institute COVID-19 Surveillance Team |
| EPI_ISL_2803851 | Lighthouse Lab in Glasgow | Wellcome Sanger Institute for the COVID-19 Genomics UK (COG-UK) Consortium | Harper VanSteenhouse, Yumi Kasai, David Gray, Carol Clugston, Anna Dominiczak and Alex Alderton, Roberto Amato, Jeffrey Barrett, Sonia Goncalves, Ewan Harrison, David K. Jackson, Ian Johnston, Dominic Kwiatkowski, Cordelia Langford, John Sillitoe on behalf of the Wellcome Sanger Institute COVID-19 Surveillance Team |
| EPI_ISL_2803852 | Lighthouse Lab in Milton Keynes | Wellcome Sanger Institute for the COVID-19 Genomics UK (COG-UK) Consortium | The Lighthouse Lab in Milton Keynes and Alex Alderton, Roberto Amato, Jeffrey Barrett, Sonia Goncalves, Ewan Harrison, David K. Jackson, Ian Johnston, Dominic Kwiatkowski, Cordelia Langford, John Sillitoe on behalf of the Wellcome Sanger Institute COVID-19 Surveillance Team |
| EPI_ISL_2803853 | Lighthouse Lab in Glasgow | Wellcome Sanger Institute for the COVID-19 Genomics UK (COG-UK) Consortium | Harper VanSteenhouse, Yumi Kasai, David Gray, Carol Clugston, Anna Dominiczak and Alex Alderton, Roberto Amato, Jeffrey Barrett, Sonia Goncalves, Ewan Harrison, David K. Jackson, Ian Johnston, Dominic Kwiatkowski, Cordelia Langford, John Sillitoe on behalf of the Wellcome Sanger Institute COVID-19 Surveillance Team |
| EPI_ISL_2803854, EPI_ISL_2803855, EPI_ISL_2803856, EPI_ISL_2803857, EPI_ISL_2803858, EPI_ISL_2803859 | Lighthouse Lab in Milton Keynes | Wellcome Sanger Institute for the COVID-19 Genomics UK (COG-UK) Consortium | The Lighthouse Lab in Milton Keynes and Alex Alderton, Roberto Amato, Jeffrey Barrett, Sonia Goncalves, Ewan Harrison, David K. Jackson, Ian Johnston, Dominic Kwiatkowski, Cordelia Langford, John Sillitoe on behalf of the Wellcome Sanger Institute COVID-19 Surveillance Team |
| EPI_ISL_2803860, EPI_ISL_2803861 | Lighthouse Lab in Glasgow | Wellcome Sanger Institute for the COVID-19 Genomics UK (COG-UK) Consortium | Harper VanSteenhouse, Yumi Kasai, David Gray, Carol Clugston, Anna Dominiczak and Alex Alderton, Roberto Amato, Jeffrey Barrett, Sonia Goncalves, Ewan Harrison, David K. Jackson, Ian Johnston, Dominic Kwiatkowski, Cordelia Langford, John Sillitoe on behalf of the Wellcome Sanger Institute COVID-19 Surveillance Team |
| EPI_ISL_2803862, EPI_ISL_2803863, | Lighthouse Lab in Milton Keynes | Wellcome Sanger Institute for the COVID-19 Genomics UK (COG-UK) Consortium | The Lighthouse Lab in Milton Keynes and Alex Alderton, Roberto Amato, Jeffrey Barrett, Sonia Goncalves, Ewan Harrison, David K. Jackson, Ian Johnston, Dominic Kwiatkowski, Cordelia Langford, John Sillitoe on behalf of the Wellcome Sanger Institute COVID-19 Surveillance Team |

| | | | |
|---|---|---|---|
| EPI_ISL_2803864 | | | |
| EPI_ISL_2803865, EPI_ISL_2803866, EPI_ISL_2803867, EPI_ISL_2803868 | Lighthouse Lab in Glasgow | Wellcome Sanger Institute for the COVID-19 Genomics UK (COG-UK) Consortium | Harper VanSteenhouse, Yumi Kasai, David Gray, Carol Clugston, Anna Dominiczak and Alex Alderton, Roberto Amato, Jeffrey Barrett, Sonia Goncalves, Ewan Harrison, David K. Jackson, Ian Johnston, Dominic Kwiatkowski, Cordelia Langford, John Sillitoe on behalf of the Wellcome Sanger Institute COVID-19 Surveillance Team |
| EPI_ISL_2803869 | Lighthouse Lab in Milton Keynes | Wellcome Sanger Institute for the COVID-19 Genomics UK (COG-UK) Consortium | The Lighthouse Lab in Milton Keynes and Alex Alderton, Roberto Amato, Jeffrey Barrett, Sonia Goncalves, Ewan Harrison, David K. Jackson, Ian Johnston, Dominic Kwiatkowski, Cordelia Langford, John Sillitoe on behalf of the Wellcome Sanger Institute COVID-19 Surveillance Team |
| EPI_ISL_2803870 | Lighthouse Lab in Glasgow | Wellcome Sanger Institute for the COVID-19 Genomics UK (COG-UK) Consortium | Harper VanSteenhouse, Yumi Kasai, David Gray, Carol Clugston, Anna Dominiczak and Alex Alderton, Roberto Amato, Jeffrey Barrett, Sonia Goncalves, Ewan Harrison, David K. Jackson, Ian Johnston, Dominic Kwiatkowski, Cordelia Langford, John Sillitoe on behalf of the Wellcome Sanger Institute COVID-19 Surveillance Team |
| EPI_ISL_2803871, EPI_ISL_2803872 | Lighthouse Lab in Milton Keynes | Wellcome Sanger Institute for the COVID-19 Genomics UK (COG-UK) Consortium | The Lighthouse Lab in Milton Keynes and Alex Alderton, Roberto Amato, Jeffrey Barrett, Sonia Goncalves, Ewan Harrison, David K. Jackson, Ian Johnston, Dominic Kwiatkowski, Cordelia Langford, John Sillitoe on behalf of the Wellcome Sanger Institute COVID-19 Surveillance Team |
| EPI_ISL_2803873, EPI_ISL_2803874, EPI_ISL_2803875 | Lighthouse Lab in Glasgow | Wellcome Sanger Institute for the COVID-19 Genomics UK (COG-UK) Consortium | Harper VanSteenhouse, Yumi Kasai, David Gray, Carol Clugston, Anna Dominiczak and Alex Alderton, Roberto Amato, Jeffrey Barrett, Sonia Goncalves, Ewan Harrison, David K. Jackson, Ian Johnston, Dominic Kwiatkowski, Cordelia Langford, John Sillitoe on behalf of the Wellcome Sanger Institute COVID-19 Surveillance Team |
| EPI_ISL_2803876 | Lighthouse Lab in Milton Keynes | Wellcome Sanger Institute for the COVID-19 Genomics UK (COG-UK) Consortium | The Lighthouse Lab in Milton Keynes and Alex Alderton, Roberto Amato, Jeffrey Barrett, Sonia Goncalves, Ewan Harrison, David K. Jackson, Ian Johnston, Dominic Kwiatkowski, Cordelia Langford, John Sillitoe on behalf of the Wellcome Sanger Institute COVID-19 Surveillance Team |
| EPI_ISL_2803877, EPI_ISL_2803878 | Lighthouse Lab in Glasgow | Wellcome Sanger Institute for the COVID-19 Genomics UK (COG-UK) Consortium | Harper VanSteenhouse, Yumi Kasai, David Gray, Carol Clugston, Anna Dominiczak and Alex Alderton, Roberto Amato, Jeffrey Barrett, Sonia Goncalves, Ewan Harrison, David K. Jackson, Ian Johnston, Dominic Kwiatkowski, Cordelia Langford, John Sillitoe on behalf of the Wellcome Sanger Institute COVID-19 Surveillance Team |
| EPI_ISL_2803879 | Lighthouse Lab in Milton Keynes | Wellcome Sanger Institute for the COVID-19 Genomics UK (COG-UK) Consortium | The Lighthouse Lab in Milton Keynes and Alex Alderton, Roberto Amato, Jeffrey Barrett, Sonia Goncalves, Ewan Harrison, David K. Jackson, Ian Johnston, Dominic Kwiatkowski, Cordelia Langford, John Sillitoe on behalf of the Wellcome Sanger Institute COVID-19 Surveillance Team |
| EPI_ISL_2803880, EPI_ISL_2803881 | Lighthouse Lab in Glasgow | Wellcome Sanger Institute for the COVID-19 Genomics UK (COG-UK) Consortium | Harper VanSteenhouse, Yumi Kasai, David Gray, Carol Clugston, Anna Dominiczak and Alex Alderton, Roberto Amato, Jeffrey Barrett, Sonia Goncalves, Ewan Harrison, David K. Jackson, Ian Johnston, Dominic Kwiatkowski, Cordelia Langford, John Sillitoe on behalf of the Wellcome Sanger Institute COVID-19 Surveillance Team |
| EPI_ISL_2803882, EPI_ISL_2803883 | Lighthouse Lab in Milton Keynes | Wellcome Sanger Institute for the COVID-19 Genomics UK (COG-UK) Consortium | The Lighthouse Lab in Milton Keynes and Alex Alderton, Roberto Amato, Jeffrey Barrett, Sonia Goncalves, Ewan Harrison, David K. Jackson, Ian Johnston, Dominic Kwiatkowski, Cordelia Langford, John Sillitoe on behalf of the Wellcome Sanger Institute COVID-19 Surveillance Team |
| EPI_ISL_2803884 | Lighthouse Lab in Glasgow | Wellcome Sanger Institute for the COVID-19 Genomics UK (COG-UK) Consortium | Harper VanSteenhouse, Yumi Kasai, David Gray, Carol Clugston, Anna Dominiczak and Alex Alderton, Roberto Amato, Jeffrey Barrett, Sonia Goncalves, Ewan Harrison, David K. Jackson, Ian Johnston, Dominic Kwiatkowski, Cordelia Langford, John Sillitoe on behalf of the Wellcome Sanger Institute COVID-19 Surveillance Team |
| EPI_ISL_2803885 | Lighthouse Lab in Milton Keynes | Wellcome Sanger Institute for the COVID-19 Genomics UK (COG-UK) Consortium | The Lighthouse Lab in Milton Keynes and Alex Alderton, Roberto Amato, Jeffrey Barrett, Sonia Goncalves, Ewan Harrison, David K. Jackson, Ian Johnston, Dominic Kwiatkowski, Cordelia Langford, John Sillitoe on behalf of the Wellcome Sanger Institute COVID-19 Surveillance Team |
| EPI_ISL_2803886 | Lighthouse Lab in Glasgow | Wellcome Sanger Institute for the COVID-19 Genomics UK (COG-UK) Consortium | Harper VanSteenhouse, Yumi Kasai, David Gray, Carol Clugston, Anna Dominiczak and Alex Alderton, Roberto Amato, Jeffrey Barrett, Sonia Goncalves, Ewan Harrison, David K. Jackson, Ian Johnston, Dominic Kwiatkowski, Cordelia Langford, John Sillitoe on behalf of the Wellcome Sanger Institute COVID-19 Surveillance Team |
| EPI_ISL_2803887 | Lighthouse Lab in Milton Keynes | Wellcome Sanger Institute for the COVID-19 Genomics UK (COG-UK) Consortium | The Lighthouse Lab in Milton Keynes and Alex Alderton, Roberto Amato, Jeffrey Barrett, Sonia Goncalves, Ewan Harrison, David K. Jackson, Ian Johnston, Dominic Kwiatkowski, Cordelia Langford, John Sillitoe on behalf of the Wellcome Sanger Institute COVID-19 Surveillance Team |
| EPI_ISL_2803888 | Lighthouse Lab in Glasgow | Wellcome Sanger Institute for the COVID-19 Genomics UK (COG-UK) Consortium | Harper VanSteenhouse, Yumi Kasai, David Gray, Carol Clugston, Anna Dominiczak and Alex Alderton, Roberto Amato, Jeffrey Barrett, Sonia Goncalves, Ewan Harrison, David K. Jackson, Ian Johnston, Dominic Kwiatkowski, Cordelia Langford, John Sillitoe on behalf of the Wellcome Sanger Institute COVID-19 Surveillance Team |
| EPI_ISL_2803889, EPI_ISL_2803890 | Lighthouse Lab in Milton Keynes | Wellcome Sanger Institute for the COVID-19 Genomics UK (COG-UK) Consortium | The Lighthouse Lab in Milton Keynes and Alex Alderton, Roberto Amato, Jeffrey Barrett, Sonia Goncalves, Ewan Harrison, David K. Jackson, Ian Johnston, Dominic Kwiatkowski, Cordelia Langford, John Sillitoe on behalf of the Wellcome Sanger Institute COVID-19 Surveillance Team |
| EPI_ISL_2803891, EPI_ISL_2803892 | Lighthouse Lab in Glasgow | Wellcome Sanger Institute for the COVID-19 Genomics UK (COG-UK) Consortium | Harper VanSteenhouse, Yumi Kasai, David Gray, Carol Clugston, Anna Dominiczak and Alex Alderton, Roberto Amato, Jeffrey Barrett, Sonia Goncalves, Ewan Harrison, David K. Jackson, Ian Johnston, Dominic Kwiatkowski, Cordelia Langford, John Sillitoe on behalf of the Wellcome Sanger Institute COVID-19 Surveillance Team |
| EPI_ISL_2803893, EPI_ISL_2803894, EPI_ISL_2803895, EPI_ISL_2803896 | Lighthouse Lab in Milton Keynes | Wellcome Sanger Institute for the COVID-19 Genomics UK (COG-UK) Consortium | The Lighthouse Lab in Milton Keynes and Alex Alderton, Roberto Amato, Jeffrey Barrett, Sonia Goncalves, Ewan Harrison, David K. Jackson, Ian Johnston, Dominic Kwiatkowski, Cordelia Langford, John Sillitoe on behalf of the Wellcome Sanger Institute COVID-19 Surveillance Team |
| EPI_ISL_2803897, EPI_ISL_2803898, EPI_ISL_2803899 | Lighthouse Lab in Glasgow | Wellcome Sanger Institute for the COVID-19 Genomics UK (COG-UK) Consortium | Harper VanSteenhouse, Yumi Kasai, David Gray, Carol Clugston, Anna Dominiczak and Alex Alderton, Roberto Amato, Jeffrey Barrett, Sonia Goncalves, Ewan Harrison, David K. Jackson, Ian Johnston, Dominic Kwiatkowski, Cordelia Langford, John Sillitoe on behalf of the Wellcome Sanger Institute COVID-19 Surveillance Team |
| EPI_ISL_2803900 | Lighthouse Lab in Milton Keynes | Wellcome Sanger Institute for the COVID-19 Genomics UK (COG-UK) Consortium | The Lighthouse Lab in Milton Keynes and Alex Alderton, Roberto Amato, Jeffrey Barrett, Sonia Goncalves, Ewan Harrison, David K. Jackson, Ian Johnston, Dominic Kwiatkowski, Cordelia Langford, John Sillitoe on behalf of the Wellcome Sanger Institute COVID-19 Surveillance Team |
| EPI_ISL_2803901 | Lighthouse Lab in Glasgow | Wellcome Sanger Institute for the COVID-19 Genomics UK (COG-UK) Consortium | Harper VanSteenhouse, Yumi Kasai, David Gray, Carol Clugston, Anna Dominiczak and Alex Alderton, Roberto Amato, Jeffrey Barrett, Sonia Goncalves, Ewan Harrison, David K. Jackson, Ian Johnston, Dominic Kwiatkowski, Cordelia Langford, John Sillitoe on behalf of the Wellcome Sanger Institute COVID-19 Surveillance Team |
| EPI_ISL_2803902, EPI_ISL_2803903, EPI_ISL_2803904, EPI_ISL_2803905, EPI_ISL_2803906 | Lighthouse Lab in Milton Keynes | Wellcome Sanger Institute for the COVID-19 Genomics UK (COG-UK) Consortium | The Lighthouse Lab in Milton Keynes and Alex Alderton, Roberto Amato, Jeffrey Barrett, Sonia Goncalves, Ewan Harrison, David K. Jackson, Ian Johnston, Dominic Kwiatkowski, Cordelia Langford, John Sillitoe on behalf of the Wellcome Sanger Institute COVID-19 Surveillance Team |
| EPI_ISL_2803907 | Lighthouse Lab in Glasgow | Wellcome Sanger Institute for the COVID-19 Genomics UK (COG-UK) Consortium | Harper VanSteenhouse, Yumi Kasai, David Gray, Carol Clugston, Anna Dominiczak and Alex Alderton, Roberto Amato, Jeffrey Barrett, Sonia Goncalves, Ewan Harrison, David K. Jackson, Ian Johnston, Dominic Kwiatkowski, Cordelia Langford, John Sillitoe on behalf of the Wellcome Sanger Institute COVID-19 Surveillance Team |
| EPI_ISL_2803908, EPI_ISL_2803910 | Lighthouse Lab in Milton Keynes | Wellcome Sanger Institute for the COVID-19 Genomics UK (COG-UK) Consortium | The Lighthouse Lab in Milton Keynes and Alex Alderton, Roberto Amato, Jeffrey Barrett, Sonia Goncalves, Ewan Harrison, David K. Jackson, Ian Johnston, Dominic Kwiatkowski, Cordelia Langford, John Sillitoe on behalf of the Wellcome Sanger Institute COVID-19 Surveillance Team |
| EPI_ISL_2803911, EPI_ISL_2803912 | Lighthouse Lab in Glasgow | Wellcome Sanger Institute for the COVID-19 Genomics UK (COG-UK) Consortium | Harper VanSteenhouse, Yumi Kasai, David Gray, Carol Clugston, Anna Dominiczak and Alex Alderton, Roberto Amato, Jeffrey Barrett, Sonia Goncalves, Ewan Harrison, David K. Jackson, Ian Johnston, Dominic Kwiatkowski, Cordelia Langford, John Sillitoe on behalf of the Wellcome Sanger Institute COVID-19 Surveillance Team |
| EPI_ISL_2803913, EPI_ISL_2803914 | Lighthouse Lab in Milton Keynes | Wellcome Sanger Institute for the COVID-19 Genomics UK (COG-UK) Consortium | The Lighthouse Lab in Milton Keynes and Alex Alderton, Roberto Amato, Jeffrey Barrett, Sonia Goncalves, Ewan Harrison, David K. Jackson, Ian Johnston, Dominic Kwiatkowski, Cordelia Langford, John Sillitoe on behalf of the Wellcome Sanger Institute COVID-19 Surveillance Team |
| EPI_ISL_2803915, EPI_ISL_2803916, EPI_ISL_2803917, EPI_ISL_2803918, EPI_ISL_2803919 | Lighthouse Lab in Glasgow | Wellcome Sanger Institute for the COVID-19 Genomics UK (COG-UK) Consortium | Harper VanSteenhouse, Yumi Kasai, David Gray, Carol Clugston, Anna Dominiczak and Alex Alderton, Roberto Amato, Jeffrey Barrett, Sonia Goncalves, Ewan Harrison, David K. Jackson, Ian Johnston, Dominic Kwiatkowski, Cordelia Langford, John Sillitoe on behalf of the Wellcome Sanger Institute COVID-19 Surveillance Team |
| EPI_ISL_2803920 | Lighthouse Lab in Milton Keynes | Wellcome Sanger Institute for the COVID-19 Genomics UK (COG-UK) Consortium | The Lighthouse Lab in Milton Keynes and Alex Alderton, Roberto Amato, Jeffrey Barrett, Sonia Goncalves, Ewan Harrison, David K. Jackson, Ian Johnston, Dominic Kwiatkowski, Cordelia Langford, John Sillitoe on behalf of the Wellcome Sanger Institute COVID-19 Surveillance Team |
| EPI_ISL_2803921 | Lighthouse Lab in Glasgow | Wellcome Sanger Institute for the COVID-19 Genomics UK (COG-UK) Consortium | Harper VanSteenhouse, Yumi Kasai, David Gray, Carol Clugston, Anna Dominiczak and Alex Alderton, Roberto Amato, Jeffrey Barrett, Sonia Goncalves, Ewan Harrison, David K. Jackson, Ian Johnston, Dominic Kwiatkowski, Cordelia Langford, John Sillitoe on behalf of the Wellcome Sanger Institute COVID-19 Surveillance Team |
| EPI_ISL_2803922, EPI_ISL_2803923 | Lighthouse Lab in Milton Keynes | Wellcome Sanger Institute for the COVID-19 Genomics UK (COG-UK) Consortium | The Lighthouse Lab in Milton Keynes and Alex Alderton, Roberto Amato, Jeffrey Barrett, Sonia Goncalves, Ewan Harrison, David K. Jackson, Ian Johnston, Dominic Kwiatkowski, Cordelia Langford, John Sillitoe on behalf of the Wellcome Sanger Institute COVID-19 Surveillance Team |
| EPI_ISL_2803924 | Lighthouse Laboratory Plymouth | Wellcome Sanger Institute for the COVID-19 Genomics UK (COG-UK) Consortium | Lighthouse Laboratory Plymouth and Alex Alderton, Roberto Amato, Jeffrey Barrett, Sonia Goncalves, Ewan Harrison, David K. Jackson, Ian Johnston, Dominic Kwiatkowski, Cordelia Langford, John Sillitoe on behalf of the Wellcome Sanger Institute COVID-19 Surveillance Team |
| EPI_ISL_2803925 | Lighthouse Lab in Alderley Park | Wellcome Sanger Institute for the COVID-19 Genomics UK (COG-UK) Consortium | Jacquelyn Wynn, Mairead Hyland, The Lighthouse Lab in Alderley Park and Alex Alderton, Roberto Amato, Jeffrey Barrett, Sonia Goncalves, Ewan Harrison, David K. Jackson, Ian Johnston, Dominic Kwiatkowski, Cordelia Langford, John Sillitoe on behalf of the Wellcome Sanger Institute COVID-19 Surveillance Team |
| EPI_ISL_2803926 | Lighthouse Laboratory Plymouth | Wellcome Sanger Institute for the COVID-19 Genomics UK (COG-UK) Consortium | Lighthouse Laboratory Plymouth and Alex Alderton, Roberto Amato, Jeffrey Barrett, Sonia Goncalves, Ewan Harrison, David K. Jackson, Ian Johnston, Dominic Kwiatkowski, Cordelia Langford, John Sillitoe on behalf of the Wellcome Sanger Institute COVID-19 Surveillance Team |
| EPI_ISL_2803927 | Lighthouse Lab in Alderley Park | Wellcome Sanger Institute for the COVID-19 Genomics UK (COG-UK) Consortium | Jacquelyn Wynn, Mairead Hyland, The Lighthouse Lab in Alderley Park and Alex Alderton, Roberto Amato, Jeffrey Barrett, Sonia Goncalves, Ewan Harrison, David K. Jackson, Ian Johnston, Dominic Kwiatkowski, Cordelia Langford, John Sillitoe on behalf of the Wellcome Sanger Institute COVID-19 Surveillance Team |
| EPI_ISL_2803928, EPI_ISL_2803929, EPI_ISL_2803930 | Lighthouse Laboratory Plymouth | Wellcome Sanger Institute for the COVID-19 Genomics UK (COG-UK) Consortium | Lighthouse Laboratory Plymouth and Alex Alderton, Roberto Amato, Jeffrey Barrett, Sonia Goncalves, Ewan Harrison, David K. Jackson, Ian Johnston, Dominic Kwiatkowski, Cordelia Langford, John Sillitoe on behalf of the Wellcome Sanger Institute COVID-19 Surveillance Team |
| EPI_ISL_2803931 | Lighthouse Lab in Milton Keynes | Wellcome Sanger Institute for the COVID-19 Genomics UK (COG-UK) Consortium | The Lighthouse Lab in Milton Keynes and Alex Alderton, Roberto Amato, Jeffrey Barrett, Sonia Goncalves, Ewan Harrison, David K. Jackson, Ian Johnston, Dominic Kwiatkowski, Cordelia Langford, John Sillitoe on behalf of the Wellcome Sanger Institute COVID-19 Surveillance Team |
| EPI_ISL_2803932 | Lighthouse Lab in Alderley Park | Wellcome Sanger Institute for the COVID-19 Genomics UK (COG-UK) Consortium | Jacquelyn Wynn, Mairead Hyland, The Lighthouse Lab in Alderley Park and Alex Alderton, Roberto Amato, Jeffrey Barrett, Sonia Goncalves, Ewan Harrison, David K. Jackson, Ian Johnston, Dominic Kwiatkowski, Cordelia Langford, John Sillitoe on behalf of the Wellcome Sanger Institute COVID-19 Surveillance Team |
| EPI_ISL_2803933 | Health Services Laboratories | Wellcome Sanger Institute for the COVID-19 Genomics UK (COG-UK) Consortium | Health Services Laboratories and Alex Alderton, Roberto Amato, Jeffrey Barrett, Sonia Goncalves, Ewan Harrison, David K. Jackson, Ian Johnston, Dominic Kwiatkowski, Cordelia Langford, John Sillitoe on behalf of the Wellcome Sanger Institute COVID-19 Surveillance Team |
| EPI_ISL_2803934 | Lighthouse Lab in Alderley Park | Wellcome Sanger Institute for the COVID-19 Genomics UK (COG-UK) Consortium | Jacquelyn Wynn, Mairead Hyland, The Lighthouse Lab in Alderley Park and Alex Alderton, Roberto Amato, Jeffrey Barrett, Sonia Goncalves, Ewan Harrison, David K. Jackson, Ian Johnston, Dominic Kwiatkowski, Cordelia Langford, John Sillitoe on behalf of the Wellcome Sanger Institute COVID-19 Surveillance Team |

| | | | |
|---|---|---|---|
| EPI_ISL_2803935 | Lighthouse Lab in Milton Keynes | Wellcome Sanger Institute for the COVID-19 Genomics UK (COG-UK) Consortium | The Lighthouse Lab in Milton Keynes and Alex Alderton, Roberto Amato, Jeffrey Barrett, Sonia Goncalves, Ewan Harrison, David K. Jackson, Ian Johnston, Dominic Kwiatkowski, Cordelia Langford, John Sillitoe on behalf of the Wellcome Sanger Institute COVID-19 Surveillance Team |
| EPI_ISL_2803936, EPI_ISL_2803937 | Lighthouse Laboratory Plymouth | Wellcome Sanger Institute for the COVID-19 Genomics UK (COG-UK) Consortium | Lighthouse Laboratory Plymouth and Alex Alderton, Roberto Amato, Jeffrey Barrett, Sonia Goncalves, Ewan Harrison, David K. Jackson, Ian Johnston, Dominic Kwiatkowski, Cordelia Langford, John Sillitoe on behalf of the Wellcome Sanger Institute COVID-19 Surveillance Team |
| EPI_ISL_2803938 | Lighthouse Lab in Alderley Park | Wellcome Sanger Institute for the COVID-19 Genomics UK (COG-UK) Consortium | Jacquelyn Wynn, Mairead Hyland, The Lighthouse Lab in Alderley Park and Alex Alderton, Roberto Amato, Jeffrey Barrett, Sonia Goncalves, Ewan Harrison, David K. Jackson, Ian Johnston, Dominic Kwiatkowski, Cordelia Langford, John Sillitoe on behalf of the Wellcome Sanger Institute COVID-19 Surveillance Team |
| EPI_ISL_2803939 | Lighthouse Lab in Milton Keynes | Wellcome Sanger Institute for the COVID-19 Genomics UK (COG-UK) Consortium | The Lighthouse Lab in Milton Keynes and Alex Alderton, Roberto Amato, Jeffrey Barrett, Sonia Goncalves, Ewan Harrison, David K. Jackson, Ian Johnston, Dominic Kwiatkowski, Cordelia Langford, John Sillitoe on behalf of the Wellcome Sanger Institute COVID-19 Surveillance Team |
| EPI_ISL_2803940, EPI_ISL_2803941, EPI_ISL_2803942, EPI_ISL_2803943, EPI_ISL_2803944, EPI_ISL_2803945, EPI_ISL_2803946, EPI_ISL_2803947 | Lighthouse Laboratory Plymouth | Wellcome Sanger Institute for the COVID-19 Genomics UK (COG-UK) Consortium | Lighthouse Laboratory Plymouth and Alex Alderton, Roberto Amato, Jeffrey Barrett, Sonia Goncalves, Ewan Harrison, David K. Jackson, Ian Johnston, Dominic Kwiatkowski, Cordelia Langford, John Sillitoe on behalf of the Wellcome Sanger Institute COVID-19 Surveillance Team |
| EPI_ISL_2803948 | Lighthouse Lab in Milton Keynes | Wellcome Sanger Institute for the COVID-19 Genomics UK (COG-UK) Consortium | The Lighthouse Lab in Milton Keynes and Alex Alderton, Roberto Amato, Jeffrey Barrett, Sonia Goncalves, Ewan Harrison, David K. Jackson, Ian Johnston, Dominic Kwiatkowski, Cordelia Langford, John Sillitoe on behalf of the Wellcome Sanger Institute COVID-19 Surveillance Team |
| EPI_ISL_2803949 | Lighthouse Lab in Alderley Park | Wellcome Sanger Institute for the COVID-19 Genomics UK (COG-UK) Consortium | Jacquelyn Wynn, Mairead Hyland, The Lighthouse Lab in Alderley Park and Alex Alderton, Roberto Amato, Jeffrey Barrett, Sonia Goncalves, Ewan Harrison, David K. Jackson, Ian Johnston, Dominic Kwiatkowski, Cordelia Langford, John Sillitoe on behalf of the Wellcome Sanger Institute COVID-19 Surveillance Team |
| EPI_ISL_2803950, EPI_ISL_2803951, EPI_ISL_2803952 | Lighthouse Laboratory Plymouth | Wellcome Sanger Institute for the COVID-19 Genomics UK (COG-UK) Consortium | Lighthouse Laboratory Plymouth and Alex Alderton, Roberto Amato, Jeffrey Barrett, Sonia Goncalves, Ewan Harrison, David K. Jackson, Ian Johnston, Dominic Kwiatkowski, Cordelia Langford, John Sillitoe on behalf of the Wellcome Sanger Institute COVID-19 Surveillance Team |
| EPI_ISL_2803953 | Lighthouse Lab in Milton Keynes | Wellcome Sanger Institute for the COVID-19 Genomics UK (COG-UK) Consortium | The Lighthouse Lab in Milton Keynes and Alex Alderton, Roberto Amato, Jeffrey Barrett, Sonia Goncalves, Ewan Harrison, David K. Jackson, Ian Johnston, Dominic Kwiatkowski, Cordelia Langford, John Sillitoe on behalf of the Wellcome Sanger Institute COVID-19 Surveillance Team |
| EPI_ISL_2803954 | Lighthouse Laboratory Plymouth | Wellcome Sanger Institute for the COVID-19 Genomics UK (COG-UK) Consortium | Lighthouse Laboratory Plymouth and Alex Alderton, Roberto Amato, Jeffrey Barrett, Sonia Goncalves, Ewan Harrison, David K. Jackson, Ian Johnston, Dominic Kwiatkowski, Cordelia Langford, John Sillitoe on behalf of the Wellcome Sanger Institute COVID-19 Surveillance Team |
| EPI_ISL_2803955 | Lighthouse Lab in Alderley Park | Wellcome Sanger Institute for the COVID-19 Genomics UK (COG-UK) Consortium | Jacquelyn Wynn, Mairead Hyland, The Lighthouse Lab in Alderley Park and Alex Alderton, Roberto Amato, Jeffrey Barrett, Sonia Goncalves, Ewan Harrison, David K. Jackson, Ian Johnston, Dominic Kwiatkowski, Cordelia Langford, John Sillitoe on behalf of the Wellcome Sanger Institute COVID-19 Surveillance Team |
| EPI_ISL_2803956, EPI_ISL_2803957, EPI_ISL_2803958 | Lighthouse Laboratory Plymouth | Wellcome Sanger Institute for the COVID-19 Genomics UK (COG-UK) Consortium | Lighthouse Laboratory Plymouth and Alex Alderton, Roberto Amato, Jeffrey Barrett, Sonia Goncalves, Ewan Harrison, David K. Jackson, Ian Johnston, Dominic Kwiatkowski, Cordelia Langford, John Sillitoe on behalf of the Wellcome Sanger Institute COVID-19 Surveillance Team |
| EPI_ISL_2803959 | Lighthouse Lab in Milton Keynes | Wellcome Sanger Institute for the COVID-19 Genomics UK (COG-UK) Consortium | The Lighthouse Lab in Milton Keynes and Alex Alderton, Roberto Amato, Jeffrey Barrett, Sonia Goncalves, Ewan Harrison, David K. Jackson, Ian Johnston, Dominic Kwiatkowski, Cordelia Langford, John Sillitoe on behalf of the Wellcome Sanger Institute COVID-19 Surveillance Team |
| EPI_ISL_2803960 | Lighthouse Laboratory Plymouth | Wellcome Sanger Institute for the COVID-19 Genomics UK (COG-UK) Consortium | Lighthouse Laboratory Plymouth and Alex Alderton, Roberto Amato, Jeffrey Barrett, Sonia Goncalves, Ewan Harrison, David K. Jackson, Ian Johnston, Dominic Kwiatkowski, Cordelia Langford, John Sillitoe on behalf of the Wellcome Sanger Institute COVID-19 Surveillance Team |
| EPI_ISL_2803961 | Lighthouse Lab in Alderley Park | Wellcome Sanger Institute for the COVID-19 Genomics UK (COG-UK) Consortium | Jacquelyn Wynn, Mairead Hyland, The Lighthouse Lab in Alderley Park and Alex Alderton, Roberto Amato, Jeffrey Barrett, Sonia Goncalves, Ewan Harrison, David K. Jackson, Ian Johnston, Dominic Kwiatkowski, Cordelia Langford, John Sillitoe on behalf of the Wellcome Sanger Institute COVID-19 Surveillance Team |
| EPI_ISL_2803962 | Health Services Laboratories | Wellcome Sanger Institute for the COVID-19 Genomics UK (COG-UK) Consortium | Health Services Laboratories and Alex Alderton, Roberto Amato, Jeffrey Barrett, Sonia Goncalves, Ewan Harrison, David K. Jackson, Ian Johnston, Dominic Kwiatkowski, Cordelia Langford, John Sillitoe on behalf of the Wellcome Sanger Institute COVID-19 Surveillance Team |
| EPI_ISL_2803963, EPI_ISL_2803964, EPI_ISL_2803965, EPI_ISL_2803966 | Lighthouse Laboratory Plymouth | Wellcome Sanger Institute for the COVID-19 Genomics UK (COG-UK) Consortium | Lighthouse Laboratory Plymouth and Alex Alderton, Roberto Amato, Jeffrey Barrett, Sonia Goncalves, Ewan Harrison, David K. Jackson, Ian Johnston, Dominic Kwiatkowski, Cordelia Langford, John Sillitoe on behalf of the Wellcome Sanger Institute COVID-19 Surveillance Team |
| EPI_ISL_2803967, EPI_ISL_2803968 | Lighthouse Lab in Alderley Park | Wellcome Sanger Institute for the COVID-19 Genomics UK (COG-UK) Consortium | Jacquelyn Wynn, Mairead Hyland, The Lighthouse Lab in Alderley Park and Alex Alderton, Roberto Amato, Jeffrey Barrett, Sonia Goncalves, Ewan Harrison, David K. Jackson, Ian Johnston, Dominic Kwiatkowski, Cordelia Langford, John Sillitoe on behalf of the Wellcome Sanger Institute COVID-19 Surveillance Team |
| EPI_ISL_2803969 | Lighthouse Lab in Milton Keynes | Wellcome Sanger Institute for the COVID-19 Genomics UK (COG-UK) Consortium | The Lighthouse Lab in Milton Keynes and Alex Alderton, Roberto Amato, Jeffrey Barrett, Sonia Goncalves, Ewan Harrison, David K. Jackson, Ian Johnston, Dominic Kwiatkowski, Cordelia Langford, John Sillitoe on behalf of the Wellcome Sanger Institute COVID-19 Surveillance Team |
| EPI_ISL_2803970, EPI_ISL_2803971 | Lighthouse Laboratory Plymouth | Wellcome Sanger Institute for the COVID-19 Genomics UK (COG-UK) Consortium | Lighthouse Laboratory Plymouth and Alex Alderton, Roberto Amato, Jeffrey Barrett, Sonia Goncalves, Ewan Harrison, David K. Jackson, Ian Johnston, Dominic Kwiatkowski, Cordelia Langford, John Sillitoe on behalf of the Wellcome Sanger Institute COVID-19 Surveillance Team |
| EPI_ISL_2803972 | Lighthouse Lab in Milton Keynes | Wellcome Sanger Institute for the COVID-19 Genomics UK (COG-UK) Consortium | The Lighthouse Lab in Milton Keynes and Alex Alderton, Roberto Amato, Jeffrey Barrett, Sonia Goncalves, Ewan Harrison, David K. Jackson, Ian Johnston, Dominic Kwiatkowski, Cordelia Langford, John Sillitoe on behalf of the Wellcome Sanger Institute COVID-19 Surveillance Team |
| EPI_ISL_2803973 | Lighthouse Lab in Alderley Park | Wellcome Sanger Institute for the COVID-19 Genomics UK (COG-UK) Consortium | Jacquelyn Wynn, Mairead Hyland, The Lighthouse Lab in Alderley Park and Alex Alderton, Roberto Amato, Jeffrey Barrett, Sonia Goncalves, Ewan Harrison, David K. Jackson, Ian Johnston, Dominic Kwiatkowski, Cordelia Langford, John Sillitoe on behalf of the Wellcome Sanger Institute COVID-19 Surveillance Team |
| EPI_ISL_2803974 | Health Services Laboratories | Wellcome Sanger Institute for the COVID-19 Genomics UK (COG-UK) Consortium | Health Services Laboratories and Alex Alderton, Roberto Amato, Jeffrey Barrett, Sonia Goncalves, Ewan Harrison, David K. Jackson, Ian Johnston, Dominic Kwiatkowski, Cordelia Langford, John Sillitoe on behalf of the Wellcome Sanger Institute COVID-19 Surveillance Team |
| EPI_ISL_2803975 | Lighthouse Lab in Milton Keynes | Wellcome Sanger Institute for the COVID-19 Genomics UK (COG-UK) Consortium | The Lighthouse Lab in Milton Keynes and Alex Alderton, Roberto Amato, Jeffrey Barrett, Sonia Goncalves, Ewan Harrison, David K. Jackson, Ian Johnston, Dominic Kwiatkowski, Cordelia Langford, John Sillitoe on behalf of the Wellcome Sanger Institute COVID-19 Surveillance Team |
| EPI_ISL_2803976, EPI_ISL_2803977 | Lighthouse Laboratory Plymouth | Wellcome Sanger Institute for the COVID-19 Genomics UK (COG-UK) Consortium | Lighthouse Laboratory Plymouth and Alex Alderton, Roberto Amato, Jeffrey Barrett, Sonia Goncalves, Ewan Harrison, David K. Jackson, Ian Johnston, Dominic Kwiatkowski, Cordelia Langford, John Sillitoe on behalf of the Wellcome Sanger Institute COVID-19 Surveillance Team |
| EPI_ISL_2803978 | Health Services Laboratories | Wellcome Sanger Institute for the COVID-19 Genomics UK (COG-UK) Consortium | Health Services Laboratories and Alex Alderton, Roberto Amato, Jeffrey Barrett, Sonia Goncalves, Ewan Harrison, David K. Jackson, Ian Johnston, Dominic Kwiatkowski, Cordelia Langford, John Sillitoe on behalf of the Wellcome Sanger Institute COVID-19 Surveillance Team |
| EPI_ISL_2803979, EPI_ISL_2803980 | Lighthouse Laboratory Plymouth | Wellcome Sanger Institute for the COVID-19 Genomics UK (COG-UK) Consortium | Lighthouse Laboratory Plymouth and Alex Alderton, Roberto Amato, Jeffrey Barrett, Sonia Goncalves, Ewan Harrison, David K. Jackson, Ian Johnston, Dominic Kwiatkowski, Cordelia Langford, John Sillitoe on behalf of the Wellcome Sanger Institute COVID-19 Surveillance Team |
| EPI_ISL_2803981, EPI_ISL_2803982 | Lighthouse Lab in Alderley Park | Wellcome Sanger Institute for the COVID-19 Genomics UK (COG-UK) Consortium | Jacquelyn Wynn, Mairead Hyland, The Lighthouse Lab in Alderley Park and Alex Alderton, Roberto Amato, Jeffrey Barrett, Sonia Goncalves, Ewan Harrison, David K. Jackson, Ian Johnston, Dominic Kwiatkowski, Cordelia Langford, John Sillitoe on behalf of the Wellcome Sanger Institute COVID-19 Surveillance Team |
| EPI_ISL_2803983 | Lighthouse Laboratory Plymouth | Wellcome Sanger Institute for the COVID-19 Genomics UK (COG-UK) Consortium | Lighthouse Laboratory Plymouth and Alex Alderton, Roberto Amato, Jeffrey Barrett, Sonia Goncalves, Ewan Harrison, David K. Jackson, Ian Johnston, Dominic Kwiatkowski, Cordelia Langford, John Sillitoe on behalf of the Wellcome Sanger Institute COVID-19 Surveillance Team |
| EPI_ISL_2803984, EPI_ISL_2803985 | Lighthouse Lab in Alderley Park | Wellcome Sanger Institute for the COVID-19 Genomics UK (COG-UK) Consortium | Jacquelyn Wynn, Mairead Hyland, The Lighthouse Lab in Alderley Park and Alex Alderton, Roberto Amato, Jeffrey Barrett, Sonia Goncalves, Ewan Harrison, David K. Jackson, Ian Johnston, Dominic Kwiatkowski, Cordelia Langford, John Sillitoe on behalf of the Wellcome Sanger Institute COVID-19 Surveillance Team |
| EPI_ISL_2803986, EPI_ISL_2803987, EPI_ISL_2803988, EPI_ISL_2803989, EPI_ISL_2803990, EPI_ISL_2803991 | Lighthouse Laboratory Plymouth | Wellcome Sanger Institute for the COVID-19 Genomics UK (COG-UK) Consortium | Lighthouse Laboratory Plymouth and Alex Alderton, Roberto Amato, Jeffrey Barrett, Sonia Goncalves, Ewan Harrison, David K. Jackson, Ian Johnston, Dominic Kwiatkowski, Cordelia Langford, John Sillitoe on behalf of the Wellcome Sanger Institute COVID-19 Surveillance Team |
| EPI_ISL_2803992 | Lighthouse Lab in Alderley Park | Wellcome Sanger Institute for the COVID-19 Genomics UK (COG-UK) Consortium | Jacquelyn Wynn, Mairead Hyland, The Lighthouse Lab in Alderley Park and Alex Alderton, Roberto Amato, Jeffrey Barrett, Sonia Goncalves, Ewan Harrison, David K. Jackson, Ian Johnston, Dominic Kwiatkowski, Cordelia Langford, John Sillitoe on behalf of the Wellcome Sanger Institute COVID-19 Surveillance Team |
| EPI_ISL_2803993 | Health Services Laboratories | Wellcome Sanger Institute for the COVID-19 Genomics UK (COG-UK) Consortium | Health Services Laboratories and Alex Alderton, Roberto Amato, Jeffrey Barrett, Sonia Goncalves, Ewan Harrison, David K. Jackson, Ian Johnston, Dominic Kwiatkowski, Cordelia Langford, John Sillitoe on behalf of the Wellcome Sanger Institute COVID-19 Surveillance Team |
| EPI_ISL_2803994, EPI_ISL_2803995 | Lighthouse Laboratory Plymouth | Wellcome Sanger Institute for the COVID-19 Genomics UK (COG-UK) Consortium | Lighthouse Laboratory Plymouth and Alex Alderton, Roberto Amato, Jeffrey Barrett, Sonia Goncalves, Ewan Harrison, David K. Jackson, Ian Johnston, Dominic Kwiatkowski, Cordelia Langford, John Sillitoe on behalf of the Wellcome Sanger Institute COVID-19 Surveillance Team |
| EPI_ISL_2803996 | Lighthouse Lab in Milton Keynes | Wellcome Sanger Institute for the COVID-19 Genomics UK (COG-UK) Consortium | The Lighthouse Lab in Milton Keynes and Alex Alderton, Roberto Amato, Jeffrey Barrett, Sonia Goncalves, Ewan Harrison, David K. Jackson, Ian Johnston, Dominic Kwiatkowski, Cordelia Langford, John Sillitoe on behalf of the Wellcome Sanger Institute COVID-19 Surveillance Team |
| EPI_ISL_2803997 | Health Services Laboratories | Wellcome Sanger Institute for the COVID-19 Genomics UK (COG-UK) Consortium | Health Services Laboratories and Alex Alderton, Roberto Amato, Jeffrey Barrett, Sonia Goncalves, Ewan Harrison, David K. Jackson, Ian Johnston, Dominic Kwiatkowski, Cordelia Langford, John Sillitoe on behalf of the Wellcome Sanger Institute COVID-19 Surveillance Team |
| EPI_ISL_2803998, EPI_ISL_2803999 | Lighthouse Lab in Milton Keynes | Wellcome Sanger Institute for the COVID-19 Genomics UK (COG-UK) Consortium | The Lighthouse Lab in Milton Keynes and Alex Alderton, Roberto Amato, Jeffrey Barrett, Sonia Goncalves, Ewan Harrison, David K. Jackson, Ian Johnston, Dominic Kwiatkowski, Cordelia Langford, John Sillitoe on behalf of the Wellcome Sanger Institute COVID-19 Surveillance Team |
| EPI_ISL_2804000 | Lighthouse Laboratory Plymouth | Wellcome Sanger Institute for the COVID-19 Genomics UK (COG-UK) Consortium | Lighthouse Laboratory Plymouth and Alex Alderton, Roberto Amato, Jeffrey Barrett, Sonia Goncalves, Ewan Harrison, David K. Jackson, Ian Johnston, Dominic Kwiatkowski, Cordelia Langford, John Sillitoe on behalf of the Wellcome Sanger Institute COVID-19 Surveillance Team |
| EPI_ISL_2804001 | Health Services Laboratories | Wellcome Sanger Institute for the COVID-19 Genomics UK (COG-UK) Consortium | Health Services Laboratories and Alex Alderton, Roberto Amato, Jeffrey Barrett, Sonia Goncalves, Ewan Harrison, David K. Jackson, Ian Johnston, Dominic Kwiatkowski, Cordelia Langford, John Sillitoe on behalf of the Wellcome Sanger Institute COVID-19 Surveillance Team |
| EPI_ISL_2804002 | Lighthouse Lab in Milton Keynes | Wellcome Sanger Institute for the COVID-19 Genomics UK (COG-UK) | The Lighthouse Lab in Milton Keynes and Alex Alderton, Roberto Amato, Jeffrey Barrett, Sonia Goncalves, Ewan Harrison, David K. Jackson, Ian Johnston, Dominic Kwiatkowski, Cordelia Langford, John Sillitoe on behalf of the Wellcome Sanger |

| | | | |
|---|---|---|---|
| | | Consortium | Institute COVID-19 Surveillance Team |
| EPI_ISL_2804003 | Lighthouse Laboratory Plymouth | Wellcome Sanger Institute for the COVID-19 Genomics UK (COG-UK) Consortium | Lighthouse Laboratory Plymouth and Alex Alderton, Roberto Amato, Jeffrey Barrett, Sonia Goncalves, Ewan Harrison, David K. Jackson, Ian Johnston, Dominic Kwiatkowski, Cordelia Langford, John Sillitoe on behalf of the Wellcome Sanger Institute COVID-19 Surveillance Team |
| EPI_ISL_2804004, EPI_ISL_2804005 | Lighthouse Lab in Alderley Park | Wellcome Sanger Institute for the COVID-19 Genomics UK (COG-UK) Consortium | Jacquelyn Wynn, Mairead Hyland, The Lighthouse Lab in Alderley Park and Alex Alderton, Roberto Amato, Jeffrey Barrett, Sonia Goncalves, Ewan Harrison, David K. Jackson, Ian Johnston, Dominic Kwiatkowski, Cordelia Langford, John Sillitoe on behalf of the Wellcome Sanger Institute COVID-19 Surveillance Team |
| EPI_ISL_2804006 | Lighthouse Laboratory Plymouth | Wellcome Sanger Institute for the COVID-19 Genomics UK (COG-UK) Consortium | Lighthouse Laboratory Plymouth and Alex Alderton, Roberto Amato, Jeffrey Barrett, Sonia Goncalves, Ewan Harrison, David K. Jackson, Ian Johnston, Dominic Kwiatkowski, Cordelia Langford, John Sillitoe on behalf of the Wellcome Sanger Institute COVID-19 Surveillance Team |
| EPI_ISL_2804007 | Lighthouse Lab in Milton Keynes | Wellcome Sanger Institute for the COVID-19 Genomics UK (COG-UK) Consortium | The Lighthouse Lab in Milton Keynes and Alex Alderton, Roberto Amato, Jeffrey Barrett, Sonia Goncalves, Ewan Harrison, David K. Jackson, Ian Johnston, Dominic Kwiatkowski, Cordelia Langford, John Sillitoe on behalf of the Wellcome Sanger Institute COVID-19 Surveillance Team |
| EPI_ISL_2804008 | Lighthouse Laboratory Plymouth | Wellcome Sanger Institute for the COVID-19 Genomics UK (COG-UK) Consortium | Lighthouse Laboratory Plymouth and Alex Alderton, Roberto Amato, Jeffrey Barrett, Sonia Goncalves, Ewan Harrison, David K. Jackson, Ian Johnston, Dominic Kwiatkowski, Cordelia Langford, John Sillitoe on behalf of the Wellcome Sanger Institute COVID-19 Surveillance Team |
| EPI_ISL_2804009 | Health Services Laboratories | Wellcome Sanger Institute for the COVID-19 Genomics UK (COG-UK) Consortium | Health Services Laboratories and Alex Alderton, Roberto Amato, Jeffrey Barrett, Sonia Goncalves, Ewan Harrison, David K. Jackson, Ian Johnston, Dominic Kwiatkowski, Cordelia Langford, John Sillitoe on behalf of the Wellcome Sanger Institute COVID-19 Surveillance Team |
| EPI_ISL_2804010, EPI_ISL_2804011 | Lighthouse Lab in Alderley Park | Wellcome Sanger Institute for the COVID-19 Genomics UK (COG-UK) Consortium | Jacquelyn Wynn, Mairead Hyland, The Lighthouse Lab in Alderley Park and Alex Alderton, Roberto Amato, Jeffrey Barrett, Sonia Goncalves, Ewan Harrison, David K. Jackson, Ian Johnston, Dominic Kwiatkowski, Cordelia Langford, John Sillitoe on behalf of the Wellcome Sanger Institute COVID-19 Surveillance Team |
| EPI_ISL_2804012, EPI_ISL_2804013 | Lighthouse Lab in Milton Keynes | Wellcome Sanger Institute for the COVID-19 Genomics UK (COG-UK) Consortium | The Lighthouse Lab in Milton Keynes and Alex Alderton, Roberto Amato, Jeffrey Barrett, Sonia Goncalves, Ewan Harrison, David K. Jackson, Ian Johnston, Dominic Kwiatkowski, Cordelia Langford, John Sillitoe on behalf of the Wellcome Sanger Institute COVID-19 Surveillance Team |
| EPI_ISL_2804014 | Lighthouse Laboratory Plymouth | Wellcome Sanger Institute for the COVID-19 Genomics UK (COG-UK) Consortium | Lighthouse Laboratory Plymouth and Alex Alderton, Roberto Amato, Jeffrey Barrett, Sonia Goncalves, Ewan Harrison, David K. Jackson, Ian Johnston, Dominic Kwiatkowski, Cordelia Langford, John Sillitoe on behalf of the Wellcome Sanger Institute COVID-19 Surveillance Team |
| EPI_ISL_2804015 | Lighthouse Lab in Alderley Park | Wellcome Sanger Institute for the COVID-19 Genomics UK (COG-UK) Consortium | Jacquelyn Wynn, Mairead Hyland, The Lighthouse Lab in Alderley Park and Alex Alderton, Roberto Amato, Jeffrey Barrett, Sonia Goncalves, Ewan Harrison, David K. Jackson, Ian Johnston, Dominic Kwiatkowski, Cordelia Langford, John Sillitoe on behalf of the Wellcome Sanger Institute COVID-19 Surveillance Team |
| EPI_ISL_2804016 | Lighthouse Laboratory Plymouth | Wellcome Sanger Institute for the COVID-19 Genomics UK (COG-UK) Consortium | Lighthouse Laboratory Plymouth and Alex Alderton, Roberto Amato, Jeffrey Barrett, Sonia Goncalves, Ewan Harrison, David K. Jackson, Ian Johnston, Dominic Kwiatkowski, Cordelia Langford, John Sillitoe on behalf of the Wellcome Sanger Institute COVID-19 Surveillance Team |
| EPI_ISL_2804017 | Lighthouse Lab in Alderley Park | Wellcome Sanger Institute for the COVID-19 Genomics UK (COG-UK) Consortium | Jacquelyn Wynn, Mairead Hyland, The Lighthouse Lab in Alderley Park and Alex Alderton, Roberto Amato, Jeffrey Barrett, Sonia Goncalves, Ewan Harrison, David K. Jackson, Ian Johnston, Dominic Kwiatkowski, Cordelia Langford, John Sillitoe on behalf of the Wellcome Sanger Institute COVID-19 Surveillance Team |
| EPI_ISL_2804018, EPI_ISL_2804019 | Lighthouse Laboratory Plymouth | Wellcome Sanger Institute for the COVID-19 Genomics UK (COG-UK) Consortium | Lighthouse Laboratory Plymouth and Alex Alderton, Roberto Amato, Jeffrey Barrett, Sonia Goncalves, Ewan Harrison, David K. Jackson, Ian Johnston, Dominic Kwiatkowski, Cordelia Langford, John Sillitoe on behalf of the Wellcome Sanger Institute COVID-19 Surveillance Team |
| EPI_ISL_2804020, EPI_ISL_2804021 | Lighthouse Lab in Alderley Park | Wellcome Sanger Institute for the COVID-19 Genomics UK (COG-UK) Consortium | Jacquelyn Wynn, Mairead Hyland, The Lighthouse Lab in Alderley Park and Alex Alderton, Roberto Amato, Jeffrey Barrett, Sonia Goncalves, Ewan Harrison, David K. Jackson, Ian Johnston, Dominic Kwiatkowski, Cordelia Langford, John Sillitoe on behalf of the Wellcome Sanger Institute COVID-19 Surveillance Team |
| EPI_ISL_2804022 | Lighthouse Lab in Milton Keynes | Wellcome Sanger Institute for the COVID-19 Genomics UK (COG-UK) Consortium | The Lighthouse Lab in Milton Keynes and Alex Alderton, Roberto Amato, Jeffrey Barrett, Sonia Goncalves, Ewan Harrison, David K. Jackson, Ian Johnston, Dominic Kwiatkowski, Cordelia Langford, John Sillitoe on behalf of the Wellcome Sanger Institute COVID-19 Surveillance Team |
| EPI_ISL_2804023, EPI_ISL_2804024, EPI_ISL_2804025, EPI_ISL_2804026, EPI_ISL_2804027, EPI_ISL_2804028, EPI_ISL_2804029, EPI_ISL_2804030 | Lighthouse Laboratory Plymouth | Wellcome Sanger Institute for the COVID-19 Genomics UK (COG-UK) Consortium | Lighthouse Laboratory Plymouth and Alex Alderton, Roberto Amato, Jeffrey Barrett, Sonia Goncalves, Ewan Harrison, David K. Jackson, Ian Johnston, Dominic Kwiatkowski, Cordelia Langford, John Sillitoe on behalf of the Wellcome Sanger Institute COVID-19 Surveillance Team |
| EPI_ISL_2804031 | Lighthouse Lab in Alderley Park | Wellcome Sanger Institute for the COVID-19 Genomics UK (COG-UK) Consortium | Jacquelyn Wynn, Mairead Hyland, The Lighthouse Lab in Alderley Park and Alex Alderton, Roberto Amato, Jeffrey Barrett, Sonia Goncalves, Ewan Harrison, David K. Jackson, Ian Johnston, Dominic Kwiatkowski, Cordelia Langford, John Sillitoe on behalf of the Wellcome Sanger Institute COVID-19 Surveillance Team |
| EPI_ISL_2804032, EPI_ISL_2804033, EPI_ISL_2804034, EPI_ISL_2804035 | Lighthouse Laboratory Plymouth | Wellcome Sanger Institute for the COVID-19 Genomics UK (COG-UK) Consortium | Lighthouse Laboratory Plymouth and Alex Alderton, Roberto Amato, Jeffrey Barrett, Sonia Goncalves, Ewan Harrison, David K. Jackson, Ian Johnston, Dominic Kwiatkowski, Cordelia Langford, John Sillitoe on behalf of the Wellcome Sanger Institute COVID-19 Surveillance Team |
| EPI_ISL_2804036 | Lighthouse Lab in Milton Keynes | Wellcome Sanger Institute for the COVID-19 Genomics UK (COG-UK) Consortium | The Lighthouse Lab in Milton Keynes and Alex Alderton, Roberto Amato, Jeffrey Barrett, Sonia Goncalves, Ewan Harrison, David K. Jackson, Ian Johnston, Dominic Kwiatkowski, Cordelia Langford, John Sillitoe on behalf of the Wellcome Sanger Institute COVID-19 Surveillance Team |
| EPI_ISL_2804037, EPI_ISL_2804038 | Lighthouse Laboratory Plymouth | Wellcome Sanger Institute for the COVID-19 Genomics UK (COG-UK) Consortium | Lighthouse Laboratory Plymouth and Alex Alderton, Roberto Amato, Jeffrey Barrett, Sonia Goncalves, Ewan Harrison, David K. Jackson, Ian Johnston, Dominic Kwiatkowski, Cordelia Langford, John Sillitoe on behalf of the Wellcome Sanger Institute COVID-19 Surveillance Team |
| EPI_ISL_2804039, EPI_ISL_2804040 | Health Services Laboratories | Wellcome Sanger Institute for the COVID-19 Genomics UK (COG-UK) Consortium | Health Services Laboratories and Alex Alderton, Roberto Amato, Jeffrey Barrett, Sonia Goncalves, Ewan Harrison, David K. Jackson, Ian Johnston, Dominic Kwiatkowski, Cordelia Langford, John Sillitoe on behalf of the Wellcome Sanger Institute COVID-19 Surveillance Team |
| EPI_ISL_2804041, EPI_ISL_2804042 | Lighthouse Laboratory Plymouth | Wellcome Sanger Institute for the COVID-19 Genomics UK (COG-UK) Consortium | Lighthouse Laboratory Plymouth and Alex Alderton, Roberto Amato, Jeffrey Barrett, Sonia Goncalves, Ewan Harrison, David K. Jackson, Ian Johnston, Dominic Kwiatkowski, Cordelia Langford, John Sillitoe on behalf of the Wellcome Sanger Institute COVID-19 Surveillance Team |
| EPI_ISL_2804043 | Lighthouse Lab in Alderley Park | Wellcome Sanger Institute for the COVID-19 Genomics UK (COG-UK) Consortium | Jacquelyn Wynn, Mairead Hyland, The Lighthouse Lab in Alderley Park and Alex Alderton, Roberto Amato, Jeffrey Barrett, Sonia Goncalves, Ewan Harrison, David K. Jackson, Ian Johnston, Dominic Kwiatkowski, Cordelia Langford, John Sillitoe on behalf of the Wellcome Sanger Institute COVID-19 Surveillance Team |
| EPI_ISL_2804044 | Lighthouse Lab in Milton Keynes | Wellcome Sanger Institute for the COVID-19 Genomics UK (COG-UK) Consortium | The Lighthouse Lab in Milton Keynes and Alex Alderton, Roberto Amato, Jeffrey Barrett, Sonia Goncalves, Ewan Harrison, David K. Jackson, Ian Johnston, Dominic Kwiatkowski, Cordelia Langford, John Sillitoe on behalf of the Wellcome Sanger Institute COVID-19 Surveillance Team |
| EPI_ISL_2804045 | Lighthouse Lab in Alderley Park | Wellcome Sanger Institute for the COVID-19 Genomics UK (COG-UK) Consortium | Jacquelyn Wynn, Mairead Hyland, The Lighthouse Lab in Alderley Park and Alex Alderton, Roberto Amato, Jeffrey Barrett, Sonia Goncalves, Ewan Harrison, David K. Jackson, Ian Johnston, Dominic Kwiatkowski, Cordelia Langford, John Sillitoe on behalf of the Wellcome Sanger Institute COVID-19 Surveillance Team |
| EPI_ISL_2804046 | Health Services Laboratories | Wellcome Sanger Institute for the COVID-19 Genomics UK (COG-UK) Consortium | Health Services Laboratories and Alex Alderton, Roberto Amato, Jeffrey Barrett, Sonia Goncalves, Ewan Harrison, David K. Jackson, Ian Johnston, Dominic Kwiatkowski, Cordelia Langford, John Sillitoe on behalf of the Wellcome Sanger Institute COVID-19 Surveillance Team |
| EPI_ISL_2804047 | Lighthouse Lab in Alderley Park | Wellcome Sanger Institute for the COVID-19 Genomics UK (COG-UK) Consortium | Jacquelyn Wynn, Mairead Hyland, The Lighthouse Lab in Alderley Park and Alex Alderton, Roberto Amato, Jeffrey Barrett, Sonia Goncalves, Ewan Harrison, David K. Jackson, Ian Johnston, Dominic Kwiatkowski, Cordelia Langford, John Sillitoe on behalf of the Wellcome Sanger Institute COVID-19 Surveillance Team |
| EPI_ISL_2804048 | Lighthouse Laboratory Plymouth | Wellcome Sanger Institute for the COVID-19 Genomics UK (COG-UK) Consortium | Lighthouse Laboratory Plymouth and Alex Alderton, Roberto Amato, Jeffrey Barrett, Sonia Goncalves, Ewan Harrison, David K. Jackson, Ian Johnston, Dominic Kwiatkowski, Cordelia Langford, John Sillitoe on behalf of the Wellcome Sanger Institute COVID-19 Surveillance Team |
| EPI_ISL_2804049 | Lighthouse Lab in Milton Keynes | Wellcome Sanger Institute for the COVID-19 Genomics UK (COG-UK) Consortium | The Lighthouse Lab in Milton Keynes and Alex Alderton, Roberto Amato, Jeffrey Barrett, Sonia Goncalves, Ewan Harrison, David K. Jackson, Ian Johnston, Dominic Kwiatkowski, Cordelia Langford, John Sillitoe on behalf of the Wellcome Sanger Institute COVID-19 Surveillance Team |
| EPI_ISL_2804050, EPI_ISL_2804051, EPI_ISL_2804052, EPI_ISL_2804053 | Lighthouse Laboratory Plymouth | Wellcome Sanger Institute for the COVID-19 Genomics UK (COG-UK) Consortium | Lighthouse Laboratory Plymouth and Alex Alderton, Roberto Amato, Jeffrey Barrett, Sonia Goncalves, Ewan Harrison, David K. Jackson, Ian Johnston, Dominic Kwiatkowski, Cordelia Langford, John Sillitoe on behalf of the Wellcome Sanger Institute COVID-19 Surveillance Team |
| EPI_ISL_2804054 | Lighthouse Lab in Milton Keynes | Wellcome Sanger Institute for the COVID-19 Genomics UK (COG-UK) Consortium | The Lighthouse Lab in Milton Keynes and Alex Alderton, Roberto Amato, Jeffrey Barrett, Sonia Goncalves, Ewan Harrison, David K. Jackson, Ian Johnston, Dominic Kwiatkowski, Cordelia Langford, John Sillitoe on behalf of the Wellcome Sanger Institute COVID-19 Surveillance Team |
| EPI_ISL_2804055, EPI_ISL_2804056, EPI_ISL_2804057 | Lighthouse Laboratory Plymouth | Wellcome Sanger Institute for the COVID-19 Genomics UK (COG-UK) Consortium | Lighthouse Laboratory Plymouth and Alex Alderton, Roberto Amato, Jeffrey Barrett, Sonia Goncalves, Ewan Harrison, David K. Jackson, Ian Johnston, Dominic Kwiatkowski, Cordelia Langford, John Sillitoe on behalf of the Wellcome Sanger Institute COVID-19 Surveillance Team |
| EPI_ISL_2804058 | Lighthouse Lab in Milton Keynes | Wellcome Sanger Institute for the COVID-19 Genomics UK (COG-UK) Consortium | The Lighthouse Lab in Milton Keynes and Alex Alderton, Roberto Amato, Jeffrey Barrett, Sonia Goncalves, Ewan Harrison, David K. Jackson, Ian Johnston, Dominic Kwiatkowski, Cordelia Langford, John Sillitoe on behalf of the Wellcome Sanger Institute COVID-19 Surveillance Team |
| EPI_ISL_2804059, EPI_ISL_2804060 | Lighthouse Lab in Alderley Park | Wellcome Sanger Institute for the COVID-19 Genomics UK (COG-UK) Consortium | Jacquelyn Wynn, Mairead Hyland, The Lighthouse Lab in Alderley Park and Alex Alderton, Roberto Amato, Jeffrey Barrett, Sonia Goncalves, Ewan Harrison, David K. Jackson, Ian Johnston, Dominic Kwiatkowski, Cordelia Langford, John Sillitoe on behalf of the Wellcome Sanger Institute COVID-19 Surveillance Team |
| EPI_ISL_2804061 | Lighthouse Laboratory Plymouth | Wellcome Sanger Institute for the COVID-19 Genomics UK (COG-UK) Consortium | Lighthouse Laboratory Plymouth and Alex Alderton, Roberto Amato, Jeffrey Barrett, Sonia Goncalves, Ewan Harrison, David K. Jackson, Ian Johnston, Dominic Kwiatkowski, Cordelia Langford, John Sillitoe on behalf of the Wellcome Sanger Institute COVID-19 Surveillance Team |
| EPI_ISL_2804062 | Health Services Laboratories | Wellcome Sanger Institute for the COVID-19 Genomics UK (COG-UK) Consortium | Health Services Laboratories and Alex Alderton, Roberto Amato, Jeffrey Barrett, Sonia Goncalves, Ewan Harrison, David K. Jackson, Ian Johnston, Dominic Kwiatkowski, Cordelia Langford, John Sillitoe on behalf of the Wellcome Sanger Institute COVID-19 Surveillance Team |
| EPI_ISL_2804064 | Lighthouse Laboratory Plymouth | Wellcome Sanger Institute for the COVID-19 Genomics UK (COG-UK) Consortium | Lighthouse Laboratory Plymouth and Alex Alderton, Roberto Amato, Jeffrey Barrett, Sonia Goncalves, Ewan Harrison, David K. Jackson, Ian Johnston, Dominic Kwiatkowski, Cordelia Langford, John Sillitoe on behalf of the Wellcome Sanger Institute COVID-19 Surveillance Team |
| EPI_ISL_2804065 | Lighthouse Lab in Milton Keynes | Wellcome Sanger Institute for the COVID-19 Genomics UK (COG-UK) Consortium | The Lighthouse Lab in Milton Keynes and Alex Alderton, Roberto Amato, Jeffrey Barrett, Sonia Goncalves, Ewan Harrison, David K. Jackson, Ian Johnston, Dominic Kwiatkowski, Cordelia Langford, John Sillitoe on behalf of the Wellcome Sanger Institute COVID-19 Surveillance Team |
| EPI_ISL_2804066 | Lighthouse Laboratory Plymouth | Wellcome Sanger Institute for the COVID-19 Genomics UK (COG-UK) Consortium | Lighthouse Laboratory Plymouth and Alex Alderton, Roberto Amato, Jeffrey Barrett, Sonia Goncalves, Ewan Harrison, David K. Jackson, Ian Johnston, Dominic Kwiatkowski, Cordelia Langford, John Sillitoe on behalf of the Wellcome Sanger Institute COVID-19 Surveillance Team |
| EPI_ISL_2804067 | Lighthouse Lab in Milton Keynes | Wellcome Sanger Institute for the COVID-19 Genomics UK (COG-UK) Consortium | The Lighthouse Lab in Milton Keynes and Alex Alderton, Roberto Amato, Jeffrey Barrett, Sonia Goncalves, Ewan Harrison, David K. Jackson, Ian Johnston, Dominic Kwiatkowski, Cordelia Langford, John Sillitoe on behalf of the Wellcome Sanger Institute COVID-19 Surveillance Team |

| | | | |
|---|---|---|---|
| EPI_ISL_2804068 | Health Services Laboratories | Wellcome Sanger Institute for the COVID-19 Genomics UK (COG-UK) Consortium | Health Services Laboratories and Alex Alderton, Roberto Amato, Jeffrey Barrett, Sonia Goncalves, Ewan Harrison, David K. Jackson, Ian Johnston, Dominic Kwiatkowski, Cordelia Langford, John Sillitoe on behalf of the Wellcome Sanger Institute COVID-19 Surveillance Team |
| EPI_ISL_2804069 | Lighthouse Laboratory Plymouth | Wellcome Sanger Institute for the COVID-19 Genomics UK (COG-UK) Consortium | Lighthouse Laboratory Plymouth and Alex Alderton, Roberto Amato, Jeffrey Barrett, Sonia Goncalves, Ewan Harrison, David K. Jackson, Ian Johnston, Dominic Kwiatkowski, Cordelia Langford, John Sillitoe on behalf of the Wellcome Sanger Institute COVID-19 Surveillance Team |
| EPI_ISL_2804070 | Lighthouse Lab in Milton Keynes | Wellcome Sanger Institute for the COVID-19 Genomics UK (COG-UK) Consortium | The Lighthouse Lab in Milton Keynes and Alex Alderton, Roberto Amato, Jeffrey Barrett, Sonia Goncalves, Ewan Harrison, David K. Jackson, Ian Johnston, Dominic Kwiatkowski, Cordelia Langford, John Sillitoe on behalf of the Wellcome Sanger Institute COVID-19 Surveillance Team |
| EPI_ISL_2804071, EPI_ISL_2804072 | Lighthouse Lab in Alderley Park | Wellcome Sanger Institute for the COVID-19 Genomics UK (COG-UK) Consortium | Jacquelyn Wynn, Mairead Hyland, The Lighthouse Lab in Alderley Park and Alex Alderton, Roberto Amato, Jeffrey Barrett, Sonia Goncalves, Ewan Harrison, David K. Jackson, Ian Johnston, Dominic Kwiatkowski, Cordelia Langford, John Sillitoe on behalf of the Wellcome Sanger Institute COVID-19 Surveillance Team |
| EPI_ISL_2804073, EPI_ISL_2804074, EPI_ISL_2804075 | Lighthouse Laboratory Plymouth | Wellcome Sanger Institute for the COVID-19 Genomics UK (COG-UK) Consortium | Lighthouse Laboratory Plymouth and Alex Alderton, Roberto Amato, Jeffrey Barrett, Sonia Goncalves, Ewan Harrison, David K. Jackson, Ian Johnston, Dominic Kwiatkowski, Cordelia Langford, John Sillitoe on behalf of the Wellcome Sanger Institute COVID-19 Surveillance Team |
| EPI_ISL_2804076, EPI_ISL_2804077 | Lighthouse Lab in Alderley Park | Wellcome Sanger Institute for the COVID-19 Genomics UK (COG-UK) Consortium | Jacquelyn Wynn, Mairead Hyland, The Lighthouse Lab in Alderley Park and Alex Alderton, Roberto Amato, Jeffrey Barrett, Sonia Goncalves, Ewan Harrison, David K. Jackson, Ian Johnston, Dominic Kwiatkowski, Cordelia Langford, John Sillitoe on behalf of the Wellcome Sanger Institute COVID-19 Surveillance Team |
| EPI_ISL_2804078, EPI_ISL_2804079, EPI_ISL_2804080 | Lighthouse Laboratory Plymouth | Wellcome Sanger Institute for the COVID-19 Genomics UK (COG-UK) Consortium | Lighthouse Laboratory Plymouth and Alex Alderton, Roberto Amato, Jeffrey Barrett, Sonia Goncalves, Ewan Harrison, David K. Jackson, Ian Johnston, Dominic Kwiatkowski, Cordelia Langford, John Sillitoe on behalf of the Wellcome Sanger Institute COVID-19 Surveillance Team |
| EPI_ISL_2804081, EPI_ISL_2804082, EPI_ISL_2804083 | Health Services Laboratories | Wellcome Sanger Institute for the COVID-19 Genomics UK (COG-UK) Consortium | Health Services Laboratories and Alex Alderton, Roberto Amato, Jeffrey Barrett, Sonia Goncalves, Ewan Harrison, David K. Jackson, Ian Johnston, Dominic Kwiatkowski, Cordelia Langford, John Sillitoe on behalf of the Wellcome Sanger Institute COVID-19 Surveillance Team |
| EPI_ISL_2804084 | Lighthouse Lab in Milton Keynes | Wellcome Sanger Institute for the COVID-19 Genomics UK (COG-UK) Consortium | The Lighthouse Lab in Milton Keynes and Alex Alderton, Roberto Amato, Jeffrey Barrett, Sonia Goncalves, Ewan Harrison, David K. Jackson, Ian Johnston, Dominic Kwiatkowski, Cordelia Langford, John Sillitoe on behalf of the Wellcome Sanger Institute COVID-19 Surveillance Team |
| EPI_ISL_2804085, EPI_ISL_2804086, EPI_ISL_2804087 | Lighthouse Lab in Alderley Park | Wellcome Sanger Institute for the COVID-19 Genomics UK (COG-UK) Consortium | Jacquelyn Wynn, Mairead Hyland, The Lighthouse Lab in Alderley Park and Alex Alderton, Roberto Amato, Jeffrey Barrett, Sonia Goncalves, Ewan Harrison, David K. Jackson, Ian Johnston, Dominic Kwiatkowski, Cordelia Langford, John Sillitoe on behalf of the Wellcome Sanger Institute COVID-19 Surveillance Team |
| EPI_ISL_2804088, EPI_ISL_2804089 | Lighthouse Laboratory Plymouth | Wellcome Sanger Institute for the COVID-19 Genomics UK (COG-UK) Consortium | Lighthouse Laboratory Plymouth and Alex Alderton, Roberto Amato, Jeffrey Barrett, Sonia Goncalves, Ewan Harrison, David K. Jackson, Ian Johnston, Dominic Kwiatkowski, Cordelia Langford, John Sillitoe on behalf of the Wellcome Sanger Institute COVID-19 Surveillance Team |
| EPI_ISL_2804090 | Health Services Laboratories | Wellcome Sanger Institute for the COVID-19 Genomics UK (COG-UK) Consortium | Health Services Laboratories and Alex Alderton, Roberto Amato, Jeffrey Barrett, Sonia Goncalves, Ewan Harrison, David K. Jackson, Ian Johnston, Dominic Kwiatkowski, Cordelia Langford, John Sillitoe on behalf of the Wellcome Sanger Institute COVID-19 Surveillance Team |
| EPI_ISL_2804091, EPI_ISL_2804092 | Lighthouse Laboratory Plymouth | Wellcome Sanger Institute for the COVID-19 Genomics UK (COG-UK) Consortium | Lighthouse Laboratory Plymouth and Alex Alderton, Roberto Amato, Jeffrey Barrett, Sonia Goncalves, Ewan Harrison, David K. Jackson, Ian Johnston, Dominic Kwiatkowski, Cordelia Langford, John Sillitoe on behalf of the Wellcome Sanger Institute COVID-19 Surveillance Team |
| EPI_ISL_2804093 | Health Services Laboratories | Wellcome Sanger Institute for the COVID-19 Genomics UK (COG-UK) Consortium | Health Services Laboratories and Alex Alderton, Roberto Amato, Jeffrey Barrett, Sonia Goncalves, Ewan Harrison, David K. Jackson, Ian Johnston, Dominic Kwiatkowski, Cordelia Langford, John Sillitoe on behalf of the Wellcome Sanger Institute COVID-19 Surveillance Team |
| EPI_ISL_2804094, EPI_ISL_2804095, EPI_ISL_2804096, EPI_ISL_2804097, EPI_ISL_2804098 | Lighthouse Laboratory Plymouth | Wellcome Sanger Institute for the COVID-19 Genomics UK (COG-UK) Consortium | Lighthouse Laboratory Plymouth and Alex Alderton, Roberto Amato, Jeffrey Barrett, Sonia Goncalves, Ewan Harrison, David K. Jackson, Ian Johnston, Dominic Kwiatkowski, Cordelia Langford, John Sillitoe on behalf of the Wellcome Sanger Institute COVID-19 Surveillance Team |
| EPI_ISL_2804099 | Lighthouse Lab in Alderley Park | Wellcome Sanger Institute for the COVID-19 Genomics UK (COG-UK) Consortium | Jacquelyn Wynn, Mairead Hyland, The Lighthouse Lab in Alderley Park and Alex Alderton, Roberto Amato, Jeffrey Barrett, Sonia Goncalves, Ewan Harrison, David K. Jackson, Ian Johnston, Dominic Kwiatkowski, Cordelia Langford, John Sillitoe on behalf of the Wellcome Sanger Institute COVID-19 Surveillance Team |
| EPI_ISL_2804100 | Lighthouse Laboratory Plymouth | Wellcome Sanger Institute for the COVID-19 Genomics UK (COG-UK) Consortium | Lighthouse Laboratory Plymouth and Alex Alderton, Roberto Amato, Jeffrey Barrett, Sonia Goncalves, Ewan Harrison, David K. Jackson, Ian Johnston, Dominic Kwiatkowski, Cordelia Langford, John Sillitoe on behalf of the Wellcome Sanger Institute COVID-19 Surveillance Team |
| EPI_ISL_2804101 | Health Services Laboratories | Wellcome Sanger Institute for the COVID-19 Genomics UK (COG-UK) Consortium | Health Services Laboratories and Alex Alderton, Roberto Amato, Jeffrey Barrett, Sonia Goncalves, Ewan Harrison, David K. Jackson, Ian Johnston, Dominic Kwiatkowski, Cordelia Langford, John Sillitoe on behalf of the Wellcome Sanger Institute COVID-19 Surveillance Team |
| EPI_ISL_2804102 | Lighthouse Lab in Alderley Park | Wellcome Sanger Institute for the COVID-19 Genomics UK (COG-UK) Consortium | Jacquelyn Wynn, Mairead Hyland, The Lighthouse Lab in Alderley Park and Alex Alderton, Roberto Amato, Jeffrey Barrett, Sonia Goncalves, Ewan Harrison, David K. Jackson, Ian Johnston, Dominic Kwiatkowski, Cordelia Langford, John Sillitoe on behalf of the Wellcome Sanger Institute COVID-19 Surveillance Team |
| EPI_ISL_2804103 | Lighthouse Laboratory Plymouth | Wellcome Sanger Institute for the COVID-19 Genomics UK (COG-UK) Consortium | Lighthouse Laboratory Plymouth and Alex Alderton, Roberto Amato, Jeffrey Barrett, Sonia Goncalves, Ewan Harrison, David K. Jackson, Ian Johnston, Dominic Kwiatkowski, Cordelia Langford, John Sillitoe on behalf of the Wellcome Sanger Institute COVID-19 Surveillance Team |
| EPI_ISL_2804104 | Lighthouse Lab in Alderley Park | Wellcome Sanger Institute for the COVID-19 Genomics UK (COG-UK) Consortium | Jacquelyn Wynn, Mairead Hyland, The Lighthouse Lab in Alderley Park and Alex Alderton, Roberto Amato, Jeffrey Barrett, Sonia Goncalves, Ewan Harrison, David K. Jackson, Ian Johnston, Dominic Kwiatkowski, Cordelia Langford, John Sillitoe on behalf of the Wellcome Sanger Institute COVID-19 Surveillance Team |
| EPI_ISL_2804105 | Lighthouse Laboratory Plymouth | Wellcome Sanger Institute for the COVID-19 Genomics UK (COG-UK) Consortium | Lighthouse Laboratory Plymouth and Alex Alderton, Roberto Amato, Jeffrey Barrett, Sonia Goncalves, Ewan Harrison, David K. Jackson, Ian Johnston, Dominic Kwiatkowski, Cordelia Langford, John Sillitoe on behalf of the Wellcome Sanger Institute COVID-19 Surveillance Team |
| EPI_ISL_2804106 | Lighthouse Lab in Alderley Park | Wellcome Sanger Institute for the COVID-19 Genomics UK (COG-UK) Consortium | Jacquelyn Wynn, Mairead Hyland, The Lighthouse Lab in Alderley Park and Alex Alderton, Roberto Amato, Jeffrey Barrett, Sonia Goncalves, Ewan Harrison, David K. Jackson, Ian Johnston, Dominic Kwiatkowski, Cordelia Langford, John Sillitoe on behalf of the Wellcome Sanger Institute COVID-19 Surveillance Team |
| EPI_ISL_2804107 | Lighthouse Lab in Milton Keynes | Wellcome Sanger Institute for the COVID-19 Genomics UK (COG-UK) Consortium | The Lighthouse Lab in Milton Keynes and Alex Alderton, Roberto Amato, Jeffrey Barrett, Sonia Goncalves, Ewan Harrison, David K. Jackson, Ian Johnston, Dominic Kwiatkowski, Cordelia Langford, John Sillitoe on behalf of the Wellcome Sanger Institute COVID-19 Surveillance Team |
| EPI_ISL_2804108 | Lighthouse Lab in Alderley Park | Wellcome Sanger Institute for the COVID-19 Genomics UK (COG-UK) Consortium | Jacquelyn Wynn, Mairead Hyland, The Lighthouse Lab in Alderley Park and Alex Alderton, Roberto Amato, Jeffrey Barrett, Sonia Goncalves, Ewan Harrison, David K. Jackson, Ian Johnston, Dominic Kwiatkowski, Cordelia Langford, John Sillitoe on behalf of the Wellcome Sanger Institute COVID-19 Surveillance Team |
| EPI_ISL_2804109, EPI_ISL_2804110 | Lighthouse Laboratory Plymouth | Wellcome Sanger Institute for the COVID-19 Genomics UK (COG-UK) Consortium | Lighthouse Laboratory Plymouth and Alex Alderton, Roberto Amato, Jeffrey Barrett, Sonia Goncalves, Ewan Harrison, David K. Jackson, Ian Johnston, Dominic Kwiatkowski, Cordelia Langford, John Sillitoe on behalf of the Wellcome Sanger Institute COVID-19 Surveillance Team |
| EPI_ISL_2804111 | Lighthouse Lab in Alderley Park | Wellcome Sanger Institute for the COVID-19 Genomics UK (COG-UK) Consortium | Jacquelyn Wynn, Mairead Hyland, The Lighthouse Lab in Alderley Park and Alex Alderton, Roberto Amato, Jeffrey Barrett, Sonia Goncalves, Ewan Harrison, David K. Jackson, Ian Johnston, Dominic Kwiatkowski, Cordelia Langford, John Sillitoe on behalf of the Wellcome Sanger Institute COVID-19 Surveillance Team |
| EPI_ISL_2804112, EPI_ISL_2804113 | Lighthouse Laboratory Plymouth | Wellcome Sanger Institute for the COVID-19 Genomics UK (COG-UK) Consortium | Lighthouse Laboratory Plymouth and Alex Alderton, Roberto Amato, Jeffrey Barrett, Sonia Goncalves, Ewan Harrison, David K. Jackson, Ian Johnston, Dominic Kwiatkowski, Cordelia Langford, John Sillitoe on behalf of the Wellcome Sanger Institute COVID-19 Surveillance Team |
| EPI_ISL_2804114 | Lighthouse Lab in Alderley Park | Wellcome Sanger Institute for the COVID-19 Genomics UK (COG-UK) Consortium | Jacquelyn Wynn, Mairead Hyland, The Lighthouse Lab in Alderley Park and Alex Alderton, Roberto Amato, Jeffrey Barrett, Sonia Goncalves, Ewan Harrison, David K. Jackson, Ian Johnston, Dominic Kwiatkowski, Cordelia Langford, John Sillitoe on behalf of the Wellcome Sanger Institute COVID-19 Surveillance Team |
| EPI_ISL_2804115 | Health Services Laboratories | Wellcome Sanger Institute for the COVID-19 Genomics UK (COG-UK) Consortium | Health Services Laboratories and Alex Alderton, Roberto Amato, Jeffrey Barrett, Sonia Goncalves, Ewan Harrison, David K. Jackson, Ian Johnston, Dominic Kwiatkowski, Cordelia Langford, John Sillitoe on behalf of the Wellcome Sanger Institute COVID-19 Surveillance Team |
| EPI_ISL_2804116, EPI_ISL_2804117 | Lighthouse Laboratory Plymouth | Wellcome Sanger Institute for the COVID-19 Genomics UK (COG-UK) Consortium | Lighthouse Laboratory Plymouth and Alex Alderton, Roberto Amato, Jeffrey Barrett, Sonia Goncalves, Ewan Harrison, David K. Jackson, Ian Johnston, Dominic Kwiatkowski, Cordelia Langford, John Sillitoe on behalf of the Wellcome Sanger Institute COVID-19 Surveillance Team |
| EPI_ISL_2804118 | Health Services Laboratories | Wellcome Sanger Institute for the COVID-19 Genomics UK (COG-UK) Consortium | Health Services Laboratories and Alex Alderton, Roberto Amato, Jeffrey Barrett, Sonia Goncalves, Ewan Harrison, David K. Jackson, Ian Johnston, Dominic Kwiatkowski, Cordelia Langford, John Sillitoe on behalf of the Wellcome Sanger Institute COVID-19 Surveillance Team |
| EPI_ISL_2804119 | Lighthouse Lab in Alderley Park | Wellcome Sanger Institute for the COVID-19 Genomics UK (COG-UK) Consortium | Jacquelyn Wynn, Mairead Hyland, The Lighthouse Lab in Alderley Park and Alex Alderton, Roberto Amato, Jeffrey Barrett, Sonia Goncalves, Ewan Harrison, David K. Jackson, Ian Johnston, Dominic Kwiatkowski, Cordelia Langford, John Sillitoe on behalf of the Wellcome Sanger Institute COVID-19 Surveillance Team |
| EPI_ISL_2804120, EPI_ISL_2804121, EPI_ISL_2804122 | Lighthouse Laboratory Plymouth | Wellcome Sanger Institute for the COVID-19 Genomics UK (COG-UK) Consortium | Lighthouse Laboratory Plymouth and Alex Alderton, Roberto Amato, Jeffrey Barrett, Sonia Goncalves, Ewan Harrison, David K. Jackson, Ian Johnston, Dominic Kwiatkowski, Cordelia Langford, John Sillitoe on behalf of the Wellcome Sanger Institute COVID-19 Surveillance Team |
| EPI_ISL_2804123 | Lighthouse Lab in Milton Keynes | Wellcome Sanger Institute for the COVID-19 Genomics UK (COG-UK) Consortium | The Lighthouse Lab in Milton Keynes and Alex Alderton, Roberto Amato, Jeffrey Barrett, Sonia Goncalves, Ewan Harrison, David K. Jackson, Ian Johnston, Dominic Kwiatkowski, Cordelia Langford, John Sillitoe on behalf of the Wellcome Sanger Institute COVID-19 Surveillance Team |
| EPI_ISL_2804124 | Lighthouse Laboratory Plymouth | Wellcome Sanger Institute for the COVID-19 Genomics UK (COG-UK) Consortium | Lighthouse Laboratory Plymouth and Alex Alderton, Roberto Amato, Jeffrey Barrett, Sonia Goncalves, Ewan Harrison, David K. Jackson, Ian Johnston, Dominic Kwiatkowski, Cordelia Langford, John Sillitoe on behalf of the Wellcome Sanger Institute COVID-19 Surveillance Team |
| EPI_ISL_2804125 | Health Services Laboratories | Wellcome Sanger Institute for the COVID-19 Genomics UK (COG-UK) Consortium | Health Services Laboratories and Alex Alderton, Roberto Amato, Jeffrey Barrett, Sonia Goncalves, Ewan Harrison, David K. Jackson, Ian Johnston, Dominic Kwiatkowski, Cordelia Langford, John Sillitoe on behalf of the Wellcome Sanger Institute COVID-19 Surveillance Team |
| EPI_ISL_2804126 | Lighthouse Lab in Alderley Park | Wellcome Sanger Institute for the COVID-19 Genomics UK (COG-UK) Consortium | Jacquelyn Wynn, Mairead Hyland, The Lighthouse Lab in Alderley Park and Alex Alderton, Roberto Amato, Jeffrey Barrett, Sonia Goncalves, Ewan Harrison, David K. Jackson, Ian Johnston, Dominic Kwiatkowski, Cordelia Langford, John Sillitoe on behalf of the Wellcome Sanger Institute COVID-19 Surveillance Team |
| EPI_ISL_2804127, EPI_ISL_2804128, EPI_ISL_2804129, EPI_ISL_2804130, EPI_ISL_2804131 | Lighthouse Laboratory Plymouth | Wellcome Sanger Institute for the COVID-19 Genomics UK (COG-UK) Consortium | Lighthouse Laboratory Plymouth and Alex Alderton, Roberto Amato, Jeffrey Barrett, Sonia Goncalves, Ewan Harrison, David K. Jackson, Ian Johnston, Dominic Kwiatkowski, Cordelia Langford, John Sillitoe on behalf of the Wellcome Sanger Institute COVID-19 Surveillance Team |
| EPI_ISL_2804132 | Lighthouse Lab in Alderley Park | Wellcome Sanger Institute for the COVID-19 Genomics UK (COG-UK) Consortium | Jacquelyn Wynn, Mairead Hyland, The Lighthouse Lab in Alderley Park and Alex Alderton, Roberto Amato, Jeffrey Barrett, Sonia Goncalves, Ewan Harrison, David K. Jackson, Ian Johnston, Dominic Kwiatkowski, Cordelia Langford, John Sillitoe on behalf of the Wellcome Sanger Institute COVID-19 Surveillance Team |
| EPI_ISL_2804133, | Lighthouse Laboratory Plymouth | Wellcome Sanger Institute for the COVID-19 Genomics UK (COG-UK) | Lighthouse Laboratory Plymouth and Alex Alderton, Roberto Amato, Jeffrey Barrett, Sonia Goncalves, Ewan Harrison, David K. Jackson, Ian Johnston, Dominic Kwiatkowski, Cordelia Langford, John Sillitoe on behalf of the Wellcome Sanger |

| | | | |
|---|---|---|---|
| EPI_ISL_2804134 | | Consortium | Institute COVID-19 Surveillance Team |
| EPI_ISL_2804135 | Health Services Laboratories | Wellcome Sanger Institute for the COVID-19 Genomics UK (COG-UK) Consortium | Health Services Laboratories and Alex Alderton, Roberto Amato, Jeffrey Barrett, Sonia Goncalves, Ewan Harrison, David K. Jackson, Ian Johnston, Dominic Kwiatkowski, Cordelia Langford, John Sillitoe on behalf of the Wellcome Sanger Institute COVID-19 Surveillance Team |
| EPI_ISL_2804136 | Lighthouse Laboratory Plymouth | Wellcome Sanger Institute for the COVID-19 Genomics UK (COG-UK) Consortium | Lighthouse Laboratory Plymouth and Alex Alderton, Roberto Amato, Jeffrey Barrett, Sonia Goncalves, Ewan Harrison, David K. Jackson, Ian Johnston, Dominic Kwiatkowski, Cordelia Langford, John Sillitoe on behalf of the Wellcome Sanger Institute COVID-19 Surveillance Team |
| EPI_ISL_2804137 | Lighthouse Lab in Alderley Park | Wellcome Sanger Institute for the COVID-19 Genomics UK (COG-UK) Consortium | Jacquelyn Wynn, Mairead Hyland, The Lighthouse Lab in Alderley Park and Alex Alderton, Roberto Amato, Jeffrey Barrett, Sonia Goncalves, Ewan Harrison, David K. Jackson, Ian Johnston, Dominic Kwiatkowski, Cordelia Langford, John Sillitoe on behalf of the Wellcome Sanger Institute COVID-19 Surveillance Team |
| EPI_ISL_2804138, EPI_ISL_2804139, EPI_ISL_2804140 | Lighthouse Laboratory Plymouth | Wellcome Sanger Institute for the COVID-19 Genomics UK (COG-UK) Consortium | Lighthouse Laboratory Plymouth and Alex Alderton, Roberto Amato, Jeffrey Barrett, Sonia Goncalves, Ewan Harrison, David K. Jackson, Ian Johnston, Dominic Kwiatkowski, Cordelia Langford, John Sillitoe on behalf of the Wellcome Sanger Institute COVID-19 Surveillance Team |
| EPI_ISL_2804141 | Lighthouse Lab in Milton Keynes | Wellcome Sanger Institute for the COVID-19 Genomics UK (COG-UK) Consortium | The Lighthouse Lab in Milton Keynes and Alex Alderton, Roberto Amato, Jeffrey Barrett, Sonia Goncalves, Ewan Harrison, David K. Jackson, Ian Johnston, Dominic Kwiatkowski, Cordelia Langford, John Sillitoe on behalf of the Wellcome Sanger Institute COVID-19 Surveillance Team |
| EPI_ISL_2804142, EPI_ISL_2804143 | Lighthouse Lab in Alderley Park | Wellcome Sanger Institute for the COVID-19 Genomics UK (COG-UK) Consortium | Jacquelyn Wynn, Mairead Hyland, The Lighthouse Lab in Alderley Park and Alex Alderton, Roberto Amato, Jeffrey Barrett, Sonia Goncalves, Ewan Harrison, David K. Jackson, Ian Johnston, Dominic Kwiatkowski, Cordelia Langford, John Sillitoe on behalf of the Wellcome Sanger Institute COVID-19 Surveillance Team |
| EPI_ISL_2804144, EPI_ISL_2804145 | Lighthouse Laboratory Plymouth | Wellcome Sanger Institute for the COVID-19 Genomics UK (COG-UK) Consortium | Lighthouse Laboratory Plymouth and Alex Alderton, Roberto Amato, Jeffrey Barrett, Sonia Goncalves, Ewan Harrison, David K. Jackson, Ian Johnston, Dominic Kwiatkowski, Cordelia Langford, John Sillitoe on behalf of the Wellcome Sanger Institute COVID-19 Surveillance Team |
| EPI_ISL_2804146 | Lighthouse Lab in Milton Keynes | Wellcome Sanger Institute for the COVID-19 Genomics UK (COG-UK) Consortium | The Lighthouse Lab in Milton Keynes and Alex Alderton, Roberto Amato, Jeffrey Barrett, Sonia Goncalves, Ewan Harrison, David K. Jackson, Ian Johnston, Dominic Kwiatkowski, Cordelia Langford, John Sillitoe on behalf of the Wellcome Sanger Institute COVID-19 Surveillance Team |
| EPI_ISL_2804147, EPI_ISL_2804148, EPI_ISL_2804149 | Lighthouse Lab in Alderley Park | Wellcome Sanger Institute for the COVID-19 Genomics UK (COG-UK) Consortium | Jacquelyn Wynn, Mairead Hyland, The Lighthouse Lab in Alderley Park and Alex Alderton, Roberto Amato, Jeffrey Barrett, Sonia Goncalves, Ewan Harrison, David K. Jackson, Ian Johnston, Dominic Kwiatkowski, Cordelia Langford, John Sillitoe on behalf of the Wellcome Sanger Institute COVID-19 Surveillance Team |
| EPI_ISL_2804150 | Lighthouse Laboratory Plymouth | Wellcome Sanger Institute for the COVID-19 Genomics UK (COG-UK) Consortium | Lighthouse Laboratory Plymouth and Alex Alderton, Roberto Amato, Jeffrey Barrett, Sonia Goncalves, Ewan Harrison, David K. Jackson, Ian Johnston, Dominic Kwiatkowski, Cordelia Langford, John Sillitoe on behalf of the Wellcome Sanger Institute COVID-19 Surveillance Team |
| EPI_ISL_2804151, EPI_ISL_2804152 | Health Services Laboratories | Wellcome Sanger Institute for the COVID-19 Genomics UK (COG-UK) Consortium | Health Services Laboratories and Alex Alderton, Roberto Amato, Jeffrey Barrett, Sonia Goncalves, Ewan Harrison, David K. Jackson, Ian Johnston, Dominic Kwiatkowski, Cordelia Langford, John Sillitoe on behalf of the Wellcome Sanger Institute COVID-19 Surveillance Team |
| EPI_ISL_2804153, EPI_ISL_2804154 | Lighthouse Laboratory Plymouth | Wellcome Sanger Institute for the COVID-19 Genomics UK (COG-UK) Consortium | Lighthouse Laboratory Plymouth and Alex Alderton, Roberto Amato, Jeffrey Barrett, Sonia Goncalves, Ewan Harrison, David K. Jackson, Ian Johnston, Dominic Kwiatkowski, Cordelia Langford, John Sillitoe on behalf of the Wellcome Sanger Institute COVID-19 Surveillance Team |
| EPI_ISL_2804155 | Health Services Laboratories | Wellcome Sanger Institute for the COVID-19 Genomics UK (COG-UK) Consortium | Health Services Laboratories and Alex Alderton, Roberto Amato, Jeffrey Barrett, Sonia Goncalves, Ewan Harrison, David K. Jackson, Ian Johnston, Dominic Kwiatkowski, Cordelia Langford, John Sillitoe on behalf of the Wellcome Sanger Institute COVID-19 Surveillance Team |
| EPI_ISL_2804156, EPI_ISL_2804157, EPI_ISL_2804158, EPI_ISL_2804159 | Lighthouse Laboratory Plymouth | Wellcome Sanger Institute for the COVID-19 Genomics UK (COG-UK) Consortium | Lighthouse Laboratory Plymouth and Alex Alderton, Roberto Amato, Jeffrey Barrett, Sonia Goncalves, Ewan Harrison, David K. Jackson, Ian Johnston, Dominic Kwiatkowski, Cordelia Langford, John Sillitoe on behalf of the Wellcome Sanger Institute COVID-19 Surveillance Team |
| EPI_ISL_2804160 | Health Services Laboratories | Wellcome Sanger Institute for the COVID-19 Genomics UK (COG-UK) Consortium | Health Services Laboratories and Alex Alderton, Roberto Amato, Jeffrey Barrett, Sonia Goncalves, Ewan Harrison, David K. Jackson, Ian Johnston, Dominic Kwiatkowski, Cordelia Langford, John Sillitoe on behalf of the Wellcome Sanger Institute COVID-19 Surveillance Team |
| EPI_ISL_2804161, EPI_ISL_2804162, EPI_ISL_2804163 | Lighthouse Laboratory Plymouth | Wellcome Sanger Institute for the COVID-19 Genomics UK (COG-UK) Consortium | Lighthouse Laboratory Plymouth and Alex Alderton, Roberto Amato, Jeffrey Barrett, Sonia Goncalves, Ewan Harrison, David K. Jackson, Ian Johnston, Dominic Kwiatkowski, Cordelia Langford, John Sillitoe on behalf of the Wellcome Sanger Institute COVID-19 Surveillance Team |
| EPI_ISL_2804164 | Health Services Laboratories | Wellcome Sanger Institute for the COVID-19 Genomics UK (COG-UK) Consortium | Health Services Laboratories and Alex Alderton, Roberto Amato, Jeffrey Barrett, Sonia Goncalves, Ewan Harrison, David K. Jackson, Ian Johnston, Dominic Kwiatkowski, Cordelia Langford, John Sillitoe on behalf of the Wellcome Sanger Institute COVID-19 Surveillance Team |
| EPI_ISL_2804165 | Lighthouse Laboratory Plymouth | Wellcome Sanger Institute for the COVID-19 Genomics UK (COG-UK) Consortium | Lighthouse Laboratory Plymouth and Alex Alderton, Roberto Amato, Jeffrey Barrett, Sonia Goncalves, Ewan Harrison, David K. Jackson, Ian Johnston, Dominic Kwiatkowski, Cordelia Langford, John Sillitoe on behalf of the Wellcome Sanger Institute COVID-19 Surveillance Team |
| EPI_ISL_2804166 | Lighthouse Lab in Alderley Park | Wellcome Sanger Institute for the COVID-19 Genomics UK (COG-UK) Consortium | Jacquelyn Wynn, Mairead Hyland, The Lighthouse Lab in Alderley Park and Alex Alderton, Roberto Amato, Jeffrey Barrett, Sonia Goncalves, Ewan Harrison, David K. Jackson, Ian Johnston, Dominic Kwiatkowski, Cordelia Langford, John Sillitoe on behalf of the Wellcome Sanger Institute COVID-19 Surveillance Team |
| EPI_ISL_2804167 | Health Services Laboratories | Wellcome Sanger Institute for the COVID-19 Genomics UK (COG-UK) Consortium | Health Services Laboratories and Alex Alderton, Roberto Amato, Jeffrey Barrett, Sonia Goncalves, Ewan Harrison, David K. Jackson, Ian Johnston, Dominic Kwiatkowski, Cordelia Langford, John Sillitoe on behalf of the Wellcome Sanger Institute COVID-19 Surveillance Team |
| EPI_ISL_2804168, EPI_ISL_2804169, EPI_ISL_2804170 | Lighthouse Laboratory Plymouth | Wellcome Sanger Institute for the COVID-19 Genomics UK (COG-UK) Consortium | Lighthouse Laboratory Plymouth and Alex Alderton, Roberto Amato, Jeffrey Barrett, Sonia Goncalves, Ewan Harrison, David K. Jackson, Ian Johnston, Dominic Kwiatkowski, Cordelia Langford, John Sillitoe on behalf of the Wellcome Sanger Institute COVID-19 Surveillance Team |
| EPI_ISL_2804171, EPI_ISL_2804172 | Lighthouse Lab in Alderley Park | Wellcome Sanger Institute for the COVID-19 Genomics UK (COG-UK) Consortium | Jacquelyn Wynn, Mairead Hyland, The Lighthouse Lab in Alderley Park and Alex Alderton, Roberto Amato, Jeffrey Barrett, Sonia Goncalves, Ewan Harrison, David K. Jackson, Ian Johnston, Dominic Kwiatkowski, Cordelia Langford, John Sillitoe on behalf of the Wellcome Sanger Institute COVID-19 Surveillance Team |
| EPI_ISL_2804173 | Lighthouse Lab in Milton Keynes | Wellcome Sanger Institute for the COVID-19 Genomics UK (COG-UK) Consortium | The Lighthouse Lab in Milton Keynes and Alex Alderton, Roberto Amato, Jeffrey Barrett, Sonia Goncalves, Ewan Harrison, David K. Jackson, Ian Johnston, Dominic Kwiatkowski, Cordelia Langford, John Sillitoe on behalf of the Wellcome Sanger Institute COVID-19 Surveillance Team |
| EPI_ISL_2804174, EPI_ISL_2804175 | Lighthouse Laboratory Plymouth | Wellcome Sanger Institute for the COVID-19 Genomics UK (COG-UK) Consortium | Lighthouse Laboratory Plymouth and Alex Alderton, Roberto Amato, Jeffrey Barrett, Sonia Goncalves, Ewan Harrison, David K. Jackson, Ian Johnston, Dominic Kwiatkowski, Cordelia Langford, John Sillitoe on behalf of the Wellcome Sanger Institute COVID-19 Surveillance Team |
| EPI_ISL_2804176, EPI_ISL_2804177 | Lighthouse Lab in Alderley Park | Wellcome Sanger Institute for the COVID-19 Genomics UK (COG-UK) Consortium | Jacquelyn Wynn, Mairead Hyland, The Lighthouse Lab in Alderley Park and Alex Alderton, Roberto Amato, Jeffrey Barrett, Sonia Goncalves, Ewan Harrison, David K. Jackson, Ian Johnston, Dominic Kwiatkowski, Cordelia Langford, John Sillitoe on behalf of the Wellcome Sanger Institute COVID-19 Surveillance Team |
| EPI_ISL_2804178, EPI_ISL_2804179, EPI_ISL_2804180 | Lighthouse Laboratory Plymouth | Wellcome Sanger Institute for the COVID-19 Genomics UK (COG-UK) Consortium | Lighthouse Laboratory Plymouth and Alex Alderton, Roberto Amato, Jeffrey Barrett, Sonia Goncalves, Ewan Harrison, David K. Jackson, Ian Johnston, Dominic Kwiatkowski, Cordelia Langford, John Sillitoe on behalf of the Wellcome Sanger Institute COVID-19 Surveillance Team |
| EPI_ISL_2804181 | Health Services Laboratories | Wellcome Sanger Institute for the COVID-19 Genomics UK (COG-UK) Consortium | Health Services Laboratories and Alex Alderton, Roberto Amato, Jeffrey Barrett, Sonia Goncalves, Ewan Harrison, David K. Jackson, Ian Johnston, Dominic Kwiatkowski, Cordelia Langford, John Sillitoe on behalf of the Wellcome Sanger Institute COVID-19 Surveillance Team |
| EPI_ISL_2804182 | Lighthouse Lab in Alderley Park | Wellcome Sanger Institute for the COVID-19 Genomics UK (COG-UK) Consortium | Jacquelyn Wynn, Mairead Hyland, The Lighthouse Lab in Alderley Park and Alex Alderton, Roberto Amato, Jeffrey Barrett, Sonia Goncalves, Ewan Harrison, David K. Jackson, Ian Johnston, Dominic Kwiatkowski, Cordelia Langford, John Sillitoe on behalf of the Wellcome Sanger Institute COVID-19 Surveillance Team |
| EPI_ISL_2804183 | Health Services Laboratories | Wellcome Sanger Institute for the COVID-19 Genomics UK (COG-UK) Consortium | Health Services Laboratories and Alex Alderton, Roberto Amato, Jeffrey Barrett, Sonia Goncalves, Ewan Harrison, David K. Jackson, Ian Johnston, Dominic Kwiatkowski, Cordelia Langford, John Sillitoe on behalf of the Wellcome Sanger Institute COVID-19 Surveillance Team |
| EPI_ISL_2804184, EPI_ISL_2804185, EPI_ISL_2804186 | Lighthouse Laboratory Plymouth | Wellcome Sanger Institute for the COVID-19 Genomics UK (COG-UK) Consortium | Lighthouse Laboratory Plymouth and Alex Alderton, Roberto Amato, Jeffrey Barrett, Sonia Goncalves, Ewan Harrison, David K. Jackson, Ian Johnston, Dominic Kwiatkowski, Cordelia Langford, John Sillitoe on behalf of the Wellcome Sanger Institute COVID-19 Surveillance Team |
| EPI_ISL_2804187 | Lighthouse Lab in Alderley Park | Wellcome Sanger Institute for the COVID-19 Genomics UK (COG-UK) Consortium | Jacquelyn Wynn, Mairead Hyland, The Lighthouse Lab in Alderley Park and Alex Alderton, Roberto Amato, Jeffrey Barrett, Sonia Goncalves, Ewan Harrison, David K. Jackson, Ian Johnston, Dominic Kwiatkowski, Cordelia Langford, John Sillitoe on behalf of the Wellcome Sanger Institute COVID-19 Surveillance Team |
| EPI_ISL_2804188, EPI_ISL_2804189, EPI_ISL_2804190 | Lighthouse Laboratory Plymouth | Wellcome Sanger Institute for the COVID-19 Genomics UK (COG-UK) Consortium | Lighthouse Laboratory Plymouth and Alex Alderton, Roberto Amato, Jeffrey Barrett, Sonia Goncalves, Ewan Harrison, David K. Jackson, Ian Johnston, Dominic Kwiatkowski, Cordelia Langford, John Sillitoe on behalf of the Wellcome Sanger Institute COVID-19 Surveillance Team |
| EPI_ISL_2804191 | Lighthouse Lab in Milton Keynes | Wellcome Sanger Institute for the COVID-19 Genomics UK (COG-UK) Consortium | The Lighthouse Lab in Milton Keynes and Alex Alderton, Roberto Amato, Jeffrey Barrett, Sonia Goncalves, Ewan Harrison, David K. Jackson, Ian Johnston, Dominic Kwiatkowski, Cordelia Langford, John Sillitoe on behalf of the Wellcome Sanger Institute COVID-19 Surveillance Team |
| EPI_ISL_2804192, EPI_ISL_2804193 | Lighthouse Laboratory Plymouth | Wellcome Sanger Institute for the COVID-19 Genomics UK (COG-UK) Consortium | Lighthouse Laboratory Plymouth and Alex Alderton, Roberto Amato, Jeffrey Barrett, Sonia Goncalves, Ewan Harrison, David K. Jackson, Ian Johnston, Dominic Kwiatkowski, Cordelia Langford, John Sillitoe on behalf of the Wellcome Sanger Institute COVID-19 Surveillance Team |
| EPI_ISL_2804194, EPI_ISL_2804195, EPI_ISL_2804196 | Lighthouse Lab in Alderley Park | Wellcome Sanger Institute for the COVID-19 Genomics UK (COG-UK) Consortium | Jacquelyn Wynn, Mairead Hyland, The Lighthouse Lab in Alderley Park and Alex Alderton, Roberto Amato, Jeffrey Barrett, Sonia Goncalves, Ewan Harrison, David K. Jackson, Ian Johnston, Dominic Kwiatkowski, Cordelia Langford, John Sillitoe on behalf of the Wellcome Sanger Institute COVID-19 Surveillance Team |
| EPI_ISL_2804197 | Lighthouse Laboratory Plymouth | Wellcome Sanger Institute for the COVID-19 Genomics UK (COG-UK) Consortium | Lighthouse Laboratory Plymouth and Alex Alderton, Roberto Amato, Jeffrey Barrett, Sonia Goncalves, Ewan Harrison, David K. Jackson, Ian Johnston, Dominic Kwiatkowski, Cordelia Langford, John Sillitoe on behalf of the Wellcome Sanger Institute COVID-19 Surveillance Team |
| EPI_ISL_2804198 | Lighthouse Lab in Alderley Park | Wellcome Sanger Institute for the COVID-19 Genomics UK (COG-UK) Consortium | Jacquelyn Wynn, Mairead Hyland, The Lighthouse Lab in Alderley Park and Alex Alderton, Roberto Amato, Jeffrey Barrett, Sonia Goncalves, Ewan Harrison, David K. Jackson, Ian Johnston, Dominic Kwiatkowski, Cordelia Langford, John Sillitoe on behalf of the Wellcome Sanger Institute COVID-19 Surveillance Team |
| EPI_ISL_2804199 | Lighthouse Lab in Milton Keynes | Wellcome Sanger Institute for the COVID-19 Genomics UK (COG-UK) Consortium | The Lighthouse Lab in Milton Keynes and Alex Alderton, Roberto Amato, Jeffrey Barrett, Sonia Goncalves, Ewan Harrison, David K. Jackson, Ian Johnston, Dominic Kwiatkowski, Cordelia Langford, John Sillitoe on behalf of the Wellcome Sanger Institute COVID-19 Surveillance Team |
| EPI_ISL_2804200, EPI_ISL_2804201 | Lighthouse Laboratory Plymouth | Wellcome Sanger Institute for the COVID-19 Genomics UK (COG-UK) Consortium | Lighthouse Laboratory Plymouth and Alex Alderton, Roberto Amato, Jeffrey Barrett, Sonia Goncalves, Ewan Harrison, David K. Jackson, Ian Johnston, Dominic Kwiatkowski, Cordelia Langford, John Sillitoe on behalf of the Wellcome Sanger Institute COVID-19 Surveillance Team |
| EPI_ISL_2804202 | Lighthouse Lab in Milton Keynes | Wellcome Sanger Institute for the COVID-19 Genomics UK (COG-UK) Consortium | The Lighthouse Lab in Milton Keynes and Alex Alderton, Roberto Amato, Jeffrey Barrett, Sonia Goncalves, Ewan Harrison, David K. Jackson, Ian Johnston, Dominic Kwiatkowski, Cordelia Langford, John Sillitoe on behalf of the Wellcome Sanger Institute COVID-19 Surveillance Team |
| EPI_ISL_2804203 | Lighthouse Laboratory Plymouth | Wellcome Sanger Institute for the COVID-19 Genomics UK (COG-UK) | Lighthouse Laboratory Plymouth and Alex Alderton, Roberto Amato, Jeffrey Barrett, Sonia Goncalves, Ewan Harrison, David K. Jackson, Ian Johnston, Dominic Kwiatkowski, Cordelia Langford, John Sillitoe on behalf of the Wellcome Sanger |

| | | | |
|---|---|---|---|
| | | Consortium | Institute COVID-19 Surveillance Team |
| EPI_ISL_2804204 | Health Services Laboratories | Wellcome Sanger Institute for the COVID-19 Genomics UK (COG-UK) Consortium | Health Services Laboratories and Alex Alderton, Roberto Amato, Jeffrey Barrett, Sonia Goncalves, Ewan Harrison, David K. Jackson, Ian Johnston, Dominic Kwiatkowski, Cordelia Langford, John Sillitoe on behalf of the Wellcome Sanger Institute COVID-19 Surveillance Team |
| EPI_ISL_2804206 | Lighthouse Laboratory Plymouth | Wellcome Sanger Institute for the COVID-19 Genomics UK (COG-UK) Consortium | Lighthouse Laboratory Plymouth and Alex Alderton, Roberto Amato, Jeffrey Barrett, Sonia Goncalves, Ewan Harrison, David K. Jackson, Ian Johnston, Dominic Kwiatkowski, Cordelia Langford, John Sillitoe on behalf of the Wellcome Sanger Institute COVID-19 Surveillance Team |
| EPI_ISL_2804207 | Lighthouse Lab in Milton Keynes | Wellcome Sanger Institute for the COVID-19 Genomics UK (COG-UK) Consortium | The Lighthouse Lab in Milton Keynes and Alex Alderton, Roberto Amato, Jeffrey Barrett, Sonia Goncalves, Ewan Harrison, David K. Jackson, Ian Johnston, Dominic Kwiatkowski, Cordelia Langford, John Sillitoe on behalf of the Wellcome Sanger Institute COVID-19 Surveillance Team |
| EPI_ISL_2804208 | Health Services Laboratories | Wellcome Sanger Institute for the COVID-19 Genomics UK (COG-UK) Consortium | Health Services Laboratories and Alex Alderton, Roberto Amato, Jeffrey Barrett, Sonia Goncalves, Ewan Harrison, David K. Jackson, Ian Johnston, Dominic Kwiatkowski, Cordelia Langford, John Sillitoe on behalf of the Wellcome Sanger Institute COVID-19 Surveillance Team |
| EPI_ISL_2804209 | Lighthouse Laboratory Plymouth | Wellcome Sanger Institute for the COVID-19 Genomics UK (COG-UK) Consortium | Lighthouse Laboratory Plymouth and Alex Alderton, Roberto Amato, Jeffrey Barrett, Sonia Goncalves, Ewan Harrison, David K. Jackson, Ian Johnston, Dominic Kwiatkowski, Cordelia Langford, John Sillitoe on behalf of the Wellcome Sanger Institute COVID-19 Surveillance Team |
| EPI_ISL_2804210 | Lighthouse Lab in Milton Keynes | Wellcome Sanger Institute for the COVID-19 Genomics UK (COG-UK) Consortium | The Lighthouse Lab in Milton Keynes and Alex Alderton, Roberto Amato, Jeffrey Barrett, Sonia Goncalves, Ewan Harrison, David K. Jackson, Ian Johnston, Dominic Kwiatkowski, Cordelia Langford, John Sillitoe on behalf of the Wellcome Sanger Institute COVID-19 Surveillance Team |
| EPI_ISL_2804211 | Lighthouse Lab in Alderley Park | Wellcome Sanger Institute for the COVID-19 Genomics UK (COG-UK) Consortium | Jacquelyn Wynn, Mairead Hyland, The Lighthouse Lab in Alderley Park and Alex Alderton, Roberto Amato, Jeffrey Barrett, Sonia Goncalves, Ewan Harrison, David K. Jackson, Ian Johnston, Dominic Kwiatkowski, Cordelia Langford, John Sillitoe on behalf of the Wellcome Sanger Institute COVID-19 Surveillance Team |
| EPI_ISL_2804212 | Health Services Laboratories | Wellcome Sanger Institute for the COVID-19 Genomics UK (COG-UK) Consortium | Health Services Laboratories and Alex Alderton, Roberto Amato, Jeffrey Barrett, Sonia Goncalves, Ewan Harrison, David K. Jackson, Ian Johnston, Dominic Kwiatkowski, Cordelia Langford, John Sillitoe on behalf of the Wellcome Sanger Institute COVID-19 Surveillance Team |
| EPI_ISL_2804213 | Lighthouse Laboratory Plymouth | Wellcome Sanger Institute for the COVID-19 Genomics UK (COG-UK) Consortium | Lighthouse Laboratory Plymouth and Alex Alderton, Roberto Amato, Jeffrey Barrett, Sonia Goncalves, Ewan Harrison, David K. Jackson, Ian Johnston, Dominic Kwiatkowski, Cordelia Langford, John Sillitoe on behalf of the Wellcome Sanger Institute COVID-19 Surveillance Team |
| EPI_ISL_2804214 | Health Services Laboratories | Wellcome Sanger Institute for the COVID-19 Genomics UK (COG-UK) Consortium | Health Services Laboratories and Alex Alderton, Roberto Amato, Jeffrey Barrett, Sonia Goncalves, Ewan Harrison, David K. Jackson, Ian Johnston, Dominic Kwiatkowski, Cordelia Langford, John Sillitoe on behalf of the Wellcome Sanger Institute COVID-19 Surveillance Team |
| EPI_ISL_2804215, EPI_ISL_2804216 | Lighthouse Lab in Alderley Park | Wellcome Sanger Institute for the COVID-19 Genomics UK (COG-UK) Consortium | Jacquelyn Wynn, Mairead Hyland, The Lighthouse Lab in Alderley Park and Alex Alderton, Roberto Amato, Jeffrey Barrett, Sonia Goncalves, Ewan Harrison, David K. Jackson, Ian Johnston, Dominic Kwiatkowski, Cordelia Langford, John Sillitoe on behalf of the Wellcome Sanger Institute COVID-19 Surveillance Team |
| EPI_ISL_2804217 | Lighthouse Laboratory Plymouth | Wellcome Sanger Institute for the COVID-19 Genomics UK (COG-UK) Consortium | Lighthouse Laboratory Plymouth and Alex Alderton, Roberto Amato, Jeffrey Barrett, Sonia Goncalves, Ewan Harrison, David K. Jackson, Ian Johnston, Dominic Kwiatkowski, Cordelia Langford, John Sillitoe on behalf of the Wellcome Sanger Institute COVID-19 Surveillance Team |
| EPI_ISL_2804218 | Health Services Laboratories | Wellcome Sanger Institute for the COVID-19 Genomics UK (COG-UK) Consortium | Health Services Laboratories and Alex Alderton, Roberto Amato, Jeffrey Barrett, Sonia Goncalves, Ewan Harrison, David K. Jackson, Ian Johnston, Dominic Kwiatkowski, Cordelia Langford, John Sillitoe on behalf of the Wellcome Sanger Institute COVID-19 Surveillance Team |
| EPI_ISL_2804219 | Lighthouse Lab in Milton Keynes | Wellcome Sanger Institute for the COVID-19 Genomics UK (COG-UK) Consortium | The Lighthouse Lab in Milton Keynes and Alex Alderton, Roberto Amato, Jeffrey Barrett, Sonia Goncalves, Ewan Harrison, David K. Jackson, Ian Johnston, Dominic Kwiatkowski, Cordelia Langford, John Sillitoe on behalf of the Wellcome Sanger Institute COVID-19 Surveillance Team |
| EPI_ISL_2804220, EPI_ISL_2804221, EPI_ISL_2804222, EPI_ISL_2804223, EPI_ISL_2804224 | Lighthouse Laboratory Plymouth | Wellcome Sanger Institute for the COVID-19 Genomics UK (COG-UK) Consortium | Lighthouse Laboratory Plymouth and Alex Alderton, Roberto Amato, Jeffrey Barrett, Sonia Goncalves, Ewan Harrison, David K. Jackson, Ian Johnston, Dominic Kwiatkowski, Cordelia Langford, John Sillitoe on behalf of the Wellcome Sanger Institute COVID-19 Surveillance Team |
| EPI_ISL_2804225 | Lighthouse Lab in Milton Keynes | Wellcome Sanger Institute for the COVID-19 Genomics UK (COG-UK) Consortium | The Lighthouse Lab in Milton Keynes and Alex Alderton, Roberto Amato, Jeffrey Barrett, Sonia Goncalves, Ewan Harrison, David K. Jackson, Ian Johnston, Dominic Kwiatkowski, Cordelia Langford, John Sillitoe on behalf of the Wellcome Sanger Institute COVID-19 Surveillance Team |
| EPI_ISL_2804226 | Health Services Laboratories | Wellcome Sanger Institute for the COVID-19 Genomics UK (COG-UK) Consortium | Health Services Laboratories and Alex Alderton, Roberto Amato, Jeffrey Barrett, Sonia Goncalves, Ewan Harrison, David K. Jackson, Ian Johnston, Dominic Kwiatkowski, Cordelia Langford, John Sillitoe on behalf of the Wellcome Sanger Institute COVID-19 Surveillance Team |
| EPI_ISL_2804227, EPI_ISL_2804228 | Lighthouse Laboratory Plymouth | Wellcome Sanger Institute for the COVID-19 Genomics UK (COG-UK) Consortium | Lighthouse Laboratory Plymouth and Alex Alderton, Roberto Amato, Jeffrey Barrett, Sonia Goncalves, Ewan Harrison, David K. Jackson, Ian Johnston, Dominic Kwiatkowski, Cordelia Langford, John Sillitoe on behalf of the Wellcome Sanger Institute COVID-19 Surveillance Team |
| EPI_ISL_2804229 | Lighthouse Lab in Milton Keynes | Wellcome Sanger Institute for the COVID-19 Genomics UK (COG-UK) Consortium | The Lighthouse Lab in Milton Keynes and Alex Alderton, Roberto Amato, Jeffrey Barrett, Sonia Goncalves, Ewan Harrison, David K. Jackson, Ian Johnston, Dominic Kwiatkowski, Cordelia Langford, John Sillitoe on behalf of the Wellcome Sanger Institute COVID-19 Surveillance Team |
| EPI_ISL_2804230 | Health Services Laboratories | Wellcome Sanger Institute for the COVID-19 Genomics UK (COG-UK) Consortium | Health Services Laboratories and Alex Alderton, Roberto Amato, Jeffrey Barrett, Sonia Goncalves, Ewan Harrison, David K. Jackson, Ian Johnston, Dominic Kwiatkowski, Cordelia Langford, John Sillitoe on behalf of the Wellcome Sanger Institute COVID-19 Surveillance Team |
| EPI_ISL_2804231 | Lighthouse Lab in Alderley Park | Wellcome Sanger Institute for the COVID-19 Genomics UK (COG-UK) Consortium | Jacquelyn Wynn, Mairead Hyland, The Lighthouse Lab in Alderley Park and Alex Alderton, Roberto Amato, Jeffrey Barrett, Sonia Goncalves, Ewan Harrison, David K. Jackson, Ian Johnston, Dominic Kwiatkowski, Cordelia Langford, John Sillitoe on behalf of the Wellcome Sanger Institute COVID-19 Surveillance Team |
| EPI_ISL_2804232, EPI_ISL_2804233 | Lighthouse Laboratory Plymouth | Wellcome Sanger Institute for the COVID-19 Genomics UK (COG-UK) Consortium | Lighthouse Laboratory Plymouth and Alex Alderton, Roberto Amato, Jeffrey Barrett, Sonia Goncalves, Ewan Harrison, David K. Jackson, Ian Johnston, Dominic Kwiatkowski, Cordelia Langford, John Sillitoe on behalf of the Wellcome Sanger Institute COVID-19 Surveillance Team |
| EPI_ISL_2804234 | Health Services Laboratories | Wellcome Sanger Institute for the COVID-19 Genomics UK (COG-UK) Consortium | Health Services Laboratories and Alex Alderton, Roberto Amato, Jeffrey Barrett, Sonia Goncalves, Ewan Harrison, David K. Jackson, Ian Johnston, Dominic Kwiatkowski, Cordelia Langford, John Sillitoe on behalf of the Wellcome Sanger Institute COVID-19 Surveillance Team |
| EPI_ISL_2804235, EPI_ISL_2804236, EPI_ISL_2804237, EPI_ISL_2804238, EPI_ISL_2804239, EPI_ISL_2804240, EPI_ISL_2804241, EPI_ISL_2804242, EPI_ISL_2804243 | Lighthouse Laboratory Plymouth | Wellcome Sanger Institute for the COVID-19 Genomics UK (COG-UK) Consortium | Lighthouse Laboratory Plymouth and Alex Alderton, Roberto Amato, Jeffrey Barrett, Sonia Goncalves, Ewan Harrison, David K. Jackson, Ian Johnston, Dominic Kwiatkowski, Cordelia Langford, John Sillitoe on behalf of the Wellcome Sanger Institute COVID-19 Surveillance Team |
| EPI_ISL_2804244 | Health Services Laboratories | Wellcome Sanger Institute for the COVID-19 Genomics UK (COG-UK) Consortium | Health Services Laboratories and Alex Alderton, Roberto Amato, Jeffrey Barrett, Sonia Goncalves, Ewan Harrison, David K. Jackson, Ian Johnston, Dominic Kwiatkowski, Cordelia Langford, John Sillitoe on behalf of the Wellcome Sanger Institute COVID-19 Surveillance Team |
| EPI_ISL_2804245 | Lighthouse Lab in Milton Keynes | Wellcome Sanger Institute for the COVID-19 Genomics UK (COG-UK) Consortium | The Lighthouse Lab in Milton Keynes and Alex Alderton, Roberto Amato, Jeffrey Barrett, Sonia Goncalves, Ewan Harrison, David K. Jackson, Ian Johnston, Dominic Kwiatkowski, Cordelia Langford, John Sillitoe on behalf of the Wellcome Sanger Institute COVID-19 Surveillance Team |
| EPI_ISL_2804246 | Lighthouse Lab in Alderley Park | Wellcome Sanger Institute for the COVID-19 Genomics UK (COG-UK) Consortium | Jacquelyn Wynn, Mairead Hyland, The Lighthouse Lab in Alderley Park and Alex Alderton, Roberto Amato, Jeffrey Barrett, Sonia Goncalves, Ewan Harrison, David K. Jackson, Ian Johnston, Dominic Kwiatkowski, Cordelia Langford, John Sillitoe on behalf of the Wellcome Sanger Institute COVID-19 Surveillance Team |
| EPI_ISL_2804247 | Lighthouse Laboratory Plymouth | Wellcome Sanger Institute for the COVID-19 Genomics UK (COG-UK) Consortium | Lighthouse Laboratory Plymouth and Alex Alderton, Roberto Amato, Jeffrey Barrett, Sonia Goncalves, Ewan Harrison, David K. Jackson, Ian Johnston, Dominic Kwiatkowski, Cordelia Langford, John Sillitoe on behalf of the Wellcome Sanger Institute COVID-19 Surveillance Team |
| EPI_ISL_2804248 | Lighthouse Lab in Alderley Park | Wellcome Sanger Institute for the COVID-19 Genomics UK (COG-UK) Consortium | Jacquelyn Wynn, Mairead Hyland, The Lighthouse Lab in Alderley Park and Alex Alderton, Roberto Amato, Jeffrey Barrett, Sonia Goncalves, Ewan Harrison, David K. Jackson, Ian Johnston, Dominic Kwiatkowski, Cordelia Langford, John Sillitoe on behalf of the Wellcome Sanger Institute COVID-19 Surveillance Team |
| EPI_ISL_2804249 | Health Services Laboratories | Wellcome Sanger Institute for the COVID-19 Genomics UK (COG-UK) Consortium | Health Services Laboratories and Alex Alderton, Roberto Amato, Jeffrey Barrett, Sonia Goncalves, Ewan Harrison, David K. Jackson, Ian Johnston, Dominic Kwiatkowski, Cordelia Langford, John Sillitoe on behalf of the Wellcome Sanger Institute COVID-19 Surveillance Team |
| EPI_ISL_2804250, EPI_ISL_2804251, EPI_ISL_2804252, EPI_ISL_2804253 | Lighthouse Laboratory Plymouth | Wellcome Sanger Institute for the COVID-19 Genomics UK (COG-UK) Consortium | Lighthouse Laboratory Plymouth and Alex Alderton, Roberto Amato, Jeffrey Barrett, Sonia Goncalves, Ewan Harrison, David K. Jackson, Ian Johnston, Dominic Kwiatkowski, Cordelia Langford, John Sillitoe on behalf of the Wellcome Sanger Institute COVID-19 Surveillance Team |
| EPI_ISL_2804254 | Lighthouse Lab in Alderley Park | Wellcome Sanger Institute for the COVID-19 Genomics UK (COG-UK) Consortium | Jacquelyn Wynn, Mairead Hyland, The Lighthouse Lab in Alderley Park and Alex Alderton, Roberto Amato, Jeffrey Barrett, Sonia Goncalves, Ewan Harrison, David K. Jackson, Ian Johnston, Dominic Kwiatkowski, Cordelia Langford, John Sillitoe on behalf of the Wellcome Sanger Institute COVID-19 Surveillance Team |
| EPI_ISL_2804255 | Lighthouse Laboratory Plymouth | Wellcome Sanger Institute for the COVID-19 Genomics UK (COG-UK) Consortium | Lighthouse Laboratory Plymouth and Alex Alderton, Roberto Amato, Jeffrey Barrett, Sonia Goncalves, Ewan Harrison, David K. Jackson, Ian Johnston, Dominic Kwiatkowski, Cordelia Langford, John Sillitoe on behalf of the Wellcome Sanger Institute COVID-19 Surveillance Team |
| EPI_ISL_2804256, EPI_ISL_2804257, EPI_ISL_2804258, EPI_ISL_2804259, EPI_ISL_2804260, EPI_ISL_2804261 | Lighthouse Lab in Alderley Park | Wellcome Sanger Institute for the COVID-19 Genomics UK (COG-UK) Consortium | Jacquelyn Wynn, Mairead Hyland, The Lighthouse Lab in Alderley Park and Alex Alderton, Roberto Amato, Jeffrey Barrett, Sonia Goncalves, Ewan Harrison, David K. Jackson, Ian Johnston, Dominic Kwiatkowski, Cordelia Langford, John Sillitoe on behalf of the Wellcome Sanger Institute COVID-19 Surveillance Team |
| EPI_ISL_2804262, EPI_ISL_2804263 | Lighthouse Laboratory Plymouth | Wellcome Sanger Institute for the COVID-19 Genomics UK (COG-UK) Consortium | Lighthouse Laboratory Plymouth and Alex Alderton, Roberto Amato, Jeffrey Barrett, Sonia Goncalves, Ewan Harrison, David K. Jackson, Ian Johnston, Dominic Kwiatkowski, Cordelia Langford, John Sillitoe on behalf of the Wellcome Sanger Institute COVID-19 Surveillance Team |
| EPI_ISL_2804264, EPI_ISL_2804265, EPI_ISL_2804266, EPI_ISL_2804267, EPI_ISL_2804268 | Lighthouse Lab in Alderley Park | Wellcome Sanger Institute for the COVID-19 Genomics UK (COG-UK) Consortium | Jacquelyn Wynn, Mairead Hyland, The Lighthouse Lab in Alderley Park and Alex Alderton, Roberto Amato, Jeffrey Barrett, Sonia Goncalves, Ewan Harrison, David K. Jackson, Ian Johnston, Dominic Kwiatkowski, Cordelia Langford, John Sillitoe on behalf of the Wellcome Sanger Institute COVID-19 Surveillance Team |
| EPI_ISL_2804269 | Lighthouse Laboratory Plymouth | Wellcome Sanger Institute for the COVID-19 Genomics UK (COG-UK) | Lighthouse Laboratory Plymouth and Alex Alderton, Roberto Amato, Jeffrey Barrett, Sonia Goncalves, Ewan Harrison, David K. Jackson, Ian Johnston, Dominic Kwiatkowski, Cordelia Langford, John Sillitoe on behalf of the Wellcome Sanger |

| | | | |
|---|---|---|---|
| | | Consortium | Institute COVID-19 Surveillance Team |
| EPI_ISL_2804270, EPI_ISL_2804271, EPI_ISL_2804272 | Lighthouse Lab in Alderley Park | Wellcome Sanger Institute for the COVID-19 Genomics UK (COG-UK) Consortium | Jacquelyn Wynn, Mairead Hyland, The Lighthouse Lab in Alderley Park and Alex Alderton, Roberto Amato, Jeffrey Barrett, Sonia Goncalves, Ewan Harrison, David K. Jackson, Ian Johnston, Dominic Kwiatkowski, Cordelia Langford, John Sillitoe on behalf of the Wellcome Sanger Institute COVID-19 Surveillance Team |
| EPI_ISL_2804273 | Lighthouse Laboratory Plymouth | Wellcome Sanger Institute for the COVID-19 Genomics UK (COG-UK) Consortium | Lighthouse Laboratory Plymouth and Alex Alderton, Roberto Amato, Jeffrey Barrett, Sonia Goncalves, Ewan Harrison, David K. Jackson, Ian Johnston, Dominic Kwiatkowski, Cordelia Langford, John Sillitoe on behalf of the Wellcome Sanger Institute COVID-19 Surveillance Team |
| EPI_ISL_2804274 | Lighthouse Lab in Alderley Park | Wellcome Sanger Institute for the COVID-19 Genomics UK (COG-UK) Consortium | Jacquelyn Wynn, Mairead Hyland, The Lighthouse Lab in Alderley Park and Alex Alderton, Roberto Amato, Jeffrey Barrett, Sonia Goncalves, Ewan Harrison, David K. Jackson, Ian Johnston, Dominic Kwiatkowski, Cordelia Langford, John Sillitoe on behalf of the Wellcome Sanger Institute COVID-19 Surveillance Team |
| EPI_ISL_2804275, EPI_ISL_2804276 | Lighthouse Laboratory Plymouth | Wellcome Sanger Institute for the COVID-19 Genomics UK (COG-UK) Consortium | Lighthouse Laboratory Plymouth and Alex Alderton, Roberto Amato, Jeffrey Barrett, Sonia Goncalves, Ewan Harrison, David K. Jackson, Ian Johnston, Dominic Kwiatkowski, Cordelia Langford, John Sillitoe on behalf of the Wellcome Sanger Institute COVID-19 Surveillance Team |
| EPI_ISL_2804277, EPI_ISL_2804278, EPI_ISL_2804279, EPI_ISL_2804280, EPI_ISL_2804281, EPI_ISL_2804282, EPI_ISL_2804283, EPI_ISL_2804284, EPI_ISL_2804285 | Lighthouse Lab in Alderley Park | Wellcome Sanger Institute for the COVID-19 Genomics UK (COG-UK) Consortium | Jacquelyn Wynn, Mairead Hyland, The Lighthouse Lab in Alderley Park and Alex Alderton, Roberto Amato, Jeffrey Barrett, Sonia Goncalves, Ewan Harrison, David K. Jackson, Ian Johnston, Dominic Kwiatkowski, Cordelia Langford, John Sillitoe on behalf of the Wellcome Sanger Institute COVID-19 Surveillance Team |
| EPI_ISL_2804286 | Lighthouse Laboratory Plymouth | Wellcome Sanger Institute for the COVID-19 Genomics UK (COG-UK) Consortium | Lighthouse Laboratory Plymouth and Alex Alderton, Roberto Amato, Jeffrey Barrett, Sonia Goncalves, Ewan Harrison, David K. Jackson, Ian Johnston, Dominic Kwiatkowski, Cordelia Langford, John Sillitoe on behalf of the Wellcome Sanger Institute COVID-19 Surveillance Team |
| EPI_ISL_2804287, EPI_ISL_2804288, EPI_ISL_2804289 | Lighthouse Lab in Alderley Park | Wellcome Sanger Institute for the COVID-19 Genomics UK (COG-UK) Consortium | Jacquelyn Wynn, Mairead Hyland, The Lighthouse Lab in Alderley Park and Alex Alderton, Roberto Amato, Jeffrey Barrett, Sonia Goncalves, Ewan Harrison, David K. Jackson, Ian Johnston, Dominic Kwiatkowski, Cordelia Langford, John Sillitoe on behalf of the Wellcome Sanger Institute COVID-19 Surveillance Team |
| EPI_ISL_2804290, EPI_ISL_2804291 | Lighthouse Laboratory Plymouth | Wellcome Sanger Institute for the COVID-19 Genomics UK (COG-UK) Consortium | Lighthouse Laboratory Plymouth and Alex Alderton, Roberto Amato, Jeffrey Barrett, Sonia Goncalves, Ewan Harrison, David K. Jackson, Ian Johnston, Dominic Kwiatkowski, Cordelia Langford, John Sillitoe on behalf of the Wellcome Sanger Institute COVID-19 Surveillance Team |
| EPI_ISL_2804292 | Lighthouse Lab in Alderley Park | Wellcome Sanger Institute for the COVID-19 Genomics UK (COG-UK) Consortium | Jacquelyn Wynn, Mairead Hyland, The Lighthouse Lab in Alderley Park and Alex Alderton, Roberto Amato, Jeffrey Barrett, Sonia Goncalves, Ewan Harrison, David K. Jackson, Ian Johnston, Dominic Kwiatkowski, Cordelia Langford, John Sillitoe on behalf of the Wellcome Sanger Institute COVID-19 Surveillance Team |
| EPI_ISL_2804293 | Lighthouse Laboratory Plymouth | Wellcome Sanger Institute for the COVID-19 Genomics UK (COG-UK) Consortium | Lighthouse Laboratory Plymouth and Alex Alderton, Roberto Amato, Jeffrey Barrett, Sonia Goncalves, Ewan Harrison, David K. Jackson, Ian Johnston, Dominic Kwiatkowski, Cordelia Langford, John Sillitoe on behalf of the Wellcome Sanger Institute COVID-19 Surveillance Team |
| EPI_ISL_2804294, EPI_ISL_2804295, EPI_ISL_2804296, EPI_ISL_2804297, EPI_ISL_2804298, EPI_ISL_2804299, EPI_ISL_2804300 | Lighthouse Lab in Alderley Park | Wellcome Sanger Institute for the COVID-19 Genomics UK (COG-UK) Consortium | Jacquelyn Wynn, Mairead Hyland, The Lighthouse Lab in Alderley Park and Alex Alderton, Roberto Amato, Jeffrey Barrett, Sonia Goncalves, Ewan Harrison, David K. Jackson, Ian Johnston, Dominic Kwiatkowski, Cordelia Langford, John Sillitoe on behalf of the Wellcome Sanger Institute COVID-19 Surveillance Team |
| EPI_ISL_2804301 | Lighthouse Laboratory Plymouth | Wellcome Sanger Institute for the COVID-19 Genomics UK (COG-UK) Consortium | Lighthouse Laboratory Plymouth and Alex Alderton, Roberto Amato, Jeffrey Barrett, Sonia Goncalves, Ewan Harrison, David K. Jackson, Ian Johnston, Dominic Kwiatkowski, Cordelia Langford, John Sillitoe on behalf of the Wellcome Sanger Institute COVID-19 Surveillance Team |
| EPI_ISL_2804302, EPI_ISL_2804303, EPI_ISL_2804304, EPI_ISL_2804305, EPI_ISL_2804306, EPI_ISL_2804307, EPI_ISL_2804308 | Lighthouse Lab in Alderley Park | Wellcome Sanger Institute for the COVID-19 Genomics UK (COG-UK) Consortium | Jacquelyn Wynn, Mairead Hyland, The Lighthouse Lab in Alderley Park and Alex Alderton, Roberto Amato, Jeffrey Barrett, Sonia Goncalves, Ewan Harrison, David K. Jackson, Ian Johnston, Dominic Kwiatkowski, Cordelia Langford, John Sillitoe on behalf of the Wellcome Sanger Institute COVID-19 Surveillance Team |
| EPI_ISL_2804309 | Health Services Laboratories | Wellcome Sanger Institute for the COVID-19 Genomics UK (COG-UK) Consortium | Health Services Laboratories and Alex Alderton, Roberto Amato, Jeffrey Barrett, Sonia Goncalves, Ewan Harrison, David K. Jackson, Ian Johnston, Dominic Kwiatkowski, Cordelia Langford, John Sillitoe on behalf of the Wellcome Sanger Institute COVID-19 Surveillance Team |
| EPI_ISL_2804310, EPI_ISL_2804311, EPI_ISL_2804312, EPI_ISL_2804313, EPI_ISL_2804314 | Lighthouse Lab in Alderley Park | Wellcome Sanger Institute for the COVID-19 Genomics UK (COG-UK) Consortium | Jacquelyn Wynn, Mairead Hyland, The Lighthouse Lab in Alderley Park and Alex Alderton, Roberto Amato, Jeffrey Barrett, Sonia Goncalves, Ewan Harrison, David K. Jackson, Ian Johnston, Dominic Kwiatkowski, Cordelia Langford, John Sillitoe on behalf of the Wellcome Sanger Institute COVID-19 Surveillance Team |
| EPI_ISL_2804315 | Lighthouse Laboratory Plymouth | Wellcome Sanger Institute for the COVID-19 Genomics UK (COG-UK) Consortium | Lighthouse Laboratory Plymouth and Alex Alderton, Roberto Amato, Jeffrey Barrett, Sonia Goncalves, Ewan Harrison, David K. Jackson, Ian Johnston, Dominic Kwiatkowski, Cordelia Langford, John Sillitoe on behalf of the Wellcome Sanger Institute COVID-19 Surveillance Team |
| EPI_ISL_2804316, EPI_ISL_2804317, EPI_ISL_2804318, EPI_ISL_2804319, EPI_ISL_2804320 | Lighthouse Lab in Alderley Park | Wellcome Sanger Institute for the COVID-19 Genomics UK (COG-UK) Consortium | Jacquelyn Wynn, Mairead Hyland, The Lighthouse Lab in Alderley Park and Alex Alderton, Roberto Amato, Jeffrey Barrett, Sonia Goncalves, Ewan Harrison, David K. Jackson, Ian Johnston, Dominic Kwiatkowski, Cordelia Langford, John Sillitoe on behalf of the Wellcome Sanger Institute COVID-19 Surveillance Team |
| EPI_ISL_2804321, EPI_ISL_2804322 | Lighthouse Laboratory Plymouth | Wellcome Sanger Institute for the COVID-19 Genomics UK (COG-UK) Consortium | Lighthouse Laboratory Plymouth and Alex Alderton, Roberto Amato, Jeffrey Barrett, Sonia Goncalves, Ewan Harrison, David K. Jackson, Ian Johnston, Dominic Kwiatkowski, Cordelia Langford, John Sillitoe on behalf of the Wellcome Sanger Institute COVID-19 Surveillance Team |
| EPI_ISL_2804323, EPI_ISL_2804324, EPI_ISL_2804325, EPI_ISL_2804326, EPI_ISL_2804327, EPI_ISL_2804328, EPI_ISL_2804329, EPI_ISL_2804330, EPI_ISL_2804331 | Lighthouse Lab in Alderley Park | Wellcome Sanger Institute for the COVID-19 Genomics UK (COG-UK) Consortium | Jacquelyn Wynn, Mairead Hyland, The Lighthouse Lab in Alderley Park and Alex Alderton, Roberto Amato, Jeffrey Barrett, Sonia Goncalves, Ewan Harrison, David K. Jackson, Ian Johnston, Dominic Kwiatkowski, Cordelia Langford, John Sillitoe on behalf of the Wellcome Sanger Institute COVID-19 Surveillance Team |
| EPI_ISL_2804332 | Lighthouse Laboratory Plymouth | Wellcome Sanger Institute for the COVID-19 Genomics UK (COG-UK) Consortium | Lighthouse Laboratory Plymouth and Alex Alderton, Roberto Amato, Jeffrey Barrett, Sonia Goncalves, Ewan Harrison, David K. Jackson, Ian Johnston, Dominic Kwiatkowski, Cordelia Langford, John Sillitoe on behalf of the Wellcome Sanger Institute COVID-19 Surveillance Team |
| EPI_ISL_2804333, EPI_ISL_2804334 | Lighthouse Lab in Alderley Park | Wellcome Sanger Institute for the COVID-19 Genomics UK (COG-UK) Consortium | Jacquelyn Wynn, Mairead Hyland, The Lighthouse Lab in Alderley Park and Alex Alderton, Roberto Amato, Jeffrey Barrett, Sonia Goncalves, Ewan Harrison, David K. Jackson, Ian Johnston, Dominic Kwiatkowski, Cordelia Langford, John Sillitoe on behalf of the Wellcome Sanger Institute COVID-19 Surveillance Team |
| EPI_ISL_2804335 | Lighthouse Laboratory Plymouth | Wellcome Sanger Institute for the COVID-19 Genomics UK (COG-UK) Consortium | Lighthouse Laboratory Plymouth and Alex Alderton, Roberto Amato, Jeffrey Barrett, Sonia Goncalves, Ewan Harrison, David K. Jackson, Ian Johnston, Dominic Kwiatkowski, Cordelia Langford, John Sillitoe on behalf of the Wellcome Sanger Institute COVID-19 Surveillance Team |
| EPI_ISL_2804336, EPI_ISL_2804337, EPI_ISL_2804338 | Lighthouse Lab in Alderley Park | Wellcome Sanger Institute for the COVID-19 Genomics UK (COG-UK) Consortium | Jacquelyn Wynn, Mairead Hyland, The Lighthouse Lab in Alderley Park and Alex Alderton, Roberto Amato, Jeffrey Barrett, Sonia Goncalves, Ewan Harrison, David K. Jackson, Ian Johnston, Dominic Kwiatkowski, Cordelia Langford, John Sillitoe on behalf of the Wellcome Sanger Institute COVID-19 Surveillance Team |
| EPI_ISL_2804339 | Lighthouse Laboratory Plymouth | Wellcome Sanger Institute for the COVID-19 Genomics UK (COG-UK) Consortium | Lighthouse Laboratory Plymouth and Alex Alderton, Roberto Amato, Jeffrey Barrett, Sonia Goncalves, Ewan Harrison, David K. Jackson, Ian Johnston, Dominic Kwiatkowski, Cordelia Langford, John Sillitoe on behalf of the Wellcome Sanger Institute COVID-19 Surveillance Team |
| EPI_ISL_2804340, EPI_ISL_2804341 | Lighthouse Lab in Alderley Park | Wellcome Sanger Institute for the COVID-19 Genomics UK (COG-UK) Consortium | Jacquelyn Wynn, Mairead Hyland, The Lighthouse Lab in Alderley Park and Alex Alderton, Roberto Amato, Jeffrey Barrett, Sonia Goncalves, Ewan Harrison, David K. Jackson, Ian Johnston, Dominic Kwiatkowski, Cordelia Langford, John Sillitoe on behalf of the Wellcome Sanger Institute COVID-19 Surveillance Team |
| EPI_ISL_2804342 | Lighthouse Laboratory Plymouth | Wellcome Sanger Institute for the COVID-19 Genomics UK (COG-UK) Consortium | Lighthouse Laboratory Plymouth and Alex Alderton, Roberto Amato, Jeffrey Barrett, Sonia Goncalves, Ewan Harrison, David K. Jackson, Ian Johnston, Dominic Kwiatkowski, Cordelia Langford, John Sillitoe on behalf of the Wellcome Sanger Institute COVID-19 Surveillance Team |
| EPI_ISL_2804343, EPI_ISL_2804344, EPI_ISL_2804345 | Lighthouse Lab in Alderley Park | Wellcome Sanger Institute for the COVID-19 Genomics UK (COG-UK) Consortium | Jacquelyn Wynn, Mairead Hyland, The Lighthouse Lab in Alderley Park and Alex Alderton, Roberto Amato, Jeffrey Barrett, Sonia Goncalves, Ewan Harrison, David K. Jackson, Ian Johnston, Dominic Kwiatkowski, Cordelia Langford, John Sillitoe on behalf of the Wellcome Sanger Institute COVID-19 Surveillance Team |
| EPI_ISL_2804346 | Health Services Laboratories | Wellcome Sanger Institute for the COVID-19 Genomics UK (COG-UK) Consortium | Health Services Laboratories and Alex Alderton, Roberto Amato, Jeffrey Barrett, Sonia Goncalves, Ewan Harrison, David K. Jackson, Ian Johnston, Dominic Kwiatkowski, Cordelia Langford, John Sillitoe on behalf of the Wellcome Sanger Institute COVID-19 Surveillance Team |
| EPI_ISL_2804347, EPI_ISL_2804348, EPI_ISL_2804349, EPI_ISL_2804350 | Lighthouse Lab in Alderley Park | Wellcome Sanger Institute for the COVID-19 Genomics UK (COG-UK) Consortium | Jacquelyn Wynn, Mairead Hyland, The Lighthouse Lab in Alderley Park and Alex Alderton, Roberto Amato, Jeffrey Barrett, Sonia Goncalves, Ewan Harrison, David K. Jackson, Ian Johnston, Dominic Kwiatkowski, Cordelia Langford, John Sillitoe on behalf of the Wellcome Sanger Institute COVID-19 Surveillance Team |
| EPI_ISL_2804351 | Health Services Laboratories | Wellcome Sanger Institute for the COVID-19 Genomics UK (COG-UK) | Health Services Laboratories and Alex Alderton, Roberto Amato, Jeffrey Barrett, Sonia Goncalves, Ewan Harrison, David K. Jackson, Ian Johnston, Dominic Kwiatkowski, Cordelia Langford, John Sillitoe on behalf of the Wellcome Sanger Institute |

| | | | |
|---|---|---|---|
| | | Consortium | COVID-19 Surveillance Team |
| EPI_ISL_2804352 | Lighthouse Lab in Alderley Park | Wellcome Sanger Institute for the COVID-19 Genomics UK (COG-UK) Consortium | Jacquelyn Wynn, Mairead Hyland, The Lighthouse Lab in Alderley Park and Alex Alderton, Roberto Amato, Jeffrey Barrett, Sonia Goncalves, Ewan Harrison, David K. Jackson, Ian Johnston, Dominic Kwiatkowski, Cordelia Langford, John Sillitoe on behalf of the Wellcome Sanger Institute COVID-19 Surveillance Team |
| EPI_ISL_2804353 | Health Services Laboratories | Wellcome Sanger Institute for the COVID-19 Genomics UK (COG-UK) Consortium | Health Services Laboratories and Alex Alderton, Roberto Amato, Jeffrey Barrett, Sonia Goncalves, Ewan Harrison, David K. Jackson, Ian Johnston, Dominic Kwiatkowski, Cordelia Langford, John Sillitoe on behalf of the Wellcome Sanger Institute COVID-19 Surveillance Team |
| EPI_ISL_2804354, EPI_ISL_2804355, EPI_ISL_2804356, EPI_ISL_2804357 | Lighthouse Lab in Alderley Park | Wellcome Sanger Institute for the COVID-19 Genomics UK (COG-UK) Consortium | Jacquelyn Wynn, Mairead Hyland, The Lighthouse Lab in Alderley Park and Alex Alderton, Roberto Amato, Jeffrey Barrett, Sonia Goncalves, Ewan Harrison, David K. Jackson, Ian Johnston, Dominic Kwiatkowski, Cordelia Langford, John Sillitoe on behalf of the Wellcome Sanger Institute COVID-19 Surveillance Team |
| EPI_ISL_2804358 | Lighthouse Laboratory Plymouth | Wellcome Sanger Institute for the COVID-19 Genomics UK (COG-UK) Consortium | Lighthouse Laboratory Plymouth and Alex Alderton, Roberto Amato, Jeffrey Barrett, Sonia Goncalves, Ewan Harrison, David K. Jackson, Ian Johnston, Dominic Kwiatkowski, Cordelia Langford, John Sillitoe on behalf of the Wellcome Sanger Institute COVID-19 Surveillance Team |
| EPI_ISL_2804359, EPI_ISL_2804360, EPI_ISL_2804361, EPI_ISL_2804362, EPI_ISL_2804363, EPI_ISL_2804364, EPI_ISL_2804365 | Lighthouse Lab in Alderley Park | Wellcome Sanger Institute for the COVID-19 Genomics UK (COG-UK) Consortium | Jacquelyn Wynn, Mairead Hyland, The Lighthouse Lab in Alderley Park and Alex Alderton, Roberto Amato, Jeffrey Barrett, Sonia Goncalves, Ewan Harrison, David K. Jackson, Ian Johnston, Dominic Kwiatkowski, Cordelia Langford, John Sillitoe on behalf of the Wellcome Sanger Institute COVID-19 Surveillance Team |
| EPI_ISL_2804366 | Lighthouse Laboratory Plymouth | Wellcome Sanger Institute for the COVID-19 Genomics UK (COG-UK) Consortium | Lighthouse Laboratory Plymouth and Alex Alderton, Roberto Amato, Jeffrey Barrett, Sonia Goncalves, Ewan Harrison, David K. Jackson, Ian Johnston, Dominic Kwiatkowski, Cordelia Langford, John Sillitoe on behalf of the Wellcome Sanger Institute COVID-19 Surveillance Team |
| EPI_ISL_2804367, EPI_ISL_2804368, EPI_ISL_2804369 | Lighthouse Lab in Alderley Park | Wellcome Sanger Institute for the COVID-19 Genomics UK (COG-UK) Consortium | Jacquelyn Wynn, Mairead Hyland, The Lighthouse Lab in Alderley Park and Alex Alderton, Roberto Amato, Jeffrey Barrett, Sonia Goncalves, Ewan Harrison, David K. Jackson, Ian Johnston, Dominic Kwiatkowski, Cordelia Langford, John Sillitoe on behalf of the Wellcome Sanger Institute COVID-19 Surveillance Team |
| EPI_ISL_2804370 | Lighthouse Laboratory Plymouth | Wellcome Sanger Institute for the COVID-19 Genomics UK (COG-UK) Consortium | Lighthouse Laboratory Plymouth and Alex Alderton, Roberto Amato, Jeffrey Barrett, Sonia Goncalves, Ewan Harrison, David K. Jackson, Ian Johnston, Dominic Kwiatkowski, Cordelia Langford, John Sillitoe on behalf of the Wellcome Sanger Institute COVID-19 Surveillance Team |
| EPI_ISL_2804371, EPI_ISL_2804372 | Lighthouse Lab in Alderley Park | Wellcome Sanger Institute for the COVID-19 Genomics UK (COG-UK) Consortium | Jacquelyn Wynn, Mairead Hyland, The Lighthouse Lab in Alderley Park and Alex Alderton, Roberto Amato, Jeffrey Barrett, Sonia Goncalves, Ewan Harrison, David K. Jackson, Ian Johnston, Dominic Kwiatkowski, Cordelia Langford, John Sillitoe on behalf of the Wellcome Sanger Institute COVID-19 Surveillance Team |
| EPI_ISL_2804373 | Lighthouse Laboratory Plymouth | Wellcome Sanger Institute for the COVID-19 Genomics UK (COG-UK) Consortium | Lighthouse Laboratory Plymouth and Alex Alderton, Roberto Amato, Jeffrey Barrett, Sonia Goncalves, Ewan Harrison, David K. Jackson, Ian Johnston, Dominic Kwiatkowski, Cordelia Langford, John Sillitoe on behalf of the Wellcome Sanger Institute COVID-19 Surveillance Team |
| EPI_ISL_2804374, EPI_ISL_2804375, EPI_ISL_2804376, EPI_ISL_2804377 | Lighthouse Lab in Alderley Park | Wellcome Sanger Institute for the COVID-19 Genomics UK (COG-UK) Consortium | Jacquelyn Wynn, Mairead Hyland, The Lighthouse Lab in Alderley Park and Alex Alderton, Roberto Amato, Jeffrey Barrett, Sonia Goncalves, Ewan Harrison, David K. Jackson, Ian Johnston, Dominic Kwiatkowski, Cordelia Langford, John Sillitoe on behalf of the Wellcome Sanger Institute COVID-19 Surveillance Team |
| EPI_ISL_2804378, EPI_ISL_2804379 | Health Services Laboratories | Wellcome Sanger Institute for the COVID-19 Genomics UK (COG-UK) Consortium | Health Services Laboratories and Alex Alderton, Roberto Amato, Jeffrey Barrett, Sonia Goncalves, Ewan Harrison, David K. Jackson, Ian Johnston, Dominic Kwiatkowski, Cordelia Langford, John Sillitoe on behalf of the Wellcome Sanger Institute COVID-19 Surveillance Team |
| EPI_ISL_2804380 | Lighthouse Laboratory Plymouth | Wellcome Sanger Institute for the COVID-19 Genomics UK (COG-UK) Consortium | Lighthouse Laboratory Plymouth and Alex Alderton, Roberto Amato, Jeffrey Barrett, Sonia Goncalves, Ewan Harrison, David K. Jackson, Ian Johnston, Dominic Kwiatkowski, Cordelia Langford, John Sillitoe on behalf of the Wellcome Sanger Institute COVID-19 Surveillance Team |
| EPI_ISL_2804381, EPI_ISL_2804382, EPI_ISL_2804383, EPI_ISL_2804384, EPI_ISL_2804385, EPI_ISL_2804386, EPI_ISL_2804387, EPI_ISL_2804388 | Lighthouse Lab in Alderley Park | Wellcome Sanger Institute for the COVID-19 Genomics UK (COG-UK) Consortium | Jacquelyn Wynn, Mairead Hyland, The Lighthouse Lab in Alderley Park and Alex Alderton, Roberto Amato, Jeffrey Barrett, Sonia Goncalves, Ewan Harrison, David K. Jackson, Ian Johnston, Dominic Kwiatkowski, Cordelia Langford, John Sillitoe on behalf of the Wellcome Sanger Institute COVID-19 Surveillance Team |
| EPI_ISL_2804389 | Lighthouse Laboratory Plymouth | Wellcome Sanger Institute for the COVID-19 Genomics UK (COG-UK) Consortium | Lighthouse Laboratory Plymouth and Alex Alderton, Roberto Amato, Jeffrey Barrett, Sonia Goncalves, Ewan Harrison, David K. Jackson, Ian Johnston, Dominic Kwiatkowski, Cordelia Langford, John Sillitoe on behalf of the Wellcome Sanger Institute COVID-19 Surveillance Team |
| EPI_ISL_2804390, EPI_ISL_2804391 | Lighthouse Lab in Alderley Park | Wellcome Sanger Institute for the COVID-19 Genomics UK (COG-UK) Consortium | Jacquelyn Wynn, Mairead Hyland, The Lighthouse Lab in Alderley Park and Alex Alderton, Roberto Amato, Jeffrey Barrett, Sonia Goncalves, Ewan Harrison, David K. Jackson, Ian Johnston, Dominic Kwiatkowski, Cordelia Langford, John Sillitoe on behalf of the Wellcome Sanger Institute COVID-19 Surveillance Team |
| EPI_ISL_2804392 | Lighthouse Laboratory Plymouth | Wellcome Sanger Institute for the COVID-19 Genomics UK (COG-UK) Consortium | Lighthouse Laboratory Plymouth and Alex Alderton, Roberto Amato, Jeffrey Barrett, Sonia Goncalves, Ewan Harrison, David K. Jackson, Ian Johnston, Dominic Kwiatkowski, Cordelia Langford, John Sillitoe on behalf of the Wellcome Sanger Institute COVID-19 Surveillance Team |
| EPI_ISL_2804393, EPI_ISL_2804394, EPI_ISL_2804395, EPI_ISL_2804396, EPI_ISL_2804397, EPI_ISL_2804398, EPI_ISL_2804399 | Lighthouse Lab in Alderley Park | Wellcome Sanger Institute for the COVID-19 Genomics UK (COG-UK) Consortium | Jacquelyn Wynn, Mairead Hyland, The Lighthouse Lab in Alderley Park and Alex Alderton, Roberto Amato, Jeffrey Barrett, Sonia Goncalves, Ewan Harrison, David K. Jackson, Ian Johnston, Dominic Kwiatkowski, Cordelia Langford, John Sillitoe on behalf of the Wellcome Sanger Institute COVID-19 Surveillance Team |
| EPI_ISL_2804400 | Lighthouse Laboratory Plymouth | Wellcome Sanger Institute for the COVID-19 Genomics UK (COG-UK) Consortium | Lighthouse Laboratory Plymouth and Alex Alderton, Roberto Amato, Jeffrey Barrett, Sonia Goncalves, Ewan Harrison, David K. Jackson, Ian Johnston, Dominic Kwiatkowski, Cordelia Langford, John Sillitoe on behalf of the Wellcome Sanger Institute COVID-19 Surveillance Team |
| EPI_ISL_2804401, EPI_ISL_2804402, EPI_ISL_2804403, EPI_ISL_2804404, EPI_ISL_2804405, EPI_ISL_2804406, EPI_ISL_2804407, EPI_ISL_2804408, EPI_ISL_2804409 | Lighthouse Lab in Alderley Park | Wellcome Sanger Institute for the COVID-19 Genomics UK (COG-UK) Consortium | Jacquelyn Wynn, Mairead Hyland, The Lighthouse Lab in Alderley Park and Alex Alderton, Roberto Amato, Jeffrey Barrett, Sonia Goncalves, Ewan Harrison, David K. Jackson, Ian Johnston, Dominic Kwiatkowski, Cordelia Langford, John Sillitoe on behalf of the Wellcome Sanger Institute COVID-19 Surveillance Team |
| EPI_ISL_2804410 | Lighthouse Laboratory Plymouth | Wellcome Sanger Institute for the COVID-19 Genomics UK (COG-UK) Consortium | Lighthouse Laboratory Plymouth and Alex Alderton, Roberto Amato, Jeffrey Barrett, Sonia Goncalves, Ewan Harrison, David K. Jackson, Ian Johnston, Dominic Kwiatkowski, Cordelia Langford, John Sillitoe on behalf of the Wellcome Sanger Institute COVID-19 Surveillance Team |
| EPI_ISL_2804411, EPI_ISL_2804412, EPI_ISL_2804413 | Lighthouse Lab in Alderley Park | Wellcome Sanger Institute for the COVID-19 Genomics UK (COG-UK) Consortium | Jacquelyn Wynn, Mairead Hyland, The Lighthouse Lab in Alderley Park and Alex Alderton, Roberto Amato, Jeffrey Barrett, Sonia Goncalves, Ewan Harrison, David K. Jackson, Ian Johnston, Dominic Kwiatkowski, Cordelia Langford, John Sillitoe on behalf of the Wellcome Sanger Institute COVID-19 Surveillance Team |
| EPI_ISL_2804414 | Lighthouse Laboratory Plymouth | Wellcome Sanger Institute for the COVID-19 Genomics UK (COG-UK) Consortium | Lighthouse Laboratory Plymouth and Alex Alderton, Roberto Amato, Jeffrey Barrett, Sonia Goncalves, Ewan Harrison, David K. Jackson, Ian Johnston, Dominic Kwiatkowski, Cordelia Langford, John Sillitoe on behalf of the Wellcome Sanger Institute COVID-19 Surveillance Team |
| EPI_ISL_2804415, EPI_ISL_2804416 | Lighthouse Lab in Alderley Park | Wellcome Sanger Institute for the COVID-19 Genomics UK (COG-UK) Consortium | Jacquelyn Wynn, Mairead Hyland, The Lighthouse Lab in Alderley Park and Alex Alderton, Roberto Amato, Jeffrey Barrett, Sonia Goncalves, Ewan Harrison, David K. Jackson, Ian Johnston, Dominic Kwiatkowski, Cordelia Langford, John Sillitoe on behalf of the Wellcome Sanger Institute COVID-19 Surveillance Team |
| EPI_ISL_2804417 | Health Services Laboratories | Wellcome Sanger Institute for the COVID-19 Genomics UK (COG-UK) Consortium | Health Services Laboratories and Alex Alderton, Roberto Amato, Jeffrey Barrett, Sonia Goncalves, Ewan Harrison, David K. Jackson, Ian Johnston, Dominic Kwiatkowski, Cordelia Langford, John Sillitoe on behalf of the Wellcome Sanger Institute COVID-19 Surveillance Team |
| EPI_ISL_2804418, EPI_ISL_2804419, EPI_ISL_2804420 | Lighthouse Lab in Alderley Park | Wellcome Sanger Institute for the COVID-19 Genomics UK (COG-UK) Consortium | Jacquelyn Wynn, Mairead Hyland, The Lighthouse Lab in Alderley Park and Alex Alderton, Roberto Amato, Jeffrey Barrett, Sonia Goncalves, Ewan Harrison, David K. Jackson, Ian Johnston, Dominic Kwiatkowski, Cordelia Langford, John Sillitoe on behalf of the Wellcome Sanger Institute COVID-19 Surveillance Team |
| EPI_ISL_2804421 | Lighthouse Laboratory Plymouth | Wellcome Sanger Institute for the COVID-19 Genomics UK (COG-UK) Consortium | Lighthouse Laboratory Plymouth and Alex Alderton, Roberto Amato, Jeffrey Barrett, Sonia Goncalves, Ewan Harrison, David K. Jackson, Ian Johnston, Dominic Kwiatkowski, Cordelia Langford, John Sillitoe on behalf of the Wellcome Sanger Institute COVID-19 Surveillance Team |
| EPI_ISL_2804422 | Lighthouse Lab in Alderley Park | Wellcome Sanger Institute for the COVID-19 Genomics UK (COG-UK) Consortium | Jacquelyn Wynn, Mairead Hyland, The Lighthouse Lab in Alderley Park and Alex Alderton, Roberto Amato, Jeffrey Barrett, Sonia Goncalves, Ewan Harrison, David K. Jackson, Ian Johnston, Dominic Kwiatkowski, Cordelia Langford, John Sillitoe on behalf of the Wellcome Sanger Institute COVID-19 Surveillance Team |
| EPI_ISL_2804423 | Lighthouse Laboratory Plymouth | Wellcome Sanger Institute for the COVID-19 Genomics UK (COG-UK) Consortium | Lighthouse Laboratory Plymouth and Alex Alderton, Roberto Amato, Jeffrey Barrett, Sonia Goncalves, Ewan Harrison, David K. Jackson, Ian Johnston, Dominic Kwiatkowski, Cordelia Langford, John Sillitoe on behalf of the Wellcome Sanger Institute COVID-19 Surveillance Team |
| EPI_ISL_2804424 | Health Services Laboratories | Wellcome Sanger Institute for the COVID-19 Genomics UK (COG-UK) Consortium | Health Services Laboratories and Alex Alderton, Roberto Amato, Jeffrey Barrett, Sonia Goncalves, Ewan Harrison, David K. Jackson, Ian Johnston, Dominic Kwiatkowski, Cordelia Langford, John Sillitoe on behalf of the Wellcome Sanger Institute COVID-19 Surveillance Team |
| EPI_ISL_2804425 | Lighthouse Laboratory Plymouth | Wellcome Sanger Institute for the COVID-19 Genomics UK (COG-UK) Consortium | Lighthouse Laboratory Plymouth and Alex Alderton, Roberto Amato, Jeffrey Barrett, Sonia Goncalves, Ewan Harrison, David K. Jackson, Ian Johnston, Dominic Kwiatkowski, Cordelia Langford, John Sillitoe on behalf of the Wellcome Sanger Institute COVID-19 Surveillance Team |
| EPI_ISL_2804426 | Lighthouse Lab in Alderley Park | Wellcome Sanger Institute for the COVID-19 Genomics UK (COG-UK) Consortium | Jacquelyn Wynn, Mairead Hyland, The Lighthouse Lab in Alderley Park and Alex Alderton, Roberto Amato, Jeffrey Barrett, Sonia Goncalves, Ewan Harrison, David K. Jackson, Ian Johnston, Dominic Kwiatkowski, Cordelia Langford, John Sillitoe on behalf of the Wellcome Sanger Institute COVID-19 Surveillance Team |
| EPI_ISL_2804427 | Lighthouse Laboratory Plymouth | Wellcome Sanger Institute for the COVID-19 Genomics UK (COG-UK) | Lighthouse Laboratory Plymouth and Alex Alderton, Roberto Amato, Jeffrey Barrett, Sonia Goncalves, Ewan Harrison, David K. Jackson, Ian Johnston, Dominic Kwiatkowski, Cordelia Langford, John Sillitoe on behalf of the Wellcome Sanger |

| | | | |
|---|---|---|---|
| | | Consortium | Institute COVID-19 Surveillance Team |
| EPI_ISL_2804428 | Lighthouse Lab in Alderley Park | Wellcome Sanger Institute for the COVID-19 Genomics UK (COG-UK) Consortium | Jacquelyn Wynn, Mairead Hyland, The Lighthouse Lab in Alderley Park and Alex Alderton, Roberto Amato, Jeffrey Barrett, Sonia Goncalves, Ewan Harrison, David K. Jackson, Ian Johnston, Dominic Kwiatkowski, Cordelia Langford, John Sillitoe on behalf of the Wellcome Sanger Institute COVID-19 Surveillance Team |
| EPI_ISL_2804429 | Lighthouse Laboratory Plymouth | Wellcome Sanger Institute for the COVID-19 Genomics UK (COG-UK) Consortium | Lighthouse Laboratory Plymouth and Alex Alderton, Roberto Amato, Jeffrey Barrett, Sonia Goncalves, Ewan Harrison, David K. Jackson, Ian Johnston, Dominic Kwiatkowski, Cordelia Langford, John Sillitoe on behalf of the Wellcome Sanger Institute COVID-19 Surveillance Team |
| EPI_ISL_2804430, EPI_ISL_2804431, EPI_ISL_2804432, EPI_ISL_2804433 | Lighthouse Lab in Alderley Park | Wellcome Sanger Institute for the COVID-19 Genomics UK (COG-UK) Consortium | Jacquelyn Wynn, Mairead Hyland, The Lighthouse Lab in Alderley Park and Alex Alderton, Roberto Amato, Jeffrey Barrett, Sonia Goncalves, Ewan Harrison, David K. Jackson, Ian Johnston, Dominic Kwiatkowski, Cordelia Langford, John Sillitoe on behalf of the Wellcome Sanger Institute COVID-19 Surveillance Team |
| EPI_ISL_2804434 | Lighthouse Laboratory Plymouth | Wellcome Sanger Institute for the COVID-19 Genomics UK (COG-UK) Consortium | Lighthouse Laboratory Plymouth and Alex Alderton, Roberto Amato, Jeffrey Barrett, Sonia Goncalves, Ewan Harrison, David K. Jackson, Ian Johnston, Dominic Kwiatkowski, Cordelia Langford, John Sillitoe on behalf of the Wellcome Sanger Institute COVID-19 Surveillance Team |
| EPI_ISL_2804435, EPI_ISL_2804436, EPI_ISL_2804437, EPI_ISL_2804438 | Lighthouse Lab in Alderley Park | Wellcome Sanger Institute for the COVID-19 Genomics UK (COG-UK) Consortium | Jacquelyn Wynn, Mairead Hyland, The Lighthouse Lab in Alderley Park and Alex Alderton, Roberto Amato, Jeffrey Barrett, Sonia Goncalves, Ewan Harrison, David K. Jackson, Ian Johnston, Dominic Kwiatkowski, Cordelia Langford, John Sillitoe on behalf of the Wellcome Sanger Institute COVID-19 Surveillance Team |
| EPI_ISL_2804439, EPI_ISL_2804440 | Lighthouse Laboratory Plymouth | Wellcome Sanger Institute for the COVID-19 Genomics UK (COG-UK) Consortium | Lighthouse Laboratory Plymouth and Alex Alderton, Roberto Amato, Jeffrey Barrett, Sonia Goncalves, Ewan Harrison, David K. Jackson, Ian Johnston, Dominic Kwiatkowski, Cordelia Langford, John Sillitoe on behalf of the Wellcome Sanger Institute COVID-19 Surveillance Team |
| EPI_ISL_2804441 | Lighthouse Lab in Alderley Park | Wellcome Sanger Institute for the COVID-19 Genomics UK (COG-UK) Consortium | Jacquelyn Wynn, Mairead Hyland, The Lighthouse Lab in Alderley Park and Alex Alderton, Roberto Amato, Jeffrey Barrett, Sonia Goncalves, Ewan Harrison, David K. Jackson, Ian Johnston, Dominic Kwiatkowski, Cordelia Langford, John Sillitoe on behalf of the Wellcome Sanger Institute COVID-19 Surveillance Team |
| EPI_ISL_2804442 | Lighthouse Laboratory Plymouth | Wellcome Sanger Institute for the COVID-19 Genomics UK (COG-UK) Consortium | Lighthouse Laboratory Plymouth and Alex Alderton, Roberto Amato, Jeffrey Barrett, Sonia Goncalves, Ewan Harrison, David K. Jackson, Ian Johnston, Dominic Kwiatkowski, Cordelia Langford, John Sillitoe on behalf of the Wellcome Sanger Institute COVID-19 Surveillance Team |
| EPI_ISL_2804443, EPI_ISL_2804444, EPI_ISL_2804445, EPI_ISL_2804446, EPI_ISL_2804447, EPI_ISL_2804448, EPI_ISL_2804449 | Lighthouse Lab in Alderley Park | Wellcome Sanger Institute for the COVID-19 Genomics UK (COG-UK) Consortium | Jacquelyn Wynn, Mairead Hyland, The Lighthouse Lab in Alderley Park and Alex Alderton, Roberto Amato, Jeffrey Barrett, Sonia Goncalves, Ewan Harrison, David K. Jackson, Ian Johnston, Dominic Kwiatkowski, Cordelia Langford, John Sillitoe on behalf of the Wellcome Sanger Institute COVID-19 Surveillance Team |
| EPI_ISL_2804450 | Health Services Laboratories | Wellcome Sanger Institute for the COVID-19 Genomics UK (COG-UK) Consortium | Health Services Laboratories and Alex Alderton, Roberto Amato, Jeffrey Barrett, Sonia Goncalves, Ewan Harrison, David K. Jackson, Ian Johnston, Dominic Kwiatkowski, Cordelia Langford, John Sillitoe on behalf of the Wellcome Sanger Institute COVID-19 Surveillance Team |
| EPI_ISL_2804451, EPI_ISL_2804452 | Lighthouse Lab in Alderley Park | Wellcome Sanger Institute for the COVID-19 Genomics UK (COG-UK) Consortium | Jacquelyn Wynn, Mairead Hyland, The Lighthouse Lab in Alderley Park and Alex Alderton, Roberto Amato, Jeffrey Barrett, Sonia Goncalves, Ewan Harrison, David K. Jackson, Ian Johnston, Dominic Kwiatkowski, Cordelia Langford, John Sillitoe on behalf of the Wellcome Sanger Institute COVID-19 Surveillance Team |
| EPI_ISL_2804453 | Lighthouse Laboratory Plymouth | Wellcome Sanger Institute for the COVID-19 Genomics UK (COG-UK) Consortium | Lighthouse Laboratory Plymouth and Alex Alderton, Roberto Amato, Jeffrey Barrett, Sonia Goncalves, Ewan Harrison, David K. Jackson, Ian Johnston, Dominic Kwiatkowski, Cordelia Langford, John Sillitoe on behalf of the Wellcome Sanger Institute COVID-19 Surveillance Team |
| EPI_ISL_2804454 | Lighthouse Lab in Alderley Park | Wellcome Sanger Institute for the COVID-19 Genomics UK (COG-UK) Consortium | Jacquelyn Wynn, Mairead Hyland, The Lighthouse Lab in Alderley Park and Alex Alderton, Roberto Amato, Jeffrey Barrett, Sonia Goncalves, Ewan Harrison, David K. Jackson, Ian Johnston, Dominic Kwiatkowski, Cordelia Langford, John Sillitoe on behalf of the Wellcome Sanger Institute COVID-19 Surveillance Team |
| EPI_ISL_2804455 | Lighthouse Laboratory Plymouth | Wellcome Sanger Institute for the COVID-19 Genomics UK (COG-UK) Consortium | Lighthouse Laboratory Plymouth and Alex Alderton, Roberto Amato, Jeffrey Barrett, Sonia Goncalves, Ewan Harrison, David K. Jackson, Ian Johnston, Dominic Kwiatkowski, Cordelia Langford, John Sillitoe on behalf of the Wellcome Sanger Institute COVID-19 Surveillance Team |
| EPI_ISL_2804456 | Health Services Laboratories | Wellcome Sanger Institute for the COVID-19 Genomics UK (COG-UK) Consortium | Health Services Laboratories and Alex Alderton, Roberto Amato, Jeffrey Barrett, Sonia Goncalves, Ewan Harrison, David K. Jackson, Ian Johnston, Dominic Kwiatkowski, Cordelia Langford, John Sillitoe on behalf of the Wellcome Sanger Institute COVID-19 Surveillance Team |
| EPI_ISL_2804457 | Lighthouse Laboratory Plymouth | Wellcome Sanger Institute for the COVID-19 Genomics UK (COG-UK) Consortium | Lighthouse Laboratory Plymouth and Alex Alderton, Roberto Amato, Jeffrey Barrett, Sonia Goncalves, Ewan Harrison, David K. Jackson, Ian Johnston, Dominic Kwiatkowski, Cordelia Langford, John Sillitoe on behalf of the Wellcome Sanger Institute COVID-19 Surveillance Team |
| EPI_ISL_2804458, EPI_ISL_2804459, EPI_ISL_2804460, EPI_ISL_2804461, EPI_ISL_2804462 | Lighthouse Lab in Alderley Park | Wellcome Sanger Institute for the COVID-19 Genomics UK (COG-UK) Consortium | Jacquelyn Wynn, Mairead Hyland, The Lighthouse Lab in Alderley Park and Alex Alderton, Roberto Amato, Jeffrey Barrett, Sonia Goncalves, Ewan Harrison, David K. Jackson, Ian Johnston, Dominic Kwiatkowski, Cordelia Langford, John Sillitoe on behalf of the Wellcome Sanger Institute COVID-19 Surveillance Team |
| EPI_ISL_2804463, EPI_ISL_2804464 | Lighthouse Laboratory Plymouth | Wellcome Sanger Institute for the COVID-19 Genomics UK (COG-UK) Consortium | Lighthouse Laboratory Plymouth and Alex Alderton, Roberto Amato, Jeffrey Barrett, Sonia Goncalves, Ewan Harrison, David K. Jackson, Ian Johnston, Dominic Kwiatkowski, Cordelia Langford, John Sillitoe on behalf of the Wellcome Sanger Institute COVID-19 Surveillance Team |
| EPI_ISL_2804465 | Lighthouse Lab in Alderley Park | Wellcome Sanger Institute for the COVID-19 Genomics UK (COG-UK) Consortium | Jacquelyn Wynn, Mairead Hyland, The Lighthouse Lab in Alderley Park and Alex Alderton, Roberto Amato, Jeffrey Barrett, Sonia Goncalves, Ewan Harrison, David K. Jackson, Ian Johnston, Dominic Kwiatkowski, Cordelia Langford, John Sillitoe on behalf of the Wellcome Sanger Institute COVID-19 Surveillance Team |
| EPI_ISL_2804466 | Lighthouse Laboratory Plymouth | Wellcome Sanger Institute for the COVID-19 Genomics UK (COG-UK) Consortium | Lighthouse Laboratory Plymouth and Alex Alderton, Roberto Amato, Jeffrey Barrett, Sonia Goncalves, Ewan Harrison, David K. Jackson, Ian Johnston, Dominic Kwiatkowski, Cordelia Langford, John Sillitoe on behalf of the Wellcome Sanger Institute COVID-19 Surveillance Team |
| EPI_ISL_2804467, EPI_ISL_2804468 | Lighthouse Lab in Alderley Park | Wellcome Sanger Institute for the COVID-19 Genomics UK (COG-UK) Consortium | Jacquelyn Wynn, Mairead Hyland, The Lighthouse Lab in Alderley Park and Alex Alderton, Roberto Amato, Jeffrey Barrett, Sonia Goncalves, Ewan Harrison, David K. Jackson, Ian Johnston, Dominic Kwiatkowski, Cordelia Langford, John Sillitoe on behalf of the Wellcome Sanger Institute COVID-19 Surveillance Team |
| EPI_ISL_2804469 | Lighthouse Laboratory Plymouth | Wellcome Sanger Institute for the COVID-19 Genomics UK (COG-UK) Consortium | Lighthouse Laboratory Plymouth and Alex Alderton, Roberto Amato, Jeffrey Barrett, Sonia Goncalves, Ewan Harrison, David K. Jackson, Ian Johnston, Dominic Kwiatkowski, Cordelia Langford, John Sillitoe on behalf of the Wellcome Sanger Institute COVID-19 Surveillance Team |
| EPI_ISL_2804470 | Lighthouse Lab in Alderley Park | Wellcome Sanger Institute for the COVID-19 Genomics UK (COG-UK) Consortium | Jacquelyn Wynn, Mairead Hyland, The Lighthouse Lab in Alderley Park and Alex Alderton, Roberto Amato, Jeffrey Barrett, Sonia Goncalves, Ewan Harrison, David K. Jackson, Ian Johnston, Dominic Kwiatkowski, Cordelia Langford, John Sillitoe on behalf of the Wellcome Sanger Institute COVID-19 Surveillance Team |
| EPI_ISL_2804471 | Lighthouse Laboratory Plymouth | Wellcome Sanger Institute for the COVID-19 Genomics UK (COG-UK) Consortium | Lighthouse Laboratory Plymouth and Alex Alderton, Roberto Amato, Jeffrey Barrett, Sonia Goncalves, Ewan Harrison, David K. Jackson, Ian Johnston, Dominic Kwiatkowski, Cordelia Langford, John Sillitoe on behalf of the Wellcome Sanger Institute COVID-19 Surveillance Team |
| EPI_ISL_2804472, EPI_ISL_2804473, EPI_ISL_2804474 | Lighthouse Lab in Alderley Park | Wellcome Sanger Institute for the COVID-19 Genomics UK (COG-UK) Consortium | Jacquelyn Wynn, Mairead Hyland, The Lighthouse Lab in Alderley Park and Alex Alderton, Roberto Amato, Jeffrey Barrett, Sonia Goncalves, Ewan Harrison, David K. Jackson, Ian Johnston, Dominic Kwiatkowski, Cordelia Langford, John Sillitoe on behalf of the Wellcome Sanger Institute COVID-19 Surveillance Team |
| EPI_ISL_2804475 | Lighthouse Laboratory Plymouth | Wellcome Sanger Institute for the COVID-19 Genomics UK (COG-UK) Consortium | Lighthouse Laboratory Plymouth and Alex Alderton, Roberto Amato, Jeffrey Barrett, Sonia Goncalves, Ewan Harrison, David K. Jackson, Ian Johnston, Dominic Kwiatkowski, Cordelia Langford, John Sillitoe on behalf of the Wellcome Sanger Institute COVID-19 Surveillance Team |
| EPI_ISL_2804476, EPI_ISL_2804477, EPI_ISL_2804478 | Lighthouse Lab in Alderley Park | Wellcome Sanger Institute for the COVID-19 Genomics UK (COG-UK) Consortium | Jacquelyn Wynn, Mairead Hyland, The Lighthouse Lab in Alderley Park and Alex Alderton, Roberto Amato, Jeffrey Barrett, Sonia Goncalves, Ewan Harrison, David K. Jackson, Ian Johnston, Dominic Kwiatkowski, Cordelia Langford, John Sillitoe on behalf of the Wellcome Sanger Institute COVID-19 Surveillance Team |
| EPI_ISL_2804479 | Lighthouse Laboratory Plymouth | Wellcome Sanger Institute for the COVID-19 Genomics UK (COG-UK) Consortium | Lighthouse Laboratory Plymouth and Alex Alderton, Roberto Amato, Jeffrey Barrett, Sonia Goncalves, Ewan Harrison, David K. Jackson, Ian Johnston, Dominic Kwiatkowski, Cordelia Langford, John Sillitoe on behalf of the Wellcome Sanger Institute COVID-19 Surveillance Team |
| EPI_ISL_2804480, EPI_ISL_2804481, EPI_ISL_2804482, EPI_ISL_2804483, EPI_ISL_2804484, EPI_ISL_2804485, EPI_ISL_2804486 | Lighthouse Lab in Alderley Park | Wellcome Sanger Institute for the COVID-19 Genomics UK (COG-UK) Consortium | Jacquelyn Wynn, Mairead Hyland, The Lighthouse Lab in Alderley Park and Alex Alderton, Roberto Amato, Jeffrey Barrett, Sonia Goncalves, Ewan Harrison, David K. Jackson, Ian Johnston, Dominic Kwiatkowski, Cordelia Langford, John Sillitoe on behalf of the Wellcome Sanger Institute COVID-19 Surveillance Team |
| EPI_ISL_2804487, EPI_ISL_2804488 | Health Services Laboratories | Wellcome Sanger Institute for the COVID-19 Genomics UK (COG-UK) Consortium | Health Services Laboratories and Alex Alderton, Roberto Amato, Jeffrey Barrett, Sonia Goncalves, Ewan Harrison, David K. Jackson, Ian Johnston, Dominic Kwiatkowski, Cordelia Langford, John Sillitoe on behalf of the Wellcome Sanger Institute COVID-19 Surveillance Team |
| EPI_ISL_2804489, EPI_ISL_2804490, EPI_ISL_2804491, EPI_ISL_2804492 | Lighthouse Lab in Alderley Park | Wellcome Sanger Institute for the COVID-19 Genomics UK (COG-UK) Consortium | Jacquelyn Wynn, Mairead Hyland, The Lighthouse Lab in Alderley Park and Alex Alderton, Roberto Amato, Jeffrey Barrett, Sonia Goncalves, Ewan Harrison, David K. Jackson, Ian Johnston, Dominic Kwiatkowski, Cordelia Langford, John Sillitoe on behalf of the Wellcome Sanger Institute COVID-19 Surveillance Team |
| EPI_ISL_2804493 | Lighthouse Laboratory Plymouth | Wellcome Sanger Institute for the COVID-19 Genomics UK (COG-UK) Consortium | Lighthouse Laboratory Plymouth and Alex Alderton, Roberto Amato, Jeffrey Barrett, Sonia Goncalves, Ewan Harrison, David K. Jackson, Ian Johnston, Dominic Kwiatkowski, Cordelia Langford, John Sillitoe on behalf of the Wellcome Sanger Institute COVID-19 Surveillance Team |
| EPI_ISL_2804494, EPI_ISL_2804495, EPI_ISL_2804496, EPI_ISL_2804497, EPI_ISL_2804498 | Lighthouse Lab in Alderley Park | Wellcome Sanger Institute for the COVID-19 Genomics UK (COG-UK) Consortium | Jacquelyn Wynn, Mairead Hyland, The Lighthouse Lab in Alderley Park and Alex Alderton, Roberto Amato, Jeffrey Barrett, Sonia Goncalves, Ewan Harrison, David K. Jackson, Ian Johnston, Dominic Kwiatkowski, Cordelia Langford, John Sillitoe on behalf of the Wellcome Sanger Institute COVID-19 Surveillance Team |
| EPI_ISL_2804499 | Health Services Laboratories | Wellcome Sanger Institute for the COVID-19 Genomics UK (COG-UK) Consortium | Health Services Laboratories and Alex Alderton, Roberto Amato, Jeffrey Barrett, Sonia Goncalves, Ewan Harrison, David K. Jackson, Ian Johnston, Dominic Kwiatkowski, Cordelia Langford, John Sillitoe on behalf of the Wellcome Sanger Institute COVID-19 Surveillance Team |
| EPI_ISL_2804500, EPI_ISL_2804501, | Lighthouse Lab in Alderley Park | Wellcome Sanger Institute for the COVID-19 Genomics UK (COG-UK) Consortium | Jacquelyn Wynn, Mairead Hyland, The Lighthouse Lab in Alderley Park and Alex Alderton, Roberto Amato, Jeffrey Barrett, Sonia Goncalves, Ewan Harrison, David K. Jackson, Ian Johnston, Dominic Kwiatkowski, Cordelia Langford, John Sillitoe on behalf of the Wellcome Sanger Institute COVID-19 Surveillance Team |

| | | | |
|---|---|---|---|
| EPI_ISL_2804502, EPI_ISL_2804503, EPI_ISL_2804504, EPI_ISL_2804505, EPI_ISL_2804506 | | | |
| EPI_ISL_2804507 | Lighthouse Laboratory Plymouth | Wellcome Sanger Institute for the COVID-19 Genomics UK (COG-UK) Consortium | Lighthouse Laboratory Plymouth and Alex Alderton, Roberto Amato, Jeffrey Barrett, Sonia Goncalves, Ewan Harrison, David K. Jackson, Ian Johnston, Dominic Kwiatkowski, Cordelia Langford, John Sillitoe on behalf of the Wellcome Sanger Institute COVID-19 Surveillance Team |
| EPI_ISL_2804508, EPI_ISL_2804509, EPI_ISL_2804510, EPI_ISL_2804511, EPI_ISL_2804512, EPI_ISL_2804513, EPI_ISL_2804514 | Lighthouse Lab in Alderley Park | Wellcome Sanger Institute for the COVID-19 Genomics UK (COG-UK) Consortium | Jacquelyn Wynn, Mairead Hyland, The Lighthouse Lab in Alderley Park and Alex Alderton, Roberto Amato, Jeffrey Barrett, Sonia Goncalves, Ewan Harrison, David K. Jackson, Ian Johnston, Dominic Kwiatkowski, Cordelia Langford, John Sillitoe on behalf of the Wellcome Sanger Institute COVID-19 Surveillance Team |
| EPI_ISL_2804515 | Lighthouse Laboratory Plymouth | Wellcome Sanger Institute for the COVID-19 Genomics UK (COG-UK) Consortium | Lighthouse Laboratory Plymouth and Alex Alderton, Roberto Amato, Jeffrey Barrett, Sonia Goncalves, Ewan Harrison, David K. Jackson, Ian Johnston, Dominic Kwiatkowski, Cordelia Langford, John Sillitoe on behalf of the Wellcome Sanger Institute COVID-19 Surveillance Team |
| EPI_ISL_2804516 | Lighthouse Lab in Alderley Park | Wellcome Sanger Institute for the COVID-19 Genomics UK (COG-UK) Consortium | Jacquelyn Wynn, Mairead Hyland, The Lighthouse Lab in Alderley Park and Alex Alderton, Roberto Amato, Jeffrey Barrett, Sonia Goncalves, Ewan Harrison, David K. Jackson, Ian Johnston, Dominic Kwiatkowski, Cordelia Langford, John Sillitoe on behalf of the Wellcome Sanger Institute COVID-19 Surveillance Team |
| EPI_ISL_2804517 | Lighthouse Laboratory Plymouth | Wellcome Sanger Institute for the COVID-19 Genomics UK (COG-UK) Consortium | Lighthouse Laboratory Plymouth and Alex Alderton, Roberto Amato, Jeffrey Barrett, Sonia Goncalves, Ewan Harrison, David K. Jackson, Ian Johnston, Dominic Kwiatkowski, Cordelia Langford, John Sillitoe on behalf of the Wellcome Sanger Institute COVID-19 Surveillance Team |
| EPI_ISL_2804518, EPI_ISL_2804519, EPI_ISL_2804520, EPI_ISL_2804521, EPI_ISL_2804522, EPI_ISL_2804523, EPI_ISL_2804524, EPI_ISL_2804525, EPI_ISL_2804526, EPI_ISL_2804527, EPI_ISL_2804528, EPI_ISL_2804529, EPI_ISL_2804530 | | | |
| see above | Lighthouse Lab in Alderley Park | Wellcome Sanger Institute for the COVID-19 Genomics UK (COG-UK) Consortium | Jacquelyn Wynn, Mairead Hyland, The Lighthouse Lab in Alderley Park and Alex Alderton, Roberto Amato, Jeffrey Barrett, Sonia Goncalves, Ewan Harrison, David K. Jackson, Ian Johnston, Dominic Kwiatkowski, Cordelia Langford, John Sillitoe on behalf of the Wellcome Sanger Institute COVID-19 Surveillance Team |
| EPI_ISL_2804531 | Health Services Laboratories | Wellcome Sanger Institute for the COVID-19 Genomics UK (COG-UK) Consortium | Health Services Laboratories and Alex Alderton, Roberto Amato, Jeffrey Barrett, Sonia Goncalves, Ewan Harrison, David K. Jackson, Ian Johnston, Dominic Kwiatkowski, Cordelia Langford, John Sillitoe on behalf of the Wellcome Sanger Institute COVID-19 Surveillance Team |
| EPI_ISL_2804532, EPI_ISL_2804533, EPI_ISL_2804534, EPI_ISL_2804535 | Lighthouse Laboratory Plymouth | Wellcome Sanger Institute for the COVID-19 Genomics UK (COG-UK) Consortium | Lighthouse Laboratory Plymouth and Alex Alderton, Roberto Amato, Jeffrey Barrett, Sonia Goncalves, Ewan Harrison, David K. Jackson, Ian Johnston, Dominic Kwiatkowski, Cordelia Langford, John Sillitoe on behalf of the Wellcome Sanger Institute COVID-19 Surveillance Team |
| EPI_ISL_2804536, EPI_ISL_2804537, EPI_ISL_2804538, EPI_ISL_2804539, EPI_ISL_2804540 | Lighthouse Lab in Alderley Park | Wellcome Sanger Institute for the COVID-19 Genomics UK (COG-UK) Consortium | Jacquelyn Wynn, Mairead Hyland, The Lighthouse Lab in Alderley Park and Alex Alderton, Roberto Amato, Jeffrey Barrett, Sonia Goncalves, Ewan Harrison, David K. Jackson, Ian Johnston, Dominic Kwiatkowski, Cordelia Langford, John Sillitoe on behalf of the Wellcome Sanger Institute COVID-19 Surveillance Team |
| EPI_ISL_2804541 | Health Services Laboratories | Wellcome Sanger Institute for the COVID-19 Genomics UK (COG-UK) Consortium | Health Services Laboratories and Alex Alderton, Roberto Amato, Jeffrey Barrett, Sonia Goncalves, Ewan Harrison, David K. Jackson, Ian Johnston, Dominic Kwiatkowski, Cordelia Langford, John Sillitoe on behalf of the Wellcome Sanger Institute COVID-19 Surveillance Team |
| EPI_ISL_2804542 | Lighthouse Lab in Alderley Park | Wellcome Sanger Institute for the COVID-19 Genomics UK (COG-UK) Consortium | Jacquelyn Wynn, Mairead Hyland, The Lighthouse Lab in Alderley Park and Alex Alderton, Roberto Amato, Jeffrey Barrett, Sonia Goncalves, Ewan Harrison, David K. Jackson, Ian Johnston, Dominic Kwiatkowski, Cordelia Langford, John Sillitoe on behalf of the Wellcome Sanger Institute COVID-19 Surveillance Team |
| EPI_ISL_2804543 | Lighthouse Laboratory Plymouth | Wellcome Sanger Institute for the COVID-19 Genomics UK (COG-UK) Consortium | Lighthouse Laboratory Plymouth and Alex Alderton, Roberto Amato, Jeffrey Barrett, Sonia Goncalves, Ewan Harrison, David K. Jackson, Ian Johnston, Dominic Kwiatkowski, Cordelia Langford, John Sillitoe on behalf of the Wellcome Sanger Institute COVID-19 Surveillance Team |
| EPI_ISL_2804544 | Lighthouse Lab in Alderley Park | Wellcome Sanger Institute for the COVID-19 Genomics UK (COG-UK) Consortium | Jacquelyn Wynn, Mairead Hyland, The Lighthouse Lab in Alderley Park and Alex Alderton, Roberto Amato, Jeffrey Barrett, Sonia Goncalves, Ewan Harrison, David K. Jackson, Ian Johnston, Dominic Kwiatkowski, Cordelia Langford, John Sillitoe on behalf of the Wellcome Sanger Institute COVID-19 Surveillance Team |
| EPI_ISL_2804545 | Lighthouse Laboratory Plymouth | Wellcome Sanger Institute for the COVID-19 Genomics UK (COG-UK) Consortium | Lighthouse Laboratory Plymouth and Alex Alderton, Roberto Amato, Jeffrey Barrett, Sonia Goncalves, Ewan Harrison, David K. Jackson, Ian Johnston, Dominic Kwiatkowski, Cordelia Langford, John Sillitoe on behalf of the Wellcome Sanger Institute COVID-19 Surveillance Team |
| EPI_ISL_2804546, EPI_ISL_2804547, EPI_ISL_2804548, EPI_ISL_2804549 | Lighthouse Lab in Alderley Park | Wellcome Sanger Institute for the COVID-19 Genomics UK (COG-UK) Consortium | Jacquelyn Wynn, Mairead Hyland, The Lighthouse Lab in Alderley Park and Alex Alderton, Roberto Amato, Jeffrey Barrett, Sonia Goncalves, Ewan Harrison, David K. Jackson, Ian Johnston, Dominic Kwiatkowski, Cordelia Langford, John Sillitoe on behalf of the Wellcome Sanger Institute COVID-19 Surveillance Team |
| EPI_ISL_2804550 | Health Services Laboratories | Wellcome Sanger Institute for the COVID-19 Genomics UK (COG-UK) Consortium | Health Services Laboratories and Alex Alderton, Roberto Amato, Jeffrey Barrett, Sonia Goncalves, Ewan Harrison, David K. Jackson, Ian Johnston, Dominic Kwiatkowski, Cordelia Langford, John Sillitoe on behalf of the Wellcome Sanger Institute COVID-19 Surveillance Team |
| EPI_ISL_2804551 | Lighthouse Laboratory Plymouth | Wellcome Sanger Institute for the COVID-19 Genomics UK (COG-UK) Consortium | Lighthouse Laboratory Plymouth and Alex Alderton, Roberto Amato, Jeffrey Barrett, Sonia Goncalves, Ewan Harrison, David K. Jackson, Ian Johnston, Dominic Kwiatkowski, Cordelia Langford, John Sillitoe on behalf of the Wellcome Sanger Institute COVID-19 Surveillance Team |
| EPI_ISL_2804552, EPI_ISL_2804553, EPI_ISL_2804554, EPI_ISL_2804555, EPI_ISL_2804556, EPI_ISL_2804557, EPI_ISL_2804558, EPI_ISL_2804559 | Lighthouse Lab in Alderley Park | Wellcome Sanger Institute for the COVID-19 Genomics UK (COG-UK) Consortium | Jacquelyn Wynn, Mairead Hyland, The Lighthouse Lab in Alderley Park and Alex Alderton, Roberto Amato, Jeffrey Barrett, Sonia Goncalves, Ewan Harrison, David K. Jackson, Ian Johnston, Dominic Kwiatkowski, Cordelia Langford, John Sillitoe on behalf of the Wellcome Sanger Institute COVID-19 Surveillance Team |
| EPI_ISL_2804560 | Lighthouse Laboratory Plymouth | Wellcome Sanger Institute for the COVID-19 Genomics UK (COG-UK) Consortium | Lighthouse Laboratory Plymouth and Alex Alderton, Roberto Amato, Jeffrey Barrett, Sonia Goncalves, Ewan Harrison, David K. Jackson, Ian Johnston, Dominic Kwiatkowski, Cordelia Langford, John Sillitoe on behalf of the Wellcome Sanger Institute COVID-19 Surveillance Team |
| EPI_ISL_2804561, EPI_ISL_2804562, EPI_ISL_2804563, EPI_ISL_2804564, EPI_ISL_2804565 | Lighthouse Lab in Alderley Park | Wellcome Sanger Institute for the COVID-19 Genomics UK (COG-UK) Consortium | Jacquelyn Wynn, Mairead Hyland, The Lighthouse Lab in Alderley Park and Alex Alderton, Roberto Amato, Jeffrey Barrett, Sonia Goncalves, Ewan Harrison, David K. Jackson, Ian Johnston, Dominic Kwiatkowski, Cordelia Langford, John Sillitoe on behalf of the Wellcome Sanger Institute COVID-19 Surveillance Team |
| EPI_ISL_2804566 | Lighthouse Laboratory Plymouth | Wellcome Sanger Institute for the COVID-19 Genomics UK (COG-UK) Consortium | Lighthouse Laboratory Plymouth and Alex Alderton, Roberto Amato, Jeffrey Barrett, Sonia Goncalves, Ewan Harrison, David K. Jackson, Ian Johnston, Dominic Kwiatkowski, Cordelia Langford, John Sillitoe on behalf of the Wellcome Sanger Institute COVID-19 Surveillance Team |
| EPI_ISL_2804567, EPI_ISL_2804568, EPI_ISL_2804569 | Lighthouse Lab in Alderley Park | Wellcome Sanger Institute for the COVID-19 Genomics UK (COG-UK) Consortium | Jacquelyn Wynn, Mairead Hyland, The Lighthouse Lab in Alderley Park and Alex Alderton, Roberto Amato, Jeffrey Barrett, Sonia Goncalves, Ewan Harrison, David K. Jackson, Ian Johnston, Dominic Kwiatkowski, Cordelia Langford, John Sillitoe on behalf of the Wellcome Sanger Institute COVID-19 Surveillance Team |
| EPI_ISL_2804570 | Lighthouse Laboratory Plymouth | Wellcome Sanger Institute for the COVID-19 Genomics UK (COG-UK) Consortium | Lighthouse Laboratory Plymouth and Alex Alderton, Roberto Amato, Jeffrey Barrett, Sonia Goncalves, Ewan Harrison, David K. Jackson, Ian Johnston, Dominic Kwiatkowski, Cordelia Langford, John Sillitoe on behalf of the Wellcome Sanger Institute COVID-19 Surveillance Team |
| EPI_ISL_2804571, EPI_ISL_2804572 | Lighthouse Lab in Alderley Park | Wellcome Sanger Institute for the COVID-19 Genomics UK (COG-UK) Consortium | Jacquelyn Wynn, Mairead Hyland, The Lighthouse Lab in Alderley Park and Alex Alderton, Roberto Amato, Jeffrey Barrett, Sonia Goncalves, Ewan Harrison, David K. Jackson, Ian Johnston, Dominic Kwiatkowski, Cordelia Langford, John Sillitoe on behalf of the Wellcome Sanger Institute COVID-19 Surveillance Team |
| EPI_ISL_2804573 | Lighthouse Laboratory Plymouth | Wellcome Sanger Institute for the COVID-19 Genomics UK (COG-UK) Consortium | Lighthouse Laboratory Plymouth and Alex Alderton, Roberto Amato, Jeffrey Barrett, Sonia Goncalves, Ewan Harrison, David K. Jackson, Ian Johnston, Dominic Kwiatkowski, Cordelia Langford, John Sillitoe on behalf of the Wellcome Sanger Institute COVID-19 Surveillance Team |
| EPI_ISL_2804574 | Health Services Laboratories | Wellcome Sanger Institute for the COVID-19 Genomics UK (COG-UK) Consortium | Health Services Laboratories and Alex Alderton, Roberto Amato, Jeffrey Barrett, Sonia Goncalves, Ewan Harrison, David K. Jackson, Ian Johnston, Dominic Kwiatkowski, Cordelia Langford, John Sillitoe on behalf of the Wellcome Sanger Institute COVID-19 Surveillance Team |
| EPI_ISL_2804575 | Lighthouse Laboratory Plymouth | Wellcome Sanger Institute for the COVID-19 Genomics UK (COG-UK) Consortium | Lighthouse Laboratory Plymouth and Alex Alderton, Roberto Amato, Jeffrey Barrett, Sonia Goncalves, Ewan Harrison, David K. Jackson, Ian Johnston, Dominic Kwiatkowski, Cordelia Langford, John Sillitoe on behalf of the Wellcome Sanger Institute COVID-19 Surveillance Team |
| EPI_ISL_2804576 | Lighthouse Lab in Alderley Park | Wellcome Sanger Institute for the COVID-19 Genomics UK (COG-UK) Consortium | Jacquelyn Wynn, Mairead Hyland, The Lighthouse Lab in Alderley Park and Alex Alderton, Roberto Amato, Jeffrey Barrett, Sonia Goncalves, Ewan Harrison, David K. Jackson, Ian Johnston, Dominic Kwiatkowski, Cordelia Langford, John Sillitoe on behalf of the Wellcome Sanger Institute COVID-19 Surveillance Team |
| EPI_ISL_2804577 | Lighthouse Laboratory Plymouth | Wellcome Sanger Institute for the COVID-19 Genomics UK (COG-UK) Consortium | Lighthouse Laboratory Plymouth and Alex Alderton, Roberto Amato, Jeffrey Barrett, Sonia Goncalves, Ewan Harrison, David K. Jackson, Ian Johnston, Dominic Kwiatkowski, Cordelia Langford, John Sillitoe on behalf of the Wellcome Sanger Institute COVID-19 Surveillance Team |
| EPI_ISL_2804578, EPI_ISL_2804579, EPI_ISL_2804580, EPI_ISL_2804581, EPI_ISL_2804582 | Lighthouse Lab in Alderley Park | Wellcome Sanger Institute for the COVID-19 Genomics UK (COG-UK) Consortium | Jacquelyn Wynn, Mairead Hyland, The Lighthouse Lab in Alderley Park and Alex Alderton, Roberto Amato, Jeffrey Barrett, Sonia Goncalves, Ewan Harrison, David K. Jackson, Ian Johnston, Dominic Kwiatkowski, Cordelia Langford, John Sillitoe on behalf of the Wellcome Sanger Institute COVID-19 Surveillance Team |
| EPI_ISL_2804583 | Lighthouse Laboratory Plymouth | Wellcome Sanger Institute for the COVID-19 Genomics UK (COG-UK) Consortium | Lighthouse Laboratory Plymouth and Alex Alderton, Roberto Amato, Jeffrey Barrett, Sonia Goncalves, Ewan Harrison, David K. Jackson, Ian Johnston, Dominic Kwiatkowski, Cordelia Langford, John Sillitoe on behalf of the Wellcome Sanger Institute COVID-19 Surveillance Team |
| EPI_ISL_2804584, EPI_ISL_2804585, | Lighthouse Lab in Alderley Park | Wellcome Sanger Institute for the COVID-19 Genomics UK (COG-UK) Consortium | Jacquelyn Wynn, Mairead Hyland, The Lighthouse Lab in Alderley Park and Alex Alderton, Roberto Amato, Jeffrey Barrett, Sonia Goncalves, Ewan Harrison, David K. Jackson, Ian Johnston, Dominic Kwiatkowski, Cordelia Langford, John Sillitoe on behalf of the Wellcome Sanger Institute COVID-19 Surveillance Team |

| | | | |
|---|---|---|---|
| EPI_ISL_2804586, EPI_ISL_2804587, EPI_ISL_2804588, EPI_ISL_2804589 | | | |
| EPI_ISL_2804590 | Lighthouse Laboratory Plymouth | Wellcome Sanger Institute for the COVID-19 Genomics UK (COG-UK) Consortium | Lighthouse Laboratory Plymouth and Alex Alderton, Roberto Amato, Jeffrey Barrett, Sonia Goncalves, Ewan Harrison, David K. Jackson, Ian Johnston, Dominic Kwiatkowski, Cordelia Langford, John Sillitoe on behalf of the Wellcome Sanger Institute COVID-19 Surveillance Team |
| EPI_ISL_2804591, EPI_ISL_2804592, EPI_ISL_2804593, EPI_ISL_2804594, EPI_ISL_2804595, EPI_ISL_2804596, EPI_ISL_2804597, EPI_ISL_2804598, EPI_ISL_2804599 | Lighthouse Lab in Alderley Park | Wellcome Sanger Institute for the COVID-19 Genomics UK (COG-UK) Consortium | Jacquelyn Wynn, Mairead Hyland, The Lighthouse Lab in Alderley Park and Alex Alderton, Roberto Amato, Jeffrey Barrett, Sonia Goncalves, Ewan Harrison, David K. Jackson, Ian Johnston, Dominic Kwiatkowski, Cordelia Langford, John Sillitoe on behalf of the Wellcome Sanger Institute COVID-19 Surveillance Team |
| EPI_ISL_2804600, EPI_ISL_2804601, EPI_ISL_2804602 | Lighthouse Laboratory Plymouth | Wellcome Sanger Institute for the COVID-19 Genomics UK (COG-UK) Consortium | Lighthouse Laboratory Plymouth and Alex Alderton, Roberto Amato, Jeffrey Barrett, Sonia Goncalves, Ewan Harrison, David K. Jackson, Ian Johnston, Dominic Kwiatkowski, Cordelia Langford, John Sillitoe on behalf of the Wellcome Sanger Institute COVID-19 Surveillance Team |
| EPI_ISL_2804603 | Lighthouse Lab in Alderley Park | Wellcome Sanger Institute for the COVID-19 Genomics UK (COG-UK) Consortium | Jacquelyn Wynn, Mairead Hyland, The Lighthouse Lab in Alderley Park and Alex Alderton, Roberto Amato, Jeffrey Barrett, Sonia Goncalves, Ewan Harrison, David K. Jackson, Ian Johnston, Dominic Kwiatkowski, Cordelia Langford, John Sillitoe on behalf of the Wellcome Sanger Institute COVID-19 Surveillance Team |
| EPI_ISL_2804604 | Health Services Laboratories | Wellcome Sanger Institute for the COVID-19 Genomics UK (COG-UK) Consortium | Health Services Laboratories and Alex Alderton, Roberto Amato, Jeffrey Barrett, Sonia Goncalves, Ewan Harrison, David K. Jackson, Ian Johnston, Dominic Kwiatkowski, Cordelia Langford, John Sillitoe on behalf of the Wellcome Sanger Institute COVID-19 Surveillance Team |
| EPI_ISL_2804605, EPI_ISL_2804606, EPI_ISL_2804607, EPI_ISL_2804608, EPI_ISL_2804609, EPI_ISL_2804610, EPI_ISL_2804611, EPI_ISL_2804612, EPI_ISL_2804613 | Lighthouse Lab in Alderley Park | Wellcome Sanger Institute for the COVID-19 Genomics UK (COG-UK) Consortium | Jacquelyn Wynn, Mairead Hyland, The Lighthouse Lab in Alderley Park and Alex Alderton, Roberto Amato, Jeffrey Barrett, Sonia Goncalves, Ewan Harrison, David K. Jackson, Ian Johnston, Dominic Kwiatkowski, Cordelia Langford, John Sillitoe on behalf of the Wellcome Sanger Institute COVID-19 Surveillance Team |
| EPI_ISL_2804614 | Lighthouse Lab in Milton Keynes | Wellcome Sanger Institute for the COVID-19 Genomics UK (COG-UK) Consortium | The Lighthouse Lab in Milton Keynes and Alex Alderton, Roberto Amato, Jeffrey Barrett, Sonia Goncalves, Ewan Harrison, David K. Jackson, Ian Johnston, Dominic Kwiatkowski, Cordelia Langford, John Sillitoe on behalf of the Wellcome Sanger Institute COVID-19 Surveillance Team |
| EPI_ISL_2804615, EPI_ISL_2804616 | Lighthouse Lab in Alderley Park | Wellcome Sanger Institute for the COVID-19 Genomics UK (COG-UK) Consortium | Jacquelyn Wynn, Mairead Hyland, The Lighthouse Lab in Alderley Park and Alex Alderton, Roberto Amato, Jeffrey Barrett, Sonia Goncalves, Ewan Harrison, David K. Jackson, Ian Johnston, Dominic Kwiatkowski, Cordelia Langford, John Sillitoe on behalf of the Wellcome Sanger Institute COVID-19 Surveillance Team |
| EPI_ISL_2804617 | Lighthouse Lab in Milton Keynes | Wellcome Sanger Institute for the COVID-19 Genomics UK (COG-UK) Consortium | The Lighthouse Lab in Milton Keynes and Alex Alderton, Roberto Amato, Jeffrey Barrett, Sonia Goncalves, Ewan Harrison, David K. Jackson, Ian Johnston, Dominic Kwiatkowski, Cordelia Langford, John Sillitoe on behalf of the Wellcome Sanger Institute COVID-19 Surveillance Team |
| EPI_ISL_2804618 | Lighthouse Lab in Alderley Park | Wellcome Sanger Institute for the COVID-19 Genomics UK (COG-UK) Consortium | Jacquelyn Wynn, Mairead Hyland, The Lighthouse Lab in Alderley Park and Alex Alderton, Roberto Amato, Jeffrey Barrett, Sonia Goncalves, Ewan Harrison, David K. Jackson, Ian Johnston, Dominic Kwiatkowski, Cordelia Langford, John Sillitoe on behalf of the Wellcome Sanger Institute COVID-19 Surveillance Team |
| EPI_ISL_2804619, EPI_ISL_2804620 | Lighthouse Lab in Glasgow | Wellcome Sanger Institute for the COVID-19 Genomics UK (COG-UK) Consortium | Harper VanSteenhouse, Yumi Kasai, David Gray, Carol Clugston, Anna Dominiczak and Alex Alderton, Roberto Amato, Jeffrey Barrett, Sonia Goncalves, Ewan Harrison, David K. Jackson, Ian Johnston, Dominic Kwiatkowski, Cordelia Langford, John Sillitoe on behalf of the Wellcome Sanger Institute COVID-19 Surveillance Team |
| EPI_ISL_2804621, EPI_ISL_2804622, EPI_ISL_2804623 | Lighthouse Lab in Milton Keynes | Wellcome Sanger Institute for the COVID-19 Genomics UK (COG-UK) Consortium | The Lighthouse Lab in Milton Keynes and Alex Alderton, Roberto Amato, Jeffrey Barrett, Sonia Goncalves, Ewan Harrison, David K. Jackson, Ian Johnston, Dominic Kwiatkowski, Cordelia Langford, John Sillitoe on behalf of the Wellcome Sanger Institute COVID-19 Surveillance Team |
| EPI_ISL_2804624 | Lighthouse Lab in Alderley Park | Wellcome Sanger Institute for the COVID-19 Genomics UK (COG-UK) Consortium | Jacquelyn Wynn, Mairead Hyland, The Lighthouse Lab in Alderley Park and Alex Alderton, Roberto Amato, Jeffrey Barrett, Sonia Goncalves, Ewan Harrison, David K. Jackson, Ian Johnston, Dominic Kwiatkowski, Cordelia Langford, John Sillitoe on behalf of the Wellcome Sanger Institute COVID-19 Surveillance Team |
| EPI_ISL_2804625 | Lighthouse Lab in Milton Keynes | Wellcome Sanger Institute for the COVID-19 Genomics UK (COG-UK) Consortium | The Lighthouse Lab in Milton Keynes and Alex Alderton, Roberto Amato, Jeffrey Barrett, Sonia Goncalves, Ewan Harrison, David K. Jackson, Ian Johnston, Dominic Kwiatkowski, Cordelia Langford, John Sillitoe on behalf of the Wellcome Sanger Institute COVID-19 Surveillance Team |
| EPI_ISL_2804626 | Lighthouse Lab in Glasgow | Wellcome Sanger Institute for the COVID-19 Genomics UK (COG-UK) Consortium | Harper VanSteenhouse, Yumi Kasai, David Gray, Carol Clugston, Anna Dominiczak and Alex Alderton, Roberto Amato, Jeffrey Barrett, Sonia Goncalves, Ewan Harrison, David K. Jackson, Ian Johnston, Dominic Kwiatkowski, Cordelia Langford, John Sillitoe on behalf of the Wellcome Sanger Institute COVID-19 Surveillance Team |
| EPI_ISL_2804627, EPI_ISL_2804628 | Lighthouse Lab in Alderley Park | Wellcome Sanger Institute for the COVID-19 Genomics UK (COG-UK) Consortium | Jacquelyn Wynn, Mairead Hyland, The Lighthouse Lab in Alderley Park and Alex Alderton, Roberto Amato, Jeffrey Barrett, Sonia Goncalves, Ewan Harrison, David K. Jackson, Ian Johnston, Dominic Kwiatkowski, Cordelia Langford, John Sillitoe on behalf of the Wellcome Sanger Institute COVID-19 Surveillance Team |
| EPI_ISL_2804629 | Lighthouse Lab in Milton Keynes | Wellcome Sanger Institute for the COVID-19 Genomics UK (COG-UK) Consortium | The Lighthouse Lab in Milton Keynes and Alex Alderton, Roberto Amato, Jeffrey Barrett, Sonia Goncalves, Ewan Harrison, David K. Jackson, Ian Johnston, Dominic Kwiatkowski, Cordelia Langford, John Sillitoe on behalf of the Wellcome Sanger Institute COVID-19 Surveillance Team |
| EPI_ISL_2804630 | Lighthouse Lab in Alderley Park | Wellcome Sanger Institute for the COVID-19 Genomics UK (COG-UK) Consortium | Jacquelyn Wynn, Mairead Hyland, The Lighthouse Lab in Alderley Park and Alex Alderton, Roberto Amato, Jeffrey Barrett, Sonia Goncalves, Ewan Harrison, David K. Jackson, Ian Johnston, Dominic Kwiatkowski, Cordelia Langford, John Sillitoe on behalf of the Wellcome Sanger Institute COVID-19 Surveillance Team |
| EPI_ISL_2804631 | Lighthouse Lab in Milton Keynes | Wellcome Sanger Institute for the COVID-19 Genomics UK (COG-UK) Consortium | The Lighthouse Lab in Milton Keynes and Alex Alderton, Roberto Amato, Jeffrey Barrett, Sonia Goncalves, Ewan Harrison, David K. Jackson, Ian Johnston, Dominic Kwiatkowski, Cordelia Langford, John Sillitoe on behalf of the Wellcome Sanger Institute COVID-19 Surveillance Team |
| EPI_ISL_2804632, EPI_ISL_2804633, EPI_ISL_2804634 | Lighthouse Lab in Alderley Park | Wellcome Sanger Institute for the COVID-19 Genomics UK (COG-UK) Consortium | Jacquelyn Wynn, Mairead Hyland, The Lighthouse Lab in Alderley Park and Alex Alderton, Roberto Amato, Jeffrey Barrett, Sonia Goncalves, Ewan Harrison, David K. Jackson, Ian Johnston, Dominic Kwiatkowski, Cordelia Langford, John Sillitoe on behalf of the Wellcome Sanger Institute COVID-19 Surveillance Team |
| EPI_ISL_2804635 | Lighthouse Lab in Glasgow | Wellcome Sanger Institute for the COVID-19 Genomics UK (COG-UK) Consortium | Harper VanSteenhouse, Yumi Kasai, David Gray, Carol Clugston, Anna Dominiczak and Alex Alderton, Roberto Amato, Jeffrey Barrett, Sonia Goncalves, Ewan Harrison, David K. Jackson, Ian Johnston, Dominic Kwiatkowski, Cordelia Langford, John Sillitoe on behalf of the Wellcome Sanger Institute COVID-19 Surveillance Team |
| EPI_ISL_2804636, EPI_ISL_2804637 | Lighthouse Lab in Alderley Park | Wellcome Sanger Institute for the COVID-19 Genomics UK (COG-UK) Consortium | Jacquelyn Wynn, Mairead Hyland, The Lighthouse Lab in Alderley Park and Alex Alderton, Roberto Amato, Jeffrey Barrett, Sonia Goncalves, Ewan Harrison, David K. Jackson, Ian Johnston, Dominic Kwiatkowski, Cordelia Langford, John Sillitoe on behalf of the Wellcome Sanger Institute COVID-19 Surveillance Team |
| EPI_ISL_2804638 | Lighthouse Lab in Glasgow | Wellcome Sanger Institute for the COVID-19 Genomics UK (COG-UK) Consortium | Harper VanSteenhouse, Yumi Kasai, David Gray, Carol Clugston, Anna Dominiczak and Alex Alderton, Roberto Amato, Jeffrey Barrett, Sonia Goncalves, Ewan Harrison, David K. Jackson, Ian Johnston, Dominic Kwiatkowski, Cordelia Langford, John Sillitoe on behalf of the Wellcome Sanger Institute COVID-19 Surveillance Team |
| EPI_ISL_2804639 | Lighthouse Lab in Milton Keynes | Wellcome Sanger Institute for the COVID-19 Genomics UK (COG-UK) Consortium | The Lighthouse Lab in Milton Keynes and Alex Alderton, Roberto Amato, Jeffrey Barrett, Sonia Goncalves, Ewan Harrison, David K. Jackson, Ian Johnston, Dominic Kwiatkowski, Cordelia Langford, John Sillitoe on behalf of the Wellcome Sanger Institute COVID-19 Surveillance Team |
| EPI_ISL_2804640 | Lighthouse Lab in Alderley Park | Wellcome Sanger Institute for the COVID-19 Genomics UK (COG-UK) Consortium | Jacquelyn Wynn, Mairead Hyland, The Lighthouse Lab in Alderley Park and Alex Alderton, Roberto Amato, Jeffrey Barrett, Sonia Goncalves, Ewan Harrison, David K. Jackson, Ian Johnston, Dominic Kwiatkowski, Cordelia Langford, John Sillitoe on behalf of the Wellcome Sanger Institute COVID-19 Surveillance Team |
| EPI_ISL_2804641 | Lighthouse Lab in Glasgow | Wellcome Sanger Institute for the COVID-19 Genomics UK (COG-UK) Consortium | Harper VanSteenhouse, Yumi Kasai, David Gray, Carol Clugston, Anna Dominiczak and Alex Alderton, Roberto Amato, Jeffrey Barrett, Sonia Goncalves, Ewan Harrison, David K. Jackson, Ian Johnston, Dominic Kwiatkowski, Cordelia Langford, John Sillitoe on behalf of the Wellcome Sanger Institute COVID-19 Surveillance Team |
| EPI_ISL_2804642, EPI_ISL_2804643, EPI_ISL_2804644 | Lighthouse Lab in Alderley Park | Wellcome Sanger Institute for the COVID-19 Genomics UK (COG-UK) Consortium | Jacquelyn Wynn, Mairead Hyland, The Lighthouse Lab in Alderley Park and Alex Alderton, Roberto Amato, Jeffrey Barrett, Sonia Goncalves, Ewan Harrison, David K. Jackson, Ian Johnston, Dominic Kwiatkowski, Cordelia Langford, John Sillitoe on behalf of the Wellcome Sanger Institute COVID-19 Surveillance Team |
| EPI_ISL_2804645 | Lighthouse Lab in Milton Keynes | Wellcome Sanger Institute for the COVID-19 Genomics UK (COG-UK) Consortium | The Lighthouse Lab in Milton Keynes and Alex Alderton, Roberto Amato, Jeffrey Barrett, Sonia Goncalves, Ewan Harrison, David K. Jackson, Ian Johnston, Dominic Kwiatkowski, Cordelia Langford, John Sillitoe on behalf of the Wellcome Sanger Institute COVID-19 Surveillance Team |
| EPI_ISL_2804646 | Lighthouse Lab in Alderley Park | Wellcome Sanger Institute for the COVID-19 Genomics UK (COG-UK) Consortium | Jacquelyn Wynn, Mairead Hyland, The Lighthouse Lab in Alderley Park and Alex Alderton, Roberto Amato, Jeffrey Barrett, Sonia Goncalves, Ewan Harrison, David K. Jackson, Ian Johnston, Dominic Kwiatkowski, Cordelia Langford, John Sillitoe on behalf of the Wellcome Sanger Institute COVID-19 Surveillance Team |
| EPI_ISL_2804647, EPI_ISL_2804648 | Lighthouse Lab in Milton Keynes | Wellcome Sanger Institute for the COVID-19 Genomics UK (COG-UK) Consortium | The Lighthouse Lab in Milton Keynes and Alex Alderton, Roberto Amato, Jeffrey Barrett, Sonia Goncalves, Ewan Harrison, David K. Jackson, Ian Johnston, Dominic Kwiatkowski, Cordelia Langford, John Sillitoe on behalf of the Wellcome Sanger Institute COVID-19 Surveillance Team |
| EPI_ISL_2804649 | Lighthouse Lab in Alderley Park | Wellcome Sanger Institute for the COVID-19 Genomics UK (COG-UK) Consortium | Jacquelyn Wynn, Mairead Hyland, The Lighthouse Lab in Alderley Park and Alex Alderton, Roberto Amato, Jeffrey Barrett, Sonia Goncalves, Ewan Harrison, David K. Jackson, Ian Johnston, Dominic Kwiatkowski, Cordelia Langford, John Sillitoe on behalf of the Wellcome Sanger Institute COVID-19 Surveillance Team |
| EPI_ISL_2804650 | Lighthouse Lab in Milton Keynes | Wellcome Sanger Institute for the COVID-19 Genomics UK (COG-UK) Consortium | The Lighthouse Lab in Milton Keynes and Alex Alderton, Roberto Amato, Jeffrey Barrett, Sonia Goncalves, Ewan Harrison, David K. Jackson, Ian Johnston, Dominic Kwiatkowski, Cordelia Langford, John Sillitoe on behalf of the Wellcome Sanger Institute COVID-19 Surveillance Team |
| EPI_ISL_2804651 | Lighthouse Lab in Glasgow | Wellcome Sanger Institute for the COVID-19 Genomics UK (COG-UK) Consortium | Harper VanSteenhouse, Yumi Kasai, David Gray, Carol Clugston, Anna Dominiczak and Alex Alderton, Roberto Amato, Jeffrey Barrett, Sonia Goncalves, Ewan Harrison, David K. Jackson, Ian Johnston, Dominic Kwiatkowski, Cordelia Langford, John Sillitoe on behalf of the Wellcome Sanger Institute COVID-19 Surveillance Team |
| EPI_ISL_2804652, EPI_ISL_2804653, EPI_ISL_2804654, EPI_ISL_2804655, EPI_ISL_2804656 | Lighthouse Lab in Alderley Park | Wellcome Sanger Institute for the COVID-19 Genomics UK (COG-UK) Consortium | Jacquelyn Wynn, Mairead Hyland, The Lighthouse Lab in Alderley Park and Alex Alderton, Roberto Amato, Jeffrey Barrett, Sonia Goncalves, Ewan Harrison, David K. Jackson, Ian Johnston, Dominic Kwiatkowski, Cordelia Langford, John Sillitoe on behalf of the Wellcome Sanger Institute COVID-19 Surveillance Team |
| EPI_ISL_2804657 | Lighthouse Lab in Milton Keynes | Wellcome Sanger Institute for the COVID-19 Genomics UK (COG-UK) Consortium | The Lighthouse Lab in Milton Keynes and Alex Alderton, Roberto Amato, Jeffrey Barrett, Sonia Goncalves, Ewan Harrison, David K. Jackson, Ian Johnston, Dominic Kwiatkowski, Cordelia Langford, John Sillitoe on behalf of the Wellcome Sanger Institute COVID-19 Surveillance Team |

| | | | |
|---|---|---|---|
| EPI_ISL_2804658, EPI_ISL_2804659, EPI_ISL_2804661, EPI_ISL_2804662 | Lighthouse Lab in Alderley Park | Wellcome Sanger Institute for the COVID-19 Genomics UK (COG-UK) Consortium | Jacquelyn Wynn, Mairead Hyland, The Lighthouse Lab in Alderley Park and Alex Alderton, Roberto Amato, Jeffrey Barrett, Sonia Goncalves, Ewan Harrison, David K. Jackson, Ian Johnston, Dominic Kwiatkowski, Cordelia Langford, John Sillitoe on behalf of the Wellcome Sanger Institute COVID-19 Surveillance Team |
| EPI_ISL_2804663 | Lighthouse Lab in Milton Keynes | Wellcome Sanger Institute for the COVID-19 Genomics UK (COG-UK) Consortium | The Lighthouse Lab in Milton Keynes and Alex Alderton, Roberto Amato, Jeffrey Barrett, Sonia Goncalves, Ewan Harrison, David K. Jackson, Ian Johnston, Dominic Kwiatkowski, Cordelia Langford, John Sillitoe on behalf of the Wellcome Sanger Institute COVID-19 Surveillance Team |
| EPI_ISL_2804664 | Lighthouse Lab in Glasgow | Wellcome Sanger Institute for the COVID-19 Genomics UK (COG-UK) Consortium | Harper VanSteenhouse, Yumi Kasai, David Gray, Carol Clugston, Anna Dominiczak and Alex Alderton, Roberto Amato, Jeffrey Barrett, Sonia Goncalves, Ewan Harrison, David K. Jackson, Ian Johnston, Dominic Kwiatkowski, Cordelia Langford, John Sillitoe on behalf of the Wellcome Sanger Institute COVID-19 Surveillance Team |
| EPI_ISL_2804665 | Lighthouse Lab in Milton Keynes | Wellcome Sanger Institute for the COVID-19 Genomics UK (COG-UK) Consortium | The Lighthouse Lab in Milton Keynes and Alex Alderton, Roberto Amato, Jeffrey Barrett, Sonia Goncalves, Ewan Harrison, David K. Jackson, Ian Johnston, Dominic Kwiatkowski, Cordelia Langford, John Sillitoe on behalf of the Wellcome Sanger Institute COVID-19 Surveillance Team |
| EPI_ISL_2804666, EPI_ISL_2804667 | Lighthouse Lab in Glasgow | Wellcome Sanger Institute for the COVID-19 Genomics UK (COG-UK) Consortium | Harper VanSteenhouse, Yumi Kasai, David Gray, Carol Clugston, Anna Dominiczak and Alex Alderton, Roberto Amato, Jeffrey Barrett, Sonia Goncalves, Ewan Harrison, David K. Jackson, Ian Johnston, Dominic Kwiatkowski, Cordelia Langford, John Sillitoe on behalf of the Wellcome Sanger Institute COVID-19 Surveillance Team |
| EPI_ISL_2804668 | Lighthouse Lab in Milton Keynes | Wellcome Sanger Institute for the COVID-19 Genomics UK (COG-UK) Consortium | The Lighthouse Lab in Milton Keynes and Alex Alderton, Roberto Amato, Jeffrey Barrett, Sonia Goncalves, Ewan Harrison, David K. Jackson, Ian Johnston, Dominic Kwiatkowski, Cordelia Langford, John Sillitoe on behalf of the Wellcome Sanger Institute COVID-19 Surveillance Team |
| EPI_ISL_2804669, EPI_ISL_2804670, EPI_ISL_2804671, EPI_ISL_2804672 | Lighthouse Lab in Alderley Park | Wellcome Sanger Institute for the COVID-19 Genomics UK (COG-UK) Consortium | Jacquelyn Wynn, Mairead Hyland, The Lighthouse Lab in Alderley Park and Alex Alderton, Roberto Amato, Jeffrey Barrett, Sonia Goncalves, Ewan Harrison, David K. Jackson, Ian Johnston, Dominic Kwiatkowski, Cordelia Langford, John Sillitoe on behalf of the Wellcome Sanger Institute COVID-19 Surveillance Team |
| EPI_ISL_2804673, EPI_ISL_2804674 | Lighthouse Lab in Milton Keynes | Wellcome Sanger Institute for the COVID-19 Genomics UK (COG-UK) Consortium | The Lighthouse Lab in Milton Keynes and Alex Alderton, Roberto Amato, Jeffrey Barrett, Sonia Goncalves, Ewan Harrison, David K. Jackson, Ian Johnston, Dominic Kwiatkowski, Cordelia Langford, John Sillitoe on behalf of the Wellcome Sanger Institute COVID-19 Surveillance Team |
| EPI_ISL_2804675, EPI_ISL_2804676, EPI_ISL_2804677, EPI_ISL_2804678, EPI_ISL_2804679 | Lighthouse Lab in Alderley Park | Wellcome Sanger Institute for the COVID-19 Genomics UK (COG-UK) Consortium | Jacquelyn Wynn, Mairead Hyland, The Lighthouse Lab in Alderley Park and Alex Alderton, Roberto Amato, Jeffrey Barrett, Sonia Goncalves, Ewan Harrison, David K. Jackson, Ian Johnston, Dominic Kwiatkowski, Cordelia Langford, John Sillitoe on behalf of the Wellcome Sanger Institute COVID-19 Surveillance Team |
| EPI_ISL_2804680 | Lighthouse Lab in Milton Keynes | Wellcome Sanger Institute for the COVID-19 Genomics UK (COG-UK) Consortium | The Lighthouse Lab in Milton Keynes and Alex Alderton, Roberto Amato, Jeffrey Barrett, Sonia Goncalves, Ewan Harrison, David K. Jackson, Ian Johnston, Dominic Kwiatkowski, Cordelia Langford, John Sillitoe on behalf of the Wellcome Sanger Institute COVID-19 Surveillance Team |
| EPI_ISL_2804681 | Lighthouse Lab in Alderley Park | Wellcome Sanger Institute for the COVID-19 Genomics UK (COG-UK) Consortium | Jacquelyn Wynn, Mairead Hyland, The Lighthouse Lab in Alderley Park and Alex Alderton, Roberto Amato, Jeffrey Barrett, Sonia Goncalves, Ewan Harrison, David K. Jackson, Ian Johnston, Dominic Kwiatkowski, Cordelia Langford, John Sillitoe on behalf of the Wellcome Sanger Institute COVID-19 Surveillance Team |
| EPI_ISL_2804682 | Lighthouse Lab in Milton Keynes | Wellcome Sanger Institute for the COVID-19 Genomics UK (COG-UK) Consortium | The Lighthouse Lab in Milton Keynes and Alex Alderton, Roberto Amato, Jeffrey Barrett, Sonia Goncalves, Ewan Harrison, David K. Jackson, Ian Johnston, Dominic Kwiatkowski, Cordelia Langford, John Sillitoe on behalf of the Wellcome Sanger Institute COVID-19 Surveillance Team |
| EPI_ISL_2804683, EPI_ISL_2804684 | Lighthouse Lab in Alderley Park | Wellcome Sanger Institute for the COVID-19 Genomics UK (COG-UK) Consortium | Jacquelyn Wynn, Mairead Hyland, The Lighthouse Lab in Alderley Park and Alex Alderton, Roberto Amato, Jeffrey Barrett, Sonia Goncalves, Ewan Harrison, David K. Jackson, Ian Johnston, Dominic Kwiatkowski, Cordelia Langford, John Sillitoe on behalf of the Wellcome Sanger Institute COVID-19 Surveillance Team |
| EPI_ISL_2804685, EPI_ISL_2804686 | Lighthouse Lab in Milton Keynes | Wellcome Sanger Institute for the COVID-19 Genomics UK (COG-UK) Consortium | The Lighthouse Lab in Milton Keynes and Alex Alderton, Roberto Amato, Jeffrey Barrett, Sonia Goncalves, Ewan Harrison, David K. Jackson, Ian Johnston, Dominic Kwiatkowski, Cordelia Langford, John Sillitoe on behalf of the Wellcome Sanger Institute COVID-19 Surveillance Team |
| EPI_ISL_2804687, EPI_ISL_2804688 | Lighthouse Lab in Alderley Park | Wellcome Sanger Institute for the COVID-19 Genomics UK (COG-UK) Consortium | Jacquelyn Wynn, Mairead Hyland, The Lighthouse Lab in Alderley Park and Alex Alderton, Roberto Amato, Jeffrey Barrett, Sonia Goncalves, Ewan Harrison, David K. Jackson, Ian Johnston, Dominic Kwiatkowski, Cordelia Langford, John Sillitoe on behalf of the Wellcome Sanger Institute COVID-19 Surveillance Team |
| EPI_ISL_2804689, EPI_ISL_2804690 | Lighthouse Lab in Milton Keynes | Wellcome Sanger Institute for the COVID-19 Genomics UK (COG-UK) Consortium | The Lighthouse Lab in Milton Keynes and Alex Alderton, Roberto Amato, Jeffrey Barrett, Sonia Goncalves, Ewan Harrison, David K. Jackson, Ian Johnston, Dominic Kwiatkowski, Cordelia Langford, John Sillitoe on behalf of the Wellcome Sanger Institute COVID-19 Surveillance Team |
| EPI_ISL_2804691 | Lighthouse Lab in Glasgow | Wellcome Sanger Institute for the COVID-19 Genomics UK (COG-UK) Consortium | Harper VanSteenhouse, Yumi Kasai, David Gray, Carol Clugston, Anna Dominiczak and Alex Alderton, Roberto Amato, Jeffrey Barrett, Sonia Goncalves, Ewan Harrison, David K. Jackson, Ian Johnston, Dominic Kwiatkowski, Cordelia Langford, John Sillitoe on behalf of the Wellcome Sanger Institute COVID-19 Surveillance Team |
| EPI_ISL_2804692, EPI_ISL_2804693, EPI_ISL_2804694 | Lighthouse Lab in Milton Keynes | Wellcome Sanger Institute for the COVID-19 Genomics UK (COG-UK) Consortium | The Lighthouse Lab in Milton Keynes and Alex Alderton, Roberto Amato, Jeffrey Barrett, Sonia Goncalves, Ewan Harrison, David K. Jackson, Ian Johnston, Dominic Kwiatkowski, Cordelia Langford, John Sillitoe on behalf of the Wellcome Sanger Institute COVID-19 Surveillance Team |
| EPI_ISL_2804695, EPI_ISL_2804696 | Lighthouse Lab in Alderley Park | Wellcome Sanger Institute for the COVID-19 Genomics UK (COG-UK) Consortium | Jacquelyn Wynn, Mairead Hyland, The Lighthouse Lab in Alderley Park and Alex Alderton, Roberto Amato, Jeffrey Barrett, Sonia Goncalves, Ewan Harrison, David K. Jackson, Ian Johnston, Dominic Kwiatkowski, Cordelia Langford, John Sillitoe on behalf of the Wellcome Sanger Institute COVID-19 Surveillance Team |
| EPI_ISL_2804697 | Lighthouse Lab in Glasgow | Wellcome Sanger Institute for the COVID-19 Genomics UK (COG-UK) Consortium | Harper VanSteenhouse, Yumi Kasai, David Gray, Carol Clugston, Anna Dominiczak and Alex Alderton, Roberto Amato, Jeffrey Barrett, Sonia Goncalves, Ewan Harrison, David K. Jackson, Ian Johnston, Dominic Kwiatkowski, Cordelia Langford, John Sillitoe on behalf of the Wellcome Sanger Institute COVID-19 Surveillance Team |
| EPI_ISL_2804698, EPI_ISL_2804699, EPI_ISL_2804700, EPI_ISL_2804701 | Lighthouse Lab in Milton Keynes | Wellcome Sanger Institute for the COVID-19 Genomics UK (COG-UK) Consortium | The Lighthouse Lab in Milton Keynes and Alex Alderton, Roberto Amato, Jeffrey Barrett, Sonia Goncalves, Ewan Harrison, David K. Jackson, Ian Johnston, Dominic Kwiatkowski, Cordelia Langford, John Sillitoe on behalf of the Wellcome Sanger Institute COVID-19 Surveillance Team |
| EPI_ISL_2804702 | Lighthouse Lab in Glasgow | Wellcome Sanger Institute for the COVID-19 Genomics UK (COG-UK) Consortium | Harper VanSteenhouse, Yumi Kasai, David Gray, Carol Clugston, Anna Dominiczak and Alex Alderton, Roberto Amato, Jeffrey Barrett, Sonia Goncalves, Ewan Harrison, David K. Jackson, Ian Johnston, Dominic Kwiatkowski, Cordelia Langford, John Sillitoe on behalf of the Wellcome Sanger Institute COVID-19 Surveillance Team |
| EPI_ISL_2804703 | Lighthouse Lab in Milton Keynes | Wellcome Sanger Institute for the COVID-19 Genomics UK (COG-UK) Consortium | The Lighthouse Lab in Milton Keynes and Alex Alderton, Roberto Amato, Jeffrey Barrett, Sonia Goncalves, Ewan Harrison, David K. Jackson, Ian Johnston, Dominic Kwiatkowski, Cordelia Langford, John Sillitoe on behalf of the Wellcome Sanger Institute COVID-19 Surveillance Team |
| EPI_ISL_2804704 | Lighthouse Lab in Alderley Park | Wellcome Sanger Institute for the COVID-19 Genomics UK (COG-UK) Consortium | Jacquelyn Wynn, Mairead Hyland, The Lighthouse Lab in Alderley Park and Alex Alderton, Roberto Amato, Jeffrey Barrett, Sonia Goncalves, Ewan Harrison, David K. Jackson, Ian Johnston, Dominic Kwiatkowski, Cordelia Langford, John Sillitoe on behalf of the Wellcome Sanger Institute COVID-19 Surveillance Team |
| EPI_ISL_2804705 | Lighthouse Lab in Milton Keynes | Wellcome Sanger Institute for the COVID-19 Genomics UK (COG-UK) Consortium | The Lighthouse Lab in Milton Keynes and Alex Alderton, Roberto Amato, Jeffrey Barrett, Sonia Goncalves, Ewan Harrison, David K. Jackson, Ian Johnston, Dominic Kwiatkowski, Cordelia Langford, John Sillitoe on behalf of the Wellcome Sanger Institute COVID-19 Surveillance Team |
| EPI_ISL_2804706, EPI_ISL_2804707, EPI_ISL_2804708 | Lighthouse Lab in Alderley Park | Wellcome Sanger Institute for the COVID-19 Genomics UK (COG-UK) Consortium | Jacquelyn Wynn, Mairead Hyland, The Lighthouse Lab in Alderley Park and Alex Alderton, Roberto Amato, Jeffrey Barrett, Sonia Goncalves, Ewan Harrison, David K. Jackson, Ian Johnston, Dominic Kwiatkowski, Cordelia Langford, John Sillitoe on behalf of the Wellcome Sanger Institute COVID-19 Surveillance Team |
| EPI_ISL_2804709 | Lighthouse Lab in Glasgow | Wellcome Sanger Institute for the COVID-19 Genomics UK (COG-UK) Consortium | Harper VanSteenhouse, Yumi Kasai, David Gray, Carol Clugston, Anna Dominiczak and Alex Alderton, Roberto Amato, Jeffrey Barrett, Sonia Goncalves, Ewan Harrison, David K. Jackson, Ian Johnston, Dominic Kwiatkowski, Cordelia Langford, John Sillitoe on behalf of the Wellcome Sanger Institute COVID-19 Surveillance Team |
| EPI_ISL_2804710, EPI_ISL_2804711, EPI_ISL_2804712 | Lighthouse Lab in Alderley Park | Wellcome Sanger Institute for the COVID-19 Genomics UK (COG-UK) Consortium | Jacquelyn Wynn, Mairead Hyland, The Lighthouse Lab in Alderley Park and Alex Alderton, Roberto Amato, Jeffrey Barrett, Sonia Goncalves, Ewan Harrison, David K. Jackson, Ian Johnston, Dominic Kwiatkowski, Cordelia Langford, John Sillitoe on behalf of the Wellcome Sanger Institute COVID-19 Surveillance Team |
| EPI_ISL_2804713 | Lighthouse Lab in Glasgow | Wellcome Sanger Institute for the COVID-19 Genomics UK (COG-UK) Consortium | Harper VanSteenhouse, Yumi Kasai, David Gray, Carol Clugston, Anna Dominiczak and Alex Alderton, Roberto Amato, Jeffrey Barrett, Sonia Goncalves, Ewan Harrison, David K. Jackson, Ian Johnston, Dominic Kwiatkowski, Cordelia Langford, John Sillitoe on behalf of the Wellcome Sanger Institute COVID-19 Surveillance Team |
| EPI_ISL_2804714, EPI_ISL_2804715 | Lighthouse Lab in Alderley Park | Wellcome Sanger Institute for the COVID-19 Genomics UK (COG-UK) Consortium | Jacquelyn Wynn, Mairead Hyland, The Lighthouse Lab in Alderley Park and Alex Alderton, Roberto Amato, Jeffrey Barrett, Sonia Goncalves, Ewan Harrison, David K. Jackson, Ian Johnston, Dominic Kwiatkowski, Cordelia Langford, John Sillitoe on behalf of the Wellcome Sanger Institute COVID-19 Surveillance Team |
| EPI_ISL_2804716, EPI_ISL_2804717, EPI_ISL_2804718 | Lighthouse Lab in Milton Keynes | Wellcome Sanger Institute for the COVID-19 Genomics UK (COG-UK) Consortium | The Lighthouse Lab in Milton Keynes and Alex Alderton, Roberto Amato, Jeffrey Barrett, Sonia Goncalves, Ewan Harrison, David K. Jackson, Ian Johnston, Dominic Kwiatkowski, Cordelia Langford, John Sillitoe on behalf of the Wellcome Sanger Institute COVID-19 Surveillance Team |
| EPI_ISL_2804719, EPI_ISL_2804720, EPI_ISL_2804721 | Lighthouse Lab in Alderley Park | Wellcome Sanger Institute for the COVID-19 Genomics UK (COG-UK) Consortium | Jacquelyn Wynn, Mairead Hyland, The Lighthouse Lab in Alderley Park and Alex Alderton, Roberto Amato, Jeffrey Barrett, Sonia Goncalves, Ewan Harrison, David K. Jackson, Ian Johnston, Dominic Kwiatkowski, Cordelia Langford, John Sillitoe on behalf of the Wellcome Sanger Institute COVID-19 Surveillance Team |
| EPI_ISL_2804722, EPI_ISL_2804723, EPI_ISL_2804724, EPI_ISL_2804725 | Lighthouse Lab in Glasgow | Wellcome Sanger Institute for the COVID-19 Genomics UK (COG-UK) Consortium | Harper VanSteenhouse, Yumi Kasai, David Gray, Carol Clugston, Anna Dominiczak and Alex Alderton, Roberto Amato, Jeffrey Barrett, Sonia Goncalves, Ewan Harrison, David K. Jackson, Ian Johnston, Dominic Kwiatkowski, Cordelia Langford, John Sillitoe on behalf of the Wellcome Sanger Institute COVID-19 Surveillance Team |
| EPI_ISL_2804726, EPI_ISL_2804727 | Lighthouse Lab in Milton Keynes | Wellcome Sanger Institute for the COVID-19 Genomics UK (COG-UK) Consortium | The Lighthouse Lab in Milton Keynes and Alex Alderton, Roberto Amato, Jeffrey Barrett, Sonia Goncalves, Ewan Harrison, David K. Jackson, Ian Johnston, Dominic Kwiatkowski, Cordelia Langford, John Sillitoe on behalf of the Wellcome Sanger Institute COVID-19 Surveillance Team |
| EPI_ISL_2804728, EPI_ISL_2804729, EPI_ISL_2804730, EPI_ISL_2804731 | Lighthouse Lab in Alderley Park | Wellcome Sanger Institute for the COVID-19 Genomics UK (COG-UK) Consortium | Jacquelyn Wynn, Mairead Hyland, The Lighthouse Lab in Alderley Park and Alex Alderton, Roberto Amato, Jeffrey Barrett, Sonia Goncalves, Ewan Harrison, David K. Jackson, Ian Johnston, Dominic Kwiatkowski, Cordelia Langford, John Sillitoe on behalf of the Wellcome Sanger Institute COVID-19 Surveillance Team |
| EPI_ISL_2804732 | Lighthouse Lab in Milton Keynes | Wellcome Sanger Institute for the COVID-19 Genomics UK (COG-UK) Consortium | The Lighthouse Lab in Milton Keynes and Alex Alderton, Roberto Amato, Jeffrey Barrett, Sonia Goncalves, Ewan Harrison, David K. Jackson, Ian Johnston, Dominic Kwiatkowski, Cordelia Langford, John Sillitoe on behalf of the Wellcome Sanger Institute COVID-19 Surveillance Team |
| EPI_ISL_2804733, EPI_ISL_2804734, EPI_ISL_2804735 | Lighthouse Lab in Alderley Park | Wellcome Sanger Institute for the COVID-19 Genomics UK (COG-UK) Consortium | Jacquelyn Wynn, Mairead Hyland, The Lighthouse Lab in Alderley Park and Alex Alderton, Roberto Amato, Jeffrey Barrett, Sonia Goncalves, Ewan Harrison, David K. Jackson, Ian Johnston, Dominic Kwiatkowski, Cordelia Langford, John Sillitoe on behalf of the Wellcome Sanger Institute COVID-19 Surveillance Team |
| EPI_ISL_2804736, | Lighthouse Lab in Milton Keynes | Wellcome Sanger Institute for the COVID-19 Genomics UK (COG-UK) | The Lighthouse Lab in Milton Keynes and Alex Alderton, Roberto Amato, Jeffrey Barrett, Sonia Goncalves, Ewan Harrison, David K. Jackson, Ian Johnston, Dominic Kwiatkowski, Cordelia Langford, John Sillitoe on behalf of the Wellcome Sanger |

| | | | |
|---|---|---|---|
| EPI_ISL_2804737 | | Consortium | Institute COVID-19 Surveillance Team |
| EPI_ISL_2804738 | Lighthouse Lab in Alderley Park | Wellcome Sanger Institute for the COVID-19 Genomics UK (COG-UK) Consortium | Jacquelyn Wynn, Mairead Hyland, The Lighthouse Lab in Alderley Park and Alex Alderton, Roberto Amato, Jeffrey Barrett, Sonia Goncalves, Ewan Harrison, David K. Jackson, Ian Johnston, Dominic Kwiatkowski, Cordelia Langford, John Sillitoe on behalf of the Wellcome Sanger Institute COVID-19 Surveillance Team |
| EPI_ISL_2804739 | Lighthouse Lab in Milton Keynes | Wellcome Sanger Institute for the COVID-19 Genomics UK (COG-UK) Consortium | The Lighthouse Lab in Milton Keynes and Alex Alderton, Roberto Amato, Jeffrey Barrett, Sonia Goncalves, Ewan Harrison, David K. Jackson, Ian Johnston, Dominic Kwiatkowski, Cordelia Langford, John Sillitoe on behalf of the Wellcome Sanger Institute COVID-19 Surveillance Team |
| EPI_ISL_2804740, EPI_ISL_2804741 | Lighthouse Lab in Alderley Park | Wellcome Sanger Institute for the COVID-19 Genomics UK (COG-UK) Consortium | Jacquelyn Wynn, Mairead Hyland, The Lighthouse Lab in Alderley Park and Alex Alderton, Roberto Amato, Jeffrey Barrett, Sonia Goncalves, Ewan Harrison, David K. Jackson, Ian Johnston, Dominic Kwiatkowski, Cordelia Langford, John Sillitoe on behalf of the Wellcome Sanger Institute COVID-19 Surveillance Team |
| EPI_ISL_2804742 | Lighthouse Lab in Glasgow | Wellcome Sanger Institute for the COVID-19 Genomics UK (COG-UK) Consortium | Harper VanSteenhouse, Yumi Kasai, David Gray, Carol Clugston, Anna Dominiczak and Alex Alderton, Roberto Amato, Jeffrey Barrett, Sonia Goncalves, Ewan Harrison, David K. Jackson, Ian Johnston, Dominic Kwiatkowski, Cordelia Langford, John Sillitoe on behalf of the Wellcome Sanger Institute COVID-19 Surveillance Team |
| EPI_ISL_2804743 | Lighthouse Lab in Alderley Park | Wellcome Sanger Institute for the COVID-19 Genomics UK (COG-UK) Consortium | Jacquelyn Wynn, Mairead Hyland, The Lighthouse Lab in Alderley Park and Alex Alderton, Roberto Amato, Jeffrey Barrett, Sonia Goncalves, Ewan Harrison, David K. Jackson, Ian Johnston, Dominic Kwiatkowski, Cordelia Langford, John Sillitoe on behalf of the Wellcome Sanger Institute COVID-19 Surveillance Team |
| EPI_ISL_2804744 | Lighthouse Lab in Milton Keynes | Wellcome Sanger Institute for the COVID-19 Genomics UK (COG-UK) Consortium | The Lighthouse Lab in Milton Keynes and Alex Alderton, Roberto Amato, Jeffrey Barrett, Sonia Goncalves, Ewan Harrison, David K. Jackson, Ian Johnston, Dominic Kwiatkowski, Cordelia Langford, John Sillitoe on behalf of the Wellcome Sanger Institute COVID-19 Surveillance Team |
| EPI_ISL_2804745 | Lighthouse Lab in Alderley Park | Wellcome Sanger Institute for the COVID-19 Genomics UK (COG-UK) Consortium | Jacquelyn Wynn, Mairead Hyland, The Lighthouse Lab in Alderley Park and Alex Alderton, Roberto Amato, Jeffrey Barrett, Sonia Goncalves, Ewan Harrison, David K. Jackson, Ian Johnston, Dominic Kwiatkowski, Cordelia Langford, John Sillitoe on behalf of the Wellcome Sanger Institute COVID-19 Surveillance Team |
| EPI_ISL_2804746 | Lighthouse Lab in Milton Keynes | Wellcome Sanger Institute for the COVID-19 Genomics UK (COG-UK) Consortium | The Lighthouse Lab in Milton Keynes and Alex Alderton, Roberto Amato, Jeffrey Barrett, Sonia Goncalves, Ewan Harrison, David K. Jackson, Ian Johnston, Dominic Kwiatkowski, Cordelia Langford, John Sillitoe on behalf of the Wellcome Sanger Institute COVID-19 Surveillance Team |
| EPI_ISL_2804747 | Lighthouse Lab in Alderley Park | Wellcome Sanger Institute for the COVID-19 Genomics UK (COG-UK) Consortium | Jacquelyn Wynn, Mairead Hyland, The Lighthouse Lab in Alderley Park and Alex Alderton, Roberto Amato, Jeffrey Barrett, Sonia Goncalves, Ewan Harrison, David K. Jackson, Ian Johnston, Dominic Kwiatkowski, Cordelia Langford, John Sillitoe on behalf of the Wellcome Sanger Institute COVID-19 Surveillance Team |
| EPI_ISL_2804748 | Lighthouse Lab in Milton Keynes | Wellcome Sanger Institute for the COVID-19 Genomics UK (COG-UK) Consortium | The Lighthouse Lab in Milton Keynes and Alex Alderton, Roberto Amato, Jeffrey Barrett, Sonia Goncalves, Ewan Harrison, David K. Jackson, Ian Johnston, Dominic Kwiatkowski, Cordelia Langford, John Sillitoe on behalf of the Wellcome Sanger Institute COVID-19 Surveillance Team |
| EPI_ISL_2804749 | Lighthouse Lab in Alderley Park | Wellcome Sanger Institute for the COVID-19 Genomics UK (COG-UK) Consortium | Jacquelyn Wynn, Mairead Hyland, The Lighthouse Lab in Alderley Park and Alex Alderton, Roberto Amato, Jeffrey Barrett, Sonia Goncalves, Ewan Harrison, David K. Jackson, Ian Johnston, Dominic Kwiatkowski, Cordelia Langford, John Sillitoe on behalf of the Wellcome Sanger Institute COVID-19 Surveillance Team |
| EPI_ISL_2804750, EPI_ISL_2804751 | Lighthouse Lab in Milton Keynes | Wellcome Sanger Institute for the COVID-19 Genomics UK (COG-UK) Consortium | The Lighthouse Lab in Milton Keynes and Alex Alderton, Roberto Amato, Jeffrey Barrett, Sonia Goncalves, Ewan Harrison, David K. Jackson, Ian Johnston, Dominic Kwiatkowski, Cordelia Langford, John Sillitoe on behalf of the Wellcome Sanger Institute COVID-19 Surveillance Team |
| EPI_ISL_2804752, EPI_ISL_2804753, EPI_ISL_2804754, EPI_ISL_2804755 | Lighthouse Lab in Alderley Park | Wellcome Sanger Institute for the COVID-19 Genomics UK (COG-UK) Consortium | Jacquelyn Wynn, Mairead Hyland, The Lighthouse Lab in Alderley Park and Alex Alderton, Roberto Amato, Jeffrey Barrett, Sonia Goncalves, Ewan Harrison, David K. Jackson, Ian Johnston, Dominic Kwiatkowski, Cordelia Langford, John Sillitoe on behalf of the Wellcome Sanger Institute COVID-19 Surveillance Team |
| EPI_ISL_2804756 | Lighthouse Lab in Milton Keynes | Wellcome Sanger Institute for the COVID-19 Genomics UK (COG-UK) Consortium | The Lighthouse Lab in Milton Keynes and Alex Alderton, Roberto Amato, Jeffrey Barrett, Sonia Goncalves, Ewan Harrison, David K. Jackson, Ian Johnston, Dominic Kwiatkowski, Cordelia Langford, John Sillitoe on behalf of the Wellcome Sanger Institute COVID-19 Surveillance Team |
| EPI_ISL_2804757 | Lighthouse Lab in Glasgow | Wellcome Sanger Institute for the COVID-19 Genomics UK (COG-UK) Consortium | Harper VanSteenhouse, Yumi Kasai, David Gray, Carol Clugston, Anna Dominiczak and Alex Alderton, Roberto Amato, Jeffrey Barrett, Sonia Goncalves, Ewan Harrison, David K. Jackson, Ian Johnston, Dominic Kwiatkowski, Cordelia Langford, John Sillitoe on behalf of the Wellcome Sanger Institute COVID-19 Surveillance Team |
| EPI_ISL_2804758 | Lighthouse Lab in Alderley Park | Wellcome Sanger Institute for the COVID-19 Genomics UK (COG-UK) Consortium | Jacquelyn Wynn, Mairead Hyland, The Lighthouse Lab in Alderley Park and Alex Alderton, Roberto Amato, Jeffrey Barrett, Sonia Goncalves, Ewan Harrison, David K. Jackson, Ian Johnston, Dominic Kwiatkowski, Cordelia Langford, John Sillitoe on behalf of the Wellcome Sanger Institute COVID-19 Surveillance Team |
| EPI_ISL_2804759 | Lighthouse Lab in Milton Keynes | Wellcome Sanger Institute for the COVID-19 Genomics UK (COG-UK) Consortium | The Lighthouse Lab in Milton Keynes and Alex Alderton, Roberto Amato, Jeffrey Barrett, Sonia Goncalves, Ewan Harrison, David K. Jackson, Ian Johnston, Dominic Kwiatkowski, Cordelia Langford, John Sillitoe on behalf of the Wellcome Sanger Institute COVID-19 Surveillance Team |
| EPI_ISL_2804760, EPI_ISL_2804761, EPI_ISL_2804762 | Lighthouse Lab in Alderley Park | Wellcome Sanger Institute for the COVID-19 Genomics UK (COG-UK) Consortium | Jacquelyn Wynn, Mairead Hyland, The Lighthouse Lab in Alderley Park and Alex Alderton, Roberto Amato, Jeffrey Barrett, Sonia Goncalves, Ewan Harrison, David K. Jackson, Ian Johnston, Dominic Kwiatkowski, Cordelia Langford, John Sillitoe on behalf of the Wellcome Sanger Institute COVID-19 Surveillance Team |
| EPI_ISL_2804763 | Lighthouse Lab in Milton Keynes | Wellcome Sanger Institute for the COVID-19 Genomics UK (COG-UK) Consortium | The Lighthouse Lab in Milton Keynes and Alex Alderton, Roberto Amato, Jeffrey Barrett, Sonia Goncalves, Ewan Harrison, David K. Jackson, Ian Johnston, Dominic Kwiatkowski, Cordelia Langford, John Sillitoe on behalf of the Wellcome Sanger Institute COVID-19 Surveillance Team |
| EPI_ISL_2804764, EPI_ISL_2804765, EPI_ISL_2804766, EPI_ISL_2804767, EPI_ISL_2804768, EPI_ISL_2804769 | Lighthouse Lab in Alderley Park | Wellcome Sanger Institute for the COVID-19 Genomics UK (COG-UK) Consortium | Jacquelyn Wynn, Mairead Hyland, The Lighthouse Lab in Alderley Park and Alex Alderton, Roberto Amato, Jeffrey Barrett, Sonia Goncalves, Ewan Harrison, David K. Jackson, Ian Johnston, Dominic Kwiatkowski, Cordelia Langford, John Sillitoe on behalf of the Wellcome Sanger Institute COVID-19 Surveillance Team |
| EPI_ISL_2804770 | Lighthouse Lab in Milton Keynes | Wellcome Sanger Institute for the COVID-19 Genomics UK (COG-UK) Consortium | The Lighthouse Lab in Milton Keynes and Alex Alderton, Roberto Amato, Jeffrey Barrett, Sonia Goncalves, Ewan Harrison, David K. Jackson, Ian Johnston, Dominic Kwiatkowski, Cordelia Langford, John Sillitoe on behalf of the Wellcome Sanger Institute COVID-19 Surveillance Team |
| EPI_ISL_2804771, EPI_ISL_2804772 | Lighthouse Lab in Alderley Park | Wellcome Sanger Institute for the COVID-19 Genomics UK (COG-UK) Consortium | Jacquelyn Wynn, Mairead Hyland, The Lighthouse Lab in Alderley Park and Alex Alderton, Roberto Amato, Jeffrey Barrett, Sonia Goncalves, Ewan Harrison, David K. Jackson, Ian Johnston, Dominic Kwiatkowski, Cordelia Langford, John Sillitoe on behalf of the Wellcome Sanger Institute COVID-19 Surveillance Team |
| EPI_ISL_2804773, EPI_ISL_2804774 | Lighthouse Lab in Milton Keynes | Wellcome Sanger Institute for the COVID-19 Genomics UK (COG-UK) Consortium | The Lighthouse Lab in Milton Keynes and Alex Alderton, Roberto Amato, Jeffrey Barrett, Sonia Goncalves, Ewan Harrison, David K. Jackson, Ian Johnston, Dominic Kwiatkowski, Cordelia Langford, John Sillitoe on behalf of the Wellcome Sanger Institute COVID-19 Surveillance Team |
| EPI_ISL_2804776, EPI_ISL_2804777 | Lighthouse Lab in Alderley Park | Wellcome Sanger Institute for the COVID-19 Genomics UK (COG-UK) Consortium | Jacquelyn Wynn, Mairead Hyland, The Lighthouse Lab in Alderley Park and Alex Alderton, Roberto Amato, Jeffrey Barrett, Sonia Goncalves, Ewan Harrison, David K. Jackson, Ian Johnston, Dominic Kwiatkowski, Cordelia Langford, John Sillitoe on behalf of the Wellcome Sanger Institute COVID-19 Surveillance Team |
| EPI_ISL_2804778 | Lighthouse Lab in Glasgow | Wellcome Sanger Institute for the COVID-19 Genomics UK (COG-UK) Consortium | Harper VanSteenhouse, Yumi Kasai, David Gray, Carol Clugston, Anna Dominiczak and Alex Alderton, Roberto Amato, Jeffrey Barrett, Sonia Goncalves, Ewan Harrison, David K. Jackson, Ian Johnston, Dominic Kwiatkowski, Cordelia Langford, John Sillitoe on behalf of the Wellcome Sanger Institute COVID-19 Surveillance Team |
| EPI_ISL_2804779, EPI_ISL_2804780, EPI_ISL_2804781, EPI_ISL_2804782, EPI_ISL_2804783, EPI_ISL_2804784, EPI_ISL_2804785, EPI_ISL_2804786, EPI_ISL_2804787 | Lighthouse Lab in Alderley Park | Wellcome Sanger Institute for the COVID-19 Genomics UK (COG-UK) Consortium | Jacquelyn Wynn, Mairead Hyland, The Lighthouse Lab in Alderley Park and Alex Alderton, Roberto Amato, Jeffrey Barrett, Sonia Goncalves, Ewan Harrison, David K. Jackson, Ian Johnston, Dominic Kwiatkowski, Cordelia Langford, John Sillitoe on behalf of the Wellcome Sanger Institute COVID-19 Surveillance Team |
| EPI_ISL_2804788 | Lighthouse Lab in Glasgow | Wellcome Sanger Institute for the COVID-19 Genomics UK (COG-UK) Consortium | Harper VanSteenhouse, Yumi Kasai, David Gray, Carol Clugston, Anna Dominiczak and Alex Alderton, Roberto Amato, Jeffrey Barrett, Sonia Goncalves, Ewan Harrison, David K. Jackson, Ian Johnston, Dominic Kwiatkowski, Cordelia Langford, John Sillitoe on behalf of the Wellcome Sanger Institute COVID-19 Surveillance Team |
| EPI_ISL_2804789 | Lighthouse Lab in Alderley Park | Wellcome Sanger Institute for the COVID-19 Genomics UK (COG-UK) Consortium | Jacquelyn Wynn, Mairead Hyland, The Lighthouse Lab in Alderley Park and Alex Alderton, Roberto Amato, Jeffrey Barrett, Sonia Goncalves, Ewan Harrison, David K. Jackson, Ian Johnston, Dominic Kwiatkowski, Cordelia Langford, John Sillitoe on behalf of the Wellcome Sanger Institute COVID-19 Surveillance Team |
| EPI_ISL_2804790 | Lighthouse Lab in Milton Keynes | Wellcome Sanger Institute for the COVID-19 Genomics UK (COG-UK) Consortium | The Lighthouse Lab in Milton Keynes and Alex Alderton, Roberto Amato, Jeffrey Barrett, Sonia Goncalves, Ewan Harrison, David K. Jackson, Ian Johnston, Dominic Kwiatkowski, Cordelia Langford, John Sillitoe on behalf of the Wellcome Sanger Institute COVID-19 Surveillance Team |
| EPI_ISL_2804791 | Lighthouse Lab in Glasgow | Wellcome Sanger Institute for the COVID-19 Genomics UK (COG-UK) Consortium | Harper VanSteenhouse, Yumi Kasai, David Gray, Carol Clugston, Anna Dominiczak and Alex Alderton, Roberto Amato, Jeffrey Barrett, Sonia Goncalves, Ewan Harrison, David K. Jackson, Ian Johnston, Dominic Kwiatkowski, Cordelia Langford, John Sillitoe on behalf of the Wellcome Sanger Institute COVID-19 Surveillance Team |
| EPI_ISL_2804792, EPI_ISL_2804793 | Lighthouse Lab in Alderley Park | Wellcome Sanger Institute for the COVID-19 Genomics UK (COG-UK) Consortium | Jacquelyn Wynn, Mairead Hyland, The Lighthouse Lab in Alderley Park and Alex Alderton, Roberto Amato, Jeffrey Barrett, Sonia Goncalves, Ewan Harrison, David K. Jackson, Ian Johnston, Dominic Kwiatkowski, Cordelia Langford, John Sillitoe on behalf of the Wellcome Sanger Institute COVID-19 Surveillance Team |
| EPI_ISL_2804794 | Lighthouse Lab in Milton Keynes | Wellcome Sanger Institute for the COVID-19 Genomics UK (COG-UK) Consortium | The Lighthouse Lab in Milton Keynes and Alex Alderton, Roberto Amato, Jeffrey Barrett, Sonia Goncalves, Ewan Harrison, David K. Jackson, Ian Johnston, Dominic Kwiatkowski, Cordelia Langford, John Sillitoe on behalf of the Wellcome Sanger Institute COVID-19 Surveillance Team |
| EPI_ISL_2804795, EPI_ISL_2804796 | Lighthouse Lab in Alderley Park | Wellcome Sanger Institute for the COVID-19 Genomics UK (COG-UK) Consortium | Jacquelyn Wynn, Mairead Hyland, The Lighthouse Lab in Alderley Park and Alex Alderton, Roberto Amato, Jeffrey Barrett, Sonia Goncalves, Ewan Harrison, David K. Jackson, Ian Johnston, Dominic Kwiatkowski, Cordelia Langford, John Sillitoe on behalf of the Wellcome Sanger Institute COVID-19 Surveillance Team |
| EPI_ISL_2804797, EPI_ISL_2804798 | Lighthouse Lab in Glasgow | Wellcome Sanger Institute for the COVID-19 Genomics UK (COG-UK) Consortium | Harper VanSteenhouse, Yumi Kasai, David Gray, Carol Clugston, Anna Dominiczak and Alex Alderton, Roberto Amato, Jeffrey Barrett, Sonia Goncalves, Ewan Harrison, David K. Jackson, Ian Johnston, Dominic Kwiatkowski, Cordelia Langford, John Sillitoe on behalf of the Wellcome Sanger Institute COVID-19 Surveillance Team |
| EPI_ISL_2804800, EPI_ISL_2804801 | Lighthouse Lab in Alderley Park | Wellcome Sanger Institute for the COVID-19 Genomics UK (COG-UK) Consortium | Jacquelyn Wynn, Mairead Hyland, The Lighthouse Lab in Alderley Park and Alex Alderton, Roberto Amato, Jeffrey Barrett, Sonia Goncalves, Ewan Harrison, David K. Jackson, Ian Johnston, Dominic Kwiatkowski, Cordelia Langford, John Sillitoe on behalf of the Wellcome Sanger Institute COVID-19 Surveillance Team |
| EPI_ISL_2804802 | Lighthouse Lab in Glasgow | Wellcome Sanger Institute for the COVID-19 Genomics UK (COG-UK) Consortium | Harper VanSteenhouse, Yumi Kasai, David Gray, Carol Clugston, Anna Dominiczak and Alex Alderton, Roberto Amato, Jeffrey Barrett, Sonia Goncalves, Ewan Harrison, David K. Jackson, Ian Johnston, Dominic Kwiatkowski, Cordelia Langford, John Sillitoe on behalf of the Wellcome Sanger Institute COVID-19 Surveillance Team |
| EPI_ISL_2804803 | Lighthouse Lab in Alderley Park | Wellcome Sanger Institute for the COVID-19 Genomics UK (COG-UK) Consortium | Jacquelyn Wynn, Mairead Hyland, The Lighthouse Lab in Alderley Park and Alex Alderton, Roberto Amato, Jeffrey Barrett, Sonia Goncalves, Ewan Harrison, David K. Jackson, Ian Johnston, Dominic Kwiatkowski, Cordelia Langford, John Sillitoe on behalf of the Wellcome Sanger Institute COVID-19 Surveillance Team |
| EPI_ISL_2804804, EPI_ISL_2804805 | Lighthouse Lab in Milton Keynes | Wellcome Sanger Institute for the COVID-19 Genomics UK (COG-UK) Consortium | The Lighthouse Lab in Milton Keynes and Alex Alderton, Roberto Amato, Jeffrey Barrett, Sonia Goncalves, Ewan Harrison, David K. Jackson, Ian Johnston, Dominic Kwiatkowski, Cordelia Langford, John Sillitoe on behalf of the Wellcome Sanger Institute COVID-19 Surveillance Team |
| EPI_ISL_2804806, EPI_ISL_2804807 | Lighthouse Lab in Glasgow | Wellcome Sanger Institute for the COVID-19 Genomics UK (COG-UK) Consortium | Harper VanSteenhouse, Yumi Kasai, David Gray, Carol Clugston, Anna Dominiczak and Alex Alderton, Roberto Amato, Jeffrey Barrett, Sonia Goncalves, Ewan Harrison, David K. Jackson, Ian Johnston, Dominic Kwiatkowski, Cordelia Langford, John Sillitoe on behalf of the Wellcome Sanger Institute COVID-19 Surveillance Team |

| | | | |
|---|---|---|---|
| EPI_ISL_2804808 | Lighthouse Lab in Milton Keynes | Wellcome Sanger Institute for the COVID-19 Genomics UK (COG-UK) Consortium | The Lighthouse Lab in Milton Keynes and Alex Alderton, Roberto Amato, Jeffrey Barrett, Sonia Goncalves, Ewan Harrison, David K. Jackson, Ian Johnston, Dominic Kwiatkowski, Cordelia Langford, John Sillitoe on behalf of the Wellcome Sanger Institute COVID-19 Surveillance Team |
| EPI_ISL_2804809 | Lighthouse Lab in Alderley Park | Wellcome Sanger Institute for the COVID-19 Genomics UK (COG-UK) Consortium | Jacquelyn Wynn, Mairead Hyland, The Lighthouse Lab in Alderley Park and Alex Alderton, Roberto Amato, Jeffrey Barrett, Sonia Goncalves, Ewan Harrison, David K. Jackson, Ian Johnston, Dominic Kwiatkowski, Cordelia Langford, John Sillitoe on behalf of the Wellcome Sanger Institute COVID-19 Surveillance Team |
| EPI_ISL_2804810 | Lighthouse Lab in Milton Keynes | Wellcome Sanger Institute for the COVID-19 Genomics UK (COG-UK) Consortium | The Lighthouse Lab in Milton Keynes and Alex Alderton, Roberto Amato, Jeffrey Barrett, Sonia Goncalves, Ewan Harrison, David K. Jackson, Ian Johnston, Dominic Kwiatkowski, Cordelia Langford, John Sillitoe on behalf of the Wellcome Sanger Institute COVID-19 Surveillance Team |
| EPI_ISL_2804811, EPI_ISL_2804812, EPI_ISL_2804813, EPI_ISL_2804814, EPI_ISL_2804815 | Lighthouse Lab in Alderley Park | Wellcome Sanger Institute for the COVID-19 Genomics UK (COG-UK) Consortium | Jacquelyn Wynn, Mairead Hyland, The Lighthouse Lab in Alderley Park and Alex Alderton, Roberto Amato, Jeffrey Barrett, Sonia Goncalves, Ewan Harrison, David K. Jackson, Ian Johnston, Dominic Kwiatkowski, Cordelia Langford, John Sillitoe on behalf of the Wellcome Sanger Institute COVID-19 Surveillance Team |
| EPI_ISL_2804816 | Lighthouse Lab in Glasgow | Wellcome Sanger Institute for the COVID-19 Genomics UK (COG-UK) Consortium | Harper VanSteenhouse, Yumi Kasai, David Gray, Carol Clugston, Anna Dominiczak and Alex Alderton, Roberto Amato, Jeffrey Barrett, Sonia Goncalves, Ewan Harrison, David K. Jackson, Ian Johnston, Dominic Kwiatkowski, Cordelia Langford, John Sillitoe on behalf of the Wellcome Sanger Institute COVID-19 Surveillance Team |
| EPI_ISL_2804817 | Lighthouse Lab in Alderley Park | Wellcome Sanger Institute for the COVID-19 Genomics UK (COG-UK) Consortium | Jacquelyn Wynn, Mairead Hyland, The Lighthouse Lab in Alderley Park and Alex Alderton, Roberto Amato, Jeffrey Barrett, Sonia Goncalves, Ewan Harrison, David K. Jackson, Ian Johnston, Dominic Kwiatkowski, Cordelia Langford, John Sillitoe on behalf of the Wellcome Sanger Institute COVID-19 Surveillance Team |
| EPI_ISL_2804818 | Lighthouse Lab in Glasgow | Wellcome Sanger Institute for the COVID-19 Genomics UK (COG-UK) Consortium | Harper VanSteenhouse, Yumi Kasai, David Gray, Carol Clugston, Anna Dominiczak and Alex Alderton, Roberto Amato, Jeffrey Barrett, Sonia Goncalves, Ewan Harrison, David K. Jackson, Ian Johnston, Dominic Kwiatkowski, Cordelia Langford, John Sillitoe on behalf of the Wellcome Sanger Institute COVID-19 Surveillance Team |
| EPI_ISL_2804819 | Lighthouse Lab in Alderley Park | Wellcome Sanger Institute for the COVID-19 Genomics UK (COG-UK) Consortium | Jacquelyn Wynn, Mairead Hyland, The Lighthouse Lab in Alderley Park and Alex Alderton, Roberto Amato, Jeffrey Barrett, Sonia Goncalves, Ewan Harrison, David K. Jackson, Ian Johnston, Dominic Kwiatkowski, Cordelia Langford, John Sillitoe on behalf of the Wellcome Sanger Institute COVID-19 Surveillance Team |
| EPI_ISL_2804820 | Lighthouse Lab in Milton Keynes | Wellcome Sanger Institute for the COVID-19 Genomics UK (COG-UK) Consortium | The Lighthouse Lab in Milton Keynes and Alex Alderton, Roberto Amato, Jeffrey Barrett, Sonia Goncalves, Ewan Harrison, David K. Jackson, Ian Johnston, Dominic Kwiatkowski, Cordelia Langford, John Sillitoe on behalf of the Wellcome Sanger Institute COVID-19 Surveillance Team |
| EPI_ISL_2804821, EPI_ISL_2804822 | Lighthouse Lab in Alderley Park | Wellcome Sanger Institute for the COVID-19 Genomics UK (COG-UK) Consortium | Jacquelyn Wynn, Mairead Hyland, The Lighthouse Lab in Alderley Park and Alex Alderton, Roberto Amato, Jeffrey Barrett, Sonia Goncalves, Ewan Harrison, David K. Jackson, Ian Johnston, Dominic Kwiatkowski, Cordelia Langford, John Sillitoe on behalf of the Wellcome Sanger Institute COVID-19 Surveillance Team |
| EPI_ISL_2804823 | Lighthouse Lab in Milton Keynes | Wellcome Sanger Institute for the COVID-19 Genomics UK (COG-UK) Consortium | The Lighthouse Lab in Milton Keynes and Alex Alderton, Roberto Amato, Jeffrey Barrett, Sonia Goncalves, Ewan Harrison, David K. Jackson, Ian Johnston, Dominic Kwiatkowski, Cordelia Langford, John Sillitoe on behalf of the Wellcome Sanger Institute COVID-19 Surveillance Team |
| EPI_ISL_2804824, EPI_ISL_2804825, EPI_ISL_2804826 | Lighthouse Lab in Alderley Park | Wellcome Sanger Institute for the COVID-19 Genomics UK (COG-UK) Consortium | Jacquelyn Wynn, Mairead Hyland, The Lighthouse Lab in Alderley Park and Alex Alderton, Roberto Amato, Jeffrey Barrett, Sonia Goncalves, Ewan Harrison, David K. Jackson, Ian Johnston, Dominic Kwiatkowski, Cordelia Langford, John Sillitoe on behalf of the Wellcome Sanger Institute COVID-19 Surveillance Team |
| EPI_ISL_2804827 | Lighthouse Lab in Glasgow | Wellcome Sanger Institute for the COVID-19 Genomics UK (COG-UK) Consortium | Harper VanSteenhouse, Yumi Kasai, David Gray, Carol Clugston, Anna Dominiczak and Alex Alderton, Roberto Amato, Jeffrey Barrett, Sonia Goncalves, Ewan Harrison, David K. Jackson, Ian Johnston, Dominic Kwiatkowski, Cordelia Langford, John Sillitoe on behalf of the Wellcome Sanger Institute COVID-19 Surveillance Team |
| EPI_ISL_2804828 | Lighthouse Lab in Alderley Park | Wellcome Sanger Institute for the COVID-19 Genomics UK (COG-UK) Consortium | Jacquelyn Wynn, Mairead Hyland, The Lighthouse Lab in Alderley Park and Alex Alderton, Roberto Amato, Jeffrey Barrett, Sonia Goncalves, Ewan Harrison, David K. Jackson, Ian Johnston, Dominic Kwiatkowski, Cordelia Langford, John Sillitoe on behalf of the Wellcome Sanger Institute COVID-19 Surveillance Team |
| EPI_ISL_2804829 | Lighthouse Lab in Milton Keynes | Wellcome Sanger Institute for the COVID-19 Genomics UK (COG-UK) Consortium | The Lighthouse Lab in Milton Keynes and Alex Alderton, Roberto Amato, Jeffrey Barrett, Sonia Goncalves, Ewan Harrison, David K. Jackson, Ian Johnston, Dominic Kwiatkowski, Cordelia Langford, John Sillitoe on behalf of the Wellcome Sanger Institute COVID-19 Surveillance Team |
| EPI_ISL_2804830 | Lighthouse Lab in Alderley Park | Wellcome Sanger Institute for the COVID-19 Genomics UK (COG-UK) Consortium | Jacquelyn Wynn, Mairead Hyland, The Lighthouse Lab in Alderley Park and Alex Alderton, Roberto Amato, Jeffrey Barrett, Sonia Goncalves, Ewan Harrison, David K. Jackson, Ian Johnston, Dominic Kwiatkowski, Cordelia Langford, John Sillitoe on behalf of the Wellcome Sanger Institute COVID-19 Surveillance Team |
| EPI_ISL_2804831 | Lighthouse Lab in Milton Keynes | Wellcome Sanger Institute for the COVID-19 Genomics UK (COG-UK) Consortium | The Lighthouse Lab in Milton Keynes and Alex Alderton, Roberto Amato, Jeffrey Barrett, Sonia Goncalves, Ewan Harrison, David K. Jackson, Ian Johnston, Dominic Kwiatkowski, Cordelia Langford, John Sillitoe on behalf of the Wellcome Sanger Institute COVID-19 Surveillance Team |
| EPI_ISL_2804832, EPI_ISL_2804833 | Lighthouse Lab in Alderley Park | Wellcome Sanger Institute for the COVID-19 Genomics UK (COG-UK) Consortium | Jacquelyn Wynn, Mairead Hyland, The Lighthouse Lab in Alderley Park and Alex Alderton, Roberto Amato, Jeffrey Barrett, Sonia Goncalves, Ewan Harrison, David K. Jackson, Ian Johnston, Dominic Kwiatkowski, Cordelia Langford, John Sillitoe on behalf of the Wellcome Sanger Institute COVID-19 Surveillance Team |
| EPI_ISL_2804834 | Lighthouse Lab in Glasgow | Wellcome Sanger Institute for the COVID-19 Genomics UK (COG-UK) Consortium | Harper VanSteenhouse, Yumi Kasai, David Gray, Carol Clugston, Anna Dominiczak and Alex Alderton, Roberto Amato, Jeffrey Barrett, Sonia Goncalves, Ewan Harrison, David K. Jackson, Ian Johnston, Dominic Kwiatkowski, Cordelia Langford, John Sillitoe on behalf of the Wellcome Sanger Institute COVID-19 Surveillance Team |
| EPI_ISL_2804835, EPI_ISL_2804836, EPI_ISL_2804837, EPI_ISL_2804838, EPI_ISL_2804840, EPI_ISL_2804841 | Lighthouse Lab in Alderley Park | Wellcome Sanger Institute for the COVID-19 Genomics UK (COG-UK) Consortium | Jacquelyn Wynn, Mairead Hyland, The Lighthouse Lab in Alderley Park and Alex Alderton, Roberto Amato, Jeffrey Barrett, Sonia Goncalves, Ewan Harrison, David K. Jackson, Ian Johnston, Dominic Kwiatkowski, Cordelia Langford, John Sillitoe on behalf of the Wellcome Sanger Institute COVID-19 Surveillance Team |
| EPI_ISL_2804842 | Lighthouse Lab in Glasgow | Wellcome Sanger Institute for the COVID-19 Genomics UK (COG-UK) Consortium | Harper VanSteenhouse, Yumi Kasai, David Gray, Carol Clugston, Anna Dominiczak and Alex Alderton, Roberto Amato, Jeffrey Barrett, Sonia Goncalves, Ewan Harrison, David K. Jackson, Ian Johnston, Dominic Kwiatkowski, Cordelia Langford, John Sillitoe on behalf of the Wellcome Sanger Institute COVID-19 Surveillance Team |
| EPI_ISL_2804843 | Lighthouse Lab in Alderley Park | Wellcome Sanger Institute for the COVID-19 Genomics UK (COG-UK) Consortium | Jacquelyn Wynn, Mairead Hyland, The Lighthouse Lab in Alderley Park and Alex Alderton, Roberto Amato, Jeffrey Barrett, Sonia Goncalves, Ewan Harrison, David K. Jackson, Ian Johnston, Dominic Kwiatkowski, Cordelia Langford, John Sillitoe on behalf of the Wellcome Sanger Institute COVID-19 Surveillance Team |
| EPI_ISL_2804844 | Lighthouse Lab in Milton Keynes | Wellcome Sanger Institute for the COVID-19 Genomics UK (COG-UK) Consortium | The Lighthouse Lab in Milton Keynes and Alex Alderton, Roberto Amato, Jeffrey Barrett, Sonia Goncalves, Ewan Harrison, David K. Jackson, Ian Johnston, Dominic Kwiatkowski, Cordelia Langford, John Sillitoe on behalf of the Wellcome Sanger Institute COVID-19 Surveillance Team |
| EPI_ISL_2804845, EPI_ISL_2804846, EPI_ISL_2804847, EPI_ISL_2804848, EPI_ISL_2804849 | Lighthouse Lab in Alderley Park | Wellcome Sanger Institute for the COVID-19 Genomics UK (COG-UK) Consortium | Jacquelyn Wynn, Mairead Hyland, The Lighthouse Lab in Alderley Park and Alex Alderton, Roberto Amato, Jeffrey Barrett, Sonia Goncalves, Ewan Harrison, David K. Jackson, Ian Johnston, Dominic Kwiatkowski, Cordelia Langford, John Sillitoe on behalf of the Wellcome Sanger Institute COVID-19 Surveillance Team |
| EPI_ISL_2804850 | Lighthouse Lab in Glasgow | Wellcome Sanger Institute for the COVID-19 Genomics UK (COG-UK) Consortium | Harper VanSteenhouse, Yumi Kasai, David Gray, Carol Clugston, Anna Dominiczak and Alex Alderton, Roberto Amato, Jeffrey Barrett, Sonia Goncalves, Ewan Harrison, David K. Jackson, Ian Johnston, Dominic Kwiatkowski, Cordelia Langford, John Sillitoe on behalf of the Wellcome Sanger Institute COVID-19 Surveillance Team |
| EPI_ISL_2804851, EPI_ISL_2804852 | Lighthouse Lab in Alderley Park | Wellcome Sanger Institute for the COVID-19 Genomics UK (COG-UK) Consortium | Jacquelyn Wynn, Mairead Hyland, The Lighthouse Lab in Alderley Park and Alex Alderton, Roberto Amato, Jeffrey Barrett, Sonia Goncalves, Ewan Harrison, David K. Jackson, Ian Johnston, Dominic Kwiatkowski, Cordelia Langford, John Sillitoe on behalf of the Wellcome Sanger Institute COVID-19 Surveillance Team |
| EPI_ISL_2804853 | Lighthouse Lab in Milton Keynes | Wellcome Sanger Institute for the COVID-19 Genomics UK (COG-UK) Consortium | The Lighthouse Lab in Milton Keynes and Alex Alderton, Roberto Amato, Jeffrey Barrett, Sonia Goncalves, Ewan Harrison, David K. Jackson, Ian Johnston, Dominic Kwiatkowski, Cordelia Langford, John Sillitoe on behalf of the Wellcome Sanger Institute COVID-19 Surveillance Team |
| EPI_ISL_2804854, EPI_ISL_2804855 | Lighthouse Lab in Alderley Park | Wellcome Sanger Institute for the COVID-19 Genomics UK (COG-UK) Consortium | Jacquelyn Wynn, Mairead Hyland, The Lighthouse Lab in Alderley Park and Alex Alderton, Roberto Amato, Jeffrey Barrett, Sonia Goncalves, Ewan Harrison, David K. Jackson, Ian Johnston, Dominic Kwiatkowski, Cordelia Langford, John Sillitoe on behalf of the Wellcome Sanger Institute COVID-19 Surveillance Team |
| EPI_ISL_2804856 | Lighthouse Lab in Milton Keynes | Wellcome Sanger Institute for the COVID-19 Genomics UK (COG-UK) Consortium | The Lighthouse Lab in Milton Keynes and Alex Alderton, Roberto Amato, Jeffrey Barrett, Sonia Goncalves, Ewan Harrison, David K. Jackson, Ian Johnston, Dominic Kwiatkowski, Cordelia Langford, John Sillitoe on behalf of the Wellcome Sanger Institute COVID-19 Surveillance Team |
| EPI_ISL_2804857, EPI_ISL_2804858, EPI_ISL_2804859, EPI_ISL_2804860, EPI_ISL_2804861, EPI_ISL_2804862, EPI_ISL_2804863, EPI_ISL_2804864 | Lighthouse Lab in Alderley Park | Wellcome Sanger Institute for the COVID-19 Genomics UK (COG-UK) Consortium | Jacquelyn Wynn, Mairead Hyland, The Lighthouse Lab in Alderley Park and Alex Alderton, Roberto Amato, Jeffrey Barrett, Sonia Goncalves, Ewan Harrison, David K. Jackson, Ian Johnston, Dominic Kwiatkowski, Cordelia Langford, John Sillitoe on behalf of the Wellcome Sanger Institute COVID-19 Surveillance Team |
| EPI_ISL_2804865 | Lighthouse Lab in Milton Keynes | Wellcome Sanger Institute for the COVID-19 Genomics UK (COG-UK) Consortium | The Lighthouse Lab in Milton Keynes and Alex Alderton, Roberto Amato, Jeffrey Barrett, Sonia Goncalves, Ewan Harrison, David K. Jackson, Ian Johnston, Dominic Kwiatkowski, Cordelia Langford, John Sillitoe on behalf of the Wellcome Sanger Institute COVID-19 Surveillance Team |
| EPI_ISL_2804866, EPI_ISL_2804867, EPI_ISL_2804868 | Lighthouse Lab in Alderley Park | Wellcome Sanger Institute for the COVID-19 Genomics UK (COG-UK) Consortium | Jacquelyn Wynn, Mairead Hyland, The Lighthouse Lab in Alderley Park and Alex Alderton, Roberto Amato, Jeffrey Barrett, Sonia Goncalves, Ewan Harrison, David K. Jackson, Ian Johnston, Dominic Kwiatkowski, Cordelia Langford, John Sillitoe on behalf of the Wellcome Sanger Institute COVID-19 Surveillance Team |
| EPI_ISL_2804869 | Lighthouse Lab in Milton Keynes | Wellcome Sanger Institute for the COVID-19 Genomics UK (COG-UK) Consortium | The Lighthouse Lab in Milton Keynes and Alex Alderton, Roberto Amato, Jeffrey Barrett, Sonia Goncalves, Ewan Harrison, David K. Jackson, Ian Johnston, Dominic Kwiatkowski, Cordelia Langford, John Sillitoe on behalf of the Wellcome Sanger Institute COVID-19 Surveillance Team |
| EPI_ISL_2804870, EPI_ISL_2804871 | Lighthouse Lab in Alderley Park | Wellcome Sanger Institute for the COVID-19 Genomics UK (COG-UK) Consortium | Jacquelyn Wynn, Mairead Hyland, The Lighthouse Lab in Alderley Park and Alex Alderton, Roberto Amato, Jeffrey Barrett, Sonia Goncalves, Ewan Harrison, David K. Jackson, Ian Johnston, Dominic Kwiatkowski, Cordelia Langford, John Sillitoe on behalf of the Wellcome Sanger Institute COVID-19 Surveillance Team |
| EPI_ISL_2804872 | Lighthouse Lab in Milton Keynes | Wellcome Sanger Institute for the COVID-19 Genomics UK (COG-UK) Consortium | The Lighthouse Lab in Milton Keynes and Alex Alderton, Roberto Amato, Jeffrey Barrett, Sonia Goncalves, Ewan Harrison, David K. Jackson, Ian Johnston, Dominic Kwiatkowski, Cordelia Langford, John Sillitoe on behalf of the Wellcome Sanger Institute COVID-19 Surveillance Team |
| EPI_ISL_2804873 | Lighthouse Lab in Glasgow | Wellcome Sanger Institute for the COVID-19 Genomics UK (COG-UK) Consortium | Harper VanSteenhouse, Yumi Kasai, David Gray, Carol Clugston, Anna Dominiczak and Alex Alderton, Roberto Amato, Jeffrey Barrett, Sonia Goncalves, Ewan Harrison, David K. Jackson, Ian Johnston, Dominic Kwiatkowski, Cordelia Langford, John Sillitoe on behalf of the Wellcome Sanger Institute COVID-19 Surveillance Team |
| EPI_ISL_2804874 | Lighthouse Lab in Alderley Park | Wellcome Sanger Institute for the COVID-19 Genomics UK (COG-UK) Consortium | Jacquelyn Wynn, Mairead Hyland, The Lighthouse Lab in Alderley Park and Alex Alderton, Roberto Amato, Jeffrey Barrett, Sonia Goncalves, Ewan Harrison, David K. Jackson, Ian Johnston, Dominic Kwiatkowski, Cordelia Langford, John Sillitoe on behalf of the Wellcome Sanger Institute COVID-19 Surveillance Team |
| EPI_ISL_2804875 | Lighthouse Lab in Glasgow | Wellcome Sanger Institute for the COVID-19 Genomics UK (COG-UK) Consortium | Harper VanSteenhouse, Yumi Kasai, David Gray, Carol Clugston, Anna Dominiczak and Alex Alderton, Roberto Amato, Jeffrey Barrett, Sonia Goncalves, Ewan Harrison, David K. Jackson, Ian Johnston, Dominic Kwiatkowski, Cordelia Langford, John Sillitoe on behalf of the Wellcome Sanger Institute COVID-19 Surveillance Team |

| | | | |
|---|---|---|---|
| EPI_ISL_2804876 | Lighthouse Lab in Milton Keynes | Wellcome Sanger Institute for the COVID-19 Genomics UK (COG-UK) Consortium | The Lighthouse Lab in Milton Keynes and Alex Alderton, Roberto Amato, Jeffrey Barrett, Sonia Goncalves, Ewan Harrison, David K. Jackson, Ian Johnston, Dominic Kwiatkowski, Cordelia Langford, John Sillitoe on behalf of the Wellcome Sanger Institute COVID-19 Surveillance Team |
| EPI_ISL_2804877 | Lighthouse Lab in Glasgow | Wellcome Sanger Institute for the COVID-19 Genomics UK (COG-UK) Consortium | Harper VanSteenhouse, Yumi Kasai, David Gray, Carol Clugston, Anna Dominiczak and Alex Alderton, Roberto Amato, Jeffrey Barrett, Sonia Goncalves, Ewan Harrison, David K. Jackson, Ian Johnston, Dominic Kwiatkowski, Cordelia Langford, John Sillitoe on behalf of the Wellcome Sanger Institute COVID-19 Surveillance Team |
| EPI_ISL_2804878, EPI_ISL_2804879, EPI_ISL_2804880, EPI_ISL_2804881, EPI_ISL_2804882, EPI_ISL_2804883, EPI_ISL_2804884 | Lighthouse Lab in Alderley Park | Wellcome Sanger Institute for the COVID-19 Genomics UK (COG-UK) Consortium | Jacquelyn Wynn, Mairead Hyland, The Lighthouse Lab in Alderley Park and Alex Alderton, Roberto Amato, Jeffrey Barrett, Sonia Goncalves, Ewan Harrison, David K. Jackson, Ian Johnston, Dominic Kwiatkowski, Cordelia Langford, John Sillitoe on behalf of the Wellcome Sanger Institute COVID-19 Surveillance Team |
| EPI_ISL_2804885 | Lighthouse Lab in Milton Keynes | Wellcome Sanger Institute for the COVID-19 Genomics UK (COG-UK) Consortium | The Lighthouse Lab in Milton Keynes and Alex Alderton, Roberto Amato, Jeffrey Barrett, Sonia Goncalves, Ewan Harrison, David K. Jackson, Ian Johnston, Dominic Kwiatkowski, Cordelia Langford, John Sillitoe on behalf of the Wellcome Sanger Institute COVID-19 Surveillance Team |
| EPI_ISL_2804886, EPI_ISL_2804887, EPI_ISL_2804888 | Lighthouse Lab in Alderley Park | Wellcome Sanger Institute for the COVID-19 Genomics UK (COG-UK) Consortium | Jacquelyn Wynn, Mairead Hyland, The Lighthouse Lab in Alderley Park and Alex Alderton, Roberto Amato, Jeffrey Barrett, Sonia Goncalves, Ewan Harrison, David K. Jackson, Ian Johnston, Dominic Kwiatkowski, Cordelia Langford, John Sillitoe on behalf of the Wellcome Sanger Institute COVID-19 Surveillance Team |
| EPI_ISL_2804889, EPI_ISL_2804890 | Lighthouse Lab in Milton Keynes | Wellcome Sanger Institute for the COVID-19 Genomics UK (COG-UK) Consortium | The Lighthouse Lab in Milton Keynes and Alex Alderton, Roberto Amato, Jeffrey Barrett, Sonia Goncalves, Ewan Harrison, David K. Jackson, Ian Johnston, Dominic Kwiatkowski, Cordelia Langford, John Sillitoe on behalf of the Wellcome Sanger Institute COVID-19 Surveillance Team |
| EPI_ISL_2804891 | Lighthouse Lab in Glasgow | Wellcome Sanger Institute for the COVID-19 Genomics UK (COG-UK) Consortium | Harper VanSteenhouse, Yumi Kasai, David Gray, Carol Clugston, Anna Dominiczak and Alex Alderton, Roberto Amato, Jeffrey Barrett, Sonia Goncalves, Ewan Harrison, David K. Jackson, Ian Johnston, Dominic Kwiatkowski, Cordelia Langford, John Sillitoe on behalf of the Wellcome Sanger Institute COVID-19 Surveillance Team |
| EPI_ISL_2804892 | Lighthouse Lab in Milton Keynes | Wellcome Sanger Institute for the COVID-19 Genomics UK (COG-UK) Consortium | The Lighthouse Lab in Milton Keynes and Alex Alderton, Roberto Amato, Jeffrey Barrett, Sonia Goncalves, Ewan Harrison, David K. Jackson, Ian Johnston, Dominic Kwiatkowski, Cordelia Langford, John Sillitoe on behalf of the Wellcome Sanger Institute COVID-19 Surveillance Team |
| EPI_ISL_2804893, EPI_ISL_2804894 | Lighthouse Lab in Alderley Park | Wellcome Sanger Institute for the COVID-19 Genomics UK (COG-UK) Consortium | Jacquelyn Wynn, Mairead Hyland, The Lighthouse Lab in Alderley Park and Alex Alderton, Roberto Amato, Jeffrey Barrett, Sonia Goncalves, Ewan Harrison, David K. Jackson, Ian Johnston, Dominic Kwiatkowski, Cordelia Langford, John Sillitoe on behalf of the Wellcome Sanger Institute COVID-19 Surveillance Team |
| EPI_ISL_2804895 | Lighthouse Lab in Glasgow | Wellcome Sanger Institute for the COVID-19 Genomics UK (COG-UK) Consortium | Harper VanSteenhouse, Yumi Kasai, David Gray, Carol Clugston, Anna Dominiczak and Alex Alderton, Roberto Amato, Jeffrey Barrett, Sonia Goncalves, Ewan Harrison, David K. Jackson, Ian Johnston, Dominic Kwiatkowski, Cordelia Langford, John Sillitoe on behalf of the Wellcome Sanger Institute COVID-19 Surveillance Team |
| EPI_ISL_2804896 | Lighthouse Lab in Milton Keynes | Wellcome Sanger Institute for the COVID-19 Genomics UK (COG-UK) Consortium | The Lighthouse Lab in Milton Keynes and Alex Alderton, Roberto Amato, Jeffrey Barrett, Sonia Goncalves, Ewan Harrison, David K. Jackson, Ian Johnston, Dominic Kwiatkowski, Cordelia Langford, John Sillitoe on behalf of the Wellcome Sanger Institute COVID-19 Surveillance Team |
| EPI_ISL_2804897, EPI_ISL_2804898 | Lighthouse Lab in Alderley Park | Wellcome Sanger Institute for the COVID-19 Genomics UK (COG-UK) Consortium | Jacquelyn Wynn, Mairead Hyland, The Lighthouse Lab in Alderley Park and Alex Alderton, Roberto Amato, Jeffrey Barrett, Sonia Goncalves, Ewan Harrison, David K. Jackson, Ian Johnston, Dominic Kwiatkowski, Cordelia Langford, John Sillitoe on behalf of the Wellcome Sanger Institute COVID-19 Surveillance Team |
| EPI_ISL_2804899 | Lighthouse Lab in Glasgow | Wellcome Sanger Institute for the COVID-19 Genomics UK (COG-UK) Consortium | Harper VanSteenhouse, Yumi Kasai, David Gray, Carol Clugston, Anna Dominiczak and Alex Alderton, Roberto Amato, Jeffrey Barrett, Sonia Goncalves, Ewan Harrison, David K. Jackson, Ian Johnston, Dominic Kwiatkowski, Cordelia Langford, John Sillitoe on behalf of the Wellcome Sanger Institute COVID-19 Surveillance Team |
| EPI_ISL_2804900 | Lighthouse Lab in Alderley Park | Wellcome Sanger Institute for the COVID-19 Genomics UK (COG-UK) Consortium | Jacquelyn Wynn, Mairead Hyland, The Lighthouse Lab in Alderley Park and Alex Alderton, Roberto Amato, Jeffrey Barrett, Sonia Goncalves, Ewan Harrison, David K. Jackson, Ian Johnston, Dominic Kwiatkowski, Cordelia Langford, John Sillitoe on behalf of the Wellcome Sanger Institute COVID-19 Surveillance Team |
| EPI_ISL_2804901, EPI_ISL_2804902 | Lighthouse Lab in Milton Keynes | Wellcome Sanger Institute for the COVID-19 Genomics UK (COG-UK) Consortium | The Lighthouse Lab in Milton Keynes and Alex Alderton, Roberto Amato, Jeffrey Barrett, Sonia Goncalves, Ewan Harrison, David K. Jackson, Ian Johnston, Dominic Kwiatkowski, Cordelia Langford, John Sillitoe on behalf of the Wellcome Sanger Institute COVID-19 Surveillance Team |
| EPI_ISL_2804903, EPI_ISL_2804904 | Lighthouse Lab in Alderley Park | Wellcome Sanger Institute for the COVID-19 Genomics UK (COG-UK) Consortium | Jacquelyn Wynn, Mairead Hyland, The Lighthouse Lab in Alderley Park and Alex Alderton, Roberto Amato, Jeffrey Barrett, Sonia Goncalves, Ewan Harrison, David K. Jackson, Ian Johnston, Dominic Kwiatkowski, Cordelia Langford, John Sillitoe on behalf of the Wellcome Sanger Institute COVID-19 Surveillance Team |
| EPI_ISL_2804905 | Lighthouse Lab in Milton Keynes | Wellcome Sanger Institute for the COVID-19 Genomics UK (COG-UK) Consortium | The Lighthouse Lab in Milton Keynes and Alex Alderton, Roberto Amato, Jeffrey Barrett, Sonia Goncalves, Ewan Harrison, David K. Jackson, Ian Johnston, Dominic Kwiatkowski, Cordelia Langford, John Sillitoe on behalf of the Wellcome Sanger Institute COVID-19 Surveillance Team |
| EPI_ISL_2804906 | Lighthouse Lab in Alderley Park | Wellcome Sanger Institute for the COVID-19 Genomics UK (COG-UK) Consortium | Jacquelyn Wynn, Mairead Hyland, The Lighthouse Lab in Alderley Park and Alex Alderton, Roberto Amato, Jeffrey Barrett, Sonia Goncalves, Ewan Harrison, David K. Jackson, Ian Johnston, Dominic Kwiatkowski, Cordelia Langford, John Sillitoe on behalf of the Wellcome Sanger Institute COVID-19 Surveillance Team |
| EPI_ISL_2804907, EPI_ISL_2804908 | Lighthouse Lab in Milton Keynes | Wellcome Sanger Institute for the COVID-19 Genomics UK (COG-UK) Consortium | The Lighthouse Lab in Milton Keynes and Alex Alderton, Roberto Amato, Jeffrey Barrett, Sonia Goncalves, Ewan Harrison, David K. Jackson, Ian Johnston, Dominic Kwiatkowski, Cordelia Langford, John Sillitoe on behalf of the Wellcome Sanger Institute COVID-19 Surveillance Team |
| EPI_ISL_2804909 | Lighthouse Lab in Glasgow | Wellcome Sanger Institute for the COVID-19 Genomics UK (COG-UK) Consortium | Harper VanSteenhouse, Yumi Kasai, David Gray, Carol Clugston, Anna Dominiczak and Alex Alderton, Roberto Amato, Jeffrey Barrett, Sonia Goncalves, Ewan Harrison, David K. Jackson, Ian Johnston, Dominic Kwiatkowski, Cordelia Langford, John Sillitoe on behalf of the Wellcome Sanger Institute COVID-19 Surveillance Team |
| EPI_ISL_2804910 | Lighthouse Lab in Milton Keynes | Wellcome Sanger Institute for the COVID-19 Genomics UK (COG-UK) Consortium | The Lighthouse Lab in Milton Keynes and Alex Alderton, Roberto Amato, Jeffrey Barrett, Sonia Goncalves, Ewan Harrison, David K. Jackson, Ian Johnston, Dominic Kwiatkowski, Cordelia Langford, John Sillitoe on behalf of the Wellcome Sanger Institute COVID-19 Surveillance Team |
| EPI_ISL_2804911 | Lighthouse Lab in Glasgow | Wellcome Sanger Institute for the COVID-19 Genomics UK (COG-UK) Consortium | Harper VanSteenhouse, Yumi Kasai, David Gray, Carol Clugston, Anna Dominiczak and Alex Alderton, Roberto Amato, Jeffrey Barrett, Sonia Goncalves, Ewan Harrison, David K. Jackson, Ian Johnston, Dominic Kwiatkowski, Cordelia Langford, John Sillitoe on behalf of the Wellcome Sanger Institute COVID-19 Surveillance Team |
| EPI_ISL_2804912, EPI_ISL_2804913 | Lighthouse Lab in Alderley Park | Wellcome Sanger Institute for the COVID-19 Genomics UK (COG-UK) Consortium | Jacquelyn Wynn, Mairead Hyland, The Lighthouse Lab in Alderley Park and Alex Alderton, Roberto Amato, Jeffrey Barrett, Sonia Goncalves, Ewan Harrison, David K. Jackson, Ian Johnston, Dominic Kwiatkowski, Cordelia Langford, John Sillitoe on behalf of the Wellcome Sanger Institute COVID-19 Surveillance Team |
| EPI_ISL_2804914 | Lighthouse Lab in Milton Keynes | Wellcome Sanger Institute for the COVID-19 Genomics UK (COG-UK) Consortium | The Lighthouse Lab in Milton Keynes and Alex Alderton, Roberto Amato, Jeffrey Barrett, Sonia Goncalves, Ewan Harrison, David K. Jackson, Ian Johnston, Dominic Kwiatkowski, Cordelia Langford, John Sillitoe on behalf of the Wellcome Sanger Institute COVID-19 Surveillance Team |
| EPI_ISL_2804915 | Lighthouse Lab in Glasgow | Wellcome Sanger Institute for the COVID-19 Genomics UK (COG-UK) Consortium | Harper VanSteenhouse, Yumi Kasai, David Gray, Carol Clugston, Anna Dominiczak and Alex Alderton, Roberto Amato, Jeffrey Barrett, Sonia Goncalves, Ewan Harrison, David K. Jackson, Ian Johnston, Dominic Kwiatkowski, Cordelia Langford, John Sillitoe on behalf of the Wellcome Sanger Institute COVID-19 Surveillance Team |
| EPI_ISL_2804916 | Lighthouse Lab in Milton Keynes | Wellcome Sanger Institute for the COVID-19 Genomics UK (COG-UK) Consortium | The Lighthouse Lab in Milton Keynes and Alex Alderton, Roberto Amato, Jeffrey Barrett, Sonia Goncalves, Ewan Harrison, David K. Jackson, Ian Johnston, Dominic Kwiatkowski, Cordelia Langford, John Sillitoe on behalf of the Wellcome Sanger Institute COVID-19 Surveillance Team |
| EPI_ISL_2804917, EPI_ISL_2804918 | Lighthouse Lab in Alderley Park | Wellcome Sanger Institute for the COVID-19 Genomics UK (COG-UK) Consortium | Jacquelyn Wynn, Mairead Hyland, The Lighthouse Lab in Alderley Park and Alex Alderton, Roberto Amato, Jeffrey Barrett, Sonia Goncalves, Ewan Harrison, David K. Jackson, Ian Johnston, Dominic Kwiatkowski, Cordelia Langford, John Sillitoe on behalf of the Wellcome Sanger Institute COVID-19 Surveillance Team |
| EPI_ISL_2804919, EPI_ISL_2804920, EPI_ISL_2804921 | Lighthouse Lab in Glasgow | Wellcome Sanger Institute for the COVID-19 Genomics UK (COG-UK) Consortium | Harper VanSteenhouse, Yumi Kasai, David Gray, Carol Clugston, Anna Dominiczak and Alex Alderton, Roberto Amato, Jeffrey Barrett, Sonia Goncalves, Ewan Harrison, David K. Jackson, Ian Johnston, Dominic Kwiatkowski, Cordelia Langford, John Sillitoe on behalf of the Wellcome Sanger Institute COVID-19 Surveillance Team |
| EPI_ISL_2804922, EPI_ISL_2804923 | Lighthouse Lab in Milton Keynes | Wellcome Sanger Institute for the COVID-19 Genomics UK (COG-UK) Consortium | The Lighthouse Lab in Milton Keynes and Alex Alderton, Roberto Amato, Jeffrey Barrett, Sonia Goncalves, Ewan Harrison, David K. Jackson, Ian Johnston, Dominic Kwiatkowski, Cordelia Langford, John Sillitoe on behalf of the Wellcome Sanger Institute COVID-19 Surveillance Team |
| EPI_ISL_2804924 | Lighthouse Lab in Alderley Park | Wellcome Sanger Institute for the COVID-19 Genomics UK (COG-UK) Consortium | Jacquelyn Wynn, Mairead Hyland, The Lighthouse Lab in Alderley Park and Alex Alderton, Roberto Amato, Jeffrey Barrett, Sonia Goncalves, Ewan Harrison, David K. Jackson, Ian Johnston, Dominic Kwiatkowski, Cordelia Langford, John Sillitoe on behalf of the Wellcome Sanger Institute COVID-19 Surveillance Team |
| EPI_ISL_2804925 | Lighthouse Lab in Milton Keynes | Wellcome Sanger Institute for the COVID-19 Genomics UK (COG-UK) Consortium | The Lighthouse Lab in Milton Keynes and Alex Alderton, Roberto Amato, Jeffrey Barrett, Sonia Goncalves, Ewan Harrison, David K. Jackson, Ian Johnston, Dominic Kwiatkowski, Cordelia Langford, John Sillitoe on behalf of the Wellcome Sanger Institute COVID-19 Surveillance Team |
| EPI_ISL_2804926 | Lighthouse Lab in Glasgow | Wellcome Sanger Institute for the COVID-19 Genomics UK (COG-UK) Consortium | Harper VanSteenhouse, Yumi Kasai, David Gray, Carol Clugston, Anna Dominiczak and Alex Alderton, Roberto Amato, Jeffrey Barrett, Sonia Goncalves, Ewan Harrison, David K. Jackson, Ian Johnston, Dominic Kwiatkowski, Cordelia Langford, John Sillitoe on behalf of the Wellcome Sanger Institute COVID-19 Surveillance Team |
| EPI_ISL_2804927, EPI_ISL_2804928 | Lighthouse Lab in Alderley Park | Wellcome Sanger Institute for the COVID-19 Genomics UK (COG-UK) Consortium | Jacquelyn Wynn, Mairead Hyland, The Lighthouse Lab in Alderley Park and Alex Alderton, Roberto Amato, Jeffrey Barrett, Sonia Goncalves, Ewan Harrison, David K. Jackson, Ian Johnston, Dominic Kwiatkowski, Cordelia Langford, John Sillitoe on behalf of the Wellcome Sanger Institute COVID-19 Surveillance Team |
| EPI_ISL_2804929 | Lighthouse Lab in Glasgow | Wellcome Sanger Institute for the COVID-19 Genomics UK (COG-UK) Consortium | Harper VanSteenhouse, Yumi Kasai, David Gray, Carol Clugston, Anna Dominiczak and Alex Alderton, Roberto Amato, Jeffrey Barrett, Sonia Goncalves, Ewan Harrison, David K. Jackson, Ian Johnston, Dominic Kwiatkowski, Cordelia Langford, John Sillitoe on behalf of the Wellcome Sanger Institute COVID-19 Surveillance Team |
| EPI_ISL_2804930 | Lighthouse Lab in Alderley Park | Wellcome Sanger Institute for the COVID-19 Genomics UK (COG-UK) Consortium | Jacquelyn Wynn, Mairead Hyland, The Lighthouse Lab in Alderley Park and Alex Alderton, Roberto Amato, Jeffrey Barrett, Sonia Goncalves, Ewan Harrison, David K. Jackson, Ian Johnston, Dominic Kwiatkowski, Cordelia Langford, John Sillitoe on behalf of the Wellcome Sanger Institute COVID-19 Surveillance Team |
| EPI_ISL_2804931 | Lighthouse Lab in Milton Keynes | Wellcome Sanger Institute for the COVID-19 Genomics UK (COG-UK) Consortium | The Lighthouse Lab in Milton Keynes and Alex Alderton, Roberto Amato, Jeffrey Barrett, Sonia Goncalves, Ewan Harrison, David K. Jackson, Ian Johnston, Dominic Kwiatkowski, Cordelia Langford, John Sillitoe on behalf of the Wellcome Sanger Institute COVID-19 Surveillance Team |
| EPI_ISL_2804932, EPI_ISL_2804933, EPI_ISL_2804934 | Lighthouse Lab in Alderley Park | Wellcome Sanger Institute for the COVID-19 Genomics UK (COG-UK) Consortium | Jacquelyn Wynn, Mairead Hyland, The Lighthouse Lab in Alderley Park and Alex Alderton, Roberto Amato, Jeffrey Barrett, Sonia Goncalves, Ewan Harrison, David K. Jackson, Ian Johnston, Dominic Kwiatkowski, Cordelia Langford, John Sillitoe on behalf of the Wellcome Sanger Institute COVID-19 Surveillance Team |
| EPI_ISL_2804935 | Lighthouse Lab in Milton Keynes | Wellcome Sanger Institute for the COVID-19 Genomics UK (COG-UK) Consortium | The Lighthouse Lab in Milton Keynes and Alex Alderton, Roberto Amato, Jeffrey Barrett, Sonia Goncalves, Ewan Harrison, David K. Jackson, Ian Johnston, Dominic Kwiatkowski, Cordelia Langford, John Sillitoe on behalf of the Wellcome Sanger Institute COVID-19 Surveillance Team |
| EPI_ISL_2804936 | Lighthouse Lab in Glasgow | Wellcome Sanger Institute for the COVID-19 Genomics UK (COG-UK) Consortium | Harper VanSteenhouse, Yumi Kasai, David Gray, Carol Clugston, Anna Dominiczak and Alex Alderton, Roberto Amato, Jeffrey Barrett, Sonia Goncalves, Ewan Harrison, David K. Jackson, Ian Johnston, Dominic Kwiatkowski, Cordelia Langford, John Sillitoe on behalf of the Wellcome Sanger Institute COVID-19 Surveillance Team |
| EPI_ISL_2804937 | Lighthouse Lab in Milton Keynes | Wellcome Sanger Institute for the COVID-19 Genomics UK (COG-UK) Consortium | The Lighthouse Lab in Milton Keynes and Alex Alderton, Roberto Amato, Jeffrey Barrett, Sonia Goncalves, Ewan Harrison, David K. Jackson, Ian Johnston, Dominic Kwiatkowski, Cordelia Langford, John Sillitoe on behalf of the Wellcome Sanger Institute COVID-19 Surveillance Team |
| EPI_ISL_2804938 | Lighthouse Lab in Alderley Park | Wellcome Sanger Institute for the COVID-19 Genomics UK (COG-UK) Consortium | Jacquelyn Wynn, Mairead Hyland, The Lighthouse Lab in Alderley Park and Alex Alderton, Roberto Amato, Jeffrey Barrett, Sonia Goncalves, Ewan Harrison, David K. Jackson, Ian Johnston, Dominic Kwiatkowski, Cordelia Langford, John Sillitoe on behalf of the Wellcome Sanger Institute COVID-19 Surveillance Team |
| EPI_ISL_2804939, EPI_ISL_2804940, | Lighthouse Lab in Glasgow | Wellcome Sanger Institute for the COVID-19 Genomics UK (COG-UK) Consortium | Harper VanSteenhouse, Yumi Kasai, David Gray, Carol Clugston, Anna Dominiczak and Alex Alderton, Roberto Amato, Jeffrey Barrett, Sonia Goncalves, Ewan Harrison, David K. Jackson, Ian Johnston, Dominic Kwiatkowski, Cordelia Langford, John Sillitoe on behalf of the Wellcome Sanger Institute COVID-19 Surveillance Team |

| | | | |
|---|---|---|---|
| EPI_ISL_2804941 | | | |
| EPI_ISL_2804942 | Lighthouse Lab in Alderley Park | Wellcome Sanger Institute for the COVID-19 Genomics UK (COG-UK) Consortium | Jacquelyn Wynn, Mairead Hyland, The Lighthouse Lab in Alderley Park and Alex Alderton, Roberto Amato, Jeffrey Barrett, Sonia Goncalves, Ewan Harrison, David K. Jackson, Ian Johnston, Dominic Kwiatkowski, Cordelia Langford, John Sillitoe on behalf of the Wellcome Sanger Institute COVID-19 Surveillance Team |
| EPI_ISL_2804943 | Lighthouse Lab in Milton Keynes | Wellcome Sanger Institute for the COVID-19 Genomics UK (COG-UK) Consortium | The Lighthouse Lab in Milton Keynes and Alex Alderton, Roberto Amato, Jeffrey Barrett, Sonia Goncalves, Ewan Harrison, David K. Jackson, Ian Johnston, Dominic Kwiatkowski, Cordelia Langford, John Sillitoe on behalf of the Wellcome Sanger Institute COVID-19 Surveillance Team |
| EPI_ISL_2804944 | Lighthouse Lab in Glasgow | Wellcome Sanger Institute for the COVID-19 Genomics UK (COG-UK) Consortium | Harper VanSteenhouse, Yumi Kasai, David Gray, Carol Clugston, Anna Dominiczak and Alex Alderton, Roberto Amato, Jeffrey Barrett, Sonia Goncalves, Ewan Harrison, David K. Jackson, Ian Johnston, Dominic Kwiatkowski, Cordelia Langford, John Sillitoe on behalf of the Wellcome Sanger Institute COVID-19 Surveillance Team |
| EPI_ISL_2804945 | Lighthouse Lab in Milton Keynes | Wellcome Sanger Institute for the COVID-19 Genomics UK (COG-UK) Consortium | The Lighthouse Lab in Milton Keynes and Alex Alderton, Roberto Amato, Jeffrey Barrett, Sonia Goncalves, Ewan Harrison, David K. Jackson, Ian Johnston, Dominic Kwiatkowski, Cordelia Langford, John Sillitoe on behalf of the Wellcome Sanger Institute COVID-19 Surveillance Team |
| EPI_ISL_2804946 | Lighthouse Lab in Glasgow | Wellcome Sanger Institute for the COVID-19 Genomics UK (COG-UK) Consortium | Harper VanSteenhouse, Yumi Kasai, David Gray, Carol Clugston, Anna Dominiczak and Alex Alderton, Roberto Amato, Jeffrey Barrett, Sonia Goncalves, Ewan Harrison, David K. Jackson, Ian Johnston, Dominic Kwiatkowski, Cordelia Langford, John Sillitoe on behalf of the Wellcome Sanger Institute COVID-19 Surveillance Team |
| EPI_ISL_2804947 | Lighthouse Lab in Alderley Park | Wellcome Sanger Institute for the COVID-19 Genomics UK (COG-UK) Consortium | Jacquelyn Wynn, Mairead Hyland, The Lighthouse Lab in Alderley Park and Alex Alderton, Roberto Amato, Jeffrey Barrett, Sonia Goncalves, Ewan Harrison, David K. Jackson, Ian Johnston, Dominic Kwiatkowski, Cordelia Langford, John Sillitoe on behalf of the Wellcome Sanger Institute COVID-19 Surveillance Team |
| EPI_ISL_2804948, EPI_ISL_2804949 | Lighthouse Lab in Milton Keynes | Wellcome Sanger Institute for the COVID-19 Genomics UK (COG-UK) Consortium | The Lighthouse Lab in Milton Keynes and Alex Alderton, Roberto Amato, Jeffrey Barrett, Sonia Goncalves, Ewan Harrison, David K. Jackson, Ian Johnston, Dominic Kwiatkowski, Cordelia Langford, John Sillitoe on behalf of the Wellcome Sanger Institute COVID-19 Surveillance Team |
| EPI_ISL_2804950 | Lighthouse Lab in Glasgow | Wellcome Sanger Institute for the COVID-19 Genomics UK (COG-UK) Consortium | Harper VanSteenhouse, Yumi Kasai, David Gray, Carol Clugston, Anna Dominiczak and Alex Alderton, Roberto Amato, Jeffrey Barrett, Sonia Goncalves, Ewan Harrison, David K. Jackson, Ian Johnston, Dominic Kwiatkowski, Cordelia Langford, John Sillitoe on behalf of the Wellcome Sanger Institute COVID-19 Surveillance Team |
| EPI_ISL_2804951, EPI_ISL_2804952 | Lighthouse Lab in Alderley Park | Wellcome Sanger Institute for the COVID-19 Genomics UK (COG-UK) Consortium | Jacquelyn Wynn, Mairead Hyland, The Lighthouse Lab in Alderley Park and Alex Alderton, Roberto Amato, Jeffrey Barrett, Sonia Goncalves, Ewan Harrison, David K. Jackson, Ian Johnston, Dominic Kwiatkowski, Cordelia Langford, John Sillitoe on behalf of the Wellcome Sanger Institute COVID-19 Surveillance Team |
| EPI_ISL_2804953, EPI_ISL_2804954 | Lighthouse Lab in Glasgow | Wellcome Sanger Institute for the COVID-19 Genomics UK (COG-UK) Consortium | Harper VanSteenhouse, Yumi Kasai, David Gray, Carol Clugston, Anna Dominiczak and Alex Alderton, Roberto Amato, Jeffrey Barrett, Sonia Goncalves, Ewan Harrison, David K. Jackson, Ian Johnston, Dominic Kwiatkowski, Cordelia Langford, John Sillitoe on behalf of the Wellcome Sanger Institute COVID-19 Surveillance Team |
| EPI_ISL_2804955 | Lighthouse Lab in Milton Keynes | Wellcome Sanger Institute for the COVID-19 Genomics UK (COG-UK) Consortium | The Lighthouse Lab in Milton Keynes and Alex Alderton, Roberto Amato, Jeffrey Barrett, Sonia Goncalves, Ewan Harrison, David K. Jackson, Ian Johnston, Dominic Kwiatkowski, Cordelia Langford, John Sillitoe on behalf of the Wellcome Sanger Institute COVID-19 Surveillance Team |
| EPI_ISL_2804956 | Lighthouse Lab in Alderley Park | Wellcome Sanger Institute for the COVID-19 Genomics UK (COG-UK) Consortium | Jacquelyn Wynn, Mairead Hyland, The Lighthouse Lab in Alderley Park and Alex Alderton, Roberto Amato, Jeffrey Barrett, Sonia Goncalves, Ewan Harrison, David K. Jackson, Ian Johnston, Dominic Kwiatkowski, Cordelia Langford, John Sillitoe on behalf of the Wellcome Sanger Institute COVID-19 Surveillance Team |
| EPI_ISL_2804957 | Lighthouse Lab in Glasgow | Wellcome Sanger Institute for the COVID-19 Genomics UK (COG-UK) Consortium | Harper VanSteenhouse, Yumi Kasai, David Gray, Carol Clugston, Anna Dominiczak and Alex Alderton, Roberto Amato, Jeffrey Barrett, Sonia Goncalves, Ewan Harrison, David K. Jackson, Ian Johnston, Dominic Kwiatkowski, Cordelia Langford, John Sillitoe on behalf of the Wellcome Sanger Institute COVID-19 Surveillance Team |
| EPI_ISL_2804958, EPI_ISL_2804959 | Lighthouse Lab in Milton Keynes | Wellcome Sanger Institute for the COVID-19 Genomics UK (COG-UK) Consortium | The Lighthouse Lab in Milton Keynes and Alex Alderton, Roberto Amato, Jeffrey Barrett, Sonia Goncalves, Ewan Harrison, David K. Jackson, Ian Johnston, Dominic Kwiatkowski, Cordelia Langford, John Sillitoe on behalf of the Wellcome Sanger Institute COVID-19 Surveillance Team |
| EPI_ISL_2804960 | Lighthouse Lab in Glasgow | Wellcome Sanger Institute for the COVID-19 Genomics UK (COG-UK) Consortium | Harper VanSteenhouse, Yumi Kasai, David Gray, Carol Clugston, Anna Dominiczak and Alex Alderton, Roberto Amato, Jeffrey Barrett, Sonia Goncalves, Ewan Harrison, David K. Jackson, Ian Johnston, Dominic Kwiatkowski, Cordelia Langford, John Sillitoe on behalf of the Wellcome Sanger Institute COVID-19 Surveillance Team |
| EPI_ISL_2804961, EPI_ISL_2804962 | Lighthouse Lab in Alderley Park | Wellcome Sanger Institute for the COVID-19 Genomics UK (COG-UK) Consortium | Jacquelyn Wynn, Mairead Hyland, The Lighthouse Lab in Alderley Park and Alex Alderton, Roberto Amato, Jeffrey Barrett, Sonia Goncalves, Ewan Harrison, David K. Jackson, Ian Johnston, Dominic Kwiatkowski, Cordelia Langford, John Sillitoe on behalf of the Wellcome Sanger Institute COVID-19 Surveillance Team |
| EPI_ISL_2804963 | Lighthouse Lab in Milton Keynes | Wellcome Sanger Institute for the COVID-19 Genomics UK (COG-UK) Consortium | The Lighthouse Lab in Milton Keynes and Alex Alderton, Roberto Amato, Jeffrey Barrett, Sonia Goncalves, Ewan Harrison, David K. Jackson, Ian Johnston, Dominic Kwiatkowski, Cordelia Langford, John Sillitoe on behalf of the Wellcome Sanger Institute COVID-19 Surveillance Team |
| EPI_ISL_2804964 | Lighthouse Lab in Alderley Park | Wellcome Sanger Institute for the COVID-19 Genomics UK (COG-UK) Consortium | Jacquelyn Wynn, Mairead Hyland, The Lighthouse Lab in Alderley Park and Alex Alderton, Roberto Amato, Jeffrey Barrett, Sonia Goncalves, Ewan Harrison, David K. Jackson, Ian Johnston, Dominic Kwiatkowski, Cordelia Langford, John Sillitoe on behalf of the Wellcome Sanger Institute COVID-19 Surveillance Team |
| EPI_ISL_2804965, EPI_ISL_2804966 | Lighthouse Lab in Milton Keynes | Wellcome Sanger Institute for the COVID-19 Genomics UK (COG-UK) Consortium | The Lighthouse Lab in Milton Keynes and Alex Alderton, Roberto Amato, Jeffrey Barrett, Sonia Goncalves, Ewan Harrison, David K. Jackson, Ian Johnston, Dominic Kwiatkowski, Cordelia Langford, John Sillitoe on behalf of the Wellcome Sanger Institute COVID-19 Surveillance Team |
| EPI_ISL_2804967, EPI_ISL_2804968 | Lighthouse Lab in Alderley Park | Wellcome Sanger Institute for the COVID-19 Genomics UK (COG-UK) Consortium | Jacquelyn Wynn, Mairead Hyland, The Lighthouse Lab in Alderley Park and Alex Alderton, Roberto Amato, Jeffrey Barrett, Sonia Goncalves, Ewan Harrison, David K. Jackson, Ian Johnston, Dominic Kwiatkowski, Cordelia Langford, John Sillitoe on behalf of the Wellcome Sanger Institute COVID-19 Surveillance Team |
| EPI_ISL_2804969 | Lighthouse Lab in Milton Keynes | Wellcome Sanger Institute for the COVID-19 Genomics UK (COG-UK) Consortium | The Lighthouse Lab in Milton Keynes and Alex Alderton, Roberto Amato, Jeffrey Barrett, Sonia Goncalves, Ewan Harrison, David K. Jackson, Ian Johnston, Dominic Kwiatkowski, Cordelia Langford, John Sillitoe on behalf of the Wellcome Sanger Institute COVID-19 Surveillance Team |
| EPI_ISL_2804970, EPI_ISL_2804971 | Lighthouse Lab in Glasgow | Wellcome Sanger Institute for the COVID-19 Genomics UK (COG-UK) Consortium | Harper VanSteenhouse, Yumi Kasai, David Gray, Carol Clugston, Anna Dominiczak and Alex Alderton, Roberto Amato, Jeffrey Barrett, Sonia Goncalves, Ewan Harrison, David K. Jackson, Ian Johnston, Dominic Kwiatkowski, Cordelia Langford, John Sillitoe on behalf of the Wellcome Sanger Institute COVID-19 Surveillance Team |
| EPI_ISL_2804972, EPI_ISL_2804973 | Lighthouse Lab in Alderley Park | Wellcome Sanger Institute for the COVID-19 Genomics UK (COG-UK) Consortium | Jacquelyn Wynn, Mairead Hyland, The Lighthouse Lab in Alderley Park and Alex Alderton, Roberto Amato, Jeffrey Barrett, Sonia Goncalves, Ewan Harrison, David K. Jackson, Ian Johnston, Dominic Kwiatkowski, Cordelia Langford, John Sillitoe on behalf of the Wellcome Sanger Institute COVID-19 Surveillance Team |
| EPI_ISL_2804974 | Lighthouse Lab in Glasgow | Wellcome Sanger Institute for the COVID-19 Genomics UK (COG-UK) Consortium | Harper VanSteenhouse, Yumi Kasai, David Gray, Carol Clugston, Anna Dominiczak and Alex Alderton, Roberto Amato, Jeffrey Barrett, Sonia Goncalves, Ewan Harrison, David K. Jackson, Ian Johnston, Dominic Kwiatkowski, Cordelia Langford, John Sillitoe on behalf of the Wellcome Sanger Institute COVID-19 Surveillance Team |
| EPI_ISL_2804975, EPI_ISL_2804976, EPI_ISL_2804977, EPI_ISL_2804978, EPI_ISL_2804979 | Lighthouse Lab in Milton Keynes | Wellcome Sanger Institute for the COVID-19 Genomics UK (COG-UK) Consortium | The Lighthouse Lab in Milton Keynes and Alex Alderton, Roberto Amato, Jeffrey Barrett, Sonia Goncalves, Ewan Harrison, David K. Jackson, Ian Johnston, Dominic Kwiatkowski, Cordelia Langford, John Sillitoe on behalf of the Wellcome Sanger Institute COVID-19 Surveillance Team |
| EPI_ISL_2804980, EPI_ISL_2804981 | Lighthouse Lab in Alderley Park | Wellcome Sanger Institute for the COVID-19 Genomics UK (COG-UK) Consortium | Jacquelyn Wynn, Mairead Hyland, The Lighthouse Lab in Alderley Park and Alex Alderton, Roberto Amato, Jeffrey Barrett, Sonia Goncalves, Ewan Harrison, David K. Jackson, Ian Johnston, Dominic Kwiatkowski, Cordelia Langford, John Sillitoe on behalf of the Wellcome Sanger Institute COVID-19 Surveillance Team |
| EPI_ISL_2804982 | Lighthouse Lab in Glasgow | Wellcome Sanger Institute for the COVID-19 Genomics UK (COG-UK) Consortium | Harper VanSteenhouse, Yumi Kasai, David Gray, Carol Clugston, Anna Dominiczak and Alex Alderton, Roberto Amato, Jeffrey Barrett, Sonia Goncalves, Ewan Harrison, David K. Jackson, Ian Johnston, Dominic Kwiatkowski, Cordelia Langford, John Sillitoe on behalf of the Wellcome Sanger Institute COVID-19 Surveillance Team |
| EPI_ISL_2804983, EPI_ISL_2804984 | Lighthouse Lab in Alderley Park | Wellcome Sanger Institute for the COVID-19 Genomics UK (COG-UK) Consortium | Jacquelyn Wynn, Mairead Hyland, The Lighthouse Lab in Alderley Park and Alex Alderton, Roberto Amato, Jeffrey Barrett, Sonia Goncalves, Ewan Harrison, David K. Jackson, Ian Johnston, Dominic Kwiatkowski, Cordelia Langford, John Sillitoe on behalf of the Wellcome Sanger Institute COVID-19 Surveillance Team |
| EPI_ISL_2804985 | Lighthouse Lab in Milton Keynes | Wellcome Sanger Institute for the COVID-19 Genomics UK (COG-UK) Consortium | The Lighthouse Lab in Milton Keynes and Alex Alderton, Roberto Amato, Jeffrey Barrett, Sonia Goncalves, Ewan Harrison, David K. Jackson, Ian Johnston, Dominic Kwiatkowski, Cordelia Langford, John Sillitoe on behalf of the Wellcome Sanger Institute COVID-19 Surveillance Team |
| EPI_ISL_2804986 | Lighthouse Lab in Alderley Park | Wellcome Sanger Institute for the COVID-19 Genomics UK (COG-UK) Consortium | Jacquelyn Wynn, Mairead Hyland, The Lighthouse Lab in Alderley Park and Alex Alderton, Roberto Amato, Jeffrey Barrett, Sonia Goncalves, Ewan Harrison, David K. Jackson, Ian Johnston, Dominic Kwiatkowski, Cordelia Langford, John Sillitoe on behalf of the Wellcome Sanger Institute COVID-19 Surveillance Team |
| EPI_ISL_2804987 | Lighthouse Lab in Glasgow | Wellcome Sanger Institute for the COVID-19 Genomics UK (COG-UK) Consortium | Harper VanSteenhouse, Yumi Kasai, David Gray, Carol Clugston, Anna Dominiczak and Alex Alderton, Roberto Amato, Jeffrey Barrett, Sonia Goncalves, Ewan Harrison, David K. Jackson, Ian Johnston, Dominic Kwiatkowski, Cordelia Langford, John Sillitoe on behalf of the Wellcome Sanger Institute COVID-19 Surveillance Team |
| EPI_ISL_2804988, EPI_ISL_2804989 | Lighthouse Lab in Alderley Park | Wellcome Sanger Institute for the COVID-19 Genomics UK (COG-UK) Consortium | Jacquelyn Wynn, Mairead Hyland, The Lighthouse Lab in Alderley Park and Alex Alderton, Roberto Amato, Jeffrey Barrett, Sonia Goncalves, Ewan Harrison, David K. Jackson, Ian Johnston, Dominic Kwiatkowski, Cordelia Langford, John Sillitoe on behalf of the Wellcome Sanger Institute COVID-19 Surveillance Team |
| EPI_ISL_2804990, EPI_ISL_2804991, EPI_ISL_2804992, EPI_ISL_2804993 | Lighthouse Lab in Glasgow | Wellcome Sanger Institute for the COVID-19 Genomics UK (COG-UK) Consortium | Harper VanSteenhouse, Yumi Kasai, David Gray, Carol Clugston, Anna Dominiczak and Alex Alderton, Roberto Amato, Jeffrey Barrett, Sonia Goncalves, Ewan Harrison, David K. Jackson, Ian Johnston, Dominic Kwiatkowski, Cordelia Langford, John Sillitoe on behalf of the Wellcome Sanger Institute COVID-19 Surveillance Team |
| EPI_ISL_2804994, EPI_ISL_2804995, EPI_ISL_2804996 | Lighthouse Lab in Alderley Park | Wellcome Sanger Institute for the COVID-19 Genomics UK (COG-UK) Consortium | Jacquelyn Wynn, Mairead Hyland, The Lighthouse Lab in Alderley Park and Alex Alderton, Roberto Amato, Jeffrey Barrett, Sonia Goncalves, Ewan Harrison, David K. Jackson, Ian Johnston, Dominic Kwiatkowski, Cordelia Langford, John Sillitoe on behalf of the Wellcome Sanger Institute COVID-19 Surveillance Team |
| EPI_ISL_2804997 | Lighthouse Lab in Glasgow | Wellcome Sanger Institute for the COVID-19 Genomics UK (COG-UK) Consortium | Harper VanSteenhouse, Yumi Kasai, David Gray, Carol Clugston, Anna Dominiczak and Alex Alderton, Roberto Amato, Jeffrey Barrett, Sonia Goncalves, Ewan Harrison, David K. Jackson, Ian Johnston, Dominic Kwiatkowski, Cordelia Langford, John Sillitoe on behalf of the Wellcome Sanger Institute COVID-19 Surveillance Team |
| EPI_ISL_2804998 | Lighthouse Lab in Alderley Park | Wellcome Sanger Institute for the COVID-19 Genomics UK (COG-UK) Consortium | Jacquelyn Wynn, Mairead Hyland, The Lighthouse Lab in Alderley Park and Alex Alderton, Roberto Amato, Jeffrey Barrett, Sonia Goncalves, Ewan Harrison, David K. Jackson, Ian Johnston, Dominic Kwiatkowski, Cordelia Langford, John Sillitoe on behalf of the Wellcome Sanger Institute COVID-19 Surveillance Team |
| EPI_ISL_2804999 | Lighthouse Lab in Glasgow | Wellcome Sanger Institute for the COVID-19 Genomics UK (COG-UK) Consortium | Harper VanSteenhouse, Yumi Kasai, David Gray, Carol Clugston, Anna Dominiczak and Alex Alderton, Roberto Amato, Jeffrey Barrett, Sonia Goncalves, Ewan Harrison, David K. Jackson, Ian Johnston, Dominic Kwiatkowski, Cordelia Langford, John Sillitoe on behalf of the Wellcome Sanger Institute COVID-19 Surveillance Team |
| EPI_ISL_2805000, EPI_ISL_2805001, EPI_ISL_2805002, EPI_ISL_2805003 | Lighthouse Lab in Milton Keynes | Wellcome Sanger Institute for the COVID-19 Genomics UK (COG-UK) Consortium | The Lighthouse Lab in Milton Keynes and Alex Alderton, Roberto Amato, Jeffrey Barrett, Sonia Goncalves, Ewan Harrison, David K. Jackson, Ian Johnston, Dominic Kwiatkowski, Cordelia Langford, John Sillitoe on behalf of the Wellcome Sanger Institute COVID-19 Surveillance Team |
| EPI_ISL_2805004 | Lighthouse Lab in Alderley Park | Wellcome Sanger Institute for the COVID-19 Genomics UK (COG-UK) Consortium | Jacquelyn Wynn, Mairead Hyland, The Lighthouse Lab in Alderley Park and Alex Alderton, Roberto Amato, Jeffrey Barrett, Sonia Goncalves, Ewan Harrison, David K. Jackson, Ian Johnston, Dominic Kwiatkowski, Cordelia Langford, John Sillitoe on behalf of the Wellcome Sanger Institute COVID-19 Surveillance Team |
| EPI_ISL_2805005 | Lighthouse Lab in Milton Keynes | Wellcome Sanger Institute for the COVID-19 Genomics UK (COG-UK) Consortium | The Lighthouse Lab in Milton Keynes and Alex Alderton, Roberto Amato, Jeffrey Barrett, Sonia Goncalves, Ewan Harrison, David K. Jackson, Ian Johnston, Dominic Kwiatkowski, Cordelia Langford, John Sillitoe on behalf of the Wellcome Sanger Institute COVID-19 Surveillance Team |
| EPI_ISL_2805006 | Lighthouse Lab in Alderley Park | Wellcome Sanger Institute for the COVID-19 Genomics UK (COG-UK) Consortium | Jacquelyn Wynn, Mairead Hyland, The Lighthouse Lab in Alderley Park and Alex Alderton, Roberto Amato, Jeffrey Barrett, Sonia Goncalves, Ewan Harrison, David K. Jackson, Ian Johnston, Dominic Kwiatkowski, Cordelia Langford, John Sillitoe on behalf of the Wellcome Sanger Institute COVID-19 Surveillance Team |

| | | | |
|---|---|---|---|
| EPI_ISL_2805007 | Lighthouse Lab in Glasgow | Wellcome Sanger Institute for the COVID-19 Genomics UK (COG-UK) Consortium | Harper VanSteenhouse, Yumi Kasai, David Gray, Carol Clugston, Anna Dominiczak and Alex Alderton, Roberto Amato, Jeffrey Barrett, Sonia Goncalves, Ewan Harrison, David K. Jackson, Ian Johnston, Dominic Kwiatkowski, Cordelia Langford, John Sillitoe on behalf of the Wellcome Sanger Institute COVID-19 Surveillance Team |
| EPI_ISL_2805008 | Lighthouse Lab in Alderley Park | Wellcome Sanger Institute for the COVID-19 Genomics UK (COG-UK) Consortium | Jacquelyn Wynn, Mairead Hyland, The Lighthouse Lab in Alderley Park and Alex Alderton, Roberto Amato, Jeffrey Barrett, Sonia Goncalves, Ewan Harrison, David K. Jackson, Ian Johnston, Dominic Kwiatkowski, Cordelia Langford, John Sillitoe on behalf of the Wellcome Sanger Institute COVID-19 Surveillance Team |
| EPI_ISL_2805009, EPI_ISL_2805010 | Lighthouse Lab in Milton Keynes | Wellcome Sanger Institute for the COVID-19 Genomics UK (COG-UK) Consortium | The Lighthouse Lab in Milton Keynes and Alex Alderton, Roberto Amato, Jeffrey Barrett, Sonia Goncalves, Ewan Harrison, David K. Jackson, Ian Johnston, Dominic Kwiatkowski, Cordelia Langford, John Sillitoe on behalf of the Wellcome Sanger Institute COVID-19 Surveillance Team |
| EPI_ISL_2805011 | Lighthouse Lab in Alderley Park | Wellcome Sanger Institute for the COVID-19 Genomics UK (COG-UK) Consortium | Jacquelyn Wynn, Mairead Hyland, The Lighthouse Lab in Alderley Park and Alex Alderton, Roberto Amato, Jeffrey Barrett, Sonia Goncalves, Ewan Harrison, David K. Jackson, Ian Johnston, Dominic Kwiatkowski, Cordelia Langford, John Sillitoe on behalf of the Wellcome Sanger Institute COVID-19 Surveillance Team |
| EPI_ISL_2805012 | Lighthouse Lab in Glasgow | Wellcome Sanger Institute for the COVID-19 Genomics UK (COG-UK) Consortium | Harper VanSteenhouse, Yumi Kasai, David Gray, Carol Clugston, Anna Dominiczak and Alex Alderton, Roberto Amato, Jeffrey Barrett, Sonia Goncalves, Ewan Harrison, David K. Jackson, Ian Johnston, Dominic Kwiatkowski, Cordelia Langford, John Sillitoe on behalf of the Wellcome Sanger Institute COVID-19 Surveillance Team |
| EPI_ISL_2805013 | Lighthouse Lab in Milton Keynes | Wellcome Sanger Institute for the COVID-19 Genomics UK (COG-UK) Consortium | The Lighthouse Lab in Milton Keynes and Alex Alderton, Roberto Amato, Jeffrey Barrett, Sonia Goncalves, Ewan Harrison, David K. Jackson, Ian Johnston, Dominic Kwiatkowski, Cordelia Langford, John Sillitoe on behalf of the Wellcome Sanger Institute COVID-19 Surveillance Team |
| EPI_ISL_2805014 | Lighthouse Lab in Alderley Park | Wellcome Sanger Institute for the COVID-19 Genomics UK (COG-UK) Consortium | Jacquelyn Wynn, Mairead Hyland, The Lighthouse Lab in Alderley Park and Alex Alderton, Roberto Amato, Jeffrey Barrett, Sonia Goncalves, Ewan Harrison, David K. Jackson, Ian Johnston, Dominic Kwiatkowski, Cordelia Langford, John Sillitoe on behalf of the Wellcome Sanger Institute COVID-19 Surveillance Team |
| EPI_ISL_2805015 | Lighthouse Lab in Glasgow | Wellcome Sanger Institute for the COVID-19 Genomics UK (COG-UK) Consortium | Harper VanSteenhouse, Yumi Kasai, David Gray, Carol Clugston, Anna Dominiczak and Alex Alderton, Roberto Amato, Jeffrey Barrett, Sonia Goncalves, Ewan Harrison, David K. Jackson, Ian Johnston, Dominic Kwiatkowski, Cordelia Langford, John Sillitoe on behalf of the Wellcome Sanger Institute COVID-19 Surveillance Team |
| EPI_ISL_2805016 | Lighthouse Lab in Milton Keynes | Wellcome Sanger Institute for the COVID-19 Genomics UK (COG-UK) Consortium | The Lighthouse Lab in Milton Keynes and Alex Alderton, Roberto Amato, Jeffrey Barrett, Sonia Goncalves, Ewan Harrison, David K. Jackson, Ian Johnston, Dominic Kwiatkowski, Cordelia Langford, John Sillitoe on behalf of the Wellcome Sanger Institute COVID-19 Surveillance Team |
| EPI_ISL_2805017, EPI_ISL_2805018 | Lighthouse Lab in Glasgow | Wellcome Sanger Institute for the COVID-19 Genomics UK (COG-UK) Consortium | Harper VanSteenhouse, Yumi Kasai, David Gray, Carol Clugston, Anna Dominiczak and Alex Alderton, Roberto Amato, Jeffrey Barrett, Sonia Goncalves, Ewan Harrison, David K. Jackson, Ian Johnston, Dominic Kwiatkowski, Cordelia Langford, John Sillitoe on behalf of the Wellcome Sanger Institute COVID-19 Surveillance Team |
| EPI_ISL_2805019, EPI_ISL_2805020, EPI_ISL_2805021 | Lighthouse Lab in Alderley Park | Wellcome Sanger Institute for the COVID-19 Genomics UK (COG-UK) Consortium | Jacquelyn Wynn, Mairead Hyland, The Lighthouse Lab in Alderley Park and Alex Alderton, Roberto Amato, Jeffrey Barrett, Sonia Goncalves, Ewan Harrison, David K. Jackson, Ian Johnston, Dominic Kwiatkowski, Cordelia Langford, John Sillitoe on behalf of the Wellcome Sanger Institute COVID-19 Surveillance Team |
| EPI_ISL_2805022 | Lighthouse Lab in Milton Keynes | Wellcome Sanger Institute for the COVID-19 Genomics UK (COG-UK) Consortium | The Lighthouse Lab in Milton Keynes and Alex Alderton, Roberto Amato, Jeffrey Barrett, Sonia Goncalves, Ewan Harrison, David K. Jackson, Ian Johnston, Dominic Kwiatkowski, Cordelia Langford, John Sillitoe on behalf of the Wellcome Sanger Institute COVID-19 Surveillance Team |
| EPI_ISL_2805023, EPI_ISL_2805024 | Lighthouse Lab in Alderley Park | Wellcome Sanger Institute for the COVID-19 Genomics UK (COG-UK) Consortium | Jacquelyn Wynn, Mairead Hyland, The Lighthouse Lab in Alderley Park and Alex Alderton, Roberto Amato, Jeffrey Barrett, Sonia Goncalves, Ewan Harrison, David K. Jackson, Ian Johnston, Dominic Kwiatkowski, Cordelia Langford, John Sillitoe on behalf of the Wellcome Sanger Institute COVID-19 Surveillance Team |
| EPI_ISL_2805025 | Lighthouse Lab in Milton Keynes | Wellcome Sanger Institute for the COVID-19 Genomics UK (COG-UK) Consortium | The Lighthouse Lab in Milton Keynes and Alex Alderton, Roberto Amato, Jeffrey Barrett, Sonia Goncalves, Ewan Harrison, David K. Jackson, Ian Johnston, Dominic Kwiatkowski, Cordelia Langford, John Sillitoe on behalf of the Wellcome Sanger Institute COVID-19 Surveillance Team |
| EPI_ISL_2805026 | Lighthouse Lab in Alderley Park | Wellcome Sanger Institute for the COVID-19 Genomics UK (COG-UK) Consortium | Jacquelyn Wynn, Mairead Hyland, The Lighthouse Lab in Alderley Park and Alex Alderton, Roberto Amato, Jeffrey Barrett, Sonia Goncalves, Ewan Harrison, David K. Jackson, Ian Johnston, Dominic Kwiatkowski, Cordelia Langford, John Sillitoe on behalf of the Wellcome Sanger Institute COVID-19 Surveillance Team |
| EPI_ISL_2805027 | Lighthouse Lab in Milton Keynes | Wellcome Sanger Institute for the COVID-19 Genomics UK (COG-UK) Consortium | The Lighthouse Lab in Milton Keynes and Alex Alderton, Roberto Amato, Jeffrey Barrett, Sonia Goncalves, Ewan Harrison, David K. Jackson, Ian Johnston, Dominic Kwiatkowski, Cordelia Langford, John Sillitoe on behalf of the Wellcome Sanger Institute COVID-19 Surveillance Team |
| EPI_ISL_2805028 | Lighthouse Lab in Alderley Park | Wellcome Sanger Institute for the COVID-19 Genomics UK (COG-UK) Consortium | Jacquelyn Wynn, Mairead Hyland, The Lighthouse Lab in Alderley Park and Alex Alderton, Roberto Amato, Jeffrey Barrett, Sonia Goncalves, Ewan Harrison, David K. Jackson, Ian Johnston, Dominic Kwiatkowski, Cordelia Langford, John Sillitoe on behalf of the Wellcome Sanger Institute COVID-19 Surveillance Team |
| EPI_ISL_2805029 | Lighthouse Lab in Glasgow | Wellcome Sanger Institute for the COVID-19 Genomics UK (COG-UK) Consortium | Harper VanSteenhouse, Yumi Kasai, David Gray, Carol Clugston, Anna Dominiczak and Alex Alderton, Roberto Amato, Jeffrey Barrett, Sonia Goncalves, Ewan Harrison, David K. Jackson, Ian Johnston, Dominic Kwiatkowski, Cordelia Langford, John Sillitoe on behalf of the Wellcome Sanger Institute COVID-19 Surveillance Team |
| EPI_ISL_2805030 | Lighthouse Lab in Alderley Park | Wellcome Sanger Institute for the COVID-19 Genomics UK (COG-UK) Consortium | Jacquelyn Wynn, Mairead Hyland, The Lighthouse Lab in Alderley Park and Alex Alderton, Roberto Amato, Jeffrey Barrett, Sonia Goncalves, Ewan Harrison, David K. Jackson, Ian Johnston, Dominic Kwiatkowski, Cordelia Langford, John Sillitoe on behalf of the Wellcome Sanger Institute COVID-19 Surveillance Team |
| EPI_ISL_2805031, EPI_ISL_2805032 | Lighthouse Lab in Glasgow | Wellcome Sanger Institute for the COVID-19 Genomics UK (COG-UK) Consortium | Harper VanSteenhouse, Yumi Kasai, David Gray, Carol Clugston, Anna Dominiczak and Alex Alderton, Roberto Amato, Jeffrey Barrett, Sonia Goncalves, Ewan Harrison, David K. Jackson, Ian Johnston, Dominic Kwiatkowski, Cordelia Langford, John Sillitoe on behalf of the Wellcome Sanger Institute COVID-19 Surveillance Team |
| EPI_ISL_2805033 | Lighthouse Lab in Alderley Park | Wellcome Sanger Institute for the COVID-19 Genomics UK (COG-UK) Consortium | Jacquelyn Wynn, Mairead Hyland, The Lighthouse Lab in Alderley Park and Alex Alderton, Roberto Amato, Jeffrey Barrett, Sonia Goncalves, Ewan Harrison, David K. Jackson, Ian Johnston, Dominic Kwiatkowski, Cordelia Langford, John Sillitoe on behalf of the Wellcome Sanger Institute COVID-19 Surveillance Team |
| EPI_ISL_2805034, EPI_ISL_2805035 | Lighthouse Lab in Milton Keynes | Wellcome Sanger Institute for the COVID-19 Genomics UK (COG-UK) Consortium | The Lighthouse Lab in Milton Keynes and Alex Alderton, Roberto Amato, Jeffrey Barrett, Sonia Goncalves, Ewan Harrison, David K. Jackson, Ian Johnston, Dominic Kwiatkowski, Cordelia Langford, John Sillitoe on behalf of the Wellcome Sanger Institute COVID-19 Surveillance Team |
| EPI_ISL_2805036 | Lighthouse Lab in Alderley Park | Wellcome Sanger Institute for the COVID-19 Genomics UK (COG-UK) Consortium | Jacquelyn Wynn, Mairead Hyland, The Lighthouse Lab in Alderley Park and Alex Alderton, Roberto Amato, Jeffrey Barrett, Sonia Goncalves, Ewan Harrison, David K. Jackson, Ian Johnston, Dominic Kwiatkowski, Cordelia Langford, John Sillitoe on behalf of the Wellcome Sanger Institute COVID-19 Surveillance Team |
| EPI_ISL_2805037 | Lighthouse Lab in Milton Keynes | Wellcome Sanger Institute for the COVID-19 Genomics UK (COG-UK) Consortium | The Lighthouse Lab in Milton Keynes and Alex Alderton, Roberto Amato, Jeffrey Barrett, Sonia Goncalves, Ewan Harrison, David K. Jackson, Ian Johnston, Dominic Kwiatkowski, Cordelia Langford, John Sillitoe on behalf of the Wellcome Sanger Institute COVID-19 Surveillance Team |
| EPI_ISL_2805038 | Lighthouse Lab in Alderley Park | Wellcome Sanger Institute for the COVID-19 Genomics UK (COG-UK) Consortium | Jacquelyn Wynn, Mairead Hyland, The Lighthouse Lab in Alderley Park and Alex Alderton, Roberto Amato, Jeffrey Barrett, Sonia Goncalves, Ewan Harrison, David K. Jackson, Ian Johnston, Dominic Kwiatkowski, Cordelia Langford, John Sillitoe on behalf of the Wellcome Sanger Institute COVID-19 Surveillance Team |
| EPI_ISL_2805039, EPI_ISL_2805040, EPI_ISL_2805041, EPI_ISL_2805042 | Lighthouse Lab in Glasgow | Wellcome Sanger Institute for the COVID-19 Genomics UK (COG-UK) Consortium | Harper VanSteenhouse, Yumi Kasai, David Gray, Carol Clugston, Anna Dominiczak and Alex Alderton, Roberto Amato, Jeffrey Barrett, Sonia Goncalves, Ewan Harrison, David K. Jackson, Ian Johnston, Dominic Kwiatkowski, Cordelia Langford, John Sillitoe on behalf of the Wellcome Sanger Institute COVID-19 Surveillance Team |
| EPI_ISL_2805043 | Lighthouse Lab in Milton Keynes | Wellcome Sanger Institute for the COVID-19 Genomics UK (COG-UK) Consortium | The Lighthouse Lab in Milton Keynes and Alex Alderton, Roberto Amato, Jeffrey Barrett, Sonia Goncalves, Ewan Harrison, David K. Jackson, Ian Johnston, Dominic Kwiatkowski, Cordelia Langford, John Sillitoe on behalf of the Wellcome Sanger Institute COVID-19 Surveillance Team |
| EPI_ISL_2805044 | Lighthouse Lab in Glasgow | Wellcome Sanger Institute for the COVID-19 Genomics UK (COG-UK) Consortium | Harper VanSteenhouse, Yumi Kasai, David Gray, Carol Clugston, Anna Dominiczak and Alex Alderton, Roberto Amato, Jeffrey Barrett, Sonia Goncalves, Ewan Harrison, David K. Jackson, Ian Johnston, Dominic Kwiatkowski, Cordelia Langford, John Sillitoe on behalf of the Wellcome Sanger Institute COVID-19 Surveillance Team |
| EPI_ISL_2805045 | Lighthouse Lab in Alderley Park | Wellcome Sanger Institute for the COVID-19 Genomics UK (COG-UK) Consortium | Jacquelyn Wynn, Mairead Hyland, The Lighthouse Lab in Alderley Park and Alex Alderton, Roberto Amato, Jeffrey Barrett, Sonia Goncalves, Ewan Harrison, David K. Jackson, Ian Johnston, Dominic Kwiatkowski, Cordelia Langford, John Sillitoe on behalf of the Wellcome Sanger Institute COVID-19 Surveillance Team |
| EPI_ISL_2805046 | Lighthouse Lab in Glasgow | Wellcome Sanger Institute for the COVID-19 Genomics UK (COG-UK) Consortium | Harper VanSteenhouse, Yumi Kasai, David Gray, Carol Clugston, Anna Dominiczak and Alex Alderton, Roberto Amato, Jeffrey Barrett, Sonia Goncalves, Ewan Harrison, David K. Jackson, Ian Johnston, Dominic Kwiatkowski, Cordelia Langford, John Sillitoe on behalf of the Wellcome Sanger Institute COVID-19 Surveillance Team |
| EPI_ISL_2805047, EPI_ISL_2805048, EPI_ISL_2805049 | Lighthouse Lab in Milton Keynes | Wellcome Sanger Institute for the COVID-19 Genomics UK (COG-UK) Consortium | The Lighthouse Lab in Milton Keynes and Alex Alderton, Roberto Amato, Jeffrey Barrett, Sonia Goncalves, Ewan Harrison, David K. Jackson, Ian Johnston, Dominic Kwiatkowski, Cordelia Langford, John Sillitoe on behalf of the Wellcome Sanger Institute COVID-19 Surveillance Team |
| EPI_ISL_2805050 | Lighthouse Lab in Glasgow | Wellcome Sanger Institute for the COVID-19 Genomics UK (COG-UK) Consortium | Harper VanSteenhouse, Yumi Kasai, David Gray, Carol Clugston, Anna Dominiczak and Alex Alderton, Roberto Amato, Jeffrey Barrett, Sonia Goncalves, Ewan Harrison, David K. Jackson, Ian Johnston, Dominic Kwiatkowski, Cordelia Langford, John Sillitoe on behalf of the Wellcome Sanger Institute COVID-19 Surveillance Team |
| EPI_ISL_2805051, EPI_ISL_2805052, EPI_ISL_2805053, EPI_ISL_2805054 | Lighthouse Lab in Alderley Park | Wellcome Sanger Institute for the COVID-19 Genomics UK (COG-UK) Consortium | Jacquelyn Wynn, Mairead Hyland, The Lighthouse Lab in Alderley Park and Alex Alderton, Roberto Amato, Jeffrey Barrett, Sonia Goncalves, Ewan Harrison, David K. Jackson, Ian Johnston, Dominic Kwiatkowski, Cordelia Langford, John Sillitoe on behalf of the Wellcome Sanger Institute COVID-19 Surveillance Team |
| EPI_ISL_2805055 | Lighthouse Lab in Milton Keynes | Wellcome Sanger Institute for the COVID-19 Genomics UK (COG-UK) Consortium | The Lighthouse Lab in Milton Keynes and Alex Alderton, Roberto Amato, Jeffrey Barrett, Sonia Goncalves, Ewan Harrison, David K. Jackson, Ian Johnston, Dominic Kwiatkowski, Cordelia Langford, John Sillitoe on behalf of the Wellcome Sanger Institute COVID-19 Surveillance Team |
| EPI_ISL_2805056, EPI_ISL_2805057 | Lighthouse Lab in Glasgow | Wellcome Sanger Institute for the COVID-19 Genomics UK (COG-UK) Consortium | Harper VanSteenhouse, Yumi Kasai, David Gray, Carol Clugston, Anna Dominiczak and Alex Alderton, Roberto Amato, Jeffrey Barrett, Sonia Goncalves, Ewan Harrison, David K. Jackson, Ian Johnston, Dominic Kwiatkowski, Cordelia Langford, John Sillitoe on behalf of the Wellcome Sanger Institute COVID-19 Surveillance Team |
| EPI_ISL_2805058 | Lighthouse Lab in Milton Keynes | Wellcome Sanger Institute for the COVID-19 Genomics UK (COG-UK) Consortium | The Lighthouse Lab in Milton Keynes and Alex Alderton, Roberto Amato, Jeffrey Barrett, Sonia Goncalves, Ewan Harrison, David K. Jackson, Ian Johnston, Dominic Kwiatkowski, Cordelia Langford, John Sillitoe on behalf of the Wellcome Sanger Institute COVID-19 Surveillance Team |
| EPI_ISL_2805059 | Lighthouse Lab in Glasgow | Wellcome Sanger Institute for the COVID-19 Genomics UK (COG-UK) Consortium | Harper VanSteenhouse, Yumi Kasai, David Gray, Carol Clugston, Anna Dominiczak and Alex Alderton, Roberto Amato, Jeffrey Barrett, Sonia Goncalves, Ewan Harrison, David K. Jackson, Ian Johnston, Dominic Kwiatkowski, Cordelia Langford, John Sillitoe on behalf of the Wellcome Sanger Institute COVID-19 Surveillance Team |
| EPI_ISL_2805060, EPI_ISL_2805061 | Lighthouse Lab in Milton Keynes | Wellcome Sanger Institute for the COVID-19 Genomics UK (COG-UK) Consortium | The Lighthouse Lab in Milton Keynes and Alex Alderton, Roberto Amato, Jeffrey Barrett, Sonia Goncalves, Ewan Harrison, David K. Jackson, Ian Johnston, Dominic Kwiatkowski, Cordelia Langford, John Sillitoe on behalf of the Wellcome Sanger Institute COVID-19 Surveillance Team |
| EPI_ISL_2805062 | Lighthouse Lab in Glasgow | Wellcome Sanger Institute for the COVID-19 Genomics UK (COG-UK) Consortium | Harper VanSteenhouse, Yumi Kasai, David Gray, Carol Clugston, Anna Dominiczak and Alex Alderton, Roberto Amato, Jeffrey Barrett, Sonia Goncalves, Ewan Harrison, David K. Jackson, Ian Johnston, Dominic Kwiatkowski, Cordelia Langford, John Sillitoe on behalf of the Wellcome Sanger Institute COVID-19 Surveillance Team |
| EPI_ISL_2805063 | Lighthouse Lab in Milton Keynes | Wellcome Sanger Institute for the COVID-19 Genomics UK (COG-UK) Consortium | The Lighthouse Lab in Milton Keynes and Alex Alderton, Roberto Amato, Jeffrey Barrett, Sonia Goncalves, Ewan Harrison, David K. Jackson, Ian Johnston, Dominic Kwiatkowski, Cordelia Langford, John Sillitoe on behalf of the Wellcome Sanger Institute COVID-19 Surveillance Team |
| EPI_ISL_2805064 | Lighthouse Lab in Glasgow | Wellcome Sanger Institute for the COVID-19 Genomics UK (COG-UK) Consortium | Harper VanSteenhouse, Yumi Kasai, David Gray, Carol Clugston, Anna Dominiczak and Alex Alderton, Roberto Amato, Jeffrey Barrett, Sonia Goncalves, Ewan Harrison, David K. Jackson, Ian Johnston, Dominic Kwiatkowski, Cordelia Langford, John Sillitoe on behalf of the Wellcome Sanger Institute COVID-19 Surveillance Team |
| EPI_ISL_2805065, EPI_ISL_2805066, EPI_ISL_2805067 | Lighthouse Lab in Alderley Park | Wellcome Sanger Institute for the COVID-19 Genomics UK (COG-UK) Consortium | Jacquelyn Wynn, Mairead Hyland, The Lighthouse Lab in Alderley Park and Alex Alderton, Roberto Amato, Jeffrey Barrett, Sonia Goncalves, Ewan Harrison, David K. Jackson, Ian Johnston, Dominic Kwiatkowski, Cordelia Langford, John Sillitoe on behalf of the Wellcome Sanger Institute COVID-19 Surveillance Team |

| | | | |
|---|---|---|---|
| EPI_ISL_2805068, EPI_ISL_2805069 | Lighthouse Lab in Milton Keynes | Wellcome Sanger Institute for the COVID-19 Genomics UK (COG-UK) Consortium | The Lighthouse Lab in Milton Keynes and Alex Alderton, Roberto Amato, Jeffrey Barrett, Sonia Goncalves, Ewan Harrison, David K. Jackson, Ian Johnston, Dominic Kwiatkowski, Cordelia Langford, John Sillitoe on behalf of the Wellcome Sanger Institute COVID-19 Surveillance Team |
| EPI_ISL_2805070 | Lighthouse Lab in Glasgow | Wellcome Sanger Institute for the COVID-19 Genomics UK (COG-UK) Consortium | Harper VanSteenhouse, Yumi Kasai, David Gray, Carol Clugston, Anna Dominiczak and Alex Alderton, Roberto Amato, Jeffrey Barrett, Sonia Goncalves, Ewan Harrison, David K. Jackson, Ian Johnston, Dominic Kwiatkowski, Cordelia Langford, John Sillitoe on behalf of the Wellcome Sanger Institute COVID-19 Surveillance Team |
| EPI_ISL_2805071, EPI_ISL_2805072, EPI_ISL_2805073, EPI_ISL_2805074, EPI_ISL_2805075 | Lighthouse Lab in Alderley Park | Wellcome Sanger Institute for the COVID-19 Genomics UK (COG-UK) Consortium | Jacquelyn Wynn, Mairead Hyland, The Lighthouse Lab in Alderley Park and Alex Alderton, Roberto Amato, Jeffrey Barrett, Sonia Goncalves, Ewan Harrison, David K. Jackson, Ian Johnston, Dominic Kwiatkowski, Cordelia Langford, John Sillitoe on behalf of the Wellcome Sanger Institute COVID-19 Surveillance Team |
| EPI_ISL_2805076 | Lighthouse Lab in Glasgow | Wellcome Sanger Institute for the COVID-19 Genomics UK (COG-UK) Consortium | Harper VanSteenhouse, Yumi Kasai, David Gray, Carol Clugston, Anna Dominiczak and Alex Alderton, Roberto Amato, Jeffrey Barrett, Sonia Goncalves, Ewan Harrison, David K. Jackson, Ian Johnston, Dominic Kwiatkowski, Cordelia Langford, John Sillitoe on behalf of the Wellcome Sanger Institute COVID-19 Surveillance Team |
| EPI_ISL_2805077 | Lighthouse Lab in Milton Keynes | Wellcome Sanger Institute for the COVID-19 Genomics UK (COG-UK) Consortium | The Lighthouse Lab in Milton Keynes and Alex Alderton, Roberto Amato, Jeffrey Barrett, Sonia Goncalves, Ewan Harrison, David K. Jackson, Ian Johnston, Dominic Kwiatkowski, Cordelia Langford, John Sillitoe on behalf of the Wellcome Sanger Institute COVID-19 Surveillance Team |
| EPI_ISL_2805078 | Lighthouse Lab in Glasgow | Wellcome Sanger Institute for the COVID-19 Genomics UK (COG-UK) Consortium | Harper VanSteenhouse, Yumi Kasai, David Gray, Carol Clugston, Anna Dominiczak and Alex Alderton, Roberto Amato, Jeffrey Barrett, Sonia Goncalves, Ewan Harrison, David K. Jackson, Ian Johnston, Dominic Kwiatkowski, Cordelia Langford, John Sillitoe on behalf of the Wellcome Sanger Institute COVID-19 Surveillance Team |
| EPI_ISL_2805079 | Lighthouse Lab in Milton Keynes | Wellcome Sanger Institute for the COVID-19 Genomics UK (COG-UK) Consortium | The Lighthouse Lab in Milton Keynes and Alex Alderton, Roberto Amato, Jeffrey Barrett, Sonia Goncalves, Ewan Harrison, David K. Jackson, Ian Johnston, Dominic Kwiatkowski, Cordelia Langford, John Sillitoe on behalf of the Wellcome Sanger Institute COVID-19 Surveillance Team |
| EPI_ISL_2805081 | Lighthouse Lab in Glasgow | Wellcome Sanger Institute for the COVID-19 Genomics UK (COG-UK) Consortium | Harper VanSteenhouse, Yumi Kasai, David Gray, Carol Clugston, Anna Dominiczak and Alex Alderton, Roberto Amato, Jeffrey Barrett, Sonia Goncalves, Ewan Harrison, David K. Jackson, Ian Johnston, Dominic Kwiatkowski, Cordelia Langford, John Sillitoe on behalf of the Wellcome Sanger Institute COVID-19 Surveillance Team |
| EPI_ISL_2805082 | Lighthouse Lab in Milton Keynes | Wellcome Sanger Institute for the COVID-19 Genomics UK (COG-UK) Consortium | The Lighthouse Lab in Milton Keynes and Alex Alderton, Roberto Amato, Jeffrey Barrett, Sonia Goncalves, Ewan Harrison, David K. Jackson, Ian Johnston, Dominic Kwiatkowski, Cordelia Langford, John Sillitoe on behalf of the Wellcome Sanger Institute COVID-19 Surveillance Team |
| EPI_ISL_2805083 | Lighthouse Lab in Alderley Park | Wellcome Sanger Institute for the COVID-19 Genomics UK (COG-UK) Consortium | Jacquelyn Wynn, Mairead Hyland, The Lighthouse Lab in Alderley Park and Alex Alderton, Roberto Amato, Jeffrey Barrett, Sonia Goncalves, Ewan Harrison, David K. Jackson, Ian Johnston, Dominic Kwiatkowski, Cordelia Langford, John Sillitoe on behalf of the Wellcome Sanger Institute COVID-19 Surveillance Team |
| EPI_ISL_2805084 | Lighthouse Lab in Glasgow | Wellcome Sanger Institute for the COVID-19 Genomics UK (COG-UK) Consortium | Harper VanSteenhouse, Yumi Kasai, David Gray, Carol Clugston, Anna Dominiczak and Alex Alderton, Roberto Amato, Jeffrey Barrett, Sonia Goncalves, Ewan Harrison, David K. Jackson, Ian Johnston, Dominic Kwiatkowski, Cordelia Langford, John Sillitoe on behalf of the Wellcome Sanger Institute COVID-19 Surveillance Team |
| EPI_ISL_2805085, EPI_ISL_2805086, EPI_ISL_2805087 | Lighthouse Lab in Milton Keynes | Wellcome Sanger Institute for the COVID-19 Genomics UK (COG-UK) Consortium | The Lighthouse Lab in Milton Keynes and Alex Alderton, Roberto Amato, Jeffrey Barrett, Sonia Goncalves, Ewan Harrison, David K. Jackson, Ian Johnston, Dominic Kwiatkowski, Cordelia Langford, John Sillitoe on behalf of the Wellcome Sanger Institute COVID-19 Surveillance Team |
| EPI_ISL_2805089 | Lighthouse Lab in Glasgow | Wellcome Sanger Institute for the COVID-19 Genomics UK (COG-UK) Consortium | Harper VanSteenhouse, Yumi Kasai, David Gray, Carol Clugston, Anna Dominiczak and Alex Alderton, Roberto Amato, Jeffrey Barrett, Sonia Goncalves, Ewan Harrison, David K. Jackson, Ian Johnston, Dominic Kwiatkowski, Cordelia Langford, John Sillitoe on behalf of the Wellcome Sanger Institute COVID-19 Surveillance Team |
| EPI_ISL_2805090 | Lighthouse Lab in Milton Keynes | Wellcome Sanger Institute for the COVID-19 Genomics UK (COG-UK) Consortium | The Lighthouse Lab in Milton Keynes and Alex Alderton, Roberto Amato, Jeffrey Barrett, Sonia Goncalves, Ewan Harrison, David K. Jackson, Ian Johnston, Dominic Kwiatkowski, Cordelia Langford, John Sillitoe on behalf of the Wellcome Sanger Institute COVID-19 Surveillance Team |
| EPI_ISL_2805091 | Lighthouse Lab in Glasgow | Wellcome Sanger Institute for the COVID-19 Genomics UK (COG-UK) Consortium | Harper VanSteenhouse, Yumi Kasai, David Gray, Carol Clugston, Anna Dominiczak and Alex Alderton, Roberto Amato, Jeffrey Barrett, Sonia Goncalves, Ewan Harrison, David K. Jackson, Ian Johnston, Dominic Kwiatkowski, Cordelia Langford, John Sillitoe on behalf of the Wellcome Sanger Institute COVID-19 Surveillance Team |
| EPI_ISL_2805092 | Lighthouse Lab in Milton Keynes | Wellcome Sanger Institute for the COVID-19 Genomics UK (COG-UK) Consortium | The Lighthouse Lab in Milton Keynes and Alex Alderton, Roberto Amato, Jeffrey Barrett, Sonia Goncalves, Ewan Harrison, David K. Jackson, Ian Johnston, Dominic Kwiatkowski, Cordelia Langford, John Sillitoe on behalf of the Wellcome Sanger Institute COVID-19 Surveillance Team |
| EPI_ISL_2805093 | Lighthouse Lab in Glasgow | Wellcome Sanger Institute for the COVID-19 Genomics UK (COG-UK) Consortium | Harper VanSteenhouse, Yumi Kasai, David Gray, Carol Clugston, Anna Dominiczak and Alex Alderton, Roberto Amato, Jeffrey Barrett, Sonia Goncalves, Ewan Harrison, David K. Jackson, Ian Johnston, Dominic Kwiatkowski, Cordelia Langford, John Sillitoe on behalf of the Wellcome Sanger Institute COVID-19 Surveillance Team |
| EPI_ISL_2805094 | Lighthouse Lab in Milton Keynes | Wellcome Sanger Institute for the COVID-19 Genomics UK (COG-UK) Consortium | The Lighthouse Lab in Milton Keynes and Alex Alderton, Roberto Amato, Jeffrey Barrett, Sonia Goncalves, Ewan Harrison, David K. Jackson, Ian Johnston, Dominic Kwiatkowski, Cordelia Langford, John Sillitoe on behalf of the Wellcome Sanger Institute COVID-19 Surveillance Team |
| EPI_ISL_2805095 | Lighthouse Lab in Glasgow | Wellcome Sanger Institute for the COVID-19 Genomics UK (COG-UK) Consortium | Harper VanSteenhouse, Yumi Kasai, David Gray, Carol Clugston, Anna Dominiczak and Alex Alderton, Roberto Amato, Jeffrey Barrett, Sonia Goncalves, Ewan Harrison, David K. Jackson, Ian Johnston, Dominic Kwiatkowski, Cordelia Langford, John Sillitoe on behalf of the Wellcome Sanger Institute COVID-19 Surveillance Team |
| EPI_ISL_2805096 | Lighthouse Lab in Alderley Park | Wellcome Sanger Institute for the COVID-19 Genomics UK (COG-UK) Consortium | Jacquelyn Wynn, Mairead Hyland, The Lighthouse Lab in Alderley Park and Alex Alderton, Roberto Amato, Jeffrey Barrett, Sonia Goncalves, Ewan Harrison, David K. Jackson, Ian Johnston, Dominic Kwiatkowski, Cordelia Langford, John Sillitoe on behalf of the Wellcome Sanger Institute COVID-19 Surveillance Team |
| EPI_ISL_2805097 | Lighthouse Lab in Glasgow | Wellcome Sanger Institute for the COVID-19 Genomics UK (COG-UK) Consortium | Harper VanSteenhouse, Yumi Kasai, David Gray, Carol Clugston, Anna Dominiczak and Alex Alderton, Roberto Amato, Jeffrey Barrett, Sonia Goncalves, Ewan Harrison, David K. Jackson, Ian Johnston, Dominic Kwiatkowski, Cordelia Langford, John Sillitoe on behalf of the Wellcome Sanger Institute COVID-19 Surveillance Team |
| EPI_ISL_2805098 | Lighthouse Lab in Alderley Park | Wellcome Sanger Institute for the COVID-19 Genomics UK (COG-UK) Consortium | Jacquelyn Wynn, Mairead Hyland, The Lighthouse Lab in Alderley Park and Alex Alderton, Roberto Amato, Jeffrey Barrett, Sonia Goncalves, Ewan Harrison, David K. Jackson, Ian Johnston, Dominic Kwiatkowski, Cordelia Langford, John Sillitoe on behalf of the Wellcome Sanger Institute COVID-19 Surveillance Team |
| EPI_ISL_2805099, EPI_ISL_2805100 | Lighthouse Lab in Milton Keynes | Wellcome Sanger Institute for the COVID-19 Genomics UK (COG-UK) Consortium | The Lighthouse Lab in Milton Keynes and Alex Alderton, Roberto Amato, Jeffrey Barrett, Sonia Goncalves, Ewan Harrison, David K. Jackson, Ian Johnston, Dominic Kwiatkowski, Cordelia Langford, John Sillitoe on behalf of the Wellcome Sanger Institute COVID-19 Surveillance Team |
| EPI_ISL_2805101 | Lighthouse Lab in Alderley Park | Wellcome Sanger Institute for the COVID-19 Genomics UK (COG-UK) Consortium | Jacquelyn Wynn, Mairead Hyland, The Lighthouse Lab in Alderley Park and Alex Alderton, Roberto Amato, Jeffrey Barrett, Sonia Goncalves, Ewan Harrison, David K. Jackson, Ian Johnston, Dominic Kwiatkowski, Cordelia Langford, John Sillitoe on behalf of the Wellcome Sanger Institute COVID-19 Surveillance Team |
| EPI_ISL_2805102, EPI_ISL_2805103 | Lighthouse Lab in Glasgow | Wellcome Sanger Institute for the COVID-19 Genomics UK (COG-UK) Consortium | Harper VanSteenhouse, Yumi Kasai, David Gray, Carol Clugston, Anna Dominiczak and Alex Alderton, Roberto Amato, Jeffrey Barrett, Sonia Goncalves, Ewan Harrison, David K. Jackson, Ian Johnston, Dominic Kwiatkowski, Cordelia Langford, John Sillitoe on behalf of the Wellcome Sanger Institute COVID-19 Surveillance Team |
| EPI_ISL_2805104 | Lighthouse Lab in Milton Keynes | Wellcome Sanger Institute for the COVID-19 Genomics UK (COG-UK) Consortium | The Lighthouse Lab in Milton Keynes and Alex Alderton, Roberto Amato, Jeffrey Barrett, Sonia Goncalves, Ewan Harrison, David K. Jackson, Ian Johnston, Dominic Kwiatkowski, Cordelia Langford, John Sillitoe on behalf of the Wellcome Sanger Institute COVID-19 Surveillance Team |
| EPI_ISL_2805105, EPI_ISL_2805106, EPI_ISL_2805107 | Lighthouse Lab in Alderley Park | Wellcome Sanger Institute for the COVID-19 Genomics UK (COG-UK) Consortium | Jacquelyn Wynn, Mairead Hyland, The Lighthouse Lab in Alderley Park and Alex Alderton, Roberto Amato, Jeffrey Barrett, Sonia Goncalves, Ewan Harrison, David K. Jackson, Ian Johnston, Dominic Kwiatkowski, Cordelia Langford, John Sillitoe on behalf of the Wellcome Sanger Institute COVID-19 Surveillance Team |
| EPI_ISL_2805108, EPI_ISL_2805109, EPI_ISL_2805110, EPI_ISL_2805111, EPI_ISL_2805112 | Lighthouse Lab in Milton Keynes | Wellcome Sanger Institute for the COVID-19 Genomics UK (COG-UK) Consortium | The Lighthouse Lab in Milton Keynes and Alex Alderton, Roberto Amato, Jeffrey Barrett, Sonia Goncalves, Ewan Harrison, David K. Jackson, Ian Johnston, Dominic Kwiatkowski, Cordelia Langford, John Sillitoe on behalf of the Wellcome Sanger Institute COVID-19 Surveillance Team |
| EPI_ISL_2805113 | Lighthouse Lab in Glasgow | Wellcome Sanger Institute for the COVID-19 Genomics UK (COG-UK) Consortium | Harper VanSteenhouse, Yumi Kasai, David Gray, Carol Clugston, Anna Dominiczak and Alex Alderton, Roberto Amato, Jeffrey Barrett, Sonia Goncalves, Ewan Harrison, David K. Jackson, Ian Johnston, Dominic Kwiatkowski, Cordelia Langford, John Sillitoe on behalf of the Wellcome Sanger Institute COVID-19 Surveillance Team |
| EPI_ISL_2805114 | Lighthouse Lab in Alderley Park | Wellcome Sanger Institute for the COVID-19 Genomics UK (COG-UK) Consortium | Jacquelyn Wynn, Mairead Hyland, The Lighthouse Lab in Alderley Park and Alex Alderton, Roberto Amato, Jeffrey Barrett, Sonia Goncalves, Ewan Harrison, David K. Jackson, Ian Johnston, Dominic Kwiatkowski, Cordelia Langford, John Sillitoe on behalf of the Wellcome Sanger Institute COVID-19 Surveillance Team |
| EPI_ISL_2805115, EPI_ISL_2805116, EPI_ISL_2805117 | Lighthouse Lab in Glasgow | Wellcome Sanger Institute for the COVID-19 Genomics UK (COG-UK) Consortium | Harper VanSteenhouse, Yumi Kasai, David Gray, Carol Clugston, Anna Dominiczak and Alex Alderton, Roberto Amato, Jeffrey Barrett, Sonia Goncalves, Ewan Harrison, David K. Jackson, Ian Johnston, Dominic Kwiatkowski, Cordelia Langford, John Sillitoe on behalf of the Wellcome Sanger Institute COVID-19 Surveillance Team |
| EPI_ISL_2805118, EPI_ISL_2805119 | Lighthouse Lab in Milton Keynes | Wellcome Sanger Institute for the COVID-19 Genomics UK (COG-UK) Consortium | The Lighthouse Lab in Milton Keynes and Alex Alderton, Roberto Amato, Jeffrey Barrett, Sonia Goncalves, Ewan Harrison, David K. Jackson, Ian Johnston, Dominic Kwiatkowski, Cordelia Langford, John Sillitoe on behalf of the Wellcome Sanger Institute COVID-19 Surveillance Team |
| EPI_ISL_2805120 | Lighthouse Lab in Glasgow | Wellcome Sanger Institute for the COVID-19 Genomics UK (COG-UK) Consortium | Harper VanSteenhouse, Yumi Kasai, David Gray, Carol Clugston, Anna Dominiczak and Alex Alderton, Roberto Amato, Jeffrey Barrett, Sonia Goncalves, Ewan Harrison, David K. Jackson, Ian Johnston, Dominic Kwiatkowski, Cordelia Langford, John Sillitoe on behalf of the Wellcome Sanger Institute COVID-19 Surveillance Team |
| EPI_ISL_2805121 | Lighthouse Lab in Alderley Park | Wellcome Sanger Institute for the COVID-19 Genomics UK (COG-UK) Consortium | Jacquelyn Wynn, Mairead Hyland, The Lighthouse Lab in Alderley Park and Alex Alderton, Roberto Amato, Jeffrey Barrett, Sonia Goncalves, Ewan Harrison, David K. Jackson, Ian Johnston, Dominic Kwiatkowski, Cordelia Langford, John Sillitoe on behalf of the Wellcome Sanger Institute COVID-19 Surveillance Team |
| EPI_ISL_2805122, EPI_ISL_2805123 | Lighthouse Lab in Glasgow | Wellcome Sanger Institute for the COVID-19 Genomics UK (COG-UK) Consortium | Harper VanSteenhouse, Yumi Kasai, David Gray, Carol Clugston, Anna Dominiczak and Alex Alderton, Roberto Amato, Jeffrey Barrett, Sonia Goncalves, Ewan Harrison, David K. Jackson, Ian Johnston, Dominic Kwiatkowski, Cordelia Langford, John Sillitoe on behalf of the Wellcome Sanger Institute COVID-19 Surveillance Team |
| EPI_ISL_2805124 | Lighthouse Lab in Alderley Park | Wellcome Sanger Institute for the COVID-19 Genomics UK (COG-UK) Consortium | Jacquelyn Wynn, Mairead Hyland, The Lighthouse Lab in Alderley Park and Alex Alderton, Roberto Amato, Jeffrey Barrett, Sonia Goncalves, Ewan Harrison, David K. Jackson, Ian Johnston, Dominic Kwiatkowski, Cordelia Langford, John Sillitoe on behalf of the Wellcome Sanger Institute COVID-19 Surveillance Team |
| EPI_ISL_2805125, EPI_ISL_2805126, EPI_ISL_2805127, EPI_ISL_2805128 | Lighthouse Lab in Milton Keynes | Wellcome Sanger Institute for the COVID-19 Genomics UK (COG-UK) Consortium | The Lighthouse Lab in Milton Keynes and Alex Alderton, Roberto Amato, Jeffrey Barrett, Sonia Goncalves, Ewan Harrison, David K. Jackson, Ian Johnston, Dominic Kwiatkowski, Cordelia Langford, John Sillitoe on behalf of the Wellcome Sanger Institute COVID-19 Surveillance Team |
| EPI_ISL_2805129 | Lighthouse Lab in Glasgow | Wellcome Sanger Institute for the COVID-19 Genomics UK (COG-UK) Consortium | Harper VanSteenhouse, Yumi Kasai, David Gray, Carol Clugston, Anna Dominiczak and Alex Alderton, Roberto Amato, Jeffrey Barrett, Sonia Goncalves, Ewan Harrison, David K. Jackson, Ian Johnston, Dominic Kwiatkowski, Cordelia Langford, John Sillitoe on behalf of the Wellcome Sanger Institute COVID-19 Surveillance Team |
| EPI_ISL_2805130 | Lighthouse Lab in Milton Keynes | Wellcome Sanger Institute for the COVID-19 Genomics UK (COG-UK) Consortium | The Lighthouse Lab in Milton Keynes and Alex Alderton, Roberto Amato, Jeffrey Barrett, Sonia Goncalves, Ewan Harrison, David K. Jackson, Ian Johnston, Dominic Kwiatkowski, Cordelia Langford, John Sillitoe on behalf of the Wellcome Sanger Institute COVID-19 Surveillance Team |
| EPI_ISL_2805131, EPI_ISL_2805132 | Lighthouse Lab in Alderley Park | Wellcome Sanger Institute for the COVID-19 Genomics UK (COG-UK) Consortium | Jacquelyn Wynn, Mairead Hyland, The Lighthouse Lab in Alderley Park and Alex Alderton, Roberto Amato, Jeffrey Barrett, Sonia Goncalves, Ewan Harrison, David K. Jackson, Ian Johnston, Dominic Kwiatkowski, Cordelia Langford, John Sillitoe on behalf of the Wellcome Sanger Institute COVID-19 Surveillance Team |
| EPI_ISL_2805133 | Lighthouse Lab in Glasgow | Wellcome Sanger Institute for the COVID-19 Genomics UK (COG-UK) | Harper VanSteenhouse, Yumi Kasai, David Gray, Carol Clugston, Anna Dominiczak and Alex Alderton, Roberto Amato, Jeffrey Barrett, Sonia Goncalves, Ewan Harrison, David K. Jackson, Ian Johnston, Dominic Kwiatkowski, Cordelia Langford, |

| | | | |
|---|---|---|---|
| | | Consortium | John Sillitoe on behalf of the Wellcome Sanger Institute COVID-19 Surveillance Team |
| EPI_ISL_2805134 | Lighthouse Lab in Milton Keynes | Wellcome Sanger Institute for the COVID-19 Genomics UK (COG-UK) Consortium | The Lighthouse Lab in Milton Keynes and Alex Alderton, Roberto Amato, Jeffrey Barrett, Sonia Goncalves, Ewan Harrison, David K. Jackson, Ian Johnston, Dominic Kwiatkowski, Cordelia Langford, John Sillitoe on behalf of the Wellcome Sanger Institute COVID-19 Surveillance Team |
| EPI_ISL_2805135 | Lighthouse Lab in Glasgow | Wellcome Sanger Institute for the COVID-19 Genomics UK (COG-UK) Consortium | Harper VanSteenhouse, Yumi Kasai, David Gray, Carol Clugston, Anna Dominiczak and Alex Alderton, Roberto Amato, Jeffrey Barrett, Sonia Goncalves, Ewan Harrison, David K. Jackson, Ian Johnston, Dominic Kwiatkowski, Cordelia Langford, John Sillitoe on behalf of the Wellcome Sanger Institute COVID-19 Surveillance Team |
| EPI_ISL_2805136 | Lighthouse Lab in Alderley Park | Wellcome Sanger Institute for the COVID-19 Genomics UK (COG-UK) Consortium | Jacquelyn Wynn, Mairead Hyland, The Lighthouse Lab in Alderley Park and Alex Alderton, Roberto Amato, Jeffrey Barrett, Sonia Goncalves, Ewan Harrison, David K. Jackson, Ian Johnston, Dominic Kwiatkowski, Cordelia Langford, John Sillitoe on behalf of the Wellcome Sanger Institute COVID-19 Surveillance Team |
| EPI_ISL_2805137, EPI_ISL_2805138, EPI_ISL_2805139, EPI_ISL_2805140 | Lighthouse Lab in Milton Keynes | Wellcome Sanger Institute for the COVID-19 Genomics UK (COG-UK) Consortium | The Lighthouse Lab in Milton Keynes and Alex Alderton, Roberto Amato, Jeffrey Barrett, Sonia Goncalves, Ewan Harrison, David K. Jackson, Ian Johnston, Dominic Kwiatkowski, Cordelia Langford, John Sillitoe on behalf of the Wellcome Sanger Institute COVID-19 Surveillance Team |
| EPI_ISL_2805141, EPI_ISL_2805142 | Lighthouse Lab in Glasgow | Wellcome Sanger Institute for the COVID-19 Genomics UK (COG-UK) Consortium | Harper VanSteenhouse, Yumi Kasai, David Gray, Carol Clugston, Anna Dominiczak and Alex Alderton, Roberto Amato, Jeffrey Barrett, Sonia Goncalves, Ewan Harrison, David K. Jackson, Ian Johnston, Dominic Kwiatkowski, Cordelia Langford, John Sillitoe on behalf of the Wellcome Sanger Institute COVID-19 Surveillance Team |
| EPI_ISL_2805143 | Lighthouse Lab in Alderley Park | Wellcome Sanger Institute for the COVID-19 Genomics UK (COG-UK) Consortium | Jacquelyn Wynn, Mairead Hyland, The Lighthouse Lab in Alderley Park and Alex Alderton, Roberto Amato, Jeffrey Barrett, Sonia Goncalves, Ewan Harrison, David K. Jackson, Ian Johnston, Dominic Kwiatkowski, Cordelia Langford, John Sillitoe on behalf of the Wellcome Sanger Institute COVID-19 Surveillance Team |
| EPI_ISL_2805144 | Lighthouse Lab in Milton Keynes | Wellcome Sanger Institute for the COVID-19 Genomics UK (COG-UK) Consortium | The Lighthouse Lab in Milton Keynes and Alex Alderton, Roberto Amato, Jeffrey Barrett, Sonia Goncalves, Ewan Harrison, David K. Jackson, Ian Johnston, Dominic Kwiatkowski, Cordelia Langford, John Sillitoe on behalf of the Wellcome Sanger Institute COVID-19 Surveillance Team |
| EPI_ISL_2805145 | Lighthouse Lab in Alderley Park | Wellcome Sanger Institute for the COVID-19 Genomics UK (COG-UK) Consortium | Jacquelyn Wynn, Mairead Hyland, The Lighthouse Lab in Alderley Park and Alex Alderton, Roberto Amato, Jeffrey Barrett, Sonia Goncalves, Ewan Harrison, David K. Jackson, Ian Johnston, Dominic Kwiatkowski, Cordelia Langford, John Sillitoe on behalf of the Wellcome Sanger Institute COVID-19 Surveillance Team |
| EPI_ISL_2805146, EPI_ISL_2805147 | Lighthouse Lab in Milton Keynes | Wellcome Sanger Institute for the COVID-19 Genomics UK (COG-UK) Consortium | The Lighthouse Lab in Milton Keynes and Alex Alderton, Roberto Amato, Jeffrey Barrett, Sonia Goncalves, Ewan Harrison, David K. Jackson, Ian Johnston, Dominic Kwiatkowski, Cordelia Langford, John Sillitoe on behalf of the Wellcome Sanger Institute COVID-19 Surveillance Team |
| EPI_ISL_2805148, EPI_ISL_2805149 | Lighthouse Lab in Alderley Park | Wellcome Sanger Institute for the COVID-19 Genomics UK (COG-UK) Consortium | Jacquelyn Wynn, Mairead Hyland, The Lighthouse Lab in Alderley Park and Alex Alderton, Roberto Amato, Jeffrey Barrett, Sonia Goncalves, Ewan Harrison, David K. Jackson, Ian Johnston, Dominic Kwiatkowski, Cordelia Langford, John Sillitoe on behalf of the Wellcome Sanger Institute COVID-19 Surveillance Team |
| EPI_ISL_2805150 | Lighthouse Lab in Glasgow | Wellcome Sanger Institute for the COVID-19 Genomics UK (COG-UK) Consortium | Harper VanSteenhouse, Yumi Kasai, David Gray, Carol Clugston, Anna Dominiczak and Alex Alderton, Roberto Amato, Jeffrey Barrett, Sonia Goncalves, Ewan Harrison, David K. Jackson, Ian Johnston, Dominic Kwiatkowski, Cordelia Langford, John Sillitoe on behalf of the Wellcome Sanger Institute COVID-19 Surveillance Team |
| EPI_ISL_2805151 | Lighthouse Lab in Alderley Park | Wellcome Sanger Institute for the COVID-19 Genomics UK (COG-UK) Consortium | Jacquelyn Wynn, Mairead Hyland, The Lighthouse Lab in Alderley Park and Alex Alderton, Roberto Amato, Jeffrey Barrett, Sonia Goncalves, Ewan Harrison, David K. Jackson, Ian Johnston, Dominic Kwiatkowski, Cordelia Langford, John Sillitoe on behalf of the Wellcome Sanger Institute COVID-19 Surveillance Team |
| EPI_ISL_2805152 | Lighthouse Lab in Milton Keynes | Wellcome Sanger Institute for the COVID-19 Genomics UK (COG-UK) Consortium | The Lighthouse Lab in Milton Keynes and Alex Alderton, Roberto Amato, Jeffrey Barrett, Sonia Goncalves, Ewan Harrison, David K. Jackson, Ian Johnston, Dominic Kwiatkowski, Cordelia Langford, John Sillitoe on behalf of the Wellcome Sanger Institute COVID-19 Surveillance Team |
| EPI_ISL_2805153, EPI_ISL_2805154 | Lighthouse Lab in Alderley Park | Wellcome Sanger Institute for the COVID-19 Genomics UK (COG-UK) Consortium | Jacquelyn Wynn, Mairead Hyland, The Lighthouse Lab in Alderley Park and Alex Alderton, Roberto Amato, Jeffrey Barrett, Sonia Goncalves, Ewan Harrison, David K. Jackson, Ian Johnston, Dominic Kwiatkowski, Cordelia Langford, John Sillitoe on behalf of the Wellcome Sanger Institute COVID-19 Surveillance Team |
| EPI_ISL_2805155 | Lighthouse Lab in Milton Keynes | Wellcome Sanger Institute for the COVID-19 Genomics UK (COG-UK) Consortium | The Lighthouse Lab in Milton Keynes and Alex Alderton, Roberto Amato, Jeffrey Barrett, Sonia Goncalves, Ewan Harrison, David K. Jackson, Ian Johnston, Dominic Kwiatkowski, Cordelia Langford, John Sillitoe on behalf of the Wellcome Sanger Institute COVID-19 Surveillance Team |
| EPI_ISL_2805156 | Lighthouse Lab in Alderley Park | Wellcome Sanger Institute for the COVID-19 Genomics UK (COG-UK) Consortium | Jacquelyn Wynn, Mairead Hyland, The Lighthouse Lab in Alderley Park and Alex Alderton, Roberto Amato, Jeffrey Barrett, Sonia Goncalves, Ewan Harrison, David K. Jackson, Ian Johnston, Dominic Kwiatkowski, Cordelia Langford, John Sillitoe on behalf of the Wellcome Sanger Institute COVID-19 Surveillance Team |
| EPI_ISL_2805157, EPI_ISL_2805158 | Lighthouse Lab in Milton Keynes | Wellcome Sanger Institute for the COVID-19 Genomics UK (COG-UK) Consortium | The Lighthouse Lab in Milton Keynes and Alex Alderton, Roberto Amato, Jeffrey Barrett, Sonia Goncalves, Ewan Harrison, David K. Jackson, Ian Johnston, Dominic Kwiatkowski, Cordelia Langford, John Sillitoe on behalf of the Wellcome Sanger Institute COVID-19 Surveillance Team |
| EPI_ISL_2805159, EPI_ISL_2805160 | Lighthouse Lab in Glasgow | Wellcome Sanger Institute for the COVID-19 Genomics UK (COG-UK) Consortium | Harper VanSteenhouse, Yumi Kasai, David Gray, Carol Clugston, Anna Dominiczak and Alex Alderton, Roberto Amato, Jeffrey Barrett, Sonia Goncalves, Ewan Harrison, David K. Jackson, Ian Johnston, Dominic Kwiatkowski, Cordelia Langford, John Sillitoe on behalf of the Wellcome Sanger Institute COVID-19 Surveillance Team |
| EPI_ISL_2805161, EPI_ISL_2805162, EPI_ISL_2805163 | Lighthouse Lab in Milton Keynes | Wellcome Sanger Institute for the COVID-19 Genomics UK (COG-UK) Consortium | The Lighthouse Lab in Milton Keynes and Alex Alderton, Roberto Amato, Jeffrey Barrett, Sonia Goncalves, Ewan Harrison, David K. Jackson, Ian Johnston, Dominic Kwiatkowski, Cordelia Langford, John Sillitoe on behalf of the Wellcome Sanger Institute COVID-19 Surveillance Team |
| EPI_ISL_2805164, EPI_ISL_2805165 | Lighthouse Lab in Alderley Park | Wellcome Sanger Institute for the COVID-19 Genomics UK (COG-UK) Consortium | Jacquelyn Wynn, Mairead Hyland, The Lighthouse Lab in Alderley Park and Alex Alderton, Roberto Amato, Jeffrey Barrett, Sonia Goncalves, Ewan Harrison, David K. Jackson, Ian Johnston, Dominic Kwiatkowski, Cordelia Langford, John Sillitoe on behalf of the Wellcome Sanger Institute COVID-19 Surveillance Team |
| EPI_ISL_2805166 | Lighthouse Lab in Milton Keynes | Wellcome Sanger Institute for the COVID-19 Genomics UK (COG-UK) Consortium | The Lighthouse Lab in Milton Keynes and Alex Alderton, Roberto Amato, Jeffrey Barrett, Sonia Goncalves, Ewan Harrison, David K. Jackson, Ian Johnston, Dominic Kwiatkowski, Cordelia Langford, John Sillitoe on behalf of the Wellcome Sanger Institute COVID-19 Surveillance Team |
| EPI_ISL_2805167 | Lighthouse Lab in Alderley Park | Wellcome Sanger Institute for the COVID-19 Genomics UK (COG-UK) Consortium | Jacquelyn Wynn, Mairead Hyland, The Lighthouse Lab in Alderley Park and Alex Alderton, Roberto Amato, Jeffrey Barrett, Sonia Goncalves, Ewan Harrison, David K. Jackson, Ian Johnston, Dominic Kwiatkowski, Cordelia Langford, John Sillitoe on behalf of the Wellcome Sanger Institute COVID-19 Surveillance Team |
| EPI_ISL_2805168, EPI_ISL_2805169, EPI_ISL_2805170 | Lighthouse Lab in Milton Keynes | Wellcome Sanger Institute for the COVID-19 Genomics UK (COG-UK) Consortium | The Lighthouse Lab in Milton Keynes and Alex Alderton, Roberto Amato, Jeffrey Barrett, Sonia Goncalves, Ewan Harrison, David K. Jackson, Ian Johnston, Dominic Kwiatkowski, Cordelia Langford, John Sillitoe on behalf of the Wellcome Sanger Institute COVID-19 Surveillance Team |
| EPI_ISL_2805171 | Lighthouse Lab in Glasgow | Wellcome Sanger Institute for the COVID-19 Genomics UK (COG-UK) Consortium | Harper VanSteenhouse, Yumi Kasai, David Gray, Carol Clugston, Anna Dominiczak and Alex Alderton, Roberto Amato, Jeffrey Barrett, Sonia Goncalves, Ewan Harrison, David K. Jackson, Ian Johnston, Dominic Kwiatkowski, Cordelia Langford, John Sillitoe on behalf of the Wellcome Sanger Institute COVID-19 Surveillance Team |
| EPI_ISL_2805172 | Lighthouse Lab in Milton Keynes | Wellcome Sanger Institute for the COVID-19 Genomics UK (COG-UK) Consortium | The Lighthouse Lab in Milton Keynes and Alex Alderton, Roberto Amato, Jeffrey Barrett, Sonia Goncalves, Ewan Harrison, David K. Jackson, Ian Johnston, Dominic Kwiatkowski, Cordelia Langford, John Sillitoe on behalf of the Wellcome Sanger Institute COVID-19 Surveillance Team |
| EPI_ISL_2805173, EPI_ISL_2805174 | Lighthouse Lab in Alderley Park | Wellcome Sanger Institute for the COVID-19 Genomics UK (COG-UK) Consortium | Jacquelyn Wynn, Mairead Hyland, The Lighthouse Lab in Alderley Park and Alex Alderton, Roberto Amato, Jeffrey Barrett, Sonia Goncalves, Ewan Harrison, David K. Jackson, Ian Johnston, Dominic Kwiatkowski, Cordelia Langford, John Sillitoe on behalf of the Wellcome Sanger Institute COVID-19 Surveillance Team |
| EPI_ISL_2805175, EPI_ISL_2805176, EPI_ISL_2805177 | Lighthouse Lab in Milton Keynes | Wellcome Sanger Institute for the COVID-19 Genomics UK (COG-UK) Consortium | The Lighthouse Lab in Milton Keynes and Alex Alderton, Roberto Amato, Jeffrey Barrett, Sonia Goncalves, Ewan Harrison, David K. Jackson, Ian Johnston, Dominic Kwiatkowski, Cordelia Langford, John Sillitoe on behalf of the Wellcome Sanger Institute COVID-19 Surveillance Team |
| EPI_ISL_2805178, EPI_ISL_2805179 | Lighthouse Lab in Alderley Park | Wellcome Sanger Institute for the COVID-19 Genomics UK (COG-UK) Consortium | Jacquelyn Wynn, Mairead Hyland, The Lighthouse Lab in Alderley Park and Alex Alderton, Roberto Amato, Jeffrey Barrett, Sonia Goncalves, Ewan Harrison, David K. Jackson, Ian Johnston, Dominic Kwiatkowski, Cordelia Langford, John Sillitoe on behalf of the Wellcome Sanger Institute COVID-19 Surveillance Team |
| EPI_ISL_2805180, EPI_ISL_2805181, EPI_ISL_2805182, EPI_ISL_2805183, EPI_ISL_2805184 | Lighthouse Lab in Milton Keynes | Wellcome Sanger Institute for the COVID-19 Genomics UK (COG-UK) Consortium | The Lighthouse Lab in Milton Keynes and Alex Alderton, Roberto Amato, Jeffrey Barrett, Sonia Goncalves, Ewan Harrison, David K. Jackson, Ian Johnston, Dominic Kwiatkowski, Cordelia Langford, John Sillitoe on behalf of the Wellcome Sanger Institute COVID-19 Surveillance Team |
| EPI_ISL_2805185 | Lighthouse Lab in Glasgow | Wellcome Sanger Institute for the COVID-19 Genomics UK (COG-UK) Consortium | Harper VanSteenhouse, Yumi Kasai, David Gray, Carol Clugston, Anna Dominiczak and Alex Alderton, Roberto Amato, Jeffrey Barrett, Sonia Goncalves, Ewan Harrison, David K. Jackson, Ian Johnston, Dominic Kwiatkowski, Cordelia Langford, John Sillitoe on behalf of the Wellcome Sanger Institute COVID-19 Surveillance Team |
| EPI_ISL_2805186 | Lighthouse Lab in Milton Keynes | Wellcome Sanger Institute for the COVID-19 Genomics UK (COG-UK) Consortium | The Lighthouse Lab in Milton Keynes and Alex Alderton, Roberto Amato, Jeffrey Barrett, Sonia Goncalves, Ewan Harrison, David K. Jackson, Ian Johnston, Dominic Kwiatkowski, Cordelia Langford, John Sillitoe on behalf of the Wellcome Sanger Institute COVID-19 Surveillance Team |
| EPI_ISL_2805187 | Lighthouse Lab in Glasgow | Wellcome Sanger Institute for the COVID-19 Genomics UK (COG-UK) Consortium | Harper VanSteenhouse, Yumi Kasai, David Gray, Carol Clugston, Anna Dominiczak and Alex Alderton, Roberto Amato, Jeffrey Barrett, Sonia Goncalves, Ewan Harrison, David K. Jackson, Ian Johnston, Dominic Kwiatkowski, Cordelia Langford, John Sillitoe on behalf of the Wellcome Sanger Institute COVID-19 Surveillance Team |
| EPI_ISL_2805188 | Lighthouse Lab in Milton Keynes | Wellcome Sanger Institute for the COVID-19 Genomics UK (COG-UK) Consortium | The Lighthouse Lab in Milton Keynes and Alex Alderton, Roberto Amato, Jeffrey Barrett, Sonia Goncalves, Ewan Harrison, David K. Jackson, Ian Johnston, Dominic Kwiatkowski, Cordelia Langford, John Sillitoe on behalf of the Wellcome Sanger Institute COVID-19 Surveillance Team |
| EPI_ISL_2805189, EPI_ISL_2805190 | Lighthouse Lab in Glasgow | Wellcome Sanger Institute for the COVID-19 Genomics UK (COG-UK) Consortium | Harper VanSteenhouse, Yumi Kasai, David Gray, Carol Clugston, Anna Dominiczak and Alex Alderton, Roberto Amato, Jeffrey Barrett, Sonia Goncalves, Ewan Harrison, David K. Jackson, Ian Johnston, Dominic Kwiatkowski, Cordelia Langford, John Sillitoe on behalf of the Wellcome Sanger Institute COVID-19 Surveillance Team |
| EPI_ISL_2805191 | Lighthouse Lab in Milton Keynes | Wellcome Sanger Institute for the COVID-19 Genomics UK (COG-UK) Consortium | The Lighthouse Lab in Milton Keynes and Alex Alderton, Roberto Amato, Jeffrey Barrett, Sonia Goncalves, Ewan Harrison, David K. Jackson, Ian Johnston, Dominic Kwiatkowski, Cordelia Langford, John Sillitoe on behalf of the Wellcome Sanger Institute COVID-19 Surveillance Team |
| EPI_ISL_2805192, EPI_ISL_2805193, EPI_ISL_2805194 | Lighthouse Lab in Alderley Park | Wellcome Sanger Institute for the COVID-19 Genomics UK (COG-UK) Consortium | Jacquelyn Wynn, Mairead Hyland, The Lighthouse Lab in Alderley Park and Alex Alderton, Roberto Amato, Jeffrey Barrett, Sonia Goncalves, Ewan Harrison, David K. Jackson, Ian Johnston, Dominic Kwiatkowski, Cordelia Langford, John Sillitoe on behalf of the Wellcome Sanger Institute COVID-19 Surveillance Team |
| EPI_ISL_2805195 | Lighthouse Lab in Milton Keynes | Wellcome Sanger Institute for the COVID-19 Genomics UK (COG-UK) Consortium | The Lighthouse Lab in Milton Keynes and Alex Alderton, Roberto Amato, Jeffrey Barrett, Sonia Goncalves, Ewan Harrison, David K. Jackson, Ian Johnston, Dominic Kwiatkowski, Cordelia Langford, John Sillitoe on behalf of the Wellcome Sanger Institute COVID-19 Surveillance Team |
| EPI_ISL_2805196, EPI_ISL_2805197 | Lighthouse Lab in Alderley Park | Wellcome Sanger Institute for the COVID-19 Genomics UK (COG-UK) Consortium | Jacquelyn Wynn, Mairead Hyland, The Lighthouse Lab in Alderley Park and Alex Alderton, Roberto Amato, Jeffrey Barrett, Sonia Goncalves, Ewan Harrison, David K. Jackson, Ian Johnston, Dominic Kwiatkowski, Cordelia Langford, John Sillitoe on behalf of the Wellcome Sanger Institute COVID-19 Surveillance Team |
| EPI_ISL_2805198, EPI_ISL_2805199 | Lighthouse Lab in Glasgow | Wellcome Sanger Institute for the COVID-19 Genomics UK (COG-UK) Consortium | Harper VanSteenhouse, Yumi Kasai, David Gray, Carol Clugston, Anna Dominiczak and Alex Alderton, Roberto Amato, Jeffrey Barrett, Sonia Goncalves, Ewan Harrison, David K. Jackson, Ian Johnston, Dominic Kwiatkowski, Cordelia Langford, John Sillitoe on behalf of the Wellcome Sanger Institute COVID-19 Surveillance Team |
| EPI_ISL_2805200, EPI_ISL_2805201, EPI_ISL_2805202 | Lighthouse Lab in Milton Keynes | Wellcome Sanger Institute for the COVID-19 Genomics UK (COG-UK) Consortium | The Lighthouse Lab in Milton Keynes and Alex Alderton, Roberto Amato, Jeffrey Barrett, Sonia Goncalves, Ewan Harrison, David K. Jackson, Ian Johnston, Dominic Kwiatkowski, Cordelia Langford, John Sillitoe on behalf of the Wellcome Sanger Institute COVID-19 Surveillance Team |
| EPI_ISL_2805203 | Lighthouse Lab in Alderley Park | Wellcome Sanger Institute for the COVID-19 Genomics UK (COG-UK) | Jacquelyn Wynn, Mairead Hyland, The Lighthouse Lab in Alderley Park and Alex Alderton, Roberto Amato, Jeffrey Barrett, Sonia Goncalves, Ewan Harrison, David K. Jackson, Ian Johnston, Dominic Kwiatkowski, Cordelia Langford, John Sillitoe on |

| | | | |
|---|---|---|---|
| | | Consortium | behalf of the Wellcome Sanger Institute COVID-19 Surveillance Team |
| EPI_ISL_2805204 | Lighthouse Lab in Glasgow | Wellcome Sanger Institute for the COVID-19 Genomics UK (COG-UK) Consortium | Harper VanSteenhouse, Yumi Kasai, David Gray, Carol Clugston, Anna Dominiczak and Alex Alderton, Roberto Amato, Jeffrey Barrett, Sonia Goncalves, Ewan Harrison, David K. Jackson, Ian Johnston, Dominic Kwiatkowski, Cordelia Langford, John Sillitoe on behalf of the Wellcome Sanger Institute COVID-19 Surveillance Team |
| EPI_ISL_2805205, EPI_ISL_2805206 | Lighthouse Lab in Alderley Park | Wellcome Sanger Institute for the COVID-19 Genomics UK (COG-UK) Consortium | Jacquelyn Wynn, Mairead Hyland, The Lighthouse Lab in Alderley Park and Alex Alderton, Roberto Amato, Jeffrey Barrett, Sonia Goncalves, Ewan Harrison, David K. Jackson, Ian Johnston, Dominic Kwiatkowski, Cordelia Langford, John Sillitoe on behalf of the Wellcome Sanger Institute COVID-19 Surveillance Team |
| EPI_ISL_2805207 | Lighthouse Lab in Milton Keynes | Wellcome Sanger Institute for the COVID-19 Genomics UK (COG-UK) Consortium | The Lighthouse Lab in Milton Keynes and Alex Alderton, Roberto Amato, Jeffrey Barrett, Sonia Goncalves, Ewan Harrison, David K. Jackson, Ian Johnston, Dominic Kwiatkowski, Cordelia Langford, John Sillitoe on behalf of the Wellcome Sanger Institute COVID-19 Surveillance Team |
| EPI_ISL_2805208 | Lighthouse Lab in Glasgow | Wellcome Sanger Institute for the COVID-19 Genomics UK (COG-UK) Consortium | Harper VanSteenhouse, Yumi Kasai, David Gray, Carol Clugston, Anna Dominiczak and Alex Alderton, Roberto Amato, Jeffrey Barrett, Sonia Goncalves, Ewan Harrison, David K. Jackson, Ian Johnston, Dominic Kwiatkowski, Cordelia Langford, John Sillitoe on behalf of the Wellcome Sanger Institute COVID-19 Surveillance Team |
| EPI_ISL_2805209 | Lighthouse Lab in Milton Keynes | Wellcome Sanger Institute for the COVID-19 Genomics UK (COG-UK) Consortium | The Lighthouse Lab in Milton Keynes and Alex Alderton, Roberto Amato, Jeffrey Barrett, Sonia Goncalves, Ewan Harrison, David K. Jackson, Ian Johnston, Dominic Kwiatkowski, Cordelia Langford, John Sillitoe on behalf of the Wellcome Sanger Institute COVID-19 Surveillance Team |
| EPI_ISL_2805210 | Lighthouse Lab in Alderley Park | Wellcome Sanger Institute for the COVID-19 Genomics UK (COG-UK) Consortium | Jacquelyn Wynn, Mairead Hyland, The Lighthouse Lab in Alderley Park and Alex Alderton, Roberto Amato, Jeffrey Barrett, Sonia Goncalves, Ewan Harrison, David K. Jackson, Ian Johnston, Dominic Kwiatkowski, Cordelia Langford, John Sillitoe on behalf of the Wellcome Sanger Institute COVID-19 Surveillance Team |
| EPI_ISL_2805211 | Lighthouse Lab in Milton Keynes | Wellcome Sanger Institute for the COVID-19 Genomics UK (COG-UK) Consortium | The Lighthouse Lab in Milton Keynes and Alex Alderton, Roberto Amato, Jeffrey Barrett, Sonia Goncalves, Ewan Harrison, David K. Jackson, Ian Johnston, Dominic Kwiatkowski, Cordelia Langford, John Sillitoe on behalf of the Wellcome Sanger Institute COVID-19 Surveillance Team |
| EPI_ISL_2805212, EPI_ISL_2805213, EPI_ISL_2805214, EPI_ISL_2805215, EPI_ISL_2805216 | Lighthouse Lab in Alderley Park | Wellcome Sanger Institute for the COVID-19 Genomics UK (COG-UK) Consortium | Jacquelyn Wynn, Mairead Hyland, The Lighthouse Lab in Alderley Park and Alex Alderton, Roberto Amato, Jeffrey Barrett, Sonia Goncalves, Ewan Harrison, David K. Jackson, Ian Johnston, Dominic Kwiatkowski, Cordelia Langford, John Sillitoe on behalf of the Wellcome Sanger Institute COVID-19 Surveillance Team |
| EPI_ISL_2805217 | Lighthouse Lab in Milton Keynes | Wellcome Sanger Institute for the COVID-19 Genomics UK (COG-UK) Consortium | The Lighthouse Lab in Milton Keynes and Alex Alderton, Roberto Amato, Jeffrey Barrett, Sonia Goncalves, Ewan Harrison, David K. Jackson, Ian Johnston, Dominic Kwiatkowski, Cordelia Langford, John Sillitoe on behalf of the Wellcome Sanger Institute COVID-19 Surveillance Team |
| EPI_ISL_2805219 | Lighthouse Lab in Glasgow | Wellcome Sanger Institute for the COVID-19 Genomics UK (COG-UK) Consortium | Harper VanSteenhouse, Yumi Kasai, David Gray, Carol Clugston, Anna Dominiczak and Alex Alderton, Roberto Amato, Jeffrey Barrett, Sonia Goncalves, Ewan Harrison, David K. Jackson, Ian Johnston, Dominic Kwiatkowski, Cordelia Langford, John Sillitoe on behalf of the Wellcome Sanger Institute COVID-19 Surveillance Team |
| EPI_ISL_2805220 | Lighthouse Lab in Alderley Park | Wellcome Sanger Institute for the COVID-19 Genomics UK (COG-UK) Consortium | Jacquelyn Wynn, Mairead Hyland, The Lighthouse Lab in Alderley Park and Alex Alderton, Roberto Amato, Jeffrey Barrett, Sonia Goncalves, Ewan Harrison, David K. Jackson, Ian Johnston, Dominic Kwiatkowski, Cordelia Langford, John Sillitoe on behalf of the Wellcome Sanger Institute COVID-19 Surveillance Team |
| EPI_ISL_2805221 | Lighthouse Lab in Milton Keynes | Wellcome Sanger Institute for the COVID-19 Genomics UK (COG-UK) Consortium | The Lighthouse Lab in Milton Keynes and Alex Alderton, Roberto Amato, Jeffrey Barrett, Sonia Goncalves, Ewan Harrison, David K. Jackson, Ian Johnston, Dominic Kwiatkowski, Cordelia Langford, John Sillitoe on behalf of the Wellcome Sanger Institute COVID-19 Surveillance Team |
| EPI_ISL_2805222 | Lighthouse Lab in Alderley Park | Wellcome Sanger Institute for the COVID-19 Genomics UK (COG-UK) Consortium | Jacquelyn Wynn, Mairead Hyland, The Lighthouse Lab in Alderley Park and Alex Alderton, Roberto Amato, Jeffrey Barrett, Sonia Goncalves, Ewan Harrison, David K. Jackson, Ian Johnston, Dominic Kwiatkowski, Cordelia Langford, John Sillitoe on behalf of the Wellcome Sanger Institute COVID-19 Surveillance Team |
| EPI_ISL_2805223 | Lighthouse Lab in Glasgow | Wellcome Sanger Institute for the COVID-19 Genomics UK (COG-UK) Consortium | Harper VanSteenhouse, Yumi Kasai, David Gray, Carol Clugston, Anna Dominiczak and Alex Alderton, Roberto Amato, Jeffrey Barrett, Sonia Goncalves, Ewan Harrison, David K. Jackson, Ian Johnston, Dominic Kwiatkowski, Cordelia Langford, John Sillitoe on behalf of the Wellcome Sanger Institute COVID-19 Surveillance Team |
| EPI_ISL_2805224, EPI_ISL_2805225 | Lighthouse Lab in Alderley Park | Wellcome Sanger Institute for the COVID-19 Genomics UK (COG-UK) Consortium | Jacquelyn Wynn, Mairead Hyland, The Lighthouse Lab in Alderley Park and Alex Alderton, Roberto Amato, Jeffrey Barrett, Sonia Goncalves, Ewan Harrison, David K. Jackson, Ian Johnston, Dominic Kwiatkowski, Cordelia Langford, John Sillitoe on behalf of the Wellcome Sanger Institute COVID-19 Surveillance Team |
| EPI_ISL_2805226, EPI_ISL_2805227 | Lighthouse Lab in Milton Keynes | Wellcome Sanger Institute for the COVID-19 Genomics UK (COG-UK) Consortium | The Lighthouse Lab in Milton Keynes and Alex Alderton, Roberto Amato, Jeffrey Barrett, Sonia Goncalves, Ewan Harrison, David K. Jackson, Ian Johnston, Dominic Kwiatkowski, Cordelia Langford, John Sillitoe on behalf of the Wellcome Sanger Institute COVID-19 Surveillance Team |
| EPI_ISL_2805228 | Lighthouse Lab in Alderley Park | Wellcome Sanger Institute for the COVID-19 Genomics UK (COG-UK) Consortium | Jacquelyn Wynn, Mairead Hyland, The Lighthouse Lab in Alderley Park and Alex Alderton, Roberto Amato, Jeffrey Barrett, Sonia Goncalves, Ewan Harrison, David K. Jackson, Ian Johnston, Dominic Kwiatkowski, Cordelia Langford, John Sillitoe on behalf of the Wellcome Sanger Institute COVID-19 Surveillance Team |
| EPI_ISL_2805229 | Lighthouse Lab in Glasgow | Wellcome Sanger Institute for the COVID-19 Genomics UK (COG-UK) Consortium | Harper VanSteenhouse, Yumi Kasai, David Gray, Carol Clugston, Anna Dominiczak and Alex Alderton, Roberto Amato, Jeffrey Barrett, Sonia Goncalves, Ewan Harrison, David K. Jackson, Ian Johnston, Dominic Kwiatkowski, Cordelia Langford, John Sillitoe on behalf of the Wellcome Sanger Institute COVID-19 Surveillance Team |
| EPI_ISL_2805230 | Lighthouse Lab in Alderley Park | Wellcome Sanger Institute for the COVID-19 Genomics UK (COG-UK) Consortium | Jacquelyn Wynn, Mairead Hyland, The Lighthouse Lab in Alderley Park and Alex Alderton, Roberto Amato, Jeffrey Barrett, Sonia Goncalves, Ewan Harrison, David K. Jackson, Ian Johnston, Dominic Kwiatkowski, Cordelia Langford, John Sillitoe on behalf of the Wellcome Sanger Institute COVID-19 Surveillance Team |
| EPI_ISL_2805231 | Lighthouse Lab in Milton Keynes | Wellcome Sanger Institute for the COVID-19 Genomics UK (COG-UK) Consortium | The Lighthouse Lab in Milton Keynes and Alex Alderton, Roberto Amato, Jeffrey Barrett, Sonia Goncalves, Ewan Harrison, David K. Jackson, Ian Johnston, Dominic Kwiatkowski, Cordelia Langford, John Sillitoe on behalf of the Wellcome Sanger Institute COVID-19 Surveillance Team |
| EPI_ISL_2805232, EPI_ISL_2805233 | Lighthouse Lab in Alderley Park | Wellcome Sanger Institute for the COVID-19 Genomics UK (COG-UK) Consortium | Jacquelyn Wynn, Mairead Hyland, The Lighthouse Lab in Alderley Park and Alex Alderton, Roberto Amato, Jeffrey Barrett, Sonia Goncalves, Ewan Harrison, David K. Jackson, Ian Johnston, Dominic Kwiatkowski, Cordelia Langford, John Sillitoe on behalf of the Wellcome Sanger Institute COVID-19 Surveillance Team |
| EPI_ISL_2805234, EPI_ISL_2805235, EPI_ISL_2805236, EPI_ISL_2805237 | Lighthouse Lab in Milton Keynes | Wellcome Sanger Institute for the COVID-19 Genomics UK (COG-UK) Consortium | The Lighthouse Lab in Milton Keynes and Alex Alderton, Roberto Amato, Jeffrey Barrett, Sonia Goncalves, Ewan Harrison, David K. Jackson, Ian Johnston, Dominic Kwiatkowski, Cordelia Langford, John Sillitoe on behalf of the Wellcome Sanger Institute COVID-19 Surveillance Team |
| EPI_ISL_2805238, EPI_ISL_2805239 | Lighthouse Lab in Alderley Park | Wellcome Sanger Institute for the COVID-19 Genomics UK (COG-UK) Consortium | Jacquelyn Wynn, Mairead Hyland, The Lighthouse Lab in Alderley Park and Alex Alderton, Roberto Amato, Jeffrey Barrett, Sonia Goncalves, Ewan Harrison, David K. Jackson, Ian Johnston, Dominic Kwiatkowski, Cordelia Langford, John Sillitoe on behalf of the Wellcome Sanger Institute COVID-19 Surveillance Team |
| EPI_ISL_2805240, EPI_ISL_2805241 | Lighthouse Lab in Milton Keynes | Wellcome Sanger Institute for the COVID-19 Genomics UK (COG-UK) Consortium | The Lighthouse Lab in Milton Keynes and Alex Alderton, Roberto Amato, Jeffrey Barrett, Sonia Goncalves, Ewan Harrison, David K. Jackson, Ian Johnston, Dominic Kwiatkowski, Cordelia Langford, John Sillitoe on behalf of the Wellcome Sanger Institute COVID-19 Surveillance Team |
| EPI_ISL_2805242 | Lighthouse Lab in Glasgow | Wellcome Sanger Institute for the COVID-19 Genomics UK (COG-UK) Consortium | Harper VanSteenhouse, Yumi Kasai, David Gray, Carol Clugston, Anna Dominiczak and Alex Alderton, Roberto Amato, Jeffrey Barrett, Sonia Goncalves, Ewan Harrison, David K. Jackson, Ian Johnston, Dominic Kwiatkowski, Cordelia Langford, John Sillitoe on behalf of the Wellcome Sanger Institute COVID-19 Surveillance Team |
| EPI_ISL_2805243 | Lighthouse Lab in Alderley Park | Wellcome Sanger Institute for the COVID-19 Genomics UK (COG-UK) Consortium | Jacquelyn Wynn, Mairead Hyland, The Lighthouse Lab in Alderley Park and Alex Alderton, Roberto Amato, Jeffrey Barrett, Sonia Goncalves, Ewan Harrison, David K. Jackson, Ian Johnston, Dominic Kwiatkowski, Cordelia Langford, John Sillitoe on behalf of the Wellcome Sanger Institute COVID-19 Surveillance Team |
| EPI_ISL_2805244 | Lighthouse Lab in Milton Keynes | Wellcome Sanger Institute for the COVID-19 Genomics UK (COG-UK) Consortium | The Lighthouse Lab in Milton Keynes and Alex Alderton, Roberto Amato, Jeffrey Barrett, Sonia Goncalves, Ewan Harrison, David K. Jackson, Ian Johnston, Dominic Kwiatkowski, Cordelia Langford, John Sillitoe on behalf of the Wellcome Sanger Institute COVID-19 Surveillance Team |
| EPI_ISL_2805245, EPI_ISL_2805246, EPI_ISL_2805247, EPI_ISL_2805248 | Lighthouse Lab in Alderley Park | Wellcome Sanger Institute for the COVID-19 Genomics UK (COG-UK) Consortium | Jacquelyn Wynn, Mairead Hyland, The Lighthouse Lab in Alderley Park and Alex Alderton, Roberto Amato, Jeffrey Barrett, Sonia Goncalves, Ewan Harrison, David K. Jackson, Ian Johnston, Dominic Kwiatkowski, Cordelia Langford, John Sillitoe on behalf of the Wellcome Sanger Institute COVID-19 Surveillance Team |
| EPI_ISL_2805249 | Lighthouse Lab in Milton Keynes | Wellcome Sanger Institute for the COVID-19 Genomics UK (COG-UK) Consortium | The Lighthouse Lab in Milton Keynes and Alex Alderton, Roberto Amato, Jeffrey Barrett, Sonia Goncalves, Ewan Harrison, David K. Jackson, Ian Johnston, Dominic Kwiatkowski, Cordelia Langford, John Sillitoe on behalf of the Wellcome Sanger Institute COVID-19 Surveillance Team |
| EPI_ISL_2805250 | Lighthouse Lab in Alderley Park | Wellcome Sanger Institute for the COVID-19 Genomics UK (COG-UK) Consortium | Jacquelyn Wynn, Mairead Hyland, The Lighthouse Lab in Alderley Park and Alex Alderton, Roberto Amato, Jeffrey Barrett, Sonia Goncalves, Ewan Harrison, David K. Jackson, Ian Johnston, Dominic Kwiatkowski, Cordelia Langford, John Sillitoe on behalf of the Wellcome Sanger Institute COVID-19 Surveillance Team |
| EPI_ISL_2805251 | Lighthouse Lab in Milton Keynes | Wellcome Sanger Institute for the COVID-19 Genomics UK (COG-UK) Consortium | The Lighthouse Lab in Milton Keynes and Alex Alderton, Roberto Amato, Jeffrey Barrett, Sonia Goncalves, Ewan Harrison, David K. Jackson, Ian Johnston, Dominic Kwiatkowski, Cordelia Langford, John Sillitoe on behalf of the Wellcome Sanger Institute COVID-19 Surveillance Team |
| EPI_ISL_2805252 | Lighthouse Lab in Alderley Park | Wellcome Sanger Institute for the COVID-19 Genomics UK (COG-UK) Consortium | Jacquelyn Wynn, Mairead Hyland, The Lighthouse Lab in Alderley Park and Alex Alderton, Roberto Amato, Jeffrey Barrett, Sonia Goncalves, Ewan Harrison, David K. Jackson, Ian Johnston, Dominic Kwiatkowski, Cordelia Langford, John Sillitoe on behalf of the Wellcome Sanger Institute COVID-19 Surveillance Team |
| EPI_ISL_2805253 | Lighthouse Lab in Milton Keynes | Wellcome Sanger Institute for the COVID-19 Genomics UK (COG-UK) Consortium | The Lighthouse Lab in Milton Keynes and Alex Alderton, Roberto Amato, Jeffrey Barrett, Sonia Goncalves, Ewan Harrison, David K. Jackson, Ian Johnston, Dominic Kwiatkowski, Cordelia Langford, John Sillitoe on behalf of the Wellcome Sanger Institute COVID-19 Surveillance Team |
| EPI_ISL_2805254 | Lighthouse Lab in Alderley Park | Wellcome Sanger Institute for the COVID-19 Genomics UK (COG-UK) Consortium | Jacquelyn Wynn, Mairead Hyland, The Lighthouse Lab in Alderley Park and Alex Alderton, Roberto Amato, Jeffrey Barrett, Sonia Goncalves, Ewan Harrison, David K. Jackson, Ian Johnston, Dominic Kwiatkowski, Cordelia Langford, John Sillitoe on behalf of the Wellcome Sanger Institute COVID-19 Surveillance Team |
| EPI_ISL_2805255 | Lighthouse Lab in Milton Keynes | Wellcome Sanger Institute for the COVID-19 Genomics UK (COG-UK) Consortium | The Lighthouse Lab in Milton Keynes and Alex Alderton, Roberto Amato, Jeffrey Barrett, Sonia Goncalves, Ewan Harrison, David K. Jackson, Ian Johnston, Dominic Kwiatkowski, Cordelia Langford, John Sillitoe on behalf of the Wellcome Sanger Institute COVID-19 Surveillance Team |
| EPI_ISL_2805256 | Lighthouse Lab in Alderley Park | Wellcome Sanger Institute for the COVID-19 Genomics UK (COG-UK) Consortium | Jacquelyn Wynn, Mairead Hyland, The Lighthouse Lab in Alderley Park and Alex Alderton, Roberto Amato, Jeffrey Barrett, Sonia Goncalves, Ewan Harrison, David K. Jackson, Ian Johnston, Dominic Kwiatkowski, Cordelia Langford, John Sillitoe on behalf of the Wellcome Sanger Institute COVID-19 Surveillance Team |
| EPI_ISL_2805257 | Lighthouse Lab in Milton Keynes | Wellcome Sanger Institute for the COVID-19 Genomics UK (COG-UK) Consortium | The Lighthouse Lab in Milton Keynes and Alex Alderton, Roberto Amato, Jeffrey Barrett, Sonia Goncalves, Ewan Harrison, David K. Jackson, Ian Johnston, Dominic Kwiatkowski, Cordelia Langford, John Sillitoe on behalf of the Wellcome Sanger Institute COVID-19 Surveillance Team |
| EPI_ISL_2805258, EPI_ISL_2805259 | Lighthouse Lab in Alderley Park | Wellcome Sanger Institute for the COVID-19 Genomics UK (COG-UK) Consortium | Jacquelyn Wynn, Mairead Hyland, The Lighthouse Lab in Alderley Park and Alex Alderton, Roberto Amato, Jeffrey Barrett, Sonia Goncalves, Ewan Harrison, David K. Jackson, Ian Johnston, Dominic Kwiatkowski, Cordelia Langford, John Sillitoe on behalf of the Wellcome Sanger Institute COVID-19 Surveillance Team |
| EPI_ISL_2805260 | Lighthouse Lab in Milton Keynes | Wellcome Sanger Institute for the COVID-19 Genomics UK (COG-UK) Consortium | The Lighthouse Lab in Milton Keynes and Alex Alderton, Roberto Amato, Jeffrey Barrett, Sonia Goncalves, Ewan Harrison, David K. Jackson, Ian Johnston, Dominic Kwiatkowski, Cordelia Langford, John Sillitoe on behalf of the Wellcome Sanger Institute COVID-19 Surveillance Team |
| EPI_ISL_2805261, EPI_ISL_2805262, EPI_ISL_2805263 | Lighthouse Lab in Alderley Park | Wellcome Sanger Institute for the COVID-19 Genomics UK (COG-UK) Consortium | Jacquelyn Wynn, Mairead Hyland, The Lighthouse Lab in Alderley Park and Alex Alderton, Roberto Amato, Jeffrey Barrett, Sonia Goncalves, Ewan Harrison, David K. Jackson, Ian Johnston, Dominic Kwiatkowski, Cordelia Langford, John Sillitoe on behalf of the Wellcome Sanger Institute COVID-19 Surveillance Team |
| EPI_ISL_2805264 | Lighthouse Lab in Glasgow | Wellcome Sanger Institute for the COVID-19 Genomics UK (COG-UK) Consortium | Harper VanSteenhouse, Yumi Kasai, David Gray, Carol Clugston, Anna Dominiczak and Alex Alderton, Roberto Amato, Jeffrey Barrett, Sonia Goncalves, Ewan Harrison, David K. Jackson, Ian Johnston, Dominic Kwiatkowski, Cordelia Langford, John Sillitoe on behalf of the Wellcome Sanger Institute COVID-19 Surveillance Team |

| | | | |
|---|---|---|---|
| EPI_ISL_2805265, EPI_ISL_2805266 | Lighthouse Lab in Milton Keynes | Wellcome Sanger Institute for the COVID-19 Genomics UK (COG-UK) Consortium | The Lighthouse Lab in Milton Keynes and Alex Alderton, Roberto Amato, Jeffrey Barrett, Sonia Goncalves, Ewan Harrison, David K. Jackson, Ian Johnston, Dominic Kwiatkowski, Cordelia Langford, John Sillitoe on behalf of the Wellcome Sanger Institute COVID-19 Surveillance Team |
| EPI_ISL_2805267 | Lighthouse Lab in Alderley Park | Wellcome Sanger Institute for the COVID-19 Genomics UK (COG-UK) Consortium | Jacquelyn Wynn, Mairead Hyland, The Lighthouse Lab in Alderley Park and Alex Alderton, Roberto Amato, Jeffrey Barrett, Sonia Goncalves, Ewan Harrison, David K. Jackson, Ian Johnston, Dominic Kwiatkowski, Cordelia Langford, John Sillitoe on behalf of the Wellcome Sanger Institute COVID-19 Surveillance Team |
| EPI_ISL_2805268 | Lighthouse Lab in Milton Keynes | Wellcome Sanger Institute for the COVID-19 Genomics UK (COG-UK) Consortium | The Lighthouse Lab in Milton Keynes and Alex Alderton, Roberto Amato, Jeffrey Barrett, Sonia Goncalves, Ewan Harrison, David K. Jackson, Ian Johnston, Dominic Kwiatkowski, Cordelia Langford, John Sillitoe on behalf of the Wellcome Sanger Institute COVID-19 Surveillance Team |
| EPI_ISL_2805269 | Lighthouse Lab in Alderley Park | Wellcome Sanger Institute for the COVID-19 Genomics UK (COG-UK) Consortium | Jacquelyn Wynn, Mairead Hyland, The Lighthouse Lab in Alderley Park and Alex Alderton, Roberto Amato, Jeffrey Barrett, Sonia Goncalves, Ewan Harrison, David K. Jackson, Ian Johnston, Dominic Kwiatkowski, Cordelia Langford, John Sillitoe on behalf of the Wellcome Sanger Institute COVID-19 Surveillance Team |
| EPI_ISL_2805270 | Lighthouse Lab in Glasgow | Wellcome Sanger Institute for the COVID-19 Genomics UK (COG-UK) Consortium | Harper VanSteenhouse, Yumi Kasai, David Gray, Carol Clugston, Anna Dominiczak and Alex Alderton, Roberto Amato, Jeffrey Barrett, Sonia Goncalves, Ewan Harrison, David K. Jackson, Ian Johnston, Dominic Kwiatkowski, Cordelia Langford, John Sillitoe on behalf of the Wellcome Sanger Institute COVID-19 Surveillance Team |
| EPI_ISL_2805271 | Lighthouse Lab in Alderley Park | Wellcome Sanger Institute for the COVID-19 Genomics UK (COG-UK) Consortium | Jacquelyn Wynn, Mairead Hyland, The Lighthouse Lab in Alderley Park and Alex Alderton, Roberto Amato, Jeffrey Barrett, Sonia Goncalves, Ewan Harrison, David K. Jackson, Ian Johnston, Dominic Kwiatkowski, Cordelia Langford, John Sillitoe on behalf of the Wellcome Sanger Institute COVID-19 Surveillance Team |
| EPI_ISL_2805272 | Lighthouse Lab in Milton Keynes | Wellcome Sanger Institute for the COVID-19 Genomics UK (COG-UK) Consortium | The Lighthouse Lab in Milton Keynes and Alex Alderton, Roberto Amato, Jeffrey Barrett, Sonia Goncalves, Ewan Harrison, David K. Jackson, Ian Johnston, Dominic Kwiatkowski, Cordelia Langford, John Sillitoe on behalf of the Wellcome Sanger Institute COVID-19 Surveillance Team |
| EPI_ISL_2805273 | Lighthouse Lab in Alderley Park | Wellcome Sanger Institute for the COVID-19 Genomics UK (COG-UK) Consortium | Jacquelyn Wynn, Mairead Hyland, The Lighthouse Lab in Alderley Park and Alex Alderton, Roberto Amato, Jeffrey Barrett, Sonia Goncalves, Ewan Harrison, David K. Jackson, Ian Johnston, Dominic Kwiatkowski, Cordelia Langford, John Sillitoe on behalf of the Wellcome Sanger Institute COVID-19 Surveillance Team |
| EPI_ISL_2805274 | Lighthouse Lab in Milton Keynes | Wellcome Sanger Institute for the COVID-19 Genomics UK (COG-UK) Consortium | The Lighthouse Lab in Milton Keynes and Alex Alderton, Roberto Amato, Jeffrey Barrett, Sonia Goncalves, Ewan Harrison, David K. Jackson, Ian Johnston, Dominic Kwiatkowski, Cordelia Langford, John Sillitoe on behalf of the Wellcome Sanger Institute COVID-19 Surveillance Team |
| EPI_ISL_2805275 | Lighthouse Lab in Glasgow | Wellcome Sanger Institute for the COVID-19 Genomics UK (COG-UK) Consortium | Harper VanSteenhouse, Yumi Kasai, David Gray, Carol Clugston, Anna Dominiczak and Alex Alderton, Roberto Amato, Jeffrey Barrett, Sonia Goncalves, Ewan Harrison, David K. Jackson, Ian Johnston, Dominic Kwiatkowski, Cordelia Langford, John Sillitoe on behalf of the Wellcome Sanger Institute COVID-19 Surveillance Team |
| EPI_ISL_2805276 | Lighthouse Lab in Milton Keynes | Wellcome Sanger Institute for the COVID-19 Genomics UK (COG-UK) Consortium | The Lighthouse Lab in Milton Keynes and Alex Alderton, Roberto Amato, Jeffrey Barrett, Sonia Goncalves, Ewan Harrison, David K. Jackson, Ian Johnston, Dominic Kwiatkowski, Cordelia Langford, John Sillitoe on behalf of the Wellcome Sanger Institute COVID-19 Surveillance Team |
| EPI_ISL_2805277, EPI_ISL_2805278, EPI_ISL_2805279 | Lighthouse Lab in Alderley Park | Wellcome Sanger Institute for the COVID-19 Genomics UK (COG-UK) Consortium | Jacquelyn Wynn, Mairead Hyland, The Lighthouse Lab in Alderley Park and Alex Alderton, Roberto Amato, Jeffrey Barrett, Sonia Goncalves, Ewan Harrison, David K. Jackson, Ian Johnston, Dominic Kwiatkowski, Cordelia Langford, John Sillitoe on behalf of the Wellcome Sanger Institute COVID-19 Surveillance Team |
| EPI_ISL_2805280 | Lighthouse Lab in Milton Keynes | Wellcome Sanger Institute for the COVID-19 Genomics UK (COG-UK) Consortium | The Lighthouse Lab in Milton Keynes and Alex Alderton, Roberto Amato, Jeffrey Barrett, Sonia Goncalves, Ewan Harrison, David K. Jackson, Ian Johnston, Dominic Kwiatkowski, Cordelia Langford, John Sillitoe on behalf of the Wellcome Sanger Institute COVID-19 Surveillance Team |
| EPI_ISL_2805281 | Lighthouse Lab in Alderley Park | Wellcome Sanger Institute for the COVID-19 Genomics UK (COG-UK) Consortium | Jacquelyn Wynn, Mairead Hyland, The Lighthouse Lab in Alderley Park and Alex Alderton, Roberto Amato, Jeffrey Barrett, Sonia Goncalves, Ewan Harrison, David K. Jackson, Ian Johnston, Dominic Kwiatkowski, Cordelia Langford, John Sillitoe on behalf of the Wellcome Sanger Institute COVID-19 Surveillance Team |
| EPI_ISL_2805282, EPI_ISL_2805283, EPI_ISL_2805284 | Lighthouse Lab in Milton Keynes | Wellcome Sanger Institute for the COVID-19 Genomics UK (COG-UK) Consortium | The Lighthouse Lab in Milton Keynes and Alex Alderton, Roberto Amato, Jeffrey Barrett, Sonia Goncalves, Ewan Harrison, David K. Jackson, Ian Johnston, Dominic Kwiatkowski, Cordelia Langford, John Sillitoe on behalf of the Wellcome Sanger Institute COVID-19 Surveillance Team |
| EPI_ISL_2805285 | Lighthouse Lab in Glasgow | Wellcome Sanger Institute for the COVID-19 Genomics UK (COG-UK) Consortium | Harper VanSteenhouse, Yumi Kasai, David Gray, Carol Clugston, Anna Dominiczak and Alex Alderton, Roberto Amato, Jeffrey Barrett, Sonia Goncalves, Ewan Harrison, David K. Jackson, Ian Johnston, Dominic Kwiatkowski, Cordelia Langford, John Sillitoe on behalf of the Wellcome Sanger Institute COVID-19 Surveillance Team |
| EPI_ISL_2805286 | Lighthouse Lab in Milton Keynes | Wellcome Sanger Institute for the COVID-19 Genomics UK (COG-UK) Consortium | The Lighthouse Lab in Milton Keynes and Alex Alderton, Roberto Amato, Jeffrey Barrett, Sonia Goncalves, Ewan Harrison, David K. Jackson, Ian Johnston, Dominic Kwiatkowski, Cordelia Langford, John Sillitoe on behalf of the Wellcome Sanger Institute COVID-19 Surveillance Team |
| EPI_ISL_2805287 | Lighthouse Lab in Alderley Park | Wellcome Sanger Institute for the COVID-19 Genomics UK (COG-UK) Consortium | Jacquelyn Wynn, Mairead Hyland, The Lighthouse Lab in Alderley Park and Alex Alderton, Roberto Amato, Jeffrey Barrett, Sonia Goncalves, Ewan Harrison, David K. Jackson, Ian Johnston, Dominic Kwiatkowski, Cordelia Langford, John Sillitoe on behalf of the Wellcome Sanger Institute COVID-19 Surveillance Team |
| EPI_ISL_2805288, EPI_ISL_2805289 | Lighthouse Lab in Milton Keynes | Wellcome Sanger Institute for the COVID-19 Genomics UK (COG-UK) Consortium | The Lighthouse Lab in Milton Keynes and Alex Alderton, Roberto Amato, Jeffrey Barrett, Sonia Goncalves, Ewan Harrison, David K. Jackson, Ian Johnston, Dominic Kwiatkowski, Cordelia Langford, John Sillitoe on behalf of the Wellcome Sanger Institute COVID-19 Surveillance Team |
| EPI_ISL_2805290 | Lighthouse Lab in Glasgow | Wellcome Sanger Institute for the COVID-19 Genomics UK (COG-UK) Consortium | Harper VanSteenhouse, Yumi Kasai, David Gray, Carol Clugston, Anna Dominiczak and Alex Alderton, Roberto Amato, Jeffrey Barrett, Sonia Goncalves, Ewan Harrison, David K. Jackson, Ian Johnston, Dominic Kwiatkowski, Cordelia Langford, John Sillitoe on behalf of the Wellcome Sanger Institute COVID-19 Surveillance Team |
| EPI_ISL_2805291 | Lighthouse Lab in Milton Keynes | Wellcome Sanger Institute for the COVID-19 Genomics UK (COG-UK) Consortium | The Lighthouse Lab in Milton Keynes and Alex Alderton, Roberto Amato, Jeffrey Barrett, Sonia Goncalves, Ewan Harrison, David K. Jackson, Ian Johnston, Dominic Kwiatkowski, Cordelia Langford, John Sillitoe on behalf of the Wellcome Sanger Institute COVID-19 Surveillance Team |
| EPI_ISL_2805292 | Lighthouse Lab in Alderley Park | Wellcome Sanger Institute for the COVID-19 Genomics UK (COG-UK) Consortium | Jacquelyn Wynn, Mairead Hyland, The Lighthouse Lab in Alderley Park and Alex Alderton, Roberto Amato, Jeffrey Barrett, Sonia Goncalves, Ewan Harrison, David K. Jackson, Ian Johnston, Dominic Kwiatkowski, Cordelia Langford, John Sillitoe on behalf of the Wellcome Sanger Institute COVID-19 Surveillance Team |
| EPI_ISL_2805293 | Lighthouse Lab in Glasgow | Wellcome Sanger Institute for the COVID-19 Genomics UK (COG-UK) Consortium | Harper VanSteenhouse, Yumi Kasai, David Gray, Carol Clugston, Anna Dominiczak and Alex Alderton, Roberto Amato, Jeffrey Barrett, Sonia Goncalves, Ewan Harrison, David K. Jackson, Ian Johnston, Dominic Kwiatkowski, Cordelia Langford, John Sillitoe on behalf of the Wellcome Sanger Institute COVID-19 Surveillance Team |
| EPI_ISL_2805294 | Lighthouse Lab in Alderley Park | Wellcome Sanger Institute for the COVID-19 Genomics UK (COG-UK) Consortium | Jacquelyn Wynn, Mairead Hyland, The Lighthouse Lab in Alderley Park and Alex Alderton, Roberto Amato, Jeffrey Barrett, Sonia Goncalves, Ewan Harrison, David K. Jackson, Ian Johnston, Dominic Kwiatkowski, Cordelia Langford, John Sillitoe on behalf of the Wellcome Sanger Institute COVID-19 Surveillance Team |
| EPI_ISL_2805295 | Lighthouse Lab in Glasgow | Wellcome Sanger Institute for the COVID-19 Genomics UK (COG-UK) Consortium | Harper VanSteenhouse, Yumi Kasai, David Gray, Carol Clugston, Anna Dominiczak and Alex Alderton, Roberto Amato, Jeffrey Barrett, Sonia Goncalves, Ewan Harrison, David K. Jackson, Ian Johnston, Dominic Kwiatkowski, Cordelia Langford, John Sillitoe on behalf of the Wellcome Sanger Institute COVID-19 Surveillance Team |
| EPI_ISL_2805296 | Lighthouse Lab in Alderley Park | Wellcome Sanger Institute for the COVID-19 Genomics UK (COG-UK) Consortium | Jacquelyn Wynn, Mairead Hyland, The Lighthouse Lab in Alderley Park and Alex Alderton, Roberto Amato, Jeffrey Barrett, Sonia Goncalves, Ewan Harrison, David K. Jackson, Ian Johnston, Dominic Kwiatkowski, Cordelia Langford, John Sillitoe on behalf of the Wellcome Sanger Institute COVID-19 Surveillance Team |
| EPI_ISL_2805297, EPI_ISL_2805298 | Lighthouse Lab in Milton Keynes | Wellcome Sanger Institute for the COVID-19 Genomics UK (COG-UK) Consortium | The Lighthouse Lab in Milton Keynes and Alex Alderton, Roberto Amato, Jeffrey Barrett, Sonia Goncalves, Ewan Harrison, David K. Jackson, Ian Johnston, Dominic Kwiatkowski, Cordelia Langford, John Sillitoe on behalf of the Wellcome Sanger Institute COVID-19 Surveillance Team |
| EPI_ISL_2805299 | Lighthouse Lab in Alderley Park | Wellcome Sanger Institute for the COVID-19 Genomics UK (COG-UK) Consortium | Jacquelyn Wynn, Mairead Hyland, The Lighthouse Lab in Alderley Park and Alex Alderton, Roberto Amato, Jeffrey Barrett, Sonia Goncalves, Ewan Harrison, David K. Jackson, Ian Johnston, Dominic Kwiatkowski, Cordelia Langford, John Sillitoe on behalf of the Wellcome Sanger Institute COVID-19 Surveillance Team |
| EPI_ISL_2805300, EPI_ISL_2805301, EPI_ISL_2805302, EPI_ISL_2805303 | Lighthouse Lab in Milton Keynes | Wellcome Sanger Institute for the COVID-19 Genomics UK (COG-UK) Consortium | The Lighthouse Lab in Milton Keynes and Alex Alderton, Roberto Amato, Jeffrey Barrett, Sonia Goncalves, Ewan Harrison, David K. Jackson, Ian Johnston, Dominic Kwiatkowski, Cordelia Langford, John Sillitoe on behalf of the Wellcome Sanger Institute COVID-19 Surveillance Team |
| EPI_ISL_2805304, EPI_ISL_2805305, EPI_ISL_2805306 | Lighthouse Lab in Alderley Park | Wellcome Sanger Institute for the COVID-19 Genomics UK (COG-UK) Consortium | Jacquelyn Wynn, Mairead Hyland, The Lighthouse Lab in Alderley Park and Alex Alderton, Roberto Amato, Jeffrey Barrett, Sonia Goncalves, Ewan Harrison, David K. Jackson, Ian Johnston, Dominic Kwiatkowski, Cordelia Langford, John Sillitoe on behalf of the Wellcome Sanger Institute COVID-19 Surveillance Team |
| EPI_ISL_2805307 | Lighthouse Lab in Milton Keynes | Wellcome Sanger Institute for the COVID-19 Genomics UK (COG-UK) Consortium | The Lighthouse Lab in Milton Keynes and Alex Alderton, Roberto Amato, Jeffrey Barrett, Sonia Goncalves, Ewan Harrison, David K. Jackson, Ian Johnston, Dominic Kwiatkowski, Cordelia Langford, John Sillitoe on behalf of the Wellcome Sanger Institute COVID-19 Surveillance Team |
| EPI_ISL_2805308, EPI_ISL_2805309, EPI_ISL_2805310, EPI_ISL_2805311 | Lighthouse Lab in Alderley Park | Wellcome Sanger Institute for the COVID-19 Genomics UK (COG-UK) Consortium | Jacquelyn Wynn, Mairead Hyland, The Lighthouse Lab in Alderley Park and Alex Alderton, Roberto Amato, Jeffrey Barrett, Sonia Goncalves, Ewan Harrison, David K. Jackson, Ian Johnston, Dominic Kwiatkowski, Cordelia Langford, John Sillitoe on behalf of the Wellcome Sanger Institute COVID-19 Surveillance Team |
| EPI_ISL_2805312 | Lighthouse Lab in Milton Keynes | Wellcome Sanger Institute for the COVID-19 Genomics UK (COG-UK) Consortium | The Lighthouse Lab in Milton Keynes and Alex Alderton, Roberto Amato, Jeffrey Barrett, Sonia Goncalves, Ewan Harrison, David K. Jackson, Ian Johnston, Dominic Kwiatkowski, Cordelia Langford, John Sillitoe on behalf of the Wellcome Sanger Institute COVID-19 Surveillance Team |
| EPI_ISL_2805313, EPI_ISL_2805314 | Lighthouse Lab in Alderley Park | Wellcome Sanger Institute for the COVID-19 Genomics UK (COG-UK) Consortium | Jacquelyn Wynn, Mairead Hyland, The Lighthouse Lab in Alderley Park and Alex Alderton, Roberto Amato, Jeffrey Barrett, Sonia Goncalves, Ewan Harrison, David K. Jackson, Ian Johnston, Dominic Kwiatkowski, Cordelia Langford, John Sillitoe on behalf of the Wellcome Sanger Institute COVID-19 Surveillance Team |
| EPI_ISL_2805315 | Lighthouse Lab in Glasgow | Wellcome Sanger Institute for the COVID-19 Genomics UK (COG-UK) Consortium | Harper VanSteenhouse, Yumi Kasai, David Gray, Carol Clugston, Anna Dominiczak and Alex Alderton, Roberto Amato, Jeffrey Barrett, Sonia Goncalves, Ewan Harrison, David K. Jackson, Ian Johnston, Dominic Kwiatkowski, Cordelia Langford, John Sillitoe on behalf of the Wellcome Sanger Institute COVID-19 Surveillance Team |
| EPI_ISL_2805316, EPI_ISL_2805317 | Lighthouse Lab in Milton Keynes | Wellcome Sanger Institute for the COVID-19 Genomics UK (COG-UK) Consortium | The Lighthouse Lab in Milton Keynes and Alex Alderton, Roberto Amato, Jeffrey Barrett, Sonia Goncalves, Ewan Harrison, David K. Jackson, Ian Johnston, Dominic Kwiatkowski, Cordelia Langford, John Sillitoe on behalf of the Wellcome Sanger Institute COVID-19 Surveillance Team |
| EPI_ISL_2805318, EPI_ISL_2805319 | Lighthouse Lab in Alderley Park | Wellcome Sanger Institute for the COVID-19 Genomics UK (COG-UK) Consortium | Jacquelyn Wynn, Mairead Hyland, The Lighthouse Lab in Alderley Park and Alex Alderton, Roberto Amato, Jeffrey Barrett, Sonia Goncalves, Ewan Harrison, David K. Jackson, Ian Johnston, Dominic Kwiatkowski, Cordelia Langford, John Sillitoe on behalf of the Wellcome Sanger Institute COVID-19 Surveillance Team |
| EPI_ISL_2805320, EPI_ISL_2805321 | Lighthouse Lab in Milton Keynes | Wellcome Sanger Institute for the COVID-19 Genomics UK (COG-UK) Consortium | The Lighthouse Lab in Milton Keynes and Alex Alderton, Roberto Amato, Jeffrey Barrett, Sonia Goncalves, Ewan Harrison, David K. Jackson, Ian Johnston, Dominic Kwiatkowski, Cordelia Langford, John Sillitoe on behalf of the Wellcome Sanger Institute COVID-19 Surveillance Team |
| EPI_ISL_2805322 | Lighthouse Lab in Glasgow | Wellcome Sanger Institute for the COVID-19 Genomics UK (COG-UK) Consortium | Harper VanSteenhouse, Yumi Kasai, David Gray, Carol Clugston, Anna Dominiczak and Alex Alderton, Roberto Amato, Jeffrey Barrett, Sonia Goncalves, Ewan Harrison, David K. Jackson, Ian Johnston, Dominic Kwiatkowski, Cordelia Langford, John Sillitoe on behalf of the Wellcome Sanger Institute COVID-19 Surveillance Team |
| EPI_ISL_2805323 | Lighthouse Lab in Milton Keynes | Wellcome Sanger Institute for the COVID-19 Genomics UK (COG-UK) Consortium | The Lighthouse Lab in Milton Keynes and Alex Alderton, Roberto Amato, Jeffrey Barrett, Sonia Goncalves, Ewan Harrison, David K. Jackson, Ian Johnston, Dominic Kwiatkowski, Cordelia Langford, John Sillitoe on behalf of the Wellcome Sanger Institute COVID-19 Surveillance Team |
| EPI_ISL_2805324 | Lighthouse Lab in Glasgow | Wellcome Sanger Institute for the COVID-19 Genomics UK (COG-UK) Consortium | Harper VanSteenhouse, Yumi Kasai, David Gray, Carol Clugston, Anna Dominiczak and Alex Alderton, Roberto Amato, Jeffrey Barrett, Sonia Goncalves, Ewan Harrison, David K. Jackson, Ian Johnston, Dominic Kwiatkowski, Cordelia Langford, John Sillitoe on behalf of the Wellcome Sanger Institute COVID-19 Surveillance Team |
| EPI_ISL_2805325 | Lighthouse Lab in Alderley Park | Wellcome Sanger Institute for the COVID-19 Genomics UK (COG-UK) Consortium | Jacquelyn Wynn, Mairead Hyland, The Lighthouse Lab in Alderley Park and Alex Alderton, Roberto Amato, Jeffrey Barrett, Sonia Goncalves, Ewan Harrison, David K. Jackson, Ian Johnston, Dominic Kwiatkowski, Cordelia Langford, John Sillitoe on behalf of the Wellcome Sanger Institute COVID-19 Surveillance Team |

| | | | |
|---|---|---|---|
| EPI_ISL_2805326, EPI_ISL_2805327 | Lighthouse Lab in Milton Keynes | Wellcome Sanger Institute for the COVID-19 Genomics UK (COG-UK) Consortium | The Lighthouse Lab in Milton Keynes and Alex Alderton, Roberto Amato, Jeffrey Barrett, Sonia Goncalves, Ewan Harrison, David K. Jackson, Ian Johnston, Dominic Kwiatkowski, Cordelia Langford, John Sillitoe on behalf of the Wellcome Sanger Institute COVID-19 Surveillance Team |
| EPI_ISL_2805328, EPI_ISL_2805329 | Lighthouse Lab in Glasgow | Wellcome Sanger Institute for the COVID-19 Genomics UK (COG-UK) Consortium | Harper VanSteenhouse, Yumi Kasai, David Gray, Carol Clugston, Anna Dominiczak and Alex Alderton, Roberto Amato, Jeffrey Barrett, Sonia Goncalves, Ewan Harrison, David K. Jackson, Ian Johnston, Dominic Kwiatkowski, Cordelia Langford, John Sillitoe on behalf of the Wellcome Sanger Institute COVID-19 Surveillance Team |
| EPI_ISL_2805330 | Lighthouse Lab in Milton Keynes | Wellcome Sanger Institute for the COVID-19 Genomics UK (COG-UK) Consortium | The Lighthouse Lab in Milton Keynes and Alex Alderton, Roberto Amato, Jeffrey Barrett, Sonia Goncalves, Ewan Harrison, David K. Jackson, Ian Johnston, Dominic Kwiatkowski, Cordelia Langford, John Sillitoe on behalf of the Wellcome Sanger Institute COVID-19 Surveillance Team |
| EPI_ISL_2805331 | Lighthouse Lab in Alderley Park | Wellcome Sanger Institute for the COVID-19 Genomics UK (COG-UK) Consortium | Jacquelyn Wynn, Mairead Hyland, The Lighthouse Lab in Alderley Park and Alex Alderton, Roberto Amato, Jeffrey Barrett, Sonia Goncalves, Ewan Harrison, David K. Jackson, Ian Johnston, Dominic Kwiatkowski, Cordelia Langford, John Sillitoe on behalf of the Wellcome Sanger Institute COVID-19 Surveillance Team |
| EPI_ISL_2805332, EPI_ISL_2805333 | Lighthouse Lab in Milton Keynes | Wellcome Sanger Institute for the COVID-19 Genomics UK (COG-UK) Consortium | The Lighthouse Lab in Milton Keynes and Alex Alderton, Roberto Amato, Jeffrey Barrett, Sonia Goncalves, Ewan Harrison, David K. Jackson, Ian Johnston, Dominic Kwiatkowski, Cordelia Langford, John Sillitoe on behalf of the Wellcome Sanger Institute COVID-19 Surveillance Team |
| EPI_ISL_2805334 | Lighthouse Lab in Alderley Park | Wellcome Sanger Institute for the COVID-19 Genomics UK (COG-UK) Consortium | Jacquelyn Wynn, Mairead Hyland, The Lighthouse Lab in Alderley Park and Alex Alderton, Roberto Amato, Jeffrey Barrett, Sonia Goncalves, Ewan Harrison, David K. Jackson, Ian Johnston, Dominic Kwiatkowski, Cordelia Langford, John Sillitoe on behalf of the Wellcome Sanger Institute COVID-19 Surveillance Team |
| EPI_ISL_2805335, EPI_ISL_2805336 | Lighthouse Lab in Milton Keynes | Wellcome Sanger Institute for the COVID-19 Genomics UK (COG-UK) Consortium | The Lighthouse Lab in Milton Keynes and Alex Alderton, Roberto Amato, Jeffrey Barrett, Sonia Goncalves, Ewan Harrison, David K. Jackson, Ian Johnston, Dominic Kwiatkowski, Cordelia Langford, John Sillitoe on behalf of the Wellcome Sanger Institute COVID-19 Surveillance Team |
| EPI_ISL_2805337 | Lighthouse Lab in Alderley Park | Wellcome Sanger Institute for the COVID-19 Genomics UK (COG-UK) Consortium | Jacquelyn Wynn, Mairead Hyland, The Lighthouse Lab in Alderley Park and Alex Alderton, Roberto Amato, Jeffrey Barrett, Sonia Goncalves, Ewan Harrison, David K. Jackson, Ian Johnston, Dominic Kwiatkowski, Cordelia Langford, John Sillitoe on behalf of the Wellcome Sanger Institute COVID-19 Surveillance Team |
| EPI_ISL_2805338 | Lighthouse Lab in Glasgow | Wellcome Sanger Institute for the COVID-19 Genomics UK (COG-UK) Consortium | Harper VanSteenhouse, Yumi Kasai, David Gray, Carol Clugston, Anna Dominiczak and Alex Alderton, Roberto Amato, Jeffrey Barrett, Sonia Goncalves, Ewan Harrison, David K. Jackson, Ian Johnston, Dominic Kwiatkowski, Cordelia Langford, John Sillitoe on behalf of the Wellcome Sanger Institute COVID-19 Surveillance Team |
| EPI_ISL_2805339 | Lighthouse Lab in Milton Keynes | Wellcome Sanger Institute for the COVID-19 Genomics UK (COG-UK) Consortium | The Lighthouse Lab in Milton Keynes and Alex Alderton, Roberto Amato, Jeffrey Barrett, Sonia Goncalves, Ewan Harrison, David K. Jackson, Ian Johnston, Dominic Kwiatkowski, Cordelia Langford, John Sillitoe on behalf of the Wellcome Sanger Institute COVID-19 Surveillance Team |
| EPI_ISL_2805340 | Lighthouse Lab in Alderley Park | Wellcome Sanger Institute for the COVID-19 Genomics UK (COG-UK) Consortium | Jacquelyn Wynn, Mairead Hyland, The Lighthouse Lab in Alderley Park and Alex Alderton, Roberto Amato, Jeffrey Barrett, Sonia Goncalves, Ewan Harrison, David K. Jackson, Ian Johnston, Dominic Kwiatkowski, Cordelia Langford, John Sillitoe on behalf of the Wellcome Sanger Institute COVID-19 Surveillance Team |
| EPI_ISL_2805341 | Lighthouse Lab in Glasgow | Wellcome Sanger Institute for the COVID-19 Genomics UK (COG-UK) Consortium | Harper VanSteenhouse, Yumi Kasai, David Gray, Carol Clugston, Anna Dominiczak and Alex Alderton, Roberto Amato, Jeffrey Barrett, Sonia Goncalves, Ewan Harrison, David K. Jackson, Ian Johnston, Dominic Kwiatkowski, Cordelia Langford, John Sillitoe on behalf of the Wellcome Sanger Institute COVID-19 Surveillance Team |
| EPI_ISL_2805342, EPI_ISL_2805343, EPI_ISL_2805344, EPI_ISL_2805345, EPI_ISL_2805346 | Lighthouse Lab in Alderley Park | Wellcome Sanger Institute for the COVID-19 Genomics UK (COG-UK) Consortium | Jacquelyn Wynn, Mairead Hyland, The Lighthouse Lab in Alderley Park and Alex Alderton, Roberto Amato, Jeffrey Barrett, Sonia Goncalves, Ewan Harrison, David K. Jackson, Ian Johnston, Dominic Kwiatkowski, Cordelia Langford, John Sillitoe on behalf of the Wellcome Sanger Institute COVID-19 Surveillance Team |
| EPI_ISL_2805347 | Lighthouse Laboratory Plymouth | Wellcome Sanger Institute for the COVID-19 Genomics UK (COG-UK) Consortium | Lighthouse Laboratory Plymouth and Alex Alderton, Roberto Amato, Jeffrey Barrett, Sonia Goncalves, Ewan Harrison, David K. Jackson, Ian Johnston, Dominic Kwiatkowski, Cordelia Langford, John Sillitoe on behalf of the Wellcome Sanger Institute COVID-19 Surveillance Team |
| EPI_ISL_2805348, EPI_ISL_2805349 | Lighthouse Lab in Alderley Park | Wellcome Sanger Institute for the COVID-19 Genomics UK (COG-UK) Consortium | Jacquelyn Wynn, Mairead Hyland, The Lighthouse Lab in Alderley Park and Alex Alderton, Roberto Amato, Jeffrey Barrett, Sonia Goncalves, Ewan Harrison, David K. Jackson, Ian Johnston, Dominic Kwiatkowski, Cordelia Langford, John Sillitoe on behalf of the Wellcome Sanger Institute COVID-19 Surveillance Team |
| EPI_ISL_2805350, EPI_ISL_2805351, EPI_ISL_2805352, EPI_ISL_2805353, EPI_ISL_2805354, EPI_ISL_2805355 | Lighthouse Laboratory Plymouth | Wellcome Sanger Institute for the COVID-19 Genomics UK (COG-UK) Consortium | Lighthouse Laboratory Plymouth and Alex Alderton, Roberto Amato, Jeffrey Barrett, Sonia Goncalves, Ewan Harrison, David K. Jackson, Ian Johnston, Dominic Kwiatkowski, Cordelia Langford, John Sillitoe on behalf of the Wellcome Sanger Institute COVID-19 Surveillance Team |
| EPI_ISL_2805356 | Lighthouse Lab in Alderley Park | Wellcome Sanger Institute for the COVID-19 Genomics UK (COG-UK) Consortium | Jacquelyn Wynn, Mairead Hyland, The Lighthouse Lab in Alderley Park and Alex Alderton, Roberto Amato, Jeffrey Barrett, Sonia Goncalves, Ewan Harrison, David K. Jackson, Ian Johnston, Dominic Kwiatkowski, Cordelia Langford, John Sillitoe on behalf of the Wellcome Sanger Institute COVID-19 Surveillance Team |
| EPI_ISL_2805357, EPI_ISL_2805358 | Lighthouse Laboratory Plymouth | Wellcome Sanger Institute for the COVID-19 Genomics UK (COG-UK) Consortium | Lighthouse Laboratory Plymouth and Alex Alderton, Roberto Amato, Jeffrey Barrett, Sonia Goncalves, Ewan Harrison, David K. Jackson, Ian Johnston, Dominic Kwiatkowski, Cordelia Langford, John Sillitoe on behalf of the Wellcome Sanger Institute COVID-19 Surveillance Team |
| EPI_ISL_2805359, EPI_ISL_2805360 | Lighthouse Lab in Alderley Park | Wellcome Sanger Institute for the COVID-19 Genomics UK (COG-UK) Consortium | Jacquelyn Wynn, Mairead Hyland, The Lighthouse Lab in Alderley Park and Alex Alderton, Roberto Amato, Jeffrey Barrett, Sonia Goncalves, Ewan Harrison, David K. Jackson, Ian Johnston, Dominic Kwiatkowski, Cordelia Langford, John Sillitoe on behalf of the Wellcome Sanger Institute COVID-19 Surveillance Team |
| EPI_ISL_2805361, EPI_ISL_2805362 | Lighthouse Laboratory Plymouth | Wellcome Sanger Institute for the COVID-19 Genomics UK (COG-UK) Consortium | Lighthouse Laboratory Plymouth and Alex Alderton, Roberto Amato, Jeffrey Barrett, Sonia Goncalves, Ewan Harrison, David K. Jackson, Ian Johnston, Dominic Kwiatkowski, Cordelia Langford, John Sillitoe on behalf of the Wellcome Sanger Institute COVID-19 Surveillance Team |
| EPI_ISL_2805363, EPI_ISL_2805364, EPI_ISL_2805365, EPI_ISL_2805366, EPI_ISL_2805367, EPI_ISL_2805368 | Lighthouse Lab in Alderley Park | Wellcome Sanger Institute for the COVID-19 Genomics UK (COG-UK) Consortium | Jacquelyn Wynn, Mairead Hyland, The Lighthouse Lab in Alderley Park and Alex Alderton, Roberto Amato, Jeffrey Barrett, Sonia Goncalves, Ewan Harrison, David K. Jackson, Ian Johnston, Dominic Kwiatkowski, Cordelia Langford, John Sillitoe on behalf of the Wellcome Sanger Institute COVID-19 Surveillance Team |
| EPI_ISL_2805369 | Lighthouse Laboratory Plymouth | Wellcome Sanger Institute for the COVID-19 Genomics UK (COG-UK) Consortium | Lighthouse Laboratory Plymouth and Alex Alderton, Roberto Amato, Jeffrey Barrett, Sonia Goncalves, Ewan Harrison, David K. Jackson, Ian Johnston, Dominic Kwiatkowski, Cordelia Langford, John Sillitoe on behalf of the Wellcome Sanger Institute COVID-19 Surveillance Team |
| EPI_ISL_2805370, EPI_ISL_2805371, EPI_ISL_2805372, EPI_ISL_2805373 | Lighthouse Lab in Alderley Park | Wellcome Sanger Institute for the COVID-19 Genomics UK (COG-UK) Consortium | Jacquelyn Wynn, Mairead Hyland, The Lighthouse Lab in Alderley Park and Alex Alderton, Roberto Amato, Jeffrey Barrett, Sonia Goncalves, Ewan Harrison, David K. Jackson, Ian Johnston, Dominic Kwiatkowski, Cordelia Langford, John Sillitoe on behalf of the Wellcome Sanger Institute COVID-19 Surveillance Team |
| EPI_ISL_2805374 | Lighthouse Laboratory Plymouth | Wellcome Sanger Institute for the COVID-19 Genomics UK (COG-UK) Consortium | Lighthouse Laboratory Plymouth and Alex Alderton, Roberto Amato, Jeffrey Barrett, Sonia Goncalves, Ewan Harrison, David K. Jackson, Ian Johnston, Dominic Kwiatkowski, Cordelia Langford, John Sillitoe on behalf of the Wellcome Sanger Institute COVID-19 Surveillance Team |
| EPI_ISL_2805375 | Lighthouse Lab in Alderley Park | Wellcome Sanger Institute for the COVID-19 Genomics UK (COG-UK) Consortium | Jacquelyn Wynn, Mairead Hyland, The Lighthouse Lab in Alderley Park and Alex Alderton, Roberto Amato, Jeffrey Barrett, Sonia Goncalves, Ewan Harrison, David K. Jackson, Ian Johnston, Dominic Kwiatkowski, Cordelia Langford, John Sillitoe on behalf of the Wellcome Sanger Institute COVID-19 Surveillance Team |
| EPI_ISL_2805376, EPI_ISL_2805377 | Lighthouse Laboratory Plymouth | Wellcome Sanger Institute for the COVID-19 Genomics UK (COG-UK) Consortium | Lighthouse Laboratory Plymouth and Alex Alderton, Roberto Amato, Jeffrey Barrett, Sonia Goncalves, Ewan Harrison, David K. Jackson, Ian Johnston, Dominic Kwiatkowski, Cordelia Langford, John Sillitoe on behalf of the Wellcome Sanger Institute COVID-19 Surveillance Team |
| EPI_ISL_2805378 | Lighthouse Lab in Alderley Park | Wellcome Sanger Institute for the COVID-19 Genomics UK (COG-UK) Consortium | Jacquelyn Wynn, Mairead Hyland, The Lighthouse Lab in Alderley Park and Alex Alderton, Roberto Amato, Jeffrey Barrett, Sonia Goncalves, Ewan Harrison, David K. Jackson, Ian Johnston, Dominic Kwiatkowski, Cordelia Langford, John Sillitoe on behalf of the Wellcome Sanger Institute COVID-19 Surveillance Team |
| EPI_ISL_2805380, EPI_ISL_2805381, EPI_ISL_2805382, EPI_ISL_2805383 | Lighthouse Laboratory Plymouth | Wellcome Sanger Institute for the COVID-19 Genomics UK (COG-UK) Consortium | Lighthouse Laboratory Plymouth and Alex Alderton, Roberto Amato, Jeffrey Barrett, Sonia Goncalves, Ewan Harrison, David K. Jackson, Ian Johnston, Dominic Kwiatkowski, Cordelia Langford, John Sillitoe on behalf of the Wellcome Sanger Institute COVID-19 Surveillance Team |
| EPI_ISL_2805384, EPI_ISL_2805385, EPI_ISL_2805386 | Lighthouse Lab in Alderley Park | Wellcome Sanger Institute for the COVID-19 Genomics UK (COG-UK) Consortium | Jacquelyn Wynn, Mairead Hyland, The Lighthouse Lab in Alderley Park and Alex Alderton, Roberto Amato, Jeffrey Barrett, Sonia Goncalves, Ewan Harrison, David K. Jackson, Ian Johnston, Dominic Kwiatkowski, Cordelia Langford, John Sillitoe on behalf of the Wellcome Sanger Institute COVID-19 Surveillance Team |
| EPI_ISL_2805387 | Lighthouse Laboratory Plymouth | Wellcome Sanger Institute for the COVID-19 Genomics UK (COG-UK) Consortium | Lighthouse Laboratory Plymouth and Alex Alderton, Roberto Amato, Jeffrey Barrett, Sonia Goncalves, Ewan Harrison, David K. Jackson, Ian Johnston, Dominic Kwiatkowski, Cordelia Langford, John Sillitoe on behalf of the Wellcome Sanger Institute COVID-19 Surveillance Team |
| EPI_ISL_2805388 | Lighthouse Lab in Alderley Park | Wellcome Sanger Institute for the COVID-19 Genomics UK (COG-UK) Consortium | Jacquelyn Wynn, Mairead Hyland, The Lighthouse Lab in Alderley Park and Alex Alderton, Roberto Amato, Jeffrey Barrett, Sonia Goncalves, Ewan Harrison, David K. Jackson, Ian Johnston, Dominic Kwiatkowski, Cordelia Langford, John Sillitoe on behalf of the Wellcome Sanger Institute COVID-19 Surveillance Team |
| EPI_ISL_2805389, EPI_ISL_2805390 | Lighthouse Laboratory Plymouth | Wellcome Sanger Institute for the COVID-19 Genomics UK (COG-UK) Consortium | Lighthouse Laboratory Plymouth and Alex Alderton, Roberto Amato, Jeffrey Barrett, Sonia Goncalves, Ewan Harrison, David K. Jackson, Ian Johnston, Dominic Kwiatkowski, Cordelia Langford, John Sillitoe on behalf of the Wellcome Sanger Institute COVID-19 Surveillance Team |
| EPI_ISL_2805391 | Lighthouse Lab in Alderley Park | Wellcome Sanger Institute for the COVID-19 Genomics UK (COG-UK) Consortium | Jacquelyn Wynn, Mairead Hyland, The Lighthouse Lab in Alderley Park and Alex Alderton, Roberto Amato, Jeffrey Barrett, Sonia Goncalves, Ewan Harrison, David K. Jackson, Ian Johnston, Dominic Kwiatkowski, Cordelia Langford, John Sillitoe on behalf of the Wellcome Sanger Institute COVID-19 Surveillance Team |
| EPI_ISL_2805392 | Lighthouse Laboratory Plymouth | Wellcome Sanger Institute for the COVID-19 Genomics UK (COG-UK) Consortium | Lighthouse Laboratory Plymouth and Alex Alderton, Roberto Amato, Jeffrey Barrett, Sonia Goncalves, Ewan Harrison, David K. Jackson, Ian Johnston, Dominic Kwiatkowski, Cordelia Langford, John Sillitoe on behalf of the Wellcome Sanger Institute COVID-19 Surveillance Team |
| EPI_ISL_2805393 | Lighthouse Lab in Alderley Park | Wellcome Sanger Institute for the COVID-19 Genomics UK (COG-UK) Consortium | Jacquelyn Wynn, Mairead Hyland, The Lighthouse Lab in Alderley Park and Alex Alderton, Roberto Amato, Jeffrey Barrett, Sonia Goncalves, Ewan Harrison, David K. Jackson, Ian Johnston, Dominic Kwiatkowski, Cordelia Langford, John Sillitoe on behalf of the Wellcome Sanger Institute COVID-19 Surveillance Team |
| EPI_ISL_2805394 | Lighthouse Laboratory Plymouth | Wellcome Sanger Institute for the COVID-19 Genomics UK (COG-UK) Consortium | Lighthouse Laboratory Plymouth and Alex Alderton, Roberto Amato, Jeffrey Barrett, Sonia Goncalves, Ewan Harrison, David K. Jackson, Ian Johnston, Dominic Kwiatkowski, Cordelia Langford, John Sillitoe on behalf of the Wellcome Sanger Institute COVID-19 Surveillance Team |
| EPI_ISL_2805395, EPI_ISL_2805396, EPI_ISL_2805397, EPI_ISL_2805398, EPI_ISL_2805399, EPI_ISL_2805400, | Lighthouse Lab in Alderley Park | Wellcome Sanger Institute for the COVID-19 Genomics UK (COG-UK) Consortium | Jacquelyn Wynn, Mairead Hyland, The Lighthouse Lab in Alderley Park and Alex Alderton, Roberto Amato, Jeffrey Barrett, Sonia Goncalves, Ewan Harrison, David K. Jackson, Ian Johnston, Dominic Kwiatkowski, Cordelia Langford, John Sillitoe on behalf of the Wellcome Sanger Institute COVID-19 Surveillance Team |

| | | | |
|---|---|---|---|
| EPI_ISL_2805401 | | | |
| EPI_ISL_2805402 | Lighthouse Laboratory Plymouth | Wellcome Sanger Institute for the COVID-19 Genomics UK (COG-UK) Consortium | Lighthouse Laboratory Plymouth and Alex Alderton, Roberto Amato, Jeffrey Barrett, Sonia Goncalves, Ewan Harrison, David K. Jackson, Ian Johnston, Dominic Kwiatkowski, Cordelia Langford, John Sillitoe on behalf of the Wellcome Sanger Institute COVID-19 Surveillance Team |
| EPI_ISL_2805403 | Lighthouse Lab in Alderley Park | Wellcome Sanger Institute for the COVID-19 Genomics UK (COG-UK) Consortium | Jacquelyn Wynn, Mairead Hyland, The Lighthouse Lab in Alderley Park and Alex Alderton, Roberto Amato, Jeffrey Barrett, Sonia Goncalves, Ewan Harrison, David K. Jackson, Ian Johnston, Dominic Kwiatkowski, Cordelia Langford, John Sillitoe on behalf of the Wellcome Sanger Institute COVID-19 Surveillance Team |
| EPI_ISL_2805404 | Lighthouse Laboratory Plymouth | Wellcome Sanger Institute for the COVID-19 Genomics UK (COG-UK) Consortium | Lighthouse Laboratory Plymouth and Alex Alderton, Roberto Amato, Jeffrey Barrett, Sonia Goncalves, Ewan Harrison, David K. Jackson, Ian Johnston, Dominic Kwiatkowski, Cordelia Langford, John Sillitoe on behalf of the Wellcome Sanger Institute COVID-19 Surveillance Team |
| EPI_ISL_2805405, EPI_ISL_2805406, EPI_ISL_2805407 | Lighthouse Lab in Alderley Park | Wellcome Sanger Institute for the COVID-19 Genomics UK (COG-UK) Consortium | Jacquelyn Wynn, Mairead Hyland, The Lighthouse Lab in Alderley Park and Alex Alderton, Roberto Amato, Jeffrey Barrett, Sonia Goncalves, Ewan Harrison, David K. Jackson, Ian Johnston, Dominic Kwiatkowski, Cordelia Langford, John Sillitoe on behalf of the Wellcome Sanger Institute COVID-19 Surveillance Team |
| EPI_ISL_2805408, EPI_ISL_2805409, EPI_ISL_2805410 | Lighthouse Laboratory Plymouth | Wellcome Sanger Institute for the COVID-19 Genomics UK (COG-UK) Consortium | Lighthouse Laboratory Plymouth and Alex Alderton, Roberto Amato, Jeffrey Barrett, Sonia Goncalves, Ewan Harrison, David K. Jackson, Ian Johnston, Dominic Kwiatkowski, Cordelia Langford, John Sillitoe on behalf of the Wellcome Sanger Institute COVID-19 Surveillance Team |
| EPI_ISL_2805411, EPI_ISL_2805412, EPI_ISL_2805413, EPI_ISL_2805414, EPI_ISL_2805415 | Lighthouse Lab in Alderley Park | Wellcome Sanger Institute for the COVID-19 Genomics UK (COG-UK) Consortium | Jacquelyn Wynn, Mairead Hyland, The Lighthouse Lab in Alderley Park and Alex Alderton, Roberto Amato, Jeffrey Barrett, Sonia Goncalves, Ewan Harrison, David K. Jackson, Ian Johnston, Dominic Kwiatkowski, Cordelia Langford, John Sillitoe on behalf of the Wellcome Sanger Institute COVID-19 Surveillance Team |
| EPI_ISL_2805416, EPI_ISL_2805417 | Lighthouse Laboratory Plymouth | Wellcome Sanger Institute for the COVID-19 Genomics UK (COG-UK) Consortium | Lighthouse Laboratory Plymouth and Alex Alderton, Roberto Amato, Jeffrey Barrett, Sonia Goncalves, Ewan Harrison, David K. Jackson, Ian Johnston, Dominic Kwiatkowski, Cordelia Langford, John Sillitoe on behalf of the Wellcome Sanger Institute COVID-19 Surveillance Team |
| EPI_ISL_2805418, EPI_ISL_2805419, EPI_ISL_2805420, EPI_ISL_2805421 | Lighthouse Lab in Alderley Park | Wellcome Sanger Institute for the COVID-19 Genomics UK (COG-UK) Consortium | Jacquelyn Wynn, Mairead Hyland, The Lighthouse Lab in Alderley Park and Alex Alderton, Roberto Amato, Jeffrey Barrett, Sonia Goncalves, Ewan Harrison, David K. Jackson, Ian Johnston, Dominic Kwiatkowski, Cordelia Langford, John Sillitoe on behalf of the Wellcome Sanger Institute COVID-19 Surveillance Team |
| EPI_ISL_2805422 | Lighthouse Laboratory Plymouth | Wellcome Sanger Institute for the COVID-19 Genomics UK (COG-UK) Consortium | Lighthouse Laboratory Plymouth and Alex Alderton, Roberto Amato, Jeffrey Barrett, Sonia Goncalves, Ewan Harrison, David K. Jackson, Ian Johnston, Dominic Kwiatkowski, Cordelia Langford, John Sillitoe on behalf of the Wellcome Sanger Institute COVID-19 Surveillance Team |
| EPI_ISL_2805423 | Lighthouse Lab in Alderley Park | Wellcome Sanger Institute for the COVID-19 Genomics UK (COG-UK) Consortium | Jacquelyn Wynn, Mairead Hyland, The Lighthouse Lab in Alderley Park and Alex Alderton, Roberto Amato, Jeffrey Barrett, Sonia Goncalves, Ewan Harrison, David K. Jackson, Ian Johnston, Dominic Kwiatkowski, Cordelia Langford, John Sillitoe on behalf of the Wellcome Sanger Institute COVID-19 Surveillance Team |
| EPI_ISL_2805424, EPI_ISL_2805425 | Lighthouse Laboratory Plymouth | Wellcome Sanger Institute for the COVID-19 Genomics UK (COG-UK) Consortium | Lighthouse Laboratory Plymouth and Alex Alderton, Roberto Amato, Jeffrey Barrett, Sonia Goncalves, Ewan Harrison, David K. Jackson, Ian Johnston, Dominic Kwiatkowski, Cordelia Langford, John Sillitoe on behalf of the Wellcome Sanger Institute COVID-19 Surveillance Team |
| EPI_ISL_2805426, EPI_ISL_2805427 | Lighthouse Lab in Alderley Park | Wellcome Sanger Institute for the COVID-19 Genomics UK (COG-UK) Consortium | Jacquelyn Wynn, Mairead Hyland, The Lighthouse Lab in Alderley Park and Alex Alderton, Roberto Amato, Jeffrey Barrett, Sonia Goncalves, Ewan Harrison, David K. Jackson, Ian Johnston, Dominic Kwiatkowski, Cordelia Langford, John Sillitoe on behalf of the Wellcome Sanger Institute COVID-19 Surveillance Team |
| EPI_ISL_2805428 | Lighthouse Laboratory Plymouth | Wellcome Sanger Institute for the COVID-19 Genomics UK (COG-UK) Consortium | Lighthouse Laboratory Plymouth and Alex Alderton, Roberto Amato, Jeffrey Barrett, Sonia Goncalves, Ewan Harrison, David K. Jackson, Ian Johnston, Dominic Kwiatkowski, Cordelia Langford, John Sillitoe on behalf of the Wellcome Sanger Institute COVID-19 Surveillance Team |
| EPI_ISL_2805429 | Lighthouse Lab in Alderley Park | Wellcome Sanger Institute for the COVID-19 Genomics UK (COG-UK) Consortium | Jacquelyn Wynn, Mairead Hyland, The Lighthouse Lab in Alderley Park and Alex Alderton, Roberto Amato, Jeffrey Barrett, Sonia Goncalves, Ewan Harrison, David K. Jackson, Ian Johnston, Dominic Kwiatkowski, Cordelia Langford, John Sillitoe on behalf of the Wellcome Sanger Institute COVID-19 Surveillance Team |
| EPI_ISL_2805430, EPI_ISL_2805431, EPI_ISL_2805432, EPI_ISL_2805433, EPI_ISL_2805434 | Lighthouse Laboratory Plymouth | Wellcome Sanger Institute for the COVID-19 Genomics UK (COG-UK) Consortium | Lighthouse Laboratory Plymouth and Alex Alderton, Roberto Amato, Jeffrey Barrett, Sonia Goncalves, Ewan Harrison, David K. Jackson, Ian Johnston, Dominic Kwiatkowski, Cordelia Langford, John Sillitoe on behalf of the Wellcome Sanger Institute COVID-19 Surveillance Team |
| EPI_ISL_2805435, EPI_ISL_2805436, EPI_ISL_2805437, EPI_ISL_2805438 | Lighthouse Lab in Alderley Park | Wellcome Sanger Institute for the COVID-19 Genomics UK (COG-UK) Consortium | Jacquelyn Wynn, Mairead Hyland, The Lighthouse Lab in Alderley Park and Alex Alderton, Roberto Amato, Jeffrey Barrett, Sonia Goncalves, Ewan Harrison, David K. Jackson, Ian Johnston, Dominic Kwiatkowski, Cordelia Langford, John Sillitoe on behalf of the Wellcome Sanger Institute COVID-19 Surveillance Team |
| EPI_ISL_2805439 | Lighthouse Laboratory Plymouth | Wellcome Sanger Institute for the COVID-19 Genomics UK (COG-UK) Consortium | Lighthouse Laboratory Plymouth and Alex Alderton, Roberto Amato, Jeffrey Barrett, Sonia Goncalves, Ewan Harrison, David K. Jackson, Ian Johnston, Dominic Kwiatkowski, Cordelia Langford, John Sillitoe on behalf of the Wellcome Sanger Institute COVID-19 Surveillance Team |
| EPI_ISL_2805440 | Lighthouse Lab in Alderley Park | Wellcome Sanger Institute for the COVID-19 Genomics UK (COG-UK) Consortium | Jacquelyn Wynn, Mairead Hyland, The Lighthouse Lab in Alderley Park and Alex Alderton, Roberto Amato, Jeffrey Barrett, Sonia Goncalves, Ewan Harrison, David K. Jackson, Ian Johnston, Dominic Kwiatkowski, Cordelia Langford, John Sillitoe on behalf of the Wellcome Sanger Institute COVID-19 Surveillance Team |
| EPI_ISL_2805441 | Lighthouse Laboratory Plymouth | Wellcome Sanger Institute for the COVID-19 Genomics UK (COG-UK) Consortium | Lighthouse Laboratory Plymouth and Alex Alderton, Roberto Amato, Jeffrey Barrett, Sonia Goncalves, Ewan Harrison, David K. Jackson, Ian Johnston, Dominic Kwiatkowski, Cordelia Langford, John Sillitoe on behalf of the Wellcome Sanger Institute COVID-19 Surveillance Team |
| EPI_ISL_2805442, EPI_ISL_2805443 | Lighthouse Lab in Alderley Park | Wellcome Sanger Institute for the COVID-19 Genomics UK (COG-UK) Consortium | Jacquelyn Wynn, Mairead Hyland, The Lighthouse Lab in Alderley Park and Alex Alderton, Roberto Amato, Jeffrey Barrett, Sonia Goncalves, Ewan Harrison, David K. Jackson, Ian Johnston, Dominic Kwiatkowski, Cordelia Langford, John Sillitoe on behalf of the Wellcome Sanger Institute COVID-19 Surveillance Team |
| EPI_ISL_2805444 | Lighthouse Laboratory Plymouth | Wellcome Sanger Institute for the COVID-19 Genomics UK (COG-UK) Consortium | Lighthouse Laboratory Plymouth and Alex Alderton, Roberto Amato, Jeffrey Barrett, Sonia Goncalves, Ewan Harrison, David K. Jackson, Ian Johnston, Dominic Kwiatkowski, Cordelia Langford, John Sillitoe on behalf of the Wellcome Sanger Institute COVID-19 Surveillance Team |
| EPI_ISL_2805445, EPI_ISL_2805446 | Lighthouse Lab in Alderley Park | Wellcome Sanger Institute for the COVID-19 Genomics UK (COG-UK) Consortium | Jacquelyn Wynn, Mairead Hyland, The Lighthouse Lab in Alderley Park and Alex Alderton, Roberto Amato, Jeffrey Barrett, Sonia Goncalves, Ewan Harrison, David K. Jackson, Ian Johnston, Dominic Kwiatkowski, Cordelia Langford, John Sillitoe on behalf of the Wellcome Sanger Institute COVID-19 Surveillance Team |
| EPI_ISL_2805447 | Lighthouse Laboratory Plymouth | Wellcome Sanger Institute for the COVID-19 Genomics UK (COG-UK) Consortium | Lighthouse Laboratory Plymouth and Alex Alderton, Roberto Amato, Jeffrey Barrett, Sonia Goncalves, Ewan Harrison, David K. Jackson, Ian Johnston, Dominic Kwiatkowski, Cordelia Langford, John Sillitoe on behalf of the Wellcome Sanger Institute COVID-19 Surveillance Team |
| EPI_ISL_2805448, EPI_ISL_2805449 | Lighthouse Lab in Alderley Park | Wellcome Sanger Institute for the COVID-19 Genomics UK (COG-UK) Consortium | Jacquelyn Wynn, Mairead Hyland, The Lighthouse Lab in Alderley Park and Alex Alderton, Roberto Amato, Jeffrey Barrett, Sonia Goncalves, Ewan Harrison, David K. Jackson, Ian Johnston, Dominic Kwiatkowski, Cordelia Langford, John Sillitoe on behalf of the Wellcome Sanger Institute COVID-19 Surveillance Team |
| EPI_ISL_2805450, EPI_ISL_2805451 | Lighthouse Laboratory Plymouth | Wellcome Sanger Institute for the COVID-19 Genomics UK (COG-UK) Consortium | Lighthouse Laboratory Plymouth and Alex Alderton, Roberto Amato, Jeffrey Barrett, Sonia Goncalves, Ewan Harrison, David K. Jackson, Ian Johnston, Dominic Kwiatkowski, Cordelia Langford, John Sillitoe on behalf of the Wellcome Sanger Institute COVID-19 Surveillance Team |
| EPI_ISL_2805452 | Lighthouse Lab in Alderley Park | Wellcome Sanger Institute for the COVID-19 Genomics UK (COG-UK) Consortium | Jacquelyn Wynn, Mairead Hyland, The Lighthouse Lab in Alderley Park and Alex Alderton, Roberto Amato, Jeffrey Barrett, Sonia Goncalves, Ewan Harrison, David K. Jackson, Ian Johnston, Dominic Kwiatkowski, Cordelia Langford, John Sillitoe on behalf of the Wellcome Sanger Institute COVID-19 Surveillance Team |
| EPI_ISL_2805453, EPI_ISL_2805454, EPI_ISL_2805455 | Lighthouse Laboratory Plymouth | Wellcome Sanger Institute for the COVID-19 Genomics UK (COG-UK) Consortium | Lighthouse Laboratory Plymouth and Alex Alderton, Roberto Amato, Jeffrey Barrett, Sonia Goncalves, Ewan Harrison, David K. Jackson, Ian Johnston, Dominic Kwiatkowski, Cordelia Langford, John Sillitoe on behalf of the Wellcome Sanger Institute COVID-19 Surveillance Team |
| EPI_ISL_2805456 | Lighthouse Lab in Alderley Park | Wellcome Sanger Institute for the COVID-19 Genomics UK (COG-UK) Consortium | Jacquelyn Wynn, Mairead Hyland, The Lighthouse Lab in Alderley Park and Alex Alderton, Roberto Amato, Jeffrey Barrett, Sonia Goncalves, Ewan Harrison, David K. Jackson, Ian Johnston, Dominic Kwiatkowski, Cordelia Langford, John Sillitoe on behalf of the Wellcome Sanger Institute COVID-19 Surveillance Team |
| EPI_ISL_2805457 | Lighthouse Laboratory Plymouth | Wellcome Sanger Institute for the COVID-19 Genomics UK (COG-UK) Consortium | Lighthouse Laboratory Plymouth and Alex Alderton, Roberto Amato, Jeffrey Barrett, Sonia Goncalves, Ewan Harrison, David K. Jackson, Ian Johnston, Dominic Kwiatkowski, Cordelia Langford, John Sillitoe on behalf of the Wellcome Sanger Institute COVID-19 Surveillance Team |
| EPI_ISL_2805458 | Lighthouse Lab in Alderley Park | Wellcome Sanger Institute for the COVID-19 Genomics UK (COG-UK) Consortium | Jacquelyn Wynn, Mairead Hyland, The Lighthouse Lab in Alderley Park and Alex Alderton, Roberto Amato, Jeffrey Barrett, Sonia Goncalves, Ewan Harrison, David K. Jackson, Ian Johnston, Dominic Kwiatkowski, Cordelia Langford, John Sillitoe on behalf of the Wellcome Sanger Institute COVID-19 Surveillance Team |
| EPI_ISL_2805459 | Lighthouse Laboratory Plymouth | Wellcome Sanger Institute for the COVID-19 Genomics UK (COG-UK) Consortium | Lighthouse Laboratory Plymouth and Alex Alderton, Roberto Amato, Jeffrey Barrett, Sonia Goncalves, Ewan Harrison, David K. Jackson, Ian Johnston, Dominic Kwiatkowski, Cordelia Langford, John Sillitoe on behalf of the Wellcome Sanger Institute COVID-19 Surveillance Team |
| EPI_ISL_2805460, EPI_ISL_2805461, EPI_ISL_2805462 | Lighthouse Lab in Alderley Park | Wellcome Sanger Institute for the COVID-19 Genomics UK (COG-UK) Consortium | Jacquelyn Wynn, Mairead Hyland, The Lighthouse Lab in Alderley Park and Alex Alderton, Roberto Amato, Jeffrey Barrett, Sonia Goncalves, Ewan Harrison, David K. Jackson, Ian Johnston, Dominic Kwiatkowski, Cordelia Langford, John Sillitoe on behalf of the Wellcome Sanger Institute COVID-19 Surveillance Team |
| EPI_ISL_2805463, EPI_ISL_2805464 | Lighthouse Laboratory Plymouth | Wellcome Sanger Institute for the COVID-19 Genomics UK (COG-UK) Consortium | Lighthouse Laboratory Plymouth and Alex Alderton, Roberto Amato, Jeffrey Barrett, Sonia Goncalves, Ewan Harrison, David K. Jackson, Ian Johnston, Dominic Kwiatkowski, Cordelia Langford, John Sillitoe on behalf of the Wellcome Sanger Institute COVID-19 Surveillance Team |
| EPI_ISL_2805465 | Lighthouse Lab in Alderley Park | Wellcome Sanger Institute for the COVID-19 Genomics UK (COG-UK) Consortium | Jacquelyn Wynn, Mairead Hyland, The Lighthouse Lab in Alderley Park and Alex Alderton, Roberto Amato, Jeffrey Barrett, Sonia Goncalves, Ewan Harrison, David K. Jackson, Ian Johnston, Dominic Kwiatkowski, Cordelia Langford, John Sillitoe on behalf of the Wellcome Sanger Institute COVID-19 Surveillance Team |
| EPI_ISL_2805466 | Lighthouse Laboratory Plymouth | Wellcome Sanger Institute for the COVID-19 Genomics UK (COG-UK) Consortium | Lighthouse Laboratory Plymouth and Alex Alderton, Roberto Amato, Jeffrey Barrett, Sonia Goncalves, Ewan Harrison, David K. Jackson, Ian Johnston, Dominic Kwiatkowski, Cordelia Langford, John Sillitoe on behalf of the Wellcome Sanger Institute COVID-19 Surveillance Team |
| EPI_ISL_2805467 | Lighthouse Lab in Alderley Park | Wellcome Sanger Institute for the COVID-19 Genomics UK (COG-UK) Consortium | Jacquelyn Wynn, Mairead Hyland, The Lighthouse Lab in Alderley Park and Alex Alderton, Roberto Amato, Jeffrey Barrett, Sonia Goncalves, Ewan Harrison, David K. Jackson, Ian Johnston, Dominic Kwiatkowski, Cordelia Langford, John Sillitoe on behalf of the Wellcome Sanger Institute COVID-19 Surveillance Team |
| EPI_ISL_2805468, EPI_ISL_2805469 | Lighthouse Laboratory Plymouth | Wellcome Sanger Institute for the COVID-19 Genomics UK (COG-UK) Consortium | Lighthouse Laboratory Plymouth and Alex Alderton, Roberto Amato, Jeffrey Barrett, Sonia Goncalves, Ewan Harrison, David K. Jackson, Ian Johnston, Dominic Kwiatkowski, Cordelia Langford, John Sillitoe on behalf of the Wellcome Sanger Institute COVID-19 Surveillance Team |
| EPI_ISL_2805470, EPI_ISL_2805471, EPI_ISL_2805472, EPI_ISL_2805473 | Lighthouse Lab in Alderley Park | Wellcome Sanger Institute for the COVID-19 Genomics UK (COG-UK) Consortium | Jacquelyn Wynn, Mairead Hyland, The Lighthouse Lab in Alderley Park and Alex Alderton, Roberto Amato, Jeffrey Barrett, Sonia Goncalves, Ewan Harrison, David K. Jackson, Ian Johnston, Dominic Kwiatkowski, Cordelia Langford, John Sillitoe on behalf of the Wellcome Sanger Institute COVID-19 Surveillance Team |

| | | | |
|---|---|---|---|
| EPI_ISL_2805474, EPI_ISL_2805475 | Lighthouse Laboratory Plymouth | Wellcome Sanger Institute for the COVID-19 Genomics UK (COG-UK) Consortium | Lighthouse Laboratory Plymouth and Alex Alderton, Roberto Amato, Jeffrey Barrett, Sonia Goncalves, Ewan Harrison, David K. Jackson, Ian Johnston, Dominic Kwiatkowski, Cordelia Langford, John Sillitoe on behalf of the Wellcome Sanger Institute COVID-19 Surveillance Team |
| EPI_ISL_2805476 | Lighthouse Lab in Alderley Park | Wellcome Sanger Institute for the COVID-19 Genomics UK (COG-UK) Consortium | Jacquelyn Wynn, Mairead Hyland, The Lighthouse Lab in Alderley Park and Alex Alderton, Roberto Amato, Jeffrey Barrett, Sonia Goncalves, Ewan Harrison, David K. Jackson, Ian Johnston, Dominic Kwiatkowski, Cordelia Langford, John Sillitoe on behalf of the Wellcome Sanger Institute COVID-19 Surveillance Team |
| EPI_ISL_2805477 | Lighthouse Laboratory Plymouth | Wellcome Sanger Institute for the COVID-19 Genomics UK (COG-UK) Consortium | Lighthouse Laboratory Plymouth and Alex Alderton, Roberto Amato, Jeffrey Barrett, Sonia Goncalves, Ewan Harrison, David K. Jackson, Ian Johnston, Dominic Kwiatkowski, Cordelia Langford, John Sillitoe on behalf of the Wellcome Sanger Institute COVID-19 Surveillance Team |
| EPI_ISL_2805478 | Lighthouse Lab in Alderley Park | Wellcome Sanger Institute for the COVID-19 Genomics UK (COG-UK) Consortium | Jacquelyn Wynn, Mairead Hyland, The Lighthouse Lab in Alderley Park and Alex Alderton, Roberto Amato, Jeffrey Barrett, Sonia Goncalves, Ewan Harrison, David K. Jackson, Ian Johnston, Dominic Kwiatkowski, Cordelia Langford, John Sillitoe on behalf of the Wellcome Sanger Institute COVID-19 Surveillance Team |
| EPI_ISL_2805479, EPI_ISL_2805480 | Lighthouse Laboratory Plymouth | Wellcome Sanger Institute for the COVID-19 Genomics UK (COG-UK) Consortium | Lighthouse Laboratory Plymouth and Alex Alderton, Roberto Amato, Jeffrey Barrett, Sonia Goncalves, Ewan Harrison, David K. Jackson, Ian Johnston, Dominic Kwiatkowski, Cordelia Langford, John Sillitoe on behalf of the Wellcome Sanger Institute COVID-19 Surveillance Team |
| EPI_ISL_2805481, EPI_ISL_2805482 | Lighthouse Lab in Alderley Park | Wellcome Sanger Institute for the COVID-19 Genomics UK (COG-UK) Consortium | Jacquelyn Wynn, Mairead Hyland, The Lighthouse Lab in Alderley Park and Alex Alderton, Roberto Amato, Jeffrey Barrett, Sonia Goncalves, Ewan Harrison, David K. Jackson, Ian Johnston, Dominic Kwiatkowski, Cordelia Langford, John Sillitoe on behalf of the Wellcome Sanger Institute COVID-19 Surveillance Team |
| EPI_ISL_2805483, EPI_ISL_2805484 | Lighthouse Laboratory Plymouth | Wellcome Sanger Institute for the COVID-19 Genomics UK (COG-UK) Consortium | Lighthouse Laboratory Plymouth and Alex Alderton, Roberto Amato, Jeffrey Barrett, Sonia Goncalves, Ewan Harrison, David K. Jackson, Ian Johnston, Dominic Kwiatkowski, Cordelia Langford, John Sillitoe on behalf of the Wellcome Sanger Institute COVID-19 Surveillance Team |
| EPI_ISL_2805485 | Lighthouse Lab in Alderley Park | Wellcome Sanger Institute for the COVID-19 Genomics UK (COG-UK) Consortium | Jacquelyn Wynn, Mairead Hyland, The Lighthouse Lab in Alderley Park and Alex Alderton, Roberto Amato, Jeffrey Barrett, Sonia Goncalves, Ewan Harrison, David K. Jackson, Ian Johnston, Dominic Kwiatkowski, Cordelia Langford, John Sillitoe on behalf of the Wellcome Sanger Institute COVID-19 Surveillance Team |
| EPI_ISL_2805486, EPI_ISL_2805487, EPI_ISL_2805488 | Lighthouse Laboratory Plymouth | Wellcome Sanger Institute for the COVID-19 Genomics UK (COG-UK) Consortium | Lighthouse Laboratory Plymouth and Alex Alderton, Roberto Amato, Jeffrey Barrett, Sonia Goncalves, Ewan Harrison, David K. Jackson, Ian Johnston, Dominic Kwiatkowski, Cordelia Langford, John Sillitoe on behalf of the Wellcome Sanger Institute COVID-19 Surveillance Team |
| EPI_ISL_2805489 | Lighthouse Lab in Alderley Park | Wellcome Sanger Institute for the COVID-19 Genomics UK (COG-UK) Consortium | Jacquelyn Wynn, Mairead Hyland, The Lighthouse Lab in Alderley Park and Alex Alderton, Roberto Amato, Jeffrey Barrett, Sonia Goncalves, Ewan Harrison, David K. Jackson, Ian Johnston, Dominic Kwiatkowski, Cordelia Langford, John Sillitoe on behalf of the Wellcome Sanger Institute COVID-19 Surveillance Team |
| EPI_ISL_2805490, EPI_ISL_2805491 | Lighthouse Laboratory Plymouth | Wellcome Sanger Institute for the COVID-19 Genomics UK (COG-UK) Consortium | Lighthouse Laboratory Plymouth and Alex Alderton, Roberto Amato, Jeffrey Barrett, Sonia Goncalves, Ewan Harrison, David K. Jackson, Ian Johnston, Dominic Kwiatkowski, Cordelia Langford, John Sillitoe on behalf of the Wellcome Sanger Institute COVID-19 Surveillance Team |
| EPI_ISL_2805492 | Lighthouse Lab in Alderley Park | Wellcome Sanger Institute for the COVID-19 Genomics UK (COG-UK) Consortium | Jacquelyn Wynn, Mairead Hyland, The Lighthouse Lab in Alderley Park and Alex Alderton, Roberto Amato, Jeffrey Barrett, Sonia Goncalves, Ewan Harrison, David K. Jackson, Ian Johnston, Dominic Kwiatkowski, Cordelia Langford, John Sillitoe on behalf of the Wellcome Sanger Institute COVID-19 Surveillance Team |
| EPI_ISL_2805493, EPI_ISL_2805494 | Lighthouse Laboratory Plymouth | Wellcome Sanger Institute for the COVID-19 Genomics UK (COG-UK) Consortium | Lighthouse Laboratory Plymouth and Alex Alderton, Roberto Amato, Jeffrey Barrett, Sonia Goncalves, Ewan Harrison, David K. Jackson, Ian Johnston, Dominic Kwiatkowski, Cordelia Langford, John Sillitoe on behalf of the Wellcome Sanger Institute COVID-19 Surveillance Team |
| EPI_ISL_2805495 | Lighthouse Lab in Alderley Park | Wellcome Sanger Institute for the COVID-19 Genomics UK (COG-UK) Consortium | Jacquelyn Wynn, Mairead Hyland, The Lighthouse Lab in Alderley Park and Alex Alderton, Roberto Amato, Jeffrey Barrett, Sonia Goncalves, Ewan Harrison, David K. Jackson, Ian Johnston, Dominic Kwiatkowski, Cordelia Langford, John Sillitoe on behalf of the Wellcome Sanger Institute COVID-19 Surveillance Team |
| EPI_ISL_2805496, EPI_ISL_2805497 | Lighthouse Laboratory Plymouth | Wellcome Sanger Institute for the COVID-19 Genomics UK (COG-UK) Consortium | Lighthouse Laboratory Plymouth and Alex Alderton, Roberto Amato, Jeffrey Barrett, Sonia Goncalves, Ewan Harrison, David K. Jackson, Ian Johnston, Dominic Kwiatkowski, Cordelia Langford, John Sillitoe on behalf of the Wellcome Sanger Institute COVID-19 Surveillance Team |
| EPI_ISL_2805498, EPI_ISL_2805499, EPI_ISL_2805500, EPI_ISL_2805501 | Lighthouse Lab in Alderley Park | Wellcome Sanger Institute for the COVID-19 Genomics UK (COG-UK) Consortium | Jacquelyn Wynn, Mairead Hyland, The Lighthouse Lab in Alderley Park and Alex Alderton, Roberto Amato, Jeffrey Barrett, Sonia Goncalves, Ewan Harrison, David K. Jackson, Ian Johnston, Dominic Kwiatkowski, Cordelia Langford, John Sillitoe on behalf of the Wellcome Sanger Institute COVID-19 Surveillance Team |
| EPI_ISL_2805502 | Lighthouse Laboratory Plymouth | Wellcome Sanger Institute for the COVID-19 Genomics UK (COG-UK) Consortium | Lighthouse Laboratory Plymouth and Alex Alderton, Roberto Amato, Jeffrey Barrett, Sonia Goncalves, Ewan Harrison, David K. Jackson, Ian Johnston, Dominic Kwiatkowski, Cordelia Langford, John Sillitoe on behalf of the Wellcome Sanger Institute COVID-19 Surveillance Team |
| EPI_ISL_2805503, EPI_ISL_2805504, EPI_ISL_2805505 | Lighthouse Lab in Alderley Park | Wellcome Sanger Institute for the COVID-19 Genomics UK (COG-UK) Consortium | Jacquelyn Wynn, Mairead Hyland, The Lighthouse Lab in Alderley Park and Alex Alderton, Roberto Amato, Jeffrey Barrett, Sonia Goncalves, Ewan Harrison, David K. Jackson, Ian Johnston, Dominic Kwiatkowski, Cordelia Langford, John Sillitoe on behalf of the Wellcome Sanger Institute COVID-19 Surveillance Team |
| EPI_ISL_2805506, EPI_ISL_2805507 | Lighthouse Laboratory Plymouth | Wellcome Sanger Institute for the COVID-19 Genomics UK (COG-UK) Consortium | Lighthouse Laboratory Plymouth and Alex Alderton, Roberto Amato, Jeffrey Barrett, Sonia Goncalves, Ewan Harrison, David K. Jackson, Ian Johnston, Dominic Kwiatkowski, Cordelia Langford, John Sillitoe on behalf of the Wellcome Sanger Institute COVID-19 Surveillance Team |
| EPI_ISL_2805508 | Lighthouse Lab in Alderley Park | Wellcome Sanger Institute for the COVID-19 Genomics UK (COG-UK) Consortium | Jacquelyn Wynn, Mairead Hyland, The Lighthouse Lab in Alderley Park and Alex Alderton, Roberto Amato, Jeffrey Barrett, Sonia Goncalves, Ewan Harrison, David K. Jackson, Ian Johnston, Dominic Kwiatkowski, Cordelia Langford, John Sillitoe on behalf of the Wellcome Sanger Institute COVID-19 Surveillance Team |
| EPI_ISL_2805509, EPI_ISL_2805510, EPI_ISL_2805511, EPI_ISL_2805512, EPI_ISL_2805513, EPI_ISL_2805514, EPI_ISL_2805515 | Lighthouse Laboratory Plymouth | Wellcome Sanger Institute for the COVID-19 Genomics UK (COG-UK) Consortium | Lighthouse Laboratory Plymouth and Alex Alderton, Roberto Amato, Jeffrey Barrett, Sonia Goncalves, Ewan Harrison, David K. Jackson, Ian Johnston, Dominic Kwiatkowski, Cordelia Langford, John Sillitoe on behalf of the Wellcome Sanger Institute COVID-19 Surveillance Team |
| EPI_ISL_2805516 | Lighthouse Lab in Alderley Park | Wellcome Sanger Institute for the COVID-19 Genomics UK (COG-UK) Consortium | Jacquelyn Wynn, Mairead Hyland, The Lighthouse Lab in Alderley Park and Alex Alderton, Roberto Amato, Jeffrey Barrett, Sonia Goncalves, Ewan Harrison, David K. Jackson, Ian Johnston, Dominic Kwiatkowski, Cordelia Langford, John Sillitoe on behalf of the Wellcome Sanger Institute COVID-19 Surveillance Team |
| EPI_ISL_2805517, EPI_ISL_2805518 | Lighthouse Laboratory Plymouth | Wellcome Sanger Institute for the COVID-19 Genomics UK (COG-UK) Consortium | Lighthouse Laboratory Plymouth and Alex Alderton, Roberto Amato, Jeffrey Barrett, Sonia Goncalves, Ewan Harrison, David K. Jackson, Ian Johnston, Dominic Kwiatkowski, Cordelia Langford, John Sillitoe on behalf of the Wellcome Sanger Institute COVID-19 Surveillance Team |
| EPI_ISL_2805519, EPI_ISL_2805520 | Lighthouse Lab in Alderley Park | Wellcome Sanger Institute for the COVID-19 Genomics UK (COG-UK) Consortium | Jacquelyn Wynn, Mairead Hyland, The Lighthouse Lab in Alderley Park and Alex Alderton, Roberto Amato, Jeffrey Barrett, Sonia Goncalves, Ewan Harrison, David K. Jackson, Ian Johnston, Dominic Kwiatkowski, Cordelia Langford, John Sillitoe on behalf of the Wellcome Sanger Institute COVID-19 Surveillance Team |
| EPI_ISL_2805521 | Lighthouse Laboratory Plymouth | Wellcome Sanger Institute for the COVID-19 Genomics UK (COG-UK) Consortium | Lighthouse Laboratory Plymouth and Alex Alderton, Roberto Amato, Jeffrey Barrett, Sonia Goncalves, Ewan Harrison, David K. Jackson, Ian Johnston, Dominic Kwiatkowski, Cordelia Langford, John Sillitoe on behalf of the Wellcome Sanger Institute COVID-19 Surveillance Team |
| EPI_ISL_2805522 | Lighthouse Lab in Alderley Park | Wellcome Sanger Institute for the COVID-19 Genomics UK (COG-UK) Consortium | Jacquelyn Wynn, Mairead Hyland, The Lighthouse Lab in Alderley Park and Alex Alderton, Roberto Amato, Jeffrey Barrett, Sonia Goncalves, Ewan Harrison, David K. Jackson, Ian Johnston, Dominic Kwiatkowski, Cordelia Langford, John Sillitoe on behalf of the Wellcome Sanger Institute COVID-19 Surveillance Team |
| EPI_ISL_2805523 | Lighthouse Laboratory Plymouth | Wellcome Sanger Institute for the COVID-19 Genomics UK (COG-UK) Consortium | Lighthouse Laboratory Plymouth and Alex Alderton, Roberto Amato, Jeffrey Barrett, Sonia Goncalves, Ewan Harrison, David K. Jackson, Ian Johnston, Dominic Kwiatkowski, Cordelia Langford, John Sillitoe on behalf of the Wellcome Sanger Institute COVID-19 Surveillance Team |
| EPI_ISL_2805524, EPI_ISL_2805525, EPI_ISL_2805526, EPI_ISL_2805527 | Lighthouse Lab in Alderley Park | Wellcome Sanger Institute for the COVID-19 Genomics UK (COG-UK) Consortium | Jacquelyn Wynn, Mairead Hyland, The Lighthouse Lab in Alderley Park and Alex Alderton, Roberto Amato, Jeffrey Barrett, Sonia Goncalves, Ewan Harrison, David K. Jackson, Ian Johnston, Dominic Kwiatkowski, Cordelia Langford, John Sillitoe on behalf of the Wellcome Sanger Institute COVID-19 Surveillance Team |
| EPI_ISL_2805528, EPI_ISL_2805529 | Lighthouse Laboratory Plymouth | Wellcome Sanger Institute for the COVID-19 Genomics UK (COG-UK) Consortium | Lighthouse Laboratory Plymouth and Alex Alderton, Roberto Amato, Jeffrey Barrett, Sonia Goncalves, Ewan Harrison, David K. Jackson, Ian Johnston, Dominic Kwiatkowski, Cordelia Langford, John Sillitoe on behalf of the Wellcome Sanger Institute COVID-19 Surveillance Team |
| EPI_ISL_2805530, EPI_ISL_2805531 | Lighthouse Lab in Alderley Park | Wellcome Sanger Institute for the COVID-19 Genomics UK (COG-UK) Consortium | Jacquelyn Wynn, Mairead Hyland, The Lighthouse Lab in Alderley Park and Alex Alderton, Roberto Amato, Jeffrey Barrett, Sonia Goncalves, Ewan Harrison, David K. Jackson, Ian Johnston, Dominic Kwiatkowski, Cordelia Langford, John Sillitoe on behalf of the Wellcome Sanger Institute COVID-19 Surveillance Team |
| EPI_ISL_2805532, EPI_ISL_2805533, EPI_ISL_2805534 | Lighthouse Laboratory Plymouth | Wellcome Sanger Institute for the COVID-19 Genomics UK (COG-UK) Consortium | Lighthouse Laboratory Plymouth and Alex Alderton, Roberto Amato, Jeffrey Barrett, Sonia Goncalves, Ewan Harrison, David K. Jackson, Ian Johnston, Dominic Kwiatkowski, Cordelia Langford, John Sillitoe on behalf of the Wellcome Sanger Institute COVID-19 Surveillance Team |
| EPI_ISL_2805535 | Lighthouse Lab in Alderley Park | Wellcome Sanger Institute for the COVID-19 Genomics UK (COG-UK) Consortium | Jacquelyn Wynn, Mairead Hyland, The Lighthouse Lab in Alderley Park and Alex Alderton, Roberto Amato, Jeffrey Barrett, Sonia Goncalves, Ewan Harrison, David K. Jackson, Ian Johnston, Dominic Kwiatkowski, Cordelia Langford, John Sillitoe on behalf of the Wellcome Sanger Institute COVID-19 Surveillance Team |
| EPI_ISL_2805536 | Lighthouse Laboratory Plymouth | Wellcome Sanger Institute for the COVID-19 Genomics UK (COG-UK) Consortium | Lighthouse Laboratory Plymouth and Alex Alderton, Roberto Amato, Jeffrey Barrett, Sonia Goncalves, Ewan Harrison, David K. Jackson, Ian Johnston, Dominic Kwiatkowski, Cordelia Langford, John Sillitoe on behalf of the Wellcome Sanger Institute COVID-19 Surveillance Team |
| EPI_ISL_2805537 | Lighthouse Lab in Alderley Park | Wellcome Sanger Institute for the COVID-19 Genomics UK (COG-UK) Consortium | Jacquelyn Wynn, Mairead Hyland, The Lighthouse Lab in Alderley Park and Alex Alderton, Roberto Amato, Jeffrey Barrett, Sonia Goncalves, Ewan Harrison, David K. Jackson, Ian Johnston, Dominic Kwiatkowski, Cordelia Langford, John Sillitoe on behalf of the Wellcome Sanger Institute COVID-19 Surveillance Team |
| EPI_ISL_2805538 | Lighthouse Laboratory Plymouth | Wellcome Sanger Institute for the COVID-19 Genomics UK (COG-UK) Consortium | Lighthouse Laboratory Plymouth and Alex Alderton, Roberto Amato, Jeffrey Barrett, Sonia Goncalves, Ewan Harrison, David K. Jackson, Ian Johnston, Dominic Kwiatkowski, Cordelia Langford, John Sillitoe on behalf of the Wellcome Sanger Institute COVID-19 Surveillance Team |
| EPI_ISL_2805539, EPI_ISL_2805540 | Lighthouse Lab in Alderley Park | Wellcome Sanger Institute for the COVID-19 Genomics UK (COG-UK) Consortium | Jacquelyn Wynn, Mairead Hyland, The Lighthouse Lab in Alderley Park and Alex Alderton, Roberto Amato, Jeffrey Barrett, Sonia Goncalves, Ewan Harrison, David K. Jackson, Ian Johnston, Dominic Kwiatkowski, Cordelia Langford, John Sillitoe on behalf of the Wellcome Sanger Institute COVID-19 Surveillance Team |
| EPI_ISL_2805541, EPI_ISL_2805542 | Lighthouse Laboratory Plymouth | Wellcome Sanger Institute for the COVID-19 Genomics UK (COG-UK) Consortium | Lighthouse Laboratory Plymouth and Alex Alderton, Roberto Amato, Jeffrey Barrett, Sonia Goncalves, Ewan Harrison, David K. Jackson, Ian Johnston, Dominic Kwiatkowski, Cordelia Langford, John Sillitoe on behalf of the Wellcome Sanger Institute COVID-19 Surveillance Team |
| EPI_ISL_2805543 | Lighthouse Lab in Alderley Park | Wellcome Sanger Institute for the COVID-19 Genomics UK (COG-UK) Consortium | Jacquelyn Wynn, Mairead Hyland, The Lighthouse Lab in Alderley Park and Alex Alderton, Roberto Amato, Jeffrey Barrett, Sonia Goncalves, Ewan Harrison, David K. Jackson, Ian Johnston, Dominic Kwiatkowski, Cordelia Langford, John Sillitoe on behalf of the Wellcome Sanger Institute COVID-19 Surveillance Team |
| EPI_ISL_2805544, EPI_ISL_2805545, EPI_ISL_2805546, EPI_ISL_2805547 | Lighthouse Laboratory Plymouth | Wellcome Sanger Institute for the COVID-19 Genomics UK (COG-UK) Consortium | Lighthouse Laboratory Plymouth and Alex Alderton, Roberto Amato, Jeffrey Barrett, Sonia Goncalves, Ewan Harrison, David K. Jackson, Ian Johnston, Dominic Kwiatkowski, Cordelia Langford, John Sillitoe on behalf of the Wellcome Sanger Institute COVID-19 Surveillance Team |
| EPI_ISL_2805548 | Lighthouse Lab in Alderley Park | Wellcome Sanger Institute for the COVID-19 Genomics UK (COG-UK) | Jacquelyn Wynn, Mairead Hyland, The Lighthouse Lab in Alderley Park and Alex Alderton, Roberto Amato, Jeffrey Barrett, Sonia Goncalves, Ewan Harrison, David K. Jackson, Ian Johnston, Dominic Kwiatkowski, Cordelia Langford, John Sillitoe on |

| | | | |
|---|---|---|---|
| | | Consortium | behalf of the Wellcome Sanger Institute COVID-19 Surveillance Team |
| EPI_ISL_2805549, EPI_ISL_2805550, EPI_ISL_2805551 | Lighthouse Laboratory Plymouth | Wellcome Sanger Institute for the COVID-19 Genomics UK (COG-UK) Consortium | Lighthouse Laboratory Plymouth and Alex Alderton, Roberto Amato, Jeffrey Barrett, Sonia Goncalves, Ewan Harrison, David K. Jackson, Ian Johnston, Dominic Kwiatkowski, Cordelia Langford, John Sillitoe on behalf of the Wellcome Sanger Institute COVID-19 Surveillance Team |
| EPI_ISL_2805552 | Lighthouse Lab in Alderley Park | Wellcome Sanger Institute for the COVID-19 Genomics UK (COG-UK) Consortium | Jacquelyn Wynn, Mairead Hyland, The Lighthouse Lab in Alderley Park and Alex Alderton, Roberto Amato, Jeffrey Barrett, Sonia Goncalves, Ewan Harrison, David K. Jackson, Ian Johnston, Dominic Kwiatkowski, Cordelia Langford, John Sillitoe on behalf of the Wellcome Sanger Institute COVID-19 Surveillance Team |
| EPI_ISL_2805553, EPI_ISL_2805554, EPI_ISL_2805555, EPI_ISL_2805556, EPI_ISL_2805557 | Lighthouse Laboratory Plymouth | Wellcome Sanger Institute for the COVID-19 Genomics UK (COG-UK) Consortium | Lighthouse Laboratory Plymouth and Alex Alderton, Roberto Amato, Jeffrey Barrett, Sonia Goncalves, Ewan Harrison, David K. Jackson, Ian Johnston, Dominic Kwiatkowski, Cordelia Langford, John Sillitoe on behalf of the Wellcome Sanger Institute COVID-19 Surveillance Team |
| EPI_ISL_2805558 | Lighthouse Lab in Alderley Park | Wellcome Sanger Institute for the COVID-19 Genomics UK (COG-UK) Consortium | Jacquelyn Wynn, Mairead Hyland, The Lighthouse Lab in Alderley Park and Alex Alderton, Roberto Amato, Jeffrey Barrett, Sonia Goncalves, Ewan Harrison, David K. Jackson, Ian Johnston, Dominic Kwiatkowski, Cordelia Langford, John Sillitoe on behalf of the Wellcome Sanger Institute COVID-19 Surveillance Team |
| EPI_ISL_2805559 | Lighthouse Laboratory Plymouth | Wellcome Sanger Institute for the COVID-19 Genomics UK (COG-UK) Consortium | Lighthouse Laboratory Plymouth and Alex Alderton, Roberto Amato, Jeffrey Barrett, Sonia Goncalves, Ewan Harrison, David K. Jackson, Ian Johnston, Dominic Kwiatkowski, Cordelia Langford, John Sillitoe on behalf of the Wellcome Sanger Institute COVID-19 Surveillance Team |
| EPI_ISL_2805560, EPI_ISL_2805561 | Lighthouse Lab in Alderley Park | Wellcome Sanger Institute for the COVID-19 Genomics UK (COG-UK) Consortium | Jacquelyn Wynn, Mairead Hyland, The Lighthouse Lab in Alderley Park and Alex Alderton, Roberto Amato, Jeffrey Barrett, Sonia Goncalves, Ewan Harrison, David K. Jackson, Ian Johnston, Dominic Kwiatkowski, Cordelia Langford, John Sillitoe on behalf of the Wellcome Sanger Institute COVID-19 Surveillance Team |
| EPI_ISL_2805562 | Lighthouse Laboratory Plymouth | Wellcome Sanger Institute for the COVID-19 Genomics UK (COG-UK) Consortium | Lighthouse Laboratory Plymouth and Alex Alderton, Roberto Amato, Jeffrey Barrett, Sonia Goncalves, Ewan Harrison, David K. Jackson, Ian Johnston, Dominic Kwiatkowski, Cordelia Langford, John Sillitoe on behalf of the Wellcome Sanger Institute COVID-19 Surveillance Team |
| EPI_ISL_2805563, EPI_ISL_2805564, EPI_ISL_2805565 | Lighthouse Lab in Alderley Park | Wellcome Sanger Institute for the COVID-19 Genomics UK (COG-UK) Consortium | Jacquelyn Wynn, Mairead Hyland, The Lighthouse Lab in Alderley Park and Alex Alderton, Roberto Amato, Jeffrey Barrett, Sonia Goncalves, Ewan Harrison, David K. Jackson, Ian Johnston, Dominic Kwiatkowski, Cordelia Langford, John Sillitoe on behalf of the Wellcome Sanger Institute COVID-19 Surveillance Team |
| EPI_ISL_2805566 | Lighthouse Laboratory Plymouth | Wellcome Sanger Institute for the COVID-19 Genomics UK (COG-UK) Consortium | Lighthouse Laboratory Plymouth and Alex Alderton, Roberto Amato, Jeffrey Barrett, Sonia Goncalves, Ewan Harrison, David K. Jackson, Ian Johnston, Dominic Kwiatkowski, Cordelia Langford, John Sillitoe on behalf of the Wellcome Sanger Institute COVID-19 Surveillance Team |
| EPI_ISL_2805567 | Lighthouse Lab in Alderley Park | Wellcome Sanger Institute for the COVID-19 Genomics UK (COG-UK) Consortium | Jacquelyn Wynn, Mairead Hyland, The Lighthouse Lab in Alderley Park and Alex Alderton, Roberto Amato, Jeffrey Barrett, Sonia Goncalves, Ewan Harrison, David K. Jackson, Ian Johnston, Dominic Kwiatkowski, Cordelia Langford, John Sillitoe on behalf of the Wellcome Sanger Institute COVID-19 Surveillance Team |
| EPI_ISL_2805568 | Lighthouse Laboratory Plymouth | Wellcome Sanger Institute for the COVID-19 Genomics UK (COG-UK) Consortium | Lighthouse Laboratory Plymouth and Alex Alderton, Roberto Amato, Jeffrey Barrett, Sonia Goncalves, Ewan Harrison, David K. Jackson, Ian Johnston, Dominic Kwiatkowski, Cordelia Langford, John Sillitoe on behalf of the Wellcome Sanger Institute COVID-19 Surveillance Team |
| EPI_ISL_2805569 | Lighthouse Lab in Alderley Park | Wellcome Sanger Institute for the COVID-19 Genomics UK (COG-UK) Consortium | Jacquelyn Wynn, Mairead Hyland, The Lighthouse Lab in Alderley Park and Alex Alderton, Roberto Amato, Jeffrey Barrett, Sonia Goncalves, Ewan Harrison, David K. Jackson, Ian Johnston, Dominic Kwiatkowski, Cordelia Langford, John Sillitoe on behalf of the Wellcome Sanger Institute COVID-19 Surveillance Team |
| EPI_ISL_2805570, EPI_ISL_2805571 | Lighthouse Laboratory Plymouth | Wellcome Sanger Institute for the COVID-19 Genomics UK (COG-UK) Consortium | Lighthouse Laboratory Plymouth and Alex Alderton, Roberto Amato, Jeffrey Barrett, Sonia Goncalves, Ewan Harrison, David K. Jackson, Ian Johnston, Dominic Kwiatkowski, Cordelia Langford, John Sillitoe on behalf of the Wellcome Sanger Institute COVID-19 Surveillance Team |
| EPI_ISL_2805572, EPI_ISL_2805573, EPI_ISL_2805574 | Lighthouse Lab in Alderley Park | Wellcome Sanger Institute for the COVID-19 Genomics UK (COG-UK) Consortium | Jacquelyn Wynn, Mairead Hyland, The Lighthouse Lab in Alderley Park and Alex Alderton, Roberto Amato, Jeffrey Barrett, Sonia Goncalves, Ewan Harrison, David K. Jackson, Ian Johnston, Dominic Kwiatkowski, Cordelia Langford, John Sillitoe on behalf of the Wellcome Sanger Institute COVID-19 Surveillance Team |
| EPI_ISL_2805575, EPI_ISL_2805576, EPI_ISL_2805577, EPI_ISL_2805578, EPI_ISL_2805579 | Lighthouse Laboratory Plymouth | Wellcome Sanger Institute for the COVID-19 Genomics UK (COG-UK) Consortium | Lighthouse Laboratory Plymouth and Alex Alderton, Roberto Amato, Jeffrey Barrett, Sonia Goncalves, Ewan Harrison, David K. Jackson, Ian Johnston, Dominic Kwiatkowski, Cordelia Langford, John Sillitoe on behalf of the Wellcome Sanger Institute COVID-19 Surveillance Team |
| EPI_ISL_2805580, EPI_ISL_2805581 | Lighthouse Lab in Alderley Park | Wellcome Sanger Institute for the COVID-19 Genomics UK (COG-UK) Consortium | Jacquelyn Wynn, Mairead Hyland, The Lighthouse Lab in Alderley Park and Alex Alderton, Roberto Amato, Jeffrey Barrett, Sonia Goncalves, Ewan Harrison, David K. Jackson, Ian Johnston, Dominic Kwiatkowski, Cordelia Langford, John Sillitoe on behalf of the Wellcome Sanger Institute COVID-19 Surveillance Team |
| EPI_ISL_2805582 | Lighthouse Laboratory Plymouth | Wellcome Sanger Institute for the COVID-19 Genomics UK (COG-UK) Consortium | Lighthouse Laboratory Plymouth and Alex Alderton, Roberto Amato, Jeffrey Barrett, Sonia Goncalves, Ewan Harrison, David K. Jackson, Ian Johnston, Dominic Kwiatkowski, Cordelia Langford, John Sillitoe on behalf of the Wellcome Sanger Institute COVID-19 Surveillance Team |
| EPI_ISL_2805583 | Lighthouse Lab in Alderley Park | Wellcome Sanger Institute for the COVID-19 Genomics UK (COG-UK) Consortium | Jacquelyn Wynn, Mairead Hyland, The Lighthouse Lab in Alderley Park and Alex Alderton, Roberto Amato, Jeffrey Barrett, Sonia Goncalves, Ewan Harrison, David K. Jackson, Ian Johnston, Dominic Kwiatkowski, Cordelia Langford, John Sillitoe on behalf of the Wellcome Sanger Institute COVID-19 Surveillance Team |
| EPI_ISL_2805584 | Lighthouse Laboratory Plymouth | Wellcome Sanger Institute for the COVID-19 Genomics UK (COG-UK) Consortium | Lighthouse Laboratory Plymouth and Alex Alderton, Roberto Amato, Jeffrey Barrett, Sonia Goncalves, Ewan Harrison, David K. Jackson, Ian Johnston, Dominic Kwiatkowski, Cordelia Langford, John Sillitoe on behalf of the Wellcome Sanger Institute COVID-19 Surveillance Team |
| EPI_ISL_2805585, EPI_ISL_2805586 | Lighthouse Lab in Alderley Park | Wellcome Sanger Institute for the COVID-19 Genomics UK (COG-UK) Consortium | Jacquelyn Wynn, Mairead Hyland, The Lighthouse Lab in Alderley Park and Alex Alderton, Roberto Amato, Jeffrey Barrett, Sonia Goncalves, Ewan Harrison, David K. Jackson, Ian Johnston, Dominic Kwiatkowski, Cordelia Langford, John Sillitoe on behalf of the Wellcome Sanger Institute COVID-19 Surveillance Team |
| EPI_ISL_2805587 | Lighthouse Laboratory Plymouth | Wellcome Sanger Institute for the COVID-19 Genomics UK (COG-UK) Consortium | Lighthouse Laboratory Plymouth and Alex Alderton, Roberto Amato, Jeffrey Barrett, Sonia Goncalves, Ewan Harrison, David K. Jackson, Ian Johnston, Dominic Kwiatkowski, Cordelia Langford, John Sillitoe on behalf of the Wellcome Sanger Institute COVID-19 Surveillance Team |
| EPI_ISL_2805588, EPI_ISL_2805589, EPI_ISL_2805590 | Lighthouse Lab in Alderley Park | Wellcome Sanger Institute for the COVID-19 Genomics UK (COG-UK) Consortium | Jacquelyn Wynn, Mairead Hyland, The Lighthouse Lab in Alderley Park and Alex Alderton, Roberto Amato, Jeffrey Barrett, Sonia Goncalves, Ewan Harrison, David K. Jackson, Ian Johnston, Dominic Kwiatkowski, Cordelia Langford, John Sillitoe on behalf of the Wellcome Sanger Institute COVID-19 Surveillance Team |
| EPI_ISL_2805591 | Lighthouse Laboratory Plymouth | Wellcome Sanger Institute for the COVID-19 Genomics UK (COG-UK) Consortium | Lighthouse Laboratory Plymouth and Alex Alderton, Roberto Amato, Jeffrey Barrett, Sonia Goncalves, Ewan Harrison, David K. Jackson, Ian Johnston, Dominic Kwiatkowski, Cordelia Langford, John Sillitoe on behalf of the Wellcome Sanger Institute COVID-19 Surveillance Team |
| EPI_ISL_2805592 | Lighthouse Lab in Alderley Park | Wellcome Sanger Institute for the COVID-19 Genomics UK (COG-UK) Consortium | Jacquelyn Wynn, Mairead Hyland, The Lighthouse Lab in Alderley Park and Alex Alderton, Roberto Amato, Jeffrey Barrett, Sonia Goncalves, Ewan Harrison, David K. Jackson, Ian Johnston, Dominic Kwiatkowski, Cordelia Langford, John Sillitoe on behalf of the Wellcome Sanger Institute COVID-19 Surveillance Team |
| EPI_ISL_2805593, EPI_ISL_2805594, EPI_ISL_2805595, EPI_ISL_2805596 | Lighthouse Laboratory Plymouth | Wellcome Sanger Institute for the COVID-19 Genomics UK (COG-UK) Consortium | Lighthouse Laboratory Plymouth and Alex Alderton, Roberto Amato, Jeffrey Barrett, Sonia Goncalves, Ewan Harrison, David K. Jackson, Ian Johnston, Dominic Kwiatkowski, Cordelia Langford, John Sillitoe on behalf of the Wellcome Sanger Institute COVID-19 Surveillance Team |
| EPI_ISL_2805597, EPI_ISL_2805598 | Lighthouse Lab in Alderley Park | Wellcome Sanger Institute for the COVID-19 Genomics UK (COG-UK) Consortium | Jacquelyn Wynn, Mairead Hyland, The Lighthouse Lab in Alderley Park and Alex Alderton, Roberto Amato, Jeffrey Barrett, Sonia Goncalves, Ewan Harrison, David K. Jackson, Ian Johnston, Dominic Kwiatkowski, Cordelia Langford, John Sillitoe on behalf of the Wellcome Sanger Institute COVID-19 Surveillance Team |
| EPI_ISL_2805599, EPI_ISL_2805600 | Lighthouse Laboratory Plymouth | Wellcome Sanger Institute for the COVID-19 Genomics UK (COG-UK) Consortium | Lighthouse Laboratory Plymouth and Alex Alderton, Roberto Amato, Jeffrey Barrett, Sonia Goncalves, Ewan Harrison, David K. Jackson, Ian Johnston, Dominic Kwiatkowski, Cordelia Langford, John Sillitoe on behalf of the Wellcome Sanger Institute COVID-19 Surveillance Team |
| EPI_ISL_2805601, EPI_ISL_2805602 | Lighthouse Lab in Alderley Park | Wellcome Sanger Institute for the COVID-19 Genomics UK (COG-UK) Consortium | Jacquelyn Wynn, Mairead Hyland, The Lighthouse Lab in Alderley Park and Alex Alderton, Roberto Amato, Jeffrey Barrett, Sonia Goncalves, Ewan Harrison, David K. Jackson, Ian Johnston, Dominic Kwiatkowski, Cordelia Langford, John Sillitoe on behalf of the Wellcome Sanger Institute COVID-19 Surveillance Team |
| EPI_ISL_2805603, EPI_ISL_2805604, EPI_ISL_2805605 | Lighthouse Laboratory Plymouth | Wellcome Sanger Institute for the COVID-19 Genomics UK (COG-UK) Consortium | Lighthouse Laboratory Plymouth and Alex Alderton, Roberto Amato, Jeffrey Barrett, Sonia Goncalves, Ewan Harrison, David K. Jackson, Ian Johnston, Dominic Kwiatkowski, Cordelia Langford, John Sillitoe on behalf of the Wellcome Sanger Institute COVID-19 Surveillance Team |
| EPI_ISL_2805606, EPI_ISL_2805607, EPI_ISL_2805608 | Lighthouse Lab in Alderley Park | Wellcome Sanger Institute for the COVID-19 Genomics UK (COG-UK) Consortium | Jacquelyn Wynn, Mairead Hyland, The Lighthouse Lab in Alderley Park and Alex Alderton, Roberto Amato, Jeffrey Barrett, Sonia Goncalves, Ewan Harrison, David K. Jackson, Ian Johnston, Dominic Kwiatkowski, Cordelia Langford, John Sillitoe on behalf of the Wellcome Sanger Institute COVID-19 Surveillance Team |
| EPI_ISL_2805609 | Lighthouse Laboratory Plymouth | Wellcome Sanger Institute for the COVID-19 Genomics UK (COG-UK) Consortium | Lighthouse Laboratory Plymouth and Alex Alderton, Roberto Amato, Jeffrey Barrett, Sonia Goncalves, Ewan Harrison, David K. Jackson, Ian Johnston, Dominic Kwiatkowski, Cordelia Langford, John Sillitoe on behalf of the Wellcome Sanger Institute COVID-19 Surveillance Team |
| EPI_ISL_2805610, EPI_ISL_2805611, EPI_ISL_2805612 | Lighthouse Lab in Alderley Park | Wellcome Sanger Institute for the COVID-19 Genomics UK (COG-UK) Consortium | Jacquelyn Wynn, Mairead Hyland, The Lighthouse Lab in Alderley Park and Alex Alderton, Roberto Amato, Jeffrey Barrett, Sonia Goncalves, Ewan Harrison, David K. Jackson, Ian Johnston, Dominic Kwiatkowski, Cordelia Langford, John Sillitoe on behalf of the Wellcome Sanger Institute COVID-19 Surveillance Team |
| EPI_ISL_2805613 | Lighthouse Laboratory Plymouth | Wellcome Sanger Institute for the COVID-19 Genomics UK (COG-UK) Consortium | Lighthouse Laboratory Plymouth and Alex Alderton, Roberto Amato, Jeffrey Barrett, Sonia Goncalves, Ewan Harrison, David K. Jackson, Ian Johnston, Dominic Kwiatkowski, Cordelia Langford, John Sillitoe on behalf of the Wellcome Sanger Institute COVID-19 Surveillance Team |
| EPI_ISL_2805614, EPI_ISL_2805615, EPI_ISL_2805616, EPI_ISL_2805617 | Lighthouse Lab in Alderley Park | Wellcome Sanger Institute for the COVID-19 Genomics UK (COG-UK) Consortium | Jacquelyn Wynn, Mairead Hyland, The Lighthouse Lab in Alderley Park and Alex Alderton, Roberto Amato, Jeffrey Barrett, Sonia Goncalves, Ewan Harrison, David K. Jackson, Ian Johnston, Dominic Kwiatkowski, Cordelia Langford, John Sillitoe on behalf of the Wellcome Sanger Institute COVID-19 Surveillance Team |
| EPI_ISL_2805618, EPI_ISL_2805619, EPI_ISL_2805620, EPI_ISL_2805621 | Lighthouse Laboratory Plymouth | Wellcome Sanger Institute for the COVID-19 Genomics UK (COG-UK) Consortium | Lighthouse Laboratory Plymouth and Alex Alderton, Roberto Amato, Jeffrey Barrett, Sonia Goncalves, Ewan Harrison, David K. Jackson, Ian Johnston, Dominic Kwiatkowski, Cordelia Langford, John Sillitoe on behalf of the Wellcome Sanger Institute COVID-19 Surveillance Team |
| EPI_ISL_2805622 | Lighthouse Lab in Alderley Park | Wellcome Sanger Institute for the COVID-19 Genomics UK (COG-UK) Consortium | Jacquelyn Wynn, Mairead Hyland, The Lighthouse Lab in Alderley Park and Alex Alderton, Roberto Amato, Jeffrey Barrett, Sonia Goncalves, Ewan Harrison, David K. Jackson, Ian Johnston, Dominic Kwiatkowski, Cordelia Langford, John Sillitoe on behalf of the Wellcome Sanger Institute COVID-19 Surveillance Team |
| EPI_ISL_2805623 | Lighthouse Laboratory Plymouth | Wellcome Sanger Institute for the COVID-19 Genomics UK (COG-UK) | Lighthouse Laboratory Plymouth and Alex Alderton, Roberto Amato, Jeffrey Barrett, Sonia Goncalves, Ewan Harrison, David K. Jackson, Ian Johnston, Dominic Kwiatkowski, Cordelia Langford, John Sillitoe on behalf of the Wellcome Sanger |

| | | | |
|---|---|---|---|
| | | Consortium | Institute COVID-19 Surveillance Team |
| EPI_ISL_2805624 | Lighthouse Lab in Alderley Park | Wellcome Sanger Institute for the COVID-19 Genomics UK (COG-UK) Consortium | Jacquelyn Wynn, Mairead Hyland, The Lighthouse Lab in Alderley Park and Alex Alderton, Roberto Amato, Jeffrey Barrett, Sonia Goncalves, Ewan Harrison, David K. Jackson, Ian Johnston, Dominic Kwiatkowski, Cordelia Langford, John Sillitoe on behalf of the Wellcome Sanger Institute COVID-19 Surveillance Team |
| EPI_ISL_2805625, EPI_ISL_2805626, EPI_ISL_2805627, EPI_ISL_2805628 | Lighthouse Laboratory Plymouth | Wellcome Sanger Institute for the COVID-19 Genomics UK (COG-UK) Consortium | Lighthouse Laboratory Plymouth and Alex Alderton, Roberto Amato, Jeffrey Barrett, Sonia Goncalves, Ewan Harrison, David K. Jackson, Ian Johnston, Dominic Kwiatkowski, Cordelia Langford, John Sillitoe on behalf of the Wellcome Sanger Institute COVID-19 Surveillance Team |
| EPI_ISL_2805629 | Lighthouse Lab in Alderley Park | Wellcome Sanger Institute for the COVID-19 Genomics UK (COG-UK) Consortium | Jacquelyn Wynn, Mairead Hyland, The Lighthouse Lab in Alderley Park and Alex Alderton, Roberto Amato, Jeffrey Barrett, Sonia Goncalves, Ewan Harrison, David K. Jackson, Ian Johnston, Dominic Kwiatkowski, Cordelia Langford, John Sillitoe on behalf of the Wellcome Sanger Institute COVID-19 Surveillance Team |
| EPI_ISL_2805630 | Lighthouse Laboratory Plymouth | Wellcome Sanger Institute for the COVID-19 Genomics UK (COG-UK) Consortium | Lighthouse Laboratory Plymouth and Alex Alderton, Roberto Amato, Jeffrey Barrett, Sonia Goncalves, Ewan Harrison, David K. Jackson, Ian Johnston, Dominic Kwiatkowski, Cordelia Langford, John Sillitoe on behalf of the Wellcome Sanger Institute COVID-19 Surveillance Team |
| EPI_ISL_2805631, EPI_ISL_2805632 | Lighthouse Lab in Alderley Park | Wellcome Sanger Institute for the COVID-19 Genomics UK (COG-UK) Consortium | Jacquelyn Wynn, Mairead Hyland, The Lighthouse Lab in Alderley Park and Alex Alderton, Roberto Amato, Jeffrey Barrett, Sonia Goncalves, Ewan Harrison, David K. Jackson, Ian Johnston, Dominic Kwiatkowski, Cordelia Langford, John Sillitoe on behalf of the Wellcome Sanger Institute COVID-19 Surveillance Team |
| EPI_ISL_2805633 | Lighthouse Laboratory Plymouth | Wellcome Sanger Institute for the COVID-19 Genomics UK (COG-UK) Consortium | Lighthouse Laboratory Plymouth and Alex Alderton, Roberto Amato, Jeffrey Barrett, Sonia Goncalves, Ewan Harrison, David K. Jackson, Ian Johnston, Dominic Kwiatkowski, Cordelia Langford, John Sillitoe on behalf of the Wellcome Sanger Institute COVID-19 Surveillance Team |
| EPI_ISL_2805634 | Lighthouse Lab in Alderley Park | Wellcome Sanger Institute for the COVID-19 Genomics UK (COG-UK) Consortium | Jacquelyn Wynn, Mairead Hyland, The Lighthouse Lab in Alderley Park and Alex Alderton, Roberto Amato, Jeffrey Barrett, Sonia Goncalves, Ewan Harrison, David K. Jackson, Ian Johnston, Dominic Kwiatkowski, Cordelia Langford, John Sillitoe on behalf of the Wellcome Sanger Institute COVID-19 Surveillance Team |
| EPI_ISL_2805635 | Lighthouse Laboratory Plymouth | Wellcome Sanger Institute for the COVID-19 Genomics UK (COG-UK) Consortium | Lighthouse Laboratory Plymouth and Alex Alderton, Roberto Amato, Jeffrey Barrett, Sonia Goncalves, Ewan Harrison, David K. Jackson, Ian Johnston, Dominic Kwiatkowski, Cordelia Langford, John Sillitoe on behalf of the Wellcome Sanger Institute COVID-19 Surveillance Team |
| EPI_ISL_2805636 | Lighthouse Lab in Alderley Park | Wellcome Sanger Institute for the COVID-19 Genomics UK (COG-UK) Consortium | Jacquelyn Wynn, Mairead Hyland, The Lighthouse Lab in Alderley Park and Alex Alderton, Roberto Amato, Jeffrey Barrett, Sonia Goncalves, Ewan Harrison, David K. Jackson, Ian Johnston, Dominic Kwiatkowski, Cordelia Langford, John Sillitoe on behalf of the Wellcome Sanger Institute COVID-19 Surveillance Team |
| EPI_ISL_2805637 | Lighthouse Laboratory Plymouth | Wellcome Sanger Institute for the COVID-19 Genomics UK (COG-UK) Consortium | Lighthouse Laboratory Plymouth and Alex Alderton, Roberto Amato, Jeffrey Barrett, Sonia Goncalves, Ewan Harrison, David K. Jackson, Ian Johnston, Dominic Kwiatkowski, Cordelia Langford, John Sillitoe on behalf of the Wellcome Sanger Institute COVID-19 Surveillance Team |
| EPI_ISL_2805638 | Lighthouse Lab in Alderley Park | Wellcome Sanger Institute for the COVID-19 Genomics UK (COG-UK) Consortium | Jacquelyn Wynn, Mairead Hyland, The Lighthouse Lab in Alderley Park and Alex Alderton, Roberto Amato, Jeffrey Barrett, Sonia Goncalves, Ewan Harrison, David K. Jackson, Ian Johnston, Dominic Kwiatkowski, Cordelia Langford, John Sillitoe on behalf of the Wellcome Sanger Institute COVID-19 Surveillance Team |
| EPI_ISL_2805639 | Lighthouse Laboratory Plymouth | Wellcome Sanger Institute for the COVID-19 Genomics UK (COG-UK) Consortium | Lighthouse Laboratory Plymouth and Alex Alderton, Roberto Amato, Jeffrey Barrett, Sonia Goncalves, Ewan Harrison, David K. Jackson, Ian Johnston, Dominic Kwiatkowski, Cordelia Langford, John Sillitoe on behalf of the Wellcome Sanger Institute COVID-19 Surveillance Team |
| EPI_ISL_2805640 | Lighthouse Lab in Alderley Park | Wellcome Sanger Institute for the COVID-19 Genomics UK (COG-UK) Consortium | Jacquelyn Wynn, Mairead Hyland, The Lighthouse Lab in Alderley Park and Alex Alderton, Roberto Amato, Jeffrey Barrett, Sonia Goncalves, Ewan Harrison, David K. Jackson, Ian Johnston, Dominic Kwiatkowski, Cordelia Langford, John Sillitoe on behalf of the Wellcome Sanger Institute COVID-19 Surveillance Team |
| EPI_ISL_2805641 | Lighthouse Laboratory Plymouth | Wellcome Sanger Institute for the COVID-19 Genomics UK (COG-UK) Consortium | Lighthouse Laboratory Plymouth and Alex Alderton, Roberto Amato, Jeffrey Barrett, Sonia Goncalves, Ewan Harrison, David K. Jackson, Ian Johnston, Dominic Kwiatkowski, Cordelia Langford, John Sillitoe on behalf of the Wellcome Sanger Institute COVID-19 Surveillance Team |
| EPI_ISL_2805642 | Lighthouse Lab in Alderley Park | Wellcome Sanger Institute for the COVID-19 Genomics UK (COG-UK) Consortium | Jacquelyn Wynn, Mairead Hyland, The Lighthouse Lab in Alderley Park and Alex Alderton, Roberto Amato, Jeffrey Barrett, Sonia Goncalves, Ewan Harrison, David K. Jackson, Ian Johnston, Dominic Kwiatkowski, Cordelia Langford, John Sillitoe on behalf of the Wellcome Sanger Institute COVID-19 Surveillance Team |
| EPI_ISL_2805643, EPI_ISL_2805644 | Lighthouse Laboratory Plymouth | Wellcome Sanger Institute for the COVID-19 Genomics UK (COG-UK) Consortium | Lighthouse Laboratory Plymouth and Alex Alderton, Roberto Amato, Jeffrey Barrett, Sonia Goncalves, Ewan Harrison, David K. Jackson, Ian Johnston, Dominic Kwiatkowski, Cordelia Langford, John Sillitoe on behalf of the Wellcome Sanger Institute COVID-19 Surveillance Team |
| EPI_ISL_2805645, EPI_ISL_2805646, EPI_ISL_2805647 | Lighthouse Lab in Alderley Park | Wellcome Sanger Institute for the COVID-19 Genomics UK (COG-UK) Consortium | Jacquelyn Wynn, Mairead Hyland, The Lighthouse Lab in Alderley Park and Alex Alderton, Roberto Amato, Jeffrey Barrett, Sonia Goncalves, Ewan Harrison, David K. Jackson, Ian Johnston, Dominic Kwiatkowski, Cordelia Langford, John Sillitoe on behalf of the Wellcome Sanger Institute COVID-19 Surveillance Team |
| EPI_ISL_2805648 | Lighthouse Laboratory Plymouth | Wellcome Sanger Institute for the COVID-19 Genomics UK (COG-UK) Consortium | Lighthouse Laboratory Plymouth and Alex Alderton, Roberto Amato, Jeffrey Barrett, Sonia Goncalves, Ewan Harrison, David K. Jackson, Ian Johnston, Dominic Kwiatkowski, Cordelia Langford, John Sillitoe on behalf of the Wellcome Sanger Institute COVID-19 Surveillance Team |
| EPI_ISL_2805649, EPI_ISL_2805650 | Lighthouse Lab in Alderley Park | Wellcome Sanger Institute for the COVID-19 Genomics UK (COG-UK) Consortium | Jacquelyn Wynn, Mairead Hyland, The Lighthouse Lab in Alderley Park and Alex Alderton, Roberto Amato, Jeffrey Barrett, Sonia Goncalves, Ewan Harrison, David K. Jackson, Ian Johnston, Dominic Kwiatkowski, Cordelia Langford, John Sillitoe on behalf of the Wellcome Sanger Institute COVID-19 Surveillance Team |
| EPI_ISL_2805651, EPI_ISL_2805652 | Lighthouse Laboratory Plymouth | Wellcome Sanger Institute for the COVID-19 Genomics UK (COG-UK) Consortium | Lighthouse Laboratory Plymouth and Alex Alderton, Roberto Amato, Jeffrey Barrett, Sonia Goncalves, Ewan Harrison, David K. Jackson, Ian Johnston, Dominic Kwiatkowski, Cordelia Langford, John Sillitoe on behalf of the Wellcome Sanger Institute COVID-19 Surveillance Team |
| EPI_ISL_2805653, EPI_ISL_2805654, EPI_ISL_2805655 | Lighthouse Lab in Alderley Park | Wellcome Sanger Institute for the COVID-19 Genomics UK (COG-UK) Consortium | Jacquelyn Wynn, Mairead Hyland, The Lighthouse Lab in Alderley Park and Alex Alderton, Roberto Amato, Jeffrey Barrett, Sonia Goncalves, Ewan Harrison, David K. Jackson, Ian Johnston, Dominic Kwiatkowski, Cordelia Langford, John Sillitoe on behalf of the Wellcome Sanger Institute COVID-19 Surveillance Team |
| EPI_ISL_2805656 | Lighthouse Laboratory Plymouth | Wellcome Sanger Institute for the COVID-19 Genomics UK (COG-UK) Consortium | Lighthouse Laboratory Plymouth and Alex Alderton, Roberto Amato, Jeffrey Barrett, Sonia Goncalves, Ewan Harrison, David K. Jackson, Ian Johnston, Dominic Kwiatkowski, Cordelia Langford, John Sillitoe on behalf of the Wellcome Sanger Institute COVID-19 Surveillance Team |
| EPI_ISL_2805657 | Lighthouse Lab in Alderley Park | Wellcome Sanger Institute for the COVID-19 Genomics UK (COG-UK) Consortium | Jacquelyn Wynn, Mairead Hyland, The Lighthouse Lab in Alderley Park and Alex Alderton, Roberto Amato, Jeffrey Barrett, Sonia Goncalves, Ewan Harrison, David K. Jackson, Ian Johnston, Dominic Kwiatkowski, Cordelia Langford, John Sillitoe on behalf of the Wellcome Sanger Institute COVID-19 Surveillance Team |
| EPI_ISL_2805658, EPI_ISL_2805659 | Lighthouse Laboratory Plymouth | Wellcome Sanger Institute for the COVID-19 Genomics UK (COG-UK) Consortium | Lighthouse Laboratory Plymouth and Alex Alderton, Roberto Amato, Jeffrey Barrett, Sonia Goncalves, Ewan Harrison, David K. Jackson, Ian Johnston, Dominic Kwiatkowski, Cordelia Langford, John Sillitoe on behalf of the Wellcome Sanger Institute COVID-19 Surveillance Team |
| EPI_ISL_2805660 | Lighthouse Lab in Alderley Park | Wellcome Sanger Institute for the COVID-19 Genomics UK (COG-UK) Consortium | Jacquelyn Wynn, Mairead Hyland, The Lighthouse Lab in Alderley Park and Alex Alderton, Roberto Amato, Jeffrey Barrett, Sonia Goncalves, Ewan Harrison, David K. Jackson, Ian Johnston, Dominic Kwiatkowski, Cordelia Langford, John Sillitoe on behalf of the Wellcome Sanger Institute COVID-19 Surveillance Team |
| EPI_ISL_2805661, EPI_ISL_2805662, EPI_ISL_2805663 | Lighthouse Laboratory Plymouth | Wellcome Sanger Institute for the COVID-19 Genomics UK (COG-UK) Consortium | Lighthouse Laboratory Plymouth and Alex Alderton, Roberto Amato, Jeffrey Barrett, Sonia Goncalves, Ewan Harrison, David K. Jackson, Ian Johnston, Dominic Kwiatkowski, Cordelia Langford, John Sillitoe on behalf of the Wellcome Sanger Institute COVID-19 Surveillance Team |
| EPI_ISL_2805664, EPI_ISL_2805665, EPI_ISL_2805666 | Lighthouse Lab in Alderley Park | Wellcome Sanger Institute for the COVID-19 Genomics UK (COG-UK) Consortium | Jacquelyn Wynn, Mairead Hyland, The Lighthouse Lab in Alderley Park and Alex Alderton, Roberto Amato, Jeffrey Barrett, Sonia Goncalves, Ewan Harrison, David K. Jackson, Ian Johnston, Dominic Kwiatkowski, Cordelia Langford, John Sillitoe on behalf of the Wellcome Sanger Institute COVID-19 Surveillance Team |
| EPI_ISL_2805667 | Lighthouse Laboratory Plymouth | Wellcome Sanger Institute for the COVID-19 Genomics UK (COG-UK) Consortium | Lighthouse Laboratory Plymouth and Alex Alderton, Roberto Amato, Jeffrey Barrett, Sonia Goncalves, Ewan Harrison, David K. Jackson, Ian Johnston, Dominic Kwiatkowski, Cordelia Langford, John Sillitoe on behalf of the Wellcome Sanger Institute COVID-19 Surveillance Team |
| EPI_ISL_2805668, EPI_ISL_2805669 | Lighthouse Lab in Alderley Park | Wellcome Sanger Institute for the COVID-19 Genomics UK (COG-UK) Consortium | Jacquelyn Wynn, Mairead Hyland, The Lighthouse Lab in Alderley Park and Alex Alderton, Roberto Amato, Jeffrey Barrett, Sonia Goncalves, Ewan Harrison, David K. Jackson, Ian Johnston, Dominic Kwiatkowski, Cordelia Langford, John Sillitoe on behalf of the Wellcome Sanger Institute COVID-19 Surveillance Team |
| EPI_ISL_2805670 | Lighthouse Laboratory Plymouth | Wellcome Sanger Institute for the COVID-19 Genomics UK (COG-UK) Consortium | Lighthouse Laboratory Plymouth and Alex Alderton, Roberto Amato, Jeffrey Barrett, Sonia Goncalves, Ewan Harrison, David K. Jackson, Ian Johnston, Dominic Kwiatkowski, Cordelia Langford, John Sillitoe on behalf of the Wellcome Sanger Institute COVID-19 Surveillance Team |
| EPI_ISL_2805671, EPI_ISL_2805672 | Lighthouse Lab in Alderley Park | Wellcome Sanger Institute for the COVID-19 Genomics UK (COG-UK) Consortium | Jacquelyn Wynn, Mairead Hyland, The Lighthouse Lab in Alderley Park and Alex Alderton, Roberto Amato, Jeffrey Barrett, Sonia Goncalves, Ewan Harrison, David K. Jackson, Ian Johnston, Dominic Kwiatkowski, Cordelia Langford, John Sillitoe on behalf of the Wellcome Sanger Institute COVID-19 Surveillance Team |
| EPI_ISL_2805673, EPI_ISL_2805674 | Lighthouse Laboratory Plymouth | Wellcome Sanger Institute for the COVID-19 Genomics UK (COG-UK) Consortium | Lighthouse Laboratory Plymouth and Alex Alderton, Roberto Amato, Jeffrey Barrett, Sonia Goncalves, Ewan Harrison, David K. Jackson, Ian Johnston, Dominic Kwiatkowski, Cordelia Langford, John Sillitoe on behalf of the Wellcome Sanger Institute COVID-19 Surveillance Team |
| EPI_ISL_2805675 | Lighthouse Lab in Alderley Park | Wellcome Sanger Institute for the COVID-19 Genomics UK (COG-UK) Consortium | Jacquelyn Wynn, Mairead Hyland, The Lighthouse Lab in Alderley Park and Alex Alderton, Roberto Amato, Jeffrey Barrett, Sonia Goncalves, Ewan Harrison, David K. Jackson, Ian Johnston, Dominic Kwiatkowski, Cordelia Langford, John Sillitoe on behalf of the Wellcome Sanger Institute COVID-19 Surveillance Team |
| EPI_ISL_2805676, EPI_ISL_2805677, EPI_ISL_2805678 | Lighthouse Laboratory Plymouth | Wellcome Sanger Institute for the COVID-19 Genomics UK (COG-UK) Consortium | Lighthouse Laboratory Plymouth and Alex Alderton, Roberto Amato, Jeffrey Barrett, Sonia Goncalves, Ewan Harrison, David K. Jackson, Ian Johnston, Dominic Kwiatkowski, Cordelia Langford, John Sillitoe on behalf of the Wellcome Sanger Institute COVID-19 Surveillance Team |
| EPI_ISL_2805679, EPI_ISL_2805680 | Lighthouse Lab in Alderley Park | Wellcome Sanger Institute for the COVID-19 Genomics UK (COG-UK) Consortium | Jacquelyn Wynn, Mairead Hyland, The Lighthouse Lab in Alderley Park and Alex Alderton, Roberto Amato, Jeffrey Barrett, Sonia Goncalves, Ewan Harrison, David K. Jackson, Ian Johnston, Dominic Kwiatkowski, Cordelia Langford, John Sillitoe on behalf of the Wellcome Sanger Institute COVID-19 Surveillance Team |
| EPI_ISL_2805681 | Lighthouse Laboratory Plymouth | Wellcome Sanger Institute for the COVID-19 Genomics UK (COG-UK) Consortium | Lighthouse Laboratory Plymouth and Alex Alderton, Roberto Amato, Jeffrey Barrett, Sonia Goncalves, Ewan Harrison, David K. Jackson, Ian Johnston, Dominic Kwiatkowski, Cordelia Langford, John Sillitoe on behalf of the Wellcome Sanger Institute COVID-19 Surveillance Team |
| EPI_ISL_2805682, EPI_ISL_2805683 | Lighthouse Lab in Alderley Park | Wellcome Sanger Institute for the COVID-19 Genomics UK (COG-UK) Consortium | Jacquelyn Wynn, Mairead Hyland, The Lighthouse Lab in Alderley Park and Alex Alderton, Roberto Amato, Jeffrey Barrett, Sonia Goncalves, Ewan Harrison, David K. Jackson, Ian Johnston, Dominic Kwiatkowski, Cordelia Langford, John Sillitoe on behalf of the Wellcome Sanger Institute COVID-19 Surveillance Team |
| EPI_ISL_2805684 | Lighthouse Laboratory Plymouth | Wellcome Sanger Institute for the COVID-19 Genomics UK (COG-UK) Consortium | Lighthouse Laboratory Plymouth and Alex Alderton, Roberto Amato, Jeffrey Barrett, Sonia Goncalves, Ewan Harrison, David K. Jackson, Ian Johnston, Dominic Kwiatkowski, Cordelia Langford, John Sillitoe on behalf of the Wellcome Sanger Institute COVID-19 Surveillance Team |
| EPI_ISL_2805685, EPI_ISL_2805686, EPI_ISL_2805687, EPI_ISL_2805688, EPI_ISL_2805689, EPI_ISL_2805690 | Lighthouse Lab in Alderley Park | Wellcome Sanger Institute for the COVID-19 Genomics UK (COG-UK) Consortium | Jacquelyn Wynn, Mairead Hyland, The Lighthouse Lab in Alderley Park and Alex Alderton, Roberto Amato, Jeffrey Barrett, Sonia Goncalves, Ewan Harrison, David K. Jackson, Ian Johnston, Dominic Kwiatkowski, Cordelia Langford, John Sillitoe on behalf of the Wellcome Sanger Institute COVID-19 Surveillance Team |
| EPI_ISL_2805691, EPI_ISL_2805692 | Lighthouse Laboratory Plymouth | Wellcome Sanger Institute for the COVID-19 Genomics UK (COG-UK) Consortium | Lighthouse Laboratory Plymouth and Alex Alderton, Roberto Amato, Jeffrey Barrett, Sonia Goncalves, Ewan Harrison, David K. Jackson, Ian Johnston, Dominic Kwiatkowski, Cordelia Langford, John Sillitoe on behalf of the Wellcome Sanger Institute COVID-19 Surveillance Team |

| | | | |
|---|---|---|---|
| EPI_ISL_2805693, EPI_ISL_2805694 | Lighthouse Lab in Alderley Park | Wellcome Sanger Institute for the COVID-19 Genomics UK (COG-UK) Consortium | Jacquelyn Wynn, Mairead Hyland, The Lighthouse Lab in Alderley Park and Alex Alderton, Roberto Amato, Jeffrey Barrett, Sonia Goncalves, Ewan Harrison, David K. Jackson, Ian Johnston, Dominic Kwiatkowski, Cordelia Langford, John Sillitoe on behalf of the Wellcome Sanger Institute COVID-19 Surveillance Team |
| EPI_ISL_2805695 | Lighthouse Lab in Milton Keynes | Wellcome Sanger Institute for the COVID-19 Genomics UK (COG-UK) Consortium | The Lighthouse Lab in Milton Keynes and Alex Alderton, Roberto Amato, Jeffrey Barrett, Sonia Goncalves, Ewan Harrison, David K. Jackson, Ian Johnston, Dominic Kwiatkowski, Cordelia Langford, John Sillitoe on behalf of the Wellcome Sanger Institute COVID-19 Surveillance Team |
| EPI_ISL_2805696 | Lighthouse Laboratory Plymouth | Wellcome Sanger Institute for the COVID-19 Genomics UK (COG-UK) Consortium | Lighthouse Laboratory Plymouth and Alex Alderton, Roberto Amato, Jeffrey Barrett, Sonia Goncalves, Ewan Harrison, David K. Jackson, Ian Johnston, Dominic Kwiatkowski, Cordelia Langford, John Sillitoe on behalf of the Wellcome Sanger Institute COVID-19 Surveillance Team |
| EPI_ISL_2805697 | Lighthouse Lab in Milton Keynes | Wellcome Sanger Institute for the COVID-19 Genomics UK (COG-UK) Consortium | The Lighthouse Lab in Milton Keynes and Alex Alderton, Roberto Amato, Jeffrey Barrett, Sonia Goncalves, Ewan Harrison, David K. Jackson, Ian Johnston, Dominic Kwiatkowski, Cordelia Langford, John Sillitoe on behalf of the Wellcome Sanger Institute COVID-19 Surveillance Team |
| EPI_ISL_2805698 | Lighthouse Lab in Glasgow | Wellcome Sanger Institute for the COVID-19 Genomics UK (COG-UK) Consortium | Harper VanSteenhouse, Yumi Kasai, David Gray, Carol Clugston, Anna Dominiczak and Alex Alderton, Roberto Amato, Jeffrey Barrett, Sonia Goncalves, Ewan Harrison, David K. Jackson, Ian Johnston, Dominic Kwiatkowski, Cordelia Langford, John Sillitoe on behalf of the Wellcome Sanger Institute COVID-19 Surveillance Team |
| EPI_ISL_2805699, EPI_ISL_2805700 | Lighthouse Lab in Milton Keynes | Wellcome Sanger Institute for the COVID-19 Genomics UK (COG-UK) Consortium | The Lighthouse Lab in Milton Keynes and Alex Alderton, Roberto Amato, Jeffrey Barrett, Sonia Goncalves, Ewan Harrison, David K. Jackson, Ian Johnston, Dominic Kwiatkowski, Cordelia Langford, John Sillitoe on behalf of the Wellcome Sanger Institute COVID-19 Surveillance Team |
| EPI_ISL_2805701 | Lighthouse Laboratory Plymouth | Wellcome Sanger Institute for the COVID-19 Genomics UK (COG-UK) Consortium | Lighthouse Laboratory Plymouth and Alex Alderton, Roberto Amato, Jeffrey Barrett, Sonia Goncalves, Ewan Harrison, David K. Jackson, Ian Johnston, Dominic Kwiatkowski, Cordelia Langford, John Sillitoe on behalf of the Wellcome Sanger Institute COVID-19 Surveillance Team |
| EPI_ISL_2805702, EPI_ISL_2805703, EPI_ISL_2805704 | Lighthouse Lab in Milton Keynes | Wellcome Sanger Institute for the COVID-19 Genomics UK (COG-UK) Consortium | The Lighthouse Lab in Milton Keynes and Alex Alderton, Roberto Amato, Jeffrey Barrett, Sonia Goncalves, Ewan Harrison, David K. Jackson, Ian Johnston, Dominic Kwiatkowski, Cordelia Langford, John Sillitoe on behalf of the Wellcome Sanger Institute COVID-19 Surveillance Team |
| EPI_ISL_2805705, EPI_ISL_2805706 | Lighthouse Laboratory Plymouth | Wellcome Sanger Institute for the COVID-19 Genomics UK (COG-UK) Consortium | Lighthouse Laboratory Plymouth and Alex Alderton, Roberto Amato, Jeffrey Barrett, Sonia Goncalves, Ewan Harrison, David K. Jackson, Ian Johnston, Dominic Kwiatkowski, Cordelia Langford, John Sillitoe on behalf of the Wellcome Sanger Institute COVID-19 Surveillance Team |
| EPI_ISL_2805707 | Lighthouse Lab in Glasgow | Wellcome Sanger Institute for the COVID-19 Genomics UK (COG-UK) Consortium | Harper VanSteenhouse, Yumi Kasai, David Gray, Carol Clugston, Anna Dominiczak and Alex Alderton, Roberto Amato, Jeffrey Barrett, Sonia Goncalves, Ewan Harrison, David K. Jackson, Ian Johnston, Dominic Kwiatkowski, Cordelia Langford, John Sillitoe on behalf of the Wellcome Sanger Institute COVID-19 Surveillance Team |
| EPI_ISL_2805708 | Lighthouse Laboratory Plymouth | Wellcome Sanger Institute for the COVID-19 Genomics UK (COG-UK) Consortium | Lighthouse Laboratory Plymouth and Alex Alderton, Roberto Amato, Jeffrey Barrett, Sonia Goncalves, Ewan Harrison, David K. Jackson, Ian Johnston, Dominic Kwiatkowski, Cordelia Langford, John Sillitoe on behalf of the Wellcome Sanger Institute COVID-19 Surveillance Team |
| EPI_ISL_2805709, EPI_ISL_2805710, EPI_ISL_2805711, EPI_ISL_2805712, EPI_ISL_2805713, EPI_ISL_2805714, EPI_ISL_2805715 | Lighthouse Lab in Milton Keynes | Wellcome Sanger Institute for the COVID-19 Genomics UK (COG-UK) Consortium | The Lighthouse Lab in Milton Keynes and Alex Alderton, Roberto Amato, Jeffrey Barrett, Sonia Goncalves, Ewan Harrison, David K. Jackson, Ian Johnston, Dominic Kwiatkowski, Cordelia Langford, John Sillitoe on behalf of the Wellcome Sanger Institute COVID-19 Surveillance Team |
| EPI_ISL_2805716 | Lighthouse Laboratory Plymouth | Wellcome Sanger Institute for the COVID-19 Genomics UK (COG-UK) Consortium | Lighthouse Laboratory Plymouth and Alex Alderton, Roberto Amato, Jeffrey Barrett, Sonia Goncalves, Ewan Harrison, David K. Jackson, Ian Johnston, Dominic Kwiatkowski, Cordelia Langford, John Sillitoe on behalf of the Wellcome Sanger Institute COVID-19 Surveillance Team |
| EPI_ISL_2805717 | Lighthouse Lab in Glasgow | Wellcome Sanger Institute for the COVID-19 Genomics UK (COG-UK) Consortium | Harper VanSteenhouse, Yumi Kasai, David Gray, Carol Clugston, Anna Dominiczak and Alex Alderton, Roberto Amato, Jeffrey Barrett, Sonia Goncalves, Ewan Harrison, David K. Jackson, Ian Johnston, Dominic Kwiatkowski, Cordelia Langford, John Sillitoe on behalf of the Wellcome Sanger Institute COVID-19 Surveillance Team |
| EPI_ISL_2805718 | Lighthouse Lab in Milton Keynes | Wellcome Sanger Institute for the COVID-19 Genomics UK (COG-UK) Consortium | The Lighthouse Lab in Milton Keynes and Alex Alderton, Roberto Amato, Jeffrey Barrett, Sonia Goncalves, Ewan Harrison, David K. Jackson, Ian Johnston, Dominic Kwiatkowski, Cordelia Langford, John Sillitoe on behalf of the Wellcome Sanger Institute COVID-19 Surveillance Team |
| EPI_ISL_2805719 | Lighthouse Laboratory Plymouth | Wellcome Sanger Institute for the COVID-19 Genomics UK (COG-UK) Consortium | Lighthouse Laboratory Plymouth and Alex Alderton, Roberto Amato, Jeffrey Barrett, Sonia Goncalves, Ewan Harrison, David K. Jackson, Ian Johnston, Dominic Kwiatkowski, Cordelia Langford, John Sillitoe on behalf of the Wellcome Sanger Institute COVID-19 Surveillance Team |
| EPI_ISL_2805720 | Lighthouse Lab in Glasgow | Wellcome Sanger Institute for the COVID-19 Genomics UK (COG-UK) Consortium | Harper VanSteenhouse, Yumi Kasai, David Gray, Carol Clugston, Anna Dominiczak and Alex Alderton, Roberto Amato, Jeffrey Barrett, Sonia Goncalves, Ewan Harrison, David K. Jackson, Ian Johnston, Dominic Kwiatkowski, Cordelia Langford, John Sillitoe on behalf of the Wellcome Sanger Institute COVID-19 Surveillance Team |
| EPI_ISL_2805721, EPI_ISL_2805722, EPI_ISL_2805723, EPI_ISL_2805724 | Lighthouse Lab in Milton Keynes | Wellcome Sanger Institute for the COVID-19 Genomics UK (COG-UK) Consortium | The Lighthouse Lab in Milton Keynes and Alex Alderton, Roberto Amato, Jeffrey Barrett, Sonia Goncalves, Ewan Harrison, David K. Jackson, Ian Johnston, Dominic Kwiatkowski, Cordelia Langford, John Sillitoe on behalf of the Wellcome Sanger Institute COVID-19 Surveillance Team |
| EPI_ISL_2805725 | Lighthouse Laboratory Plymouth | Wellcome Sanger Institute for the COVID-19 Genomics UK (COG-UK) Consortium | Lighthouse Laboratory Plymouth and Alex Alderton, Roberto Amato, Jeffrey Barrett, Sonia Goncalves, Ewan Harrison, David K. Jackson, Ian Johnston, Dominic Kwiatkowski, Cordelia Langford, John Sillitoe on behalf of the Wellcome Sanger Institute COVID-19 Surveillance Team |
| EPI_ISL_2805726, EPI_ISL_2805727, EPI_ISL_2805728, EPI_ISL_2805729, EPI_ISL_2805730, EPI_ISL_2805731, EPI_ISL_2805732, EPI_ISL_2805733 | Lighthouse Lab in Milton Keynes | Wellcome Sanger Institute for the COVID-19 Genomics UK (COG-UK) Consortium | The Lighthouse Lab in Milton Keynes and Alex Alderton, Roberto Amato, Jeffrey Barrett, Sonia Goncalves, Ewan Harrison, David K. Jackson, Ian Johnston, Dominic Kwiatkowski, Cordelia Langford, John Sillitoe on behalf of the Wellcome Sanger Institute COVID-19 Surveillance Team |
| EPI_ISL_2805734, EPI_ISL_2805735 | Lighthouse Laboratory Plymouth | Wellcome Sanger Institute for the COVID-19 Genomics UK (COG-UK) Consortium | Lighthouse Laboratory Plymouth and Alex Alderton, Roberto Amato, Jeffrey Barrett, Sonia Goncalves, Ewan Harrison, David K. Jackson, Ian Johnston, Dominic Kwiatkowski, Cordelia Langford, John Sillitoe on behalf of the Wellcome Sanger Institute COVID-19 Surveillance Team |
| EPI_ISL_2805736, EPI_ISL_2805737, EPI_ISL_2805738 | Lighthouse Lab in Milton Keynes | Wellcome Sanger Institute for the COVID-19 Genomics UK (COG-UK) Consortium | The Lighthouse Lab in Milton Keynes and Alex Alderton, Roberto Amato, Jeffrey Barrett, Sonia Goncalves, Ewan Harrison, David K. Jackson, Ian Johnston, Dominic Kwiatkowski, Cordelia Langford, John Sillitoe on behalf of the Wellcome Sanger Institute COVID-19 Surveillance Team |
| EPI_ISL_2805739, EPI_ISL_2805741 | Lighthouse Laboratory Plymouth | Wellcome Sanger Institute for the COVID-19 Genomics UK (COG-UK) Consortium | Lighthouse Laboratory Plymouth and Alex Alderton, Roberto Amato, Jeffrey Barrett, Sonia Goncalves, Ewan Harrison, David K. Jackson, Ian Johnston, Dominic Kwiatkowski, Cordelia Langford, John Sillitoe on behalf of the Wellcome Sanger Institute COVID-19 Surveillance Team |
| EPI_ISL_2805742, EPI_ISL_2805743 | Lighthouse Lab in Milton Keynes | Wellcome Sanger Institute for the COVID-19 Genomics UK (COG-UK) Consortium | The Lighthouse Lab in Milton Keynes and Alex Alderton, Roberto Amato, Jeffrey Barrett, Sonia Goncalves, Ewan Harrison, David K. Jackson, Ian Johnston, Dominic Kwiatkowski, Cordelia Langford, John Sillitoe on behalf of the Wellcome Sanger Institute COVID-19 Surveillance Team |
| EPI_ISL_2805744 | Lighthouse Laboratory Plymouth | Wellcome Sanger Institute for the COVID-19 Genomics UK (COG-UK) Consortium | Lighthouse Laboratory Plymouth and Alex Alderton, Roberto Amato, Jeffrey Barrett, Sonia Goncalves, Ewan Harrison, David K. Jackson, Ian Johnston, Dominic Kwiatkowski, Cordelia Langford, John Sillitoe on behalf of the Wellcome Sanger Institute COVID-19 Surveillance Team |
| EPI_ISL_2805745 | Lighthouse Lab in Milton Keynes | Wellcome Sanger Institute for the COVID-19 Genomics UK (COG-UK) Consortium | The Lighthouse Lab in Milton Keynes and Alex Alderton, Roberto Amato, Jeffrey Barrett, Sonia Goncalves, Ewan Harrison, David K. Jackson, Ian Johnston, Dominic Kwiatkowski, Cordelia Langford, John Sillitoe on behalf of the Wellcome Sanger Institute COVID-19 Surveillance Team |
| EPI_ISL_2805746, EPI_ISL_2805747 | Lighthouse Laboratory Plymouth | Wellcome Sanger Institute for the COVID-19 Genomics UK (COG-UK) Consortium | Lighthouse Laboratory Plymouth and Alex Alderton, Roberto Amato, Jeffrey Barrett, Sonia Goncalves, Ewan Harrison, David K. Jackson, Ian Johnston, Dominic Kwiatkowski, Cordelia Langford, John Sillitoe on behalf of the Wellcome Sanger Institute COVID-19 Surveillance Team |
| EPI_ISL_2805748, EPI_ISL_2805749 | Lighthouse Lab in Milton Keynes | Wellcome Sanger Institute for the COVID-19 Genomics UK (COG-UK) Consortium | The Lighthouse Lab in Milton Keynes and Alex Alderton, Roberto Amato, Jeffrey Barrett, Sonia Goncalves, Ewan Harrison, David K. Jackson, Ian Johnston, Dominic Kwiatkowski, Cordelia Langford, John Sillitoe on behalf of the Wellcome Sanger Institute COVID-19 Surveillance Team |
| EPI_ISL_2805750, EPI_ISL_2805751, EPI_ISL_2805752 | Lighthouse Laboratory Plymouth | Wellcome Sanger Institute for the COVID-19 Genomics UK (COG-UK) Consortium | Lighthouse Laboratory Plymouth and Alex Alderton, Roberto Amato, Jeffrey Barrett, Sonia Goncalves, Ewan Harrison, David K. Jackson, Ian Johnston, Dominic Kwiatkowski, Cordelia Langford, John Sillitoe on behalf of the Wellcome Sanger Institute COVID-19 Surveillance Team |
| EPI_ISL_2805753 | Lighthouse Lab in Milton Keynes | Wellcome Sanger Institute for the COVID-19 Genomics UK (COG-UK) Consortium | The Lighthouse Lab in Milton Keynes and Alex Alderton, Roberto Amato, Jeffrey Barrett, Sonia Goncalves, Ewan Harrison, David K. Jackson, Ian Johnston, Dominic Kwiatkowski, Cordelia Langford, John Sillitoe on behalf of the Wellcome Sanger Institute COVID-19 Surveillance Team |
| EPI_ISL_2805754 | Lighthouse Lab in Glasgow | Wellcome Sanger Institute for the COVID-19 Genomics UK (COG-UK) Consortium | Harper VanSteenhouse, Yumi Kasai, David Gray, Carol Clugston, Anna Dominiczak and Alex Alderton, Roberto Amato, Jeffrey Barrett, Sonia Goncalves, Ewan Harrison, David K. Jackson, Ian Johnston, Dominic Kwiatkowski, Cordelia Langford, John Sillitoe on behalf of the Wellcome Sanger Institute COVID-19 Surveillance Team |
| EPI_ISL_2805755 | Lighthouse Lab in Milton Keynes | Wellcome Sanger Institute for the COVID-19 Genomics UK (COG-UK) Consortium | The Lighthouse Lab in Milton Keynes and Alex Alderton, Roberto Amato, Jeffrey Barrett, Sonia Goncalves, Ewan Harrison, David K. Jackson, Ian Johnston, Dominic Kwiatkowski, Cordelia Langford, John Sillitoe on behalf of the Wellcome Sanger Institute COVID-19 Surveillance Team |
| EPI_ISL_2805756, EPI_ISL_2805757 | Lighthouse Laboratory Plymouth | Wellcome Sanger Institute for the COVID-19 Genomics UK (COG-UK) Consortium | Lighthouse Laboratory Plymouth and Alex Alderton, Roberto Amato, Jeffrey Barrett, Sonia Goncalves, Ewan Harrison, David K. Jackson, Ian Johnston, Dominic Kwiatkowski, Cordelia Langford, John Sillitoe on behalf of the Wellcome Sanger Institute COVID-19 Surveillance Team |
| EPI_ISL_2805758, EPI_ISL_2805759, EPI_ISL_2805760, EPI_ISL_2805761 | Lighthouse Lab in Milton Keynes | Wellcome Sanger Institute for the COVID-19 Genomics UK (COG-UK) Consortium | The Lighthouse Lab in Milton Keynes and Alex Alderton, Roberto Amato, Jeffrey Barrett, Sonia Goncalves, Ewan Harrison, David K. Jackson, Ian Johnston, Dominic Kwiatkowski, Cordelia Langford, John Sillitoe on behalf of the Wellcome Sanger Institute COVID-19 Surveillance Team |
| EPI_ISL_2805762 | Lighthouse Laboratory Plymouth | Wellcome Sanger Institute for the COVID-19 Genomics UK (COG-UK) Consortium | Lighthouse Laboratory Plymouth and Alex Alderton, Roberto Amato, Jeffrey Barrett, Sonia Goncalves, Ewan Harrison, David K. Jackson, Ian Johnston, Dominic Kwiatkowski, Cordelia Langford, John Sillitoe on behalf of the Wellcome Sanger Institute COVID-19 Surveillance Team |
| EPI_ISL_2805763, EPI_ISL_2805764, EPI_ISL_2805765, EPI_ISL_2805766, EPI_ISL_2805767, EPI_ISL_2805768 | Lighthouse Lab in Milton Keynes | Wellcome Sanger Institute for the COVID-19 Genomics UK (COG-UK) Consortium | The Lighthouse Lab in Milton Keynes and Alex Alderton, Roberto Amato, Jeffrey Barrett, Sonia Goncalves, Ewan Harrison, David K. Jackson, Ian Johnston, Dominic Kwiatkowski, Cordelia Langford, John Sillitoe on behalf of the Wellcome Sanger Institute COVID-19 Surveillance Team |

| | | | |
|---|---|---|---|
| EPI_ISL_2805769 | Lighthouse Lab in Glasgow | Wellcome Sanger Institute for the COVID-19 Genomics UK (COG-UK) Consortium | Harper VanSteenhouse, Yumi Kasai, David Gray, Carol Clugston, Anna Dominiczak and Alex Alderton, Roberto Amato, Jeffrey Barrett, Sonia Goncalves, Ewan Harrison, David K. Jackson, Ian Johnston, Dominic Kwiatkowski, Cordelia Langford, John Sillitoe on behalf of the Wellcome Sanger Institute COVID-19 Surveillance Team |
| EPI_ISL_2805770 | Lighthouse Laboratory Plymouth | Wellcome Sanger Institute for the COVID-19 Genomics UK (COG-UK) Consortium | Lighthouse Laboratory Plymouth and Alex Alderton, Roberto Amato, Jeffrey Barrett, Sonia Goncalves, Ewan Harrison, David K. Jackson, Ian Johnston, Dominic Kwiatkowski, Cordelia Langford, John Sillitoe on behalf of the Wellcome Sanger Institute COVID-19 Surveillance Team |
| EPI_ISL_2805771 | Lighthouse Lab in Milton Keynes | Wellcome Sanger Institute for the COVID-19 Genomics UK (COG-UK) Consortium | The Lighthouse Lab in Milton Keynes and Alex Alderton, Roberto Amato, Jeffrey Barrett, Sonia Goncalves, Ewan Harrison, David K. Jackson, Ian Johnston, Dominic Kwiatkowski, Cordelia Langford, John Sillitoe on behalf of the Wellcome Sanger Institute COVID-19 Surveillance Team |
| EPI_ISL_2805772 | Lighthouse Laboratory Plymouth | Wellcome Sanger Institute for the COVID-19 Genomics UK (COG-UK) Consortium | Lighthouse Laboratory Plymouth and Alex Alderton, Roberto Amato, Jeffrey Barrett, Sonia Goncalves, Ewan Harrison, David K. Jackson, Ian Johnston, Dominic Kwiatkowski, Cordelia Langford, John Sillitoe on behalf of the Wellcome Sanger Institute COVID-19 Surveillance Team |
| EPI_ISL_2805773 | Lighthouse Lab in Milton Keynes | Wellcome Sanger Institute for the COVID-19 Genomics UK (COG-UK) Consortium | The Lighthouse Lab in Milton Keynes and Alex Alderton, Roberto Amato, Jeffrey Barrett, Sonia Goncalves, Ewan Harrison, David K. Jackson, Ian Johnston, Dominic Kwiatkowski, Cordelia Langford, John Sillitoe on behalf of the Wellcome Sanger Institute COVID-19 Surveillance Team |
| EPI_ISL_2805774 | Lighthouse Laboratory Plymouth | Wellcome Sanger Institute for the COVID-19 Genomics UK (COG-UK) Consortium | Lighthouse Laboratory Plymouth and Alex Alderton, Roberto Amato, Jeffrey Barrett, Sonia Goncalves, Ewan Harrison, David K. Jackson, Ian Johnston, Dominic Kwiatkowski, Cordelia Langford, John Sillitoe on behalf of the Wellcome Sanger Institute COVID-19 Surveillance Team |
| EPI_ISL_2805775 | Lighthouse Lab in Glasgow | Wellcome Sanger Institute for the COVID-19 Genomics UK (COG-UK) Consortium | Harper VanSteenhouse, Yumi Kasai, David Gray, Carol Clugston, Anna Dominiczak and Alex Alderton, Roberto Amato, Jeffrey Barrett, Sonia Goncalves, Ewan Harrison, David K. Jackson, Ian Johnston, Dominic Kwiatkowski, Cordelia Langford, John Sillitoe on behalf of the Wellcome Sanger Institute COVID-19 Surveillance Team |
| EPI_ISL_2805776, EPI_ISL_2805777 | Lighthouse Lab in Milton Keynes | Wellcome Sanger Institute for the COVID-19 Genomics UK (COG-UK) Consortium | The Lighthouse Lab in Milton Keynes and Alex Alderton, Roberto Amato, Jeffrey Barrett, Sonia Goncalves, Ewan Harrison, David K. Jackson, Ian Johnston, Dominic Kwiatkowski, Cordelia Langford, John Sillitoe on behalf of the Wellcome Sanger Institute COVID-19 Surveillance Team |
| EPI_ISL_2805778 | Lighthouse Lab in Glasgow | Wellcome Sanger Institute for the COVID-19 Genomics UK (COG-UK) Consortium | Harper VanSteenhouse, Yumi Kasai, David Gray, Carol Clugston, Anna Dominiczak and Alex Alderton, Roberto Amato, Jeffrey Barrett, Sonia Goncalves, Ewan Harrison, David K. Jackson, Ian Johnston, Dominic Kwiatkowski, Cordelia Langford, John Sillitoe on behalf of the Wellcome Sanger Institute COVID-19 Surveillance Team |
| EPI_ISL_2805779, EPI_ISL_2805780 | Lighthouse Lab in Milton Keynes | Wellcome Sanger Institute for the COVID-19 Genomics UK (COG-UK) Consortium | The Lighthouse Lab in Milton Keynes and Alex Alderton, Roberto Amato, Jeffrey Barrett, Sonia Goncalves, Ewan Harrison, David K. Jackson, Ian Johnston, Dominic Kwiatkowski, Cordelia Langford, John Sillitoe on behalf of the Wellcome Sanger Institute COVID-19 Surveillance Team |
| EPI_ISL_2805781 | Lighthouse Laboratory Plymouth | Wellcome Sanger Institute for the COVID-19 Genomics UK (COG-UK) Consortium | Lighthouse Laboratory Plymouth and Alex Alderton, Roberto Amato, Jeffrey Barrett, Sonia Goncalves, Ewan Harrison, David K. Jackson, Ian Johnston, Dominic Kwiatkowski, Cordelia Langford, John Sillitoe on behalf of the Wellcome Sanger Institute COVID-19 Surveillance Team |
| EPI_ISL_2805782, EPI_ISL_2805783 | Lighthouse Lab in Milton Keynes | Wellcome Sanger Institute for the COVID-19 Genomics UK (COG-UK) Consortium | The Lighthouse Lab in Milton Keynes and Alex Alderton, Roberto Amato, Jeffrey Barrett, Sonia Goncalves, Ewan Harrison, David K. Jackson, Ian Johnston, Dominic Kwiatkowski, Cordelia Langford, John Sillitoe on behalf of the Wellcome Sanger Institute COVID-19 Surveillance Team |
| EPI_ISL_2805784 | Lighthouse Lab in Glasgow | Wellcome Sanger Institute for the COVID-19 Genomics UK (COG-UK) Consortium | Harper VanSteenhouse, Yumi Kasai, David Gray, Carol Clugston, Anna Dominiczak and Alex Alderton, Roberto Amato, Jeffrey Barrett, Sonia Goncalves, Ewan Harrison, David K. Jackson, Ian Johnston, Dominic Kwiatkowski, Cordelia Langford, John Sillitoe on behalf of the Wellcome Sanger Institute COVID-19 Surveillance Team |
| EPI_ISL_2805785, EPI_ISL_2805786 | Lighthouse Laboratory Plymouth | Wellcome Sanger Institute for the COVID-19 Genomics UK (COG-UK) Consortium | Lighthouse Laboratory Plymouth and Alex Alderton, Roberto Amato, Jeffrey Barrett, Sonia Goncalves, Ewan Harrison, David K. Jackson, Ian Johnston, Dominic Kwiatkowski, Cordelia Langford, John Sillitoe on behalf of the Wellcome Sanger Institute COVID-19 Surveillance Team |
| EPI_ISL_2805787, EPI_ISL_2805788 | Lighthouse Lab in Milton Keynes | Wellcome Sanger Institute for the COVID-19 Genomics UK (COG-UK) Consortium | The Lighthouse Lab in Milton Keynes and Alex Alderton, Roberto Amato, Jeffrey Barrett, Sonia Goncalves, Ewan Harrison, David K. Jackson, Ian Johnston, Dominic Kwiatkowski, Cordelia Langford, John Sillitoe on behalf of the Wellcome Sanger Institute COVID-19 Surveillance Team |
| EPI_ISL_2805789, EPI_ISL_2805790 | Lighthouse Laboratory Plymouth | Wellcome Sanger Institute for the COVID-19 Genomics UK (COG-UK) Consortium | Lighthouse Laboratory Plymouth and Alex Alderton, Roberto Amato, Jeffrey Barrett, Sonia Goncalves, Ewan Harrison, David K. Jackson, Ian Johnston, Dominic Kwiatkowski, Cordelia Langford, John Sillitoe on behalf of the Wellcome Sanger Institute COVID-19 Surveillance Team |
| EPI_ISL_2805791 | Lighthouse Lab in Glasgow | Wellcome Sanger Institute for the COVID-19 Genomics UK (COG-UK) Consortium | Harper VanSteenhouse, Yumi Kasai, David Gray, Carol Clugston, Anna Dominiczak and Alex Alderton, Roberto Amato, Jeffrey Barrett, Sonia Goncalves, Ewan Harrison, David K. Jackson, Ian Johnston, Dominic Kwiatkowski, Cordelia Langford, John Sillitoe on behalf of the Wellcome Sanger Institute COVID-19 Surveillance Team |
| EPI_ISL_2805792, EPI_ISL_2805793, EPI_ISL_2805794, EPI_ISL_2805795, EPI_ISL_2805796, EPI_ISL_2805797, EPI_ISL_2805798, EPI_ISL_2805799 | Lighthouse Lab in Milton Keynes | Wellcome Sanger Institute for the COVID-19 Genomics UK (COG-UK) Consortium | The Lighthouse Lab in Milton Keynes and Alex Alderton, Roberto Amato, Jeffrey Barrett, Sonia Goncalves, Ewan Harrison, David K. Jackson, Ian Johnston, Dominic Kwiatkowski, Cordelia Langford, John Sillitoe on behalf of the Wellcome Sanger Institute COVID-19 Surveillance Team |
| EPI_ISL_2805800 | Lighthouse Laboratory Plymouth | Wellcome Sanger Institute for the COVID-19 Genomics UK (COG-UK) Consortium | Lighthouse Laboratory Plymouth and Alex Alderton, Roberto Amato, Jeffrey Barrett, Sonia Goncalves, Ewan Harrison, David K. Jackson, Ian Johnston, Dominic Kwiatkowski, Cordelia Langford, John Sillitoe on behalf of the Wellcome Sanger Institute COVID-19 Surveillance Team |
| EPI_ISL_2805801, EPI_ISL_2805802, EPI_ISL_2805803, EPI_ISL_2805804 | Lighthouse Lab in Milton Keynes | Wellcome Sanger Institute for the COVID-19 Genomics UK (COG-UK) Consortium | The Lighthouse Lab in Milton Keynes and Alex Alderton, Roberto Amato, Jeffrey Barrett, Sonia Goncalves, Ewan Harrison, David K. Jackson, Ian Johnston, Dominic Kwiatkowski, Cordelia Langford, John Sillitoe on behalf of the Wellcome Sanger Institute COVID-19 Surveillance Team |
| EPI_ISL_2805805 | Lighthouse Laboratory Plymouth | Wellcome Sanger Institute for the COVID-19 Genomics UK (COG-UK) Consortium | Lighthouse Laboratory Plymouth and Alex Alderton, Roberto Amato, Jeffrey Barrett, Sonia Goncalves, Ewan Harrison, David K. Jackson, Ian Johnston, Dominic Kwiatkowski, Cordelia Langford, John Sillitoe on behalf of the Wellcome Sanger Institute COVID-19 Surveillance Team |
| EPI_ISL_2805806 | Lighthouse Lab in Milton Keynes | Wellcome Sanger Institute for the COVID-19 Genomics UK (COG-UK) Consortium | The Lighthouse Lab in Milton Keynes and Alex Alderton, Roberto Amato, Jeffrey Barrett, Sonia Goncalves, Ewan Harrison, David K. Jackson, Ian Johnston, Dominic Kwiatkowski, Cordelia Langford, John Sillitoe on behalf of the Wellcome Sanger Institute COVID-19 Surveillance Team |
| EPI_ISL_2805807 | Lighthouse Laboratory Plymouth | Wellcome Sanger Institute for the COVID-19 Genomics UK (COG-UK) Consortium | Lighthouse Laboratory Plymouth and Alex Alderton, Roberto Amato, Jeffrey Barrett, Sonia Goncalves, Ewan Harrison, David K. Jackson, Ian Johnston, Dominic Kwiatkowski, Cordelia Langford, John Sillitoe on behalf of the Wellcome Sanger Institute COVID-19 Surveillance Team |
| EPI_ISL_2805808 | Lighthouse Lab in Glasgow | Wellcome Sanger Institute for the COVID-19 Genomics UK (COG-UK) Consortium | Harper VanSteenhouse, Yumi Kasai, David Gray, Carol Clugston, Anna Dominiczak and Alex Alderton, Roberto Amato, Jeffrey Barrett, Sonia Goncalves, Ewan Harrison, David K. Jackson, Ian Johnston, Dominic Kwiatkowski, Cordelia Langford, John Sillitoe on behalf of the Wellcome Sanger Institute COVID-19 Surveillance Team |
| EPI_ISL_2805809, EPI_ISL_2805810 | Lighthouse Lab in Milton Keynes | Wellcome Sanger Institute for the COVID-19 Genomics UK (COG-UK) Consortium | The Lighthouse Lab in Milton Keynes and Alex Alderton, Roberto Amato, Jeffrey Barrett, Sonia Goncalves, Ewan Harrison, David K. Jackson, Ian Johnston, Dominic Kwiatkowski, Cordelia Langford, John Sillitoe on behalf of the Wellcome Sanger Institute COVID-19 Surveillance Team |
| EPI_ISL_2805811 | Lighthouse Lab in Glasgow | Wellcome Sanger Institute for the COVID-19 Genomics UK (COG-UK) Consortium | Harper VanSteenhouse, Yumi Kasai, David Gray, Carol Clugston, Anna Dominiczak and Alex Alderton, Roberto Amato, Jeffrey Barrett, Sonia Goncalves, Ewan Harrison, David K. Jackson, Ian Johnston, Dominic Kwiatkowski, Cordelia Langford, John Sillitoe on behalf of the Wellcome Sanger Institute COVID-19 Surveillance Team |
| EPI_ISL_2805812 | Lighthouse Lab in Milton Keynes | Wellcome Sanger Institute for the COVID-19 Genomics UK (COG-UK) Consortium | The Lighthouse Lab in Milton Keynes and Alex Alderton, Roberto Amato, Jeffrey Barrett, Sonia Goncalves, Ewan Harrison, David K. Jackson, Ian Johnston, Dominic Kwiatkowski, Cordelia Langford, John Sillitoe on behalf of the Wellcome Sanger Institute COVID-19 Surveillance Team |
| EPI_ISL_2805813, EPI_ISL_2805814 | Lighthouse Lab in Glasgow | Wellcome Sanger Institute for the COVID-19 Genomics UK (COG-UK) Consortium | Harper VanSteenhouse, Yumi Kasai, David Gray, Carol Clugston, Anna Dominiczak and Alex Alderton, Roberto Amato, Jeffrey Barrett, Sonia Goncalves, Ewan Harrison, David K. Jackson, Ian Johnston, Dominic Kwiatkowski, Cordelia Langford, John Sillitoe on behalf of the Wellcome Sanger Institute COVID-19 Surveillance Team |
| EPI_ISL_2805815 | Lighthouse Lab in Milton Keynes | Wellcome Sanger Institute for the COVID-19 Genomics UK (COG-UK) Consortium | The Lighthouse Lab in Milton Keynes and Alex Alderton, Roberto Amato, Jeffrey Barrett, Sonia Goncalves, Ewan Harrison, David K. Jackson, Ian Johnston, Dominic Kwiatkowski, Cordelia Langford, John Sillitoe on behalf of the Wellcome Sanger Institute COVID-19 Surveillance Team |
| EPI_ISL_2805816 | Lighthouse Lab in Glasgow | Wellcome Sanger Institute for the COVID-19 Genomics UK (COG-UK) Consortium | Harper VanSteenhouse, Yumi Kasai, David Gray, Carol Clugston, Anna Dominiczak and Alex Alderton, Roberto Amato, Jeffrey Barrett, Sonia Goncalves, Ewan Harrison, David K. Jackson, Ian Johnston, Dominic Kwiatkowski, Cordelia Langford, John Sillitoe on behalf of the Wellcome Sanger Institute COVID-19 Surveillance Team |
| EPI_ISL_2805817 | Lighthouse Laboratory Plymouth | Wellcome Sanger Institute for the COVID-19 Genomics UK (COG-UK) Consortium | Lighthouse Laboratory Plymouth and Alex Alderton, Roberto Amato, Jeffrey Barrett, Sonia Goncalves, Ewan Harrison, David K. Jackson, Ian Johnston, Dominic Kwiatkowski, Cordelia Langford, John Sillitoe on behalf of the Wellcome Sanger Institute COVID-19 Surveillance Team |
| EPI_ISL_2805818, EPI_ISL_2805819, EPI_ISL_2805820, EPI_ISL_2805821, EPI_ISL_2805822 | Lighthouse Lab in Milton Keynes | Wellcome Sanger Institute for the COVID-19 Genomics UK (COG-UK) Consortium | The Lighthouse Lab in Milton Keynes and Alex Alderton, Roberto Amato, Jeffrey Barrett, Sonia Goncalves, Ewan Harrison, David K. Jackson, Ian Johnston, Dominic Kwiatkowski, Cordelia Langford, John Sillitoe on behalf of the Wellcome Sanger Institute COVID-19 Surveillance Team |
| EPI_ISL_2805823, EPI_ISL_2805824 | Lighthouse Laboratory Plymouth | Wellcome Sanger Institute for the COVID-19 Genomics UK (COG-UK) Consortium | Lighthouse Laboratory Plymouth and Alex Alderton, Roberto Amato, Jeffrey Barrett, Sonia Goncalves, Ewan Harrison, David K. Jackson, Ian Johnston, Dominic Kwiatkowski, Cordelia Langford, John Sillitoe on behalf of the Wellcome Sanger Institute COVID-19 Surveillance Team |
| EPI_ISL_2805825, EPI_ISL_2805826, EPI_ISL_2805827 | Lighthouse Lab in Milton Keynes | Wellcome Sanger Institute for the COVID-19 Genomics UK (COG-UK) Consortium | The Lighthouse Lab in Milton Keynes and Alex Alderton, Roberto Amato, Jeffrey Barrett, Sonia Goncalves, Ewan Harrison, David K. Jackson, Ian Johnston, Dominic Kwiatkowski, Cordelia Langford, John Sillitoe on behalf of the Wellcome Sanger Institute COVID-19 Surveillance Team |
| EPI_ISL_2805828 | Lighthouse Laboratory Plymouth | Wellcome Sanger Institute for the COVID-19 Genomics UK (COG-UK) Consortium | Lighthouse Laboratory Plymouth and Alex Alderton, Roberto Amato, Jeffrey Barrett, Sonia Goncalves, Ewan Harrison, David K. Jackson, Ian Johnston, Dominic Kwiatkowski, Cordelia Langford, John Sillitoe on behalf of the Wellcome Sanger Institute COVID-19 Surveillance Team |
| EPI_ISL_2805829, EPI_ISL_2805830, EPI_ISL_2805831, EPI_ISL_2805832, EPI_ISL_2805833, EPI_ISL_2805834, EPI_ISL_2805835 | Lighthouse Lab in Milton Keynes | Wellcome Sanger Institute for the COVID-19 Genomics UK (COG-UK) Consortium | The Lighthouse Lab in Milton Keynes and Alex Alderton, Roberto Amato, Jeffrey Barrett, Sonia Goncalves, Ewan Harrison, David K. Jackson, Ian Johnston, Dominic Kwiatkowski, Cordelia Langford, John Sillitoe on behalf of the Wellcome Sanger Institute COVID-19 Surveillance Team |
| EPI_ISL_2805836 | Lighthouse Lab in Glasgow | Wellcome Sanger Institute for the COVID-19 Genomics UK (COG-UK) Consortium | Harper VanSteenhouse, Yumi Kasai, David Gray, Carol Clugston, Anna Dominiczak and Alex Alderton, Roberto Amato, Jeffrey Barrett, Sonia Goncalves, Ewan Harrison, David K. Jackson, Ian Johnston, Dominic Kwiatkowski, Cordelia Langford, John Sillitoe on behalf of the Wellcome Sanger Institute COVID-19 Surveillance Team |
| EPI_ISL_2805837 | Lighthouse Lab in Milton Keynes | Wellcome Sanger Institute for the COVID-19 Genomics UK (COG-UK) Consortium | The Lighthouse Lab in Milton Keynes and Alex Alderton, Roberto Amato, Jeffrey Barrett, Sonia Goncalves, Ewan Harrison, David K. Jackson, Ian Johnston, Dominic Kwiatkowski, Cordelia Langford, John Sillitoe on behalf of the Wellcome Sanger Institute COVID-19 Surveillance Team |

| | | | |
|---|---|---|---|
| EPI_ISL_2805838 | Lighthouse Lab in Glasgow | Wellcome Sanger Institute for the COVID-19 Genomics UK (COG-UK) Consortium | Harper VanSteenhouse, Yumi Kasai, David Gray, Carol Clugston, Anna Dominiczak and Alex Alderton, Roberto Amato, Jeffrey Barrett, Sonia Goncalves, Ewan Harrison, David K. Jackson, Ian Johnston, Dominic Kwiatkowski, Cordelia Langford, John Sillitoe on behalf of the Wellcome Sanger Institute COVID-19 Surveillance Team |
| EPI_ISL_2805839 | Lighthouse Lab in Milton Keynes | Wellcome Sanger Institute for the COVID-19 Genomics UK (COG-UK) Consortium | The Lighthouse Lab in Milton Keynes and Alex Alderton, Roberto Amato, Jeffrey Barrett, Sonia Goncalves, Ewan Harrison, David K. Jackson, Ian Johnston, Dominic Kwiatkowski, Cordelia Langford, John Sillitoe on behalf of the Wellcome Sanger Institute COVID-19 Surveillance Team |
| EPI_ISL_2805840 | Lighthouse Lab in Glasgow | Wellcome Sanger Institute for the COVID-19 Genomics UK (COG-UK) Consortium | Harper VanSteenhouse, Yumi Kasai, David Gray, Carol Clugston, Anna Dominiczak and Alex Alderton, Roberto Amato, Jeffrey Barrett, Sonia Goncalves, Ewan Harrison, David K. Jackson, Ian Johnston, Dominic Kwiatkowski, Cordelia Langford, John Sillitoe on behalf of the Wellcome Sanger Institute COVID-19 Surveillance Team |
| EPI_ISL_2805841 | Lighthouse Lab in Milton Keynes | Wellcome Sanger Institute for the COVID-19 Genomics UK (COG-UK) Consortium | The Lighthouse Lab in Milton Keynes and Alex Alderton, Roberto Amato, Jeffrey Barrett, Sonia Goncalves, Ewan Harrison, David K. Jackson, Ian Johnston, Dominic Kwiatkowski, Cordelia Langford, John Sillitoe on behalf of the Wellcome Sanger Institute COVID-19 Surveillance Team |
| EPI_ISL_2805842 | Lighthouse Laboratory Plymouth | Wellcome Sanger Institute for the COVID-19 Genomics UK (COG-UK) Consortium | Lighthouse Laboratory Plymouth and Alex Alderton, Roberto Amato, Jeffrey Barrett, Sonia Goncalves, Ewan Harrison, David K. Jackson, Ian Johnston, Dominic Kwiatkowski, Cordelia Langford, John Sillitoe on behalf of the Wellcome Sanger Institute COVID-19 Surveillance Team |
| EPI_ISL_2805843 | Lighthouse Lab in Milton Keynes | Wellcome Sanger Institute for the COVID-19 Genomics UK (COG-UK) Consortium | The Lighthouse Lab in Milton Keynes and Alex Alderton, Roberto Amato, Jeffrey Barrett, Sonia Goncalves, Ewan Harrison, David K. Jackson, Ian Johnston, Dominic Kwiatkowski, Cordelia Langford, John Sillitoe on behalf of the Wellcome Sanger Institute COVID-19 Surveillance Team |
| EPI_ISL_2805844 | Lighthouse Laboratory Plymouth | Wellcome Sanger Institute for the COVID-19 Genomics UK (COG-UK) Consortium | Lighthouse Laboratory Plymouth and Alex Alderton, Roberto Amato, Jeffrey Barrett, Sonia Goncalves, Ewan Harrison, David K. Jackson, Ian Johnston, Dominic Kwiatkowski, Cordelia Langford, John Sillitoe on behalf of the Wellcome Sanger Institute COVID-19 Surveillance Team |
| EPI_ISL_2805845 | Lighthouse Lab in Glasgow | Wellcome Sanger Institute for the COVID-19 Genomics UK (COG-UK) Consortium | Harper VanSteenhouse, Yumi Kasai, David Gray, Carol Clugston, Anna Dominiczak and Alex Alderton, Roberto Amato, Jeffrey Barrett, Sonia Goncalves, Ewan Harrison, David K. Jackson, Ian Johnston, Dominic Kwiatkowski, Cordelia Langford, John Sillitoe on behalf of the Wellcome Sanger Institute COVID-19 Surveillance Team |
| EPI_ISL_2805846, EPI_ISL_2805847 | Lighthouse Lab in Milton Keynes | Wellcome Sanger Institute for the COVID-19 Genomics UK (COG-UK) Consortium | The Lighthouse Lab in Milton Keynes and Alex Alderton, Roberto Amato, Jeffrey Barrett, Sonia Goncalves, Ewan Harrison, David K. Jackson, Ian Johnston, Dominic Kwiatkowski, Cordelia Langford, John Sillitoe on behalf of the Wellcome Sanger Institute COVID-19 Surveillance Team |
| EPI_ISL_2805848 | Lighthouse Laboratory Plymouth | Wellcome Sanger Institute for the COVID-19 Genomics UK (COG-UK) Consortium | Lighthouse Laboratory Plymouth and Alex Alderton, Roberto Amato, Jeffrey Barrett, Sonia Goncalves, Ewan Harrison, David K. Jackson, Ian Johnston, Dominic Kwiatkowski, Cordelia Langford, John Sillitoe on behalf of the Wellcome Sanger Institute COVID-19 Surveillance Team |
| EPI_ISL_2805849, EPI_ISL_2805850 | Lighthouse Lab in Milton Keynes | Wellcome Sanger Institute for the COVID-19 Genomics UK (COG-UK) Consortium | The Lighthouse Lab in Milton Keynes and Alex Alderton, Roberto Amato, Jeffrey Barrett, Sonia Goncalves, Ewan Harrison, David K. Jackson, Ian Johnston, Dominic Kwiatkowski, Cordelia Langford, John Sillitoe on behalf of the Wellcome Sanger Institute COVID-19 Surveillance Team |
| EPI_ISL_2805851, EPI_ISL_2805852, EPI_ISL_2805853 | Lighthouse Laboratory Plymouth | Wellcome Sanger Institute for the COVID-19 Genomics UK (COG-UK) Consortium | Lighthouse Laboratory Plymouth and Alex Alderton, Roberto Amato, Jeffrey Barrett, Sonia Goncalves, Ewan Harrison, David K. Jackson, Ian Johnston, Dominic Kwiatkowski, Cordelia Langford, John Sillitoe on behalf of the Wellcome Sanger Institute COVID-19 Surveillance Team |
| EPI_ISL_2805854 | Lighthouse Lab in Milton Keynes | Wellcome Sanger Institute for the COVID-19 Genomics UK (COG-UK) Consortium | The Lighthouse Lab in Milton Keynes and Alex Alderton, Roberto Amato, Jeffrey Barrett, Sonia Goncalves, Ewan Harrison, David K. Jackson, Ian Johnston, Dominic Kwiatkowski, Cordelia Langford, John Sillitoe on behalf of the Wellcome Sanger Institute COVID-19 Surveillance Team |
| EPI_ISL_2805855, EPI_ISL_2805856 | Lighthouse Laboratory Plymouth | Wellcome Sanger Institute for the COVID-19 Genomics UK (COG-UK) Consortium | Lighthouse Laboratory Plymouth and Alex Alderton, Roberto Amato, Jeffrey Barrett, Sonia Goncalves, Ewan Harrison, David K. Jackson, Ian Johnston, Dominic Kwiatkowski, Cordelia Langford, John Sillitoe on behalf of the Wellcome Sanger Institute COVID-19 Surveillance Team |
| EPI_ISL_2805857, EPI_ISL_2805858, EPI_ISL_2805859, EPI_ISL_2805860 | Lighthouse Lab in Milton Keynes | Wellcome Sanger Institute for the COVID-19 Genomics UK (COG-UK) Consortium | The Lighthouse Lab in Milton Keynes and Alex Alderton, Roberto Amato, Jeffrey Barrett, Sonia Goncalves, Ewan Harrison, David K. Jackson, Ian Johnston, Dominic Kwiatkowski, Cordelia Langford, John Sillitoe on behalf of the Wellcome Sanger Institute COVID-19 Surveillance Team |
| EPI_ISL_2805861 | Lighthouse Lab in Glasgow | Wellcome Sanger Institute for the COVID-19 Genomics UK (COG-UK) Consortium | Harper VanSteenhouse, Yumi Kasai, David Gray, Carol Clugston, Anna Dominiczak and Alex Alderton, Roberto Amato, Jeffrey Barrett, Sonia Goncalves, Ewan Harrison, David K. Jackson, Ian Johnston, Dominic Kwiatkowski, Cordelia Langford, John Sillitoe on behalf of the Wellcome Sanger Institute COVID-19 Surveillance Team |
| EPI_ISL_2805862, EPI_ISL_2805863, EPI_ISL_2805864 | Lighthouse Lab in Milton Keynes | Wellcome Sanger Institute for the COVID-19 Genomics UK (COG-UK) Consortium | The Lighthouse Lab in Milton Keynes and Alex Alderton, Roberto Amato, Jeffrey Barrett, Sonia Goncalves, Ewan Harrison, David K. Jackson, Ian Johnston, Dominic Kwiatkowski, Cordelia Langford, John Sillitoe on behalf of the Wellcome Sanger Institute COVID-19 Surveillance Team |
| EPI_ISL_2805865 | Lighthouse Laboratory Plymouth | Wellcome Sanger Institute for the COVID-19 Genomics UK (COG-UK) Consortium | Lighthouse Laboratory Plymouth and Alex Alderton, Roberto Amato, Jeffrey Barrett, Sonia Goncalves, Ewan Harrison, David K. Jackson, Ian Johnston, Dominic Kwiatkowski, Cordelia Langford, John Sillitoe on behalf of the Wellcome Sanger Institute COVID-19 Surveillance Team |
| EPI_ISL_2805866, EPI_ISL_2805867 | Lighthouse Lab in Milton Keynes | Wellcome Sanger Institute for the COVID-19 Genomics UK (COG-UK) Consortium | The Lighthouse Lab in Milton Keynes and Alex Alderton, Roberto Amato, Jeffrey Barrett, Sonia Goncalves, Ewan Harrison, David K. Jackson, Ian Johnston, Dominic Kwiatkowski, Cordelia Langford, John Sillitoe on behalf of the Wellcome Sanger Institute COVID-19 Surveillance Team |
| EPI_ISL_2805868, EPI_ISL_2805869, EPI_ISL_2805870 | Lighthouse Laboratory Plymouth | Wellcome Sanger Institute for the COVID-19 Genomics UK (COG-UK) Consortium | Lighthouse Laboratory Plymouth and Alex Alderton, Roberto Amato, Jeffrey Barrett, Sonia Goncalves, Ewan Harrison, David K. Jackson, Ian Johnston, Dominic Kwiatkowski, Cordelia Langford, John Sillitoe on behalf of the Wellcome Sanger Institute COVID-19 Surveillance Team |
| EPI_ISL_2805871, EPI_ISL_2805872 | Lighthouse Lab in Milton Keynes | Wellcome Sanger Institute for the COVID-19 Genomics UK (COG-UK) Consortium | The Lighthouse Lab in Milton Keynes and Alex Alderton, Roberto Amato, Jeffrey Barrett, Sonia Goncalves, Ewan Harrison, David K. Jackson, Ian Johnston, Dominic Kwiatkowski, Cordelia Langford, John Sillitoe on behalf of the Wellcome Sanger Institute COVID-19 Surveillance Team |
| EPI_ISL_2805873, EPI_ISL_2805874 | Lighthouse Laboratory Plymouth | Wellcome Sanger Institute for the COVID-19 Genomics UK (COG-UK) Consortium | Lighthouse Laboratory Plymouth and Alex Alderton, Roberto Amato, Jeffrey Barrett, Sonia Goncalves, Ewan Harrison, David K. Jackson, Ian Johnston, Dominic Kwiatkowski, Cordelia Langford, John Sillitoe on behalf of the Wellcome Sanger Institute COVID-19 Surveillance Team |
| EPI_ISL_2805875 | Lighthouse Lab in Glasgow | Wellcome Sanger Institute for the COVID-19 Genomics UK (COG-UK) Consortium | Harper VanSteenhouse, Yumi Kasai, David Gray, Carol Clugston, Anna Dominiczak and Alex Alderton, Roberto Amato, Jeffrey Barrett, Sonia Goncalves, Ewan Harrison, David K. Jackson, Ian Johnston, Dominic Kwiatkowski, Cordelia Langford, John Sillitoe on behalf of the Wellcome Sanger Institute COVID-19 Surveillance Team |
| EPI_ISL_2805876, EPI_ISL_2805877, EPI_ISL_2805878, EPI_ISL_2805879, EPI_ISL_2805880 | Lighthouse Lab in Milton Keynes | Wellcome Sanger Institute for the COVID-19 Genomics UK (COG-UK) Consortium | The Lighthouse Lab in Milton Keynes and Alex Alderton, Roberto Amato, Jeffrey Barrett, Sonia Goncalves, Ewan Harrison, David K. Jackson, Ian Johnston, Dominic Kwiatkowski, Cordelia Langford, John Sillitoe on behalf of the Wellcome Sanger Institute COVID-19 Surveillance Team |
| EPI_ISL_2805881, EPI_ISL_2805882 | Lighthouse Laboratory Plymouth | Wellcome Sanger Institute for the COVID-19 Genomics UK (COG-UK) Consortium | Lighthouse Laboratory Plymouth and Alex Alderton, Roberto Amato, Jeffrey Barrett, Sonia Goncalves, Ewan Harrison, David K. Jackson, Ian Johnston, Dominic Kwiatkowski, Cordelia Langford, John Sillitoe on behalf of the Wellcome Sanger Institute COVID-19 Surveillance Team |
| EPI_ISL_2805883, EPI_ISL_2805884 | Lighthouse Lab in Milton Keynes | Wellcome Sanger Institute for the COVID-19 Genomics UK (COG-UK) Consortium | The Lighthouse Lab in Milton Keynes and Alex Alderton, Roberto Amato, Jeffrey Barrett, Sonia Goncalves, Ewan Harrison, David K. Jackson, Ian Johnston, Dominic Kwiatkowski, Cordelia Langford, John Sillitoe on behalf of the Wellcome Sanger Institute COVID-19 Surveillance Team |
| EPI_ISL_2805885 | Lighthouse Lab in Glasgow | Wellcome Sanger Institute for the COVID-19 Genomics UK (COG-UK) Consortium | Harper VanSteenhouse, Yumi Kasai, David Gray, Carol Clugston, Anna Dominiczak and Alex Alderton, Roberto Amato, Jeffrey Barrett, Sonia Goncalves, Ewan Harrison, David K. Jackson, Ian Johnston, Dominic Kwiatkowski, Cordelia Langford, John Sillitoe on behalf of the Wellcome Sanger Institute COVID-19 Surveillance Team |
| EPI_ISL_2805886 | Lighthouse Laboratory Plymouth | Wellcome Sanger Institute for the COVID-19 Genomics UK (COG-UK) Consortium | Lighthouse Laboratory Plymouth and Alex Alderton, Roberto Amato, Jeffrey Barrett, Sonia Goncalves, Ewan Harrison, David K. Jackson, Ian Johnston, Dominic Kwiatkowski, Cordelia Langford, John Sillitoe on behalf of the Wellcome Sanger Institute COVID-19 Surveillance Team |
| EPI_ISL_2805887 | Lighthouse Lab in Glasgow | Wellcome Sanger Institute for the COVID-19 Genomics UK (COG-UK) Consortium | Harper VanSteenhouse, Yumi Kasai, David Gray, Carol Clugston, Anna Dominiczak and Alex Alderton, Roberto Amato, Jeffrey Barrett, Sonia Goncalves, Ewan Harrison, David K. Jackson, Ian Johnston, Dominic Kwiatkowski, Cordelia Langford, John Sillitoe on behalf of the Wellcome Sanger Institute COVID-19 Surveillance Team |
| EPI_ISL_2805888 | Lighthouse Lab in Milton Keynes | Wellcome Sanger Institute for the COVID-19 Genomics UK (COG-UK) Consortium | The Lighthouse Lab in Milton Keynes and Alex Alderton, Roberto Amato, Jeffrey Barrett, Sonia Goncalves, Ewan Harrison, David K. Jackson, Ian Johnston, Dominic Kwiatkowski, Cordelia Langford, John Sillitoe on behalf of the Wellcome Sanger Institute COVID-19 Surveillance Team |
| EPI_ISL_2805889, EPI_ISL_2805890, EPI_ISL_2805891, EPI_ISL_2805892, EPI_ISL_2805893 | Lighthouse Laboratory Plymouth | Wellcome Sanger Institute for the COVID-19 Genomics UK (COG-UK) Consortium | Lighthouse Laboratory Plymouth and Alex Alderton, Roberto Amato, Jeffrey Barrett, Sonia Goncalves, Ewan Harrison, David K. Jackson, Ian Johnston, Dominic Kwiatkowski, Cordelia Langford, John Sillitoe on behalf of the Wellcome Sanger Institute COVID-19 Surveillance Team |
| EPI_ISL_2805894, EPI_ISL_2805895, EPI_ISL_2805896, EPI_ISL_2805897 | Lighthouse Lab in Milton Keynes | Wellcome Sanger Institute for the COVID-19 Genomics UK (COG-UK) Consortium | The Lighthouse Lab in Milton Keynes and Alex Alderton, Roberto Amato, Jeffrey Barrett, Sonia Goncalves, Ewan Harrison, David K. Jackson, Ian Johnston, Dominic Kwiatkowski, Cordelia Langford, John Sillitoe on behalf of the Wellcome Sanger Institute COVID-19 Surveillance Team |
| EPI_ISL_2805898, EPI_ISL_2805899 | Lighthouse Lab in Glasgow | Wellcome Sanger Institute for the COVID-19 Genomics UK (COG-UK) Consortium | Harper VanSteenhouse, Yumi Kasai, David Gray, Carol Clugston, Anna Dominiczak and Alex Alderton, Roberto Amato, Jeffrey Barrett, Sonia Goncalves, Ewan Harrison, David K. Jackson, Ian Johnston, Dominic Kwiatkowski, Cordelia Langford, John Sillitoe on behalf of the Wellcome Sanger Institute COVID-19 Surveillance Team |
| EPI_ISL_2805900, EPI_ISL_2805901 | Lighthouse Lab in Milton Keynes | Wellcome Sanger Institute for the COVID-19 Genomics UK (COG-UK) Consortium | The Lighthouse Lab in Milton Keynes and Alex Alderton, Roberto Amato, Jeffrey Barrett, Sonia Goncalves, Ewan Harrison, David K. Jackson, Ian Johnston, Dominic Kwiatkowski, Cordelia Langford, John Sillitoe on behalf of the Wellcome Sanger Institute COVID-19 Surveillance Team |
| EPI_ISL_2805902, EPI_ISL_2805903 | Lighthouse Laboratory Plymouth | Wellcome Sanger Institute for the COVID-19 Genomics UK (COG-UK) Consortium | Lighthouse Laboratory Plymouth and Alex Alderton, Roberto Amato, Jeffrey Barrett, Sonia Goncalves, Ewan Harrison, David K. Jackson, Ian Johnston, Dominic Kwiatkowski, Cordelia Langford, John Sillitoe on behalf of the Wellcome Sanger Institute COVID-19 Surveillance Team |
| EPI_ISL_2805904, EPI_ISL_2805905 | Lighthouse Lab in Milton Keynes | Wellcome Sanger Institute for the COVID-19 Genomics UK (COG-UK) Consortium | The Lighthouse Lab in Milton Keynes and Alex Alderton, Roberto Amato, Jeffrey Barrett, Sonia Goncalves, Ewan Harrison, David K. Jackson, Ian Johnston, Dominic Kwiatkowski, Cordelia Langford, John Sillitoe on behalf of the Wellcome Sanger Institute COVID-19 Surveillance Team |
| EPI_ISL_2805906 | Lighthouse Lab in Glasgow | Wellcome Sanger Institute for the COVID-19 Genomics UK (COG-UK) Consortium | Harper VanSteenhouse, Yumi Kasai, David Gray, Carol Clugston, Anna Dominiczak and Alex Alderton, Roberto Amato, Jeffrey Barrett, Sonia Goncalves, Ewan Harrison, David K. Jackson, Ian Johnston, Dominic Kwiatkowski, Cordelia Langford, John Sillitoe on behalf of the Wellcome Sanger Institute COVID-19 Surveillance Team |
| EPI_ISL_2805907 | Lighthouse Lab in Milton Keynes | Wellcome Sanger Institute for the COVID-19 Genomics UK (COG-UK) Consortium | The Lighthouse Lab in Milton Keynes and Alex Alderton, Roberto Amato, Jeffrey Barrett, Sonia Goncalves, Ewan Harrison, David K. Jackson, Ian Johnston, Dominic Kwiatkowski, Cordelia Langford, John Sillitoe on behalf of the Wellcome Sanger Institute COVID-19 Surveillance Team |
| EPI_ISL_2805908 | Lighthouse Laboratory Plymouth | Wellcome Sanger Institute for the COVID-19 Genomics UK (COG-UK) Consortium | Lighthouse Laboratory Plymouth and Alex Alderton, Roberto Amato, Jeffrey Barrett, Sonia Goncalves, Ewan Harrison, David K. Jackson, Ian Johnston, Dominic Kwiatkowski, Cordelia Langford, John Sillitoe on behalf of the Wellcome Sanger Institute COVID-19 Surveillance Team |
| EPI_ISL_2805909, EPI_ISL_2805910, | Lighthouse Lab in Milton Keynes | Wellcome Sanger Institute for the COVID-19 Genomics UK (COG-UK) Consortium | The Lighthouse Lab in Milton Keynes and Alex Alderton, Roberto Amato, Jeffrey Barrett, Sonia Goncalves, Ewan Harrison, David K. Jackson, Ian Johnston, Dominic Kwiatkowski, Cordelia Langford, John Sillitoe on behalf of the Wellcome Sanger Institute COVID-19 Surveillance Team |

| | | | |
|---|---|---|---|
| EPI_ISL_2805911 | | | |
| EPI_ISL_2805912 | Lighthouse Laboratory Plymouth | Wellcome Sanger Institute for the COVID-19 Genomics UK (COG-UK) Consortium | Lighthouse Laboratory Plymouth and Alex Alderton, Roberto Amato, Jeffrey Barrett, Sonia Goncalves, Ewan Harrison, David K. Jackson, Ian Johnston, Dominic Kwiatkowski, Cordelia Langford, John Sillitoe on behalf of the Wellcome Sanger Institute COVID-19 Surveillance Team |
| EPI_ISL_2805913 | Lighthouse Lab in Glasgow | Wellcome Sanger Institute for the COVID-19 Genomics UK (COG-UK) Consortium | Harper VanSteenhouse, Yumi Kasai, David Gray, Carol Clugston, Anna Dominiczak and Alex Alderton, Roberto Amato, Jeffrey Barrett, Sonia Goncalves, Ewan Harrison, David K. Jackson, Ian Johnston, Dominic Kwiatkowski, Cordelia Langford, John Sillitoe on behalf of the Wellcome Sanger Institute COVID-19 Surveillance Team |
| EPI_ISL_2805914, EPI_ISL_2805915, EPI_ISL_2805916 | Lighthouse Lab in Milton Keynes | Wellcome Sanger Institute for the COVID-19 Genomics UK (COG-UK) Consortium | The Lighthouse Lab in Milton Keynes and Alex Alderton, Roberto Amato, Jeffrey Barrett, Sonia Goncalves, Ewan Harrison, David K. Jackson, Ian Johnston, Dominic Kwiatkowski, Cordelia Langford, John Sillitoe on behalf of the Wellcome Sanger Institute COVID-19 Surveillance Team |
| EPI_ISL_2805917 | Lighthouse Laboratory Plymouth | Wellcome Sanger Institute for the COVID-19 Genomics UK (COG-UK) Consortium | Lighthouse Laboratory Plymouth and Alex Alderton, Roberto Amato, Jeffrey Barrett, Sonia Goncalves, Ewan Harrison, David K. Jackson, Ian Johnston, Dominic Kwiatkowski, Cordelia Langford, John Sillitoe on behalf of the Wellcome Sanger Institute COVID-19 Surveillance Team |
| EPI_ISL_2805919 | Lighthouse Lab in Milton Keynes | Wellcome Sanger Institute for the COVID-19 Genomics UK (COG-UK) Consortium | The Lighthouse Lab in Milton Keynes and Alex Alderton, Roberto Amato, Jeffrey Barrett, Sonia Goncalves, Ewan Harrison, David K. Jackson, Ian Johnston, Dominic Kwiatkowski, Cordelia Langford, John Sillitoe on behalf of the Wellcome Sanger Institute COVID-19 Surveillance Team |
| EPI_ISL_2805920 | Lighthouse Laboratory Plymouth | Wellcome Sanger Institute for the COVID-19 Genomics UK (COG-UK) Consortium | Lighthouse Laboratory Plymouth and Alex Alderton, Roberto Amato, Jeffrey Barrett, Sonia Goncalves, Ewan Harrison, David K. Jackson, Ian Johnston, Dominic Kwiatkowski, Cordelia Langford, John Sillitoe on behalf of the Wellcome Sanger Institute COVID-19 Surveillance Team |
| EPI_ISL_2805921 | Lighthouse Lab in Milton Keynes | Wellcome Sanger Institute for the COVID-19 Genomics UK (COG-UK) Consortium | The Lighthouse Lab in Milton Keynes and Alex Alderton, Roberto Amato, Jeffrey Barrett, Sonia Goncalves, Ewan Harrison, David K. Jackson, Ian Johnston, Dominic Kwiatkowski, Cordelia Langford, John Sillitoe on behalf of the Wellcome Sanger Institute COVID-19 Surveillance Team |
| EPI_ISL_2805922 | Lighthouse Lab in Glasgow | Wellcome Sanger Institute for the COVID-19 Genomics UK (COG-UK) Consortium | Harper VanSteenhouse, Yumi Kasai, David Gray, Carol Clugston, Anna Dominiczak and Alex Alderton, Roberto Amato, Jeffrey Barrett, Sonia Goncalves, Ewan Harrison, David K. Jackson, Ian Johnston, Dominic Kwiatkowski, Cordelia Langford, John Sillitoe on behalf of the Wellcome Sanger Institute COVID-19 Surveillance Team |
| EPI_ISL_2805923, EPI_ISL_2805924 | Lighthouse Lab in Milton Keynes | Wellcome Sanger Institute for the COVID-19 Genomics UK (COG-UK) Consortium | The Lighthouse Lab in Milton Keynes and Alex Alderton, Roberto Amato, Jeffrey Barrett, Sonia Goncalves, Ewan Harrison, David K. Jackson, Ian Johnston, Dominic Kwiatkowski, Cordelia Langford, John Sillitoe on behalf of the Wellcome Sanger Institute COVID-19 Surveillance Team |
| EPI_ISL_2805925, EPI_ISL_2805926 | Lighthouse Laboratory Plymouth | Wellcome Sanger Institute for the COVID-19 Genomics UK (COG-UK) Consortium | Lighthouse Laboratory Plymouth and Alex Alderton, Roberto Amato, Jeffrey Barrett, Sonia Goncalves, Ewan Harrison, David K. Jackson, Ian Johnston, Dominic Kwiatkowski, Cordelia Langford, John Sillitoe on behalf of the Wellcome Sanger Institute COVID-19 Surveillance Team |
| EPI_ISL_2805927, EPI_ISL_2805928, EPI_ISL_2805929 | Lighthouse Lab in Milton Keynes | Wellcome Sanger Institute for the COVID-19 Genomics UK (COG-UK) Consortium | The Lighthouse Lab in Milton Keynes and Alex Alderton, Roberto Amato, Jeffrey Barrett, Sonia Goncalves, Ewan Harrison, David K. Jackson, Ian Johnston, Dominic Kwiatkowski, Cordelia Langford, John Sillitoe on behalf of the Wellcome Sanger Institute COVID-19 Surveillance Team |
| EPI_ISL_2805930 | Lighthouse Lab in Glasgow | Wellcome Sanger Institute for the COVID-19 Genomics UK (COG-UK) Consortium | Harper VanSteenhouse, Yumi Kasai, David Gray, Carol Clugston, Anna Dominiczak and Alex Alderton, Roberto Amato, Jeffrey Barrett, Sonia Goncalves, Ewan Harrison, David K. Jackson, Ian Johnston, Dominic Kwiatkowski, Cordelia Langford, John Sillitoe on behalf of the Wellcome Sanger Institute COVID-19 Surveillance Team |
| EPI_ISL_2805931, EPI_ISL_2805932, EPI_ISL_2805933 | Lighthouse Lab in Milton Keynes | Wellcome Sanger Institute for the COVID-19 Genomics UK (COG-UK) Consortium | The Lighthouse Lab in Milton Keynes and Alex Alderton, Roberto Amato, Jeffrey Barrett, Sonia Goncalves, Ewan Harrison, David K. Jackson, Ian Johnston, Dominic Kwiatkowski, Cordelia Langford, John Sillitoe on behalf of the Wellcome Sanger Institute COVID-19 Surveillance Team |
| EPI_ISL_2805934 | Lighthouse Laboratory Plymouth | Wellcome Sanger Institute for the COVID-19 Genomics UK (COG-UK) Consortium | Lighthouse Laboratory Plymouth and Alex Alderton, Roberto Amato, Jeffrey Barrett, Sonia Goncalves, Ewan Harrison, David K. Jackson, Ian Johnston, Dominic Kwiatkowski, Cordelia Langford, John Sillitoe on behalf of the Wellcome Sanger Institute COVID-19 Surveillance Team |
| EPI_ISL_2805935, EPI_ISL_2805936, EPI_ISL_2805937 | Lighthouse Lab in Milton Keynes | Wellcome Sanger Institute for the COVID-19 Genomics UK (COG-UK) Consortium | The Lighthouse Lab in Milton Keynes and Alex Alderton, Roberto Amato, Jeffrey Barrett, Sonia Goncalves, Ewan Harrison, David K. Jackson, Ian Johnston, Dominic Kwiatkowski, Cordelia Langford, John Sillitoe on behalf of the Wellcome Sanger Institute COVID-19 Surveillance Team |
| EPI_ISL_2805938 | Lighthouse Laboratory Plymouth | Wellcome Sanger Institute for the COVID-19 Genomics UK (COG-UK) Consortium | Lighthouse Laboratory Plymouth and Alex Alderton, Roberto Amato, Jeffrey Barrett, Sonia Goncalves, Ewan Harrison, David K. Jackson, Ian Johnston, Dominic Kwiatkowski, Cordelia Langford, John Sillitoe on behalf of the Wellcome Sanger Institute COVID-19 Surveillance Team |
| EPI_ISL_2805939, EPI_ISL_2805940 | Lighthouse Lab in Milton Keynes | Wellcome Sanger Institute for the COVID-19 Genomics UK (COG-UK) Consortium | The Lighthouse Lab in Milton Keynes and Alex Alderton, Roberto Amato, Jeffrey Barrett, Sonia Goncalves, Ewan Harrison, David K. Jackson, Ian Johnston, Dominic Kwiatkowski, Cordelia Langford, John Sillitoe on behalf of the Wellcome Sanger Institute COVID-19 Surveillance Team |
| EPI_ISL_2805941 | Lighthouse Laboratory Plymouth | Wellcome Sanger Institute for the COVID-19 Genomics UK (COG-UK) Consortium | Lighthouse Laboratory Plymouth and Alex Alderton, Roberto Amato, Jeffrey Barrett, Sonia Goncalves, Ewan Harrison, David K. Jackson, Ian Johnston, Dominic Kwiatkowski, Cordelia Langford, John Sillitoe on behalf of the Wellcome Sanger Institute COVID-19 Surveillance Team |
| EPI_ISL_2805942, EPI_ISL_2805943 | Lighthouse Lab in Milton Keynes | Wellcome Sanger Institute for the COVID-19 Genomics UK (COG-UK) Consortium | The Lighthouse Lab in Milton Keynes and Alex Alderton, Roberto Amato, Jeffrey Barrett, Sonia Goncalves, Ewan Harrison, David K. Jackson, Ian Johnston, Dominic Kwiatkowski, Cordelia Langford, John Sillitoe on behalf of the Wellcome Sanger Institute COVID-19 Surveillance Team |
| EPI_ISL_2805944 | Lighthouse Laboratory Plymouth | Wellcome Sanger Institute for the COVID-19 Genomics UK (COG-UK) Consortium | Lighthouse Laboratory Plymouth and Alex Alderton, Roberto Amato, Jeffrey Barrett, Sonia Goncalves, Ewan Harrison, David K. Jackson, Ian Johnston, Dominic Kwiatkowski, Cordelia Langford, John Sillitoe on behalf of the Wellcome Sanger Institute COVID-19 Surveillance Team |
| EPI_ISL_2805945 | Lighthouse Lab in Glasgow | Wellcome Sanger Institute for the COVID-19 Genomics UK (COG-UK) Consortium | Harper VanSteenhouse, Yumi Kasai, David Gray, Carol Clugston, Anna Dominiczak and Alex Alderton, Roberto Amato, Jeffrey Barrett, Sonia Goncalves, Ewan Harrison, David K. Jackson, Ian Johnston, Dominic Kwiatkowski, Cordelia Langford, John Sillitoe on behalf of the Wellcome Sanger Institute COVID-19 Surveillance Team |
| EPI_ISL_2805946 | Lighthouse Laboratory Plymouth | Wellcome Sanger Institute for the COVID-19 Genomics UK (COG-UK) Consortium | Lighthouse Laboratory Plymouth and Alex Alderton, Roberto Amato, Jeffrey Barrett, Sonia Goncalves, Ewan Harrison, David K. Jackson, Ian Johnston, Dominic Kwiatkowski, Cordelia Langford, John Sillitoe on behalf of the Wellcome Sanger Institute COVID-19 Surveillance Team |
| EPI_ISL_2805947 | Lighthouse Lab in Milton Keynes | Wellcome Sanger Institute for the COVID-19 Genomics UK (COG-UK) Consortium | The Lighthouse Lab in Milton Keynes and Alex Alderton, Roberto Amato, Jeffrey Barrett, Sonia Goncalves, Ewan Harrison, David K. Jackson, Ian Johnston, Dominic Kwiatkowski, Cordelia Langford, John Sillitoe on behalf of the Wellcome Sanger Institute COVID-19 Surveillance Team |
| EPI_ISL_2805948, EPI_ISL_2805949 | Lighthouse Lab in Glasgow | Wellcome Sanger Institute for the COVID-19 Genomics UK (COG-UK) Consortium | Harper VanSteenhouse, Yumi Kasai, David Gray, Carol Clugston, Anna Dominiczak and Alex Alderton, Roberto Amato, Jeffrey Barrett, Sonia Goncalves, Ewan Harrison, David K. Jackson, Ian Johnston, Dominic Kwiatkowski, Cordelia Langford, John Sillitoe on behalf of the Wellcome Sanger Institute COVID-19 Surveillance Team |
| EPI_ISL_2805950, EPI_ISL_2805951, EPI_ISL_2805952, EPI_ISL_2805953 | Lighthouse Lab in Milton Keynes | Wellcome Sanger Institute for the COVID-19 Genomics UK (COG-UK) Consortium | The Lighthouse Lab in Milton Keynes and Alex Alderton, Roberto Amato, Jeffrey Barrett, Sonia Goncalves, Ewan Harrison, David K. Jackson, Ian Johnston, Dominic Kwiatkowski, Cordelia Langford, John Sillitoe on behalf of the Wellcome Sanger Institute COVID-19 Surveillance Team |
| EPI_ISL_2805954, EPI_ISL_2805955 | Lighthouse Laboratory Plymouth | Wellcome Sanger Institute for the COVID-19 Genomics UK (COG-UK) Consortium | Lighthouse Laboratory Plymouth and Alex Alderton, Roberto Amato, Jeffrey Barrett, Sonia Goncalves, Ewan Harrison, David K. Jackson, Ian Johnston, Dominic Kwiatkowski, Cordelia Langford, John Sillitoe on behalf of the Wellcome Sanger Institute COVID-19 Surveillance Team |
| EPI_ISL_2805956 | Lighthouse Lab in Glasgow | Wellcome Sanger Institute for the COVID-19 Genomics UK (COG-UK) Consortium | Harper VanSteenhouse, Yumi Kasai, David Gray, Carol Clugston, Anna Dominiczak and Alex Alderton, Roberto Amato, Jeffrey Barrett, Sonia Goncalves, Ewan Harrison, David K. Jackson, Ian Johnston, Dominic Kwiatkowski, Cordelia Langford, John Sillitoe on behalf of the Wellcome Sanger Institute COVID-19 Surveillance Team |
| EPI_ISL_2805957, EPI_ISL_2805958 | Lighthouse Lab in Milton Keynes | Wellcome Sanger Institute for the COVID-19 Genomics UK (COG-UK) Consortium | The Lighthouse Lab in Milton Keynes and Alex Alderton, Roberto Amato, Jeffrey Barrett, Sonia Goncalves, Ewan Harrison, David K. Jackson, Ian Johnston, Dominic Kwiatkowski, Cordelia Langford, John Sillitoe on behalf of the Wellcome Sanger Institute COVID-19 Surveillance Team |
| EPI_ISL_2805959 | Lighthouse Lab in Glasgow | Wellcome Sanger Institute for the COVID-19 Genomics UK (COG-UK) Consortium | Harper VanSteenhouse, Yumi Kasai, David Gray, Carol Clugston, Anna Dominiczak and Alex Alderton, Roberto Amato, Jeffrey Barrett, Sonia Goncalves, Ewan Harrison, David K. Jackson, Ian Johnston, Dominic Kwiatkowski, Cordelia Langford, John Sillitoe on behalf of the Wellcome Sanger Institute COVID-19 Surveillance Team |
| EPI_ISL_2805960, EPI_ISL_2805961, EPI_ISL_2805962 | Lighthouse Lab in Milton Keynes | Wellcome Sanger Institute for the COVID-19 Genomics UK (COG-UK) Consortium | The Lighthouse Lab in Milton Keynes and Alex Alderton, Roberto Amato, Jeffrey Barrett, Sonia Goncalves, Ewan Harrison, David K. Jackson, Ian Johnston, Dominic Kwiatkowski, Cordelia Langford, John Sillitoe on behalf of the Wellcome Sanger Institute COVID-19 Surveillance Team |
| EPI_ISL_2805963 | Lighthouse Lab in Glasgow | Wellcome Sanger Institute for the COVID-19 Genomics UK (COG-UK) Consortium | Harper VanSteenhouse, Yumi Kasai, David Gray, Carol Clugston, Anna Dominiczak and Alex Alderton, Roberto Amato, Jeffrey Barrett, Sonia Goncalves, Ewan Harrison, David K. Jackson, Ian Johnston, Dominic Kwiatkowski, Cordelia Langford, John Sillitoe on behalf of the Wellcome Sanger Institute COVID-19 Surveillance Team |
| EPI_ISL_2805964 | Lighthouse Lab in Milton Keynes | Wellcome Sanger Institute for the COVID-19 Genomics UK (COG-UK) Consortium | The Lighthouse Lab in Milton Keynes and Alex Alderton, Roberto Amato, Jeffrey Barrett, Sonia Goncalves, Ewan Harrison, David K. Jackson, Ian Johnston, Dominic Kwiatkowski, Cordelia Langford, John Sillitoe on behalf of the Wellcome Sanger Institute COVID-19 Surveillance Team |
| EPI_ISL_2805965 | Lighthouse Laboratory Plymouth | Wellcome Sanger Institute for the COVID-19 Genomics UK (COG-UK) Consortium | Lighthouse Laboratory Plymouth and Alex Alderton, Roberto Amato, Jeffrey Barrett, Sonia Goncalves, Ewan Harrison, David K. Jackson, Ian Johnston, Dominic Kwiatkowski, Cordelia Langford, John Sillitoe on behalf of the Wellcome Sanger Institute COVID-19 Surveillance Team |
| EPI_ISL_2805966, EPI_ISL_2805967 | Lighthouse Lab in Milton Keynes | Wellcome Sanger Institute for the COVID-19 Genomics UK (COG-UK) Consortium | The Lighthouse Lab in Milton Keynes and Alex Alderton, Roberto Amato, Jeffrey Barrett, Sonia Goncalves, Ewan Harrison, David K. Jackson, Ian Johnston, Dominic Kwiatkowski, Cordelia Langford, John Sillitoe on behalf of the Wellcome Sanger Institute COVID-19 Surveillance Team |
| EPI_ISL_2805968 | Lighthouse Lab in Glasgow | Wellcome Sanger Institute for the COVID-19 Genomics UK (COG-UK) Consortium | Harper VanSteenhouse, Yumi Kasai, David Gray, Carol Clugston, Anna Dominiczak and Alex Alderton, Roberto Amato, Jeffrey Barrett, Sonia Goncalves, Ewan Harrison, David K. Jackson, Ian Johnston, Dominic Kwiatkowski, Cordelia Langford, John Sillitoe on behalf of the Wellcome Sanger Institute COVID-19 Surveillance Team |
| EPI_ISL_2805969, EPI_ISL_2805970 | Lighthouse Lab in Milton Keynes | Wellcome Sanger Institute for the COVID-19 Genomics UK (COG-UK) Consortium | The Lighthouse Lab in Milton Keynes and Alex Alderton, Roberto Amato, Jeffrey Barrett, Sonia Goncalves, Ewan Harrison, David K. Jackson, Ian Johnston, Dominic Kwiatkowski, Cordelia Langford, John Sillitoe on behalf of the Wellcome Sanger Institute COVID-19 Surveillance Team |
| EPI_ISL_2805971, EPI_ISL_2805972, EPI_ISL_2805973 | Lighthouse Lab in Glasgow | Wellcome Sanger Institute for the COVID-19 Genomics UK (COG-UK) Consortium | Harper VanSteenhouse, Yumi Kasai, David Gray, Carol Clugston, Anna Dominiczak and Alex Alderton, Roberto Amato, Jeffrey Barrett, Sonia Goncalves, Ewan Harrison, David K. Jackson, Ian Johnston, Dominic Kwiatkowski, Cordelia Langford, John Sillitoe on behalf of the Wellcome Sanger Institute COVID-19 Surveillance Team |
| EPI_ISL_2805974 | Lighthouse Lab in Milton Keynes | Wellcome Sanger Institute for the COVID-19 Genomics UK (COG-UK) Consortium | The Lighthouse Lab in Milton Keynes and Alex Alderton, Roberto Amato, Jeffrey Barrett, Sonia Goncalves, Ewan Harrison, David K. Jackson, Ian Johnston, Dominic Kwiatkowski, Cordelia Langford, John Sillitoe on behalf of the Wellcome Sanger Institute COVID-19 Surveillance Team |
| EPI_ISL_2805975 | Lighthouse Laboratory Plymouth | Wellcome Sanger Institute for the COVID-19 Genomics UK (COG-UK) Consortium | Lighthouse Laboratory Plymouth and Alex Alderton, Roberto Amato, Jeffrey Barrett, Sonia Goncalves, Ewan Harrison, David K. Jackson, Ian Johnston, Dominic Kwiatkowski, Cordelia Langford, John Sillitoe on behalf of the Wellcome Sanger Institute COVID-19 Surveillance Team |
| EPI_ISL_2805976, EPI_ISL_2805977 | Lighthouse Lab in Milton Keynes | Wellcome Sanger Institute for the COVID-19 Genomics UK (COG-UK) Consortium | The Lighthouse Lab in Milton Keynes and Alex Alderton, Roberto Amato, Jeffrey Barrett, Sonia Goncalves, Ewan Harrison, David K. Jackson, Ian Johnston, Dominic Kwiatkowski, Cordelia Langford, John Sillitoe on behalf of the Wellcome Sanger Institute COVID-19 Surveillance Team |
| EPI_ISL_2805978 | Lighthouse Lab in Glasgow | Wellcome Sanger Institute for the COVID-19 Genomics UK (COG-UK) Consortium | Harper VanSteenhouse, Yumi Kasai, David Gray, Carol Clugston, Anna Dominiczak and Alex Alderton, Roberto Amato, Jeffrey Barrett, Sonia Goncalves, Ewan Harrison, David K. Jackson, Ian Johnston, Dominic Kwiatkowski, Cordelia Langford, John Sillitoe on behalf of the Wellcome Sanger Institute COVID-19 Surveillance Team |

| | | | |
|---|---|---|---|
| EPI_ISL_2805979, EPI_ISL_2805980 | Lighthouse Lab in Milton Keynes | Wellcome Sanger Institute for the COVID-19 Genomics UK (COG-UK) Consortium | The Lighthouse Lab in Milton Keynes and Alex Alderton, Roberto Amato, Jeffrey Barrett, Sonia Goncalves, Ewan Harrison, David K. Jackson, Ian Johnston, Dominic Kwiatkowski, Cordelia Langford, John Sillitoe on behalf of the Wellcome Sanger Institute COVID-19 Surveillance Team |
| EPI_ISL_2805981, EPI_ISL_2805982 | Lighthouse Laboratory Plymouth | Wellcome Sanger Institute for the COVID-19 Genomics UK (COG-UK) Consortium | Lighthouse Laboratory Plymouth and Alex Alderton, Roberto Amato, Jeffrey Barrett, Sonia Goncalves, Ewan Harrison, David K. Jackson, Ian Johnston, Dominic Kwiatkowski, Cordelia Langford, John Sillitoe on behalf of the Wellcome Sanger Institute COVID-19 Surveillance Team |
| EPI_ISL_2805983 | Lighthouse Lab in Glasgow | Wellcome Sanger Institute for the COVID-19 Genomics UK (COG-UK) Consortium | Harper VanSteenhouse, Yumi Kasai, David Gray, Carol Clugston, Anna Dominiczak and Alex Alderton, Roberto Amato, Jeffrey Barrett, Sonia Goncalves, Ewan Harrison, David K. Jackson, Ian Johnston, Dominic Kwiatkowski, Cordelia Langford, John Sillitoe on behalf of the Wellcome Sanger Institute COVID-19 Surveillance Team |
| EPI_ISL_2805984 | Lighthouse Laboratory Plymouth | Wellcome Sanger Institute for the COVID-19 Genomics UK (COG-UK) Consortium | Lighthouse Laboratory Plymouth and Alex Alderton, Roberto Amato, Jeffrey Barrett, Sonia Goncalves, Ewan Harrison, David K. Jackson, Ian Johnston, Dominic Kwiatkowski, Cordelia Langford, John Sillitoe on behalf of the Wellcome Sanger Institute COVID-19 Surveillance Team |
| EPI_ISL_2805985 | Lighthouse Lab in Glasgow | Wellcome Sanger Institute for the COVID-19 Genomics UK (COG-UK) Consortium | Harper VanSteenhouse, Yumi Kasai, David Gray, Carol Clugston, Anna Dominiczak and Alex Alderton, Roberto Amato, Jeffrey Barrett, Sonia Goncalves, Ewan Harrison, David K. Jackson, Ian Johnston, Dominic Kwiatkowski, Cordelia Langford, John Sillitoe on behalf of the Wellcome Sanger Institute COVID-19 Surveillance Team |
| EPI_ISL_2805986 | Lighthouse Laboratory Plymouth | Wellcome Sanger Institute for the COVID-19 Genomics UK (COG-UK) Consortium | Lighthouse Laboratory Plymouth and Alex Alderton, Roberto Amato, Jeffrey Barrett, Sonia Goncalves, Ewan Harrison, David K. Jackson, Ian Johnston, Dominic Kwiatkowski, Cordelia Langford, John Sillitoe on behalf of the Wellcome Sanger Institute COVID-19 Surveillance Team |
| EPI_ISL_2805987 | Lighthouse Lab in Milton Keynes | Wellcome Sanger Institute for the COVID-19 Genomics UK (COG-UK) Consortium | The Lighthouse Lab in Milton Keynes and Alex Alderton, Roberto Amato, Jeffrey Barrett, Sonia Goncalves, Ewan Harrison, David K. Jackson, Ian Johnston, Dominic Kwiatkowski, Cordelia Langford, John Sillitoe on behalf of the Wellcome Sanger Institute COVID-19 Surveillance Team |
| EPI_ISL_2805988 | Lighthouse Lab in Glasgow | Wellcome Sanger Institute for the COVID-19 Genomics UK (COG-UK) Consortium | Harper VanSteenhouse, Yumi Kasai, David Gray, Carol Clugston, Anna Dominiczak and Alex Alderton, Roberto Amato, Jeffrey Barrett, Sonia Goncalves, Ewan Harrison, David K. Jackson, Ian Johnston, Dominic Kwiatkowski, Cordelia Langford, John Sillitoe on behalf of the Wellcome Sanger Institute COVID-19 Surveillance Team |
| EPI_ISL_2805989 | Lighthouse Lab in Milton Keynes | Wellcome Sanger Institute for the COVID-19 Genomics UK (COG-UK) Consortium | The Lighthouse Lab in Milton Keynes and Alex Alderton, Roberto Amato, Jeffrey Barrett, Sonia Goncalves, Ewan Harrison, David K. Jackson, Ian Johnston, Dominic Kwiatkowski, Cordelia Langford, John Sillitoe on behalf of the Wellcome Sanger Institute COVID-19 Surveillance Team |
| EPI_ISL_2805990 | Lighthouse Laboratory Plymouth | Wellcome Sanger Institute for the COVID-19 Genomics UK (COG-UK) Consortium | Lighthouse Laboratory Plymouth and Alex Alderton, Roberto Amato, Jeffrey Barrett, Sonia Goncalves, Ewan Harrison, David K. Jackson, Ian Johnston, Dominic Kwiatkowski, Cordelia Langford, John Sillitoe on behalf of the Wellcome Sanger Institute COVID-19 Surveillance Team |
| EPI_ISL_2805991 | Lighthouse Lab in Milton Keynes | Wellcome Sanger Institute for the COVID-19 Genomics UK (COG-UK) Consortium | The Lighthouse Lab in Milton Keynes and Alex Alderton, Roberto Amato, Jeffrey Barrett, Sonia Goncalves, Ewan Harrison, David K. Jackson, Ian Johnston, Dominic Kwiatkowski, Cordelia Langford, John Sillitoe on behalf of the Wellcome Sanger Institute COVID-19 Surveillance Team |
| EPI_ISL_2805992, EPI_ISL_2805993, EPI_ISL_2805994, EPI_ISL_2805995, EPI_ISL_2805996 | Lighthouse Lab in Alderley Park | Wellcome Sanger Institute for the COVID-19 Genomics UK (COG-UK) Consortium | Jacquelyn Wynn, Mairead Hyland, The Lighthouse Lab in Alderley Park and Alex Alderton, Roberto Amato, Jeffrey Barrett, Sonia Goncalves, Ewan Harrison, David K. Jackson, Ian Johnston, Dominic Kwiatkowski, Cordelia Langford, John Sillitoe on behalf of the Wellcome Sanger Institute COVID-19 Surveillance Team |
| EPI_ISL_2805997 | Lighthouse Lab in Glasgow | Wellcome Sanger Institute for the COVID-19 Genomics UK (COG-UK) Consortium | Harper VanSteenhouse, Yumi Kasai, David Gray, Carol Clugston, Anna Dominiczak and Alex Alderton, Roberto Amato, Jeffrey Barrett, Sonia Goncalves, Ewan Harrison, David K. Jackson, Ian Johnston, Dominic Kwiatkowski, Cordelia Langford, John Sillitoe on behalf of the Wellcome Sanger Institute COVID-19 Surveillance Team |
| EPI_ISL_2805998, EPI_ISL_2805999, EPI_ISL_2806000, EPI_ISL_2806001, EPI_ISL_2806002, EPI_ISL_2806003, EPI_ISL_2806004, EPI_ISL_2806005, EPI_ISL_2806006, EPI_ISL_2806007 | Lighthouse Lab in Alderley Park | Wellcome Sanger Institute for the COVID-19 Genomics UK (COG-UK) Consortium | Jacquelyn Wynn, Mairead Hyland, The Lighthouse Lab in Alderley Park and Alex Alderton, Roberto Amato, Jeffrey Barrett, Sonia Goncalves, Ewan Harrison, David K. Jackson, Ian Johnston, Dominic Kwiatkowski, Cordelia Langford, John Sillitoe on behalf of the Wellcome Sanger Institute COVID-19 Surveillance Team |
| EPI_ISL_2806008 | Lighthouse Lab in Glasgow | Wellcome Sanger Institute for the COVID-19 Genomics UK (COG-UK) Consortium | Harper VanSteenhouse, Yumi Kasai, David Gray, Carol Clugston, Anna Dominiczak and Alex Alderton, Roberto Amato, Jeffrey Barrett, Sonia Goncalves, Ewan Harrison, David K. Jackson, Ian Johnston, Dominic Kwiatkowski, Cordelia Langford, John Sillitoe on behalf of the Wellcome Sanger Institute COVID-19 Surveillance Team |
| EPI_ISL_2806009, EPI_ISL_2806010, EPI_ISL_2806011, EPI_ISL_2806012 | Lighthouse Lab in Alderley Park | Wellcome Sanger Institute for the COVID-19 Genomics UK (COG-UK) Consortium | Jacquelyn Wynn, Mairead Hyland, The Lighthouse Lab in Alderley Park and Alex Alderton, Roberto Amato, Jeffrey Barrett, Sonia Goncalves, Ewan Harrison, David K. Jackson, Ian Johnston, Dominic Kwiatkowski, Cordelia Langford, John Sillitoe on behalf of the Wellcome Sanger Institute COVID-19 Surveillance Team |
| EPI_ISL_2806013 | Lighthouse Lab in Glasgow | Wellcome Sanger Institute for the COVID-19 Genomics UK (COG-UK) Consortium | Harper VanSteenhouse, Yumi Kasai, David Gray, Carol Clugston, Anna Dominiczak and Alex Alderton, Roberto Amato, Jeffrey Barrett, Sonia Goncalves, Ewan Harrison, David K. Jackson, Ian Johnston, Dominic Kwiatkowski, Cordelia Langford, John Sillitoe on behalf of the Wellcome Sanger Institute COVID-19 Surveillance Team |
| EPI_ISL_2806014, EPI_ISL_2806015, EPI_ISL_2806016, EPI_ISL_2806017 | Lighthouse Lab in Alderley Park | Wellcome Sanger Institute for the COVID-19 Genomics UK (COG-UK) Consortium | Jacquelyn Wynn, Mairead Hyland, The Lighthouse Lab in Alderley Park and Alex Alderton, Roberto Amato, Jeffrey Barrett, Sonia Goncalves, Ewan Harrison, David K. Jackson, Ian Johnston, Dominic Kwiatkowski, Cordelia Langford, John Sillitoe on behalf of the Wellcome Sanger Institute COVID-19 Surveillance Team |
| EPI_ISL_2806018, EPI_ISL_2806019 | Lighthouse Lab in Glasgow | Wellcome Sanger Institute for the COVID-19 Genomics UK (COG-UK) Consortium | Harper VanSteenhouse, Yumi Kasai, David Gray, Carol Clugston, Anna Dominiczak and Alex Alderton, Roberto Amato, Jeffrey Barrett, Sonia Goncalves, Ewan Harrison, David K. Jackson, Ian Johnston, Dominic Kwiatkowski, Cordelia Langford, John Sillitoe on behalf of the Wellcome Sanger Institute COVID-19 Surveillance Team |
| EPI_ISL_2806020, EPI_ISL_2806021, EPI_ISL_2806022, EPI_ISL_2806023 | Lighthouse Lab in Alderley Park | Wellcome Sanger Institute for the COVID-19 Genomics UK (COG-UK) Consortium | Jacquelyn Wynn, Mairead Hyland, The Lighthouse Lab in Alderley Park and Alex Alderton, Roberto Amato, Jeffrey Barrett, Sonia Goncalves, Ewan Harrison, David K. Jackson, Ian Johnston, Dominic Kwiatkowski, Cordelia Langford, John Sillitoe on behalf of the Wellcome Sanger Institute COVID-19 Surveillance Team |
| EPI_ISL_2806024, EPI_ISL_2806025 | Lighthouse Lab in Glasgow | Wellcome Sanger Institute for the COVID-19 Genomics UK (COG-UK) Consortium | Harper VanSteenhouse, Yumi Kasai, David Gray, Carol Clugston, Anna Dominiczak and Alex Alderton, Roberto Amato, Jeffrey Barrett, Sonia Goncalves, Ewan Harrison, David K. Jackson, Ian Johnston, Dominic Kwiatkowski, Cordelia Langford, John Sillitoe on behalf of the Wellcome Sanger Institute COVID-19 Surveillance Team |
| EPI_ISL_2806026, EPI_ISL_2806027, EPI_ISL_2806028, EPI_ISL_2806029, EPI_ISL_2806030 | Lighthouse Lab in Alderley Park | Wellcome Sanger Institute for the COVID-19 Genomics UK (COG-UK) Consortium | Jacquelyn Wynn, Mairead Hyland, The Lighthouse Lab in Alderley Park and Alex Alderton, Roberto Amato, Jeffrey Barrett, Sonia Goncalves, Ewan Harrison, David K. Jackson, Ian Johnston, Dominic Kwiatkowski, Cordelia Langford, John Sillitoe on behalf of the Wellcome Sanger Institute COVID-19 Surveillance Team |
| EPI_ISL_2806031 | Lighthouse Lab in Glasgow | Wellcome Sanger Institute for the COVID-19 Genomics UK (COG-UK) Consortium | Harper VanSteenhouse, Yumi Kasai, David Gray, Carol Clugston, Anna Dominiczak and Alex Alderton, Roberto Amato, Jeffrey Barrett, Sonia Goncalves, Ewan Harrison, David K. Jackson, Ian Johnston, Dominic Kwiatkowski, Cordelia Langford, John Sillitoe on behalf of the Wellcome Sanger Institute COVID-19 Surveillance Team |
| EPI_ISL_2806032, EPI_ISL_2806033 | Lighthouse Lab in Alderley Park | Wellcome Sanger Institute for the COVID-19 Genomics UK (COG-UK) Consortium | Jacquelyn Wynn, Mairead Hyland, The Lighthouse Lab in Alderley Park and Alex Alderton, Roberto Amato, Jeffrey Barrett, Sonia Goncalves, Ewan Harrison, David K. Jackson, Ian Johnston, Dominic Kwiatkowski, Cordelia Langford, John Sillitoe on behalf of the Wellcome Sanger Institute COVID-19 Surveillance Team |
| EPI_ISL_2806034 | Lighthouse Lab in Glasgow | Wellcome Sanger Institute for the COVID-19 Genomics UK (COG-UK) Consortium | Harper VanSteenhouse, Yumi Kasai, David Gray, Carol Clugston, Anna Dominiczak and Alex Alderton, Roberto Amato, Jeffrey Barrett, Sonia Goncalves, Ewan Harrison, David K. Jackson, Ian Johnston, Dominic Kwiatkowski, Cordelia Langford, John Sillitoe on behalf of the Wellcome Sanger Institute COVID-19 Surveillance Team |
| EPI_ISL_2806035, EPI_ISL_2806036 | Lighthouse Lab in Alderley Park | Wellcome Sanger Institute for the COVID-19 Genomics UK (COG-UK) Consortium | Jacquelyn Wynn, Mairead Hyland, The Lighthouse Lab in Alderley Park and Alex Alderton, Roberto Amato, Jeffrey Barrett, Sonia Goncalves, Ewan Harrison, David K. Jackson, Ian Johnston, Dominic Kwiatkowski, Cordelia Langford, John Sillitoe on behalf of the Wellcome Sanger Institute COVID-19 Surveillance Team |
| EPI_ISL_2806037 | Lighthouse Lab in Glasgow | Wellcome Sanger Institute for the COVID-19 Genomics UK (COG-UK) Consortium | Harper VanSteenhouse, Yumi Kasai, David Gray, Carol Clugston, Anna Dominiczak and Alex Alderton, Roberto Amato, Jeffrey Barrett, Sonia Goncalves, Ewan Harrison, David K. Jackson, Ian Johnston, Dominic Kwiatkowski, Cordelia Langford, John Sillitoe on behalf of the Wellcome Sanger Institute COVID-19 Surveillance Team |
| EPI_ISL_2806038 | Lighthouse Lab in Alderley Park | Wellcome Sanger Institute for the COVID-19 Genomics UK (COG-UK) Consortium | Jacquelyn Wynn, Mairead Hyland, The Lighthouse Lab in Alderley Park and Alex Alderton, Roberto Amato, Jeffrey Barrett, Sonia Goncalves, Ewan Harrison, David K. Jackson, Ian Johnston, Dominic Kwiatkowski, Cordelia Langford, John Sillitoe on behalf of the Wellcome Sanger Institute COVID-19 Surveillance Team |
| EPI_ISL_2806039 | Lighthouse Lab in Glasgow | Wellcome Sanger Institute for the COVID-19 Genomics UK (COG-UK) Consortium | Harper VanSteenhouse, Yumi Kasai, David Gray, Carol Clugston, Anna Dominiczak and Alex Alderton, Roberto Amato, Jeffrey Barrett, Sonia Goncalves, Ewan Harrison, David K. Jackson, Ian Johnston, Dominic Kwiatkowski, Cordelia Langford, John Sillitoe on behalf of the Wellcome Sanger Institute COVID-19 Surveillance Team |
| EPI_ISL_2806040, EPI_ISL_2806041, EPI_ISL_2806042 | Lighthouse Lab in Alderley Park | Wellcome Sanger Institute for the COVID-19 Genomics UK (COG-UK) Consortium | Jacquelyn Wynn, Mairead Hyland, The Lighthouse Lab in Alderley Park and Alex Alderton, Roberto Amato, Jeffrey Barrett, Sonia Goncalves, Ewan Harrison, David K. Jackson, Ian Johnston, Dominic Kwiatkowski, Cordelia Langford, John Sillitoe on behalf of the Wellcome Sanger Institute COVID-19 Surveillance Team |
| EPI_ISL_2806043 | Lighthouse Lab in Glasgow | Wellcome Sanger Institute for the COVID-19 Genomics UK (COG-UK) Consortium | Harper VanSteenhouse, Yumi Kasai, David Gray, Carol Clugston, Anna Dominiczak and Alex Alderton, Roberto Amato, Jeffrey Barrett, Sonia Goncalves, Ewan Harrison, David K. Jackson, Ian Johnston, Dominic Kwiatkowski, Cordelia Langford, John Sillitoe on behalf of the Wellcome Sanger Institute COVID-19 Surveillance Team |
| EPI_ISL_2806044 | Lighthouse Lab in Alderley Park | Wellcome Sanger Institute for the COVID-19 Genomics UK (COG-UK) Consortium | Jacquelyn Wynn, Mairead Hyland, The Lighthouse Lab in Alderley Park and Alex Alderton, Roberto Amato, Jeffrey Barrett, Sonia Goncalves, Ewan Harrison, David K. Jackson, Ian Johnston, Dominic Kwiatkowski, Cordelia Langford, John Sillitoe on behalf of the Wellcome Sanger Institute COVID-19 Surveillance Team |
| EPI_ISL_2806045 | Lighthouse Lab in Glasgow | Wellcome Sanger Institute for the COVID-19 Genomics UK (COG-UK) Consortium | Harper VanSteenhouse, Yumi Kasai, David Gray, Carol Clugston, Anna Dominiczak and Alex Alderton, Roberto Amato, Jeffrey Barrett, Sonia Goncalves, Ewan Harrison, David K. Jackson, Ian Johnston, Dominic Kwiatkowski, Cordelia Langford, John Sillitoe on behalf of the Wellcome Sanger Institute COVID-19 Surveillance Team |
| EPI_ISL_2806046, EPI_ISL_2806047, EPI_ISL_2806048, EPI_ISL_2806049, EPI_ISL_2806050, EPI_ISL_2806051, EPI_ISL_2806052, EPI_ISL_2806053, EPI_ISL_2806054, EPI_ISL_2806055, EPI_ISL_2806056 | | | |
| see above | Lighthouse Lab in Alderley Park | Wellcome Sanger Institute for the COVID-19 Genomics UK (COG-UK) Consortium | Jacquelyn Wynn, Mairead Hyland, The Lighthouse Lab in Alderley Park and Alex Alderton, Roberto Amato, Jeffrey Barrett, Sonia Goncalves, Ewan Harrison, David K. Jackson, Ian Johnston, Dominic Kwiatkowski, Cordelia Langford, John Sillitoe on behalf of the Wellcome Sanger Institute COVID-19 Surveillance Team |
| EPI_ISL_2806057, EPI_ISL_2806058 | Lighthouse Lab in Glasgow | Wellcome Sanger Institute for the COVID-19 Genomics UK (COG-UK) Consortium | Harper VanSteenhouse, Yumi Kasai, David Gray, Carol Clugston, Anna Dominiczak and Alex Alderton, Roberto Amato, Jeffrey Barrett, Sonia Goncalves, Ewan Harrison, David K. Jackson, Ian Johnston, Dominic Kwiatkowski, Cordelia Langford, John Sillitoe on behalf of the Wellcome Sanger Institute COVID-19 Surveillance Team |
| EPI_ISL_2806059, EPI_ISL_2806060, | Lighthouse Lab in Alderley Park | Wellcome Sanger Institute for the COVID-19 Genomics UK (COG-UK) Consortium | Jacquelyn Wynn, Mairead Hyland, The Lighthouse Lab in Alderley Park and Alex Alderton, Roberto Amato, Jeffrey Barrett, Sonia Goncalves, Ewan Harrison, David K. Jackson, Ian Johnston, Dominic Kwiatkowski, Cordelia Langford, John Sillitoe on behalf of the Wellcome Sanger Institute COVID-19 Surveillance Team |

| | | | |
|---|---|---|---|
| EPI_ISL_2806062, EPI_ISL_2806063 | | | |
| EPI_ISL_2806064 | Lighthouse Lab in Glasgow | Wellcome Sanger Institute for the COVID-19 Genomics UK (COG-UK) Consortium | Harper VanSteenhouse, Yumi Kasai, David Gray, Carol Clugston, Anna Dominiczak and Alex Alderton, Roberto Amato, Jeffrey Barrett, Sonia Goncalves, Ewan Harrison, David K. Jackson, Ian Johnston, Dominic Kwiatkowski, Cordelia Langford, John Sillitoe on behalf of the Wellcome Sanger Institute COVID-19 Surveillance Team |
| EPI_ISL_2806065, EPI_ISL_2806066, EPI_ISL_2806067 | Lighthouse Lab in Alderley Park | Wellcome Sanger Institute for the COVID-19 Genomics UK (COG-UK) Consortium | Jacquelyn Wynn, Mairead Hyland, The Lighthouse Lab in Alderley Park and Alex Alderton, Roberto Amato, Jeffrey Barrett, Sonia Goncalves, Ewan Harrison, David K. Jackson, Ian Johnston, Dominic Kwiatkowski, Cordelia Langford, John Sillitoe on behalf of the Wellcome Sanger Institute COVID-19 Surveillance Team |
| EPI_ISL_2806068 | Lighthouse Lab in Glasgow | Wellcome Sanger Institute for the COVID-19 Genomics UK (COG-UK) Consortium | Harper VanSteenhouse, Yumi Kasai, David Gray, Carol Clugston, Anna Dominiczak and Alex Alderton, Roberto Amato, Jeffrey Barrett, Sonia Goncalves, Ewan Harrison, David K. Jackson, Ian Johnston, Dominic Kwiatkowski, Cordelia Langford, John Sillitoe on behalf of the Wellcome Sanger Institute COVID-19 Surveillance Team |
| EPI_ISL_2806069, EPI_ISL_2806070, EPI_ISL_2806071, EPI_ISL_2806072, EPI_ISL_2806073 | Lighthouse Lab in Alderley Park | Wellcome Sanger Institute for the COVID-19 Genomics UK (COG-UK) Consortium | Jacquelyn Wynn, Mairead Hyland, The Lighthouse Lab in Alderley Park and Alex Alderton, Roberto Amato, Jeffrey Barrett, Sonia Goncalves, Ewan Harrison, David K. Jackson, Ian Johnston, Dominic Kwiatkowski, Cordelia Langford, John Sillitoe on behalf of the Wellcome Sanger Institute COVID-19 Surveillance Team |
| EPI_ISL_2806074 | Lighthouse Lab in Glasgow | Wellcome Sanger Institute for the COVID-19 Genomics UK (COG-UK) Consortium | Harper VanSteenhouse, Yumi Kasai, David Gray, Carol Clugston, Anna Dominiczak and Alex Alderton, Roberto Amato, Jeffrey Barrett, Sonia Goncalves, Ewan Harrison, David K. Jackson, Ian Johnston, Dominic Kwiatkowski, Cordelia Langford, John Sillitoe on behalf of the Wellcome Sanger Institute COVID-19 Surveillance Team |
| EPI_ISL_2806075 | Lighthouse Lab in Alderley Park | Wellcome Sanger Institute for the COVID-19 Genomics UK (COG-UK) Consortium | Jacquelyn Wynn, Mairead Hyland, The Lighthouse Lab in Alderley Park and Alex Alderton, Roberto Amato, Jeffrey Barrett, Sonia Goncalves, Ewan Harrison, David K. Jackson, Ian Johnston, Dominic Kwiatkowski, Cordelia Langford, John Sillitoe on behalf of the Wellcome Sanger Institute COVID-19 Surveillance Team |
| EPI_ISL_2806076 | Lighthouse Lab in Glasgow | Wellcome Sanger Institute for the COVID-19 Genomics UK (COG-UK) Consortium | Harper VanSteenhouse, Yumi Kasai, David Gray, Carol Clugston, Anna Dominiczak and Alex Alderton, Roberto Amato, Jeffrey Barrett, Sonia Goncalves, Ewan Harrison, David K. Jackson, Ian Johnston, Dominic Kwiatkowski, Cordelia Langford, John Sillitoe on behalf of the Wellcome Sanger Institute COVID-19 Surveillance Team |
| EPI_ISL_2806077, EPI_ISL_2806078, EPI_ISL_2806079, EPI_ISL_2806080, EPI_ISL_2806081, EPI_ISL_2806082 | Lighthouse Lab in Alderley Park | Wellcome Sanger Institute for the COVID-19 Genomics UK (COG-UK) Consortium | Jacquelyn Wynn, Mairead Hyland, The Lighthouse Lab in Alderley Park and Alex Alderton, Roberto Amato, Jeffrey Barrett, Sonia Goncalves, Ewan Harrison, David K. Jackson, Ian Johnston, Dominic Kwiatkowski, Cordelia Langford, John Sillitoe on behalf of the Wellcome Sanger Institute COVID-19 Surveillance Team |
| EPI_ISL_2806083 | Lighthouse Lab in Glasgow | Wellcome Sanger Institute for the COVID-19 Genomics UK (COG-UK) Consortium | Harper VanSteenhouse, Yumi Kasai, David Gray, Carol Clugston, Anna Dominiczak and Alex Alderton, Roberto Amato, Jeffrey Barrett, Sonia Goncalves, Ewan Harrison, David K. Jackson, Ian Johnston, Dominic Kwiatkowski, Cordelia Langford, John Sillitoe on behalf of the Wellcome Sanger Institute COVID-19 Surveillance Team |
| EPI_ISL_2806084 | Lighthouse Lab in Alderley Park | Wellcome Sanger Institute for the COVID-19 Genomics UK (COG-UK) Consortium | Jacquelyn Wynn, Mairead Hyland, The Lighthouse Lab in Alderley Park and Alex Alderton, Roberto Amato, Jeffrey Barrett, Sonia Goncalves, Ewan Harrison, David K. Jackson, Ian Johnston, Dominic Kwiatkowski, Cordelia Langford, John Sillitoe on behalf of the Wellcome Sanger Institute COVID-19 Surveillance Team |
| EPI_ISL_2806085 | Lighthouse Lab in Glasgow | Wellcome Sanger Institute for the COVID-19 Genomics UK (COG-UK) Consortium | Harper VanSteenhouse, Yumi Kasai, David Gray, Carol Clugston, Anna Dominiczak and Alex Alderton, Roberto Amato, Jeffrey Barrett, Sonia Goncalves, Ewan Harrison, David K. Jackson, Ian Johnston, Dominic Kwiatkowski, Cordelia Langford, John Sillitoe on behalf of the Wellcome Sanger Institute COVID-19 Surveillance Team |
| EPI_ISL_2806086, EPI_ISL_2806087, EPI_ISL_2806088, EPI_ISL_2806089, EPI_ISL_2806090, EPI_ISL_2806091, EPI_ISL_2806092, EPI_ISL_2806093 | Lighthouse Lab in Alderley Park | Wellcome Sanger Institute for the COVID-19 Genomics UK (COG-UK) Consortium | Jacquelyn Wynn, Mairead Hyland, The Lighthouse Lab in Alderley Park and Alex Alderton, Roberto Amato, Jeffrey Barrett, Sonia Goncalves, Ewan Harrison, David K. Jackson, Ian Johnston, Dominic Kwiatkowski, Cordelia Langford, John Sillitoe on behalf of the Wellcome Sanger Institute COVID-19 Surveillance Team |
| EPI_ISL_2806094, EPI_ISL_2806096 | Lighthouse Lab in Glasgow | Wellcome Sanger Institute for the COVID-19 Genomics UK (COG-UK) Consortium | Harper VanSteenhouse, Yumi Kasai, David Gray, Carol Clugston, Anna Dominiczak and Alex Alderton, Roberto Amato, Jeffrey Barrett, Sonia Goncalves, Ewan Harrison, David K. Jackson, Ian Johnston, Dominic Kwiatkowski, Cordelia Langford, John Sillitoe on behalf of the Wellcome Sanger Institute COVID-19 Surveillance Team |
| EPI_ISL_2806097 | Lighthouse Lab in Alderley Park | Wellcome Sanger Institute for the COVID-19 Genomics UK (COG-UK) Consortium | Jacquelyn Wynn, Mairead Hyland, The Lighthouse Lab in Alderley Park and Alex Alderton, Roberto Amato, Jeffrey Barrett, Sonia Goncalves, Ewan Harrison, David K. Jackson, Ian Johnston, Dominic Kwiatkowski, Cordelia Langford, John Sillitoe on behalf of the Wellcome Sanger Institute COVID-19 Surveillance Team |
| EPI_ISL_2806098 | Lighthouse Lab in Glasgow | Wellcome Sanger Institute for the COVID-19 Genomics UK (COG-UK) Consortium | Harper VanSteenhouse, Yumi Kasai, David Gray, Carol Clugston, Anna Dominiczak and Alex Alderton, Roberto Amato, Jeffrey Barrett, Sonia Goncalves, Ewan Harrison, David K. Jackson, Ian Johnston, Dominic Kwiatkowski, Cordelia Langford, John Sillitoe on behalf of the Wellcome Sanger Institute COVID-19 Surveillance Team |
| EPI_ISL_2806099 | Lighthouse Lab in Alderley Park | Wellcome Sanger Institute for the COVID-19 Genomics UK (COG-UK) Consortium | Jacquelyn Wynn, Mairead Hyland, The Lighthouse Lab in Alderley Park and Alex Alderton, Roberto Amato, Jeffrey Barrett, Sonia Goncalves, Ewan Harrison, David K. Jackson, Ian Johnston, Dominic Kwiatkowski, Cordelia Langford, John Sillitoe on behalf of the Wellcome Sanger Institute COVID-19 Surveillance Team |
| EPI_ISL_2806100 | Lighthouse Lab in Glasgow | Wellcome Sanger Institute for the COVID-19 Genomics UK (COG-UK) Consortium | Harper VanSteenhouse, Yumi Kasai, David Gray, Carol Clugston, Anna Dominiczak and Alex Alderton, Roberto Amato, Jeffrey Barrett, Sonia Goncalves, Ewan Harrison, David K. Jackson, Ian Johnston, Dominic Kwiatkowski, Cordelia Langford, John Sillitoe on behalf of the Wellcome Sanger Institute COVID-19 Surveillance Team |
| EPI_ISL_2806101 | Lighthouse Lab in Alderley Park | Wellcome Sanger Institute for the COVID-19 Genomics UK (COG-UK) Consortium | Jacquelyn Wynn, Mairead Hyland, The Lighthouse Lab in Alderley Park and Alex Alderton, Roberto Amato, Jeffrey Barrett, Sonia Goncalves, Ewan Harrison, David K. Jackson, Ian Johnston, Dominic Kwiatkowski, Cordelia Langford, John Sillitoe on behalf of the Wellcome Sanger Institute COVID-19 Surveillance Team |
| EPI_ISL_2806102, EPI_ISL_2806103, EPI_ISL_2806104 | Lighthouse Lab in Glasgow | Wellcome Sanger Institute for the COVID-19 Genomics UK (COG-UK) Consortium | Harper VanSteenhouse, Yumi Kasai, David Gray, Carol Clugston, Anna Dominiczak and Alex Alderton, Roberto Amato, Jeffrey Barrett, Sonia Goncalves, Ewan Harrison, David K. Jackson, Ian Johnston, Dominic Kwiatkowski, Cordelia Langford, John Sillitoe on behalf of the Wellcome Sanger Institute COVID-19 Surveillance Team |
| EPI_ISL_2806105, EPI_ISL_2806107 | Lighthouse Lab in Alderley Park | Wellcome Sanger Institute for the COVID-19 Genomics UK (COG-UK) Consortium | Jacquelyn Wynn, Mairead Hyland, The Lighthouse Lab in Alderley Park and Alex Alderton, Roberto Amato, Jeffrey Barrett, Sonia Goncalves, Ewan Harrison, David K. Jackson, Ian Johnston, Dominic Kwiatkowski, Cordelia Langford, John Sillitoe on behalf of the Wellcome Sanger Institute COVID-19 Surveillance Team |
| EPI_ISL_2806108 | Lighthouse Lab in Glasgow | Wellcome Sanger Institute for the COVID-19 Genomics UK (COG-UK) Consortium | Harper VanSteenhouse, Yumi Kasai, David Gray, Carol Clugston, Anna Dominiczak and Alex Alderton, Roberto Amato, Jeffrey Barrett, Sonia Goncalves, Ewan Harrison, David K. Jackson, Ian Johnston, Dominic Kwiatkowski, Cordelia Langford, John Sillitoe on behalf of the Wellcome Sanger Institute COVID-19 Surveillance Team |
| EPI_ISL_2806109 | Lighthouse Lab in Alderley Park | Wellcome Sanger Institute for the COVID-19 Genomics UK (COG-UK) Consortium | Jacquelyn Wynn, Mairead Hyland, The Lighthouse Lab in Alderley Park and Alex Alderton, Roberto Amato, Jeffrey Barrett, Sonia Goncalves, Ewan Harrison, David K. Jackson, Ian Johnston, Dominic Kwiatkowski, Cordelia Langford, John Sillitoe on behalf of the Wellcome Sanger Institute COVID-19 Surveillance Team |
| EPI_ISL_2806110, EPI_ISL_2806111 | Lighthouse Lab in Glasgow | Wellcome Sanger Institute for the COVID-19 Genomics UK (COG-UK) Consortium | Harper VanSteenhouse, Yumi Kasai, David Gray, Carol Clugston, Anna Dominiczak and Alex Alderton, Roberto Amato, Jeffrey Barrett, Sonia Goncalves, Ewan Harrison, David K. Jackson, Ian Johnston, Dominic Kwiatkowski, Cordelia Langford, John Sillitoe on behalf of the Wellcome Sanger Institute COVID-19 Surveillance Team |
| EPI_ISL_2806112, EPI_ISL_2806113, EPI_ISL_2806114, EPI_ISL_2806115 | Lighthouse Lab in Alderley Park | Wellcome Sanger Institute for the COVID-19 Genomics UK (COG-UK) Consortium | Jacquelyn Wynn, Mairead Hyland, The Lighthouse Lab in Alderley Park and Alex Alderton, Roberto Amato, Jeffrey Barrett, Sonia Goncalves, Ewan Harrison, David K. Jackson, Ian Johnston, Dominic Kwiatkowski, Cordelia Langford, John Sillitoe on behalf of the Wellcome Sanger Institute COVID-19 Surveillance Team |
| EPI_ISL_2806116, EPI_ISL_2806117 | Lighthouse Lab in Glasgow | Wellcome Sanger Institute for the COVID-19 Genomics UK (COG-UK) Consortium | Harper VanSteenhouse, Yumi Kasai, David Gray, Carol Clugston, Anna Dominiczak and Alex Alderton, Roberto Amato, Jeffrey Barrett, Sonia Goncalves, Ewan Harrison, David K. Jackson, Ian Johnston, Dominic Kwiatkowski, Cordelia Langford, John Sillitoe on behalf of the Wellcome Sanger Institute COVID-19 Surveillance Team |
| EPI_ISL_2806118, EPI_ISL_2806119 | Lighthouse Lab in Alderley Park | Wellcome Sanger Institute for the COVID-19 Genomics UK (COG-UK) Consortium | Jacquelyn Wynn, Mairead Hyland, The Lighthouse Lab in Alderley Park and Alex Alderton, Roberto Amato, Jeffrey Barrett, Sonia Goncalves, Ewan Harrison, David K. Jackson, Ian Johnston, Dominic Kwiatkowski, Cordelia Langford, John Sillitoe on behalf of the Wellcome Sanger Institute COVID-19 Surveillance Team |
| EPI_ISL_2806120 | Lighthouse Lab in Glasgow | Wellcome Sanger Institute for the COVID-19 Genomics UK (COG-UK) Consortium | Harper VanSteenhouse, Yumi Kasai, David Gray, Carol Clugston, Anna Dominiczak and Alex Alderton, Roberto Amato, Jeffrey Barrett, Sonia Goncalves, Ewan Harrison, David K. Jackson, Ian Johnston, Dominic Kwiatkowski, Cordelia Langford, John Sillitoe on behalf of the Wellcome Sanger Institute COVID-19 Surveillance Team |
| EPI_ISL_2806121 | Lighthouse Lab in Alderley Park | Wellcome Sanger Institute for the COVID-19 Genomics UK (COG-UK) Consortium | Jacquelyn Wynn, Mairead Hyland, The Lighthouse Lab in Alderley Park and Alex Alderton, Roberto Amato, Jeffrey Barrett, Sonia Goncalves, Ewan Harrison, David K. Jackson, Ian Johnston, Dominic Kwiatkowski, Cordelia Langford, John Sillitoe on behalf of the Wellcome Sanger Institute COVID-19 Surveillance Team |
| EPI_ISL_2806122 | Lighthouse Lab in Glasgow | Wellcome Sanger Institute for the COVID-19 Genomics UK (COG-UK) Consortium | Harper VanSteenhouse, Yumi Kasai, David Gray, Carol Clugston, Anna Dominiczak and Alex Alderton, Roberto Amato, Jeffrey Barrett, Sonia Goncalves, Ewan Harrison, David K. Jackson, Ian Johnston, Dominic Kwiatkowski, Cordelia Langford, John Sillitoe on behalf of the Wellcome Sanger Institute COVID-19 Surveillance Team |
| EPI_ISL_2806123 | Lighthouse Lab in Alderley Park | Wellcome Sanger Institute for the COVID-19 Genomics UK (COG-UK) Consortium | Jacquelyn Wynn, Mairead Hyland, The Lighthouse Lab in Alderley Park and Alex Alderton, Roberto Amato, Jeffrey Barrett, Sonia Goncalves, Ewan Harrison, David K. Jackson, Ian Johnston, Dominic Kwiatkowski, Cordelia Langford, John Sillitoe on behalf of the Wellcome Sanger Institute COVID-19 Surveillance Team |
| EPI_ISL_2806124, EPI_ISL_2806125 | Lighthouse Lab in Glasgow | Wellcome Sanger Institute for the COVID-19 Genomics UK (COG-UK) Consortium | Harper VanSteenhouse, Yumi Kasai, David Gray, Carol Clugston, Anna Dominiczak and Alex Alderton, Roberto Amato, Jeffrey Barrett, Sonia Goncalves, Ewan Harrison, David K. Jackson, Ian Johnston, Dominic Kwiatkowski, Cordelia Langford, John Sillitoe on behalf of the Wellcome Sanger Institute COVID-19 Surveillance Team |
| EPI_ISL_2806126, EPI_ISL_2806127, EPI_ISL_2806128, EPI_ISL_2806129, EPI_ISL_2806130, EPI_ISL_2806131, EPI_ISL_2806132, EPI_ISL_2806133, EPI_ISL_2806134, EPI_ISL_2806135, EPI_ISL_2806136, EPI_ISL_2806137, EPI_ISL_2806138, EPI_ISL_2806139, EPI_ISL_2806140, EPI_ISL_2806141, EPI_ISL_2806142, EPI_ISL_2806143 | | | |
| see above | Lighthouse Lab in Alderley Park | Wellcome Sanger Institute for the COVID-19 Genomics UK (COG-UK) Consortium | Jacquelyn Wynn, Mairead Hyland, The Lighthouse Lab in Alderley Park and Alex Alderton, Roberto Amato, Jeffrey Barrett, Sonia Goncalves, Ewan Harrison, David K. Jackson, Ian Johnston, Dominic Kwiatkowski, Cordelia Langford, John Sillitoe on behalf of the Wellcome Sanger Institute COVID-19 Surveillance Team |
| EPI_ISL_2806144 | Lighthouse Lab in Glasgow | Wellcome Sanger Institute for the COVID-19 Genomics UK (COG-UK) Consortium | Harper VanSteenhouse, Yumi Kasai, David Gray, Carol Clugston, Anna Dominiczak and Alex Alderton, Roberto Amato, Jeffrey Barrett, Sonia Goncalves, Ewan Harrison, David K. Jackson, Ian Johnston, Dominic Kwiatkowski, Cordelia Langford, John Sillitoe on behalf of the Wellcome Sanger Institute COVID-19 Surveillance Team |
| EPI_ISL_2806145, EPI_ISL_2806146, EPI_ISL_2806147, EPI_ISL_2806148, EPI_ISL_2806149, EPI_ISL_2806150, EPI_ISL_2806151 | Lighthouse Lab in Alderley Park | Wellcome Sanger Institute for the COVID-19 Genomics UK (COG-UK) Consortium | Jacquelyn Wynn, Mairead Hyland, The Lighthouse Lab in Alderley Park and Alex Alderton, Roberto Amato, Jeffrey Barrett, Sonia Goncalves, Ewan Harrison, David K. Jackson, Ian Johnston, Dominic Kwiatkowski, Cordelia Langford, John Sillitoe on behalf of the Wellcome Sanger Institute COVID-19 Surveillance Team |
| EPI_ISL_2806152 | Lighthouse Lab in Glasgow | Wellcome Sanger Institute for the COVID-19 Genomics UK (COG-UK) | Harper VanSteenhouse, Yumi Kasai, David Gray, Carol Clugston, Anna Dominiczak and Alex Alderton, Roberto Amato, Jeffrey Barrett, Sonia Goncalves, Ewan Harrison, David K. Jackson, Ian Johnston, Dominic Kwiatkowski, Cordelia Langford, |

| | | | |
|---|---|---|---|
| | | Consortium | John Sillitoe on behalf of the Wellcome Sanger Institute COVID-19 Surveillance Team |
| EPI_ISL_2806153 | Lighthouse Lab in Alderley Park | Wellcome Sanger Institute for the COVID-19 Genomics UK (COG-UK) Consortium | Jacquelyn Wynn, Mairead Hyland, The Lighthouse Lab in Alderley Park and Alex Alderton, Roberto Amato, Jeffrey Barrett, Sonia Goncalves, Ewan Harrison, David K. Jackson, Ian Johnston, Dominic Kwiatkowski, Cordelia Langford, John Sillitoe on behalf of the Wellcome Sanger Institute COVID-19 Surveillance Team |
| EPI_ISL_2806154, EPI_ISL_2806155 | Lighthouse Lab in Glasgow | Wellcome Sanger Institute for the COVID-19 Genomics UK (COG-UK) Consortium | Harper VanSteenhouse, Yumi Kasai, David Gray, Carol Clugston, Anna Dominiczak and Alex Alderton, Roberto Amato, Jeffrey Barrett, Sonia Goncalves, Ewan Harrison, David K. Jackson, Ian Johnston, Dominic Kwiatkowski, Cordelia Langford, John Sillitoe on behalf of the Wellcome Sanger Institute COVID-19 Surveillance Team |
| EPI_ISL_2806156, EPI_ISL_2806157, EPI_ISL_2806158, EPI_ISL_2806159, EPI_ISL_2806160, EPI_ISL_2806161 | Lighthouse Lab in Alderley Park | Wellcome Sanger Institute for the COVID-19 Genomics UK (COG-UK) Consortium | Jacquelyn Wynn, Mairead Hyland, The Lighthouse Lab in Alderley Park and Alex Alderton, Roberto Amato, Jeffrey Barrett, Sonia Goncalves, Ewan Harrison, David K. Jackson, Ian Johnston, Dominic Kwiatkowski, Cordelia Langford, John Sillitoe on behalf of the Wellcome Sanger Institute COVID-19 Surveillance Team |
| EPI_ISL_2806162, EPI_ISL_2806163 | Lighthouse Lab in Glasgow | Wellcome Sanger Institute for the COVID-19 Genomics UK (COG-UK) Consortium | Harper VanSteenhouse, Yumi Kasai, David Gray, Carol Clugston, Anna Dominiczak and Alex Alderton, Roberto Amato, Jeffrey Barrett, Sonia Goncalves, Ewan Harrison, David K. Jackson, Ian Johnston, Dominic Kwiatkowski, Cordelia Langford, John Sillitoe on behalf of the Wellcome Sanger Institute COVID-19 Surveillance Team |
| EPI_ISL_2806164, EPI_ISL_2806165, EPI_ISL_2806166, EPI_ISL_2806167, EPI_ISL_2806168, EPI_ISL_2806169, EPI_ISL_2806170 | Lighthouse Lab in Alderley Park | Wellcome Sanger Institute for the COVID-19 Genomics UK (COG-UK) Consortium | Jacquelyn Wynn, Mairead Hyland, The Lighthouse Lab in Alderley Park and Alex Alderton, Roberto Amato, Jeffrey Barrett, Sonia Goncalves, Ewan Harrison, David K. Jackson, Ian Johnston, Dominic Kwiatkowski, Cordelia Langford, John Sillitoe on behalf of the Wellcome Sanger Institute COVID-19 Surveillance Team |
| EPI_ISL_2806171 | Lighthouse Lab in Glasgow | Wellcome Sanger Institute for the COVID-19 Genomics UK (COG-UK) Consortium | Harper VanSteenhouse, Yumi Kasai, David Gray, Carol Clugston, Anna Dominiczak and Alex Alderton, Roberto Amato, Jeffrey Barrett, Sonia Goncalves, Ewan Harrison, David K. Jackson, Ian Johnston, Dominic Kwiatkowski, Cordelia Langford, John Sillitoe on behalf of the Wellcome Sanger Institute COVID-19 Surveillance Team |
| EPI_ISL_2806172, EPI_ISL_2806173, EPI_ISL_2806174, EPI_ISL_2806175 | Lighthouse Lab in Alderley Park | Wellcome Sanger Institute for the COVID-19 Genomics UK (COG-UK) Consortium | Jacquelyn Wynn, Mairead Hyland, The Lighthouse Lab in Alderley Park and Alex Alderton, Roberto Amato, Jeffrey Barrett, Sonia Goncalves, Ewan Harrison, David K. Jackson, Ian Johnston, Dominic Kwiatkowski, Cordelia Langford, John Sillitoe on behalf of the Wellcome Sanger Institute COVID-19 Surveillance Team |
| EPI_ISL_2806176 | Lighthouse Lab in Glasgow | Wellcome Sanger Institute for the COVID-19 Genomics UK (COG-UK) Consortium | Harper VanSteenhouse, Yumi Kasai, David Gray, Carol Clugston, Anna Dominiczak and Alex Alderton, Roberto Amato, Jeffrey Barrett, Sonia Goncalves, Ewan Harrison, David K. Jackson, Ian Johnston, Dominic Kwiatkowski, Cordelia Langford, John Sillitoe on behalf of the Wellcome Sanger Institute COVID-19 Surveillance Team |
| EPI_ISL_2806177 | Lighthouse Lab in Alderley Park | Wellcome Sanger Institute for the COVID-19 Genomics UK (COG-UK) Consortium | Jacquelyn Wynn, Mairead Hyland, The Lighthouse Lab in Alderley Park and Alex Alderton, Roberto Amato, Jeffrey Barrett, Sonia Goncalves, Ewan Harrison, David K. Jackson, Ian Johnston, Dominic Kwiatkowski, Cordelia Langford, John Sillitoe on behalf of the Wellcome Sanger Institute COVID-19 Surveillance Team |
| EPI_ISL_2806179 | Lighthouse Lab in Glasgow | Wellcome Sanger Institute for the COVID-19 Genomics UK (COG-UK) Consortium | Harper VanSteenhouse, Yumi Kasai, David Gray, Carol Clugston, Anna Dominiczak and Alex Alderton, Roberto Amato, Jeffrey Barrett, Sonia Goncalves, Ewan Harrison, David K. Jackson, Ian Johnston, Dominic Kwiatkowski, Cordelia Langford, John Sillitoe on behalf of the Wellcome Sanger Institute COVID-19 Surveillance Team |
| EPI_ISL_2806180 | Lighthouse Lab in Alderley Park | Wellcome Sanger Institute for the COVID-19 Genomics UK (COG-UK) Consortium | Jacquelyn Wynn, Mairead Hyland, The Lighthouse Lab in Alderley Park and Alex Alderton, Roberto Amato, Jeffrey Barrett, Sonia Goncalves, Ewan Harrison, David K. Jackson, Ian Johnston, Dominic Kwiatkowski, Cordelia Langford, John Sillitoe on behalf of the Wellcome Sanger Institute COVID-19 Surveillance Team |
| EPI_ISL_2806181 | Lighthouse Lab in Glasgow | Wellcome Sanger Institute for the COVID-19 Genomics UK (COG-UK) Consortium | Harper VanSteenhouse, Yumi Kasai, David Gray, Carol Clugston, Anna Dominiczak and Alex Alderton, Roberto Amato, Jeffrey Barrett, Sonia Goncalves, Ewan Harrison, David K. Jackson, Ian Johnston, Dominic Kwiatkowski, Cordelia Langford, John Sillitoe on behalf of the Wellcome Sanger Institute COVID-19 Surveillance Team |
| EPI_ISL_2806182, EPI_ISL_2806183, EPI_ISL_2806184, EPI_ISL_2806185, EPI_ISL_2806186 | Lighthouse Lab in Alderley Park | Wellcome Sanger Institute for the COVID-19 Genomics UK (COG-UK) Consortium | Jacquelyn Wynn, Mairead Hyland, The Lighthouse Lab in Alderley Park and Alex Alderton, Roberto Amato, Jeffrey Barrett, Sonia Goncalves, Ewan Harrison, David K. Jackson, Ian Johnston, Dominic Kwiatkowski, Cordelia Langford, John Sillitoe on behalf of the Wellcome Sanger Institute COVID-19 Surveillance Team |
| EPI_ISL_2806187 | Lighthouse Lab in Glasgow | Wellcome Sanger Institute for the COVID-19 Genomics UK (COG-UK) Consortium | Harper VanSteenhouse, Yumi Kasai, David Gray, Carol Clugston, Anna Dominiczak and Alex Alderton, Roberto Amato, Jeffrey Barrett, Sonia Goncalves, Ewan Harrison, David K. Jackson, Ian Johnston, Dominic Kwiatkowski, Cordelia Langford, John Sillitoe on behalf of the Wellcome Sanger Institute COVID-19 Surveillance Team |
| EPI_ISL_2806188 | Lighthouse Lab in Alderley Park | Wellcome Sanger Institute for the COVID-19 Genomics UK (COG-UK) Consortium | Jacquelyn Wynn, Mairead Hyland, The Lighthouse Lab in Alderley Park and Alex Alderton, Roberto Amato, Jeffrey Barrett, Sonia Goncalves, Ewan Harrison, David K. Jackson, Ian Johnston, Dominic Kwiatkowski, Cordelia Langford, John Sillitoe on behalf of the Wellcome Sanger Institute COVID-19 Surveillance Team |
| EPI_ISL_2806189, EPI_ISL_2806190 | Lighthouse Lab in Glasgow | Wellcome Sanger Institute for the COVID-19 Genomics UK (COG-UK) Consortium | Harper VanSteenhouse, Yumi Kasai, David Gray, Carol Clugston, Anna Dominiczak and Alex Alderton, Roberto Amato, Jeffrey Barrett, Sonia Goncalves, Ewan Harrison, David K. Jackson, Ian Johnston, Dominic Kwiatkowski, Cordelia Langford, John Sillitoe on behalf of the Wellcome Sanger Institute COVID-19 Surveillance Team |
| EPI_ISL_2806191 | Lighthouse Lab in Alderley Park | Wellcome Sanger Institute for the COVID-19 Genomics UK (COG-UK) Consortium | Jacquelyn Wynn, Mairead Hyland, The Lighthouse Lab in Alderley Park and Alex Alderton, Roberto Amato, Jeffrey Barrett, Sonia Goncalves, Ewan Harrison, David K. Jackson, Ian Johnston, Dominic Kwiatkowski, Cordelia Langford, John Sillitoe on behalf of the Wellcome Sanger Institute COVID-19 Surveillance Team |
| EPI_ISL_2806192 | Lighthouse Lab in Glasgow | Wellcome Sanger Institute for the COVID-19 Genomics UK (COG-UK) Consortium | Harper VanSteenhouse, Yumi Kasai, David Gray, Carol Clugston, Anna Dominiczak and Alex Alderton, Roberto Amato, Jeffrey Barrett, Sonia Goncalves, Ewan Harrison, David K. Jackson, Ian Johnston, Dominic Kwiatkowski, Cordelia Langford, John Sillitoe on behalf of the Wellcome Sanger Institute COVID-19 Surveillance Team |
| EPI_ISL_2806193 | Lighthouse Lab in Alderley Park | Wellcome Sanger Institute for the COVID-19 Genomics UK (COG-UK) Consortium | Jacquelyn Wynn, Mairead Hyland, The Lighthouse Lab in Alderley Park and Alex Alderton, Roberto Amato, Jeffrey Barrett, Sonia Goncalves, Ewan Harrison, David K. Jackson, Ian Johnston, Dominic Kwiatkowski, Cordelia Langford, John Sillitoe on behalf of the Wellcome Sanger Institute COVID-19 Surveillance Team |
| EPI_ISL_2806194 | Lighthouse Lab in Glasgow | Wellcome Sanger Institute for the COVID-19 Genomics UK (COG-UK) Consortium | Harper VanSteenhouse, Yumi Kasai, David Gray, Carol Clugston, Anna Dominiczak and Alex Alderton, Roberto Amato, Jeffrey Barrett, Sonia Goncalves, Ewan Harrison, David K. Jackson, Ian Johnston, Dominic Kwiatkowski, Cordelia Langford, John Sillitoe on behalf of the Wellcome Sanger Institute COVID-19 Surveillance Team |
| EPI_ISL_2806195, EPI_ISL_2806196, EPI_ISL_2806197 | Lighthouse Lab in Alderley Park | Wellcome Sanger Institute for the COVID-19 Genomics UK (COG-UK) Consortium | Jacquelyn Wynn, Mairead Hyland, The Lighthouse Lab in Alderley Park and Alex Alderton, Roberto Amato, Jeffrey Barrett, Sonia Goncalves, Ewan Harrison, David K. Jackson, Ian Johnston, Dominic Kwiatkowski, Cordelia Langford, John Sillitoe on behalf of the Wellcome Sanger Institute COVID-19 Surveillance Team |
| EPI_ISL_2806198 | Lighthouse Lab in Glasgow | Wellcome Sanger Institute for the COVID-19 Genomics UK (COG-UK) Consortium | Harper VanSteenhouse, Yumi Kasai, David Gray, Carol Clugston, Anna Dominiczak and Alex Alderton, Roberto Amato, Jeffrey Barrett, Sonia Goncalves, Ewan Harrison, David K. Jackson, Ian Johnston, Dominic Kwiatkowski, Cordelia Langford, John Sillitoe on behalf of the Wellcome Sanger Institute COVID-19 Surveillance Team |
| EPI_ISL_2806199, EPI_ISL_2806200 | Lighthouse Lab in Alderley Park | Wellcome Sanger Institute for the COVID-19 Genomics UK (COG-UK) Consortium | Jacquelyn Wynn, Mairead Hyland, The Lighthouse Lab in Alderley Park and Alex Alderton, Roberto Amato, Jeffrey Barrett, Sonia Goncalves, Ewan Harrison, David K. Jackson, Ian Johnston, Dominic Kwiatkowski, Cordelia Langford, John Sillitoe on behalf of the Wellcome Sanger Institute COVID-19 Surveillance Team |
| EPI_ISL_2806201 | Lighthouse Lab in Glasgow | Wellcome Sanger Institute for the COVID-19 Genomics UK (COG-UK) Consortium | Harper VanSteenhouse, Yumi Kasai, David Gray, Carol Clugston, Anna Dominiczak and Alex Alderton, Roberto Amato, Jeffrey Barrett, Sonia Goncalves, Ewan Harrison, David K. Jackson, Ian Johnston, Dominic Kwiatkowski, Cordelia Langford, John Sillitoe on behalf of the Wellcome Sanger Institute COVID-19 Surveillance Team |
| EPI_ISL_2806202 | Lighthouse Lab in Alderley Park | Wellcome Sanger Institute for the COVID-19 Genomics UK (COG-UK) Consortium | Jacquelyn Wynn, Mairead Hyland, The Lighthouse Lab in Alderley Park and Alex Alderton, Roberto Amato, Jeffrey Barrett, Sonia Goncalves, Ewan Harrison, David K. Jackson, Ian Johnston, Dominic Kwiatkowski, Cordelia Langford, John Sillitoe on behalf of the Wellcome Sanger Institute COVID-19 Surveillance Team |
| EPI_ISL_2806203 | Lighthouse Lab in Glasgow | Wellcome Sanger Institute for the COVID-19 Genomics UK (COG-UK) Consortium | Harper VanSteenhouse, Yumi Kasai, David Gray, Carol Clugston, Anna Dominiczak and Alex Alderton, Roberto Amato, Jeffrey Barrett, Sonia Goncalves, Ewan Harrison, David K. Jackson, Ian Johnston, Dominic Kwiatkowski, Cordelia Langford, John Sillitoe on behalf of the Wellcome Sanger Institute COVID-19 Surveillance Team |
| EPI_ISL_2806204, EPI_ISL_2806205, EPI_ISL_2806206 | Lighthouse Lab in Alderley Park | Wellcome Sanger Institute for the COVID-19 Genomics UK (COG-UK) Consortium | Jacquelyn Wynn, Mairead Hyland, The Lighthouse Lab in Alderley Park and Alex Alderton, Roberto Amato, Jeffrey Barrett, Sonia Goncalves, Ewan Harrison, David K. Jackson, Ian Johnston, Dominic Kwiatkowski, Cordelia Langford, John Sillitoe on behalf of the Wellcome Sanger Institute COVID-19 Surveillance Team |
| EPI_ISL_2806207 | Lighthouse Lab in Glasgow | Wellcome Sanger Institute for the COVID-19 Genomics UK (COG-UK) Consortium | Harper VanSteenhouse, Yumi Kasai, David Gray, Carol Clugston, Anna Dominiczak and Alex Alderton, Roberto Amato, Jeffrey Barrett, Sonia Goncalves, Ewan Harrison, David K. Jackson, Ian Johnston, Dominic Kwiatkowski, Cordelia Langford, John Sillitoe on behalf of the Wellcome Sanger Institute COVID-19 Surveillance Team |
| EPI_ISL_2806208 | Lighthouse Lab in Alderley Park | Wellcome Sanger Institute for the COVID-19 Genomics UK (COG-UK) Consortium | Jacquelyn Wynn, Mairead Hyland, The Lighthouse Lab in Alderley Park and Alex Alderton, Roberto Amato, Jeffrey Barrett, Sonia Goncalves, Ewan Harrison, David K. Jackson, Ian Johnston, Dominic Kwiatkowski, Cordelia Langford, John Sillitoe on behalf of the Wellcome Sanger Institute COVID-19 Surveillance Team |
| EPI_ISL_2806209 | Health Services Laboratories | Wellcome Sanger Institute for the COVID-19 Genomics UK (COG-UK) Consortium | Health Services Laboratories and Alex Alderton, Roberto Amato, Jeffrey Barrett, Sonia Goncalves, Ewan Harrison, David K. Jackson, Ian Johnston, Dominic Kwiatkowski, Cordelia Langford, John Sillitoe on behalf of the Wellcome Sanger Institute COVID-19 Surveillance Team |
| EPI_ISL_2806210 | Lighthouse Lab in Glasgow | Wellcome Sanger Institute for the COVID-19 Genomics UK (COG-UK) Consortium | Harper VanSteenhouse, Yumi Kasai, David Gray, Carol Clugston, Anna Dominiczak and Alex Alderton, Roberto Amato, Jeffrey Barrett, Sonia Goncalves, Ewan Harrison, David K. Jackson, Ian Johnston, Dominic Kwiatkowski, Cordelia Langford, John Sillitoe on behalf of the Wellcome Sanger Institute COVID-19 Surveillance Team |
| EPI_ISL_2806211 | Health Services Laboratories | Wellcome Sanger Institute for the COVID-19 Genomics UK (COG-UK) Consortium | Health Services Laboratories and Alex Alderton, Roberto Amato, Jeffrey Barrett, Sonia Goncalves, Ewan Harrison, David K. Jackson, Ian Johnston, Dominic Kwiatkowski, Cordelia Langford, John Sillitoe on behalf of the Wellcome Sanger Institute COVID-19 Surveillance Team |
| EPI_ISL_2806212 | Lighthouse Lab in Glasgow | Wellcome Sanger Institute for the COVID-19 Genomics UK (COG-UK) Consortium | Harper VanSteenhouse, Yumi Kasai, David Gray, Carol Clugston, Anna Dominiczak and Alex Alderton, Roberto Amato, Jeffrey Barrett, Sonia Goncalves, Ewan Harrison, David K. Jackson, Ian Johnston, Dominic Kwiatkowski, Cordelia Langford, John Sillitoe on behalf of the Wellcome Sanger Institute COVID-19 Surveillance Team |
| EPI_ISL_2806213 | Health Services Laboratories | Wellcome Sanger Institute for the COVID-19 Genomics UK (COG-UK) Consortium | Health Services Laboratories and Alex Alderton, Roberto Amato, Jeffrey Barrett, Sonia Goncalves, Ewan Harrison, David K. Jackson, Ian Johnston, Dominic Kwiatkowski, Cordelia Langford, John Sillitoe on behalf of the Wellcome Sanger Institute COVID-19 Surveillance Team |
| EPI_ISL_2806214, EPI_ISL_2806215, EPI_ISL_2806216, EPI_ISL_2806217 | Lighthouse Lab in Alderley Park | Wellcome Sanger Institute for the COVID-19 Genomics UK (COG-UK) Consortium | Jacquelyn Wynn, Mairead Hyland, The Lighthouse Lab in Alderley Park and Alex Alderton, Roberto Amato, Jeffrey Barrett, Sonia Goncalves, Ewan Harrison, David K. Jackson, Ian Johnston, Dominic Kwiatkowski, Cordelia Langford, John Sillitoe on behalf of the Wellcome Sanger Institute COVID-19 Surveillance Team |
| EPI_ISL_2806218, EPI_ISL_2806219 | Health Services Laboratories | Wellcome Sanger Institute for the COVID-19 Genomics UK (COG-UK) Consortium | Health Services Laboratories and Alex Alderton, Roberto Amato, Jeffrey Barrett, Sonia Goncalves, Ewan Harrison, David K. Jackson, Ian Johnston, Dominic Kwiatkowski, Cordelia Langford, John Sillitoe on behalf of the Wellcome Sanger Institute COVID-19 Surveillance Team |
| EPI_ISL_2806220 | Lighthouse Lab in Alderley Park | Wellcome Sanger Institute for the COVID-19 Genomics UK (COG-UK) Consortium | Jacquelyn Wynn, Mairead Hyland, The Lighthouse Lab in Alderley Park and Alex Alderton, Roberto Amato, Jeffrey Barrett, Sonia Goncalves, Ewan Harrison, David K. Jackson, Ian Johnston, Dominic Kwiatkowski, Cordelia Langford, John Sillitoe on behalf of the Wellcome Sanger Institute COVID-19 Surveillance Team |

| | | | |
|---|---|---|---|
| EPI_ISL_2806221 | Lighthouse Lab in Glasgow | Wellcome Sanger Institute for the COVID-19 Genomics UK (COG-UK) Consortium | Harper VanSteenhouse, Yumi Kasai, David Gray, Carol Clugston, Anna Dominiczak and Alex Alderton, Roberto Amato, Jeffrey Barrett, Sonia Goncalves, Ewan Harrison, David K. Jackson, Ian Johnston, Dominic Kwiatkowski, Cordelia Langford, John Sillitoe on behalf of the Wellcome Sanger Institute COVID-19 Surveillance Team |
| EPI_ISL_2806222, EPI_ISL_2806223 | Health Services Laboratories | Wellcome Sanger Institute for the COVID-19 Genomics UK (COG-UK) Consortium | Health Services Laboratories and Alex Alderton, Roberto Amato, Jeffrey Barrett, Sonia Goncalves, Ewan Harrison, David K. Jackson, Ian Johnston, Dominic Kwiatkowski, Cordelia Langford, John Sillitoe on behalf of the Wellcome Sanger Institute COVID-19 Surveillance Team |
| EPI_ISL_2806224 | Lighthouse Lab in Glasgow | Wellcome Sanger Institute for the COVID-19 Genomics UK (COG-UK) Consortium | Harper VanSteenhouse, Yumi Kasai, David Gray, Carol Clugston, Anna Dominiczak and Alex Alderton, Roberto Amato, Jeffrey Barrett, Sonia Goncalves, Ewan Harrison, David K. Jackson, Ian Johnston, Dominic Kwiatkowski, Cordelia Langford, John Sillitoe on behalf of the Wellcome Sanger Institute COVID-19 Surveillance Team |
| EPI_ISL_2806225 | Lighthouse Lab in Alderley Park | Wellcome Sanger Institute for the COVID-19 Genomics UK (COG-UK) Consortium | Jacquelyn Wynn, Mairead Hyland, The Lighthouse Lab in Alderley Park and Alex Alderton, Roberto Amato, Jeffrey Barrett, Sonia Goncalves, Ewan Harrison, David K. Jackson, Ian Johnston, Dominic Kwiatkowski, Cordelia Langford, John Sillitoe on behalf of the Wellcome Sanger Institute COVID-19 Surveillance Team |
| EPI_ISL_2806226 | Lighthouse Lab in Glasgow | Wellcome Sanger Institute for the COVID-19 Genomics UK (COG-UK) Consortium | Harper VanSteenhouse, Yumi Kasai, David Gray, Carol Clugston, Anna Dominiczak and Alex Alderton, Roberto Amato, Jeffrey Barrett, Sonia Goncalves, Ewan Harrison, David K. Jackson, Ian Johnston, Dominic Kwiatkowski, Cordelia Langford, John Sillitoe on behalf of the Wellcome Sanger Institute COVID-19 Surveillance Team |
| EPI_ISL_2806227 | Lighthouse Lab in Alderley Park | Wellcome Sanger Institute for the COVID-19 Genomics UK (COG-UK) Consortium | Jacquelyn Wynn, Mairead Hyland, The Lighthouse Lab in Alderley Park and Alex Alderton, Roberto Amato, Jeffrey Barrett, Sonia Goncalves, Ewan Harrison, David K. Jackson, Ian Johnston, Dominic Kwiatkowski, Cordelia Langford, John Sillitoe on behalf of the Wellcome Sanger Institute COVID-19 Surveillance Team |
| EPI_ISL_2806229 | Health Services Laboratories | Wellcome Sanger Institute for the COVID-19 Genomics UK (COG-UK) Consortium | Health Services Laboratories and Alex Alderton, Roberto Amato, Jeffrey Barrett, Sonia Goncalves, Ewan Harrison, David K. Jackson, Ian Johnston, Dominic Kwiatkowski, Cordelia Langford, John Sillitoe on behalf of the Wellcome Sanger Institute COVID-19 Surveillance Team |
| EPI_ISL_2806230 | Lighthouse Lab in Alderley Park | Wellcome Sanger Institute for the COVID-19 Genomics UK (COG-UK) Consortium | Jacquelyn Wynn, Mairead Hyland, The Lighthouse Lab in Alderley Park and Alex Alderton, Roberto Amato, Jeffrey Barrett, Sonia Goncalves, Ewan Harrison, David K. Jackson, Ian Johnston, Dominic Kwiatkowski, Cordelia Langford, John Sillitoe on behalf of the Wellcome Sanger Institute COVID-19 Surveillance Team |
| EPI_ISL_2806231 | Health Services Laboratories | Wellcome Sanger Institute for the COVID-19 Genomics UK (COG-UK) Consortium | Health Services Laboratories and Alex Alderton, Roberto Amato, Jeffrey Barrett, Sonia Goncalves, Ewan Harrison, David K. Jackson, Ian Johnston, Dominic Kwiatkowski, Cordelia Langford, John Sillitoe on behalf of the Wellcome Sanger Institute COVID-19 Surveillance Team |
| EPI_ISL_2806232, EPI_ISL_2806233, EPI_ISL_2806234 | Lighthouse Lab in Alderley Park | Wellcome Sanger Institute for the COVID-19 Genomics UK (COG-UK) Consortium | Jacquelyn Wynn, Mairead Hyland, The Lighthouse Lab in Alderley Park and Alex Alderton, Roberto Amato, Jeffrey Barrett, Sonia Goncalves, Ewan Harrison, David K. Jackson, Ian Johnston, Dominic Kwiatkowski, Cordelia Langford, John Sillitoe on behalf of the Wellcome Sanger Institute COVID-19 Surveillance Team |
| EPI_ISL_2806235, EPI_ISL_2806236 | Health Services Laboratories | Wellcome Sanger Institute for the COVID-19 Genomics UK (COG-UK) Consortium | Health Services Laboratories and Alex Alderton, Roberto Amato, Jeffrey Barrett, Sonia Goncalves, Ewan Harrison, David K. Jackson, Ian Johnston, Dominic Kwiatkowski, Cordelia Langford, John Sillitoe on behalf of the Wellcome Sanger Institute COVID-19 Surveillance Team |
| EPI_ISL_2806237 | Lighthouse Lab in Alderley Park | Wellcome Sanger Institute for the COVID-19 Genomics UK (COG-UK) Consortium | Jacquelyn Wynn, Mairead Hyland, The Lighthouse Lab in Alderley Park and Alex Alderton, Roberto Amato, Jeffrey Barrett, Sonia Goncalves, Ewan Harrison, David K. Jackson, Ian Johnston, Dominic Kwiatkowski, Cordelia Langford, John Sillitoe on behalf of the Wellcome Sanger Institute COVID-19 Surveillance Team |
| EPI_ISL_2806238 | Health Services Laboratories | Wellcome Sanger Institute for the COVID-19 Genomics UK (COG-UK) Consortium | Health Services Laboratories and Alex Alderton, Roberto Amato, Jeffrey Barrett, Sonia Goncalves, Ewan Harrison, David K. Jackson, Ian Johnston, Dominic Kwiatkowski, Cordelia Langford, John Sillitoe on behalf of the Wellcome Sanger Institute COVID-19 Surveillance Team |
| EPI_ISL_2806239 | Lighthouse Lab in Glasgow | Wellcome Sanger Institute for the COVID-19 Genomics UK (COG-UK) Consortium | Harper VanSteenhouse, Yumi Kasai, David Gray, Carol Clugston, Anna Dominiczak and Alex Alderton, Roberto Amato, Jeffrey Barrett, Sonia Goncalves, Ewan Harrison, David K. Jackson, Ian Johnston, Dominic Kwiatkowski, Cordelia Langford, John Sillitoe on behalf of the Wellcome Sanger Institute COVID-19 Surveillance Team |
| EPI_ISL_2806240 | Lighthouse Lab in Alderley Park | Wellcome Sanger Institute for the COVID-19 Genomics UK (COG-UK) Consortium | Jacquelyn Wynn, Mairead Hyland, The Lighthouse Lab in Alderley Park and Alex Alderton, Roberto Amato, Jeffrey Barrett, Sonia Goncalves, Ewan Harrison, David K. Jackson, Ian Johnston, Dominic Kwiatkowski, Cordelia Langford, John Sillitoe on behalf of the Wellcome Sanger Institute COVID-19 Surveillance Team |
| EPI_ISL_2806241, EPI_ISL_2806242, EPI_ISL_2806243, EPI_ISL_2806244 | Lighthouse Lab in Glasgow | Wellcome Sanger Institute for the COVID-19 Genomics UK (COG-UK) Consortium | Harper VanSteenhouse, Yumi Kasai, David Gray, Carol Clugston, Anna Dominiczak and Alex Alderton, Roberto Amato, Jeffrey Barrett, Sonia Goncalves, Ewan Harrison, David K. Jackson, Ian Johnston, Dominic Kwiatkowski, Cordelia Langford, John Sillitoe on behalf of the Wellcome Sanger Institute COVID-19 Surveillance Team |
| EPI_ISL_2806245, EPI_ISL_2806246 | Health Services Laboratories | Wellcome Sanger Institute for the COVID-19 Genomics UK (COG-UK) Consortium | Health Services Laboratories and Alex Alderton, Roberto Amato, Jeffrey Barrett, Sonia Goncalves, Ewan Harrison, David K. Jackson, Ian Johnston, Dominic Kwiatkowski, Cordelia Langford, John Sillitoe on behalf of the Wellcome Sanger Institute COVID-19 Surveillance Team |
| EPI_ISL_2806247 | Lighthouse Lab in Alderley Park | Wellcome Sanger Institute for the COVID-19 Genomics UK (COG-UK) Consortium | Jacquelyn Wynn, Mairead Hyland, The Lighthouse Lab in Alderley Park and Alex Alderton, Roberto Amato, Jeffrey Barrett, Sonia Goncalves, Ewan Harrison, David K. Jackson, Ian Johnston, Dominic Kwiatkowski, Cordelia Langford, John Sillitoe on behalf of the Wellcome Sanger Institute COVID-19 Surveillance Team |
| EPI_ISL_2806248 | Health Services Laboratories | Wellcome Sanger Institute for the COVID-19 Genomics UK (COG-UK) Consortium | Health Services Laboratories and Alex Alderton, Roberto Amato, Jeffrey Barrett, Sonia Goncalves, Ewan Harrison, David K. Jackson, Ian Johnston, Dominic Kwiatkowski, Cordelia Langford, John Sillitoe on behalf of the Wellcome Sanger Institute COVID-19 Surveillance Team |
| EPI_ISL_2806249 | Lighthouse Lab in Alderley Park | Wellcome Sanger Institute for the COVID-19 Genomics UK (COG-UK) Consortium | Jacquelyn Wynn, Mairead Hyland, The Lighthouse Lab in Alderley Park and Alex Alderton, Roberto Amato, Jeffrey Barrett, Sonia Goncalves, Ewan Harrison, David K. Jackson, Ian Johnston, Dominic Kwiatkowski, Cordelia Langford, John Sillitoe on behalf of the Wellcome Sanger Institute COVID-19 Surveillance Team |
| EPI_ISL_2806250, EPI_ISL_2806251 | Lighthouse Lab in Glasgow | Wellcome Sanger Institute for the COVID-19 Genomics UK (COG-UK) Consortium | Harper VanSteenhouse, Yumi Kasai, David Gray, Carol Clugston, Anna Dominiczak and Alex Alderton, Roberto Amato, Jeffrey Barrett, Sonia Goncalves, Ewan Harrison, David K. Jackson, Ian Johnston, Dominic Kwiatkowski, Cordelia Langford, John Sillitoe on behalf of the Wellcome Sanger Institute COVID-19 Surveillance Team |
| EPI_ISL_2806252, EPI_ISL_2806253, EPI_ISL_2806254, EPI_ISL_2806255 | Lighthouse Lab in Alderley Park | Wellcome Sanger Institute for the COVID-19 Genomics UK (COG-UK) Consortium | Jacquelyn Wynn, Mairead Hyland, The Lighthouse Lab in Alderley Park and Alex Alderton, Roberto Amato, Jeffrey Barrett, Sonia Goncalves, Ewan Harrison, David K. Jackson, Ian Johnston, Dominic Kwiatkowski, Cordelia Langford, John Sillitoe on behalf of the Wellcome Sanger Institute COVID-19 Surveillance Team |
| EPI_ISL_2806256 | Health Services Laboratories | Wellcome Sanger Institute for the COVID-19 Genomics UK (COG-UK) Consortium | Health Services Laboratories and Alex Alderton, Roberto Amato, Jeffrey Barrett, Sonia Goncalves, Ewan Harrison, David K. Jackson, Ian Johnston, Dominic Kwiatkowski, Cordelia Langford, John Sillitoe on behalf of the Wellcome Sanger Institute COVID-19 Surveillance Team |
| EPI_ISL_2806257 | Lighthouse Lab in Alderley Park | Wellcome Sanger Institute for the COVID-19 Genomics UK (COG-UK) Consortium | Jacquelyn Wynn, Mairead Hyland, The Lighthouse Lab in Alderley Park and Alex Alderton, Roberto Amato, Jeffrey Barrett, Sonia Goncalves, Ewan Harrison, David K. Jackson, Ian Johnston, Dominic Kwiatkowski, Cordelia Langford, John Sillitoe on behalf of the Wellcome Sanger Institute COVID-19 Surveillance Team |
| EPI_ISL_2806258 | Health Services Laboratories | Wellcome Sanger Institute for the COVID-19 Genomics UK (COG-UK) Consortium | Health Services Laboratories and Alex Alderton, Roberto Amato, Jeffrey Barrett, Sonia Goncalves, Ewan Harrison, David K. Jackson, Ian Johnston, Dominic Kwiatkowski, Cordelia Langford, John Sillitoe on behalf of the Wellcome Sanger Institute COVID-19 Surveillance Team |
| EPI_ISL_2806259 | Lighthouse Lab in Glasgow | Wellcome Sanger Institute for the COVID-19 Genomics UK (COG-UK) Consortium | Harper VanSteenhouse, Yumi Kasai, David Gray, Carol Clugston, Anna Dominiczak and Alex Alderton, Roberto Amato, Jeffrey Barrett, Sonia Goncalves, Ewan Harrison, David K. Jackson, Ian Johnston, Dominic Kwiatkowski, Cordelia Langford, John Sillitoe on behalf of the Wellcome Sanger Institute COVID-19 Surveillance Team |
| EPI_ISL_2806260, EPI_ISL_2806261 | Health Services Laboratories | Wellcome Sanger Institute for the COVID-19 Genomics UK (COG-UK) Consortium | Health Services Laboratories and Alex Alderton, Roberto Amato, Jeffrey Barrett, Sonia Goncalves, Ewan Harrison, David K. Jackson, Ian Johnston, Dominic Kwiatkowski, Cordelia Langford, John Sillitoe on behalf of the Wellcome Sanger Institute COVID-19 Surveillance Team |
| EPI_ISL_2806262 | Lighthouse Lab in Alderley Park | Wellcome Sanger Institute for the COVID-19 Genomics UK (COG-UK) Consortium | Jacquelyn Wynn, Mairead Hyland, The Lighthouse Lab in Alderley Park and Alex Alderton, Roberto Amato, Jeffrey Barrett, Sonia Goncalves, Ewan Harrison, David K. Jackson, Ian Johnston, Dominic Kwiatkowski, Cordelia Langford, John Sillitoe on behalf of the Wellcome Sanger Institute COVID-19 Surveillance Team |
| EPI_ISL_2806263 | Health Services Laboratories | Wellcome Sanger Institute for the COVID-19 Genomics UK (COG-UK) Consortium | Health Services Laboratories and Alex Alderton, Roberto Amato, Jeffrey Barrett, Sonia Goncalves, Ewan Harrison, David K. Jackson, Ian Johnston, Dominic Kwiatkowski, Cordelia Langford, John Sillitoe on behalf of the Wellcome Sanger Institute COVID-19 Surveillance Team |
| EPI_ISL_2806264 | Lighthouse Lab in Alderley Park | Wellcome Sanger Institute for the COVID-19 Genomics UK (COG-UK) Consortium | Jacquelyn Wynn, Mairead Hyland, The Lighthouse Lab in Alderley Park and Alex Alderton, Roberto Amato, Jeffrey Barrett, Sonia Goncalves, Ewan Harrison, David K. Jackson, Ian Johnston, Dominic Kwiatkowski, Cordelia Langford, John Sillitoe on behalf of the Wellcome Sanger Institute COVID-19 Surveillance Team |
| EPI_ISL_2806265 | Lighthouse Lab in Glasgow | Wellcome Sanger Institute for the COVID-19 Genomics UK (COG-UK) Consortium | Harper VanSteenhouse, Yumi Kasai, David Gray, Carol Clugston, Anna Dominiczak and Alex Alderton, Roberto Amato, Jeffrey Barrett, Sonia Goncalves, Ewan Harrison, David K. Jackson, Ian Johnston, Dominic Kwiatkowski, Cordelia Langford, John Sillitoe on behalf of the Wellcome Sanger Institute COVID-19 Surveillance Team |
| EPI_ISL_2806266 | Lighthouse Lab in Alderley Park | Wellcome Sanger Institute for the COVID-19 Genomics UK (COG-UK) Consortium | Jacquelyn Wynn, Mairead Hyland, The Lighthouse Lab in Alderley Park and Alex Alderton, Roberto Amato, Jeffrey Barrett, Sonia Goncalves, Ewan Harrison, David K. Jackson, Ian Johnston, Dominic Kwiatkowski, Cordelia Langford, John Sillitoe on behalf of the Wellcome Sanger Institute COVID-19 Surveillance Team |
| EPI_ISL_2806267 | Health Services Laboratories | Wellcome Sanger Institute for the COVID-19 Genomics UK (COG-UK) Consortium | Health Services Laboratories and Alex Alderton, Roberto Amato, Jeffrey Barrett, Sonia Goncalves, Ewan Harrison, David K. Jackson, Ian Johnston, Dominic Kwiatkowski, Cordelia Langford, John Sillitoe on behalf of the Wellcome Sanger Institute COVID-19 Surveillance Team |
| EPI_ISL_2806268 | Lighthouse Lab in Alderley Park | Wellcome Sanger Institute for the COVID-19 Genomics UK (COG-UK) Consortium | Jacquelyn Wynn, Mairead Hyland, The Lighthouse Lab in Alderley Park and Alex Alderton, Roberto Amato, Jeffrey Barrett, Sonia Goncalves, Ewan Harrison, David K. Jackson, Ian Johnston, Dominic Kwiatkowski, Cordelia Langford, John Sillitoe on behalf of the Wellcome Sanger Institute COVID-19 Surveillance Team |
| EPI_ISL_2806269, EPI_ISL_2806270, EPI_ISL_2806271 | Health Services Laboratories | Wellcome Sanger Institute for the COVID-19 Genomics UK (COG-UK) Consortium | Health Services Laboratories and Alex Alderton, Roberto Amato, Jeffrey Barrett, Sonia Goncalves, Ewan Harrison, David K. Jackson, Ian Johnston, Dominic Kwiatkowski, Cordelia Langford, John Sillitoe on behalf of the Wellcome Sanger Institute COVID-19 Surveillance Team |
| EPI_ISL_2806272 | Lighthouse Lab in Glasgow | Wellcome Sanger Institute for the COVID-19 Genomics UK (COG-UK) Consortium | Harper VanSteenhouse, Yumi Kasai, David Gray, Carol Clugston, Anna Dominiczak and Alex Alderton, Roberto Amato, Jeffrey Barrett, Sonia Goncalves, Ewan Harrison, David K. Jackson, Ian Johnston, Dominic Kwiatkowski, Cordelia Langford, John Sillitoe on behalf of the Wellcome Sanger Institute COVID-19 Surveillance Team |
| EPI_ISL_2806273 | Lighthouse Lab in Alderley Park | Wellcome Sanger Institute for the COVID-19 Genomics UK (COG-UK) Consortium | Jacquelyn Wynn, Mairead Hyland, The Lighthouse Lab in Alderley Park and Alex Alderton, Roberto Amato, Jeffrey Barrett, Sonia Goncalves, Ewan Harrison, David K. Jackson, Ian Johnston, Dominic Kwiatkowski, Cordelia Langford, John Sillitoe on behalf of the Wellcome Sanger Institute COVID-19 Surveillance Team |
| EPI_ISL_2806274 | Health Services Laboratories | Wellcome Sanger Institute for the COVID-19 Genomics UK (COG-UK) Consortium | Health Services Laboratories and Alex Alderton, Roberto Amato, Jeffrey Barrett, Sonia Goncalves, Ewan Harrison, David K. Jackson, Ian Johnston, Dominic Kwiatkowski, Cordelia Langford, John Sillitoe on behalf of the Wellcome Sanger Institute COVID-19 Surveillance Team |
| EPI_ISL_2806275 | Lighthouse Lab in Alderley Park | Wellcome Sanger Institute for the COVID-19 Genomics UK (COG-UK) Consortium | Jacquelyn Wynn, Mairead Hyland, The Lighthouse Lab in Alderley Park and Alex Alderton, Roberto Amato, Jeffrey Barrett, Sonia Goncalves, Ewan Harrison, David K. Jackson, Ian Johnston, Dominic Kwiatkowski, Cordelia Langford, John Sillitoe on behalf of the Wellcome Sanger Institute COVID-19 Surveillance Team |
| EPI_ISL_2806276 | Health Services Laboratories | Wellcome Sanger Institute for the COVID-19 Genomics UK (COG-UK) Consortium | Health Services Laboratories and Alex Alderton, Roberto Amato, Jeffrey Barrett, Sonia Goncalves, Ewan Harrison, David K. Jackson, Ian Johnston, Dominic Kwiatkowski, Cordelia Langford, John Sillitoe on behalf of the Wellcome Sanger Institute COVID-19 Surveillance Team |
| EPI_ISL_2806277 | Lighthouse Lab in Alderley Park | Wellcome Sanger Institute for the COVID-19 Genomics UK (COG-UK) Consortium | Jacquelyn Wynn, Mairead Hyland, The Lighthouse Lab in Alderley Park and Alex Alderton, Roberto Amato, Jeffrey Barrett, Sonia Goncalves, Ewan Harrison, David K. Jackson, Ian Johnston, Dominic Kwiatkowski, Cordelia Langford, John Sillitoe on behalf of the Wellcome Sanger Institute COVID-19 Surveillance Team |
| EPI_ISL_2806278, EPI_ISL_2806279 | Health Services Laboratories | Wellcome Sanger Institute for the COVID-19 Genomics UK (COG-UK) Consortium | Health Services Laboratories and Alex Alderton, Roberto Amato, Jeffrey Barrett, Sonia Goncalves, Ewan Harrison, David K. Jackson, Ian Johnston, Dominic Kwiatkowski, Cordelia Langford, John Sillitoe on behalf of the Wellcome Sanger Institute COVID-19 Surveillance Team |
| EPI_ISL_2806280 | Lighthouse Lab in Glasgow | Wellcome Sanger Institute for the COVID-19 Genomics UK (COG-UK) Consortium | Harper VanSteenhouse, Yumi Kasai, David Gray, Carol Clugston, Anna Dominiczak and Alex Alderton, Roberto Amato, Jeffrey Barrett, Sonia Goncalves, Ewan Harrison, David K. Jackson, Ian Johnston, Dominic Kwiatkowski, Cordelia Langford, John Sillitoe on behalf of the Wellcome Sanger Institute COVID-19 Surveillance Team |

| | | | |
|---|---|---|---|
| EPI_ISL_2806281, EPI_ISL_2806282, EPI_ISL_2806283, EPI_ISL_2806284, EPI_ISL_2806285 | Lighthouse Lab in Alderley Park | Wellcome Sanger Institute for the COVID-19 Genomics UK (COG-UK) Consortium | Jacquelyn Wynn, Mairead Hyland, The Lighthouse Lab in Alderley Park and Alex Alderton, Roberto Amato, Jeffrey Barrett, Sonia Goncalves, Ewan Harrison, David K. Jackson, Ian Johnston, Dominic Kwiatkowski, Cordelia Langford, John Sillitoe on behalf of the Wellcome Sanger Institute COVID-19 Surveillance Team |
| EPI_ISL_2806286 | Health Services Laboratories | Wellcome Sanger Institute for the COVID-19 Genomics UK (COG-UK) Consortium | Health Services Laboratories and Alex Alderton, Roberto Amato, Jeffrey Barrett, Sonia Goncalves, Ewan Harrison, David K. Jackson, Ian Johnston, Dominic Kwiatkowski, Cordelia Langford, John Sillitoe on behalf of the Wellcome Sanger Institute COVID-19 Surveillance Team |
| EPI_ISL_2806287 | Lighthouse Lab in Alderley Park | Wellcome Sanger Institute for the COVID-19 Genomics UK (COG-UK) Consortium | Jacquelyn Wynn, Mairead Hyland, The Lighthouse Lab in Alderley Park and Alex Alderton, Roberto Amato, Jeffrey Barrett, Sonia Goncalves, Ewan Harrison, David K. Jackson, Ian Johnston, Dominic Kwiatkowski, Cordelia Langford, John Sillitoe on behalf of the Wellcome Sanger Institute COVID-19 Surveillance Team |
| EPI_ISL_2806288 | Health Services Laboratories | Wellcome Sanger Institute for the COVID-19 Genomics UK (COG-UK) Consortium | Health Services Laboratories and Alex Alderton, Roberto Amato, Jeffrey Barrett, Sonia Goncalves, Ewan Harrison, David K. Jackson, Ian Johnston, Dominic Kwiatkowski, Cordelia Langford, John Sillitoe on behalf of the Wellcome Sanger Institute COVID-19 Surveillance Team |
| EPI_ISL_2806289, EPI_ISL_2806290 | Lighthouse Lab in Glasgow | Wellcome Sanger Institute for the COVID-19 Genomics UK (COG-UK) Consortium | Harper VanSteenhouse, Yumi Kasai, David Gray, Carol Clugston, Anna Dominiczak and Alex Alderton, Roberto Amato, Jeffrey Barrett, Sonia Goncalves, Ewan Harrison, David K. Jackson, Ian Johnston, Dominic Kwiatkowski, Cordelia Langford, John Sillitoe on behalf of the Wellcome Sanger Institute COVID-19 Surveillance Team |
| EPI_ISL_2806291 | Lighthouse Lab in Alderley Park | Wellcome Sanger Institute for the COVID-19 Genomics UK (COG-UK) Consortium | Jacquelyn Wynn, Mairead Hyland, The Lighthouse Lab in Alderley Park and Alex Alderton, Roberto Amato, Jeffrey Barrett, Sonia Goncalves, Ewan Harrison, David K. Jackson, Ian Johnston, Dominic Kwiatkowski, Cordelia Langford, John Sillitoe on behalf of the Wellcome Sanger Institute COVID-19 Surveillance Team |
| EPI_ISL_2806292 | Lighthouse Lab in Glasgow | Wellcome Sanger Institute for the COVID-19 Genomics UK (COG-UK) Consortium | Harper VanSteenhouse, Yumi Kasai, David Gray, Carol Clugston, Anna Dominiczak and Alex Alderton, Roberto Amato, Jeffrey Barrett, Sonia Goncalves, Ewan Harrison, David K. Jackson, Ian Johnston, Dominic Kwiatkowski, Cordelia Langford, John Sillitoe on behalf of the Wellcome Sanger Institute COVID-19 Surveillance Team |
| EPI_ISL_2806293, EPI_ISL_2806294 | Health Services Laboratories | Wellcome Sanger Institute for the COVID-19 Genomics UK (COG-UK) Consortium | Health Services Laboratories and Alex Alderton, Roberto Amato, Jeffrey Barrett, Sonia Goncalves, Ewan Harrison, David K. Jackson, Ian Johnston, Dominic Kwiatkowski, Cordelia Langford, John Sillitoe on behalf of the Wellcome Sanger Institute COVID-19 Surveillance Team |
| EPI_ISL_2806295 | Lighthouse Lab in Glasgow | Wellcome Sanger Institute for the COVID-19 Genomics UK (COG-UK) Consortium | Harper VanSteenhouse, Yumi Kasai, David Gray, Carol Clugston, Anna Dominiczak and Alex Alderton, Roberto Amato, Jeffrey Barrett, Sonia Goncalves, Ewan Harrison, David K. Jackson, Ian Johnston, Dominic Kwiatkowski, Cordelia Langford, John Sillitoe on behalf of the Wellcome Sanger Institute COVID-19 Surveillance Team |
| EPI_ISL_2806296, EPI_ISL_2806297 | Health Services Laboratories | Wellcome Sanger Institute for the COVID-19 Genomics UK (COG-UK) Consortium | Health Services Laboratories and Alex Alderton, Roberto Amato, Jeffrey Barrett, Sonia Goncalves, Ewan Harrison, David K. Jackson, Ian Johnston, Dominic Kwiatkowski, Cordelia Langford, John Sillitoe on behalf of the Wellcome Sanger Institute COVID-19 Surveillance Team |
| EPI_ISL_2806298 | Lighthouse Lab in Glasgow | Wellcome Sanger Institute for the COVID-19 Genomics UK (COG-UK) Consortium | Harper VanSteenhouse, Yumi Kasai, David Gray, Carol Clugston, Anna Dominiczak and Alex Alderton, Roberto Amato, Jeffrey Barrett, Sonia Goncalves, Ewan Harrison, David K. Jackson, Ian Johnston, Dominic Kwiatkowski, Cordelia Langford, John Sillitoe on behalf of the Wellcome Sanger Institute COVID-19 Surveillance Team |
| EPI_ISL_2806299 | Lighthouse Lab in Alderley Park | Wellcome Sanger Institute for the COVID-19 Genomics UK (COG-UK) Consortium | Jacquelyn Wynn, Mairead Hyland, The Lighthouse Lab in Alderley Park and Alex Alderton, Roberto Amato, Jeffrey Barrett, Sonia Goncalves, Ewan Harrison, David K. Jackson, Ian Johnston, Dominic Kwiatkowski, Cordelia Langford, John Sillitoe on behalf of the Wellcome Sanger Institute COVID-19 Surveillance Team |
| EPI_ISL_2806300 | Lighthouse Lab in Glasgow | Wellcome Sanger Institute for the COVID-19 Genomics UK (COG-UK) Consortium | Harper VanSteenhouse, Yumi Kasai, David Gray, Carol Clugston, Anna Dominiczak and Alex Alderton, Roberto Amato, Jeffrey Barrett, Sonia Goncalves, Ewan Harrison, David K. Jackson, Ian Johnston, Dominic Kwiatkowski, Cordelia Langford, John Sillitoe on behalf of the Wellcome Sanger Institute COVID-19 Surveillance Team |
| EPI_ISL_2806301, EPI_ISL_2806302 | Health Services Laboratories | Wellcome Sanger Institute for the COVID-19 Genomics UK (COG-UK) Consortium | Health Services Laboratories and Alex Alderton, Roberto Amato, Jeffrey Barrett, Sonia Goncalves, Ewan Harrison, David K. Jackson, Ian Johnston, Dominic Kwiatkowski, Cordelia Langford, John Sillitoe on behalf of the Wellcome Sanger Institute COVID-19 Surveillance Team |
| EPI_ISL_2806303, EPI_ISL_2806304 | Lighthouse Lab in Alderley Park | Wellcome Sanger Institute for the COVID-19 Genomics UK (COG-UK) Consortium | Jacquelyn Wynn, Mairead Hyland, The Lighthouse Lab in Alderley Park and Alex Alderton, Roberto Amato, Jeffrey Barrett, Sonia Goncalves, Ewan Harrison, David K. Jackson, Ian Johnston, Dominic Kwiatkowski, Cordelia Langford, John Sillitoe on behalf of the Wellcome Sanger Institute COVID-19 Surveillance Team |
| EPI_ISL_2806305, EPI_ISL_2806306 | Health Services Laboratories | Wellcome Sanger Institute for the COVID-19 Genomics UK (COG-UK) Consortium | Health Services Laboratories and Alex Alderton, Roberto Amato, Jeffrey Barrett, Sonia Goncalves, Ewan Harrison, David K. Jackson, Ian Johnston, Dominic Kwiatkowski, Cordelia Langford, John Sillitoe on behalf of the Wellcome Sanger Institute COVID-19 Surveillance Team |
| EPI_ISL_2806307 | Lighthouse Lab in Alderley Park | Wellcome Sanger Institute for the COVID-19 Genomics UK (COG-UK) Consortium | Jacquelyn Wynn, Mairead Hyland, The Lighthouse Lab in Alderley Park and Alex Alderton, Roberto Amato, Jeffrey Barrett, Sonia Goncalves, Ewan Harrison, David K. Jackson, Ian Johnston, Dominic Kwiatkowski, Cordelia Langford, John Sillitoe on behalf of the Wellcome Sanger Institute COVID-19 Surveillance Team |
| EPI_ISL_2806308 | Health Services Laboratories | Wellcome Sanger Institute for the COVID-19 Genomics UK (COG-UK) Consortium | Health Services Laboratories and Alex Alderton, Roberto Amato, Jeffrey Barrett, Sonia Goncalves, Ewan Harrison, David K. Jackson, Ian Johnston, Dominic Kwiatkowski, Cordelia Langford, John Sillitoe on behalf of the Wellcome Sanger Institute COVID-19 Surveillance Team |
| EPI_ISL_2806309, EPI_ISL_2806310 | Lighthouse Lab in Glasgow | Wellcome Sanger Institute for the COVID-19 Genomics UK (COG-UK) Consortium | Harper VanSteenhouse, Yumi Kasai, David Gray, Carol Clugston, Anna Dominiczak and Alex Alderton, Roberto Amato, Jeffrey Barrett, Sonia Goncalves, Ewan Harrison, David K. Jackson, Ian Johnston, Dominic Kwiatkowski, Cordelia Langford, John Sillitoe on behalf of the Wellcome Sanger Institute COVID-19 Surveillance Team |
| EPI_ISL_2806311, EPI_ISL_2806312 | Lighthouse Lab in Alderley Park | Wellcome Sanger Institute for the COVID-19 Genomics UK (COG-UK) Consortium | Jacquelyn Wynn, Mairead Hyland, The Lighthouse Lab in Alderley Park and Alex Alderton, Roberto Amato, Jeffrey Barrett, Sonia Goncalves, Ewan Harrison, David K. Jackson, Ian Johnston, Dominic Kwiatkowski, Cordelia Langford, John Sillitoe on behalf of the Wellcome Sanger Institute COVID-19 Surveillance Team |
| EPI_ISL_2806313 | Health Services Laboratories | Wellcome Sanger Institute for the COVID-19 Genomics UK (COG-UK) Consortium | Health Services Laboratories and Alex Alderton, Roberto Amato, Jeffrey Barrett, Sonia Goncalves, Ewan Harrison, David K. Jackson, Ian Johnston, Dominic Kwiatkowski, Cordelia Langford, John Sillitoe on behalf of the Wellcome Sanger Institute COVID-19 Surveillance Team |
| EPI_ISL_2806314, EPI_ISL_2806315, EPI_ISL_2806316 | Lighthouse Lab in Glasgow | Wellcome Sanger Institute for the COVID-19 Genomics UK (COG-UK) Consortium | Harper VanSteenhouse, Yumi Kasai, David Gray, Carol Clugston, Anna Dominiczak and Alex Alderton, Roberto Amato, Jeffrey Barrett, Sonia Goncalves, Ewan Harrison, David K. Jackson, Ian Johnston, Dominic Kwiatkowski, Cordelia Langford, John Sillitoe on behalf of the Wellcome Sanger Institute COVID-19 Surveillance Team |
| EPI_ISL_2806317 | Health Services Laboratories | Wellcome Sanger Institute for the COVID-19 Genomics UK (COG-UK) Consortium | Health Services Laboratories and Alex Alderton, Roberto Amato, Jeffrey Barrett, Sonia Goncalves, Ewan Harrison, David K. Jackson, Ian Johnston, Dominic Kwiatkowski, Cordelia Langford, John Sillitoe on behalf of the Wellcome Sanger Institute COVID-19 Surveillance Team |
| EPI_ISL_2806318 | Lighthouse Lab in Glasgow | Wellcome Sanger Institute for the COVID-19 Genomics UK (COG-UK) Consortium | Harper VanSteenhouse, Yumi Kasai, David Gray, Carol Clugston, Anna Dominiczak and Alex Alderton, Roberto Amato, Jeffrey Barrett, Sonia Goncalves, Ewan Harrison, David K. Jackson, Ian Johnston, Dominic Kwiatkowski, Cordelia Langford, John Sillitoe on behalf of the Wellcome Sanger Institute COVID-19 Surveillance Team |
| EPI_ISL_2806319 | Lighthouse Lab in Alderley Park | Wellcome Sanger Institute for the COVID-19 Genomics UK (COG-UK) Consortium | Jacquelyn Wynn, Mairead Hyland, The Lighthouse Lab in Alderley Park and Alex Alderton, Roberto Amato, Jeffrey Barrett, Sonia Goncalves, Ewan Harrison, David K. Jackson, Ian Johnston, Dominic Kwiatkowski, Cordelia Langford, John Sillitoe on behalf of the Wellcome Sanger Institute COVID-19 Surveillance Team |
| EPI_ISL_2806320 | Lighthouse Lab in Glasgow | Wellcome Sanger Institute for the COVID-19 Genomics UK (COG-UK) Consortium | Harper VanSteenhouse, Yumi Kasai, David Gray, Carol Clugston, Anna Dominiczak and Alex Alderton, Roberto Amato, Jeffrey Barrett, Sonia Goncalves, Ewan Harrison, David K. Jackson, Ian Johnston, Dominic Kwiatkowski, Cordelia Langford, John Sillitoe on behalf of the Wellcome Sanger Institute COVID-19 Surveillance Team |
| EPI_ISL_2806321 | Health Services Laboratories | Wellcome Sanger Institute for the COVID-19 Genomics UK (COG-UK) Consortium | Health Services Laboratories and Alex Alderton, Roberto Amato, Jeffrey Barrett, Sonia Goncalves, Ewan Harrison, David K. Jackson, Ian Johnston, Dominic Kwiatkowski, Cordelia Langford, John Sillitoe on behalf of the Wellcome Sanger Institute COVID-19 Surveillance Team |
| EPI_ISL_2806323 | Lighthouse Lab in Glasgow | Wellcome Sanger Institute for the COVID-19 Genomics UK (COG-UK) Consortium | Harper VanSteenhouse, Yumi Kasai, David Gray, Carol Clugston, Anna Dominiczak and Alex Alderton, Roberto Amato, Jeffrey Barrett, Sonia Goncalves, Ewan Harrison, David K. Jackson, Ian Johnston, Dominic Kwiatkowski, Cordelia Langford, John Sillitoe on behalf of the Wellcome Sanger Institute COVID-19 Surveillance Team |
| EPI_ISL_2806324 | Lighthouse Lab in Alderley Park | Wellcome Sanger Institute for the COVID-19 Genomics UK (COG-UK) Consortium | Jacquelyn Wynn, Mairead Hyland, The Lighthouse Lab in Alderley Park and Alex Alderton, Roberto Amato, Jeffrey Barrett, Sonia Goncalves, Ewan Harrison, David K. Jackson, Ian Johnston, Dominic Kwiatkowski, Cordelia Langford, John Sillitoe on behalf of the Wellcome Sanger Institute COVID-19 Surveillance Team |
| EPI_ISL_2806325, EPI_ISL_2806326 | Health Services Laboratories | Wellcome Sanger Institute for the COVID-19 Genomics UK (COG-UK) Consortium | Health Services Laboratories and Alex Alderton, Roberto Amato, Jeffrey Barrett, Sonia Goncalves, Ewan Harrison, David K. Jackson, Ian Johnston, Dominic Kwiatkowski, Cordelia Langford, John Sillitoe on behalf of the Wellcome Sanger Institute COVID-19 Surveillance Team |
| EPI_ISL_2806327 | Lighthouse Lab in Alderley Park | Wellcome Sanger Institute for the COVID-19 Genomics UK (COG-UK) Consortium | Jacquelyn Wynn, Mairead Hyland, The Lighthouse Lab in Alderley Park and Alex Alderton, Roberto Amato, Jeffrey Barrett, Sonia Goncalves, Ewan Harrison, David K. Jackson, Ian Johnston, Dominic Kwiatkowski, Cordelia Langford, John Sillitoe on behalf of the Wellcome Sanger Institute COVID-19 Surveillance Team |
| EPI_ISL_2806328 | Health Services Laboratories | Wellcome Sanger Institute for the COVID-19 Genomics UK (COG-UK) Consortium | Health Services Laboratories and Alex Alderton, Roberto Amato, Jeffrey Barrett, Sonia Goncalves, Ewan Harrison, David K. Jackson, Ian Johnston, Dominic Kwiatkowski, Cordelia Langford, John Sillitoe on behalf of the Wellcome Sanger Institute COVID-19 Surveillance Team |
| EPI_ISL_2806329, EPI_ISL_2806330 | Lighthouse Lab in Glasgow | Wellcome Sanger Institute for the COVID-19 Genomics UK (COG-UK) Consortium | Harper VanSteenhouse, Yumi Kasai, David Gray, Carol Clugston, Anna Dominiczak and Alex Alderton, Roberto Amato, Jeffrey Barrett, Sonia Goncalves, Ewan Harrison, David K. Jackson, Ian Johnston, Dominic Kwiatkowski, Cordelia Langford, John Sillitoe on behalf of the Wellcome Sanger Institute COVID-19 Surveillance Team |
| EPI_ISL_2806331, EPI_ISL_2806332 | Health Services Laboratories | Wellcome Sanger Institute for the COVID-19 Genomics UK (COG-UK) Consortium | Health Services Laboratories and Alex Alderton, Roberto Amato, Jeffrey Barrett, Sonia Goncalves, Ewan Harrison, David K. Jackson, Ian Johnston, Dominic Kwiatkowski, Cordelia Langford, John Sillitoe on behalf of the Wellcome Sanger Institute COVID-19 Surveillance Team |
| EPI_ISL_2806333, EPI_ISL_2806334 | Lighthouse Lab in Alderley Park | Wellcome Sanger Institute for the COVID-19 Genomics UK (COG-UK) Consortium | Jacquelyn Wynn, Mairead Hyland, The Lighthouse Lab in Alderley Park and Alex Alderton, Roberto Amato, Jeffrey Barrett, Sonia Goncalves, Ewan Harrison, David K. Jackson, Ian Johnston, Dominic Kwiatkowski, Cordelia Langford, John Sillitoe on behalf of the Wellcome Sanger Institute COVID-19 Surveillance Team |
| EPI_ISL_2806335 | Lighthouse Lab in Glasgow | Wellcome Sanger Institute for the COVID-19 Genomics UK (COG-UK) Consortium | Harper VanSteenhouse, Yumi Kasai, David Gray, Carol Clugston, Anna Dominiczak and Alex Alderton, Roberto Amato, Jeffrey Barrett, Sonia Goncalves, Ewan Harrison, David K. Jackson, Ian Johnston, Dominic Kwiatkowski, Cordelia Langford, John Sillitoe on behalf of the Wellcome Sanger Institute COVID-19 Surveillance Team |
| EPI_ISL_2806336 | Health Services Laboratories | Wellcome Sanger Institute for the COVID-19 Genomics UK (COG-UK) Consortium | Health Services Laboratories and Alex Alderton, Roberto Amato, Jeffrey Barrett, Sonia Goncalves, Ewan Harrison, David K. Jackson, Ian Johnston, Dominic Kwiatkowski, Cordelia Langford, John Sillitoe on behalf of the Wellcome Sanger Institute COVID-19 Surveillance Team |
| EPI_ISL_2806337, EPI_ISL_2806338 | Lighthouse Lab in Glasgow | Wellcome Sanger Institute for the COVID-19 Genomics UK (COG-UK) Consortium | Harper VanSteenhouse, Yumi Kasai, David Gray, Carol Clugston, Anna Dominiczak and Alex Alderton, Roberto Amato, Jeffrey Barrett, Sonia Goncalves, Ewan Harrison, David K. Jackson, Ian Johnston, Dominic Kwiatkowski, Cordelia Langford, John Sillitoe on behalf of the Wellcome Sanger Institute COVID-19 Surveillance Team |
| EPI_ISL_2806339, EPI_ISL_2806340 | Health Services Laboratories | Wellcome Sanger Institute for the COVID-19 Genomics UK (COG-UK) Consortium | Health Services Laboratories and Alex Alderton, Roberto Amato, Jeffrey Barrett, Sonia Goncalves, Ewan Harrison, David K. Jackson, Ian Johnston, Dominic Kwiatkowski, Cordelia Langford, John Sillitoe on behalf of the Wellcome Sanger Institute COVID-19 Surveillance Team |
| EPI_ISL_2806341 | Lighthouse Lab in Alderley Park | Wellcome Sanger Institute for the COVID-19 Genomics UK (COG-UK) Consortium | Jacquelyn Wynn, Mairead Hyland, The Lighthouse Lab in Alderley Park and Alex Alderton, Roberto Amato, Jeffrey Barrett, Sonia Goncalves, Ewan Harrison, David K. Jackson, Ian Johnston, Dominic Kwiatkowski, Cordelia Langford, John Sillitoe on behalf of the Wellcome Sanger Institute COVID-19 Surveillance Team |
| EPI_ISL_2806342, EPI_ISL_2806343 | Lighthouse Lab in Glasgow | Wellcome Sanger Institute for the COVID-19 Genomics UK (COG-UK) Consortium | Harper VanSteenhouse, Yumi Kasai, David Gray, Carol Clugston, Anna Dominiczak and Alex Alderton, Roberto Amato, Jeffrey Barrett, Sonia Goncalves, Ewan Harrison, David K. Jackson, Ian Johnston, Dominic Kwiatkowski, Cordelia Langford, John Sillitoe on behalf of the Wellcome Sanger Institute COVID-19 Surveillance Team |
| EPI_ISL_2806344 | Health Services Laboratories | Wellcome Sanger Institute for the COVID-19 Genomics UK (COG-UK) Consortium | Health Services Laboratories and Alex Alderton, Roberto Amato, Jeffrey Barrett, Sonia Goncalves, Ewan Harrison, David K. Jackson, Ian Johnston, Dominic Kwiatkowski, Cordelia Langford, John Sillitoe on behalf of the Wellcome Sanger Institute COVID-19 Surveillance Team |
| EPI_ISL_2806345, EPI_ISL_2806346 | Lighthouse Lab in Alderley Park | Wellcome Sanger Institute for the COVID-19 Genomics UK (COG-UK) Consortium | Jacquelyn Wynn, Mairead Hyland, The Lighthouse Lab in Alderley Park and Alex Alderton, Roberto Amato, Jeffrey Barrett, Sonia Goncalves, Ewan Harrison, David K. Jackson, Ian Johnston, Dominic Kwiatkowski, Cordelia Langford, John Sillitoe on behalf of the Wellcome Sanger Institute COVID-19 Surveillance Team |
| EPI_ISL_2806347, | Health Services Laboratories | Wellcome Sanger Institute for the COVID-19 Genomics UK (COG-UK) | Health Services Laboratories and Alex Alderton, Roberto Amato, Jeffrey Barrett, Sonia Goncalves, Ewan Harrison, David K. Jackson, Ian Johnston, Dominic Kwiatkowski, Cordelia Langford, John Sillitoe on behalf of the Wellcome Sanger Institute |

| | | | |
|---|---|---|---|
| EPI_ISL_2806348 | | Consortium | COVID-19 Surveillance Team |
| EPI_ISL_2806349 | Lighthouse Lab in Alderley Park | Wellcome Sanger Institute for the COVID-19 Genomics UK (COG-UK) Consortium | Jacquelyn Wynn, Mairead Hyland, The Lighthouse Lab in Alderley Park and Alex Alderton, Roberto Amato, Jeffrey Barrett, Sonia Goncalves, Ewan Harrison, David K. Jackson, Ian Johnston, Dominic Kwiatkowski, Cordelia Langford, John Sillitoe on behalf of the Wellcome Sanger Institute COVID-19 Surveillance Team |
| EPI_ISL_2806350 | Lighthouse Lab in Glasgow | Wellcome Sanger Institute for the COVID-19 Genomics UK (COG-UK) Consortium | Harper VanSteenhouse, Yumi Kasai, David Gray, Carol Clugston, Anna Dominiczak and Alex Alderton, Roberto Amato, Jeffrey Barrett, Sonia Goncalves, Ewan Harrison, David K. Jackson, Ian Johnston, Dominic Kwiatkowski, Cordelia Langford, John Sillitoe on behalf of the Wellcome Sanger Institute COVID-19 Surveillance Team |
| EPI_ISL_2806351, EPI_ISL_2806352, EPI_ISL_2806353, EPI_ISL_2806354, EPI_ISL_2806355 | Lighthouse Lab in Alderley Park | Wellcome Sanger Institute for the COVID-19 Genomics UK (COG-UK) Consortium | Jacquelyn Wynn, Mairead Hyland, The Lighthouse Lab in Alderley Park and Alex Alderton, Roberto Amato, Jeffrey Barrett, Sonia Goncalves, Ewan Harrison, David K. Jackson, Ian Johnston, Dominic Kwiatkowski, Cordelia Langford, John Sillitoe on behalf of the Wellcome Sanger Institute COVID-19 Surveillance Team |
| EPI_ISL_2806356 | Health Services Laboratories | Wellcome Sanger Institute for the COVID-19 Genomics UK (COG-UK) Consortium | Health Services Laboratories and Alex Alderton, Roberto Amato, Jeffrey Barrett, Sonia Goncalves, Ewan Harrison, David K. Jackson, Ian Johnston, Dominic Kwiatkowski, Cordelia Langford, John Sillitoe on behalf of the Wellcome Sanger Institute COVID-19 Surveillance Team |
| EPI_ISL_2806357, EPI_ISL_2806358 | Lighthouse Lab in Alderley Park | Wellcome Sanger Institute for the COVID-19 Genomics UK (COG-UK) Consortium | Jacquelyn Wynn, Mairead Hyland, The Lighthouse Lab in Alderley Park and Alex Alderton, Roberto Amato, Jeffrey Barrett, Sonia Goncalves, Ewan Harrison, David K. Jackson, Ian Johnston, Dominic Kwiatkowski, Cordelia Langford, John Sillitoe on behalf of the Wellcome Sanger Institute COVID-19 Surveillance Team |
| EPI_ISL_2806359 | Health Services Laboratories | Wellcome Sanger Institute for the COVID-19 Genomics UK (COG-UK) Consortium | Health Services Laboratories and Alex Alderton, Roberto Amato, Jeffrey Barrett, Sonia Goncalves, Ewan Harrison, David K. Jackson, Ian Johnston, Dominic Kwiatkowski, Cordelia Langford, John Sillitoe on behalf of the Wellcome Sanger Institute COVID-19 Surveillance Team |
| EPI_ISL_2806360, EPI_ISL_2806361 | Lighthouse Lab in Alderley Park | Wellcome Sanger Institute for the COVID-19 Genomics UK (COG-UK) Consortium | Jacquelyn Wynn, Mairead Hyland, The Lighthouse Lab in Alderley Park and Alex Alderton, Roberto Amato, Jeffrey Barrett, Sonia Goncalves, Ewan Harrison, David K. Jackson, Ian Johnston, Dominic Kwiatkowski, Cordelia Langford, John Sillitoe on behalf of the Wellcome Sanger Institute COVID-19 Surveillance Team |
| EPI_ISL_2806362 | Health Services Laboratories | Wellcome Sanger Institute for the COVID-19 Genomics UK (COG-UK) Consortium | Health Services Laboratories and Alex Alderton, Roberto Amato, Jeffrey Barrett, Sonia Goncalves, Ewan Harrison, David K. Jackson, Ian Johnston, Dominic Kwiatkowski, Cordelia Langford, John Sillitoe on behalf of the Wellcome Sanger Institute COVID-19 Surveillance Team |
| EPI_ISL_2806363 | Lighthouse Lab in Glasgow | Wellcome Sanger Institute for the COVID-19 Genomics UK (COG-UK) Consortium | Harper VanSteenhouse, Yumi Kasai, David Gray, Carol Clugston, Anna Dominiczak and Alex Alderton, Roberto Amato, Jeffrey Barrett, Sonia Goncalves, Ewan Harrison, David K. Jackson, Ian Johnston, Dominic Kwiatkowski, Cordelia Langford, John Sillitoe on behalf of the Wellcome Sanger Institute COVID-19 Surveillance Team |
| EPI_ISL_2806364, EPI_ISL_2806365 | Health Services Laboratories | Wellcome Sanger Institute for the COVID-19 Genomics UK (COG-UK) Consortium | Health Services Laboratories and Alex Alderton, Roberto Amato, Jeffrey Barrett, Sonia Goncalves, Ewan Harrison, David K. Jackson, Ian Johnston, Dominic Kwiatkowski, Cordelia Langford, John Sillitoe on behalf of the Wellcome Sanger Institute COVID-19 Surveillance Team |
| EPI_ISL_2806366 | Lighthouse Lab in Alderley Park | Wellcome Sanger Institute for the COVID-19 Genomics UK (COG-UK) Consortium | Jacquelyn Wynn, Mairead Hyland, The Lighthouse Lab in Alderley Park and Alex Alderton, Roberto Amato, Jeffrey Barrett, Sonia Goncalves, Ewan Harrison, David K. Jackson, Ian Johnston, Dominic Kwiatkowski, Cordelia Langford, John Sillitoe on behalf of the Wellcome Sanger Institute COVID-19 Surveillance Team |
| EPI_ISL_2806367 | Lighthouse Lab in Glasgow | Wellcome Sanger Institute for the COVID-19 Genomics UK (COG-UK) Consortium | Harper VanSteenhouse, Yumi Kasai, David Gray, Carol Clugston, Anna Dominiczak and Alex Alderton, Roberto Amato, Jeffrey Barrett, Sonia Goncalves, Ewan Harrison, David K. Jackson, Ian Johnston, Dominic Kwiatkowski, Cordelia Langford, John Sillitoe on behalf of the Wellcome Sanger Institute COVID-19 Surveillance Team |
| EPI_ISL_2806368 | Health Services Laboratories | Wellcome Sanger Institute for the COVID-19 Genomics UK (COG-UK) Consortium | Health Services Laboratories and Alex Alderton, Roberto Amato, Jeffrey Barrett, Sonia Goncalves, Ewan Harrison, David K. Jackson, Ian Johnston, Dominic Kwiatkowski, Cordelia Langford, John Sillitoe on behalf of the Wellcome Sanger Institute COVID-19 Surveillance Team |
| EPI_ISL_2806369 | Lighthouse Lab in Glasgow | Wellcome Sanger Institute for the COVID-19 Genomics UK (COG-UK) Consortium | Harper VanSteenhouse, Yumi Kasai, David Gray, Carol Clugston, Anna Dominiczak and Alex Alderton, Roberto Amato, Jeffrey Barrett, Sonia Goncalves, Ewan Harrison, David K. Jackson, Ian Johnston, Dominic Kwiatkowski, Cordelia Langford, John Sillitoe on behalf of the Wellcome Sanger Institute COVID-19 Surveillance Team |
| EPI_ISL_2806370 | Health Services Laboratories | Wellcome Sanger Institute for the COVID-19 Genomics UK (COG-UK) Consortium | Health Services Laboratories and Alex Alderton, Roberto Amato, Jeffrey Barrett, Sonia Goncalves, Ewan Harrison, David K. Jackson, Ian Johnston, Dominic Kwiatkowski, Cordelia Langford, John Sillitoe on behalf of the Wellcome Sanger Institute COVID-19 Surveillance Team |
| EPI_ISL_2806371, EPI_ISL_2806372 | Lighthouse Lab in Alderley Park | Wellcome Sanger Institute for the COVID-19 Genomics UK (COG-UK) Consortium | Jacquelyn Wynn, Mairead Hyland, The Lighthouse Lab in Alderley Park and Alex Alderton, Roberto Amato, Jeffrey Barrett, Sonia Goncalves, Ewan Harrison, David K. Jackson, Ian Johnston, Dominic Kwiatkowski, Cordelia Langford, John Sillitoe on behalf of the Wellcome Sanger Institute COVID-19 Surveillance Team |
| EPI_ISL_2806373 | Health Services Laboratories | Wellcome Sanger Institute for the COVID-19 Genomics UK (COG-UK) Consortium | Health Services Laboratories and Alex Alderton, Roberto Amato, Jeffrey Barrett, Sonia Goncalves, Ewan Harrison, David K. Jackson, Ian Johnston, Dominic Kwiatkowski, Cordelia Langford, John Sillitoe on behalf of the Wellcome Sanger Institute COVID-19 Surveillance Team |
| EPI_ISL_2806374 | Lighthouse Lab in Glasgow | Wellcome Sanger Institute for the COVID-19 Genomics UK (COG-UK) Consortium | Harper VanSteenhouse, Yumi Kasai, David Gray, Carol Clugston, Anna Dominiczak and Alex Alderton, Roberto Amato, Jeffrey Barrett, Sonia Goncalves, Ewan Harrison, David K. Jackson, Ian Johnston, Dominic Kwiatkowski, Cordelia Langford, John Sillitoe on behalf of the Wellcome Sanger Institute COVID-19 Surveillance Team |
| EPI_ISL_2806375, EPI_ISL_2806376, EPI_ISL_2806377, EPI_ISL_2806378, EPI_ISL_2806379 | Health Services Laboratories | Wellcome Sanger Institute for the COVID-19 Genomics UK (COG-UK) Consortium | Health Services Laboratories and Alex Alderton, Roberto Amato, Jeffrey Barrett, Sonia Goncalves, Ewan Harrison, David K. Jackson, Ian Johnston, Dominic Kwiatkowski, Cordelia Langford, John Sillitoe on behalf of the Wellcome Sanger Institute COVID-19 Surveillance Team |
| EPI_ISL_2806380 | Lighthouse Lab in Glasgow | Wellcome Sanger Institute for the COVID-19 Genomics UK (COG-UK) Consortium | Harper VanSteenhouse, Yumi Kasai, David Gray, Carol Clugston, Anna Dominiczak and Alex Alderton, Roberto Amato, Jeffrey Barrett, Sonia Goncalves, Ewan Harrison, David K. Jackson, Ian Johnston, Dominic Kwiatkowski, Cordelia Langford, John Sillitoe on behalf of the Wellcome Sanger Institute COVID-19 Surveillance Team |
| EPI_ISL_2806381 | Lighthouse Lab in Alderley Park | Wellcome Sanger Institute for the COVID-19 Genomics UK (COG-UK) Consortium | Jacquelyn Wynn, Mairead Hyland, The Lighthouse Lab in Alderley Park and Alex Alderton, Roberto Amato, Jeffrey Barrett, Sonia Goncalves, Ewan Harrison, David K. Jackson, Ian Johnston, Dominic Kwiatkowski, Cordelia Langford, John Sillitoe on behalf of the Wellcome Sanger Institute COVID-19 Surveillance Team |
| EPI_ISL_2806382, EPI_ISL_2806383, EPI_ISL_2806384, EPI_ISL_2806385 | Health Services Laboratories | Wellcome Sanger Institute for the COVID-19 Genomics UK (COG-UK) Consortium | Health Services Laboratories and Alex Alderton, Roberto Amato, Jeffrey Barrett, Sonia Goncalves, Ewan Harrison, David K. Jackson, Ian Johnston, Dominic Kwiatkowski, Cordelia Langford, John Sillitoe on behalf of the Wellcome Sanger Institute COVID-19 Surveillance Team |
| EPI_ISL_2806386 | Lighthouse Lab in Glasgow | Wellcome Sanger Institute for the COVID-19 Genomics UK (COG-UK) Consortium | Harper VanSteenhouse, Yumi Kasai, David Gray, Carol Clugston, Anna Dominiczak and Alex Alderton, Roberto Amato, Jeffrey Barrett, Sonia Goncalves, Ewan Harrison, David K. Jackson, Ian Johnston, Dominic Kwiatkowski, Cordelia Langford, John Sillitoe on behalf of the Wellcome Sanger Institute COVID-19 Surveillance Team |
| EPI_ISL_2806387 | Lighthouse Lab in Alderley Park | Wellcome Sanger Institute for the COVID-19 Genomics UK (COG-UK) Consortium | Jacquelyn Wynn, Mairead Hyland, The Lighthouse Lab in Alderley Park and Alex Alderton, Roberto Amato, Jeffrey Barrett, Sonia Goncalves, Ewan Harrison, David K. Jackson, Ian Johnston, Dominic Kwiatkowski, Cordelia Langford, John Sillitoe on behalf of the Wellcome Sanger Institute COVID-19 Surveillance Team |
| EPI_ISL_2806388, EPI_ISL_2806389, EPI_ISL_2806390 | Health Services Laboratories | Wellcome Sanger Institute for the COVID-19 Genomics UK (COG-UK) Consortium | Health Services Laboratories and Alex Alderton, Roberto Amato, Jeffrey Barrett, Sonia Goncalves, Ewan Harrison, David K. Jackson, Ian Johnston, Dominic Kwiatkowski, Cordelia Langford, John Sillitoe on behalf of the Wellcome Sanger Institute COVID-19 Surveillance Team |
| EPI_ISL_2806391, EPI_ISL_2806392 | Lighthouse Lab in Alderley Park | Wellcome Sanger Institute for the COVID-19 Genomics UK (COG-UK) Consortium | Jacquelyn Wynn, Mairead Hyland, The Lighthouse Lab in Alderley Park and Alex Alderton, Roberto Amato, Jeffrey Barrett, Sonia Goncalves, Ewan Harrison, David K. Jackson, Ian Johnston, Dominic Kwiatkowski, Cordelia Langford, John Sillitoe on behalf of the Wellcome Sanger Institute COVID-19 Surveillance Team |
| EPI_ISL_2806393 | Lighthouse Lab in Glasgow | Wellcome Sanger Institute for the COVID-19 Genomics UK (COG-UK) Consortium | Harper VanSteenhouse, Yumi Kasai, David Gray, Carol Clugston, Anna Dominiczak and Alex Alderton, Roberto Amato, Jeffrey Barrett, Sonia Goncalves, Ewan Harrison, David K. Jackson, Ian Johnston, Dominic Kwiatkowski, Cordelia Langford, John Sillitoe on behalf of the Wellcome Sanger Institute COVID-19 Surveillance Team |
| EPI_ISL_2806394 | Lighthouse Lab in Alderley Park | Wellcome Sanger Institute for the COVID-19 Genomics UK (COG-UK) Consortium | Jacquelyn Wynn, Mairead Hyland, The Lighthouse Lab in Alderley Park and Alex Alderton, Roberto Amato, Jeffrey Barrett, Sonia Goncalves, Ewan Harrison, David K. Jackson, Ian Johnston, Dominic Kwiatkowski, Cordelia Langford, John Sillitoe on behalf of the Wellcome Sanger Institute COVID-19 Surveillance Team |
| EPI_ISL_2806395, EPI_ISL_2806396, EPI_ISL_2806397, EPI_ISL_2806398, EPI_ISL_2806399, EPI_ISL_2806400, EPI_ISL_2806401 | Lighthouse Lab in Glasgow | Wellcome Sanger Institute for the COVID-19 Genomics UK (COG-UK) Consortium | Harper VanSteenhouse, Yumi Kasai, David Gray, Carol Clugston, Anna Dominiczak and Alex Alderton, Roberto Amato, Jeffrey Barrett, Sonia Goncalves, Ewan Harrison, David K. Jackson, Ian Johnston, Dominic Kwiatkowski, Cordelia Langford, John Sillitoe on behalf of the Wellcome Sanger Institute COVID-19 Surveillance Team |
| EPI_ISL_2806402 | Lighthouse Lab in Alderley Park | Wellcome Sanger Institute for the COVID-19 Genomics UK (COG-UK) Consortium | Jacquelyn Wynn, Mairead Hyland, The Lighthouse Lab in Alderley Park and Alex Alderton, Roberto Amato, Jeffrey Barrett, Sonia Goncalves, Ewan Harrison, David K. Jackson, Ian Johnston, Dominic Kwiatkowski, Cordelia Langford, John Sillitoe on behalf of the Wellcome Sanger Institute COVID-19 Surveillance Team |
| EPI_ISL_2806403, EPI_ISL_2806404, EPI_ISL_2806405 | Lighthouse Lab in Glasgow | Wellcome Sanger Institute for the COVID-19 Genomics UK (COG-UK) Consortium | Harper VanSteenhouse, Yumi Kasai, David Gray, Carol Clugston, Anna Dominiczak and Alex Alderton, Roberto Amato, Jeffrey Barrett, Sonia Goncalves, Ewan Harrison, David K. Jackson, Ian Johnston, Dominic Kwiatkowski, Cordelia Langford, John Sillitoe on behalf of the Wellcome Sanger Institute COVID-19 Surveillance Team |
| EPI_ISL_2806406, EPI_ISL_2806407, EPI_ISL_2806408 | Lighthouse Lab in Alderley Park | Wellcome Sanger Institute for the COVID-19 Genomics UK (COG-UK) Consortium | Jacquelyn Wynn, Mairead Hyland, The Lighthouse Lab in Alderley Park and Alex Alderton, Roberto Amato, Jeffrey Barrett, Sonia Goncalves, Ewan Harrison, David K. Jackson, Ian Johnston, Dominic Kwiatkowski, Cordelia Langford, John Sillitoe on behalf of the Wellcome Sanger Institute COVID-19 Surveillance Team |
| EPI_ISL_2806409, EPI_ISL_2806410, EPI_ISL_2806411 | Health Services Laboratories | Wellcome Sanger Institute for the COVID-19 Genomics UK (COG-UK) Consortium | Health Services Laboratories and Alex Alderton, Roberto Amato, Jeffrey Barrett, Sonia Goncalves, Ewan Harrison, David K. Jackson, Ian Johnston, Dominic Kwiatkowski, Cordelia Langford, John Sillitoe on behalf of the Wellcome Sanger Institute COVID-19 Surveillance Team |
| EPI_ISL_2806412, EPI_ISL_2806413, EPI_ISL_2806414 | Lighthouse Lab in Alderley Park | Wellcome Sanger Institute for the COVID-19 Genomics UK (COG-UK) Consortium | Jacquelyn Wynn, Mairead Hyland, The Lighthouse Lab in Alderley Park and Alex Alderton, Roberto Amato, Jeffrey Barrett, Sonia Goncalves, Ewan Harrison, David K. Jackson, Ian Johnston, Dominic Kwiatkowski, Cordelia Langford, John Sillitoe on behalf of the Wellcome Sanger Institute COVID-19 Surveillance Team |
| EPI_ISL_2806415, EPI_ISL_2806416 | Lighthouse Lab in Glasgow | Wellcome Sanger Institute for the COVID-19 Genomics UK (COG-UK) Consortium | Harper VanSteenhouse, Yumi Kasai, David Gray, Carol Clugston, Anna Dominiczak and Alex Alderton, Roberto Amato, Jeffrey Barrett, Sonia Goncalves, Ewan Harrison, David K. Jackson, Ian Johnston, Dominic Kwiatkowski, Cordelia Langford, John Sillitoe on behalf of the Wellcome Sanger Institute COVID-19 Surveillance Team |
| EPI_ISL_2806417 | Lighthouse Lab in Alderley Park | Wellcome Sanger Institute for the COVID-19 Genomics UK (COG-UK) Consortium | Jacquelyn Wynn, Mairead Hyland, The Lighthouse Lab in Alderley Park and Alex Alderton, Roberto Amato, Jeffrey Barrett, Sonia Goncalves, Ewan Harrison, David K. Jackson, Ian Johnston, Dominic Kwiatkowski, Cordelia Langford, John Sillitoe on behalf of the Wellcome Sanger Institute COVID-19 Surveillance Team |
| EPI_ISL_2806418, EPI_ISL_2806419, | Health Services Laboratories | Wellcome Sanger Institute for the COVID-19 Genomics UK (COG-UK) Consortium | Health Services Laboratories and Alex Alderton, Roberto Amato, Jeffrey Barrett, Sonia Goncalves, Ewan Harrison, David K. Jackson, Ian Johnston, Dominic Kwiatkowski, Cordelia Langford, John Sillitoe on behalf of the Wellcome Sanger Institute COVID-19 Surveillance Team |

| | | | |
|---|---|---|---|
| EPI_ISL_2806420 | | | |
| EPI_ISL_2806421 | Lighthouse Lab in Glasgow | Wellcome Sanger Institute for the COVID-19 Genomics UK (COG-UK) Consortium | Harper VanSteenhouse, Yumi Kasai, David Gray, Carol Clugston, Anna Dominiczak and Alex Alderton, Roberto Amato, Jeffrey Barrett, Sonia Goncalves, Ewan Harrison, David K. Jackson, Ian Johnston, Dominic Kwiatkowski, Cordelia Langford, John Sillitoe on behalf of the Wellcome Sanger Institute COVID-19 Surveillance Team |
| EPI_ISL_2806422 | Lighthouse Lab in Alderley Park | Wellcome Sanger Institute for the COVID-19 Genomics UK (COG-UK) Consortium | Jacquelyn Wynn, Mairead Hyland, The Lighthouse Lab in Alderley Park and Alex Alderton, Roberto Amato, Jeffrey Barrett, Sonia Goncalves, Ewan Harrison, David K. Jackson, Ian Johnston, Dominic Kwiatkowski, Cordelia Langford, John Sillitoe on behalf of the Wellcome Sanger Institute COVID-19 Surveillance Team |
| EPI_ISL_2806423, EPI_ISL_2806424 | Health Services Laboratories | Wellcome Sanger Institute for the COVID-19 Genomics UK (COG-UK) Consortium | Health Services Laboratories and Alex Alderton, Roberto Amato, Jeffrey Barrett, Sonia Goncalves, Ewan Harrison, David K. Jackson, Ian Johnston, Dominic Kwiatkowski, Cordelia Langford, John Sillitoe on behalf of the Wellcome Sanger Institute COVID-19 Surveillance Team |
| EPI_ISL_2806425 | Lighthouse Lab in Glasgow | Wellcome Sanger Institute for the COVID-19 Genomics UK (COG-UK) Consortium | Harper VanSteenhouse, Yumi Kasai, David Gray, Carol Clugston, Anna Dominiczak and Alex Alderton, Roberto Amato, Jeffrey Barrett, Sonia Goncalves, Ewan Harrison, David K. Jackson, Ian Johnston, Dominic Kwiatkowski, Cordelia Langford, John Sillitoe on behalf of the Wellcome Sanger Institute COVID-19 Surveillance Team |
| EPI_ISL_2806426, EPI_ISL_2806427, EPI_ISL_2806428 | Health Services Laboratories | Wellcome Sanger Institute for the COVID-19 Genomics UK (COG-UK) Consortium | Health Services Laboratories and Alex Alderton, Roberto Amato, Jeffrey Barrett, Sonia Goncalves, Ewan Harrison, David K. Jackson, Ian Johnston, Dominic Kwiatkowski, Cordelia Langford, John Sillitoe on behalf of the Wellcome Sanger Institute COVID-19 Surveillance Team |
| EPI_ISL_2806429, EPI_ISL_2806430 | Lighthouse Lab in Alderley Park | Wellcome Sanger Institute for the COVID-19 Genomics UK (COG-UK) Consortium | Jacquelyn Wynn, Mairead Hyland, The Lighthouse Lab in Alderley Park and Alex Alderton, Roberto Amato, Jeffrey Barrett, Sonia Goncalves, Ewan Harrison, David K. Jackson, Ian Johnston, Dominic Kwiatkowski, Cordelia Langford, John Sillitoe on behalf of the Wellcome Sanger Institute COVID-19 Surveillance Team |
| EPI_ISL_2806431 | Health Services Laboratories | Wellcome Sanger Institute for the COVID-19 Genomics UK (COG-UK) Consortium | Health Services Laboratories and Alex Alderton, Roberto Amato, Jeffrey Barrett, Sonia Goncalves, Ewan Harrison, David K. Jackson, Ian Johnston, Dominic Kwiatkowski, Cordelia Langford, John Sillitoe on behalf of the Wellcome Sanger Institute COVID-19 Surveillance Team |
| EPI_ISL_2806432, EPI_ISL_2806433 | Lighthouse Lab in Alderley Park | Wellcome Sanger Institute for the COVID-19 Genomics UK (COG-UK) Consortium | Jacquelyn Wynn, Mairead Hyland, The Lighthouse Lab in Alderley Park and Alex Alderton, Roberto Amato, Jeffrey Barrett, Sonia Goncalves, Ewan Harrison, David K. Jackson, Ian Johnston, Dominic Kwiatkowski, Cordelia Langford, John Sillitoe on behalf of the Wellcome Sanger Institute COVID-19 Surveillance Team |
| EPI_ISL_2806434, EPI_ISL_2806435 | Health Services Laboratories | Wellcome Sanger Institute for the COVID-19 Genomics UK (COG-UK) Consortium | Health Services Laboratories and Alex Alderton, Roberto Amato, Jeffrey Barrett, Sonia Goncalves, Ewan Harrison, David K. Jackson, Ian Johnston, Dominic Kwiatkowski, Cordelia Langford, John Sillitoe on behalf of the Wellcome Sanger Institute COVID-19 Surveillance Team |
| EPI_ISL_2806436, EPI_ISL_2806437, EPI_ISL_2806438 | Lighthouse Lab in Alderley Park | Wellcome Sanger Institute for the COVID-19 Genomics UK (COG-UK) Consortium | Jacquelyn Wynn, Mairead Hyland, The Lighthouse Lab in Alderley Park and Alex Alderton, Roberto Amato, Jeffrey Barrett, Sonia Goncalves, Ewan Harrison, David K. Jackson, Ian Johnston, Dominic Kwiatkowski, Cordelia Langford, John Sillitoe on behalf of the Wellcome Sanger Institute COVID-19 Surveillance Team |
| EPI_ISL_2806439, EPI_ISL_2806440 | Health Services Laboratories | Wellcome Sanger Institute for the COVID-19 Genomics UK (COG-UK) Consortium | Health Services Laboratories and Alex Alderton, Roberto Amato, Jeffrey Barrett, Sonia Goncalves, Ewan Harrison, David K. Jackson, Ian Johnston, Dominic Kwiatkowski, Cordelia Langford, John Sillitoe on behalf of the Wellcome Sanger Institute COVID-19 Surveillance Team |
| EPI_ISL_2806441 | Lighthouse Lab in Alderley Park | Wellcome Sanger Institute for the COVID-19 Genomics UK (COG-UK) Consortium | Jacquelyn Wynn, Mairead Hyland, The Lighthouse Lab in Alderley Park and Alex Alderton, Roberto Amato, Jeffrey Barrett, Sonia Goncalves, Ewan Harrison, David K. Jackson, Ian Johnston, Dominic Kwiatkowski, Cordelia Langford, John Sillitoe on behalf of the Wellcome Sanger Institute COVID-19 Surveillance Team |
| EPI_ISL_2806442 | Lighthouse Lab in Glasgow | Wellcome Sanger Institute for the COVID-19 Genomics UK (COG-UK) Consortium | Harper VanSteenhouse, Yumi Kasai, David Gray, Carol Clugston, Anna Dominiczak and Alex Alderton, Roberto Amato, Jeffrey Barrett, Sonia Goncalves, Ewan Harrison, David K. Jackson, Ian Johnston, Dominic Kwiatkowski, Cordelia Langford, John Sillitoe on behalf of the Wellcome Sanger Institute COVID-19 Surveillance Team |
| EPI_ISL_2806443 | Health Services Laboratories | Wellcome Sanger Institute for the COVID-19 Genomics UK (COG-UK) Consortium | Health Services Laboratories and Alex Alderton, Roberto Amato, Jeffrey Barrett, Sonia Goncalves, Ewan Harrison, David K. Jackson, Ian Johnston, Dominic Kwiatkowski, Cordelia Langford, John Sillitoe on behalf of the Wellcome Sanger Institute COVID-19 Surveillance Team |
| EPI_ISL_2806444 | Lighthouse Laboratory Plymouth | Wellcome Sanger Institute for the COVID-19 Genomics UK (COG-UK) Consortium | Lighthouse Laboratory Plymouth and Alex Alderton, Roberto Amato, Jeffrey Barrett, Sonia Goncalves, Ewan Harrison, David K. Jackson, Ian Johnston, Dominic Kwiatkowski, Cordelia Langford, John Sillitoe on behalf of the Wellcome Sanger Institute COVID-19 Surveillance Team |
| EPI_ISL_2806445, EPI_ISL_2806446 | Lighthouse Lab in Milton Keynes | Wellcome Sanger Institute for the COVID-19 Genomics UK (COG-UK) Consortium | The Lighthouse Lab in Milton Keynes and Alex Alderton, Roberto Amato, Jeffrey Barrett, Sonia Goncalves, Ewan Harrison, David K. Jackson, Ian Johnston, Dominic Kwiatkowski, Cordelia Langford, John Sillitoe on behalf of the Wellcome Sanger Institute COVID-19 Surveillance Team |
| EPI_ISL_2806447, EPI_ISL_2806448, EPI_ISL_2806449, EPI_ISL_2806450 | Health Services Laboratories | Wellcome Sanger Institute for the COVID-19 Genomics UK (COG-UK) Consortium | Health Services Laboratories and Alex Alderton, Roberto Amato, Jeffrey Barrett, Sonia Goncalves, Ewan Harrison, David K. Jackson, Ian Johnston, Dominic Kwiatkowski, Cordelia Langford, John Sillitoe on behalf of the Wellcome Sanger Institute COVID-19 Surveillance Team |
| EPI_ISL_2806451 | Lighthouse Lab in Milton Keynes | Wellcome Sanger Institute for the COVID-19 Genomics UK (COG-UK) Consortium | The Lighthouse Lab in Milton Keynes and Alex Alderton, Roberto Amato, Jeffrey Barrett, Sonia Goncalves, Ewan Harrison, David K. Jackson, Ian Johnston, Dominic Kwiatkowski, Cordelia Langford, John Sillitoe on behalf of the Wellcome Sanger Institute COVID-19 Surveillance Team |
| EPI_ISL_2806452, EPI_ISL_2806453, EPI_ISL_2806454 | Health Services Laboratories | Wellcome Sanger Institute for the COVID-19 Genomics UK (COG-UK) Consortium | Health Services Laboratories and Alex Alderton, Roberto Amato, Jeffrey Barrett, Sonia Goncalves, Ewan Harrison, David K. Jackson, Ian Johnston, Dominic Kwiatkowski, Cordelia Langford, John Sillitoe on behalf of the Wellcome Sanger Institute COVID-19 Surveillance Team |
| EPI_ISL_2806455 | Lighthouse Lab in Milton Keynes | Wellcome Sanger Institute for the COVID-19 Genomics UK (COG-UK) Consortium | The Lighthouse Lab in Milton Keynes and Alex Alderton, Roberto Amato, Jeffrey Barrett, Sonia Goncalves, Ewan Harrison, David K. Jackson, Ian Johnston, Dominic Kwiatkowski, Cordelia Langford, John Sillitoe on behalf of the Wellcome Sanger Institute COVID-19 Surveillance Team |
| EPI_ISL_2806456 | Lighthouse Laboratory Plymouth | Wellcome Sanger Institute for the COVID-19 Genomics UK (COG-UK) Consortium | Lighthouse Laboratory Plymouth and Alex Alderton, Roberto Amato, Jeffrey Barrett, Sonia Goncalves, Ewan Harrison, David K. Jackson, Ian Johnston, Dominic Kwiatkowski, Cordelia Langford, John Sillitoe on behalf of the Wellcome Sanger Institute COVID-19 Surveillance Team |
| EPI_ISL_2806457, EPI_ISL_2806458 | Health Services Laboratories | Wellcome Sanger Institute for the COVID-19 Genomics UK (COG-UK) Consortium | Health Services Laboratories and Alex Alderton, Roberto Amato, Jeffrey Barrett, Sonia Goncalves, Ewan Harrison, David K. Jackson, Ian Johnston, Dominic Kwiatkowski, Cordelia Langford, John Sillitoe on behalf of the Wellcome Sanger Institute COVID-19 Surveillance Team |
| EPI_ISL_2806459 | Lighthouse Laboratory Plymouth | Wellcome Sanger Institute for the COVID-19 Genomics UK (COG-UK) Consortium | Lighthouse Laboratory Plymouth and Alex Alderton, Roberto Amato, Jeffrey Barrett, Sonia Goncalves, Ewan Harrison, David K. Jackson, Ian Johnston, Dominic Kwiatkowski, Cordelia Langford, John Sillitoe on behalf of the Wellcome Sanger Institute COVID-19 Surveillance Team |
| EPI_ISL_2806460 | Health Services Laboratories | Wellcome Sanger Institute for the COVID-19 Genomics UK (COG-UK) Consortium | Health Services Laboratories and Alex Alderton, Roberto Amato, Jeffrey Barrett, Sonia Goncalves, Ewan Harrison, David K. Jackson, Ian Johnston, Dominic Kwiatkowski, Cordelia Langford, John Sillitoe on behalf of the Wellcome Sanger Institute COVID-19 Surveillance Team |
| EPI_ISL_2806461, EPI_ISL_2806462 | Lighthouse Laboratory Plymouth | Wellcome Sanger Institute for the COVID-19 Genomics UK (COG-UK) Consortium | Lighthouse Laboratory Plymouth and Alex Alderton, Roberto Amato, Jeffrey Barrett, Sonia Goncalves, Ewan Harrison, David K. Jackson, Ian Johnston, Dominic Kwiatkowski, Cordelia Langford, John Sillitoe on behalf of the Wellcome Sanger Institute COVID-19 Surveillance Team |
| EPI_ISL_2806463, EPI_ISL_2806464 | Lighthouse Lab in Milton Keynes | Wellcome Sanger Institute for the COVID-19 Genomics UK (COG-UK) Consortium | The Lighthouse Lab in Milton Keynes and Alex Alderton, Roberto Amato, Jeffrey Barrett, Sonia Goncalves, Ewan Harrison, David K. Jackson, Ian Johnston, Dominic Kwiatkowski, Cordelia Langford, John Sillitoe on behalf of the Wellcome Sanger Institute COVID-19 Surveillance Team |
| EPI_ISL_2806465 | Lighthouse Laboratory Plymouth | Wellcome Sanger Institute for the COVID-19 Genomics UK (COG-UK) Consortium | Lighthouse Laboratory Plymouth and Alex Alderton, Roberto Amato, Jeffrey Barrett, Sonia Goncalves, Ewan Harrison, David K. Jackson, Ian Johnston, Dominic Kwiatkowski, Cordelia Langford, John Sillitoe on behalf of the Wellcome Sanger Institute COVID-19 Surveillance Team |
| EPI_ISL_2806466 | Lighthouse Lab in Milton Keynes | Wellcome Sanger Institute for the COVID-19 Genomics UK (COG-UK) Consortium | The Lighthouse Lab in Milton Keynes and Alex Alderton, Roberto Amato, Jeffrey Barrett, Sonia Goncalves, Ewan Harrison, David K. Jackson, Ian Johnston, Dominic Kwiatkowski, Cordelia Langford, John Sillitoe on behalf of the Wellcome Sanger Institute COVID-19 Surveillance Team |
| EPI_ISL_2806467 | Lighthouse Laboratory Plymouth | Wellcome Sanger Institute for the COVID-19 Genomics UK (COG-UK) Consortium | Lighthouse Laboratory Plymouth and Alex Alderton, Roberto Amato, Jeffrey Barrett, Sonia Goncalves, Ewan Harrison, David K. Jackson, Ian Johnston, Dominic Kwiatkowski, Cordelia Langford, John Sillitoe on behalf of the Wellcome Sanger Institute COVID-19 Surveillance Team |
| EPI_ISL_2806468, EPI_ISL_2806469 | Health Services Laboratories | Wellcome Sanger Institute for the COVID-19 Genomics UK (COG-UK) Consortium | Health Services Laboratories and Alex Alderton, Roberto Amato, Jeffrey Barrett, Sonia Goncalves, Ewan Harrison, David K. Jackson, Ian Johnston, Dominic Kwiatkowski, Cordelia Langford, John Sillitoe on behalf of the Wellcome Sanger Institute COVID-19 Surveillance Team |
| EPI_ISL_2806470 | Lighthouse Laboratory Plymouth | Wellcome Sanger Institute for the COVID-19 Genomics UK (COG-UK) Consortium | Lighthouse Laboratory Plymouth and Alex Alderton, Roberto Amato, Jeffrey Barrett, Sonia Goncalves, Ewan Harrison, David K. Jackson, Ian Johnston, Dominic Kwiatkowski, Cordelia Langford, John Sillitoe on behalf of the Wellcome Sanger Institute COVID-19 Surveillance Team |
| EPI_ISL_2806471 | Health Services Laboratories | Wellcome Sanger Institute for the COVID-19 Genomics UK (COG-UK) Consortium | Health Services Laboratories and Alex Alderton, Roberto Amato, Jeffrey Barrett, Sonia Goncalves, Ewan Harrison, David K. Jackson, Ian Johnston, Dominic Kwiatkowski, Cordelia Langford, John Sillitoe on behalf of the Wellcome Sanger Institute COVID-19 Surveillance Team |
| EPI_ISL_2806472 | Lighthouse Laboratory Plymouth | Wellcome Sanger Institute for the COVID-19 Genomics UK (COG-UK) Consortium | Lighthouse Laboratory Plymouth and Alex Alderton, Roberto Amato, Jeffrey Barrett, Sonia Goncalves, Ewan Harrison, David K. Jackson, Ian Johnston, Dominic Kwiatkowski, Cordelia Langford, John Sillitoe on behalf of the Wellcome Sanger Institute COVID-19 Surveillance Team |
| EPI_ISL_2806473, EPI_ISL_2806474, EPI_ISL_2806475 | Health Services Laboratories | Wellcome Sanger Institute for the COVID-19 Genomics UK (COG-UK) Consortium | Health Services Laboratories and Alex Alderton, Roberto Amato, Jeffrey Barrett, Sonia Goncalves, Ewan Harrison, David K. Jackson, Ian Johnston, Dominic Kwiatkowski, Cordelia Langford, John Sillitoe on behalf of the Wellcome Sanger Institute COVID-19 Surveillance Team |
| EPI_ISL_2806476 | Lighthouse Lab in Milton Keynes | Wellcome Sanger Institute for the COVID-19 Genomics UK (COG-UK) Consortium | The Lighthouse Lab in Milton Keynes and Alex Alderton, Roberto Amato, Jeffrey Barrett, Sonia Goncalves, Ewan Harrison, David K. Jackson, Ian Johnston, Dominic Kwiatkowski, Cordelia Langford, John Sillitoe on behalf of the Wellcome Sanger Institute COVID-19 Surveillance Team |
| EPI_ISL_2806477 | Health Services Laboratories | Wellcome Sanger Institute for the COVID-19 Genomics UK (COG-UK) Consortium | Health Services Laboratories and Alex Alderton, Roberto Amato, Jeffrey Barrett, Sonia Goncalves, Ewan Harrison, David K. Jackson, Ian Johnston, Dominic Kwiatkowski, Cordelia Langford, John Sillitoe on behalf of the Wellcome Sanger Institute COVID-19 Surveillance Team |
| EPI_ISL_2806478 | Lighthouse Lab in Milton Keynes | Wellcome Sanger Institute for the COVID-19 Genomics UK (COG-UK) Consortium | The Lighthouse Lab in Milton Keynes and Alex Alderton, Roberto Amato, Jeffrey Barrett, Sonia Goncalves, Ewan Harrison, David K. Jackson, Ian Johnston, Dominic Kwiatkowski, Cordelia Langford, John Sillitoe on behalf of the Wellcome Sanger Institute COVID-19 Surveillance Team |
| EPI_ISL_2806479 | Lighthouse Laboratory Plymouth | Wellcome Sanger Institute for the COVID-19 Genomics UK (COG-UK) Consortium | Lighthouse Laboratory Plymouth and Alex Alderton, Roberto Amato, Jeffrey Barrett, Sonia Goncalves, Ewan Harrison, David K. Jackson, Ian Johnston, Dominic Kwiatkowski, Cordelia Langford, John Sillitoe on behalf of the Wellcome Sanger Institute COVID-19 Surveillance Team |
| EPI_ISL_2806480 | Health Services Laboratories | Wellcome Sanger Institute for the COVID-19 Genomics UK (COG-UK) Consortium | Health Services Laboratories and Alex Alderton, Roberto Amato, Jeffrey Barrett, Sonia Goncalves, Ewan Harrison, David K. Jackson, Ian Johnston, Dominic Kwiatkowski, Cordelia Langford, John Sillitoe on behalf of the Wellcome Sanger Institute COVID-19 Surveillance Team |
| EPI_ISL_2806481, EPI_ISL_2806482 | Lighthouse Laboratory Plymouth | Wellcome Sanger Institute for the COVID-19 Genomics UK (COG-UK) Consortium | Lighthouse Laboratory Plymouth and Alex Alderton, Roberto Amato, Jeffrey Barrett, Sonia Goncalves, Ewan Harrison, David K. Jackson, Ian Johnston, Dominic Kwiatkowski, Cordelia Langford, John Sillitoe on behalf of the Wellcome Sanger Institute COVID-19 Surveillance Team |
| EPI_ISL_2806483, EPI_ISL_2806484, EPI_ISL_2806485 | Health Services Laboratories | Wellcome Sanger Institute for the COVID-19 Genomics UK (COG-UK) Consortium | Health Services Laboratories and Alex Alderton, Roberto Amato, Jeffrey Barrett, Sonia Goncalves, Ewan Harrison, David K. Jackson, Ian Johnston, Dominic Kwiatkowski, Cordelia Langford, John Sillitoe on behalf of the Wellcome Sanger Institute COVID-19 Surveillance Team |
| EPI_ISL_2806486 | Lighthouse Laboratory Plymouth | Wellcome Sanger Institute for the COVID-19 Genomics UK (COG-UK) | Lighthouse Laboratory Plymouth and Alex Alderton, Roberto Amato, Jeffrey Barrett, Sonia Goncalves, Ewan Harrison, David K. Jackson, Ian Johnston, Dominic Kwiatkowski, Cordelia Langford, John Sillitoe on behalf of the Wellcome Sanger |

| | | | |
|---|---|---|---|
| | | Consortium | Institute COVID-19 Surveillance Team |
| EPI_ISL_2806488 | Lighthouse Lab in Milton Keynes | Wellcome Sanger Institute for the COVID-19 Genomics UK (COG-UK) Consortium | The Lighthouse Lab in Milton Keynes and Alex Alderton, Roberto Amato, Jeffrey Barrett, Sonia Goncalves, Ewan Harrison, David K. Jackson, Ian Johnston, Dominic Kwiatkowski, Cordelia Langford, John Sillitoe on behalf of the Wellcome Sanger Institute COVID-19 Surveillance Team |
| EPI_ISL_2806489 | Health Services Laboratories | Wellcome Sanger Institute for the COVID-19 Genomics UK (COG-UK) Consortium | Health Services Laboratories and Alex Alderton, Roberto Amato, Jeffrey Barrett, Sonia Goncalves, Ewan Harrison, David K. Jackson, Ian Johnston, Dominic Kwiatkowski, Cordelia Langford, John Sillitoe on behalf of the Wellcome Sanger Institute COVID-19 Surveillance Team |
| EPI_ISL_2806490 | Lighthouse Lab in Milton Keynes | Wellcome Sanger Institute for the COVID-19 Genomics UK (COG-UK) Consortium | The Lighthouse Lab in Milton Keynes and Alex Alderton, Roberto Amato, Jeffrey Barrett, Sonia Goncalves, Ewan Harrison, David K. Jackson, Ian Johnston, Dominic Kwiatkowski, Cordelia Langford, John Sillitoe on behalf of the Wellcome Sanger Institute COVID-19 Surveillance Team |
| EPI_ISL_2806491 | Health Services Laboratories | Wellcome Sanger Institute for the COVID-19 Genomics UK (COG-UK) Consortium | Health Services Laboratories and Alex Alderton, Roberto Amato, Jeffrey Barrett, Sonia Goncalves, Ewan Harrison, David K. Jackson, Ian Johnston, Dominic Kwiatkowski, Cordelia Langford, John Sillitoe on behalf of the Wellcome Sanger Institute COVID-19 Surveillance Team |
| EPI_ISL_2806492 | Lighthouse Laboratory Plymouth | Wellcome Sanger Institute for the COVID-19 Genomics UK (COG-UK) Consortium | Lighthouse Laboratory Plymouth and Alex Alderton, Roberto Amato, Jeffrey Barrett, Sonia Goncalves, Ewan Harrison, David K. Jackson, Ian Johnston, Dominic Kwiatkowski, Cordelia Langford, John Sillitoe on behalf of the Wellcome Sanger Institute COVID-19 Surveillance Team |
| EPI_ISL_2806493 | Lighthouse Lab in Milton Keynes | Wellcome Sanger Institute for the COVID-19 Genomics UK (COG-UK) Consortium | The Lighthouse Lab in Milton Keynes and Alex Alderton, Roberto Amato, Jeffrey Barrett, Sonia Goncalves, Ewan Harrison, David K. Jackson, Ian Johnston, Dominic Kwiatkowski, Cordelia Langford, John Sillitoe on behalf of the Wellcome Sanger Institute COVID-19 Surveillance Team |
| EPI_ISL_2806494 | Health Services Laboratories | Wellcome Sanger Institute for the COVID-19 Genomics UK (COG-UK) Consortium | Health Services Laboratories and Alex Alderton, Roberto Amato, Jeffrey Barrett, Sonia Goncalves, Ewan Harrison, David K. Jackson, Ian Johnston, Dominic Kwiatkowski, Cordelia Langford, John Sillitoe on behalf of the Wellcome Sanger Institute COVID-19 Surveillance Team |
| EPI_ISL_2806495 | Lighthouse Lab in Milton Keynes | Wellcome Sanger Institute for the COVID-19 Genomics UK (COG-UK) Consortium | The Lighthouse Lab in Milton Keynes and Alex Alderton, Roberto Amato, Jeffrey Barrett, Sonia Goncalves, Ewan Harrison, David K. Jackson, Ian Johnston, Dominic Kwiatkowski, Cordelia Langford, John Sillitoe on behalf of the Wellcome Sanger Institute COVID-19 Surveillance Team |
| EPI_ISL_2806496, EPI_ISL_2806497, EPI_ISL_2806498 | Health Services Laboratories | Wellcome Sanger Institute for the COVID-19 Genomics UK (COG-UK) Consortium | Health Services Laboratories and Alex Alderton, Roberto Amato, Jeffrey Barrett, Sonia Goncalves, Ewan Harrison, David K. Jackson, Ian Johnston, Dominic Kwiatkowski, Cordelia Langford, John Sillitoe on behalf of the Wellcome Sanger Institute COVID-19 Surveillance Team |
| EPI_ISL_2806499 | Lighthouse Lab in Milton Keynes | Wellcome Sanger Institute for the COVID-19 Genomics UK (COG-UK) Consortium | The Lighthouse Lab in Milton Keynes and Alex Alderton, Roberto Amato, Jeffrey Barrett, Sonia Goncalves, Ewan Harrison, David K. Jackson, Ian Johnston, Dominic Kwiatkowski, Cordelia Langford, John Sillitoe on behalf of the Wellcome Sanger Institute COVID-19 Surveillance Team |
| EPI_ISL_2806500, EPI_ISL_2806501 | Health Services Laboratories | Wellcome Sanger Institute for the COVID-19 Genomics UK (COG-UK) Consortium | Health Services Laboratories and Alex Alderton, Roberto Amato, Jeffrey Barrett, Sonia Goncalves, Ewan Harrison, David K. Jackson, Ian Johnston, Dominic Kwiatkowski, Cordelia Langford, John Sillitoe on behalf of the Wellcome Sanger Institute COVID-19 Surveillance Team |
| EPI_ISL_2806502 | Lighthouse Laboratory Plymouth | Wellcome Sanger Institute for the COVID-19 Genomics UK (COG-UK) Consortium | Lighthouse Laboratory Plymouth and Alex Alderton, Roberto Amato, Jeffrey Barrett, Sonia Goncalves, Ewan Harrison, David K. Jackson, Ian Johnston, Dominic Kwiatkowski, Cordelia Langford, John Sillitoe on behalf of the Wellcome Sanger Institute COVID-19 Surveillance Team |
| EPI_ISL_2806503, EPI_ISL_2806505 | Health Services Laboratories | Wellcome Sanger Institute for the COVID-19 Genomics UK (COG-UK) Consortium | Health Services Laboratories and Alex Alderton, Roberto Amato, Jeffrey Barrett, Sonia Goncalves, Ewan Harrison, David K. Jackson, Ian Johnston, Dominic Kwiatkowski, Cordelia Langford, John Sillitoe on behalf of the Wellcome Sanger Institute COVID-19 Surveillance Team |
| EPI_ISL_2806506, EPI_ISL_2806507 | Lighthouse Lab in Milton Keynes | Wellcome Sanger Institute for the COVID-19 Genomics UK (COG-UK) Consortium | The Lighthouse Lab in Milton Keynes and Alex Alderton, Roberto Amato, Jeffrey Barrett, Sonia Goncalves, Ewan Harrison, David K. Jackson, Ian Johnston, Dominic Kwiatkowski, Cordelia Langford, John Sillitoe on behalf of the Wellcome Sanger Institute COVID-19 Surveillance Team |
| EPI_ISL_2806508, EPI_ISL_2806509 | Health Services Laboratories | Wellcome Sanger Institute for the COVID-19 Genomics UK (COG-UK) Consortium | Health Services Laboratories and Alex Alderton, Roberto Amato, Jeffrey Barrett, Sonia Goncalves, Ewan Harrison, David K. Jackson, Ian Johnston, Dominic Kwiatkowski, Cordelia Langford, John Sillitoe on behalf of the Wellcome Sanger Institute COVID-19 Surveillance Team |
| EPI_ISL_2806510 | Lighthouse Laboratory Plymouth | Wellcome Sanger Institute for the COVID-19 Genomics UK (COG-UK) Consortium | Lighthouse Laboratory Plymouth and Alex Alderton, Roberto Amato, Jeffrey Barrett, Sonia Goncalves, Ewan Harrison, David K. Jackson, Ian Johnston, Dominic Kwiatkowski, Cordelia Langford, John Sillitoe on behalf of the Wellcome Sanger Institute COVID-19 Surveillance Team |
| EPI_ISL_2806511, EPI_ISL_2806512, EPI_ISL_2806513, EPI_ISL_2806514 | Health Services Laboratories | Wellcome Sanger Institute for the COVID-19 Genomics UK (COG-UK) Consortium | Health Services Laboratories and Alex Alderton, Roberto Amato, Jeffrey Barrett, Sonia Goncalves, Ewan Harrison, David K. Jackson, Ian Johnston, Dominic Kwiatkowski, Cordelia Langford, John Sillitoe on behalf of the Wellcome Sanger Institute COVID-19 Surveillance Team |
| EPI_ISL_2806515, EPI_ISL_2806516 | Lighthouse Lab in Milton Keynes | Wellcome Sanger Institute for the COVID-19 Genomics UK (COG-UK) Consortium | The Lighthouse Lab in Milton Keynes and Alex Alderton, Roberto Amato, Jeffrey Barrett, Sonia Goncalves, Ewan Harrison, David K. Jackson, Ian Johnston, Dominic Kwiatkowski, Cordelia Langford, John Sillitoe on behalf of the Wellcome Sanger Institute COVID-19 Surveillance Team |
| EPI_ISL_2806517 | Lighthouse Laboratory Plymouth | Wellcome Sanger Institute for the COVID-19 Genomics UK (COG-UK) Consortium | Lighthouse Laboratory Plymouth and Alex Alderton, Roberto Amato, Jeffrey Barrett, Sonia Goncalves, Ewan Harrison, David K. Jackson, Ian Johnston, Dominic Kwiatkowski, Cordelia Langford, John Sillitoe on behalf of the Wellcome Sanger Institute COVID-19 Surveillance Team |
| EPI_ISL_2806518 | Health Services Laboratories | Wellcome Sanger Institute for the COVID-19 Genomics UK (COG-UK) Consortium | Health Services Laboratories and Alex Alderton, Roberto Amato, Jeffrey Barrett, Sonia Goncalves, Ewan Harrison, David K. Jackson, Ian Johnston, Dominic Kwiatkowski, Cordelia Langford, John Sillitoe on behalf of the Wellcome Sanger Institute COVID-19 Surveillance Team |
| EPI_ISL_2806519 | Lighthouse Lab in Milton Keynes | Wellcome Sanger Institute for the COVID-19 Genomics UK (COG-UK) Consortium | The Lighthouse Lab in Milton Keynes and Alex Alderton, Roberto Amato, Jeffrey Barrett, Sonia Goncalves, Ewan Harrison, David K. Jackson, Ian Johnston, Dominic Kwiatkowski, Cordelia Langford, John Sillitoe on behalf of the Wellcome Sanger Institute COVID-19 Surveillance Team |
| EPI_ISL_2806520 | Health Services Laboratories | Wellcome Sanger Institute for the COVID-19 Genomics UK (COG-UK) Consortium | Health Services Laboratories and Alex Alderton, Roberto Amato, Jeffrey Barrett, Sonia Goncalves, Ewan Harrison, David K. Jackson, Ian Johnston, Dominic Kwiatkowski, Cordelia Langford, John Sillitoe on behalf of the Wellcome Sanger Institute COVID-19 Surveillance Team |
| EPI_ISL_2806521 | Lighthouse Laboratory Plymouth | Wellcome Sanger Institute for the COVID-19 Genomics UK (COG-UK) Consortium | Lighthouse Laboratory Plymouth and Alex Alderton, Roberto Amato, Jeffrey Barrett, Sonia Goncalves, Ewan Harrison, David K. Jackson, Ian Johnston, Dominic Kwiatkowski, Cordelia Langford, John Sillitoe on behalf of the Wellcome Sanger Institute COVID-19 Surveillance Team |
| EPI_ISL_2806522 | Lighthouse Lab in Milton Keynes | Wellcome Sanger Institute for the COVID-19 Genomics UK (COG-UK) Consortium | The Lighthouse Lab in Milton Keynes and Alex Alderton, Roberto Amato, Jeffrey Barrett, Sonia Goncalves, Ewan Harrison, David K. Jackson, Ian Johnston, Dominic Kwiatkowski, Cordelia Langford, John Sillitoe on behalf of the Wellcome Sanger Institute COVID-19 Surveillance Team |
| EPI_ISL_2806523, EPI_ISL_2806524 | Lighthouse Laboratory Plymouth | Wellcome Sanger Institute for the COVID-19 Genomics UK (COG-UK) Consortium | Lighthouse Laboratory Plymouth and Alex Alderton, Roberto Amato, Jeffrey Barrett, Sonia Goncalves, Ewan Harrison, David K. Jackson, Ian Johnston, Dominic Kwiatkowski, Cordelia Langford, John Sillitoe on behalf of the Wellcome Sanger Institute COVID-19 Surveillance Team |
| EPI_ISL_2806525 | Health Services Laboratories | Wellcome Sanger Institute for the COVID-19 Genomics UK (COG-UK) Consortium | Health Services Laboratories and Alex Alderton, Roberto Amato, Jeffrey Barrett, Sonia Goncalves, Ewan Harrison, David K. Jackson, Ian Johnston, Dominic Kwiatkowski, Cordelia Langford, John Sillitoe on behalf of the Wellcome Sanger Institute COVID-19 Surveillance Team |
| EPI_ISL_2806526 | Lighthouse Lab in Milton Keynes | Wellcome Sanger Institute for the COVID-19 Genomics UK (COG-UK) Consortium | The Lighthouse Lab in Milton Keynes and Alex Alderton, Roberto Amato, Jeffrey Barrett, Sonia Goncalves, Ewan Harrison, David K. Jackson, Ian Johnston, Dominic Kwiatkowski, Cordelia Langford, John Sillitoe on behalf of the Wellcome Sanger Institute COVID-19 Surveillance Team |
| EPI_ISL_2806527, EPI_ISL_2806528 | Health Services Laboratories | Wellcome Sanger Institute for the COVID-19 Genomics UK (COG-UK) Consortium | Health Services Laboratories and Alex Alderton, Roberto Amato, Jeffrey Barrett, Sonia Goncalves, Ewan Harrison, David K. Jackson, Ian Johnston, Dominic Kwiatkowski, Cordelia Langford, John Sillitoe on behalf of the Wellcome Sanger Institute COVID-19 Surveillance Team |
| EPI_ISL_2806529 | Lighthouse Lab in Milton Keynes | Wellcome Sanger Institute for the COVID-19 Genomics UK (COG-UK) Consortium | The Lighthouse Lab in Milton Keynes and Alex Alderton, Roberto Amato, Jeffrey Barrett, Sonia Goncalves, Ewan Harrison, David K. Jackson, Ian Johnston, Dominic Kwiatkowski, Cordelia Langford, John Sillitoe on behalf of the Wellcome Sanger Institute COVID-19 Surveillance Team |
| EPI_ISL_2806530 | Health Services Laboratories | Wellcome Sanger Institute for the COVID-19 Genomics UK (COG-UK) Consortium | Health Services Laboratories and Alex Alderton, Roberto Amato, Jeffrey Barrett, Sonia Goncalves, Ewan Harrison, David K. Jackson, Ian Johnston, Dominic Kwiatkowski, Cordelia Langford, John Sillitoe on behalf of the Wellcome Sanger Institute COVID-19 Surveillance Team |
| EPI_ISL_2806531 | Lighthouse Laboratory Plymouth | Wellcome Sanger Institute for the COVID-19 Genomics UK (COG-UK) Consortium | Lighthouse Laboratory Plymouth and Alex Alderton, Roberto Amato, Jeffrey Barrett, Sonia Goncalves, Ewan Harrison, David K. Jackson, Ian Johnston, Dominic Kwiatkowski, Cordelia Langford, John Sillitoe on behalf of the Wellcome Sanger Institute COVID-19 Surveillance Team |
| EPI_ISL_2806532, EPI_ISL_2806533 | Lighthouse Lab in Milton Keynes | Wellcome Sanger Institute for the COVID-19 Genomics UK (COG-UK) Consortium | The Lighthouse Lab in Milton Keynes and Alex Alderton, Roberto Amato, Jeffrey Barrett, Sonia Goncalves, Ewan Harrison, David K. Jackson, Ian Johnston, Dominic Kwiatkowski, Cordelia Langford, John Sillitoe on behalf of the Wellcome Sanger Institute COVID-19 Surveillance Team |
| EPI_ISL_2806534 | Lighthouse Laboratory Plymouth | Wellcome Sanger Institute for the COVID-19 Genomics UK (COG-UK) Consortium | Lighthouse Laboratory Plymouth and Alex Alderton, Roberto Amato, Jeffrey Barrett, Sonia Goncalves, Ewan Harrison, David K. Jackson, Ian Johnston, Dominic Kwiatkowski, Cordelia Langford, John Sillitoe on behalf of the Wellcome Sanger Institute COVID-19 Surveillance Team |
| EPI_ISL_2806535, EPI_ISL_2806536, EPI_ISL_2806537, EPI_ISL_2806538, EPI_ISL_2806539, EPI_ISL_2806540 | Health Services Laboratories | Wellcome Sanger Institute for the COVID-19 Genomics UK (COG-UK) Consortium | Health Services Laboratories and Alex Alderton, Roberto Amato, Jeffrey Barrett, Sonia Goncalves, Ewan Harrison, David K. Jackson, Ian Johnston, Dominic Kwiatkowski, Cordelia Langford, John Sillitoe on behalf of the Wellcome Sanger Institute COVID-19 Surveillance Team |
| EPI_ISL_2806541 | Lighthouse Laboratory Plymouth | Wellcome Sanger Institute for the COVID-19 Genomics UK (COG-UK) Consortium | Lighthouse Laboratory Plymouth and Alex Alderton, Roberto Amato, Jeffrey Barrett, Sonia Goncalves, Ewan Harrison, David K. Jackson, Ian Johnston, Dominic Kwiatkowski, Cordelia Langford, John Sillitoe on behalf of the Wellcome Sanger Institute COVID-19 Surveillance Team |
| EPI_ISL_2806542 | Lighthouse Lab in Milton Keynes | Wellcome Sanger Institute for the COVID-19 Genomics UK (COG-UK) Consortium | The Lighthouse Lab in Milton Keynes and Alex Alderton, Roberto Amato, Jeffrey Barrett, Sonia Goncalves, Ewan Harrison, David K. Jackson, Ian Johnston, Dominic Kwiatkowski, Cordelia Langford, John Sillitoe on behalf of the Wellcome Sanger Institute COVID-19 Surveillance Team |
| EPI_ISL_2806543, EPI_ISL_2806544 | Lighthouse Laboratory Plymouth | Wellcome Sanger Institute for the COVID-19 Genomics UK (COG-UK) Consortium | Lighthouse Laboratory Plymouth and Alex Alderton, Roberto Amato, Jeffrey Barrett, Sonia Goncalves, Ewan Harrison, David K. Jackson, Ian Johnston, Dominic Kwiatkowski, Cordelia Langford, John Sillitoe on behalf of the Wellcome Sanger Institute COVID-19 Surveillance Team |
| EPI_ISL_2806545 | Health Services Laboratories | Wellcome Sanger Institute for the COVID-19 Genomics UK (COG-UK) Consortium | Health Services Laboratories and Alex Alderton, Roberto Amato, Jeffrey Barrett, Sonia Goncalves, Ewan Harrison, David K. Jackson, Ian Johnston, Dominic Kwiatkowski, Cordelia Langford, John Sillitoe on behalf of the Wellcome Sanger Institute COVID-19 Surveillance Team |
| EPI_ISL_2806546 | Lighthouse Lab in Milton Keynes | Wellcome Sanger Institute for the COVID-19 Genomics UK (COG-UK) Consortium | The Lighthouse Lab in Milton Keynes and Alex Alderton, Roberto Amato, Jeffrey Barrett, Sonia Goncalves, Ewan Harrison, David K. Jackson, Ian Johnston, Dominic Kwiatkowski, Cordelia Langford, John Sillitoe on behalf of the Wellcome Sanger Institute COVID-19 Surveillance Team |
| EPI_ISL_2806547, EPI_ISL_2806548 | Lighthouse Laboratory Plymouth | Wellcome Sanger Institute for the COVID-19 Genomics UK (COG-UK) Consortium | Lighthouse Laboratory Plymouth and Alex Alderton, Roberto Amato, Jeffrey Barrett, Sonia Goncalves, Ewan Harrison, David K. Jackson, Ian Johnston, Dominic Kwiatkowski, Cordelia Langford, John Sillitoe on behalf of the Wellcome Sanger Institute COVID-19 Surveillance Team |
| EPI_ISL_2806549 | Health Services Laboratories | Wellcome Sanger Institute for the COVID-19 Genomics UK (COG-UK) Consortium | Health Services Laboratories and Alex Alderton, Roberto Amato, Jeffrey Barrett, Sonia Goncalves, Ewan Harrison, David K. Jackson, Ian Johnston, Dominic Kwiatkowski, Cordelia Langford, John Sillitoe on behalf of the Wellcome Sanger Institute COVID-19 Surveillance Team |
| EPI_ISL_2806550 | Lighthouse Lab in Milton Keynes | Wellcome Sanger Institute for the COVID-19 Genomics UK (COG-UK) Consortium | The Lighthouse Lab in Milton Keynes and Alex Alderton, Roberto Amato, Jeffrey Barrett, Sonia Goncalves, Ewan Harrison, David K. Jackson, Ian Johnston, Dominic Kwiatkowski, Cordelia Langford, John Sillitoe on behalf of the Wellcome Sanger Institute COVID-19 Surveillance Team |

| | | | |
|---|---|---|---|
| EPI_ISL_2806551, EPI_ISL_2806552, EPI_ISL_2806553 | Health Services Laboratories | Wellcome Sanger Institute for the COVID-19 Genomics UK (COG-UK) Consortium | Health Services Laboratories and Alex Alderton, Roberto Amato, Jeffrey Barrett, Sonia Goncalves, Ewan Harrison, David K. Jackson, Ian Johnston, Dominic Kwiatkowski, Cordelia Langford, John Sillitoe on behalf of the Wellcome Sanger Institute COVID-19 Surveillance Team |
| EPI_ISL_2806554 | Lighthouse Laboratory Plymouth | Wellcome Sanger Institute for the COVID-19 Genomics UK (COG-UK) Consortium | Lighthouse Laboratory Plymouth and Alex Alderton, Roberto Amato, Jeffrey Barrett, Sonia Goncalves, Ewan Harrison, David K. Jackson, Ian Johnston, Dominic Kwiatkowski, Cordelia Langford, John Sillitoe on behalf of the Wellcome Sanger Institute COVID-19 Surveillance Team |
| EPI_ISL_2806555 | Lighthouse Lab in Milton Keynes | Wellcome Sanger Institute for the COVID-19 Genomics UK (COG-UK) Consortium | The Lighthouse Lab in Milton Keynes and Alex Alderton, Roberto Amato, Jeffrey Barrett, Sonia Goncalves, Ewan Harrison, David K. Jackson, Ian Johnston, Dominic Kwiatkowski, Cordelia Langford, John Sillitoe on behalf of the Wellcome Sanger Institute COVID-19 Surveillance Team |
| EPI_ISL_2806556, EPI_ISL_2806557, EPI_ISL_2806558 | Health Services Laboratories | Wellcome Sanger Institute for the COVID-19 Genomics UK (COG-UK) Consortium | Health Services Laboratories and Alex Alderton, Roberto Amato, Jeffrey Barrett, Sonia Goncalves, Ewan Harrison, David K. Jackson, Ian Johnston, Dominic Kwiatkowski, Cordelia Langford, John Sillitoe on behalf of the Wellcome Sanger Institute COVID-19 Surveillance Team |
| EPI_ISL_2806559 | Lighthouse Laboratory Plymouth | Wellcome Sanger Institute for the COVID-19 Genomics UK (COG-UK) Consortium | Lighthouse Laboratory Plymouth and Alex Alderton, Roberto Amato, Jeffrey Barrett, Sonia Goncalves, Ewan Harrison, David K. Jackson, Ian Johnston, Dominic Kwiatkowski, Cordelia Langford, John Sillitoe on behalf of the Wellcome Sanger Institute COVID-19 Surveillance Team |
| EPI_ISL_2806560 | Lighthouse Lab in Milton Keynes | Wellcome Sanger Institute for the COVID-19 Genomics UK (COG-UK) Consortium | The Lighthouse Lab in Milton Keynes and Alex Alderton, Roberto Amato, Jeffrey Barrett, Sonia Goncalves, Ewan Harrison, David K. Jackson, Ian Johnston, Dominic Kwiatkowski, Cordelia Langford, John Sillitoe on behalf of the Wellcome Sanger Institute COVID-19 Surveillance Team |
| EPI_ISL_2806561, EPI_ISL_2806562 | Lighthouse Laboratory Plymouth | Wellcome Sanger Institute for the COVID-19 Genomics UK (COG-UK) Consortium | Lighthouse Laboratory Plymouth and Alex Alderton, Roberto Amato, Jeffrey Barrett, Sonia Goncalves, Ewan Harrison, David K. Jackson, Ian Johnston, Dominic Kwiatkowski, Cordelia Langford, John Sillitoe on behalf of the Wellcome Sanger Institute COVID-19 Surveillance Team |
| EPI_ISL_2806563 | Lighthouse Lab in Milton Keynes | Wellcome Sanger Institute for the COVID-19 Genomics UK (COG-UK) Consortium | The Lighthouse Lab in Milton Keynes and Alex Alderton, Roberto Amato, Jeffrey Barrett, Sonia Goncalves, Ewan Harrison, David K. Jackson, Ian Johnston, Dominic Kwiatkowski, Cordelia Langford, John Sillitoe on behalf of the Wellcome Sanger Institute COVID-19 Surveillance Team |
| EPI_ISL_2806564 | Lighthouse Laboratory Plymouth | Wellcome Sanger Institute for the COVID-19 Genomics UK (COG-UK) Consortium | Lighthouse Laboratory Plymouth and Alex Alderton, Roberto Amato, Jeffrey Barrett, Sonia Goncalves, Ewan Harrison, David K. Jackson, Ian Johnston, Dominic Kwiatkowski, Cordelia Langford, John Sillitoe on behalf of the Wellcome Sanger Institute COVID-19 Surveillance Team |
| EPI_ISL_2806565 | Health Services Laboratories | Wellcome Sanger Institute for the COVID-19 Genomics UK (COG-UK) Consortium | Health Services Laboratories and Alex Alderton, Roberto Amato, Jeffrey Barrett, Sonia Goncalves, Ewan Harrison, David K. Jackson, Ian Johnston, Dominic Kwiatkowski, Cordelia Langford, John Sillitoe on behalf of the Wellcome Sanger Institute COVID-19 Surveillance Team |
| EPI_ISL_2806566 | Lighthouse Lab in Milton Keynes | Wellcome Sanger Institute for the COVID-19 Genomics UK (COG-UK) Consortium | The Lighthouse Lab in Milton Keynes and Alex Alderton, Roberto Amato, Jeffrey Barrett, Sonia Goncalves, Ewan Harrison, David K. Jackson, Ian Johnston, Dominic Kwiatkowski, Cordelia Langford, John Sillitoe on behalf of the Wellcome Sanger Institute COVID-19 Surveillance Team |
| EPI_ISL_2806567, EPI_ISL_2806568, EPI_ISL_2806569 | Health Services Laboratories | Wellcome Sanger Institute for the COVID-19 Genomics UK (COG-UK) Consortium | Health Services Laboratories and Alex Alderton, Roberto Amato, Jeffrey Barrett, Sonia Goncalves, Ewan Harrison, David K. Jackson, Ian Johnston, Dominic Kwiatkowski, Cordelia Langford, John Sillitoe on behalf of the Wellcome Sanger Institute COVID-19 Surveillance Team |
| EPI_ISL_2806570 | Lighthouse Laboratory Plymouth | Wellcome Sanger Institute for the COVID-19 Genomics UK (COG-UK) Consortium | Lighthouse Laboratory Plymouth and Alex Alderton, Roberto Amato, Jeffrey Barrett, Sonia Goncalves, Ewan Harrison, David K. Jackson, Ian Johnston, Dominic Kwiatkowski, Cordelia Langford, John Sillitoe on behalf of the Wellcome Sanger Institute COVID-19 Surveillance Team |
| EPI_ISL_2806571, EPI_ISL_2806572 | Health Services Laboratories | Wellcome Sanger Institute for the COVID-19 Genomics UK (COG-UK) Consortium | Health Services Laboratories and Alex Alderton, Roberto Amato, Jeffrey Barrett, Sonia Goncalves, Ewan Harrison, David K. Jackson, Ian Johnston, Dominic Kwiatkowski, Cordelia Langford, John Sillitoe on behalf of the Wellcome Sanger Institute COVID-19 Surveillance Team |
| EPI_ISL_2806573 | Lighthouse Lab in Milton Keynes | Wellcome Sanger Institute for the COVID-19 Genomics UK (COG-UK) Consortium | The Lighthouse Lab in Milton Keynes and Alex Alderton, Roberto Amato, Jeffrey Barrett, Sonia Goncalves, Ewan Harrison, David K. Jackson, Ian Johnston, Dominic Kwiatkowski, Cordelia Langford, John Sillitoe on behalf of the Wellcome Sanger Institute COVID-19 Surveillance Team |
| EPI_ISL_2806574 | Health Services Laboratories | Wellcome Sanger Institute for the COVID-19 Genomics UK (COG-UK) Consortium | Health Services Laboratories and Alex Alderton, Roberto Amato, Jeffrey Barrett, Sonia Goncalves, Ewan Harrison, David K. Jackson, Ian Johnston, Dominic Kwiatkowski, Cordelia Langford, John Sillitoe on behalf of the Wellcome Sanger Institute COVID-19 Surveillance Team |
| EPI_ISL_2806575 | Lighthouse Laboratory Plymouth | Wellcome Sanger Institute for the COVID-19 Genomics UK (COG-UK) Consortium | Lighthouse Laboratory Plymouth and Alex Alderton, Roberto Amato, Jeffrey Barrett, Sonia Goncalves, Ewan Harrison, David K. Jackson, Ian Johnston, Dominic Kwiatkowski, Cordelia Langford, John Sillitoe on behalf of the Wellcome Sanger Institute COVID-19 Surveillance Team |
| EPI_ISL_2806576 | Health Services Laboratories | Wellcome Sanger Institute for the COVID-19 Genomics UK (COG-UK) Consortium | Health Services Laboratories and Alex Alderton, Roberto Amato, Jeffrey Barrett, Sonia Goncalves, Ewan Harrison, David K. Jackson, Ian Johnston, Dominic Kwiatkowski, Cordelia Langford, John Sillitoe on behalf of the Wellcome Sanger Institute COVID-19 Surveillance Team |
| EPI_ISL_2806577 | Lighthouse Laboratory Plymouth | Wellcome Sanger Institute for the COVID-19 Genomics UK (COG-UK) Consortium | Lighthouse Laboratory Plymouth and Alex Alderton, Roberto Amato, Jeffrey Barrett, Sonia Goncalves, Ewan Harrison, David K. Jackson, Ian Johnston, Dominic Kwiatkowski, Cordelia Langford, John Sillitoe on behalf of the Wellcome Sanger Institute COVID-19 Surveillance Team |
| EPI_ISL_2806578, EPI_ISL_2806579, EPI_ISL_2806580 | Health Services Laboratories | Wellcome Sanger Institute for the COVID-19 Genomics UK (COG-UK) Consortium | Health Services Laboratories and Alex Alderton, Roberto Amato, Jeffrey Barrett, Sonia Goncalves, Ewan Harrison, David K. Jackson, Ian Johnston, Dominic Kwiatkowski, Cordelia Langford, John Sillitoe on behalf of the Wellcome Sanger Institute COVID-19 Surveillance Team |
| EPI_ISL_2806581, EPI_ISL_2806582 | Lighthouse Lab in Milton Keynes | Wellcome Sanger Institute for the COVID-19 Genomics UK (COG-UK) Consortium | The Lighthouse Lab in Milton Keynes and Alex Alderton, Roberto Amato, Jeffrey Barrett, Sonia Goncalves, Ewan Harrison, David K. Jackson, Ian Johnston, Dominic Kwiatkowski, Cordelia Langford, John Sillitoe on behalf of the Wellcome Sanger Institute COVID-19 Surveillance Team |
| EPI_ISL_2806583 | Lighthouse Laboratory Plymouth | Wellcome Sanger Institute for the COVID-19 Genomics UK (COG-UK) Consortium | Lighthouse Laboratory Plymouth and Alex Alderton, Roberto Amato, Jeffrey Barrett, Sonia Goncalves, Ewan Harrison, David K. Jackson, Ian Johnston, Dominic Kwiatkowski, Cordelia Langford, John Sillitoe on behalf of the Wellcome Sanger Institute COVID-19 Surveillance Team |
| EPI_ISL_2806584, EPI_ISL_2806585 | Health Services Laboratories | Wellcome Sanger Institute for the COVID-19 Genomics UK (COG-UK) Consortium | Health Services Laboratories and Alex Alderton, Roberto Amato, Jeffrey Barrett, Sonia Goncalves, Ewan Harrison, David K. Jackson, Ian Johnston, Dominic Kwiatkowski, Cordelia Langford, John Sillitoe on behalf of the Wellcome Sanger Institute COVID-19 Surveillance Team |
| EPI_ISL_2806586 | Lighthouse Laboratory Plymouth | Wellcome Sanger Institute for the COVID-19 Genomics UK (COG-UK) Consortium | Lighthouse Laboratory Plymouth and Alex Alderton, Roberto Amato, Jeffrey Barrett, Sonia Goncalves, Ewan Harrison, David K. Jackson, Ian Johnston, Dominic Kwiatkowski, Cordelia Langford, John Sillitoe on behalf of the Wellcome Sanger Institute COVID-19 Surveillance Team |
| EPI_ISL_2806587 | Lighthouse Lab in Milton Keynes | Wellcome Sanger Institute for the COVID-19 Genomics UK (COG-UK) Consortium | The Lighthouse Lab in Milton Keynes and Alex Alderton, Roberto Amato, Jeffrey Barrett, Sonia Goncalves, Ewan Harrison, David K. Jackson, Ian Johnston, Dominic Kwiatkowski, Cordelia Langford, John Sillitoe on behalf of the Wellcome Sanger Institute COVID-19 Surveillance Team |
| EPI_ISL_2806588 | Health Services Laboratories | Wellcome Sanger Institute for the COVID-19 Genomics UK (COG-UK) Consortium | Health Services Laboratories and Alex Alderton, Roberto Amato, Jeffrey Barrett, Sonia Goncalves, Ewan Harrison, David K. Jackson, Ian Johnston, Dominic Kwiatkowski, Cordelia Langford, John Sillitoe on behalf of the Wellcome Sanger Institute COVID-19 Surveillance Team |
| EPI_ISL_2806589 | Lighthouse Laboratory Plymouth | Wellcome Sanger Institute for the COVID-19 Genomics UK (COG-UK) Consortium | Lighthouse Laboratory Plymouth and Alex Alderton, Roberto Amato, Jeffrey Barrett, Sonia Goncalves, Ewan Harrison, David K. Jackson, Ian Johnston, Dominic Kwiatkowski, Cordelia Langford, John Sillitoe on behalf of the Wellcome Sanger Institute COVID-19 Surveillance Team |
| EPI_ISL_2806590 | Health Services Laboratories | Wellcome Sanger Institute for the COVID-19 Genomics UK (COG-UK) Consortium | Health Services Laboratories and Alex Alderton, Roberto Amato, Jeffrey Barrett, Sonia Goncalves, Ewan Harrison, David K. Jackson, Ian Johnston, Dominic Kwiatkowski, Cordelia Langford, John Sillitoe on behalf of the Wellcome Sanger Institute COVID-19 Surveillance Team |
| EPI_ISL_2806591, EPI_ISL_2806592 | Lighthouse Lab in Milton Keynes | Wellcome Sanger Institute for the COVID-19 Genomics UK (COG-UK) Consortium | The Lighthouse Lab in Milton Keynes and Alex Alderton, Roberto Amato, Jeffrey Barrett, Sonia Goncalves, Ewan Harrison, David K. Jackson, Ian Johnston, Dominic Kwiatkowski, Cordelia Langford, John Sillitoe on behalf of the Wellcome Sanger Institute COVID-19 Surveillance Team |
| EPI_ISL_2806593, EPI_ISL_2806594, EPI_ISL_2806595 | Lighthouse Laboratory Plymouth | Wellcome Sanger Institute for the COVID-19 Genomics UK (COG-UK) Consortium | Lighthouse Laboratory Plymouth and Alex Alderton, Roberto Amato, Jeffrey Barrett, Sonia Goncalves, Ewan Harrison, David K. Jackson, Ian Johnston, Dominic Kwiatkowski, Cordelia Langford, John Sillitoe on behalf of the Wellcome Sanger Institute COVID-19 Surveillance Team |
| EPI_ISL_2806596, EPI_ISL_2806597, EPI_ISL_2806598 | Lighthouse Lab in Milton Keynes | Wellcome Sanger Institute for the COVID-19 Genomics UK (COG-UK) Consortium | The Lighthouse Lab in Milton Keynes and Alex Alderton, Roberto Amato, Jeffrey Barrett, Sonia Goncalves, Ewan Harrison, David K. Jackson, Ian Johnston, Dominic Kwiatkowski, Cordelia Langford, John Sillitoe on behalf of the Wellcome Sanger Institute COVID-19 Surveillance Team |
| EPI_ISL_2806599 | Health Services Laboratories | Wellcome Sanger Institute for the COVID-19 Genomics UK (COG-UK) Consortium | Health Services Laboratories and Alex Alderton, Roberto Amato, Jeffrey Barrett, Sonia Goncalves, Ewan Harrison, David K. Jackson, Ian Johnston, Dominic Kwiatkowski, Cordelia Langford, John Sillitoe on behalf of the Wellcome Sanger Institute COVID-19 Surveillance Team |
| EPI_ISL_2806600 | Lighthouse Laboratory Plymouth | Wellcome Sanger Institute for the COVID-19 Genomics UK (COG-UK) Consortium | Lighthouse Laboratory Plymouth and Alex Alderton, Roberto Amato, Jeffrey Barrett, Sonia Goncalves, Ewan Harrison, David K. Jackson, Ian Johnston, Dominic Kwiatkowski, Cordelia Langford, John Sillitoe on behalf of the Wellcome Sanger Institute COVID-19 Surveillance Team |
| EPI_ISL_2806601, EPI_ISL_2806602 | Health Services Laboratories | Wellcome Sanger Institute for the COVID-19 Genomics UK (COG-UK) Consortium | Health Services Laboratories and Alex Alderton, Roberto Amato, Jeffrey Barrett, Sonia Goncalves, Ewan Harrison, David K. Jackson, Ian Johnston, Dominic Kwiatkowski, Cordelia Langford, John Sillitoe on behalf of the Wellcome Sanger Institute COVID-19 Surveillance Team |
| EPI_ISL_2806603, EPI_ISL_2806604 | Lighthouse Lab in Milton Keynes | Wellcome Sanger Institute for the COVID-19 Genomics UK (COG-UK) Consortium | The Lighthouse Lab in Milton Keynes and Alex Alderton, Roberto Amato, Jeffrey Barrett, Sonia Goncalves, Ewan Harrison, David K. Jackson, Ian Johnston, Dominic Kwiatkowski, Cordelia Langford, John Sillitoe on behalf of the Wellcome Sanger Institute COVID-19 Surveillance Team |
| EPI_ISL_2806605 | Health Services Laboratories | Wellcome Sanger Institute for the COVID-19 Genomics UK (COG-UK) Consortium | Health Services Laboratories and Alex Alderton, Roberto Amato, Jeffrey Barrett, Sonia Goncalves, Ewan Harrison, David K. Jackson, Ian Johnston, Dominic Kwiatkowski, Cordelia Langford, John Sillitoe on behalf of the Wellcome Sanger Institute COVID-19 Surveillance Team |
| EPI_ISL_2806606 | Lighthouse Laboratory Plymouth | Wellcome Sanger Institute for the COVID-19 Genomics UK (COG-UK) Consortium | Lighthouse Laboratory Plymouth and Alex Alderton, Roberto Amato, Jeffrey Barrett, Sonia Goncalves, Ewan Harrison, David K. Jackson, Ian Johnston, Dominic Kwiatkowski, Cordelia Langford, John Sillitoe on behalf of the Wellcome Sanger Institute COVID-19 Surveillance Team |
| EPI_ISL_2806607 | Lighthouse Lab in Milton Keynes | Wellcome Sanger Institute for the COVID-19 Genomics UK (COG-UK) Consortium | The Lighthouse Lab in Milton Keynes and Alex Alderton, Roberto Amato, Jeffrey Barrett, Sonia Goncalves, Ewan Harrison, David K. Jackson, Ian Johnston, Dominic Kwiatkowski, Cordelia Langford, John Sillitoe on behalf of the Wellcome Sanger Institute COVID-19 Surveillance Team |
| EPI_ISL_2806608, EPI_ISL_2806609 | Health Services Laboratories | Wellcome Sanger Institute for the COVID-19 Genomics UK (COG-UK) Consortium | Health Services Laboratories and Alex Alderton, Roberto Amato, Jeffrey Barrett, Sonia Goncalves, Ewan Harrison, David K. Jackson, Ian Johnston, Dominic Kwiatkowski, Cordelia Langford, John Sillitoe on behalf of the Wellcome Sanger Institute COVID-19 Surveillance Team |
| EPI_ISL_2806610 | Lighthouse Lab in Milton Keynes | Wellcome Sanger Institute for the COVID-19 Genomics UK (COG-UK) Consortium | The Lighthouse Lab in Milton Keynes and Alex Alderton, Roberto Amato, Jeffrey Barrett, Sonia Goncalves, Ewan Harrison, David K. Jackson, Ian Johnston, Dominic Kwiatkowski, Cordelia Langford, John Sillitoe on behalf of the Wellcome Sanger Institute COVID-19 Surveillance Team |
| EPI_ISL_2806611 | Health Services Laboratories | Wellcome Sanger Institute for the COVID-19 Genomics UK (COG-UK) Consortium | Health Services Laboratories and Alex Alderton, Roberto Amato, Jeffrey Barrett, Sonia Goncalves, Ewan Harrison, David K. Jackson, Ian Johnston, Dominic Kwiatkowski, Cordelia Langford, John Sillitoe on behalf of the Wellcome Sanger Institute COVID-19 Surveillance Team |
| EPI_ISL_2806612, EPI_ISL_2806613 | Lighthouse Laboratory Plymouth | Wellcome Sanger Institute for the COVID-19 Genomics UK (COG-UK) Consortium | Lighthouse Laboratory Plymouth and Alex Alderton, Roberto Amato, Jeffrey Barrett, Sonia Goncalves, Ewan Harrison, David K. Jackson, Ian Johnston, Dominic Kwiatkowski, Cordelia Langford, John Sillitoe on behalf of the Wellcome Sanger Institute COVID-19 Surveillance Team |
| EPI_ISL_2806614 | Health Services Laboratories | Wellcome Sanger Institute for the COVID-19 Genomics UK (COG-UK) | Health Services Laboratories and Alex Alderton, Roberto Amato, Jeffrey Barrett, Sonia Goncalves, Ewan Harrison, David K. Jackson, Ian Johnston, Dominic Kwiatkowski, Cordelia Langford, John Sillitoe on behalf of the Wellcome Sanger Institute |

| | | | |
|---|---|---|---|
| | | Consortium | COVID-19 Surveillance Team |
| EPI_ISL_2806615, EPI_ISL_2806616 | Lighthouse Lab in Milton Keynes | Wellcome Sanger Institute for the COVID-19 Genomics UK (COG-UK) Consortium | The Lighthouse Lab in Milton Keynes and Alex Alderton, Roberto Amato, Jeffrey Barrett, Sonia Goncalves, Ewan Harrison, David K. Jackson, Ian Johnston, Dominic Kwiatkowski, Cordelia Langford, John Sillitoe on behalf of the Wellcome Sanger Institute COVID-19 Surveillance Team |
| EPI_ISL_2806617 | Health Services Laboratories | Wellcome Sanger Institute for the COVID-19 Genomics UK (COG-UK) Consortium | Health Services Laboratories and Alex Alderton, Roberto Amato, Jeffrey Barrett, Sonia Goncalves, Ewan Harrison, David K. Jackson, Ian Johnston, Dominic Kwiatkowski, Cordelia Langford, John Sillitoe on behalf of the Wellcome Sanger Institute COVID-19 Surveillance Team |
| EPI_ISL_2806618 | Lighthouse Laboratory Plymouth | Wellcome Sanger Institute for the COVID-19 Genomics UK (COG-UK) Consortium | Lighthouse Laboratory Plymouth and Alex Alderton, Roberto Amato, Jeffrey Barrett, Sonia Goncalves, Ewan Harrison, David K. Jackson, Ian Johnston, Dominic Kwiatkowski, Cordelia Langford, John Sillitoe on behalf of the Wellcome Sanger Institute COVID-19 Surveillance Team |
| EPI_ISL_2806619 | Health Services Laboratories | Wellcome Sanger Institute for the COVID-19 Genomics UK (COG-UK) Consortium | Health Services Laboratories and Alex Alderton, Roberto Amato, Jeffrey Barrett, Sonia Goncalves, Ewan Harrison, David K. Jackson, Ian Johnston, Dominic Kwiatkowski, Cordelia Langford, John Sillitoe on behalf of the Wellcome Sanger Institute COVID-19 Surveillance Team |
| EPI_ISL_2806620 | Lighthouse Lab in Milton Keynes | Wellcome Sanger Institute for the COVID-19 Genomics UK (COG-UK) Consortium | The Lighthouse Lab in Milton Keynes and Alex Alderton, Roberto Amato, Jeffrey Barrett, Sonia Goncalves, Ewan Harrison, David K. Jackson, Ian Johnston, Dominic Kwiatkowski, Cordelia Langford, John Sillitoe on behalf of the Wellcome Sanger Institute COVID-19 Surveillance Team |
| EPI_ISL_2806621, EPI_ISL_2806622, EPI_ISL_2806623 | Health Services Laboratories | Wellcome Sanger Institute for the COVID-19 Genomics UK (COG-UK) Consortium | Health Services Laboratories and Alex Alderton, Roberto Amato, Jeffrey Barrett, Sonia Goncalves, Ewan Harrison, David K. Jackson, Ian Johnston, Dominic Kwiatkowski, Cordelia Langford, John Sillitoe on behalf of the Wellcome Sanger Institute COVID-19 Surveillance Team |
| EPI_ISL_2806624 | Lighthouse Laboratory Plymouth | Wellcome Sanger Institute for the COVID-19 Genomics UK (COG-UK) Consortium | Lighthouse Laboratory Plymouth and Alex Alderton, Roberto Amato, Jeffrey Barrett, Sonia Goncalves, Ewan Harrison, David K. Jackson, Ian Johnston, Dominic Kwiatkowski, Cordelia Langford, John Sillitoe on behalf of the Wellcome Sanger Institute COVID-19 Surveillance Team |
| EPI_ISL_2806625, EPI_ISL_2806626, EPI_ISL_2806627 | Health Services Laboratories | Wellcome Sanger Institute for the COVID-19 Genomics UK (COG-UK) Consortium | Health Services Laboratories and Alex Alderton, Roberto Amato, Jeffrey Barrett, Sonia Goncalves, Ewan Harrison, David K. Jackson, Ian Johnston, Dominic Kwiatkowski, Cordelia Langford, John Sillitoe on behalf of the Wellcome Sanger Institute COVID-19 Surveillance Team |
| EPI_ISL_2806628 | Lighthouse Laboratory Plymouth | Wellcome Sanger Institute for the COVID-19 Genomics UK (COG-UK) Consortium | Lighthouse Laboratory Plymouth and Alex Alderton, Roberto Amato, Jeffrey Barrett, Sonia Goncalves, Ewan Harrison, David K. Jackson, Ian Johnston, Dominic Kwiatkowski, Cordelia Langford, John Sillitoe on behalf of the Wellcome Sanger Institute COVID-19 Surveillance Team |
| EPI_ISL_2806629, EPI_ISL_2806630, EPI_ISL_2806631, EPI_ISL_2806632, EPI_ISL_2806633, EPI_ISL_2806634 | Health Services Laboratories | Wellcome Sanger Institute for the COVID-19 Genomics UK (COG-UK) Consortium | Health Services Laboratories and Alex Alderton, Roberto Amato, Jeffrey Barrett, Sonia Goncalves, Ewan Harrison, David K. Jackson, Ian Johnston, Dominic Kwiatkowski, Cordelia Langford, John Sillitoe on behalf of the Wellcome Sanger Institute COVID-19 Surveillance Team |
| EPI_ISL_2806635 | Lighthouse Lab in Milton Keynes | Wellcome Sanger Institute for the COVID-19 Genomics UK (COG-UK) Consortium | The Lighthouse Lab in Milton Keynes and Alex Alderton, Roberto Amato, Jeffrey Barrett, Sonia Goncalves, Ewan Harrison, David K. Jackson, Ian Johnston, Dominic Kwiatkowski, Cordelia Langford, John Sillitoe on behalf of the Wellcome Sanger Institute COVID-19 Surveillance Team |
| EPI_ISL_2806636 | Lighthouse Laboratory Plymouth | Wellcome Sanger Institute for the COVID-19 Genomics UK (COG-UK) Consortium | Lighthouse Laboratory Plymouth and Alex Alderton, Roberto Amato, Jeffrey Barrett, Sonia Goncalves, Ewan Harrison, David K. Jackson, Ian Johnston, Dominic Kwiatkowski, Cordelia Langford, John Sillitoe on behalf of the Wellcome Sanger Institute COVID-19 Surveillance Team |
| EPI_ISL_2806637 | Lighthouse Lab in Milton Keynes | Wellcome Sanger Institute for the COVID-19 Genomics UK (COG-UK) Consortium | The Lighthouse Lab in Milton Keynes and Alex Alderton, Roberto Amato, Jeffrey Barrett, Sonia Goncalves, Ewan Harrison, David K. Jackson, Ian Johnston, Dominic Kwiatkowski, Cordelia Langford, John Sillitoe on behalf of the Wellcome Sanger Institute COVID-19 Surveillance Team |
| EPI_ISL_2806638 | Health Services Laboratories | Wellcome Sanger Institute for the COVID-19 Genomics UK (COG-UK) Consortium | Health Services Laboratories and Alex Alderton, Roberto Amato, Jeffrey Barrett, Sonia Goncalves, Ewan Harrison, David K. Jackson, Ian Johnston, Dominic Kwiatkowski, Cordelia Langford, John Sillitoe on behalf of the Wellcome Sanger Institute COVID-19 Surveillance Team |
| EPI_ISL_2806639 | Lighthouse Laboratory Plymouth | Wellcome Sanger Institute for the COVID-19 Genomics UK (COG-UK) Consortium | Lighthouse Laboratory Plymouth and Alex Alderton, Roberto Amato, Jeffrey Barrett, Sonia Goncalves, Ewan Harrison, David K. Jackson, Ian Johnston, Dominic Kwiatkowski, Cordelia Langford, John Sillitoe on behalf of the Wellcome Sanger Institute COVID-19 Surveillance Team |
| EPI_ISL_2806640 | Health Services Laboratories | Wellcome Sanger Institute for the COVID-19 Genomics UK (COG-UK) Consortium | Health Services Laboratories and Alex Alderton, Roberto Amato, Jeffrey Barrett, Sonia Goncalves, Ewan Harrison, David K. Jackson, Ian Johnston, Dominic Kwiatkowski, Cordelia Langford, John Sillitoe on behalf of the Wellcome Sanger Institute COVID-19 Surveillance Team |
| EPI_ISL_2806641, EPI_ISL_2806642 | Lighthouse Lab in Milton Keynes | Wellcome Sanger Institute for the COVID-19 Genomics UK (COG-UK) Consortium | The Lighthouse Lab in Milton Keynes and Alex Alderton, Roberto Amato, Jeffrey Barrett, Sonia Goncalves, Ewan Harrison, David K. Jackson, Ian Johnston, Dominic Kwiatkowski, Cordelia Langford, John Sillitoe on behalf of the Wellcome Sanger Institute COVID-19 Surveillance Team |
| EPI_ISL_2806643 | Health Services Laboratories | Wellcome Sanger Institute for the COVID-19 Genomics UK (COG-UK) Consortium | Health Services Laboratories and Alex Alderton, Roberto Amato, Jeffrey Barrett, Sonia Goncalves, Ewan Harrison, David K. Jackson, Ian Johnston, Dominic Kwiatkowski, Cordelia Langford, John Sillitoe on behalf of the Wellcome Sanger Institute COVID-19 Surveillance Team |
| EPI_ISL_2806644 | Lighthouse Laboratory Plymouth | Wellcome Sanger Institute for the COVID-19 Genomics UK (COG-UK) Consortium | Lighthouse Laboratory Plymouth and Alex Alderton, Roberto Amato, Jeffrey Barrett, Sonia Goncalves, Ewan Harrison, David K. Jackson, Ian Johnston, Dominic Kwiatkowski, Cordelia Langford, John Sillitoe on behalf of the Wellcome Sanger Institute COVID-19 Surveillance Team |
| EPI_ISL_2806645 | Health Services Laboratories | Wellcome Sanger Institute for the COVID-19 Genomics UK (COG-UK) Consortium | Health Services Laboratories and Alex Alderton, Roberto Amato, Jeffrey Barrett, Sonia Goncalves, Ewan Harrison, David K. Jackson, Ian Johnston, Dominic Kwiatkowski, Cordelia Langford, John Sillitoe on behalf of the Wellcome Sanger Institute COVID-19 Surveillance Team |
| EPI_ISL_2806646 | Lighthouse Lab in Milton Keynes | Wellcome Sanger Institute for the COVID-19 Genomics UK (COG-UK) Consortium | The Lighthouse Lab in Milton Keynes and Alex Alderton, Roberto Amato, Jeffrey Barrett, Sonia Goncalves, Ewan Harrison, David K. Jackson, Ian Johnston, Dominic Kwiatkowski, Cordelia Langford, John Sillitoe on behalf of the Wellcome Sanger Institute COVID-19 Surveillance Team |
| EPI_ISL_2806647, EPI_ISL_2806648, EPI_ISL_2806649 | Health Services Laboratories | Wellcome Sanger Institute for the COVID-19 Genomics UK (COG-UK) Consortium | Health Services Laboratories and Alex Alderton, Roberto Amato, Jeffrey Barrett, Sonia Goncalves, Ewan Harrison, David K. Jackson, Ian Johnston, Dominic Kwiatkowski, Cordelia Langford, John Sillitoe on behalf of the Wellcome Sanger Institute COVID-19 Surveillance Team |
| EPI_ISL_2806650 | Lighthouse Lab in Milton Keynes | Wellcome Sanger Institute for the COVID-19 Genomics UK (COG-UK) Consortium | The Lighthouse Lab in Milton Keynes and Alex Alderton, Roberto Amato, Jeffrey Barrett, Sonia Goncalves, Ewan Harrison, David K. Jackson, Ian Johnston, Dominic Kwiatkowski, Cordelia Langford, John Sillitoe on behalf of the Wellcome Sanger Institute COVID-19 Surveillance Team |
| EPI_ISL_2806651, EPI_ISL_2806652 | Lighthouse Laboratory Plymouth | Wellcome Sanger Institute for the COVID-19 Genomics UK (COG-UK) Consortium | Lighthouse Laboratory Plymouth and Alex Alderton, Roberto Amato, Jeffrey Barrett, Sonia Goncalves, Ewan Harrison, David K. Jackson, Ian Johnston, Dominic Kwiatkowski, Cordelia Langford, John Sillitoe on behalf of the Wellcome Sanger Institute COVID-19 Surveillance Team |
| EPI_ISL_2806653 | Health Services Laboratories | Wellcome Sanger Institute for the COVID-19 Genomics UK (COG-UK) Consortium | Health Services Laboratories and Alex Alderton, Roberto Amato, Jeffrey Barrett, Sonia Goncalves, Ewan Harrison, David K. Jackson, Ian Johnston, Dominic Kwiatkowski, Cordelia Langford, John Sillitoe on behalf of the Wellcome Sanger Institute COVID-19 Surveillance Team |
| EPI_ISL_2806654 | Lighthouse Lab in Milton Keynes | Wellcome Sanger Institute for the COVID-19 Genomics UK (COG-UK) Consortium | The Lighthouse Lab in Milton Keynes and Alex Alderton, Roberto Amato, Jeffrey Barrett, Sonia Goncalves, Ewan Harrison, David K. Jackson, Ian Johnston, Dominic Kwiatkowski, Cordelia Langford, John Sillitoe on behalf of the Wellcome Sanger Institute COVID-19 Surveillance Team |
| EPI_ISL_2806655 | Health Services Laboratories | Wellcome Sanger Institute for the COVID-19 Genomics UK (COG-UK) Consortium | Health Services Laboratories and Alex Alderton, Roberto Amato, Jeffrey Barrett, Sonia Goncalves, Ewan Harrison, David K. Jackson, Ian Johnston, Dominic Kwiatkowski, Cordelia Langford, John Sillitoe on behalf of the Wellcome Sanger Institute COVID-19 Surveillance Team |
| EPI_ISL_2806656 | Lighthouse Lab in Milton Keynes | Wellcome Sanger Institute for the COVID-19 Genomics UK (COG-UK) Consortium | The Lighthouse Lab in Milton Keynes and Alex Alderton, Roberto Amato, Jeffrey Barrett, Sonia Goncalves, Ewan Harrison, David K. Jackson, Ian Johnston, Dominic Kwiatkowski, Cordelia Langford, John Sillitoe on behalf of the Wellcome Sanger Institute COVID-19 Surveillance Team |
| EPI_ISL_2806657, EPI_ISL_2806658, EPI_ISL_2806659 | Lighthouse Laboratory Plymouth | Wellcome Sanger Institute for the COVID-19 Genomics UK (COG-UK) Consortium | Lighthouse Laboratory Plymouth and Alex Alderton, Roberto Amato, Jeffrey Barrett, Sonia Goncalves, Ewan Harrison, David K. Jackson, Ian Johnston, Dominic Kwiatkowski, Cordelia Langford, John Sillitoe on behalf of the Wellcome Sanger Institute COVID-19 Surveillance Team |
| EPI_ISL_2806660, EPI_ISL_2806661 | Health Services Laboratories | Wellcome Sanger Institute for the COVID-19 Genomics UK (COG-UK) Consortium | Health Services Laboratories and Alex Alderton, Roberto Amato, Jeffrey Barrett, Sonia Goncalves, Ewan Harrison, David K. Jackson, Ian Johnston, Dominic Kwiatkowski, Cordelia Langford, John Sillitoe on behalf of the Wellcome Sanger Institute COVID-19 Surveillance Team |
| EPI_ISL_2806662, EPI_ISL_2806663 | Lighthouse Laboratory Plymouth | Wellcome Sanger Institute for the COVID-19 Genomics UK (COG-UK) Consortium | Lighthouse Laboratory Plymouth and Alex Alderton, Roberto Amato, Jeffrey Barrett, Sonia Goncalves, Ewan Harrison, David K. Jackson, Ian Johnston, Dominic Kwiatkowski, Cordelia Langford, John Sillitoe on behalf of the Wellcome Sanger Institute COVID-19 Surveillance Team |
| EPI_ISL_2806664 | Health Services Laboratories | Wellcome Sanger Institute for the COVID-19 Genomics UK (COG-UK) Consortium | Health Services Laboratories and Alex Alderton, Roberto Amato, Jeffrey Barrett, Sonia Goncalves, Ewan Harrison, David K. Jackson, Ian Johnston, Dominic Kwiatkowski, Cordelia Langford, John Sillitoe on behalf of the Wellcome Sanger Institute COVID-19 Surveillance Team |
| EPI_ISL_2806665 | Lighthouse Laboratory Plymouth | Wellcome Sanger Institute for the COVID-19 Genomics UK (COG-UK) Consortium | Lighthouse Laboratory Plymouth and Alex Alderton, Roberto Amato, Jeffrey Barrett, Sonia Goncalves, Ewan Harrison, David K. Jackson, Ian Johnston, Dominic Kwiatkowski, Cordelia Langford, John Sillitoe on behalf of the Wellcome Sanger Institute COVID-19 Surveillance Team |
| EPI_ISL_2806666, EPI_ISL_2806667, EPI_ISL_2806668, EPI_ISL_2806669, EPI_ISL_2806670 | Health Services Laboratories | Wellcome Sanger Institute for the COVID-19 Genomics UK (COG-UK) Consortium | Health Services Laboratories and Alex Alderton, Roberto Amato, Jeffrey Barrett, Sonia Goncalves, Ewan Harrison, David K. Jackson, Ian Johnston, Dominic Kwiatkowski, Cordelia Langford, John Sillitoe on behalf of the Wellcome Sanger Institute COVID-19 Surveillance Team |
| EPI_ISL_2806671 | Lighthouse Laboratory Plymouth | Wellcome Sanger Institute for the COVID-19 Genomics UK (COG-UK) Consortium | Lighthouse Laboratory Plymouth and Alex Alderton, Roberto Amato, Jeffrey Barrett, Sonia Goncalves, Ewan Harrison, David K. Jackson, Ian Johnston, Dominic Kwiatkowski, Cordelia Langford, John Sillitoe on behalf of the Wellcome Sanger Institute COVID-19 Surveillance Team |
| EPI_ISL_2806672, EPI_ISL_2806673 | Health Services Laboratories | Wellcome Sanger Institute for the COVID-19 Genomics UK (COG-UK) Consortium | Health Services Laboratories and Alex Alderton, Roberto Amato, Jeffrey Barrett, Sonia Goncalves, Ewan Harrison, David K. Jackson, Ian Johnston, Dominic Kwiatkowski, Cordelia Langford, John Sillitoe on behalf of the Wellcome Sanger Institute COVID-19 Surveillance Team |
| EPI_ISL_2806674 | Lighthouse Lab in Milton Keynes | Wellcome Sanger Institute for the COVID-19 Genomics UK (COG-UK) Consortium | The Lighthouse Lab in Milton Keynes and Alex Alderton, Roberto Amato, Jeffrey Barrett, Sonia Goncalves, Ewan Harrison, David K. Jackson, Ian Johnston, Dominic Kwiatkowski, Cordelia Langford, John Sillitoe on behalf of the Wellcome Sanger Institute COVID-19 Surveillance Team |
| EPI_ISL_2806675, EPI_ISL_2806676 | Lighthouse Laboratory Plymouth | Wellcome Sanger Institute for the COVID-19 Genomics UK (COG-UK) Consortium | Lighthouse Laboratory Plymouth and Alex Alderton, Roberto Amato, Jeffrey Barrett, Sonia Goncalves, Ewan Harrison, David K. Jackson, Ian Johnston, Dominic Kwiatkowski, Cordelia Langford, John Sillitoe on behalf of the Wellcome Sanger Institute COVID-19 Surveillance Team |
| EPI_ISL_2806677, EPI_ISL_2806678 | Health Services Laboratories | Wellcome Sanger Institute for the COVID-19 Genomics UK (COG-UK) Consortium | Health Services Laboratories and Alex Alderton, Roberto Amato, Jeffrey Barrett, Sonia Goncalves, Ewan Harrison, David K. Jackson, Ian Johnston, Dominic Kwiatkowski, Cordelia Langford, John Sillitoe on behalf of the Wellcome Sanger Institute COVID-19 Surveillance Team |
| EPI_ISL_2806679 | Lighthouse Lab in Milton Keynes | Wellcome Sanger Institute for the COVID-19 Genomics UK (COG-UK) Consortium | The Lighthouse Lab in Milton Keynes and Alex Alderton, Roberto Amato, Jeffrey Barrett, Sonia Goncalves, Ewan Harrison, David K. Jackson, Ian Johnston, Dominic Kwiatkowski, Cordelia Langford, John Sillitoe on behalf of the Wellcome Sanger Institute COVID-19 Surveillance Team |

| | | | |
|---|---|---|---|
| EPI_ISL_2806680 | Health Services Laboratories | Wellcome Sanger Institute for the COVID-19 Genomics UK (COG-UK) Consortium | Health Services Laboratories and Alex Alderton, Roberto Amato, Jeffrey Barrett, Sonia Goncalves, Ewan Harrison, David K. Jackson, Ian Johnston, Dominic Kwiatkowski, Cordelia Langford, John Sillitoe on behalf of the Wellcome Sanger Institute COVID-19 Surveillance Team |
| EPI_ISL_2806681, EPI_ISL_2806682 | Lighthouse Lab in Milton Keynes | Wellcome Sanger Institute for the COVID-19 Genomics UK (COG-UK) Consortium | The Lighthouse Lab in Milton Keynes and Alex Alderton, Roberto Amato, Jeffrey Barrett, Sonia Goncalves, Ewan Harrison, David K. Jackson, Ian Johnston, Dominic Kwiatkowski, Cordelia Langford, John Sillitoe on behalf of the Wellcome Sanger Institute COVID-19 Surveillance Team |
| EPI_ISL_2806683 | Health Services Laboratories | Wellcome Sanger Institute for the COVID-19 Genomics UK (COG-UK) Consortium | Health Services Laboratories and Alex Alderton, Roberto Amato, Jeffrey Barrett, Sonia Goncalves, Ewan Harrison, David K. Jackson, Ian Johnston, Dominic Kwiatkowski, Cordelia Langford, John Sillitoe on behalf of the Wellcome Sanger Institute COVID-19 Surveillance Team |
| EPI_ISL_2806684 | Lighthouse Lab in Milton Keynes | Wellcome Sanger Institute for the COVID-19 Genomics UK (COG-UK) Consortium | The Lighthouse Lab in Milton Keynes and Alex Alderton, Roberto Amato, Jeffrey Barrett, Sonia Goncalves, Ewan Harrison, David K. Jackson, Ian Johnston, Dominic Kwiatkowski, Cordelia Langford, John Sillitoe on behalf of the Wellcome Sanger Institute COVID-19 Surveillance Team |
| EPI_ISL_2806685 | Health Services Laboratories | Wellcome Sanger Institute for the COVID-19 Genomics UK (COG-UK) Consortium | Health Services Laboratories and Alex Alderton, Roberto Amato, Jeffrey Barrett, Sonia Goncalves, Ewan Harrison, David K. Jackson, Ian Johnston, Dominic Kwiatkowski, Cordelia Langford, John Sillitoe on behalf of the Wellcome Sanger Institute COVID-19 Surveillance Team |
| EPI_ISL_2806686, EPI_ISL_2806687 | Lighthouse Laboratory Plymouth | Wellcome Sanger Institute for the COVID-19 Genomics UK (COG-UK) Consortium | Lighthouse Laboratory Plymouth and Alex Alderton, Roberto Amato, Jeffrey Barrett, Sonia Goncalves, Ewan Harrison, David K. Jackson, Ian Johnston, Dominic Kwiatkowski, Cordelia Langford, John Sillitoe on behalf of the Wellcome Sanger Institute COVID-19 Surveillance Team |
| EPI_ISL_2806688, EPI_ISL_2806689 | Health Services Laboratories | Wellcome Sanger Institute for the COVID-19 Genomics UK (COG-UK) Consortium | Health Services Laboratories and Alex Alderton, Roberto Amato, Jeffrey Barrett, Sonia Goncalves, Ewan Harrison, David K. Jackson, Ian Johnston, Dominic Kwiatkowski, Cordelia Langford, John Sillitoe on behalf of the Wellcome Sanger Institute COVID-19 Surveillance Team |
| EPI_ISL_2806690 | Lighthouse Laboratory Plymouth | Wellcome Sanger Institute for the COVID-19 Genomics UK (COG-UK) Consortium | Lighthouse Laboratory Plymouth and Alex Alderton, Roberto Amato, Jeffrey Barrett, Sonia Goncalves, Ewan Harrison, David K. Jackson, Ian Johnston, Dominic Kwiatkowski, Cordelia Langford, John Sillitoe on behalf of the Wellcome Sanger Institute COVID-19 Surveillance Team |
| EPI_ISL_2806691, EPI_ISL_2806692 | Lighthouse Lab in Milton Keynes | Wellcome Sanger Institute for the COVID-19 Genomics UK (COG-UK) Consortium | The Lighthouse Lab in Milton Keynes and Alex Alderton, Roberto Amato, Jeffrey Barrett, Sonia Goncalves, Ewan Harrison, David K. Jackson, Ian Johnston, Dominic Kwiatkowski, Cordelia Langford, John Sillitoe on behalf of the Wellcome Sanger Institute COVID-19 Surveillance Team |
| EPI_ISL_2806693, EPI_ISL_2806694, EPI_ISL_2806695, EPI_ISL_2806696 | Health Services Laboratories | Wellcome Sanger Institute for the COVID-19 Genomics UK (COG-UK) Consortium | Health Services Laboratories and Alex Alderton, Roberto Amato, Jeffrey Barrett, Sonia Goncalves, Ewan Harrison, David K. Jackson, Ian Johnston, Dominic Kwiatkowski, Cordelia Langford, John Sillitoe on behalf of the Wellcome Sanger Institute COVID-19 Surveillance Team |
| EPI_ISL_2806697, EPI_ISL_2806698, EPI_ISL_2806699 | Lighthouse Lab in Milton Keynes | Wellcome Sanger Institute for the COVID-19 Genomics UK (COG-UK) Consortium | The Lighthouse Lab in Milton Keynes and Alex Alderton, Roberto Amato, Jeffrey Barrett, Sonia Goncalves, Ewan Harrison, David K. Jackson, Ian Johnston, Dominic Kwiatkowski, Cordelia Langford, John Sillitoe on behalf of the Wellcome Sanger Institute COVID-19 Surveillance Team |
| EPI_ISL_2806700 | Health Services Laboratories | Wellcome Sanger Institute for the COVID-19 Genomics UK (COG-UK) Consortium | Health Services Laboratories and Alex Alderton, Roberto Amato, Jeffrey Barrett, Sonia Goncalves, Ewan Harrison, David K. Jackson, Ian Johnston, Dominic Kwiatkowski, Cordelia Langford, John Sillitoe on behalf of the Wellcome Sanger Institute COVID-19 Surveillance Team |
| EPI_ISL_2806701, EPI_ISL_2806702 | Lighthouse Lab in Milton Keynes | Wellcome Sanger Institute for the COVID-19 Genomics UK (COG-UK) Consortium | The Lighthouse Lab in Milton Keynes and Alex Alderton, Roberto Amato, Jeffrey Barrett, Sonia Goncalves, Ewan Harrison, David K. Jackson, Ian Johnston, Dominic Kwiatkowski, Cordelia Langford, John Sillitoe on behalf of the Wellcome Sanger Institute COVID-19 Surveillance Team |
| EPI_ISL_2806703 | Health Services Laboratories | Wellcome Sanger Institute for the COVID-19 Genomics UK (COG-UK) Consortium | Health Services Laboratories and Alex Alderton, Roberto Amato, Jeffrey Barrett, Sonia Goncalves, Ewan Harrison, David K. Jackson, Ian Johnston, Dominic Kwiatkowski, Cordelia Langford, John Sillitoe on behalf of the Wellcome Sanger Institute COVID-19 Surveillance Team |
| EPI_ISL_2806704 | Lighthouse Lab in Milton Keynes | Wellcome Sanger Institute for the COVID-19 Genomics UK (COG-UK) Consortium | The Lighthouse Lab in Milton Keynes and Alex Alderton, Roberto Amato, Jeffrey Barrett, Sonia Goncalves, Ewan Harrison, David K. Jackson, Ian Johnston, Dominic Kwiatkowski, Cordelia Langford, John Sillitoe on behalf of the Wellcome Sanger Institute COVID-19 Surveillance Team |
| EPI_ISL_2806705, EPI_ISL_2806706 | Health Services Laboratories | Wellcome Sanger Institute for the COVID-19 Genomics UK (COG-UK) Consortium | Health Services Laboratories and Alex Alderton, Roberto Amato, Jeffrey Barrett, Sonia Goncalves, Ewan Harrison, David K. Jackson, Ian Johnston, Dominic Kwiatkowski, Cordelia Langford, John Sillitoe on behalf of the Wellcome Sanger Institute COVID-19 Surveillance Team |
| EPI_ISL_2806707 | Lighthouse Laboratory Plymouth | Wellcome Sanger Institute for the COVID-19 Genomics UK (COG-UK) Consortium | Lighthouse Laboratory Plymouth and Alex Alderton, Roberto Amato, Jeffrey Barrett, Sonia Goncalves, Ewan Harrison, David K. Jackson, Ian Johnston, Dominic Kwiatkowski, Cordelia Langford, John Sillitoe on behalf of the Wellcome Sanger Institute COVID-19 Surveillance Team |
| EPI_ISL_2806708 | Health Services Laboratories | Wellcome Sanger Institute for the COVID-19 Genomics UK (COG-UK) Consortium | Health Services Laboratories and Alex Alderton, Roberto Amato, Jeffrey Barrett, Sonia Goncalves, Ewan Harrison, David K. Jackson, Ian Johnston, Dominic Kwiatkowski, Cordelia Langford, John Sillitoe on behalf of the Wellcome Sanger Institute COVID-19 Surveillance Team |
| EPI_ISL_2806709, EPI_ISL_2806710 | Lighthouse Laboratory Plymouth | Wellcome Sanger Institute for the COVID-19 Genomics UK (COG-UK) Consortium | Lighthouse Laboratory Plymouth and Alex Alderton, Roberto Amato, Jeffrey Barrett, Sonia Goncalves, Ewan Harrison, David K. Jackson, Ian Johnston, Dominic Kwiatkowski, Cordelia Langford, John Sillitoe on behalf of the Wellcome Sanger Institute COVID-19 Surveillance Team |
| EPI_ISL_2806711, EPI_ISL_2806712 | Health Services Laboratories | Wellcome Sanger Institute for the COVID-19 Genomics UK (COG-UK) Consortium | Health Services Laboratories and Alex Alderton, Roberto Amato, Jeffrey Barrett, Sonia Goncalves, Ewan Harrison, David K. Jackson, Ian Johnston, Dominic Kwiatkowski, Cordelia Langford, John Sillitoe on behalf of the Wellcome Sanger Institute COVID-19 Surveillance Team |
| EPI_ISL_2806713 | Lighthouse Laboratory Plymouth | Wellcome Sanger Institute for the COVID-19 Genomics UK (COG-UK) Consortium | Lighthouse Laboratory Plymouth and Alex Alderton, Roberto Amato, Jeffrey Barrett, Sonia Goncalves, Ewan Harrison, David K. Jackson, Ian Johnston, Dominic Kwiatkowski, Cordelia Langford, John Sillitoe on behalf of the Wellcome Sanger Institute COVID-19 Surveillance Team |
| EPI_ISL_2806714 | Lighthouse Lab in Milton Keynes | Wellcome Sanger Institute for the COVID-19 Genomics UK (COG-UK) Consortium | The Lighthouse Lab in Milton Keynes and Alex Alderton, Roberto Amato, Jeffrey Barrett, Sonia Goncalves, Ewan Harrison, David K. Jackson, Ian Johnston, Dominic Kwiatkowski, Cordelia Langford, John Sillitoe on behalf of the Wellcome Sanger Institute COVID-19 Surveillance Team |
| EPI_ISL_2806715 | Health Services Laboratories | Wellcome Sanger Institute for the COVID-19 Genomics UK (COG-UK) Consortium | Health Services Laboratories and Alex Alderton, Roberto Amato, Jeffrey Barrett, Sonia Goncalves, Ewan Harrison, David K. Jackson, Ian Johnston, Dominic Kwiatkowski, Cordelia Langford, John Sillitoe on behalf of the Wellcome Sanger Institute COVID-19 Surveillance Team |
| EPI_ISL_2806716, EPI_ISL_2806717, EPI_ISL_2806718, EPI_ISL_2806719 | Lighthouse Lab in Milton Keynes | Wellcome Sanger Institute for the COVID-19 Genomics UK (COG-UK) Consortium | The Lighthouse Lab in Milton Keynes and Alex Alderton, Roberto Amato, Jeffrey Barrett, Sonia Goncalves, Ewan Harrison, David K. Jackson, Ian Johnston, Dominic Kwiatkowski, Cordelia Langford, John Sillitoe on behalf of the Wellcome Sanger Institute COVID-19 Surveillance Team |
| EPI_ISL_2806720 | Lighthouse Laboratory Plymouth | Wellcome Sanger Institute for the COVID-19 Genomics UK (COG-UK) Consortium | Lighthouse Laboratory Plymouth and Alex Alderton, Roberto Amato, Jeffrey Barrett, Sonia Goncalves, Ewan Harrison, David K. Jackson, Ian Johnston, Dominic Kwiatkowski, Cordelia Langford, John Sillitoe on behalf of the Wellcome Sanger Institute COVID-19 Surveillance Team |
| EPI_ISL_2806721 | Lighthouse Lab in Milton Keynes | Wellcome Sanger Institute for the COVID-19 Genomics UK (COG-UK) Consortium | The Lighthouse Lab in Milton Keynes and Alex Alderton, Roberto Amato, Jeffrey Barrett, Sonia Goncalves, Ewan Harrison, David K. Jackson, Ian Johnston, Dominic Kwiatkowski, Cordelia Langford, John Sillitoe on behalf of the Wellcome Sanger Institute COVID-19 Surveillance Team |
| EPI_ISL_2806722, EPI_ISL_2806723 | Health Services Laboratories | Wellcome Sanger Institute for the COVID-19 Genomics UK (COG-UK) Consortium | Health Services Laboratories and Alex Alderton, Roberto Amato, Jeffrey Barrett, Sonia Goncalves, Ewan Harrison, David K. Jackson, Ian Johnston, Dominic Kwiatkowski, Cordelia Langford, John Sillitoe on behalf of the Wellcome Sanger Institute COVID-19 Surveillance Team |
| EPI_ISL_2806724 | Lighthouse Laboratory Plymouth | Wellcome Sanger Institute for the COVID-19 Genomics UK (COG-UK) Consortium | Lighthouse Laboratory Plymouth and Alex Alderton, Roberto Amato, Jeffrey Barrett, Sonia Goncalves, Ewan Harrison, David K. Jackson, Ian Johnston, Dominic Kwiatkowski, Cordelia Langford, John Sillitoe on behalf of the Wellcome Sanger Institute COVID-19 Surveillance Team |
| EPI_ISL_2806725 | Health Services Laboratories | Wellcome Sanger Institute for the COVID-19 Genomics UK (COG-UK) Consortium | Health Services Laboratories and Alex Alderton, Roberto Amato, Jeffrey Barrett, Sonia Goncalves, Ewan Harrison, David K. Jackson, Ian Johnston, Dominic Kwiatkowski, Cordelia Langford, John Sillitoe on behalf of the Wellcome Sanger Institute COVID-19 Surveillance Team |
| EPI_ISL_2806726 | Lighthouse Lab in Milton Keynes | Wellcome Sanger Institute for the COVID-19 Genomics UK (COG-UK) Consortium | The Lighthouse Lab in Milton Keynes and Alex Alderton, Roberto Amato, Jeffrey Barrett, Sonia Goncalves, Ewan Harrison, David K. Jackson, Ian Johnston, Dominic Kwiatkowski, Cordelia Langford, John Sillitoe on behalf of the Wellcome Sanger Institute COVID-19 Surveillance Team |
| EPI_ISL_2806727 | Health Services Laboratories | Wellcome Sanger Institute for the COVID-19 Genomics UK (COG-UK) Consortium | Health Services Laboratories and Alex Alderton, Roberto Amato, Jeffrey Barrett, Sonia Goncalves, Ewan Harrison, David K. Jackson, Ian Johnston, Dominic Kwiatkowski, Cordelia Langford, John Sillitoe on behalf of the Wellcome Sanger Institute COVID-19 Surveillance Team |
| EPI_ISL_2806728 | Lighthouse Laboratory Plymouth | Wellcome Sanger Institute for the COVID-19 Genomics UK (COG-UK) Consortium | Lighthouse Laboratory Plymouth and Alex Alderton, Roberto Amato, Jeffrey Barrett, Sonia Goncalves, Ewan Harrison, David K. Jackson, Ian Johnston, Dominic Kwiatkowski, Cordelia Langford, John Sillitoe on behalf of the Wellcome Sanger Institute COVID-19 Surveillance Team |
| EPI_ISL_2806729, EPI_ISL_2806730, EPI_ISL_2806731 | Health Services Laboratories | Wellcome Sanger Institute for the COVID-19 Genomics UK (COG-UK) Consortium | Health Services Laboratories and Alex Alderton, Roberto Amato, Jeffrey Barrett, Sonia Goncalves, Ewan Harrison, David K. Jackson, Ian Johnston, Dominic Kwiatkowski, Cordelia Langford, John Sillitoe on behalf of the Wellcome Sanger Institute COVID-19 Surveillance Team |
| EPI_ISL_2806732 | Lighthouse Laboratory Plymouth | Wellcome Sanger Institute for the COVID-19 Genomics UK (COG-UK) Consortium | Lighthouse Laboratory Plymouth and Alex Alderton, Roberto Amato, Jeffrey Barrett, Sonia Goncalves, Ewan Harrison, David K. Jackson, Ian Johnston, Dominic Kwiatkowski, Cordelia Langford, John Sillitoe on behalf of the Wellcome Sanger Institute COVID-19 Surveillance Team |
| EPI_ISL_2806733, EPI_ISL_2806735, EPI_ISL_2806736 | Health Services Laboratories | Wellcome Sanger Institute for the COVID-19 Genomics UK (COG-UK) Consortium | Health Services Laboratories and Alex Alderton, Roberto Amato, Jeffrey Barrett, Sonia Goncalves, Ewan Harrison, David K. Jackson, Ian Johnston, Dominic Kwiatkowski, Cordelia Langford, John Sillitoe on behalf of the Wellcome Sanger Institute COVID-19 Surveillance Team |
| EPI_ISL_2806737 | Lighthouse Laboratory Plymouth | Wellcome Sanger Institute for the COVID-19 Genomics UK (COG-UK) Consortium | Lighthouse Laboratory Plymouth and Alex Alderton, Roberto Amato, Jeffrey Barrett, Sonia Goncalves, Ewan Harrison, David K. Jackson, Ian Johnston, Dominic Kwiatkowski, Cordelia Langford, John Sillitoe on behalf of the Wellcome Sanger Institute COVID-19 Surveillance Team |
| EPI_ISL_2806738 | Lighthouse Lab in Milton Keynes | Wellcome Sanger Institute for the COVID-19 Genomics UK (COG-UK) Consortium | The Lighthouse Lab in Milton Keynes and Alex Alderton, Roberto Amato, Jeffrey Barrett, Sonia Goncalves, Ewan Harrison, David K. Jackson, Ian Johnston, Dominic Kwiatkowski, Cordelia Langford, John Sillitoe on behalf of the Wellcome Sanger Institute COVID-19 Surveillance Team |
| EPI_ISL_2806739, EPI_ISL_2806740, EPI_ISL_2806741 | Health Services Laboratories | Wellcome Sanger Institute for the COVID-19 Genomics UK (COG-UK) Consortium | Health Services Laboratories and Alex Alderton, Roberto Amato, Jeffrey Barrett, Sonia Goncalves, Ewan Harrison, David K. Jackson, Ian Johnston, Dominic Kwiatkowski, Cordelia Langford, John Sillitoe on behalf of the Wellcome Sanger Institute COVID-19 Surveillance Team |
| EPI_ISL_2806742 | Lighthouse Laboratory Plymouth | Wellcome Sanger Institute for the COVID-19 Genomics UK (COG-UK) Consortium | Lighthouse Laboratory Plymouth and Alex Alderton, Roberto Amato, Jeffrey Barrett, Sonia Goncalves, Ewan Harrison, David K. Jackson, Ian Johnston, Dominic Kwiatkowski, Cordelia Langford, John Sillitoe on behalf of the Wellcome Sanger Institute COVID-19 Surveillance Team |
| EPI_ISL_2806743 | Health Services Laboratories | Wellcome Sanger Institute for the COVID-19 Genomics UK (COG-UK) Consortium | Health Services Laboratories and Alex Alderton, Roberto Amato, Jeffrey Barrett, Sonia Goncalves, Ewan Harrison, David K. Jackson, Ian Johnston, Dominic Kwiatkowski, Cordelia Langford, John Sillitoe on behalf of the Wellcome Sanger Institute COVID-19 Surveillance Team |
| EPI_ISL_2806744 | Lighthouse Lab in Milton Keynes | Wellcome Sanger Institute for the COVID-19 Genomics UK (COG-UK) Consortium | The Lighthouse Lab in Milton Keynes and Alex Alderton, Roberto Amato, Jeffrey Barrett, Sonia Goncalves, Ewan Harrison, David K. Jackson, Ian Johnston, Dominic Kwiatkowski, Cordelia Langford, John Sillitoe on behalf of the Wellcome Sanger Institute COVID-19 Surveillance Team |
| EPI_ISL_2806745, EPI_ISL_2806746, | Lighthouse Laboratory Plymouth | Wellcome Sanger Institute for the COVID-19 Genomics UK (COG-UK) Consortium | Lighthouse Laboratory Plymouth and Alex Alderton, Roberto Amato, Jeffrey Barrett, Sonia Goncalves, Ewan Harrison, David K. Jackson, Ian Johnston, Dominic Kwiatkowski, Cordelia Langford, John Sillitoe on behalf of the Wellcome Sanger Institute COVID-19 Surveillance Team |

| | | | |
|---|---|---|---|
| EPI_ISL_2806747 | | | |
| EPI_ISL_2806748 | Health Services Laboratories | Wellcome Sanger Institute for the COVID-19 Genomics UK (COG-UK) Consortium | Health Services Laboratories and Alex Alderton, Roberto Amato, Jeffrey Barrett, Sonia Goncalves, Ewan Harrison, David K. Jackson, Ian Johnston, Dominic Kwiatkowski, Cordelia Langford, John Sillitoe on behalf of the Wellcome Sanger Institute COVID-19 Surveillance Team |
| EPI_ISL_2806749 | Lighthouse Laboratory Plymouth | Wellcome Sanger Institute for the COVID-19 Genomics UK (COG-UK) Consortium | Lighthouse Laboratory Plymouth and Alex Alderton, Roberto Amato, Jeffrey Barrett, Sonia Goncalves, Ewan Harrison, David K. Jackson, Ian Johnston, Dominic Kwiatkowski, Cordelia Langford, John Sillitoe on behalf of the Wellcome Sanger Institute COVID-19 Surveillance Team |
| EPI_ISL_2806750 | Health Services Laboratories | Wellcome Sanger Institute for the COVID-19 Genomics UK (COG-UK) Consortium | Health Services Laboratories and Alex Alderton, Roberto Amato, Jeffrey Barrett, Sonia Goncalves, Ewan Harrison, David K. Jackson, Ian Johnston, Dominic Kwiatkowski, Cordelia Langford, John Sillitoe on behalf of the Wellcome Sanger Institute COVID-19 Surveillance Team |
| EPI_ISL_2806751 | Lighthouse Laboratory Plymouth | Wellcome Sanger Institute for the COVID-19 Genomics UK (COG-UK) Consortium | Lighthouse Laboratory Plymouth and Alex Alderton, Roberto Amato, Jeffrey Barrett, Sonia Goncalves, Ewan Harrison, David K. Jackson, Ian Johnston, Dominic Kwiatkowski, Cordelia Langford, John Sillitoe on behalf of the Wellcome Sanger Institute COVID-19 Surveillance Team |
| EPI_ISL_2806752 | Lighthouse Lab in Milton Keynes | Wellcome Sanger Institute for the COVID-19 Genomics UK (COG-UK) Consortium | The Lighthouse Lab in Milton Keynes and Alex Alderton, Roberto Amato, Jeffrey Barrett, Sonia Goncalves, Ewan Harrison, David K. Jackson, Ian Johnston, Dominic Kwiatkowski, Cordelia Langford, John Sillitoe on behalf of the Wellcome Sanger Institute COVID-19 Surveillance Team |
| EPI_ISL_2806753 | Lighthouse Laboratory Plymouth | Wellcome Sanger Institute for the COVID-19 Genomics UK (COG-UK) Consortium | Lighthouse Laboratory Plymouth and Alex Alderton, Roberto Amato, Jeffrey Barrett, Sonia Goncalves, Ewan Harrison, David K. Jackson, Ian Johnston, Dominic Kwiatkowski, Cordelia Langford, John Sillitoe on behalf of the Wellcome Sanger Institute COVID-19 Surveillance Team |
| EPI_ISL_2806754, EPI_ISL_2806755 | Health Services Laboratories | Wellcome Sanger Institute for the COVID-19 Genomics UK (COG-UK) Consortium | Health Services Laboratories and Alex Alderton, Roberto Amato, Jeffrey Barrett, Sonia Goncalves, Ewan Harrison, David K. Jackson, Ian Johnston, Dominic Kwiatkowski, Cordelia Langford, John Sillitoe on behalf of the Wellcome Sanger Institute COVID-19 Surveillance Team |
| EPI_ISL_2806756, EPI_ISL_2806757 | Lighthouse Laboratory Plymouth | Wellcome Sanger Institute for the COVID-19 Genomics UK (COG-UK) Consortium | Lighthouse Laboratory Plymouth and Alex Alderton, Roberto Amato, Jeffrey Barrett, Sonia Goncalves, Ewan Harrison, David K. Jackson, Ian Johnston, Dominic Kwiatkowski, Cordelia Langford, John Sillitoe on behalf of the Wellcome Sanger Institute COVID-19 Surveillance Team |
| EPI_ISL_2806758, EPI_ISL_2806759, EPI_ISL_2806760 | Health Services Laboratories | Wellcome Sanger Institute for the COVID-19 Genomics UK (COG-UK) Consortium | Health Services Laboratories and Alex Alderton, Roberto Amato, Jeffrey Barrett, Sonia Goncalves, Ewan Harrison, David K. Jackson, Ian Johnston, Dominic Kwiatkowski, Cordelia Langford, John Sillitoe on behalf of the Wellcome Sanger Institute COVID-19 Surveillance Team |
| EPI_ISL_2806761, EPI_ISL_2806762 | Lighthouse Lab in Milton Keynes | Wellcome Sanger Institute for the COVID-19 Genomics UK (COG-UK) Consortium | The Lighthouse Lab in Milton Keynes and Alex Alderton, Roberto Amato, Jeffrey Barrett, Sonia Goncalves, Ewan Harrison, David K. Jackson, Ian Johnston, Dominic Kwiatkowski, Cordelia Langford, John Sillitoe on behalf of the Wellcome Sanger Institute COVID-19 Surveillance Team |
| EPI_ISL_2806763 | Health Services Laboratories | Wellcome Sanger Institute for the COVID-19 Genomics UK (COG-UK) Consortium | Health Services Laboratories and Alex Alderton, Roberto Amato, Jeffrey Barrett, Sonia Goncalves, Ewan Harrison, David K. Jackson, Ian Johnston, Dominic Kwiatkowski, Cordelia Langford, John Sillitoe on behalf of the Wellcome Sanger Institute COVID-19 Surveillance Team |
| EPI_ISL_2806764 | Lighthouse Laboratory Plymouth | Wellcome Sanger Institute for the COVID-19 Genomics UK (COG-UK) Consortium | Lighthouse Laboratory Plymouth and Alex Alderton, Roberto Amato, Jeffrey Barrett, Sonia Goncalves, Ewan Harrison, David K. Jackson, Ian Johnston, Dominic Kwiatkowski, Cordelia Langford, John Sillitoe on behalf of the Wellcome Sanger Institute COVID-19 Surveillance Team |
| EPI_ISL_2806765 | Health Services Laboratories | Wellcome Sanger Institute for the COVID-19 Genomics UK (COG-UK) Consortium | Health Services Laboratories and Alex Alderton, Roberto Amato, Jeffrey Barrett, Sonia Goncalves, Ewan Harrison, David K. Jackson, Ian Johnston, Dominic Kwiatkowski, Cordelia Langford, John Sillitoe on behalf of the Wellcome Sanger Institute COVID-19 Surveillance Team |
| EPI_ISL_2806766 | Lighthouse Laboratory Plymouth | Wellcome Sanger Institute for the COVID-19 Genomics UK (COG-UK) Consortium | Lighthouse Laboratory Plymouth and Alex Alderton, Roberto Amato, Jeffrey Barrett, Sonia Goncalves, Ewan Harrison, David K. Jackson, Ian Johnston, Dominic Kwiatkowski, Cordelia Langford, John Sillitoe on behalf of the Wellcome Sanger Institute COVID-19 Surveillance Team |
| EPI_ISL_2806767, EPI_ISL_2806768 | Health Services Laboratories | Wellcome Sanger Institute for the COVID-19 Genomics UK (COG-UK) Consortium | Health Services Laboratories and Alex Alderton, Roberto Amato, Jeffrey Barrett, Sonia Goncalves, Ewan Harrison, David K. Jackson, Ian Johnston, Dominic Kwiatkowski, Cordelia Langford, John Sillitoe on behalf of the Wellcome Sanger Institute COVID-19 Surveillance Team |
| EPI_ISL_2806769 | Lighthouse Laboratory Plymouth | Wellcome Sanger Institute for the COVID-19 Genomics UK (COG-UK) Consortium | Lighthouse Laboratory Plymouth and Alex Alderton, Roberto Amato, Jeffrey Barrett, Sonia Goncalves, Ewan Harrison, David K. Jackson, Ian Johnston, Dominic Kwiatkowski, Cordelia Langford, John Sillitoe on behalf of the Wellcome Sanger Institute COVID-19 Surveillance Team |
| EPI_ISL_2806770 | Health Services Laboratories | Wellcome Sanger Institute for the COVID-19 Genomics UK (COG-UK) Consortium | Health Services Laboratories and Alex Alderton, Roberto Amato, Jeffrey Barrett, Sonia Goncalves, Ewan Harrison, David K. Jackson, Ian Johnston, Dominic Kwiatkowski, Cordelia Langford, John Sillitoe on behalf of the Wellcome Sanger Institute COVID-19 Surveillance Team |
| EPI_ISL_2806771 | Lighthouse Laboratory Plymouth | Wellcome Sanger Institute for the COVID-19 Genomics UK (COG-UK) Consortium | Lighthouse Laboratory Plymouth and Alex Alderton, Roberto Amato, Jeffrey Barrett, Sonia Goncalves, Ewan Harrison, David K. Jackson, Ian Johnston, Dominic Kwiatkowski, Cordelia Langford, John Sillitoe on behalf of the Wellcome Sanger Institute COVID-19 Surveillance Team |
| EPI_ISL_2806772 | Health Services Laboratories | Wellcome Sanger Institute for the COVID-19 Genomics UK (COG-UK) Consortium | Health Services Laboratories and Alex Alderton, Roberto Amato, Jeffrey Barrett, Sonia Goncalves, Ewan Harrison, David K. Jackson, Ian Johnston, Dominic Kwiatkowski, Cordelia Langford, John Sillitoe on behalf of the Wellcome Sanger Institute COVID-19 Surveillance Team |
| EPI_ISL_2806773 | Lighthouse Laboratory Plymouth | Wellcome Sanger Institute for the COVID-19 Genomics UK (COG-UK) Consortium | Lighthouse Laboratory Plymouth and Alex Alderton, Roberto Amato, Jeffrey Barrett, Sonia Goncalves, Ewan Harrison, David K. Jackson, Ian Johnston, Dominic Kwiatkowski, Cordelia Langford, John Sillitoe on behalf of the Wellcome Sanger Institute COVID-19 Surveillance Team |
| EPI_ISL_2806774 | Health Services Laboratories | Wellcome Sanger Institute for the COVID-19 Genomics UK (COG-UK) Consortium | Health Services Laboratories and Alex Alderton, Roberto Amato, Jeffrey Barrett, Sonia Goncalves, Ewan Harrison, David K. Jackson, Ian Johnston, Dominic Kwiatkowski, Cordelia Langford, John Sillitoe on behalf of the Wellcome Sanger Institute COVID-19 Surveillance Team |
| EPI_ISL_2806775, EPI_ISL_2806776 | Lighthouse Lab in Milton Keynes | Wellcome Sanger Institute for the COVID-19 Genomics UK (COG-UK) Consortium | The Lighthouse Lab in Milton Keynes and Alex Alderton, Roberto Amato, Jeffrey Barrett, Sonia Goncalves, Ewan Harrison, David K. Jackson, Ian Johnston, Dominic Kwiatkowski, Cordelia Langford, John Sillitoe on behalf of the Wellcome Sanger Institute COVID-19 Surveillance Team |
| EPI_ISL_2806777 | Health Services Laboratories | Wellcome Sanger Institute for the COVID-19 Genomics UK (COG-UK) Consortium | Health Services Laboratories and Alex Alderton, Roberto Amato, Jeffrey Barrett, Sonia Goncalves, Ewan Harrison, David K. Jackson, Ian Johnston, Dominic Kwiatkowski, Cordelia Langford, John Sillitoe on behalf of the Wellcome Sanger Institute COVID-19 Surveillance Team |
| EPI_ISL_2806778 | Lighthouse Lab in Milton Keynes | Wellcome Sanger Institute for the COVID-19 Genomics UK (COG-UK) Consortium | The Lighthouse Lab in Milton Keynes and Alex Alderton, Roberto Amato, Jeffrey Barrett, Sonia Goncalves, Ewan Harrison, David K. Jackson, Ian Johnston, Dominic Kwiatkowski, Cordelia Langford, John Sillitoe on behalf of the Wellcome Sanger Institute COVID-19 Surveillance Team |
| EPI_ISL_2806779, EPI_ISL_2806780, EPI_ISL_2806781, EPI_ISL_2806782 | Lighthouse Laboratory Plymouth | Wellcome Sanger Institute for the COVID-19 Genomics UK (COG-UK) Consortium | Lighthouse Laboratory Plymouth and Alex Alderton, Roberto Amato, Jeffrey Barrett, Sonia Goncalves, Ewan Harrison, David K. Jackson, Ian Johnston, Dominic Kwiatkowski, Cordelia Langford, John Sillitoe on behalf of the Wellcome Sanger Institute COVID-19 Surveillance Team |
| EPI_ISL_2806783 | Lighthouse Lab in Milton Keynes | Wellcome Sanger Institute for the COVID-19 Genomics UK (COG-UK) Consortium | The Lighthouse Lab in Milton Keynes and Alex Alderton, Roberto Amato, Jeffrey Barrett, Sonia Goncalves, Ewan Harrison, David K. Jackson, Ian Johnston, Dominic Kwiatkowski, Cordelia Langford, John Sillitoe on behalf of the Wellcome Sanger Institute COVID-19 Surveillance Team |
| EPI_ISL_2806784, EPI_ISL_2806785 | Lighthouse Laboratory Plymouth | Wellcome Sanger Institute for the COVID-19 Genomics UK (COG-UK) Consortium | Lighthouse Laboratory Plymouth and Alex Alderton, Roberto Amato, Jeffrey Barrett, Sonia Goncalves, Ewan Harrison, David K. Jackson, Ian Johnston, Dominic Kwiatkowski, Cordelia Langford, John Sillitoe on behalf of the Wellcome Sanger Institute COVID-19 Surveillance Team |
| EPI_ISL_2806786 | Health Services Laboratories | Wellcome Sanger Institute for the COVID-19 Genomics UK (COG-UK) Consortium | Health Services Laboratories and Alex Alderton, Roberto Amato, Jeffrey Barrett, Sonia Goncalves, Ewan Harrison, David K. Jackson, Ian Johnston, Dominic Kwiatkowski, Cordelia Langford, John Sillitoe on behalf of the Wellcome Sanger Institute COVID-19 Surveillance Team |
| EPI_ISL_2806787 | Lighthouse Lab in Milton Keynes | Wellcome Sanger Institute for the COVID-19 Genomics UK (COG-UK) Consortium | The Lighthouse Lab in Milton Keynes and Alex Alderton, Roberto Amato, Jeffrey Barrett, Sonia Goncalves, Ewan Harrison, David K. Jackson, Ian Johnston, Dominic Kwiatkowski, Cordelia Langford, John Sillitoe on behalf of the Wellcome Sanger Institute COVID-19 Surveillance Team |
| EPI_ISL_2806788 | Lighthouse Laboratory Plymouth | Wellcome Sanger Institute for the COVID-19 Genomics UK (COG-UK) Consortium | Lighthouse Laboratory Plymouth and Alex Alderton, Roberto Amato, Jeffrey Barrett, Sonia Goncalves, Ewan Harrison, David K. Jackson, Ian Johnston, Dominic Kwiatkowski, Cordelia Langford, John Sillitoe on behalf of the Wellcome Sanger Institute COVID-19 Surveillance Team |
| EPI_ISL_2806789, EPI_ISL_2806790, EPI_ISL_2806791, EPI_ISL_2806792, EPI_ISL_2806793, EPI_ISL_2806794, EPI_ISL_2806795 | Health Services Laboratories | Wellcome Sanger Institute for the COVID-19 Genomics UK (COG-UK) Consortium | Health Services Laboratories and Alex Alderton, Roberto Amato, Jeffrey Barrett, Sonia Goncalves, Ewan Harrison, David K. Jackson, Ian Johnston, Dominic Kwiatkowski, Cordelia Langford, John Sillitoe on behalf of the Wellcome Sanger Institute COVID-19 Surveillance Team |
| EPI_ISL_2806796, EPI_ISL_2806797 | Lighthouse Lab in Milton Keynes | Wellcome Sanger Institute for the COVID-19 Genomics UK (COG-UK) Consortium | The Lighthouse Lab in Milton Keynes and Alex Alderton, Roberto Amato, Jeffrey Barrett, Sonia Goncalves, Ewan Harrison, David K. Jackson, Ian Johnston, Dominic Kwiatkowski, Cordelia Langford, John Sillitoe on behalf of the Wellcome Sanger Institute COVID-19 Surveillance Team |
| EPI_ISL_2806798, EPI_ISL_2806799 | Lighthouse Laboratory Plymouth | Wellcome Sanger Institute for the COVID-19 Genomics UK (COG-UK) Consortium | Lighthouse Laboratory Plymouth and Alex Alderton, Roberto Amato, Jeffrey Barrett, Sonia Goncalves, Ewan Harrison, David K. Jackson, Ian Johnston, Dominic Kwiatkowski, Cordelia Langford, John Sillitoe on behalf of the Wellcome Sanger Institute COVID-19 Surveillance Team |
| EPI_ISL_2806800, EPI_ISL_2806801 | Lighthouse Lab in Milton Keynes | Wellcome Sanger Institute for the COVID-19 Genomics UK (COG-UK) Consortium | The Lighthouse Lab in Milton Keynes and Alex Alderton, Roberto Amato, Jeffrey Barrett, Sonia Goncalves, Ewan Harrison, David K. Jackson, Ian Johnston, Dominic Kwiatkowski, Cordelia Langford, John Sillitoe on behalf of the Wellcome Sanger Institute COVID-19 Surveillance Team |
| EPI_ISL_2806802 | Lighthouse Laboratory Plymouth | Wellcome Sanger Institute for the COVID-19 Genomics UK (COG-UK) Consortium | Lighthouse Laboratory Plymouth and Alex Alderton, Roberto Amato, Jeffrey Barrett, Sonia Goncalves, Ewan Harrison, David K. Jackson, Ian Johnston, Dominic Kwiatkowski, Cordelia Langford, John Sillitoe on behalf of the Wellcome Sanger Institute COVID-19 Surveillance Team |
| EPI_ISL_2806803, EPI_ISL_2806804, EPI_ISL_2806805 | Lighthouse Lab in Milton Keynes | Wellcome Sanger Institute for the COVID-19 Genomics UK (COG-UK) Consortium | The Lighthouse Lab in Milton Keynes and Alex Alderton, Roberto Amato, Jeffrey Barrett, Sonia Goncalves, Ewan Harrison, David K. Jackson, Ian Johnston, Dominic Kwiatkowski, Cordelia Langford, John Sillitoe on behalf of the Wellcome Sanger Institute COVID-19 Surveillance Team |
| EPI_ISL_2806806, EPI_ISL_2806807, EPI_ISL_2806808 | Lighthouse Laboratory Plymouth | Wellcome Sanger Institute for the COVID-19 Genomics UK (COG-UK) Consortium | Lighthouse Laboratory Plymouth and Alex Alderton, Roberto Amato, Jeffrey Barrett, Sonia Goncalves, Ewan Harrison, David K. Jackson, Ian Johnston, Dominic Kwiatkowski, Cordelia Langford, John Sillitoe on behalf of the Wellcome Sanger Institute COVID-19 Surveillance Team |
| EPI_ISL_2806809, EPI_ISL_2806810 | Lighthouse Lab in Milton Keynes | Wellcome Sanger Institute for the COVID-19 Genomics UK (COG-UK) Consortium | The Lighthouse Lab in Milton Keynes and Alex Alderton, Roberto Amato, Jeffrey Barrett, Sonia Goncalves, Ewan Harrison, David K. Jackson, Ian Johnston, Dominic Kwiatkowski, Cordelia Langford, John Sillitoe on behalf of the Wellcome Sanger Institute COVID-19 Surveillance Team |
| EPI_ISL_2806811, EPI_ISL_2806812, EPI_ISL_2806813, EPI_ISL_2806814 | Lighthouse Laboratory Plymouth | Wellcome Sanger Institute for the COVID-19 Genomics UK (COG-UK) Consortium | Lighthouse Laboratory Plymouth and Alex Alderton, Roberto Amato, Jeffrey Barrett, Sonia Goncalves, Ewan Harrison, David K. Jackson, Ian Johnston, Dominic Kwiatkowski, Cordelia Langford, John Sillitoe on behalf of the Wellcome Sanger Institute COVID-19 Surveillance Team |
| EPI_ISL_2806815, | Lighthouse Lab in Milton Keynes | Wellcome Sanger Institute for the COVID-19 Genomics UK (COG-UK) | The Lighthouse Lab in Milton Keynes and Alex Alderton, Roberto Amato, Jeffrey Barrett, Sonia Goncalves, Ewan Harrison, David K. Jackson, Ian Johnston, Dominic Kwiatkowski, Cordelia Langford, John Sillitoe on behalf of the Wellcome Sanger |

| | | | |
|---|---|---|---|
| EPI_ISL_2806816, EPI_ISL_2806817 | | Consortium | Institute COVID-19 Surveillance Team |
| EPI_ISL_2806818, EPI_ISL_2806819 | Lighthouse Laboratory Plymouth | Wellcome Sanger Institute for the COVID-19 Genomics UK (COG-UK) Consortium | Lighthouse Laboratory Plymouth and Alex Alderton, Roberto Amato, Jeffrey Barrett, Sonia Goncalves, Ewan Harrison, David K. Jackson, Ian Johnston, Dominic Kwiatkowski, Cordelia Langford, John Sillitoe on behalf of the Wellcome Sanger Institute COVID-19 Surveillance Team |
| EPI_ISL_2806820, EPI_ISL_2806821 | Lighthouse Lab in Milton Keynes | Wellcome Sanger Institute for the COVID-19 Genomics UK (COG-UK) Consortium | The Lighthouse Lab in Milton Keynes and Alex Alderton, Roberto Amato, Jeffrey Barrett, Sonia Goncalves, Ewan Harrison, David K. Jackson, Ian Johnston, Dominic Kwiatkowski, Cordelia Langford, John Sillitoe on behalf of the Wellcome Sanger Institute COVID-19 Surveillance Team |
| EPI_ISL_2806822 | Lighthouse Laboratory Plymouth | Wellcome Sanger Institute for the COVID-19 Genomics UK (COG-UK) Consortium | Lighthouse Laboratory Plymouth and Alex Alderton, Roberto Amato, Jeffrey Barrett, Sonia Goncalves, Ewan Harrison, David K. Jackson, Ian Johnston, Dominic Kwiatkowski, Cordelia Langford, John Sillitoe on behalf of the Wellcome Sanger Institute COVID-19 Surveillance Team |
| EPI_ISL_2806823, EPI_ISL_2806824 | Lighthouse Lab in Milton Keynes | Wellcome Sanger Institute for the COVID-19 Genomics UK (COG-UK) Consortium | The Lighthouse Lab in Milton Keynes and Alex Alderton, Roberto Amato, Jeffrey Barrett, Sonia Goncalves, Ewan Harrison, David K. Jackson, Ian Johnston, Dominic Kwiatkowski, Cordelia Langford, John Sillitoe on behalf of the Wellcome Sanger Institute COVID-19 Surveillance Team |
| EPI_ISL_2806825 | Lighthouse Laboratory Plymouth | Wellcome Sanger Institute for the COVID-19 Genomics UK (COG-UK) Consortium | Lighthouse Laboratory Plymouth and Alex Alderton, Roberto Amato, Jeffrey Barrett, Sonia Goncalves, Ewan Harrison, David K. Jackson, Ian Johnston, Dominic Kwiatkowski, Cordelia Langford, John Sillitoe on behalf of the Wellcome Sanger Institute COVID-19 Surveillance Team |
| EPI_ISL_2806826 | Lighthouse Lab in Milton Keynes | Wellcome Sanger Institute for the COVID-19 Genomics UK (COG-UK) Consortium | The Lighthouse Lab in Milton Keynes and Alex Alderton, Roberto Amato, Jeffrey Barrett, Sonia Goncalves, Ewan Harrison, David K. Jackson, Ian Johnston, Dominic Kwiatkowski, Cordelia Langford, John Sillitoe on behalf of the Wellcome Sanger Institute COVID-19 Surveillance Team |
| EPI_ISL_2806827 | Lighthouse Laboratory Plymouth | Wellcome Sanger Institute for the COVID-19 Genomics UK (COG-UK) Consortium | Lighthouse Laboratory Plymouth and Alex Alderton, Roberto Amato, Jeffrey Barrett, Sonia Goncalves, Ewan Harrison, David K. Jackson, Ian Johnston, Dominic Kwiatkowski, Cordelia Langford, John Sillitoe on behalf of the Wellcome Sanger Institute COVID-19 Surveillance Team |
| EPI_ISL_2806828 | Lighthouse Lab in Milton Keynes | Wellcome Sanger Institute for the COVID-19 Genomics UK (COG-UK) Consortium | The Lighthouse Lab in Milton Keynes and Alex Alderton, Roberto Amato, Jeffrey Barrett, Sonia Goncalves, Ewan Harrison, David K. Jackson, Ian Johnston, Dominic Kwiatkowski, Cordelia Langford, John Sillitoe on behalf of the Wellcome Sanger Institute COVID-19 Surveillance Team |
| EPI_ISL_2806829 | Lighthouse Laboratory Plymouth | Wellcome Sanger Institute for the COVID-19 Genomics UK (COG-UK) Consortium | Lighthouse Laboratory Plymouth and Alex Alderton, Roberto Amato, Jeffrey Barrett, Sonia Goncalves, Ewan Harrison, David K. Jackson, Ian Johnston, Dominic Kwiatkowski, Cordelia Langford, John Sillitoe on behalf of the Wellcome Sanger Institute COVID-19 Surveillance Team |
| EPI_ISL_2806830 | Lighthouse Lab in Milton Keynes | Wellcome Sanger Institute for the COVID-19 Genomics UK (COG-UK) Consortium | The Lighthouse Lab in Milton Keynes and Alex Alderton, Roberto Amato, Jeffrey Barrett, Sonia Goncalves, Ewan Harrison, David K. Jackson, Ian Johnston, Dominic Kwiatkowski, Cordelia Langford, John Sillitoe on behalf of the Wellcome Sanger Institute COVID-19 Surveillance Team |
| EPI_ISL_2806831, EPI_ISL_2806832 | Lighthouse Laboratory Plymouth | Wellcome Sanger Institute for the COVID-19 Genomics UK (COG-UK) Consortium | Lighthouse Laboratory Plymouth and Alex Alderton, Roberto Amato, Jeffrey Barrett, Sonia Goncalves, Ewan Harrison, David K. Jackson, Ian Johnston, Dominic Kwiatkowski, Cordelia Langford, John Sillitoe on behalf of the Wellcome Sanger Institute COVID-19 Surveillance Team |
| EPI_ISL_2806833, EPI_ISL_2806834 | Lighthouse Lab in Milton Keynes | Wellcome Sanger Institute for the COVID-19 Genomics UK (COG-UK) Consortium | The Lighthouse Lab in Milton Keynes and Alex Alderton, Roberto Amato, Jeffrey Barrett, Sonia Goncalves, Ewan Harrison, David K. Jackson, Ian Johnston, Dominic Kwiatkowski, Cordelia Langford, John Sillitoe on behalf of the Wellcome Sanger Institute COVID-19 Surveillance Team |
| EPI_ISL_2806835, EPI_ISL_2806836, EPI_ISL_2806837 | Lighthouse Laboratory Plymouth | Wellcome Sanger Institute for the COVID-19 Genomics UK (COG-UK) Consortium | Lighthouse Laboratory Plymouth and Alex Alderton, Roberto Amato, Jeffrey Barrett, Sonia Goncalves, Ewan Harrison, David K. Jackson, Ian Johnston, Dominic Kwiatkowski, Cordelia Langford, John Sillitoe on behalf of the Wellcome Sanger Institute COVID-19 Surveillance Team |
| EPI_ISL_2806838, EPI_ISL_2806839, EPI_ISL_2806840 | Lighthouse Lab in Milton Keynes | Wellcome Sanger Institute for the COVID-19 Genomics UK (COG-UK) Consortium | The Lighthouse Lab in Milton Keynes and Alex Alderton, Roberto Amato, Jeffrey Barrett, Sonia Goncalves, Ewan Harrison, David K. Jackson, Ian Johnston, Dominic Kwiatkowski, Cordelia Langford, John Sillitoe on behalf of the Wellcome Sanger Institute COVID-19 Surveillance Team |
| EPI_ISL_2806841, EPI_ISL_2806842, EPI_ISL_2806843 | Lighthouse Laboratory Plymouth | Wellcome Sanger Institute for the COVID-19 Genomics UK (COG-UK) Consortium | Lighthouse Laboratory Plymouth and Alex Alderton, Roberto Amato, Jeffrey Barrett, Sonia Goncalves, Ewan Harrison, David K. Jackson, Ian Johnston, Dominic Kwiatkowski, Cordelia Langford, John Sillitoe on behalf of the Wellcome Sanger Institute COVID-19 Surveillance Team |
| EPI_ISL_2806844, EPI_ISL_2806845 | Lighthouse Lab in Milton Keynes | Wellcome Sanger Institute for the COVID-19 Genomics UK (COG-UK) Consortium | The Lighthouse Lab in Milton Keynes and Alex Alderton, Roberto Amato, Jeffrey Barrett, Sonia Goncalves, Ewan Harrison, David K. Jackson, Ian Johnston, Dominic Kwiatkowski, Cordelia Langford, John Sillitoe on behalf of the Wellcome Sanger Institute COVID-19 Surveillance Team |
| EPI_ISL_2806846 | Lighthouse Laboratory Plymouth | Wellcome Sanger Institute for the COVID-19 Genomics UK (COG-UK) Consortium | Lighthouse Laboratory Plymouth and Alex Alderton, Roberto Amato, Jeffrey Barrett, Sonia Goncalves, Ewan Harrison, David K. Jackson, Ian Johnston, Dominic Kwiatkowski, Cordelia Langford, John Sillitoe on behalf of the Wellcome Sanger Institute COVID-19 Surveillance Team |
| EPI_ISL_2806847 | Lighthouse Lab in Milton Keynes | Wellcome Sanger Institute for the COVID-19 Genomics UK (COG-UK) Consortium | The Lighthouse Lab in Milton Keynes and Alex Alderton, Roberto Amato, Jeffrey Barrett, Sonia Goncalves, Ewan Harrison, David K. Jackson, Ian Johnston, Dominic Kwiatkowski, Cordelia Langford, John Sillitoe on behalf of the Wellcome Sanger Institute COVID-19 Surveillance Team |
| EPI_ISL_2806848 | Lighthouse Laboratory Plymouth | Wellcome Sanger Institute for the COVID-19 Genomics UK (COG-UK) Consortium | Lighthouse Laboratory Plymouth and Alex Alderton, Roberto Amato, Jeffrey Barrett, Sonia Goncalves, Ewan Harrison, David K. Jackson, Ian Johnston, Dominic Kwiatkowski, Cordelia Langford, John Sillitoe on behalf of the Wellcome Sanger Institute COVID-19 Surveillance Team |
| EPI_ISL_2806849, EPI_ISL_2806850, EPI_ISL_2806851, EPI_ISL_2806852 | Lighthouse Lab in Milton Keynes | Wellcome Sanger Institute for the COVID-19 Genomics UK (COG-UK) Consortium | The Lighthouse Lab in Milton Keynes and Alex Alderton, Roberto Amato, Jeffrey Barrett, Sonia Goncalves, Ewan Harrison, David K. Jackson, Ian Johnston, Dominic Kwiatkowski, Cordelia Langford, John Sillitoe on behalf of the Wellcome Sanger Institute COVID-19 Surveillance Team |
| EPI_ISL_2806853 | Lighthouse Laboratory Plymouth | Wellcome Sanger Institute for the COVID-19 Genomics UK (COG-UK) Consortium | Lighthouse Laboratory Plymouth and Alex Alderton, Roberto Amato, Jeffrey Barrett, Sonia Goncalves, Ewan Harrison, David K. Jackson, Ian Johnston, Dominic Kwiatkowski, Cordelia Langford, John Sillitoe on behalf of the Wellcome Sanger Institute COVID-19 Surveillance Team |
| EPI_ISL_2806854, EPI_ISL_2806855, EPI_ISL_2806856, EPI_ISL_2806857, EPI_ISL_2806858 | Lighthouse Lab in Milton Keynes | Wellcome Sanger Institute for the COVID-19 Genomics UK (COG-UK) Consortium | The Lighthouse Lab in Milton Keynes and Alex Alderton, Roberto Amato, Jeffrey Barrett, Sonia Goncalves, Ewan Harrison, David K. Jackson, Ian Johnston, Dominic Kwiatkowski, Cordelia Langford, John Sillitoe on behalf of the Wellcome Sanger Institute COVID-19 Surveillance Team |
| EPI_ISL_2806859 | Lighthouse Laboratory Plymouth | Wellcome Sanger Institute for the COVID-19 Genomics UK (COG-UK) Consortium | Lighthouse Laboratory Plymouth and Alex Alderton, Roberto Amato, Jeffrey Barrett, Sonia Goncalves, Ewan Harrison, David K. Jackson, Ian Johnston, Dominic Kwiatkowski, Cordelia Langford, John Sillitoe on behalf of the Wellcome Sanger Institute COVID-19 Surveillance Team |
| EPI_ISL_2806860 | Lighthouse Lab in Milton Keynes | Wellcome Sanger Institute for the COVID-19 Genomics UK (COG-UK) Consortium | The Lighthouse Lab in Milton Keynes and Alex Alderton, Roberto Amato, Jeffrey Barrett, Sonia Goncalves, Ewan Harrison, David K. Jackson, Ian Johnston, Dominic Kwiatkowski, Cordelia Langford, John Sillitoe on behalf of the Wellcome Sanger Institute COVID-19 Surveillance Team |
| EPI_ISL_2806861, EPI_ISL_2806862, EPI_ISL_2806863 | Lighthouse Laboratory Plymouth | Wellcome Sanger Institute for the COVID-19 Genomics UK (COG-UK) Consortium | Lighthouse Laboratory Plymouth and Alex Alderton, Roberto Amato, Jeffrey Barrett, Sonia Goncalves, Ewan Harrison, David K. Jackson, Ian Johnston, Dominic Kwiatkowski, Cordelia Langford, John Sillitoe on behalf of the Wellcome Sanger Institute COVID-19 Surveillance Team |
| EPI_ISL_2806864 | Lighthouse Lab in Milton Keynes | Wellcome Sanger Institute for the COVID-19 Genomics UK (COG-UK) Consortium | The Lighthouse Lab in Milton Keynes and Alex Alderton, Roberto Amato, Jeffrey Barrett, Sonia Goncalves, Ewan Harrison, David K. Jackson, Ian Johnston, Dominic Kwiatkowski, Cordelia Langford, John Sillitoe on behalf of the Wellcome Sanger Institute COVID-19 Surveillance Team |
| EPI_ISL_2806865, EPI_ISL_2806866, EPI_ISL_2806867 | Lighthouse Laboratory Plymouth | Wellcome Sanger Institute for the COVID-19 Genomics UK (COG-UK) Consortium | Lighthouse Laboratory Plymouth and Alex Alderton, Roberto Amato, Jeffrey Barrett, Sonia Goncalves, Ewan Harrison, David K. Jackson, Ian Johnston, Dominic Kwiatkowski, Cordelia Langford, John Sillitoe on behalf of the Wellcome Sanger Institute COVID-19 Surveillance Team |
| EPI_ISL_2806868 | Lighthouse Lab in Milton Keynes | Wellcome Sanger Institute for the COVID-19 Genomics UK (COG-UK) Consortium | The Lighthouse Lab in Milton Keynes and Alex Alderton, Roberto Amato, Jeffrey Barrett, Sonia Goncalves, Ewan Harrison, David K. Jackson, Ian Johnston, Dominic Kwiatkowski, Cordelia Langford, John Sillitoe on behalf of the Wellcome Sanger Institute COVID-19 Surveillance Team |
| EPI_ISL_2806869 | Lighthouse Laboratory Plymouth | Wellcome Sanger Institute for the COVID-19 Genomics UK (COG-UK) Consortium | Lighthouse Laboratory Plymouth and Alex Alderton, Roberto Amato, Jeffrey Barrett, Sonia Goncalves, Ewan Harrison, David K. Jackson, Ian Johnston, Dominic Kwiatkowski, Cordelia Langford, John Sillitoe on behalf of the Wellcome Sanger Institute COVID-19 Surveillance Team |
| EPI_ISL_2806870 | Lighthouse Lab in Milton Keynes | Wellcome Sanger Institute for the COVID-19 Genomics UK (COG-UK) Consortium | The Lighthouse Lab in Milton Keynes and Alex Alderton, Roberto Amato, Jeffrey Barrett, Sonia Goncalves, Ewan Harrison, David K. Jackson, Ian Johnston, Dominic Kwiatkowski, Cordelia Langford, John Sillitoe on behalf of the Wellcome Sanger Institute COVID-19 Surveillance Team |
| EPI_ISL_2806871, EPI_ISL_2806872 | Lighthouse Laboratory Plymouth | Wellcome Sanger Institute for the COVID-19 Genomics UK (COG-UK) Consortium | Lighthouse Laboratory Plymouth and Alex Alderton, Roberto Amato, Jeffrey Barrett, Sonia Goncalves, Ewan Harrison, David K. Jackson, Ian Johnston, Dominic Kwiatkowski, Cordelia Langford, John Sillitoe on behalf of the Wellcome Sanger Institute COVID-19 Surveillance Team |
| EPI_ISL_2806873, EPI_ISL_2806874, EPI_ISL_2806875, EPI_ISL_2806876 | Lighthouse Lab in Milton Keynes | Wellcome Sanger Institute for the COVID-19 Genomics UK (COG-UK) Consortium | The Lighthouse Lab in Milton Keynes and Alex Alderton, Roberto Amato, Jeffrey Barrett, Sonia Goncalves, Ewan Harrison, David K. Jackson, Ian Johnston, Dominic Kwiatkowski, Cordelia Langford, John Sillitoe on behalf of the Wellcome Sanger Institute COVID-19 Surveillance Team |
| EPI_ISL_2806877 | Lighthouse Laboratory Plymouth | Wellcome Sanger Institute for the COVID-19 Genomics UK (COG-UK) Consortium | Lighthouse Laboratory Plymouth and Alex Alderton, Roberto Amato, Jeffrey Barrett, Sonia Goncalves, Ewan Harrison, David K. Jackson, Ian Johnston, Dominic Kwiatkowski, Cordelia Langford, John Sillitoe on behalf of the Wellcome Sanger Institute COVID-19 Surveillance Team |
| EPI_ISL_2806878 | Lighthouse Lab in Milton Keynes | Wellcome Sanger Institute for the COVID-19 Genomics UK (COG-UK) Consortium | The Lighthouse Lab in Milton Keynes and Alex Alderton, Roberto Amato, Jeffrey Barrett, Sonia Goncalves, Ewan Harrison, David K. Jackson, Ian Johnston, Dominic Kwiatkowski, Cordelia Langford, John Sillitoe on behalf of the Wellcome Sanger Institute COVID-19 Surveillance Team |
| EPI_ISL_2806879 | Lighthouse Laboratory Plymouth | Wellcome Sanger Institute for the COVID-19 Genomics UK (COG-UK) Consortium | Lighthouse Laboratory Plymouth and Alex Alderton, Roberto Amato, Jeffrey Barrett, Sonia Goncalves, Ewan Harrison, David K. Jackson, Ian Johnston, Dominic Kwiatkowski, Cordelia Langford, John Sillitoe on behalf of the Wellcome Sanger Institute COVID-19 Surveillance Team |
| EPI_ISL_2806880 | Lighthouse Lab in Milton Keynes | Wellcome Sanger Institute for the COVID-19 Genomics UK (COG-UK) Consortium | The Lighthouse Lab in Milton Keynes and Alex Alderton, Roberto Amato, Jeffrey Barrett, Sonia Goncalves, Ewan Harrison, David K. Jackson, Ian Johnston, Dominic Kwiatkowski, Cordelia Langford, John Sillitoe on behalf of the Wellcome Sanger Institute COVID-19 Surveillance Team |
| EPI_ISL_2806881, EPI_ISL_2806882, EPI_ISL_2806883, EPI_ISL_2806884 | Lighthouse Laboratory Plymouth | Wellcome Sanger Institute for the COVID-19 Genomics UK (COG-UK) Consortium | Lighthouse Laboratory Plymouth and Alex Alderton, Roberto Amato, Jeffrey Barrett, Sonia Goncalves, Ewan Harrison, David K. Jackson, Ian Johnston, Dominic Kwiatkowski, Cordelia Langford, John Sillitoe on behalf of the Wellcome Sanger Institute COVID-19 Surveillance Team |
| EPI_ISL_2806885 | Lighthouse Lab in Milton Keynes | Wellcome Sanger Institute for the COVID-19 Genomics UK (COG-UK) Consortium | The Lighthouse Lab in Milton Keynes and Alex Alderton, Roberto Amato, Jeffrey Barrett, Sonia Goncalves, Ewan Harrison, David K. Jackson, Ian Johnston, Dominic Kwiatkowski, Cordelia Langford, John Sillitoe on behalf of the Wellcome Sanger Institute COVID-19 Surveillance Team |

| | | | |
|---|---|---|---|
| EPI_ISL_2806886 | Lighthouse Laboratory Plymouth | Wellcome Sanger Institute for the COVID-19 Genomics UK (COG-UK) Consortium | Lighthouse Laboratory Plymouth and Alex Alderton, Roberto Amato, Jeffrey Barrett, Sonia Goncalves, Ewan Harrison, David K. Jackson, Ian Johnston, Dominic Kwiatkowski, Cordelia Langford, John Sillitoe on behalf of the Wellcome Sanger Institute COVID-19 Surveillance Team |
| EPI_ISL_2806887, EPI_ISL_2806888, EPI_ISL_2806889 | Lighthouse Lab in Milton Keynes | Wellcome Sanger Institute for the COVID-19 Genomics UK (COG-UK) Consortium | The Lighthouse Lab in Milton Keynes and Alex Alderton, Roberto Amato, Jeffrey Barrett, Sonia Goncalves, Ewan Harrison, David K. Jackson, Ian Johnston, Dominic Kwiatkowski, Cordelia Langford, John Sillitoe on behalf of the Wellcome Sanger Institute COVID-19 Surveillance Team |
| EPI_ISL_2806890, EPI_ISL_2806891 | Lighthouse Laboratory Plymouth | Wellcome Sanger Institute for the COVID-19 Genomics UK (COG-UK) Consortium | Lighthouse Laboratory Plymouth and Alex Alderton, Roberto Amato, Jeffrey Barrett, Sonia Goncalves, Ewan Harrison, David K. Jackson, Ian Johnston, Dominic Kwiatkowski, Cordelia Langford, John Sillitoe on behalf of the Wellcome Sanger Institute COVID-19 Surveillance Team |
| EPI_ISL_2806892, EPI_ISL_2806893, EPI_ISL_2806894 | Lighthouse Lab in Milton Keynes | Wellcome Sanger Institute for the COVID-19 Genomics UK (COG-UK) Consortium | The Lighthouse Lab in Milton Keynes and Alex Alderton, Roberto Amato, Jeffrey Barrett, Sonia Goncalves, Ewan Harrison, David K. Jackson, Ian Johnston, Dominic Kwiatkowski, Cordelia Langford, John Sillitoe on behalf of the Wellcome Sanger Institute COVID-19 Surveillance Team |
| EPI_ISL_2806895 | Lighthouse Laboratory Plymouth | Wellcome Sanger Institute for the COVID-19 Genomics UK (COG-UK) Consortium | Lighthouse Laboratory Plymouth and Alex Alderton, Roberto Amato, Jeffrey Barrett, Sonia Goncalves, Ewan Harrison, David K. Jackson, Ian Johnston, Dominic Kwiatkowski, Cordelia Langford, John Sillitoe on behalf of the Wellcome Sanger Institute COVID-19 Surveillance Team |
| EPI_ISL_2806896, EPI_ISL_2806897 | Lighthouse Lab in Milton Keynes | Wellcome Sanger Institute for the COVID-19 Genomics UK (COG-UK) Consortium | The Lighthouse Lab in Milton Keynes and Alex Alderton, Roberto Amato, Jeffrey Barrett, Sonia Goncalves, Ewan Harrison, David K. Jackson, Ian Johnston, Dominic Kwiatkowski, Cordelia Langford, John Sillitoe on behalf of the Wellcome Sanger Institute COVID-19 Surveillance Team |
| EPI_ISL_2806898 | Lighthouse Laboratory Plymouth | Wellcome Sanger Institute for the COVID-19 Genomics UK (COG-UK) Consortium | Lighthouse Laboratory Plymouth and Alex Alderton, Roberto Amato, Jeffrey Barrett, Sonia Goncalves, Ewan Harrison, David K. Jackson, Ian Johnston, Dominic Kwiatkowski, Cordelia Langford, John Sillitoe on behalf of the Wellcome Sanger Institute COVID-19 Surveillance Team |
| EPI_ISL_2806899 | Lighthouse Lab in Milton Keynes | Wellcome Sanger Institute for the COVID-19 Genomics UK (COG-UK) Consortium | The Lighthouse Lab in Milton Keynes and Alex Alderton, Roberto Amato, Jeffrey Barrett, Sonia Goncalves, Ewan Harrison, David K. Jackson, Ian Johnston, Dominic Kwiatkowski, Cordelia Langford, John Sillitoe on behalf of the Wellcome Sanger Institute COVID-19 Surveillance Team |
| EPI_ISL_2806900 | Lighthouse Laboratory Plymouth | Wellcome Sanger Institute for the COVID-19 Genomics UK (COG-UK) Consortium | Lighthouse Laboratory Plymouth and Alex Alderton, Roberto Amato, Jeffrey Barrett, Sonia Goncalves, Ewan Harrison, David K. Jackson, Ian Johnston, Dominic Kwiatkowski, Cordelia Langford, John Sillitoe on behalf of the Wellcome Sanger Institute COVID-19 Surveillance Team |
| EPI_ISL_2806901, EPI_ISL_2806902 | Lighthouse Lab in Milton Keynes | Wellcome Sanger Institute for the COVID-19 Genomics UK (COG-UK) Consortium | The Lighthouse Lab in Milton Keynes and Alex Alderton, Roberto Amato, Jeffrey Barrett, Sonia Goncalves, Ewan Harrison, David K. Jackson, Ian Johnston, Dominic Kwiatkowski, Cordelia Langford, John Sillitoe on behalf of the Wellcome Sanger Institute COVID-19 Surveillance Team |
| EPI_ISL_2806903 | Lighthouse Laboratory Plymouth | Wellcome Sanger Institute for the COVID-19 Genomics UK (COG-UK) Consortium | Lighthouse Laboratory Plymouth and Alex Alderton, Roberto Amato, Jeffrey Barrett, Sonia Goncalves, Ewan Harrison, David K. Jackson, Ian Johnston, Dominic Kwiatkowski, Cordelia Langford, John Sillitoe on behalf of the Wellcome Sanger Institute COVID-19 Surveillance Team |
| EPI_ISL_2806904 | Lighthouse Lab in Milton Keynes | Wellcome Sanger Institute for the COVID-19 Genomics UK (COG-UK) Consortium | The Lighthouse Lab in Milton Keynes and Alex Alderton, Roberto Amato, Jeffrey Barrett, Sonia Goncalves, Ewan Harrison, David K. Jackson, Ian Johnston, Dominic Kwiatkowski, Cordelia Langford, John Sillitoe on behalf of the Wellcome Sanger Institute COVID-19 Surveillance Team |
| EPI_ISL_2806906 | Lighthouse Laboratory Plymouth | Wellcome Sanger Institute for the COVID-19 Genomics UK (COG-UK) Consortium | Lighthouse Laboratory Plymouth and Alex Alderton, Roberto Amato, Jeffrey Barrett, Sonia Goncalves, Ewan Harrison, David K. Jackson, Ian Johnston, Dominic Kwiatkowski, Cordelia Langford, John Sillitoe on behalf of the Wellcome Sanger Institute COVID-19 Surveillance Team |
| EPI_ISL_2806907, EPI_ISL_2806908 | Lighthouse Lab in Milton Keynes | Wellcome Sanger Institute for the COVID-19 Genomics UK (COG-UK) Consortium | The Lighthouse Lab in Milton Keynes and Alex Alderton, Roberto Amato, Jeffrey Barrett, Sonia Goncalves, Ewan Harrison, David K. Jackson, Ian Johnston, Dominic Kwiatkowski, Cordelia Langford, John Sillitoe on behalf of the Wellcome Sanger Institute COVID-19 Surveillance Team |
| EPI_ISL_2806909 | Lighthouse Laboratory Plymouth | Wellcome Sanger Institute for the COVID-19 Genomics UK (COG-UK) Consortium | Lighthouse Laboratory Plymouth and Alex Alderton, Roberto Amato, Jeffrey Barrett, Sonia Goncalves, Ewan Harrison, David K. Jackson, Ian Johnston, Dominic Kwiatkowski, Cordelia Langford, John Sillitoe on behalf of the Wellcome Sanger Institute COVID-19 Surveillance Team |
| EPI_ISL_2806910 | Lighthouse Lab in Milton Keynes | Wellcome Sanger Institute for the COVID-19 Genomics UK (COG-UK) Consortium | The Lighthouse Lab in Milton Keynes and Alex Alderton, Roberto Amato, Jeffrey Barrett, Sonia Goncalves, Ewan Harrison, David K. Jackson, Ian Johnston, Dominic Kwiatkowski, Cordelia Langford, John Sillitoe on behalf of the Wellcome Sanger Institute COVID-19 Surveillance Team |
| EPI_ISL_2806911 | Lighthouse Laboratory Plymouth | Wellcome Sanger Institute for the COVID-19 Genomics UK (COG-UK) Consortium | Lighthouse Laboratory Plymouth and Alex Alderton, Roberto Amato, Jeffrey Barrett, Sonia Goncalves, Ewan Harrison, David K. Jackson, Ian Johnston, Dominic Kwiatkowski, Cordelia Langford, John Sillitoe on behalf of the Wellcome Sanger Institute COVID-19 Surveillance Team |
| EPI_ISL_2806912 | Lighthouse Lab in Milton Keynes | Wellcome Sanger Institute for the COVID-19 Genomics UK (COG-UK) Consortium | The Lighthouse Lab in Milton Keynes and Alex Alderton, Roberto Amato, Jeffrey Barrett, Sonia Goncalves, Ewan Harrison, David K. Jackson, Ian Johnston, Dominic Kwiatkowski, Cordelia Langford, John Sillitoe on behalf of the Wellcome Sanger Institute COVID-19 Surveillance Team |
| EPI_ISL_2806913, EPI_ISL_2806914 | Lighthouse Laboratory Plymouth | Wellcome Sanger Institute for the COVID-19 Genomics UK (COG-UK) Consortium | Lighthouse Laboratory Plymouth and Alex Alderton, Roberto Amato, Jeffrey Barrett, Sonia Goncalves, Ewan Harrison, David K. Jackson, Ian Johnston, Dominic Kwiatkowski, Cordelia Langford, John Sillitoe on behalf of the Wellcome Sanger Institute COVID-19 Surveillance Team |
| EPI_ISL_2806915, EPI_ISL_2806916 | Lighthouse Lab in Milton Keynes | Wellcome Sanger Institute for the COVID-19 Genomics UK (COG-UK) Consortium | The Lighthouse Lab in Milton Keynes and Alex Alderton, Roberto Amato, Jeffrey Barrett, Sonia Goncalves, Ewan Harrison, David K. Jackson, Ian Johnston, Dominic Kwiatkowski, Cordelia Langford, John Sillitoe on behalf of the Wellcome Sanger Institute COVID-19 Surveillance Team |
| EPI_ISL_2806917 | Lighthouse Laboratory Plymouth | Wellcome Sanger Institute for the COVID-19 Genomics UK (COG-UK) Consortium | Lighthouse Laboratory Plymouth and Alex Alderton, Roberto Amato, Jeffrey Barrett, Sonia Goncalves, Ewan Harrison, David K. Jackson, Ian Johnston, Dominic Kwiatkowski, Cordelia Langford, John Sillitoe on behalf of the Wellcome Sanger Institute COVID-19 Surveillance Team |
| EPI_ISL_2806918 | Lighthouse Lab in Milton Keynes | Wellcome Sanger Institute for the COVID-19 Genomics UK (COG-UK) Consortium | The Lighthouse Lab in Milton Keynes and Alex Alderton, Roberto Amato, Jeffrey Barrett, Sonia Goncalves, Ewan Harrison, David K. Jackson, Ian Johnston, Dominic Kwiatkowski, Cordelia Langford, John Sillitoe on behalf of the Wellcome Sanger Institute COVID-19 Surveillance Team |
| EPI_ISL_2806919 | Lighthouse Laboratory Plymouth | Wellcome Sanger Institute for the COVID-19 Genomics UK (COG-UK) Consortium | Lighthouse Laboratory Plymouth and Alex Alderton, Roberto Amato, Jeffrey Barrett, Sonia Goncalves, Ewan Harrison, David K. Jackson, Ian Johnston, Dominic Kwiatkowski, Cordelia Langford, John Sillitoe on behalf of the Wellcome Sanger Institute COVID-19 Surveillance Team |
| EPI_ISL_2806920, EPI_ISL_2806921, EPI_ISL_2806922 | Lighthouse Lab in Milton Keynes | Wellcome Sanger Institute for the COVID-19 Genomics UK (COG-UK) Consortium | The Lighthouse Lab in Milton Keynes and Alex Alderton, Roberto Amato, Jeffrey Barrett, Sonia Goncalves, Ewan Harrison, David K. Jackson, Ian Johnston, Dominic Kwiatkowski, Cordelia Langford, John Sillitoe on behalf of the Wellcome Sanger Institute COVID-19 Surveillance Team |
| EPI_ISL_2806923, EPI_ISL_2806924 | Lighthouse Laboratory Plymouth | Wellcome Sanger Institute for the COVID-19 Genomics UK (COG-UK) Consortium | Lighthouse Laboratory Plymouth and Alex Alderton, Roberto Amato, Jeffrey Barrett, Sonia Goncalves, Ewan Harrison, David K. Jackson, Ian Johnston, Dominic Kwiatkowski, Cordelia Langford, John Sillitoe on behalf of the Wellcome Sanger Institute COVID-19 Surveillance Team |
| EPI_ISL_2806925, EPI_ISL_2806926, EPI_ISL_2806927, EPI_ISL_2806928 | Lighthouse Lab in Milton Keynes | Wellcome Sanger Institute for the COVID-19 Genomics UK (COG-UK) Consortium | The Lighthouse Lab in Milton Keynes and Alex Alderton, Roberto Amato, Jeffrey Barrett, Sonia Goncalves, Ewan Harrison, David K. Jackson, Ian Johnston, Dominic Kwiatkowski, Cordelia Langford, John Sillitoe on behalf of the Wellcome Sanger Institute COVID-19 Surveillance Team |
| EPI_ISL_2806929, EPI_ISL_2806930 | Lighthouse Laboratory Plymouth | Wellcome Sanger Institute for the COVID-19 Genomics UK (COG-UK) Consortium | Lighthouse Laboratory Plymouth and Alex Alderton, Roberto Amato, Jeffrey Barrett, Sonia Goncalves, Ewan Harrison, David K. Jackson, Ian Johnston, Dominic Kwiatkowski, Cordelia Langford, John Sillitoe on behalf of the Wellcome Sanger Institute COVID-19 Surveillance Team |
| EPI_ISL_2806931 | Lighthouse Lab in Milton Keynes | Wellcome Sanger Institute for the COVID-19 Genomics UK (COG-UK) Consortium | The Lighthouse Lab in Milton Keynes and Alex Alderton, Roberto Amato, Jeffrey Barrett, Sonia Goncalves, Ewan Harrison, David K. Jackson, Ian Johnston, Dominic Kwiatkowski, Cordelia Langford, John Sillitoe on behalf of the Wellcome Sanger Institute COVID-19 Surveillance Team |
| EPI_ISL_2806932 | Lighthouse Laboratory Plymouth | Wellcome Sanger Institute for the COVID-19 Genomics UK (COG-UK) Consortium | Lighthouse Laboratory Plymouth and Alex Alderton, Roberto Amato, Jeffrey Barrett, Sonia Goncalves, Ewan Harrison, David K. Jackson, Ian Johnston, Dominic Kwiatkowski, Cordelia Langford, John Sillitoe on behalf of the Wellcome Sanger Institute COVID-19 Surveillance Team |
| EPI_ISL_2806933 | Lighthouse Lab in Milton Keynes | Wellcome Sanger Institute for the COVID-19 Genomics UK (COG-UK) Consortium | The Lighthouse Lab in Milton Keynes and Alex Alderton, Roberto Amato, Jeffrey Barrett, Sonia Goncalves, Ewan Harrison, David K. Jackson, Ian Johnston, Dominic Kwiatkowski, Cordelia Langford, John Sillitoe on behalf of the Wellcome Sanger Institute COVID-19 Surveillance Team |
| EPI_ISL_2806934, EPI_ISL_2806935, EPI_ISL_2806936 | Lighthouse Laboratory Plymouth | Wellcome Sanger Institute for the COVID-19 Genomics UK (COG-UK) Consortium | Lighthouse Laboratory Plymouth and Alex Alderton, Roberto Amato, Jeffrey Barrett, Sonia Goncalves, Ewan Harrison, David K. Jackson, Ian Johnston, Dominic Kwiatkowski, Cordelia Langford, John Sillitoe on behalf of the Wellcome Sanger Institute COVID-19 Surveillance Team |
| EPI_ISL_2806937, EPI_ISL_2806938, EPI_ISL_2806939, EPI_ISL_2806940 | Lighthouse Lab in Milton Keynes | Wellcome Sanger Institute for the COVID-19 Genomics UK (COG-UK) Consortium | The Lighthouse Lab in Milton Keynes and Alex Alderton, Roberto Amato, Jeffrey Barrett, Sonia Goncalves, Ewan Harrison, David K. Jackson, Ian Johnston, Dominic Kwiatkowski, Cordelia Langford, John Sillitoe on behalf of the Wellcome Sanger Institute COVID-19 Surveillance Team |
| EPI_ISL_2806941, EPI_ISL_2806942, EPI_ISL_2806943 | Lighthouse Laboratory Plymouth | Wellcome Sanger Institute for the COVID-19 Genomics UK (COG-UK) Consortium | Lighthouse Laboratory Plymouth and Alex Alderton, Roberto Amato, Jeffrey Barrett, Sonia Goncalves, Ewan Harrison, David K. Jackson, Ian Johnston, Dominic Kwiatkowski, Cordelia Langford, John Sillitoe on behalf of the Wellcome Sanger Institute COVID-19 Surveillance Team |
| EPI_ISL_2806944, EPI_ISL_2806945, EPI_ISL_2806946, EPI_ISL_2806947, EPI_ISL_2806948, EPI_ISL_2806949, EPI_ISL_2806950, EPI_ISL_2806951 | Lighthouse Lab in Milton Keynes | Wellcome Sanger Institute for the COVID-19 Genomics UK (COG-UK) Consortium | The Lighthouse Lab in Milton Keynes and Alex Alderton, Roberto Amato, Jeffrey Barrett, Sonia Goncalves, Ewan Harrison, David K. Jackson, Ian Johnston, Dominic Kwiatkowski, Cordelia Langford, John Sillitoe on behalf of the Wellcome Sanger Institute COVID-19 Surveillance Team |
| EPI_ISL_2806952, EPI_ISL_2806953, EPI_ISL_2806954 | Lighthouse Laboratory Plymouth | Wellcome Sanger Institute for the COVID-19 Genomics UK (COG-UK) Consortium | Lighthouse Laboratory Plymouth and Alex Alderton, Roberto Amato, Jeffrey Barrett, Sonia Goncalves, Ewan Harrison, David K. Jackson, Ian Johnston, Dominic Kwiatkowski, Cordelia Langford, John Sillitoe on behalf of the Wellcome Sanger Institute COVID-19 Surveillance Team |
| EPI_ISL_2806955 | Lighthouse Lab in Milton Keynes | Wellcome Sanger Institute for the COVID-19 Genomics UK (COG-UK) Consortium | The Lighthouse Lab in Milton Keynes and Alex Alderton, Roberto Amato, Jeffrey Barrett, Sonia Goncalves, Ewan Harrison, David K. Jackson, Ian Johnston, Dominic Kwiatkowski, Cordelia Langford, John Sillitoe on behalf of the Wellcome Sanger Institute COVID-19 Surveillance Team |
| EPI_ISL_2806956, EPI_ISL_2806957 | Lighthouse Laboratory Plymouth | Wellcome Sanger Institute for the COVID-19 Genomics UK (COG-UK) Consortium | Lighthouse Laboratory Plymouth and Alex Alderton, Roberto Amato, Jeffrey Barrett, Sonia Goncalves, Ewan Harrison, David K. Jackson, Ian Johnston, Dominic Kwiatkowski, Cordelia Langford, John Sillitoe on behalf of the Wellcome Sanger Institute COVID-19 Surveillance Team |
| EPI_ISL_2806958, EPI_ISL_2806959, EPI_ISL_2806960, | Lighthouse Lab in Milton Keynes | Wellcome Sanger Institute for the COVID-19 Genomics UK (COG-UK) Consortium | The Lighthouse Lab in Milton Keynes and Alex Alderton, Roberto Amato, Jeffrey Barrett, Sonia Goncalves, Ewan Harrison, David K. Jackson, Ian Johnston, Dominic Kwiatkowski, Cordelia Langford, John Sillitoe on behalf of the Wellcome Sanger Institute COVID-19 Surveillance Team |

| | | | |
|---|---|---|---|
| EPI_ISL_2806961, EPI_ISL_2806962 | | | |
| EPI_ISL_2806963, EPI_ISL_2806964, EPI_ISL_2806965 | Lighthouse Laboratory Plymouth | Wellcome Sanger Institute for the COVID-19 Genomics UK (COG-UK) Consortium | Lighthouse Laboratory Plymouth and Alex Alderton, Roberto Amato, Jeffrey Barrett, Sonia Goncalves, Ewan Harrison, David K. Jackson, Ian Johnston, Dominic Kwiatkowski, Cordelia Langford, John Sillitoe on behalf of the Wellcome Sanger Institute COVID-19 Surveillance Team |
| EPI_ISL_2806966, EPI_ISL_2806967, EPI_ISL_2806968 | Lighthouse Lab in Milton Keynes | Wellcome Sanger Institute for the COVID-19 Genomics UK (COG-UK) Consortium | The Lighthouse Lab in Milton Keynes and Alex Alderton, Roberto Amato, Jeffrey Barrett, Sonia Goncalves, Ewan Harrison, David K. Jackson, Ian Johnston, Dominic Kwiatkowski, Cordelia Langford, John Sillitoe on behalf of the Wellcome Sanger Institute COVID-19 Surveillance Team |
| EPI_ISL_2806969 | Lighthouse Laboratory Plymouth | Wellcome Sanger Institute for the COVID-19 Genomics UK (COG-UK) Consortium | Lighthouse Laboratory Plymouth and Alex Alderton, Roberto Amato, Jeffrey Barrett, Sonia Goncalves, Ewan Harrison, David K. Jackson, Ian Johnston, Dominic Kwiatkowski, Cordelia Langford, John Sillitoe on behalf of the Wellcome Sanger Institute COVID-19 Surveillance Team |
| EPI_ISL_2806970 | Lighthouse Lab in Milton Keynes | Wellcome Sanger Institute for the COVID-19 Genomics UK (COG-UK) Consortium | The Lighthouse Lab in Milton Keynes and Alex Alderton, Roberto Amato, Jeffrey Barrett, Sonia Goncalves, Ewan Harrison, David K. Jackson, Ian Johnston, Dominic Kwiatkowski, Cordelia Langford, John Sillitoe on behalf of the Wellcome Sanger Institute COVID-19 Surveillance Team |
| EPI_ISL_2806971, EPI_ISL_2806972, EPI_ISL_2806973, EPI_ISL_2806974, EPI_ISL_2806975 | Lighthouse Laboratory Plymouth | Wellcome Sanger Institute for the COVID-19 Genomics UK (COG-UK) Consortium | Lighthouse Laboratory Plymouth and Alex Alderton, Roberto Amato, Jeffrey Barrett, Sonia Goncalves, Ewan Harrison, David K. Jackson, Ian Johnston, Dominic Kwiatkowski, Cordelia Langford, John Sillitoe on behalf of the Wellcome Sanger Institute COVID-19 Surveillance Team |
| EPI_ISL_2806976 | Lighthouse Lab in Milton Keynes | Wellcome Sanger Institute for the COVID-19 Genomics UK (COG-UK) Consortium | The Lighthouse Lab in Milton Keynes and Alex Alderton, Roberto Amato, Jeffrey Barrett, Sonia Goncalves, Ewan Harrison, David K. Jackson, Ian Johnston, Dominic Kwiatkowski, Cordelia Langford, John Sillitoe on behalf of the Wellcome Sanger Institute COVID-19 Surveillance Team |
| EPI_ISL_2806977 | Lighthouse Laboratory Plymouth | Wellcome Sanger Institute for the COVID-19 Genomics UK (COG-UK) Consortium | Lighthouse Laboratory Plymouth and Alex Alderton, Roberto Amato, Jeffrey Barrett, Sonia Goncalves, Ewan Harrison, David K. Jackson, Ian Johnston, Dominic Kwiatkowski, Cordelia Langford, John Sillitoe on behalf of the Wellcome Sanger Institute COVID-19 Surveillance Team |
| EPI_ISL_2806978 | Lighthouse Lab in Milton Keynes | Wellcome Sanger Institute for the COVID-19 Genomics UK (COG-UK) Consortium | The Lighthouse Lab in Milton Keynes and Alex Alderton, Roberto Amato, Jeffrey Barrett, Sonia Goncalves, Ewan Harrison, David K. Jackson, Ian Johnston, Dominic Kwiatkowski, Cordelia Langford, John Sillitoe on behalf of the Wellcome Sanger Institute COVID-19 Surveillance Team |
| EPI_ISL_2806979 | Lighthouse Laboratory Plymouth | Wellcome Sanger Institute for the COVID-19 Genomics UK (COG-UK) Consortium | Lighthouse Laboratory Plymouth and Alex Alderton, Roberto Amato, Jeffrey Barrett, Sonia Goncalves, Ewan Harrison, David K. Jackson, Ian Johnston, Dominic Kwiatkowski, Cordelia Langford, John Sillitoe on behalf of the Wellcome Sanger Institute COVID-19 Surveillance Team |
| EPI_ISL_2806980, EPI_ISL_2806981 | Lighthouse Lab in Milton Keynes | Wellcome Sanger Institute for the COVID-19 Genomics UK (COG-UK) Consortium | The Lighthouse Lab in Milton Keynes and Alex Alderton, Roberto Amato, Jeffrey Barrett, Sonia Goncalves, Ewan Harrison, David K. Jackson, Ian Johnston, Dominic Kwiatkowski, Cordelia Langford, John Sillitoe on behalf of the Wellcome Sanger Institute COVID-19 Surveillance Team |
| EPI_ISL_2806982 | Lighthouse Laboratory Plymouth | Wellcome Sanger Institute for the COVID-19 Genomics UK (COG-UK) Consortium | Lighthouse Laboratory Plymouth and Alex Alderton, Roberto Amato, Jeffrey Barrett, Sonia Goncalves, Ewan Harrison, David K. Jackson, Ian Johnston, Dominic Kwiatkowski, Cordelia Langford, John Sillitoe on behalf of the Wellcome Sanger Institute COVID-19 Surveillance Team |
| EPI_ISL_2806983 | Lighthouse Lab in Milton Keynes | Wellcome Sanger Institute for the COVID-19 Genomics UK (COG-UK) Consortium | The Lighthouse Lab in Milton Keynes and Alex Alderton, Roberto Amato, Jeffrey Barrett, Sonia Goncalves, Ewan Harrison, David K. Jackson, Ian Johnston, Dominic Kwiatkowski, Cordelia Langford, John Sillitoe on behalf of the Wellcome Sanger Institute COVID-19 Surveillance Team |
| EPI_ISL_2806984, EPI_ISL_2806985, EPI_ISL_2806986 | Lighthouse Laboratory Plymouth | Wellcome Sanger Institute for the COVID-19 Genomics UK (COG-UK) Consortium | Lighthouse Laboratory Plymouth and Alex Alderton, Roberto Amato, Jeffrey Barrett, Sonia Goncalves, Ewan Harrison, David K. Jackson, Ian Johnston, Dominic Kwiatkowski, Cordelia Langford, John Sillitoe on behalf of the Wellcome Sanger Institute COVID-19 Surveillance Team |
| EPI_ISL_2806987, EPI_ISL_2806988, EPI_ISL_2806989 | Lighthouse Lab in Milton Keynes | Wellcome Sanger Institute for the COVID-19 Genomics UK (COG-UK) Consortium | The Lighthouse Lab in Milton Keynes and Alex Alderton, Roberto Amato, Jeffrey Barrett, Sonia Goncalves, Ewan Harrison, David K. Jackson, Ian Johnston, Dominic Kwiatkowski, Cordelia Langford, John Sillitoe on behalf of the Wellcome Sanger Institute COVID-19 Surveillance Team |
| EPI_ISL_2806990, EPI_ISL_2806991 | Lighthouse Laboratory Plymouth | Wellcome Sanger Institute for the COVID-19 Genomics UK (COG-UK) Consortium | Lighthouse Laboratory Plymouth and Alex Alderton, Roberto Amato, Jeffrey Barrett, Sonia Goncalves, Ewan Harrison, David K. Jackson, Ian Johnston, Dominic Kwiatkowski, Cordelia Langford, John Sillitoe on behalf of the Wellcome Sanger Institute COVID-19 Surveillance Team |
| EPI_ISL_2806992, EPI_ISL_2806993, EPI_ISL_2806994, EPI_ISL_2806995, EPI_ISL_2806996 | Lighthouse Lab in Milton Keynes | Wellcome Sanger Institute for the COVID-19 Genomics UK (COG-UK) Consortium | The Lighthouse Lab in Milton Keynes and Alex Alderton, Roberto Amato, Jeffrey Barrett, Sonia Goncalves, Ewan Harrison, David K. Jackson, Ian Johnston, Dominic Kwiatkowski, Cordelia Langford, John Sillitoe on behalf of the Wellcome Sanger Institute COVID-19 Surveillance Team |
| EPI_ISL_2806997, EPI_ISL_2806998 | Lighthouse Laboratory Plymouth | Wellcome Sanger Institute for the COVID-19 Genomics UK (COG-UK) Consortium | Lighthouse Laboratory Plymouth and Alex Alderton, Roberto Amato, Jeffrey Barrett, Sonia Goncalves, Ewan Harrison, David K. Jackson, Ian Johnston, Dominic Kwiatkowski, Cordelia Langford, John Sillitoe on behalf of the Wellcome Sanger Institute COVID-19 Surveillance Team |
| EPI_ISL_2806999 | Lighthouse Lab in Milton Keynes | Wellcome Sanger Institute for the COVID-19 Genomics UK (COG-UK) Consortium | The Lighthouse Lab in Milton Keynes and Alex Alderton, Roberto Amato, Jeffrey Barrett, Sonia Goncalves, Ewan Harrison, David K. Jackson, Ian Johnston, Dominic Kwiatkowski, Cordelia Langford, John Sillitoe on behalf of the Wellcome Sanger Institute COVID-19 Surveillance Team |
| EPI_ISL_2807000 | Lighthouse Laboratory Plymouth | Wellcome Sanger Institute for the COVID-19 Genomics UK (COG-UK) Consortium | Lighthouse Laboratory Plymouth and Alex Alderton, Roberto Amato, Jeffrey Barrett, Sonia Goncalves, Ewan Harrison, David K. Jackson, Ian Johnston, Dominic Kwiatkowski, Cordelia Langford, John Sillitoe on behalf of the Wellcome Sanger Institute COVID-19 Surveillance Team |
| EPI_ISL_2807001, EPI_ISL_2807002, EPI_ISL_2807003 | Lighthouse Lab in Milton Keynes | Wellcome Sanger Institute for the COVID-19 Genomics UK (COG-UK) Consortium | The Lighthouse Lab in Milton Keynes and Alex Alderton, Roberto Amato, Jeffrey Barrett, Sonia Goncalves, Ewan Harrison, David K. Jackson, Ian Johnston, Dominic Kwiatkowski, Cordelia Langford, John Sillitoe on behalf of the Wellcome Sanger Institute COVID-19 Surveillance Team |
| EPI_ISL_2807004, EPI_ISL_2807005, EPI_ISL_2807006, EPI_ISL_2807007, EPI_ISL_2807008, EPI_ISL_2807009 | Lighthouse Laboratory Plymouth | Wellcome Sanger Institute for the COVID-19 Genomics UK (COG-UK) Consortium | Lighthouse Laboratory Plymouth and Alex Alderton, Roberto Amato, Jeffrey Barrett, Sonia Goncalves, Ewan Harrison, David K. Jackson, Ian Johnston, Dominic Kwiatkowski, Cordelia Langford, John Sillitoe on behalf of the Wellcome Sanger Institute COVID-19 Surveillance Team |
| EPI_ISL_2807010, EPI_ISL_2807011, EPI_ISL_2807012, EPI_ISL_2807013, EPI_ISL_2807014, EPI_ISL_2807015 | Lighthouse Lab in Milton Keynes | Wellcome Sanger Institute for the COVID-19 Genomics UK (COG-UK) Consortium | The Lighthouse Lab in Milton Keynes and Alex Alderton, Roberto Amato, Jeffrey Barrett, Sonia Goncalves, Ewan Harrison, David K. Jackson, Ian Johnston, Dominic Kwiatkowski, Cordelia Langford, John Sillitoe on behalf of the Wellcome Sanger Institute COVID-19 Surveillance Team |
| EPI_ISL_2807016, EPI_ISL_2807017 | Lighthouse Laboratory Plymouth | Wellcome Sanger Institute for the COVID-19 Genomics UK (COG-UK) Consortium | Lighthouse Laboratory Plymouth and Alex Alderton, Roberto Amato, Jeffrey Barrett, Sonia Goncalves, Ewan Harrison, David K. Jackson, Ian Johnston, Dominic Kwiatkowski, Cordelia Langford, John Sillitoe on behalf of the Wellcome Sanger Institute COVID-19 Surveillance Team |
| EPI_ISL_2807018 | Lighthouse Lab in Milton Keynes | Wellcome Sanger Institute for the COVID-19 Genomics UK (COG-UK) Consortium | The Lighthouse Lab in Milton Keynes and Alex Alderton, Roberto Amato, Jeffrey Barrett, Sonia Goncalves, Ewan Harrison, David K. Jackson, Ian Johnston, Dominic Kwiatkowski, Cordelia Langford, John Sillitoe on behalf of the Wellcome Sanger Institute COVID-19 Surveillance Team |
| EPI_ISL_2807019 | Lighthouse Laboratory Plymouth | Wellcome Sanger Institute for the COVID-19 Genomics UK (COG-UK) Consortium | Lighthouse Laboratory Plymouth and Alex Alderton, Roberto Amato, Jeffrey Barrett, Sonia Goncalves, Ewan Harrison, David K. Jackson, Ian Johnston, Dominic Kwiatkowski, Cordelia Langford, John Sillitoe on behalf of the Wellcome Sanger Institute COVID-19 Surveillance Team |
| EPI_ISL_2807020 | Lighthouse Lab in Milton Keynes | Wellcome Sanger Institute for the COVID-19 Genomics UK (COG-UK) Consortium | The Lighthouse Lab in Milton Keynes and Alex Alderton, Roberto Amato, Jeffrey Barrett, Sonia Goncalves, Ewan Harrison, David K. Jackson, Ian Johnston, Dominic Kwiatkowski, Cordelia Langford, John Sillitoe on behalf of the Wellcome Sanger Institute COVID-19 Surveillance Team |
| EPI_ISL_2807021 | Lighthouse Laboratory Plymouth | Wellcome Sanger Institute for the COVID-19 Genomics UK (COG-UK) Consortium | Lighthouse Laboratory Plymouth and Alex Alderton, Roberto Amato, Jeffrey Barrett, Sonia Goncalves, Ewan Harrison, David K. Jackson, Ian Johnston, Dominic Kwiatkowski, Cordelia Langford, John Sillitoe on behalf of the Wellcome Sanger Institute COVID-19 Surveillance Team |
| EPI_ISL_2807022 | Lighthouse Lab in Milton Keynes | Wellcome Sanger Institute for the COVID-19 Genomics UK (COG-UK) Consortium | The Lighthouse Lab in Milton Keynes and Alex Alderton, Roberto Amato, Jeffrey Barrett, Sonia Goncalves, Ewan Harrison, David K. Jackson, Ian Johnston, Dominic Kwiatkowski, Cordelia Langford, John Sillitoe on behalf of the Wellcome Sanger Institute COVID-19 Surveillance Team |
| EPI_ISL_2807023 | Lighthouse Laboratory Plymouth | Wellcome Sanger Institute for the COVID-19 Genomics UK (COG-UK) Consortium | Lighthouse Laboratory Plymouth and Alex Alderton, Roberto Amato, Jeffrey Barrett, Sonia Goncalves, Ewan Harrison, David K. Jackson, Ian Johnston, Dominic Kwiatkowski, Cordelia Langford, John Sillitoe on behalf of the Wellcome Sanger Institute COVID-19 Surveillance Team |
| EPI_ISL_2807024 | Lighthouse Lab in Milton Keynes | Wellcome Sanger Institute for the COVID-19 Genomics UK (COG-UK) Consortium | The Lighthouse Lab in Milton Keynes and Alex Alderton, Roberto Amato, Jeffrey Barrett, Sonia Goncalves, Ewan Harrison, David K. Jackson, Ian Johnston, Dominic Kwiatkowski, Cordelia Langford, John Sillitoe on behalf of the Wellcome Sanger Institute COVID-19 Surveillance Team |
| EPI_ISL_2807025, EPI_ISL_2807026, EPI_ISL_2807027, EPI_ISL_2807028, EPI_ISL_2807029, EPI_ISL_2807030 | Lighthouse Laboratory Plymouth | Wellcome Sanger Institute for the COVID-19 Genomics UK (COG-UK) Consortium | Lighthouse Laboratory Plymouth and Alex Alderton, Roberto Amato, Jeffrey Barrett, Sonia Goncalves, Ewan Harrison, David K. Jackson, Ian Johnston, Dominic Kwiatkowski, Cordelia Langford, John Sillitoe on behalf of the Wellcome Sanger Institute COVID-19 Surveillance Team |
| EPI_ISL_2807031 | Lighthouse Lab in Milton Keynes | Wellcome Sanger Institute for the COVID-19 Genomics UK (COG-UK) Consortium | The Lighthouse Lab in Milton Keynes and Alex Alderton, Roberto Amato, Jeffrey Barrett, Sonia Goncalves, Ewan Harrison, David K. Jackson, Ian Johnston, Dominic Kwiatkowski, Cordelia Langford, John Sillitoe on behalf of the Wellcome Sanger Institute COVID-19 Surveillance Team |
| EPI_ISL_2807032 | Lighthouse Laboratory Plymouth | Wellcome Sanger Institute for the COVID-19 Genomics UK (COG-UK) Consortium | Lighthouse Laboratory Plymouth and Alex Alderton, Roberto Amato, Jeffrey Barrett, Sonia Goncalves, Ewan Harrison, David K. Jackson, Ian Johnston, Dominic Kwiatkowski, Cordelia Langford, John Sillitoe on behalf of the Wellcome Sanger Institute COVID-19 Surveillance Team |
| EPI_ISL_2807033 | Lighthouse Lab in Milton Keynes | Wellcome Sanger Institute for the COVID-19 Genomics UK (COG-UK) Consortium | The Lighthouse Lab in Milton Keynes and Alex Alderton, Roberto Amato, Jeffrey Barrett, Sonia Goncalves, Ewan Harrison, David K. Jackson, Ian Johnston, Dominic Kwiatkowski, Cordelia Langford, John Sillitoe on behalf of the Wellcome Sanger Institute COVID-19 Surveillance Team |
| EPI_ISL_2807034, | Lighthouse Laboratory Plymouth | Wellcome Sanger Institute for the COVID-19 Genomics UK (COG-UK) | Lighthouse Laboratory Plymouth and Alex Alderton, Roberto Amato, Jeffrey Barrett, Sonia Goncalves, Ewan Harrison, David K. Jackson, Ian Johnston, Dominic Kwiatkowski, Cordelia Langford, John Sillitoe on behalf of the Wellcome Sanger |

| | | | |
|---|---|---|---|
| EPI_ISL_2807035, EPI_ISL_2807036 | | Consortium | Institute COVID-19 Surveillance Team |
| EPI_ISL_2807037, EPI_ISL_2807038 | Lighthouse Lab in Milton Keynes | Wellcome Sanger Institute for the COVID-19 Genomics UK (COG-UK) Consortium | The Lighthouse Lab in Milton Keynes and Alex Alderton, Roberto Amato, Jeffrey Barrett, Sonia Goncalves, Ewan Harrison, David K. Jackson, Ian Johnston, Dominic Kwiatkowski, Cordelia Langford, John Sillitoe on behalf of the Wellcome Sanger Institute COVID-19 Surveillance Team |
| EPI_ISL_2807039, EPI_ISL_2807040, EPI_ISL_2807041, EPI_ISL_2807042, EPI_ISL_2807043 | Lighthouse Laboratory Plymouth | Wellcome Sanger Institute for the COVID-19 Genomics UK (COG-UK) Consortium | Lighthouse Laboratory Plymouth and Alex Alderton, Roberto Amato, Jeffrey Barrett, Sonia Goncalves, Ewan Harrison, David K. Jackson, Ian Johnston, Dominic Kwiatkowski, Cordelia Langford, John Sillitoe on behalf of the Wellcome Sanger Institute COVID-19 Surveillance Team |
| EPI_ISL_2807044 | Lighthouse Lab in Milton Keynes | Wellcome Sanger Institute for the COVID-19 Genomics UK (COG-UK) Consortium | The Lighthouse Lab in Milton Keynes and Alex Alderton, Roberto Amato, Jeffrey Barrett, Sonia Goncalves, Ewan Harrison, David K. Jackson, Ian Johnston, Dominic Kwiatkowski, Cordelia Langford, John Sillitoe on behalf of the Wellcome Sanger Institute COVID-19 Surveillance Team |
| EPI_ISL_2807045, EPI_ISL_2807046 | Lighthouse Laboratory Plymouth | Wellcome Sanger Institute for the COVID-19 Genomics UK (COG-UK) Consortium | Lighthouse Laboratory Plymouth and Alex Alderton, Roberto Amato, Jeffrey Barrett, Sonia Goncalves, Ewan Harrison, David K. Jackson, Ian Johnston, Dominic Kwiatkowski, Cordelia Langford, John Sillitoe on behalf of the Wellcome Sanger Institute COVID-19 Surveillance Team |
| EPI_ISL_2807047, EPI_ISL_2807048, EPI_ISL_2807049 | Lighthouse Lab in Milton Keynes | Wellcome Sanger Institute for the COVID-19 Genomics UK (COG-UK) Consortium | The Lighthouse Lab in Milton Keynes and Alex Alderton, Roberto Amato, Jeffrey Barrett, Sonia Goncalves, Ewan Harrison, David K. Jackson, Ian Johnston, Dominic Kwiatkowski, Cordelia Langford, John Sillitoe on behalf of the Wellcome Sanger Institute COVID-19 Surveillance Team |
| EPI_ISL_2807051, EPI_ISL_2807052, EPI_ISL_2807053 | Lighthouse Laboratory Plymouth | Wellcome Sanger Institute for the COVID-19 Genomics UK (COG-UK) Consortium | Lighthouse Laboratory Plymouth and Alex Alderton, Roberto Amato, Jeffrey Barrett, Sonia Goncalves, Ewan Harrison, David K. Jackson, Ian Johnston, Dominic Kwiatkowski, Cordelia Langford, John Sillitoe on behalf of the Wellcome Sanger Institute COVID-19 Surveillance Team |
| EPI_ISL_2807054 | Lighthouse Lab in Milton Keynes | Wellcome Sanger Institute for the COVID-19 Genomics UK (COG-UK) Consortium | The Lighthouse Lab in Milton Keynes and Alex Alderton, Roberto Amato, Jeffrey Barrett, Sonia Goncalves, Ewan Harrison, David K. Jackson, Ian Johnston, Dominic Kwiatkowski, Cordelia Langford, John Sillitoe on behalf of the Wellcome Sanger Institute COVID-19 Surveillance Team |
| EPI_ISL_2807055, EPI_ISL_2807056, EPI_ISL_2807057, EPI_ISL_2807058 | Lighthouse Laboratory Plymouth | Wellcome Sanger Institute for the COVID-19 Genomics UK (COG-UK) Consortium | Lighthouse Laboratory Plymouth and Alex Alderton, Roberto Amato, Jeffrey Barrett, Sonia Goncalves, Ewan Harrison, David K. Jackson, Ian Johnston, Dominic Kwiatkowski, Cordelia Langford, John Sillitoe on behalf of the Wellcome Sanger Institute COVID-19 Surveillance Team |
| EPI_ISL_2807059 | Lighthouse Lab in Milton Keynes | Wellcome Sanger Institute for the COVID-19 Genomics UK (COG-UK) Consortium | The Lighthouse Lab in Milton Keynes and Alex Alderton, Roberto Amato, Jeffrey Barrett, Sonia Goncalves, Ewan Harrison, David K. Jackson, Ian Johnston, Dominic Kwiatkowski, Cordelia Langford, John Sillitoe on behalf of the Wellcome Sanger Institute COVID-19 Surveillance Team |
| EPI_ISL_2807060 | Lighthouse Laboratory Plymouth | Wellcome Sanger Institute for the COVID-19 Genomics UK (COG-UK) Consortium | Lighthouse Laboratory Plymouth and Alex Alderton, Roberto Amato, Jeffrey Barrett, Sonia Goncalves, Ewan Harrison, David K. Jackson, Ian Johnston, Dominic Kwiatkowski, Cordelia Langford, John Sillitoe on behalf of the Wellcome Sanger Institute COVID-19 Surveillance Team |
| EPI_ISL_2807061, EPI_ISL_2807062, EPI_ISL_2807063 | Lighthouse Lab in Milton Keynes | Wellcome Sanger Institute for the COVID-19 Genomics UK (COG-UK) Consortium | The Lighthouse Lab in Milton Keynes and Alex Alderton, Roberto Amato, Jeffrey Barrett, Sonia Goncalves, Ewan Harrison, David K. Jackson, Ian Johnston, Dominic Kwiatkowski, Cordelia Langford, John Sillitoe on behalf of the Wellcome Sanger Institute COVID-19 Surveillance Team |
| EPI_ISL_2807064, EPI_ISL_2807065, EPI_ISL_2807066, EPI_ISL_2807067 | Lighthouse Laboratory Plymouth | Wellcome Sanger Institute for the COVID-19 Genomics UK (COG-UK) Consortium | Lighthouse Laboratory Plymouth and Alex Alderton, Roberto Amato, Jeffrey Barrett, Sonia Goncalves, Ewan Harrison, David K. Jackson, Ian Johnston, Dominic Kwiatkowski, Cordelia Langford, John Sillitoe on behalf of the Wellcome Sanger Institute COVID-19 Surveillance Team |
| EPI_ISL_2807068 | Lighthouse Lab in Milton Keynes | Wellcome Sanger Institute for the COVID-19 Genomics UK (COG-UK) Consortium | The Lighthouse Lab in Milton Keynes and Alex Alderton, Roberto Amato, Jeffrey Barrett, Sonia Goncalves, Ewan Harrison, David K. Jackson, Ian Johnston, Dominic Kwiatkowski, Cordelia Langford, John Sillitoe on behalf of the Wellcome Sanger Institute COVID-19 Surveillance Team |
| EPI_ISL_2807069 | Lighthouse Laboratory Plymouth | Wellcome Sanger Institute for the COVID-19 Genomics UK (COG-UK) Consortium | Lighthouse Laboratory Plymouth and Alex Alderton, Roberto Amato, Jeffrey Barrett, Sonia Goncalves, Ewan Harrison, David K. Jackson, Ian Johnston, Dominic Kwiatkowski, Cordelia Langford, John Sillitoe on behalf of the Wellcome Sanger Institute COVID-19 Surveillance Team |
| EPI_ISL_2807070, EPI_ISL_2807071, EPI_ISL_2807072 | Lighthouse Lab in Milton Keynes | Wellcome Sanger Institute for the COVID-19 Genomics UK (COG-UK) Consortium | The Lighthouse Lab in Milton Keynes and Alex Alderton, Roberto Amato, Jeffrey Barrett, Sonia Goncalves, Ewan Harrison, David K. Jackson, Ian Johnston, Dominic Kwiatkowski, Cordelia Langford, John Sillitoe on behalf of the Wellcome Sanger Institute COVID-19 Surveillance Team |
| EPI_ISL_2807073 | Lighthouse Laboratory Plymouth | Wellcome Sanger Institute for the COVID-19 Genomics UK (COG-UK) Consortium | Lighthouse Laboratory Plymouth and Alex Alderton, Roberto Amato, Jeffrey Barrett, Sonia Goncalves, Ewan Harrison, David K. Jackson, Ian Johnston, Dominic Kwiatkowski, Cordelia Langford, John Sillitoe on behalf of the Wellcome Sanger Institute COVID-19 Surveillance Team |
| EPI_ISL_2807074 | Lighthouse Lab in Milton Keynes | Wellcome Sanger Institute for the COVID-19 Genomics UK (COG-UK) Consortium | The Lighthouse Lab in Milton Keynes and Alex Alderton, Roberto Amato, Jeffrey Barrett, Sonia Goncalves, Ewan Harrison, David K. Jackson, Ian Johnston, Dominic Kwiatkowski, Cordelia Langford, John Sillitoe on behalf of the Wellcome Sanger Institute COVID-19 Surveillance Team |
| EPI_ISL_2807075 | Lighthouse Laboratory Plymouth | Wellcome Sanger Institute for the COVID-19 Genomics UK (COG-UK) Consortium | Lighthouse Laboratory Plymouth and Alex Alderton, Roberto Amato, Jeffrey Barrett, Sonia Goncalves, Ewan Harrison, David K. Jackson, Ian Johnston, Dominic Kwiatkowski, Cordelia Langford, John Sillitoe on behalf of the Wellcome Sanger Institute COVID-19 Surveillance Team |
| EPI_ISL_2807076, EPI_ISL_2807077, EPI_ISL_2807078, EPI_ISL_2807079 | Lighthouse Lab in Milton Keynes | Wellcome Sanger Institute for the COVID-19 Genomics UK (COG-UK) Consortium | The Lighthouse Lab in Milton Keynes and Alex Alderton, Roberto Amato, Jeffrey Barrett, Sonia Goncalves, Ewan Harrison, David K. Jackson, Ian Johnston, Dominic Kwiatkowski, Cordelia Langford, John Sillitoe on behalf of the Wellcome Sanger Institute COVID-19 Surveillance Team |
| EPI_ISL_2807080, EPI_ISL_2807081, EPI_ISL_2807082, EPI_ISL_2807083 | Lighthouse Laboratory Plymouth | Wellcome Sanger Institute for the COVID-19 Genomics UK (COG-UK) Consortium | Lighthouse Laboratory Plymouth and Alex Alderton, Roberto Amato, Jeffrey Barrett, Sonia Goncalves, Ewan Harrison, David K. Jackson, Ian Johnston, Dominic Kwiatkowski, Cordelia Langford, John Sillitoe on behalf of the Wellcome Sanger Institute COVID-19 Surveillance Team |
| EPI_ISL_2807084, EPI_ISL_2807085, EPI_ISL_2807086 | Lighthouse Lab in Milton Keynes | Wellcome Sanger Institute for the COVID-19 Genomics UK (COG-UK) Consortium | The Lighthouse Lab in Milton Keynes and Alex Alderton, Roberto Amato, Jeffrey Barrett, Sonia Goncalves, Ewan Harrison, David K. Jackson, Ian Johnston, Dominic Kwiatkowski, Cordelia Langford, John Sillitoe on behalf of the Wellcome Sanger Institute COVID-19 Surveillance Team |
| EPI_ISL_2807087 | Lighthouse Laboratory Plymouth | Wellcome Sanger Institute for the COVID-19 Genomics UK (COG-UK) Consortium | Lighthouse Laboratory Plymouth and Alex Alderton, Roberto Amato, Jeffrey Barrett, Sonia Goncalves, Ewan Harrison, David K. Jackson, Ian Johnston, Dominic Kwiatkowski, Cordelia Langford, John Sillitoe on behalf of the Wellcome Sanger Institute COVID-19 Surveillance Team |
| EPI_ISL_2807088, EPI_ISL_2807089 | Lighthouse Lab in Milton Keynes | Wellcome Sanger Institute for the COVID-19 Genomics UK (COG-UK) Consortium | The Lighthouse Lab in Milton Keynes and Alex Alderton, Roberto Amato, Jeffrey Barrett, Sonia Goncalves, Ewan Harrison, David K. Jackson, Ian Johnston, Dominic Kwiatkowski, Cordelia Langford, John Sillitoe on behalf of the Wellcome Sanger Institute COVID-19 Surveillance Team |
| EPI_ISL_2807090, EPI_ISL_2807091, EPI_ISL_2807092 | Lighthouse Laboratory Plymouth | Wellcome Sanger Institute for the COVID-19 Genomics UK (COG-UK) Consortium | Lighthouse Laboratory Plymouth and Alex Alderton, Roberto Amato, Jeffrey Barrett, Sonia Goncalves, Ewan Harrison, David K. Jackson, Ian Johnston, Dominic Kwiatkowski, Cordelia Langford, John Sillitoe on behalf of the Wellcome Sanger Institute COVID-19 Surveillance Team |
| EPI_ISL_2807093 | Lighthouse Lab in Milton Keynes | Wellcome Sanger Institute for the COVID-19 Genomics UK (COG-UK) Consortium | The Lighthouse Lab in Milton Keynes and Alex Alderton, Roberto Amato, Jeffrey Barrett, Sonia Goncalves, Ewan Harrison, David K. Jackson, Ian Johnston, Dominic Kwiatkowski, Cordelia Langford, John Sillitoe on behalf of the Wellcome Sanger Institute COVID-19 Surveillance Team |
| EPI_ISL_2807094, EPI_ISL_2807095 | Lighthouse Laboratory Plymouth | Wellcome Sanger Institute for the COVID-19 Genomics UK (COG-UK) Consortium | Lighthouse Laboratory Plymouth and Alex Alderton, Roberto Amato, Jeffrey Barrett, Sonia Goncalves, Ewan Harrison, David K. Jackson, Ian Johnston, Dominic Kwiatkowski, Cordelia Langford, John Sillitoe on behalf of the Wellcome Sanger Institute COVID-19 Surveillance Team |
| EPI_ISL_2807096, EPI_ISL_2807097, EPI_ISL_2807098 | Lighthouse Lab in Milton Keynes | Wellcome Sanger Institute for the COVID-19 Genomics UK (COG-UK) Consortium | The Lighthouse Lab in Milton Keynes and Alex Alderton, Roberto Amato, Jeffrey Barrett, Sonia Goncalves, Ewan Harrison, David K. Jackson, Ian Johnston, Dominic Kwiatkowski, Cordelia Langford, John Sillitoe on behalf of the Wellcome Sanger Institute COVID-19 Surveillance Team |
| EPI_ISL_2807099 | Lighthouse Laboratory Plymouth | Wellcome Sanger Institute for the COVID-19 Genomics UK (COG-UK) Consortium | Lighthouse Laboratory Plymouth and Alex Alderton, Roberto Amato, Jeffrey Barrett, Sonia Goncalves, Ewan Harrison, David K. Jackson, Ian Johnston, Dominic Kwiatkowski, Cordelia Langford, John Sillitoe on behalf of the Wellcome Sanger Institute COVID-19 Surveillance Team |
| EPI_ISL_2807100 | Lighthouse Lab in Milton Keynes | Wellcome Sanger Institute for the COVID-19 Genomics UK (COG-UK) Consortium | The Lighthouse Lab in Milton Keynes and Alex Alderton, Roberto Amato, Jeffrey Barrett, Sonia Goncalves, Ewan Harrison, David K. Jackson, Ian Johnston, Dominic Kwiatkowski, Cordelia Langford, John Sillitoe on behalf of the Wellcome Sanger Institute COVID-19 Surveillance Team |
| EPI_ISL_2807101 | Lighthouse Laboratory Plymouth | Wellcome Sanger Institute for the COVID-19 Genomics UK (COG-UK) Consortium | Lighthouse Laboratory Plymouth and Alex Alderton, Roberto Amato, Jeffrey Barrett, Sonia Goncalves, Ewan Harrison, David K. Jackson, Ian Johnston, Dominic Kwiatkowski, Cordelia Langford, John Sillitoe on behalf of the Wellcome Sanger Institute COVID-19 Surveillance Team |
| EPI_ISL_2807102 | Lighthouse Lab in Milton Keynes | Wellcome Sanger Institute for the COVID-19 Genomics UK (COG-UK) Consortium | The Lighthouse Lab in Milton Keynes and Alex Alderton, Roberto Amato, Jeffrey Barrett, Sonia Goncalves, Ewan Harrison, David K. Jackson, Ian Johnston, Dominic Kwiatkowski, Cordelia Langford, John Sillitoe on behalf of the Wellcome Sanger Institute COVID-19 Surveillance Team |
| EPI_ISL_2807103 | Lighthouse Laboratory Plymouth | Wellcome Sanger Institute for the COVID-19 Genomics UK (COG-UK) Consortium | Lighthouse Laboratory Plymouth and Alex Alderton, Roberto Amato, Jeffrey Barrett, Sonia Goncalves, Ewan Harrison, David K. Jackson, Ian Johnston, Dominic Kwiatkowski, Cordelia Langford, John Sillitoe on behalf of the Wellcome Sanger Institute COVID-19 Surveillance Team |
| EPI_ISL_2807104, EPI_ISL_2807105 | Lighthouse Lab in Milton Keynes | Wellcome Sanger Institute for the COVID-19 Genomics UK (COG-UK) Consortium | The Lighthouse Lab in Milton Keynes and Alex Alderton, Roberto Amato, Jeffrey Barrett, Sonia Goncalves, Ewan Harrison, David K. Jackson, Ian Johnston, Dominic Kwiatkowski, Cordelia Langford, John Sillitoe on behalf of the Wellcome Sanger Institute COVID-19 Surveillance Team |
| EPI_ISL_2807106 | Lighthouse Laboratory Plymouth | Wellcome Sanger Institute for the COVID-19 Genomics UK (COG-UK) Consortium | Lighthouse Laboratory Plymouth and Alex Alderton, Roberto Amato, Jeffrey Barrett, Sonia Goncalves, Ewan Harrison, David K. Jackson, Ian Johnston, Dominic Kwiatkowski, Cordelia Langford, John Sillitoe on behalf of the Wellcome Sanger Institute COVID-19 Surveillance Team |
| EPI_ISL_2807107, EPI_ISL_2807108, EPI_ISL_2807109, EPI_ISL_2807110 | Lighthouse Lab in Milton Keynes | Wellcome Sanger Institute for the COVID-19 Genomics UK (COG-UK) Consortium | The Lighthouse Lab in Milton Keynes and Alex Alderton, Roberto Amato, Jeffrey Barrett, Sonia Goncalves, Ewan Harrison, David K. Jackson, Ian Johnston, Dominic Kwiatkowski, Cordelia Langford, John Sillitoe on behalf of the Wellcome Sanger Institute COVID-19 Surveillance Team |
| EPI_ISL_2807111 | Lighthouse Laboratory Plymouth | Wellcome Sanger Institute for the COVID-19 Genomics UK (COG-UK) | Lighthouse Laboratory Plymouth and Alex Alderton, Roberto Amato, Jeffrey Barrett, Sonia Goncalves, Ewan Harrison, David K. Jackson, Ian Johnston, Dominic Kwiatkowski, Cordelia Langford, John Sillitoe on behalf of the Wellcome Sanger |

| | | | |
|---|---|---|---|
| | | Consortium | Institute COVID-19 Surveillance Team |
| EPI_ISL_2807112, EPI_ISL_2807113 | Lighthouse Lab in Milton Keynes | Wellcome Sanger Institute for the COVID-19 Genomics UK (COG-UK) Consortium | The Lighthouse Lab in Milton Keynes and Alex Alderton, Roberto Amato, Jeffrey Barrett, Sonia Goncalves, Ewan Harrison, David K. Jackson, Ian Johnston, Dominic Kwiatkowski, Cordelia Langford, John Sillitoe on behalf of the Wellcome Sanger Institute COVID-19 Surveillance Team |
| EPI_ISL_2807114, EPI_ISL_2807115 | Lighthouse Laboratory Plymouth | Wellcome Sanger Institute for the COVID-19 Genomics UK (COG-UK) Consortium | Lighthouse Laboratory Plymouth and Alex Alderton, Roberto Amato, Jeffrey Barrett, Sonia Goncalves, Ewan Harrison, David K. Jackson, Ian Johnston, Dominic Kwiatkowski, Cordelia Langford, John Sillitoe on behalf of the Wellcome Sanger Institute COVID-19 Surveillance Team |
| EPI_ISL_2807116 | Lighthouse Lab in Milton Keynes | Wellcome Sanger Institute for the COVID-19 Genomics UK (COG-UK) Consortium | The Lighthouse Lab in Milton Keynes and Alex Alderton, Roberto Amato, Jeffrey Barrett, Sonia Goncalves, Ewan Harrison, David K. Jackson, Ian Johnston, Dominic Kwiatkowski, Cordelia Langford, John Sillitoe on behalf of the Wellcome Sanger Institute COVID-19 Surveillance Team |
| EPI_ISL_2807117 | Lighthouse Laboratory Plymouth | Wellcome Sanger Institute for the COVID-19 Genomics UK (COG-UK) Consortium | Lighthouse Laboratory Plymouth and Alex Alderton, Roberto Amato, Jeffrey Barrett, Sonia Goncalves, Ewan Harrison, David K. Jackson, Ian Johnston, Dominic Kwiatkowski, Cordelia Langford, John Sillitoe on behalf of the Wellcome Sanger Institute COVID-19 Surveillance Team |
| EPI_ISL_2807118, EPI_ISL_2807119 | Lighthouse Lab in Milton Keynes | Wellcome Sanger Institute for the COVID-19 Genomics UK (COG-UK) Consortium | The Lighthouse Lab in Milton Keynes and Alex Alderton, Roberto Amato, Jeffrey Barrett, Sonia Goncalves, Ewan Harrison, David K. Jackson, Ian Johnston, Dominic Kwiatkowski, Cordelia Langford, John Sillitoe on behalf of the Wellcome Sanger Institute COVID-19 Surveillance Team |
| EPI_ISL_2807120, EPI_ISL_2807121 | Lighthouse Laboratory Plymouth | Wellcome Sanger Institute for the COVID-19 Genomics UK (COG-UK) Consortium | Lighthouse Laboratory Plymouth and Alex Alderton, Roberto Amato, Jeffrey Barrett, Sonia Goncalves, Ewan Harrison, David K. Jackson, Ian Johnston, Dominic Kwiatkowski, Cordelia Langford, John Sillitoe on behalf of the Wellcome Sanger Institute COVID-19 Surveillance Team |
| EPI_ISL_2807122, EPI_ISL_2807123 | Lighthouse Lab in Milton Keynes | Wellcome Sanger Institute for the COVID-19 Genomics UK (COG-UK) Consortium | The Lighthouse Lab in Milton Keynes and Alex Alderton, Roberto Amato, Jeffrey Barrett, Sonia Goncalves, Ewan Harrison, David K. Jackson, Ian Johnston, Dominic Kwiatkowski, Cordelia Langford, John Sillitoe on behalf of the Wellcome Sanger Institute COVID-19 Surveillance Team |
| EPI_ISL_2807124, EPI_ISL_2807125, EPI_ISL_2807126 | Lighthouse Laboratory Plymouth | Wellcome Sanger Institute for the COVID-19 Genomics UK (COG-UK) Consortium | Lighthouse Laboratory Plymouth and Alex Alderton, Roberto Amato, Jeffrey Barrett, Sonia Goncalves, Ewan Harrison, David K. Jackson, Ian Johnston, Dominic Kwiatkowski, Cordelia Langford, John Sillitoe on behalf of the Wellcome Sanger Institute COVID-19 Surveillance Team |
| EPI_ISL_2807127 | Lighthouse Lab in Milton Keynes | Wellcome Sanger Institute for the COVID-19 Genomics UK (COG-UK) Consortium | The Lighthouse Lab in Milton Keynes and Alex Alderton, Roberto Amato, Jeffrey Barrett, Sonia Goncalves, Ewan Harrison, David K. Jackson, Ian Johnston, Dominic Kwiatkowski, Cordelia Langford, John Sillitoe on behalf of the Wellcome Sanger Institute COVID-19 Surveillance Team |
| EPI_ISL_2807128 | Lighthouse Laboratory Plymouth | Wellcome Sanger Institute for the COVID-19 Genomics UK (COG-UK) Consortium | Lighthouse Laboratory Plymouth and Alex Alderton, Roberto Amato, Jeffrey Barrett, Sonia Goncalves, Ewan Harrison, David K. Jackson, Ian Johnston, Dominic Kwiatkowski, Cordelia Langford, John Sillitoe on behalf of the Wellcome Sanger Institute COVID-19 Surveillance Team |
| EPI_ISL_2807129 | Lighthouse Lab in Milton Keynes | Wellcome Sanger Institute for the COVID-19 Genomics UK (COG-UK) Consortium | The Lighthouse Lab in Milton Keynes and Alex Alderton, Roberto Amato, Jeffrey Barrett, Sonia Goncalves, Ewan Harrison, David K. Jackson, Ian Johnston, Dominic Kwiatkowski, Cordelia Langford, John Sillitoe on behalf of the Wellcome Sanger Institute COVID-19 Surveillance Team |
| EPI_ISL_2807130 | Lighthouse Laboratory Plymouth | Wellcome Sanger Institute for the COVID-19 Genomics UK (COG-UK) Consortium | Lighthouse Laboratory Plymouth and Alex Alderton, Roberto Amato, Jeffrey Barrett, Sonia Goncalves, Ewan Harrison, David K. Jackson, Ian Johnston, Dominic Kwiatkowski, Cordelia Langford, John Sillitoe on behalf of the Wellcome Sanger Institute COVID-19 Surveillance Team |
| EPI_ISL_2807131, EPI_ISL_2807132 | Lighthouse Lab in Milton Keynes | Wellcome Sanger Institute for the COVID-19 Genomics UK (COG-UK) Consortium | The Lighthouse Lab in Milton Keynes and Alex Alderton, Roberto Amato, Jeffrey Barrett, Sonia Goncalves, Ewan Harrison, David K. Jackson, Ian Johnston, Dominic Kwiatkowski, Cordelia Langford, John Sillitoe on behalf of the Wellcome Sanger Institute COVID-19 Surveillance Team |
| EPI_ISL_2807133 | Lighthouse Laboratory Plymouth | Wellcome Sanger Institute for the COVID-19 Genomics UK (COG-UK) Consortium | Lighthouse Laboratory Plymouth and Alex Alderton, Roberto Amato, Jeffrey Barrett, Sonia Goncalves, Ewan Harrison, David K. Jackson, Ian Johnston, Dominic Kwiatkowski, Cordelia Langford, John Sillitoe on behalf of the Wellcome Sanger Institute COVID-19 Surveillance Team |
| EPI_ISL_2807134 | Lighthouse Lab in Milton Keynes | Wellcome Sanger Institute for the COVID-19 Genomics UK (COG-UK) Consortium | The Lighthouse Lab in Milton Keynes and Alex Alderton, Roberto Amato, Jeffrey Barrett, Sonia Goncalves, Ewan Harrison, David K. Jackson, Ian Johnston, Dominic Kwiatkowski, Cordelia Langford, John Sillitoe on behalf of the Wellcome Sanger Institute COVID-19 Surveillance Team |
| EPI_ISL_2807135, EPI_ISL_2807136 | Lighthouse Laboratory Plymouth | Wellcome Sanger Institute for the COVID-19 Genomics UK (COG-UK) Consortium | Lighthouse Laboratory Plymouth and Alex Alderton, Roberto Amato, Jeffrey Barrett, Sonia Goncalves, Ewan Harrison, David K. Jackson, Ian Johnston, Dominic Kwiatkowski, Cordelia Langford, John Sillitoe on behalf of the Wellcome Sanger Institute COVID-19 Surveillance Team |
| EPI_ISL_2807137, EPI_ISL_2807138, EPI_ISL_2807139 | Lighthouse Lab in Milton Keynes | Wellcome Sanger Institute for the COVID-19 Genomics UK (COG-UK) Consortium | The Lighthouse Lab in Milton Keynes and Alex Alderton, Roberto Amato, Jeffrey Barrett, Sonia Goncalves, Ewan Harrison, David K. Jackson, Ian Johnston, Dominic Kwiatkowski, Cordelia Langford, John Sillitoe on behalf of the Wellcome Sanger Institute COVID-19 Surveillance Team |
| EPI_ISL_2807140 | Lighthouse Laboratory Plymouth | Wellcome Sanger Institute for the COVID-19 Genomics UK (COG-UK) Consortium | Lighthouse Laboratory Plymouth and Alex Alderton, Roberto Amato, Jeffrey Barrett, Sonia Goncalves, Ewan Harrison, David K. Jackson, Ian Johnston, Dominic Kwiatkowski, Cordelia Langford, John Sillitoe on behalf of the Wellcome Sanger Institute COVID-19 Surveillance Team |
| EPI_ISL_2807141, EPI_ISL_2807143, EPI_ISL_2807144, EPI_ISL_2807145, EPI_ISL_2807146, EPI_ISL_2807147, EPI_ISL_2807148, EPI_ISL_2807149, EPI_ISL_2807150, EPI_ISL_2807151, EPI_ISL_2807152, EPI_ISL_2807153, EPI_ISL_2807154, EPI_ISL_2807155, EPI_ISL_2807156, EPI_ISL_2807157, EPI_ISL_2807158, EPI_ISL_2807159, EPI_ISL_2807160, EPI_ISL_2807161, EPI_ISL_2807162, EPI_ISL_2807163, EPI_ISL_2807164, EPI_ISL_2807165, EPI_ISL_2807166, EPI_ISL_2807167, EPI_ISL_2807168, EPI_ISL_2807169, EPI_ISL_2807170, EPI_ISL_2807171, EPI_ISL_2807172, EPI_ISL_2807174, EPI_ISL_2807175, EPI_ISL_2807176, EPI_ISL_2807177, EPI_ISL_2807178, EPI_ISL_2807179, EPI_ISL_2807180, EPI_ISL_2807181, EPI_ISL_2807182, EPI_ISL_2807183, EPI_ISL_2807184, EPI_ISL_2807185, EPI_ISL_2807186, EPI_ISL_2807187, EPI_ISL_2807188, EPI_ISL_2807189, EPI_ISL_2807190, EPI_ISL_2807191, EPI_ISL_2807192, EPI_ISL_2807193, EPI_ISL_2807194, EPI_ISL_2807195, EPI_ISL_2807196, EPI_ISL_2807197, EPI_ISL_2807198, EPI_ISL_2807199, EPI_ISL_2807200, EPI_ISL_2807201, EPI_ISL_2807202, EPI_ISL_2807203, EPI_ISL_2807204, EPI_ISL_2807205, EPI_ISL_2807206, EPI_ISL_2807207, EPI_ISL_2807208, EPI_ISL_2807209, EPI_ISL_2807210, EPI_ISL_2807211, EPI_ISL_2807212, EPI_ISL_2807213, EPI_ISL_2807214, EPI_ISL_2807215, EPI_ISL_2807216, EPI_ISL_2807217, EPI_ISL_2807218, EPI_ISL_2807219, EPI_ISL_2807220, EPI_ISL_2807221, EPI_ISL_2807223, EPI_ISL_2807224, EPI_ISL_2807225, EPI_ISL_2807226, EPI_ISL_2807227, EPI_ISL_2807228, EPI_ISL_2807229, EPI_ISL_2807230, EPI_ISL_2807231, EPI_ISL_2807232, EPI_ISL_2807233, EPI_ISL_2807234, EPI_ISL_2807235, EPI_ISL_2807236, EPI_ISL_2807237, EPI_ISL_2807238, EPI_ISL_2807239, EPI_ISL_2807240, EPI_ISL_2807241, EPI_ISL_2807242, EPI_ISL_2807243, EPI_ISL_2807244, EPI_ISL_2807245, EPI_ISL_2807246, EPI_ISL_2807247, EPI_ISL_2807248, EPI_ISL_2807249, EPI_ISL_2807250, EPI_ISL_2807251, EPI_ISL_2807252, EPI_ISL_2807253, EPI_ISL_2807254, EPI_ISL_2807255, EPI_ISL_2807256, EPI_ISL_2807257, EPI_ISL_2807258, EPI_ISL_2807259, EPI_ISL_2807260, EPI_ISL_2807261, EPI_ISL_2807262, EPI_ISL_2807263, EPI_ISL_2807264, EPI_ISL_2807265, EPI_ISL_2807266, EPI_ISL_2807267, EPI_ISL_2807268, EPI_ISL_2807269, EPI_ISL_2807270, EPI_ISL_2807271, EPI_ISL_2807272, EPI_ISL_2807273, EPI_ISL_2807274, EPI_ISL_2807275, EPI_ISL_2807276, EPI_ISL_2807277, EPI_ISL_2807280, EPI_ISL_2807281, EPI_ISL_2807282, EPI_ISL_2807283, EPI_ISL_2807284, EPI_ISL_2807285, EPI_ISL_2807286, EPI_ISL_2807287, EPI_ISL_2807288, EPI_ISL_2807290, EPI_ISL_2807291, EPI_ISL_2807292, EPI_ISL_2807293, EPI_ISL_2807294, EPI_ISL_2807295, EPI_ISL_2807296, EPI_ISL_2807297, EPI_ISL_2807298, EPI_ISL_2807299, EPI_ISL_2807300, EPI_ISL_2807301, EPI_ISL_2807302, EPI_ISL_2807303, EPI_ISL_2807305, EPI_ISL_2807306, EPI_ISL_2807307, EPI_ISL_2807308, EPI_ISL_2807309, EPI_ISL_2807310, EPI_ISL_2807311, EPI_ISL_2807312, EPI_ISL_2807313, EPI_ISL_2807314, EPI_ISL_2807315, EPI_ISL_2807316, EPI_ISL_2807317, EPI_ISL_2807318, EPI_ISL_2807319, EPI_ISL_2807320, EPI_ISL_2807321, EPI_ISL_2807322, EPI_ISL_2807323, EPI_ISL_2807324, EPI_ISL_2807325, EPI_ISL_2807326, EPI_ISL_2807327, EPI_ISL_2807328, EPI_ISL_2807329, EPI_ISL_2807330, EPI_ISL_2807331, EPI_ISL_2807332, EPI_ISL_2807333, EPI_ISL_2807334, EPI_ISL_2807336, EPI_ISL_2807338, EPI_ISL_2807339, EPI_ISL_2807340, EPI_ISL_2807341, EPI_ISL_2807342, EPI_ISL_2807343, EPI_ISL_2807344, EPI_ISL_2807345, EPI_ISL_2807346, EPI_ISL_2807347, EPI_ISL_2807348, EPI_ISL_2807349, EPI_ISL_2807350, EPI_ISL_2807351, EPI_ISL_2807352, EPI_ISL_2807353 | | | |
| see above | Lighthouse Lab in Glasgow | Wellcome Sanger Institute for the COVID-19 Genomics UK (COG-UK) Consortium | Harper VanSteenhouse, Yumi Kasai, David Gray, Carol Clugston, Anna Dominiczak and Alex Alderton, Roberto Amato, Jeffrey Barrett, Sonia Goncalves, Ewan Harrison, David K. Jackson, Ian Johnston, Dominic Kwiatkowski, Cordelia Langford, John Sillitoe on behalf of the Wellcome Sanger Institute COVID-19 Surveillance Team |
| EPI_ISL_2807354, EPI_ISL_2807355 | Lighthouse Lab in Milton Keynes | Wellcome Sanger Institute for the COVID-19 Genomics UK (COG-UK) Consortium | The Lighthouse Lab in Milton Keynes and Alex Alderton, Roberto Amato, Jeffrey Barrett, Sonia Goncalves, Ewan Harrison, David K. Jackson, Ian Johnston, Dominic Kwiatkowski, Cordelia Langford, John Sillitoe on behalf of the Wellcome Sanger Institute COVID-19 Surveillance Team |
| EPI_ISL_2807356, EPI_ISL_2807357 | Lighthouse Lab in Alderley Park | Wellcome Sanger Institute for the COVID-19 Genomics UK (COG-UK) Consortium | Jacquelyn Wynn, Mairead Hyland, The Lighthouse Lab in Alderley Park and Alex Alderton, Roberto Amato, Jeffrey Barrett, Sonia Goncalves, Ewan Harrison, David K. Jackson, Ian Johnston, Dominic Kwiatkowski, Cordelia Langford, John Sillitoe on behalf of the Wellcome Sanger Institute COVID-19 Surveillance Team |
| EPI_ISL_2807358 | Lighthouse Lab in Milton Keynes | Wellcome Sanger Institute for the COVID-19 Genomics UK (COG-UK) Consortium | The Lighthouse Lab in Milton Keynes and Alex Alderton, Roberto Amato, Jeffrey Barrett, Sonia Goncalves, Ewan Harrison, David K. Jackson, Ian Johnston, Dominic Kwiatkowski, Cordelia Langford, John Sillitoe on behalf of the Wellcome Sanger Institute COVID-19 Surveillance Team |
| EPI_ISL_2807359, EPI_ISL_2807360 | Lighthouse Lab in Glasgow | Wellcome Sanger Institute for the COVID-19 Genomics UK (COG-UK) Consortium | Harper VanSteenhouse, Yumi Kasai, David Gray, Carol Clugston, Anna Dominiczak and Alex Alderton, Roberto Amato, Jeffrey Barrett, Sonia Goncalves, Ewan Harrison, David K. Jackson, Ian Johnston, Dominic Kwiatkowski, Cordelia Langford, John Sillitoe on behalf of the Wellcome Sanger Institute COVID-19 Surveillance Team |
| EPI_ISL_2807361, EPI_ISL_2807362 | Lighthouse Lab in Milton Keynes | Wellcome Sanger Institute for the COVID-19 Genomics UK (COG-UK) Consortium | The Lighthouse Lab in Milton Keynes and Alex Alderton, Roberto Amato, Jeffrey Barrett, Sonia Goncalves, Ewan Harrison, David K. Jackson, Ian Johnston, Dominic Kwiatkowski, Cordelia Langford, John Sillitoe on behalf of the Wellcome Sanger Institute COVID-19 Surveillance Team |
| EPI_ISL_2807363 | Lighthouse Lab in Glasgow | Wellcome Sanger Institute for the COVID-19 Genomics UK (COG-UK) Consortium | Harper VanSteenhouse, Yumi Kasai, David Gray, Carol Clugston, Anna Dominiczak and Alex Alderton, Roberto Amato, Jeffrey Barrett, Sonia Goncalves, Ewan Harrison, David K. Jackson, Ian Johnston, Dominic Kwiatkowski, Cordelia Langford, John Sillitoe on behalf of the Wellcome Sanger Institute COVID-19 Surveillance Team |
| EPI_ISL_2807364 | Lighthouse Lab in Alderley Park | Wellcome Sanger Institute for the COVID-19 Genomics UK (COG-UK) Consortium | Jacquelyn Wynn, Mairead Hyland, The Lighthouse Lab in Alderley Park and Alex Alderton, Roberto Amato, Jeffrey Barrett, Sonia Goncalves, Ewan Harrison, David K. Jackson, Ian Johnston, Dominic Kwiatkowski, Cordelia Langford, John Sillitoe on behalf of the Wellcome Sanger Institute COVID-19 Surveillance Team |
| EPI_ISL_2807365 | Lighthouse Lab in Glasgow | Wellcome Sanger Institute for the COVID-19 Genomics UK (COG-UK) Consortium | Harper VanSteenhouse, Yumi Kasai, David Gray, Carol Clugston, Anna Dominiczak and Alex Alderton, Roberto Amato, Jeffrey Barrett, Sonia Goncalves, Ewan Harrison, David K. Jackson, Ian Johnston, Dominic Kwiatkowski, Cordelia Langford, John Sillitoe on behalf of the Wellcome Sanger Institute COVID-19 Surveillance Team |
| EPI_ISL_2807366, EPI_ISL_2807367 | Lighthouse Lab in Milton Keynes | Wellcome Sanger Institute for the COVID-19 Genomics UK (COG-UK) Consortium | The Lighthouse Lab in Milton Keynes and Alex Alderton, Roberto Amato, Jeffrey Barrett, Sonia Goncalves, Ewan Harrison, David K. Jackson, Ian Johnston, Dominic Kwiatkowski, Cordelia Langford, John Sillitoe on behalf of the Wellcome Sanger Institute COVID-19 Surveillance Team |
| EPI_ISL_2807368, EPI_ISL_2807369 | Lighthouse Lab in Glasgow | Wellcome Sanger Institute for the COVID-19 Genomics UK (COG-UK) Consortium | Harper VanSteenhouse, Yumi Kasai, David Gray, Carol Clugston, Anna Dominiczak and Alex Alderton, Roberto Amato, Jeffrey Barrett, Sonia Goncalves, Ewan Harrison, David K. Jackson, Ian Johnston, Dominic Kwiatkowski, Cordelia Langford, John Sillitoe on behalf of the Wellcome Sanger Institute COVID-19 Surveillance Team |
| EPI_ISL_2807370 | Lighthouse Lab in Milton Keynes | Wellcome Sanger Institute for the COVID-19 Genomics UK (COG-UK) Consortium | The Lighthouse Lab in Milton Keynes and Alex Alderton, Roberto Amato, Jeffrey Barrett, Sonia Goncalves, Ewan Harrison, David K. Jackson, Ian Johnston, Dominic Kwiatkowski, Cordelia Langford, John Sillitoe on behalf of the Wellcome Sanger Institute COVID-19 Surveillance Team |
| EPI_ISL_2807371 | Lighthouse Lab in Alderley Park | Wellcome Sanger Institute for the COVID-19 Genomics UK (COG-UK) Consortium | Jacquelyn Wynn, Mairead Hyland, The Lighthouse Lab in Alderley Park and Alex Alderton, Roberto Amato, Jeffrey Barrett, Sonia Goncalves, Ewan Harrison, David K. Jackson, Ian Johnston, Dominic Kwiatkowski, Cordelia Langford, John Sillitoe on behalf of the Wellcome Sanger Institute COVID-19 Surveillance Team |
| EPI_ISL_2807372 | Lighthouse Lab in Glasgow | Wellcome Sanger Institute for the COVID-19 Genomics UK (COG-UK) Consortium | Harper VanSteenhouse, Yumi Kasai, David Gray, Carol Clugston, Anna Dominiczak and Alex Alderton, Roberto Amato, Jeffrey Barrett, Sonia Goncalves, Ewan Harrison, David K. Jackson, Ian Johnston, Dominic Kwiatkowski, Cordelia Langford, John Sillitoe on behalf of the Wellcome Sanger Institute COVID-19 Surveillance Team |
| EPI_ISL_2807373, EPI_ISL_2807374 | Lighthouse Lab in Alderley Park | Wellcome Sanger Institute for the COVID-19 Genomics UK (COG-UK) Consortium | Jacquelyn Wynn, Mairead Hyland, The Lighthouse Lab in Alderley Park and Alex Alderton, Roberto Amato, Jeffrey Barrett, Sonia Goncalves, Ewan Harrison, David K. Jackson, Ian Johnston, Dominic Kwiatkowski, Cordelia Langford, John Sillitoe on behalf of the Wellcome Sanger Institute COVID-19 Surveillance Team |
| EPI_ISL_2807375 | Lighthouse Lab in Glasgow | Wellcome Sanger Institute for the COVID-19 Genomics UK (COG-UK) Consortium | Harper VanSteenhouse, Yumi Kasai, David Gray, Carol Clugston, Anna Dominiczak and Alex Alderton, Roberto Amato, Jeffrey Barrett, Sonia Goncalves, Ewan Harrison, David K. Jackson, Ian Johnston, Dominic Kwiatkowski, Cordelia Langford, John Sillitoe on behalf of the Wellcome Sanger Institute COVID-19 Surveillance Team |
| EPI_ISL_2807376, EPI_ISL_2807377, EPI_ISL_2807378 | Lighthouse Lab in Milton Keynes | Wellcome Sanger Institute for the COVID-19 Genomics UK (COG-UK) Consortium | The Lighthouse Lab in Milton Keynes and Alex Alderton, Roberto Amato, Jeffrey Barrett, Sonia Goncalves, Ewan Harrison, David K. Jackson, Ian Johnston, Dominic Kwiatkowski, Cordelia Langford, John Sillitoe on behalf of the Wellcome Sanger Institute COVID-19 Surveillance Team |
| EPI_ISL_2807379, EPI_ISL_2807380, EPI_ISL_2807381 | Lighthouse Lab in Glasgow | Wellcome Sanger Institute for the COVID-19 Genomics UK (COG-UK) Consortium | Harper VanSteenhouse, Yumi Kasai, David Gray, Carol Clugston, Anna Dominiczak and Alex Alderton, Roberto Amato, Jeffrey Barrett, Sonia Goncalves, Ewan Harrison, David K. Jackson, Ian Johnston, Dominic Kwiatkowski, Cordelia Langford, John Sillitoe on behalf of the Wellcome Sanger Institute COVID-19 Surveillance Team |
| EPI_ISL_2807382, EPI_ISL_2807383, EPI_ISL_2807384 | Lighthouse Lab in Milton Keynes | Wellcome Sanger Institute for the COVID-19 Genomics UK (COG-UK) Consortium | The Lighthouse Lab in Milton Keynes and Alex Alderton, Roberto Amato, Jeffrey Barrett, Sonia Goncalves, Ewan Harrison, David K. Jackson, Ian Johnston, Dominic Kwiatkowski, Cordelia Langford, John Sillitoe on behalf of the Wellcome Sanger Institute COVID-19 Surveillance Team |
| EPI_ISL_2807385 | Lighthouse Lab in Glasgow | Wellcome Sanger Institute for the COVID-19 Genomics UK (COG-UK) Consortium | Harper VanSteenhouse, Yumi Kasai, David Gray, Carol Clugston, Anna Dominiczak and Alex Alderton, Roberto Amato, Jeffrey Barrett, Sonia Goncalves, Ewan Harrison, David K. Jackson, Ian Johnston, Dominic Kwiatkowski, Cordelia Langford, John Sillitoe on behalf of the Wellcome Sanger Institute COVID-19 Surveillance Team |

| | | | |
|---|---|---|---|
| EPI_ISL_2807386 | Lighthouse Lab in Alderley Park | Wellcome Sanger Institute for the COVID-19 Genomics UK (COG-UK) Consortium | Jacquelyn Wynn, Mairead Hyland, The Lighthouse Lab in Alderley Park and Alex Alderton, Roberto Amato, Jeffrey Barrett, Sonia Goncalves, Ewan Harrison, David K. Jackson, Ian Johnston, Dominic Kwiatkowski, Cordelia Langford, John Sillitoe on behalf of the Wellcome Sanger Institute COVID-19 Surveillance Team |
| EPI_ISL_2807387, EPI_ISL_2807388 | Lighthouse Lab in Milton Keynes | Wellcome Sanger Institute for the COVID-19 Genomics UK (COG-UK) Consortium | The Lighthouse Lab in Milton Keynes and Alex Alderton, Roberto Amato, Jeffrey Barrett, Sonia Goncalves, Ewan Harrison, David K. Jackson, Ian Johnston, Dominic Kwiatkowski, Cordelia Langford, John Sillitoe on behalf of the Wellcome Sanger Institute COVID-19 Surveillance Team |
| EPI_ISL_2807389 | Lighthouse Lab in Glasgow | Wellcome Sanger Institute for the COVID-19 Genomics UK (COG-UK) Consortium | Harper VanSteenhouse, Yumi Kasai, David Gray, Carol Clugston, Anna Dominiczak and Alex Alderton, Roberto Amato, Jeffrey Barrett, Sonia Goncalves, Ewan Harrison, David K. Jackson, Ian Johnston, Dominic Kwiatkowski, Cordelia Langford, John Sillitoe on behalf of the Wellcome Sanger Institute COVID-19 Surveillance Team |
| EPI_ISL_2807390 | Lighthouse Lab in Alderley Park | Wellcome Sanger Institute for the COVID-19 Genomics UK (COG-UK) Consortium | Jacquelyn Wynn, Mairead Hyland, The Lighthouse Lab in Alderley Park and Alex Alderton, Roberto Amato, Jeffrey Barrett, Sonia Goncalves, Ewan Harrison, David K. Jackson, Ian Johnston, Dominic Kwiatkowski, Cordelia Langford, John Sillitoe on behalf of the Wellcome Sanger Institute COVID-19 Surveillance Team |
| EPI_ISL_2807391 | Lighthouse Lab in Milton Keynes | Wellcome Sanger Institute for the COVID-19 Genomics UK (COG-UK) Consortium | The Lighthouse Lab in Milton Keynes and Alex Alderton, Roberto Amato, Jeffrey Barrett, Sonia Goncalves, Ewan Harrison, David K. Jackson, Ian Johnston, Dominic Kwiatkowski, Cordelia Langford, John Sillitoe on behalf of the Wellcome Sanger Institute COVID-19 Surveillance Team |
| EPI_ISL_2807392 | Lighthouse Lab in Glasgow | Wellcome Sanger Institute for the COVID-19 Genomics UK (COG-UK) Consortium | Harper VanSteenhouse, Yumi Kasai, David Gray, Carol Clugston, Anna Dominiczak and Alex Alderton, Roberto Amato, Jeffrey Barrett, Sonia Goncalves, Ewan Harrison, David K. Jackson, Ian Johnston, Dominic Kwiatkowski, Cordelia Langford, John Sillitoe on behalf of the Wellcome Sanger Institute COVID-19 Surveillance Team |
| EPI_ISL_2807394 | Lighthouse Lab in Alderley Park | Wellcome Sanger Institute for the COVID-19 Genomics UK (COG-UK) Consortium | Jacquelyn Wynn, Mairead Hyland, The Lighthouse Lab in Alderley Park and Alex Alderton, Roberto Amato, Jeffrey Barrett, Sonia Goncalves, Ewan Harrison, David K. Jackson, Ian Johnston, Dominic Kwiatkowski, Cordelia Langford, John Sillitoe on behalf of the Wellcome Sanger Institute COVID-19 Surveillance Team |
| EPI_ISL_2807395 | Lighthouse Lab in Milton Keynes | Wellcome Sanger Institute for the COVID-19 Genomics UK (COG-UK) Consortium | The Lighthouse Lab in Milton Keynes and Alex Alderton, Roberto Amato, Jeffrey Barrett, Sonia Goncalves, Ewan Harrison, David K. Jackson, Ian Johnston, Dominic Kwiatkowski, Cordelia Langford, John Sillitoe on behalf of the Wellcome Sanger Institute COVID-19 Surveillance Team |
| EPI_ISL_2807396, EPI_ISL_2807397, EPI_ISL_2807398 | Lighthouse Lab in Alderley Park | Wellcome Sanger Institute for the COVID-19 Genomics UK (COG-UK) Consortium | Jacquelyn Wynn, Mairead Hyland, The Lighthouse Lab in Alderley Park and Alex Alderton, Roberto Amato, Jeffrey Barrett, Sonia Goncalves, Ewan Harrison, David K. Jackson, Ian Johnston, Dominic Kwiatkowski, Cordelia Langford, John Sillitoe on behalf of the Wellcome Sanger Institute COVID-19 Surveillance Team |
| EPI_ISL_2807399 | Lighthouse Lab in Glasgow | Wellcome Sanger Institute for the COVID-19 Genomics UK (COG-UK) Consortium | Harper VanSteenhouse, Yumi Kasai, David Gray, Carol Clugston, Anna Dominiczak and Alex Alderton, Roberto Amato, Jeffrey Barrett, Sonia Goncalves, Ewan Harrison, David K. Jackson, Ian Johnston, Dominic Kwiatkowski, Cordelia Langford, John Sillitoe on behalf of the Wellcome Sanger Institute COVID-19 Surveillance Team |
| EPI_ISL_2807400, EPI_ISL_2807401, EPI_ISL_2807402 | Lighthouse Lab in Milton Keynes | Wellcome Sanger Institute for the COVID-19 Genomics UK (COG-UK) Consortium | The Lighthouse Lab in Milton Keynes and Alex Alderton, Roberto Amato, Jeffrey Barrett, Sonia Goncalves, Ewan Harrison, David K. Jackson, Ian Johnston, Dominic Kwiatkowski, Cordelia Langford, John Sillitoe on behalf of the Wellcome Sanger Institute COVID-19 Surveillance Team |
| EPI_ISL_2807403 | Lighthouse Lab in Glasgow | Wellcome Sanger Institute for the COVID-19 Genomics UK (COG-UK) Consortium | Harper VanSteenhouse, Yumi Kasai, David Gray, Carol Clugston, Anna Dominiczak and Alex Alderton, Roberto Amato, Jeffrey Barrett, Sonia Goncalves, Ewan Harrison, David K. Jackson, Ian Johnston, Dominic Kwiatkowski, Cordelia Langford, John Sillitoe on behalf of the Wellcome Sanger Institute COVID-19 Surveillance Team |
| EPI_ISL_2807404, EPI_ISL_2807405 | Lighthouse Lab in Milton Keynes | Wellcome Sanger Institute for the COVID-19 Genomics UK (COG-UK) Consortium | The Lighthouse Lab in Milton Keynes and Alex Alderton, Roberto Amato, Jeffrey Barrett, Sonia Goncalves, Ewan Harrison, David K. Jackson, Ian Johnston, Dominic Kwiatkowski, Cordelia Langford, John Sillitoe on behalf of the Wellcome Sanger Institute COVID-19 Surveillance Team |
| EPI_ISL_2807406 | Lighthouse Lab in Alderley Park | Wellcome Sanger Institute for the COVID-19 Genomics UK (COG-UK) Consortium | Jacquelyn Wynn, Mairead Hyland, The Lighthouse Lab in Alderley Park and Alex Alderton, Roberto Amato, Jeffrey Barrett, Sonia Goncalves, Ewan Harrison, David K. Jackson, Ian Johnston, Dominic Kwiatkowski, Cordelia Langford, John Sillitoe on behalf of the Wellcome Sanger Institute COVID-19 Surveillance Team |
| EPI_ISL_2807407, EPI_ISL_2807408, EPI_ISL_2807409, EPI_ISL_2807410 | Lighthouse Lab in Milton Keynes | Wellcome Sanger Institute for the COVID-19 Genomics UK (COG-UK) Consortium | The Lighthouse Lab in Milton Keynes and Alex Alderton, Roberto Amato, Jeffrey Barrett, Sonia Goncalves, Ewan Harrison, David K. Jackson, Ian Johnston, Dominic Kwiatkowski, Cordelia Langford, John Sillitoe on behalf of the Wellcome Sanger Institute COVID-19 Surveillance Team |
| EPI_ISL_2807411, EPI_ISL_2807412, EPI_ISL_2807413 | Lighthouse Lab in Glasgow | Wellcome Sanger Institute for the COVID-19 Genomics UK (COG-UK) Consortium | Harper VanSteenhouse, Yumi Kasai, David Gray, Carol Clugston, Anna Dominiczak and Alex Alderton, Roberto Amato, Jeffrey Barrett, Sonia Goncalves, Ewan Harrison, David K. Jackson, Ian Johnston, Dominic Kwiatkowski, Cordelia Langford, John Sillitoe on behalf of the Wellcome Sanger Institute COVID-19 Surveillance Team |
| EPI_ISL_2807414, EPI_ISL_2807415 | Lighthouse Lab in Alderley Park | Wellcome Sanger Institute for the COVID-19 Genomics UK (COG-UK) Consortium | Jacquelyn Wynn, Mairead Hyland, The Lighthouse Lab in Alderley Park and Alex Alderton, Roberto Amato, Jeffrey Barrett, Sonia Goncalves, Ewan Harrison, David K. Jackson, Ian Johnston, Dominic Kwiatkowski, Cordelia Langford, John Sillitoe on behalf of the Wellcome Sanger Institute COVID-19 Surveillance Team |
| EPI_ISL_2807416 | Lighthouse Lab in Milton Keynes | Wellcome Sanger Institute for the COVID-19 Genomics UK (COG-UK) Consortium | The Lighthouse Lab in Milton Keynes and Alex Alderton, Roberto Amato, Jeffrey Barrett, Sonia Goncalves, Ewan Harrison, David K. Jackson, Ian Johnston, Dominic Kwiatkowski, Cordelia Langford, John Sillitoe on behalf of the Wellcome Sanger Institute COVID-19 Surveillance Team |
| EPI_ISL_2807417, EPI_ISL_2807418, EPI_ISL_2807419 | Lighthouse Lab in Alderley Park | Wellcome Sanger Institute for the COVID-19 Genomics UK (COG-UK) Consortium | Jacquelyn Wynn, Mairead Hyland, The Lighthouse Lab in Alderley Park and Alex Alderton, Roberto Amato, Jeffrey Barrett, Sonia Goncalves, Ewan Harrison, David K. Jackson, Ian Johnston, Dominic Kwiatkowski, Cordelia Langford, John Sillitoe on behalf of the Wellcome Sanger Institute COVID-19 Surveillance Team |
| EPI_ISL_2807420, EPI_ISL_2807421 | Lighthouse Lab in Milton Keynes | Wellcome Sanger Institute for the COVID-19 Genomics UK (COG-UK) Consortium | The Lighthouse Lab in Milton Keynes and Alex Alderton, Roberto Amato, Jeffrey Barrett, Sonia Goncalves, Ewan Harrison, David K. Jackson, Ian Johnston, Dominic Kwiatkowski, Cordelia Langford, John Sillitoe on behalf of the Wellcome Sanger Institute COVID-19 Surveillance Team |
| EPI_ISL_2807422 | Lighthouse Lab in Alderley Park | Wellcome Sanger Institute for the COVID-19 Genomics UK (COG-UK) Consortium | Jacquelyn Wynn, Mairead Hyland, The Lighthouse Lab in Alderley Park and Alex Alderton, Roberto Amato, Jeffrey Barrett, Sonia Goncalves, Ewan Harrison, David K. Jackson, Ian Johnston, Dominic Kwiatkowski, Cordelia Langford, John Sillitoe on behalf of the Wellcome Sanger Institute COVID-19 Surveillance Team |
| EPI_ISL_2807423, EPI_ISL_2807424, EPI_ISL_2807426 | Lighthouse Lab in Milton Keynes | Wellcome Sanger Institute for the COVID-19 Genomics UK (COG-UK) Consortium | The Lighthouse Lab in Milton Keynes and Alex Alderton, Roberto Amato, Jeffrey Barrett, Sonia Goncalves, Ewan Harrison, David K. Jackson, Ian Johnston, Dominic Kwiatkowski, Cordelia Langford, John Sillitoe on behalf of the Wellcome Sanger Institute COVID-19 Surveillance Team |
| EPI_ISL_2807427 | Lighthouse Lab in Alderley Park | Wellcome Sanger Institute for the COVID-19 Genomics UK (COG-UK) Consortium | Jacquelyn Wynn, Mairead Hyland, The Lighthouse Lab in Alderley Park and Alex Alderton, Roberto Amato, Jeffrey Barrett, Sonia Goncalves, Ewan Harrison, David K. Jackson, Ian Johnston, Dominic Kwiatkowski, Cordelia Langford, John Sillitoe on behalf of the Wellcome Sanger Institute COVID-19 Surveillance Team |
| EPI_ISL_2807428 | Lighthouse Lab in Milton Keynes | Wellcome Sanger Institute for the COVID-19 Genomics UK (COG-UK) Consortium | The Lighthouse Lab in Milton Keynes and Alex Alderton, Roberto Amato, Jeffrey Barrett, Sonia Goncalves, Ewan Harrison, David K. Jackson, Ian Johnston, Dominic Kwiatkowski, Cordelia Langford, John Sillitoe on behalf of the Wellcome Sanger Institute COVID-19 Surveillance Team |
| EPI_ISL_2807429 | Lighthouse Lab in Glasgow | Wellcome Sanger Institute for the COVID-19 Genomics UK (COG-UK) Consortium | Harper VanSteenhouse, Yumi Kasai, David Gray, Carol Clugston, Anna Dominiczak and Alex Alderton, Roberto Amato, Jeffrey Barrett, Sonia Goncalves, Ewan Harrison, David K. Jackson, Ian Johnston, Dominic Kwiatkowski, Cordelia Langford, John Sillitoe on behalf of the Wellcome Sanger Institute COVID-19 Surveillance Team |
| EPI_ISL_2807430, EPI_ISL_2807431, EPI_ISL_2807432 | Lighthouse Lab in Milton Keynes | Wellcome Sanger Institute for the COVID-19 Genomics UK (COG-UK) Consortium | The Lighthouse Lab in Milton Keynes and Alex Alderton, Roberto Amato, Jeffrey Barrett, Sonia Goncalves, Ewan Harrison, David K. Jackson, Ian Johnston, Dominic Kwiatkowski, Cordelia Langford, John Sillitoe on behalf of the Wellcome Sanger Institute COVID-19 Surveillance Team |
| EPI_ISL_2807433, EPI_ISL_2807434 | Lighthouse Lab in Glasgow | Wellcome Sanger Institute for the COVID-19 Genomics UK (COG-UK) Consortium | Harper VanSteenhouse, Yumi Kasai, David Gray, Carol Clugston, Anna Dominiczak and Alex Alderton, Roberto Amato, Jeffrey Barrett, Sonia Goncalves, Ewan Harrison, David K. Jackson, Ian Johnston, Dominic Kwiatkowski, Cordelia Langford, John Sillitoe on behalf of the Wellcome Sanger Institute COVID-19 Surveillance Team |
| EPI_ISL_2807435, EPI_ISL_2807436 | Lighthouse Lab in Alderley Park | Wellcome Sanger Institute for the COVID-19 Genomics UK (COG-UK) Consortium | Jacquelyn Wynn, Mairead Hyland, The Lighthouse Lab in Alderley Park and Alex Alderton, Roberto Amato, Jeffrey Barrett, Sonia Goncalves, Ewan Harrison, David K. Jackson, Ian Johnston, Dominic Kwiatkowski, Cordelia Langford, John Sillitoe on behalf of the Wellcome Sanger Institute COVID-19 Surveillance Team |
| EPI_ISL_2807437 | Lighthouse Lab in Glasgow | Wellcome Sanger Institute for the COVID-19 Genomics UK (COG-UK) Consortium | Harper VanSteenhouse, Yumi Kasai, David Gray, Carol Clugston, Anna Dominiczak and Alex Alderton, Roberto Amato, Jeffrey Barrett, Sonia Goncalves, Ewan Harrison, David K. Jackson, Ian Johnston, Dominic Kwiatkowski, Cordelia Langford, John Sillitoe on behalf of the Wellcome Sanger Institute COVID-19 Surveillance Team |
| EPI_ISL_2807438, EPI_ISL_2807439, EPI_ISL_2807440, EPI_ISL_2807441 | Lighthouse Lab in Alderley Park | Wellcome Sanger Institute for the COVID-19 Genomics UK (COG-UK) Consortium | Jacquelyn Wynn, Mairead Hyland, The Lighthouse Lab in Alderley Park and Alex Alderton, Roberto Amato, Jeffrey Barrett, Sonia Goncalves, Ewan Harrison, David K. Jackson, Ian Johnston, Dominic Kwiatkowski, Cordelia Langford, John Sillitoe on behalf of the Wellcome Sanger Institute COVID-19 Surveillance Team |
| EPI_ISL_2807442 | Lighthouse Lab in Milton Keynes | Wellcome Sanger Institute for the COVID-19 Genomics UK (COG-UK) Consortium | The Lighthouse Lab in Milton Keynes and Alex Alderton, Roberto Amato, Jeffrey Barrett, Sonia Goncalves, Ewan Harrison, David K. Jackson, Ian Johnston, Dominic Kwiatkowski, Cordelia Langford, John Sillitoe on behalf of the Wellcome Sanger Institute COVID-19 Surveillance Team |
| EPI_ISL_2807443, EPI_ISL_2807444 | Lighthouse Lab in Glasgow | Wellcome Sanger Institute for the COVID-19 Genomics UK (COG-UK) Consortium | Harper VanSteenhouse, Yumi Kasai, David Gray, Carol Clugston, Anna Dominiczak and Alex Alderton, Roberto Amato, Jeffrey Barrett, Sonia Goncalves, Ewan Harrison, David K. Jackson, Ian Johnston, Dominic Kwiatkowski, Cordelia Langford, John Sillitoe on behalf of the Wellcome Sanger Institute COVID-19 Surveillance Team |
| EPI_ISL_2807445, EPI_ISL_2807446 | Lighthouse Lab in Milton Keynes | Wellcome Sanger Institute for the COVID-19 Genomics UK (COG-UK) Consortium | The Lighthouse Lab in Milton Keynes and Alex Alderton, Roberto Amato, Jeffrey Barrett, Sonia Goncalves, Ewan Harrison, David K. Jackson, Ian Johnston, Dominic Kwiatkowski, Cordelia Langford, John Sillitoe on behalf of the Wellcome Sanger Institute COVID-19 Surveillance Team |
| EPI_ISL_2807447 | Lighthouse Lab in Alderley Park | Wellcome Sanger Institute for the COVID-19 Genomics UK (COG-UK) Consortium | Jacquelyn Wynn, Mairead Hyland, The Lighthouse Lab in Alderley Park and Alex Alderton, Roberto Amato, Jeffrey Barrett, Sonia Goncalves, Ewan Harrison, David K. Jackson, Ian Johnston, Dominic Kwiatkowski, Cordelia Langford, John Sillitoe on behalf of the Wellcome Sanger Institute COVID-19 Surveillance Team |
| EPI_ISL_2807448 | Lighthouse Lab in Milton Keynes | Wellcome Sanger Institute for the COVID-19 Genomics UK (COG-UK) Consortium | The Lighthouse Lab in Milton Keynes and Alex Alderton, Roberto Amato, Jeffrey Barrett, Sonia Goncalves, Ewan Harrison, David K. Jackson, Ian Johnston, Dominic Kwiatkowski, Cordelia Langford, John Sillitoe on behalf of the Wellcome Sanger Institute COVID-19 Surveillance Team |
| EPI_ISL_2807449, EPI_ISL_2807450, EPI_ISL_2807451, EPI_ISL_2807452, EPI_ISL_2807453 | Lighthouse Lab in Alderley Park | Wellcome Sanger Institute for the COVID-19 Genomics UK (COG-UK) Consortium | Jacquelyn Wynn, Mairead Hyland, The Lighthouse Lab in Alderley Park and Alex Alderton, Roberto Amato, Jeffrey Barrett, Sonia Goncalves, Ewan Harrison, David K. Jackson, Ian Johnston, Dominic Kwiatkowski, Cordelia Langford, John Sillitoe on behalf of the Wellcome Sanger Institute COVID-19 Surveillance Team |
| EPI_ISL_2807454, EPI_ISL_2807455 | Lighthouse Lab in Glasgow | Wellcome Sanger Institute for the COVID-19 Genomics UK (COG-UK) Consortium | Harper VanSteenhouse, Yumi Kasai, David Gray, Carol Clugston, Anna Dominiczak and Alex Alderton, Roberto Amato, Jeffrey Barrett, Sonia Goncalves, Ewan Harrison, David K. Jackson, Ian Johnston, Dominic Kwiatkowski, Cordelia Langford, John Sillitoe on behalf of the Wellcome Sanger Institute COVID-19 Surveillance Team |
| EPI_ISL_2807456 | Lighthouse Lab in Milton Keynes | Wellcome Sanger Institute for the COVID-19 Genomics UK (COG-UK) Consortium | The Lighthouse Lab in Milton Keynes and Alex Alderton, Roberto Amato, Jeffrey Barrett, Sonia Goncalves, Ewan Harrison, David K. Jackson, Ian Johnston, Dominic Kwiatkowski, Cordelia Langford, John Sillitoe on behalf of the Wellcome Sanger Institute COVID-19 Surveillance Team |
| EPI_ISL_2807457 | Lighthouse Lab in Glasgow | Wellcome Sanger Institute for the COVID-19 Genomics UK (COG-UK) Consortium | Harper VanSteenhouse, Yumi Kasai, David Gray, Carol Clugston, Anna Dominiczak and Alex Alderton, Roberto Amato, Jeffrey Barrett, Sonia Goncalves, Ewan Harrison, David K. Jackson, Ian Johnston, Dominic Kwiatkowski, Cordelia Langford, John Sillitoe on behalf of the Wellcome Sanger Institute COVID-19 Surveillance Team |
| EPI_ISL_2807458, | Lighthouse Lab in Milton Keynes | Wellcome Sanger Institute for the COVID-19 Genomics UK (COG-UK) | The Lighthouse Lab in Milton Keynes and Alex Alderton, Roberto Amato, Jeffrey Barrett, Sonia Goncalves, Ewan Harrison, David K. Jackson, Ian Johnston, Dominic Kwiatkowski, Cordelia Langford, John Sillitoe on behalf of the Wellcome Sanger |

| | | | |
|---|---|---|---|
| EPI_ISL_2807459, EPI_ISL_2807460, EPI_ISL_2807461 | | Consortium | Institute COVID-19 Surveillance Team |
| EPI_ISL_2807462 | Lighthouse Lab in Alderley Park | Wellcome Sanger Institute for the COVID-19 Genomics UK (COG-UK) Consortium | Jacquelyn Wynn, Mairead Hyland, The Lighthouse Lab in Alderley Park and Alex Alderton, Roberto Amato, Jeffrey Barrett, Sonia Goncalves, Ewan Harrison, David K. Jackson, Ian Johnston, Dominic Kwiatkowski, Cordelia Langford, John Sillitoe on behalf of the Wellcome Sanger Institute COVID-19 Surveillance Team |
| EPI_ISL_2807463 | Lighthouse Lab in Milton Keynes | Wellcome Sanger Institute for the COVID-19 Genomics UK (COG-UK) Consortium | The Lighthouse Lab in Milton Keynes and Alex Alderton, Roberto Amato, Jeffrey Barrett, Sonia Goncalves, Ewan Harrison, David K. Jackson, Ian Johnston, Dominic Kwiatkowski, Cordelia Langford, John Sillitoe on behalf of the Wellcome Sanger Institute COVID-19 Surveillance Team |
| EPI_ISL_2807464, EPI_ISL_2807465 | Lighthouse Lab in Glasgow | Wellcome Sanger Institute for the COVID-19 Genomics UK (COG-UK) Consortium | Harper VanSteenhouse, Yumi Kasai, David Gray, Carol Clugston, Anna Dominiczak and Alex Alderton, Roberto Amato, Jeffrey Barrett, Sonia Goncalves, Ewan Harrison, David K. Jackson, Ian Johnston, Dominic Kwiatkowski, Cordelia Langford, John Sillitoe on behalf of the Wellcome Sanger Institute COVID-19 Surveillance Team |
| EPI_ISL_2807467 | Lighthouse Lab in Alderley Park | Wellcome Sanger Institute for the COVID-19 Genomics UK (COG-UK) Consortium | Jacquelyn Wynn, Mairead Hyland, The Lighthouse Lab in Alderley Park and Alex Alderton, Roberto Amato, Jeffrey Barrett, Sonia Goncalves, Ewan Harrison, David K. Jackson, Ian Johnston, Dominic Kwiatkowski, Cordelia Langford, John Sillitoe on behalf of the Wellcome Sanger Institute COVID-19 Surveillance Team |
| EPI_ISL_2807468, EPI_ISL_2807469, EPI_ISL_2807470 | Lighthouse Lab in Glasgow | Wellcome Sanger Institute for the COVID-19 Genomics UK (COG-UK) Consortium | Harper VanSteenhouse, Yumi Kasai, David Gray, Carol Clugston, Anna Dominiczak and Alex Alderton, Roberto Amato, Jeffrey Barrett, Sonia Goncalves, Ewan Harrison, David K. Jackson, Ian Johnston, Dominic Kwiatkowski, Cordelia Langford, John Sillitoe on behalf of the Wellcome Sanger Institute COVID-19 Surveillance Team |
| EPI_ISL_2807471, EPI_ISL_2807472 | Lighthouse Lab in Alderley Park | Wellcome Sanger Institute for the COVID-19 Genomics UK (COG-UK) Consortium | Jacquelyn Wynn, Mairead Hyland, The Lighthouse Lab in Alderley Park and Alex Alderton, Roberto Amato, Jeffrey Barrett, Sonia Goncalves, Ewan Harrison, David K. Jackson, Ian Johnston, Dominic Kwiatkowski, Cordelia Langford, John Sillitoe on behalf of the Wellcome Sanger Institute COVID-19 Surveillance Team |
| EPI_ISL_2807473 | Lighthouse Lab in Milton Keynes | Wellcome Sanger Institute for the COVID-19 Genomics UK (COG-UK) Consortium | The Lighthouse Lab in Milton Keynes and Alex Alderton, Roberto Amato, Jeffrey Barrett, Sonia Goncalves, Ewan Harrison, David K. Jackson, Ian Johnston, Dominic Kwiatkowski, Cordelia Langford, John Sillitoe on behalf of the Wellcome Sanger Institute COVID-19 Surveillance Team |
| EPI_ISL_2807474 | Lighthouse Lab in Glasgow | Wellcome Sanger Institute for the COVID-19 Genomics UK (COG-UK) Consortium | Harper VanSteenhouse, Yumi Kasai, David Gray, Carol Clugston, Anna Dominiczak and Alex Alderton, Roberto Amato, Jeffrey Barrett, Sonia Goncalves, Ewan Harrison, David K. Jackson, Ian Johnston, Dominic Kwiatkowski, Cordelia Langford, John Sillitoe on behalf of the Wellcome Sanger Institute COVID-19 Surveillance Team |
| EPI_ISL_2807476, EPI_ISL_2807477, EPI_ISL_2807478 | Lighthouse Lab in Milton Keynes | Wellcome Sanger Institute for the COVID-19 Genomics UK (COG-UK) Consortium | The Lighthouse Lab in Milton Keynes and Alex Alderton, Roberto Amato, Jeffrey Barrett, Sonia Goncalves, Ewan Harrison, David K. Jackson, Ian Johnston, Dominic Kwiatkowski, Cordelia Langford, John Sillitoe on behalf of the Wellcome Sanger Institute COVID-19 Surveillance Team |
| EPI_ISL_2807479 | Lighthouse Lab in Alderley Park | Wellcome Sanger Institute for the COVID-19 Genomics UK (COG-UK) Consortium | Jacquelyn Wynn, Mairead Hyland, The Lighthouse Lab in Alderley Park and Alex Alderton, Roberto Amato, Jeffrey Barrett, Sonia Goncalves, Ewan Harrison, David K. Jackson, Ian Johnston, Dominic Kwiatkowski, Cordelia Langford, John Sillitoe on behalf of the Wellcome Sanger Institute COVID-19 Surveillance Team |
| EPI_ISL_2807480 | Lighthouse Lab in Milton Keynes | Wellcome Sanger Institute for the COVID-19 Genomics UK (COG-UK) Consortium | The Lighthouse Lab in Milton Keynes and Alex Alderton, Roberto Amato, Jeffrey Barrett, Sonia Goncalves, Ewan Harrison, David K. Jackson, Ian Johnston, Dominic Kwiatkowski, Cordelia Langford, John Sillitoe on behalf of the Wellcome Sanger Institute COVID-19 Surveillance Team |
| EPI_ISL_2807481 | Lighthouse Lab in Glasgow | Wellcome Sanger Institute for the COVID-19 Genomics UK (COG-UK) Consortium | Harper VanSteenhouse, Yumi Kasai, David Gray, Carol Clugston, Anna Dominiczak and Alex Alderton, Roberto Amato, Jeffrey Barrett, Sonia Goncalves, Ewan Harrison, David K. Jackson, Ian Johnston, Dominic Kwiatkowski, Cordelia Langford, John Sillitoe on behalf of the Wellcome Sanger Institute COVID-19 Surveillance Team |
| EPI_ISL_2807482, EPI_ISL_2807483, EPI_ISL_2807484 | Lighthouse Lab in Milton Keynes | Wellcome Sanger Institute for the COVID-19 Genomics UK (COG-UK) Consortium | The Lighthouse Lab in Milton Keynes and Alex Alderton, Roberto Amato, Jeffrey Barrett, Sonia Goncalves, Ewan Harrison, David K. Jackson, Ian Johnston, Dominic Kwiatkowski, Cordelia Langford, John Sillitoe on behalf of the Wellcome Sanger Institute COVID-19 Surveillance Team |
| EPI_ISL_2807485, EPI_ISL_2807486 | Lighthouse Lab in Alderley Park | Wellcome Sanger Institute for the COVID-19 Genomics UK (COG-UK) Consortium | Jacquelyn Wynn, Mairead Hyland, The Lighthouse Lab in Alderley Park and Alex Alderton, Roberto Amato, Jeffrey Barrett, Sonia Goncalves, Ewan Harrison, David K. Jackson, Ian Johnston, Dominic Kwiatkowski, Cordelia Langford, John Sillitoe on behalf of the Wellcome Sanger Institute COVID-19 Surveillance Team |
| EPI_ISL_2807487 | Lighthouse Lab in Milton Keynes | Wellcome Sanger Institute for the COVID-19 Genomics UK (COG-UK) Consortium | The Lighthouse Lab in Milton Keynes and Alex Alderton, Roberto Amato, Jeffrey Barrett, Sonia Goncalves, Ewan Harrison, David K. Jackson, Ian Johnston, Dominic Kwiatkowski, Cordelia Langford, John Sillitoe on behalf of the Wellcome Sanger Institute COVID-19 Surveillance Team |
| EPI_ISL_2807488 | Lighthouse Lab in Alderley Park | Wellcome Sanger Institute for the COVID-19 Genomics UK (COG-UK) Consortium | Jacquelyn Wynn, Mairead Hyland, The Lighthouse Lab in Alderley Park and Alex Alderton, Roberto Amato, Jeffrey Barrett, Sonia Goncalves, Ewan Harrison, David K. Jackson, Ian Johnston, Dominic Kwiatkowski, Cordelia Langford, John Sillitoe on behalf of the Wellcome Sanger Institute COVID-19 Surveillance Team |
| EPI_ISL_2807489 | Lighthouse Lab in Glasgow | Wellcome Sanger Institute for the COVID-19 Genomics UK (COG-UK) Consortium | Harper VanSteenhouse, Yumi Kasai, David Gray, Carol Clugston, Anna Dominiczak and Alex Alderton, Roberto Amato, Jeffrey Barrett, Sonia Goncalves, Ewan Harrison, David K. Jackson, Ian Johnston, Dominic Kwiatkowski, Cordelia Langford, John Sillitoe on behalf of the Wellcome Sanger Institute COVID-19 Surveillance Team |
| EPI_ISL_2807490, EPI_ISL_2807491, EPI_ISL_2807492 | Lighthouse Lab in Alderley Park | Wellcome Sanger Institute for the COVID-19 Genomics UK (COG-UK) Consortium | Jacquelyn Wynn, Mairead Hyland, The Lighthouse Lab in Alderley Park and Alex Alderton, Roberto Amato, Jeffrey Barrett, Sonia Goncalves, Ewan Harrison, David K. Jackson, Ian Johnston, Dominic Kwiatkowski, Cordelia Langford, John Sillitoe on behalf of the Wellcome Sanger Institute COVID-19 Surveillance Team |
| EPI_ISL_2807493 | Lighthouse Lab in Glasgow | Wellcome Sanger Institute for the COVID-19 Genomics UK (COG-UK) Consortium | Harper VanSteenhouse, Yumi Kasai, David Gray, Carol Clugston, Anna Dominiczak and Alex Alderton, Roberto Amato, Jeffrey Barrett, Sonia Goncalves, Ewan Harrison, David K. Jackson, Ian Johnston, Dominic Kwiatkowski, Cordelia Langford, John Sillitoe on behalf of the Wellcome Sanger Institute COVID-19 Surveillance Team |
| EPI_ISL_2807494, EPI_ISL_2807495, EPI_ISL_2807496 | Lighthouse Lab in Milton Keynes | Wellcome Sanger Institute for the COVID-19 Genomics UK (COG-UK) Consortium | The Lighthouse Lab in Milton Keynes and Alex Alderton, Roberto Amato, Jeffrey Barrett, Sonia Goncalves, Ewan Harrison, David K. Jackson, Ian Johnston, Dominic Kwiatkowski, Cordelia Langford, John Sillitoe on behalf of the Wellcome Sanger Institute COVID-19 Surveillance Team |
| EPI_ISL_2807497, EPI_ISL_2807498 | Lighthouse Lab in Alderley Park | Wellcome Sanger Institute for the COVID-19 Genomics UK (COG-UK) Consortium | Jacquelyn Wynn, Mairead Hyland, The Lighthouse Lab in Alderley Park and Alex Alderton, Roberto Amato, Jeffrey Barrett, Sonia Goncalves, Ewan Harrison, David K. Jackson, Ian Johnston, Dominic Kwiatkowski, Cordelia Langford, John Sillitoe on behalf of the Wellcome Sanger Institute COVID-19 Surveillance Team |
| EPI_ISL_2807499 | Lighthouse Lab in Milton Keynes | Wellcome Sanger Institute for the COVID-19 Genomics UK (COG-UK) Consortium | The Lighthouse Lab in Milton Keynes and Alex Alderton, Roberto Amato, Jeffrey Barrett, Sonia Goncalves, Ewan Harrison, David K. Jackson, Ian Johnston, Dominic Kwiatkowski, Cordelia Langford, John Sillitoe on behalf of the Wellcome Sanger Institute COVID-19 Surveillance Team |
| EPI_ISL_2807500, EPI_ISL_2807501 | Lighthouse Lab in Glasgow | Wellcome Sanger Institute for the COVID-19 Genomics UK (COG-UK) Consortium | Harper VanSteenhouse, Yumi Kasai, David Gray, Carol Clugston, Anna Dominiczak and Alex Alderton, Roberto Amato, Jeffrey Barrett, Sonia Goncalves, Ewan Harrison, David K. Jackson, Ian Johnston, Dominic Kwiatkowski, Cordelia Langford, John Sillitoe on behalf of the Wellcome Sanger Institute COVID-19 Surveillance Team |
| EPI_ISL_2807502, EPI_ISL_2807503 | Lighthouse Lab in Milton Keynes | Wellcome Sanger Institute for the COVID-19 Genomics UK (COG-UK) Consortium | The Lighthouse Lab in Milton Keynes and Alex Alderton, Roberto Amato, Jeffrey Barrett, Sonia Goncalves, Ewan Harrison, David K. Jackson, Ian Johnston, Dominic Kwiatkowski, Cordelia Langford, John Sillitoe on behalf of the Wellcome Sanger Institute COVID-19 Surveillance Team |
| EPI_ISL_2807504 | Lighthouse Lab in Alderley Park | Wellcome Sanger Institute for the COVID-19 Genomics UK (COG-UK) Consortium | Jacquelyn Wynn, Mairead Hyland, The Lighthouse Lab in Alderley Park and Alex Alderton, Roberto Amato, Jeffrey Barrett, Sonia Goncalves, Ewan Harrison, David K. Jackson, Ian Johnston, Dominic Kwiatkowski, Cordelia Langford, John Sillitoe on behalf of the Wellcome Sanger Institute COVID-19 Surveillance Team |
| EPI_ISL_2807505 | Lighthouse Lab in Glasgow | Wellcome Sanger Institute for the COVID-19 Genomics UK (COG-UK) Consortium | Harper VanSteenhouse, Yumi Kasai, David Gray, Carol Clugston, Anna Dominiczak and Alex Alderton, Roberto Amato, Jeffrey Barrett, Sonia Goncalves, Ewan Harrison, David K. Jackson, Ian Johnston, Dominic Kwiatkowski, Cordelia Langford, John Sillitoe on behalf of the Wellcome Sanger Institute COVID-19 Surveillance Team |
| EPI_ISL_2807506, EPI_ISL_2807507 | Lighthouse Lab in Milton Keynes | Wellcome Sanger Institute for the COVID-19 Genomics UK (COG-UK) Consortium | The Lighthouse Lab in Milton Keynes and Alex Alderton, Roberto Amato, Jeffrey Barrett, Sonia Goncalves, Ewan Harrison, David K. Jackson, Ian Johnston, Dominic Kwiatkowski, Cordelia Langford, John Sillitoe on behalf of the Wellcome Sanger Institute COVID-19 Surveillance Team |
| EPI_ISL_2807508 | Lighthouse Lab in Alderley Park | Wellcome Sanger Institute for the COVID-19 Genomics UK (COG-UK) Consortium | Jacquelyn Wynn, Mairead Hyland, The Lighthouse Lab in Alderley Park and Alex Alderton, Roberto Amato, Jeffrey Barrett, Sonia Goncalves, Ewan Harrison, David K. Jackson, Ian Johnston, Dominic Kwiatkowski, Cordelia Langford, John Sillitoe on behalf of the Wellcome Sanger Institute COVID-19 Surveillance Team |
| EPI_ISL_2807509, EPI_ISL_2807510, EPI_ISL_2807511, EPI_ISL_2807512 | Lighthouse Lab in Milton Keynes | Wellcome Sanger Institute for the COVID-19 Genomics UK (COG-UK) Consortium | The Lighthouse Lab in Milton Keynes and Alex Alderton, Roberto Amato, Jeffrey Barrett, Sonia Goncalves, Ewan Harrison, David K. Jackson, Ian Johnston, Dominic Kwiatkowski, Cordelia Langford, John Sillitoe on behalf of the Wellcome Sanger Institute COVID-19 Surveillance Team |
| EPI_ISL_2807513 | Lighthouse Lab in Alderley Park | Wellcome Sanger Institute for the COVID-19 Genomics UK (COG-UK) Consortium | Jacquelyn Wynn, Mairead Hyland, The Lighthouse Lab in Alderley Park and Alex Alderton, Roberto Amato, Jeffrey Barrett, Sonia Goncalves, Ewan Harrison, David K. Jackson, Ian Johnston, Dominic Kwiatkowski, Cordelia Langford, John Sillitoe on behalf of the Wellcome Sanger Institute COVID-19 Surveillance Team |
| EPI_ISL_2807514, EPI_ISL_2807515, EPI_ISL_2807516, EPI_ISL_2807517, EPI_ISL_2807518, EPI_ISL_2807519 | Lighthouse Lab in Milton Keynes | Wellcome Sanger Institute for the COVID-19 Genomics UK (COG-UK) Consortium | The Lighthouse Lab in Milton Keynes and Alex Alderton, Roberto Amato, Jeffrey Barrett, Sonia Goncalves, Ewan Harrison, David K. Jackson, Ian Johnston, Dominic Kwiatkowski, Cordelia Langford, John Sillitoe on behalf of the Wellcome Sanger Institute COVID-19 Surveillance Team |
| EPI_ISL_2807520 | Lighthouse Lab in Glasgow | Wellcome Sanger Institute for the COVID-19 Genomics UK (COG-UK) Consortium | Harper VanSteenhouse, Yumi Kasai, David Gray, Carol Clugston, Anna Dominiczak and Alex Alderton, Roberto Amato, Jeffrey Barrett, Sonia Goncalves, Ewan Harrison, David K. Jackson, Ian Johnston, Dominic Kwiatkowski, Cordelia Langford, John Sillitoe on behalf of the Wellcome Sanger Institute COVID-19 Surveillance Team |
| EPI_ISL_2807521, EPI_ISL_2807522, EPI_ISL_2807523 | Lighthouse Lab in Milton Keynes | Wellcome Sanger Institute for the COVID-19 Genomics UK (COG-UK) Consortium | The Lighthouse Lab in Milton Keynes and Alex Alderton, Roberto Amato, Jeffrey Barrett, Sonia Goncalves, Ewan Harrison, David K. Jackson, Ian Johnston, Dominic Kwiatkowski, Cordelia Langford, John Sillitoe on behalf of the Wellcome Sanger Institute COVID-19 Surveillance Team |
| EPI_ISL_2807524 | Lighthouse Lab in Alderley Park | Wellcome Sanger Institute for the COVID-19 Genomics UK (COG-UK) Consortium | Jacquelyn Wynn, Mairead Hyland, The Lighthouse Lab in Alderley Park and Alex Alderton, Roberto Amato, Jeffrey Barrett, Sonia Goncalves, Ewan Harrison, David K. Jackson, Ian Johnston, Dominic Kwiatkowski, Cordelia Langford, John Sillitoe on behalf of the Wellcome Sanger Institute COVID-19 Surveillance Team |
| EPI_ISL_2807525, EPI_ISL_2807526 | Lighthouse Lab in Milton Keynes | Wellcome Sanger Institute for the COVID-19 Genomics UK (COG-UK) Consortium | The Lighthouse Lab in Milton Keynes and Alex Alderton, Roberto Amato, Jeffrey Barrett, Sonia Goncalves, Ewan Harrison, David K. Jackson, Ian Johnston, Dominic Kwiatkowski, Cordelia Langford, John Sillitoe on behalf of the Wellcome Sanger Institute COVID-19 Surveillance Team |
| EPI_ISL_2807527 | Lighthouse Lab in Glasgow | Wellcome Sanger Institute for the COVID-19 Genomics UK (COG-UK) Consortium | Harper VanSteenhouse, Yumi Kasai, David Gray, Carol Clugston, Anna Dominiczak and Alex Alderton, Roberto Amato, Jeffrey Barrett, Sonia Goncalves, Ewan Harrison, David K. Jackson, Ian Johnston, Dominic Kwiatkowski, Cordelia Langford, John Sillitoe on behalf of the Wellcome Sanger Institute COVID-19 Surveillance Team |
| EPI_ISL_2807528, EPI_ISL_2807529 | Lighthouse Lab in Milton Keynes | Wellcome Sanger Institute for the COVID-19 Genomics UK (COG-UK) Consortium | The Lighthouse Lab in Milton Keynes and Alex Alderton, Roberto Amato, Jeffrey Barrett, Sonia Goncalves, Ewan Harrison, David K. Jackson, Ian Johnston, Dominic Kwiatkowski, Cordelia Langford, John Sillitoe on behalf of the Wellcome Sanger Institute COVID-19 Surveillance Team |
| EPI_ISL_2807530, EPI_ISL_2807532, EPI_ISL_2807533 | Lighthouse Lab in Glasgow | Wellcome Sanger Institute for the COVID-19 Genomics UK (COG-UK) Consortium | Harper VanSteenhouse, Yumi Kasai, David Gray, Carol Clugston, Anna Dominiczak and Alex Alderton, Roberto Amato, Jeffrey Barrett, Sonia Goncalves, Ewan Harrison, David K. Jackson, Ian Johnston, Dominic Kwiatkowski, Cordelia Langford, John Sillitoe on behalf of the Wellcome Sanger Institute COVID-19 Surveillance Team |

| | | | |
|---|---|---|---|
| EPI_ISL_2807534, EPI_ISL_2807537 | Lighthouse Lab in Milton Keynes | Wellcome Sanger Institute for the COVID-19 Genomics UK (COG-UK) Consortium | The Lighthouse Lab in Milton Keynes and Alex Alderton, Roberto Amato, Jeffrey Barrett, Sonia Goncalves, Ewan Harrison, David K. Jackson, Ian Johnston, Dominic Kwiatkowski, Cordelia Langford, John Sillitoe on behalf of the Wellcome Sanger Institute COVID-19 Surveillance Team |
| EPI_ISL_2807538 | Lighthouse Lab in Glasgow | Wellcome Sanger Institute for the COVID-19 Genomics UK (COG-UK) Consortium | Harper VanSteenhouse, Yumi Kasai, David Gray, Carol Clugston, Anna Dominiczak and Alex Alderton, Roberto Amato, Jeffrey Barrett, Sonia Goncalves, Ewan Harrison, David K. Jackson, Ian Johnston, Dominic Kwiatkowski, Cordelia Langford, John Sillitoe on behalf of the Wellcome Sanger Institute COVID-19 Surveillance Team |
| EPI_ISL_2807539 | Lighthouse Lab in Alderley Park | Wellcome Sanger Institute for the COVID-19 Genomics UK (COG-UK) Consortium | Jacquelyn Wynn, Mairead Hyland, The Lighthouse Lab in Alderley Park and Alex Alderton, Roberto Amato, Jeffrey Barrett, Sonia Goncalves, Ewan Harrison, David K. Jackson, Ian Johnston, Dominic Kwiatkowski, Cordelia Langford, John Sillitoe on behalf of the Wellcome Sanger Institute COVID-19 Surveillance Team |
| EPI_ISL_2807540 | Lighthouse Lab in Glasgow | Wellcome Sanger Institute for the COVID-19 Genomics UK (COG-UK) Consortium | Harper VanSteenhouse, Yumi Kasai, David Gray, Carol Clugston, Anna Dominiczak and Alex Alderton, Roberto Amato, Jeffrey Barrett, Sonia Goncalves, Ewan Harrison, David K. Jackson, Ian Johnston, Dominic Kwiatkowski, Cordelia Langford, John Sillitoe on behalf of the Wellcome Sanger Institute COVID-19 Surveillance Team |
| EPI_ISL_2807541 | Lighthouse Lab in Milton Keynes | Wellcome Sanger Institute for the COVID-19 Genomics UK (COG-UK) Consortium | The Lighthouse Lab in Milton Keynes and Alex Alderton, Roberto Amato, Jeffrey Barrett, Sonia Goncalves, Ewan Harrison, David K. Jackson, Ian Johnston, Dominic Kwiatkowski, Cordelia Langford, John Sillitoe on behalf of the Wellcome Sanger Institute COVID-19 Surveillance Team |
| EPI_ISL_2807542 | Lighthouse Lab in Glasgow | Wellcome Sanger Institute for the COVID-19 Genomics UK (COG-UK) Consortium | Harper VanSteenhouse, Yumi Kasai, David Gray, Carol Clugston, Anna Dominiczak and Alex Alderton, Roberto Amato, Jeffrey Barrett, Sonia Goncalves, Ewan Harrison, David K. Jackson, Ian Johnston, Dominic Kwiatkowski, Cordelia Langford, John Sillitoe on behalf of the Wellcome Sanger Institute COVID-19 Surveillance Team |
| EPI_ISL_2807543 | Lighthouse Lab in Alderley Park | Wellcome Sanger Institute for the COVID-19 Genomics UK (COG-UK) Consortium | Jacquelyn Wynn, Mairead Hyland, The Lighthouse Lab in Alderley Park and Alex Alderton, Roberto Amato, Jeffrey Barrett, Sonia Goncalves, Ewan Harrison, David K. Jackson, Ian Johnston, Dominic Kwiatkowski, Cordelia Langford, John Sillitoe on behalf of the Wellcome Sanger Institute COVID-19 Surveillance Team |
| EPI_ISL_2807544 | Lighthouse Lab in Milton Keynes | Wellcome Sanger Institute for the COVID-19 Genomics UK (COG-UK) Consortium | The Lighthouse Lab in Milton Keynes and Alex Alderton, Roberto Amato, Jeffrey Barrett, Sonia Goncalves, Ewan Harrison, David K. Jackson, Ian Johnston, Dominic Kwiatkowski, Cordelia Langford, John Sillitoe on behalf of the Wellcome Sanger Institute COVID-19 Surveillance Team |
| EPI_ISL_2807545 | Lighthouse Lab in Glasgow | Wellcome Sanger Institute for the COVID-19 Genomics UK (COG-UK) Consortium | Harper VanSteenhouse, Yumi Kasai, David Gray, Carol Clugston, Anna Dominiczak and Alex Alderton, Roberto Amato, Jeffrey Barrett, Sonia Goncalves, Ewan Harrison, David K. Jackson, Ian Johnston, Dominic Kwiatkowski, Cordelia Langford, John Sillitoe on behalf of the Wellcome Sanger Institute COVID-19 Surveillance Team |
| EPI_ISL_2807546 | Lighthouse Lab in Alderley Park | Wellcome Sanger Institute for the COVID-19 Genomics UK (COG-UK) Consortium | Jacquelyn Wynn, Mairead Hyland, The Lighthouse Lab in Alderley Park and Alex Alderton, Roberto Amato, Jeffrey Barrett, Sonia Goncalves, Ewan Harrison, David K. Jackson, Ian Johnston, Dominic Kwiatkowski, Cordelia Langford, John Sillitoe on behalf of the Wellcome Sanger Institute COVID-19 Surveillance Team |
| EPI_ISL_2807547 | Lighthouse Lab in Glasgow | Wellcome Sanger Institute for the COVID-19 Genomics UK (COG-UK) Consortium | Harper VanSteenhouse, Yumi Kasai, David Gray, Carol Clugston, Anna Dominiczak and Alex Alderton, Roberto Amato, Jeffrey Barrett, Sonia Goncalves, Ewan Harrison, David K. Jackson, Ian Johnston, Dominic Kwiatkowski, Cordelia Langford, John Sillitoe on behalf of the Wellcome Sanger Institute COVID-19 Surveillance Team |
| EPI_ISL_2807548, EPI_ISL_2807549, EPI_ISL_2807550 | Lighthouse Lab in Milton Keynes | Wellcome Sanger Institute for the COVID-19 Genomics UK (COG-UK) Consortium | The Lighthouse Lab in Milton Keynes and Alex Alderton, Roberto Amato, Jeffrey Barrett, Sonia Goncalves, Ewan Harrison, David K. Jackson, Ian Johnston, Dominic Kwiatkowski, Cordelia Langford, John Sillitoe on behalf of the Wellcome Sanger Institute COVID-19 Surveillance Team |
| EPI_ISL_2807551 | Lighthouse Lab in Glasgow | Wellcome Sanger Institute for the COVID-19 Genomics UK (COG-UK) Consortium | Harper VanSteenhouse, Yumi Kasai, David Gray, Carol Clugston, Anna Dominiczak and Alex Alderton, Roberto Amato, Jeffrey Barrett, Sonia Goncalves, Ewan Harrison, David K. Jackson, Ian Johnston, Dominic Kwiatkowski, Cordelia Langford, John Sillitoe on behalf of the Wellcome Sanger Institute COVID-19 Surveillance Team |
| EPI_ISL_2807552 | Lighthouse Lab in Milton Keynes | Wellcome Sanger Institute for the COVID-19 Genomics UK (COG-UK) Consortium | The Lighthouse Lab in Milton Keynes and Alex Alderton, Roberto Amato, Jeffrey Barrett, Sonia Goncalves, Ewan Harrison, David K. Jackson, Ian Johnston, Dominic Kwiatkowski, Cordelia Langford, John Sillitoe on behalf of the Wellcome Sanger Institute COVID-19 Surveillance Team |
| EPI_ISL_2807553 | Lighthouse Lab in Alderley Park | Wellcome Sanger Institute for the COVID-19 Genomics UK (COG-UK) Consortium | Jacquelyn Wynn, Mairead Hyland, The Lighthouse Lab in Alderley Park and Alex Alderton, Roberto Amato, Jeffrey Barrett, Sonia Goncalves, Ewan Harrison, David K. Jackson, Ian Johnston, Dominic Kwiatkowski, Cordelia Langford, John Sillitoe on behalf of the Wellcome Sanger Institute COVID-19 Surveillance Team |
| EPI_ISL_2807554, EPI_ISL_2807555, EPI_ISL_2807556, EPI_ISL_2807557, EPI_ISL_2807558 | Lighthouse Lab in Milton Keynes | Wellcome Sanger Institute for the COVID-19 Genomics UK (COG-UK) Consortium | The Lighthouse Lab in Milton Keynes and Alex Alderton, Roberto Amato, Jeffrey Barrett, Sonia Goncalves, Ewan Harrison, David K. Jackson, Ian Johnston, Dominic Kwiatkowski, Cordelia Langford, John Sillitoe on behalf of the Wellcome Sanger Institute COVID-19 Surveillance Team |
| EPI_ISL_2807559 | Lighthouse Lab in Glasgow | Wellcome Sanger Institute for the COVID-19 Genomics UK (COG-UK) Consortium | Harper VanSteenhouse, Yumi Kasai, David Gray, Carol Clugston, Anna Dominiczak and Alex Alderton, Roberto Amato, Jeffrey Barrett, Sonia Goncalves, Ewan Harrison, David K. Jackson, Ian Johnston, Dominic Kwiatkowski, Cordelia Langford, John Sillitoe on behalf of the Wellcome Sanger Institute COVID-19 Surveillance Team |
| EPI_ISL_2807560 | Lighthouse Lab in Alderley Park | Wellcome Sanger Institute for the COVID-19 Genomics UK (COG-UK) Consortium | Jacquelyn Wynn, Mairead Hyland, The Lighthouse Lab in Alderley Park and Alex Alderton, Roberto Amato, Jeffrey Barrett, Sonia Goncalves, Ewan Harrison, David K. Jackson, Ian Johnston, Dominic Kwiatkowski, Cordelia Langford, John Sillitoe on behalf of the Wellcome Sanger Institute COVID-19 Surveillance Team |
| EPI_ISL_2807561 | Lighthouse Lab in Milton Keynes | Wellcome Sanger Institute for the COVID-19 Genomics UK (COG-UK) Consortium | The Lighthouse Lab in Milton Keynes and Alex Alderton, Roberto Amato, Jeffrey Barrett, Sonia Goncalves, Ewan Harrison, David K. Jackson, Ian Johnston, Dominic Kwiatkowski, Cordelia Langford, John Sillitoe on behalf of the Wellcome Sanger Institute COVID-19 Surveillance Team |
| EPI_ISL_2807562 | Lighthouse Lab in Glasgow | Wellcome Sanger Institute for the COVID-19 Genomics UK (COG-UK) Consortium | Harper VanSteenhouse, Yumi Kasai, David Gray, Carol Clugston, Anna Dominiczak and Alex Alderton, Roberto Amato, Jeffrey Barrett, Sonia Goncalves, Ewan Harrison, David K. Jackson, Ian Johnston, Dominic Kwiatkowski, Cordelia Langford, John Sillitoe on behalf of the Wellcome Sanger Institute COVID-19 Surveillance Team |
| EPI_ISL_2807563 | Lighthouse Lab in Milton Keynes | Wellcome Sanger Institute for the COVID-19 Genomics UK (COG-UK) Consortium | The Lighthouse Lab in Milton Keynes and Alex Alderton, Roberto Amato, Jeffrey Barrett, Sonia Goncalves, Ewan Harrison, David K. Jackson, Ian Johnston, Dominic Kwiatkowski, Cordelia Langford, John Sillitoe on behalf of the Wellcome Sanger Institute COVID-19 Surveillance Team |
| EPI_ISL_2807564, EPI_ISL_2807565 | Lighthouse Lab in Glasgow | Wellcome Sanger Institute for the COVID-19 Genomics UK (COG-UK) Consortium | Harper VanSteenhouse, Yumi Kasai, David Gray, Carol Clugston, Anna Dominiczak and Alex Alderton, Roberto Amato, Jeffrey Barrett, Sonia Goncalves, Ewan Harrison, David K. Jackson, Ian Johnston, Dominic Kwiatkowski, Cordelia Langford, John Sillitoe on behalf of the Wellcome Sanger Institute COVID-19 Surveillance Team |
| EPI_ISL_2807566 | Lighthouse Lab in Milton Keynes | Wellcome Sanger Institute for the COVID-19 Genomics UK (COG-UK) Consortium | The Lighthouse Lab in Milton Keynes and Alex Alderton, Roberto Amato, Jeffrey Barrett, Sonia Goncalves, Ewan Harrison, David K. Jackson, Ian Johnston, Dominic Kwiatkowski, Cordelia Langford, John Sillitoe on behalf of the Wellcome Sanger Institute COVID-19 Surveillance Team |
| EPI_ISL_2807567 | Lighthouse Lab in Glasgow | Wellcome Sanger Institute for the COVID-19 Genomics UK (COG-UK) Consortium | Harper VanSteenhouse, Yumi Kasai, David Gray, Carol Clugston, Anna Dominiczak and Alex Alderton, Roberto Amato, Jeffrey Barrett, Sonia Goncalves, Ewan Harrison, David K. Jackson, Ian Johnston, Dominic Kwiatkowski, Cordelia Langford, John Sillitoe on behalf of the Wellcome Sanger Institute COVID-19 Surveillance Team |
| EPI_ISL_2807568, EPI_ISL_2807569 | Lighthouse Lab in Milton Keynes | Wellcome Sanger Institute for the COVID-19 Genomics UK (COG-UK) Consortium | The Lighthouse Lab in Milton Keynes and Alex Alderton, Roberto Amato, Jeffrey Barrett, Sonia Goncalves, Ewan Harrison, David K. Jackson, Ian Johnston, Dominic Kwiatkowski, Cordelia Langford, John Sillitoe on behalf of the Wellcome Sanger Institute COVID-19 Surveillance Team |
| EPI_ISL_2807570 | Lighthouse Lab in Alderley Park | Wellcome Sanger Institute for the COVID-19 Genomics UK (COG-UK) Consortium | Jacquelyn Wynn, Mairead Hyland, The Lighthouse Lab in Alderley Park and Alex Alderton, Roberto Amato, Jeffrey Barrett, Sonia Goncalves, Ewan Harrison, David K. Jackson, Ian Johnston, Dominic Kwiatkowski, Cordelia Langford, John Sillitoe on behalf of the Wellcome Sanger Institute COVID-19 Surveillance Team |
| EPI_ISL_2807571, EPI_ISL_2807572, EPI_ISL_2807573, EPI_ISL_2807574, EPI_ISL_2807575 | Lighthouse Lab in Milton Keynes | Wellcome Sanger Institute for the COVID-19 Genomics UK (COG-UK) Consortium | The Lighthouse Lab in Milton Keynes and Alex Alderton, Roberto Amato, Jeffrey Barrett, Sonia Goncalves, Ewan Harrison, David K. Jackson, Ian Johnston, Dominic Kwiatkowski, Cordelia Langford, John Sillitoe on behalf of the Wellcome Sanger Institute COVID-19 Surveillance Team |
| EPI_ISL_2807576 | Lighthouse Lab in Alderley Park | Wellcome Sanger Institute for the COVID-19 Genomics UK (COG-UK) Consortium | Jacquelyn Wynn, Mairead Hyland, The Lighthouse Lab in Alderley Park and Alex Alderton, Roberto Amato, Jeffrey Barrett, Sonia Goncalves, Ewan Harrison, David K. Jackson, Ian Johnston, Dominic Kwiatkowski, Cordelia Langford, John Sillitoe on behalf of the Wellcome Sanger Institute COVID-19 Surveillance Team |
| EPI_ISL_2807577, EPI_ISL_2807578 | Lighthouse Lab in Milton Keynes | Wellcome Sanger Institute for the COVID-19 Genomics UK (COG-UK) Consortium | The Lighthouse Lab in Milton Keynes and Alex Alderton, Roberto Amato, Jeffrey Barrett, Sonia Goncalves, Ewan Harrison, David K. Jackson, Ian Johnston, Dominic Kwiatkowski, Cordelia Langford, John Sillitoe on behalf of the Wellcome Sanger Institute COVID-19 Surveillance Team |
| EPI_ISL_2807579 | Lighthouse Lab in Alderley Park | Wellcome Sanger Institute for the COVID-19 Genomics UK (COG-UK) Consortium | Jacquelyn Wynn, Mairead Hyland, The Lighthouse Lab in Alderley Park and Alex Alderton, Roberto Amato, Jeffrey Barrett, Sonia Goncalves, Ewan Harrison, David K. Jackson, Ian Johnston, Dominic Kwiatkowski, Cordelia Langford, John Sillitoe on behalf of the Wellcome Sanger Institute COVID-19 Surveillance Team |
| EPI_ISL_2807580, EPI_ISL_2807581, EPI_ISL_2807582 | Lighthouse Lab in Milton Keynes | Wellcome Sanger Institute for the COVID-19 Genomics UK (COG-UK) Consortium | The Lighthouse Lab in Milton Keynes and Alex Alderton, Roberto Amato, Jeffrey Barrett, Sonia Goncalves, Ewan Harrison, David K. Jackson, Ian Johnston, Dominic Kwiatkowski, Cordelia Langford, John Sillitoe on behalf of the Wellcome Sanger Institute COVID-19 Surveillance Team |
| EPI_ISL_2807583 | Lighthouse Lab in Alderley Park | Wellcome Sanger Institute for the COVID-19 Genomics UK (COG-UK) Consortium | Jacquelyn Wynn, Mairead Hyland, The Lighthouse Lab in Alderley Park and Alex Alderton, Roberto Amato, Jeffrey Barrett, Sonia Goncalves, Ewan Harrison, David K. Jackson, Ian Johnston, Dominic Kwiatkowski, Cordelia Langford, John Sillitoe on behalf of the Wellcome Sanger Institute COVID-19 Surveillance Team |
| EPI_ISL_2807584, EPI_ISL_2807585 | Lighthouse Lab in Glasgow | Wellcome Sanger Institute for the COVID-19 Genomics UK (COG-UK) Consortium | Harper VanSteenhouse, Yumi Kasai, David Gray, Carol Clugston, Anna Dominiczak and Alex Alderton, Roberto Amato, Jeffrey Barrett, Sonia Goncalves, Ewan Harrison, David K. Jackson, Ian Johnston, Dominic Kwiatkowski, Cordelia Langford, John Sillitoe on behalf of the Wellcome Sanger Institute COVID-19 Surveillance Team |
| EPI_ISL_2807586, EPI_ISL_2807587, EPI_ISL_2807588 | Lighthouse Lab in Alderley Park | Wellcome Sanger Institute for the COVID-19 Genomics UK (COG-UK) Consortium | Jacquelyn Wynn, Mairead Hyland, The Lighthouse Lab in Alderley Park and Alex Alderton, Roberto Amato, Jeffrey Barrett, Sonia Goncalves, Ewan Harrison, David K. Jackson, Ian Johnston, Dominic Kwiatkowski, Cordelia Langford, John Sillitoe on behalf of the Wellcome Sanger Institute COVID-19 Surveillance Team |
| EPI_ISL_2807589, EPI_ISL_2807590 | Lighthouse Lab in Milton Keynes | Wellcome Sanger Institute for the COVID-19 Genomics UK (COG-UK) Consortium | The Lighthouse Lab in Milton Keynes and Alex Alderton, Roberto Amato, Jeffrey Barrett, Sonia Goncalves, Ewan Harrison, David K. Jackson, Ian Johnston, Dominic Kwiatkowski, Cordelia Langford, John Sillitoe on behalf of the Wellcome Sanger Institute COVID-19 Surveillance Team |
| EPI_ISL_2807591, EPI_ISL_2807592 | Lighthouse Lab in Alderley Park | Wellcome Sanger Institute for the COVID-19 Genomics UK (COG-UK) Consortium | Jacquelyn Wynn, Mairead Hyland, The Lighthouse Lab in Alderley Park and Alex Alderton, Roberto Amato, Jeffrey Barrett, Sonia Goncalves, Ewan Harrison, David K. Jackson, Ian Johnston, Dominic Kwiatkowski, Cordelia Langford, John Sillitoe on behalf of the Wellcome Sanger Institute COVID-19 Surveillance Team |
| EPI_ISL_2807593 | Lighthouse Lab in Milton Keynes | Wellcome Sanger Institute for the COVID-19 Genomics UK (COG-UK) Consortium | The Lighthouse Lab in Milton Keynes and Alex Alderton, Roberto Amato, Jeffrey Barrett, Sonia Goncalves, Ewan Harrison, David K. Jackson, Ian Johnston, Dominic Kwiatkowski, Cordelia Langford, John Sillitoe on behalf of the Wellcome Sanger Institute COVID-19 Surveillance Team |
| EPI_ISL_2807594 | Lighthouse Lab in Alderley Park | Wellcome Sanger Institute for the COVID-19 Genomics UK (COG-UK) Consortium | Jacquelyn Wynn, Mairead Hyland, The Lighthouse Lab in Alderley Park and Alex Alderton, Roberto Amato, Jeffrey Barrett, Sonia Goncalves, Ewan Harrison, David K. Jackson, Ian Johnston, Dominic Kwiatkowski, Cordelia Langford, John Sillitoe on behalf of the Wellcome Sanger Institute COVID-19 Surveillance Team |
| EPI_ISL_2807595 | Lighthouse Lab in Milton Keynes | Wellcome Sanger Institute for the COVID-19 Genomics UK (COG-UK) Consortium | The Lighthouse Lab in Milton Keynes and Alex Alderton, Roberto Amato, Jeffrey Barrett, Sonia Goncalves, Ewan Harrison, David K. Jackson, Ian Johnston, Dominic Kwiatkowski, Cordelia Langford, John Sillitoe on behalf of the Wellcome Sanger Institute COVID-19 Surveillance Team |
| EPI_ISL_2807596, EPI_ISL_2807597 | Lighthouse Lab in Alderley Park | Wellcome Sanger Institute for the COVID-19 Genomics UK (COG-UK) Consortium | Jacquelyn Wynn, Mairead Hyland, The Lighthouse Lab in Alderley Park and Alex Alderton, Roberto Amato, Jeffrey Barrett, Sonia Goncalves, Ewan Harrison, David K. Jackson, Ian Johnston, Dominic Kwiatkowski, Cordelia Langford, John Sillitoe on behalf of the Wellcome Sanger Institute COVID-19 Surveillance Team |
| EPI_ISL_2807598, EPI_ISL_2807599 | Lighthouse Lab in Milton Keynes | Wellcome Sanger Institute for the COVID-19 Genomics UK (COG-UK) Consortium | The Lighthouse Lab in Milton Keynes and Alex Alderton, Roberto Amato, Jeffrey Barrett, Sonia Goncalves, Ewan Harrison, David K. Jackson, Ian Johnston, Dominic Kwiatkowski, Cordelia Langford, John Sillitoe on behalf of the Wellcome Sanger Institute COVID-19 Surveillance Team |

| | | | |
|---|---|---|---|
| EPI_ISL_2807600 | Lighthouse Lab in Glasgow | Wellcome Sanger Institute for the COVID-19 Genomics UK (COG-UK) Consortium | Harper VanSteenhouse, Yumi Kasai, David Gray, Carol Clugston, Anna Dominiczak and Alex Alderton, Roberto Amato, Jeffrey Barrett, Sonia Goncalves, Ewan Harrison, David K. Jackson, Ian Johnston, Dominic Kwiatkowski, Cordelia Langford, John Sillitoe on behalf of the Wellcome Sanger Institute COVID-19 Surveillance Team |
| EPI_ISL_2807601, EPI_ISL_2807602 | Lighthouse Lab in Milton Keynes | Wellcome Sanger Institute for the COVID-19 Genomics UK (COG-UK) Consortium | The Lighthouse Lab in Milton Keynes and Alex Alderton, Roberto Amato, Jeffrey Barrett, Sonia Goncalves, Ewan Harrison, David K. Jackson, Ian Johnston, Dominic Kwiatkowski, Cordelia Langford, John Sillitoe on behalf of the Wellcome Sanger Institute COVID-19 Surveillance Team |
| EPI_ISL_2807603 | Lighthouse Lab in Glasgow | Wellcome Sanger Institute for the COVID-19 Genomics UK (COG-UK) Consortium | Harper VanSteenhouse, Yumi Kasai, David Gray, Carol Clugston, Anna Dominiczak and Alex Alderton, Roberto Amato, Jeffrey Barrett, Sonia Goncalves, Ewan Harrison, David K. Jackson, Ian Johnston, Dominic Kwiatkowski, Cordelia Langford, John Sillitoe on behalf of the Wellcome Sanger Institute COVID-19 Surveillance Team |
| EPI_ISL_2807604 | Lighthouse Lab in Alderley Park | Wellcome Sanger Institute for the COVID-19 Genomics UK (COG-UK) Consortium | Jacquelyn Wynn, Mairead Hyland, The Lighthouse Lab in Alderley Park and Alex Alderton, Roberto Amato, Jeffrey Barrett, Sonia Goncalves, Ewan Harrison, David K. Jackson, Ian Johnston, Dominic Kwiatkowski, Cordelia Langford, John Sillitoe on behalf of the Wellcome Sanger Institute COVID-19 Surveillance Team |
| EPI_ISL_2807605, EPI_ISL_2807606, EPI_ISL_2807607, EPI_ISL_2807608 | Lighthouse Lab in Milton Keynes | Wellcome Sanger Institute for the COVID-19 Genomics UK (COG-UK) Consortium | The Lighthouse Lab in Milton Keynes and Alex Alderton, Roberto Amato, Jeffrey Barrett, Sonia Goncalves, Ewan Harrison, David K. Jackson, Ian Johnston, Dominic Kwiatkowski, Cordelia Langford, John Sillitoe on behalf of the Wellcome Sanger Institute COVID-19 Surveillance Team |
| EPI_ISL_2807609 | Lighthouse Lab in Alderley Park | Wellcome Sanger Institute for the COVID-19 Genomics UK (COG-UK) Consortium | Jacquelyn Wynn, Mairead Hyland, The Lighthouse Lab in Alderley Park and Alex Alderton, Roberto Amato, Jeffrey Barrett, Sonia Goncalves, Ewan Harrison, David K. Jackson, Ian Johnston, Dominic Kwiatkowski, Cordelia Langford, John Sillitoe on behalf of the Wellcome Sanger Institute COVID-19 Surveillance Team |
| EPI_ISL_2807610 | Lighthouse Lab in Milton Keynes | Wellcome Sanger Institute for the COVID-19 Genomics UK (COG-UK) Consortium | The Lighthouse Lab in Milton Keynes and Alex Alderton, Roberto Amato, Jeffrey Barrett, Sonia Goncalves, Ewan Harrison, David K. Jackson, Ian Johnston, Dominic Kwiatkowski, Cordelia Langford, John Sillitoe on behalf of the Wellcome Sanger Institute COVID-19 Surveillance Team |
| EPI_ISL_2807611 | Lighthouse Lab in Alderley Park | Wellcome Sanger Institute for the COVID-19 Genomics UK (COG-UK) Consortium | Jacquelyn Wynn, Mairead Hyland, The Lighthouse Lab in Alderley Park and Alex Alderton, Roberto Amato, Jeffrey Barrett, Sonia Goncalves, Ewan Harrison, David K. Jackson, Ian Johnston, Dominic Kwiatkowski, Cordelia Langford, John Sillitoe on behalf of the Wellcome Sanger Institute COVID-19 Surveillance Team |
| EPI_ISL_2807612 | Lighthouse Lab in Milton Keynes | Wellcome Sanger Institute for the COVID-19 Genomics UK (COG-UK) Consortium | The Lighthouse Lab in Milton Keynes and Alex Alderton, Roberto Amato, Jeffrey Barrett, Sonia Goncalves, Ewan Harrison, David K. Jackson, Ian Johnston, Dominic Kwiatkowski, Cordelia Langford, John Sillitoe on behalf of the Wellcome Sanger Institute COVID-19 Surveillance Team |
| EPI_ISL_2807613 | Lighthouse Lab in Glasgow | Wellcome Sanger Institute for the COVID-19 Genomics UK (COG-UK) Consortium | Harper VanSteenhouse, Yumi Kasai, David Gray, Carol Clugston, Anna Dominiczak and Alex Alderton, Roberto Amato, Jeffrey Barrett, Sonia Goncalves, Ewan Harrison, David K. Jackson, Ian Johnston, Dominic Kwiatkowski, Cordelia Langford, John Sillitoe on behalf of the Wellcome Sanger Institute COVID-19 Surveillance Team |
| EPI_ISL_2807614 | Lighthouse Lab in Alderley Park | Wellcome Sanger Institute for the COVID-19 Genomics UK (COG-UK) Consortium | Jacquelyn Wynn, Mairead Hyland, The Lighthouse Lab in Alderley Park and Alex Alderton, Roberto Amato, Jeffrey Barrett, Sonia Goncalves, Ewan Harrison, David K. Jackson, Ian Johnston, Dominic Kwiatkowski, Cordelia Langford, John Sillitoe on behalf of the Wellcome Sanger Institute COVID-19 Surveillance Team |
| EPI_ISL_2807615, EPI_ISL_2807616, EPI_ISL_2807617, EPI_ISL_2807618 | Lighthouse Lab in Glasgow | Wellcome Sanger Institute for the COVID-19 Genomics UK (COG-UK) Consortium | Harper VanSteenhouse, Yumi Kasai, David Gray, Carol Clugston, Anna Dominiczak and Alex Alderton, Roberto Amato, Jeffrey Barrett, Sonia Goncalves, Ewan Harrison, David K. Jackson, Ian Johnston, Dominic Kwiatkowski, Cordelia Langford, John Sillitoe on behalf of the Wellcome Sanger Institute COVID-19 Surveillance Team |
| EPI_ISL_2807619 | Lighthouse Lab in Milton Keynes | Wellcome Sanger Institute for the COVID-19 Genomics UK (COG-UK) Consortium | The Lighthouse Lab in Milton Keynes and Alex Alderton, Roberto Amato, Jeffrey Barrett, Sonia Goncalves, Ewan Harrison, David K. Jackson, Ian Johnston, Dominic Kwiatkowski, Cordelia Langford, John Sillitoe on behalf of the Wellcome Sanger Institute COVID-19 Surveillance Team |
| EPI_ISL_2807620 | Lighthouse Lab in Glasgow | Wellcome Sanger Institute for the COVID-19 Genomics UK (COG-UK) Consortium | Harper VanSteenhouse, Yumi Kasai, David Gray, Carol Clugston, Anna Dominiczak and Alex Alderton, Roberto Amato, Jeffrey Barrett, Sonia Goncalves, Ewan Harrison, David K. Jackson, Ian Johnston, Dominic Kwiatkowski, Cordelia Langford, John Sillitoe on behalf of the Wellcome Sanger Institute COVID-19 Surveillance Team |
| EPI_ISL_2807621 | Lighthouse Lab in Milton Keynes | Wellcome Sanger Institute for the COVID-19 Genomics UK (COG-UK) Consortium | The Lighthouse Lab in Milton Keynes and Alex Alderton, Roberto Amato, Jeffrey Barrett, Sonia Goncalves, Ewan Harrison, David K. Jackson, Ian Johnston, Dominic Kwiatkowski, Cordelia Langford, John Sillitoe on behalf of the Wellcome Sanger Institute COVID-19 Surveillance Team |
| EPI_ISL_2807622 | Lighthouse Lab in Glasgow | Wellcome Sanger Institute for the COVID-19 Genomics UK (COG-UK) Consortium | Harper VanSteenhouse, Yumi Kasai, David Gray, Carol Clugston, Anna Dominiczak and Alex Alderton, Roberto Amato, Jeffrey Barrett, Sonia Goncalves, Ewan Harrison, David K. Jackson, Ian Johnston, Dominic Kwiatkowski, Cordelia Langford, John Sillitoe on behalf of the Wellcome Sanger Institute COVID-19 Surveillance Team |
| EPI_ISL_2807623, EPI_ISL_2807624 | Lighthouse Lab in Alderley Park | Wellcome Sanger Institute for the COVID-19 Genomics UK (COG-UK) Consortium | Jacquelyn Wynn, Mairead Hyland, The Lighthouse Lab in Alderley Park and Alex Alderton, Roberto Amato, Jeffrey Barrett, Sonia Goncalves, Ewan Harrison, David K. Jackson, Ian Johnston, Dominic Kwiatkowski, Cordelia Langford, John Sillitoe on behalf of the Wellcome Sanger Institute COVID-19 Surveillance Team |
| EPI_ISL_2807625 | Lighthouse Lab in Glasgow | Wellcome Sanger Institute for the COVID-19 Genomics UK (COG-UK) Consortium | Harper VanSteenhouse, Yumi Kasai, David Gray, Carol Clugston, Anna Dominiczak and Alex Alderton, Roberto Amato, Jeffrey Barrett, Sonia Goncalves, Ewan Harrison, David K. Jackson, Ian Johnston, Dominic Kwiatkowski, Cordelia Langford, John Sillitoe on behalf of the Wellcome Sanger Institute COVID-19 Surveillance Team |
| EPI_ISL_2807626 | Lighthouse Lab in Milton Keynes | Wellcome Sanger Institute for the COVID-19 Genomics UK (COG-UK) Consortium | The Lighthouse Lab in Milton Keynes and Alex Alderton, Roberto Amato, Jeffrey Barrett, Sonia Goncalves, Ewan Harrison, David K. Jackson, Ian Johnston, Dominic Kwiatkowski, Cordelia Langford, John Sillitoe on behalf of the Wellcome Sanger Institute COVID-19 Surveillance Team |
| EPI_ISL_2807627 | Lighthouse Lab in Glasgow | Wellcome Sanger Institute for the COVID-19 Genomics UK (COG-UK) Consortium | Harper VanSteenhouse, Yumi Kasai, David Gray, Carol Clugston, Anna Dominiczak and Alex Alderton, Roberto Amato, Jeffrey Barrett, Sonia Goncalves, Ewan Harrison, David K. Jackson, Ian Johnston, Dominic Kwiatkowski, Cordelia Langford, John Sillitoe on behalf of the Wellcome Sanger Institute COVID-19 Surveillance Team |
| EPI_ISL_2807628 | Lighthouse Lab in Milton Keynes | Wellcome Sanger Institute for the COVID-19 Genomics UK (COG-UK) Consortium | The Lighthouse Lab in Milton Keynes and Alex Alderton, Roberto Amato, Jeffrey Barrett, Sonia Goncalves, Ewan Harrison, David K. Jackson, Ian Johnston, Dominic Kwiatkowski, Cordelia Langford, John Sillitoe on behalf of the Wellcome Sanger Institute COVID-19 Surveillance Team |
| EPI_ISL_2807629 | Lighthouse Lab in Glasgow | Wellcome Sanger Institute for the COVID-19 Genomics UK (COG-UK) Consortium | Harper VanSteenhouse, Yumi Kasai, David Gray, Carol Clugston, Anna Dominiczak and Alex Alderton, Roberto Amato, Jeffrey Barrett, Sonia Goncalves, Ewan Harrison, David K. Jackson, Ian Johnston, Dominic Kwiatkowski, Cordelia Langford, John Sillitoe on behalf of the Wellcome Sanger Institute COVID-19 Surveillance Team |
| EPI_ISL_2807630 | Lighthouse Lab in Milton Keynes | Wellcome Sanger Institute for the COVID-19 Genomics UK (COG-UK) Consortium | The Lighthouse Lab in Milton Keynes and Alex Alderton, Roberto Amato, Jeffrey Barrett, Sonia Goncalves, Ewan Harrison, David K. Jackson, Ian Johnston, Dominic Kwiatkowski, Cordelia Langford, John Sillitoe on behalf of the Wellcome Sanger Institute COVID-19 Surveillance Team |
| EPI_ISL_2807631, EPI_ISL_2807632 | Lighthouse Lab in Glasgow | Wellcome Sanger Institute for the COVID-19 Genomics UK (COG-UK) Consortium | Harper VanSteenhouse, Yumi Kasai, David Gray, Carol Clugston, Anna Dominiczak and Alex Alderton, Roberto Amato, Jeffrey Barrett, Sonia Goncalves, Ewan Harrison, David K. Jackson, Ian Johnston, Dominic Kwiatkowski, Cordelia Langford, John Sillitoe on behalf of the Wellcome Sanger Institute COVID-19 Surveillance Team |
| EPI_ISL_2807633 | Lighthouse Lab in Alderley Park | Wellcome Sanger Institute for the COVID-19 Genomics UK (COG-UK) Consortium | Jacquelyn Wynn, Mairead Hyland, The Lighthouse Lab in Alderley Park and Alex Alderton, Roberto Amato, Jeffrey Barrett, Sonia Goncalves, Ewan Harrison, David K. Jackson, Ian Johnston, Dominic Kwiatkowski, Cordelia Langford, John Sillitoe on behalf of the Wellcome Sanger Institute COVID-19 Surveillance Team |
| EPI_ISL_2807634 | Lighthouse Lab in Milton Keynes | Wellcome Sanger Institute for the COVID-19 Genomics UK (COG-UK) Consortium | The Lighthouse Lab in Milton Keynes and Alex Alderton, Roberto Amato, Jeffrey Barrett, Sonia Goncalves, Ewan Harrison, David K. Jackson, Ian Johnston, Dominic Kwiatkowski, Cordelia Langford, John Sillitoe on behalf of the Wellcome Sanger Institute COVID-19 Surveillance Team |
| EPI_ISL_2807635 | Lighthouse Lab in Alderley Park | Wellcome Sanger Institute for the COVID-19 Genomics UK (COG-UK) Consortium | Jacquelyn Wynn, Mairead Hyland, The Lighthouse Lab in Alderley Park and Alex Alderton, Roberto Amato, Jeffrey Barrett, Sonia Goncalves, Ewan Harrison, David K. Jackson, Ian Johnston, Dominic Kwiatkowski, Cordelia Langford, John Sillitoe on behalf of the Wellcome Sanger Institute COVID-19 Surveillance Team |
| EPI_ISL_2807636, EPI_ISL_2807637 | Lighthouse Lab in Milton Keynes | Wellcome Sanger Institute for the COVID-19 Genomics UK (COG-UK) Consortium | The Lighthouse Lab in Milton Keynes and Alex Alderton, Roberto Amato, Jeffrey Barrett, Sonia Goncalves, Ewan Harrison, David K. Jackson, Ian Johnston, Dominic Kwiatkowski, Cordelia Langford, John Sillitoe on behalf of the Wellcome Sanger Institute COVID-19 Surveillance Team |
| EPI_ISL_2807638 | Lighthouse Lab in Glasgow | Wellcome Sanger Institute for the COVID-19 Genomics UK (COG-UK) Consortium | Harper VanSteenhouse, Yumi Kasai, David Gray, Carol Clugston, Anna Dominiczak and Alex Alderton, Roberto Amato, Jeffrey Barrett, Sonia Goncalves, Ewan Harrison, David K. Jackson, Ian Johnston, Dominic Kwiatkowski, Cordelia Langford, John Sillitoe on behalf of the Wellcome Sanger Institute COVID-19 Surveillance Team |
| EPI_ISL_2807639, EPI_ISL_2807640 | Lighthouse Lab in Milton Keynes | Wellcome Sanger Institute for the COVID-19 Genomics UK (COG-UK) Consortium | The Lighthouse Lab in Milton Keynes and Alex Alderton, Roberto Amato, Jeffrey Barrett, Sonia Goncalves, Ewan Harrison, David K. Jackson, Ian Johnston, Dominic Kwiatkowski, Cordelia Langford, John Sillitoe on behalf of the Wellcome Sanger Institute COVID-19 Surveillance Team |
| EPI_ISL_2807641 | Lighthouse Lab in Glasgow | Wellcome Sanger Institute for the COVID-19 Genomics UK (COG-UK) Consortium | Harper VanSteenhouse, Yumi Kasai, David Gray, Carol Clugston, Anna Dominiczak and Alex Alderton, Roberto Amato, Jeffrey Barrett, Sonia Goncalves, Ewan Harrison, David K. Jackson, Ian Johnston, Dominic Kwiatkowski, Cordelia Langford, John Sillitoe on behalf of the Wellcome Sanger Institute COVID-19 Surveillance Team |
| EPI_ISL_2807642, EPI_ISL_2807643 | Lighthouse Lab in Milton Keynes | Wellcome Sanger Institute for the COVID-19 Genomics UK (COG-UK) Consortium | The Lighthouse Lab in Milton Keynes and Alex Alderton, Roberto Amato, Jeffrey Barrett, Sonia Goncalves, Ewan Harrison, David K. Jackson, Ian Johnston, Dominic Kwiatkowski, Cordelia Langford, John Sillitoe on behalf of the Wellcome Sanger Institute COVID-19 Surveillance Team |
| EPI_ISL_2807644 | Lighthouse Lab in Glasgow | Wellcome Sanger Institute for the COVID-19 Genomics UK (COG-UK) Consortium | Harper VanSteenhouse, Yumi Kasai, David Gray, Carol Clugston, Anna Dominiczak and Alex Alderton, Roberto Amato, Jeffrey Barrett, Sonia Goncalves, Ewan Harrison, David K. Jackson, Ian Johnston, Dominic Kwiatkowski, Cordelia Langford, John Sillitoe on behalf of the Wellcome Sanger Institute COVID-19 Surveillance Team |
| EPI_ISL_2807645 | Lighthouse Lab in Milton Keynes | Wellcome Sanger Institute for the COVID-19 Genomics UK (COG-UK) Consortium | The Lighthouse Lab in Milton Keynes and Alex Alderton, Roberto Amato, Jeffrey Barrett, Sonia Goncalves, Ewan Harrison, David K. Jackson, Ian Johnston, Dominic Kwiatkowski, Cordelia Langford, John Sillitoe on behalf of the Wellcome Sanger Institute COVID-19 Surveillance Team |
| EPI_ISL_2807646 | Lighthouse Lab in Glasgow | Wellcome Sanger Institute for the COVID-19 Genomics UK (COG-UK) Consortium | Harper VanSteenhouse, Yumi Kasai, David Gray, Carol Clugston, Anna Dominiczak and Alex Alderton, Roberto Amato, Jeffrey Barrett, Sonia Goncalves, Ewan Harrison, David K. Jackson, Ian Johnston, Dominic Kwiatkowski, Cordelia Langford, John Sillitoe on behalf of the Wellcome Sanger Institute COVID-19 Surveillance Team |
| EPI_ISL_2807647 | Lighthouse Lab in Alderley Park | Wellcome Sanger Institute for the COVID-19 Genomics UK (COG-UK) Consortium | Jacquelyn Wynn, Mairead Hyland, The Lighthouse Lab in Alderley Park and Alex Alderton, Roberto Amato, Jeffrey Barrett, Sonia Goncalves, Ewan Harrison, David K. Jackson, Ian Johnston, Dominic Kwiatkowski, Cordelia Langford, John Sillitoe on behalf of the Wellcome Sanger Institute COVID-19 Surveillance Team |
| EPI_ISL_2807648 | Lighthouse Lab in Glasgow | Wellcome Sanger Institute for the COVID-19 Genomics UK (COG-UK) Consortium | Harper VanSteenhouse, Yumi Kasai, David Gray, Carol Clugston, Anna Dominiczak and Alex Alderton, Roberto Amato, Jeffrey Barrett, Sonia Goncalves, Ewan Harrison, David K. Jackson, Ian Johnston, Dominic Kwiatkowski, Cordelia Langford, John Sillitoe on behalf of the Wellcome Sanger Institute COVID-19 Surveillance Team |
| EPI_ISL_2807649 | Lighthouse Lab in Milton Keynes | Wellcome Sanger Institute for the COVID-19 Genomics UK (COG-UK) Consortium | The Lighthouse Lab in Milton Keynes and Alex Alderton, Roberto Amato, Jeffrey Barrett, Sonia Goncalves, Ewan Harrison, David K. Jackson, Ian Johnston, Dominic Kwiatkowski, Cordelia Langford, John Sillitoe on behalf of the Wellcome Sanger Institute COVID-19 Surveillance Team |
| EPI_ISL_2807650 | Lighthouse Lab in Glasgow | Wellcome Sanger Institute for the COVID-19 Genomics UK (COG-UK) Consortium | Harper VanSteenhouse, Yumi Kasai, David Gray, Carol Clugston, Anna Dominiczak and Alex Alderton, Roberto Amato, Jeffrey Barrett, Sonia Goncalves, Ewan Harrison, David K. Jackson, Ian Johnston, Dominic Kwiatkowski, Cordelia Langford, John Sillitoe on behalf of the Wellcome Sanger Institute COVID-19 Surveillance Team |
| EPI_ISL_2807651, EPI_ISL_2807652, EPI_ISL_2807653, EPI_ISL_2807654 | Lighthouse Lab in Milton Keynes | Wellcome Sanger Institute for the COVID-19 Genomics UK (COG-UK) Consortium | The Lighthouse Lab in Milton Keynes and Alex Alderton, Roberto Amato, Jeffrey Barrett, Sonia Goncalves, Ewan Harrison, David K. Jackson, Ian Johnston, Dominic Kwiatkowski, Cordelia Langford, John Sillitoe on behalf of the Wellcome Sanger Institute COVID-19 Surveillance Team |
| EPI_ISL_2807655 | Lighthouse Lab in Glasgow | Wellcome Sanger Institute for the COVID-19 Genomics UK (COG-UK) Consortium | Harper VanSteenhouse, Yumi Kasai, David Gray, Carol Clugston, Anna Dominiczak and Alex Alderton, Roberto Amato, Jeffrey Barrett, Sonia Goncalves, Ewan Harrison, David K. Jackson, Ian Johnston, Dominic Kwiatkowski, Cordelia Langford, John Sillitoe on behalf of the Wellcome Sanger Institute COVID-19 Surveillance Team |
| EPI_ISL_2807656 | Lighthouse Lab in Alderley Park | Wellcome Sanger Institute for the COVID-19 Genomics UK (COG-UK) Consortium | Jacquelyn Wynn, Mairead Hyland, The Lighthouse Lab in Alderley Park and Alex Alderton, Roberto Amato, Jeffrey Barrett, Sonia Goncalves, Ewan Harrison, David K. Jackson, Ian Johnston, Dominic Kwiatkowski, Cordelia Langford, John Sillitoe on behalf of the Wellcome Sanger Institute COVID-19 Surveillance Team |
| EPI_ISL_2807657 | Lighthouse Lab in Glasgow | Wellcome Sanger Institute for the COVID-19 Genomics UK (COG-UK) | Harper VanSteenhouse, Yumi Kasai, David Gray, Carol Clugston, Anna Dominiczak and Alex Alderton, Roberto Amato, Jeffrey Barrett, Sonia Goncalves, Ewan Harrison, David K. Jackson, Ian Johnston, Dominic Kwiatkowski, Cordelia Langford, |

| | | | |
|---|---|---|---|
| | | Consortium | John Sillitoe on behalf of the Wellcome Sanger Institute COVID-19 Surveillance Team |
| EPI_ISL_2807658 | Lighthouse Lab in Milton Keynes | Wellcome Sanger Institute for the COVID-19 Genomics UK (COG-UK) Consortium | The Lighthouse Lab in Milton Keynes and Alex Alderton, Roberto Amato, Jeffrey Barrett, Sonia Goncalves, Ewan Harrison, David K. Jackson, Ian Johnston, Dominic Kwiatkowski, Cordelia Langford, John Sillitoe on behalf of the Wellcome Sanger Institute COVID-19 Surveillance Team |
| EPI_ISL_2807659 | Lighthouse Lab in Alderley Park | Wellcome Sanger Institute for the COVID-19 Genomics UK (COG-UK) Consortium | Jacquelyn Wynn, Mairead Hyland, The Lighthouse Lab in Alderley Park and Alex Alderton, Roberto Amato, Jeffrey Barrett, Sonia Goncalves, Ewan Harrison, David K. Jackson, Ian Johnston, Dominic Kwiatkowski, Cordelia Langford, John Sillitoe on behalf of the Wellcome Sanger Institute COVID-19 Surveillance Team |
| EPI_ISL_2807660, EPI_ISL_2807661, EPI_ISL_2807662, EPI_ISL_2807663 | Lighthouse Lab in Milton Keynes | Wellcome Sanger Institute for the COVID-19 Genomics UK (COG-UK) Consortium | The Lighthouse Lab in Milton Keynes and Alex Alderton, Roberto Amato, Jeffrey Barrett, Sonia Goncalves, Ewan Harrison, David K. Jackson, Ian Johnston, Dominic Kwiatkowski, Cordelia Langford, John Sillitoe on behalf of the Wellcome Sanger Institute COVID-19 Surveillance Team |
| EPI_ISL_2807664, EPI_ISL_2807665 | Lighthouse Lab in Alderley Park | Wellcome Sanger Institute for the COVID-19 Genomics UK (COG-UK) Consortium | Jacquelyn Wynn, Mairead Hyland, The Lighthouse Lab in Alderley Park and Alex Alderton, Roberto Amato, Jeffrey Barrett, Sonia Goncalves, Ewan Harrison, David K. Jackson, Ian Johnston, Dominic Kwiatkowski, Cordelia Langford, John Sillitoe on behalf of the Wellcome Sanger Institute COVID-19 Surveillance Team |
| EPI_ISL_2807666 | Lighthouse Lab in Milton Keynes | Wellcome Sanger Institute for the COVID-19 Genomics UK (COG-UK) Consortium | The Lighthouse Lab in Milton Keynes and Alex Alderton, Roberto Amato, Jeffrey Barrett, Sonia Goncalves, Ewan Harrison, David K. Jackson, Ian Johnston, Dominic Kwiatkowski, Cordelia Langford, John Sillitoe on behalf of the Wellcome Sanger Institute COVID-19 Surveillance Team |
| EPI_ISL_2807667 | Lighthouse Lab in Alderley Park | Wellcome Sanger Institute for the COVID-19 Genomics UK (COG-UK) Consortium | Jacquelyn Wynn, Mairead Hyland, The Lighthouse Lab in Alderley Park and Alex Alderton, Roberto Amato, Jeffrey Barrett, Sonia Goncalves, Ewan Harrison, David K. Jackson, Ian Johnston, Dominic Kwiatkowski, Cordelia Langford, John Sillitoe on behalf of the Wellcome Sanger Institute COVID-19 Surveillance Team |
| EPI_ISL_2807668, EPI_ISL_2807669, EPI_ISL_2807670, EPI_ISL_2807671 | Lighthouse Lab in Milton Keynes | Wellcome Sanger Institute for the COVID-19 Genomics UK (COG-UK) Consortium | The Lighthouse Lab in Milton Keynes and Alex Alderton, Roberto Amato, Jeffrey Barrett, Sonia Goncalves, Ewan Harrison, David K. Jackson, Ian Johnston, Dominic Kwiatkowski, Cordelia Langford, John Sillitoe on behalf of the Wellcome Sanger Institute COVID-19 Surveillance Team |
| EPI_ISL_2807672 | Lighthouse Lab in Alderley Park | Wellcome Sanger Institute for the COVID-19 Genomics UK (COG-UK) Consortium | Jacquelyn Wynn, Mairead Hyland, The Lighthouse Lab in Alderley Park and Alex Alderton, Roberto Amato, Jeffrey Barrett, Sonia Goncalves, Ewan Harrison, David K. Jackson, Ian Johnston, Dominic Kwiatkowski, Cordelia Langford, John Sillitoe on behalf of the Wellcome Sanger Institute COVID-19 Surveillance Team |
| EPI_ISL_2807673 | Lighthouse Lab in Milton Keynes | Wellcome Sanger Institute for the COVID-19 Genomics UK (COG-UK) Consortium | The Lighthouse Lab in Milton Keynes and Alex Alderton, Roberto Amato, Jeffrey Barrett, Sonia Goncalves, Ewan Harrison, David K. Jackson, Ian Johnston, Dominic Kwiatkowski, Cordelia Langford, John Sillitoe on behalf of the Wellcome Sanger Institute COVID-19 Surveillance Team |
| EPI_ISL_2807674 | Lighthouse Lab in Glasgow | Wellcome Sanger Institute for the COVID-19 Genomics UK (COG-UK) Consortium | Harper VanSteenhouse, Yumi Kasai, David Gray, Carol Clugston, Anna Dominiczak and Alex Alderton, Roberto Amato, Jeffrey Barrett, Sonia Goncalves, Ewan Harrison, David K. Jackson, Ian Johnston, Dominic Kwiatkowski, Cordelia Langford, John Sillitoe on behalf of the Wellcome Sanger Institute COVID-19 Surveillance Team |
| EPI_ISL_2807675 | Lighthouse Lab in Alderley Park | Wellcome Sanger Institute for the COVID-19 Genomics UK (COG-UK) Consortium | Jacquelyn Wynn, Mairead Hyland, The Lighthouse Lab in Alderley Park and Alex Alderton, Roberto Amato, Jeffrey Barrett, Sonia Goncalves, Ewan Harrison, David K. Jackson, Ian Johnston, Dominic Kwiatkowski, Cordelia Langford, John Sillitoe on behalf of the Wellcome Sanger Institute COVID-19 Surveillance Team |
| EPI_ISL_2807676 | Lighthouse Lab in Glasgow | Wellcome Sanger Institute for the COVID-19 Genomics UK (COG-UK) Consortium | Harper VanSteenhouse, Yumi Kasai, David Gray, Carol Clugston, Anna Dominiczak and Alex Alderton, Roberto Amato, Jeffrey Barrett, Sonia Goncalves, Ewan Harrison, David K. Jackson, Ian Johnston, Dominic Kwiatkowski, Cordelia Langford, John Sillitoe on behalf of the Wellcome Sanger Institute COVID-19 Surveillance Team |
| EPI_ISL_2807677 | Lighthouse Lab in Alderley Park | Wellcome Sanger Institute for the COVID-19 Genomics UK (COG-UK) Consortium | Jacquelyn Wynn, Mairead Hyland, The Lighthouse Lab in Alderley Park and Alex Alderton, Roberto Amato, Jeffrey Barrett, Sonia Goncalves, Ewan Harrison, David K. Jackson, Ian Johnston, Dominic Kwiatkowski, Cordelia Langford, John Sillitoe on behalf of the Wellcome Sanger Institute COVID-19 Surveillance Team |
| EPI_ISL_2807678, EPI_ISL_2807679, EPI_ISL_2807680 | Lighthouse Lab in Milton Keynes | Wellcome Sanger Institute for the COVID-19 Genomics UK (COG-UK) Consortium | The Lighthouse Lab in Milton Keynes and Alex Alderton, Roberto Amato, Jeffrey Barrett, Sonia Goncalves, Ewan Harrison, David K. Jackson, Ian Johnston, Dominic Kwiatkowski, Cordelia Langford, John Sillitoe on behalf of the Wellcome Sanger Institute COVID-19 Surveillance Team |
| EPI_ISL_2807681 | Lighthouse Lab in Alderley Park | Wellcome Sanger Institute for the COVID-19 Genomics UK (COG-UK) Consortium | Jacquelyn Wynn, Mairead Hyland, The Lighthouse Lab in Alderley Park and Alex Alderton, Roberto Amato, Jeffrey Barrett, Sonia Goncalves, Ewan Harrison, David K. Jackson, Ian Johnston, Dominic Kwiatkowski, Cordelia Langford, John Sillitoe on behalf of the Wellcome Sanger Institute COVID-19 Surveillance Team |
| EPI_ISL_2807682 | Lighthouse Lab in Glasgow | Wellcome Sanger Institute for the COVID-19 Genomics UK (COG-UK) Consortium | Harper VanSteenhouse, Yumi Kasai, David Gray, Carol Clugston, Anna Dominiczak and Alex Alderton, Roberto Amato, Jeffrey Barrett, Sonia Goncalves, Ewan Harrison, David K. Jackson, Ian Johnston, Dominic Kwiatkowski, Cordelia Langford, John Sillitoe on behalf of the Wellcome Sanger Institute COVID-19 Surveillance Team |
| EPI_ISL_2807683 | Lighthouse Lab in Alderley Park | Wellcome Sanger Institute for the COVID-19 Genomics UK (COG-UK) Consortium | Jacquelyn Wynn, Mairead Hyland, The Lighthouse Lab in Alderley Park and Alex Alderton, Roberto Amato, Jeffrey Barrett, Sonia Goncalves, Ewan Harrison, David K. Jackson, Ian Johnston, Dominic Kwiatkowski, Cordelia Langford, John Sillitoe on behalf of the Wellcome Sanger Institute COVID-19 Surveillance Team |
| EPI_ISL_2807684 | Lighthouse Lab in Milton Keynes | Wellcome Sanger Institute for the COVID-19 Genomics UK (COG-UK) Consortium | The Lighthouse Lab in Milton Keynes and Alex Alderton, Roberto Amato, Jeffrey Barrett, Sonia Goncalves, Ewan Harrison, David K. Jackson, Ian Johnston, Dominic Kwiatkowski, Cordelia Langford, John Sillitoe on behalf of the Wellcome Sanger Institute COVID-19 Surveillance Team |
| EPI_ISL_2807685, EPI_ISL_2807686 | Lighthouse Lab in Glasgow | Wellcome Sanger Institute for the COVID-19 Genomics UK (COG-UK) Consortium | Harper VanSteenhouse, Yumi Kasai, David Gray, Carol Clugston, Anna Dominiczak and Alex Alderton, Roberto Amato, Jeffrey Barrett, Sonia Goncalves, Ewan Harrison, David K. Jackson, Ian Johnston, Dominic Kwiatkowski, Cordelia Langford, John Sillitoe on behalf of the Wellcome Sanger Institute COVID-19 Surveillance Team |
| EPI_ISL_2807687 | Lighthouse Lab in Milton Keynes | Wellcome Sanger Institute for the COVID-19 Genomics UK (COG-UK) Consortium | The Lighthouse Lab in Milton Keynes and Alex Alderton, Roberto Amato, Jeffrey Barrett, Sonia Goncalves, Ewan Harrison, David K. Jackson, Ian Johnston, Dominic Kwiatkowski, Cordelia Langford, John Sillitoe on behalf of the Wellcome Sanger Institute COVID-19 Surveillance Team |
| EPI_ISL_2807688 | Lighthouse Lab in Glasgow | Wellcome Sanger Institute for the COVID-19 Genomics UK (COG-UK) Consortium | Harper VanSteenhouse, Yumi Kasai, David Gray, Carol Clugston, Anna Dominiczak and Alex Alderton, Roberto Amato, Jeffrey Barrett, Sonia Goncalves, Ewan Harrison, David K. Jackson, Ian Johnston, Dominic Kwiatkowski, Cordelia Langford, John Sillitoe on behalf of the Wellcome Sanger Institute COVID-19 Surveillance Team |
| EPI_ISL_2807689, EPI_ISL_2807690 | Lighthouse Lab in Milton Keynes | Wellcome Sanger Institute for the COVID-19 Genomics UK (COG-UK) Consortium | The Lighthouse Lab in Milton Keynes and Alex Alderton, Roberto Amato, Jeffrey Barrett, Sonia Goncalves, Ewan Harrison, David K. Jackson, Ian Johnston, Dominic Kwiatkowski, Cordelia Langford, John Sillitoe on behalf of the Wellcome Sanger Institute COVID-19 Surveillance Team |
| EPI_ISL_2807691 | Lighthouse Lab in Alderley Park | Wellcome Sanger Institute for the COVID-19 Genomics UK (COG-UK) Consortium | Jacquelyn Wynn, Mairead Hyland, The Lighthouse Lab in Alderley Park and Alex Alderton, Roberto Amato, Jeffrey Barrett, Sonia Goncalves, Ewan Harrison, David K. Jackson, Ian Johnston, Dominic Kwiatkowski, Cordelia Langford, John Sillitoe on behalf of the Wellcome Sanger Institute COVID-19 Surveillance Team |
| EPI_ISL_2807692, EPI_ISL_2807693, EPI_ISL_2807694, EPI_ISL_2807695 | Lighthouse Lab in Milton Keynes | Wellcome Sanger Institute for the COVID-19 Genomics UK (COG-UK) Consortium | The Lighthouse Lab in Milton Keynes and Alex Alderton, Roberto Amato, Jeffrey Barrett, Sonia Goncalves, Ewan Harrison, David K. Jackson, Ian Johnston, Dominic Kwiatkowski, Cordelia Langford, John Sillitoe on behalf of the Wellcome Sanger Institute COVID-19 Surveillance Team |
| EPI_ISL_2807696 | Lighthouse Lab in Alderley Park | Wellcome Sanger Institute for the COVID-19 Genomics UK (COG-UK) Consortium | Jacquelyn Wynn, Mairead Hyland, The Lighthouse Lab in Alderley Park and Alex Alderton, Roberto Amato, Jeffrey Barrett, Sonia Goncalves, Ewan Harrison, David K. Jackson, Ian Johnston, Dominic Kwiatkowski, Cordelia Langford, John Sillitoe on behalf of the Wellcome Sanger Institute COVID-19 Surveillance Team |
| EPI_ISL_2807697, EPI_ISL_2807698 | Lighthouse Lab in Milton Keynes | Wellcome Sanger Institute for the COVID-19 Genomics UK (COG-UK) Consortium | The Lighthouse Lab in Milton Keynes and Alex Alderton, Roberto Amato, Jeffrey Barrett, Sonia Goncalves, Ewan Harrison, David K. Jackson, Ian Johnston, Dominic Kwiatkowski, Cordelia Langford, John Sillitoe on behalf of the Wellcome Sanger Institute COVID-19 Surveillance Team |
| EPI_ISL_2807700, EPI_ISL_2807701 | Lighthouse Lab in Alderley Park | Wellcome Sanger Institute for the COVID-19 Genomics UK (COG-UK) Consortium | Jacquelyn Wynn, Mairead Hyland, The Lighthouse Lab in Alderley Park and Alex Alderton, Roberto Amato, Jeffrey Barrett, Sonia Goncalves, Ewan Harrison, David K. Jackson, Ian Johnston, Dominic Kwiatkowski, Cordelia Langford, John Sillitoe on behalf of the Wellcome Sanger Institute COVID-19 Surveillance Team |
| EPI_ISL_2807702 | Lighthouse Lab in Milton Keynes | Wellcome Sanger Institute for the COVID-19 Genomics UK (COG-UK) Consortium | The Lighthouse Lab in Milton Keynes and Alex Alderton, Roberto Amato, Jeffrey Barrett, Sonia Goncalves, Ewan Harrison, David K. Jackson, Ian Johnston, Dominic Kwiatkowski, Cordelia Langford, John Sillitoe on behalf of the Wellcome Sanger Institute COVID-19 Surveillance Team |
| EPI_ISL_2807703 | Lighthouse Lab in Alderley Park | Wellcome Sanger Institute for the COVID-19 Genomics UK (COG-UK) Consortium | Jacquelyn Wynn, Mairead Hyland, The Lighthouse Lab in Alderley Park and Alex Alderton, Roberto Amato, Jeffrey Barrett, Sonia Goncalves, Ewan Harrison, David K. Jackson, Ian Johnston, Dominic Kwiatkowski, Cordelia Langford, John Sillitoe on behalf of the Wellcome Sanger Institute COVID-19 Surveillance Team |
| EPI_ISL_2807704, EPI_ISL_2807705, EPI_ISL_2807706, EPI_ISL_2807707, EPI_ISL_2807708, EPI_ISL_2807709, EPI_ISL_2807710, EPI_ISL_2807711 | Lighthouse Lab in Milton Keynes | Wellcome Sanger Institute for the COVID-19 Genomics UK (COG-UK) Consortium | The Lighthouse Lab in Milton Keynes and Alex Alderton, Roberto Amato, Jeffrey Barrett, Sonia Goncalves, Ewan Harrison, David K. Jackson, Ian Johnston, Dominic Kwiatkowski, Cordelia Langford, John Sillitoe on behalf of the Wellcome Sanger Institute COVID-19 Surveillance Team |
| EPI_ISL_2807712 | Lighthouse Lab in Glasgow | Wellcome Sanger Institute for the COVID-19 Genomics UK (COG-UK) Consortium | Harper VanSteenhouse, Yumi Kasai, David Gray, Carol Clugston, Anna Dominiczak and Alex Alderton, Roberto Amato, Jeffrey Barrett, Sonia Goncalves, Ewan Harrison, David K. Jackson, Ian Johnston, Dominic Kwiatkowski, Cordelia Langford, John Sillitoe on behalf of the Wellcome Sanger Institute COVID-19 Surveillance Team |
| EPI_ISL_2807713 | Lighthouse Lab in Milton Keynes | Wellcome Sanger Institute for the COVID-19 Genomics UK (COG-UK) Consortium | The Lighthouse Lab in Milton Keynes and Alex Alderton, Roberto Amato, Jeffrey Barrett, Sonia Goncalves, Ewan Harrison, David K. Jackson, Ian Johnston, Dominic Kwiatkowski, Cordelia Langford, John Sillitoe on behalf of the Wellcome Sanger Institute COVID-19 Surveillance Team |
| EPI_ISL_2807714 | Lighthouse Lab in Glasgow | Wellcome Sanger Institute for the COVID-19 Genomics UK (COG-UK) Consortium | Harper VanSteenhouse, Yumi Kasai, David Gray, Carol Clugston, Anna Dominiczak and Alex Alderton, Roberto Amato, Jeffrey Barrett, Sonia Goncalves, Ewan Harrison, David K. Jackson, Ian Johnston, Dominic Kwiatkowski, Cordelia Langford, John Sillitoe on behalf of the Wellcome Sanger Institute COVID-19 Surveillance Team |
| EPI_ISL_2807715 | Lighthouse Lab in Milton Keynes | Wellcome Sanger Institute for the COVID-19 Genomics UK (COG-UK) Consortium | The Lighthouse Lab in Milton Keynes and Alex Alderton, Roberto Amato, Jeffrey Barrett, Sonia Goncalves, Ewan Harrison, David K. Jackson, Ian Johnston, Dominic Kwiatkowski, Cordelia Langford, John Sillitoe on behalf of the Wellcome Sanger Institute COVID-19 Surveillance Team |
| EPI_ISL_2807716, EPI_ISL_2807717, EPI_ISL_2807718, EPI_ISL_2807719, EPI_ISL_2807720, EPI_ISL_2807721, EPI_ISL_2807722, EPI_ISL_2807723, EPI_ISL_2807724, EPI_ISL_2807725, EPI_ISL_2807726 | | | |
| see above | Lighthouse Lab in Glasgow | Wellcome Sanger Institute for the COVID-19 Genomics UK (COG-UK) Consortium | Harper VanSteenhouse, Yumi Kasai, David Gray, Carol Clugston, Anna Dominiczak and Alex Alderton, Roberto Amato, Jeffrey Barrett, Sonia Goncalves, Ewan Harrison, David K. Jackson, Ian Johnston, Dominic Kwiatkowski, Cordelia Langford, John Sillitoe on behalf of the Wellcome Sanger Institute COVID-19 Surveillance Team |
| EPI_ISL_2807727, EPI_ISL_2807728 | Lighthouse Lab in Milton Keynes | Wellcome Sanger Institute for the COVID-19 Genomics UK (COG-UK) Consortium | The Lighthouse Lab in Milton Keynes and Alex Alderton, Roberto Amato, Jeffrey Barrett, Sonia Goncalves, Ewan Harrison, David K. Jackson, Ian Johnston, Dominic Kwiatkowski, Cordelia Langford, John Sillitoe on behalf of the Wellcome Sanger Institute COVID-19 Surveillance Team |
| EPI_ISL_2807729, EPI_ISL_2807730, EPI_ISL_2807731, EPI_ISL_2807732 | Lighthouse Lab in Glasgow | Wellcome Sanger Institute for the COVID-19 Genomics UK (COG-UK) Consortium | Harper VanSteenhouse, Yumi Kasai, David Gray, Carol Clugston, Anna Dominiczak and Alex Alderton, Roberto Amato, Jeffrey Barrett, Sonia Goncalves, Ewan Harrison, David K. Jackson, Ian Johnston, Dominic Kwiatkowski, Cordelia Langford, John Sillitoe on behalf of the Wellcome Sanger Institute COVID-19 Surveillance Team |
| EPI_ISL_2807733, EPI_ISL_2807734, | Lighthouse Lab in Milton Keynes | Wellcome Sanger Institute for the COVID-19 Genomics UK (COG-UK) Consortium | The Lighthouse Lab in Milton Keynes and Alex Alderton, Roberto Amato, Jeffrey Barrett, Sonia Goncalves, Ewan Harrison, David K. Jackson, Ian Johnston, Dominic Kwiatkowski, Cordelia Langford, John Sillitoe on behalf of the Wellcome Sanger Institute COVID-19 Surveillance Team |

| | | | |
|---|---|---|---|
| EPI_ISL_2807735, EPI_ISL_2807736, EPI_ISL_2807738 | | | |
| EPI_ISL_2807739 | Lighthouse Lab in Glasgow | Wellcome Sanger Institute for the COVID-19 Genomics UK (COG-UK) Consortium | Harper VanSteenhouse, Yumi Kasai, David Gray, Carol Clugston, Anna Dominiczak and Alex Alderton, Roberto Amato, Jeffrey Barrett, Sonia Goncalves, Ewan Harrison, David K. Jackson, Ian Johnston, Dominic Kwiatkowski, Cordelia Langford, John Sillitoe on behalf of the Wellcome Sanger Institute COVID-19 Surveillance Team |
| EPI_ISL_2807740 | Lighthouse Lab in Milton Keynes | Wellcome Sanger Institute for the COVID-19 Genomics UK (COG-UK) Consortium | The Lighthouse Lab in Milton Keynes and Alex Alderton, Roberto Amato, Jeffrey Barrett, Sonia Goncalves, Ewan Harrison, David K. Jackson, Ian Johnston, Dominic Kwiatkowski, Cordelia Langford, John Sillitoe on behalf of the Wellcome Sanger Institute COVID-19 Surveillance Team |
| EPI_ISL_2807741, EPI_ISL_2807742 | Lighthouse Lab in Glasgow | Wellcome Sanger Institute for the COVID-19 Genomics UK (COG-UK) Consortium | Harper VanSteenhouse, Yumi Kasai, David Gray, Carol Clugston, Anna Dominiczak and Alex Alderton, Roberto Amato, Jeffrey Barrett, Sonia Goncalves, Ewan Harrison, David K. Jackson, Ian Johnston, Dominic Kwiatkowski, Cordelia Langford, John Sillitoe on behalf of the Wellcome Sanger Institute COVID-19 Surveillance Team |
| EPI_ISL_2807743, EPI_ISL_2807744, EPI_ISL_2807745 | Lighthouse Lab in Milton Keynes | Wellcome Sanger Institute for the COVID-19 Genomics UK (COG-UK) Consortium | The Lighthouse Lab in Milton Keynes and Alex Alderton, Roberto Amato, Jeffrey Barrett, Sonia Goncalves, Ewan Harrison, David K. Jackson, Ian Johnston, Dominic Kwiatkowski, Cordelia Langford, John Sillitoe on behalf of the Wellcome Sanger Institute COVID-19 Surveillance Team |
| EPI_ISL_2807746, EPI_ISL_2807747 | Lighthouse Lab in Glasgow | Wellcome Sanger Institute for the COVID-19 Genomics UK (COG-UK) Consortium | Harper VanSteenhouse, Yumi Kasai, David Gray, Carol Clugston, Anna Dominiczak and Alex Alderton, Roberto Amato, Jeffrey Barrett, Sonia Goncalves, Ewan Harrison, David K. Jackson, Ian Johnston, Dominic Kwiatkowski, Cordelia Langford, John Sillitoe on behalf of the Wellcome Sanger Institute COVID-19 Surveillance Team |
| EPI_ISL_2807748, EPI_ISL_2807749 | Lighthouse Lab in Milton Keynes | Wellcome Sanger Institute for the COVID-19 Genomics UK (COG-UK) Consortium | The Lighthouse Lab in Milton Keynes and Alex Alderton, Roberto Amato, Jeffrey Barrett, Sonia Goncalves, Ewan Harrison, David K. Jackson, Ian Johnston, Dominic Kwiatkowski, Cordelia Langford, John Sillitoe on behalf of the Wellcome Sanger Institute COVID-19 Surveillance Team |
| EPI_ISL_2807750, EPI_ISL_2807751 | Lighthouse Lab in Glasgow | Wellcome Sanger Institute for the COVID-19 Genomics UK (COG-UK) Consortium | Harper VanSteenhouse, Yumi Kasai, David Gray, Carol Clugston, Anna Dominiczak and Alex Alderton, Roberto Amato, Jeffrey Barrett, Sonia Goncalves, Ewan Harrison, David K. Jackson, Ian Johnston, Dominic Kwiatkowski, Cordelia Langford, John Sillitoe on behalf of the Wellcome Sanger Institute COVID-19 Surveillance Team |
| EPI_ISL_2807752 | Lighthouse Lab in Milton Keynes | Wellcome Sanger Institute for the COVID-19 Genomics UK (COG-UK) Consortium | The Lighthouse Lab in Milton Keynes and Alex Alderton, Roberto Amato, Jeffrey Barrett, Sonia Goncalves, Ewan Harrison, David K. Jackson, Ian Johnston, Dominic Kwiatkowski, Cordelia Langford, John Sillitoe on behalf of the Wellcome Sanger Institute COVID-19 Surveillance Team |
| EPI_ISL_2807753 | Lighthouse Lab in Glasgow | Wellcome Sanger Institute for the COVID-19 Genomics UK (COG-UK) Consortium | Harper VanSteenhouse, Yumi Kasai, David Gray, Carol Clugston, Anna Dominiczak and Alex Alderton, Roberto Amato, Jeffrey Barrett, Sonia Goncalves, Ewan Harrison, David K. Jackson, Ian Johnston, Dominic Kwiatkowski, Cordelia Langford, John Sillitoe on behalf of the Wellcome Sanger Institute COVID-19 Surveillance Team |
| EPI_ISL_2807754 | Lighthouse Lab in Milton Keynes | Wellcome Sanger Institute for the COVID-19 Genomics UK (COG-UK) Consortium | The Lighthouse Lab in Milton Keynes and Alex Alderton, Roberto Amato, Jeffrey Barrett, Sonia Goncalves, Ewan Harrison, David K. Jackson, Ian Johnston, Dominic Kwiatkowski, Cordelia Langford, John Sillitoe on behalf of the Wellcome Sanger Institute COVID-19 Surveillance Team |
| EPI_ISL_2807755 | Lighthouse Lab in Glasgow | Wellcome Sanger Institute for the COVID-19 Genomics UK (COG-UK) Consortium | Harper VanSteenhouse, Yumi Kasai, David Gray, Carol Clugston, Anna Dominiczak and Alex Alderton, Roberto Amato, Jeffrey Barrett, Sonia Goncalves, Ewan Harrison, David K. Jackson, Ian Johnston, Dominic Kwiatkowski, Cordelia Langford, John Sillitoe on behalf of the Wellcome Sanger Institute COVID-19 Surveillance Team |
| EPI_ISL_2807756 | Lighthouse Lab in Milton Keynes | Wellcome Sanger Institute for the COVID-19 Genomics UK (COG-UK) Consortium | The Lighthouse Lab in Milton Keynes and Alex Alderton, Roberto Amato, Jeffrey Barrett, Sonia Goncalves, Ewan Harrison, David K. Jackson, Ian Johnston, Dominic Kwiatkowski, Cordelia Langford, John Sillitoe on behalf of the Wellcome Sanger Institute COVID-19 Surveillance Team |
| EPI_ISL_2807757 | Lighthouse Lab in Glasgow | Wellcome Sanger Institute for the COVID-19 Genomics UK (COG-UK) Consortium | Harper VanSteenhouse, Yumi Kasai, David Gray, Carol Clugston, Anna Dominiczak and Alex Alderton, Roberto Amato, Jeffrey Barrett, Sonia Goncalves, Ewan Harrison, David K. Jackson, Ian Johnston, Dominic Kwiatkowski, Cordelia Langford, John Sillitoe on behalf of the Wellcome Sanger Institute COVID-19 Surveillance Team |
| EPI_ISL_2807758 | Lighthouse Lab in Milton Keynes | Wellcome Sanger Institute for the COVID-19 Genomics UK (COG-UK) Consortium | The Lighthouse Lab in Milton Keynes and Alex Alderton, Roberto Amato, Jeffrey Barrett, Sonia Goncalves, Ewan Harrison, David K. Jackson, Ian Johnston, Dominic Kwiatkowski, Cordelia Langford, John Sillitoe on behalf of the Wellcome Sanger Institute COVID-19 Surveillance Team |
| EPI_ISL_2807759, EPI_ISL_2807760, EPI_ISL_2807761 | Lighthouse Lab in Glasgow | Wellcome Sanger Institute for the COVID-19 Genomics UK (COG-UK) Consortium | Harper VanSteenhouse, Yumi Kasai, David Gray, Carol Clugston, Anna Dominiczak and Alex Alderton, Roberto Amato, Jeffrey Barrett, Sonia Goncalves, Ewan Harrison, David K. Jackson, Ian Johnston, Dominic Kwiatkowski, Cordelia Langford, John Sillitoe on behalf of the Wellcome Sanger Institute COVID-19 Surveillance Team |
| EPI_ISL_2807762, EPI_ISL_2807763, EPI_ISL_2807764 | Lighthouse Lab in Milton Keynes | Wellcome Sanger Institute for the COVID-19 Genomics UK (COG-UK) Consortium | The Lighthouse Lab in Milton Keynes and Alex Alderton, Roberto Amato, Jeffrey Barrett, Sonia Goncalves, Ewan Harrison, David K. Jackson, Ian Johnston, Dominic Kwiatkowski, Cordelia Langford, John Sillitoe on behalf of the Wellcome Sanger Institute COVID-19 Surveillance Team |
| EPI_ISL_2807765 | Lighthouse Lab in Glasgow | Wellcome Sanger Institute for the COVID-19 Genomics UK (COG-UK) Consortium | Harper VanSteenhouse, Yumi Kasai, David Gray, Carol Clugston, Anna Dominiczak and Alex Alderton, Roberto Amato, Jeffrey Barrett, Sonia Goncalves, Ewan Harrison, David K. Jackson, Ian Johnston, Dominic Kwiatkowski, Cordelia Langford, John Sillitoe on behalf of the Wellcome Sanger Institute COVID-19 Surveillance Team |
| EPI_ISL_2807766 | Lighthouse Lab in Milton Keynes | Wellcome Sanger Institute for the COVID-19 Genomics UK (COG-UK) Consortium | The Lighthouse Lab in Milton Keynes and Alex Alderton, Roberto Amato, Jeffrey Barrett, Sonia Goncalves, Ewan Harrison, David K. Jackson, Ian Johnston, Dominic Kwiatkowski, Cordelia Langford, John Sillitoe on behalf of the Wellcome Sanger Institute COVID-19 Surveillance Team |
| EPI_ISL_2807767, EPI_ISL_2807768, EPI_ISL_2807769 | Lighthouse Lab in Glasgow | Wellcome Sanger Institute for the COVID-19 Genomics UK (COG-UK) Consortium | Harper VanSteenhouse, Yumi Kasai, David Gray, Carol Clugston, Anna Dominiczak and Alex Alderton, Roberto Amato, Jeffrey Barrett, Sonia Goncalves, Ewan Harrison, David K. Jackson, Ian Johnston, Dominic Kwiatkowski, Cordelia Langford, John Sillitoe on behalf of the Wellcome Sanger Institute COVID-19 Surveillance Team |
| EPI_ISL_2807770 | Lighthouse Lab in Milton Keynes | Wellcome Sanger Institute for the COVID-19 Genomics UK (COG-UK) Consortium | The Lighthouse Lab in Milton Keynes and Alex Alderton, Roberto Amato, Jeffrey Barrett, Sonia Goncalves, Ewan Harrison, David K. Jackson, Ian Johnston, Dominic Kwiatkowski, Cordelia Langford, John Sillitoe on behalf of the Wellcome Sanger Institute COVID-19 Surveillance Team |
| EPI_ISL_2807771, EPI_ISL_2807772, EPI_ISL_2807773 | Lighthouse Lab in Glasgow | Wellcome Sanger Institute for the COVID-19 Genomics UK (COG-UK) Consortium | Harper VanSteenhouse, Yumi Kasai, David Gray, Carol Clugston, Anna Dominiczak and Alex Alderton, Roberto Amato, Jeffrey Barrett, Sonia Goncalves, Ewan Harrison, David K. Jackson, Ian Johnston, Dominic Kwiatkowski, Cordelia Langford, John Sillitoe on behalf of the Wellcome Sanger Institute COVID-19 Surveillance Team |
| EPI_ISL_2807774, EPI_ISL_2807775 | Lighthouse Lab in Milton Keynes | Wellcome Sanger Institute for the COVID-19 Genomics UK (COG-UK) Consortium | The Lighthouse Lab in Milton Keynes and Alex Alderton, Roberto Amato, Jeffrey Barrett, Sonia Goncalves, Ewan Harrison, David K. Jackson, Ian Johnston, Dominic Kwiatkowski, Cordelia Langford, John Sillitoe on behalf of the Wellcome Sanger Institute COVID-19 Surveillance Team |
| EPI_ISL_2807776, EPI_ISL_2807777, EPI_ISL_2807778 | Lighthouse Lab in Glasgow | Wellcome Sanger Institute for the COVID-19 Genomics UK (COG-UK) Consortium | Harper VanSteenhouse, Yumi Kasai, David Gray, Carol Clugston, Anna Dominiczak and Alex Alderton, Roberto Amato, Jeffrey Barrett, Sonia Goncalves, Ewan Harrison, David K. Jackson, Ian Johnston, Dominic Kwiatkowski, Cordelia Langford, John Sillitoe on behalf of the Wellcome Sanger Institute COVID-19 Surveillance Team |
| EPI_ISL_2807779, EPI_ISL_2807780 | Lighthouse Lab in Milton Keynes | Wellcome Sanger Institute for the COVID-19 Genomics UK (COG-UK) Consortium | The Lighthouse Lab in Milton Keynes and Alex Alderton, Roberto Amato, Jeffrey Barrett, Sonia Goncalves, Ewan Harrison, David K. Jackson, Ian Johnston, Dominic Kwiatkowski, Cordelia Langford, John Sillitoe on behalf of the Wellcome Sanger Institute COVID-19 Surveillance Team |
| EPI_ISL_2807781, EPI_ISL_2807782, EPI_ISL_2807783, EPI_ISL_2807784, EPI_ISL_2807785 | Lighthouse Lab in Glasgow | Wellcome Sanger Institute for the COVID-19 Genomics UK (COG-UK) Consortium | Harper VanSteenhouse, Yumi Kasai, David Gray, Carol Clugston, Anna Dominiczak and Alex Alderton, Roberto Amato, Jeffrey Barrett, Sonia Goncalves, Ewan Harrison, David K. Jackson, Ian Johnston, Dominic Kwiatkowski, Cordelia Langford, John Sillitoe on behalf of the Wellcome Sanger Institute COVID-19 Surveillance Team |
| EPI_ISL_2807786 | Lighthouse Lab in Milton Keynes | Wellcome Sanger Institute for the COVID-19 Genomics UK (COG-UK) Consortium | The Lighthouse Lab in Milton Keynes and Alex Alderton, Roberto Amato, Jeffrey Barrett, Sonia Goncalves, Ewan Harrison, David K. Jackson, Ian Johnston, Dominic Kwiatkowski, Cordelia Langford, John Sillitoe on behalf of the Wellcome Sanger Institute COVID-19 Surveillance Team |
| EPI_ISL_2807787, EPI_ISL_2807788 | Lighthouse Lab in Glasgow | Wellcome Sanger Institute for the COVID-19 Genomics UK (COG-UK) Consortium | Harper VanSteenhouse, Yumi Kasai, David Gray, Carol Clugston, Anna Dominiczak and Alex Alderton, Roberto Amato, Jeffrey Barrett, Sonia Goncalves, Ewan Harrison, David K. Jackson, Ian Johnston, Dominic Kwiatkowski, Cordelia Langford, John Sillitoe on behalf of the Wellcome Sanger Institute COVID-19 Surveillance Team |
| EPI_ISL_2807789 | Lighthouse Lab in Milton Keynes | Wellcome Sanger Institute for the COVID-19 Genomics UK (COG-UK) Consortium | The Lighthouse Lab in Milton Keynes and Alex Alderton, Roberto Amato, Jeffrey Barrett, Sonia Goncalves, Ewan Harrison, David K. Jackson, Ian Johnston, Dominic Kwiatkowski, Cordelia Langford, John Sillitoe on behalf of the Wellcome Sanger Institute COVID-19 Surveillance Team |
| EPI_ISL_2807790, EPI_ISL_2807791, EPI_ISL_2807792, EPI_ISL_2807793 | Lighthouse Lab in Glasgow | Wellcome Sanger Institute for the COVID-19 Genomics UK (COG-UK) Consortium | Harper VanSteenhouse, Yumi Kasai, David Gray, Carol Clugston, Anna Dominiczak and Alex Alderton, Roberto Amato, Jeffrey Barrett, Sonia Goncalves, Ewan Harrison, David K. Jackson, Ian Johnston, Dominic Kwiatkowski, Cordelia Langford, John Sillitoe on behalf of the Wellcome Sanger Institute COVID-19 Surveillance Team |
| EPI_ISL_2807794, EPI_ISL_2807795, EPI_ISL_2807796 | Lighthouse Lab in Milton Keynes | Wellcome Sanger Institute for the COVID-19 Genomics UK (COG-UK) Consortium | The Lighthouse Lab in Milton Keynes and Alex Alderton, Roberto Amato, Jeffrey Barrett, Sonia Goncalves, Ewan Harrison, David K. Jackson, Ian Johnston, Dominic Kwiatkowski, Cordelia Langford, John Sillitoe on behalf of the Wellcome Sanger Institute COVID-19 Surveillance Team |
| EPI_ISL_2807797 | Lighthouse Lab in Glasgow | Wellcome Sanger Institute for the COVID-19 Genomics UK (COG-UK) Consortium | Harper VanSteenhouse, Yumi Kasai, David Gray, Carol Clugston, Anna Dominiczak and Alex Alderton, Roberto Amato, Jeffrey Barrett, Sonia Goncalves, Ewan Harrison, David K. Jackson, Ian Johnston, Dominic Kwiatkowski, Cordelia Langford, John Sillitoe on behalf of the Wellcome Sanger Institute COVID-19 Surveillance Team |
| EPI_ISL_2807798, EPI_ISL_2807799 | Lighthouse Lab in Milton Keynes | Wellcome Sanger Institute for the COVID-19 Genomics UK (COG-UK) Consortium | The Lighthouse Lab in Milton Keynes and Alex Alderton, Roberto Amato, Jeffrey Barrett, Sonia Goncalves, Ewan Harrison, David K. Jackson, Ian Johnston, Dominic Kwiatkowski, Cordelia Langford, John Sillitoe on behalf of the Wellcome Sanger Institute COVID-19 Surveillance Team |
| EPI_ISL_2807800 | Lighthouse Lab in Glasgow | Wellcome Sanger Institute for the COVID-19 Genomics UK (COG-UK) Consortium | Harper VanSteenhouse, Yumi Kasai, David Gray, Carol Clugston, Anna Dominiczak and Alex Alderton, Roberto Amato, Jeffrey Barrett, Sonia Goncalves, Ewan Harrison, David K. Jackson, Ian Johnston, Dominic Kwiatkowski, Cordelia Langford, John Sillitoe on behalf of the Wellcome Sanger Institute COVID-19 Surveillance Team |
| EPI_ISL_2807801 | Lighthouse Lab in Milton Keynes | Wellcome Sanger Institute for the COVID-19 Genomics UK (COG-UK) Consortium | The Lighthouse Lab in Milton Keynes and Alex Alderton, Roberto Amato, Jeffrey Barrett, Sonia Goncalves, Ewan Harrison, David K. Jackson, Ian Johnston, Dominic Kwiatkowski, Cordelia Langford, John Sillitoe on behalf of the Wellcome Sanger Institute COVID-19 Surveillance Team |
| EPI_ISL_2807802, EPI_ISL_2807803 | Lighthouse Lab in Glasgow | Wellcome Sanger Institute for the COVID-19 Genomics UK (COG-UK) Consortium | Harper VanSteenhouse, Yumi Kasai, David Gray, Carol Clugston, Anna Dominiczak and Alex Alderton, Roberto Amato, Jeffrey Barrett, Sonia Goncalves, Ewan Harrison, David K. Jackson, Ian Johnston, Dominic Kwiatkowski, Cordelia Langford, John Sillitoe on behalf of the Wellcome Sanger Institute COVID-19 Surveillance Team |
| EPI_ISL_2807804, EPI_ISL_2807805 | Lighthouse Lab in Milton Keynes | Wellcome Sanger Institute for the COVID-19 Genomics UK (COG-UK) Consortium | The Lighthouse Lab in Milton Keynes and Alex Alderton, Roberto Amato, Jeffrey Barrett, Sonia Goncalves, Ewan Harrison, David K. Jackson, Ian Johnston, Dominic Kwiatkowski, Cordelia Langford, John Sillitoe on behalf of the Wellcome Sanger Institute COVID-19 Surveillance Team |
| EPI_ISL_2807806 | Lighthouse Lab in Glasgow | Wellcome Sanger Institute for the COVID-19 Genomics UK (COG-UK) Consortium | Harper VanSteenhouse, Yumi Kasai, David Gray, Carol Clugston, Anna Dominiczak and Alex Alderton, Roberto Amato, Jeffrey Barrett, Sonia Goncalves, Ewan Harrison, David K. Jackson, Ian Johnston, Dominic Kwiatkowski, Cordelia Langford, John Sillitoe on behalf of the Wellcome Sanger Institute COVID-19 Surveillance Team |
| EPI_ISL_2807807 | Lighthouse Lab in Milton Keynes | Wellcome Sanger Institute for the COVID-19 Genomics UK (COG-UK) Consortium | The Lighthouse Lab in Milton Keynes and Alex Alderton, Roberto Amato, Jeffrey Barrett, Sonia Goncalves, Ewan Harrison, David K. Jackson, Ian Johnston, Dominic Kwiatkowski, Cordelia Langford, John Sillitoe on behalf of the Wellcome Sanger Institute COVID-19 Surveillance Team |
| EPI_ISL_2807808 | Lighthouse Lab in Glasgow | Wellcome Sanger Institute for the COVID-19 Genomics UK (COG-UK) | Harper VanSteenhouse, Yumi Kasai, David Gray, Carol Clugston, Anna Dominiczak and Alex Alderton, Roberto Amato, Jeffrey Barrett, Sonia Goncalves, Ewan Harrison, David K. Jackson, Ian Johnston, Dominic Kwiatkowski, Cordelia Langford, |

| | | | |
|---|---|---|---|
| | | Consortium | John Sillitoe on behalf of the Wellcome Sanger Institute COVID-19 Surveillance Team |
| EPI_ISL_2807809 | Lighthouse Lab in Milton Keynes | Wellcome Sanger Institute for the COVID-19 Genomics UK (COG-UK) Consortium | The Lighthouse Lab in Milton Keynes and Alex Alderton, Roberto Amato, Jeffrey Barrett, Sonia Goncalves, Ewan Harrison, David K. Jackson, Ian Johnston, Dominic Kwiatkowski, Cordelia Langford, John Sillitoe on behalf of the Wellcome Sanger Institute COVID-19 Surveillance Team |
| EPI_ISL_2807810 | Lighthouse Lab in Glasgow | Wellcome Sanger Institute for the COVID-19 Genomics UK (COG-UK) Consortium | Harper VanSteenhouse, Yumi Kasai, David Gray, Carol Clugston, Anna Dominiczak and Alex Alderton, Roberto Amato, Jeffrey Barrett, Sonia Goncalves, Ewan Harrison, David K. Jackson, Ian Johnston, Dominic Kwiatkowski, Cordelia Langford, John Sillitoe on behalf of the Wellcome Sanger Institute COVID-19 Surveillance Team |
| EPI_ISL_2807811, EPI_ISL_2807812 | Lighthouse Lab in Milton Keynes | Wellcome Sanger Institute for the COVID-19 Genomics UK (COG-UK) Consortium | The Lighthouse Lab in Milton Keynes and Alex Alderton, Roberto Amato, Jeffrey Barrett, Sonia Goncalves, Ewan Harrison, David K. Jackson, Ian Johnston, Dominic Kwiatkowski, Cordelia Langford, John Sillitoe on behalf of the Wellcome Sanger Institute COVID-19 Surveillance Team |
| EPI_ISL_2807813 | Lighthouse Lab in Glasgow | Wellcome Sanger Institute for the COVID-19 Genomics UK (COG-UK) Consortium | Harper VanSteenhouse, Yumi Kasai, David Gray, Carol Clugston, Anna Dominiczak and Alex Alderton, Roberto Amato, Jeffrey Barrett, Sonia Goncalves, Ewan Harrison, David K. Jackson, Ian Johnston, Dominic Kwiatkowski, Cordelia Langford, John Sillitoe on behalf of the Wellcome Sanger Institute COVID-19 Surveillance Team |
| EPI_ISL_2807814, EPI_ISL_2807815, EPI_ISL_2807816 | Lighthouse Lab in Milton Keynes | Wellcome Sanger Institute for the COVID-19 Genomics UK (COG-UK) Consortium | The Lighthouse Lab in Milton Keynes and Alex Alderton, Roberto Amato, Jeffrey Barrett, Sonia Goncalves, Ewan Harrison, David K. Jackson, Ian Johnston, Dominic Kwiatkowski, Cordelia Langford, John Sillitoe on behalf of the Wellcome Sanger Institute COVID-19 Surveillance Team |
| EPI_ISL_2807817 | Lighthouse Lab in Glasgow | Wellcome Sanger Institute for the COVID-19 Genomics UK (COG-UK) Consortium | Harper VanSteenhouse, Yumi Kasai, David Gray, Carol Clugston, Anna Dominiczak and Alex Alderton, Roberto Amato, Jeffrey Barrett, Sonia Goncalves, Ewan Harrison, David K. Jackson, Ian Johnston, Dominic Kwiatkowski, Cordelia Langford, John Sillitoe on behalf of the Wellcome Sanger Institute COVID-19 Surveillance Team |
| EPI_ISL_2807818 | Lighthouse Lab in Milton Keynes | Wellcome Sanger Institute for the COVID-19 Genomics UK (COG-UK) Consortium | The Lighthouse Lab in Milton Keynes and Alex Alderton, Roberto Amato, Jeffrey Barrett, Sonia Goncalves, Ewan Harrison, David K. Jackson, Ian Johnston, Dominic Kwiatkowski, Cordelia Langford, John Sillitoe on behalf of the Wellcome Sanger Institute COVID-19 Surveillance Team |
| EPI_ISL_2807819 | Lighthouse Lab in Glasgow | Wellcome Sanger Institute for the COVID-19 Genomics UK (COG-UK) Consortium | Harper VanSteenhouse, Yumi Kasai, David Gray, Carol Clugston, Anna Dominiczak and Alex Alderton, Roberto Amato, Jeffrey Barrett, Sonia Goncalves, Ewan Harrison, David K. Jackson, Ian Johnston, Dominic Kwiatkowski, Cordelia Langford, John Sillitoe on behalf of the Wellcome Sanger Institute COVID-19 Surveillance Team |
| EPI_ISL_2807820, EPI_ISL_2807821 | Lighthouse Lab in Milton Keynes | Wellcome Sanger Institute for the COVID-19 Genomics UK (COG-UK) Consortium | The Lighthouse Lab in Milton Keynes and Alex Alderton, Roberto Amato, Jeffrey Barrett, Sonia Goncalves, Ewan Harrison, David K. Jackson, Ian Johnston, Dominic Kwiatkowski, Cordelia Langford, John Sillitoe on behalf of the Wellcome Sanger Institute COVID-19 Surveillance Team |
| EPI_ISL_2807822, EPI_ISL_2807823 | Lighthouse Lab in Glasgow | Wellcome Sanger Institute for the COVID-19 Genomics UK (COG-UK) Consortium | Harper VanSteenhouse, Yumi Kasai, David Gray, Carol Clugston, Anna Dominiczak and Alex Alderton, Roberto Amato, Jeffrey Barrett, Sonia Goncalves, Ewan Harrison, David K. Jackson, Ian Johnston, Dominic Kwiatkowski, Cordelia Langford, John Sillitoe on behalf of the Wellcome Sanger Institute COVID-19 Surveillance Team |
| EPI_ISL_2807824 | Lighthouse Lab in Milton Keynes | Wellcome Sanger Institute for the COVID-19 Genomics UK (COG-UK) Consortium | The Lighthouse Lab in Milton Keynes and Alex Alderton, Roberto Amato, Jeffrey Barrett, Sonia Goncalves, Ewan Harrison, David K. Jackson, Ian Johnston, Dominic Kwiatkowski, Cordelia Langford, John Sillitoe on behalf of the Wellcome Sanger Institute COVID-19 Surveillance Team |
| EPI_ISL_2807825 | Lighthouse Lab in Glasgow | Wellcome Sanger Institute for the COVID-19 Genomics UK (COG-UK) Consortium | Harper VanSteenhouse, Yumi Kasai, David Gray, Carol Clugston, Anna Dominiczak and Alex Alderton, Roberto Amato, Jeffrey Barrett, Sonia Goncalves, Ewan Harrison, David K. Jackson, Ian Johnston, Dominic Kwiatkowski, Cordelia Langford, John Sillitoe on behalf of the Wellcome Sanger Institute COVID-19 Surveillance Team |
| EPI_ISL_2807826 | Lighthouse Lab in Milton Keynes | Wellcome Sanger Institute for the COVID-19 Genomics UK (COG-UK) Consortium | The Lighthouse Lab in Milton Keynes and Alex Alderton, Roberto Amato, Jeffrey Barrett, Sonia Goncalves, Ewan Harrison, David K. Jackson, Ian Johnston, Dominic Kwiatkowski, Cordelia Langford, John Sillitoe on behalf of the Wellcome Sanger Institute COVID-19 Surveillance Team |
| EPI_ISL_2807827 | Lighthouse Lab in Glasgow | Wellcome Sanger Institute for the COVID-19 Genomics UK (COG-UK) Consortium | Harper VanSteenhouse, Yumi Kasai, David Gray, Carol Clugston, Anna Dominiczak and Alex Alderton, Roberto Amato, Jeffrey Barrett, Sonia Goncalves, Ewan Harrison, David K. Jackson, Ian Johnston, Dominic Kwiatkowski, Cordelia Langford, John Sillitoe on behalf of the Wellcome Sanger Institute COVID-19 Surveillance Team |
| EPI_ISL_2807828 | Lighthouse Lab in Milton Keynes | Wellcome Sanger Institute for the COVID-19 Genomics UK (COG-UK) Consortium | The Lighthouse Lab in Milton Keynes and Alex Alderton, Roberto Amato, Jeffrey Barrett, Sonia Goncalves, Ewan Harrison, David K. Jackson, Ian Johnston, Dominic Kwiatkowski, Cordelia Langford, John Sillitoe on behalf of the Wellcome Sanger Institute COVID-19 Surveillance Team |
| EPI_ISL_2807829, EPI_ISL_2807830 | Lighthouse Lab in Glasgow | Wellcome Sanger Institute for the COVID-19 Genomics UK (COG-UK) Consortium | Harper VanSteenhouse, Yumi Kasai, David Gray, Carol Clugston, Anna Dominiczak and Alex Alderton, Roberto Amato, Jeffrey Barrett, Sonia Goncalves, Ewan Harrison, David K. Jackson, Ian Johnston, Dominic Kwiatkowski, Cordelia Langford, John Sillitoe on behalf of the Wellcome Sanger Institute COVID-19 Surveillance Team |
| EPI_ISL_2807831 | Lighthouse Lab in Milton Keynes | Wellcome Sanger Institute for the COVID-19 Genomics UK (COG-UK) Consortium | The Lighthouse Lab in Milton Keynes and Alex Alderton, Roberto Amato, Jeffrey Barrett, Sonia Goncalves, Ewan Harrison, David K. Jackson, Ian Johnston, Dominic Kwiatkowski, Cordelia Langford, John Sillitoe on behalf of the Wellcome Sanger Institute COVID-19 Surveillance Team |
| EPI_ISL_2807832, EPI_ISL_2807833 | Lighthouse Lab in Glasgow | Wellcome Sanger Institute for the COVID-19 Genomics UK (COG-UK) Consortium | Harper VanSteenhouse, Yumi Kasai, David Gray, Carol Clugston, Anna Dominiczak and Alex Alderton, Roberto Amato, Jeffrey Barrett, Sonia Goncalves, Ewan Harrison, David K. Jackson, Ian Johnston, Dominic Kwiatkowski, Cordelia Langford, John Sillitoe on behalf of the Wellcome Sanger Institute COVID-19 Surveillance Team |
| EPI_ISL_2807834, EPI_ISL_2807835, EPI_ISL_2807836 | Lighthouse Lab in Milton Keynes | Wellcome Sanger Institute for the COVID-19 Genomics UK (COG-UK) Consortium | The Lighthouse Lab in Milton Keynes and Alex Alderton, Roberto Amato, Jeffrey Barrett, Sonia Goncalves, Ewan Harrison, David K. Jackson, Ian Johnston, Dominic Kwiatkowski, Cordelia Langford, John Sillitoe on behalf of the Wellcome Sanger Institute COVID-19 Surveillance Team |
| EPI_ISL_2807837, EPI_ISL_2807838, EPI_ISL_2807839 | Lighthouse Lab in Glasgow | Wellcome Sanger Institute for the COVID-19 Genomics UK (COG-UK) Consortium | Harper VanSteenhouse, Yumi Kasai, David Gray, Carol Clugston, Anna Dominiczak and Alex Alderton, Roberto Amato, Jeffrey Barrett, Sonia Goncalves, Ewan Harrison, David K. Jackson, Ian Johnston, Dominic Kwiatkowski, Cordelia Langford, John Sillitoe on behalf of the Wellcome Sanger Institute COVID-19 Surveillance Team |
| EPI_ISL_2807840, EPI_ISL_2807841 | Lighthouse Lab in Milton Keynes | Wellcome Sanger Institute for the COVID-19 Genomics UK (COG-UK) Consortium | The Lighthouse Lab in Milton Keynes and Alex Alderton, Roberto Amato, Jeffrey Barrett, Sonia Goncalves, Ewan Harrison, David K. Jackson, Ian Johnston, Dominic Kwiatkowski, Cordelia Langford, John Sillitoe on behalf of the Wellcome Sanger Institute COVID-19 Surveillance Team |
| EPI_ISL_2807842 | Lighthouse Lab in Glasgow | Wellcome Sanger Institute for the COVID-19 Genomics UK (COG-UK) Consortium | Harper VanSteenhouse, Yumi Kasai, David Gray, Carol Clugston, Anna Dominiczak and Alex Alderton, Roberto Amato, Jeffrey Barrett, Sonia Goncalves, Ewan Harrison, David K. Jackson, Ian Johnston, Dominic Kwiatkowski, Cordelia Langford, John Sillitoe on behalf of the Wellcome Sanger Institute COVID-19 Surveillance Team |
| EPI_ISL_2807843, EPI_ISL_2807844, EPI_ISL_2807845, EPI_ISL_2807846, EPI_ISL_2807847, EPI_ISL_2807848, EPI_ISL_2807849 | Lighthouse Lab in Milton Keynes | Wellcome Sanger Institute for the COVID-19 Genomics UK (COG-UK) Consortium | The Lighthouse Lab in Milton Keynes and Alex Alderton, Roberto Amato, Jeffrey Barrett, Sonia Goncalves, Ewan Harrison, David K. Jackson, Ian Johnston, Dominic Kwiatkowski, Cordelia Langford, John Sillitoe on behalf of the Wellcome Sanger Institute COVID-19 Surveillance Team |
| EPI_ISL_2807850 | Lighthouse Lab in Glasgow | Wellcome Sanger Institute for the COVID-19 Genomics UK (COG-UK) Consortium | Harper VanSteenhouse, Yumi Kasai, David Gray, Carol Clugston, Anna Dominiczak and Alex Alderton, Roberto Amato, Jeffrey Barrett, Sonia Goncalves, Ewan Harrison, David K. Jackson, Ian Johnston, Dominic Kwiatkowski, Cordelia Langford, John Sillitoe on behalf of the Wellcome Sanger Institute COVID-19 Surveillance Team |
| EPI_ISL_2807851 | Lighthouse Lab in Milton Keynes | Wellcome Sanger Institute for the COVID-19 Genomics UK (COG-UK) Consortium | The Lighthouse Lab in Milton Keynes and Alex Alderton, Roberto Amato, Jeffrey Barrett, Sonia Goncalves, Ewan Harrison, David K. Jackson, Ian Johnston, Dominic Kwiatkowski, Cordelia Langford, John Sillitoe on behalf of the Wellcome Sanger Institute COVID-19 Surveillance Team |
| EPI_ISL_2807852 | Lighthouse Lab in Glasgow | Wellcome Sanger Institute for the COVID-19 Genomics UK (COG-UK) Consortium | Harper VanSteenhouse, Yumi Kasai, David Gray, Carol Clugston, Anna Dominiczak and Alex Alderton, Roberto Amato, Jeffrey Barrett, Sonia Goncalves, Ewan Harrison, David K. Jackson, Ian Johnston, Dominic Kwiatkowski, Cordelia Langford, John Sillitoe on behalf of the Wellcome Sanger Institute COVID-19 Surveillance Team |
| EPI_ISL_2807853 | Lighthouse Lab in Milton Keynes | Wellcome Sanger Institute for the COVID-19 Genomics UK (COG-UK) Consortium | The Lighthouse Lab in Milton Keynes and Alex Alderton, Roberto Amato, Jeffrey Barrett, Sonia Goncalves, Ewan Harrison, David K. Jackson, Ian Johnston, Dominic Kwiatkowski, Cordelia Langford, John Sillitoe on behalf of the Wellcome Sanger Institute COVID-19 Surveillance Team |
| EPI_ISL_2807854, EPI_ISL_2807855 | Lighthouse Lab in Glasgow | Wellcome Sanger Institute for the COVID-19 Genomics UK (COG-UK) Consortium | Harper VanSteenhouse, Yumi Kasai, David Gray, Carol Clugston, Anna Dominiczak and Alex Alderton, Roberto Amato, Jeffrey Barrett, Sonia Goncalves, Ewan Harrison, David K. Jackson, Ian Johnston, Dominic Kwiatkowski, Cordelia Langford, John Sillitoe on behalf of the Wellcome Sanger Institute COVID-19 Surveillance Team |
| EPI_ISL_2807856, EPI_ISL_2807857 | Lighthouse Lab in Milton Keynes | Wellcome Sanger Institute for the COVID-19 Genomics UK (COG-UK) Consortium | The Lighthouse Lab in Milton Keynes and Alex Alderton, Roberto Amato, Jeffrey Barrett, Sonia Goncalves, Ewan Harrison, David K. Jackson, Ian Johnston, Dominic Kwiatkowski, Cordelia Langford, John Sillitoe on behalf of the Wellcome Sanger Institute COVID-19 Surveillance Team |
| EPI_ISL_2807858 | Lighthouse Lab in Glasgow | Wellcome Sanger Institute for the COVID-19 Genomics UK (COG-UK) Consortium | Harper VanSteenhouse, Yumi Kasai, David Gray, Carol Clugston, Anna Dominiczak and Alex Alderton, Roberto Amato, Jeffrey Barrett, Sonia Goncalves, Ewan Harrison, David K. Jackson, Ian Johnston, Dominic Kwiatkowski, Cordelia Langford, John Sillitoe on behalf of the Wellcome Sanger Institute COVID-19 Surveillance Team |
| EPI_ISL_2807859, EPI_ISL_2807860 | Lighthouse Lab in Milton Keynes | Wellcome Sanger Institute for the COVID-19 Genomics UK (COG-UK) Consortium | The Lighthouse Lab in Milton Keynes and Alex Alderton, Roberto Amato, Jeffrey Barrett, Sonia Goncalves, Ewan Harrison, David K. Jackson, Ian Johnston, Dominic Kwiatkowski, Cordelia Langford, John Sillitoe on behalf of the Wellcome Sanger Institute COVID-19 Surveillance Team |
| EPI_ISL_2807861, EPI_ISL_2807862, EPI_ISL_2807863 | Lighthouse Lab in Glasgow | Wellcome Sanger Institute for the COVID-19 Genomics UK (COG-UK) Consortium | Harper VanSteenhouse, Yumi Kasai, David Gray, Carol Clugston, Anna Dominiczak and Alex Alderton, Roberto Amato, Jeffrey Barrett, Sonia Goncalves, Ewan Harrison, David K. Jackson, Ian Johnston, Dominic Kwiatkowski, Cordelia Langford, John Sillitoe on behalf of the Wellcome Sanger Institute COVID-19 Surveillance Team |
| EPI_ISL_2807864, EPI_ISL_2807865 | Lighthouse Lab in Milton Keynes | Wellcome Sanger Institute for the COVID-19 Genomics UK (COG-UK) Consortium | The Lighthouse Lab in Milton Keynes and Alex Alderton, Roberto Amato, Jeffrey Barrett, Sonia Goncalves, Ewan Harrison, David K. Jackson, Ian Johnston, Dominic Kwiatkowski, Cordelia Langford, John Sillitoe on behalf of the Wellcome Sanger Institute COVID-19 Surveillance Team |
| EPI_ISL_2807866 | Lighthouse Lab in Glasgow | Wellcome Sanger Institute for the COVID-19 Genomics UK (COG-UK) Consortium | Harper VanSteenhouse, Yumi Kasai, David Gray, Carol Clugston, Anna Dominiczak and Alex Alderton, Roberto Amato, Jeffrey Barrett, Sonia Goncalves, Ewan Harrison, David K. Jackson, Ian Johnston, Dominic Kwiatkowski, Cordelia Langford, John Sillitoe on behalf of the Wellcome Sanger Institute COVID-19 Surveillance Team |
| EPI_ISL_2807867 | Lighthouse Lab in Milton Keynes | Wellcome Sanger Institute for the COVID-19 Genomics UK (COG-UK) Consortium | The Lighthouse Lab in Milton Keynes and Alex Alderton, Roberto Amato, Jeffrey Barrett, Sonia Goncalves, Ewan Harrison, David K. Jackson, Ian Johnston, Dominic Kwiatkowski, Cordelia Langford, John Sillitoe on behalf of the Wellcome Sanger Institute COVID-19 Surveillance Team |
| EPI_ISL_2807868, EPI_ISL_2807869 | Lighthouse Lab in Glasgow | Wellcome Sanger Institute for the COVID-19 Genomics UK (COG-UK) Consortium | Harper VanSteenhouse, Yumi Kasai, David Gray, Carol Clugston, Anna Dominiczak and Alex Alderton, Roberto Amato, Jeffrey Barrett, Sonia Goncalves, Ewan Harrison, David K. Jackson, Ian Johnston, Dominic Kwiatkowski, Cordelia Langford, John Sillitoe on behalf of the Wellcome Sanger Institute COVID-19 Surveillance Team |
| EPI_ISL_2807870 | Lighthouse Lab in Milton Keynes | Wellcome Sanger Institute for the COVID-19 Genomics UK (COG-UK) Consortium | The Lighthouse Lab in Milton Keynes and Alex Alderton, Roberto Amato, Jeffrey Barrett, Sonia Goncalves, Ewan Harrison, David K. Jackson, Ian Johnston, Dominic Kwiatkowski, Cordelia Langford, John Sillitoe on behalf of the Wellcome Sanger Institute COVID-19 Surveillance Team |
| EPI_ISL_2807871 | Lighthouse Lab in Glasgow | Wellcome Sanger Institute for the COVID-19 Genomics UK (COG-UK) Consortium | Harper VanSteenhouse, Yumi Kasai, David Gray, Carol Clugston, Anna Dominiczak and Alex Alderton, Roberto Amato, Jeffrey Barrett, Sonia Goncalves, Ewan Harrison, David K. Jackson, Ian Johnston, Dominic Kwiatkowski, Cordelia Langford, John Sillitoe on behalf of the Wellcome Sanger Institute COVID-19 Surveillance Team |
| EPI_ISL_2807872, EPI_ISL_2807873, EPI_ISL_2807874, EPI_ISL_2807875, EPI_ISL_2807876 | Lighthouse Lab in Milton Keynes | Wellcome Sanger Institute for the COVID-19 Genomics UK (COG-UK) Consortium | The Lighthouse Lab in Milton Keynes and Alex Alderton, Roberto Amato, Jeffrey Barrett, Sonia Goncalves, Ewan Harrison, David K. Jackson, Ian Johnston, Dominic Kwiatkowski, Cordelia Langford, John Sillitoe on behalf of the Wellcome Sanger Institute COVID-19 Surveillance Team |
| EPI_ISL_2807877, | Lighthouse Lab in Glasgow | Wellcome Sanger Institute for the COVID-19 Genomics UK (COG-UK) | Harper VanSteenhouse, Yumi Kasai, David Gray, Carol Clugston, Anna Dominiczak and Alex Alderton, Roberto Amato, Jeffrey Barrett, Sonia Goncalves, Ewan Harrison, David K. Jackson, Ian Johnston, Dominic Kwiatkowski, Cordelia Langford, |

| | | | |
|---|---|---|---|
| EPI_ISL_2807878 | | Consortium | John Sillitoe on behalf of the Wellcome Sanger Institute COVID-19 Surveillance Team |
| EPI_ISL_2807879 | Lighthouse Lab in Milton Keynes | Wellcome Sanger Institute for the COVID-19 Genomics UK (COG-UK) Consortium | The Lighthouse Lab in Milton Keynes and Alex Alderton, Roberto Amato, Jeffrey Barrett, Sonia Goncalves, Ewan Harrison, David K. Jackson, Ian Johnston, Dominic Kwiatkowski, Cordelia Langford, John Sillitoe on behalf of the Wellcome Sanger Institute COVID-19 Surveillance Team |
| EPI_ISL_2807880 | Lighthouse Lab in Glasgow | Wellcome Sanger Institute for the COVID-19 Genomics UK (COG-UK) Consortium | Harper VanSteenhouse, Yumi Kasai, David Gray, Carol Clugston, Anna Dominiczak and Alex Alderton, Roberto Amato, Jeffrey Barrett, Sonia Goncalves, Ewan Harrison, David K. Jackson, Ian Johnston, Dominic Kwiatkowski, Cordelia Langford, John Sillitoe on behalf of the Wellcome Sanger Institute COVID-19 Surveillance Team |
| EPI_ISL_2807881 | Lighthouse Lab in Milton Keynes | Wellcome Sanger Institute for the COVID-19 Genomics UK (COG-UK) Consortium | The Lighthouse Lab in Milton Keynes and Alex Alderton, Roberto Amato, Jeffrey Barrett, Sonia Goncalves, Ewan Harrison, David K. Jackson, Ian Johnston, Dominic Kwiatkowski, Cordelia Langford, John Sillitoe on behalf of the Wellcome Sanger Institute COVID-19 Surveillance Team |
| EPI_ISL_2807882 | Lighthouse Lab in Glasgow | Wellcome Sanger Institute for the COVID-19 Genomics UK (COG-UK) Consortium | Harper VanSteenhouse, Yumi Kasai, David Gray, Carol Clugston, Anna Dominiczak and Alex Alderton, Roberto Amato, Jeffrey Barrett, Sonia Goncalves, Ewan Harrison, David K. Jackson, Ian Johnston, Dominic Kwiatkowski, Cordelia Langford, John Sillitoe on behalf of the Wellcome Sanger Institute COVID-19 Surveillance Team |
| EPI_ISL_2807883, EPI_ISL_2807884, EPI_ISL_2807885, EPI_ISL_2807886 | Lighthouse Lab in Milton Keynes | Wellcome Sanger Institute for the COVID-19 Genomics UK (COG-UK) Consortium | The Lighthouse Lab in Milton Keynes and Alex Alderton, Roberto Amato, Jeffrey Barrett, Sonia Goncalves, Ewan Harrison, David K. Jackson, Ian Johnston, Dominic Kwiatkowski, Cordelia Langford, John Sillitoe on behalf of the Wellcome Sanger Institute COVID-19 Surveillance Team |
| EPI_ISL_2807887, EPI_ISL_2807888 | Lighthouse Lab in Glasgow | Wellcome Sanger Institute for the COVID-19 Genomics UK (COG-UK) Consortium | Harper VanSteenhouse, Yumi Kasai, David Gray, Carol Clugston, Anna Dominiczak and Alex Alderton, Roberto Amato, Jeffrey Barrett, Sonia Goncalves, Ewan Harrison, David K. Jackson, Ian Johnston, Dominic Kwiatkowski, Cordelia Langford, John Sillitoe on behalf of the Wellcome Sanger Institute COVID-19 Surveillance Team |
| EPI_ISL_2807889 | Lighthouse Lab in Milton Keynes | Wellcome Sanger Institute for the COVID-19 Genomics UK (COG-UK) Consortium | The Lighthouse Lab in Milton Keynes and Alex Alderton, Roberto Amato, Jeffrey Barrett, Sonia Goncalves, Ewan Harrison, David K. Jackson, Ian Johnston, Dominic Kwiatkowski, Cordelia Langford, John Sillitoe on behalf of the Wellcome Sanger Institute COVID-19 Surveillance Team |
| EPI_ISL_2807890, EPI_ISL_2807891 | Lighthouse Lab in Glasgow | Wellcome Sanger Institute for the COVID-19 Genomics UK (COG-UK) Consortium | Harper VanSteenhouse, Yumi Kasai, David Gray, Carol Clugston, Anna Dominiczak and Alex Alderton, Roberto Amato, Jeffrey Barrett, Sonia Goncalves, Ewan Harrison, David K. Jackson, Ian Johnston, Dominic Kwiatkowski, Cordelia Langford, John Sillitoe on behalf of the Wellcome Sanger Institute COVID-19 Surveillance Team |
| EPI_ISL_2807892 | Lighthouse Lab in Milton Keynes | Wellcome Sanger Institute for the COVID-19 Genomics UK (COG-UK) Consortium | The Lighthouse Lab in Milton Keynes and Alex Alderton, Roberto Amato, Jeffrey Barrett, Sonia Goncalves, Ewan Harrison, David K. Jackson, Ian Johnston, Dominic Kwiatkowski, Cordelia Langford, John Sillitoe on behalf of the Wellcome Sanger Institute COVID-19 Surveillance Team |
| EPI_ISL_2807893, EPI_ISL_2807894 | Lighthouse Lab in Glasgow | Wellcome Sanger Institute for the COVID-19 Genomics UK (COG-UK) Consortium | Harper VanSteenhouse, Yumi Kasai, David Gray, Carol Clugston, Anna Dominiczak and Alex Alderton, Roberto Amato, Jeffrey Barrett, Sonia Goncalves, Ewan Harrison, David K. Jackson, Ian Johnston, Dominic Kwiatkowski, Cordelia Langford, John Sillitoe on behalf of the Wellcome Sanger Institute COVID-19 Surveillance Team |
| EPI_ISL_2807895, EPI_ISL_2807896, EPI_ISL_2807897 | Lighthouse Lab in Milton Keynes | Wellcome Sanger Institute for the COVID-19 Genomics UK (COG-UK) Consortium | The Lighthouse Lab in Milton Keynes and Alex Alderton, Roberto Amato, Jeffrey Barrett, Sonia Goncalves, Ewan Harrison, David K. Jackson, Ian Johnston, Dominic Kwiatkowski, Cordelia Langford, John Sillitoe on behalf of the Wellcome Sanger Institute COVID-19 Surveillance Team |
| EPI_ISL_2807898 | Lighthouse Lab in Glasgow | Wellcome Sanger Institute for the COVID-19 Genomics UK (COG-UK) Consortium | Harper VanSteenhouse, Yumi Kasai, David Gray, Carol Clugston, Anna Dominiczak and Alex Alderton, Roberto Amato, Jeffrey Barrett, Sonia Goncalves, Ewan Harrison, David K. Jackson, Ian Johnston, Dominic Kwiatkowski, Cordelia Langford, John Sillitoe on behalf of the Wellcome Sanger Institute COVID-19 Surveillance Team |
| EPI_ISL_2807899, EPI_ISL_2807900, EPI_ISL_2807901 | Lighthouse Lab in Milton Keynes | Wellcome Sanger Institute for the COVID-19 Genomics UK (COG-UK) Consortium | The Lighthouse Lab in Milton Keynes and Alex Alderton, Roberto Amato, Jeffrey Barrett, Sonia Goncalves, Ewan Harrison, David K. Jackson, Ian Johnston, Dominic Kwiatkowski, Cordelia Langford, John Sillitoe on behalf of the Wellcome Sanger Institute COVID-19 Surveillance Team |
| EPI_ISL_2807902 | Lighthouse Lab in Glasgow | Wellcome Sanger Institute for the COVID-19 Genomics UK (COG-UK) Consortium | Harper VanSteenhouse, Yumi Kasai, David Gray, Carol Clugston, Anna Dominiczak and Alex Alderton, Roberto Amato, Jeffrey Barrett, Sonia Goncalves, Ewan Harrison, David K. Jackson, Ian Johnston, Dominic Kwiatkowski, Cordelia Langford, John Sillitoe on behalf of the Wellcome Sanger Institute COVID-19 Surveillance Team |
| EPI_ISL_2807903 | Lighthouse Lab in Milton Keynes | Wellcome Sanger Institute for the COVID-19 Genomics UK (COG-UK) Consortium | The Lighthouse Lab in Milton Keynes and Alex Alderton, Roberto Amato, Jeffrey Barrett, Sonia Goncalves, Ewan Harrison, David K. Jackson, Ian Johnston, Dominic Kwiatkowski, Cordelia Langford, John Sillitoe on behalf of the Wellcome Sanger Institute COVID-19 Surveillance Team |
| EPI_ISL_2807904, EPI_ISL_2807905, EPI_ISL_2807906, EPI_ISL_2807907, EPI_ISL_2807908, EPI_ISL_2807909 | Lighthouse Lab in Glasgow | Wellcome Sanger Institute for the COVID-19 Genomics UK (COG-UK) Consortium | Harper VanSteenhouse, Yumi Kasai, David Gray, Carol Clugston, Anna Dominiczak and Alex Alderton, Roberto Amato, Jeffrey Barrett, Sonia Goncalves, Ewan Harrison, David K. Jackson, Ian Johnston, Dominic Kwiatkowski, Cordelia Langford, John Sillitoe on behalf of the Wellcome Sanger Institute COVID-19 Surveillance Team |
| EPI_ISL_2807910, EPI_ISL_2807911 | Lighthouse Lab in Milton Keynes | Wellcome Sanger Institute for the COVID-19 Genomics UK (COG-UK) Consortium | The Lighthouse Lab in Milton Keynes and Alex Alderton, Roberto Amato, Jeffrey Barrett, Sonia Goncalves, Ewan Harrison, David K. Jackson, Ian Johnston, Dominic Kwiatkowski, Cordelia Langford, John Sillitoe on behalf of the Wellcome Sanger Institute COVID-19 Surveillance Team |
| EPI_ISL_2807912 | Lighthouse Lab in Glasgow | Wellcome Sanger Institute for the COVID-19 Genomics UK (COG-UK) Consortium | Harper VanSteenhouse, Yumi Kasai, David Gray, Carol Clugston, Anna Dominiczak and Alex Alderton, Roberto Amato, Jeffrey Barrett, Sonia Goncalves, Ewan Harrison, David K. Jackson, Ian Johnston, Dominic Kwiatkowski, Cordelia Langford, John Sillitoe on behalf of the Wellcome Sanger Institute COVID-19 Surveillance Team |
| EPI_ISL_2807913 | Lighthouse Lab in Milton Keynes | Wellcome Sanger Institute for the COVID-19 Genomics UK (COG-UK) Consortium | The Lighthouse Lab in Milton Keynes and Alex Alderton, Roberto Amato, Jeffrey Barrett, Sonia Goncalves, Ewan Harrison, David K. Jackson, Ian Johnston, Dominic Kwiatkowski, Cordelia Langford, John Sillitoe on behalf of the Wellcome Sanger Institute COVID-19 Surveillance Team |
| EPI_ISL_2807914, EPI_ISL_2807915, EPI_ISL_2807916, EPI_ISL_2807917 | Lighthouse Lab in Glasgow | Wellcome Sanger Institute for the COVID-19 Genomics UK (COG-UK) Consortium | Harper VanSteenhouse, Yumi Kasai, David Gray, Carol Clugston, Anna Dominiczak and Alex Alderton, Roberto Amato, Jeffrey Barrett, Sonia Goncalves, Ewan Harrison, David K. Jackson, Ian Johnston, Dominic Kwiatkowski, Cordelia Langford, John Sillitoe on behalf of the Wellcome Sanger Institute COVID-19 Surveillance Team |
| EPI_ISL_2807918, EPI_ISL_2807919 | Lighthouse Lab in Milton Keynes | Wellcome Sanger Institute for the COVID-19 Genomics UK (COG-UK) Consortium | The Lighthouse Lab in Milton Keynes and Alex Alderton, Roberto Amato, Jeffrey Barrett, Sonia Goncalves, Ewan Harrison, David K. Jackson, Ian Johnston, Dominic Kwiatkowski, Cordelia Langford, John Sillitoe on behalf of the Wellcome Sanger Institute COVID-19 Surveillance Team |
| EPI_ISL_2807920, EPI_ISL_2807921 | Lighthouse Lab in Glasgow | Wellcome Sanger Institute for the COVID-19 Genomics UK (COG-UK) Consortium | Harper VanSteenhouse, Yumi Kasai, David Gray, Carol Clugston, Anna Dominiczak and Alex Alderton, Roberto Amato, Jeffrey Barrett, Sonia Goncalves, Ewan Harrison, David K. Jackson, Ian Johnston, Dominic Kwiatkowski, Cordelia Langford, John Sillitoe on behalf of the Wellcome Sanger Institute COVID-19 Surveillance Team |
| EPI_ISL_2807922, EPI_ISL_2807923 | Lighthouse Lab in Milton Keynes | Wellcome Sanger Institute for the COVID-19 Genomics UK (COG-UK) Consortium | The Lighthouse Lab in Milton Keynes and Alex Alderton, Roberto Amato, Jeffrey Barrett, Sonia Goncalves, Ewan Harrison, David K. Jackson, Ian Johnston, Dominic Kwiatkowski, Cordelia Langford, John Sillitoe on behalf of the Wellcome Sanger Institute COVID-19 Surveillance Team |
| EPI_ISL_2807924, EPI_ISL_2807925, EPI_ISL_2807926 | Lighthouse Lab in Glasgow | Wellcome Sanger Institute for the COVID-19 Genomics UK (COG-UK) Consortium | Harper VanSteenhouse, Yumi Kasai, David Gray, Carol Clugston, Anna Dominiczak and Alex Alderton, Roberto Amato, Jeffrey Barrett, Sonia Goncalves, Ewan Harrison, David K. Jackson, Ian Johnston, Dominic Kwiatkowski, Cordelia Langford, John Sillitoe on behalf of the Wellcome Sanger Institute COVID-19 Surveillance Team |
| EPI_ISL_2807927, EPI_ISL_2807928, EPI_ISL_2807929, EPI_ISL_2807930 | Lighthouse Lab in Milton Keynes | Wellcome Sanger Institute for the COVID-19 Genomics UK (COG-UK) Consortium | The Lighthouse Lab in Milton Keynes and Alex Alderton, Roberto Amato, Jeffrey Barrett, Sonia Goncalves, Ewan Harrison, David K. Jackson, Ian Johnston, Dominic Kwiatkowski, Cordelia Langford, John Sillitoe on behalf of the Wellcome Sanger Institute COVID-19 Surveillance Team |
| EPI_ISL_2807931, EPI_ISL_2807932, EPI_ISL_2807933 | Lighthouse Lab in Glasgow | Wellcome Sanger Institute for the COVID-19 Genomics UK (COG-UK) Consortium | Harper VanSteenhouse, Yumi Kasai, David Gray, Carol Clugston, Anna Dominiczak and Alex Alderton, Roberto Amato, Jeffrey Barrett, Sonia Goncalves, Ewan Harrison, David K. Jackson, Ian Johnston, Dominic Kwiatkowski, Cordelia Langford, John Sillitoe on behalf of the Wellcome Sanger Institute COVID-19 Surveillance Team |
| EPI_ISL_2807934 | Lighthouse Lab in Milton Keynes | Wellcome Sanger Institute for the COVID-19 Genomics UK (COG-UK) Consortium | The Lighthouse Lab in Milton Keynes and Alex Alderton, Roberto Amato, Jeffrey Barrett, Sonia Goncalves, Ewan Harrison, David K. Jackson, Ian Johnston, Dominic Kwiatkowski, Cordelia Langford, John Sillitoe on behalf of the Wellcome Sanger Institute COVID-19 Surveillance Team |
| EPI_ISL_2807935 | Lighthouse Lab in Glasgow | Wellcome Sanger Institute for the COVID-19 Genomics UK (COG-UK) Consortium | Harper VanSteenhouse, Yumi Kasai, David Gray, Carol Clugston, Anna Dominiczak and Alex Alderton, Roberto Amato, Jeffrey Barrett, Sonia Goncalves, Ewan Harrison, David K. Jackson, Ian Johnston, Dominic Kwiatkowski, Cordelia Langford, John Sillitoe on behalf of the Wellcome Sanger Institute COVID-19 Surveillance Team |
| EPI_ISL_2807936 | Lighthouse Lab in Milton Keynes | Wellcome Sanger Institute for the COVID-19 Genomics UK (COG-UK) Consortium | The Lighthouse Lab in Milton Keynes and Alex Alderton, Roberto Amato, Jeffrey Barrett, Sonia Goncalves, Ewan Harrison, David K. Jackson, Ian Johnston, Dominic Kwiatkowski, Cordelia Langford, John Sillitoe on behalf of the Wellcome Sanger Institute COVID-19 Surveillance Team |
| EPI_ISL_2807937, EPI_ISL_2807938 | Lighthouse Lab in Glasgow | Wellcome Sanger Institute for the COVID-19 Genomics UK (COG-UK) Consortium | Harper VanSteenhouse, Yumi Kasai, David Gray, Carol Clugston, Anna Dominiczak and Alex Alderton, Roberto Amato, Jeffrey Barrett, Sonia Goncalves, Ewan Harrison, David K. Jackson, Ian Johnston, Dominic Kwiatkowski, Cordelia Langford, John Sillitoe on behalf of the Wellcome Sanger Institute COVID-19 Surveillance Team |
| EPI_ISL_2807939, EPI_ISL_2807940, EPI_ISL_2807941 | Lighthouse Lab in Milton Keynes | Wellcome Sanger Institute for the COVID-19 Genomics UK (COG-UK) Consortium | The Lighthouse Lab in Milton Keynes and Alex Alderton, Roberto Amato, Jeffrey Barrett, Sonia Goncalves, Ewan Harrison, David K. Jackson, Ian Johnston, Dominic Kwiatkowski, Cordelia Langford, John Sillitoe on behalf of the Wellcome Sanger Institute COVID-19 Surveillance Team |
| EPI_ISL_2807942, EPI_ISL_2807943 | Lighthouse Lab in Glasgow | Wellcome Sanger Institute for the COVID-19 Genomics UK (COG-UK) Consortium | Harper VanSteenhouse, Yumi Kasai, David Gray, Carol Clugston, Anna Dominiczak and Alex Alderton, Roberto Amato, Jeffrey Barrett, Sonia Goncalves, Ewan Harrison, David K. Jackson, Ian Johnston, Dominic Kwiatkowski, Cordelia Langford, John Sillitoe on behalf of the Wellcome Sanger Institute COVID-19 Surveillance Team |
| EPI_ISL_2807944, EPI_ISL_2807945, EPI_ISL_2807946, EPI_ISL_2807947, EPI_ISL_2807948 | Lighthouse Lab in Milton Keynes | Wellcome Sanger Institute for the COVID-19 Genomics UK (COG-UK) Consortium | The Lighthouse Lab in Milton Keynes and Alex Alderton, Roberto Amato, Jeffrey Barrett, Sonia Goncalves, Ewan Harrison, David K. Jackson, Ian Johnston, Dominic Kwiatkowski, Cordelia Langford, John Sillitoe on behalf of the Wellcome Sanger Institute COVID-19 Surveillance Team |
| EPI_ISL_2807949, EPI_ISL_2807950, EPI_ISL_2807951, EPI_ISL_2807952, EPI_ISL_2807953, EPI_ISL_2807954 | Lighthouse Lab in Glasgow | Wellcome Sanger Institute for the COVID-19 Genomics UK (COG-UK) Consortium | Harper VanSteenhouse, Yumi Kasai, David Gray, Carol Clugston, Anna Dominiczak and Alex Alderton, Roberto Amato, Jeffrey Barrett, Sonia Goncalves, Ewan Harrison, David K. Jackson, Ian Johnston, Dominic Kwiatkowski, Cordelia Langford, John Sillitoe on behalf of the Wellcome Sanger Institute COVID-19 Surveillance Team |
| EPI_ISL_2807955 | Lighthouse Lab in Milton Keynes | Wellcome Sanger Institute for the COVID-19 Genomics UK (COG-UK) Consortium | The Lighthouse Lab in Milton Keynes and Alex Alderton, Roberto Amato, Jeffrey Barrett, Sonia Goncalves, Ewan Harrison, David K. Jackson, Ian Johnston, Dominic Kwiatkowski, Cordelia Langford, John Sillitoe on behalf of the Wellcome Sanger Institute COVID-19 Surveillance Team |
| EPI_ISL_2807956 | Lighthouse Lab in Glasgow | Wellcome Sanger Institute for the COVID-19 Genomics UK (COG-UK) | Harper VanSteenhouse, Yumi Kasai, David Gray, Carol Clugston, Anna Dominiczak and Alex Alderton, Roberto Amato, Jeffrey Barrett, Sonia Goncalves, Ewan Harrison, David K. Jackson, Ian Johnston, Dominic Kwiatkowski, Cordelia Langford, |

| | | | |
|---|---|---|---|
| | | Consortium | John Sillitoe on behalf of the Wellcome Sanger Institute COVID-19 Surveillance Team |
| EPI_ISL_2807957, EPI_ISL_2807958 | Lighthouse Lab in Milton Keynes | Wellcome Sanger Institute for the COVID-19 Genomics UK (COG-UK) Consortium | The Lighthouse Lab in Milton Keynes and Alex Alderton, Roberto Amato, Jeffrey Barrett, Sonia Goncalves, Ewan Harrison, David K. Jackson, Ian Johnston, Dominic Kwiatkowski, Cordelia Langford, John Sillitoe on behalf of the Wellcome Sanger Institute COVID-19 Surveillance Team |
| EPI_ISL_2807959, EPI_ISL_2807960, EPI_ISL_2807961 | Lighthouse Lab in Glasgow | Wellcome Sanger Institute for the COVID-19 Genomics UK (COG-UK) Consortium | Harper VanSteenhouse, Yumi Kasai, David Gray, Carol Clugston, Anna Dominiczak and Alex Alderton, Roberto Amato, Jeffrey Barrett, Sonia Goncalves, Ewan Harrison, David K. Jackson, Ian Johnston, Dominic Kwiatkowski, Cordelia Langford, John Sillitoe on behalf of the Wellcome Sanger Institute COVID-19 Surveillance Team |
| EPI_ISL_2807962, EPI_ISL_2807963 | Lighthouse Lab in Milton Keynes | Wellcome Sanger Institute for the COVID-19 Genomics UK (COG-UK) Consortium | The Lighthouse Lab in Milton Keynes and Alex Alderton, Roberto Amato, Jeffrey Barrett, Sonia Goncalves, Ewan Harrison, David K. Jackson, Ian Johnston, Dominic Kwiatkowski, Cordelia Langford, John Sillitoe on behalf of the Wellcome Sanger Institute COVID-19 Surveillance Team |
| EPI_ISL_2807964 | Lighthouse Lab in Glasgow | Wellcome Sanger Institute for the COVID-19 Genomics UK (COG-UK) Consortium | Harper VanSteenhouse, Yumi Kasai, David Gray, Carol Clugston, Anna Dominiczak and Alex Alderton, Roberto Amato, Jeffrey Barrett, Sonia Goncalves, Ewan Harrison, David K. Jackson, Ian Johnston, Dominic Kwiatkowski, Cordelia Langford, John Sillitoe on behalf of the Wellcome Sanger Institute COVID-19 Surveillance Team |
| EPI_ISL_2807965, EPI_ISL_2807966, EPI_ISL_2807967 | Lighthouse Lab in Milton Keynes | Wellcome Sanger Institute for the COVID-19 Genomics UK (COG-UK) Consortium | The Lighthouse Lab in Milton Keynes and Alex Alderton, Roberto Amato, Jeffrey Barrett, Sonia Goncalves, Ewan Harrison, David K. Jackson, Ian Johnston, Dominic Kwiatkowski, Cordelia Langford, John Sillitoe on behalf of the Wellcome Sanger Institute COVID-19 Surveillance Team |
| EPI_ISL_2807968 | Lighthouse Lab in Glasgow | Wellcome Sanger Institute for the COVID-19 Genomics UK (COG-UK) Consortium | Harper VanSteenhouse, Yumi Kasai, David Gray, Carol Clugston, Anna Dominiczak and Alex Alderton, Roberto Amato, Jeffrey Barrett, Sonia Goncalves, Ewan Harrison, David K. Jackson, Ian Johnston, Dominic Kwiatkowski, Cordelia Langford, John Sillitoe on behalf of the Wellcome Sanger Institute COVID-19 Surveillance Team |
| EPI_ISL_2807969, EPI_ISL_2807970, EPI_ISL_2807971 | Lighthouse Lab in Milton Keynes | Wellcome Sanger Institute for the COVID-19 Genomics UK (COG-UK) Consortium | The Lighthouse Lab in Milton Keynes and Alex Alderton, Roberto Amato, Jeffrey Barrett, Sonia Goncalves, Ewan Harrison, David K. Jackson, Ian Johnston, Dominic Kwiatkowski, Cordelia Langford, John Sillitoe on behalf of the Wellcome Sanger Institute COVID-19 Surveillance Team |
| EPI_ISL_2807972, EPI_ISL_2807973 | Lighthouse Lab in Glasgow | Wellcome Sanger Institute for the COVID-19 Genomics UK (COG-UK) Consortium | Harper VanSteenhouse, Yumi Kasai, David Gray, Carol Clugston, Anna Dominiczak and Alex Alderton, Roberto Amato, Jeffrey Barrett, Sonia Goncalves, Ewan Harrison, David K. Jackson, Ian Johnston, Dominic Kwiatkowski, Cordelia Langford, John Sillitoe on behalf of the Wellcome Sanger Institute COVID-19 Surveillance Team |
| EPI_ISL_2807974, EPI_ISL_2807975, EPI_ISL_2807976, EPI_ISL_2807977 | Lighthouse Lab in Milton Keynes | Wellcome Sanger Institute for the COVID-19 Genomics UK (COG-UK) Consortium | The Lighthouse Lab in Milton Keynes and Alex Alderton, Roberto Amato, Jeffrey Barrett, Sonia Goncalves, Ewan Harrison, David K. Jackson, Ian Johnston, Dominic Kwiatkowski, Cordelia Langford, John Sillitoe on behalf of the Wellcome Sanger Institute COVID-19 Surveillance Team |
| EPI_ISL_2807978, EPI_ISL_2807979, EPI_ISL_2807980, EPI_ISL_2807981 | Lighthouse Lab in Glasgow | Wellcome Sanger Institute for the COVID-19 Genomics UK (COG-UK) Consortium | Harper VanSteenhouse, Yumi Kasai, David Gray, Carol Clugston, Anna Dominiczak and Alex Alderton, Roberto Amato, Jeffrey Barrett, Sonia Goncalves, Ewan Harrison, David K. Jackson, Ian Johnston, Dominic Kwiatkowski, Cordelia Langford, John Sillitoe on behalf of the Wellcome Sanger Institute COVID-19 Surveillance Team |
| EPI_ISL_2807982, EPI_ISL_2807983, EPI_ISL_2807984, EPI_ISL_2807985, EPI_ISL_2807986 | Lighthouse Lab in Milton Keynes | Wellcome Sanger Institute for the COVID-19 Genomics UK (COG-UK) Consortium | The Lighthouse Lab in Milton Keynes and Alex Alderton, Roberto Amato, Jeffrey Barrett, Sonia Goncalves, Ewan Harrison, David K. Jackson, Ian Johnston, Dominic Kwiatkowski, Cordelia Langford, John Sillitoe on behalf of the Wellcome Sanger Institute COVID-19 Surveillance Team |
| EPI_ISL_2807987 | Lighthouse Lab in Glasgow | Wellcome Sanger Institute for the COVID-19 Genomics UK (COG-UK) Consortium | Harper VanSteenhouse, Yumi Kasai, David Gray, Carol Clugston, Anna Dominiczak and Alex Alderton, Roberto Amato, Jeffrey Barrett, Sonia Goncalves, Ewan Harrison, David K. Jackson, Ian Johnston, Dominic Kwiatkowski, Cordelia Langford, John Sillitoe on behalf of the Wellcome Sanger Institute COVID-19 Surveillance Team |
| EPI_ISL_2807988 | Lighthouse Lab in Milton Keynes | Wellcome Sanger Institute for the COVID-19 Genomics UK (COG-UK) Consortium | The Lighthouse Lab in Milton Keynes and Alex Alderton, Roberto Amato, Jeffrey Barrett, Sonia Goncalves, Ewan Harrison, David K. Jackson, Ian Johnston, Dominic Kwiatkowski, Cordelia Langford, John Sillitoe on behalf of the Wellcome Sanger Institute COVID-19 Surveillance Team |
| EPI_ISL_2807989 | Lighthouse Lab in Glasgow | Wellcome Sanger Institute for the COVID-19 Genomics UK (COG-UK) Consortium | Harper VanSteenhouse, Yumi Kasai, David Gray, Carol Clugston, Anna Dominiczak and Alex Alderton, Roberto Amato, Jeffrey Barrett, Sonia Goncalves, Ewan Harrison, David K. Jackson, Ian Johnston, Dominic Kwiatkowski, Cordelia Langford, John Sillitoe on behalf of the Wellcome Sanger Institute COVID-19 Surveillance Team |
| EPI_ISL_2807990, EPI_ISL_2807991, EPI_ISL_2807992, EPI_ISL_2807993 | Lighthouse Lab in Milton Keynes | Wellcome Sanger Institute for the COVID-19 Genomics UK (COG-UK) Consortium | The Lighthouse Lab in Milton Keynes and Alex Alderton, Roberto Amato, Jeffrey Barrett, Sonia Goncalves, Ewan Harrison, David K. Jackson, Ian Johnston, Dominic Kwiatkowski, Cordelia Langford, John Sillitoe on behalf of the Wellcome Sanger Institute COVID-19 Surveillance Team |
| EPI_ISL_2807994, EPI_ISL_2807995 | Lighthouse Lab in Glasgow | Wellcome Sanger Institute for the COVID-19 Genomics UK (COG-UK) Consortium | Harper VanSteenhouse, Yumi Kasai, David Gray, Carol Clugston, Anna Dominiczak and Alex Alderton, Roberto Amato, Jeffrey Barrett, Sonia Goncalves, Ewan Harrison, David K. Jackson, Ian Johnston, Dominic Kwiatkowski, Cordelia Langford, John Sillitoe on behalf of the Wellcome Sanger Institute COVID-19 Surveillance Team |
| EPI_ISL_2807996 | Lighthouse Lab in Milton Keynes | Wellcome Sanger Institute for the COVID-19 Genomics UK (COG-UK) Consortium | The Lighthouse Lab in Milton Keynes and Alex Alderton, Roberto Amato, Jeffrey Barrett, Sonia Goncalves, Ewan Harrison, David K. Jackson, Ian Johnston, Dominic Kwiatkowski, Cordelia Langford, John Sillitoe on behalf of the Wellcome Sanger Institute COVID-19 Surveillance Team |
| EPI_ISL_2807997, EPI_ISL_2807998, EPI_ISL_2807999, EPI_ISL_2808000 | Lighthouse Lab in Glasgow | Wellcome Sanger Institute for the COVID-19 Genomics UK (COG-UK) Consortium | Harper VanSteenhouse, Yumi Kasai, David Gray, Carol Clugston, Anna Dominiczak and Alex Alderton, Roberto Amato, Jeffrey Barrett, Sonia Goncalves, Ewan Harrison, David K. Jackson, Ian Johnston, Dominic Kwiatkowski, Cordelia Langford, John Sillitoe on behalf of the Wellcome Sanger Institute COVID-19 Surveillance Team |
| EPI_ISL_2808001, EPI_ISL_2808002 | Lighthouse Lab in Milton Keynes | Wellcome Sanger Institute for the COVID-19 Genomics UK (COG-UK) Consortium | The Lighthouse Lab in Milton Keynes and Alex Alderton, Roberto Amato, Jeffrey Barrett, Sonia Goncalves, Ewan Harrison, David K. Jackson, Ian Johnston, Dominic Kwiatkowski, Cordelia Langford, John Sillitoe on behalf of the Wellcome Sanger Institute COVID-19 Surveillance Team |
| EPI_ISL_2808003, EPI_ISL_2808004 | Lighthouse Lab in Glasgow | Wellcome Sanger Institute for the COVID-19 Genomics UK (COG-UK) Consortium | Harper VanSteenhouse, Yumi Kasai, David Gray, Carol Clugston, Anna Dominiczak and Alex Alderton, Roberto Amato, Jeffrey Barrett, Sonia Goncalves, Ewan Harrison, David K. Jackson, Ian Johnston, Dominic Kwiatkowski, Cordelia Langford, John Sillitoe on behalf of the Wellcome Sanger Institute COVID-19 Surveillance Team |
| EPI_ISL_2808005, EPI_ISL_2808006 | Lighthouse Lab in Milton Keynes | Wellcome Sanger Institute for the COVID-19 Genomics UK (COG-UK) Consortium | The Lighthouse Lab in Milton Keynes and Alex Alderton, Roberto Amato, Jeffrey Barrett, Sonia Goncalves, Ewan Harrison, David K. Jackson, Ian Johnston, Dominic Kwiatkowski, Cordelia Langford, John Sillitoe on behalf of the Wellcome Sanger Institute COVID-19 Surveillance Team |
| EPI_ISL_2808007, EPI_ISL_2808008, EPI_ISL_2808009, EPI_ISL_2808010, EPI_ISL_2808011, EPI_ISL_2808012, EPI_ISL_2808013, EPI_ISL_2808014 | Lighthouse Lab in Glasgow | Wellcome Sanger Institute for the COVID-19 Genomics UK (COG-UK) Consortium | Harper VanSteenhouse, Yumi Kasai, David Gray, Carol Clugston, Anna Dominiczak and Alex Alderton, Roberto Amato, Jeffrey Barrett, Sonia Goncalves, Ewan Harrison, David K. Jackson, Ian Johnston, Dominic Kwiatkowski, Cordelia Langford, John Sillitoe on behalf of the Wellcome Sanger Institute COVID-19 Surveillance Team |
| EPI_ISL_2808015, EPI_ISL_2808017, EPI_ISL_2808018 | Lighthouse Lab in Milton Keynes | Wellcome Sanger Institute for the COVID-19 Genomics UK (COG-UK) Consortium | The Lighthouse Lab in Milton Keynes and Alex Alderton, Roberto Amato, Jeffrey Barrett, Sonia Goncalves, Ewan Harrison, David K. Jackson, Ian Johnston, Dominic Kwiatkowski, Cordelia Langford, John Sillitoe on behalf of the Wellcome Sanger Institute COVID-19 Surveillance Team |
| EPI_ISL_2808019 | Lighthouse Lab in Glasgow | Wellcome Sanger Institute for the COVID-19 Genomics UK (COG-UK) Consortium | Harper VanSteenhouse, Yumi Kasai, David Gray, Carol Clugston, Anna Dominiczak and Alex Alderton, Roberto Amato, Jeffrey Barrett, Sonia Goncalves, Ewan Harrison, David K. Jackson, Ian Johnston, Dominic Kwiatkowski, Cordelia Langford, John Sillitoe on behalf of the Wellcome Sanger Institute COVID-19 Surveillance Team |
| EPI_ISL_2808020, EPI_ISL_2808021 | Lighthouse Lab in Milton Keynes | Wellcome Sanger Institute for the COVID-19 Genomics UK (COG-UK) Consortium | The Lighthouse Lab in Milton Keynes and Alex Alderton, Roberto Amato, Jeffrey Barrett, Sonia Goncalves, Ewan Harrison, David K. Jackson, Ian Johnston, Dominic Kwiatkowski, Cordelia Langford, John Sillitoe on behalf of the Wellcome Sanger Institute COVID-19 Surveillance Team |
| EPI_ISL_2808022, EPI_ISL_2808023 | Lighthouse Lab in Glasgow | Wellcome Sanger Institute for the COVID-19 Genomics UK (COG-UK) Consortium | Harper VanSteenhouse, Yumi Kasai, David Gray, Carol Clugston, Anna Dominiczak and Alex Alderton, Roberto Amato, Jeffrey Barrett, Sonia Goncalves, Ewan Harrison, David K. Jackson, Ian Johnston, Dominic Kwiatkowski, Cordelia Langford, John Sillitoe on behalf of the Wellcome Sanger Institute COVID-19 Surveillance Team |
| EPI_ISL_2808024, EPI_ISL_2808025 | Lighthouse Lab in Milton Keynes | Wellcome Sanger Institute for the COVID-19 Genomics UK (COG-UK) Consortium | The Lighthouse Lab in Milton Keynes and Alex Alderton, Roberto Amato, Jeffrey Barrett, Sonia Goncalves, Ewan Harrison, David K. Jackson, Ian Johnston, Dominic Kwiatkowski, Cordelia Langford, John Sillitoe on behalf of the Wellcome Sanger Institute COVID-19 Surveillance Team |
| EPI_ISL_2808026 | Lighthouse Lab in Glasgow | Wellcome Sanger Institute for the COVID-19 Genomics UK (COG-UK) Consortium | Harper VanSteenhouse, Yumi Kasai, David Gray, Carol Clugston, Anna Dominiczak and Alex Alderton, Roberto Amato, Jeffrey Barrett, Sonia Goncalves, Ewan Harrison, David K. Jackson, Ian Johnston, Dominic Kwiatkowski, Cordelia Langford, John Sillitoe on behalf of the Wellcome Sanger Institute COVID-19 Surveillance Team |
| EPI_ISL_2808027, EPI_ISL_2808028 | Lighthouse Lab in Milton Keynes | Wellcome Sanger Institute for the COVID-19 Genomics UK (COG-UK) Consortium | The Lighthouse Lab in Milton Keynes and Alex Alderton, Roberto Amato, Jeffrey Barrett, Sonia Goncalves, Ewan Harrison, David K. Jackson, Ian Johnston, Dominic Kwiatkowski, Cordelia Langford, John Sillitoe on behalf of the Wellcome Sanger Institute COVID-19 Surveillance Team |
| EPI_ISL_2808029, EPI_ISL_2808030 | Lighthouse Lab in Glasgow | Wellcome Sanger Institute for the COVID-19 Genomics UK (COG-UK) Consortium | Harper VanSteenhouse, Yumi Kasai, David Gray, Carol Clugston, Anna Dominiczak and Alex Alderton, Roberto Amato, Jeffrey Barrett, Sonia Goncalves, Ewan Harrison, David K. Jackson, Ian Johnston, Dominic Kwiatkowski, Cordelia Langford, John Sillitoe on behalf of the Wellcome Sanger Institute COVID-19 Surveillance Team |
| EPI_ISL_2808031 | Lighthouse Lab in Milton Keynes | Wellcome Sanger Institute for the COVID-19 Genomics UK (COG-UK) Consortium | The Lighthouse Lab in Milton Keynes and Alex Alderton, Roberto Amato, Jeffrey Barrett, Sonia Goncalves, Ewan Harrison, David K. Jackson, Ian Johnston, Dominic Kwiatkowski, Cordelia Langford, John Sillitoe on behalf of the Wellcome Sanger Institute COVID-19 Surveillance Team |
| EPI_ISL_2808032 | Lighthouse Lab in Glasgow | Wellcome Sanger Institute for the COVID-19 Genomics UK (COG-UK) Consortium | Harper VanSteenhouse, Yumi Kasai, David Gray, Carol Clugston, Anna Dominiczak and Alex Alderton, Roberto Amato, Jeffrey Barrett, Sonia Goncalves, Ewan Harrison, David K. Jackson, Ian Johnston, Dominic Kwiatkowski, Cordelia Langford, John Sillitoe on behalf of the Wellcome Sanger Institute COVID-19 Surveillance Team |
| EPI_ISL_2808033, EPI_ISL_2808034 | Lighthouse Lab in Milton Keynes | Wellcome Sanger Institute for the COVID-19 Genomics UK (COG-UK) Consortium | The Lighthouse Lab in Milton Keynes and Alex Alderton, Roberto Amato, Jeffrey Barrett, Sonia Goncalves, Ewan Harrison, David K. Jackson, Ian Johnston, Dominic Kwiatkowski, Cordelia Langford, John Sillitoe on behalf of the Wellcome Sanger Institute COVID-19 Surveillance Team |
| EPI_ISL_2808035 | Lighthouse Lab in Glasgow | Wellcome Sanger Institute for the COVID-19 Genomics UK (COG-UK) Consortium | Harper VanSteenhouse, Yumi Kasai, David Gray, Carol Clugston, Anna Dominiczak and Alex Alderton, Roberto Amato, Jeffrey Barrett, Sonia Goncalves, Ewan Harrison, David K. Jackson, Ian Johnston, Dominic Kwiatkowski, Cordelia Langford, John Sillitoe on behalf of the Wellcome Sanger Institute COVID-19 Surveillance Team |
| EPI_ISL_2808036, EPI_ISL_2808037 | Lighthouse Lab in Milton Keynes | Wellcome Sanger Institute for the COVID-19 Genomics UK (COG-UK) Consortium | The Lighthouse Lab in Milton Keynes and Alex Alderton, Roberto Amato, Jeffrey Barrett, Sonia Goncalves, Ewan Harrison, David K. Jackson, Ian Johnston, Dominic Kwiatkowski, Cordelia Langford, John Sillitoe on behalf of the Wellcome Sanger Institute COVID-19 Surveillance Team |
| EPI_ISL_2808038 | Lighthouse Lab in Glasgow | Wellcome Sanger Institute for the COVID-19 Genomics UK (COG-UK) Consortium | Harper VanSteenhouse, Yumi Kasai, David Gray, Carol Clugston, Anna Dominiczak and Alex Alderton, Roberto Amato, Jeffrey Barrett, Sonia Goncalves, Ewan Harrison, David K. Jackson, Ian Johnston, Dominic Kwiatkowski, Cordelia Langford, John Sillitoe on behalf of the Wellcome Sanger Institute COVID-19 Surveillance Team |

| | | | |
|---|---|---|---|
| EPI_ISL_2808039 | Lighthouse Lab in Milton Keynes | Wellcome Sanger Institute for the COVID-19 Genomics UK (COG-UK) Consortium | The Lighthouse Lab in Milton Keynes and Alex Alderton, Roberto Amato, Jeffrey Barrett, Sonia Goncalves, Ewan Harrison, David K. Jackson, Ian Johnston, Dominic Kwiatkowski, Cordelia Langford, John Sillitoe on behalf of the Wellcome Sanger Institute COVID-19 Surveillance Team |
| EPI_ISL_2808040, EPI_ISL_2808041 | Lighthouse Lab in Glasgow | Wellcome Sanger Institute for the COVID-19 Genomics UK (COG-UK) Consortium | Harper VanSteenhouse, Yumi Kasai, David Gray, Carol Clugston, Anna Dominiczak and Alex Alderton, Roberto Amato, Jeffrey Barrett, Sonia Goncalves, Ewan Harrison, David K. Jackson, Ian Johnston, Dominic Kwiatkowski, Cordelia Langford, John Sillitoe on behalf of the Wellcome Sanger Institute COVID-19 Surveillance Team |
| EPI_ISL_2808042 | Lighthouse Lab in Milton Keynes | Wellcome Sanger Institute for the COVID-19 Genomics UK (COG-UK) Consortium | The Lighthouse Lab in Milton Keynes and Alex Alderton, Roberto Amato, Jeffrey Barrett, Sonia Goncalves, Ewan Harrison, David K. Jackson, Ian Johnston, Dominic Kwiatkowski, Cordelia Langford, John Sillitoe on behalf of the Wellcome Sanger Institute COVID-19 Surveillance Team |
| EPI_ISL_2808043, EPI_ISL_2808044 | Lighthouse Lab in Glasgow | Wellcome Sanger Institute for the COVID-19 Genomics UK (COG-UK) Consortium | Harper VanSteenhouse, Yumi Kasai, David Gray, Carol Clugston, Anna Dominiczak and Alex Alderton, Roberto Amato, Jeffrey Barrett, Sonia Goncalves, Ewan Harrison, David K. Jackson, Ian Johnston, Dominic Kwiatkowski, Cordelia Langford, John Sillitoe on behalf of the Wellcome Sanger Institute COVID-19 Surveillance Team |
| EPI_ISL_2808045 | Lighthouse Lab in Milton Keynes | Wellcome Sanger Institute for the COVID-19 Genomics UK (COG-UK) Consortium | The Lighthouse Lab in Milton Keynes and Alex Alderton, Roberto Amato, Jeffrey Barrett, Sonia Goncalves, Ewan Harrison, David K. Jackson, Ian Johnston, Dominic Kwiatkowski, Cordelia Langford, John Sillitoe on behalf of the Wellcome Sanger Institute COVID-19 Surveillance Team |
| EPI_ISL_2808046, EPI_ISL_2808047, EPI_ISL_2808048, EPI_ISL_2808049, EPI_ISL_2808050 | Lighthouse Lab in Glasgow | Wellcome Sanger Institute for the COVID-19 Genomics UK (COG-UK) Consortium | Harper VanSteenhouse, Yumi Kasai, David Gray, Carol Clugston, Anna Dominiczak and Alex Alderton, Roberto Amato, Jeffrey Barrett, Sonia Goncalves, Ewan Harrison, David K. Jackson, Ian Johnston, Dominic Kwiatkowski, Cordelia Langford, John Sillitoe on behalf of the Wellcome Sanger Institute COVID-19 Surveillance Team |
| EPI_ISL_2808051, EPI_ISL_2808052, EPI_ISL_2808053 | Lighthouse Lab in Milton Keynes | Wellcome Sanger Institute for the COVID-19 Genomics UK (COG-UK) Consortium | The Lighthouse Lab in Milton Keynes and Alex Alderton, Roberto Amato, Jeffrey Barrett, Sonia Goncalves, Ewan Harrison, David K. Jackson, Ian Johnston, Dominic Kwiatkowski, Cordelia Langford, John Sillitoe on behalf of the Wellcome Sanger Institute COVID-19 Surveillance Team |
| EPI_ISL_2808054 | Lighthouse Lab in Glasgow | Wellcome Sanger Institute for the COVID-19 Genomics UK (COG-UK) Consortium | Harper VanSteenhouse, Yumi Kasai, David Gray, Carol Clugston, Anna Dominiczak and Alex Alderton, Roberto Amato, Jeffrey Barrett, Sonia Goncalves, Ewan Harrison, David K. Jackson, Ian Johnston, Dominic Kwiatkowski, Cordelia Langford, John Sillitoe on behalf of the Wellcome Sanger Institute COVID-19 Surveillance Team |
| EPI_ISL_2808055 | Lighthouse Lab in Milton Keynes | Wellcome Sanger Institute for the COVID-19 Genomics UK (COG-UK) Consortium | The Lighthouse Lab in Milton Keynes and Alex Alderton, Roberto Amato, Jeffrey Barrett, Sonia Goncalves, Ewan Harrison, David K. Jackson, Ian Johnston, Dominic Kwiatkowski, Cordelia Langford, John Sillitoe on behalf of the Wellcome Sanger Institute COVID-19 Surveillance Team |
| EPI_ISL_2808056 | Lighthouse Lab in Glasgow | Wellcome Sanger Institute for the COVID-19 Genomics UK (COG-UK) Consortium | Harper VanSteenhouse, Yumi Kasai, David Gray, Carol Clugston, Anna Dominiczak and Alex Alderton, Roberto Amato, Jeffrey Barrett, Sonia Goncalves, Ewan Harrison, David K. Jackson, Ian Johnston, Dominic Kwiatkowski, Cordelia Langford, John Sillitoe on behalf of the Wellcome Sanger Institute COVID-19 Surveillance Team |
| EPI_ISL_2808057, EPI_ISL_2808058, EPI_ISL_2808059 | Lighthouse Lab in Milton Keynes | Wellcome Sanger Institute for the COVID-19 Genomics UK (COG-UK) Consortium | The Lighthouse Lab in Milton Keynes and Alex Alderton, Roberto Amato, Jeffrey Barrett, Sonia Goncalves, Ewan Harrison, David K. Jackson, Ian Johnston, Dominic Kwiatkowski, Cordelia Langford, John Sillitoe on behalf of the Wellcome Sanger Institute COVID-19 Surveillance Team |
| EPI_ISL_2808060 | Lighthouse Lab in Glasgow | Wellcome Sanger Institute for the COVID-19 Genomics UK (COG-UK) Consortium | Harper VanSteenhouse, Yumi Kasai, David Gray, Carol Clugston, Anna Dominiczak and Alex Alderton, Roberto Amato, Jeffrey Barrett, Sonia Goncalves, Ewan Harrison, David K. Jackson, Ian Johnston, Dominic Kwiatkowski, Cordelia Langford, John Sillitoe on behalf of the Wellcome Sanger Institute COVID-19 Surveillance Team |
| EPI_ISL_2808061 | Lighthouse Lab in Milton Keynes | Wellcome Sanger Institute for the COVID-19 Genomics UK (COG-UK) Consortium | The Lighthouse Lab in Milton Keynes and Alex Alderton, Roberto Amato, Jeffrey Barrett, Sonia Goncalves, Ewan Harrison, David K. Jackson, Ian Johnston, Dominic Kwiatkowski, Cordelia Langford, John Sillitoe on behalf of the Wellcome Sanger Institute COVID-19 Surveillance Team |
| EPI_ISL_2808062, EPI_ISL_2808063, EPI_ISL_2808064 | Lighthouse Lab in Glasgow | Wellcome Sanger Institute for the COVID-19 Genomics UK (COG-UK) Consortium | Harper VanSteenhouse, Yumi Kasai, David Gray, Carol Clugston, Anna Dominiczak and Alex Alderton, Roberto Amato, Jeffrey Barrett, Sonia Goncalves, Ewan Harrison, David K. Jackson, Ian Johnston, Dominic Kwiatkowski, Cordelia Langford, John Sillitoe on behalf of the Wellcome Sanger Institute COVID-19 Surveillance Team |
| EPI_ISL_2808065 | Lighthouse Lab in Milton Keynes | Wellcome Sanger Institute for the COVID-19 Genomics UK (COG-UK) Consortium | The Lighthouse Lab in Milton Keynes and Alex Alderton, Roberto Amato, Jeffrey Barrett, Sonia Goncalves, Ewan Harrison, David K. Jackson, Ian Johnston, Dominic Kwiatkowski, Cordelia Langford, John Sillitoe on behalf of the Wellcome Sanger Institute COVID-19 Surveillance Team |
| EPI_ISL_2808066, EPI_ISL_2808067 | Lighthouse Lab in Glasgow | Wellcome Sanger Institute for the COVID-19 Genomics UK (COG-UK) Consortium | Harper VanSteenhouse, Yumi Kasai, David Gray, Carol Clugston, Anna Dominiczak and Alex Alderton, Roberto Amato, Jeffrey Barrett, Sonia Goncalves, Ewan Harrison, David K. Jackson, Ian Johnston, Dominic Kwiatkowski, Cordelia Langford, John Sillitoe on behalf of the Wellcome Sanger Institute COVID-19 Surveillance Team |
| EPI_ISL_2808068, EPI_ISL_2808069, EPI_ISL_2808070, EPI_ISL_2808071 | Lighthouse Lab in Milton Keynes | Wellcome Sanger Institute for the COVID-19 Genomics UK (COG-UK) Consortium | The Lighthouse Lab in Milton Keynes and Alex Alderton, Roberto Amato, Jeffrey Barrett, Sonia Goncalves, Ewan Harrison, David K. Jackson, Ian Johnston, Dominic Kwiatkowski, Cordelia Langford, John Sillitoe on behalf of the Wellcome Sanger Institute COVID-19 Surveillance Team |
| EPI_ISL_2808072, EPI_ISL_2808073 | Health Services Laboratories | Wellcome Sanger Institute for the COVID-19 Genomics UK (COG-UK) Consortium | Health Services Laboratories and Alex Alderton, Roberto Amato, Jeffrey Barrett, Sonia Goncalves, Ewan Harrison, David K. Jackson, Ian Johnston, Dominic Kwiatkowski, Cordelia Langford, John Sillitoe on behalf of the Wellcome Sanger Institute COVID-19 Surveillance Team |
| EPI_ISL_2808074 | Lighthouse Lab in Milton Keynes | Wellcome Sanger Institute for the COVID-19 Genomics UK (COG-UK) Consortium | The Lighthouse Lab in Milton Keynes and Alex Alderton, Roberto Amato, Jeffrey Barrett, Sonia Goncalves, Ewan Harrison, David K. Jackson, Ian Johnston, Dominic Kwiatkowski, Cordelia Langford, John Sillitoe on behalf of the Wellcome Sanger Institute COVID-19 Surveillance Team |
| EPI_ISL_2808075, EPI_ISL_2808076 | Lighthouse Laboratory Plymouth | Wellcome Sanger Institute for the COVID-19 Genomics UK (COG-UK) Consortium | Lighthouse Laboratory Plymouth and Alex Alderton, Roberto Amato, Jeffrey Barrett, Sonia Goncalves, Ewan Harrison, David K. Jackson, Ian Johnston, Dominic Kwiatkowski, Cordelia Langford, John Sillitoe on behalf of the Wellcome Sanger Institute COVID-19 Surveillance Team |
| EPI_ISL_2808077 | Health Services Laboratories | Wellcome Sanger Institute for the COVID-19 Genomics UK (COG-UK) Consortium | Health Services Laboratories and Alex Alderton, Roberto Amato, Jeffrey Barrett, Sonia Goncalves, Ewan Harrison, David K. Jackson, Ian Johnston, Dominic Kwiatkowski, Cordelia Langford, John Sillitoe on behalf of the Wellcome Sanger Institute COVID-19 Surveillance Team |
| EPI_ISL_2808078 | Lighthouse Laboratory Plymouth | Wellcome Sanger Institute for the COVID-19 Genomics UK (COG-UK) Consortium | Lighthouse Laboratory Plymouth and Alex Alderton, Roberto Amato, Jeffrey Barrett, Sonia Goncalves, Ewan Harrison, David K. Jackson, Ian Johnston, Dominic Kwiatkowski, Cordelia Langford, John Sillitoe on behalf of the Wellcome Sanger Institute COVID-19 Surveillance Team |
| EPI_ISL_2808079, EPI_ISL_2808080 | Health Services Laboratories | Wellcome Sanger Institute for the COVID-19 Genomics UK (COG-UK) Consortium | Health Services Laboratories and Alex Alderton, Roberto Amato, Jeffrey Barrett, Sonia Goncalves, Ewan Harrison, David K. Jackson, Ian Johnston, Dominic Kwiatkowski, Cordelia Langford, John Sillitoe on behalf of the Wellcome Sanger Institute COVID-19 Surveillance Team |
| EPI_ISL_2808081, EPI_ISL_2808082, EPI_ISL_2808083 | Lighthouse Laboratory Plymouth | Wellcome Sanger Institute for the COVID-19 Genomics UK (COG-UK) Consortium | Lighthouse Laboratory Plymouth and Alex Alderton, Roberto Amato, Jeffrey Barrett, Sonia Goncalves, Ewan Harrison, David K. Jackson, Ian Johnston, Dominic Kwiatkowski, Cordelia Langford, John Sillitoe on behalf of the Wellcome Sanger Institute COVID-19 Surveillance Team |
| EPI_ISL_2808084 | Health Services Laboratories | Wellcome Sanger Institute for the COVID-19 Genomics UK (COG-UK) Consortium | Health Services Laboratories and Alex Alderton, Roberto Amato, Jeffrey Barrett, Sonia Goncalves, Ewan Harrison, David K. Jackson, Ian Johnston, Dominic Kwiatkowski, Cordelia Langford, John Sillitoe on behalf of the Wellcome Sanger Institute COVID-19 Surveillance Team |
| EPI_ISL_2808085, EPI_ISL_2808086, EPI_ISL_2808087, EPI_ISL_2808088 | Lighthouse Laboratory Plymouth | Wellcome Sanger Institute for the COVID-19 Genomics UK (COG-UK) Consortium | Lighthouse Laboratory Plymouth and Alex Alderton, Roberto Amato, Jeffrey Barrett, Sonia Goncalves, Ewan Harrison, David K. Jackson, Ian Johnston, Dominic Kwiatkowski, Cordelia Langford, John Sillitoe on behalf of the Wellcome Sanger Institute COVID-19 Surveillance Team |
| EPI_ISL_2808089 | Health Services Laboratories | Wellcome Sanger Institute for the COVID-19 Genomics UK (COG-UK) Consortium | Health Services Laboratories and Alex Alderton, Roberto Amato, Jeffrey Barrett, Sonia Goncalves, Ewan Harrison, David K. Jackson, Ian Johnston, Dominic Kwiatkowski, Cordelia Langford, John Sillitoe on behalf of the Wellcome Sanger Institute COVID-19 Surveillance Team |
| EPI_ISL_2808090 | Lighthouse Lab in Milton Keynes | Wellcome Sanger Institute for the COVID-19 Genomics UK (COG-UK) Consortium | The Lighthouse Lab in Milton Keynes and Alex Alderton, Roberto Amato, Jeffrey Barrett, Sonia Goncalves, Ewan Harrison, David K. Jackson, Ian Johnston, Dominic Kwiatkowski, Cordelia Langford, John Sillitoe on behalf of the Wellcome Sanger Institute COVID-19 Surveillance Team |
| EPI_ISL_2808091 | Lighthouse Laboratory Plymouth | Wellcome Sanger Institute for the COVID-19 Genomics UK (COG-UK) Consortium | Lighthouse Laboratory Plymouth and Alex Alderton, Roberto Amato, Jeffrey Barrett, Sonia Goncalves, Ewan Harrison, David K. Jackson, Ian Johnston, Dominic Kwiatkowski, Cordelia Langford, John Sillitoe on behalf of the Wellcome Sanger Institute COVID-19 Surveillance Team |
| EPI_ISL_2808092 | Lighthouse Lab in Milton Keynes | Wellcome Sanger Institute for the COVID-19 Genomics UK (COG-UK) Consortium | The Lighthouse Lab in Milton Keynes and Alex Alderton, Roberto Amato, Jeffrey Barrett, Sonia Goncalves, Ewan Harrison, David K. Jackson, Ian Johnston, Dominic Kwiatkowski, Cordelia Langford, John Sillitoe on behalf of the Wellcome Sanger Institute COVID-19 Surveillance Team |
| EPI_ISL_2808093 | Health Services Laboratories | Wellcome Sanger Institute for the COVID-19 Genomics UK (COG-UK) Consortium | Health Services Laboratories and Alex Alderton, Roberto Amato, Jeffrey Barrett, Sonia Goncalves, Ewan Harrison, David K. Jackson, Ian Johnston, Dominic Kwiatkowski, Cordelia Langford, John Sillitoe on behalf of the Wellcome Sanger Institute COVID-19 Surveillance Team |
| EPI_ISL_2808094, EPI_ISL_2808095, EPI_ISL_2808096, EPI_ISL_2808097 | Lighthouse Laboratory Plymouth | Wellcome Sanger Institute for the COVID-19 Genomics UK (COG-UK) Consortium | Lighthouse Laboratory Plymouth and Alex Alderton, Roberto Amato, Jeffrey Barrett, Sonia Goncalves, Ewan Harrison, David K. Jackson, Ian Johnston, Dominic Kwiatkowski, Cordelia Langford, John Sillitoe on behalf of the Wellcome Sanger Institute COVID-19 Surveillance Team |
| EPI_ISL_2808099, EPI_ISL_2808100 | Health Services Laboratories | Wellcome Sanger Institute for the COVID-19 Genomics UK (COG-UK) Consortium | Health Services Laboratories and Alex Alderton, Roberto Amato, Jeffrey Barrett, Sonia Goncalves, Ewan Harrison, David K. Jackson, Ian Johnston, Dominic Kwiatkowski, Cordelia Langford, John Sillitoe on behalf of the Wellcome Sanger Institute COVID-19 Surveillance Team |
| EPI_ISL_2808101, EPI_ISL_2808102, EPI_ISL_2808103 | Lighthouse Laboratory Plymouth | Wellcome Sanger Institute for the COVID-19 Genomics UK (COG-UK) Consortium | Lighthouse Laboratory Plymouth and Alex Alderton, Roberto Amato, Jeffrey Barrett, Sonia Goncalves, Ewan Harrison, David K. Jackson, Ian Johnston, Dominic Kwiatkowski, Cordelia Langford, John Sillitoe on behalf of the Wellcome Sanger Institute COVID-19 Surveillance Team |
| EPI_ISL_2808104 | Health Services Laboratories | Wellcome Sanger Institute for the COVID-19 Genomics UK (COG-UK) Consortium | Health Services Laboratories and Alex Alderton, Roberto Amato, Jeffrey Barrett, Sonia Goncalves, Ewan Harrison, David K. Jackson, Ian Johnston, Dominic Kwiatkowski, Cordelia Langford, John Sillitoe on behalf of the Wellcome Sanger Institute COVID-19 Surveillance Team |
| EPI_ISL_2808105, EPI_ISL_2808106, EPI_ISL_2808107, EPI_ISL_2808108, EPI_ISL_2808109, EPI_ISL_2808110, EPI_ISL_2808111 | Lighthouse Laboratory Plymouth | Wellcome Sanger Institute for the COVID-19 Genomics UK (COG-UK) Consortium | Lighthouse Laboratory Plymouth and Alex Alderton, Roberto Amato, Jeffrey Barrett, Sonia Goncalves, Ewan Harrison, David K. Jackson, Ian Johnston, Dominic Kwiatkowski, Cordelia Langford, John Sillitoe on behalf of the Wellcome Sanger Institute COVID-19 Surveillance Team |
| EPI_ISL_2808112 | Lighthouse Lab in Milton Keynes | Wellcome Sanger Institute for the COVID-19 Genomics UK (COG-UK) Consortium | The Lighthouse Lab in Milton Keynes and Alex Alderton, Roberto Amato, Jeffrey Barrett, Sonia Goncalves, Ewan Harrison, David K. Jackson, Ian Johnston, Dominic Kwiatkowski, Cordelia Langford, John Sillitoe on behalf of the Wellcome Sanger Institute COVID-19 Surveillance Team |

| | | | |
|---|---|---|---|
| EPI_ISL_2808113, EPI_ISL_2808114, EPI_ISL_2808115 | Lighthouse Laboratory Plymouth | Wellcome Sanger Institute for the COVID-19 Genomics UK (COG-UK) Consortium | Lighthouse Laboratory Plymouth and Alex Alderton, Roberto Amato, Jeffrey Barrett, Sonia Goncalves, Ewan Harrison, David K. Jackson, Ian Johnston, Dominic Kwiatkowski, Cordelia Langford, John Sillitoe on behalf of the Wellcome Sanger Institute COVID-19 Surveillance Team |
| EPI_ISL_2808116 | Lighthouse Lab in Milton Keynes | Wellcome Sanger Institute for the COVID-19 Genomics UK (COG-UK) Consortium | The Lighthouse Lab in Milton Keynes and Alex Alderton, Roberto Amato, Jeffrey Barrett, Sonia Goncalves, Ewan Harrison, David K. Jackson, Ian Johnston, Dominic Kwiatkowski, Cordelia Langford, John Sillitoe on behalf of the Wellcome Sanger Institute COVID-19 Surveillance Team |
| EPI_ISL_2808117, EPI_ISL_2808118, EPI_ISL_2808119 | Health Services Laboratories | Wellcome Sanger Institute for the COVID-19 Genomics UK (COG-UK) Consortium | Health Services Laboratories and Alex Alderton, Roberto Amato, Jeffrey Barrett, Sonia Goncalves, Ewan Harrison, David K. Jackson, Ian Johnston, Dominic Kwiatkowski, Cordelia Langford, John Sillitoe on behalf of the Wellcome Sanger Institute COVID-19 Surveillance Team |
| EPI_ISL_2808120 | Lighthouse Laboratory Plymouth | Wellcome Sanger Institute for the COVID-19 Genomics UK (COG-UK) Consortium | Lighthouse Laboratory Plymouth and Alex Alderton, Roberto Amato, Jeffrey Barrett, Sonia Goncalves, Ewan Harrison, David K. Jackson, Ian Johnston, Dominic Kwiatkowski, Cordelia Langford, John Sillitoe on behalf of the Wellcome Sanger Institute COVID-19 Surveillance Team |
| EPI_ISL_2808121 | Health Services Laboratories | Wellcome Sanger Institute for the COVID-19 Genomics UK (COG-UK) Consortium | Health Services Laboratories and Alex Alderton, Roberto Amato, Jeffrey Barrett, Sonia Goncalves, Ewan Harrison, David K. Jackson, Ian Johnston, Dominic Kwiatkowski, Cordelia Langford, John Sillitoe on behalf of the Wellcome Sanger Institute COVID-19 Surveillance Team |
| EPI_ISL_2808122 | Lighthouse Laboratory Plymouth | Wellcome Sanger Institute for the COVID-19 Genomics UK (COG-UK) Consortium | Lighthouse Laboratory Plymouth and Alex Alderton, Roberto Amato, Jeffrey Barrett, Sonia Goncalves, Ewan Harrison, David K. Jackson, Ian Johnston, Dominic Kwiatkowski, Cordelia Langford, John Sillitoe on behalf of the Wellcome Sanger Institute COVID-19 Surveillance Team |
| EPI_ISL_2808123, EPI_ISL_2808124, EPI_ISL_2808125, EPI_ISL_2808126 | Health Services Laboratories | Wellcome Sanger Institute for the COVID-19 Genomics UK (COG-UK) Consortium | Health Services Laboratories and Alex Alderton, Roberto Amato, Jeffrey Barrett, Sonia Goncalves, Ewan Harrison, David K. Jackson, Ian Johnston, Dominic Kwiatkowski, Cordelia Langford, John Sillitoe on behalf of the Wellcome Sanger Institute COVID-19 Surveillance Team |
| EPI_ISL_2808127, EPI_ISL_2808128 | Lighthouse Laboratory Plymouth | Wellcome Sanger Institute for the COVID-19 Genomics UK (COG-UK) Consortium | Lighthouse Laboratory Plymouth and Alex Alderton, Roberto Amato, Jeffrey Barrett, Sonia Goncalves, Ewan Harrison, David K. Jackson, Ian Johnston, Dominic Kwiatkowski, Cordelia Langford, John Sillitoe on behalf of the Wellcome Sanger Institute COVID-19 Surveillance Team |
| EPI_ISL_2808129 | Health Services Laboratories | Wellcome Sanger Institute for the COVID-19 Genomics UK (COG-UK) Consortium | Health Services Laboratories and Alex Alderton, Roberto Amato, Jeffrey Barrett, Sonia Goncalves, Ewan Harrison, David K. Jackson, Ian Johnston, Dominic Kwiatkowski, Cordelia Langford, John Sillitoe on behalf of the Wellcome Sanger Institute COVID-19 Surveillance Team |
| EPI_ISL_2808130, EPI_ISL_2808131, EPI_ISL_2808132 | Lighthouse Laboratory Plymouth | Wellcome Sanger Institute for the COVID-19 Genomics UK (COG-UK) Consortium | Lighthouse Laboratory Plymouth and Alex Alderton, Roberto Amato, Jeffrey Barrett, Sonia Goncalves, Ewan Harrison, David K. Jackson, Ian Johnston, Dominic Kwiatkowski, Cordelia Langford, John Sillitoe on behalf of the Wellcome Sanger Institute COVID-19 Surveillance Team |
| EPI_ISL_2808133 | Lighthouse Lab in Milton Keynes | Wellcome Sanger Institute for the COVID-19 Genomics UK (COG-UK) Consortium | The Lighthouse Lab in Milton Keynes and Alex Alderton, Roberto Amato, Jeffrey Barrett, Sonia Goncalves, Ewan Harrison, David K. Jackson, Ian Johnston, Dominic Kwiatkowski, Cordelia Langford, John Sillitoe on behalf of the Wellcome Sanger Institute COVID-19 Surveillance Team |
| EPI_ISL_2808134, EPI_ISL_2808135 | Lighthouse Laboratory Plymouth | Wellcome Sanger Institute for the COVID-19 Genomics UK (COG-UK) Consortium | Lighthouse Laboratory Plymouth and Alex Alderton, Roberto Amato, Jeffrey Barrett, Sonia Goncalves, Ewan Harrison, David K. Jackson, Ian Johnston, Dominic Kwiatkowski, Cordelia Langford, John Sillitoe on behalf of the Wellcome Sanger Institute COVID-19 Surveillance Team |
| EPI_ISL_2808136 | Lighthouse Lab in Milton Keynes | Wellcome Sanger Institute for the COVID-19 Genomics UK (COG-UK) Consortium | The Lighthouse Lab in Milton Keynes and Alex Alderton, Roberto Amato, Jeffrey Barrett, Sonia Goncalves, Ewan Harrison, David K. Jackson, Ian Johnston, Dominic Kwiatkowski, Cordelia Langford, John Sillitoe on behalf of the Wellcome Sanger Institute COVID-19 Surveillance Team |
| EPI_ISL_2808137, EPI_ISL_2808138, EPI_ISL_2808139, EPI_ISL_2808140, EPI_ISL_2808141 | Lighthouse Laboratory Plymouth | Wellcome Sanger Institute for the COVID-19 Genomics UK (COG-UK) Consortium | Lighthouse Laboratory Plymouth and Alex Alderton, Roberto Amato, Jeffrey Barrett, Sonia Goncalves, Ewan Harrison, David K. Jackson, Ian Johnston, Dominic Kwiatkowski, Cordelia Langford, John Sillitoe on behalf of the Wellcome Sanger Institute COVID-19 Surveillance Team |
| EPI_ISL_2808142 | Lighthouse Lab in Milton Keynes | Wellcome Sanger Institute for the COVID-19 Genomics UK (COG-UK) Consortium | The Lighthouse Lab in Milton Keynes and Alex Alderton, Roberto Amato, Jeffrey Barrett, Sonia Goncalves, Ewan Harrison, David K. Jackson, Ian Johnston, Dominic Kwiatkowski, Cordelia Langford, John Sillitoe on behalf of the Wellcome Sanger Institute COVID-19 Surveillance Team |
| EPI_ISL_2808143 | Lighthouse Laboratory Plymouth | Wellcome Sanger Institute for the COVID-19 Genomics UK (COG-UK) Consortium | Lighthouse Laboratory Plymouth and Alex Alderton, Roberto Amato, Jeffrey Barrett, Sonia Goncalves, Ewan Harrison, David K. Jackson, Ian Johnston, Dominic Kwiatkowski, Cordelia Langford, John Sillitoe on behalf of the Wellcome Sanger Institute COVID-19 Surveillance Team |
| EPI_ISL_2808144 | Health Services Laboratories | Wellcome Sanger Institute for the COVID-19 Genomics UK (COG-UK) Consortium | Health Services Laboratories and Alex Alderton, Roberto Amato, Jeffrey Barrett, Sonia Goncalves, Ewan Harrison, David K. Jackson, Ian Johnston, Dominic Kwiatkowski, Cordelia Langford, John Sillitoe on behalf of the Wellcome Sanger Institute COVID-19 Surveillance Team |
| EPI_ISL_2808145, EPI_ISL_2808146, EPI_ISL_2808147 | Lighthouse Laboratory Plymouth | Wellcome Sanger Institute for the COVID-19 Genomics UK (COG-UK) Consortium | Lighthouse Laboratory Plymouth and Alex Alderton, Roberto Amato, Jeffrey Barrett, Sonia Goncalves, Ewan Harrison, David K. Jackson, Ian Johnston, Dominic Kwiatkowski, Cordelia Langford, John Sillitoe on behalf of the Wellcome Sanger Institute COVID-19 Surveillance Team |
| EPI_ISL_2808148 | Health Services Laboratories | Wellcome Sanger Institute for the COVID-19 Genomics UK (COG-UK) Consortium | Health Services Laboratories and Alex Alderton, Roberto Amato, Jeffrey Barrett, Sonia Goncalves, Ewan Harrison, David K. Jackson, Ian Johnston, Dominic Kwiatkowski, Cordelia Langford, John Sillitoe on behalf of the Wellcome Sanger Institute COVID-19 Surveillance Team |
| EPI_ISL_2808149 | Lighthouse Laboratory Plymouth | Wellcome Sanger Institute for the COVID-19 Genomics UK (COG-UK) Consortium | Lighthouse Laboratory Plymouth and Alex Alderton, Roberto Amato, Jeffrey Barrett, Sonia Goncalves, Ewan Harrison, David K. Jackson, Ian Johnston, Dominic Kwiatkowski, Cordelia Langford, John Sillitoe on behalf of the Wellcome Sanger Institute COVID-19 Surveillance Team |
| EPI_ISL_2808150 | Health Services Laboratories | Wellcome Sanger Institute for the COVID-19 Genomics UK (COG-UK) Consortium | Health Services Laboratories and Alex Alderton, Roberto Amato, Jeffrey Barrett, Sonia Goncalves, Ewan Harrison, David K. Jackson, Ian Johnston, Dominic Kwiatkowski, Cordelia Langford, John Sillitoe on behalf of the Wellcome Sanger Institute COVID-19 Surveillance Team |
| EPI_ISL_2808151, EPI_ISL_2808152, EPI_ISL_2808153, EPI_ISL_2808154 | Lighthouse Laboratory Plymouth | Wellcome Sanger Institute for the COVID-19 Genomics UK (COG-UK) Consortium | Lighthouse Laboratory Plymouth and Alex Alderton, Roberto Amato, Jeffrey Barrett, Sonia Goncalves, Ewan Harrison, David K. Jackson, Ian Johnston, Dominic Kwiatkowski, Cordelia Langford, John Sillitoe on behalf of the Wellcome Sanger Institute COVID-19 Surveillance Team |
| EPI_ISL_2808155 | Lighthouse Lab in Milton Keynes | Wellcome Sanger Institute for the COVID-19 Genomics UK (COG-UK) Consortium | The Lighthouse Lab in Milton Keynes and Alex Alderton, Roberto Amato, Jeffrey Barrett, Sonia Goncalves, Ewan Harrison, David K. Jackson, Ian Johnston, Dominic Kwiatkowski, Cordelia Langford, John Sillitoe on behalf of the Wellcome Sanger Institute COVID-19 Surveillance Team |
| EPI_ISL_2808156, EPI_ISL_2808158 | Health Services Laboratories | Wellcome Sanger Institute for the COVID-19 Genomics UK (COG-UK) Consortium | Health Services Laboratories and Alex Alderton, Roberto Amato, Jeffrey Barrett, Sonia Goncalves, Ewan Harrison, David K. Jackson, Ian Johnston, Dominic Kwiatkowski, Cordelia Langford, John Sillitoe on behalf of the Wellcome Sanger Institute COVID-19 Surveillance Team |
| EPI_ISL_2808159, EPI_ISL_2808161, EPI_ISL_2808162, EPI_ISL_2808163, EPI_ISL_2808164, EPI_ISL_2808165, EPI_ISL_2808166, EPI_ISL_2808167 | Lighthouse Laboratory Plymouth | Wellcome Sanger Institute for the COVID-19 Genomics UK (COG-UK) Consortium | Lighthouse Laboratory Plymouth and Alex Alderton, Roberto Amato, Jeffrey Barrett, Sonia Goncalves, Ewan Harrison, David K. Jackson, Ian Johnston, Dominic Kwiatkowski, Cordelia Langford, John Sillitoe on behalf of the Wellcome Sanger Institute COVID-19 Surveillance Team |
| EPI_ISL_2808168 | Health Services Laboratories | Wellcome Sanger Institute for the COVID-19 Genomics UK (COG-UK) Consortium | Health Services Laboratories and Alex Alderton, Roberto Amato, Jeffrey Barrett, Sonia Goncalves, Ewan Harrison, David K. Jackson, Ian Johnston, Dominic Kwiatkowski, Cordelia Langford, John Sillitoe on behalf of the Wellcome Sanger Institute COVID-19 Surveillance Team |
| EPI_ISL_2808169, EPI_ISL_2808170 | Lighthouse Laboratory Plymouth | Wellcome Sanger Institute for the COVID-19 Genomics UK (COG-UK) Consortium | Lighthouse Laboratory Plymouth and Alex Alderton, Roberto Amato, Jeffrey Barrett, Sonia Goncalves, Ewan Harrison, David K. Jackson, Ian Johnston, Dominic Kwiatkowski, Cordelia Langford, John Sillitoe on behalf of the Wellcome Sanger Institute COVID-19 Surveillance Team |
| EPI_ISL_2808171 | Lighthouse Lab in Milton Keynes | Wellcome Sanger Institute for the COVID-19 Genomics UK (COG-UK) Consortium | The Lighthouse Lab in Milton Keynes and Alex Alderton, Roberto Amato, Jeffrey Barrett, Sonia Goncalves, Ewan Harrison, David K. Jackson, Ian Johnston, Dominic Kwiatkowski, Cordelia Langford, John Sillitoe on behalf of the Wellcome Sanger Institute COVID-19 Surveillance Team |
| EPI_ISL_2808172 | Health Services Laboratories | Wellcome Sanger Institute for the COVID-19 Genomics UK (COG-UK) Consortium | Health Services Laboratories and Alex Alderton, Roberto Amato, Jeffrey Barrett, Sonia Goncalves, Ewan Harrison, David K. Jackson, Ian Johnston, Dominic Kwiatkowski, Cordelia Langford, John Sillitoe on behalf of the Wellcome Sanger Institute COVID-19 Surveillance Team |
| EPI_ISL_2808173 | Lighthouse Laboratory Plymouth | Wellcome Sanger Institute for the COVID-19 Genomics UK (COG-UK) Consortium | Lighthouse Laboratory Plymouth and Alex Alderton, Roberto Amato, Jeffrey Barrett, Sonia Goncalves, Ewan Harrison, David K. Jackson, Ian Johnston, Dominic Kwiatkowski, Cordelia Langford, John Sillitoe on behalf of the Wellcome Sanger Institute COVID-19 Surveillance Team |
| EPI_ISL_2808174 | Health Services Laboratories | Wellcome Sanger Institute for the COVID-19 Genomics UK (COG-UK) Consortium | Health Services Laboratories and Alex Alderton, Roberto Amato, Jeffrey Barrett, Sonia Goncalves, Ewan Harrison, David K. Jackson, Ian Johnston, Dominic Kwiatkowski, Cordelia Langford, John Sillitoe on behalf of the Wellcome Sanger Institute COVID-19 Surveillance Team |
| EPI_ISL_2808175 | Lighthouse Laboratory Plymouth | Wellcome Sanger Institute for the COVID-19 Genomics UK (COG-UK) Consortium | Lighthouse Laboratory Plymouth and Alex Alderton, Roberto Amato, Jeffrey Barrett, Sonia Goncalves, Ewan Harrison, David K. Jackson, Ian Johnston, Dominic Kwiatkowski, Cordelia Langford, John Sillitoe on behalf of the Wellcome Sanger Institute COVID-19 Surveillance Team |
| EPI_ISL_2808177 | Lighthouse Lab in Milton Keynes | Wellcome Sanger Institute for the COVID-19 Genomics UK (COG-UK) Consortium | The Lighthouse Lab in Milton Keynes and Alex Alderton, Roberto Amato, Jeffrey Barrett, Sonia Goncalves, Ewan Harrison, David K. Jackson, Ian Johnston, Dominic Kwiatkowski, Cordelia Langford, John Sillitoe on behalf of the Wellcome Sanger Institute COVID-19 Surveillance Team |
| EPI_ISL_2808178 | Health Services Laboratories | Wellcome Sanger Institute for the COVID-19 Genomics UK (COG-UK) Consortium | Health Services Laboratories and Alex Alderton, Roberto Amato, Jeffrey Barrett, Sonia Goncalves, Ewan Harrison, David K. Jackson, Ian Johnston, Dominic Kwiatkowski, Cordelia Langford, John Sillitoe on behalf of the Wellcome Sanger Institute COVID-19 Surveillance Team |
| EPI_ISL_2808179, EPI_ISL_2808180, EPI_ISL_2808181 | Lighthouse Laboratory Plymouth | Wellcome Sanger Institute for the COVID-19 Genomics UK (COG-UK) Consortium | Lighthouse Laboratory Plymouth and Alex Alderton, Roberto Amato, Jeffrey Barrett, Sonia Goncalves, Ewan Harrison, David K. Jackson, Ian Johnston, Dominic Kwiatkowski, Cordelia Langford, John Sillitoe on behalf of the Wellcome Sanger Institute COVID-19 Surveillance Team |
| EPI_ISL_2808182, EPI_ISL_2808183 | Lighthouse Lab in Milton Keynes | Wellcome Sanger Institute for the COVID-19 Genomics UK (COG-UK) Consortium | The Lighthouse Lab in Milton Keynes and Alex Alderton, Roberto Amato, Jeffrey Barrett, Sonia Goncalves, Ewan Harrison, David K. Jackson, Ian Johnston, Dominic Kwiatkowski, Cordelia Langford, John Sillitoe on behalf of the Wellcome Sanger Institute COVID-19 Surveillance Team |
| EPI_ISL_2808184, EPI_ISL_2808185, EPI_ISL_2808186 | Lighthouse Laboratory Plymouth | Wellcome Sanger Institute for the COVID-19 Genomics UK (COG-UK) Consortium | Lighthouse Laboratory Plymouth and Alex Alderton, Roberto Amato, Jeffrey Barrett, Sonia Goncalves, Ewan Harrison, David K. Jackson, Ian Johnston, Dominic Kwiatkowski, Cordelia Langford, John Sillitoe on behalf of the Wellcome Sanger Institute COVID-19 Surveillance Team |
| EPI_ISL_2808187 | Health Services Laboratories | Wellcome Sanger Institute for the COVID-19 Genomics UK (COG-UK) | Health Services Laboratories and Alex Alderton, Roberto Amato, Jeffrey Barrett, Sonia Goncalves, Ewan Harrison, David K. Jackson, Ian Johnston, Dominic Kwiatkowski, Cordelia Langford, John Sillitoe on behalf of the Wellcome Sanger Institute |

| | | | |
|---|---|---|---|
| | | Consortium | COVID-19 Surveillance Team |
| EPI_ISL_2808188 | Lighthouse Laboratory Plymouth | Wellcome Sanger Institute for the COVID-19 Genomics UK (COG-UK) Consortium | Lighthouse Laboratory Plymouth and Alex Alderton, Roberto Amato, Jeffrey Barrett, Sonia Goncalves, Ewan Harrison, David K. Jackson, Ian Johnston, Dominic Kwiatkowski, Cordelia Langford, John Sillitoe on behalf of the Wellcome Sanger Institute COVID-19 Surveillance Team |
| EPI_ISL_2808189 | Health Services Laboratories | Wellcome Sanger Institute for the COVID-19 Genomics UK (COG-UK) Consortium | Health Services Laboratories and Alex Alderton, Roberto Amato, Jeffrey Barrett, Sonia Goncalves, Ewan Harrison, David K. Jackson, Ian Johnston, Dominic Kwiatkowski, Cordelia Langford, John Sillitoe on behalf of the Wellcome Sanger Institute COVID-19 Surveillance Team |
| EPI_ISL_2808190, EPI_ISL_2808191, EPI_ISL_2808192, EPI_ISL_2808194, EPI_ISL_2808195, EPI_ISL_2808196, EPI_ISL_2808197 | Lighthouse Laboratory Plymouth | Wellcome Sanger Institute for the COVID-19 Genomics UK (COG-UK) Consortium | Lighthouse Laboratory Plymouth and Alex Alderton, Roberto Amato, Jeffrey Barrett, Sonia Goncalves, Ewan Harrison, David K. Jackson, Ian Johnston, Dominic Kwiatkowski, Cordelia Langford, John Sillitoe on behalf of the Wellcome Sanger Institute COVID-19 Surveillance Team |
| EPI_ISL_2808198 | Health Services Laboratories | Wellcome Sanger Institute for the COVID-19 Genomics UK (COG-UK) Consortium | Health Services Laboratories and Alex Alderton, Roberto Amato, Jeffrey Barrett, Sonia Goncalves, Ewan Harrison, David K. Jackson, Ian Johnston, Dominic Kwiatkowski, Cordelia Langford, John Sillitoe on behalf of the Wellcome Sanger Institute COVID-19 Surveillance Team |
| EPI_ISL_2808199 | Lighthouse Laboratory Plymouth | Wellcome Sanger Institute for the COVID-19 Genomics UK (COG-UK) Consortium | Lighthouse Laboratory Plymouth and Alex Alderton, Roberto Amato, Jeffrey Barrett, Sonia Goncalves, Ewan Harrison, David K. Jackson, Ian Johnston, Dominic Kwiatkowski, Cordelia Langford, John Sillitoe on behalf of the Wellcome Sanger Institute COVID-19 Surveillance Team |
| EPI_ISL_2808200 | Health Services Laboratories | Wellcome Sanger Institute for the COVID-19 Genomics UK (COG-UK) Consortium | Health Services Laboratories and Alex Alderton, Roberto Amato, Jeffrey Barrett, Sonia Goncalves, Ewan Harrison, David K. Jackson, Ian Johnston, Dominic Kwiatkowski, Cordelia Langford, John Sillitoe on behalf of the Wellcome Sanger Institute COVID-19 Surveillance Team |
| EPI_ISL_2808201, EPI_ISL_2808202, EPI_ISL_2808203, EPI_ISL_2808204, EPI_ISL_2808205 | Lighthouse Laboratory Plymouth | Wellcome Sanger Institute for the COVID-19 Genomics UK (COG-UK) Consortium | Lighthouse Laboratory Plymouth and Alex Alderton, Roberto Amato, Jeffrey Barrett, Sonia Goncalves, Ewan Harrison, David K. Jackson, Ian Johnston, Dominic Kwiatkowski, Cordelia Langford, John Sillitoe on behalf of the Wellcome Sanger Institute COVID-19 Surveillance Team |
| EPI_ISL_2808206 | Lighthouse Lab in Milton Keynes | Wellcome Sanger Institute for the COVID-19 Genomics UK (COG-UK) Consortium | The Lighthouse Lab in Milton Keynes and Alex Alderton, Roberto Amato, Jeffrey Barrett, Sonia Goncalves, Ewan Harrison, David K. Jackson, Ian Johnston, Dominic Kwiatkowski, Cordelia Langford, John Sillitoe on behalf of the Wellcome Sanger Institute COVID-19 Surveillance Team |
| EPI_ISL_2808207, EPI_ISL_2808208, EPI_ISL_2808209 | Lighthouse Laboratory Plymouth | Wellcome Sanger Institute for the COVID-19 Genomics UK (COG-UK) Consortium | Lighthouse Laboratory Plymouth and Alex Alderton, Roberto Amato, Jeffrey Barrett, Sonia Goncalves, Ewan Harrison, David K. Jackson, Ian Johnston, Dominic Kwiatkowski, Cordelia Langford, John Sillitoe on behalf of the Wellcome Sanger Institute COVID-19 Surveillance Team |
| EPI_ISL_2808210 | Health Services Laboratories | Wellcome Sanger Institute for the COVID-19 Genomics UK (COG-UK) Consortium | Health Services Laboratories and Alex Alderton, Roberto Amato, Jeffrey Barrett, Sonia Goncalves, Ewan Harrison, David K. Jackson, Ian Johnston, Dominic Kwiatkowski, Cordelia Langford, John Sillitoe on behalf of the Wellcome Sanger Institute COVID-19 Surveillance Team |
| EPI_ISL_2808211, EPI_ISL_2808212, EPI_ISL_2808213, EPI_ISL_2808214, EPI_ISL_2808215 | Lighthouse Laboratory Plymouth | Wellcome Sanger Institute for the COVID-19 Genomics UK (COG-UK) Consortium | Lighthouse Laboratory Plymouth and Alex Alderton, Roberto Amato, Jeffrey Barrett, Sonia Goncalves, Ewan Harrison, David K. Jackson, Ian Johnston, Dominic Kwiatkowski, Cordelia Langford, John Sillitoe on behalf of the Wellcome Sanger Institute COVID-19 Surveillance Team |
| EPI_ISL_2808216 | Health Services Laboratories | Wellcome Sanger Institute for the COVID-19 Genomics UK (COG-UK) Consortium | Health Services Laboratories and Alex Alderton, Roberto Amato, Jeffrey Barrett, Sonia Goncalves, Ewan Harrison, David K. Jackson, Ian Johnston, Dominic Kwiatkowski, Cordelia Langford, John Sillitoe on behalf of the Wellcome Sanger Institute COVID-19 Surveillance Team |
| EPI_ISL_2808217, EPI_ISL_2808218, EPI_ISL_2808219 | Lighthouse Laboratory Plymouth | Wellcome Sanger Institute for the COVID-19 Genomics UK (COG-UK) Consortium | Lighthouse Laboratory Plymouth and Alex Alderton, Roberto Amato, Jeffrey Barrett, Sonia Goncalves, Ewan Harrison, David K. Jackson, Ian Johnston, Dominic Kwiatkowski, Cordelia Langford, John Sillitoe on behalf of the Wellcome Sanger Institute COVID-19 Surveillance Team |
| EPI_ISL_2808220 | Health Services Laboratories | Wellcome Sanger Institute for the COVID-19 Genomics UK (COG-UK) Consortium | Health Services Laboratories and Alex Alderton, Roberto Amato, Jeffrey Barrett, Sonia Goncalves, Ewan Harrison, David K. Jackson, Ian Johnston, Dominic Kwiatkowski, Cordelia Langford, John Sillitoe on behalf of the Wellcome Sanger Institute COVID-19 Surveillance Team |
| EPI_ISL_2808221, EPI_ISL_2808222, EPI_ISL_2808223 | Lighthouse Laboratory Plymouth | Wellcome Sanger Institute for the COVID-19 Genomics UK (COG-UK) Consortium | Lighthouse Laboratory Plymouth and Alex Alderton, Roberto Amato, Jeffrey Barrett, Sonia Goncalves, Ewan Harrison, David K. Jackson, Ian Johnston, Dominic Kwiatkowski, Cordelia Langford, John Sillitoe on behalf of the Wellcome Sanger Institute COVID-19 Surveillance Team |
| EPI_ISL_2808224, EPI_ISL_2808225 | Health Services Laboratories | Wellcome Sanger Institute for the COVID-19 Genomics UK (COG-UK) Consortium | Health Services Laboratories and Alex Alderton, Roberto Amato, Jeffrey Barrett, Sonia Goncalves, Ewan Harrison, David K. Jackson, Ian Johnston, Dominic Kwiatkowski, Cordelia Langford, John Sillitoe on behalf of the Wellcome Sanger Institute COVID-19 Surveillance Team |
| EPI_ISL_2808226, EPI_ISL_2808227, EPI_ISL_2808228, EPI_ISL_2808229, EPI_ISL_2808230, EPI_ISL_2808231, EPI_ISL_2808232 | Lighthouse Laboratory Plymouth | Wellcome Sanger Institute for the COVID-19 Genomics UK (COG-UK) Consortium | Lighthouse Laboratory Plymouth and Alex Alderton, Roberto Amato, Jeffrey Barrett, Sonia Goncalves, Ewan Harrison, David K. Jackson, Ian Johnston, Dominic Kwiatkowski, Cordelia Langford, John Sillitoe on behalf of the Wellcome Sanger Institute COVID-19 Surveillance Team |
| EPI_ISL_2808233 | Health Services Laboratories | Wellcome Sanger Institute for the COVID-19 Genomics UK (COG-UK) Consortium | Health Services Laboratories and Alex Alderton, Roberto Amato, Jeffrey Barrett, Sonia Goncalves, Ewan Harrison, David K. Jackson, Ian Johnston, Dominic Kwiatkowski, Cordelia Langford, John Sillitoe on behalf of the Wellcome Sanger Institute COVID-19 Surveillance Team |
| EPI_ISL_2808234, EPI_ISL_2808235, EPI_ISL_2808236 | Lighthouse Laboratory Plymouth | Wellcome Sanger Institute for the COVID-19 Genomics UK (COG-UK) Consortium | Lighthouse Laboratory Plymouth and Alex Alderton, Roberto Amato, Jeffrey Barrett, Sonia Goncalves, Ewan Harrison, David K. Jackson, Ian Johnston, Dominic Kwiatkowski, Cordelia Langford, John Sillitoe on behalf of the Wellcome Sanger Institute COVID-19 Surveillance Team |
| EPI_ISL_2808237, EPI_ISL_2808238 | Health Services Laboratories | Wellcome Sanger Institute for the COVID-19 Genomics UK (COG-UK) Consortium | Health Services Laboratories and Alex Alderton, Roberto Amato, Jeffrey Barrett, Sonia Goncalves, Ewan Harrison, David K. Jackson, Ian Johnston, Dominic Kwiatkowski, Cordelia Langford, John Sillitoe on behalf of the Wellcome Sanger Institute COVID-19 Surveillance Team |
| EPI_ISL_2808239 | Lighthouse Laboratory Plymouth | Wellcome Sanger Institute for the COVID-19 Genomics UK (COG-UK) Consortium | Lighthouse Laboratory Plymouth and Alex Alderton, Roberto Amato, Jeffrey Barrett, Sonia Goncalves, Ewan Harrison, David K. Jackson, Ian Johnston, Dominic Kwiatkowski, Cordelia Langford, John Sillitoe on behalf of the Wellcome Sanger Institute COVID-19 Surveillance Team |
| EPI_ISL_2808240, EPI_ISL_2808241 | Health Services Laboratories | Wellcome Sanger Institute for the COVID-19 Genomics UK (COG-UK) Consortium | Health Services Laboratories and Alex Alderton, Roberto Amato, Jeffrey Barrett, Sonia Goncalves, Ewan Harrison, David K. Jackson, Ian Johnston, Dominic Kwiatkowski, Cordelia Langford, John Sillitoe on behalf of the Wellcome Sanger Institute COVID-19 Surveillance Team |
| EPI_ISL_2808242, EPI_ISL_2808243, EPI_ISL_2808244, EPI_ISL_2808245, EPI_ISL_2808246, EPI_ISL_2808247, EPI_ISL_2808249, EPI_ISL_2808250, EPI_ISL_2808251, EPI_ISL_2808252, EPI_ISL_2808253, EPI_ISL_2808254, EPI_ISL_2808255 | | | |
| see above | Lighthouse Laboratory Plymouth | Wellcome Sanger Institute for the COVID-19 Genomics UK (COG-UK) Consortium | Lighthouse Laboratory Plymouth and Alex Alderton, Roberto Amato, Jeffrey Barrett, Sonia Goncalves, Ewan Harrison, David K. Jackson, Ian Johnston, Dominic Kwiatkowski, Cordelia Langford, John Sillitoe on behalf of the Wellcome Sanger Institute COVID-19 Surveillance Team |
| EPI_ISL_2808256 | Health Services Laboratories | Wellcome Sanger Institute for the COVID-19 Genomics UK (COG-UK) Consortium | Health Services Laboratories and Alex Alderton, Roberto Amato, Jeffrey Barrett, Sonia Goncalves, Ewan Harrison, David K. Jackson, Ian Johnston, Dominic Kwiatkowski, Cordelia Langford, John Sillitoe on behalf of the Wellcome Sanger Institute COVID-19 Surveillance Team |
| EPI_ISL_2808257, EPI_ISL_2808258 | Lighthouse Laboratory Plymouth | Wellcome Sanger Institute for the COVID-19 Genomics UK (COG-UK) Consortium | Lighthouse Laboratory Plymouth and Alex Alderton, Roberto Amato, Jeffrey Barrett, Sonia Goncalves, Ewan Harrison, David K. Jackson, Ian Johnston, Dominic Kwiatkowski, Cordelia Langford, John Sillitoe on behalf of the Wellcome Sanger Institute COVID-19 Surveillance Team |
| EPI_ISL_2808259 | Lighthouse Lab in Milton Keynes | Wellcome Sanger Institute for the COVID-19 Genomics UK (COG-UK) Consortium | The Lighthouse Lab in Milton Keynes and Alex Alderton, Roberto Amato, Jeffrey Barrett, Sonia Goncalves, Ewan Harrison, David K. Jackson, Ian Johnston, Dominic Kwiatkowski, Cordelia Langford, John Sillitoe on behalf of the Wellcome Sanger Institute COVID-19 Surveillance Team |
| EPI_ISL_2808260 | Lighthouse Laboratory Plymouth | Wellcome Sanger Institute for the COVID-19 Genomics UK (COG-UK) Consortium | Lighthouse Laboratory Plymouth and Alex Alderton, Roberto Amato, Jeffrey Barrett, Sonia Goncalves, Ewan Harrison, David K. Jackson, Ian Johnston, Dominic Kwiatkowski, Cordelia Langford, John Sillitoe on behalf of the Wellcome Sanger Institute COVID-19 Surveillance Team |
| EPI_ISL_2808261 | Health Services Laboratories | Wellcome Sanger Institute for the COVID-19 Genomics UK (COG-UK) Consortium | Health Services Laboratories and Alex Alderton, Roberto Amato, Jeffrey Barrett, Sonia Goncalves, Ewan Harrison, David K. Jackson, Ian Johnston, Dominic Kwiatkowski, Cordelia Langford, John Sillitoe on behalf of the Wellcome Sanger Institute COVID-19 Surveillance Team |
| EPI_ISL_2808262, EPI_ISL_2808263 | Lighthouse Laboratory Plymouth | Wellcome Sanger Institute for the COVID-19 Genomics UK (COG-UK) Consortium | Lighthouse Laboratory Plymouth and Alex Alderton, Roberto Amato, Jeffrey Barrett, Sonia Goncalves, Ewan Harrison, David K. Jackson, Ian Johnston, Dominic Kwiatkowski, Cordelia Langford, John Sillitoe on behalf of the Wellcome Sanger Institute COVID-19 Surveillance Team |
| EPI_ISL_2808264 | Health Services Laboratories | Wellcome Sanger Institute for the COVID-19 Genomics UK (COG-UK) Consortium | Health Services Laboratories and Alex Alderton, Roberto Amato, Jeffrey Barrett, Sonia Goncalves, Ewan Harrison, David K. Jackson, Ian Johnston, Dominic Kwiatkowski, Cordelia Langford, John Sillitoe on behalf of the Wellcome Sanger Institute COVID-19 Surveillance Team |
| EPI_ISL_2808265 | Lighthouse Laboratory Plymouth | Wellcome Sanger Institute for the COVID-19 Genomics UK (COG-UK) Consortium | Lighthouse Laboratory Plymouth and Alex Alderton, Roberto Amato, Jeffrey Barrett, Sonia Goncalves, Ewan Harrison, David K. Jackson, Ian Johnston, Dominic Kwiatkowski, Cordelia Langford, John Sillitoe on behalf of the Wellcome Sanger Institute COVID-19 Surveillance Team |
| EPI_ISL_2808266 | Health Services Laboratories | Wellcome Sanger Institute for the COVID-19 Genomics UK (COG-UK) Consortium | Health Services Laboratories and Alex Alderton, Roberto Amato, Jeffrey Barrett, Sonia Goncalves, Ewan Harrison, David K. Jackson, Ian Johnston, Dominic Kwiatkowski, Cordelia Langford, John Sillitoe on behalf of the Wellcome Sanger Institute COVID-19 Surveillance Team |
| EPI_ISL_2808267 | Lighthouse Lab in Milton Keynes | Wellcome Sanger Institute for the COVID-19 Genomics UK (COG-UK) Consortium | The Lighthouse Lab in Milton Keynes and Alex Alderton, Roberto Amato, Jeffrey Barrett, Sonia Goncalves, Ewan Harrison, David K. Jackson, Ian Johnston, Dominic Kwiatkowski, Cordelia Langford, John Sillitoe on behalf of the Wellcome Sanger Institute COVID-19 Surveillance Team |
| EPI_ISL_2808268, EPI_ISL_2808269, EPI_ISL_2808270, EPI_ISL_2808271 | Lighthouse Laboratory Plymouth | Wellcome Sanger Institute for the COVID-19 Genomics UK (COG-UK) Consortium | Lighthouse Laboratory Plymouth and Alex Alderton, Roberto Amato, Jeffrey Barrett, Sonia Goncalves, Ewan Harrison, David K. Jackson, Ian Johnston, Dominic Kwiatkowski, Cordelia Langford, John Sillitoe on behalf of the Wellcome Sanger Institute COVID-19 Surveillance Team |
| EPI_ISL_2808272 | Health Services Laboratories | Wellcome Sanger Institute for the COVID-19 Genomics UK (COG-UK) Consortium | Health Services Laboratories and Alex Alderton, Roberto Amato, Jeffrey Barrett, Sonia Goncalves, Ewan Harrison, David K. Jackson, Ian Johnston, Dominic Kwiatkowski, Cordelia Langford, John Sillitoe on behalf of the Wellcome Sanger Institute COVID-19 Surveillance Team |

| | | | |
|---|---|---|---|
| EPI_ISL_2808273, EPI_ISL_2808274, EPI_ISL_2808275, EPI_ISL_2808276 | Lighthouse Laboratory Plymouth | Wellcome Sanger Institute for the COVID-19 Genomics UK (COG-UK) Consortium | Lighthouse Laboratory Plymouth and Alex Alderton, Roberto Amato, Jeffrey Barrett, Sonia Goncalves, Ewan Harrison, David K. Jackson, Ian Johnston, Dominic Kwiatkowski, Cordelia Langford, John Sillitoe on behalf of the Wellcome Sanger Institute COVID-19 Surveillance Team |
| EPI_ISL_2808277 | Health Services Laboratories | Wellcome Sanger Institute for the COVID-19 Genomics UK (COG-UK) Consortium | Health Services Laboratories and Alex Alderton, Roberto Amato, Jeffrey Barrett, Sonia Goncalves, Ewan Harrison, David K. Jackson, Ian Johnston, Dominic Kwiatkowski, Cordelia Langford, John Sillitoe on behalf of the Wellcome Sanger Institute COVID-19 Surveillance Team |
| EPI_ISL_2808278, EPI_ISL_2808279, EPI_ISL_2808280 | Lighthouse Laboratory Plymouth | Wellcome Sanger Institute for the COVID-19 Genomics UK (COG-UK) Consortium | Lighthouse Laboratory Plymouth and Alex Alderton, Roberto Amato, Jeffrey Barrett, Sonia Goncalves, Ewan Harrison, David K. Jackson, Ian Johnston, Dominic Kwiatkowski, Cordelia Langford, John Sillitoe on behalf of the Wellcome Sanger Institute COVID-19 Surveillance Team |
| EPI_ISL_2808281 | Lighthouse Lab in Milton Keynes | Wellcome Sanger Institute for the COVID-19 Genomics UK (COG-UK) Consortium | The Lighthouse Lab in Milton Keynes and Alex Alderton, Roberto Amato, Jeffrey Barrett, Sonia Goncalves, Ewan Harrison, David K. Jackson, Ian Johnston, Dominic Kwiatkowski, Cordelia Langford, John Sillitoe on behalf of the Wellcome Sanger Institute COVID-19 Surveillance Team |
| EPI_ISL_2808282, EPI_ISL_2808283, EPI_ISL_2808285, EPI_ISL_2808286, EPI_ISL_2808287, EPI_ISL_2808288, EPI_ISL_2808289 | Lighthouse Laboratory Plymouth | Wellcome Sanger Institute for the COVID-19 Genomics UK (COG-UK) Consortium | Lighthouse Laboratory Plymouth and Alex Alderton, Roberto Amato, Jeffrey Barrett, Sonia Goncalves, Ewan Harrison, David K. Jackson, Ian Johnston, Dominic Kwiatkowski, Cordelia Langford, John Sillitoe on behalf of the Wellcome Sanger Institute COVID-19 Surveillance Team |
| EPI_ISL_2808290 | Health Services Laboratories | Wellcome Sanger Institute for the COVID-19 Genomics UK (COG-UK) Consortium | Health Services Laboratories and Alex Alderton, Roberto Amato, Jeffrey Barrett, Sonia Goncalves, Ewan Harrison, David K. Jackson, Ian Johnston, Dominic Kwiatkowski, Cordelia Langford, John Sillitoe on behalf of the Wellcome Sanger Institute COVID-19 Surveillance Team |
| EPI_ISL_2808291, EPI_ISL_2808292 | Lighthouse Laboratory Plymouth | Wellcome Sanger Institute for the COVID-19 Genomics UK (COG-UK) Consortium | Lighthouse Laboratory Plymouth and Alex Alderton, Roberto Amato, Jeffrey Barrett, Sonia Goncalves, Ewan Harrison, David K. Jackson, Ian Johnston, Dominic Kwiatkowski, Cordelia Langford, John Sillitoe on behalf of the Wellcome Sanger Institute COVID-19 Surveillance Team |
| EPI_ISL_2808293 | Health Services Laboratories | Wellcome Sanger Institute for the COVID-19 Genomics UK (COG-UK) Consortium | Health Services Laboratories and Alex Alderton, Roberto Amato, Jeffrey Barrett, Sonia Goncalves, Ewan Harrison, David K. Jackson, Ian Johnston, Dominic Kwiatkowski, Cordelia Langford, John Sillitoe on behalf of the Wellcome Sanger Institute COVID-19 Surveillance Team |
| EPI_ISL_2808294, EPI_ISL_2808295, EPI_ISL_2808296, EPI_ISL_2808297, EPI_ISL_2808298, EPI_ISL_2808299, EPI_ISL_2808300 | Lighthouse Laboratory Plymouth | Wellcome Sanger Institute for the COVID-19 Genomics UK (COG-UK) Consortium | Lighthouse Laboratory Plymouth and Alex Alderton, Roberto Amato, Jeffrey Barrett, Sonia Goncalves, Ewan Harrison, David K. Jackson, Ian Johnston, Dominic Kwiatkowski, Cordelia Langford, John Sillitoe on behalf of the Wellcome Sanger Institute COVID-19 Surveillance Team |
| EPI_ISL_2808301 | Health Services Laboratories | Wellcome Sanger Institute for the COVID-19 Genomics UK (COG-UK) Consortium | Health Services Laboratories and Alex Alderton, Roberto Amato, Jeffrey Barrett, Sonia Goncalves, Ewan Harrison, David K. Jackson, Ian Johnston, Dominic Kwiatkowski, Cordelia Langford, John Sillitoe on behalf of the Wellcome Sanger Institute COVID-19 Surveillance Team |
| EPI_ISL_2808302, EPI_ISL_2808303, EPI_ISL_2808304 | Lighthouse Lab in Milton Keynes | Wellcome Sanger Institute for the COVID-19 Genomics UK (COG-UK) Consortium | The Lighthouse Lab in Milton Keynes and Alex Alderton, Roberto Amato, Jeffrey Barrett, Sonia Goncalves, Ewan Harrison, David K. Jackson, Ian Johnston, Dominic Kwiatkowski, Cordelia Langford, John Sillitoe on behalf of the Wellcome Sanger Institute COVID-19 Surveillance Team |
| EPI_ISL_2808305, EPI_ISL_2808306, EPI_ISL_2808307, EPI_ISL_2808308, EPI_ISL_2808309 | Lighthouse Laboratory Plymouth | Wellcome Sanger Institute for the COVID-19 Genomics UK (COG-UK) Consortium | Lighthouse Laboratory Plymouth and Alex Alderton, Roberto Amato, Jeffrey Barrett, Sonia Goncalves, Ewan Harrison, David K. Jackson, Ian Johnston, Dominic Kwiatkowski, Cordelia Langford, John Sillitoe on behalf of the Wellcome Sanger Institute COVID-19 Surveillance Team |
| EPI_ISL_2808310 | Health Services Laboratories | Wellcome Sanger Institute for the COVID-19 Genomics UK (COG-UK) Consortium | Health Services Laboratories and Alex Alderton, Roberto Amato, Jeffrey Barrett, Sonia Goncalves, Ewan Harrison, David K. Jackson, Ian Johnston, Dominic Kwiatkowski, Cordelia Langford, John Sillitoe on behalf of the Wellcome Sanger Institute COVID-19 Surveillance Team |
| EPI_ISL_2808311, EPI_ISL_2808312, EPI_ISL_2808313, EPI_ISL_2808314 | Lighthouse Laboratory Plymouth | Wellcome Sanger Institute for the COVID-19 Genomics UK (COG-UK) Consortium | Lighthouse Laboratory Plymouth and Alex Alderton, Roberto Amato, Jeffrey Barrett, Sonia Goncalves, Ewan Harrison, David K. Jackson, Ian Johnston, Dominic Kwiatkowski, Cordelia Langford, John Sillitoe on behalf of the Wellcome Sanger Institute COVID-19 Surveillance Team |
| EPI_ISL_2808315 | Health Services Laboratories | Wellcome Sanger Institute for the COVID-19 Genomics UK (COG-UK) Consortium | Health Services Laboratories and Alex Alderton, Roberto Amato, Jeffrey Barrett, Sonia Goncalves, Ewan Harrison, David K. Jackson, Ian Johnston, Dominic Kwiatkowski, Cordelia Langford, John Sillitoe on behalf of the Wellcome Sanger Institute COVID-19 Surveillance Team |
| EPI_ISL_2808316, EPI_ISL_2808317, EPI_ISL_2808318 | Lighthouse Laboratory Plymouth | Wellcome Sanger Institute for the COVID-19 Genomics UK (COG-UK) Consortium | Lighthouse Laboratory Plymouth and Alex Alderton, Roberto Amato, Jeffrey Barrett, Sonia Goncalves, Ewan Harrison, David K. Jackson, Ian Johnston, Dominic Kwiatkowski, Cordelia Langford, John Sillitoe on behalf of the Wellcome Sanger Institute COVID-19 Surveillance Team |
| EPI_ISL_2808319 | Lighthouse Lab in Milton Keynes | Wellcome Sanger Institute for the COVID-19 Genomics UK (COG-UK) Consortium | The Lighthouse Lab in Milton Keynes and Alex Alderton, Roberto Amato, Jeffrey Barrett, Sonia Goncalves, Ewan Harrison, David K. Jackson, Ian Johnston, Dominic Kwiatkowski, Cordelia Langford, John Sillitoe on behalf of the Wellcome Sanger Institute COVID-19 Surveillance Team |
| EPI_ISL_2808320, EPI_ISL_2808321, EPI_ISL_2808322 | Lighthouse Laboratory Plymouth | Wellcome Sanger Institute for the COVID-19 Genomics UK (COG-UK) Consortium | Lighthouse Laboratory Plymouth and Alex Alderton, Roberto Amato, Jeffrey Barrett, Sonia Goncalves, Ewan Harrison, David K. Jackson, Ian Johnston, Dominic Kwiatkowski, Cordelia Langford, John Sillitoe on behalf of the Wellcome Sanger Institute COVID-19 Surveillance Team |
| EPI_ISL_2808323, EPI_ISL_2808324 | Lighthouse Lab in Milton Keynes | Wellcome Sanger Institute for the COVID-19 Genomics UK (COG-UK) Consortium | The Lighthouse Lab in Milton Keynes and Alex Alderton, Roberto Amato, Jeffrey Barrett, Sonia Goncalves, Ewan Harrison, David K. Jackson, Ian Johnston, Dominic Kwiatkowski, Cordelia Langford, John Sillitoe on behalf of the Wellcome Sanger Institute COVID-19 Surveillance Team |
| EPI_ISL_2808325 | Lighthouse Laboratory Plymouth | Wellcome Sanger Institute for the COVID-19 Genomics UK (COG-UK) Consortium | Lighthouse Laboratory Plymouth and Alex Alderton, Roberto Amato, Jeffrey Barrett, Sonia Goncalves, Ewan Harrison, David K. Jackson, Ian Johnston, Dominic Kwiatkowski, Cordelia Langford, John Sillitoe on behalf of the Wellcome Sanger Institute COVID-19 Surveillance Team |
| EPI_ISL_2808326 | Health Services Laboratories | Wellcome Sanger Institute for the COVID-19 Genomics UK (COG-UK) Consortium | Health Services Laboratories and Alex Alderton, Roberto Amato, Jeffrey Barrett, Sonia Goncalves, Ewan Harrison, David K. Jackson, Ian Johnston, Dominic Kwiatkowski, Cordelia Langford, John Sillitoe on behalf of the Wellcome Sanger Institute COVID-19 Surveillance Team |
| EPI_ISL_2808327 | Lighthouse Laboratory Plymouth | Wellcome Sanger Institute for the COVID-19 Genomics UK (COG-UK) Consortium | Lighthouse Laboratory Plymouth and Alex Alderton, Roberto Amato, Jeffrey Barrett, Sonia Goncalves, Ewan Harrison, David K. Jackson, Ian Johnston, Dominic Kwiatkowski, Cordelia Langford, John Sillitoe on behalf of the Wellcome Sanger Institute COVID-19 Surveillance Team |
| EPI_ISL_2808328 | Lighthouse Lab in Milton Keynes | Wellcome Sanger Institute for the COVID-19 Genomics UK (COG-UK) Consortium | The Lighthouse Lab in Milton Keynes and Alex Alderton, Roberto Amato, Jeffrey Barrett, Sonia Goncalves, Ewan Harrison, David K. Jackson, Ian Johnston, Dominic Kwiatkowski, Cordelia Langford, John Sillitoe on behalf of the Wellcome Sanger Institute COVID-19 Surveillance Team |
| EPI_ISL_2808329 | Health Services Laboratories | Wellcome Sanger Institute for the COVID-19 Genomics UK (COG-UK) Consortium | Health Services Laboratories and Alex Alderton, Roberto Amato, Jeffrey Barrett, Sonia Goncalves, Ewan Harrison, David K. Jackson, Ian Johnston, Dominic Kwiatkowski, Cordelia Langford, John Sillitoe on behalf of the Wellcome Sanger Institute COVID-19 Surveillance Team |
| EPI_ISL_2808330, EPI_ISL_2808331 | Lighthouse Laboratory Plymouth | Wellcome Sanger Institute for the COVID-19 Genomics UK (COG-UK) Consortium | Lighthouse Laboratory Plymouth and Alex Alderton, Roberto Amato, Jeffrey Barrett, Sonia Goncalves, Ewan Harrison, David K. Jackson, Ian Johnston, Dominic Kwiatkowski, Cordelia Langford, John Sillitoe on behalf of the Wellcome Sanger Institute COVID-19 Surveillance Team |
| EPI_ISL_2808332 | Health Services Laboratories | Wellcome Sanger Institute for the COVID-19 Genomics UK (COG-UK) Consortium | Health Services Laboratories and Alex Alderton, Roberto Amato, Jeffrey Barrett, Sonia Goncalves, Ewan Harrison, David K. Jackson, Ian Johnston, Dominic Kwiatkowski, Cordelia Langford, John Sillitoe on behalf of the Wellcome Sanger Institute COVID-19 Surveillance Team |
| EPI_ISL_2808333, EPI_ISL_2808334, EPI_ISL_2808335, EPI_ISL_2808336, EPI_ISL_2808339, EPI_ISL_2808340 | Lighthouse Laboratory Plymouth | Wellcome Sanger Institute for the COVID-19 Genomics UK (COG-UK) Consortium | Lighthouse Laboratory Plymouth and Alex Alderton, Roberto Amato, Jeffrey Barrett, Sonia Goncalves, Ewan Harrison, David K. Jackson, Ian Johnston, Dominic Kwiatkowski, Cordelia Langford, John Sillitoe on behalf of the Wellcome Sanger Institute COVID-19 Surveillance Team |
| EPI_ISL_2808341 | Health Services Laboratories | Wellcome Sanger Institute for the COVID-19 Genomics UK (COG-UK) Consortium | Health Services Laboratories and Alex Alderton, Roberto Amato, Jeffrey Barrett, Sonia Goncalves, Ewan Harrison, David K. Jackson, Ian Johnston, Dominic Kwiatkowski, Cordelia Langford, John Sillitoe on behalf of the Wellcome Sanger Institute COVID-19 Surveillance Team |
| EPI_ISL_2808342, EPI_ISL_2808343 | Lighthouse Laboratory Plymouth | Wellcome Sanger Institute for the COVID-19 Genomics UK (COG-UK) Consortium | Lighthouse Laboratory Plymouth and Alex Alderton, Roberto Amato, Jeffrey Barrett, Sonia Goncalves, Ewan Harrison, David K. Jackson, Ian Johnston, Dominic Kwiatkowski, Cordelia Langford, John Sillitoe on behalf of the Wellcome Sanger Institute COVID-19 Surveillance Team |
| EPI_ISL_2808344 | Health Services Laboratories | Wellcome Sanger Institute for the COVID-19 Genomics UK (COG-UK) Consortium | Health Services Laboratories and Alex Alderton, Roberto Amato, Jeffrey Barrett, Sonia Goncalves, Ewan Harrison, David K. Jackson, Ian Johnston, Dominic Kwiatkowski, Cordelia Langford, John Sillitoe on behalf of the Wellcome Sanger Institute COVID-19 Surveillance Team |
| EPI_ISL_2808345, EPI_ISL_2808346 | Lighthouse Laboratory Plymouth | Wellcome Sanger Institute for the COVID-19 Genomics UK (COG-UK) Consortium | Lighthouse Laboratory Plymouth and Alex Alderton, Roberto Amato, Jeffrey Barrett, Sonia Goncalves, Ewan Harrison, David K. Jackson, Ian Johnston, Dominic Kwiatkowski, Cordelia Langford, John Sillitoe on behalf of the Wellcome Sanger Institute COVID-19 Surveillance Team |
| EPI_ISL_2808347 | Lighthouse Lab in Milton Keynes | Wellcome Sanger Institute for the COVID-19 Genomics UK (COG-UK) Consortium | The Lighthouse Lab in Milton Keynes and Alex Alderton, Roberto Amato, Jeffrey Barrett, Sonia Goncalves, Ewan Harrison, David K. Jackson, Ian Johnston, Dominic Kwiatkowski, Cordelia Langford, John Sillitoe on behalf of the Wellcome Sanger Institute COVID-19 Surveillance Team |
| EPI_ISL_2808348, EPI_ISL_2808349 | Lighthouse Laboratory Plymouth | Wellcome Sanger Institute for the COVID-19 Genomics UK (COG-UK) Consortium | Lighthouse Laboratory Plymouth and Alex Alderton, Roberto Amato, Jeffrey Barrett, Sonia Goncalves, Ewan Harrison, David K. Jackson, Ian Johnston, Dominic Kwiatkowski, Cordelia Langford, John Sillitoe on behalf of the Wellcome Sanger Institute COVID-19 Surveillance Team |
| EPI_ISL_2808350 | Health Services Laboratories | Wellcome Sanger Institute for the COVID-19 Genomics UK (COG-UK) Consortium | Health Services Laboratories and Alex Alderton, Roberto Amato, Jeffrey Barrett, Sonia Goncalves, Ewan Harrison, David K. Jackson, Ian Johnston, Dominic Kwiatkowski, Cordelia Langford, John Sillitoe on behalf of the Wellcome Sanger Institute COVID-19 Surveillance Team |
| EPI_ISL_2808351 | Lighthouse Lab in Milton Keynes | Wellcome Sanger Institute for the COVID-19 Genomics UK (COG-UK) | The Lighthouse Lab in Milton Keynes and Alex Alderton, Roberto Amato, Jeffrey Barrett, Sonia Goncalves, Ewan Harrison, David K. Jackson, Ian Johnston, Dominic Kwiatkowski, Cordelia Langford, John Sillitoe on behalf of the Wellcome Sanger |

| | | | |
|---|---|---|---|
| | | Consortium | Institute COVID-19 Surveillance Team |
| EPI_ISL_2808352, EPI_ISL_2808353 | Health Services Laboratories | Wellcome Sanger Institute for the COVID-19 Genomics UK (COG-UK) Consortium | Health Services Laboratories and Alex Alderton, Roberto Amato, Jeffrey Barrett, Sonia Goncalves, Ewan Harrison, David K. Jackson, Ian Johnston, Dominic Kwiatkowski, Cordelia Langford, John Sillitoe on behalf of the Wellcome Sanger Institute COVID-19 Surveillance Team |
| EPI_ISL_2808354, EPI_ISL_2808355, EPI_ISL_2808356, EPI_ISL_2808357, EPI_ISL_2808358 | Lighthouse Laboratory Plymouth | Wellcome Sanger Institute for the COVID-19 Genomics UK (COG-UK) Consortium | Lighthouse Laboratory Plymouth and Alex Alderton, Roberto Amato, Jeffrey Barrett, Sonia Goncalves, Ewan Harrison, David K. Jackson, Ian Johnston, Dominic Kwiatkowski, Cordelia Langford, John Sillitoe on behalf of the Wellcome Sanger Institute COVID-19 Surveillance Team |
| EPI_ISL_2808359, EPI_ISL_2808360, EPI_ISL_2808361 | Health Services Laboratories | Wellcome Sanger Institute for the COVID-19 Genomics UK (COG-UK) Consortium | Health Services Laboratories and Alex Alderton, Roberto Amato, Jeffrey Barrett, Sonia Goncalves, Ewan Harrison, David K. Jackson, Ian Johnston, Dominic Kwiatkowski, Cordelia Langford, John Sillitoe on behalf of the Wellcome Sanger Institute COVID-19 Surveillance Team |
| EPI_ISL_2808362, EPI_ISL_2808363, EPI_ISL_2808364, EPI_ISL_2808365, EPI_ISL_2808366, EPI_ISL_2808367, EPI_ISL_2808368, EPI_ISL_2808369 | Lighthouse Laboratory Plymouth | Wellcome Sanger Institute for the COVID-19 Genomics UK (COG-UK) Consortium | Lighthouse Laboratory Plymouth and Alex Alderton, Roberto Amato, Jeffrey Barrett, Sonia Goncalves, Ewan Harrison, David K. Jackson, Ian Johnston, Dominic Kwiatkowski, Cordelia Langford, John Sillitoe on behalf of the Wellcome Sanger Institute COVID-19 Surveillance Team |
| EPI_ISL_2808370 | Lighthouse Lab in Milton Keynes | Wellcome Sanger Institute for the COVID-19 Genomics UK (COG-UK) Consortium | The Lighthouse Lab in Milton Keynes and Alex Alderton, Roberto Amato, Jeffrey Barrett, Sonia Goncalves, Ewan Harrison, David K. Jackson, Ian Johnston, Dominic Kwiatkowski, Cordelia Langford, John Sillitoe on behalf of the Wellcome Sanger Institute COVID-19 Surveillance Team |
| EPI_ISL_2808371, EPI_ISL_2808372, EPI_ISL_2808373, EPI_ISL_2808374 | Lighthouse Laboratory Plymouth | Wellcome Sanger Institute for the COVID-19 Genomics UK (COG-UK) Consortium | Lighthouse Laboratory Plymouth and Alex Alderton, Roberto Amato, Jeffrey Barrett, Sonia Goncalves, Ewan Harrison, David K. Jackson, Ian Johnston, Dominic Kwiatkowski, Cordelia Langford, John Sillitoe on behalf of the Wellcome Sanger Institute COVID-19 Surveillance Team |
| EPI_ISL_2808375 | Lighthouse Lab in Milton Keynes | Wellcome Sanger Institute for the COVID-19 Genomics UK (COG-UK) Consortium | The Lighthouse Lab in Milton Keynes and Alex Alderton, Roberto Amato, Jeffrey Barrett, Sonia Goncalves, Ewan Harrison, David K. Jackson, Ian Johnston, Dominic Kwiatkowski, Cordelia Langford, John Sillitoe on behalf of the Wellcome Sanger Institute COVID-19 Surveillance Team |
| EPI_ISL_2808376, EPI_ISL_2808377, EPI_ISL_2808378 | Lighthouse Laboratory Plymouth | Wellcome Sanger Institute for the COVID-19 Genomics UK (COG-UK) Consortium | Lighthouse Laboratory Plymouth and Alex Alderton, Roberto Amato, Jeffrey Barrett, Sonia Goncalves, Ewan Harrison, David K. Jackson, Ian Johnston, Dominic Kwiatkowski, Cordelia Langford, John Sillitoe on behalf of the Wellcome Sanger Institute COVID-19 Surveillance Team |
| EPI_ISL_2808379 | Lighthouse Lab in Milton Keynes | Wellcome Sanger Institute for the COVID-19 Genomics UK (COG-UK) Consortium | The Lighthouse Lab in Milton Keynes and Alex Alderton, Roberto Amato, Jeffrey Barrett, Sonia Goncalves, Ewan Harrison, David K. Jackson, Ian Johnston, Dominic Kwiatkowski, Cordelia Langford, John Sillitoe on behalf of the Wellcome Sanger Institute COVID-19 Surveillance Team |
| EPI_ISL_2808380 | Lighthouse Laboratory Plymouth | Wellcome Sanger Institute for the COVID-19 Genomics UK (COG-UK) Consortium | Lighthouse Laboratory Plymouth and Alex Alderton, Roberto Amato, Jeffrey Barrett, Sonia Goncalves, Ewan Harrison, David K. Jackson, Ian Johnston, Dominic Kwiatkowski, Cordelia Langford, John Sillitoe on behalf of the Wellcome Sanger Institute COVID-19 Surveillance Team |
| EPI_ISL_2808382 | Health Services Laboratories | Wellcome Sanger Institute for the COVID-19 Genomics UK (COG-UK) Consortium | Health Services Laboratories and Alex Alderton, Roberto Amato, Jeffrey Barrett, Sonia Goncalves, Ewan Harrison, David K. Jackson, Ian Johnston, Dominic Kwiatkowski, Cordelia Langford, John Sillitoe on behalf of the Wellcome Sanger Institute COVID-19 Surveillance Team |
| EPI_ISL_2808383, EPI_ISL_2808384 | Lighthouse Laboratory Plymouth | Wellcome Sanger Institute for the COVID-19 Genomics UK (COG-UK) Consortium | Lighthouse Laboratory Plymouth and Alex Alderton, Roberto Amato, Jeffrey Barrett, Sonia Goncalves, Ewan Harrison, David K. Jackson, Ian Johnston, Dominic Kwiatkowski, Cordelia Langford, John Sillitoe on behalf of the Wellcome Sanger Institute COVID-19 Surveillance Team |
| EPI_ISL_2808385 | Lighthouse Lab in Milton Keynes | Wellcome Sanger Institute for the COVID-19 Genomics UK (COG-UK) Consortium | The Lighthouse Lab in Milton Keynes and Alex Alderton, Roberto Amato, Jeffrey Barrett, Sonia Goncalves, Ewan Harrison, David K. Jackson, Ian Johnston, Dominic Kwiatkowski, Cordelia Langford, John Sillitoe on behalf of the Wellcome Sanger Institute COVID-19 Surveillance Team |
| EPI_ISL_2808386, EPI_ISL_2808387 | Lighthouse Laboratory Plymouth | Wellcome Sanger Institute for the COVID-19 Genomics UK (COG-UK) Consortium | Lighthouse Laboratory Plymouth and Alex Alderton, Roberto Amato, Jeffrey Barrett, Sonia Goncalves, Ewan Harrison, David K. Jackson, Ian Johnston, Dominic Kwiatkowski, Cordelia Langford, John Sillitoe on behalf of the Wellcome Sanger Institute COVID-19 Surveillance Team |
| EPI_ISL_2808388 | Lighthouse Lab in Milton Keynes | Wellcome Sanger Institute for the COVID-19 Genomics UK (COG-UK) Consortium | The Lighthouse Lab in Milton Keynes and Alex Alderton, Roberto Amato, Jeffrey Barrett, Sonia Goncalves, Ewan Harrison, David K. Jackson, Ian Johnston, Dominic Kwiatkowski, Cordelia Langford, John Sillitoe on behalf of the Wellcome Sanger Institute COVID-19 Surveillance Team |
| EPI_ISL_2808389 | Lighthouse Laboratory Plymouth | Wellcome Sanger Institute for the COVID-19 Genomics UK (COG-UK) Consortium | Lighthouse Laboratory Plymouth and Alex Alderton, Roberto Amato, Jeffrey Barrett, Sonia Goncalves, Ewan Harrison, David K. Jackson, Ian Johnston, Dominic Kwiatkowski, Cordelia Langford, John Sillitoe on behalf of the Wellcome Sanger Institute COVID-19 Surveillance Team |
| EPI_ISL_2808390 | Lighthouse Lab in Milton Keynes | Wellcome Sanger Institute for the COVID-19 Genomics UK (COG-UK) Consortium | The Lighthouse Lab in Milton Keynes and Alex Alderton, Roberto Amato, Jeffrey Barrett, Sonia Goncalves, Ewan Harrison, David K. Jackson, Ian Johnston, Dominic Kwiatkowski, Cordelia Langford, John Sillitoe on behalf of the Wellcome Sanger Institute COVID-19 Surveillance Team |
| EPI_ISL_2808391 | Health Services Laboratories | Wellcome Sanger Institute for the COVID-19 Genomics UK (COG-UK) Consortium | Health Services Laboratories and Alex Alderton, Roberto Amato, Jeffrey Barrett, Sonia Goncalves, Ewan Harrison, David K. Jackson, Ian Johnston, Dominic Kwiatkowski, Cordelia Langford, John Sillitoe on behalf of the Wellcome Sanger Institute COVID-19 Surveillance Team |
| EPI_ISL_2808392, EPI_ISL_2808393, EPI_ISL_2808394, EPI_ISL_2808396 | Lighthouse Laboratory Plymouth | Wellcome Sanger Institute for the COVID-19 Genomics UK (COG-UK) Consortium | Lighthouse Laboratory Plymouth and Alex Alderton, Roberto Amato, Jeffrey Barrett, Sonia Goncalves, Ewan Harrison, David K. Jackson, Ian Johnston, Dominic Kwiatkowski, Cordelia Langford, John Sillitoe on behalf of the Wellcome Sanger Institute COVID-19 Surveillance Team |
| EPI_ISL_2808397 | Health Services Laboratories | Wellcome Sanger Institute for the COVID-19 Genomics UK (COG-UK) Consortium | Health Services Laboratories and Alex Alderton, Roberto Amato, Jeffrey Barrett, Sonia Goncalves, Ewan Harrison, David K. Jackson, Ian Johnston, Dominic Kwiatkowski, Cordelia Langford, John Sillitoe on behalf of the Wellcome Sanger Institute COVID-19 Surveillance Team |
| EPI_ISL_2808398 | Lighthouse Laboratory Plymouth | Wellcome Sanger Institute for the COVID-19 Genomics UK (COG-UK) Consortium | Lighthouse Laboratory Plymouth and Alex Alderton, Roberto Amato, Jeffrey Barrett, Sonia Goncalves, Ewan Harrison, David K. Jackson, Ian Johnston, Dominic Kwiatkowski, Cordelia Langford, John Sillitoe on behalf of the Wellcome Sanger Institute COVID-19 Surveillance Team |
| EPI_ISL_2808399, EPI_ISL_2808400 | Health Services Laboratories | Wellcome Sanger Institute for the COVID-19 Genomics UK (COG-UK) Consortium | Health Services Laboratories and Alex Alderton, Roberto Amato, Jeffrey Barrett, Sonia Goncalves, Ewan Harrison, David K. Jackson, Ian Johnston, Dominic Kwiatkowski, Cordelia Langford, John Sillitoe on behalf of the Wellcome Sanger Institute COVID-19 Surveillance Team |
| EPI_ISL_2808401 | Lighthouse Laboratory Plymouth | Wellcome Sanger Institute for the COVID-19 Genomics UK (COG-UK) Consortium | Lighthouse Laboratory Plymouth and Alex Alderton, Roberto Amato, Jeffrey Barrett, Sonia Goncalves, Ewan Harrison, David K. Jackson, Ian Johnston, Dominic Kwiatkowski, Cordelia Langford, John Sillitoe on behalf of the Wellcome Sanger Institute COVID-19 Surveillance Team |
| EPI_ISL_2808403, EPI_ISL_2808404 | Lighthouse Lab in Milton Keynes | Wellcome Sanger Institute for the COVID-19 Genomics UK (COG-UK) Consortium | The Lighthouse Lab in Milton Keynes and Alex Alderton, Roberto Amato, Jeffrey Barrett, Sonia Goncalves, Ewan Harrison, David K. Jackson, Ian Johnston, Dominic Kwiatkowski, Cordelia Langford, John Sillitoe on behalf of the Wellcome Sanger Institute COVID-19 Surveillance Team |
| EPI_ISL_2808405, EPI_ISL_2808406 | Lighthouse Laboratory Plymouth | Wellcome Sanger Institute for the COVID-19 Genomics UK (COG-UK) Consortium | Lighthouse Laboratory Plymouth and Alex Alderton, Roberto Amato, Jeffrey Barrett, Sonia Goncalves, Ewan Harrison, David K. Jackson, Ian Johnston, Dominic Kwiatkowski, Cordelia Langford, John Sillitoe on behalf of the Wellcome Sanger Institute COVID-19 Surveillance Team |
| EPI_ISL_2808407 | Lighthouse Lab in Milton Keynes | Wellcome Sanger Institute for the COVID-19 Genomics UK (COG-UK) Consortium | The Lighthouse Lab in Milton Keynes and Alex Alderton, Roberto Amato, Jeffrey Barrett, Sonia Goncalves, Ewan Harrison, David K. Jackson, Ian Johnston, Dominic Kwiatkowski, Cordelia Langford, John Sillitoe on behalf of the Wellcome Sanger Institute COVID-19 Surveillance Team |
| EPI_ISL_2808409, EPI_ISL_2808410, EPI_ISL_2808411, EPI_ISL_2808412, EPI_ISL_2808413 | Lighthouse Laboratory Plymouth | Wellcome Sanger Institute for the COVID-19 Genomics UK (COG-UK) Consortium | Lighthouse Laboratory Plymouth and Alex Alderton, Roberto Amato, Jeffrey Barrett, Sonia Goncalves, Ewan Harrison, David K. Jackson, Ian Johnston, Dominic Kwiatkowski, Cordelia Langford, John Sillitoe on behalf of the Wellcome Sanger Institute COVID-19 Surveillance Team |
| EPI_ISL_2808414 | Lighthouse Lab in Milton Keynes | Wellcome Sanger Institute for the COVID-19 Genomics UK (COG-UK) Consortium | The Lighthouse Lab in Milton Keynes and Alex Alderton, Roberto Amato, Jeffrey Barrett, Sonia Goncalves, Ewan Harrison, David K. Jackson, Ian Johnston, Dominic Kwiatkowski, Cordelia Langford, John Sillitoe on behalf of the Wellcome Sanger Institute COVID-19 Surveillance Team |
| EPI_ISL_2808415, EPI_ISL_2808416, EPI_ISL_2808417, EPI_ISL_2808418, EPI_ISL_2808419, EPI_ISL_2808420, EPI_ISL_2808421, EPI_ISL_2808422 | Lighthouse Laboratory Plymouth | Wellcome Sanger Institute for the COVID-19 Genomics UK (COG-UK) Consortium | Lighthouse Laboratory Plymouth and Alex Alderton, Roberto Amato, Jeffrey Barrett, Sonia Goncalves, Ewan Harrison, David K. Jackson, Ian Johnston, Dominic Kwiatkowski, Cordelia Langford, John Sillitoe on behalf of the Wellcome Sanger Institute COVID-19 Surveillance Team |
| EPI_ISL_2808423 | Health Services Laboratories | Wellcome Sanger Institute for the COVID-19 Genomics UK (COG-UK) Consortium | Health Services Laboratories and Alex Alderton, Roberto Amato, Jeffrey Barrett, Sonia Goncalves, Ewan Harrison, David K. Jackson, Ian Johnston, Dominic Kwiatkowski, Cordelia Langford, John Sillitoe on behalf of the Wellcome Sanger Institute COVID-19 Surveillance Team |
| EPI_ISL_2808424, EPI_ISL_2808425 | Lighthouse Laboratory Plymouth | Wellcome Sanger Institute for the COVID-19 Genomics UK (COG-UK) Consortium | Lighthouse Laboratory Plymouth and Alex Alderton, Roberto Amato, Jeffrey Barrett, Sonia Goncalves, Ewan Harrison, David K. Jackson, Ian Johnston, Dominic Kwiatkowski, Cordelia Langford, John Sillitoe on behalf of the Wellcome Sanger Institute COVID-19 Surveillance Team |
| EPI_ISL_2808426 | Lighthouse Lab in Milton Keynes | Wellcome Sanger Institute for the COVID-19 Genomics UK (COG-UK) Consortium | The Lighthouse Lab in Milton Keynes and Alex Alderton, Roberto Amato, Jeffrey Barrett, Sonia Goncalves, Ewan Harrison, David K. Jackson, Ian Johnston, Dominic Kwiatkowski, Cordelia Langford, John Sillitoe on behalf of the Wellcome Sanger Institute COVID-19 Surveillance Team |
| EPI_ISL_2808427, EPI_ISL_2808428 | Lighthouse Laboratory Plymouth | Wellcome Sanger Institute for the COVID-19 Genomics UK (COG-UK) Consortium | Lighthouse Laboratory Plymouth and Alex Alderton, Roberto Amato, Jeffrey Barrett, Sonia Goncalves, Ewan Harrison, David K. Jackson, Ian Johnston, Dominic Kwiatkowski, Cordelia Langford, John Sillitoe on behalf of the Wellcome Sanger Institute COVID-19 Surveillance Team |
| EPI_ISL_2808429 | Lighthouse Lab in Milton Keynes | Wellcome Sanger Institute for the COVID-19 Genomics UK (COG-UK) Consortium | The Lighthouse Lab in Milton Keynes and Alex Alderton, Roberto Amato, Jeffrey Barrett, Sonia Goncalves, Ewan Harrison, David K. Jackson, Ian Johnston, Dominic Kwiatkowski, Cordelia Langford, John Sillitoe on behalf of the Wellcome Sanger Institute COVID-19 Surveillance Team |
| EPI_ISL_2808430 | Lighthouse Lab in Glasgow | Wellcome Sanger Institute for the COVID-19 Genomics UK (COG-UK) | Harper VanSteenhouse, Yumi Kasai, David Gray, Carol Clugston, Anna Dominiczak and Alex Alderton, Roberto Amato, Jeffrey Barrett, Sonia Goncalves, Ewan Harrison, David K. Jackson, Ian Johnston, Dominic Kwiatkowski, Cordelia Langford, |

| | | | |
|---|---|---|---|
| | | Consortium | John Sillitoe on behalf of the Wellcome Sanger Institute COVID-19 Surveillance Team |
| EPI_ISL_2808431 | Lighthouse Laboratory Plymouth | Wellcome Sanger Institute for the COVID-19 Genomics UK (COG-UK) Consortium | Lighthouse Laboratory Plymouth and Alex Alderton, Roberto Amato, Jeffrey Barrett, Sonia Goncalves, Ewan Harrison, David K. Jackson, Ian Johnston, Dominic Kwiatkowski, Cordelia Langford, John Sillitoe on behalf of the Wellcome Sanger Institute COVID-19 Surveillance Team |
| EPI_ISL_2808432 | Lighthouse Lab in Glasgow | Wellcome Sanger Institute for the COVID-19 Genomics UK (COG-UK) Consortium | Harper VanSteenhouse, Yumi Kasai, David Gray, Carol Clugston, Anna Dominiczak and Alex Alderton, Roberto Amato, Jeffrey Barrett, Sonia Goncalves, Ewan Harrison, David K. Jackson, Ian Johnston, Dominic Kwiatkowski, Cordelia Langford, John Sillitoe on behalf of the Wellcome Sanger Institute COVID-19 Surveillance Team |
| EPI_ISL_2808433 | Lighthouse Lab in Milton Keynes | Wellcome Sanger Institute for the COVID-19 Genomics UK (COG-UK) Consortium | The Lighthouse Lab in Milton Keynes and Alex Alderton, Roberto Amato, Jeffrey Barrett, Sonia Goncalves, Ewan Harrison, David K. Jackson, Ian Johnston, Dominic Kwiatkowski, Cordelia Langford, John Sillitoe on behalf of the Wellcome Sanger Institute COVID-19 Surveillance Team |
| EPI_ISL_2808434, EPI_ISL_2808435 | Lighthouse Lab in Glasgow | Wellcome Sanger Institute for the COVID-19 Genomics UK (COG-UK) Consortium | Harper VanSteenhouse, Yumi Kasai, David Gray, Carol Clugston, Anna Dominiczak and Alex Alderton, Roberto Amato, Jeffrey Barrett, Sonia Goncalves, Ewan Harrison, David K. Jackson, Ian Johnston, Dominic Kwiatkowski, Cordelia Langford, John Sillitoe on behalf of the Wellcome Sanger Institute COVID-19 Surveillance Team |
| EPI_ISL_2808436, EPI_ISL_2808437, EPI_ISL_2808438 | Lighthouse Laboratory Plymouth | Wellcome Sanger Institute for the COVID-19 Genomics UK (COG-UK) Consortium | Lighthouse Laboratory Plymouth and Alex Alderton, Roberto Amato, Jeffrey Barrett, Sonia Goncalves, Ewan Harrison, David K. Jackson, Ian Johnston, Dominic Kwiatkowski, Cordelia Langford, John Sillitoe on behalf of the Wellcome Sanger Institute COVID-19 Surveillance Team |
| EPI_ISL_2808439 | Lighthouse Lab in Glasgow | Wellcome Sanger Institute for the COVID-19 Genomics UK (COG-UK) Consortium | Harper VanSteenhouse, Yumi Kasai, David Gray, Carol Clugston, Anna Dominiczak and Alex Alderton, Roberto Amato, Jeffrey Barrett, Sonia Goncalves, Ewan Harrison, David K. Jackson, Ian Johnston, Dominic Kwiatkowski, Cordelia Langford, John Sillitoe on behalf of the Wellcome Sanger Institute COVID-19 Surveillance Team |
| EPI_ISL_2808440 | Lighthouse Lab in Milton Keynes | Wellcome Sanger Institute for the COVID-19 Genomics UK (COG-UK) Consortium | The Lighthouse Lab in Milton Keynes and Alex Alderton, Roberto Amato, Jeffrey Barrett, Sonia Goncalves, Ewan Harrison, David K. Jackson, Ian Johnston, Dominic Kwiatkowski, Cordelia Langford, John Sillitoe on behalf of the Wellcome Sanger Institute COVID-19 Surveillance Team |
| EPI_ISL_2808441 | Lighthouse Laboratory Plymouth | Wellcome Sanger Institute for the COVID-19 Genomics UK (COG-UK) Consortium | Lighthouse Laboratory Plymouth and Alex Alderton, Roberto Amato, Jeffrey Barrett, Sonia Goncalves, Ewan Harrison, David K. Jackson, Ian Johnston, Dominic Kwiatkowski, Cordelia Langford, John Sillitoe on behalf of the Wellcome Sanger Institute COVID-19 Surveillance Team |
| EPI_ISL_2808442 | Lighthouse Lab in Glasgow | Wellcome Sanger Institute for the COVID-19 Genomics UK (COG-UK) Consortium | Harper VanSteenhouse, Yumi Kasai, David Gray, Carol Clugston, Anna Dominiczak and Alex Alderton, Roberto Amato, Jeffrey Barrett, Sonia Goncalves, Ewan Harrison, David K. Jackson, Ian Johnston, Dominic Kwiatkowski, Cordelia Langford, John Sillitoe on behalf of the Wellcome Sanger Institute COVID-19 Surveillance Team |
| EPI_ISL_2808443 | Lighthouse Lab in Milton Keynes | Wellcome Sanger Institute for the COVID-19 Genomics UK (COG-UK) Consortium | The Lighthouse Lab in Milton Keynes and Alex Alderton, Roberto Amato, Jeffrey Barrett, Sonia Goncalves, Ewan Harrison, David K. Jackson, Ian Johnston, Dominic Kwiatkowski, Cordelia Langford, John Sillitoe on behalf of the Wellcome Sanger Institute COVID-19 Surveillance Team |
| EPI_ISL_2808444, EPI_ISL_2808445, EPI_ISL_2808446, EPI_ISL_2808447, EPI_ISL_2808448, EPI_ISL_2808449 | Lighthouse Laboratory Plymouth | Wellcome Sanger Institute for the COVID-19 Genomics UK (COG-UK) Consortium | Lighthouse Laboratory Plymouth and Alex Alderton, Roberto Amato, Jeffrey Barrett, Sonia Goncalves, Ewan Harrison, David K. Jackson, Ian Johnston, Dominic Kwiatkowski, Cordelia Langford, John Sillitoe on behalf of the Wellcome Sanger Institute COVID-19 Surveillance Team |
| EPI_ISL_2808450 | Lighthouse Lab in Milton Keynes | Wellcome Sanger Institute for the COVID-19 Genomics UK (COG-UK) Consortium | The Lighthouse Lab in Milton Keynes and Alex Alderton, Roberto Amato, Jeffrey Barrett, Sonia Goncalves, Ewan Harrison, David K. Jackson, Ian Johnston, Dominic Kwiatkowski, Cordelia Langford, John Sillitoe on behalf of the Wellcome Sanger Institute COVID-19 Surveillance Team |
| EPI_ISL_2808451 | Lighthouse Laboratory Plymouth | Wellcome Sanger Institute for the COVID-19 Genomics UK (COG-UK) Consortium | Lighthouse Laboratory Plymouth and Alex Alderton, Roberto Amato, Jeffrey Barrett, Sonia Goncalves, Ewan Harrison, David K. Jackson, Ian Johnston, Dominic Kwiatkowski, Cordelia Langford, John Sillitoe on behalf of the Wellcome Sanger Institute COVID-19 Surveillance Team |
| EPI_ISL_2808452 | Lighthouse Lab in Glasgow | Wellcome Sanger Institute for the COVID-19 Genomics UK (COG-UK) Consortium | Harper VanSteenhouse, Yumi Kasai, David Gray, Carol Clugston, Anna Dominiczak and Alex Alderton, Roberto Amato, Jeffrey Barrett, Sonia Goncalves, Ewan Harrison, David K. Jackson, Ian Johnston, Dominic Kwiatkowski, Cordelia Langford, John Sillitoe on behalf of the Wellcome Sanger Institute COVID-19 Surveillance Team |
| EPI_ISL_2808453, EPI_ISL_2808454, EPI_ISL_2808455, EPI_ISL_2808456, EPI_ISL_2808457, EPI_ISL_2808458, EPI_ISL_2808459 | Lighthouse Laboratory Plymouth | Wellcome Sanger Institute for the COVID-19 Genomics UK (COG-UK) Consortium | Lighthouse Laboratory Plymouth and Alex Alderton, Roberto Amato, Jeffrey Barrett, Sonia Goncalves, Ewan Harrison, David K. Jackson, Ian Johnston, Dominic Kwiatkowski, Cordelia Langford, John Sillitoe on behalf of the Wellcome Sanger Institute COVID-19 Surveillance Team |
| EPI_ISL_2808460 | Lighthouse Lab in Glasgow | Wellcome Sanger Institute for the COVID-19 Genomics UK (COG-UK) Consortium | Harper VanSteenhouse, Yumi Kasai, David Gray, Carol Clugston, Anna Dominiczak and Alex Alderton, Roberto Amato, Jeffrey Barrett, Sonia Goncalves, Ewan Harrison, David K. Jackson, Ian Johnston, Dominic Kwiatkowski, Cordelia Langford, John Sillitoe on behalf of the Wellcome Sanger Institute COVID-19 Surveillance Team |
| EPI_ISL_2808461, EPI_ISL_2808462 | Lighthouse Laboratory Plymouth | Wellcome Sanger Institute for the COVID-19 Genomics UK (COG-UK) Consortium | Lighthouse Laboratory Plymouth and Alex Alderton, Roberto Amato, Jeffrey Barrett, Sonia Goncalves, Ewan Harrison, David K. Jackson, Ian Johnston, Dominic Kwiatkowski, Cordelia Langford, John Sillitoe on behalf of the Wellcome Sanger Institute COVID-19 Surveillance Team |
| EPI_ISL_2808463 | Health Services Laboratories | Wellcome Sanger Institute for the COVID-19 Genomics UK (COG-UK) Consortium | Health Services Laboratories and Alex Alderton, Roberto Amato, Jeffrey Barrett, Sonia Goncalves, Ewan Harrison, David K. Jackson, Ian Johnston, Dominic Kwiatkowski, Cordelia Langford, John Sillitoe on behalf of the Wellcome Sanger Institute COVID-19 Surveillance Team |
| EPI_ISL_2808464 | Lighthouse Lab in Milton Keynes | Wellcome Sanger Institute for the COVID-19 Genomics UK (COG-UK) Consortium | The Lighthouse Lab in Milton Keynes and Alex Alderton, Roberto Amato, Jeffrey Barrett, Sonia Goncalves, Ewan Harrison, David K. Jackson, Ian Johnston, Dominic Kwiatkowski, Cordelia Langford, John Sillitoe on behalf of the Wellcome Sanger Institute COVID-19 Surveillance Team |
| EPI_ISL_2808465, EPI_ISL_2808466 | Lighthouse Laboratory Plymouth | Wellcome Sanger Institute for the COVID-19 Genomics UK (COG-UK) Consortium | Lighthouse Laboratory Plymouth and Alex Alderton, Roberto Amato, Jeffrey Barrett, Sonia Goncalves, Ewan Harrison, David K. Jackson, Ian Johnston, Dominic Kwiatkowski, Cordelia Langford, John Sillitoe on behalf of the Wellcome Sanger Institute COVID-19 Surveillance Team |
| EPI_ISL_2808467 | Lighthouse Lab in Glasgow | Wellcome Sanger Institute for the COVID-19 Genomics UK (COG-UK) Consortium | Harper VanSteenhouse, Yumi Kasai, David Gray, Carol Clugston, Anna Dominiczak and Alex Alderton, Roberto Amato, Jeffrey Barrett, Sonia Goncalves, Ewan Harrison, David K. Jackson, Ian Johnston, Dominic Kwiatkowski, Cordelia Langford, John Sillitoe on behalf of the Wellcome Sanger Institute COVID-19 Surveillance Team |
| EPI_ISL_2808468, EPI_ISL_2808469, EPI_ISL_2808470 | Lighthouse Laboratory Plymouth | Wellcome Sanger Institute for the COVID-19 Genomics UK (COG-UK) Consortium | Lighthouse Laboratory Plymouth and Alex Alderton, Roberto Amato, Jeffrey Barrett, Sonia Goncalves, Ewan Harrison, David K. Jackson, Ian Johnston, Dominic Kwiatkowski, Cordelia Langford, John Sillitoe on behalf of the Wellcome Sanger Institute COVID-19 Surveillance Team |
| EPI_ISL_2808471 | Lighthouse Lab in Milton Keynes | Wellcome Sanger Institute for the COVID-19 Genomics UK (COG-UK) Consortium | The Lighthouse Lab in Milton Keynes and Alex Alderton, Roberto Amato, Jeffrey Barrett, Sonia Goncalves, Ewan Harrison, David K. Jackson, Ian Johnston, Dominic Kwiatkowski, Cordelia Langford, John Sillitoe on behalf of the Wellcome Sanger Institute COVID-19 Surveillance Team |
| EPI_ISL_2808472, EPI_ISL_2808473 | Lighthouse Lab in Glasgow | Wellcome Sanger Institute for the COVID-19 Genomics UK (COG-UK) Consortium | Harper VanSteenhouse, Yumi Kasai, David Gray, Carol Clugston, Anna Dominiczak and Alex Alderton, Roberto Amato, Jeffrey Barrett, Sonia Goncalves, Ewan Harrison, David K. Jackson, Ian Johnston, Dominic Kwiatkowski, Cordelia Langford, John Sillitoe on behalf of the Wellcome Sanger Institute COVID-19 Surveillance Team |
| EPI_ISL_2808474, EPI_ISL_2808475, EPI_ISL_2808476 | Lighthouse Lab in Milton Keynes | Wellcome Sanger Institute for the COVID-19 Genomics UK (COG-UK) Consortium | The Lighthouse Lab in Milton Keynes and Alex Alderton, Roberto Amato, Jeffrey Barrett, Sonia Goncalves, Ewan Harrison, David K. Jackson, Ian Johnston, Dominic Kwiatkowski, Cordelia Langford, John Sillitoe on behalf of the Wellcome Sanger Institute COVID-19 Surveillance Team |
| EPI_ISL_2808477 | Health Services Laboratories | Wellcome Sanger Institute for the COVID-19 Genomics UK (COG-UK) Consortium | Health Services Laboratories and Alex Alderton, Roberto Amato, Jeffrey Barrett, Sonia Goncalves, Ewan Harrison, David K. Jackson, Ian Johnston, Dominic Kwiatkowski, Cordelia Langford, John Sillitoe on behalf of the Wellcome Sanger Institute COVID-19 Surveillance Team |
| EPI_ISL_2808478, EPI_ISL_2808479 | Lighthouse Laboratory Plymouth | Wellcome Sanger Institute for the COVID-19 Genomics UK (COG-UK) Consortium | Lighthouse Laboratory Plymouth and Alex Alderton, Roberto Amato, Jeffrey Barrett, Sonia Goncalves, Ewan Harrison, David K. Jackson, Ian Johnston, Dominic Kwiatkowski, Cordelia Langford, John Sillitoe on behalf of the Wellcome Sanger Institute COVID-19 Surveillance Team |
| EPI_ISL_2808480 | Lighthouse Lab in Milton Keynes | Wellcome Sanger Institute for the COVID-19 Genomics UK (COG-UK) Consortium | The Lighthouse Lab in Milton Keynes and Alex Alderton, Roberto Amato, Jeffrey Barrett, Sonia Goncalves, Ewan Harrison, David K. Jackson, Ian Johnston, Dominic Kwiatkowski, Cordelia Langford, John Sillitoe on behalf of the Wellcome Sanger Institute COVID-19 Surveillance Team |
| EPI_ISL_2808481, EPI_ISL_2808482, EPI_ISL_2808483 | Lighthouse Lab in Glasgow | Wellcome Sanger Institute for the COVID-19 Genomics UK (COG-UK) Consortium | Harper VanSteenhouse, Yumi Kasai, David Gray, Carol Clugston, Anna Dominiczak and Alex Alderton, Roberto Amato, Jeffrey Barrett, Sonia Goncalves, Ewan Harrison, David K. Jackson, Ian Johnston, Dominic Kwiatkowski, Cordelia Langford, John Sillitoe on behalf of the Wellcome Sanger Institute COVID-19 Surveillance Team |
| EPI_ISL_2808484, EPI_ISL_2808485, EPI_ISL_2808486 | Lighthouse Lab in Milton Keynes | Wellcome Sanger Institute for the COVID-19 Genomics UK (COG-UK) Consortium | The Lighthouse Lab in Milton Keynes and Alex Alderton, Roberto Amato, Jeffrey Barrett, Sonia Goncalves, Ewan Harrison, David K. Jackson, Ian Johnston, Dominic Kwiatkowski, Cordelia Langford, John Sillitoe on behalf of the Wellcome Sanger Institute COVID-19 Surveillance Team |
| EPI_ISL_2808487 | Lighthouse Laboratory Plymouth | Wellcome Sanger Institute for the COVID-19 Genomics UK (COG-UK) Consortium | Lighthouse Laboratory Plymouth and Alex Alderton, Roberto Amato, Jeffrey Barrett, Sonia Goncalves, Ewan Harrison, David K. Jackson, Ian Johnston, Dominic Kwiatkowski, Cordelia Langford, John Sillitoe on behalf of the Wellcome Sanger Institute COVID-19 Surveillance Team |
| EPI_ISL_2808488 | Lighthouse Lab in Milton Keynes | Wellcome Sanger Institute for the COVID-19 Genomics UK (COG-UK) Consortium | The Lighthouse Lab in Milton Keynes and Alex Alderton, Roberto Amato, Jeffrey Barrett, Sonia Goncalves, Ewan Harrison, David K. Jackson, Ian Johnston, Dominic Kwiatkowski, Cordelia Langford, John Sillitoe on behalf of the Wellcome Sanger Institute COVID-19 Surveillance Team |
| EPI_ISL_2808489 | Lighthouse Laboratory Plymouth | Wellcome Sanger Institute for the COVID-19 Genomics UK (COG-UK) Consortium | Lighthouse Laboratory Plymouth and Alex Alderton, Roberto Amato, Jeffrey Barrett, Sonia Goncalves, Ewan Harrison, David K. Jackson, Ian Johnston, Dominic Kwiatkowski, Cordelia Langford, John Sillitoe on behalf of the Wellcome Sanger Institute COVID-19 Surveillance Team |
| EPI_ISL_2808490 | Lighthouse Lab in Milton Keynes | Wellcome Sanger Institute for the COVID-19 Genomics UK (COG-UK) Consortium | The Lighthouse Lab in Milton Keynes and Alex Alderton, Roberto Amato, Jeffrey Barrett, Sonia Goncalves, Ewan Harrison, David K. Jackson, Ian Johnston, Dominic Kwiatkowski, Cordelia Langford, John Sillitoe on behalf of the Wellcome Sanger Institute COVID-19 Surveillance Team |
| EPI_ISL_2808491 | Lighthouse Laboratory Plymouth | Wellcome Sanger Institute for the COVID-19 Genomics UK (COG-UK) Consortium | Lighthouse Laboratory Plymouth and Alex Alderton, Roberto Amato, Jeffrey Barrett, Sonia Goncalves, Ewan Harrison, David K. Jackson, Ian Johnston, Dominic Kwiatkowski, Cordelia Langford, John Sillitoe on behalf of the Wellcome Sanger Institute COVID-19 Surveillance Team |
| EPI_ISL_2808492 | Lighthouse Lab in Milton Keynes | Wellcome Sanger Institute for the COVID-19 Genomics UK (COG-UK) Consortium | The Lighthouse Lab in Milton Keynes and Alex Alderton, Roberto Amato, Jeffrey Barrett, Sonia Goncalves, Ewan Harrison, David K. Jackson, Ian Johnston, Dominic Kwiatkowski, Cordelia Langford, John Sillitoe on behalf of the Wellcome Sanger Institute COVID-19 Surveillance Team |
| EPI_ISL_2808493 | Lighthouse Laboratory Plymouth | Wellcome Sanger Institute for the COVID-19 Genomics UK (COG-UK) Consortium | Lighthouse Laboratory Plymouth and Alex Alderton, Roberto Amato, Jeffrey Barrett, Sonia Goncalves, Ewan Harrison, David K. Jackson, Ian Johnston, Dominic Kwiatkowski, Cordelia Langford, John Sillitoe on behalf of the Wellcome Sanger Institute COVID-19 Surveillance Team |
| EPI_ISL_2808494, EPI_ISL_2808495 | Lighthouse Lab in Milton Keynes | Wellcome Sanger Institute for the COVID-19 Genomics UK (COG-UK) Consortium | The Lighthouse Lab in Milton Keynes and Alex Alderton, Roberto Amato, Jeffrey Barrett, Sonia Goncalves, Ewan Harrison, David K. Jackson, Ian Johnston, Dominic Kwiatkowski, Cordelia Langford, John Sillitoe on behalf of the Wellcome Sanger Institute COVID-19 Surveillance Team |
| EPI_ISL_2808496, EPI_ISL_2808497 | Lighthouse Laboratory Plymouth | Wellcome Sanger Institute for the COVID-19 Genomics UK (COG-UK) Consortium | Lighthouse Laboratory Plymouth and Alex Alderton, Roberto Amato, Jeffrey Barrett, Sonia Goncalves, Ewan Harrison, David K. Jackson, Ian Johnston, Dominic Kwiatkowski, Cordelia Langford, John Sillitoe on behalf of the Wellcome Sanger Institute COVID-19 Surveillance Team |

| | | | |
|---|---|---|---|
| EPI_ISL_2808498 | Lighthouse Lab in Milton Keynes | Wellcome Sanger Institute for the COVID-19 Genomics UK (COG-UK) Consortium | The Lighthouse Lab in Milton Keynes and Alex Alderton, Roberto Amato, Jeffrey Barrett, Sonia Goncalves, Ewan Harrison, David K. Jackson, Ian Johnston, Dominic Kwiatkowski, Cordelia Langford, John Sillitoe on behalf of the Wellcome Sanger Institute COVID-19 Surveillance Team |
| EPI_ISL_2808499 | Lighthouse Laboratory Plymouth | Wellcome Sanger Institute for the COVID-19 Genomics UK (COG-UK) Consortium | Lighthouse Laboratory Plymouth and Alex Alderton, Roberto Amato, Jeffrey Barrett, Sonia Goncalves, Ewan Harrison, David K. Jackson, Ian Johnston, Dominic Kwiatkowski, Cordelia Langford, John Sillitoe on behalf of the Wellcome Sanger Institute COVID-19 Surveillance Team |
| EPI_ISL_2808500 | Lighthouse Lab in Glasgow | Wellcome Sanger Institute for the COVID-19 Genomics UK (COG-UK) Consortium | Harper VanSteenhouse, Yumi Kasai, David Gray, Carol Clugston, Anna Dominiczak and Alex Alderton, Roberto Amato, Jeffrey Barrett, Sonia Goncalves, Ewan Harrison, David K. Jackson, Ian Johnston, Dominic Kwiatkowski, Cordelia Langford, John Sillitoe on behalf of the Wellcome Sanger Institute COVID-19 Surveillance Team |
| EPI_ISL_2808501 | Lighthouse Laboratory Plymouth | Wellcome Sanger Institute for the COVID-19 Genomics UK (COG-UK) Consortium | Lighthouse Laboratory Plymouth and Alex Alderton, Roberto Amato, Jeffrey Barrett, Sonia Goncalves, Ewan Harrison, David K. Jackson, Ian Johnston, Dominic Kwiatkowski, Cordelia Langford, John Sillitoe on behalf of the Wellcome Sanger Institute COVID-19 Surveillance Team |
| EPI_ISL_2808502, EPI_ISL_2808503 | Lighthouse Lab in Milton Keynes | Wellcome Sanger Institute for the COVID-19 Genomics UK (COG-UK) Consortium | The Lighthouse Lab in Milton Keynes and Alex Alderton, Roberto Amato, Jeffrey Barrett, Sonia Goncalves, Ewan Harrison, David K. Jackson, Ian Johnston, Dominic Kwiatkowski, Cordelia Langford, John Sillitoe on behalf of the Wellcome Sanger Institute COVID-19 Surveillance Team |
| EPI_ISL_2808504, EPI_ISL_2808505 | Lighthouse Lab in Glasgow | Wellcome Sanger Institute for the COVID-19 Genomics UK (COG-UK) Consortium | Harper VanSteenhouse, Yumi Kasai, David Gray, Carol Clugston, Anna Dominiczak and Alex Alderton, Roberto Amato, Jeffrey Barrett, Sonia Goncalves, Ewan Harrison, David K. Jackson, Ian Johnston, Dominic Kwiatkowski, Cordelia Langford, John Sillitoe on behalf of the Wellcome Sanger Institute COVID-19 Surveillance Team |
| EPI_ISL_2808506, EPI_ISL_2808507 | Lighthouse Lab in Milton Keynes | Wellcome Sanger Institute for the COVID-19 Genomics UK (COG-UK) Consortium | The Lighthouse Lab in Milton Keynes and Alex Alderton, Roberto Amato, Jeffrey Barrett, Sonia Goncalves, Ewan Harrison, David K. Jackson, Ian Johnston, Dominic Kwiatkowski, Cordelia Langford, John Sillitoe on behalf of the Wellcome Sanger Institute COVID-19 Surveillance Team |
| EPI_ISL_2808508 | Lighthouse Lab in Glasgow | Wellcome Sanger Institute for the COVID-19 Genomics UK (COG-UK) Consortium | Harper VanSteenhouse, Yumi Kasai, David Gray, Carol Clugston, Anna Dominiczak and Alex Alderton, Roberto Amato, Jeffrey Barrett, Sonia Goncalves, Ewan Harrison, David K. Jackson, Ian Johnston, Dominic Kwiatkowski, Cordelia Langford, John Sillitoe on behalf of the Wellcome Sanger Institute COVID-19 Surveillance Team |
| EPI_ISL_2808509 | Lighthouse Laboratory Plymouth | Wellcome Sanger Institute for the COVID-19 Genomics UK (COG-UK) Consortium | Lighthouse Laboratory Plymouth and Alex Alderton, Roberto Amato, Jeffrey Barrett, Sonia Goncalves, Ewan Harrison, David K. Jackson, Ian Johnston, Dominic Kwiatkowski, Cordelia Langford, John Sillitoe on behalf of the Wellcome Sanger Institute COVID-19 Surveillance Team |
| EPI_ISL_2808510 | Lighthouse Lab in Glasgow | Wellcome Sanger Institute for the COVID-19 Genomics UK (COG-UK) Consortium | Harper VanSteenhouse, Yumi Kasai, David Gray, Carol Clugston, Anna Dominiczak and Alex Alderton, Roberto Amato, Jeffrey Barrett, Sonia Goncalves, Ewan Harrison, David K. Jackson, Ian Johnston, Dominic Kwiatkowski, Cordelia Langford, John Sillitoe on behalf of the Wellcome Sanger Institute COVID-19 Surveillance Team |
| EPI_ISL_2808511, EPI_ISL_2808512, EPI_ISL_2808513 | Lighthouse Lab in Milton Keynes | Wellcome Sanger Institute for the COVID-19 Genomics UK (COG-UK) Consortium | The Lighthouse Lab in Milton Keynes and Alex Alderton, Roberto Amato, Jeffrey Barrett, Sonia Goncalves, Ewan Harrison, David K. Jackson, Ian Johnston, Dominic Kwiatkowski, Cordelia Langford, John Sillitoe on behalf of the Wellcome Sanger Institute COVID-19 Surveillance Team |
| EPI_ISL_2808514, EPI_ISL_2808515 | Lighthouse Laboratory Plymouth | Wellcome Sanger Institute for the COVID-19 Genomics UK (COG-UK) Consortium | Lighthouse Laboratory Plymouth and Alex Alderton, Roberto Amato, Jeffrey Barrett, Sonia Goncalves, Ewan Harrison, David K. Jackson, Ian Johnston, Dominic Kwiatkowski, Cordelia Langford, John Sillitoe on behalf of the Wellcome Sanger Institute COVID-19 Surveillance Team |
| EPI_ISL_2808516 | Lighthouse Lab in Milton Keynes | Wellcome Sanger Institute for the COVID-19 Genomics UK (COG-UK) Consortium | The Lighthouse Lab in Milton Keynes and Alex Alderton, Roberto Amato, Jeffrey Barrett, Sonia Goncalves, Ewan Harrison, David K. Jackson, Ian Johnston, Dominic Kwiatkowski, Cordelia Langford, John Sillitoe on behalf of the Wellcome Sanger Institute COVID-19 Surveillance Team |
| EPI_ISL_2808517 | Health Services Laboratories | Wellcome Sanger Institute for the COVID-19 Genomics UK (COG-UK) Consortium | Health Services Laboratories and Alex Alderton, Roberto Amato, Jeffrey Barrett, Sonia Goncalves, Ewan Harrison, David K. Jackson, Ian Johnston, Dominic Kwiatkowski, Cordelia Langford, John Sillitoe on behalf of the Wellcome Sanger Institute COVID-19 Surveillance Team |
| EPI_ISL_2808518, EPI_ISL_2808519 | Lighthouse Lab in Milton Keynes | Wellcome Sanger Institute for the COVID-19 Genomics UK (COG-UK) Consortium | The Lighthouse Lab in Milton Keynes and Alex Alderton, Roberto Amato, Jeffrey Barrett, Sonia Goncalves, Ewan Harrison, David K. Jackson, Ian Johnston, Dominic Kwiatkowski, Cordelia Langford, John Sillitoe on behalf of the Wellcome Sanger Institute COVID-19 Surveillance Team |
| EPI_ISL_2808520, EPI_ISL_2808521 | Lighthouse Laboratory Plymouth | Wellcome Sanger Institute for the COVID-19 Genomics UK (COG-UK) Consortium | Lighthouse Laboratory Plymouth and Alex Alderton, Roberto Amato, Jeffrey Barrett, Sonia Goncalves, Ewan Harrison, David K. Jackson, Ian Johnston, Dominic Kwiatkowski, Cordelia Langford, John Sillitoe on behalf of the Wellcome Sanger Institute COVID-19 Surveillance Team |
| EPI_ISL_2808522 | Health Services Laboratories | Wellcome Sanger Institute for the COVID-19 Genomics UK (COG-UK) Consortium | Health Services Laboratories and Alex Alderton, Roberto Amato, Jeffrey Barrett, Sonia Goncalves, Ewan Harrison, David K. Jackson, Ian Johnston, Dominic Kwiatkowski, Cordelia Langford, John Sillitoe on behalf of the Wellcome Sanger Institute COVID-19 Surveillance Team |
| EPI_ISL_2808523 | Lighthouse Laboratory Plymouth | Wellcome Sanger Institute for the COVID-19 Genomics UK (COG-UK) Consortium | Lighthouse Laboratory Plymouth and Alex Alderton, Roberto Amato, Jeffrey Barrett, Sonia Goncalves, Ewan Harrison, David K. Jackson, Ian Johnston, Dominic Kwiatkowski, Cordelia Langford, John Sillitoe on behalf of the Wellcome Sanger Institute COVID-19 Surveillance Team |
| EPI_ISL_2808524 | Lighthouse Lab in Glasgow | Wellcome Sanger Institute for the COVID-19 Genomics UK (COG-UK) Consortium | Harper VanSteenhouse, Yumi Kasai, David Gray, Carol Clugston, Anna Dominiczak and Alex Alderton, Roberto Amato, Jeffrey Barrett, Sonia Goncalves, Ewan Harrison, David K. Jackson, Ian Johnston, Dominic Kwiatkowski, Cordelia Langford, John Sillitoe on behalf of the Wellcome Sanger Institute COVID-19 Surveillance Team |
| EPI_ISL_2808525, EPI_ISL_2808526, EPI_ISL_2808527 | Lighthouse Laboratory Plymouth | Wellcome Sanger Institute for the COVID-19 Genomics UK (COG-UK) Consortium | Lighthouse Laboratory Plymouth and Alex Alderton, Roberto Amato, Jeffrey Barrett, Sonia Goncalves, Ewan Harrison, David K. Jackson, Ian Johnston, Dominic Kwiatkowski, Cordelia Langford, John Sillitoe on behalf of the Wellcome Sanger Institute COVID-19 Surveillance Team |
| EPI_ISL_2808528 | Health Services Laboratories | Wellcome Sanger Institute for the COVID-19 Genomics UK (COG-UK) Consortium | Health Services Laboratories and Alex Alderton, Roberto Amato, Jeffrey Barrett, Sonia Goncalves, Ewan Harrison, David K. Jackson, Ian Johnston, Dominic Kwiatkowski, Cordelia Langford, John Sillitoe on behalf of the Wellcome Sanger Institute COVID-19 Surveillance Team |
| EPI_ISL_2808529 | Lighthouse Laboratory Plymouth | Wellcome Sanger Institute for the COVID-19 Genomics UK (COG-UK) Consortium | Lighthouse Laboratory Plymouth and Alex Alderton, Roberto Amato, Jeffrey Barrett, Sonia Goncalves, Ewan Harrison, David K. Jackson, Ian Johnston, Dominic Kwiatkowski, Cordelia Langford, John Sillitoe on behalf of the Wellcome Sanger Institute COVID-19 Surveillance Team |
| EPI_ISL_2808530, EPI_ISL_2808531 | Lighthouse Lab in Milton Keynes | Wellcome Sanger Institute for the COVID-19 Genomics UK (COG-UK) Consortium | The Lighthouse Lab in Milton Keynes and Alex Alderton, Roberto Amato, Jeffrey Barrett, Sonia Goncalves, Ewan Harrison, David K. Jackson, Ian Johnston, Dominic Kwiatkowski, Cordelia Langford, John Sillitoe on behalf of the Wellcome Sanger Institute COVID-19 Surveillance Team |
| EPI_ISL_2808532, EPI_ISL_2808533, EPI_ISL_2808534 | Lighthouse Laboratory Plymouth | Wellcome Sanger Institute for the COVID-19 Genomics UK (COG-UK) Consortium | Lighthouse Laboratory Plymouth and Alex Alderton, Roberto Amato, Jeffrey Barrett, Sonia Goncalves, Ewan Harrison, David K. Jackson, Ian Johnston, Dominic Kwiatkowski, Cordelia Langford, John Sillitoe on behalf of the Wellcome Sanger Institute COVID-19 Surveillance Team |
| EPI_ISL_2808535 | Lighthouse Lab in Milton Keynes | Wellcome Sanger Institute for the COVID-19 Genomics UK (COG-UK) Consortium | The Lighthouse Lab in Milton Keynes and Alex Alderton, Roberto Amato, Jeffrey Barrett, Sonia Goncalves, Ewan Harrison, David K. Jackson, Ian Johnston, Dominic Kwiatkowski, Cordelia Langford, John Sillitoe on behalf of the Wellcome Sanger Institute COVID-19 Surveillance Team |
| EPI_ISL_2808536 | Lighthouse Laboratory Plymouth | Wellcome Sanger Institute for the COVID-19 Genomics UK (COG-UK) Consortium | Lighthouse Laboratory Plymouth and Alex Alderton, Roberto Amato, Jeffrey Barrett, Sonia Goncalves, Ewan Harrison, David K. Jackson, Ian Johnston, Dominic Kwiatkowski, Cordelia Langford, John Sillitoe on behalf of the Wellcome Sanger Institute COVID-19 Surveillance Team |
| EPI_ISL_2808537 | Lighthouse Lab in Milton Keynes | Wellcome Sanger Institute for the COVID-19 Genomics UK (COG-UK) Consortium | The Lighthouse Lab in Milton Keynes and Alex Alderton, Roberto Amato, Jeffrey Barrett, Sonia Goncalves, Ewan Harrison, David K. Jackson, Ian Johnston, Dominic Kwiatkowski, Cordelia Langford, John Sillitoe on behalf of the Wellcome Sanger Institute COVID-19 Surveillance Team |
| EPI_ISL_2808538 | Lighthouse Lab in Glasgow | Wellcome Sanger Institute for the COVID-19 Genomics UK (COG-UK) Consortium | Harper VanSteenhouse, Yumi Kasai, David Gray, Carol Clugston, Anna Dominiczak and Alex Alderton, Roberto Amato, Jeffrey Barrett, Sonia Goncalves, Ewan Harrison, David K. Jackson, Ian Johnston, Dominic Kwiatkowski, Cordelia Langford, John Sillitoe on behalf of the Wellcome Sanger Institute COVID-19 Surveillance Team |
| EPI_ISL_2808539 | Lighthouse Laboratory Plymouth | Wellcome Sanger Institute for the COVID-19 Genomics UK (COG-UK) Consortium | Lighthouse Laboratory Plymouth and Alex Alderton, Roberto Amato, Jeffrey Barrett, Sonia Goncalves, Ewan Harrison, David K. Jackson, Ian Johnston, Dominic Kwiatkowski, Cordelia Langford, John Sillitoe on behalf of the Wellcome Sanger Institute COVID-19 Surveillance Team |
| EPI_ISL_2808540 | Lighthouse Lab in Milton Keynes | Wellcome Sanger Institute for the COVID-19 Genomics UK (COG-UK) Consortium | The Lighthouse Lab in Milton Keynes and Alex Alderton, Roberto Amato, Jeffrey Barrett, Sonia Goncalves, Ewan Harrison, David K. Jackson, Ian Johnston, Dominic Kwiatkowski, Cordelia Langford, John Sillitoe on behalf of the Wellcome Sanger Institute COVID-19 Surveillance Team |
| EPI_ISL_2808541, EPI_ISL_2808542 | Lighthouse Laboratory Plymouth | Wellcome Sanger Institute for the COVID-19 Genomics UK (COG-UK) Consortium | Lighthouse Laboratory Plymouth and Alex Alderton, Roberto Amato, Jeffrey Barrett, Sonia Goncalves, Ewan Harrison, David K. Jackson, Ian Johnston, Dominic Kwiatkowski, Cordelia Langford, John Sillitoe on behalf of the Wellcome Sanger Institute COVID-19 Surveillance Team |
| EPI_ISL_2808543 | Lighthouse Lab in Milton Keynes | Wellcome Sanger Institute for the COVID-19 Genomics UK (COG-UK) Consortium | The Lighthouse Lab in Milton Keynes and Alex Alderton, Roberto Amato, Jeffrey Barrett, Sonia Goncalves, Ewan Harrison, David K. Jackson, Ian Johnston, Dominic Kwiatkowski, Cordelia Langford, John Sillitoe on behalf of the Wellcome Sanger Institute COVID-19 Surveillance Team |
| EPI_ISL_2808544 | Lighthouse Lab in Glasgow | Wellcome Sanger Institute for the COVID-19 Genomics UK (COG-UK) Consortium | Harper VanSteenhouse, Yumi Kasai, David Gray, Carol Clugston, Anna Dominiczak and Alex Alderton, Roberto Amato, Jeffrey Barrett, Sonia Goncalves, Ewan Harrison, David K. Jackson, Ian Johnston, Dominic Kwiatkowski, Cordelia Langford, John Sillitoe on behalf of the Wellcome Sanger Institute COVID-19 Surveillance Team |
| EPI_ISL_2808545, EPI_ISL_2808546, EPI_ISL_2808547, EPI_ISL_2808548, EPI_ISL_2808549 | Lighthouse Laboratory Plymouth | Wellcome Sanger Institute for the COVID-19 Genomics UK (COG-UK) Consortium | Lighthouse Laboratory Plymouth and Alex Alderton, Roberto Amato, Jeffrey Barrett, Sonia Goncalves, Ewan Harrison, David K. Jackson, Ian Johnston, Dominic Kwiatkowski, Cordelia Langford, John Sillitoe on behalf of the Wellcome Sanger Institute COVID-19 Surveillance Team |
| EPI_ISL_2808550 | Lighthouse Lab in Milton Keynes | Wellcome Sanger Institute for the COVID-19 Genomics UK (COG-UK) Consortium | The Lighthouse Lab in Milton Keynes and Alex Alderton, Roberto Amato, Jeffrey Barrett, Sonia Goncalves, Ewan Harrison, David K. Jackson, Ian Johnston, Dominic Kwiatkowski, Cordelia Langford, John Sillitoe on behalf of the Wellcome Sanger Institute COVID-19 Surveillance Team |
| EPI_ISL_2808551 | Lighthouse Lab in Glasgow | Wellcome Sanger Institute for the COVID-19 Genomics UK (COG-UK) Consortium | Harper VanSteenhouse, Yumi Kasai, David Gray, Carol Clugston, Anna Dominiczak and Alex Alderton, Roberto Amato, Jeffrey Barrett, Sonia Goncalves, Ewan Harrison, David K. Jackson, Ian Johnston, Dominic Kwiatkowski, Cordelia Langford, John Sillitoe on behalf of the Wellcome Sanger Institute COVID-19 Surveillance Team |
| EPI_ISL_2808552 | Lighthouse Laboratory Plymouth | Wellcome Sanger Institute for the COVID-19 Genomics UK (COG-UK) Consortium | Lighthouse Laboratory Plymouth and Alex Alderton, Roberto Amato, Jeffrey Barrett, Sonia Goncalves, Ewan Harrison, David K. Jackson, Ian Johnston, Dominic Kwiatkowski, Cordelia Langford, John Sillitoe on behalf of the Wellcome Sanger Institute COVID-19 Surveillance Team |
| EPI_ISL_2808553 | Lighthouse Lab in Glasgow | Wellcome Sanger Institute for the COVID-19 Genomics UK (COG-UK) Consortium | Harper VanSteenhouse, Yumi Kasai, David Gray, Carol Clugston, Anna Dominiczak and Alex Alderton, Roberto Amato, Jeffrey Barrett, Sonia Goncalves, Ewan Harrison, David K. Jackson, Ian Johnston, Dominic Kwiatkowski, Cordelia Langford, John Sillitoe on behalf of the Wellcome Sanger Institute COVID-19 Surveillance Team |
| EPI_ISL_2808554 | Lighthouse Lab in Milton Keynes | Wellcome Sanger Institute for the COVID-19 Genomics UK (COG-UK) Consortium | The Lighthouse Lab in Milton Keynes and Alex Alderton, Roberto Amato, Jeffrey Barrett, Sonia Goncalves, Ewan Harrison, David K. Jackson, Ian Johnston, Dominic Kwiatkowski, Cordelia Langford, John Sillitoe on behalf of the Wellcome Sanger Institute COVID-19 Surveillance Team |
| EPI_ISL_2808555 | Lighthouse Laboratory Plymouth | Wellcome Sanger Institute for the COVID-19 Genomics UK (COG-UK) Consortium | Lighthouse Laboratory Plymouth and Alex Alderton, Roberto Amato, Jeffrey Barrett, Sonia Goncalves, Ewan Harrison, David K. Jackson, Ian Johnston, Dominic Kwiatkowski, Cordelia Langford, John Sillitoe on behalf of the Wellcome Sanger Institute COVID-19 Surveillance Team |
| EPI_ISL_2808556 | Lighthouse Lab in Milton Keynes | Wellcome Sanger Institute for the COVID-19 Genomics UK (COG-UK) Consortium | The Lighthouse Lab in Milton Keynes and Alex Alderton, Roberto Amato, Jeffrey Barrett, Sonia Goncalves, Ewan Harrison, David K. Jackson, Ian Johnston, Dominic Kwiatkowski, Cordelia Langford, John Sillitoe on behalf of the Wellcome Sanger Institute COVID-19 Surveillance Team |
| EPI_ISL_2808557 | Lighthouse Laboratory Plymouth | Wellcome Sanger Institute for the COVID-19 Genomics UK (COG-UK) Consortium | Lighthouse Laboratory Plymouth and Alex Alderton, Roberto Amato, Jeffrey Barrett, Sonia Goncalves, Ewan Harrison, David K. Jackson, Ian Johnston, Dominic Kwiatkowski, Cordelia Langford, John Sillitoe on behalf of the Wellcome Sanger Institute COVID-19 Surveillance Team |
| EPI_ISL_2808558, | Lighthouse Lab in Glasgow | Wellcome Sanger Institute for the COVID-19 Genomics UK (COG-UK) | Harper VanSteenhouse, Yumi Kasai, David Gray, Carol Clugston, Anna Dominiczak and Alex Alderton, Roberto Amato, Jeffrey Barrett, Sonia Goncalves, Ewan Harrison, David K. Jackson, Ian Johnston, Dominic Kwiatkowski, Cordelia Langford, |

| | | | |
|---|---|---|---|
| EPI_ISL_2808559 | | Consortium | John Sillitoe on behalf of the Wellcome Sanger Institute COVID-19 Surveillance Team |
| EPI_ISL_2808560, EPI_ISL_2808561 | Lighthouse Laboratory Plymouth | Wellcome Sanger Institute for the COVID-19 Genomics UK (COG-UK) Consortium | Lighthouse Laboratory Plymouth and Alex Alderton, Roberto Amato, Jeffrey Barrett, Sonia Goncalves, Ewan Harrison, David K. Jackson, Ian Johnston, Dominic Kwiatkowski, Cordelia Langford, John Sillitoe on behalf of the Wellcome Sanger Institute COVID-19 Surveillance Team |
| EPI_ISL_2808562 | Health Services Laboratories | Wellcome Sanger Institute for the COVID-19 Genomics UK (COG-UK) Consortium | Health Services Laboratories and Alex Alderton, Roberto Amato, Jeffrey Barrett, Sonia Goncalves, Ewan Harrison, David K. Jackson, Ian Johnston, Dominic Kwiatkowski, Cordelia Langford, John Sillitoe on behalf of the Wellcome Sanger Institute COVID-19 Surveillance Team |
| EPI_ISL_2808563, EPI_ISL_2808564, EPI_ISL_2808565, EPI_ISL_2808566 | Lighthouse Laboratory Plymouth | Wellcome Sanger Institute for the COVID-19 Genomics UK (COG-UK) Consortium | Lighthouse Laboratory Plymouth and Alex Alderton, Roberto Amato, Jeffrey Barrett, Sonia Goncalves, Ewan Harrison, David K. Jackson, Ian Johnston, Dominic Kwiatkowski, Cordelia Langford, John Sillitoe on behalf of the Wellcome Sanger Institute COVID-19 Surveillance Team |
| EPI_ISL_2808567 | Health Services Laboratories | Wellcome Sanger Institute for the COVID-19 Genomics UK (COG-UK) Consortium | Health Services Laboratories and Alex Alderton, Roberto Amato, Jeffrey Barrett, Sonia Goncalves, Ewan Harrison, David K. Jackson, Ian Johnston, Dominic Kwiatkowski, Cordelia Langford, John Sillitoe on behalf of the Wellcome Sanger Institute COVID-19 Surveillance Team |
| EPI_ISL_2808568 | Lighthouse Lab in Glasgow | Wellcome Sanger Institute for the COVID-19 Genomics UK (COG-UK) Consortium | Harper VanSteenhouse, Yumi Kasai, David Gray, Carol Clugston, Anna Dominiczak and Alex Alderton, Roberto Amato, Jeffrey Barrett, Sonia Goncalves, Ewan Harrison, David K. Jackson, Ian Johnston, Dominic Kwiatkowski, Cordelia Langford, John Sillitoe on behalf of the Wellcome Sanger Institute COVID-19 Surveillance Team |
| EPI_ISL_2808569 | Lighthouse Laboratory Plymouth | Wellcome Sanger Institute for the COVID-19 Genomics UK (COG-UK) Consortium | Lighthouse Laboratory Plymouth and Alex Alderton, Roberto Amato, Jeffrey Barrett, Sonia Goncalves, Ewan Harrison, David K. Jackson, Ian Johnston, Dominic Kwiatkowski, Cordelia Langford, John Sillitoe on behalf of the Wellcome Sanger Institute COVID-19 Surveillance Team |
| EPI_ISL_2808570 | Health Services Laboratories | Wellcome Sanger Institute for the COVID-19 Genomics UK (COG-UK) Consortium | Health Services Laboratories and Alex Alderton, Roberto Amato, Jeffrey Barrett, Sonia Goncalves, Ewan Harrison, David K. Jackson, Ian Johnston, Dominic Kwiatkowski, Cordelia Langford, John Sillitoe on behalf of the Wellcome Sanger Institute COVID-19 Surveillance Team |
| EPI_ISL_2808571 | Lighthouse Lab in Glasgow | Wellcome Sanger Institute for the COVID-19 Genomics UK (COG-UK) Consortium | Harper VanSteenhouse, Yumi Kasai, David Gray, Carol Clugston, Anna Dominiczak and Alex Alderton, Roberto Amato, Jeffrey Barrett, Sonia Goncalves, Ewan Harrison, David K. Jackson, Ian Johnston, Dominic Kwiatkowski, Cordelia Langford, John Sillitoe on behalf of the Wellcome Sanger Institute COVID-19 Surveillance Team |
| EPI_ISL_2808572, EPI_ISL_2808573 | Lighthouse Laboratory Plymouth | Wellcome Sanger Institute for the COVID-19 Genomics UK (COG-UK) Consortium | Lighthouse Laboratory Plymouth and Alex Alderton, Roberto Amato, Jeffrey Barrett, Sonia Goncalves, Ewan Harrison, David K. Jackson, Ian Johnston, Dominic Kwiatkowski, Cordelia Langford, John Sillitoe on behalf of the Wellcome Sanger Institute COVID-19 Surveillance Team |
| EPI_ISL_2808574, EPI_ISL_2808575 | Lighthouse Lab in Milton Keynes | Wellcome Sanger Institute for the COVID-19 Genomics UK (COG-UK) Consortium | The Lighthouse Lab in Milton Keynes and Alex Alderton, Roberto Amato, Jeffrey Barrett, Sonia Goncalves, Ewan Harrison, David K. Jackson, Ian Johnston, Dominic Kwiatkowski, Cordelia Langford, John Sillitoe on behalf of the Wellcome Sanger Institute COVID-19 Surveillance Team |
| EPI_ISL_2808576, EPI_ISL_2808577, EPI_ISL_2808578, EPI_ISL_2808579, EPI_ISL_2808580 | Lighthouse Laboratory Plymouth | Wellcome Sanger Institute for the COVID-19 Genomics UK (COG-UK) Consortium | Lighthouse Laboratory Plymouth and Alex Alderton, Roberto Amato, Jeffrey Barrett, Sonia Goncalves, Ewan Harrison, David K. Jackson, Ian Johnston, Dominic Kwiatkowski, Cordelia Langford, John Sillitoe on behalf of the Wellcome Sanger Institute COVID-19 Surveillance Team |
| EPI_ISL_2808581, EPI_ISL_2808582, EPI_ISL_2808583 | Lighthouse Lab in Milton Keynes | Wellcome Sanger Institute for the COVID-19 Genomics UK (COG-UK) Consortium | The Lighthouse Lab in Milton Keynes and Alex Alderton, Roberto Amato, Jeffrey Barrett, Sonia Goncalves, Ewan Harrison, David K. Jackson, Ian Johnston, Dominic Kwiatkowski, Cordelia Langford, John Sillitoe on behalf of the Wellcome Sanger Institute COVID-19 Surveillance Team |
| EPI_ISL_2808584 | Lighthouse Lab in Glasgow | Wellcome Sanger Institute for the COVID-19 Genomics UK (COG-UK) Consortium | Harper VanSteenhouse, Yumi Kasai, David Gray, Carol Clugston, Anna Dominiczak and Alex Alderton, Roberto Amato, Jeffrey Barrett, Sonia Goncalves, Ewan Harrison, David K. Jackson, Ian Johnston, Dominic Kwiatkowski, Cordelia Langford, John Sillitoe on behalf of the Wellcome Sanger Institute COVID-19 Surveillance Team |
| EPI_ISL_2808585 | Lighthouse Laboratory Plymouth | Wellcome Sanger Institute for the COVID-19 Genomics UK (COG-UK) Consortium | Lighthouse Laboratory Plymouth and Alex Alderton, Roberto Amato, Jeffrey Barrett, Sonia Goncalves, Ewan Harrison, David K. Jackson, Ian Johnston, Dominic Kwiatkowski, Cordelia Langford, John Sillitoe on behalf of the Wellcome Sanger Institute COVID-19 Surveillance Team |
| EPI_ISL_2808586 | Health Services Laboratories | Wellcome Sanger Institute for the COVID-19 Genomics UK (COG-UK) Consortium | Health Services Laboratories and Alex Alderton, Roberto Amato, Jeffrey Barrett, Sonia Goncalves, Ewan Harrison, David K. Jackson, Ian Johnston, Dominic Kwiatkowski, Cordelia Langford, John Sillitoe on behalf of the Wellcome Sanger Institute COVID-19 Surveillance Team |
| EPI_ISL_2808587 | Lighthouse Laboratory Plymouth | Wellcome Sanger Institute for the COVID-19 Genomics UK (COG-UK) Consortium | Lighthouse Laboratory Plymouth and Alex Alderton, Roberto Amato, Jeffrey Barrett, Sonia Goncalves, Ewan Harrison, David K. Jackson, Ian Johnston, Dominic Kwiatkowski, Cordelia Langford, John Sillitoe on behalf of the Wellcome Sanger Institute COVID-19 Surveillance Team |
| EPI_ISL_2808588 | Lighthouse Lab in Milton Keynes | Wellcome Sanger Institute for the COVID-19 Genomics UK (COG-UK) Consortium | The Lighthouse Lab in Milton Keynes and Alex Alderton, Roberto Amato, Jeffrey Barrett, Sonia Goncalves, Ewan Harrison, David K. Jackson, Ian Johnston, Dominic Kwiatkowski, Cordelia Langford, John Sillitoe on behalf of the Wellcome Sanger Institute COVID-19 Surveillance Team |
| EPI_ISL_2808589, EPI_ISL_2808590 | Lighthouse Laboratory Plymouth | Wellcome Sanger Institute for the COVID-19 Genomics UK (COG-UK) Consortium | Lighthouse Laboratory Plymouth and Alex Alderton, Roberto Amato, Jeffrey Barrett, Sonia Goncalves, Ewan Harrison, David K. Jackson, Ian Johnston, Dominic Kwiatkowski, Cordelia Langford, John Sillitoe on behalf of the Wellcome Sanger Institute COVID-19 Surveillance Team |
| EPI_ISL_2808591 | Lighthouse Lab in Glasgow | Wellcome Sanger Institute for the COVID-19 Genomics UK (COG-UK) Consortium | Harper VanSteenhouse, Yumi Kasai, David Gray, Carol Clugston, Anna Dominiczak and Alex Alderton, Roberto Amato, Jeffrey Barrett, Sonia Goncalves, Ewan Harrison, David K. Jackson, Ian Johnston, Dominic Kwiatkowski, Cordelia Langford, John Sillitoe on behalf of the Wellcome Sanger Institute COVID-19 Surveillance Team |
| EPI_ISL_2808592, EPI_ISL_2808593, EPI_ISL_2808594 | Lighthouse Laboratory Plymouth | Wellcome Sanger Institute for the COVID-19 Genomics UK (COG-UK) Consortium | Lighthouse Laboratory Plymouth and Alex Alderton, Roberto Amato, Jeffrey Barrett, Sonia Goncalves, Ewan Harrison, David K. Jackson, Ian Johnston, Dominic Kwiatkowski, Cordelia Langford, John Sillitoe on behalf of the Wellcome Sanger Institute COVID-19 Surveillance Team |
| EPI_ISL_2808595 | Lighthouse Lab in Glasgow | Wellcome Sanger Institute for the COVID-19 Genomics UK (COG-UK) Consortium | Harper VanSteenhouse, Yumi Kasai, David Gray, Carol Clugston, Anna Dominiczak and Alex Alderton, Roberto Amato, Jeffrey Barrett, Sonia Goncalves, Ewan Harrison, David K. Jackson, Ian Johnston, Dominic Kwiatkowski, Cordelia Langford, John Sillitoe on behalf of the Wellcome Sanger Institute COVID-19 Surveillance Team |
| EPI_ISL_2808596 | Lighthouse Laboratory Plymouth | Wellcome Sanger Institute for the COVID-19 Genomics UK (COG-UK) Consortium | Lighthouse Laboratory Plymouth and Alex Alderton, Roberto Amato, Jeffrey Barrett, Sonia Goncalves, Ewan Harrison, David K. Jackson, Ian Johnston, Dominic Kwiatkowski, Cordelia Langford, John Sillitoe on behalf of the Wellcome Sanger Institute COVID-19 Surveillance Team |
| EPI_ISL_2808597, EPI_ISL_2808598 | Lighthouse Lab in Milton Keynes | Wellcome Sanger Institute for the COVID-19 Genomics UK (COG-UK) Consortium | The Lighthouse Lab in Milton Keynes and Alex Alderton, Roberto Amato, Jeffrey Barrett, Sonia Goncalves, Ewan Harrison, David K. Jackson, Ian Johnston, Dominic Kwiatkowski, Cordelia Langford, John Sillitoe on behalf of the Wellcome Sanger Institute COVID-19 Surveillance Team |
| EPI_ISL_2808599 | Lighthouse Laboratory Plymouth | Wellcome Sanger Institute for the COVID-19 Genomics UK (COG-UK) Consortium | Lighthouse Laboratory Plymouth and Alex Alderton, Roberto Amato, Jeffrey Barrett, Sonia Goncalves, Ewan Harrison, David K. Jackson, Ian Johnston, Dominic Kwiatkowski, Cordelia Langford, John Sillitoe on behalf of the Wellcome Sanger Institute COVID-19 Surveillance Team |
| EPI_ISL_2808600 | Health Services Laboratories | Wellcome Sanger Institute for the COVID-19 Genomics UK (COG-UK) Consortium | Health Services Laboratories and Alex Alderton, Roberto Amato, Jeffrey Barrett, Sonia Goncalves, Ewan Harrison, David K. Jackson, Ian Johnston, Dominic Kwiatkowski, Cordelia Langford, John Sillitoe on behalf of the Wellcome Sanger Institute COVID-19 Surveillance Team |
| EPI_ISL_2808601 | Lighthouse Lab in Milton Keynes | Wellcome Sanger Institute for the COVID-19 Genomics UK (COG-UK) Consortium | The Lighthouse Lab in Milton Keynes and Alex Alderton, Roberto Amato, Jeffrey Barrett, Sonia Goncalves, Ewan Harrison, David K. Jackson, Ian Johnston, Dominic Kwiatkowski, Cordelia Langford, John Sillitoe on behalf of the Wellcome Sanger Institute COVID-19 Surveillance Team |
| EPI_ISL_2808602, EPI_ISL_2808603, EPI_ISL_2808604 | Lighthouse Laboratory Plymouth | Wellcome Sanger Institute for the COVID-19 Genomics UK (COG-UK) Consortium | Lighthouse Laboratory Plymouth and Alex Alderton, Roberto Amato, Jeffrey Barrett, Sonia Goncalves, Ewan Harrison, David K. Jackson, Ian Johnston, Dominic Kwiatkowski, Cordelia Langford, John Sillitoe on behalf of the Wellcome Sanger Institute COVID-19 Surveillance Team |
| EPI_ISL_2808605 | Lighthouse Lab in Milton Keynes | Wellcome Sanger Institute for the COVID-19 Genomics UK (COG-UK) Consortium | The Lighthouse Lab in Milton Keynes and Alex Alderton, Roberto Amato, Jeffrey Barrett, Sonia Goncalves, Ewan Harrison, David K. Jackson, Ian Johnston, Dominic Kwiatkowski, Cordelia Langford, John Sillitoe on behalf of the Wellcome Sanger Institute COVID-19 Surveillance Team |
| EPI_ISL_2808606 | Lighthouse Laboratory Plymouth | Wellcome Sanger Institute for the COVID-19 Genomics UK (COG-UK) Consortium | Lighthouse Laboratory Plymouth and Alex Alderton, Roberto Amato, Jeffrey Barrett, Sonia Goncalves, Ewan Harrison, David K. Jackson, Ian Johnston, Dominic Kwiatkowski, Cordelia Langford, John Sillitoe on behalf of the Wellcome Sanger Institute COVID-19 Surveillance Team |
| EPI_ISL_2808607 | Lighthouse Lab in Glasgow | Wellcome Sanger Institute for the COVID-19 Genomics UK (COG-UK) Consortium | Harper VanSteenhouse, Yumi Kasai, David Gray, Carol Clugston, Anna Dominiczak and Alex Alderton, Roberto Amato, Jeffrey Barrett, Sonia Goncalves, Ewan Harrison, David K. Jackson, Ian Johnston, Dominic Kwiatkowski, Cordelia Langford, John Sillitoe on behalf of the Wellcome Sanger Institute COVID-19 Surveillance Team |
| EPI_ISL_2808608, EPI_ISL_2808609, EPI_ISL_2808610, EPI_ISL_2808611, EPI_ISL_2808612 | Lighthouse Laboratory Plymouth | Wellcome Sanger Institute for the COVID-19 Genomics UK (COG-UK) Consortium | Lighthouse Laboratory Plymouth and Alex Alderton, Roberto Amato, Jeffrey Barrett, Sonia Goncalves, Ewan Harrison, David K. Jackson, Ian Johnston, Dominic Kwiatkowski, Cordelia Langford, John Sillitoe on behalf of the Wellcome Sanger Institute COVID-19 Surveillance Team |
| EPI_ISL_2808613 | Health Services Laboratories | Wellcome Sanger Institute for the COVID-19 Genomics UK (COG-UK) Consortium | Health Services Laboratories and Alex Alderton, Roberto Amato, Jeffrey Barrett, Sonia Goncalves, Ewan Harrison, David K. Jackson, Ian Johnston, Dominic Kwiatkowski, Cordelia Langford, John Sillitoe on behalf of the Wellcome Sanger Institute COVID-19 Surveillance Team |
| EPI_ISL_2808614 | Lighthouse Lab in Glasgow | Wellcome Sanger Institute for the COVID-19 Genomics UK (COG-UK) Consortium | Harper VanSteenhouse, Yumi Kasai, David Gray, Carol Clugston, Anna Dominiczak and Alex Alderton, Roberto Amato, Jeffrey Barrett, Sonia Goncalves, Ewan Harrison, David K. Jackson, Ian Johnston, Dominic Kwiatkowski, Cordelia Langford, John Sillitoe on behalf of the Wellcome Sanger Institute COVID-19 Surveillance Team |
| EPI_ISL_2808615 | Lighthouse Laboratory Plymouth | Wellcome Sanger Institute for the COVID-19 Genomics UK (COG-UK) Consortium | Lighthouse Laboratory Plymouth and Alex Alderton, Roberto Amato, Jeffrey Barrett, Sonia Goncalves, Ewan Harrison, David K. Jackson, Ian Johnston, Dominic Kwiatkowski, Cordelia Langford, John Sillitoe on behalf of the Wellcome Sanger Institute COVID-19 Surveillance Team |
| EPI_ISL_2808616 | Lighthouse Lab in Milton Keynes | Wellcome Sanger Institute for the COVID-19 Genomics UK (COG-UK) Consortium | The Lighthouse Lab in Milton Keynes and Alex Alderton, Roberto Amato, Jeffrey Barrett, Sonia Goncalves, Ewan Harrison, David K. Jackson, Ian Johnston, Dominic Kwiatkowski, Cordelia Langford, John Sillitoe on behalf of the Wellcome Sanger Institute COVID-19 Surveillance Team |
| EPI_ISL_2808617 | Health Services Laboratories | Wellcome Sanger Institute for the COVID-19 Genomics UK (COG-UK) Consortium | Health Services Laboratories and Alex Alderton, Roberto Amato, Jeffrey Barrett, Sonia Goncalves, Ewan Harrison, David K. Jackson, Ian Johnston, Dominic Kwiatkowski, Cordelia Langford, John Sillitoe on behalf of the Wellcome Sanger Institute COVID-19 Surveillance Team |
| EPI_ISL_2808618 | Lighthouse Lab in Glasgow | Wellcome Sanger Institute for the COVID-19 Genomics UK (COG-UK) Consortium | Harper VanSteenhouse, Yumi Kasai, David Gray, Carol Clugston, Anna Dominiczak and Alex Alderton, Roberto Amato, Jeffrey Barrett, Sonia Goncalves, Ewan Harrison, David K. Jackson, Ian Johnston, Dominic Kwiatkowski, Cordelia Langford, John Sillitoe on behalf of the Wellcome Sanger Institute COVID-19 Surveillance Team |
| EPI_ISL_2808619 | Lighthouse Laboratory Plymouth | Wellcome Sanger Institute for the COVID-19 Genomics UK (COG-UK) Consortium | Lighthouse Laboratory Plymouth and Alex Alderton, Roberto Amato, Jeffrey Barrett, Sonia Goncalves, Ewan Harrison, David K. Jackson, Ian Johnston, Dominic Kwiatkowski, Cordelia Langford, John Sillitoe on behalf of the Wellcome Sanger Institute COVID-19 Surveillance Team |
| EPI_ISL_2808620, EPI_ISL_2808621 | Lighthouse Lab in Milton Keynes | Wellcome Sanger Institute for the COVID-19 Genomics UK (COG-UK) Consortium | The Lighthouse Lab in Milton Keynes and Alex Alderton, Roberto Amato, Jeffrey Barrett, Sonia Goncalves, Ewan Harrison, David K. Jackson, Ian Johnston, Dominic Kwiatkowski, Cordelia Langford, John Sillitoe on behalf of the Wellcome Sanger Institute COVID-19 Surveillance Team |
| EPI_ISL_2808622 | Lighthouse Laboratory Plymouth | Wellcome Sanger Institute for the COVID-19 Genomics UK (COG-UK) Consortium | Lighthouse Laboratory Plymouth and Alex Alderton, Roberto Amato, Jeffrey Barrett, Sonia Goncalves, Ewan Harrison, David K. Jackson, Ian Johnston, Dominic Kwiatkowski, Cordelia Langford, John Sillitoe on behalf of the Wellcome Sanger Institute COVID-19 Surveillance Team |

| | | | |
|---|---|---|---|
| EPI_ISL_2808623 | Health Services Laboratories | Wellcome Sanger Institute for the COVID-19 Genomics UK (COG-UK) Consortium | Health Services Laboratories and Alex Alderton, Roberto Amato, Jeffrey Barrett, Sonia Goncalves, Ewan Harrison, David K. Jackson, Ian Johnston, Dominic Kwiatkowski, Cordelia Langford, John Sillitoe on behalf of the Wellcome Sanger Institute COVID-19 Surveillance Team |
| EPI_ISL_2808624 | Lighthouse Laboratory Plymouth | Wellcome Sanger Institute for the COVID-19 Genomics UK (COG-UK) Consortium | Lighthouse Laboratory Plymouth and Alex Alderton, Roberto Amato, Jeffrey Barrett, Sonia Goncalves, Ewan Harrison, David K. Jackson, Ian Johnston, Dominic Kwiatkowski, Cordelia Langford, John Sillitoe on behalf of the Wellcome Sanger Institute COVID-19 Surveillance Team |
| EPI_ISL_2808625 | Lighthouse Lab in Milton Keynes | Wellcome Sanger Institute for the COVID-19 Genomics UK (COG-UK) Consortium | The Lighthouse Lab in Milton Keynes and Alex Alderton, Roberto Amato, Jeffrey Barrett, Sonia Goncalves, Ewan Harrison, David K. Jackson, Ian Johnston, Dominic Kwiatkowski, Cordelia Langford, John Sillitoe on behalf of the Wellcome Sanger Institute COVID-19 Surveillance Team |
| EPI_ISL_2808626 | Lighthouse Lab in Glasgow | Wellcome Sanger Institute for the COVID-19 Genomics UK (COG-UK) Consortium | Harper VanSteenhouse, Yumi Kasai, David Gray, Carol Clugston, Anna Dominiczak and Alex Alderton, Roberto Amato, Jeffrey Barrett, Sonia Goncalves, Ewan Harrison, David K. Jackson, Ian Johnston, Dominic Kwiatkowski, Cordelia Langford, John Sillitoe on behalf of the Wellcome Sanger Institute COVID-19 Surveillance Team |
| EPI_ISL_2808627 | Lighthouse Lab in Milton Keynes | Wellcome Sanger Institute for the COVID-19 Genomics UK (COG-UK) Consortium | The Lighthouse Lab in Milton Keynes and Alex Alderton, Roberto Amato, Jeffrey Barrett, Sonia Goncalves, Ewan Harrison, David K. Jackson, Ian Johnston, Dominic Kwiatkowski, Cordelia Langford, John Sillitoe on behalf of the Wellcome Sanger Institute COVID-19 Surveillance Team |
| EPI_ISL_2808628, EPI_ISL_2808629 | Lighthouse Laboratory Plymouth | Wellcome Sanger Institute for the COVID-19 Genomics UK (COG-UK) Consortium | Lighthouse Laboratory Plymouth and Alex Alderton, Roberto Amato, Jeffrey Barrett, Sonia Goncalves, Ewan Harrison, David K. Jackson, Ian Johnston, Dominic Kwiatkowski, Cordelia Langford, John Sillitoe on behalf of the Wellcome Sanger Institute COVID-19 Surveillance Team |
| EPI_ISL_2808630 | Health Services Laboratories | Wellcome Sanger Institute for the COVID-19 Genomics UK (COG-UK) Consortium | Health Services Laboratories and Alex Alderton, Roberto Amato, Jeffrey Barrett, Sonia Goncalves, Ewan Harrison, David K. Jackson, Ian Johnston, Dominic Kwiatkowski, Cordelia Langford, John Sillitoe on behalf of the Wellcome Sanger Institute COVID-19 Surveillance Team |
| EPI_ISL_2808631 | Lighthouse Laboratory Plymouth | Wellcome Sanger Institute for the COVID-19 Genomics UK (COG-UK) Consortium | Lighthouse Laboratory Plymouth and Alex Alderton, Roberto Amato, Jeffrey Barrett, Sonia Goncalves, Ewan Harrison, David K. Jackson, Ian Johnston, Dominic Kwiatkowski, Cordelia Langford, John Sillitoe on behalf of the Wellcome Sanger Institute COVID-19 Surveillance Team |
| EPI_ISL_2808632, EPI_ISL_2808633 | Lighthouse Lab in Milton Keynes | Wellcome Sanger Institute for the COVID-19 Genomics UK (COG-UK) Consortium | The Lighthouse Lab in Milton Keynes and Alex Alderton, Roberto Amato, Jeffrey Barrett, Sonia Goncalves, Ewan Harrison, David K. Jackson, Ian Johnston, Dominic Kwiatkowski, Cordelia Langford, John Sillitoe on behalf of the Wellcome Sanger Institute COVID-19 Surveillance Team |
| EPI_ISL_2808634, EPI_ISL_2808635 | Lighthouse Laboratory Plymouth | Wellcome Sanger Institute for the COVID-19 Genomics UK (COG-UK) Consortium | Lighthouse Laboratory Plymouth and Alex Alderton, Roberto Amato, Jeffrey Barrett, Sonia Goncalves, Ewan Harrison, David K. Jackson, Ian Johnston, Dominic Kwiatkowski, Cordelia Langford, John Sillitoe on behalf of the Wellcome Sanger Institute COVID-19 Surveillance Team |
| EPI_ISL_2808636, EPI_ISL_2808637 | Lighthouse Lab in Milton Keynes | Wellcome Sanger Institute for the COVID-19 Genomics UK (COG-UK) Consortium | The Lighthouse Lab in Milton Keynes and Alex Alderton, Roberto Amato, Jeffrey Barrett, Sonia Goncalves, Ewan Harrison, David K. Jackson, Ian Johnston, Dominic Kwiatkowski, Cordelia Langford, John Sillitoe on behalf of the Wellcome Sanger Institute COVID-19 Surveillance Team |
| EPI_ISL_2808638, EPI_ISL_2808639, EPI_ISL_2808640 | Lighthouse Laboratory Plymouth | Wellcome Sanger Institute for the COVID-19 Genomics UK (COG-UK) Consortium | Lighthouse Laboratory Plymouth and Alex Alderton, Roberto Amato, Jeffrey Barrett, Sonia Goncalves, Ewan Harrison, David K. Jackson, Ian Johnston, Dominic Kwiatkowski, Cordelia Langford, John Sillitoe on behalf of the Wellcome Sanger Institute COVID-19 Surveillance Team |
| EPI_ISL_2808641 | Lighthouse Lab in Milton Keynes | Wellcome Sanger Institute for the COVID-19 Genomics UK (COG-UK) Consortium | The Lighthouse Lab in Milton Keynes and Alex Alderton, Roberto Amato, Jeffrey Barrett, Sonia Goncalves, Ewan Harrison, David K. Jackson, Ian Johnston, Dominic Kwiatkowski, Cordelia Langford, John Sillitoe on behalf of the Wellcome Sanger Institute COVID-19 Surveillance Team |
| EPI_ISL_2808642 | Lighthouse Laboratory Plymouth | Wellcome Sanger Institute for the COVID-19 Genomics UK (COG-UK) Consortium | Lighthouse Laboratory Plymouth and Alex Alderton, Roberto Amato, Jeffrey Barrett, Sonia Goncalves, Ewan Harrison, David K. Jackson, Ian Johnston, Dominic Kwiatkowski, Cordelia Langford, John Sillitoe on behalf of the Wellcome Sanger Institute COVID-19 Surveillance Team |
| EPI_ISL_2808643 | Lighthouse Lab in Glasgow | Wellcome Sanger Institute for the COVID-19 Genomics UK (COG-UK) Consortium | Harper VanSteenhouse, Yumi Kasai, David Gray, Carol Clugston, Anna Dominiczak and Alex Alderton, Roberto Amato, Jeffrey Barrett, Sonia Goncalves, Ewan Harrison, David K. Jackson, Ian Johnston, Dominic Kwiatkowski, Cordelia Langford, John Sillitoe on behalf of the Wellcome Sanger Institute COVID-19 Surveillance Team |
| EPI_ISL_2808644, EPI_ISL_2808645 | Lighthouse Laboratory Plymouth | Wellcome Sanger Institute for the COVID-19 Genomics UK (COG-UK) Consortium | Lighthouse Laboratory Plymouth and Alex Alderton, Roberto Amato, Jeffrey Barrett, Sonia Goncalves, Ewan Harrison, David K. Jackson, Ian Johnston, Dominic Kwiatkowski, Cordelia Langford, John Sillitoe on behalf of the Wellcome Sanger Institute COVID-19 Surveillance Team |
| EPI_ISL_2808646 | Lighthouse Lab in Glasgow | Wellcome Sanger Institute for the COVID-19 Genomics UK (COG-UK) Consortium | Harper VanSteenhouse, Yumi Kasai, David Gray, Carol Clugston, Anna Dominiczak and Alex Alderton, Roberto Amato, Jeffrey Barrett, Sonia Goncalves, Ewan Harrison, David K. Jackson, Ian Johnston, Dominic Kwiatkowski, Cordelia Langford, John Sillitoe on behalf of the Wellcome Sanger Institute COVID-19 Surveillance Team |
| EPI_ISL_2808647 | Lighthouse Lab in Milton Keynes | Wellcome Sanger Institute for the COVID-19 Genomics UK (COG-UK) Consortium | The Lighthouse Lab in Milton Keynes and Alex Alderton, Roberto Amato, Jeffrey Barrett, Sonia Goncalves, Ewan Harrison, David K. Jackson, Ian Johnston, Dominic Kwiatkowski, Cordelia Langford, John Sillitoe on behalf of the Wellcome Sanger Institute COVID-19 Surveillance Team |
| EPI_ISL_2808648 | Health Services Laboratories | Wellcome Sanger Institute for the COVID-19 Genomics UK (COG-UK) Consortium | Health Services Laboratories and Alex Alderton, Roberto Amato, Jeffrey Barrett, Sonia Goncalves, Ewan Harrison, David K. Jackson, Ian Johnston, Dominic Kwiatkowski, Cordelia Langford, John Sillitoe on behalf of the Wellcome Sanger Institute COVID-19 Surveillance Team |
| EPI_ISL_2808649 | Lighthouse Lab in Milton Keynes | Wellcome Sanger Institute for the COVID-19 Genomics UK (COG-UK) Consortium | The Lighthouse Lab in Milton Keynes and Alex Alderton, Roberto Amato, Jeffrey Barrett, Sonia Goncalves, Ewan Harrison, David K. Jackson, Ian Johnston, Dominic Kwiatkowski, Cordelia Langford, John Sillitoe on behalf of the Wellcome Sanger Institute COVID-19 Surveillance Team |
| EPI_ISL_2808650, EPI_ISL_2808651 | Lighthouse Laboratory Plymouth | Wellcome Sanger Institute for the COVID-19 Genomics UK (COG-UK) Consortium | Lighthouse Laboratory Plymouth and Alex Alderton, Roberto Amato, Jeffrey Barrett, Sonia Goncalves, Ewan Harrison, David K. Jackson, Ian Johnston, Dominic Kwiatkowski, Cordelia Langford, John Sillitoe on behalf of the Wellcome Sanger Institute COVID-19 Surveillance Team |
| EPI_ISL_2808652, EPI_ISL_2808653 | Lighthouse Lab in Milton Keynes | Wellcome Sanger Institute for the COVID-19 Genomics UK (COG-UK) Consortium | The Lighthouse Lab in Milton Keynes and Alex Alderton, Roberto Amato, Jeffrey Barrett, Sonia Goncalves, Ewan Harrison, David K. Jackson, Ian Johnston, Dominic Kwiatkowski, Cordelia Langford, John Sillitoe on behalf of the Wellcome Sanger Institute COVID-19 Surveillance Team |
| EPI_ISL_2808654 | Health Services Laboratories | Wellcome Sanger Institute for the COVID-19 Genomics UK (COG-UK) Consortium | Health Services Laboratories and Alex Alderton, Roberto Amato, Jeffrey Barrett, Sonia Goncalves, Ewan Harrison, David K. Jackson, Ian Johnston, Dominic Kwiatkowski, Cordelia Langford, John Sillitoe on behalf of the Wellcome Sanger Institute COVID-19 Surveillance Team |
| EPI_ISL_2808655 | Lighthouse Laboratory Plymouth | Wellcome Sanger Institute for the COVID-19 Genomics UK (COG-UK) Consortium | Lighthouse Laboratory Plymouth and Alex Alderton, Roberto Amato, Jeffrey Barrett, Sonia Goncalves, Ewan Harrison, David K. Jackson, Ian Johnston, Dominic Kwiatkowski, Cordelia Langford, John Sillitoe on behalf of the Wellcome Sanger Institute COVID-19 Surveillance Team |
| EPI_ISL_2808656, EPI_ISL_2808657 | Lighthouse Lab in Glasgow | Wellcome Sanger Institute for the COVID-19 Genomics UK (COG-UK) Consortium | Harper VanSteenhouse, Yumi Kasai, David Gray, Carol Clugston, Anna Dominiczak and Alex Alderton, Roberto Amato, Jeffrey Barrett, Sonia Goncalves, Ewan Harrison, David K. Jackson, Ian Johnston, Dominic Kwiatkowski, Cordelia Langford, John Sillitoe on behalf of the Wellcome Sanger Institute COVID-19 Surveillance Team |
| EPI_ISL_2808658, EPI_ISL_2808659 | Lighthouse Laboratory Plymouth | Wellcome Sanger Institute for the COVID-19 Genomics UK (COG-UK) Consortium | Lighthouse Laboratory Plymouth and Alex Alderton, Roberto Amato, Jeffrey Barrett, Sonia Goncalves, Ewan Harrison, David K. Jackson, Ian Johnston, Dominic Kwiatkowski, Cordelia Langford, John Sillitoe on behalf of the Wellcome Sanger Institute COVID-19 Surveillance Team |
| EPI_ISL_2808660 | Lighthouse Lab in Glasgow | Wellcome Sanger Institute for the COVID-19 Genomics UK (COG-UK) Consortium | Harper VanSteenhouse, Yumi Kasai, David Gray, Carol Clugston, Anna Dominiczak and Alex Alderton, Roberto Amato, Jeffrey Barrett, Sonia Goncalves, Ewan Harrison, David K. Jackson, Ian Johnston, Dominic Kwiatkowski, Cordelia Langford, John Sillitoe on behalf of the Wellcome Sanger Institute COVID-19 Surveillance Team |
| EPI_ISL_2808661 | Health Services Laboratories | Wellcome Sanger Institute for the COVID-19 Genomics UK (COG-UK) Consortium | Health Services Laboratories and Alex Alderton, Roberto Amato, Jeffrey Barrett, Sonia Goncalves, Ewan Harrison, David K. Jackson, Ian Johnston, Dominic Kwiatkowski, Cordelia Langford, John Sillitoe on behalf of the Wellcome Sanger Institute COVID-19 Surveillance Team |
| EPI_ISL_2808662 | Lighthouse Lab in Milton Keynes | Wellcome Sanger Institute for the COVID-19 Genomics UK (COG-UK) Consortium | The Lighthouse Lab in Milton Keynes and Alex Alderton, Roberto Amato, Jeffrey Barrett, Sonia Goncalves, Ewan Harrison, David K. Jackson, Ian Johnston, Dominic Kwiatkowski, Cordelia Langford, John Sillitoe on behalf of the Wellcome Sanger Institute COVID-19 Surveillance Team |
| EPI_ISL_2808663, EPI_ISL_2808664 | Lighthouse Laboratory Plymouth | Wellcome Sanger Institute for the COVID-19 Genomics UK (COG-UK) Consortium | Lighthouse Laboratory Plymouth and Alex Alderton, Roberto Amato, Jeffrey Barrett, Sonia Goncalves, Ewan Harrison, David K. Jackson, Ian Johnston, Dominic Kwiatkowski, Cordelia Langford, John Sillitoe on behalf of the Wellcome Sanger Institute COVID-19 Surveillance Team |
| EPI_ISL_2808665 | Health Services Laboratories | Wellcome Sanger Institute for the COVID-19 Genomics UK (COG-UK) Consortium | Health Services Laboratories and Alex Alderton, Roberto Amato, Jeffrey Barrett, Sonia Goncalves, Ewan Harrison, David K. Jackson, Ian Johnston, Dominic Kwiatkowski, Cordelia Langford, John Sillitoe on behalf of the Wellcome Sanger Institute COVID-19 Surveillance Team |
| EPI_ISL_2808666, EPI_ISL_2808667 | Lighthouse Laboratory Plymouth | Wellcome Sanger Institute for the COVID-19 Genomics UK (COG-UK) Consortium | Lighthouse Laboratory Plymouth and Alex Alderton, Roberto Amato, Jeffrey Barrett, Sonia Goncalves, Ewan Harrison, David K. Jackson, Ian Johnston, Dominic Kwiatkowski, Cordelia Langford, John Sillitoe on behalf of the Wellcome Sanger Institute COVID-19 Surveillance Team |
| EPI_ISL_2808668, EPI_ISL_2808669 | Lighthouse Lab in Milton Keynes | Wellcome Sanger Institute for the COVID-19 Genomics UK (COG-UK) Consortium | The Lighthouse Lab in Milton Keynes and Alex Alderton, Roberto Amato, Jeffrey Barrett, Sonia Goncalves, Ewan Harrison, David K. Jackson, Ian Johnston, Dominic Kwiatkowski, Cordelia Langford, John Sillitoe on behalf of the Wellcome Sanger Institute COVID-19 Surveillance Team |
| EPI_ISL_2808670, EPI_ISL_2808671 | Lighthouse Laboratory Plymouth | Wellcome Sanger Institute for the COVID-19 Genomics UK (COG-UK) Consortium | Lighthouse Laboratory Plymouth and Alex Alderton, Roberto Amato, Jeffrey Barrett, Sonia Goncalves, Ewan Harrison, David K. Jackson, Ian Johnston, Dominic Kwiatkowski, Cordelia Langford, John Sillitoe on behalf of the Wellcome Sanger Institute COVID-19 Surveillance Team |
| EPI_ISL_2808672 | Lighthouse Lab in Glasgow | Wellcome Sanger Institute for the COVID-19 Genomics UK (COG-UK) Consortium | Harper VanSteenhouse, Yumi Kasai, David Gray, Carol Clugston, Anna Dominiczak and Alex Alderton, Roberto Amato, Jeffrey Barrett, Sonia Goncalves, Ewan Harrison, David K. Jackson, Ian Johnston, Dominic Kwiatkowski, Cordelia Langford, John Sillitoe on behalf of the Wellcome Sanger Institute COVID-19 Surveillance Team |
| EPI_ISL_2808673 | Lighthouse Lab in Milton Keynes | Wellcome Sanger Institute for the COVID-19 Genomics UK (COG-UK) Consortium | The Lighthouse Lab in Milton Keynes and Alex Alderton, Roberto Amato, Jeffrey Barrett, Sonia Goncalves, Ewan Harrison, David K. Jackson, Ian Johnston, Dominic Kwiatkowski, Cordelia Langford, John Sillitoe on behalf of the Wellcome Sanger Institute COVID-19 Surveillance Team |
| EPI_ISL_2808674 | Lighthouse Laboratory Plymouth | Wellcome Sanger Institute for the COVID-19 Genomics UK (COG-UK) Consortium | Lighthouse Laboratory Plymouth and Alex Alderton, Roberto Amato, Jeffrey Barrett, Sonia Goncalves, Ewan Harrison, David K. Jackson, Ian Johnston, Dominic Kwiatkowski, Cordelia Langford, John Sillitoe on behalf of the Wellcome Sanger Institute COVID-19 Surveillance Team |
| EPI_ISL_2808675 | Lighthouse Lab in Milton Keynes | Wellcome Sanger Institute for the COVID-19 Genomics UK (COG-UK) Consortium | The Lighthouse Lab in Milton Keynes and Alex Alderton, Roberto Amato, Jeffrey Barrett, Sonia Goncalves, Ewan Harrison, David K. Jackson, Ian Johnston, Dominic Kwiatkowski, Cordelia Langford, John Sillitoe on behalf of the Wellcome Sanger Institute COVID-19 Surveillance Team |
| EPI_ISL_2808676 | Lighthouse Lab in Glasgow | Wellcome Sanger Institute for the COVID-19 Genomics UK (COG-UK) Consortium | Harper VanSteenhouse, Yumi Kasai, David Gray, Carol Clugston, Anna Dominiczak and Alex Alderton, Roberto Amato, Jeffrey Barrett, Sonia Goncalves, Ewan Harrison, David K. Jackson, Ian Johnston, Dominic Kwiatkowski, Cordelia Langford, John Sillitoe on behalf of the Wellcome Sanger Institute COVID-19 Surveillance Team |
| EPI_ISL_2808677 | Lighthouse Lab in Milton Keynes | Wellcome Sanger Institute for the COVID-19 Genomics UK (COG-UK) Consortium | The Lighthouse Lab in Milton Keynes and Alex Alderton, Roberto Amato, Jeffrey Barrett, Sonia Goncalves, Ewan Harrison, David K. Jackson, Ian Johnston, Dominic Kwiatkowski, Cordelia Langford, John Sillitoe on behalf of the Wellcome Sanger Institute COVID-19 Surveillance Team |
| EPI_ISL_2808678 | Lighthouse Laboratory Plymouth | Wellcome Sanger Institute for the COVID-19 Genomics UK (COG-UK) Consortium | Lighthouse Laboratory Plymouth and Alex Alderton, Roberto Amato, Jeffrey Barrett, Sonia Goncalves, Ewan Harrison, David K. Jackson, Ian Johnston, Dominic Kwiatkowski, Cordelia Langford, John Sillitoe on behalf of the Wellcome Sanger Institute COVID-19 Surveillance Team |
| EPI_ISL_2808679, EPI_ISL_2808680 | Lighthouse Lab in Milton Keynes | Wellcome Sanger Institute for the COVID-19 Genomics UK (COG-UK) Consortium | The Lighthouse Lab in Milton Keynes and Alex Alderton, Roberto Amato, Jeffrey Barrett, Sonia Goncalves, Ewan Harrison, David K. Jackson, Ian Johnston, Dominic Kwiatkowski, Cordelia Langford, John Sillitoe on behalf of the Wellcome Sanger Institute COVID-19 Surveillance Team |
| EPI_ISL_2808681 | Lighthouse Laboratory Plymouth | Wellcome Sanger Institute for the COVID-19 Genomics UK (COG-UK) Consortium | Lighthouse Laboratory Plymouth and Alex Alderton, Roberto Amato, Jeffrey Barrett, Sonia Goncalves, Ewan Harrison, David K. Jackson, Ian Johnston, Dominic Kwiatkowski, Cordelia Langford, John Sillitoe on behalf of the Wellcome Sanger Institute COVID-19 Surveillance Team |
| EPI_ISL_2808682 | Lighthouse Lab in Glasgow | Wellcome Sanger Institute for the COVID-19 Genomics UK (COG-UK) Consortium | Harper VanSteenhouse, Yumi Kasai, David Gray, Carol Clugston, Anna Dominiczak and Alex Alderton, Roberto Amato, Jeffrey Barrett, Sonia Goncalves, Ewan Harrison, David K. Jackson, Ian Johnston, Dominic Kwiatkowski, Cordelia Langford, John Sillitoe on behalf of the Wellcome Sanger Institute COVID-19 Surveillance Team |
| EPI_ISL_2808683 | Lighthouse Laboratory Plymouth | Wellcome Sanger Institute for the COVID-19 Genomics UK (COG-UK) Consortium | Lighthouse Laboratory Plymouth and Alex Alderton, Roberto Amato, Jeffrey Barrett, Sonia Goncalves, Ewan Harrison, David K. Jackson, Ian Johnston, Dominic Kwiatkowski, Cordelia Langford, John Sillitoe on behalf of the Wellcome Sanger Institute COVID-19 Surveillance Team |

| | | | |
|---|---|---|---|
| EPI_ISL_2808684 | Health Services Laboratories | Wellcome Sanger Institute for the COVID-19 Genomics UK (COG-UK) Consortium | Health Services Laboratories and Alex Alderton, Roberto Amato, Jeffrey Barrett, Sonia Goncalves, Ewan Harrison, David K. Jackson, Ian Johnston, Dominic Kwiatkowski, Cordelia Langford, John Sillitoe on behalf of the Wellcome Sanger Institute COVID-19 Surveillance Team |
| EPI_ISL_2808685, EPI_ISL_2808686, EPI_ISL_2808687, EPI_ISL_2808688 | Lighthouse Lab in Milton Keynes | Wellcome Sanger Institute for the COVID-19 Genomics UK (COG-UK) Consortium | The Lighthouse Lab in Milton Keynes and Alex Alderton, Roberto Amato, Jeffrey Barrett, Sonia Goncalves, Ewan Harrison, David K. Jackson, Ian Johnston, Dominic Kwiatkowski, Cordelia Langford, John Sillitoe on behalf of the Wellcome Sanger Institute COVID-19 Surveillance Team |
| EPI_ISL_2808689 | Lighthouse Laboratory Plymouth | Wellcome Sanger Institute for the COVID-19 Genomics UK (COG-UK) Consortium | Lighthouse Laboratory Plymouth and Alex Alderton, Roberto Amato, Jeffrey Barrett, Sonia Goncalves, Ewan Harrison, David K. Jackson, Ian Johnston, Dominic Kwiatkowski, Cordelia Langford, John Sillitoe on behalf of the Wellcome Sanger Institute COVID-19 Surveillance Team |
| EPI_ISL_2808690 | Lighthouse Lab in Milton Keynes | Wellcome Sanger Institute for the COVID-19 Genomics UK (COG-UK) Consortium | The Lighthouse Lab in Milton Keynes and Alex Alderton, Roberto Amato, Jeffrey Barrett, Sonia Goncalves, Ewan Harrison, David K. Jackson, Ian Johnston, Dominic Kwiatkowski, Cordelia Langford, John Sillitoe on behalf of the Wellcome Sanger Institute COVID-19 Surveillance Team |
| EPI_ISL_2808691 | Lighthouse Laboratory Plymouth | Wellcome Sanger Institute for the COVID-19 Genomics UK (COG-UK) Consortium | Lighthouse Laboratory Plymouth and Alex Alderton, Roberto Amato, Jeffrey Barrett, Sonia Goncalves, Ewan Harrison, David K. Jackson, Ian Johnston, Dominic Kwiatkowski, Cordelia Langford, John Sillitoe on behalf of the Wellcome Sanger Institute COVID-19 Surveillance Team |
| EPI_ISL_2808692 | Lighthouse Lab in Milton Keynes | Wellcome Sanger Institute for the COVID-19 Genomics UK (COG-UK) Consortium | The Lighthouse Lab in Milton Keynes and Alex Alderton, Roberto Amato, Jeffrey Barrett, Sonia Goncalves, Ewan Harrison, David K. Jackson, Ian Johnston, Dominic Kwiatkowski, Cordelia Langford, John Sillitoe on behalf of the Wellcome Sanger Institute COVID-19 Surveillance Team |
| EPI_ISL_2808693, EPI_ISL_2808694 | Lighthouse Laboratory Plymouth | Wellcome Sanger Institute for the COVID-19 Genomics UK (COG-UK) Consortium | Lighthouse Laboratory Plymouth and Alex Alderton, Roberto Amato, Jeffrey Barrett, Sonia Goncalves, Ewan Harrison, David K. Jackson, Ian Johnston, Dominic Kwiatkowski, Cordelia Langford, John Sillitoe on behalf of the Wellcome Sanger Institute COVID-19 Surveillance Team |
| EPI_ISL_2808695 | Lighthouse Lab in Milton Keynes | Wellcome Sanger Institute for the COVID-19 Genomics UK (COG-UK) Consortium | The Lighthouse Lab in Milton Keynes and Alex Alderton, Roberto Amato, Jeffrey Barrett, Sonia Goncalves, Ewan Harrison, David K. Jackson, Ian Johnston, Dominic Kwiatkowski, Cordelia Langford, John Sillitoe on behalf of the Wellcome Sanger Institute COVID-19 Surveillance Team |
| EPI_ISL_2808696, EPI_ISL_2808697 | Lighthouse Laboratory Plymouth | Wellcome Sanger Institute for the COVID-19 Genomics UK (COG-UK) Consortium | Lighthouse Laboratory Plymouth and Alex Alderton, Roberto Amato, Jeffrey Barrett, Sonia Goncalves, Ewan Harrison, David K. Jackson, Ian Johnston, Dominic Kwiatkowski, Cordelia Langford, John Sillitoe on behalf of the Wellcome Sanger Institute COVID-19 Surveillance Team |
| EPI_ISL_2808698 | Lighthouse Lab in Milton Keynes | Wellcome Sanger Institute for the COVID-19 Genomics UK (COG-UK) Consortium | The Lighthouse Lab in Milton Keynes and Alex Alderton, Roberto Amato, Jeffrey Barrett, Sonia Goncalves, Ewan Harrison, David K. Jackson, Ian Johnston, Dominic Kwiatkowski, Cordelia Langford, John Sillitoe on behalf of the Wellcome Sanger Institute COVID-19 Surveillance Team |
| EPI_ISL_2808699, EPI_ISL_2808700, EPI_ISL_2808701 | Lighthouse Laboratory Plymouth | Wellcome Sanger Institute for the COVID-19 Genomics UK (COG-UK) Consortium | Lighthouse Laboratory Plymouth and Alex Alderton, Roberto Amato, Jeffrey Barrett, Sonia Goncalves, Ewan Harrison, David K. Jackson, Ian Johnston, Dominic Kwiatkowski, Cordelia Langford, John Sillitoe on behalf of the Wellcome Sanger Institute COVID-19 Surveillance Team |
| EPI_ISL_2808702 | Lighthouse Lab in Glasgow | Wellcome Sanger Institute for the COVID-19 Genomics UK (COG-UK) Consortium | Harper VanSteenhouse, Yumi Kasai, David Gray, Carol Clugston, Anna Dominiczak and Alex Alderton, Roberto Amato, Jeffrey Barrett, Sonia Goncalves, Ewan Harrison, David K. Jackson, Ian Johnston, Dominic Kwiatkowski, Cordelia Langford, John Sillitoe on behalf of the Wellcome Sanger Institute COVID-19 Surveillance Team |
| EPI_ISL_2808703 | Lighthouse Laboratory Plymouth | Wellcome Sanger Institute for the COVID-19 Genomics UK (COG-UK) Consortium | Lighthouse Laboratory Plymouth and Alex Alderton, Roberto Amato, Jeffrey Barrett, Sonia Goncalves, Ewan Harrison, David K. Jackson, Ian Johnston, Dominic Kwiatkowski, Cordelia Langford, John Sillitoe on behalf of the Wellcome Sanger Institute COVID-19 Surveillance Team |
| EPI_ISL_2808704 | Health Services Laboratories | Wellcome Sanger Institute for the COVID-19 Genomics UK (COG-UK) Consortium | Health Services Laboratories and Alex Alderton, Roberto Amato, Jeffrey Barrett, Sonia Goncalves, Ewan Harrison, David K. Jackson, Ian Johnston, Dominic Kwiatkowski, Cordelia Langford, John Sillitoe on behalf of the Wellcome Sanger Institute COVID-19 Surveillance Team |
| EPI_ISL_2808705, EPI_ISL_2808706 | Lighthouse Laboratory Plymouth | Wellcome Sanger Institute for the COVID-19 Genomics UK (COG-UK) Consortium | Lighthouse Laboratory Plymouth and Alex Alderton, Roberto Amato, Jeffrey Barrett, Sonia Goncalves, Ewan Harrison, David K. Jackson, Ian Johnston, Dominic Kwiatkowski, Cordelia Langford, John Sillitoe on behalf of the Wellcome Sanger Institute COVID-19 Surveillance Team |
| EPI_ISL_2808707 | Lighthouse Lab in Glasgow | Wellcome Sanger Institute for the COVID-19 Genomics UK (COG-UK) Consortium | Harper VanSteenhouse, Yumi Kasai, David Gray, Carol Clugston, Anna Dominiczak and Alex Alderton, Roberto Amato, Jeffrey Barrett, Sonia Goncalves, Ewan Harrison, David K. Jackson, Ian Johnston, Dominic Kwiatkowski, Cordelia Langford, John Sillitoe on behalf of the Wellcome Sanger Institute COVID-19 Surveillance Team |
| EPI_ISL_2808708 | Lighthouse Laboratory Plymouth | Wellcome Sanger Institute for the COVID-19 Genomics UK (COG-UK) Consortium | Lighthouse Laboratory Plymouth and Alex Alderton, Roberto Amato, Jeffrey Barrett, Sonia Goncalves, Ewan Harrison, David K. Jackson, Ian Johnston, Dominic Kwiatkowski, Cordelia Langford, John Sillitoe on behalf of the Wellcome Sanger Institute COVID-19 Surveillance Team |
| EPI_ISL_2808709 | Lighthouse Lab in Milton Keynes | Wellcome Sanger Institute for the COVID-19 Genomics UK (COG-UK) Consortium | The Lighthouse Lab in Milton Keynes and Alex Alderton, Roberto Amato, Jeffrey Barrett, Sonia Goncalves, Ewan Harrison, David K. Jackson, Ian Johnston, Dominic Kwiatkowski, Cordelia Langford, John Sillitoe on behalf of the Wellcome Sanger Institute COVID-19 Surveillance Team |
| EPI_ISL_2808710 | Lighthouse Laboratory Plymouth | Wellcome Sanger Institute for the COVID-19 Genomics UK (COG-UK) Consortium | Lighthouse Laboratory Plymouth and Alex Alderton, Roberto Amato, Jeffrey Barrett, Sonia Goncalves, Ewan Harrison, David K. Jackson, Ian Johnston, Dominic Kwiatkowski, Cordelia Langford, John Sillitoe on behalf of the Wellcome Sanger Institute COVID-19 Surveillance Team |
| EPI_ISL_2808711, EPI_ISL_2808712, EPI_ISL_2808713 | Lighthouse Lab in Milton Keynes | Wellcome Sanger Institute for the COVID-19 Genomics UK (COG-UK) Consortium | The Lighthouse Lab in Milton Keynes and Alex Alderton, Roberto Amato, Jeffrey Barrett, Sonia Goncalves, Ewan Harrison, David K. Jackson, Ian Johnston, Dominic Kwiatkowski, Cordelia Langford, John Sillitoe on behalf of the Wellcome Sanger Institute COVID-19 Surveillance Team |
| EPI_ISL_2808714 | Lighthouse Laboratory Plymouth | Wellcome Sanger Institute for the COVID-19 Genomics UK (COG-UK) Consortium | Lighthouse Laboratory Plymouth and Alex Alderton, Roberto Amato, Jeffrey Barrett, Sonia Goncalves, Ewan Harrison, David K. Jackson, Ian Johnston, Dominic Kwiatkowski, Cordelia Langford, John Sillitoe on behalf of the Wellcome Sanger Institute COVID-19 Surveillance Team |
| EPI_ISL_2808715 | Lighthouse Lab in Milton Keynes | Wellcome Sanger Institute for the COVID-19 Genomics UK (COG-UK) Consortium | The Lighthouse Lab in Milton Keynes and Alex Alderton, Roberto Amato, Jeffrey Barrett, Sonia Goncalves, Ewan Harrison, David K. Jackson, Ian Johnston, Dominic Kwiatkowski, Cordelia Langford, John Sillitoe on behalf of the Wellcome Sanger Institute COVID-19 Surveillance Team |
| EPI_ISL_2808716 | Lighthouse Lab in Glasgow | Wellcome Sanger Institute for the COVID-19 Genomics UK (COG-UK) Consortium | Harper VanSteenhouse, Yumi Kasai, David Gray, Carol Clugston, Anna Dominiczak and Alex Alderton, Roberto Amato, Jeffrey Barrett, Sonia Goncalves, Ewan Harrison, David K. Jackson, Ian Johnston, Dominic Kwiatkowski, Cordelia Langford, John Sillitoe on behalf of the Wellcome Sanger Institute COVID-19 Surveillance Team |
| EPI_ISL_2808717 | Health Services Laboratories | Wellcome Sanger Institute for the COVID-19 Genomics UK (COG-UK) Consortium | Health Services Laboratories and Alex Alderton, Roberto Amato, Jeffrey Barrett, Sonia Goncalves, Ewan Harrison, David K. Jackson, Ian Johnston, Dominic Kwiatkowski, Cordelia Langford, John Sillitoe on behalf of the Wellcome Sanger Institute COVID-19 Surveillance Team |
| EPI_ISL_2808718, EPI_ISL_2808719 | Lighthouse Laboratory Plymouth | Wellcome Sanger Institute for the COVID-19 Genomics UK (COG-UK) Consortium | Lighthouse Laboratory Plymouth and Alex Alderton, Roberto Amato, Jeffrey Barrett, Sonia Goncalves, Ewan Harrison, David K. Jackson, Ian Johnston, Dominic Kwiatkowski, Cordelia Langford, John Sillitoe on behalf of the Wellcome Sanger Institute COVID-19 Surveillance Team |
| EPI_ISL_2808720, EPI_ISL_2808721, EPI_ISL_2808722, EPI_ISL_2808723 | Lighthouse Lab in Milton Keynes | Wellcome Sanger Institute for the COVID-19 Genomics UK (COG-UK) Consortium | The Lighthouse Lab in Milton Keynes and Alex Alderton, Roberto Amato, Jeffrey Barrett, Sonia Goncalves, Ewan Harrison, David K. Jackson, Ian Johnston, Dominic Kwiatkowski, Cordelia Langford, John Sillitoe on behalf of the Wellcome Sanger Institute COVID-19 Surveillance Team |
| EPI_ISL_2808724 | Lighthouse Lab in Glasgow | Wellcome Sanger Institute for the COVID-19 Genomics UK (COG-UK) Consortium | Harper VanSteenhouse, Yumi Kasai, David Gray, Carol Clugston, Anna Dominiczak and Alex Alderton, Roberto Amato, Jeffrey Barrett, Sonia Goncalves, Ewan Harrison, David K. Jackson, Ian Johnston, Dominic Kwiatkowski, Cordelia Langford, John Sillitoe on behalf of the Wellcome Sanger Institute COVID-19 Surveillance Team |
| EPI_ISL_2808725 | Lighthouse Lab in Milton Keynes | Wellcome Sanger Institute for the COVID-19 Genomics UK (COG-UK) Consortium | The Lighthouse Lab in Milton Keynes and Alex Alderton, Roberto Amato, Jeffrey Barrett, Sonia Goncalves, Ewan Harrison, David K. Jackson, Ian Johnston, Dominic Kwiatkowski, Cordelia Langford, John Sillitoe on behalf of the Wellcome Sanger Institute COVID-19 Surveillance Team |
| EPI_ISL_2808726, EPI_ISL_2808727 | Lighthouse Lab in Glasgow | Wellcome Sanger Institute for the COVID-19 Genomics UK (COG-UK) Consortium | Harper VanSteenhouse, Yumi Kasai, David Gray, Carol Clugston, Anna Dominiczak and Alex Alderton, Roberto Amato, Jeffrey Barrett, Sonia Goncalves, Ewan Harrison, David K. Jackson, Ian Johnston, Dominic Kwiatkowski, Cordelia Langford, John Sillitoe on behalf of the Wellcome Sanger Institute COVID-19 Surveillance Team |
| EPI_ISL_2808728 | Lighthouse Laboratory Plymouth | Wellcome Sanger Institute for the COVID-19 Genomics UK (COG-UK) Consortium | Lighthouse Laboratory Plymouth and Alex Alderton, Roberto Amato, Jeffrey Barrett, Sonia Goncalves, Ewan Harrison, David K. Jackson, Ian Johnston, Dominic Kwiatkowski, Cordelia Langford, John Sillitoe on behalf of the Wellcome Sanger Institute COVID-19 Surveillance Team |
| EPI_ISL_2808729, EPI_ISL_2808730 | Lighthouse Lab in Milton Keynes | Wellcome Sanger Institute for the COVID-19 Genomics UK (COG-UK) Consortium | The Lighthouse Lab in Milton Keynes and Alex Alderton, Roberto Amato, Jeffrey Barrett, Sonia Goncalves, Ewan Harrison, David K. Jackson, Ian Johnston, Dominic Kwiatkowski, Cordelia Langford, John Sillitoe on behalf of the Wellcome Sanger Institute COVID-19 Surveillance Team |
| EPI_ISL_2808731, EPI_ISL_2808732 | Lighthouse Laboratory Plymouth | Wellcome Sanger Institute for the COVID-19 Genomics UK (COG-UK) Consortium | Lighthouse Laboratory Plymouth and Alex Alderton, Roberto Amato, Jeffrey Barrett, Sonia Goncalves, Ewan Harrison, David K. Jackson, Ian Johnston, Dominic Kwiatkowski, Cordelia Langford, John Sillitoe on behalf of the Wellcome Sanger Institute COVID-19 Surveillance Team |
| EPI_ISL_2808733, EPI_ISL_2808734, EPI_ISL_2808735, EPI_ISL_2808736, EPI_ISL_2808737, EPI_ISL_2808738, EPI_ISL_2808739, EPI_ISL_2808740, EPI_ISL_2808741, EPI_ISL_2808742, EPI_ISL_2808743, EPI_ISL_2808744, EPI_ISL_2808745, EPI_ISL_2808746, EPI_ISL_2808747, EPI_ISL_2808748, EPI_ISL_2808749, EPI_ISL_2808750, EPI_ISL_2808751, EPI_ISL_2808752, EPI_ISL_2808753, EPI_ISL_2808754, EPI_ISL_2808755, EPI_ISL_2808756, EPI_ISL_2808757, EPI_ISL_2808758, EPI_ISL_2808759, EPI_ISL_2808760, EPI_ISL_2808761 | | | |
| see above | Lighthouse Lab in Glasgow | Wellcome Sanger Institute for the COVID-19 Genomics UK (COG-UK) Consortium | Harper VanSteenhouse, Yumi Kasai, David Gray, Carol Clugston, Anna Dominiczak and Alex Alderton, Roberto Amato, Jeffrey Barrett, Sonia Goncalves, Ewan Harrison, David K. Jackson, Ian Johnston, Dominic Kwiatkowski, Cordelia Langford, John Sillitoe on behalf of the Wellcome Sanger Institute COVID-19 Surveillance Team |
| EPI_ISL_2808763 | Lighthouse Lab in Milton Keynes | Wellcome Sanger Institute for the COVID-19 Genomics UK (COG-UK) Consortium | The Lighthouse Lab in Milton Keynes and Alex Alderton, Roberto Amato, Jeffrey Barrett, Sonia Goncalves, Ewan Harrison, David K. Jackson, Ian Johnston, Dominic Kwiatkowski, Cordelia Langford, John Sillitoe on behalf of the Wellcome Sanger Institute COVID-19 Surveillance Team |
| EPI_ISL_2808764, EPI_ISL_2808765, EPI_ISL_2808766, EPI_ISL_2808767, EPI_ISL_2808768, EPI_ISL_2808769, EPI_ISL_2808770, EPI_ISL_2808771, EPI_ISL_2808772, EPI_ISL_2808773, EPI_ISL_2808774, EPI_ISL_2808775, EPI_ISL_2808776, EPI_ISL_2808777, EPI_ISL_2808778, EPI_ISL_2808779, EPI_ISL_2808780, EPI_ISL_2808781, EPI_ISL_2808782, EPI_ISL_2808783, EPI_ISL_2808784, EPI_ISL_2808785, EPI_ISL_2808786, EPI_ISL_2808787, EPI_ISL_2808788, EPI_ISL_2808789, EPI_ISL_2808790, EPI_ISL_2808791, EPI_ISL_2808792 | | | |
| see above | Lighthouse Lab in Glasgow | Wellcome Sanger Institute for the COVID-19 Genomics UK (COG-UK) Consortium | Harper VanSteenhouse, Yumi Kasai, David Gray, Carol Clugston, Anna Dominiczak and Alex Alderton, Roberto Amato, Jeffrey Barrett, Sonia Goncalves, Ewan Harrison, David K. Jackson, Ian Johnston, Dominic Kwiatkowski, Cordelia Langford, John Sillitoe on behalf of the Wellcome Sanger Institute COVID-19 Surveillance Team |
| EPI_ISL_2808793 | Lighthouse Lab in Milton Keynes | Wellcome Sanger Institute for the COVID-19 Genomics UK (COG-UK) Consortium | The Lighthouse Lab in Milton Keynes and Alex Alderton, Roberto Amato, Jeffrey Barrett, Sonia Goncalves, Ewan Harrison, David K. Jackson, Ian Johnston, Dominic Kwiatkowski, Cordelia Langford, John Sillitoe on behalf of the Wellcome Sanger Institute COVID-19 Surveillance Team |
| EPI_ISL_2808794 | Lighthouse Lab in Alderley Park | Wellcome Sanger Institute for the COVID-19 Genomics UK (COG-UK) Consortium | Jacquelyn Wynn, Mairead Hyland, The Lighthouse Lab in Alderley Park and Alex Alderton, Roberto Amato, Jeffrey Barrett, Sonia Goncalves, Ewan Harrison, David K. Jackson, Ian Johnston, Dominic Kwiatkowski, Cordelia Langford, John Sillitoe on behalf of the Wellcome Sanger Institute COVID-19 Surveillance Team |
| EPI_ISL_2808795, EPI_ISL_2808796 | Health Services Laboratories | Wellcome Sanger Institute for the COVID-19 Genomics UK (COG-UK) Consortium | Health Services Laboratories and Alex Alderton, Roberto Amato, Jeffrey Barrett, Sonia Goncalves, Ewan Harrison, David K. Jackson, Ian Johnston, Dominic Kwiatkowski, Cordelia Langford, John Sillitoe on behalf of the Wellcome Sanger Institute COVID-19 Surveillance Team |
| EPI_ISL_2808797 | Lighthouse Lab in Alderley Park | Wellcome Sanger Institute for the COVID-19 Genomics UK (COG-UK) Consortium | Jacquelyn Wynn, Mairead Hyland, The Lighthouse Lab in Alderley Park and Alex Alderton, Roberto Amato, Jeffrey Barrett, Sonia Goncalves, Ewan Harrison, David K. Jackson, Ian Johnston, Dominic Kwiatkowski, Cordelia Langford, John Sillitoe on behalf of the Wellcome Sanger Institute COVID-19 Surveillance Team |
| EPI_ISL_2808798, EPI_ISL_2808799, EPI_ISL_2808800 | Lighthouse Laboratory Plymouth | Wellcome Sanger Institute for the COVID-19 Genomics UK (COG-UK) Consortium | Lighthouse Laboratory Plymouth and Alex Alderton, Roberto Amato, Jeffrey Barrett, Sonia Goncalves, Ewan Harrison, David K. Jackson, Ian Johnston, Dominic Kwiatkowski, Cordelia Langford, John Sillitoe on behalf of the Wellcome Sanger Institute COVID-19 Surveillance Team |

| | | | |
|---|---|---|---|
| EPI_ISL_2808801 | Health Services Laboratories | Wellcome Sanger Institute for the COVID-19 Genomics UK (COG-UK) Consortium | Health Services Laboratories and Alex Alderton, Roberto Amato, Jeffrey Barrett, Sonia Goncalves, Ewan Harrison, David K. Jackson, Ian Johnston, Dominic Kwiatkowski, Cordelia Langford, John Sillitoe on behalf of the Wellcome Sanger Institute COVID-19 Surveillance Team |
| EPI_ISL_2808802, EPI_ISL_2808803 | Lighthouse Lab in Milton Keynes | Wellcome Sanger Institute for the COVID-19 Genomics UK (COG-UK) Consortium | The Lighthouse Lab in Milton Keynes and Alex Alderton, Roberto Amato, Jeffrey Barrett, Sonia Goncalves, Ewan Harrison, David K. Jackson, Ian Johnston, Dominic Kwiatkowski, Cordelia Langford, John Sillitoe on behalf of the Wellcome Sanger Institute COVID-19 Surveillance Team |
| EPI_ISL_2808804 | Lighthouse Lab in Alderley Park | Wellcome Sanger Institute for the COVID-19 Genomics UK (COG-UK) Consortium | Jacquelyn Wynn, Mairead Hyland, The Lighthouse Lab in Alderley Park and Alex Alderton, Roberto Amato, Jeffrey Barrett, Sonia Goncalves, Ewan Harrison, David K. Jackson, Ian Johnston, Dominic Kwiatkowski, Cordelia Langford, John Sillitoe on behalf of the Wellcome Sanger Institute COVID-19 Surveillance Team |
| EPI_ISL_2808805 | Lighthouse Laboratory Plymouth | Wellcome Sanger Institute for the COVID-19 Genomics UK (COG-UK) Consortium | Lighthouse Laboratory Plymouth and Alex Alderton, Roberto Amato, Jeffrey Barrett, Sonia Goncalves, Ewan Harrison, David K. Jackson, Ian Johnston, Dominic Kwiatkowski, Cordelia Langford, John Sillitoe on behalf of the Wellcome Sanger Institute COVID-19 Surveillance Team |
| EPI_ISL_2808806 | Lighthouse Lab in Alderley Park | Wellcome Sanger Institute for the COVID-19 Genomics UK (COG-UK) Consortium | Jacquelyn Wynn, Mairead Hyland, The Lighthouse Lab in Alderley Park and Alex Alderton, Roberto Amato, Jeffrey Barrett, Sonia Goncalves, Ewan Harrison, David K. Jackson, Ian Johnston, Dominic Kwiatkowski, Cordelia Langford, John Sillitoe on behalf of the Wellcome Sanger Institute COVID-19 Surveillance Team |
| EPI_ISL_2808807, EPI_ISL_2808808, EPI_ISL_2808809 | Lighthouse Lab in Glasgow | Wellcome Sanger Institute for the COVID-19 Genomics UK (COG-UK) Consortium | Harper VanSteenhouse, Yumi Kasai, David Gray, Carol Clugston, Anna Dominiczak and Alex Alderton, Roberto Amato, Jeffrey Barrett, Sonia Goncalves, Ewan Harrison, David K. Jackson, Ian Johnston, Dominic Kwiatkowski, Cordelia Langford, John Sillitoe on behalf of the Wellcome Sanger Institute COVID-19 Surveillance Team |
| EPI_ISL_2808810 | Lighthouse Lab in Milton Keynes | Wellcome Sanger Institute for the COVID-19 Genomics UK (COG-UK) Consortium | The Lighthouse Lab in Milton Keynes and Alex Alderton, Roberto Amato, Jeffrey Barrett, Sonia Goncalves, Ewan Harrison, David K. Jackson, Ian Johnston, Dominic Kwiatkowski, Cordelia Langford, John Sillitoe on behalf of the Wellcome Sanger Institute COVID-19 Surveillance Team |
| EPI_ISL_2808811, EPI_ISL_2808812, EPI_ISL_2808813, EPI_ISL_2808814, EPI_ISL_2808815, EPI_ISL_2808816, EPI_ISL_2808817, EPI_ISL_2808818, EPI_ISL_2808819, EPI_ISL_2808820 | Lighthouse Lab in Glasgow | Wellcome Sanger Institute for the COVID-19 Genomics UK (COG-UK) Consortium | Harper VanSteenhouse, Yumi Kasai, David Gray, Carol Clugston, Anna Dominiczak and Alex Alderton, Roberto Amato, Jeffrey Barrett, Sonia Goncalves, Ewan Harrison, David K. Jackson, Ian Johnston, Dominic Kwiatkowski, Cordelia Langford, John Sillitoe on behalf of the Wellcome Sanger Institute COVID-19 Surveillance Team |
| EPI_ISL_2808821 | Lighthouse Lab in Alderley Park | Wellcome Sanger Institute for the COVID-19 Genomics UK (COG-UK) Consortium | Jacquelyn Wynn, Mairead Hyland, The Lighthouse Lab in Alderley Park and Alex Alderton, Roberto Amato, Jeffrey Barrett, Sonia Goncalves, Ewan Harrison, David K. Jackson, Ian Johnston, Dominic Kwiatkowski, Cordelia Langford, John Sillitoe on behalf of the Wellcome Sanger Institute COVID-19 Surveillance Team |
| EPI_ISL_2808822, EPI_ISL_2808823, EPI_ISL_2808824, EPI_ISL_2808825, EPI_ISL_2808826, EPI_ISL_2808827, EPI_ISL_2808828, EPI_ISL_2808829, EPI_ISL_2808830, EPI_ISL_2808831, EPI_ISL_2808832, EPI_ISL_2808833, EPI_ISL_2808834, EPI_ISL_2808835, EPI_ISL_2808836, EPI_ISL_2808837, EPI_ISL_2808838, EPI_ISL_2808839, EPI_ISL_2808840, EPI_ISL_2808841, EPI_ISL_2808842, EPI_ISL_2808843, EPI_ISL_2808844, EPI_ISL_2808846, EPI_ISL_2808847, EPI_ISL_2808848, EPI_ISL_2808849, EPI_ISL_2808850, EPI_ISL_2808851, EPI_ISL_2808852, EPI_ISL_2808853, EPI_ISL_2808854, EPI_ISL_2808855, EPI_ISL_2808856, EPI_ISL_2808857, EPI_ISL_2808858, EPI_ISL_2808859, EPI_ISL_2808860, EPI_ISL_2808861, EPI_ISL_2808862, EPI_ISL_2808863, EPI_ISL_2808864, EPI_ISL_2808865, EPI_ISL_2808866, EPI_ISL_2808867, EPI_ISL_2808868, EPI_ISL_2808869, EPI_ISL_2808870, EPI_ISL_2808871, EPI_ISL_2808872, EPI_ISL_2808873, EPI_ISL_2808874, EPI_ISL_2808875, EPI_ISL_2808876, EPI_ISL_2808877, EPI_ISL_2808878, EPI_ISL_2808879, EPI_ISL_2808880, EPI_ISL_2808881, EPI_ISL_2808882, EPI_ISL_2808883, EPI_ISL_2808884, EPI_ISL_2808885, EPI_ISL_2808886, EPI_ISL_2808887, EPI_ISL_2808888, EPI_ISL_2808889, EPI_ISL_2808890, EPI_ISL_2808891, EPI_ISL_2808893, EPI_ISL_2808894, EPI_ISL_2808895, EPI_ISL_2808896, EPI_ISL_2808897, EPI_ISL_2808898, EPI_ISL_2808899, EPI_ISL_2808900 | | | |
| see above | Lighthouse Lab in Glasgow | Wellcome Sanger Institute for the COVID-19 Genomics UK (COG-UK) Consortium | Harper VanSteenhouse, Yumi Kasai, David Gray, Carol Clugston, Anna Dominiczak and Alex Alderton, Roberto Amato, Jeffrey Barrett, Sonia Goncalves, Ewan Harrison, David K. Jackson, Ian Johnston, Dominic Kwiatkowski, Cordelia Langford, John Sillitoe on behalf of the Wellcome Sanger Institute COVID-19 Surveillance Team |
| EPI_ISL_2808901 | Lighthouse Lab in Milton Keynes | Wellcome Sanger Institute for the COVID-19 Genomics UK (COG-UK) Consortium | The Lighthouse Lab in Milton Keynes and Alex Alderton, Roberto Amato, Jeffrey Barrett, Sonia Goncalves, Ewan Harrison, David K. Jackson, Ian Johnston, Dominic Kwiatkowski, Cordelia Langford, John Sillitoe on behalf of the Wellcome Sanger Institute COVID-19 Surveillance Team |
| EPI_ISL_2808902, EPI_ISL_2808903, EPI_ISL_2808904, EPI_ISL_2808905, EPI_ISL_2808906, EPI_ISL_2808907, EPI_ISL_2808908, EPI_ISL_2808909, EPI_ISL_2808910, EPI_ISL_2808911, EPI_ISL_2808912, EPI_ISL_2808913, EPI_ISL_2808914, EPI_ISL_2808915, EPI_ISL_2808916, EPI_ISL_2808917, EPI_ISL_2808918, EPI_ISL_2808919, EPI_ISL_2808920, EPI_ISL_2808921, EPI_ISL_2808922, EPI_ISL_2808923, EPI_ISL_2808924, EPI_ISL_2808925, EPI_ISL_2808926, EPI_ISL_2808927, EPI_ISL_2808928, EPI_ISL_2808929, EPI_ISL_2808930, EPI_ISL_2808931, EPI_ISL_2808932, EPI_ISL_2808933, EPI_ISL_2808934, EPI_ISL_2808935, EPI_ISL_2808936, EPI_ISL_2808937, EPI_ISL_2808938, EPI_ISL_2808939, EPI_ISL_2808940, EPI_ISL_2808941, EPI_ISL_2808942, EPI_ISL_2808943, EPI_ISL_2808945, EPI_ISL_2808946, EPI_ISL_2808947, EPI_ISL_2808948, EPI_ISL_2808949, EPI_ISL_2808950, EPI_ISL_2808951, EPI_ISL_2808952, EPI_ISL_2808954, EPI_ISL_2808955, EPI_ISL_2808956, EPI_ISL_2808957, EPI_ISL_2808958, EPI_ISL_2808959, EPI_ISL_2808960, EPI_ISL_2808961, EPI_ISL_2808962, EPI_ISL_2808963, EPI_ISL_2808964, EPI_ISL_2808965, EPI_ISL_2808966, EPI_ISL_2808967, EPI_ISL_2808968, EPI_ISL_2808969, EPI_ISL_2808970, EPI_ISL_2808971, EPI_ISL_2808972, EPI_ISL_2808973, EPI_ISL_2808974, EPI_ISL_2808975, EPI_ISL_2808976, EPI_ISL_2808977, EPI_ISL_2808978, EPI_ISL_2808979, EPI_ISL_2808980, EPI_ISL_2808981, EPI_ISL_2808982, EPI_ISL_2808983, EPI_ISL_2808984, EPI_ISL_2808985, EPI_ISL_2808986, EPI_ISL_2808987 | | | |
| see above | Lighthouse Lab in Glasgow | Wellcome Sanger Institute for the COVID-19 Genomics UK (COG-UK) Consortium | Harper VanSteenhouse, Yumi Kasai, David Gray, Carol Clugston, Anna Dominiczak and Alex Alderton, Roberto Amato, Jeffrey Barrett, Sonia Goncalves, Ewan Harrison, David K. Jackson, Ian Johnston, Dominic Kwiatkowski, Cordelia Langford, John Sillitoe on behalf of the Wellcome Sanger Institute COVID-19 Surveillance Team |
| EPI_ISL_2808988 | Lighthouse Lab in Alderley Park | Wellcome Sanger Institute for the COVID-19 Genomics UK (COG-UK) Consortium | Jacquelyn Wynn, Mairead Hyland, The Lighthouse Lab in Alderley Park and Alex Alderton, Roberto Amato, Jeffrey Barrett, Sonia Goncalves, Ewan Harrison, David K. Jackson, Ian Johnston, Dominic Kwiatkowski, Cordelia Langford, John Sillitoe on behalf of the Wellcome Sanger Institute COVID-19 Surveillance Team |
| EPI_ISL_2808989, EPI_ISL_2808990, EPI_ISL_2808991, EPI_ISL_2808992, EPI_ISL_2808993, EPI_ISL_2808994, EPI_ISL_2808995, EPI_ISL_2808996, EPI_ISL_2808997, EPI_ISL_2808998, EPI_ISL_2808999, EPI_ISL_2809000, EPI_ISL_2809001, EPI_ISL_2809002, EPI_ISL_2809003, EPI_ISL_2809004, EPI_ISL_2809005, EPI_ISL_2809008, EPI_ISL_2809009, EPI_ISL_2809010, EPI_ISL_2809011, EPI_ISL_2809012, EPI_ISL_2809013, EPI_ISL_2809014, EPI_ISL_2809015, EPI_ISL_2809016, EPI_ISL_2809017, EPI_ISL_2809018, EPI_ISL_2809019, EPI_ISL_2809020, EPI_ISL_2809021, EPI_ISL_2809022, EPI_ISL_2809023, EPI_ISL_2809024, EPI_ISL_2809025, EPI_ISL_2809026, EPI_ISL_2809027, EPI_ISL_2809029, EPI_ISL_2809030, EPI_ISL_2809031, EPI_ISL_2809032, EPI_ISL_2809033, EPI_ISL_2809034, EPI_ISL_2809035, EPI_ISL_2809036, EPI_ISL_2809037, EPI_ISL_2809038, EPI_ISL_2809039, EPI_ISL_2809040, EPI_ISL_2809041, EPI_ISL_2809042, EPI_ISL_2809043, EPI_ISL_2809044, EPI_ISL_2809045, EPI_ISL_2809046 | | | |
| see above | Lighthouse Lab in Glasgow | Wellcome Sanger Institute for the COVID-19 Genomics UK (COG-UK) Consortium | Harper VanSteenhouse, Yumi Kasai, David Gray, Carol Clugston, Anna Dominiczak and Alex Alderton, Roberto Amato, Jeffrey Barrett, Sonia Goncalves, Ewan Harrison, David K. Jackson, Ian Johnston, Dominic Kwiatkowski, Cordelia Langford, John Sillitoe on behalf of the Wellcome Sanger Institute COVID-19 Surveillance Team |
| EPI_ISL_2809047 | Lighthouse Lab in Alderley Park | Wellcome Sanger Institute for the COVID-19 Genomics UK (COG-UK) Consortium | Jacquelyn Wynn, Mairead Hyland, The Lighthouse Lab in Alderley Park and Alex Alderton, Roberto Amato, Jeffrey Barrett, Sonia Goncalves, Ewan Harrison, David K. Jackson, Ian Johnston, Dominic Kwiatkowski, Cordelia Langford, John Sillitoe on behalf of the Wellcome Sanger Institute COVID-19 Surveillance Team |
| EPI_ISL_2809048, EPI_ISL_2809049, EPI_ISL_2809050, EPI_ISL_2809051, EPI_ISL_2809052, EPI_ISL_2809053, EPI_ISL_2809054, EPI_ISL_2809055, EPI_ISL_2809056, EPI_ISL_2809057, EPI_ISL_2809058, EPI_ISL_2809059, EPI_ISL_2809060, EPI_ISL_2809061, EPI_ISL_2809062, EPI_ISL_2809063, EPI_ISL_2809064, EPI_ISL_2809065, EPI_ISL_2809066, EPI_ISL_2809067, EPI_ISL_2809068, EPI_ISL_2809069, EPI_ISL_2809070, EPI_ISL_2809071, EPI_ISL_2809072, EPI_ISL_2809073, EPI_ISL_2809074 | | | |
| see above | Lighthouse Lab in Glasgow | Wellcome Sanger Institute for the COVID-19 Genomics UK (COG-UK) Consortium | Harper VanSteenhouse, Yumi Kasai, David Gray, Carol Clugston, Anna Dominiczak and Alex Alderton, Roberto Amato, Jeffrey Barrett, Sonia Goncalves, Ewan Harrison, David K. Jackson, Ian Johnston, Dominic Kwiatkowski, Cordelia Langford, John Sillitoe on behalf of the Wellcome Sanger Institute COVID-19 Surveillance Team |
| EPI_ISL_2809075 | Lighthouse Laboratory Plymouth | Wellcome Sanger Institute for the COVID-19 Genomics UK (COG-UK) Consortium | Lighthouse Laboratory Plymouth and Alex Alderton, Roberto Amato, Jeffrey Barrett, Sonia Goncalves, Ewan Harrison, David K. Jackson, Ian Johnston, Dominic Kwiatkowski, Cordelia Langford, John Sillitoe on behalf of the Wellcome Sanger Institute COVID-19 Surveillance Team |
| EPI_ISL_2809076, EPI_ISL_2809077, EPI_ISL_2809078, EPI_ISL_2809079, EPI_ISL_2809080, EPI_ISL_2809081, EPI_ISL_2809082, EPI_ISL_2809083, EPI_ISL_2809084, EPI_ISL_2809085, EPI_ISL_2809086, EPI_ISL_2809087, EPI_ISL_2809088, EPI_ISL_2809089, EPI_ISL_2809090, EPI_ISL_2809091, EPI_ISL_2809092, EPI_ISL_2809093, EPI_ISL_2809094 | | | |
| see above | Lighthouse Lab in Glasgow | Wellcome Sanger Institute for the COVID-19 Genomics UK (COG-UK) Consortium | Harper VanSteenhouse, Yumi Kasai, David Gray, Carol Clugston, Anna Dominiczak and Alex Alderton, Roberto Amato, Jeffrey Barrett, Sonia Goncalves, Ewan Harrison, David K. Jackson, Ian Johnston, Dominic Kwiatkowski, Cordelia Langford, John Sillitoe on behalf of the Wellcome Sanger Institute COVID-19 Surveillance Team |
| EPI_ISL_2809095 | Lighthouse Lab in Milton Keynes | Wellcome Sanger Institute for the COVID-19 Genomics UK (COG-UK) Consortium | The Lighthouse Lab in Milton Keynes and Alex Alderton, Roberto Amato, Jeffrey Barrett, Sonia Goncalves, Ewan Harrison, David K. Jackson, Ian Johnston, Dominic Kwiatkowski, Cordelia Langford, John Sillitoe on behalf of the Wellcome Sanger Institute COVID-19 Surveillance Team |
| EPI_ISL_2809096 | Lighthouse Lab in Alderley Park | Wellcome Sanger Institute for the COVID-19 Genomics UK (COG-UK) Consortium | Jacquelyn Wynn, Mairead Hyland, The Lighthouse Lab in Alderley Park and Alex Alderton, Roberto Amato, Jeffrey Barrett, Sonia Goncalves, Ewan Harrison, David K. Jackson, Ian Johnston, Dominic Kwiatkowski, Cordelia Langford, John Sillitoe on behalf of the Wellcome Sanger Institute COVID-19 Surveillance Team |
| EPI_ISL_2809097 | Health Services Laboratories | Wellcome Sanger Institute for the COVID-19 Genomics UK (COG-UK) Consortium | Health Services Laboratories and Alex Alderton, Roberto Amato, Jeffrey Barrett, Sonia Goncalves, Ewan Harrison, David K. Jackson, Ian Johnston, Dominic Kwiatkowski, Cordelia Langford, John Sillitoe on behalf of the Wellcome Sanger Institute COVID-19 Surveillance Team |
| EPI_ISL_2809098 | Lighthouse Lab in Glasgow | Wellcome Sanger Institute for the COVID-19 Genomics UK (COG-UK) Consortium | Harper VanSteenhouse, Yumi Kasai, David Gray, Carol Clugston, Anna Dominiczak and Alex Alderton, Roberto Amato, Jeffrey Barrett, Sonia Goncalves, Ewan Harrison, David K. Jackson, Ian Johnston, Dominic Kwiatkowski, Cordelia Langford, John Sillitoe on behalf of the Wellcome Sanger Institute COVID-19 Surveillance Team |
| EPI_ISL_2809099, EPI_ISL_2809100 | Lighthouse Lab in Alderley Park | Wellcome Sanger Institute for the COVID-19 Genomics UK (COG-UK) Consortium | Jacquelyn Wynn, Mairead Hyland, The Lighthouse Lab in Alderley Park and Alex Alderton, Roberto Amato, Jeffrey Barrett, Sonia Goncalves, Ewan Harrison, David K. Jackson, Ian Johnston, Dominic Kwiatkowski, Cordelia Langford, John Sillitoe on behalf of the Wellcome Sanger Institute COVID-19 Surveillance Team |
| EPI_ISL_2809101, EPI_ISL_2809102 | Lighthouse Lab in Milton Keynes | Wellcome Sanger Institute for the COVID-19 Genomics UK (COG-UK) Consortium | The Lighthouse Lab in Milton Keynes and Alex Alderton, Roberto Amato, Jeffrey Barrett, Sonia Goncalves, Ewan Harrison, David K. Jackson, Ian Johnston, Dominic Kwiatkowski, Cordelia Langford, John Sillitoe on behalf of the Wellcome Sanger Institute COVID-19 Surveillance Team |
| EPI_ISL_2809103 | Health Services Laboratories | Wellcome Sanger Institute for the COVID-19 Genomics UK (COG-UK) Consortium | Health Services Laboratories and Alex Alderton, Roberto Amato, Jeffrey Barrett, Sonia Goncalves, Ewan Harrison, David K. Jackson, Ian Johnston, Dominic Kwiatkowski, Cordelia Langford, John Sillitoe on behalf of the Wellcome Sanger Institute COVID-19 Surveillance Team |
| EPI_ISL_2809104 | Lighthouse Lab in Milton Keynes | Wellcome Sanger Institute for the COVID-19 Genomics UK (COG-UK) Consortium | The Lighthouse Lab in Milton Keynes and Alex Alderton, Roberto Amato, Jeffrey Barrett, Sonia Goncalves, Ewan Harrison, David K. Jackson, Ian Johnston, Dominic Kwiatkowski, Cordelia Langford, John Sillitoe on behalf of the Wellcome Sanger Institute COVID-19 Surveillance Team |
| EPI_ISL_2809105, EPI_ISL_2809106 | Lighthouse Lab in Alderley Park | Wellcome Sanger Institute for the COVID-19 Genomics UK (COG-UK) Consortium | Jacquelyn Wynn, Mairead Hyland, The Lighthouse Lab in Alderley Park and Alex Alderton, Roberto Amato, Jeffrey Barrett, Sonia Goncalves, Ewan Harrison, David K. Jackson, Ian Johnston, Dominic Kwiatkowski, Cordelia Langford, John Sillitoe on behalf of the Wellcome Sanger Institute COVID-19 Surveillance Team |
| EPI_ISL_2809107 | Lighthouse Lab in Milton Keynes | Wellcome Sanger Institute for the COVID-19 Genomics UK (COG-UK) Consortium | The Lighthouse Lab in Milton Keynes and Alex Alderton, Roberto Amato, Jeffrey Barrett, Sonia Goncalves, Ewan Harrison, David K. Jackson, Ian Johnston, Dominic Kwiatkowski, Cordelia Langford, John Sillitoe on behalf of the Wellcome Sanger Institute COVID-19 Surveillance Team |
| EPI_ISL_2809108 | Lighthouse Lab in Alderley Park | Wellcome Sanger Institute for the COVID-19 Genomics UK (COG-UK) Consortium | Jacquelyn Wynn, Mairead Hyland, The Lighthouse Lab in Alderley Park and Alex Alderton, Roberto Amato, Jeffrey Barrett, Sonia Goncalves, Ewan Harrison, David K. Jackson, Ian Johnston, Dominic Kwiatkowski, Cordelia Langford, John Sillitoe on behalf of the Wellcome Sanger Institute COVID-19 Surveillance Team |
| EPI_ISL_2809109 | Health Services Laboratories | Wellcome Sanger Institute for the COVID-19 Genomics UK (COG-UK) Consortium | Health Services Laboratories and Alex Alderton, Roberto Amato, Jeffrey Barrett, Sonia Goncalves, Ewan Harrison, David K. Jackson, Ian Johnston, Dominic Kwiatkowski, Cordelia Langford, John Sillitoe on behalf of the Wellcome Sanger Institute COVID-19 Surveillance Team |
| EPI_ISL_2809110, EPI_ISL_2809111, EPI_ISL_2809113, EPI_ISL_2809114 | Lighthouse Lab in Alderley Park | Wellcome Sanger Institute for the COVID-19 Genomics UK (COG-UK) Consortium | Jacquelyn Wynn, Mairead Hyland, The Lighthouse Lab in Alderley Park and Alex Alderton, Roberto Amato, Jeffrey Barrett, Sonia Goncalves, Ewan Harrison, David K. Jackson, Ian Johnston, Dominic Kwiatkowski, Cordelia Langford, John Sillitoe on behalf of the Wellcome Sanger Institute COVID-19 Surveillance Team |
| EPI_ISL_2809115 | Health Services Laboratories | Wellcome Sanger Institute for the COVID-19 Genomics UK (COG-UK) Consortium | Health Services Laboratories and Alex Alderton, Roberto Amato, Jeffrey Barrett, Sonia Goncalves, Ewan Harrison, David K. Jackson, Ian Johnston, Dominic Kwiatkowski, Cordelia Langford, John Sillitoe on behalf of the Wellcome Sanger Institute COVID-19 Surveillance Team |
| EPI_ISL_2809116, EPI_ISL_2809117, EPI_ISL_2809118 | Lighthouse Lab in Alderley Park | Wellcome Sanger Institute for the COVID-19 Genomics UK (COG-UK) Consortium | Jacquelyn Wynn, Mairead Hyland, The Lighthouse Lab in Alderley Park and Alex Alderton, Roberto Amato, Jeffrey Barrett, Sonia Goncalves, Ewan Harrison, David K. Jackson, Ian Johnston, Dominic Kwiatkowski, Cordelia Langford, John Sillitoe on behalf of the Wellcome Sanger Institute COVID-19 Surveillance Team |
| EPI_ISL_2809119 | Lighthouse Lab in Milton Keynes | Wellcome Sanger Institute for the COVID-19 Genomics UK (COG-UK) | The Lighthouse Lab in Milton Keynes and Alex Alderton, Roberto Amato, Jeffrey Barrett, Sonia Goncalves, Ewan Harrison, David K. Jackson, Ian Johnston, Dominic Kwiatkowski, Cordelia Langford, John Sillitoe on behalf of the Wellcome Sanger |

| | | | |
|---|---|---|---|
| | | Consortium | Institute COVID-19 Surveillance Team |
| EPI_ISL_2809120, EPI_ISL_2809121 | Lighthouse Lab in Alderley Park | Wellcome Sanger Institute for the COVID-19 Genomics UK (COG-UK) Consortium | Jacquelyn Wynn, Mairead Hyland, The Lighthouse Lab in Alderley Park and Alex Alderton, Roberto Amato, Jeffrey Barrett, Sonia Goncalves, Ewan Harrison, David K. Jackson, Ian Johnston, Dominic Kwiatkowski, Cordelia Langford, John Sillitoe on behalf of the Wellcome Sanger Institute COVID-19 Surveillance Team |
| EPI_ISL_2809122 | Health Services Laboratories | Wellcome Sanger Institute for the COVID-19 Genomics UK (COG-UK) Consortium | Health Services Laboratories and Alex Alderton, Roberto Amato, Jeffrey Barrett, Sonia Goncalves, Ewan Harrison, David K. Jackson, Ian Johnston, Dominic Kwiatkowski, Cordelia Langford, John Sillitoe on behalf of the Wellcome Sanger Institute COVID-19 Surveillance Team |
| EPI_ISL_2809123 | Lighthouse Lab in Alderley Park | Wellcome Sanger Institute for the COVID-19 Genomics UK (COG-UK) Consortium | Jacquelyn Wynn, Mairead Hyland, The Lighthouse Lab in Alderley Park and Alex Alderton, Roberto Amato, Jeffrey Barrett, Sonia Goncalves, Ewan Harrison, David K. Jackson, Ian Johnston, Dominic Kwiatkowski, Cordelia Langford, John Sillitoe on behalf of the Wellcome Sanger Institute COVID-19 Surveillance Team |
| EPI_ISL_2809124 | Lighthouse Lab in Milton Keynes | Wellcome Sanger Institute for the COVID-19 Genomics UK (COG-UK) Consortium | The Lighthouse Lab in Milton Keynes and Alex Alderton, Roberto Amato, Jeffrey Barrett, Sonia Goncalves, Ewan Harrison, David K. Jackson, Ian Johnston, Dominic Kwiatkowski, Cordelia Langford, John Sillitoe on behalf of the Wellcome Sanger Institute COVID-19 Surveillance Team |
| EPI_ISL_2809125 | Health Services Laboratories | Wellcome Sanger Institute for the COVID-19 Genomics UK (COG-UK) Consortium | Health Services Laboratories and Alex Alderton, Roberto Amato, Jeffrey Barrett, Sonia Goncalves, Ewan Harrison, David K. Jackson, Ian Johnston, Dominic Kwiatkowski, Cordelia Langford, John Sillitoe on behalf of the Wellcome Sanger Institute COVID-19 Surveillance Team |
| EPI_ISL_2809126 | Lighthouse Lab in Alderley Park | Wellcome Sanger Institute for the COVID-19 Genomics UK (COG-UK) Consortium | Jacquelyn Wynn, Mairead Hyland, The Lighthouse Lab in Alderley Park and Alex Alderton, Roberto Amato, Jeffrey Barrett, Sonia Goncalves, Ewan Harrison, David K. Jackson, Ian Johnston, Dominic Kwiatkowski, Cordelia Langford, John Sillitoe on behalf of the Wellcome Sanger Institute COVID-19 Surveillance Team |
| EPI_ISL_2809127 | Lighthouse Lab in Milton Keynes | Wellcome Sanger Institute for the COVID-19 Genomics UK (COG-UK) Consortium | The Lighthouse Lab in Milton Keynes and Alex Alderton, Roberto Amato, Jeffrey Barrett, Sonia Goncalves, Ewan Harrison, David K. Jackson, Ian Johnston, Dominic Kwiatkowski, Cordelia Langford, John Sillitoe on behalf of the Wellcome Sanger Institute COVID-19 Surveillance Team |
| EPI_ISL_2809128, EPI_ISL_2809129 | Lighthouse Lab in Alderley Park | Wellcome Sanger Institute for the COVID-19 Genomics UK (COG-UK) Consortium | Jacquelyn Wynn, Mairead Hyland, The Lighthouse Lab in Alderley Park and Alex Alderton, Roberto Amato, Jeffrey Barrett, Sonia Goncalves, Ewan Harrison, David K. Jackson, Ian Johnston, Dominic Kwiatkowski, Cordelia Langford, John Sillitoe on behalf of the Wellcome Sanger Institute COVID-19 Surveillance Team |
| EPI_ISL_2809130 | Health Services Laboratories | Wellcome Sanger Institute for the COVID-19 Genomics UK (COG-UK) Consortium | Health Services Laboratories and Alex Alderton, Roberto Amato, Jeffrey Barrett, Sonia Goncalves, Ewan Harrison, David K. Jackson, Ian Johnston, Dominic Kwiatkowski, Cordelia Langford, John Sillitoe on behalf of the Wellcome Sanger Institute COVID-19 Surveillance Team |
| EPI_ISL_2809131 | Lighthouse Lab in Alderley Park | Wellcome Sanger Institute for the COVID-19 Genomics UK (COG-UK) Consortium | Jacquelyn Wynn, Mairead Hyland, The Lighthouse Lab in Alderley Park and Alex Alderton, Roberto Amato, Jeffrey Barrett, Sonia Goncalves, Ewan Harrison, David K. Jackson, Ian Johnston, Dominic Kwiatkowski, Cordelia Langford, John Sillitoe on behalf of the Wellcome Sanger Institute COVID-19 Surveillance Team |
| EPI_ISL_2809132 | Health Services Laboratories | Wellcome Sanger Institute for the COVID-19 Genomics UK (COG-UK) Consortium | Health Services Laboratories and Alex Alderton, Roberto Amato, Jeffrey Barrett, Sonia Goncalves, Ewan Harrison, David K. Jackson, Ian Johnston, Dominic Kwiatkowski, Cordelia Langford, John Sillitoe on behalf of the Wellcome Sanger Institute COVID-19 Surveillance Team |
| EPI_ISL_2809133 | Lighthouse Lab in Alderley Park | Wellcome Sanger Institute for the COVID-19 Genomics UK (COG-UK) Consortium | Jacquelyn Wynn, Mairead Hyland, The Lighthouse Lab in Alderley Park and Alex Alderton, Roberto Amato, Jeffrey Barrett, Sonia Goncalves, Ewan Harrison, David K. Jackson, Ian Johnston, Dominic Kwiatkowski, Cordelia Langford, John Sillitoe on behalf of the Wellcome Sanger Institute COVID-19 Surveillance Team |
| EPI_ISL_2809134 | Lighthouse Lab in Milton Keynes | Wellcome Sanger Institute for the COVID-19 Genomics UK (COG-UK) Consortium | The Lighthouse Lab in Milton Keynes and Alex Alderton, Roberto Amato, Jeffrey Barrett, Sonia Goncalves, Ewan Harrison, David K. Jackson, Ian Johnston, Dominic Kwiatkowski, Cordelia Langford, John Sillitoe on behalf of the Wellcome Sanger Institute COVID-19 Surveillance Team |
| EPI_ISL_2809135, EPI_ISL_2809136 | Lighthouse Lab in Alderley Park | Wellcome Sanger Institute for the COVID-19 Genomics UK (COG-UK) Consortium | Jacquelyn Wynn, Mairead Hyland, The Lighthouse Lab in Alderley Park and Alex Alderton, Roberto Amato, Jeffrey Barrett, Sonia Goncalves, Ewan Harrison, David K. Jackson, Ian Johnston, Dominic Kwiatkowski, Cordelia Langford, John Sillitoe on behalf of the Wellcome Sanger Institute COVID-19 Surveillance Team |
| EPI_ISL_2809137 | Lighthouse Lab in Milton Keynes | Wellcome Sanger Institute for the COVID-19 Genomics UK (COG-UK) Consortium | The Lighthouse Lab in Milton Keynes and Alex Alderton, Roberto Amato, Jeffrey Barrett, Sonia Goncalves, Ewan Harrison, David K. Jackson, Ian Johnston, Dominic Kwiatkowski, Cordelia Langford, John Sillitoe on behalf of the Wellcome Sanger Institute COVID-19 Surveillance Team |
| EPI_ISL_2809138 | Lighthouse Lab in Alderley Park | Wellcome Sanger Institute for the COVID-19 Genomics UK (COG-UK) Consortium | Jacquelyn Wynn, Mairead Hyland, The Lighthouse Lab in Alderley Park and Alex Alderton, Roberto Amato, Jeffrey Barrett, Sonia Goncalves, Ewan Harrison, David K. Jackson, Ian Johnston, Dominic Kwiatkowski, Cordelia Langford, John Sillitoe on behalf of the Wellcome Sanger Institute COVID-19 Surveillance Team |
| EPI_ISL_2809139 | Lighthouse Lab in Milton Keynes | Wellcome Sanger Institute for the COVID-19 Genomics UK (COG-UK) Consortium | The Lighthouse Lab in Milton Keynes and Alex Alderton, Roberto Amato, Jeffrey Barrett, Sonia Goncalves, Ewan Harrison, David K. Jackson, Ian Johnston, Dominic Kwiatkowski, Cordelia Langford, John Sillitoe on behalf of the Wellcome Sanger Institute COVID-19 Surveillance Team |
| EPI_ISL_2809140 | Lighthouse Lab in Alderley Park | Wellcome Sanger Institute for the COVID-19 Genomics UK (COG-UK) Consortium | Jacquelyn Wynn, Mairead Hyland, The Lighthouse Lab in Alderley Park and Alex Alderton, Roberto Amato, Jeffrey Barrett, Sonia Goncalves, Ewan Harrison, David K. Jackson, Ian Johnston, Dominic Kwiatkowski, Cordelia Langford, John Sillitoe on behalf of the Wellcome Sanger Institute COVID-19 Surveillance Team |
| EPI_ISL_2809141 | Health Services Laboratories | Wellcome Sanger Institute for the COVID-19 Genomics UK (COG-UK) Consortium | Health Services Laboratories and Alex Alderton, Roberto Amato, Jeffrey Barrett, Sonia Goncalves, Ewan Harrison, David K. Jackson, Ian Johnston, Dominic Kwiatkowski, Cordelia Langford, John Sillitoe on behalf of the Wellcome Sanger Institute COVID-19 Surveillance Team |
| EPI_ISL_2809142, EPI_ISL_2809143, EPI_ISL_2809144, EPI_ISL_2809145 | Lighthouse Lab in Alderley Park | Wellcome Sanger Institute for the COVID-19 Genomics UK (COG-UK) Consortium | Jacquelyn Wynn, Mairead Hyland, The Lighthouse Lab in Alderley Park and Alex Alderton, Roberto Amato, Jeffrey Barrett, Sonia Goncalves, Ewan Harrison, David K. Jackson, Ian Johnston, Dominic Kwiatkowski, Cordelia Langford, John Sillitoe on behalf of the Wellcome Sanger Institute COVID-19 Surveillance Team |
| EPI_ISL_2809146 | Health Services Laboratories | Wellcome Sanger Institute for the COVID-19 Genomics UK (COG-UK) Consortium | Health Services Laboratories and Alex Alderton, Roberto Amato, Jeffrey Barrett, Sonia Goncalves, Ewan Harrison, David K. Jackson, Ian Johnston, Dominic Kwiatkowski, Cordelia Langford, John Sillitoe on behalf of the Wellcome Sanger Institute COVID-19 Surveillance Team |
| EPI_ISL_2809147 | Lighthouse Lab in Alderley Park | Wellcome Sanger Institute for the COVID-19 Genomics UK (COG-UK) Consortium | Jacquelyn Wynn, Mairead Hyland, The Lighthouse Lab in Alderley Park and Alex Alderton, Roberto Amato, Jeffrey Barrett, Sonia Goncalves, Ewan Harrison, David K. Jackson, Ian Johnston, Dominic Kwiatkowski, Cordelia Langford, John Sillitoe on behalf of the Wellcome Sanger Institute COVID-19 Surveillance Team |
| EPI_ISL_2809148 | Lighthouse Lab in Milton Keynes | Wellcome Sanger Institute for the COVID-19 Genomics UK (COG-UK) Consortium | The Lighthouse Lab in Milton Keynes and Alex Alderton, Roberto Amato, Jeffrey Barrett, Sonia Goncalves, Ewan Harrison, David K. Jackson, Ian Johnston, Dominic Kwiatkowski, Cordelia Langford, John Sillitoe on behalf of the Wellcome Sanger Institute COVID-19 Surveillance Team |
| EPI_ISL_2809149 | Lighthouse Lab in Alderley Park | Wellcome Sanger Institute for the COVID-19 Genomics UK (COG-UK) Consortium | Jacquelyn Wynn, Mairead Hyland, The Lighthouse Lab in Alderley Park and Alex Alderton, Roberto Amato, Jeffrey Barrett, Sonia Goncalves, Ewan Harrison, David K. Jackson, Ian Johnston, Dominic Kwiatkowski, Cordelia Langford, John Sillitoe on behalf of the Wellcome Sanger Institute COVID-19 Surveillance Team |
| EPI_ISL_2809150 | Health Services Laboratories | Wellcome Sanger Institute for the COVID-19 Genomics UK (COG-UK) Consortium | Health Services Laboratories and Alex Alderton, Roberto Amato, Jeffrey Barrett, Sonia Goncalves, Ewan Harrison, David K. Jackson, Ian Johnston, Dominic Kwiatkowski, Cordelia Langford, John Sillitoe on behalf of the Wellcome Sanger Institute COVID-19 Surveillance Team |
| EPI_ISL_2809151 | Lighthouse Lab in Milton Keynes | Wellcome Sanger Institute for the COVID-19 Genomics UK (COG-UK) Consortium | The Lighthouse Lab in Milton Keynes and Alex Alderton, Roberto Amato, Jeffrey Barrett, Sonia Goncalves, Ewan Harrison, David K. Jackson, Ian Johnston, Dominic Kwiatkowski, Cordelia Langford, John Sillitoe on behalf of the Wellcome Sanger Institute COVID-19 Surveillance Team |
| EPI_ISL_2809152 | Health Services Laboratories | Wellcome Sanger Institute for the COVID-19 Genomics UK (COG-UK) Consortium | Health Services Laboratories and Alex Alderton, Roberto Amato, Jeffrey Barrett, Sonia Goncalves, Ewan Harrison, David K. Jackson, Ian Johnston, Dominic Kwiatkowski, Cordelia Langford, John Sillitoe on behalf of the Wellcome Sanger Institute COVID-19 Surveillance Team |
| EPI_ISL_2809153 | Lighthouse Lab in Alderley Park | Wellcome Sanger Institute for the COVID-19 Genomics UK (COG-UK) Consortium | Jacquelyn Wynn, Mairead Hyland, The Lighthouse Lab in Alderley Park and Alex Alderton, Roberto Amato, Jeffrey Barrett, Sonia Goncalves, Ewan Harrison, David K. Jackson, Ian Johnston, Dominic Kwiatkowski, Cordelia Langford, John Sillitoe on behalf of the Wellcome Sanger Institute COVID-19 Surveillance Team |
| EPI_ISL_2809154 | Lighthouse Lab in Milton Keynes | Wellcome Sanger Institute for the COVID-19 Genomics UK (COG-UK) Consortium | The Lighthouse Lab in Milton Keynes and Alex Alderton, Roberto Amato, Jeffrey Barrett, Sonia Goncalves, Ewan Harrison, David K. Jackson, Ian Johnston, Dominic Kwiatkowski, Cordelia Langford, John Sillitoe on behalf of the Wellcome Sanger Institute COVID-19 Surveillance Team |
| EPI_ISL_2809155 | Lighthouse Lab in Alderley Park | Wellcome Sanger Institute for the COVID-19 Genomics UK (COG-UK) Consortium | Jacquelyn Wynn, Mairead Hyland, The Lighthouse Lab in Alderley Park and Alex Alderton, Roberto Amato, Jeffrey Barrett, Sonia Goncalves, Ewan Harrison, David K. Jackson, Ian Johnston, Dominic Kwiatkowski, Cordelia Langford, John Sillitoe on behalf of the Wellcome Sanger Institute COVID-19 Surveillance Team |
| EPI_ISL_2809156, EPI_ISL_2809157 | Lighthouse Lab in Milton Keynes | Wellcome Sanger Institute for the COVID-19 Genomics UK (COG-UK) Consortium | The Lighthouse Lab in Milton Keynes and Alex Alderton, Roberto Amato, Jeffrey Barrett, Sonia Goncalves, Ewan Harrison, David K. Jackson, Ian Johnston, Dominic Kwiatkowski, Cordelia Langford, John Sillitoe on behalf of the Wellcome Sanger Institute COVID-19 Surveillance Team |
| EPI_ISL_2809158 | Lighthouse Lab in Alderley Park | Wellcome Sanger Institute for the COVID-19 Genomics UK (COG-UK) Consortium | Jacquelyn Wynn, Mairead Hyland, The Lighthouse Lab in Alderley Park and Alex Alderton, Roberto Amato, Jeffrey Barrett, Sonia Goncalves, Ewan Harrison, David K. Jackson, Ian Johnston, Dominic Kwiatkowski, Cordelia Langford, John Sillitoe on behalf of the Wellcome Sanger Institute COVID-19 Surveillance Team |
| EPI_ISL_2809159 | Health Services Laboratories | Wellcome Sanger Institute for the COVID-19 Genomics UK (COG-UK) Consortium | Health Services Laboratories and Alex Alderton, Roberto Amato, Jeffrey Barrett, Sonia Goncalves, Ewan Harrison, David K. Jackson, Ian Johnston, Dominic Kwiatkowski, Cordelia Langford, John Sillitoe on behalf of the Wellcome Sanger Institute COVID-19 Surveillance Team |
| EPI_ISL_2809160, EPI_ISL_2809161 | Lighthouse Lab in Alderley Park | Wellcome Sanger Institute for the COVID-19 Genomics UK (COG-UK) Consortium | Jacquelyn Wynn, Mairead Hyland, The Lighthouse Lab in Alderley Park and Alex Alderton, Roberto Amato, Jeffrey Barrett, Sonia Goncalves, Ewan Harrison, David K. Jackson, Ian Johnston, Dominic Kwiatkowski, Cordelia Langford, John Sillitoe on behalf of the Wellcome Sanger Institute COVID-19 Surveillance Team |
| EPI_ISL_2809162 | Health Services Laboratories | Wellcome Sanger Institute for the COVID-19 Genomics UK (COG-UK) Consortium | Health Services Laboratories and Alex Alderton, Roberto Amato, Jeffrey Barrett, Sonia Goncalves, Ewan Harrison, David K. Jackson, Ian Johnston, Dominic Kwiatkowski, Cordelia Langford, John Sillitoe on behalf of the Wellcome Sanger Institute COVID-19 Surveillance Team |
| EPI_ISL_2809163 | Lighthouse Lab in Alderley Park | Wellcome Sanger Institute for the COVID-19 Genomics UK (COG-UK) Consortium | Jacquelyn Wynn, Mairead Hyland, The Lighthouse Lab in Alderley Park and Alex Alderton, Roberto Amato, Jeffrey Barrett, Sonia Goncalves, Ewan Harrison, David K. Jackson, Ian Johnston, Dominic Kwiatkowski, Cordelia Langford, John Sillitoe on behalf of the Wellcome Sanger Institute COVID-19 Surveillance Team |
| EPI_ISL_2809164 | Health Services Laboratories | Wellcome Sanger Institute for the COVID-19 Genomics UK (COG-UK) Consortium | Health Services Laboratories and Alex Alderton, Roberto Amato, Jeffrey Barrett, Sonia Goncalves, Ewan Harrison, David K. Jackson, Ian Johnston, Dominic Kwiatkowski, Cordelia Langford, John Sillitoe on behalf of the Wellcome Sanger Institute COVID-19 Surveillance Team |
| EPI_ISL_2809165, EPI_ISL_2809166, EPI_ISL_2809167 | Lighthouse Lab in Alderley Park | Wellcome Sanger Institute for the COVID-19 Genomics UK (COG-UK) Consortium | Jacquelyn Wynn, Mairead Hyland, The Lighthouse Lab in Alderley Park and Alex Alderton, Roberto Amato, Jeffrey Barrett, Sonia Goncalves, Ewan Harrison, David K. Jackson, Ian Johnston, Dominic Kwiatkowski, Cordelia Langford, John Sillitoe on behalf of the Wellcome Sanger Institute COVID-19 Surveillance Team |
| EPI_ISL_2809168 | Health Services Laboratories | Wellcome Sanger Institute for the COVID-19 Genomics UK (COG-UK) Consortium | Health Services Laboratories and Alex Alderton, Roberto Amato, Jeffrey Barrett, Sonia Goncalves, Ewan Harrison, David K. Jackson, Ian Johnston, Dominic Kwiatkowski, Cordelia Langford, John Sillitoe on behalf of the Wellcome Sanger Institute COVID-19 Surveillance Team |
| EPI_ISL_2809169 | Lighthouse Lab in Milton Keynes | Wellcome Sanger Institute for the COVID-19 Genomics UK (COG-UK) Consortium | The Lighthouse Lab in Milton Keynes and Alex Alderton, Roberto Amato, Jeffrey Barrett, Sonia Goncalves, Ewan Harrison, David K. Jackson, Ian Johnston, Dominic Kwiatkowski, Cordelia Langford, John Sillitoe on behalf of the Wellcome Sanger Institute COVID-19 Surveillance Team |
| EPI_ISL_2809170 | Lighthouse Lab in Alderley Park | Wellcome Sanger Institute for the COVID-19 Genomics UK (COG-UK) Consortium | Jacquelyn Wynn, Mairead Hyland, The Lighthouse Lab in Alderley Park and Alex Alderton, Roberto Amato, Jeffrey Barrett, Sonia Goncalves, Ewan Harrison, David K. Jackson, Ian Johnston, Dominic Kwiatkowski, Cordelia Langford, John Sillitoe on behalf of the Wellcome Sanger Institute COVID-19 Surveillance Team |
| EPI_ISL_2809171, EPI_ISL_2809172 | Health Services Laboratories | Wellcome Sanger Institute for the COVID-19 Genomics UK (COG-UK) Consortium | Health Services Laboratories and Alex Alderton, Roberto Amato, Jeffrey Barrett, Sonia Goncalves, Ewan Harrison, David K. Jackson, Ian Johnston, Dominic Kwiatkowski, Cordelia Langford, John Sillitoe on behalf of the Wellcome Sanger Institute COVID-19 Surveillance Team |
| EPI_ISL_2809173 | Lighthouse Lab in Milton Keynes | Wellcome Sanger Institute for the COVID-19 Genomics UK (COG-UK) Consortium | The Lighthouse Lab in Milton Keynes and Alex Alderton, Roberto Amato, Jeffrey Barrett, Sonia Goncalves, Ewan Harrison, David K. Jackson, Ian Johnston, Dominic Kwiatkowski, Cordelia Langford, John Sillitoe on behalf of the Wellcome Sanger Institute COVID-19 Surveillance Team |
| EPI_ISL_2809174, | Lighthouse Lab in Alderley Park | Wellcome Sanger Institute for the COVID-19 Genomics UK (COG-UK) | Jacquelyn Wynn, Mairead Hyland, The Lighthouse Lab in Alderley Park and Alex Alderton, Roberto Amato, Jeffrey Barrett, Sonia Goncalves, Ewan Harrison, David K. Jackson, Ian Johnston, Dominic Kwiatkowski, Cordelia Langford, John Sillitoe on |

| | | | |
|---|---|---|---|
| EPI_ISL_2809175 | | Consortium | behalf of the Wellcome Sanger Institute COVID-19 Surveillance Team |
| EPI_ISL_2809176 | Health Services Laboratories | Wellcome Sanger Institute for the COVID-19 Genomics UK (COG-UK) Consortium | Health Services Laboratories and Alex Alderton, Roberto Amato, Jeffrey Barrett, Sonia Goncalves, Ewan Harrison, David K. Jackson, Ian Johnston, Dominic Kwiatkowski, Cordelia Langford, John Sillitoe on behalf of the Wellcome Sanger Institute COVID-19 Surveillance Team |
| EPI_ISL_2809177 | Lighthouse Lab in Alderley Park | Wellcome Sanger Institute for the COVID-19 Genomics UK (COG-UK) Consortium | Jacquelyn Wynn, Mairead Hyland, The Lighthouse Lab in Alderley Park and Alex Alderton, Roberto Amato, Jeffrey Barrett, Sonia Goncalves, Ewan Harrison, David K. Jackson, Ian Johnston, Dominic Kwiatkowski, Cordelia Langford, John Sillitoe on behalf of the Wellcome Sanger Institute COVID-19 Surveillance Team |
| EPI_ISL_2809178 | Health Services Laboratories | Wellcome Sanger Institute for the COVID-19 Genomics UK (COG-UK) Consortium | Health Services Laboratories and Alex Alderton, Roberto Amato, Jeffrey Barrett, Sonia Goncalves, Ewan Harrison, David K. Jackson, Ian Johnston, Dominic Kwiatkowski, Cordelia Langford, John Sillitoe on behalf of the Wellcome Sanger Institute COVID-19 Surveillance Team |
| EPI_ISL_2809179 | Lighthouse Lab in Alderley Park | Wellcome Sanger Institute for the COVID-19 Genomics UK (COG-UK) Consortium | Jacquelyn Wynn, Mairead Hyland, The Lighthouse Lab in Alderley Park and Alex Alderton, Roberto Amato, Jeffrey Barrett, Sonia Goncalves, Ewan Harrison, David K. Jackson, Ian Johnston, Dominic Kwiatkowski, Cordelia Langford, John Sillitoe on behalf of the Wellcome Sanger Institute COVID-19 Surveillance Team |
| EPI_ISL_2809180 | Health Services Laboratories | Wellcome Sanger Institute for the COVID-19 Genomics UK (COG-UK) Consortium | Health Services Laboratories and Alex Alderton, Roberto Amato, Jeffrey Barrett, Sonia Goncalves, Ewan Harrison, David K. Jackson, Ian Johnston, Dominic Kwiatkowski, Cordelia Langford, John Sillitoe on behalf of the Wellcome Sanger Institute COVID-19 Surveillance Team |
| EPI_ISL_2809181 | Lighthouse Lab in Milton Keynes | Wellcome Sanger Institute for the COVID-19 Genomics UK (COG-UK) Consortium | The Lighthouse Lab in Milton Keynes and Alex Alderton, Roberto Amato, Jeffrey Barrett, Sonia Goncalves, Ewan Harrison, David K. Jackson, Ian Johnston, Dominic Kwiatkowski, Cordelia Langford, John Sillitoe on behalf of the Wellcome Sanger Institute COVID-19 Surveillance Team |
| EPI_ISL_2809182, EPI_ISL_2809183 | Health Services Laboratories | Wellcome Sanger Institute for the COVID-19 Genomics UK (COG-UK) Consortium | Health Services Laboratories and Alex Alderton, Roberto Amato, Jeffrey Barrett, Sonia Goncalves, Ewan Harrison, David K. Jackson, Ian Johnston, Dominic Kwiatkowski, Cordelia Langford, John Sillitoe on behalf of the Wellcome Sanger Institute COVID-19 Surveillance Team |
| EPI_ISL_2809184, EPI_ISL_2809185 | Lighthouse Lab in Alderley Park | Wellcome Sanger Institute for the COVID-19 Genomics UK (COG-UK) Consortium | Jacquelyn Wynn, Mairead Hyland, The Lighthouse Lab in Alderley Park and Alex Alderton, Roberto Amato, Jeffrey Barrett, Sonia Goncalves, Ewan Harrison, David K. Jackson, Ian Johnston, Dominic Kwiatkowski, Cordelia Langford, John Sillitoe on behalf of the Wellcome Sanger Institute COVID-19 Surveillance Team |
| EPI_ISL_2809186 | Health Services Laboratories | Wellcome Sanger Institute for the COVID-19 Genomics UK (COG-UK) Consortium | Health Services Laboratories and Alex Alderton, Roberto Amato, Jeffrey Barrett, Sonia Goncalves, Ewan Harrison, David K. Jackson, Ian Johnston, Dominic Kwiatkowski, Cordelia Langford, John Sillitoe on behalf of the Wellcome Sanger Institute COVID-19 Surveillance Team |
| EPI_ISL_2809187 | Lighthouse Lab in Milton Keynes | Wellcome Sanger Institute for the COVID-19 Genomics UK (COG-UK) Consortium | The Lighthouse Lab in Milton Keynes and Alex Alderton, Roberto Amato, Jeffrey Barrett, Sonia Goncalves, Ewan Harrison, David K. Jackson, Ian Johnston, Dominic Kwiatkowski, Cordelia Langford, John Sillitoe on behalf of the Wellcome Sanger Institute COVID-19 Surveillance Team |
| EPI_ISL_2809188 | Lighthouse Lab in Alderley Park | Wellcome Sanger Institute for the COVID-19 Genomics UK (COG-UK) Consortium | Jacquelyn Wynn, Mairead Hyland, The Lighthouse Lab in Alderley Park and Alex Alderton, Roberto Amato, Jeffrey Barrett, Sonia Goncalves, Ewan Harrison, David K. Jackson, Ian Johnston, Dominic Kwiatkowski, Cordelia Langford, John Sillitoe on behalf of the Wellcome Sanger Institute COVID-19 Surveillance Team |
| EPI_ISL_2809189 | Lighthouse Lab in Milton Keynes | Wellcome Sanger Institute for the COVID-19 Genomics UK (COG-UK) Consortium | The Lighthouse Lab in Milton Keynes and Alex Alderton, Roberto Amato, Jeffrey Barrett, Sonia Goncalves, Ewan Harrison, David K. Jackson, Ian Johnston, Dominic Kwiatkowski, Cordelia Langford, John Sillitoe on behalf of the Wellcome Sanger Institute COVID-19 Surveillance Team |
| EPI_ISL_2809190 | Lighthouse Lab in Alderley Park | Wellcome Sanger Institute for the COVID-19 Genomics UK (COG-UK) Consortium | Jacquelyn Wynn, Mairead Hyland, The Lighthouse Lab in Alderley Park and Alex Alderton, Roberto Amato, Jeffrey Barrett, Sonia Goncalves, Ewan Harrison, David K. Jackson, Ian Johnston, Dominic Kwiatkowski, Cordelia Langford, John Sillitoe on behalf of the Wellcome Sanger Institute COVID-19 Surveillance Team |
| EPI_ISL_2809191 | Lighthouse Lab in Milton Keynes | Wellcome Sanger Institute for the COVID-19 Genomics UK (COG-UK) Consortium | The Lighthouse Lab in Milton Keynes and Alex Alderton, Roberto Amato, Jeffrey Barrett, Sonia Goncalves, Ewan Harrison, David K. Jackson, Ian Johnston, Dominic Kwiatkowski, Cordelia Langford, John Sillitoe on behalf of the Wellcome Sanger Institute COVID-19 Surveillance Team |
| EPI_ISL_2809192 | Health Services Laboratories | Wellcome Sanger Institute for the COVID-19 Genomics UK (COG-UK) Consortium | Health Services Laboratories and Alex Alderton, Roberto Amato, Jeffrey Barrett, Sonia Goncalves, Ewan Harrison, David K. Jackson, Ian Johnston, Dominic Kwiatkowski, Cordelia Langford, John Sillitoe on behalf of the Wellcome Sanger Institute COVID-19 Surveillance Team |
| EPI_ISL_2809193 | Lighthouse Lab in Alderley Park | Wellcome Sanger Institute for the COVID-19 Genomics UK (COG-UK) Consortium | Jacquelyn Wynn, Mairead Hyland, The Lighthouse Lab in Alderley Park and Alex Alderton, Roberto Amato, Jeffrey Barrett, Sonia Goncalves, Ewan Harrison, David K. Jackson, Ian Johnston, Dominic Kwiatkowski, Cordelia Langford, John Sillitoe on behalf of the Wellcome Sanger Institute COVID-19 Surveillance Team |
| EPI_ISL_2809194 | Lighthouse Lab in Milton Keynes | Wellcome Sanger Institute for the COVID-19 Genomics UK (COG-UK) Consortium | The Lighthouse Lab in Milton Keynes and Alex Alderton, Roberto Amato, Jeffrey Barrett, Sonia Goncalves, Ewan Harrison, David K. Jackson, Ian Johnston, Dominic Kwiatkowski, Cordelia Langford, John Sillitoe on behalf of the Wellcome Sanger Institute COVID-19 Surveillance Team |
| EPI_ISL_2809195, EPI_ISL_2809196 | Health Services Laboratories | Wellcome Sanger Institute for the COVID-19 Genomics UK (COG-UK) Consortium | Health Services Laboratories and Alex Alderton, Roberto Amato, Jeffrey Barrett, Sonia Goncalves, Ewan Harrison, David K. Jackson, Ian Johnston, Dominic Kwiatkowski, Cordelia Langford, John Sillitoe on behalf of the Wellcome Sanger Institute COVID-19 Surveillance Team |
| EPI_ISL_2809197 | Lighthouse Lab in Alderley Park | Wellcome Sanger Institute for the COVID-19 Genomics UK (COG-UK) Consortium | Jacquelyn Wynn, Mairead Hyland, The Lighthouse Lab in Alderley Park and Alex Alderton, Roberto Amato, Jeffrey Barrett, Sonia Goncalves, Ewan Harrison, David K. Jackson, Ian Johnston, Dominic Kwiatkowski, Cordelia Langford, John Sillitoe on behalf of the Wellcome Sanger Institute COVID-19 Surveillance Team |
| EPI_ISL_2809198 | Health Services Laboratories | Wellcome Sanger Institute for the COVID-19 Genomics UK (COG-UK) Consortium | Health Services Laboratories and Alex Alderton, Roberto Amato, Jeffrey Barrett, Sonia Goncalves, Ewan Harrison, David K. Jackson, Ian Johnston, Dominic Kwiatkowski, Cordelia Langford, John Sillitoe on behalf of the Wellcome Sanger Institute COVID-19 Surveillance Team |
| EPI_ISL_2809199 | Lighthouse Lab in Milton Keynes | Wellcome Sanger Institute for the COVID-19 Genomics UK (COG-UK) Consortium | The Lighthouse Lab in Milton Keynes and Alex Alderton, Roberto Amato, Jeffrey Barrett, Sonia Goncalves, Ewan Harrison, David K. Jackson, Ian Johnston, Dominic Kwiatkowski, Cordelia Langford, John Sillitoe on behalf of the Wellcome Sanger Institute COVID-19 Surveillance Team |
| EPI_ISL_2809200, EPI_ISL_2809201, EPI_ISL_2809202 | Lighthouse Lab in Alderley Park | Wellcome Sanger Institute for the COVID-19 Genomics UK (COG-UK) Consortium | Jacquelyn Wynn, Mairead Hyland, The Lighthouse Lab in Alderley Park and Alex Alderton, Roberto Amato, Jeffrey Barrett, Sonia Goncalves, Ewan Harrison, David K. Jackson, Ian Johnston, Dominic Kwiatkowski, Cordelia Langford, John Sillitoe on behalf of the Wellcome Sanger Institute COVID-19 Surveillance Team |
| EPI_ISL_2809203 | Health Services Laboratories | Wellcome Sanger Institute for the COVID-19 Genomics UK (COG-UK) Consortium | Health Services Laboratories and Alex Alderton, Roberto Amato, Jeffrey Barrett, Sonia Goncalves, Ewan Harrison, David K. Jackson, Ian Johnston, Dominic Kwiatkowski, Cordelia Langford, John Sillitoe on behalf of the Wellcome Sanger Institute COVID-19 Surveillance Team |
| EPI_ISL_2809204 | Lighthouse Lab in Alderley Park | Wellcome Sanger Institute for the COVID-19 Genomics UK (COG-UK) Consortium | Jacquelyn Wynn, Mairead Hyland, The Lighthouse Lab in Alderley Park and Alex Alderton, Roberto Amato, Jeffrey Barrett, Sonia Goncalves, Ewan Harrison, David K. Jackson, Ian Johnston, Dominic Kwiatkowski, Cordelia Langford, John Sillitoe on behalf of the Wellcome Sanger Institute COVID-19 Surveillance Team |
| EPI_ISL_2809205, EPI_ISL_2809206 | Health Services Laboratories | Wellcome Sanger Institute for the COVID-19 Genomics UK (COG-UK) Consortium | Health Services Laboratories and Alex Alderton, Roberto Amato, Jeffrey Barrett, Sonia Goncalves, Ewan Harrison, David K. Jackson, Ian Johnston, Dominic Kwiatkowski, Cordelia Langford, John Sillitoe on behalf of the Wellcome Sanger Institute COVID-19 Surveillance Team |
| EPI_ISL_2809207 | Lighthouse Lab in Milton Keynes | Wellcome Sanger Institute for the COVID-19 Genomics UK (COG-UK) Consortium | The Lighthouse Lab in Milton Keynes and Alex Alderton, Roberto Amato, Jeffrey Barrett, Sonia Goncalves, Ewan Harrison, David K. Jackson, Ian Johnston, Dominic Kwiatkowski, Cordelia Langford, John Sillitoe on behalf of the Wellcome Sanger Institute COVID-19 Surveillance Team |
| EPI_ISL_2809208, EPI_ISL_2809209, EPI_ISL_2809210, EPI_ISL_2809211 | Lighthouse Lab in Alderley Park | Wellcome Sanger Institute for the COVID-19 Genomics UK (COG-UK) Consortium | Jacquelyn Wynn, Mairead Hyland, The Lighthouse Lab in Alderley Park and Alex Alderton, Roberto Amato, Jeffrey Barrett, Sonia Goncalves, Ewan Harrison, David K. Jackson, Ian Johnston, Dominic Kwiatkowski, Cordelia Langford, John Sillitoe on behalf of the Wellcome Sanger Institute COVID-19 Surveillance Team |
| EPI_ISL_2809212 | Health Services Laboratories | Wellcome Sanger Institute for the COVID-19 Genomics UK (COG-UK) Consortium | Health Services Laboratories and Alex Alderton, Roberto Amato, Jeffrey Barrett, Sonia Goncalves, Ewan Harrison, David K. Jackson, Ian Johnston, Dominic Kwiatkowski, Cordelia Langford, John Sillitoe on behalf of the Wellcome Sanger Institute COVID-19 Surveillance Team |
| EPI_ISL_2809213, EPI_ISL_2809214, EPI_ISL_2809215, EPI_ISL_2809216, EPI_ISL_2809217 | Lighthouse Lab in Alderley Park | Wellcome Sanger Institute for the COVID-19 Genomics UK (COG-UK) Consortium | Jacquelyn Wynn, Mairead Hyland, The Lighthouse Lab in Alderley Park and Alex Alderton, Roberto Amato, Jeffrey Barrett, Sonia Goncalves, Ewan Harrison, David K. Jackson, Ian Johnston, Dominic Kwiatkowski, Cordelia Langford, John Sillitoe on behalf of the Wellcome Sanger Institute COVID-19 Surveillance Team |
| EPI_ISL_2809218 | Lighthouse Lab in Milton Keynes | Wellcome Sanger Institute for the COVID-19 Genomics UK (COG-UK) Consortium | The Lighthouse Lab in Milton Keynes and Alex Alderton, Roberto Amato, Jeffrey Barrett, Sonia Goncalves, Ewan Harrison, David K. Jackson, Ian Johnston, Dominic Kwiatkowski, Cordelia Langford, John Sillitoe on behalf of the Wellcome Sanger Institute COVID-19 Surveillance Team |
| EPI_ISL_2809219 | Lighthouse Lab in Alderley Park | Wellcome Sanger Institute for the COVID-19 Genomics UK (COG-UK) Consortium | Jacquelyn Wynn, Mairead Hyland, The Lighthouse Lab in Alderley Park and Alex Alderton, Roberto Amato, Jeffrey Barrett, Sonia Goncalves, Ewan Harrison, David K. Jackson, Ian Johnston, Dominic Kwiatkowski, Cordelia Langford, John Sillitoe on behalf of the Wellcome Sanger Institute COVID-19 Surveillance Team |
| EPI_ISL_2809220 | Health Services Laboratories | Wellcome Sanger Institute for the COVID-19 Genomics UK (COG-UK) Consortium | Health Services Laboratories and Alex Alderton, Roberto Amato, Jeffrey Barrett, Sonia Goncalves, Ewan Harrison, David K. Jackson, Ian Johnston, Dominic Kwiatkowski, Cordelia Langford, John Sillitoe on behalf of the Wellcome Sanger Institute COVID-19 Surveillance Team |
| EPI_ISL_2809221, EPI_ISL_2809222, EPI_ISL_2809223, EPI_ISL_2809224 | Lighthouse Lab in Alderley Park | Wellcome Sanger Institute for the COVID-19 Genomics UK (COG-UK) Consortium | Jacquelyn Wynn, Mairead Hyland, The Lighthouse Lab in Alderley Park and Alex Alderton, Roberto Amato, Jeffrey Barrett, Sonia Goncalves, Ewan Harrison, David K. Jackson, Ian Johnston, Dominic Kwiatkowski, Cordelia Langford, John Sillitoe on behalf of the Wellcome Sanger Institute COVID-19 Surveillance Team |
| EPI_ISL_2809225, EPI_ISL_2809226, EPI_ISL_2809227 | Health Services Laboratories | Wellcome Sanger Institute for the COVID-19 Genomics UK (COG-UK) Consortium | Health Services Laboratories and Alex Alderton, Roberto Amato, Jeffrey Barrett, Sonia Goncalves, Ewan Harrison, David K. Jackson, Ian Johnston, Dominic Kwiatkowski, Cordelia Langford, John Sillitoe on behalf of the Wellcome Sanger Institute COVID-19 Surveillance Team |
| EPI_ISL_2809228, EPI_ISL_2809229, EPI_ISL_2809231, EPI_ISL_2809232, EPI_ISL_2809233 | Lighthouse Lab in Alderley Park | Wellcome Sanger Institute for the COVID-19 Genomics UK (COG-UK) Consortium | Jacquelyn Wynn, Mairead Hyland, The Lighthouse Lab in Alderley Park and Alex Alderton, Roberto Amato, Jeffrey Barrett, Sonia Goncalves, Ewan Harrison, David K. Jackson, Ian Johnston, Dominic Kwiatkowski, Cordelia Langford, John Sillitoe on behalf of the Wellcome Sanger Institute COVID-19 Surveillance Team |
| EPI_ISL_2809234 | Lighthouse Lab in Milton Keynes | Wellcome Sanger Institute for the COVID-19 Genomics UK (COG-UK) Consortium | The Lighthouse Lab in Milton Keynes and Alex Alderton, Roberto Amato, Jeffrey Barrett, Sonia Goncalves, Ewan Harrison, David K. Jackson, Ian Johnston, Dominic Kwiatkowski, Cordelia Langford, John Sillitoe on behalf of the Wellcome Sanger Institute COVID-19 Surveillance Team |
| EPI_ISL_2809235, EPI_ISL_2809236, EPI_ISL_2809237, EPI_ISL_2809238, EPI_ISL_2809239 | Lighthouse Lab in Alderley Park | Wellcome Sanger Institute for the COVID-19 Genomics UK (COG-UK) Consortium | Jacquelyn Wynn, Mairead Hyland, The Lighthouse Lab in Alderley Park and Alex Alderton, Roberto Amato, Jeffrey Barrett, Sonia Goncalves, Ewan Harrison, David K. Jackson, Ian Johnston, Dominic Kwiatkowski, Cordelia Langford, John Sillitoe on behalf of the Wellcome Sanger Institute COVID-19 Surveillance Team |
| EPI_ISL_2809240 | Health Services Laboratories | Wellcome Sanger Institute for the COVID-19 Genomics UK (COG-UK) Consortium | Health Services Laboratories and Alex Alderton, Roberto Amato, Jeffrey Barrett, Sonia Goncalves, Ewan Harrison, David K. Jackson, Ian Johnston, Dominic Kwiatkowski, Cordelia Langford, John Sillitoe on behalf of the Wellcome Sanger Institute COVID-19 Surveillance Team |
| EPI_ISL_2809241 | Lighthouse Lab in Alderley Park | Wellcome Sanger Institute for the COVID-19 Genomics UK (COG-UK) | Jacquelyn Wynn, Mairead Hyland, The Lighthouse Lab in Alderley Park and Alex Alderton, Roberto Amato, Jeffrey Barrett, Sonia Goncalves, Ewan Harrison, David K. Jackson, Ian Johnston, Dominic Kwiatkowski, Cordelia Langford, John Sillitoe on |

| | | | |
|---|---|---|---|
| | | Consortium | behalf of the Wellcome Sanger Institute COVID-19 Surveillance Team |
| EPI_ISL_2809242 | Health Services Laboratories | Wellcome Sanger Institute for the COVID-19 Genomics UK (COG-UK) Consortium | Health Services Laboratories and Alex Alderton, Roberto Amato, Jeffrey Barrett, Sonia Goncalves, Ewan Harrison, David K. Jackson, Ian Johnston, Dominic Kwiatkowski, Cordelia Langford, John Sillitoe on behalf of the Wellcome Sanger Institute COVID-19 Surveillance Team |
| EPI_ISL_2809243 | Lighthouse Lab in Milton Keynes | Wellcome Sanger Institute for the COVID-19 Genomics UK (COG-UK) Consortium | The Lighthouse Lab in Milton Keynes and Alex Alderton, Roberto Amato, Jeffrey Barrett, Sonia Goncalves, Ewan Harrison, David K. Jackson, Ian Johnston, Dominic Kwiatkowski, Cordelia Langford, John Sillitoe on behalf of the Wellcome Sanger Institute COVID-19 Surveillance Team |
| EPI_ISL_2809244 | Health Services Laboratories | Wellcome Sanger Institute for the COVID-19 Genomics UK (COG-UK) Consortium | Health Services Laboratories and Alex Alderton, Roberto Amato, Jeffrey Barrett, Sonia Goncalves, Ewan Harrison, David K. Jackson, Ian Johnston, Dominic Kwiatkowski, Cordelia Langford, John Sillitoe on behalf of the Wellcome Sanger Institute COVID-19 Surveillance Team |
| EPI_ISL_2809245, EPI_ISL_2809246 | Lighthouse Lab in Milton Keynes | Wellcome Sanger Institute for the COVID-19 Genomics UK (COG-UK) Consortium | The Lighthouse Lab in Milton Keynes and Alex Alderton, Roberto Amato, Jeffrey Barrett, Sonia Goncalves, Ewan Harrison, David K. Jackson, Ian Johnston, Dominic Kwiatkowski, Cordelia Langford, John Sillitoe on behalf of the Wellcome Sanger Institute COVID-19 Surveillance Team |
| EPI_ISL_2809247, EPI_ISL_2809248, EPI_ISL_2809249 | Lighthouse Lab in Alderley Park | Wellcome Sanger Institute for the COVID-19 Genomics UK (COG-UK) Consortium | Jacquelyn Wynn, Mairead Hyland, The Lighthouse Lab in Alderley Park and Alex Alderton, Roberto Amato, Jeffrey Barrett, Sonia Goncalves, Ewan Harrison, David K. Jackson, Ian Johnston, Dominic Kwiatkowski, Cordelia Langford, John Sillitoe on behalf of the Wellcome Sanger Institute COVID-19 Surveillance Team |
| EPI_ISL_2809250 | Health Services Laboratories | Wellcome Sanger Institute for the COVID-19 Genomics UK (COG-UK) Consortium | Health Services Laboratories and Alex Alderton, Roberto Amato, Jeffrey Barrett, Sonia Goncalves, Ewan Harrison, David K. Jackson, Ian Johnston, Dominic Kwiatkowski, Cordelia Langford, John Sillitoe on behalf of the Wellcome Sanger Institute COVID-19 Surveillance Team |
| EPI_ISL_2809251 | Lighthouse Lab in Alderley Park | Wellcome Sanger Institute for the COVID-19 Genomics UK (COG-UK) Consortium | Jacquelyn Wynn, Mairead Hyland, The Lighthouse Lab in Alderley Park and Alex Alderton, Roberto Amato, Jeffrey Barrett, Sonia Goncalves, Ewan Harrison, David K. Jackson, Ian Johnston, Dominic Kwiatkowski, Cordelia Langford, John Sillitoe on behalf of the Wellcome Sanger Institute COVID-19 Surveillance Team |
| EPI_ISL_2809252 | Lighthouse Lab in Milton Keynes | Wellcome Sanger Institute for the COVID-19 Genomics UK (COG-UK) Consortium | The Lighthouse Lab in Milton Keynes and Alex Alderton, Roberto Amato, Jeffrey Barrett, Sonia Goncalves, Ewan Harrison, David K. Jackson, Ian Johnston, Dominic Kwiatkowski, Cordelia Langford, John Sillitoe on behalf of the Wellcome Sanger Institute COVID-19 Surveillance Team |
| EPI_ISL_2809253 | Lighthouse Lab in Alderley Park | Wellcome Sanger Institute for the COVID-19 Genomics UK (COG-UK) Consortium | Jacquelyn Wynn, Mairead Hyland, The Lighthouse Lab in Alderley Park and Alex Alderton, Roberto Amato, Jeffrey Barrett, Sonia Goncalves, Ewan Harrison, David K. Jackson, Ian Johnston, Dominic Kwiatkowski, Cordelia Langford, John Sillitoe on behalf of the Wellcome Sanger Institute COVID-19 Surveillance Team |
| EPI_ISL_2809254, EPI_ISL_2809255 | Lighthouse Lab in Milton Keynes | Wellcome Sanger Institute for the COVID-19 Genomics UK (COG-UK) Consortium | The Lighthouse Lab in Milton Keynes and Alex Alderton, Roberto Amato, Jeffrey Barrett, Sonia Goncalves, Ewan Harrison, David K. Jackson, Ian Johnston, Dominic Kwiatkowski, Cordelia Langford, John Sillitoe on behalf of the Wellcome Sanger Institute COVID-19 Surveillance Team |
| EPI_ISL_2809256, EPI_ISL_2809257 | Lighthouse Lab in Alderley Park | Wellcome Sanger Institute for the COVID-19 Genomics UK (COG-UK) Consortium | Jacquelyn Wynn, Mairead Hyland, The Lighthouse Lab in Alderley Park and Alex Alderton, Roberto Amato, Jeffrey Barrett, Sonia Goncalves, Ewan Harrison, David K. Jackson, Ian Johnston, Dominic Kwiatkowski, Cordelia Langford, John Sillitoe on behalf of the Wellcome Sanger Institute COVID-19 Surveillance Team |
| EPI_ISL_2809258 | Lighthouse Lab in Glasgow | Wellcome Sanger Institute for the COVID-19 Genomics UK (COG-UK) Consortium | Harper VanSteenhouse, Yumi Kasai, David Gray, Carol Clugston, Anna Dominiczak and Alex Alderton, Roberto Amato, Jeffrey Barrett, Sonia Goncalves, Ewan Harrison, David K. Jackson, Ian Johnston, Dominic Kwiatkowski, Cordelia Langford, John Sillitoe on behalf of the Wellcome Sanger Institute COVID-19 Surveillance Team |
| EPI_ISL_2809259 | Lighthouse Lab in Milton Keynes | Wellcome Sanger Institute for the COVID-19 Genomics UK (COG-UK) Consortium | The Lighthouse Lab in Milton Keynes and Alex Alderton, Roberto Amato, Jeffrey Barrett, Sonia Goncalves, Ewan Harrison, David K. Jackson, Ian Johnston, Dominic Kwiatkowski, Cordelia Langford, John Sillitoe on behalf of the Wellcome Sanger Institute COVID-19 Surveillance Team |
| EPI_ISL_2809260 | Lighthouse Lab in Alderley Park | Wellcome Sanger Institute for the COVID-19 Genomics UK (COG-UK) Consortium | Jacquelyn Wynn, Mairead Hyland, The Lighthouse Lab in Alderley Park and Alex Alderton, Roberto Amato, Jeffrey Barrett, Sonia Goncalves, Ewan Harrison, David K. Jackson, Ian Johnston, Dominic Kwiatkowski, Cordelia Langford, John Sillitoe on behalf of the Wellcome Sanger Institute COVID-19 Surveillance Team |
| EPI_ISL_2809261 | Health Services Laboratories | Wellcome Sanger Institute for the COVID-19 Genomics UK (COG-UK) Consortium | Health Services Laboratories and Alex Alderton, Roberto Amato, Jeffrey Barrett, Sonia Goncalves, Ewan Harrison, David K. Jackson, Ian Johnston, Dominic Kwiatkowski, Cordelia Langford, John Sillitoe on behalf of the Wellcome Sanger Institute COVID-19 Surveillance Team |
| EPI_ISL_2809262 | Lighthouse Lab in Alderley Park | Wellcome Sanger Institute for the COVID-19 Genomics UK (COG-UK) Consortium | Jacquelyn Wynn, Mairead Hyland, The Lighthouse Lab in Alderley Park and Alex Alderton, Roberto Amato, Jeffrey Barrett, Sonia Goncalves, Ewan Harrison, David K. Jackson, Ian Johnston, Dominic Kwiatkowski, Cordelia Langford, John Sillitoe on behalf of the Wellcome Sanger Institute COVID-19 Surveillance Team |
| EPI_ISL_2809263, EPI_ISL_2809264 | Lighthouse Lab in Milton Keynes | Wellcome Sanger Institute for the COVID-19 Genomics UK (COG-UK) Consortium | The Lighthouse Lab in Milton Keynes and Alex Alderton, Roberto Amato, Jeffrey Barrett, Sonia Goncalves, Ewan Harrison, David K. Jackson, Ian Johnston, Dominic Kwiatkowski, Cordelia Langford, John Sillitoe on behalf of the Wellcome Sanger Institute COVID-19 Surveillance Team |
| EPI_ISL_2809265 | Health Services Laboratories | Wellcome Sanger Institute for the COVID-19 Genomics UK (COG-UK) Consortium | Health Services Laboratories and Alex Alderton, Roberto Amato, Jeffrey Barrett, Sonia Goncalves, Ewan Harrison, David K. Jackson, Ian Johnston, Dominic Kwiatkowski, Cordelia Langford, John Sillitoe on behalf of the Wellcome Sanger Institute COVID-19 Surveillance Team |
| EPI_ISL_2809266 | Lighthouse Lab in Milton Keynes | Wellcome Sanger Institute for the COVID-19 Genomics UK (COG-UK) Consortium | The Lighthouse Lab in Milton Keynes and Alex Alderton, Roberto Amato, Jeffrey Barrett, Sonia Goncalves, Ewan Harrison, David K. Jackson, Ian Johnston, Dominic Kwiatkowski, Cordelia Langford, John Sillitoe on behalf of the Wellcome Sanger Institute COVID-19 Surveillance Team |
| EPI_ISL_2809267, EPI_ISL_2809268 | Lighthouse Lab in Alderley Park | Wellcome Sanger Institute for the COVID-19 Genomics UK (COG-UK) Consortium | Jacquelyn Wynn, Mairead Hyland, The Lighthouse Lab in Alderley Park and Alex Alderton, Roberto Amato, Jeffrey Barrett, Sonia Goncalves, Ewan Harrison, David K. Jackson, Ian Johnston, Dominic Kwiatkowski, Cordelia Langford, John Sillitoe on behalf of the Wellcome Sanger Institute COVID-19 Surveillance Team |
| EPI_ISL_2809269 | Health Services Laboratories | Wellcome Sanger Institute for the COVID-19 Genomics UK (COG-UK) Consortium | Health Services Laboratories and Alex Alderton, Roberto Amato, Jeffrey Barrett, Sonia Goncalves, Ewan Harrison, David K. Jackson, Ian Johnston, Dominic Kwiatkowski, Cordelia Langford, John Sillitoe on behalf of the Wellcome Sanger Institute COVID-19 Surveillance Team |
| EPI_ISL_2809270, EPI_ISL_2809271 | Lighthouse Lab in Milton Keynes | Wellcome Sanger Institute for the COVID-19 Genomics UK (COG-UK) Consortium | The Lighthouse Lab in Milton Keynes and Alex Alderton, Roberto Amato, Jeffrey Barrett, Sonia Goncalves, Ewan Harrison, David K. Jackson, Ian Johnston, Dominic Kwiatkowski, Cordelia Langford, John Sillitoe on behalf of the Wellcome Sanger Institute COVID-19 Surveillance Team |
| EPI_ISL_2809272 | Lighthouse Lab in Glasgow | Wellcome Sanger Institute for the COVID-19 Genomics UK (COG-UK) Consortium | Harper VanSteenhouse, Yumi Kasai, David Gray, Carol Clugston, Anna Dominiczak and Alex Alderton, Roberto Amato, Jeffrey Barrett, Sonia Goncalves, Ewan Harrison, David K. Jackson, Ian Johnston, Dominic Kwiatkowski, Cordelia Langford, John Sillitoe on behalf of the Wellcome Sanger Institute COVID-19 Surveillance Team |
| EPI_ISL_2809273 | Health Services Laboratories | Wellcome Sanger Institute for the COVID-19 Genomics UK (COG-UK) Consortium | Health Services Laboratories and Alex Alderton, Roberto Amato, Jeffrey Barrett, Sonia Goncalves, Ewan Harrison, David K. Jackson, Ian Johnston, Dominic Kwiatkowski, Cordelia Langford, John Sillitoe on behalf of the Wellcome Sanger Institute COVID-19 Surveillance Team |
| EPI_ISL_2809274 | Lighthouse Lab in Milton Keynes | Wellcome Sanger Institute for the COVID-19 Genomics UK (COG-UK) Consortium | The Lighthouse Lab in Milton Keynes and Alex Alderton, Roberto Amato, Jeffrey Barrett, Sonia Goncalves, Ewan Harrison, David K. Jackson, Ian Johnston, Dominic Kwiatkowski, Cordelia Langford, John Sillitoe on behalf of the Wellcome Sanger Institute COVID-19 Surveillance Team |
| EPI_ISL_2809275 | Lighthouse Lab in Alderley Park | Wellcome Sanger Institute for the COVID-19 Genomics UK (COG-UK) Consortium | Jacquelyn Wynn, Mairead Hyland, The Lighthouse Lab in Alderley Park and Alex Alderton, Roberto Amato, Jeffrey Barrett, Sonia Goncalves, Ewan Harrison, David K. Jackson, Ian Johnston, Dominic Kwiatkowski, Cordelia Langford, John Sillitoe on behalf of the Wellcome Sanger Institute COVID-19 Surveillance Team |
| EPI_ISL_2809276 | Lighthouse Lab in Milton Keynes | Wellcome Sanger Institute for the COVID-19 Genomics UK (COG-UK) Consortium | The Lighthouse Lab in Milton Keynes and Alex Alderton, Roberto Amato, Jeffrey Barrett, Sonia Goncalves, Ewan Harrison, David K. Jackson, Ian Johnston, Dominic Kwiatkowski, Cordelia Langford, John Sillitoe on behalf of the Wellcome Sanger Institute COVID-19 Surveillance Team |
| EPI_ISL_2809277 | Lighthouse Lab in Alderley Park | Wellcome Sanger Institute for the COVID-19 Genomics UK (COG-UK) Consortium | Jacquelyn Wynn, Mairead Hyland, The Lighthouse Lab in Alderley Park and Alex Alderton, Roberto Amato, Jeffrey Barrett, Sonia Goncalves, Ewan Harrison, David K. Jackson, Ian Johnston, Dominic Kwiatkowski, Cordelia Langford, John Sillitoe on behalf of the Wellcome Sanger Institute COVID-19 Surveillance Team |
| EPI_ISL_2809278, EPI_ISL_2809279, EPI_ISL_2809280 | Lighthouse Lab in Milton Keynes | Wellcome Sanger Institute for the COVID-19 Genomics UK (COG-UK) Consortium | The Lighthouse Lab in Milton Keynes and Alex Alderton, Roberto Amato, Jeffrey Barrett, Sonia Goncalves, Ewan Harrison, David K. Jackson, Ian Johnston, Dominic Kwiatkowski, Cordelia Langford, John Sillitoe on behalf of the Wellcome Sanger Institute COVID-19 Surveillance Team |
| EPI_ISL_2809281 | Lighthouse Lab in Alderley Park | Wellcome Sanger Institute for the COVID-19 Genomics UK (COG-UK) Consortium | Jacquelyn Wynn, Mairead Hyland, The Lighthouse Lab in Alderley Park and Alex Alderton, Roberto Amato, Jeffrey Barrett, Sonia Goncalves, Ewan Harrison, David K. Jackson, Ian Johnston, Dominic Kwiatkowski, Cordelia Langford, John Sillitoe on behalf of the Wellcome Sanger Institute COVID-19 Surveillance Team |
| EPI_ISL_2809282 | Health Services Laboratories | Wellcome Sanger Institute for the COVID-19 Genomics UK (COG-UK) Consortium | Health Services Laboratories and Alex Alderton, Roberto Amato, Jeffrey Barrett, Sonia Goncalves, Ewan Harrison, David K. Jackson, Ian Johnston, Dominic Kwiatkowski, Cordelia Langford, John Sillitoe on behalf of the Wellcome Sanger Institute COVID-19 Surveillance Team |
| EPI_ISL_2809283 | Lighthouse Lab in Alderley Park | Wellcome Sanger Institute for the COVID-19 Genomics UK (COG-UK) Consortium | Jacquelyn Wynn, Mairead Hyland, The Lighthouse Lab in Alderley Park and Alex Alderton, Roberto Amato, Jeffrey Barrett, Sonia Goncalves, Ewan Harrison, David K. Jackson, Ian Johnston, Dominic Kwiatkowski, Cordelia Langford, John Sillitoe on behalf of the Wellcome Sanger Institute COVID-19 Surveillance Team |
| EPI_ISL_2809284, EPI_ISL_2809285 | Lighthouse Lab in Milton Keynes | Wellcome Sanger Institute for the COVID-19 Genomics UK (COG-UK) Consortium | The Lighthouse Lab in Milton Keynes and Alex Alderton, Roberto Amato, Jeffrey Barrett, Sonia Goncalves, Ewan Harrison, David K. Jackson, Ian Johnston, Dominic Kwiatkowski, Cordelia Langford, John Sillitoe on behalf of the Wellcome Sanger Institute COVID-19 Surveillance Team |
| EPI_ISL_2809286 | Health Services Laboratories | Wellcome Sanger Institute for the COVID-19 Genomics UK (COG-UK) Consortium | Health Services Laboratories and Alex Alderton, Roberto Amato, Jeffrey Barrett, Sonia Goncalves, Ewan Harrison, David K. Jackson, Ian Johnston, Dominic Kwiatkowski, Cordelia Langford, John Sillitoe on behalf of the Wellcome Sanger Institute COVID-19 Surveillance Team |
| EPI_ISL_2809287 | Lighthouse Lab in Glasgow | Wellcome Sanger Institute for the COVID-19 Genomics UK (COG-UK) Consortium | Harper VanSteenhouse, Yumi Kasai, David Gray, Carol Clugston, Anna Dominiczak and Alex Alderton, Roberto Amato, Jeffrey Barrett, Sonia Goncalves, Ewan Harrison, David K. Jackson, Ian Johnston, Dominic Kwiatkowski, Cordelia Langford, John Sillitoe on behalf of the Wellcome Sanger Institute COVID-19 Surveillance Team |
| EPI_ISL_2809288 | Lighthouse Lab in Milton Keynes | Wellcome Sanger Institute for the COVID-19 Genomics UK (COG-UK) Consortium | The Lighthouse Lab in Milton Keynes and Alex Alderton, Roberto Amato, Jeffrey Barrett, Sonia Goncalves, Ewan Harrison, David K. Jackson, Ian Johnston, Dominic Kwiatkowski, Cordelia Langford, John Sillitoe on behalf of the Wellcome Sanger Institute COVID-19 Surveillance Team |
| EPI_ISL_2809289 | Lighthouse Lab in Alderley Park | Wellcome Sanger Institute for the COVID-19 Genomics UK (COG-UK) Consortium | Jacquelyn Wynn, Mairead Hyland, The Lighthouse Lab in Alderley Park and Alex Alderton, Roberto Amato, Jeffrey Barrett, Sonia Goncalves, Ewan Harrison, David K. Jackson, Ian Johnston, Dominic Kwiatkowski, Cordelia Langford, John Sillitoe on behalf of the Wellcome Sanger Institute COVID-19 Surveillance Team |
| EPI_ISL_2809290 | Health Services Laboratories | Wellcome Sanger Institute for the COVID-19 Genomics UK (COG-UK) Consortium | Health Services Laboratories and Alex Alderton, Roberto Amato, Jeffrey Barrett, Sonia Goncalves, Ewan Harrison, David K. Jackson, Ian Johnston, Dominic Kwiatkowski, Cordelia Langford, John Sillitoe on behalf of the Wellcome Sanger Institute COVID-19 Surveillance Team |
| EPI_ISL_2809291 | Lighthouse Lab in Alderley Park | Wellcome Sanger Institute for the COVID-19 Genomics UK (COG-UK) Consortium | Jacquelyn Wynn, Mairead Hyland, The Lighthouse Lab in Alderley Park and Alex Alderton, Roberto Amato, Jeffrey Barrett, Sonia Goncalves, Ewan Harrison, David K. Jackson, Ian Johnston, Dominic Kwiatkowski, Cordelia Langford, John Sillitoe on behalf of the Wellcome Sanger Institute COVID-19 Surveillance Team |
| EPI_ISL_2809292 | Lighthouse Lab in Milton Keynes | Wellcome Sanger Institute for the COVID-19 Genomics UK (COG-UK) Consortium | The Lighthouse Lab in Milton Keynes and Alex Alderton, Roberto Amato, Jeffrey Barrett, Sonia Goncalves, Ewan Harrison, David K. Jackson, Ian Johnston, Dominic Kwiatkowski, Cordelia Langford, John Sillitoe on behalf of the Wellcome Sanger Institute COVID-19 Surveillance Team |
| EPI_ISL_2809293 | Lighthouse Lab in Alderley Park | Wellcome Sanger Institute for the COVID-19 Genomics UK (COG-UK) Consortium | Jacquelyn Wynn, Mairead Hyland, The Lighthouse Lab in Alderley Park and Alex Alderton, Roberto Amato, Jeffrey Barrett, Sonia Goncalves, Ewan Harrison, David K. Jackson, Ian Johnston, Dominic Kwiatkowski, Cordelia Langford, John Sillitoe on behalf of the Wellcome Sanger Institute COVID-19 Surveillance Team |
| EPI_ISL_2809294, EPI_ISL_2809295, EPI_ISL_2809296, EPI_ISL_2809297 | Health Services Laboratories | Wellcome Sanger Institute for the COVID-19 Genomics UK (COG-UK) Consortium | Health Services Laboratories and Alex Alderton, Roberto Amato, Jeffrey Barrett, Sonia Goncalves, Ewan Harrison, David K. Jackson, Ian Johnston, Dominic Kwiatkowski, Cordelia Langford, John Sillitoe on behalf of the Wellcome Sanger Institute COVID-19 Surveillance Team |
| EPI_ISL_2809298, EPI_ISL_2809299 | Lighthouse Lab in Alderley Park | Wellcome Sanger Institute for the COVID-19 Genomics UK (COG-UK) Consortium | Jacquelyn Wynn, Mairead Hyland, The Lighthouse Lab in Alderley Park and Alex Alderton, Roberto Amato, Jeffrey Barrett, Sonia Goncalves, Ewan Harrison, David K. Jackson, Ian Johnston, Dominic Kwiatkowski, Cordelia Langford, John Sillitoe on behalf of the Wellcome Sanger Institute COVID-19 Surveillance Team |

| | | | |
|---|---|---|---|
| EPI_ISL_2809300 | Lighthouse Lab in Glasgow | Wellcome Sanger Institute for the COVID-19 Genomics UK (COG-UK) Consortium | Harper VanSteenhouse, Yumi Kasai, David Gray, Carol Clugston, Anna Dominiczak and Alex Alderton, Roberto Amato, Jeffrey Barrett, Sonia Goncalves, Ewan Harrison, David K. Jackson, Ian Johnston, Dominic Kwiatkowski, Cordelia Langford, John Sillitoe on behalf of the Wellcome Sanger Institute COVID-19 Surveillance Team |
| EPI_ISL_2809301, EPI_ISL_2809302, EPI_ISL_2809303 | Health Services Laboratories | Wellcome Sanger Institute for the COVID-19 Genomics UK (COG-UK) Consortium | Health Services Laboratories and Alex Alderton, Roberto Amato, Jeffrey Barrett, Sonia Goncalves, Ewan Harrison, David K. Jackson, Ian Johnston, Dominic Kwiatkowski, Cordelia Langford, John Sillitoe on behalf of the Wellcome Sanger Institute COVID-19 Surveillance Team |
| EPI_ISL_2809304 | Lighthouse Lab in Alderley Park | Wellcome Sanger Institute for the COVID-19 Genomics UK (COG-UK) Consortium | Jacquelyn Wynn, Mairead Hyland, The Lighthouse Lab in Alderley Park and Alex Alderton, Roberto Amato, Jeffrey Barrett, Sonia Goncalves, Ewan Harrison, David K. Jackson, Ian Johnston, Dominic Kwiatkowski, Cordelia Langford, John Sillitoe on behalf of the Wellcome Sanger Institute COVID-19 Surveillance Team |
| EPI_ISL_2809305 | Health Services Laboratories | Wellcome Sanger Institute for the COVID-19 Genomics UK (COG-UK) Consortium | Health Services Laboratories and Alex Alderton, Roberto Amato, Jeffrey Barrett, Sonia Goncalves, Ewan Harrison, David K. Jackson, Ian Johnston, Dominic Kwiatkowski, Cordelia Langford, John Sillitoe on behalf of the Wellcome Sanger Institute COVID-19 Surveillance Team |
| EPI_ISL_2809306 | Lighthouse Lab in Alderley Park | Wellcome Sanger Institute for the COVID-19 Genomics UK (COG-UK) Consortium | Jacquelyn Wynn, Mairead Hyland, The Lighthouse Lab in Alderley Park and Alex Alderton, Roberto Amato, Jeffrey Barrett, Sonia Goncalves, Ewan Harrison, David K. Jackson, Ian Johnston, Dominic Kwiatkowski, Cordelia Langford, John Sillitoe on behalf of the Wellcome Sanger Institute COVID-19 Surveillance Team |
| EPI_ISL_2809307, EPI_ISL_2809308, EPI_ISL_2809309 | Lighthouse Lab in Milton Keynes | Wellcome Sanger Institute for the COVID-19 Genomics UK (COG-UK) Consortium | The Lighthouse Lab in Milton Keynes and Alex Alderton, Roberto Amato, Jeffrey Barrett, Sonia Goncalves, Ewan Harrison, David K. Jackson, Ian Johnston, Dominic Kwiatkowski, Cordelia Langford, John Sillitoe on behalf of the Wellcome Sanger Institute COVID-19 Surveillance Team |
| EPI_ISL_2809310 | Lighthouse Lab in Alderley Park | Wellcome Sanger Institute for the COVID-19 Genomics UK (COG-UK) Consortium | Jacquelyn Wynn, Mairead Hyland, The Lighthouse Lab in Alderley Park and Alex Alderton, Roberto Amato, Jeffrey Barrett, Sonia Goncalves, Ewan Harrison, David K. Jackson, Ian Johnston, Dominic Kwiatkowski, Cordelia Langford, John Sillitoe on behalf of the Wellcome Sanger Institute COVID-19 Surveillance Team |
| EPI_ISL_2809311 | Health Services Laboratories | Wellcome Sanger Institute for the COVID-19 Genomics UK (COG-UK) Consortium | Health Services Laboratories and Alex Alderton, Roberto Amato, Jeffrey Barrett, Sonia Goncalves, Ewan Harrison, David K. Jackson, Ian Johnston, Dominic Kwiatkowski, Cordelia Langford, John Sillitoe on behalf of the Wellcome Sanger Institute COVID-19 Surveillance Team |
| EPI_ISL_2809312 | Lighthouse Lab in Milton Keynes | Wellcome Sanger Institute for the COVID-19 Genomics UK (COG-UK) Consortium | The Lighthouse Lab in Milton Keynes and Alex Alderton, Roberto Amato, Jeffrey Barrett, Sonia Goncalves, Ewan Harrison, David K. Jackson, Ian Johnston, Dominic Kwiatkowski, Cordelia Langford, John Sillitoe on behalf of the Wellcome Sanger Institute COVID-19 Surveillance Team |
| EPI_ISL_2809313 | Lighthouse Lab in Alderley Park | Wellcome Sanger Institute for the COVID-19 Genomics UK (COG-UK) Consortium | Jacquelyn Wynn, Mairead Hyland, The Lighthouse Lab in Alderley Park and Alex Alderton, Roberto Amato, Jeffrey Barrett, Sonia Goncalves, Ewan Harrison, David K. Jackson, Ian Johnston, Dominic Kwiatkowski, Cordelia Langford, John Sillitoe on behalf of the Wellcome Sanger Institute COVID-19 Surveillance Team |
| EPI_ISL_2809314 | Health Services Laboratories | Wellcome Sanger Institute for the COVID-19 Genomics UK (COG-UK) Consortium | Health Services Laboratories and Alex Alderton, Roberto Amato, Jeffrey Barrett, Sonia Goncalves, Ewan Harrison, David K. Jackson, Ian Johnston, Dominic Kwiatkowski, Cordelia Langford, John Sillitoe on behalf of the Wellcome Sanger Institute COVID-19 Surveillance Team |
| EPI_ISL_2809315, EPI_ISL_2809316, EPI_ISL_2809317 | Lighthouse Lab in Alderley Park | Wellcome Sanger Institute for the COVID-19 Genomics UK (COG-UK) Consortium | Jacquelyn Wynn, Mairead Hyland, The Lighthouse Lab in Alderley Park and Alex Alderton, Roberto Amato, Jeffrey Barrett, Sonia Goncalves, Ewan Harrison, David K. Jackson, Ian Johnston, Dominic Kwiatkowski, Cordelia Langford, John Sillitoe on behalf of the Wellcome Sanger Institute COVID-19 Surveillance Team |
| EPI_ISL_2809318 | Health Services Laboratories | Wellcome Sanger Institute for the COVID-19 Genomics UK (COG-UK) Consortium | Health Services Laboratories and Alex Alderton, Roberto Amato, Jeffrey Barrett, Sonia Goncalves, Ewan Harrison, David K. Jackson, Ian Johnston, Dominic Kwiatkowski, Cordelia Langford, John Sillitoe on behalf of the Wellcome Sanger Institute COVID-19 Surveillance Team |
| EPI_ISL_2809319 | Lighthouse Lab in Alderley Park | Wellcome Sanger Institute for the COVID-19 Genomics UK (COG-UK) Consortium | Jacquelyn Wynn, Mairead Hyland, The Lighthouse Lab in Alderley Park and Alex Alderton, Roberto Amato, Jeffrey Barrett, Sonia Goncalves, Ewan Harrison, David K. Jackson, Ian Johnston, Dominic Kwiatkowski, Cordelia Langford, John Sillitoe on behalf of the Wellcome Sanger Institute COVID-19 Surveillance Team |
| EPI_ISL_2809320 | Lighthouse Lab in Milton Keynes | Wellcome Sanger Institute for the COVID-19 Genomics UK (COG-UK) Consortium | The Lighthouse Lab in Milton Keynes and Alex Alderton, Roberto Amato, Jeffrey Barrett, Sonia Goncalves, Ewan Harrison, David K. Jackson, Ian Johnston, Dominic Kwiatkowski, Cordelia Langford, John Sillitoe on behalf of the Wellcome Sanger Institute COVID-19 Surveillance Team |
| EPI_ISL_2809321, EPI_ISL_2809322 | Lighthouse Lab in Alderley Park | Wellcome Sanger Institute for the COVID-19 Genomics UK (COG-UK) Consortium | Jacquelyn Wynn, Mairead Hyland, The Lighthouse Lab in Alderley Park and Alex Alderton, Roberto Amato, Jeffrey Barrett, Sonia Goncalves, Ewan Harrison, David K. Jackson, Ian Johnston, Dominic Kwiatkowski, Cordelia Langford, John Sillitoe on behalf of the Wellcome Sanger Institute COVID-19 Surveillance Team |
| EPI_ISL_2809323 | Health Services Laboratories | Wellcome Sanger Institute for the COVID-19 Genomics UK (COG-UK) Consortium | Health Services Laboratories and Alex Alderton, Roberto Amato, Jeffrey Barrett, Sonia Goncalves, Ewan Harrison, David K. Jackson, Ian Johnston, Dominic Kwiatkowski, Cordelia Langford, John Sillitoe on behalf of the Wellcome Sanger Institute COVID-19 Surveillance Team |
| EPI_ISL_2809324 | Lighthouse Lab in Milton Keynes | Wellcome Sanger Institute for the COVID-19 Genomics UK (COG-UK) Consortium | The Lighthouse Lab in Milton Keynes and Alex Alderton, Roberto Amato, Jeffrey Barrett, Sonia Goncalves, Ewan Harrison, David K. Jackson, Ian Johnston, Dominic Kwiatkowski, Cordelia Langford, John Sillitoe on behalf of the Wellcome Sanger Institute COVID-19 Surveillance Team |
| EPI_ISL_2809325 | Lighthouse Lab in Alderley Park | Wellcome Sanger Institute for the COVID-19 Genomics UK (COG-UK) Consortium | Jacquelyn Wynn, Mairead Hyland, The Lighthouse Lab in Alderley Park and Alex Alderton, Roberto Amato, Jeffrey Barrett, Sonia Goncalves, Ewan Harrison, David K. Jackson, Ian Johnston, Dominic Kwiatkowski, Cordelia Langford, John Sillitoe on behalf of the Wellcome Sanger Institute COVID-19 Surveillance Team |
| EPI_ISL_2809326 | Lighthouse Lab in Glasgow | Wellcome Sanger Institute for the COVID-19 Genomics UK (COG-UK) Consortium | Harper VanSteenhouse, Yumi Kasai, David Gray, Carol Clugston, Anna Dominiczak and Alex Alderton, Roberto Amato, Jeffrey Barrett, Sonia Goncalves, Ewan Harrison, David K. Jackson, Ian Johnston, Dominic Kwiatkowski, Cordelia Langford, John Sillitoe on behalf of the Wellcome Sanger Institute COVID-19 Surveillance Team |
| EPI_ISL_2809327, EPI_ISL_2809328 | Health Services Laboratories | Wellcome Sanger Institute for the COVID-19 Genomics UK (COG-UK) Consortium | Health Services Laboratories and Alex Alderton, Roberto Amato, Jeffrey Barrett, Sonia Goncalves, Ewan Harrison, David K. Jackson, Ian Johnston, Dominic Kwiatkowski, Cordelia Langford, John Sillitoe on behalf of the Wellcome Sanger Institute COVID-19 Surveillance Team |
| EPI_ISL_2809329 | Lighthouse Lab in Alderley Park | Wellcome Sanger Institute for the COVID-19 Genomics UK (COG-UK) Consortium | Jacquelyn Wynn, Mairead Hyland, The Lighthouse Lab in Alderley Park and Alex Alderton, Roberto Amato, Jeffrey Barrett, Sonia Goncalves, Ewan Harrison, David K. Jackson, Ian Johnston, Dominic Kwiatkowski, Cordelia Langford, John Sillitoe on behalf of the Wellcome Sanger Institute COVID-19 Surveillance Team |
| EPI_ISL_2809330 | Lighthouse Lab in Milton Keynes | Wellcome Sanger Institute for the COVID-19 Genomics UK (COG-UK) Consortium | The Lighthouse Lab in Milton Keynes and Alex Alderton, Roberto Amato, Jeffrey Barrett, Sonia Goncalves, Ewan Harrison, David K. Jackson, Ian Johnston, Dominic Kwiatkowski, Cordelia Langford, John Sillitoe on behalf of the Wellcome Sanger Institute COVID-19 Surveillance Team |
| EPI_ISL_2809331 | Health Services Laboratories | Wellcome Sanger Institute for the COVID-19 Genomics UK (COG-UK) Consortium | Health Services Laboratories and Alex Alderton, Roberto Amato, Jeffrey Barrett, Sonia Goncalves, Ewan Harrison, David K. Jackson, Ian Johnston, Dominic Kwiatkowski, Cordelia Langford, John Sillitoe on behalf of the Wellcome Sanger Institute COVID-19 Surveillance Team |
| EPI_ISL_2809332 | Lighthouse Lab in Milton Keynes | Wellcome Sanger Institute for the COVID-19 Genomics UK (COG-UK) Consortium | The Lighthouse Lab in Milton Keynes and Alex Alderton, Roberto Amato, Jeffrey Barrett, Sonia Goncalves, Ewan Harrison, David K. Jackson, Ian Johnston, Dominic Kwiatkowski, Cordelia Langford, John Sillitoe on behalf of the Wellcome Sanger Institute COVID-19 Surveillance Team |
| EPI_ISL_2809333 | Lighthouse Lab in Alderley Park | Wellcome Sanger Institute for the COVID-19 Genomics UK (COG-UK) Consortium | Jacquelyn Wynn, Mairead Hyland, The Lighthouse Lab in Alderley Park and Alex Alderton, Roberto Amato, Jeffrey Barrett, Sonia Goncalves, Ewan Harrison, David K. Jackson, Ian Johnston, Dominic Kwiatkowski, Cordelia Langford, John Sillitoe on behalf of the Wellcome Sanger Institute COVID-19 Surveillance Team |
| EPI_ISL_2809334 | Health Services Laboratories | Wellcome Sanger Institute for the COVID-19 Genomics UK (COG-UK) Consortium | Health Services Laboratories and Alex Alderton, Roberto Amato, Jeffrey Barrett, Sonia Goncalves, Ewan Harrison, David K. Jackson, Ian Johnston, Dominic Kwiatkowski, Cordelia Langford, John Sillitoe on behalf of the Wellcome Sanger Institute COVID-19 Surveillance Team |
| EPI_ISL_2809335 | Lighthouse Lab in Milton Keynes | Wellcome Sanger Institute for the COVID-19 Genomics UK (COG-UK) Consortium | The Lighthouse Lab in Milton Keynes and Alex Alderton, Roberto Amato, Jeffrey Barrett, Sonia Goncalves, Ewan Harrison, David K. Jackson, Ian Johnston, Dominic Kwiatkowski, Cordelia Langford, John Sillitoe on behalf of the Wellcome Sanger Institute COVID-19 Surveillance Team |
| EPI_ISL_2809336 | Health Services Laboratories | Wellcome Sanger Institute for the COVID-19 Genomics UK (COG-UK) Consortium | Health Services Laboratories and Alex Alderton, Roberto Amato, Jeffrey Barrett, Sonia Goncalves, Ewan Harrison, David K. Jackson, Ian Johnston, Dominic Kwiatkowski, Cordelia Langford, John Sillitoe on behalf of the Wellcome Sanger Institute COVID-19 Surveillance Team |
| EPI_ISL_2809337 | Lighthouse Lab in Alderley Park | Wellcome Sanger Institute for the COVID-19 Genomics UK (COG-UK) Consortium | Jacquelyn Wynn, Mairead Hyland, The Lighthouse Lab in Alderley Park and Alex Alderton, Roberto Amato, Jeffrey Barrett, Sonia Goncalves, Ewan Harrison, David K. Jackson, Ian Johnston, Dominic Kwiatkowski, Cordelia Langford, John Sillitoe on behalf of the Wellcome Sanger Institute COVID-19 Surveillance Team |
| EPI_ISL_2809338 | Lighthouse Lab in Milton Keynes | Wellcome Sanger Institute for the COVID-19 Genomics UK (COG-UK) Consortium | The Lighthouse Lab in Milton Keynes and Alex Alderton, Roberto Amato, Jeffrey Barrett, Sonia Goncalves, Ewan Harrison, David K. Jackson, Ian Johnston, Dominic Kwiatkowski, Cordelia Langford, John Sillitoe on behalf of the Wellcome Sanger Institute COVID-19 Surveillance Team |
| EPI_ISL_2809339, EPI_ISL_2809341 | Health Services Laboratories | Wellcome Sanger Institute for the COVID-19 Genomics UK (COG-UK) Consortium | Health Services Laboratories and Alex Alderton, Roberto Amato, Jeffrey Barrett, Sonia Goncalves, Ewan Harrison, David K. Jackson, Ian Johnston, Dominic Kwiatkowski, Cordelia Langford, John Sillitoe on behalf of the Wellcome Sanger Institute COVID-19 Surveillance Team |
| EPI_ISL_2809342 | Lighthouse Lab in Glasgow | Wellcome Sanger Institute for the COVID-19 Genomics UK (COG-UK) Consortium | Harper VanSteenhouse, Yumi Kasai, David Gray, Carol Clugston, Anna Dominiczak and Alex Alderton, Roberto Amato, Jeffrey Barrett, Sonia Goncalves, Ewan Harrison, David K. Jackson, Ian Johnston, Dominic Kwiatkowski, Cordelia Langford, John Sillitoe on behalf of the Wellcome Sanger Institute COVID-19 Surveillance Team |
| EPI_ISL_2809343 | Lighthouse Lab in Alderley Park | Wellcome Sanger Institute for the COVID-19 Genomics UK (COG-UK) Consortium | Jacquelyn Wynn, Mairead Hyland, The Lighthouse Lab in Alderley Park and Alex Alderton, Roberto Amato, Jeffrey Barrett, Sonia Goncalves, Ewan Harrison, David K. Jackson, Ian Johnston, Dominic Kwiatkowski, Cordelia Langford, John Sillitoe on behalf of the Wellcome Sanger Institute COVID-19 Surveillance Team |
| EPI_ISL_2809344, EPI_ISL_2809345, EPI_ISL_2809346, EPI_ISL_2809347 | Health Services Laboratories | Wellcome Sanger Institute for the COVID-19 Genomics UK (COG-UK) Consortium | Health Services Laboratories and Alex Alderton, Roberto Amato, Jeffrey Barrett, Sonia Goncalves, Ewan Harrison, David K. Jackson, Ian Johnston, Dominic Kwiatkowski, Cordelia Langford, John Sillitoe on behalf of the Wellcome Sanger Institute COVID-19 Surveillance Team |
| EPI_ISL_2809348 | Lighthouse Lab in Milton Keynes | Wellcome Sanger Institute for the COVID-19 Genomics UK (COG-UK) Consortium | The Lighthouse Lab in Milton Keynes and Alex Alderton, Roberto Amato, Jeffrey Barrett, Sonia Goncalves, Ewan Harrison, David K. Jackson, Ian Johnston, Dominic Kwiatkowski, Cordelia Langford, John Sillitoe on behalf of the Wellcome Sanger Institute COVID-19 Surveillance Team |
| EPI_ISL_2809349 | Lighthouse Lab in Alderley Park | Wellcome Sanger Institute for the COVID-19 Genomics UK (COG-UK) Consortium | Jacquelyn Wynn, Mairead Hyland, The Lighthouse Lab in Alderley Park and Alex Alderton, Roberto Amato, Jeffrey Barrett, Sonia Goncalves, Ewan Harrison, David K. Jackson, Ian Johnston, Dominic Kwiatkowski, Cordelia Langford, John Sillitoe on behalf of the Wellcome Sanger Institute COVID-19 Surveillance Team |
| EPI_ISL_2809350 | Health Services Laboratories | Wellcome Sanger Institute for the COVID-19 Genomics UK (COG-UK) Consortium | Health Services Laboratories and Alex Alderton, Roberto Amato, Jeffrey Barrett, Sonia Goncalves, Ewan Harrison, David K. Jackson, Ian Johnston, Dominic Kwiatkowski, Cordelia Langford, John Sillitoe on behalf of the Wellcome Sanger Institute COVID-19 Surveillance Team |
| EPI_ISL_2809351 | Lighthouse Lab in Milton Keynes | Wellcome Sanger Institute for the COVID-19 Genomics UK (COG-UK) Consortium | The Lighthouse Lab in Milton Keynes and Alex Alderton, Roberto Amato, Jeffrey Barrett, Sonia Goncalves, Ewan Harrison, David K. Jackson, Ian Johnston, Dominic Kwiatkowski, Cordelia Langford, John Sillitoe on behalf of the Wellcome Sanger Institute COVID-19 Surveillance Team |
| EPI_ISL_2809352, EPI_ISL_2809353 | Lighthouse Lab in Alderley Park | Wellcome Sanger Institute for the COVID-19 Genomics UK (COG-UK) Consortium | Jacquelyn Wynn, Mairead Hyland, The Lighthouse Lab in Alderley Park and Alex Alderton, Roberto Amato, Jeffrey Barrett, Sonia Goncalves, Ewan Harrison, David K. Jackson, Ian Johnston, Dominic Kwiatkowski, Cordelia Langford, John Sillitoe on behalf of the Wellcome Sanger Institute COVID-19 Surveillance Team |
| EPI_ISL_2809354 | Health Services Laboratories | Wellcome Sanger Institute for the COVID-19 Genomics UK (COG-UK) Consortium | Health Services Laboratories and Alex Alderton, Roberto Amato, Jeffrey Barrett, Sonia Goncalves, Ewan Harrison, David K. Jackson, Ian Johnston, Dominic Kwiatkowski, Cordelia Langford, John Sillitoe on behalf of the Wellcome Sanger Institute COVID-19 Surveillance Team |
| EPI_ISL_2809355 | Lighthouse Lab in Milton Keynes | Wellcome Sanger Institute for the COVID-19 Genomics UK (COG-UK) Consortium | The Lighthouse Lab in Milton Keynes and Alex Alderton, Roberto Amato, Jeffrey Barrett, Sonia Goncalves, Ewan Harrison, David K. Jackson, Ian Johnston, Dominic Kwiatkowski, Cordelia Langford, John Sillitoe on behalf of the Wellcome Sanger Institute COVID-19 Surveillance Team |
| EPI_ISL_2809356 | Health Services Laboratories | Wellcome Sanger Institute for the COVID-19 Genomics UK (COG-UK) Consortium | Health Services Laboratories and Alex Alderton, Roberto Amato, Jeffrey Barrett, Sonia Goncalves, Ewan Harrison, David K. Jackson, Ian Johnston, Dominic Kwiatkowski, Cordelia Langford, John Sillitoe on behalf of the Wellcome Sanger Institute COVID-19 Surveillance Team |

| | | | |
|---|---|---|---|
| EPI_ISL_2809357 | Lighthouse Lab in Alderley Park | Wellcome Sanger Institute for the COVID-19 Genomics UK (COG-UK) Consortium | Jacquelyn Wynn, Mairead Hyland, The Lighthouse Lab in Alderley Park and Alex Alderton, Roberto Amato, Jeffrey Barrett, Sonia Goncalves, Ewan Harrison, David K. Jackson, Ian Johnston, Dominic Kwiatkowski, Cordelia Langford, John Sillitoe on behalf of the Wellcome Sanger Institute COVID-19 Surveillance Team |
| EPI_ISL_2809358 | Lighthouse Lab in Milton Keynes | Wellcome Sanger Institute for the COVID-19 Genomics UK (COG-UK) Consortium | The Lighthouse Lab in Milton Keynes and Alex Alderton, Roberto Amato, Jeffrey Barrett, Sonia Goncalves, Ewan Harrison, David K. Jackson, Ian Johnston, Dominic Kwiatkowski, Cordelia Langford, John Sillitoe on behalf of the Wellcome Sanger Institute COVID-19 Surveillance Team |
| EPI_ISL_2809359 | Lighthouse Lab in Alderley Park | Wellcome Sanger Institute for the COVID-19 Genomics UK (COG-UK) Consortium | Jacquelyn Wynn, Mairead Hyland, The Lighthouse Lab in Alderley Park and Alex Alderton, Roberto Amato, Jeffrey Barrett, Sonia Goncalves, Ewan Harrison, David K. Jackson, Ian Johnston, Dominic Kwiatkowski, Cordelia Langford, John Sillitoe on behalf of the Wellcome Sanger Institute COVID-19 Surveillance Team |
| EPI_ISL_2809360, EPI_ISL_2809361 | Lighthouse Lab in Milton Keynes | Wellcome Sanger Institute for the COVID-19 Genomics UK (COG-UK) Consortium | The Lighthouse Lab in Milton Keynes and Alex Alderton, Roberto Amato, Jeffrey Barrett, Sonia Goncalves, Ewan Harrison, David K. Jackson, Ian Johnston, Dominic Kwiatkowski, Cordelia Langford, John Sillitoe on behalf of the Wellcome Sanger Institute COVID-19 Surveillance Team |
| EPI_ISL_2809362 | Health Services Laboratories | Wellcome Sanger Institute for the COVID-19 Genomics UK (COG-UK) Consortium | Health Services Laboratories and Alex Alderton, Roberto Amato, Jeffrey Barrett, Sonia Goncalves, Ewan Harrison, David K. Jackson, Ian Johnston, Dominic Kwiatkowski, Cordelia Langford, John Sillitoe on behalf of the Wellcome Sanger Institute COVID-19 Surveillance Team |
| EPI_ISL_2809363, EPI_ISL_2809364 | Lighthouse Lab in Alderley Park | Wellcome Sanger Institute for the COVID-19 Genomics UK (COG-UK) Consortium | Jacquelyn Wynn, Mairead Hyland, The Lighthouse Lab in Alderley Park and Alex Alderton, Roberto Amato, Jeffrey Barrett, Sonia Goncalves, Ewan Harrison, David K. Jackson, Ian Johnston, Dominic Kwiatkowski, Cordelia Langford, John Sillitoe on behalf of the Wellcome Sanger Institute COVID-19 Surveillance Team |
| EPI_ISL_2809365 | Health Services Laboratories | Wellcome Sanger Institute for the COVID-19 Genomics UK (COG-UK) Consortium | Health Services Laboratories and Alex Alderton, Roberto Amato, Jeffrey Barrett, Sonia Goncalves, Ewan Harrison, David K. Jackson, Ian Johnston, Dominic Kwiatkowski, Cordelia Langford, John Sillitoe on behalf of the Wellcome Sanger Institute COVID-19 Surveillance Team |
| EPI_ISL_2809366 | Lighthouse Lab in Alderley Park | Wellcome Sanger Institute for the COVID-19 Genomics UK (COG-UK) Consortium | Jacquelyn Wynn, Mairead Hyland, The Lighthouse Lab in Alderley Park and Alex Alderton, Roberto Amato, Jeffrey Barrett, Sonia Goncalves, Ewan Harrison, David K. Jackson, Ian Johnston, Dominic Kwiatkowski, Cordelia Langford, John Sillitoe on behalf of the Wellcome Sanger Institute COVID-19 Surveillance Team |
| EPI_ISL_2809367, EPI_ISL_2809368 | Lighthouse Lab in Milton Keynes | Wellcome Sanger Institute for the COVID-19 Genomics UK (COG-UK) Consortium | The Lighthouse Lab in Milton Keynes and Alex Alderton, Roberto Amato, Jeffrey Barrett, Sonia Goncalves, Ewan Harrison, David K. Jackson, Ian Johnston, Dominic Kwiatkowski, Cordelia Langford, John Sillitoe on behalf of the Wellcome Sanger Institute COVID-19 Surveillance Team |
| EPI_ISL_2809369, EPI_ISL_2809370 | Lighthouse Lab in Alderley Park | Wellcome Sanger Institute for the COVID-19 Genomics UK (COG-UK) Consortium | Jacquelyn Wynn, Mairead Hyland, The Lighthouse Lab in Alderley Park and Alex Alderton, Roberto Amato, Jeffrey Barrett, Sonia Goncalves, Ewan Harrison, David K. Jackson, Ian Johnston, Dominic Kwiatkowski, Cordelia Langford, John Sillitoe on behalf of the Wellcome Sanger Institute COVID-19 Surveillance Team |
| EPI_ISL_2809371 | Lighthouse Lab in Milton Keynes | Wellcome Sanger Institute for the COVID-19 Genomics UK (COG-UK) Consortium | The Lighthouse Lab in Milton Keynes and Alex Alderton, Roberto Amato, Jeffrey Barrett, Sonia Goncalves, Ewan Harrison, David K. Jackson, Ian Johnston, Dominic Kwiatkowski, Cordelia Langford, John Sillitoe on behalf of the Wellcome Sanger Institute COVID-19 Surveillance Team |
| EPI_ISL_2809372 | Health Services Laboratories | Wellcome Sanger Institute for the COVID-19 Genomics UK (COG-UK) Consortium | Health Services Laboratories and Alex Alderton, Roberto Amato, Jeffrey Barrett, Sonia Goncalves, Ewan Harrison, David K. Jackson, Ian Johnston, Dominic Kwiatkowski, Cordelia Langford, John Sillitoe on behalf of the Wellcome Sanger Institute COVID-19 Surveillance Team |
| EPI_ISL_2809373 | Lighthouse Lab in Alderley Park | Wellcome Sanger Institute for the COVID-19 Genomics UK (COG-UK) Consortium | Jacquelyn Wynn, Mairead Hyland, The Lighthouse Lab in Alderley Park and Alex Alderton, Roberto Amato, Jeffrey Barrett, Sonia Goncalves, Ewan Harrison, David K. Jackson, Ian Johnston, Dominic Kwiatkowski, Cordelia Langford, John Sillitoe on behalf of the Wellcome Sanger Institute COVID-19 Surveillance Team |
| EPI_ISL_2809374 | Health Services Laboratories | Wellcome Sanger Institute for the COVID-19 Genomics UK (COG-UK) Consortium | Health Services Laboratories and Alex Alderton, Roberto Amato, Jeffrey Barrett, Sonia Goncalves, Ewan Harrison, David K. Jackson, Ian Johnston, Dominic Kwiatkowski, Cordelia Langford, John Sillitoe on behalf of the Wellcome Sanger Institute COVID-19 Surveillance Team |
| EPI_ISL_2809375 | Lighthouse Lab in Alderley Park | Wellcome Sanger Institute for the COVID-19 Genomics UK (COG-UK) Consortium | Jacquelyn Wynn, Mairead Hyland, The Lighthouse Lab in Alderley Park and Alex Alderton, Roberto Amato, Jeffrey Barrett, Sonia Goncalves, Ewan Harrison, David K. Jackson, Ian Johnston, Dominic Kwiatkowski, Cordelia Langford, John Sillitoe on behalf of the Wellcome Sanger Institute COVID-19 Surveillance Team |
| EPI_ISL_2809376, EPI_ISL_2809377 | Lighthouse Lab in Milton Keynes | Wellcome Sanger Institute for the COVID-19 Genomics UK (COG-UK) Consortium | The Lighthouse Lab in Milton Keynes and Alex Alderton, Roberto Amato, Jeffrey Barrett, Sonia Goncalves, Ewan Harrison, David K. Jackson, Ian Johnston, Dominic Kwiatkowski, Cordelia Langford, John Sillitoe on behalf of the Wellcome Sanger Institute COVID-19 Surveillance Team |
| EPI_ISL_2809378 | Lighthouse Lab in Alderley Park | Wellcome Sanger Institute for the COVID-19 Genomics UK (COG-UK) Consortium | Jacquelyn Wynn, Mairead Hyland, The Lighthouse Lab in Alderley Park and Alex Alderton, Roberto Amato, Jeffrey Barrett, Sonia Goncalves, Ewan Harrison, David K. Jackson, Ian Johnston, Dominic Kwiatkowski, Cordelia Langford, John Sillitoe on behalf of the Wellcome Sanger Institute COVID-19 Surveillance Team |
| EPI_ISL_2809379 | Lighthouse Lab in Milton Keynes | Wellcome Sanger Institute for the COVID-19 Genomics UK (COG-UK) Consortium | The Lighthouse Lab in Milton Keynes and Alex Alderton, Roberto Amato, Jeffrey Barrett, Sonia Goncalves, Ewan Harrison, David K. Jackson, Ian Johnston, Dominic Kwiatkowski, Cordelia Langford, John Sillitoe on behalf of the Wellcome Sanger Institute COVID-19 Surveillance Team |
| EPI_ISL_2809380 | Lighthouse Lab in Alderley Park | Wellcome Sanger Institute for the COVID-19 Genomics UK (COG-UK) Consortium | Jacquelyn Wynn, Mairead Hyland, The Lighthouse Lab in Alderley Park and Alex Alderton, Roberto Amato, Jeffrey Barrett, Sonia Goncalves, Ewan Harrison, David K. Jackson, Ian Johnston, Dominic Kwiatkowski, Cordelia Langford, John Sillitoe on behalf of the Wellcome Sanger Institute COVID-19 Surveillance Team |
| EPI_ISL_2809381, EPI_ISL_2809382 | Lighthouse Lab in Milton Keynes | Wellcome Sanger Institute for the COVID-19 Genomics UK (COG-UK) Consortium | The Lighthouse Lab in Milton Keynes and Alex Alderton, Roberto Amato, Jeffrey Barrett, Sonia Goncalves, Ewan Harrison, David K. Jackson, Ian Johnston, Dominic Kwiatkowski, Cordelia Langford, John Sillitoe on behalf of the Wellcome Sanger Institute COVID-19 Surveillance Team |
| EPI_ISL_2809383, EPI_ISL_2809384, EPI_ISL_2809385, EPI_ISL_2809386, EPI_ISL_2809387 | Lighthouse Lab in Alderley Park | Wellcome Sanger Institute for the COVID-19 Genomics UK (COG-UK) Consortium | Jacquelyn Wynn, Mairead Hyland, The Lighthouse Lab in Alderley Park and Alex Alderton, Roberto Amato, Jeffrey Barrett, Sonia Goncalves, Ewan Harrison, David K. Jackson, Ian Johnston, Dominic Kwiatkowski, Cordelia Langford, John Sillitoe on behalf of the Wellcome Sanger Institute COVID-19 Surveillance Team |
| EPI_ISL_2809388 | Health Services Laboratories | Wellcome Sanger Institute for the COVID-19 Genomics UK (COG-UK) Consortium | Health Services Laboratories and Alex Alderton, Roberto Amato, Jeffrey Barrett, Sonia Goncalves, Ewan Harrison, David K. Jackson, Ian Johnston, Dominic Kwiatkowski, Cordelia Langford, John Sillitoe on behalf of the Wellcome Sanger Institute COVID-19 Surveillance Team |
| EPI_ISL_2809389 | Lighthouse Lab in Alderley Park | Wellcome Sanger Institute for the COVID-19 Genomics UK (COG-UK) Consortium | Jacquelyn Wynn, Mairead Hyland, The Lighthouse Lab in Alderley Park and Alex Alderton, Roberto Amato, Jeffrey Barrett, Sonia Goncalves, Ewan Harrison, David K. Jackson, Ian Johnston, Dominic Kwiatkowski, Cordelia Langford, John Sillitoe on behalf of the Wellcome Sanger Institute COVID-19 Surveillance Team |
| EPI_ISL_2809390 | Health Services Laboratories | Wellcome Sanger Institute for the COVID-19 Genomics UK (COG-UK) Consortium | Health Services Laboratories and Alex Alderton, Roberto Amato, Jeffrey Barrett, Sonia Goncalves, Ewan Harrison, David K. Jackson, Ian Johnston, Dominic Kwiatkowski, Cordelia Langford, John Sillitoe on behalf of the Wellcome Sanger Institute COVID-19 Surveillance Team |
| EPI_ISL_2809391 | Lighthouse Lab in Alderley Park | Wellcome Sanger Institute for the COVID-19 Genomics UK (COG-UK) Consortium | Jacquelyn Wynn, Mairead Hyland, The Lighthouse Lab in Alderley Park and Alex Alderton, Roberto Amato, Jeffrey Barrett, Sonia Goncalves, Ewan Harrison, David K. Jackson, Ian Johnston, Dominic Kwiatkowski, Cordelia Langford, John Sillitoe on behalf of the Wellcome Sanger Institute COVID-19 Surveillance Team |
| EPI_ISL_2809392, EPI_ISL_2809393 | Health Services Laboratories | Wellcome Sanger Institute for the COVID-19 Genomics UK (COG-UK) Consortium | Health Services Laboratories and Alex Alderton, Roberto Amato, Jeffrey Barrett, Sonia Goncalves, Ewan Harrison, David K. Jackson, Ian Johnston, Dominic Kwiatkowski, Cordelia Langford, John Sillitoe on behalf of the Wellcome Sanger Institute COVID-19 Surveillance Team |
| EPI_ISL_2809394 | Lighthouse Lab in Alderley Park | Wellcome Sanger Institute for the COVID-19 Genomics UK (COG-UK) Consortium | Jacquelyn Wynn, Mairead Hyland, The Lighthouse Lab in Alderley Park and Alex Alderton, Roberto Amato, Jeffrey Barrett, Sonia Goncalves, Ewan Harrison, David K. Jackson, Ian Johnston, Dominic Kwiatkowski, Cordelia Langford, John Sillitoe on behalf of the Wellcome Sanger Institute COVID-19 Surveillance Team |
| EPI_ISL_2809395 | Health Services Laboratories | Wellcome Sanger Institute for the COVID-19 Genomics UK (COG-UK) Consortium | Health Services Laboratories and Alex Alderton, Roberto Amato, Jeffrey Barrett, Sonia Goncalves, Ewan Harrison, David K. Jackson, Ian Johnston, Dominic Kwiatkowski, Cordelia Langford, John Sillitoe on behalf of the Wellcome Sanger Institute COVID-19 Surveillance Team |
| EPI_ISL_2809396, EPI_ISL_2809397, EPI_ISL_2809398, EPI_ISL_2809399 | Lighthouse Lab in Alderley Park | Wellcome Sanger Institute for the COVID-19 Genomics UK (COG-UK) Consortium | Jacquelyn Wynn, Mairead Hyland, The Lighthouse Lab in Alderley Park and Alex Alderton, Roberto Amato, Jeffrey Barrett, Sonia Goncalves, Ewan Harrison, David K. Jackson, Ian Johnston, Dominic Kwiatkowski, Cordelia Langford, John Sillitoe on behalf of the Wellcome Sanger Institute COVID-19 Surveillance Team |
| EPI_ISL_2809400 | Lighthouse Lab in Milton Keynes | Wellcome Sanger Institute for the COVID-19 Genomics UK (COG-UK) Consortium | The Lighthouse Lab in Milton Keynes and Alex Alderton, Roberto Amato, Jeffrey Barrett, Sonia Goncalves, Ewan Harrison, David K. Jackson, Ian Johnston, Dominic Kwiatkowski, Cordelia Langford, John Sillitoe on behalf of the Wellcome Sanger Institute COVID-19 Surveillance Team |
| EPI_ISL_2809401, EPI_ISL_2809402 | Health Services Laboratories | Wellcome Sanger Institute for the COVID-19 Genomics UK (COG-UK) Consortium | Health Services Laboratories and Alex Alderton, Roberto Amato, Jeffrey Barrett, Sonia Goncalves, Ewan Harrison, David K. Jackson, Ian Johnston, Dominic Kwiatkowski, Cordelia Langford, John Sillitoe on behalf of the Wellcome Sanger Institute COVID-19 Surveillance Team |
| EPI_ISL_2809403 | Lighthouse Lab in Milton Keynes | Wellcome Sanger Institute for the COVID-19 Genomics UK (COG-UK) Consortium | The Lighthouse Lab in Milton Keynes and Alex Alderton, Roberto Amato, Jeffrey Barrett, Sonia Goncalves, Ewan Harrison, David K. Jackson, Ian Johnston, Dominic Kwiatkowski, Cordelia Langford, John Sillitoe on behalf of the Wellcome Sanger Institute COVID-19 Surveillance Team |
| EPI_ISL_2809404 | Lighthouse Lab in Alderley Park | Wellcome Sanger Institute for the COVID-19 Genomics UK (COG-UK) Consortium | Jacquelyn Wynn, Mairead Hyland, The Lighthouse Lab in Alderley Park and Alex Alderton, Roberto Amato, Jeffrey Barrett, Sonia Goncalves, Ewan Harrison, David K. Jackson, Ian Johnston, Dominic Kwiatkowski, Cordelia Langford, John Sillitoe on behalf of the Wellcome Sanger Institute COVID-19 Surveillance Team |
| EPI_ISL_2809405 | Lighthouse Lab in Milton Keynes | Wellcome Sanger Institute for the COVID-19 Genomics UK (COG-UK) Consortium | The Lighthouse Lab in Milton Keynes and Alex Alderton, Roberto Amato, Jeffrey Barrett, Sonia Goncalves, Ewan Harrison, David K. Jackson, Ian Johnston, Dominic Kwiatkowski, Cordelia Langford, John Sillitoe on behalf of the Wellcome Sanger Institute COVID-19 Surveillance Team |
| EPI_ISL_2809406 | Lighthouse Lab in Alderley Park | Wellcome Sanger Institute for the COVID-19 Genomics UK (COG-UK) Consortium | Jacquelyn Wynn, Mairead Hyland, The Lighthouse Lab in Alderley Park and Alex Alderton, Roberto Amato, Jeffrey Barrett, Sonia Goncalves, Ewan Harrison, David K. Jackson, Ian Johnston, Dominic Kwiatkowski, Cordelia Langford, John Sillitoe on behalf of the Wellcome Sanger Institute COVID-19 Surveillance Team |
| EPI_ISL_2809407, EPI_ISL_2809408 | Health Services Laboratories | Wellcome Sanger Institute for the COVID-19 Genomics UK (COG-UK) Consortium | Health Services Laboratories and Alex Alderton, Roberto Amato, Jeffrey Barrett, Sonia Goncalves, Ewan Harrison, David K. Jackson, Ian Johnston, Dominic Kwiatkowski, Cordelia Langford, John Sillitoe on behalf of the Wellcome Sanger Institute COVID-19 Surveillance Team |
| EPI_ISL_2809409 | Lighthouse Lab in Alderley Park | Wellcome Sanger Institute for the COVID-19 Genomics UK (COG-UK) Consortium | Jacquelyn Wynn, Mairead Hyland, The Lighthouse Lab in Alderley Park and Alex Alderton, Roberto Amato, Jeffrey Barrett, Sonia Goncalves, Ewan Harrison, David K. Jackson, Ian Johnston, Dominic Kwiatkowski, Cordelia Langford, John Sillitoe on behalf of the Wellcome Sanger Institute COVID-19 Surveillance Team |
| EPI_ISL_2809410 | Lighthouse Lab in Milton Keynes | Wellcome Sanger Institute for the COVID-19 Genomics UK (COG-UK) Consortium | The Lighthouse Lab in Milton Keynes and Alex Alderton, Roberto Amato, Jeffrey Barrett, Sonia Goncalves, Ewan Harrison, David K. Jackson, Ian Johnston, Dominic Kwiatkowski, Cordelia Langford, John Sillitoe on behalf of the Wellcome Sanger Institute COVID-19 Surveillance Team |
| EPI_ISL_2809411 | Lighthouse Lab in Alderley Park | Wellcome Sanger Institute for the COVID-19 Genomics UK (COG-UK) Consortium | Jacquelyn Wynn, Mairead Hyland, The Lighthouse Lab in Alderley Park and Alex Alderton, Roberto Amato, Jeffrey Barrett, Sonia Goncalves, Ewan Harrison, David K. Jackson, Ian Johnston, Dominic Kwiatkowski, Cordelia Langford, John Sillitoe on behalf of the Wellcome Sanger Institute COVID-19 Surveillance Team |
| EPI_ISL_2809412 | Lighthouse Lab in Milton Keynes | Wellcome Sanger Institute for the COVID-19 Genomics UK (COG-UK) Consortium | The Lighthouse Lab in Milton Keynes and Alex Alderton, Roberto Amato, Jeffrey Barrett, Sonia Goncalves, Ewan Harrison, David K. Jackson, Ian Johnston, Dominic Kwiatkowski, Cordelia Langford, John Sillitoe on behalf of the Wellcome Sanger Institute COVID-19 Surveillance Team |
| EPI_ISL_2809413 | Health Services Laboratories | Wellcome Sanger Institute for the COVID-19 Genomics UK (COG-UK) Consortium | Health Services Laboratories and Alex Alderton, Roberto Amato, Jeffrey Barrett, Sonia Goncalves, Ewan Harrison, David K. Jackson, Ian Johnston, Dominic Kwiatkowski, Cordelia Langford, John Sillitoe on behalf of the Wellcome Sanger Institute COVID-19 Surveillance Team |
| EPI_ISL_2809414 | Lighthouse Lab in Milton Keynes | Wellcome Sanger Institute for the COVID-19 Genomics UK (COG-UK) Consortium | The Lighthouse Lab in Milton Keynes and Alex Alderton, Roberto Amato, Jeffrey Barrett, Sonia Goncalves, Ewan Harrison, David K. Jackson, Ian Johnston, Dominic Kwiatkowski, Cordelia Langford, John Sillitoe on behalf of the Wellcome Sanger Institute COVID-19 Surveillance Team |
| EPI_ISL_2809415 | Lighthouse Lab in Alderley Park | Wellcome Sanger Institute for the COVID-19 Genomics UK (COG-UK) Consortium | Jacquelyn Wynn, Mairead Hyland, The Lighthouse Lab in Alderley Park and Alex Alderton, Roberto Amato, Jeffrey Barrett, Sonia Goncalves, Ewan Harrison, David K. Jackson, Ian Johnston, Dominic Kwiatkowski, Cordelia Langford, John Sillitoe on behalf of the Wellcome Sanger Institute COVID-19 Surveillance Team |

| | | | |
|---|---|---|---|
| EPI_ISL_2809416 | Lighthouse Lab in Milton Keynes | Wellcome Sanger Institute for the COVID-19 Genomics UK (COG-UK) Consortium | The Lighthouse Lab in Milton Keynes and Alex Alderton, Roberto Amato, Jeffrey Barrett, Sonia Goncalves, Ewan Harrison, David K. Jackson, Ian Johnston, Dominic Kwiatkowski, Cordelia Langford, John Sillitoe on behalf of the Wellcome Sanger Institute COVID-19 Surveillance Team |
| EPI_ISL_2809417 | Health Services Laboratories | Wellcome Sanger Institute for the COVID-19 Genomics UK (COG-UK) Consortium | Health Services Laboratories and Alex Alderton, Roberto Amato, Jeffrey Barrett, Sonia Goncalves, Ewan Harrison, David K. Jackson, Ian Johnston, Dominic Kwiatkowski, Cordelia Langford, John Sillitoe on behalf of the Wellcome Sanger Institute COVID-19 Surveillance Team |
| EPI_ISL_2809418, EPI_ISL_2809419 | Lighthouse Lab in Alderley Park | Wellcome Sanger Institute for the COVID-19 Genomics UK (COG-UK) Consortium | Jacquelyn Wynn, Mairead Hyland, The Lighthouse Lab in Alderley Park and Alex Alderton, Roberto Amato, Jeffrey Barrett, Sonia Goncalves, Ewan Harrison, David K. Jackson, Ian Johnston, Dominic Kwiatkowski, Cordelia Langford, John Sillitoe on behalf of the Wellcome Sanger Institute COVID-19 Surveillance Team |
| EPI_ISL_2809420 | Health Services Laboratories | Wellcome Sanger Institute for the COVID-19 Genomics UK (COG-UK) Consortium | Health Services Laboratories and Alex Alderton, Roberto Amato, Jeffrey Barrett, Sonia Goncalves, Ewan Harrison, David K. Jackson, Ian Johnston, Dominic Kwiatkowski, Cordelia Langford, John Sillitoe on behalf of the Wellcome Sanger Institute COVID-19 Surveillance Team |
| EPI_ISL_2809421 | Lighthouse Lab in Alderley Park | Wellcome Sanger Institute for the COVID-19 Genomics UK (COG-UK) Consortium | Jacquelyn Wynn, Mairead Hyland, The Lighthouse Lab in Alderley Park and Alex Alderton, Roberto Amato, Jeffrey Barrett, Sonia Goncalves, Ewan Harrison, David K. Jackson, Ian Johnston, Dominic Kwiatkowski, Cordelia Langford, John Sillitoe on behalf of the Wellcome Sanger Institute COVID-19 Surveillance Team |
| EPI_ISL_2809422, EPI_ISL_2809423, EPI_ISL_2809424, EPI_ISL_2809425, EPI_ISL_2809427, EPI_ISL_2809428, EPI_ISL_2809429, EPI_ISL_2809430, EPI_ISL_2809431, EPI_ISL_2809432, EPI_ISL_2809433, EPI_ISL_2809434 | | | |
| see above | Lighthouse Lab in Glasgow | Wellcome Sanger Institute for the COVID-19 Genomics UK (COG-UK) Consortium | Harper VanSteenhouse, Yumi Kasai, David Gray, Carol Clugston, Anna Dominiczak and Alex Alderton, Roberto Amato, Jeffrey Barrett, Sonia Goncalves, Ewan Harrison, David K. Jackson, Ian Johnston, Dominic Kwiatkowski, Cordelia Langford, John Sillitoe on behalf of the Wellcome Sanger Institute COVID-19 Surveillance Team |
| EPI_ISL_2809435 | Lighthouse Lab in Milton Keynes | Wellcome Sanger Institute for the COVID-19 Genomics UK (COG-UK) Consortium | The Lighthouse Lab in Milton Keynes and Alex Alderton, Roberto Amato, Jeffrey Barrett, Sonia Goncalves, Ewan Harrison, David K. Jackson, Ian Johnston, Dominic Kwiatkowski, Cordelia Langford, John Sillitoe on behalf of the Wellcome Sanger Institute COVID-19 Surveillance Team |
| EPI_ISL_2809436, EPI_ISL_2809437, EPI_ISL_2809438, EPI_ISL_2809439, EPI_ISL_2809440, EPI_ISL_2809441, EPI_ISL_2809442, EPI_ISL_2809443, EPI_ISL_2809444, EPI_ISL_2809445, EPI_ISL_2809446, EPI_ISL_2809447, EPI_ISL_2809448, EPI_ISL_2809449, EPI_ISL_2809450, EPI_ISL_2809451, EPI_ISL_2809452, EPI_ISL_2809453, EPI_ISL_2809454, EPI_ISL_2809455, EPI_ISL_2809456, EPI_ISL_2809457, EPI_ISL_2809458, EPI_ISL_2809459, EPI_ISL_2809460 | | | |
| see above | Lighthouse Lab in Glasgow | Wellcome Sanger Institute for the COVID-19 Genomics UK (COG-UK) Consortium | Harper VanSteenhouse, Yumi Kasai, David Gray, Carol Clugston, Anna Dominiczak and Alex Alderton, Roberto Amato, Jeffrey Barrett, Sonia Goncalves, Ewan Harrison, David K. Jackson, Ian Johnston, Dominic Kwiatkowski, Cordelia Langford, John Sillitoe on behalf of the Wellcome Sanger Institute COVID-19 Surveillance Team |
| EPI_ISL_2809461 | Lighthouse Lab in Alderley Park | Wellcome Sanger Institute for the COVID-19 Genomics UK (COG-UK) Consortium | Jacquelyn Wynn, Mairead Hyland, The Lighthouse Lab in Alderley Park and Alex Alderton, Roberto Amato, Jeffrey Barrett, Sonia Goncalves, Ewan Harrison, David K. Jackson, Ian Johnston, Dominic Kwiatkowski, Cordelia Langford, John Sillitoe on behalf of the Wellcome Sanger Institute COVID-19 Surveillance Team |
| EPI_ISL_2809462, EPI_ISL_2809463, EPI_ISL_2809464, EPI_ISL_2809465, EPI_ISL_2809466, EPI_ISL_2809467, EPI_ISL_2809468, EPI_ISL_2809469, EPI_ISL_2809470, EPI_ISL_2809471, EPI_ISL_2809472, EPI_ISL_2809473, EPI_ISL_2809474, EPI_ISL_2809475, EPI_ISL_2809476, EPI_ISL_2809477, EPI_ISL_2809479, EPI_ISL_2809480, EPI_ISL_2809481, EPI_ISL_2809482, EPI_ISL_2809483, EPI_ISL_2809484, EPI_ISL_2809485, EPI_ISL_2809486, EPI_ISL_2809487 | | | |
| see above | Lighthouse Lab in Glasgow | Wellcome Sanger Institute for the COVID-19 Genomics UK (COG-UK) Consortium | Harper VanSteenhouse, Yumi Kasai, David Gray, Carol Clugston, Anna Dominiczak and Alex Alderton, Roberto Amato, Jeffrey Barrett, Sonia Goncalves, Ewan Harrison, David K. Jackson, Ian Johnston, Dominic Kwiatkowski, Cordelia Langford, John Sillitoe on behalf of the Wellcome Sanger Institute COVID-19 Surveillance Team |
| EPI_ISL_2809488 | Lighthouse Lab in Alderley Park | Wellcome Sanger Institute for the COVID-19 Genomics UK (COG-UK) Consortium | Jacquelyn Wynn, Mairead Hyland, The Lighthouse Lab in Alderley Park and Alex Alderton, Roberto Amato, Jeffrey Barrett, Sonia Goncalves, Ewan Harrison, David K. Jackson, Ian Johnston, Dominic Kwiatkowski, Cordelia Langford, John Sillitoe on behalf of the Wellcome Sanger Institute COVID-19 Surveillance Team |
| EPI_ISL_2809489, EPI_ISL_2809490, EPI_ISL_2809492, EPI_ISL_2809493, EPI_ISL_2809494, EPI_ISL_2809495, EPI_ISL_2809496, EPI_ISL_2809497, EPI_ISL_2809498, EPI_ISL_2809499, EPI_ISL_2809500, EPI_ISL_2809501, EPI_ISL_2809502, EPI_ISL_2809503, EPI_ISL_2809504, EPI_ISL_2809505, EPI_ISL_2809506, EPI_ISL_2809507, EPI_ISL_2809508, EPI_ISL_2809509, EPI_ISL_2809510, EPI_ISL_2809511, EPI_ISL_2809512, EPI_ISL_2809513, EPI_ISL_2809514, EPI_ISL_2809515, EPI_ISL_2809516, EPI_ISL_2809517, EPI_ISL_2809518, EPI_ISL_2809519, EPI_ISL_2809520, EPI_ISL_2809521, EPI_ISL_2809522 | | | |
| see above | Lighthouse Lab in Glasgow | Wellcome Sanger Institute for the COVID-19 Genomics UK (COG-UK) Consortium | Harper VanSteenhouse, Yumi Kasai, David Gray, Carol Clugston, Anna Dominiczak and Alex Alderton, Roberto Amato, Jeffrey Barrett, Sonia Goncalves, Ewan Harrison, David K. Jackson, Ian Johnston, Dominic Kwiatkowski, Cordelia Langford, John Sillitoe on behalf of the Wellcome Sanger Institute COVID-19 Surveillance Team |
| EPI_ISL_2809523 | Lighthouse Lab in Milton Keynes | Wellcome Sanger Institute for the COVID-19 Genomics UK (COG-UK) Consortium | The Lighthouse Lab in Milton Keynes and Alex Alderton, Roberto Amato, Jeffrey Barrett, Sonia Goncalves, Ewan Harrison, David K. Jackson, Ian Johnston, Dominic Kwiatkowski, Cordelia Langford, John Sillitoe on behalf of the Wellcome Sanger Institute COVID-19 Surveillance Team |
| EPI_ISL_2809524, EPI_ISL_2809525, EPI_ISL_2809526, EPI_ISL_2809527, EPI_ISL_2809528, EPI_ISL_2809529, EPI_ISL_2809530, EPI_ISL_2809531, EPI_ISL_2809532, EPI_ISL_2809533, EPI_ISL_2809534, EPI_ISL_2809535, EPI_ISL_2809536, EPI_ISL_2809537, EPI_ISL_2809538, EPI_ISL_2809539, EPI_ISL_2809540, EPI_ISL_2809541, EPI_ISL_2809542, EPI_ISL_2809543, EPI_ISL_2809544 | | | |
| see above | Lighthouse Lab in Glasgow | Wellcome Sanger Institute for the COVID-19 Genomics UK (COG-UK) Consortium | Harper VanSteenhouse, Yumi Kasai, David Gray, Carol Clugston, Anna Dominiczak and Alex Alderton, Roberto Amato, Jeffrey Barrett, Sonia Goncalves, Ewan Harrison, David K. Jackson, Ian Johnston, Dominic Kwiatkowski, Cordelia Langford, John Sillitoe on behalf of the Wellcome Sanger Institute COVID-19 Surveillance Team |
| EPI_ISL_2809545 | Lighthouse Lab in Alderley Park | Wellcome Sanger Institute for the COVID-19 Genomics UK (COG-UK) Consortium | Jacquelyn Wynn, Mairead Hyland, The Lighthouse Lab in Alderley Park and Alex Alderton, Roberto Amato, Jeffrey Barrett, Sonia Goncalves, Ewan Harrison, David K. Jackson, Ian Johnston, Dominic Kwiatkowski, Cordelia Langford, John Sillitoe on behalf of the Wellcome Sanger Institute COVID-19 Surveillance Team |
| EPI_ISL_2809546, EPI_ISL_2809547, EPI_ISL_2809548, EPI_ISL_2809549, EPI_ISL_2809550, EPI_ISL_2809551, EPI_ISL_2809552, EPI_ISL_2809553, EPI_ISL_2809554, EPI_ISL_2809555, EPI_ISL_2809556, EPI_ISL_2809557, EPI_ISL_2809558 | | | |
| see above | Lighthouse Lab in Glasgow | Wellcome Sanger Institute for the COVID-19 Genomics UK (COG-UK) Consortium | Harper VanSteenhouse, Yumi Kasai, David Gray, Carol Clugston, Anna Dominiczak and Alex Alderton, Roberto Amato, Jeffrey Barrett, Sonia Goncalves, Ewan Harrison, David K. Jackson, Ian Johnston, Dominic Kwiatkowski, Cordelia Langford, John Sillitoe on behalf of the Wellcome Sanger Institute COVID-19 Surveillance Team |
| EPI_ISL_2809559 | Lighthouse Laboratory Plymouth | Wellcome Sanger Institute for the COVID-19 Genomics UK (COG-UK) Consortium | Lighthouse Laboratory Plymouth and Alex Alderton, Roberto Amato, Jeffrey Barrett, Sonia Goncalves, Ewan Harrison, David K. Jackson, Ian Johnston, Dominic Kwiatkowski, Cordelia Langford, John Sillitoe on behalf of the Wellcome Sanger Institute COVID-19 Surveillance Team |
| EPI_ISL_2809560, EPI_ISL_2809561, EPI_ISL_2809562, EPI_ISL_2809563, EPI_ISL_2809564, EPI_ISL_2809565, EPI_ISL_2809566, EPI_ISL_2809567, EPI_ISL_2809568, EPI_ISL_2809569, EPI_ISL_2809570, EPI_ISL_2809571, EPI_ISL_2809572 | | | |
| see above | Lighthouse Lab in Glasgow | Wellcome Sanger Institute for the COVID-19 Genomics UK (COG-UK) Consortium | Harper VanSteenhouse, Yumi Kasai, David Gray, Carol Clugston, Anna Dominiczak and Alex Alderton, Roberto Amato, Jeffrey Barrett, Sonia Goncalves, Ewan Harrison, David K. Jackson, Ian Johnston, Dominic Kwiatkowski, Cordelia Langford, John Sillitoe on behalf of the Wellcome Sanger Institute COVID-19 Surveillance Team |
| EPI_ISL_2809573 | Lighthouse Lab in Milton Keynes | Wellcome Sanger Institute for the COVID-19 Genomics UK (COG-UK) Consortium | The Lighthouse Lab in Milton Keynes and Alex Alderton, Roberto Amato, Jeffrey Barrett, Sonia Goncalves, Ewan Harrison, David K. Jackson, Ian Johnston, Dominic Kwiatkowski, Cordelia Langford, John Sillitoe on behalf of the Wellcome Sanger Institute COVID-19 Surveillance Team |
| EPI_ISL_2809574, EPI_ISL_2809575, EPI_ISL_2809576, EPI_ISL_2809577, EPI_ISL_2809578, EPI_ISL_2809579, EPI_ISL_2809580, EPI_ISL_2809581, EPI_ISL_2809582, EPI_ISL_2809583, EPI_ISL_2809584, EPI_ISL_2809585, EPI_ISL_2809586, EPI_ISL_2809587, EPI_ISL_2809588, EPI_ISL_2809589, EPI_ISL_2809590, EPI_ISL_2809591, EPI_ISL_2809592, EPI_ISL_2809593, EPI_ISL_2809594, EPI_ISL_2809595, EPI_ISL_2809596, EPI_ISL_2809597, EPI_ISL_2809598, EPI_ISL_2809599, EPI_ISL_2809600, EPI_ISL_2809601, EPI_ISL_2809602, EPI_ISL_2809603, EPI_ISL_2809604, EPI_ISL_2809605, EPI_ISL_2809607, EPI_ISL_2809608, EPI_ISL_2809609, EPI_ISL_2809610, EPI_ISL_2809611, EPI_ISL_2809612, EPI_ISL_2809613, EPI_ISL_2809614, EPI_ISL_2809615, EPI_ISL_2809616, EPI_ISL_2809617, EPI_ISL_2809618, EPI_ISL_2809619, EPI_ISL_2809620, EPI_ISL_2809621, EPI_ISL_2809622, EPI_ISL_2809623, EPI_ISL_2809624, EPI_ISL_2809625, EPI_ISL_2809626, EPI_ISL_2809627, EPI_ISL_2809628, EPI_ISL_2809629, EPI_ISL_2809630, EPI_ISL_2809631, EPI_ISL_2809632, EPI_ISL_2809633, EPI_ISL_2809634, EPI_ISL_2809635, EPI_ISL_2809636, EPI_ISL_2809637, EPI_ISL_2809639, EPI_ISL_2809640, EPI_ISL_2809641, EPI_ISL_2809642, EPI_ISL_2809643, EPI_ISL_2809644, EPI_ISL_2809645, EPI_ISL_2809647, EPI_ISL_2809648, EPI_ISL_2809649, EPI_ISL_2809650, EPI_ISL_2809651, EPI_ISL_2809652, EPI_ISL_2809653, EPI_ISL_2809654, EPI_ISL_2809655, EPI_ISL_2809656, EPI_ISL_2809657, EPI_ISL_2809658, EPI_ISL_2809659, EPI_ISL_2809660, EPI_ISL_2809661, EPI_ISL_2809662, EPI_ISL_2809663, EPI_ISL_2809664, EPI_ISL_2809665, EPI_ISL_2809666, EPI_ISL_2809667, EPI_ISL_2809668, EPI_ISL_2809669, EPI_ISL_2809670, EPI_ISL_2809671, EPI_ISL_2809672, EPI_ISL_2809673, EPI_ISL_2809674, EPI_ISL_2809675, EPI_ISL_2809676, EPI_ISL_2809677, EPI_ISL_2809678, EPI_ISL_2809679, EPI_ISL_2809680, EPI_ISL_2809681, EPI_ISL_2809682, EPI_ISL_2809683, EPI_ISL_2809684, EPI_ISL_2809685, EPI_ISL_2809686, EPI_ISL_2809687, EPI_ISL_2809688, EPI_ISL_2809689, EPI_ISL_2809690, EPI_ISL_2809691, EPI_ISL_2809692, EPI_ISL_2809693, EPI_ISL_2809694, EPI_ISL_2809695, EPI_ISL_2809696, EPI_ISL_2809697, EPI_ISL_2809698, EPI_ISL_2809699, EPI_ISL_2809700, EPI_ISL_2809701, EPI_ISL_2809702, EPI_ISL_2809703, EPI_ISL_2809704, EPI_ISL_2809705, EPI_ISL_2809706, EPI_ISL_2809707, EPI_ISL_2809708 | | | |
| see above | Lighthouse Lab in Glasgow | Wellcome Sanger Institute for the COVID-19 Genomics UK (COG-UK) Consortium | Harper VanSteenhouse, Yumi Kasai, David Gray, Carol Clugston, Anna Dominiczak and Alex Alderton, Roberto Amato, Jeffrey Barrett, Sonia Goncalves, Ewan Harrison, David K. Jackson, Ian Johnston, Dominic Kwiatkowski, Cordelia Langford, John Sillitoe on behalf of the Wellcome Sanger Institute COVID-19 Surveillance Team |
| EPI_ISL_2809709 | Lighthouse Lab in Alderley Park | Wellcome Sanger Institute for the COVID-19 Genomics UK (COG-UK) Consortium | Jacquelyn Wynn, Mairead Hyland, The Lighthouse Lab in Alderley Park and Alex Alderton, Roberto Amato, Jeffrey Barrett, Sonia Goncalves, Ewan Harrison, David K. Jackson, Ian Johnston, Dominic Kwiatkowski, Cordelia Langford, John Sillitoe on behalf of the Wellcome Sanger Institute COVID-19 Surveillance Team |
| EPI_ISL_2809710, EPI_ISL_2809711, EPI_ISL_2809712, EPI_ISL_2809713, EPI_ISL_2809714, EPI_ISL_2809715, EPI_ISL_2809716, EPI_ISL_2809717, EPI_ISL_2809718, EPI_ISL_2809719, EPI_ISL_2809720, EPI_ISL_2809721, EPI_ISL_2809722, EPI_ISL_2809723, EPI_ISL_2809724, EPI_ISL_2809725, EPI_ISL_2809726, EPI_ISL_2809727, EPI_ISL_2809728, EPI_ISL_2809729, EPI_ISL_2809730, EPI_ISL_2809731, EPI_ISL_2809732, EPI_ISL_2809733, EPI_ISL_2809734, EPI_ISL_2809735, EPI_ISL_2809736, EPI_ISL_2809737, EPI_ISL_2809738, EPI_ISL_2809739, EPI_ISL_2809740, EPI_ISL_2809741, EPI_ISL_2809742, EPI_ISL_2809743, EPI_ISL_2809744, EPI_ISL_2809745, EPI_ISL_2809746, EPI_ISL_2809747, EPI_ISL_2809748, EPI_ISL_2809749, EPI_ISL_2809750, EPI_ISL_2809751, EPI_ISL_2809752, EPI_ISL_2809753, EPI_ISL_2809754, EPI_ISL_2809755, EPI_ISL_2809756, EPI_ISL_2809757, EPI_ISL_2809758, EPI_ISL_2809759, EPI_ISL_2809760, EPI_ISL_2809761, EPI_ISL_2809762, EPI_ISL_2809763, EPI_ISL_2809764, EPI_ISL_2809765, EPI_ISL_2809766, EPI_ISL_2809767, EPI_ISL_2809768, EPI_ISL_2809770, EPI_ISL_2809771, EPI_ISL_2809772, EPI_ISL_2809773, EPI_ISL_2809774, EPI_ISL_2809775, EPI_ISL_2809776, EPI_ISL_2809777, EPI_ISL_2809778, EPI_ISL_2809779, EPI_ISL_2809780, EPI_ISL_2809781, EPI_ISL_2809782, EPI_ISL_2809783, EPI_ISL_2809785, EPI_ISL_2809786, EPI_ISL_2809787, EPI_ISL_2809788, EPI_ISL_2809789, EPI_ISL_2809790, EPI_ISL_2809791, EPI_ISL_2809793, EPI_ISL_2809794, EPI_ISL_2809795, EPI_ISL_2809796, EPI_ISL_2809797, EPI_ISL_2809799, EPI_ISL_2809800, EPI_ISL_2809801, EPI_ISL_2809802, EPI_ISL_2809803, EPI_ISL_2809804, EPI_ISL_2809805, EPI_ISL_2809806, EPI_ISL_2809807, EPI_ISL_2809808, EPI_ISL_2809809, EPI_ISL_2809810, EPI_ISL_2809811, EPI_ISL_2809812, EPI_ISL_2809813, EPI_ISL_2809814, EPI_ISL_2809815, EPI_ISL_2809816, EPI_ISL_2809817, EPI_ISL_2809818, EPI_ISL_2809819, EPI_ISL_2809820, EPI_ISL_2809821, EPI_ISL_2809822, EPI_ISL_2809823, EPI_ISL_2809824, EPI_ISL_2809825, EPI_ISL_2809826, EPI_ISL_2809827, EPI_ISL_2809828, EPI_ISL_2809829, EPI_ISL_2809830, EPI_ISL_2809831, EPI_ISL_2809832, EPI_ISL_2809833, EPI_ISL_2809834, EPI_ISL_2809835, EPI_ISL_2809836, EPI_ISL_2809837, EPI_ISL_2809838, EPI_ISL_2809839, EPI_ISL_2809840, EPI_ISL_2809841, EPI_ISL_2809842, EPI_ISL_2809843, EPI_ISL_2809844, EPI_ISL_2809845, EPI_ISL_2809846, EPI_ISL_2809847, EPI_ISL_2809848, EPI_ISL_2809849, EPI_ISL_2809850, EPI_ISL_2809851, EPI_ISL_2809852, EPI_ISL_2809853, EPI_ISL_2809854 | | | |
| see above | Lighthouse Lab in Glasgow | Wellcome Sanger Institute for the COVID-19 Genomics UK (COG-UK) Consortium | Harper VanSteenhouse, Yumi Kasai, David Gray, Carol Clugston, Anna Dominiczak and Alex Alderton, Roberto Amato, Jeffrey Barrett, Sonia Goncalves, Ewan Harrison, David K. Jackson, Ian Johnston, Dominic Kwiatkowski, Cordelia Langford, John Sillitoe on behalf of the Wellcome Sanger Institute COVID-19 Surveillance Team |
| EPI_ISL_2809855 | Lighthouse Laboratory Plymouth | Wellcome Sanger Institute for the COVID-19 Genomics UK (COG-UK) Consortium | Lighthouse Laboratory Plymouth and Alex Alderton, Roberto Amato, Jeffrey Barrett, Sonia Goncalves, Ewan Harrison, David K. Jackson, Ian Johnston, Dominic Kwiatkowski, Cordelia Langford, John Sillitoe on behalf of the Wellcome Sanger Institute COVID-19 Surveillance Team |
| EPI_ISL_2809856, EPI_ISL_2809857, EPI_ISL_2809858, EPI_ISL_2809859, EPI_ISL_2809860, EPI_ISL_2809861, EPI_ISL_2809862, EPI_ISL_2809863, EPI_ISL_2809864, EPI_ISL_2809865, EPI_ISL_2809866, EPI_ISL_2809867, EPI_ISL_2809868, EPI_ISL_2809869, EPI_ISL_2809870, EPI_ISL_2809871, EPI_ISL_2809872, EPI_ISL_2809873, EPI_ISL_2809874, EPI_ISL_2809875, EPI_ISL_2809876, EPI_ISL_2809877, EPI_ISL_2809878, EPI_ISL_2809879, EPI_ISL_2809880, EPI_ISL_2809881 | | | |
| see above | Lighthouse Lab in Glasgow | Wellcome Sanger Institute for the COVID-19 Genomics UK (COG-UK) Consortium | Harper VanSteenhouse, Yumi Kasai, David Gray, Carol Clugston, Anna Dominiczak and Alex Alderton, Roberto Amato, Jeffrey Barrett, Sonia Goncalves, Ewan Harrison, David K. Jackson, Ian Johnston, Dominic Kwiatkowski, Cordelia Langford, John Sillitoe on behalf of the Wellcome Sanger Institute COVID-19 Surveillance Team |
| EPI_ISL_2809882, EPI_ISL_2809883, EPI_ISL_2809884 | Lighthouse Lab in Milton Keynes | Wellcome Sanger Institute for the COVID-19 Genomics UK (COG-UK) Consortium | The Lighthouse Lab in Milton Keynes and Alex Alderton, Roberto Amato, Jeffrey Barrett, Sonia Goncalves, Ewan Harrison, David K. Jackson, Ian Johnston, Dominic Kwiatkowski, Cordelia Langford, John Sillitoe on behalf of the Wellcome Sanger Institute COVID-19 Surveillance Team |
| EPI_ISL_2809885 | Lighthouse Lab in Glasgow | Wellcome Sanger Institute for the COVID-19 Genomics UK (COG-UK) Consortium | Harper VanSteenhouse, Yumi Kasai, David Gray, Carol Clugston, Anna Dominiczak and Alex Alderton, Roberto Amato, Jeffrey Barrett, Sonia Goncalves, Ewan Harrison, David K. Jackson, Ian Johnston, Dominic Kwiatkowski, Cordelia Langford, John Sillitoe on behalf of the Wellcome Sanger Institute COVID-19 Surveillance Team |
| EPI_ISL_2809886, EPI_ISL_2809887, EPI_ISL_2809888, EPI_ISL_2809889, EPI_ISL_2809890, EPI_ISL_2809891, EPI_ISL_2809892 | Lighthouse Lab in Milton Keynes | Wellcome Sanger Institute for the COVID-19 Genomics UK (COG-UK) Consortium | The Lighthouse Lab in Milton Keynes and Alex Alderton, Roberto Amato, Jeffrey Barrett, Sonia Goncalves, Ewan Harrison, David K. Jackson, Ian Johnston, Dominic Kwiatkowski, Cordelia Langford, John Sillitoe on behalf of the Wellcome Sanger Institute COVID-19 Surveillance Team |
| EPI_ISL_2809893 | Lighthouse Lab in Glasgow | Wellcome Sanger Institute for the COVID-19 Genomics UK (COG-UK) Consortium | Harper VanSteenhouse, Yumi Kasai, David Gray, Carol Clugston, Anna Dominiczak and Alex Alderton, Roberto Amato, Jeffrey Barrett, Sonia Goncalves, Ewan Harrison, David K. Jackson, Ian Johnston, Dominic Kwiatkowski, Cordelia Langford, John Sillitoe on behalf of the Wellcome Sanger Institute COVID-19 Surveillance Team |
| EPI_ISL_2809894 | Lighthouse Lab in Milton Keynes | Wellcome Sanger Institute for the COVID-19 Genomics UK (COG-UK) Consortium | The Lighthouse Lab in Milton Keynes and Alex Alderton, Roberto Amato, Jeffrey Barrett, Sonia Goncalves, Ewan Harrison, David K. Jackson, Ian Johnston, Dominic Kwiatkowski, Cordelia Langford, John Sillitoe on behalf of the Wellcome Sanger Institute COVID-19 Surveillance Team |
| EPI_ISL_2809895 | Lighthouse Lab in Glasgow | Wellcome Sanger Institute for the COVID-19 Genomics UK (COG-UK) Consortium | Harper VanSteenhouse, Yumi Kasai, David Gray, Carol Clugston, Anna Dominiczak and Alex Alderton, Roberto Amato, Jeffrey Barrett, Sonia Goncalves, Ewan Harrison, David K. Jackson, Ian Johnston, Dominic Kwiatkowski, Cordelia Langford, John Sillitoe on behalf of the Wellcome Sanger Institute COVID-19 Surveillance Team |
| EPI_ISL_2809896, | Lighthouse Lab in Milton Keynes | Wellcome Sanger Institute for the COVID-19 Genomics UK (COG-UK) Consortium | The Lighthouse Lab in Milton Keynes and Alex Alderton, Roberto Amato, Jeffrey Barrett, Sonia Goncalves, Ewan Harrison, David K. Jackson, Ian Johnston, Dominic Kwiatkowski, Cordelia Langford, John Sillitoe on behalf of the Wellcome Sanger |

| | | | |
|---|---|---|---|
| EPI_ISL_2809897, EPI_ISL_2809898, EPI_ISL_2809899, EPI_ISL_2809900, EPI_ISL_2809901, EPI_ISL_2809902, EPI_ISL_2809903, EPI_ISL_2809904, EPI_ISL_2809905 | | Consortium | Institute COVID-19 Surveillance Team |
| EPI_ISL_2809906 | Lighthouse Lab in Glasgow | Wellcome Sanger Institute for the COVID-19 Genomics UK (COG-UK) Consortium | Harper VanSteenhouse, Yumi Kasai, David Gray, Carol Clugston, Anna Dominiczak and Alex Alderton, Roberto Amato, Jeffrey Barrett, Sonia Goncalves, Ewan Harrison, David K. Jackson, Ian Johnston, Dominic Kwiatkowski, Cordelia Langford, John Sillitoe on behalf of the Wellcome Sanger Institute COVID-19 Surveillance Team |
| EPI_ISL_2809907, EPI_ISL_2809908 | Lighthouse Lab in Milton Keynes | Wellcome Sanger Institute for the COVID-19 Genomics UK (COG-UK) Consortium | The Lighthouse Lab in Milton Keynes and Alex Alderton, Roberto Amato, Jeffrey Barrett, Sonia Goncalves, Ewan Harrison, David K. Jackson, Ian Johnston, Dominic Kwiatkowski, Cordelia Langford, John Sillitoe on behalf of the Wellcome Sanger Institute COVID-19 Surveillance Team |
| EPI_ISL_2809909 | Lighthouse Lab in Glasgow | Wellcome Sanger Institute for the COVID-19 Genomics UK (COG-UK) Consortium | Harper VanSteenhouse, Yumi Kasai, David Gray, Carol Clugston, Anna Dominiczak and Alex Alderton, Roberto Amato, Jeffrey Barrett, Sonia Goncalves, Ewan Harrison, David K. Jackson, Ian Johnston, Dominic Kwiatkowski, Cordelia Langford, John Sillitoe on behalf of the Wellcome Sanger Institute COVID-19 Surveillance Team |
| EPI_ISL_2809910, EPI_ISL_2809911, EPI_ISL_2809912 | Lighthouse Lab in Milton Keynes | Wellcome Sanger Institute for the COVID-19 Genomics UK (COG-UK) Consortium | The Lighthouse Lab in Milton Keynes and Alex Alderton, Roberto Amato, Jeffrey Barrett, Sonia Goncalves, Ewan Harrison, David K. Jackson, Ian Johnston, Dominic Kwiatkowski, Cordelia Langford, John Sillitoe on behalf of the Wellcome Sanger Institute COVID-19 Surveillance Team |
| EPI_ISL_2809913 | Lighthouse Lab in Glasgow | Wellcome Sanger Institute for the COVID-19 Genomics UK (COG-UK) Consortium | Harper VanSteenhouse, Yumi Kasai, David Gray, Carol Clugston, Anna Dominiczak and Alex Alderton, Roberto Amato, Jeffrey Barrett, Sonia Goncalves, Ewan Harrison, David K. Jackson, Ian Johnston, Dominic Kwiatkowski, Cordelia Langford, John Sillitoe on behalf of the Wellcome Sanger Institute COVID-19 Surveillance Team |
| EPI_ISL_2809914, EPI_ISL_2809915 | Lighthouse Lab in Milton Keynes | Wellcome Sanger Institute for the COVID-19 Genomics UK (COG-UK) Consortium | The Lighthouse Lab in Milton Keynes and Alex Alderton, Roberto Amato, Jeffrey Barrett, Sonia Goncalves, Ewan Harrison, David K. Jackson, Ian Johnston, Dominic Kwiatkowski, Cordelia Langford, John Sillitoe on behalf of the Wellcome Sanger Institute COVID-19 Surveillance Team |
| EPI_ISL_2809916 | Lighthouse Lab in Glasgow | Wellcome Sanger Institute for the COVID-19 Genomics UK (COG-UK) Consortium | Harper VanSteenhouse, Yumi Kasai, David Gray, Carol Clugston, Anna Dominiczak and Alex Alderton, Roberto Amato, Jeffrey Barrett, Sonia Goncalves, Ewan Harrison, David K. Jackson, Ian Johnston, Dominic Kwiatkowski, Cordelia Langford, John Sillitoe on behalf of the Wellcome Sanger Institute COVID-19 Surveillance Team |
| EPI_ISL_2809917, EPI_ISL_2809918, EPI_ISL_2809919, EPI_ISL_2809920, EPI_ISL_2809921 | Lighthouse Lab in Milton Keynes | Wellcome Sanger Institute for the COVID-19 Genomics UK (COG-UK) Consortium | The Lighthouse Lab in Milton Keynes and Alex Alderton, Roberto Amato, Jeffrey Barrett, Sonia Goncalves, Ewan Harrison, David K. Jackson, Ian Johnston, Dominic Kwiatkowski, Cordelia Langford, John Sillitoe on behalf of the Wellcome Sanger Institute COVID-19 Surveillance Team |
| EPI_ISL_2809922, EPI_ISL_2809923, EPI_ISL_2809924 | Lighthouse Lab in Glasgow | Wellcome Sanger Institute for the COVID-19 Genomics UK (COG-UK) Consortium | Harper VanSteenhouse, Yumi Kasai, David Gray, Carol Clugston, Anna Dominiczak and Alex Alderton, Roberto Amato, Jeffrey Barrett, Sonia Goncalves, Ewan Harrison, David K. Jackson, Ian Johnston, Dominic Kwiatkowski, Cordelia Langford, John Sillitoe on behalf of the Wellcome Sanger Institute COVID-19 Surveillance Team |
| EPI_ISL_2809926, EPI_ISL_2809927 | Lighthouse Lab in Milton Keynes | Wellcome Sanger Institute for the COVID-19 Genomics UK (COG-UK) Consortium | The Lighthouse Lab in Milton Keynes and Alex Alderton, Roberto Amato, Jeffrey Barrett, Sonia Goncalves, Ewan Harrison, David K. Jackson, Ian Johnston, Dominic Kwiatkowski, Cordelia Langford, John Sillitoe on behalf of the Wellcome Sanger Institute COVID-19 Surveillance Team |
| EPI_ISL_2809928 | Lighthouse Lab in Glasgow | Wellcome Sanger Institute for the COVID-19 Genomics UK (COG-UK) Consortium | Harper VanSteenhouse, Yumi Kasai, David Gray, Carol Clugston, Anna Dominiczak and Alex Alderton, Roberto Amato, Jeffrey Barrett, Sonia Goncalves, Ewan Harrison, David K. Jackson, Ian Johnston, Dominic Kwiatkowski, Cordelia Langford, John Sillitoe on behalf of the Wellcome Sanger Institute COVID-19 Surveillance Team |
| EPI_ISL_2809929, EPI_ISL_2809930, EPI_ISL_2809931, EPI_ISL_2809932 | Lighthouse Lab in Milton Keynes | Wellcome Sanger Institute for the COVID-19 Genomics UK (COG-UK) Consortium | The Lighthouse Lab in Milton Keynes and Alex Alderton, Roberto Amato, Jeffrey Barrett, Sonia Goncalves, Ewan Harrison, David K. Jackson, Ian Johnston, Dominic Kwiatkowski, Cordelia Langford, John Sillitoe on behalf of the Wellcome Sanger Institute COVID-19 Surveillance Team |
| EPI_ISL_2809933, EPI_ISL_2809934 | Lighthouse Lab in Glasgow | Wellcome Sanger Institute for the COVID-19 Genomics UK (COG-UK) Consortium | Harper VanSteenhouse, Yumi Kasai, David Gray, Carol Clugston, Anna Dominiczak and Alex Alderton, Roberto Amato, Jeffrey Barrett, Sonia Goncalves, Ewan Harrison, David K. Jackson, Ian Johnston, Dominic Kwiatkowski, Cordelia Langford, John Sillitoe on behalf of the Wellcome Sanger Institute COVID-19 Surveillance Team |
| EPI_ISL_2809935 | Lighthouse Lab in Milton Keynes | Wellcome Sanger Institute for the COVID-19 Genomics UK (COG-UK) Consortium | The Lighthouse Lab in Milton Keynes and Alex Alderton, Roberto Amato, Jeffrey Barrett, Sonia Goncalves, Ewan Harrison, David K. Jackson, Ian Johnston, Dominic Kwiatkowski, Cordelia Langford, John Sillitoe on behalf of the Wellcome Sanger Institute COVID-19 Surveillance Team |
| EPI_ISL_2809936, EPI_ISL_2809937 | Lighthouse Lab in Glasgow | Wellcome Sanger Institute for the COVID-19 Genomics UK (COG-UK) Consortium | Harper VanSteenhouse, Yumi Kasai, David Gray, Carol Clugston, Anna Dominiczak and Alex Alderton, Roberto Amato, Jeffrey Barrett, Sonia Goncalves, Ewan Harrison, David K. Jackson, Ian Johnston, Dominic Kwiatkowski, Cordelia Langford, John Sillitoe on behalf of the Wellcome Sanger Institute COVID-19 Surveillance Team |
| EPI_ISL_2809938 | Lighthouse Lab in Milton Keynes | Wellcome Sanger Institute for the COVID-19 Genomics UK (COG-UK) Consortium | The Lighthouse Lab in Milton Keynes and Alex Alderton, Roberto Amato, Jeffrey Barrett, Sonia Goncalves, Ewan Harrison, David K. Jackson, Ian Johnston, Dominic Kwiatkowski, Cordelia Langford, John Sillitoe on behalf of the Wellcome Sanger Institute COVID-19 Surveillance Team |
| EPI_ISL_2809939 | Lighthouse Lab in Glasgow | Wellcome Sanger Institute for the COVID-19 Genomics UK (COG-UK) Consortium | Harper VanSteenhouse, Yumi Kasai, David Gray, Carol Clugston, Anna Dominiczak and Alex Alderton, Roberto Amato, Jeffrey Barrett, Sonia Goncalves, Ewan Harrison, David K. Jackson, Ian Johnston, Dominic Kwiatkowski, Cordelia Langford, John Sillitoe on behalf of the Wellcome Sanger Institute COVID-19 Surveillance Team |
| EPI_ISL_2809940, EPI_ISL_2809941, EPI_ISL_2809942, EPI_ISL_2809943, EPI_ISL_2809944, EPI_ISL_2809945, EPI_ISL_2809946, EPI_ISL_2809947, EPI_ISL_2809948 | Lighthouse Lab in Milton Keynes | Wellcome Sanger Institute for the COVID-19 Genomics UK (COG-UK) Consortium | The Lighthouse Lab in Milton Keynes and Alex Alderton, Roberto Amato, Jeffrey Barrett, Sonia Goncalves, Ewan Harrison, David K. Jackson, Ian Johnston, Dominic Kwiatkowski, Cordelia Langford, John Sillitoe on behalf of the Wellcome Sanger Institute COVID-19 Surveillance Team |
| EPI_ISL_2809949 | Lighthouse Lab in Glasgow | Wellcome Sanger Institute for the COVID-19 Genomics UK (COG-UK) Consortium | Harper VanSteenhouse, Yumi Kasai, David Gray, Carol Clugston, Anna Dominiczak and Alex Alderton, Roberto Amato, Jeffrey Barrett, Sonia Goncalves, Ewan Harrison, David K. Jackson, Ian Johnston, Dominic Kwiatkowski, Cordelia Langford, John Sillitoe on behalf of the Wellcome Sanger Institute COVID-19 Surveillance Team |
| EPI_ISL_2809950, EPI_ISL_2809951, EPI_ISL_2809952 | Lighthouse Lab in Milton Keynes | Wellcome Sanger Institute for the COVID-19 Genomics UK (COG-UK) Consortium | The Lighthouse Lab in Milton Keynes and Alex Alderton, Roberto Amato, Jeffrey Barrett, Sonia Goncalves, Ewan Harrison, David K. Jackson, Ian Johnston, Dominic Kwiatkowski, Cordelia Langford, John Sillitoe on behalf of the Wellcome Sanger Institute COVID-19 Surveillance Team |
| EPI_ISL_2809953 | Lighthouse Lab in Glasgow | Wellcome Sanger Institute for the COVID-19 Genomics UK (COG-UK) Consortium | Harper VanSteenhouse, Yumi Kasai, David Gray, Carol Clugston, Anna Dominiczak and Alex Alderton, Roberto Amato, Jeffrey Barrett, Sonia Goncalves, Ewan Harrison, David K. Jackson, Ian Johnston, Dominic Kwiatkowski, Cordelia Langford, John Sillitoe on behalf of the Wellcome Sanger Institute COVID-19 Surveillance Team |
| EPI_ISL_2809954 | Lighthouse Lab in Milton Keynes | Wellcome Sanger Institute for the COVID-19 Genomics UK (COG-UK) Consortium | The Lighthouse Lab in Milton Keynes and Alex Alderton, Roberto Amato, Jeffrey Barrett, Sonia Goncalves, Ewan Harrison, David K. Jackson, Ian Johnston, Dominic Kwiatkowski, Cordelia Langford, John Sillitoe on behalf of the Wellcome Sanger Institute COVID-19 Surveillance Team |
| EPI_ISL_2809955, EPI_ISL_2809956, EPI_ISL_2809957, EPI_ISL_2809958 | Lighthouse Lab in Glasgow | Wellcome Sanger Institute for the COVID-19 Genomics UK (COG-UK) Consortium | Harper VanSteenhouse, Yumi Kasai, David Gray, Carol Clugston, Anna Dominiczak and Alex Alderton, Roberto Amato, Jeffrey Barrett, Sonia Goncalves, Ewan Harrison, David K. Jackson, Ian Johnston, Dominic Kwiatkowski, Cordelia Langford, John Sillitoe on behalf of the Wellcome Sanger Institute COVID-19 Surveillance Team |
| EPI_ISL_2809959, EPI_ISL_2809960, EPI_ISL_2809961, EPI_ISL_2809962 | Lighthouse Lab in Milton Keynes | Wellcome Sanger Institute for the COVID-19 Genomics UK (COG-UK) Consortium | The Lighthouse Lab in Milton Keynes and Alex Alderton, Roberto Amato, Jeffrey Barrett, Sonia Goncalves, Ewan Harrison, David K. Jackson, Ian Johnston, Dominic Kwiatkowski, Cordelia Langford, John Sillitoe on behalf of the Wellcome Sanger Institute COVID-19 Surveillance Team |
| EPI_ISL_2809963 | Lighthouse Lab in Glasgow | Wellcome Sanger Institute for the COVID-19 Genomics UK (COG-UK) Consortium | Harper VanSteenhouse, Yumi Kasai, David Gray, Carol Clugston, Anna Dominiczak and Alex Alderton, Roberto Amato, Jeffrey Barrett, Sonia Goncalves, Ewan Harrison, David K. Jackson, Ian Johnston, Dominic Kwiatkowski, Cordelia Langford, John Sillitoe on behalf of the Wellcome Sanger Institute COVID-19 Surveillance Team |
| EPI_ISL_2809964, EPI_ISL_2809965 | Lighthouse Lab in Milton Keynes | Wellcome Sanger Institute for the COVID-19 Genomics UK (COG-UK) Consortium | The Lighthouse Lab in Milton Keynes and Alex Alderton, Roberto Amato, Jeffrey Barrett, Sonia Goncalves, Ewan Harrison, David K. Jackson, Ian Johnston, Dominic Kwiatkowski, Cordelia Langford, John Sillitoe on behalf of the Wellcome Sanger Institute COVID-19 Surveillance Team |
| EPI_ISL_2809966 | Lighthouse Lab in Glasgow | Wellcome Sanger Institute for the COVID-19 Genomics UK (COG-UK) Consortium | Harper VanSteenhouse, Yumi Kasai, David Gray, Carol Clugston, Anna Dominiczak and Alex Alderton, Roberto Amato, Jeffrey Barrett, Sonia Goncalves, Ewan Harrison, David K. Jackson, Ian Johnston, Dominic Kwiatkowski, Cordelia Langford, John Sillitoe on behalf of the Wellcome Sanger Institute COVID-19 Surveillance Team |
| EPI_ISL_2809967, EPI_ISL_2809968 | Lighthouse Lab in Milton Keynes | Wellcome Sanger Institute for the COVID-19 Genomics UK (COG-UK) Consortium | The Lighthouse Lab in Milton Keynes and Alex Alderton, Roberto Amato, Jeffrey Barrett, Sonia Goncalves, Ewan Harrison, David K. Jackson, Ian Johnston, Dominic Kwiatkowski, Cordelia Langford, John Sillitoe on behalf of the Wellcome Sanger Institute COVID-19 Surveillance Team |
| EPI_ISL_2809969 | Lighthouse Lab in Glasgow | Wellcome Sanger Institute for the COVID-19 Genomics UK (COG-UK) Consortium | Harper VanSteenhouse, Yumi Kasai, David Gray, Carol Clugston, Anna Dominiczak and Alex Alderton, Roberto Amato, Jeffrey Barrett, Sonia Goncalves, Ewan Harrison, David K. Jackson, Ian Johnston, Dominic Kwiatkowski, Cordelia Langford, John Sillitoe on behalf of the Wellcome Sanger Institute COVID-19 Surveillance Team |
| EPI_ISL_2809970, EPI_ISL_2809971 | Lighthouse Lab in Milton Keynes | Wellcome Sanger Institute for the COVID-19 Genomics UK (COG-UK) Consortium | The Lighthouse Lab in Milton Keynes and Alex Alderton, Roberto Amato, Jeffrey Barrett, Sonia Goncalves, Ewan Harrison, David K. Jackson, Ian Johnston, Dominic Kwiatkowski, Cordelia Langford, John Sillitoe on behalf of the Wellcome Sanger Institute COVID-19 Surveillance Team |
| EPI_ISL_2809972, EPI_ISL_2809973 | Lighthouse Lab in Glasgow | Wellcome Sanger Institute for the COVID-19 Genomics UK (COG-UK) Consortium | Harper VanSteenhouse, Yumi Kasai, David Gray, Carol Clugston, Anna Dominiczak and Alex Alderton, Roberto Amato, Jeffrey Barrett, Sonia Goncalves, Ewan Harrison, David K. Jackson, Ian Johnston, Dominic Kwiatkowski, Cordelia Langford, John Sillitoe on behalf of the Wellcome Sanger Institute COVID-19 Surveillance Team |
| EPI_ISL_2809974 | Lighthouse Lab in Milton Keynes | Wellcome Sanger Institute for the COVID-19 Genomics UK (COG-UK) Consortium | The Lighthouse Lab in Milton Keynes and Alex Alderton, Roberto Amato, Jeffrey Barrett, Sonia Goncalves, Ewan Harrison, David K. Jackson, Ian Johnston, Dominic Kwiatkowski, Cordelia Langford, John Sillitoe on behalf of the Wellcome Sanger Institute COVID-19 Surveillance Team |
| EPI_ISL_2809975 | Lighthouse Lab in Glasgow | Wellcome Sanger Institute for the COVID-19 Genomics UK (COG-UK) Consortium | Harper VanSteenhouse, Yumi Kasai, David Gray, Carol Clugston, Anna Dominiczak and Alex Alderton, Roberto Amato, Jeffrey Barrett, Sonia Goncalves, Ewan Harrison, David K. Jackson, Ian Johnston, Dominic Kwiatkowski, Cordelia Langford, John Sillitoe on behalf of the Wellcome Sanger Institute COVID-19 Surveillance Team |

| | | | |
|---|---|---|---|
| EPI_ISL_2809976, EPI_ISL_2809977 | Lighthouse Lab in Milton Keynes | Wellcome Sanger Institute for the COVID-19 Genomics UK (COG-UK) Consortium | The Lighthouse Lab in Milton Keynes and Alex Alderton, Roberto Amato, Jeffrey Barrett, Sonia Goncalves, Ewan Harrison, David K. Jackson, Ian Johnston, Dominic Kwiatkowski, Cordelia Langford, John Sillitoe on behalf of the Wellcome Sanger Institute COVID-19 Surveillance Team |
| EPI_ISL_2809978, EPI_ISL_2809979 | Lighthouse Lab in Glasgow | Wellcome Sanger Institute for the COVID-19 Genomics UK (COG-UK) Consortium | Harper VanSteenhouse, Yumi Kasai, David Gray, Carol Clugston, Anna Dominiczak and Alex Alderton, Roberto Amato, Jeffrey Barrett, Sonia Goncalves, Ewan Harrison, David K. Jackson, Ian Johnston, Dominic Kwiatkowski, Cordelia Langford, John Sillitoe on behalf of the Wellcome Sanger Institute COVID-19 Surveillance Team |
| EPI_ISL_2809980, EPI_ISL_2809981, EPI_ISL_2809982 | Lighthouse Lab in Milton Keynes | Wellcome Sanger Institute for the COVID-19 Genomics UK (COG-UK) Consortium | The Lighthouse Lab in Milton Keynes and Alex Alderton, Roberto Amato, Jeffrey Barrett, Sonia Goncalves, Ewan Harrison, David K. Jackson, Ian Johnston, Dominic Kwiatkowski, Cordelia Langford, John Sillitoe on behalf of the Wellcome Sanger Institute COVID-19 Surveillance Team |
| EPI_ISL_2809983, EPI_ISL_2809984, EPI_ISL_2809985, EPI_ISL_2809986 | Lighthouse Lab in Glasgow | Wellcome Sanger Institute for the COVID-19 Genomics UK (COG-UK) Consortium | Harper VanSteenhouse, Yumi Kasai, David Gray, Carol Clugston, Anna Dominiczak and Alex Alderton, Roberto Amato, Jeffrey Barrett, Sonia Goncalves, Ewan Harrison, David K. Jackson, Ian Johnston, Dominic Kwiatkowski, Cordelia Langford, John Sillitoe on behalf of the Wellcome Sanger Institute COVID-19 Surveillance Team |
| EPI_ISL_2809987, EPI_ISL_2809988, EPI_ISL_2809989 | Lighthouse Lab in Milton Keynes | Wellcome Sanger Institute for the COVID-19 Genomics UK (COG-UK) Consortium | The Lighthouse Lab in Milton Keynes and Alex Alderton, Roberto Amato, Jeffrey Barrett, Sonia Goncalves, Ewan Harrison, David K. Jackson, Ian Johnston, Dominic Kwiatkowski, Cordelia Langford, John Sillitoe on behalf of the Wellcome Sanger Institute COVID-19 Surveillance Team |
| EPI_ISL_2809990 | Lighthouse Lab in Glasgow | Wellcome Sanger Institute for the COVID-19 Genomics UK (COG-UK) Consortium | Harper VanSteenhouse, Yumi Kasai, David Gray, Carol Clugston, Anna Dominiczak and Alex Alderton, Roberto Amato, Jeffrey Barrett, Sonia Goncalves, Ewan Harrison, David K. Jackson, Ian Johnston, Dominic Kwiatkowski, Cordelia Langford, John Sillitoe on behalf of the Wellcome Sanger Institute COVID-19 Surveillance Team |
| EPI_ISL_2809991 | Lighthouse Lab in Milton Keynes | Wellcome Sanger Institute for the COVID-19 Genomics UK (COG-UK) Consortium | The Lighthouse Lab in Milton Keynes and Alex Alderton, Roberto Amato, Jeffrey Barrett, Sonia Goncalves, Ewan Harrison, David K. Jackson, Ian Johnston, Dominic Kwiatkowski, Cordelia Langford, John Sillitoe on behalf of the Wellcome Sanger Institute COVID-19 Surveillance Team |
| EPI_ISL_2809992 | Lighthouse Lab in Glasgow | Wellcome Sanger Institute for the COVID-19 Genomics UK (COG-UK) Consortium | Harper VanSteenhouse, Yumi Kasai, David Gray, Carol Clugston, Anna Dominiczak and Alex Alderton, Roberto Amato, Jeffrey Barrett, Sonia Goncalves, Ewan Harrison, David K. Jackson, Ian Johnston, Dominic Kwiatkowski, Cordelia Langford, John Sillitoe on behalf of the Wellcome Sanger Institute COVID-19 Surveillance Team |
| EPI_ISL_2809993, EPI_ISL_2809994, EPI_ISL_2809995 | Lighthouse Lab in Milton Keynes | Wellcome Sanger Institute for the COVID-19 Genomics UK (COG-UK) Consortium | The Lighthouse Lab in Milton Keynes and Alex Alderton, Roberto Amato, Jeffrey Barrett, Sonia Goncalves, Ewan Harrison, David K. Jackson, Ian Johnston, Dominic Kwiatkowski, Cordelia Langford, John Sillitoe on behalf of the Wellcome Sanger Institute COVID-19 Surveillance Team |
| EPI_ISL_2809996 | Lighthouse Lab in Glasgow | Wellcome Sanger Institute for the COVID-19 Genomics UK (COG-UK) Consortium | Harper VanSteenhouse, Yumi Kasai, David Gray, Carol Clugston, Anna Dominiczak and Alex Alderton, Roberto Amato, Jeffrey Barrett, Sonia Goncalves, Ewan Harrison, David K. Jackson, Ian Johnston, Dominic Kwiatkowski, Cordelia Langford, John Sillitoe on behalf of the Wellcome Sanger Institute COVID-19 Surveillance Team |
| EPI_ISL_2809997 | Lighthouse Lab in Milton Keynes | Wellcome Sanger Institute for the COVID-19 Genomics UK (COG-UK) Consortium | The Lighthouse Lab in Milton Keynes and Alex Alderton, Roberto Amato, Jeffrey Barrett, Sonia Goncalves, Ewan Harrison, David K. Jackson, Ian Johnston, Dominic Kwiatkowski, Cordelia Langford, John Sillitoe on behalf of the Wellcome Sanger Institute COVID-19 Surveillance Team |
| EPI_ISL_2809998, EPI_ISL_2809999 | Lighthouse Lab in Glasgow | Wellcome Sanger Institute for the COVID-19 Genomics UK (COG-UK) Consortium | Harper VanSteenhouse, Yumi Kasai, David Gray, Carol Clugston, Anna Dominiczak and Alex Alderton, Roberto Amato, Jeffrey Barrett, Sonia Goncalves, Ewan Harrison, David K. Jackson, Ian Johnston, Dominic Kwiatkowski, Cordelia Langford, John Sillitoe on behalf of the Wellcome Sanger Institute COVID-19 Surveillance Team |
| EPI_ISL_2810000, EPI_ISL_2810001, EPI_ISL_2810002 | Lighthouse Lab in Milton Keynes | Wellcome Sanger Institute for the COVID-19 Genomics UK (COG-UK) Consortium | The Lighthouse Lab in Milton Keynes and Alex Alderton, Roberto Amato, Jeffrey Barrett, Sonia Goncalves, Ewan Harrison, David K. Jackson, Ian Johnston, Dominic Kwiatkowski, Cordelia Langford, John Sillitoe on behalf of the Wellcome Sanger Institute COVID-19 Surveillance Team |
| EPI_ISL_2810003, EPI_ISL_2810004 | Lighthouse Lab in Glasgow | Wellcome Sanger Institute for the COVID-19 Genomics UK (COG-UK) Consortium | Harper VanSteenhouse, Yumi Kasai, David Gray, Carol Clugston, Anna Dominiczak and Alex Alderton, Roberto Amato, Jeffrey Barrett, Sonia Goncalves, Ewan Harrison, David K. Jackson, Ian Johnston, Dominic Kwiatkowski, Cordelia Langford, John Sillitoe on behalf of the Wellcome Sanger Institute COVID-19 Surveillance Team |
| EPI_ISL_2810005, EPI_ISL_2810006 | Lighthouse Lab in Milton Keynes | Wellcome Sanger Institute for the COVID-19 Genomics UK (COG-UK) Consortium | The Lighthouse Lab in Milton Keynes and Alex Alderton, Roberto Amato, Jeffrey Barrett, Sonia Goncalves, Ewan Harrison, David K. Jackson, Ian Johnston, Dominic Kwiatkowski, Cordelia Langford, John Sillitoe on behalf of the Wellcome Sanger Institute COVID-19 Surveillance Team |
| EPI_ISL_2810007, EPI_ISL_2810008 | Lighthouse Lab in Glasgow | Wellcome Sanger Institute for the COVID-19 Genomics UK (COG-UK) Consortium | Harper VanSteenhouse, Yumi Kasai, David Gray, Carol Clugston, Anna Dominiczak and Alex Alderton, Roberto Amato, Jeffrey Barrett, Sonia Goncalves, Ewan Harrison, David K. Jackson, Ian Johnston, Dominic Kwiatkowski, Cordelia Langford, John Sillitoe on behalf of the Wellcome Sanger Institute COVID-19 Surveillance Team |
| EPI_ISL_2810009, EPI_ISL_2810010, EPI_ISL_2810011 | Lighthouse Lab in Milton Keynes | Wellcome Sanger Institute for the COVID-19 Genomics UK (COG-UK) Consortium | The Lighthouse Lab in Milton Keynes and Alex Alderton, Roberto Amato, Jeffrey Barrett, Sonia Goncalves, Ewan Harrison, David K. Jackson, Ian Johnston, Dominic Kwiatkowski, Cordelia Langford, John Sillitoe on behalf of the Wellcome Sanger Institute COVID-19 Surveillance Team |
| EPI_ISL_2810012 | Lighthouse Lab in Glasgow | Wellcome Sanger Institute for the COVID-19 Genomics UK (COG-UK) Consortium | Harper VanSteenhouse, Yumi Kasai, David Gray, Carol Clugston, Anna Dominiczak and Alex Alderton, Roberto Amato, Jeffrey Barrett, Sonia Goncalves, Ewan Harrison, David K. Jackson, Ian Johnston, Dominic Kwiatkowski, Cordelia Langford, John Sillitoe on behalf of the Wellcome Sanger Institute COVID-19 Surveillance Team |
| EPI_ISL_2810013, EPI_ISL_2810014, EPI_ISL_2810015 | Lighthouse Lab in Milton Keynes | Wellcome Sanger Institute for the COVID-19 Genomics UK (COG-UK) Consortium | The Lighthouse Lab in Milton Keynes and Alex Alderton, Roberto Amato, Jeffrey Barrett, Sonia Goncalves, Ewan Harrison, David K. Jackson, Ian Johnston, Dominic Kwiatkowski, Cordelia Langford, John Sillitoe on behalf of the Wellcome Sanger Institute COVID-19 Surveillance Team |
| EPI_ISL_2810016, EPI_ISL_2810017 | Lighthouse Lab in Glasgow | Wellcome Sanger Institute for the COVID-19 Genomics UK (COG-UK) Consortium | Harper VanSteenhouse, Yumi Kasai, David Gray, Carol Clugston, Anna Dominiczak and Alex Alderton, Roberto Amato, Jeffrey Barrett, Sonia Goncalves, Ewan Harrison, David K. Jackson, Ian Johnston, Dominic Kwiatkowski, Cordelia Langford, John Sillitoe on behalf of the Wellcome Sanger Institute COVID-19 Surveillance Team |
| EPI_ISL_2810018 | Lighthouse Lab in Milton Keynes | Wellcome Sanger Institute for the COVID-19 Genomics UK (COG-UK) Consortium | The Lighthouse Lab in Milton Keynes and Alex Alderton, Roberto Amato, Jeffrey Barrett, Sonia Goncalves, Ewan Harrison, David K. Jackson, Ian Johnston, Dominic Kwiatkowski, Cordelia Langford, John Sillitoe on behalf of the Wellcome Sanger Institute COVID-19 Surveillance Team |
| EPI_ISL_2810019 | Lighthouse Lab in Glasgow | Wellcome Sanger Institute for the COVID-19 Genomics UK (COG-UK) Consortium | Harper VanSteenhouse, Yumi Kasai, David Gray, Carol Clugston, Anna Dominiczak and Alex Alderton, Roberto Amato, Jeffrey Barrett, Sonia Goncalves, Ewan Harrison, David K. Jackson, Ian Johnston, Dominic Kwiatkowski, Cordelia Langford, John Sillitoe on behalf of the Wellcome Sanger Institute COVID-19 Surveillance Team |
| EPI_ISL_2810020, EPI_ISL_2810021 | Lighthouse Lab in Milton Keynes | Wellcome Sanger Institute for the COVID-19 Genomics UK (COG-UK) Consortium | The Lighthouse Lab in Milton Keynes and Alex Alderton, Roberto Amato, Jeffrey Barrett, Sonia Goncalves, Ewan Harrison, David K. Jackson, Ian Johnston, Dominic Kwiatkowski, Cordelia Langford, John Sillitoe on behalf of the Wellcome Sanger Institute COVID-19 Surveillance Team |
| EPI_ISL_2810031 | Instituto Nacional de Medicina Genomica | Instituto Nacional de Medicina Genomica | Cedro-Tanda A, Hidalgo-Miranda A, Mendoza-Vargas A, Reyes-Grajeda JP, Gonzalez-Barrera D, Rangel-DeLeon D, Munguia-Garza P, Ramirez-Vega O, Escobar-Arrazola, M, Herrera-Montalvo LA. |
| EPI_ISL_2810036, EPI_ISL_2810037, EPI_ISL_2810038, EPI_ISL_2810039 | Salud Digna | Instituto Nacional de Medicina Genomica | Cedro-Tanda A, Hidalgo-Miranda A, Mendoza-Vargas A, Reyes-Grajeda JP, Abraham Campos-Romero, Moreno-Camacho José Luis, Rodriguez-Gallegos Jorge, Luna-Ruiz Marco, Gonzalez-Barrera D, Rangel-DeLeon D, Munguia-Garza P, Ramirez-Vega O, Escobar-Arrazola, M, Herrera-Montalvo LA. |
| EPI_ISL_2810041 | Instituto Nacional de Medicina Genomica | Instituto Nacional de Medicina Genomica | Cedro-Tanda A, Hidalgo-Miranda A, Mendoza-Vargas A, Reyes-Grajeda JP, Gonzalez-Barrera D, Rangel-DeLeon D, Munguia-Garza P, Ramirez-Vega O, Escobar-Arrazola, M, Herrera-Montalvo LA. |
| EPI_ISL_2810042, EPI_ISL_2810043, EPI_ISL_2810045 | Salud Digna | Instituto Nacional de Medicina Genomica | Cedro-Tanda A, Hidalgo-Miranda A, Mendoza-Vargas A, Reyes-Grajeda JP, Abraham Campos-Romero, Moreno-Camacho José Luis, Rodriguez-Gallegos Jorge, Luna-Ruiz Marco, Gonzalez-Barrera D, Rangel-DeLeon D, Munguia-Garza P, Ramirez-Vega O, Escobar-Arrazola, M, Herrera-Montalvo LA. |
| EPI_ISL_2810046 | Instituto Nacional de Medicina Genomica | Instituto Nacional de Medicina Genomica | Cedro-Tanda A, Hidalgo-Miranda A, Mendoza-Vargas A, Reyes-Grajeda JP, Gonzalez-Barrera D, Rangel-DeLeon D, Munguia-Garza P, Ramirez-Vega O, Escobar-Arrazola, M, Herrera-Montalvo LA. |
| EPI_ISL_2810047 | Salud Digna | Instituto Nacional de Medicina Genomica | Cedro-Tanda A, Hidalgo-Miranda A, Mendoza-Vargas A, Reyes-Grajeda JP, Abraham Campos-Romero, Moreno-Camacho José Luis, Rodriguez-Gallegos Jorge, Luna-Ruiz Marco, Gonzalez-Barrera D, Rangel-DeLeon D, Munguia-Garza P, Ramirez-Vega O, Escobar-Arrazola, M, Herrera-Montalvo LA. |
| EPI_ISL_2810048 | Instituto Nacional de Medicina Genomica | Instituto Nacional de Medicina Genomica | PENDIENTE |
| EPI_ISL_2810059, EPI_ISL_2810061 | Salud Digna | Instituto Nacional de Medicina Genomica | Cedro-Tanda A, Hidalgo-Miranda A, Mendoza-Vargas A, Reyes-Grajeda JP, Abraham Campos-Romero, Moreno-Camacho José Luis, Rodriguez-Gallegos Jorge, Luna-Ruiz Marco, Gonzalez-Barrera D, Rangel-DeLeon D, Munguia-Garza P, Ramirez-Vega O, Escobar-Arrazola, M, Herrera-Montalvo LA. |
| EPI_ISL_2810062 | Salud Digna | Instituto Nacional de Medicina Genomica | Cedro-Tanda A, Hidalgo-Miranda A, Mendoza-Vargas A, Reyes-Grajeda JP, Gonzalez-Barrera D, Rangel-DeLeon D, Munguia-Garza P, Ramirez-Vega O, Escobar-Arrazola, M, Herrera-Montalvo LA. |
| EPI_ISL_2810064, EPI_ISL_2810066, EPI_ISL_2810071, EPI_ISL_2810075, EPI_ISL_2810076, EPI_ISL_2810077, EPI_ISL_2810083, EPI_ISL_2810084, EPI_ISL_2810085, EPI_ISL_2810087, EPI_ISL_2810089, EPI_ISL_2810090, EPI_ISL_2810092, EPI_ISL_2810093, EPI_ISL_2810095, EPI_ISL_2810097, EPI_ISL_2810099, EPI_ISL_2810106 | | | |
| see above | Salud Digna | Instituto Nacional de Medicina Genomica | Cedro-Tanda A, Hidalgo-Miranda A, Mendoza-Vargas A, Reyes-Grajeda JP, Abraham Campos-Romero, Moreno-Camacho José Luis, Rodriguez-Gallegos Jorge, Luna-Ruiz Marco, Gonzalez-Barrera D, Rangel-DeLeon D, Munguia-Garza P, Ramirez-Vega O, Escobar-Arrazola, M, Herrera-Montalvo LA. |
| EPI_ISL_2810110 | Arión Genética | Instituto Nacional de Medicina Genomica | Cedro-Tanda A, Hidalgo-Miranda A, Mendoza-Vargas A, Reyes-Grajeda JP, Gonzalez-Barrera D, Rangel-DeLeon D, Munguia-Garza P, Ramirez-Vega O, Escobar-Arrazola M, Uribe-Figueroa Laura, Roldan-Castillo Magaly, Verea Jazmin, Díaz-Quiñonez José Alberto, Herrera-Montalvo LA. |
| EPI_ISL_2810111 | Salud Digna | Instituto Nacional de Medicina Genomica | Cedro-Tanda A, Hidalgo-Miranda A, Mendoza-Vargas A, Reyes-Grajeda JP, Abraham Campos-Romero, Moreno-Camacho José Luis, Rodriguez-Gallegos Jorge, Luna-Ruiz Marco, Gonzalez-Barrera D, Rangel-DeLeon D, Munguia-Garza P, Ramirez-Vega O, Escobar-Arrazola, M, Herrera-Montalvo LA. |
| EPI_ISL_2810112, EPI_ISL_2810113 | Arión Genética | Instituto Nacional de Medicina Genomica | Cedro-Tanda A, Hidalgo-Miranda A, Mendoza-Vargas A, Reyes-Grajeda JP, Gonzalez-Barrera D, Rangel-DeLeon D, Munguia-Garza P, Ramirez-Vega O, Escobar-Arrazola M, Uribe-Figueroa Laura, Roldan-Castillo Magaly, Verea Jazmin, Díaz-Quiñonez José Alberto, Herrera-Montalvo LA. |
| EPI_ISL_2810116 | Salud Digna | Instituto Nacional de Medicina Genomica | Cedro-Tanda A, Hidalgo-Miranda A, Mendoza-Vargas A, Reyes-Grajeda JP, Abraham Campos-Romero, Moreno-Camacho José Luis, Rodriguez-Gallegos Jorge, Luna-Ruiz Marco, Gonzalez-Barrera D, Rangel-DeLeon D, Munguia-Garza P, Ramirez-Vega O, Escobar-Arrazola, M, Herrera-Montalvo LA. |
| EPI_ISL_2810117 | Arión Genética | Instituto Nacional de Medicina Genomica | Cedro-Tanda A, Hidalgo-Miranda A, Mendoza-Vargas A, Reyes-Grajeda JP, Gonzalez-Barrera D, Rangel-DeLeon D, Munguia-Garza P, Ramirez-Vega O, Escobar-Arrazola M, Uribe-Figueroa Laura, Roldan-Castillo Magaly, Verea Jazmin, Díaz-Quiñonez José Alberto, Herrera-Montalvo LA. |
| EPI_ISL_2810118, EPI_ISL_2810119 | Salud Digna | Instituto Nacional de Medicina Genomica | Cedro-Tanda A, Hidalgo-Miranda A, Mendoza-Vargas A, Reyes-Grajeda JP, Abraham Campos-Romero, Moreno-Camacho José Luis, Rodriguez-Gallegos Jorge, Luna-Ruiz Marco, Gonzalez-Barrera D, Rangel-DeLeon D, Munguia-Garza P, Ramirez-Vega O, Escobar-Arrazola, M, Herrera-Montalvo LA. |
| EPI_ISL_2810120 | Arión Genética | Instituto Nacional de Medicina Genomica | Cedro-Tanda A, Hidalgo-Miranda A, Mendoza-Vargas A, Reyes-Grajeda JP, Gonzalez-Barrera D, Rangel-DeLeon D, Munguia-Garza P, Ramirez-Vega O, Escobar-Arrazola M, Uribe-Figueroa Laura, Roldan-Castillo Magaly, Verea Jazmin, Díaz-Quiñonez José Alberto, Herrera-Montalvo LA. |
| EPI_ISL_2810123, EPI_ISL_2810124, EPI_ISL_2810125, EPI_ISL_2810129, EPI_ISL_2810130, | Salud Digna | Instituto Nacional de Medicina Genomica | Cedro-Tanda A, Hidalgo-Miranda A, Mendoza-Vargas A, Reyes-Grajeda JP, Abraham Campos-Romero, Moreno-Camacho José Luis, Rodriguez-Gallegos Jorge, Luna-Ruiz Marco, Gonzalez-Barrera D, Rangel-DeLeon D, Munguia-Garza P, Ramirez-Vega O, Escobar-Arrazola, M, Herrera-Montalvo LA. |

| | | | |
|---|---|---|---|
| EPI_ISL_2810131 | | | |
| EPI_ISL_2810132 | Arión Genética | Instituto Nacional de Medicina Genomica | Cedro-Tanda A, Hidalgo-Miranda A, Mendoza-Vargas A, Reyes-Grajeda JP, Gonzalez-Barrera D, Rangel-DeLeon D, Munguia-Garza P, Ramirez-Vega O, Escobar-Arrazola M, Uribe-Figueroa Laura, Roldan-Castillo Magaly, Verea Jazmin, Díaz-Quiñonez José Alberto, Herrera-Montalvo LA. |
| EPI_ISL_2810134, EPI_ISL_2810138, EPI_ISL_2810142, EPI_ISL_2810144 | Salud Digna | Instituto Nacional de Medicina Genomica | Cedro-Tanda A, Hidalgo-Miranda A, Mendoza-Vargas A, Reyes-Grajeda JP, Abraham Campos-Romero, Moreno-Camacho José Luis, Rodriguez-Gallegos Jorge, Luna-Ruiz Marco, Gonzalez-Barrera D, Rangel-DeLeon D, Munguia-Garza P, Ramirez-Vega O, Escobar-Arrazola, M, Herrera-Montalvo LA. |
| EPI_ISL_2810150, EPI_ISL_2810151, EPI_ISL_2810152 | Arión Genética | Instituto Nacional de Medicina Genomica | Cedro-Tanda A, Hidalgo-Miranda A, Mendoza-Vargas A, Reyes-Grajeda JP, Gonzalez-Barrera D, Rangel-DeLeon D, Munguia-Garza P, Ramirez-Vega O, Escobar-Arrazola M, Uribe-Figueroa Laura, Roldan-Castillo Magaly, Verea Jazmin, Díaz-Quiñonez José Alberto, Herrera-Montalvo LA. |
| EPI_ISL_2810153 | Salud Digna | Instituto Nacional de Medicina Genomica | Cedro-Tanda A, Hidalgo-Miranda A, Mendoza-Vargas A, Reyes-Grajeda JP, Abraham Campos-Romero, Moreno-Camacho José Luis, Rodriguez-Gallegos Jorge, Luna-Ruiz Marco, Gonzalez-Barrera D, Rangel-DeLeon D, Munguia-Garza P, Ramirez-Vega O, Escobar-Arrazola, M, Herrera-Montalvo LA. |
| EPI_ISL_2810154 | Arión Genética | Instituto Nacional de Medicina Genomica | Cedro-Tanda A, Hidalgo-Miranda A, Mendoza-Vargas A, Reyes-Grajeda JP, Gonzalez-Barrera D, Rangel-DeLeon D, Munguia-Garza P, Ramirez-Vega O, Escobar-Arrazola M, Uribe-Figueroa Laura, Roldan-Castillo Magaly, Verea Jazmin, Díaz-Quiñonez José Alberto, Herrera-Montalvo LA. |
| EPI_ISL_2810155, EPI_ISL_2810158 | Salud Digna | Instituto Nacional de Medicina Genomica | Cedro-Tanda A, Hidalgo-Miranda A, Mendoza-Vargas A, Reyes-Grajeda JP, Abraham Campos-Romero, Moreno-Camacho José Luis, Rodriguez-Gallegos Jorge, Luna-Ruiz Marco, Gonzalez-Barrera D, Rangel-DeLeon D, Munguia-Garza P, Ramirez-Vega O, Escobar-Arrazola, M, Herrera-Montalvo LA. |
| EPI_ISL_2810159 | Arión Genética | Instituto Nacional de Medicina Genomica | Cedro-Tanda A, Hidalgo-Miranda A, Mendoza-Vargas A, Reyes-Grajeda JP, Gonzalez-Barrera D, Rangel-DeLeon D, Munguia-Garza P, Ramirez-Vega O, Escobar-Arrazola M, Uribe-Figueroa Laura, Roldan-Castillo Magaly, Verea Jazmin, Díaz-Quiñonez José Alberto, Herrera-Montalvo LA. |
| EPI_ISL_2810160, EPI_ISL_2810162 | Salud Digna | Instituto Nacional de Medicina Genomica | Cedro-Tanda A, Hidalgo-Miranda A, Mendoza-Vargas A, Reyes-Grajeda JP, Abraham Campos-Romero, Moreno-Camacho José Luis, Rodriguez-Gallegos Jorge, Luna-Ruiz Marco, Gonzalez-Barrera D, Rangel-DeLeon D, Munguia-Garza P, Ramirez-Vega O, Escobar-Arrazola, M, Herrera-Montalvo LA. |
| EPI_ISL_2810163 | Arión Genética | Instituto Nacional de Medicina Genomica | Cedro-Tanda A, Hidalgo-Miranda A, Mendoza-Vargas A, Reyes-Grajeda JP, Gonzalez-Barrera D, Rangel-DeLeon D, Munguia-Garza P, Ramirez-Vega O, Escobar-Arrazola M, Uribe-Figueroa Laura, Roldan-Castillo Magaly, Verea Jazmin, Díaz-Quiñonez José Alberto, Herrera-Montalvo LA. |
| EPI_ISL_2810164 | Salud Digna | Instituto Nacional de Medicina Genomica | Cedro-Tanda A, Hidalgo-Miranda A, Mendoza-Vargas A, Reyes-Grajeda JP, Abraham Campos-Romero, Moreno-Camacho José Luis, Rodriguez-Gallegos Jorge, Luna-Ruiz Marco, Gonzalez-Barrera D, Rangel-DeLeon D, Munguia-Garza P, Ramirez-Vega O, Escobar-Arrazola, M, Herrera-Montalvo LA. |
| EPI_ISL_2810165 | Arión Genética | Instituto Nacional de Medicina Genomica | Cedro-Tanda A, Hidalgo-Miranda A, Mendoza-Vargas A, Reyes-Grajeda JP, Gonzalez-Barrera D, Rangel-DeLeon D, Munguia-Garza P, Ramirez-Vega O, Escobar-Arrazola M, Uribe-Figueroa Laura, Roldan-Castillo Magaly, Verea Jazmin, Díaz-Quiñonez José Alberto, Herrera-Montalvo LA. |
| EPI_ISL_2810166, EPI_ISL_2810172, EPI_ISL_2810173, EPI_ISL_2810175, EPI_ISL_2810177, EPI_ISL_2810178, EPI_ISL_2810180, EPI_ISL_2810181, EPI_ISL_2810182, EPI_ISL_2810184, EPI_ISL_2810186, EPI_ISL_2810187, EPI_ISL_2810189, EPI_ISL_2810190, EPI_ISL_2810191, EPI_ISL_2810192, EPI_ISL_2810193, EPI_ISL_2810194, EPI_ISL_2810195, EPI_ISL_2810196, EPI_ISL_2810197, EPI_ISL_2810198, EPI_ISL_2810199, EPI_ISL_2810200, EPI_ISL_2810201, EPI_ISL_2810202, EPI_ISL_2810203, EPI_ISL_2810204, EPI_ISL_2810205, EPI_ISL_2810206, EPI_ISL_2810207, EPI_ISL_2810209, EPI_ISL_2810210, EPI_ISL_2810211, EPI_ISL_2810212, EPI_ISL_2810213, EPI_ISL_2810214, EPI_ISL_2810215, EPI_ISL_2810216, EPI_ISL_2810217, EPI_ISL_2810219, EPI_ISL_2810220, EPI_ISL_2810221, EPI_ISL_2810222, EPI_ISL_2810223, EPI_ISL_2810224, EPI_ISL_2810225, EPI_ISL_2810226, EPI_ISL_2810227, EPI_ISL_2810228, EPI_ISL_2810229, EPI_ISL_2810230, EPI_ISL_2810231, EPI_ISL_2810233, EPI_ISL_2810234, EPI_ISL_2810235, EPI_ISL_2810236, EPI_ISL_2810237, EPI_ISL_2810238, EPI_ISL_2810239, EPI_ISL_2810240, EPI_ISL_2810241, EPI_ISL_2810242, EPI_ISL_2810243, EPI_ISL_2810244, EPI_ISL_2810245, EPI_ISL_2810248, EPI_ISL_2810249, EPI_ISL_2810252, EPI_ISL_2810253, EPI_ISL_2810254, EPI_ISL_2810255, EPI_ISL_2810257, EPI_ISL_2810260, EPI_ISL_2810261, EPI_ISL_2810266, EPI_ISL_2810267, EPI_ISL_2810268, EPI_ISL_2810269, EPI_ISL_2810272, EPI_ISL_2810273, EPI_ISL_2810274, EPI_ISL_2810276, EPI_ISL_2810280, EPI_ISL_2810281, EPI_ISL_2810282, EPI_ISL_2810283, EPI_ISL_2810287, EPI_ISL_2810288, EPI_ISL_2810289, EPI_ISL_2810290, EPI_ISL_2810291, EPI_ISL_2810297, EPI_ISL_2810300, EPI_ISL_2810302, EPI_ISL_2810303, EPI_ISL_2810307, EPI_ISL_2810308, EPI_ISL_2810309, EPI_ISL_2810315, EPI_ISL_2810316, EPI_ISL_2810317, EPI_ISL_2810321, EPI_ISL_2810323, EPI_ISL_2810324 | | | |
| see above | Salud Digna | Instituto Nacional de Medicina Genomica | Cedro-Tanda A, Hidalgo-Miranda A, Mendoza-Vargas A, Reyes-Grajeda JP, Abraham Campos-Romero, Moreno-Camacho José Luis, Rodriguez-Gallegos Jorge, Luna-Ruiz Marco, Gonzalez-Barrera D, Rangel-DeLeon D, Munguia-Garza P, Ramirez-Vega O, Escobar-Arrazola, M, Herrera-Montalvo LA. |
| EPI_ISL_2810328, EPI_ISL_2810329, EPI_ISL_2810330, EPI_ISL_2810332, EPI_ISL_2810333, EPI_ISL_2810334, EPI_ISL_2810335, EPI_ISL_2810336, EPI_ISL_2810338, EPI_ISL_2810339, EPI_ISL_2810340, EPI_ISL_2810341, EPI_ISL_2810342, EPI_ISL_2810343, EPI_ISL_2810344, EPI_ISL_2810345, EPI_ISL_2810346, EPI_ISL_2810347, EPI_ISL_2810348, EPI_ISL_2810349, EPI_ISL_2810350, EPI_ISL_2810351, EPI_ISL_2810352, EPI_ISL_2810353, EPI_ISL_2810354, EPI_ISL_2810356, EPI_ISL_2810357, EPI_ISL_2810359, EPI_ISL_2810360, EPI_ISL_2810361, EPI_ISL_2810362, EPI_ISL_2810363, EPI_ISL_2810364, EPI_ISL_2810365, EPI_ISL_2810366, EPI_ISL_2810367, EPI_ISL_2810368, EPI_ISL_2810369, EPI_ISL_2810370, EPI_ISL_2810371, EPI_ISL_2810372, EPI_ISL_2810373, EPI_ISL_2810374, EPI_ISL_2810375, EPI_ISL_2810377, EPI_ISL_2810378, EPI_ISL_2810379, EPI_ISL_2810380, EPI_ISL_2810381, EPI_ISL_2810382, EPI_ISL_2810383, EPI_ISL_2810384, EPI_ISL_2810385, EPI_ISL_2810386, EPI_ISL_2810387, EPI_ISL_2810388, EPI_ISL_2810390, EPI_ISL_2810393, EPI_ISL_2810394, EPI_ISL_2810395, EPI_ISL_2810396, EPI_ISL_2810397, EPI_ISL_2810399, EPI_ISL_2810400, EPI_ISL_2810401, EPI_ISL_2810402, EPI_ISL_2810403, EPI_ISL_2810404, EPI_ISL_2810405, EPI_ISL_2810406, EPI_ISL_2810407, EPI_ISL_2810408, EPI_ISL_2810409, EPI_ISL_2810410, EPI_ISL_2810411, EPI_ISL_2810412, EPI_ISL_2810413, EPI_ISL_2810414, EPI_ISL_2810415, EPI_ISL_2810416, EPI_ISL_2810417, EPI_ISL_2810418, EPI_ISL_2810419, EPI_ISL_2810420, EPI_ISL_2810421, EPI_ISL_2810422, EPI_ISL_2810423, EPI_ISL_2810424, EPI_ISL_2810425, EPI_ISL_2810426, EPI_ISL_2810427, EPI_ISL_2810428, EPI_ISL_2810429, EPI_ISL_2810430, EPI_ISL_2810431, EPI_ISL_2810433, EPI_ISL_2810434, EPI_ISL_2810435, EPI_ISL_2810436, EPI_ISL_2810437, EPI_ISL_2810438, EPI_ISL_2810439, EPI_ISL_2810441, EPI_ISL_2810443, EPI_ISL_2810444, EPI_ISL_2810445, EPI_ISL_2810446, EPI_ISL_2810447, EPI_ISL_2810448, EPI_ISL_2810450, EPI_ISL_2810451, EPI_ISL_2810452, EPI_ISL_2810453, EPI_ISL_2810454, EPI_ISL_2810455, EPI_ISL_2810457, EPI_ISL_2810458, EPI_ISL_2810460, EPI_ISL_2810461, EPI_ISL_2810462, EPI_ISL_2810463, EPI_ISL_2810464, EPI_ISL_2810465, EPI_ISL_2810466, EPI_ISL_2810467, EPI_ISL_2810468, EPI_ISL_2810469, EPI_ISL_2810470, EPI_ISL_2810471, EPI_ISL_2810472 | | | |
| see above | UNILABS | Instituto Nacional de Saude (INSA) and BioSystems & Integrative Sciences Institute (BioISI) Genomics Unit, FCUL | Borges et al |
| EPI_ISL_2810473, EPI_ISL_2810474, EPI_ISL_2810475, EPI_ISL_2810476, EPI_ISL_2810477, EPI_ISL_2810478 | ARS Algarve - Laboratorio Laura Ayres | Instituto Nacional de Saude (INSA) | Borges et al |
| EPI_ISL_2810479 | CHLC | Instituto Nacional de Saude (INSA) | Borges et al |
| EPI_ISL_2810480, EPI_ISL_2810481, EPI_ISL_2810482 | CH Setubal | Instituto Nacional de Saude (INSA) | Borges et al |
| EPI_ISL_2810485 | Labocentro | Instituto Nacional de Saude (INSA) | Borges et al |
| EPI_ISL_2810486, EPI_ISL_2810487, EPI_ISL_2810488, EPI_ISL_2810489, EPI_ISL_2810490 | SYNLAB | Instituto Nacional de Saude (INSA) | Borges et al |
| EPI_ISL_2810491, EPI_ISL_2810492, EPI_ISL_2810493, EPI_ISL_2810494, EPI_ISL_2810495, EPI_ISL_2810496, EPI_ISL_2810499, EPI_ISL_2810500, EPI_ISL_2810501, EPI_ISL_2810502, EPI_ISL_2810503, EPI_ISL_2810504, EPI_ISL_2810505, EPI_ISL_2810507, EPI_ISL_2810510, EPI_ISL_2810511, EPI_ISL_2810512, EPI_ISL_2810513, EPI_ISL_2810515, EPI_ISL_2810517, EPI_ISL_2810518, EPI_ISL_2810519, EPI_ISL_2810520, EPI_ISL_2810521, EPI_ISL_2810522, EPI_ISL_2810524, EPI_ISL_2810525, EPI_ISL_2810526, EPI_ISL_2810527, EPI_ISL_2810528, EPI_ISL_2810529, EPI_ISL_2810531, EPI_ISL_2810532, EPI_ISL_2810533, EPI_ISL_2810534, EPI_ISL_2810535, EPI_ISL_2810536, EPI_ISL_2810537, EPI_ISL_2810538, EPI_ISL_2810539, EPI_ISL_2810540, EPI_ISL_2810541, EPI_ISL_2810542, EPI_ISL_2810543, EPI_ISL_2810544, EPI_ISL_2810545, EPI_ISL_2810546, EPI_ISL_2810547, EPI_ISL_2810548, EPI_ISL_2810549, EPI_ISL_2810550, EPI_ISL_2810551, EPI_ISL_2810552, EPI_ISL_2810553, EPI_ISL_2810554, EPI_ISL_2810556, EPI_ISL_2810557, EPI_ISL_2810558, EPI_ISL_2810559, EPI_ISL_2810560, EPI_ISL_2810561, EPI_ISL_2810562, EPI_ISL_2810563, EPI_ISL_2810564, EPI_ISL_2810566, EPI_ISL_2810567, EPI_ISL_2810569, EPI_ISL_2810570, EPI_ISL_2810571, EPI_ISL_2810572, EPI_ISL_2810574, EPI_ISL_2810576, EPI_ISL_2810577, EPI_ISL_2810578, EPI_ISL_2810579, EPI_ISL_2810580, EPI_ISL_2810581, EPI_ISL_2810582, EPI_ISL_2810583, EPI_ISL_2810584, EPI_ISL_2810585, EPI_ISL_2810586, EPI_ISL_2810587, EPI_ISL_2810588, EPI_ISL_2810589, EPI_ISL_2810590, EPI_ISL_2810591, EPI_ISL_2810592, EPI_ISL_2810593, EPI_ISL_2810594, EPI_ISL_2810595, EPI_ISL_2810596, EPI_ISL_2810597, EPI_ISL_2810598, EPI_ISL_2810599, EPI_ISL_2810600, EPI_ISL_2810601, EPI_ISL_2810602, EPI_ISL_2810603, EPI_ISL_2810604, EPI_ISL_2810605, EPI_ISL_2810606, EPI_ISL_2810607, EPI_ISL_2810608, EPI_ISL_2810609, EPI_ISL_2810610, EPI_ISL_2810611, EPI_ISL_2810612, EPI_ISL_2810613, EPI_ISL_2810614, EPI_ISL_2810615, EPI_ISL_2810616, EPI_ISL_2810617, EPI_ISL_2810618, EPI_ISL_2810619, EPI_ISL_2810620, EPI_ISL_2810621, EPI_ISL_2810623, EPI_ISL_2810624, EPI_ISL_2810626, EPI_ISL_2810627, EPI_ISL_2810628, EPI_ISL_2810629, EPI_ISL_2810630, EPI_ISL_2810631, EPI_ISL_2810632, EPI_ISL_2810633, EPI_ISL_2810634, EPI_ISL_2810636, EPI_ISL_2810637, EPI_ISL_2810638, EPI_ISL_2810639, EPI_ISL_2810640, EPI_ISL_2810641, EPI_ISL_2810642, EPI_ISL_2810644, EPI_ISL_2810645, EPI_ISL_2810646, EPI_ISL_2810648, EPI_ISL_2810650, EPI_ISL_2810651, EPI_ISL_2810652, EPI_ISL_2810653, EPI_ISL_2810654, EPI_ISL_2810655, EPI_ISL_2810656, EPI_ISL_2810657, EPI_ISL_2810658, EPI_ISL_2810659, EPI_ISL_2810660, EPI_ISL_2810661, EPI_ISL_2810662, EPI_ISL_2810663, EPI_ISL_2810664, EPI_ISL_2810665, EPI_ISL_2810667, EPI_ISL_2810668, EPI_ISL_2810670, EPI_ISL_2810671, EPI_ISL_2810672, EPI_ISL_2810673, EPI_ISL_2810674, EPI_ISL_2810675, EPI_ISL_2810676, EPI_ISL_2810677, EPI_ISL_2810678, EPI_ISL_2810679, EPI_ISL_2810680, EPI_ISL_2810681, EPI_ISL_2810682, EPI_ISL_2810683, EPI_ISL_2810684, EPI_ISL_2810685, EPI_ISL_2810686, EPI_ISL_2810687, EPI_ISL_2810688, EPI_ISL_2810689, EPI_ISL_2810691, EPI_ISL_2810692, EPI_ISL_2810694, EPI_ISL_2810695, EPI_ISL_2810696, EPI_ISL_2810698, EPI_ISL_2810699, EPI_ISL_2810700, EPI_ISL_2810701, EPI_ISL_2810702, EPI_ISL_2810703, EPI_ISL_2810704, EPI_ISL_2810706, EPI_ISL_2810707, EPI_ISL_2810708, EPI_ISL_2810709, EPI_ISL_2810710, EPI_ISL_2810711, EPI_ISL_2810713, EPI_ISL_2810714, EPI_ISL_2810715, EPI_ISL_2810716, EPI_ISL_2810717, EPI_ISL_2810718, EPI_ISL_2810719, EPI_ISL_2810720, EPI_ISL_2810721, EPI_ISL_2810722, EPI_ISL_2810723, EPI_ISL_2810724, EPI_ISL_2810726, EPI_ISL_2810727, EPI_ISL_2810728, EPI_ISL_2810729, EPI_ISL_2810730, EPI_ISL_2810731, EPI_ISL_2810732, EPI_ISL_2810733, EPI_ISL_2810734, EPI_ISL_2810735, EPI_ISL_2810736, EPI_ISL_2810737, EPI_ISL_2810738, EPI_ISL_2810739, EPI_ISL_2810740, EPI_ISL_2810741, EPI_ISL_2810742, EPI_ISL_2810743, EPI_ISL_2810744, EPI_ISL_2810745, EPI_ISL_2810746, EPI_ISL_2810747 | | | |
| see above | UNILABS | Instituto Nacional de Saude (INSA) | Borges et al |
| EPI_ISL_2811394 | MAX Hospital, Delhi | CSIR-IGIB | Rajesh Pandey, Priti Devi, Priyanka Mehta, Jayanthi Shastri, Swapneil Parikh, Veena Aggarwal, Sachee Agrawal, Nirjhar Chaterjee, Rajit Shah |
| EPI_ISL_2811395 | MAX Hospital, Delhi | CSIR-IGIB | Rajesh Pandey, Jayanthi Shastri, Swapneil Parikh, Veena Aggarwal, Sachee Agrawal, Nirjhar Chaterjee, Rajit Shah, Priti Devi, Priyanka Mehta |
| EPI_ISL_2811411 | Genome Analysis Center, Yamanashi Central Hospital | Genome Analysis Center, Yamanashi Central Hospital | Yosuke Hirotsu |
| EPI_ISL_2811412, EPI_ISL_2811413, EPI_ISL_2811414, EPI_ISL_2811415, EPI_ISL_2811416, EPI_ISL_2811417, EPI_ISL_2811418, EPI_ISL_2811419, EPI_ISL_2811420, EPI_ISL_2811421, EPI_ISL_2811422, EPI_ISL_2811423, EPI_ISL_2811424, EPI_ISL_2811425, EPI_ISL_2811426, EPI_ISL_2811427, EPI_ISL_2811428, EPI_ISL_2811429, EPI_ISL_2811430, EPI_ISL_2811431, EPI_ISL_2811432, EPI_ISL_2811433, EPI_ISL_2811434, EPI_ISL_2811435, EPI_ISL_2811436, EPI_ISL_2811437, EPI_ISL_2811438, EPI_ISL_2811440, EPI_ISL_2811441, EPI_ISL_2811443, EPI_ISL_2811444, EPI_ISL_2811445, EPI_ISL_2811446, EPI_ISL_2811447, EPI_ISL_2811448, EPI_ISL_2811449, EPI_ISL_2811450, EPI_ISL_2811451, EPI_ISL_2811452, EPI_ISL_2811453, EPI_ISL_2811454, EPI_ISL_2811455, EPI_ISL_2811456, EPI_ISL_2811457, EPI_ISL_2811458, EPI_ISL_2811459, EPI_ISL_2811460, EPI_ISL_2811462, EPI_ISL_2811463, EPI_ISL_2811464, EPI_ISL_2811465, EPI_ISL_2811466, EPI_ISL_2811467, EPI_ISL_2811468, EPI_ISL_2811469, EPI_ISL_2811470, EPI_ISL_2811471, EPI_ISL_2811472, EPI_ISL_2811473, EPI_ISL_2811474, EPI_ISL_2811475, EPI_ISL_2811476, EPI_ISL_2811477, EPI_ISL_2811478, EPI_ISL_2811479, EPI_ISL_2811480, EPI_ISL_2811481, EPI_ISL_2811482, EPI_ISL_2811483, EPI_ISL_2811484, EPI_ISL_2811485, EPI_ISL_2811486, EPI_ISL_2811487, EPI_ISL_2811488, EPI_ISL_2811489, EPI_ISL_2811490, EPI_ISL_2811491, EPI_ISL_2811493, EPI_ISL_2811495, EPI_ISL_2811496, EPI_ISL_2811497, EPI_ISL_2811498, EPI_ISL_2811499, EPI_ISL_2811500, EPI_ISL_2811501, EPI_ISL_2811502, EPI_ISL_2811503, EPI_ISL_2811504, EPI_ISL_2811505, EPI_ISL_2811506, EPI_ISL_2811508, EPI_ISL_2811510, EPI_ISL_2811511, EPI_ISL_2811513, EPI_ISL_2811514, EPI_ISL_2811515, EPI_ISL_2811516, EPI_ISL_2811517, EPI_ISL_2811518, EPI_ISL_2811519, EPI_ISL_2811520, EPI_ISL_2811521, EPI_ISL_2811522, EPI_ISL_2811524, EPI_ISL_2811525, EPI_ISL_2811526, EPI_ISL_2811527, EPI_ISL_2811528, EPI_ISL_2811529, EPI_ISL_2811530, EPI_ISL_2811532, EPI_ISL_2811533, EPI_ISL_2811534, EPI_ISL_2811535, EPI_ISL_2811536, EPI_ISL_2811537, EPI_ISL_2811538, EPI_ISL_2811539, EPI_ISL_2811540, EPI_ISL_2811541, EPI_ISL_2811542, EPI_ISL_2811543, EPI_ISL_2811544, EPI_ISL_2811545, EPI_ISL_2811546, EPI_ISL_2811547, EPI_ISL_2811548, EPI_ISL_2811549, EPI_ISL_2811550, EPI_ISL_2811551, EPI_ISL_2811552, EPI_ISL_2811553, EPI_ISL_2811556, EPI_ISL_2811558, EPI_ISL_2811559, EPI_ISL_2811560, EPI_ISL_2811561, EPI_ISL_2811562, EPI_ISL_2811564, EPI_ISL_2811565, EPI_ISL_2811566, EPI_ISL_2811567, EPI_ISL_2811568, EPI_ISL_2811569, EPI_ISL_2811571, EPI_ISL_2811572, EPI_ISL_2811573, EPI_ISL_2811574, EPI_ISL_2811576, EPI_ISL_2811577, EPI_ISL_2811578, EPI_ISL_2811579, EPI_ISL_2811580, EPI_ISL_2811581, EPI_ISL_2811582, EPI_ISL_2811583, EPI_ISL_2811584, EPI_ISL_2811586, EPI_ISL_2811587, EPI_ISL_2811588, EPI_ISL_2811589, EPI_ISL_2811590, EPI_ISL_2811591, EPI_ISL_2811592, EPI_ISL_2811593, EPI_ISL_2811594, EPI_ISL_2811596, EPI_ISL_2811597, EPI_ISL_2811598, EPI_ISL_2811599, EPI_ISL_2811600, EPI_ISL_2811601, EPI_ISL_2811602, EPI_ISL_2811603, EPI_ISL_2811604, EPI_ISL_2811605, EPI_ISL_2811606, EPI_ISL_2811607, EPI_ISL_2811608, EPI_ISL_2811609, EPI_ISL_2811610, EPI_ISL_2811612, EPI_ISL_2811614, EPI_ISL_2811615, EPI_ISL_2811616, EPI_ISL_2811617, EPI_ISL_2811618, EPI_ISL_2811619, EPI_ISL_2811620, EPI_ISL_2811621, EPI_ISL_2811622, EPI_ISL_2811623, EPI_ISL_2811624, EPI_ISL_2811625, EPI_ISL_2811626, EPI_ISL_2811627, EPI_ISL_2811628, EPI_ISL_2811629, EPI_ISL_2811630, EPI_ISL_2811631, EPI_ISL_2811632, EPI_ISL_2811633, EPI_ISL_2811634, EPI_ISL_2811635, EPI_ISL_2811636, EPI_ISL_2811637, EPI_ISL_2811638, EPI_ISL_2811639, EPI_ISL_2811641, EPI_ISL_2811642, EPI_ISL_2811643, EPI_ISL_2811644, EPI_ISL_2811645, EPI_ISL_2811646, EPI_ISL_2811647, EPI_ISL_2811648, EPI_ISL_2811649, EPI_ISL_2811650, EPI_ISL_2811651, EPI_ISL_2811652, EPI_ISL_2811653, EPI_ISL_2811654, EPI_ISL_2811655, EPI_ISL_2811656, EPI_ISL_2811657, EPI_ISL_2811658, EPI_ISL_2811659, EPI_ISL_2811660, EPI_ISL_2811661, EPI_ISL_2811662, EPI_ISL_2811664, EPI_ISL_2811665, EPI_ISL_2811666, EPI_ISL_2811667, EPI_ISL_2811668, EPI_ISL_2811669, EPI_ISL_2811670, EPI_ISL_2811671, EPI_ISL_2811672, EPI_ISL_2811673, EPI_ISL_2811674, EPI_ISL_2811675, EPI_ISL_2811676, EPI_ISL_2811677, EPI_ISL_2811678, EPI_ISL_2811679, EPI_ISL_2811680, EPI_ISL_2811681, EPI_ISL_2811682, EPI_ISL_2811683, EPI_ISL_2811685, EPI_ISL_2811687, EPI_ISL_2811688, EPI_ISL_2811689, EPI_ISL_2811690, EPI_ISL_2811691, EPI_ISL_2811692, EPI_ISL_2811693, EPI_ISL_2811694, EPI_ISL_2811695, EPI_ISL_2811696, EPI_ISL_2811697, EPI_ISL_2811698, EPI_ISL_2811700, EPI_ISL_2811701, EPI_ISL_2811702, EPI_ISL_2811703, EPI_ISL_2811704, EPI_ISL_2811705, EPI_ISL_2811706, EPI_ISL_2811707, EPI_ISL_2811709, EPI_ISL_2811710, EPI_ISL_2811711, EPI_ISL_2811712, EPI_ISL_2811713, EPI_ISL_2811714, EPI_ISL_2811715 | | | |
| see above | Shamir Medical Center (Asaf Harofe) | Shamir Medical Center (Asaf Harofe) | Nir Rainy,Chen Weiner,Reut Sorek Abramovich,Yevgeni Yegorov,Anna Vishnevsky,Patricia Benveniste-Lekovitz,Abu Hamad Ramzia,Adina Bar Chaim |
| EPI_ISL_2811720, EPI_ISL_2811721, EPI_ISL_2811722, EPI_ISL_2811723, EPI_ISL_2811724, EPI_ISL_2811725, EPI_ISL_2811726, EPI_ISL_2811728 | Nucleic Acid Testing, National Reference Laboratory | GIGA Medical Genomics | Yvan Butera, Keith Durkin, Maria Artesi, Bouchra Boujemla, Robert Rutayisire, Patrick Tuyisenge, Esperence Umumararungu, Sébastien Bontems, Marie-Pierre Hayette, Nathalie Renotte, Swaibu Gatare, Jacob Souopgui, Sabin Nsanzimana, Vincent Bours, Léon Mutesa |
| EPI_ISL_2811796, EPI_ISL_2811797, EPI_ISL_2811798, EPI_ISL_2811799, EPI_ISL_2811800, EPI_ISL_2811801, EPI_ISL_2811802, EPI_ISL_2811803, EPI_ISL_2811804, EPI_ISL_2811806, EPI_ISL_2811807, EPI_ISL_2811808, EPI_ISL_2811809, EPI_ISL_2811810, EPI_ISL_2811811, EPI_ISL_2811812, EPI_ISL_2811813, EPI_ISL_2811814, EPI_ISL_2811815, EPI_ISL_2811816, EPI_ISL_2811817, EPI_ISL_2811818, EPI_ISL_2811819, EPI_ISL_2811820, EPI_ISL_2811821, EPI_ISL_2811822, EPI_ISL_2811823, EPI_ISL_2811824, EPI_ISL_2811825 | | | |

| | | | |
|---|---|---|---|
| see above | New South Wales Health Pathology Royal Prince Alfred Hospital | Microbiology RPAH | Foster, C.; Au, J.; Ruiz Silva, M.; Deveson, I.; Bull, R.; Van Hal, S.; Rawlinson, W. |
| EPI_ISL_2811830 | RS OMNI Pulomas | Eijkman Institute for Molecular Biology, National Research and Innovation Agency | Edison Johar, Frilasita A Yudhaputri, Muhammad Rezki Rasyak, Willy Agustine, Hidayat Trimarsanto, Lydia V. Panggalo, Iskandar Adnan, Sukma Oktavianthi, Lidwina Priliani, Safarina G Malik, Khin Saw Myint, Amin Soebandrio |
| EPI_ISL_2811831 | RS OMNI Pulomas | Eijkman Institute for Molecular Biology, National Research and Innovation Agency | Hidayat Trimarsanto, Lydia V. Panggalo, Iskandar Adnan, Sukma Oktavianthi, Lidwina Priliani, Edison Johar, Frilasita A Yudhaputri, Muhammad Rezki Rasyak, Willy Agustine, Safarina G Malik, Khin Saw Myint, Amin Soebandrio |
| EPI_ISL_2811832 | RS OMNI Pulomas | Eijkman Institute for Molecular Biology, National Research and Innovation Agency | Willy Agustine, Hidayat Trimarsanto, Lydia V. Panggalo, Iskandar Adnan, Sukma Oktavianthi, Lidwina Priliani, Edison Johar, Frilasita A Yudhaputri, Muhammad Rezki Rasyak, Safarina G Malik, Khin Saw Myint, Amin Soebandrio |
| EPI_ISL_2811834 | RS Pondok Indah Pondok Indah | Eijkman Institute for Molecular Biology, National Research and Innovation Agency | Frilasita A Yudhaputri, Muhammad Rezki Rasyak, Willy Agustine, Hidayat Trimarsanto, Lydia V. Panggalo, Iskandar Adnan, Sukma Oktavianthi, Lidwina Priliani, Edison Johar, Safarina G Malik, Khin Saw Myint, Amin Soebandrio |
| EPI_ISL_2811835 | RS Pondok Indah Pondok Indah | Eijkman Institute for Molecular Biology, National Research and Innovation Agency | Edison Johar, Frilasita A Yudhaputri, Muhammad Rezki Rasyak, Willy Agustine, Hidayat Trimarsanto, Lydia V. Panggalo, Iskandar Adnan, Sukma Oktavianthi, Lidwina Priliani, Safarina G Malik, Khin Saw Myint, Amin Soebandrio |
| EPI_ISL_2811836 | RS Pondok Indah Pondok Indah | Eijkman Institute for Molecular Biology, National Research and Innovation Agency | Iskandar Adnan, Sukma Oktavianthi, Lidwina Priliani, Edison Johar, Frilasita A Yudhaputri, Muhammad Rezki Rasyak, Willy Agustine, Hidayat Trimarsanto, Lydia V. Panggalo, Safarina G Malik, Khin Saw Myint, Amin Soebandrio |
| EPI_ISL_2811837 | BMC Mayapada Hospital | Eijkman Institute for Molecular Biology, National Research and Innovation Agency | Sukma Oktavianthi, Lidwina Priliani, Edison Johar, Frilasita A Yudhaputri, Muhammad Rezki Rasyak, Willy Agustine, Hidayat Trimarsanto, Lydia V. Panggalo, Iskandar Adnan, Safarina G Malik, Khin Saw Myint, Amin Soebandrio |
| EPI_ISL_2811888 | National Centre For Cell Science | National Centre For Cell Science – INSACOG | Dhiraj Paul, Girish R Nair, Kartik Jayant Deopujari, Shivang P. Bhanushali, Mitali Inamdar, Harshada Ghode,Kalpesh Chandrakant Bavkar, Manoj Kumar Bhat, Ajay Pillai, INSACOG Consortium team, Yogesh Shouche |
| EPI_ISL_2811890, EPI_ISL_2811891, EPI_ISL_2811894, EPI_ISL_2811897, EPI_ISL_2811898, EPI_ISL_2811901, EPI_ISL_2811903 | Queensland Health Forensic and Scientific Services | Queensland Health Forensic and Scientific Services | Son Nguyen |
| EPI_ISL_2811920 | UCLA Clinical Micro Lab | Los Angeles County Public Health Laboratories | P. Hemarajata et al. |
| EPI_ISL_2811922, EPI_ISL_2811939, EPI_ISL_2811940 | Los Angeles County Public Health Laboratories | Los Angeles County Public Health Laboratories | P. Hemarajata et al. |
| EPI_ISL_2811943 | USC Clinical Lab | Los Angeles County Public Health Laboratories | P. Hemarajata et al. |
| EPI_ISL_2811944, EPI_ISL_2811945 | Middlemore Hospital | Institute of Environmental Science and Research (ESR) | Rachel Boyle, SallyAnn Harbison, Olivia Stroeven, Xiaoyun Ren, Matt Storey, Nikki Freed, Muhammad Faisal, Jing Wang, Hermes Perez, Anja Werno, Antje van der Linden, Arlo Upton, Chris Mansell, David Hammer, Dragana Drinkovic, Gary McAuliffe, Hana Sofia Andersson, James Ussher, Jill Sherwood, Josh Freeman, Julia Howard, Juliet Elvy, Mary DeAlmeida, Matt Blakiston, Matthew Rogers, Max Bloomfield, Michael Addidle, Michelle Balm, Sally Roberts, Sarah Jefferies, Sharmini Muttaiyah, Susan Morpeth, Susan Taylor, Timothy Blackmore, Vani Sathyendran, Veronica Playle, Virginia Hope, Erasmus Smit, Lauren Jelly, Olin Silander, Joep de Ligt |
| EPI_ISL_2811947 | Canterbury Health Laboratories | Institute of Environmental Science and Research (ESR) | Paula scholes, Susan Lin, Xiaoyun Ren, Matt Storey, Nikki Freed, Muhammad Faisal, Jing Wang, Hermes Perez, Anja Werno, Antje van der Linden, Arlo Upton, Chris Mansell, David Hammer, Dragana Drinkovic, Gary McAuliffe, Hana Sofia Andersson, James Ussher, Jill Sherwood, Josh Freeman, Julia Howard, Juliet Elvy, Mary DeAlmeida, Matt Blakiston, Matthew Rogers, Max Bloomfield, Michael Addidle, Michelle Balm, Sally Roberts, Sarah Jefferies, Sharmini Muttaiyah, Susan Morpeth, Susan Taylor, Timothy Blackmore, Vani Sathyendran, Veronica Playle, Virginia Hope, Erasmus Smit, Lauren Jelly, Olin Silander, Joep de Ligt |
| EPI_ISL_2811949 | LabPLUS | Institute of Environmental Science and Research (ESR) | Rachel Boyle, SallyAnn Harbison, Olivia Stroeven, Xiaoyun Ren, Matt Storey, Nikki Freed, Muhammad Faisal, Jing Wang, Hermes Perez, Anja Werno, Antje van der Linden, Arlo Upton, Chris Mansell, David Hammer, Dragana Drinkovic, Gary McAuliffe, Hana Sofia Andersson, James Ussher, Jill Sherwood, Josh Freeman, Julia Howard, Juliet Elvy, Mary DeAlmeida, Matt Blakiston, Matthew Rogers, Max Bloomfield, Michael Addidle, Michelle Balm, Sally Roberts, Sarah Jefferies, Sharmini Muttaiyah, Susan Morpeth, Susan Taylor, Timothy Blackmore, Vani Sathyendran, Veronica Playle, Virginia Hope, Erasmus Smit, Lauren Jelly, Olin Silander, Joep de Ligt |
| EPI_ISL_2811954 | Middlemore Hospital | Institute of Environmental Science and Research (ESR) | Rachel Boyle, SallyAnn Harbison, Olivia Stroeven, Xiaoyun Ren, Matt Storey, Nikki Freed, Muhammad Faisal, Jing Wang, Hermes Perez, Anja Werno, Antje van der Linden, Arlo Upton, Chris Mansell, David Hammer, Dragana Drinkovic, Gary McAuliffe, Hana Sofia Andersson, James Ussher, Jill Sherwood, Josh Freeman, Julia Howard, Juliet Elvy, Mary DeAlmeida, Matt Blakiston, Matthew Rogers, Max Bloomfield, Michael Addidle, Michelle Balm, Sally Roberts, Sarah Jefferies, Sharmini Muttaiyah, Susan Morpeth, Susan Taylor, Timothy Blackmore, Vani Sathyendran, Veronica Playle, Virginia Hope, Erasmus Smit, Lauren Jelly, Olin Silander, Joep de Ligt |
| EPI_ISL_2811955, EPI_ISL_2811957 | LabPLUS | Institute of Environmental Science and Research (ESR) | Rachel Boyle, SallyAnn Harbison, Olivia Stroeven, Xiaoyun Ren, Matt Storey, Nikki Freed, Muhammad Faisal, Jing Wang, Hermes Perez, Anja Werno, Antje van der Linden, Arlo Upton, Chris Mansell, David Hammer, Dragana Drinkovic, Gary McAuliffe, Hana Sofia Andersson, James Ussher, Jill Sherwood, Josh Freeman, Julia Howard, Juliet Elvy, Mary DeAlmeida, Matt Blakiston, Matthew Rogers, Max Bloomfield, Michael Addidle, Michelle Balm, Sally Roberts, Sarah Jefferies, Sharmini Muttaiyah, Susan Morpeth, Susan Taylor, Timothy Blackmore, Vani Sathyendran, Veronica Playle, Virginia Hope, Erasmus Smit, Lauren Jelly, Olin Silander, Joep de Ligt |
| EPI_ISL_2811958 | Canterbury Health Laboratories | Institute of Environmental Science and Research (ESR) | Paula scholes, Susan Lin, Xiaoyun Ren, Matt Storey, Nikki Freed, Muhammad Faisal, Jing Wang, Hermes Perez, Anja Werno, Antje van der Linden, Arlo Upton, Chris Mansell, David Hammer, Dragana Drinkovic, Gary McAuliffe, Hana Sofia Andersson, James Ussher, Jill Sherwood, Josh Freeman, Julia Howard, Juliet Elvy, Mary DeAlmeida, Matt Blakiston, Matthew Rogers, Max Bloomfield, Michael Addidle, Michelle Balm, Sally Roberts, Sarah Jefferies, Sharmini Muttaiyah, Susan Morpeth, Susan Taylor, Timothy Blackmore, Vani Sathyendran, Veronica Playle, Virginia Hope, Erasmus Smit, Lauren Jelly, Olin Silander, Joep de Ligt |
| EPI_ISL_2811961 | Hollywood Presbyterian Medical Center | Los Angeles County Public Health Laboratories | P. Hemarajata et al. |
| EPI_ISL_2811966 | USC Clinical Lab | Los Angeles County Public Health Laboratories | P. Hemarajata et al. |
| EPI_ISL_2811967, EPI_ISL_2811968, EPI_ISL_2811969, EPI_ISL_2811970, EPI_ISL_2811971, EPI_ISL_2811972, EPI_ISL_2811974, EPI_ISL_2811975, EPI_ISL_2811976, EPI_ISL_2811977, EPI_ISL_2811978, EPI_ISL_2811979 | | | |
| see above | Palmdale Regional Medical Center | Los Angeles County Public Health Laboratories | P. Hemarajata et al. |
| EPI_ISL_2811984 | Pekan Tajau Health Clinic | Institute for Medical Research, Infectious Disease Research Centre, National Institutes of Health, Ministry of Health Malaysia | Suppiah J, Kamel K, Mohd Zawawi Z, Azizan MA, Ramly N, Robert F, Thayan R |
| EPI_ISL_2811985 | St Vincent's Pathology (SydPath) | NSW Health Pathology - Institute of Clinical Pathology and Medical Research; Westmead Hospital; University of Sydney | CIDM-PH et al. |
| EPI_ISL_2811986, EPI_ISL_2811987, EPI_ISL_2811988 | Sydney South West Pathology Service (SSWPS) - Liverpool Hospital - NSW Health Pathology | NSW Health Pathology - Institute of Clinical Pathology and Medical Research; Westmead Hospital; University of Sydney | CIDM-PH et al. |
| EPI_ISL_2811989 | Laverty Pathology | NSW Health Pathology - Institute of Clinical Pathology and Medical Research; Westmead Hospital; University of Sydney | CIDM-PH et al. |
| EPI_ISL_2811990 | 4Cyte Pathology | NSW Health Pathology - Institute of Clinical Pathology and Medical Research; Westmead Hospital; University of Sydney | CIDM-PH et al. |
| EPI_ISL_2811991, EPI_ISL_2811992 | Histopath | NSW Health Pathology - Institute of Clinical Pathology and Medical Research; Westmead Hospital; University of Sydney | CIDM-PH et al. |
| EPI_ISL_2811993, EPI_ISL_2811994 | 4Cyte Pathology | NSW Health Pathology - Institute of Clinical Pathology and Medical Research; Westmead Hospital; University of Sydney | CIDM-PH et al. |
| EPI_ISL_2811995 | Laverty Pathology | NSW Health Pathology - Institute of Clinical Pathology and Medical Research; Westmead Hospital; University of Sydney | CIDM-PH et al. |
| EPI_ISL_2811996 | St Vincent's Pathology (SydPath) | NSW Health Pathology - Institute of Clinical Pathology and Medical Research; Westmead Hospital; University of Sydney | CIDM-PH et al. |
| EPI_ISL_2811997 | Austech Medical Laboratories | NSW Health Pathology - Institute of Clinical Pathology and Medical Research; Westmead Hospital; University of Sydney | CIDM-PH et al. |
| EPI_ISL_2812002 | Flow Diagnostics Lab | Los Angeles County Public Health Laboratories | P. Hemarajata et al. |
| EPI_ISL_2812005 | Saint Francis Medical Center | Los Angeles County Public Health Laboratories | P. Hemarajata et al. |
| EPI_ISL_2812006, EPI_ISL_2812007, EPI_ISL_2812008, EPI_ISL_2812009, EPI_ISL_2812010, EPI_ISL_2812011, EPI_ISL_2812012, EPI_ISL_2812013, EPI_ISL_2812014 | Palmdale Regional Medical Center | Los Angeles County Public Health Laboratories | P. Hemarajata et al. |
| EPI_ISL_2812026, EPI_ISL_2812027, EPI_ISL_2812028 | Rembau Health Clinic | Institute for Medical Research, Infectious Disease Research Centre, National Institutes of Health, Ministry of Health Malaysia | Suppiah J, Kamel K, Mohd Zawawi Z, Azizan MA, Ramly N, Robert F, Thayan R |
| EPI_ISL_2812145 | Saitama Prefectural Institute of Public Health | Saitama Prefectural Institute of Public Health | Tsuyoshi Sekizuka, Kentaro Itokawa, Rina Tanaka, Masanori Hashino, Makoto Kuroda,Satoshi Daisaki,Eri Aonuma,Hayato Ehara |
| EPI_ISL_2812146, EPI_ISL_2812147, EPI_ISL_2812148, EPI_ISL_2812149, EPI_ISL_2812150, EPI_ISL_2812151, EPI_ISL_2812152, EPI_ISL_2812153, EPI_ISL_2812154, EPI_ISL_2812155, EPI_ISL_2812156 | | | |
| see above | Singapore General Hospital | Department of Microbiology | Nurdyana Abdul Rahman, Kenneth Xin Long Chan, Chayaporn Suphavilai, Kun Lee Lim, Sui Sin Goh, Kian Sing Chan, Lynette Oon, Kern Rei Chng, James Sim Heng Chiak, Niranjan Nagarajan, Karrie Ko |
| EPI_ISL_2812180, EPI_ISL_2812182, EPI_ISL_2812183, EPI_ISL_2812184 | Institute for Infectious Diseases | Institute for Infectious Diseases | Stefan Neuenschwander, Christian Baumann, Miguel A Terrazos Miani, Cora SÃ¤gesser, Pascal Bittel, Peter Keller, Franziska Suter-Riniker, Stephen L Leib, Alban Ramette |

| | | | |
|---|---|---|---|
| EPI_ISL_2812256, EPI_ISL_2812258, EPI_ISL_2812259, EPI_ISL_2812260, EPI_ISL_2812261, EPI_ISL_2812264, EPI_ISL_2812265, EPI_ISL_2812266, EPI_ISL_2812267, EPI_ISL_2812268, EPI_ISL_2812269 | | | |
| see above | Microvida | Microvida | Suzan D. Pas, Jaco J. Verweij, Joep J. J. M. Stohr |
| EPI_ISL_2812369, EPI_ISL_2812374 | UFS Virology | KRISP, KZn Research Innovation and Sequencing Platform | Giandhari Jennifer, Pillay Sureshnee, Yajna Ramphal, Naidoo Yeshnee, Tshabuila Derek,Tegally Houriiyah, San James, Wilkinson Eduan, de Oliveira Tulio |
| EPI_ISL_2812444 | Microbiologia CATLAB | Can Ruti SARS-CoV-2 Sequencing Hub (HUGTiP/IrsiCaixa/IGTP) | Marc Noguera-Julian, Pilar Armengol, Ignacio Blanco, Antoni E Bordoy, Francesc Catala-Moll, Pere-Joan Cardona, Maria Casadellà, Cristina Casañ, Gemma Clara, Bonaventura Clotet, Montserrat Giménez, Mercedes Guerrero, Anna Not, David Panisello, Roger Paredes, Mariona Parera, Alexia París, Verónica Saludes, Alba Sánchez, Laia Soler, Lauro Sumoy, and Elisa Martró on behalf of the Can Ruti SARS-CoV-2 Sequencing Hub |
| EPI_ISL_2812446, EPI_ISL_2812447, EPI_ISL_2812451, EPI_ISL_2812454 | Microbiology Department, Laboratori Clínic Metropolitana Nord. Hospital Universitari Germans Trias i Pujol | Can Ruti SARS-CoV-2 Sequencing Hub (HUGTiP/IrsiCaixa/IGTP) | Marc Noguera-Julian, Pilar Armengol, Ignacio Blanco, Antoni E Bordoy, Francesc Catala-Moll, Pere-Joan Cardona, Maria Casadellà, Cristina Casañ, Gemma Clara, Bonaventura Clotet, Montserrat Giménez, Mercedes Guerrero, Anna Not, David Panisello, Roger Paredes, Mariona Parera, Alexia París, Verónica Saludes, Alba Sánchez, Laia Soler, Lauro Sumoy, and Elisa Martró on behalf of the Can Ruti SARS-CoV-2 Sequencing Hub |
| EPI_ISL_2812457 | Microbiologia CATLAB | Can Ruti SARS-CoV-2 Sequencing Hub (HUGTiP/IrsiCaixa/IGTP) | Marc Noguera-Julian, Pilar Armengol, Ignacio Blanco, Antoni E Bordoy, Francesc Catala-Moll, Pere-Joan Cardona, Maria Casadellà, Cristina Casañ, Gemma Clara, Bonaventura Clotet, Montserrat Giménez, Mercedes Guerrero, Anna Not, David Panisello, Roger Paredes, Mariona Parera, Alexia París, Verónica Saludes, Alba Sánchez, Laia Soler, Lauro Sumoy, and Elisa Martró on behalf of the Can Ruti SARS-CoV-2 Sequencing Hub |
| EPI_ISL_2812458, EPI_ISL_2812459, EPI_ISL_2812461, EPI_ISL_2812466, EPI_ISL_2812467, EPI_ISL_2812470, EPI_ISL_2812473, EPI_ISL_2812474, EPI_ISL_2812476 | Microbiology Department, Laboratori Clínic Metropolitana Nord. Hospital Universitari Germans Trias i Pujol | Can Ruti SARS-CoV-2 Sequencing Hub (HUGTiP/IrsiCaixa/IGTP) | Marc Noguera-Julian, Pilar Armengol, Ignacio Blanco, Antoni E Bordoy, Francesc Catala-Moll, Pere-Joan Cardona, Maria Casadellà, Cristina Casañ, Gemma Clara, Bonaventura Clotet, Montserrat Giménez, Mercedes Guerrero, Anna Not, David Panisello, Roger Paredes, Mariona Parera, Alexia París, Verónica Saludes, Alba Sánchez, Laia Soler, Lauro Sumoy, and Elisa Martró on behalf of the Can Ruti SARS-CoV-2 Sequencing Hub |
| EPI_ISL_2812480 | Microbiologia CATLAB | Can Ruti SARS-CoV-2 Sequencing Hub (HUGTiP/IrsiCaixa/IGTP) | Marc Noguera-Julian, Pilar Armengol, Ignacio Blanco, Antoni E Bordoy, Francesc Catala-Moll, Pere-Joan Cardona, Maria Casadellà, Cristina Casañ, Gemma Clara, Bonaventura Clotet, Montserrat Giménez, Mercedes Guerrero, Anna Not, David Panisello, Roger Paredes, Mariona Parera, Alexia París, Verónica Saludes, Alba Sánchez, Laia Soler, Lauro Sumoy, and Elisa Martró on behalf of the Can Ruti SARS-CoV-2 Sequencing Hub |
| EPI_ISL_2812481, EPI_ISL_2812482, EPI_ISL_2812483, EPI_ISL_2812484, EPI_ISL_2812490, EPI_ISL_2812494, EPI_ISL_2812497, EPI_ISL_2812500 | Microbiology Department, Laboratori Clínic Metropolitana Nord. Hospital Universitari Germans Trias i Pujol | Can Ruti SARS-CoV-2 Sequencing Hub (HUGTiP/IrsiCaixa/IGTP) | Marc Noguera-Julian, Pilar Armengol, Ignacio Blanco, Antoni E Bordoy, Francesc Catala-Moll, Pere-Joan Cardona, Maria Casadellà, Cristina Casañ, Gemma Clara, Bonaventura Clotet, Montserrat Giménez, Mercedes Guerrero, Anna Not, David Panisello, Roger Paredes, Mariona Parera, Alexia París, Verónica Saludes, Alba Sánchez, Laia Soler, Lauro Sumoy, and Elisa Martró on behalf of the Can Ruti SARS-CoV-2 Sequencing Hub |
| EPI_ISL_2812502 | Parc Taulí Hospital Universitari | Can Ruti SARS-CoV-2 Sequencing Hub (HUGTiP/IrsiCaixa/IGTP) | Marc Noguera-Julian, Pilar Armengol, Ignacio Blanco, Antoni E Bordoy, Francesc Catala-Moll, Pere-Joan Cardona, Maria Casadellà, Cristina Casañ, Gemma Clara, Bonaventura Clotet, Montserrat Giménez, Mercedes Guerrero, Anna Not, David Panisello, Roger Paredes, Mariona Parera, Alexia París, Verónica Saludes, Alba Sánchez, Laia Soler, Lauro Sumoy, and Elisa Martró on behalf of the Can Ruti SARS-CoV-2 Sequencing Hub |
| EPI_ISL_2812505, EPI_ISL_2812508, EPI_ISL_2812509, EPI_ISL_2812510, EPI_ISL_2812512, EPI_ISL_2812513 | Microbiology Department, Laboratori Clínic Metropolitana Nord. Hospital Universitari Germans Trias i Pujol | Can Ruti SARS-CoV-2 Sequencing Hub (HUGTiP/IrsiCaixa/IGTP) | Marc Noguera-Julian, Pilar Armengol, Ignacio Blanco, Antoni E Bordoy, Francesc Catala-Moll, Pere-Joan Cardona, Maria Casadellà, Cristina Casañ, Gemma Clara, Bonaventura Clotet, Montserrat Giménez, Mercedes Guerrero, Anna Not, David Panisello, Roger Paredes, Mariona Parera, Alexia París, Verónica Saludes, Alba Sánchez, Laia Soler, Lauro Sumoy, and Elisa Martró on behalf of the Can Ruti SARS-CoV-2 Sequencing Hub |
| EPI_ISL_2812514 | Parc Taulí Hospital Universitari | Can Ruti SARS-CoV-2 Sequencing Hub (HUGTiP/IrsiCaixa/IGTP) | Marc Noguera-Julian, Pilar Armengol, Ignacio Blanco, Antoni E Bordoy, Francesc Catala-Moll, Pere-Joan Cardona, Maria Casadellà, Cristina Casañ, Gemma Clara, Bonaventura Clotet, Montserrat Giménez, Mercedes Guerrero, Anna Not, David Panisello, Roger Paredes, Mariona Parera, Alexia París, Verónica Saludes, Alba Sánchez, Laia Soler, Lauro Sumoy, and Elisa Martró on behalf of the Can Ruti SARS-CoV-2 Sequencing Hub |
| EPI_ISL_2812515 | Microbiologia CATLAB | Can Ruti SARS-CoV-2 Sequencing Hub (HUGTiP/IrsiCaixa/IGTP) | Marc Noguera-Julian, Pilar Armengol, Ignacio Blanco, Antoni E Bordoy, Francesc Catala-Moll, Pere-Joan Cardona, Maria Casadellà, Cristina Casañ, Gemma Clara, Bonaventura Clotet, Montserrat Giménez, Mercedes Guerrero, Anna Not, David Panisello, Roger Paredes, Mariona Parera, Alexia París, Verónica Saludes, Alba Sánchez, Laia Soler, Lauro Sumoy, and Elisa Martró on behalf of the Can Ruti SARS-CoV-2 Sequencing Hub |
| EPI_ISL_2812516, EPI_ISL_2812517, EPI_ISL_2812518, EPI_ISL_2812519, EPI_ISL_2812521, EPI_ISL_2812523 | Microbiology Department, Laboratori Clínic Metropolitana Nord. Hospital Universitari Germans Trias i Pujol | Can Ruti SARS-CoV-2 Sequencing Hub (HUGTiP/IrsiCaixa/IGTP) | Marc Noguera-Julian, Pilar Armengol, Ignacio Blanco, Antoni E Bordoy, Francesc Catala-Moll, Pere-Joan Cardona, Maria Casadellà, Cristina Casañ, Gemma Clara, Bonaventura Clotet, Montserrat Giménez, Mercedes Guerrero, Anna Not, David Panisello, Roger Paredes, Mariona Parera, Alexia París, Verónica Saludes, Alba Sánchez, Laia Soler, Lauro Sumoy, and Elisa Martró on behalf of the Can Ruti SARS-CoV-2 Sequencing Hub |
| EPI_ISL_2812524, EPI_ISL_2812526 | Microbiologia CATLAB | Can Ruti SARS-CoV-2 Sequencing Hub (HUGTiP/IrsiCaixa/IGTP) | Marc Noguera-Julian, Pilar Armengol, Ignacio Blanco, Antoni E Bordoy, Francesc Catala-Moll, Pere-Joan Cardona, Maria Casadellà, Cristina Casañ, Gemma Clara, Bonaventura Clotet, Montserrat Giménez, Mercedes Guerrero, Anna Not, David Panisello, Roger Paredes, Mariona Parera, Alexia París, Verónica Saludes, Alba Sánchez, Laia Soler, Lauro Sumoy, and Elisa Martró on behalf of the Can Ruti SARS-CoV-2 Sequencing Hub |
| EPI_ISL_2812533 | Microbiology Department, Laboratori Clínic Metropolitana Nord. Hospital Universitari Germans Trias i Pujol | Can Ruti SARS-CoV-2 Sequencing Hub (HUGTiP/IrsiCaixa/IGTP) | Marc Noguera-Julian, Pilar Armengol, Ignacio Blanco, Antoni E Bordoy, Francesc Catala-Moll, Pere-Joan Cardona, Maria Casadellà, Cristina Casañ, Gemma Clara, Bonaventura Clotet, Montserrat Giménez, Mercedes Guerrero, Anna Not, David Panisello, Roger Paredes, Mariona Parera, Alexia París, Verónica Saludes, Alba Sánchez, Laia Soler, Lauro Sumoy, and Elisa Martró on behalf of the Can Ruti SARS-CoV-2 Sequencing Hub |
| EPI_ISL_2812579, EPI_ISL_2812580, EPI_ISL_2812581, EPI_ISL_2812582, EPI_ISL_2812583, EPI_ISL_2812584, EPI_ISL_2812585, EPI_ISL_2812586, EPI_ISL_2812587, EPI_ISL_2812588, EPI_ISL_2812589, EPI_ISL_2812590, EPI_ISL_2812591, EPI_ISL_2812592, EPI_ISL_2812593, EPI_ISL_2812594, EPI_ISL_2812595, EPI_ISL_2812596, EPI_ISL_2812597, EPI_ISL_2812598, EPI_ISL_2812599, EPI_ISL_2812600, EPI_ISL_2812601, EPI_ISL_2812602, EPI_ISL_2812603, EPI_ISL_2812604, EPI_ISL_2812605, EPI_ISL_2812606, EPI_ISL_2812607, EPI_ISL_2812608, EPI_ISL_2812609, EPI_ISL_2812610, EPI_ISL_2812611, EPI_ISL_2812612, EPI_ISL_2812613, EPI_ISL_2812614, EPI_ISL_2812615, EPI_ISL_2812616, EPI_ISL_2812617, EPI_ISL_2812618, EPI_ISL_2812619, EPI_ISL_2812620, EPI_ISL_2812621, EPI_ISL_2812622, EPI_ISL_2812623, EPI_ISL_2812624, EPI_ISL_2812625, EPI_ISL_2812626, EPI_ISL_2812627, EPI_ISL_2812628, EPI_ISL_2812629, EPI_ISL_2812630, EPI_ISL_2812631, EPI_ISL_2812632, EPI_ISL_2812633, EPI_ISL_2812634, EPI_ISL_2812635, EPI_ISL_2812636, EPI_ISL_2812637, EPI_ISL_2812638, EPI_ISL_2812639, EPI_ISL_2812640, EPI_ISL_2812641, EPI_ISL_2812642, EPI_ISL_2812643, EPI_ISL_2812644, EPI_ISL_2812645, EPI_ISL_2812646 | | | |
| see above | INSACOG-MANIPUR | National Institute of Biomedical Genomics – INSACOG | Arindam Maitra, Nanaocha Sharma, Nidhan Kumar Biswas, R K Manglem Singh, Sreedhar Chinnaswamy, Khumanthem Ranjana, Saumitra Das |
| EPI_ISL_2812651 | Labuan Hospital | Institute for Medical Research, Infectious Disease Research Centre, National Institutes of Health, Ministry of Health Malaysia | Suppiah J, Kamel K, Mohd Zawawi Z, Azizan MA, Ramly N, Robert F, Thayan R |
| EPI_ISL_2812656, EPI_ISL_2812660, EPI_ISL_2812663, EPI_ISL_2812664, EPI_ISL_2812665, EPI_ISL_2812666, EPI_ISL_2812673, EPI_ISL_2812674, EPI_ISL_2812675, EPI_ISL_2812677, EPI_ISL_2812678, EPI_ISL_2812687, EPI_ISL_2812689, EPI_ISL_2812690, EPI_ISL_2812694, EPI_ISL_2812695, EPI_ISL_2812696, EPI_ISL_2812697, EPI_ISL_2812698, EPI_ISL_2812699, EPI_ISL_2812700, EPI_ISL_2812701, EPI_ISL_2812703 | | | |
| see above | ASL Napoli 1 Centro | AMES Centro Polidiagnostico Strumentale S.r.l. | "Giovanni Savarese, Raffaella Ruggiero, Eloisa Evangelista, Antonella Di Carlo, Luisa Circelli, Luigi D'Amore, Roberto Sirica, Monica Ianniello, Nadia Petrillo, Maurizio D'Amora, Antonio Fico" |
| EPI_ISL_2812706 | Sultan Haji Ahmad Shah Hospital | Institute for Medical Research, Infectious Disease Research Centre, National Institutes of Health, Ministry of Health Malaysia | Suppiah J, Kamel K, Mohd Zawawi Z, Azizan MA, Ramly N, Robert F, Thayan R |
| EPI_ISL_2812710, EPI_ISL_2812712, EPI_ISL_2812715, EPI_ISL_2812718, EPI_ISL_2812721, EPI_ISL_2812723, EPI_ISL_2812724, EPI_ISL_2812725, EPI_ISL_2812726 | HOSPITAL UNIVERSITARIO SON ESPASES | HOSPITAL UNIVERSITARIO SON ESPASES | Servicio de Microbiología, Hospital Universitario Son Espases |
| EPI_ISL_2812727 | HOSPITAL CAN MISSES | HOSPITAL UNIVERSITARIO SON ESPASES | Servicio de Microbiología, Hospital Universitario Son Espases |
| EPI_ISL_2812729, EPI_ISL_2812730 | HOSPITAL UNIVERSITARIO SON ESPASES | HOSPITAL UNIVERSITARIO SON ESPASES | Servicio de Microbiología, Hospital Universitario Son Espases |
| EPI_ISL_2812731 | HOSPITAL MANACOR | HOSPITAL UNIVERSITARIO SON ESPASES | Servicio de Microbiología, Hospital Universitario Son Espases |
| EPI_ISL_2812733, EPI_ISL_2812734, EPI_ISL_2812735, EPI_ISL_2812741, EPI_ISL_2812742 | HOSPITAL UNIVERSITARIO SON ESPASES | HOSPITAL UNIVERSITARIO SON ESPASES | Servicio de Microbiología, Hospital Universitario Son Espases |
| EPI_ISL_2812744 | HOSPITAL UNIVERSITARIO SON LLATZER | HOSPITAL UNIVERSITARIO SON ESPASES | Servicio de Microbiología, Hospital Universitario Son Espases |
| EPI_ISL_2812748, EPI_ISL_2812749, EPI_ISL_2812752 | HOSPITAL UNIVERSITARIO SON ESPASES | HOSPITAL UNIVERSITARIO SON ESPASES | Servicio de Microbiología, Hospital Universitario Son Espases |
| EPI_ISL_2812754 | HOSPITAL CAN MISSES | HOSPITAL UNIVERSITARIO SON ESPASES | Servicio de Microbiología, Hospital Universitario Son Espases |
| EPI_ISL_2812755 | HOSPITAL UNIVERSITARIO SON LLATZER | HOSPITAL UNIVERSITARIO SON ESPASES | Servicio de Microbiología, Hospital Universitario Son Espases |
| EPI_ISL_2812756, EPI_ISL_2812757 | HOSPITAL UNIVERSITARIO SON ESPASES | HOSPITAL UNIVERSITARIO SON ESPASES | Servicio de Microbiología, Hospital Universitario Son Espases |
| EPI_ISL_2812758 | HOSPITAL MATEU ORFILA | HOSPITAL UNIVERSITARIO SON ESPASES | Servicio de Microbiología, Hospital Universitario Son Espases |
| EPI_ISL_2812760, EPI_ISL_2812761, EPI_ISL_2812762, | HOSPITAL UNIVERSITARIO SON ESPASES | HOSPITAL UNIVERSITARIO SON ESPASES | Servicio de Microbiología, Hospital Universitario Son Espases |

| | | | |
|---|---|---|---|
| EPI_ISL_2812764 | | | |
| EPI_ISL_2812766 | HOSPITAL MANACOR | HOSPITAL UNIVERSITARIO SON ESPASES | Servicio de Microbiología, Hospital Universitario Son Espases |
| EPI_ISL_2812768 | HOSPITAL UNIVERSITARIO SON LLATZER | HOSPITAL UNIVERSITARIO SON ESPASES | Servicio de Microbiología, Hospital Universitario Son Espases |
| EPI_ISL_2812769 | HOSPITAL MATEU ORFILA | HOSPITAL UNIVERSITARIO SON ESPASES | Servicio de Microbiología, Hospital Universitario Son Espases |
| EPI_ISL_2812770, EPI_ISL_2812771, EPI_ISL_2812772, EPI_ISL_2812773, EPI_ISL_2812774, EPI_ISL_2812775 | HOSPITAL UNIVERSITARIO SON ESPASES | HOSPITAL UNIVERSITARIO SON ESPASES | Servicio de Microbiología, Hospital Universitario Son Espases |
| EPI_ISL_2812777 | HOSPITAL MANACOR | HOSPITAL UNIVERSITARIO SON ESPASES | Servicio de Microbiología, Hospital Universitario Son Espases |
| EPI_ISL_2812778 | HOSPITAL CAN MISSES | HOSPITAL UNIVERSITARIO SON ESPASES | Servicio de Microbiología, Hospital Universitario Son Espases |
| EPI_ISL_2812779, EPI_ISL_2812782, EPI_ISL_2812783, EPI_ISL_2812784, EPI_ISL_2812785 | HOSPITAL UNIVERSITARIO SON ESPASES | HOSPITAL UNIVERSITARIO SON ESPASES | Servicio de Microbiología, Hospital Universitario Son Espases |
| EPI_ISL_2812789, EPI_ISL_2812790 | Ifo Regina Elena | National Institute for Infectious Diseases (INMI) L. Spallanzani I.R.C.C.S | E Giombini, B Bartolini, M Rueca, G Bonfiglio, O Butera, CEM Gruber, F Santini, F Messina, A Di Caro, MR Capobianchi |
| EPI_ISL_2812791 | INMI Lazzaro Spallanzani IRCCS | National Institute for Infectious Diseases (INMI) L. Spallanzani I.R.C.C.S | E Giombini, B Bartolini, M Rueca, G Bonfiglio, O Butera, CEM Gruber, F Santini, F Messina, A Di Caro, MR Capobianchi |
| EPI_ISL_2812792 | Ospedale di Genzano - ASL RM 6 | National Institute for Infectious Diseases (INMI) L. Spallanzani I.R.C.C.S | E Giombini, B Bartolini, M Rueca, G Bonfiglio, O Butera, CEM Gruber, F Santini, F Messina, A Di Caro, MR Capobianchi |
| EPI_ISL_2812793, EPI_ISL_2812795 | RM1 | National Institute for Infectious Diseases (INMI) L. Spallanzani I.R.C.C.S | E Giombini, B Bartolini, M Rueca, G Bonfiglio, O Butera, CEM Gruber, F Santini, F Messina, A Di Caro, MR Capobianchi |
| EPI_ISL_2812796 | Ospedale di Genzano - ASL RM 6 | National Institute for Infectious Diseases (INMI) L. Spallanzani I.R.C.C.S | E Giombini, B Bartolini, M Rueca, G Bonfiglio, O Butera, CEM Gruber, F Santini, F Messina, A Di Caro, MR Capobianchi |
| EPI_ISL_2812797 | RM1 | National Institute for Infectious Diseases (INMI) L. Spallanzani I.R.C.C.S | E Giombini, B Bartolini, M Rueca, G Bonfiglio, O Butera, CEM Gruber, F Santini, F Messina, A Di Caro, MR Capobianchi |
| EPI_ISL_2812798 | Ospedale di Genzano - ASL RM 6 | National Institute for Infectious Diseases (INMI) L. Spallanzani I.R.C.C.S | E Giombini, B Bartolini, M Rueca, G Bonfiglio, O Butera, CEM Gruber, F Santini, F Messina, A Di Caro, MR Capobianchi |
| EPI_ISL_2812799 | RM1 | National Institute for Infectious Diseases (INMI) L. Spallanzani I.R.C.C.S | E Giombini, B Bartolini, M Rueca, G Bonfiglio, O Butera, CEM Gruber, F Santini, F Messina, A Di Caro, MR Capobianchi |
| EPI_ISL_2812801, EPI_ISL_2812802, EPI_ISL_2812803, EPI_ISL_2812804, EPI_ISL_2812805, EPI_ISL_2812806, EPI_ISL_2812807 | Ifo Regina Elena | National Institute for Infectious Diseases (INMI) L. Spallanzani I.R.C.C.S | E Giombini, B Bartolini, M Rueca, G Bonfiglio, O Butera, CEM Gruber, F Santini, F Messina, A Di Caro, MR Capobianchi |
| EPI_ISL_2812811 | IFO Regina Elena | National Institute for Infectious Diseases (INMI) L. Spallanzani I.R.C.C.S | E Giombini, B Bartolini, M Rueca, G Bonfiglio, O Butera, CEM Gruber, F Santini, F Messina, A Di Caro, MR Capobianchi |
| EPI_ISL_2812812, EPI_ISL_2812813 | Genzano | National Institute for Infectious Diseases (INMI) L. Spallanzani I.R.C.C.S | E Giombini, B Bartolini, M Rueca, G Bonfiglio, O Butera, CEM Gruber, F Santini, F Messina, A Di Caro, MR Capobianchi |
| EPI_ISL_2812814 | Southern IML Pathology | NSW Health Pathology - Institute of Clinical Pathology and Medical Research; Westmead Hospital; University of Sydney | CIDM-PH et al. |
| EPI_ISL_2812815 | Histopath | NSW Health Pathology - Institute of Clinical Pathology and Medical Research; Westmead Hospital; University of Sydney | CIDM-PH et al. |
| EPI_ISL_2812816 | Laverty Pathology | NSW Health Pathology - Institute of Clinical Pathology and Medical Research; Westmead Hospital; University of Sydney | CIDM-PH et al. |
| EPI_ISL_2812817 | Pathology North - Royal North Shore Hospital - NSW Health Pathology | NSW Health Pathology - Institute of Clinical Pathology and Medical Research; Westmead Hospital; University of Sydney | CIDM-PH et al. |
| EPI_ISL_2812818 | 4Cyte Pathology | NSW Health Pathology - Institute of Clinical Pathology and Medical Research; Westmead Hospital; University of Sydney | CIDM-PH et al. |
| EPI_ISL_2812819 | St Vincent's Pathology (SydPath) | NSW Health Pathology - Institute of Clinical Pathology and Medical Research; Westmead Hospital; University of Sydney | CIDM-PH et al. |
| EPI_ISL_2812820 | Laverty Pathology | NSW Health Pathology - Institute of Clinical Pathology and Medical Research; Westmead Hospital; University of Sydney | CIDM-PH et al. |
| EPI_ISL_2812821, EPI_ISL_2812822 | Sydney South West Pathology Service (SSWPS) - Liverpool Hospital - NSW Health Pathology | NSW Health Pathology - Institute of Clinical Pathology and Medical Research; Westmead Hospital; University of Sydney | CIDM-PH et al. |
| EPI_ISL_2812823 | Pathology West - NSW Health Pathology | NSW Health Pathology - Institute of Clinical Pathology and Medical Research; Westmead Hospital; University of Sydney | CIDM-PH et al. |
| EPI_ISL_2812824 | Histopath | NSW Health Pathology - Institute of Clinical Pathology and Medical Research; Westmead Hospital; University of Sydney | CIDM-PH et al. |
| EPI_ISL_2812825 | Australian Clinical Labs | NSW Health Pathology - Institute of Clinical Pathology and Medical Research; Westmead Hospital; University of Sydney | CIDM-PH et al. |
| EPI_ISL_2812826 | Southern. IML Pathology | NSW Health Pathology - Institute of Clinical Pathology and Medical Research; Westmead Hospital; University of Sydney | CIDM-PH et al. |
| EPI_ISL_2812827 | Sydney South West Pathology Service (SSWPS) - Liverpool Hospital - NSW Health Pathology | NSW Health Pathology - Institute of Clinical Pathology and Medical Research; Westmead Hospital; University of Sydney | CIDM-PH et al. |
| EPI_ISL_2812828, EPI_ISL_2812829, EPI_ISL_2812830 | Histopath | NSW Health Pathology - Institute of Clinical Pathology and Medical Research; Westmead Hospital; University of Sydney | CIDM-PH et al. |
| EPI_ISL_2812831, EPI_ISL_2812832, EPI_ISL_2812833, EPI_ISL_2812834, EPI_ISL_2812835, EPI_ISL_2812836, EPI_ISL_2812837, EPI_ISL_2812838, EPI_ISL_2812839, EPI_ISL_2812840, EPI_ISL_2812841 | | | |
| see above | Sydney South West Pathology Service (SSWPS) - Liverpool Hospital - NSW Health Pathology | NSW Health Pathology - Institute of Clinical Pathology and Medical Research; Westmead Hospital; University of Sydney | CIDM-PH et al. |
| EPI_ISL_2812842, EPI_ISL_2812843, EPI_ISL_2812844, EPI_ISL_2812845 | St Vincent's Pathology (SydPath) | NSW Health Pathology - Institute of Clinical Pathology and Medical Research; Westmead Hospital; University of Sydney | CIDM-PH et al. |
| EPI_ISL_2812846 | Pathology West - NSW Health Pathology | NSW Health Pathology - Institute of Clinical Pathology and Medical Research; Westmead Hospital; University of Sydney | CIDM-PH et al. |
| EPI_ISL_2812847, EPI_ISL_2812848, EPI_ISL_2812849, EPI_ISL_2812850, EPI_ISL_2812851 | Histopath | NSW Health Pathology - Institute of Clinical Pathology and Medical Research; Westmead Hospital; University of Sydney | CIDM-PH et al. |
| EPI_ISL_2812852 | 4Cyte Pathology | NSW Health Pathology - Institute of Clinical Pathology and Medical Research; Westmead Hospital; University of Sydney | CIDM-PH et al. |
| EPI_ISL_2812853, EPI_ISL_2812854 | Laverty Pathology | NSW Health Pathology - Institute of Clinical Pathology and Medical Research; Westmead Hospital; University of Sydney | CIDM-PH et al. |
| EPI_ISL_2812855 | Sydney South West Pathology Service (SSWPS) - Liverpool Hospital - NSW Health Pathology | NSW Health Pathology - Institute of Clinical Pathology and Medical Research; Westmead Hospital; University of Sydney | CIDM-PH et al. |

| | | | |
|---|---|---|---|
| EPI_ISL_2812856 | Histopath | NSW Health Pathology - Institute of Clinical Pathology and Medical Research; Westmead Hospital; University of Sydney | CIDM-PH et al. |
| EPI_ISL_2812857 | Sydney South West Pathology Service (SSWPS) - Liverpool Hospital - NSW Health Pathology | NSW Health Pathology - Institute of Clinical Pathology and Medical Research; Westmead Hospital; University of Sydney | CIDM-PH et al. |
| EPI_ISL_2812858 | 4Cyte Pathology | NSW Health Pathology - Institute of Clinical Pathology and Medical Research; Westmead Hospital; University of Sydney | CIDM-PH et al. |
| EPI_ISL_2812859, EPI_ISL_2812860, EPI_ISL_2812861, EPI_ISL_2812862, EPI_ISL_2812863, EPI_ISL_2812864, EPI_ISL_2812865, EPI_ISL_2812866, EPI_ISL_2812867, EPI_ISL_2812868, EPI_ISL_2812869 | | | |
| see above | St Vincent's Pathology (SydPath) | NSW Health Pathology - Institute of Clinical Pathology and Medical Research; Westmead Hospital; University of Sydney | CIDM-PH et al. |
| EPI_ISL_2812870 | Douglass Hanly Moir Pathology | NSW Health Pathology - Institute of Clinical Pathology and Medical Research; Westmead Hospital; University of Sydney | CIDM-PH et al. |
| EPI_ISL_2812871 | Histopath | NSW Health Pathology - Institute of Clinical Pathology and Medical Research; Westmead Hospital; University of Sydney | CIDM-PH et al. |
| EPI_ISL_2812872 | Sydney South West Pathology Service (SSWPS) - Liverpool Hospital - NSW Health Pathology | NSW Health Pathology - Institute of Clinical Pathology and Medical Research; Westmead Hospital; University of Sydney | CIDM-PH et al. |
| EPI_ISL_2812873 | 4Cyte Pathology | NSW Health Pathology - Institute of Clinical Pathology and Medical Research; Westmead Hospital; University of Sydney | CIDM-PH et al. |
| EPI_ISL_2812874, EPI_ISL_2812875, EPI_ISL_2812876 | Laverty Pathology | NSW Health Pathology - Institute of Clinical Pathology and Medical Research; Westmead Hospital; University of Sydney | CIDM-PH et al. |
| EPI_ISL_2812877 | Sydney South West Pathology Service (SSWPS) - Liverpool Hospital - NSW Health Pathology | NSW Health Pathology - Institute of Clinical Pathology and Medical Research; Westmead Hospital; University of Sydney | CIDM-PH et al. |
| EPI_ISL_2812878, EPI_ISL_2812879 | St Vincent's Pathology (SydPath) | NSW Health Pathology - Institute of Clinical Pathology and Medical Research; Westmead Hospital; University of Sydney | CIDM-PH et al. |
| EPI_ISL_2812880, EPI_ISL_2812881 | Laverty Pathology | NSW Health Pathology - Institute of Clinical Pathology and Medical Research; Westmead Hospital; University of Sydney | CIDM-PH et al. |
| EPI_ISL_2812882, EPI_ISL_2812883 | Histopath | NSW Health Pathology - Institute of Clinical Pathology and Medical Research; Westmead Hospital; University of Sydney | CIDM-PH et al. |
| EPI_ISL_2812884 | Medlab Pathology | NSW Health Pathology - Institute of Clinical Pathology and Medical Research; Westmead Hospital; University of Sydney | CIDM-PH et al. |
| EPI_ISL_2812885 | St Vincent's Pathology (SydPath) | NSW Health Pathology - Institute of Clinical Pathology and Medical Research; Westmead Hospital; University of Sydney | CIDM-PH et al. |
| EPI_ISL_2812886, EPI_ISL_2812887, EPI_ISL_2812888 | Histopath | NSW Health Pathology - Institute of Clinical Pathology and Medical Research; Westmead Hospital; University of Sydney | CIDM-PH et al. |
| EPI_ISL_2812889, EPI_ISL_2812890 | Laverty Pathology | NSW Health Pathology - Institute of Clinical Pathology and Medical Research; Westmead Hospital; University of Sydney | CIDM-PH et al. |
| EPI_ISL_2812891 | St Vincent's Pathology (SydPath) | NSW Health Pathology - Institute of Clinical Pathology and Medical Research; Westmead Hospital; University of Sydney | CIDM-PH et al. |
| EPI_ISL_2812892, EPI_ISL_2812893, EPI_ISL_2812894, EPI_ISL_2812895 | Pathology West - NSW Health Pathology | NSW Health Pathology - Institute of Clinical Pathology and Medical Research; Westmead Hospital; University of Sydney | CIDM-PH et al. |
| EPI_ISL_2812896, EPI_ISL_2812897 | Sydney South West Pathology Service (SSWPS) - Liverpool Hospital - NSW Health Pathology | NSW Health Pathology - Institute of Clinical Pathology and Medical Research; Westmead Hospital; University of Sydney | CIDM-PH et al. |
| EPI_ISL_2812898, EPI_ISL_2812899 | Laverty Pathology | NSW Health Pathology - Institute of Clinical Pathology and Medical Research; Westmead Hospital; University of Sydney | CIDM-PH et al. |
| EPI_ISL_2812900 | 4Cyte Pathology | NSW Health Pathology - Institute of Clinical Pathology and Medical Research; Westmead Hospital; University of Sydney | CIDM-PH et al. |
| EPI_ISL_2812901 | Australian Clinical Labs (formerly Healthscope Pathology) | NSW Health Pathology - Institute of Clinical Pathology and Medical Research; Westmead Hospital; University of Sydney | CIDM-PH et al. |
| EPI_ISL_2812902 | Pathology West - NSW Health Pathology | NSW Health Pathology - Institute of Clinical Pathology and Medical Research; Westmead Hospital; University of Sydney | CIDM-PH et al. |
| EPI_ISL_2812903 | Medlab Pathology | NSW Health Pathology - Institute of Clinical Pathology and Medical Research; Westmead Hospital; University of Sydney | CIDM-PH et al. |
| EPI_ISL_2812904, EPI_ISL_2812905, EPI_ISL_2812906, EPI_ISL_2812907, EPI_ISL_2812908, EPI_ISL_2812909, EPI_ISL_2812910 | Histopath | NSW Health Pathology - Institute of Clinical Pathology and Medical Research; Westmead Hospital; University of Sydney | CIDM-PH et al. |
| EPI_ISL_2812911 | 4Cyte Pathology | NSW Health Pathology - Institute of Clinical Pathology and Medical Research; Westmead Hospital; University of Sydney | CIDM-PH et al. |
| EPI_ISL_2812912 | Australian Clinical Labs (formerly Healthscope Pathology) | NSW Health Pathology - Institute of Clinical Pathology and Medical Research; Westmead Hospital; University of Sydney | CIDM-PH et al. |
| EPI_ISL_2812913, EPI_ISL_2812914, EPI_ISL_2812915, EPI_ISL_2812916, EPI_ISL_2812917 | Laverty Pathology | NSW Health Pathology - Institute of Clinical Pathology and Medical Research; Westmead Hospital; University of Sydney | CIDM-PH et al. |
| EPI_ISL_2812918, EPI_ISL_2812919 | St Vincent's Pathology (SydPath) | NSW Health Pathology - Institute of Clinical Pathology and Medical Research; Westmead Hospital; University of Sydney | CIDM-PH et al. |
| EPI_ISL_2812958, EPI_ISL_2812959, EPI_ISL_2812960, EPI_ISL_2812961, EPI_ISL_2812962, EPI_ISL_2812963, EPI_ISL_2812964, EPI_ISL_2812965, EPI_ISL_2812966, EPI_ISL_2812967, EPI_ISL_2812968, EPI_ISL_2812969 | | | |
| see above | Department of Health Technology and Informatics, The Hong Kong Polytechnic University | Department of Health Technology and Informatics, The Hong Kong Polytechnic University | Gilman Kit-Hang Siu, Lam-Kwong Lee, Kenneth Siu-Sing Leung, Jake Siu-Lun Leung, Wing-Hei Lo, Timothy Ting-Leung Ng, Chloe Toi-Mei Chan, Hoi-Ching Jim, Kingsley King-Gee Tam, Hiu-Yin Lao, Denise Sze-Hang Wong, Alan Ka-Lun Wu, Miranda Chong-Yee Yau, Yvette Wai-Man Lai, Kitty Sau-Chun Fung, Sandy Ka-Yee Chau, Barry Kin-Chung Wong, Wing-Kin To, Kristine Luk, Alex Yat-Man Ho, Tak-Lun Que, Kam-Tong Yip, Wing Cheong Yam, David Ho-Keung Shum, Shea Ping Yip |
| EPI_ISL_2812971 | Labuan Hospital | Institute for Medical Research, Infectious Disease Research Centre, National Institutes of Health, Ministry of Health Malaysia | Suppiah J, Kamel K, Mohd Zawawi Z, Azizan MA, Ramly N, Robert F, Thayan R |
| EPI_ISL_2812972, EPI_ISL_2812978 | Dianalabs SA | Genesupport | Tanguy ARAUD, Geraldine Jost, Katia Jaton, Nadia Liassine |
| EPI_ISL_2812984, EPI_ISL_2812985, EPI_ISL_2812986, EPI_ISL_2812987, EPI_ISL_2812988, EPI_ISL_2812989, EPI_ISL_2812990, EPI_ISL_2812991, EPI_ISL_2812992, EPI_ISL_2812993, EPI_ISL_2812994, EPI_ISL_2812995, EPI_ISL_2812996, EPI_ISL_2812997, EPI_ISL_2812998, EPI_ISL_2812999, EPI_ISL_2813000, EPI_ISL_2813001, EPI_ISL_2813002, EPI_ISL_2813003, EPI_ISL_2813004, EPI_ISL_2813005, EPI_ISL_2813006, EPI_ISL_2813007, EPI_ISL_2813008, EPI_ISL_2813009, EPI_ISL_2813010, EPI_ISL_2813011, EPI_ISL_2813012, EPI_ISL_2813013, EPI_ISL_2813014, EPI_ISL_2813015, EPI_ISL_2813016, EPI_ISL_2813017, EPI_ISL_2813018, EPI_ISL_2813019, EPI_ISL_2813020, EPI_ISL_2813021, EPI_ISL_2813022, EPI_ISL_2813023, EPI_ISL_2813024, EPI_ISL_2813025, EPI_ISL_2813026, EPI_ISL_2813027, EPI_ISL_2813028, EPI_ISL_2813029, EPI_ISL_2813030, EPI_ISL_2813031, EPI_ISL_2813032, EPI_ISL_2813033, EPI_ISL_2813034, EPI_ISL_2813035, EPI_ISL_2813036, EPI_ISL_2813037, EPI_ISL_2813038, EPI_ISL_2813039, EPI_ISL_2813040, EPI_ISL_2813041, EPI_ISL_2813042, EPI_ISL_2813043, EPI_ISL_2813044, EPI_ISL_2813045, EPI_ISL_2813046, EPI_ISL_2813047, EPI_ISL_2813048, EPI_ISL_2813049, EPI_ISL_2813050, EPI_ISL_2813051, EPI_ISL_2813052, EPI_ISL_2813053, EPI_ISL_2813054, EPI_ISL_2813055, EPI_ISL_2813057, EPI_ISL_2813058, EPI_ISL_2813059, EPI_ISL_2813060, EPI_ISL_2813061, EPI_ISL_2813062, EPI_ISL_2813063, EPI_ISL_2813064, EPI_ISL_2813065, EPI_ISL_2813066, EPI_ISL_2813067, EPI_ISL_2813068, EPI_ISL_2813069, EPI_ISL_2813070, EPI_ISL_2813071, EPI_ISL_2813072, EPI_ISL_2813073, EPI_ISL_2813074, EPI_ISL_2813075, EPI_ISL_2813076, EPI_ISL_2813077, EPI_ISL_2813078, EPI_ISL_2813079, EPI_ISL_2813080, EPI_ISL_2813081, EPI_ISL_2813082, EPI_ISL_2813083, EPI_ISL_2813084, EPI_ISL_2813085, EPI_ISL_2813086, EPI_ISL_2813087, EPI_ISL_2813088, EPI_ISL_2813089, EPI_ISL_2813090, EPI_ISL_2813091, EPI_ISL_2813092, EPI_ISL_2813093, EPI_ISL_2813094, EPI_ISL_2813095, EPI_ISL_2813096, EPI_ISL_2813097, EPI_ISL_2813098, EPI_ISL_2813099, EPI_ISL_2813100, EPI_ISL_2813101, EPI_ISL_2813102, EPI_ISL_2813103, EPI_ISL_2813104, EPI_ISL_2813105, EPI_ISL_2813106, EPI_ISL_2813107, EPI_ISL_2813108, EPI_ISL_2813109, EPI_ISL_2813110, EPI_ISL_2813111, EPI_ISL_2813112, EPI_ISL_2813113, EPI_ISL_2813114, EPI_ISL_2813115, EPI_ISL_2813116, EPI_ISL_2813117, EPI_ISL_2813118, EPI_ISL_2813119, EPI_ISL_2813120, EPI_ISL_2813121, EPI_ISL_2813122, EPI_ISL_2813123, EPI_ISL_2813124, EPI_ISL_2813125, EPI_ISL_2813126, EPI_ISL_2813127, EPI_ISL_2813128, EPI_ISL_2813129, EPI_ISL_2813130, EPI_ISL_2813131, EPI_ISL_2813132, EPI_ISL_2813133, EPI_ISL_2813134, EPI_ISL_2813135, EPI_ISL_2813136, EPI_ISL_2813137, EPI_ISL_2813138, EPI_ISL_2813139, EPI_ISL_2813140, EPI_ISL_2813141, EPI_ISL_2813142, EPI_ISL_2813143, EPI_ISL_2813144, EPI_ISL_2813145, EPI_ISL_2813146, EPI_ISL_2813147, EPI_ISL_2813148, EPI_ISL_2813149, EPI_ISL_2813150, EPI_ISL_2813151, EPI_ISL_2813152, EPI_ISL_2813153, EPI_ISL_2813154, EPI_ISL_2813155, EPI_ISL_2813156, EPI_ISL_2813157, EPI_ISL_2813158, EPI_ISL_2813159, EPI_ISL_2813160, EPI_ISL_2813161, EPI_ISL_2813162, EPI_ISL_2813163, EPI_ISL_2813164, EPI_ISL_2813165, EPI_ISL_2813166, EPI_ISL_2813167, EPI_ISL_2813168, EPI_ISL_2813169, EPI_ISL_2813170, EPI_ISL_2813171, EPI_ISL_2813172, EPI_ISL_2813173, EPI_ISL_2813174, EPI_ISL_2813175, EPI_ISL_2813176, EPI_ISL_2813177, EPI_ISL_2813178, EPI_ISL_2813179, EPI_ISL_2813180, EPI_ISL_2813181, EPI_ISL_2813182, EPI_ISL_2813183, EPI_ISL_2813184, EPI_ISL_2813185, EPI_ISL_2813186, EPI_ISL_2813187, EPI_ISL_2813188, EPI_ISL_2813189, EPI_ISL_2813190, EPI_ISL_2813191, EPI_ISL_2813192, EPI_ISL_2813193, EPI_ISL_2813194, EPI_ISL_2813195, EPI_ISL_2813196, EPI_ISL_2813197, EPI_ISL_2813198, EPI_ISL_2813199, EPI_ISL_2813200, EPI_ISL_2813201, EPI_ISL_2813202, EPI_ISL_2813203, EPI_ISL_2813204, EPI_ISL_2813205, EPI_ISL_2813206, EPI_ISL_2813207, EPI_ISL_2813208, EPI_ISL_2813209, EPI_ISL_2813210, EPI_ISL_2813211, EPI_ISL_2813212, EPI_ISL_2813213, EPI_ISL_2813214, EPI_ISL_2813215, EPI_ISL_2813216, EPI_ISL_2813217, EPI_ISL_2813218, EPI_ISL_2813219, EPI_ISL_2813220, EPI_ISL_2813221, EPI_ISL_2813222, EPI_ISL_2813223, EPI_ISL_2813224, EPI_ISL_2813225, EPI_ISL_2813226, EPI_ISL_2813227, EPI_ISL_2813228, EPI_ISL_2813229, EPI_ISL_2813230, EPI_ISL_2813231, EPI_ISL_2813232, EPI_ISL_2813233, EPI_ISL_2813234, EPI_ISL_2813235, EPI_ISL_2813236, EPI_ISL_2813237, EPI_ISL_2813238, EPI_ISL_2813239, EPI_ISL_2813240, EPI_ISL_2813241, EPI_ISL_2813242, EPI_ISL_2813243, EPI_ISL_2813244, EPI_ISL_2813245, EPI_ISL_2813246, EPI_ISL_2813247, EPI_ISL_2813248, EPI_ISL_2813249, EPI_ISL_2813250, EPI_ISL_2813251, EPI_ISL_2813252, EPI_ISL_2813253, EPI_ISL_2813254, EPI_ISL_2813255, EPI_ISL_2813256, EPI_ISL_2813257, EPI_ISL_2813258, EPI_ISL_2813259, EPI_ISL_2813260, EPI_ISL_2813261, EPI_ISL_2813262, EPI_ISL_2813263, EPI_ISL_2813264, EPI_ISL_2813265, EPI_ISL_2813266, EPI_ISL_2813267, EPI_ISL_2813269, EPI_ISL_2813270, EPI_ISL_2813271, EPI_ISL_2813272, EPI_ISL_2813273, EPI_ISL_2813274, EPI_ISL_2813275, EPI_ISL_2813276, EPI_ISL_2813277, EPI_ISL_2813278, EPI_ISL_2813279, EPI_ISL_2813280, EPI_ISL_2813281, EPI_ISL_2813282, EPI_ISL_2813283, EPI_ISL_2813284, EPI_ISL_2813285, EPI_ISL_2813286, EPI_ISL_2813287, EPI_ISL_2813288, EPI_ISL_2813289, EPI_ISL_2813290, EPI_ISL_2813291, EPI_ISL_2813292, EPI_ISL_2813293, | | | |

| | | | |
|---|---|---|---|
| EPI_ISL_2813294, EPI_ISL_2813295, EPI_ISL_2813296, EPI_ISL_2813297, EPI_ISL_2813298, EPI_ISL_2813299, EPI_ISL_2813300, EPI_ISL_2813301, EPI_ISL_2813302, EPI_ISL_2813303, EPI_ISL_2813304, EPI_ISL_2813305, EPI_ISL_2813306, EPI_ISL_2813307, EPI_ISL_2813308, EPI_ISL_2813309, EPI_ISL_2813310, EPI_ISL_2813311, EPI_ISL_2813312, EPI_ISL_2813313, EPI_ISL_2813314, EPI_ISL_2813315, EPI_ISL_2813316, EPI_ISL_2813317, EPI_ISL_2813318, EPI_ISL_2813319 | | | |
| see above | Lighthouse Lab in Milton Keynes | Wellcome Sanger Institute for the COVID-19 Genomics UK (COG-UK) Consortium | The Lighthouse Lab in Milton Keynes and Alex Alderton, Roberto Amato, Jeffrey Barrett, Sonia Goncalves, Ewan Harrison, David K. Jackson, Ian Johnston, Dominic Kwiatkowski, Cordelia Langford, John Sillitoe on behalf of the Wellcome Sanger Institute COVID-19 Surveillance Team |
| EPI_ISL_2813320, EPI_ISL_2813321, EPI_ISL_2813322 | Lighthouse Laboratory Plymouth | Wellcome Sanger Institute for the COVID-19 Genomics UK (COG-UK) Consortium | Lighthouse Laboratory Plymouth and Alex Alderton, Roberto Amato, Jeffrey Barrett, Sonia Goncalves, Ewan Harrison, David K. Jackson, Ian Johnston, Dominic Kwiatkowski, Cordelia Langford, John Sillitoe on behalf of the Wellcome Sanger Institute COVID-19 Surveillance Team |
| EPI_ISL_2813323 | Lighthouse Lab in Milton Keynes | Wellcome Sanger Institute for the COVID-19 Genomics UK (COG-UK) Consortium | The Lighthouse Lab in Milton Keynes and Alex Alderton, Roberto Amato, Jeffrey Barrett, Sonia Goncalves, Ewan Harrison, David K. Jackson, Ian Johnston, Dominic Kwiatkowski, Cordelia Langford, John Sillitoe on behalf of the Wellcome Sanger Institute COVID-19 Surveillance Team |
| EPI_ISL_2813324, EPI_ISL_2813325, EPI_ISL_2813326, EPI_ISL_2813327, EPI_ISL_2813328, EPI_ISL_2813329, EPI_ISL_2813330, EPI_ISL_2813331 | Lighthouse Laboratory Plymouth | Wellcome Sanger Institute for the COVID-19 Genomics UK (COG-UK) Consortium | Lighthouse Laboratory Plymouth and Alex Alderton, Roberto Amato, Jeffrey Barrett, Sonia Goncalves, Ewan Harrison, David K. Jackson, Ian Johnston, Dominic Kwiatkowski, Cordelia Langford, John Sillitoe on behalf of the Wellcome Sanger Institute COVID-19 Surveillance Team |
| EPI_ISL_2813332 | Lighthouse Lab in Milton Keynes | Wellcome Sanger Institute for the COVID-19 Genomics UK (COG-UK) Consortium | The Lighthouse Lab in Milton Keynes and Alex Alderton, Roberto Amato, Jeffrey Barrett, Sonia Goncalves, Ewan Harrison, David K. Jackson, Ian Johnston, Dominic Kwiatkowski, Cordelia Langford, John Sillitoe on behalf of the Wellcome Sanger Institute COVID-19 Surveillance Team |
| EPI_ISL_2813333, EPI_ISL_2813334, EPI_ISL_2813335, EPI_ISL_2813336, EPI_ISL_2813337 | Lighthouse Laboratory Plymouth | Wellcome Sanger Institute for the COVID-19 Genomics UK (COG-UK) Consortium | Lighthouse Laboratory Plymouth and Alex Alderton, Roberto Amato, Jeffrey Barrett, Sonia Goncalves, Ewan Harrison, David K. Jackson, Ian Johnston, Dominic Kwiatkowski, Cordelia Langford, John Sillitoe on behalf of the Wellcome Sanger Institute COVID-19 Surveillance Team |
| EPI_ISL_2813338 | Lighthouse Lab in Milton Keynes | Wellcome Sanger Institute for the COVID-19 Genomics UK (COG-UK) Consortium | The Lighthouse Lab in Milton Keynes and Alex Alderton, Roberto Amato, Jeffrey Barrett, Sonia Goncalves, Ewan Harrison, David K. Jackson, Ian Johnston, Dominic Kwiatkowski, Cordelia Langford, John Sillitoe on behalf of the Wellcome Sanger Institute COVID-19 Surveillance Team |
| EPI_ISL_2813339, EPI_ISL_2813340, EPI_ISL_2813341 | Lighthouse Laboratory Plymouth | Wellcome Sanger Institute for the COVID-19 Genomics UK (COG-UK) Consortium | Lighthouse Laboratory Plymouth and Alex Alderton, Roberto Amato, Jeffrey Barrett, Sonia Goncalves, Ewan Harrison, David K. Jackson, Ian Johnston, Dominic Kwiatkowski, Cordelia Langford, John Sillitoe on behalf of the Wellcome Sanger Institute COVID-19 Surveillance Team |
| EPI_ISL_2813342 | Lighthouse Lab in Milton Keynes | Wellcome Sanger Institute for the COVID-19 Genomics UK (COG-UK) Consortium | The Lighthouse Lab in Milton Keynes and Alex Alderton, Roberto Amato, Jeffrey Barrett, Sonia Goncalves, Ewan Harrison, David K. Jackson, Ian Johnston, Dominic Kwiatkowski, Cordelia Langford, John Sillitoe on behalf of the Wellcome Sanger Institute COVID-19 Surveillance Team |
| EPI_ISL_2813343, EPI_ISL_2813344, EPI_ISL_2813345, EPI_ISL_2813346, EPI_ISL_2813347, EPI_ISL_2813348, EPI_ISL_2813349, EPI_ISL_2813350, EPI_ISL_2813351 | Lighthouse Laboratory Plymouth | Wellcome Sanger Institute for the COVID-19 Genomics UK (COG-UK) Consortium | Lighthouse Laboratory Plymouth and Alex Alderton, Roberto Amato, Jeffrey Barrett, Sonia Goncalves, Ewan Harrison, David K. Jackson, Ian Johnston, Dominic Kwiatkowski, Cordelia Langford, John Sillitoe on behalf of the Wellcome Sanger Institute COVID-19 Surveillance Team |
| EPI_ISL_2813352 | Lighthouse Lab in Milton Keynes | Wellcome Sanger Institute for the COVID-19 Genomics UK (COG-UK) Consortium | The Lighthouse Lab in Milton Keynes and Alex Alderton, Roberto Amato, Jeffrey Barrett, Sonia Goncalves, Ewan Harrison, David K. Jackson, Ian Johnston, Dominic Kwiatkowski, Cordelia Langford, John Sillitoe on behalf of the Wellcome Sanger Institute COVID-19 Surveillance Team |
| EPI_ISL_2813353, EPI_ISL_2813354, EPI_ISL_2813355, EPI_ISL_2813356 | Lighthouse Laboratory Plymouth | Wellcome Sanger Institute for the COVID-19 Genomics UK (COG-UK) Consortium | Lighthouse Laboratory Plymouth and Alex Alderton, Roberto Amato, Jeffrey Barrett, Sonia Goncalves, Ewan Harrison, David K. Jackson, Ian Johnston, Dominic Kwiatkowski, Cordelia Langford, John Sillitoe on behalf of the Wellcome Sanger Institute COVID-19 Surveillance Team |
| EPI_ISL_2813357 | Lighthouse Lab in Milton Keynes | Wellcome Sanger Institute for the COVID-19 Genomics UK (COG-UK) Consortium | The Lighthouse Lab in Milton Keynes and Alex Alderton, Roberto Amato, Jeffrey Barrett, Sonia Goncalves, Ewan Harrison, David K. Jackson, Ian Johnston, Dominic Kwiatkowski, Cordelia Langford, John Sillitoe on behalf of the Wellcome Sanger Institute COVID-19 Surveillance Team |
| EPI_ISL_2813358 | Lighthouse Laboratory Plymouth | Wellcome Sanger Institute for the COVID-19 Genomics UK (COG-UK) Consortium | Lighthouse Laboratory Plymouth and Alex Alderton, Roberto Amato, Jeffrey Barrett, Sonia Goncalves, Ewan Harrison, David K. Jackson, Ian Johnston, Dominic Kwiatkowski, Cordelia Langford, John Sillitoe on behalf of the Wellcome Sanger Institute COVID-19 Surveillance Team |
| EPI_ISL_2813359, EPI_ISL_2813360 | Lighthouse Lab in Milton Keynes | Wellcome Sanger Institute for the COVID-19 Genomics UK (COG-UK) Consortium | The Lighthouse Lab in Milton Keynes and Alex Alderton, Roberto Amato, Jeffrey Barrett, Sonia Goncalves, Ewan Harrison, David K. Jackson, Ian Johnston, Dominic Kwiatkowski, Cordelia Langford, John Sillitoe on behalf of the Wellcome Sanger Institute COVID-19 Surveillance Team |
| EPI_ISL_2813361 | Lighthouse Laboratory Plymouth | Wellcome Sanger Institute for the COVID-19 Genomics UK (COG-UK) Consortium | Lighthouse Laboratory Plymouth and Alex Alderton, Roberto Amato, Jeffrey Barrett, Sonia Goncalves, Ewan Harrison, David K. Jackson, Ian Johnston, Dominic Kwiatkowski, Cordelia Langford, John Sillitoe on behalf of the Wellcome Sanger Institute COVID-19 Surveillance Team |
| EPI_ISL_2813362, EPI_ISL_2813363 | Lighthouse Lab in Milton Keynes | Wellcome Sanger Institute for the COVID-19 Genomics UK (COG-UK) Consortium | The Lighthouse Lab in Milton Keynes and Alex Alderton, Roberto Amato, Jeffrey Barrett, Sonia Goncalves, Ewan Harrison, David K. Jackson, Ian Johnston, Dominic Kwiatkowski, Cordelia Langford, John Sillitoe on behalf of the Wellcome Sanger Institute COVID-19 Surveillance Team |
| EPI_ISL_2813364, EPI_ISL_2813365, EPI_ISL_2813366 | Lighthouse Laboratory Plymouth | Wellcome Sanger Institute for the COVID-19 Genomics UK (COG-UK) Consortium | Lighthouse Laboratory Plymouth and Alex Alderton, Roberto Amato, Jeffrey Barrett, Sonia Goncalves, Ewan Harrison, David K. Jackson, Ian Johnston, Dominic Kwiatkowski, Cordelia Langford, John Sillitoe on behalf of the Wellcome Sanger Institute COVID-19 Surveillance Team |
| EPI_ISL_2813367 | Lighthouse Lab in Milton Keynes | Wellcome Sanger Institute for the COVID-19 Genomics UK (COG-UK) Consortium | The Lighthouse Lab in Milton Keynes and Alex Alderton, Roberto Amato, Jeffrey Barrett, Sonia Goncalves, Ewan Harrison, David K. Jackson, Ian Johnston, Dominic Kwiatkowski, Cordelia Langford, John Sillitoe on behalf of the Wellcome Sanger Institute COVID-19 Surveillance Team |
| EPI_ISL_2813368, EPI_ISL_2813369, EPI_ISL_2813370, EPI_ISL_2813371, EPI_ISL_2813372, EPI_ISL_2813373, EPI_ISL_2813374, EPI_ISL_2813375, EPI_ISL_2813376, EPI_ISL_2813377, EPI_ISL_2813378, EPI_ISL_2813379, EPI_ISL_2813380, EPI_ISL_2813381 | | | |
| see above | Lighthouse Laboratory Plymouth | Wellcome Sanger Institute for the COVID-19 Genomics UK (COG-UK) Consortium | Lighthouse Laboratory Plymouth and Alex Alderton, Roberto Amato, Jeffrey Barrett, Sonia Goncalves, Ewan Harrison, David K. Jackson, Ian Johnston, Dominic Kwiatkowski, Cordelia Langford, John Sillitoe on behalf of the Wellcome Sanger Institute COVID-19 Surveillance Team |
| EPI_ISL_2813382 | Lighthouse Lab in Milton Keynes | Wellcome Sanger Institute for the COVID-19 Genomics UK (COG-UK) Consortium | The Lighthouse Lab in Milton Keynes and Alex Alderton, Roberto Amato, Jeffrey Barrett, Sonia Goncalves, Ewan Harrison, David K. Jackson, Ian Johnston, Dominic Kwiatkowski, Cordelia Langford, John Sillitoe on behalf of the Wellcome Sanger Institute COVID-19 Surveillance Team |
| EPI_ISL_2813383 | Lighthouse Laboratory Plymouth | Wellcome Sanger Institute for the COVID-19 Genomics UK (COG-UK) Consortium | Lighthouse Laboratory Plymouth and Alex Alderton, Roberto Amato, Jeffrey Barrett, Sonia Goncalves, Ewan Harrison, David K. Jackson, Ian Johnston, Dominic Kwiatkowski, Cordelia Langford, John Sillitoe on behalf of the Wellcome Sanger Institute COVID-19 Surveillance Team |
| EPI_ISL_2813384 | Lighthouse Lab in Milton Keynes | Wellcome Sanger Institute for the COVID-19 Genomics UK (COG-UK) Consortium | The Lighthouse Lab in Milton Keynes and Alex Alderton, Roberto Amato, Jeffrey Barrett, Sonia Goncalves, Ewan Harrison, David K. Jackson, Ian Johnston, Dominic Kwiatkowski, Cordelia Langford, John Sillitoe on behalf of the Wellcome Sanger Institute COVID-19 Surveillance Team |
| EPI_ISL_2813385, EPI_ISL_2813386, EPI_ISL_2813387, EPI_ISL_2813388, EPI_ISL_2813389, EPI_ISL_2813390 | Lighthouse Laboratory Plymouth | Wellcome Sanger Institute for the COVID-19 Genomics UK (COG-UK) Consortium | Lighthouse Laboratory Plymouth and Alex Alderton, Roberto Amato, Jeffrey Barrett, Sonia Goncalves, Ewan Harrison, David K. Jackson, Ian Johnston, Dominic Kwiatkowski, Cordelia Langford, John Sillitoe on behalf of the Wellcome Sanger Institute COVID-19 Surveillance Team |
| EPI_ISL_2813391 | Lighthouse Lab in Milton Keynes | Wellcome Sanger Institute for the COVID-19 Genomics UK (COG-UK) Consortium | The Lighthouse Lab in Milton Keynes and Alex Alderton, Roberto Amato, Jeffrey Barrett, Sonia Goncalves, Ewan Harrison, David K. Jackson, Ian Johnston, Dominic Kwiatkowski, Cordelia Langford, John Sillitoe on behalf of the Wellcome Sanger Institute COVID-19 Surveillance Team |
| EPI_ISL_2813392, EPI_ISL_2813393, EPI_ISL_2813394, EPI_ISL_2813395, EPI_ISL_2813396 | Lighthouse Laboratory Plymouth | Wellcome Sanger Institute for the COVID-19 Genomics UK (COG-UK) Consortium | Lighthouse Laboratory Plymouth and Alex Alderton, Roberto Amato, Jeffrey Barrett, Sonia Goncalves, Ewan Harrison, David K. Jackson, Ian Johnston, Dominic Kwiatkowski, Cordelia Langford, John Sillitoe on behalf of the Wellcome Sanger Institute COVID-19 Surveillance Team |
| EPI_ISL_2813397 | Lighthouse Lab in Milton Keynes | Wellcome Sanger Institute for the COVID-19 Genomics UK (COG-UK) Consortium | The Lighthouse Lab in Milton Keynes and Alex Alderton, Roberto Amato, Jeffrey Barrett, Sonia Goncalves, Ewan Harrison, David K. Jackson, Ian Johnston, Dominic Kwiatkowski, Cordelia Langford, John Sillitoe on behalf of the Wellcome Sanger Institute COVID-19 Surveillance Team |
| EPI_ISL_2813398, EPI_ISL_2813399 | Lighthouse Laboratory Plymouth | Wellcome Sanger Institute for the COVID-19 Genomics UK (COG-UK) Consortium | Lighthouse Laboratory Plymouth and Alex Alderton, Roberto Amato, Jeffrey Barrett, Sonia Goncalves, Ewan Harrison, David K. Jackson, Ian Johnston, Dominic Kwiatkowski, Cordelia Langford, John Sillitoe on behalf of the Wellcome Sanger Institute COVID-19 Surveillance Team |
| EPI_ISL_2813400, EPI_ISL_2813401 | Lighthouse Lab in Milton Keynes | Wellcome Sanger Institute for the COVID-19 Genomics UK (COG-UK) Consortium | The Lighthouse Lab in Milton Keynes and Alex Alderton, Roberto Amato, Jeffrey Barrett, Sonia Goncalves, Ewan Harrison, David K. Jackson, Ian Johnston, Dominic Kwiatkowski, Cordelia Langford, John Sillitoe on behalf of the Wellcome Sanger Institute COVID-19 Surveillance Team |
| EPI_ISL_2813402 | Lighthouse Laboratory Plymouth | Wellcome Sanger Institute for the COVID-19 Genomics UK (COG-UK) Consortium | Lighthouse Laboratory Plymouth and Alex Alderton, Roberto Amato, Jeffrey Barrett, Sonia Goncalves, Ewan Harrison, David K. Jackson, Ian Johnston, Dominic Kwiatkowski, Cordelia Langford, John Sillitoe on behalf of the Wellcome Sanger Institute COVID-19 Surveillance Team |
| EPI_ISL_2813403 | Lighthouse Lab in Milton Keynes | Wellcome Sanger Institute for the COVID-19 Genomics UK (COG-UK) Consortium | The Lighthouse Lab in Milton Keynes and Alex Alderton, Roberto Amato, Jeffrey Barrett, Sonia Goncalves, Ewan Harrison, David K. Jackson, Ian Johnston, Dominic Kwiatkowski, Cordelia Langford, John Sillitoe on behalf of the Wellcome Sanger Institute COVID-19 Surveillance Team |
| EPI_ISL_2813404 | Lighthouse Laboratory Plymouth | Wellcome Sanger Institute for the COVID-19 Genomics UK (COG-UK) Consortium | Lighthouse Laboratory Plymouth and Alex Alderton, Roberto Amato, Jeffrey Barrett, Sonia Goncalves, Ewan Harrison, David K. Jackson, Ian Johnston, Dominic Kwiatkowski, Cordelia Langford, John Sillitoe on behalf of the Wellcome Sanger Institute COVID-19 Surveillance Team |

| | | | |
|---|---|---|---|
| EPI_ISL_2813405 | Lighthouse Lab in Milton Keynes | Wellcome Sanger Institute for the COVID-19 Genomics UK (COG-UK) Consortium | The Lighthouse Lab in Milton Keynes and Alex Alderton, Roberto Amato, Jeffrey Barrett, Sonia Goncalves, Ewan Harrison, David K. Jackson, Ian Johnston, Dominic Kwiatkowski, Cordelia Langford, John Sillitoe on behalf of the Wellcome Sanger Institute COVID-19 Surveillance Team |
| EPI_ISL_2813406 | Lighthouse Laboratory Plymouth | Wellcome Sanger Institute for the COVID-19 Genomics UK (COG-UK) Consortium | Lighthouse Laboratory Plymouth and Alex Alderton, Roberto Amato, Jeffrey Barrett, Sonia Goncalves, Ewan Harrison, David K. Jackson, Ian Johnston, Dominic Kwiatkowski, Cordelia Langford, John Sillitoe on behalf of the Wellcome Sanger Institute COVID-19 Surveillance Team |
| EPI_ISL_2813407, EPI_ISL_2813408 | Lighthouse Lab in Milton Keynes | Wellcome Sanger Institute for the COVID-19 Genomics UK (COG-UK) Consortium | The Lighthouse Lab in Milton Keynes and Alex Alderton, Roberto Amato, Jeffrey Barrett, Sonia Goncalves, Ewan Harrison, David K. Jackson, Ian Johnston, Dominic Kwiatkowski, Cordelia Langford, John Sillitoe on behalf of the Wellcome Sanger Institute COVID-19 Surveillance Team |
| EPI_ISL_2813409, EPI_ISL_2813410 | Lighthouse Laboratory Plymouth | Wellcome Sanger Institute for the COVID-19 Genomics UK (COG-UK) Consortium | Lighthouse Laboratory Plymouth and Alex Alderton, Roberto Amato, Jeffrey Barrett, Sonia Goncalves, Ewan Harrison, David K. Jackson, Ian Johnston, Dominic Kwiatkowski, Cordelia Langford, John Sillitoe on behalf of the Wellcome Sanger Institute COVID-19 Surveillance Team |
| EPI_ISL_2813411 | Lighthouse Lab in Milton Keynes | Wellcome Sanger Institute for the COVID-19 Genomics UK (COG-UK) Consortium | The Lighthouse Lab in Milton Keynes and Alex Alderton, Roberto Amato, Jeffrey Barrett, Sonia Goncalves, Ewan Harrison, David K. Jackson, Ian Johnston, Dominic Kwiatkowski, Cordelia Langford, John Sillitoe on behalf of the Wellcome Sanger Institute COVID-19 Surveillance Team |
| EPI_ISL_2813412, EPI_ISL_2813413, EPI_ISL_2813414, EPI_ISL_2813415, EPI_ISL_2813416 | Lighthouse Laboratory Plymouth | Wellcome Sanger Institute for the COVID-19 Genomics UK (COG-UK) Consortium | Lighthouse Laboratory Plymouth and Alex Alderton, Roberto Amato, Jeffrey Barrett, Sonia Goncalves, Ewan Harrison, David K. Jackson, Ian Johnston, Dominic Kwiatkowski, Cordelia Langford, John Sillitoe on behalf of the Wellcome Sanger Institute COVID-19 Surveillance Team |
| EPI_ISL_2813417, EPI_ISL_2813418 | Lighthouse Lab in Milton Keynes | Wellcome Sanger Institute for the COVID-19 Genomics UK (COG-UK) Consortium | The Lighthouse Lab in Milton Keynes and Alex Alderton, Roberto Amato, Jeffrey Barrett, Sonia Goncalves, Ewan Harrison, David K. Jackson, Ian Johnston, Dominic Kwiatkowski, Cordelia Langford, John Sillitoe on behalf of the Wellcome Sanger Institute COVID-19 Surveillance Team |
| EPI_ISL_2813419, EPI_ISL_2813420, EPI_ISL_2813421, EPI_ISL_2813422 | Lighthouse Laboratory Plymouth | Wellcome Sanger Institute for the COVID-19 Genomics UK (COG-UK) Consortium | Lighthouse Laboratory Plymouth and Alex Alderton, Roberto Amato, Jeffrey Barrett, Sonia Goncalves, Ewan Harrison, David K. Jackson, Ian Johnston, Dominic Kwiatkowski, Cordelia Langford, John Sillitoe on behalf of the Wellcome Sanger Institute COVID-19 Surveillance Team |
| EPI_ISL_2813423 | Lighthouse Lab in Milton Keynes | Wellcome Sanger Institute for the COVID-19 Genomics UK (COG-UK) Consortium | The Lighthouse Lab in Milton Keynes and Alex Alderton, Roberto Amato, Jeffrey Barrett, Sonia Goncalves, Ewan Harrison, David K. Jackson, Ian Johnston, Dominic Kwiatkowski, Cordelia Langford, John Sillitoe on behalf of the Wellcome Sanger Institute COVID-19 Surveillance Team |
| EPI_ISL_2813424, EPI_ISL_2813425 | Lighthouse Laboratory Plymouth | Wellcome Sanger Institute for the COVID-19 Genomics UK (COG-UK) Consortium | Lighthouse Laboratory Plymouth and Alex Alderton, Roberto Amato, Jeffrey Barrett, Sonia Goncalves, Ewan Harrison, David K. Jackson, Ian Johnston, Dominic Kwiatkowski, Cordelia Langford, John Sillitoe on behalf of the Wellcome Sanger Institute COVID-19 Surveillance Team |
| EPI_ISL_2813426 | Lighthouse Lab in Milton Keynes | Wellcome Sanger Institute for the COVID-19 Genomics UK (COG-UK) Consortium | The Lighthouse Lab in Milton Keynes and Alex Alderton, Roberto Amato, Jeffrey Barrett, Sonia Goncalves, Ewan Harrison, David K. Jackson, Ian Johnston, Dominic Kwiatkowski, Cordelia Langford, John Sillitoe on behalf of the Wellcome Sanger Institute COVID-19 Surveillance Team |
| EPI_ISL_2813427 | Lighthouse Laboratory Plymouth | Wellcome Sanger Institute for the COVID-19 Genomics UK (COG-UK) Consortium | Lighthouse Laboratory Plymouth and Alex Alderton, Roberto Amato, Jeffrey Barrett, Sonia Goncalves, Ewan Harrison, David K. Jackson, Ian Johnston, Dominic Kwiatkowski, Cordelia Langford, John Sillitoe on behalf of the Wellcome Sanger Institute COVID-19 Surveillance Team |
| EPI_ISL_2813428 | Lighthouse Lab in Milton Keynes | Wellcome Sanger Institute for the COVID-19 Genomics UK (COG-UK) Consortium | The Lighthouse Lab in Milton Keynes and Alex Alderton, Roberto Amato, Jeffrey Barrett, Sonia Goncalves, Ewan Harrison, David K. Jackson, Ian Johnston, Dominic Kwiatkowski, Cordelia Langford, John Sillitoe on behalf of the Wellcome Sanger Institute COVID-19 Surveillance Team |
| EPI_ISL_2813429, EPI_ISL_2813430, EPI_ISL_2813432, EPI_ISL_2813434 | Lighthouse Laboratory Plymouth | Wellcome Sanger Institute for the COVID-19 Genomics UK (COG-UK) Consortium | Lighthouse Laboratory Plymouth and Alex Alderton, Roberto Amato, Jeffrey Barrett, Sonia Goncalves, Ewan Harrison, David K. Jackson, Ian Johnston, Dominic Kwiatkowski, Cordelia Langford, John Sillitoe on behalf of the Wellcome Sanger Institute COVID-19 Surveillance Team |
| EPI_ISL_2813435 | Lighthouse Lab in Milton Keynes | Wellcome Sanger Institute for the COVID-19 Genomics UK (COG-UK) Consortium | The Lighthouse Lab in Milton Keynes and Alex Alderton, Roberto Amato, Jeffrey Barrett, Sonia Goncalves, Ewan Harrison, David K. Jackson, Ian Johnston, Dominic Kwiatkowski, Cordelia Langford, John Sillitoe on behalf of the Wellcome Sanger Institute COVID-19 Surveillance Team |
| EPI_ISL_2813436, EPI_ISL_2813437, EPI_ISL_2813438 | Lighthouse Laboratory Plymouth | Wellcome Sanger Institute for the COVID-19 Genomics UK (COG-UK) Consortium | Lighthouse Laboratory Plymouth and Alex Alderton, Roberto Amato, Jeffrey Barrett, Sonia Goncalves, Ewan Harrison, David K. Jackson, Ian Johnston, Dominic Kwiatkowski, Cordelia Langford, John Sillitoe on behalf of the Wellcome Sanger Institute COVID-19 Surveillance Team |
| EPI_ISL_2813439 | Lighthouse Lab in Milton Keynes | Wellcome Sanger Institute for the COVID-19 Genomics UK (COG-UK) Consortium | The Lighthouse Lab in Milton Keynes and Alex Alderton, Roberto Amato, Jeffrey Barrett, Sonia Goncalves, Ewan Harrison, David K. Jackson, Ian Johnston, Dominic Kwiatkowski, Cordelia Langford, John Sillitoe on behalf of the Wellcome Sanger Institute COVID-19 Surveillance Team |
| EPI_ISL_2813440, EPI_ISL_2813441, EPI_ISL_2813442, EPI_ISL_2813443, EPI_ISL_2813444, EPI_ISL_2813445, EPI_ISL_2813446 | Lighthouse Laboratory Plymouth | Wellcome Sanger Institute for the COVID-19 Genomics UK (COG-UK) Consortium | Lighthouse Laboratory Plymouth and Alex Alderton, Roberto Amato, Jeffrey Barrett, Sonia Goncalves, Ewan Harrison, David K. Jackson, Ian Johnston, Dominic Kwiatkowski, Cordelia Langford, John Sillitoe on behalf of the Wellcome Sanger Institute COVID-19 Surveillance Team |
| EPI_ISL_2813447 | Lighthouse Lab in Milton Keynes | Wellcome Sanger Institute for the COVID-19 Genomics UK (COG-UK) Consortium | The Lighthouse Lab in Milton Keynes and Alex Alderton, Roberto Amato, Jeffrey Barrett, Sonia Goncalves, Ewan Harrison, David K. Jackson, Ian Johnston, Dominic Kwiatkowski, Cordelia Langford, John Sillitoe on behalf of the Wellcome Sanger Institute COVID-19 Surveillance Team |
| EPI_ISL_2813448, EPI_ISL_2813449, EPI_ISL_2813450, EPI_ISL_2813451, EPI_ISL_2813452 | Lighthouse Laboratory Plymouth | Wellcome Sanger Institute for the COVID-19 Genomics UK (COG-UK) Consortium | Lighthouse Laboratory Plymouth and Alex Alderton, Roberto Amato, Jeffrey Barrett, Sonia Goncalves, Ewan Harrison, David K. Jackson, Ian Johnston, Dominic Kwiatkowski, Cordelia Langford, John Sillitoe on behalf of the Wellcome Sanger Institute COVID-19 Surveillance Team |
| EPI_ISL_2813453, EPI_ISL_2813454 | Lighthouse Lab in Milton Keynes | Wellcome Sanger Institute for the COVID-19 Genomics UK (COG-UK) Consortium | The Lighthouse Lab in Milton Keynes and Alex Alderton, Roberto Amato, Jeffrey Barrett, Sonia Goncalves, Ewan Harrison, David K. Jackson, Ian Johnston, Dominic Kwiatkowski, Cordelia Langford, John Sillitoe on behalf of the Wellcome Sanger Institute COVID-19 Surveillance Team |
| EPI_ISL_2813455 | Lighthouse Laboratory Plymouth | Wellcome Sanger Institute for the COVID-19 Genomics UK (COG-UK) Consortium | Lighthouse Laboratory Plymouth and Alex Alderton, Roberto Amato, Jeffrey Barrett, Sonia Goncalves, Ewan Harrison, David K. Jackson, Ian Johnston, Dominic Kwiatkowski, Cordelia Langford, John Sillitoe on behalf of the Wellcome Sanger Institute COVID-19 Surveillance Team |
| EPI_ISL_2813456 | Lighthouse Lab in Milton Keynes | Wellcome Sanger Institute for the COVID-19 Genomics UK (COG-UK) Consortium | The Lighthouse Lab in Milton Keynes and Alex Alderton, Roberto Amato, Jeffrey Barrett, Sonia Goncalves, Ewan Harrison, David K. Jackson, Ian Johnston, Dominic Kwiatkowski, Cordelia Langford, John Sillitoe on behalf of the Wellcome Sanger Institute COVID-19 Surveillance Team |
| EPI_ISL_2813457, EPI_ISL_2813458, EPI_ISL_2813459, EPI_ISL_2813460, EPI_ISL_2813461, EPI_ISL_2813462, EPI_ISL_2813463, EPI_ISL_2813464 | Lighthouse Laboratory Plymouth | Wellcome Sanger Institute for the COVID-19 Genomics UK (COG-UK) Consortium | Lighthouse Laboratory Plymouth and Alex Alderton, Roberto Amato, Jeffrey Barrett, Sonia Goncalves, Ewan Harrison, David K. Jackson, Ian Johnston, Dominic Kwiatkowski, Cordelia Langford, John Sillitoe on behalf of the Wellcome Sanger Institute COVID-19 Surveillance Team |
| EPI_ISL_2813465 | Lighthouse Lab in Milton Keynes | Wellcome Sanger Institute for the COVID-19 Genomics UK (COG-UK) Consortium | The Lighthouse Lab in Milton Keynes and Alex Alderton, Roberto Amato, Jeffrey Barrett, Sonia Goncalves, Ewan Harrison, David K. Jackson, Ian Johnston, Dominic Kwiatkowski, Cordelia Langford, John Sillitoe on behalf of the Wellcome Sanger Institute COVID-19 Surveillance Team |
| EPI_ISL_2813466, EPI_ISL_2813467, EPI_ISL_2813468, EPI_ISL_2813469, EPI_ISL_2813470, EPI_ISL_2813471, EPI_ISL_2813472 | Lighthouse Laboratory Plymouth | Wellcome Sanger Institute for the COVID-19 Genomics UK (COG-UK) Consortium | Lighthouse Laboratory Plymouth and Alex Alderton, Roberto Amato, Jeffrey Barrett, Sonia Goncalves, Ewan Harrison, David K. Jackson, Ian Johnston, Dominic Kwiatkowski, Cordelia Langford, John Sillitoe on behalf of the Wellcome Sanger Institute COVID-19 Surveillance Team |
| EPI_ISL_2813473 | Lighthouse Lab in Milton Keynes | Wellcome Sanger Institute for the COVID-19 Genomics UK (COG-UK) Consortium | The Lighthouse Lab in Milton Keynes and Alex Alderton, Roberto Amato, Jeffrey Barrett, Sonia Goncalves, Ewan Harrison, David K. Jackson, Ian Johnston, Dominic Kwiatkowski, Cordelia Langford, John Sillitoe on behalf of the Wellcome Sanger Institute COVID-19 Surveillance Team |
| EPI_ISL_2813474, EPI_ISL_2813475 | Lighthouse Laboratory Plymouth | Wellcome Sanger Institute for the COVID-19 Genomics UK (COG-UK) Consortium | Lighthouse Laboratory Plymouth and Alex Alderton, Roberto Amato, Jeffrey Barrett, Sonia Goncalves, Ewan Harrison, David K. Jackson, Ian Johnston, Dominic Kwiatkowski, Cordelia Langford, John Sillitoe on behalf of the Wellcome Sanger Institute COVID-19 Surveillance Team |
| EPI_ISL_2813476, EPI_ISL_2813477 | Lighthouse Lab in Milton Keynes | Wellcome Sanger Institute for the COVID-19 Genomics UK (COG-UK) Consortium | The Lighthouse Lab in Milton Keynes and Alex Alderton, Roberto Amato, Jeffrey Barrett, Sonia Goncalves, Ewan Harrison, David K. Jackson, Ian Johnston, Dominic Kwiatkowski, Cordelia Langford, John Sillitoe on behalf of the Wellcome Sanger Institute COVID-19 Surveillance Team |
| EPI_ISL_2813478, EPI_ISL_2813479, EPI_ISL_2813480, EPI_ISL_2813481, EPI_ISL_2813482, EPI_ISL_2813483, EPI_ISL_2813484, EPI_ISL_2813485, EPI_ISL_2813486, EPI_ISL_2813487, EPI_ISL_2813488, EPI_ISL_2813489, EPI_ISL_2813490, EPI_ISL_2813491, EPI_ISL_2813492 | | | |
| see above | Lighthouse Laboratory Plymouth | Wellcome Sanger Institute for the COVID-19 Genomics UK (COG-UK) Consortium | Lighthouse Laboratory Plymouth and Alex Alderton, Roberto Amato, Jeffrey Barrett, Sonia Goncalves, Ewan Harrison, David K. Jackson, Ian Johnston, Dominic Kwiatkowski, Cordelia Langford, John Sillitoe on behalf of the Wellcome Sanger Institute COVID-19 Surveillance Team |
| EPI_ISL_2813493 | Lighthouse Lab in Milton Keynes | Wellcome Sanger Institute for the COVID-19 Genomics UK (COG-UK) Consortium | The Lighthouse Lab in Milton Keynes and Alex Alderton, Roberto Amato, Jeffrey Barrett, Sonia Goncalves, Ewan Harrison, David K. Jackson, Ian Johnston, Dominic Kwiatkowski, Cordelia Langford, John Sillitoe on behalf of the Wellcome Sanger Institute COVID-19 Surveillance Team |
| EPI_ISL_2813494 | Lighthouse Laboratory Plymouth | Wellcome Sanger Institute for the COVID-19 Genomics UK (COG-UK) Consortium | Lighthouse Laboratory Plymouth and Alex Alderton, Roberto Amato, Jeffrey Barrett, Sonia Goncalves, Ewan Harrison, David K. Jackson, Ian Johnston, Dominic Kwiatkowski, Cordelia Langford, John Sillitoe on behalf of the Wellcome Sanger Institute COVID-19 Surveillance Team |
| EPI_ISL_2813495, EPI_ISL_2813496, EPI_ISL_2813497 | Lighthouse Lab in Milton Keynes | Wellcome Sanger Institute for the COVID-19 Genomics UK (COG-UK) Consortium | The Lighthouse Lab in Milton Keynes and Alex Alderton, Roberto Amato, Jeffrey Barrett, Sonia Goncalves, Ewan Harrison, David K. Jackson, Ian Johnston, Dominic Kwiatkowski, Cordelia Langford, John Sillitoe on behalf of the Wellcome Sanger Institute COVID-19 Surveillance Team |

| | | | |
|---|---|---|---|
| EPI_ISL_2813498, EPI_ISL_2813499 | Lighthouse Laboratory Plymouth | Wellcome Sanger Institute for the COVID-19 Genomics UK (COG-UK) Consortium | Lighthouse Laboratory Plymouth and Alex Alderton, Roberto Amato, Jeffrey Barrett, Sonia Goncalves, Ewan Harrison, David K. Jackson, Ian Johnston, Dominic Kwiatkowski, Cordelia Langford, John Sillitoe on behalf of the Wellcome Sanger Institute COVID-19 Surveillance Team |
| EPI_ISL_2813500 | Lighthouse Lab in Milton Keynes | Wellcome Sanger Institute for the COVID-19 Genomics UK (COG-UK) Consortium | The Lighthouse Lab in Milton Keynes and Alex Alderton, Roberto Amato, Jeffrey Barrett, Sonia Goncalves, Ewan Harrison, David K. Jackson, Ian Johnston, Dominic Kwiatkowski, Cordelia Langford, John Sillitoe on behalf of the Wellcome Sanger Institute COVID-19 Surveillance Team |
| EPI_ISL_2813501, EPI_ISL_2813502, EPI_ISL_2813503, EPI_ISL_2813504, EPI_ISL_2813505, EPI_ISL_2813506 | Lighthouse Laboratory Plymouth | Wellcome Sanger Institute for the COVID-19 Genomics UK (COG-UK) Consortium | Lighthouse Laboratory Plymouth and Alex Alderton, Roberto Amato, Jeffrey Barrett, Sonia Goncalves, Ewan Harrison, David K. Jackson, Ian Johnston, Dominic Kwiatkowski, Cordelia Langford, John Sillitoe on behalf of the Wellcome Sanger Institute COVID-19 Surveillance Team |
| EPI_ISL_2813507 | Lighthouse Lab in Milton Keynes | Wellcome Sanger Institute for the COVID-19 Genomics UK (COG-UK) Consortium | The Lighthouse Lab in Milton Keynes and Alex Alderton, Roberto Amato, Jeffrey Barrett, Sonia Goncalves, Ewan Harrison, David K. Jackson, Ian Johnston, Dominic Kwiatkowski, Cordelia Langford, John Sillitoe on behalf of the Wellcome Sanger Institute COVID-19 Surveillance Team |
| EPI_ISL_2813508, EPI_ISL_2813509 | Lighthouse Laboratory Plymouth | Wellcome Sanger Institute for the COVID-19 Genomics UK (COG-UK) Consortium | Lighthouse Laboratory Plymouth and Alex Alderton, Roberto Amato, Jeffrey Barrett, Sonia Goncalves, Ewan Harrison, David K. Jackson, Ian Johnston, Dominic Kwiatkowski, Cordelia Langford, John Sillitoe on behalf of the Wellcome Sanger Institute COVID-19 Surveillance Team |
| EPI_ISL_2813510, EPI_ISL_2813511 | Lighthouse Lab in Milton Keynes | Wellcome Sanger Institute for the COVID-19 Genomics UK (COG-UK) Consortium | The Lighthouse Lab in Milton Keynes and Alex Alderton, Roberto Amato, Jeffrey Barrett, Sonia Goncalves, Ewan Harrison, David K. Jackson, Ian Johnston, Dominic Kwiatkowski, Cordelia Langford, John Sillitoe on behalf of the Wellcome Sanger Institute COVID-19 Surveillance Team |
| EPI_ISL_2813513, EPI_ISL_2813514 | Lighthouse Laboratory Plymouth | Wellcome Sanger Institute for the COVID-19 Genomics UK (COG-UK) Consortium | Lighthouse Laboratory Plymouth and Alex Alderton, Roberto Amato, Jeffrey Barrett, Sonia Goncalves, Ewan Harrison, David K. Jackson, Ian Johnston, Dominic Kwiatkowski, Cordelia Langford, John Sillitoe on behalf of the Wellcome Sanger Institute COVID-19 Surveillance Team |
| EPI_ISL_2813516 | Lighthouse Lab in Milton Keynes | Wellcome Sanger Institute for the COVID-19 Genomics UK (COG-UK) Consortium | The Lighthouse Lab in Milton Keynes and Alex Alderton, Roberto Amato, Jeffrey Barrett, Sonia Goncalves, Ewan Harrison, David K. Jackson, Ian Johnston, Dominic Kwiatkowski, Cordelia Langford, John Sillitoe on behalf of the Wellcome Sanger Institute COVID-19 Surveillance Team |
| EPI_ISL_2813517, EPI_ISL_2813518, EPI_ISL_2813519, EPI_ISL_2813520, EPI_ISL_2813521, EPI_ISL_2813522, EPI_ISL_2813523, EPI_ISL_2813524, EPI_ISL_2813525, EPI_ISL_2813526 | Lighthouse Laboratory Plymouth | Wellcome Sanger Institute for the COVID-19 Genomics UK (COG-UK) Consortium | Lighthouse Laboratory Plymouth and Alex Alderton, Roberto Amato, Jeffrey Barrett, Sonia Goncalves, Ewan Harrison, David K. Jackson, Ian Johnston, Dominic Kwiatkowski, Cordelia Langford, John Sillitoe on behalf of the Wellcome Sanger Institute COVID-19 Surveillance Team |
| EPI_ISL_2813527 | Lighthouse Lab in Milton Keynes | Wellcome Sanger Institute for the COVID-19 Genomics UK (COG-UK) Consortium | The Lighthouse Lab in Milton Keynes and Alex Alderton, Roberto Amato, Jeffrey Barrett, Sonia Goncalves, Ewan Harrison, David K. Jackson, Ian Johnston, Dominic Kwiatkowski, Cordelia Langford, John Sillitoe on behalf of the Wellcome Sanger Institute COVID-19 Surveillance Team |
| EPI_ISL_2813528, EPI_ISL_2813529 | Lighthouse Laboratory Plymouth | Wellcome Sanger Institute for the COVID-19 Genomics UK (COG-UK) Consortium | Lighthouse Laboratory Plymouth and Alex Alderton, Roberto Amato, Jeffrey Barrett, Sonia Goncalves, Ewan Harrison, David K. Jackson, Ian Johnston, Dominic Kwiatkowski, Cordelia Langford, John Sillitoe on behalf of the Wellcome Sanger Institute COVID-19 Surveillance Team |
| EPI_ISL_2813530, EPI_ISL_2813531 | Lighthouse Lab in Milton Keynes | Wellcome Sanger Institute for the COVID-19 Genomics UK (COG-UK) Consortium | The Lighthouse Lab in Milton Keynes and Alex Alderton, Roberto Amato, Jeffrey Barrett, Sonia Goncalves, Ewan Harrison, David K. Jackson, Ian Johnston, Dominic Kwiatkowski, Cordelia Langford, John Sillitoe on behalf of the Wellcome Sanger Institute COVID-19 Surveillance Team |
| EPI_ISL_2813532, EPI_ISL_2813533, EPI_ISL_2813534, EPI_ISL_2813536, EPI_ISL_2813537, EPI_ISL_2813538, EPI_ISL_2813539, EPI_ISL_2813540, EPI_ISL_2813541, EPI_ISL_2813542, EPI_ISL_2813543, EPI_ISL_2813544, EPI_ISL_2813545 | | | |
| see above | Lighthouse Laboratory Plymouth | Wellcome Sanger Institute for the COVID-19 Genomics UK (COG-UK) Consortium | Lighthouse Laboratory Plymouth and Alex Alderton, Roberto Amato, Jeffrey Barrett, Sonia Goncalves, Ewan Harrison, David K. Jackson, Ian Johnston, Dominic Kwiatkowski, Cordelia Langford, John Sillitoe on behalf of the Wellcome Sanger Institute COVID-19 Surveillance Team |
| EPI_ISL_2813546, EPI_ISL_2813547 | Lighthouse Lab in Milton Keynes | Wellcome Sanger Institute for the COVID-19 Genomics UK (COG-UK) Consortium | The Lighthouse Lab in Milton Keynes and Alex Alderton, Roberto Amato, Jeffrey Barrett, Sonia Goncalves, Ewan Harrison, David K. Jackson, Ian Johnston, Dominic Kwiatkowski, Cordelia Langford, John Sillitoe on behalf of the Wellcome Sanger Institute COVID-19 Surveillance Team |
| EPI_ISL_2813548, EPI_ISL_2813549, EPI_ISL_2813550, EPI_ISL_2813551 | Lighthouse Laboratory Plymouth | Wellcome Sanger Institute for the COVID-19 Genomics UK (COG-UK) Consortium | Lighthouse Laboratory Plymouth and Alex Alderton, Roberto Amato, Jeffrey Barrett, Sonia Goncalves, Ewan Harrison, David K. Jackson, Ian Johnston, Dominic Kwiatkowski, Cordelia Langford, John Sillitoe on behalf of the Wellcome Sanger Institute COVID-19 Surveillance Team |
| EPI_ISL_2813552, EPI_ISL_2813553 | Lighthouse Lab in Milton Keynes | Wellcome Sanger Institute for the COVID-19 Genomics UK (COG-UK) Consortium | The Lighthouse Lab in Milton Keynes and Alex Alderton, Roberto Amato, Jeffrey Barrett, Sonia Goncalves, Ewan Harrison, David K. Jackson, Ian Johnston, Dominic Kwiatkowski, Cordelia Langford, John Sillitoe on behalf of the Wellcome Sanger Institute COVID-19 Surveillance Team |
| EPI_ISL_2813554, EPI_ISL_2813555 | Lighthouse Laboratory Plymouth | Wellcome Sanger Institute for the COVID-19 Genomics UK (COG-UK) Consortium | Lighthouse Laboratory Plymouth and Alex Alderton, Roberto Amato, Jeffrey Barrett, Sonia Goncalves, Ewan Harrison, David K. Jackson, Ian Johnston, Dominic Kwiatkowski, Cordelia Langford, John Sillitoe on behalf of the Wellcome Sanger Institute COVID-19 Surveillance Team |
| EPI_ISL_2813556, EPI_ISL_2813557 | Lighthouse Lab in Milton Keynes | Wellcome Sanger Institute for the COVID-19 Genomics UK (COG-UK) Consortium | The Lighthouse Lab in Milton Keynes and Alex Alderton, Roberto Amato, Jeffrey Barrett, Sonia Goncalves, Ewan Harrison, David K. Jackson, Ian Johnston, Dominic Kwiatkowski, Cordelia Langford, John Sillitoe on behalf of the Wellcome Sanger Institute COVID-19 Surveillance Team |
| EPI_ISL_2813558, EPI_ISL_2813559, EPI_ISL_2813560, EPI_ISL_2813561, EPI_ISL_2813562, EPI_ISL_2813563, EPI_ISL_2813564, EPI_ISL_2813565 | Lighthouse Laboratory Plymouth | Wellcome Sanger Institute for the COVID-19 Genomics UK (COG-UK) Consortium | Lighthouse Laboratory Plymouth and Alex Alderton, Roberto Amato, Jeffrey Barrett, Sonia Goncalves, Ewan Harrison, David K. Jackson, Ian Johnston, Dominic Kwiatkowski, Cordelia Langford, John Sillitoe on behalf of the Wellcome Sanger Institute COVID-19 Surveillance Team |
| EPI_ISL_2813566 | Lighthouse Lab in Milton Keynes | Wellcome Sanger Institute for the COVID-19 Genomics UK (COG-UK) Consortium | The Lighthouse Lab in Milton Keynes and Alex Alderton, Roberto Amato, Jeffrey Barrett, Sonia Goncalves, Ewan Harrison, David K. Jackson, Ian Johnston, Dominic Kwiatkowski, Cordelia Langford, John Sillitoe on behalf of the Wellcome Sanger Institute COVID-19 Surveillance Team |
| EPI_ISL_2813567, EPI_ISL_2813568, EPI_ISL_2813569, EPI_ISL_2813570, EPI_ISL_2813571, EPI_ISL_2813572 | Lighthouse Laboratory Plymouth | Wellcome Sanger Institute for the COVID-19 Genomics UK (COG-UK) Consortium | Lighthouse Laboratory Plymouth and Alex Alderton, Roberto Amato, Jeffrey Barrett, Sonia Goncalves, Ewan Harrison, David K. Jackson, Ian Johnston, Dominic Kwiatkowski, Cordelia Langford, John Sillitoe on behalf of the Wellcome Sanger Institute COVID-19 Surveillance Team |
| EPI_ISL_2813573 | Lighthouse Lab in Milton Keynes | Wellcome Sanger Institute for the COVID-19 Genomics UK (COG-UK) Consortium | The Lighthouse Lab in Milton Keynes and Alex Alderton, Roberto Amato, Jeffrey Barrett, Sonia Goncalves, Ewan Harrison, David K. Jackson, Ian Johnston, Dominic Kwiatkowski, Cordelia Langford, John Sillitoe on behalf of the Wellcome Sanger Institute COVID-19 Surveillance Team |
| EPI_ISL_2813574 | Lighthouse Laboratory Plymouth | Wellcome Sanger Institute for the COVID-19 Genomics UK (COG-UK) Consortium | Lighthouse Laboratory Plymouth and Alex Alderton, Roberto Amato, Jeffrey Barrett, Sonia Goncalves, Ewan Harrison, David K. Jackson, Ian Johnston, Dominic Kwiatkowski, Cordelia Langford, John Sillitoe on behalf of the Wellcome Sanger Institute COVID-19 Surveillance Team |
| EPI_ISL_2813575 | Lighthouse Lab in Milton Keynes | Wellcome Sanger Institute for the COVID-19 Genomics UK (COG-UK) Consortium | The Lighthouse Lab in Milton Keynes and Alex Alderton, Roberto Amato, Jeffrey Barrett, Sonia Goncalves, Ewan Harrison, David K. Jackson, Ian Johnston, Dominic Kwiatkowski, Cordelia Langford, John Sillitoe on behalf of the Wellcome Sanger Institute COVID-19 Surveillance Team |
| EPI_ISL_2813576, EPI_ISL_2813577, EPI_ISL_2813578, EPI_ISL_2813579, EPI_ISL_2813580 | Lighthouse Laboratory Plymouth | Wellcome Sanger Institute for the COVID-19 Genomics UK (COG-UK) Consortium | Lighthouse Laboratory Plymouth and Alex Alderton, Roberto Amato, Jeffrey Barrett, Sonia Goncalves, Ewan Harrison, David K. Jackson, Ian Johnston, Dominic Kwiatkowski, Cordelia Langford, John Sillitoe on behalf of the Wellcome Sanger Institute COVID-19 Surveillance Team |
| EPI_ISL_2813581 | Lighthouse Lab in Milton Keynes | Wellcome Sanger Institute for the COVID-19 Genomics UK (COG-UK) Consortium | The Lighthouse Lab in Milton Keynes and Alex Alderton, Roberto Amato, Jeffrey Barrett, Sonia Goncalves, Ewan Harrison, David K. Jackson, Ian Johnston, Dominic Kwiatkowski, Cordelia Langford, John Sillitoe on behalf of the Wellcome Sanger Institute COVID-19 Surveillance Team |
| EPI_ISL_2813582, EPI_ISL_2813583 | Lighthouse Laboratory Plymouth | Wellcome Sanger Institute for the COVID-19 Genomics UK (COG-UK) Consortium | Lighthouse Laboratory Plymouth and Alex Alderton, Roberto Amato, Jeffrey Barrett, Sonia Goncalves, Ewan Harrison, David K. Jackson, Ian Johnston, Dominic Kwiatkowski, Cordelia Langford, John Sillitoe on behalf of the Wellcome Sanger Institute COVID-19 Surveillance Team |
| EPI_ISL_2813584 | Lighthouse Lab in Milton Keynes | Wellcome Sanger Institute for the COVID-19 Genomics UK (COG-UK) Consortium | The Lighthouse Lab in Milton Keynes and Alex Alderton, Roberto Amato, Jeffrey Barrett, Sonia Goncalves, Ewan Harrison, David K. Jackson, Ian Johnston, Dominic Kwiatkowski, Cordelia Langford, John Sillitoe on behalf of the Wellcome Sanger Institute COVID-19 Surveillance Team |
| EPI_ISL_2813585, EPI_ISL_2813586 | Lighthouse Laboratory Plymouth | Wellcome Sanger Institute for the COVID-19 Genomics UK (COG-UK) Consortium | Lighthouse Laboratory Plymouth and Alex Alderton, Roberto Amato, Jeffrey Barrett, Sonia Goncalves, Ewan Harrison, David K. Jackson, Ian Johnston, Dominic Kwiatkowski, Cordelia Langford, John Sillitoe on behalf of the Wellcome Sanger Institute COVID-19 Surveillance Team |
| EPI_ISL_2813587, EPI_ISL_2813588 | Lighthouse Lab in Milton Keynes | Wellcome Sanger Institute for the COVID-19 Genomics UK (COG-UK) Consortium | The Lighthouse Lab in Milton Keynes and Alex Alderton, Roberto Amato, Jeffrey Barrett, Sonia Goncalves, Ewan Harrison, David K. Jackson, Ian Johnston, Dominic Kwiatkowski, Cordelia Langford, John Sillitoe on behalf of the Wellcome Sanger Institute COVID-19 Surveillance Team |
| EPI_ISL_2813589 | Lighthouse Laboratory Plymouth | Wellcome Sanger Institute for the COVID-19 Genomics UK (COG-UK) Consortium | Lighthouse Laboratory Plymouth and Alex Alderton, Roberto Amato, Jeffrey Barrett, Sonia Goncalves, Ewan Harrison, David K. Jackson, Ian Johnston, Dominic Kwiatkowski, Cordelia Langford, John Sillitoe on behalf of the Wellcome Sanger Institute COVID-19 Surveillance Team |
| EPI_ISL_2813590 | Lighthouse Lab in Milton Keynes | Wellcome Sanger Institute for the COVID-19 Genomics UK (COG-UK) Consortium | The Lighthouse Lab in Milton Keynes and Alex Alderton, Roberto Amato, Jeffrey Barrett, Sonia Goncalves, Ewan Harrison, David K. Jackson, Ian Johnston, Dominic Kwiatkowski, Cordelia Langford, John Sillitoe on behalf of the Wellcome Sanger Institute COVID-19 Surveillance Team |
| EPI_ISL_2813591, EPI_ISL_2813592, EPI_ISL_2813593, EPI_ISL_2813594, EPI_ISL_2813595, EPI_ISL_2813596, EPI_ISL_2813597, EPI_ISL_2813598, EPI_ISL_2813599, EPI_ISL_2813600, EPI_ISL_2813601, EPI_ISL_2813602, EPI_ISL_2813603, EPI_ISL_2813604 | | | |
| see above | Lighthouse Laboratory Plymouth | Wellcome Sanger Institute for the COVID-19 Genomics UK (COG-UK) Consortium | Lighthouse Laboratory Plymouth and Alex Alderton, Roberto Amato, Jeffrey Barrett, Sonia Goncalves, Ewan Harrison, David K. Jackson, Ian Johnston, Dominic Kwiatkowski, Cordelia Langford, John Sillitoe on behalf of the Wellcome Sanger Institute COVID-19 Surveillance Team |

| | | | |
|---|---|---|---|
| EPI_ISL_2813605, EPI_ISL_2813606, EPI_ISL_2813607 | Lighthouse Lab in Milton Keynes | Wellcome Sanger Institute for the COVID-19 Genomics UK (COG-UK) Consortium | The Lighthouse Lab in Milton Keynes and Alex Alderton, Roberto Amato, Jeffrey Barrett, Sonia Goncalves, Ewan Harrison, David K. Jackson, Ian Johnston, Dominic Kwiatkowski, Cordelia Langford, John Sillitoe on behalf of the Wellcome Sanger Institute COVID-19 Surveillance Team |
| EPI_ISL_2813608, EPI_ISL_2813609, EPI_ISL_2813610, EPI_ISL_2813611, EPI_ISL_2813612, EPI_ISL_2813613, EPI_ISL_2813614, EPI_ISL_2813615, EPI_ISL_2813616, EPI_ISL_2813617 | Lighthouse Laboratory Plymouth | Wellcome Sanger Institute for the COVID-19 Genomics UK (COG-UK) Consortium | Lighthouse Laboratory Plymouth and Alex Alderton, Roberto Amato, Jeffrey Barrett, Sonia Goncalves, Ewan Harrison, David K. Jackson, Ian Johnston, Dominic Kwiatkowski, Cordelia Langford, John Sillitoe on behalf of the Wellcome Sanger Institute COVID-19 Surveillance Team |
| EPI_ISL_2813618 | Lighthouse Lab in Milton Keynes | Wellcome Sanger Institute for the COVID-19 Genomics UK (COG-UK) Consortium | The Lighthouse Lab in Milton Keynes and Alex Alderton, Roberto Amato, Jeffrey Barrett, Sonia Goncalves, Ewan Harrison, David K. Jackson, Ian Johnston, Dominic Kwiatkowski, Cordelia Langford, John Sillitoe on behalf of the Wellcome Sanger Institute COVID-19 Surveillance Team |
| EPI_ISL_2813619 | Lighthouse Laboratory Plymouth | Wellcome Sanger Institute for the COVID-19 Genomics UK (COG-UK) Consortium | Lighthouse Laboratory Plymouth and Alex Alderton, Roberto Amato, Jeffrey Barrett, Sonia Goncalves, Ewan Harrison, David K. Jackson, Ian Johnston, Dominic Kwiatkowski, Cordelia Langford, John Sillitoe on behalf of the Wellcome Sanger Institute COVID-19 Surveillance Team |
| EPI_ISL_2813620 | Lighthouse Lab in Milton Keynes | Wellcome Sanger Institute for the COVID-19 Genomics UK (COG-UK) Consortium | The Lighthouse Lab in Milton Keynes and Alex Alderton, Roberto Amato, Jeffrey Barrett, Sonia Goncalves, Ewan Harrison, David K. Jackson, Ian Johnston, Dominic Kwiatkowski, Cordelia Langford, John Sillitoe on behalf of the Wellcome Sanger Institute COVID-19 Surveillance Team |
| EPI_ISL_2813621, EPI_ISL_2813622, EPI_ISL_2813623 | Lighthouse Laboratory Plymouth | Wellcome Sanger Institute for the COVID-19 Genomics UK (COG-UK) Consortium | Lighthouse Laboratory Plymouth and Alex Alderton, Roberto Amato, Jeffrey Barrett, Sonia Goncalves, Ewan Harrison, David K. Jackson, Ian Johnston, Dominic Kwiatkowski, Cordelia Langford, John Sillitoe on behalf of the Wellcome Sanger Institute COVID-19 Surveillance Team |
| EPI_ISL_2813624 | Lighthouse Lab in Milton Keynes | Wellcome Sanger Institute for the COVID-19 Genomics UK (COG-UK) Consortium | The Lighthouse Lab in Milton Keynes and Alex Alderton, Roberto Amato, Jeffrey Barrett, Sonia Goncalves, Ewan Harrison, David K. Jackson, Ian Johnston, Dominic Kwiatkowski, Cordelia Langford, John Sillitoe on behalf of the Wellcome Sanger Institute COVID-19 Surveillance Team |
| EPI_ISL_2813625 | Lighthouse Laboratory Plymouth | Wellcome Sanger Institute for the COVID-19 Genomics UK (COG-UK) Consortium | Lighthouse Laboratory Plymouth and Alex Alderton, Roberto Amato, Jeffrey Barrett, Sonia Goncalves, Ewan Harrison, David K. Jackson, Ian Johnston, Dominic Kwiatkowski, Cordelia Langford, John Sillitoe on behalf of the Wellcome Sanger Institute COVID-19 Surveillance Team |
| EPI_ISL_2813626 | Lighthouse Lab in Milton Keynes | Wellcome Sanger Institute for the COVID-19 Genomics UK (COG-UK) Consortium | The Lighthouse Lab in Milton Keynes and Alex Alderton, Roberto Amato, Jeffrey Barrett, Sonia Goncalves, Ewan Harrison, David K. Jackson, Ian Johnston, Dominic Kwiatkowski, Cordelia Langford, John Sillitoe on behalf of the Wellcome Sanger Institute COVID-19 Surveillance Team |
| EPI_ISL_2813627, EPI_ISL_2813628, EPI_ISL_2813629, EPI_ISL_2813630, EPI_ISL_2813631, EPI_ISL_2813632, EPI_ISL_2813633, EPI_ISL_2813634, EPI_ISL_2813635, EPI_ISL_2813636 | Lighthouse Laboratory Plymouth | Wellcome Sanger Institute for the COVID-19 Genomics UK (COG-UK) Consortium | Lighthouse Laboratory Plymouth and Alex Alderton, Roberto Amato, Jeffrey Barrett, Sonia Goncalves, Ewan Harrison, David K. Jackson, Ian Johnston, Dominic Kwiatkowski, Cordelia Langford, John Sillitoe on behalf of the Wellcome Sanger Institute COVID-19 Surveillance Team |
| EPI_ISL_2813637 | Lighthouse Lab in Milton Keynes | Wellcome Sanger Institute for the COVID-19 Genomics UK (COG-UK) Consortium | The Lighthouse Lab in Milton Keynes and Alex Alderton, Roberto Amato, Jeffrey Barrett, Sonia Goncalves, Ewan Harrison, David K. Jackson, Ian Johnston, Dominic Kwiatkowski, Cordelia Langford, John Sillitoe on behalf of the Wellcome Sanger Institute COVID-19 Surveillance Team |
| EPI_ISL_2813638, EPI_ISL_2813639, EPI_ISL_2813640, EPI_ISL_2813641 | Lighthouse Laboratory Plymouth | Wellcome Sanger Institute for the COVID-19 Genomics UK (COG-UK) Consortium | Lighthouse Laboratory Plymouth and Alex Alderton, Roberto Amato, Jeffrey Barrett, Sonia Goncalves, Ewan Harrison, David K. Jackson, Ian Johnston, Dominic Kwiatkowski, Cordelia Langford, John Sillitoe on behalf of the Wellcome Sanger Institute COVID-19 Surveillance Team |
| EPI_ISL_2813642, EPI_ISL_2813643, EPI_ISL_2813644 | Lighthouse Lab in Milton Keynes | Wellcome Sanger Institute for the COVID-19 Genomics UK (COG-UK) Consortium | The Lighthouse Lab in Milton Keynes and Alex Alderton, Roberto Amato, Jeffrey Barrett, Sonia Goncalves, Ewan Harrison, David K. Jackson, Ian Johnston, Dominic Kwiatkowski, Cordelia Langford, John Sillitoe on behalf of the Wellcome Sanger Institute COVID-19 Surveillance Team |
| EPI_ISL_2813645 | Lighthouse Laboratory Plymouth | Wellcome Sanger Institute for the COVID-19 Genomics UK (COG-UK) Consortium | Lighthouse Laboratory Plymouth and Alex Alderton, Roberto Amato, Jeffrey Barrett, Sonia Goncalves, Ewan Harrison, David K. Jackson, Ian Johnston, Dominic Kwiatkowski, Cordelia Langford, John Sillitoe on behalf of the Wellcome Sanger Institute COVID-19 Surveillance Team |
| EPI_ISL_2813646 | Lighthouse Lab in Milton Keynes | Wellcome Sanger Institute for the COVID-19 Genomics UK (COG-UK) Consortium | The Lighthouse Lab in Milton Keynes and Alex Alderton, Roberto Amato, Jeffrey Barrett, Sonia Goncalves, Ewan Harrison, David K. Jackson, Ian Johnston, Dominic Kwiatkowski, Cordelia Langford, John Sillitoe on behalf of the Wellcome Sanger Institute COVID-19 Surveillance Team |
| EPI_ISL_2813647, EPI_ISL_2813648, EPI_ISL_2813649, EPI_ISL_2813650, EPI_ISL_2813651, EPI_ISL_2813652, EPI_ISL_2813653, EPI_ISL_2813654, EPI_ISL_2813655 | Lighthouse Laboratory Plymouth | Wellcome Sanger Institute for the COVID-19 Genomics UK (COG-UK) Consortium | Lighthouse Laboratory Plymouth and Alex Alderton, Roberto Amato, Jeffrey Barrett, Sonia Goncalves, Ewan Harrison, David K. Jackson, Ian Johnston, Dominic Kwiatkowski, Cordelia Langford, John Sillitoe on behalf of the Wellcome Sanger Institute COVID-19 Surveillance Team |
| EPI_ISL_2813656 | Lighthouse Lab in Milton Keynes | Wellcome Sanger Institute for the COVID-19 Genomics UK (COG-UK) Consortium | The Lighthouse Lab in Milton Keynes and Alex Alderton, Roberto Amato, Jeffrey Barrett, Sonia Goncalves, Ewan Harrison, David K. Jackson, Ian Johnston, Dominic Kwiatkowski, Cordelia Langford, John Sillitoe on behalf of the Wellcome Sanger Institute COVID-19 Surveillance Team |
| EPI_ISL_2813657, EPI_ISL_2813658, EPI_ISL_2813659 | Lighthouse Laboratory Plymouth | Wellcome Sanger Institute for the COVID-19 Genomics UK (COG-UK) Consortium | Lighthouse Laboratory Plymouth and Alex Alderton, Roberto Amato, Jeffrey Barrett, Sonia Goncalves, Ewan Harrison, David K. Jackson, Ian Johnston, Dominic Kwiatkowski, Cordelia Langford, John Sillitoe on behalf of the Wellcome Sanger Institute COVID-19 Surveillance Team |
| EPI_ISL_2813660 | Lighthouse Lab in Alderley Park | Wellcome Sanger Institute for the COVID-19 Genomics UK (COG-UK) Consortium | Jacquelyn Wynn, Mairead Hyland, The Lighthouse Lab in Alderley Park and Alex Alderton, Roberto Amato, Jeffrey Barrett, Sonia Goncalves, Ewan Harrison, David K. Jackson, Ian Johnston, Dominic Kwiatkowski, Cordelia Langford, John Sillitoe on behalf of the Wellcome Sanger Institute COVID-19 Surveillance Team |
| EPI_ISL_2813661, EPI_ISL_2813662, EPI_ISL_2813663, EPI_ISL_2813664, EPI_ISL_2813665, EPI_ISL_2813666, EPI_ISL_2813667, EPI_ISL_2813668, EPI_ISL_2813669 | Lighthouse Lab in Milton Keynes | Wellcome Sanger Institute for the COVID-19 Genomics UK (COG-UK) Consortium | The Lighthouse Lab in Milton Keynes and Alex Alderton, Roberto Amato, Jeffrey Barrett, Sonia Goncalves, Ewan Harrison, David K. Jackson, Ian Johnston, Dominic Kwiatkowski, Cordelia Langford, John Sillitoe on behalf of the Wellcome Sanger Institute COVID-19 Surveillance Team |
| EPI_ISL_2813670 | Lighthouse Lab in Alderley Park | Wellcome Sanger Institute for the COVID-19 Genomics UK (COG-UK) Consortium | Jacquelyn Wynn, Mairead Hyland, The Lighthouse Lab in Alderley Park and Alex Alderton, Roberto Amato, Jeffrey Barrett, Sonia Goncalves, Ewan Harrison, David K. Jackson, Ian Johnston, Dominic Kwiatkowski, Cordelia Langford, John Sillitoe on behalf of the Wellcome Sanger Institute COVID-19 Surveillance Team |
| EPI_ISL_2813671 | Lighthouse Lab in Milton Keynes | Wellcome Sanger Institute for the COVID-19 Genomics UK (COG-UK) Consortium | The Lighthouse Lab in Milton Keynes and Alex Alderton, Roberto Amato, Jeffrey Barrett, Sonia Goncalves, Ewan Harrison, David K. Jackson, Ian Johnston, Dominic Kwiatkowski, Cordelia Langford, John Sillitoe on behalf of the Wellcome Sanger Institute COVID-19 Surveillance Team |
| EPI_ISL_2813672 | Lighthouse Lab in Alderley Park | Wellcome Sanger Institute for the COVID-19 Genomics UK (COG-UK) Consortium | Jacquelyn Wynn, Mairead Hyland, The Lighthouse Lab in Alderley Park and Alex Alderton, Roberto Amato, Jeffrey Barrett, Sonia Goncalves, Ewan Harrison, David K. Jackson, Ian Johnston, Dominic Kwiatkowski, Cordelia Langford, John Sillitoe on behalf of the Wellcome Sanger Institute COVID-19 Surveillance Team |
| EPI_ISL_2813673, EPI_ISL_2813674, EPI_ISL_2813675, EPI_ISL_2813676, EPI_ISL_2813677, EPI_ISL_2813678, EPI_ISL_2813679, EPI_ISL_2813680, EPI_ISL_2813681, EPI_ISL_2813682, EPI_ISL_2813683 | | | |
| see above | Lighthouse Lab in Milton Keynes | Wellcome Sanger Institute for the COVID-19 Genomics UK (COG-UK) Consortium | The Lighthouse Lab in Milton Keynes and Alex Alderton, Roberto Amato, Jeffrey Barrett, Sonia Goncalves, Ewan Harrison, David K. Jackson, Ian Johnston, Dominic Kwiatkowski, Cordelia Langford, John Sillitoe on behalf of the Wellcome Sanger Institute COVID-19 Surveillance Team |
| EPI_ISL_2813684, EPI_ISL_2813685 | Lighthouse Lab in Alderley Park | Wellcome Sanger Institute for the COVID-19 Genomics UK (COG-UK) Consortium | Jacquelyn Wynn, Mairead Hyland, The Lighthouse Lab in Alderley Park and Alex Alderton, Roberto Amato, Jeffrey Barrett, Sonia Goncalves, Ewan Harrison, David K. Jackson, Ian Johnston, Dominic Kwiatkowski, Cordelia Langford, John Sillitoe on behalf of the Wellcome Sanger Institute COVID-19 Surveillance Team |
| EPI_ISL_2813686, EPI_ISL_2813687, EPI_ISL_2813688 | Lighthouse Lab in Milton Keynes | Wellcome Sanger Institute for the COVID-19 Genomics UK (COG-UK) Consortium | The Lighthouse Lab in Milton Keynes and Alex Alderton, Roberto Amato, Jeffrey Barrett, Sonia Goncalves, Ewan Harrison, David K. Jackson, Ian Johnston, Dominic Kwiatkowski, Cordelia Langford, John Sillitoe on behalf of the Wellcome Sanger Institute COVID-19 Surveillance Team |
| EPI_ISL_2813689 | Lighthouse Lab in Alderley Park | Wellcome Sanger Institute for the COVID-19 Genomics UK (COG-UK) Consortium | Jacquelyn Wynn, Mairead Hyland, The Lighthouse Lab in Alderley Park and Alex Alderton, Roberto Amato, Jeffrey Barrett, Sonia Goncalves, Ewan Harrison, David K. Jackson, Ian Johnston, Dominic Kwiatkowski, Cordelia Langford, John Sillitoe on behalf of the Wellcome Sanger Institute COVID-19 Surveillance Team |
| EPI_ISL_2813690, EPI_ISL_2813691 | Lighthouse Lab in Milton Keynes | Wellcome Sanger Institute for the COVID-19 Genomics UK (COG-UK) Consortium | The Lighthouse Lab in Milton Keynes and Alex Alderton, Roberto Amato, Jeffrey Barrett, Sonia Goncalves, Ewan Harrison, David K. Jackson, Ian Johnston, Dominic Kwiatkowski, Cordelia Langford, John Sillitoe on behalf of the Wellcome Sanger Institute COVID-19 Surveillance Team |
| EPI_ISL_2813692, EPI_ISL_2813693 | Lighthouse Lab in Alderley Park | Wellcome Sanger Institute for the COVID-19 Genomics UK (COG-UK) Consortium | Jacquelyn Wynn, Mairead Hyland, The Lighthouse Lab in Alderley Park and Alex Alderton, Roberto Amato, Jeffrey Barrett, Sonia Goncalves, Ewan Harrison, David K. Jackson, Ian Johnston, Dominic Kwiatkowski, Cordelia Langford, John Sillitoe on behalf of the Wellcome Sanger Institute COVID-19 Surveillance Team |

| | | | |
|---|---|---|---|
| EPI_ISL_2813694, EPI_ISL_2813695, EPI_ISL_2813696, EPI_ISL_2813697 | Lighthouse Lab in Milton Keynes | Wellcome Sanger Institute for the COVID-19 Genomics UK (COG-UK) Consortium | The Lighthouse Lab in Milton Keynes and Alex Alderton, Roberto Amato, Jeffrey Barrett, Sonia Goncalves, Ewan Harrison, David K. Jackson, Ian Johnston, Dominic Kwiatkowski, Cordelia Langford, John Sillitoe on behalf of the Wellcome Sanger Institute COVID-19 Surveillance Team |
| EPI_ISL_2813698 | Lighthouse Lab in Alderley Park | Wellcome Sanger Institute for the COVID-19 Genomics UK (COG-UK) Consortium | Jacquelyn Wynn, Mairead Hyland, The Lighthouse Lab in Alderley Park and Alex Alderton, Roberto Amato, Jeffrey Barrett, Sonia Goncalves, Ewan Harrison, David K. Jackson, Ian Johnston, Dominic Kwiatkowski, Cordelia Langford, John Sillitoe on behalf of the Wellcome Sanger Institute COVID-19 Surveillance Team |
| EPI_ISL_2813699, EPI_ISL_2813700 | Lighthouse Lab in Milton Keynes | Wellcome Sanger Institute for the COVID-19 Genomics UK (COG-UK) Consortium | The Lighthouse Lab in Milton Keynes and Alex Alderton, Roberto Amato, Jeffrey Barrett, Sonia Goncalves, Ewan Harrison, David K. Jackson, Ian Johnston, Dominic Kwiatkowski, Cordelia Langford, John Sillitoe on behalf of the Wellcome Sanger Institute COVID-19 Surveillance Team |
| EPI_ISL_2813701 | Lighthouse Lab in Alderley Park | Wellcome Sanger Institute for the COVID-19 Genomics UK (COG-UK) Consortium | Jacquelyn Wynn, Mairead Hyland, The Lighthouse Lab in Alderley Park and Alex Alderton, Roberto Amato, Jeffrey Barrett, Sonia Goncalves, Ewan Harrison, David K. Jackson, Ian Johnston, Dominic Kwiatkowski, Cordelia Langford, John Sillitoe on behalf of the Wellcome Sanger Institute COVID-19 Surveillance Team |
| EPI_ISL_2813702 | Lighthouse Lab in Milton Keynes | Wellcome Sanger Institute for the COVID-19 Genomics UK (COG-UK) Consortium | The Lighthouse Lab in Milton Keynes and Alex Alderton, Roberto Amato, Jeffrey Barrett, Sonia Goncalves, Ewan Harrison, David K. Jackson, Ian Johnston, Dominic Kwiatkowski, Cordelia Langford, John Sillitoe on behalf of the Wellcome Sanger Institute COVID-19 Surveillance Team |
| EPI_ISL_2813703, EPI_ISL_2813704 | Lighthouse Lab in Alderley Park | Wellcome Sanger Institute for the COVID-19 Genomics UK (COG-UK) Consortium | Jacquelyn Wynn, Mairead Hyland, The Lighthouse Lab in Alderley Park and Alex Alderton, Roberto Amato, Jeffrey Barrett, Sonia Goncalves, Ewan Harrison, David K. Jackson, Ian Johnston, Dominic Kwiatkowski, Cordelia Langford, John Sillitoe on behalf of the Wellcome Sanger Institute COVID-19 Surveillance Team |
| EPI_ISL_2813705 | Lighthouse Lab in Milton Keynes | Wellcome Sanger Institute for the COVID-19 Genomics UK (COG-UK) Consortium | The Lighthouse Lab in Milton Keynes and Alex Alderton, Roberto Amato, Jeffrey Barrett, Sonia Goncalves, Ewan Harrison, David K. Jackson, Ian Johnston, Dominic Kwiatkowski, Cordelia Langford, John Sillitoe on behalf of the Wellcome Sanger Institute COVID-19 Surveillance Team |
| EPI_ISL_2813706 | Lighthouse Lab in Alderley Park | Wellcome Sanger Institute for the COVID-19 Genomics UK (COG-UK) Consortium | Jacquelyn Wynn, Mairead Hyland, The Lighthouse Lab in Alderley Park and Alex Alderton, Roberto Amato, Jeffrey Barrett, Sonia Goncalves, Ewan Harrison, David K. Jackson, Ian Johnston, Dominic Kwiatkowski, Cordelia Langford, John Sillitoe on behalf of the Wellcome Sanger Institute COVID-19 Surveillance Team |
| EPI_ISL_2813707 | Lighthouse Lab in Milton Keynes | Wellcome Sanger Institute for the COVID-19 Genomics UK (COG-UK) Consortium | The Lighthouse Lab in Milton Keynes and Alex Alderton, Roberto Amato, Jeffrey Barrett, Sonia Goncalves, Ewan Harrison, David K. Jackson, Ian Johnston, Dominic Kwiatkowski, Cordelia Langford, John Sillitoe on behalf of the Wellcome Sanger Institute COVID-19 Surveillance Team |
| EPI_ISL_2813708 | Lighthouse Lab in Alderley Park | Wellcome Sanger Institute for the COVID-19 Genomics UK (COG-UK) Consortium | Jacquelyn Wynn, Mairead Hyland, The Lighthouse Lab in Alderley Park and Alex Alderton, Roberto Amato, Jeffrey Barrett, Sonia Goncalves, Ewan Harrison, David K. Jackson, Ian Johnston, Dominic Kwiatkowski, Cordelia Langford, John Sillitoe on behalf of the Wellcome Sanger Institute COVID-19 Surveillance Team |
| EPI_ISL_2813709, EPI_ISL_2813710, EPI_ISL_2813711, EPI_ISL_2813712, EPI_ISL_2813713 | Lighthouse Lab in Milton Keynes | Wellcome Sanger Institute for the COVID-19 Genomics UK (COG-UK) Consortium | The Lighthouse Lab in Milton Keynes and Alex Alderton, Roberto Amato, Jeffrey Barrett, Sonia Goncalves, Ewan Harrison, David K. Jackson, Ian Johnston, Dominic Kwiatkowski, Cordelia Langford, John Sillitoe on behalf of the Wellcome Sanger Institute COVID-19 Surveillance Team |
| EPI_ISL_2813714, EPI_ISL_2813715, EPI_ISL_2813716 | Lighthouse Lab in Alderley Park | Wellcome Sanger Institute for the COVID-19 Genomics UK (COG-UK) Consortium | Jacquelyn Wynn, Mairead Hyland, The Lighthouse Lab in Alderley Park and Alex Alderton, Roberto Amato, Jeffrey Barrett, Sonia Goncalves, Ewan Harrison, David K. Jackson, Ian Johnston, Dominic Kwiatkowski, Cordelia Langford, John Sillitoe on behalf of the Wellcome Sanger Institute COVID-19 Surveillance Team |
| EPI_ISL_2813717, EPI_ISL_2813718, EPI_ISL_2813719 | Lighthouse Lab in Milton Keynes | Wellcome Sanger Institute for the COVID-19 Genomics UK (COG-UK) Consortium | The Lighthouse Lab in Milton Keynes and Alex Alderton, Roberto Amato, Jeffrey Barrett, Sonia Goncalves, Ewan Harrison, David K. Jackson, Ian Johnston, Dominic Kwiatkowski, Cordelia Langford, John Sillitoe on behalf of the Wellcome Sanger Institute COVID-19 Surveillance Team |
| EPI_ISL_2813720, EPI_ISL_2813721 | Lighthouse Lab in Alderley Park | Wellcome Sanger Institute for the COVID-19 Genomics UK (COG-UK) Consortium | Jacquelyn Wynn, Mairead Hyland, The Lighthouse Lab in Alderley Park and Alex Alderton, Roberto Amato, Jeffrey Barrett, Sonia Goncalves, Ewan Harrison, David K. Jackson, Ian Johnston, Dominic Kwiatkowski, Cordelia Langford, John Sillitoe on behalf of the Wellcome Sanger Institute COVID-19 Surveillance Team |
| EPI_ISL_2813722, EPI_ISL_2813723 | Lighthouse Lab in Milton Keynes | Wellcome Sanger Institute for the COVID-19 Genomics UK (COG-UK) Consortium | The Lighthouse Lab in Milton Keynes and Alex Alderton, Roberto Amato, Jeffrey Barrett, Sonia Goncalves, Ewan Harrison, David K. Jackson, Ian Johnston, Dominic Kwiatkowski, Cordelia Langford, John Sillitoe on behalf of the Wellcome Sanger Institute COVID-19 Surveillance Team |
| EPI_ISL_2813724 | Lighthouse Lab in Alderley Park | Wellcome Sanger Institute for the COVID-19 Genomics UK (COG-UK) Consortium | Jacquelyn Wynn, Mairead Hyland, The Lighthouse Lab in Alderley Park and Alex Alderton, Roberto Amato, Jeffrey Barrett, Sonia Goncalves, Ewan Harrison, David K. Jackson, Ian Johnston, Dominic Kwiatkowski, Cordelia Langford, John Sillitoe on behalf of the Wellcome Sanger Institute COVID-19 Surveillance Team |
| EPI_ISL_2813725, EPI_ISL_2813726, EPI_ISL_2813727 | Lighthouse Lab in Milton Keynes | Wellcome Sanger Institute for the COVID-19 Genomics UK (COG-UK) Consortium | The Lighthouse Lab in Milton Keynes and Alex Alderton, Roberto Amato, Jeffrey Barrett, Sonia Goncalves, Ewan Harrison, David K. Jackson, Ian Johnston, Dominic Kwiatkowski, Cordelia Langford, John Sillitoe on behalf of the Wellcome Sanger Institute COVID-19 Surveillance Team |
| EPI_ISL_2813728 | Lighthouse Lab in Alderley Park | Wellcome Sanger Institute for the COVID-19 Genomics UK (COG-UK) Consortium | Jacquelyn Wynn, Mairead Hyland, The Lighthouse Lab in Alderley Park and Alex Alderton, Roberto Amato, Jeffrey Barrett, Sonia Goncalves, Ewan Harrison, David K. Jackson, Ian Johnston, Dominic Kwiatkowski, Cordelia Langford, John Sillitoe on behalf of the Wellcome Sanger Institute COVID-19 Surveillance Team |
| EPI_ISL_2813729 | Lighthouse Lab in Milton Keynes | Wellcome Sanger Institute for the COVID-19 Genomics UK (COG-UK) Consortium | The Lighthouse Lab in Milton Keynes and Alex Alderton, Roberto Amato, Jeffrey Barrett, Sonia Goncalves, Ewan Harrison, David K. Jackson, Ian Johnston, Dominic Kwiatkowski, Cordelia Langford, John Sillitoe on behalf of the Wellcome Sanger Institute COVID-19 Surveillance Team |
| EPI_ISL_2813730 | Lighthouse Lab in Alderley Park | Wellcome Sanger Institute for the COVID-19 Genomics UK (COG-UK) Consortium | Jacquelyn Wynn, Mairead Hyland, The Lighthouse Lab in Alderley Park and Alex Alderton, Roberto Amato, Jeffrey Barrett, Sonia Goncalves, Ewan Harrison, David K. Jackson, Ian Johnston, Dominic Kwiatkowski, Cordelia Langford, John Sillitoe on behalf of the Wellcome Sanger Institute COVID-19 Surveillance Team |
| EPI_ISL_2813731 | Lighthouse Lab in Milton Keynes | Wellcome Sanger Institute for the COVID-19 Genomics UK (COG-UK) Consortium | The Lighthouse Lab in Milton Keynes and Alex Alderton, Roberto Amato, Jeffrey Barrett, Sonia Goncalves, Ewan Harrison, David K. Jackson, Ian Johnston, Dominic Kwiatkowski, Cordelia Langford, John Sillitoe on behalf of the Wellcome Sanger Institute COVID-19 Surveillance Team |
| EPI_ISL_2813732, EPI_ISL_2813733 | Lighthouse Lab in Alderley Park | Wellcome Sanger Institute for the COVID-19 Genomics UK (COG-UK) Consortium | Jacquelyn Wynn, Mairead Hyland, The Lighthouse Lab in Alderley Park and Alex Alderton, Roberto Amato, Jeffrey Barrett, Sonia Goncalves, Ewan Harrison, David K. Jackson, Ian Johnston, Dominic Kwiatkowski, Cordelia Langford, John Sillitoe on behalf of the Wellcome Sanger Institute COVID-19 Surveillance Team |
| EPI_ISL_2813734, EPI_ISL_2813735, EPI_ISL_2813736 | Lighthouse Lab in Milton Keynes | Wellcome Sanger Institute for the COVID-19 Genomics UK (COG-UK) Consortium | The Lighthouse Lab in Milton Keynes and Alex Alderton, Roberto Amato, Jeffrey Barrett, Sonia Goncalves, Ewan Harrison, David K. Jackson, Ian Johnston, Dominic Kwiatkowski, Cordelia Langford, John Sillitoe on behalf of the Wellcome Sanger Institute COVID-19 Surveillance Team |
| EPI_ISL_2813737 | Lighthouse Lab in Alderley Park | Wellcome Sanger Institute for the COVID-19 Genomics UK (COG-UK) Consortium | Jacquelyn Wynn, Mairead Hyland, The Lighthouse Lab in Alderley Park and Alex Alderton, Roberto Amato, Jeffrey Barrett, Sonia Goncalves, Ewan Harrison, David K. Jackson, Ian Johnston, Dominic Kwiatkowski, Cordelia Langford, John Sillitoe on behalf of the Wellcome Sanger Institute COVID-19 Surveillance Team |
| EPI_ISL_2813738, EPI_ISL_2813739 | Lighthouse Lab in Milton Keynes | Wellcome Sanger Institute for the COVID-19 Genomics UK (COG-UK) Consortium | The Lighthouse Lab in Milton Keynes and Alex Alderton, Roberto Amato, Jeffrey Barrett, Sonia Goncalves, Ewan Harrison, David K. Jackson, Ian Johnston, Dominic Kwiatkowski, Cordelia Langford, John Sillitoe on behalf of the Wellcome Sanger Institute COVID-19 Surveillance Team |
| EPI_ISL_2813740 | Lighthouse Lab in Alderley Park | Wellcome Sanger Institute for the COVID-19 Genomics UK (COG-UK) Consortium | Jacquelyn Wynn, Mairead Hyland, The Lighthouse Lab in Alderley Park and Alex Alderton, Roberto Amato, Jeffrey Barrett, Sonia Goncalves, Ewan Harrison, David K. Jackson, Ian Johnston, Dominic Kwiatkowski, Cordelia Langford, John Sillitoe on behalf of the Wellcome Sanger Institute COVID-19 Surveillance Team |
| EPI_ISL_2813741, EPI_ISL_2813742, EPI_ISL_2813743, EPI_ISL_2813744, EPI_ISL_2813745, EPI_ISL_2813746, EPI_ISL_2813747, EPI_ISL_2813748, EPI_ISL_2813749, EPI_ISL_2813750 | Lighthouse Lab in Milton Keynes | Wellcome Sanger Institute for the COVID-19 Genomics UK (COG-UK) Consortium | The Lighthouse Lab in Milton Keynes and Alex Alderton, Roberto Amato, Jeffrey Barrett, Sonia Goncalves, Ewan Harrison, David K. Jackson, Ian Johnston, Dominic Kwiatkowski, Cordelia Langford, John Sillitoe on behalf of the Wellcome Sanger Institute COVID-19 Surveillance Team |
| EPI_ISL_2813751 | Lighthouse Lab in Alderley Park | Wellcome Sanger Institute for the COVID-19 Genomics UK (COG-UK) Consortium | Jacquelyn Wynn, Mairead Hyland, The Lighthouse Lab in Alderley Park and Alex Alderton, Roberto Amato, Jeffrey Barrett, Sonia Goncalves, Ewan Harrison, David K. Jackson, Ian Johnston, Dominic Kwiatkowski, Cordelia Langford, John Sillitoe on behalf of the Wellcome Sanger Institute COVID-19 Surveillance Team |
| EPI_ISL_2813752, EPI_ISL_2813753, EPI_ISL_2813754 | Lighthouse Lab in Milton Keynes | Wellcome Sanger Institute for the COVID-19 Genomics UK (COG-UK) Consortium | The Lighthouse Lab in Milton Keynes and Alex Alderton, Roberto Amato, Jeffrey Barrett, Sonia Goncalves, Ewan Harrison, David K. Jackson, Ian Johnston, Dominic Kwiatkowski, Cordelia Langford, John Sillitoe on behalf of the Wellcome Sanger Institute COVID-19 Surveillance Team |
| EPI_ISL_2813755 | Lighthouse Lab in Alderley Park | Wellcome Sanger Institute for the COVID-19 Genomics UK (COG-UK) Consortium | Jacquelyn Wynn, Mairead Hyland, The Lighthouse Lab in Alderley Park and Alex Alderton, Roberto Amato, Jeffrey Barrett, Sonia Goncalves, Ewan Harrison, David K. Jackson, Ian Johnston, Dominic Kwiatkowski, Cordelia Langford, John Sillitoe on behalf of the Wellcome Sanger Institute COVID-19 Surveillance Team |
| EPI_ISL_2813756 | Lighthouse Lab in Milton Keynes | Wellcome Sanger Institute for the COVID-19 Genomics UK (COG-UK) Consortium | The Lighthouse Lab in Milton Keynes and Alex Alderton, Roberto Amato, Jeffrey Barrett, Sonia Goncalves, Ewan Harrison, David K. Jackson, Ian Johnston, Dominic Kwiatkowski, Cordelia Langford, John Sillitoe on behalf of the Wellcome Sanger Institute COVID-19 Surveillance Team |
| EPI_ISL_2813757 | Lighthouse Lab in Alderley Park | Wellcome Sanger Institute for the COVID-19 Genomics UK (COG-UK) Consortium | Jacquelyn Wynn, Mairead Hyland, The Lighthouse Lab in Alderley Park and Alex Alderton, Roberto Amato, Jeffrey Barrett, Sonia Goncalves, Ewan Harrison, David K. Jackson, Ian Johnston, Dominic Kwiatkowski, Cordelia Langford, John Sillitoe on behalf of the Wellcome Sanger Institute COVID-19 Surveillance Team |
| EPI_ISL_2813758 | Lighthouse Lab in Milton Keynes | Wellcome Sanger Institute for the COVID-19 Genomics UK (COG-UK) Consortium | The Lighthouse Lab in Milton Keynes and Alex Alderton, Roberto Amato, Jeffrey Barrett, Sonia Goncalves, Ewan Harrison, David K. Jackson, Ian Johnston, Dominic Kwiatkowski, Cordelia Langford, John Sillitoe on behalf of the Wellcome Sanger Institute COVID-19 Surveillance Team |
| EPI_ISL_2813759 | Lighthouse Lab in Alderley Park | Wellcome Sanger Institute for the COVID-19 Genomics UK (COG-UK) Consortium | Jacquelyn Wynn, Mairead Hyland, The Lighthouse Lab in Alderley Park and Alex Alderton, Roberto Amato, Jeffrey Barrett, Sonia Goncalves, Ewan Harrison, David K. Jackson, Ian Johnston, Dominic Kwiatkowski, Cordelia Langford, John Sillitoe on behalf of the Wellcome Sanger Institute COVID-19 Surveillance Team |
| EPI_ISL_2813760 | Lighthouse Lab in Milton Keynes | Wellcome Sanger Institute for the COVID-19 Genomics UK (COG-UK) Consortium | The Lighthouse Lab in Milton Keynes and Alex Alderton, Roberto Amato, Jeffrey Barrett, Sonia Goncalves, Ewan Harrison, David K. Jackson, Ian Johnston, Dominic Kwiatkowski, Cordelia Langford, John Sillitoe on behalf of the Wellcome Sanger Institute COVID-19 Surveillance Team |
| EPI_ISL_2813761 | Lighthouse Lab in Alderley Park | Wellcome Sanger Institute for the COVID-19 Genomics UK (COG-UK) Consortium | Jacquelyn Wynn, Mairead Hyland, The Lighthouse Lab in Alderley Park and Alex Alderton, Roberto Amato, Jeffrey Barrett, Sonia Goncalves, Ewan Harrison, David K. Jackson, Ian Johnston, Dominic Kwiatkowski, Cordelia Langford, John Sillitoe on behalf of the Wellcome Sanger Institute COVID-19 Surveillance Team |
| EPI_ISL_2813762, EPI_ISL_2813763, EPI_ISL_2813764, EPI_ISL_2813765, | Lighthouse Lab in Milton Keynes | Wellcome Sanger Institute for the COVID-19 Genomics UK (COG-UK) Consortium | The Lighthouse Lab in Milton Keynes and Alex Alderton, Roberto Amato, Jeffrey Barrett, Sonia Goncalves, Ewan Harrison, David K. Jackson, Ian Johnston, Dominic Kwiatkowski, Cordelia Langford, John Sillitoe on behalf of the Wellcome Sanger Institute COVID-19 Surveillance Team |

| | | | |
|---|---|---|---|
| EPI_ISL_2813766, EPI_ISL_2813767, EPI_ISL_2813768 | | | |
| EPI_ISL_2813769, EPI_ISL_2813770 | Lighthouse Lab in Alderley Park | Wellcome Sanger Institute for the COVID-19 Genomics UK (COG-UK) Consortium | Jacquelyn Wynn, Mairead Hyland, The Lighthouse Lab in Alderley Park and Alex Alderton, Roberto Amato, Jeffrey Barrett, Sonia Goncalves, Ewan Harrison, David K. Jackson, Ian Johnston, Dominic Kwiatkowski, Cordelia Langford, John Sillitoe on behalf of the Wellcome Sanger Institute COVID-19 Surveillance Team |
| EPI_ISL_2813771 | Lighthouse Lab in Milton Keynes | Wellcome Sanger Institute for the COVID-19 Genomics UK (COG-UK) Consortium | The Lighthouse Lab in Milton Keynes and Alex Alderton, Roberto Amato, Jeffrey Barrett, Sonia Goncalves, Ewan Harrison, David K. Jackson, Ian Johnston, Dominic Kwiatkowski, Cordelia Langford, John Sillitoe on behalf of the Wellcome Sanger Institute COVID-19 Surveillance Team |
| EPI_ISL_2813772, EPI_ISL_2813773 | Lighthouse Lab in Alderley Park | Wellcome Sanger Institute for the COVID-19 Genomics UK (COG-UK) Consortium | Jacquelyn Wynn, Mairead Hyland, The Lighthouse Lab in Alderley Park and Alex Alderton, Roberto Amato, Jeffrey Barrett, Sonia Goncalves, Ewan Harrison, David K. Jackson, Ian Johnston, Dominic Kwiatkowski, Cordelia Langford, John Sillitoe on behalf of the Wellcome Sanger Institute COVID-19 Surveillance Team |
| EPI_ISL_2813774, EPI_ISL_2813775, EPI_ISL_2813776, EPI_ISL_2813777, EPI_ISL_2813778, EPI_ISL_2813779, EPI_ISL_2813780, EPI_ISL_2813781, EPI_ISL_2813782, EPI_ISL_2813783, EPI_ISL_2813784, EPI_ISL_2813785, EPI_ISL_2813786, EPI_ISL_2813787, EPI_ISL_2813788 | | | |
| see above | Lighthouse Lab in Milton Keynes | Wellcome Sanger Institute for the COVID-19 Genomics UK (COG-UK) Consortium | The Lighthouse Lab in Milton Keynes and Alex Alderton, Roberto Amato, Jeffrey Barrett, Sonia Goncalves, Ewan Harrison, David K. Jackson, Ian Johnston, Dominic Kwiatkowski, Cordelia Langford, John Sillitoe on behalf of the Wellcome Sanger Institute COVID-19 Surveillance Team |
| EPI_ISL_2813789, EPI_ISL_2813790, EPI_ISL_2813791 | Lighthouse Lab in Alderley Park | Wellcome Sanger Institute for the COVID-19 Genomics UK (COG-UK) Consortium | Jacquelyn Wynn, Mairead Hyland, The Lighthouse Lab in Alderley Park and Alex Alderton, Roberto Amato, Jeffrey Barrett, Sonia Goncalves, Ewan Harrison, David K. Jackson, Ian Johnston, Dominic Kwiatkowski, Cordelia Langford, John Sillitoe on behalf of the Wellcome Sanger Institute COVID-19 Surveillance Team |
| EPI_ISL_2813792 | Lighthouse Lab in Milton Keynes | Wellcome Sanger Institute for the COVID-19 Genomics UK (COG-UK) Consortium | The Lighthouse Lab in Milton Keynes and Alex Alderton, Roberto Amato, Jeffrey Barrett, Sonia Goncalves, Ewan Harrison, David K. Jackson, Ian Johnston, Dominic Kwiatkowski, Cordelia Langford, John Sillitoe on behalf of the Wellcome Sanger Institute COVID-19 Surveillance Team |
| EPI_ISL_2813793 | Lighthouse Lab in Alderley Park | Wellcome Sanger Institute for the COVID-19 Genomics UK (COG-UK) Consortium | Jacquelyn Wynn, Mairead Hyland, The Lighthouse Lab in Alderley Park and Alex Alderton, Roberto Amato, Jeffrey Barrett, Sonia Goncalves, Ewan Harrison, David K. Jackson, Ian Johnston, Dominic Kwiatkowski, Cordelia Langford, John Sillitoe on behalf of the Wellcome Sanger Institute COVID-19 Surveillance Team |
| EPI_ISL_2813794 | Lighthouse Lab in Milton Keynes | Wellcome Sanger Institute for the COVID-19 Genomics UK (COG-UK) Consortium | The Lighthouse Lab in Milton Keynes and Alex Alderton, Roberto Amato, Jeffrey Barrett, Sonia Goncalves, Ewan Harrison, David K. Jackson, Ian Johnston, Dominic Kwiatkowski, Cordelia Langford, John Sillitoe on behalf of the Wellcome Sanger Institute COVID-19 Surveillance Team |
| EPI_ISL_2813795 | Lighthouse Lab in Alderley Park | Wellcome Sanger Institute for the COVID-19 Genomics UK (COG-UK) Consortium | Jacquelyn Wynn, Mairead Hyland, The Lighthouse Lab in Alderley Park and Alex Alderton, Roberto Amato, Jeffrey Barrett, Sonia Goncalves, Ewan Harrison, David K. Jackson, Ian Johnston, Dominic Kwiatkowski, Cordelia Langford, John Sillitoe on behalf of the Wellcome Sanger Institute COVID-19 Surveillance Team |
| EPI_ISL_2813796 | Lighthouse Lab in Milton Keynes | Wellcome Sanger Institute for the COVID-19 Genomics UK (COG-UK) Consortium | The Lighthouse Lab in Milton Keynes and Alex Alderton, Roberto Amato, Jeffrey Barrett, Sonia Goncalves, Ewan Harrison, David K. Jackson, Ian Johnston, Dominic Kwiatkowski, Cordelia Langford, John Sillitoe on behalf of the Wellcome Sanger Institute COVID-19 Surveillance Team |
| EPI_ISL_2813797 | Lighthouse Lab in Alderley Park | Wellcome Sanger Institute for the COVID-19 Genomics UK (COG-UK) Consortium | Jacquelyn Wynn, Mairead Hyland, The Lighthouse Lab in Alderley Park and Alex Alderton, Roberto Amato, Jeffrey Barrett, Sonia Goncalves, Ewan Harrison, David K. Jackson, Ian Johnston, Dominic Kwiatkowski, Cordelia Langford, John Sillitoe on behalf of the Wellcome Sanger Institute COVID-19 Surveillance Team |
| EPI_ISL_2813798, EPI_ISL_2813799, EPI_ISL_2813800 | Lighthouse Lab in Milton Keynes | Wellcome Sanger Institute for the COVID-19 Genomics UK (COG-UK) Consortium | The Lighthouse Lab in Milton Keynes and Alex Alderton, Roberto Amato, Jeffrey Barrett, Sonia Goncalves, Ewan Harrison, David K. Jackson, Ian Johnston, Dominic Kwiatkowski, Cordelia Langford, John Sillitoe on behalf of the Wellcome Sanger Institute COVID-19 Surveillance Team |
| EPI_ISL_2813801, EPI_ISL_2813802, EPI_ISL_2813803 | Lighthouse Lab in Alderley Park | Wellcome Sanger Institute for the COVID-19 Genomics UK (COG-UK) Consortium | Jacquelyn Wynn, Mairead Hyland, The Lighthouse Lab in Alderley Park and Alex Alderton, Roberto Amato, Jeffrey Barrett, Sonia Goncalves, Ewan Harrison, David K. Jackson, Ian Johnston, Dominic Kwiatkowski, Cordelia Langford, John Sillitoe on behalf of the Wellcome Sanger Institute COVID-19 Surveillance Team |
| EPI_ISL_2813804, EPI_ISL_2813805, EPI_ISL_2813806, EPI_ISL_2813807, EPI_ISL_2813808 | Lighthouse Lab in Milton Keynes | Wellcome Sanger Institute for the COVID-19 Genomics UK (COG-UK) Consortium | The Lighthouse Lab in Milton Keynes and Alex Alderton, Roberto Amato, Jeffrey Barrett, Sonia Goncalves, Ewan Harrison, David K. Jackson, Ian Johnston, Dominic Kwiatkowski, Cordelia Langford, John Sillitoe on behalf of the Wellcome Sanger Institute COVID-19 Surveillance Team |
| EPI_ISL_2813809 | Lighthouse Lab in Alderley Park | Wellcome Sanger Institute for the COVID-19 Genomics UK (COG-UK) Consortium | Jacquelyn Wynn, Mairead Hyland, The Lighthouse Lab in Alderley Park and Alex Alderton, Roberto Amato, Jeffrey Barrett, Sonia Goncalves, Ewan Harrison, David K. Jackson, Ian Johnston, Dominic Kwiatkowski, Cordelia Langford, John Sillitoe on behalf of the Wellcome Sanger Institute COVID-19 Surveillance Team |
| EPI_ISL_2813810 | Lighthouse Lab in Milton Keynes | Wellcome Sanger Institute for the COVID-19 Genomics UK (COG-UK) Consortium | The Lighthouse Lab in Milton Keynes and Alex Alderton, Roberto Amato, Jeffrey Barrett, Sonia Goncalves, Ewan Harrison, David K. Jackson, Ian Johnston, Dominic Kwiatkowski, Cordelia Langford, John Sillitoe on behalf of the Wellcome Sanger Institute COVID-19 Surveillance Team |
| EPI_ISL_2813811 | Lighthouse Lab in Alderley Park | Wellcome Sanger Institute for the COVID-19 Genomics UK (COG-UK) Consortium | Jacquelyn Wynn, Mairead Hyland, The Lighthouse Lab in Alderley Park and Alex Alderton, Roberto Amato, Jeffrey Barrett, Sonia Goncalves, Ewan Harrison, David K. Jackson, Ian Johnston, Dominic Kwiatkowski, Cordelia Langford, John Sillitoe on behalf of the Wellcome Sanger Institute COVID-19 Surveillance Team |
| EPI_ISL_2813812, EPI_ISL_2813813, EPI_ISL_2813814, EPI_ISL_2813815, EPI_ISL_2813816, EPI_ISL_2813817, EPI_ISL_2813818, EPI_ISL_2813819, EPI_ISL_2813820, EPI_ISL_2813821, EPI_ISL_2813822, EPI_ISL_2813823, EPI_ISL_2813824 | | | |
| see above | Lighthouse Lab in Milton Keynes | Wellcome Sanger Institute for the COVID-19 Genomics UK (COG-UK) Consortium | The Lighthouse Lab in Milton Keynes and Alex Alderton, Roberto Amato, Jeffrey Barrett, Sonia Goncalves, Ewan Harrison, David K. Jackson, Ian Johnston, Dominic Kwiatkowski, Cordelia Langford, John Sillitoe on behalf of the Wellcome Sanger Institute COVID-19 Surveillance Team |
| EPI_ISL_2813825 | Lighthouse Lab in Alderley Park | Wellcome Sanger Institute for the COVID-19 Genomics UK (COG-UK) Consortium | Jacquelyn Wynn, Mairead Hyland, The Lighthouse Lab in Alderley Park and Alex Alderton, Roberto Amato, Jeffrey Barrett, Sonia Goncalves, Ewan Harrison, David K. Jackson, Ian Johnston, Dominic Kwiatkowski, Cordelia Langford, John Sillitoe on behalf of the Wellcome Sanger Institute COVID-19 Surveillance Team |
| EPI_ISL_2813826, EPI_ISL_2813827, EPI_ISL_2813828, EPI_ISL_2813829, EPI_ISL_2813830 | Lighthouse Lab in Milton Keynes | Wellcome Sanger Institute for the COVID-19 Genomics UK (COG-UK) Consortium | The Lighthouse Lab in Milton Keynes and Alex Alderton, Roberto Amato, Jeffrey Barrett, Sonia Goncalves, Ewan Harrison, David K. Jackson, Ian Johnston, Dominic Kwiatkowski, Cordelia Langford, John Sillitoe on behalf of the Wellcome Sanger Institute COVID-19 Surveillance Team |
| EPI_ISL_2813831 | Lighthouse Lab in Alderley Park | Wellcome Sanger Institute for the COVID-19 Genomics UK (COG-UK) Consortium | Jacquelyn Wynn, Mairead Hyland, The Lighthouse Lab in Alderley Park and Alex Alderton, Roberto Amato, Jeffrey Barrett, Sonia Goncalves, Ewan Harrison, David K. Jackson, Ian Johnston, Dominic Kwiatkowski, Cordelia Langford, John Sillitoe on behalf of the Wellcome Sanger Institute COVID-19 Surveillance Team |
| EPI_ISL_2813832 | Lighthouse Lab in Milton Keynes | Wellcome Sanger Institute for the COVID-19 Genomics UK (COG-UK) Consortium | The Lighthouse Lab in Milton Keynes and Alex Alderton, Roberto Amato, Jeffrey Barrett, Sonia Goncalves, Ewan Harrison, David K. Jackson, Ian Johnston, Dominic Kwiatkowski, Cordelia Langford, John Sillitoe on behalf of the Wellcome Sanger Institute COVID-19 Surveillance Team |
| EPI_ISL_2813833 | Lighthouse Lab in Alderley Park | Wellcome Sanger Institute for the COVID-19 Genomics UK (COG-UK) Consortium | Jacquelyn Wynn, Mairead Hyland, The Lighthouse Lab in Alderley Park and Alex Alderton, Roberto Amato, Jeffrey Barrett, Sonia Goncalves, Ewan Harrison, David K. Jackson, Ian Johnston, Dominic Kwiatkowski, Cordelia Langford, John Sillitoe on behalf of the Wellcome Sanger Institute COVID-19 Surveillance Team |
| EPI_ISL_2813834, EPI_ISL_2813835, EPI_ISL_2813836, EPI_ISL_2813837 | Lighthouse Lab in Milton Keynes | Wellcome Sanger Institute for the COVID-19 Genomics UK (COG-UK) Consortium | The Lighthouse Lab in Milton Keynes and Alex Alderton, Roberto Amato, Jeffrey Barrett, Sonia Goncalves, Ewan Harrison, David K. Jackson, Ian Johnston, Dominic Kwiatkowski, Cordelia Langford, John Sillitoe on behalf of the Wellcome Sanger Institute COVID-19 Surveillance Team |
| EPI_ISL_2813838, EPI_ISL_2813839 | Lighthouse Lab in Alderley Park | Wellcome Sanger Institute for the COVID-19 Genomics UK (COG-UK) Consortium | Jacquelyn Wynn, Mairead Hyland, The Lighthouse Lab in Alderley Park and Alex Alderton, Roberto Amato, Jeffrey Barrett, Sonia Goncalves, Ewan Harrison, David K. Jackson, Ian Johnston, Dominic Kwiatkowski, Cordelia Langford, John Sillitoe on behalf of the Wellcome Sanger Institute COVID-19 Surveillance Team |
| EPI_ISL_2813840, EPI_ISL_2813841, EPI_ISL_2813842, EPI_ISL_2813843, EPI_ISL_2813844 | Lighthouse Lab in Milton Keynes | Wellcome Sanger Institute for the COVID-19 Genomics UK (COG-UK) Consortium | The Lighthouse Lab in Milton Keynes and Alex Alderton, Roberto Amato, Jeffrey Barrett, Sonia Goncalves, Ewan Harrison, David K. Jackson, Ian Johnston, Dominic Kwiatkowski, Cordelia Langford, John Sillitoe on behalf of the Wellcome Sanger Institute COVID-19 Surveillance Team |
| EPI_ISL_2813845 | Lighthouse Lab in Alderley Park | Wellcome Sanger Institute for the COVID-19 Genomics UK (COG-UK) Consortium | Jacquelyn Wynn, Mairead Hyland, The Lighthouse Lab in Alderley Park and Alex Alderton, Roberto Amato, Jeffrey Barrett, Sonia Goncalves, Ewan Harrison, David K. Jackson, Ian Johnston, Dominic Kwiatkowski, Cordelia Langford, John Sillitoe on behalf of the Wellcome Sanger Institute COVID-19 Surveillance Team |
| EPI_ISL_2813846, EPI_ISL_2813847 | Lighthouse Lab in Milton Keynes | Wellcome Sanger Institute for the COVID-19 Genomics UK (COG-UK) Consortium | The Lighthouse Lab in Milton Keynes and Alex Alderton, Roberto Amato, Jeffrey Barrett, Sonia Goncalves, Ewan Harrison, David K. Jackson, Ian Johnston, Dominic Kwiatkowski, Cordelia Langford, John Sillitoe on behalf of the Wellcome Sanger Institute COVID-19 Surveillance Team |
| EPI_ISL_2813848 | Lighthouse Lab in Alderley Park | Wellcome Sanger Institute for the COVID-19 Genomics UK (COG-UK) Consortium | Jacquelyn Wynn, Mairead Hyland, The Lighthouse Lab in Alderley Park and Alex Alderton, Roberto Amato, Jeffrey Barrett, Sonia Goncalves, Ewan Harrison, David K. Jackson, Ian Johnston, Dominic Kwiatkowski, Cordelia Langford, John Sillitoe on behalf of the Wellcome Sanger Institute COVID-19 Surveillance Team |
| EPI_ISL_2813849 | Lighthouse Lab in Milton Keynes | Wellcome Sanger Institute for the COVID-19 Genomics UK (COG-UK) Consortium | The Lighthouse Lab in Milton Keynes and Alex Alderton, Roberto Amato, Jeffrey Barrett, Sonia Goncalves, Ewan Harrison, David K. Jackson, Ian Johnston, Dominic Kwiatkowski, Cordelia Langford, John Sillitoe on behalf of the Wellcome Sanger Institute COVID-19 Surveillance Team |
| EPI_ISL_2813850, EPI_ISL_2813851 | Lighthouse Lab in Alderley Park | Wellcome Sanger Institute for the COVID-19 Genomics UK (COG-UK) Consortium | Jacquelyn Wynn, Mairead Hyland, The Lighthouse Lab in Alderley Park and Alex Alderton, Roberto Amato, Jeffrey Barrett, Sonia Goncalves, Ewan Harrison, David K. Jackson, Ian Johnston, Dominic Kwiatkowski, Cordelia Langford, John Sillitoe on behalf of the Wellcome Sanger Institute COVID-19 Surveillance Team |
| EPI_ISL_2813852, EPI_ISL_2813853, EPI_ISL_2813854, EPI_ISL_2813855, EPI_ISL_2813856 | Lighthouse Lab in Milton Keynes | Wellcome Sanger Institute for the COVID-19 Genomics UK (COG-UK) Consortium | The Lighthouse Lab in Milton Keynes and Alex Alderton, Roberto Amato, Jeffrey Barrett, Sonia Goncalves, Ewan Harrison, David K. Jackson, Ian Johnston, Dominic Kwiatkowski, Cordelia Langford, John Sillitoe on behalf of the Wellcome Sanger Institute COVID-19 Surveillance Team |
| EPI_ISL_2813857 | Lighthouse Lab in Alderley Park | Wellcome Sanger Institute for the COVID-19 Genomics UK (COG-UK) Consortium | Jacquelyn Wynn, Mairead Hyland, The Lighthouse Lab in Alderley Park and Alex Alderton, Roberto Amato, Jeffrey Barrett, Sonia Goncalves, Ewan Harrison, David K. Jackson, Ian Johnston, Dominic Kwiatkowski, Cordelia Langford, John Sillitoe on behalf of the Wellcome Sanger Institute COVID-19 Surveillance Team |
| EPI_ISL_2813858, EPI_ISL_2813859, EPI_ISL_2813860 | Lighthouse Lab in Milton Keynes | Wellcome Sanger Institute for the COVID-19 Genomics UK (COG-UK) Consortium | The Lighthouse Lab in Milton Keynes and Alex Alderton, Roberto Amato, Jeffrey Barrett, Sonia Goncalves, Ewan Harrison, David K. Jackson, Ian Johnston, Dominic Kwiatkowski, Cordelia Langford, John Sillitoe on behalf of the Wellcome Sanger Institute COVID-19 Surveillance Team |
| EPI_ISL_2813861 | Lighthouse Lab in Alderley Park | Wellcome Sanger Institute for the COVID-19 Genomics UK (COG-UK) Consortium | Jacquelyn Wynn, Mairead Hyland, The Lighthouse Lab in Alderley Park and Alex Alderton, Roberto Amato, Jeffrey Barrett, Sonia Goncalves, Ewan Harrison, David K. Jackson, Ian Johnston, Dominic Kwiatkowski, Cordelia Langford, John Sillitoe on behalf of the Wellcome Sanger Institute COVID-19 Surveillance Team |

| | | | |
|---|---|---|---|
| EPI_ISL_2813862, EPI_ISL_2813863, EPI_ISL_2813864, EPI_ISL_2813865, EPI_ISL_2813866, EPI_ISL_2813867, EPI_ISL_2813868, EPI_ISL_2813869, EPI_ISL_2813870 | Lighthouse Lab in Milton Keynes | Wellcome Sanger Institute for the COVID-19 Genomics UK (COG-UK) Consortium | The Lighthouse Lab in Milton Keynes and Alex Alderton, Roberto Amato, Jeffrey Barrett, Sonia Goncalves, Ewan Harrison, David K. Jackson, Ian Johnston, Dominic Kwiatkowski, Cordelia Langford, John Sillitoe on behalf of the Wellcome Sanger Institute COVID-19 Surveillance Team |
| EPI_ISL_2813871, EPI_ISL_2813872 | Lighthouse Lab in Alderley Park | Wellcome Sanger Institute for the COVID-19 Genomics UK (COG-UK) Consortium | Jacquelyn Wynn, Mairead Hyland, The Lighthouse Lab in Alderley Park and Alex Alderton, Roberto Amato, Jeffrey Barrett, Sonia Goncalves, Ewan Harrison, David K. Jackson, Ian Johnston, Dominic Kwiatkowski, Cordelia Langford, John Sillitoe on behalf of the Wellcome Sanger Institute COVID-19 Surveillance Team |
| EPI_ISL_2813873, EPI_ISL_2813874, EPI_ISL_2813876, EPI_ISL_2813877, EPI_ISL_2813878, EPI_ISL_2813879, EPI_ISL_2813880, EPI_ISL_2813881, EPI_ISL_2813882, EPI_ISL_2813883 | Lighthouse Lab in Milton Keynes | Wellcome Sanger Institute for the COVID-19 Genomics UK (COG-UK) Consortium | The Lighthouse Lab in Milton Keynes and Alex Alderton, Roberto Amato, Jeffrey Barrett, Sonia Goncalves, Ewan Harrison, David K. Jackson, Ian Johnston, Dominic Kwiatkowski, Cordelia Langford, John Sillitoe on behalf of the Wellcome Sanger Institute COVID-19 Surveillance Team |
| EPI_ISL_2813884 | Lighthouse Lab in Alderley Park | Wellcome Sanger Institute for the COVID-19 Genomics UK (COG-UK) Consortium | Jacquelyn Wynn, Mairead Hyland, The Lighthouse Lab in Alderley Park and Alex Alderton, Roberto Amato, Jeffrey Barrett, Sonia Goncalves, Ewan Harrison, David K. Jackson, Ian Johnston, Dominic Kwiatkowski, Cordelia Langford, John Sillitoe on behalf of the Wellcome Sanger Institute COVID-19 Surveillance Team |
| EPI_ISL_2813885, EPI_ISL_2813886, EPI_ISL_2813887 | Lighthouse Lab in Milton Keynes | Wellcome Sanger Institute for the COVID-19 Genomics UK (COG-UK) Consortium | The Lighthouse Lab in Milton Keynes and Alex Alderton, Roberto Amato, Jeffrey Barrett, Sonia Goncalves, Ewan Harrison, David K. Jackson, Ian Johnston, Dominic Kwiatkowski, Cordelia Langford, John Sillitoe on behalf of the Wellcome Sanger Institute COVID-19 Surveillance Team |
| EPI_ISL_2813888 | Lighthouse Lab in Alderley Park | Wellcome Sanger Institute for the COVID-19 Genomics UK (COG-UK) Consortium | Jacquelyn Wynn, Mairead Hyland, The Lighthouse Lab in Alderley Park and Alex Alderton, Roberto Amato, Jeffrey Barrett, Sonia Goncalves, Ewan Harrison, David K. Jackson, Ian Johnston, Dominic Kwiatkowski, Cordelia Langford, John Sillitoe on behalf of the Wellcome Sanger Institute COVID-19 Surveillance Team |
| EPI_ISL_2813889, EPI_ISL_2813890, EPI_ISL_2813891, EPI_ISL_2813892, EPI_ISL_2813893 | Lighthouse Lab in Milton Keynes | Wellcome Sanger Institute for the COVID-19 Genomics UK (COG-UK) Consortium | The Lighthouse Lab in Milton Keynes and Alex Alderton, Roberto Amato, Jeffrey Barrett, Sonia Goncalves, Ewan Harrison, David K. Jackson, Ian Johnston, Dominic Kwiatkowski, Cordelia Langford, John Sillitoe on behalf of the Wellcome Sanger Institute COVID-19 Surveillance Team |
| EPI_ISL_2813894, EPI_ISL_2813895 | Lighthouse Lab in Alderley Park | Wellcome Sanger Institute for the COVID-19 Genomics UK (COG-UK) Consortium | Jacquelyn Wynn, Mairead Hyland, The Lighthouse Lab in Alderley Park and Alex Alderton, Roberto Amato, Jeffrey Barrett, Sonia Goncalves, Ewan Harrison, David K. Jackson, Ian Johnston, Dominic Kwiatkowski, Cordelia Langford, John Sillitoe on behalf of the Wellcome Sanger Institute COVID-19 Surveillance Team |
| EPI_ISL_2813896, EPI_ISL_2813897, EPI_ISL_2813898 | Lighthouse Lab in Milton Keynes | Wellcome Sanger Institute for the COVID-19 Genomics UK (COG-UK) Consortium | The Lighthouse Lab in Milton Keynes and Alex Alderton, Roberto Amato, Jeffrey Barrett, Sonia Goncalves, Ewan Harrison, David K. Jackson, Ian Johnston, Dominic Kwiatkowski, Cordelia Langford, John Sillitoe on behalf of the Wellcome Sanger Institute COVID-19 Surveillance Team |
| EPI_ISL_2813899 | Lighthouse Lab in Alderley Park | Wellcome Sanger Institute for the COVID-19 Genomics UK (COG-UK) Consortium | Jacquelyn Wynn, Mairead Hyland, The Lighthouse Lab in Alderley Park and Alex Alderton, Roberto Amato, Jeffrey Barrett, Sonia Goncalves, Ewan Harrison, David K. Jackson, Ian Johnston, Dominic Kwiatkowski, Cordelia Langford, John Sillitoe on behalf of the Wellcome Sanger Institute COVID-19 Surveillance Team |
| EPI_ISL_2813900, EPI_ISL_2813901, EPI_ISL_2813902, EPI_ISL_2813903, EPI_ISL_2813904, EPI_ISL_2813905 | Lighthouse Lab in Milton Keynes | Wellcome Sanger Institute for the COVID-19 Genomics UK (COG-UK) Consortium | The Lighthouse Lab in Milton Keynes and Alex Alderton, Roberto Amato, Jeffrey Barrett, Sonia Goncalves, Ewan Harrison, David K. Jackson, Ian Johnston, Dominic Kwiatkowski, Cordelia Langford, John Sillitoe on behalf of the Wellcome Sanger Institute COVID-19 Surveillance Team |
| EPI_ISL_2813906 | Lighthouse Lab in Alderley Park | Wellcome Sanger Institute for the COVID-19 Genomics UK (COG-UK) Consortium | Jacquelyn Wynn, Mairead Hyland, The Lighthouse Lab in Alderley Park and Alex Alderton, Roberto Amato, Jeffrey Barrett, Sonia Goncalves, Ewan Harrison, David K. Jackson, Ian Johnston, Dominic Kwiatkowski, Cordelia Langford, John Sillitoe on behalf of the Wellcome Sanger Institute COVID-19 Surveillance Team |
| EPI_ISL_2813907, EPI_ISL_2813908, EPI_ISL_2813909 | Lighthouse Lab in Milton Keynes | Wellcome Sanger Institute for the COVID-19 Genomics UK (COG-UK) Consortium | The Lighthouse Lab in Milton Keynes and Alex Alderton, Roberto Amato, Jeffrey Barrett, Sonia Goncalves, Ewan Harrison, David K. Jackson, Ian Johnston, Dominic Kwiatkowski, Cordelia Langford, John Sillitoe on behalf of the Wellcome Sanger Institute COVID-19 Surveillance Team |
| EPI_ISL_2813910 | Lighthouse Lab in Alderley Park | Wellcome Sanger Institute for the COVID-19 Genomics UK (COG-UK) Consortium | Jacquelyn Wynn, Mairead Hyland, The Lighthouse Lab in Alderley Park and Alex Alderton, Roberto Amato, Jeffrey Barrett, Sonia Goncalves, Ewan Harrison, David K. Jackson, Ian Johnston, Dominic Kwiatkowski, Cordelia Langford, John Sillitoe on behalf of the Wellcome Sanger Institute COVID-19 Surveillance Team |
| EPI_ISL_2813911, EPI_ISL_2813912, EPI_ISL_2813913, EPI_ISL_2813914, EPI_ISL_2813915, EPI_ISL_2813916, EPI_ISL_2813917, EPI_ISL_2813918, EPI_ISL_2813919, EPI_ISL_2813920, EPI_ISL_2813921, EPI_ISL_2813922, EPI_ISL_2813923, EPI_ISL_2813924, EPI_ISL_2813925, EPI_ISL_2813926, EPI_ISL_2813927 | | | |
| see above | Lighthouse Lab in Milton Keynes | Wellcome Sanger Institute for the COVID-19 Genomics UK (COG-UK) Consortium | The Lighthouse Lab in Milton Keynes and Alex Alderton, Roberto Amato, Jeffrey Barrett, Sonia Goncalves, Ewan Harrison, David K. Jackson, Ian Johnston, Dominic Kwiatkowski, Cordelia Langford, John Sillitoe on behalf of the Wellcome Sanger Institute COVID-19 Surveillance Team |
| EPI_ISL_2813928 | Lighthouse Lab in Alderley Park | Wellcome Sanger Institute for the COVID-19 Genomics UK (COG-UK) Consortium | Jacquelyn Wynn, Mairead Hyland, The Lighthouse Lab in Alderley Park and Alex Alderton, Roberto Amato, Jeffrey Barrett, Sonia Goncalves, Ewan Harrison, David K. Jackson, Ian Johnston, Dominic Kwiatkowski, Cordelia Langford, John Sillitoe on behalf of the Wellcome Sanger Institute COVID-19 Surveillance Team |
| EPI_ISL_2813929, EPI_ISL_2813930, EPI_ISL_2813931, EPI_ISL_2813932, EPI_ISL_2813933, EPI_ISL_2813934, EPI_ISL_2813935, EPI_ISL_2813936 | Lighthouse Lab in Milton Keynes | Wellcome Sanger Institute for the COVID-19 Genomics UK (COG-UK) Consortium | The Lighthouse Lab in Milton Keynes and Alex Alderton, Roberto Amato, Jeffrey Barrett, Sonia Goncalves, Ewan Harrison, David K. Jackson, Ian Johnston, Dominic Kwiatkowski, Cordelia Langford, John Sillitoe on behalf of the Wellcome Sanger Institute COVID-19 Surveillance Team |
| EPI_ISL_2813937 | Lighthouse Lab in Alderley Park | Wellcome Sanger Institute for the COVID-19 Genomics UK (COG-UK) Consortium | Jacquelyn Wynn, Mairead Hyland, The Lighthouse Lab in Alderley Park and Alex Alderton, Roberto Amato, Jeffrey Barrett, Sonia Goncalves, Ewan Harrison, David K. Jackson, Ian Johnston, Dominic Kwiatkowski, Cordelia Langford, John Sillitoe on behalf of the Wellcome Sanger Institute COVID-19 Surveillance Team |
| EPI_ISL_2813938, EPI_ISL_2813939, EPI_ISL_2813940 | Lighthouse Lab in Milton Keynes | Wellcome Sanger Institute for the COVID-19 Genomics UK (COG-UK) Consortium | The Lighthouse Lab in Milton Keynes and Alex Alderton, Roberto Amato, Jeffrey Barrett, Sonia Goncalves, Ewan Harrison, David K. Jackson, Ian Johnston, Dominic Kwiatkowski, Cordelia Langford, John Sillitoe on behalf of the Wellcome Sanger Institute COVID-19 Surveillance Team |
| EPI_ISL_2813941 | Lighthouse Lab in Alderley Park | Wellcome Sanger Institute for the COVID-19 Genomics UK (COG-UK) Consortium | Jacquelyn Wynn, Mairead Hyland, The Lighthouse Lab in Alderley Park and Alex Alderton, Roberto Amato, Jeffrey Barrett, Sonia Goncalves, Ewan Harrison, David K. Jackson, Ian Johnston, Dominic Kwiatkowski, Cordelia Langford, John Sillitoe on behalf of the Wellcome Sanger Institute COVID-19 Surveillance Team |
| EPI_ISL_2813942, EPI_ISL_2813943, EPI_ISL_2813944, EPI_ISL_2813945 | Lighthouse Lab in Milton Keynes | Wellcome Sanger Institute for the COVID-19 Genomics UK (COG-UK) Consortium | The Lighthouse Lab in Milton Keynes and Alex Alderton, Roberto Amato, Jeffrey Barrett, Sonia Goncalves, Ewan Harrison, David K. Jackson, Ian Johnston, Dominic Kwiatkowski, Cordelia Langford, John Sillitoe on behalf of the Wellcome Sanger Institute COVID-19 Surveillance Team |
| EPI_ISL_2813946 | Lighthouse Lab in Alderley Park | Wellcome Sanger Institute for the COVID-19 Genomics UK (COG-UK) Consortium | Jacquelyn Wynn, Mairead Hyland, The Lighthouse Lab in Alderley Park and Alex Alderton, Roberto Amato, Jeffrey Barrett, Sonia Goncalves, Ewan Harrison, David K. Jackson, Ian Johnston, Dominic Kwiatkowski, Cordelia Langford, John Sillitoe on behalf of the Wellcome Sanger Institute COVID-19 Surveillance Team |
| EPI_ISL_2813947 | Lighthouse Lab in Milton Keynes | Wellcome Sanger Institute for the COVID-19 Genomics UK (COG-UK) Consortium | The Lighthouse Lab in Milton Keynes and Alex Alderton, Roberto Amato, Jeffrey Barrett, Sonia Goncalves, Ewan Harrison, David K. Jackson, Ian Johnston, Dominic Kwiatkowski, Cordelia Langford, John Sillitoe on behalf of the Wellcome Sanger Institute COVID-19 Surveillance Team |
| EPI_ISL_2813948, EPI_ISL_2813949 | Lighthouse Lab in Alderley Park | Wellcome Sanger Institute for the COVID-19 Genomics UK (COG-UK) Consortium | Jacquelyn Wynn, Mairead Hyland, The Lighthouse Lab in Alderley Park and Alex Alderton, Roberto Amato, Jeffrey Barrett, Sonia Goncalves, Ewan Harrison, David K. Jackson, Ian Johnston, Dominic Kwiatkowski, Cordelia Langford, John Sillitoe on behalf of the Wellcome Sanger Institute COVID-19 Surveillance Team |
| EPI_ISL_2813950, EPI_ISL_2813951, EPI_ISL_2813952 | Lighthouse Lab in Milton Keynes | Wellcome Sanger Institute for the COVID-19 Genomics UK (COG-UK) Consortium | The Lighthouse Lab in Milton Keynes and Alex Alderton, Roberto Amato, Jeffrey Barrett, Sonia Goncalves, Ewan Harrison, David K. Jackson, Ian Johnston, Dominic Kwiatkowski, Cordelia Langford, John Sillitoe on behalf of the Wellcome Sanger Institute COVID-19 Surveillance Team |
| EPI_ISL_2813953 | Lighthouse Lab in Alderley Park | Wellcome Sanger Institute for the COVID-19 Genomics UK (COG-UK) Consortium | Jacquelyn Wynn, Mairead Hyland, The Lighthouse Lab in Alderley Park and Alex Alderton, Roberto Amato, Jeffrey Barrett, Sonia Goncalves, Ewan Harrison, David K. Jackson, Ian Johnston, Dominic Kwiatkowski, Cordelia Langford, John Sillitoe on behalf of the Wellcome Sanger Institute COVID-19 Surveillance Team |
| EPI_ISL_2813954 | Lighthouse Lab in Milton Keynes | Wellcome Sanger Institute for the COVID-19 Genomics UK (COG-UK) Consortium | The Lighthouse Lab in Milton Keynes and Alex Alderton, Roberto Amato, Jeffrey Barrett, Sonia Goncalves, Ewan Harrison, David K. Jackson, Ian Johnston, Dominic Kwiatkowski, Cordelia Langford, John Sillitoe on behalf of the Wellcome Sanger Institute COVID-19 Surveillance Team |
| EPI_ISL_2813955 | Lighthouse Lab in Alderley Park | Wellcome Sanger Institute for the COVID-19 Genomics UK (COG-UK) Consortium | Jacquelyn Wynn, Mairead Hyland, The Lighthouse Lab in Alderley Park and Alex Alderton, Roberto Amato, Jeffrey Barrett, Sonia Goncalves, Ewan Harrison, David K. Jackson, Ian Johnston, Dominic Kwiatkowski, Cordelia Langford, John Sillitoe on behalf of the Wellcome Sanger Institute COVID-19 Surveillance Team |
| EPI_ISL_2813956, EPI_ISL_2813957, EPI_ISL_2813958, EPI_ISL_2813959, EPI_ISL_2813960, EPI_ISL_2813961, EPI_ISL_2813962, EPI_ISL_2813963, EPI_ISL_2813964, EPI_ISL_2813965, EPI_ISL_2813966, EPI_ISL_2813967, EPI_ISL_2813968, EPI_ISL_2813969, EPI_ISL_2813970, EPI_ISL_2813971, EPI_ISL_2813972, EPI_ISL_2813973, EPI_ISL_2813974, EPI_ISL_2813975, EPI_ISL_2813976 | | | |
| see above | Lighthouse Lab in Milton Keynes | Wellcome Sanger Institute for the COVID-19 Genomics UK (COG-UK) Consortium | The Lighthouse Lab in Milton Keynes and Alex Alderton, Roberto Amato, Jeffrey Barrett, Sonia Goncalves, Ewan Harrison, David K. Jackson, Ian Johnston, Dominic Kwiatkowski, Cordelia Langford, John Sillitoe on behalf of the Wellcome Sanger Institute COVID-19 Surveillance Team |
| EPI_ISL_2813977, EPI_ISL_2813978, | Lighthouse Lab in Alderley Park | Wellcome Sanger Institute for the COVID-19 Genomics UK (COG-UK) Consortium | Jacquelyn Wynn, Mairead Hyland, The Lighthouse Lab in Alderley Park and Alex Alderton, Roberto Amato, Jeffrey Barrett, Sonia Goncalves, Ewan Harrison, David K. Jackson, Ian Johnston, Dominic Kwiatkowski, Cordelia Langford, John Sillitoe on behalf of the Wellcome Sanger Institute COVID-19 Surveillance Team |

| | | | |
|---|---|---|---|
| EPI_ISL_2813979, EPI_ISL_2813980 | | | |
| EPI_ISL_2813981, EPI_ISL_2813982, EPI_ISL_2813983, EPI_ISL_2813984, EPI_ISL_2813985, EPI_ISL_2813986, EPI_ISL_2813987, EPI_ISL_2813988, EPI_ISL_2813989, EPI_ISL_2813990, EPI_ISL_2813991, EPI_ISL_2813992, EPI_ISL_2813993, EPI_ISL_2813994, EPI_ISL_2813995, EPI_ISL_2813996, EPI_ISL_2813997, EPI_ISL_2813998, EPI_ISL_2813999 | | | |
| see above | Lighthouse Lab in Milton Keynes | Wellcome Sanger Institute for the COVID-19 Genomics UK (COG-UK) Consortium | The Lighthouse Lab in Milton Keynes and Alex Alderton, Roberto Amato, Jeffrey Barrett, Sonia Goncalves, Ewan Harrison, David K. Jackson, Ian Johnston, Dominic Kwiatkowski, Cordelia Langford, John Sillitoe on behalf of the Wellcome Sanger Institute COVID-19 Surveillance Team |
| EPI_ISL_2814000, EPI_ISL_2814001 | Lighthouse Lab in Alderley Park | Wellcome Sanger Institute for the COVID-19 Genomics UK (COG-UK) Consortium | Jacquelyn Wynn, Mairead Hyland, The Lighthouse Lab in Alderley Park and Alex Alderton, Roberto Amato, Jeffrey Barrett, Sonia Goncalves, Ewan Harrison, David K. Jackson, Ian Johnston, Dominic Kwiatkowski, Cordelia Langford, John Sillitoe on behalf of the Wellcome Sanger Institute COVID-19 Surveillance Team |
| EPI_ISL_2814002 | Lighthouse Lab in Milton Keynes | Wellcome Sanger Institute for the COVID-19 Genomics UK (COG-UK) Consortium | The Lighthouse Lab in Milton Keynes and Alex Alderton, Roberto Amato, Jeffrey Barrett, Sonia Goncalves, Ewan Harrison, David K. Jackson, Ian Johnston, Dominic Kwiatkowski, Cordelia Langford, John Sillitoe on behalf of the Wellcome Sanger Institute COVID-19 Surveillance Team |
| EPI_ISL_2814003, EPI_ISL_2814004 | Lighthouse Lab in Alderley Park | Wellcome Sanger Institute for the COVID-19 Genomics UK (COG-UK) Consortium | Jacquelyn Wynn, Mairead Hyland, The Lighthouse Lab in Alderley Park and Alex Alderton, Roberto Amato, Jeffrey Barrett, Sonia Goncalves, Ewan Harrison, David K. Jackson, Ian Johnston, Dominic Kwiatkowski, Cordelia Langford, John Sillitoe on behalf of the Wellcome Sanger Institute COVID-19 Surveillance Team |
| EPI_ISL_2814005, EPI_ISL_2814006 | Lighthouse Lab in Milton Keynes | Wellcome Sanger Institute for the COVID-19 Genomics UK (COG-UK) Consortium | The Lighthouse Lab in Milton Keynes and Alex Alderton, Roberto Amato, Jeffrey Barrett, Sonia Goncalves, Ewan Harrison, David K. Jackson, Ian Johnston, Dominic Kwiatkowski, Cordelia Langford, John Sillitoe on behalf of the Wellcome Sanger Institute COVID-19 Surveillance Team |
| EPI_ISL_2814007, EPI_ISL_2814008, EPI_ISL_2814009, EPI_ISL_2814010 | Lighthouse Lab in Alderley Park | Wellcome Sanger Institute for the COVID-19 Genomics UK (COG-UK) Consortium | Jacquelyn Wynn, Mairead Hyland, The Lighthouse Lab in Alderley Park and Alex Alderton, Roberto Amato, Jeffrey Barrett, Sonia Goncalves, Ewan Harrison, David K. Jackson, Ian Johnston, Dominic Kwiatkowski, Cordelia Langford, John Sillitoe on behalf of the Wellcome Sanger Institute COVID-19 Surveillance Team |
| EPI_ISL_2814011 | Lighthouse Lab in Milton Keynes | Wellcome Sanger Institute for the COVID-19 Genomics UK (COG-UK) Consortium | The Lighthouse Lab in Milton Keynes and Alex Alderton, Roberto Amato, Jeffrey Barrett, Sonia Goncalves, Ewan Harrison, David K. Jackson, Ian Johnston, Dominic Kwiatkowski, Cordelia Langford, John Sillitoe on behalf of the Wellcome Sanger Institute COVID-19 Surveillance Team |
| EPI_ISL_2814012, EPI_ISL_2814013, EPI_ISL_2814014, EPI_ISL_2814015, EPI_ISL_2814016 | Lighthouse Lab in Alderley Park | Wellcome Sanger Institute for the COVID-19 Genomics UK (COG-UK) Consortium | Jacquelyn Wynn, Mairead Hyland, The Lighthouse Lab in Alderley Park and Alex Alderton, Roberto Amato, Jeffrey Barrett, Sonia Goncalves, Ewan Harrison, David K. Jackson, Ian Johnston, Dominic Kwiatkowski, Cordelia Langford, John Sillitoe on behalf of the Wellcome Sanger Institute COVID-19 Surveillance Team |
| EPI_ISL_2814017, EPI_ISL_2814018 | Lighthouse Lab in Milton Keynes | Wellcome Sanger Institute for the COVID-19 Genomics UK (COG-UK) Consortium | The Lighthouse Lab in Milton Keynes and Alex Alderton, Roberto Amato, Jeffrey Barrett, Sonia Goncalves, Ewan Harrison, David K. Jackson, Ian Johnston, Dominic Kwiatkowski, Cordelia Langford, John Sillitoe on behalf of the Wellcome Sanger Institute COVID-19 Surveillance Team |
| EPI_ISL_2814019 | Lighthouse Lab in Glasgow | Wellcome Sanger Institute for the COVID-19 Genomics UK (COG-UK) Consortium | Harper VanSteenhouse, Yumi Kasai, David Gray, Carol Clugston, Anna Dominiczak and Alex Alderton, Roberto Amato, Jeffrey Barrett, Sonia Goncalves, Ewan Harrison, David K. Jackson, Ian Johnston, Dominic Kwiatkowski, Cordelia Langford, John Sillitoe on behalf of the Wellcome Sanger Institute COVID-19 Surveillance Team |
| EPI_ISL_2814020 | Lighthouse Lab in Alderley Park | Wellcome Sanger Institute for the COVID-19 Genomics UK (COG-UK) Consortium | Jacquelyn Wynn, Mairead Hyland, The Lighthouse Lab in Alderley Park and Alex Alderton, Roberto Amato, Jeffrey Barrett, Sonia Goncalves, Ewan Harrison, David K. Jackson, Ian Johnston, Dominic Kwiatkowski, Cordelia Langford, John Sillitoe on behalf of the Wellcome Sanger Institute COVID-19 Surveillance Team |
| EPI_ISL_2814021, EPI_ISL_2814022 | Lighthouse Lab in Milton Keynes | Wellcome Sanger Institute for the COVID-19 Genomics UK (COG-UK) Consortium | The Lighthouse Lab in Milton Keynes and Alex Alderton, Roberto Amato, Jeffrey Barrett, Sonia Goncalves, Ewan Harrison, David K. Jackson, Ian Johnston, Dominic Kwiatkowski, Cordelia Langford, John Sillitoe on behalf of the Wellcome Sanger Institute COVID-19 Surveillance Team |
| EPI_ISL_2814023 | Lighthouse Lab in Glasgow | Wellcome Sanger Institute for the COVID-19 Genomics UK (COG-UK) Consortium | Harper VanSteenhouse, Yumi Kasai, David Gray, Carol Clugston, Anna Dominiczak and Alex Alderton, Roberto Amato, Jeffrey Barrett, Sonia Goncalves, Ewan Harrison, David K. Jackson, Ian Johnston, Dominic Kwiatkowski, Cordelia Langford, John Sillitoe on behalf of the Wellcome Sanger Institute COVID-19 Surveillance Team |
| EPI_ISL_2814024, EPI_ISL_2814025, EPI_ISL_2814026, EPI_ISL_2814027 | Lighthouse Lab in Milton Keynes | Wellcome Sanger Institute for the COVID-19 Genomics UK (COG-UK) Consortium | The Lighthouse Lab in Milton Keynes and Alex Alderton, Roberto Amato, Jeffrey Barrett, Sonia Goncalves, Ewan Harrison, David K. Jackson, Ian Johnston, Dominic Kwiatkowski, Cordelia Langford, John Sillitoe on behalf of the Wellcome Sanger Institute COVID-19 Surveillance Team |
| EPI_ISL_2814028 | Lighthouse Lab in Alderley Park | Wellcome Sanger Institute for the COVID-19 Genomics UK (COG-UK) Consortium | Jacquelyn Wynn, Mairead Hyland, The Lighthouse Lab in Alderley Park and Alex Alderton, Roberto Amato, Jeffrey Barrett, Sonia Goncalves, Ewan Harrison, David K. Jackson, Ian Johnston, Dominic Kwiatkowski, Cordelia Langford, John Sillitoe on behalf of the Wellcome Sanger Institute COVID-19 Surveillance Team |
| EPI_ISL_2814029 | Lighthouse Lab in Glasgow | Wellcome Sanger Institute for the COVID-19 Genomics UK (COG-UK) Consortium | Harper VanSteenhouse, Yumi Kasai, David Gray, Carol Clugston, Anna Dominiczak and Alex Alderton, Roberto Amato, Jeffrey Barrett, Sonia Goncalves, Ewan Harrison, David K. Jackson, Ian Johnston, Dominic Kwiatkowski, Cordelia Langford, John Sillitoe on behalf of the Wellcome Sanger Institute COVID-19 Surveillance Team |
| EPI_ISL_2814030 | Lighthouse Lab in Milton Keynes | Wellcome Sanger Institute for the COVID-19 Genomics UK (COG-UK) Consortium | The Lighthouse Lab in Milton Keynes and Alex Alderton, Roberto Amato, Jeffrey Barrett, Sonia Goncalves, Ewan Harrison, David K. Jackson, Ian Johnston, Dominic Kwiatkowski, Cordelia Langford, John Sillitoe on behalf of the Wellcome Sanger Institute COVID-19 Surveillance Team |
| EPI_ISL_2814031 | Lighthouse Lab in Alderley Park | Wellcome Sanger Institute for the COVID-19 Genomics UK (COG-UK) Consortium | Jacquelyn Wynn, Mairead Hyland, The Lighthouse Lab in Alderley Park and Alex Alderton, Roberto Amato, Jeffrey Barrett, Sonia Goncalves, Ewan Harrison, David K. Jackson, Ian Johnston, Dominic Kwiatkowski, Cordelia Langford, John Sillitoe on behalf of the Wellcome Sanger Institute COVID-19 Surveillance Team |
| EPI_ISL_2814032 | Lighthouse Lab in Glasgow | Wellcome Sanger Institute for the COVID-19 Genomics UK (COG-UK) Consortium | Harper VanSteenhouse, Yumi Kasai, David Gray, Carol Clugston, Anna Dominiczak and Alex Alderton, Roberto Amato, Jeffrey Barrett, Sonia Goncalves, Ewan Harrison, David K. Jackson, Ian Johnston, Dominic Kwiatkowski, Cordelia Langford, John Sillitoe on behalf of the Wellcome Sanger Institute COVID-19 Surveillance Team |
| EPI_ISL_2814033 | Lighthouse Lab in Milton Keynes | Wellcome Sanger Institute for the COVID-19 Genomics UK (COG-UK) Consortium | The Lighthouse Lab in Milton Keynes and Alex Alderton, Roberto Amato, Jeffrey Barrett, Sonia Goncalves, Ewan Harrison, David K. Jackson, Ian Johnston, Dominic Kwiatkowski, Cordelia Langford, John Sillitoe on behalf of the Wellcome Sanger Institute COVID-19 Surveillance Team |
| EPI_ISL_2814034 | Lighthouse Lab in Alderley Park | Wellcome Sanger Institute for the COVID-19 Genomics UK (COG-UK) Consortium | Jacquelyn Wynn, Mairead Hyland, The Lighthouse Lab in Alderley Park and Alex Alderton, Roberto Amato, Jeffrey Barrett, Sonia Goncalves, Ewan Harrison, David K. Jackson, Ian Johnston, Dominic Kwiatkowski, Cordelia Langford, John Sillitoe on behalf of the Wellcome Sanger Institute COVID-19 Surveillance Team |
| EPI_ISL_2814035 | Lighthouse Lab in Milton Keynes | Wellcome Sanger Institute for the COVID-19 Genomics UK (COG-UK) Consortium | The Lighthouse Lab in Milton Keynes and Alex Alderton, Roberto Amato, Jeffrey Barrett, Sonia Goncalves, Ewan Harrison, David K. Jackson, Ian Johnston, Dominic Kwiatkowski, Cordelia Langford, John Sillitoe on behalf of the Wellcome Sanger Institute COVID-19 Surveillance Team |
| EPI_ISL_2814036, EPI_ISL_2814037, EPI_ISL_2814038 | Lighthouse Lab in Alderley Park | Wellcome Sanger Institute for the COVID-19 Genomics UK (COG-UK) Consortium | Jacquelyn Wynn, Mairead Hyland, The Lighthouse Lab in Alderley Park and Alex Alderton, Roberto Amato, Jeffrey Barrett, Sonia Goncalves, Ewan Harrison, David K. Jackson, Ian Johnston, Dominic Kwiatkowski, Cordelia Langford, John Sillitoe on behalf of the Wellcome Sanger Institute COVID-19 Surveillance Team |
| EPI_ISL_2814039 | Lighthouse Lab in Milton Keynes | Wellcome Sanger Institute for the COVID-19 Genomics UK (COG-UK) Consortium | The Lighthouse Lab in Milton Keynes and Alex Alderton, Roberto Amato, Jeffrey Barrett, Sonia Goncalves, Ewan Harrison, David K. Jackson, Ian Johnston, Dominic Kwiatkowski, Cordelia Langford, John Sillitoe on behalf of the Wellcome Sanger Institute COVID-19 Surveillance Team |
| EPI_ISL_2814040 | Lighthouse Lab in Alderley Park | Wellcome Sanger Institute for the COVID-19 Genomics UK (COG-UK) Consortium | Jacquelyn Wynn, Mairead Hyland, The Lighthouse Lab in Alderley Park and Alex Alderton, Roberto Amato, Jeffrey Barrett, Sonia Goncalves, Ewan Harrison, David K. Jackson, Ian Johnston, Dominic Kwiatkowski, Cordelia Langford, John Sillitoe on behalf of the Wellcome Sanger Institute COVID-19 Surveillance Team |
| EPI_ISL_2814041, EPI_ISL_2814042 | Lighthouse Lab in Milton Keynes | Wellcome Sanger Institute for the COVID-19 Genomics UK (COG-UK) Consortium | The Lighthouse Lab in Milton Keynes and Alex Alderton, Roberto Amato, Jeffrey Barrett, Sonia Goncalves, Ewan Harrison, David K. Jackson, Ian Johnston, Dominic Kwiatkowski, Cordelia Langford, John Sillitoe on behalf of the Wellcome Sanger Institute COVID-19 Surveillance Team |
| EPI_ISL_2814043 | Lighthouse Lab in Alderley Park | Wellcome Sanger Institute for the COVID-19 Genomics UK (COG-UK) Consortium | Jacquelyn Wynn, Mairead Hyland, The Lighthouse Lab in Alderley Park and Alex Alderton, Roberto Amato, Jeffrey Barrett, Sonia Goncalves, Ewan Harrison, David K. Jackson, Ian Johnston, Dominic Kwiatkowski, Cordelia Langford, John Sillitoe on behalf of the Wellcome Sanger Institute COVID-19 Surveillance Team |
| EPI_ISL_2814044, EPI_ISL_2814045 | Lighthouse Lab in Milton Keynes | Wellcome Sanger Institute for the COVID-19 Genomics UK (COG-UK) Consortium | The Lighthouse Lab in Milton Keynes and Alex Alderton, Roberto Amato, Jeffrey Barrett, Sonia Goncalves, Ewan Harrison, David K. Jackson, Ian Johnston, Dominic Kwiatkowski, Cordelia Langford, John Sillitoe on behalf of the Wellcome Sanger Institute COVID-19 Surveillance Team |
| EPI_ISL_2814046 | Lighthouse Lab in Alderley Park | Wellcome Sanger Institute for the COVID-19 Genomics UK (COG-UK) Consortium | Jacquelyn Wynn, Mairead Hyland, The Lighthouse Lab in Alderley Park and Alex Alderton, Roberto Amato, Jeffrey Barrett, Sonia Goncalves, Ewan Harrison, David K. Jackson, Ian Johnston, Dominic Kwiatkowski, Cordelia Langford, John Sillitoe on behalf of the Wellcome Sanger Institute COVID-19 Surveillance Team |
| EPI_ISL_2814047 | Lighthouse Lab in Milton Keynes | Wellcome Sanger Institute for the COVID-19 Genomics UK (COG-UK) Consortium | The Lighthouse Lab in Milton Keynes and Alex Alderton, Roberto Amato, Jeffrey Barrett, Sonia Goncalves, Ewan Harrison, David K. Jackson, Ian Johnston, Dominic Kwiatkowski, Cordelia Langford, John Sillitoe on behalf of the Wellcome Sanger Institute COVID-19 Surveillance Team |
| EPI_ISL_2814048, EPI_ISL_2814049 | Lighthouse Lab in Alderley Park | Wellcome Sanger Institute for the COVID-19 Genomics UK (COG-UK) Consortium | Jacquelyn Wynn, Mairead Hyland, The Lighthouse Lab in Alderley Park and Alex Alderton, Roberto Amato, Jeffrey Barrett, Sonia Goncalves, Ewan Harrison, David K. Jackson, Ian Johnston, Dominic Kwiatkowski, Cordelia Langford, John Sillitoe on behalf of the Wellcome Sanger Institute COVID-19 Surveillance Team |
| EPI_ISL_2814050 | Lighthouse Lab in Glasgow | Wellcome Sanger Institute for the COVID-19 Genomics UK (COG-UK) Consortium | Harper VanSteenhouse, Yumi Kasai, David Gray, Carol Clugston, Anna Dominiczak and Alex Alderton, Roberto Amato, Jeffrey Barrett, Sonia Goncalves, Ewan Harrison, David K. Jackson, Ian Johnston, Dominic Kwiatkowski, Cordelia Langford, John Sillitoe on behalf of the Wellcome Sanger Institute COVID-19 Surveillance Team |
| EPI_ISL_2814051 | Lighthouse Lab in Milton Keynes | Wellcome Sanger Institute for the COVID-19 Genomics UK (COG-UK) Consortium | The Lighthouse Lab in Milton Keynes and Alex Alderton, Roberto Amato, Jeffrey Barrett, Sonia Goncalves, Ewan Harrison, David K. Jackson, Ian Johnston, Dominic Kwiatkowski, Cordelia Langford, John Sillitoe on behalf of the Wellcome Sanger Institute COVID-19 Surveillance Team |
| EPI_ISL_2814052, EPI_ISL_2814053 | Lighthouse Lab in Alderley Park | Wellcome Sanger Institute for the COVID-19 Genomics UK (COG-UK) Consortium | Jacquelyn Wynn, Mairead Hyland, The Lighthouse Lab in Alderley Park and Alex Alderton, Roberto Amato, Jeffrey Barrett, Sonia Goncalves, Ewan Harrison, David K. Jackson, Ian Johnston, Dominic Kwiatkowski, Cordelia Langford, John Sillitoe on behalf of the Wellcome Sanger Institute COVID-19 Surveillance Team |
| EPI_ISL_2814054 | Lighthouse Lab in Milton Keynes | Wellcome Sanger Institute for the COVID-19 Genomics UK (COG-UK) Consortium | The Lighthouse Lab in Milton Keynes and Alex Alderton, Roberto Amato, Jeffrey Barrett, Sonia Goncalves, Ewan Harrison, David K. Jackson, Ian Johnston, Dominic Kwiatkowski, Cordelia Langford, John Sillitoe on behalf of the Wellcome Sanger Institute COVID-19 Surveillance Team |
| EPI_ISL_2814055 | Lighthouse Lab in Alderley Park | Wellcome Sanger Institute for the COVID-19 Genomics UK (COG-UK) Consortium | Jacquelyn Wynn, Mairead Hyland, The Lighthouse Lab in Alderley Park and Alex Alderton, Roberto Amato, Jeffrey Barrett, Sonia Goncalves, Ewan Harrison, David K. Jackson, Ian Johnston, Dominic Kwiatkowski, Cordelia Langford, John Sillitoe on behalf of the Wellcome Sanger Institute COVID-19 Surveillance Team |
| EPI_ISL_2814056, EPI_ISL_2814057, EPI_ISL_2814058, EPI_ISL_2814059 | Lighthouse Lab in Milton Keynes | Wellcome Sanger Institute for the COVID-19 Genomics UK (COG-UK) Consortium | The Lighthouse Lab in Milton Keynes and Alex Alderton, Roberto Amato, Jeffrey Barrett, Sonia Goncalves, Ewan Harrison, David K. Jackson, Ian Johnston, Dominic Kwiatkowski, Cordelia Langford, John Sillitoe on behalf of the Wellcome Sanger Institute COVID-19 Surveillance Team |
| EPI_ISL_2814060 | Lighthouse Lab in Glasgow | Wellcome Sanger Institute for the COVID-19 Genomics UK (COG-UK) Consortium | Harper VanSteenhouse, Yumi Kasai, David Gray, Carol Clugston, Anna Dominiczak and Alex Alderton, Roberto Amato, Jeffrey Barrett, Sonia Goncalves, Ewan Harrison, David K. Jackson, Ian Johnston, Dominic Kwiatkowski, Cordelia Langford, John Sillitoe on behalf of the Wellcome Sanger Institute COVID-19 Surveillance Team |
| EPI_ISL_2814061 | Lighthouse Lab in Milton Keynes | Wellcome Sanger Institute for the COVID-19 Genomics UK (COG-UK) Consortium | The Lighthouse Lab in Milton Keynes and Alex Alderton, Roberto Amato, Jeffrey Barrett, Sonia Goncalves, Ewan Harrison, David K. Jackson, Ian Johnston, Dominic Kwiatkowski, Cordelia Langford, John Sillitoe on behalf of the Wellcome Sanger Institute COVID-19 Surveillance Team |
| EPI_ISL_2814062 | Lighthouse Lab in Glasgow | Wellcome Sanger Institute for the COVID-19 Genomics UK (COG-UK) | Harper VanSteenhouse, Yumi Kasai, David Gray, Carol Clugston, Anna Dominiczak and Alex Alderton, Roberto Amato, Jeffrey Barrett, Sonia Goncalves, Ewan Harrison, David K. Jackson, Ian Johnston, Dominic Kwiatkowski, Cordelia Langford, |

| | | | |
|---|---|---|---|
| | | Consortium | John Sillitoe on behalf of the Wellcome Sanger Institute COVID-19 Surveillance Team |
| EPI_ISL_2814063, EPI_ISL_2814064, EPI_ISL_2814065 | Lighthouse Lab in Alderley Park | Wellcome Sanger Institute for the COVID-19 Genomics UK (COG-UK) Consortium | Jacquelyn Wynn, Mairead Hyland, The Lighthouse Lab in Alderley Park and Alex Alderton, Roberto Amato, Jeffrey Barrett, Sonia Goncalves, Ewan Harrison, David K. Jackson, Ian Johnston, Dominic Kwiatkowski, Cordelia Langford, John Sillitoe on behalf of the Wellcome Sanger Institute COVID-19 Surveillance Team |
| EPI_ISL_2814066, EPI_ISL_2814067 | Lighthouse Lab in Milton Keynes | Wellcome Sanger Institute for the COVID-19 Genomics UK (COG-UK) Consortium | The Lighthouse Lab in Milton Keynes and Alex Alderton, Roberto Amato, Jeffrey Barrett, Sonia Goncalves, Ewan Harrison, David K. Jackson, Ian Johnston, Dominic Kwiatkowski, Cordelia Langford, John Sillitoe on behalf of the Wellcome Sanger Institute COVID-19 Surveillance Team |
| EPI_ISL_2814068 | Lighthouse Lab in Alderley Park | Wellcome Sanger Institute for the COVID-19 Genomics UK (COG-UK) Consortium | Jacquelyn Wynn, Mairead Hyland, The Lighthouse Lab in Alderley Park and Alex Alderton, Roberto Amato, Jeffrey Barrett, Sonia Goncalves, Ewan Harrison, David K. Jackson, Ian Johnston, Dominic Kwiatkowski, Cordelia Langford, John Sillitoe on behalf of the Wellcome Sanger Institute COVID-19 Surveillance Team |
| EPI_ISL_2814069 | Lighthouse Lab in Milton Keynes | Wellcome Sanger Institute for the COVID-19 Genomics UK (COG-UK) Consortium | The Lighthouse Lab in Milton Keynes and Alex Alderton, Roberto Amato, Jeffrey Barrett, Sonia Goncalves, Ewan Harrison, David K. Jackson, Ian Johnston, Dominic Kwiatkowski, Cordelia Langford, John Sillitoe on behalf of the Wellcome Sanger Institute COVID-19 Surveillance Team |
| EPI_ISL_2814070 | Lighthouse Lab in Alderley Park | Wellcome Sanger Institute for the COVID-19 Genomics UK (COG-UK) Consortium | Jacquelyn Wynn, Mairead Hyland, The Lighthouse Lab in Alderley Park and Alex Alderton, Roberto Amato, Jeffrey Barrett, Sonia Goncalves, Ewan Harrison, David K. Jackson, Ian Johnston, Dominic Kwiatkowski, Cordelia Langford, John Sillitoe on behalf of the Wellcome Sanger Institute COVID-19 Surveillance Team |
| EPI_ISL_2814071, EPI_ISL_2814072, EPI_ISL_2814073 | Lighthouse Lab in Milton Keynes | Wellcome Sanger Institute for the COVID-19 Genomics UK (COG-UK) Consortium | The Lighthouse Lab in Milton Keynes and Alex Alderton, Roberto Amato, Jeffrey Barrett, Sonia Goncalves, Ewan Harrison, David K. Jackson, Ian Johnston, Dominic Kwiatkowski, Cordelia Langford, John Sillitoe on behalf of the Wellcome Sanger Institute COVID-19 Surveillance Team |
| EPI_ISL_2814074, EPI_ISL_2814075, EPI_ISL_2814076 | Lighthouse Lab in Alderley Park | Wellcome Sanger Institute for the COVID-19 Genomics UK (COG-UK) Consortium | Jacquelyn Wynn, Mairead Hyland, The Lighthouse Lab in Alderley Park and Alex Alderton, Roberto Amato, Jeffrey Barrett, Sonia Goncalves, Ewan Harrison, David K. Jackson, Ian Johnston, Dominic Kwiatkowski, Cordelia Langford, John Sillitoe on behalf of the Wellcome Sanger Institute COVID-19 Surveillance Team |
| EPI_ISL_2814077, EPI_ISL_2814078, EPI_ISL_2814079 | Lighthouse Lab in Milton Keynes | Wellcome Sanger Institute for the COVID-19 Genomics UK (COG-UK) Consortium | The Lighthouse Lab in Milton Keynes and Alex Alderton, Roberto Amato, Jeffrey Barrett, Sonia Goncalves, Ewan Harrison, David K. Jackson, Ian Johnston, Dominic Kwiatkowski, Cordelia Langford, John Sillitoe on behalf of the Wellcome Sanger Institute COVID-19 Surveillance Team |
| EPI_ISL_2814080 | Lighthouse Lab in Alderley Park | Wellcome Sanger Institute for the COVID-19 Genomics UK (COG-UK) Consortium | Jacquelyn Wynn, Mairead Hyland, The Lighthouse Lab in Alderley Park and Alex Alderton, Roberto Amato, Jeffrey Barrett, Sonia Goncalves, Ewan Harrison, David K. Jackson, Ian Johnston, Dominic Kwiatkowski, Cordelia Langford, John Sillitoe on behalf of the Wellcome Sanger Institute COVID-19 Surveillance Team |
| EPI_ISL_2814081, EPI_ISL_2814082, EPI_ISL_2814083, EPI_ISL_2814084 | Lighthouse Lab in Milton Keynes | Wellcome Sanger Institute for the COVID-19 Genomics UK (COG-UK) Consortium | The Lighthouse Lab in Milton Keynes and Alex Alderton, Roberto Amato, Jeffrey Barrett, Sonia Goncalves, Ewan Harrison, David K. Jackson, Ian Johnston, Dominic Kwiatkowski, Cordelia Langford, John Sillitoe on behalf of the Wellcome Sanger Institute COVID-19 Surveillance Team |
| EPI_ISL_2814085 | Lighthouse Lab in Glasgow | Wellcome Sanger Institute for the COVID-19 Genomics UK (COG-UK) Consortium | Harper VanSteenhouse, Yumi Kasai, David Gray, Carol Clugston, Anna Dominiczak and Alex Alderton, Roberto Amato, Jeffrey Barrett, Sonia Goncalves, Ewan Harrison, David K. Jackson, Ian Johnston, Dominic Kwiatkowski, Cordelia Langford, John Sillitoe on behalf of the Wellcome Sanger Institute COVID-19 Surveillance Team |
| EPI_ISL_2814086, EPI_ISL_2814087 | Lighthouse Lab in Milton Keynes | Wellcome Sanger Institute for the COVID-19 Genomics UK (COG-UK) Consortium | The Lighthouse Lab in Milton Keynes and Alex Alderton, Roberto Amato, Jeffrey Barrett, Sonia Goncalves, Ewan Harrison, David K. Jackson, Ian Johnston, Dominic Kwiatkowski, Cordelia Langford, John Sillitoe on behalf of the Wellcome Sanger Institute COVID-19 Surveillance Team |
| EPI_ISL_2814088, EPI_ISL_2814089 | Lighthouse Lab in Alderley Park | Wellcome Sanger Institute for the COVID-19 Genomics UK (COG-UK) Consortium | Jacquelyn Wynn, Mairead Hyland, The Lighthouse Lab in Alderley Park and Alex Alderton, Roberto Amato, Jeffrey Barrett, Sonia Goncalves, Ewan Harrison, David K. Jackson, Ian Johnston, Dominic Kwiatkowski, Cordelia Langford, John Sillitoe on behalf of the Wellcome Sanger Institute COVID-19 Surveillance Team |
| EPI_ISL_2814090, EPI_ISL_2814091 | Lighthouse Lab in Milton Keynes | Wellcome Sanger Institute for the COVID-19 Genomics UK (COG-UK) Consortium | The Lighthouse Lab in Milton Keynes and Alex Alderton, Roberto Amato, Jeffrey Barrett, Sonia Goncalves, Ewan Harrison, David K. Jackson, Ian Johnston, Dominic Kwiatkowski, Cordelia Langford, John Sillitoe on behalf of the Wellcome Sanger Institute COVID-19 Surveillance Team |
| EPI_ISL_2814092 | Lighthouse Lab in Alderley Park | Wellcome Sanger Institute for the COVID-19 Genomics UK (COG-UK) Consortium | Jacquelyn Wynn, Mairead Hyland, The Lighthouse Lab in Alderley Park and Alex Alderton, Roberto Amato, Jeffrey Barrett, Sonia Goncalves, Ewan Harrison, David K. Jackson, Ian Johnston, Dominic Kwiatkowski, Cordelia Langford, John Sillitoe on behalf of the Wellcome Sanger Institute COVID-19 Surveillance Team |
| EPI_ISL_2814093, EPI_ISL_2814094, EPI_ISL_2814095 | Lighthouse Lab in Milton Keynes | Wellcome Sanger Institute for the COVID-19 Genomics UK (COG-UK) Consortium | The Lighthouse Lab in Milton Keynes and Alex Alderton, Roberto Amato, Jeffrey Barrett, Sonia Goncalves, Ewan Harrison, David K. Jackson, Ian Johnston, Dominic Kwiatkowski, Cordelia Langford, John Sillitoe on behalf of the Wellcome Sanger Institute COVID-19 Surveillance Team |
| EPI_ISL_2814096, EPI_ISL_2814097, EPI_ISL_2814098, EPI_ISL_2814099, EPI_ISL_2814100, EPI_ISL_2814101 | Lighthouse Lab in Alderley Park | Wellcome Sanger Institute for the COVID-19 Genomics UK (COG-UK) Consortium | Jacquelyn Wynn, Mairead Hyland, The Lighthouse Lab in Alderley Park and Alex Alderton, Roberto Amato, Jeffrey Barrett, Sonia Goncalves, Ewan Harrison, David K. Jackson, Ian Johnston, Dominic Kwiatkowski, Cordelia Langford, John Sillitoe on behalf of the Wellcome Sanger Institute COVID-19 Surveillance Team |
| EPI_ISL_2814102, EPI_ISL_2814103 | Lighthouse Lab in Glasgow | Wellcome Sanger Institute for the COVID-19 Genomics UK (COG-UK) Consortium | Harper VanSteenhouse, Yumi Kasai, David Gray, Carol Clugston, Anna Dominiczak and Alex Alderton, Roberto Amato, Jeffrey Barrett, Sonia Goncalves, Ewan Harrison, David K. Jackson, Ian Johnston, Dominic Kwiatkowski, Cordelia Langford, John Sillitoe on behalf of the Wellcome Sanger Institute COVID-19 Surveillance Team |
| EPI_ISL_2814104 | Lighthouse Lab in Alderley Park | Wellcome Sanger Institute for the COVID-19 Genomics UK (COG-UK) Consortium | Jacquelyn Wynn, Mairead Hyland, The Lighthouse Lab in Alderley Park and Alex Alderton, Roberto Amato, Jeffrey Barrett, Sonia Goncalves, Ewan Harrison, David K. Jackson, Ian Johnston, Dominic Kwiatkowski, Cordelia Langford, John Sillitoe on behalf of the Wellcome Sanger Institute COVID-19 Surveillance Team |
| EPI_ISL_2814105, EPI_ISL_2814106, EPI_ISL_2814107, EPI_ISL_2814108, EPI_ISL_2814109, EPI_ISL_2814110 | Lighthouse Lab in Milton Keynes | Wellcome Sanger Institute for the COVID-19 Genomics UK (COG-UK) Consortium | The Lighthouse Lab in Milton Keynes and Alex Alderton, Roberto Amato, Jeffrey Barrett, Sonia Goncalves, Ewan Harrison, David K. Jackson, Ian Johnston, Dominic Kwiatkowski, Cordelia Langford, John Sillitoe on behalf of the Wellcome Sanger Institute COVID-19 Surveillance Team |
| EPI_ISL_2814111, EPI_ISL_2814112 | Lighthouse Lab in Alderley Park | Wellcome Sanger Institute for the COVID-19 Genomics UK (COG-UK) Consortium | Jacquelyn Wynn, Mairead Hyland, The Lighthouse Lab in Alderley Park and Alex Alderton, Roberto Amato, Jeffrey Barrett, Sonia Goncalves, Ewan Harrison, David K. Jackson, Ian Johnston, Dominic Kwiatkowski, Cordelia Langford, John Sillitoe on behalf of the Wellcome Sanger Institute COVID-19 Surveillance Team |
| EPI_ISL_2814113, EPI_ISL_2814114 | Lighthouse Lab in Milton Keynes | Wellcome Sanger Institute for the COVID-19 Genomics UK (COG-UK) Consortium | The Lighthouse Lab in Milton Keynes and Alex Alderton, Roberto Amato, Jeffrey Barrett, Sonia Goncalves, Ewan Harrison, David K. Jackson, Ian Johnston, Dominic Kwiatkowski, Cordelia Langford, John Sillitoe on behalf of the Wellcome Sanger Institute COVID-19 Surveillance Team |
| EPI_ISL_2814115, EPI_ISL_2814116, EPI_ISL_2814117, EPI_ISL_2814118, EPI_ISL_2814119, EPI_ISL_2814120 | Lighthouse Lab in Alderley Park | Wellcome Sanger Institute for the COVID-19 Genomics UK (COG-UK) Consortium | Jacquelyn Wynn, Mairead Hyland, The Lighthouse Lab in Alderley Park and Alex Alderton, Roberto Amato, Jeffrey Barrett, Sonia Goncalves, Ewan Harrison, David K. Jackson, Ian Johnston, Dominic Kwiatkowski, Cordelia Langford, John Sillitoe on behalf of the Wellcome Sanger Institute COVID-19 Surveillance Team |
| EPI_ISL_2814121, EPI_ISL_2814122, EPI_ISL_2814123 | Lighthouse Lab in Milton Keynes | Wellcome Sanger Institute for the COVID-19 Genomics UK (COG-UK) Consortium | The Lighthouse Lab in Milton Keynes and Alex Alderton, Roberto Amato, Jeffrey Barrett, Sonia Goncalves, Ewan Harrison, David K. Jackson, Ian Johnston, Dominic Kwiatkowski, Cordelia Langford, John Sillitoe on behalf of the Wellcome Sanger Institute COVID-19 Surveillance Team |
| EPI_ISL_2814124 | Lighthouse Lab in Glasgow | Wellcome Sanger Institute for the COVID-19 Genomics UK (COG-UK) Consortium | Harper VanSteenhouse, Yumi Kasai, David Gray, Carol Clugston, Anna Dominiczak and Alex Alderton, Roberto Amato, Jeffrey Barrett, Sonia Goncalves, Ewan Harrison, David K. Jackson, Ian Johnston, Dominic Kwiatkowski, Cordelia Langford, John Sillitoe on behalf of the Wellcome Sanger Institute COVID-19 Surveillance Team |
| EPI_ISL_2814125, EPI_ISL_2814126 | Lighthouse Lab in Alderley Park | Wellcome Sanger Institute for the COVID-19 Genomics UK (COG-UK) Consortium | Jacquelyn Wynn, Mairead Hyland, The Lighthouse Lab in Alderley Park and Alex Alderton, Roberto Amato, Jeffrey Barrett, Sonia Goncalves, Ewan Harrison, David K. Jackson, Ian Johnston, Dominic Kwiatkowski, Cordelia Langford, John Sillitoe on behalf of the Wellcome Sanger Institute COVID-19 Surveillance Team |
| EPI_ISL_2814127, EPI_ISL_2814128 | Lighthouse Lab in Milton Keynes | Wellcome Sanger Institute for the COVID-19 Genomics UK (COG-UK) Consortium | The Lighthouse Lab in Milton Keynes and Alex Alderton, Roberto Amato, Jeffrey Barrett, Sonia Goncalves, Ewan Harrison, David K. Jackson, Ian Johnston, Dominic Kwiatkowski, Cordelia Langford, John Sillitoe on behalf of the Wellcome Sanger Institute COVID-19 Surveillance Team |
| EPI_ISL_2814129, EPI_ISL_2814130, EPI_ISL_2814131 | Lighthouse Lab in Glasgow | Wellcome Sanger Institute for the COVID-19 Genomics UK (COG-UK) Consortium | Harper VanSteenhouse, Yumi Kasai, David Gray, Carol Clugston, Anna Dominiczak and Alex Alderton, Roberto Amato, Jeffrey Barrett, Sonia Goncalves, Ewan Harrison, David K. Jackson, Ian Johnston, Dominic Kwiatkowski, Cordelia Langford, John Sillitoe on behalf of the Wellcome Sanger Institute COVID-19 Surveillance Team |
| EPI_ISL_2814132 | Lighthouse Lab in Alderley Park | Wellcome Sanger Institute for the COVID-19 Genomics UK (COG-UK) Consortium | Jacquelyn Wynn, Mairead Hyland, The Lighthouse Lab in Alderley Park and Alex Alderton, Roberto Amato, Jeffrey Barrett, Sonia Goncalves, Ewan Harrison, David K. Jackson, Ian Johnston, Dominic Kwiatkowski, Cordelia Langford, John Sillitoe on behalf of the Wellcome Sanger Institute COVID-19 Surveillance Team |
| EPI_ISL_2814133 | Lighthouse Lab in Glasgow | Wellcome Sanger Institute for the COVID-19 Genomics UK (COG-UK) Consortium | Harper VanSteenhouse, Yumi Kasai, David Gray, Carol Clugston, Anna Dominiczak and Alex Alderton, Roberto Amato, Jeffrey Barrett, Sonia Goncalves, Ewan Harrison, David K. Jackson, Ian Johnston, Dominic Kwiatkowski, Cordelia Langford, John Sillitoe on behalf of the Wellcome Sanger Institute COVID-19 Surveillance Team |
| EPI_ISL_2814134 | Lighthouse Lab in Alderley Park | Wellcome Sanger Institute for the COVID-19 Genomics UK (COG-UK) Consortium | Jacquelyn Wynn, Mairead Hyland, The Lighthouse Lab in Alderley Park and Alex Alderton, Roberto Amato, Jeffrey Barrett, Sonia Goncalves, Ewan Harrison, David K. Jackson, Ian Johnston, Dominic Kwiatkowski, Cordelia Langford, John Sillitoe on behalf of the Wellcome Sanger Institute COVID-19 Surveillance Team |
| EPI_ISL_2814135, EPI_ISL_2814136 | Lighthouse Lab in Milton Keynes | Wellcome Sanger Institute for the COVID-19 Genomics UK (COG-UK) Consortium | The Lighthouse Lab in Milton Keynes and Alex Alderton, Roberto Amato, Jeffrey Barrett, Sonia Goncalves, Ewan Harrison, David K. Jackson, Ian Johnston, Dominic Kwiatkowski, Cordelia Langford, John Sillitoe on behalf of the Wellcome Sanger Institute COVID-19 Surveillance Team |
| EPI_ISL_2814137 | Lighthouse Lab in Alderley Park | Wellcome Sanger Institute for the COVID-19 Genomics UK (COG-UK) Consortium | Jacquelyn Wynn, Mairead Hyland, The Lighthouse Lab in Alderley Park and Alex Alderton, Roberto Amato, Jeffrey Barrett, Sonia Goncalves, Ewan Harrison, David K. Jackson, Ian Johnston, Dominic Kwiatkowski, Cordelia Langford, John Sillitoe on behalf of the Wellcome Sanger Institute COVID-19 Surveillance Team |
| EPI_ISL_2814138 | Lighthouse Lab in Milton Keynes | Wellcome Sanger Institute for the COVID-19 Genomics UK (COG-UK) Consortium | The Lighthouse Lab in Milton Keynes and Alex Alderton, Roberto Amato, Jeffrey Barrett, Sonia Goncalves, Ewan Harrison, David K. Jackson, Ian Johnston, Dominic Kwiatkowski, Cordelia Langford, John Sillitoe on behalf of the Wellcome Sanger Institute COVID-19 Surveillance Team |
| EPI_ISL_2814139 | Lighthouse Lab in Glasgow | Wellcome Sanger Institute for the COVID-19 Genomics UK (COG-UK) Consortium | Harper VanSteenhouse, Yumi Kasai, David Gray, Carol Clugston, Anna Dominiczak and Alex Alderton, Roberto Amato, Jeffrey Barrett, Sonia Goncalves, Ewan Harrison, David K. Jackson, Ian Johnston, Dominic Kwiatkowski, Cordelia Langford, John Sillitoe on behalf of the Wellcome Sanger Institute COVID-19 Surveillance Team |
| EPI_ISL_2814140 | Lighthouse Lab in Milton Keynes | Wellcome Sanger Institute for the COVID-19 Genomics UK (COG-UK) Consortium | The Lighthouse Lab in Milton Keynes and Alex Alderton, Roberto Amato, Jeffrey Barrett, Sonia Goncalves, Ewan Harrison, David K. Jackson, Ian Johnston, Dominic Kwiatkowski, Cordelia Langford, John Sillitoe on behalf of the Wellcome Sanger Institute COVID-19 Surveillance Team |

| | | | |
|---|---|---|---|
| EPI_ISL_2814141 | Lighthouse Lab in Alderley Park | Wellcome Sanger Institute for the COVID-19 Genomics UK (COG-UK) Consortium | Jacquelyn Wynn, Mairead Hyland, The Lighthouse Lab in Alderley Park and Alex Alderton, Roberto Amato, Jeffrey Barrett, Sonia Goncalves, Ewan Harrison, David K. Jackson, Ian Johnston, Dominic Kwiatkowski, Cordelia Langford, John Sillitoe on behalf of the Wellcome Sanger Institute COVID-19 Surveillance Team |
| EPI_ISL_2814142 | Lighthouse Lab in Glasgow | Wellcome Sanger Institute for the COVID-19 Genomics UK (COG-UK) Consortium | Harper VanSteenhouse, Yumi Kasai, David Gray, Carol Clugston, Anna Dominiczak and Alex Alderton, Roberto Amato, Jeffrey Barrett, Sonia Goncalves, Ewan Harrison, David K. Jackson, Ian Johnston, Dominic Kwiatkowski, Cordelia Langford, John Sillitoe on behalf of the Wellcome Sanger Institute COVID-19 Surveillance Team |
| EPI_ISL_2814143, EPI_ISL_2814144 | Lighthouse Lab in Alderley Park | Wellcome Sanger Institute for the COVID-19 Genomics UK (COG-UK) Consortium | Jacquelyn Wynn, Mairead Hyland, The Lighthouse Lab in Alderley Park and Alex Alderton, Roberto Amato, Jeffrey Barrett, Sonia Goncalves, Ewan Harrison, David K. Jackson, Ian Johnston, Dominic Kwiatkowski, Cordelia Langford, John Sillitoe on behalf of the Wellcome Sanger Institute COVID-19 Surveillance Team |
| EPI_ISL_2814145 | Lighthouse Lab in Milton Keynes | Wellcome Sanger Institute for the COVID-19 Genomics UK (COG-UK) Consortium | The Lighthouse Lab in Milton Keynes and Alex Alderton, Roberto Amato, Jeffrey Barrett, Sonia Goncalves, Ewan Harrison, David K. Jackson, Ian Johnston, Dominic Kwiatkowski, Cordelia Langford, John Sillitoe on behalf of the Wellcome Sanger Institute COVID-19 Surveillance Team |
| EPI_ISL_2814146 | Lighthouse Lab in Glasgow | Wellcome Sanger Institute for the COVID-19 Genomics UK (COG-UK) Consortium | Harper VanSteenhouse, Yumi Kasai, David Gray, Carol Clugston, Anna Dominiczak and Alex Alderton, Roberto Amato, Jeffrey Barrett, Sonia Goncalves, Ewan Harrison, David K. Jackson, Ian Johnston, Dominic Kwiatkowski, Cordelia Langford, John Sillitoe on behalf of the Wellcome Sanger Institute COVID-19 Surveillance Team |
| EPI_ISL_2814147 | Lighthouse Lab in Alderley Park | Wellcome Sanger Institute for the COVID-19 Genomics UK (COG-UK) Consortium | Jacquelyn Wynn, Mairead Hyland, The Lighthouse Lab in Alderley Park and Alex Alderton, Roberto Amato, Jeffrey Barrett, Sonia Goncalves, Ewan Harrison, David K. Jackson, Ian Johnston, Dominic Kwiatkowski, Cordelia Langford, John Sillitoe on behalf of the Wellcome Sanger Institute COVID-19 Surveillance Team |
| EPI_ISL_2814148 | Lighthouse Lab in Glasgow | Wellcome Sanger Institute for the COVID-19 Genomics UK (COG-UK) Consortium | Harper VanSteenhouse, Yumi Kasai, David Gray, Carol Clugston, Anna Dominiczak and Alex Alderton, Roberto Amato, Jeffrey Barrett, Sonia Goncalves, Ewan Harrison, David K. Jackson, Ian Johnston, Dominic Kwiatkowski, Cordelia Langford, John Sillitoe on behalf of the Wellcome Sanger Institute COVID-19 Surveillance Team |
| EPI_ISL_2814149, EPI_ISL_2814150 | Lighthouse Lab in Alderley Park | Wellcome Sanger Institute for the COVID-19 Genomics UK (COG-UK) Consortium | Jacquelyn Wynn, Mairead Hyland, The Lighthouse Lab in Alderley Park and Alex Alderton, Roberto Amato, Jeffrey Barrett, Sonia Goncalves, Ewan Harrison, David K. Jackson, Ian Johnston, Dominic Kwiatkowski, Cordelia Langford, John Sillitoe on behalf of the Wellcome Sanger Institute COVID-19 Surveillance Team |
| EPI_ISL_2814151 | Lighthouse Lab in Milton Keynes | Wellcome Sanger Institute for the COVID-19 Genomics UK (COG-UK) Consortium | The Lighthouse Lab in Milton Keynes and Alex Alderton, Roberto Amato, Jeffrey Barrett, Sonia Goncalves, Ewan Harrison, David K. Jackson, Ian Johnston, Dominic Kwiatkowski, Cordelia Langford, John Sillitoe on behalf of the Wellcome Sanger Institute COVID-19 Surveillance Team |
| EPI_ISL_2814152 | Lighthouse Lab in Alderley Park | Wellcome Sanger Institute for the COVID-19 Genomics UK (COG-UK) Consortium | Jacquelyn Wynn, Mairead Hyland, The Lighthouse Lab in Alderley Park and Alex Alderton, Roberto Amato, Jeffrey Barrett, Sonia Goncalves, Ewan Harrison, David K. Jackson, Ian Johnston, Dominic Kwiatkowski, Cordelia Langford, John Sillitoe on behalf of the Wellcome Sanger Institute COVID-19 Surveillance Team |
| EPI_ISL_2814153 | Lighthouse Lab in Glasgow | Wellcome Sanger Institute for the COVID-19 Genomics UK (COG-UK) Consortium | Harper VanSteenhouse, Yumi Kasai, David Gray, Carol Clugston, Anna Dominiczak and Alex Alderton, Roberto Amato, Jeffrey Barrett, Sonia Goncalves, Ewan Harrison, David K. Jackson, Ian Johnston, Dominic Kwiatkowski, Cordelia Langford, John Sillitoe on behalf of the Wellcome Sanger Institute COVID-19 Surveillance Team |
| EPI_ISL_2814154 | Lighthouse Lab in Alderley Park | Wellcome Sanger Institute for the COVID-19 Genomics UK (COG-UK) Consortium | Jacquelyn Wynn, Mairead Hyland, The Lighthouse Lab in Alderley Park and Alex Alderton, Roberto Amato, Jeffrey Barrett, Sonia Goncalves, Ewan Harrison, David K. Jackson, Ian Johnston, Dominic Kwiatkowski, Cordelia Langford, John Sillitoe on behalf of the Wellcome Sanger Institute COVID-19 Surveillance Team |
| EPI_ISL_2814155 | Lighthouse Lab in Milton Keynes | Wellcome Sanger Institute for the COVID-19 Genomics UK (COG-UK) Consortium | The Lighthouse Lab in Milton Keynes and Alex Alderton, Roberto Amato, Jeffrey Barrett, Sonia Goncalves, Ewan Harrison, David K. Jackson, Ian Johnston, Dominic Kwiatkowski, Cordelia Langford, John Sillitoe on behalf of the Wellcome Sanger Institute COVID-19 Surveillance Team |
| EPI_ISL_2814156 | Lighthouse Lab in Glasgow | Wellcome Sanger Institute for the COVID-19 Genomics UK (COG-UK) Consortium | Harper VanSteenhouse, Yumi Kasai, David Gray, Carol Clugston, Anna Dominiczak and Alex Alderton, Roberto Amato, Jeffrey Barrett, Sonia Goncalves, Ewan Harrison, David K. Jackson, Ian Johnston, Dominic Kwiatkowski, Cordelia Langford, John Sillitoe on behalf of the Wellcome Sanger Institute COVID-19 Surveillance Team |
| EPI_ISL_2814157 | Lighthouse Lab in Alderley Park | Wellcome Sanger Institute for the COVID-19 Genomics UK (COG-UK) Consortium | Jacquelyn Wynn, Mairead Hyland, The Lighthouse Lab in Alderley Park and Alex Alderton, Roberto Amato, Jeffrey Barrett, Sonia Goncalves, Ewan Harrison, David K. Jackson, Ian Johnston, Dominic Kwiatkowski, Cordelia Langford, John Sillitoe on behalf of the Wellcome Sanger Institute COVID-19 Surveillance Team |
| EPI_ISL_2814158, EPI_ISL_2814159 | Lighthouse Lab in Milton Keynes | Wellcome Sanger Institute for the COVID-19 Genomics UK (COG-UK) Consortium | The Lighthouse Lab in Milton Keynes and Alex Alderton, Roberto Amato, Jeffrey Barrett, Sonia Goncalves, Ewan Harrison, David K. Jackson, Ian Johnston, Dominic Kwiatkowski, Cordelia Langford, John Sillitoe on behalf of the Wellcome Sanger Institute COVID-19 Surveillance Team |
| EPI_ISL_2814160, EPI_ISL_2814161, EPI_ISL_2814162 | Lighthouse Lab in Alderley Park | Wellcome Sanger Institute for the COVID-19 Genomics UK (COG-UK) Consortium | Jacquelyn Wynn, Mairead Hyland, The Lighthouse Lab in Alderley Park and Alex Alderton, Roberto Amato, Jeffrey Barrett, Sonia Goncalves, Ewan Harrison, David K. Jackson, Ian Johnston, Dominic Kwiatkowski, Cordelia Langford, John Sillitoe on behalf of the Wellcome Sanger Institute COVID-19 Surveillance Team |
| EPI_ISL_2814163, EPI_ISL_2814164, EPI_ISL_2814165 | Lighthouse Lab in Milton Keynes | Wellcome Sanger Institute for the COVID-19 Genomics UK (COG-UK) Consortium | The Lighthouse Lab in Milton Keynes and Alex Alderton, Roberto Amato, Jeffrey Barrett, Sonia Goncalves, Ewan Harrison, David K. Jackson, Ian Johnston, Dominic Kwiatkowski, Cordelia Langford, John Sillitoe on behalf of the Wellcome Sanger Institute COVID-19 Surveillance Team |
| EPI_ISL_2814166, EPI_ISL_2814167, EPI_ISL_2814169 | Lighthouse Lab in Alderley Park | Wellcome Sanger Institute for the COVID-19 Genomics UK (COG-UK) Consortium | Jacquelyn Wynn, Mairead Hyland, The Lighthouse Lab in Alderley Park and Alex Alderton, Roberto Amato, Jeffrey Barrett, Sonia Goncalves, Ewan Harrison, David K. Jackson, Ian Johnston, Dominic Kwiatkowski, Cordelia Langford, John Sillitoe on behalf of the Wellcome Sanger Institute COVID-19 Surveillance Team |
| EPI_ISL_2814170, EPI_ISL_2814171, EPI_ISL_2814172 | Lighthouse Lab in Milton Keynes | Wellcome Sanger Institute for the COVID-19 Genomics UK (COG-UK) Consortium | The Lighthouse Lab in Milton Keynes and Alex Alderton, Roberto Amato, Jeffrey Barrett, Sonia Goncalves, Ewan Harrison, David K. Jackson, Ian Johnston, Dominic Kwiatkowski, Cordelia Langford, John Sillitoe on behalf of the Wellcome Sanger Institute COVID-19 Surveillance Team |
| EPI_ISL_2814173, EPI_ISL_2814174 | Lighthouse Lab in Alderley Park | Wellcome Sanger Institute for the COVID-19 Genomics UK (COG-UK) Consortium | Jacquelyn Wynn, Mairead Hyland, The Lighthouse Lab in Alderley Park and Alex Alderton, Roberto Amato, Jeffrey Barrett, Sonia Goncalves, Ewan Harrison, David K. Jackson, Ian Johnston, Dominic Kwiatkowski, Cordelia Langford, John Sillitoe on behalf of the Wellcome Sanger Institute COVID-19 Surveillance Team |
| EPI_ISL_2814175 | Lighthouse Lab in Glasgow | Wellcome Sanger Institute for the COVID-19 Genomics UK (COG-UK) Consortium | Harper VanSteenhouse, Yumi Kasai, David Gray, Carol Clugston, Anna Dominiczak and Alex Alderton, Roberto Amato, Jeffrey Barrett, Sonia Goncalves, Ewan Harrison, David K. Jackson, Ian Johnston, Dominic Kwiatkowski, Cordelia Langford, John Sillitoe on behalf of the Wellcome Sanger Institute COVID-19 Surveillance Team |
| EPI_ISL_2814176 | Lighthouse Lab in Milton Keynes | Wellcome Sanger Institute for the COVID-19 Genomics UK (COG-UK) Consortium | The Lighthouse Lab in Milton Keynes and Alex Alderton, Roberto Amato, Jeffrey Barrett, Sonia Goncalves, Ewan Harrison, David K. Jackson, Ian Johnston, Dominic Kwiatkowski, Cordelia Langford, John Sillitoe on behalf of the Wellcome Sanger Institute COVID-19 Surveillance Team |
| EPI_ISL_2814177, EPI_ISL_2814178 | Lighthouse Lab in Glasgow | Wellcome Sanger Institute for the COVID-19 Genomics UK (COG-UK) Consortium | Harper VanSteenhouse, Yumi Kasai, David Gray, Carol Clugston, Anna Dominiczak and Alex Alderton, Roberto Amato, Jeffrey Barrett, Sonia Goncalves, Ewan Harrison, David K. Jackson, Ian Johnston, Dominic Kwiatkowski, Cordelia Langford, John Sillitoe on behalf of the Wellcome Sanger Institute COVID-19 Surveillance Team |
| EPI_ISL_2814179 | Lighthouse Lab in Milton Keynes | Wellcome Sanger Institute for the COVID-19 Genomics UK (COG-UK) Consortium | The Lighthouse Lab in Milton Keynes and Alex Alderton, Roberto Amato, Jeffrey Barrett, Sonia Goncalves, Ewan Harrison, David K. Jackson, Ian Johnston, Dominic Kwiatkowski, Cordelia Langford, John Sillitoe on behalf of the Wellcome Sanger Institute COVID-19 Surveillance Team |
| EPI_ISL_2814180, EPI_ISL_2814181 | Lighthouse Lab in Alderley Park | Wellcome Sanger Institute for the COVID-19 Genomics UK (COG-UK) Consortium | Jacquelyn Wynn, Mairead Hyland, The Lighthouse Lab in Alderley Park and Alex Alderton, Roberto Amato, Jeffrey Barrett, Sonia Goncalves, Ewan Harrison, David K. Jackson, Ian Johnston, Dominic Kwiatkowski, Cordelia Langford, John Sillitoe on behalf of the Wellcome Sanger Institute COVID-19 Surveillance Team |
| EPI_ISL_2814182, EPI_ISL_2814183 | Lighthouse Lab in Glasgow | Wellcome Sanger Institute for the COVID-19 Genomics UK (COG-UK) Consortium | Harper VanSteenhouse, Yumi Kasai, David Gray, Carol Clugston, Anna Dominiczak and Alex Alderton, Roberto Amato, Jeffrey Barrett, Sonia Goncalves, Ewan Harrison, David K. Jackson, Ian Johnston, Dominic Kwiatkowski, Cordelia Langford, John Sillitoe on behalf of the Wellcome Sanger Institute COVID-19 Surveillance Team |
| EPI_ISL_2814184 | Lighthouse Lab in Milton Keynes | Wellcome Sanger Institute for the COVID-19 Genomics UK (COG-UK) Consortium | The Lighthouse Lab in Milton Keynes and Alex Alderton, Roberto Amato, Jeffrey Barrett, Sonia Goncalves, Ewan Harrison, David K. Jackson, Ian Johnston, Dominic Kwiatkowski, Cordelia Langford, John Sillitoe on behalf of the Wellcome Sanger Institute COVID-19 Surveillance Team |
| EPI_ISL_2814185, EPI_ISL_2814186, EPI_ISL_2814187, EPI_ISL_2814188, EPI_ISL_2814189 | Lighthouse Lab in Alderley Park | Wellcome Sanger Institute for the COVID-19 Genomics UK (COG-UK) Consortium | Jacquelyn Wynn, Mairead Hyland, The Lighthouse Lab in Alderley Park and Alex Alderton, Roberto Amato, Jeffrey Barrett, Sonia Goncalves, Ewan Harrison, David K. Jackson, Ian Johnston, Dominic Kwiatkowski, Cordelia Langford, John Sillitoe on behalf of the Wellcome Sanger Institute COVID-19 Surveillance Team |
| EPI_ISL_2814190, EPI_ISL_2814191 | Lighthouse Lab in Milton Keynes | Wellcome Sanger Institute for the COVID-19 Genomics UK (COG-UK) Consortium | The Lighthouse Lab in Milton Keynes and Alex Alderton, Roberto Amato, Jeffrey Barrett, Sonia Goncalves, Ewan Harrison, David K. Jackson, Ian Johnston, Dominic Kwiatkowski, Cordelia Langford, John Sillitoe on behalf of the Wellcome Sanger Institute COVID-19 Surveillance Team |
| EPI_ISL_2814192, EPI_ISL_2814193, EPI_ISL_2814194 | Lighthouse Lab in Alderley Park | Wellcome Sanger Institute for the COVID-19 Genomics UK (COG-UK) Consortium | Jacquelyn Wynn, Mairead Hyland, The Lighthouse Lab in Alderley Park and Alex Alderton, Roberto Amato, Jeffrey Barrett, Sonia Goncalves, Ewan Harrison, David K. Jackson, Ian Johnston, Dominic Kwiatkowski, Cordelia Langford, John Sillitoe on behalf of the Wellcome Sanger Institute COVID-19 Surveillance Team |
| EPI_ISL_2814195, EPI_ISL_2814196, EPI_ISL_2814197, EPI_ISL_2814198, EPI_ISL_2814199 | Lighthouse Lab in Milton Keynes | Wellcome Sanger Institute for the COVID-19 Genomics UK (COG-UK) Consortium | The Lighthouse Lab in Milton Keynes and Alex Alderton, Roberto Amato, Jeffrey Barrett, Sonia Goncalves, Ewan Harrison, David K. Jackson, Ian Johnston, Dominic Kwiatkowski, Cordelia Langford, John Sillitoe on behalf of the Wellcome Sanger Institute COVID-19 Surveillance Team |
| EPI_ISL_2814200 | Lighthouse Lab in Alderley Park | Wellcome Sanger Institute for the COVID-19 Genomics UK (COG-UK) Consortium | Jacquelyn Wynn, Mairead Hyland, The Lighthouse Lab in Alderley Park and Alex Alderton, Roberto Amato, Jeffrey Barrett, Sonia Goncalves, Ewan Harrison, David K. Jackson, Ian Johnston, Dominic Kwiatkowski, Cordelia Langford, John Sillitoe on behalf of the Wellcome Sanger Institute COVID-19 Surveillance Team |
| EPI_ISL_2814201 | Lighthouse Lab in Milton Keynes | Wellcome Sanger Institute for the COVID-19 Genomics UK (COG-UK) Consortium | The Lighthouse Lab in Milton Keynes and Alex Alderton, Roberto Amato, Jeffrey Barrett, Sonia Goncalves, Ewan Harrison, David K. Jackson, Ian Johnston, Dominic Kwiatkowski, Cordelia Langford, John Sillitoe on behalf of the Wellcome Sanger Institute COVID-19 Surveillance Team |
| EPI_ISL_2814202, EPI_ISL_2814203 | Lighthouse Lab in Alderley Park | Wellcome Sanger Institute for the COVID-19 Genomics UK (COG-UK) Consortium | Jacquelyn Wynn, Mairead Hyland, The Lighthouse Lab in Alderley Park and Alex Alderton, Roberto Amato, Jeffrey Barrett, Sonia Goncalves, Ewan Harrison, David K. Jackson, Ian Johnston, Dominic Kwiatkowski, Cordelia Langford, John Sillitoe on behalf of the Wellcome Sanger Institute COVID-19 Surveillance Team |
| EPI_ISL_2814204, EPI_ISL_2814205, EPI_ISL_2814206 | Lighthouse Lab in Glasgow | Wellcome Sanger Institute for the COVID-19 Genomics UK (COG-UK) Consortium | Harper VanSteenhouse, Yumi Kasai, David Gray, Carol Clugston, Anna Dominiczak and Alex Alderton, Roberto Amato, Jeffrey Barrett, Sonia Goncalves, Ewan Harrison, David K. Jackson, Ian Johnston, Dominic Kwiatkowski, Cordelia Langford, John Sillitoe on behalf of the Wellcome Sanger Institute COVID-19 Surveillance Team |
| EPI_ISL_2814207 | Lighthouse Lab in Alderley Park | Wellcome Sanger Institute for the COVID-19 Genomics UK (COG-UK) Consortium | Jacquelyn Wynn, Mairead Hyland, The Lighthouse Lab in Alderley Park and Alex Alderton, Roberto Amato, Jeffrey Barrett, Sonia Goncalves, Ewan Harrison, David K. Jackson, Ian Johnston, Dominic Kwiatkowski, Cordelia Langford, John Sillitoe on behalf of the Wellcome Sanger Institute COVID-19 Surveillance Team |
| EPI_ISL_2814208, EPI_ISL_2814209 | Lighthouse Lab in Milton Keynes | Wellcome Sanger Institute for the COVID-19 Genomics UK (COG-UK) Consortium | The Lighthouse Lab in Milton Keynes and Alex Alderton, Roberto Amato, Jeffrey Barrett, Sonia Goncalves, Ewan Harrison, David K. Jackson, Ian Johnston, Dominic Kwiatkowski, Cordelia Langford, John Sillitoe on behalf of the Wellcome Sanger Institute COVID-19 Surveillance Team |
| EPI_ISL_2814210, EPI_ISL_2814211 | Lighthouse Lab in Alderley Park | Wellcome Sanger Institute for the COVID-19 Genomics UK (COG-UK) Consortium | Jacquelyn Wynn, Mairead Hyland, The Lighthouse Lab in Alderley Park and Alex Alderton, Roberto Amato, Jeffrey Barrett, Sonia Goncalves, Ewan Harrison, David K. Jackson, Ian Johnston, Dominic Kwiatkowski, Cordelia Langford, John Sillitoe on behalf of the Wellcome Sanger Institute COVID-19 Surveillance Team |
| EPI_ISL_2814212 | Lighthouse Lab in Milton Keynes | Wellcome Sanger Institute for the COVID-19 Genomics UK (COG-UK) Consortium | The Lighthouse Lab in Milton Keynes and Alex Alderton, Roberto Amato, Jeffrey Barrett, Sonia Goncalves, Ewan Harrison, David K. Jackson, Ian Johnston, Dominic Kwiatkowski, Cordelia Langford, John Sillitoe on behalf of the Wellcome Sanger Institute COVID-19 Surveillance Team |

| | | | |
|---|---|---|---|
| EPI_ISL_2814213 | Lighthouse Lab in Glasgow | Wellcome Sanger Institute for the COVID-19 Genomics UK (COG-UK) Consortium | Harper VanSteenhouse, Yumi Kasai, David Gray, Carol Clugston, Anna Dominiczak and Alex Alderton, Roberto Amato, Jeffrey Barrett, Sonia Goncalves, Ewan Harrison, David K. Jackson, Ian Johnston, Dominic Kwiatkowski, Cordelia Langford, John Sillitoe on behalf of the Wellcome Sanger Institute COVID-19 Surveillance Team |
| EPI_ISL_2814214 | Lighthouse Lab in Milton Keynes | Wellcome Sanger Institute for the COVID-19 Genomics UK (COG-UK) Consortium | The Lighthouse Lab in Milton Keynes and Alex Alderton, Roberto Amato, Jeffrey Barrett, Sonia Goncalves, Ewan Harrison, David K. Jackson, Ian Johnston, Dominic Kwiatkowski, Cordelia Langford, John Sillitoe on behalf of the Wellcome Sanger Institute COVID-19 Surveillance Team |
| EPI_ISL_2814215, EPI_ISL_2814216 | Lighthouse Lab in Alderley Park | Wellcome Sanger Institute for the COVID-19 Genomics UK (COG-UK) Consortium | Jacquelyn Wynn, Mairead Hyland, The Lighthouse Lab in Alderley Park and Alex Alderton, Roberto Amato, Jeffrey Barrett, Sonia Goncalves, Ewan Harrison, David K. Jackson, Ian Johnston, Dominic Kwiatkowski, Cordelia Langford, John Sillitoe on behalf of the Wellcome Sanger Institute COVID-19 Surveillance Team |
| EPI_ISL_2814217 | Lighthouse Lab in Milton Keynes | Wellcome Sanger Institute for the COVID-19 Genomics UK (COG-UK) Consortium | The Lighthouse Lab in Milton Keynes and Alex Alderton, Roberto Amato, Jeffrey Barrett, Sonia Goncalves, Ewan Harrison, David K. Jackson, Ian Johnston, Dominic Kwiatkowski, Cordelia Langford, John Sillitoe on behalf of the Wellcome Sanger Institute COVID-19 Surveillance Team |
| EPI_ISL_2814218 | Lighthouse Lab in Glasgow | Wellcome Sanger Institute for the COVID-19 Genomics UK (COG-UK) Consortium | Harper VanSteenhouse, Yumi Kasai, David Gray, Carol Clugston, Anna Dominiczak and Alex Alderton, Roberto Amato, Jeffrey Barrett, Sonia Goncalves, Ewan Harrison, David K. Jackson, Ian Johnston, Dominic Kwiatkowski, Cordelia Langford, John Sillitoe on behalf of the Wellcome Sanger Institute COVID-19 Surveillance Team |
| EPI_ISL_2814219 | Lighthouse Lab in Alderley Park | Wellcome Sanger Institute for the COVID-19 Genomics UK (COG-UK) Consortium | Jacquelyn Wynn, Mairead Hyland, The Lighthouse Lab in Alderley Park and Alex Alderton, Roberto Amato, Jeffrey Barrett, Sonia Goncalves, Ewan Harrison, David K. Jackson, Ian Johnston, Dominic Kwiatkowski, Cordelia Langford, John Sillitoe on behalf of the Wellcome Sanger Institute COVID-19 Surveillance Team |
| EPI_ISL_2814220 | Lighthouse Lab in Glasgow | Wellcome Sanger Institute for the COVID-19 Genomics UK (COG-UK) Consortium | Harper VanSteenhouse, Yumi Kasai, David Gray, Carol Clugston, Anna Dominiczak and Alex Alderton, Roberto Amato, Jeffrey Barrett, Sonia Goncalves, Ewan Harrison, David K. Jackson, Ian Johnston, Dominic Kwiatkowski, Cordelia Langford, John Sillitoe on behalf of the Wellcome Sanger Institute COVID-19 Surveillance Team |
| EPI_ISL_2814221, EPI_ISL_2814222, EPI_ISL_2814223 | Lighthouse Lab in Milton Keynes | Wellcome Sanger Institute for the COVID-19 Genomics UK (COG-UK) Consortium | The Lighthouse Lab in Milton Keynes and Alex Alderton, Roberto Amato, Jeffrey Barrett, Sonia Goncalves, Ewan Harrison, David K. Jackson, Ian Johnston, Dominic Kwiatkowski, Cordelia Langford, John Sillitoe on behalf of the Wellcome Sanger Institute COVID-19 Surveillance Team |
| EPI_ISL_2814224, EPI_ISL_2814225 | Lighthouse Lab in Alderley Park | Wellcome Sanger Institute for the COVID-19 Genomics UK (COG-UK) Consortium | Jacquelyn Wynn, Mairead Hyland, The Lighthouse Lab in Alderley Park and Alex Alderton, Roberto Amato, Jeffrey Barrett, Sonia Goncalves, Ewan Harrison, David K. Jackson, Ian Johnston, Dominic Kwiatkowski, Cordelia Langford, John Sillitoe on behalf of the Wellcome Sanger Institute COVID-19 Surveillance Team |
| EPI_ISL_2814226 | Lighthouse Lab in Milton Keynes | Wellcome Sanger Institute for the COVID-19 Genomics UK (COG-UK) Consortium | The Lighthouse Lab in Milton Keynes and Alex Alderton, Roberto Amato, Jeffrey Barrett, Sonia Goncalves, Ewan Harrison, David K. Jackson, Ian Johnston, Dominic Kwiatkowski, Cordelia Langford, John Sillitoe on behalf of the Wellcome Sanger Institute COVID-19 Surveillance Team |
| EPI_ISL_2814227, EPI_ISL_2814228, EPI_ISL_2814229, EPI_ISL_2814230, EPI_ISL_2814231, EPI_ISL_2814232, EPI_ISL_2814233, EPI_ISL_2814234, EPI_ISL_2814235, EPI_ISL_2814236, EPI_ISL_2814237, EPI_ISL_2814238, EPI_ISL_2814239, EPI_ISL_2814240, EPI_ISL_2814241, EPI_ISL_2814242, EPI_ISL_2814243, EPI_ISL_2814244, EPI_ISL_2814245, EPI_ISL_2814246, EPI_ISL_2814247, EPI_ISL_2814248, EPI_ISL_2814249, EPI_ISL_2814250, EPI_ISL_2814251, EPI_ISL_2814252, EPI_ISL_2814253, EPI_ISL_2814254, EPI_ISL_2814255, EPI_ISL_2814256, EPI_ISL_2814257, EPI_ISL_2814258, EPI_ISL_2814259, EPI_ISL_2814260, EPI_ISL_2814261, EPI_ISL_2814262, EPI_ISL_2814263, EPI_ISL_2814264, EPI_ISL_2814265, EPI_ISL_2814266, EPI_ISL_2814267, EPI_ISL_2814268, EPI_ISL_2814269, EPI_ISL_2814270, EPI_ISL_2814271, EPI_ISL_2814272, EPI_ISL_2814273, EPI_ISL_2814274, EPI_ISL_2814275, EPI_ISL_2814276, EPI_ISL_2814277, EPI_ISL_2814278, EPI_ISL_2814279, EPI_ISL_2814280, EPI_ISL_2814281, EPI_ISL_2814282, EPI_ISL_2814283, EPI_ISL_2814284, EPI_ISL_2814285, EPI_ISL_2814286, EPI_ISL_2814287, EPI_ISL_2814288, EPI_ISL_2814289, EPI_ISL_2814290, EPI_ISL_2814291, EPI_ISL_2814292, EPI_ISL_2814293, EPI_ISL_2814294, EPI_ISL_2814295, EPI_ISL_2814296, EPI_ISL_2814297, EPI_ISL_2814298, EPI_ISL_2814299, EPI_ISL_2814300, EPI_ISL_2814301, EPI_ISL_2814302, EPI_ISL_2814303, EPI_ISL_2814304, EPI_ISL_2814305, EPI_ISL_2814306, EPI_ISL_2814307, EPI_ISL_2814308, EPI_ISL_2814309, EPI_ISL_2814310, EPI_ISL_2814311, EPI_ISL_2814312, EPI_ISL_2814313, EPI_ISL_2814314, EPI_ISL_2814315, EPI_ISL_2814316, EPI_ISL_2814317, EPI_ISL_2814318, EPI_ISL_2814319, EPI_ISL_2814320, EPI_ISL_2814321, EPI_ISL_2814323, EPI_ISL_2814324, EPI_ISL_2814325, EPI_ISL_2814326, EPI_ISL_2814327, EPI_ISL_2814328, EPI_ISL_2814329, EPI_ISL_2814330, EPI_ISL_2814331, EPI_ISL_2814332, EPI_ISL_2814333, EPI_ISL_2814334, EPI_ISL_2814336, EPI_ISL_2814337, EPI_ISL_2814338, EPI_ISL_2814339, EPI_ISL_2814340, EPI_ISL_2814341, EPI_ISL_2814342, EPI_ISL_2814343, EPI_ISL_2814344, EPI_ISL_2814345, EPI_ISL_2814346, EPI_ISL_2814347, EPI_ISL_2814348, EPI_ISL_2814349, EPI_ISL_2814350, EPI_ISL_2814351, EPI_ISL_2814352, EPI_ISL_2814353, EPI_ISL_2814354, EPI_ISL_2814355, EPI_ISL_2814356, EPI_ISL_2814357, EPI_ISL_2814358, EPI_ISL_2814359, EPI_ISL_2814360, EPI_ISL_2814361, EPI_ISL_2814362, EPI_ISL_2814363, EPI_ISL_2814364, EPI_ISL_2814365, EPI_ISL_2814366, EPI_ISL_2814367, EPI_ISL_2814368, EPI_ISL_2814369, EPI_ISL_2814370, EPI_ISL_2814372, EPI_ISL_2814373, EPI_ISL_2814374, EPI_ISL_2814375, EPI_ISL_2814376, EPI_ISL_2814377, EPI_ISL_2814378, EPI_ISL_2814379, EPI_ISL_2814380, EPI_ISL_2814381, EPI_ISL_2814382, EPI_ISL_2814383, EPI_ISL_2814384, EPI_ISL_2814385, EPI_ISL_2814386, EPI_ISL_2814387, EPI_ISL_2814388, EPI_ISL_2814389, EPI_ISL_2814390, EPI_ISL_2814391, EPI_ISL_2814392, EPI_ISL_2814393, EPI_ISL_2814394, EPI_ISL_2814395, EPI_ISL_2814396, EPI_ISL_2814397, EPI_ISL_2814398, EPI_ISL_2814399, EPI_ISL_2814400, EPI_ISL_2814401, EPI_ISL_2814402, EPI_ISL_2814403, EPI_ISL_2814404, EPI_ISL_2814405, EPI_ISL_2814406, EPI_ISL_2814407, EPI_ISL_2814408, EPI_ISL_2814409, EPI_ISL_2814410, EPI_ISL_2814411, EPI_ISL_2814412, EPI_ISL_2814413, EPI_ISL_2814414, EPI_ISL_2814416, EPI_ISL_2814417, EPI_ISL_2814418, EPI_ISL_2814419, EPI_ISL_2814420, EPI_ISL_2814421, EPI_ISL_2814422, EPI_ISL_2814423 | | | |
| see above | Lighthouse Lab in Glasgow | Wellcome Sanger Institute for the COVID-19 Genomics UK (COG-UK) Consortium | Harper VanSteenhouse, Yumi Kasai, David Gray, Carol Clugston, Anna Dominiczak and Alex Alderton, Roberto Amato, Jeffrey Barrett, Sonia Goncalves, Ewan Harrison, David K. Jackson, Ian Johnston, Dominic Kwiatkowski, Cordelia Langford, John Sillitoe on behalf of the Wellcome Sanger Institute COVID-19 Surveillance Team |
| EPI_ISL_2814424 | Lighthouse Laboratory Plymouth | Wellcome Sanger Institute for the COVID-19 Genomics UK (COG-UK) Consortium | Lighthouse Laboratory Plymouth and Alex Alderton, Roberto Amato, Jeffrey Barrett, Sonia Goncalves, Ewan Harrison, David K. Jackson, Ian Johnston, Dominic Kwiatkowski, Cordelia Langford, John Sillitoe on behalf of the Wellcome Sanger Institute COVID-19 Surveillance Team |
| EPI_ISL_2814425, EPI_ISL_2814426, EPI_ISL_2814427, EPI_ISL_2814428, EPI_ISL_2814429, EPI_ISL_2814430, EPI_ISL_2814431, EPI_ISL_2814432, EPI_ISL_2814433, EPI_ISL_2814434, EPI_ISL_2814435, EPI_ISL_2814436, EPI_ISL_2814437, EPI_ISL_2814438, EPI_ISL_2814439, EPI_ISL_2814440, EPI_ISL_2814441, EPI_ISL_2814442, EPI_ISL_2814443, EPI_ISL_2814444, EPI_ISL_2814445, EPI_ISL_2814446, EPI_ISL_2814447, EPI_ISL_2814448, EPI_ISL_2814449, EPI_ISL_2814450, EPI_ISL_2814451, EPI_ISL_2814452, EPI_ISL_2814453, EPI_ISL_2814454, EPI_ISL_2814455, EPI_ISL_2814456, EPI_ISL_2814457, EPI_ISL_2814458, EPI_ISL_2814459, EPI_ISL_2814460, EPI_ISL_2814461, EPI_ISL_2814462, EPI_ISL_2814463, EPI_ISL_2814464, EPI_ISL_2814465, EPI_ISL_2814466, EPI_ISL_2814467, EPI_ISL_2814468, EPI_ISL_2814469, EPI_ISL_2814470, EPI_ISL_2814471, EPI_ISL_2814472, EPI_ISL_2814473, EPI_ISL_2814474, EPI_ISL_2814475, EPI_ISL_2814476, EPI_ISL_2814477, EPI_ISL_2814478, EPI_ISL_2814479, EPI_ISL_2814480, EPI_ISL_2814481, EPI_ISL_2814483, EPI_ISL_2814484, EPI_ISL_2814485, EPI_ISL_2814486, EPI_ISL_2814487, EPI_ISL_2814488, EPI_ISL_2814489, EPI_ISL_2814490, EPI_ISL_2814491, EPI_ISL_2814492, EPI_ISL_2814493, EPI_ISL_2814494, EPI_ISL_2814495, EPI_ISL_2814496, EPI_ISL_2814497, EPI_ISL_2814498, EPI_ISL_2814499, EPI_ISL_2814500, EPI_ISL_2814501, EPI_ISL_2814502, EPI_ISL_2814503, EPI_ISL_2814504, EPI_ISL_2814505, EPI_ISL_2814506, EPI_ISL_2814507, EPI_ISL_2814508, EPI_ISL_2814509, EPI_ISL_2814510, EPI_ISL_2814511, EPI_ISL_2814512, EPI_ISL_2814513, EPI_ISL_2814514, EPI_ISL_2814515, EPI_ISL_2814516, EPI_ISL_2814517, EPI_ISL_2814518, EPI_ISL_2814519, EPI_ISL_2814520, EPI_ISL_2814521, EPI_ISL_2814522, EPI_ISL_2814523, EPI_ISL_2814524, EPI_ISL_2814525, EPI_ISL_2814526, EPI_ISL_2814527, EPI_ISL_2814528, EPI_ISL_2814529, EPI_ISL_2814530, EPI_ISL_2814531, EPI_ISL_2814532, EPI_ISL_2814533, EPI_ISL_2814534, EPI_ISL_2814535, EPI_ISL_2814536, EPI_ISL_2814537, EPI_ISL_2814538, EPI_ISL_2814539, EPI_ISL_2814540, EPI_ISL_2814541, EPI_ISL_2814542, EPI_ISL_2814543, EPI_ISL_2814544, EPI_ISL_2814545, EPI_ISL_2814546, EPI_ISL_2814547, EPI_ISL_2814548, EPI_ISL_2814549, EPI_ISL_2814551, EPI_ISL_2814552, EPI_ISL_2814553, EPI_ISL_2814554, EPI_ISL_2814555, EPI_ISL_2814557, EPI_ISL_2814558, EPI_ISL_2814559, EPI_ISL_2814560, EPI_ISL_2814561, EPI_ISL_2814562, EPI_ISL_2814563, EPI_ISL_2814564, EPI_ISL_2814565, EPI_ISL_2814566, EPI_ISL_2814567, EPI_ISL_2814568, EPI_ISL_2814569, EPI_ISL_2814570, EPI_ISL_2814571, EPI_ISL_2814572, EPI_ISL_2814573, EPI_ISL_2814574, EPI_ISL_2814575, EPI_ISL_2814576, EPI_ISL_2814577, EPI_ISL_2814578, EPI_ISL_2814579, EPI_ISL_2814580, EPI_ISL_2814581, EPI_ISL_2814582, EPI_ISL_2814583, EPI_ISL_2814584, EPI_ISL_2814585, EPI_ISL_2814586, EPI_ISL_2814587, EPI_ISL_2814588, EPI_ISL_2814589, EPI_ISL_2814590, EPI_ISL_2814591, EPI_ISL_2814592, EPI_ISL_2814593, EPI_ISL_2814594, EPI_ISL_2814595, EPI_ISL_2814596, EPI_ISL_2814597, EPI_ISL_2814598, EPI_ISL_2814599, EPI_ISL_2814600, EPI_ISL_2814601, EPI_ISL_2814602, EPI_ISL_2814603, EPI_ISL_2814604, EPI_ISL_2814605, EPI_ISL_2814606, EPI_ISL_2814607, EPI_ISL_2814608, EPI_ISL_2814609, EPI_ISL_2814610, EPI_ISL_2814611, EPI_ISL_2814612, EPI_ISL_2814613, EPI_ISL_2814614, EPI_ISL_2814615, EPI_ISL_2814616, EPI_ISL_2814617, EPI_ISL_2814618, EPI_ISL_2814619, EPI_ISL_2814620, EPI_ISL_2814621, EPI_ISL_2814622, EPI_ISL_2814623, EPI_ISL_2814624, EPI_ISL_2814625, EPI_ISL_2814626, EPI_ISL_2814627, EPI_ISL_2814628, EPI_ISL_2814629, EPI_ISL_2814630, EPI_ISL_2814631, EPI_ISL_2814632, EPI_ISL_2814633, EPI_ISL_2814635, EPI_ISL_2814637, EPI_ISL_2814638, EPI_ISL_2814639, EPI_ISL_2814640, EPI_ISL_2814641, EPI_ISL_2814642, EPI_ISL_2814643, EPI_ISL_2814644, EPI_ISL_2814645, EPI_ISL_2814646, EPI_ISL_2814647, EPI_ISL_2814648, EPI_ISL_2814649, EPI_ISL_2814650, EPI_ISL_2814651, EPI_ISL_2814652, EPI_ISL_2814653, EPI_ISL_2814654, EPI_ISL_2814655, EPI_ISL_2814656, EPI_ISL_2814657, EPI_ISL_2814658, EPI_ISL_2814659, EPI_ISL_2814660, EPI_ISL_2814661, EPI_ISL_2814662, EPI_ISL_2814663, EPI_ISL_2814664, EPI_ISL_2814665, EPI_ISL_2814666, EPI_ISL_2814667, EPI_ISL_2814668, EPI_ISL_2814669, EPI_ISL_2814671, EPI_ISL_2814672, EPI_ISL_2814673, EPI_ISL_2814674, EPI_ISL_2814675, EPI_ISL_2814676, EPI_ISL_2814677, EPI_ISL_2814678, EPI_ISL_2814679, EPI_ISL_2814680, EPI_ISL_2814681, EPI_ISL_2814682, EPI_ISL_2814683, EPI_ISL_2814684, EPI_ISL_2814685, EPI_ISL_2814686, EPI_ISL_2814687, EPI_ISL_2814688, EPI_ISL_2814689, EPI_ISL_2814690, EPI_ISL_2814691, EPI_ISL_2814692, EPI_ISL_2814693, EPI_ISL_2814694, EPI_ISL_2814695, EPI_ISL_2814696, EPI_ISL_2814697, EPI_ISL_2814698, EPI_ISL_2814699, EPI_ISL_2814700, EPI_ISL_2814701, EPI_ISL_2814702, EPI_ISL_2814703, EPI_ISL_2814704, EPI_ISL_2814705, EPI_ISL_2814706, EPI_ISL_2814707, EPI_ISL_2814708, EPI_ISL_2814709, EPI_ISL_2814710, EPI_ISL_2814711, EPI_ISL_2814712, EPI_ISL_2814713, EPI_ISL_2814714, EPI_ISL_2814715, EPI_ISL_2814716, EPI_ISL_2814717, EPI_ISL_2814718, EPI_ISL_2814719, EPI_ISL_2814720, EPI_ISL_2814721, EPI_ISL_2814722, EPI_ISL_2814723, EPI_ISL_2814724, EPI_ISL_2814725, EPI_ISL_2814726, EPI_ISL_2814727, EPI_ISL_2814728, EPI_ISL_2814729 | | | |
| see above | Lighthouse Lab in Glasgow | Wellcome Sanger Institute for the COVID-19 Genomics UK (COG-UK) Consortium | Harper VanSteenhouse, Yumi Kasai, David Gray, Carol Clugston, Anna Dominiczak and Alex Alderton, Roberto Amato, Jeffrey Barrett, Sonia Goncalves, Ewan Harrison, David K. Jackson, Ian Johnston, Dominic Kwiatkowski, Cordelia Langford, John Sillitoe on behalf of the Wellcome Sanger Institute COVID-19 Surveillance Team |
| EPI_ISL_2814730 | Lighthouse Lab in Alderley Park | Wellcome Sanger Institute for the COVID-19 Genomics UK (COG-UK) Consortium | Jacquelyn Wynn, Mairead Hyland, The Lighthouse Lab in Alderley Park and Alex Alderton, Roberto Amato, Jeffrey Barrett, Sonia Goncalves, Ewan Harrison, David K. Jackson, Ian Johnston, Dominic Kwiatkowski, Cordelia Langford, John Sillitoe on behalf of the Wellcome Sanger Institute COVID-19 Surveillance Team |
| EPI_ISL_2814731, EPI_ISL_2814732, EPI_ISL_2814733, EPI_ISL_2814734, EPI_ISL_2814735, EPI_ISL_2814736, EPI_ISL_2814737, EPI_ISL_2814738, EPI_ISL_2814739, EPI_ISL_2814740, EPI_ISL_2814741, EPI_ISL_2814742, EPI_ISL_2814743, EPI_ISL_2814744, EPI_ISL_2814745, EPI_ISL_2814746, EPI_ISL_2814747, EPI_ISL_2814748, EPI_ISL_2814749, EPI_ISL_2814750, EPI_ISL_2814751, EPI_ISL_2814752, EPI_ISL_2814753, EPI_ISL_2814755, EPI_ISL_2814756, EPI_ISL_2814757, EPI_ISL_2814758, EPI_ISL_2814759, EPI_ISL_2814760, EPI_ISL_2814761, EPI_ISL_2814762, EPI_ISL_2814763, EPI_ISL_2814764, EPI_ISL_2814765, EPI_ISL_2814766, EPI_ISL_2814767 | | | |
| see above | Lighthouse Lab in Glasgow | Wellcome Sanger Institute for the COVID-19 Genomics UK (COG-UK) Consortium | Harper VanSteenhouse, Yumi Kasai, David Gray, Carol Clugston, Anna Dominiczak and Alex Alderton, Roberto Amato, Jeffrey Barrett, Sonia Goncalves, Ewan Harrison, David K. Jackson, Ian Johnston, Dominic Kwiatkowski, Cordelia Langford, John Sillitoe on behalf of the Wellcome Sanger Institute COVID-19 Surveillance Team |
| EPI_ISL_2814768 | Lighthouse Lab in Milton Keynes | Wellcome Sanger Institute for the COVID-19 Genomics UK (COG-UK) Consortium | The Lighthouse Lab in Milton Keynes and Alex Alderton, Roberto Amato, Jeffrey Barrett, Sonia Goncalves, Ewan Harrison, David K. Jackson, Ian Johnston, Dominic Kwiatkowski, Cordelia Langford, John Sillitoe on behalf of the Wellcome Sanger Institute COVID-19 Surveillance Team |
| EPI_ISL_2814769, EPI_ISL_2814770, EPI_ISL_2814771, EPI_ISL_2814772, EPI_ISL_2814774, EPI_ISL_2814775, EPI_ISL_2814776, EPI_ISL_2814777, EPI_ISL_2814778, EPI_ISL_2814779, EPI_ISL_2814780, EPI_ISL_2814781, EPI_ISL_2814782, EPI_ISL_2814783, EPI_ISL_2814784, EPI_ISL_2814785, EPI_ISL_2814786, EPI_ISL_2814787, EPI_ISL_2814788, EPI_ISL_2814789, EPI_ISL_2814790 | | | |
| see above | Lighthouse Lab in Glasgow | Wellcome Sanger Institute for the COVID-19 Genomics UK (COG-UK) Consortium | Harper VanSteenhouse, Yumi Kasai, David Gray, Carol Clugston, Anna Dominiczak and Alex Alderton, Roberto Amato, Jeffrey Barrett, Sonia Goncalves, Ewan Harrison, David K. Jackson, Ian Johnston, Dominic Kwiatkowski, Cordelia Langford, John Sillitoe on behalf of the Wellcome Sanger Institute COVID-19 Surveillance Team |
| EPI_ISL_2814791 | Lighthouse Lab in Alderley Park | Wellcome Sanger Institute for the COVID-19 Genomics UK (COG-UK) Consortium | Jacquelyn Wynn, Mairead Hyland, The Lighthouse Lab in Alderley Park and Alex Alderton, Roberto Amato, Jeffrey Barrett, Sonia Goncalves, Ewan Harrison, David K. Jackson, Ian Johnston, Dominic Kwiatkowski, Cordelia Langford, John Sillitoe on behalf of the Wellcome Sanger Institute COVID-19 Surveillance Team |
| EPI_ISL_2814792, EPI_ISL_2814793, EPI_ISL_2814794, EPI_ISL_2814795, EPI_ISL_2814796, EPI_ISL_2814797, EPI_ISL_2814798, EPI_ISL_2814799, EPI_ISL_2814800, EPI_ISL_2814801 | Lighthouse Lab in Glasgow | Wellcome Sanger Institute for the COVID-19 Genomics UK (COG-UK) Consortium | Harper VanSteenhouse, Yumi Kasai, David Gray, Carol Clugston, Anna Dominiczak and Alex Alderton, Roberto Amato, Jeffrey Barrett, Sonia Goncalves, Ewan Harrison, David K. Jackson, Ian Johnston, Dominic Kwiatkowski, Cordelia Langford, John Sillitoe on behalf of the Wellcome Sanger Institute COVID-19 Surveillance Team |
| EPI_ISL_2814802 | Lighthouse Lab in Alderley Park | Wellcome Sanger Institute for the COVID-19 Genomics UK (COG-UK) Consortium | Jacquelyn Wynn, Mairead Hyland, The Lighthouse Lab in Alderley Park and Alex Alderton, Roberto Amato, Jeffrey Barrett, Sonia Goncalves, Ewan Harrison, David K. Jackson, Ian Johnston, Dominic Kwiatkowski, Cordelia Langford, John Sillitoe on behalf of the Wellcome Sanger Institute COVID-19 Surveillance Team |
| EPI_ISL_2814803, EPI_ISL_2814804, EPI_ISL_2814805, EPI_ISL_2814806, EPI_ISL_2814807, EPI_ISL_2814808, EPI_ISL_2814809, EPI_ISL_2814810 | Lighthouse Lab in Glasgow | Wellcome Sanger Institute for the COVID-19 Genomics UK (COG-UK) Consortium | Harper VanSteenhouse, Yumi Kasai, David Gray, Carol Clugston, Anna Dominiczak and Alex Alderton, Roberto Amato, Jeffrey Barrett, Sonia Goncalves, Ewan Harrison, David K. Jackson, Ian Johnston, Dominic Kwiatkowski, Cordelia Langford, John Sillitoe on behalf of the Wellcome Sanger Institute COVID-19 Surveillance Team |
| EPI_ISL_2814811 | Lighthouse Lab in Alderley Park | Wellcome Sanger Institute for the COVID-19 Genomics UK (COG-UK) Consortium | Jacquelyn Wynn, Mairead Hyland, The Lighthouse Lab in Alderley Park and Alex Alderton, Roberto Amato, Jeffrey Barrett, Sonia Goncalves, Ewan Harrison, David K. Jackson, Ian Johnston, Dominic Kwiatkowski, Cordelia Langford, John Sillitoe on behalf of the Wellcome Sanger Institute COVID-19 Surveillance Team |
| EPI_ISL_2814812, EPI_ISL_2814813 | Lighthouse Lab in Glasgow | Wellcome Sanger Institute for the COVID-19 Genomics UK (COG-UK) Consortium | Harper VanSteenhouse, Yumi Kasai, David Gray, Carol Clugston, Anna Dominiczak and Alex Alderton, Roberto Amato, Jeffrey Barrett, Sonia Goncalves, Ewan Harrison, David K. Jackson, Ian Johnston, Dominic Kwiatkowski, Cordelia Langford, John Sillitoe on behalf of the Wellcome Sanger Institute COVID-19 Surveillance Team |
| EPI_ISL_2814814 | Lighthouse Lab in Alderley Park | Wellcome Sanger Institute for the COVID-19 Genomics UK (COG-UK) Consortium | Jacquelyn Wynn, Mairead Hyland, The Lighthouse Lab in Alderley Park and Alex Alderton, Roberto Amato, Jeffrey Barrett, Sonia Goncalves, Ewan Harrison, David K. Jackson, Ian Johnston, Dominic Kwiatkowski, Cordelia Langford, John Sillitoe on behalf of the Wellcome Sanger Institute COVID-19 Surveillance Team |
| EPI_ISL_2814815 | Lighthouse Lab in Glasgow | Wellcome Sanger Institute for the COVID-19 Genomics UK (COG-UK) Consortium | Harper VanSteenhouse, Yumi Kasai, David Gray, Carol Clugston, Anna Dominiczak and Alex Alderton, Roberto Amato, Jeffrey Barrett, Sonia Goncalves, Ewan Harrison, David K. Jackson, Ian Johnston, Dominic Kwiatkowski, Cordelia Langford, John Sillitoe on behalf of the Wellcome Sanger Institute COVID-19 Surveillance Team |
| EPI_ISL_2814816 | Lighthouse Lab in Alderley Park | Wellcome Sanger Institute for the COVID-19 Genomics UK (COG-UK) Consortium | Jacquelyn Wynn, Mairead Hyland, The Lighthouse Lab in Alderley Park and Alex Alderton, Roberto Amato, Jeffrey Barrett, Sonia Goncalves, Ewan Harrison, David K. Jackson, Ian Johnston, Dominic Kwiatkowski, Cordelia Langford, John Sillitoe on behalf of the Wellcome Sanger Institute COVID-19 Surveillance Team |
| EPI_ISL_2814817, EPI_ISL_2814818, EPI_ISL_2814819, EPI_ISL_2814820, EPI_ISL_2814821, EPI_ISL_2814822, EPI_ISL_2814823, EPI_ISL_2814824, EPI_ISL_2814825, EPI_ISL_2814826, EPI_ISL_2814827, EPI_ISL_2814828, EPI_ISL_2814829, EPI_ISL_2814830, EPI_ISL_2814831, EPI_ISL_2814832, EPI_ISL_2814833 | | | |
| see above | Lighthouse Lab in Glasgow | Wellcome Sanger Institute for the COVID-19 Genomics UK (COG-UK) | Harper VanSteenhouse, Yumi Kasai, David Gray, Carol Clugston, Anna Dominiczak and Alex Alderton, Roberto Amato, Jeffrey Barrett, Sonia Goncalves, Ewan Harrison, David K. Jackson, Ian Johnston, Dominic Kwiatkowski, Cordelia Langford, |

| | | | |
|---|---|---|---|
| | | Consortium | John Sillitoe on behalf of the Wellcome Sanger Institute COVID-19 Surveillance Team |
| EPI_ISL_2814834 | Lighthouse Lab in Alderley Park | Wellcome Sanger Institute for the COVID-19 Genomics UK (COG-UK) Consortium | Jacquelyn Wynn, Mairead Hyland, The Lighthouse Lab in Alderley Park and Alex Alderton, Roberto Amato, Jeffrey Barrett, Sonia Goncalves, Ewan Harrison, David K. Jackson, Ian Johnston, Dominic Kwiatkowski, Cordelia Langford, John Sillitoe on behalf of the Wellcome Sanger Institute COVID-19 Surveillance Team |
| EPI_ISL_2814835, EPI_ISL_2814836, EPI_ISL_2814837, EPI_ISL_2814838, EPI_ISL_2814839, EPI_ISL_2814840, EPI_ISL_2814841, EPI_ISL_2814842 | Lighthouse Lab in Glasgow | Wellcome Sanger Institute for the COVID-19 Genomics UK (COG-UK) Consortium | Harper VanSteenhouse, Yumi Kasai, David Gray, Carol Clugston, Anna Dominiczak and Alex Alderton, Roberto Amato, Jeffrey Barrett, Sonia Goncalves, Ewan Harrison, David K. Jackson, Ian Johnston, Dominic Kwiatkowski, Cordelia Langford, John Sillitoe on behalf of the Wellcome Sanger Institute COVID-19 Surveillance Team |
| EPI_ISL_2814843 | Lighthouse Lab in Alderley Park | Wellcome Sanger Institute for the COVID-19 Genomics UK (COG-UK) Consortium | Jacquelyn Wynn, Mairead Hyland, The Lighthouse Lab in Alderley Park and Alex Alderton, Roberto Amato, Jeffrey Barrett, Sonia Goncalves, Ewan Harrison, David K. Jackson, Ian Johnston, Dominic Kwiatkowski, Cordelia Langford, John Sillitoe on behalf of the Wellcome Sanger Institute COVID-19 Surveillance Team |
| EPI_ISL_2814844 | Lighthouse Lab in Glasgow | Wellcome Sanger Institute for the COVID-19 Genomics UK (COG-UK) Consortium | Harper VanSteenhouse, Yumi Kasai, David Gray, Carol Clugston, Anna Dominiczak and Alex Alderton, Roberto Amato, Jeffrey Barrett, Sonia Goncalves, Ewan Harrison, David K. Jackson, Ian Johnston, Dominic Kwiatkowski, Cordelia Langford, John Sillitoe on behalf of the Wellcome Sanger Institute COVID-19 Surveillance Team |
| EPI_ISL_2814845 | Lighthouse Lab in Alderley Park | Wellcome Sanger Institute for the COVID-19 Genomics UK (COG-UK) Consortium | Jacquelyn Wynn, Mairead Hyland, The Lighthouse Lab in Alderley Park and Alex Alderton, Roberto Amato, Jeffrey Barrett, Sonia Goncalves, Ewan Harrison, David K. Jackson, Ian Johnston, Dominic Kwiatkowski, Cordelia Langford, John Sillitoe on behalf of the Wellcome Sanger Institute COVID-19 Surveillance Team |
| EPI_ISL_2814846, EPI_ISL_2814847, EPI_ISL_2814848, EPI_ISL_2814849, EPI_ISL_2814850, EPI_ISL_2814851, EPI_ISL_2814852, EPI_ISL_2814853, EPI_ISL_2814854, EPI_ISL_2814856, EPI_ISL_2814857, EPI_ISL_2814858, EPI_ISL_2814859, EPI_ISL_2814860, EPI_ISL_2814861, EPI_ISL_2814862, EPI_ISL_2814863, EPI_ISL_2814864, EPI_ISL_2814865, EPI_ISL_2814866, EPI_ISL_2814867, EPI_ISL_2814868, EPI_ISL_2814869, EPI_ISL_2814870, EPI_ISL_2814871, EPI_ISL_2814872, EPI_ISL_2814873, EPI_ISL_2814874, EPI_ISL_2814875, EPI_ISL_2814876 | | | |
| see above | Lighthouse Lab in Glasgow | Wellcome Sanger Institute for the COVID-19 Genomics UK (COG-UK) Consortium | Harper VanSteenhouse, Yumi Kasai, David Gray, Carol Clugston, Anna Dominiczak and Alex Alderton, Roberto Amato, Jeffrey Barrett, Sonia Goncalves, Ewan Harrison, David K. Jackson, Ian Johnston, Dominic Kwiatkowski, Cordelia Langford, John Sillitoe on behalf of the Wellcome Sanger Institute COVID-19 Surveillance Team |
| EPI_ISL_2814877 | Lighthouse Lab in Alderley Park | Wellcome Sanger Institute for the COVID-19 Genomics UK (COG-UK) Consortium | Jacquelyn Wynn, Mairead Hyland, The Lighthouse Lab in Alderley Park and Alex Alderton, Roberto Amato, Jeffrey Barrett, Sonia Goncalves, Ewan Harrison, David K. Jackson, Ian Johnston, Dominic Kwiatkowski, Cordelia Langford, John Sillitoe on behalf of the Wellcome Sanger Institute COVID-19 Surveillance Team |
| EPI_ISL_2814878 | Lighthouse Lab in Glasgow | Wellcome Sanger Institute for the COVID-19 Genomics UK (COG-UK) Consortium | Harper VanSteenhouse, Yumi Kasai, David Gray, Carol Clugston, Anna Dominiczak and Alex Alderton, Roberto Amato, Jeffrey Barrett, Sonia Goncalves, Ewan Harrison, David K. Jackson, Ian Johnston, Dominic Kwiatkowski, Cordelia Langford, John Sillitoe on behalf of the Wellcome Sanger Institute COVID-19 Surveillance Team |
| EPI_ISL_2814879 | Lighthouse Lab in Alderley Park | Wellcome Sanger Institute for the COVID-19 Genomics UK (COG-UK) Consortium | Jacquelyn Wynn, Mairead Hyland, The Lighthouse Lab in Alderley Park and Alex Alderton, Roberto Amato, Jeffrey Barrett, Sonia Goncalves, Ewan Harrison, David K. Jackson, Ian Johnston, Dominic Kwiatkowski, Cordelia Langford, John Sillitoe on behalf of the Wellcome Sanger Institute COVID-19 Surveillance Team |
| EPI_ISL_2814880, EPI_ISL_2814881, EPI_ISL_2814882, EPI_ISL_2814883, EPI_ISL_2814884, EPI_ISL_2814885 | Lighthouse Lab in Glasgow | Wellcome Sanger Institute for the COVID-19 Genomics UK (COG-UK) Consortium | Harper VanSteenhouse, Yumi Kasai, David Gray, Carol Clugston, Anna Dominiczak and Alex Alderton, Roberto Amato, Jeffrey Barrett, Sonia Goncalves, Ewan Harrison, David K. Jackson, Ian Johnston, Dominic Kwiatkowski, Cordelia Langford, John Sillitoe on behalf of the Wellcome Sanger Institute COVID-19 Surveillance Team |
| EPI_ISL_2814886 | Lighthouse Lab in Alderley Park | Wellcome Sanger Institute for the COVID-19 Genomics UK (COG-UK) Consortium | Jacquelyn Wynn, Mairead Hyland, The Lighthouse Lab in Alderley Park and Alex Alderton, Roberto Amato, Jeffrey Barrett, Sonia Goncalves, Ewan Harrison, David K. Jackson, Ian Johnston, Dominic Kwiatkowski, Cordelia Langford, John Sillitoe on behalf of the Wellcome Sanger Institute COVID-19 Surveillance Team |
| EPI_ISL_2814887, EPI_ISL_2814888, EPI_ISL_2814889, EPI_ISL_2814890, EPI_ISL_2814891, EPI_ISL_2814892, EPI_ISL_2814893, EPI_ISL_2814894, EPI_ISL_2814895, EPI_ISL_2814896, EPI_ISL_2814897, EPI_ISL_2814898, EPI_ISL_2814899, EPI_ISL_2814900, EPI_ISL_2814901, EPI_ISL_2814902, EPI_ISL_2814903, EPI_ISL_2814904, EPI_ISL_2814905, EPI_ISL_2814906, EPI_ISL_2814907, EPI_ISL_2814908, EPI_ISL_2814909, EPI_ISL_2814910, EPI_ISL_2814911, EPI_ISL_2814912, EPI_ISL_2814913, EPI_ISL_2814914, EPI_ISL_2814915, EPI_ISL_2814917, EPI_ISL_2814918, EPI_ISL_2814919, EPI_ISL_2814920, EPI_ISL_2814921, EPI_ISL_2814922, EPI_ISL_2814923, EPI_ISL_2814924, EPI_ISL_2814925, EPI_ISL_2814926, EPI_ISL_2814927, EPI_ISL_2814928, EPI_ISL_2814929, EPI_ISL_2814930, EPI_ISL_2814931, EPI_ISL_2814932, EPI_ISL_2814933, EPI_ISL_2814934, EPI_ISL_2814935, EPI_ISL_2814936, EPI_ISL_2814938, EPI_ISL_2814939, EPI_ISL_2814940, EPI_ISL_2814941, EPI_ISL_2814942, EPI_ISL_2814943, EPI_ISL_2814944, EPI_ISL_2814945, EPI_ISL_2814946, EPI_ISL_2814947, EPI_ISL_2814948, EPI_ISL_2814949, EPI_ISL_2814950, EPI_ISL_2814951, EPI_ISL_2814952, EPI_ISL_2814953, EPI_ISL_2814954, EPI_ISL_2814955, EPI_ISL_2814956, EPI_ISL_2814957, EPI_ISL_2814958, EPI_ISL_2814959, EPI_ISL_2814960, EPI_ISL_2814961, EPI_ISL_2814962, EPI_ISL_2814963, EPI_ISL_2814964, EPI_ISL_2814965, EPI_ISL_2814966, EPI_ISL_2814967, EPI_ISL_2814968, EPI_ISL_2814969, EPI_ISL_2814970, EPI_ISL_2814971, EPI_ISL_2814972, EPI_ISL_2814973, EPI_ISL_2814974, EPI_ISL_2814975, EPI_ISL_2814976, EPI_ISL_2814977, EPI_ISL_2814978, EPI_ISL_2814979, EPI_ISL_2814980, EPI_ISL_2814981, EPI_ISL_2814982, EPI_ISL_2814983, EPI_ISL_2814984, EPI_ISL_2814985, EPI_ISL_2814986, EPI_ISL_2814987, EPI_ISL_2814988, EPI_ISL_2814989, EPI_ISL_2814990, EPI_ISL_2814991, EPI_ISL_2814992, EPI_ISL_2814993, EPI_ISL_2814994, EPI_ISL_2814995, EPI_ISL_2814996, EPI_ISL_2814997, EPI_ISL_2814998, EPI_ISL_2814999, EPI_ISL_2815000, EPI_ISL_2815001, EPI_ISL_2815002, EPI_ISL_2815003, EPI_ISL_2815004, EPI_ISL_2815005, EPI_ISL_2815006, EPI_ISL_2815007, EPI_ISL_2815008, EPI_ISL_2815009, EPI_ISL_2815010, EPI_ISL_2815011, EPI_ISL_2815012, EPI_ISL_2815013, EPI_ISL_2815014, EPI_ISL_2815015, EPI_ISL_2815016, EPI_ISL_2815017, EPI_ISL_2815018, EPI_ISL_2815019, EPI_ISL_2815020, EPI_ISL_2815021, EPI_ISL_2815022, EPI_ISL_2815023, EPI_ISL_2815024, EPI_ISL_2815025, EPI_ISL_2815026, EPI_ISL_2815027, EPI_ISL_2815028, EPI_ISL_2815029, EPI_ISL_2815030, EPI_ISL_2815031, EPI_ISL_2815032, EPI_ISL_2815033, EPI_ISL_2815034, EPI_ISL_2815035, EPI_ISL_2815037, EPI_ISL_2815038 | | | |
| see above | Lighthouse Lab in Glasgow | Wellcome Sanger Institute for the COVID-19 Genomics UK (COG-UK) Consortium | Harper VanSteenhouse, Yumi Kasai, David Gray, Carol Clugston, Anna Dominiczak and Alex Alderton, Roberto Amato, Jeffrey Barrett, Sonia Goncalves, Ewan Harrison, David K. Jackson, Ian Johnston, Dominic Kwiatkowski, Cordelia Langford, John Sillitoe on behalf of the Wellcome Sanger Institute COVID-19 Surveillance Team |
| EPI_ISL_2815039, EPI_ISL_2815040 | Lighthouse Lab in Alderley Park | Wellcome Sanger Institute for the COVID-19 Genomics UK (COG-UK) Consortium | Jacquelyn Wynn, Mairead Hyland, The Lighthouse Lab in Alderley Park and Alex Alderton, Roberto Amato, Jeffrey Barrett, Sonia Goncalves, Ewan Harrison, David K. Jackson, Ian Johnston, Dominic Kwiatkowski, Cordelia Langford, John Sillitoe on behalf of the Wellcome Sanger Institute COVID-19 Surveillance Team |
| EPI_ISL_2815041 | Lighthouse Lab in Glasgow | Wellcome Sanger Institute for the COVID-19 Genomics UK (COG-UK) Consortium | Harper VanSteenhouse, Yumi Kasai, David Gray, Carol Clugston, Anna Dominiczak and Alex Alderton, Roberto Amato, Jeffrey Barrett, Sonia Goncalves, Ewan Harrison, David K. Jackson, Ian Johnston, Dominic Kwiatkowski, Cordelia Langford, John Sillitoe on behalf of the Wellcome Sanger Institute COVID-19 Surveillance Team |
| EPI_ISL_2815042 | Lighthouse Laboratory Plymouth | Wellcome Sanger Institute for the COVID-19 Genomics UK (COG-UK) Consortium | Lighthouse Laboratory Plymouth and Alex Alderton, Roberto Amato, Jeffrey Barrett, Sonia Goncalves, Ewan Harrison, David K. Jackson, Ian Johnston, Dominic Kwiatkowski, Cordelia Langford, John Sillitoe on behalf of the Wellcome Sanger Institute COVID-19 Surveillance Team |
| EPI_ISL_2815043, EPI_ISL_2815044, EPI_ISL_2815045, EPI_ISL_2815046, EPI_ISL_2815047, EPI_ISL_2815048, EPI_ISL_2815049, EPI_ISL_2815050, EPI_ISL_2815051, EPI_ISL_2815052, EPI_ISL_2815053, EPI_ISL_2815054, EPI_ISL_2815055, EPI_ISL_2815056, EPI_ISL_2815057, EPI_ISL_2815058, EPI_ISL_2815059, EPI_ISL_2815060, EPI_ISL_2815061, EPI_ISL_2815062, EPI_ISL_2815063, EPI_ISL_2815064, EPI_ISL_2815065, EPI_ISL_2815066, EPI_ISL_2815067, EPI_ISL_2815068, EPI_ISL_2815069, EPI_ISL_2815070, EPI_ISL_2815071, EPI_ISL_2815072, EPI_ISL_2815073, EPI_ISL_2815074, EPI_ISL_2815075, EPI_ISL_2815076, EPI_ISL_2815078, EPI_ISL_2815079, EPI_ISL_2815080, EPI_ISL_2815081 | | | |
| see above | Lighthouse Lab in Glasgow | Wellcome Sanger Institute for the COVID-19 Genomics UK (COG-UK) Consortium | Harper VanSteenhouse, Yumi Kasai, David Gray, Carol Clugston, Anna Dominiczak and Alex Alderton, Roberto Amato, Jeffrey Barrett, Sonia Goncalves, Ewan Harrison, David K. Jackson, Ian Johnston, Dominic Kwiatkowski, Cordelia Langford, John Sillitoe on behalf of the Wellcome Sanger Institute COVID-19 Surveillance Team |
| EPI_ISL_2815082 | Health Services Laboratories | Wellcome Sanger Institute for the COVID-19 Genomics UK (COG-UK) Consortium | Health Services Laboratories and Alex Alderton, Roberto Amato, Jeffrey Barrett, Sonia Goncalves, Ewan Harrison, David K. Jackson, Ian Johnston, Dominic Kwiatkowski, Cordelia Langford, John Sillitoe on behalf of the Wellcome Sanger Institute COVID-19 Surveillance Team |
| EPI_ISL_2815083, EPI_ISL_2815084, EPI_ISL_2815086, EPI_ISL_2815087, EPI_ISL_2815089, EPI_ISL_2815090 | Lighthouse Lab in Glasgow | Wellcome Sanger Institute for the COVID-19 Genomics UK (COG-UK) Consortium | Harper VanSteenhouse, Yumi Kasai, David Gray, Carol Clugston, Anna Dominiczak and Alex Alderton, Roberto Amato, Jeffrey Barrett, Sonia Goncalves, Ewan Harrison, David K. Jackson, Ian Johnston, Dominic Kwiatkowski, Cordelia Langford, John Sillitoe on behalf of the Wellcome Sanger Institute COVID-19 Surveillance Team |
| EPI_ISL_2815091 | Health Services Laboratories | Wellcome Sanger Institute for the COVID-19 Genomics UK (COG-UK) Consortium | Health Services Laboratories and Alex Alderton, Roberto Amato, Jeffrey Barrett, Sonia Goncalves, Ewan Harrison, David K. Jackson, Ian Johnston, Dominic Kwiatkowski, Cordelia Langford, John Sillitoe on behalf of the Wellcome Sanger Institute COVID-19 Surveillance Team |
| EPI_ISL_2815092, EPI_ISL_2815093 | Lighthouse Lab in Glasgow | Wellcome Sanger Institute for the COVID-19 Genomics UK (COG-UK) Consortium | Harper VanSteenhouse, Yumi Kasai, David Gray, Carol Clugston, Anna Dominiczak and Alex Alderton, Roberto Amato, Jeffrey Barrett, Sonia Goncalves, Ewan Harrison, David K. Jackson, Ian Johnston, Dominic Kwiatkowski, Cordelia Langford, John Sillitoe on behalf of the Wellcome Sanger Institute COVID-19 Surveillance Team |
| EPI_ISL_2815094, EPI_ISL_2815095 | Health Services Laboratories | Wellcome Sanger Institute for the COVID-19 Genomics UK (COG-UK) Consortium | Health Services Laboratories and Alex Alderton, Roberto Amato, Jeffrey Barrett, Sonia Goncalves, Ewan Harrison, David K. Jackson, Ian Johnston, Dominic Kwiatkowski, Cordelia Langford, John Sillitoe on behalf of the Wellcome Sanger Institute COVID-19 Surveillance Team |
| EPI_ISL_2815096, EPI_ISL_2815097, EPI_ISL_2815098, EPI_ISL_2815099, EPI_ISL_2815100, EPI_ISL_2815101, EPI_ISL_2815102, EPI_ISL_2815103, EPI_ISL_2815104, EPI_ISL_2815105, EPI_ISL_2815106, EPI_ISL_2815107, EPI_ISL_2815108, EPI_ISL_2815109, EPI_ISL_2815110, EPI_ISL_2815111, EPI_ISL_2815112, EPI_ISL_2815114, EPI_ISL_2815115, EPI_ISL_2815116, EPI_ISL_2815117, EPI_ISL_2815118, EPI_ISL_2815119, EPI_ISL_2815120, EPI_ISL_2815121, EPI_ISL_2815122, EPI_ISL_2815123, EPI_ISL_2815124, EPI_ISL_2815125, EPI_ISL_2815126, EPI_ISL_2815127, EPI_ISL_2815128, EPI_ISL_2815129, EPI_ISL_2815130, EPI_ISL_2815131, EPI_ISL_2815132, EPI_ISL_2815133, EPI_ISL_2815134, EPI_ISL_2815135, EPI_ISL_2815136, EPI_ISL_2815137, EPI_ISL_2815138, EPI_ISL_2815139, EPI_ISL_2815140, EPI_ISL_2815141, EPI_ISL_2815142, EPI_ISL_2815143, EPI_ISL_2815144, EPI_ISL_2815145, EPI_ISL_2815146, EPI_ISL_2815147, EPI_ISL_2815148, EPI_ISL_2815149, EPI_ISL_2815150, EPI_ISL_2815151, EPI_ISL_2815152, EPI_ISL_2815153, EPI_ISL_2815154, EPI_ISL_2815155, EPI_ISL_2815156, EPI_ISL_2815157, EPI_ISL_2815158, EPI_ISL_2815159, EPI_ISL_2815160, EPI_ISL_2815161, EPI_ISL_2815162, EPI_ISL_2815163, EPI_ISL_2815164, EPI_ISL_2815165, EPI_ISL_2815166, EPI_ISL_2815167, EPI_ISL_2815168, EPI_ISL_2815169, EPI_ISL_2815170, EPI_ISL_2815171, EPI_ISL_2815172, EPI_ISL_2815173, EPI_ISL_2815174, EPI_ISL_2815175, EPI_ISL_2815176, EPI_ISL_2815177, EPI_ISL_2815178, EPI_ISL_2815179, EPI_ISL_2815180, EPI_ISL_2815181, EPI_ISL_2815182, EPI_ISL_2815183, EPI_ISL_2815184, EPI_ISL_2815185, EPI_ISL_2815186, EPI_ISL_2815187, EPI_ISL_2815188, EPI_ISL_2815189, EPI_ISL_2815190, EPI_ISL_2815191, EPI_ISL_2815192, EPI_ISL_2815193, EPI_ISL_2815194, EPI_ISL_2815195, EPI_ISL_2815196, EPI_ISL_2815197, EPI_ISL_2815198, EPI_ISL_2815199, EPI_ISL_2815200, EPI_ISL_2815201, EPI_ISL_2815202, EPI_ISL_2815203, EPI_ISL_2815204, EPI_ISL_2815205, EPI_ISL_2815206, EPI_ISL_2815207, EPI_ISL_2815208, EPI_ISL_2815209, EPI_ISL_2815210, EPI_ISL_2815211, EPI_ISL_2815212, EPI_ISL_2815213, EPI_ISL_2815214, EPI_ISL_2815215, EPI_ISL_2815216, EPI_ISL_2815217, EPI_ISL_2815218, EPI_ISL_2815219, EPI_ISL_2815220, EPI_ISL_2815221, EPI_ISL_2815222, EPI_ISL_2815223, EPI_ISL_2815224, EPI_ISL_2815225, EPI_ISL_2815226, EPI_ISL_2815227, EPI_ISL_2815228, EPI_ISL_2815229, EPI_ISL_2815230, EPI_ISL_2815231, EPI_ISL_2815232, EPI_ISL_2815233, EPI_ISL_2815234, EPI_ISL_2815235, EPI_ISL_2815236, EPI_ISL_2815237, EPI_ISL_2815238, EPI_ISL_2815239, EPI_ISL_2815240, EPI_ISL_2815241, EPI_ISL_2815242, EPI_ISL_2815243, EPI_ISL_2815244, EPI_ISL_2815245, EPI_ISL_2815246, EPI_ISL_2815247, EPI_ISL_2815248, EPI_ISL_2815249, EPI_ISL_2815250, EPI_ISL_2815251, EPI_ISL_2815252, EPI_ISL_2815253, EPI_ISL_2815254, EPI_ISL_2815255, EPI_ISL_2815256, EPI_ISL_2815257, EPI_ISL_2815258, EPI_ISL_2815259, EPI_ISL_2815260, EPI_ISL_2815261, EPI_ISL_2815262, EPI_ISL_2815263, EPI_ISL_2815264, EPI_ISL_2815265, EPI_ISL_2815266, EPI_ISL_2815267, EPI_ISL_2815268, EPI_ISL_2815269, EPI_ISL_2815270, EPI_ISL_2815271, EPI_ISL_2815272, EPI_ISL_2815273, EPI_ISL_2815274, EPI_ISL_2815275, EPI_ISL_2815276, EPI_ISL_2815278, EPI_ISL_2815279, EPI_ISL_2815280, EPI_ISL_2815281, EPI_ISL_2815282, EPI_ISL_2815283, EPI_ISL_2815284, EPI_ISL_2815285, EPI_ISL_2815286, EPI_ISL_2815287, EPI_ISL_2815288, EPI_ISL_2815289, EPI_ISL_2815290, EPI_ISL_2815291, EPI_ISL_2815292, EPI_ISL_2815293, EPI_ISL_2815294, EPI_ISL_2815295, EPI_ISL_2815296, EPI_ISL_2815297, EPI_ISL_2815298, EPI_ISL_2815299, EPI_ISL_2815300, EPI_ISL_2815301, EPI_ISL_2815302, EPI_ISL_2815303, EPI_ISL_2815304, EPI_ISL_2815305, EPI_ISL_2815306, EPI_ISL_2815307, EPI_ISL_2815308, EPI_ISL_2815309, EPI_ISL_2815310, EPI_ISL_2815311, EPI_ISL_2815312, EPI_ISL_2815313, EPI_ISL_2815314, EPI_ISL_2815315, EPI_ISL_2815316, EPI_ISL_2815317, EPI_ISL_2815318, EPI_ISL_2815319, EPI_ISL_2815320, EPI_ISL_2815321, EPI_ISL_2815322, EPI_ISL_2815323, EPI_ISL_2815324, EPI_ISL_2815325, EPI_ISL_2815326 | | | |
| see above | Lighthouse Lab in Glasgow | Wellcome Sanger Institute for the COVID-19 Genomics UK (COG-UK) Consortium | Harper VanSteenhouse, Yumi Kasai, David Gray, Carol Clugston, Anna Dominiczak and Alex Alderton, Roberto Amato, Jeffrey Barrett, Sonia Goncalves, Ewan Harrison, David K. Jackson, Ian Johnston, Dominic Kwiatkowski, Cordelia Langford, John Sillitoe on behalf of the Wellcome Sanger Institute COVID-19 Surveillance Team |
| EPI_ISL_2815327, EPI_ISL_2815328 | Melaka Hospital | Institute for Medical Research, Infectious Disease Research Centre, National Institutes of Health, Ministry of Health Malaysia | Suppiah J, Kamel K, Mohd Zawawi Z, Azizan MA, Ramly N, Robert F, Thayan R |
| EPI_ISL_2815329, EPI_ISL_2815330 | Ipoh Public Health Laboratory (MKAI), Ministry of Health Malaysia | Institute for Medical Research, Infectious Disease Research Centre, National Institutes of Health, Ministry of Health Malaysia | Suppiah J, Kamel K, Mohd Zawawi Z, Azizan MA, Ramly N, Robert F, Thayan R |
| EPI_ISL_2815331 | Kuala Lumpur General Hospital | Institute for Medical Research, Infectious Disease Research Centre, National Institutes of Health, Ministry of Health Malaysia | Suppiah J, Kamel K, Mohd Zawawi Z, Azizan MA, Ramly N, Robert F, Thayan R |
| EPI_ISL_2815332, EPI_ISL_2815333, EPI_ISL_2815334, EPI_ISL_2815335, EPI_ISL_2815337, EPI_ISL_2815339, EPI_ISL_2815340, EPI_ISL_2815342 | ASL BARI | University of Bari Biomedical Sciences and Human Oncology | Maria Chironna |
| EPI_ISL_2815343, EPI_ISL_2815344 | ASL BARI | University of Bari Biomedical Sciences and Human Oncology | Chironna M., Sallustio A., Loconsole D., Accogli M. |

| | | | |
|---|---|---|---|
| EPI_ISL_2815345, EPI_ISL_2815349, EPI_ISL_2815350 | ASL BARI | University of Bari Biomedical Sciences and Human Oncology | Maria Chironna |
| EPI_ISL_2815378, EPI_ISL_2815379, EPI_ISL_2815380, EPI_ISL_2815381, EPI_ISL_2815382, EPI_ISL_2815383, EPI_ISL_2815385, EPI_ISL_2815386, EPI_ISL_2815387, EPI_ISL_2815388, EPI_ISL_2815389, EPI_ISL_2815390, EPI_ISL_2815391, EPI_ISL_2815392, EPI_ISL_2815393, EPI_ISL_2815394, EPI_ISL_2815395, EPI_ISL_2815396, EPI_ISL_2815397, EPI_ISL_2815398, EPI_ISL_2815399, EPI_ISL_2815400, EPI_ISL_2815401, EPI_ISL_2815402, EPI_ISL_2815403, EPI_ISL_2815404, EPI_ISL_2815405, EPI_ISL_2815406, EPI_ISL_2815407, EPI_ISL_2815408, EPI_ISL_2815409, EPI_ISL_2815410, EPI_ISL_2815411, EPI_ISL_2815412, EPI_ISL_2815413, EPI_ISL_2815414, EPI_ISL_2815415, EPI_ISL_2815416, EPI_ISL_2815417, EPI_ISL_2815418, EPI_ISL_2815419, EPI_ISL_2815420, EPI_ISL_2815421, EPI_ISL_2815422, EPI_ISL_2815423, EPI_ISL_2815424, EPI_ISL_2815425, EPI_ISL_2815426, EPI_ISL_2815427, EPI_ISL_2815428, EPI_ISL_2815429, EPI_ISL_2815430, EPI_ISL_2815431, EPI_ISL_2815432, EPI_ISL_2815433, EPI_ISL_2815434, EPI_ISL_2815435, EPI_ISL_2815436, EPI_ISL_2815437, EPI_ISL_2815438, EPI_ISL_2815439, EPI_ISL_2815440, EPI_ISL_2815441, EPI_ISL_2815442, EPI_ISL_2815443, EPI_ISL_2815444, EPI_ISL_2815445, EPI_ISL_2815446, EPI_ISL_2815447, EPI_ISL_2815450, EPI_ISL_2815451, EPI_ISL_2815452, EPI_ISL_2815453, EPI_ISL_2815454, EPI_ISL_2815455, EPI_ISL_2815456, EPI_ISL_2815457, EPI_ISL_2815458, EPI_ISL_2815459, EPI_ISL_2815460, EPI_ISL_2815461, EPI_ISL_2815462, EPI_ISL_2815463, EPI_ISL_2815464, EPI_ISL_2815465, EPI_ISL_2815466, EPI_ISL_2815467, EPI_ISL_2815468, EPI_ISL_2815469, EPI_ISL_2815471, EPI_ISL_2815472, EPI_ISL_2815473, EPI_ISL_2815474, EPI_ISL_2815475, EPI_ISL_2815476, EPI_ISL_2815477, EPI_ISL_2815478, EPI_ISL_2815479, EPI_ISL_2815480, EPI_ISL_2815481, EPI_ISL_2815482, EPI_ISL_2815483, EPI_ISL_2815484, EPI_ISL_2815485, EPI_ISL_2815486, EPI_ISL_2815487, EPI_ISL_2815488, EPI_ISL_2815489, EPI_ISL_2815490, EPI_ISL_2815491, EPI_ISL_2815492, EPI_ISL_2815493, EPI_ISL_2815494, EPI_ISL_2815495, EPI_ISL_2815496, EPI_ISL_2815497, EPI_ISL_2815498, EPI_ISL_2815499, EPI_ISL_2815500, EPI_ISL_2815501, EPI_ISL_2815502, EPI_ISL_2815503, EPI_ISL_2815504, EPI_ISL_2815505, EPI_ISL_2815506, EPI_ISL_2815507, EPI_ISL_2815508, EPI_ISL_2815509, EPI_ISL_2815510, EPI_ISL_2815511, EPI_ISL_2815512, EPI_ISL_2815513, EPI_ISL_2815514, EPI_ISL_2815515, EPI_ISL_2815517, EPI_ISL_2815518, EPI_ISL_2815519, EPI_ISL_2815520, EPI_ISL_2815521, EPI_ISL_2815522, EPI_ISL_2815523, EPI_ISL_2815524, EPI_ISL_2815525, EPI_ISL_2815526, EPI_ISL_2815527, EPI_ISL_2815528, EPI_ISL_2815529, EPI_ISL_2815530, EPI_ISL_2815531, EPI_ISL_2815532, EPI_ISL_2815533, EPI_ISL_2815534, EPI_ISL_2815535, EPI_ISL_2815536, EPI_ISL_2815537, EPI_ISL_2815538, EPI_ISL_2815539, EPI_ISL_2815540, EPI_ISL_2815541, EPI_ISL_2815542, EPI_ISL_2815543, EPI_ISL_2815544, EPI_ISL_2815545, EPI_ISL_2815546, EPI_ISL_2815547, EPI_ISL_2815548 | | | |
| see above | University of Birmingham | COVID-19 Genomics UK (COG-UK) Consortium | Institute of Microbiology, University of Birmingham: Claire McMurray, Joanne Stockton, Samuel Nicholls, Radoslaw Poplawski, Will Rowe, Josh Quick, Nicholas Loman. University of Birmingham Testing Laboratory: Celina M Whalley, Andrew Bosworth, Charlotte Poxon, Kasun Wanigasooriya, Oliver Pickles, Mike Kidd, Alex Richter, Andrew D Beggs PHE Heartlands Lab: Husam Osman, Andrew Bosworth. Queen Elizabeth Hospital: Anna Casey |
| EPI_ISL_2815549, EPI_ISL_2815550, EPI_ISL_2815551, EPI_ISL_2815552, EPI_ISL_2815553, EPI_ISL_2815554, EPI_ISL_2815555, EPI_ISL_2815556, EPI_ISL_2815557, EPI_ISL_2815558, EPI_ISL_2815559, EPI_ISL_2815560, EPI_ISL_2815561, EPI_ISL_2815562, EPI_ISL_2815563, EPI_ISL_2815564, EPI_ISL_2815565, EPI_ISL_2815566, EPI_ISL_2815567, EPI_ISL_2815568, EPI_ISL_2815569, EPI_ISL_2815570, EPI_ISL_2815571, EPI_ISL_2815572, EPI_ISL_2815573, EPI_ISL_2815574, EPI_ISL_2815575, EPI_ISL_2815576, EPI_ISL_2815577, EPI_ISL_2815578, EPI_ISL_2815579, EPI_ISL_2815580, EPI_ISL_2815581, EPI_ISL_2815582, EPI_ISL_2815583, EPI_ISL_2815584, EPI_ISL_2815585, EPI_ISL_2815586, EPI_ISL_2815587, EPI_ISL_2815588, EPI_ISL_2815589, EPI_ISL_2815590, EPI_ISL_2815591, EPI_ISL_2815592, EPI_ISL_2815593, EPI_ISL_2815594, EPI_ISL_2815595, EPI_ISL_2815596, EPI_ISL_2815597, EPI_ISL_2815598, EPI_ISL_2815599, EPI_ISL_2815600, EPI_ISL_2815601, EPI_ISL_2815602, EPI_ISL_2815603, EPI_ISL_2815604, EPI_ISL_2815605, EPI_ISL_2815606, EPI_ISL_2815607, EPI_ISL_2815608, EPI_ISL_2815609, EPI_ISL_2815610, EPI_ISL_2815611, EPI_ISL_2815612, EPI_ISL_2815613, EPI_ISL_2815614, EPI_ISL_2815615, EPI_ISL_2815616, EPI_ISL_2815617, EPI_ISL_2815618, EPI_ISL_2815619, EPI_ISL_2815620, EPI_ISL_2815621, EPI_ISL_2815622, EPI_ISL_2815623, EPI_ISL_2815624, EPI_ISL_2815625, EPI_ISL_2815626, EPI_ISL_2815627, EPI_ISL_2815628, EPI_ISL_2815629, EPI_ISL_2815630, EPI_ISL_2815631, EPI_ISL_2815632, EPI_ISL_2815633, EPI_ISL_2815634, EPI_ISL_2815635, EPI_ISL_2815636, EPI_ISL_2815637, EPI_ISL_2815638, EPI_ISL_2815639, EPI_ISL_2815640, EPI_ISL_2815641, EPI_ISL_2815642, EPI_ISL_2815643, EPI_ISL_2815644, EPI_ISL_2815645, EPI_ISL_2815646, EPI_ISL_2815647, EPI_ISL_2815648, EPI_ISL_2815649, EPI_ISL_2815650, EPI_ISL_2815651, EPI_ISL_2815652, EPI_ISL_2815653, EPI_ISL_2815654, EPI_ISL_2815655, EPI_ISL_2815657, EPI_ISL_2815658, EPI_ISL_2815659, EPI_ISL_2815660, EPI_ISL_2815661, EPI_ISL_2815662, EPI_ISL_2815663, EPI_ISL_2815664, EPI_ISL_2815665, EPI_ISL_2815666, EPI_ISL_2815667, EPI_ISL_2815668, EPI_ISL_2815669, EPI_ISL_2815670, EPI_ISL_2815671, EPI_ISL_2815672, EPI_ISL_2815673, EPI_ISL_2815674, EPI_ISL_2815675, EPI_ISL_2815676, EPI_ISL_2815677, EPI_ISL_2815678, EPI_ISL_2815679, EPI_ISL_2815680, EPI_ISL_2815681, EPI_ISL_2815682, EPI_ISL_2815683, EPI_ISL_2815684, EPI_ISL_2815685, EPI_ISL_2815686, EPI_ISL_2815687, EPI_ISL_2815688, EPI_ISL_2815689, EPI_ISL_2815690, EPI_ISL_2815691, EPI_ISL_2815692, EPI_ISL_2815693, EPI_ISL_2815694, EPI_ISL_2815695, EPI_ISL_2815696, EPI_ISL_2815697, EPI_ISL_2815698, EPI_ISL_2815699, EPI_ISL_2815700, EPI_ISL_2815701, EPI_ISL_2815702, EPI_ISL_2815703, EPI_ISL_2815704, EPI_ISL_2815705, EPI_ISL_2815706, EPI_ISL_2815707, EPI_ISL_2815708, EPI_ISL_2815709, EPI_ISL_2815710, EPI_ISL_2815711, EPI_ISL_2815712, EPI_ISL_2815713, EPI_ISL_2815714, EPI_ISL_2815715, EPI_ISL_2815716, EPI_ISL_2815717, EPI_ISL_2815718, EPI_ISL_2815719, EPI_ISL_2815720, EPI_ISL_2815721, EPI_ISL_2815722, EPI_ISL_2815723, EPI_ISL_2815724, EPI_ISL_2815725, EPI_ISL_2815726, EPI_ISL_2815727, EPI_ISL_2815728, EPI_ISL_2815729, EPI_ISL_2815730, EPI_ISL_2815731, EPI_ISL_2815732, EPI_ISL_2815733, EPI_ISL_2815734, EPI_ISL_2815735, EPI_ISL_2815736, EPI_ISL_2815737, EPI_ISL_2815738, EPI_ISL_2815739, EPI_ISL_2815740, EPI_ISL_2815741, EPI_ISL_2815742, EPI_ISL_2815743, EPI_ISL_2815744, EPI_ISL_2815745, EPI_ISL_2815746, EPI_ISL_2815747, EPI_ISL_2815748, EPI_ISL_2815749, EPI_ISL_2815750, EPI_ISL_2815751, EPI_ISL_2815752, EPI_ISL_2815753, EPI_ISL_2815754, EPI_ISL_2815755, EPI_ISL_2815756, EPI_ISL_2815757, EPI_ISL_2815758, EPI_ISL_2815759, EPI_ISL_2815760, EPI_ISL_2815761, EPI_ISL_2815762, EPI_ISL_2815763, EPI_ISL_2815764, EPI_ISL_2815765, EPI_ISL_2815766, EPI_ISL_2815767, EPI_ISL_2815768, EPI_ISL_2815769, EPI_ISL_2815770, EPI_ISL_2815771, EPI_ISL_2815772, EPI_ISL_2815773, EPI_ISL_2815774, EPI_ISL_2815775 | | | |
| see above | Department of Pathology, University of Cambridge | COVID-19 Genomics UK (COG-UK) Consortium | Aminu S. Jahun, Yasmin Chaudhry, Iliana Georgana, Myra Hosmillo, Rhys Izuagbe, William L. Hamilton, Martin D. Curran, Surendra Parmar, Ian Goodfellow |
| EPI_ISL_2815776 | ASL BARI | University of Bari Biomedical Sciences and Human Oncology | Maria Chironna |
| EPI_ISL_2815777 | Department of Pathology, University of Cambridge | COVID-19 Genomics UK (COG-UK) Consortium | Aminu S. Jahun, Yasmin Chaudhry, Iliana Georgana, Myra Hosmillo, Rhys Izuagbe, William L. Hamilton, Martin D. Curran, Surendra Parmar, Ian Goodfellow |
| EPI_ISL_2815778 | ASL BARI | University of Bari Biomedical Sciences and Human Oncology | Maria Chironna |
| EPI_ISL_2815779, EPI_ISL_2815780, EPI_ISL_2815781, EPI_ISL_2815782, EPI_ISL_2815783, EPI_ISL_2815784, EPI_ISL_2815785, EPI_ISL_2815786, EPI_ISL_2815787, EPI_ISL_2815788, EPI_ISL_2815789, EPI_ISL_2815790, EPI_ISL_2815791, EPI_ISL_2815792, EPI_ISL_2815793, EPI_ISL_2815794, EPI_ISL_2815795, EPI_ISL_2815796, EPI_ISL_2815797, EPI_ISL_2815798, EPI_ISL_2815799, EPI_ISL_2815800, EPI_ISL_2815801, EPI_ISL_2815802, EPI_ISL_2815803, EPI_ISL_2815804, EPI_ISL_2815805, EPI_ISL_2815806, EPI_ISL_2815807, EPI_ISL_2815808, EPI_ISL_2815809, EPI_ISL_2815810, EPI_ISL_2815811, EPI_ISL_2815812, EPI_ISL_2815813, EPI_ISL_2815814, EPI_ISL_2815815, EPI_ISL_2815816, EPI_ISL_2815818, EPI_ISL_2815819, EPI_ISL_2815820, EPI_ISL_2815821, EPI_ISL_2815822, EPI_ISL_2815823, EPI_ISL_2815824, EPI_ISL_2815825, EPI_ISL_2815826, EPI_ISL_2815827, EPI_ISL_2815828, EPI_ISL_2815829, EPI_ISL_2815830, EPI_ISL_2815831, EPI_ISL_2815832, EPI_ISL_2815833, EPI_ISL_2815834, EPI_ISL_2815835, EPI_ISL_2815836, EPI_ISL_2815838, EPI_ISL_2815839, EPI_ISL_2815840, EPI_ISL_2815841, EPI_ISL_2815842, EPI_ISL_2815843, EPI_ISL_2815844, EPI_ISL_2815845, EPI_ISL_2815846, EPI_ISL_2815847, EPI_ISL_2815848, EPI_ISL_2815849, EPI_ISL_2815850, EPI_ISL_2815851, EPI_ISL_2815852, EPI_ISL_2815853, EPI_ISL_2815854, EPI_ISL_2815855, EPI_ISL_2815856, EPI_ISL_2815857, EPI_ISL_2815859, EPI_ISL_2815860, EPI_ISL_2815861, EPI_ISL_2815862, EPI_ISL_2815863, EPI_ISL_2815864, EPI_ISL_2815865, EPI_ISL_2815866, EPI_ISL_2815867, EPI_ISL_2815868, EPI_ISL_2815869, EPI_ISL_2815870, EPI_ISL_2815871, EPI_ISL_2815872, EPI_ISL_2815873, EPI_ISL_2815874, EPI_ISL_2815875, EPI_ISL_2815876, EPI_ISL_2815877, EPI_ISL_2815878, EPI_ISL_2815879, EPI_ISL_2815880, EPI_ISL_2815881, EPI_ISL_2815882, EPI_ISL_2815883, EPI_ISL_2815884, EPI_ISL_2815885, EPI_ISL_2815886, EPI_ISL_2815887, EPI_ISL_2815888, EPI_ISL_2815889, EPI_ISL_2815890, EPI_ISL_2815891, EPI_ISL_2815892, EPI_ISL_2815893, EPI_ISL_2815894 | | | |
| see above | Department of Pathology, University of Cambridge | COVID-19 Genomics UK (COG-UK) Consortium | Aminu S. Jahun, Yasmin Chaudhry, Iliana Georgana, Myra Hosmillo, Rhys Izuagbe, William L. Hamilton, Martin D. Curran, Surendra Parmar, Ian Goodfellow |
| EPI_ISL_2815895, EPI_ISL_2815896, EPI_ISL_2815899, EPI_ISL_2815900, EPI_ISL_2815901, EPI_ISL_2815902, EPI_ISL_2815903, EPI_ISL_2815904, EPI_ISL_2815905, EPI_ISL_2815907, EPI_ISL_2815908, EPI_ISL_2815909, EPI_ISL_2815910, EPI_ISL_2815912, EPI_ISL_2815914, EPI_ISL_2815915, EPI_ISL_2815916, EPI_ISL_2815918, EPI_ISL_2815920, EPI_ISL_2815921, EPI_ISL_2815922, EPI_ISL_2815923, EPI_ISL_2815924, EPI_ISL_2815925, EPI_ISL_2815926, EPI_ISL_2815927, EPI_ISL_2815928, EPI_ISL_2815929, EPI_ISL_2815930, EPI_ISL_2815931, EPI_ISL_2815933, EPI_ISL_2815934, EPI_ISL_2815935, EPI_ISL_2815936, EPI_ISL_2815937, EPI_ISL_2815938, EPI_ISL_2815939, EPI_ISL_2815940, EPI_ISL_2815941, EPI_ISL_2815942, EPI_ISL_2815943, EPI_ISL_2815944, EPI_ISL_2815945, EPI_ISL_2815946, EPI_ISL_2815947, EPI_ISL_2815948, EPI_ISL_2815949, EPI_ISL_2815950, EPI_ISL_2815951, EPI_ISL_2815952, EPI_ISL_2815953, EPI_ISL_2815954, EPI_ISL_2815955, EPI_ISL_2815956, EPI_ISL_2815957, EPI_ISL_2815958, EPI_ISL_2815959, EPI_ISL_2815960, EPI_ISL_2815961, EPI_ISL_2815962, EPI_ISL_2815963, EPI_ISL_2815964, EPI_ISL_2815965, EPI_ISL_2815966, EPI_ISL_2815967 | | | |
| see above | West of Scotland Specialist Virology Centre, NHSGGC / MRC-University of Glasgow Centre for Virus Research | COVID-19 Genomics UK (COG-UK) Consortium | Ana da Silva Filipe, Natasha Johnson, Kathy Smollett, Daniel Mair, Stephen Carmichael, Alice Broos, Lily Tong, Jenna Nichols, Kyriaki Nomikou; Sarah McDonald; Ioulia Tsatsani, Richard Orton, Joseph Hughes, Sreenu Vattipally, David L Robertson; Alasdair MacLean, Rory Gunson; Sharif Shaaban, Matthew Holden; Rachel Blacow, Guy Mollett, Kathy Li, James Shepherd, Antonia Ho, Emma Thomson |
| EPI_ISL_2815970, EPI_ISL_2815972, EPI_ISL_2815973, EPI_ISL_2815975, EPI_ISL_2815976, EPI_ISL_2815977, EPI_ISL_2815978, EPI_ISL_2815979, EPI_ISL_2815980, EPI_ISL_2815981, EPI_ISL_2815982, EPI_ISL_2815983, EPI_ISL_2815984, EPI_ISL_2815985, EPI_ISL_2815988, EPI_ISL_2815989, EPI_ISL_2815990, EPI_ISL_2815991, EPI_ISL_2815992, EPI_ISL_2815993, EPI_ISL_2815994, EPI_ISL_2815995, EPI_ISL_2815996, EPI_ISL_2815997, EPI_ISL_2815998, EPI_ISL_2815999, EPI_ISL_2816000, EPI_ISL_2816001, EPI_ISL_2816002, EPI_ISL_2816003, EPI_ISL_2816004, EPI_ISL_2816005, EPI_ISL_2816006, EPI_ISL_2816007, EPI_ISL_2816008, EPI_ISL_2816009, EPI_ISL_2816010, EPI_ISL_2816011, EPI_ISL_2816012, EPI_ISL_2816013, EPI_ISL_2816014, EPI_ISL_2816015, EPI_ISL_2816016, EPI_ISL_2816017, EPI_ISL_2816018, EPI_ISL_2816019, EPI_ISL_2816020, EPI_ISL_2816021, EPI_ISL_2816022, EPI_ISL_2816023, EPI_ISL_2816024, EPI_ISL_2816025, EPI_ISL_2816026, EPI_ISL_2816027, EPI_ISL_2816028, EPI_ISL_2816030, EPI_ISL_2816031, EPI_ISL_2816032, EPI_ISL_2816033, EPI_ISL_2816034, EPI_ISL_2816035, EPI_ISL_2816036, EPI_ISL_2816037, EPI_ISL_2816038, EPI_ISL_2816039, EPI_ISL_2816040, EPI_ISL_2816041, EPI_ISL_2816042, EPI_ISL_2816043, EPI_ISL_2816044, EPI_ISL_2816045, EPI_ISL_2816046, EPI_ISL_2816047, EPI_ISL_2816048, EPI_ISL_2816049, EPI_ISL_2816050, EPI_ISL_2816051, EPI_ISL_2816052, EPI_ISL_2816053, EPI_ISL_2816054, EPI_ISL_2816056, EPI_ISL_2816057, EPI_ISL_2816058, EPI_ISL_2816059, EPI_ISL_2816060, EPI_ISL_2816061, EPI_ISL_2816062, EPI_ISL_2816063, EPI_ISL_2816064, EPI_ISL_2816065, EPI_ISL_2816066, EPI_ISL_2816067, EPI_ISL_2816068, EPI_ISL_2816069, EPI_ISL_2816070, EPI_ISL_2816071, EPI_ISL_2816072, EPI_ISL_2816074, EPI_ISL_2816075, EPI_ISL_2816076, EPI_ISL_2816077, EPI_ISL_2816078, EPI_ISL_2816079, EPI_ISL_2816080, EPI_ISL_2816081, EPI_ISL_2816082, EPI_ISL_2816083, EPI_ISL_2816084, EPI_ISL_2816085, EPI_ISL_2816086, EPI_ISL_2816087, EPI_ISL_2816088, EPI_ISL_2816089, EPI_ISL_2816091 | | | |
| see above | Virology Department, Royal Infirmary of Edinburgh, NHS Lothian / School of Biological Sciences, University of Edinburgh | COVID-19 Genomics UK (COG-UK) Consortium | McHugh M, Dewar R, Cotton S, Gallagher A, Maloney D, Rooke S, O'Toole Á, Scher E, Hill V, McCrone JT, Colquhoun R, Yu X, Jackson B, Rambaut A, Templeton K |
| EPI_ISL_2816092, EPI_ISL_2816094, EPI_ISL_2816095, EPI_ISL_2816096, EPI_ISL_2816097, EPI_ISL_2816098, EPI_ISL_2816099, EPI_ISL_2816100, EPI_ISL_2816101, EPI_ISL_2816102, EPI_ISL_2816103, EPI_ISL_2816104, EPI_ISL_2816105, EPI_ISL_2816106, EPI_ISL_2816107, EPI_ISL_2816108, EPI_ISL_2816109, EPI_ISL_2816110, EPI_ISL_2816111, EPI_ISL_2816113, EPI_ISL_2816114, EPI_ISL_2816115, EPI_ISL_2816116, EPI_ISL_2816117, EPI_ISL_2816118, EPI_ISL_2816119, EPI_ISL_2816120, EPI_ISL_2816121, EPI_ISL_2816122, EPI_ISL_2816123, EPI_ISL_2816124, EPI_ISL_2816125, EPI_ISL_2816126, EPI_ISL_2816127, EPI_ISL_2816128, EPI_ISL_2816129, EPI_ISL_2816130, EPI_ISL_2816131, EPI_ISL_2816132, EPI_ISL_2816133, EPI_ISL_2816134, EPI_ISL_2816135, EPI_ISL_2816136, EPI_ISL_2816137, EPI_ISL_2816138, EPI_ISL_2816139, EPI_ISL_2816140, EPI_ISL_2816141, EPI_ISL_2816142, EPI_ISL_2816143 | | | |
| see above | Liverpool Clinical Laboratories | COVID-19 Genomics UK (COG-UK) Consortium | Sam Haldenby, Alistair Darby, Steve Paterson, Anita Lucaci, Julian Hiscox, M Almsaud, A Alrezaihi, Muhannad Alruwaili, Stuart D Armstrong, Jones Benjamin, Eleanor G Bentley, Anu Chawla, Jordan J Clark, Angela Cowell, Richard Eccles, Isabel García-Dorival, Matthew Gemmell, Alessandro Gerada, PKF Gilmore, Richard Gregory, Ximeng Han, Catherine Hartley, Margaret Hughes, Miren Iturriza-Gomara, James Johnson, L Luu, Jenifer Manson, Charlotte Nelson, Elaine O'Toole, Cassie Olateju, Rebekah Penrice-Randal , Lucille Rainbow, N.P Randle, Trevor Ian Robinson, Parul Sharma, Ghada T Shawli, James P Stewart, Neil Swainston, Ecaterina Vamos, Joanne Watts, Mark Whitehead, Hermione Webster |
| EPI_ISL_2816144 | WHO National Influenza Centre Russian Federation | WHO National Influenza Centre Russian Federation | Andrey Komissarov, Artem Fadeev, Kseniya Komissarova, Oula Mansour, Kirill Varchenko, Mikhail Bakaev, Tamila Musaeva, Veronika Eder, Maria Pisareva, Nikita Yolshin, Daria Danilenko, Ksenia Safina, Elena Nabieva, Georgii Bazykin, Dmitry Lioznov |
| EPI_ISL_2816145 | Liverpool Clinical Laboratories | COVID-19 Genomics UK (COG-UK) Consortium | Sam Haldenby, Alistair Darby, Steve Paterson, Anita Lucaci, Julian Hiscox, M Almsaud, A Alrezaihi, Muhannad Alruwaili, Stuart D Armstrong, Jones Benjamin, Eleanor G Bentley, Anu Chawla, Jordan J Clark, Angela Cowell, Richard Eccles, Isabel García-Dorival, Matthew Gemmell, Alessandro Gerada, PKF Gilmore, Richard Gregory, Ximeng Han, Catherine Hartley, Margaret Hughes, Miren Iturriza-Gomara, James Johnson, L Luu, Jenifer Manson, Charlotte Nelson, Elaine O'Toole, Cassie Olateju, Rebekah Penrice-Randal , Lucille Rainbow, N.P Randle, Trevor Ian Robinson, Parul Sharma, Ghada T Shawli, James P Stewart, Neil Swainston, Ecaterina Vamos, Joanne Watts, Mark Whitehead, Hermione Webster |
| EPI_ISL_2816146 | WHO National Influenza Centre Russian Federation | WHO National Influenza Centre Russian Federation | Andrey Komissarov, Artem Fadeev, Kseniya Komissarova, Oula Mansour, Kirill Varchenko, Mikhail Bakaev, Tamila Musaeva, Veronika Eder, Maria Pisareva, Nikita Yolshin, Daria Danilenko, Ksenia Safina, Elena Nabieva, Georgii Bazykin, Dmitry Lioznov |
| EPI_ISL_2816147, EPI_ISL_2816148 | Liverpool Clinical Laboratories | COVID-19 Genomics UK (COG-UK) Consortium | Sam Haldenby, Alistair Darby, Steve Paterson, Anita Lucaci, Julian Hiscox, M Almsaud, A Alrezaihi, Muhannad Alruwaili, Stuart D Armstrong, Jones Benjamin, Eleanor G Bentley, Anu Chawla, Jordan J Clark, Angela Cowell, Richard Eccles, Isabel García-Dorival, Matthew Gemmell, Alessandro Gerada, PKF Gilmore, Richard Gregory, Ximeng Han, Catherine Hartley, Margaret Hughes, Miren Iturriza-Gomara, James Johnson, L Luu, Jenifer Manson, Charlotte Nelson, Elaine O'Toole, Cassie Olateju, Rebekah Penrice-Randal , Lucille Rainbow, N.P Randle, Trevor Ian Robinson, Parul Sharma, Ghada T Shawli, James P Stewart, Neil Swainston, Ecaterina Vamos, Joanne Watts, Mark Whitehead, Hermione Webster |
| EPI_ISL_2816149, EPI_ISL_2816150, EPI_ISL_2816151, EPI_ISL_2816152, EPI_ISL_2816153, EPI_ISL_2816154, EPI_ISL_2816155, EPI_ISL_2816156, EPI_ISL_2816157, EPI_ISL_2816158, EPI_ISL_2816159, EPI_ISL_2816160, EPI_ISL_2816161, EPI_ISL_2816163, EPI_ISL_2816164, EPI_ISL_2816165, EPI_ISL_2816167, EPI_ISL_2816168, EPI_ISL_2816169, EPI_ISL_2816170, EPI_ISL_2816171, EPI_ISL_2816172, EPI_ISL_2816173, EPI_ISL_2816174, EPI_ISL_2816175, EPI_ISL_2816176, EPI_ISL_2816177, EPI_ISL_2816178, EPI_ISL_2816179, EPI_ISL_2816180, EPI_ISL_2816181, EPI_ISL_2816182, EPI_ISL_2816183, EPI_ISL_2816184, EPI_ISL_2816185, EPI_ISL_2816186, EPI_ISL_2816187, EPI_ISL_2816188, EPI_ISL_2816189, EPI_ISL_2816190, EPI_ISL_2816191, EPI_ISL_2816192, EPI_ISL_2816193, EPI_ISL_2816194 | | | |
| see above | WHO National Influenza Centre Russian Federation | WHO National Influenza Centre Russian Federation | Andrey Komissarov, Artem Fadeev, Kseniya Komissarova, Oula Mansour, Kirill Varchenko, Mikhail Bakaev, Tamila Musaeva, Veronika Eder, Maria Pisareva, Nikita Yolshin, Daria Danilenko, Ksenia Safina, Elena Nabieva, Georgii Bazykin, Dmitry Lioznov |
| EPI_ISL_2816197, EPI_ISL_2816198, EPI_ISL_2816199, EPI_ISL_2816200, EPI_ISL_2816201, EPI_ISL_2816202, EPI_ISL_2816203, EPI_ISL_2816204, EPI_ISL_2816205, EPI_ISL_2816206, EPI_ISL_2816207, EPI_ISL_2816209, EPI_ISL_2816210, EPI_ISL_2816211, EPI_ISL_2816212, EPI_ISL_2816213, EPI_ISL_2816214, EPI_ISL_2816215, EPI_ISL_2816216, EPI_ISL_2816217, EPI_ISL_2816218, EPI_ISL_2816219, EPI_ISL_2816220, EPI_ISL_2816221, EPI_ISL_2816222 | | | |
| see above | Sklifosovsky Research Institute of Emergency Medicine | WHO National Influenza Centre Russian Federation | Andrey Komissarov, Artem Fadeev, Kseniya Komissarova, Oula Mansour, Kirill Varchenko, Mikhail Bakaev, Tamila Musaeva, Veronika Eder, Maria Pisareva, Nikita Yolshin, Daria Danilenko, Ksenia Safina, Elena Nabieva, Georgii Bazykin, Dmitry Lioznov |
| EPI_ISL_2816224, EPI_ISL_2816225, EPI_ISL_2816226, EPI_ISL_2816227, EPI_ISL_2816228, EPI_ISL_2816229, EPI_ISL_2816230, EPI_ISL_2816231, EPI_ISL_2816232, EPI_ISL_2816233, EPI_ISL_2816234, EPI_ISL_2816235, EPI_ISL_2816236, EPI_ISL_2816237, EPI_ISL_2816238, EPI_ISL_2816240, EPI_ISL_2816241, EPI_ISL_2816242 | | | |
| see above | Infectious Diseases Hospital No 2 | WHO National Influenza Centre Russian Federation | Andrey Komissarov, Artem Fadeev, Kseniya Komissarova, Oula Mansour, Kirill Varchenko, Mikhail Bakaev, Tamila Musaeva, Veronika Eder, Maria Pisareva, Nikita Yolshin, Daria Danilenko, Ksenia Safina, Elena Nabieva, Georgii Bazykin, Dmitry Lioznov |
| EPI_ISL_2816243, EPI_ISL_2816244, EPI_ISL_2816245, EPI_ISL_2816246, EPI_ISL_2816247, EPI_ISL_2816249, EPI_ISL_2816250, EPI_ISL_2816251, EPI_ISL_2816252, EPI_ISL_2816253, EPI_ISL_2816255, EPI_ISL_2816257, EPI_ISL_2816259, EPI_ISL_2816260, EPI_ISL_2816261, EPI_ISL_2816262, EPI_ISL_2816263, EPI_ISL_2816264, EPI_ISL_2816265, EPI_ISL_2816266, EPI_ISL_2816267, EPI_ISL_2816268, EPI_ISL_2816269, EPI_ISL_2816270, EPI_ISL_2816272, EPI_ISL_2816273, EPI_ISL_2816274, EPI_ISL_2816275, EPI_ISL_2816276, EPI_ISL_2816277, EPI_ISL_2816278, EPI_ISL_2816279, EPI_ISL_2816280, EPI_ISL_2816281, EPI_ISL_2816282, EPI_ISL_2816283 | | | |
| see above | N.F. Gamaleya Research Center for Epidemiology and Microbiology | WHO National Influenza Centre Russian Federation | Andrey Komissarov, Artem Fadeev, Kseniya Komissarova, Oula Mansour, Kirill Varchenko, Mikhail Bakaev, Tamila Musaeva, Veronika Eder, Maria Pisareva, Nikita Yolshin, Daria Danilenko,Elizaveta Divisenko, Alina Odintsova, Elena Shidlovskaya, Ludmila Vasilchenko, Nadezhda Kuznetsova, Maria Nikiforova, Alexey Shchetinin, Artem Tkachuk, Andrei Pochtovyi, Andrei Siniavin, Denis Kleymenov, Evgeniia Bykonia, Liubov Popova, Elena Mazunina, Inna Dolzhikova, Anna Kovyrshina, Andrei Botikov, Dmitry Shcheblyakov, Vladimir Gushchin, Denis Logunov, Alexander Gintsburg, Ksenia Safina, Elena Nabieva, Georgii Bazykin, Dmitry Lioznov |
| EPI_ISL_2816284 | University College London, Great Ormond Street Hospital for Children NHS Foundation Trust, Imperial College Healthcare NHS Trust | COVID-19 Genomics UK (COG-UK) Consortium | Sergi Castellano, Rachel Williams, Mark Kristiansen, Sunando Roy, Charlotte Williams, Nadua Bayzid, Marius Cotic, Helena Tutill, Patricia Dyal, Judith Breuer |
| EPI_ISL_2816285 | N.F. Gamaleya Research Center for Epidemiology and Microbiology | WHO National Influenza Centre Russian Federation | Andrey Komissarov, Artem Fadeev, Kseniya Komissarova, Oula Mansour, Kirill Varchenko, Mikhail Bakaev, Tamila Musaeva, Veronika Eder, Maria Pisareva, Nikita Yolshin, Daria Danilenko,Elizaveta Divisenko, Alina Odintsova, Elena Shidlovskaya, Ludmila Vasilchenko, Nadezhda Kuznetsova, Maria Nikiforova, Alexey Shchetinin, Artem Tkachuk, Andrei Pochtovyi, Andrei Siniavin, Denis Kleymenov, Evgeniia Bykonia, Liubov Popova, Elena Mazunina, Inna Dolzhikova, Anna Kovyrshina, Andrei Botikov, Dmitry Shcheblyakov, Vladimir Gushchin, Denis Logunov, Alexander Gintsburg, Ksenia Safina, Elena Nabieva, Georgii Bazykin, Dmitry Lioznov |
| EPI_ISL_2816286, EPI_ISL_2816287 | University College London, Great Ormond Street Hospital for Children NHS Foundation Trust, Imperial College Healthcare NHS Trust | COVID-19 Genomics UK (COG-UK) Consortium | Sergi Castellano, Rachel Williams, Mark Kristiansen, Sunando Roy, Charlotte Williams, Nadua Bayzid, Marius Cotic, Helena Tutill, Patricia Dyal, Judith Breuer |
| EPI_ISL_2816288 | N.F. Gamaleya Research Center for Epidemiology and Microbiology | WHO National Influenza Centre Russian Federation | Andrey Komissarov, Artem Fadeev, Kseniya Komissarova, Oula Mansour, Kirill Varchenko, Mikhail Bakaev, Tamila Musaeva, Veronika Eder, Maria Pisareva, Nikita Yolshin, Daria Danilenko,Elizaveta Divisenko, Alina Odintsova, Elena Shidlovskaya, Ludmila Vasilchenko, Nadezhda Kuznetsova, Maria Nikiforova, Alexey Shchetinin, Artem Tkachuk, Andrei Pochtovyi, Andrei Siniavin, Denis Kleymenov, Evgeniia Bykonia, Liubov Popova, Elena Mazunina, Inna Dolzhikova, Anna Kovyrshina, Andrei Botikov, Dmitry Shcheblyakov, Vladimir Gushchin, Denis Logunov, Alexander Gintsburg, Ksenia Safina, Elena Nabieva, Georgii Bazykin, Dmitry Lioznov |
| EPI_ISL_2816289 | University College London, Great Ormond Street Hospital for Children NHS Foundation Trust, Imperial College Healthcare NHS Trust | COVID-19 Genomics UK (COG-UK) Consortium | Sergi Castellano, Rachel Williams, Mark Kristiansen, Sunando Roy, Charlotte Williams, Nadua Bayzid, Marius Cotic, Helena Tutill, Patricia Dyal, Judith Breuer |

| | | | |
|---|---|---|---|
| EPI_ISL_2816290 | N.F. Gamaleya Research Center for Epidemiology and Microbiology | WHO National Influenza Centre Russian Federation | Andrey Komissarov, Artem Fadeev, Kseniya Komissarova, Oula Mansour, Kirill Varchenko, Mikhail Bakaev, Tamila Musaeva, Veronika Eder, Maria Pisareva, Nikita Yolshin, Daria Danilenko,Elizaveta Divisenko, Alina Odintsova, Elena Shidlovskaya, Ludmila Vasilchenko, Nadezhda Kuznetsova, Maria Nikiforova, Alexey Shchetinin, Artem Tkachuk, Andrei Pochtovyi, Andrei Siniavin, Denis Kleymenov, Evgeniia Bykonia, Liubov Popova, Elena Mazunina, Inna Dolzhikova, Anna Kovyrshina, Andrei Botikov, Dmitry Shcheblyakov, Vladimir Gushchin, Denis Logunov, Alexander Gintsburg, Ksenia Safina, Elena Nabieva, Georgii Bazykin, Dmitry Liozno |
| EPI_ISL_2816291 | University College London, Great Ormond Street Hospital for Children NHS Foundation Trust, Imperial College Healthcare NHS Trust | COVID-19 Genomics UK (COG-UK) Consortium | Sergi Castellano, Rachel Williams, Mark Kristiansen, Sunando Roy, Charlotte Williams, Nadua Bayzid, Marius Cotic, Helena Tutill, Patricia Dyal, Judith Breuer |
| EPI_ISL_2816292 | N.F. Gamaleya Research Center for Epidemiology and Microbiology | WHO National Influenza Centre Russian Federation | Andrey Komissarov, Artem Fadeev, Kseniya Komissarova, Oula Mansour, Kirill Varchenko, Mikhail Bakaev, Tamila Musaeva, Veronika Eder, Maria Pisareva, Nikita Yolshin, Daria Danilenko,Elizaveta Divisenko, Alina Odintsova, Elena Shidlovskaya, Ludmila Vasilchenko, Nadezhda Kuznetsova, Maria Nikiforova, Alexey Shchetinin, Artem Tkachuk, Andrei Pochtovyi, Andrei Siniavin, Denis Kleymenov, Evgeniia Bykonia, Liubov Popova, Elena Mazunina, Inna Dolzhikova, Anna Kovyrshina, Andrei Botikov, Dmitry Shcheblyakov, Vladimir Gushchin, Denis Logunov, Alexander Gintsburg, Ksenia Safina, Elena Nabieva, Georgii Bazykin, Dmitry Liozno |
| EPI_ISL_2816293, EPI_ISL_2816295, EPI_ISL_2816296, EPI_ISL_2816297 | University College London, Great Ormond Street Hospital for Children NHS Foundation Trust, Imperial College Healthcare NHS Trust | COVID-19 Genomics UK (COG-UK) Consortium | Sergi Castellano, Rachel Williams, Mark Kristiansen, Sunando Roy, Charlotte Williams, Nadua Bayzid, Marius Cotic, Helena Tutill, Patricia Dyal, Judith Breuer |
| EPI_ISL_2816298 | N.F. Gamaleya Research Center for Epidemiology and Microbiology | WHO National Influenza Centre Russian Federation | Andrey Komissarov, Artem Fadeev, Kseniya Komissarova, Oula Mansour, Kirill Varchenko, Mikhail Bakaev, Tamila Musaeva, Veronika Eder, Maria Pisareva, Nikita Yolshin, Daria Danilenko,Elizaveta Divisenko, Alina Odintsova, Elena Shidlovskaya, Ludmila Vasilchenko, Nadezhda Kuznetsova, Maria Nikiforova, Alexey Shchetinin, Artem Tkachuk, Andrei Pochtovyi, Andrei Siniavin, Denis Kleymenov, Evgeniia Bykonia, Liubov Popova, Elena Mazunina, Inna Dolzhikova, Anna Kovyrshina, Andrei Botikov, Dmitry Shcheblyakov, Vladimir Gushchin, Denis Logunov, Alexander Gintsburg, Ksenia Safina, Elena Nabieva, Georgii Bazykin, Dmitry Liozno |
| EPI_ISL_2816299, EPI_ISL_2816300 | University College London, Great Ormond Street Hospital for Children NHS Foundation Trust, Imperial College Healthcare NHS Trust | COVID-19 Genomics UK (COG-UK) Consortium | Sergi Castellano, Rachel Williams, Mark Kristiansen, Sunando Roy, Charlotte Williams, Nadua Bayzid, Marius Cotic, Helena Tutill, Patricia Dyal, Judith Breuer |
| EPI_ISL_2816301 | N.F. Gamaleya Research Center for Epidemiology and Microbiology | WHO National Influenza Centre Russian Federation | Andrey Komissarov, Artem Fadeev, Kseniya Komissarova, Oula Mansour, Kirill Varchenko, Mikhail Bakaev, Tamila Musaeva, Veronika Eder, Maria Pisareva, Nikita Yolshin, Daria Danilenko,Elizaveta Divisenko, Alina Odintsova, Elena Shidlovskaya, Ludmila Vasilchenko, Nadezhda Kuznetsova, Maria Nikiforova, Alexey Shchetinin, Artem Tkachuk, Andrei Pochtovyi, Andrei Siniavin, Denis Kleymenov, Evgeniia Bykonia, Liubov Popova, Elena Mazunina, Inna Dolzhikova, Anna Kovyrshina, Andrei Botikov, Dmitry Shcheblyakov, Vladimir Gushchin, Denis Logunov, Alexander Gintsburg, Ksenia Safina, Elena Nabieva, Georgii Bazykin, Dmitry Liozno |
| EPI_ISL_2816302, EPI_ISL_2816303 | University College London, Great Ormond Street Hospital for Children NHS Foundation Trust, Imperial College Healthcare NHS Trust | COVID-19 Genomics UK (COG-UK) Consortium | Sergi Castellano, Rachel Williams, Mark Kristiansen, Sunando Roy, Charlotte Williams, Nadua Bayzid, Marius Cotic, Helena Tutill, Patricia Dyal, Judith Breuer |
| EPI_ISL_2816304 | N.F. Gamaleya Research Center for Epidemiology and Microbiology | WHO National Influenza Centre Russian Federation | Andrey Komissarov, Artem Fadeev, Kseniya Komissarova, Oula Mansour, Kirill Varchenko, Mikhail Bakaev, Tamila Musaeva, Veronika Eder, Maria Pisareva, Nikita Yolshin, Daria Danilenko,Elizaveta Divisenko, Alina Odintsova, Elena Shidlovskaya, Ludmila Vasilchenko, Nadezhda Kuznetsova, Maria Nikiforova, Alexey Shchetinin, Artem Tkachuk, Andrei Pochtovyi, Andrei Siniavin, Denis Kleymenov, Evgeniia Bykonia, Liubov Popova, Elena Mazunina, Inna Dolzhikova, Anna Kovyrshina, Andrei Botikov, Dmitry Shcheblyakov, Vladimir Gushchin, Denis Logunov, Alexander Gintsburg, Ksenia Safina, Elena Nabieva, Georgii Bazykin, Dmitry Liozno |
| EPI_ISL_2816305 | University College London, Great Ormond Street Hospital for Children NHS Foundation Trust, Imperial College Healthcare NHS Trust | COVID-19 Genomics UK (COG-UK) Consortium | Sergi Castellano, Rachel Williams, Mark Kristiansen, Sunando Roy, Charlotte Williams, Nadua Bayzid, Marius Cotic, Helena Tutill, Patricia Dyal, Judith Breuer |
| EPI_ISL_2816306 | N.F. Gamaleya Research Center for Epidemiology and Microbiology | WHO National Influenza Centre Russian Federation | Andrey Komissarov, Artem Fadeev, Kseniya Komissarova, Oula Mansour, Kirill Varchenko, Mikhail Bakaev, Tamila Musaeva, Veronika Eder, Maria Pisareva, Nikita Yolshin, Daria Danilenko,Elizaveta Divisenko, Alina Odintsova, Elena Shidlovskaya, Ludmila Vasilchenko, Nadezhda Kuznetsova, Maria Nikiforova, Alexey Shchetinin, Artem Tkachuk, Andrei Pochtovyi, Andrei Siniavin, Denis Kleymenov, Evgeniia Bykonia, Liubov Popova, Elena Mazunina, Inna Dolzhikova, Anna Kovyrshina, Andrei Botikov, Dmitry Shcheblyakov, Vladimir Gushchin, Denis Logunov, Alexander Gintsburg, Ksenia Safina, Elena Nabieva, Georgii Bazykin, Dmitry Liozno |
| EPI_ISL_2816307, EPI_ISL_2816308 | University College London, Great Ormond Street Hospital for Children NHS Foundation Trust, Imperial College Healthcare NHS Trust | COVID-19 Genomics UK (COG-UK) Consortium | Sergi Castellano, Rachel Williams, Mark Kristiansen, Sunando Roy, Charlotte Williams, Nadua Bayzid, Marius Cotic, Helena Tutill, Patricia Dyal, Judith Breuer |
| EPI_ISL_2816309 | N.F. Gamaleya Research Center for Epidemiology and Microbiology | WHO National Influenza Centre Russian Federation | Andrey Komissarov, Artem Fadeev, Kseniya Komissarova, Oula Mansour, Kirill Varchenko, Mikhail Bakaev, Tamila Musaeva, Veronika Eder, Maria Pisareva, Nikita Yolshin, Daria Danilenko,Elizaveta Divisenko, Alina Odintsova, Elena Shidlovskaya, Ludmila Vasilchenko, Nadezhda Kuznetsova, Maria Nikiforova, Alexey Shchetinin, Artem Tkachuk, Andrei Pochtovyi, Andrei Siniavin, Denis Kleymenov, Evgeniia Bykonia, Liubov Popova, Elena Mazunina, Inna Dolzhikova, Anna Kovyrshina, Andrei Botikov, Dmitry Shcheblyakov, Vladimir Gushchin, Denis Logunov, Alexander Gintsburg, Ksenia Safina, Elena Nabieva, Georgii Bazykin, Dmitry Liozno |
| EPI_ISL_2816310 | University College London, Great Ormond Street Hospital for Children NHS Foundation Trust, Imperial College Healthcare NHS Trust | COVID-19 Genomics UK (COG-UK) Consortium | Sergi Castellano, Rachel Williams, Mark Kristiansen, Sunando Roy, Charlotte Williams, Nadua Bayzid, Marius Cotic, Helena Tutill, Patricia Dyal, Judith Breuer |
| EPI_ISL_2816311 | N.F. Gamaleya Research Center for Epidemiology and Microbiology | WHO National Influenza Centre Russian Federation | Andrey Komissarov, Artem Fadeev, Kseniya Komissarova, Oula Mansour, Kirill Varchenko, Mikhail Bakaev, Tamila Musaeva, Veronika Eder, Maria Pisareva, Nikita Yolshin, Daria Danilenko,Elizaveta Divisenko, Alina Odintsova, Elena Shidlovskaya, Ludmila Vasilchenko, Nadezhda Kuznetsova, Maria Nikiforova, Alexey Shchetinin, Artem Tkachuk, Andrei Pochtovyi, Andrei Siniavin, Denis Kleymenov, Evgeniia Bykonia, Liubov Popova, Elena Mazunina, Inna Dolzhikova, Anna Kovyrshina, Andrei Botikov, Dmitry Shcheblyakov, Vladimir Gushchin, Denis Logunov, Alexander Gintsburg, Ksenia Safina, Elena Nabieva, Georgii Bazykin, Dmitry Liozno |
| EPI_ISL_2816312, EPI_ISL_2816313, EPI_ISL_2816315 | University College London, Great Ormond Street Hospital for Children NHS Foundation Trust, Imperial College Healthcare NHS Trust | COVID-19 Genomics UK (COG-UK) Consortium | Sergi Castellano, Rachel Williams, Mark Kristiansen, Sunando Roy, Charlotte Williams, Nadua Bayzid, Marius Cotic, Helena Tutill, Patricia Dyal, Judith Breuer |
| EPI_ISL_2816316 | N.F. Gamaleya Research Center for Epidemiology and Microbiology | WHO National Influenza Centre Russian Federation | Andrey Komissarov, Artem Fadeev, Kseniya Komissarova, Oula Mansour, Kirill Varchenko, Mikhail Bakaev, Tamila Musaeva, Veronika Eder, Maria Pisareva, Nikita Yolshin, Daria Danilenko,Elizaveta Divisenko, Alina Odintsova, Elena Shidlovskaya, Ludmila Vasilchenko, Nadezhda Kuznetsova, Maria Nikiforova, Alexey Shchetinin, Artem Tkachuk, Andrei Pochtovyi, Andrei Siniavin, Denis Kleymenov, Evgeniia Bykonia, Liubov Popova, Elena Mazunina, Inna Dolzhikova, Anna Kovyrshina, Andrei Botikov, Dmitry Shcheblyakov, Vladimir Gushchin, Denis Logunov, Alexander Gintsburg, Ksenia Safina, Elena Nabieva, Georgii Bazykin, Dmitry Liozno |
| EPI_ISL_2816317 | University College London, Great Ormond Street Hospital for Children NHS Foundation Trust, Imperial College Healthcare NHS Trust | COVID-19 Genomics UK (COG-UK) Consortium | Sergi Castellano, Rachel Williams, Mark Kristiansen, Sunando Roy, Charlotte Williams, Nadua Bayzid, Marius Cotic, Helena Tutill, Patricia Dyal, Judith Breuer |
| EPI_ISL_2816318 | N.F. Gamaleya Research Center for Epidemiology and Microbiology | WHO National Influenza Centre Russian Federation | Andrey Komissarov, Artem Fadeev, Kseniya Komissarova, Oula Mansour, Kirill Varchenko, Mikhail Bakaev, Tamila Musaeva, Veronika Eder, Maria Pisareva, Nikita Yolshin, Daria Danilenko,Elizaveta Divisenko, Alina Odintsova, Elena Shidlovskaya, Ludmila Vasilchenko, Nadezhda Kuznetsova, Maria Nikiforova, Alexey Shchetinin, Artem Tkachuk, Andrei Pochtovyi, Andrei Siniavin, Denis Kleymenov, Evgeniia Bykonia, Liubov Popova, Elena Mazunina, Inna Dolzhikova, Anna Kovyrshina, Andrei Botikov, Dmitry Shcheblyakov, Vladimir Gushchin, Denis Logunov, Alexander Gintsburg, Ksenia Safina, Elena Nabieva, Georgii Bazykin, Dmitry Liozno |
| EPI_ISL_2816319, EPI_ISL_2816320 | University College London, Great Ormond Street Hospital for Children NHS Foundation Trust, Imperial College Healthcare NHS Trust | COVID-19 Genomics UK (COG-UK) Consortium | Sergi Castellano, Rachel Williams, Mark Kristiansen, Sunando Roy, Charlotte Williams, Nadua Bayzid, Marius Cotic, Helena Tutill, Patricia Dyal, Judith Breuer |
| EPI_ISL_2816321 | N.F. Gamaleya Research Center for Epidemiology and Microbiology | WHO National Influenza Centre Russian Federation | Andrey Komissarov, Artem Fadeev, Kseniya Komissarova, Oula Mansour, Kirill Varchenko, Mikhail Bakaev, Tamila Musaeva, Veronika Eder, Maria Pisareva, Nikita Yolshin, Daria Danilenko,Elizaveta Divisenko, Alina Odintsova, Elena Shidlovskaya, Ludmila Vasilchenko, Nadezhda Kuznetsova, Maria Nikiforova, Alexey Shchetinin, Artem Tkachuk, Andrei Pochtovyi, Andrei Siniavin, Denis Kleymenov, Evgeniia Bykonia, Liubov Popova, Elena Mazunina, Inna Dolzhikova, Anna Kovyrshina, Andrei Botikov, Dmitry Shcheblyakov, Vladimir Gushchin, Denis Logunov, Alexander Gintsburg, Ksenia Safina, Elena Nabieva, Georgii Bazykin, Dmitry Liozno |
| EPI_ISL_2816322 | University College London, Great Ormond Street Hospital for Children NHS Foundation Trust, Imperial College Healthcare NHS Trust | COVID-19 Genomics UK (COG-UK) Consortium | Sergi Castellano, Rachel Williams, Mark Kristiansen, Sunando Roy, Charlotte Williams, Nadua Bayzid, Marius Cotic, Helena Tutill, Patricia Dyal, Judith Breuer |
| EPI_ISL_2816323 | N.F. Gamaleya Research Center for Epidemiology and Microbiology | WHO National Influenza Centre Russian Federation | Andrey Komissarov, Artem Fadeev, Kseniya Komissarova, Oula Mansour, Kirill Varchenko, Mikhail Bakaev, Tamila Musaeva, Veronika Eder, Maria Pisareva, Nikita Yolshin, Daria Danilenko,Elizaveta Divisenko, Alina Odintsova, Elena Shidlovskaya, Ludmila Vasilchenko, Nadezhda Kuznetsova, Maria Nikiforova, Alexey Shchetinin, Artem Tkachuk, Andrei Pochtovyi, Andrei Siniavin, Denis Kleymenov, Evgeniia Bykonia, Liubov Popova, Elena Mazunina, Inna Dolzhikova, Anna Kovyrshina, Andrei Botikov, Dmitry Shcheblyakov, Vladimir Gushchin, Denis Logunov, Alexander Gintsburg, Ksenia Safina, Elena Nabieva, Georgii Bazykin, Dmitry Liozno |
| EPI_ISL_2816324, EPI_ISL_2816325 | University College London, Great Ormond Street Hospital for Children NHS Foundation Trust, Imperial College Healthcare NHS Trust | COVID-19 Genomics UK (COG-UK) Consortium | Sergi Castellano, Rachel Williams, Mark Kristiansen, Sunando Roy, Charlotte Williams, Nadua Bayzid, Marius Cotic, Helena Tutill, Patricia Dyal, Judith Breuer |
| EPI_ISL_2816326 | N.F. Gamaleya Research Center for Epidemiology and Microbiology | WHO National Influenza Centre Russian Federation | Andrey Komissarov, Artem Fadeev, Kseniya Komissarova, Oula Mansour, Kirill Varchenko, Mikhail Bakaev, Tamila Musaeva, Veronika Eder, Maria Pisareva, Nikita Yolshin, Daria Danilenko,Elizaveta Divisenko, Alina Odintsova, Elena Shidlovskaya, Ludmila Vasilchenko, Nadezhda Kuznetsova, Maria Nikiforova, Alexey Shchetinin, Artem Tkachuk, Andrei Pochtovyi, Andrei Siniavin, Denis Kleymenov, Evgeniia Bykonia, Liubov Popova, Elena Mazunina, Inna Dolzhikova, Anna Kovyrshina, Andrei Botikov, Dmitry Shcheblyakov, Vladimir Gushchin, Denis Logunov, Alexander Gintsburg, Ksenia Safina, Elena Nabieva, Georgii Bazykin, Dmitry Liozno |
| EPI_ISL_2816327, EPI_ISL_2816329, EPI_ISL_2816330 | University College London, Great Ormond Street Hospital for Children NHS Foundation Trust, Imperial College Healthcare NHS Trust | COVID-19 Genomics UK (COG-UK) Consortium | Sergi Castellano, Rachel Williams, Mark Kristiansen, Sunando Roy, Charlotte Williams, Nadua Bayzid, Marius Cotic, Helena Tutill, Patricia Dyal, Judith Breuer |
| EPI_ISL_2816331 | N.F. Gamaleya Research Center for Epidemiology and Microbiology | WHO National Influenza Centre Russian Federation | Andrey Komissarov, Artem Fadeev, Kseniya Komissarova, Oula Mansour, Kirill Varchenko, Mikhail Bakaev, Tamila Musaeva, Veronika Eder, Maria Pisareva, Nikita Yolshin, Daria Danilenko,Elizaveta Divisenko, Alina Odintsova, Elena Shidlovskaya, Ludmila Vasilchenko, Nadezhda Kuznetsova, Maria Nikiforova, Alexey Shchetinin, Artem Tkachuk, Andrei Pochtovyi, Andrei Siniavin, Denis Kleymenov, Evgeniia Bykonia, Liubov Popova, Elena Mazunina, Inna Dolzhikova, Anna Kovyrshina, Andrei Botikov, Dmitry Shcheblyakov, Vladimir Gushchin, Denis Logunov, Alexander Gintsburg, Ksenia Safina, Elena Nabieva, Georgii Bazykin, Dmitry Liozno |
| EPI_ISL_2816332 | University College London, Great Ormond Street Hospital for Children NHS Foundation Trust, Imperial College Healthcare NHS Trust | COVID-19 Genomics UK (COG-UK) Consortium | Sergi Castellano, Rachel Williams, Mark Kristiansen, Sunando Roy, Charlotte Williams, Nadua Bayzid, Marius Cotic, Helena Tutill, Patricia Dyal, Judith Breuer |
| EPI_ISL_2816334 | N.F. Gamaleya Research Center for Epidemiology and Microbiology | WHO National Influenza Centre Russian Federation | Andrey Komissarov, Artem Fadeev, Kseniya Komissarova, Oula Mansour, Kirill Varchenko, Mikhail Bakaev, Tamila Musaeva, Veronika Eder, Maria Pisareva, Nikita Yolshin, Daria Danilenko,Elizaveta Divisenko, Alina Odintsova, Elena Shidlovskaya, Ludmila Vasilchenko, Nadezhda Kuznetsova, Maria Nikiforova, Alexey Shchetinin, Artem Tkachuk, Andrei Pochtovyi, Andrei Siniavin, Denis Kleymenov, Evgeniia Bykonia, Liubov Popova, Elena Mazunina, Inna Dolzhikova, Anna Kovyrshina, Andrei Botikov, Dmitry Shcheblyakov, Vladimir Gushchin, Denis Logunov, Alexander Gintsburg, Ksenia Safina, Elena Nabieva, Georgii Bazykin, Dmitry Liozno |
| EPI_ISL_2816335 | University College London, Great Ormond Street Hospital for Children NHS Foundation Trust, Imperial College Healthcare NHS Trust | COVID-19 Genomics UK (COG-UK) Consortium | Sergi Castellano, Rachel Williams, Mark Kristiansen, Sunando Roy, Charlotte Williams, Nadua Bayzid, Marius Cotic, Helena Tutill, Patricia Dyal, Judith Breuer |
| EPI_ISL_2816336 | N.F. Gamaleya Research Center for Epidemiology and Microbiology | WHO National Influenza Centre Russian Federation | Andrey Komissarov, Artem Fadeev, Kseniya Komissarova, Oula Mansour, Kirill Varchenko, Mikhail Bakaev, Tamila Musaeva, Veronika Eder, Maria Pisareva, Nikita Yolshin, Daria Danilenko,Elizaveta Divisenko, Alina Odintsova, Elena Shidlovskaya, Ludmila Vasilchenko, Nadezhda Kuznetsova, Maria Nikiforova, Alexey Shchetinin, Artem Tkachuk, Andrei Pochtovyi, Andrei Siniavin, Denis Kleymenov, Evgeniia Bykonia, Liubov Popova, Elena Mazunina, Inna Dolzhikova, Anna Kovyrshina, Andrei Botikov, Dmitry Shcheblyakov, Vladimir Gushchin, Denis Logunov, Alexander Gintsburg, Ksenia Safina, Elena Nabieva, Georgii Bazykin, Dmitry Liozno |
| EPI_ISL_2816337, EPI_ISL_2816338 | University College London, Great Ormond Street Hospital for Children NHS Foundation Trust, Imperial College Healthcare NHS | COVID-19 Genomics UK (COG-UK) Consortium | Sergi Castellano, Rachel Williams, Mark Kristiansen, Sunando Roy, Charlotte Williams, Nadua Bayzid, Marius Cotic, Helena Tutill, Patricia Dyal, Judith Breuer |

| | | | |
|---|---|---|---|
| | Trust | | |
| EPI_ISL_2816339 | N.F. Gamaleya Research Center for Epidemiology and Microbiology | WHO National Influenza Centre Russian Federation | Andrey Komissarov, Artem Fadeev, Kseniya Komissarova, Oula Mansour, Kirill Varchenko, Mikhail Bakaev, Tamila Musaeva, Veronika Eder, Maria Pisareva, Nikita Yolshin, Daria Danilenko,Elizaveta Divisenko, Alina Odintsova, Elena Shidlovskaya, Ludmila Vasilchenko, Nadezhda Kuznetsova, Maria Nikiforova, Alexey Shchetinin, Artem Tkachuk, Andrei Pochtovyi, Andrei Siniavin, Denis Kleymenov, Evgeniia Bykonia, Liubov Popova, Elena Mazunina, Inna Dolzhikova, Anna Kovyrshina, Andrei Botikov, Dmitry Shcheblyakov, Vladimir Gushchin, Denis Logunov, Alexander Gintsburg, Ksenia Safina, Elena Nabieva, Georgii Bazykin, Dmitry Lioznov |
| EPI_ISL_2816340 | University College London, Great Ormond Street Hospital for Children NHS Foundation Trust, Imperial College Healthcare NHS Trust | COVID-19 Genomics UK (COG-UK) Consortium | Sergi Castellano, Rachel Williams, Mark Kristiansen, Sunando Roy, Charlotte Williams, Nadua Bayzid, Marius Cotic, Helena Tutill, Patricia Dyal, Judith Breuer |
| EPI_ISL_2816341 | N.F. Gamaleya Research Center for Epidemiology and Microbiology | WHO National Influenza Centre Russian Federation | Andrey Komissarov, Artem Fadeev, Kseniya Komissarova, Oula Mansour, Kirill Varchenko, Mikhail Bakaev, Tamila Musaeva, Veronika Eder, Maria Pisareva, Nikita Yolshin, Daria Danilenko,Elizaveta Divisenko, Alina Odintsova, Elena Shidlovskaya, Ludmila Vasilchenko, Nadezhda Kuznetsova, Maria Nikiforova, Alexey Shchetinin, Artem Tkachuk, Andrei Pochtovyi, Andrei Siniavin, Denis Kleymenov, Evgeniia Bykonia, Liubov Popova, Elena Mazunina, Inna Dolzhikova, Anna Kovyrshina, Andrei Botikov, Dmitry Shcheblyakov, Vladimir Gushchin, Denis Logunov, Alexander Gintsburg, Ksenia Safina, Elena Nabieva, Georgii Bazykin, Dmitry Lioznov |
| EPI_ISL_2816342, EPI_ISL_2816343 | University College London, Great Ormond Street Hospital for Children NHS Foundation Trust, Imperial College Healthcare NHS Trust | COVID-19 Genomics UK (COG-UK) Consortium | Sergi Castellano, Rachel Williams, Mark Kristiansen, Sunando Roy, Charlotte Williams, Nadua Bayzid, Marius Cotic, Helena Tutill, Patricia Dyal, Judith Breuer |
| EPI_ISL_2816344 | N.F. Gamaleya Research Center for Epidemiology and Microbiology | WHO National Influenza Centre Russian Federation | Andrey Komissarov, Artem Fadeev, Kseniya Komissarova, Oula Mansour, Kirill Varchenko, Mikhail Bakaev, Tamila Musaeva, Veronika Eder, Maria Pisareva, Nikita Yolshin, Daria Danilenko,Elizaveta Divisenko, Alina Odintsova, Elena Shidlovskaya, Ludmila Vasilchenko, Nadezhda Kuznetsova, Maria Nikiforova, Alexey Shchetinin, Artem Tkachuk, Andrei Pochtovyi, Andrei Siniavin, Denis Kleymenov, Evgeniia Bykonia, Liubov Popova, Elena Mazunina, Inna Dolzhikova, Anna Kovyrshina, Andrei Botikov, Dmitry Shcheblyakov, Vladimir Gushchin, Denis Logunov, Alexander Gintsburg, Ksenia Safina, Elena Nabieva, Georgii Bazykin, Dmitry Lioznov |
| EPI_ISL_2816345 | University College London, Great Ormond Street Hospital for Children NHS Foundation Trust, Imperial College Healthcare NHS Trust | COVID-19 Genomics UK (COG-UK) Consortium | Sergi Castellano, Rachel Williams, Mark Kristiansen, Sunando Roy, Charlotte Williams, Nadua Bayzid, Marius Cotic, Helena Tutill, Patricia Dyal, Judith Breuer |
| EPI_ISL_2816346 | N.F. Gamaleya Research Center for Epidemiology and Microbiology | WHO National Influenza Centre Russian Federation | Andrey Komissarov, Artem Fadeev, Kseniya Komissarova, Oula Mansour, Kirill Varchenko, Mikhail Bakaev, Tamila Musaeva, Veronika Eder, Maria Pisareva, Nikita Yolshin, Daria Danilenko,Elizaveta Divisenko, Alina Odintsova, Elena Shidlovskaya, Ludmila Vasilchenko, Nadezhda Kuznetsova, Maria Nikiforova, Alexey Shchetinin, Artem Tkachuk, Andrei Pochtovyi, Andrei Siniavin, Denis Kleymenov, Evgeniia Bykonia, Liubov Popova, Elena Mazunina, Inna Dolzhikova, Anna Kovyrshina, Andrei Botikov, Dmitry Shcheblyakov, Vladimir Gushchin, Denis Logunov, Alexander Gintsburg, Ksenia Safina, Elena Nabieva, Georgii Bazykin, Dmitry Lioznov |
| EPI_ISL_2816347, EPI_ISL_2816348 | University College London, Great Ormond Street Hospital for Children NHS Foundation Trust, Imperial College Healthcare NHS Trust | COVID-19 Genomics UK (COG-UK) Consortium | Sergi Castellano, Rachel Williams, Mark Kristiansen, Sunando Roy, Charlotte Williams, Nadua Bayzid, Marius Cotic, Helena Tutill, Patricia Dyal, Judith Breuer |
| EPI_ISL_2816349 | N.F. Gamaleya Research Center for Epidemiology and Microbiology | WHO National Influenza Centre Russian Federation | Andrey Komissarov, Artem Fadeev, Kseniya Komissarova, Oula Mansour, Kirill Varchenko, Mikhail Bakaev, Tamila Musaeva, Veronika Eder, Maria Pisareva, Nikita Yolshin, Daria Danilenko,Elizaveta Divisenko, Alina Odintsova, Elena Shidlovskaya, Ludmila Vasilchenko, Nadezhda Kuznetsova, Maria Nikiforova, Alexey Shchetinin, Artem Tkachuk, Andrei Pochtovyi, Andrei Siniavin, Denis Kleymenov, Evgeniia Bykonia, Liubov Popova, Elena Mazunina, Inna Dolzhikova, Anna Kovyrshina, Andrei Botikov, Dmitry Shcheblyakov, Vladimir Gushchin, Denis Logunov, Alexander Gintsburg, Ksenia Safina, Elena Nabieva, Georgii Bazykin, Dmitry Lioznov |
| EPI_ISL_2816350 | University College London, Great Ormond Street Hospital for Children NHS Foundation Trust, Imperial College Healthcare NHS Trust | COVID-19 Genomics UK (COG-UK) Consortium | Sergi Castellano, Rachel Williams, Mark Kristiansen, Sunando Roy, Charlotte Williams, Nadua Bayzid, Marius Cotic, Helena Tutill, Patricia Dyal, Judith Breuer |
| EPI_ISL_2816351 | N.F. Gamaleya Research Center for Epidemiology and Microbiology | WHO National Influenza Centre Russian Federation | Andrey Komissarov, Artem Fadeev, Kseniya Komissarova, Oula Mansour, Kirill Varchenko, Mikhail Bakaev, Tamila Musaeva, Veronika Eder, Maria Pisareva, Nikita Yolshin, Daria Danilenko,Elizaveta Divisenko, Alina Odintsova, Elena Shidlovskaya, Ludmila Vasilchenko, Nadezhda Kuznetsova, Maria Nikiforova, Alexey Shchetinin, Artem Tkachuk, Andrei Pochtovyi, Andrei Siniavin, Denis Kleymenov, Evgeniia Bykonia, Liubov Popova, Elena Mazunina, Inna Dolzhikova, Anna Kovyrshina, Andrei Botikov, Dmitry Shcheblyakov, Vladimir Gushchin, Denis Logunov, Alexander Gintsburg, Ksenia Safina, Elena Nabieva, Georgii Bazykin, Dmitry Lioznov |
| EPI_ISL_2816352, EPI_ISL_2816353 | University College London, Great Ormond Street Hospital for Children NHS Foundation Trust, Imperial College Healthcare NHS Trust | COVID-19 Genomics UK (COG-UK) Consortium | Sergi Castellano, Rachel Williams, Mark Kristiansen, Sunando Roy, Charlotte Williams, Nadua Bayzid, Marius Cotic, Helena Tutill, Patricia Dyal, Judith Breuer |
| EPI_ISL_2816354 | N.F. Gamaleya Research Center for Epidemiology and Microbiology | WHO National Influenza Centre Russian Federation | Andrey Komissarov, Artem Fadeev, Kseniya Komissarova, Oula Mansour, Kirill Varchenko, Mikhail Bakaev, Tamila Musaeva, Veronika Eder, Maria Pisareva, Nikita Yolshin, Daria Danilenko,Elizaveta Divisenko, Alina Odintsova, Elena Shidlovskaya, Ludmila Vasilchenko, Nadezhda Kuznetsova, Maria Nikiforova, Alexey Shchetinin, Artem Tkachuk, Andrei Pochtovyi, Andrei Siniavin, Denis Kleymenov, Evgeniia Bykonia, Liubov Popova, Elena Mazunina, Inna Dolzhikova, Anna Kovyrshina, Andrei Botikov, Dmitry Shcheblyakov, Vladimir Gushchin, Denis Logunov, Alexander Gintsburg, Ksenia Safina, Elena Nabieva, Georgii Bazykin, Dmitry Lioznov |
| EPI_ISL_2816355 | University College London, Great Ormond Street Hospital for Children NHS Foundation Trust, Imperial College Healthcare NHS Trust | COVID-19 Genomics UK (COG-UK) Consortium | Sergi Castellano, Rachel Williams, Mark Kristiansen, Sunando Roy, Charlotte Williams, Nadua Bayzid, Marius Cotic, Helena Tutill, Patricia Dyal, Judith Breuer |
| EPI_ISL_2816356 | N.F. Gamaleya Research Center for Epidemiology and Microbiology | WHO National Influenza Centre Russian Federation | Andrey Komissarov, Artem Fadeev, Kseniya Komissarova, Oula Mansour, Kirill Varchenko, Mikhail Bakaev, Tamila Musaeva, Veronika Eder, Maria Pisareva, Nikita Yolshin, Daria Danilenko,Elizaveta Divisenko, Alina Odintsova, Elena Shidlovskaya, Ludmila Vasilchenko, Nadezhda Kuznetsova, Maria Nikiforova, Alexey Shchetinin, Artem Tkachuk, Andrei Pochtovyi, Andrei Siniavin, Denis Kleymenov, Evgeniia Bykonia, Liubov Popova, Elena Mazunina, Inna Dolzhikova, Anna Kovyrshina, Andrei Botikov, Dmitry Shcheblyakov, Vladimir Gushchin, Denis Logunov, Alexander Gintsburg, Ksenia Safina, Elena Nabieva, Georgii Bazykin, Dmitry Lioznov |
| EPI_ISL_2816357, EPI_ISL_2816358 | University College London, Great Ormond Street Hospital for Children NHS Foundation Trust, Imperial College Healthcare NHS Trust | COVID-19 Genomics UK (COG-UK) Consortium | Sergi Castellano, Rachel Williams, Mark Kristiansen, Sunando Roy, Charlotte Williams, Nadua Bayzid, Marius Cotic, Helena Tutill, Patricia Dyal, Judith Breuer |
| EPI_ISL_2816359 | N.F. Gamaleya Research Center for Epidemiology and Microbiology | WHO National Influenza Centre Russian Federation | Andrey Komissarov, Artem Fadeev, Kseniya Komissarova, Oula Mansour, Kirill Varchenko, Mikhail Bakaev, Tamila Musaeva, Veronika Eder, Maria Pisareva, Nikita Yolshin, Daria Danilenko,Elizaveta Divisenko, Alina Odintsova, Elena Shidlovskaya, Ludmila Vasilchenko, Nadezhda Kuznetsova, Maria Nikiforova, Alexey Shchetinin, Artem Tkachuk, Andrei Pochtovyi, Andrei Siniavin, Denis Kleymenov, Evgeniia Bykonia, Liubov Popova, Elena Mazunina, Inna Dolzhikova, Anna Kovyrshina, Andrei Botikov, Dmitry Shcheblyakov, Vladimir Gushchin, Denis Logunov, Alexander Gintsburg, Ksenia Safina, Elena Nabieva, Georgii Bazykin, Dmitry Lioznov |
| EPI_ISL_2816360 | University College London, Great Ormond Street Hospital for Children NHS Foundation Trust, Imperial College Healthcare NHS Trust | COVID-19 Genomics UK (COG-UK) Consortium | Sergi Castellano, Rachel Williams, Mark Kristiansen, Sunando Roy, Charlotte Williams, Nadua Bayzid, Marius Cotic, Helena Tutill, Patricia Dyal, Judith Breuer |
| EPI_ISL_2816361 | N.F. Gamaleya Research Center for Epidemiology and Microbiology | WHO National Influenza Centre Russian Federation | Andrey Komissarov, Artem Fadeev, Kseniya Komissarova, Oula Mansour, Kirill Varchenko, Mikhail Bakaev, Tamila Musaeva, Veronika Eder, Maria Pisareva, Nikita Yolshin, Daria Danilenko,Elizaveta Divisenko, Alina Odintsova, Elena Shidlovskaya, Ludmila Vasilchenko, Nadezhda Kuznetsova, Maria Nikiforova, Alexey Shchetinin, Artem Tkachuk, Andrei Pochtovyi, Andrei Siniavin, Denis Kleymenov, Evgeniia Bykonia, Liubov Popova, Elena Mazunina, Inna Dolzhikova, Anna Kovyrshina, Andrei Botikov, Dmitry Shcheblyakov, Vladimir Gushchin, Denis Logunov, Alexander Gintsburg, Ksenia Safina, Elena Nabieva, Georgii Bazykin, Dmitry Lioznov |
| EPI_ISL_2816362 | University College London, Great Ormond Street Hospital for Children NHS Foundation Trust, Imperial College Healthcare NHS Trust | COVID-19 Genomics UK (COG-UK) Consortium | Sergi Castellano, Rachel Williams, Mark Kristiansen, Sunando Roy, Charlotte Williams, Nadua Bayzid, Marius Cotic, Helena Tutill, Patricia Dyal, Judith Breuer |
| EPI_ISL_2816363 | N.F. Gamaleya Research Center for Epidemiology and Microbiology | WHO National Influenza Centre Russian Federation | Andrey Komissarov, Artem Fadeev, Kseniya Komissarova, Oula Mansour, Kirill Varchenko, Mikhail Bakaev, Tamila Musaeva, Veronika Eder, Maria Pisareva, Nikita Yolshin, Daria Danilenko,Elizaveta Divisenko, Alina Odintsova, Elena Shidlovskaya, Ludmila Vasilchenko, Nadezhda Kuznetsova, Maria Nikiforova, Alexey Shchetinin, Artem Tkachuk, Andrei Pochtovyi, Andrei Siniavin, Denis Kleymenov, Evgeniia Bykonia, Liubov Popova, Elena Mazunina, Inna Dolzhikova, Anna Kovyrshina, Andrei Botikov, Dmitry Shcheblyakov, Vladimir Gushchin, Denis Logunov, Alexander Gintsburg, Ksenia Safina, Elena Nabieva, Georgii Bazykin, Dmitry Lioznov |
| EPI_ISL_2816364, EPI_ISL_2816365 | University College London, Great Ormond Street Hospital for Children NHS Foundation Trust, Imperial College Healthcare NHS Trust | COVID-19 Genomics UK (COG-UK) Consortium | Sergi Castellano, Rachel Williams, Mark Kristiansen, Sunando Roy, Charlotte Williams, Nadua Bayzid, Marius Cotic, Helena Tutill, Patricia Dyal, Judith Breuer |
| EPI_ISL_2816366 | N.F. Gamaleya Research Center for Epidemiology and Microbiology | WHO National Influenza Centre Russian Federation | Andrey Komissarov, Artem Fadeev, Kseniya Komissarova, Oula Mansour, Kirill Varchenko, Mikhail Bakaev, Tamila Musaeva, Veronika Eder, Maria Pisareva, Nikita Yolshin, Daria Danilenko,Elizaveta Divisenko, Alina Odintsova, Elena Shidlovskaya, Ludmila Vasilchenko, Nadezhda Kuznetsova, Maria Nikiforova, Alexey Shchetinin, Artem Tkachuk, Andrei Pochtovyi, Andrei Siniavin, Denis Kleymenov, Evgeniia Bykonia, Liubov Popova, Elena Mazunina, Inna Dolzhikova, Anna Kovyrshina, Andrei Botikov, Dmitry Shcheblyakov, Vladimir Gushchin, Denis Logunov, Alexander Gintsburg, Ksenia Safina, Elena Nabieva, Georgii Bazykin, Dmitry Lioznov |
| EPI_ISL_2816367 | University College London, Great Ormond Street Hospital for Children NHS Foundation Trust, Imperial College Healthcare NHS Trust | COVID-19 Genomics UK (COG-UK) Consortium | Sergi Castellano, Rachel Williams, Mark Kristiansen, Sunando Roy, Charlotte Williams, Nadua Bayzid, Marius Cotic, Helena Tutill, Patricia Dyal, Judith Breuer |
| EPI_ISL_2816368 | N.F. Gamaleya Research Center for Epidemiology and Microbiology | WHO National Influenza Centre Russian Federation | Andrey Komissarov, Artem Fadeev, Kseniya Komissarova, Oula Mansour, Kirill Varchenko, Mikhail Bakaev, Tamila Musaeva, Veronika Eder, Maria Pisareva, Nikita Yolshin, Daria Danilenko,Elizaveta Divisenko, Alina Odintsova, Elena Shidlovskaya, Ludmila Vasilchenko, Nadezhda Kuznetsova, Maria Nikiforova, Alexey Shchetinin, Artem Tkachuk, Andrei Pochtovyi, Andrei Siniavin, Denis Kleymenov, Evgeniia Bykonia, Liubov Popova, Elena Mazunina, Inna Dolzhikova, Anna Kovyrshina, Andrei Botikov, Dmitry Shcheblyakov, Vladimir Gushchin, Denis Logunov, Alexander Gintsburg, Ksenia Safina, Elena Nabieva, Georgii Bazykin, Dmitry Lioznov |
| EPI_ISL_2816369, EPI_ISL_2816370 | University College London, Great Ormond Street Hospital for Children NHS Foundation Trust, Imperial College Healthcare NHS Trust | COVID-19 Genomics UK (COG-UK) Consortium | Sergi Castellano, Rachel Williams, Mark Kristiansen, Sunando Roy, Charlotte Williams, Nadua Bayzid, Marius Cotic, Helena Tutill, Patricia Dyal, Judith Breuer |
| EPI_ISL_2816371 | N.F. Gamaleya Research Center for Epidemiology and Microbiology | WHO National Influenza Centre Russian Federation | Andrey Komissarov, Artem Fadeev, Kseniya Komissarova, Oula Mansour, Kirill Varchenko, Mikhail Bakaev, Tamila Musaeva, Veronika Eder, Maria Pisareva, Nikita Yolshin, Daria Danilenko,Elizaveta Divisenko, Alina Odintsova, Elena Shidlovskaya, Ludmila Vasilchenko, Nadezhda Kuznetsova, Maria Nikiforova, Alexey Shchetinin, Artem Tkachuk, Andrei Pochtovyi, Andrei Siniavin, Denis Kleymenov, Evgeniia Bykonia, Liubov Popova, Elena Mazunina, Inna Dolzhikova, Anna Kovyrshina, Andrei Botikov, Dmitry Shcheblyakov, Vladimir Gushchin, Denis Logunov, Alexander Gintsburg, Ksenia Safina, Elena Nabieva, Georgii Bazykin, Dmitry Lioznov |
| EPI_ISL_2816372 | University College London, Great Ormond Street Hospital for Children NHS Foundation Trust, Imperial College Healthcare NHS Trust | COVID-19 Genomics UK (COG-UK) Consortium | Sergi Castellano, Rachel Williams, Mark Kristiansen, Sunando Roy, Charlotte Williams, Nadua Bayzid, Marius Cotic, Helena Tutill, Patricia Dyal, Judith Breuer |
| EPI_ISL_2816373 | N.F. Gamaleya Research Center for Epidemiology and Microbiology | WHO National Influenza Centre Russian Federation | Andrey Komissarov, Artem Fadeev, Kseniya Komissarova, Oula Mansour, Kirill Varchenko, Mikhail Bakaev, Tamila Musaeva, Veronika Eder, Maria Pisareva, Nikita Yolshin, Daria Danilenko,Elizaveta Divisenko, Alina Odintsova, Elena Shidlovskaya, Ludmila Vasilchenko, Nadezhda Kuznetsova, Maria Nikiforova, Alexey Shchetinin, Artem Tkachuk, Andrei Pochtovyi, Andrei Siniavin, Denis Kleymenov, Evgeniia Bykonia, Liubov Popova, Elena Mazunina, Inna Dolzhikova, Anna Kovyrshina, Andrei Botikov, Dmitry Shcheblyakov, Vladimir Gushchin, Denis Logunov, Alexander Gintsburg, Ksenia Safina, Elena Nabieva, Georgii Bazykin, Dmitry Lioznov |
| EPI_ISL_2816374, EPI_ISL_2816375 | University College London, Great Ormond Street Hospital for Children NHS Foundation Trust, Imperial College Healthcare NHS Trust | COVID-19 Genomics UK (COG-UK) Consortium | Sergi Castellano, Rachel Williams, Mark Kristiansen, Sunando Roy, Charlotte Williams, Nadua Bayzid, Marius Cotic, Helena Tutill, Patricia Dyal, Judith Breuer |
| EPI_ISL_2816376 | N.F. Gamaleya Research Center for Epidemiology and Microbiology | WHO National Influenza Centre Russian Federation | Andrey Komissarov, Artem Fadeev, Kseniya Komissarova, Oula Mansour, Kirill Varchenko, Mikhail Bakaev, Tamila Musaeva, Veronika Eder, Maria Pisareva, Nikita Yolshin, Daria Danilenko,Elizaveta Divisenko, Alina Odintsova, Elena Shidlovskaya, Ludmila Vasilchenko, Nadezhda Kuznetsova, Maria Nikiforova, Alexey Shchetinin, Artem Tkachuk, Andrei Pochtovyi, Andrei Siniavin, Denis Kleymenov, Evgeniia Bykonia, Liubov Popova, Elena Mazunina, Inna Dolzhikova, Anna Kovyrshina, Andrei Botikov, Dmitry Shcheblyakov, Vladimir Gushchin, Denis Logunov, Alexander Gintsburg, Ksenia Safina, Elena Nabieva, Georgii Bazykin, Dmitry Lioznov |
| EPI_ISL_2816377 | University College London, Great Ormond Street Hospital for Children NHS Foundation Trust, Imperial College Healthcare NHS | COVID-19 Genomics UK (COG-UK) Consortium | Sergi Castellano, Rachel Williams, Mark Kristiansen, Sunando Roy, Charlotte Williams, Nadua Bayzid, Marius Cotic, Helena Tutill, Patricia Dyal, Judith Breuer |

| | | | |
|---|---|---|---|
| | Trust | | |
| EPI_ISL_2816378 | N.F. Gamaleya Research Center for Epidemiology and Microbiology | WHO National Influenza Centre Russian Federation | Andrey Komissarov, Artem Fadeev, Kseniya Komissarova, Oula Mansour, Kirill Varchenko, Mikhail Bakaev, Tamila Musaeva, Veronika Eder, Maria Pisareva, Nikita Yolshin, Daria Danilenko,Elizaveta Divisenko, Alina Odintsova, Elena Shidlovskaya, Ludmila Vasilchenko, Nadezhda Kuznetsova, Maria Nikiforova, Alexey Shchetinin, Artem Tkachuk, Andrei Pochtovyi, Andrei Siniavin, Denis Kleymenov, Evgeniia Bykonia, Liubov Popova, Elena Mazunina, Inna Dolzhikova, Anna Kovyrshina, Andrei Botikov, Dmitry Shcheblyakov, Vladimir Gushchin, Denis Logunov, Alexander Gintsburg, Ksenia Safina, Elena Nabieva, Georgii Bazykin, Dmitry Lioznov |
| EPI_ISL_2816379 | University College London, Great Ormond Street Hospital for Children NHS Foundation Trust, Imperial College Healthcare NHS Trust | COVID-19 Genomics UK (COG-UK) Consortium | Sergi Castellano, Rachel Williams, Mark Kristiansen, Sunando Roy, Charlotte Williams, Nadua Bayzid, Marius Cotic, Helena Tutill, Patricia Dyal, Judith Breuer |
| EPI_ISL_2816380 | N.F. Gamaleya Research Center for Epidemiology and Microbiology | WHO National Influenza Centre Russian Federation | Andrey Komissarov, Artem Fadeev, Kseniya Komissarova, Oula Mansour, Kirill Varchenko, Mikhail Bakaev, Tamila Musaeva, Veronika Eder, Maria Pisareva, Nikita Yolshin, Daria Danilenko,Elizaveta Divisenko, Alina Odintsova, Elena Shidlovskaya, Ludmila Vasilchenko, Nadezhda Kuznetsova, Maria Nikiforova, Alexey Shchetinin, Artem Tkachuk, Andrei Pochtovyi, Andrei Siniavin, Denis Kleymenov, Evgeniia Bykonia, Liubov Popova, Elena Mazunina, Inna Dolzhikova, Anna Kovyrshina, Andrei Botikov, Dmitry Shcheblyakov, Vladimir Gushchin, Denis Logunov, Alexander Gintsburg, Ksenia Safina, Elena Nabieva, Georgii Bazykin, Dmitry Lioznov |
| EPI_ISL_2816381, EPI_ISL_2816382 | University College London, Great Ormond Street Hospital for Children NHS Foundation Trust, Imperial College Healthcare NHS Trust | COVID-19 Genomics UK (COG-UK) Consortium | Sergi Castellano, Rachel Williams, Mark Kristiansen, Sunando Roy, Charlotte Williams, Nadua Bayzid, Marius Cotic, Helena Tutill, Patricia Dyal, Judith Breuer |
| EPI_ISL_2816383 | N.F. Gamaleya Research Center for Epidemiology and Microbiology | WHO National Influenza Centre Russian Federation | Andrey Komissarov, Artem Fadeev, Kseniya Komissarova, Oula Mansour, Kirill Varchenko, Mikhail Bakaev, Tamila Musaeva, Veronika Eder, Maria Pisareva, Nikita Yolshin, Daria Danilenko,Elizaveta Divisenko, Alina Odintsova, Elena Shidlovskaya, Ludmila Vasilchenko, Nadezhda Kuznetsova, Maria Nikiforova, Alexey Shchetinin, Artem Tkachuk, Andrei Pochtovyi, Andrei Siniavin, Denis Kleymenov, Evgeniia Bykonia, Liubov Popova, Elena Mazunina, Inna Dolzhikova, Anna Kovyrshina, Andrei Botikov, Dmitry Shcheblyakov, Vladimir Gushchin, Denis Logunov, Alexander Gintsburg, Ksenia Safina, Elena Nabieva, Georgii Bazykin, Dmitry Lioznov |
| EPI_ISL_2816384, EPI_ISL_2816386, EPI_ISL_2816387 | University College London, Great Ormond Street Hospital for Children NHS Foundation Trust, Imperial College Healthcare NHS Trust | COVID-19 Genomics UK (COG-UK) Consortium | Sergi Castellano, Rachel Williams, Mark Kristiansen, Sunando Roy, Charlotte Williams, Nadua Bayzid, Marius Cotic, Helena Tutill, Patricia Dyal, Judith Breuer |
| EPI_ISL_2816388 | N.F. Gamaleya Research Center for Epidemiology and Microbiology | WHO National Influenza Centre Russian Federation | Andrey Komissarov, Artem Fadeev, Kseniya Komissarova, Oula Mansour, Kirill Varchenko, Mikhail Bakaev, Tamila Musaeva, Veronika Eder, Maria Pisareva, Nikita Yolshin, Daria Danilenko,Elizaveta Divisenko, Alina Odintsova, Elena Shidlovskaya, Ludmila Vasilchenko, Nadezhda Kuznetsova, Maria Nikiforova, Alexey Shchetinin, Artem Tkachuk, Andrei Pochtovyi, Andrei Siniavin, Denis Kleymenov, Evgeniia Bykonia, Liubov Popova, Elena Mazunina, Inna Dolzhikova, Anna Kovyrshina, Andrei Botikov, Dmitry Shcheblyakov, Vladimir Gushchin, Denis Logunov, Alexander Gintsburg, Ksenia Safina, Elena Nabieva, Georgii Bazykin, Dmitry Lioznov |
| EPI_ISL_2816389 | University College London, Great Ormond Street Hospital for Children NHS Foundation Trust, Imperial College Healthcare NHS Trust | COVID-19 Genomics UK (COG-UK) Consortium | Sergi Castellano, Rachel Williams, Mark Kristiansen, Sunando Roy, Charlotte Williams, Nadua Bayzid, Marius Cotic, Helena Tutill, Patricia Dyal, Judith Breuer |
| EPI_ISL_2816390 | N.F. Gamaleya Research Center for Epidemiology and Microbiology | WHO National Influenza Centre Russian Federation | Andrey Komissarov, Artem Fadeev, Kseniya Komissarova, Oula Mansour, Kirill Varchenko, Mikhail Bakaev, Tamila Musaeva, Veronika Eder, Maria Pisareva, Nikita Yolshin, Daria Danilenko,Elizaveta Divisenko, Alina Odintsova, Elena Shidlovskaya, Ludmila Vasilchenko, Nadezhda Kuznetsova, Maria Nikiforova, Alexey Shchetinin, Artem Tkachuk, Andrei Pochtovyi, Andrei Siniavin, Denis Kleymenov, Evgeniia Bykonia, Liubov Popova, Elena Mazunina, Inna Dolzhikova, Anna Kovyrshina, Andrei Botikov, Dmitry Shcheblyakov, Vladimir Gushchin, Denis Logunov, Alexander Gintsburg, Ksenia Safina, Elena Nabieva, Georgii Bazykin, Dmitry Lioznov |
| EPI_ISL_2816391, EPI_ISL_2816392 | University College London, Great Ormond Street Hospital for Children NHS Foundation Trust, Imperial College Healthcare NHS Trust | COVID-19 Genomics UK (COG-UK) Consortium | Sergi Castellano, Rachel Williams, Mark Kristiansen, Sunando Roy, Charlotte Williams, Nadua Bayzid, Marius Cotic, Helena Tutill, Patricia Dyal, Judith Breuer |
| EPI_ISL_2816393 | N.F. Gamaleya Research Center for Epidemiology and Microbiology | WHO National Influenza Centre Russian Federation | Andrey Komissarov, Artem Fadeev, Kseniya Komissarova, Oula Mansour, Kirill Varchenko, Mikhail Bakaev, Tamila Musaeva, Veronika Eder, Maria Pisareva, Nikita Yolshin, Daria Danilenko,Elizaveta Divisenko, Alina Odintsova, Elena Shidlovskaya, Ludmila Vasilchenko, Nadezhda Kuznetsova, Maria Nikiforova, Alexey Shchetinin, Artem Tkachuk, Andrei Pochtovyi, Andrei Siniavin, Denis Kleymenov, Evgeniia Bykonia, Liubov Popova, Elena Mazunina, Inna Dolzhikova, Anna Kovyrshina, Andrei Botikov, Dmitry Shcheblyakov, Vladimir Gushchin, Denis Logunov, Alexander Gintsburg, Ksenia Safina, Elena Nabieva, Georgii Bazykin, Dmitry Lioznov |
| EPI_ISL_2816394 | University College London, Great Ormond Street Hospital for Children NHS Foundation Trust, Imperial College Healthcare NHS Trust | COVID-19 Genomics UK (COG-UK) Consortium | Sergi Castellano, Rachel Williams, Mark Kristiansen, Sunando Roy, Charlotte Williams, Nadua Bayzid, Marius Cotic, Helena Tutill, Patricia Dyal, Judith Breuer |
| EPI_ISL_2816395 | N.F. Gamaleya Research Center for Epidemiology and Microbiology | WHO National Influenza Centre Russian Federation | Andrey Komissarov, Artem Fadeev, Kseniya Komissarova, Oula Mansour, Kirill Varchenko, Mikhail Bakaev, Tamila Musaeva, Veronika Eder, Maria Pisareva, Nikita Yolshin, Daria Danilenko,Elizaveta Divisenko, Alina Odintsova, Elena Shidlovskaya, Ludmila Vasilchenko, Nadezhda Kuznetsova, Maria Nikiforova, Alexey Shchetinin, Artem Tkachuk, Andrei Pochtovyi, Andrei Siniavin, Denis Kleymenov, Evgeniia Bykonia, Liubov Popova, Elena Mazunina, Inna Dolzhikova, Anna Kovyrshina, Andrei Botikov, Dmitry Shcheblyakov, Vladimir Gushchin, Denis Logunov, Alexander Gintsburg, Ksenia Safina, Elena Nabieva, Georgii Bazykin, Dmitry Lioznov |
| EPI_ISL_2816396, EPI_ISL_2816397, EPI_ISL_2816399, EPI_ISL_2816400 | University College London, Great Ormond Street Hospital for Children NHS Foundation Trust, Imperial College Healthcare NHS Trust | COVID-19 Genomics UK (COG-UK) Consortium | Sergi Castellano, Rachel Williams, Mark Kristiansen, Sunando Roy, Charlotte Williams, Nadua Bayzid, Marius Cotic, Helena Tutill, Patricia Dyal, Judith Breuer |
| EPI_ISL_2816401 | N.F. Gamaleya Research Center for Epidemiology and Microbiology | WHO National Influenza Centre Russian Federation | Andrey Komissarov, Artem Fadeev, Kseniya Komissarova, Oula Mansour, Kirill Varchenko, Mikhail Bakaev, Tamila Musaeva, Veronika Eder, Maria Pisareva, Nikita Yolshin, Daria Danilenko,Elizaveta Divisenko, Alina Odintsova, Elena Shidlovskaya, Ludmila Vasilchenko, Nadezhda Kuznetsova, Maria Nikiforova, Alexey Shchetinin, Artem Tkachuk, Andrei Pochtovyi, Andrei Siniavin, Denis Kleymenov, Evgeniia Bykonia, Liubov Popova, Elena Mazunina, Inna Dolzhikova, Anna Kovyrshina, Andrei Botikov, Dmitry Shcheblyakov, Vladimir Gushchin, Denis Logunov, Alexander Gintsburg, Ksenia Safina, Elena Nabieva, Georgii Bazykin, Dmitry Lioznov |
| EPI_ISL_2816402 | University College London, Great Ormond Street Hospital for Children NHS Foundation Trust, Imperial College Healthcare NHS Trust | COVID-19 Genomics UK (COG-UK) Consortium | Sergi Castellano, Rachel Williams, Mark Kristiansen, Sunando Roy, Charlotte Williams, Nadua Bayzid, Marius Cotic, Helena Tutill, Patricia Dyal, Judith Breuer |
| EPI_ISL_2816403 | N.F. Gamaleya Research Center for Epidemiology and Microbiology | WHO National Influenza Centre Russian Federation | Andrey Komissarov, Artem Fadeev, Kseniya Komissarova, Oula Mansour, Kirill Varchenko, Mikhail Bakaev, Tamila Musaeva, Veronika Eder, Maria Pisareva, Nikita Yolshin, Daria Danilenko,Elizaveta Divisenko, Alina Odintsova, Elena Shidlovskaya, Ludmila Vasilchenko, Nadezhda Kuznetsova, Maria Nikiforova, Alexey Shchetinin, Artem Tkachuk, Andrei Pochtovyi, Andrei Siniavin, Denis Kleymenov, Evgeniia Bykonia, Liubov Popova, Elena Mazunina, Inna Dolzhikova, Anna Kovyrshina, Andrei Botikov, Dmitry Shcheblyakov, Vladimir Gushchin, Denis Logunov, Alexander Gintsburg, Ksenia Safina, Elena Nabieva, Georgii Bazykin, Dmitry Lioznov |
| EPI_ISL_2816404, EPI_ISL_2816405 | University College London, Great Ormond Street Hospital for Children NHS Foundation Trust, Imperial College Healthcare NHS Trust | COVID-19 Genomics UK (COG-UK) Consortium | Sergi Castellano, Rachel Williams, Mark Kristiansen, Sunando Roy, Charlotte Williams, Nadua Bayzid, Marius Cotic, Helena Tutill, Patricia Dyal, Judith Breuer |
| EPI_ISL_2816406 | N.F. Gamaleya Research Center for Epidemiology and Microbiology | WHO National Influenza Centre Russian Federation | Andrey Komissarov, Artem Fadeev, Kseniya Komissarova, Oula Mansour, Kirill Varchenko, Mikhail Bakaev, Tamila Musaeva, Veronika Eder, Maria Pisareva, Nikita Yolshin, Daria Danilenko,Elizaveta Divisenko, Alina Odintsova, Elena Shidlovskaya, Ludmila Vasilchenko, Nadezhda Kuznetsova, Maria Nikiforova, Alexey Shchetinin, Artem Tkachuk, Andrei Pochtovyi, Andrei Siniavin, Denis Kleymenov, Evgeniia Bykonia, Liubov Popova, Elena Mazunina, Inna Dolzhikova, Anna Kovyrshina, Andrei Botikov, Dmitry Shcheblyakov, Vladimir Gushchin, Denis Logunov, Alexander Gintsburg, Ksenia Safina, Elena Nabieva, Georgii Bazykin, Dmitry Lioznov |
| EPI_ISL_2816407 | University College London, Great Ormond Street Hospital for Children NHS Foundation Trust, Imperial College Healthcare NHS Trust | COVID-19 Genomics UK (COG-UK) Consortium | Sergi Castellano, Rachel Williams, Mark Kristiansen, Sunando Roy, Charlotte Williams, Nadua Bayzid, Marius Cotic, Helena Tutill, Patricia Dyal, Judith Breuer |
| EPI_ISL_2816408 | N.F. Gamaleya Research Center for Epidemiology and Microbiology | WHO National Influenza Centre Russian Federation | Andrey Komissarov, Artem Fadeev, Kseniya Komissarova, Oula Mansour, Kirill Varchenko, Mikhail Bakaev, Tamila Musaeva, Veronika Eder, Maria Pisareva, Nikita Yolshin, Daria Danilenko,Elizaveta Divisenko, Alina Odintsova, Elena Shidlovskaya, Ludmila Vasilchenko, Nadezhda Kuznetsova, Maria Nikiforova, Alexey Shchetinin, Artem Tkachuk, Andrei Pochtovyi, Andrei Siniavin, Denis Kleymenov, Evgeniia Bykonia, Liubov Popova, Elena Mazunina, Inna Dolzhikova, Anna Kovyrshina, Andrei Botikov, Dmitry Shcheblyakov, Vladimir Gushchin, Denis Logunov, Alexander Gintsburg, Ksenia Safina, Elena Nabieva, Georgii Bazykin, Dmitry Lioznov |
| EPI_ISL_2816409 | University College London, Great Ormond Street Hospital for Children NHS Foundation Trust, Imperial College Healthcare NHS Trust | COVID-19 Genomics UK (COG-UK) Consortium | Sergi Castellano, Rachel Williams, Mark Kristiansen, Sunando Roy, Charlotte Williams, Nadua Bayzid, Marius Cotic, Helena Tutill, Patricia Dyal, Judith Breuer |
| EPI_ISL_2816410 | N.F. Gamaleya Research Center for Epidemiology and Microbiology | WHO National Influenza Centre Russian Federation | Andrey Komissarov, Artem Fadeev, Kseniya Komissarova, Oula Mansour, Kirill Varchenko, Mikhail Bakaev, Tamila Musaeva, Veronika Eder, Maria Pisareva, Nikita Yolshin, Daria Danilenko,Elizaveta Divisenko, Alina Odintsova, Elena Shidlovskaya, Ludmila Vasilchenko, Nadezhda Kuznetsova, Maria Nikiforova, Alexey Shchetinin, Artem Tkachuk, Andrei Pochtovyi, Andrei Siniavin, Denis Kleymenov, Evgeniia Bykonia, Liubov Popova, Elena Mazunina, Inna Dolzhikova, Anna Kovyrshina, Andrei Botikov, Dmitry Shcheblyakov, Vladimir Gushchin, Denis Logunov, Alexander Gintsburg, Ksenia Safina, Elena Nabieva, Georgii Bazykin, Dmitry Lioznov |
| EPI_ISL_2816411, EPI_ISL_2816412, EPI_ISL_2816413, EPI_ISL_2816414, EPI_ISL_2816415, EPI_ISL_2816416, EPI_ISL_2816417, EPI_ISL_2816418, EPI_ISL_2816419, EPI_ISL_2816420, EPI_ISL_2816421, EPI_ISL_2816422, EPI_ISL_2816423, EPI_ISL_2816424, EPI_ISL_2816425, EPI_ISL_2816426, EPI_ISL_2816427, EPI_ISL_2816428, EPI_ISL_2816429, EPI_ISL_2816430, EPI_ISL_2816431, EPI_ISL_2816432, EPI_ISL_2816433, EPI_ISL_2816434, EPI_ISL_2816435, EPI_ISL_2816436, EPI_ISL_2816437, EPI_ISL_2816438, EPI_ISL_2816439, EPI_ISL_2816440, EPI_ISL_2816441, EPI_ISL_2816442, EPI_ISL_2816443, EPI_ISL_2816444, EPI_ISL_2816445, EPI_ISL_2816446, EPI_ISL_2816447, EPI_ISL_2816448, EPI_ISL_2816449, EPI_ISL_2816450, EPI_ISL_2816451, EPI_ISL_2816452, EPI_ISL_2816453, EPI_ISL_2816454, EPI_ISL_2816455, EPI_ISL_2816456, EPI_ISL_2816457, EPI_ISL_2816458, EPI_ISL_2816459, EPI_ISL_2816460, EPI_ISL_2816461, EPI_ISL_2816462, EPI_ISL_2816463, EPI_ISL_2816464, EPI_ISL_2816465, EPI_ISL_2816466, EPI_ISL_2816467, EPI_ISL_2816468, EPI_ISL_2816469, EPI_ISL_2816470, EPI_ISL_2816471, EPI_ISL_2816472, EPI_ISL_2816473, EPI_ISL_2816474, EPI_ISL_2816475, EPI_ISL_2816476, EPI_ISL_2816477, EPI_ISL_2816478, EPI_ISL_2816479, EPI_ISL_2816480, EPI_ISL_2816481, EPI_ISL_2816482, EPI_ISL_2816483, EPI_ISL_2816484, EPI_ISL_2816485, EPI_ISL_2816486, EPI_ISL_2816487, EPI_ISL_2816488, EPI_ISL_2816489, EPI_ISL_2816490, EPI_ISL_2816491, EPI_ISL_2816492, EPI_ISL_2816493, EPI_ISL_2816494, EPI_ISL_2816495, EPI_ISL_2816496, EPI_ISL_2816497, EPI_ISL_2816498, EPI_ISL_2816499, EPI_ISL_2816500, EPI_ISL_2816501, EPI_ISL_2816502, EPI_ISL_2816503, EPI_ISL_2816504, EPI_ISL_2816505, EPI_ISL_2816506, EPI_ISL_2816507, EPI_ISL_2816508, EPI_ISL_2816509, EPI_ISL_2816510, EPI_ISL_2816511, EPI_ISL_2816512, EPI_ISL_2816513, EPI_ISL_2816514, EPI_ISL_2816515, EPI_ISL_2816516, EPI_ISL_2816517, EPI_ISL_2816518, EPI_ISL_2816519, EPI_ISL_2816520, EPI_ISL_2816521, EPI_ISL_2816522, EPI_ISL_2816523, EPI_ISL_2816524, EPI_ISL_2816525, EPI_ISL_2816526, EPI_ISL_2816527, EPI_ISL_2816528, EPI_ISL_2816529, EPI_ISL_2816530, EPI_ISL_2816531, EPI_ISL_2816532, EPI_ISL_2816533, EPI_ISL_2816534, EPI_ISL_2816535, EPI_ISL_2816536, EPI_ISL_2816537, EPI_ISL_2816538, EPI_ISL_2816539, EPI_ISL_2816540, EPI_ISL_2816541, EPI_ISL_2816542, EPI_ISL_2816543, EPI_ISL_2816544, EPI_ISL_2816545, EPI_ISL_2816546, EPI_ISL_2816547, EPI_ISL_2816548, EPI_ISL_2816549, EPI_ISL_2816550, EPI_ISL_2816551, EPI_ISL_2816552, EPI_ISL_2816553, EPI_ISL_2816554, EPI_ISL_2816555, EPI_ISL_2816556, EPI_ISL_2816557, EPI_ISL_2816558, EPI_ISL_2816559, EPI_ISL_2816560, EPI_ISL_2816561, EPI_ISL_2816562, EPI_ISL_2816563 | | | |
| see above | University College London, Great Ormond Street Hospital for Children NHS Foundation Trust, Imperial College Healthcare NHS Trust | COVID-19 Genomics UK (COG-UK) Consortium | Sergi Castellano, Rachel Williams, Mark Kristiansen, Sunando Roy, Charlotte Williams, Nadua Bayzid, Marius Cotic, Helena Tutill, Patricia Dyal, Judith Breuer |
| EPI_ISL_2816564, EPI_ISL_2816565, EPI_ISL_2816566, EPI_ISL_2816567, EPI_ISL_2816568, EPI_ISL_2816569, EPI_ISL_2816570, EPI_ISL_2816571, EPI_ISL_2816572, EPI_ISL_2816573, EPI_ISL_2816574, EPI_ISL_2816575, EPI_ISL_2816576, EPI_ISL_2816577, EPI_ISL_2816578, EPI_ISL_2816579, EPI_ISL_2816580, EPI_ISL_2816581, EPI_ISL_2816582, EPI_ISL_2816583, EPI_ISL_2816584, EPI_ISL_2816585, EPI_ISL_2816586, EPI_ISL_2816587, EPI_ISL_2816588, EPI_ISL_2816589, EPI_ISL_2816590, EPI_ISL_2816591, EPI_ISL_2816592, EPI_ISL_2816593, EPI_ISL_2816594, EPI_ISL_2816595, EPI_ISL_2816596, EPI_ISL_2816597, EPI_ISL_2816598, EPI_ISL_2816599, EPI_ISL_2816600, EPI_ISL_2816601, EPI_ISL_2816602, EPI_ISL_2816603, EPI_ISL_2816604, EPI_ISL_2816605, EPI_ISL_2816606, EPI_ISL_2816607, EPI_ISL_2816608, EPI_ISL_2816609, EPI_ISL_2816610, EPI_ISL_2816611, EPI_ISL_2816612, EPI_ISL_2816613, EPI_ISL_2816614, EPI_ISL_2816615, EPI_ISL_2816616, EPI_ISL_2816617, EPI_ISL_2816618, EPI_ISL_2816619, EPI_ISL_2816620, EPI_ISL_2816621, EPI_ISL_2816622, EPI_ISL_2816623, EPI_ISL_2816624, EPI_ISL_2816625, EPI_ISL_2816626, EPI_ISL_2816627, EPI_ISL_2816628, EPI_ISL_2816629, EPI_ISL_2816630, EPI_ISL_2816631, EPI_ISL_2816632, EPI_ISL_2816633, EPI_ISL_2816634, EPI_ISL_2816635, EPI_ISL_2816636, EPI_ISL_2816637, EPI_ISL_2816638, EPI_ISL_2816639, EPI_ISL_2816640, EPI_ISL_2816641, EPI_ISL_2816642, EPI_ISL_2816643, EPI_ISL_2816644, EPI_ISL_2816645, EPI_ISL_2816646, EPI_ISL_2816647, EPI_ISL_2816648, EPI_ISL_2816649, EPI_ISL_2816650, EPI_ISL_2816651, EPI_ISL_2816652, EPI_ISL_2816653, EPI_ISL_2816654, EPI_ISL_2816655, EPI_ISL_2816656, EPI_ISL_2816657, EPI_ISL_2816658, EPI_ISL_2816659, EPI_ISL_2816660, EPI_ISL_2816661, EPI_ISL_2816662, EPI_ISL_2816663, EPI_ISL_2816664, EPI_ISL_2816665, EPI_ISL_2816666, EPI_ISL_2816667, EPI_ISL_2816668, EPI_ISL_2816669, EPI_ISL_2816670, EPI_ISL_2816671, EPI_ISL_2816672, EPI_ISL_2816673, EPI_ISL_2816674, EPI_ISL_2816675, EPI_ISL_2816676, EPI_ISL_2816677, EPI_ISL_2816678, EPI_ISL_2816679, EPI_ISL_2816680, EPI_ISL_2816681, EPI_ISL_2816682, EPI_ISL_2816683, EPI_ISL_2816684, EPI_ISL_2816685, EPI_ISL_2816686, EPI_ISL_2816687, EPI_ISL_2816690, EPI_ISL_2816691, EPI_ISL_2816692, EPI_ISL_2816693, EPI_ISL_2816694, EPI_ISL_2816695, EPI_ISL_2816696, EPI_ISL_2816697, EPI_ISL_2816698, EPI_ISL_2816699, EPI_ISL_2816700, EPI_ISL_2816701, EPI_ISL_2816702, EPI_ISL_2816703, EPI_ISL_2816704, EPI_ISL_2816705, EPI_ISL_2816706, EPI_ISL_2816707, EPI_ISL_2816708, EPI_ISL_2816709, EPI_ISL_2816710, EPI_ISL_2816711, EPI_ISL_2816712, EPI_ISL_2816713, EPI_ISL_2816714, EPI_ISL_2816715, EPI_ISL_2816716, EPI_ISL_2816717, EPI_ISL_2816718, EPI_ISL_2816719, EPI_ISL_2816720, EPI_ISL_2816721, EPI_ISL_2816722, EPI_ISL_2816723, EPI_ISL_2816724, EPI_ISL_2816725, EPI_ISL_2816726, EPI_ISL_2816727, EPI_ISL_2816728, EPI_ISL_2816729, EPI_ISL_2816730, EPI_ISL_2816731, EPI_ISL_2816732, EPI_ISL_2816733, EPI_ISL_2816734, EPI_ISL_2816735, EPI_ISL_2816736, EPI_ISL_2816737, EPI_ISL_2816738, EPI_ISL_2816739, EPI_ISL_2816740, EPI_ISL_2816741, EPI_ISL_2816742, EPI_ISL_2816743, EPI_ISL_2816744, EPI_ISL_2816745, EPI_ISL_2816746, EPI_ISL_2816747, EPI_ISL_2816748, EPI_ISL_2816749, EPI_ISL_2816750, EPI_ISL_2816751, EPI_ISL_2816752, EPI_ISL_2816753, EPI_ISL_2816754, EPI_ISL_2816755, EPI_ISL_2816756, EPI_ISL_2816757, EPI_ISL_2816758, EPI_ISL_2816759, EPI_ISL_2816760, EPI_ISL_2816761, EPI_ISL_2816762, EPI_ISL_2816763, EPI_ISL_2816764, EPI_ISL_2816765, EPI_ISL_2816766, EPI_ISL_2816767, EPI_ISL_2816768, EPI_ISL_2816769, EPI_ISL_2816770, EPI_ISL_2816771, EPI_ISL_2816772, EPI_ISL_2816773, EPI_ISL_2816774, EPI_ISL_2816775, EPI_ISL_2816776, EPI_ISL_2816777, EPI_ISL_2816778, EPI_ISL_2816779, EPI_ISL_2816780, EPI_ISL_2816781, EPI_ISL_2816782, EPI_ISL_2816783, EPI_ISL_2816784, EPI_ISL_2816785, EPI_ISL_2816786, EPI_ISL_2816787, EPI_ISL_2816788, EPI_ISL_2816789, EPI_ISL_2816790, EPI_ISL_2816791, EPI_ISL_2816792, EPI_ISL_2816793, EPI_ISL_2816794, EPI_ISL_2816795, EPI_ISL_2816796, EPI_ISL_2816797, EPI_ISL_2816798, EPI_ISL_2816799, EPI_ISL_2816800, EPI_ISL_2816801, EPI_ISL_2816802, EPI_ISL_2816803, EPI_ISL_2816804, EPI_ISL_2816805, EPI_ISL_2816806, EPI_ISL_2816807, EPI_ISL_2816808, EPI_ISL_2816809, EPI_ISL_2816810, EPI_ISL_2816811, EPI_ISL_2816812, EPI_ISL_2816813, EPI_ISL_2816814, EPI_ISL_2816815, EPI_ISL_2816816, EPI_ISL_2816817, EPI_ISL_2816818, EPI_ISL_2816819, EPI_ISL_2816820, EPI_ISL_2816821, EPI_ISL_2816822, EPI_ISL_2816823, EPI_ISL_2816824, EPI_ISL_2816825, EPI_ISL_2816826, EPI_ISL_2816827, EPI_ISL_2816828, EPI_ISL_2816829, EPI_ISL_2816830, EPI_ISL_2816831, EPI_ISL_2816832, EPI_ISL_2816833, EPI_ISL_2816834, EPI_ISL_2816835, EPI_ISL_2816836, EPI_ISL_2816837, EPI_ISL_2816838, EPI_ISL_2816839, EPI_ISL_2816840, EPI_ISL_2816841, EPI_ISL_2816842, EPI_ISL_2816843, EPI_ISL_2816844, EPI_ISL_2816845, EPI_ISL_2816846, EPI_ISL_2816847, EPI_ISL_2816848, EPI_ISL_2816849, EPI_ISL_2816850, EPI_ISL_2816851, EPI_ISL_2816852, EPI_ISL_2816853, EPI_ISL_2816854, EPI_ISL_2816855, EPI_ISL_2816856, EPI_ISL_2816857, EPI_ISL_2816858, EPI_ISL_2816859, EPI_ISL_2816860, EPI_ISL_2816861, EPI_ISL_2816862, EPI_ISL_2816863, EPI_ISL_2816864, EPI_ISL_2816865, EPI_ISL_2816866, EPI_ISL_2816867, EPI_ISL_2816868, EPI_ISL_2816869, EPI_ISL_2816870, EPI_ISL_2816871, EPI_ISL_2816872, EPI_ISL_2816873, EPI_ISL_2816874, EPI_ISL_2816875, EPI_ISL_2816876, EPI_ISL_2816877, EPI_ISL_2816878, EPI_ISL_2816879, EPI_ISL_2816880, EPI_ISL_2816881, EPI_ISL_2816882 | | | |
| see above | Northumbria University / South Tees Hospitals NHS Foundation Trust / North Cumbria Integrated Care NHS Foundation Trust / | COVID-19 Genomics UK (COG-UK) Consortium | Darren L Smith,Andrew Nelson,Matthew Bashton,Greg R Young,Joshua Loh,John Allan,Mohammad A Tariq,Giles S Holt,Gary Black,Wen C Yew,Lynn Dover,Paul Baker,Steve Liggett,Sarah Essex,Jane Greenaway,Debra Padgett,Clive Graham,Garren Scott,Edward Barton,Emma Swindells,Brendan Payne,Jennifer Collins,Yusri Taha,Gary Eltringham |

North Tees and Hartlepool NHS Foundation Trust / Newcastle Hospitals NHS Foundation Trust

EPI_ISL_2816883, EPI_ISL_2816884, EPI_ISL_2816885, EPI_ISL_2816886, EPI_ISL_2816887, EPI_ISL_2816888, EPI_ISL_2816889, EPI_ISL_2816890, EPI_ISL_2816891, EPI_ISL_2816892, EPI_ISL_2816893, EPI_ISL_2816894, EPI_ISL_2816895, EPI_ISL_2816896, EPI_ISL_2816897, EPI_ISL_2816898, EPI_ISL_2816899, EPI_ISL_2816900, EPI_ISL_2816901, EPI_ISL_2816902, EPI_ISL_2816903, EPI_ISL_2816904, EPI_ISL_2816905, EPI_ISL_2816906, EPI_ISL_2816907, EPI_ISL_2816908, EPI_ISL_2816909, EPI_ISL_2816910, EPI_ISL_2816911, EPI_ISL_2816912, EPI_ISL_2816913, EPI_ISL_2816914, EPI_ISL_2816915, EPI_ISL_2816916, EPI_ISL_2816917, EPI_ISL_2816918, EPI_ISL_2816919, EPI_ISL_2816920, EPI_ISL_2816921, EPI_ISL_2816922, EPI_ISL_2816923, EPI_ISL_2816924, EPI_ISL_2816925, EPI_ISL_2816926, EPI_ISL_2816927, EPI_ISL_2816928, EPI_ISL_2816929, EPI_ISL_2816930, EPI_ISL_2816931, EPI_ISL_2816932, EPI_ISL_2816933, EPI_ISL_2816934, EPI_ISL_2816935, EPI_ISL_2816936, EPI_ISL_2816937, EPI_ISL_2816938, EPI_ISL_2816939, EPI_ISL_2816940, EPI_ISL_2816941, EPI_ISL_2816942, EPI_ISL_2816943, EPI_ISL_2816944, EPI_ISL_2816945, EPI_ISL_2816946, EPI_ISL_2816947, EPI_ISL_2816948, EPI_ISL_2816949, EPI_ISL_2816950, EPI_ISL_2816951, EPI_ISL_2816952, EPI_ISL_2816953, EPI_ISL_2816954, EPI_ISL_2816955, EPI_ISL_2816956, EPI_ISL_2816957, EPI_ISL_2816958, EPI_ISL_2816959, EPI_ISL_2816960, EPI_ISL_2816961, EPI_ISL_2816962, EPI_ISL_2816963, EPI_ISL_2816964, EPI_ISL_2816965, EPI_ISL_2816966, EPI_ISL_2816967, EPI_ISL_2816969, EPI_ISL_2816970, EPI_ISL_2816971, EPI_ISL_2816972, EPI_ISL_2816973, EPI_ISL_2816974, EPI_ISL_2816975, EPI_ISL_2816976, EPI_ISL_2816977, EPI_ISL_2816978, EPI_ISL_2816979, EPI_ISL_2816980, EPI_ISL_2816981, EPI_ISL_2816982, EPI_ISL_2816983, EPI_ISL_2816984, EPI_ISL_2816985, EPI_ISL_2816986, EPI_ISL_2816987, EPI_ISL_2816988, EPI_ISL_2816989, EPI_ISL_2816990, EPI_ISL_2816991, EPI_ISL_2816992, EPI_ISL_2816993, EPI_ISL_2816994, EPI_ISL_2816995, EPI_ISL_2816996, EPI_ISL_2816997, EPI_ISL_2816998, EPI_ISL_2816999, EPI_ISL_2817000, EPI_ISL_2817001, EPI_ISL_2817002, EPI_ISL_2817003, EPI_ISL_2817004, EPI_ISL_2817005, EPI_ISL_2817006, EPI_ISL_2817007, EPI_ISL_2817008, EPI_ISL_2817009, EPI_ISL_2817010, EPI_ISL_2817011, EPI_ISL_2817012, EPI_ISL_2817013, EPI_ISL_2817014, EPI_ISL_2817015, EPI_ISL_2817016, EPI_ISL_2817017, EPI_ISL_2817018, EPI_ISL_2817019, EPI_ISL_2817020, EPI_ISL_2817021, EPI_ISL_2817022, EPI_ISL_2817023, EPI_ISL_2817024, EPI_ISL_2817025, EPI_ISL_2817026, EPI_ISL_2817027, EPI_ISL_2817028, EPI_ISL_2817029, EPI_ISL_2817030, EPI_ISL_2817031, EPI_ISL_2817032, EPI_ISL_2817033, EPI_ISL_2817034, EPI_ISL_2817035, EPI_ISL_2817036, EPI_ISL_2817037, EPI_ISL_2817039, EPI_ISL_2817040, EPI_ISL_2817041, EPI_ISL_2817042, EPI_ISL_2817043, EPI_ISL_2817044, EPI_ISL_2817045, EPI_ISL_2817046, EPI_ISL_2817047, EPI_ISL_2817048, EPI_ISL_2817049, EPI_ISL_2817050, EPI_ISL_2817051, EPI_ISL_2817052, EPI_ISL_2817053, EPI_ISL_2817054, EPI_ISL_2817055, EPI_ISL_2817056, EPI_ISL_2817057, EPI_ISL_2817058, EPI_ISL_2817059, EPI_ISL_2817060, EPI_ISL_2817061, EPI_ISL_2817062, EPI_ISL_2817063, EPI_ISL_2817064, EPI_ISL_2817065, EPI_ISL_2817066, EPI_ISL_2817067, EPI_ISL_2817068, EPI_ISL_2817069, EPI_ISL_2817070, EPI_ISL_2817071, EPI_ISL_2817072, EPI_ISL_2817073, EPI_ISL_2817074, EPI_ISL_2817075, EPI_ISL_2817076, EPI_ISL_2817077, EPI_ISL_2817078, EPI_ISL_2817079, EPI_ISL_2817080, EPI_ISL_2817081, EPI_ISL_2817082, EPI_ISL_2817083, EPI_ISL_2817084, EPI_ISL_2817085, EPI_ISL_2817086, EPI_ISL_2817087, EPI_ISL_2817089, EPI_ISL_2817090, EPI_ISL_2817091, EPI_ISL_2817092, EPI_ISL_2817093, EPI_ISL_2817094, EPI_ISL_2817095, EPI_ISL_2817096, EPI_ISL_2817097, EPI_ISL_2817098, EPI_ISL_2817099, EPI_ISL_2817100, EPI_ISL_2817101, EPI_ISL_2817102, EPI_ISL_2817103, EPI_ISL_2817104, EPI_ISL_2817105, EPI_ISL_2817106, EPI_ISL_2817107, EPI_ISL_2817108, EPI_ISL_2817109, EPI_ISL_2817110, EPI_ISL_2817111, EPI_ISL_2817112, EPI_ISL_2817113, EPI_ISL_2817114, EPI_ISL_2817115, EPI_ISL_2817116, EPI_ISL_2817117, EPI_ISL_2817118, EPI_ISL_2817119, EPI_ISL_2817120, EPI_ISL_2817121, EPI_ISL_2817122, EPI_ISL_2817123, EPI_ISL_2817124, EPI_ISL_2817125, EPI_ISL_2817126, EPI_ISL_2817127, EPI_ISL_2817128, EPI_ISL_2817129, EPI_ISL_2817130, EPI_ISL_2817131, EPI_ISL_2817132, EPI_ISL_2817133, EPI_ISL_2817134, EPI_ISL_2817135, EPI_ISL_2817136, EPI_ISL_2817137, EPI_ISL_2817138, EPI_ISL_2817139, EPI_ISL_2817140, EPI_ISL_2817141, EPI_ISL_2817142, EPI_ISL_2817143, EPI_ISL_2817144, EPI_ISL_2817145, EPI_ISL_2817146, EPI_ISL_2817147, EPI_ISL_2817148, EPI_ISL_2817149, EPI_ISL_2817150, EPI_ISL_2817151, EPI_ISL_2817152, EPI_ISL_2817153, EPI_ISL_2817154, EPI_ISL_2817155, EPI_ISL_2817156, EPI_ISL_2817157, EPI_ISL_2817158, EPI_ISL_2817159, EPI_ISL_2817160, EPI_ISL_2817161, EPI_ISL_2817162, EPI_ISL_2817163, EPI_ISL_2817164, EPI_ISL_2817165, EPI_ISL_2817166, EPI_ISL_2817167, EPI_ISL_2817168, EPI_ISL_2817169, EPI_ISL_2817170, EPI_ISL_2817171, EPI_ISL_2817172

see above | Quadram Institute Bioscience | COVID-19 Genomics UK (COG-UK) Consortium | Dave J. Baker, Gemma L. Kay, Alp Aydin, Thanh Le-Viet, Steven Rudder, Ana P. Tedim, Anastasia Kolyva, Maria Diaz, Leonardo de Oliveira Martins, Nabil-Fareed Alikhan, Lizzie Meadows, Rachael Stanley, Ngozi Elumogo, Muhammed Yasir, Nicholas M. Thomson, Alexander J Trotter, Rachel Gilroy, Samuel Bloomfield, Claire Stuart, Andrew Bell, Reenesh Prakash, Samir Dervisevic, Alison E. Mather, John Wain, Mark Webber, Andrew J. Page, Justin O’Grady

EPI_ISL_2817173, EPI_ISL_2817174, EPI_ISL_2817176, EPI_ISL_2817177, EPI_ISL_2817178, EPI_ISL_2817179, EPI_ISL_2817180, EPI_ISL_2817181, EPI_ISL_2817182, EPI_ISL_2817183, EPI_ISL_2817184, EPI_ISL_2817185, EPI_ISL_2817186, EPI_ISL_2817187, EPI_ISL_2817188, EPI_ISL_2817189, EPI_ISL_2817190, EPI_ISL_2817191, EPI_ISL_2817192, EPI_ISL_2817193, EPI_ISL_2817194, EPI_ISL_2817195, EPI_ISL_2817196, EPI_ISL_2817197, EPI_ISL_2817198, EPI_ISL_2817199, EPI_ISL_2817200, EPI_ISL_2817201, EPI_ISL_2817202, EPI_ISL_2817203, EPI_ISL_2817204, EPI_ISL_2817205, EPI_ISL_2817206, EPI_ISL_2817207, EPI_ISL_2817208, EPI_ISL_2817209, EPI_ISL_2817210, EPI_ISL_2817211, EPI_ISL_2817212, EPI_ISL_2817213, EPI_ISL_2817214, EPI_ISL_2817215, EPI_ISL_2817216, EPI_ISL_2817217, EPI_ISL_2817218, EPI_ISL_2817219, EPI_ISL_2817220, EPI_ISL_2817221, EPI_ISL_2817222, EPI_ISL_2817223, EPI_ISL_2817224, EPI_ISL_2817225, EPI_ISL_2817226, EPI_ISL_2817227, EPI_ISL_2817228, EPI_ISL_2817229, EPI_ISL_2817230, EPI_ISL_2817231, EPI_ISL_2817232, EPI_ISL_2817233, EPI_ISL_2817234, EPI_ISL_2817235, EPI_ISL_2817236, EPI_ISL_2817237, EPI_ISL_2817238, EPI_ISL_2817239, EPI_ISL_2817240, EPI_ISL_2817241, EPI_ISL_2817242, EPI_ISL_2817243, EPI_ISL_2817244, EPI_ISL_2817245, EPI_ISL_2817246, EPI_ISL_2817247, EPI_ISL_2817248, EPI_ISL_2817249, EPI_ISL_2817250, EPI_ISL_2817251, EPI_ISL_2817252, EPI_ISL_2817253, EPI_ISL_2817254, EPI_ISL_2817255, EPI_ISL_2817256, EPI_ISL_2817257, EPI_ISL_2817258, EPI_ISL_2817259, EPI_ISL_2817260, EPI_ISL_2817261, EPI_ISL_2817262, EPI_ISL_2817263, EPI_ISL_2817264, EPI_ISL_2817265, EPI_ISL_2817266, EPI_ISL_2817267, EPI_ISL_2817268, EPI_ISL_2817269, EPI_ISL_2817270, EPI_ISL_2817271, EPI_ISL_2817272, EPI_ISL_2817273, EPI_ISL_2817274, EPI_ISL_2817275, EPI_ISL_2817276, EPI_ISL_2817277, EPI_ISL_2817278, EPI_ISL_2817279, EPI_ISL_2817280, EPI_ISL_2817281

see above | Queens Medical Centre, Clinical Microbiology Department / DeepSeq Nottingham | COVID-19 Genomics UK (COG-UK) Consortium | Gemma Clark, Wendy Smith, Manjinder Khakh, Vicki M Fleming, Michelle M Lister, Hannah Howson-Wells, Jonathan Ball, Timothy Byaruhanga, Jayasree Dey, Emily Park, Jack Hill, Patrick McClure, Joseph Chappell, Theocharis Tsoleridis, Nadine Holmes, Matthew Carlisle, Christopher Moore, Fei Sang, Johnny Debebe, Victoria Wright, Matthew Loose

EPI_ISL_2817282, EPI_ISL_2817283, EPI_ISL_2817284, EPI_ISL_2817285, EPI_ISL_2817286, EPI_ISL_2817287, EPI_ISL_2817288, EPI_ISL_2817289, EPI_ISL_2817290, EPI_ISL_2817291, EPI_ISL_2817292, EPI_ISL_2817293, EPI_ISL_2817294, EPI_ISL_2817295, EPI_ISL_2817296, EPI_ISL_2817297, EPI_ISL_2817298, EPI_ISL_2817299, EPI_ISL_2817300, EPI_ISL_2817301, EPI_ISL_2817302, EPI_ISL_2817303, EPI_ISL_2817304, EPI_ISL_2817305, EPI_ISL_2817306, EPI_ISL_2817307, EPI_ISL_2817308, EPI_ISL_2817309, EPI_ISL_2817310, EPI_ISL_2817311, EPI_ISL_2817312, EPI_ISL_2817313, EPI_ISL_2817314, EPI_ISL_2817315, EPI_ISL_2817316, EPI_ISL_2817318, EPI_ISL_2817319, EPI_ISL_2817320, EPI_ISL_2817321, EPI_ISL_2817323, EPI_ISL_2817325, EPI_ISL_2817326, EPI_ISL_2817327, EPI_ISL_2817328, EPI_ISL_2817329, EPI_ISL_2817330, EPI_ISL_2817331, EPI_ISL_2817332, EPI_ISL_2817333, EPI_ISL_2817334, EPI_ISL_2817335, EPI_ISL_2817336, EPI_ISL_2817337, EPI_ISL_2817338, EPI_ISL_2817339, EPI_ISL_2817340, EPI_ISL_2817341, EPI_ISL_2817342, EPI_ISL_2817343, EPI_ISL_2817344, EPI_ISL_2817345, EPI_ISL_2817346, EPI_ISL_2817347, EPI_ISL_2817348, EPI_ISL_2817349, EPI_ISL_2817350, EPI_ISL_2817351, EPI_ISL_2817352, EPI_ISL_2817353, EPI_ISL_2817354, EPI_ISL_2817355, EPI_ISL_2817356, EPI_ISL_2817358, EPI_ISL_2817359, EPI_ISL_2817360, EPI_ISL_2817361, EPI_ISL_2817362, EPI_ISL_2817363, EPI_ISL_2817364, EPI_ISL_2817365, EPI_ISL_2817366, EPI_ISL_2817367, EPI_ISL_2817368, EPI_ISL_2817369, EPI_ISL_2817370, EPI_ISL_2817371, EPI_ISL_2817372, EPI_ISL_2817373, EPI_ISL_2817374, EPI_ISL_2817375, EPI_ISL_2817376, EPI_ISL_2817377, EPI_ISL_2817379, EPI_ISL_2817380, EPI_ISL_2817381, EPI_ISL_2817382, EPI_ISL_2817383, EPI_ISL_2817384, EPI_ISL_2817385, EPI_ISL_2817386, EPI_ISL_2817387, EPI_ISL_2817388, EPI_ISL_2817389, EPI_ISL_2817390, EPI_ISL_2817391, EPI_ISL_2817392, EPI_ISL_2817393, EPI_ISL_2817394, EPI_ISL_2817395, EPI_ISL_2817396, EPI_ISL_2817397, EPI_ISL_2817398, EPI_ISL_2817399, EPI_ISL_2817400, EPI_ISL_2817401, EPI_ISL_2817402, EPI_ISL_2817403, EPI_ISL_2817404, EPI_ISL_2817405, EPI_ISL_2817406, EPI_ISL_2817407, EPI_ISL_2817408, EPI_ISL_2817409, EPI_ISL_2817410, EPI_ISL_2817411, EPI_ISL_2817412, EPI_ISL_2817413, EPI_ISL_2817414, EPI_ISL_2817415, EPI_ISL_2817416, EPI_ISL_2817417, EPI_ISL_2817418, EPI_ISL_2817419, EPI_ISL_2817420, EPI_ISL_2817421, EPI_ISL_2817422, EPI_ISL_2817423, EPI_ISL_2817424, EPI_ISL_2817425, EPI_ISL_2817426, EPI_ISL_2817427, EPI_ISL_2817428, EPI_ISL_2817429, EPI_ISL_2817430, EPI_ISL_2817431, EPI_ISL_2817432, EPI_ISL_2817433, EPI_ISL_2817434, EPI_ISL_2817435, EPI_ISL_2817436, EPI_ISL_2817437, EPI_ISL_2817438, EPI_ISL_2817439, EPI_ISL_2817440, EPI_ISL_2817441, EPI_ISL_2817442, EPI_ISL_2817443, EPI_ISL_2817444, EPI_ISL_2817445, EPI_ISL_2817446, EPI_ISL_2817447, EPI_ISL_2817448, EPI_ISL_2817449, EPI_ISL_2817450, EPI_ISL_2817451, EPI_ISL_2817452, EPI_ISL_2817453, EPI_ISL_2817454, EPI_ISL_2817455, EPI_ISL_2817456, EPI_ISL_2817457, EPI_ISL_2817458, EPI_ISL_2817459, EPI_ISL_2817461, EPI_ISL_2817462, EPI_ISL_2817463, EPI_ISL_2817464, EPI_ISL_2817465, EPI_ISL_2817466, EPI_ISL_2817467, EPI_ISL_2817468, EPI_ISL_2817469, EPI_ISL_2817470, EPI_ISL_2817471, EPI_ISL_2817472, EPI_ISL_2817473, EPI_ISL_2817474, EPI_ISL_2817475, EPI_ISL_2817476, EPI_ISL_2817477, EPI_ISL_2817478, EPI_ISL_2817479, EPI_ISL_2817480, EPI_ISL_2817481, EPI_ISL_2817482, EPI_ISL_2817483, EPI_ISL_2817484, EPI_ISL_2817485, EPI_ISL_2817486, EPI_ISL_2817487, EPI_ISL_2817488, EPI_ISL_2817489, EPI_ISL_2817490, EPI_ISL_2817491, EPI_ISL_2817492, EPI_ISL_2817493, EPI_ISL_2817494, EPI_ISL_2817495, EPI_ISL_2817496, EPI_ISL_2817497, EPI_ISL_2817498, EPI_ISL_2817499, EPI_ISL_2817500, EPI_ISL_2817501, EPI_ISL_2817502, EPI_ISL_2817503, EPI_ISL_2817505, EPI_ISL_2817506, EPI_ISL_2817507, EPI_ISL_2817508, EPI_ISL_2817509, EPI_ISL_2817510, EPI_ISL_2817511, EPI_ISL_2817512, EPI_ISL_2817513, EPI_ISL_2817514, EPI_ISL_2817515, EPI_ISL_2817516, EPI_ISL_2817517, EPI_ISL_2817518, EPI_ISL_2817519, EPI_ISL_2817520, EPI_ISL_2817521, EPI_ISL_2817522, EPI_ISL_2817524, EPI_ISL_2817525, EPI_ISL_2817526, EPI_ISL_2817527, EPI_ISL_2817528, EPI_ISL_2817529, EPI_ISL_2817530, EPI_ISL_2817531, EPI_ISL_2817532, EPI_ISL_2817533, EPI_ISL_2817534, EPI_ISL_2817535, EPI_ISL_2817536, EPI_ISL_2817537, EPI_ISL_2817538, EPI_ISL_2817539, EPI_ISL_2817540, EPI_ISL_2817541, EPI_ISL_2817542, EPI_ISL_2817543, EPI_ISL_2817544, EPI_ISL_2817545, EPI_ISL_2817546, EPI_ISL_2817547, EPI_ISL_2817548, EPI_ISL_2817549, EPI_ISL_2817550, EPI_ISL_2817551, EPI_ISL_2817552, EPI_ISL_2817553, EPI_ISL_2817554, EPI_ISL_2817555, EPI_ISL_2817556, EPI_ISL_2817557, EPI_ISL_2817558, EPI_ISL_2817559, EPI_ISL_2817560, EPI_ISL_2817561, EPI_ISL_2817562, EPI_ISL_2817563, EPI_ISL_2817564, EPI_ISL_2817565, EPI_ISL_2817566, EPI_ISL_2817567, EPI_ISL_2817568, EPI_ISL_2817569, EPI_ISL_2817570, EPI_ISL_2817571, EPI_ISL_2817572, EPI_ISL_2817573, EPI_ISL_2817574, EPI_ISL_2817575, EPI_ISL_2817576, EPI_ISL_2817577, EPI_ISL_2817578, EPI_ISL_2817579, EPI_ISL_2817580, EPI_ISL_2817581, EPI_ISL_2817582, EPI_ISL_2817583, EPI_ISL_2817584, EPI_ISL_2817585, EPI_ISL_2817586, EPI_ISL_2817587, EPI_ISL_2817588, EPI_ISL_2817589, EPI_ISL_2817590, EPI_ISL_2817591, EPI_ISL_2817592, EPI_ISL_2817593, EPI_ISL_2817594, EPI_ISL_2817595, EPI_ISL_2817596, EPI_ISL_2817597, EPI_ISL_2817598, EPI_ISL_2817599, EPI_ISL_2817600, EPI_ISL_2817601, EPI_ISL_2817602, EPI_ISL_2817603, EPI_ISL_2817604, EPI_ISL_2817605, EPI_ISL_2817606, EPI_ISL_2817607, EPI_ISL_2817608, EPI_ISL_2817609, EPI_ISL_2817610, EPI_ISL_2817611, EPI_ISL_2817612, EPI_ISL_2817613, EPI_ISL_2817614, EPI_ISL_2817615, EPI_ISL_2817616, EPI_ISL_2817617, EPI_ISL_2817618, EPI_ISL_2817619, EPI_ISL_2817620, EPI_ISL_2817621, EPI_ISL_2817622, EPI_ISL_2817623, EPI_ISL_2817624, EPI_ISL_2817625, EPI_ISL_2817626, EPI_ISL_2817627, EPI_ISL_2817628, EPI_ISL_2817629, EPI_ISL_2817630, EPI_ISL_2817631, EPI_ISL_2817632, EPI_ISL_2817633, EPI_ISL_2817634, EPI_ISL_2817635, EPI_ISL_2817636, EPI_ISL_2817637, EPI_ISL_2817638, EPI_ISL_2817639, EPI_ISL_2817640, EPI_ISL_2817641, EPI_ISL_2817642, EPI_ISL_2817643, EPI_ISL_2817644, EPI_ISL_2817645, EPI_ISL_2817646, EPI_ISL_2817647, EPI_ISL_2817648, EPI_ISL_2817649, EPI_ISL_2817650, EPI_ISL_2817651, EPI_ISL_2817652, EPI_ISL_2817653, EPI_ISL_2817654, EPI_ISL_2817655, EPI_ISL_2817656, EPI_ISL_2817657, EPI_ISL_2817658, EPI_ISL_2817659, EPI_ISL_2817660, EPI_ISL_2817661, EPI_ISL_2817662, EPI_ISL_2817663, EPI_ISL_2817664, EPI_ISL_2817665, EPI_ISL_2817666, EPI_ISL_2817667, EPI_ISL_2817668, EPI_ISL_2817669, EPI_ISL_2817670, EPI_ISL_2817671, EPI_ISL_2817672, EPI_ISL_2817673, EPI_ISL_2817674, EPI_ISL_2817675, EPI_ISL_2817676, EPI_ISL_2817677, EPI_ISL_2817678, EPI_ISL_2817679, EPI_ISL_2817680, EPI_ISL_2817681, EPI_ISL_2817682, EPI_ISL_2817683, EPI_ISL_2817684, EPI_ISL_2817685, EPI_ISL_2817686, EPI_ISL_2817687, EPI_ISL_2817688, EPI_ISL_2817689, EPI_ISL_2817690, EPI_ISL_2817691, EPI_ISL_2817692, EPI_ISL_2817693, EPI_ISL_2817694, EPI_ISL_2817695, EPI_ISL_2817696, EPI_ISL_2817697, EPI_ISL_2817698, EPI_ISL_2817699, EPI_ISL_2817700, EPI_ISL_2817701, EPI_ISL_2817702, EPI_ISL_2817703, EPI_ISL_2817704, EPI_ISL_2817705, EPI_ISL_2817706, EPI_ISL_2817707, EPI_ISL_2817708, EPI_ISL_2817709, EPI_ISL_2817710, EPI_ISL_2817711, EPI_ISL_2817712, EPI_ISL_2817713, EPI_ISL_2817714, EPI_ISL_2817715, EPI_ISL_2817716, EPI_ISL_2817717, EPI_ISL_2817718, EPI_ISL_2817719, EPI_ISL_2817720, EPI_ISL_2817721, EPI_ISL_2817722, EPI_ISL_2817723, EPI_ISL_2817724, EPI_ISL_2817725, EPI_ISL_2817726, EPI_ISL_2817727, EPI_ISL_2817728, EPI_ISL_2817729, EPI_ISL_2817730, EPI_ISL_2817731, EPI_ISL_2817732, EPI_ISL_2817733, EPI_ISL_2817734, EPI_ISL_2817735, EPI_ISL_2817736, EPI_ISL_2817737, EPI_ISL_2817738, EPI_ISL_2817739, EPI_ISL_2817740, EPI_ISL_2817741, EPI_ISL_2817742, EPI_ISL_2817744, EPI_ISL_2817745, EPI_ISL_2817746, EPI_ISL_2817747, EPI_ISL_2817748, EPI_ISL_2817749, EPI_ISL_2817750, EPI_ISL_2817751, EPI_ISL_2817752, EPI_ISL_2817753, EPI_ISL_2817754, EPI_ISL_2817755, EPI_ISL_2817756, EPI_ISL_2817758, EPI_ISL_2817759, EPI_ISL_2817760, EPI_ISL_2817761, EPI_ISL_2817762, EPI_ISL_2817763, EPI_ISL_2817764, EPI_ISL_2817765, EPI_ISL_2817766, EPI_ISL_2817767, EPI_ISL_2817768, EPI_ISL_2817770, EPI_ISL_2817771, EPI_ISL_2817772, EPI_ISL_2817773, EPI_ISL_2817774, EPI_ISL_2817776, EPI_ISL_2817777, EPI_ISL_2817778, EPI_ISL_2817780, EPI_ISL_2817781, EPI_ISL_2817782, EPI_ISL_2817784, EPI_ISL_2817785, EPI_ISL_2817786, EPI_ISL_2817787, EPI_ISL_2817788, EPI_ISL_2817789, EPI_ISL_2817790, EPI_ISL_2817791, EPI_ISL_2817792, EPI_ISL_2817793, EPI_ISL_2817794, EPI_ISL_2817795, EPI_ISL_2817796, EPI_ISL_2817797, EPI_ISL_2817798, EPI_ISL_2817799, EPI_ISL_2817800, EPI_ISL_2817801, EPI_ISL_2817802, EPI_ISL_2817803, EPI_ISL_2817804, EPI_ISL_2817805, EPI_ISL_2817806, EPI_ISL_2817807, EPI_ISL_2817808, EPI_ISL_2817809, EPI_ISL_2817810, EPI_ISL_2817811, EPI_ISL_2817812, EPI_ISL_2817813, EPI_ISL_2817814, EPI_ISL_2817815, EPI_ISL_2817816, EPI_ISL_2817817, EPI_ISL_2817818, EPI_ISL_2817819, EPI_ISL_2817820, EPI_ISL_2817821, EPI_ISL_2817822, EPI_ISL_2817823, EPI_ISL_2817824, EPI_ISL_2817825, EPI_ISL_2817826, EPI_ISL_2817827, EPI_ISL_2817828, EPI_ISL_2817829, EPI_ISL_2817830, EPI_ISL_2817831, EPI_ISL_2817832, EPI_ISL_2817833, EPI_ISL_2817834, EPI_ISL_2817835, EPI_ISL_2817836, EPI_ISL_2817837, EPI_ISL_2817838, EPI_ISL_2817839, EPI_ISL_2817840, EPI_ISL_2817841, EPI_ISL_2817842, EPI_ISL_2817843, EPI_ISL_2817844, EPI_ISL_2817845, EPI_ISL_2817846, EPI_ISL_2817847, EPI_ISL_2817848, EPI_ISL_2817849, EPI_ISL_2817850, EPI_ISL_2817851, EPI_ISL_2817852, EPI_ISL_2817853, EPI_ISL_2817854, EPI_ISL_2817855, EPI_ISL_2817856, EPI_ISL_2817857, EPI_ISL_2817858, EPI_ISL_2817859, EPI_ISL_2817860, EPI_ISL_2817861, EPI_ISL_2817862, EPI_ISL_2817863, EPI_ISL_2817864, EPI_ISL_2817865, EPI_ISL_2817866, EPI_ISL_2817867, EPI_ISL_2817868, EPI_ISL_2817869, EPI_ISL_2817870, EPI_ISL_2817871, EPI_ISL_2817872, EPI_ISL_2817873, EPI_ISL_2817874, EPI_ISL_2817875, EPI_ISL_2817876, EPI_ISL_2817877, EPI_ISL_2817878, EPI_ISL_2817879, EPI_ISL_2817880, EPI_ISL_2817881, EPI_ISL_2817882, EPI_ISL_2817883, EPI_ISL_2817884, EPI_ISL_2817885, EPI_ISL_2817886, EPI_ISL_2817887, EPI_ISL_2817888, EPI_ISL_2817889, EPI_ISL_2817890, EPI_ISL_2817891, EPI_ISL_2817892, EPI_ISL_2817893, EPI_ISL_2817895, EPI_ISL_2817896, EPI_ISL_2817897, EPI_ISL_2817898, EPI_ISL_2817899, EPI_ISL_2817900, EPI_ISL_2817901, EPI_ISL_2817902, EPI_ISL_2817903, EPI_ISL_2817905, EPI_ISL_2817906, EPI_ISL_2817907, EPI_ISL_2817908, EPI_ISL_2817909, EPI_ISL_2817910, EPI_ISL_2817911, EPI_ISL_2817912, EPI_ISL_2817913, EPI_ISL_2817914, EPI_ISL_2817915, EPI_ISL_2817916, EPI_ISL_2817917, EPI_ISL_2817918, EPI_ISL_2817919, EPI_ISL_2817920, EPI_ISL_2817921, EPI_ISL_2817922, EPI_ISL_2817923, EPI_ISL_2817924, EPI_ISL_2817925, EPI_ISL_2817926, EPI_ISL_2817927, EPI_ISL_2817928, EPI_ISL_2817929, EPI_ISL_2817930, EPI_ISL_2817931, EPI_ISL_2817932, EPI_ISL_2817933, EPI_ISL_2817935, EPI_ISL_2817936, EPI_ISL_2817937, EPI_ISL_2817938, EPI_ISL_2817939, EPI_ISL_2817940, EPI_ISL_2817941, EPI_ISL_2817942, EPI_ISL_2817943, EPI_ISL_2817944, EPI_ISL_2817945, EPI_ISL_2817946, EPI_ISL_2817947, EPI_ISL_2817948, EPI_ISL_2817949, EPI_ISL_2817950, EPI_ISL_2817951, EPI_ISL_2817952, EPI_ISL_2817953, EPI_ISL_2817954, EPI_ISL_2817955, EPI_ISL_2817956, EPI_ISL_2817957, EPI_ISL_2817958, EPI_ISL_2817959, EPI_ISL_2817960, EPI_ISL_2817961, EPI_ISL_2817962, EPI_ISL_2817963, EPI_ISL_2817964, EPI_ISL_2817965, EPI_ISL_2817966, EPI_ISL_2817967, EPI_ISL_2817968, EPI_ISL_2817969, EPI_ISL_2817970, EPI_ISL_2817971, EPI_ISL_2817972, EPI_ISL_2817973, EPI_ISL_2817974, EPI_ISL_2817975, EPI_ISL_2817976, EPI_ISL_2817977, EPI_ISL_2817978, EPI_ISL_2817979, EPI_ISL_2817980, EPI_ISL_2817981, EPI_ISL_2817982, EPI_ISL_2817983, EPI_ISL_2817984, EPI_ISL_2817985, EPI_ISL_2817986, EPI_ISL_2817987, EPI_ISL_2817988, EPI_ISL_2817989, EPI_ISL_2817990, EPI_ISL_2817991, EPI_ISL_2817992, EPI_ISL_2817993, EPI_ISL_2817994, EPI_ISL_2817995, EPI_ISL_2817996, EPI_ISL_2817997, EPI_ISL_2817998, EPI_ISL_2817999, EPI_ISL_2818000, EPI_ISL_2818001, EPI_ISL_2818002, EPI_ISL_2818003, EPI_ISL_2818004, EPI_ISL_2818005, EPI_ISL_2818006, EPI_ISL_2818007, EPI_ISL_2818008, EPI_ISL_2818009, EPI_ISL_2818010, EPI_ISL_2818011, EPI_ISL_2818012, EPI_ISL_2818013, EPI_ISL_2818014, EPI_ISL_2818015, EPI_ISL_2818016, EPI_ISL_2818017, EPI_ISL_2818018, EPI_ISL_2818019, EPI_ISL_2818020, EPI_ISL_2818021, EPI_ISL_2818022, EPI_ISL_2818023, EPI_ISL_2818024, EPI_ISL_2818025, EPI_ISL_2818026, EPI_ISL_2818027, EPI_ISL_2818028, EPI_ISL_2818029, EPI_ISL_2818030, EPI_ISL_2818031, EPI_ISL_2818032, EPI_ISL_2818033, EPI_ISL_2818034, EPI_ISL_2818035, EPI_ISL_2818036, EPI_ISL_2818037, EPI_ISL_2818038, EPI_ISL_2818039, EPI_ISL_2818040, EPI_ISL_2818041, EPI_ISL_2818042, EPI_ISL_2818043, EPI_ISL_2818044, EPI_ISL_2818045, EPI_ISL_2818046, EPI_ISL_2818047, EPI_ISL_2818048, EPI_ISL_2818049, EPI_ISL_2818050, EPI_ISL_2818051, EPI_ISL_2818052, EPI_ISL_2818053, EPI_ISL_2818054, EPI_ISL_2818055, EPI_ISL_2818056, EPI_ISL_2818057, EPI_ISL_2818058, EPI_ISL_2818059, EPI_ISL_2818060, EPI_ISL_2818061, EPI_ISL_2818062, EPI_ISL_2818063, EPI_ISL_2818064, EPI_ISL_2818065, EPI_ISL_2818066, EPI_ISL_2818067, EPI_ISL_2818068, EPI_ISL_2818069, EPI_ISL_2818070, EPI_ISL_2818071, EPI_ISL_2818072, EPI_ISL_2818073, EPI_ISL_2818074, EPI_ISL_2818075, EPI_ISL_2818077, EPI_ISL_2818078, EPI_ISL_2818079, EPI_ISL_2818081, EPI_ISL_2818082, EPI_ISL_2818083, EPI_ISL_2818084, EPI_ISL_2818085, EPI_ISL_2818086, EPI_ISL_2818087, EPI_ISL_2818088, EPI_ISL_2818089, EPI_ISL_2818091, EPI_ISL_2818092, EPI_ISL_2818093, EPI_ISL_2818094, EPI_ISL_2818095, EPI_ISL_2818096, EPI_ISL_2818098, EPI_ISL_2818100, EPI_ISL_2818101, EPI_ISL_2818102, EPI_ISL_2818103, EPI_ISL_2818104, EPI_ISL_2818105, EPI_ISL_2818106, EPI_ISL_2818107, EPI_ISL_2818108, EPI_ISL_2818109, EPI_ISL_2818110, EPI_ISL_2818114, EPI_ISL_2818115, EPI_ISL_2818116, EPI_ISL_2818118, EPI_ISL_2818119, EPI_ISL_2818120, EPI_ISL_2818121, EPI_ISL_2818122, EPI_ISL_2818123, EPI_ISL_2818124, EPI_ISL_2818125, EPI_ISL_2818126, EPI_ISL_2818127, EPI_ISL_2818128, EPI_ISL_2818129, EPI_ISL_2818130, EPI_ISL_2818131, EPI_ISL_2818132, EPI_ISL_2818134, EPI_ISL_2818136, EPI_ISL_2818137, EPI_ISL_2818139, EPI_ISL_2818140, EPI_ISL_2818141, EPI_ISL_2818142, EPI_ISL_2818144, EPI_ISL_2818146, EPI_ISL_2818147, EPI_ISL_2818148, EPI_ISL_2818149, EPI_ISL_2818150, EPI_ISL_2818151, EPI_ISL_2818152, EPI_ISL_2818153, EPI_ISL_2818154, EPI_ISL_2818155, EPI_ISL_2818156, EPI_ISL_2818157, EPI_ISL_2818158, EPI_ISL_2818159, EPI_ISL_2818160, EPI_ISL_2818161, EPI_ISL_2818162, EPI_ISL_2818163, EPI_ISL_2818164, EPI_ISL_2818165, EPI_ISL_2818166, EPI_ISL_2818167, EPI_ISL_2818168, EPI_ISL_2818169, EPI_ISL_2818172, EPI_ISL_2818173, EPI_ISL_2818174, EPI_ISL_2818175, EPI_ISL_2818176, EPI_ISL_2818178, EPI_ISL_2818180, EPI_ISL_2818181, EPI_ISL_2818182, EPI_ISL_2818183, EPI_ISL_2818184, EPI_ISL_2818185, EPI_ISL_2818186, EPI_ISL_2818187, EPI_ISL_2818188, EPI_ISL_2818189, EPI_ISL_2818191, EPI_ISL_2818192, EPI_ISL_2818193, EPI_ISL_2818195, EPI_ISL_2818196, EPI_ISL_2818198, EPI_ISL_2818199, EPI_ISL_2818200, EPI_ISL_2818201, EPI_ISL_2818202, EPI_ISL_2818203, EPI_ISL_2818204, EPI_ISL_2818205, EPI_ISL_2818206, EPI_ISL_2818207, EPI_ISL_2818209, EPI_ISL_2818210, EPI_ISL_2818211, EPI_ISL_2818212, EPI_ISL_2818213, EPI_ISL_2818214, EPI_ISL_2818216, EPI_ISL_2818217, EPI_ISL_2818218, EPI_ISL_2818219, EPI_ISL_2818220, EPI_ISL_2818221, EPI_ISL_2818222, EPI_ISL_2818223, EPI_ISL_2818224, EPI_ISL_2818226, EPI_ISL_2818227, EPI_ISL_2818228, EPI_ISL_2818229, EPI_ISL_2818230, EPI_ISL_2818231, EPI_ISL_2818232, EPI_ISL_2818233, EPI_ISL_2818234, EPI_ISL_2818235, EPI_ISL_2818236, EPI_ISL_2818237, EPI_ISL_2818238, EPI_ISL_2818239, EPI_ISL_2818240, EPI_ISL_2818241, EPI_ISL_2818242, EPI_ISL_2818243, EPI_ISL_2818244, EPI_ISL_2818245, EPI_ISL_2818246, EPI_ISL_2818247, EPI_ISL_2818248, EPI_ISL_2818249, EPI_ISL_2818250, EPI_ISL_2818251, EPI_ISL_2818252, EPI_ISL_2818253, EPI_ISL_2818254, EPI_ISL_2818255, EPI_ISL_2818256, EPI_ISL_2818257, EPI_ISL_2818258, EPI_ISL_2818259, EPI_ISL_2818260, EPI_ISL_2818261, EPI_ISL_2818262, EPI_ISL_2818263, EPI_ISL_2818264, EPI_ISL_2818265, EPI_ISL_2818266, EPI_ISL_2818267, EPI_ISL_2818268, EPI_ISL_2818269, EPI_ISL_2818270, EPI_ISL_2818271, EPI_ISL_2818272, EPI_ISL_2818273, EPI_ISL_2818274, EPI_ISL_2818275, EPI_ISL_2818276, EPI_ISL_2818277, EPI_ISL_2818278, EPI_ISL_2818279, EPI_ISL_2818280, EPI_ISL_2818281, EPI_ISL_2818282, EPI_ISL_2818283, EPI_ISL_2818284, EPI_ISL_2818285, EPI_ISL_2818286, EPI_ISL_2818287, EPI_ISL_2818288, EPI_ISL_2818289, EPI_ISL_2818290, EPI_ISL_2818291, EPI_ISL_2818292, EPI_ISL_2818293, EPI_ISL_2818295, EPI_ISL_2818296, EPI_ISL_2818297, EPI_ISL_2818298, EPI_ISL_2818302, EPI_ISL_2818303, EPI_ISL_2818305, EPI_ISL_2818309, EPI_ISL_2818310, EPI_ISL_2818311, EPI_ISL_2818313, EPI_ISL_2818316, EPI_ISL_2818317, EPI_ISL_2818318, EPI_ISL_2818319, EPI_ISL_2818320, EPI_ISL_2818321, EPI_ISL_2818322, EPI_ISL_2818323, EPI_ISL_2818324

see above | Respiratory Virus Unit, Microbiology Services Colindale, Public Health England | COVID-19 Genomics UK (COG-UK) Consortium | PHE Covid Sequencing Team

EPI_ISL_2818325, EPI_ISL_2818326, EPI_ISL_2818327, EPI_ISL_2818328, EPI_ISL_2818329, EPI_ISL_2818330, EPI_ISL_2818331, EPI_ISL_2818332, EPI_ISL_2818333, EPI_ISL_2818334, EPI_ISL_2818335, EPI_ISL_2818336, EPI_ISL_2818337, EPI_ISL_2818338, EPI_ISL_2818339, EPI_ISL_2818340, EPI_ISL_2818341, EPI_ISL_2818342, EPI_ISL_2818343, EPI_ISL_2818344, EPI_ISL_2818345, EPI_ISL_2818346, EPI_ISL_2818347, EPI_ISL_2818348, EPI_ISL_2818349, EPI_ISL_2818350, EPI_ISL_2818351, EPI_ISL_2818352, EPI_ISL_2818353, EPI_ISL_2818354, EPI_ISL_2818355, EPI_ISL_2818356, EPI_ISL_2818357, EPI_ISL_2818358, EPI_ISL_2818359, EPI_ISL_2818360, EPI_ISL_2818361, EPI_ISL_2818362, EPI_ISL_2818363, EPI_ISL_2818364, EPI_ISL_2818365, EPI_ISL_2818366, EPI_ISL_2818367, EPI_ISL_2818368, EPI_ISL_2818369, EPI_ISL_2818370, EPI_ISL_2818371, EPI_ISL_2818372, EPI_ISL_2818373, EPI_ISL_2818374, EPI_ISL_2818375, EPI_ISL_2818376, EPI_ISL_2818377, EPI_ISL_2818378, EPI_ISL_2818379, EPI_ISL_2818380, EPI_ISL_2818381, EPI_ISL_2818382, EPI_ISL_2818383, EPI_ISL_2818384, EPI_ISL_2818385, EPI_ISL_2818386, EPI_ISL_2818387, EPI_ISL_2818388, EPI_ISL_2818389, EPI_ISL_2818390, EPI_ISL_2818391, EPI_ISL_2818392, EPI_ISL_2818393, EPI_ISL_2818394, EPI_ISL_2818395, EPI_ISL_2818396, EPI_ISL_2818397, EPI_ISL_2818398, EPI_ISL_2818399, EPI_ISL_2818400, EPI_ISL_2818401, EPI_ISL_2818402, EPI_ISL_2818403, EPI_ISL_2818404, EPI_ISL_2818405, EPI_ISL_2818406, EPI_ISL_2818407, EPI_ISL_2818408, EPI_ISL_2818409, EPI_ISL_2818410, EPI_ISL_2818411, EPI_ISL_2818412, EPI_ISL_2818413, EPI_ISL_2818414, EPI_ISL_2818415, EPI_ISL_2818416, EPI_ISL_2818417, EPI_ISL_2818418, EPI_ISL_2818419, EPI_ISL_2818420, EPI_ISL_2818421, EPI_ISL_2818422, EPI_ISL_2818423, EPI_ISL_2818424, EPI_ISL_2818425, EPI_ISL_2818426, EPI_ISL_2818427, EPI_ISL_2818428, EPI_ISL_2818429, EPI_ISL_2818430, EPI_ISL_2818431, EPI_ISL_2818432, EPI_ISL_2818433, EPI_ISL_2818434, EPI_ISL_2818435, EPI_ISL_2818436, EPI_ISL_2818437, EPI_ISL_2818438, EPI_ISL_2818439, EPI_ISL_2818440, EPI_ISL_2818441, EPI_ISL_2818442, EPI_ISL_2818443, EPI_ISL_2818444, EPI_ISL_2818445, EPI_ISL_2818446, EPI_ISL_2818447, EPI_ISL_2818448, EPI_ISL_2818449, EPI_ISL_2818451, EPI_ISL_2818452, EPI_ISL_2818453, EPI_ISL_2818454, EPI_ISL_2818455, EPI_ISL_2818456, EPI_ISL_2818457, EPI_ISL_2818458, EPI_ISL_2818459, EPI_ISL_2818460, EPI_ISL_2818461, EPI_ISL_2818462, EPI_ISL_2818463, EPI_ISL_2818464, EPI_ISL_2818465, EPI_ISL_2818466, EPI_ISL_2818467, EPI_ISL_2818468, EPI_ISL_2818469, EPI_ISL_2818470, EPI_ISL_2818471, EPI_ISL_2818472, EPI_ISL_2818473, EPI_ISL_2818474, EPI_ISL_2818475, EPI_ISL_2818476, EPI_ISL_2818477, EPI_ISL_2818478, EPI_ISL_2818479, EPI_ISL_2818480, EPI_ISL_2818481, EPI_ISL_2818482, EPI_ISL_2818483, EPI_ISL_2818484, EPI_ISL_2818485, EPI_ISL_2818486, EPI_ISL_2818487, EPI_ISL_2818488, EPI_ISL_2818489, EPI_ISL_2818490, EPI_ISL_2818491, EPI_ISL_2818492, EPI_ISL_2818493, EPI_ISL_2818494, EPI_ISL_2818495, EPI_ISL_2818496, EPI_ISL_2818497, EPI_ISL_2818498, EPI_ISL_2818499, EPI_ISL_2818500, EPI_ISL_2818501, EPI_ISL_2818502, EPI_ISL_2818503, EPI_ISL_2818504, EPI_ISL_2818505, EPI_ISL_2818506, EPI_ISL_2818507, EPI_ISL_2818508, EPI_ISL_2818509, EPI_ISL_2818511, EPI_ISL_2818512, EPI_ISL_2818513, EPI_ISL_2818514, EPI_ISL_2818515, EPI_ISL_2818516, EPI_ISL_2818517, EPI_ISL_2818518, EPI_ISL_2818519, EPI_ISL_2818520, EPI_ISL_2818521, EPI_ISL_2818522, EPI_ISL_2818523, EPI_ISL_2818524, EPI_ISL_2818526, EPI_ISL_2818527, EPI_ISL_2818528, EPI_ISL_2818529, EPI_ISL_2818530, EPI_ISL_2818531, EPI_ISL_2818532, EPI_ISL_2818533, EPI_ISL_2818534, EPI_ISL_2818535, EPI_ISL_2818536, EPI_ISL_2818537, EPI_ISL_2818538, EPI_ISL_2818539, EPI_ISL_2818540, EPI_ISL_2818541, EPI_ISL_2818542, EPI_ISL_2818543, EPI_ISL_2818544, EPI_ISL_2818545, EPI_ISL_2818546, EPI_ISL_2818547, EPI_ISL_2818548, EPI_ISL_2818549, EPI_ISL_2818550, EPI_ISL_2818551, EPI_ISL_2818552, EPI_ISL_2818553, EPI_ISL_2818554, EPI_ISL_2818555, EPI_ISL_2818556, EPI_ISL_2818557, EPI_ISL_2818558, EPI_ISL_2818559, EPI_ISL_2818560, EPI_ISL_2818561, EPI_ISL_2818562, EPI_ISL_2818563, EPI_ISL_2818564, EPI_ISL_2818565, EPI_ISL_2818566, EPI_ISL_2818567, EPI_ISL_2818568, EPI_ISL_2818569, EPI_ISL_2818570, EPI_ISL_2818571, EPI_ISL_2818572, EPI_ISL_2818573, EPI_ISL_2818574, EPI_ISL_2818575, EPI_ISL_2818576, EPI_ISL_2818577, EPI_ISL_2818578, EPI_ISL_2818579, EPI_ISL_2818580, EPI_ISL_2818581, EPI_ISL_2818582, EPI_ISL_2818583, EPI_ISL_2818584, EPI_ISL_2818585, EPI_ISL_2818586, EPI_ISL_2818587, EPI_ISL_2818588, EPI_ISL_2818589, EPI_ISL_2818590, EPI_ISL_2818591, EPI_ISL_2818592, EPI_ISL_2818593, EPI_ISL_2818594, EPI_ISL_2818595, EPI_ISL_2818596, EPI_ISL_2818597, EPI_ISL_2818598, EPI_ISL_2818599, EPI_ISL_2818600, EPI_ISL_2818601, EPI_ISL_2818602, EPI_ISL_2818603, EPI_ISL_2818604, EPI_ISL_2818605, EPI_ISL_2818606, EPI_ISL_2818607, EPI_ISL_2818608, EPI_ISL_2818609, EPI_ISL_2818610, EPI_ISL_2818611, EPI_ISL_2818612, EPI_ISL_2818613, EPI_ISL_2818614, EPI_ISL_2818615, EPI_ISL_2818616, EPI_ISL_2818617, EPI_ISL_2818618, EPI_ISL_2818619, EPI_ISL_2818620, EPI_ISL_2818621, EPI_ISL_2818622, EPI_ISL_2818623, EPI_ISL_2818624, EPI_ISL_2818625, EPI_ISL_2818626, EPI_ISL_2818627, EPI_ISL_2818628, EPI_ISL_2818629, EPI_ISL_2818630, EPI_ISL_2818631, EPI_ISL_2818632, EPI_ISL_2818633, EPI_ISL_2818634, EPI_ISL_2818635, EPI_ISL_2818636, EPI_ISL_2818637, EPI_ISL_2818638, EPI_ISL_2818639, EPI_ISL_2818640, EPI_ISL_2818641, EPI_ISL_2818642, EPI_ISL_2818643, EPI_ISL_2818644, EPI_ISL_2818645, EPI_ISL_2818646, EPI_ISL_2818647, EPI_ISL_2818648, EPI_ISL_2818649, EPI_ISL_2818650, EPI_ISL_2818651, EPI_ISL_2818652, EPI_ISL_2818653, EPI_ISL_2818654, EPI_ISL_2818655, EPI_ISL_2818656, EPI_ISL_2818657, EPI_ISL_2818658, EPI_ISL_2818659, EPI_ISL_2818660, EPI_ISL_2818661, EPI_ISL_2818662, EPI_ISL_2818663, EPI_ISL_2818665, EPI_ISL_2818666, EPI_ISL_2818667, EPI_ISL_2818669, EPI_ISL_2818670, EPI_ISL_2818671, EPI_ISL_2818672, EPI_ISL_2818673, EPI_ISL_2818674, EPI_ISL_2818675, EPI_ISL_2818676, EPI_ISL_2818677, EPI_ISL_2818678, EPI_ISL_2818679, EPI_ISL_2818680, EPI_ISL_2818681, EPI_ISL_2818683, EPI_ISL_2818684, EPI_ISL_2818685, EPI_ISL_2818686, EPI_ISL_2818687, EPI_ISL_2818688, EPI_ISL_2818689, EPI_ISL_2818691, EPI_ISL_2818692, EPI_ISL_2818693, EPI_ISL_2818694, EPI_ISL_2818695, EPI_ISL_2818696, EPI_ISL_2818697, EPI_ISL_2818698, EPI_ISL_2818699, EPI_ISL_2818700, EPI_ISL_2818701, EPI_ISL_2818702, EPI_ISL_2818703, EPI_ISL_2818704, EPI_ISL_2818705, EPI_ISL_2818706, EPI_ISL_2818707, EPI_ISL_2818709, EPI_ISL_2818710, EPI_ISL_2818711, EPI_ISL_2818712, EPI_ISL_2818713, EPI_ISL_2818714, EPI_ISL_2818715, EPI_ISL_2818716, EPI_ISL_2818717, EPI_ISL_2818718, EPI_ISL_2818719, EPI_ISL_2818720, EPI_ISL_2818721, EPI_ISL_2818722, EPI_ISL_2818723, EPI_ISL_2818724, EPI_ISL_2818725, EPI_ISL_2818726, EPI_ISL_2818727, EPI_ISL_2818728, EPI_ISL_2818729, EPI_ISL_2818730, EPI_ISL_2818731, EPI_ISL_2818732, EPI_ISL_2818733, EPI_ISL_2818734, EPI_ISL_2818735, EPI_ISL_2818736, EPI_ISL_2818737, EPI_ISL_2818738, EPI_ISL_2818739, EPI_ISL_2818740, EPI_ISL_2818741, EPI_ISL_2818742, EPI_ISL_2818743, EPI_ISL_2818744, EPI_ISL_2818745, EPI_ISL_2818746, EPI_ISL_2818747, EPI_ISL_2818748, EPI_ISL_2818749, EPI_ISL_2818750, EPI_ISL_2818751, EPI_ISL_2818752, EPI_ISL_2818753, EPI_ISL_2818754, EPI_ISL_2818755, EPI_ISL_2818756, EPI_ISL_2818757, EPI_ISL_2818758, EPI_ISL_2818759, EPI_ISL_2818760, EPI_ISL_2818761, EPI_ISL_2818762, EPI_ISL_2818763, EPI_ISL_2818764, EPI_ISL_2818765, EPI_ISL_2818766, EPI_ISL_2818767, EPI_ISL_2818768, EPI_ISL_2818769, EPI_ISL_2818770, EPI_ISL_2818771, EPI_ISL_2818772, EPI_ISL_2818773, EPI_ISL_2818774, EPI_ISL_2818775, EPI_ISL_2818776, EPI_ISL_2818777, EPI_ISL_2818778, EPI_ISL_2818779, EPI_ISL_2818780, EPI_ISL_2818781, EPI_ISL_2818782, EPI_ISL_2818783, EPI_ISL_2818784, EPI_ISL_2818785, EPI_ISL_2818786, EPI_ISL_2818787, EPI_ISL_2818788, EPI_ISL_2818789, EPI_ISL_2818790, EPI_ISL_2818791, EPI_ISL_2818792, EPI_ISL_2818793, EPI_ISL_2818794, EPI_ISL_2818795, EPI_ISL_2818796, EPI_ISL_2818797, EPI_ISL_2818798, EPI_ISL_2818799, EPI_ISL_2818800, EPI_ISL_2818801, EPI_ISL_2818802, EPI_ISL_2818803, EPI_ISL_2818804, EPI_ISL_2818805, EPI_ISL_2818806, EPI_ISL_2818807, EPI_ISL_2818808, EPI_ISL_2818809, EPI_ISL_2818810, EPI_ISL_2818811, EPI_ISL_2818812, EPI_ISL_2818813, EPI_ISL_2818814, EPI_ISL_2818815, EPI_ISL_2818816, EPI_ISL_2818817, EPI_ISL_2818818, EPI_ISL_2818819, EPI_ISL_2818820, EPI_ISL_2818821, EPI_ISL_2818822, EPI_ISL_2818823, EPI_ISL_2818824, EPI_ISL_2818825, EPI_ISL_2818826, EPI_ISL_2818827, EPI_ISL_2818828, EPI_ISL_2818829, EPI_ISL_2818830, EPI_ISL_2818831, EPI_ISL_2818832, EPI_ISL_2818833, EPI_ISL_2818834, EPI_ISL_2818835, EPI_ISL_2818836, EPI_ISL_2818837, EPI_ISL_2818839, EPI_ISL_2818840, EPI_ISL_2818841, EPI_ISL_2818842, EPI_ISL_2818843, EPI_ISL_2818844, EPI_ISL_2818845, EPI_ISL_2818846, EPI_ISL_2818847, EPI_ISL_2818848, EPI_ISL_2818849, EPI_ISL_2818850, EPI_ISL_2818852, EPI_ISL_2818853, EPI_ISL_2818854, EPI_ISL_2818855, EPI_ISL_2818856, EPI_ISL_2818857, EPI_ISL_2818858, EPI_ISL_2818859, EPI_ISL_2818860, EPI_ISL_2818861, EPI_ISL_2818862, EPI_ISL_2818863, EPI_ISL_2818864, EPI_ISL_2818865, EPI_ISL_2818866, EPI_ISL_2818867, EPI_ISL_2818868, EPI_ISL_2818869, EPI_ISL_2818870, EPI_ISL_2818871, EPI_ISL_2818872, EPI_ISL_2818873, EPI_ISL_2818874, EPI_ISL_2818875, EPI_ISL_2818876, EPI_ISL_2818877, EPI_ISL_2818878, EPI_ISL_2818879, EPI_ISL_2818880, EPI_ISL_2818881, EPI_ISL_2818882, EPI_ISL_2818884, EPI_ISL_2818885, EPI_ISL_2818886, EPI_ISL_2818887, EPI_ISL_2818888, EPI_ISL_2818889, EPI_ISL_2818890, EPI_ISL_2818891, EPI_ISL_2818892, EPI_ISL_2818893, EPI_ISL_2818894, EPI_ISL_2818895, EPI_ISL_2818896, EPI_ISL_2818897, EPI_ISL_2818898, EPI_ISL_2818899, EPI_ISL_2818900, EPI_ISL_2818901, EPI_ISL_2818902, EPI_ISL_2818903, EPI_ISL_2818904, EPI_ISL_2818905, EPI_ISL_2818906, EPI_ISL_2818907, EPI_ISL_2818908, EPI_ISL_2818909, EPI_ISL_2818910, EPI_ISL_2818911, EPI_ISL_2818912, EPI_ISL_2818913, EPI_ISL_2818914, EPI_ISL_2818915, EPI_ISL_2818916, EPI_ISL_2818917, EPI_ISL_2818918, EPI_ISL_2818919, EPI_ISL_2818920, EPI_ISL_2818921, EPI_ISL_2818922, EPI_ISL_2818923, EPI_ISL_2818924, EPI_ISL_2818925, EPI_ISL_2818926, EPI_ISL_2818927, EPI_ISL_2818928, EPI_ISL_2818929, EPI_ISL_2818930, EPI_ISL_2818931, EPI_ISL_2818932, EPI_ISL_2818933, EPI_ISL_2818934, EPI_ISL_2818935, EPI_ISL_2818936, EPI_ISL_2818937, EPI_ISL_2818938, EPI_ISL_2818939, EPI_ISL_2818940, EPI_ISL_2818941, EPI_ISL_2818942, EPI_ISL_2818943, EPI_ISL_2818944, EPI_ISL_2818945, EPI_ISL_2818946, EPI_ISL_2818947, EPI_ISL_2818948, EPI_ISL_2818949, EPI_ISL_2818950, EPI_ISL_2818951, EPI_ISL_2818952, EPI_ISL_2818953, EPI_ISL_2818954, EPI_ISL_2818955, EPI_ISL_2818956, EPI_ISL_2818957, EPI_ISL_2818958, EPI_ISL_2818959, EPI_ISL_2818960, EPI_ISL_2818961, EPI_ISL_2818962, EPI_ISL_2818963, EPI_ISL_2818964, EPI_ISL_2818965, EPI_ISL_2818966, EPI_ISL_2818967, EPI_ISL_2818968, EPI_ISL_2818969, EPI_ISL_2818970, EPI_ISL_2818971, EPI_ISL_2818972, EPI_ISL_2818973, EPI_ISL_2818974, EPI_ISL_2818975, EPI_ISL_2818976, EPI_ISL_2818977, EPI_ISL_2818978, EPI_ISL_2818979, EPI_ISL_2818980, EPI_ISL_2818981, EPI_ISL_2818982, EPI_ISL_2818983, EPI_ISL_2818984, EPI_ISL_2818985, EPI_ISL_2818986, EPI_ISL_2818987, EPI_ISL_2818988, EPI_ISL_2818989, EPI_ISL_2818990, EPI_ISL_2818991, EPI_ISL_2818992, EPI_ISL_2818993, EPI_ISL_2818994, EPI_ISL_2818995, EPI_ISL_2818996, EPI_ISL_2818997, EPI_ISL_2818998, EPI_ISL_2818999, EPI_ISL_2819000, EPI_ISL_2819001, EPI_ISL_2819002, EPI_ISL_2819003, EPI_ISL_2819004, EPI_ISL_2819005, EPI_ISL_2819006, EPI_ISL_2819007, EPI_ISL_2819008, EPI_ISL_2819009, EPI_ISL_2819010, EPI_ISL_2819011, EPI_ISL_2819012, EPI_ISL_2819013, EPI_ISL_2819014, EPI_ISL_2819015, EPI_ISL_2819016, EPI_ISL_2819017, EPI_ISL_2819018, EPI_ISL_2819019, EPI_ISL_2819020, EPI_ISL_2819021, EPI_ISL_2819022, EPI_ISL_2819023

see above | Originating lab: Wales Specialist Virology Centre Sequencing lab: Pathogen Genomics Unit | Public Health Wales Microbiology Cardiff Wales Specialist Virology Centre | Catherine Moore, Johnathan Evans, Laura Gifford, Malorie Perry, Simon Cottrell, Angela Marchbank, Alec Birchley, Alexander Adams, Amy Gaskin, Bree Gatica-Wilcox, Jason Coombes, Joel Southgate, Lauren Gilbert, Lee Graham, Nicole Pacchiarini, Sara Kumziene-Summerhayes, Sarah Taylor, Sophie Jones, Sara Rey, Matthew Bull, Joanne Watkins, Sally Corden, Tom Connor

| | | | |
|---|---|---|---|
| EPI_ISL_2819024 | Institute for Medical Research, Infectious Disease Research Centre, National Institutes of Health, Ministry of Health Malaysia | Institute for Medical Research, Infectious Disease Research Centre, National Institutes of Health, Ministry of Health Malaysia | Suppiah J, Kamel K, Mohd Zawawi Z, Azizan MA, Ramly N, Robert F, Thayan R |
| EPI_ISL_2819025, EPI_ISL_2819026, EPI_ISL_2819027, EPI_ISL_2819028, EPI_ISL_2819029, EPI_ISL_2819030, EPI_ISL_2819031, EPI_ISL_2819032, EPI_ISL_2819033, EPI_ISL_2819034, EPI_ISL_2819035, EPI_ISL_2819036, EPI_ISL_2819037, EPI_ISL_2819038, EPI_ISL_2819039, EPI_ISL_2819040, EPI_ISL_2819041, EPI_ISL_2819042, EPI_ISL_2819043, EPI_ISL_2819044 | | | |
| see above | Originating lab: Wales Specialist Virology Centre Sequencing lab: Pathogen Genomics Unit | Public Health Wales Microbiology Cardiff Wales Specialist Virology Centre | Catherine Moore, Johnathan Evans, Laura Gifford, Malorie Perry, Simon Cottrell, Angela Marchbank, Alec Birchley, Alexander Adams, Amy Gaskin, Bree Gatica-Wilcox, Jason Coombes, Joel Southgate, Lauren Gilbert, Lee Graham, Nicole Pacchiarini, Sara Kumziene-Summerhayes, Sarah Taylor, Sophie Jones, Sara Rey, Matthew Bull, Joanne Watkins, Sally Corden, Tom Connor |
| EPI_ISL_2819045 | Department of Bacteria, Parasites and Fungi, Statens Serum Institut, Copenhagen, Denmark | Statens Serum Institut Bioinformatics and Microbial Genomics | Danish Covid-19 Genome Consortium |
| EPI_ISL_2819046 | Originating lab: Wales Specialist Virology Centre Sequencing lab: Pathogen Genomics Unit | Public Health Wales Microbiology Cardiff Wales Specialist Virology Centre | Catherine Moore, Johnathan Evans, Laura Gifford, Malorie Perry, Simon Cottrell, Angela Marchbank, Alec Birchley, Alexander Adams, Amy Gaskin, Bree Gatica-Wilcox, Jason Coombes, Joel Southgate, Lauren Gilbert, Lee Graham, Nicole Pacchiarini, Sara Kumziene-Summerhayes, Sarah Taylor, Sophie Jones, Sara Rey, Matthew Bull, Joanne Watkins, Sally Corden, Tom Connor |
| EPI_ISL_2819047 | Department of Bacteria, Parasites and Fungi, Statens Serum Institut, Copenhagen, Denmark | Statens Serum Institut Bioinformatics and Microbial Genomics | Danish Covid-19 Genome Consortium |
| EPI_ISL_2819048, EPI_ISL_2819049 | Originating lab: Wales Specialist Virology Centre Sequencing lab: Pathogen Genomics Unit | Public Health Wales Microbiology Cardiff Wales Specialist Virology Centre | Catherine Moore, Johnathan Evans, Laura Gifford, Malorie Perry, Simon Cottrell, Angela Marchbank, Alec Birchley, Alexander Adams, Amy Gaskin, Bree Gatica-Wilcox, Jason Coombes, Joel Southgate, Lauren Gilbert, Lee Graham, Nicole Pacchiarini, Sara Kumziene-Summerhayes, Sarah Taylor, Sophie Jones, Sara Rey, Matthew Bull, Joanne Watkins, Sally Corden, Tom Connor |
| EPI_ISL_2819050 | Department of Bacteria, Parasites and Fungi, Statens Serum Institut, Copenhagen, Denmark | Statens Serum Institut Bioinformatics and Microbial Genomics | Danish Covid-19 Genome Consortium |
| EPI_ISL_2819051 | Originating lab: Wales Specialist Virology Centre Sequencing lab: Pathogen Genomics Unit | Public Health Wales Microbiology Cardiff Wales Specialist Virology Centre | Catherine Moore, Johnathan Evans, Laura Gifford, Malorie Perry, Simon Cottrell, Angela Marchbank, Alec Birchley, Alexander Adams, Amy Gaskin, Bree Gatica-Wilcox, Jason Coombes, Joel Southgate, Lauren Gilbert, Lee Graham, Nicole Pacchiarini, Sara Kumziene-Summerhayes, Sarah Taylor, Sophie Jones, Sara Rey, Matthew Bull, Joanne Watkins, Sally Corden, Tom Connor |
| EPI_ISL_2819052 | Department of Bacteria, Parasites and Fungi, Statens Serum Institut, Copenhagen, Denmark | Statens Serum Institut Bioinformatics and Microbial Genomics | Danish Covid-19 Genome Consortium |
| EPI_ISL_2819053 | Originating lab: Wales Specialist Virology Centre Sequencing lab: Pathogen Genomics Unit | Public Health Wales Microbiology Cardiff Wales Specialist Virology Centre | Catherine Moore, Johnathan Evans, Laura Gifford, Malorie Perry, Simon Cottrell, Angela Marchbank, Alec Birchley, Alexander Adams, Amy Gaskin, Bree Gatica-Wilcox, Jason Coombes, Joel Southgate, Lauren Gilbert, Lee Graham, Nicole Pacchiarini, Sara Kumziene-Summerhayes, Sarah Taylor, Sophie Jones, Sara Rey, Matthew Bull, Joanne Watkins, Sally Corden, Tom Connor |
| EPI_ISL_2819054 | Department of Bacteria, Parasites and Fungi, Statens Serum Institut, Copenhagen, Denmark | Statens Serum Institut Bioinformatics and Microbial Genomics | Danish Covid-19 Genome Consortium |
| EPI_ISL_2819055, EPI_ISL_2819057, EPI_ISL_2819059 | Originating lab: Wales Specialist Virology Centre Sequencing lab: Pathogen Genomics Unit | Public Health Wales Microbiology Cardiff Wales Specialist Virology Centre | Catherine Moore, Johnathan Evans, Laura Gifford, Malorie Perry, Simon Cottrell, Angela Marchbank, Alec Birchley, Alexander Adams, Amy Gaskin, Bree Gatica-Wilcox, Jason Coombes, Joel Southgate, Lauren Gilbert, Lee Graham, Nicole Pacchiarini, Sara Kumziene-Summerhayes, Sarah Taylor, Sophie Jones, Sara Rey, Matthew Bull, Joanne Watkins, Sally Corden, Tom Connor |
| EPI_ISL_2819060 | Department of Bacteria, Parasites and Fungi, Statens Serum Institut, Copenhagen, Denmark | Statens Serum Institut Bioinformatics and Microbial Genomics | Danish Covid-19 Genome Consortium |
| EPI_ISL_2819061 | Originating lab: Wales Specialist Virology Centre Sequencing lab: Pathogen Genomics Unit | Public Health Wales Microbiology Cardiff Wales Specialist Virology Centre | Catherine Moore, Johnathan Evans, Laura Gifford, Malorie Perry, Simon Cottrell, Angela Marchbank, Alec Birchley, Alexander Adams, Amy Gaskin, Bree Gatica-Wilcox, Jason Coombes, Joel Southgate, Lauren Gilbert, Lee Graham, Nicole Pacchiarini, Sara Kumziene-Summerhayes, Sarah Taylor, Sophie Jones, Sara Rey, Matthew Bull, Joanne Watkins, Sally Corden, Tom Connor |
| EPI_ISL_2819062 | Department of Bacteria, Parasites and Fungi, Statens Serum Institut, Copenhagen, Denmark | Statens Serum Institut Bioinformatics and Microbial Genomics | Danish Covid-19 Genome Consortium |
| EPI_ISL_2819063 | Originating lab: Wales Specialist Virology Centre Sequencing lab: Pathogen Genomics Unit | Public Health Wales Microbiology Cardiff Wales Specialist Virology Centre | Catherine Moore, Johnathan Evans, Laura Gifford, Malorie Perry, Simon Cottrell, Angela Marchbank, Alec Birchley, Alexander Adams, Amy Gaskin, Bree Gatica-Wilcox, Jason Coombes, Joel Southgate, Lauren Gilbert, Lee Graham, Nicole Pacchiarini, Sara Kumziene-Summerhayes, Sarah Taylor, Sophie Jones, Sara Rey, Matthew Bull, Joanne Watkins, Sally Corden, Tom Connor |
| EPI_ISL_2819064 | Department of Bacteria, Parasites and Fungi, Statens Serum Institut, Copenhagen, Denmark | Statens Serum Institut Bioinformatics and Microbial Genomics | Danish Covid-19 Genome Consortium |
| EPI_ISL_2819065, EPI_ISL_2819066 | Originating lab: Wales Specialist Virology Centre Sequencing lab: Pathogen Genomics Unit | Public Health Wales Microbiology Cardiff Wales Specialist Virology Centre | Catherine Moore, Johnathan Evans, Laura Gifford, Malorie Perry, Simon Cottrell, Angela Marchbank, Alec Birchley, Alexander Adams, Amy Gaskin, Bree Gatica-Wilcox, Jason Coombes, Joel Southgate, Lauren Gilbert, Lee Graham, Nicole Pacchiarini, Sara Kumziene-Summerhayes, Sarah Taylor, Sophie Jones, Sara Rey, Matthew Bull, Joanne Watkins, Sally Corden, Tom Connor |
| EPI_ISL_2819067 | Department of Bacteria, Parasites and Fungi, Statens Serum Institut, Copenhagen, Denmark | Statens Serum Institut Bioinformatics and Microbial Genomics | Danish Covid-19 Genome Consortium |
| EPI_ISL_2819068 | Originating lab: Wales Specialist Virology Centre Sequencing lab: Pathogen Genomics Unit | Public Health Wales Microbiology Cardiff Wales Specialist Virology Centre | Catherine Moore, Johnathan Evans, Laura Gifford, Malorie Perry, Simon Cottrell, Angela Marchbank, Alec Birchley, Alexander Adams, Amy Gaskin, Bree Gatica-Wilcox, Jason Coombes, Joel Southgate, Lauren Gilbert, Lee Graham, Nicole Pacchiarini, Sara Kumziene-Summerhayes, Sarah Taylor, Sophie Jones, Sara Rey, Matthew Bull, Joanne Watkins, Sally Corden, Tom Connor |
| EPI_ISL_2819069 | Department of Bacteria, Parasites and Fungi, Statens Serum Institut, Copenhagen, Denmark | Statens Serum Institut Bioinformatics and Microbial Genomics | Danish Covid-19 Genome Consortium |
| EPI_ISL_2819070, EPI_ISL_2819072 | Originating lab: Wales Specialist Virology Centre Sequencing lab: Pathogen Genomics Unit | Public Health Wales Microbiology Cardiff Wales Specialist Virology Centre | Catherine Moore, Johnathan Evans, Laura Gifford, Malorie Perry, Simon Cottrell, Angela Marchbank, Alec Birchley, Alexander Adams, Amy Gaskin, Bree Gatica-Wilcox, Jason Coombes, Joel Southgate, Lauren Gilbert, Lee Graham, Nicole Pacchiarini, Sara Kumziene-Summerhayes, Sarah Taylor, Sophie Jones, Sara Rey, Matthew Bull, Joanne Watkins, Sally Corden, Tom Connor |
| EPI_ISL_2819073 | Department of Bacteria, Parasites and Fungi, Statens Serum Institut, Copenhagen, Denmark | Statens Serum Institut Bioinformatics and Microbial Genomics | Danish Covid-19 Genome Consortium |
| EPI_ISL_2819074 | Originating lab: Wales Specialist Virology Centre Sequencing lab: Pathogen Genomics Unit | Public Health Wales Microbiology Cardiff Wales Specialist Virology Centre | Catherine Moore, Johnathan Evans, Laura Gifford, Malorie Perry, Simon Cottrell, Angela Marchbank, Alec Birchley, Alexander Adams, Amy Gaskin, Bree Gatica-Wilcox, Jason Coombes, Joel Southgate, Lauren Gilbert, Lee Graham, Nicole Pacchiarini, Sara Kumziene-Summerhayes, Sarah Taylor, Sophie Jones, Sara Rey, Matthew Bull, Joanne Watkins, Sally Corden, Tom Connor |
| EPI_ISL_2819075 | Department of Bacteria, Parasites and Fungi, Statens Serum Institut, Copenhagen, Denmark | Statens Serum Institut Bioinformatics and Microbial Genomics | Danish Covid-19 Genome Consortium |
| EPI_ISL_2819076 | Originating lab: Wales Specialist Virology Centre Sequencing lab: Pathogen Genomics Unit | Public Health Wales Microbiology Cardiff Wales Specialist Virology Centre | Catherine Moore, Johnathan Evans, Laura Gifford, Malorie Perry, Simon Cottrell, Angela Marchbank, Alec Birchley, Alexander Adams, Amy Gaskin, Bree Gatica-Wilcox, Jason Coombes, Joel Southgate, Lauren Gilbert, Lee Graham, Nicole Pacchiarini, Sara Kumziene-Summerhayes, Sarah Taylor, Sophie Jones, Sara Rey, Matthew Bull, Joanne Watkins, Sally Corden, Tom Connor |
| EPI_ISL_2819077 | Department of Bacteria, Parasites and Fungi, Statens Serum Institut, Copenhagen, Denmark | Statens Serum Institut Bioinformatics and Microbial Genomics | Danish Covid-19 Genome Consortium |
| EPI_ISL_2819078, EPI_ISL_2819079 | Originating lab: Wales Specialist Virology Centre Sequencing lab: Pathogen Genomics Unit | Public Health Wales Microbiology Cardiff Wales Specialist Virology Centre | Catherine Moore, Johnathan Evans, Laura Gifford, Malorie Perry, Simon Cottrell, Angela Marchbank, Alec Birchley, Alexander Adams, Amy Gaskin, Bree Gatica-Wilcox, Jason Coombes, Joel Southgate, Lauren Gilbert, Lee Graham, Nicole Pacchiarini, Sara Kumziene-Summerhayes, Sarah Taylor, Sophie Jones, Sara Rey, Matthew Bull, Joanne Watkins, Sally Corden, Tom Connor |
| EPI_ISL_2819080 | Department of Bacteria, Parasites and Fungi, Statens Serum Institut, Copenhagen, Denmark | Statens Serum Institut Bioinformatics and Microbial Genomics | Danish Covid-19 Genome Consortium |
| EPI_ISL_2819081 | Originating lab: Wales Specialist Virology Centre Sequencing lab: Pathogen Genomics Unit | Public Health Wales Microbiology Cardiff Wales Specialist Virology Centre | Catherine Moore, Johnathan Evans, Laura Gifford, Malorie Perry, Simon Cottrell, Angela Marchbank, Alec Birchley, Alexander Adams, Amy Gaskin, Bree Gatica-Wilcox, Jason Coombes, Joel Southgate, Lauren Gilbert, Lee Graham, Nicole Pacchiarini, Sara Kumziene-Summerhayes, Sarah Taylor, Sophie Jones, Sara Rey, Matthew Bull, Joanne Watkins, Sally Corden, Tom Connor |
| EPI_ISL_2819082, EPI_ISL_2819083 | Department of Bacteria, Parasites and Fungi, Statens Serum Institut, Copenhagen, Denmark | Statens Serum Institut Bioinformatics and Microbial Genomics | Danish Covid-19 Genome Consortium |
| EPI_ISL_2819084, EPI_ISL_2819085 | Originating lab: Wales Specialist Virology Centre Sequencing lab: Pathogen Genomics Unit | Public Health Wales Microbiology Cardiff Wales Specialist Virology Centre | Catherine Moore, Johnathan Evans, Laura Gifford, Malorie Perry, Simon Cottrell, Angela Marchbank, Alec Birchley, Alexander Adams, Amy Gaskin, Bree Gatica-Wilcox, Jason Coombes, Joel Southgate, Lauren Gilbert, Lee Graham, Nicole Pacchiarini, Sara Kumziene-Summerhayes, Sarah Taylor, Sophie Jones, Sara Rey, Matthew Bull, Joanne Watkins, Sally Corden, Tom Connor |
| EPI_ISL_2819086 | Department of Bacteria, Parasites and Fungi, Statens Serum Institut, Copenhagen, Denmark | Statens Serum Institut Bioinformatics and Microbial Genomics | Danish Covid-19 Genome Consortium |
| EPI_ISL_2819087 | Originating lab: Wales Specialist Virology Centre Sequencing lab: Pathogen Genomics Unit | Public Health Wales Microbiology Cardiff Wales Specialist Virology Centre | Catherine Moore, Johnathan Evans, Laura Gifford, Malorie Perry, Simon Cottrell, Angela Marchbank, Alec Birchley, Alexander Adams, Amy Gaskin, Bree Gatica-Wilcox, Jason Coombes, Joel Southgate, Lauren Gilbert, Lee Graham, Nicole Pacchiarini, Sara Kumziene-Summerhayes, Sarah Taylor, Sophie Jones, Sara Rey, Matthew Bull, Joanne Watkins, Sally Corden, Tom Connor |
| EPI_ISL_2819088 | Department of Bacteria, Parasites and Fungi, Statens Serum Institut, Copenhagen, Denmark | Statens Serum Institut Bioinformatics and Microbial Genomics | Danish Covid-19 Genome Consortium |
| EPI_ISL_2819089 | Originating lab: Wales Specialist Virology Centre Sequencing lab: Pathogen Genomics Unit | Public Health Wales Microbiology Cardiff Wales Specialist Virology Centre | Catherine Moore, Johnathan Evans, Laura Gifford, Malorie Perry, Simon Cottrell, Angela Marchbank, Alec Birchley, Alexander Adams, Amy Gaskin, Bree Gatica-Wilcox, Jason Coombes, Joel Southgate, Lauren Gilbert, Lee Graham, Nicole Pacchiarini, Sara Kumziene-Summerhayes, Sarah Taylor, Sophie Jones, Sara Rey, Matthew Bull, Joanne Watkins, Sally Corden, Tom Connor |
| EPI_ISL_2819090 | Department of Bacteria, Parasites and Fungi, Statens Serum Institut, Copenhagen, Denmark | Statens Serum Institut Bioinformatics and Microbial Genomics | Danish Covid-19 Genome Consortium |
| EPI_ISL_2819091 | Originating lab: Wales Specialist Virology Centre Sequencing lab: Pathogen Genomics Unit | Public Health Wales Microbiology Cardiff Wales Specialist Virology Centre | Catherine Moore, Johnathan Evans, Laura Gifford, Malorie Perry, Simon Cottrell, Angela Marchbank, Alec Birchley, Alexander Adams, Amy Gaskin, Bree Gatica-Wilcox, Jason Coombes, Joel Southgate, Lauren Gilbert, Lee Graham, Nicole Pacchiarini, Sara Kumziene-Summerhayes, Sarah Taylor, Sophie Jones, Sara Rey, Matthew Bull, Joanne Watkins, Sally Corden, Tom Connor |
| EPI_ISL_2819092 | Department of Bacteria, Parasites and Fungi, Statens Serum Institut, Copenhagen, Denmark | Statens Serum Institut Bioinformatics and Microbial Genomics | Danish Covid-19 Genome Consortium |
| EPI_ISL_2819093 | Originating lab: Wales Specialist Virology Centre Sequencing lab: Pathogen Genomics Unit | Public Health Wales Microbiology Cardiff Wales Specialist Virology Centre | Catherine Moore, Johnathan Evans, Laura Gifford, Malorie Perry, Simon Cottrell, Angela Marchbank, Alec Birchley, Alexander Adams, Amy Gaskin, Bree Gatica-Wilcox, Jason Coombes, Joel Southgate, Lauren Gilbert, Lee Graham, Nicole Pacchiarini, Sara Kumziene-Summerhayes, Sarah Taylor, Sophie Jones, Sara Rey, Matthew Bull, Joanne Watkins, Sally Corden, Tom Connor |
| EPI_ISL_2819094 | Department of Bacteria, Parasites and Fungi, Statens Serum Institut, Copenhagen, Denmark | Statens Serum Institut Bioinformatics and Microbial Genomics | Danish Covid-19 Genome Consortium |
| EPI_ISL_2819095, EPI_ISL_2819097, EPI_ISL_2819099, EPI_ISL_2819101, EPI_ISL_2819103, EPI_ISL_2819105 | Originating lab: Wales Specialist Virology Centre Sequencing lab: Pathogen Genomics Unit | Public Health Wales Microbiology Cardiff Wales Specialist Virology Centre | Catherine Moore, Johnathan Evans, Laura Gifford, Malorie Perry, Simon Cottrell, Angela Marchbank, Alec Birchley, Alexander Adams, Amy Gaskin, Bree Gatica-Wilcox, Jason Coombes, Joel Southgate, Lauren Gilbert, Lee Graham, Nicole Pacchiarini, Sara Kumziene-Summerhayes, Sarah Taylor, Sophie Jones, Sara Rey, Matthew Bull, Joanne Watkins, Sally Corden, Tom Connor |
| EPI_ISL_2819106 | Department of Bacteria, Parasites and Fungi, Statens Serum | Statens Serum Institut Bioinformatics and Microbial Genomics | Danish Covid-19 Genome Consortium |

| | | | |
|---|---|---|---|
| | Institut, Copenhagen, Denmark | | |
| EPI_ISL_2819107, EPI_ISL_2819109, EPI_ISL_2819111 | Originating lab: Wales Specialist Virology Centre Sequencing lab: Pathogen Genomics Unit | Public Health Wales Microbiology Cardiff Wales Specialist Virology Centre | Catherine Moore, Johnathan Evans, Laura Gifford, Malorie Perry, Simon Cottrell, Angela Marchbank, Alec Birchley, Alexander Adams, Amy Gaskin, Bree Gatica-Wilcox, Jason Coombes, Joel Southgate, Lauren Gilbert, Lee Graham, Nicole Pacchiarini, Sara Kumziene-Summerhayes, Sarah Taylor, Sophie Jones, Sara Rey, Matthew Bull, Joanne Watkins, Sally Corden, Tom Connor |
| EPI_ISL_2819112 | Department of Bacteria, Parasites and Fungi, Statens Serum Institut, Copenhagen, Denmark | Statens Serum Institut Bioinformatics and Microbial Genomics | Danish Covid-19 Genome Consortium |
| EPI_ISL_2819113 | Originating lab: Wales Specialist Virology Centre Sequencing lab: Pathogen Genomics Unit | Public Health Wales Microbiology Cardiff Wales Specialist Virology Centre | Catherine Moore, Johnathan Evans, Laura Gifford, Malorie Perry, Simon Cottrell, Angela Marchbank, Alec Birchley, Alexander Adams, Amy Gaskin, Bree Gatica-Wilcox, Jason Coombes, Joel Southgate, Lauren Gilbert, Lee Graham, Nicole Pacchiarini, Sara Kumziene-Summerhayes, Sarah Taylor, Sophie Jones, Sara Rey, Matthew Bull, Joanne Watkins, Sally Corden, Tom Connor |
| EPI_ISL_2819114 | Department of Bacteria, Parasites and Fungi, Statens Serum Institut, Copenhagen, Denmark | Statens Serum Institut Bioinformatics and Microbial Genomics | Danish Covid-19 Genome Consortium |
| EPI_ISL_2819115 | Originating lab: Wales Specialist Virology Centre Sequencing lab: Pathogen Genomics Unit | Public Health Wales Microbiology Cardiff Wales Specialist Virology Centre | Catherine Moore, Johnathan Evans, Laura Gifford, Malorie Perry, Simon Cottrell, Angela Marchbank, Alec Birchley, Alexander Adams, Amy Gaskin, Bree Gatica-Wilcox, Jason Coombes, Joel Southgate, Lauren Gilbert, Lee Graham, Nicole Pacchiarini, Sara Kumziene-Summerhayes, Sarah Taylor, Sophie Jones, Sara Rey, Matthew Bull, Joanne Watkins, Sally Corden, Tom Connor |
| EPI_ISL_2819116 | Department of Bacteria, Parasites and Fungi, Statens Serum Institut, Copenhagen, Denmark | Statens Serum Institut Bioinformatics and Microbial Genomics | Danish Covid-19 Genome Consortium |
| EPI_ISL_2819117, EPI_ISL_2819120, EPI_ISL_2819121 | Originating lab: Wales Specialist Virology Centre Sequencing lab: Pathogen Genomics Unit | Public Health Wales Microbiology Cardiff Wales Specialist Virology Centre | Catherine Moore, Johnathan Evans, Laura Gifford, Malorie Perry, Simon Cottrell, Angela Marchbank, Alec Birchley, Alexander Adams, Amy Gaskin, Bree Gatica-Wilcox, Jason Coombes, Joel Southgate, Lauren Gilbert, Lee Graham, Nicole Pacchiarini, Sara Kumziene-Summerhayes, Sarah Taylor, Sophie Jones, Sara Rey, Matthew Bull, Joanne Watkins, Sally Corden, Tom Connor |
| EPI_ISL_2819122 | Department of Bacteria, Parasites and Fungi, Statens Serum Institut, Copenhagen, Denmark | Statens Serum Institut Bioinformatics and Microbial Genomics | Danish Covid-19 Genome Consortium |
| EPI_ISL_2819123 | Originating lab: Wales Specialist Virology Centre Sequencing lab: Pathogen Genomics Unit | Public Health Wales Microbiology Cardiff Wales Specialist Virology Centre | Catherine Moore, Johnathan Evans, Laura Gifford, Malorie Perry, Simon Cottrell, Angela Marchbank, Alec Birchley, Alexander Adams, Amy Gaskin, Bree Gatica-Wilcox, Jason Coombes, Joel Southgate, Lauren Gilbert, Lee Graham, Nicole Pacchiarini, Sara Kumziene-Summerhayes, Sarah Taylor, Sophie Jones, Sara Rey, Matthew Bull, Joanne Watkins, Sally Corden, Tom Connor |
| EPI_ISL_2819124 | Department of Bacteria, Parasites and Fungi, Statens Serum Institut, Copenhagen, Denmark | Statens Serum Institut Bioinformatics and Microbial Genomics | Danish Covid-19 Genome Consortium |
| EPI_ISL_2819125, EPI_ISL_2819127 | Originating lab: Wales Specialist Virology Centre Sequencing lab: Pathogen Genomics Unit | Public Health Wales Microbiology Cardiff Wales Specialist Virology Centre | Catherine Moore, Johnathan Evans, Laura Gifford, Malorie Perry, Simon Cottrell, Angela Marchbank, Alec Birchley, Alexander Adams, Amy Gaskin, Bree Gatica-Wilcox, Jason Coombes, Joel Southgate, Lauren Gilbert, Lee Graham, Nicole Pacchiarini, Sara Kumziene-Summerhayes, Sarah Taylor, Sophie Jones, Sara Rey, Matthew Bull, Joanne Watkins, Sally Corden, Tom Connor |
| EPI_ISL_2819128 | Department of Bacteria, Parasites and Fungi, Statens Serum Institut, Copenhagen, Denmark | Statens Serum Institut Bioinformatics and Microbial Genomics | Danish Covid-19 Genome Consortium |
| EPI_ISL_2819129 | Originating lab: Wales Specialist Virology Centre Sequencing lab: Pathogen Genomics Unit | Public Health Wales Microbiology Cardiff Wales Specialist Virology Centre | Catherine Moore, Johnathan Evans, Laura Gifford, Malorie Perry, Simon Cottrell, Angela Marchbank, Alec Birchley, Alexander Adams, Amy Gaskin, Bree Gatica-Wilcox, Jason Coombes, Joel Southgate, Lauren Gilbert, Lee Graham, Nicole Pacchiarini, Sara Kumziene-Summerhayes, Sarah Taylor, Sophie Jones, Sara Rey, Matthew Bull, Joanne Watkins, Sally Corden, Tom Connor |
| EPI_ISL_2819130 | Department of Bacteria, Parasites and Fungi, Statens Serum Institut, Copenhagen, Denmark | Statens Serum Institut Bioinformatics and Microbial Genomics | Danish Covid-19 Genome Consortium |
| EPI_ISL_2819131 | Originating lab: Wales Specialist Virology Centre Sequencing lab: Pathogen Genomics Unit | Public Health Wales Microbiology Cardiff Wales Specialist Virology Centre | Catherine Moore, Johnathan Evans, Laura Gifford, Malorie Perry, Simon Cottrell, Angela Marchbank, Alec Birchley, Alexander Adams, Amy Gaskin, Bree Gatica-Wilcox, Jason Coombes, Joel Southgate, Lauren Gilbert, Lee Graham, Nicole Pacchiarini, Sara Kumziene-Summerhayes, Sarah Taylor, Sophie Jones, Sara Rey, Matthew Bull, Joanne Watkins, Sally Corden, Tom Connor |
| EPI_ISL_2819132 | Department of Bacteria, Parasites and Fungi, Statens Serum Institut, Copenhagen, Denmark | Statens Serum Institut Bioinformatics and Microbial Genomics | Danish Covid-19 Genome Consortium |
| EPI_ISL_2819133 | Originating lab: Wales Specialist Virology Centre Sequencing lab: Pathogen Genomics Unit | Public Health Wales Microbiology Cardiff Wales Specialist Virology Centre | Catherine Moore, Johnathan Evans, Laura Gifford, Malorie Perry, Simon Cottrell, Angela Marchbank, Alec Birchley, Alexander Adams, Amy Gaskin, Bree Gatica-Wilcox, Jason Coombes, Joel Southgate, Lauren Gilbert, Lee Graham, Nicole Pacchiarini, Sara Kumziene-Summerhayes, Sarah Taylor, Sophie Jones, Sara Rey, Matthew Bull, Joanne Watkins, Sally Corden, Tom Connor |
| EPI_ISL_2819134 | Department of Bacteria, Parasites and Fungi, Statens Serum Institut, Copenhagen, Denmark | Statens Serum Institut Bioinformatics and Microbial Genomics | Danish Covid-19 Genome Consortium |
| EPI_ISL_2819135, EPI_ISL_2819137, EPI_ISL_2819139 | Originating lab: Wales Specialist Virology Centre Sequencing lab: Pathogen Genomics Unit | Public Health Wales Microbiology Cardiff Wales Specialist Virology Centre | Catherine Moore, Johnathan Evans, Laura Gifford, Malorie Perry, Simon Cottrell, Angela Marchbank, Alec Birchley, Alexander Adams, Amy Gaskin, Bree Gatica-Wilcox, Jason Coombes, Joel Southgate, Lauren Gilbert, Lee Graham, Nicole Pacchiarini, Sara Kumziene-Summerhayes, Sarah Taylor, Sophie Jones, Sara Rey, Matthew Bull, Joanne Watkins, Sally Corden, Tom Connor |
| EPI_ISL_2819140 | Department of Bacteria, Parasites and Fungi, Statens Serum Institut, Copenhagen, Denmark | Statens Serum Institut Bioinformatics and Microbial Genomics | Danish Covid-19 Genome Consortium |
| EPI_ISL_2819141 | Originating lab: Wales Specialist Virology Centre Sequencing lab: Pathogen Genomics Unit | Public Health Wales Microbiology Cardiff Wales Specialist Virology Centre | Catherine Moore, Johnathan Evans, Laura Gifford, Malorie Perry, Simon Cottrell, Angela Marchbank, Alec Birchley, Alexander Adams, Amy Gaskin, Bree Gatica-Wilcox, Jason Coombes, Joel Southgate, Lauren Gilbert, Lee Graham, Nicole Pacchiarini, Sara Kumziene-Summerhayes, Sarah Taylor, Sophie Jones, Sara Rey, Matthew Bull, Joanne Watkins, Sally Corden, Tom Connor |
| EPI_ISL_2819142 | Department of Bacteria, Parasites and Fungi, Statens Serum Institut, Copenhagen, Denmark | Statens Serum Institut Bioinformatics and Microbial Genomics | Danish Covid-19 Genome Consortium |
| EPI_ISL_2819143 | Originating lab: Wales Specialist Virology Centre Sequencing lab: Pathogen Genomics Unit | Public Health Wales Microbiology Cardiff Wales Specialist Virology Centre | Catherine Moore, Johnathan Evans, Laura Gifford, Malorie Perry, Simon Cottrell, Angela Marchbank, Alec Birchley, Alexander Adams, Amy Gaskin, Bree Gatica-Wilcox, Jason Coombes, Joel Southgate, Lauren Gilbert, Lee Graham, Nicole Pacchiarini, Sara Kumziene-Summerhayes, Sarah Taylor, Sophie Jones, Sara Rey, Matthew Bull, Joanne Watkins, Sally Corden, Tom Connor |
| EPI_ISL_2819144 | Department of Bacteria, Parasites and Fungi, Statens Serum Institut, Copenhagen, Denmark | Statens Serum Institut Bioinformatics and Microbial Genomics | Danish Covid-19 Genome Consortium |
| EPI_ISL_2819145 | Originating lab: Wales Specialist Virology Centre Sequencing lab: Pathogen Genomics Unit | Public Health Wales Microbiology Cardiff Wales Specialist Virology Centre | Catherine Moore, Johnathan Evans, Laura Gifford, Malorie Perry, Simon Cottrell, Angela Marchbank, Alec Birchley, Alexander Adams, Amy Gaskin, Bree Gatica-Wilcox, Jason Coombes, Joel Southgate, Lauren Gilbert, Lee Graham, Nicole Pacchiarini, Sara Kumziene-Summerhayes, Sarah Taylor, Sophie Jones, Sara Rey, Matthew Bull, Joanne Watkins, Sally Corden, Tom Connor |
| EPI_ISL_2819146 | Department of Bacteria, Parasites and Fungi, Statens Serum Institut, Copenhagen, Denmark | Statens Serum Institut Bioinformatics and Microbial Genomics | Danish Covid-19 Genome Consortium |
| EPI_ISL_2819147 | Originating lab: Wales Specialist Virology Centre Sequencing lab: Pathogen Genomics Unit | Public Health Wales Microbiology Cardiff Wales Specialist Virology Centre | Catherine Moore, Johnathan Evans, Laura Gifford, Malorie Perry, Simon Cottrell, Angela Marchbank, Alec Birchley, Alexander Adams, Amy Gaskin, Bree Gatica-Wilcox, Jason Coombes, Joel Southgate, Lauren Gilbert, Lee Graham, Nicole Pacchiarini, Sara Kumziene-Summerhayes, Sarah Taylor, Sophie Jones, Sara Rey, Matthew Bull, Joanne Watkins, Sally Corden, Tom Connor |
| EPI_ISL_2819148 | Department of Bacteria, Parasites and Fungi, Statens Serum Institut, Copenhagen, Denmark | Statens Serum Institut Bioinformatics and Microbial Genomics | Danish Covid-19 Genome Consortium |
| EPI_ISL_2819149 | Originating lab: Wales Specialist Virology Centre Sequencing lab: Pathogen Genomics Unit | Public Health Wales Microbiology Cardiff Wales Specialist Virology Centre | Catherine Moore, Johnathan Evans, Laura Gifford, Malorie Perry, Simon Cottrell, Angela Marchbank, Alec Birchley, Alexander Adams, Amy Gaskin, Bree Gatica-Wilcox, Jason Coombes, Joel Southgate, Lauren Gilbert, Lee Graham, Nicole Pacchiarini, Sara Kumziene-Summerhayes, Sarah Taylor, Sophie Jones, Sara Rey, Matthew Bull, Joanne Watkins, Sally Corden, Tom Connor |
| EPI_ISL_2819150 | Department of Bacteria, Parasites and Fungi, Statens Serum Institut, Copenhagen, Denmark | Statens Serum Institut Bioinformatics and Microbial Genomics | Danish Covid-19 Genome Consortium |
| EPI_ISL_2819151 | Originating lab: Wales Specialist Virology Centre Sequencing lab: Pathogen Genomics Unit | Public Health Wales Microbiology Cardiff Wales Specialist Virology Centre | Catherine Moore, Johnathan Evans, Laura Gifford, Malorie Perry, Simon Cottrell, Angela Marchbank, Alec Birchley, Alexander Adams, Amy Gaskin, Bree Gatica-Wilcox, Jason Coombes, Joel Southgate, Lauren Gilbert, Lee Graham, Nicole Pacchiarini, Sara Kumziene-Summerhayes, Sarah Taylor, Sophie Jones, Sara Rey, Matthew Bull, Joanne Watkins, Sally Corden, Tom Connor |
| EPI_ISL_2819152 | Department of Bacteria, Parasites and Fungi, Statens Serum Institut, Copenhagen, Denmark | Statens Serum Institut Bioinformatics and Microbial Genomics | Danish Covid-19 Genome Consortium |
| EPI_ISL_2819153 | Originating lab: Wales Specialist Virology Centre Sequencing lab: Pathogen Genomics Unit | Public Health Wales Microbiology Cardiff Wales Specialist Virology Centre | Catherine Moore, Johnathan Evans, Laura Gifford, Malorie Perry, Simon Cottrell, Angela Marchbank, Alec Birchley, Alexander Adams, Amy Gaskin, Bree Gatica-Wilcox, Jason Coombes, Joel Southgate, Lauren Gilbert, Lee Graham, Nicole Pacchiarini, Sara Kumziene-Summerhayes, Sarah Taylor, Sophie Jones, Sara Rey, Matthew Bull, Joanne Watkins, Sally Corden, Tom Connor |
| EPI_ISL_2819154 | Department of Bacteria, Parasites and Fungi, Statens Serum Institut, Copenhagen, Denmark | Statens Serum Institut Bioinformatics and Microbial Genomics | Danish Covid-19 Genome Consortium |
| EPI_ISL_2819155, EPI_ISL_2819156 | Originating lab: Wales Specialist Virology Centre Sequencing lab: Pathogen Genomics Unit | Public Health Wales Microbiology Cardiff Wales Specialist Virology Centre | Catherine Moore, Johnathan Evans, Laura Gifford, Malorie Perry, Simon Cottrell, Angela Marchbank, Alec Birchley, Alexander Adams, Amy Gaskin, Bree Gatica-Wilcox, Jason Coombes, Joel Southgate, Lauren Gilbert, Lee Graham, Nicole Pacchiarini, Sara Kumziene-Summerhayes, Sarah Taylor, Sophie Jones, Sara Rey, Matthew Bull, Joanne Watkins, Sally Corden, Tom Connor |
| EPI_ISL_2819157 | Department of Bacteria, Parasites and Fungi, Statens Serum Institut, Copenhagen, Denmark | Statens Serum Institut Bioinformatics and Microbial Genomics | Danish Covid-19 Genome Consortium |
| EPI_ISL_2819158 | Originating lab: Wales Specialist Virology Centre Sequencing lab: Pathogen Genomics Unit | Public Health Wales Microbiology Cardiff Wales Specialist Virology Centre | Catherine Moore, Johnathan Evans, Laura Gifford, Malorie Perry, Simon Cottrell, Angela Marchbank, Alec Birchley, Alexander Adams, Amy Gaskin, Bree Gatica-Wilcox, Jason Coombes, Joel Southgate, Lauren Gilbert, Lee Graham, Nicole Pacchiarini, Sara Kumziene-Summerhayes, Sarah Taylor, Sophie Jones, Sara Rey, Matthew Bull, Joanne Watkins, Sally Corden, Tom Connor |
| EPI_ISL_2819159 | Department of Bacteria, Parasites and Fungi, Statens Serum Institut, Copenhagen, Denmark | Statens Serum Institut Bioinformatics and Microbial Genomics | Danish Covid-19 Genome Consortium |
| EPI_ISL_2819160 | Originating lab: Wales Specialist Virology Centre Sequencing lab: Pathogen Genomics Unit | Public Health Wales Microbiology Cardiff Wales Specialist Virology Centre | Catherine Moore, Johnathan Evans, Laura Gifford, Malorie Perry, Simon Cottrell, Angela Marchbank, Alec Birchley, Alexander Adams, Amy Gaskin, Bree Gatica-Wilcox, Jason Coombes, Joel Southgate, Lauren Gilbert, Lee Graham, Nicole Pacchiarini, Sara Kumziene-Summerhayes, Sarah Taylor, Sophie Jones, Sara Rey, Matthew Bull, Joanne Watkins, Sally Corden, Tom Connor |
| EPI_ISL_2819161 | Department of Bacteria, Parasites and Fungi, Statens Serum Institut, Copenhagen, Denmark | Statens Serum Institut Bioinformatics and Microbial Genomics | Danish Covid-19 Genome Consortium |
| EPI_ISL_2819162 | Originating lab: Wales Specialist Virology Centre Sequencing lab: Pathogen Genomics Unit | Public Health Wales Microbiology Cardiff Wales Specialist Virology Centre | Catherine Moore, Johnathan Evans, Laura Gifford, Malorie Perry, Simon Cottrell, Angela Marchbank, Alec Birchley, Alexander Adams, Amy Gaskin, Bree Gatica-Wilcox, Jason Coombes, Joel Southgate, Lauren Gilbert, Lee Graham, Nicole Pacchiarini, Sara Kumziene-Summerhayes, Sarah Taylor, Sophie Jones, Sara Rey, Matthew Bull, Joanne Watkins, Sally Corden, Tom Connor |
| EPI_ISL_2819163 | Department of Bacteria, Parasites and Fungi, Statens Serum Institut, Copenhagen, Denmark | Statens Serum Institut Bioinformatics and Microbial Genomics | Danish Covid-19 Genome Consortium |
| EPI_ISL_2819164, EPI_ISL_2819166 | Originating lab: Wales Specialist Virology Centre Sequencing lab: Pathogen Genomics Unit | Public Health Wales Microbiology Cardiff Wales Specialist Virology Centre | Catherine Moore, Johnathan Evans, Laura Gifford, Malorie Perry, Simon Cottrell, Angela Marchbank, Alec Birchley, Alexander Adams, Amy Gaskin, Bree Gatica-Wilcox, Jason Coombes, Joel Southgate, Lauren Gilbert, Lee Graham, Nicole Pacchiarini, Sara Kumziene-Summerhayes, Sarah Taylor, Sophie Jones, Sara Rey, Matthew Bull, Joanne Watkins, Sally Corden, Tom Connor |

| | | | |
|---|---|---|---|
| EPI_ISL_2819167 | Department of Bacteria, Parasites and Fungi, Statens Serum Institut, Copenhagen, Denmark | Statens Serum Institut Bioinformatics and Microbial Genomics | Danish Covid-19 Genome Consortium |
| EPI_ISL_2819168 | Originating lab: Wales Specialist Virology Centre Sequencing lab: Pathogen Genomics Unit | Public Health Wales Microbiology Cardiff Wales Specialist Virology Centre | Catherine Moore, Johnathan Evans, Laura Gifford, Malorie Perry, Simon Cottrell, Angela Marchbank, Alec Birchley, Alexander Adams, Amy Gaskin, Bree Gatica-Wilcox, Jason Coombes, Joel Southgate, Lauren Gilbert, Lee Graham, Nicole Pacchiarini, Sara Kumziene-Summerhayes, Sarah Taylor, Sophie Jones, Sara Rey, Matthew Bull, Joanne Watkins, Sally Corden, Tom Connor |
| EPI_ISL_2819169 | Department of Bacteria, Parasites and Fungi, Statens Serum Institut, Copenhagen, Denmark | Statens Serum Institut Bioinformatics and Microbial Genomics | Danish Covid-19 Genome Consortium |
| EPI_ISL_2819170, EPI_ISL_2819172 | Originating lab: Wales Specialist Virology Centre Sequencing lab: Pathogen Genomics Unit | Public Health Wales Microbiology Cardiff Wales Specialist Virology Centre | Catherine Moore, Johnathan Evans, Laura Gifford, Malorie Perry, Simon Cottrell, Angela Marchbank, Alec Birchley, Alexander Adams, Amy Gaskin, Bree Gatica-Wilcox, Jason Coombes, Joel Southgate, Lauren Gilbert, Lee Graham, Nicole Pacchiarini, Sara Kumziene-Summerhayes, Sarah Taylor, Sophie Jones, Sara Rey, Matthew Bull, Joanne Watkins, Sally Corden, Tom Connor |
| EPI_ISL_2819173 | Department of Bacteria, Parasites and Fungi, Statens Serum Institut, Copenhagen, Denmark | Statens Serum Institut Bioinformatics and Microbial Genomics | Danish Covid-19 Genome Consortium |
| EPI_ISL_2819174, EPI_ISL_2819176 | Originating lab: Wales Specialist Virology Centre Sequencing lab: Pathogen Genomics Unit | Public Health Wales Microbiology Cardiff Wales Specialist Virology Centre | Catherine Moore, Johnathan Evans, Laura Gifford, Malorie Perry, Simon Cottrell, Angela Marchbank, Alec Birchley, Alexander Adams, Amy Gaskin, Bree Gatica-Wilcox, Jason Coombes, Joel Southgate, Lauren Gilbert, Lee Graham, Nicole Pacchiarini, Sara Kumziene-Summerhayes, Sarah Taylor, Sophie Jones, Sara Rey, Matthew Bull, Joanne Watkins, Sally Corden, Tom Connor |
| EPI_ISL_2819177 | Department of Bacteria, Parasites and Fungi, Statens Serum Institut, Copenhagen, Denmark | Statens Serum Institut Bioinformatics and Microbial Genomics | Danish Covid-19 Genome Consortium |
| EPI_ISL_2819178 | Originating lab: Wales Specialist Virology Centre Sequencing lab: Pathogen Genomics Unit | Public Health Wales Microbiology Cardiff Wales Specialist Virology Centre | Catherine Moore, Johnathan Evans, Laura Gifford, Malorie Perry, Simon Cottrell, Angela Marchbank, Alec Birchley, Alexander Adams, Amy Gaskin, Bree Gatica-Wilcox, Jason Coombes, Joel Southgate, Lauren Gilbert, Lee Graham, Nicole Pacchiarini, Sara Kumziene-Summerhayes, Sarah Taylor, Sophie Jones, Sara Rey, Matthew Bull, Joanne Watkins, Sally Corden, Tom Connor |
| EPI_ISL_2819179 | Department of Bacteria, Parasites and Fungi, Statens Serum Institut, Copenhagen, Denmark | Statens Serum Institut Bioinformatics and Microbial Genomics | Danish Covid-19 Genome Consortium |
| EPI_ISL_2819180 | Originating lab: Wales Specialist Virology Centre Sequencing lab: Pathogen Genomics Unit | Public Health Wales Microbiology Cardiff Wales Specialist Virology Centre | Catherine Moore, Johnathan Evans, Laura Gifford, Malorie Perry, Simon Cottrell, Angela Marchbank, Alec Birchley, Alexander Adams, Amy Gaskin, Bree Gatica-Wilcox, Jason Coombes, Joel Southgate, Lauren Gilbert, Lee Graham, Nicole Pacchiarini, Sara Kumziene-Summerhayes, Sarah Taylor, Sophie Jones, Sara Rey, Matthew Bull, Joanne Watkins, Sally Corden, Tom Connor |
| EPI_ISL_2819181 | Department of Bacteria, Parasites and Fungi, Statens Serum Institut, Copenhagen, Denmark | Statens Serum Institut Bioinformatics and Microbial Genomics | Danish Covid-19 Genome Consortium |
| EPI_ISL_2819182 | Originating lab: Wales Specialist Virology Centre Sequencing lab: Pathogen Genomics Unit | Public Health Wales Microbiology Cardiff Wales Specialist Virology Centre | Catherine Moore, Johnathan Evans, Laura Gifford, Malorie Perry, Simon Cottrell, Angela Marchbank, Alec Birchley, Alexander Adams, Amy Gaskin, Bree Gatica-Wilcox, Jason Coombes, Joel Southgate, Lauren Gilbert, Lee Graham, Nicole Pacchiarini, Sara Kumziene-Summerhayes, Sarah Taylor, Sophie Jones, Sara Rey, Matthew Bull, Joanne Watkins, Sally Corden, Tom Connor |
| EPI_ISL_2819183 | Department of Bacteria, Parasites and Fungi, Statens Serum Institut, Copenhagen, Denmark | Statens Serum Institut Bioinformatics and Microbial Genomics | Danish Covid-19 Genome Consortium |
| EPI_ISL_2819184, EPI_ISL_2819185 | Originating lab: Wales Specialist Virology Centre Sequencing lab: Pathogen Genomics Unit | Public Health Wales Microbiology Cardiff Wales Specialist Virology Centre | Catherine Moore, Johnathan Evans, Laura Gifford, Malorie Perry, Simon Cottrell, Angela Marchbank, Alec Birchley, Alexander Adams, Amy Gaskin, Bree Gatica-Wilcox, Jason Coombes, Joel Southgate, Lauren Gilbert, Lee Graham, Nicole Pacchiarini, Sara Kumziene-Summerhayes, Sarah Taylor, Sophie Jones, Sara Rey, Matthew Bull, Joanne Watkins, Sally Corden, Tom Connor |
| EPI_ISL_2819186 | Department of Bacteria, Parasites and Fungi, Statens Serum Institut, Copenhagen, Denmark | Statens Serum Institut Bioinformatics and Microbial Genomics | Danish Covid-19 Genome Consortium |
| EPI_ISL_2819187 | Originating lab: Wales Specialist Virology Centre Sequencing lab: Pathogen Genomics Unit | Public Health Wales Microbiology Cardiff Wales Specialist Virology Centre | Catherine Moore, Johnathan Evans, Laura Gifford, Malorie Perry, Simon Cottrell, Angela Marchbank, Alec Birchley, Alexander Adams, Amy Gaskin, Bree Gatica-Wilcox, Jason Coombes, Joel Southgate, Lauren Gilbert, Lee Graham, Nicole Pacchiarini, Sara Kumziene-Summerhayes, Sarah Taylor, Sophie Jones, Sara Rey, Matthew Bull, Joanne Watkins, Sally Corden, Tom Connor |
| EPI_ISL_2819188 | Department of Bacteria, Parasites and Fungi, Statens Serum Institut, Copenhagen, Denmark | Statens Serum Institut Bioinformatics and Microbial Genomics | Danish Covid-19 Genome Consortium |
| EPI_ISL_2819189 | Originating lab: Wales Specialist Virology Centre Sequencing lab: Pathogen Genomics Unit | Public Health Wales Microbiology Cardiff Wales Specialist Virology Centre | Catherine Moore, Johnathan Evans, Laura Gifford, Malorie Perry, Simon Cottrell, Angela Marchbank, Alec Birchley, Alexander Adams, Amy Gaskin, Bree Gatica-Wilcox, Jason Coombes, Joel Southgate, Lauren Gilbert, Lee Graham, Nicole Pacchiarini, Sara Kumziene-Summerhayes, Sarah Taylor, Sophie Jones, Sara Rey, Matthew Bull, Joanne Watkins, Sally Corden, Tom Connor |
| EPI_ISL_2819190 | Department of Bacteria, Parasites and Fungi, Statens Serum Institut, Copenhagen, Denmark | Statens Serum Institut Bioinformatics and Microbial Genomics | Danish Covid-19 Genome Consortium |
| EPI_ISL_2819191, EPI_ISL_2819193 | Originating lab: Wales Specialist Virology Centre Sequencing lab: Pathogen Genomics Unit | Public Health Wales Microbiology Cardiff Wales Specialist Virology Centre | Catherine Moore, Johnathan Evans, Laura Gifford, Malorie Perry, Simon Cottrell, Angela Marchbank, Alec Birchley, Alexander Adams, Amy Gaskin, Bree Gatica-Wilcox, Jason Coombes, Joel Southgate, Lauren Gilbert, Lee Graham, Nicole Pacchiarini, Sara Kumziene-Summerhayes, Sarah Taylor, Sophie Jones, Sara Rey, Matthew Bull, Joanne Watkins, Sally Corden, Tom Connor |
| EPI_ISL_2819194 | Department of Bacteria, Parasites and Fungi, Statens Serum Institut, Copenhagen, Denmark | Statens Serum Institut Bioinformatics and Microbial Genomics | Danish Covid-19 Genome Consortium |
| EPI_ISL_2819195 | Originating lab: Wales Specialist Virology Centre Sequencing lab: Pathogen Genomics Unit | Public Health Wales Microbiology Cardiff Wales Specialist Virology Centre | Catherine Moore, Johnathan Evans, Laura Gifford, Malorie Perry, Simon Cottrell, Angela Marchbank, Alec Birchley, Alexander Adams, Amy Gaskin, Bree Gatica-Wilcox, Jason Coombes, Joel Southgate, Lauren Gilbert, Lee Graham, Nicole Pacchiarini, Sara Kumziene-Summerhayes, Sarah Taylor, Sophie Jones, Sara Rey, Matthew Bull, Joanne Watkins, Sally Corden, Tom Connor |
| EPI_ISL_2819196 | Department of Bacteria, Parasites and Fungi, Statens Serum Institut, Copenhagen, Denmark | Statens Serum Institut Bioinformatics and Microbial Genomics | Danish Covid-19 Genome Consortium |
| EPI_ISL_2819197 | Originating lab: Wales Specialist Virology Centre Sequencing lab: Pathogen Genomics Unit | Public Health Wales Microbiology Cardiff Wales Specialist Virology Centre | Catherine Moore, Johnathan Evans, Laura Gifford, Malorie Perry, Simon Cottrell, Angela Marchbank, Alec Birchley, Alexander Adams, Amy Gaskin, Bree Gatica-Wilcox, Jason Coombes, Joel Southgate, Lauren Gilbert, Lee Graham, Nicole Pacchiarini, Sara Kumziene-Summerhayes, Sarah Taylor, Sophie Jones, Sara Rey, Matthew Bull, Joanne Watkins, Sally Corden, Tom Connor |
| EPI_ISL_2819198 | Department of Bacteria, Parasites and Fungi, Statens Serum Institut, Copenhagen, Denmark | Statens Serum Institut Bioinformatics and Microbial Genomics | Danish Covid-19 Genome Consortium |
| EPI_ISL_2819199 | Originating lab: Wales Specialist Virology Centre Sequencing lab: Pathogen Genomics Unit | Public Health Wales Microbiology Cardiff Wales Specialist Virology Centre | Catherine Moore, Johnathan Evans, Laura Gifford, Malorie Perry, Simon Cottrell, Angela Marchbank, Alec Birchley, Alexander Adams, Amy Gaskin, Bree Gatica-Wilcox, Jason Coombes, Joel Southgate, Lauren Gilbert, Lee Graham, Nicole Pacchiarini, Sara Kumziene-Summerhayes, Sarah Taylor, Sophie Jones, Sara Rey, Matthew Bull, Joanne Watkins, Sally Corden, Tom Connor |
| EPI_ISL_2819200 | Department of Bacteria, Parasites and Fungi, Statens Serum Institut, Copenhagen, Denmark | Statens Serum Institut Bioinformatics and Microbial Genomics | Danish Covid-19 Genome Consortium |
| EPI_ISL_2819201, EPI_ISL_2819203 | Originating lab: Wales Specialist Virology Centre Sequencing lab: Pathogen Genomics Unit | Public Health Wales Microbiology Cardiff Wales Specialist Virology Centre | Catherine Moore, Johnathan Evans, Laura Gifford, Malorie Perry, Simon Cottrell, Angela Marchbank, Alec Birchley, Alexander Adams, Amy Gaskin, Bree Gatica-Wilcox, Jason Coombes, Joel Southgate, Lauren Gilbert, Lee Graham, Nicole Pacchiarini, Sara Kumziene-Summerhayes, Sarah Taylor, Sophie Jones, Sara Rey, Matthew Bull, Joanne Watkins, Sally Corden, Tom Connor |
| EPI_ISL_2819204 | Department of Bacteria, Parasites and Fungi, Statens Serum Institut, Copenhagen, Denmark | Statens Serum Institut Bioinformatics and Microbial Genomics | Danish Covid-19 Genome Consortium |
| EPI_ISL_2819205 | Originating lab: Wales Specialist Virology Centre Sequencing lab: Pathogen Genomics Unit | Public Health Wales Microbiology Cardiff Wales Specialist Virology Centre | Catherine Moore, Johnathan Evans, Laura Gifford, Malorie Perry, Simon Cottrell, Angela Marchbank, Alec Birchley, Alexander Adams, Amy Gaskin, Bree Gatica-Wilcox, Jason Coombes, Joel Southgate, Lauren Gilbert, Lee Graham, Nicole Pacchiarini, Sara Kumziene-Summerhayes, Sarah Taylor, Sophie Jones, Sara Rey, Matthew Bull, Joanne Watkins, Sally Corden, Tom Connor |
| EPI_ISL_2819206 | Department of Bacteria, Parasites and Fungi, Statens Serum Institut, Copenhagen, Denmark | Statens Serum Institut Bioinformatics and Microbial Genomics | Danish Covid-19 Genome Consortium |
| EPI_ISL_2819207, EPI_ISL_2819209 | Originating lab: Wales Specialist Virology Centre Sequencing lab: Pathogen Genomics Unit | Public Health Wales Microbiology Cardiff Wales Specialist Virology Centre | Catherine Moore, Johnathan Evans, Laura Gifford, Malorie Perry, Simon Cottrell, Angela Marchbank, Alec Birchley, Alexander Adams, Amy Gaskin, Bree Gatica-Wilcox, Jason Coombes, Joel Southgate, Lauren Gilbert, Lee Graham, Nicole Pacchiarini, Sara Kumziene-Summerhayes, Sarah Taylor, Sophie Jones, Sara Rey, Matthew Bull, Joanne Watkins, Sally Corden, Tom Connor |
| EPI_ISL_2819210 | Department of Bacteria, Parasites and Fungi, Statens Serum Institut, Copenhagen, Denmark | Statens Serum Institut Bioinformatics and Microbial Genomics | Danish Covid-19 Genome Consortium |
| EPI_ISL_2819211, EPI_ISL_2819212 | Originating lab: Wales Specialist Virology Centre Sequencing lab: Pathogen Genomics Unit | Public Health Wales Microbiology Cardiff Wales Specialist Virology Centre | Catherine Moore, Johnathan Evans, Laura Gifford, Malorie Perry, Simon Cottrell, Angela Marchbank, Alec Birchley, Alexander Adams, Amy Gaskin, Bree Gatica-Wilcox, Jason Coombes, Joel Southgate, Lauren Gilbert, Lee Graham, Nicole Pacchiarini, Sara Kumziene-Summerhayes, Sarah Taylor, Sophie Jones, Sara Rey, Matthew Bull, Joanne Watkins, Sally Corden, Tom Connor |
| EPI_ISL_2819213 | Department of Bacteria, Parasites and Fungi, Statens Serum Institut, Copenhagen, Denmark | Statens Serum Institut Bioinformatics and Microbial Genomics | Danish Covid-19 Genome Consortium |
| EPI_ISL_2819214 | Originating lab: Wales Specialist Virology Centre Sequencing lab: Pathogen Genomics Unit | Public Health Wales Microbiology Cardiff Wales Specialist Virology Centre | Catherine Moore, Johnathan Evans, Laura Gifford, Malorie Perry, Simon Cottrell, Angela Marchbank, Alec Birchley, Alexander Adams, Amy Gaskin, Bree Gatica-Wilcox, Jason Coombes, Joel Southgate, Lauren Gilbert, Lee Graham, Nicole Pacchiarini, Sara Kumziene-Summerhayes, Sarah Taylor, Sophie Jones, Sara Rey, Matthew Bull, Joanne Watkins, Sally Corden, Tom Connor |
| EPI_ISL_2819215 | Department of Bacteria, Parasites and Fungi, Statens Serum Institut, Copenhagen, Denmark | Statens Serum Institut Bioinformatics and Microbial Genomics | Danish Covid-19 Genome Consortium |
| EPI_ISL_2819216 | Originating lab: Wales Specialist Virology Centre Sequencing lab: Pathogen Genomics Unit | Public Health Wales Microbiology Cardiff Wales Specialist Virology Centre | Catherine Moore, Johnathan Evans, Laura Gifford, Malorie Perry, Simon Cottrell, Angela Marchbank, Alec Birchley, Alexander Adams, Amy Gaskin, Bree Gatica-Wilcox, Jason Coombes, Joel Southgate, Lauren Gilbert, Lee Graham, Nicole Pacchiarini, Sara Kumziene-Summerhayes, Sarah Taylor, Sophie Jones, Sara Rey, Matthew Bull, Joanne Watkins, Sally Corden, Tom Connor |
| EPI_ISL_2819217 | Department of Bacteria, Parasites and Fungi, Statens Serum Institut, Copenhagen, Denmark | Statens Serum Institut Bioinformatics and Microbial Genomics | Danish Covid-19 Genome Consortium |
| EPI_ISL_2819218 | Originating lab: Wales Specialist Virology Centre Sequencing lab: Pathogen Genomics Unit | Public Health Wales Microbiology Cardiff Wales Specialist Virology Centre | Catherine Moore, Johnathan Evans, Laura Gifford, Malorie Perry, Simon Cottrell, Angela Marchbank, Alec Birchley, Alexander Adams, Amy Gaskin, Bree Gatica-Wilcox, Jason Coombes, Joel Southgate, Lauren Gilbert, Lee Graham, Nicole Pacchiarini, Sara Kumziene-Summerhayes, Sarah Taylor, Sophie Jones, Sara Rey, Matthew Bull, Joanne Watkins, Sally Corden, Tom Connor |
| EPI_ISL_2819219 | Department of Bacteria, Parasites and Fungi, Statens Serum Institut, Copenhagen, Denmark | Statens Serum Institut Bioinformatics and Microbial Genomics | Danish Covid-19 Genome Consortium |
| EPI_ISL_2819220 | Originating lab: Wales Specialist Virology Centre Sequencing lab: Pathogen Genomics Unit | Public Health Wales Microbiology Cardiff Wales Specialist Virology Centre | Catherine Moore, Johnathan Evans, Laura Gifford, Malorie Perry, Simon Cottrell, Angela Marchbank, Alec Birchley, Alexander Adams, Amy Gaskin, Bree Gatica-Wilcox, Jason Coombes, Joel Southgate, Lauren Gilbert, Lee Graham, Nicole Pacchiarini, Sara Kumziene-Summerhayes, Sarah Taylor, Sophie Jones, Sara Rey, Matthew Bull, Joanne Watkins, Sally Corden, Tom Connor |
| EPI_ISL_2819221 | Department of Bacteria, Parasites and Fungi, Statens Serum Institut, Copenhagen, Denmark | Statens Serum Institut Bioinformatics and Microbial Genomics | Danish Covid-19 Genome Consortium |
| EPI_ISL_2819222, EPI_ISL_2819224 | Originating lab: Wales Specialist Virology Centre Sequencing lab: Pathogen Genomics Unit | Public Health Wales Microbiology Cardiff Wales Specialist Virology Centre | Catherine Moore, Johnathan Evans, Laura Gifford, Malorie Perry, Simon Cottrell, Angela Marchbank, Alec Birchley, Alexander Adams, Amy Gaskin, Bree Gatica-Wilcox, Jason Coombes, Joel Southgate, Lauren Gilbert, Lee Graham, Nicole Pacchiarini, Sara Kumziene-Summerhayes, Sarah Taylor, Sophie Jones, Sara Rey, Matthew Bull, Joanne Watkins, Sally Corden, Tom Connor |
| EPI_ISL_2819225 | Department of Bacteria, Parasites and Fungi, Statens Serum Institut, Copenhagen, Denmark | Statens Serum Institut Bioinformatics and Microbial Genomics | Danish Covid-19 Genome Consortium |

| | | | |
|---|---|---|---|
| EPI_ISL_2819226, EPI_ISL_2819228 | Originating lab: Wales Specialist Virology Centre Sequencing lab: Pathogen Genomics Unit | Public Health Wales Microbiology Cardiff Wales Specialist Virology Centre | Catherine Moore, Johnathan Evans, Laura Gifford, Malorie Perry, Simon Cottrell, Angela Marchbank, Alec Birchley, Alexander Adams, Amy Gaskin, Bree Gatica-Wilcox, Jason Coombes, Joel Southgate, Lauren Gilbert, Lee Graham, Nicole Pacchiarini, Sara Kumziene-Summerhayes, Sarah Taylor, Sophie Jones, Sara Rey, Matthew Bull, Joanne Watkins, Sally Corden, Tom Connor |
| EPI_ISL_2819229 | Department of Bacteria, Parasites and Fungi, Statens Serum Institut, Copenhagen, Denmark | Statens Serum Institut Bioinformatics and Microbial Genomics | Danish Covid-19 Genome Consortium |
| EPI_ISL_2819230 | Originating lab: Wales Specialist Virology Centre Sequencing lab: Pathogen Genomics Unit | Public Health Wales Microbiology Cardiff Wales Specialist Virology Centre | Catherine Moore, Johnathan Evans, Laura Gifford, Malorie Perry, Simon Cottrell, Angela Marchbank, Alec Birchley, Alexander Adams, Amy Gaskin, Bree Gatica-Wilcox, Jason Coombes, Joel Southgate, Lauren Gilbert, Lee Graham, Nicole Pacchiarini, Sara Kumziene-Summerhayes, Sarah Taylor, Sophie Jones, Sara Rey, Matthew Bull, Joanne Watkins, Sally Corden, Tom Connor |
| EPI_ISL_2819231 | Department of Bacteria, Parasites and Fungi, Statens Serum Institut, Copenhagen, Denmark | Statens Serum Institut Bioinformatics and Microbial Genomics | Danish Covid-19 Genome Consortium |
| EPI_ISL_2819232, EPI_ISL_2819233, EPI_ISL_2819235, EPI_ISL_2819237 | Originating lab: Wales Specialist Virology Centre Sequencing lab: Pathogen Genomics Unit | Public Health Wales Microbiology Cardiff Wales Specialist Virology Centre | Catherine Moore, Johnathan Evans, Laura Gifford, Malorie Perry, Simon Cottrell, Angela Marchbank, Alec Birchley, Alexander Adams, Amy Gaskin, Bree Gatica-Wilcox, Jason Coombes, Joel Southgate, Lauren Gilbert, Lee Graham, Nicole Pacchiarini, Sara Kumziene-Summerhayes, Sarah Taylor, Sophie Jones, Sara Rey, Matthew Bull, Joanne Watkins, Sally Corden, Tom Connor |
| EPI_ISL_2819238 | Department of Bacteria, Parasites and Fungi, Statens Serum Institut, Copenhagen, Denmark | Statens Serum Institut Bioinformatics and Microbial Genomics | Danish Covid-19 Genome Consortium |
| EPI_ISL_2819239 | Originating lab: Wales Specialist Virology Centre Sequencing lab: Pathogen Genomics Unit | Public Health Wales Microbiology Cardiff Wales Specialist Virology Centre | Catherine Moore, Johnathan Evans, Laura Gifford, Malorie Perry, Simon Cottrell, Angela Marchbank, Alec Birchley, Alexander Adams, Amy Gaskin, Bree Gatica-Wilcox, Jason Coombes, Joel Southgate, Lauren Gilbert, Lee Graham, Nicole Pacchiarini, Sara Kumziene-Summerhayes, Sarah Taylor, Sophie Jones, Sara Rey, Matthew Bull, Joanne Watkins, Sally Corden, Tom Connor |
| EPI_ISL_2819240 | Department of Bacteria, Parasites and Fungi, Statens Serum Institut, Copenhagen, Denmark | Statens Serum Institut Bioinformatics and Microbial Genomics | Danish Covid-19 Genome Consortium |
| EPI_ISL_2819242, EPI_ISL_2819244, EPI_ISL_2819245, EPI_ISL_2819247 | Originating lab: Wales Specialist Virology Centre Sequencing lab: Pathogen Genomics Unit | Public Health Wales Microbiology Cardiff Wales Specialist Virology Centre | Catherine Moore, Johnathan Evans, Laura Gifford, Malorie Perry, Simon Cottrell, Angela Marchbank, Alec Birchley, Alexander Adams, Amy Gaskin, Bree Gatica-Wilcox, Jason Coombes, Joel Southgate, Lauren Gilbert, Lee Graham, Nicole Pacchiarini, Sara Kumziene-Summerhayes, Sarah Taylor, Sophie Jones, Sara Rey, Matthew Bull, Joanne Watkins, Sally Corden, Tom Connor |
| EPI_ISL_2819248 | Department of Bacteria, Parasites and Fungi, Statens Serum Institut, Copenhagen, Denmark | Statens Serum Institut Bioinformatics and Microbial Genomics | Danish Covid-19 Genome Consortium |
| EPI_ISL_2819249 | Originating lab: Wales Specialist Virology Centre Sequencing lab: Pathogen Genomics Unit | Public Health Wales Microbiology Cardiff Wales Specialist Virology Centre | Catherine Moore, Johnathan Evans, Laura Gifford, Malorie Perry, Simon Cottrell, Angela Marchbank, Alec Birchley, Alexander Adams, Amy Gaskin, Bree Gatica-Wilcox, Jason Coombes, Joel Southgate, Lauren Gilbert, Lee Graham, Nicole Pacchiarini, Sara Kumziene-Summerhayes, Sarah Taylor, Sophie Jones, Sara Rey, Matthew Bull, Joanne Watkins, Sally Corden, Tom Connor |
| EPI_ISL_2819250 | Department of Bacteria, Parasites and Fungi, Statens Serum Institut, Copenhagen, Denmark | Statens Serum Institut Bioinformatics and Microbial Genomics | Danish Covid-19 Genome Consortium |
| EPI_ISL_2819251 | Originating lab: Wales Specialist Virology Centre Sequencing lab: Pathogen Genomics Unit | Public Health Wales Microbiology Cardiff Wales Specialist Virology Centre | Catherine Moore, Johnathan Evans, Laura Gifford, Malorie Perry, Simon Cottrell, Angela Marchbank, Alec Birchley, Alexander Adams, Amy Gaskin, Bree Gatica-Wilcox, Jason Coombes, Joel Southgate, Lauren Gilbert, Lee Graham, Nicole Pacchiarini, Sara Kumziene-Summerhayes, Sarah Taylor, Sophie Jones, Sara Rey, Matthew Bull, Joanne Watkins, Sally Corden, Tom Connor |
| EPI_ISL_2819252 | Department of Bacteria, Parasites and Fungi, Statens Serum Institut, Copenhagen, Denmark | Statens Serum Institut Bioinformatics and Microbial Genomics | Danish Covid-19 Genome Consortium |
| EPI_ISL_2819253, EPI_ISL_2819255 | Originating lab: Wales Specialist Virology Centre Sequencing lab: Pathogen Genomics Unit | Public Health Wales Microbiology Cardiff Wales Specialist Virology Centre | Catherine Moore, Johnathan Evans, Laura Gifford, Malorie Perry, Simon Cottrell, Angela Marchbank, Alec Birchley, Alexander Adams, Amy Gaskin, Bree Gatica-Wilcox, Jason Coombes, Joel Southgate, Lauren Gilbert, Lee Graham, Nicole Pacchiarini, Sara Kumziene-Summerhayes, Sarah Taylor, Sophie Jones, Sara Rey, Matthew Bull, Joanne Watkins, Sally Corden, Tom Connor |
| EPI_ISL_2819256 | Department of Bacteria, Parasites and Fungi, Statens Serum Institut, Copenhagen, Denmark | Statens Serum Institut Bioinformatics and Microbial Genomics | Danish Covid-19 Genome Consortium |
| EPI_ISL_2819257 | Originating lab: Wales Specialist Virology Centre Sequencing lab: Pathogen Genomics Unit | Public Health Wales Microbiology Cardiff Wales Specialist Virology Centre | Catherine Moore, Johnathan Evans, Laura Gifford, Malorie Perry, Simon Cottrell, Angela Marchbank, Alec Birchley, Alexander Adams, Amy Gaskin, Bree Gatica-Wilcox, Jason Coombes, Joel Southgate, Lauren Gilbert, Lee Graham, Nicole Pacchiarini, Sara Kumziene-Summerhayes, Sarah Taylor, Sophie Jones, Sara Rey, Matthew Bull, Joanne Watkins, Sally Corden, Tom Connor |
| EPI_ISL_2819258 | Department of Bacteria, Parasites and Fungi, Statens Serum Institut, Copenhagen, Denmark | Statens Serum Institut Bioinformatics and Microbial Genomics | Danish Covid-19 Genome Consortium |
| EPI_ISL_2819259, EPI_ISL_2819260 | Originating lab: Wales Specialist Virology Centre Sequencing lab: Pathogen Genomics Unit | Public Health Wales Microbiology Cardiff Wales Specialist Virology Centre | Catherine Moore, Johnathan Evans, Laura Gifford, Malorie Perry, Simon Cottrell, Angela Marchbank, Alec Birchley, Alexander Adams, Amy Gaskin, Bree Gatica-Wilcox, Jason Coombes, Joel Southgate, Lauren Gilbert, Lee Graham, Nicole Pacchiarini, Sara Kumziene-Summerhayes, Sarah Taylor, Sophie Jones, Sara Rey, Matthew Bull, Joanne Watkins, Sally Corden, Tom Connor |
| EPI_ISL_2819261 | Department of Bacteria, Parasites and Fungi, Statens Serum Institut, Copenhagen, Denmark | Statens Serum Institut Bioinformatics and Microbial Genomics | Danish Covid-19 Genome Consortium |
| EPI_ISL_2819262 | Originating lab: Wales Specialist Virology Centre Sequencing lab: Pathogen Genomics Unit | Public Health Wales Microbiology Cardiff Wales Specialist Virology Centre | Catherine Moore, Johnathan Evans, Laura Gifford, Malorie Perry, Simon Cottrell, Angela Marchbank, Alec Birchley, Alexander Adams, Amy Gaskin, Bree Gatica-Wilcox, Jason Coombes, Joel Southgate, Lauren Gilbert, Lee Graham, Nicole Pacchiarini, Sara Kumziene-Summerhayes, Sarah Taylor, Sophie Jones, Sara Rey, Matthew Bull, Joanne Watkins, Sally Corden, Tom Connor |
| EPI_ISL_2819263 | Department of Bacteria, Parasites and Fungi, Statens Serum Institut, Copenhagen, Denmark | Statens Serum Institut Bioinformatics and Microbial Genomics | Danish Covid-19 Genome Consortium |
| EPI_ISL_2819264 | Originating lab: Wales Specialist Virology Centre Sequencing lab: Pathogen Genomics Unit | Public Health Wales Microbiology Cardiff Wales Specialist Virology Centre | Catherine Moore, Johnathan Evans, Laura Gifford, Malorie Perry, Simon Cottrell, Angela Marchbank, Alec Birchley, Alexander Adams, Amy Gaskin, Bree Gatica-Wilcox, Jason Coombes, Joel Southgate, Lauren Gilbert, Lee Graham, Nicole Pacchiarini, Sara Kumziene-Summerhayes, Sarah Taylor, Sophie Jones, Sara Rey, Matthew Bull, Joanne Watkins, Sally Corden, Tom Connor |
| EPI_ISL_2819265 | Department of Bacteria, Parasites and Fungi, Statens Serum Institut, Copenhagen, Denmark | Statens Serum Institut Bioinformatics and Microbial Genomics | Danish Covid-19 Genome Consortium |
| EPI_ISL_2819266 | Originating lab: Wales Specialist Virology Centre Sequencing lab: Pathogen Genomics Unit | Public Health Wales Microbiology Cardiff Wales Specialist Virology Centre | Catherine Moore, Johnathan Evans, Laura Gifford, Malorie Perry, Simon Cottrell, Angela Marchbank, Alec Birchley, Alexander Adams, Amy Gaskin, Bree Gatica-Wilcox, Jason Coombes, Joel Southgate, Lauren Gilbert, Lee Graham, Nicole Pacchiarini, Sara Kumziene-Summerhayes, Sarah Taylor, Sophie Jones, Sara Rey, Matthew Bull, Joanne Watkins, Sally Corden, Tom Connor |
| EPI_ISL_2819267 | Department of Bacteria, Parasites and Fungi, Statens Serum Institut, Copenhagen, Denmark | Statens Serum Institut Bioinformatics and Microbial Genomics | Danish Covid-19 Genome Consortium |
| EPI_ISL_2819268, EPI_ISL_2819270, EPI_ISL_2819271 | Originating lab: Wales Specialist Virology Centre Sequencing lab: Pathogen Genomics Unit | Public Health Wales Microbiology Cardiff Wales Specialist Virology Centre | Catherine Moore, Johnathan Evans, Laura Gifford, Malorie Perry, Simon Cottrell, Angela Marchbank, Alec Birchley, Alexander Adams, Amy Gaskin, Bree Gatica-Wilcox, Jason Coombes, Joel Southgate, Lauren Gilbert, Lee Graham, Nicole Pacchiarini, Sara Kumziene-Summerhayes, Sarah Taylor, Sophie Jones, Sara Rey, Matthew Bull, Joanne Watkins, Sally Corden, Tom Connor |
| EPI_ISL_2819272 | Department of Bacteria, Parasites and Fungi, Statens Serum Institut, Copenhagen, Denmark | Statens Serum Institut Bioinformatics and Microbial Genomics | Danish Covid-19 Genome Consortium |
| EPI_ISL_2819273 | Originating lab: Wales Specialist Virology Centre Sequencing lab: Pathogen Genomics Unit | Public Health Wales Microbiology Cardiff Wales Specialist Virology Centre | Catherine Moore, Johnathan Evans, Laura Gifford, Malorie Perry, Simon Cottrell, Angela Marchbank, Alec Birchley, Alexander Adams, Amy Gaskin, Bree Gatica-Wilcox, Jason Coombes, Joel Southgate, Lauren Gilbert, Lee Graham, Nicole Pacchiarini, Sara Kumziene-Summerhayes, Sarah Taylor, Sophie Jones, Sara Rey, Matthew Bull, Joanne Watkins, Sally Corden, Tom Connor |
| EPI_ISL_2819274 | Department of Bacteria, Parasites and Fungi, Statens Serum Institut, Copenhagen, Denmark | Statens Serum Institut Bioinformatics and Microbial Genomics | Danish Covid-19 Genome Consortium |
| EPI_ISL_2819275 | Originating lab: Wales Specialist Virology Centre Sequencing lab: Pathogen Genomics Unit | Public Health Wales Microbiology Cardiff Wales Specialist Virology Centre | Catherine Moore, Johnathan Evans, Laura Gifford, Malorie Perry, Simon Cottrell, Angela Marchbank, Alec Birchley, Alexander Adams, Amy Gaskin, Bree Gatica-Wilcox, Jason Coombes, Joel Southgate, Lauren Gilbert, Lee Graham, Nicole Pacchiarini, Sara Kumziene-Summerhayes, Sarah Taylor, Sophie Jones, Sara Rey, Matthew Bull, Joanne Watkins, Sally Corden, Tom Connor |
| EPI_ISL_2819276 | Department of Bacteria, Parasites and Fungi, Statens Serum Institut, Copenhagen, Denmark | Statens Serum Institut Bioinformatics and Microbial Genomics | Danish Covid-19 Genome Consortium |
| EPI_ISL_2819277, EPI_ISL_2819279, EPI_ISL_2819280, EPI_ISL_2819282, EPI_ISL_2819284 | Originating lab: Wales Specialist Virology Centre Sequencing lab: Pathogen Genomics Unit | Public Health Wales Microbiology Cardiff Wales Specialist Virology Centre | Catherine Moore, Johnathan Evans, Laura Gifford, Malorie Perry, Simon Cottrell, Angela Marchbank, Alec Birchley, Alexander Adams, Amy Gaskin, Bree Gatica-Wilcox, Jason Coombes, Joel Southgate, Lauren Gilbert, Lee Graham, Nicole Pacchiarini, Sara Kumziene-Summerhayes, Sarah Taylor, Sophie Jones, Sara Rey, Matthew Bull, Joanne Watkins, Sally Corden, Tom Connor |
| EPI_ISL_2819285 | Department of Bacteria, Parasites and Fungi, Statens Serum Institut, Copenhagen, Denmark | Statens Serum Institut Bioinformatics and Microbial Genomics | Danish Covid-19 Genome Consortium |
| EPI_ISL_2819286 | Originating lab: Wales Specialist Virology Centre Sequencing lab: Pathogen Genomics Unit | Public Health Wales Microbiology Cardiff Wales Specialist Virology Centre | Catherine Moore, Johnathan Evans, Laura Gifford, Malorie Perry, Simon Cottrell, Angela Marchbank, Alec Birchley, Alexander Adams, Amy Gaskin, Bree Gatica-Wilcox, Jason Coombes, Joel Southgate, Lauren Gilbert, Lee Graham, Nicole Pacchiarini, Sara Kumziene-Summerhayes, Sarah Taylor, Sophie Jones, Sara Rey, Matthew Bull, Joanne Watkins, Sally Corden, Tom Connor |
| EPI_ISL_2819287 | Department of Bacteria, Parasites and Fungi, Statens Serum Institut, Copenhagen, Denmark | Statens Serum Institut Bioinformatics and Microbial Genomics | Danish Covid-19 Genome Consortium |
| EPI_ISL_2819288 | Originating lab: Wales Specialist Virology Centre Sequencing lab: Pathogen Genomics Unit | Public Health Wales Microbiology Cardiff Wales Specialist Virology Centre | Catherine Moore, Johnathan Evans, Laura Gifford, Malorie Perry, Simon Cottrell, Angela Marchbank, Alec Birchley, Alexander Adams, Amy Gaskin, Bree Gatica-Wilcox, Jason Coombes, Joel Southgate, Lauren Gilbert, Lee Graham, Nicole Pacchiarini, Sara Kumziene-Summerhayes, Sarah Taylor, Sophie Jones, Sara Rey, Matthew Bull, Joanne Watkins, Sally Corden, Tom Connor |
| EPI_ISL_2819289 | Department of Bacteria, Parasites and Fungi, Statens Serum Institut, Copenhagen, Denmark | Statens Serum Institut Bioinformatics and Microbial Genomics | Danish Covid-19 Genome Consortium |
| EPI_ISL_2819290 | Originating lab: Wales Specialist Virology Centre Sequencing lab: Pathogen Genomics Unit | Public Health Wales Microbiology Cardiff Wales Specialist Virology Centre | Catherine Moore, Johnathan Evans, Laura Gifford, Malorie Perry, Simon Cottrell, Angela Marchbank, Alec Birchley, Alexander Adams, Amy Gaskin, Bree Gatica-Wilcox, Jason Coombes, Joel Southgate, Lauren Gilbert, Lee Graham, Nicole Pacchiarini, Sara Kumziene-Summerhayes, Sarah Taylor, Sophie Jones, Sara Rey, Matthew Bull, Joanne Watkins, Sally Corden, Tom Connor |
| EPI_ISL_2819291 | Department of Bacteria, Parasites and Fungi, Statens Serum Institut, Copenhagen, Denmark | Statens Serum Institut Bioinformatics and Microbial Genomics | Danish Covid-19 Genome Consortium |
| EPI_ISL_2819292 | Originating lab: Wales Specialist Virology Centre Sequencing lab: Pathogen Genomics Unit | Public Health Wales Microbiology Cardiff Wales Specialist Virology Centre | Catherine Moore, Johnathan Evans, Laura Gifford, Malorie Perry, Simon Cottrell, Angela Marchbank, Alec Birchley, Alexander Adams, Amy Gaskin, Bree Gatica-Wilcox, Jason Coombes, Joel Southgate, Lauren Gilbert, Lee Graham, Nicole Pacchiarini, Sara Kumziene-Summerhayes, Sarah Taylor, Sophie Jones, Sara Rey, Matthew Bull, Joanne Watkins, Sally Corden, Tom Connor |
| EPI_ISL_2819293 | Department of Bacteria, Parasites and Fungi, Statens Serum Institut, Copenhagen, Denmark | Statens Serum Institut Bioinformatics and Microbial Genomics | Danish Covid-19 Genome Consortium |

| | | | |
|---|---|---|---|
| EPI_ISL_2819294 | Originating lab: Wales Specialist Virology Centre Sequencing lab: Pathogen Genomics Unit | Public Health Wales Microbiology Cardiff Wales Specialist Virology Centre | Catherine Moore, Johnathan Evans, Laura Gifford, Malorie Perry, Simon Cottrell, Angela Marchbank, Alec Birchley, Alexander Adams, Amy Gaskin, Bree Gatica-Wilcox, Jason Coombes, Joel Southgate, Lauren Gilbert, Lee Graham, Nicole Pacchiarini, Sara Kumziene-Summerhayes, Sarah Taylor, Sophie Jones, Sara Rey, Matthew Bull, Joanne Watkins, Sally Corden, Tom Connor |
| EPI_ISL_2819295 | Department of Bacteria, Parasites and Fungi, Statens Serum Institut, Copenhagen, Denmark | Statens Serum Institut Bioinformatics and Microbial Genomics | Danish Covid-19 Genome Consortium |
| EPI_ISL_2819296 | Originating lab: Wales Specialist Virology Centre Sequencing lab: Pathogen Genomics Unit | Public Health Wales Microbiology Cardiff Wales Specialist Virology Centre | Catherine Moore, Johnathan Evans, Laura Gifford, Malorie Perry, Simon Cottrell, Angela Marchbank, Alec Birchley, Alexander Adams, Amy Gaskin, Bree Gatica-Wilcox, Jason Coombes, Joel Southgate, Lauren Gilbert, Lee Graham, Nicole Pacchiarini, Sara Kumziene-Summerhayes, Sarah Taylor, Sophie Jones, Sara Rey, Matthew Bull, Joanne Watkins, Sally Corden, Tom Connor |
| EPI_ISL_2819297 | Department of Bacteria, Parasites and Fungi, Statens Serum Institut, Copenhagen, Denmark | Statens Serum Institut Bioinformatics and Microbial Genomics | Danish Covid-19 Genome Consortium |
| EPI_ISL_2819298, EPI_ISL_2819300, EPI_ISL_2819301, EPI_ISL_2819303 | Originating lab: Wales Specialist Virology Centre Sequencing lab: Pathogen Genomics Unit | Public Health Wales Microbiology Cardiff Wales Specialist Virology Centre | Catherine Moore, Johnathan Evans, Laura Gifford, Malorie Perry, Simon Cottrell, Angela Marchbank, Alec Birchley, Alexander Adams, Amy Gaskin, Bree Gatica-Wilcox, Jason Coombes, Joel Southgate, Lauren Gilbert, Lee Graham, Nicole Pacchiarini, Sara Kumziene-Summerhayes, Sarah Taylor, Sophie Jones, Sara Rey, Matthew Bull, Joanne Watkins, Sally Corden, Tom Connor |
| EPI_ISL_2819304 | Department of Bacteria, Parasites and Fungi, Statens Serum Institut, Copenhagen, Denmark | Statens Serum Institut Bioinformatics and Microbial Genomics | Danish Covid-19 Genome Consortium |
| EPI_ISL_2819305 | Originating lab: Wales Specialist Virology Centre Sequencing lab: Pathogen Genomics Unit | Public Health Wales Microbiology Cardiff Wales Specialist Virology Centre | Catherine Moore, Johnathan Evans, Laura Gifford, Malorie Perry, Simon Cottrell, Angela Marchbank, Alec Birchley, Alexander Adams, Amy Gaskin, Bree Gatica-Wilcox, Jason Coombes, Joel Southgate, Lauren Gilbert, Lee Graham, Nicole Pacchiarini, Sara Kumziene-Summerhayes, Sarah Taylor, Sophie Jones, Sara Rey, Matthew Bull, Joanne Watkins, Sally Corden, Tom Connor |
| EPI_ISL_2819306 | Department of Bacteria, Parasites and Fungi, Statens Serum Institut, Copenhagen, Denmark | Statens Serum Institut Bioinformatics and Microbial Genomics | Danish Covid-19 Genome Consortium |
| EPI_ISL_2819307 | Originating lab: Wales Specialist Virology Centre Sequencing lab: Pathogen Genomics Unit | Public Health Wales Microbiology Cardiff Wales Specialist Virology Centre | Catherine Moore, Johnathan Evans, Laura Gifford, Malorie Perry, Simon Cottrell, Angela Marchbank, Alec Birchley, Alexander Adams, Amy Gaskin, Bree Gatica-Wilcox, Jason Coombes, Joel Southgate, Lauren Gilbert, Lee Graham, Nicole Pacchiarini, Sara Kumziene-Summerhayes, Sarah Taylor, Sophie Jones, Sara Rey, Matthew Bull, Joanne Watkins, Sally Corden, Tom Connor |
| EPI_ISL_2819308 | Department of Bacteria, Parasites and Fungi, Statens Serum Institut, Copenhagen, Denmark | Statens Serum Institut Bioinformatics and Microbial Genomics | Danish Covid-19 Genome Consortium |
| EPI_ISL_2819309, EPI_ISL_2819310, EPI_ISL_2819312, EPI_ISL_2819314, EPI_ISL_2819316 | Originating lab: Wales Specialist Virology Centre Sequencing lab: Pathogen Genomics Unit | Public Health Wales Microbiology Cardiff Wales Specialist Virology Centre | Catherine Moore, Johnathan Evans, Laura Gifford, Malorie Perry, Simon Cottrell, Angela Marchbank, Alec Birchley, Alexander Adams, Amy Gaskin, Bree Gatica-Wilcox, Jason Coombes, Joel Southgate, Lauren Gilbert, Lee Graham, Nicole Pacchiarini, Sara Kumziene-Summerhayes, Sarah Taylor, Sophie Jones, Sara Rey, Matthew Bull, Joanne Watkins, Sally Corden, Tom Connor |
| EPI_ISL_2819317 | Department of Bacteria, Parasites and Fungi, Statens Serum Institut, Copenhagen, Denmark | Statens Serum Institut Bioinformatics and Microbial Genomics | Danish Covid-19 Genome Consortium |
| EPI_ISL_2819318 | Originating lab: Wales Specialist Virology Centre Sequencing lab: Pathogen Genomics Unit | Public Health Wales Microbiology Cardiff Wales Specialist Virology Centre | Catherine Moore, Johnathan Evans, Laura Gifford, Malorie Perry, Simon Cottrell, Angela Marchbank, Alec Birchley, Alexander Adams, Amy Gaskin, Bree Gatica-Wilcox, Jason Coombes, Joel Southgate, Lauren Gilbert, Lee Graham, Nicole Pacchiarini, Sara Kumziene-Summerhayes, Sarah Taylor, Sophie Jones, Sara Rey, Matthew Bull, Joanne Watkins, Sally Corden, Tom Connor |
| EPI_ISL_2819319 | Department of Bacteria, Parasites and Fungi, Statens Serum Institut, Copenhagen, Denmark | Statens Serum Institut Bioinformatics and Microbial Genomics | Danish Covid-19 Genome Consortium |
| EPI_ISL_2819320, EPI_ISL_2819322, EPI_ISL_2819323 | Originating lab: Wales Specialist Virology Centre Sequencing lab: Pathogen Genomics Unit | Public Health Wales Microbiology Cardiff Wales Specialist Virology Centre | Catherine Moore, Johnathan Evans, Laura Gifford, Malorie Perry, Simon Cottrell, Angela Marchbank, Alec Birchley, Alexander Adams, Amy Gaskin, Bree Gatica-Wilcox, Jason Coombes, Joel Southgate, Lauren Gilbert, Lee Graham, Nicole Pacchiarini, Sara Kumziene-Summerhayes, Sarah Taylor, Sophie Jones, Sara Rey, Matthew Bull, Joanne Watkins, Sally Corden, Tom Connor |
| EPI_ISL_2819324 | Department of Bacteria, Parasites and Fungi, Statens Serum Institut, Copenhagen, Denmark | Statens Serum Institut Bioinformatics and Microbial Genomics | Danish Covid-19 Genome Consortium |
| EPI_ISL_2819325 | Originating lab: Wales Specialist Virology Centre Sequencing lab: Pathogen Genomics Unit | Public Health Wales Microbiology Cardiff Wales Specialist Virology Centre | Catherine Moore, Johnathan Evans, Laura Gifford, Malorie Perry, Simon Cottrell, Angela Marchbank, Alec Birchley, Alexander Adams, Amy Gaskin, Bree Gatica-Wilcox, Jason Coombes, Joel Southgate, Lauren Gilbert, Lee Graham, Nicole Pacchiarini, Sara Kumziene-Summerhayes, Sarah Taylor, Sophie Jones, Sara Rey, Matthew Bull, Joanne Watkins, Sally Corden, Tom Connor |
| EPI_ISL_2819326 | Department of Bacteria, Parasites and Fungi, Statens Serum Institut, Copenhagen, Denmark | Statens Serum Institut Bioinformatics and Microbial Genomics | Danish Covid-19 Genome Consortium |
| EPI_ISL_2819327 | Originating lab: Wales Specialist Virology Centre Sequencing lab: Pathogen Genomics Unit | Public Health Wales Microbiology Cardiff Wales Specialist Virology Centre | Catherine Moore, Johnathan Evans, Laura Gifford, Malorie Perry, Simon Cottrell, Angela Marchbank, Alec Birchley, Alexander Adams, Amy Gaskin, Bree Gatica-Wilcox, Jason Coombes, Joel Southgate, Lauren Gilbert, Lee Graham, Nicole Pacchiarini, Sara Kumziene-Summerhayes, Sarah Taylor, Sophie Jones, Sara Rey, Matthew Bull, Joanne Watkins, Sally Corden, Tom Connor |
| EPI_ISL_2819328 | Department of Bacteria, Parasites and Fungi, Statens Serum Institut, Copenhagen, Denmark | Statens Serum Institut Bioinformatics and Microbial Genomics | Danish Covid-19 Genome Consortium |
| EPI_ISL_2819329 | Originating lab: Wales Specialist Virology Centre Sequencing lab: Pathogen Genomics Unit | Public Health Wales Microbiology Cardiff Wales Specialist Virology Centre | Catherine Moore, Johnathan Evans, Laura Gifford, Malorie Perry, Simon Cottrell, Angela Marchbank, Alec Birchley, Alexander Adams, Amy Gaskin, Bree Gatica-Wilcox, Jason Coombes, Joel Southgate, Lauren Gilbert, Lee Graham, Nicole Pacchiarini, Sara Kumziene-Summerhayes, Sarah Taylor, Sophie Jones, Sara Rey, Matthew Bull, Joanne Watkins, Sally Corden, Tom Connor |
| EPI_ISL_2819330, EPI_ISL_2819332 | Department of Bacteria, Parasites and Fungi, Statens Serum Institut, Copenhagen, Denmark | Statens Serum Institut Bioinformatics and Microbial Genomics | Danish Covid-19 Genome Consortium |
| EPI_ISL_2819333, EPI_ISL_2819335 | Originating lab: Wales Specialist Virology Centre Sequencing lab: Pathogen Genomics Unit | Public Health Wales Microbiology Cardiff Wales Specialist Virology Centre | Catherine Moore, Johnathan Evans, Laura Gifford, Malorie Perry, Simon Cottrell, Angela Marchbank, Alec Birchley, Alexander Adams, Amy Gaskin, Bree Gatica-Wilcox, Jason Coombes, Joel Southgate, Lauren Gilbert, Lee Graham, Nicole Pacchiarini, Sara Kumziene-Summerhayes, Sarah Taylor, Sophie Jones, Sara Rey, Matthew Bull, Joanne Watkins, Sally Corden, Tom Connor |
| EPI_ISL_2819336 | Department of Bacteria, Parasites and Fungi, Statens Serum Institut, Copenhagen, Denmark | Statens Serum Institut Bioinformatics and Microbial Genomics | Danish Covid-19 Genome Consortium |
| EPI_ISL_2819337 | Originating lab: Wales Specialist Virology Centre Sequencing lab: Pathogen Genomics Unit | Public Health Wales Microbiology Cardiff Wales Specialist Virology Centre | Catherine Moore, Johnathan Evans, Laura Gifford, Malorie Perry, Simon Cottrell, Angela Marchbank, Alec Birchley, Alexander Adams, Amy Gaskin, Bree Gatica-Wilcox, Jason Coombes, Joel Southgate, Lauren Gilbert, Lee Graham, Nicole Pacchiarini, Sara Kumziene-Summerhayes, Sarah Taylor, Sophie Jones, Sara Rey, Matthew Bull, Joanne Watkins, Sally Corden, Tom Connor |
| EPI_ISL_2819338 | Department of Bacteria, Parasites and Fungi, Statens Serum Institut, Copenhagen, Denmark | Statens Serum Institut Bioinformatics and Microbial Genomics | Danish Covid-19 Genome Consortium |
| EPI_ISL_2819339 | Originating lab: Wales Specialist Virology Centre Sequencing lab: Pathogen Genomics Unit | Public Health Wales Microbiology Cardiff Wales Specialist Virology Centre | Catherine Moore, Johnathan Evans, Laura Gifford, Malorie Perry, Simon Cottrell, Angela Marchbank, Alec Birchley, Alexander Adams, Amy Gaskin, Bree Gatica-Wilcox, Jason Coombes, Joel Southgate, Lauren Gilbert, Lee Graham, Nicole Pacchiarini, Sara Kumziene-Summerhayes, Sarah Taylor, Sophie Jones, Sara Rey, Matthew Bull, Joanne Watkins, Sally Corden, Tom Connor |
| EPI_ISL_2819340 | Department of Bacteria, Parasites and Fungi, Statens Serum Institut, Copenhagen, Denmark | Statens Serum Institut Bioinformatics and Microbial Genomics | Danish Covid-19 Genome Consortium |
| EPI_ISL_2819341 | Originating lab: Wales Specialist Virology Centre Sequencing lab: Pathogen Genomics Unit | Public Health Wales Microbiology Cardiff Wales Specialist Virology Centre | Catherine Moore, Johnathan Evans, Laura Gifford, Malorie Perry, Simon Cottrell, Angela Marchbank, Alec Birchley, Alexander Adams, Amy Gaskin, Bree Gatica-Wilcox, Jason Coombes, Joel Southgate, Lauren Gilbert, Lee Graham, Nicole Pacchiarini, Sara Kumziene-Summerhayes, Sarah Taylor, Sophie Jones, Sara Rey, Matthew Bull, Joanne Watkins, Sally Corden, Tom Connor |
| EPI_ISL_2819342 | Department of Bacteria, Parasites and Fungi, Statens Serum Institut, Copenhagen, Denmark | Statens Serum Institut Bioinformatics and Microbial Genomics | Danish Covid-19 Genome Consortium |
| EPI_ISL_2819343, EPI_ISL_2819344 | Originating lab: Wales Specialist Virology Centre Sequencing lab: Pathogen Genomics Unit | Public Health Wales Microbiology Cardiff Wales Specialist Virology Centre | Catherine Moore, Johnathan Evans, Laura Gifford, Malorie Perry, Simon Cottrell, Angela Marchbank, Alec Birchley, Alexander Adams, Amy Gaskin, Bree Gatica-Wilcox, Jason Coombes, Joel Southgate, Lauren Gilbert, Lee Graham, Nicole Pacchiarini, Sara Kumziene-Summerhayes, Sarah Taylor, Sophie Jones, Sara Rey, Matthew Bull, Joanne Watkins, Sally Corden, Tom Connor |
| EPI_ISL_2819345 | Department of Bacteria, Parasites and Fungi, Statens Serum Institut, Copenhagen, Denmark | Statens Serum Institut Bioinformatics and Microbial Genomics | Danish Covid-19 Genome Consortium |
| EPI_ISL_2819346 | Originating lab: Wales Specialist Virology Centre Sequencing lab: Pathogen Genomics Unit | Public Health Wales Microbiology Cardiff Wales Specialist Virology Centre | Catherine Moore, Johnathan Evans, Laura Gifford, Malorie Perry, Simon Cottrell, Angela Marchbank, Alec Birchley, Alexander Adams, Amy Gaskin, Bree Gatica-Wilcox, Jason Coombes, Joel Southgate, Lauren Gilbert, Lee Graham, Nicole Pacchiarini, Sara Kumziene-Summerhayes, Sarah Taylor, Sophie Jones, Sara Rey, Matthew Bull, Joanne Watkins, Sally Corden, Tom Connor |
| EPI_ISL_2819347 | Department of Bacteria, Parasites and Fungi, Statens Serum Institut, Copenhagen, Denmark | Statens Serum Institut Bioinformatics and Microbial Genomics | Danish Covid-19 Genome Consortium |
| EPI_ISL_2819348 | Originating lab: Wales Specialist Virology Centre Sequencing lab: Pathogen Genomics Unit | Public Health Wales Microbiology Cardiff Wales Specialist Virology Centre | Catherine Moore, Johnathan Evans, Laura Gifford, Malorie Perry, Simon Cottrell, Angela Marchbank, Alec Birchley, Alexander Adams, Amy Gaskin, Bree Gatica-Wilcox, Jason Coombes, Joel Southgate, Lauren Gilbert, Lee Graham, Nicole Pacchiarini, Sara Kumziene-Summerhayes, Sarah Taylor, Sophie Jones, Sara Rey, Matthew Bull, Joanne Watkins, Sally Corden, Tom Connor |
| EPI_ISL_2819349 | Department of Bacteria, Parasites and Fungi, Statens Serum Institut, Copenhagen, Denmark | Statens Serum Institut Bioinformatics and Microbial Genomics | Danish Covid-19 Genome Consortium |
| EPI_ISL_2819350, EPI_ISL_2819352, EPI_ISL_2819354 | Originating lab: Wales Specialist Virology Centre Sequencing lab: Pathogen Genomics Unit | Public Health Wales Microbiology Cardiff Wales Specialist Virology Centre | Catherine Moore, Johnathan Evans, Laura Gifford, Malorie Perry, Simon Cottrell, Angela Marchbank, Alec Birchley, Alexander Adams, Amy Gaskin, Bree Gatica-Wilcox, Jason Coombes, Joel Southgate, Lauren Gilbert, Lee Graham, Nicole Pacchiarini, Sara Kumziene-Summerhayes, Sarah Taylor, Sophie Jones, Sara Rey, Matthew Bull, Joanne Watkins, Sally Corden, Tom Connor |
| EPI_ISL_2819355 | Department of Bacteria, Parasites and Fungi, Statens Serum Institut, Copenhagen, Denmark | Statens Serum Institut Bioinformatics and Microbial Genomics | Danish Covid-19 Genome Consortium |
| EPI_ISL_2819356 | Originating lab: Wales Specialist Virology Centre Sequencing lab: Pathogen Genomics Unit | Public Health Wales Microbiology Cardiff Wales Specialist Virology Centre | Catherine Moore, Johnathan Evans, Laura Gifford, Malorie Perry, Simon Cottrell, Angela Marchbank, Alec Birchley, Alexander Adams, Amy Gaskin, Bree Gatica-Wilcox, Jason Coombes, Joel Southgate, Lauren Gilbert, Lee Graham, Nicole Pacchiarini, Sara Kumziene-Summerhayes, Sarah Taylor, Sophie Jones, Sara Rey, Matthew Bull, Joanne Watkins, Sally Corden, Tom Connor |
| EPI_ISL_2819357 | Department of Bacteria, Parasites and Fungi, Statens Serum Institut, Copenhagen, Denmark | Statens Serum Institut Bioinformatics and Microbial Genomics | Danish Covid-19 Genome Consortium |
| EPI_ISL_2819358, EPI_ISL_2819359 | Originating lab: Wales Specialist Virology Centre Sequencing lab: Pathogen Genomics Unit | Public Health Wales Microbiology Cardiff Wales Specialist Virology Centre | Catherine Moore, Johnathan Evans, Laura Gifford, Malorie Perry, Simon Cottrell, Angela Marchbank, Alec Birchley, Alexander Adams, Amy Gaskin, Bree Gatica-Wilcox, Jason Coombes, Joel Southgate, Lauren Gilbert, Lee Graham, Nicole Pacchiarini, Sara Kumziene-Summerhayes, Sarah Taylor, Sophie Jones, Sara Rey, Matthew Bull, Joanne Watkins, Sally Corden, Tom Connor |
| EPI_ISL_2819360 | Department of Bacteria, Parasites and Fungi, Statens Serum Institut, Copenhagen, Denmark | Statens Serum Institut Bioinformatics and Microbial Genomics | Danish Covid-19 Genome Consortium |

| | | | |
|---|---|---|---|
| EPI_ISL_2819361 | Originating lab: Wales Specialist Virology Centre Sequencing lab: Pathogen Genomics Unit | Public Health Wales Microbiology Cardiff Wales Specialist Virology Centre | Catherine Moore, Johnathan Evans, Laura Gifford, Malorie Perry, Simon Cottrell, Angela Marchbank, Alec Birchley, Alexander Adams, Amy Gaskin, Bree Gatica-Wilcox, Jason Coombes, Joel Southgate, Lauren Gilbert, Lee Graham, Nicole Pacchiarini, Sara Kumziene-Summerhayes, Sarah Taylor, Sophie Jones, Sara Rey, Matthew Bull, Joanne Watkins, Sally Corden, Tom Connor |
| EPI_ISL_2819362 | Department of Bacteria, Parasites and Fungi, Statens Serum Institut, Copenhagen, Denmark | Statens Serum Institut Bioinformatics and Microbial Genomics | Danish Covid-19 Genome Consortium |
| EPI_ISL_2819363, EPI_ISL_2819365 | Originating lab: Wales Specialist Virology Centre Sequencing lab: Pathogen Genomics Unit | Public Health Wales Microbiology Cardiff Wales Specialist Virology Centre | Catherine Moore, Johnathan Evans, Laura Gifford, Malorie Perry, Simon Cottrell, Angela Marchbank, Alec Birchley, Alexander Adams, Amy Gaskin, Bree Gatica-Wilcox, Jason Coombes, Joel Southgate, Lauren Gilbert, Lee Graham, Nicole Pacchiarini, Sara Kumziene-Summerhayes, Sarah Taylor, Sophie Jones, Sara Rey, Matthew Bull, Joanne Watkins, Sally Corden, Tom Connor |
| EPI_ISL_2819366 | Department of Bacteria, Parasites and Fungi, Statens Serum Institut, Copenhagen, Denmark | Statens Serum Institut Bioinformatics and Microbial Genomics | Danish Covid-19 Genome Consortium |
| EPI_ISL_2819367, EPI_ISL_2819369 | Originating lab: Wales Specialist Virology Centre Sequencing lab: Pathogen Genomics Unit | Public Health Wales Microbiology Cardiff Wales Specialist Virology Centre | Catherine Moore, Johnathan Evans, Laura Gifford, Malorie Perry, Simon Cottrell, Angela Marchbank, Alec Birchley, Alexander Adams, Amy Gaskin, Bree Gatica-Wilcox, Jason Coombes, Joel Southgate, Lauren Gilbert, Lee Graham, Nicole Pacchiarini, Sara Kumziene-Summerhayes, Sarah Taylor, Sophie Jones, Sara Rey, Matthew Bull, Joanne Watkins, Sally Corden, Tom Connor |
| EPI_ISL_2819370 | Department of Bacteria, Parasites and Fungi, Statens Serum Institut, Copenhagen, Denmark | Statens Serum Institut Bioinformatics and Microbial Genomics | Danish Covid-19 Genome Consortium |
| EPI_ISL_2819371 | Originating lab: Wales Specialist Virology Centre Sequencing lab: Pathogen Genomics Unit | Public Health Wales Microbiology Cardiff Wales Specialist Virology Centre | Catherine Moore, Johnathan Evans, Laura Gifford, Malorie Perry, Simon Cottrell, Angela Marchbank, Alec Birchley, Alexander Adams, Amy Gaskin, Bree Gatica-Wilcox, Jason Coombes, Joel Southgate, Lauren Gilbert, Lee Graham, Nicole Pacchiarini, Sara Kumziene-Summerhayes, Sarah Taylor, Sophie Jones, Sara Rey, Matthew Bull, Joanne Watkins, Sally Corden, Tom Connor |
| EPI_ISL_2819372 | Department of Bacteria, Parasites and Fungi, Statens Serum Institut, Copenhagen, Denmark | Statens Serum Institut Bioinformatics and Microbial Genomics | Danish Covid-19 Genome Consortium |
| EPI_ISL_2819373 | Originating lab: Wales Specialist Virology Centre Sequencing lab: Pathogen Genomics Unit | Public Health Wales Microbiology Cardiff Wales Specialist Virology Centre | Catherine Moore, Johnathan Evans, Laura Gifford, Malorie Perry, Simon Cottrell, Angela Marchbank, Alec Birchley, Alexander Adams, Amy Gaskin, Bree Gatica-Wilcox, Jason Coombes, Joel Southgate, Lauren Gilbert, Lee Graham, Nicole Pacchiarini, Sara Kumziene-Summerhayes, Sarah Taylor, Sophie Jones, Sara Rey, Matthew Bull, Joanne Watkins, Sally Corden, Tom Connor |
| EPI_ISL_2819374 | Department of Bacteria, Parasites and Fungi, Statens Serum Institut, Copenhagen, Denmark | Statens Serum Institut Bioinformatics and Microbial Genomics | Danish Covid-19 Genome Consortium |
| EPI_ISL_2819375 | Originating lab: Wales Specialist Virology Centre Sequencing lab: Pathogen Genomics Unit | Public Health Wales Microbiology Cardiff Wales Specialist Virology Centre | Catherine Moore, Johnathan Evans, Laura Gifford, Malorie Perry, Simon Cottrell, Angela Marchbank, Alec Birchley, Alexander Adams, Amy Gaskin, Bree Gatica-Wilcox, Jason Coombes, Joel Southgate, Lauren Gilbert, Lee Graham, Nicole Pacchiarini, Sara Kumziene-Summerhayes, Sarah Taylor, Sophie Jones, Sara Rey, Matthew Bull, Joanne Watkins, Sally Corden, Tom Connor |
| EPI_ISL_2819376 | Department of Bacteria, Parasites and Fungi, Statens Serum Institut, Copenhagen, Denmark | Statens Serum Institut Bioinformatics and Microbial Genomics | Danish Covid-19 Genome Consortium |
| EPI_ISL_2819377 | Originating lab: Wales Specialist Virology Centre Sequencing lab: Pathogen Genomics Unit | Public Health Wales Microbiology Cardiff Wales Specialist Virology Centre | Catherine Moore, Johnathan Evans, Laura Gifford, Malorie Perry, Simon Cottrell, Angela Marchbank, Alec Birchley, Alexander Adams, Amy Gaskin, Bree Gatica-Wilcox, Jason Coombes, Joel Southgate, Lauren Gilbert, Lee Graham, Nicole Pacchiarini, Sara Kumziene-Summerhayes, Sarah Taylor, Sophie Jones, Sara Rey, Matthew Bull, Joanne Watkins, Sally Corden, Tom Connor |
| EPI_ISL_2819378 | Department of Bacteria, Parasites and Fungi, Statens Serum Institut, Copenhagen, Denmark | Statens Serum Institut Bioinformatics and Microbial Genomics | Danish Covid-19 Genome Consortium |
| EPI_ISL_2819379, EPI_ISL_2819381 | Originating lab: Wales Specialist Virology Centre Sequencing lab: Pathogen Genomics Unit | Public Health Wales Microbiology Cardiff Wales Specialist Virology Centre | Catherine Moore, Johnathan Evans, Laura Gifford, Malorie Perry, Simon Cottrell, Angela Marchbank, Alec Birchley, Alexander Adams, Amy Gaskin, Bree Gatica-Wilcox, Jason Coombes, Joel Southgate, Lauren Gilbert, Lee Graham, Nicole Pacchiarini, Sara Kumziene-Summerhayes, Sarah Taylor, Sophie Jones, Sara Rey, Matthew Bull, Joanne Watkins, Sally Corden, Tom Connor |
| EPI_ISL_2819383 | Department of Bacteria, Parasites and Fungi, Statens Serum Institut, Copenhagen, Denmark | Statens Serum Institut Bioinformatics and Microbial Genomics | Danish Covid-19 Genome Consortium |
| EPI_ISL_2819384 | Originating lab: Wales Specialist Virology Centre Sequencing lab: Pathogen Genomics Unit | Public Health Wales Microbiology Cardiff Wales Specialist Virology Centre | Catherine Moore, Johnathan Evans, Laura Gifford, Malorie Perry, Simon Cottrell, Angela Marchbank, Alec Birchley, Alexander Adams, Amy Gaskin, Bree Gatica-Wilcox, Jason Coombes, Joel Southgate, Lauren Gilbert, Lee Graham, Nicole Pacchiarini, Sara Kumziene-Summerhayes, Sarah Taylor, Sophie Jones, Sara Rey, Matthew Bull, Joanne Watkins, Sally Corden, Tom Connor |
| EPI_ISL_2819385, EPI_ISL_2819387 | Department of Bacteria, Parasites and Fungi, Statens Serum Institut, Copenhagen, Denmark | Statens Serum Institut Bioinformatics and Microbial Genomics | Danish Covid-19 Genome Consortium |
| EPI_ISL_2819388, EPI_ISL_2819390 | Originating lab: Wales Specialist Virology Centre Sequencing lab: Pathogen Genomics Unit | Public Health Wales Microbiology Cardiff Wales Specialist Virology Centre | Catherine Moore, Johnathan Evans, Laura Gifford, Malorie Perry, Simon Cottrell, Angela Marchbank, Alec Birchley, Alexander Adams, Amy Gaskin, Bree Gatica-Wilcox, Jason Coombes, Joel Southgate, Lauren Gilbert, Lee Graham, Nicole Pacchiarini, Sara Kumziene-Summerhayes, Sarah Taylor, Sophie Jones, Sara Rey, Matthew Bull, Joanne Watkins, Sally Corden, Tom Connor |
| EPI_ISL_2819391 | Department of Bacteria, Parasites and Fungi, Statens Serum Institut, Copenhagen, Denmark | Statens Serum Institut Bioinformatics and Microbial Genomics | Danish Covid-19 Genome Consortium |
| EPI_ISL_2819392 | Originating lab: Wales Specialist Virology Centre Sequencing lab: Pathogen Genomics Unit | Public Health Wales Microbiology Cardiff Wales Specialist Virology Centre | Catherine Moore, Johnathan Evans, Laura Gifford, Malorie Perry, Simon Cottrell, Angela Marchbank, Alec Birchley, Alexander Adams, Amy Gaskin, Bree Gatica-Wilcox, Jason Coombes, Joel Southgate, Lauren Gilbert, Lee Graham, Nicole Pacchiarini, Sara Kumziene-Summerhayes, Sarah Taylor, Sophie Jones, Sara Rey, Matthew Bull, Joanne Watkins, Sally Corden, Tom Connor |
| EPI_ISL_2819393 | Department of Bacteria, Parasites and Fungi, Statens Serum Institut, Copenhagen, Denmark | Statens Serum Institut Bioinformatics and Microbial Genomics | Danish Covid-19 Genome Consortium |
| EPI_ISL_2819394 | Originating lab: Wales Specialist Virology Centre Sequencing lab: Pathogen Genomics Unit | Public Health Wales Microbiology Cardiff Wales Specialist Virology Centre | Catherine Moore, Johnathan Evans, Laura Gifford, Malorie Perry, Simon Cottrell, Angela Marchbank, Alec Birchley, Alexander Adams, Amy Gaskin, Bree Gatica-Wilcox, Jason Coombes, Joel Southgate, Lauren Gilbert, Lee Graham, Nicole Pacchiarini, Sara Kumziene-Summerhayes, Sarah Taylor, Sophie Jones, Sara Rey, Matthew Bull, Joanne Watkins, Sally Corden, Tom Connor |
| EPI_ISL_2819395 | Department of Bacteria, Parasites and Fungi, Statens Serum Institut, Copenhagen, Denmark | Statens Serum Institut Bioinformatics and Microbial Genomics | Danish Covid-19 Genome Consortium |
| EPI_ISL_2819396, EPI_ISL_2819398, EPI_ISL_2819400, EPI_ISL_2819401, EPI_ISL_2819403 | Originating lab: Wales Specialist Virology Centre Sequencing lab: Pathogen Genomics Unit | Public Health Wales Microbiology Cardiff Wales Specialist Virology Centre | Catherine Moore, Johnathan Evans, Laura Gifford, Malorie Perry, Simon Cottrell, Angela Marchbank, Alec Birchley, Alexander Adams, Amy Gaskin, Bree Gatica-Wilcox, Jason Coombes, Joel Southgate, Lauren Gilbert, Lee Graham, Nicole Pacchiarini, Sara Kumziene-Summerhayes, Sarah Taylor, Sophie Jones, Sara Rey, Matthew Bull, Joanne Watkins, Sally Corden, Tom Connor |
| EPI_ISL_2819404 | Department of Bacteria, Parasites and Fungi, Statens Serum Institut, Copenhagen, Denmark | Statens Serum Institut Bioinformatics and Microbial Genomics | Danish Covid-19 Genome Consortium |
| EPI_ISL_2819405, EPI_ISL_2819407 | Originating lab: Wales Specialist Virology Centre Sequencing lab: Pathogen Genomics Unit | Public Health Wales Microbiology Cardiff Wales Specialist Virology Centre | Catherine Moore, Johnathan Evans, Laura Gifford, Malorie Perry, Simon Cottrell, Angela Marchbank, Alec Birchley, Alexander Adams, Amy Gaskin, Bree Gatica-Wilcox, Jason Coombes, Joel Southgate, Lauren Gilbert, Lee Graham, Nicole Pacchiarini, Sara Kumziene-Summerhayes, Sarah Taylor, Sophie Jones, Sara Rey, Matthew Bull, Joanne Watkins, Sally Corden, Tom Connor |
| EPI_ISL_2819408 | Department of Bacteria, Parasites and Fungi, Statens Serum Institut, Copenhagen, Denmark | Statens Serum Institut Bioinformatics and Microbial Genomics | Danish Covid-19 Genome Consortium |
| EPI_ISL_2819409 | Originating lab: Wales Specialist Virology Centre Sequencing lab: Pathogen Genomics Unit | Public Health Wales Microbiology Cardiff Wales Specialist Virology Centre | Catherine Moore, Johnathan Evans, Laura Gifford, Malorie Perry, Simon Cottrell, Angela Marchbank, Alec Birchley, Alexander Adams, Amy Gaskin, Bree Gatica-Wilcox, Jason Coombes, Joel Southgate, Lauren Gilbert, Lee Graham, Nicole Pacchiarini, Sara Kumziene-Summerhayes, Sarah Taylor, Sophie Jones, Sara Rey, Matthew Bull, Joanne Watkins, Sally Corden, Tom Connor |
| EPI_ISL_2819410 | Department of Bacteria, Parasites and Fungi, Statens Serum Institut, Copenhagen, Denmark | Statens Serum Institut Bioinformatics and Microbial Genomics | Danish Covid-19 Genome Consortium |
| EPI_ISL_2819411 | Originating lab: Wales Specialist Virology Centre Sequencing lab: Pathogen Genomics Unit | Public Health Wales Microbiology Cardiff Wales Specialist Virology Centre | Catherine Moore, Johnathan Evans, Laura Gifford, Malorie Perry, Simon Cottrell, Angela Marchbank, Alec Birchley, Alexander Adams, Amy Gaskin, Bree Gatica-Wilcox, Jason Coombes, Joel Southgate, Lauren Gilbert, Lee Graham, Nicole Pacchiarini, Sara Kumziene-Summerhayes, Sarah Taylor, Sophie Jones, Sara Rey, Matthew Bull, Joanne Watkins, Sally Corden, Tom Connor |
| EPI_ISL_2819412 | Department of Bacteria, Parasites and Fungi, Statens Serum Institut, Copenhagen, Denmark | Statens Serum Institut Bioinformatics and Microbial Genomics | Danish Covid-19 Genome Consortium |
| EPI_ISL_2819413, EPI_ISL_2819414 | Originating lab: Wales Specialist Virology Centre Sequencing lab: Pathogen Genomics Unit | Public Health Wales Microbiology Cardiff Wales Specialist Virology Centre | Catherine Moore, Johnathan Evans, Laura Gifford, Malorie Perry, Simon Cottrell, Angela Marchbank, Alec Birchley, Alexander Adams, Amy Gaskin, Bree Gatica-Wilcox, Jason Coombes, Joel Southgate, Lauren Gilbert, Lee Graham, Nicole Pacchiarini, Sara Kumziene-Summerhayes, Sarah Taylor, Sophie Jones, Sara Rey, Matthew Bull, Joanne Watkins, Sally Corden, Tom Connor |
| EPI_ISL_2819415 | Department of Bacteria, Parasites and Fungi, Statens Serum Institut, Copenhagen, Denmark | Statens Serum Institut Bioinformatics and Microbial Genomics | Danish Covid-19 Genome Consortium |
| EPI_ISL_2819416 | Originating lab: Wales Specialist Virology Centre Sequencing lab: Pathogen Genomics Unit | Public Health Wales Microbiology Cardiff Wales Specialist Virology Centre | Catherine Moore, Johnathan Evans, Laura Gifford, Malorie Perry, Simon Cottrell, Angela Marchbank, Alec Birchley, Alexander Adams, Amy Gaskin, Bree Gatica-Wilcox, Jason Coombes, Joel Southgate, Lauren Gilbert, Lee Graham, Nicole Pacchiarini, Sara Kumziene-Summerhayes, Sarah Taylor, Sophie Jones, Sara Rey, Matthew Bull, Joanne Watkins, Sally Corden, Tom Connor |
| EPI_ISL_2819417 | Department of Bacteria, Parasites and Fungi, Statens Serum Institut, Copenhagen, Denmark | Statens Serum Institut Bioinformatics and Microbial Genomics | Danish Covid-19 Genome Consortium |
| EPI_ISL_2819418 | Originating lab: Wales Specialist Virology Centre Sequencing lab: Pathogen Genomics Unit | Public Health Wales Microbiology Cardiff Wales Specialist Virology Centre | Catherine Moore, Johnathan Evans, Laura Gifford, Malorie Perry, Simon Cottrell, Angela Marchbank, Alec Birchley, Alexander Adams, Amy Gaskin, Bree Gatica-Wilcox, Jason Coombes, Joel Southgate, Lauren Gilbert, Lee Graham, Nicole Pacchiarini, Sara Kumziene-Summerhayes, Sarah Taylor, Sophie Jones, Sara Rey, Matthew Bull, Joanne Watkins, Sally Corden, Tom Connor |
| EPI_ISL_2819419 | Department of Bacteria, Parasites and Fungi, Statens Serum Institut, Copenhagen, Denmark | Statens Serum Institut Bioinformatics and Microbial Genomics | Danish Covid-19 Genome Consortium |
| EPI_ISL_2819420 | Originating lab: Wales Specialist Virology Centre Sequencing lab: Pathogen Genomics Unit | Public Health Wales Microbiology Cardiff Wales Specialist Virology Centre | Catherine Moore, Johnathan Evans, Laura Gifford, Malorie Perry, Simon Cottrell, Angela Marchbank, Alec Birchley, Alexander Adams, Amy Gaskin, Bree Gatica-Wilcox, Jason Coombes, Joel Southgate, Lauren Gilbert, Lee Graham, Nicole Pacchiarini, Sara Kumziene-Summerhayes, Sarah Taylor, Sophie Jones, Sara Rey, Matthew Bull, Joanne Watkins, Sally Corden, Tom Connor |
| EPI_ISL_2819421 | Department of Bacteria, Parasites and Fungi, Statens Serum Institut, Copenhagen, Denmark | Statens Serum Institut Bioinformatics and Microbial Genomics | Danish Covid-19 Genome Consortium |
| EPI_ISL_2819422, EPI_ISL_2819423 | Originating lab: Wales Specialist Virology Centre Sequencing lab: Pathogen Genomics Unit | Public Health Wales Microbiology Cardiff Wales Specialist Virology Centre | Catherine Moore, Johnathan Evans, Laura Gifford, Malorie Perry, Simon Cottrell, Angela Marchbank, Alec Birchley, Alexander Adams, Amy Gaskin, Bree Gatica-Wilcox, Jason Coombes, Joel Southgate, Lauren Gilbert, Lee Graham, Nicole Pacchiarini, Sara Kumziene-Summerhayes, Sarah Taylor, Sophie Jones, Sara Rey, Matthew Bull, Joanne Watkins, Sally Corden, Tom Connor |
| EPI_ISL_2819424 | Department of Bacteria, Parasites and Fungi, Statens Serum Institut, Copenhagen, Denmark | Statens Serum Institut Bioinformatics and Microbial Genomics | Danish Covid-19 Genome Consortium |
| EPI_ISL_2819425 | Originating lab: Wales Specialist Virology Centre Sequencing lab: Pathogen Genomics Unit | Public Health Wales Microbiology Cardiff Wales Specialist Virology Centre | Catherine Moore, Johnathan Evans, Laura Gifford, Malorie Perry, Simon Cottrell, Angela Marchbank, Alec Birchley, Alexander Adams, Amy Gaskin, Bree Gatica-Wilcox, Jason Coombes, Joel Southgate, Lauren Gilbert, Lee Graham, Nicole Pacchiarini, Sara Kumziene-Summerhayes, Sarah Taylor, Sophie Jones, Sara Rey, Matthew Bull, Joanne Watkins, Sally Corden, Tom Connor |
| EPI_ISL_2819426 | Department of Bacteria, Parasites and Fungi, Statens Serum Institut, Copenhagen, Denmark | Statens Serum Institut Bioinformatics and Microbial Genomics | Danish Covid-19 Genome Consortium |

| | | | |
|---|---|---|---|
| EPI_ISL_2819427 | Originating lab: Wales Specialist Virology Centre Sequencing lab: Pathogen Genomics Unit | Public Health Wales Microbiology Cardiff Wales Specialist Virology Centre | Catherine Moore, Johnathan Evans, Laura Gifford, Malorie Perry, Simon Cottrell, Angela Marchbank, Alec Birchley, Alexander Adams, Amy Gaskin, Bree Gatica-Wilcox, Jason Coombes, Joel Southgate, Lauren Gilbert, Lee Graham, Nicole Pacchiarini, Sara Kumziene-Summerhayes, Sarah Taylor, Sophie Jones, Sara Rey, Matthew Bull, Joanne Watkins, Sally Corden, Tom Connor |
| EPI_ISL_2819428 | Department of Bacteria, Parasites and Fungi, Statens Serum Institut, Copenhagen, Denmark | Statens Serum Institut Bioinformatics and Microbial Genomics | Danish Covid-19 Genome Consortium |
| EPI_ISL_2819429 | Originating lab: Wales Specialist Virology Centre Sequencing lab: Pathogen Genomics Unit | Public Health Wales Microbiology Cardiff Wales Specialist Virology Centre | Catherine Moore, Johnathan Evans, Laura Gifford, Malorie Perry, Simon Cottrell, Angela Marchbank, Alec Birchley, Alexander Adams, Amy Gaskin, Bree Gatica-Wilcox, Jason Coombes, Joel Southgate, Lauren Gilbert, Lee Graham, Nicole Pacchiarini, Sara Kumziene-Summerhayes, Sarah Taylor, Sophie Jones, Sara Rey, Matthew Bull, Joanne Watkins, Sally Corden, Tom Connor |
| EPI_ISL_2819430 | Department of Bacteria, Parasites and Fungi, Statens Serum Institut, Copenhagen, Denmark | Statens Serum Institut Bioinformatics and Microbial Genomics | Danish Covid-19 Genome Consortium |
| EPI_ISL_2819431 | Originating lab: Wales Specialist Virology Centre Sequencing lab: Pathogen Genomics Unit | Public Health Wales Microbiology Cardiff Wales Specialist Virology Centre | Catherine Moore, Johnathan Evans, Laura Gifford, Malorie Perry, Simon Cottrell, Angela Marchbank, Alec Birchley, Alexander Adams, Amy Gaskin, Bree Gatica-Wilcox, Jason Coombes, Joel Southgate, Lauren Gilbert, Lee Graham, Nicole Pacchiarini, Sara Kumziene-Summerhayes, Sarah Taylor, Sophie Jones, Sara Rey, Matthew Bull, Joanne Watkins, Sally Corden, Tom Connor |
| EPI_ISL_2819432 | Department of Bacteria, Parasites and Fungi, Statens Serum Institut, Copenhagen, Denmark | Statens Serum Institut Bioinformatics and Microbial Genomics | Danish Covid-19 Genome Consortium |
| EPI_ISL_2819433, EPI_ISL_2819435 | Originating lab: Wales Specialist Virology Centre Sequencing lab: Pathogen Genomics Unit | Public Health Wales Microbiology Cardiff Wales Specialist Virology Centre | Catherine Moore, Johnathan Evans, Laura Gifford, Malorie Perry, Simon Cottrell, Angela Marchbank, Alec Birchley, Alexander Adams, Amy Gaskin, Bree Gatica-Wilcox, Jason Coombes, Joel Southgate, Lauren Gilbert, Lee Graham, Nicole Pacchiarini, Sara Kumziene-Summerhayes, Sarah Taylor, Sophie Jones, Sara Rey, Matthew Bull, Joanne Watkins, Sally Corden, Tom Connor |
| EPI_ISL_2819436 | Department of Bacteria, Parasites and Fungi, Statens Serum Institut, Copenhagen, Denmark | Statens Serum Institut Bioinformatics and Microbial Genomics | Danish Covid-19 Genome Consortium |
| EPI_ISL_2819437 | Originating lab: Wales Specialist Virology Centre Sequencing lab: Pathogen Genomics Unit | Public Health Wales Microbiology Cardiff Wales Specialist Virology Centre | Catherine Moore, Johnathan Evans, Laura Gifford, Malorie Perry, Simon Cottrell, Angela Marchbank, Alec Birchley, Alexander Adams, Amy Gaskin, Bree Gatica-Wilcox, Jason Coombes, Joel Southgate, Lauren Gilbert, Lee Graham, Nicole Pacchiarini, Sara Kumziene-Summerhayes, Sarah Taylor, Sophie Jones, Sara Rey, Matthew Bull, Joanne Watkins, Sally Corden, Tom Connor |
| EPI_ISL_2819438 | Department of Bacteria, Parasites and Fungi, Statens Serum Institut, Copenhagen, Denmark | Statens Serum Institut Bioinformatics and Microbial Genomics | Danish Covid-19 Genome Consortium |
| EPI_ISL_2819439, EPI_ISL_2819441, EPI_ISL_2819442 | Originating lab: Wales Specialist Virology Centre Sequencing lab: Pathogen Genomics Unit | Public Health Wales Microbiology Cardiff Wales Specialist Virology Centre | Catherine Moore, Johnathan Evans, Laura Gifford, Malorie Perry, Simon Cottrell, Angela Marchbank, Alec Birchley, Alexander Adams, Amy Gaskin, Bree Gatica-Wilcox, Jason Coombes, Joel Southgate, Lauren Gilbert, Lee Graham, Nicole Pacchiarini, Sara Kumziene-Summerhayes, Sarah Taylor, Sophie Jones, Sara Rey, Matthew Bull, Joanne Watkins, Sally Corden, Tom Connor |
| EPI_ISL_2819443 | Department of Bacteria, Parasites and Fungi, Statens Serum Institut, Copenhagen, Denmark | Statens Serum Institut Bioinformatics and Microbial Genomics | Danish Covid-19 Genome Consortium |
| EPI_ISL_2819444, EPI_ISL_2819446, EPI_ISL_2819447 | Originating lab: Wales Specialist Virology Centre Sequencing lab: Pathogen Genomics Unit | Public Health Wales Microbiology Cardiff Wales Specialist Virology Centre | Catherine Moore, Johnathan Evans, Laura Gifford, Malorie Perry, Simon Cottrell, Angela Marchbank, Alec Birchley, Alexander Adams, Amy Gaskin, Bree Gatica-Wilcox, Jason Coombes, Joel Southgate, Lauren Gilbert, Lee Graham, Nicole Pacchiarini, Sara Kumziene-Summerhayes, Sarah Taylor, Sophie Jones, Sara Rey, Matthew Bull, Joanne Watkins, Sally Corden, Tom Connor |
| EPI_ISL_2819448 | Department of Bacteria, Parasites and Fungi, Statens Serum Institut, Copenhagen, Denmark | Statens Serum Institut Bioinformatics and Microbial Genomics | Danish Covid-19 Genome Consortium |
| EPI_ISL_2819449 | Originating lab: Wales Specialist Virology Centre Sequencing lab: Pathogen Genomics Unit | Public Health Wales Microbiology Cardiff Wales Specialist Virology Centre | Catherine Moore, Johnathan Evans, Laura Gifford, Malorie Perry, Simon Cottrell, Angela Marchbank, Alec Birchley, Alexander Adams, Amy Gaskin, Bree Gatica-Wilcox, Jason Coombes, Joel Southgate, Lauren Gilbert, Lee Graham, Nicole Pacchiarini, Sara Kumziene-Summerhayes, Sarah Taylor, Sophie Jones, Sara Rey, Matthew Bull, Joanne Watkins, Sally Corden, Tom Connor |
| EPI_ISL_2819450 | Department of Bacteria, Parasites and Fungi, Statens Serum Institut, Copenhagen, Denmark | Statens Serum Institut Bioinformatics and Microbial Genomics | Danish Covid-19 Genome Consortium |
| EPI_ISL_2819451 | Originating lab: Wales Specialist Virology Centre Sequencing lab: Pathogen Genomics Unit | Public Health Wales Microbiology Cardiff Wales Specialist Virology Centre | Catherine Moore, Johnathan Evans, Laura Gifford, Malorie Perry, Simon Cottrell, Angela Marchbank, Alec Birchley, Alexander Adams, Amy Gaskin, Bree Gatica-Wilcox, Jason Coombes, Joel Southgate, Lauren Gilbert, Lee Graham, Nicole Pacchiarini, Sara Kumziene-Summerhayes, Sarah Taylor, Sophie Jones, Sara Rey, Matthew Bull, Joanne Watkins, Sally Corden, Tom Connor |
| EPI_ISL_2819452 | Department of Bacteria, Parasites and Fungi, Statens Serum Institut, Copenhagen, Denmark | Statens Serum Institut Bioinformatics and Microbial Genomics | Danish Covid-19 Genome Consortium |
| EPI_ISL_2819453 | Originating lab: Wales Specialist Virology Centre Sequencing lab: Pathogen Genomics Unit | Public Health Wales Microbiology Cardiff Wales Specialist Virology Centre | Catherine Moore, Johnathan Evans, Laura Gifford, Malorie Perry, Simon Cottrell, Angela Marchbank, Alec Birchley, Alexander Adams, Amy Gaskin, Bree Gatica-Wilcox, Jason Coombes, Joel Southgate, Lauren Gilbert, Lee Graham, Nicole Pacchiarini, Sara Kumziene-Summerhayes, Sarah Taylor, Sophie Jones, Sara Rey, Matthew Bull, Joanne Watkins, Sally Corden, Tom Connor |
| EPI_ISL_2819454 | Department of Bacteria, Parasites and Fungi, Statens Serum Institut, Copenhagen, Denmark | Statens Serum Institut Bioinformatics and Microbial Genomics | Danish Covid-19 Genome Consortium |
| EPI_ISL_2819455, EPI_ISL_2819457, EPI_ISL_2819458 | Originating lab: Wales Specialist Virology Centre Sequencing lab: Pathogen Genomics Unit | Public Health Wales Microbiology Cardiff Wales Specialist Virology Centre | Catherine Moore, Johnathan Evans, Laura Gifford, Malorie Perry, Simon Cottrell, Angela Marchbank, Alec Birchley, Alexander Adams, Amy Gaskin, Bree Gatica-Wilcox, Jason Coombes, Joel Southgate, Lauren Gilbert, Lee Graham, Nicole Pacchiarini, Sara Kumziene-Summerhayes, Sarah Taylor, Sophie Jones, Sara Rey, Matthew Bull, Joanne Watkins, Sally Corden, Tom Connor |
| EPI_ISL_2819459 | Department of Bacteria, Parasites and Fungi, Statens Serum Institut, Copenhagen, Denmark | Statens Serum Institut Bioinformatics and Microbial Genomics | Danish Covid-19 Genome Consortium |
| EPI_ISL_2819460 | Originating lab: Wales Specialist Virology Centre Sequencing lab: Pathogen Genomics Unit | Public Health Wales Microbiology Cardiff Wales Specialist Virology Centre | Catherine Moore, Johnathan Evans, Laura Gifford, Malorie Perry, Simon Cottrell, Angela Marchbank, Alec Birchley, Alexander Adams, Amy Gaskin, Bree Gatica-Wilcox, Jason Coombes, Joel Southgate, Lauren Gilbert, Lee Graham, Nicole Pacchiarini, Sara Kumziene-Summerhayes, Sarah Taylor, Sophie Jones, Sara Rey, Matthew Bull, Joanne Watkins, Sally Corden, Tom Connor |
| EPI_ISL_2819461 | Department of Bacteria, Parasites and Fungi, Statens Serum Institut, Copenhagen, Denmark | Statens Serum Institut Bioinformatics and Microbial Genomics | Danish Covid-19 Genome Consortium |
| EPI_ISL_2819462 | Originating lab: Wales Specialist Virology Centre Sequencing lab: Pathogen Genomics Unit | Public Health Wales Microbiology Cardiff Wales Specialist Virology Centre | Catherine Moore, Johnathan Evans, Laura Gifford, Malorie Perry, Simon Cottrell, Angela Marchbank, Alec Birchley, Alexander Adams, Amy Gaskin, Bree Gatica-Wilcox, Jason Coombes, Joel Southgate, Lauren Gilbert, Lee Graham, Nicole Pacchiarini, Sara Kumziene-Summerhayes, Sarah Taylor, Sophie Jones, Sara Rey, Matthew Bull, Joanne Watkins, Sally Corden, Tom Connor |
| EPI_ISL_2819463 | Department of Bacteria, Parasites and Fungi, Statens Serum Institut, Copenhagen, Denmark | Statens Serum Institut Bioinformatics and Microbial Genomics | Danish Covid-19 Genome Consortium |
| EPI_ISL_2819464 | Originating lab: Wales Specialist Virology Centre Sequencing lab: Pathogen Genomics Unit | Public Health Wales Microbiology Cardiff Wales Specialist Virology Centre | Catherine Moore, Johnathan Evans, Laura Gifford, Malorie Perry, Simon Cottrell, Angela Marchbank, Alec Birchley, Alexander Adams, Amy Gaskin, Bree Gatica-Wilcox, Jason Coombes, Joel Southgate, Lauren Gilbert, Lee Graham, Nicole Pacchiarini, Sara Kumziene-Summerhayes, Sarah Taylor, Sophie Jones, Sara Rey, Matthew Bull, Joanne Watkins, Sally Corden, Tom Connor |
| EPI_ISL_2819465 | Department of Bacteria, Parasites and Fungi, Statens Serum Institut, Copenhagen, Denmark | Statens Serum Institut Bioinformatics and Microbial Genomics | Danish Covid-19 Genome Consortium |
| EPI_ISL_2819466, EPI_ISL_2819468, EPI_ISL_2819469 | Originating lab: Wales Specialist Virology Centre Sequencing lab: Pathogen Genomics Unit | Public Health Wales Microbiology Cardiff Wales Specialist Virology Centre | Catherine Moore, Johnathan Evans, Laura Gifford, Malorie Perry, Simon Cottrell, Angela Marchbank, Alec Birchley, Alexander Adams, Amy Gaskin, Bree Gatica-Wilcox, Jason Coombes, Joel Southgate, Lauren Gilbert, Lee Graham, Nicole Pacchiarini, Sara Kumziene-Summerhayes, Sarah Taylor, Sophie Jones, Sara Rey, Matthew Bull, Joanne Watkins, Sally Corden, Tom Connor |
| EPI_ISL_2819470 | Department of Bacteria, Parasites and Fungi, Statens Serum Institut, Copenhagen, Denmark | Statens Serum Institut Bioinformatics and Microbial Genomics | Danish Covid-19 Genome Consortium |
| EPI_ISL_2819471 | Originating lab: Wales Specialist Virology Centre Sequencing lab: Pathogen Genomics Unit | Public Health Wales Microbiology Cardiff Wales Specialist Virology Centre | Catherine Moore, Johnathan Evans, Laura Gifford, Malorie Perry, Simon Cottrell, Angela Marchbank, Alec Birchley, Alexander Adams, Amy Gaskin, Bree Gatica-Wilcox, Jason Coombes, Joel Southgate, Lauren Gilbert, Lee Graham, Nicole Pacchiarini, Sara Kumziene-Summerhayes, Sarah Taylor, Sophie Jones, Sara Rey, Matthew Bull, Joanne Watkins, Sally Corden, Tom Connor |
| EPI_ISL_2819472 | Department of Bacteria, Parasites and Fungi, Statens Serum Institut, Copenhagen, Denmark | Statens Serum Institut Bioinformatics and Microbial Genomics | Danish Covid-19 Genome Consortium |
| EPI_ISL_2819473 | Originating lab: Wales Specialist Virology Centre Sequencing lab: Pathogen Genomics Unit | Public Health Wales Microbiology Cardiff Wales Specialist Virology Centre | Catherine Moore, Johnathan Evans, Laura Gifford, Malorie Perry, Simon Cottrell, Angela Marchbank, Alec Birchley, Alexander Adams, Amy Gaskin, Bree Gatica-Wilcox, Jason Coombes, Joel Southgate, Lauren Gilbert, Lee Graham, Nicole Pacchiarini, Sara Kumziene-Summerhayes, Sarah Taylor, Sophie Jones, Sara Rey, Matthew Bull, Joanne Watkins, Sally Corden, Tom Connor |
| EPI_ISL_2819474 | Department of Bacteria, Parasites and Fungi, Statens Serum Institut, Copenhagen, Denmark | Statens Serum Institut Bioinformatics and Microbial Genomics | Danish Covid-19 Genome Consortium |
| EPI_ISL_2819475 | Originating lab: Wales Specialist Virology Centre Sequencing lab: Pathogen Genomics Unit | Public Health Wales Microbiology Cardiff Wales Specialist Virology Centre | Catherine Moore, Johnathan Evans, Laura Gifford, Malorie Perry, Simon Cottrell, Angela Marchbank, Alec Birchley, Alexander Adams, Amy Gaskin, Bree Gatica-Wilcox, Jason Coombes, Joel Southgate, Lauren Gilbert, Lee Graham, Nicole Pacchiarini, Sara Kumziene-Summerhayes, Sarah Taylor, Sophie Jones, Sara Rey, Matthew Bull, Joanne Watkins, Sally Corden, Tom Connor |
| EPI_ISL_2819476 | Department of Bacteria, Parasites and Fungi, Statens Serum Institut, Copenhagen, Denmark | Statens Serum Institut Bioinformatics and Microbial Genomics | Danish Covid-19 Genome Consortium |
| EPI_ISL_2819477 | Originating lab: Wales Specialist Virology Centre Sequencing lab: Pathogen Genomics Unit | Public Health Wales Microbiology Cardiff Wales Specialist Virology Centre | Catherine Moore, Johnathan Evans, Laura Gifford, Malorie Perry, Simon Cottrell, Angela Marchbank, Alec Birchley, Alexander Adams, Amy Gaskin, Bree Gatica-Wilcox, Jason Coombes, Joel Southgate, Lauren Gilbert, Lee Graham, Nicole Pacchiarini, Sara Kumziene-Summerhayes, Sarah Taylor, Sophie Jones, Sara Rey, Matthew Bull, Joanne Watkins, Sally Corden, Tom Connor |
| EPI_ISL_2819478 | Department of Bacteria, Parasites and Fungi, Statens Serum Institut, Copenhagen, Denmark | Statens Serum Institut Bioinformatics and Microbial Genomics | Danish Covid-19 Genome Consortium |
| EPI_ISL_2819479 | Originating lab: Wales Specialist Virology Centre Sequencing lab: Pathogen Genomics Unit | Public Health Wales Microbiology Cardiff Wales Specialist Virology Centre | Catherine Moore, Johnathan Evans, Laura Gifford, Malorie Perry, Simon Cottrell, Angela Marchbank, Alec Birchley, Alexander Adams, Amy Gaskin, Bree Gatica-Wilcox, Jason Coombes, Joel Southgate, Lauren Gilbert, Lee Graham, Nicole Pacchiarini, Sara Kumziene-Summerhayes, Sarah Taylor, Sophie Jones, Sara Rey, Matthew Bull, Joanne Watkins, Sally Corden, Tom Connor |
| EPI_ISL_2819480 | Department of Bacteria, Parasites and Fungi, Statens Serum Institut, Copenhagen, Denmark | Statens Serum Institut Bioinformatics and Microbial Genomics | Danish Covid-19 Genome Consortium |
| EPI_ISL_2819481, EPI_ISL_2819482 | Originating lab: Wales Specialist Virology Centre Sequencing lab: Pathogen Genomics Unit | Public Health Wales Microbiology Cardiff Wales Specialist Virology Centre | Catherine Moore, Johnathan Evans, Laura Gifford, Malorie Perry, Simon Cottrell, Angela Marchbank, Alec Birchley, Alexander Adams, Amy Gaskin, Bree Gatica-Wilcox, Jason Coombes, Joel Southgate, Lauren Gilbert, Lee Graham, Nicole Pacchiarini, Sara Kumziene-Summerhayes, Sarah Taylor, Sophie Jones, Sara Rey, Matthew Bull, Joanne Watkins, Sally Corden, Tom Connor |
| EPI_ISL_2819483 | Department of Bacteria, Parasites and Fungi, Statens Serum Institut, Copenhagen, Denmark | Statens Serum Institut Bioinformatics and Microbial Genomics | Danish Covid-19 Genome Consortium |
| EPI_ISL_2819484 | Originating lab: Wales Specialist Virology Centre Sequencing lab: Pathogen Genomics Unit | Public Health Wales Microbiology Cardiff Wales Specialist Virology Centre | Catherine Moore, Johnathan Evans, Laura Gifford, Malorie Perry, Simon Cottrell, Angela Marchbank, Alec Birchley, Alexander Adams, Amy Gaskin, Bree Gatica-Wilcox, Jason Coombes, Joel Southgate, Lauren Gilbert, Lee Graham, Nicole Pacchiarini, Sara Kumziene-Summerhayes, Sarah Taylor, Sophie Jones, Sara Rey, Matthew Bull, Joanne Watkins, Sally Corden, Tom Connor |
| EPI_ISL_2819485 | Department of Bacteria, Parasites and Fungi, Statens Serum | Statens Serum Institut Bioinformatics and Microbial Genomics | Danish Covid-19 Genome Consortium |

| | | | |
|---|---|---|---|
| | Institut, Copenhagen, Denmark | | |
| EPI_ISL_2819486 | Originating lab: Wales Specialist Virology Centre Sequencing lab: Pathogen Genomics Unit | Public Health Wales Microbiology Cardiff Wales Specialist Virology Centre | Catherine Moore, Johnathan Evans, Laura Gifford, Malorie Perry, Simon Cottrell, Angela Marchbank, Alec Birchley, Alexander Adams, Amy Gaskin, Bree Gatica-Wilcox, Jason Coombes, Joel Southgate, Lauren Gilbert, Lee Graham, Nicole Pacchiarini, Sara Kumziene-Summerhayes, Sarah Taylor, Sophie Jones, Sara Rey, Matthew Bull, Joanne Watkins, Sally Corden, Tom Connor |
| EPI_ISL_2819487 | Department of Bacteria, Parasites and Fungi, Statens Serum Institut, Copenhagen, Denmark | Statens Serum Institut Bioinformatics and Microbial Genomics | Danish Covid-19 Genome Consortium |
| EPI_ISL_2819488 | Originating lab: Wales Specialist Virology Centre Sequencing lab: Pathogen Genomics Unit | Public Health Wales Microbiology Cardiff Wales Specialist Virology Centre | Catherine Moore, Johnathan Evans, Laura Gifford, Malorie Perry, Simon Cottrell, Angela Marchbank, Alec Birchley, Alexander Adams, Amy Gaskin, Bree Gatica-Wilcox, Jason Coombes, Joel Southgate, Lauren Gilbert, Lee Graham, Nicole Pacchiarini, Sara Kumziene-Summerhayes, Sarah Taylor, Sophie Jones, Sara Rey, Matthew Bull, Joanne Watkins, Sally Corden, Tom Connor |
| EPI_ISL_2819489 | Department of Clinical Microbiology and Center for Genomic Medicine, Rigshospitalet, Copenhagen, Denmark | Statens Serum Institut Bioinformatics and Microbial Genomics | Danish Covid-19 Genome Consortium |
| EPI_ISL_2819490 | Department of Bacteria, Parasites and Fungi, Statens Serum Institut, Copenhagen, Denmark | Statens Serum Institut Bioinformatics and Microbial Genomics | Danish Covid-19 Genome Consortium |
| EPI_ISL_2819491 | Originating lab: Wales Specialist Virology Centre Sequencing lab: Pathogen Genomics Unit | Public Health Wales Microbiology Cardiff Wales Specialist Virology Centre | Catherine Moore, Johnathan Evans, Laura Gifford, Malorie Perry, Simon Cottrell, Angela Marchbank, Alec Birchley, Alexander Adams, Amy Gaskin, Bree Gatica-Wilcox, Jason Coombes, Joel Southgate, Lauren Gilbert, Lee Graham, Nicole Pacchiarini, Sara Kumziene-Summerhayes, Sarah Taylor, Sophie Jones, Sara Rey, Matthew Bull, Joanne Watkins, Sally Corden, Tom Connor |
| EPI_ISL_2819492 | Department of Bacteria, Parasites and Fungi, Statens Serum Institut, Copenhagen, Denmark | Statens Serum Institut Bioinformatics and Microbial Genomics | Danish Covid-19 Genome Consortium |
| EPI_ISL_2819493 | Originating lab: Wales Specialist Virology Centre Sequencing lab: Pathogen Genomics Unit | Public Health Wales Microbiology Cardiff Wales Specialist Virology Centre | Catherine Moore, Johnathan Evans, Laura Gifford, Malorie Perry, Simon Cottrell, Angela Marchbank, Alec Birchley, Alexander Adams, Amy Gaskin, Bree Gatica-Wilcox, Jason Coombes, Joel Southgate, Lauren Gilbert, Lee Graham, Nicole Pacchiarini, Sara Kumziene-Summerhayes, Sarah Taylor, Sophie Jones, Sara Rey, Matthew Bull, Joanne Watkins, Sally Corden, Tom Connor |
| EPI_ISL_2819494 | Department of Bacteria, Parasites and Fungi, Statens Serum Institut, Copenhagen, Denmark | Statens Serum Institut Bioinformatics and Microbial Genomics | Danish Covid-19 Genome Consortium |
| EPI_ISL_2819495 | Originating lab: Wales Specialist Virology Centre Sequencing lab: Pathogen Genomics Unit | Public Health Wales Microbiology Cardiff Wales Specialist Virology Centre | Catherine Moore, Johnathan Evans, Laura Gifford, Malorie Perry, Simon Cottrell, Angela Marchbank, Alec Birchley, Alexander Adams, Amy Gaskin, Bree Gatica-Wilcox, Jason Coombes, Joel Southgate, Lauren Gilbert, Lee Graham, Nicole Pacchiarini, Sara Kumziene-Summerhayes, Sarah Taylor, Sophie Jones, Sara Rey, Matthew Bull, Joanne Watkins, Sally Corden, Tom Connor |
| EPI_ISL_2819496 | Department of Bacteria, Parasites and Fungi, Statens Serum Institut, Copenhagen, Denmark | Statens Serum Institut Bioinformatics and Microbial Genomics | Danish Covid-19 Genome Consortium |
| EPI_ISL_2819497, EPI_ISL_2819499, EPI_ISL_2819500 | Originating lab: Wales Specialist Virology Centre Sequencing lab: Pathogen Genomics Unit | Public Health Wales Microbiology Cardiff Wales Specialist Virology Centre | Catherine Moore, Johnathan Evans, Laura Gifford, Malorie Perry, Simon Cottrell, Angela Marchbank, Alec Birchley, Alexander Adams, Amy Gaskin, Bree Gatica-Wilcox, Jason Coombes, Joel Southgate, Lauren Gilbert, Lee Graham, Nicole Pacchiarini, Sara Kumziene-Summerhayes, Sarah Taylor, Sophie Jones, Sara Rey, Matthew Bull, Joanne Watkins, Sally Corden, Tom Connor |
| EPI_ISL_2819501, EPI_ISL_2819502 | Department of Bacteria, Parasites and Fungi, Statens Serum Institut, Copenhagen, Denmark | Statens Serum Institut Bioinformatics and Microbial Genomics | Danish Covid-19 Genome Consortium |
| EPI_ISL_2819503 | Originating lab: Wales Specialist Virology Centre Sequencing lab: Pathogen Genomics Unit | Public Health Wales Microbiology Cardiff Wales Specialist Virology Centre | Catherine Moore, Johnathan Evans, Laura Gifford, Malorie Perry, Simon Cottrell, Angela Marchbank, Alec Birchley, Alexander Adams, Amy Gaskin, Bree Gatica-Wilcox, Jason Coombes, Joel Southgate, Lauren Gilbert, Lee Graham, Nicole Pacchiarini, Sara Kumziene-Summerhayes, Sarah Taylor, Sophie Jones, Sara Rey, Matthew Bull, Joanne Watkins, Sally Corden, Tom Connor |
| EPI_ISL_2819504 | Department of Bacteria, Parasites and Fungi, Statens Serum Institut, Copenhagen, Denmark | Statens Serum Institut Bioinformatics and Microbial Genomics | Danish Covid-19 Genome Consortium |
| EPI_ISL_2819505, EPI_ISL_2819506 | Originating lab: Wales Specialist Virology Centre Sequencing lab: Pathogen Genomics Unit | Public Health Wales Microbiology Cardiff Wales Specialist Virology Centre | Catherine Moore, Johnathan Evans, Laura Gifford, Malorie Perry, Simon Cottrell, Angela Marchbank, Alec Birchley, Alexander Adams, Amy Gaskin, Bree Gatica-Wilcox, Jason Coombes, Joel Southgate, Lauren Gilbert, Lee Graham, Nicole Pacchiarini, Sara Kumziene-Summerhayes, Sarah Taylor, Sophie Jones, Sara Rey, Matthew Bull, Joanne Watkins, Sally Corden, Tom Connor |
| EPI_ISL_2819507 | Department of Bacteria, Parasites and Fungi, Statens Serum Institut, Copenhagen, Denmark | Statens Serum Institut Bioinformatics and Microbial Genomics | Danish Covid-19 Genome Consortium |
| EPI_ISL_2819508, EPI_ISL_2819510 | Originating lab: Wales Specialist Virology Centre Sequencing lab: Pathogen Genomics Unit | Public Health Wales Microbiology Cardiff Wales Specialist Virology Centre | Catherine Moore, Johnathan Evans, Laura Gifford, Malorie Perry, Simon Cottrell, Angela Marchbank, Alec Birchley, Alexander Adams, Amy Gaskin, Bree Gatica-Wilcox, Jason Coombes, Joel Southgate, Lauren Gilbert, Lee Graham, Nicole Pacchiarini, Sara Kumziene-Summerhayes, Sarah Taylor, Sophie Jones, Sara Rey, Matthew Bull, Joanne Watkins, Sally Corden, Tom Connor |
| EPI_ISL_2819511 | Department of Bacteria, Parasites and Fungi, Statens Serum Institut, Copenhagen, Denmark | Statens Serum Institut Bioinformatics and Microbial Genomics | Danish Covid-19 Genome Consortium |
| EPI_ISL_2819512 | Originating lab: Wales Specialist Virology Centre Sequencing lab: Pathogen Genomics Unit | Public Health Wales Microbiology Cardiff Wales Specialist Virology Centre | Catherine Moore, Johnathan Evans, Laura Gifford, Malorie Perry, Simon Cottrell, Angela Marchbank, Alec Birchley, Alexander Adams, Amy Gaskin, Bree Gatica-Wilcox, Jason Coombes, Joel Southgate, Lauren Gilbert, Lee Graham, Nicole Pacchiarini, Sara Kumziene-Summerhayes, Sarah Taylor, Sophie Jones, Sara Rey, Matthew Bull, Joanne Watkins, Sally Corden, Tom Connor |
| EPI_ISL_2819513 | Department of Bacteria, Parasites and Fungi, Statens Serum Institut, Copenhagen, Denmark | Statens Serum Institut Bioinformatics and Microbial Genomics | Danish Covid-19 Genome Consortium |
| EPI_ISL_2819514 | Originating lab: Wales Specialist Virology Centre Sequencing lab: Pathogen Genomics Unit | Public Health Wales Microbiology Cardiff Wales Specialist Virology Centre | Catherine Moore, Johnathan Evans, Laura Gifford, Malorie Perry, Simon Cottrell, Angela Marchbank, Alec Birchley, Alexander Adams, Amy Gaskin, Bree Gatica-Wilcox, Jason Coombes, Joel Southgate, Lauren Gilbert, Lee Graham, Nicole Pacchiarini, Sara Kumziene-Summerhayes, Sarah Taylor, Sophie Jones, Sara Rey, Matthew Bull, Joanne Watkins, Sally Corden, Tom Connor |
| EPI_ISL_2819515 | Department of Bacteria, Parasites and Fungi, Statens Serum Institut, Copenhagen, Denmark | Statens Serum Institut Bioinformatics and Microbial Genomics | Danish Covid-19 Genome Consortium |
| EPI_ISL_2819516 | Originating lab: Wales Specialist Virology Centre Sequencing lab: Pathogen Genomics Unit | Public Health Wales Microbiology Cardiff Wales Specialist Virology Centre | Catherine Moore, Johnathan Evans, Laura Gifford, Malorie Perry, Simon Cottrell, Angela Marchbank, Alec Birchley, Alexander Adams, Amy Gaskin, Bree Gatica-Wilcox, Jason Coombes, Joel Southgate, Lauren Gilbert, Lee Graham, Nicole Pacchiarini, Sara Kumziene-Summerhayes, Sarah Taylor, Sophie Jones, Sara Rey, Matthew Bull, Joanne Watkins, Sally Corden, Tom Connor |
| EPI_ISL_2819517 | Department of Bacteria, Parasites and Fungi, Statens Serum Institut, Copenhagen, Denmark | Statens Serum Institut Bioinformatics and Microbial Genomics | Danish Covid-19 Genome Consortium |
| EPI_ISL_2819518, EPI_ISL_2819520, EPI_ISL_2819522, EPI_ISL_2819524 | Originating lab: Wales Specialist Virology Centre Sequencing lab: Pathogen Genomics Unit | Public Health Wales Microbiology Cardiff Wales Specialist Virology Centre | Catherine Moore, Johnathan Evans, Laura Gifford, Malorie Perry, Simon Cottrell, Angela Marchbank, Alec Birchley, Alexander Adams, Amy Gaskin, Bree Gatica-Wilcox, Jason Coombes, Joel Southgate, Lauren Gilbert, Lee Graham, Nicole Pacchiarini, Sara Kumziene-Summerhayes, Sarah Taylor, Sophie Jones, Sara Rey, Matthew Bull, Joanne Watkins, Sally Corden, Tom Connor |
| EPI_ISL_2819525 | Department of Bacteria, Parasites and Fungi, Statens Serum Institut, Copenhagen, Denmark | Statens Serum Institut Bioinformatics and Microbial Genomics | Danish Covid-19 Genome Consortium |
| EPI_ISL_2819526, EPI_ISL_2819528, EPI_ISL_2819529, EPI_ISL_2819531 | Originating lab: Wales Specialist Virology Centre Sequencing lab: Pathogen Genomics Unit | Public Health Wales Microbiology Cardiff Wales Specialist Virology Centre | Catherine Moore, Johnathan Evans, Laura Gifford, Malorie Perry, Simon Cottrell, Angela Marchbank, Alec Birchley, Alexander Adams, Amy Gaskin, Bree Gatica-Wilcox, Jason Coombes, Joel Southgate, Lauren Gilbert, Lee Graham, Nicole Pacchiarini, Sara Kumziene-Summerhayes, Sarah Taylor, Sophie Jones, Sara Rey, Matthew Bull, Joanne Watkins, Sally Corden, Tom Connor |
| EPI_ISL_2819532 | Department of Bacteria, Parasites and Fungi, Statens Serum Institut, Copenhagen, Denmark | Statens Serum Institut Bioinformatics and Microbial Genomics | Danish Covid-19 Genome Consortium |
| EPI_ISL_2819533, EPI_ISL_2819534 | Originating lab: Wales Specialist Virology Centre Sequencing lab: Pathogen Genomics Unit | Public Health Wales Microbiology Cardiff Wales Specialist Virology Centre | Catherine Moore, Johnathan Evans, Laura Gifford, Malorie Perry, Simon Cottrell, Angela Marchbank, Alec Birchley, Alexander Adams, Amy Gaskin, Bree Gatica-Wilcox, Jason Coombes, Joel Southgate, Lauren Gilbert, Lee Graham, Nicole Pacchiarini, Sara Kumziene-Summerhayes, Sarah Taylor, Sophie Jones, Sara Rey, Matthew Bull, Joanne Watkins, Sally Corden, Tom Connor |
| EPI_ISL_2819535 | Department of Bacteria, Parasites and Fungi, Statens Serum Institut, Copenhagen, Denmark | Statens Serum Institut Bioinformatics and Microbial Genomics | Danish Covid-19 Genome Consortium |
| EPI_ISL_2819536 | Originating lab: Wales Specialist Virology Centre Sequencing lab: Pathogen Genomics Unit | Public Health Wales Microbiology Cardiff Wales Specialist Virology Centre | Catherine Moore, Johnathan Evans, Laura Gifford, Malorie Perry, Simon Cottrell, Angela Marchbank, Alec Birchley, Alexander Adams, Amy Gaskin, Bree Gatica-Wilcox, Jason Coombes, Joel Southgate, Lauren Gilbert, Lee Graham, Nicole Pacchiarini, Sara Kumziene-Summerhayes, Sarah Taylor, Sophie Jones, Sara Rey, Matthew Bull, Joanne Watkins, Sally Corden, Tom Connor |
| EPI_ISL_2819537 | Department of Bacteria, Parasites and Fungi, Statens Serum Institut, Copenhagen, Denmark | Statens Serum Institut Bioinformatics and Microbial Genomics | Danish Covid-19 Genome Consortium |
| EPI_ISL_2819538 | Originating lab: Wales Specialist Virology Centre Sequencing lab: Pathogen Genomics Unit | Public Health Wales Microbiology Cardiff Wales Specialist Virology Centre | Catherine Moore, Johnathan Evans, Laura Gifford, Malorie Perry, Simon Cottrell, Angela Marchbank, Alec Birchley, Alexander Adams, Amy Gaskin, Bree Gatica-Wilcox, Jason Coombes, Joel Southgate, Lauren Gilbert, Lee Graham, Nicole Pacchiarini, Sara Kumziene-Summerhayes, Sarah Taylor, Sophie Jones, Sara Rey, Matthew Bull, Joanne Watkins, Sally Corden, Tom Connor |
| EPI_ISL_2819539 | Department of Bacteria, Parasites and Fungi, Statens Serum Institut, Copenhagen, Denmark | Statens Serum Institut Bioinformatics and Microbial Genomics | Danish Covid-19 Genome Consortium |
| EPI_ISL_2819540 | Originating lab: Wales Specialist Virology Centre Sequencing lab: Pathogen Genomics Unit | Public Health Wales Microbiology Cardiff Wales Specialist Virology Centre | Catherine Moore, Johnathan Evans, Laura Gifford, Malorie Perry, Simon Cottrell, Angela Marchbank, Alec Birchley, Alexander Adams, Amy Gaskin, Bree Gatica-Wilcox, Jason Coombes, Joel Southgate, Lauren Gilbert, Lee Graham, Nicole Pacchiarini, Sara Kumziene-Summerhayes, Sarah Taylor, Sophie Jones, Sara Rey, Matthew Bull, Joanne Watkins, Sally Corden, Tom Connor |
| EPI_ISL_2819541 | Department of Bacteria, Parasites and Fungi, Statens Serum Institut, Copenhagen, Denmark | Statens Serum Institut Bioinformatics and Microbial Genomics | Danish Covid-19 Genome Consortium |
| EPI_ISL_2819542 | Originating lab: Wales Specialist Virology Centre Sequencing lab: Pathogen Genomics Unit | Public Health Wales Microbiology Cardiff Wales Specialist Virology Centre | Catherine Moore, Johnathan Evans, Laura Gifford, Malorie Perry, Simon Cottrell, Angela Marchbank, Alec Birchley, Alexander Adams, Amy Gaskin, Bree Gatica-Wilcox, Jason Coombes, Joel Southgate, Lauren Gilbert, Lee Graham, Nicole Pacchiarini, Sara Kumziene-Summerhayes, Sarah Taylor, Sophie Jones, Sara Rey, Matthew Bull, Joanne Watkins, Sally Corden, Tom Connor |
| EPI_ISL_2819543 | Department of Bacteria, Parasites and Fungi, Statens Serum Institut, Copenhagen, Denmark | Statens Serum Institut Bioinformatics and Microbial Genomics | Danish Covid-19 Genome Consortium |
| EPI_ISL_2819544, EPI_ISL_2819546 | Originating lab: Wales Specialist Virology Centre Sequencing lab: Pathogen Genomics Unit | Public Health Wales Microbiology Cardiff Wales Specialist Virology Centre | Catherine Moore, Johnathan Evans, Laura Gifford, Malorie Perry, Simon Cottrell, Angela Marchbank, Alec Birchley, Alexander Adams, Amy Gaskin, Bree Gatica-Wilcox, Jason Coombes, Joel Southgate, Lauren Gilbert, Lee Graham, Nicole Pacchiarini, Sara Kumziene-Summerhayes, Sarah Taylor, Sophie Jones, Sara Rey, Matthew Bull, Joanne Watkins, Sally Corden, Tom Connor |
| EPI_ISL_2819547 | Department of Bacteria, Parasites and Fungi, Statens Serum Institut, Copenhagen, Denmark | Statens Serum Institut Bioinformatics and Microbial Genomics | Danish Covid-19 Genome Consortium |
| EPI_ISL_2819548 | Originating lab: Wales Specialist Virology Centre Sequencing lab: Pathogen Genomics Unit | Public Health Wales Microbiology Cardiff Wales Specialist Virology Centre | Catherine Moore, Johnathan Evans, Laura Gifford, Malorie Perry, Simon Cottrell, Angela Marchbank, Alec Birchley, Alexander Adams, Amy Gaskin, Bree Gatica-Wilcox, Jason Coombes, Joel Southgate, Lauren Gilbert, Lee Graham, Nicole Pacchiarini, Sara Kumziene-Summerhayes, Sarah Taylor, Sophie Jones, Sara Rey, Matthew Bull, Joanne Watkins, Sally Corden, Tom Connor |
| EPI_ISL_2819549 | Department of Bacteria, Parasites and Fungi, Statens Serum | Statens Serum Institut Bioinformatics and Microbial Genomics | Danish Covid-19 Genome Consortium |

| | | | |
|---|---|---|---|
| | Institut, Copenhagen, Denmark | | |
| EPI_ISL_2819550 | Originating lab: Wales Specialist Virology Centre Sequencing lab: Pathogen Genomics Unit | Public Health Wales Microbiology Cardiff Wales Specialist Virology Centre | Catherine Moore, Johnathan Evans, Laura Gifford, Malorie Perry, Simon Cottrell, Angela Marchbank, Alec Birchley, Alexander Adams, Amy Gaskin, Bree Gatica-Wilcox, Jason Coombes, Joel Southgate, Lauren Gilbert, Lee Graham, Nicole Pacchiarini, Sara Kumziene-Summerhayes, Sarah Taylor, Sophie Jones, Sara Rey, Matthew Bull, Joanne Watkins, Sally Corden, Tom Connor |
| EPI_ISL_2819551 | Department of Bacteria, Parasites and Fungi, Statens Serum Institut, Copenhagen, Denmark | Statens Serum Institut Bioinformatics and Microbial Genomics | Danish Covid-19 Genome Consortium |
| EPI_ISL_2819552 | Originating lab: Wales Specialist Virology Centre Sequencing lab: Pathogen Genomics Unit | Public Health Wales Microbiology Cardiff Wales Specialist Virology Centre | Catherine Moore, Johnathan Evans, Laura Gifford, Malorie Perry, Simon Cottrell, Angela Marchbank, Alec Birchley, Alexander Adams, Amy Gaskin, Bree Gatica-Wilcox, Jason Coombes, Joel Southgate, Lauren Gilbert, Lee Graham, Nicole Pacchiarini, Sara Kumziene-Summerhayes, Sarah Taylor, Sophie Jones, Sara Rey, Matthew Bull, Joanne Watkins, Sally Corden, Tom Connor |
| EPI_ISL_2819553 | Department of Bacteria, Parasites and Fungi, Statens Serum Institut, Copenhagen, Denmark | Statens Serum Institut Bioinformatics and Microbial Genomics | Danish Covid-19 Genome Consortium |
| EPI_ISL_2819554 | Originating lab: Wales Specialist Virology Centre Sequencing lab: Pathogen Genomics Unit | Public Health Wales Microbiology Cardiff Wales Specialist Virology Centre | Catherine Moore, Johnathan Evans, Laura Gifford, Malorie Perry, Simon Cottrell, Angela Marchbank, Alec Birchley, Alexander Adams, Amy Gaskin, Bree Gatica-Wilcox, Jason Coombes, Joel Southgate, Lauren Gilbert, Lee Graham, Nicole Pacchiarini, Sara Kumziene-Summerhayes, Sarah Taylor, Sophie Jones, Sara Rey, Matthew Bull, Joanne Watkins, Sally Corden, Tom Connor |
| EPI_ISL_2819555 | Department of Bacteria, Parasites and Fungi, Statens Serum Institut, Copenhagen, Denmark | Statens Serum Institut Bioinformatics and Microbial Genomics | Danish Covid-19 Genome Consortium |
| EPI_ISL_2819556 | Originating lab: Wales Specialist Virology Centre Sequencing lab: Pathogen Genomics Unit | Public Health Wales Microbiology Cardiff Wales Specialist Virology Centre | Catherine Moore, Johnathan Evans, Laura Gifford, Malorie Perry, Simon Cottrell, Angela Marchbank, Alec Birchley, Alexander Adams, Amy Gaskin, Bree Gatica-Wilcox, Jason Coombes, Joel Southgate, Lauren Gilbert, Lee Graham, Nicole Pacchiarini, Sara Kumziene-Summerhayes, Sarah Taylor, Sophie Jones, Sara Rey, Matthew Bull, Joanne Watkins, Sally Corden, Tom Connor |
| EPI_ISL_2819557 | Department of Bacteria, Parasites and Fungi, Statens Serum Institut, Copenhagen, Denmark | Statens Serum Institut Bioinformatics and Microbial Genomics | Danish Covid-19 Genome Consortium |
| EPI_ISL_2819558, EPI_ISL_2819559 | Originating lab: Wales Specialist Virology Centre Sequencing lab: Pathogen Genomics Unit | Public Health Wales Microbiology Cardiff Wales Specialist Virology Centre | Catherine Moore, Johnathan Evans, Laura Gifford, Malorie Perry, Simon Cottrell, Angela Marchbank, Alec Birchley, Alexander Adams, Amy Gaskin, Bree Gatica-Wilcox, Jason Coombes, Joel Southgate, Lauren Gilbert, Lee Graham, Nicole Pacchiarini, Sara Kumziene-Summerhayes, Sarah Taylor, Sophie Jones, Sara Rey, Matthew Bull, Joanne Watkins, Sally Corden, Tom Connor |
| EPI_ISL_2819560 | Department of Bacteria, Parasites and Fungi, Statens Serum Institut, Copenhagen, Denmark | Statens Serum Institut Bioinformatics and Microbial Genomics | Danish Covid-19 Genome Consortium |
| EPI_ISL_2819561, EPI_ISL_2819563, EPI_ISL_2819565, EPI_ISL_2819567, EPI_ISL_2819569 | Originating lab: Wales Specialist Virology Centre Sequencing lab: Pathogen Genomics Unit | Public Health Wales Microbiology Cardiff Wales Specialist Virology Centre | Catherine Moore, Johnathan Evans, Laura Gifford, Malorie Perry, Simon Cottrell, Angela Marchbank, Alec Birchley, Alexander Adams, Amy Gaskin, Bree Gatica-Wilcox, Jason Coombes, Joel Southgate, Lauren Gilbert, Lee Graham, Nicole Pacchiarini, Sara Kumziene-Summerhayes, Sarah Taylor, Sophie Jones, Sara Rey, Matthew Bull, Joanne Watkins, Sally Corden, Tom Connor |
| EPI_ISL_2819570 | Department of Bacteria, Parasites and Fungi, Statens Serum Institut, Copenhagen, Denmark | Statens Serum Institut Bioinformatics and Microbial Genomics | Danish Covid-19 Genome Consortium |
| EPI_ISL_2819571 | Institute for Medical Research, Infectious Disease Research Centre, National Institutes of Health, Ministry of Health Malaysia | Institute for Medical Research, Infectious Disease Research Centre, National Institutes of Health, Ministry of Health Malaysia | Suppiah J, Kamel K, Mohd Zawawi Z, Azizan MA, Ramly N, Robert F, Thayan R |
| EPI_ISL_2819572, EPI_ISL_2819574 | Originating lab: Wales Specialist Virology Centre Sequencing lab: Pathogen Genomics Unit | Public Health Wales Microbiology Cardiff Wales Specialist Virology Centre | Catherine Moore, Johnathan Evans, Laura Gifford, Malorie Perry, Simon Cottrell, Angela Marchbank, Alec Birchley, Alexander Adams, Amy Gaskin, Bree Gatica-Wilcox, Jason Coombes, Joel Southgate, Lauren Gilbert, Lee Graham, Nicole Pacchiarini, Sara Kumziene-Summerhayes, Sarah Taylor, Sophie Jones, Sara Rey, Matthew Bull, Joanne Watkins, Sally Corden, Tom Connor |
| EPI_ISL_2819575 | Department of Bacteria, Parasites and Fungi, Statens Serum Institut, Copenhagen, Denmark | Statens Serum Institut Bioinformatics and Microbial Genomics | Danish Covid-19 Genome Consortium |
| EPI_ISL_2819576 | Originating lab: Wales Specialist Virology Centre Sequencing lab: Pathogen Genomics Unit | Public Health Wales Microbiology Cardiff Wales Specialist Virology Centre | Catherine Moore, Johnathan Evans, Laura Gifford, Malorie Perry, Simon Cottrell, Angela Marchbank, Alec Birchley, Alexander Adams, Amy Gaskin, Bree Gatica-Wilcox, Jason Coombes, Joel Southgate, Lauren Gilbert, Lee Graham, Nicole Pacchiarini, Sara Kumziene-Summerhayes, Sarah Taylor, Sophie Jones, Sara Rey, Matthew Bull, Joanne Watkins, Sally Corden, Tom Connor |
| EPI_ISL_2819577 | Department of Bacteria, Parasites and Fungi, Statens Serum Institut, Copenhagen, Denmark | Statens Serum Institut Bioinformatics and Microbial Genomics | Danish Covid-19 Genome Consortium |
| EPI_ISL_2819578 | Originating lab: Wales Specialist Virology Centre Sequencing lab: Pathogen Genomics Unit | Public Health Wales Microbiology Cardiff Wales Specialist Virology Centre | Catherine Moore, Johnathan Evans, Laura Gifford, Malorie Perry, Simon Cottrell, Angela Marchbank, Alec Birchley, Alexander Adams, Amy Gaskin, Bree Gatica-Wilcox, Jason Coombes, Joel Southgate, Lauren Gilbert, Lee Graham, Nicole Pacchiarini, Sara Kumziene-Summerhayes, Sarah Taylor, Sophie Jones, Sara Rey, Matthew Bull, Joanne Watkins, Sally Corden, Tom Connor |
| EPI_ISL_2819579 | Department of Bacteria, Parasites and Fungi, Statens Serum Institut, Copenhagen, Denmark | Statens Serum Institut Bioinformatics and Microbial Genomics | Danish Covid-19 Genome Consortium |
| EPI_ISL_2819580, EPI_ISL_2819581 | Originating lab: Wales Specialist Virology Centre Sequencing lab: Pathogen Genomics Unit | Public Health Wales Microbiology Cardiff Wales Specialist Virology Centre | Catherine Moore, Johnathan Evans, Laura Gifford, Malorie Perry, Simon Cottrell, Angela Marchbank, Alec Birchley, Alexander Adams, Amy Gaskin, Bree Gatica-Wilcox, Jason Coombes, Joel Southgate, Lauren Gilbert, Lee Graham, Nicole Pacchiarini, Sara Kumziene-Summerhayes, Sarah Taylor, Sophie Jones, Sara Rey, Matthew Bull, Joanne Watkins, Sally Corden, Tom Connor |
| EPI_ISL_2819583 | Department of Bacteria, Parasites and Fungi, Statens Serum Institut, Copenhagen, Denmark | Statens Serum Institut Bioinformatics and Microbial Genomics | Danish Covid-19 Genome Consortium |
| EPI_ISL_2819584 | Originating lab: Wales Specialist Virology Centre Sequencing lab: Pathogen Genomics Unit | Public Health Wales Microbiology Cardiff Wales Specialist Virology Centre | Catherine Moore, Johnathan Evans, Laura Gifford, Malorie Perry, Simon Cottrell, Angela Marchbank, Alec Birchley, Alexander Adams, Amy Gaskin, Bree Gatica-Wilcox, Jason Coombes, Joel Southgate, Lauren Gilbert, Lee Graham, Nicole Pacchiarini, Sara Kumziene-Summerhayes, Sarah Taylor, Sophie Jones, Sara Rey, Matthew Bull, Joanne Watkins, Sally Corden, Tom Connor |
| EPI_ISL_2819585 | Department of Bacteria, Parasites and Fungi, Statens Serum Institut, Copenhagen, Denmark | Statens Serum Institut Bioinformatics and Microbial Genomics | Danish Covid-19 Genome Consortium |
| EPI_ISL_2819586 | Originating lab: Wales Specialist Virology Centre Sequencing lab: Pathogen Genomics Unit | Public Health Wales Microbiology Cardiff Wales Specialist Virology Centre | Catherine Moore, Johnathan Evans, Laura Gifford, Malorie Perry, Simon Cottrell, Angela Marchbank, Alec Birchley, Alexander Adams, Amy Gaskin, Bree Gatica-Wilcox, Jason Coombes, Joel Southgate, Lauren Gilbert, Lee Graham, Nicole Pacchiarini, Sara Kumziene-Summerhayes, Sarah Taylor, Sophie Jones, Sara Rey, Matthew Bull, Joanne Watkins, Sally Corden, Tom Connor |
| EPI_ISL_2819587 | Department of Bacteria, Parasites and Fungi, Statens Serum Institut, Copenhagen, Denmark | Statens Serum Institut Bioinformatics and Microbial Genomics | Danish Covid-19 Genome Consortium |
| EPI_ISL_2819588, EPI_ISL_2819590 | Originating lab: Wales Specialist Virology Centre Sequencing lab: Pathogen Genomics Unit | Public Health Wales Microbiology Cardiff Wales Specialist Virology Centre | Catherine Moore, Johnathan Evans, Laura Gifford, Malorie Perry, Simon Cottrell, Angela Marchbank, Alec Birchley, Alexander Adams, Amy Gaskin, Bree Gatica-Wilcox, Jason Coombes, Joel Southgate, Lauren Gilbert, Lee Graham, Nicole Pacchiarini, Sara Kumziene-Summerhayes, Sarah Taylor, Sophie Jones, Sara Rey, Matthew Bull, Joanne Watkins, Sally Corden, Tom Connor |
| EPI_ISL_2819591 | Department of Bacteria, Parasites and Fungi, Statens Serum Institut, Copenhagen, Denmark | Statens Serum Institut Bioinformatics and Microbial Genomics | Danish Covid-19 Genome Consortium |
| EPI_ISL_2819592 | Originating lab: Wales Specialist Virology Centre Sequencing lab: Pathogen Genomics Unit | Public Health Wales Microbiology Cardiff Wales Specialist Virology Centre | Catherine Moore, Johnathan Evans, Laura Gifford, Malorie Perry, Simon Cottrell, Angela Marchbank, Alec Birchley, Alexander Adams, Amy Gaskin, Bree Gatica-Wilcox, Jason Coombes, Joel Southgate, Lauren Gilbert, Lee Graham, Nicole Pacchiarini, Sara Kumziene-Summerhayes, Sarah Taylor, Sophie Jones, Sara Rey, Matthew Bull, Joanne Watkins, Sally Corden, Tom Connor |
| EPI_ISL_2819593 | Department of Bacteria, Parasites and Fungi, Statens Serum Institut, Copenhagen, Denmark | Statens Serum Institut Bioinformatics and Microbial Genomics | Danish Covid-19 Genome Consortium |
| EPI_ISL_2819594, EPI_ISL_2819595 | Originating lab: Wales Specialist Virology Centre Sequencing lab: Pathogen Genomics Unit | Public Health Wales Microbiology Cardiff Wales Specialist Virology Centre | Catherine Moore, Johnathan Evans, Laura Gifford, Malorie Perry, Simon Cottrell, Angela Marchbank, Alec Birchley, Alexander Adams, Amy Gaskin, Bree Gatica-Wilcox, Jason Coombes, Joel Southgate, Lauren Gilbert, Lee Graham, Nicole Pacchiarini, Sara Kumziene-Summerhayes, Sarah Taylor, Sophie Jones, Sara Rey, Matthew Bull, Joanne Watkins, Sally Corden, Tom Connor |
| EPI_ISL_2819596 | Department of Bacteria, Parasites and Fungi, Statens Serum Institut, Copenhagen, Denmark | Statens Serum Institut Bioinformatics and Microbial Genomics | Danish Covid-19 Genome Consortium |
| EPI_ISL_2819597 | Originating lab: Wales Specialist Virology Centre Sequencing lab: Pathogen Genomics Unit | Public Health Wales Microbiology Cardiff Wales Specialist Virology Centre | Catherine Moore, Johnathan Evans, Laura Gifford, Malorie Perry, Simon Cottrell, Angela Marchbank, Alec Birchley, Alexander Adams, Amy Gaskin, Bree Gatica-Wilcox, Jason Coombes, Joel Southgate, Lauren Gilbert, Lee Graham, Nicole Pacchiarini, Sara Kumziene-Summerhayes, Sarah Taylor, Sophie Jones, Sara Rey, Matthew Bull, Joanne Watkins, Sally Corden, Tom Connor |
| EPI_ISL_2819598 | Department of Bacteria, Parasites and Fungi, Statens Serum Institut, Copenhagen, Denmark | Statens Serum Institut Bioinformatics and Microbial Genomics | Danish Covid-19 Genome Consortium |
| EPI_ISL_2819599, EPI_ISL_2819601 | Originating lab: Wales Specialist Virology Centre Sequencing lab: Pathogen Genomics Unit | Public Health Wales Microbiology Cardiff Wales Specialist Virology Centre | Catherine Moore, Johnathan Evans, Laura Gifford, Malorie Perry, Simon Cottrell, Angela Marchbank, Alec Birchley, Alexander Adams, Amy Gaskin, Bree Gatica-Wilcox, Jason Coombes, Joel Southgate, Lauren Gilbert, Lee Graham, Nicole Pacchiarini, Sara Kumziene-Summerhayes, Sarah Taylor, Sophie Jones, Sara Rey, Matthew Bull, Joanne Watkins, Sally Corden, Tom Connor |
| EPI_ISL_2819602 | Department of Bacteria, Parasites and Fungi, Statens Serum Institut, Copenhagen, Denmark | Statens Serum Institut Bioinformatics and Microbial Genomics | Danish Covid-19 Genome Consortium |
| EPI_ISL_2819603 | Originating lab: Wales Specialist Virology Centre Sequencing lab: Pathogen Genomics Unit | Public Health Wales Microbiology Cardiff Wales Specialist Virology Centre | Catherine Moore, Johnathan Evans, Laura Gifford, Malorie Perry, Simon Cottrell, Angela Marchbank, Alec Birchley, Alexander Adams, Amy Gaskin, Bree Gatica-Wilcox, Jason Coombes, Joel Southgate, Lauren Gilbert, Lee Graham, Nicole Pacchiarini, Sara Kumziene-Summerhayes, Sarah Taylor, Sophie Jones, Sara Rey, Matthew Bull, Joanne Watkins, Sally Corden, Tom Connor |
| EPI_ISL_2819604 | Department of Bacteria, Parasites and Fungi, Statens Serum Institut, Copenhagen, Denmark | Statens Serum Institut Bioinformatics and Microbial Genomics | Danish Covid-19 Genome Consortium |
| EPI_ISL_2819605 | Originating lab: Wales Specialist Virology Centre Sequencing lab: Pathogen Genomics Unit | Public Health Wales Microbiology Cardiff Wales Specialist Virology Centre | Catherine Moore, Johnathan Evans, Laura Gifford, Malorie Perry, Simon Cottrell, Angela Marchbank, Alec Birchley, Alexander Adams, Amy Gaskin, Bree Gatica-Wilcox, Jason Coombes, Joel Southgate, Lauren Gilbert, Lee Graham, Nicole Pacchiarini, Sara Kumziene-Summerhayes, Sarah Taylor, Sophie Jones, Sara Rey, Matthew Bull, Joanne Watkins, Sally Corden, Tom Connor |
| EPI_ISL_2819606 | Department of Bacteria, Parasites and Fungi, Statens Serum Institut, Copenhagen, Denmark | Statens Serum Institut Bioinformatics and Microbial Genomics | Danish Covid-19 Genome Consortium |
| EPI_ISL_2819607 | Originating lab: Wales Specialist Virology Centre Sequencing lab: Pathogen Genomics Unit | Public Health Wales Microbiology Cardiff Wales Specialist Virology Centre | Catherine Moore, Johnathan Evans, Laura Gifford, Malorie Perry, Simon Cottrell, Angela Marchbank, Alec Birchley, Alexander Adams, Amy Gaskin, Bree Gatica-Wilcox, Jason Coombes, Joel Southgate, Lauren Gilbert, Lee Graham, Nicole Pacchiarini, Sara Kumziene-Summerhayes, Sarah Taylor, Sophie Jones, Sara Rey, Matthew Bull, Joanne Watkins, Sally Corden, Tom Connor |
| EPI_ISL_2819608 | Department of Bacteria, Parasites and Fungi, Statens Serum Institut, Copenhagen, Denmark | Statens Serum Institut Bioinformatics and Microbial Genomics | Danish Covid-19 Genome Consortium |
| EPI_ISL_2819609 | Originating lab: Wales Specialist Virology Centre Sequencing lab: Pathogen Genomics Unit | Public Health Wales Microbiology Cardiff Wales Specialist Virology Centre | Catherine Moore, Johnathan Evans, Laura Gifford, Malorie Perry, Simon Cottrell, Angela Marchbank, Alec Birchley, Alexander Adams, Amy Gaskin, Bree Gatica-Wilcox, Jason Coombes, Joel Southgate, Lauren Gilbert, Lee Graham, Nicole Pacchiarini, Sara Kumziene-Summerhayes, Sarah Taylor, Sophie Jones, Sara Rey, Matthew Bull, Joanne Watkins, Sally Corden, Tom Connor |
| EPI_ISL_2819610 | Department of Bacteria, Parasites and Fungi, Statens Serum Institut, Copenhagen, Denmark | Statens Serum Institut Bioinformatics and Microbial Genomics | Danish Covid-19 Genome Consortium |

| | | | |
|---|---|---|---|
| EPI_ISL_2819611 | Originating lab: Wales Specialist Virology Centre Sequencing lab: Pathogen Genomics Unit | Public Health Wales Microbiology Cardiff Wales Specialist Virology Centre | Catherine Moore, Johnathan Evans, Laura Gifford, Malorie Perry, Simon Cottrell, Angela Marchbank, Alec Birchley, Alexander Adams, Amy Gaskin, Bree Gatica-Wilcox, Jason Coombes, Joel Southgate, Lauren Gilbert, Lee Graham, Nicole Pacchiarini, Sara Kumziene-Summerhayes, Sarah Taylor, Sophie Jones, Sara Rey, Matthew Bull, Joanne Watkins, Sally Corden, Tom Connor |
| EPI_ISL_2819613 | Department of Bacteria, Parasites and Fungi, Statens Serum Institut, Copenhagen, Denmark | Statens Serum Institut Bioinformatics and Microbial Genomics | Danish Covid-19 Genome Consortium |
| EPI_ISL_2819614 | Originating lab: Wales Specialist Virology Centre Sequencing lab: Pathogen Genomics Unit | Public Health Wales Microbiology Cardiff Wales Specialist Virology Centre | Catherine Moore, Johnathan Evans, Laura Gifford, Malorie Perry, Simon Cottrell, Angela Marchbank, Alec Birchley, Alexander Adams, Amy Gaskin, Bree Gatica-Wilcox, Jason Coombes, Joel Southgate, Lauren Gilbert, Lee Graham, Nicole Pacchiarini, Sara Kumziene-Summerhayes, Sarah Taylor, Sophie Jones, Sara Rey, Matthew Bull, Joanne Watkins, Sally Corden, Tom Connor |
| EPI_ISL_2819615 | Department of Bacteria, Parasites and Fungi, Statens Serum Institut, Copenhagen, Denmark | Statens Serum Institut Bioinformatics and Microbial Genomics | Danish Covid-19 Genome Consortium |
| EPI_ISL_2819616, EPI_ISL_2819618 | Originating lab: Wales Specialist Virology Centre Sequencing lab: Pathogen Genomics Unit | Public Health Wales Microbiology Cardiff Wales Specialist Virology Centre | Catherine Moore, Johnathan Evans, Laura Gifford, Malorie Perry, Simon Cottrell, Angela Marchbank, Alec Birchley, Alexander Adams, Amy Gaskin, Bree Gatica-Wilcox, Jason Coombes, Joel Southgate, Lauren Gilbert, Lee Graham, Nicole Pacchiarini, Sara Kumziene-Summerhayes, Sarah Taylor, Sophie Jones, Sara Rey, Matthew Bull, Joanne Watkins, Sally Corden, Tom Connor |
| EPI_ISL_2819619 | Department of Bacteria, Parasites and Fungi, Statens Serum Institut, Copenhagen, Denmark | Statens Serum Institut Bioinformatics and Microbial Genomics | Danish Covid-19 Genome Consortium |
| EPI_ISL_2819620, EPI_ISL_2819622 | Originating lab: Wales Specialist Virology Centre Sequencing lab: Pathogen Genomics Unit | Public Health Wales Microbiology Cardiff Wales Specialist Virology Centre | Catherine Moore, Johnathan Evans, Laura Gifford, Malorie Perry, Simon Cottrell, Angela Marchbank, Alec Birchley, Alexander Adams, Amy Gaskin, Bree Gatica-Wilcox, Jason Coombes, Joel Southgate, Lauren Gilbert, Lee Graham, Nicole Pacchiarini, Sara Kumziene-Summerhayes, Sarah Taylor, Sophie Jones, Sara Rey, Matthew Bull, Joanne Watkins, Sally Corden, Tom Connor |
| EPI_ISL_2819623 | Department of Bacteria, Parasites and Fungi, Statens Serum Institut, Copenhagen, Denmark | Statens Serum Institut Bioinformatics and Microbial Genomics | Danish Covid-19 Genome Consortium |
| EPI_ISL_2819624, EPI_ISL_2819625 | Originating lab: Wales Specialist Virology Centre Sequencing lab: Pathogen Genomics Unit | Public Health Wales Microbiology Cardiff Wales Specialist Virology Centre | Catherine Moore, Johnathan Evans, Laura Gifford, Malorie Perry, Simon Cottrell, Angela Marchbank, Alec Birchley, Alexander Adams, Amy Gaskin, Bree Gatica-Wilcox, Jason Coombes, Joel Southgate, Lauren Gilbert, Lee Graham, Nicole Pacchiarini, Sara Kumziene-Summerhayes, Sarah Taylor, Sophie Jones, Sara Rey, Matthew Bull, Joanne Watkins, Sally Corden, Tom Connor |
| EPI_ISL_2819626 | Department of Bacteria, Parasites and Fungi, Statens Serum Institut, Copenhagen, Denmark | Statens Serum Institut Bioinformatics and Microbial Genomics | Danish Covid-19 Genome Consortium |
| EPI_ISL_2819627, EPI_ISL_2819629, EPI_ISL_2819631 | Originating lab: Wales Specialist Virology Centre Sequencing lab: Pathogen Genomics Unit | Public Health Wales Microbiology Cardiff Wales Specialist Virology Centre | Catherine Moore, Johnathan Evans, Laura Gifford, Malorie Perry, Simon Cottrell, Angela Marchbank, Alec Birchley, Alexander Adams, Amy Gaskin, Bree Gatica-Wilcox, Jason Coombes, Joel Southgate, Lauren Gilbert, Lee Graham, Nicole Pacchiarini, Sara Kumziene-Summerhayes, Sarah Taylor, Sophie Jones, Sara Rey, Matthew Bull, Joanne Watkins, Sally Corden, Tom Connor |
| EPI_ISL_2819632 | Department of Bacteria, Parasites and Fungi, Statens Serum Institut, Copenhagen, Denmark | Statens Serum Institut Bioinformatics and Microbial Genomics | Danish Covid-19 Genome Consortium |
| EPI_ISL_2819633 | Originating lab: Wales Specialist Virology Centre Sequencing lab: Pathogen Genomics Unit | Public Health Wales Microbiology Cardiff Wales Specialist Virology Centre | Catherine Moore, Johnathan Evans, Laura Gifford, Malorie Perry, Simon Cottrell, Angela Marchbank, Alec Birchley, Alexander Adams, Amy Gaskin, Bree Gatica-Wilcox, Jason Coombes, Joel Southgate, Lauren Gilbert, Lee Graham, Nicole Pacchiarini, Sara Kumziene-Summerhayes, Sarah Taylor, Sophie Jones, Sara Rey, Matthew Bull, Joanne Watkins, Sally Corden, Tom Connor |
| EPI_ISL_2819634, EPI_ISL_2819635, EPI_ISL_2819637 | Department of Bacteria, Parasites and Fungi, Statens Serum Institut, Copenhagen, Denmark | Statens Serum Institut Bioinformatics and Microbial Genomics | Danish Covid-19 Genome Consortium |
| EPI_ISL_2819638 | Originating lab: Wales Specialist Virology Centre Sequencing lab: Pathogen Genomics Unit | Public Health Wales Microbiology Cardiff Wales Specialist Virology Centre | Catherine Moore, Johnathan Evans, Laura Gifford, Malorie Perry, Simon Cottrell, Angela Marchbank, Alec Birchley, Alexander Adams, Amy Gaskin, Bree Gatica-Wilcox, Jason Coombes, Joel Southgate, Lauren Gilbert, Lee Graham, Nicole Pacchiarini, Sara Kumziene-Summerhayes, Sarah Taylor, Sophie Jones, Sara Rey, Matthew Bull, Joanne Watkins, Sally Corden, Tom Connor |
| EPI_ISL_2819639 | Department of Bacteria, Parasites and Fungi, Statens Serum Institut, Copenhagen, Denmark | Statens Serum Institut Bioinformatics and Microbial Genomics | Danish Covid-19 Genome Consortium |
| EPI_ISL_2819640, EPI_ISL_2819641 | Originating lab: Wales Specialist Virology Centre Sequencing lab: Pathogen Genomics Unit | Public Health Wales Microbiology Cardiff Wales Specialist Virology Centre | Catherine Moore, Johnathan Evans, Laura Gifford, Malorie Perry, Simon Cottrell, Angela Marchbank, Alec Birchley, Alexander Adams, Amy Gaskin, Bree Gatica-Wilcox, Jason Coombes, Joel Southgate, Lauren Gilbert, Lee Graham, Nicole Pacchiarini, Sara Kumziene-Summerhayes, Sarah Taylor, Sophie Jones, Sara Rey, Matthew Bull, Joanne Watkins, Sally Corden, Tom Connor |
| EPI_ISL_2819642 | Department of Bacteria, Parasites and Fungi, Statens Serum Institut, Copenhagen, Denmark | Statens Serum Institut Bioinformatics and Microbial Genomics | Danish Covid-19 Genome Consortium |
| EPI_ISL_2819643 | Originating lab: Wales Specialist Virology Centre Sequencing lab: Pathogen Genomics Unit | Public Health Wales Microbiology Cardiff Wales Specialist Virology Centre | Catherine Moore, Johnathan Evans, Laura Gifford, Malorie Perry, Simon Cottrell, Angela Marchbank, Alec Birchley, Alexander Adams, Amy Gaskin, Bree Gatica-Wilcox, Jason Coombes, Joel Southgate, Lauren Gilbert, Lee Graham, Nicole Pacchiarini, Sara Kumziene-Summerhayes, Sarah Taylor, Sophie Jones, Sara Rey, Matthew Bull, Joanne Watkins, Sally Corden, Tom Connor |
| EPI_ISL_2819644 | Department of Clinical Microbiology, Odense University Hospital, Odense, Denmark | Statens Serum Institut Bioinformatics and Microbial Genomics | Danish Covid-19 Genome Consortium |
| EPI_ISL_2819645 | Originating lab: Wales Specialist Virology Centre Sequencing lab: Pathogen Genomics Unit | Public Health Wales Microbiology Cardiff Wales Specialist Virology Centre | Catherine Moore, Johnathan Evans, Laura Gifford, Malorie Perry, Simon Cottrell, Angela Marchbank, Alec Birchley, Alexander Adams, Amy Gaskin, Bree Gatica-Wilcox, Jason Coombes, Joel Southgate, Lauren Gilbert, Lee Graham, Nicole Pacchiarini, Sara Kumziene-Summerhayes, Sarah Taylor, Sophie Jones, Sara Rey, Matthew Bull, Joanne Watkins, Sally Corden, Tom Connor |
| EPI_ISL_2819646 | Department of Bacteria, Parasites and Fungi, Statens Serum Institut, Copenhagen, Denmark | Statens Serum Institut Bioinformatics and Microbial Genomics | Danish Covid-19 Genome Consortium |
| EPI_ISL_2819647, EPI_ISL_2819649 | Originating lab: Wales Specialist Virology Centre Sequencing lab: Pathogen Genomics Unit | Public Health Wales Microbiology Cardiff Wales Specialist Virology Centre | Catherine Moore, Johnathan Evans, Laura Gifford, Malorie Perry, Simon Cottrell, Angela Marchbank, Alec Birchley, Alexander Adams, Amy Gaskin, Bree Gatica-Wilcox, Jason Coombes, Joel Southgate, Lauren Gilbert, Lee Graham, Nicole Pacchiarini, Sara Kumziene-Summerhayes, Sarah Taylor, Sophie Jones, Sara Rey, Matthew Bull, Joanne Watkins, Sally Corden, Tom Connor |
| EPI_ISL_2819650 | Department of Bacteria, Parasites and Fungi, Statens Serum Institut, Copenhagen, Denmark | Statens Serum Institut Bioinformatics and Microbial Genomics | Danish Covid-19 Genome Consortium |
| EPI_ISL_2819651, EPI_ISL_2819653 | Originating lab: Wales Specialist Virology Centre Sequencing lab: Pathogen Genomics Unit | Public Health Wales Microbiology Cardiff Wales Specialist Virology Centre | Catherine Moore, Johnathan Evans, Laura Gifford, Malorie Perry, Simon Cottrell, Angela Marchbank, Alec Birchley, Alexander Adams, Amy Gaskin, Bree Gatica-Wilcox, Jason Coombes, Joel Southgate, Lauren Gilbert, Lee Graham, Nicole Pacchiarini, Sara Kumziene-Summerhayes, Sarah Taylor, Sophie Jones, Sara Rey, Matthew Bull, Joanne Watkins, Sally Corden, Tom Connor |
| EPI_ISL_2819654 | Department of Bacteria, Parasites and Fungi, Statens Serum Institut, Copenhagen, Denmark | Statens Serum Institut Bioinformatics and Microbial Genomics | Danish Covid-19 Genome Consortium |
| EPI_ISL_2819655, EPI_ISL_2819657 | Originating lab: Wales Specialist Virology Centre Sequencing lab: Pathogen Genomics Unit | Public Health Wales Microbiology Cardiff Wales Specialist Virology Centre | Catherine Moore, Johnathan Evans, Laura Gifford, Malorie Perry, Simon Cottrell, Angela Marchbank, Alec Birchley, Alexander Adams, Amy Gaskin, Bree Gatica-Wilcox, Jason Coombes, Joel Southgate, Lauren Gilbert, Lee Graham, Nicole Pacchiarini, Sara Kumziene-Summerhayes, Sarah Taylor, Sophie Jones, Sara Rey, Matthew Bull, Joanne Watkins, Sally Corden, Tom Connor |
| EPI_ISL_2819658 | Department of Bacteria, Parasites and Fungi, Statens Serum Institut, Copenhagen, Denmark | Statens Serum Institut Bioinformatics and Microbial Genomics | Danish Covid-19 Genome Consortium |
| EPI_ISL_2819659, EPI_ISL_2819661 | Originating lab: Wales Specialist Virology Centre Sequencing lab: Pathogen Genomics Unit | Public Health Wales Microbiology Cardiff Wales Specialist Virology Centre | Catherine Moore, Johnathan Evans, Laura Gifford, Malorie Perry, Simon Cottrell, Angela Marchbank, Alec Birchley, Alexander Adams, Amy Gaskin, Bree Gatica-Wilcox, Jason Coombes, Joel Southgate, Lauren Gilbert, Lee Graham, Nicole Pacchiarini, Sara Kumziene-Summerhayes, Sarah Taylor, Sophie Jones, Sara Rey, Matthew Bull, Joanne Watkins, Sally Corden, Tom Connor |
| EPI_ISL_2819662 | Department of Bacteria, Parasites and Fungi, Statens Serum Institut, Copenhagen, Denmark | Statens Serum Institut Bioinformatics and Microbial Genomics | Danish Covid-19 Genome Consortium |
| EPI_ISL_2819663 | Originating lab: Wales Specialist Virology Centre Sequencing lab: Pathogen Genomics Unit | Public Health Wales Microbiology Cardiff Wales Specialist Virology Centre | Catherine Moore, Johnathan Evans, Laura Gifford, Malorie Perry, Simon Cottrell, Angela Marchbank, Alec Birchley, Alexander Adams, Amy Gaskin, Bree Gatica-Wilcox, Jason Coombes, Joel Southgate, Lauren Gilbert, Lee Graham, Nicole Pacchiarini, Sara Kumziene-Summerhayes, Sarah Taylor, Sophie Jones, Sara Rey, Matthew Bull, Joanne Watkins, Sally Corden, Tom Connor |
| EPI_ISL_2819664 | Department of Bacteria, Parasites and Fungi, Statens Serum Institut, Copenhagen, Denmark | Statens Serum Institut Bioinformatics and Microbial Genomics | Danish Covid-19 Genome Consortium |
| EPI_ISL_2819665 | Originating lab: Wales Specialist Virology Centre Sequencing lab: Pathogen Genomics Unit | Public Health Wales Microbiology Cardiff Wales Specialist Virology Centre | Catherine Moore, Johnathan Evans, Laura Gifford, Malorie Perry, Simon Cottrell, Angela Marchbank, Alec Birchley, Alexander Adams, Amy Gaskin, Bree Gatica-Wilcox, Jason Coombes, Joel Southgate, Lauren Gilbert, Lee Graham, Nicole Pacchiarini, Sara Kumziene-Summerhayes, Sarah Taylor, Sophie Jones, Sara Rey, Matthew Bull, Joanne Watkins, Sally Corden, Tom Connor |
| EPI_ISL_2819666 | Department of Bacteria, Parasites and Fungi, Statens Serum Institut, Copenhagen, Denmark | Statens Serum Institut Bioinformatics and Microbial Genomics | Danish Covid-19 Genome Consortium |
| EPI_ISL_2819667 | Originating lab: Wales Specialist Virology Centre Sequencing lab: Pathogen Genomics Unit | Public Health Wales Microbiology Cardiff Wales Specialist Virology Centre | Catherine Moore, Johnathan Evans, Laura Gifford, Malorie Perry, Simon Cottrell, Angela Marchbank, Alec Birchley, Alexander Adams, Amy Gaskin, Bree Gatica-Wilcox, Jason Coombes, Joel Southgate, Lauren Gilbert, Lee Graham, Nicole Pacchiarini, Sara Kumziene-Summerhayes, Sarah Taylor, Sophie Jones, Sara Rey, Matthew Bull, Joanne Watkins, Sally Corden, Tom Connor |
| EPI_ISL_2819668 | Department of Bacteria, Parasites and Fungi, Statens Serum Institut, Copenhagen, Denmark | Statens Serum Institut Bioinformatics and Microbial Genomics | Danish Covid-19 Genome Consortium |
| EPI_ISL_2819669 | Originating lab: Wales Specialist Virology Centre Sequencing lab: Pathogen Genomics Unit | Public Health Wales Microbiology Cardiff Wales Specialist Virology Centre | Catherine Moore, Johnathan Evans, Laura Gifford, Malorie Perry, Simon Cottrell, Angela Marchbank, Alec Birchley, Alexander Adams, Amy Gaskin, Bree Gatica-Wilcox, Jason Coombes, Joel Southgate, Lauren Gilbert, Lee Graham, Nicole Pacchiarini, Sara Kumziene-Summerhayes, Sarah Taylor, Sophie Jones, Sara Rey, Matthew Bull, Joanne Watkins, Sally Corden, Tom Connor |
| EPI_ISL_2819670 | Department of Bacteria, Parasites and Fungi, Statens Serum Institut, Copenhagen, Denmark | Statens Serum Institut Bioinformatics and Microbial Genomics | Danish Covid-19 Genome Consortium |
| EPI_ISL_2819671 | Originating lab: Wales Specialist Virology Centre Sequencing lab: Pathogen Genomics Unit | Public Health Wales Microbiology Cardiff Wales Specialist Virology Centre | Catherine Moore, Johnathan Evans, Laura Gifford, Malorie Perry, Simon Cottrell, Angela Marchbank, Alec Birchley, Alexander Adams, Amy Gaskin, Bree Gatica-Wilcox, Jason Coombes, Joel Southgate, Lauren Gilbert, Lee Graham, Nicole Pacchiarini, Sara Kumziene-Summerhayes, Sarah Taylor, Sophie Jones, Sara Rey, Matthew Bull, Joanne Watkins, Sally Corden, Tom Connor |
| EPI_ISL_2819672 | Department of Bacteria, Parasites and Fungi, Statens Serum Institut, Copenhagen, Denmark | Statens Serum Institut Bioinformatics and Microbial Genomics | Danish Covid-19 Genome Consortium |
| EPI_ISL_2819673, EPI_ISL_2819674 | Originating lab: Wales Specialist Virology Centre Sequencing lab: Pathogen Genomics Unit | Public Health Wales Microbiology Cardiff Wales Specialist Virology Centre | Catherine Moore, Johnathan Evans, Laura Gifford, Malorie Perry, Simon Cottrell, Angela Marchbank, Alec Birchley, Alexander Adams, Amy Gaskin, Bree Gatica-Wilcox, Jason Coombes, Joel Southgate, Lauren Gilbert, Lee Graham, Nicole Pacchiarini, Sara Kumziene-Summerhayes, Sarah Taylor, Sophie Jones, Sara Rey, Matthew Bull, Joanne Watkins, Sally Corden, Tom Connor |
| EPI_ISL_2819675 | Department of Bacteria, Parasites and Fungi, Statens Serum Institut, Copenhagen, Denmark | Statens Serum Institut Bioinformatics and Microbial Genomics | Danish Covid-19 Genome Consortium |
| EPI_ISL_2819676, EPI_ISL_2819678 | Originating lab: Wales Specialist Virology Centre Sequencing lab: Pathogen Genomics Unit | Public Health Wales Microbiology Cardiff Wales Specialist Virology Centre | Catherine Moore, Johnathan Evans, Laura Gifford, Malorie Perry, Simon Cottrell, Angela Marchbank, Alec Birchley, Alexander Adams, Amy Gaskin, Bree Gatica-Wilcox, Jason Coombes, Joel Southgate, Lauren Gilbert, Lee Graham, Nicole Pacchiarini, Sara Kumziene-Summerhayes, Sarah Taylor, Sophie Jones, Sara Rey, Matthew Bull, Joanne Watkins, Sally Corden, Tom Connor |
| EPI_ISL_2819679 | Department of Bacteria, Parasites and Fungi, Statens Serum Institut, Copenhagen, Denmark | Statens Serum Institut Bioinformatics and Microbial Genomics | Danish Covid-19 Genome Consortium |

| | | | |
|---|---|---|---|
| EPI_ISL_2819680, EPI_ISL_2819682 | Originating lab: Wales Specialist Virology Centre Sequencing lab: Pathogen Genomics Unit | Public Health Wales Microbiology Cardiff Wales Specialist Virology Centre | Catherine Moore, Johnathan Evans, Laura Gifford, Malorie Perry, Simon Cottrell, Angela Marchbank, Alec Birchley, Alexander Adams, Amy Gaskin, Bree Gatica-Wilcox, Jason Coombes, Joel Southgate, Lauren Gilbert, Lee Graham, Nicole Pacchiarini, Sara Kumziene-Summerhayes, Sarah Taylor, Sophie Jones, Sara Rey, Matthew Bull, Joanne Watkins, Sally Corden, Tom Connor |
| EPI_ISL_2819683 | Department of Bacteria, Parasites and Fungi, Statens Serum Institut, Copenhagen, Denmark | Statens Serum Institut Bioinformatics and Microbial Genomics | Danish Covid-19 Genome Consortium |
| EPI_ISL_2819684 | Originating lab: Wales Specialist Virology Centre Sequencing lab: Pathogen Genomics Unit | Public Health Wales Microbiology Cardiff Wales Specialist Virology Centre | Catherine Moore, Johnathan Evans, Laura Gifford, Malorie Perry, Simon Cottrell, Angela Marchbank, Alec Birchley, Alexander Adams, Amy Gaskin, Bree Gatica-Wilcox, Jason Coombes, Joel Southgate, Lauren Gilbert, Lee Graham, Nicole Pacchiarini, Sara Kumziene-Summerhayes, Sarah Taylor, Sophie Jones, Sara Rey, Matthew Bull, Joanne Watkins, Sally Corden, Tom Connor |
| EPI_ISL_2819685 | Department of Bacteria, Parasites and Fungi, Statens Serum Institut, Copenhagen, Denmark | Statens Serum Institut Bioinformatics and Microbial Genomics | Danish Covid-19 Genome Consortium |
| EPI_ISL_2819686 | Originating lab: Wales Specialist Virology Centre Sequencing lab: Pathogen Genomics Unit | Public Health Wales Microbiology Cardiff Wales Specialist Virology Centre | Catherine Moore, Johnathan Evans, Laura Gifford, Malorie Perry, Simon Cottrell, Angela Marchbank, Alec Birchley, Alexander Adams, Amy Gaskin, Bree Gatica-Wilcox, Jason Coombes, Joel Southgate, Lauren Gilbert, Lee Graham, Nicole Pacchiarini, Sara Kumziene-Summerhayes, Sarah Taylor, Sophie Jones, Sara Rey, Matthew Bull, Joanne Watkins, Sally Corden, Tom Connor |
| EPI_ISL_2819687 | Department of Bacteria, Parasites and Fungi, Statens Serum Institut, Copenhagen, Denmark | Statens Serum Institut Bioinformatics and Microbial Genomics | Danish Covid-19 Genome Consortium |
| EPI_ISL_2819688, EPI_ISL_2819689 | Originating lab: Wales Specialist Virology Centre Sequencing lab: Pathogen Genomics Unit | Public Health Wales Microbiology Cardiff Wales Specialist Virology Centre | Catherine Moore, Johnathan Evans, Laura Gifford, Malorie Perry, Simon Cottrell, Angela Marchbank, Alec Birchley, Alexander Adams, Amy Gaskin, Bree Gatica-Wilcox, Jason Coombes, Joel Southgate, Lauren Gilbert, Lee Graham, Nicole Pacchiarini, Sara Kumziene-Summerhayes, Sarah Taylor, Sophie Jones, Sara Rey, Matthew Bull, Joanne Watkins, Sally Corden, Tom Connor |
| EPI_ISL_2819690 | Department of Bacteria, Parasites and Fungi, Statens Serum Institut, Copenhagen, Denmark | Statens Serum Institut Bioinformatics and Microbial Genomics | Danish Covid-19 Genome Consortium |
| EPI_ISL_2819691, EPI_ISL_2819692, EPI_ISL_2819693, EPI_ISL_2819694, EPI_ISL_2819695, EPI_ISL_2819696, EPI_ISL_2819697, EPI_ISL_2819698, EPI_ISL_2819699, EPI_ISL_2819700, EPI_ISL_2819701, EPI_ISL_2819702, EPI_ISL_2819703, EPI_ISL_2819704, EPI_ISL_2819705, EPI_ISL_2819706, EPI_ISL_2819707, EPI_ISL_2819708, EPI_ISL_2819709, EPI_ISL_2819710, EPI_ISL_2819711, EPI_ISL_2819712, EPI_ISL_2819713, EPI_ISL_2819714, EPI_ISL_2819715, EPI_ISL_2819716, EPI_ISL_2819717, EPI_ISL_2819718, EPI_ISL_2819719, EPI_ISL_2819720, EPI_ISL_2819721, EPI_ISL_2819722, EPI_ISL_2819723, EPI_ISL_2819724, EPI_ISL_2819725, EPI_ISL_2819726, EPI_ISL_2819727, EPI_ISL_2819728, EPI_ISL_2819729, EPI_ISL_2819730, EPI_ISL_2819731, EPI_ISL_2819732, EPI_ISL_2819733, EPI_ISL_2819734, EPI_ISL_2819735, EPI_ISL_2819736, EPI_ISL_2819737, EPI_ISL_2819738, EPI_ISL_2819739, EPI_ISL_2819740, EPI_ISL_2819741, EPI_ISL_2819742, EPI_ISL_2819743, EPI_ISL_2819744, EPI_ISL_2819745, EPI_ISL_2819746, EPI_ISL_2819747, EPI_ISL_2819748, EPI_ISL_2819749, EPI_ISL_2819750, EPI_ISL_2819751, EPI_ISL_2819752, EPI_ISL_2819753, EPI_ISL_2819754, EPI_ISL_2819755, EPI_ISL_2819757, EPI_ISL_2819758, EPI_ISL_2819759, EPI_ISL_2819760, EPI_ISL_2819761, EPI_ISL_2819762, EPI_ISL_2819763, EPI_ISL_2819764, EPI_ISL_2819765, EPI_ISL_2819766, EPI_ISL_2819767, EPI_ISL_2819768, EPI_ISL_2819769, EPI_ISL_2819770, EPI_ISL_2819771, EPI_ISL_2819772, EPI_ISL_2819773, EPI_ISL_2819774, EPI_ISL_2819775, EPI_ISL_2819776, EPI_ISL_2819777, EPI_ISL_2819778, EPI_ISL_2819779, EPI_ISL_2819780, EPI_ISL_2819781, EPI_ISL_2819782, EPI_ISL_2819783, EPI_ISL_2819784, EPI_ISL_2819785, EPI_ISL_2819786, EPI_ISL_2819787, EPI_ISL_2819788, EPI_ISL_2819790, EPI_ISL_2819791, EPI_ISL_2819792, EPI_ISL_2819793, EPI_ISL_2819794, EPI_ISL_2819795, EPI_ISL_2819796, EPI_ISL_2819797, EPI_ISL_2819798, EPI_ISL_2819799, EPI_ISL_2819800, EPI_ISL_2819801, EPI_ISL_2819802, EPI_ISL_2819803, EPI_ISL_2819804, EPI_ISL_2819805, EPI_ISL_2819806, EPI_ISL_2819807, EPI_ISL_2819808, EPI_ISL_2819809, EPI_ISL_2819810, EPI_ISL_2819811, EPI_ISL_2819812, EPI_ISL_2819813, EPI_ISL_2819814, EPI_ISL_2819815, EPI_ISL_2819816, EPI_ISL_2819817, EPI_ISL_2819818, EPI_ISL_2819819, EPI_ISL_2819820, EPI_ISL_2819821, EPI_ISL_2819822, EPI_ISL_2819823, EPI_ISL_2819824, EPI_ISL_2819825, EPI_ISL_2819826, EPI_ISL_2819827, EPI_ISL_2819828, EPI_ISL_2819829, EPI_ISL_2819830, EPI_ISL_2819831, EPI_ISL_2819832, EPI_ISL_2819833, EPI_ISL_2819834, EPI_ISL_2819835, EPI_ISL_2819836, EPI_ISL_2819837, EPI_ISL_2819838, EPI_ISL_2819839, EPI_ISL_2819840, EPI_ISL_2819841, EPI_ISL_2819842, EPI_ISL_2819843, EPI_ISL_2819844, EPI_ISL_2819845, EPI_ISL_2819846, EPI_ISL_2819847, EPI_ISL_2819848, EPI_ISL_2819849 | | | |
| see above | Originating lab: Wales Specialist Virology Centre Sequencing lab: Pathogen Genomics Unit | Public Health Wales Microbiology Cardiff Wales Specialist Virology Centre | Catherine Moore, Johnathan Evans, Laura Gifford, Malorie Perry, Simon Cottrell, Angela Marchbank, Alec Birchley, Alexander Adams, Amy Gaskin, Bree Gatica-Wilcox, Jason Coombes, Joel Southgate, Lauren Gilbert, Lee Graham, Nicole Pacchiarini, Sara Kumziene-Summerhayes, Sarah Taylor, Sophie Jones, Sara Rey, Matthew Bull, Joanne Watkins, Sally Corden, Tom Connor |
| EPI_ISL_2819850, EPI_ISL_2819851, EPI_ISL_2819852, EPI_ISL_2819853, EPI_ISL_2819854, EPI_ISL_2819855, EPI_ISL_2819856, EPI_ISL_2819857, EPI_ISL_2819859, EPI_ISL_2819860, EPI_ISL_2819861, EPI_ISL_2819862, EPI_ISL_2819863, EPI_ISL_2819864, EPI_ISL_2819865, EPI_ISL_2819870, EPI_ISL_2819871, EPI_ISL_2819872, EPI_ISL_2819873, EPI_ISL_2819874, EPI_ISL_2819875, EPI_ISL_2819876, EPI_ISL_2819877, EPI_ISL_2819878, EPI_ISL_2819879, EPI_ISL_2819880, EPI_ISL_2819881, EPI_ISL_2819882, EPI_ISL_2819883, EPI_ISL_2819884, EPI_ISL_2819885, EPI_ISL_2819886, EPI_ISL_2819887, EPI_ISL_2819888, EPI_ISL_2819889, EPI_ISL_2819890, EPI_ISL_2819891, EPI_ISL_2819892, EPI_ISL_2819893, EPI_ISL_2819894, EPI_ISL_2819895, EPI_ISL_2819896, EPI_ISL_2819897, EPI_ISL_2819898, EPI_ISL_2819899, EPI_ISL_2819900, EPI_ISL_2819901, EPI_ISL_2819902, EPI_ISL_2819903, EPI_ISL_2819904, EPI_ISL_2819905, EPI_ISL_2819906, EPI_ISL_2819907, EPI_ISL_2819908, EPI_ISL_2819909, EPI_ISL_2819910, EPI_ISL_2819911, EPI_ISL_2819912, EPI_ISL_2819913, EPI_ISL_2819914, EPI_ISL_2819915, EPI_ISL_2819916, EPI_ISL_2819917, EPI_ISL_2819918 | | | |
| see above | Centre for Enzyme Innovation, University of Portsmouth / Translational Research Laboratory, Portsmouth Hospitals NHS Trust / Hampshire Hospitals NHS Foundation Trust / Bournemouth University / University Hospitals Dorset NHS Foundation Trust / Brighton and Sussex University Hospitals NHS Trust / Dartford and Gravesham NHS Trust / Isle of Wight NHS Trust / Maidstone and Tunbridge Wells NHS Trust / University Hospital Southampton NHS Foundation Trust | COVID-19 Genomics UK (COG-UK) Consortium | Samuel Robson, Angela Beckett, Salman Goudarzi, Christopher Fearn, Kate Cook, Katie Loveson, Sharon Glaysher, Scott Elliott, Kelly Bicknell, Sarah Wyllie, Allyson Lloyd, Robert Impey, Anoop Chauhan, Stephen Kidd, Nathan Moore, Nick Cortes, Claire Thomas, Anna Mantzaouratou, Sarah Buchan, Magdalena Barrow, Andrew Butt, Liz Sheridan, Jonnie Seymour, Dorian Crudgington, Ben Macklin, Mohammed Hassan-Ibrahim, Cassandra Malone, Benjamin Cogger, Kevin Tucker, Samirakhon Raupova, Rachael Jeremiah, Anibolina Castigador, Emily Macnaughton, Karen Withell, Kordo Saeed, Jacqui Prieto, Adhyana Mahanama, Buddhini Samaraweera, Siona Silvieri, Emanuela Pelosi, Eleri Wilson-Davies, Sarah Jeremiah, Helen Wheeler, Matthew Harvey, Thea Sass, Helen Umpleby, Stephen Aplin |
| EPI_ISL_2819919 | Institute for Medical Research, Infectious Disease Research Centre, National Institutes of Health, Ministry of Health Malaysia | Institute for Medical Research, Infectious Disease Research Centre, National Institutes of Health, Ministry of Health Malaysia | Suppiah J, Kamel K, Mohd Zawawi Z, Azizan MA, Ramly N, Robert F, Thayan R |
| EPI_ISL_2819920, EPI_ISL_2819921, EPI_ISL_2819922, EPI_ISL_2819923, EPI_ISL_2819924, EPI_ISL_2819925, EPI_ISL_2819926, EPI_ISL_2819927, EPI_ISL_2819928, EPI_ISL_2819929, EPI_ISL_2819930, EPI_ISL_2819931, EPI_ISL_2819932, EPI_ISL_2819933, EPI_ISL_2819934, EPI_ISL_2819935, EPI_ISL_2819936, EPI_ISL_2819937, EPI_ISL_2819938, EPI_ISL_2819939, EPI_ISL_2819940, EPI_ISL_2819941, EPI_ISL_2819942, EPI_ISL_2819943, EPI_ISL_2819944, EPI_ISL_2819945, EPI_ISL_2819946, EPI_ISL_2819947, EPI_ISL_2819948, EPI_ISL_2819949, EPI_ISL_2819950, EPI_ISL_2819951, EPI_ISL_2819952, EPI_ISL_2819953, EPI_ISL_2819954, EPI_ISL_2819955, EPI_ISL_2819956, EPI_ISL_2819957, EPI_ISL_2819958, EPI_ISL_2819959, EPI_ISL_2819960, EPI_ISL_2819961, EPI_ISL_2819962, EPI_ISL_2819964, EPI_ISL_2819965, EPI_ISL_2819966, EPI_ISL_2819967, EPI_ISL_2819968, EPI_ISL_2819969, EPI_ISL_2819970, EPI_ISL_2819971, EPI_ISL_2819972, EPI_ISL_2819973, EPI_ISL_2819974, EPI_ISL_2819975, EPI_ISL_2819976, EPI_ISL_2819977, EPI_ISL_2819978, EPI_ISL_2819979, EPI_ISL_2819980, EPI_ISL_2819981, EPI_ISL_2819982, EPI_ISL_2819983, EPI_ISL_2819984, EPI_ISL_2819985, EPI_ISL_2819986, EPI_ISL_2819987, EPI_ISL_2819988, EPI_ISL_2819989, EPI_ISL_2819990, EPI_ISL_2819991, EPI_ISL_2819992, EPI_ISL_2819993, EPI_ISL_2819994, EPI_ISL_2819995, EPI_ISL_2819996, EPI_ISL_2819997, EPI_ISL_2819998, EPI_ISL_2819999, EPI_ISL_2820000, EPI_ISL_2820001, EPI_ISL_2820002, EPI_ISL_2820003, EPI_ISL_2820004, EPI_ISL_2820005, EPI_ISL_2820006, EPI_ISL_2820007, EPI_ISL_2820008, EPI_ISL_2820009, EPI_ISL_2820010, EPI_ISL_2820011, EPI_ISL_2820012, EPI_ISL_2820013, EPI_ISL_2820014, EPI_ISL_2820015, EPI_ISL_2820016, EPI_ISL_2820017, EPI_ISL_2820018, EPI_ISL_2820019, EPI_ISL_2820020, EPI_ISL_2820021, EPI_ISL_2820022, EPI_ISL_2820023, EPI_ISL_2820024, EPI_ISL_2820025, EPI_ISL_2820026, EPI_ISL_2820027, EPI_ISL_2820028, EPI_ISL_2820029, EPI_ISL_2820030, EPI_ISL_2820031, EPI_ISL_2820032, EPI_ISL_2820033, EPI_ISL_2820034, EPI_ISL_2820035, EPI_ISL_2820036, EPI_ISL_2820037, EPI_ISL_2820039, EPI_ISL_2820040, EPI_ISL_2820041, EPI_ISL_2820042, EPI_ISL_2820043, EPI_ISL_2820045, EPI_ISL_2820046, EPI_ISL_2820047, EPI_ISL_2820048, EPI_ISL_2820049, EPI_ISL_2820050, EPI_ISL_2820051, EPI_ISL_2820052, EPI_ISL_2820053, EPI_ISL_2820054, EPI_ISL_2820055, EPI_ISL_2820056, EPI_ISL_2820057, EPI_ISL_2820058, EPI_ISL_2820059, EPI_ISL_2820060, EPI_ISL_2820061, EPI_ISL_2820062, EPI_ISL_2820063, EPI_ISL_2820064, EPI_ISL_2820065, EPI_ISL_2820066, EPI_ISL_2820067, EPI_ISL_2820068, EPI_ISL_2820069, EPI_ISL_2820070, EPI_ISL_2820071, EPI_ISL_2820072, EPI_ISL_2820073, EPI_ISL_2820074, EPI_ISL_2820075, EPI_ISL_2820076, EPI_ISL_2820077, EPI_ISL_2820078, EPI_ISL_2820079, EPI_ISL_2820080, EPI_ISL_2820081, EPI_ISL_2820082, EPI_ISL_2820083, EPI_ISL_2820084, EPI_ISL_2820085, EPI_ISL_2820086, EPI_ISL_2820087, EPI_ISL_2820088, EPI_ISL_2820089, EPI_ISL_2820090, EPI_ISL_2820091, EPI_ISL_2820092, EPI_ISL_2820093, EPI_ISL_2820094, EPI_ISL_2820095, EPI_ISL_2820096, EPI_ISL_2820097, EPI_ISL_2820098, EPI_ISL_2820099, EPI_ISL_2820100, EPI_ISL_2820101, EPI_ISL_2820102, EPI_ISL_2820103, EPI_ISL_2820104, EPI_ISL_2820105, EPI_ISL_2820106, EPI_ISL_2820107, EPI_ISL_2820108, EPI_ISL_2820109, EPI_ISL_2820110, EPI_ISL_2820111, EPI_ISL_2820112, EPI_ISL_2820113, EPI_ISL_2820114, EPI_ISL_2820115, EPI_ISL_2820116, EPI_ISL_2820117, EPI_ISL_2820118, EPI_ISL_2820119, EPI_ISL_2820120, EPI_ISL_2820121, EPI_ISL_2820122, EPI_ISL_2820123, EPI_ISL_2820125, EPI_ISL_2820126, EPI_ISL_2820127, EPI_ISL_2820128, EPI_ISL_2820129, EPI_ISL_2820130, EPI_ISL_2820131, EPI_ISL_2820132, EPI_ISL_2820133, EPI_ISL_2820134, EPI_ISL_2820135, EPI_ISL_2820136, EPI_ISL_2820137, EPI_ISL_2820138, EPI_ISL_2820139, EPI_ISL_2820140, EPI_ISL_2820141, EPI_ISL_2820142, EPI_ISL_2820143, EPI_ISL_2820144, EPI_ISL_2820145, EPI_ISL_2820146, EPI_ISL_2820147, EPI_ISL_2820148, EPI_ISL_2820149, EPI_ISL_2820150, EPI_ISL_2820151, EPI_ISL_2820152, EPI_ISL_2820153, EPI_ISL_2820154, EPI_ISL_2820155, EPI_ISL_2820156, EPI_ISL_2820157, EPI_ISL_2820158, EPI_ISL_2820159, EPI_ISL_2820160, EPI_ISL_2820161, EPI_ISL_2820162, EPI_ISL_2820163, EPI_ISL_2820164, EPI_ISL_2820165, EPI_ISL_2820166, EPI_ISL_2820167, EPI_ISL_2820168, EPI_ISL_2820169, EPI_ISL_2820170, EPI_ISL_2820171, EPI_ISL_2820172, EPI_ISL_2820173, EPI_ISL_2820174, EPI_ISL_2820175, EPI_ISL_2820176, EPI_ISL_2820177, EPI_ISL_2820178, EPI_ISL_2820179, EPI_ISL_2820180, EPI_ISL_2820181, EPI_ISL_2820182, EPI_ISL_2820183, EPI_ISL_2820184, EPI_ISL_2820185, EPI_ISL_2820186, EPI_ISL_2820187, EPI_ISL_2820188, EPI_ISL_2820189, EPI_ISL_2820190, EPI_ISL_2820191, EPI_ISL_2820192, EPI_ISL_2820193, EPI_ISL_2820194, EPI_ISL_2820195, EPI_ISL_2820196, EPI_ISL_2820197, EPI_ISL_2820198, EPI_ISL_2820199, EPI_ISL_2820200, EPI_ISL_2820201, EPI_ISL_2820202, EPI_ISL_2820203, EPI_ISL_2820204, EPI_ISL_2820205, EPI_ISL_2820206, EPI_ISL_2820207, EPI_ISL_2820208, EPI_ISL_2820209, EPI_ISL_2820210, EPI_ISL_2820211, EPI_ISL_2820212, EPI_ISL_2820213, EPI_ISL_2820214, EPI_ISL_2820215, EPI_ISL_2820216, EPI_ISL_2820217, EPI_ISL_2820218, EPI_ISL_2820219, EPI_ISL_2820220, EPI_ISL_2820221, EPI_ISL_2820222 | | | |
| see above | Virology Department, Sheffield Teaching Hospitals NHS Foundation Trust/Department of Infection, Immunity and Cardiovascular Disease, The Medical School, University of Sheffield | COVID-19 Genomics UK (COG-UK) Consortium | Thushan de Silva, Matthew Parker, Nikki Smith, Adri Angyal, Rebecca Brown, Luke Green, Rachel Tucker, Paul Parsons, Danielle Groves, Katie Johnson, Laura Carrilero, Alex Keeley, Dave Partridge, Matthew Wyles, Benjamin Lindsey, Mehmet Yavuz, Mohammad Raza, Cariad Evans |
| EPI_ISL_2820237, EPI_ISL_2820240, EPI_ISL_2820241 | Dianalabs SA | Genesupport | Tanguy ARAUD, Geraldine Jost, Katia Jaton, Nadia Liassine |
| EPI_ISL_2820243, EPI_ISL_2820244, EPI_ISL_2820245, EPI_ISL_2820246, EPI_ISL_2820248, EPI_ISL_2820249 | Hospital Universitari Joan XXIII | Hospital Universitari Joan XXIII | Gemma Recio, Cristina Gutiérrez, Clara Benavent, Carla Martín, Margarida Terrón, Ester Picó, Natalia Bastón |
| EPI_ISL_2820251 | ASL BARI | University of Bari Biomedical Sciences and Human Oncology | Maria Chironna |
| EPI_ISL_2820263, EPI_ISL_2820264, EPI_ISL_2820265, EPI_ISL_2820266, EPI_ISL_2820267, EPI_ISL_2820268, EPI_ISL_2820269, EPI_ISL_2820270, EPI_ISL_2820271, EPI_ISL_2820272, EPI_ISL_2820273, EPI_ISL_2820274, EPI_ISL_2820275, EPI_ISL_2820276, EPI_ISL_2820277, EPI_ISL_2820278, EPI_ISL_2820279, EPI_ISL_2820280, EPI_ISL_2820281, EPI_ISL_2820282, EPI_ISL_2820283, EPI_ISL_2820284, EPI_ISL_2820285, EPI_ISL_2820286 | | | |
| see above | National Public Health Laboratory, National Centre for Infectious Diseases | National Public Health Laboratory, National Centre for Infectious Diseases | Tze Minn Mak, Katherine Ching, Zhenyang Zhou, Royce Ang, Lin Cui, Raymond Tzer Pin Lin |
| EPI_ISL_2820291, EPI_ISL_2820292, EPI_ISL_2820294, EPI_ISL_2820295, EPI_ISL_2820296, EPI_ISL_2820297, EPI_ISL_2820298, EPI_ISL_2820299, EPI_ISL_2820300, EPI_ISL_2820302, EPI_ISL_2820303, EPI_ISL_2820304, EPI_ISL_2820305, EPI_ISL_2820306, EPI_ISL_2820307, EPI_ISL_2820308, EPI_ISL_2820309, EPI_ISL_2820311, EPI_ISL_2820312, EPI_ISL_2820313, EPI_ISL_2820314, EPI_ISL_2820316, EPI_ISL_2820317, EPI_ISL_2820318, EPI_ISL_2820319, EPI_ISL_2820320, EPI_ISL_2820321, EPI_ISL_2820322, EPI_ISL_2820325, EPI_ISL_2820326, EPI_ISL_2820327, EPI_ISL_2820328, EPI_ISL_2820329, EPI_ISL_2820330, EPI_ISL_2820331, EPI_ISL_2820332, EPI_ISL_2820333, EPI_ISL_2820334, EPI_ISL_2820335, EPI_ISL_2820336 | | | |
| see above | Kasane Primary Hospital Laboratory | Botswana Harvard HIV Reference Laboratory | Sikhulile Moyo, Wonderful T. Choga, Dorcas Maruapula, Agnes Karutwaeng, Rose Munyere, Keoratile Ntshambiwa, Thongbotho Mphoyakgosi, Boitumelo Zuze, Botshelo Radibe, Legodile Kooepile, Ontlametse T. Bareng, Roger Shapiro, Shahin Lockman, Joseph Makhema, Madisa Mine, Mosepele Mosepele, Simani Gaseitsiwe |
| EPI_ISL_2820337, EPI_ISL_2820338, EPI_ISL_2820339, EPI_ISL_2820340, EPI_ISL_2820341, EPI_ISL_2820342, EPI_ISL_2820343, EPI_ISL_2820344, EPI_ISL_2820345, EPI_ISL_2820346, EPI_ISL_2820347, EPI_ISL_2820348, EPI_ISL_2820349, EPI_ISL_2820350, EPI_ISL_2820353, EPI_ISL_2820354, EPI_ISL_2820355, EPI_ISL_2820356, EPI_ISL_2820357, EPI_ISL_2820358, EPI_ISL_2820364, EPI_ISL_2820365, EPI_ISL_2820366, EPI_ISL_2820367, EPI_ISL_2820368, EPI_ISL_2820369, EPI_ISL_2820370, EPI_ISL_2820371, EPI_ISL_2820373, EPI_ISL_2820374 | | | |
| see above | Botswana Harvard HIV Reference Laboratory | Botswana Harvard HIV Reference Laboratory | Sikhulile Moyo, Wonderful T. Choga, Dorcas Maruapula, Thongbotho Mphoyakgosi, Boitumelo Zuze, Botshelo Radibe, Legodile Kooepile, Ontlametse T. Bareng, Motlalepule L. Pone, Godfrey Simoonga, Kereng Mphoyakgosi, Keoratile Ntshambiwa, Modisa Motswaledi, Madisa Mine, Joseph Makhema, Roger Shapiro, Shahin Lockman, Mosepele Mosepele, Simani Gaseitsiwe |
| EPI_ISL_2820375, EPI_ISL_2820376 | Botswana Harvard HIV Reference Laboratory | Botswana Harvard HIV Reference Laboratory | Sikhulile Moyo, Wonderful T. Choga, Dorcas Maruapula, Thongbotho Mphoyakgosi, Boitumelo Zuze, Botshelo Radibe, Legodile Kooepile, Ontlametse T. Bareng, Keoratile Ntshambiwa, Modisa Motswaledi, Madisa Mine, Joseph Makhema, Roger Shapiro, Shahin Lockman, Mosepele Mosepele, Simani Gaseitsiwe |
| EPI_ISL_2820377, EPI_ISL_2820378, EPI_ISL_2820381, EPI_ISL_2820382, EPI_ISL_2820383, EPI_ISL_2820384, EPI_ISL_2820385, EPI_ISL_2820386, EPI_ISL_2820387, EPI_ISL_2820390, EPI_ISL_2820391, EPI_ISL_2820392, EPI_ISL_2820393, EPI_ISL_2820394, EPI_ISL_2820397, EPI_ISL_2820398, EPI_ISL_2820400, EPI_ISL_2820401, EPI_ISL_2820402, EPI_ISL_2820404, EPI_ISL_2820405, EPI_ISL_2820407, EPI_ISL_2820408, EPI_ISL_2820409, EPI_ISL_2820410, EPI_ISL_2820411, EPI_ISL_2820413, EPI_ISL_2820414, EPI_ISL_2820415, EPI_ISL_2820417 | | | |
| see above | Kasane Primary Hospital Laboratory | Botswana Harvard HIV Reference Laboratory | Sikhulile Moyo, Wonderful T. Choga, Dorcas Maruapula, Agnes Karutwaeng, Rose Munyere, Keoratile Ntshambiwa, Thongbotho Mphoyakgosi, Boitumelo Zuze, Botshelo Radibe, Legodile Kooepile, Ontlametse T. Bareng, Roger Shapiro, Shahin Lockman, Joseph Makhema, Madisa Mine, Mosepele Mosepele, Simani Gaseitsiwe |
| EPI_ISL_2820419, EPI_ISL_2820420 | Botswana Harvard HIV Reference Laboratory | Botswana Harvard HIV Reference Laboratory | Sikhulile Moyo, Wonderful T. Choga, Dorcas Maruapula, Thongbotho Mphoyakgosi, Boitumelo Zuze, Botshelo Radibe, Legodile Kooepile, Ontlametse T. Bareng, Motlalepule L. Pone, Godfrey Simoonga, Kereng Mphoyakgosi, Keoratile Ntshambiwa, Modisa Motswaledi, Madisa Mine, Joseph Makhema, Roger Shapiro, Shahin Lockman, Mosepele Mosepele, Simani Gaseitsiwe |
| EPI_ISL_2820424, EPI_ISL_2820425, EPI_ISL_2820427, EPI_ISL_2820428, EPI_ISL_2820430, EPI_ISL_2820432, EPI_ISL_2820433, EPI_ISL_2820434 | Botswana Harvard HIV Reference Laboratory | Botswana Harvard HIV Reference Laboratory | Sikhulile Moyo, Wonderful T. Choga, Dorcas Maruapula, Thongbotho Mphoyakgosi, Boitumelo Zuze, Botshelo Radibe, Legodile Kooepile, Ontlametse T. Bareng, Keoratile Ntshambiwa, Modisa Motswaledi, Madisa Mine, Joseph Makhema, Roger Shapiro, Shahin Lockman, Mosepele Mosepele, Simani Gaseitsiwe |
| EPI_ISL_2820439, EPI_ISL_2820440, | Botswana Harvard HIV Reference Laboratory | Botswana Harvard HIV Reference Laboratory | Sikhulile Moyo, Wonderful T. Choga, Dorcas Maruapula, Thongbotho Mphoyakgosi, Boitumelo Zuze, Botshelo Radibe, Legodile Kooepile, Ontlametse T. Bareng, Motlalepule L. Pone, Godfrey Simoonga, Kereng Mphoyakgosi, Keoratile Ntshambiwa, Modisa Motswaledi, Madisa Mine, Joseph Makhema, Roger Shapiro, Shahin Lockman, Mosepele Mosepele, Simani Gaseitsiwe |

| | | | |
|---|---|---|---|
| EPI_ISL_2820441, EPI_ISL_2820442, EPI_ISL_2820444, EPI_ISL_2820445 | | | |
| EPI_ISL_2820447, EPI_ISL_2820452, EPI_ISL_2820454, EPI_ISL_2820455, EPI_ISL_2820456, EPI_ISL_2820458, EPI_ISL_2820459, EPI_ISL_2820461, EPI_ISL_2820462, EPI_ISL_2820463 | Botswana Harvard HIV Reference Laboratory | Botswana Harvard HIV Reference Laboratory | Sikhulile Moyo, Wonderful T. Choga, Dorcas Maruapula, Thongbotho Mphoyakgosi, Boitumelo Zuze, Botshelo Radibe, Legodile Kooepile, Ontlametse T. Bareng, Keoratile Ntshambiwa, Modisa Motswaledi, Madisa Mine, Joseph Makhema, Roger Shapiro, Shahin Lockman, Mosepele Mosepele, Simani Gaseitsiwe |
| EPI_ISL_2820531 | LA SOURCE | Laboratory of genomics and metagenomics | Trestan Pillonel, Damien Jacot, Sébastien Aeby, Gilbert Greub, Claire Bertelli |
| EPI_ISL_2820533 | CHUV | Laboratory of genomics and metagenomics | Trestan Pillonel, Damien Jacot, Sébastien Aeby, Gilbert Greub, Claire Bertelli |
| EPI_ISL_2820534 | UNILABS - LAUSANNE | Laboratory of genomics and metagenomics | Trestan Pillonel, Damien Jacot, Sébastien Aeby, Gilbert Greub, Claire Bertelli |
| EPI_ISL_2820536 | CHUV | Laboratory of genomics and metagenomics | Trestan Pillonel, Damien Jacot, Sébastien Aeby, Gilbert Greub, Claire Bertelli |
| EPI_ISL_2820537 | LA SOURCE | Laboratory of genomics and metagenomics | Trestan Pillonel, Damien Jacot, Sébastien Aeby, Gilbert Greub, Claire Bertelli |
| EPI_ISL_2820540 | EHNV | Laboratory of genomics and metagenomics | Trestan Pillonel, Damien Jacot, Sébastien Aeby, Gilbert Greub, Claire Bertelli |
| EPI_ISL_2820555, EPI_ISL_2820562, EPI_ISL_2820565, EPI_ISL_2820571, EPI_ISL_2820577, EPI_ISL_2820583, EPI_ISL_2820584, EPI_ISL_2820585, EPI_ISL_2820589, EPI_ISL_2820590, EPI_ISL_2820592, EPI_ISL_2820594, EPI_ISL_2820596, EPI_ISL_2820600 | | | |
| see above | Viollier AG | Department of Biosystems Science and Engineering, ETH Zürich | Christian Beisel, Sarah Nadeau, Chaoran Chen, Ivan Topolsky, Kim Philipp Jablonski, Lara Fuhrmann, Rebecca Denes, Mirjam Feldkamp, Ina Nissen, Natascha Santacroce, Elodie Burcklen, Christiane Beckmann, Maurice Redondo, Olivier Kobel, Christoph Noppen, Niko Beerenwinkel, Tanja Stadler |
| EPI_ISL_2820624, EPI_ISL_2820625 | Omega Diagnostics at Mounes | Omega Diagnostics at Mounes | Maria Echeverria, Latira Haynes-Jacob, Cynthia Corley, Vivek Khare MD |
| EPI_ISL_2820629, EPI_ISL_2820630 | Servicio de Microbiología. Hospital Clínico Universitario de Valencia | SeqCOVID-SPAIN consortium/IBV(CSIC) | David Navarro Ortega, Eliseo Albert Vicent, Ignacio Torres and SeqCOVID-SPAIN consortium |
| EPI_ISL_2820632, EPI_ISL_2820637, EPI_ISL_2820639, EPI_ISL_2820640, EPI_ISL_2820643, EPI_ISL_2820647, EPI_ISL_2820649 | Servicio de Microbiología. Hospital Universitario Doctor Peset | SeqCOVID-SPAIN consortium/IBV(CSIC) | Juan Alberola Enguídanos, Juan José Camarena Miñana, Rosa González Pellicer, José Miguel Nogueira Coito and SeqCOVID-SPAIN consortium |
| EPI_ISL_2820653, EPI_ISL_2820666, EPI_ISL_2820667, EPI_ISL_2820668, EPI_ISL_2820669 | Hospital Universitari i Politècnic La Fe de València | SeqCOVID-SPAIN consortium/IBV(CSIC) | María Dolores Gómez Ruiz, Eva González Barberá, Ana Gil Brusola, Salvador Giner Almaraz, José Luis López Hontangas and SeqCOVID-SPAIN consortium |
| EPI_ISL_2820680 | Omega Diagnostics at Mounes | Omega Diagnostics at Mounes | Maria Echeverria, Latira Haynes-Jacob, Cynthia Corley, Vivek Khare MD |
| EPI_ISL_2820697, EPI_ISL_2820699 | MRCG at LSHTM Genomics lab | MRCG at LSHTM Genomics lab | Abdul Karim sesay, Abdoulie Kanteh, Jarra Manneh, Mariama Kujabi, Bakary Sanyang,Dabiri Damilari |
| EPI_ISL_2820702 | San Gallicano Dermatological Institute IFO | INMI Lazzaro Spallanzani IRCCS | CEM Gruber, F Santini, B Bartolini, E Giombini, F Messina, M Rueca, G Bonfiglio, O Butera, F Pimpinelli, F Ensoli, A Mastrofrancesco, A Di Caro, MR Capobianchi |
| EPI_ISL_2820703 | Ospedale S.M. Goretti di Latina - ASL Latina presidio ospedaliero Nord | INMI Lazzaro Spallanzani IRCCS | E Giombini, F Messina, M Rueca, G Bonfiglio, O Butera, CEM Gruber, F Santini, B Bartolini, A Lucci, L Corso, L Di Biase, MR Capobianchi, A Di Caro |
| EPI_ISL_2820708, EPI_ISL_2820709, EPI_ISL_2820710, EPI_ISL_2820711, EPI_ISL_2820713 | Presidio di Brindisi Di Summa - Perrino | Istituto Zooprofilattico Sperimentale della Puglia e della Basilicata | Parisi A., Bianco A., Capozzi L., Del Sambro L., Simone D., Difato L., Santoro A., Ridolfi D., Giannico A., Manzari C., Pesole G. |
| EPI_ISL_2820716 | I.R.C.C.S. "S. De Bellis" - Ente Ospedaliero | Istituto Zooprofilattico Sperimentale della Puglia e della Basilicata | Parisi A., Bianco A., Capozzi L., Del Sambro L., Simone D., Difato L., Notarnicola M., Lippolis A., Manzari C., Pesole G. |
| EPI_ISL_2820719, EPI_ISL_2820722 | Ospedale Santa Caterina Novella | Istituto Zooprofilattico Sperimentale della Puglia e della Basilicata | Parisi A., Bianco A., Capozzi L., Del Sambro L., Simone D., Difato L., Bruno A. R., Manzari C., Pesole G. |
| EPI_ISL_2820725, EPI_ISL_2820726, EPI_ISL_2820727 | Ospedale Vito Fazzi | Istituto Zooprofilattico Sperimentale della Puglia e della Basilicata | Parisi A., Bianco A., Capozzi L., Del Sambro L., Simone D., Difato L., Lobreglio G., Manzari C., Pesole G. |
| EPI_ISL_2820730, EPI_ISL_2820733, EPI_ISL_2820734, EPI_ISL_2820736, EPI_ISL_2820738, EPI_ISL_2820739, EPI_ISL_2820741, EPI_ISL_2820742, EPI_ISL_2820743 | Presidio di Brindisi Di Summa - Perrino | Istituto Zooprofilattico Sperimentale della Puglia e della Basilicata | Parisi A., Bianco A., Capozzi L., Del Sambro L., Simone D., Difato L., Santoro A., Ridolfi D., Giannico A., Manzari C., Pesole G. |
| EPI_ISL_2820745, EPI_ISL_2820754 | Ospedale Santissima Annunziata | Istituto Zooprofilattico Sperimentale della Puglia e della Basilicata | Parisi A., Bianco A., Capozzi L., Del Sambro L., Simone D., Difato L., Ridolfi D., Giannico A, Tampoia M., Manzari C., Pesole G. |
| EPI_ISL_2820773, EPI_ISL_2820776, EPI_ISL_2820778, EPI_ISL_2820779, EPI_ISL_2820780, EPI_ISL_2820784, EPI_ISL_2820785, EPI_ISL_2820786, EPI_ISL_2820793, EPI_ISL_2820794, EPI_ISL_2820795, EPI_ISL_2820796, EPI_ISL_2820797, EPI_ISL_2820798, EPI_ISL_2820799, EPI_ISL_2820800, EPI_ISL_2820801, EPI_ISL_2820802, EPI_ISL_2820808, EPI_ISL_2820811, EPI_ISL_2820812, EPI_ISL_2820817 | | | |
| see above | Swedish national genomic surveillance program of SARS-CoV-2 | The Public Health Agency of Sweden | Maximilian Riess, Maria Lind Karlberg, Alma Brolund, Swedish national genomic surveillance program of SARS-CoV-2 |
| EPI_ISL_2820835, EPI_ISL_2820838, EPI_ISL_2820840 | Furst Medical Laboratory | Norwegian Institute of Public Health, Department of Virology | Kathrine Stene-Johansen, Kamilla Heddeland Instefjord, Hilde Elshaug, Garcia Llorente Ignacio, Jon Bråte, Engebretsen Serina Beate,Pedersen Benedikte Nevjen, Line Victoria Moen, Debech Nadia, Atiya R Ali,Marie Paulsen Madsen, Rasmus Riis Kopperud, Hilde Vollan, Karoline Bragstad, Olav Hungnes |
| EPI_ISL_2820841 | Innlandet Hospital Trust, Division Lillehammer, Department for Medical Microbiology | Norwegian Institute of Public Health, Department of Virology | Kathrine Stene-Johansen, Kamilla Heddeland Instefjord, Hilde Elshaug, Garcia Llorente Ignacio, Jon Bråte, Engebretsen Serina Beate,Pedersen Benedikte Nevjen, Line Victoria Moen, Debech Nadia, Atiya R Ali,Marie Paulsen Madsen, Rasmus Riis Kopperud, Hilde Vollan, Karoline Bragstad, Olav Hungnes |
| EPI_ISL_2820844 | Furst Medical Laboratory | Norwegian Institute of Public Health, Department of Virology | Kathrine Stene-Johansen, Kamilla Heddeland Instefjord, Hilde Elshaug, Garcia Llorente Ignacio, Jon Bråte, Engebretsen Serina Beate,Pedersen Benedikte Nevjen, Line Victoria Moen, Debech Nadia, Atiya R Ali,Marie Paulsen Madsen, Rasmus Riis Kopperud, Hilde Vollan, Karoline Bragstad, Olav Hungnes |
| EPI_ISL_2820845 | Akershus University Hospital, Department for Microbiology and Infectious Disease Control | Norwegian Institute of Public Health, Department of Virology | Kathrine Stene-Johansen, Kamilla Heddeland Instefjord, Hilde Elshaug, Garcia Llorente Ignacio, Jon Bråte, Engebretsen Serina Beate,Pedersen Benedikte Nevjen, Line Victoria Moen, Debech Nadia, Atiya R Ali,Marie Paulsen Madsen, Rasmus Riis Kopperud, Hilde Vollan, Karoline Bragstad, Olav Hungnes |
| EPI_ISL_2820850 | Innlandet Hospital Trust, Division Lillehammer, Department for Medical Microbiology | Norwegian Institute of Public Health, Department of Virology | Kathrine Stene-Johansen, Kamilla Heddeland Instefjord, Hilde Elshaug, Garcia Llorente Ignacio, Jon Bråte, Engebretsen Serina Beate,Pedersen Benedikte Nevjen, Line Victoria Moen, Debech Nadia, Atiya R Ali,Marie Paulsen Madsen, Rasmus Riis Kopperud, Hilde Vollan, Karoline Bragstad, Olav Hungnes |
| EPI_ISL_2820852 | Dept. of Medical Microbiology, Stavanger University Hospital, Helse Stavanger HF | Norwegian Institute of Public Health, Department of Virology | Kathrine Stene-Johansen, Kamilla Heddeland Instefjord, Hilde Elshaug, Garcia Llorente Ignacio, Jon Bråte, Engebretsen Serina Beate,Pedersen Benedikte Nevjen, Line Victoria Moen, Debech Nadia, Atiya R Ali,Marie Paulsen Madsen, Rasmus Riis Kopperud, Hilde Vollan, Karoline Bragstad, Olav Hungnes |
| EPI_ISL_2820867 | Unilabs Laboratory Medicine | Norwegian Institute of Public Health, Department of Virology | Kathrine Stene-Johansen, Kamilla Heddeland Instefjord, Hilde Elshaug, Garcia Llorente Ignacio, Jon Bråte, Engebretsen Serina Beate,Pedersen Benedikte Nevjen, Line Victoria Moen, Debech Nadia, Atiya R Ali,Marie Paulsen Madsen, Rasmus Riis Kopperud, Hilde Vollan, Karoline Bragstad, Olav Hungnes |
| EPI_ISL_2820868 | Furst Medical Laboratory | Norwegian Institute of Public Health, Department of Virology | Kathrine Stene-Johansen, Kamilla Heddeland Instefjord, Hilde Elshaug, Garcia Llorente Ignacio, Jon Bråte, Engebretsen Serina Beate,Pedersen Benedikte Nevjen, Line Victoria Moen, Debech Nadia, Atiya R Ali,Marie Paulsen Madsen, Rasmus Riis Kopperud, Hilde Vollan, Karoline Bragstad, Olav Hungnes |
| EPI_ISL_2820869, EPI_ISL_2820870, EPI_ISL_2820871 | Medical Microbiology Unit, Department for Laboratory Medicine, Drammen Hospital, Vestre Viken Health Trust, | Norwegian Institute of Public Health, Department of Virology | Kathrine Stene-Johansen, Kamilla Heddeland Instefjord, Hilde Elshaug, Garcia Llorente Ignacio, Jon Bråte, Engebretsen Serina Beate,Pedersen Benedikte Nevjen, Line Victoria Moen, Debech Nadia, Atiya R Ali,Marie Paulsen Madsen, Rasmus Riis Kopperud, Hilde Vollan, Karoline Bragstad, Olav Hungnes |
| EPI_ISL_2820872, EPI_ISL_2820873 | Innlandet Hospital Trust, Division Lillehammer, Department for Medical Microbiology | Norwegian Institute of Public Health, Department of Virology | Kathrine Stene-Johansen, Kamilla Heddeland Instefjord, Hilde Elshaug, Garcia Llorente Ignacio, Jon Bråte, Engebretsen Serina Beate,Pedersen Benedikte Nevjen, Line Victoria Moen, Debech Nadia, Atiya R Ali,Marie Paulsen Madsen, Rasmus Riis Kopperud, Hilde Vollan, Karoline Bragstad, Olav Hungnes |
| EPI_ISL_2820878 | Furst Medical Laboratory | Norwegian Institute of Public Health, Department of Virology | Kathrine Stene-Johansen, Kamilla Heddeland Instefjord, Hilde Elshaug, Garcia Llorente Ignacio, Jon Bråte, Engebretsen Serina Beate,Pedersen Benedikte Nevjen, Line Victoria Moen, Debech Nadia, Atiya R Ali,Marie Paulsen Madsen, Rasmus Riis Kopperud, Hilde Vollan, Karoline Bragstad, Olav Hungnes |
| EPI_ISL_2820880 | Unilabs Laboratory Medicine | Norwegian Institute of Public Health, Department of Virology | Kathrine Stene-Johansen, Kamilla Heddeland Instefjord, Hilde Elshaug, Garcia Llorente Ignacio, Jon Bråte, Engebretsen Serina Beate,Pedersen Benedikte Nevjen, Line Victoria Moen, Debech Nadia, Atiya R Ali,Marie Paulsen Madsen, Rasmus Riis |

| | | | |
|---|---|---|---|
| | | | Kopperud, Hilde Vollan, Karoline Bragstad, Olav Hungnes |
| EPI_ISL_2820888 | Medical Microbiology Unit, Department for Laboratory Medicine, Drammen Hospital, Vestre Viken Health Trust, | Norwegian Institute of Public Health, Department of Virology | Kathrine Stene-Johansen, Kamilla Heddeland Instefjord, Hilde Elshaug, Garcia Llorente Ignacio, Jon Bråte, Engebretsen Serina Beate,Pedersen Benedikte Nevjen, Line Victoria Moen, Debech Nadia, Atiya R Ali,Marie Paulsen Madsen, Rasmus Riis Kopperud, Hilde Vollan, Karoline Bragstad, Olav Hungnes |
| EPI_ISL_2820895 | Vestfold Hospital, Toensberg Department of Microbiology | Norwegian Institute of Public Health, Department of Virology | Kathrine Stene-Johansen, Kamilla Heddeland Instefjord, Hilde Elshaug, Garcia Llorente Ignacio, Jon Bråte, Engebretsen Serina Beate,Pedersen Benedikte Nevjen, Line Victoria Moen, Debech Nadia, Atiya R Ali,Marie Paulsen Madsen, Rasmus Riis Kopperud, Hilde Vollan, Karoline Bragstad, Olav Hungnes |
| EPI_ISL_2820911 | Akershus University Hospital, Department for Microbiology and Infectious Disease Control | Norwegian Institute of Public Health, Department of Virology | Kathrine Stene-Johansen, Kamilla Heddeland Instefjord, Hilde Elshaug, Garcia Llorente Ignacio, Jon Bråte, Engebretsen Serina Beate,Pedersen Benedikte Nevjen, Line Victoria Moen, Debech Nadia, Atiya R Ali,Marie Paulsen Madsen, Rasmus Riis Kopperud, Hilde Vollan, Karoline Bragstad, Olav Hungnes |
| EPI_ISL_2820913, EPI_ISL_2820917, EPI_ISL_2820919 | Medical Microbiology Unit, Department for Laboratory Medicine, Drammen Hospital, Vestre Viken Health Trust, | Norwegian Institute of Public Health, Department of Virology | Kathrine Stene-Johansen, Kamilla Heddeland Instefjord, Hilde Elshaug, Garcia Llorente Ignacio, Jon Bråte, Engebretsen Serina Beate,Pedersen Benedikte Nevjen, Line Victoria Moen, Debech Nadia, Atiya R Ali,Marie Paulsen Madsen, Rasmus Riis Kopperud, Hilde Vollan, Karoline Bragstad, Olav Hungnes |
| EPI_ISL_2820923, EPI_ISL_2820925 | Akershus University Hospital, Department for Microbiology and Infectious Disease Control | Norwegian Institute of Public Health, Department of Virology | Kathrine Stene-Johansen, Kamilla Heddeland Instefjord, Hilde Elshaug, Garcia Llorente Ignacio, Jon Bråte, Engebretsen Serina Beate,Pedersen Benedikte Nevjen, Line Victoria Moen, Debech Nadia, Atiya R Ali,Marie Paulsen Madsen, Rasmus Riis Kopperud, Hilde Vollan, Karoline Bragstad, Olav Hungnes |
| EPI_ISL_2820932 | Vestfold Hospital, Toensberg Department of Microbiology | Norwegian Institute of Public Health, Department of Virology | Kathrine Stene-Johansen, Kamilla Heddeland Instefjord, Hilde Elshaug, Garcia Llorente Ignacio, Jon Bråte, Engebretsen Serina Beate,Pedersen Benedikte Nevjen, Line Victoria Moen, Debech Nadia, Atiya R Ali,Marie Paulsen Madsen, Rasmus Riis Kopperud, Hilde Vollan, Karoline Bragstad, Olav Hungnes |
| EPI_ISL_2820933 | Medical Microbiology Unit, Department for Laboratory Medicine, Drammen Hospital, Vestre Viken Health Trust, | Norwegian Institute of Public Health, Department of Virology | Kathrine Stene-Johansen, Kamilla Heddeland Instefjord, Hilde Elshaug, Garcia Llorente Ignacio, Jon Bråte, Engebretsen Serina Beate,Pedersen Benedikte Nevjen, Line Victoria Moen, Debech Nadia, Atiya R Ali,Marie Paulsen Madsen, Rasmus Riis Kopperud, Hilde Vollan, Karoline Bragstad, Olav Hungnes |
| EPI_ISL_2820952 | INSIDE DIAGNÓSTICOS | Instituto Butantan | Dimas Tadeu Covas, Antonio Jorge Martins, Claudia Renata dos Santos Barros, David Schlesinger, Debora Botequio Moretti, Elaine Cristina Marqueze, Elaine Vieira Santos, Evandra Strazza Rodrigues, Heidge Fukumasu, Jayme Augusto de Souza-Neto, José Salvatore Leister Patané, Luiz Alcantara, Luiz Lehmann Coutinho, Maria Carolina Elias, Maurício Lacerda Nogueira, Rafael dos Santos Bezerra, Raul Machado Neto, Rejane Maria Tommasini Grotto, Ricardo Haddad, Sandra Coccuzzo Sampaio Vessoni, Simone Kashima, Svetoslav Nanev Slavov, Vincent Louis Viala |
| EPI_ISL_2820959, EPI_ISL_2820962 | UOC Microbiologia e Virologia, Azienda Ospedaliera Universitaria Senese, Siena, Italy | Dipartimento di Biotecnologie Mediche | Maria Grazia Cusi, David Pinzauti, Claudia Gandolfo, Gabriele Anichini, Gianni Pozzi, Gianni Gori Savellini, Francesco Santoro |
| EPI_ISL_2820995, EPI_ISL_2821003, EPI_ISL_2821007, EPI_ISL_2821009, EPI_ISL_2821010, EPI_ISL_2821011, EPI_ISL_2821024, EPI_ISL_2821025, EPI_ISL_2821028, EPI_ISL_2821030, EPI_ISL_2821031, EPI_ISL_2821032, EPI_ISL_2821033, EPI_ISL_2821034, EPI_ISL_2821037, EPI_ISL_2821038, EPI_ISL_2821043, EPI_ISL_2821044, EPI_ISL_2821045, EPI_ISL_2821048 | | | |
| see above | Azienda Ospedaliero - Universitaria di Modena Policlinico - Virologia e Microbiologia Molecolare | Istituto Zooprofilattico Sperimentale della Lombardia e dell'Emilia Romagna (IZSLER), Risk Analysis and Genomic Epidemiology Unit | Monica Pecorari, William Gennari, Giulia Fregni Serpini, Marina Morganti, Ilaria Menozzi, Erika Scaltriti, Stefano Pongolini |
| EPI_ISL_2821066, EPI_ISL_2821069 | Diagnostyka. Laboratoria Medyczne. | 1. ViroGenetics - BSL3 Laboratory of Virology, Małopolska Centre of Biotechnology, Jagiellonian University; 2. Diagtron Laboratoria Łukasz Rąbalski | Rabalski L., Gromowski,T., Rogalska-Kupiec M., Swadzba J., Wydmanski,W., Mazur-Panasiuk,N., Kowalski,M., Maciej Kosinski, Natalia Derewonko, Szulc,P., Sylwia Januszczak, Labaj,P.P., Pyrc,K. |
| EPI_ISL_2821077, EPI_ISL_2821078, EPI_ISL_2821079, EPI_ISL_2821080 | National Institute of High Security Animal Diseases | National Institute of High Security Animal Diseases | Anamika Mishra, Ashutosh Aasdev, Sandeep Bhatia, Naveen Kumar, Pradeep N Gandhale, Chandan K Dubey, Meghna Tripathi, Sushil Tripathi, Dipesh K Nayak, Ashwin A Raut, Vijendra P Singh |
| EPI_ISL_2821095, EPI_ISL_2821105, EPI_ISL_2821109, EPI_ISL_2821135, EPI_ISL_2821140, EPI_ISL_2821153, EPI_ISL_2821161, EPI_ISL_2821177, EPI_ISL_2821178, EPI_ISL_2821179, EPI_ISL_2821180, EPI_ISL_2821189, EPI_ISL_2821190, EPI_ISL_2821191, EPI_ISL_2821192, EPI_ISL_2821200, EPI_ISL_2821201, EPI_ISL_2821202, EPI_ISL_2821203, EPI_ISL_2821212, EPI_ISL_2821213, EPI_ISL_2821214, EPI_ISL_2821215, EPI_ISL_2821223, EPI_ISL_2821225, EPI_ISL_2821233, EPI_ISL_2821235, EPI_ISL_2821236, EPI_ISL_2821244, EPI_ISL_2821245, EPI_ISL_2821246, EPI_ISL_2821247, EPI_ISL_2821252, EPI_ISL_2821253, EPI_ISL_2821254, EPI_ISL_2821255 | | | |
| see above | National Virus Reference Laboratory | National Virus Reference Laboratory | Fiona Crispie, Calum Walsh, Matthew McCabe, Zoe Yandle, Charlene Bennet, Gabriel Gonzalez, Michael Carr, Jonathan Dean, Paul Cotter, Cillian F De Gascun |
| EPI_ISL_2822332, EPI_ISL_2822337, EPI_ISL_2822339, EPI_ISL_2822346, EPI_ISL_2822418, EPI_ISL_2822421, EPI_ISL_2822427, EPI_ISL_2822435, EPI_ISL_2822436, EPI_ISL_2822437, EPI_ISL_2822438, EPI_ISL_2822439, EPI_ISL_2822440, EPI_ISL_2822441, EPI_ISL_2822442, EPI_ISL_2822443, EPI_ISL_2822444, EPI_ISL_2822445, EPI_ISL_2822450, EPI_ISL_2822452, EPI_ISL_2822455, EPI_ISL_2822456, EPI_ISL_2822457, EPI_ISL_2822465, EPI_ISL_2822467, EPI_ISL_2822468, EPI_ISL_2822471, EPI_ISL_2822477, EPI_ISL_2822478, EPI_ISL_2822479, EPI_ISL_2822480, EPI_ISL_2822481, EPI_ISL_2822482, EPI_ISL_2822483, EPI_ISL_2822488, EPI_ISL_2822503, EPI_ISL_2822507, EPI_ISL_2822508, EPI_ISL_2822510, EPI_ISL_2822511, EPI_ISL_2822512, EPI_ISL_2822513, EPI_ISL_2822515, EPI_ISL_2822516, EPI_ISL_2822517, EPI_ISL_2822518, EPI_ISL_2822519, EPI_ISL_2822520, EPI_ISL_2822521, EPI_ISL_2822522, EPI_ISL_2822523, EPI_ISL_2822524, EPI_ISL_2822525, EPI_ISL_2822526, EPI_ISL_2822528, EPI_ISL_2822529, EPI_ISL_2822530, EPI_ISL_2822532, EPI_ISL_2822533, EPI_ISL_2822534, EPI_ISL_2822535, EPI_ISL_2822536, EPI_ISL_2822537, EPI_ISL_2822540, EPI_ISL_2822541, EPI_ISL_2822542, EPI_ISL_2822543, EPI_ISL_2822544, EPI_ISL_2822545, EPI_ISL_2822546, EPI_ISL_2822547, EPI_ISL_2822549, EPI_ISL_2822550, EPI_ISL_2822551, EPI_ISL_2822552, EPI_ISL_2822553, EPI_ISL_2822554, EPI_ISL_2822556, EPI_ISL_2822557, EPI_ISL_2822558, EPI_ISL_2822559, EPI_ISL_2822560, EPI_ISL_2822562, EPI_ISL_2822563, EPI_ISL_2822564, EPI_ISL_2822565, EPI_ISL_2822566, EPI_ISL_2822567, EPI_ISL_2822568, EPI_ISL_2822569, EPI_ISL_2822570, EPI_ISL_2822571, EPI_ISL_2822572, EPI_ISL_2822575, EPI_ISL_2822577, EPI_ISL_2822578, EPI_ISL_2822579, EPI_ISL_2822580, EPI_ISL_2822581, EPI_ISL_2822586, EPI_ISL_2822587, EPI_ISL_2822588, EPI_ISL_2822591, EPI_ISL_2822597, EPI_ISL_2822605, EPI_ISL_2822606, EPI_ISL_2822607, EPI_ISL_2822608, EPI_ISL_2822609, EPI_ISL_2822611, EPI_ISL_2822612, EPI_ISL_2822614, EPI_ISL_2822615, EPI_ISL_2822616, EPI_ISL_2822617, EPI_ISL_2822619, EPI_ISL_2822621, EPI_ISL_2822622, EPI_ISL_2822623, EPI_ISL_2822624, EPI_ISL_2822626, EPI_ISL_2822627, EPI_ISL_2822628, EPI_ISL_2822629, EPI_ISL_2822630, EPI_ISL_2822631, EPI_ISL_2822632, EPI_ISL_2822633, EPI_ISL_2822635, EPI_ISL_2822636, EPI_ISL_2822637, EPI_ISL_2822651, EPI_ISL_2822653, EPI_ISL_2822655, EPI_ISL_2822659, EPI_ISL_2822663, EPI_ISL_2822665, EPI_ISL_2822668, EPI_ISL_2822670, EPI_ISL_2822671, EPI_ISL_2822673, EPI_ISL_2822674, EPI_ISL_2822675, EPI_ISL_2822676, EPI_ISL_2822677, EPI_ISL_2822678, EPI_ISL_2822680, EPI_ISL_2822681, EPI_ISL_2822687, EPI_ISL_2822689, EPI_ISL_2822703, EPI_ISL_2822704, EPI_ISL_2822706, EPI_ISL_2822707, EPI_ISL_2822717, EPI_ISL_2822719, EPI_ISL_2822723, EPI_ISL_2822732, EPI_ISL_2822735, EPI_ISL_2822742, EPI_ISL_2822743, EPI_ISL_2822745, EPI_ISL_2822746, EPI_ISL_2822748, EPI_ISL_2822749, EPI_ISL_2822750, EPI_ISL_2822752, EPI_ISL_2822753, EPI_ISL_2822754, EPI_ISL_2822755, EPI_ISL_2822756, EPI_ISL_2822760, EPI_ISL_2822767, EPI_ISL_2822768, EPI_ISL_2822769, EPI_ISL_2822770, EPI_ISL_2822771, EPI_ISL_2822772, EPI_ISL_2822773, EPI_ISL_2822775, EPI_ISL_2822776, EPI_ISL_2822777, EPI_ISL_2822778, EPI_ISL_2822779, EPI_ISL_2822782, EPI_ISL_2822783, EPI_ISL_2822784, EPI_ISL_2822785, EPI_ISL_2822786, EPI_ISL_2822787, EPI_ISL_2822788, EPI_ISL_2822789, EPI_ISL_2822790, EPI_ISL_2822791, EPI_ISL_2822792, EPI_ISL_2822793, EPI_ISL_2822794, EPI_ISL_2822795, EPI_ISL_2822796, EPI_ISL_2822799, EPI_ISL_2822802, EPI_ISL_2822807, EPI_ISL_2822808, EPI_ISL_2822809, EPI_ISL_2822811, EPI_ISL_2822817, EPI_ISL_2822818, EPI_ISL_2822822, EPI_ISL_2822823, EPI_ISL_2822824, EPI_ISL_2822829, EPI_ISL_2822831, EPI_ISL_2822832, EPI_ISL_2822836, EPI_ISL_2822843, EPI_ISL_2822848, EPI_ISL_2822849, EPI_ISL_2822853, EPI_ISL_2822854, EPI_ISL_2822855, EPI_ISL_2822856, EPI_ISL_2822865, EPI_ISL_2822869, EPI_ISL_2822870, EPI_ISL_2822878, EPI_ISL_2822880, EPI_ISL_2822882, EPI_ISL_2822891, EPI_ISL_2822895, EPI_ISL_2822896, EPI_ISL_2822898, EPI_ISL_2822899, EPI_ISL_2822900, EPI_ISL_2822901, EPI_ISL_2822910, EPI_ISL_2822911, EPI_ISL_2822947, EPI_ISL_2822949, EPI_ISL_2822981, EPI_ISL_2822984, EPI_ISL_2822985, EPI_ISL_2822986, EPI_ISL_2822989, EPI_ISL_2822990, EPI_ISL_2822996, EPI_ISL_2822997, EPI_ISL_2822998, EPI_ISL_2823000, EPI_ISL_2823002, EPI_ISL_2823007, EPI_ISL_2823008, EPI_ISL_2823009, EPI_ISL_2823014, EPI_ISL_2823018, EPI_ISL_2823025, EPI_ISL_2823026, EPI_ISL_2823027, EPI_ISL_2823028, EPI_ISL_2823029, EPI_ISL_2823031, EPI_ISL_2823032, EPI_ISL_2823036, EPI_ISL_2823038, EPI_ISL_2823041, EPI_ISL_2823043, EPI_ISL_2823045, EPI_ISL_2823047, EPI_ISL_2823048, EPI_ISL_2823054, EPI_ISL_2823055, EPI_ISL_2823056, EPI_ISL_2823058, EPI_ISL_2823059, EPI_ISL_2823060, EPI_ISL_2823061, EPI_ISL_2823062, EPI_ISL_2823063, EPI_ISL_2823064, EPI_ISL_2823065, EPI_ISL_2823066, EPI_ISL_2823067, EPI_ISL_2823068, EPI_ISL_2823069, EPI_ISL_2823070, EPI_ISL_2823071, EPI_ISL_2823072, EPI_ISL_2823073, EPI_ISL_2823075, EPI_ISL_2823076, EPI_ISL_2823077, EPI_ISL_2823078, EPI_ISL_2823079, EPI_ISL_2823080, EPI_ISL_2823082, EPI_ISL_2823083, EPI_ISL_2823084, EPI_ISL_2823085, EPI_ISL_2823086, EPI_ISL_2823088, EPI_ISL_2823089, EPI_ISL_2823090, EPI_ISL_2823091, EPI_ISL_2823093, EPI_ISL_2823095, EPI_ISL_2823096, EPI_ISL_2823097, EPI_ISL_2823099, EPI_ISL_2823100, EPI_ISL_2823101, EPI_ISL_2823102, EPI_ISL_2823103, EPI_ISL_2823104, EPI_ISL_2823105, EPI_ISL_2823106, EPI_ISL_2823107, EPI_ISL_2823108, EPI_ISL_2823109, EPI_ISL_2823110, EPI_ISL_2823111, EPI_ISL_2823112, EPI_ISL_2823113, EPI_ISL_2823114, EPI_ISL_2823115, EPI_ISL_2823116, EPI_ISL_2823117, EPI_ISL_2823118, EPI_ISL_2823119, EPI_ISL_2823120, EPI_ISL_2823121, EPI_ISL_2823122, EPI_ISL_2823123, EPI_ISL_2823124, EPI_ISL_2823126, EPI_ISL_2823127, EPI_ISL_2823128, EPI_ISL_2823129, EPI_ISL_2823130, EPI_ISL_2823131, EPI_ISL_2823132, EPI_ISL_2823133, EPI_ISL_2823134, EPI_ISL_2823137, EPI_ISL_2823138, EPI_ISL_2823140, EPI_ISL_2823144, EPI_ISL_2823145, EPI_ISL_2823146, EPI_ISL_2823147, EPI_ISL_2823155, EPI_ISL_2823157, EPI_ISL_2823160, EPI_ISL_2823163, EPI_ISL_2823165, EPI_ISL_2823176, EPI_ISL_2823178, EPI_ISL_2823179, EPI_ISL_2823184, EPI_ISL_2823185, EPI_ISL_2823186, EPI_ISL_2823189, EPI_ISL_2823192, EPI_ISL_2823194, EPI_ISL_2823195, EPI_ISL_2823199, EPI_ISL_2823200, EPI_ISL_2823201, EPI_ISL_2823202, EPI_ISL_2823203, EPI_ISL_2823204, EPI_ISL_2823205, EPI_ISL_2823206, EPI_ISL_2823207, EPI_ISL_2823208, EPI_ISL_2823209, EPI_ISL_2823210, EPI_ISL_2823211, EPI_ISL_2823212, EPI_ISL_2823213, EPI_ISL_2823214, EPI_ISL_2823215, EPI_ISL_2823216, EPI_ISL_2823217, EPI_ISL_2823218, EPI_ISL_2823219, EPI_ISL_2823222, EPI_ISL_2823233, EPI_ISL_2823240, EPI_ISL_2823241, EPI_ISL_2823243, EPI_ISL_2823245, EPI_ISL_2823246, EPI_ISL_2823247, EPI_ISL_2823248, EPI_ISL_2823249, EPI_ISL_2823251, EPI_ISL_2823255, EPI_ISL_2823259, EPI_ISL_2823263, EPI_ISL_2823264, EPI_ISL_2823265, EPI_ISL_2823266, EPI_ISL_2823269, EPI_ISL_2823270, EPI_ISL_2823271, EPI_ISL_2823273, EPI_ISL_2823274, EPI_ISL_2823276, EPI_ISL_2823279, EPI_ISL_2823289, EPI_ISL_2823294, EPI_ISL_2823299, EPI_ISL_2823301, EPI_ISL_2823303, EPI_ISL_2823304, EPI_ISL_2823306, EPI_ISL_2823309, EPI_ISL_2823315, EPI_ISL_2823371, EPI_ISL_2823375, EPI_ISL_2823378, EPI_ISL_2823384, EPI_ISL_2823385, EPI_ISL_2823386, EPI_ISL_2823387, EPI_ISL_2823388, EPI_ISL_2823389, EPI_ISL_2823390, EPI_ISL_2823391, EPI_ISL_2823392, EPI_ISL_2823393, EPI_ISL_2823394, EPI_ISL_2823395, EPI_ISL_2823396, EPI_ISL_2823397, EPI_ISL_2823398, EPI_ISL_2823399, EPI_ISL_2823400, EPI_ISL_2823401, EPI_ISL_2823402, EPI_ISL_2823403, EPI_ISL_2823404, EPI_ISL_2823408, EPI_ISL_2823410, EPI_ISL_2823413, EPI_ISL_2823417, EPI_ISL_2823418, EPI_ISL_2823419, EPI_ISL_2823420, EPI_ISL_2823421, EPI_ISL_2823422, EPI_ISL_2823423, EPI_ISL_2823424, EPI_ISL_2823425, EPI_ISL_2823426, EPI_ISL_2823427, EPI_ISL_2823428, EPI_ISL_2823429, EPI_ISL_2823430, EPI_ISL_2823431, EPI_ISL_2823432, EPI_ISL_2823433, EPI_ISL_2823434, EPI_ISL_2823435, EPI_ISL_2823437, EPI_ISL_2823438, EPI_ISL_2823439, EPI_ISL_2823451, EPI_ISL_2823452, EPI_ISL_2823453, EPI_ISL_2823459, EPI_ISL_2823460, EPI_ISL_2823461, EPI_ISL_2823462, EPI_ISL_2823463, EPI_ISL_2823464, EPI_ISL_2823465, EPI_ISL_2823466, EPI_ISL_2823467, EPI_ISL_2823468, EPI_ISL_2823469, EPI_ISL_2823470, EPI_ISL_2823474, EPI_ISL_2823475, EPI_ISL_2823476, EPI_ISL_2823477, EPI_ISL_2823483, EPI_ISL_2823484, EPI_ISL_2823485, EPI_ISL_2823486, EPI_ISL_2823487, EPI_ISL_2823488, EPI_ISL_2823489, EPI_ISL_2823490, EPI_ISL_2823491, EPI_ISL_2823492, EPI_ISL_2823493, EPI_ISL_2823494, EPI_ISL_2823495, EPI_ISL_2823496, EPI_ISL_2823497, EPI_ISL_2823498, EPI_ISL_2823500, EPI_ISL_2823501, EPI_ISL_2823502, EPI_ISL_2823503, EPI_ISL_2823504, EPI_ISL_2823505, EPI_ISL_2823506, EPI_ISL_2823507, EPI_ISL_2823508, EPI_ISL_2823509, EPI_ISL_2823510, EPI_ISL_2823511, EPI_ISL_2823512, EPI_ISL_2823513, EPI_ISL_2823514, EPI_ISL_2823516, EPI_ISL_2823517, EPI_ISL_2823518, EPI_ISL_2823519, EPI_ISL_2823521, EPI_ISL_2823522, EPI_ISL_2823523, EPI_ISL_2823524, EPI_ISL_2823525, EPI_ISL_2823526 | | | |
| see above | Public Health Ontario Laboratory | Public Health Ontario Laboratory | Vanessa G Allen, Philip Banh, Yao Chen, Richard de Borja, Alireza Eshaghi, Nahuel Fittipaldi, Christine Frantz, Jonathan B Gubbay, Jennifer L Guthrie, Lawrence Heisler, Esha Joshi, Michael Laszloffy, Aimin Li, Michael CY Li, Dean Maxwell, Sandeep Nagra, Samir N Patel, Jared Simpson, Karthikeyan Sivaraman, Ashleigh Sullivan, Yogi Sundaravadanam, Sarah Teatero, Andre Villegas, Matthew Watson, Sandra Zittermann |
| EPI_ISL_2824953, EPI_ISL_2824956, EPI_ISL_2824958, EPI_ISL_2824959, EPI_ISL_2824960, EPI_ISL_2824961 | Hospital Universitari Joan XXIII | Hospital Universitari Joan XXIII | Gemma Recio, Cristina Gutiérrez, Clara Benavent, Carla Martín, Margarida Terrón, Ester Picó, Natalia Bastón |
| EPI_ISL_2824965 | Laboratoire Hôpital Sainte Musse | Laboratoire Hôpital Sainte Musse | Lionel Chollet |
| EPI_ISL_2824969 | ABC Algarve | Instituto Nacional de Saude (INSA) and Faculdade de Medicina da Universidade de Coimbra | Borges et al |
| EPI_ISL_2824974, EPI_ISL_2824975, EPI_ISL_2824976, EPI_ISL_2824977, EPI_ISL_2824978 | ULS Norte Alentejano | Instituto Nacional de Saude (INSA) and Faculdade de Medicina da Universidade de Coimbra | Borges et al |
| EPI_ISL_2824979, EPI_ISL_2824980, EPI_ISL_2824982, EPI_ISL_2824983 | CEDOC | Instituto Nacional de Saude (INSA) and Faculdade de Medicina da Universidade de Coimbra | Borges et al |
| EPI_ISL_2824984 | UNILABS | Instituto Nacional de Saude (INSA) | Borges et al |
| EPI_ISL_2824985, EPI_ISL_2824986 | UNILABS | Instituto Nacional de Saude (INSA) and BioSystems & Integrative Sciences Institute (BioISI) Genomics Unit, FCUL | Borges et al |
| EPI_ISL_2824988, EPI_ISL_2824989, EPI_ISL_2824990, EPI_ISL_2824991, EPI_ISL_2824992, EPI_ISL_2824993, EPI_ISL_2824994, EPI_ISL_2824995 | RSUP Dr. Hasan Sadikin Bandung | Genomik Solidaritas Indonesia Laboratorium | Reinhart Gabriel, Anna Christina Brazia, Vania Gavrila Wikasa, Carissa Sintca Wijaya, Dhahlia Agustina Cahyono, Alfin Mohammad Abdillah, Normastuti Adhini Tantyo, Annisa Muthiah Sukirman, Gracia Felias Enos Korompis, Louisa Markus, Basti Andriyoko, Delita Prihatni, Dewi Kartika Turbawati, Vivi Setyawati, Anuraj Shankar, Meutia Ayuputeri Kumaheri, Ariel Pradipta |
| EPI_ISL_2824996, EPI_ISL_2824997, EPI_ISL_2824998, EPI_ISL_2824999, EPI_ISL_2825000, EPI_ISL_2825001, EPI_ISL_2825002, EPI_ISL_2825003, EPI_ISL_2825004 | RSUP Dr. Hasan Sadikin Bandung | Genomik Solidaritas Indonesia Laboratorium | Anna Christina Brazia, Reinhart Gabriel, Vania Gavrila Wikasa, Carissa Sintca Wijaya, Dhahlia Agustina Cahyono, Alfin Mohammad Abdillah, Normastuti Adhini Tantyo, Annisa Muthiah Sukirman, Gracia Felias Enos Korompis, Louisa Markus, Basti Andriyoko, Delita Prihatni, Dewi Kartika Turbawati, Vivi Setyawati, Anuraj Shankar, Meutia Ayuputeri Kumaheri, Ariel Pradipta |

| | | | |
|---|---|---|---|
| EPI_ISL_2825006, EPI_ISL_2825007, EPI_ISL_2825008, EPI_ISL_2825009, EPI_ISL_2825010 | RSUP Dr. Hasan Sadikin Bandung | Genomik Solidaritas Indonesia Laboratorium | Vania Gavrila Wikasa, Carissa Sintca Wijaya, Anna Christina Brazia, Reinhart Gabriel, Dhahlia Agustina Cahyono, Alfin Mohammad Abdillah, Normastuti Adhini Tantyo, Annisa Muthiah Sukirman, Gracia Felias Enos Korompis, Louisa Markus, Basti Andriyoko, Delita Prihatni, Dewi Kartika Turbawati, Vivi Setyawati, Anuraj Shankar, Meutia Ayuputeri Kumaheri, Ariel Pradipta |
| EPI_ISL_2825013, EPI_ISL_2825014, EPI_ISL_2825015, EPI_ISL_2825016, EPI_ISL_2825017, EPI_ISL_2825018, EPI_ISL_2825019, EPI_ISL_2825020 | RSUP Dr. Hasan Sadikin Bandung | Genomik Solidaritas Indonesia Laboratorium | Carissa Sintca Wijaya, Vania Gavrila Wikasa, Anna Christina Brazia, Reinhart Gabriel, Dhahlia Agustina Cahyono, Alfin Mohammad Abdillah, Normastuti Adhini Tantyo, Annisa Muthiah Sukirman, Gracia Felias Enos Korompis, Louisa Markus, Basti Andriyoko, Delita Prihatni, Dewi Kartika Turbawati, Vivi Setyawati, Anuraj Shankar, Meutia Ayuputeri Kumaheri, Ariel Pradipta |
| EPI_ISL_2825021 | RSUP Dr. Hasan Sadikin Bandung | Genomik Solidaritas Indonesia Laboratorium | Vania Gavrila Wikasa, Carissa Sintca Wijaya, Anna Christina Brazia, Reinhart Gabriel, Dhahlia Agustina Cahyono, Alfin Mohammad Abdillah, Normastuti Adhini Tantyo, Annisa Muthiah Sukirman, Gracia Felias Enos Korompis, Louisa Markus, Basti Andriyoko, Delita Prihatni, Dewi Kartika Turbawati, Vivi Setyawati, Anuraj Shankar, Meutia Ayuputeri Kumaheri, Ariel Pradipta |
| EPI_ISL_2825022 | RSUP Dr. Hasan Sadikin Bandung | Genomik Solidaritas Indonesia Laboratorium | Carissa Sintca Wijaya, Vania Gavrila Wikasa, Anna Christina Brazia, Reinhart Gabriel, Dhahlia Agustina Cahyono, Alfin Mohammad Abdillah, Normastuti Adhini Tantyo, Annisa Muthiah Sukirman, Gracia Felias Enos Korompis, Louisa Markus, Basti Andriyoko, Delita Prihatni, Dewi Kartika Turbawati, Vivi Setyawati, Anuraj Shankar, Meutia Ayuputeri Kumaheri, Ariel Pradipta |
| EPI_ISL_2825796, EPI_ISL_2825877, EPI_ISL_2825878, EPI_ISL_2825904, EPI_ISL_2825929, EPI_ISL_2825971, EPI_ISL_2825999 | CDPH VBL | California Department of Public Health | CDPH-COVIDNet,UCLA Technology Center for Genomics & Bioinformatics |
| EPI_ISL_2827047, EPI_ISL_2827048, EPI_ISL_2827049, EPI_ISL_2827051, EPI_ISL_2827059, EPI_ISL_2827066 | TAMBO MEMORIAL LABORATORY | National Institute for Communicable Diseases of the National Health Laboratory Service | Amoako DG, Everatt J, Scheepers C, Mohale T, Ntuli N, Mahlangu B, Mnguni A, Ismail A, Bhiman JN |
| EPI_ISL_2827071 | MANKWENG LABORATORY | National Institute for Communicable Diseases of the National Health Laboratory Service | Amoako DG, Everatt J, Scheepers C, Mohale T, Ntuli N, Mahlangu B, Mnguni A, Ismail A, Bhiman JN |
| EPI_ISL_2827073, EPI_ISL_2827078, EPI_ISL_2827079, EPI_ISL_2827080, EPI_ISL_2827083, EPI_ISL_2827087, EPI_ISL_2827088, EPI_ISL_2827092, EPI_ISL_2827094, EPI_ISL_2827096, EPI_ISL_2827097, EPI_ISL_2827099, EPI_ISL_2827100, EPI_ISL_2827102, EPI_ISL_2827106, EPI_ISL_2827108, EPI_ISL_2827110, EPI_ISL_2827114, EPI_ISL_2827115, EPI_ISL_2827116, EPI_ISL_2827117, EPI_ISL_2827120, EPI_ISL_2827121, EPI_ISL_2827122, EPI_ISL_2827124, EPI_ISL_2827128, EPI_ISL_2827130, EPI_ISL_2827132 | | | |
| see above | HELEN JOSEPH LABORATORY | National Institute for Communicable Diseases of the National Health Laboratory Service | Amoako DG, Everatt J, Scheepers C, Mohale T, Ntuli N, Mahlangu B, Mnguni A, Ismail A, Bhiman JN |
| EPI_ISL_2827133, EPI_ISL_2827134, EPI_ISL_2827135, EPI_ISL_2827138, EPI_ISL_2827141 | MANKWENG LABORATORY | National Institute for Communicable Diseases of the National Health Laboratory Service | Amoako DG, Everatt J, Scheepers C, Mohale T, Ntuli N, Mahlangu B, Mnguni A, Ismail A, Bhiman JN |
| EPI_ISL_2827155 | TAMBO MEMORIAL LABORATORY | National Institute for Communicable Diseases of the National Health Laboratory Service | Amoako DG, Everatt J, Scheepers C, Mohale T, Ntuli N, Mahlangu B, Mnguni A, Ismail A, Bhiman JN |
| EPI_ISL_2827156, EPI_ISL_2827157 | CHRIS HANI BARAGWANATH LABORATORY | National Institute for Communicable Diseases of the National Health Laboratory Service | Amoako DG, Everatt J, Scheepers C, Mohale T, Ntuli N, Mahlangu B, Mnguni A, Ismail A, Bhiman JN |
| EPI_ISL_2827161, EPI_ISL_2827163, EPI_ISL_2827164 | MANKWENG LABORATORY | National Institute for Communicable Diseases of the National Health Laboratory Service | Amoako DG, Everatt J, Scheepers C, Mohale T, Ntuli N, Mahlangu B, Mnguni A, Ismail A, Bhiman JN |
| EPI_ISL_2827474, EPI_ISL_2827476, EPI_ISL_2827478, EPI_ISL_2827480, EPI_ISL_2827481, EPI_ISL_2827482, EPI_ISL_2827483, EPI_ISL_2827484, EPI_ISL_2827493, EPI_ISL_2827494, EPI_ISL_2827495, EPI_ISL_2827496, EPI_ISL_2827497, EPI_ISL_2827499, EPI_ISL_2827502, EPI_ISL_2827504, EPI_ISL_2827505, EPI_ISL_2827506, EPI_ISL_2827507, EPI_ISL_2827508, EPI_ISL_2827509, EPI_ISL_2827510, EPI_ISL_2827512, EPI_ISL_2827514, EPI_ISL_2827516, EPI_ISL_2827520, EPI_ISL_2827521 | | | |
| see above | Pandemic Response Lab - NYC | Pandemic Response Lab, R&D | Henry Lee, Michael Hammerling, Melissa Hopkins, Cybill del Castillo, Shinyoung Clair Kang, William Ward, Pradeep Bugga, Sol Rey, Dylan Law, Isabel Fernandez Escapa, Haiping Hao, Jon Laurent |
| EPI_ISL_2827868, EPI_ISL_2827869, EPI_ISL_2827870, EPI_ISL_2827871, EPI_ISL_2827872, EPI_ISL_2827874, EPI_ISL_2827875, EPI_ISL_2827876, EPI_ISL_2827878, EPI_ISL_2827879, EPI_ISL_2827881, EPI_ISL_2827883, EPI_ISL_2827884, EPI_ISL_2827887, EPI_ISL_2827888, EPI_ISL_2827889, EPI_ISL_2827890, EPI_ISL_2827891, EPI_ISL_2827892, EPI_ISL_2827893, EPI_ISL_2827894, EPI_ISL_2827895, EPI_ISL_2827896, EPI_ISL_2827897, EPI_ISL_2827898 | | | |
| see above | RUSTENBURG LABORATORY | National Institute for Communicable Diseases of the National Health Laboratory Service | Amoako DG, Everatt J, Scheepers C, Mohale T, Ntuli N, Mahlangu B, Mnguni A, Ismail A, Bhiman JN |
| EPI_ISL_2827899, EPI_ISL_2827901, EPI_ISL_2827902, EPI_ISL_2827904 | PORT ELIZABETH LABORATORY | National Institute for Communicable Diseases of the National Health Laboratory Service | Amoako DG, Everatt J, Scheepers C, Mohale T, Ntuli N, Mahlangu B, Mnguni A, Ismail A, Bhiman JN |
| EPI_ISL_2827906, EPI_ISL_2827907, EPI_ISL_2827913, EPI_ISL_2827914, EPI_ISL_2827915, EPI_ISL_2827916 | TSHEPONG LABORATORY | National Institute for Communicable Diseases of the National Health Laboratory Service | Amoako DG, Everatt J, Scheepers C, Mohale T, Ntuli N, Mahlangu B, Mnguni A, Ismail A, Bhiman JN |
| EPI_ISL_2827918, EPI_ISL_2827920 | POTCHEFSTROOM LABORATORY | National Institute for Communicable Diseases of the National Health Laboratory Service | Amoako DG, Everatt J, Scheepers C, Mohale T, Ntuli N, Mahlangu B, Mnguni A, Ismail A, Bhiman JN |
| EPI_ISL_2827925 | TSHEPONG LABORATORY | National Institute for Communicable Diseases of the National Health Laboratory Service | Amoako DG, Everatt J, Scheepers C, Mohale T, Ntuli N, Mahlangu B, Mnguni A, Ismail A, Bhiman JN |
| EPI_ISL_2827927, EPI_ISL_2827928, EPI_ISL_2827931, EPI_ISL_2827932, EPI_ISL_2827933, EPI_ISL_2827935, EPI_ISL_2827936, EPI_ISL_2827938, EPI_ISL_2827939, EPI_ISL_2827940, EPI_ISL_2827941, EPI_ISL_2827942, EPI_ISL_2827943, EPI_ISL_2827945, EPI_ISL_2827946, EPI_ISL_2827947, EPI_ISL_2827948, EPI_ISL_2827949, EPI_ISL_2827950, EPI_ISL_2827951 | | | |
| see above | MANKWENG LABORATORY | National Institute for Communicable Diseases of the National Health Laboratory Service | Amoako DG, Everatt J, Scheepers C, Mohale T, Ntuli N, Mahlangu B, Mnguni A, Ismail A, Bhiman JN |
| EPI_ISL_2827952, EPI_ISL_2827953, EPI_ISL_2827954, EPI_ISL_2827955, EPI_ISL_2827956, EPI_ISL_2827957, EPI_ISL_2827958, EPI_ISL_2827959, EPI_ISL_2827960 | New South Wales Health Pathology Royal Prince Alfred Hospital | Microbiology RPAH | Foster, C.; Au, J.; Ruiz Silva, M.; Deveson, I.; Bull, R.; Van Hal, S.; Rawlinson, W. |
| EPI_ISL_2828017, EPI_ISL_2828018 | South Eastern Area Laboratory Services (SEALS) | NSW Health Pathology - Institute of Clinical Pathology and Medical Research; Westmead Hospital; University of Sydney | CIDM-PH et al. |
| EPI_ISL_2828057 | Histopath | NSW Health Pathology - Institute of Clinical Pathology and Medical Research; Westmead Hospital; University of Sydney | CIDM-PH et al. |
| EPI_ISL_2828058 | Pathology West - NSW Health Pathology | NSW Health Pathology - Institute of Clinical Pathology and Medical Research; Westmead Hospital; University of Sydney | CIDM-PH et al. |
| EPI_ISL_2828059, EPI_ISL_2828060, EPI_ISL_2828061, EPI_ISL_2828062, EPI_ISL_2828063, EPI_ISL_2828064, EPI_ISL_2828065, EPI_ISL_2828066, EPI_ISL_2828067, EPI_ISL_2828068 | Sydney South West Pathology Service (SSWPS) - Royal Prince Alfred Hospital - NSW Health Pathology | NSW Health Pathology - Institute of Clinical Pathology and Medical Research; Westmead Hospital; University of Sydney | CIDM-PH et al. |

| | | | |
|---|---|---|---|
| EPI_ISL_2828069, EPI_ISL_2828070, EPI_ISL_2828071, EPI_ISL_2828072, EPI_ISL_2828073, EPI_ISL_2828074 | South Eastern Area Laboratory Services (SEALS) | NSW Health Pathology - Institute of Clinical Pathology and Medical Research; Westmead Hospital; University of Sydney | CIDM-PH et al. |
| EPI_ISL_2828075 | Laverty | NSW Health Pathology - Institute of Clinical Pathology and Medical Research; Westmead Hospital; University of Sydney | CIDM-PH et al. |
| EPI_ISL_2828076 | 4Cyte Pathology | NSW Health Pathology - Institute of Clinical Pathology and Medical Research; Westmead Hospital; University of Sydney | CIDM-PH et al. |
| EPI_ISL_2828077, EPI_ISL_2828078 | Laverty | NSW Health Pathology - Institute of Clinical Pathology and Medical Research; Westmead Hospital; University of Sydney | CIDM-PH et al. |
| EPI_ISL_2828079 | Sydney South West Pathology Service (SSWPS) - Liverpool Hospital - NSW Health Pathology | NSW Health Pathology - Institute of Clinical Pathology and Medical Research; Westmead Hospital; University of Sydney | CIDM-PH et al. |
| EPI_ISL_2828080, EPI_ISL_2828081, EPI_ISL_2828082, EPI_ISL_2828083, EPI_ISL_2828084 | Histopath | NSW Health Pathology - Institute of Clinical Pathology and Medical Research; Westmead Hospital; University of Sydney | CIDM-PH et al. |
| EPI_ISL_2828085 | Sydney South West Pathology Service (SSWPS) - Liverpool Hospital - NSW Health Pathology | NSW Health Pathology - Institute of Clinical Pathology and Medical Research; Westmead Hospital; University of Sydney | CIDM-PH et al. |
| EPI_ISL_2828086 | Laverty | NSW Health Pathology - Institute of Clinical Pathology and Medical Research; Westmead Hospital; University of Sydney | CIDM-PH et al. |
| EPI_ISL_2828087, EPI_ISL_2828088, EPI_ISL_2828089 | Sydney South West Pathology Service (SSWPS) - Liverpool Hospital - NSW Health Pathology | NSW Health Pathology - Institute of Clinical Pathology and Medical Research; Westmead Hospital; University of Sydney | CIDM-PH et al. |
| EPI_ISL_2828090, EPI_ISL_2828091 | Histopath | NSW Health Pathology - Institute of Clinical Pathology and Medical Research; Westmead Hospital; University of Sydney | CIDM-PH et al. |
| EPI_ISL_2828106, EPI_ISL_2828108, EPI_ISL_2828110, EPI_ISL_2828112, EPI_ISL_2828113, EPI_ISL_2828117, EPI_ISL_2828118, EPI_ISL_2828122, EPI_ISL_2828123, EPI_ISL_2828124, EPI_ISL_2828125, EPI_ISL_2828126, EPI_ISL_2828127, EPI_ISL_2828128, EPI_ISL_2828129, EPI_ISL_2828133, EPI_ISL_2828136, EPI_ISL_2828137, EPI_ISL_2828139, EPI_ISL_2828140, EPI_ISL_2828141, EPI_ISL_2828143, EPI_ISL_2828144, EPI_ISL_2828146, EPI_ISL_2828147, EPI_ISL_2828150, EPI_ISL_2828153, EPI_ISL_2828154, EPI_ISL_2828155, EPI_ISL_2828157, EPI_ISL_2828158, EPI_ISL_2828159, EPI_ISL_2828161, EPI_ISL_2828162, EPI_ISL_2828163, EPI_ISL_2828164, EPI_ISL_2828165, EPI_ISL_2828166, EPI_ISL_2828167, EPI_ISL_2828168, EPI_ISL_2828169, EPI_ISL_2828173, EPI_ISL_2828174, EPI_ISL_2828175, EPI_ISL_2828176, EPI_ISL_2828177, EPI_ISL_2828178, EPI_ISL_2828179, EPI_ISL_2828180, EPI_ISL_2828181, EPI_ISL_2828182, EPI_ISL_2828183, EPI_ISL_2828184, EPI_ISL_2828185, EPI_ISL_2828187, EPI_ISL_2828188, EPI_ISL_2828189, EPI_ISL_2828190, EPI_ISL_2828191, EPI_ISL_2828192, EPI_ISL_2828193, EPI_ISL_2828194, EPI_ISL_2828195, EPI_ISL_2828197, EPI_ISL_2828199, EPI_ISL_2828201, EPI_ISL_2828202, EPI_ISL_2828203, EPI_ISL_2828204, EPI_ISL_2828205, EPI_ISL_2828206, EPI_ISL_2828207, EPI_ISL_2828208, EPI_ISL_2828209, EPI_ISL_2828210, EPI_ISL_2828211, EPI_ISL_2828212, EPI_ISL_2828216, EPI_ISL_2828218, EPI_ISL_2828221 | | | |
| see above | Hospital Universitari Vall d'Hebron - Vall d'Hebron Institut de Recerca | Hospital Universitari Vall d'Hebron - Vall d'Hebron Institut de Recerca | Cristina Andrés, Maria Piñana, Alejandra González-Sánchez, Damir Garcia-Cehic, Ariadna Rando, Juliana Esperalba, Maria Gema Codina, Carla Castillo, Maria Carmen Martin, Tomàs Pumarola, Josep Quer, Andrés Antón |
| EPI_ISL_2828228, EPI_ISL_2828229, EPI_ISL_2828234, EPI_ISL_2828235, EPI_ISL_2828236, EPI_ISL_2828237, EPI_ISL_2828238, EPI_ISL_2828239, EPI_ISL_2828240, EPI_ISL_2828241, EPI_ISL_2828242, EPI_ISL_2828243, EPI_ISL_2828244, EPI_ISL_2828245, EPI_ISL_2828246, EPI_ISL_2828247, EPI_ISL_2828248, EPI_ISL_2828249, EPI_ISL_2828250 | | | |
| see above | Hospital Universitari Arnau de Vilanova | Hospital Universitari Vall d'Hebron - Vall d'Hebron Institut de Recerca | Cristina Andrés, Maria Piñana, Alejandra González-Sánchez, Damir Garcia-Cehic, Ariadna Rando, Juliana Esperalba, Maria Gema Codina, Carla Castillo, Maria Carmen Martin, Tomàs Pumarola, Josep Quer, Andrés Antón |
| EPI_ISL_2828255, EPI_ISL_2828256, EPI_ISL_2828257, EPI_ISL_2828259, EPI_ISL_2828260, EPI_ISL_2828261, EPI_ISL_2828262, EPI_ISL_2828263, EPI_ISL_2828264, EPI_ISL_2828265, EPI_ISL_2828267, EPI_ISL_2828268, EPI_ISL_2828269, EPI_ISL_2828270, EPI_ISL_2828271, EPI_ISL_2828272, EPI_ISL_2828273, EPI_ISL_2828274, EPI_ISL_2828275, EPI_ISL_2828276, EPI_ISL_2828278, EPI_ISL_2828279, EPI_ISL_2828280, EPI_ISL_2828281, EPI_ISL_2828282, EPI_ISL_2828284, EPI_ISL_2828288, EPI_ISL_2828289, EPI_ISL_2828290, EPI_ISL_2828291, EPI_ISL_2828293, EPI_ISL_2828294, EPI_ISL_2828295, EPI_ISL_2828299, EPI_ISL_2828300, EPI_ISL_2828301, EPI_ISL_2828302, EPI_ISL_2828303, EPI_ISL_2828305, EPI_ISL_2828306, EPI_ISL_2828307, EPI_ISL_2828308, EPI_ISL_2828309, EPI_ISL_2828310, EPI_ISL_2828311, EPI_ISL_2828313, EPI_ISL_2828314, EPI_ISL_2828315, EPI_ISL_2828316, EPI_ISL_2828317, EPI_ISL_2828318, EPI_ISL_2828319, EPI_ISL_2828323, EPI_ISL_2828325 | | | |
| see above | SARS-CoV-2 testing team, National Institute of Infectious Diseases | Pathogen Genomics Center, National Institute of Infectious Diseases | Tsuyoshi Sekizuka, Kentaro Itokawa, Rina Tanaka, Masanori Hashino, Nozomu Hanaoka, Masumichi Saito, Naomi Nojiri, Hazuka Y Furihata, Hiromizu Takahashi, Tsuguto Fujimoto, Makoto Kuroda |
| EPI_ISL_2828333, EPI_ISL_2828334, EPI_ISL_2828335, EPI_ISL_2828341, EPI_ISL_2828342, EPI_ISL_2828347, EPI_ISL_2828350, EPI_ISL_2828352, EPI_ISL_2828356, EPI_ISL_2828357, EPI_ISL_2828358, EPI_ISL_2828359, EPI_ISL_2828360, EPI_ISL_2828361, EPI_ISL_2828362, EPI_ISL_2828363, EPI_ISL_2828364, EPI_ISL_2828365, EPI_ISL_2828368, EPI_ISL_2828369, EPI_ISL_2828370, EPI_ISL_2828373, EPI_ISL_2828375, EPI_ISL_2828376, EPI_ISL_2828379, EPI_ISL_2828380, EPI_ISL_2828381 | | | |
| see above | Dept. of Microbiology and Infection Control, Akershus University Hospital HF | Dept. of Microbiology and Infection Control, Akershus University Hospital HF | Hege Vangstein Aamot, Alexander Hesselberg Løvestad |
| EPI_ISL_2828385, EPI_ISL_2828386, EPI_ISL_2828392, EPI_ISL_2828393, EPI_ISL_2828394, EPI_ISL_2828396, EPI_ISL_2828398, EPI_ISL_2828399, EPI_ISL_2828400, EPI_ISL_2828401, EPI_ISL_2828402, EPI_ISL_2828403, EPI_ISL_2828406, EPI_ISL_2828408, EPI_ISL_2828441 | | | |
| see above | SC (UCO) Igiene e Sanità Pubblica, ASUGI, Trieste | ARGO Laboratorio Genomica ed Epigenomica | Licastro D, Dal Monego S, Degasperi M, Marcello A, Segat L, Piscianz E, Busetti M, Fontana F, Breda C, Barbone F, D'Agaro P |
| EPI_ISL_2828454, EPI_ISL_2828455, EPI_ISL_2828457 | SC (UCO) Igiene e Sanità Pubblica, ASUGI, Trieste and SC Microbiologia e Virologia, ASFO, Pordenone | ARGO Laboratorio Genomica ed Epigenomica | Licastro D, Dal Monego S, Degasperi M, Marcello A, Segat L, De Rosa R, Breda C, Barbone F, D'Agaro P |
| EPI_ISL_2828459, EPI_ISL_2828460, EPI_ISL_2828461, EPI_ISL_2828462, EPI_ISL_2828464, EPI_ISL_2828465, EPI_ISL_2828466, EPI_ISL_2828467, EPI_ISL_2828468 | SC (UCO) Igiene e Sanità Pubblica, ASUGI, Trieste | ARGO Laboratorio Genomica ed Epigenomica | Licastro D, Dal Monego S, Degasperi M, Marcello A, Segat L, Piscianz E, Busetti M, Fontana F, Breda C, Barbone F, D'Agaro P |
| EPI_ISL_2828472, EPI_ISL_2828477, EPI_ISL_2828479, EPI_ISL_2828480, EPI_ISL_2828481, EPI_ISL_2828482, EPI_ISL_2828483, EPI_ISL_2828484, EPI_ISL_2828485, EPI_ISL_2828486, EPI_ISL_2828487, EPI_ISL_2828488, EPI_ISL_2828489, EPI_ISL_2828490, EPI_ISL_2828491, EPI_ISL_2828492, EPI_ISL_2828493, EPI_ISL_2828494, EPI_ISL_2828495, EPI_ISL_2828497, EPI_ISL_2828499, EPI_ISL_2828500, EPI_ISL_2828505, EPI_ISL_2828506, EPI_ISL_2828507, EPI_ISL_2828508, EPI_ISL_2828509, EPI_ISL_2828510, EPI_ISL_2828511, EPI_ISL_2828512, EPI_ISL_2828513, EPI_ISL_2828514, EPI_ISL_2828515 | | | |
| see above | Nucleic Acid Testing, National Reference Laboratory | GIGA Medical Genomics | Yvan Butera, Keith Durkin, Maria Artesi, Bouchra Boujemla, Robert Rutayisire, Reuben Sindayiheba, Patrick Tuyisenge, Esperence Umumararungu, Sébastien Bontems, Marie-Pierre Hayette, Nathalie Renotte, Swaibu Gatare, Jacob Souopgui, Sabin Nsanzimana, Vincent Bours, Léon Mutesa |
| EPI_ISL_2828577 | CHUGA-IBP-laboratoire de Virologie | IBP-laboratoire de virologie | Sylvie Larrat; Patrice Morand; Pascal Poignard; Raphaële Germi; Julien Lupo; Anne Signori -Schmuck; Anne-Karen Faure; Benjamin Nemoz; Aurélie Truffot; Léa Ponderand; Julien Andréani; Alban Caporossi; Hugo Jardin |
| EPI_ISL_2828629, EPI_ISL_2828635, EPI_ISL_2828637, EPI_ISL_2828638, EPI_ISL_2828640, EPI_ISL_2828642 | Utah Public Health Laboratory, Utah Public Health Laboratory Infectious Disease submission group | Utah Public Health Laboratory, Utah Public Health Laboratory Infectious Disease submission group | Gallagher,T., Young,E.L., Oakeson,K.F. |
| EPI_ISL_2828727 | SISP Lecce | University of Bari Biomedical Sciences and Human Oncology | Maria Chironna |
| EPI_ISL_2828732 | Lancet | National Institute for Communicable Diseases of the National Health Laboratory Service | Amoako DG, Everatt J, Scheepers C, Mohale T, Ntuli N, Mahlangu B, Mnguni A, Ismail A, Bhiman JN |
| EPI_ISL_2828735, EPI_ISL_2828736, EPI_ISL_2828738, EPI_ISL_2828739, EPI_ISL_2828742, EPI_ISL_2828744, EPI_ISL_2828745, EPI_ISL_2828747, EPI_ISL_2828751, EPI_ISL_2828752, EPI_ISL_2828753, EPI_ISL_2828755, EPI_ISL_2828756, EPI_ISL_2828757, EPI_ISL_2828758 | | | |
| see above | Pathcare Vaal | National Institute for Communicable Diseases of the National Health Laboratory Service | Amoako DG, Everatt J, Scheepers C, Mohale T, Ntuli N, Mahlangu B, Mnguni A, Ismail A, Bhiman JN |
| EPI_ISL_2828759 | Lighthouse Lab in Glasgow | Wellcome Sanger Institute for the COVID-19 Genomics UK (COG-UK) Consortium | Harper VanSteenhouse, Yumi Kasai, David Gray, Carol Clugston, Anna Dominiczak and Alex Alderton, Roberto Amato, Jeffrey Barrett, Sonia Goncalves, Ewan Harrison, David K. Jackson, Ian Johnston, Dominic Kwiatkowski, Cordelia Langford, John Sillitoe on behalf of the Wellcome Sanger Institute COVID-19 Surveillance Team |
| EPI_ISL_2828760 | Lighthouse Lab in Alderley Park | Wellcome Sanger Institute for the COVID-19 Genomics UK (COG-UK) Consortium | Jacquelyn Wynn, Mairead Hyland, The Lighthouse Lab in Alderley Park and Alex Alderton, Roberto Amato, Jeffrey Barrett, Sonia Goncalves, Ewan Harrison, David K. Jackson, Ian Johnston, Dominic Kwiatkowski, Cordelia Langford, John Sillitoe on behalf of the Wellcome Sanger Institute COVID-19 Surveillance Team |
| EPI_ISL_2828761 | Lighthouse Laboratory Plymouth | Wellcome Sanger Institute for the COVID-19 Genomics UK (COG-UK) Consortium | Lighthouse Laboratory Plymouth and Alex Alderton, Roberto Amato, Jeffrey Barrett, Sonia Goncalves, Ewan Harrison, David K. Jackson, Ian Johnston, Dominic Kwiatkowski, Cordelia Langford, John Sillitoe on behalf of the Wellcome Sanger Institute COVID-19 Surveillance Team |
| EPI_ISL_2828762, EPI_ISL_2828763, EPI_ISL_2828764, EPI_ISL_2828765 | Lighthouse Lab in Milton Keynes | Wellcome Sanger Institute for the COVID-19 Genomics UK (COG-UK) Consortium | The Lighthouse Lab in Milton Keynes and Alex Alderton, Roberto Amato, Jeffrey Barrett, Sonia Goncalves, Ewan Harrison, David K. Jackson, Ian Johnston, Dominic Kwiatkowski, Cordelia Langford, John Sillitoe on behalf of the Wellcome Sanger Institute COVID-19 Surveillance Team |
| EPI_ISL_2828766 | Lighthouse Laboratory Plymouth | Wellcome Sanger Institute for the COVID-19 Genomics UK (COG-UK) Consortium | Lighthouse Laboratory Plymouth and Alex Alderton, Roberto Amato, Jeffrey Barrett, Sonia Goncalves, Ewan Harrison, David K. Jackson, Ian Johnston, Dominic Kwiatkowski, Cordelia Langford, John Sillitoe on behalf of the Wellcome Sanger Institute COVID-19 Surveillance Team |
| EPI_ISL_2828767, EPI_ISL_2828768 | Lighthouse Lab in Alderley Park | Wellcome Sanger Institute for the COVID-19 Genomics UK (COG-UK) Consortium | Jacquelyn Wynn, Mairead Hyland, The Lighthouse Lab in Alderley Park and Alex Alderton, Roberto Amato, Jeffrey Barrett, Sonia Goncalves, Ewan Harrison, David K. Jackson, Ian Johnston, Dominic Kwiatkowski, Cordelia Langford, John Sillitoe on behalf of the Wellcome Sanger Institute COVID-19 Surveillance Team |
| EPI_ISL_2828769, EPI_ISL_2828770, EPI_ISL_2828771, | Lighthouse Laboratory Plymouth | Wellcome Sanger Institute for the COVID-19 Genomics UK (COG-UK) Consortium | Lighthouse Laboratory Plymouth and Alex Alderton, Roberto Amato, Jeffrey Barrett, Sonia Goncalves, Ewan Harrison, David K. Jackson, Ian Johnston, Dominic Kwiatkowski, Cordelia Langford, John Sillitoe on behalf of the Wellcome Sanger Institute COVID-19 Surveillance Team |

| | | | |
|---|---|---|---|
| EPI_ISL_2828772 | | | |
| EPI_ISL_2828773 | Lighthouse Lab in Milton Keynes | Wellcome Sanger Institute for the COVID-19 Genomics UK (COG-UK) Consortium | The Lighthouse Lab in Milton Keynes and Alex Alderton, Roberto Amato, Jeffrey Barrett, Sonia Goncalves, Ewan Harrison, David K. Jackson, Ian Johnston, Dominic Kwiatkowski, Cordelia Langford, John Sillitoe on behalf of the Wellcome Sanger Institute COVID-19 Surveillance Team |
| EPI_ISL_2828774 | Lighthouse Laboratory Plymouth | Wellcome Sanger Institute for the COVID-19 Genomics UK (COG-UK) Consortium | Lighthouse Laboratory Plymouth and Alex Alderton, Roberto Amato, Jeffrey Barrett, Sonia Goncalves, Ewan Harrison, David K. Jackson, Ian Johnston, Dominic Kwiatkowski, Cordelia Langford, John Sillitoe on behalf of the Wellcome Sanger Institute COVID-19 Surveillance Team |
| EPI_ISL_2828775 | Lighthouse Lab in Alderley Park | Wellcome Sanger Institute for the COVID-19 Genomics UK (COG-UK) Consortium | Jacquelyn Wynn, Mairead Hyland, The Lighthouse Lab in Alderley Park and Alex Alderton, Roberto Amato, Jeffrey Barrett, Sonia Goncalves, Ewan Harrison, David K. Jackson, Ian Johnston, Dominic Kwiatkowski, Cordelia Langford, John Sillitoe on behalf of the Wellcome Sanger Institute COVID-19 Surveillance Team |
| EPI_ISL_2828776 | Lighthouse Laboratory Plymouth | Wellcome Sanger Institute for the COVID-19 Genomics UK (COG-UK) Consortium | Lighthouse Laboratory Plymouth and Alex Alderton, Roberto Amato, Jeffrey Barrett, Sonia Goncalves, Ewan Harrison, David K. Jackson, Ian Johnston, Dominic Kwiatkowski, Cordelia Langford, John Sillitoe on behalf of the Wellcome Sanger Institute COVID-19 Surveillance Team |
| EPI_ISL_2828777 | Lighthouse Lab in Milton Keynes | Wellcome Sanger Institute for the COVID-19 Genomics UK (COG-UK) Consortium | The Lighthouse Lab in Milton Keynes and Alex Alderton, Roberto Amato, Jeffrey Barrett, Sonia Goncalves, Ewan Harrison, David K. Jackson, Ian Johnston, Dominic Kwiatkowski, Cordelia Langford, John Sillitoe on behalf of the Wellcome Sanger Institute COVID-19 Surveillance Team |
| EPI_ISL_2828778, EPI_ISL_2828779 | Lighthouse Lab in Alderley Park | Wellcome Sanger Institute for the COVID-19 Genomics UK (COG-UK) Consortium | Jacquelyn Wynn, Mairead Hyland, The Lighthouse Lab in Alderley Park and Alex Alderton, Roberto Amato, Jeffrey Barrett, Sonia Goncalves, Ewan Harrison, David K. Jackson, Ian Johnston, Dominic Kwiatkowski, Cordelia Langford, John Sillitoe on behalf of the Wellcome Sanger Institute COVID-19 Surveillance Team |
| EPI_ISL_2828780 | Lighthouse Laboratory Plymouth | Wellcome Sanger Institute for the COVID-19 Genomics UK (COG-UK) Consortium | Lighthouse Laboratory Plymouth and Alex Alderton, Roberto Amato, Jeffrey Barrett, Sonia Goncalves, Ewan Harrison, David K. Jackson, Ian Johnston, Dominic Kwiatkowski, Cordelia Langford, John Sillitoe on behalf of the Wellcome Sanger Institute COVID-19 Surveillance Team |
| EPI_ISL_2828781, EPI_ISL_2828782 | Lighthouse Lab in Milton Keynes | Wellcome Sanger Institute for the COVID-19 Genomics UK (COG-UK) Consortium | The Lighthouse Lab in Milton Keynes and Alex Alderton, Roberto Amato, Jeffrey Barrett, Sonia Goncalves, Ewan Harrison, David K. Jackson, Ian Johnston, Dominic Kwiatkowski, Cordelia Langford, John Sillitoe on behalf of the Wellcome Sanger Institute COVID-19 Surveillance Team |
| EPI_ISL_2828783, EPI_ISL_2828784, EPI_ISL_2828785, EPI_ISL_2828786, EPI_ISL_2828787 | Lighthouse Laboratory Plymouth | Wellcome Sanger Institute for the COVID-19 Genomics UK (COG-UK) Consortium | Lighthouse Laboratory Plymouth and Alex Alderton, Roberto Amato, Jeffrey Barrett, Sonia Goncalves, Ewan Harrison, David K. Jackson, Ian Johnston, Dominic Kwiatkowski, Cordelia Langford, John Sillitoe on behalf of the Wellcome Sanger Institute COVID-19 Surveillance Team |
| EPI_ISL_2828788, EPI_ISL_2828789, EPI_ISL_2828790 | Lighthouse Lab in Alderley Park | Wellcome Sanger Institute for the COVID-19 Genomics UK (COG-UK) Consortium | Jacquelyn Wynn, Mairead Hyland, The Lighthouse Lab in Alderley Park and Alex Alderton, Roberto Amato, Jeffrey Barrett, Sonia Goncalves, Ewan Harrison, David K. Jackson, Ian Johnston, Dominic Kwiatkowski, Cordelia Langford, John Sillitoe on behalf of the Wellcome Sanger Institute COVID-19 Surveillance Team |
| EPI_ISL_2828791 | Lighthouse Laboratory Plymouth | Wellcome Sanger Institute for the COVID-19 Genomics UK (COG-UK) Consortium | Lighthouse Laboratory Plymouth and Alex Alderton, Roberto Amato, Jeffrey Barrett, Sonia Goncalves, Ewan Harrison, David K. Jackson, Ian Johnston, Dominic Kwiatkowski, Cordelia Langford, John Sillitoe on behalf of the Wellcome Sanger Institute COVID-19 Surveillance Team |
| EPI_ISL_2828792, EPI_ISL_2828793, EPI_ISL_2828794, EPI_ISL_2828795 | Lighthouse Lab in Milton Keynes | Wellcome Sanger Institute for the COVID-19 Genomics UK (COG-UK) Consortium | The Lighthouse Lab in Milton Keynes and Alex Alderton, Roberto Amato, Jeffrey Barrett, Sonia Goncalves, Ewan Harrison, David K. Jackson, Ian Johnston, Dominic Kwiatkowski, Cordelia Langford, John Sillitoe on behalf of the Wellcome Sanger Institute COVID-19 Surveillance Team |
| EPI_ISL_2828796 | Lighthouse Lab in Alderley Park | Wellcome Sanger Institute for the COVID-19 Genomics UK (COG-UK) Consortium | Jacquelyn Wynn, Mairead Hyland, The Lighthouse Lab in Alderley Park and Alex Alderton, Roberto Amato, Jeffrey Barrett, Sonia Goncalves, Ewan Harrison, David K. Jackson, Ian Johnston, Dominic Kwiatkowski, Cordelia Langford, John Sillitoe on behalf of the Wellcome Sanger Institute COVID-19 Surveillance Team |
| EPI_ISL_2828797 | Lighthouse Lab in Milton Keynes | Wellcome Sanger Institute for the COVID-19 Genomics UK (COG-UK) Consortium | The Lighthouse Lab in Milton Keynes and Alex Alderton, Roberto Amato, Jeffrey Barrett, Sonia Goncalves, Ewan Harrison, David K. Jackson, Ian Johnston, Dominic Kwiatkowski, Cordelia Langford, John Sillitoe on behalf of the Wellcome Sanger Institute COVID-19 Surveillance Team |
| EPI_ISL_2828798 | Lighthouse Lab in Alderley Park | Wellcome Sanger Institute for the COVID-19 Genomics UK (COG-UK) Consortium | Jacquelyn Wynn, Mairead Hyland, The Lighthouse Lab in Alderley Park and Alex Alderton, Roberto Amato, Jeffrey Barrett, Sonia Goncalves, Ewan Harrison, David K. Jackson, Ian Johnston, Dominic Kwiatkowski, Cordelia Langford, John Sillitoe on behalf of the Wellcome Sanger Institute COVID-19 Surveillance Team |
| EPI_ISL_2828799 | Lighthouse Laboratory Plymouth | Wellcome Sanger Institute for the COVID-19 Genomics UK (COG-UK) Consortium | Lighthouse Laboratory Plymouth and Alex Alderton, Roberto Amato, Jeffrey Barrett, Sonia Goncalves, Ewan Harrison, David K. Jackson, Ian Johnston, Dominic Kwiatkowski, Cordelia Langford, John Sillitoe on behalf of the Wellcome Sanger Institute COVID-19 Surveillance Team |
| EPI_ISL_2828800 | Lighthouse Lab in Milton Keynes | Wellcome Sanger Institute for the COVID-19 Genomics UK (COG-UK) Consortium | The Lighthouse Lab in Milton Keynes and Alex Alderton, Roberto Amato, Jeffrey Barrett, Sonia Goncalves, Ewan Harrison, David K. Jackson, Ian Johnston, Dominic Kwiatkowski, Cordelia Langford, John Sillitoe on behalf of the Wellcome Sanger Institute COVID-19 Surveillance Team |
| EPI_ISL_2828801 | Lighthouse Lab in Alderley Park | Wellcome Sanger Institute for the COVID-19 Genomics UK (COG-UK) Consortium | Jacquelyn Wynn, Mairead Hyland, The Lighthouse Lab in Alderley Park and Alex Alderton, Roberto Amato, Jeffrey Barrett, Sonia Goncalves, Ewan Harrison, David K. Jackson, Ian Johnston, Dominic Kwiatkowski, Cordelia Langford, John Sillitoe on behalf of the Wellcome Sanger Institute COVID-19 Surveillance Team |
| EPI_ISL_2828802 | Lighthouse Laboratory Plymouth | Wellcome Sanger Institute for the COVID-19 Genomics UK (COG-UK) Consortium | Lighthouse Laboratory Plymouth and Alex Alderton, Roberto Amato, Jeffrey Barrett, Sonia Goncalves, Ewan Harrison, David K. Jackson, Ian Johnston, Dominic Kwiatkowski, Cordelia Langford, John Sillitoe on behalf of the Wellcome Sanger Institute COVID-19 Surveillance Team |
| EPI_ISL_2828803 | Lighthouse Lab in Milton Keynes | Wellcome Sanger Institute for the COVID-19 Genomics UK (COG-UK) Consortium | The Lighthouse Lab in Milton Keynes and Alex Alderton, Roberto Amato, Jeffrey Barrett, Sonia Goncalves, Ewan Harrison, David K. Jackson, Ian Johnston, Dominic Kwiatkowski, Cordelia Langford, John Sillitoe on behalf of the Wellcome Sanger Institute COVID-19 Surveillance Team |
| EPI_ISL_2828804, EPI_ISL_2828805 | Lighthouse Laboratory Plymouth | Wellcome Sanger Institute for the COVID-19 Genomics UK (COG-UK) Consortium | Lighthouse Laboratory Plymouth and Alex Alderton, Roberto Amato, Jeffrey Barrett, Sonia Goncalves, Ewan Harrison, David K. Jackson, Ian Johnston, Dominic Kwiatkowski, Cordelia Langford, John Sillitoe on behalf of the Wellcome Sanger Institute COVID-19 Surveillance Team |
| EPI_ISL_2828806, EPI_ISL_2828807 | Lighthouse Lab in Alderley Park | Wellcome Sanger Institute for the COVID-19 Genomics UK (COG-UK) Consortium | Jacquelyn Wynn, Mairead Hyland, The Lighthouse Lab in Alderley Park and Alex Alderton, Roberto Amato, Jeffrey Barrett, Sonia Goncalves, Ewan Harrison, David K. Jackson, Ian Johnston, Dominic Kwiatkowski, Cordelia Langford, John Sillitoe on behalf of the Wellcome Sanger Institute COVID-19 Surveillance Team |
| EPI_ISL_2828808 | Lighthouse Laboratory Plymouth | Wellcome Sanger Institute for the COVID-19 Genomics UK (COG-UK) Consortium | Lighthouse Laboratory Plymouth and Alex Alderton, Roberto Amato, Jeffrey Barrett, Sonia Goncalves, Ewan Harrison, David K. Jackson, Ian Johnston, Dominic Kwiatkowski, Cordelia Langford, John Sillitoe on behalf of the Wellcome Sanger Institute COVID-19 Surveillance Team |
| EPI_ISL_2828809 | Lighthouse Lab in Milton Keynes | Wellcome Sanger Institute for the COVID-19 Genomics UK (COG-UK) Consortium | The Lighthouse Lab in Milton Keynes and Alex Alderton, Roberto Amato, Jeffrey Barrett, Sonia Goncalves, Ewan Harrison, David K. Jackson, Ian Johnston, Dominic Kwiatkowski, Cordelia Langford, John Sillitoe on behalf of the Wellcome Sanger Institute COVID-19 Surveillance Team |
| EPI_ISL_2828810 | Lighthouse Laboratory Plymouth | Wellcome Sanger Institute for the COVID-19 Genomics UK (COG-UK) Consortium | Lighthouse Laboratory Plymouth and Alex Alderton, Roberto Amato, Jeffrey Barrett, Sonia Goncalves, Ewan Harrison, David K. Jackson, Ian Johnston, Dominic Kwiatkowski, Cordelia Langford, John Sillitoe on behalf of the Wellcome Sanger Institute COVID-19 Surveillance Team |
| EPI_ISL_2828811 | Lighthouse Lab in Milton Keynes | Wellcome Sanger Institute for the COVID-19 Genomics UK (COG-UK) Consortium | The Lighthouse Lab in Milton Keynes and Alex Alderton, Roberto Amato, Jeffrey Barrett, Sonia Goncalves, Ewan Harrison, David K. Jackson, Ian Johnston, Dominic Kwiatkowski, Cordelia Langford, John Sillitoe on behalf of the Wellcome Sanger Institute COVID-19 Surveillance Team |
| EPI_ISL_2828812 | Lighthouse Lab in Alderley Park | Wellcome Sanger Institute for the COVID-19 Genomics UK (COG-UK) Consortium | Jacquelyn Wynn, Mairead Hyland, The Lighthouse Lab in Alderley Park and Alex Alderton, Roberto Amato, Jeffrey Barrett, Sonia Goncalves, Ewan Harrison, David K. Jackson, Ian Johnston, Dominic Kwiatkowski, Cordelia Langford, John Sillitoe on behalf of the Wellcome Sanger Institute COVID-19 Surveillance Team |
| EPI_ISL_2828813 | Lighthouse Lab in Milton Keynes | Wellcome Sanger Institute for the COVID-19 Genomics UK (COG-UK) Consortium | The Lighthouse Lab in Milton Keynes and Alex Alderton, Roberto Amato, Jeffrey Barrett, Sonia Goncalves, Ewan Harrison, David K. Jackson, Ian Johnston, Dominic Kwiatkowski, Cordelia Langford, John Sillitoe on behalf of the Wellcome Sanger Institute COVID-19 Surveillance Team |
| EPI_ISL_2828814 | Lighthouse Lab in Alderley Park | Wellcome Sanger Institute for the COVID-19 Genomics UK (COG-UK) Consortium | Jacquelyn Wynn, Mairead Hyland, The Lighthouse Lab in Alderley Park and Alex Alderton, Roberto Amato, Jeffrey Barrett, Sonia Goncalves, Ewan Harrison, David K. Jackson, Ian Johnston, Dominic Kwiatkowski, Cordelia Langford, John Sillitoe on behalf of the Wellcome Sanger Institute COVID-19 Surveillance Team |
| EPI_ISL_2828815 | Lighthouse Laboratory Plymouth | Wellcome Sanger Institute for the COVID-19 Genomics UK (COG-UK) Consortium | Lighthouse Laboratory Plymouth and Alex Alderton, Roberto Amato, Jeffrey Barrett, Sonia Goncalves, Ewan Harrison, David K. Jackson, Ian Johnston, Dominic Kwiatkowski, Cordelia Langford, John Sillitoe on behalf of the Wellcome Sanger Institute COVID-19 Surveillance Team |
| EPI_ISL_2828816 | Lighthouse Lab in Alderley Park | Wellcome Sanger Institute for the COVID-19 Genomics UK (COG-UK) Consortium | Jacquelyn Wynn, Mairead Hyland, The Lighthouse Lab in Alderley Park and Alex Alderton, Roberto Amato, Jeffrey Barrett, Sonia Goncalves, Ewan Harrison, David K. Jackson, Ian Johnston, Dominic Kwiatkowski, Cordelia Langford, John Sillitoe on behalf of the Wellcome Sanger Institute COVID-19 Surveillance Team |
| EPI_ISL_2828817 | Lighthouse Lab in Milton Keynes | Wellcome Sanger Institute for the COVID-19 Genomics UK (COG-UK) Consortium | The Lighthouse Lab in Milton Keynes and Alex Alderton, Roberto Amato, Jeffrey Barrett, Sonia Goncalves, Ewan Harrison, David K. Jackson, Ian Johnston, Dominic Kwiatkowski, Cordelia Langford, John Sillitoe on behalf of the Wellcome Sanger Institute COVID-19 Surveillance Team |
| EPI_ISL_2828818 | Lighthouse Lab in Alderley Park | Wellcome Sanger Institute for the COVID-19 Genomics UK (COG-UK) Consortium | Jacquelyn Wynn, Mairead Hyland, The Lighthouse Lab in Alderley Park and Alex Alderton, Roberto Amato, Jeffrey Barrett, Sonia Goncalves, Ewan Harrison, David K. Jackson, Ian Johnston, Dominic Kwiatkowski, Cordelia Langford, John Sillitoe on behalf of the Wellcome Sanger Institute COVID-19 Surveillance Team |
| EPI_ISL_2828819, EPI_ISL_2828820, EPI_ISL_2828821 | Lighthouse Lab in Milton Keynes | Wellcome Sanger Institute for the COVID-19 Genomics UK (COG-UK) Consortium | The Lighthouse Lab in Milton Keynes and Alex Alderton, Roberto Amato, Jeffrey Barrett, Sonia Goncalves, Ewan Harrison, David K. Jackson, Ian Johnston, Dominic Kwiatkowski, Cordelia Langford, John Sillitoe on behalf of the Wellcome Sanger Institute COVID-19 Surveillance Team |
| EPI_ISL_2828822 | Lighthouse Lab in Alderley Park | Wellcome Sanger Institute for the COVID-19 Genomics UK (COG-UK) Consortium | Jacquelyn Wynn, Mairead Hyland, The Lighthouse Lab in Alderley Park and Alex Alderton, Roberto Amato, Jeffrey Barrett, Sonia Goncalves, Ewan Harrison, David K. Jackson, Ian Johnston, Dominic Kwiatkowski, Cordelia Langford, John Sillitoe on behalf of the Wellcome Sanger Institute COVID-19 Surveillance Team |
| EPI_ISL_2828823 | Lighthouse Lab in Milton Keynes | Wellcome Sanger Institute for the COVID-19 Genomics UK (COG-UK) Consortium | The Lighthouse Lab in Milton Keynes and Alex Alderton, Roberto Amato, Jeffrey Barrett, Sonia Goncalves, Ewan Harrison, David K. Jackson, Ian Johnston, Dominic Kwiatkowski, Cordelia Langford, John Sillitoe on behalf of the Wellcome Sanger Institute COVID-19 Surveillance Team |
| EPI_ISL_2828824 | Lighthouse Laboratory Plymouth | Wellcome Sanger Institute for the COVID-19 Genomics UK (COG-UK) Consortium | Lighthouse Laboratory Plymouth and Alex Alderton, Roberto Amato, Jeffrey Barrett, Sonia Goncalves, Ewan Harrison, David K. Jackson, Ian Johnston, Dominic Kwiatkowski, Cordelia Langford, John Sillitoe on behalf of the Wellcome Sanger Institute COVID-19 Surveillance Team |
| EPI_ISL_2828825, EPI_ISL_2828826, EPI_ISL_2828827 | Lighthouse Lab in Milton Keynes | Wellcome Sanger Institute for the COVID-19 Genomics UK (COG-UK) Consortium | The Lighthouse Lab in Milton Keynes and Alex Alderton, Roberto Amato, Jeffrey Barrett, Sonia Goncalves, Ewan Harrison, David K. Jackson, Ian Johnston, Dominic Kwiatkowski, Cordelia Langford, John Sillitoe on behalf of the Wellcome Sanger Institute COVID-19 Surveillance Team |
| EPI_ISL_2828828, EPI_ISL_2828829 | Lighthouse Lab in Alderley Park | Wellcome Sanger Institute for the COVID-19 Genomics UK (COG-UK) Consortium | Jacquelyn Wynn, Mairead Hyland, The Lighthouse Lab in Alderley Park and Alex Alderton, Roberto Amato, Jeffrey Barrett, Sonia Goncalves, Ewan Harrison, David K. Jackson, Ian Johnston, Dominic Kwiatkowski, Cordelia Langford, John Sillitoe on behalf of the Wellcome Sanger Institute COVID-19 Surveillance Team |
| EPI_ISL_2828830 | Lighthouse Laboratory Plymouth | Wellcome Sanger Institute for the COVID-19 Genomics UK (COG-UK) Consortium | Lighthouse Laboratory Plymouth and Alex Alderton, Roberto Amato, Jeffrey Barrett, Sonia Goncalves, Ewan Harrison, David K. Jackson, Ian Johnston, Dominic Kwiatkowski, Cordelia Langford, John Sillitoe on behalf of the Wellcome Sanger Institute COVID-19 Surveillance Team |
| EPI_ISL_2828831 | Lighthouse Lab in Milton Keynes | Wellcome Sanger Institute for the COVID-19 Genomics UK (COG-UK) Consortium | The Lighthouse Lab in Milton Keynes and Alex Alderton, Roberto Amato, Jeffrey Barrett, Sonia Goncalves, Ewan Harrison, David K. Jackson, Ian Johnston, Dominic Kwiatkowski, Cordelia Langford, John Sillitoe on behalf of the Wellcome Sanger Institute COVID-19 Surveillance Team |
| EPI_ISL_2828832 | Lighthouse Laboratory Plymouth | Wellcome Sanger Institute for the COVID-19 Genomics UK (COG-UK) | Lighthouse Laboratory Plymouth and Alex Alderton, Roberto Amato, Jeffrey Barrett, Sonia Goncalves, Ewan Harrison, David K. Jackson, Ian Johnston, Dominic Kwiatkowski, Cordelia Langford, John Sillitoe on behalf of the Wellcome Sanger |

| | | | |
|---|---|---|---|
| | | Consortium | Institute COVID-19 Surveillance Team |
| EPI_ISL_2828833 | Lighthouse Lab in Milton Keynes | Wellcome Sanger Institute for the COVID-19 Genomics UK (COG-UK) Consortium | The Lighthouse Lab in Milton Keynes and Alex Alderton, Roberto Amato, Jeffrey Barrett, Sonia Goncalves, Ewan Harrison, David K. Jackson, Ian Johnston, Dominic Kwiatkowski, Cordelia Langford, John Sillitoe on behalf of the Wellcome Sanger Institute COVID-19 Surveillance Team |
| EPI_ISL_2828834 | Lighthouse Lab in Alderley Park | Wellcome Sanger Institute for the COVID-19 Genomics UK (COG-UK) Consortium | Jacquelyn Wynn, Mairead Hyland, The Lighthouse Lab in Alderley Park and Alex Alderton, Roberto Amato, Jeffrey Barrett, Sonia Goncalves, Ewan Harrison, David K. Jackson, Ian Johnston, Dominic Kwiatkowski, Cordelia Langford, John Sillitoe on behalf of the Wellcome Sanger Institute COVID-19 Surveillance Team |
| EPI_ISL_2828835 | Lighthouse Lab in Milton Keynes | Wellcome Sanger Institute for the COVID-19 Genomics UK (COG-UK) Consortium | The Lighthouse Lab in Milton Keynes and Alex Alderton, Roberto Amato, Jeffrey Barrett, Sonia Goncalves, Ewan Harrison, David K. Jackson, Ian Johnston, Dominic Kwiatkowski, Cordelia Langford, John Sillitoe on behalf of the Wellcome Sanger Institute COVID-19 Surveillance Team |
| EPI_ISL_2828836, EPI_ISL_2828837 | Lighthouse Laboratory Plymouth | Wellcome Sanger Institute for the COVID-19 Genomics UK (COG-UK) Consortium | Lighthouse Laboratory Plymouth and Alex Alderton, Roberto Amato, Jeffrey Barrett, Sonia Goncalves, Ewan Harrison, David K. Jackson, Ian Johnston, Dominic Kwiatkowski, Cordelia Langford, John Sillitoe on behalf of the Wellcome Sanger Institute COVID-19 Surveillance Team |
| EPI_ISL_2828838, EPI_ISL_2828839, EPI_ISL_2828840 | Lighthouse Lab in Alderley Park | Wellcome Sanger Institute for the COVID-19 Genomics UK (COG-UK) Consortium | Jacquelyn Wynn, Mairead Hyland, The Lighthouse Lab in Alderley Park and Alex Alderton, Roberto Amato, Jeffrey Barrett, Sonia Goncalves, Ewan Harrison, David K. Jackson, Ian Johnston, Dominic Kwiatkowski, Cordelia Langford, John Sillitoe on behalf of the Wellcome Sanger Institute COVID-19 Surveillance Team |
| EPI_ISL_2828841 | Lighthouse Laboratory Plymouth | Wellcome Sanger Institute for the COVID-19 Genomics UK (COG-UK) Consortium | Lighthouse Laboratory Plymouth and Alex Alderton, Roberto Amato, Jeffrey Barrett, Sonia Goncalves, Ewan Harrison, David K. Jackson, Ian Johnston, Dominic Kwiatkowski, Cordelia Langford, John Sillitoe on behalf of the Wellcome Sanger Institute COVID-19 Surveillance Team |
| EPI_ISL_2828842, EPI_ISL_2828843 | Lighthouse Lab in Alderley Park | Wellcome Sanger Institute for the COVID-19 Genomics UK (COG-UK) Consortium | Jacquelyn Wynn, Mairead Hyland, The Lighthouse Lab in Alderley Park and Alex Alderton, Roberto Amato, Jeffrey Barrett, Sonia Goncalves, Ewan Harrison, David K. Jackson, Ian Johnston, Dominic Kwiatkowski, Cordelia Langford, John Sillitoe on behalf of the Wellcome Sanger Institute COVID-19 Surveillance Team |
| EPI_ISL_2828844 | Lighthouse Laboratory Plymouth | Wellcome Sanger Institute for the COVID-19 Genomics UK (COG-UK) Consortium | Lighthouse Laboratory Plymouth and Alex Alderton, Roberto Amato, Jeffrey Barrett, Sonia Goncalves, Ewan Harrison, David K. Jackson, Ian Johnston, Dominic Kwiatkowski, Cordelia Langford, John Sillitoe on behalf of the Wellcome Sanger Institute COVID-19 Surveillance Team |
| EPI_ISL_2828845 | Lighthouse Lab in Milton Keynes | Wellcome Sanger Institute for the COVID-19 Genomics UK (COG-UK) Consortium | The Lighthouse Lab in Milton Keynes and Alex Alderton, Roberto Amato, Jeffrey Barrett, Sonia Goncalves, Ewan Harrison, David K. Jackson, Ian Johnston, Dominic Kwiatkowski, Cordelia Langford, John Sillitoe on behalf of the Wellcome Sanger Institute COVID-19 Surveillance Team |
| EPI_ISL_2828846 | Lighthouse Laboratory Plymouth | Wellcome Sanger Institute for the COVID-19 Genomics UK (COG-UK) Consortium | Lighthouse Laboratory Plymouth and Alex Alderton, Roberto Amato, Jeffrey Barrett, Sonia Goncalves, Ewan Harrison, David K. Jackson, Ian Johnston, Dominic Kwiatkowski, Cordelia Langford, John Sillitoe on behalf of the Wellcome Sanger Institute COVID-19 Surveillance Team |
| EPI_ISL_2828847 | Lighthouse Lab in Milton Keynes | Wellcome Sanger Institute for the COVID-19 Genomics UK (COG-UK) Consortium | The Lighthouse Lab in Milton Keynes and Alex Alderton, Roberto Amato, Jeffrey Barrett, Sonia Goncalves, Ewan Harrison, David K. Jackson, Ian Johnston, Dominic Kwiatkowski, Cordelia Langford, John Sillitoe on behalf of the Wellcome Sanger Institute COVID-19 Surveillance Team |
| EPI_ISL_2828848, EPI_ISL_2828849 | Lighthouse Lab in Alderley Park | Wellcome Sanger Institute for the COVID-19 Genomics UK (COG-UK) Consortium | Jacquelyn Wynn, Mairead Hyland, The Lighthouse Lab in Alderley Park and Alex Alderton, Roberto Amato, Jeffrey Barrett, Sonia Goncalves, Ewan Harrison, David K. Jackson, Ian Johnston, Dominic Kwiatkowski, Cordelia Langford, John Sillitoe on behalf of the Wellcome Sanger Institute COVID-19 Surveillance Team |
| EPI_ISL_2828850 | Lighthouse Lab in Milton Keynes | Wellcome Sanger Institute for the COVID-19 Genomics UK (COG-UK) Consortium | The Lighthouse Lab in Milton Keynes and Alex Alderton, Roberto Amato, Jeffrey Barrett, Sonia Goncalves, Ewan Harrison, David K. Jackson, Ian Johnston, Dominic Kwiatkowski, Cordelia Langford, John Sillitoe on behalf of the Wellcome Sanger Institute COVID-19 Surveillance Team |
| EPI_ISL_2828851, EPI_ISL_2828852 | Lighthouse Lab in Alderley Park | Wellcome Sanger Institute for the COVID-19 Genomics UK (COG-UK) Consortium | Jacquelyn Wynn, Mairead Hyland, The Lighthouse Lab in Alderley Park and Alex Alderton, Roberto Amato, Jeffrey Barrett, Sonia Goncalves, Ewan Harrison, David K. Jackson, Ian Johnston, Dominic Kwiatkowski, Cordelia Langford, John Sillitoe on behalf of the Wellcome Sanger Institute COVID-19 Surveillance Team |
| EPI_ISL_2828853 | Lighthouse Laboratory Plymouth | Wellcome Sanger Institute for the COVID-19 Genomics UK (COG-UK) Consortium | Lighthouse Laboratory Plymouth and Alex Alderton, Roberto Amato, Jeffrey Barrett, Sonia Goncalves, Ewan Harrison, David K. Jackson, Ian Johnston, Dominic Kwiatkowski, Cordelia Langford, John Sillitoe on behalf of the Wellcome Sanger Institute COVID-19 Surveillance Team |
| EPI_ISL_2828854 | Lighthouse Lab in Milton Keynes | Wellcome Sanger Institute for the COVID-19 Genomics UK (COG-UK) Consortium | The Lighthouse Lab in Milton Keynes and Alex Alderton, Roberto Amato, Jeffrey Barrett, Sonia Goncalves, Ewan Harrison, David K. Jackson, Ian Johnston, Dominic Kwiatkowski, Cordelia Langford, John Sillitoe on behalf of the Wellcome Sanger Institute COVID-19 Surveillance Team |
| EPI_ISL_2828855 | Lighthouse Lab in Alderley Park | Wellcome Sanger Institute for the COVID-19 Genomics UK (COG-UK) Consortium | Jacquelyn Wynn, Mairead Hyland, The Lighthouse Lab in Alderley Park and Alex Alderton, Roberto Amato, Jeffrey Barrett, Sonia Goncalves, Ewan Harrison, David K. Jackson, Ian Johnston, Dominic Kwiatkowski, Cordelia Langford, John Sillitoe on behalf of the Wellcome Sanger Institute COVID-19 Surveillance Team |
| EPI_ISL_2828856, EPI_ISL_2828857 | Lighthouse Lab in Milton Keynes | Wellcome Sanger Institute for the COVID-19 Genomics UK (COG-UK) Consortium | The Lighthouse Lab in Milton Keynes and Alex Alderton, Roberto Amato, Jeffrey Barrett, Sonia Goncalves, Ewan Harrison, David K. Jackson, Ian Johnston, Dominic Kwiatkowski, Cordelia Langford, John Sillitoe on behalf of the Wellcome Sanger Institute COVID-19 Surveillance Team |
| EPI_ISL_2828858 | Lighthouse Laboratory Plymouth | Wellcome Sanger Institute for the COVID-19 Genomics UK (COG-UK) Consortium | Lighthouse Laboratory Plymouth and Alex Alderton, Roberto Amato, Jeffrey Barrett, Sonia Goncalves, Ewan Harrison, David K. Jackson, Ian Johnston, Dominic Kwiatkowski, Cordelia Langford, John Sillitoe on behalf of the Wellcome Sanger Institute COVID-19 Surveillance Team |
| EPI_ISL_2828859, EPI_ISL_2828860, EPI_ISL_2828861 | Lighthouse Lab in Alderley Park | Wellcome Sanger Institute for the COVID-19 Genomics UK (COG-UK) Consortium | Jacquelyn Wynn, Mairead Hyland, The Lighthouse Lab in Alderley Park and Alex Alderton, Roberto Amato, Jeffrey Barrett, Sonia Goncalves, Ewan Harrison, David K. Jackson, Ian Johnston, Dominic Kwiatkowski, Cordelia Langford, John Sillitoe on behalf of the Wellcome Sanger Institute COVID-19 Surveillance Team |
| EPI_ISL_2828862 | Lighthouse Laboratory Plymouth | Wellcome Sanger Institute for the COVID-19 Genomics UK (COG-UK) Consortium | Lighthouse Laboratory Plymouth and Alex Alderton, Roberto Amato, Jeffrey Barrett, Sonia Goncalves, Ewan Harrison, David K. Jackson, Ian Johnston, Dominic Kwiatkowski, Cordelia Langford, John Sillitoe on behalf of the Wellcome Sanger Institute COVID-19 Surveillance Team |
| EPI_ISL_2828863 | Lighthouse Lab in Milton Keynes | Wellcome Sanger Institute for the COVID-19 Genomics UK (COG-UK) Consortium | The Lighthouse Lab in Milton Keynes and Alex Alderton, Roberto Amato, Jeffrey Barrett, Sonia Goncalves, Ewan Harrison, David K. Jackson, Ian Johnston, Dominic Kwiatkowski, Cordelia Langford, John Sillitoe on behalf of the Wellcome Sanger Institute COVID-19 Surveillance Team |
| EPI_ISL_2828864 | Lighthouse Lab in Alderley Park | Wellcome Sanger Institute for the COVID-19 Genomics UK (COG-UK) Consortium | Jacquelyn Wynn, Mairead Hyland, The Lighthouse Lab in Alderley Park and Alex Alderton, Roberto Amato, Jeffrey Barrett, Sonia Goncalves, Ewan Harrison, David K. Jackson, Ian Johnston, Dominic Kwiatkowski, Cordelia Langford, John Sillitoe on behalf of the Wellcome Sanger Institute COVID-19 Surveillance Team |
| EPI_ISL_2828865, EPI_ISL_2828866 | Lighthouse Laboratory Plymouth | Wellcome Sanger Institute for the COVID-19 Genomics UK (COG-UK) Consortium | Lighthouse Laboratory Plymouth and Alex Alderton, Roberto Amato, Jeffrey Barrett, Sonia Goncalves, Ewan Harrison, David K. Jackson, Ian Johnston, Dominic Kwiatkowski, Cordelia Langford, John Sillitoe on behalf of the Wellcome Sanger Institute COVID-19 Surveillance Team |
| EPI_ISL_2828867 | Lighthouse Lab in Alderley Park | Wellcome Sanger Institute for the COVID-19 Genomics UK (COG-UK) Consortium | Jacquelyn Wynn, Mairead Hyland, The Lighthouse Lab in Alderley Park and Alex Alderton, Roberto Amato, Jeffrey Barrett, Sonia Goncalves, Ewan Harrison, David K. Jackson, Ian Johnston, Dominic Kwiatkowski, Cordelia Langford, John Sillitoe on behalf of the Wellcome Sanger Institute COVID-19 Surveillance Team |
| EPI_ISL_2828868 | Lighthouse Lab in Milton Keynes | Wellcome Sanger Institute for the COVID-19 Genomics UK (COG-UK) Consortium | The Lighthouse Lab in Milton Keynes and Alex Alderton, Roberto Amato, Jeffrey Barrett, Sonia Goncalves, Ewan Harrison, David K. Jackson, Ian Johnston, Dominic Kwiatkowski, Cordelia Langford, John Sillitoe on behalf of the Wellcome Sanger Institute COVID-19 Surveillance Team |
| EPI_ISL_2828869, EPI_ISL_2828870 | Lighthouse Laboratory Plymouth | Wellcome Sanger Institute for the COVID-19 Genomics UK (COG-UK) Consortium | Lighthouse Laboratory Plymouth and Alex Alderton, Roberto Amato, Jeffrey Barrett, Sonia Goncalves, Ewan Harrison, David K. Jackson, Ian Johnston, Dominic Kwiatkowski, Cordelia Langford, John Sillitoe on behalf of the Wellcome Sanger Institute COVID-19 Surveillance Team |
| EPI_ISL_2828871, EPI_ISL_2828872 | Lighthouse Lab in Alderley Park | Wellcome Sanger Institute for the COVID-19 Genomics UK (COG-UK) Consortium | Jacquelyn Wynn, Mairead Hyland, The Lighthouse Lab in Alderley Park and Alex Alderton, Roberto Amato, Jeffrey Barrett, Sonia Goncalves, Ewan Harrison, David K. Jackson, Ian Johnston, Dominic Kwiatkowski, Cordelia Langford, John Sillitoe on behalf of the Wellcome Sanger Institute COVID-19 Surveillance Team |
| EPI_ISL_2828873, EPI_ISL_2828874, EPI_ISL_2828875 | Lighthouse Lab in Milton Keynes | Wellcome Sanger Institute for the COVID-19 Genomics UK (COG-UK) Consortium | The Lighthouse Lab in Milton Keynes and Alex Alderton, Roberto Amato, Jeffrey Barrett, Sonia Goncalves, Ewan Harrison, David K. Jackson, Ian Johnston, Dominic Kwiatkowski, Cordelia Langford, John Sillitoe on behalf of the Wellcome Sanger Institute COVID-19 Surveillance Team |
| EPI_ISL_2828876 | Lighthouse Lab in Alderley Park | Wellcome Sanger Institute for the COVID-19 Genomics UK (COG-UK) Consortium | Jacquelyn Wynn, Mairead Hyland, The Lighthouse Lab in Alderley Park and Alex Alderton, Roberto Amato, Jeffrey Barrett, Sonia Goncalves, Ewan Harrison, David K. Jackson, Ian Johnston, Dominic Kwiatkowski, Cordelia Langford, John Sillitoe on behalf of the Wellcome Sanger Institute COVID-19 Surveillance Team |
| EPI_ISL_2828877 | Lighthouse Laboratory Plymouth | Wellcome Sanger Institute for the COVID-19 Genomics UK (COG-UK) Consortium | Lighthouse Laboratory Plymouth and Alex Alderton, Roberto Amato, Jeffrey Barrett, Sonia Goncalves, Ewan Harrison, David K. Jackson, Ian Johnston, Dominic Kwiatkowski, Cordelia Langford, John Sillitoe on behalf of the Wellcome Sanger Institute COVID-19 Surveillance Team |
| EPI_ISL_2828878 | Lighthouse Lab in Milton Keynes | Wellcome Sanger Institute for the COVID-19 Genomics UK (COG-UK) Consortium | The Lighthouse Lab in Milton Keynes and Alex Alderton, Roberto Amato, Jeffrey Barrett, Sonia Goncalves, Ewan Harrison, David K. Jackson, Ian Johnston, Dominic Kwiatkowski, Cordelia Langford, John Sillitoe on behalf of the Wellcome Sanger Institute COVID-19 Surveillance Team |
| EPI_ISL_2828879, EPI_ISL_2828880, EPI_ISL_2828881 | Lighthouse Laboratory Plymouth | Wellcome Sanger Institute for the COVID-19 Genomics UK (COG-UK) Consortium | Lighthouse Laboratory Plymouth and Alex Alderton, Roberto Amato, Jeffrey Barrett, Sonia Goncalves, Ewan Harrison, David K. Jackson, Ian Johnston, Dominic Kwiatkowski, Cordelia Langford, John Sillitoe on behalf of the Wellcome Sanger Institute COVID-19 Surveillance Team |
| EPI_ISL_2828882 | Lighthouse Lab in Alderley Park | Wellcome Sanger Institute for the COVID-19 Genomics UK (COG-UK) Consortium | Jacquelyn Wynn, Mairead Hyland, The Lighthouse Lab in Alderley Park and Alex Alderton, Roberto Amato, Jeffrey Barrett, Sonia Goncalves, Ewan Harrison, David K. Jackson, Ian Johnston, Dominic Kwiatkowski, Cordelia Langford, John Sillitoe on behalf of the Wellcome Sanger Institute COVID-19 Surveillance Team |
| EPI_ISL_2828883, EPI_ISL_2828884 | Lighthouse Lab in Milton Keynes | Wellcome Sanger Institute for the COVID-19 Genomics UK (COG-UK) Consortium | The Lighthouse Lab in Milton Keynes and Alex Alderton, Roberto Amato, Jeffrey Barrett, Sonia Goncalves, Ewan Harrison, David K. Jackson, Ian Johnston, Dominic Kwiatkowski, Cordelia Langford, John Sillitoe on behalf of the Wellcome Sanger Institute COVID-19 Surveillance Team |
| EPI_ISL_2828885 | Lighthouse Laboratory Plymouth | Wellcome Sanger Institute for the COVID-19 Genomics UK (COG-UK) Consortium | Lighthouse Laboratory Plymouth and Alex Alderton, Roberto Amato, Jeffrey Barrett, Sonia Goncalves, Ewan Harrison, David K. Jackson, Ian Johnston, Dominic Kwiatkowski, Cordelia Langford, John Sillitoe on behalf of the Wellcome Sanger Institute COVID-19 Surveillance Team |
| EPI_ISL_2828886, EPI_ISL_2828887, EPI_ISL_2828888 | Lighthouse Lab in Alderley Park | Wellcome Sanger Institute for the COVID-19 Genomics UK (COG-UK) Consortium | Jacquelyn Wynn, Mairead Hyland, The Lighthouse Lab in Alderley Park and Alex Alderton, Roberto Amato, Jeffrey Barrett, Sonia Goncalves, Ewan Harrison, David K. Jackson, Ian Johnston, Dominic Kwiatkowski, Cordelia Langford, John Sillitoe on behalf of the Wellcome Sanger Institute COVID-19 Surveillance Team |
| EPI_ISL_2828889 | Lighthouse Lab in Milton Keynes | Wellcome Sanger Institute for the COVID-19 Genomics UK (COG-UK) Consortium | The Lighthouse Lab in Milton Keynes and Alex Alderton, Roberto Amato, Jeffrey Barrett, Sonia Goncalves, Ewan Harrison, David K. Jackson, Ian Johnston, Dominic Kwiatkowski, Cordelia Langford, John Sillitoe on behalf of the Wellcome Sanger Institute COVID-19 Surveillance Team |
| EPI_ISL_2828890, EPI_ISL_2828891 | Lighthouse Lab in Alderley Park | Wellcome Sanger Institute for the COVID-19 Genomics UK (COG-UK) Consortium | Jacquelyn Wynn, Mairead Hyland, The Lighthouse Lab in Alderley Park and Alex Alderton, Roberto Amato, Jeffrey Barrett, Sonia Goncalves, Ewan Harrison, David K. Jackson, Ian Johnston, Dominic Kwiatkowski, Cordelia Langford, John Sillitoe on behalf of the Wellcome Sanger Institute COVID-19 Surveillance Team |
| EPI_ISL_2828892 | Lighthouse Laboratory Plymouth | Wellcome Sanger Institute for the COVID-19 Genomics UK (COG-UK) Consortium | Lighthouse Laboratory Plymouth and Alex Alderton, Roberto Amato, Jeffrey Barrett, Sonia Goncalves, Ewan Harrison, David K. Jackson, Ian Johnston, Dominic Kwiatkowski, Cordelia Langford, John Sillitoe on behalf of the Wellcome Sanger Institute COVID-19 Surveillance Team |
| EPI_ISL_2828893, EPI_ISL_2828894 | Lighthouse Lab in Milton Keynes | Wellcome Sanger Institute for the COVID-19 Genomics UK (COG-UK) Consortium | The Lighthouse Lab in Milton Keynes and Alex Alderton, Roberto Amato, Jeffrey Barrett, Sonia Goncalves, Ewan Harrison, David K. Jackson, Ian Johnston, Dominic Kwiatkowski, Cordelia Langford, John Sillitoe on behalf of the Wellcome Sanger Institute COVID-19 Surveillance Team |
| EPI_ISL_2828895 | Lighthouse Laboratory Plymouth | Wellcome Sanger Institute for the COVID-19 Genomics UK (COG-UK) Consortium | Lighthouse Laboratory Plymouth and Alex Alderton, Roberto Amato, Jeffrey Barrett, Sonia Goncalves, Ewan Harrison, David K. Jackson, Ian Johnston, Dominic Kwiatkowski, Cordelia Langford, John Sillitoe on behalf of the Wellcome Sanger Institute COVID-19 Surveillance Team |
| EPI_ISL_2828896 | Lighthouse Lab in Alderley Park | Wellcome Sanger Institute for the COVID-19 Genomics UK (COG-UK) | Jacquelyn Wynn, Mairead Hyland, The Lighthouse Lab in Alderley Park and Alex Alderton, Roberto Amato, Jeffrey Barrett, Sonia Goncalves, Ewan Harrison, David K. Jackson, Ian Johnston, Dominic Kwiatkowski, Cordelia Langford, John Sillitoe on |

| | | | |
|---|---|---|---|
| | | Consortium | behalf of the Wellcome Sanger Institute COVID-19 Surveillance Team |
| EPI_ISL_2828897, EPI_ISL_2828898, EPI_ISL_2828899 | Lighthouse Laboratory Plymouth | Wellcome Sanger Institute for the COVID-19 Genomics UK (COG-UK) Consortium | Lighthouse Laboratory Plymouth and Alex Alderton, Roberto Amato, Jeffrey Barrett, Sonia Goncalves, Ewan Harrison, David K. Jackson, Ian Johnston, Dominic Kwiatkowski, Cordelia Langford, John Sillitoe on behalf of the Wellcome Sanger Institute COVID-19 Surveillance Team |
| EPI_ISL_2828900, EPI_ISL_2828901, EPI_ISL_2828902 | Lighthouse Lab in Milton Keynes | Wellcome Sanger Institute for the COVID-19 Genomics UK (COG-UK) Consortium | The Lighthouse Lab in Milton Keynes and Alex Alderton, Roberto Amato, Jeffrey Barrett, Sonia Goncalves, Ewan Harrison, David K. Jackson, Ian Johnston, Dominic Kwiatkowski, Cordelia Langford, John Sillitoe on behalf of the Wellcome Sanger Institute COVID-19 Surveillance Team |
| EPI_ISL_2828903 | Lighthouse Laboratory Plymouth | Wellcome Sanger Institute for the COVID-19 Genomics UK (COG-UK) Consortium | Lighthouse Laboratory Plymouth and Alex Alderton, Roberto Amato, Jeffrey Barrett, Sonia Goncalves, Ewan Harrison, David K. Jackson, Ian Johnston, Dominic Kwiatkowski, Cordelia Langford, John Sillitoe on behalf of the Wellcome Sanger Institute COVID-19 Surveillance Team |
| EPI_ISL_2828904, EPI_ISL_2828905 | Lighthouse Lab in Milton Keynes | Wellcome Sanger Institute for the COVID-19 Genomics UK (COG-UK) Consortium | The Lighthouse Lab in Milton Keynes and Alex Alderton, Roberto Amato, Jeffrey Barrett, Sonia Goncalves, Ewan Harrison, David K. Jackson, Ian Johnston, Dominic Kwiatkowski, Cordelia Langford, John Sillitoe on behalf of the Wellcome Sanger Institute COVID-19 Surveillance Team |
| EPI_ISL_2828906 | Lighthouse Lab in Alderley Park | Wellcome Sanger Institute for the COVID-19 Genomics UK (COG-UK) Consortium | Jacquelyn Wynn, Mairead Hyland, The Lighthouse Lab in Alderley Park and Alex Alderton, Roberto Amato, Jeffrey Barrett, Sonia Goncalves, Ewan Harrison, David K. Jackson, Ian Johnston, Dominic Kwiatkowski, Cordelia Langford, John Sillitoe on behalf of the Wellcome Sanger Institute COVID-19 Surveillance Team |
| EPI_ISL_2828907, EPI_ISL_2828908 | Lighthouse Lab in Milton Keynes | Wellcome Sanger Institute for the COVID-19 Genomics UK (COG-UK) Consortium | The Lighthouse Lab in Milton Keynes and Alex Alderton, Roberto Amato, Jeffrey Barrett, Sonia Goncalves, Ewan Harrison, David K. Jackson, Ian Johnston, Dominic Kwiatkowski, Cordelia Langford, John Sillitoe on behalf of the Wellcome Sanger Institute COVID-19 Surveillance Team |
| EPI_ISL_2828909, EPI_ISL_2828910, EPI_ISL_2828911 | Lighthouse Laboratory Plymouth | Wellcome Sanger Institute for the COVID-19 Genomics UK (COG-UK) Consortium | Lighthouse Laboratory Plymouth and Alex Alderton, Roberto Amato, Jeffrey Barrett, Sonia Goncalves, Ewan Harrison, David K. Jackson, Ian Johnston, Dominic Kwiatkowski, Cordelia Langford, John Sillitoe on behalf of the Wellcome Sanger Institute COVID-19 Surveillance Team |
| EPI_ISL_2828912, EPI_ISL_2828913 | Lighthouse Lab in Alderley Park | Wellcome Sanger Institute for the COVID-19 Genomics UK (COG-UK) Consortium | Jacquelyn Wynn, Mairead Hyland, The Lighthouse Lab in Alderley Park and Alex Alderton, Roberto Amato, Jeffrey Barrett, Sonia Goncalves, Ewan Harrison, David K. Jackson, Ian Johnston, Dominic Kwiatkowski, Cordelia Langford, John Sillitoe on behalf of the Wellcome Sanger Institute COVID-19 Surveillance Team |
| EPI_ISL_2828914 | Lighthouse Lab in Milton Keynes | Wellcome Sanger Institute for the COVID-19 Genomics UK (COG-UK) Consortium | The Lighthouse Lab in Milton Keynes and Alex Alderton, Roberto Amato, Jeffrey Barrett, Sonia Goncalves, Ewan Harrison, David K. Jackson, Ian Johnston, Dominic Kwiatkowski, Cordelia Langford, John Sillitoe on behalf of the Wellcome Sanger Institute COVID-19 Surveillance Team |
| EPI_ISL_2828915 | Lighthouse Laboratory Plymouth | Wellcome Sanger Institute for the COVID-19 Genomics UK (COG-UK) Consortium | Lighthouse Laboratory Plymouth and Alex Alderton, Roberto Amato, Jeffrey Barrett, Sonia Goncalves, Ewan Harrison, David K. Jackson, Ian Johnston, Dominic Kwiatkowski, Cordelia Langford, John Sillitoe on behalf of the Wellcome Sanger Institute COVID-19 Surveillance Team |
| EPI_ISL_2828916, EPI_ISL_2828917, EPI_ISL_2828918 | Lighthouse Lab in Alderley Park | Wellcome Sanger Institute for the COVID-19 Genomics UK (COG-UK) Consortium | Jacquelyn Wynn, Mairead Hyland, The Lighthouse Lab in Alderley Park and Alex Alderton, Roberto Amato, Jeffrey Barrett, Sonia Goncalves, Ewan Harrison, David K. Jackson, Ian Johnston, Dominic Kwiatkowski, Cordelia Langford, John Sillitoe on behalf of the Wellcome Sanger Institute COVID-19 Surveillance Team |
| EPI_ISL_2828919, EPI_ISL_2828920, EPI_ISL_2828921, EPI_ISL_2828922 | Lighthouse Lab in Milton Keynes | Wellcome Sanger Institute for the COVID-19 Genomics UK (COG-UK) Consortium | The Lighthouse Lab in Milton Keynes and Alex Alderton, Roberto Amato, Jeffrey Barrett, Sonia Goncalves, Ewan Harrison, David K. Jackson, Ian Johnston, Dominic Kwiatkowski, Cordelia Langford, John Sillitoe on behalf of the Wellcome Sanger Institute COVID-19 Surveillance Team |
| EPI_ISL_2828923 | Lighthouse Laboratory Plymouth | Wellcome Sanger Institute for the COVID-19 Genomics UK (COG-UK) Consortium | Lighthouse Laboratory Plymouth and Alex Alderton, Roberto Amato, Jeffrey Barrett, Sonia Goncalves, Ewan Harrison, David K. Jackson, Ian Johnston, Dominic Kwiatkowski, Cordelia Langford, John Sillitoe on behalf of the Wellcome Sanger Institute COVID-19 Surveillance Team |
| EPI_ISL_2828924, EPI_ISL_2828925, EPI_ISL_2828926, EPI_ISL_2828927 | Lighthouse Lab in Milton Keynes | Wellcome Sanger Institute for the COVID-19 Genomics UK (COG-UK) Consortium | The Lighthouse Lab in Milton Keynes and Alex Alderton, Roberto Amato, Jeffrey Barrett, Sonia Goncalves, Ewan Harrison, David K. Jackson, Ian Johnston, Dominic Kwiatkowski, Cordelia Langford, John Sillitoe on behalf of the Wellcome Sanger Institute COVID-19 Surveillance Team |
| EPI_ISL_2828928, EPI_ISL_2828929, EPI_ISL_2828930 | Lighthouse Laboratory Plymouth | Wellcome Sanger Institute for the COVID-19 Genomics UK (COG-UK) Consortium | Lighthouse Laboratory Plymouth and Alex Alderton, Roberto Amato, Jeffrey Barrett, Sonia Goncalves, Ewan Harrison, David K. Jackson, Ian Johnston, Dominic Kwiatkowski, Cordelia Langford, John Sillitoe on behalf of the Wellcome Sanger Institute COVID-19 Surveillance Team |
| EPI_ISL_2828931, EPI_ISL_2828932 | Lighthouse Lab in Alderley Park | Wellcome Sanger Institute for the COVID-19 Genomics UK (COG-UK) Consortium | Jacquelyn Wynn, Mairead Hyland, The Lighthouse Lab in Alderley Park and Alex Alderton, Roberto Amato, Jeffrey Barrett, Sonia Goncalves, Ewan Harrison, David K. Jackson, Ian Johnston, Dominic Kwiatkowski, Cordelia Langford, John Sillitoe on behalf of the Wellcome Sanger Institute COVID-19 Surveillance Team |
| EPI_ISL_2828933 | Lighthouse Laboratory Plymouth | Wellcome Sanger Institute for the COVID-19 Genomics UK (COG-UK) Consortium | Lighthouse Laboratory Plymouth and Alex Alderton, Roberto Amato, Jeffrey Barrett, Sonia Goncalves, Ewan Harrison, David K. Jackson, Ian Johnston, Dominic Kwiatkowski, Cordelia Langford, John Sillitoe on behalf of the Wellcome Sanger Institute COVID-19 Surveillance Team |
| EPI_ISL_2828934 | Lighthouse Lab in Alderley Park | Wellcome Sanger Institute for the COVID-19 Genomics UK (COG-UK) Consortium | Jacquelyn Wynn, Mairead Hyland, The Lighthouse Lab in Alderley Park and Alex Alderton, Roberto Amato, Jeffrey Barrett, Sonia Goncalves, Ewan Harrison, David K. Jackson, Ian Johnston, Dominic Kwiatkowski, Cordelia Langford, John Sillitoe on behalf of the Wellcome Sanger Institute COVID-19 Surveillance Team |
| EPI_ISL_2828935, EPI_ISL_2828936 | Lighthouse Lab in Milton Keynes | Wellcome Sanger Institute for the COVID-19 Genomics UK (COG-UK) Consortium | The Lighthouse Lab in Milton Keynes and Alex Alderton, Roberto Amato, Jeffrey Barrett, Sonia Goncalves, Ewan Harrison, David K. Jackson, Ian Johnston, Dominic Kwiatkowski, Cordelia Langford, John Sillitoe on behalf of the Wellcome Sanger Institute COVID-19 Surveillance Team |
| EPI_ISL_2828937 | Lighthouse Laboratory Plymouth | Wellcome Sanger Institute for the COVID-19 Genomics UK (COG-UK) Consortium | Lighthouse Laboratory Plymouth and Alex Alderton, Roberto Amato, Jeffrey Barrett, Sonia Goncalves, Ewan Harrison, David K. Jackson, Ian Johnston, Dominic Kwiatkowski, Cordelia Langford, John Sillitoe on behalf of the Wellcome Sanger Institute COVID-19 Surveillance Team |
| EPI_ISL_2828938 | Lighthouse Lab in Alderley Park | Wellcome Sanger Institute for the COVID-19 Genomics UK (COG-UK) Consortium | Jacquelyn Wynn, Mairead Hyland, The Lighthouse Lab in Alderley Park and Alex Alderton, Roberto Amato, Jeffrey Barrett, Sonia Goncalves, Ewan Harrison, David K. Jackson, Ian Johnston, Dominic Kwiatkowski, Cordelia Langford, John Sillitoe on behalf of the Wellcome Sanger Institute COVID-19 Surveillance Team |
| EPI_ISL_2828939 | Lighthouse Laboratory Plymouth | Wellcome Sanger Institute for the COVID-19 Genomics UK (COG-UK) Consortium | Lighthouse Laboratory Plymouth and Alex Alderton, Roberto Amato, Jeffrey Barrett, Sonia Goncalves, Ewan Harrison, David K. Jackson, Ian Johnston, Dominic Kwiatkowski, Cordelia Langford, John Sillitoe on behalf of the Wellcome Sanger Institute COVID-19 Surveillance Team |
| EPI_ISL_2828940 | Lighthouse Lab in Milton Keynes | Wellcome Sanger Institute for the COVID-19 Genomics UK (COG-UK) Consortium | The Lighthouse Lab in Milton Keynes and Alex Alderton, Roberto Amato, Jeffrey Barrett, Sonia Goncalves, Ewan Harrison, David K. Jackson, Ian Johnston, Dominic Kwiatkowski, Cordelia Langford, John Sillitoe on behalf of the Wellcome Sanger Institute COVID-19 Surveillance Team |
| EPI_ISL_2828941, EPI_ISL_2828942 | Lighthouse Laboratory Plymouth | Wellcome Sanger Institute for the COVID-19 Genomics UK (COG-UK) Consortium | Lighthouse Laboratory Plymouth and Alex Alderton, Roberto Amato, Jeffrey Barrett, Sonia Goncalves, Ewan Harrison, David K. Jackson, Ian Johnston, Dominic Kwiatkowski, Cordelia Langford, John Sillitoe on behalf of the Wellcome Sanger Institute COVID-19 Surveillance Team |
| EPI_ISL_2828943 | Lighthouse Lab in Milton Keynes | Wellcome Sanger Institute for the COVID-19 Genomics UK (COG-UK) Consortium | The Lighthouse Lab in Milton Keynes and Alex Alderton, Roberto Amato, Jeffrey Barrett, Sonia Goncalves, Ewan Harrison, David K. Jackson, Ian Johnston, Dominic Kwiatkowski, Cordelia Langford, John Sillitoe on behalf of the Wellcome Sanger Institute COVID-19 Surveillance Team |
| EPI_ISL_2828944 | Lighthouse Laboratory Plymouth | Wellcome Sanger Institute for the COVID-19 Genomics UK (COG-UK) Consortium | Lighthouse Laboratory Plymouth and Alex Alderton, Roberto Amato, Jeffrey Barrett, Sonia Goncalves, Ewan Harrison, David K. Jackson, Ian Johnston, Dominic Kwiatkowski, Cordelia Langford, John Sillitoe on behalf of the Wellcome Sanger Institute COVID-19 Surveillance Team |
| EPI_ISL_2828945 | Lighthouse Lab in Alderley Park | Wellcome Sanger Institute for the COVID-19 Genomics UK (COG-UK) Consortium | Jacquelyn Wynn, Mairead Hyland, The Lighthouse Lab in Alderley Park and Alex Alderton, Roberto Amato, Jeffrey Barrett, Sonia Goncalves, Ewan Harrison, David K. Jackson, Ian Johnston, Dominic Kwiatkowski, Cordelia Langford, John Sillitoe on behalf of the Wellcome Sanger Institute COVID-19 Surveillance Team |
| EPI_ISL_2828946, EPI_ISL_2828947 | Lighthouse Lab in Milton Keynes | Wellcome Sanger Institute for the COVID-19 Genomics UK (COG-UK) Consortium | The Lighthouse Lab in Milton Keynes and Alex Alderton, Roberto Amato, Jeffrey Barrett, Sonia Goncalves, Ewan Harrison, David K. Jackson, Ian Johnston, Dominic Kwiatkowski, Cordelia Langford, John Sillitoe on behalf of the Wellcome Sanger Institute COVID-19 Surveillance Team |
| EPI_ISL_2828948 | Lighthouse Lab in Alderley Park | Wellcome Sanger Institute for the COVID-19 Genomics UK (COG-UK) Consortium | Jacquelyn Wynn, Mairead Hyland, The Lighthouse Lab in Alderley Park and Alex Alderton, Roberto Amato, Jeffrey Barrett, Sonia Goncalves, Ewan Harrison, David K. Jackson, Ian Johnston, Dominic Kwiatkowski, Cordelia Langford, John Sillitoe on behalf of the Wellcome Sanger Institute COVID-19 Surveillance Team |
| EPI_ISL_2828949 | Lighthouse Laboratory Plymouth | Wellcome Sanger Institute for the COVID-19 Genomics UK (COG-UK) Consortium | Lighthouse Laboratory Plymouth and Alex Alderton, Roberto Amato, Jeffrey Barrett, Sonia Goncalves, Ewan Harrison, David K. Jackson, Ian Johnston, Dominic Kwiatkowski, Cordelia Langford, John Sillitoe on behalf of the Wellcome Sanger Institute COVID-19 Surveillance Team |
| EPI_ISL_2828950 | Lighthouse Lab in Milton Keynes | Wellcome Sanger Institute for the COVID-19 Genomics UK (COG-UK) Consortium | The Lighthouse Lab in Milton Keynes and Alex Alderton, Roberto Amato, Jeffrey Barrett, Sonia Goncalves, Ewan Harrison, David K. Jackson, Ian Johnston, Dominic Kwiatkowski, Cordelia Langford, John Sillitoe on behalf of the Wellcome Sanger Institute COVID-19 Surveillance Team |
| EPI_ISL_2828951, EPI_ISL_2828952 | Lighthouse Lab in Alderley Park | Wellcome Sanger Institute for the COVID-19 Genomics UK (COG-UK) Consortium | Jacquelyn Wynn, Mairead Hyland, The Lighthouse Lab in Alderley Park and Alex Alderton, Roberto Amato, Jeffrey Barrett, Sonia Goncalves, Ewan Harrison, David K. Jackson, Ian Johnston, Dominic Kwiatkowski, Cordelia Langford, John Sillitoe on behalf of the Wellcome Sanger Institute COVID-19 Surveillance Team |
| EPI_ISL_2828953 | Lighthouse Lab in Milton Keynes | Wellcome Sanger Institute for the COVID-19 Genomics UK (COG-UK) Consortium | The Lighthouse Lab in Milton Keynes and Alex Alderton, Roberto Amato, Jeffrey Barrett, Sonia Goncalves, Ewan Harrison, David K. Jackson, Ian Johnston, Dominic Kwiatkowski, Cordelia Langford, John Sillitoe on behalf of the Wellcome Sanger Institute COVID-19 Surveillance Team |
| EPI_ISL_2828954, EPI_ISL_2828955 | Lighthouse Lab in Alderley Park | Wellcome Sanger Institute for the COVID-19 Genomics UK (COG-UK) Consortium | Jacquelyn Wynn, Mairead Hyland, The Lighthouse Lab in Alderley Park and Alex Alderton, Roberto Amato, Jeffrey Barrett, Sonia Goncalves, Ewan Harrison, David K. Jackson, Ian Johnston, Dominic Kwiatkowski, Cordelia Langford, John Sillitoe on behalf of the Wellcome Sanger Institute COVID-19 Surveillance Team |
| EPI_ISL_2828956, EPI_ISL_2828957 | Lighthouse Lab in Milton Keynes | Wellcome Sanger Institute for the COVID-19 Genomics UK (COG-UK) Consortium | The Lighthouse Lab in Milton Keynes and Alex Alderton, Roberto Amato, Jeffrey Barrett, Sonia Goncalves, Ewan Harrison, David K. Jackson, Ian Johnston, Dominic Kwiatkowski, Cordelia Langford, John Sillitoe on behalf of the Wellcome Sanger Institute COVID-19 Surveillance Team |
| EPI_ISL_2828958 | Lighthouse Laboratory Plymouth | Wellcome Sanger Institute for the COVID-19 Genomics UK (COG-UK) Consortium | Lighthouse Laboratory Plymouth and Alex Alderton, Roberto Amato, Jeffrey Barrett, Sonia Goncalves, Ewan Harrison, David K. Jackson, Ian Johnston, Dominic Kwiatkowski, Cordelia Langford, John Sillitoe on behalf of the Wellcome Sanger Institute COVID-19 Surveillance Team |
| EPI_ISL_2828959 | Lighthouse Lab in Alderley Park | Wellcome Sanger Institute for the COVID-19 Genomics UK (COG-UK) Consortium | Jacquelyn Wynn, Mairead Hyland, The Lighthouse Lab in Alderley Park and Alex Alderton, Roberto Amato, Jeffrey Barrett, Sonia Goncalves, Ewan Harrison, David K. Jackson, Ian Johnston, Dominic Kwiatkowski, Cordelia Langford, John Sillitoe on behalf of the Wellcome Sanger Institute COVID-19 Surveillance Team |
| EPI_ISL_2828960 | Lighthouse Laboratory Plymouth | Wellcome Sanger Institute for the COVID-19 Genomics UK (COG-UK) Consortium | Lighthouse Laboratory Plymouth and Alex Alderton, Roberto Amato, Jeffrey Barrett, Sonia Goncalves, Ewan Harrison, David K. Jackson, Ian Johnston, Dominic Kwiatkowski, Cordelia Langford, John Sillitoe on behalf of the Wellcome Sanger Institute COVID-19 Surveillance Team |
| EPI_ISL_2828961 | Lighthouse Lab in Alderley Park | Wellcome Sanger Institute for the COVID-19 Genomics UK (COG-UK) Consortium | Jacquelyn Wynn, Mairead Hyland, The Lighthouse Lab in Alderley Park and Alex Alderton, Roberto Amato, Jeffrey Barrett, Sonia Goncalves, Ewan Harrison, David K. Jackson, Ian Johnston, Dominic Kwiatkowski, Cordelia Langford, John Sillitoe on behalf of the Wellcome Sanger Institute COVID-19 Surveillance Team |
| EPI_ISL_2828962 | Lighthouse Laboratory Plymouth | Wellcome Sanger Institute for the COVID-19 Genomics UK (COG-UK) Consortium | Lighthouse Laboratory Plymouth and Alex Alderton, Roberto Amato, Jeffrey Barrett, Sonia Goncalves, Ewan Harrison, David K. Jackson, Ian Johnston, Dominic Kwiatkowski, Cordelia Langford, John Sillitoe on behalf of the Wellcome Sanger Institute COVID-19 Surveillance Team |
| EPI_ISL_2828963 | Lighthouse Lab in Alderley Park | Wellcome Sanger Institute for the COVID-19 Genomics UK (COG-UK) Consortium | Jacquelyn Wynn, Mairead Hyland, The Lighthouse Lab in Alderley Park and Alex Alderton, Roberto Amato, Jeffrey Barrett, Sonia Goncalves, Ewan Harrison, David K. Jackson, Ian Johnston, Dominic Kwiatkowski, Cordelia Langford, John Sillitoe on behalf of the Wellcome Sanger Institute COVID-19 Surveillance Team |

| | | | |
|---|---|---|---|
| EPI_ISL_2828964 | Lighthouse Lab in Milton Keynes | Wellcome Sanger Institute for the COVID-19 Genomics UK (COG-UK) Consortium | The Lighthouse Lab in Milton Keynes and Alex Alderton, Roberto Amato, Jeffrey Barrett, Sonia Goncalves, Ewan Harrison, David K. Jackson, Ian Johnston, Dominic Kwiatkowski, Cordelia Langford, John Sillitoe on behalf of the Wellcome Sanger Institute COVID-19 Surveillance Team |
| EPI_ISL_2828965, EPI_ISL_2828966, EPI_ISL_2828967 | Lighthouse Lab in Alderley Park | Wellcome Sanger Institute for the COVID-19 Genomics UK (COG-UK) Consortium | Jacquelyn Wynn, Mairead Hyland, The Lighthouse Lab in Alderley Park and Alex Alderton, Roberto Amato, Jeffrey Barrett, Sonia Goncalves, Ewan Harrison, David K. Jackson, Ian Johnston, Dominic Kwiatkowski, Cordelia Langford, John Sillitoe on behalf of the Wellcome Sanger Institute COVID-19 Surveillance Team |
| EPI_ISL_2828968 | Lighthouse Lab in Milton Keynes | Wellcome Sanger Institute for the COVID-19 Genomics UK (COG-UK) Consortium | The Lighthouse Lab in Milton Keynes and Alex Alderton, Roberto Amato, Jeffrey Barrett, Sonia Goncalves, Ewan Harrison, David K. Jackson, Ian Johnston, Dominic Kwiatkowski, Cordelia Langford, John Sillitoe on behalf of the Wellcome Sanger Institute COVID-19 Surveillance Team |
| EPI_ISL_2828969 | Lighthouse Lab in Alderley Park | Wellcome Sanger Institute for the COVID-19 Genomics UK (COG-UK) Consortium | Jacquelyn Wynn, Mairead Hyland, The Lighthouse Lab in Alderley Park and Alex Alderton, Roberto Amato, Jeffrey Barrett, Sonia Goncalves, Ewan Harrison, David K. Jackson, Ian Johnston, Dominic Kwiatkowski, Cordelia Langford, John Sillitoe on behalf of the Wellcome Sanger Institute COVID-19 Surveillance Team |
| EPI_ISL_2828970 | Lighthouse Lab in Milton Keynes | Wellcome Sanger Institute for the COVID-19 Genomics UK (COG-UK) Consortium | The Lighthouse Lab in Milton Keynes and Alex Alderton, Roberto Amato, Jeffrey Barrett, Sonia Goncalves, Ewan Harrison, David K. Jackson, Ian Johnston, Dominic Kwiatkowski, Cordelia Langford, John Sillitoe on behalf of the Wellcome Sanger Institute COVID-19 Surveillance Team |
| EPI_ISL_2828971 | Lighthouse Lab in Alderley Park | Wellcome Sanger Institute for the COVID-19 Genomics UK (COG-UK) Consortium | Jacquelyn Wynn, Mairead Hyland, The Lighthouse Lab in Alderley Park and Alex Alderton, Roberto Amato, Jeffrey Barrett, Sonia Goncalves, Ewan Harrison, David K. Jackson, Ian Johnston, Dominic Kwiatkowski, Cordelia Langford, John Sillitoe on behalf of the Wellcome Sanger Institute COVID-19 Surveillance Team |
| EPI_ISL_2828972 | Lighthouse Laboratory Plymouth | Wellcome Sanger Institute for the COVID-19 Genomics UK (COG-UK) Consortium | Lighthouse Laboratory Plymouth and Alex Alderton, Roberto Amato, Jeffrey Barrett, Sonia Goncalves, Ewan Harrison, David K. Jackson, Ian Johnston, Dominic Kwiatkowski, Cordelia Langford, John Sillitoe on behalf of the Wellcome Sanger Institute COVID-19 Surveillance Team |
| EPI_ISL_2828973 | Lighthouse Lab in Alderley Park | Wellcome Sanger Institute for the COVID-19 Genomics UK (COG-UK) Consortium | Jacquelyn Wynn, Mairead Hyland, The Lighthouse Lab in Alderley Park and Alex Alderton, Roberto Amato, Jeffrey Barrett, Sonia Goncalves, Ewan Harrison, David K. Jackson, Ian Johnston, Dominic Kwiatkowski, Cordelia Langford, John Sillitoe on behalf of the Wellcome Sanger Institute COVID-19 Surveillance Team |
| EPI_ISL_2828974 | Lighthouse Laboratory Plymouth | Wellcome Sanger Institute for the COVID-19 Genomics UK (COG-UK) Consortium | Lighthouse Laboratory Plymouth and Alex Alderton, Roberto Amato, Jeffrey Barrett, Sonia Goncalves, Ewan Harrison, David K. Jackson, Ian Johnston, Dominic Kwiatkowski, Cordelia Langford, John Sillitoe on behalf of the Wellcome Sanger Institute COVID-19 Surveillance Team |
| EPI_ISL_2828975 | Lighthouse Lab in Alderley Park | Wellcome Sanger Institute for the COVID-19 Genomics UK (COG-UK) Consortium | Jacquelyn Wynn, Mairead Hyland, The Lighthouse Lab in Alderley Park and Alex Alderton, Roberto Amato, Jeffrey Barrett, Sonia Goncalves, Ewan Harrison, David K. Jackson, Ian Johnston, Dominic Kwiatkowski, Cordelia Langford, John Sillitoe on behalf of the Wellcome Sanger Institute COVID-19 Surveillance Team |
| EPI_ISL_2828976 | Lighthouse Lab in Milton Keynes | Wellcome Sanger Institute for the COVID-19 Genomics UK (COG-UK) Consortium | The Lighthouse Lab in Milton Keynes and Alex Alderton, Roberto Amato, Jeffrey Barrett, Sonia Goncalves, Ewan Harrison, David K. Jackson, Ian Johnston, Dominic Kwiatkowski, Cordelia Langford, John Sillitoe on behalf of the Wellcome Sanger Institute COVID-19 Surveillance Team |
| EPI_ISL_2828977 | Lighthouse Lab in Alderley Park | Wellcome Sanger Institute for the COVID-19 Genomics UK (COG-UK) Consortium | Jacquelyn Wynn, Mairead Hyland, The Lighthouse Lab in Alderley Park and Alex Alderton, Roberto Amato, Jeffrey Barrett, Sonia Goncalves, Ewan Harrison, David K. Jackson, Ian Johnston, Dominic Kwiatkowski, Cordelia Langford, John Sillitoe on behalf of the Wellcome Sanger Institute COVID-19 Surveillance Team |
| EPI_ISL_2828978 | Lighthouse Lab in Milton Keynes | Wellcome Sanger Institute for the COVID-19 Genomics UK (COG-UK) Consortium | The Lighthouse Lab in Milton Keynes and Alex Alderton, Roberto Amato, Jeffrey Barrett, Sonia Goncalves, Ewan Harrison, David K. Jackson, Ian Johnston, Dominic Kwiatkowski, Cordelia Langford, John Sillitoe on behalf of the Wellcome Sanger Institute COVID-19 Surveillance Team |
| EPI_ISL_2828979 | Lighthouse Laboratory Plymouth | Wellcome Sanger Institute for the COVID-19 Genomics UK (COG-UK) Consortium | Lighthouse Laboratory Plymouth and Alex Alderton, Roberto Amato, Jeffrey Barrett, Sonia Goncalves, Ewan Harrison, David K. Jackson, Ian Johnston, Dominic Kwiatkowski, Cordelia Langford, John Sillitoe on behalf of the Wellcome Sanger Institute COVID-19 Surveillance Team |
| EPI_ISL_2828980 | Lighthouse Lab in Milton Keynes | Wellcome Sanger Institute for the COVID-19 Genomics UK (COG-UK) Consortium | The Lighthouse Lab in Milton Keynes and Alex Alderton, Roberto Amato, Jeffrey Barrett, Sonia Goncalves, Ewan Harrison, David K. Jackson, Ian Johnston, Dominic Kwiatkowski, Cordelia Langford, John Sillitoe on behalf of the Wellcome Sanger Institute COVID-19 Surveillance Team |
| EPI_ISL_2828981 | Lighthouse Lab in Alderley Park | Wellcome Sanger Institute for the COVID-19 Genomics UK (COG-UK) Consortium | Jacquelyn Wynn, Mairead Hyland, The Lighthouse Lab in Alderley Park and Alex Alderton, Roberto Amato, Jeffrey Barrett, Sonia Goncalves, Ewan Harrison, David K. Jackson, Ian Johnston, Dominic Kwiatkowski, Cordelia Langford, John Sillitoe on behalf of the Wellcome Sanger Institute COVID-19 Surveillance Team |
| EPI_ISL_2828982 | Lighthouse Lab in Milton Keynes | Wellcome Sanger Institute for the COVID-19 Genomics UK (COG-UK) Consortium | The Lighthouse Lab in Milton Keynes and Alex Alderton, Roberto Amato, Jeffrey Barrett, Sonia Goncalves, Ewan Harrison, David K. Jackson, Ian Johnston, Dominic Kwiatkowski, Cordelia Langford, John Sillitoe on behalf of the Wellcome Sanger Institute COVID-19 Surveillance Team |
| EPI_ISL_2828983, EPI_ISL_2828984 | Lighthouse Lab in Alderley Park | Wellcome Sanger Institute for the COVID-19 Genomics UK (COG-UK) Consortium | Jacquelyn Wynn, Mairead Hyland, The Lighthouse Lab in Alderley Park and Alex Alderton, Roberto Amato, Jeffrey Barrett, Sonia Goncalves, Ewan Harrison, David K. Jackson, Ian Johnston, Dominic Kwiatkowski, Cordelia Langford, John Sillitoe on behalf of the Wellcome Sanger Institute COVID-19 Surveillance Team |
| EPI_ISL_2828985 | Lighthouse Laboratory Plymouth | Wellcome Sanger Institute for the COVID-19 Genomics UK (COG-UK) Consortium | Lighthouse Laboratory Plymouth and Alex Alderton, Roberto Amato, Jeffrey Barrett, Sonia Goncalves, Ewan Harrison, David K. Jackson, Ian Johnston, Dominic Kwiatkowski, Cordelia Langford, John Sillitoe on behalf of the Wellcome Sanger Institute COVID-19 Surveillance Team |
| EPI_ISL_2828986 | Lighthouse Lab in Milton Keynes | Wellcome Sanger Institute for the COVID-19 Genomics UK (COG-UK) Consortium | The Lighthouse Lab in Milton Keynes and Alex Alderton, Roberto Amato, Jeffrey Barrett, Sonia Goncalves, Ewan Harrison, David K. Jackson, Ian Johnston, Dominic Kwiatkowski, Cordelia Langford, John Sillitoe on behalf of the Wellcome Sanger Institute COVID-19 Surveillance Team |
| EPI_ISL_2828987, EPI_ISL_2828988 | Lighthouse Laboratory Plymouth | Wellcome Sanger Institute for the COVID-19 Genomics UK (COG-UK) Consortium | Lighthouse Laboratory Plymouth and Alex Alderton, Roberto Amato, Jeffrey Barrett, Sonia Goncalves, Ewan Harrison, David K. Jackson, Ian Johnston, Dominic Kwiatkowski, Cordelia Langford, John Sillitoe on behalf of the Wellcome Sanger Institute COVID-19 Surveillance Team |
| EPI_ISL_2828989, EPI_ISL_2828990 | Lighthouse Lab in Alderley Park | Wellcome Sanger Institute for the COVID-19 Genomics UK (COG-UK) Consortium | Jacquelyn Wynn, Mairead Hyland, The Lighthouse Lab in Alderley Park and Alex Alderton, Roberto Amato, Jeffrey Barrett, Sonia Goncalves, Ewan Harrison, David K. Jackson, Ian Johnston, Dominic Kwiatkowski, Cordelia Langford, John Sillitoe on behalf of the Wellcome Sanger Institute COVID-19 Surveillance Team |
| EPI_ISL_2828991 | Lighthouse Laboratory Plymouth | Wellcome Sanger Institute for the COVID-19 Genomics UK (COG-UK) Consortium | Lighthouse Laboratory Plymouth and Alex Alderton, Roberto Amato, Jeffrey Barrett, Sonia Goncalves, Ewan Harrison, David K. Jackson, Ian Johnston, Dominic Kwiatkowski, Cordelia Langford, John Sillitoe on behalf of the Wellcome Sanger Institute COVID-19 Surveillance Team |
| EPI_ISL_2828992 | Lighthouse Lab in Alderley Park | Wellcome Sanger Institute for the COVID-19 Genomics UK (COG-UK) Consortium | Jacquelyn Wynn, Mairead Hyland, The Lighthouse Lab in Alderley Park and Alex Alderton, Roberto Amato, Jeffrey Barrett, Sonia Goncalves, Ewan Harrison, David K. Jackson, Ian Johnston, Dominic Kwiatkowski, Cordelia Langford, John Sillitoe on behalf of the Wellcome Sanger Institute COVID-19 Surveillance Team |
| EPI_ISL_2828993 | Lighthouse Lab in Milton Keynes | Wellcome Sanger Institute for the COVID-19 Genomics UK (COG-UK) Consortium | The Lighthouse Lab in Milton Keynes and Alex Alderton, Roberto Amato, Jeffrey Barrett, Sonia Goncalves, Ewan Harrison, David K. Jackson, Ian Johnston, Dominic Kwiatkowski, Cordelia Langford, John Sillitoe on behalf of the Wellcome Sanger Institute COVID-19 Surveillance Team |
| EPI_ISL_2828994 | Lighthouse Laboratory Plymouth | Wellcome Sanger Institute for the COVID-19 Genomics UK (COG-UK) Consortium | Lighthouse Laboratory Plymouth and Alex Alderton, Roberto Amato, Jeffrey Barrett, Sonia Goncalves, Ewan Harrison, David K. Jackson, Ian Johnston, Dominic Kwiatkowski, Cordelia Langford, John Sillitoe on behalf of the Wellcome Sanger Institute COVID-19 Surveillance Team |
| EPI_ISL_2828995 | Lighthouse Lab in Alderley Park | Wellcome Sanger Institute for the COVID-19 Genomics UK (COG-UK) Consortium | Jacquelyn Wynn, Mairead Hyland, The Lighthouse Lab in Alderley Park and Alex Alderton, Roberto Amato, Jeffrey Barrett, Sonia Goncalves, Ewan Harrison, David K. Jackson, Ian Johnston, Dominic Kwiatkowski, Cordelia Langford, John Sillitoe on behalf of the Wellcome Sanger Institute COVID-19 Surveillance Team |
| EPI_ISL_2828996 | Lighthouse Lab in Milton Keynes | Wellcome Sanger Institute for the COVID-19 Genomics UK (COG-UK) Consortium | The Lighthouse Lab in Milton Keynes and Alex Alderton, Roberto Amato, Jeffrey Barrett, Sonia Goncalves, Ewan Harrison, David K. Jackson, Ian Johnston, Dominic Kwiatkowski, Cordelia Langford, John Sillitoe on behalf of the Wellcome Sanger Institute COVID-19 Surveillance Team |
| EPI_ISL_2828997 | Lighthouse Laboratory Plymouth | Wellcome Sanger Institute for the COVID-19 Genomics UK (COG-UK) Consortium | Lighthouse Laboratory Plymouth and Alex Alderton, Roberto Amato, Jeffrey Barrett, Sonia Goncalves, Ewan Harrison, David K. Jackson, Ian Johnston, Dominic Kwiatkowski, Cordelia Langford, John Sillitoe on behalf of the Wellcome Sanger Institute COVID-19 Surveillance Team |
| EPI_ISL_2828998 | Lighthouse Lab in Alderley Park | Wellcome Sanger Institute for the COVID-19 Genomics UK (COG-UK) Consortium | Jacquelyn Wynn, Mairead Hyland, The Lighthouse Lab in Alderley Park and Alex Alderton, Roberto Amato, Jeffrey Barrett, Sonia Goncalves, Ewan Harrison, David K. Jackson, Ian Johnston, Dominic Kwiatkowski, Cordelia Langford, John Sillitoe on behalf of the Wellcome Sanger Institute COVID-19 Surveillance Team |
| EPI_ISL_2828999, EPI_ISL_2829000 | Lighthouse Laboratory Plymouth | Wellcome Sanger Institute for the COVID-19 Genomics UK (COG-UK) Consortium | Lighthouse Laboratory Plymouth and Alex Alderton, Roberto Amato, Jeffrey Barrett, Sonia Goncalves, Ewan Harrison, David K. Jackson, Ian Johnston, Dominic Kwiatkowski, Cordelia Langford, John Sillitoe on behalf of the Wellcome Sanger Institute COVID-19 Surveillance Team |
| EPI_ISL_2829001 | Lighthouse Lab in Alderley Park | Wellcome Sanger Institute for the COVID-19 Genomics UK (COG-UK) Consortium | Jacquelyn Wynn, Mairead Hyland, The Lighthouse Lab in Alderley Park and Alex Alderton, Roberto Amato, Jeffrey Barrett, Sonia Goncalves, Ewan Harrison, David K. Jackson, Ian Johnston, Dominic Kwiatkowski, Cordelia Langford, John Sillitoe on behalf of the Wellcome Sanger Institute COVID-19 Surveillance Team |
| EPI_ISL_2829002, EPI_ISL_2829003, EPI_ISL_2829004 | Lighthouse Lab in Milton Keynes | Wellcome Sanger Institute for the COVID-19 Genomics UK (COG-UK) Consortium | The Lighthouse Lab in Milton Keynes and Alex Alderton, Roberto Amato, Jeffrey Barrett, Sonia Goncalves, Ewan Harrison, David K. Jackson, Ian Johnston, Dominic Kwiatkowski, Cordelia Langford, John Sillitoe on behalf of the Wellcome Sanger Institute COVID-19 Surveillance Team |
| EPI_ISL_2829005 | Lighthouse Laboratory Plymouth | Wellcome Sanger Institute for the COVID-19 Genomics UK (COG-UK) Consortium | Lighthouse Laboratory Plymouth and Alex Alderton, Roberto Amato, Jeffrey Barrett, Sonia Goncalves, Ewan Harrison, David K. Jackson, Ian Johnston, Dominic Kwiatkowski, Cordelia Langford, John Sillitoe on behalf of the Wellcome Sanger Institute COVID-19 Surveillance Team |
| EPI_ISL_2829006, EPI_ISL_2829007 | Lighthouse Lab in Alderley Park | Wellcome Sanger Institute for the COVID-19 Genomics UK (COG-UK) Consortium | Jacquelyn Wynn, Mairead Hyland, The Lighthouse Lab in Alderley Park and Alex Alderton, Roberto Amato, Jeffrey Barrett, Sonia Goncalves, Ewan Harrison, David K. Jackson, Ian Johnston, Dominic Kwiatkowski, Cordelia Langford, John Sillitoe on behalf of the Wellcome Sanger Institute COVID-19 Surveillance Team |
| EPI_ISL_2829008 | Lighthouse Laboratory Plymouth | Wellcome Sanger Institute for the COVID-19 Genomics UK (COG-UK) Consortium | Lighthouse Laboratory Plymouth and Alex Alderton, Roberto Amato, Jeffrey Barrett, Sonia Goncalves, Ewan Harrison, David K. Jackson, Ian Johnston, Dominic Kwiatkowski, Cordelia Langford, John Sillitoe on behalf of the Wellcome Sanger Institute COVID-19 Surveillance Team |
| EPI_ISL_2829009 | Lighthouse Lab in Alderley Park | Wellcome Sanger Institute for the COVID-19 Genomics UK (COG-UK) Consortium | Jacquelyn Wynn, Mairead Hyland, The Lighthouse Lab in Alderley Park and Alex Alderton, Roberto Amato, Jeffrey Barrett, Sonia Goncalves, Ewan Harrison, David K. Jackson, Ian Johnston, Dominic Kwiatkowski, Cordelia Langford, John Sillitoe on behalf of the Wellcome Sanger Institute COVID-19 Surveillance Team |
| EPI_ISL_2829010, EPI_ISL_2829011 | Lighthouse Laboratory Plymouth | Wellcome Sanger Institute for the COVID-19 Genomics UK (COG-UK) Consortium | Lighthouse Laboratory Plymouth and Alex Alderton, Roberto Amato, Jeffrey Barrett, Sonia Goncalves, Ewan Harrison, David K. Jackson, Ian Johnston, Dominic Kwiatkowski, Cordelia Langford, John Sillitoe on behalf of the Wellcome Sanger Institute COVID-19 Surveillance Team |
| EPI_ISL_2829012, EPI_ISL_2829013 | Lighthouse Lab in Milton Keynes | Wellcome Sanger Institute for the COVID-19 Genomics UK (COG-UK) Consortium | The Lighthouse Lab in Milton Keynes and Alex Alderton, Roberto Amato, Jeffrey Barrett, Sonia Goncalves, Ewan Harrison, David K. Jackson, Ian Johnston, Dominic Kwiatkowski, Cordelia Langford, John Sillitoe on behalf of the Wellcome Sanger Institute COVID-19 Surveillance Team |
| EPI_ISL_2829014, EPI_ISL_2829015, EPI_ISL_2829016, EPI_ISL_2829017 | Lighthouse Laboratory Plymouth | Wellcome Sanger Institute for the COVID-19 Genomics UK (COG-UK) Consortium | Lighthouse Laboratory Plymouth and Alex Alderton, Roberto Amato, Jeffrey Barrett, Sonia Goncalves, Ewan Harrison, David K. Jackson, Ian Johnston, Dominic Kwiatkowski, Cordelia Langford, John Sillitoe on behalf of the Wellcome Sanger Institute COVID-19 Surveillance Team |
| EPI_ISL_2829018 | Lighthouse Lab in Milton Keynes | Wellcome Sanger Institute for the COVID-19 Genomics UK (COG-UK) Consortium | The Lighthouse Lab in Milton Keynes and Alex Alderton, Roberto Amato, Jeffrey Barrett, Sonia Goncalves, Ewan Harrison, David K. Jackson, Ian Johnston, Dominic Kwiatkowski, Cordelia Langford, John Sillitoe on behalf of the Wellcome Sanger Institute COVID-19 Surveillance Team |
| EPI_ISL_2829019 | Lighthouse Lab in Alderley Park | Wellcome Sanger Institute for the COVID-19 Genomics UK (COG-UK) Consortium | Jacquelyn Wynn, Mairead Hyland, The Lighthouse Lab in Alderley Park and Alex Alderton, Roberto Amato, Jeffrey Barrett, Sonia Goncalves, Ewan Harrison, David K. Jackson, Ian Johnston, Dominic Kwiatkowski, Cordelia Langford, John Sillitoe on behalf of the Wellcome Sanger Institute COVID-19 Surveillance Team |
| EPI_ISL_2829020, EPI_ISL_2829021 | Lighthouse Laboratory Plymouth | Wellcome Sanger Institute for the COVID-19 Genomics UK (COG-UK) Consortium | Lighthouse Laboratory Plymouth and Alex Alderton, Roberto Amato, Jeffrey Barrett, Sonia Goncalves, Ewan Harrison, David K. Jackson, Ian Johnston, Dominic Kwiatkowski, Cordelia Langford, John Sillitoe on behalf of the Wellcome Sanger Institute COVID-19 Surveillance Team |

| | | | |
|---|---|---|---|
| EPI_ISL_2829022 | Lighthouse Lab in Milton Keynes | Wellcome Sanger Institute for the COVID-19 Genomics UK (COG-UK) Consortium | The Lighthouse Lab in Milton Keynes and Alex Alderton, Roberto Amato, Jeffrey Barrett, Sonia Goncalves, Ewan Harrison, David K. Jackson, Ian Johnston, Dominic Kwiatkowski, Cordelia Langford, John Sillitoe on behalf of the Wellcome Sanger Institute COVID-19 Surveillance Team |
| EPI_ISL_2829023 | Lighthouse Lab in Alderley Park | Wellcome Sanger Institute for the COVID-19 Genomics UK (COG-UK) Consortium | Jacquelyn Wynn, Mairead Hyland, The Lighthouse Lab in Alderley Park and Alex Alderton, Roberto Amato, Jeffrey Barrett, Sonia Goncalves, Ewan Harrison, David K. Jackson, Ian Johnston, Dominic Kwiatkowski, Cordelia Langford, John Sillitoe on behalf of the Wellcome Sanger Institute COVID-19 Surveillance Team |
| EPI_ISL_2829024, EPI_ISL_2829025 | Lighthouse Lab in Milton Keynes | Wellcome Sanger Institute for the COVID-19 Genomics UK (COG-UK) Consortium | The Lighthouse Lab in Milton Keynes and Alex Alderton, Roberto Amato, Jeffrey Barrett, Sonia Goncalves, Ewan Harrison, David K. Jackson, Ian Johnston, Dominic Kwiatkowski, Cordelia Langford, John Sillitoe on behalf of the Wellcome Sanger Institute COVID-19 Surveillance Team |
| EPI_ISL_2829026 | Lighthouse Lab in Alderley Park | Wellcome Sanger Institute for the COVID-19 Genomics UK (COG-UK) Consortium | Jacquelyn Wynn, Mairead Hyland, The Lighthouse Lab in Alderley Park and Alex Alderton, Roberto Amato, Jeffrey Barrett, Sonia Goncalves, Ewan Harrison, David K. Jackson, Ian Johnston, Dominic Kwiatkowski, Cordelia Langford, John Sillitoe on behalf of the Wellcome Sanger Institute COVID-19 Surveillance Team |
| EPI_ISL_2829027 | Health Services Laboratories | Wellcome Sanger Institute for the COVID-19 Genomics UK (COG-UK) Consortium | Health Services Laboratories and Alex Alderton, Roberto Amato, Jeffrey Barrett, Sonia Goncalves, Ewan Harrison, David K. Jackson, Ian Johnston, Dominic Kwiatkowski, Cordelia Langford, John Sillitoe on behalf of the Wellcome Sanger Institute COVID-19 Surveillance Team |
| EPI_ISL_2829028 | Lighthouse Lab in Alderley Park | Wellcome Sanger Institute for the COVID-19 Genomics UK (COG-UK) Consortium | Jacquelyn Wynn, Mairead Hyland, The Lighthouse Lab in Alderley Park and Alex Alderton, Roberto Amato, Jeffrey Barrett, Sonia Goncalves, Ewan Harrison, David K. Jackson, Ian Johnston, Dominic Kwiatkowski, Cordelia Langford, John Sillitoe on behalf of the Wellcome Sanger Institute COVID-19 Surveillance Team |
| EPI_ISL_2829029 | Lighthouse Laboratory Plymouth | Wellcome Sanger Institute for the COVID-19 Genomics UK (COG-UK) Consortium | Lighthouse Laboratory Plymouth and Alex Alderton, Roberto Amato, Jeffrey Barrett, Sonia Goncalves, Ewan Harrison, David K. Jackson, Ian Johnston, Dominic Kwiatkowski, Cordelia Langford, John Sillitoe on behalf of the Wellcome Sanger Institute COVID-19 Surveillance Team |
| EPI_ISL_2829030 | Health Services Laboratories | Wellcome Sanger Institute for the COVID-19 Genomics UK (COG-UK) Consortium | Health Services Laboratories and Alex Alderton, Roberto Amato, Jeffrey Barrett, Sonia Goncalves, Ewan Harrison, David K. Jackson, Ian Johnston, Dominic Kwiatkowski, Cordelia Langford, John Sillitoe on behalf of the Wellcome Sanger Institute COVID-19 Surveillance Team |
| EPI_ISL_2829031 | Lighthouse Lab in Milton Keynes | Wellcome Sanger Institute for the COVID-19 Genomics UK (COG-UK) Consortium | The Lighthouse Lab in Milton Keynes and Alex Alderton, Roberto Amato, Jeffrey Barrett, Sonia Goncalves, Ewan Harrison, David K. Jackson, Ian Johnston, Dominic Kwiatkowski, Cordelia Langford, John Sillitoe on behalf of the Wellcome Sanger Institute COVID-19 Surveillance Team |
| EPI_ISL_2829032 | Health Services Laboratories | Wellcome Sanger Institute for the COVID-19 Genomics UK (COG-UK) Consortium | Health Services Laboratories and Alex Alderton, Roberto Amato, Jeffrey Barrett, Sonia Goncalves, Ewan Harrison, David K. Jackson, Ian Johnston, Dominic Kwiatkowski, Cordelia Langford, John Sillitoe on behalf of the Wellcome Sanger Institute COVID-19 Surveillance Team |
| EPI_ISL_2829033, EPI_ISL_2829034 | Lighthouse Lab in Milton Keynes | Wellcome Sanger Institute for the COVID-19 Genomics UK (COG-UK) Consortium | The Lighthouse Lab in Milton Keynes and Alex Alderton, Roberto Amato, Jeffrey Barrett, Sonia Goncalves, Ewan Harrison, David K. Jackson, Ian Johnston, Dominic Kwiatkowski, Cordelia Langford, John Sillitoe on behalf of the Wellcome Sanger Institute COVID-19 Surveillance Team |
| EPI_ISL_2829035 | Health Services Laboratories | Wellcome Sanger Institute for the COVID-19 Genomics UK (COG-UK) Consortium | Health Services Laboratories and Alex Alderton, Roberto Amato, Jeffrey Barrett, Sonia Goncalves, Ewan Harrison, David K. Jackson, Ian Johnston, Dominic Kwiatkowski, Cordelia Langford, John Sillitoe on behalf of the Wellcome Sanger Institute COVID-19 Surveillance Team |
| EPI_ISL_2829036 | Lighthouse Lab in Alderley Park | Wellcome Sanger Institute for the COVID-19 Genomics UK (COG-UK) Consortium | Jacquelyn Wynn, Mairead Hyland, The Lighthouse Lab in Alderley Park and Alex Alderton, Roberto Amato, Jeffrey Barrett, Sonia Goncalves, Ewan Harrison, David K. Jackson, Ian Johnston, Dominic Kwiatkowski, Cordelia Langford, John Sillitoe on behalf of the Wellcome Sanger Institute COVID-19 Surveillance Team |
| EPI_ISL_2829037 | Health Services Laboratories | Wellcome Sanger Institute for the COVID-19 Genomics UK (COG-UK) Consortium | Health Services Laboratories and Alex Alderton, Roberto Amato, Jeffrey Barrett, Sonia Goncalves, Ewan Harrison, David K. Jackson, Ian Johnston, Dominic Kwiatkowski, Cordelia Langford, John Sillitoe on behalf of the Wellcome Sanger Institute COVID-19 Surveillance Team |
| EPI_ISL_2829038 | Lighthouse Lab in Alderley Park | Wellcome Sanger Institute for the COVID-19 Genomics UK (COG-UK) Consortium | Jacquelyn Wynn, Mairead Hyland, The Lighthouse Lab in Alderley Park and Alex Alderton, Roberto Amato, Jeffrey Barrett, Sonia Goncalves, Ewan Harrison, David K. Jackson, Ian Johnston, Dominic Kwiatkowski, Cordelia Langford, John Sillitoe on behalf of the Wellcome Sanger Institute COVID-19 Surveillance Team |
| EPI_ISL_2829039 | Health Services Laboratories | Wellcome Sanger Institute for the COVID-19 Genomics UK (COG-UK) Consortium | Health Services Laboratories and Alex Alderton, Roberto Amato, Jeffrey Barrett, Sonia Goncalves, Ewan Harrison, David K. Jackson, Ian Johnston, Dominic Kwiatkowski, Cordelia Langford, John Sillitoe on behalf of the Wellcome Sanger Institute COVID-19 Surveillance Team |
| EPI_ISL_2829040 | Lighthouse Lab in Alderley Park | Wellcome Sanger Institute for the COVID-19 Genomics UK (COG-UK) Consortium | Jacquelyn Wynn, Mairead Hyland, The Lighthouse Lab in Alderley Park and Alex Alderton, Roberto Amato, Jeffrey Barrett, Sonia Goncalves, Ewan Harrison, David K. Jackson, Ian Johnston, Dominic Kwiatkowski, Cordelia Langford, John Sillitoe on behalf of the Wellcome Sanger Institute COVID-19 Surveillance Team |
| EPI_ISL_2829041 | Health Services Laboratories | Wellcome Sanger Institute for the COVID-19 Genomics UK (COG-UK) Consortium | Health Services Laboratories and Alex Alderton, Roberto Amato, Jeffrey Barrett, Sonia Goncalves, Ewan Harrison, David K. Jackson, Ian Johnston, Dominic Kwiatkowski, Cordelia Langford, John Sillitoe on behalf of the Wellcome Sanger Institute COVID-19 Surveillance Team |
| EPI_ISL_2829042, EPI_ISL_2829043, EPI_ISL_2829044 | Lighthouse Lab in Alderley Park | Wellcome Sanger Institute for the COVID-19 Genomics UK (COG-UK) Consortium | Jacquelyn Wynn, Mairead Hyland, The Lighthouse Lab in Alderley Park and Alex Alderton, Roberto Amato, Jeffrey Barrett, Sonia Goncalves, Ewan Harrison, David K. Jackson, Ian Johnston, Dominic Kwiatkowski, Cordelia Langford, John Sillitoe on behalf of the Wellcome Sanger Institute COVID-19 Surveillance Team |
| EPI_ISL_2829045 | Lighthouse Lab in Milton Keynes | Wellcome Sanger Institute for the COVID-19 Genomics UK (COG-UK) Consortium | The Lighthouse Lab in Milton Keynes and Alex Alderton, Roberto Amato, Jeffrey Barrett, Sonia Goncalves, Ewan Harrison, David K. Jackson, Ian Johnston, Dominic Kwiatkowski, Cordelia Langford, John Sillitoe on behalf of the Wellcome Sanger Institute COVID-19 Surveillance Team |
| EPI_ISL_2829046 | Lighthouse Laboratory Plymouth | Wellcome Sanger Institute for the COVID-19 Genomics UK (COG-UK) Consortium | Lighthouse Laboratory Plymouth and Alex Alderton, Roberto Amato, Jeffrey Barrett, Sonia Goncalves, Ewan Harrison, David K. Jackson, Ian Johnston, Dominic Kwiatkowski, Cordelia Langford, John Sillitoe on behalf of the Wellcome Sanger Institute COVID-19 Surveillance Team |
| EPI_ISL_2829047 | Lighthouse Lab in Alderley Park | Wellcome Sanger Institute for the COVID-19 Genomics UK (COG-UK) Consortium | Jacquelyn Wynn, Mairead Hyland, The Lighthouse Lab in Alderley Park and Alex Alderton, Roberto Amato, Jeffrey Barrett, Sonia Goncalves, Ewan Harrison, David K. Jackson, Ian Johnston, Dominic Kwiatkowski, Cordelia Langford, John Sillitoe on behalf of the Wellcome Sanger Institute COVID-19 Surveillance Team |
| EPI_ISL_2829048, EPI_ISL_2829049, EPI_ISL_2829050 | Lighthouse Lab in Milton Keynes | Wellcome Sanger Institute for the COVID-19 Genomics UK (COG-UK) Consortium | The Lighthouse Lab in Milton Keynes and Alex Alderton, Roberto Amato, Jeffrey Barrett, Sonia Goncalves, Ewan Harrison, David K. Jackson, Ian Johnston, Dominic Kwiatkowski, Cordelia Langford, John Sillitoe on behalf of the Wellcome Sanger Institute COVID-19 Surveillance Team |
| EPI_ISL_2829051 | Lighthouse Lab in Alderley Park | Wellcome Sanger Institute for the COVID-19 Genomics UK (COG-UK) Consortium | Jacquelyn Wynn, Mairead Hyland, The Lighthouse Lab in Alderley Park and Alex Alderton, Roberto Amato, Jeffrey Barrett, Sonia Goncalves, Ewan Harrison, David K. Jackson, Ian Johnston, Dominic Kwiatkowski, Cordelia Langford, John Sillitoe on behalf of the Wellcome Sanger Institute COVID-19 Surveillance Team |
| EPI_ISL_2829052 | Lighthouse Lab in Milton Keynes | Wellcome Sanger Institute for the COVID-19 Genomics UK (COG-UK) Consortium | The Lighthouse Lab in Milton Keynes and Alex Alderton, Roberto Amato, Jeffrey Barrett, Sonia Goncalves, Ewan Harrison, David K. Jackson, Ian Johnston, Dominic Kwiatkowski, Cordelia Langford, John Sillitoe on behalf of the Wellcome Sanger Institute COVID-19 Surveillance Team |
| EPI_ISL_2829053 | Lighthouse Lab in Alderley Park | Wellcome Sanger Institute for the COVID-19 Genomics UK (COG-UK) Consortium | Jacquelyn Wynn, Mairead Hyland, The Lighthouse Lab in Alderley Park and Alex Alderton, Roberto Amato, Jeffrey Barrett, Sonia Goncalves, Ewan Harrison, David K. Jackson, Ian Johnston, Dominic Kwiatkowski, Cordelia Langford, John Sillitoe on behalf of the Wellcome Sanger Institute COVID-19 Surveillance Team |
| EPI_ISL_2829054, EPI_ISL_2829055 | Lighthouse Lab in Milton Keynes | Wellcome Sanger Institute for the COVID-19 Genomics UK (COG-UK) Consortium | The Lighthouse Lab in Milton Keynes and Alex Alderton, Roberto Amato, Jeffrey Barrett, Sonia Goncalves, Ewan Harrison, David K. Jackson, Ian Johnston, Dominic Kwiatkowski, Cordelia Langford, John Sillitoe on behalf of the Wellcome Sanger Institute COVID-19 Surveillance Team |
| EPI_ISL_2829056 | Lighthouse Laboratory Plymouth | Wellcome Sanger Institute for the COVID-19 Genomics UK (COG-UK) Consortium | Lighthouse Laboratory Plymouth and Alex Alderton, Roberto Amato, Jeffrey Barrett, Sonia Goncalves, Ewan Harrison, David K. Jackson, Ian Johnston, Dominic Kwiatkowski, Cordelia Langford, John Sillitoe on behalf of the Wellcome Sanger Institute COVID-19 Surveillance Team |
| EPI_ISL_2829057 | Health Services Laboratories | Wellcome Sanger Institute for the COVID-19 Genomics UK (COG-UK) Consortium | Health Services Laboratories and Alex Alderton, Roberto Amato, Jeffrey Barrett, Sonia Goncalves, Ewan Harrison, David K. Jackson, Ian Johnston, Dominic Kwiatkowski, Cordelia Langford, John Sillitoe on behalf of the Wellcome Sanger Institute COVID-19 Surveillance Team |
| EPI_ISL_2829058 | Lighthouse Lab in Alderley Park | Wellcome Sanger Institute for the COVID-19 Genomics UK (COG-UK) Consortium | Jacquelyn Wynn, Mairead Hyland, The Lighthouse Lab in Alderley Park and Alex Alderton, Roberto Amato, Jeffrey Barrett, Sonia Goncalves, Ewan Harrison, David K. Jackson, Ian Johnston, Dominic Kwiatkowski, Cordelia Langford, John Sillitoe on behalf of the Wellcome Sanger Institute COVID-19 Surveillance Team |
| EPI_ISL_2829059 | Lighthouse Lab in Milton Keynes | Wellcome Sanger Institute for the COVID-19 Genomics UK (COG-UK) Consortium | The Lighthouse Lab in Milton Keynes and Alex Alderton, Roberto Amato, Jeffrey Barrett, Sonia Goncalves, Ewan Harrison, David K. Jackson, Ian Johnston, Dominic Kwiatkowski, Cordelia Langford, John Sillitoe on behalf of the Wellcome Sanger Institute COVID-19 Surveillance Team |
| EPI_ISL_2829060, EPI_ISL_2829061 | Lighthouse Lab in Alderley Park | Wellcome Sanger Institute for the COVID-19 Genomics UK (COG-UK) Consortium | Jacquelyn Wynn, Mairead Hyland, The Lighthouse Lab in Alderley Park and Alex Alderton, Roberto Amato, Jeffrey Barrett, Sonia Goncalves, Ewan Harrison, David K. Jackson, Ian Johnston, Dominic Kwiatkowski, Cordelia Langford, John Sillitoe on behalf of the Wellcome Sanger Institute COVID-19 Surveillance Team |
| EPI_ISL_2829062 | Health Services Laboratories | Wellcome Sanger Institute for the COVID-19 Genomics UK (COG-UK) Consortium | Health Services Laboratories and Alex Alderton, Roberto Amato, Jeffrey Barrett, Sonia Goncalves, Ewan Harrison, David K. Jackson, Ian Johnston, Dominic Kwiatkowski, Cordelia Langford, John Sillitoe on behalf of the Wellcome Sanger Institute COVID-19 Surveillance Team |
| EPI_ISL_2829063 | Lighthouse Lab in Alderley Park | Wellcome Sanger Institute for the COVID-19 Genomics UK (COG-UK) Consortium | Jacquelyn Wynn, Mairead Hyland, The Lighthouse Lab in Alderley Park and Alex Alderton, Roberto Amato, Jeffrey Barrett, Sonia Goncalves, Ewan Harrison, David K. Jackson, Ian Johnston, Dominic Kwiatkowski, Cordelia Langford, John Sillitoe on behalf of the Wellcome Sanger Institute COVID-19 Surveillance Team |
| EPI_ISL_2829064, EPI_ISL_2829065 | Lighthouse Lab in Milton Keynes | Wellcome Sanger Institute for the COVID-19 Genomics UK (COG-UK) Consortium | The Lighthouse Lab in Milton Keynes and Alex Alderton, Roberto Amato, Jeffrey Barrett, Sonia Goncalves, Ewan Harrison, David K. Jackson, Ian Johnston, Dominic Kwiatkowski, Cordelia Langford, John Sillitoe on behalf of the Wellcome Sanger Institute COVID-19 Surveillance Team |
| EPI_ISL_2829067 | Health Services Laboratories | Wellcome Sanger Institute for the COVID-19 Genomics UK (COG-UK) Consortium | Health Services Laboratories and Alex Alderton, Roberto Amato, Jeffrey Barrett, Sonia Goncalves, Ewan Harrison, David K. Jackson, Ian Johnston, Dominic Kwiatkowski, Cordelia Langford, John Sillitoe on behalf of the Wellcome Sanger Institute COVID-19 Surveillance Team |
| EPI_ISL_2829068 | Lighthouse Lab in Alderley Park | Wellcome Sanger Institute for the COVID-19 Genomics UK (COG-UK) Consortium | Jacquelyn Wynn, Mairead Hyland, The Lighthouse Lab in Alderley Park and Alex Alderton, Roberto Amato, Jeffrey Barrett, Sonia Goncalves, Ewan Harrison, David K. Jackson, Ian Johnston, Dominic Kwiatkowski, Cordelia Langford, John Sillitoe on behalf of the Wellcome Sanger Institute COVID-19 Surveillance Team |
| EPI_ISL_2829069, EPI_ISL_2829070 | Lighthouse Lab in Milton Keynes | Wellcome Sanger Institute for the COVID-19 Genomics UK (COG-UK) Consortium | The Lighthouse Lab in Milton Keynes and Alex Alderton, Roberto Amato, Jeffrey Barrett, Sonia Goncalves, Ewan Harrison, David K. Jackson, Ian Johnston, Dominic Kwiatkowski, Cordelia Langford, John Sillitoe on behalf of the Wellcome Sanger Institute COVID-19 Surveillance Team |
| EPI_ISL_2829071 | Health Services Laboratories | Wellcome Sanger Institute for the COVID-19 Genomics UK (COG-UK) Consortium | Health Services Laboratories and Alex Alderton, Roberto Amato, Jeffrey Barrett, Sonia Goncalves, Ewan Harrison, David K. Jackson, Ian Johnston, Dominic Kwiatkowski, Cordelia Langford, John Sillitoe on behalf of the Wellcome Sanger Institute COVID-19 Surveillance Team |
| EPI_ISL_2829072 | Lighthouse Lab in Alderley Park | Wellcome Sanger Institute for the COVID-19 Genomics UK (COG-UK) Consortium | Jacquelyn Wynn, Mairead Hyland, The Lighthouse Lab in Alderley Park and Alex Alderton, Roberto Amato, Jeffrey Barrett, Sonia Goncalves, Ewan Harrison, David K. Jackson, Ian Johnston, Dominic Kwiatkowski, Cordelia Langford, John Sillitoe on behalf of the Wellcome Sanger Institute COVID-19 Surveillance Team |
| EPI_ISL_2829073 | Lighthouse Lab in Milton Keynes | Wellcome Sanger Institute for the COVID-19 Genomics UK (COG-UK) Consortium | The Lighthouse Lab in Milton Keynes and Alex Alderton, Roberto Amato, Jeffrey Barrett, Sonia Goncalves, Ewan Harrison, David K. Jackson, Ian Johnston, Dominic Kwiatkowski, Cordelia Langford, John Sillitoe on behalf of the Wellcome Sanger Institute COVID-19 Surveillance Team |
| EPI_ISL_2829074, EPI_ISL_2829075 | Health Services Laboratories | Wellcome Sanger Institute for the COVID-19 Genomics UK (COG-UK) Consortium | Health Services Laboratories and Alex Alderton, Roberto Amato, Jeffrey Barrett, Sonia Goncalves, Ewan Harrison, David K. Jackson, Ian Johnston, Dominic Kwiatkowski, Cordelia Langford, John Sillitoe on behalf of the Wellcome Sanger Institute COVID-19 Surveillance Team |
| EPI_ISL_2829076 | Lighthouse Lab in Alderley Park | Wellcome Sanger Institute for the COVID-19 Genomics UK (COG-UK) Consortium | Jacquelyn Wynn, Mairead Hyland, The Lighthouse Lab in Alderley Park and Alex Alderton, Roberto Amato, Jeffrey Barrett, Sonia Goncalves, Ewan Harrison, David K. Jackson, Ian Johnston, Dominic Kwiatkowski, Cordelia Langford, John Sillitoe on behalf of the Wellcome Sanger Institute COVID-19 Surveillance Team |
| EPI_ISL_2829077 | Lighthouse Lab in Milton Keynes | Wellcome Sanger Institute for the COVID-19 Genomics UK (COG-UK) Consortium | The Lighthouse Lab in Milton Keynes and Alex Alderton, Roberto Amato, Jeffrey Barrett, Sonia Goncalves, Ewan Harrison, David K. Jackson, Ian Johnston, Dominic Kwiatkowski, Cordelia Langford, John Sillitoe on behalf of the Wellcome Sanger Institute COVID-19 Surveillance Team |
| EPI_ISL_2829078 | Lighthouse Laboratory Plymouth | Wellcome Sanger Institute for the COVID-19 Genomics UK (COG-UK) | Lighthouse Laboratory Plymouth and Alex Alderton, Roberto Amato, Jeffrey Barrett, Sonia Goncalves, Ewan Harrison, David K. Jackson, Ian Johnston, Dominic Kwiatkowski, Cordelia Langford, John Sillitoe on behalf of the Wellcome Sanger |

| | | | |
|---|---|---|---|
| | | Consortium | Institute COVID-19 Surveillance Team |
| EPI_ISL_2829079 | Lighthouse Lab in Milton Keynes | Wellcome Sanger Institute for the COVID-19 Genomics UK (COG-UK) Consortium | The Lighthouse Lab in Milton Keynes and Alex Alderton, Roberto Amato, Jeffrey Barrett, Sonia Goncalves, Ewan Harrison, David K. Jackson, Ian Johnston, Dominic Kwiatkowski, Cordelia Langford, John Sillitoe on behalf of the Wellcome Sanger Institute COVID-19 Surveillance Team |
| EPI_ISL_2829080, EPI_ISL_2829081 | Lighthouse Lab in Alderley Park | Wellcome Sanger Institute for the COVID-19 Genomics UK (COG-UK) Consortium | Jacquelyn Wynn, Mairead Hyland, The Lighthouse Lab in Alderley Park and Alex Alderton, Roberto Amato, Jeffrey Barrett, Sonia Goncalves, Ewan Harrison, David K. Jackson, Ian Johnston, Dominic Kwiatkowski, Cordelia Langford, John Sillitoe on behalf of the Wellcome Sanger Institute COVID-19 Surveillance Team |
| EPI_ISL_2829082 | Lighthouse Laboratory Plymouth | Wellcome Sanger Institute for the COVID-19 Genomics UK (COG-UK) Consortium | Lighthouse Laboratory Plymouth and Alex Alderton, Roberto Amato, Jeffrey Barrett, Sonia Goncalves, Ewan Harrison, David K. Jackson, Ian Johnston, Dominic Kwiatkowski, Cordelia Langford, John Sillitoe on behalf of the Wellcome Sanger Institute COVID-19 Surveillance Team |
| EPI_ISL_2829083, EPI_ISL_2829084 | Lighthouse Lab in Alderley Park | Wellcome Sanger Institute for the COVID-19 Genomics UK (COG-UK) Consortium | Jacquelyn Wynn, Mairead Hyland, The Lighthouse Lab in Alderley Park and Alex Alderton, Roberto Amato, Jeffrey Barrett, Sonia Goncalves, Ewan Harrison, David K. Jackson, Ian Johnston, Dominic Kwiatkowski, Cordelia Langford, John Sillitoe on behalf of the Wellcome Sanger Institute COVID-19 Surveillance Team |
| EPI_ISL_2829085 | Lighthouse Lab in Milton Keynes | Wellcome Sanger Institute for the COVID-19 Genomics UK (COG-UK) Consortium | The Lighthouse Lab in Milton Keynes and Alex Alderton, Roberto Amato, Jeffrey Barrett, Sonia Goncalves, Ewan Harrison, David K. Jackson, Ian Johnston, Dominic Kwiatkowski, Cordelia Langford, John Sillitoe on behalf of the Wellcome Sanger Institute COVID-19 Surveillance Team |
| EPI_ISL_2829086 | Health Services Laboratories | Wellcome Sanger Institute for the COVID-19 Genomics UK (COG-UK) Consortium | Health Services Laboratories and Alex Alderton, Roberto Amato, Jeffrey Barrett, Sonia Goncalves, Ewan Harrison, David K. Jackson, Ian Johnston, Dominic Kwiatkowski, Cordelia Langford, John Sillitoe on behalf of the Wellcome Sanger Institute COVID-19 Surveillance Team |
| EPI_ISL_2829087 | Lighthouse Lab in Milton Keynes | Wellcome Sanger Institute for the COVID-19 Genomics UK (COG-UK) Consortium | The Lighthouse Lab in Milton Keynes and Alex Alderton, Roberto Amato, Jeffrey Barrett, Sonia Goncalves, Ewan Harrison, David K. Jackson, Ian Johnston, Dominic Kwiatkowski, Cordelia Langford, John Sillitoe on behalf of the Wellcome Sanger Institute COVID-19 Surveillance Team |
| EPI_ISL_2829088 | Lighthouse Lab in Alderley Park | Wellcome Sanger Institute for the COVID-19 Genomics UK (COG-UK) Consortium | Jacquelyn Wynn, Mairead Hyland, The Lighthouse Lab in Alderley Park and Alex Alderton, Roberto Amato, Jeffrey Barrett, Sonia Goncalves, Ewan Harrison, David K. Jackson, Ian Johnston, Dominic Kwiatkowski, Cordelia Langford, John Sillitoe on behalf of the Wellcome Sanger Institute COVID-19 Surveillance Team |
| EPI_ISL_2829089 | Lighthouse Lab in Milton Keynes | Wellcome Sanger Institute for the COVID-19 Genomics UK (COG-UK) Consortium | The Lighthouse Lab in Milton Keynes and Alex Alderton, Roberto Amato, Jeffrey Barrett, Sonia Goncalves, Ewan Harrison, David K. Jackson, Ian Johnston, Dominic Kwiatkowski, Cordelia Langford, John Sillitoe on behalf of the Wellcome Sanger Institute COVID-19 Surveillance Team |
| EPI_ISL_2829090 | Lighthouse Lab in Glasgow | Wellcome Sanger Institute for the COVID-19 Genomics UK (COG-UK) Consortium | Harper VanSteenhouse, Yumi Kasai, David Gray, Carol Clugston, Anna Dominiczak and Alex Alderton, Roberto Amato, Jeffrey Barrett, Sonia Goncalves, Ewan Harrison, David K. Jackson, Ian Johnston, Dominic Kwiatkowski, Cordelia Langford, John Sillitoe on behalf of the Wellcome Sanger Institute COVID-19 Surveillance Team |
| EPI_ISL_2829091 | Health Services Laboratories | Wellcome Sanger Institute for the COVID-19 Genomics UK (COG-UK) Consortium | Health Services Laboratories and Alex Alderton, Roberto Amato, Jeffrey Barrett, Sonia Goncalves, Ewan Harrison, David K. Jackson, Ian Johnston, Dominic Kwiatkowski, Cordelia Langford, John Sillitoe on behalf of the Wellcome Sanger Institute COVID-19 Surveillance Team |
| EPI_ISL_2829092, EPI_ISL_2829093 | Lighthouse Lab in Alderley Park | Wellcome Sanger Institute for the COVID-19 Genomics UK (COG-UK) Consortium | Jacquelyn Wynn, Mairead Hyland, The Lighthouse Lab in Alderley Park and Alex Alderton, Roberto Amato, Jeffrey Barrett, Sonia Goncalves, Ewan Harrison, David K. Jackson, Ian Johnston, Dominic Kwiatkowski, Cordelia Langford, John Sillitoe on behalf of the Wellcome Sanger Institute COVID-19 Surveillance Team |
| EPI_ISL_2829094 | Lighthouse Lab in Milton Keynes | Wellcome Sanger Institute for the COVID-19 Genomics UK (COG-UK) Consortium | The Lighthouse Lab in Milton Keynes and Alex Alderton, Roberto Amato, Jeffrey Barrett, Sonia Goncalves, Ewan Harrison, David K. Jackson, Ian Johnston, Dominic Kwiatkowski, Cordelia Langford, John Sillitoe on behalf of the Wellcome Sanger Institute COVID-19 Surveillance Team |
| EPI_ISL_2829095 | Lighthouse Lab in Alderley Park | Wellcome Sanger Institute for the COVID-19 Genomics UK (COG-UK) Consortium | Jacquelyn Wynn, Mairead Hyland, The Lighthouse Lab in Alderley Park and Alex Alderton, Roberto Amato, Jeffrey Barrett, Sonia Goncalves, Ewan Harrison, David K. Jackson, Ian Johnston, Dominic Kwiatkowski, Cordelia Langford, John Sillitoe on behalf of the Wellcome Sanger Institute COVID-19 Surveillance Team |
| EPI_ISL_2829096 | Lighthouse Laboratory Plymouth | Wellcome Sanger Institute for the COVID-19 Genomics UK (COG-UK) Consortium | Lighthouse Laboratory Plymouth and Alex Alderton, Roberto Amato, Jeffrey Barrett, Sonia Goncalves, Ewan Harrison, David K. Jackson, Ian Johnston, Dominic Kwiatkowski, Cordelia Langford, John Sillitoe on behalf of the Wellcome Sanger Institute COVID-19 Surveillance Team |
| EPI_ISL_2829097 | Lighthouse Lab in Milton Keynes | Wellcome Sanger Institute for the COVID-19 Genomics UK (COG-UK) Consortium | The Lighthouse Lab in Milton Keynes and Alex Alderton, Roberto Amato, Jeffrey Barrett, Sonia Goncalves, Ewan Harrison, David K. Jackson, Ian Johnston, Dominic Kwiatkowski, Cordelia Langford, John Sillitoe on behalf of the Wellcome Sanger Institute COVID-19 Surveillance Team |
| EPI_ISL_2829098 | Lighthouse Lab in Alderley Park | Wellcome Sanger Institute for the COVID-19 Genomics UK (COG-UK) Consortium | Jacquelyn Wynn, Mairead Hyland, The Lighthouse Lab in Alderley Park and Alex Alderton, Roberto Amato, Jeffrey Barrett, Sonia Goncalves, Ewan Harrison, David K. Jackson, Ian Johnston, Dominic Kwiatkowski, Cordelia Langford, John Sillitoe on behalf of the Wellcome Sanger Institute COVID-19 Surveillance Team |
| EPI_ISL_2829099 | Lighthouse Laboratory Plymouth | Wellcome Sanger Institute for the COVID-19 Genomics UK (COG-UK) Consortium | Lighthouse Laboratory Plymouth and Alex Alderton, Roberto Amato, Jeffrey Barrett, Sonia Goncalves, Ewan Harrison, David K. Jackson, Ian Johnston, Dominic Kwiatkowski, Cordelia Langford, John Sillitoe on behalf of the Wellcome Sanger Institute COVID-19 Surveillance Team |
| EPI_ISL_2829100 | Lighthouse Lab in Milton Keynes | Wellcome Sanger Institute for the COVID-19 Genomics UK (COG-UK) Consortium | The Lighthouse Lab in Milton Keynes and Alex Alderton, Roberto Amato, Jeffrey Barrett, Sonia Goncalves, Ewan Harrison, David K. Jackson, Ian Johnston, Dominic Kwiatkowski, Cordelia Langford, John Sillitoe on behalf of the Wellcome Sanger Institute COVID-19 Surveillance Team |
| EPI_ISL_2829101 | Lighthouse Lab in Alderley Park | Wellcome Sanger Institute for the COVID-19 Genomics UK (COG-UK) Consortium | Jacquelyn Wynn, Mairead Hyland, The Lighthouse Lab in Alderley Park and Alex Alderton, Roberto Amato, Jeffrey Barrett, Sonia Goncalves, Ewan Harrison, David K. Jackson, Ian Johnston, Dominic Kwiatkowski, Cordelia Langford, John Sillitoe on behalf of the Wellcome Sanger Institute COVID-19 Surveillance Team |
| EPI_ISL_2829102, EPI_ISL_2829103 | Lighthouse Lab in Milton Keynes | Wellcome Sanger Institute for the COVID-19 Genomics UK (COG-UK) Consortium | The Lighthouse Lab in Milton Keynes and Alex Alderton, Roberto Amato, Jeffrey Barrett, Sonia Goncalves, Ewan Harrison, David K. Jackson, Ian Johnston, Dominic Kwiatkowski, Cordelia Langford, John Sillitoe on behalf of the Wellcome Sanger Institute COVID-19 Surveillance Team |
| EPI_ISL_2829104, EPI_ISL_2829105 | Health Services Laboratories | Wellcome Sanger Institute for the COVID-19 Genomics UK (COG-UK) Consortium | Health Services Laboratories and Alex Alderton, Roberto Amato, Jeffrey Barrett, Sonia Goncalves, Ewan Harrison, David K. Jackson, Ian Johnston, Dominic Kwiatkowski, Cordelia Langford, John Sillitoe on behalf of the Wellcome Sanger Institute COVID-19 Surveillance Team |
| EPI_ISL_2829106 | Lighthouse Lab in Alderley Park | Wellcome Sanger Institute for the COVID-19 Genomics UK (COG-UK) Consortium | Jacquelyn Wynn, Mairead Hyland, The Lighthouse Lab in Alderley Park and Alex Alderton, Roberto Amato, Jeffrey Barrett, Sonia Goncalves, Ewan Harrison, David K. Jackson, Ian Johnston, Dominic Kwiatkowski, Cordelia Langford, John Sillitoe on behalf of the Wellcome Sanger Institute COVID-19 Surveillance Team |
| EPI_ISL_2829107 | Health Services Laboratories | Wellcome Sanger Institute for the COVID-19 Genomics UK (COG-UK) Consortium | Health Services Laboratories and Alex Alderton, Roberto Amato, Jeffrey Barrett, Sonia Goncalves, Ewan Harrison, David K. Jackson, Ian Johnston, Dominic Kwiatkowski, Cordelia Langford, John Sillitoe on behalf of the Wellcome Sanger Institute COVID-19 Surveillance Team |
| EPI_ISL_2829108 | Lighthouse Laboratory Plymouth | Wellcome Sanger Institute for the COVID-19 Genomics UK (COG-UK) Consortium | Lighthouse Laboratory Plymouth and Alex Alderton, Roberto Amato, Jeffrey Barrett, Sonia Goncalves, Ewan Harrison, David K. Jackson, Ian Johnston, Dominic Kwiatkowski, Cordelia Langford, John Sillitoe on behalf of the Wellcome Sanger Institute COVID-19 Surveillance Team |
| EPI_ISL_2829109 | Lighthouse Lab in Alderley Park | Wellcome Sanger Institute for the COVID-19 Genomics UK (COG-UK) Consortium | Jacquelyn Wynn, Mairead Hyland, The Lighthouse Lab in Alderley Park and Alex Alderton, Roberto Amato, Jeffrey Barrett, Sonia Goncalves, Ewan Harrison, David K. Jackson, Ian Johnston, Dominic Kwiatkowski, Cordelia Langford, John Sillitoe on behalf of the Wellcome Sanger Institute COVID-19 Surveillance Team |
| EPI_ISL_2829110 | Lighthouse Lab in Milton Keynes | Wellcome Sanger Institute for the COVID-19 Genomics UK (COG-UK) Consortium | The Lighthouse Lab in Milton Keynes and Alex Alderton, Roberto Amato, Jeffrey Barrett, Sonia Goncalves, Ewan Harrison, David K. Jackson, Ian Johnston, Dominic Kwiatkowski, Cordelia Langford, John Sillitoe on behalf of the Wellcome Sanger Institute COVID-19 Surveillance Team |
| EPI_ISL_2829111, EPI_ISL_2829112 | Lighthouse Laboratory Plymouth | Wellcome Sanger Institute for the COVID-19 Genomics UK (COG-UK) Consortium | Lighthouse Laboratory Plymouth and Alex Alderton, Roberto Amato, Jeffrey Barrett, Sonia Goncalves, Ewan Harrison, David K. Jackson, Ian Johnston, Dominic Kwiatkowski, Cordelia Langford, John Sillitoe on behalf of the Wellcome Sanger Institute COVID-19 Surveillance Team |
| EPI_ISL_2829113, EPI_ISL_2829114 | Lighthouse Lab in Milton Keynes | Wellcome Sanger Institute for the COVID-19 Genomics UK (COG-UK) Consortium | The Lighthouse Lab in Milton Keynes and Alex Alderton, Roberto Amato, Jeffrey Barrett, Sonia Goncalves, Ewan Harrison, David K. Jackson, Ian Johnston, Dominic Kwiatkowski, Cordelia Langford, John Sillitoe on behalf of the Wellcome Sanger Institute COVID-19 Surveillance Team |
| EPI_ISL_2829115 | Lighthouse Lab in Alderley Park | Wellcome Sanger Institute for the COVID-19 Genomics UK (COG-UK) Consortium | Jacquelyn Wynn, Mairead Hyland, The Lighthouse Lab in Alderley Park and Alex Alderton, Roberto Amato, Jeffrey Barrett, Sonia Goncalves, Ewan Harrison, David K. Jackson, Ian Johnston, Dominic Kwiatkowski, Cordelia Langford, John Sillitoe on behalf of the Wellcome Sanger Institute COVID-19 Surveillance Team |
| EPI_ISL_2829116 | Health Services Laboratories | Wellcome Sanger Institute for the COVID-19 Genomics UK (COG-UK) Consortium | Health Services Laboratories and Alex Alderton, Roberto Amato, Jeffrey Barrett, Sonia Goncalves, Ewan Harrison, David K. Jackson, Ian Johnston, Dominic Kwiatkowski, Cordelia Langford, John Sillitoe on behalf of the Wellcome Sanger Institute COVID-19 Surveillance Team |
| EPI_ISL_2829117, EPI_ISL_2829118 | Lighthouse Laboratory Plymouth | Wellcome Sanger Institute for the COVID-19 Genomics UK (COG-UK) Consortium | Lighthouse Laboratory Plymouth and Alex Alderton, Roberto Amato, Jeffrey Barrett, Sonia Goncalves, Ewan Harrison, David K. Jackson, Ian Johnston, Dominic Kwiatkowski, Cordelia Langford, John Sillitoe on behalf of the Wellcome Sanger Institute COVID-19 Surveillance Team |
| EPI_ISL_2829119, EPI_ISL_2829120, EPI_ISL_2829121 | Lighthouse Lab in Alderley Park | Wellcome Sanger Institute for the COVID-19 Genomics UK (COG-UK) Consortium | Jacquelyn Wynn, Mairead Hyland, The Lighthouse Lab in Alderley Park and Alex Alderton, Roberto Amato, Jeffrey Barrett, Sonia Goncalves, Ewan Harrison, David K. Jackson, Ian Johnston, Dominic Kwiatkowski, Cordelia Langford, John Sillitoe on behalf of the Wellcome Sanger Institute COVID-19 Surveillance Team |
| EPI_ISL_2829122 | Lighthouse Lab in Milton Keynes | Wellcome Sanger Institute for the COVID-19 Genomics UK (COG-UK) Consortium | The Lighthouse Lab in Milton Keynes and Alex Alderton, Roberto Amato, Jeffrey Barrett, Sonia Goncalves, Ewan Harrison, David K. Jackson, Ian Johnston, Dominic Kwiatkowski, Cordelia Langford, John Sillitoe on behalf of the Wellcome Sanger Institute COVID-19 Surveillance Team |
| EPI_ISL_2829123, EPI_ISL_2829124 | Lighthouse Lab in Alderley Park | Wellcome Sanger Institute for the COVID-19 Genomics UK (COG-UK) Consortium | Jacquelyn Wynn, Mairead Hyland, The Lighthouse Lab in Alderley Park and Alex Alderton, Roberto Amato, Jeffrey Barrett, Sonia Goncalves, Ewan Harrison, David K. Jackson, Ian Johnston, Dominic Kwiatkowski, Cordelia Langford, John Sillitoe on behalf of the Wellcome Sanger Institute COVID-19 Surveillance Team |
| EPI_ISL_2829125 | Lighthouse Lab in Milton Keynes | Wellcome Sanger Institute for the COVID-19 Genomics UK (COG-UK) Consortium | The Lighthouse Lab in Milton Keynes and Alex Alderton, Roberto Amato, Jeffrey Barrett, Sonia Goncalves, Ewan Harrison, David K. Jackson, Ian Johnston, Dominic Kwiatkowski, Cordelia Langford, John Sillitoe on behalf of the Wellcome Sanger Institute COVID-19 Surveillance Team |
| EPI_ISL_2829126, EPI_ISL_2829127, EPI_ISL_2829128, EPI_ISL_2829129 | Lighthouse Lab in Alderley Park | Wellcome Sanger Institute for the COVID-19 Genomics UK (COG-UK) Consortium | Jacquelyn Wynn, Mairead Hyland, The Lighthouse Lab in Alderley Park and Alex Alderton, Roberto Amato, Jeffrey Barrett, Sonia Goncalves, Ewan Harrison, David K. Jackson, Ian Johnston, Dominic Kwiatkowski, Cordelia Langford, John Sillitoe on behalf of the Wellcome Sanger Institute COVID-19 Surveillance Team |
| EPI_ISL_2829130 | Lighthouse Lab in Milton Keynes | Wellcome Sanger Institute for the COVID-19 Genomics UK (COG-UK) Consortium | The Lighthouse Lab in Milton Keynes and Alex Alderton, Roberto Amato, Jeffrey Barrett, Sonia Goncalves, Ewan Harrison, David K. Jackson, Ian Johnston, Dominic Kwiatkowski, Cordelia Langford, John Sillitoe on behalf of the Wellcome Sanger Institute COVID-19 Surveillance Team |
| EPI_ISL_2829131 | Lighthouse Laboratory Plymouth | Wellcome Sanger Institute for the COVID-19 Genomics UK (COG-UK) Consortium | Lighthouse Laboratory Plymouth and Alex Alderton, Roberto Amato, Jeffrey Barrett, Sonia Goncalves, Ewan Harrison, David K. Jackson, Ian Johnston, Dominic Kwiatkowski, Cordelia Langford, John Sillitoe on behalf of the Wellcome Sanger Institute COVID-19 Surveillance Team |
| EPI_ISL_2829132 | Lighthouse Lab in Milton Keynes | Wellcome Sanger Institute for the COVID-19 Genomics UK (COG-UK) Consortium | The Lighthouse Lab in Milton Keynes and Alex Alderton, Roberto Amato, Jeffrey Barrett, Sonia Goncalves, Ewan Harrison, David K. Jackson, Ian Johnston, Dominic Kwiatkowski, Cordelia Langford, John Sillitoe on behalf of the Wellcome Sanger Institute COVID-19 Surveillance Team |
| EPI_ISL_2829133 | Lighthouse Lab in Alderley Park | Wellcome Sanger Institute for the COVID-19 Genomics UK (COG-UK) Consortium | Jacquelyn Wynn, Mairead Hyland, The Lighthouse Lab in Alderley Park and Alex Alderton, Roberto Amato, Jeffrey Barrett, Sonia Goncalves, Ewan Harrison, David K. Jackson, Ian Johnston, Dominic Kwiatkowski, Cordelia Langford, John Sillitoe on behalf of the Wellcome Sanger Institute COVID-19 Surveillance Team |
| EPI_ISL_2829134 | Lighthouse Lab in Milton Keynes | Wellcome Sanger Institute for the COVID-19 Genomics UK (COG-UK) Consortium | The Lighthouse Lab in Milton Keynes and Alex Alderton, Roberto Amato, Jeffrey Barrett, Sonia Goncalves, Ewan Harrison, David K. Jackson, Ian Johnston, Dominic Kwiatkowski, Cordelia Langford, John Sillitoe on behalf of the Wellcome Sanger Institute COVID-19 Surveillance Team |
| EPI_ISL_2829135, EPI_ISL_2829136, EPI_ISL_2829137 | Lighthouse Lab in Alderley Park | Wellcome Sanger Institute for the COVID-19 Genomics UK (COG-UK) Consortium | Jacquelyn Wynn, Mairead Hyland, The Lighthouse Lab in Alderley Park and Alex Alderton, Roberto Amato, Jeffrey Barrett, Sonia Goncalves, Ewan Harrison, David K. Jackson, Ian Johnston, Dominic Kwiatkowski, Cordelia Langford, John Sillitoe on behalf of the Wellcome Sanger Institute COVID-19 Surveillance Team |

| | | | |
|---|---|---|---|
| EPI_ISL_2829138, EPI_ISL_2829139, EPI_ISL_2829140, EPI_ISL_2829141 | Lighthouse Lab in Milton Keynes | Wellcome Sanger Institute for the COVID-19 Genomics UK (COG-UK) Consortium | The Lighthouse Lab in Milton Keynes and Alex Alderton, Roberto Amato, Jeffrey Barrett, Sonia Goncalves, Ewan Harrison, David K. Jackson, Ian Johnston, Dominic Kwiatkowski, Cordelia Langford, John Sillitoe on behalf of the Wellcome Sanger Institute COVID-19 Surveillance Team |
| EPI_ISL_2829142, EPI_ISL_2829143 | Health Services Laboratories | Wellcome Sanger Institute for the COVID-19 Genomics UK (COG-UK) Consortium | Health Services Laboratories and Alex Alderton, Roberto Amato, Jeffrey Barrett, Sonia Goncalves, Ewan Harrison, David K. Jackson, Ian Johnston, Dominic Kwiatkowski, Cordelia Langford, John Sillitoe on behalf of the Wellcome Sanger Institute COVID-19 Surveillance Team |
| EPI_ISL_2829144, EPI_ISL_2829145 | Lighthouse Laboratory Plymouth | Wellcome Sanger Institute for the COVID-19 Genomics UK (COG-UK) Consortium | Lighthouse Laboratory Plymouth and Alex Alderton, Roberto Amato, Jeffrey Barrett, Sonia Goncalves, Ewan Harrison, David K. Jackson, Ian Johnston, Dominic Kwiatkowski, Cordelia Langford, John Sillitoe on behalf of the Wellcome Sanger Institute COVID-19 Surveillance Team |
| EPI_ISL_2829146 | Lighthouse Lab in Alderley Park | Wellcome Sanger Institute for the COVID-19 Genomics UK (COG-UK) Consortium | Jacquelyn Wynn, Mairead Hyland, The Lighthouse Lab in Alderley Park and Alex Alderton, Roberto Amato, Jeffrey Barrett, Sonia Goncalves, Ewan Harrison, David K. Jackson, Ian Johnston, Dominic Kwiatkowski, Cordelia Langford, John Sillitoe on behalf of the Wellcome Sanger Institute COVID-19 Surveillance Team |
| EPI_ISL_2829147, EPI_ISL_2829148, EPI_ISL_2829149, EPI_ISL_2829150, EPI_ISL_2829151 | Lighthouse Lab in Milton Keynes | Wellcome Sanger Institute for the COVID-19 Genomics UK (COG-UK) Consortium | The Lighthouse Lab in Milton Keynes and Alex Alderton, Roberto Amato, Jeffrey Barrett, Sonia Goncalves, Ewan Harrison, David K. Jackson, Ian Johnston, Dominic Kwiatkowski, Cordelia Langford, John Sillitoe on behalf of the Wellcome Sanger Institute COVID-19 Surveillance Team |
| EPI_ISL_2829152 | Health Services Laboratories | Wellcome Sanger Institute for the COVID-19 Genomics UK (COG-UK) Consortium | Health Services Laboratories and Alex Alderton, Roberto Amato, Jeffrey Barrett, Sonia Goncalves, Ewan Harrison, David K. Jackson, Ian Johnston, Dominic Kwiatkowski, Cordelia Langford, John Sillitoe on behalf of the Wellcome Sanger Institute COVID-19 Surveillance Team |
| EPI_ISL_2829153 | Lighthouse Lab in Alderley Park | Wellcome Sanger Institute for the COVID-19 Genomics UK (COG-UK) Consortium | Jacquelyn Wynn, Mairead Hyland, The Lighthouse Lab in Alderley Park and Alex Alderton, Roberto Amato, Jeffrey Barrett, Sonia Goncalves, Ewan Harrison, David K. Jackson, Ian Johnston, Dominic Kwiatkowski, Cordelia Langford, John Sillitoe on behalf of the Wellcome Sanger Institute COVID-19 Surveillance Team |
| EPI_ISL_2829154, EPI_ISL_2829155, EPI_ISL_2829156 | Lighthouse Lab in Milton Keynes | Wellcome Sanger Institute for the COVID-19 Genomics UK (COG-UK) Consortium | The Lighthouse Lab in Milton Keynes and Alex Alderton, Roberto Amato, Jeffrey Barrett, Sonia Goncalves, Ewan Harrison, David K. Jackson, Ian Johnston, Dominic Kwiatkowski, Cordelia Langford, John Sillitoe on behalf of the Wellcome Sanger Institute COVID-19 Surveillance Team |
| EPI_ISL_2829157, EPI_ISL_2829158, EPI_ISL_2829159 | Lighthouse Laboratory Plymouth | Wellcome Sanger Institute for the COVID-19 Genomics UK (COG-UK) Consortium | Lighthouse Laboratory Plymouth and Alex Alderton, Roberto Amato, Jeffrey Barrett, Sonia Goncalves, Ewan Harrison, David K. Jackson, Ian Johnston, Dominic Kwiatkowski, Cordelia Langford, John Sillitoe on behalf of the Wellcome Sanger Institute COVID-19 Surveillance Team |
| EPI_ISL_2829160 | Lighthouse Lab in Milton Keynes | Wellcome Sanger Institute for the COVID-19 Genomics UK (COG-UK) Consortium | The Lighthouse Lab in Milton Keynes and Alex Alderton, Roberto Amato, Jeffrey Barrett, Sonia Goncalves, Ewan Harrison, David K. Jackson, Ian Johnston, Dominic Kwiatkowski, Cordelia Langford, John Sillitoe on behalf of the Wellcome Sanger Institute COVID-19 Surveillance Team |
| EPI_ISL_2829161 | Lighthouse Lab in Alderley Park | Wellcome Sanger Institute for the COVID-19 Genomics UK (COG-UK) Consortium | Jacquelyn Wynn, Mairead Hyland, The Lighthouse Lab in Alderley Park and Alex Alderton, Roberto Amato, Jeffrey Barrett, Sonia Goncalves, Ewan Harrison, David K. Jackson, Ian Johnston, Dominic Kwiatkowski, Cordelia Langford, John Sillitoe on behalf of the Wellcome Sanger Institute COVID-19 Surveillance Team |
| EPI_ISL_2829162 | Lighthouse Laboratory Plymouth | Wellcome Sanger Institute for the COVID-19 Genomics UK (COG-UK) Consortium | Lighthouse Laboratory Plymouth and Alex Alderton, Roberto Amato, Jeffrey Barrett, Sonia Goncalves, Ewan Harrison, David K. Jackson, Ian Johnston, Dominic Kwiatkowski, Cordelia Langford, John Sillitoe on behalf of the Wellcome Sanger Institute COVID-19 Surveillance Team |
| EPI_ISL_2829163 | Lighthouse Lab in Milton Keynes | Wellcome Sanger Institute for the COVID-19 Genomics UK (COG-UK) Consortium | The Lighthouse Lab in Milton Keynes and Alex Alderton, Roberto Amato, Jeffrey Barrett, Sonia Goncalves, Ewan Harrison, David K. Jackson, Ian Johnston, Dominic Kwiatkowski, Cordelia Langford, John Sillitoe on behalf of the Wellcome Sanger Institute COVID-19 Surveillance Team |
| EPI_ISL_2829164 | Lighthouse Lab in Alderley Park | Wellcome Sanger Institute for the COVID-19 Genomics UK (COG-UK) Consortium | Jacquelyn Wynn, Mairead Hyland, The Lighthouse Lab in Alderley Park and Alex Alderton, Roberto Amato, Jeffrey Barrett, Sonia Goncalves, Ewan Harrison, David K. Jackson, Ian Johnston, Dominic Kwiatkowski, Cordelia Langford, John Sillitoe on behalf of the Wellcome Sanger Institute COVID-19 Surveillance Team |
| EPI_ISL_2829165, EPI_ISL_2829166, EPI_ISL_2829167 | Lighthouse Lab in Milton Keynes | Wellcome Sanger Institute for the COVID-19 Genomics UK (COG-UK) Consortium | The Lighthouse Lab in Milton Keynes and Alex Alderton, Roberto Amato, Jeffrey Barrett, Sonia Goncalves, Ewan Harrison, David K. Jackson, Ian Johnston, Dominic Kwiatkowski, Cordelia Langford, John Sillitoe on behalf of the Wellcome Sanger Institute COVID-19 Surveillance Team |
| EPI_ISL_2829168 | Health Services Laboratories | Wellcome Sanger Institute for the COVID-19 Genomics UK (COG-UK) Consortium | Health Services Laboratories and Alex Alderton, Roberto Amato, Jeffrey Barrett, Sonia Goncalves, Ewan Harrison, David K. Jackson, Ian Johnston, Dominic Kwiatkowski, Cordelia Langford, John Sillitoe on behalf of the Wellcome Sanger Institute COVID-19 Surveillance Team |
| EPI_ISL_2829169 | Lighthouse Lab in Glasgow | Wellcome Sanger Institute for the COVID-19 Genomics UK (COG-UK) Consortium | Harper VanSteenhouse, Yumi Kasai, David Gray, Carol Clugston, Anna Dominiczak and Alex Alderton, Roberto Amato, Jeffrey Barrett, Sonia Goncalves, Ewan Harrison, David K. Jackson, Ian Johnston, Dominic Kwiatkowski, Cordelia Langford, John Sillitoe on behalf of the Wellcome Sanger Institute COVID-19 Surveillance Team |
| EPI_ISL_2829170, EPI_ISL_2829171, EPI_ISL_2829172, EPI_ISL_2829173 | Lighthouse Lab in Alderley Park | Wellcome Sanger Institute for the COVID-19 Genomics UK (COG-UK) Consortium | Jacquelyn Wynn, Mairead Hyland, The Lighthouse Lab in Alderley Park and Alex Alderton, Roberto Amato, Jeffrey Barrett, Sonia Goncalves, Ewan Harrison, David K. Jackson, Ian Johnston, Dominic Kwiatkowski, Cordelia Langford, John Sillitoe on behalf of the Wellcome Sanger Institute COVID-19 Surveillance Team |
| EPI_ISL_2829174 | Lighthouse Laboratory Plymouth | Wellcome Sanger Institute for the COVID-19 Genomics UK (COG-UK) Consortium | Lighthouse Laboratory Plymouth and Alex Alderton, Roberto Amato, Jeffrey Barrett, Sonia Goncalves, Ewan Harrison, David K. Jackson, Ian Johnston, Dominic Kwiatkowski, Cordelia Langford, John Sillitoe on behalf of the Wellcome Sanger Institute COVID-19 Surveillance Team |
| EPI_ISL_2829175 | Lighthouse Lab in Milton Keynes | Wellcome Sanger Institute for the COVID-19 Genomics UK (COG-UK) Consortium | The Lighthouse Lab in Milton Keynes and Alex Alderton, Roberto Amato, Jeffrey Barrett, Sonia Goncalves, Ewan Harrison, David K. Jackson, Ian Johnston, Dominic Kwiatkowski, Cordelia Langford, John Sillitoe on behalf of the Wellcome Sanger Institute COVID-19 Surveillance Team |
| EPI_ISL_2829176 | Lighthouse Laboratory Plymouth | Wellcome Sanger Institute for the COVID-19 Genomics UK (COG-UK) Consortium | Lighthouse Laboratory Plymouth and Alex Alderton, Roberto Amato, Jeffrey Barrett, Sonia Goncalves, Ewan Harrison, David K. Jackson, Ian Johnston, Dominic Kwiatkowski, Cordelia Langford, John Sillitoe on behalf of the Wellcome Sanger Institute COVID-19 Surveillance Team |
| EPI_ISL_2829177, EPI_ISL_2829178 | Lighthouse Lab in Milton Keynes | Wellcome Sanger Institute for the COVID-19 Genomics UK (COG-UK) Consortium | The Lighthouse Lab in Milton Keynes and Alex Alderton, Roberto Amato, Jeffrey Barrett, Sonia Goncalves, Ewan Harrison, David K. Jackson, Ian Johnston, Dominic Kwiatkowski, Cordelia Langford, John Sillitoe on behalf of the Wellcome Sanger Institute COVID-19 Surveillance Team |
| EPI_ISL_2829179 | Lighthouse Lab in Alderley Park | Wellcome Sanger Institute for the COVID-19 Genomics UK (COG-UK) Consortium | Jacquelyn Wynn, Mairead Hyland, The Lighthouse Lab in Alderley Park and Alex Alderton, Roberto Amato, Jeffrey Barrett, Sonia Goncalves, Ewan Harrison, David K. Jackson, Ian Johnston, Dominic Kwiatkowski, Cordelia Langford, John Sillitoe on behalf of the Wellcome Sanger Institute COVID-19 Surveillance Team |
| EPI_ISL_2829180, EPI_ISL_2829181 | Lighthouse Lab in Milton Keynes | Wellcome Sanger Institute for the COVID-19 Genomics UK (COG-UK) Consortium | The Lighthouse Lab in Milton Keynes and Alex Alderton, Roberto Amato, Jeffrey Barrett, Sonia Goncalves, Ewan Harrison, David K. Jackson, Ian Johnston, Dominic Kwiatkowski, Cordelia Langford, John Sillitoe on behalf of the Wellcome Sanger Institute COVID-19 Surveillance Team |
| EPI_ISL_2829182 | Lighthouse Laboratory Plymouth | Wellcome Sanger Institute for the COVID-19 Genomics UK (COG-UK) Consortium | Lighthouse Laboratory Plymouth and Alex Alderton, Roberto Amato, Jeffrey Barrett, Sonia Goncalves, Ewan Harrison, David K. Jackson, Ian Johnston, Dominic Kwiatkowski, Cordelia Langford, John Sillitoe on behalf of the Wellcome Sanger Institute COVID-19 Surveillance Team |
| EPI_ISL_2829183 | Health Services Laboratories | Wellcome Sanger Institute for the COVID-19 Genomics UK (COG-UK) Consortium | Health Services Laboratories and Alex Alderton, Roberto Amato, Jeffrey Barrett, Sonia Goncalves, Ewan Harrison, David K. Jackson, Ian Johnston, Dominic Kwiatkowski, Cordelia Langford, John Sillitoe on behalf of the Wellcome Sanger Institute COVID-19 Surveillance Team |
| EPI_ISL_2829184, EPI_ISL_2829185 | Lighthouse Lab in Alderley Park | Wellcome Sanger Institute for the COVID-19 Genomics UK (COG-UK) Consortium | Jacquelyn Wynn, Mairead Hyland, The Lighthouse Lab in Alderley Park and Alex Alderton, Roberto Amato, Jeffrey Barrett, Sonia Goncalves, Ewan Harrison, David K. Jackson, Ian Johnston, Dominic Kwiatkowski, Cordelia Langford, John Sillitoe on behalf of the Wellcome Sanger Institute COVID-19 Surveillance Team |
| EPI_ISL_2829186 | Health Services Laboratories | Wellcome Sanger Institute for the COVID-19 Genomics UK (COG-UK) Consortium | Health Services Laboratories and Alex Alderton, Roberto Amato, Jeffrey Barrett, Sonia Goncalves, Ewan Harrison, David K. Jackson, Ian Johnston, Dominic Kwiatkowski, Cordelia Langford, John Sillitoe on behalf of the Wellcome Sanger Institute COVID-19 Surveillance Team |
| EPI_ISL_2829187, EPI_ISL_2829188 | Lighthouse Lab in Alderley Park | Wellcome Sanger Institute for the COVID-19 Genomics UK (COG-UK) Consortium | Jacquelyn Wynn, Mairead Hyland, The Lighthouse Lab in Alderley Park and Alex Alderton, Roberto Amato, Jeffrey Barrett, Sonia Goncalves, Ewan Harrison, David K. Jackson, Ian Johnston, Dominic Kwiatkowski, Cordelia Langford, John Sillitoe on behalf of the Wellcome Sanger Institute COVID-19 Surveillance Team |
| EPI_ISL_2829189 | Lighthouse Laboratory Plymouth | Wellcome Sanger Institute for the COVID-19 Genomics UK (COG-UK) Consortium | Lighthouse Laboratory Plymouth and Alex Alderton, Roberto Amato, Jeffrey Barrett, Sonia Goncalves, Ewan Harrison, David K. Jackson, Ian Johnston, Dominic Kwiatkowski, Cordelia Langford, John Sillitoe on behalf of the Wellcome Sanger Institute COVID-19 Surveillance Team |
| EPI_ISL_2829190 | Lighthouse Lab in Alderley Park | Wellcome Sanger Institute for the COVID-19 Genomics UK (COG-UK) Consortium | Jacquelyn Wynn, Mairead Hyland, The Lighthouse Lab in Alderley Park and Alex Alderton, Roberto Amato, Jeffrey Barrett, Sonia Goncalves, Ewan Harrison, David K. Jackson, Ian Johnston, Dominic Kwiatkowski, Cordelia Langford, John Sillitoe on behalf of the Wellcome Sanger Institute COVID-19 Surveillance Team |
| EPI_ISL_2829191 | Lighthouse Laboratory Plymouth | Wellcome Sanger Institute for the COVID-19 Genomics UK (COG-UK) Consortium | Lighthouse Laboratory Plymouth and Alex Alderton, Roberto Amato, Jeffrey Barrett, Sonia Goncalves, Ewan Harrison, David K. Jackson, Ian Johnston, Dominic Kwiatkowski, Cordelia Langford, John Sillitoe on behalf of the Wellcome Sanger Institute COVID-19 Surveillance Team |
| EPI_ISL_2829192 | Health Services Laboratories | Wellcome Sanger Institute for the COVID-19 Genomics UK (COG-UK) Consortium | Health Services Laboratories and Alex Alderton, Roberto Amato, Jeffrey Barrett, Sonia Goncalves, Ewan Harrison, David K. Jackson, Ian Johnston, Dominic Kwiatkowski, Cordelia Langford, John Sillitoe on behalf of the Wellcome Sanger Institute COVID-19 Surveillance Team |
| EPI_ISL_2829193 | Lighthouse Lab in Alderley Park | Wellcome Sanger Institute for the COVID-19 Genomics UK (COG-UK) Consortium | Jacquelyn Wynn, Mairead Hyland, The Lighthouse Lab in Alderley Park and Alex Alderton, Roberto Amato, Jeffrey Barrett, Sonia Goncalves, Ewan Harrison, David K. Jackson, Ian Johnston, Dominic Kwiatkowski, Cordelia Langford, John Sillitoe on behalf of the Wellcome Sanger Institute COVID-19 Surveillance Team |
| EPI_ISL_2829194, EPI_ISL_2829195, EPI_ISL_2829196, EPI_ISL_2829197 | Health Services Laboratories | Wellcome Sanger Institute for the COVID-19 Genomics UK (COG-UK) Consortium | Health Services Laboratories and Alex Alderton, Roberto Amato, Jeffrey Barrett, Sonia Goncalves, Ewan Harrison, David K. Jackson, Ian Johnston, Dominic Kwiatkowski, Cordelia Langford, John Sillitoe on behalf of the Wellcome Sanger Institute COVID-19 Surveillance Team |
| EPI_ISL_2829198 | Lighthouse Lab in Milton Keynes | Wellcome Sanger Institute for the COVID-19 Genomics UK (COG-UK) Consortium | The Lighthouse Lab in Milton Keynes and Alex Alderton, Roberto Amato, Jeffrey Barrett, Sonia Goncalves, Ewan Harrison, David K. Jackson, Ian Johnston, Dominic Kwiatkowski, Cordelia Langford, John Sillitoe on behalf of the Wellcome Sanger Institute COVID-19 Surveillance Team |
| EPI_ISL_2829199, EPI_ISL_2829200 | Lighthouse Lab in Alderley Park | Wellcome Sanger Institute for the COVID-19 Genomics UK (COG-UK) Consortium | Jacquelyn Wynn, Mairead Hyland, The Lighthouse Lab in Alderley Park and Alex Alderton, Roberto Amato, Jeffrey Barrett, Sonia Goncalves, Ewan Harrison, David K. Jackson, Ian Johnston, Dominic Kwiatkowski, Cordelia Langford, John Sillitoe on behalf of the Wellcome Sanger Institute COVID-19 Surveillance Team |
| EPI_ISL_2829201, EPI_ISL_2829202 | Health Services Laboratories | Wellcome Sanger Institute for the COVID-19 Genomics UK (COG-UK) Consortium | Health Services Laboratories and Alex Alderton, Roberto Amato, Jeffrey Barrett, Sonia Goncalves, Ewan Harrison, David K. Jackson, Ian Johnston, Dominic Kwiatkowski, Cordelia Langford, John Sillitoe on behalf of the Wellcome Sanger Institute COVID-19 Surveillance Team |
| EPI_ISL_2829203 | Lighthouse Lab in Milton Keynes | Wellcome Sanger Institute for the COVID-19 Genomics UK (COG-UK) Consortium | The Lighthouse Lab in Milton Keynes and Alex Alderton, Roberto Amato, Jeffrey Barrett, Sonia Goncalves, Ewan Harrison, David K. Jackson, Ian Johnston, Dominic Kwiatkowski, Cordelia Langford, John Sillitoe on behalf of the Wellcome Sanger Institute COVID-19 Surveillance Team |
| EPI_ISL_2829204, EPI_ISL_2829205 | Lighthouse Laboratory Plymouth | Wellcome Sanger Institute for the COVID-19 Genomics UK (COG-UK) Consortium | Lighthouse Laboratory Plymouth and Alex Alderton, Roberto Amato, Jeffrey Barrett, Sonia Goncalves, Ewan Harrison, David K. Jackson, Ian Johnston, Dominic Kwiatkowski, Cordelia Langford, John Sillitoe on behalf of the Wellcome Sanger Institute COVID-19 Surveillance Team |

| | | | |
|---|---|---|---|
| EPI_ISL_2829206, EPI_ISL_2829207, EPI_ISL_2829208, EPI_ISL_2829209 | Lighthouse Lab in Milton Keynes | Wellcome Sanger Institute for the COVID-19 Genomics UK (COG-UK) Consortium | The Lighthouse Lab in Milton Keynes and Alex Alderton, Roberto Amato, Jeffrey Barrett, Sonia Goncalves, Ewan Harrison, David K. Jackson, Ian Johnston, Dominic Kwiatkowski, Cordelia Langford, John Sillitoe on behalf of the Wellcome Sanger Institute COVID-19 Surveillance Team |
| EPI_ISL_2829210 | Lighthouse Lab in Alderley Park | Wellcome Sanger Institute for the COVID-19 Genomics UK (COG-UK) Consortium | Jacquelyn Wynn, Mairead Hyland, The Lighthouse Lab in Alderley Park and Alex Alderton, Roberto Amato, Jeffrey Barrett, Sonia Goncalves, Ewan Harrison, David K. Jackson, Ian Johnston, Dominic Kwiatkowski, Cordelia Langford, John Sillitoe on behalf of the Wellcome Sanger Institute COVID-19 Surveillance Team |
| EPI_ISL_2829211 | Health Services Laboratories | Wellcome Sanger Institute for the COVID-19 Genomics UK (COG-UK) Consortium | Health Services Laboratories and Alex Alderton, Roberto Amato, Jeffrey Barrett, Sonia Goncalves, Ewan Harrison, David K. Jackson, Ian Johnston, Dominic Kwiatkowski, Cordelia Langford, John Sillitoe on behalf of the Wellcome Sanger Institute COVID-19 Surveillance Team |
| EPI_ISL_2829212 | Lighthouse Lab in Alderley Park | Wellcome Sanger Institute for the COVID-19 Genomics UK (COG-UK) Consortium | Jacquelyn Wynn, Mairead Hyland, The Lighthouse Lab in Alderley Park and Alex Alderton, Roberto Amato, Jeffrey Barrett, Sonia Goncalves, Ewan Harrison, David K. Jackson, Ian Johnston, Dominic Kwiatkowski, Cordelia Langford, John Sillitoe on behalf of the Wellcome Sanger Institute COVID-19 Surveillance Team |
| EPI_ISL_2829213 | Lighthouse Laboratory Plymouth | Wellcome Sanger Institute for the COVID-19 Genomics UK (COG-UK) Consortium | Lighthouse Laboratory Plymouth and Alex Alderton, Roberto Amato, Jeffrey Barrett, Sonia Goncalves, Ewan Harrison, David K. Jackson, Ian Johnston, Dominic Kwiatkowski, Cordelia Langford, John Sillitoe on behalf of the Wellcome Sanger Institute COVID-19 Surveillance Team |
| EPI_ISL_2829214 | Lighthouse Lab in Milton Keynes | Wellcome Sanger Institute for the COVID-19 Genomics UK (COG-UK) Consortium | The Lighthouse Lab in Milton Keynes and Alex Alderton, Roberto Amato, Jeffrey Barrett, Sonia Goncalves, Ewan Harrison, David K. Jackson, Ian Johnston, Dominic Kwiatkowski, Cordelia Langford, John Sillitoe on behalf of the Wellcome Sanger Institute COVID-19 Surveillance Team |
| EPI_ISL_2829215 | Lighthouse Lab in Alderley Park | Wellcome Sanger Institute for the COVID-19 Genomics UK (COG-UK) Consortium | Jacquelyn Wynn, Mairead Hyland, The Lighthouse Lab in Alderley Park and Alex Alderton, Roberto Amato, Jeffrey Barrett, Sonia Goncalves, Ewan Harrison, David K. Jackson, Ian Johnston, Dominic Kwiatkowski, Cordelia Langford, John Sillitoe on behalf of the Wellcome Sanger Institute COVID-19 Surveillance Team |
| EPI_ISL_2829216 | Lighthouse Laboratory Plymouth | Wellcome Sanger Institute for the COVID-19 Genomics UK (COG-UK) Consortium | Lighthouse Laboratory Plymouth and Alex Alderton, Roberto Amato, Jeffrey Barrett, Sonia Goncalves, Ewan Harrison, David K. Jackson, Ian Johnston, Dominic Kwiatkowski, Cordelia Langford, John Sillitoe on behalf of the Wellcome Sanger Institute COVID-19 Surveillance Team |
| EPI_ISL_2829217 | Health Services Laboratories | Wellcome Sanger Institute for the COVID-19 Genomics UK (COG-UK) Consortium | Health Services Laboratories and Alex Alderton, Roberto Amato, Jeffrey Barrett, Sonia Goncalves, Ewan Harrison, David K. Jackson, Ian Johnston, Dominic Kwiatkowski, Cordelia Langford, John Sillitoe on behalf of the Wellcome Sanger Institute COVID-19 Surveillance Team |
| EPI_ISL_2829218, EPI_ISL_2829219 | Lighthouse Lab in Milton Keynes | Wellcome Sanger Institute for the COVID-19 Genomics UK (COG-UK) Consortium | The Lighthouse Lab in Milton Keynes and Alex Alderton, Roberto Amato, Jeffrey Barrett, Sonia Goncalves, Ewan Harrison, David K. Jackson, Ian Johnston, Dominic Kwiatkowski, Cordelia Langford, John Sillitoe on behalf of the Wellcome Sanger Institute COVID-19 Surveillance Team |
| EPI_ISL_2829220 | Lighthouse Laboratory Plymouth | Wellcome Sanger Institute for the COVID-19 Genomics UK (COG-UK) Consortium | Lighthouse Laboratory Plymouth and Alex Alderton, Roberto Amato, Jeffrey Barrett, Sonia Goncalves, Ewan Harrison, David K. Jackson, Ian Johnston, Dominic Kwiatkowski, Cordelia Langford, John Sillitoe on behalf of the Wellcome Sanger Institute COVID-19 Surveillance Team |
| EPI_ISL_2829221 | Lighthouse Lab in Alderley Park | Wellcome Sanger Institute for the COVID-19 Genomics UK (COG-UK) Consortium | Jacquelyn Wynn, Mairead Hyland, The Lighthouse Lab in Alderley Park and Alex Alderton, Roberto Amato, Jeffrey Barrett, Sonia Goncalves, Ewan Harrison, David K. Jackson, Ian Johnston, Dominic Kwiatkowski, Cordelia Langford, John Sillitoe on behalf of the Wellcome Sanger Institute COVID-19 Surveillance Team |
| EPI_ISL_2829222 | Lighthouse Lab in Milton Keynes | Wellcome Sanger Institute for the COVID-19 Genomics UK (COG-UK) Consortium | The Lighthouse Lab in Milton Keynes and Alex Alderton, Roberto Amato, Jeffrey Barrett, Sonia Goncalves, Ewan Harrison, David K. Jackson, Ian Johnston, Dominic Kwiatkowski, Cordelia Langford, John Sillitoe on behalf of the Wellcome Sanger Institute COVID-19 Surveillance Team |
| EPI_ISL_2829223 | Lighthouse Lab in Alderley Park | Wellcome Sanger Institute for the COVID-19 Genomics UK (COG-UK) Consortium | Jacquelyn Wynn, Mairead Hyland, The Lighthouse Lab in Alderley Park and Alex Alderton, Roberto Amato, Jeffrey Barrett, Sonia Goncalves, Ewan Harrison, David K. Jackson, Ian Johnston, Dominic Kwiatkowski, Cordelia Langford, John Sillitoe on behalf of the Wellcome Sanger Institute COVID-19 Surveillance Team |
| EPI_ISL_2829224 | Lighthouse Lab in Milton Keynes | Wellcome Sanger Institute for the COVID-19 Genomics UK (COG-UK) Consortium | The Lighthouse Lab in Milton Keynes and Alex Alderton, Roberto Amato, Jeffrey Barrett, Sonia Goncalves, Ewan Harrison, David K. Jackson, Ian Johnston, Dominic Kwiatkowski, Cordelia Langford, John Sillitoe on behalf of the Wellcome Sanger Institute COVID-19 Surveillance Team |
| EPI_ISL_2829226, EPI_ISL_2829227 | Lighthouse Lab in Alderley Park | Wellcome Sanger Institute for the COVID-19 Genomics UK (COG-UK) Consortium | Jacquelyn Wynn, Mairead Hyland, The Lighthouse Lab in Alderley Park and Alex Alderton, Roberto Amato, Jeffrey Barrett, Sonia Goncalves, Ewan Harrison, David K. Jackson, Ian Johnston, Dominic Kwiatkowski, Cordelia Langford, John Sillitoe on behalf of the Wellcome Sanger Institute COVID-19 Surveillance Team |
| EPI_ISL_2829228 | Health Services Laboratories | Wellcome Sanger Institute for the COVID-19 Genomics UK (COG-UK) Consortium | Health Services Laboratories and Alex Alderton, Roberto Amato, Jeffrey Barrett, Sonia Goncalves, Ewan Harrison, David K. Jackson, Ian Johnston, Dominic Kwiatkowski, Cordelia Langford, John Sillitoe on behalf of the Wellcome Sanger Institute COVID-19 Surveillance Team |
| EPI_ISL_2829229 | Lighthouse Lab in Alderley Park | Wellcome Sanger Institute for the COVID-19 Genomics UK (COG-UK) Consortium | Jacquelyn Wynn, Mairead Hyland, The Lighthouse Lab in Alderley Park and Alex Alderton, Roberto Amato, Jeffrey Barrett, Sonia Goncalves, Ewan Harrison, David K. Jackson, Ian Johnston, Dominic Kwiatkowski, Cordelia Langford, John Sillitoe on behalf of the Wellcome Sanger Institute COVID-19 Surveillance Team |
| EPI_ISL_2829230, EPI_ISL_2829231, EPI_ISL_2829232, EPI_ISL_2829233 | Lighthouse Lab in Milton Keynes | Wellcome Sanger Institute for the COVID-19 Genomics UK (COG-UK) Consortium | The Lighthouse Lab in Milton Keynes and Alex Alderton, Roberto Amato, Jeffrey Barrett, Sonia Goncalves, Ewan Harrison, David K. Jackson, Ian Johnston, Dominic Kwiatkowski, Cordelia Langford, John Sillitoe on behalf of the Wellcome Sanger Institute COVID-19 Surveillance Team |
| EPI_ISL_2829234 | Lighthouse Lab in Alderley Park | Wellcome Sanger Institute for the COVID-19 Genomics UK (COG-UK) Consortium | Jacquelyn Wynn, Mairead Hyland, The Lighthouse Lab in Alderley Park and Alex Alderton, Roberto Amato, Jeffrey Barrett, Sonia Goncalves, Ewan Harrison, David K. Jackson, Ian Johnston, Dominic Kwiatkowski, Cordelia Langford, John Sillitoe on behalf of the Wellcome Sanger Institute COVID-19 Surveillance Team |
| EPI_ISL_2829235 | Health Services Laboratories | Wellcome Sanger Institute for the COVID-19 Genomics UK (COG-UK) Consortium | Health Services Laboratories and Alex Alderton, Roberto Amato, Jeffrey Barrett, Sonia Goncalves, Ewan Harrison, David K. Jackson, Ian Johnston, Dominic Kwiatkowski, Cordelia Langford, John Sillitoe on behalf of the Wellcome Sanger Institute COVID-19 Surveillance Team |
| EPI_ISL_2829236 | Lighthouse Lab in Alderley Park | Wellcome Sanger Institute for the COVID-19 Genomics UK (COG-UK) Consortium | Jacquelyn Wynn, Mairead Hyland, The Lighthouse Lab in Alderley Park and Alex Alderton, Roberto Amato, Jeffrey Barrett, Sonia Goncalves, Ewan Harrison, David K. Jackson, Ian Johnston, Dominic Kwiatkowski, Cordelia Langford, John Sillitoe on behalf of the Wellcome Sanger Institute COVID-19 Surveillance Team |
| EPI_ISL_2829237 | Lighthouse Lab in Milton Keynes | Wellcome Sanger Institute for the COVID-19 Genomics UK (COG-UK) Consortium | The Lighthouse Lab in Milton Keynes and Alex Alderton, Roberto Amato, Jeffrey Barrett, Sonia Goncalves, Ewan Harrison, David K. Jackson, Ian Johnston, Dominic Kwiatkowski, Cordelia Langford, John Sillitoe on behalf of the Wellcome Sanger Institute COVID-19 Surveillance Team |
| EPI_ISL_2829238 | Health Services Laboratories | Wellcome Sanger Institute for the COVID-19 Genomics UK (COG-UK) Consortium | Health Services Laboratories and Alex Alderton, Roberto Amato, Jeffrey Barrett, Sonia Goncalves, Ewan Harrison, David K. Jackson, Ian Johnston, Dominic Kwiatkowski, Cordelia Langford, John Sillitoe on behalf of the Wellcome Sanger Institute COVID-19 Surveillance Team |
| EPI_ISL_2829239 | Lighthouse Lab in Milton Keynes | Wellcome Sanger Institute for the COVID-19 Genomics UK (COG-UK) Consortium | The Lighthouse Lab in Milton Keynes and Alex Alderton, Roberto Amato, Jeffrey Barrett, Sonia Goncalves, Ewan Harrison, David K. Jackson, Ian Johnston, Dominic Kwiatkowski, Cordelia Langford, John Sillitoe on behalf of the Wellcome Sanger Institute COVID-19 Surveillance Team |
| EPI_ISL_2829240 | Lighthouse Lab in Alderley Park | Wellcome Sanger Institute for the COVID-19 Genomics UK (COG-UK) Consortium | Jacquelyn Wynn, Mairead Hyland, The Lighthouse Lab in Alderley Park and Alex Alderton, Roberto Amato, Jeffrey Barrett, Sonia Goncalves, Ewan Harrison, David K. Jackson, Ian Johnston, Dominic Kwiatkowski, Cordelia Langford, John Sillitoe on behalf of the Wellcome Sanger Institute COVID-19 Surveillance Team |
| EPI_ISL_2829241 | Health Services Laboratories | Wellcome Sanger Institute for the COVID-19 Genomics UK (COG-UK) Consortium | Health Services Laboratories and Alex Alderton, Roberto Amato, Jeffrey Barrett, Sonia Goncalves, Ewan Harrison, David K. Jackson, Ian Johnston, Dominic Kwiatkowski, Cordelia Langford, John Sillitoe on behalf of the Wellcome Sanger Institute COVID-19 Surveillance Team |
| EPI_ISL_2829242 | Lighthouse Lab in Milton Keynes | Wellcome Sanger Institute for the COVID-19 Genomics UK (COG-UK) Consortium | The Lighthouse Lab in Milton Keynes and Alex Alderton, Roberto Amato, Jeffrey Barrett, Sonia Goncalves, Ewan Harrison, David K. Jackson, Ian Johnston, Dominic Kwiatkowski, Cordelia Langford, John Sillitoe on behalf of the Wellcome Sanger Institute COVID-19 Surveillance Team |
| EPI_ISL_2829243 | Lighthouse Laboratory Plymouth | Wellcome Sanger Institute for the COVID-19 Genomics UK (COG-UK) Consortium | Lighthouse Laboratory Plymouth and Alex Alderton, Roberto Amato, Jeffrey Barrett, Sonia Goncalves, Ewan Harrison, David K. Jackson, Ian Johnston, Dominic Kwiatkowski, Cordelia Langford, John Sillitoe on behalf of the Wellcome Sanger Institute COVID-19 Surveillance Team |
| EPI_ISL_2829244, EPI_ISL_2829245, EPI_ISL_2829246 | Lighthouse Lab in Milton Keynes | Wellcome Sanger Institute for the COVID-19 Genomics UK (COG-UK) Consortium | The Lighthouse Lab in Milton Keynes and Alex Alderton, Roberto Amato, Jeffrey Barrett, Sonia Goncalves, Ewan Harrison, David K. Jackson, Ian Johnston, Dominic Kwiatkowski, Cordelia Langford, John Sillitoe on behalf of the Wellcome Sanger Institute COVID-19 Surveillance Team |
| EPI_ISL_2829247 | Health Services Laboratories | Wellcome Sanger Institute for the COVID-19 Genomics UK (COG-UK) Consortium | Health Services Laboratories and Alex Alderton, Roberto Amato, Jeffrey Barrett, Sonia Goncalves, Ewan Harrison, David K. Jackson, Ian Johnston, Dominic Kwiatkowski, Cordelia Langford, John Sillitoe on behalf of the Wellcome Sanger Institute COVID-19 Surveillance Team |
| EPI_ISL_2829248 | Lighthouse Lab in Milton Keynes | Wellcome Sanger Institute for the COVID-19 Genomics UK (COG-UK) Consortium | The Lighthouse Lab in Milton Keynes and Alex Alderton, Roberto Amato, Jeffrey Barrett, Sonia Goncalves, Ewan Harrison, David K. Jackson, Ian Johnston, Dominic Kwiatkowski, Cordelia Langford, John Sillitoe on behalf of the Wellcome Sanger Institute COVID-19 Surveillance Team |
| EPI_ISL_2829249 | Health Services Laboratories | Wellcome Sanger Institute for the COVID-19 Genomics UK (COG-UK) Consortium | Health Services Laboratories and Alex Alderton, Roberto Amato, Jeffrey Barrett, Sonia Goncalves, Ewan Harrison, David K. Jackson, Ian Johnston, Dominic Kwiatkowski, Cordelia Langford, John Sillitoe on behalf of the Wellcome Sanger Institute COVID-19 Surveillance Team |
| EPI_ISL_2829250 | Lighthouse Laboratory Plymouth | Wellcome Sanger Institute for the COVID-19 Genomics UK (COG-UK) Consortium | Lighthouse Laboratory Plymouth and Alex Alderton, Roberto Amato, Jeffrey Barrett, Sonia Goncalves, Ewan Harrison, David K. Jackson, Ian Johnston, Dominic Kwiatkowski, Cordelia Langford, John Sillitoe on behalf of the Wellcome Sanger Institute COVID-19 Surveillance Team |
| EPI_ISL_2829251 | Lighthouse Lab in Milton Keynes | Wellcome Sanger Institute for the COVID-19 Genomics UK (COG-UK) Consortium | The Lighthouse Lab in Milton Keynes and Alex Alderton, Roberto Amato, Jeffrey Barrett, Sonia Goncalves, Ewan Harrison, David K. Jackson, Ian Johnston, Dominic Kwiatkowski, Cordelia Langford, John Sillitoe on behalf of the Wellcome Sanger Institute COVID-19 Surveillance Team |
| EPI_ISL_2829252, EPI_ISL_2829253, EPI_ISL_2829254, EPI_ISL_2829255 | Lighthouse Lab in Alderley Park | Wellcome Sanger Institute for the COVID-19 Genomics UK (COG-UK) Consortium | Jacquelyn Wynn, Mairead Hyland, The Lighthouse Lab in Alderley Park and Alex Alderton, Roberto Amato, Jeffrey Barrett, Sonia Goncalves, Ewan Harrison, David K. Jackson, Ian Johnston, Dominic Kwiatkowski, Cordelia Langford, John Sillitoe on behalf of the Wellcome Sanger Institute COVID-19 Surveillance Team |
| EPI_ISL_2829256 | Health Services Laboratories | Wellcome Sanger Institute for the COVID-19 Genomics UK (COG-UK) Consortium | Health Services Laboratories and Alex Alderton, Roberto Amato, Jeffrey Barrett, Sonia Goncalves, Ewan Harrison, David K. Jackson, Ian Johnston, Dominic Kwiatkowski, Cordelia Langford, John Sillitoe on behalf of the Wellcome Sanger Institute COVID-19 Surveillance Team |
| EPI_ISL_2829257 | Lighthouse Laboratory Plymouth | Wellcome Sanger Institute for the COVID-19 Genomics UK (COG-UK) Consortium | Lighthouse Laboratory Plymouth and Alex Alderton, Roberto Amato, Jeffrey Barrett, Sonia Goncalves, Ewan Harrison, David K. Jackson, Ian Johnston, Dominic Kwiatkowski, Cordelia Langford, John Sillitoe on behalf of the Wellcome Sanger Institute COVID-19 Surveillance Team |
| EPI_ISL_2829258 | Lighthouse Lab in Milton Keynes | Wellcome Sanger Institute for the COVID-19 Genomics UK (COG-UK) Consortium | The Lighthouse Lab in Milton Keynes and Alex Alderton, Roberto Amato, Jeffrey Barrett, Sonia Goncalves, Ewan Harrison, David K. Jackson, Ian Johnston, Dominic Kwiatkowski, Cordelia Langford, John Sillitoe on behalf of the Wellcome Sanger Institute COVID-19 Surveillance Team |
| EPI_ISL_2829259 | Health Services Laboratories | Wellcome Sanger Institute for the COVID-19 Genomics UK (COG-UK) Consortium | Health Services Laboratories and Alex Alderton, Roberto Amato, Jeffrey Barrett, Sonia Goncalves, Ewan Harrison, David K. Jackson, Ian Johnston, Dominic Kwiatkowski, Cordelia Langford, John Sillitoe on behalf of the Wellcome Sanger Institute COVID-19 Surveillance Team |
| EPI_ISL_2829260 | Lighthouse Lab in Glasgow | Wellcome Sanger Institute for the COVID-19 Genomics UK (COG-UK) Consortium | Harper VanSteenhouse, Yumi Kasai, David Gray, Carol Clugston, Anna Dominiczak and Alex Alderton, Roberto Amato, Jeffrey Barrett, Sonia Goncalves, Ewan Harrison, David K. Jackson, Ian Johnston, Dominic Kwiatkowski, Cordelia Langford, John Sillitoe on behalf of the Wellcome Sanger Institute COVID-19 Surveillance Team |
| EPI_ISL_2829261 | Lighthouse Lab in Alderley Park | Wellcome Sanger Institute for the COVID-19 Genomics UK (COG-UK) Consortium | Jacquelyn Wynn, Mairead Hyland, The Lighthouse Lab in Alderley Park and Alex Alderton, Roberto Amato, Jeffrey Barrett, Sonia Goncalves, Ewan Harrison, David K. Jackson, Ian Johnston, Dominic Kwiatkowski, Cordelia Langford, John Sillitoe on behalf of the Wellcome Sanger Institute COVID-19 Surveillance Team |

| | | | |
|---|---|---|---|
| EPI_ISL_2829262 | Lighthouse Lab in Milton Keynes | Wellcome Sanger Institute for the COVID-19 Genomics UK (COG-UK) Consortium | The Lighthouse Lab in Milton Keynes and Alex Alderton, Roberto Amato, Jeffrey Barrett, Sonia Goncalves, Ewan Harrison, David K. Jackson, Ian Johnston, Dominic Kwiatkowski, Cordelia Langford, John Sillitoe on behalf of the Wellcome Sanger Institute COVID-19 Surveillance Team |
| EPI_ISL_2829263 | Lighthouse Laboratory Plymouth | Wellcome Sanger Institute for the COVID-19 Genomics UK (COG-UK) Consortium | Lighthouse Laboratory Plymouth and Alex Alderton, Roberto Amato, Jeffrey Barrett, Sonia Goncalves, Ewan Harrison, David K. Jackson, Ian Johnston, Dominic Kwiatkowski, Cordelia Langford, John Sillitoe on behalf of the Wellcome Sanger Institute COVID-19 Surveillance Team |
| EPI_ISL_2829264 | Lighthouse Lab in Alderley Park | Wellcome Sanger Institute for the COVID-19 Genomics UK (COG-UK) Consortium | Jacquelyn Wynn, Mairead Hyland, The Lighthouse Lab in Alderley Park and Alex Alderton, Roberto Amato, Jeffrey Barrett, Sonia Goncalves, Ewan Harrison, David K. Jackson, Ian Johnston, Dominic Kwiatkowski, Cordelia Langford, John Sillitoe on behalf of the Wellcome Sanger Institute COVID-19 Surveillance Team |
| EPI_ISL_2829265 | Health Services Laboratories | Wellcome Sanger Institute for the COVID-19 Genomics UK (COG-UK) Consortium | Health Services Laboratories and Alex Alderton, Roberto Amato, Jeffrey Barrett, Sonia Goncalves, Ewan Harrison, David K. Jackson, Ian Johnston, Dominic Kwiatkowski, Cordelia Langford, John Sillitoe on behalf of the Wellcome Sanger Institute COVID-19 Surveillance Team |
| EPI_ISL_2829266 | Lighthouse Lab in Glasgow | Wellcome Sanger Institute for the COVID-19 Genomics UK (COG-UK) Consortium | Harper VanSteenhouse, Yumi Kasai, David Gray, Carol Clugston, Anna Dominiczak and Alex Alderton, Roberto Amato, Jeffrey Barrett, Sonia Goncalves, Ewan Harrison, David K. Jackson, Ian Johnston, Dominic Kwiatkowski, Cordelia Langford, John Sillitoe on behalf of the Wellcome Sanger Institute COVID-19 Surveillance Team |
| EPI_ISL_2829267 | Health Services Laboratories | Wellcome Sanger Institute for the COVID-19 Genomics UK (COG-UK) Consortium | Health Services Laboratories and Alex Alderton, Roberto Amato, Jeffrey Barrett, Sonia Goncalves, Ewan Harrison, David K. Jackson, Ian Johnston, Dominic Kwiatkowski, Cordelia Langford, John Sillitoe on behalf of the Wellcome Sanger Institute COVID-19 Surveillance Team |
| EPI_ISL_2829268 | Lighthouse Lab in Alderley Park | Wellcome Sanger Institute for the COVID-19 Genomics UK (COG-UK) Consortium | Jacquelyn Wynn, Mairead Hyland, The Lighthouse Lab in Alderley Park and Alex Alderton, Roberto Amato, Jeffrey Barrett, Sonia Goncalves, Ewan Harrison, David K. Jackson, Ian Johnston, Dominic Kwiatkowski, Cordelia Langford, John Sillitoe on behalf of the Wellcome Sanger Institute COVID-19 Surveillance Team |
| EPI_ISL_2829269 | Lighthouse Lab in Milton Keynes | Wellcome Sanger Institute for the COVID-19 Genomics UK (COG-UK) Consortium | The Lighthouse Lab in Milton Keynes and Alex Alderton, Roberto Amato, Jeffrey Barrett, Sonia Goncalves, Ewan Harrison, David K. Jackson, Ian Johnston, Dominic Kwiatkowski, Cordelia Langford, John Sillitoe on behalf of the Wellcome Sanger Institute COVID-19 Surveillance Team |
| EPI_ISL_2829270 | Health Services Laboratories | Wellcome Sanger Institute for the COVID-19 Genomics UK (COG-UK) Consortium | Health Services Laboratories and Alex Alderton, Roberto Amato, Jeffrey Barrett, Sonia Goncalves, Ewan Harrison, David K. Jackson, Ian Johnston, Dominic Kwiatkowski, Cordelia Langford, John Sillitoe on behalf of the Wellcome Sanger Institute COVID-19 Surveillance Team |
| EPI_ISL_2829271 | Lighthouse Laboratory Plymouth | Wellcome Sanger Institute for the COVID-19 Genomics UK (COG-UK) Consortium | Lighthouse Laboratory Plymouth and Alex Alderton, Roberto Amato, Jeffrey Barrett, Sonia Goncalves, Ewan Harrison, David K. Jackson, Ian Johnston, Dominic Kwiatkowski, Cordelia Langford, John Sillitoe on behalf of the Wellcome Sanger Institute COVID-19 Surveillance Team |
| EPI_ISL_2829272 | Health Services Laboratories | Wellcome Sanger Institute for the COVID-19 Genomics UK (COG-UK) Consortium | Health Services Laboratories and Alex Alderton, Roberto Amato, Jeffrey Barrett, Sonia Goncalves, Ewan Harrison, David K. Jackson, Ian Johnston, Dominic Kwiatkowski, Cordelia Langford, John Sillitoe on behalf of the Wellcome Sanger Institute COVID-19 Surveillance Team |
| EPI_ISL_2829273 | Lighthouse Lab in Milton Keynes | Wellcome Sanger Institute for the COVID-19 Genomics UK (COG-UK) Consortium | The Lighthouse Lab in Milton Keynes and Alex Alderton, Roberto Amato, Jeffrey Barrett, Sonia Goncalves, Ewan Harrison, David K. Jackson, Ian Johnston, Dominic Kwiatkowski, Cordelia Langford, John Sillitoe on behalf of the Wellcome Sanger Institute COVID-19 Surveillance Team |
| EPI_ISL_2829274, EPI_ISL_2829275, EPI_ISL_2829276 | Lighthouse Lab in Alderley Park | Wellcome Sanger Institute for the COVID-19 Genomics UK (COG-UK) Consortium | Jacquelyn Wynn, Mairead Hyland, The Lighthouse Lab in Alderley Park and Alex Alderton, Roberto Amato, Jeffrey Barrett, Sonia Goncalves, Ewan Harrison, David K. Jackson, Ian Johnston, Dominic Kwiatkowski, Cordelia Langford, John Sillitoe on behalf of the Wellcome Sanger Institute COVID-19 Surveillance Team |
| EPI_ISL_2829277, EPI_ISL_2829278 | Lighthouse Laboratory Plymouth | Wellcome Sanger Institute for the COVID-19 Genomics UK (COG-UK) Consortium | Lighthouse Laboratory Plymouth and Alex Alderton, Roberto Amato, Jeffrey Barrett, Sonia Goncalves, Ewan Harrison, David K. Jackson, Ian Johnston, Dominic Kwiatkowski, Cordelia Langford, John Sillitoe on behalf of the Wellcome Sanger Institute COVID-19 Surveillance Team |
| EPI_ISL_2829279 | Health Services Laboratories | Wellcome Sanger Institute for the COVID-19 Genomics UK (COG-UK) Consortium | Health Services Laboratories and Alex Alderton, Roberto Amato, Jeffrey Barrett, Sonia Goncalves, Ewan Harrison, David K. Jackson, Ian Johnston, Dominic Kwiatkowski, Cordelia Langford, John Sillitoe on behalf of the Wellcome Sanger Institute COVID-19 Surveillance Team |
| EPI_ISL_2829280 | Lighthouse Lab in Glasgow | Wellcome Sanger Institute for the COVID-19 Genomics UK (COG-UK) Consortium | Harper VanSteenhouse, Yumi Kasai, David Gray, Carol Clugston, Anna Dominiczak and Alex Alderton, Roberto Amato, Jeffrey Barrett, Sonia Goncalves, Ewan Harrison, David K. Jackson, Ian Johnston, Dominic Kwiatkowski, Cordelia Langford, John Sillitoe on behalf of the Wellcome Sanger Institute COVID-19 Surveillance Team |
| EPI_ISL_2829281 | Lighthouse Lab in Milton Keynes | Wellcome Sanger Institute for the COVID-19 Genomics UK (COG-UK) Consortium | The Lighthouse Lab in Milton Keynes and Alex Alderton, Roberto Amato, Jeffrey Barrett, Sonia Goncalves, Ewan Harrison, David K. Jackson, Ian Johnston, Dominic Kwiatkowski, Cordelia Langford, John Sillitoe on behalf of the Wellcome Sanger Institute COVID-19 Surveillance Team |
| EPI_ISL_2829282 | Lighthouse Laboratory Plymouth | Wellcome Sanger Institute for the COVID-19 Genomics UK (COG-UK) Consortium | Lighthouse Laboratory Plymouth and Alex Alderton, Roberto Amato, Jeffrey Barrett, Sonia Goncalves, Ewan Harrison, David K. Jackson, Ian Johnston, Dominic Kwiatkowski, Cordelia Langford, John Sillitoe on behalf of the Wellcome Sanger Institute COVID-19 Surveillance Team |
| EPI_ISL_2829283 | Health Services Laboratories | Wellcome Sanger Institute for the COVID-19 Genomics UK (COG-UK) Consortium | Health Services Laboratories and Alex Alderton, Roberto Amato, Jeffrey Barrett, Sonia Goncalves, Ewan Harrison, David K. Jackson, Ian Johnston, Dominic Kwiatkowski, Cordelia Langford, John Sillitoe on behalf of the Wellcome Sanger Institute COVID-19 Surveillance Team |
| EPI_ISL_2829284 | Lighthouse Lab in Milton Keynes | Wellcome Sanger Institute for the COVID-19 Genomics UK (COG-UK) Consortium | The Lighthouse Lab in Milton Keynes and Alex Alderton, Roberto Amato, Jeffrey Barrett, Sonia Goncalves, Ewan Harrison, David K. Jackson, Ian Johnston, Dominic Kwiatkowski, Cordelia Langford, John Sillitoe on behalf of the Wellcome Sanger Institute COVID-19 Surveillance Team |
| EPI_ISL_2829285 | Lighthouse Lab in Alderley Park | Wellcome Sanger Institute for the COVID-19 Genomics UK (COG-UK) Consortium | Jacquelyn Wynn, Mairead Hyland, The Lighthouse Lab in Alderley Park and Alex Alderton, Roberto Amato, Jeffrey Barrett, Sonia Goncalves, Ewan Harrison, David K. Jackson, Ian Johnston, Dominic Kwiatkowski, Cordelia Langford, John Sillitoe on behalf of the Wellcome Sanger Institute COVID-19 Surveillance Team |
| EPI_ISL_2829286, EPI_ISL_2829287 | Lighthouse Lab in Milton Keynes | Wellcome Sanger Institute for the COVID-19 Genomics UK (COG-UK) Consortium | The Lighthouse Lab in Milton Keynes and Alex Alderton, Roberto Amato, Jeffrey Barrett, Sonia Goncalves, Ewan Harrison, David K. Jackson, Ian Johnston, Dominic Kwiatkowski, Cordelia Langford, John Sillitoe on behalf of the Wellcome Sanger Institute COVID-19 Surveillance Team |
| EPI_ISL_2829288, EPI_ISL_2829289 | Lighthouse Lab in Alderley Park | Wellcome Sanger Institute for the COVID-19 Genomics UK (COG-UK) Consortium | Jacquelyn Wynn, Mairead Hyland, The Lighthouse Lab in Alderley Park and Alex Alderton, Roberto Amato, Jeffrey Barrett, Sonia Goncalves, Ewan Harrison, David K. Jackson, Ian Johnston, Dominic Kwiatkowski, Cordelia Langford, John Sillitoe on behalf of the Wellcome Sanger Institute COVID-19 Surveillance Team |
| EPI_ISL_2829290 | Lighthouse Lab in Milton Keynes | Wellcome Sanger Institute for the COVID-19 Genomics UK (COG-UK) Consortium | The Lighthouse Lab in Milton Keynes and Alex Alderton, Roberto Amato, Jeffrey Barrett, Sonia Goncalves, Ewan Harrison, David K. Jackson, Ian Johnston, Dominic Kwiatkowski, Cordelia Langford, John Sillitoe on behalf of the Wellcome Sanger Institute COVID-19 Surveillance Team |
| EPI_ISL_2829291 | Health Services Laboratories | Wellcome Sanger Institute for the COVID-19 Genomics UK (COG-UK) Consortium | Health Services Laboratories and Alex Alderton, Roberto Amato, Jeffrey Barrett, Sonia Goncalves, Ewan Harrison, David K. Jackson, Ian Johnston, Dominic Kwiatkowski, Cordelia Langford, John Sillitoe on behalf of the Wellcome Sanger Institute COVID-19 Surveillance Team |
| EPI_ISL_2829292 | Lighthouse Laboratory Plymouth | Wellcome Sanger Institute for the COVID-19 Genomics UK (COG-UK) Consortium | Lighthouse Laboratory Plymouth and Alex Alderton, Roberto Amato, Jeffrey Barrett, Sonia Goncalves, Ewan Harrison, David K. Jackson, Ian Johnston, Dominic Kwiatkowski, Cordelia Langford, John Sillitoe on behalf of the Wellcome Sanger Institute COVID-19 Surveillance Team |
| EPI_ISL_2829293 | Lighthouse Lab in Milton Keynes | Wellcome Sanger Institute for the COVID-19 Genomics UK (COG-UK) Consortium | The Lighthouse Lab in Milton Keynes and Alex Alderton, Roberto Amato, Jeffrey Barrett, Sonia Goncalves, Ewan Harrison, David K. Jackson, Ian Johnston, Dominic Kwiatkowski, Cordelia Langford, John Sillitoe on behalf of the Wellcome Sanger Institute COVID-19 Surveillance Team |
| EPI_ISL_2829294 | Lighthouse Lab in Alderley Park | Wellcome Sanger Institute for the COVID-19 Genomics UK (COG-UK) Consortium | Jacquelyn Wynn, Mairead Hyland, The Lighthouse Lab in Alderley Park and Alex Alderton, Roberto Amato, Jeffrey Barrett, Sonia Goncalves, Ewan Harrison, David K. Jackson, Ian Johnston, Dominic Kwiatkowski, Cordelia Langford, John Sillitoe on behalf of the Wellcome Sanger Institute COVID-19 Surveillance Team |
| EPI_ISL_2829295 | Lighthouse Lab in Milton Keynes | Wellcome Sanger Institute for the COVID-19 Genomics UK (COG-UK) Consortium | The Lighthouse Lab in Milton Keynes and Alex Alderton, Roberto Amato, Jeffrey Barrett, Sonia Goncalves, Ewan Harrison, David K. Jackson, Ian Johnston, Dominic Kwiatkowski, Cordelia Langford, John Sillitoe on behalf of the Wellcome Sanger Institute COVID-19 Surveillance Team |
| EPI_ISL_2829296 | Health Services Laboratories | Wellcome Sanger Institute for the COVID-19 Genomics UK (COG-UK) Consortium | Health Services Laboratories and Alex Alderton, Roberto Amato, Jeffrey Barrett, Sonia Goncalves, Ewan Harrison, David K. Jackson, Ian Johnston, Dominic Kwiatkowski, Cordelia Langford, John Sillitoe on behalf of the Wellcome Sanger Institute COVID-19 Surveillance Team |
| EPI_ISL_2829297 | Lighthouse Lab in Alderley Park | Wellcome Sanger Institute for the COVID-19 Genomics UK (COG-UK) Consortium | Jacquelyn Wynn, Mairead Hyland, The Lighthouse Lab in Alderley Park and Alex Alderton, Roberto Amato, Jeffrey Barrett, Sonia Goncalves, Ewan Harrison, David K. Jackson, Ian Johnston, Dominic Kwiatkowski, Cordelia Langford, John Sillitoe on behalf of the Wellcome Sanger Institute COVID-19 Surveillance Team |
| EPI_ISL_2829298 | Lighthouse Laboratory Plymouth | Wellcome Sanger Institute for the COVID-19 Genomics UK (COG-UK) Consortium | Lighthouse Laboratory Plymouth and Alex Alderton, Roberto Amato, Jeffrey Barrett, Sonia Goncalves, Ewan Harrison, David K. Jackson, Ian Johnston, Dominic Kwiatkowski, Cordelia Langford, John Sillitoe on behalf of the Wellcome Sanger Institute COVID-19 Surveillance Team |
| EPI_ISL_2829299, EPI_ISL_2829300, EPI_ISL_2829301 | Lighthouse Lab in Alderley Park | Wellcome Sanger Institute for the COVID-19 Genomics UK (COG-UK) Consortium | Jacquelyn Wynn, Mairead Hyland, The Lighthouse Lab in Alderley Park and Alex Alderton, Roberto Amato, Jeffrey Barrett, Sonia Goncalves, Ewan Harrison, David K. Jackson, Ian Johnston, Dominic Kwiatkowski, Cordelia Langford, John Sillitoe on behalf of the Wellcome Sanger Institute COVID-19 Surveillance Team |
| EPI_ISL_2829302 | Lighthouse Lab in Glasgow | Wellcome Sanger Institute for the COVID-19 Genomics UK (COG-UK) Consortium | Harper VanSteenhouse, Yumi Kasai, David Gray, Carol Clugston, Anna Dominiczak and Alex Alderton, Roberto Amato, Jeffrey Barrett, Sonia Goncalves, Ewan Harrison, David K. Jackson, Ian Johnston, Dominic Kwiatkowski, Cordelia Langford, John Sillitoe on behalf of the Wellcome Sanger Institute COVID-19 Surveillance Team |
| EPI_ISL_2829303, EPI_ISL_2829304, EPI_ISL_2829305 | Lighthouse Lab in Alderley Park | Wellcome Sanger Institute for the COVID-19 Genomics UK (COG-UK) Consortium | Jacquelyn Wynn, Mairead Hyland, The Lighthouse Lab in Alderley Park and Alex Alderton, Roberto Amato, Jeffrey Barrett, Sonia Goncalves, Ewan Harrison, David K. Jackson, Ian Johnston, Dominic Kwiatkowski, Cordelia Langford, John Sillitoe on behalf of the Wellcome Sanger Institute COVID-19 Surveillance Team |
| EPI_ISL_2829306 | Lighthouse Lab in Glasgow | Wellcome Sanger Institute for the COVID-19 Genomics UK (COG-UK) Consortium | Harper VanSteenhouse, Yumi Kasai, David Gray, Carol Clugston, Anna Dominiczak and Alex Alderton, Roberto Amato, Jeffrey Barrett, Sonia Goncalves, Ewan Harrison, David K. Jackson, Ian Johnston, Dominic Kwiatkowski, Cordelia Langford, John Sillitoe on behalf of the Wellcome Sanger Institute COVID-19 Surveillance Team |
| EPI_ISL_2829307, EPI_ISL_2829308 | Lighthouse Lab in Alderley Park | Wellcome Sanger Institute for the COVID-19 Genomics UK (COG-UK) Consortium | Jacquelyn Wynn, Mairead Hyland, The Lighthouse Lab in Alderley Park and Alex Alderton, Roberto Amato, Jeffrey Barrett, Sonia Goncalves, Ewan Harrison, David K. Jackson, Ian Johnston, Dominic Kwiatkowski, Cordelia Langford, John Sillitoe on behalf of the Wellcome Sanger Institute COVID-19 Surveillance Team |
| EPI_ISL_2829309 | Lighthouse Lab in Glasgow | Wellcome Sanger Institute for the COVID-19 Genomics UK (COG-UK) Consortium | Harper VanSteenhouse, Yumi Kasai, David Gray, Carol Clugston, Anna Dominiczak and Alex Alderton, Roberto Amato, Jeffrey Barrett, Sonia Goncalves, Ewan Harrison, David K. Jackson, Ian Johnston, Dominic Kwiatkowski, Cordelia Langford, John Sillitoe on behalf of the Wellcome Sanger Institute COVID-19 Surveillance Team |
| EPI_ISL_2829310 | Lighthouse Lab in Alderley Park | Wellcome Sanger Institute for the COVID-19 Genomics UK (COG-UK) Consortium | Jacquelyn Wynn, Mairead Hyland, The Lighthouse Lab in Alderley Park and Alex Alderton, Roberto Amato, Jeffrey Barrett, Sonia Goncalves, Ewan Harrison, David K. Jackson, Ian Johnston, Dominic Kwiatkowski, Cordelia Langford, John Sillitoe on behalf of the Wellcome Sanger Institute COVID-19 Surveillance Team |
| EPI_ISL_2829311 | Lighthouse Lab in Glasgow | Wellcome Sanger Institute for the COVID-19 Genomics UK (COG-UK) Consortium | Harper VanSteenhouse, Yumi Kasai, David Gray, Carol Clugston, Anna Dominiczak and Alex Alderton, Roberto Amato, Jeffrey Barrett, Sonia Goncalves, Ewan Harrison, David K. Jackson, Ian Johnston, Dominic Kwiatkowski, Cordelia Langford, John Sillitoe on behalf of the Wellcome Sanger Institute COVID-19 Surveillance Team |
| EPI_ISL_2829312, EPI_ISL_2829313, EPI_ISL_2829314 | Lighthouse Lab in Alderley Park | Wellcome Sanger Institute for the COVID-19 Genomics UK (COG-UK) Consortium | Jacquelyn Wynn, Mairead Hyland, The Lighthouse Lab in Alderley Park and Alex Alderton, Roberto Amato, Jeffrey Barrett, Sonia Goncalves, Ewan Harrison, David K. Jackson, Ian Johnston, Dominic Kwiatkowski, Cordelia Langford, John Sillitoe on behalf of the Wellcome Sanger Institute COVID-19 Surveillance Team |
| EPI_ISL_2829315 | Lighthouse Laboratory Plymouth | Wellcome Sanger Institute for the COVID-19 Genomics UK (COG-UK) Consortium | Lighthouse Laboratory Plymouth and Alex Alderton, Roberto Amato, Jeffrey Barrett, Sonia Goncalves, Ewan Harrison, David K. Jackson, Ian Johnston, Dominic Kwiatkowski, Cordelia Langford, John Sillitoe on behalf of the Wellcome Sanger Institute COVID-19 Surveillance Team |
| EPI_ISL_2829316 | Lighthouse Lab in Alderley Park | Wellcome Sanger Institute for the COVID-19 Genomics UK (COG-UK) Consortium | Jacquelyn Wynn, Mairead Hyland, The Lighthouse Lab in Alderley Park and Alex Alderton, Roberto Amato, Jeffrey Barrett, Sonia Goncalves, Ewan Harrison, David K. Jackson, Ian Johnston, Dominic Kwiatkowski, Cordelia Langford, John Sillitoe on behalf of the Wellcome Sanger Institute COVID-19 Surveillance Team |
| EPI_ISL_2829317 | Lighthouse Laboratory Plymouth | Wellcome Sanger Institute for the COVID-19 Genomics UK (COG-UK) | Lighthouse Laboratory Plymouth and Alex Alderton, Roberto Amato, Jeffrey Barrett, Sonia Goncalves, Ewan Harrison, David K. Jackson, Ian Johnston, Dominic Kwiatkowski, Cordelia Langford, John Sillitoe on behalf of the Wellcome Sanger |

| | | | |
|---|---|---|---|
| | | Consortium | Institute COVID-19 Surveillance Team |
| EPI_ISL_2829318 | Lighthouse Lab in Alderley Park | Wellcome Sanger Institute for the COVID-19 Genomics UK (COG-UK) Consortium | Jacquelyn Wynn, Mairead Hyland, The Lighthouse Lab in Alderley Park and Alex Alderton, Roberto Amato, Jeffrey Barrett, Sonia Goncalves, Ewan Harrison, David K. Jackson, Ian Johnston, Dominic Kwiatkowski, Cordelia Langford, John Sillitoe on behalf of the Wellcome Sanger Institute COVID-19 Surveillance Team |
| EPI_ISL_2829319 | Lighthouse Laboratory Plymouth | Wellcome Sanger Institute for the COVID-19 Genomics UK (COG-UK) Consortium | Lighthouse Laboratory Plymouth and Alex Alderton, Roberto Amato, Jeffrey Barrett, Sonia Goncalves, Ewan Harrison, David K. Jackson, Ian Johnston, Dominic Kwiatkowski, Cordelia Langford, John Sillitoe on behalf of the Wellcome Sanger Institute COVID-19 Surveillance Team |
| EPI_ISL_2829320 | Lighthouse Lab in Alderley Park | Wellcome Sanger Institute for the COVID-19 Genomics UK (COG-UK) Consortium | Jacquelyn Wynn, Mairead Hyland, The Lighthouse Lab in Alderley Park and Alex Alderton, Roberto Amato, Jeffrey Barrett, Sonia Goncalves, Ewan Harrison, David K. Jackson, Ian Johnston, Dominic Kwiatkowski, Cordelia Langford, John Sillitoe on behalf of the Wellcome Sanger Institute COVID-19 Surveillance Team |
| EPI_ISL_2829321 | Lighthouse Laboratory Plymouth | Wellcome Sanger Institute for the COVID-19 Genomics UK (COG-UK) Consortium | Lighthouse Laboratory Plymouth and Alex Alderton, Roberto Amato, Jeffrey Barrett, Sonia Goncalves, Ewan Harrison, David K. Jackson, Ian Johnston, Dominic Kwiatkowski, Cordelia Langford, John Sillitoe on behalf of the Wellcome Sanger Institute COVID-19 Surveillance Team |
| EPI_ISL_2829322, EPI_ISL_2829323, EPI_ISL_2829324 | Lighthouse Lab in Alderley Park | Wellcome Sanger Institute for the COVID-19 Genomics UK (COG-UK) Consortium | Jacquelyn Wynn, Mairead Hyland, The Lighthouse Lab in Alderley Park and Alex Alderton, Roberto Amato, Jeffrey Barrett, Sonia Goncalves, Ewan Harrison, David K. Jackson, Ian Johnston, Dominic Kwiatkowski, Cordelia Langford, John Sillitoe on behalf of the Wellcome Sanger Institute COVID-19 Surveillance Team |
| EPI_ISL_2829325 | Lighthouse Laboratory Plymouth | Wellcome Sanger Institute for the COVID-19 Genomics UK (COG-UK) Consortium | Lighthouse Laboratory Plymouth and Alex Alderton, Roberto Amato, Jeffrey Barrett, Sonia Goncalves, Ewan Harrison, David K. Jackson, Ian Johnston, Dominic Kwiatkowski, Cordelia Langford, John Sillitoe on behalf of the Wellcome Sanger Institute COVID-19 Surveillance Team |
| EPI_ISL_2829326, EPI_ISL_2829327, EPI_ISL_2829328 | Lighthouse Lab in Alderley Park | Wellcome Sanger Institute for the COVID-19 Genomics UK (COG-UK) Consortium | Jacquelyn Wynn, Mairead Hyland, The Lighthouse Lab in Alderley Park and Alex Alderton, Roberto Amato, Jeffrey Barrett, Sonia Goncalves, Ewan Harrison, David K. Jackson, Ian Johnston, Dominic Kwiatkowski, Cordelia Langford, John Sillitoe on behalf of the Wellcome Sanger Institute COVID-19 Surveillance Team |
| EPI_ISL_2829329, EPI_ISL_2829330 | Lighthouse Lab in Glasgow | Wellcome Sanger Institute for the COVID-19 Genomics UK (COG-UK) Consortium | Harper VanSteenhouse, Yumi Kasai, David Gray, Carol Clugston, Anna Dominiczak and Alex Alderton, Roberto Amato, Jeffrey Barrett, Sonia Goncalves, Ewan Harrison, David K. Jackson, Ian Johnston, Dominic Kwiatkowski, Cordelia Langford, John Sillitoe on behalf of the Wellcome Sanger Institute COVID-19 Surveillance Team |
| EPI_ISL_2829332 | Lighthouse Laboratory Plymouth | Wellcome Sanger Institute for the COVID-19 Genomics UK (COG-UK) Consortium | Lighthouse Laboratory Plymouth and Alex Alderton, Roberto Amato, Jeffrey Barrett, Sonia Goncalves, Ewan Harrison, David K. Jackson, Ian Johnston, Dominic Kwiatkowski, Cordelia Langford, John Sillitoe on behalf of the Wellcome Sanger Institute COVID-19 Surveillance Team |
| EPI_ISL_2829333 | Lighthouse Lab in Alderley Park | Wellcome Sanger Institute for the COVID-19 Genomics UK (COG-UK) Consortium | Jacquelyn Wynn, Mairead Hyland, The Lighthouse Lab in Alderley Park and Alex Alderton, Roberto Amato, Jeffrey Barrett, Sonia Goncalves, Ewan Harrison, David K. Jackson, Ian Johnston, Dominic Kwiatkowski, Cordelia Langford, John Sillitoe on behalf of the Wellcome Sanger Institute COVID-19 Surveillance Team |
| EPI_ISL_2829334 | Lighthouse Lab in Glasgow | Wellcome Sanger Institute for the COVID-19 Genomics UK (COG-UK) Consortium | Harper VanSteenhouse, Yumi Kasai, David Gray, Carol Clugston, Anna Dominiczak and Alex Alderton, Roberto Amato, Jeffrey Barrett, Sonia Goncalves, Ewan Harrison, David K. Jackson, Ian Johnston, Dominic Kwiatkowski, Cordelia Langford, John Sillitoe on behalf of the Wellcome Sanger Institute COVID-19 Surveillance Team |
| EPI_ISL_2829335 | Lighthouse Lab in Alderley Park | Wellcome Sanger Institute for the COVID-19 Genomics UK (COG-UK) Consortium | Jacquelyn Wynn, Mairead Hyland, The Lighthouse Lab in Alderley Park and Alex Alderton, Roberto Amato, Jeffrey Barrett, Sonia Goncalves, Ewan Harrison, David K. Jackson, Ian Johnston, Dominic Kwiatkowski, Cordelia Langford, John Sillitoe on behalf of the Wellcome Sanger Institute COVID-19 Surveillance Team |
| EPI_ISL_2829336 | Lighthouse Laboratory Plymouth | Wellcome Sanger Institute for the COVID-19 Genomics UK (COG-UK) Consortium | Lighthouse Laboratory Plymouth and Alex Alderton, Roberto Amato, Jeffrey Barrett, Sonia Goncalves, Ewan Harrison, David K. Jackson, Ian Johnston, Dominic Kwiatkowski, Cordelia Langford, John Sillitoe on behalf of the Wellcome Sanger Institute COVID-19 Surveillance Team |
| EPI_ISL_2829337 | Lighthouse Lab in Alderley Park | Wellcome Sanger Institute for the COVID-19 Genomics UK (COG-UK) Consortium | Jacquelyn Wynn, Mairead Hyland, The Lighthouse Lab in Alderley Park and Alex Alderton, Roberto Amato, Jeffrey Barrett, Sonia Goncalves, Ewan Harrison, David K. Jackson, Ian Johnston, Dominic Kwiatkowski, Cordelia Langford, John Sillitoe on behalf of the Wellcome Sanger Institute COVID-19 Surveillance Team |
| EPI_ISL_2829338 | Lighthouse Laboratory Plymouth | Wellcome Sanger Institute for the COVID-19 Genomics UK (COG-UK) Consortium | Lighthouse Laboratory Plymouth and Alex Alderton, Roberto Amato, Jeffrey Barrett, Sonia Goncalves, Ewan Harrison, David K. Jackson, Ian Johnston, Dominic Kwiatkowski, Cordelia Langford, John Sillitoe on behalf of the Wellcome Sanger Institute COVID-19 Surveillance Team |
| EPI_ISL_2829339 | Lighthouse Lab in Alderley Park | Wellcome Sanger Institute for the COVID-19 Genomics UK (COG-UK) Consortium | Jacquelyn Wynn, Mairead Hyland, The Lighthouse Lab in Alderley Park and Alex Alderton, Roberto Amato, Jeffrey Barrett, Sonia Goncalves, Ewan Harrison, David K. Jackson, Ian Johnston, Dominic Kwiatkowski, Cordelia Langford, John Sillitoe on behalf of the Wellcome Sanger Institute COVID-19 Surveillance Team |
| EPI_ISL_2829340 | Lighthouse Laboratory Plymouth | Wellcome Sanger Institute for the COVID-19 Genomics UK (COG-UK) Consortium | Lighthouse Laboratory Plymouth and Alex Alderton, Roberto Amato, Jeffrey Barrett, Sonia Goncalves, Ewan Harrison, David K. Jackson, Ian Johnston, Dominic Kwiatkowski, Cordelia Langford, John Sillitoe on behalf of the Wellcome Sanger Institute COVID-19 Surveillance Team |
| EPI_ISL_2829341 | Lighthouse Lab in Alderley Park | Wellcome Sanger Institute for the COVID-19 Genomics UK (COG-UK) Consortium | Jacquelyn Wynn, Mairead Hyland, The Lighthouse Lab in Alderley Park and Alex Alderton, Roberto Amato, Jeffrey Barrett, Sonia Goncalves, Ewan Harrison, David K. Jackson, Ian Johnston, Dominic Kwiatkowski, Cordelia Langford, John Sillitoe on behalf of the Wellcome Sanger Institute COVID-19 Surveillance Team |
| EPI_ISL_2829342 | Lighthouse Laboratory Plymouth | Wellcome Sanger Institute for the COVID-19 Genomics UK (COG-UK) Consortium | Lighthouse Laboratory Plymouth and Alex Alderton, Roberto Amato, Jeffrey Barrett, Sonia Goncalves, Ewan Harrison, David K. Jackson, Ian Johnston, Dominic Kwiatkowski, Cordelia Langford, John Sillitoe on behalf of the Wellcome Sanger Institute COVID-19 Surveillance Team |
| EPI_ISL_2829343, EPI_ISL_2829344 | Lighthouse Lab in Alderley Park | Wellcome Sanger Institute for the COVID-19 Genomics UK (COG-UK) Consortium | Jacquelyn Wynn, Mairead Hyland, The Lighthouse Lab in Alderley Park and Alex Alderton, Roberto Amato, Jeffrey Barrett, Sonia Goncalves, Ewan Harrison, David K. Jackson, Ian Johnston, Dominic Kwiatkowski, Cordelia Langford, John Sillitoe on behalf of the Wellcome Sanger Institute COVID-19 Surveillance Team |
| EPI_ISL_2829345, EPI_ISL_2829346, EPI_ISL_2829347 | Lighthouse Laboratory Plymouth | Wellcome Sanger Institute for the COVID-19 Genomics UK (COG-UK) Consortium | Lighthouse Laboratory Plymouth and Alex Alderton, Roberto Amato, Jeffrey Barrett, Sonia Goncalves, Ewan Harrison, David K. Jackson, Ian Johnston, Dominic Kwiatkowski, Cordelia Langford, John Sillitoe on behalf of the Wellcome Sanger Institute COVID-19 Surveillance Team |
| EPI_ISL_2829348, EPI_ISL_2829349, EPI_ISL_2829350 | Lighthouse Lab in Alderley Park | Wellcome Sanger Institute for the COVID-19 Genomics UK (COG-UK) Consortium | Jacquelyn Wynn, Mairead Hyland, The Lighthouse Lab in Alderley Park and Alex Alderton, Roberto Amato, Jeffrey Barrett, Sonia Goncalves, Ewan Harrison, David K. Jackson, Ian Johnston, Dominic Kwiatkowski, Cordelia Langford, John Sillitoe on behalf of the Wellcome Sanger Institute COVID-19 Surveillance Team |
| EPI_ISL_2829351 | Lighthouse Laboratory Plymouth | Wellcome Sanger Institute for the COVID-19 Genomics UK (COG-UK) Consortium | Lighthouse Laboratory Plymouth and Alex Alderton, Roberto Amato, Jeffrey Barrett, Sonia Goncalves, Ewan Harrison, David K. Jackson, Ian Johnston, Dominic Kwiatkowski, Cordelia Langford, John Sillitoe on behalf of the Wellcome Sanger Institute COVID-19 Surveillance Team |
| EPI_ISL_2829352, EPI_ISL_2829353 | Lighthouse Lab in Alderley Park | Wellcome Sanger Institute for the COVID-19 Genomics UK (COG-UK) Consortium | Jacquelyn Wynn, Mairead Hyland, The Lighthouse Lab in Alderley Park and Alex Alderton, Roberto Amato, Jeffrey Barrett, Sonia Goncalves, Ewan Harrison, David K. Jackson, Ian Johnston, Dominic Kwiatkowski, Cordelia Langford, John Sillitoe on behalf of the Wellcome Sanger Institute COVID-19 Surveillance Team |
| EPI_ISL_2829354 | Lighthouse Laboratory Plymouth | Wellcome Sanger Institute for the COVID-19 Genomics UK (COG-UK) Consortium | Lighthouse Laboratory Plymouth and Alex Alderton, Roberto Amato, Jeffrey Barrett, Sonia Goncalves, Ewan Harrison, David K. Jackson, Ian Johnston, Dominic Kwiatkowski, Cordelia Langford, John Sillitoe on behalf of the Wellcome Sanger Institute COVID-19 Surveillance Team |
| EPI_ISL_2829355, EPI_ISL_2829356 | Lighthouse Lab in Alderley Park | Wellcome Sanger Institute for the COVID-19 Genomics UK (COG-UK) Consortium | Jacquelyn Wynn, Mairead Hyland, The Lighthouse Lab in Alderley Park and Alex Alderton, Roberto Amato, Jeffrey Barrett, Sonia Goncalves, Ewan Harrison, David K. Jackson, Ian Johnston, Dominic Kwiatkowski, Cordelia Langford, John Sillitoe on behalf of the Wellcome Sanger Institute COVID-19 Surveillance Team |
| EPI_ISL_2829357 | Lighthouse Laboratory Plymouth | Wellcome Sanger Institute for the COVID-19 Genomics UK (COG-UK) Consortium | Lighthouse Laboratory Plymouth and Alex Alderton, Roberto Amato, Jeffrey Barrett, Sonia Goncalves, Ewan Harrison, David K. Jackson, Ian Johnston, Dominic Kwiatkowski, Cordelia Langford, John Sillitoe on behalf of the Wellcome Sanger Institute COVID-19 Surveillance Team |
| EPI_ISL_2829358 | Lighthouse Lab in Glasgow | Wellcome Sanger Institute for the COVID-19 Genomics UK (COG-UK) Consortium | Harper VanSteenhouse, Yumi Kasai, David Gray, Carol Clugston, Anna Dominiczak and Alex Alderton, Roberto Amato, Jeffrey Barrett, Sonia Goncalves, Ewan Harrison, David K. Jackson, Ian Johnston, Dominic Kwiatkowski, Cordelia Langford, John Sillitoe on behalf of the Wellcome Sanger Institute COVID-19 Surveillance Team |
| EPI_ISL_2829359, EPI_ISL_2829360, EPI_ISL_2829361 | Lighthouse Laboratory Plymouth | Wellcome Sanger Institute for the COVID-19 Genomics UK (COG-UK) Consortium | Lighthouse Laboratory Plymouth and Alex Alderton, Roberto Amato, Jeffrey Barrett, Sonia Goncalves, Ewan Harrison, David K. Jackson, Ian Johnston, Dominic Kwiatkowski, Cordelia Langford, John Sillitoe on behalf of the Wellcome Sanger Institute COVID-19 Surveillance Team |
| EPI_ISL_2829362 | Lighthouse Lab in Alderley Park | Wellcome Sanger Institute for the COVID-19 Genomics UK (COG-UK) Consortium | Jacquelyn Wynn, Mairead Hyland, The Lighthouse Lab in Alderley Park and Alex Alderton, Roberto Amato, Jeffrey Barrett, Sonia Goncalves, Ewan Harrison, David K. Jackson, Ian Johnston, Dominic Kwiatkowski, Cordelia Langford, John Sillitoe on behalf of the Wellcome Sanger Institute COVID-19 Surveillance Team |
| EPI_ISL_2829363 | Lighthouse Laboratory Plymouth | Wellcome Sanger Institute for the COVID-19 Genomics UK (COG-UK) Consortium | Lighthouse Laboratory Plymouth and Alex Alderton, Roberto Amato, Jeffrey Barrett, Sonia Goncalves, Ewan Harrison, David K. Jackson, Ian Johnston, Dominic Kwiatkowski, Cordelia Langford, John Sillitoe on behalf of the Wellcome Sanger Institute COVID-19 Surveillance Team |
| EPI_ISL_2829364 | Lighthouse Lab in Alderley Park | Wellcome Sanger Institute for the COVID-19 Genomics UK (COG-UK) Consortium | Jacquelyn Wynn, Mairead Hyland, The Lighthouse Lab in Alderley Park and Alex Alderton, Roberto Amato, Jeffrey Barrett, Sonia Goncalves, Ewan Harrison, David K. Jackson, Ian Johnston, Dominic Kwiatkowski, Cordelia Langford, John Sillitoe on behalf of the Wellcome Sanger Institute COVID-19 Surveillance Team |
| EPI_ISL_2829365 | Lighthouse Lab in Glasgow | Wellcome Sanger Institute for the COVID-19 Genomics UK (COG-UK) Consortium | Harper VanSteenhouse, Yumi Kasai, David Gray, Carol Clugston, Anna Dominiczak and Alex Alderton, Roberto Amato, Jeffrey Barrett, Sonia Goncalves, Ewan Harrison, David K. Jackson, Ian Johnston, Dominic Kwiatkowski, Cordelia Langford, John Sillitoe on behalf of the Wellcome Sanger Institute COVID-19 Surveillance Team |
| EPI_ISL_2829366, EPI_ISL_2829367, EPI_ISL_2829368, EPI_ISL_2829369, EPI_ISL_2829370 | Lighthouse Lab in Alderley Park | Wellcome Sanger Institute for the COVID-19 Genomics UK (COG-UK) Consortium | Jacquelyn Wynn, Mairead Hyland, The Lighthouse Lab in Alderley Park and Alex Alderton, Roberto Amato, Jeffrey Barrett, Sonia Goncalves, Ewan Harrison, David K. Jackson, Ian Johnston, Dominic Kwiatkowski, Cordelia Langford, John Sillitoe on behalf of the Wellcome Sanger Institute COVID-19 Surveillance Team |
| EPI_ISL_2829371 | Lighthouse Laboratory Plymouth | Wellcome Sanger Institute for the COVID-19 Genomics UK (COG-UK) Consortium | Lighthouse Laboratory Plymouth and Alex Alderton, Roberto Amato, Jeffrey Barrett, Sonia Goncalves, Ewan Harrison, David K. Jackson, Ian Johnston, Dominic Kwiatkowski, Cordelia Langford, John Sillitoe on behalf of the Wellcome Sanger Institute COVID-19 Surveillance Team |
| EPI_ISL_2829372 | Lighthouse Lab in Alderley Park | Wellcome Sanger Institute for the COVID-19 Genomics UK (COG-UK) Consortium | Jacquelyn Wynn, Mairead Hyland, The Lighthouse Lab in Alderley Park and Alex Alderton, Roberto Amato, Jeffrey Barrett, Sonia Goncalves, Ewan Harrison, David K. Jackson, Ian Johnston, Dominic Kwiatkowski, Cordelia Langford, John Sillitoe on behalf of the Wellcome Sanger Institute COVID-19 Surveillance Team |
| EPI_ISL_2829373, EPI_ISL_2829374 | Lighthouse Laboratory Plymouth | Wellcome Sanger Institute for the COVID-19 Genomics UK (COG-UK) Consortium | Lighthouse Laboratory Plymouth and Alex Alderton, Roberto Amato, Jeffrey Barrett, Sonia Goncalves, Ewan Harrison, David K. Jackson, Ian Johnston, Dominic Kwiatkowski, Cordelia Langford, John Sillitoe on behalf of the Wellcome Sanger Institute COVID-19 Surveillance Team |
| EPI_ISL_2829375 | Lighthouse Lab in Alderley Park | Wellcome Sanger Institute for the COVID-19 Genomics UK (COG-UK) Consortium | Jacquelyn Wynn, Mairead Hyland, The Lighthouse Lab in Alderley Park and Alex Alderton, Roberto Amato, Jeffrey Barrett, Sonia Goncalves, Ewan Harrison, David K. Jackson, Ian Johnston, Dominic Kwiatkowski, Cordelia Langford, John Sillitoe on behalf of the Wellcome Sanger Institute COVID-19 Surveillance Team |
| EPI_ISL_2829376 | Lighthouse Laboratory Plymouth | Wellcome Sanger Institute for the COVID-19 Genomics UK (COG-UK) Consortium | Lighthouse Laboratory Plymouth and Alex Alderton, Roberto Amato, Jeffrey Barrett, Sonia Goncalves, Ewan Harrison, David K. Jackson, Ian Johnston, Dominic Kwiatkowski, Cordelia Langford, John Sillitoe on behalf of the Wellcome Sanger Institute COVID-19 Surveillance Team |
| EPI_ISL_2829377, EPI_ISL_2829378 | Lighthouse Lab in Alderley Park | Wellcome Sanger Institute for the COVID-19 Genomics UK (COG-UK) Consortium | Jacquelyn Wynn, Mairead Hyland, The Lighthouse Lab in Alderley Park and Alex Alderton, Roberto Amato, Jeffrey Barrett, Sonia Goncalves, Ewan Harrison, David K. Jackson, Ian Johnston, Dominic Kwiatkowski, Cordelia Langford, John Sillitoe on behalf of the Wellcome Sanger Institute COVID-19 Surveillance Team |
| EPI_ISL_2829379, EPI_ISL_2829380, EPI_ISL_2829381 | Lighthouse Laboratory Plymouth | Wellcome Sanger Institute for the COVID-19 Genomics UK (COG-UK) Consortium | Lighthouse Laboratory Plymouth and Alex Alderton, Roberto Amato, Jeffrey Barrett, Sonia Goncalves, Ewan Harrison, David K. Jackson, Ian Johnston, Dominic Kwiatkowski, Cordelia Langford, John Sillitoe on behalf of the Wellcome Sanger Institute COVID-19 Surveillance Team |

| | | | |
|---|---|---|---|
| EPI_ISL_2829382 | Lighthouse Lab in Glasgow | Wellcome Sanger Institute for the COVID-19 Genomics UK (COG-UK) Consortium | Harper VanSteenhouse, Yumi Kasai, David Gray, Carol Clugston, Anna Dominiczak and Alex Alderton, Roberto Amato, Jeffrey Barrett, Sonia Goncalves, Ewan Harrison, David K. Jackson, Ian Johnston, Dominic Kwiatkowski, Cordelia Langford, John Sillitoe on behalf of the Wellcome Sanger Institute COVID-19 Surveillance Team |
| EPI_ISL_2829383 | Lighthouse Laboratory Plymouth | Wellcome Sanger Institute for the COVID-19 Genomics UK (COG-UK) Consortium | Lighthouse Laboratory Plymouth and Alex Alderton, Roberto Amato, Jeffrey Barrett, Sonia Goncalves, Ewan Harrison, David K. Jackson, Ian Johnston, Dominic Kwiatkowski, Cordelia Langford, John Sillitoe on behalf of the Wellcome Sanger Institute COVID-19 Surveillance Team |
| EPI_ISL_2829384, EPI_ISL_2829385, EPI_ISL_2829386, EPI_ISL_2829387, EPI_ISL_2829389 | Lighthouse Lab in Alderley Park | Wellcome Sanger Institute for the COVID-19 Genomics UK (COG-UK) Consortium | Jacquelyn Wynn, Mairead Hyland, The Lighthouse Lab in Alderley Park and Alex Alderton, Roberto Amato, Jeffrey Barrett, Sonia Goncalves, Ewan Harrison, David K. Jackson, Ian Johnston, Dominic Kwiatkowski, Cordelia Langford, John Sillitoe on behalf of the Wellcome Sanger Institute COVID-19 Surveillance Team |
| EPI_ISL_2829390 | Lighthouse Laboratory Plymouth | Wellcome Sanger Institute for the COVID-19 Genomics UK (COG-UK) Consortium | Lighthouse Laboratory Plymouth and Alex Alderton, Roberto Amato, Jeffrey Barrett, Sonia Goncalves, Ewan Harrison, David K. Jackson, Ian Johnston, Dominic Kwiatkowski, Cordelia Langford, John Sillitoe on behalf of the Wellcome Sanger Institute COVID-19 Surveillance Team |
| EPI_ISL_2829391 | Lighthouse Lab in Alderley Park | Wellcome Sanger Institute for the COVID-19 Genomics UK (COG-UK) Consortium | Jacquelyn Wynn, Mairead Hyland, The Lighthouse Lab in Alderley Park and Alex Alderton, Roberto Amato, Jeffrey Barrett, Sonia Goncalves, Ewan Harrison, David K. Jackson, Ian Johnston, Dominic Kwiatkowski, Cordelia Langford, John Sillitoe on behalf of the Wellcome Sanger Institute COVID-19 Surveillance Team |
| EPI_ISL_2829392 | Lighthouse Laboratory Plymouth | Wellcome Sanger Institute for the COVID-19 Genomics UK (COG-UK) Consortium | Lighthouse Laboratory Plymouth and Alex Alderton, Roberto Amato, Jeffrey Barrett, Sonia Goncalves, Ewan Harrison, David K. Jackson, Ian Johnston, Dominic Kwiatkowski, Cordelia Langford, John Sillitoe on behalf of the Wellcome Sanger Institute COVID-19 Surveillance Team |
| EPI_ISL_2829393 | Lighthouse Lab in Alderley Park | Wellcome Sanger Institute for the COVID-19 Genomics UK (COG-UK) Consortium | Jacquelyn Wynn, Mairead Hyland, The Lighthouse Lab in Alderley Park and Alex Alderton, Roberto Amato, Jeffrey Barrett, Sonia Goncalves, Ewan Harrison, David K. Jackson, Ian Johnston, Dominic Kwiatkowski, Cordelia Langford, John Sillitoe on behalf of the Wellcome Sanger Institute COVID-19 Surveillance Team |
| EPI_ISL_2829394 | Lighthouse Laboratory Plymouth | Wellcome Sanger Institute for the COVID-19 Genomics UK (COG-UK) Consortium | Lighthouse Laboratory Plymouth and Alex Alderton, Roberto Amato, Jeffrey Barrett, Sonia Goncalves, Ewan Harrison, David K. Jackson, Ian Johnston, Dominic Kwiatkowski, Cordelia Langford, John Sillitoe on behalf of the Wellcome Sanger Institute COVID-19 Surveillance Team |
| EPI_ISL_2829395, EPI_ISL_2829396, EPI_ISL_2829397 | Lighthouse Lab in Alderley Park | Wellcome Sanger Institute for the COVID-19 Genomics UK (COG-UK) Consortium | Jacquelyn Wynn, Mairead Hyland, The Lighthouse Lab in Alderley Park and Alex Alderton, Roberto Amato, Jeffrey Barrett, Sonia Goncalves, Ewan Harrison, David K. Jackson, Ian Johnston, Dominic Kwiatkowski, Cordelia Langford, John Sillitoe on behalf of the Wellcome Sanger Institute COVID-19 Surveillance Team |
| EPI_ISL_2829398 | Lighthouse Laboratory Plymouth | Wellcome Sanger Institute for the COVID-19 Genomics UK (COG-UK) Consortium | Lighthouse Laboratory Plymouth and Alex Alderton, Roberto Amato, Jeffrey Barrett, Sonia Goncalves, Ewan Harrison, David K. Jackson, Ian Johnston, Dominic Kwiatkowski, Cordelia Langford, John Sillitoe on behalf of the Wellcome Sanger Institute COVID-19 Surveillance Team |
| EPI_ISL_2829399 | Lighthouse Lab in Glasgow | Wellcome Sanger Institute for the COVID-19 Genomics UK (COG-UK) Consortium | Harper VanSteenhouse, Yumi Kasai, David Gray, Carol Clugston, Anna Dominiczak and Alex Alderton, Roberto Amato, Jeffrey Barrett, Sonia Goncalves, Ewan Harrison, David K. Jackson, Ian Johnston, Dominic Kwiatkowski, Cordelia Langford, John Sillitoe on behalf of the Wellcome Sanger Institute COVID-19 Surveillance Team |
| EPI_ISL_2829400 | Lighthouse Lab in Alderley Park | Wellcome Sanger Institute for the COVID-19 Genomics UK (COG-UK) Consortium | Jacquelyn Wynn, Mairead Hyland, The Lighthouse Lab in Alderley Park and Alex Alderton, Roberto Amato, Jeffrey Barrett, Sonia Goncalves, Ewan Harrison, David K. Jackson, Ian Johnston, Dominic Kwiatkowski, Cordelia Langford, John Sillitoe on behalf of the Wellcome Sanger Institute COVID-19 Surveillance Team |
| EPI_ISL_2829401, EPI_ISL_2829402 | Lighthouse Laboratory Plymouth | Wellcome Sanger Institute for the COVID-19 Genomics UK (COG-UK) Consortium | Lighthouse Laboratory Plymouth and Alex Alderton, Roberto Amato, Jeffrey Barrett, Sonia Goncalves, Ewan Harrison, David K. Jackson, Ian Johnston, Dominic Kwiatkowski, Cordelia Langford, John Sillitoe on behalf of the Wellcome Sanger Institute COVID-19 Surveillance Team |
| EPI_ISL_2829403 | Lighthouse Lab in Alderley Park | Wellcome Sanger Institute for the COVID-19 Genomics UK (COG-UK) Consortium | Jacquelyn Wynn, Mairead Hyland, The Lighthouse Lab in Alderley Park and Alex Alderton, Roberto Amato, Jeffrey Barrett, Sonia Goncalves, Ewan Harrison, David K. Jackson, Ian Johnston, Dominic Kwiatkowski, Cordelia Langford, John Sillitoe on behalf of the Wellcome Sanger Institute COVID-19 Surveillance Team |
| EPI_ISL_2829404 | Lighthouse Laboratory Plymouth | Wellcome Sanger Institute for the COVID-19 Genomics UK (COG-UK) Consortium | Lighthouse Laboratory Plymouth and Alex Alderton, Roberto Amato, Jeffrey Barrett, Sonia Goncalves, Ewan Harrison, David K. Jackson, Ian Johnston, Dominic Kwiatkowski, Cordelia Langford, John Sillitoe on behalf of the Wellcome Sanger Institute COVID-19 Surveillance Team |
| EPI_ISL_2829405 | Lighthouse Lab in Alderley Park | Wellcome Sanger Institute for the COVID-19 Genomics UK (COG-UK) Consortium | Jacquelyn Wynn, Mairead Hyland, The Lighthouse Lab in Alderley Park and Alex Alderton, Roberto Amato, Jeffrey Barrett, Sonia Goncalves, Ewan Harrison, David K. Jackson, Ian Johnston, Dominic Kwiatkowski, Cordelia Langford, John Sillitoe on behalf of the Wellcome Sanger Institute COVID-19 Surveillance Team |
| EPI_ISL_2829406, EPI_ISL_2829407 | Lighthouse Lab in Glasgow | Wellcome Sanger Institute for the COVID-19 Genomics UK (COG-UK) Consortium | Harper VanSteenhouse, Yumi Kasai, David Gray, Carol Clugston, Anna Dominiczak and Alex Alderton, Roberto Amato, Jeffrey Barrett, Sonia Goncalves, Ewan Harrison, David K. Jackson, Ian Johnston, Dominic Kwiatkowski, Cordelia Langford, John Sillitoe on behalf of the Wellcome Sanger Institute COVID-19 Surveillance Team |
| EPI_ISL_2829408, EPI_ISL_2829409, EPI_ISL_2829410, EPI_ISL_2829411 | Lighthouse Lab in Alderley Park | Wellcome Sanger Institute for the COVID-19 Genomics UK (COG-UK) Consortium | Jacquelyn Wynn, Mairead Hyland, The Lighthouse Lab in Alderley Park and Alex Alderton, Roberto Amato, Jeffrey Barrett, Sonia Goncalves, Ewan Harrison, David K. Jackson, Ian Johnston, Dominic Kwiatkowski, Cordelia Langford, John Sillitoe on behalf of the Wellcome Sanger Institute COVID-19 Surveillance Team |
| EPI_ISL_2829412 | Lighthouse Laboratory Plymouth | Wellcome Sanger Institute for the COVID-19 Genomics UK (COG-UK) Consortium | Lighthouse Laboratory Plymouth and Alex Alderton, Roberto Amato, Jeffrey Barrett, Sonia Goncalves, Ewan Harrison, David K. Jackson, Ian Johnston, Dominic Kwiatkowski, Cordelia Langford, John Sillitoe on behalf of the Wellcome Sanger Institute COVID-19 Surveillance Team |
| EPI_ISL_2829413, EPI_ISL_2829414 | Lighthouse Lab in Alderley Park | Wellcome Sanger Institute for the COVID-19 Genomics UK (COG-UK) Consortium | Jacquelyn Wynn, Mairead Hyland, The Lighthouse Lab in Alderley Park and Alex Alderton, Roberto Amato, Jeffrey Barrett, Sonia Goncalves, Ewan Harrison, David K. Jackson, Ian Johnston, Dominic Kwiatkowski, Cordelia Langford, John Sillitoe on behalf of the Wellcome Sanger Institute COVID-19 Surveillance Team |
| EPI_ISL_2829415 | Lighthouse Laboratory Plymouth | Wellcome Sanger Institute for the COVID-19 Genomics UK (COG-UK) Consortium | Lighthouse Laboratory Plymouth and Alex Alderton, Roberto Amato, Jeffrey Barrett, Sonia Goncalves, Ewan Harrison, David K. Jackson, Ian Johnston, Dominic Kwiatkowski, Cordelia Langford, John Sillitoe on behalf of the Wellcome Sanger Institute COVID-19 Surveillance Team |
| EPI_ISL_2829416 | Lighthouse Lab in Alderley Park | Wellcome Sanger Institute for the COVID-19 Genomics UK (COG-UK) Consortium | Jacquelyn Wynn, Mairead Hyland, The Lighthouse Lab in Alderley Park and Alex Alderton, Roberto Amato, Jeffrey Barrett, Sonia Goncalves, Ewan Harrison, David K. Jackson, Ian Johnston, Dominic Kwiatkowski, Cordelia Langford, John Sillitoe on behalf of the Wellcome Sanger Institute COVID-19 Surveillance Team |
| EPI_ISL_2829417 | Lighthouse Lab in Glasgow | Wellcome Sanger Institute for the COVID-19 Genomics UK (COG-UK) Consortium | Harper VanSteenhouse, Yumi Kasai, David Gray, Carol Clugston, Anna Dominiczak and Alex Alderton, Roberto Amato, Jeffrey Barrett, Sonia Goncalves, Ewan Harrison, David K. Jackson, Ian Johnston, Dominic Kwiatkowski, Cordelia Langford, John Sillitoe on behalf of the Wellcome Sanger Institute COVID-19 Surveillance Team |
| EPI_ISL_2829418, EPI_ISL_2829419, EPI_ISL_2829420 | Lighthouse Lab in Alderley Park | Wellcome Sanger Institute for the COVID-19 Genomics UK (COG-UK) Consortium | Jacquelyn Wynn, Mairead Hyland, The Lighthouse Lab in Alderley Park and Alex Alderton, Roberto Amato, Jeffrey Barrett, Sonia Goncalves, Ewan Harrison, David K. Jackson, Ian Johnston, Dominic Kwiatkowski, Cordelia Langford, John Sillitoe on behalf of the Wellcome Sanger Institute COVID-19 Surveillance Team |
| EPI_ISL_2829421 | Lighthouse Laboratory Plymouth | Wellcome Sanger Institute for the COVID-19 Genomics UK (COG-UK) Consortium | Lighthouse Laboratory Plymouth and Alex Alderton, Roberto Amato, Jeffrey Barrett, Sonia Goncalves, Ewan Harrison, David K. Jackson, Ian Johnston, Dominic Kwiatkowski, Cordelia Langford, John Sillitoe on behalf of the Wellcome Sanger Institute COVID-19 Surveillance Team |
| EPI_ISL_2829422, EPI_ISL_2829423, EPI_ISL_2829424 | Lighthouse Lab in Alderley Park | Wellcome Sanger Institute for the COVID-19 Genomics UK (COG-UK) Consortium | Jacquelyn Wynn, Mairead Hyland, The Lighthouse Lab in Alderley Park and Alex Alderton, Roberto Amato, Jeffrey Barrett, Sonia Goncalves, Ewan Harrison, David K. Jackson, Ian Johnston, Dominic Kwiatkowski, Cordelia Langford, John Sillitoe on behalf of the Wellcome Sanger Institute COVID-19 Surveillance Team |
| EPI_ISL_2829425, EPI_ISL_2829426 | Lighthouse Laboratory Plymouth | Wellcome Sanger Institute for the COVID-19 Genomics UK (COG-UK) Consortium | Lighthouse Laboratory Plymouth and Alex Alderton, Roberto Amato, Jeffrey Barrett, Sonia Goncalves, Ewan Harrison, David K. Jackson, Ian Johnston, Dominic Kwiatkowski, Cordelia Langford, John Sillitoe on behalf of the Wellcome Sanger Institute COVID-19 Surveillance Team |
| EPI_ISL_2829427 | Lighthouse Lab in Alderley Park | Wellcome Sanger Institute for the COVID-19 Genomics UK (COG-UK) Consortium | Jacquelyn Wynn, Mairead Hyland, The Lighthouse Lab in Alderley Park and Alex Alderton, Roberto Amato, Jeffrey Barrett, Sonia Goncalves, Ewan Harrison, David K. Jackson, Ian Johnston, Dominic Kwiatkowski, Cordelia Langford, John Sillitoe on behalf of the Wellcome Sanger Institute COVID-19 Surveillance Team |
| EPI_ISL_2829428 | Lighthouse Laboratory Plymouth | Wellcome Sanger Institute for the COVID-19 Genomics UK (COG-UK) Consortium | Lighthouse Laboratory Plymouth and Alex Alderton, Roberto Amato, Jeffrey Barrett, Sonia Goncalves, Ewan Harrison, David K. Jackson, Ian Johnston, Dominic Kwiatkowski, Cordelia Langford, John Sillitoe on behalf of the Wellcome Sanger Institute COVID-19 Surveillance Team |
| EPI_ISL_2829429, EPI_ISL_2829430, EPI_ISL_2829431 | Lighthouse Lab in Alderley Park | Wellcome Sanger Institute for the COVID-19 Genomics UK (COG-UK) Consortium | Jacquelyn Wynn, Mairead Hyland, The Lighthouse Lab in Alderley Park and Alex Alderton, Roberto Amato, Jeffrey Barrett, Sonia Goncalves, Ewan Harrison, David K. Jackson, Ian Johnston, Dominic Kwiatkowski, Cordelia Langford, John Sillitoe on behalf of the Wellcome Sanger Institute COVID-19 Surveillance Team |
| EPI_ISL_2829432, EPI_ISL_2829433 | Lighthouse Laboratory Plymouth | Wellcome Sanger Institute for the COVID-19 Genomics UK (COG-UK) Consortium | Lighthouse Laboratory Plymouth and Alex Alderton, Roberto Amato, Jeffrey Barrett, Sonia Goncalves, Ewan Harrison, David K. Jackson, Ian Johnston, Dominic Kwiatkowski, Cordelia Langford, John Sillitoe on behalf of the Wellcome Sanger Institute COVID-19 Surveillance Team |
| EPI_ISL_2829434, EPI_ISL_2829435, EPI_ISL_2829436, EPI_ISL_2829437 | Lighthouse Lab in Alderley Park | Wellcome Sanger Institute for the COVID-19 Genomics UK (COG-UK) Consortium | Jacquelyn Wynn, Mairead Hyland, The Lighthouse Lab in Alderley Park and Alex Alderton, Roberto Amato, Jeffrey Barrett, Sonia Goncalves, Ewan Harrison, David K. Jackson, Ian Johnston, Dominic Kwiatkowski, Cordelia Langford, John Sillitoe on behalf of the Wellcome Sanger Institute COVID-19 Surveillance Team |
| EPI_ISL_2829438 | Lighthouse Laboratory Plymouth | Wellcome Sanger Institute for the COVID-19 Genomics UK (COG-UK) Consortium | Lighthouse Laboratory Plymouth and Alex Alderton, Roberto Amato, Jeffrey Barrett, Sonia Goncalves, Ewan Harrison, David K. Jackson, Ian Johnston, Dominic Kwiatkowski, Cordelia Langford, John Sillitoe on behalf of the Wellcome Sanger Institute COVID-19 Surveillance Team |
| EPI_ISL_2829439, EPI_ISL_2829440, EPI_ISL_2829441, EPI_ISL_2829442, EPI_ISL_2829443, EPI_ISL_2829444 | Lighthouse Lab in Alderley Park | Wellcome Sanger Institute for the COVID-19 Genomics UK (COG-UK) Consortium | Jacquelyn Wynn, Mairead Hyland, The Lighthouse Lab in Alderley Park and Alex Alderton, Roberto Amato, Jeffrey Barrett, Sonia Goncalves, Ewan Harrison, David K. Jackson, Ian Johnston, Dominic Kwiatkowski, Cordelia Langford, John Sillitoe on behalf of the Wellcome Sanger Institute COVID-19 Surveillance Team |
| EPI_ISL_2829445 | Lighthouse Laboratory Plymouth | Wellcome Sanger Institute for the COVID-19 Genomics UK (COG-UK) Consortium | Lighthouse Laboratory Plymouth and Alex Alderton, Roberto Amato, Jeffrey Barrett, Sonia Goncalves, Ewan Harrison, David K. Jackson, Ian Johnston, Dominic Kwiatkowski, Cordelia Langford, John Sillitoe on behalf of the Wellcome Sanger Institute COVID-19 Surveillance Team |
| EPI_ISL_2829446 | Lighthouse Lab in Glasgow | Wellcome Sanger Institute for the COVID-19 Genomics UK (COG-UK) Consortium | Harper VanSteenhouse, Yumi Kasai, David Gray, Carol Clugston, Anna Dominiczak and Alex Alderton, Roberto Amato, Jeffrey Barrett, Sonia Goncalves, Ewan Harrison, David K. Jackson, Ian Johnston, Dominic Kwiatkowski, Cordelia Langford, John Sillitoe on behalf of the Wellcome Sanger Institute COVID-19 Surveillance Team |
| EPI_ISL_2829447 | Lighthouse Laboratory Plymouth | Wellcome Sanger Institute for the COVID-19 Genomics UK (COG-UK) Consortium | Lighthouse Laboratory Plymouth and Alex Alderton, Roberto Amato, Jeffrey Barrett, Sonia Goncalves, Ewan Harrison, David K. Jackson, Ian Johnston, Dominic Kwiatkowski, Cordelia Langford, John Sillitoe on behalf of the Wellcome Sanger Institute COVID-19 Surveillance Team |
| EPI_ISL_2829448, EPI_ISL_2829449 | Lighthouse Lab in Alderley Park | Wellcome Sanger Institute for the COVID-19 Genomics UK (COG-UK) Consortium | Jacquelyn Wynn, Mairead Hyland, The Lighthouse Lab in Alderley Park and Alex Alderton, Roberto Amato, Jeffrey Barrett, Sonia Goncalves, Ewan Harrison, David K. Jackson, Ian Johnston, Dominic Kwiatkowski, Cordelia Langford, John Sillitoe on behalf of the Wellcome Sanger Institute COVID-19 Surveillance Team |
| EPI_ISL_2829450, EPI_ISL_2829451 | Lighthouse Laboratory Plymouth | Wellcome Sanger Institute for the COVID-19 Genomics UK (COG-UK) Consortium | Lighthouse Laboratory Plymouth and Alex Alderton, Roberto Amato, Jeffrey Barrett, Sonia Goncalves, Ewan Harrison, David K. Jackson, Ian Johnston, Dominic Kwiatkowski, Cordelia Langford, John Sillitoe on behalf of the Wellcome Sanger Institute COVID-19 Surveillance Team |

| | | | |
|---|---|---|---|
| EPI_ISL_2829452 | Lighthouse Lab in Alderley Park | Wellcome Sanger Institute for the COVID-19 Genomics UK (COG-UK) Consortium | Jacquelyn Wynn, Mairead Hyland, The Lighthouse Lab in Alderley Park and Alex Alderton, Roberto Amato, Jeffrey Barrett, Sonia Goncalves, Ewan Harrison, David K. Jackson, Ian Johnston, Dominic Kwiatkowski, Cordelia Langford, John Sillitoe on behalf of the Wellcome Sanger Institute COVID-19 Surveillance Team |
| EPI_ISL_2829453, EPI_ISL_2829454 | Lighthouse Laboratory Plymouth | Wellcome Sanger Institute for the COVID-19 Genomics UK (COG-UK) Consortium | Lighthouse Laboratory Plymouth and Alex Alderton, Roberto Amato, Jeffrey Barrett, Sonia Goncalves, Ewan Harrison, David K. Jackson, Ian Johnston, Dominic Kwiatkowski, Cordelia Langford, John Sillitoe on behalf of the Wellcome Sanger Institute COVID-19 Surveillance Team |
| EPI_ISL_2829455, EPI_ISL_2829456 | Lighthouse Lab in Alderley Park | Wellcome Sanger Institute for the COVID-19 Genomics UK (COG-UK) Consortium | Jacquelyn Wynn, Mairead Hyland, The Lighthouse Lab in Alderley Park and Alex Alderton, Roberto Amato, Jeffrey Barrett, Sonia Goncalves, Ewan Harrison, David K. Jackson, Ian Johnston, Dominic Kwiatkowski, Cordelia Langford, John Sillitoe on behalf of the Wellcome Sanger Institute COVID-19 Surveillance Team |
| EPI_ISL_2829457 | Lighthouse Lab in Glasgow | Wellcome Sanger Institute for the COVID-19 Genomics UK (COG-UK) Consortium | Harper VanSteenhouse, Yumi Kasai, David Gray, Carol Clugston, Anna Dominiczak and Alex Alderton, Roberto Amato, Jeffrey Barrett, Sonia Goncalves, Ewan Harrison, David K. Jackson, Ian Johnston, Dominic Kwiatkowski, Cordelia Langford, John Sillitoe on behalf of the Wellcome Sanger Institute COVID-19 Surveillance Team |
| EPI_ISL_2829458 | Lighthouse Lab in Alderley Park | Wellcome Sanger Institute for the COVID-19 Genomics UK (COG-UK) Consortium | Jacquelyn Wynn, Mairead Hyland, The Lighthouse Lab in Alderley Park and Alex Alderton, Roberto Amato, Jeffrey Barrett, Sonia Goncalves, Ewan Harrison, David K. Jackson, Ian Johnston, Dominic Kwiatkowski, Cordelia Langford, John Sillitoe on behalf of the Wellcome Sanger Institute COVID-19 Surveillance Team |
| EPI_ISL_2829459 | Lighthouse Laboratory Plymouth | Wellcome Sanger Institute for the COVID-19 Genomics UK (COG-UK) Consortium | Lighthouse Laboratory Plymouth and Alex Alderton, Roberto Amato, Jeffrey Barrett, Sonia Goncalves, Ewan Harrison, David K. Jackson, Ian Johnston, Dominic Kwiatkowski, Cordelia Langford, John Sillitoe on behalf of the Wellcome Sanger Institute COVID-19 Surveillance Team |
| EPI_ISL_2829460 | Lighthouse Lab in Alderley Park | Wellcome Sanger Institute for the COVID-19 Genomics UK (COG-UK) Consortium | Jacquelyn Wynn, Mairead Hyland, The Lighthouse Lab in Alderley Park and Alex Alderton, Roberto Amato, Jeffrey Barrett, Sonia Goncalves, Ewan Harrison, David K. Jackson, Ian Johnston, Dominic Kwiatkowski, Cordelia Langford, John Sillitoe on behalf of the Wellcome Sanger Institute COVID-19 Surveillance Team |
| EPI_ISL_2829461 | Lighthouse Laboratory Plymouth | Wellcome Sanger Institute for the COVID-19 Genomics UK (COG-UK) Consortium | Lighthouse Laboratory Plymouth and Alex Alderton, Roberto Amato, Jeffrey Barrett, Sonia Goncalves, Ewan Harrison, David K. Jackson, Ian Johnston, Dominic Kwiatkowski, Cordelia Langford, John Sillitoe on behalf of the Wellcome Sanger Institute COVID-19 Surveillance Team |
| EPI_ISL_2829462 | Lighthouse Lab in Alderley Park | Wellcome Sanger Institute for the COVID-19 Genomics UK (COG-UK) Consortium | Jacquelyn Wynn, Mairead Hyland, The Lighthouse Lab in Alderley Park and Alex Alderton, Roberto Amato, Jeffrey Barrett, Sonia Goncalves, Ewan Harrison, David K. Jackson, Ian Johnston, Dominic Kwiatkowski, Cordelia Langford, John Sillitoe on behalf of the Wellcome Sanger Institute COVID-19 Surveillance Team |
| EPI_ISL_2829463 | Lighthouse Laboratory Plymouth | Wellcome Sanger Institute for the COVID-19 Genomics UK (COG-UK) Consortium | Lighthouse Laboratory Plymouth and Alex Alderton, Roberto Amato, Jeffrey Barrett, Sonia Goncalves, Ewan Harrison, David K. Jackson, Ian Johnston, Dominic Kwiatkowski, Cordelia Langford, John Sillitoe on behalf of the Wellcome Sanger Institute COVID-19 Surveillance Team |
| EPI_ISL_2829464 | Lighthouse Lab in Alderley Park | Wellcome Sanger Institute for the COVID-19 Genomics UK (COG-UK) Consortium | Jacquelyn Wynn, Mairead Hyland, The Lighthouse Lab in Alderley Park and Alex Alderton, Roberto Amato, Jeffrey Barrett, Sonia Goncalves, Ewan Harrison, David K. Jackson, Ian Johnston, Dominic Kwiatkowski, Cordelia Langford, John Sillitoe on behalf of the Wellcome Sanger Institute COVID-19 Surveillance Team |
| EPI_ISL_2829465 | Lighthouse Laboratory Plymouth | Wellcome Sanger Institute for the COVID-19 Genomics UK (COG-UK) Consortium | Lighthouse Laboratory Plymouth and Alex Alderton, Roberto Amato, Jeffrey Barrett, Sonia Goncalves, Ewan Harrison, David K. Jackson, Ian Johnston, Dominic Kwiatkowski, Cordelia Langford, John Sillitoe on behalf of the Wellcome Sanger Institute COVID-19 Surveillance Team |
| EPI_ISL_2829466, EPI_ISL_2829467 | Lighthouse Lab in Glasgow | Wellcome Sanger Institute for the COVID-19 Genomics UK (COG-UK) Consortium | Harper VanSteenhouse, Yumi Kasai, David Gray, Carol Clugston, Anna Dominiczak and Alex Alderton, Roberto Amato, Jeffrey Barrett, Sonia Goncalves, Ewan Harrison, David K. Jackson, Ian Johnston, Dominic Kwiatkowski, Cordelia Langford, John Sillitoe on behalf of the Wellcome Sanger Institute COVID-19 Surveillance Team |
| EPI_ISL_2829468, EPI_ISL_2829469, EPI_ISL_2829470, EPI_ISL_2829471, EPI_ISL_2829472 | Lighthouse Lab in Alderley Park | Wellcome Sanger Institute for the COVID-19 Genomics UK (COG-UK) Consortium | Jacquelyn Wynn, Mairead Hyland, The Lighthouse Lab in Alderley Park and Alex Alderton, Roberto Amato, Jeffrey Barrett, Sonia Goncalves, Ewan Harrison, David K. Jackson, Ian Johnston, Dominic Kwiatkowski, Cordelia Langford, John Sillitoe on behalf of the Wellcome Sanger Institute COVID-19 Surveillance Team |
| EPI_ISL_2829473 | Lighthouse Laboratory Plymouth | Wellcome Sanger Institute for the COVID-19 Genomics UK (COG-UK) Consortium | Lighthouse Laboratory Plymouth and Alex Alderton, Roberto Amato, Jeffrey Barrett, Sonia Goncalves, Ewan Harrison, David K. Jackson, Ian Johnston, Dominic Kwiatkowski, Cordelia Langford, John Sillitoe on behalf of the Wellcome Sanger Institute COVID-19 Surveillance Team |
| EPI_ISL_2829474 | Lighthouse Lab in Alderley Park | Wellcome Sanger Institute for the COVID-19 Genomics UK (COG-UK) Consortium | Jacquelyn Wynn, Mairead Hyland, The Lighthouse Lab in Alderley Park and Alex Alderton, Roberto Amato, Jeffrey Barrett, Sonia Goncalves, Ewan Harrison, David K. Jackson, Ian Johnston, Dominic Kwiatkowski, Cordelia Langford, John Sillitoe on behalf of the Wellcome Sanger Institute COVID-19 Surveillance Team |
| EPI_ISL_2829475, EPI_ISL_2829476 | Lighthouse Laboratory Plymouth | Wellcome Sanger Institute for the COVID-19 Genomics UK (COG-UK) Consortium | Lighthouse Laboratory Plymouth and Alex Alderton, Roberto Amato, Jeffrey Barrett, Sonia Goncalves, Ewan Harrison, David K. Jackson, Ian Johnston, Dominic Kwiatkowski, Cordelia Langford, John Sillitoe on behalf of the Wellcome Sanger Institute COVID-19 Surveillance Team |
| EPI_ISL_2829477 | Lighthouse Lab in Alderley Park | Wellcome Sanger Institute for the COVID-19 Genomics UK (COG-UK) Consortium | Jacquelyn Wynn, Mairead Hyland, The Lighthouse Lab in Alderley Park and Alex Alderton, Roberto Amato, Jeffrey Barrett, Sonia Goncalves, Ewan Harrison, David K. Jackson, Ian Johnston, Dominic Kwiatkowski, Cordelia Langford, John Sillitoe on behalf of the Wellcome Sanger Institute COVID-19 Surveillance Team |
| EPI_ISL_2829478 | Lighthouse Laboratory Plymouth | Wellcome Sanger Institute for the COVID-19 Genomics UK (COG-UK) Consortium | Lighthouse Laboratory Plymouth and Alex Alderton, Roberto Amato, Jeffrey Barrett, Sonia Goncalves, Ewan Harrison, David K. Jackson, Ian Johnston, Dominic Kwiatkowski, Cordelia Langford, John Sillitoe on behalf of the Wellcome Sanger Institute COVID-19 Surveillance Team |
| EPI_ISL_2829479, EPI_ISL_2829480, EPI_ISL_2829482, EPI_ISL_2829483, EPI_ISL_2829484, EPI_ISL_2829485 | Lighthouse Lab in Alderley Park | Wellcome Sanger Institute for the COVID-19 Genomics UK (COG-UK) Consortium | Jacquelyn Wynn, Mairead Hyland, The Lighthouse Lab in Alderley Park and Alex Alderton, Roberto Amato, Jeffrey Barrett, Sonia Goncalves, Ewan Harrison, David K. Jackson, Ian Johnston, Dominic Kwiatkowski, Cordelia Langford, John Sillitoe on behalf of the Wellcome Sanger Institute COVID-19 Surveillance Team |
| EPI_ISL_2829486 | Lighthouse Laboratory Plymouth | Wellcome Sanger Institute for the COVID-19 Genomics UK (COG-UK) Consortium | Lighthouse Laboratory Plymouth and Alex Alderton, Roberto Amato, Jeffrey Barrett, Sonia Goncalves, Ewan Harrison, David K. Jackson, Ian Johnston, Dominic Kwiatkowski, Cordelia Langford, John Sillitoe on behalf of the Wellcome Sanger Institute COVID-19 Surveillance Team |
| EPI_ISL_2829487, EPI_ISL_2829488 | Lighthouse Lab in Alderley Park | Wellcome Sanger Institute for the COVID-19 Genomics UK (COG-UK) Consortium | Jacquelyn Wynn, Mairead Hyland, The Lighthouse Lab in Alderley Park and Alex Alderton, Roberto Amato, Jeffrey Barrett, Sonia Goncalves, Ewan Harrison, David K. Jackson, Ian Johnston, Dominic Kwiatkowski, Cordelia Langford, John Sillitoe on behalf of the Wellcome Sanger Institute COVID-19 Surveillance Team |
| EPI_ISL_2829489 | Lighthouse Laboratory Plymouth | Wellcome Sanger Institute for the COVID-19 Genomics UK (COG-UK) Consortium | Lighthouse Laboratory Plymouth and Alex Alderton, Roberto Amato, Jeffrey Barrett, Sonia Goncalves, Ewan Harrison, David K. Jackson, Ian Johnston, Dominic Kwiatkowski, Cordelia Langford, John Sillitoe on behalf of the Wellcome Sanger Institute COVID-19 Surveillance Team |
| EPI_ISL_2829490 | Lighthouse Lab in Alderley Park | Wellcome Sanger Institute for the COVID-19 Genomics UK (COG-UK) Consortium | Jacquelyn Wynn, Mairead Hyland, The Lighthouse Lab in Alderley Park and Alex Alderton, Roberto Amato, Jeffrey Barrett, Sonia Goncalves, Ewan Harrison, David K. Jackson, Ian Johnston, Dominic Kwiatkowski, Cordelia Langford, John Sillitoe on behalf of the Wellcome Sanger Institute COVID-19 Surveillance Team |
| EPI_ISL_2829491 | Lighthouse Lab in Glasgow | Wellcome Sanger Institute for the COVID-19 Genomics UK (COG-UK) Consortium | Harper VanSteenhouse, Yumi Kasai, David Gray, Carol Clugston, Anna Dominiczak and Alex Alderton, Roberto Amato, Jeffrey Barrett, Sonia Goncalves, Ewan Harrison, David K. Jackson, Ian Johnston, Dominic Kwiatkowski, Cordelia Langford, John Sillitoe on behalf of the Wellcome Sanger Institute COVID-19 Surveillance Team |
| EPI_ISL_2829492 | Lighthouse Lab in Alderley Park | Wellcome Sanger Institute for the COVID-19 Genomics UK (COG-UK) Consortium | Jacquelyn Wynn, Mairead Hyland, The Lighthouse Lab in Alderley Park and Alex Alderton, Roberto Amato, Jeffrey Barrett, Sonia Goncalves, Ewan Harrison, David K. Jackson, Ian Johnston, Dominic Kwiatkowski, Cordelia Langford, John Sillitoe on behalf of the Wellcome Sanger Institute COVID-19 Surveillance Team |
| EPI_ISL_2829493 | Lighthouse Laboratory Plymouth | Wellcome Sanger Institute for the COVID-19 Genomics UK (COG-UK) Consortium | Lighthouse Laboratory Plymouth and Alex Alderton, Roberto Amato, Jeffrey Barrett, Sonia Goncalves, Ewan Harrison, David K. Jackson, Ian Johnston, Dominic Kwiatkowski, Cordelia Langford, John Sillitoe on behalf of the Wellcome Sanger Institute COVID-19 Surveillance Team |
| EPI_ISL_2829494, EPI_ISL_2829495, EPI_ISL_2829496 | Lighthouse Lab in Alderley Park | Wellcome Sanger Institute for the COVID-19 Genomics UK (COG-UK) Consortium | Jacquelyn Wynn, Mairead Hyland, The Lighthouse Lab in Alderley Park and Alex Alderton, Roberto Amato, Jeffrey Barrett, Sonia Goncalves, Ewan Harrison, David K. Jackson, Ian Johnston, Dominic Kwiatkowski, Cordelia Langford, John Sillitoe on behalf of the Wellcome Sanger Institute COVID-19 Surveillance Team |
| EPI_ISL_2829497 | Lighthouse Laboratory Plymouth | Wellcome Sanger Institute for the COVID-19 Genomics UK (COG-UK) Consortium | Lighthouse Laboratory Plymouth and Alex Alderton, Roberto Amato, Jeffrey Barrett, Sonia Goncalves, Ewan Harrison, David K. Jackson, Ian Johnston, Dominic Kwiatkowski, Cordelia Langford, John Sillitoe on behalf of the Wellcome Sanger Institute COVID-19 Surveillance Team |
| EPI_ISL_2829498 | Lighthouse Lab in Alderley Park | Wellcome Sanger Institute for the COVID-19 Genomics UK (COG-UK) Consortium | Jacquelyn Wynn, Mairead Hyland, The Lighthouse Lab in Alderley Park and Alex Alderton, Roberto Amato, Jeffrey Barrett, Sonia Goncalves, Ewan Harrison, David K. Jackson, Ian Johnston, Dominic Kwiatkowski, Cordelia Langford, John Sillitoe on behalf of the Wellcome Sanger Institute COVID-19 Surveillance Team |
| EPI_ISL_2829499 | Lighthouse Lab in Glasgow | Wellcome Sanger Institute for the COVID-19 Genomics UK (COG-UK) Consortium | Harper VanSteenhouse, Yumi Kasai, David Gray, Carol Clugston, Anna Dominiczak and Alex Alderton, Roberto Amato, Jeffrey Barrett, Sonia Goncalves, Ewan Harrison, David K. Jackson, Ian Johnston, Dominic Kwiatkowski, Cordelia Langford, John Sillitoe on behalf of the Wellcome Sanger Institute COVID-19 Surveillance Team |
| EPI_ISL_2829500, EPI_ISL_2829501 | Lighthouse Lab in Alderley Park | Wellcome Sanger Institute for the COVID-19 Genomics UK (COG-UK) Consortium | Jacquelyn Wynn, Mairead Hyland, The Lighthouse Lab in Alderley Park and Alex Alderton, Roberto Amato, Jeffrey Barrett, Sonia Goncalves, Ewan Harrison, David K. Jackson, Ian Johnston, Dominic Kwiatkowski, Cordelia Langford, John Sillitoe on behalf of the Wellcome Sanger Institute COVID-19 Surveillance Team |
| EPI_ISL_2829502 | Lighthouse Laboratory Plymouth | Wellcome Sanger Institute for the COVID-19 Genomics UK (COG-UK) Consortium | Lighthouse Laboratory Plymouth and Alex Alderton, Roberto Amato, Jeffrey Barrett, Sonia Goncalves, Ewan Harrison, David K. Jackson, Ian Johnston, Dominic Kwiatkowski, Cordelia Langford, John Sillitoe on behalf of the Wellcome Sanger Institute COVID-19 Surveillance Team |
| EPI_ISL_2829503, EPI_ISL_2829504, EPI_ISL_2829505, EPI_ISL_2829506, EPI_ISL_2829507, EPI_ISL_2829508, EPI_ISL_2829509, EPI_ISL_2829510, EPI_ISL_2829511, EPI_ISL_2829512, EPI_ISL_2829513, EPI_ISL_2829514 | | | |
| see above | Lighthouse Lab in Alderley Park | Wellcome Sanger Institute for the COVID-19 Genomics UK (COG-UK) Consortium | Jacquelyn Wynn, Mairead Hyland, The Lighthouse Lab in Alderley Park and Alex Alderton, Roberto Amato, Jeffrey Barrett, Sonia Goncalves, Ewan Harrison, David K. Jackson, Ian Johnston, Dominic Kwiatkowski, Cordelia Langford, John Sillitoe on behalf of the Wellcome Sanger Institute COVID-19 Surveillance Team |
| EPI_ISL_2829515 | Lighthouse Lab in Glasgow | Wellcome Sanger Institute for the COVID-19 Genomics UK (COG-UK) Consortium | Harper VanSteenhouse, Yumi Kasai, David Gray, Carol Clugston, Anna Dominiczak and Alex Alderton, Roberto Amato, Jeffrey Barrett, Sonia Goncalves, Ewan Harrison, David K. Jackson, Ian Johnston, Dominic Kwiatkowski, Cordelia Langford, John Sillitoe on behalf of the Wellcome Sanger Institute COVID-19 Surveillance Team |
| EPI_ISL_2829516, EPI_ISL_2829517 | Lighthouse Lab in Alderley Park | Wellcome Sanger Institute for the COVID-19 Genomics UK (COG-UK) Consortium | Jacquelyn Wynn, Mairead Hyland, The Lighthouse Lab in Alderley Park and Alex Alderton, Roberto Amato, Jeffrey Barrett, Sonia Goncalves, Ewan Harrison, David K. Jackson, Ian Johnston, Dominic Kwiatkowski, Cordelia Langford, John Sillitoe on behalf of the Wellcome Sanger Institute COVID-19 Surveillance Team |
| EPI_ISL_2829518 | Lighthouse Lab in Glasgow | Wellcome Sanger Institute for the COVID-19 Genomics UK (COG-UK) Consortium | Harper VanSteenhouse, Yumi Kasai, David Gray, Carol Clugston, Anna Dominiczak and Alex Alderton, Roberto Amato, Jeffrey Barrett, Sonia Goncalves, Ewan Harrison, David K. Jackson, Ian Johnston, Dominic Kwiatkowski, Cordelia Langford, John Sillitoe on behalf of the Wellcome Sanger Institute COVID-19 Surveillance Team |
| EPI_ISL_2829519 | Lighthouse Laboratory Plymouth | Wellcome Sanger Institute for the COVID-19 Genomics UK (COG-UK) Consortium | Lighthouse Laboratory Plymouth and Alex Alderton, Roberto Amato, Jeffrey Barrett, Sonia Goncalves, Ewan Harrison, David K. Jackson, Ian Johnston, Dominic Kwiatkowski, Cordelia Langford, John Sillitoe on behalf of the Wellcome Sanger Institute COVID-19 Surveillance Team |
| EPI_ISL_2829520 | Lighthouse Lab in Alderley Park | Wellcome Sanger Institute for the COVID-19 Genomics UK (COG-UK) Consortium | Jacquelyn Wynn, Mairead Hyland, The Lighthouse Lab in Alderley Park and Alex Alderton, Roberto Amato, Jeffrey Barrett, Sonia Goncalves, Ewan Harrison, David K. Jackson, Ian Johnston, Dominic Kwiatkowski, Cordelia Langford, John Sillitoe on behalf of the Wellcome Sanger Institute COVID-19 Surveillance Team |
| EPI_ISL_2829521, EPI_ISL_2829522 | Lighthouse Laboratory Plymouth | Wellcome Sanger Institute for the COVID-19 Genomics UK (COG-UK) Consortium | Lighthouse Laboratory Plymouth and Alex Alderton, Roberto Amato, Jeffrey Barrett, Sonia Goncalves, Ewan Harrison, David K. Jackson, Ian Johnston, Dominic Kwiatkowski, Cordelia Langford, John Sillitoe on behalf of the Wellcome Sanger Institute COVID-19 Surveillance Team |
| EPI_ISL_2829523, EPI_ISL_2829524, EPI_ISL_2829525, | Lighthouse Lab in Alderley Park | Wellcome Sanger Institute for the COVID-19 Genomics UK (COG-UK) Consortium | Jacquelyn Wynn, Mairead Hyland, The Lighthouse Lab in Alderley Park and Alex Alderton, Roberto Amato, Jeffrey Barrett, Sonia Goncalves, Ewan Harrison, David K. Jackson, Ian Johnston, Dominic Kwiatkowski, Cordelia Langford, John Sillitoe on behalf of the Wellcome Sanger Institute COVID-19 Surveillance Team |

| | | | |
|---|---|---|---|
| EPI_ISL_2829526 | | | |
| EPI_ISL_2829527 | Lighthouse Lab in Glasgow | Wellcome Sanger Institute for the COVID-19 Genomics UK (COG-UK) Consortium | Harper VanSteenhouse, Yumi Kasai, David Gray, Carol Clugston, Anna Dominiczak and Alex Alderton, Roberto Amato, Jeffrey Barrett, Sonia Goncalves, Ewan Harrison, David K. Jackson, Ian Johnston, Dominic Kwiatkowski, Cordelia Langford, John Sillitoe on behalf of the Wellcome Sanger Institute COVID-19 Surveillance Team |
| EPI_ISL_2829528, EPI_ISL_2829529, EPI_ISL_2829530 | Lighthouse Lab in Alderley Park | Wellcome Sanger Institute for the COVID-19 Genomics UK (COG-UK) Consortium | Jacquelyn Wynn, Mairead Hyland, The Lighthouse Lab in Alderley Park and Alex Alderton, Roberto Amato, Jeffrey Barrett, Sonia Goncalves, Ewan Harrison, David K. Jackson, Ian Johnston, Dominic Kwiatkowski, Cordelia Langford, John Sillitoe on behalf of the Wellcome Sanger Institute COVID-19 Surveillance Team |
| EPI_ISL_2829531 | Lighthouse Laboratory Plymouth | Wellcome Sanger Institute for the COVID-19 Genomics UK (COG-UK) Consortium | Lighthouse Laboratory Plymouth and Alex Alderton, Roberto Amato, Jeffrey Barrett, Sonia Goncalves, Ewan Harrison, David K. Jackson, Ian Johnston, Dominic Kwiatkowski, Cordelia Langford, John Sillitoe on behalf of the Wellcome Sanger Institute COVID-19 Surveillance Team |
| EPI_ISL_2829532 | Lighthouse Lab in Alderley Park | Wellcome Sanger Institute for the COVID-19 Genomics UK (COG-UK) Consortium | Jacquelyn Wynn, Mairead Hyland, The Lighthouse Lab in Alderley Park and Alex Alderton, Roberto Amato, Jeffrey Barrett, Sonia Goncalves, Ewan Harrison, David K. Jackson, Ian Johnston, Dominic Kwiatkowski, Cordelia Langford, John Sillitoe on behalf of the Wellcome Sanger Institute COVID-19 Surveillance Team |
| EPI_ISL_2829533 | Lighthouse Laboratory Plymouth | Wellcome Sanger Institute for the COVID-19 Genomics UK (COG-UK) Consortium | Lighthouse Laboratory Plymouth and Alex Alderton, Roberto Amato, Jeffrey Barrett, Sonia Goncalves, Ewan Harrison, David K. Jackson, Ian Johnston, Dominic Kwiatkowski, Cordelia Langford, John Sillitoe on behalf of the Wellcome Sanger Institute COVID-19 Surveillance Team |
| EPI_ISL_2829534 | Lighthouse Lab in Glasgow | Wellcome Sanger Institute for the COVID-19 Genomics UK (COG-UK) Consortium | Harper VanSteenhouse, Yumi Kasai, David Gray, Carol Clugston, Anna Dominiczak and Alex Alderton, Roberto Amato, Jeffrey Barrett, Sonia Goncalves, Ewan Harrison, David K. Jackson, Ian Johnston, Dominic Kwiatkowski, Cordelia Langford, John Sillitoe on behalf of the Wellcome Sanger Institute COVID-19 Surveillance Team |
| EPI_ISL_2829535, EPI_ISL_2829536 | Lighthouse Lab in Alderley Park | Wellcome Sanger Institute for the COVID-19 Genomics UK (COG-UK) Consortium | Jacquelyn Wynn, Mairead Hyland, The Lighthouse Lab in Alderley Park and Alex Alderton, Roberto Amato, Jeffrey Barrett, Sonia Goncalves, Ewan Harrison, David K. Jackson, Ian Johnston, Dominic Kwiatkowski, Cordelia Langford, John Sillitoe on behalf of the Wellcome Sanger Institute COVID-19 Surveillance Team |
| EPI_ISL_2829537 | Lighthouse Laboratory Plymouth | Wellcome Sanger Institute for the COVID-19 Genomics UK (COG-UK) Consortium | Lighthouse Laboratory Plymouth and Alex Alderton, Roberto Amato, Jeffrey Barrett, Sonia Goncalves, Ewan Harrison, David K. Jackson, Ian Johnston, Dominic Kwiatkowski, Cordelia Langford, John Sillitoe on behalf of the Wellcome Sanger Institute COVID-19 Surveillance Team |
| EPI_ISL_2829538, EPI_ISL_2829539, EPI_ISL_2829540, EPI_ISL_2829541, EPI_ISL_2829542, EPI_ISL_2829543, EPI_ISL_2829544 | Lighthouse Lab in Alderley Park | Wellcome Sanger Institute for the COVID-19 Genomics UK (COG-UK) Consortium | Jacquelyn Wynn, Mairead Hyland, The Lighthouse Lab in Alderley Park and Alex Alderton, Roberto Amato, Jeffrey Barrett, Sonia Goncalves, Ewan Harrison, David K. Jackson, Ian Johnston, Dominic Kwiatkowski, Cordelia Langford, John Sillitoe on behalf of the Wellcome Sanger Institute COVID-19 Surveillance Team |
| EPI_ISL_2829545, EPI_ISL_2829546 | Lighthouse Laboratory Plymouth | Wellcome Sanger Institute for the COVID-19 Genomics UK (COG-UK) Consortium | Lighthouse Laboratory Plymouth and Alex Alderton, Roberto Amato, Jeffrey Barrett, Sonia Goncalves, Ewan Harrison, David K. Jackson, Ian Johnston, Dominic Kwiatkowski, Cordelia Langford, John Sillitoe on behalf of the Wellcome Sanger Institute COVID-19 Surveillance Team |
| EPI_ISL_2829547 | Lighthouse Lab in Alderley Park | Wellcome Sanger Institute for the COVID-19 Genomics UK (COG-UK) Consortium | Jacquelyn Wynn, Mairead Hyland, The Lighthouse Lab in Alderley Park and Alex Alderton, Roberto Amato, Jeffrey Barrett, Sonia Goncalves, Ewan Harrison, David K. Jackson, Ian Johnston, Dominic Kwiatkowski, Cordelia Langford, John Sillitoe on behalf of the Wellcome Sanger Institute COVID-19 Surveillance Team |
| EPI_ISL_2829548, EPI_ISL_2829549 | Lighthouse Laboratory Plymouth | Wellcome Sanger Institute for the COVID-19 Genomics UK (COG-UK) Consortium | Lighthouse Laboratory Plymouth and Alex Alderton, Roberto Amato, Jeffrey Barrett, Sonia Goncalves, Ewan Harrison, David K. Jackson, Ian Johnston, Dominic Kwiatkowski, Cordelia Langford, John Sillitoe on behalf of the Wellcome Sanger Institute COVID-19 Surveillance Team |
| EPI_ISL_2829550 | Lighthouse Lab in Glasgow | Wellcome Sanger Institute for the COVID-19 Genomics UK (COG-UK) Consortium | Harper VanSteenhouse, Yumi Kasai, David Gray, Carol Clugston, Anna Dominiczak and Alex Alderton, Roberto Amato, Jeffrey Barrett, Sonia Goncalves, Ewan Harrison, David K. Jackson, Ian Johnston, Dominic Kwiatkowski, Cordelia Langford, John Sillitoe on behalf of the Wellcome Sanger Institute COVID-19 Surveillance Team |
| EPI_ISL_2829551 | Lighthouse Laboratory Plymouth | Wellcome Sanger Institute for the COVID-19 Genomics UK (COG-UK) Consortium | Lighthouse Laboratory Plymouth and Alex Alderton, Roberto Amato, Jeffrey Barrett, Sonia Goncalves, Ewan Harrison, David K. Jackson, Ian Johnston, Dominic Kwiatkowski, Cordelia Langford, John Sillitoe on behalf of the Wellcome Sanger Institute COVID-19 Surveillance Team |
| EPI_ISL_2829552, EPI_ISL_2829553, EPI_ISL_2829554, EPI_ISL_2829555 | Lighthouse Lab in Alderley Park | Wellcome Sanger Institute for the COVID-19 Genomics UK (COG-UK) Consortium | Jacquelyn Wynn, Mairead Hyland, The Lighthouse Lab in Alderley Park and Alex Alderton, Roberto Amato, Jeffrey Barrett, Sonia Goncalves, Ewan Harrison, David K. Jackson, Ian Johnston, Dominic Kwiatkowski, Cordelia Langford, John Sillitoe on behalf of the Wellcome Sanger Institute COVID-19 Surveillance Team |
| EPI_ISL_2829556 | Lighthouse Laboratory Plymouth | Wellcome Sanger Institute for the COVID-19 Genomics UK (COG-UK) Consortium | Lighthouse Laboratory Plymouth and Alex Alderton, Roberto Amato, Jeffrey Barrett, Sonia Goncalves, Ewan Harrison, David K. Jackson, Ian Johnston, Dominic Kwiatkowski, Cordelia Langford, John Sillitoe on behalf of the Wellcome Sanger Institute COVID-19 Surveillance Team |
| EPI_ISL_2829557 | Lighthouse Lab in Alderley Park | Wellcome Sanger Institute for the COVID-19 Genomics UK (COG-UK) Consortium | Jacquelyn Wynn, Mairead Hyland, The Lighthouse Lab in Alderley Park and Alex Alderton, Roberto Amato, Jeffrey Barrett, Sonia Goncalves, Ewan Harrison, David K. Jackson, Ian Johnston, Dominic Kwiatkowski, Cordelia Langford, John Sillitoe on behalf of the Wellcome Sanger Institute COVID-19 Surveillance Team |
| EPI_ISL_2829558 | Lighthouse Lab in Glasgow | Wellcome Sanger Institute for the COVID-19 Genomics UK (COG-UK) Consortium | Harper VanSteenhouse, Yumi Kasai, David Gray, Carol Clugston, Anna Dominiczak and Alex Alderton, Roberto Amato, Jeffrey Barrett, Sonia Goncalves, Ewan Harrison, David K. Jackson, Ian Johnston, Dominic Kwiatkowski, Cordelia Langford, John Sillitoe on behalf of the Wellcome Sanger Institute COVID-19 Surveillance Team |
| EPI_ISL_2829559, EPI_ISL_2829560, EPI_ISL_2829561, EPI_ISL_2829562 | Lighthouse Lab in Alderley Park | Wellcome Sanger Institute for the COVID-19 Genomics UK (COG-UK) Consortium | Jacquelyn Wynn, Mairead Hyland, The Lighthouse Lab in Alderley Park and Alex Alderton, Roberto Amato, Jeffrey Barrett, Sonia Goncalves, Ewan Harrison, David K. Jackson, Ian Johnston, Dominic Kwiatkowski, Cordelia Langford, John Sillitoe on behalf of the Wellcome Sanger Institute COVID-19 Surveillance Team |
| EPI_ISL_2829563 | Lighthouse Lab in Glasgow | Wellcome Sanger Institute for the COVID-19 Genomics UK (COG-UK) Consortium | Harper VanSteenhouse, Yumi Kasai, David Gray, Carol Clugston, Anna Dominiczak and Alex Alderton, Roberto Amato, Jeffrey Barrett, Sonia Goncalves, Ewan Harrison, David K. Jackson, Ian Johnston, Dominic Kwiatkowski, Cordelia Langford, John Sillitoe on behalf of the Wellcome Sanger Institute COVID-19 Surveillance Team |
| EPI_ISL_2829564 | Lighthouse Lab in Alderley Park | Wellcome Sanger Institute for the COVID-19 Genomics UK (COG-UK) Consortium | Jacquelyn Wynn, Mairead Hyland, The Lighthouse Lab in Alderley Park and Alex Alderton, Roberto Amato, Jeffrey Barrett, Sonia Goncalves, Ewan Harrison, David K. Jackson, Ian Johnston, Dominic Kwiatkowski, Cordelia Langford, John Sillitoe on behalf of the Wellcome Sanger Institute COVID-19 Surveillance Team |
| EPI_ISL_2829565 | Lighthouse Laboratory Plymouth | Wellcome Sanger Institute for the COVID-19 Genomics UK (COG-UK) Consortium | Lighthouse Laboratory Plymouth and Alex Alderton, Roberto Amato, Jeffrey Barrett, Sonia Goncalves, Ewan Harrison, David K. Jackson, Ian Johnston, Dominic Kwiatkowski, Cordelia Langford, John Sillitoe on behalf of the Wellcome Sanger Institute COVID-19 Surveillance Team |
| EPI_ISL_2829566 | Lighthouse Lab in Glasgow | Wellcome Sanger Institute for the COVID-19 Genomics UK (COG-UK) Consortium | Harper VanSteenhouse, Yumi Kasai, David Gray, Carol Clugston, Anna Dominiczak and Alex Alderton, Roberto Amato, Jeffrey Barrett, Sonia Goncalves, Ewan Harrison, David K. Jackson, Ian Johnston, Dominic Kwiatkowski, Cordelia Langford, John Sillitoe on behalf of the Wellcome Sanger Institute COVID-19 Surveillance Team |
| EPI_ISL_2829567 | Lighthouse Laboratory Plymouth | Wellcome Sanger Institute for the COVID-19 Genomics UK (COG-UK) Consortium | Lighthouse Laboratory Plymouth and Alex Alderton, Roberto Amato, Jeffrey Barrett, Sonia Goncalves, Ewan Harrison, David K. Jackson, Ian Johnston, Dominic Kwiatkowski, Cordelia Langford, John Sillitoe on behalf of the Wellcome Sanger Institute COVID-19 Surveillance Team |
| EPI_ISL_2829568, EPI_ISL_2829569, EPI_ISL_2829570, EPI_ISL_2829571 | Lighthouse Lab in Alderley Park | Wellcome Sanger Institute for the COVID-19 Genomics UK (COG-UK) Consortium | Jacquelyn Wynn, Mairead Hyland, The Lighthouse Lab in Alderley Park and Alex Alderton, Roberto Amato, Jeffrey Barrett, Sonia Goncalves, Ewan Harrison, David K. Jackson, Ian Johnston, Dominic Kwiatkowski, Cordelia Langford, John Sillitoe on behalf of the Wellcome Sanger Institute COVID-19 Surveillance Team |
| EPI_ISL_2829572 | Lighthouse Laboratory Plymouth | Wellcome Sanger Institute for the COVID-19 Genomics UK (COG-UK) Consortium | Lighthouse Laboratory Plymouth and Alex Alderton, Roberto Amato, Jeffrey Barrett, Sonia Goncalves, Ewan Harrison, David K. Jackson, Ian Johnston, Dominic Kwiatkowski, Cordelia Langford, John Sillitoe on behalf of the Wellcome Sanger Institute COVID-19 Surveillance Team |
| EPI_ISL_2829573, EPI_ISL_2829574, EPI_ISL_2829575, EPI_ISL_2829576, EPI_ISL_2829577, EPI_ISL_2829578, EPI_ISL_2829579, EPI_ISL_2829580, EPI_ISL_2829581, EPI_ISL_2829582 | Lighthouse Lab in Alderley Park | Wellcome Sanger Institute for the COVID-19 Genomics UK (COG-UK) Consortium | Jacquelyn Wynn, Mairead Hyland, The Lighthouse Lab in Alderley Park and Alex Alderton, Roberto Amato, Jeffrey Barrett, Sonia Goncalves, Ewan Harrison, David K. Jackson, Ian Johnston, Dominic Kwiatkowski, Cordelia Langford, John Sillitoe on behalf of the Wellcome Sanger Institute COVID-19 Surveillance Team |
| EPI_ISL_2829583, EPI_ISL_2829584 | Lighthouse Laboratory Plymouth | Wellcome Sanger Institute for the COVID-19 Genomics UK (COG-UK) Consortium | Lighthouse Laboratory Plymouth and Alex Alderton, Roberto Amato, Jeffrey Barrett, Sonia Goncalves, Ewan Harrison, David K. Jackson, Ian Johnston, Dominic Kwiatkowski, Cordelia Langford, John Sillitoe on behalf of the Wellcome Sanger Institute COVID-19 Surveillance Team |
| EPI_ISL_2829585 | Lighthouse Lab in Alderley Park | Wellcome Sanger Institute for the COVID-19 Genomics UK (COG-UK) Consortium | Jacquelyn Wynn, Mairead Hyland, The Lighthouse Lab in Alderley Park and Alex Alderton, Roberto Amato, Jeffrey Barrett, Sonia Goncalves, Ewan Harrison, David K. Jackson, Ian Johnston, Dominic Kwiatkowski, Cordelia Langford, John Sillitoe on behalf of the Wellcome Sanger Institute COVID-19 Surveillance Team |
| EPI_ISL_2829586 | Lighthouse Laboratory Plymouth | Wellcome Sanger Institute for the COVID-19 Genomics UK (COG-UK) Consortium | Lighthouse Laboratory Plymouth and Alex Alderton, Roberto Amato, Jeffrey Barrett, Sonia Goncalves, Ewan Harrison, David K. Jackson, Ian Johnston, Dominic Kwiatkowski, Cordelia Langford, John Sillitoe on behalf of the Wellcome Sanger Institute COVID-19 Surveillance Team |
| EPI_ISL_2829587 | Lighthouse Lab in Alderley Park | Wellcome Sanger Institute for the COVID-19 Genomics UK (COG-UK) Consortium | Jacquelyn Wynn, Mairead Hyland, The Lighthouse Lab in Alderley Park and Alex Alderton, Roberto Amato, Jeffrey Barrett, Sonia Goncalves, Ewan Harrison, David K. Jackson, Ian Johnston, Dominic Kwiatkowski, Cordelia Langford, John Sillitoe on behalf of the Wellcome Sanger Institute COVID-19 Surveillance Team |
| EPI_ISL_2829588, EPI_ISL_2829589 | Lighthouse Laboratory Plymouth | Wellcome Sanger Institute for the COVID-19 Genomics UK (COG-UK) Consortium | Lighthouse Laboratory Plymouth and Alex Alderton, Roberto Amato, Jeffrey Barrett, Sonia Goncalves, Ewan Harrison, David K. Jackson, Ian Johnston, Dominic Kwiatkowski, Cordelia Langford, John Sillitoe on behalf of the Wellcome Sanger Institute COVID-19 Surveillance Team |
| EPI_ISL_2829590 | Lighthouse Lab in Alderley Park | Wellcome Sanger Institute for the COVID-19 Genomics UK (COG-UK) Consortium | Jacquelyn Wynn, Mairead Hyland, The Lighthouse Lab in Alderley Park and Alex Alderton, Roberto Amato, Jeffrey Barrett, Sonia Goncalves, Ewan Harrison, David K. Jackson, Ian Johnston, Dominic Kwiatkowski, Cordelia Langford, John Sillitoe on behalf of the Wellcome Sanger Institute COVID-19 Surveillance Team |
| EPI_ISL_2829591, EPI_ISL_2829592 | Health Services Laboratories | Wellcome Sanger Institute for the COVID-19 Genomics UK (COG-UK) Consortium | Health Services Laboratories and Alex Alderton, Roberto Amato, Jeffrey Barrett, Sonia Goncalves, Ewan Harrison, David K. Jackson, Ian Johnston, Dominic Kwiatkowski, Cordelia Langford, John Sillitoe on behalf of the Wellcome Sanger Institute COVID-19 Surveillance Team |
| EPI_ISL_2829593 | Lighthouse Lab in Milton Keynes | Wellcome Sanger Institute for the COVID-19 Genomics UK (COG-UK) Consortium | The Lighthouse Lab in Milton Keynes and Alex Alderton, Roberto Amato, Jeffrey Barrett, Sonia Goncalves, Ewan Harrison, David K. Jackson, Ian Johnston, Dominic Kwiatkowski, Cordelia Langford, John Sillitoe on behalf of the Wellcome Sanger Institute COVID-19 Surveillance Team |
| EPI_ISL_2829594 | Health Services Laboratories | Wellcome Sanger Institute for the COVID-19 Genomics UK (COG-UK) Consortium | Health Services Laboratories and Alex Alderton, Roberto Amato, Jeffrey Barrett, Sonia Goncalves, Ewan Harrison, David K. Jackson, Ian Johnston, Dominic Kwiatkowski, Cordelia Langford, John Sillitoe on behalf of the Wellcome Sanger Institute COVID-19 Surveillance Team |
| EPI_ISL_2829595, | Lighthouse Lab in Milton Keynes | Wellcome Sanger Institute for the COVID-19 Genomics UK (COG-UK) | The Lighthouse Lab in Milton Keynes and Alex Alderton, Roberto Amato, Jeffrey Barrett, Sonia Goncalves, Ewan Harrison, David K. Jackson, Ian Johnston, Dominic Kwiatkowski, Cordelia Langford, John Sillitoe on behalf of the Wellcome Sanger |

| | | | |
|---|---|---|---|
| EPI_ISL_2829596 | | Consortium | Institute COVID-19 Surveillance Team |
| EPI_ISL_2829597 | Health Services Laboratories | Wellcome Sanger Institute for the COVID-19 Genomics UK (COG-UK) Consortium | Health Services Laboratories and Alex Alderton, Roberto Amato, Jeffrey Barrett, Sonia Goncalves, Ewan Harrison, David K. Jackson, Ian Johnston, Dominic Kwiatkowski, Cordelia Langford, John Sillitoe on behalf of the Wellcome Sanger Institute COVID-19 Surveillance Team |
| EPI_ISL_2829598, EPI_ISL_2829599, EPI_ISL_2829600 | Lighthouse Lab in Alderley Park | Wellcome Sanger Institute for the COVID-19 Genomics UK (COG-UK) Consortium | Jacquelyn Wynn, Mairead Hyland, The Lighthouse Lab in Alderley Park and Alex Alderton, Roberto Amato, Jeffrey Barrett, Sonia Goncalves, Ewan Harrison, David K. Jackson, Ian Johnston, Dominic Kwiatkowski, Cordelia Langford, John Sillitoe on behalf of the Wellcome Sanger Institute COVID-19 Surveillance Team |
| EPI_ISL_2829601 | Lighthouse Lab in Milton Keynes | Wellcome Sanger Institute for the COVID-19 Genomics UK (COG-UK) Consortium | The Lighthouse Lab in Milton Keynes and Alex Alderton, Roberto Amato, Jeffrey Barrett, Sonia Goncalves, Ewan Harrison, David K. Jackson, Ian Johnston, Dominic Kwiatkowski, Cordelia Langford, John Sillitoe on behalf of the Wellcome Sanger Institute COVID-19 Surveillance Team |
| EPI_ISL_2829602 | Health Services Laboratories | Wellcome Sanger Institute for the COVID-19 Genomics UK (COG-UK) Consortium | Health Services Laboratories and Alex Alderton, Roberto Amato, Jeffrey Barrett, Sonia Goncalves, Ewan Harrison, David K. Jackson, Ian Johnston, Dominic Kwiatkowski, Cordelia Langford, John Sillitoe on behalf of the Wellcome Sanger Institute COVID-19 Surveillance Team |
| EPI_ISL_2829603, EPI_ISL_2829604 | Lighthouse Lab in Milton Keynes | Wellcome Sanger Institute for the COVID-19 Genomics UK (COG-UK) Consortium | The Lighthouse Lab in Milton Keynes and Alex Alderton, Roberto Amato, Jeffrey Barrett, Sonia Goncalves, Ewan Harrison, David K. Jackson, Ian Johnston, Dominic Kwiatkowski, Cordelia Langford, John Sillitoe on behalf of the Wellcome Sanger Institute COVID-19 Surveillance Team |
| EPI_ISL_2829605 | Lighthouse Lab in Alderley Park | Wellcome Sanger Institute for the COVID-19 Genomics UK (COG-UK) Consortium | Jacquelyn Wynn, Mairead Hyland, The Lighthouse Lab in Alderley Park and Alex Alderton, Roberto Amato, Jeffrey Barrett, Sonia Goncalves, Ewan Harrison, David K. Jackson, Ian Johnston, Dominic Kwiatkowski, Cordelia Langford, John Sillitoe on behalf of the Wellcome Sanger Institute COVID-19 Surveillance Team |
| EPI_ISL_2829606 | Lighthouse Lab in Milton Keynes | Wellcome Sanger Institute for the COVID-19 Genomics UK (COG-UK) Consortium | The Lighthouse Lab in Milton Keynes and Alex Alderton, Roberto Amato, Jeffrey Barrett, Sonia Goncalves, Ewan Harrison, David K. Jackson, Ian Johnston, Dominic Kwiatkowski, Cordelia Langford, John Sillitoe on behalf of the Wellcome Sanger Institute COVID-19 Surveillance Team |
| EPI_ISL_2829607, EPI_ISL_2829608, EPI_ISL_2829609 | Health Services Laboratories | Wellcome Sanger Institute for the COVID-19 Genomics UK (COG-UK) Consortium | Health Services Laboratories and Alex Alderton, Roberto Amato, Jeffrey Barrett, Sonia Goncalves, Ewan Harrison, David K. Jackson, Ian Johnston, Dominic Kwiatkowski, Cordelia Langford, John Sillitoe on behalf of the Wellcome Sanger Institute COVID-19 Surveillance Team |
| EPI_ISL_2829610 | Lighthouse Lab in Milton Keynes | Wellcome Sanger Institute for the COVID-19 Genomics UK (COG-UK) Consortium | The Lighthouse Lab in Milton Keynes and Alex Alderton, Roberto Amato, Jeffrey Barrett, Sonia Goncalves, Ewan Harrison, David K. Jackson, Ian Johnston, Dominic Kwiatkowski, Cordelia Langford, John Sillitoe on behalf of the Wellcome Sanger Institute COVID-19 Surveillance Team |
| EPI_ISL_2829611, EPI_ISL_2829612 | Health Services Laboratories | Wellcome Sanger Institute for the COVID-19 Genomics UK (COG-UK) Consortium | Health Services Laboratories and Alex Alderton, Roberto Amato, Jeffrey Barrett, Sonia Goncalves, Ewan Harrison, David K. Jackson, Ian Johnston, Dominic Kwiatkowski, Cordelia Langford, John Sillitoe on behalf of the Wellcome Sanger Institute COVID-19 Surveillance Team |
| EPI_ISL_2829613, EPI_ISL_2829614 | Lighthouse Lab in Alderley Park | Wellcome Sanger Institute for the COVID-19 Genomics UK (COG-UK) Consortium | Jacquelyn Wynn, Mairead Hyland, The Lighthouse Lab in Alderley Park and Alex Alderton, Roberto Amato, Jeffrey Barrett, Sonia Goncalves, Ewan Harrison, David K. Jackson, Ian Johnston, Dominic Kwiatkowski, Cordelia Langford, John Sillitoe on behalf of the Wellcome Sanger Institute COVID-19 Surveillance Team |
| EPI_ISL_2829615, EPI_ISL_2829616, EPI_ISL_2829617, EPI_ISL_2829618 | Health Services Laboratories | Wellcome Sanger Institute for the COVID-19 Genomics UK (COG-UK) Consortium | Health Services Laboratories and Alex Alderton, Roberto Amato, Jeffrey Barrett, Sonia Goncalves, Ewan Harrison, David K. Jackson, Ian Johnston, Dominic Kwiatkowski, Cordelia Langford, John Sillitoe on behalf of the Wellcome Sanger Institute COVID-19 Surveillance Team |
| EPI_ISL_2829619 | Lighthouse Lab in Alderley Park | Wellcome Sanger Institute for the COVID-19 Genomics UK (COG-UK) Consortium | Jacquelyn Wynn, Mairead Hyland, The Lighthouse Lab in Alderley Park and Alex Alderton, Roberto Amato, Jeffrey Barrett, Sonia Goncalves, Ewan Harrison, David K. Jackson, Ian Johnston, Dominic Kwiatkowski, Cordelia Langford, John Sillitoe on behalf of the Wellcome Sanger Institute COVID-19 Surveillance Team |
| EPI_ISL_2829620 | Health Services Laboratories | Wellcome Sanger Institute for the COVID-19 Genomics UK (COG-UK) Consortium | Health Services Laboratories and Alex Alderton, Roberto Amato, Jeffrey Barrett, Sonia Goncalves, Ewan Harrison, David K. Jackson, Ian Johnston, Dominic Kwiatkowski, Cordelia Langford, John Sillitoe on behalf of the Wellcome Sanger Institute COVID-19 Surveillance Team |
| EPI_ISL_2829621 | Lighthouse Lab in Milton Keynes | Wellcome Sanger Institute for the COVID-19 Genomics UK (COG-UK) Consortium | The Lighthouse Lab in Milton Keynes and Alex Alderton, Roberto Amato, Jeffrey Barrett, Sonia Goncalves, Ewan Harrison, David K. Jackson, Ian Johnston, Dominic Kwiatkowski, Cordelia Langford, John Sillitoe on behalf of the Wellcome Sanger Institute COVID-19 Surveillance Team |
| EPI_ISL_2829622 | Health Services Laboratories | Wellcome Sanger Institute for the COVID-19 Genomics UK (COG-UK) Consortium | Health Services Laboratories and Alex Alderton, Roberto Amato, Jeffrey Barrett, Sonia Goncalves, Ewan Harrison, David K. Jackson, Ian Johnston, Dominic Kwiatkowski, Cordelia Langford, John Sillitoe on behalf of the Wellcome Sanger Institute COVID-19 Surveillance Team |
| EPI_ISL_2829623 | Lighthouse Lab in Alderley Park | Wellcome Sanger Institute for the COVID-19 Genomics UK (COG-UK) Consortium | Jacquelyn Wynn, Mairead Hyland, The Lighthouse Lab in Alderley Park and Alex Alderton, Roberto Amato, Jeffrey Barrett, Sonia Goncalves, Ewan Harrison, David K. Jackson, Ian Johnston, Dominic Kwiatkowski, Cordelia Langford, John Sillitoe on behalf of the Wellcome Sanger Institute COVID-19 Surveillance Team |
| EPI_ISL_2829624, EPI_ISL_2829625 | Health Services Laboratories | Wellcome Sanger Institute for the COVID-19 Genomics UK (COG-UK) Consortium | Health Services Laboratories and Alex Alderton, Roberto Amato, Jeffrey Barrett, Sonia Goncalves, Ewan Harrison, David K. Jackson, Ian Johnston, Dominic Kwiatkowski, Cordelia Langford, John Sillitoe on behalf of the Wellcome Sanger Institute COVID-19 Surveillance Team |
| EPI_ISL_2829626, EPI_ISL_2829627 | Lighthouse Lab in Milton Keynes | Wellcome Sanger Institute for the COVID-19 Genomics UK (COG-UK) Consortium | The Lighthouse Lab in Milton Keynes and Alex Alderton, Roberto Amato, Jeffrey Barrett, Sonia Goncalves, Ewan Harrison, David K. Jackson, Ian Johnston, Dominic Kwiatkowski, Cordelia Langford, John Sillitoe on behalf of the Wellcome Sanger Institute COVID-19 Surveillance Team |
| EPI_ISL_2829628 | Lighthouse Lab in Alderley Park | Wellcome Sanger Institute for the COVID-19 Genomics UK (COG-UK) Consortium | Jacquelyn Wynn, Mairead Hyland, The Lighthouse Lab in Alderley Park and Alex Alderton, Roberto Amato, Jeffrey Barrett, Sonia Goncalves, Ewan Harrison, David K. Jackson, Ian Johnston, Dominic Kwiatkowski, Cordelia Langford, John Sillitoe on behalf of the Wellcome Sanger Institute COVID-19 Surveillance Team |
| EPI_ISL_2829629 | Lighthouse Lab in Milton Keynes | Wellcome Sanger Institute for the COVID-19 Genomics UK (COG-UK) Consortium | The Lighthouse Lab in Milton Keynes and Alex Alderton, Roberto Amato, Jeffrey Barrett, Sonia Goncalves, Ewan Harrison, David K. Jackson, Ian Johnston, Dominic Kwiatkowski, Cordelia Langford, John Sillitoe on behalf of the Wellcome Sanger Institute COVID-19 Surveillance Team |
| EPI_ISL_2829630 | Health Services Laboratories | Wellcome Sanger Institute for the COVID-19 Genomics UK (COG-UK) Consortium | Health Services Laboratories and Alex Alderton, Roberto Amato, Jeffrey Barrett, Sonia Goncalves, Ewan Harrison, David K. Jackson, Ian Johnston, Dominic Kwiatkowski, Cordelia Langford, John Sillitoe on behalf of the Wellcome Sanger Institute COVID-19 Surveillance Team |
| EPI_ISL_2829631 | Lighthouse Lab in Milton Keynes | Wellcome Sanger Institute for the COVID-19 Genomics UK (COG-UK) Consortium | The Lighthouse Lab in Milton Keynes and Alex Alderton, Roberto Amato, Jeffrey Barrett, Sonia Goncalves, Ewan Harrison, David K. Jackson, Ian Johnston, Dominic Kwiatkowski, Cordelia Langford, John Sillitoe on behalf of the Wellcome Sanger Institute COVID-19 Surveillance Team |
| EPI_ISL_2829632, EPI_ISL_2829633 | Lighthouse Lab in Alderley Park | Wellcome Sanger Institute for the COVID-19 Genomics UK (COG-UK) Consortium | Jacquelyn Wynn, Mairead Hyland, The Lighthouse Lab in Alderley Park and Alex Alderton, Roberto Amato, Jeffrey Barrett, Sonia Goncalves, Ewan Harrison, David K. Jackson, Ian Johnston, Dominic Kwiatkowski, Cordelia Langford, John Sillitoe on behalf of the Wellcome Sanger Institute COVID-19 Surveillance Team |
| EPI_ISL_2829634, EPI_ISL_2829635 | Lighthouse Lab in Milton Keynes | Wellcome Sanger Institute for the COVID-19 Genomics UK (COG-UK) Consortium | The Lighthouse Lab in Milton Keynes and Alex Alderton, Roberto Amato, Jeffrey Barrett, Sonia Goncalves, Ewan Harrison, David K. Jackson, Ian Johnston, Dominic Kwiatkowski, Cordelia Langford, John Sillitoe on behalf of the Wellcome Sanger Institute COVID-19 Surveillance Team |
| EPI_ISL_2829636 | Health Services Laboratories | Wellcome Sanger Institute for the COVID-19 Genomics UK (COG-UK) Consortium | Health Services Laboratories and Alex Alderton, Roberto Amato, Jeffrey Barrett, Sonia Goncalves, Ewan Harrison, David K. Jackson, Ian Johnston, Dominic Kwiatkowski, Cordelia Langford, John Sillitoe on behalf of the Wellcome Sanger Institute COVID-19 Surveillance Team |
| EPI_ISL_2829637, EPI_ISL_2829638, EPI_ISL_2829639, EPI_ISL_2829640 | Lighthouse Lab in Milton Keynes | Wellcome Sanger Institute for the COVID-19 Genomics UK (COG-UK) Consortium | The Lighthouse Lab in Milton Keynes and Alex Alderton, Roberto Amato, Jeffrey Barrett, Sonia Goncalves, Ewan Harrison, David K. Jackson, Ian Johnston, Dominic Kwiatkowski, Cordelia Langford, John Sillitoe on behalf of the Wellcome Sanger Institute COVID-19 Surveillance Team |
| EPI_ISL_2829641, EPI_ISL_2829642 | Lighthouse Lab in Alderley Park | Wellcome Sanger Institute for the COVID-19 Genomics UK (COG-UK) Consortium | Jacquelyn Wynn, Mairead Hyland, The Lighthouse Lab in Alderley Park and Alex Alderton, Roberto Amato, Jeffrey Barrett, Sonia Goncalves, Ewan Harrison, David K. Jackson, Ian Johnston, Dominic Kwiatkowski, Cordelia Langford, John Sillitoe on behalf of the Wellcome Sanger Institute COVID-19 Surveillance Team |
| EPI_ISL_2829643, EPI_ISL_2829644, EPI_ISL_2829645 | Health Services Laboratories | Wellcome Sanger Institute for the COVID-19 Genomics UK (COG-UK) Consortium | Health Services Laboratories and Alex Alderton, Roberto Amato, Jeffrey Barrett, Sonia Goncalves, Ewan Harrison, David K. Jackson, Ian Johnston, Dominic Kwiatkowski, Cordelia Langford, John Sillitoe on behalf of the Wellcome Sanger Institute COVID-19 Surveillance Team |
| EPI_ISL_2829646 | Lighthouse Lab in Alderley Park | Wellcome Sanger Institute for the COVID-19 Genomics UK (COG-UK) Consortium | Jacquelyn Wynn, Mairead Hyland, The Lighthouse Lab in Alderley Park and Alex Alderton, Roberto Amato, Jeffrey Barrett, Sonia Goncalves, Ewan Harrison, David K. Jackson, Ian Johnston, Dominic Kwiatkowski, Cordelia Langford, John Sillitoe on behalf of the Wellcome Sanger Institute COVID-19 Surveillance Team |
| EPI_ISL_2829647, EPI_ISL_2829648 | Lighthouse Lab in Milton Keynes | Wellcome Sanger Institute for the COVID-19 Genomics UK (COG-UK) Consortium | The Lighthouse Lab in Milton Keynes and Alex Alderton, Roberto Amato, Jeffrey Barrett, Sonia Goncalves, Ewan Harrison, David K. Jackson, Ian Johnston, Dominic Kwiatkowski, Cordelia Langford, John Sillitoe on behalf of the Wellcome Sanger Institute COVID-19 Surveillance Team |
| EPI_ISL_2829649, EPI_ISL_2829650 | Health Services Laboratories | Wellcome Sanger Institute for the COVID-19 Genomics UK (COG-UK) Consortium | Health Services Laboratories and Alex Alderton, Roberto Amato, Jeffrey Barrett, Sonia Goncalves, Ewan Harrison, David K. Jackson, Ian Johnston, Dominic Kwiatkowski, Cordelia Langford, John Sillitoe on behalf of the Wellcome Sanger Institute COVID-19 Surveillance Team |
| EPI_ISL_2829651 | Lighthouse Lab in Milton Keynes | Wellcome Sanger Institute for the COVID-19 Genomics UK (COG-UK) Consortium | The Lighthouse Lab in Milton Keynes and Alex Alderton, Roberto Amato, Jeffrey Barrett, Sonia Goncalves, Ewan Harrison, David K. Jackson, Ian Johnston, Dominic Kwiatkowski, Cordelia Langford, John Sillitoe on behalf of the Wellcome Sanger Institute COVID-19 Surveillance Team |
| EPI_ISL_2829652 | Health Services Laboratories | Wellcome Sanger Institute for the COVID-19 Genomics UK (COG-UK) Consortium | Health Services Laboratories and Alex Alderton, Roberto Amato, Jeffrey Barrett, Sonia Goncalves, Ewan Harrison, David K. Jackson, Ian Johnston, Dominic Kwiatkowski, Cordelia Langford, John Sillitoe on behalf of the Wellcome Sanger Institute COVID-19 Surveillance Team |
| EPI_ISL_2829653, EPI_ISL_2829654 | Lighthouse Lab in Milton Keynes | Wellcome Sanger Institute for the COVID-19 Genomics UK (COG-UK) Consortium | The Lighthouse Lab in Milton Keynes and Alex Alderton, Roberto Amato, Jeffrey Barrett, Sonia Goncalves, Ewan Harrison, David K. Jackson, Ian Johnston, Dominic Kwiatkowski, Cordelia Langford, John Sillitoe on behalf of the Wellcome Sanger Institute COVID-19 Surveillance Team |
| EPI_ISL_2829655 | Lighthouse Lab in Alderley Park | Wellcome Sanger Institute for the COVID-19 Genomics UK (COG-UK) Consortium | Jacquelyn Wynn, Mairead Hyland, The Lighthouse Lab in Alderley Park and Alex Alderton, Roberto Amato, Jeffrey Barrett, Sonia Goncalves, Ewan Harrison, David K. Jackson, Ian Johnston, Dominic Kwiatkowski, Cordelia Langford, John Sillitoe on behalf of the Wellcome Sanger Institute COVID-19 Surveillance Team |
| EPI_ISL_2829656 | Lighthouse Lab in Milton Keynes | Wellcome Sanger Institute for the COVID-19 Genomics UK (COG-UK) Consortium | The Lighthouse Lab in Milton Keynes and Alex Alderton, Roberto Amato, Jeffrey Barrett, Sonia Goncalves, Ewan Harrison, David K. Jackson, Ian Johnston, Dominic Kwiatkowski, Cordelia Langford, John Sillitoe on behalf of the Wellcome Sanger Institute COVID-19 Surveillance Team |
| EPI_ISL_2829657 | Lighthouse Lab in Alderley Park | Wellcome Sanger Institute for the COVID-19 Genomics UK (COG-UK) Consortium | Jacquelyn Wynn, Mairead Hyland, The Lighthouse Lab in Alderley Park and Alex Alderton, Roberto Amato, Jeffrey Barrett, Sonia Goncalves, Ewan Harrison, David K. Jackson, Ian Johnston, Dominic Kwiatkowski, Cordelia Langford, John Sillitoe on behalf of the Wellcome Sanger Institute COVID-19 Surveillance Team |
| EPI_ISL_2829658, EPI_ISL_2829659, EPI_ISL_2829660, EPI_ISL_2829661 | Lighthouse Lab in Milton Keynes | Wellcome Sanger Institute for the COVID-19 Genomics UK (COG-UK) Consortium | The Lighthouse Lab in Milton Keynes and Alex Alderton, Roberto Amato, Jeffrey Barrett, Sonia Goncalves, Ewan Harrison, David K. Jackson, Ian Johnston, Dominic Kwiatkowski, Cordelia Langford, John Sillitoe on behalf of the Wellcome Sanger Institute COVID-19 Surveillance Team |
| EPI_ISL_2829662, EPI_ISL_2829663, EPI_ISL_2829664, EPI_ISL_2829665 | Lighthouse Lab in Alderley Park | Wellcome Sanger Institute for the COVID-19 Genomics UK (COG-UK) Consortium | Jacquelyn Wynn, Mairead Hyland, The Lighthouse Lab in Alderley Park and Alex Alderton, Roberto Amato, Jeffrey Barrett, Sonia Goncalves, Ewan Harrison, David K. Jackson, Ian Johnston, Dominic Kwiatkowski, Cordelia Langford, John Sillitoe on behalf of the Wellcome Sanger Institute COVID-19 Surveillance Team |

| | | | |
|---|---|---|---|
| EPI_ISL_2829666 | Lighthouse Lab in Milton Keynes | Wellcome Sanger Institute for the COVID-19 Genomics UK (COG-UK) Consortium | The Lighthouse Lab in Milton Keynes and Alex Alderton, Roberto Amato, Jeffrey Barrett, Sonia Goncalves, Ewan Harrison, David K. Jackson, Ian Johnston, Dominic Kwiatkowski, Cordelia Langford, John Sillitoe on behalf of the Wellcome Sanger Institute COVID-19 Surveillance Team |
| EPI_ISL_2829667 | Health Services Laboratories | Wellcome Sanger Institute for the COVID-19 Genomics UK (COG-UK) Consortium | Health Services Laboratories and Alex Alderton, Roberto Amato, Jeffrey Barrett, Sonia Goncalves, Ewan Harrison, David K. Jackson, Ian Johnston, Dominic Kwiatkowski, Cordelia Langford, John Sillitoe on behalf of the Wellcome Sanger Institute COVID-19 Surveillance Team |
| EPI_ISL_2829668 | Lighthouse Lab in Milton Keynes | Wellcome Sanger Institute for the COVID-19 Genomics UK (COG-UK) Consortium | The Lighthouse Lab in Milton Keynes and Alex Alderton, Roberto Amato, Jeffrey Barrett, Sonia Goncalves, Ewan Harrison, David K. Jackson, Ian Johnston, Dominic Kwiatkowski, Cordelia Langford, John Sillitoe on behalf of the Wellcome Sanger Institute COVID-19 Surveillance Team |
| EPI_ISL_2829669 | Health Services Laboratories | Wellcome Sanger Institute for the COVID-19 Genomics UK (COG-UK) Consortium | Health Services Laboratories and Alex Alderton, Roberto Amato, Jeffrey Barrett, Sonia Goncalves, Ewan Harrison, David K. Jackson, Ian Johnston, Dominic Kwiatkowski, Cordelia Langford, John Sillitoe on behalf of the Wellcome Sanger Institute COVID-19 Surveillance Team |
| EPI_ISL_2829670, EPI_ISL_2829671 | Lighthouse Lab in Milton Keynes | Wellcome Sanger Institute for the COVID-19 Genomics UK (COG-UK) Consortium | The Lighthouse Lab in Milton Keynes and Alex Alderton, Roberto Amato, Jeffrey Barrett, Sonia Goncalves, Ewan Harrison, David K. Jackson, Ian Johnston, Dominic Kwiatkowski, Cordelia Langford, John Sillitoe on behalf of the Wellcome Sanger Institute COVID-19 Surveillance Team |
| EPI_ISL_2829672 | Lighthouse Lab in Alderley Park | Wellcome Sanger Institute for the COVID-19 Genomics UK (COG-UK) Consortium | Jacquelyn Wynn, Mairead Hyland, The Lighthouse Lab in Alderley Park and Alex Alderton, Roberto Amato, Jeffrey Barrett, Sonia Goncalves, Ewan Harrison, David K. Jackson, Ian Johnston, Dominic Kwiatkowski, Cordelia Langford, John Sillitoe on behalf of the Wellcome Sanger Institute COVID-19 Surveillance Team |
| EPI_ISL_2829673 | Health Services Laboratories | Wellcome Sanger Institute for the COVID-19 Genomics UK (COG-UK) Consortium | Health Services Laboratories and Alex Alderton, Roberto Amato, Jeffrey Barrett, Sonia Goncalves, Ewan Harrison, David K. Jackson, Ian Johnston, Dominic Kwiatkowski, Cordelia Langford, John Sillitoe on behalf of the Wellcome Sanger Institute COVID-19 Surveillance Team |
| EPI_ISL_2829674, EPI_ISL_2829675 | Lighthouse Lab in Milton Keynes | Wellcome Sanger Institute for the COVID-19 Genomics UK (COG-UK) Consortium | The Lighthouse Lab in Milton Keynes and Alex Alderton, Roberto Amato, Jeffrey Barrett, Sonia Goncalves, Ewan Harrison, David K. Jackson, Ian Johnston, Dominic Kwiatkowski, Cordelia Langford, John Sillitoe on behalf of the Wellcome Sanger Institute COVID-19 Surveillance Team |
| EPI_ISL_2829676, EPI_ISL_2829677 | Lighthouse Lab in Alderley Park | Wellcome Sanger Institute for the COVID-19 Genomics UK (COG-UK) Consortium | Jacquelyn Wynn, Mairead Hyland, The Lighthouse Lab in Alderley Park and Alex Alderton, Roberto Amato, Jeffrey Barrett, Sonia Goncalves, Ewan Harrison, David K. Jackson, Ian Johnston, Dominic Kwiatkowski, Cordelia Langford, John Sillitoe on behalf of the Wellcome Sanger Institute COVID-19 Surveillance Team |
| EPI_ISL_2829678, EPI_ISL_2829679 | Lighthouse Lab in Milton Keynes | Wellcome Sanger Institute for the COVID-19 Genomics UK (COG-UK) Consortium | The Lighthouse Lab in Milton Keynes and Alex Alderton, Roberto Amato, Jeffrey Barrett, Sonia Goncalves, Ewan Harrison, David K. Jackson, Ian Johnston, Dominic Kwiatkowski, Cordelia Langford, John Sillitoe on behalf of the Wellcome Sanger Institute COVID-19 Surveillance Team |
| EPI_ISL_2829680 | Lighthouse Lab in Alderley Park | Wellcome Sanger Institute for the COVID-19 Genomics UK (COG-UK) Consortium | Jacquelyn Wynn, Mairead Hyland, The Lighthouse Lab in Alderley Park and Alex Alderton, Roberto Amato, Jeffrey Barrett, Sonia Goncalves, Ewan Harrison, David K. Jackson, Ian Johnston, Dominic Kwiatkowski, Cordelia Langford, John Sillitoe on behalf of the Wellcome Sanger Institute COVID-19 Surveillance Team |
| EPI_ISL_2829681, EPI_ISL_2829682 | Lighthouse Lab in Milton Keynes | Wellcome Sanger Institute for the COVID-19 Genomics UK (COG-UK) Consortium | The Lighthouse Lab in Milton Keynes and Alex Alderton, Roberto Amato, Jeffrey Barrett, Sonia Goncalves, Ewan Harrison, David K. Jackson, Ian Johnston, Dominic Kwiatkowski, Cordelia Langford, John Sillitoe on behalf of the Wellcome Sanger Institute COVID-19 Surveillance Team |
| EPI_ISL_2829683 | Health Services Laboratories | Wellcome Sanger Institute for the COVID-19 Genomics UK (COG-UK) Consortium | Health Services Laboratories and Alex Alderton, Roberto Amato, Jeffrey Barrett, Sonia Goncalves, Ewan Harrison, David K. Jackson, Ian Johnston, Dominic Kwiatkowski, Cordelia Langford, John Sillitoe on behalf of the Wellcome Sanger Institute COVID-19 Surveillance Team |
| EPI_ISL_2829684, EPI_ISL_2829685 | Lighthouse Lab in Milton Keynes | Wellcome Sanger Institute for the COVID-19 Genomics UK (COG-UK) Consortium | The Lighthouse Lab in Milton Keynes and Alex Alderton, Roberto Amato, Jeffrey Barrett, Sonia Goncalves, Ewan Harrison, David K. Jackson, Ian Johnston, Dominic Kwiatkowski, Cordelia Langford, John Sillitoe on behalf of the Wellcome Sanger Institute COVID-19 Surveillance Team |
| EPI_ISL_2829686 | Health Services Laboratories | Wellcome Sanger Institute for the COVID-19 Genomics UK (COG-UK) Consortium | Health Services Laboratories and Alex Alderton, Roberto Amato, Jeffrey Barrett, Sonia Goncalves, Ewan Harrison, David K. Jackson, Ian Johnston, Dominic Kwiatkowski, Cordelia Langford, John Sillitoe on behalf of the Wellcome Sanger Institute COVID-19 Surveillance Team |
| EPI_ISL_2829687 | Lighthouse Lab in Milton Keynes | Wellcome Sanger Institute for the COVID-19 Genomics UK (COG-UK) Consortium | The Lighthouse Lab in Milton Keynes and Alex Alderton, Roberto Amato, Jeffrey Barrett, Sonia Goncalves, Ewan Harrison, David K. Jackson, Ian Johnston, Dominic Kwiatkowski, Cordelia Langford, John Sillitoe on behalf of the Wellcome Sanger Institute COVID-19 Surveillance Team |
| EPI_ISL_2829688 | Lighthouse Lab in Alderley Park | Wellcome Sanger Institute for the COVID-19 Genomics UK (COG-UK) Consortium | Jacquelyn Wynn, Mairead Hyland, The Lighthouse Lab in Alderley Park and Alex Alderton, Roberto Amato, Jeffrey Barrett, Sonia Goncalves, Ewan Harrison, David K. Jackson, Ian Johnston, Dominic Kwiatkowski, Cordelia Langford, John Sillitoe on behalf of the Wellcome Sanger Institute COVID-19 Surveillance Team |
| EPI_ISL_2829689, EPI_ISL_2829690 | Lighthouse Lab in Milton Keynes | Wellcome Sanger Institute for the COVID-19 Genomics UK (COG-UK) Consortium | The Lighthouse Lab in Milton Keynes and Alex Alderton, Roberto Amato, Jeffrey Barrett, Sonia Goncalves, Ewan Harrison, David K. Jackson, Ian Johnston, Dominic Kwiatkowski, Cordelia Langford, John Sillitoe on behalf of the Wellcome Sanger Institute COVID-19 Surveillance Team |
| EPI_ISL_2829691 | Lighthouse Lab in Alderley Park | Wellcome Sanger Institute for the COVID-19 Genomics UK (COG-UK) Consortium | Jacquelyn Wynn, Mairead Hyland, The Lighthouse Lab in Alderley Park and Alex Alderton, Roberto Amato, Jeffrey Barrett, Sonia Goncalves, Ewan Harrison, David K. Jackson, Ian Johnston, Dominic Kwiatkowski, Cordelia Langford, John Sillitoe on behalf of the Wellcome Sanger Institute COVID-19 Surveillance Team |
| EPI_ISL_2829692 | Lighthouse Lab in Milton Keynes | Wellcome Sanger Institute for the COVID-19 Genomics UK (COG-UK) Consortium | The Lighthouse Lab in Milton Keynes and Alex Alderton, Roberto Amato, Jeffrey Barrett, Sonia Goncalves, Ewan Harrison, David K. Jackson, Ian Johnston, Dominic Kwiatkowski, Cordelia Langford, John Sillitoe on behalf of the Wellcome Sanger Institute COVID-19 Surveillance Team |
| EPI_ISL_2829693, EPI_ISL_2829694, EPI_ISL_2829695 | Health Services Laboratories | Wellcome Sanger Institute for the COVID-19 Genomics UK (COG-UK) Consortium | Health Services Laboratories and Alex Alderton, Roberto Amato, Jeffrey Barrett, Sonia Goncalves, Ewan Harrison, David K. Jackson, Ian Johnston, Dominic Kwiatkowski, Cordelia Langford, John Sillitoe on behalf of the Wellcome Sanger Institute COVID-19 Surveillance Team |
| EPI_ISL_2829696, EPI_ISL_2829697, EPI_ISL_2829698, EPI_ISL_2829699 | Lighthouse Lab in Milton Keynes | Wellcome Sanger Institute for the COVID-19 Genomics UK (COG-UK) Consortium | The Lighthouse Lab in Milton Keynes and Alex Alderton, Roberto Amato, Jeffrey Barrett, Sonia Goncalves, Ewan Harrison, David K. Jackson, Ian Johnston, Dominic Kwiatkowski, Cordelia Langford, John Sillitoe on behalf of the Wellcome Sanger Institute COVID-19 Surveillance Team |
| EPI_ISL_2829700 | Health Services Laboratories | Wellcome Sanger Institute for the COVID-19 Genomics UK (COG-UK) Consortium | Health Services Laboratories and Alex Alderton, Roberto Amato, Jeffrey Barrett, Sonia Goncalves, Ewan Harrison, David K. Jackson, Ian Johnston, Dominic Kwiatkowski, Cordelia Langford, John Sillitoe on behalf of the Wellcome Sanger Institute COVID-19 Surveillance Team |
| EPI_ISL_2829701 | Lighthouse Lab in Alderley Park | Wellcome Sanger Institute for the COVID-19 Genomics UK (COG-UK) Consortium | Jacquelyn Wynn, Mairead Hyland, The Lighthouse Lab in Alderley Park and Alex Alderton, Roberto Amato, Jeffrey Barrett, Sonia Goncalves, Ewan Harrison, David K. Jackson, Ian Johnston, Dominic Kwiatkowski, Cordelia Langford, John Sillitoe on behalf of the Wellcome Sanger Institute COVID-19 Surveillance Team |
| EPI_ISL_2829702 | Lighthouse Lab in Milton Keynes | Wellcome Sanger Institute for the COVID-19 Genomics UK (COG-UK) Consortium | The Lighthouse Lab in Milton Keynes and Alex Alderton, Roberto Amato, Jeffrey Barrett, Sonia Goncalves, Ewan Harrison, David K. Jackson, Ian Johnston, Dominic Kwiatkowski, Cordelia Langford, John Sillitoe on behalf of the Wellcome Sanger Institute COVID-19 Surveillance Team |
| EPI_ISL_2829703 | Lighthouse Lab in Alderley Park | Wellcome Sanger Institute for the COVID-19 Genomics UK (COG-UK) Consortium | Jacquelyn Wynn, Mairead Hyland, The Lighthouse Lab in Alderley Park and Alex Alderton, Roberto Amato, Jeffrey Barrett, Sonia Goncalves, Ewan Harrison, David K. Jackson, Ian Johnston, Dominic Kwiatkowski, Cordelia Langford, John Sillitoe on behalf of the Wellcome Sanger Institute COVID-19 Surveillance Team |
| EPI_ISL_2829704 | Lighthouse Lab in Milton Keynes | Wellcome Sanger Institute for the COVID-19 Genomics UK (COG-UK) Consortium | The Lighthouse Lab in Milton Keynes and Alex Alderton, Roberto Amato, Jeffrey Barrett, Sonia Goncalves, Ewan Harrison, David K. Jackson, Ian Johnston, Dominic Kwiatkowski, Cordelia Langford, John Sillitoe on behalf of the Wellcome Sanger Institute COVID-19 Surveillance Team |
| EPI_ISL_2829705 | Health Services Laboratories | Wellcome Sanger Institute for the COVID-19 Genomics UK (COG-UK) Consortium | Health Services Laboratories and Alex Alderton, Roberto Amato, Jeffrey Barrett, Sonia Goncalves, Ewan Harrison, David K. Jackson, Ian Johnston, Dominic Kwiatkowski, Cordelia Langford, John Sillitoe on behalf of the Wellcome Sanger Institute COVID-19 Surveillance Team |
| EPI_ISL_2829706 | Lighthouse Lab in Milton Keynes | Wellcome Sanger Institute for the COVID-19 Genomics UK (COG-UK) Consortium | The Lighthouse Lab in Milton Keynes and Alex Alderton, Roberto Amato, Jeffrey Barrett, Sonia Goncalves, Ewan Harrison, David K. Jackson, Ian Johnston, Dominic Kwiatkowski, Cordelia Langford, John Sillitoe on behalf of the Wellcome Sanger Institute COVID-19 Surveillance Team |
| EPI_ISL_2829707 | Lighthouse Lab in Alderley Park | Wellcome Sanger Institute for the COVID-19 Genomics UK (COG-UK) Consortium | Jacquelyn Wynn, Mairead Hyland, The Lighthouse Lab in Alderley Park and Alex Alderton, Roberto Amato, Jeffrey Barrett, Sonia Goncalves, Ewan Harrison, David K. Jackson, Ian Johnston, Dominic Kwiatkowski, Cordelia Langford, John Sillitoe on behalf of the Wellcome Sanger Institute COVID-19 Surveillance Team |
| EPI_ISL_2829708, EPI_ISL_2829709, EPI_ISL_2829710, EPI_ISL_2829711, EPI_ISL_2829712 | Lighthouse Lab in Milton Keynes | Wellcome Sanger Institute for the COVID-19 Genomics UK (COG-UK) Consortium | The Lighthouse Lab in Milton Keynes and Alex Alderton, Roberto Amato, Jeffrey Barrett, Sonia Goncalves, Ewan Harrison, David K. Jackson, Ian Johnston, Dominic Kwiatkowski, Cordelia Langford, John Sillitoe on behalf of the Wellcome Sanger Institute COVID-19 Surveillance Team |
| EPI_ISL_2829713, EPI_ISL_2829714 | Health Services Laboratories | Wellcome Sanger Institute for the COVID-19 Genomics UK (COG-UK) Consortium | Health Services Laboratories and Alex Alderton, Roberto Amato, Jeffrey Barrett, Sonia Goncalves, Ewan Harrison, David K. Jackson, Ian Johnston, Dominic Kwiatkowski, Cordelia Langford, John Sillitoe on behalf of the Wellcome Sanger Institute COVID-19 Surveillance Team |
| EPI_ISL_2829715 | Lighthouse Lab in Milton Keynes | Wellcome Sanger Institute for the COVID-19 Genomics UK (COG-UK) Consortium | The Lighthouse Lab in Milton Keynes and Alex Alderton, Roberto Amato, Jeffrey Barrett, Sonia Goncalves, Ewan Harrison, David K. Jackson, Ian Johnston, Dominic Kwiatkowski, Cordelia Langford, John Sillitoe on behalf of the Wellcome Sanger Institute COVID-19 Surveillance Team |
| EPI_ISL_2829716 | Health Services Laboratories | Wellcome Sanger Institute for the COVID-19 Genomics UK (COG-UK) Consortium | Health Services Laboratories and Alex Alderton, Roberto Amato, Jeffrey Barrett, Sonia Goncalves, Ewan Harrison, David K. Jackson, Ian Johnston, Dominic Kwiatkowski, Cordelia Langford, John Sillitoe on behalf of the Wellcome Sanger Institute COVID-19 Surveillance Team |
| EPI_ISL_2829717, EPI_ISL_2829718 | Lighthouse Lab in Alderley Park | Wellcome Sanger Institute for the COVID-19 Genomics UK (COG-UK) Consortium | Jacquelyn Wynn, Mairead Hyland, The Lighthouse Lab in Alderley Park and Alex Alderton, Roberto Amato, Jeffrey Barrett, Sonia Goncalves, Ewan Harrison, David K. Jackson, Ian Johnston, Dominic Kwiatkowski, Cordelia Langford, John Sillitoe on behalf of the Wellcome Sanger Institute COVID-19 Surveillance Team |
| EPI_ISL_2829719 | Health Services Laboratories | Wellcome Sanger Institute for the COVID-19 Genomics UK (COG-UK) Consortium | Health Services Laboratories and Alex Alderton, Roberto Amato, Jeffrey Barrett, Sonia Goncalves, Ewan Harrison, David K. Jackson, Ian Johnston, Dominic Kwiatkowski, Cordelia Langford, John Sillitoe on behalf of the Wellcome Sanger Institute COVID-19 Surveillance Team |
| EPI_ISL_2829720 | Lighthouse Lab in Alderley Park | Wellcome Sanger Institute for the COVID-19 Genomics UK (COG-UK) Consortium | Jacquelyn Wynn, Mairead Hyland, The Lighthouse Lab in Alderley Park and Alex Alderton, Roberto Amato, Jeffrey Barrett, Sonia Goncalves, Ewan Harrison, David K. Jackson, Ian Johnston, Dominic Kwiatkowski, Cordelia Langford, John Sillitoe on behalf of the Wellcome Sanger Institute COVID-19 Surveillance Team |
| EPI_ISL_2829721 | Lighthouse Lab in Milton Keynes | Wellcome Sanger Institute for the COVID-19 Genomics UK (COG-UK) Consortium | The Lighthouse Lab in Milton Keynes and Alex Alderton, Roberto Amato, Jeffrey Barrett, Sonia Goncalves, Ewan Harrison, David K. Jackson, Ian Johnston, Dominic Kwiatkowski, Cordelia Langford, John Sillitoe on behalf of the Wellcome Sanger Institute COVID-19 Surveillance Team |
| EPI_ISL_2829722 | Health Services Laboratories | Wellcome Sanger Institute for the COVID-19 Genomics UK (COG-UK) Consortium | Health Services Laboratories and Alex Alderton, Roberto Amato, Jeffrey Barrett, Sonia Goncalves, Ewan Harrison, David K. Jackson, Ian Johnston, Dominic Kwiatkowski, Cordelia Langford, John Sillitoe on behalf of the Wellcome Sanger Institute COVID-19 Surveillance Team |
| EPI_ISL_2829723, EPI_ISL_2829724 | Lighthouse Lab in Milton Keynes | Wellcome Sanger Institute for the COVID-19 Genomics UK (COG-UK) Consortium | The Lighthouse Lab in Milton Keynes and Alex Alderton, Roberto Amato, Jeffrey Barrett, Sonia Goncalves, Ewan Harrison, David K. Jackson, Ian Johnston, Dominic Kwiatkowski, Cordelia Langford, John Sillitoe on behalf of the Wellcome Sanger Institute COVID-19 Surveillance Team |
| EPI_ISL_2829725 | Health Services Laboratories | Wellcome Sanger Institute for the COVID-19 Genomics UK (COG-UK) Consortium | Health Services Laboratories and Alex Alderton, Roberto Amato, Jeffrey Barrett, Sonia Goncalves, Ewan Harrison, David K. Jackson, Ian Johnston, Dominic Kwiatkowski, Cordelia Langford, John Sillitoe on behalf of the Wellcome Sanger Institute COVID-19 Surveillance Team |
| EPI_ISL_2829726 | Lighthouse Lab in Alderley Park | Wellcome Sanger Institute for the COVID-19 Genomics UK (COG-UK) Consortium | Jacquelyn Wynn, Mairead Hyland, The Lighthouse Lab in Alderley Park and Alex Alderton, Roberto Amato, Jeffrey Barrett, Sonia Goncalves, Ewan Harrison, David K. Jackson, Ian Johnston, Dominic Kwiatkowski, Cordelia Langford, John Sillitoe on behalf of the Wellcome Sanger Institute COVID-19 Surveillance Team |
| EPI_ISL_2829727, | Lighthouse Lab in Milton Keynes | Wellcome Sanger Institute for the COVID-19 Genomics UK (COG-UK) | The Lighthouse Lab in Milton Keynes and Alex Alderton, Roberto Amato, Jeffrey Barrett, Sonia Goncalves, Ewan Harrison, David K. Jackson, Ian Johnston, Dominic Kwiatkowski, Cordelia Langford, John Sillitoe on behalf of the Wellcome Sanger |

| | | | |
|---|---|---|---|
| EPI_ISL_2829728, EPI_ISL_2829729 | | Consortium | Institute COVID-19 Surveillance Team |
| EPI_ISL_2829730 | Health Services Laboratories | Wellcome Sanger Institute for the COVID-19 Genomics UK (COG-UK) Consortium | Health Services Laboratories and Alex Alderton, Roberto Amato, Jeffrey Barrett, Sonia Goncalves, Ewan Harrison, David K. Jackson, Ian Johnston, Dominic Kwiatkowski, Cordelia Langford, John Sillitoe on behalf of the Wellcome Sanger Institute COVID-19 Surveillance Team |
| EPI_ISL_2829731 | Lighthouse Lab in Alderley Park | Wellcome Sanger Institute for the COVID-19 Genomics UK (COG-UK) Consortium | Jacquelyn Wynn, Mairead Hyland, The Lighthouse Lab in Alderley Park and Alex Alderton, Roberto Amato, Jeffrey Barrett, Sonia Goncalves, Ewan Harrison, David K. Jackson, Ian Johnston, Dominic Kwiatkowski, Cordelia Langford, John Sillitoe on behalf of the Wellcome Sanger Institute COVID-19 Surveillance Team |
| EPI_ISL_2829732 | Lighthouse Lab in Milton Keynes | Wellcome Sanger Institute for the COVID-19 Genomics UK (COG-UK) Consortium | The Lighthouse Lab in Milton Keynes and Alex Alderton, Roberto Amato, Jeffrey Barrett, Sonia Goncalves, Ewan Harrison, David K. Jackson, Ian Johnston, Dominic Kwiatkowski, Cordelia Langford, John Sillitoe on behalf of the Wellcome Sanger Institute COVID-19 Surveillance Team |
| EPI_ISL_2829733 | Lighthouse Lab in Alderley Park | Wellcome Sanger Institute for the COVID-19 Genomics UK (COG-UK) Consortium | Jacquelyn Wynn, Mairead Hyland, The Lighthouse Lab in Alderley Park and Alex Alderton, Roberto Amato, Jeffrey Barrett, Sonia Goncalves, Ewan Harrison, David K. Jackson, Ian Johnston, Dominic Kwiatkowski, Cordelia Langford, John Sillitoe on behalf of the Wellcome Sanger Institute COVID-19 Surveillance Team |
| EPI_ISL_2829734 | Lighthouse Lab in Milton Keynes | Wellcome Sanger Institute for the COVID-19 Genomics UK (COG-UK) Consortium | The Lighthouse Lab in Milton Keynes and Alex Alderton, Roberto Amato, Jeffrey Barrett, Sonia Goncalves, Ewan Harrison, David K. Jackson, Ian Johnston, Dominic Kwiatkowski, Cordelia Langford, John Sillitoe on behalf of the Wellcome Sanger Institute COVID-19 Surveillance Team |
| EPI_ISL_2829735 | Health Services Laboratories | Wellcome Sanger Institute for the COVID-19 Genomics UK (COG-UK) Consortium | Health Services Laboratories and Alex Alderton, Roberto Amato, Jeffrey Barrett, Sonia Goncalves, Ewan Harrison, David K. Jackson, Ian Johnston, Dominic Kwiatkowski, Cordelia Langford, John Sillitoe on behalf of the Wellcome Sanger Institute COVID-19 Surveillance Team |
| EPI_ISL_2829736, EPI_ISL_2829737 | Lighthouse Lab in Milton Keynes | Wellcome Sanger Institute for the COVID-19 Genomics UK (COG-UK) Consortium | The Lighthouse Lab in Milton Keynes and Alex Alderton, Roberto Amato, Jeffrey Barrett, Sonia Goncalves, Ewan Harrison, David K. Jackson, Ian Johnston, Dominic Kwiatkowski, Cordelia Langford, John Sillitoe on behalf of the Wellcome Sanger Institute COVID-19 Surveillance Team |
| EPI_ISL_2829738 | Lighthouse Lab in Alderley Park | Wellcome Sanger Institute for the COVID-19 Genomics UK (COG-UK) Consortium | Jacquelyn Wynn, Mairead Hyland, The Lighthouse Lab in Alderley Park and Alex Alderton, Roberto Amato, Jeffrey Barrett, Sonia Goncalves, Ewan Harrison, David K. Jackson, Ian Johnston, Dominic Kwiatkowski, Cordelia Langford, John Sillitoe on behalf of the Wellcome Sanger Institute COVID-19 Surveillance Team |
| EPI_ISL_2829739 | Health Services Laboratories | Wellcome Sanger Institute for the COVID-19 Genomics UK (COG-UK) Consortium | Health Services Laboratories and Alex Alderton, Roberto Amato, Jeffrey Barrett, Sonia Goncalves, Ewan Harrison, David K. Jackson, Ian Johnston, Dominic Kwiatkowski, Cordelia Langford, John Sillitoe on behalf of the Wellcome Sanger Institute COVID-19 Surveillance Team |
| EPI_ISL_2829740 | Lighthouse Lab in Alderley Park | Wellcome Sanger Institute for the COVID-19 Genomics UK (COG-UK) Consortium | Jacquelyn Wynn, Mairead Hyland, The Lighthouse Lab in Alderley Park and Alex Alderton, Roberto Amato, Jeffrey Barrett, Sonia Goncalves, Ewan Harrison, David K. Jackson, Ian Johnston, Dominic Kwiatkowski, Cordelia Langford, John Sillitoe on behalf of the Wellcome Sanger Institute COVID-19 Surveillance Team |
| EPI_ISL_2829741 | Lighthouse Lab in Milton Keynes | Wellcome Sanger Institute for the COVID-19 Genomics UK (COG-UK) Consortium | The Lighthouse Lab in Milton Keynes and Alex Alderton, Roberto Amato, Jeffrey Barrett, Sonia Goncalves, Ewan Harrison, David K. Jackson, Ian Johnston, Dominic Kwiatkowski, Cordelia Langford, John Sillitoe on behalf of the Wellcome Sanger Institute COVID-19 Surveillance Team |
| EPI_ISL_2829742 | Lighthouse Lab in Alderley Park | Wellcome Sanger Institute for the COVID-19 Genomics UK (COG-UK) Consortium | Jacquelyn Wynn, Mairead Hyland, The Lighthouse Lab in Alderley Park and Alex Alderton, Roberto Amato, Jeffrey Barrett, Sonia Goncalves, Ewan Harrison, David K. Jackson, Ian Johnston, Dominic Kwiatkowski, Cordelia Langford, John Sillitoe on behalf of the Wellcome Sanger Institute COVID-19 Surveillance Team |
| EPI_ISL_2829743 | Lighthouse Lab in Milton Keynes | Wellcome Sanger Institute for the COVID-19 Genomics UK (COG-UK) Consortium | The Lighthouse Lab in Milton Keynes and Alex Alderton, Roberto Amato, Jeffrey Barrett, Sonia Goncalves, Ewan Harrison, David K. Jackson, Ian Johnston, Dominic Kwiatkowski, Cordelia Langford, John Sillitoe on behalf of the Wellcome Sanger Institute COVID-19 Surveillance Team |
| EPI_ISL_2829744 | Lighthouse Lab in Alderley Park | Wellcome Sanger Institute for the COVID-19 Genomics UK (COG-UK) Consortium | Jacquelyn Wynn, Mairead Hyland, The Lighthouse Lab in Alderley Park and Alex Alderton, Roberto Amato, Jeffrey Barrett, Sonia Goncalves, Ewan Harrison, David K. Jackson, Ian Johnston, Dominic Kwiatkowski, Cordelia Langford, John Sillitoe on behalf of the Wellcome Sanger Institute COVID-19 Surveillance Team |
| EPI_ISL_2829745 | Health Services Laboratories | Wellcome Sanger Institute for the COVID-19 Genomics UK (COG-UK) Consortium | Health Services Laboratories and Alex Alderton, Roberto Amato, Jeffrey Barrett, Sonia Goncalves, Ewan Harrison, David K. Jackson, Ian Johnston, Dominic Kwiatkowski, Cordelia Langford, John Sillitoe on behalf of the Wellcome Sanger Institute COVID-19 Surveillance Team |
| EPI_ISL_2829746 | Lighthouse Lab in Alderley Park | Wellcome Sanger Institute for the COVID-19 Genomics UK (COG-UK) Consortium | Jacquelyn Wynn, Mairead Hyland, The Lighthouse Lab in Alderley Park and Alex Alderton, Roberto Amato, Jeffrey Barrett, Sonia Goncalves, Ewan Harrison, David K. Jackson, Ian Johnston, Dominic Kwiatkowski, Cordelia Langford, John Sillitoe on behalf of the Wellcome Sanger Institute COVID-19 Surveillance Team |
| EPI_ISL_2829747 | Health Services Laboratories | Wellcome Sanger Institute for the COVID-19 Genomics UK (COG-UK) Consortium | Health Services Laboratories and Alex Alderton, Roberto Amato, Jeffrey Barrett, Sonia Goncalves, Ewan Harrison, David K. Jackson, Ian Johnston, Dominic Kwiatkowski, Cordelia Langford, John Sillitoe on behalf of the Wellcome Sanger Institute COVID-19 Surveillance Team |
| EPI_ISL_2829748, EPI_ISL_2829749 | Lighthouse Lab in Milton Keynes | Wellcome Sanger Institute for the COVID-19 Genomics UK (COG-UK) Consortium | The Lighthouse Lab in Milton Keynes and Alex Alderton, Roberto Amato, Jeffrey Barrett, Sonia Goncalves, Ewan Harrison, David K. Jackson, Ian Johnston, Dominic Kwiatkowski, Cordelia Langford, John Sillitoe on behalf of the Wellcome Sanger Institute COVID-19 Surveillance Team |
| EPI_ISL_2829750, EPI_ISL_2829751 | Lighthouse Lab in Alderley Park | Wellcome Sanger Institute for the COVID-19 Genomics UK (COG-UK) Consortium | Jacquelyn Wynn, Mairead Hyland, The Lighthouse Lab in Alderley Park and Alex Alderton, Roberto Amato, Jeffrey Barrett, Sonia Goncalves, Ewan Harrison, David K. Jackson, Ian Johnston, Dominic Kwiatkowski, Cordelia Langford, John Sillitoe on behalf of the Wellcome Sanger Institute COVID-19 Surveillance Team |
| EPI_ISL_2829752, EPI_ISL_2829753, EPI_ISL_2829754 | Health Services Laboratories | Wellcome Sanger Institute for the COVID-19 Genomics UK (COG-UK) Consortium | Health Services Laboratories and Alex Alderton, Roberto Amato, Jeffrey Barrett, Sonia Goncalves, Ewan Harrison, David K. Jackson, Ian Johnston, Dominic Kwiatkowski, Cordelia Langford, John Sillitoe on behalf of the Wellcome Sanger Institute COVID-19 Surveillance Team |
| EPI_ISL_2829755 | Lighthouse Lab in Milton Keynes | Wellcome Sanger Institute for the COVID-19 Genomics UK (COG-UK) Consortium | The Lighthouse Lab in Milton Keynes and Alex Alderton, Roberto Amato, Jeffrey Barrett, Sonia Goncalves, Ewan Harrison, David K. Jackson, Ian Johnston, Dominic Kwiatkowski, Cordelia Langford, John Sillitoe on behalf of the Wellcome Sanger Institute COVID-19 Surveillance Team |
| EPI_ISL_2829756, EPI_ISL_2829757 | Health Services Laboratories | Wellcome Sanger Institute for the COVID-19 Genomics UK (COG-UK) Consortium | Health Services Laboratories and Alex Alderton, Roberto Amato, Jeffrey Barrett, Sonia Goncalves, Ewan Harrison, David K. Jackson, Ian Johnston, Dominic Kwiatkowski, Cordelia Langford, John Sillitoe on behalf of the Wellcome Sanger Institute COVID-19 Surveillance Team |
| EPI_ISL_2829758 | Lighthouse Lab in Milton Keynes | Wellcome Sanger Institute for the COVID-19 Genomics UK (COG-UK) Consortium | The Lighthouse Lab in Milton Keynes and Alex Alderton, Roberto Amato, Jeffrey Barrett, Sonia Goncalves, Ewan Harrison, David K. Jackson, Ian Johnston, Dominic Kwiatkowski, Cordelia Langford, John Sillitoe on behalf of the Wellcome Sanger Institute COVID-19 Surveillance Team |
| EPI_ISL_2829759, EPI_ISL_2829760 | Health Services Laboratories | Wellcome Sanger Institute for the COVID-19 Genomics UK (COG-UK) Consortium | Health Services Laboratories and Alex Alderton, Roberto Amato, Jeffrey Barrett, Sonia Goncalves, Ewan Harrison, David K. Jackson, Ian Johnston, Dominic Kwiatkowski, Cordelia Langford, John Sillitoe on behalf of the Wellcome Sanger Institute COVID-19 Surveillance Team |
| EPI_ISL_2829761 | Lighthouse Lab in Milton Keynes | Wellcome Sanger Institute for the COVID-19 Genomics UK (COG-UK) Consortium | The Lighthouse Lab in Milton Keynes and Alex Alderton, Roberto Amato, Jeffrey Barrett, Sonia Goncalves, Ewan Harrison, David K. Jackson, Ian Johnston, Dominic Kwiatkowski, Cordelia Langford, John Sillitoe on behalf of the Wellcome Sanger Institute COVID-19 Surveillance Team |
| EPI_ISL_2829762 | Lighthouse Lab in Alderley Park | Wellcome Sanger Institute for the COVID-19 Genomics UK (COG-UK) Consortium | Jacquelyn Wynn, Mairead Hyland, The Lighthouse Lab in Alderley Park and Alex Alderton, Roberto Amato, Jeffrey Barrett, Sonia Goncalves, Ewan Harrison, David K. Jackson, Ian Johnston, Dominic Kwiatkowski, Cordelia Langford, John Sillitoe on behalf of the Wellcome Sanger Institute COVID-19 Surveillance Team |
| EPI_ISL_2829763 | Health Services Laboratories | Wellcome Sanger Institute for the COVID-19 Genomics UK (COG-UK) Consortium | Health Services Laboratories and Alex Alderton, Roberto Amato, Jeffrey Barrett, Sonia Goncalves, Ewan Harrison, David K. Jackson, Ian Johnston, Dominic Kwiatkowski, Cordelia Langford, John Sillitoe on behalf of the Wellcome Sanger Institute COVID-19 Surveillance Team |
| EPI_ISL_2829764 | Lighthouse Lab in Milton Keynes | Wellcome Sanger Institute for the COVID-19 Genomics UK (COG-UK) Consortium | The Lighthouse Lab in Milton Keynes and Alex Alderton, Roberto Amato, Jeffrey Barrett, Sonia Goncalves, Ewan Harrison, David K. Jackson, Ian Johnston, Dominic Kwiatkowski, Cordelia Langford, John Sillitoe on behalf of the Wellcome Sanger Institute COVID-19 Surveillance Team |
| EPI_ISL_2829765 | Lighthouse Lab in Alderley Park | Wellcome Sanger Institute for the COVID-19 Genomics UK (COG-UK) Consortium | Jacquelyn Wynn, Mairead Hyland, The Lighthouse Lab in Alderley Park and Alex Alderton, Roberto Amato, Jeffrey Barrett, Sonia Goncalves, Ewan Harrison, David K. Jackson, Ian Johnston, Dominic Kwiatkowski, Cordelia Langford, John Sillitoe on behalf of the Wellcome Sanger Institute COVID-19 Surveillance Team |
| EPI_ISL_2829766, EPI_ISL_2829767, EPI_ISL_2829768 | Lighthouse Lab in Milton Keynes | Wellcome Sanger Institute for the COVID-19 Genomics UK (COG-UK) Consortium | The Lighthouse Lab in Milton Keynes and Alex Alderton, Roberto Amato, Jeffrey Barrett, Sonia Goncalves, Ewan Harrison, David K. Jackson, Ian Johnston, Dominic Kwiatkowski, Cordelia Langford, John Sillitoe on behalf of the Wellcome Sanger Institute COVID-19 Surveillance Team |
| EPI_ISL_2829769 | Lighthouse Lab in Alderley Park | Wellcome Sanger Institute for the COVID-19 Genomics UK (COG-UK) Consortium | Jacquelyn Wynn, Mairead Hyland, The Lighthouse Lab in Alderley Park and Alex Alderton, Roberto Amato, Jeffrey Barrett, Sonia Goncalves, Ewan Harrison, David K. Jackson, Ian Johnston, Dominic Kwiatkowski, Cordelia Langford, John Sillitoe on behalf of the Wellcome Sanger Institute COVID-19 Surveillance Team |
| EPI_ISL_2829770, EPI_ISL_2829771, EPI_ISL_2829772 | Lighthouse Lab in Milton Keynes | Wellcome Sanger Institute for the COVID-19 Genomics UK (COG-UK) Consortium | The Lighthouse Lab in Milton Keynes and Alex Alderton, Roberto Amato, Jeffrey Barrett, Sonia Goncalves, Ewan Harrison, David K. Jackson, Ian Johnston, Dominic Kwiatkowski, Cordelia Langford, John Sillitoe on behalf of the Wellcome Sanger Institute COVID-19 Surveillance Team |
| EPI_ISL_2829773 | Lighthouse Lab in Alderley Park | Wellcome Sanger Institute for the COVID-19 Genomics UK (COG-UK) Consortium | Jacquelyn Wynn, Mairead Hyland, The Lighthouse Lab in Alderley Park and Alex Alderton, Roberto Amato, Jeffrey Barrett, Sonia Goncalves, Ewan Harrison, David K. Jackson, Ian Johnston, Dominic Kwiatkowski, Cordelia Langford, John Sillitoe on behalf of the Wellcome Sanger Institute COVID-19 Surveillance Team |
| EPI_ISL_2829774, EPI_ISL_2829775 | Lighthouse Lab in Milton Keynes | Wellcome Sanger Institute for the COVID-19 Genomics UK (COG-UK) Consortium | The Lighthouse Lab in Milton Keynes and Alex Alderton, Roberto Amato, Jeffrey Barrett, Sonia Goncalves, Ewan Harrison, David K. Jackson, Ian Johnston, Dominic Kwiatkowski, Cordelia Langford, John Sillitoe on behalf of the Wellcome Sanger Institute COVID-19 Surveillance Team |
| EPI_ISL_2829776 | Lighthouse Lab in Alderley Park | Wellcome Sanger Institute for the COVID-19 Genomics UK (COG-UK) Consortium | Jacquelyn Wynn, Mairead Hyland, The Lighthouse Lab in Alderley Park and Alex Alderton, Roberto Amato, Jeffrey Barrett, Sonia Goncalves, Ewan Harrison, David K. Jackson, Ian Johnston, Dominic Kwiatkowski, Cordelia Langford, John Sillitoe on behalf of the Wellcome Sanger Institute COVID-19 Surveillance Team |
| EPI_ISL_2829777 | Lighthouse Lab in Milton Keynes | Wellcome Sanger Institute for the COVID-19 Genomics UK (COG-UK) Consortium | The Lighthouse Lab in Milton Keynes and Alex Alderton, Roberto Amato, Jeffrey Barrett, Sonia Goncalves, Ewan Harrison, David K. Jackson, Ian Johnston, Dominic Kwiatkowski, Cordelia Langford, John Sillitoe on behalf of the Wellcome Sanger Institute COVID-19 Surveillance Team |
| EPI_ISL_2829778, EPI_ISL_2829779, EPI_ISL_2829780 | Health Services Laboratories | Wellcome Sanger Institute for the COVID-19 Genomics UK (COG-UK) Consortium | Health Services Laboratories and Alex Alderton, Roberto Amato, Jeffrey Barrett, Sonia Goncalves, Ewan Harrison, David K. Jackson, Ian Johnston, Dominic Kwiatkowski, Cordelia Langford, John Sillitoe on behalf of the Wellcome Sanger Institute COVID-19 Surveillance Team |
| EPI_ISL_2829781 | Lighthouse Lab in Milton Keynes | Wellcome Sanger Institute for the COVID-19 Genomics UK (COG-UK) Consortium | The Lighthouse Lab in Milton Keynes and Alex Alderton, Roberto Amato, Jeffrey Barrett, Sonia Goncalves, Ewan Harrison, David K. Jackson, Ian Johnston, Dominic Kwiatkowski, Cordelia Langford, John Sillitoe on behalf of the Wellcome Sanger Institute COVID-19 Surveillance Team |
| EPI_ISL_2829782 | Lighthouse Lab in Alderley Park | Wellcome Sanger Institute for the COVID-19 Genomics UK (COG-UK) Consortium | Jacquelyn Wynn, Mairead Hyland, The Lighthouse Lab in Alderley Park and Alex Alderton, Roberto Amato, Jeffrey Barrett, Sonia Goncalves, Ewan Harrison, David K. Jackson, Ian Johnston, Dominic Kwiatkowski, Cordelia Langford, John Sillitoe on behalf of the Wellcome Sanger Institute COVID-19 Surveillance Team |
| EPI_ISL_2829783 | Lighthouse Lab in Milton Keynes | Wellcome Sanger Institute for the COVID-19 Genomics UK (COG-UK) Consortium | The Lighthouse Lab in Milton Keynes and Alex Alderton, Roberto Amato, Jeffrey Barrett, Sonia Goncalves, Ewan Harrison, David K. Jackson, Ian Johnston, Dominic Kwiatkowski, Cordelia Langford, John Sillitoe on behalf of the Wellcome Sanger Institute COVID-19 Surveillance Team |
| EPI_ISL_2829784 | Lighthouse Lab in Alderley Park | Wellcome Sanger Institute for the COVID-19 Genomics UK (COG-UK) Consortium | Jacquelyn Wynn, Mairead Hyland, The Lighthouse Lab in Alderley Park and Alex Alderton, Roberto Amato, Jeffrey Barrett, Sonia Goncalves, Ewan Harrison, David K. Jackson, Ian Johnston, Dominic Kwiatkowski, Cordelia Langford, John Sillitoe on behalf of the Wellcome Sanger Institute COVID-19 Surveillance Team |
| EPI_ISL_2829785, EPI_ISL_2829786 | Lighthouse Lab in Milton Keynes | Wellcome Sanger Institute for the COVID-19 Genomics UK (COG-UK) Consortium | The Lighthouse Lab in Milton Keynes and Alex Alderton, Roberto Amato, Jeffrey Barrett, Sonia Goncalves, Ewan Harrison, David K. Jackson, Ian Johnston, Dominic Kwiatkowski, Cordelia Langford, John Sillitoe on behalf of the Wellcome Sanger Institute COVID-19 Surveillance Team |
| EPI_ISL_2829787 | Health Services Laboratories | Wellcome Sanger Institute for the COVID-19 Genomics UK (COG-UK) | Health Services Laboratories and Alex Alderton, Roberto Amato, Jeffrey Barrett, Sonia Goncalves, Ewan Harrison, David K. Jackson, Ian Johnston, Dominic Kwiatkowski, Cordelia Langford, John Sillitoe on behalf of the Wellcome Sanger Institute |

| | | | |
|---|---|---|---|
| | | Consortium | COVID-19 Surveillance Team |
| EPI_ISL_2829788 | Lighthouse Lab in Milton Keynes | Wellcome Sanger Institute for the COVID-19 Genomics UK (COG-UK) Consortium | The Lighthouse Lab in Milton Keynes and Alex Alderton, Roberto Amato, Jeffrey Barrett, Sonia Goncalves, Ewan Harrison, David K. Jackson, Ian Johnston, Dominic Kwiatkowski, Cordelia Langford, John Sillitoe on behalf of the Wellcome Sanger Institute COVID-19 Surveillance Team |
| EPI_ISL_2829789 | Health Services Laboratories | Wellcome Sanger Institute for the COVID-19 Genomics UK (COG-UK) Consortium | Health Services Laboratories and Alex Alderton, Roberto Amato, Jeffrey Barrett, Sonia Goncalves, Ewan Harrison, David K. Jackson, Ian Johnston, Dominic Kwiatkowski, Cordelia Langford, John Sillitoe on behalf of the Wellcome Sanger Institute COVID-19 Surveillance Team |
| EPI_ISL_2829790 | Lighthouse Lab in Milton Keynes | Wellcome Sanger Institute for the COVID-19 Genomics UK (COG-UK) Consortium | The Lighthouse Lab in Milton Keynes and Alex Alderton, Roberto Amato, Jeffrey Barrett, Sonia Goncalves, Ewan Harrison, David K. Jackson, Ian Johnston, Dominic Kwiatkowski, Cordelia Langford, John Sillitoe on behalf of the Wellcome Sanger Institute COVID-19 Surveillance Team |
| EPI_ISL_2829791 | Lighthouse Lab in Alderley Park | Wellcome Sanger Institute for the COVID-19 Genomics UK (COG-UK) Consortium | Jacquelyn Wynn, Mairead Hyland, The Lighthouse Lab in Alderley Park and Alex Alderton, Roberto Amato, Jeffrey Barrett, Sonia Goncalves, Ewan Harrison, David K. Jackson, Ian Johnston, Dominic Kwiatkowski, Cordelia Langford, John Sillitoe on behalf of the Wellcome Sanger Institute COVID-19 Surveillance Team |
| EPI_ISL_2829792, EPI_ISL_2829793 | Lighthouse Lab in Milton Keynes | Wellcome Sanger Institute for the COVID-19 Genomics UK (COG-UK) Consortium | The Lighthouse Lab in Milton Keynes and Alex Alderton, Roberto Amato, Jeffrey Barrett, Sonia Goncalves, Ewan Harrison, David K. Jackson, Ian Johnston, Dominic Kwiatkowski, Cordelia Langford, John Sillitoe on behalf of the Wellcome Sanger Institute COVID-19 Surveillance Team |
| EPI_ISL_2829794 | Lighthouse Lab in Alderley Park | Wellcome Sanger Institute for the COVID-19 Genomics UK (COG-UK) Consortium | Jacquelyn Wynn, Mairead Hyland, The Lighthouse Lab in Alderley Park and Alex Alderton, Roberto Amato, Jeffrey Barrett, Sonia Goncalves, Ewan Harrison, David K. Jackson, Ian Johnston, Dominic Kwiatkowski, Cordelia Langford, John Sillitoe on behalf of the Wellcome Sanger Institute COVID-19 Surveillance Team |
| EPI_ISL_2829795 | Lighthouse Lab in Milton Keynes | Wellcome Sanger Institute for the COVID-19 Genomics UK (COG-UK) Consortium | The Lighthouse Lab in Milton Keynes and Alex Alderton, Roberto Amato, Jeffrey Barrett, Sonia Goncalves, Ewan Harrison, David K. Jackson, Ian Johnston, Dominic Kwiatkowski, Cordelia Langford, John Sillitoe on behalf of the Wellcome Sanger Institute COVID-19 Surveillance Team |
| EPI_ISL_2829796 | Health Services Laboratories | Wellcome Sanger Institute for the COVID-19 Genomics UK (COG-UK) Consortium | Health Services Laboratories and Alex Alderton, Roberto Amato, Jeffrey Barrett, Sonia Goncalves, Ewan Harrison, David K. Jackson, Ian Johnston, Dominic Kwiatkowski, Cordelia Langford, John Sillitoe on behalf of the Wellcome Sanger Institute COVID-19 Surveillance Team |
| EPI_ISL_2829797 | Lighthouse Lab in Milton Keynes | Wellcome Sanger Institute for the COVID-19 Genomics UK (COG-UK) Consortium | The Lighthouse Lab in Milton Keynes and Alex Alderton, Roberto Amato, Jeffrey Barrett, Sonia Goncalves, Ewan Harrison, David K. Jackson, Ian Johnston, Dominic Kwiatkowski, Cordelia Langford, John Sillitoe on behalf of the Wellcome Sanger Institute COVID-19 Surveillance Team |
| EPI_ISL_2829798, EPI_ISL_2829799 | Lighthouse Lab in Alderley Park | Wellcome Sanger Institute for the COVID-19 Genomics UK (COG-UK) Consortium | Jacquelyn Wynn, Mairead Hyland, The Lighthouse Lab in Alderley Park and Alex Alderton, Roberto Amato, Jeffrey Barrett, Sonia Goncalves, Ewan Harrison, David K. Jackson, Ian Johnston, Dominic Kwiatkowski, Cordelia Langford, John Sillitoe on behalf of the Wellcome Sanger Institute COVID-19 Surveillance Team |
| EPI_ISL_2829800, EPI_ISL_2829801, EPI_ISL_2829802, EPI_ISL_2829803, EPI_ISL_2829804 | Lighthouse Lab in Milton Keynes | Wellcome Sanger Institute for the COVID-19 Genomics UK (COG-UK) Consortium | The Lighthouse Lab in Milton Keynes and Alex Alderton, Roberto Amato, Jeffrey Barrett, Sonia Goncalves, Ewan Harrison, David K. Jackson, Ian Johnston, Dominic Kwiatkowski, Cordelia Langford, John Sillitoe on behalf of the Wellcome Sanger Institute COVID-19 Surveillance Team |
| EPI_ISL_2829805 | Health Services Laboratories | Wellcome Sanger Institute for the COVID-19 Genomics UK (COG-UK) Consortium | Health Services Laboratories and Alex Alderton, Roberto Amato, Jeffrey Barrett, Sonia Goncalves, Ewan Harrison, David K. Jackson, Ian Johnston, Dominic Kwiatkowski, Cordelia Langford, John Sillitoe on behalf of the Wellcome Sanger Institute COVID-19 Surveillance Team |
| EPI_ISL_2829806 | Lighthouse Lab in Milton Keynes | Wellcome Sanger Institute for the COVID-19 Genomics UK (COG-UK) Consortium | The Lighthouse Lab in Milton Keynes and Alex Alderton, Roberto Amato, Jeffrey Barrett, Sonia Goncalves, Ewan Harrison, David K. Jackson, Ian Johnston, Dominic Kwiatkowski, Cordelia Langford, John Sillitoe on behalf of the Wellcome Sanger Institute COVID-19 Surveillance Team |
| EPI_ISL_2829807 | Lighthouse Lab in Alderley Park | Wellcome Sanger Institute for the COVID-19 Genomics UK (COG-UK) Consortium | Jacquelyn Wynn, Mairead Hyland, The Lighthouse Lab in Alderley Park and Alex Alderton, Roberto Amato, Jeffrey Barrett, Sonia Goncalves, Ewan Harrison, David K. Jackson, Ian Johnston, Dominic Kwiatkowski, Cordelia Langford, John Sillitoe on behalf of the Wellcome Sanger Institute COVID-19 Surveillance Team |
| EPI_ISL_2829808, EPI_ISL_2829809 | Lighthouse Lab in Milton Keynes | Wellcome Sanger Institute for the COVID-19 Genomics UK (COG-UK) Consortium | The Lighthouse Lab in Milton Keynes and Alex Alderton, Roberto Amato, Jeffrey Barrett, Sonia Goncalves, Ewan Harrison, David K. Jackson, Ian Johnston, Dominic Kwiatkowski, Cordelia Langford, John Sillitoe on behalf of the Wellcome Sanger Institute COVID-19 Surveillance Team |
| EPI_ISL_2829810 | Lighthouse Lab in Alderley Park | Wellcome Sanger Institute for the COVID-19 Genomics UK (COG-UK) Consortium | Jacquelyn Wynn, Mairead Hyland, The Lighthouse Lab in Alderley Park and Alex Alderton, Roberto Amato, Jeffrey Barrett, Sonia Goncalves, Ewan Harrison, David K. Jackson, Ian Johnston, Dominic Kwiatkowski, Cordelia Langford, John Sillitoe on behalf of the Wellcome Sanger Institute COVID-19 Surveillance Team |
| EPI_ISL_2829811 | Lighthouse Lab in Milton Keynes | Wellcome Sanger Institute for the COVID-19 Genomics UK (COG-UK) Consortium | The Lighthouse Lab in Milton Keynes and Alex Alderton, Roberto Amato, Jeffrey Barrett, Sonia Goncalves, Ewan Harrison, David K. Jackson, Ian Johnston, Dominic Kwiatkowski, Cordelia Langford, John Sillitoe on behalf of the Wellcome Sanger Institute COVID-19 Surveillance Team |
| EPI_ISL_2829812 | Lighthouse Lab in Alderley Park | Wellcome Sanger Institute for the COVID-19 Genomics UK (COG-UK) Consortium | Jacquelyn Wynn, Mairead Hyland, The Lighthouse Lab in Alderley Park and Alex Alderton, Roberto Amato, Jeffrey Barrett, Sonia Goncalves, Ewan Harrison, David K. Jackson, Ian Johnston, Dominic Kwiatkowski, Cordelia Langford, John Sillitoe on behalf of the Wellcome Sanger Institute COVID-19 Surveillance Team |
| EPI_ISL_2829813, EPI_ISL_2829814, EPI_ISL_2829815, EPI_ISL_2829816, EPI_ISL_2829817, EPI_ISL_2829818 | Lighthouse Lab in Milton Keynes | Wellcome Sanger Institute for the COVID-19 Genomics UK (COG-UK) Consortium | The Lighthouse Lab in Milton Keynes and Alex Alderton, Roberto Amato, Jeffrey Barrett, Sonia Goncalves, Ewan Harrison, David K. Jackson, Ian Johnston, Dominic Kwiatkowski, Cordelia Langford, John Sillitoe on behalf of the Wellcome Sanger Institute COVID-19 Surveillance Team |
| EPI_ISL_2829819 | Lighthouse Lab in Alderley Park | Wellcome Sanger Institute for the COVID-19 Genomics UK (COG-UK) Consortium | Jacquelyn Wynn, Mairead Hyland, The Lighthouse Lab in Alderley Park and Alex Alderton, Roberto Amato, Jeffrey Barrett, Sonia Goncalves, Ewan Harrison, David K. Jackson, Ian Johnston, Dominic Kwiatkowski, Cordelia Langford, John Sillitoe on behalf of the Wellcome Sanger Institute COVID-19 Surveillance Team |
| EPI_ISL_2829820 | Health Services Laboratories | Wellcome Sanger Institute for the COVID-19 Genomics UK (COG-UK) Consortium | Health Services Laboratories and Alex Alderton, Roberto Amato, Jeffrey Barrett, Sonia Goncalves, Ewan Harrison, David K. Jackson, Ian Johnston, Dominic Kwiatkowski, Cordelia Langford, John Sillitoe on behalf of the Wellcome Sanger Institute COVID-19 Surveillance Team |
| EPI_ISL_2829821, EPI_ISL_2829822 | Lighthouse Lab in Milton Keynes | Wellcome Sanger Institute for the COVID-19 Genomics UK (COG-UK) Consortium | The Lighthouse Lab in Milton Keynes and Alex Alderton, Roberto Amato, Jeffrey Barrett, Sonia Goncalves, Ewan Harrison, David K. Jackson, Ian Johnston, Dominic Kwiatkowski, Cordelia Langford, John Sillitoe on behalf of the Wellcome Sanger Institute COVID-19 Surveillance Team |
| EPI_ISL_2829823, EPI_ISL_2829824 | Health Services Laboratories | Wellcome Sanger Institute for the COVID-19 Genomics UK (COG-UK) Consortium | Health Services Laboratories and Alex Alderton, Roberto Amato, Jeffrey Barrett, Sonia Goncalves, Ewan Harrison, David K. Jackson, Ian Johnston, Dominic Kwiatkowski, Cordelia Langford, John Sillitoe on behalf of the Wellcome Sanger Institute COVID-19 Surveillance Team |
| EPI_ISL_2829825, EPI_ISL_2829826, EPI_ISL_2829827, EPI_ISL_2829828, EPI_ISL_2829829, EPI_ISL_2829830 | Lighthouse Lab in Milton Keynes | Wellcome Sanger Institute for the COVID-19 Genomics UK (COG-UK) Consortium | The Lighthouse Lab in Milton Keynes and Alex Alderton, Roberto Amato, Jeffrey Barrett, Sonia Goncalves, Ewan Harrison, David K. Jackson, Ian Johnston, Dominic Kwiatkowski, Cordelia Langford, John Sillitoe on behalf of the Wellcome Sanger Institute COVID-19 Surveillance Team |
| EPI_ISL_2829831 | Lighthouse Lab in Alderley Park | Wellcome Sanger Institute for the COVID-19 Genomics UK (COG-UK) Consortium | Jacquelyn Wynn, Mairead Hyland, The Lighthouse Lab in Alderley Park and Alex Alderton, Roberto Amato, Jeffrey Barrett, Sonia Goncalves, Ewan Harrison, David K. Jackson, Ian Johnston, Dominic Kwiatkowski, Cordelia Langford, John Sillitoe on behalf of the Wellcome Sanger Institute COVID-19 Surveillance Team |
| EPI_ISL_2829832 | Health Services Laboratories | Wellcome Sanger Institute for the COVID-19 Genomics UK (COG-UK) Consortium | Health Services Laboratories and Alex Alderton, Roberto Amato, Jeffrey Barrett, Sonia Goncalves, Ewan Harrison, David K. Jackson, Ian Johnston, Dominic Kwiatkowski, Cordelia Langford, John Sillitoe on behalf of the Wellcome Sanger Institute COVID-19 Surveillance Team |
| EPI_ISL_2829833 | Lighthouse Lab in Alderley Park | Wellcome Sanger Institute for the COVID-19 Genomics UK (COG-UK) Consortium | Jacquelyn Wynn, Mairead Hyland, The Lighthouse Lab in Alderley Park and Alex Alderton, Roberto Amato, Jeffrey Barrett, Sonia Goncalves, Ewan Harrison, David K. Jackson, Ian Johnston, Dominic Kwiatkowski, Cordelia Langford, John Sillitoe on behalf of the Wellcome Sanger Institute COVID-19 Surveillance Team |
| EPI_ISL_2829834, EPI_ISL_2829835, EPI_ISL_2829836 | Lighthouse Lab in Milton Keynes | Wellcome Sanger Institute for the COVID-19 Genomics UK (COG-UK) Consortium | The Lighthouse Lab in Milton Keynes and Alex Alderton, Roberto Amato, Jeffrey Barrett, Sonia Goncalves, Ewan Harrison, David K. Jackson, Ian Johnston, Dominic Kwiatkowski, Cordelia Langford, John Sillitoe on behalf of the Wellcome Sanger Institute COVID-19 Surveillance Team |
| EPI_ISL_2829837 | Health Services Laboratories | Wellcome Sanger Institute for the COVID-19 Genomics UK (COG-UK) Consortium | Health Services Laboratories and Alex Alderton, Roberto Amato, Jeffrey Barrett, Sonia Goncalves, Ewan Harrison, David K. Jackson, Ian Johnston, Dominic Kwiatkowski, Cordelia Langford, John Sillitoe on behalf of the Wellcome Sanger Institute COVID-19 Surveillance Team |
| EPI_ISL_2829838, EPI_ISL_2829839 | Lighthouse Lab in Milton Keynes | Wellcome Sanger Institute for the COVID-19 Genomics UK (COG-UK) Consortium | The Lighthouse Lab in Milton Keynes and Alex Alderton, Roberto Amato, Jeffrey Barrett, Sonia Goncalves, Ewan Harrison, David K. Jackson, Ian Johnston, Dominic Kwiatkowski, Cordelia Langford, John Sillitoe on behalf of the Wellcome Sanger Institute COVID-19 Surveillance Team |
| EPI_ISL_2829840, EPI_ISL_2829841 | Lighthouse Lab in Alderley Park | Wellcome Sanger Institute for the COVID-19 Genomics UK (COG-UK) Consortium | Jacquelyn Wynn, Mairead Hyland, The Lighthouse Lab in Alderley Park and Alex Alderton, Roberto Amato, Jeffrey Barrett, Sonia Goncalves, Ewan Harrison, David K. Jackson, Ian Johnston, Dominic Kwiatkowski, Cordelia Langford, John Sillitoe on behalf of the Wellcome Sanger Institute COVID-19 Surveillance Team |
| EPI_ISL_2829842, EPI_ISL_2829843 | Health Services Laboratories | Wellcome Sanger Institute for the COVID-19 Genomics UK (COG-UK) Consortium | Health Services Laboratories and Alex Alderton, Roberto Amato, Jeffrey Barrett, Sonia Goncalves, Ewan Harrison, David K. Jackson, Ian Johnston, Dominic Kwiatkowski, Cordelia Langford, John Sillitoe on behalf of the Wellcome Sanger Institute COVID-19 Surveillance Team |
| EPI_ISL_2829844, EPI_ISL_2829845 | Lighthouse Lab in Milton Keynes | Wellcome Sanger Institute for the COVID-19 Genomics UK (COG-UK) Consortium | The Lighthouse Lab in Milton Keynes and Alex Alderton, Roberto Amato, Jeffrey Barrett, Sonia Goncalves, Ewan Harrison, David K. Jackson, Ian Johnston, Dominic Kwiatkowski, Cordelia Langford, John Sillitoe on behalf of the Wellcome Sanger Institute COVID-19 Surveillance Team |
| EPI_ISL_2829846 | Lighthouse Lab in Alderley Park | Wellcome Sanger Institute for the COVID-19 Genomics UK (COG-UK) Consortium | Jacquelyn Wynn, Mairead Hyland, The Lighthouse Lab in Alderley Park and Alex Alderton, Roberto Amato, Jeffrey Barrett, Sonia Goncalves, Ewan Harrison, David K. Jackson, Ian Johnston, Dominic Kwiatkowski, Cordelia Langford, John Sillitoe on behalf of the Wellcome Sanger Institute COVID-19 Surveillance Team |
| EPI_ISL_2829847, EPI_ISL_2829848, EPI_ISL_2829849, EPI_ISL_2829850, EPI_ISL_2829851 | Lighthouse Lab in Milton Keynes | Wellcome Sanger Institute for the COVID-19 Genomics UK (COG-UK) Consortium | The Lighthouse Lab in Milton Keynes and Alex Alderton, Roberto Amato, Jeffrey Barrett, Sonia Goncalves, Ewan Harrison, David K. Jackson, Ian Johnston, Dominic Kwiatkowski, Cordelia Langford, John Sillitoe on behalf of the Wellcome Sanger Institute COVID-19 Surveillance Team |
| EPI_ISL_2829852 | Lighthouse Lab in Alderley Park | Wellcome Sanger Institute for the COVID-19 Genomics UK (COG-UK) Consortium | Jacquelyn Wynn, Mairead Hyland, The Lighthouse Lab in Alderley Park and Alex Alderton, Roberto Amato, Jeffrey Barrett, Sonia Goncalves, Ewan Harrison, David K. Jackson, Ian Johnston, Dominic Kwiatkowski, Cordelia Langford, John Sillitoe on behalf of the Wellcome Sanger Institute COVID-19 Surveillance Team |
| EPI_ISL_2829853 | Health Services Laboratories | Wellcome Sanger Institute for the COVID-19 Genomics UK (COG-UK) Consortium | Health Services Laboratories and Alex Alderton, Roberto Amato, Jeffrey Barrett, Sonia Goncalves, Ewan Harrison, David K. Jackson, Ian Johnston, Dominic Kwiatkowski, Cordelia Langford, John Sillitoe on behalf of the Wellcome Sanger Institute COVID-19 Surveillance Team |
| EPI_ISL_2829854 | Lighthouse Lab in Milton Keynes | Wellcome Sanger Institute for the COVID-19 Genomics UK (COG-UK) Consortium | The Lighthouse Lab in Milton Keynes and Alex Alderton, Roberto Amato, Jeffrey Barrett, Sonia Goncalves, Ewan Harrison, David K. Jackson, Ian Johnston, Dominic Kwiatkowski, Cordelia Langford, John Sillitoe on behalf of the Wellcome Sanger Institute COVID-19 Surveillance Team |

| | | | |
|---|---|---|---|
| EPI_ISL_2829855 | Lighthouse Lab in Alderley Park | Wellcome Sanger Institute for the COVID-19 Genomics UK (COG-UK) Consortium | Jacquelyn Wynn, Mairead Hyland, The Lighthouse Lab in Alderley Park and Alex Alderton, Roberto Amato, Jeffrey Barrett, Sonia Goncalves, Ewan Harrison, David K. Jackson, Ian Johnston, Dominic Kwiatkowski, Cordelia Langford, John Sillitoe on behalf of the Wellcome Sanger Institute COVID-19 Surveillance Team |
| EPI_ISL_2829856 | Health Services Laboratories | Wellcome Sanger Institute for the COVID-19 Genomics UK (COG-UK) Consortium | Health Services Laboratories and Alex Alderton, Roberto Amato, Jeffrey Barrett, Sonia Goncalves, Ewan Harrison, David K. Jackson, Ian Johnston, Dominic Kwiatkowski, Cordelia Langford, John Sillitoe on behalf of the Wellcome Sanger Institute COVID-19 Surveillance Team |
| EPI_ISL_2829857 | Lighthouse Lab in Milton Keynes | Wellcome Sanger Institute for the COVID-19 Genomics UK (COG-UK) Consortium | The Lighthouse Lab in Milton Keynes and Alex Alderton, Roberto Amato, Jeffrey Barrett, Sonia Goncalves, Ewan Harrison, David K. Jackson, Ian Johnston, Dominic Kwiatkowski, Cordelia Langford, John Sillitoe on behalf of the Wellcome Sanger Institute COVID-19 Surveillance Team |
| EPI_ISL_2829858 | Lighthouse Lab in Alderley Park | Wellcome Sanger Institute for the COVID-19 Genomics UK (COG-UK) Consortium | Jacquelyn Wynn, Mairead Hyland, The Lighthouse Lab in Alderley Park and Alex Alderton, Roberto Amato, Jeffrey Barrett, Sonia Goncalves, Ewan Harrison, David K. Jackson, Ian Johnston, Dominic Kwiatkowski, Cordelia Langford, John Sillitoe on behalf of the Wellcome Sanger Institute COVID-19 Surveillance Team |
| EPI_ISL_2829859 | Lighthouse Lab in Milton Keynes | Wellcome Sanger Institute for the COVID-19 Genomics UK (COG-UK) Consortium | The Lighthouse Lab in Milton Keynes and Alex Alderton, Roberto Amato, Jeffrey Barrett, Sonia Goncalves, Ewan Harrison, David K. Jackson, Ian Johnston, Dominic Kwiatkowski, Cordelia Langford, John Sillitoe on behalf of the Wellcome Sanger Institute COVID-19 Surveillance Team |
| EPI_ISL_2829860, EPI_ISL_2829861 | Health Services Laboratories | Wellcome Sanger Institute for the COVID-19 Genomics UK (COG-UK) Consortium | Health Services Laboratories and Alex Alderton, Roberto Amato, Jeffrey Barrett, Sonia Goncalves, Ewan Harrison, David K. Jackson, Ian Johnston, Dominic Kwiatkowski, Cordelia Langford, John Sillitoe on behalf of the Wellcome Sanger Institute COVID-19 Surveillance Team |
| EPI_ISL_2829862 | Lighthouse Lab in Alderley Park | Wellcome Sanger Institute for the COVID-19 Genomics UK (COG-UK) Consortium | Jacquelyn Wynn, Mairead Hyland, The Lighthouse Lab in Alderley Park and Alex Alderton, Roberto Amato, Jeffrey Barrett, Sonia Goncalves, Ewan Harrison, David K. Jackson, Ian Johnston, Dominic Kwiatkowski, Cordelia Langford, John Sillitoe on behalf of the Wellcome Sanger Institute COVID-19 Surveillance Team |
| EPI_ISL_2829863 | Lighthouse Lab in Milton Keynes | Wellcome Sanger Institute for the COVID-19 Genomics UK (COG-UK) Consortium | The Lighthouse Lab in Milton Keynes and Alex Alderton, Roberto Amato, Jeffrey Barrett, Sonia Goncalves, Ewan Harrison, David K. Jackson, Ian Johnston, Dominic Kwiatkowski, Cordelia Langford, John Sillitoe on behalf of the Wellcome Sanger Institute COVID-19 Surveillance Team |
| EPI_ISL_2829864 | Health Services Laboratories | Wellcome Sanger Institute for the COVID-19 Genomics UK (COG-UK) Consortium | Health Services Laboratories and Alex Alderton, Roberto Amato, Jeffrey Barrett, Sonia Goncalves, Ewan Harrison, David K. Jackson, Ian Johnston, Dominic Kwiatkowski, Cordelia Langford, John Sillitoe on behalf of the Wellcome Sanger Institute COVID-19 Surveillance Team |
| EPI_ISL_2829865 | Lighthouse Lab in Alderley Park | Wellcome Sanger Institute for the COVID-19 Genomics UK (COG-UK) Consortium | Jacquelyn Wynn, Mairead Hyland, The Lighthouse Lab in Alderley Park and Alex Alderton, Roberto Amato, Jeffrey Barrett, Sonia Goncalves, Ewan Harrison, David K. Jackson, Ian Johnston, Dominic Kwiatkowski, Cordelia Langford, John Sillitoe on behalf of the Wellcome Sanger Institute COVID-19 Surveillance Team |
| EPI_ISL_2829866 | Lighthouse Lab in Milton Keynes | Wellcome Sanger Institute for the COVID-19 Genomics UK (COG-UK) Consortium | The Lighthouse Lab in Milton Keynes and Alex Alderton, Roberto Amato, Jeffrey Barrett, Sonia Goncalves, Ewan Harrison, David K. Jackson, Ian Johnston, Dominic Kwiatkowski, Cordelia Langford, John Sillitoe on behalf of the Wellcome Sanger Institute COVID-19 Surveillance Team |
| EPI_ISL_2829867 | Health Services Laboratories | Wellcome Sanger Institute for the COVID-19 Genomics UK (COG-UK) Consortium | Health Services Laboratories and Alex Alderton, Roberto Amato, Jeffrey Barrett, Sonia Goncalves, Ewan Harrison, David K. Jackson, Ian Johnston, Dominic Kwiatkowski, Cordelia Langford, John Sillitoe on behalf of the Wellcome Sanger Institute COVID-19 Surveillance Team |
| EPI_ISL_2829868, EPI_ISL_2829869 | Lighthouse Lab in Alderley Park | Wellcome Sanger Institute for the COVID-19 Genomics UK (COG-UK) Consortium | Jacquelyn Wynn, Mairead Hyland, The Lighthouse Lab in Alderley Park and Alex Alderton, Roberto Amato, Jeffrey Barrett, Sonia Goncalves, Ewan Harrison, David K. Jackson, Ian Johnston, Dominic Kwiatkowski, Cordelia Langford, John Sillitoe on behalf of the Wellcome Sanger Institute COVID-19 Surveillance Team |
| EPI_ISL_2829870, EPI_ISL_2829871, EPI_ISL_2829872 | Lighthouse Lab in Milton Keynes | Wellcome Sanger Institute for the COVID-19 Genomics UK (COG-UK) Consortium | The Lighthouse Lab in Milton Keynes and Alex Alderton, Roberto Amato, Jeffrey Barrett, Sonia Goncalves, Ewan Harrison, David K. Jackson, Ian Johnston, Dominic Kwiatkowski, Cordelia Langford, John Sillitoe on behalf of the Wellcome Sanger Institute COVID-19 Surveillance Team |
| EPI_ISL_2829873 | Health Services Laboratories | Wellcome Sanger Institute for the COVID-19 Genomics UK (COG-UK) Consortium | Health Services Laboratories and Alex Alderton, Roberto Amato, Jeffrey Barrett, Sonia Goncalves, Ewan Harrison, David K. Jackson, Ian Johnston, Dominic Kwiatkowski, Cordelia Langford, John Sillitoe on behalf of the Wellcome Sanger Institute COVID-19 Surveillance Team |
| EPI_ISL_2829874 | Lighthouse Lab in Milton Keynes | Wellcome Sanger Institute for the COVID-19 Genomics UK (COG-UK) Consortium | The Lighthouse Lab in Milton Keynes and Alex Alderton, Roberto Amato, Jeffrey Barrett, Sonia Goncalves, Ewan Harrison, David K. Jackson, Ian Johnston, Dominic Kwiatkowski, Cordelia Langford, John Sillitoe on behalf of the Wellcome Sanger Institute COVID-19 Surveillance Team |
| EPI_ISL_2829875 | Lighthouse Lab in Alderley Park | Wellcome Sanger Institute for the COVID-19 Genomics UK (COG-UK) Consortium | Jacquelyn Wynn, Mairead Hyland, The Lighthouse Lab in Alderley Park and Alex Alderton, Roberto Amato, Jeffrey Barrett, Sonia Goncalves, Ewan Harrison, David K. Jackson, Ian Johnston, Dominic Kwiatkowski, Cordelia Langford, John Sillitoe on behalf of the Wellcome Sanger Institute COVID-19 Surveillance Team |
| EPI_ISL_2829876 | Lighthouse Lab in Milton Keynes | Wellcome Sanger Institute for the COVID-19 Genomics UK (COG-UK) Consortium | The Lighthouse Lab in Milton Keynes and Alex Alderton, Roberto Amato, Jeffrey Barrett, Sonia Goncalves, Ewan Harrison, David K. Jackson, Ian Johnston, Dominic Kwiatkowski, Cordelia Langford, John Sillitoe on behalf of the Wellcome Sanger Institute COVID-19 Surveillance Team |
| EPI_ISL_2829877 | Health Services Laboratories | Wellcome Sanger Institute for the COVID-19 Genomics UK (COG-UK) Consortium | Health Services Laboratories and Alex Alderton, Roberto Amato, Jeffrey Barrett, Sonia Goncalves, Ewan Harrison, David K. Jackson, Ian Johnston, Dominic Kwiatkowski, Cordelia Langford, John Sillitoe on behalf of the Wellcome Sanger Institute COVID-19 Surveillance Team |
| EPI_ISL_2829878 | Lighthouse Lab in Alderley Park | Wellcome Sanger Institute for the COVID-19 Genomics UK (COG-UK) Consortium | Jacquelyn Wynn, Mairead Hyland, The Lighthouse Lab in Alderley Park and Alex Alderton, Roberto Amato, Jeffrey Barrett, Sonia Goncalves, Ewan Harrison, David K. Jackson, Ian Johnston, Dominic Kwiatkowski, Cordelia Langford, John Sillitoe on behalf of the Wellcome Sanger Institute COVID-19 Surveillance Team |
| EPI_ISL_2829879 | Lighthouse Lab in Milton Keynes | Wellcome Sanger Institute for the COVID-19 Genomics UK (COG-UK) Consortium | The Lighthouse Lab in Milton Keynes and Alex Alderton, Roberto Amato, Jeffrey Barrett, Sonia Goncalves, Ewan Harrison, David K. Jackson, Ian Johnston, Dominic Kwiatkowski, Cordelia Langford, John Sillitoe on behalf of the Wellcome Sanger Institute COVID-19 Surveillance Team |
| EPI_ISL_2829880 | Health Services Laboratories | Wellcome Sanger Institute for the COVID-19 Genomics UK (COG-UK) Consortium | Health Services Laboratories and Alex Alderton, Roberto Amato, Jeffrey Barrett, Sonia Goncalves, Ewan Harrison, David K. Jackson, Ian Johnston, Dominic Kwiatkowski, Cordelia Langford, John Sillitoe on behalf of the Wellcome Sanger Institute COVID-19 Surveillance Team |
| EPI_ISL_2829881, EPI_ISL_2829882 | Lighthouse Lab in Milton Keynes | Wellcome Sanger Institute for the COVID-19 Genomics UK (COG-UK) Consortium | The Lighthouse Lab in Milton Keynes and Alex Alderton, Roberto Amato, Jeffrey Barrett, Sonia Goncalves, Ewan Harrison, David K. Jackson, Ian Johnston, Dominic Kwiatkowski, Cordelia Langford, John Sillitoe on behalf of the Wellcome Sanger Institute COVID-19 Surveillance Team |
| EPI_ISL_2829883 | Lighthouse Lab in Alderley Park | Wellcome Sanger Institute for the COVID-19 Genomics UK (COG-UK) Consortium | Jacquelyn Wynn, Mairead Hyland, The Lighthouse Lab in Alderley Park and Alex Alderton, Roberto Amato, Jeffrey Barrett, Sonia Goncalves, Ewan Harrison, David K. Jackson, Ian Johnston, Dominic Kwiatkowski, Cordelia Langford, John Sillitoe on behalf of the Wellcome Sanger Institute COVID-19 Surveillance Team |
| EPI_ISL_2829884 | Health Services Laboratories | Wellcome Sanger Institute for the COVID-19 Genomics UK (COG-UK) Consortium | Health Services Laboratories and Alex Alderton, Roberto Amato, Jeffrey Barrett, Sonia Goncalves, Ewan Harrison, David K. Jackson, Ian Johnston, Dominic Kwiatkowski, Cordelia Langford, John Sillitoe on behalf of the Wellcome Sanger Institute COVID-19 Surveillance Team |
| EPI_ISL_2829885, EPI_ISL_2829886, EPI_ISL_2829887, EPI_ISL_2829888, EPI_ISL_2829889, EPI_ISL_2829890, EPI_ISL_2829891 | Lighthouse Lab in Milton Keynes | Wellcome Sanger Institute for the COVID-19 Genomics UK (COG-UK) Consortium | The Lighthouse Lab in Milton Keynes and Alex Alderton, Roberto Amato, Jeffrey Barrett, Sonia Goncalves, Ewan Harrison, David K. Jackson, Ian Johnston, Dominic Kwiatkowski, Cordelia Langford, John Sillitoe on behalf of the Wellcome Sanger Institute COVID-19 Surveillance Team |
| EPI_ISL_2829892 | Lighthouse Lab in Alderley Park | Wellcome Sanger Institute for the COVID-19 Genomics UK (COG-UK) Consortium | Jacquelyn Wynn, Mairead Hyland, The Lighthouse Lab in Alderley Park and Alex Alderton, Roberto Amato, Jeffrey Barrett, Sonia Goncalves, Ewan Harrison, David K. Jackson, Ian Johnston, Dominic Kwiatkowski, Cordelia Langford, John Sillitoe on behalf of the Wellcome Sanger Institute COVID-19 Surveillance Team |
| EPI_ISL_2829893 | Health Services Laboratories | Wellcome Sanger Institute for the COVID-19 Genomics UK (COG-UK) Consortium | Health Services Laboratories and Alex Alderton, Roberto Amato, Jeffrey Barrett, Sonia Goncalves, Ewan Harrison, David K. Jackson, Ian Johnston, Dominic Kwiatkowski, Cordelia Langford, John Sillitoe on behalf of the Wellcome Sanger Institute COVID-19 Surveillance Team |
| EPI_ISL_2829894, EPI_ISL_2829895, EPI_ISL_2829896, EPI_ISL_2829897, EPI_ISL_2829898, EPI_ISL_2829899 | Lighthouse Lab in Milton Keynes | Wellcome Sanger Institute for the COVID-19 Genomics UK (COG-UK) Consortium | The Lighthouse Lab in Milton Keynes and Alex Alderton, Roberto Amato, Jeffrey Barrett, Sonia Goncalves, Ewan Harrison, David K. Jackson, Ian Johnston, Dominic Kwiatkowski, Cordelia Langford, John Sillitoe on behalf of the Wellcome Sanger Institute COVID-19 Surveillance Team |
| EPI_ISL_2829900 | Lighthouse Lab in Alderley Park | Wellcome Sanger Institute for the COVID-19 Genomics UK (COG-UK) Consortium | Jacquelyn Wynn, Mairead Hyland, The Lighthouse Lab in Alderley Park and Alex Alderton, Roberto Amato, Jeffrey Barrett, Sonia Goncalves, Ewan Harrison, David K. Jackson, Ian Johnston, Dominic Kwiatkowski, Cordelia Langford, John Sillitoe on behalf of the Wellcome Sanger Institute COVID-19 Surveillance Team |
| EPI_ISL_2829901, EPI_ISL_2829902, EPI_ISL_2829903, EPI_ISL_2829904 | Lighthouse Lab in Milton Keynes | Wellcome Sanger Institute for the COVID-19 Genomics UK (COG-UK) Consortium | The Lighthouse Lab in Milton Keynes and Alex Alderton, Roberto Amato, Jeffrey Barrett, Sonia Goncalves, Ewan Harrison, David K. Jackson, Ian Johnston, Dominic Kwiatkowski, Cordelia Langford, John Sillitoe on behalf of the Wellcome Sanger Institute COVID-19 Surveillance Team |
| EPI_ISL_2829905, EPI_ISL_2829906, EPI_ISL_2829907 | Health Services Laboratories | Wellcome Sanger Institute for the COVID-19 Genomics UK (COG-UK) Consortium | Health Services Laboratories and Alex Alderton, Roberto Amato, Jeffrey Barrett, Sonia Goncalves, Ewan Harrison, David K. Jackson, Ian Johnston, Dominic Kwiatkowski, Cordelia Langford, John Sillitoe on behalf of the Wellcome Sanger Institute COVID-19 Surveillance Team |
| EPI_ISL_2829908 | Lighthouse Lab in Alderley Park | Wellcome Sanger Institute for the COVID-19 Genomics UK (COG-UK) Consortium | Jacquelyn Wynn, Mairead Hyland, The Lighthouse Lab in Alderley Park and Alex Alderton, Roberto Amato, Jeffrey Barrett, Sonia Goncalves, Ewan Harrison, David K. Jackson, Ian Johnston, Dominic Kwiatkowski, Cordelia Langford, John Sillitoe on behalf of the Wellcome Sanger Institute COVID-19 Surveillance Team |
| EPI_ISL_2829909 | Health Services Laboratories | Wellcome Sanger Institute for the COVID-19 Genomics UK (COG-UK) Consortium | Health Services Laboratories and Alex Alderton, Roberto Amato, Jeffrey Barrett, Sonia Goncalves, Ewan Harrison, David K. Jackson, Ian Johnston, Dominic Kwiatkowski, Cordelia Langford, John Sillitoe on behalf of the Wellcome Sanger Institute COVID-19 Surveillance Team |
| EPI_ISL_2829910, EPI_ISL_2829911, EPI_ISL_2829912 | Lighthouse Lab in Milton Keynes | Wellcome Sanger Institute for the COVID-19 Genomics UK (COG-UK) Consortium | The Lighthouse Lab in Milton Keynes and Alex Alderton, Roberto Amato, Jeffrey Barrett, Sonia Goncalves, Ewan Harrison, David K. Jackson, Ian Johnston, Dominic Kwiatkowski, Cordelia Langford, John Sillitoe on behalf of the Wellcome Sanger Institute COVID-19 Surveillance Team |
| EPI_ISL_2829913 | Health Services Laboratories | Wellcome Sanger Institute for the COVID-19 Genomics UK (COG-UK) Consortium | Health Services Laboratories and Alex Alderton, Roberto Amato, Jeffrey Barrett, Sonia Goncalves, Ewan Harrison, David K. Jackson, Ian Johnston, Dominic Kwiatkowski, Cordelia Langford, John Sillitoe on behalf of the Wellcome Sanger Institute COVID-19 Surveillance Team |
| EPI_ISL_2829914, EPI_ISL_2829915 | Lighthouse Lab in Alderley Park | Wellcome Sanger Institute for the COVID-19 Genomics UK (COG-UK) Consortium | Jacquelyn Wynn, Mairead Hyland, The Lighthouse Lab in Alderley Park and Alex Alderton, Roberto Amato, Jeffrey Barrett, Sonia Goncalves, Ewan Harrison, David K. Jackson, Ian Johnston, Dominic Kwiatkowski, Cordelia Langford, John Sillitoe on behalf of the Wellcome Sanger Institute COVID-19 Surveillance Team |
| EPI_ISL_2829916, EPI_ISL_2829917 | Health Services Laboratories | Wellcome Sanger Institute for the COVID-19 Genomics UK (COG-UK) Consortium | Health Services Laboratories and Alex Alderton, Roberto Amato, Jeffrey Barrett, Sonia Goncalves, Ewan Harrison, David K. Jackson, Ian Johnston, Dominic Kwiatkowski, Cordelia Langford, John Sillitoe on behalf of the Wellcome Sanger Institute COVID-19 Surveillance Team |
| EPI_ISL_2829918 | Lighthouse Lab in Milton Keynes | Wellcome Sanger Institute for the COVID-19 Genomics UK (COG-UK) Consortium | The Lighthouse Lab in Milton Keynes and Alex Alderton, Roberto Amato, Jeffrey Barrett, Sonia Goncalves, Ewan Harrison, David K. Jackson, Ian Johnston, Dominic Kwiatkowski, Cordelia Langford, John Sillitoe on behalf of the Wellcome Sanger Institute COVID-19 Surveillance Team |
| EPI_ISL_2829919, | Lighthouse Lab in Alderley Park | Wellcome Sanger Institute for the COVID-19 Genomics UK (COG-UK) | Jacquelyn Wynn, Mairead Hyland, The Lighthouse Lab in Alderley Park and Alex Alderton, Roberto Amato, Jeffrey Barrett, Sonia Goncalves, Ewan Harrison, David K. Jackson, Ian Johnston, Dominic Kwiatkowski, Cordelia Langford, John Sillitoe on |

| | | | |
|---|---|---|---|
| EPI_ISL_2829920, EPI_ISL_2829921 | | Consortium | behalf of the Wellcome Sanger Institute COVID-19 Surveillance Team |
| EPI_ISL_2829922 | Lighthouse Lab in Milton Keynes | Wellcome Sanger Institute for the COVID-19 Genomics UK (COG-UK) Consortium | The Lighthouse Lab in Milton Keynes and Alex Alderton, Roberto Amato, Jeffrey Barrett, Sonia Goncalves, Ewan Harrison, David K. Jackson, Ian Johnston, Dominic Kwiatkowski, Cordelia Langford, John Sillitoe on behalf of the Wellcome Sanger Institute COVID-19 Surveillance Team |
| EPI_ISL_2829923 | Health Services Laboratories | Wellcome Sanger Institute for the COVID-19 Genomics UK (COG-UK) Consortium | Health Services Laboratories and Alex Alderton, Roberto Amato, Jeffrey Barrett, Sonia Goncalves, Ewan Harrison, David K. Jackson, Ian Johnston, Dominic Kwiatkowski, Cordelia Langford, John Sillitoe on behalf of the Wellcome Sanger Institute COVID-19 Surveillance Team |
| EPI_ISL_2829924 | Lighthouse Lab in Milton Keynes | Wellcome Sanger Institute for the COVID-19 Genomics UK (COG-UK) Consortium | The Lighthouse Lab in Milton Keynes and Alex Alderton, Roberto Amato, Jeffrey Barrett, Sonia Goncalves, Ewan Harrison, David K. Jackson, Ian Johnston, Dominic Kwiatkowski, Cordelia Langford, John Sillitoe on behalf of the Wellcome Sanger Institute COVID-19 Surveillance Team |
| EPI_ISL_2829925 | Lighthouse Lab in Alderley Park | Wellcome Sanger Institute for the COVID-19 Genomics UK (COG-UK) Consortium | Jacquelyn Wynn, Mairead Hyland, The Lighthouse Lab in Alderley Park and Alex Alderton, Roberto Amato, Jeffrey Barrett, Sonia Goncalves, Ewan Harrison, David K. Jackson, Ian Johnston, Dominic Kwiatkowski, Cordelia Langford, John Sillitoe on behalf of the Wellcome Sanger Institute COVID-19 Surveillance Team |
| EPI_ISL_2829926 | Lighthouse Lab in Milton Keynes | Wellcome Sanger Institute for the COVID-19 Genomics UK (COG-UK) Consortium | The Lighthouse Lab in Milton Keynes and Alex Alderton, Roberto Amato, Jeffrey Barrett, Sonia Goncalves, Ewan Harrison, David K. Jackson, Ian Johnston, Dominic Kwiatkowski, Cordelia Langford, John Sillitoe on behalf of the Wellcome Sanger Institute COVID-19 Surveillance Team |
| EPI_ISL_2829927, EPI_ISL_2829928 | Lighthouse Lab in Alderley Park | Wellcome Sanger Institute for the COVID-19 Genomics UK (COG-UK) Consortium | Jacquelyn Wynn, Mairead Hyland, The Lighthouse Lab in Alderley Park and Alex Alderton, Roberto Amato, Jeffrey Barrett, Sonia Goncalves, Ewan Harrison, David K. Jackson, Ian Johnston, Dominic Kwiatkowski, Cordelia Langford, John Sillitoe on behalf of the Wellcome Sanger Institute COVID-19 Surveillance Team |
| EPI_ISL_2829929 | Health Services Laboratories | Wellcome Sanger Institute for the COVID-19 Genomics UK (COG-UK) Consortium | Health Services Laboratories and Alex Alderton, Roberto Amato, Jeffrey Barrett, Sonia Goncalves, Ewan Harrison, David K. Jackson, Ian Johnston, Dominic Kwiatkowski, Cordelia Langford, John Sillitoe on behalf of the Wellcome Sanger Institute COVID-19 Surveillance Team |
| EPI_ISL_2829930, EPI_ISL_2829931, EPI_ISL_2829932, EPI_ISL_2829933 | Lighthouse Lab in Milton Keynes | Wellcome Sanger Institute for the COVID-19 Genomics UK (COG-UK) Consortium | The Lighthouse Lab in Milton Keynes and Alex Alderton, Roberto Amato, Jeffrey Barrett, Sonia Goncalves, Ewan Harrison, David K. Jackson, Ian Johnston, Dominic Kwiatkowski, Cordelia Langford, John Sillitoe on behalf of the Wellcome Sanger Institute COVID-19 Surveillance Team |
| EPI_ISL_2829934 | Health Services Laboratories | Wellcome Sanger Institute for the COVID-19 Genomics UK (COG-UK) Consortium | Health Services Laboratories and Alex Alderton, Roberto Amato, Jeffrey Barrett, Sonia Goncalves, Ewan Harrison, David K. Jackson, Ian Johnston, Dominic Kwiatkowski, Cordelia Langford, John Sillitoe on behalf of the Wellcome Sanger Institute COVID-19 Surveillance Team |
| EPI_ISL_2829935 | Lighthouse Lab in Milton Keynes | Wellcome Sanger Institute for the COVID-19 Genomics UK (COG-UK) Consortium | The Lighthouse Lab in Milton Keynes and Alex Alderton, Roberto Amato, Jeffrey Barrett, Sonia Goncalves, Ewan Harrison, David K. Jackson, Ian Johnston, Dominic Kwiatkowski, Cordelia Langford, John Sillitoe on behalf of the Wellcome Sanger Institute COVID-19 Surveillance Team |
| EPI_ISL_2829936 | Health Services Laboratories | Wellcome Sanger Institute for the COVID-19 Genomics UK (COG-UK) Consortium | Health Services Laboratories and Alex Alderton, Roberto Amato, Jeffrey Barrett, Sonia Goncalves, Ewan Harrison, David K. Jackson, Ian Johnston, Dominic Kwiatkowski, Cordelia Langford, John Sillitoe on behalf of the Wellcome Sanger Institute COVID-19 Surveillance Team |
| EPI_ISL_2829937 | Lighthouse Lab in Milton Keynes | Wellcome Sanger Institute for the COVID-19 Genomics UK (COG-UK) Consortium | The Lighthouse Lab in Milton Keynes and Alex Alderton, Roberto Amato, Jeffrey Barrett, Sonia Goncalves, Ewan Harrison, David K. Jackson, Ian Johnston, Dominic Kwiatkowski, Cordelia Langford, John Sillitoe on behalf of the Wellcome Sanger Institute COVID-19 Surveillance Team |
| EPI_ISL_2829938 | Lighthouse Lab in Alderley Park | Wellcome Sanger Institute for the COVID-19 Genomics UK (COG-UK) Consortium | Jacquelyn Wynn, Mairead Hyland, The Lighthouse Lab in Alderley Park and Alex Alderton, Roberto Amato, Jeffrey Barrett, Sonia Goncalves, Ewan Harrison, David K. Jackson, Ian Johnston, Dominic Kwiatkowski, Cordelia Langford, John Sillitoe on behalf of the Wellcome Sanger Institute COVID-19 Surveillance Team |
| EPI_ISL_2829939 | Lighthouse Lab in Milton Keynes | Wellcome Sanger Institute for the COVID-19 Genomics UK (COG-UK) Consortium | The Lighthouse Lab in Milton Keynes and Alex Alderton, Roberto Amato, Jeffrey Barrett, Sonia Goncalves, Ewan Harrison, David K. Jackson, Ian Johnston, Dominic Kwiatkowski, Cordelia Langford, John Sillitoe on behalf of the Wellcome Sanger Institute COVID-19 Surveillance Team |
| EPI_ISL_2829940 | Lighthouse Lab in Alderley Park | Wellcome Sanger Institute for the COVID-19 Genomics UK (COG-UK) Consortium | Jacquelyn Wynn, Mairead Hyland, The Lighthouse Lab in Alderley Park and Alex Alderton, Roberto Amato, Jeffrey Barrett, Sonia Goncalves, Ewan Harrison, David K. Jackson, Ian Johnston, Dominic Kwiatkowski, Cordelia Langford, John Sillitoe on behalf of the Wellcome Sanger Institute COVID-19 Surveillance Team |
| EPI_ISL_2829941 | Health Services Laboratories | Wellcome Sanger Institute for the COVID-19 Genomics UK (COG-UK) Consortium | Health Services Laboratories and Alex Alderton, Roberto Amato, Jeffrey Barrett, Sonia Goncalves, Ewan Harrison, David K. Jackson, Ian Johnston, Dominic Kwiatkowski, Cordelia Langford, John Sillitoe on behalf of the Wellcome Sanger Institute COVID-19 Surveillance Team |
| EPI_ISL_2829942, EPI_ISL_2829943, EPI_ISL_2829944, EPI_ISL_2829945 | Lighthouse Lab in Milton Keynes | Wellcome Sanger Institute for the COVID-19 Genomics UK (COG-UK) Consortium | The Lighthouse Lab in Milton Keynes and Alex Alderton, Roberto Amato, Jeffrey Barrett, Sonia Goncalves, Ewan Harrison, David K. Jackson, Ian Johnston, Dominic Kwiatkowski, Cordelia Langford, John Sillitoe on behalf of the Wellcome Sanger Institute COVID-19 Surveillance Team |
| EPI_ISL_2829946 | Lighthouse Lab in Alderley Park | Wellcome Sanger Institute for the COVID-19 Genomics UK (COG-UK) Consortium | Jacquelyn Wynn, Mairead Hyland, The Lighthouse Lab in Alderley Park and Alex Alderton, Roberto Amato, Jeffrey Barrett, Sonia Goncalves, Ewan Harrison, David K. Jackson, Ian Johnston, Dominic Kwiatkowski, Cordelia Langford, John Sillitoe on behalf of the Wellcome Sanger Institute COVID-19 Surveillance Team |
| EPI_ISL_2829947 | Health Services Laboratories | Wellcome Sanger Institute for the COVID-19 Genomics UK (COG-UK) Consortium | Health Services Laboratories and Alex Alderton, Roberto Amato, Jeffrey Barrett, Sonia Goncalves, Ewan Harrison, David K. Jackson, Ian Johnston, Dominic Kwiatkowski, Cordelia Langford, John Sillitoe on behalf of the Wellcome Sanger Institute COVID-19 Surveillance Team |
| EPI_ISL_2829948 | Lighthouse Lab in Alderley Park | Wellcome Sanger Institute for the COVID-19 Genomics UK (COG-UK) Consortium | Jacquelyn Wynn, Mairead Hyland, The Lighthouse Lab in Alderley Park and Alex Alderton, Roberto Amato, Jeffrey Barrett, Sonia Goncalves, Ewan Harrison, David K. Jackson, Ian Johnston, Dominic Kwiatkowski, Cordelia Langford, John Sillitoe on behalf of the Wellcome Sanger Institute COVID-19 Surveillance Team |
| EPI_ISL_2829949, EPI_ISL_2829950, EPI_ISL_2829951 | Lighthouse Lab in Milton Keynes | Wellcome Sanger Institute for the COVID-19 Genomics UK (COG-UK) Consortium | The Lighthouse Lab in Milton Keynes and Alex Alderton, Roberto Amato, Jeffrey Barrett, Sonia Goncalves, Ewan Harrison, David K. Jackson, Ian Johnston, Dominic Kwiatkowski, Cordelia Langford, John Sillitoe on behalf of the Wellcome Sanger Institute COVID-19 Surveillance Team |
| EPI_ISL_2829952 | Health Services Laboratories | Wellcome Sanger Institute for the COVID-19 Genomics UK (COG-UK) Consortium | Health Services Laboratories and Alex Alderton, Roberto Amato, Jeffrey Barrett, Sonia Goncalves, Ewan Harrison, David K. Jackson, Ian Johnston, Dominic Kwiatkowski, Cordelia Langford, John Sillitoe on behalf of the Wellcome Sanger Institute COVID-19 Surveillance Team |
| EPI_ISL_2829953, EPI_ISL_2829954, EPI_ISL_2829955 | Lighthouse Lab in Milton Keynes | Wellcome Sanger Institute for the COVID-19 Genomics UK (COG-UK) Consortium | The Lighthouse Lab in Milton Keynes and Alex Alderton, Roberto Amato, Jeffrey Barrett, Sonia Goncalves, Ewan Harrison, David K. Jackson, Ian Johnston, Dominic Kwiatkowski, Cordelia Langford, John Sillitoe on behalf of the Wellcome Sanger Institute COVID-19 Surveillance Team |
| EPI_ISL_2829956 | Lighthouse Lab in Glasgow | Wellcome Sanger Institute for the COVID-19 Genomics UK (COG-UK) Consortium | Harper VanSteenhouse, Yumi Kasai, David Gray, Carol Clugston, Anna Dominiczak and Alex Alderton, Roberto Amato, Jeffrey Barrett, Sonia Goncalves, Ewan Harrison, David K. Jackson, Ian Johnston, Dominic Kwiatkowski, Cordelia Langford, John Sillitoe on behalf of the Wellcome Sanger Institute COVID-19 Surveillance Team |
| EPI_ISL_2829957 | Lighthouse Lab in Alderley Park | Wellcome Sanger Institute for the COVID-19 Genomics UK (COG-UK) Consortium | Jacquelyn Wynn, Mairead Hyland, The Lighthouse Lab in Alderley Park and Alex Alderton, Roberto Amato, Jeffrey Barrett, Sonia Goncalves, Ewan Harrison, David K. Jackson, Ian Johnston, Dominic Kwiatkowski, Cordelia Langford, John Sillitoe on behalf of the Wellcome Sanger Institute COVID-19 Surveillance Team |
| EPI_ISL_2829958 | Lighthouse Lab in Milton Keynes | Wellcome Sanger Institute for the COVID-19 Genomics UK (COG-UK) Consortium | The Lighthouse Lab in Milton Keynes and Alex Alderton, Roberto Amato, Jeffrey Barrett, Sonia Goncalves, Ewan Harrison, David K. Jackson, Ian Johnston, Dominic Kwiatkowski, Cordelia Langford, John Sillitoe on behalf of the Wellcome Sanger Institute COVID-19 Surveillance Team |
| EPI_ISL_2829959 | Lighthouse Lab in Alderley Park | Wellcome Sanger Institute for the COVID-19 Genomics UK (COG-UK) Consortium | Jacquelyn Wynn, Mairead Hyland, The Lighthouse Lab in Alderley Park and Alex Alderton, Roberto Amato, Jeffrey Barrett, Sonia Goncalves, Ewan Harrison, David K. Jackson, Ian Johnston, Dominic Kwiatkowski, Cordelia Langford, John Sillitoe on behalf of the Wellcome Sanger Institute COVID-19 Surveillance Team |
| EPI_ISL_2829960, EPI_ISL_2829961 | Lighthouse Lab in Milton Keynes | Wellcome Sanger Institute for the COVID-19 Genomics UK (COG-UK) Consortium | The Lighthouse Lab in Milton Keynes and Alex Alderton, Roberto Amato, Jeffrey Barrett, Sonia Goncalves, Ewan Harrison, David K. Jackson, Ian Johnston, Dominic Kwiatkowski, Cordelia Langford, John Sillitoe on behalf of the Wellcome Sanger Institute COVID-19 Surveillance Team |
| EPI_ISL_2829962, EPI_ISL_2829963, EPI_ISL_2829964, EPI_ISL_2829965 | Lighthouse Lab in Alderley Park | Wellcome Sanger Institute for the COVID-19 Genomics UK (COG-UK) Consortium | Jacquelyn Wynn, Mairead Hyland, The Lighthouse Lab in Alderley Park and Alex Alderton, Roberto Amato, Jeffrey Barrett, Sonia Goncalves, Ewan Harrison, David K. Jackson, Ian Johnston, Dominic Kwiatkowski, Cordelia Langford, John Sillitoe on behalf of the Wellcome Sanger Institute COVID-19 Surveillance Team |
| EPI_ISL_2829966 | Lighthouse Lab in Glasgow | Wellcome Sanger Institute for the COVID-19 Genomics UK (COG-UK) Consortium | Harper VanSteenhouse, Yumi Kasai, David Gray, Carol Clugston, Anna Dominiczak and Alex Alderton, Roberto Amato, Jeffrey Barrett, Sonia Goncalves, Ewan Harrison, David K. Jackson, Ian Johnston, Dominic Kwiatkowski, Cordelia Langford, John Sillitoe on behalf of the Wellcome Sanger Institute COVID-19 Surveillance Team |
| EPI_ISL_2829967 | Lighthouse Lab in Milton Keynes | Wellcome Sanger Institute for the COVID-19 Genomics UK (COG-UK) Consortium | The Lighthouse Lab in Milton Keynes and Alex Alderton, Roberto Amato, Jeffrey Barrett, Sonia Goncalves, Ewan Harrison, David K. Jackson, Ian Johnston, Dominic Kwiatkowski, Cordelia Langford, John Sillitoe on behalf of the Wellcome Sanger Institute COVID-19 Surveillance Team |
| EPI_ISL_2829968, EPI_ISL_2829969, EPI_ISL_2829970, EPI_ISL_2829971 | Lighthouse Lab in Alderley Park | Wellcome Sanger Institute for the COVID-19 Genomics UK (COG-UK) Consortium | Jacquelyn Wynn, Mairead Hyland, The Lighthouse Lab in Alderley Park and Alex Alderton, Roberto Amato, Jeffrey Barrett, Sonia Goncalves, Ewan Harrison, David K. Jackson, Ian Johnston, Dominic Kwiatkowski, Cordelia Langford, John Sillitoe on behalf of the Wellcome Sanger Institute COVID-19 Surveillance Team |
| EPI_ISL_2829972 | Lighthouse Lab in Milton Keynes | Wellcome Sanger Institute for the COVID-19 Genomics UK (COG-UK) Consortium | The Lighthouse Lab in Milton Keynes and Alex Alderton, Roberto Amato, Jeffrey Barrett, Sonia Goncalves, Ewan Harrison, David K. Jackson, Ian Johnston, Dominic Kwiatkowski, Cordelia Langford, John Sillitoe on behalf of the Wellcome Sanger Institute COVID-19 Surveillance Team |
| EPI_ISL_2829973 | Lighthouse Lab in Glasgow | Wellcome Sanger Institute for the COVID-19 Genomics UK (COG-UK) Consortium | Harper VanSteenhouse, Yumi Kasai, David Gray, Carol Clugston, Anna Dominiczak and Alex Alderton, Roberto Amato, Jeffrey Barrett, Sonia Goncalves, Ewan Harrison, David K. Jackson, Ian Johnston, Dominic Kwiatkowski, Cordelia Langford, John Sillitoe on behalf of the Wellcome Sanger Institute COVID-19 Surveillance Team |
| EPI_ISL_2829974, EPI_ISL_2829975, EPI_ISL_2829976, EPI_ISL_2829977, EPI_ISL_2829978, EPI_ISL_2829979, EPI_ISL_2829980, EPI_ISL_2829981, EPI_ISL_2829982, EPI_ISL_2829983, EPI_ISL_2829984 | | | |
| see above | Lighthouse Lab in Milton Keynes | Wellcome Sanger Institute for the COVID-19 Genomics UK (COG-UK) Consortium | The Lighthouse Lab in Milton Keynes and Alex Alderton, Roberto Amato, Jeffrey Barrett, Sonia Goncalves, Ewan Harrison, David K. Jackson, Ian Johnston, Dominic Kwiatkowski, Cordelia Langford, John Sillitoe on behalf of the Wellcome Sanger Institute COVID-19 Surveillance Team |
| EPI_ISL_2829985, EPI_ISL_2829986 | Lighthouse Lab in Alderley Park | Wellcome Sanger Institute for the COVID-19 Genomics UK (COG-UK) Consortium | Jacquelyn Wynn, Mairead Hyland, The Lighthouse Lab in Alderley Park and Alex Alderton, Roberto Amato, Jeffrey Barrett, Sonia Goncalves, Ewan Harrison, David K. Jackson, Ian Johnston, Dominic Kwiatkowski, Cordelia Langford, John Sillitoe on behalf of the Wellcome Sanger Institute COVID-19 Surveillance Team |
| EPI_ISL_2829987, EPI_ISL_2829988, EPI_ISL_2829989 | Lighthouse Lab in Milton Keynes | Wellcome Sanger Institute for the COVID-19 Genomics UK (COG-UK) Consortium | The Lighthouse Lab in Milton Keynes and Alex Alderton, Roberto Amato, Jeffrey Barrett, Sonia Goncalves, Ewan Harrison, David K. Jackson, Ian Johnston, Dominic Kwiatkowski, Cordelia Langford, John Sillitoe on behalf of the Wellcome Sanger Institute COVID-19 Surveillance Team |
| EPI_ISL_2829990, EPI_ISL_2829991 | Lighthouse Lab in Alderley Park | Wellcome Sanger Institute for the COVID-19 Genomics UK (COG-UK) Consortium | Jacquelyn Wynn, Mairead Hyland, The Lighthouse Lab in Alderley Park and Alex Alderton, Roberto Amato, Jeffrey Barrett, Sonia Goncalves, Ewan Harrison, David K. Jackson, Ian Johnston, Dominic Kwiatkowski, Cordelia Langford, John Sillitoe on behalf of the Wellcome Sanger Institute COVID-19 Surveillance Team |
| EPI_ISL_2829992 | Lighthouse Lab in Milton Keynes | Wellcome Sanger Institute for the COVID-19 Genomics UK (COG-UK) Consortium | The Lighthouse Lab in Milton Keynes and Alex Alderton, Roberto Amato, Jeffrey Barrett, Sonia Goncalves, Ewan Harrison, David K. Jackson, Ian Johnston, Dominic Kwiatkowski, Cordelia Langford, John Sillitoe on behalf of the Wellcome Sanger Institute COVID-19 Surveillance Team |

| | | | |
|---|---|---|---|
| EPI_ISL_2829993 | Lighthouse Lab in Glasgow | Wellcome Sanger Institute for the COVID-19 Genomics UK (COG-UK) Consortium | Harper VanSteenhouse, Yumi Kasai, David Gray, Carol Clugston, Anna Dominiczak and Alex Alderton, Roberto Amato, Jeffrey Barrett, Sonia Goncalves, Ewan Harrison, David K. Jackson, Ian Johnston, Dominic Kwiatkowski, Cordelia Langford, John Sillitoe on behalf of the Wellcome Sanger Institute COVID-19 Surveillance Team |
| EPI_ISL_2829994 | Lighthouse Lab in Milton Keynes | Wellcome Sanger Institute for the COVID-19 Genomics UK (COG-UK) Consortium | The Lighthouse Lab in Milton Keynes and Alex Alderton, Roberto Amato, Jeffrey Barrett, Sonia Goncalves, Ewan Harrison, David K. Jackson, Ian Johnston, Dominic Kwiatkowski, Cordelia Langford, John Sillitoe on behalf of the Wellcome Sanger Institute COVID-19 Surveillance Team |
| EPI_ISL_2829996 | Lighthouse Lab in Alderley Park | Wellcome Sanger Institute for the COVID-19 Genomics UK (COG-UK) Consortium | Jacquelyn Wynn, Mairead Hyland, The Lighthouse Lab in Alderley Park and Alex Alderton, Roberto Amato, Jeffrey Barrett, Sonia Goncalves, Ewan Harrison, David K. Jackson, Ian Johnston, Dominic Kwiatkowski, Cordelia Langford, John Sillitoe on behalf of the Wellcome Sanger Institute COVID-19 Surveillance Team |
| EPI_ISL_2829997 | Lighthouse Lab in Milton Keynes | Wellcome Sanger Institute for the COVID-19 Genomics UK (COG-UK) Consortium | The Lighthouse Lab in Milton Keynes and Alex Alderton, Roberto Amato, Jeffrey Barrett, Sonia Goncalves, Ewan Harrison, David K. Jackson, Ian Johnston, Dominic Kwiatkowski, Cordelia Langford, John Sillitoe on behalf of the Wellcome Sanger Institute COVID-19 Surveillance Team |
| EPI_ISL_2829998, EPI_ISL_2829999 | Lighthouse Lab in Alderley Park | Wellcome Sanger Institute for the COVID-19 Genomics UK (COG-UK) Consortium | Jacquelyn Wynn, Mairead Hyland, The Lighthouse Lab in Alderley Park and Alex Alderton, Roberto Amato, Jeffrey Barrett, Sonia Goncalves, Ewan Harrison, David K. Jackson, Ian Johnston, Dominic Kwiatkowski, Cordelia Langford, John Sillitoe on behalf of the Wellcome Sanger Institute COVID-19 Surveillance Team |
| EPI_ISL_2830000 | Lighthouse Lab in Milton Keynes | Wellcome Sanger Institute for the COVID-19 Genomics UK (COG-UK) Consortium | The Lighthouse Lab in Milton Keynes and Alex Alderton, Roberto Amato, Jeffrey Barrett, Sonia Goncalves, Ewan Harrison, David K. Jackson, Ian Johnston, Dominic Kwiatkowski, Cordelia Langford, John Sillitoe on behalf of the Wellcome Sanger Institute COVID-19 Surveillance Team |
| EPI_ISL_2830001 | Lighthouse Lab in Glasgow | Wellcome Sanger Institute for the COVID-19 Genomics UK (COG-UK) Consortium | Harper VanSteenhouse, Yumi Kasai, David Gray, Carol Clugston, Anna Dominiczak and Alex Alderton, Roberto Amato, Jeffrey Barrett, Sonia Goncalves, Ewan Harrison, David K. Jackson, Ian Johnston, Dominic Kwiatkowski, Cordelia Langford, John Sillitoe on behalf of the Wellcome Sanger Institute COVID-19 Surveillance Team |
| EPI_ISL_2830002, EPI_ISL_2830003 | Lighthouse Lab in Milton Keynes | Wellcome Sanger Institute for the COVID-19 Genomics UK (COG-UK) Consortium | The Lighthouse Lab in Milton Keynes and Alex Alderton, Roberto Amato, Jeffrey Barrett, Sonia Goncalves, Ewan Harrison, David K. Jackson, Ian Johnston, Dominic Kwiatkowski, Cordelia Langford, John Sillitoe on behalf of the Wellcome Sanger Institute COVID-19 Surveillance Team |
| EPI_ISL_2830004, EPI_ISL_2830005 | Lighthouse Lab in Glasgow | Wellcome Sanger Institute for the COVID-19 Genomics UK (COG-UK) Consortium | Harper VanSteenhouse, Yumi Kasai, David Gray, Carol Clugston, Anna Dominiczak and Alex Alderton, Roberto Amato, Jeffrey Barrett, Sonia Goncalves, Ewan Harrison, David K. Jackson, Ian Johnston, Dominic Kwiatkowski, Cordelia Langford, John Sillitoe on behalf of the Wellcome Sanger Institute COVID-19 Surveillance Team |
| EPI_ISL_2830006 | Lighthouse Lab in Alderley Park | Wellcome Sanger Institute for the COVID-19 Genomics UK (COG-UK) Consortium | Jacquelyn Wynn, Mairead Hyland, The Lighthouse Lab in Alderley Park and Alex Alderton, Roberto Amato, Jeffrey Barrett, Sonia Goncalves, Ewan Harrison, David K. Jackson, Ian Johnston, Dominic Kwiatkowski, Cordelia Langford, John Sillitoe on behalf of the Wellcome Sanger Institute COVID-19 Surveillance Team |
| EPI_ISL_2830007 | Lighthouse Lab in Glasgow | Wellcome Sanger Institute for the COVID-19 Genomics UK (COG-UK) Consortium | Harper VanSteenhouse, Yumi Kasai, David Gray, Carol Clugston, Anna Dominiczak and Alex Alderton, Roberto Amato, Jeffrey Barrett, Sonia Goncalves, Ewan Harrison, David K. Jackson, Ian Johnston, Dominic Kwiatkowski, Cordelia Langford, John Sillitoe on behalf of the Wellcome Sanger Institute COVID-19 Surveillance Team |
| EPI_ISL_2830008 | Lighthouse Lab in Alderley Park | Wellcome Sanger Institute for the COVID-19 Genomics UK (COG-UK) Consortium | Jacquelyn Wynn, Mairead Hyland, The Lighthouse Lab in Alderley Park and Alex Alderton, Roberto Amato, Jeffrey Barrett, Sonia Goncalves, Ewan Harrison, David K. Jackson, Ian Johnston, Dominic Kwiatkowski, Cordelia Langford, John Sillitoe on behalf of the Wellcome Sanger Institute COVID-19 Surveillance Team |
| EPI_ISL_2830009 | Lighthouse Lab in Glasgow | Wellcome Sanger Institute for the COVID-19 Genomics UK (COG-UK) Consortium | Harper VanSteenhouse, Yumi Kasai, David Gray, Carol Clugston, Anna Dominiczak and Alex Alderton, Roberto Amato, Jeffrey Barrett, Sonia Goncalves, Ewan Harrison, David K. Jackson, Ian Johnston, Dominic Kwiatkowski, Cordelia Langford, John Sillitoe on behalf of the Wellcome Sanger Institute COVID-19 Surveillance Team |
| EPI_ISL_2830010 | Lighthouse Lab in Alderley Park | Wellcome Sanger Institute for the COVID-19 Genomics UK (COG-UK) Consortium | Jacquelyn Wynn, Mairead Hyland, The Lighthouse Lab in Alderley Park and Alex Alderton, Roberto Amato, Jeffrey Barrett, Sonia Goncalves, Ewan Harrison, David K. Jackson, Ian Johnston, Dominic Kwiatkowski, Cordelia Langford, John Sillitoe on behalf of the Wellcome Sanger Institute COVID-19 Surveillance Team |
| EPI_ISL_2830011 | Lighthouse Lab in Milton Keynes | Wellcome Sanger Institute for the COVID-19 Genomics UK (COG-UK) Consortium | The Lighthouse Lab in Milton Keynes and Alex Alderton, Roberto Amato, Jeffrey Barrett, Sonia Goncalves, Ewan Harrison, David K. Jackson, Ian Johnston, Dominic Kwiatkowski, Cordelia Langford, John Sillitoe on behalf of the Wellcome Sanger Institute COVID-19 Surveillance Team |
| EPI_ISL_2830012 | Lighthouse Lab in Alderley Park | Wellcome Sanger Institute for the COVID-19 Genomics UK (COG-UK) Consortium | Jacquelyn Wynn, Mairead Hyland, The Lighthouse Lab in Alderley Park and Alex Alderton, Roberto Amato, Jeffrey Barrett, Sonia Goncalves, Ewan Harrison, David K. Jackson, Ian Johnston, Dominic Kwiatkowski, Cordelia Langford, John Sillitoe on behalf of the Wellcome Sanger Institute COVID-19 Surveillance Team |
| EPI_ISL_2830013 | Lighthouse Lab in Glasgow | Wellcome Sanger Institute for the COVID-19 Genomics UK (COG-UK) Consortium | Harper VanSteenhouse, Yumi Kasai, David Gray, Carol Clugston, Anna Dominiczak and Alex Alderton, Roberto Amato, Jeffrey Barrett, Sonia Goncalves, Ewan Harrison, David K. Jackson, Ian Johnston, Dominic Kwiatkowski, Cordelia Langford, John Sillitoe on behalf of the Wellcome Sanger Institute COVID-19 Surveillance Team |
| EPI_ISL_2830014 | Lighthouse Lab in Alderley Park | Wellcome Sanger Institute for the COVID-19 Genomics UK (COG-UK) Consortium | Jacquelyn Wynn, Mairead Hyland, The Lighthouse Lab in Alderley Park and Alex Alderton, Roberto Amato, Jeffrey Barrett, Sonia Goncalves, Ewan Harrison, David K. Jackson, Ian Johnston, Dominic Kwiatkowski, Cordelia Langford, John Sillitoe on behalf of the Wellcome Sanger Institute COVID-19 Surveillance Team |
| EPI_ISL_2830015, EPI_ISL_2830016 | Lighthouse Lab in Milton Keynes | Wellcome Sanger Institute for the COVID-19 Genomics UK (COG-UK) Consortium | The Lighthouse Lab in Milton Keynes and Alex Alderton, Roberto Amato, Jeffrey Barrett, Sonia Goncalves, Ewan Harrison, David K. Jackson, Ian Johnston, Dominic Kwiatkowski, Cordelia Langford, John Sillitoe on behalf of the Wellcome Sanger Institute COVID-19 Surveillance Team |
| EPI_ISL_2830017 | Lighthouse Lab in Glasgow | Wellcome Sanger Institute for the COVID-19 Genomics UK (COG-UK) Consortium | Harper VanSteenhouse, Yumi Kasai, David Gray, Carol Clugston, Anna Dominiczak and Alex Alderton, Roberto Amato, Jeffrey Barrett, Sonia Goncalves, Ewan Harrison, David K. Jackson, Ian Johnston, Dominic Kwiatkowski, Cordelia Langford, John Sillitoe on behalf of the Wellcome Sanger Institute COVID-19 Surveillance Team |
| EPI_ISL_2830018, EPI_ISL_2830019 | Lighthouse Lab in Milton Keynes | Wellcome Sanger Institute for the COVID-19 Genomics UK (COG-UK) Consortium | The Lighthouse Lab in Milton Keynes and Alex Alderton, Roberto Amato, Jeffrey Barrett, Sonia Goncalves, Ewan Harrison, David K. Jackson, Ian Johnston, Dominic Kwiatkowski, Cordelia Langford, John Sillitoe on behalf of the Wellcome Sanger Institute COVID-19 Surveillance Team |
| EPI_ISL_2830020, EPI_ISL_2830021 | Lighthouse Lab in Alderley Park | Wellcome Sanger Institute for the COVID-19 Genomics UK (COG-UK) Consortium | Jacquelyn Wynn, Mairead Hyland, The Lighthouse Lab in Alderley Park and Alex Alderton, Roberto Amato, Jeffrey Barrett, Sonia Goncalves, Ewan Harrison, David K. Jackson, Ian Johnston, Dominic Kwiatkowski, Cordelia Langford, John Sillitoe on behalf of the Wellcome Sanger Institute COVID-19 Surveillance Team |
| EPI_ISL_2830022, EPI_ISL_2830023, EPI_ISL_2830024, EPI_ISL_2830025, EPI_ISL_2830026 | Lighthouse Lab in Glasgow | Wellcome Sanger Institute for the COVID-19 Genomics UK (COG-UK) Consortium | Harper VanSteenhouse, Yumi Kasai, David Gray, Carol Clugston, Anna Dominiczak and Alex Alderton, Roberto Amato, Jeffrey Barrett, Sonia Goncalves, Ewan Harrison, David K. Jackson, Ian Johnston, Dominic Kwiatkowski, Cordelia Langford, John Sillitoe on behalf of the Wellcome Sanger Institute COVID-19 Surveillance Team |
| EPI_ISL_2830027 | Lighthouse Lab in Milton Keynes | Wellcome Sanger Institute for the COVID-19 Genomics UK (COG-UK) Consortium | The Lighthouse Lab in Milton Keynes and Alex Alderton, Roberto Amato, Jeffrey Barrett, Sonia Goncalves, Ewan Harrison, David K. Jackson, Ian Johnston, Dominic Kwiatkowski, Cordelia Langford, John Sillitoe on behalf of the Wellcome Sanger Institute COVID-19 Surveillance Team |
| EPI_ISL_2830028, EPI_ISL_2830029 | Lighthouse Lab in Alderley Park | Wellcome Sanger Institute for the COVID-19 Genomics UK (COG-UK) Consortium | Jacquelyn Wynn, Mairead Hyland, The Lighthouse Lab in Alderley Park and Alex Alderton, Roberto Amato, Jeffrey Barrett, Sonia Goncalves, Ewan Harrison, David K. Jackson, Ian Johnston, Dominic Kwiatkowski, Cordelia Langford, John Sillitoe on behalf of the Wellcome Sanger Institute COVID-19 Surveillance Team |
| EPI_ISL_2830030 | Lighthouse Lab in Glasgow | Wellcome Sanger Institute for the COVID-19 Genomics UK (COG-UK) Consortium | Harper VanSteenhouse, Yumi Kasai, David Gray, Carol Clugston, Anna Dominiczak and Alex Alderton, Roberto Amato, Jeffrey Barrett, Sonia Goncalves, Ewan Harrison, David K. Jackson, Ian Johnston, Dominic Kwiatkowski, Cordelia Langford, John Sillitoe on behalf of the Wellcome Sanger Institute COVID-19 Surveillance Team |
| EPI_ISL_2830031 | Lighthouse Lab in Milton Keynes | Wellcome Sanger Institute for the COVID-19 Genomics UK (COG-UK) Consortium | The Lighthouse Lab in Milton Keynes and Alex Alderton, Roberto Amato, Jeffrey Barrett, Sonia Goncalves, Ewan Harrison, David K. Jackson, Ian Johnston, Dominic Kwiatkowski, Cordelia Langford, John Sillitoe on behalf of the Wellcome Sanger Institute COVID-19 Surveillance Team |
| EPI_ISL_2830032 | Lighthouse Lab in Glasgow | Wellcome Sanger Institute for the COVID-19 Genomics UK (COG-UK) Consortium | Harper VanSteenhouse, Yumi Kasai, David Gray, Carol Clugston, Anna Dominiczak and Alex Alderton, Roberto Amato, Jeffrey Barrett, Sonia Goncalves, Ewan Harrison, David K. Jackson, Ian Johnston, Dominic Kwiatkowski, Cordelia Langford, John Sillitoe on behalf of the Wellcome Sanger Institute COVID-19 Surveillance Team |
| EPI_ISL_2830033 | Lighthouse Lab in Alderley Park | Wellcome Sanger Institute for the COVID-19 Genomics UK (COG-UK) Consortium | Jacquelyn Wynn, Mairead Hyland, The Lighthouse Lab in Alderley Park and Alex Alderton, Roberto Amato, Jeffrey Barrett, Sonia Goncalves, Ewan Harrison, David K. Jackson, Ian Johnston, Dominic Kwiatkowski, Cordelia Langford, John Sillitoe on behalf of the Wellcome Sanger Institute COVID-19 Surveillance Team |
| EPI_ISL_2830034 | Lighthouse Lab in Glasgow | Wellcome Sanger Institute for the COVID-19 Genomics UK (COG-UK) Consortium | Harper VanSteenhouse, Yumi Kasai, David Gray, Carol Clugston, Anna Dominiczak and Alex Alderton, Roberto Amato, Jeffrey Barrett, Sonia Goncalves, Ewan Harrison, David K. Jackson, Ian Johnston, Dominic Kwiatkowski, Cordelia Langford, John Sillitoe on behalf of the Wellcome Sanger Institute COVID-19 Surveillance Team |
| EPI_ISL_2830035 | Lighthouse Lab in Alderley Park | Wellcome Sanger Institute for the COVID-19 Genomics UK (COG-UK) Consortium | Jacquelyn Wynn, Mairead Hyland, The Lighthouse Lab in Alderley Park and Alex Alderton, Roberto Amato, Jeffrey Barrett, Sonia Goncalves, Ewan Harrison, David K. Jackson, Ian Johnston, Dominic Kwiatkowski, Cordelia Langford, John Sillitoe on behalf of the Wellcome Sanger Institute COVID-19 Surveillance Team |
| EPI_ISL_2830036, EPI_ISL_2830037 | Lighthouse Lab in Milton Keynes | Wellcome Sanger Institute for the COVID-19 Genomics UK (COG-UK) Consortium | The Lighthouse Lab in Milton Keynes and Alex Alderton, Roberto Amato, Jeffrey Barrett, Sonia Goncalves, Ewan Harrison, David K. Jackson, Ian Johnston, Dominic Kwiatkowski, Cordelia Langford, John Sillitoe on behalf of the Wellcome Sanger Institute COVID-19 Surveillance Team |
| EPI_ISL_2830038 | Lighthouse Lab in Alderley Park | Wellcome Sanger Institute for the COVID-19 Genomics UK (COG-UK) Consortium | Jacquelyn Wynn, Mairead Hyland, The Lighthouse Lab in Alderley Park and Alex Alderton, Roberto Amato, Jeffrey Barrett, Sonia Goncalves, Ewan Harrison, David K. Jackson, Ian Johnston, Dominic Kwiatkowski, Cordelia Langford, John Sillitoe on behalf of the Wellcome Sanger Institute COVID-19 Surveillance Team |
| EPI_ISL_2830039, EPI_ISL_2830040 | Lighthouse Lab in Milton Keynes | Wellcome Sanger Institute for the COVID-19 Genomics UK (COG-UK) Consortium | The Lighthouse Lab in Milton Keynes and Alex Alderton, Roberto Amato, Jeffrey Barrett, Sonia Goncalves, Ewan Harrison, David K. Jackson, Ian Johnston, Dominic Kwiatkowski, Cordelia Langford, John Sillitoe on behalf of the Wellcome Sanger Institute COVID-19 Surveillance Team |
| EPI_ISL_2830041 | Lighthouse Lab in Alderley Park | Wellcome Sanger Institute for the COVID-19 Genomics UK (COG-UK) Consortium | Jacquelyn Wynn, Mairead Hyland, The Lighthouse Lab in Alderley Park and Alex Alderton, Roberto Amato, Jeffrey Barrett, Sonia Goncalves, Ewan Harrison, David K. Jackson, Ian Johnston, Dominic Kwiatkowski, Cordelia Langford, John Sillitoe on behalf of the Wellcome Sanger Institute COVID-19 Surveillance Team |
| EPI_ISL_2830042, EPI_ISL_2830043 | Lighthouse Lab in Milton Keynes | Wellcome Sanger Institute for the COVID-19 Genomics UK (COG-UK) Consortium | The Lighthouse Lab in Milton Keynes and Alex Alderton, Roberto Amato, Jeffrey Barrett, Sonia Goncalves, Ewan Harrison, David K. Jackson, Ian Johnston, Dominic Kwiatkowski, Cordelia Langford, John Sillitoe on behalf of the Wellcome Sanger Institute COVID-19 Surveillance Team |
| EPI_ISL_2830044 | Lighthouse Lab in Glasgow | Wellcome Sanger Institute for the COVID-19 Genomics UK (COG-UK) Consortium | Harper VanSteenhouse, Yumi Kasai, David Gray, Carol Clugston, Anna Dominiczak and Alex Alderton, Roberto Amato, Jeffrey Barrett, Sonia Goncalves, Ewan Harrison, David K. Jackson, Ian Johnston, Dominic Kwiatkowski, Cordelia Langford, John Sillitoe on behalf of the Wellcome Sanger Institute COVID-19 Surveillance Team |
| EPI_ISL_2830045 | Lighthouse Lab in Alderley Park | Wellcome Sanger Institute for the COVID-19 Genomics UK (COG-UK) Consortium | Jacquelyn Wynn, Mairead Hyland, The Lighthouse Lab in Alderley Park and Alex Alderton, Roberto Amato, Jeffrey Barrett, Sonia Goncalves, Ewan Harrison, David K. Jackson, Ian Johnston, Dominic Kwiatkowski, Cordelia Langford, John Sillitoe on behalf of the Wellcome Sanger Institute COVID-19 Surveillance Team |
| EPI_ISL_2830046 | Lighthouse Lab in Milton Keynes | Wellcome Sanger Institute for the COVID-19 Genomics UK (COG-UK) Consortium | The Lighthouse Lab in Milton Keynes and Alex Alderton, Roberto Amato, Jeffrey Barrett, Sonia Goncalves, Ewan Harrison, David K. Jackson, Ian Johnston, Dominic Kwiatkowski, Cordelia Langford, John Sillitoe on behalf of the Wellcome Sanger Institute COVID-19 Surveillance Team |
| EPI_ISL_2830047, EPI_ISL_2830048, EPI_ISL_2830049 | Lighthouse Lab in Alderley Park | Wellcome Sanger Institute for the COVID-19 Genomics UK (COG-UK) Consortium | Jacquelyn Wynn, Mairead Hyland, The Lighthouse Lab in Alderley Park and Alex Alderton, Roberto Amato, Jeffrey Barrett, Sonia Goncalves, Ewan Harrison, David K. Jackson, Ian Johnston, Dominic Kwiatkowski, Cordelia Langford, John Sillitoe on behalf of the Wellcome Sanger Institute COVID-19 Surveillance Team |
| EPI_ISL_2830050 | Lighthouse Lab in Milton Keynes | Wellcome Sanger Institute for the COVID-19 Genomics UK (COG-UK) Consortium | The Lighthouse Lab in Milton Keynes and Alex Alderton, Roberto Amato, Jeffrey Barrett, Sonia Goncalves, Ewan Harrison, David K. Jackson, Ian Johnston, Dominic Kwiatkowski, Cordelia Langford, John Sillitoe on behalf of the Wellcome Sanger Institute COVID-19 Surveillance Team |
| EPI_ISL_2830051 | Lighthouse Lab in Glasgow | Wellcome Sanger Institute for the COVID-19 Genomics UK (COG-UK) Consortium | Harper VanSteenhouse, Yumi Kasai, David Gray, Carol Clugston, Anna Dominiczak and Alex Alderton, Roberto Amato, Jeffrey Barrett, Sonia Goncalves, Ewan Harrison, David K. Jackson, Ian Johnston, Dominic Kwiatkowski, Cordelia Langford, John Sillitoe on behalf of the Wellcome Sanger Institute COVID-19 Surveillance Team |
| EPI_ISL_2830052, EPI_ISL_2830053 | Lighthouse Lab in Milton Keynes | Wellcome Sanger Institute for the COVID-19 Genomics UK (COG-UK) Consortium | The Lighthouse Lab in Milton Keynes and Alex Alderton, Roberto Amato, Jeffrey Barrett, Sonia Goncalves, Ewan Harrison, David K. Jackson, Ian Johnston, Dominic Kwiatkowski, Cordelia Langford, John Sillitoe on behalf of the Wellcome Sanger Institute COVID-19 Surveillance Team |

| | | | |
|---|---|---|---|
| EPI_ISL_2830054 | Lighthouse Lab in Alderley Park | Wellcome Sanger Institute for the COVID-19 Genomics UK (COG-UK) Consortium | Jacquelyn Wynn, Mairead Hyland, The Lighthouse Lab in Alderley Park and Alex Alderton, Roberto Amato, Jeffrey Barrett, Sonia Goncalves, Ewan Harrison, David K. Jackson, Ian Johnston, Dominic Kwiatkowski, Cordelia Langford, John Sillitoe on behalf of the Wellcome Sanger Institute COVID-19 Surveillance Team |
| EPI_ISL_2830055 | Lighthouse Lab in Glasgow | Wellcome Sanger Institute for the COVID-19 Genomics UK (COG-UK) Consortium | Harper VanSteenhouse, Yumi Kasai, David Gray, Carol Clugston, Anna Dominiczak and Alex Alderton, Roberto Amato, Jeffrey Barrett, Sonia Goncalves, Ewan Harrison, David K. Jackson, Ian Johnston, Dominic Kwiatkowski, Cordelia Langford, John Sillitoe on behalf of the Wellcome Sanger Institute COVID-19 Surveillance Team |
| EPI_ISL_2830056 | Lighthouse Lab in Alderley Park | Wellcome Sanger Institute for the COVID-19 Genomics UK (COG-UK) Consortium | Jacquelyn Wynn, Mairead Hyland, The Lighthouse Lab in Alderley Park and Alex Alderton, Roberto Amato, Jeffrey Barrett, Sonia Goncalves, Ewan Harrison, David K. Jackson, Ian Johnston, Dominic Kwiatkowski, Cordelia Langford, John Sillitoe on behalf of the Wellcome Sanger Institute COVID-19 Surveillance Team |
| EPI_ISL_2830057, EPI_ISL_2830058 | Lighthouse Lab in Milton Keynes | Wellcome Sanger Institute for the COVID-19 Genomics UK (COG-UK) Consortium | The Lighthouse Lab in Milton Keynes and Alex Alderton, Roberto Amato, Jeffrey Barrett, Sonia Goncalves, Ewan Harrison, David K. Jackson, Ian Johnston, Dominic Kwiatkowski, Cordelia Langford, John Sillitoe on behalf of the Wellcome Sanger Institute COVID-19 Surveillance Team |
| EPI_ISL_2830059, EPI_ISL_2830060 | Lighthouse Lab in Alderley Park | Wellcome Sanger Institute for the COVID-19 Genomics UK (COG-UK) Consortium | Jacquelyn Wynn, Mairead Hyland, The Lighthouse Lab in Alderley Park and Alex Alderton, Roberto Amato, Jeffrey Barrett, Sonia Goncalves, Ewan Harrison, David K. Jackson, Ian Johnston, Dominic Kwiatkowski, Cordelia Langford, John Sillitoe on behalf of the Wellcome Sanger Institute COVID-19 Surveillance Team |
| EPI_ISL_2830061 | Lighthouse Lab in Milton Keynes | Wellcome Sanger Institute for the COVID-19 Genomics UK (COG-UK) Consortium | The Lighthouse Lab in Milton Keynes and Alex Alderton, Roberto Amato, Jeffrey Barrett, Sonia Goncalves, Ewan Harrison, David K. Jackson, Ian Johnston, Dominic Kwiatkowski, Cordelia Langford, John Sillitoe on behalf of the Wellcome Sanger Institute COVID-19 Surveillance Team |
| EPI_ISL_2830062 | Lighthouse Lab in Alderley Park | Wellcome Sanger Institute for the COVID-19 Genomics UK (COG-UK) Consortium | Jacquelyn Wynn, Mairead Hyland, The Lighthouse Lab in Alderley Park and Alex Alderton, Roberto Amato, Jeffrey Barrett, Sonia Goncalves, Ewan Harrison, David K. Jackson, Ian Johnston, Dominic Kwiatkowski, Cordelia Langford, John Sillitoe on behalf of the Wellcome Sanger Institute COVID-19 Surveillance Team |
| EPI_ISL_2830063 | Lighthouse Lab in Milton Keynes | Wellcome Sanger Institute for the COVID-19 Genomics UK (COG-UK) Consortium | The Lighthouse Lab in Milton Keynes and Alex Alderton, Roberto Amato, Jeffrey Barrett, Sonia Goncalves, Ewan Harrison, David K. Jackson, Ian Johnston, Dominic Kwiatkowski, Cordelia Langford, John Sillitoe on behalf of the Wellcome Sanger Institute COVID-19 Surveillance Team |
| EPI_ISL_2830064, EPI_ISL_2830065 | Lighthouse Lab in Alderley Park | Wellcome Sanger Institute for the COVID-19 Genomics UK (COG-UK) Consortium | Jacquelyn Wynn, Mairead Hyland, The Lighthouse Lab in Alderley Park and Alex Alderton, Roberto Amato, Jeffrey Barrett, Sonia Goncalves, Ewan Harrison, David K. Jackson, Ian Johnston, Dominic Kwiatkowski, Cordelia Langford, John Sillitoe on behalf of the Wellcome Sanger Institute COVID-19 Surveillance Team |
| EPI_ISL_2830066 | Lighthouse Lab in Milton Keynes | Wellcome Sanger Institute for the COVID-19 Genomics UK (COG-UK) Consortium | The Lighthouse Lab in Milton Keynes and Alex Alderton, Roberto Amato, Jeffrey Barrett, Sonia Goncalves, Ewan Harrison, David K. Jackson, Ian Johnston, Dominic Kwiatkowski, Cordelia Langford, John Sillitoe on behalf of the Wellcome Sanger Institute COVID-19 Surveillance Team |
| EPI_ISL_2830067, EPI_ISL_2830068 | Lighthouse Lab in Alderley Park | Wellcome Sanger Institute for the COVID-19 Genomics UK (COG-UK) Consortium | Jacquelyn Wynn, Mairead Hyland, The Lighthouse Lab in Alderley Park and Alex Alderton, Roberto Amato, Jeffrey Barrett, Sonia Goncalves, Ewan Harrison, David K. Jackson, Ian Johnston, Dominic Kwiatkowski, Cordelia Langford, John Sillitoe on behalf of the Wellcome Sanger Institute COVID-19 Surveillance Team |
| EPI_ISL_2830069, EPI_ISL_2830070 | Lighthouse Lab in Milton Keynes | Wellcome Sanger Institute for the COVID-19 Genomics UK (COG-UK) Consortium | The Lighthouse Lab in Milton Keynes and Alex Alderton, Roberto Amato, Jeffrey Barrett, Sonia Goncalves, Ewan Harrison, David K. Jackson, Ian Johnston, Dominic Kwiatkowski, Cordelia Langford, John Sillitoe on behalf of the Wellcome Sanger Institute COVID-19 Surveillance Team |
| EPI_ISL_2830071 | Lighthouse Lab in Glasgow | Wellcome Sanger Institute for the COVID-19 Genomics UK (COG-UK) Consortium | Harper VanSteenhouse, Yumi Kasai, David Gray, Carol Clugston, Anna Dominiczak and Alex Alderton, Roberto Amato, Jeffrey Barrett, Sonia Goncalves, Ewan Harrison, David K. Jackson, Ian Johnston, Dominic Kwiatkowski, Cordelia Langford, John Sillitoe on behalf of the Wellcome Sanger Institute COVID-19 Surveillance Team |
| EPI_ISL_2830072, EPI_ISL_2830073, EPI_ISL_2830074, EPI_ISL_2830075 | Lighthouse Lab in Milton Keynes | Wellcome Sanger Institute for the COVID-19 Genomics UK (COG-UK) Consortium | The Lighthouse Lab in Milton Keynes and Alex Alderton, Roberto Amato, Jeffrey Barrett, Sonia Goncalves, Ewan Harrison, David K. Jackson, Ian Johnston, Dominic Kwiatkowski, Cordelia Langford, John Sillitoe on behalf of the Wellcome Sanger Institute COVID-19 Surveillance Team |
| EPI_ISL_2830076 | Lighthouse Lab in Glasgow | Wellcome Sanger Institute for the COVID-19 Genomics UK (COG-UK) Consortium | Harper VanSteenhouse, Yumi Kasai, David Gray, Carol Clugston, Anna Dominiczak and Alex Alderton, Roberto Amato, Jeffrey Barrett, Sonia Goncalves, Ewan Harrison, David K. Jackson, Ian Johnston, Dominic Kwiatkowski, Cordelia Langford, John Sillitoe on behalf of the Wellcome Sanger Institute COVID-19 Surveillance Team |
| EPI_ISL_2830077, EPI_ISL_2830078, EPI_ISL_2830079 | Lighthouse Lab in Alderley Park | Wellcome Sanger Institute for the COVID-19 Genomics UK (COG-UK) Consortium | Jacquelyn Wynn, Mairead Hyland, The Lighthouse Lab in Alderley Park and Alex Alderton, Roberto Amato, Jeffrey Barrett, Sonia Goncalves, Ewan Harrison, David K. Jackson, Ian Johnston, Dominic Kwiatkowski, Cordelia Langford, John Sillitoe on behalf of the Wellcome Sanger Institute COVID-19 Surveillance Team |
| EPI_ISL_2830080 | Lighthouse Lab in Milton Keynes | Wellcome Sanger Institute for the COVID-19 Genomics UK (COG-UK) Consortium | The Lighthouse Lab in Milton Keynes and Alex Alderton, Roberto Amato, Jeffrey Barrett, Sonia Goncalves, Ewan Harrison, David K. Jackson, Ian Johnston, Dominic Kwiatkowski, Cordelia Langford, John Sillitoe on behalf of the Wellcome Sanger Institute COVID-19 Surveillance Team |
| EPI_ISL_2830081, EPI_ISL_2830082 | Lighthouse Lab in Alderley Park | Wellcome Sanger Institute for the COVID-19 Genomics UK (COG-UK) Consortium | Jacquelyn Wynn, Mairead Hyland, The Lighthouse Lab in Alderley Park and Alex Alderton, Roberto Amato, Jeffrey Barrett, Sonia Goncalves, Ewan Harrison, David K. Jackson, Ian Johnston, Dominic Kwiatkowski, Cordelia Langford, John Sillitoe on behalf of the Wellcome Sanger Institute COVID-19 Surveillance Team |
| EPI_ISL_2830083 | Lighthouse Lab in Glasgow | Wellcome Sanger Institute for the COVID-19 Genomics UK (COG-UK) Consortium | Harper VanSteenhouse, Yumi Kasai, David Gray, Carol Clugston, Anna Dominiczak and Alex Alderton, Roberto Amato, Jeffrey Barrett, Sonia Goncalves, Ewan Harrison, David K. Jackson, Ian Johnston, Dominic Kwiatkowski, Cordelia Langford, John Sillitoe on behalf of the Wellcome Sanger Institute COVID-19 Surveillance Team |
| EPI_ISL_2830084 | Lighthouse Lab in Alderley Park | Wellcome Sanger Institute for the COVID-19 Genomics UK (COG-UK) Consortium | Jacquelyn Wynn, Mairead Hyland, The Lighthouse Lab in Alderley Park and Alex Alderton, Roberto Amato, Jeffrey Barrett, Sonia Goncalves, Ewan Harrison, David K. Jackson, Ian Johnston, Dominic Kwiatkowski, Cordelia Langford, John Sillitoe on behalf of the Wellcome Sanger Institute COVID-19 Surveillance Team |
| EPI_ISL_2830085 | Lighthouse Lab in Milton Keynes | Wellcome Sanger Institute for the COVID-19 Genomics UK (COG-UK) Consortium | The Lighthouse Lab in Milton Keynes and Alex Alderton, Roberto Amato, Jeffrey Barrett, Sonia Goncalves, Ewan Harrison, David K. Jackson, Ian Johnston, Dominic Kwiatkowski, Cordelia Langford, John Sillitoe on behalf of the Wellcome Sanger Institute COVID-19 Surveillance Team |
| EPI_ISL_2830086 | Lighthouse Lab in Glasgow | Wellcome Sanger Institute for the COVID-19 Genomics UK (COG-UK) Consortium | Harper VanSteenhouse, Yumi Kasai, David Gray, Carol Clugston, Anna Dominiczak and Alex Alderton, Roberto Amato, Jeffrey Barrett, Sonia Goncalves, Ewan Harrison, David K. Jackson, Ian Johnston, Dominic Kwiatkowski, Cordelia Langford, John Sillitoe on behalf of the Wellcome Sanger Institute COVID-19 Surveillance Team |
| EPI_ISL_2830087, EPI_ISL_2830088, EPI_ISL_2830089, EPI_ISL_2830090 | Lighthouse Lab in Alderley Park | Wellcome Sanger Institute for the COVID-19 Genomics UK (COG-UK) Consortium | Jacquelyn Wynn, Mairead Hyland, The Lighthouse Lab in Alderley Park and Alex Alderton, Roberto Amato, Jeffrey Barrett, Sonia Goncalves, Ewan Harrison, David K. Jackson, Ian Johnston, Dominic Kwiatkowski, Cordelia Langford, John Sillitoe on behalf of the Wellcome Sanger Institute COVID-19 Surveillance Team |
| EPI_ISL_2830091, EPI_ISL_2830092, EPI_ISL_2830093 | Lighthouse Lab in Milton Keynes | Wellcome Sanger Institute for the COVID-19 Genomics UK (COG-UK) Consortium | The Lighthouse Lab in Milton Keynes and Alex Alderton, Roberto Amato, Jeffrey Barrett, Sonia Goncalves, Ewan Harrison, David K. Jackson, Ian Johnston, Dominic Kwiatkowski, Cordelia Langford, John Sillitoe on behalf of the Wellcome Sanger Institute COVID-19 Surveillance Team |
| EPI_ISL_2830094 | Lighthouse Lab in Glasgow | Wellcome Sanger Institute for the COVID-19 Genomics UK (COG-UK) Consortium | Harper VanSteenhouse, Yumi Kasai, David Gray, Carol Clugston, Anna Dominiczak and Alex Alderton, Roberto Amato, Jeffrey Barrett, Sonia Goncalves, Ewan Harrison, David K. Jackson, Ian Johnston, Dominic Kwiatkowski, Cordelia Langford, John Sillitoe on behalf of the Wellcome Sanger Institute COVID-19 Surveillance Team |
| EPI_ISL_2830095, EPI_ISL_2830096 | Lighthouse Lab in Milton Keynes | Wellcome Sanger Institute for the COVID-19 Genomics UK (COG-UK) Consortium | The Lighthouse Lab in Milton Keynes and Alex Alderton, Roberto Amato, Jeffrey Barrett, Sonia Goncalves, Ewan Harrison, David K. Jackson, Ian Johnston, Dominic Kwiatkowski, Cordelia Langford, John Sillitoe on behalf of the Wellcome Sanger Institute COVID-19 Surveillance Team |
| EPI_ISL_2830097 | Lighthouse Lab in Alderley Park | Wellcome Sanger Institute for the COVID-19 Genomics UK (COG-UK) Consortium | Jacquelyn Wynn, Mairead Hyland, The Lighthouse Lab in Alderley Park and Alex Alderton, Roberto Amato, Jeffrey Barrett, Sonia Goncalves, Ewan Harrison, David K. Jackson, Ian Johnston, Dominic Kwiatkowski, Cordelia Langford, John Sillitoe on behalf of the Wellcome Sanger Institute COVID-19 Surveillance Team |
| EPI_ISL_2830098 | Lighthouse Lab in Milton Keynes | Wellcome Sanger Institute for the COVID-19 Genomics UK (COG-UK) Consortium | The Lighthouse Lab in Milton Keynes and Alex Alderton, Roberto Amato, Jeffrey Barrett, Sonia Goncalves, Ewan Harrison, David K. Jackson, Ian Johnston, Dominic Kwiatkowski, Cordelia Langford, John Sillitoe on behalf of the Wellcome Sanger Institute COVID-19 Surveillance Team |
| EPI_ISL_2830099, EPI_ISL_2830100 | Lighthouse Lab in Alderley Park | Wellcome Sanger Institute for the COVID-19 Genomics UK (COG-UK) Consortium | Jacquelyn Wynn, Mairead Hyland, The Lighthouse Lab in Alderley Park and Alex Alderton, Roberto Amato, Jeffrey Barrett, Sonia Goncalves, Ewan Harrison, David K. Jackson, Ian Johnston, Dominic Kwiatkowski, Cordelia Langford, John Sillitoe on behalf of the Wellcome Sanger Institute COVID-19 Surveillance Team |
| EPI_ISL_2830101, EPI_ISL_2830102 | Lighthouse Lab in Glasgow | Wellcome Sanger Institute for the COVID-19 Genomics UK (COG-UK) Consortium | Harper VanSteenhouse, Yumi Kasai, David Gray, Carol Clugston, Anna Dominiczak and Alex Alderton, Roberto Amato, Jeffrey Barrett, Sonia Goncalves, Ewan Harrison, David K. Jackson, Ian Johnston, Dominic Kwiatkowski, Cordelia Langford, John Sillitoe on behalf of the Wellcome Sanger Institute COVID-19 Surveillance Team |
| EPI_ISL_2830103 | Lighthouse Lab in Milton Keynes | Wellcome Sanger Institute for the COVID-19 Genomics UK (COG-UK) Consortium | The Lighthouse Lab in Milton Keynes and Alex Alderton, Roberto Amato, Jeffrey Barrett, Sonia Goncalves, Ewan Harrison, David K. Jackson, Ian Johnston, Dominic Kwiatkowski, Cordelia Langford, John Sillitoe on behalf of the Wellcome Sanger Institute COVID-19 Surveillance Team |
| EPI_ISL_2830104, EPI_ISL_2830105 | Lighthouse Lab in Alderley Park | Wellcome Sanger Institute for the COVID-19 Genomics UK (COG-UK) Consortium | Jacquelyn Wynn, Mairead Hyland, The Lighthouse Lab in Alderley Park and Alex Alderton, Roberto Amato, Jeffrey Barrett, Sonia Goncalves, Ewan Harrison, David K. Jackson, Ian Johnston, Dominic Kwiatkowski, Cordelia Langford, John Sillitoe on behalf of the Wellcome Sanger Institute COVID-19 Surveillance Team |
| EPI_ISL_2830106 | Lighthouse Lab in Glasgow | Wellcome Sanger Institute for the COVID-19 Genomics UK (COG-UK) Consortium | Harper VanSteenhouse, Yumi Kasai, David Gray, Carol Clugston, Anna Dominiczak and Alex Alderton, Roberto Amato, Jeffrey Barrett, Sonia Goncalves, Ewan Harrison, David K. Jackson, Ian Johnston, Dominic Kwiatkowski, Cordelia Langford, John Sillitoe on behalf of the Wellcome Sanger Institute COVID-19 Surveillance Team |
| EPI_ISL_2830107, EPI_ISL_2830108, EPI_ISL_2830109 | Lighthouse Lab in Alderley Park | Wellcome Sanger Institute for the COVID-19 Genomics UK (COG-UK) Consortium | Jacquelyn Wynn, Mairead Hyland, The Lighthouse Lab in Alderley Park and Alex Alderton, Roberto Amato, Jeffrey Barrett, Sonia Goncalves, Ewan Harrison, David K. Jackson, Ian Johnston, Dominic Kwiatkowski, Cordelia Langford, John Sillitoe on behalf of the Wellcome Sanger Institute COVID-19 Surveillance Team |
| EPI_ISL_2830110, EPI_ISL_2830111, EPI_ISL_2830112 | Lighthouse Lab in Milton Keynes | Wellcome Sanger Institute for the COVID-19 Genomics UK (COG-UK) Consortium | The Lighthouse Lab in Milton Keynes and Alex Alderton, Roberto Amato, Jeffrey Barrett, Sonia Goncalves, Ewan Harrison, David K. Jackson, Ian Johnston, Dominic Kwiatkowski, Cordelia Langford, John Sillitoe on behalf of the Wellcome Sanger Institute COVID-19 Surveillance Team |
| EPI_ISL_2830113 | Lighthouse Lab in Glasgow | Wellcome Sanger Institute for the COVID-19 Genomics UK (COG-UK) Consortium | Harper VanSteenhouse, Yumi Kasai, David Gray, Carol Clugston, Anna Dominiczak and Alex Alderton, Roberto Amato, Jeffrey Barrett, Sonia Goncalves, Ewan Harrison, David K. Jackson, Ian Johnston, Dominic Kwiatkowski, Cordelia Langford, John Sillitoe on behalf of the Wellcome Sanger Institute COVID-19 Surveillance Team |
| EPI_ISL_2830114, EPI_ISL_2830115, EPI_ISL_2830116 | Lighthouse Lab in Alderley Park | Wellcome Sanger Institute for the COVID-19 Genomics UK (COG-UK) Consortium | Jacquelyn Wynn, Mairead Hyland, The Lighthouse Lab in Alderley Park and Alex Alderton, Roberto Amato, Jeffrey Barrett, Sonia Goncalves, Ewan Harrison, David K. Jackson, Ian Johnston, Dominic Kwiatkowski, Cordelia Langford, John Sillitoe on behalf of the Wellcome Sanger Institute COVID-19 Surveillance Team |
| EPI_ISL_2830117 | Lighthouse Lab in Milton Keynes | Wellcome Sanger Institute for the COVID-19 Genomics UK (COG-UK) Consortium | The Lighthouse Lab in Milton Keynes and Alex Alderton, Roberto Amato, Jeffrey Barrett, Sonia Goncalves, Ewan Harrison, David K. Jackson, Ian Johnston, Dominic Kwiatkowski, Cordelia Langford, John Sillitoe on behalf of the Wellcome Sanger Institute COVID-19 Surveillance Team |
| EPI_ISL_2830118, EPI_ISL_2830119 | Lighthouse Lab in Alderley Park | Wellcome Sanger Institute for the COVID-19 Genomics UK (COG-UK) Consortium | Jacquelyn Wynn, Mairead Hyland, The Lighthouse Lab in Alderley Park and Alex Alderton, Roberto Amato, Jeffrey Barrett, Sonia Goncalves, Ewan Harrison, David K. Jackson, Ian Johnston, Dominic Kwiatkowski, Cordelia Langford, John Sillitoe on behalf of the Wellcome Sanger Institute COVID-19 Surveillance Team |
| EPI_ISL_2830120 | Lighthouse Lab in Milton Keynes | Wellcome Sanger Institute for the COVID-19 Genomics UK (COG-UK) Consortium | The Lighthouse Lab in Milton Keynes and Alex Alderton, Roberto Amato, Jeffrey Barrett, Sonia Goncalves, Ewan Harrison, David K. Jackson, Ian Johnston, Dominic Kwiatkowski, Cordelia Langford, John Sillitoe on behalf of the Wellcome Sanger Institute COVID-19 Surveillance Team |
| EPI_ISL_2830121 | Lighthouse Lab in Glasgow | Wellcome Sanger Institute for the COVID-19 Genomics UK (COG-UK) Consortium | Harper VanSteenhouse, Yumi Kasai, David Gray, Carol Clugston, Anna Dominiczak and Alex Alderton, Roberto Amato, Jeffrey Barrett, Sonia Goncalves, Ewan Harrison, David K. Jackson, Ian Johnston, Dominic Kwiatkowski, Cordelia Langford, John Sillitoe on behalf of the Wellcome Sanger Institute COVID-19 Surveillance Team |

| | | | |
|---|---|---|---|
| EPI_ISL_2830122 | Lighthouse Lab in Milton Keynes | Wellcome Sanger Institute for the COVID-19 Genomics UK (COG-UK) Consortium | The Lighthouse Lab in Milton Keynes and Alex Alderton, Roberto Amato, Jeffrey Barrett, Sonia Goncalves, Ewan Harrison, David K. Jackson, Ian Johnston, Dominic Kwiatkowski, Cordelia Langford, John Sillitoe on behalf of the Wellcome Sanger Institute COVID-19 Surveillance Team |
| EPI_ISL_2830123 | Lighthouse Lab in Alderley Park | Wellcome Sanger Institute for the COVID-19 Genomics UK (COG-UK) Consortium | Jacquelyn Wynn, Mairead Hyland, The Lighthouse Lab in Alderley Park and Alex Alderton, Roberto Amato, Jeffrey Barrett, Sonia Goncalves, Ewan Harrison, David K. Jackson, Ian Johnston, Dominic Kwiatkowski, Cordelia Langford, John Sillitoe on behalf of the Wellcome Sanger Institute COVID-19 Surveillance Team |
| EPI_ISL_2830124 | Lighthouse Lab in Milton Keynes | Wellcome Sanger Institute for the COVID-19 Genomics UK (COG-UK) Consortium | The Lighthouse Lab in Milton Keynes and Alex Alderton, Roberto Amato, Jeffrey Barrett, Sonia Goncalves, Ewan Harrison, David K. Jackson, Ian Johnston, Dominic Kwiatkowski, Cordelia Langford, John Sillitoe on behalf of the Wellcome Sanger Institute COVID-19 Surveillance Team |
| EPI_ISL_2830125 | Lighthouse Lab in Alderley Park | Wellcome Sanger Institute for the COVID-19 Genomics UK (COG-UK) Consortium | Jacquelyn Wynn, Mairead Hyland, The Lighthouse Lab in Alderley Park and Alex Alderton, Roberto Amato, Jeffrey Barrett, Sonia Goncalves, Ewan Harrison, David K. Jackson, Ian Johnston, Dominic Kwiatkowski, Cordelia Langford, John Sillitoe on behalf of the Wellcome Sanger Institute COVID-19 Surveillance Team |
| EPI_ISL_2830126, EPI_ISL_2830127 | Lighthouse Lab in Milton Keynes | Wellcome Sanger Institute for the COVID-19 Genomics UK (COG-UK) Consortium | The Lighthouse Lab in Milton Keynes and Alex Alderton, Roberto Amato, Jeffrey Barrett, Sonia Goncalves, Ewan Harrison, David K. Jackson, Ian Johnston, Dominic Kwiatkowski, Cordelia Langford, John Sillitoe on behalf of the Wellcome Sanger Institute COVID-19 Surveillance Team |
| EPI_ISL_2830128 | Lighthouse Lab in Alderley Park | Wellcome Sanger Institute for the COVID-19 Genomics UK (COG-UK) Consortium | Jacquelyn Wynn, Mairead Hyland, The Lighthouse Lab in Alderley Park and Alex Alderton, Roberto Amato, Jeffrey Barrett, Sonia Goncalves, Ewan Harrison, David K. Jackson, Ian Johnston, Dominic Kwiatkowski, Cordelia Langford, John Sillitoe on behalf of the Wellcome Sanger Institute COVID-19 Surveillance Team |
| EPI_ISL_2830129 | Lighthouse Lab in Glasgow | Wellcome Sanger Institute for the COVID-19 Genomics UK (COG-UK) Consortium | Harper VanSteenhouse, Yumi Kasai, David Gray, Carol Clugston, Anna Dominiczak and Alex Alderton, Roberto Amato, Jeffrey Barrett, Sonia Goncalves, Ewan Harrison, David K. Jackson, Ian Johnston, Dominic Kwiatkowski, Cordelia Langford, John Sillitoe on behalf of the Wellcome Sanger Institute COVID-19 Surveillance Team |
| EPI_ISL_2830130 | Lighthouse Lab in Milton Keynes | Wellcome Sanger Institute for the COVID-19 Genomics UK (COG-UK) Consortium | The Lighthouse Lab in Milton Keynes and Alex Alderton, Roberto Amato, Jeffrey Barrett, Sonia Goncalves, Ewan Harrison, David K. Jackson, Ian Johnston, Dominic Kwiatkowski, Cordelia Langford, John Sillitoe on behalf of the Wellcome Sanger Institute COVID-19 Surveillance Team |
| EPI_ISL_2830131 | Lighthouse Lab in Glasgow | Wellcome Sanger Institute for the COVID-19 Genomics UK (COG-UK) Consortium | Harper VanSteenhouse, Yumi Kasai, David Gray, Carol Clugston, Anna Dominiczak and Alex Alderton, Roberto Amato, Jeffrey Barrett, Sonia Goncalves, Ewan Harrison, David K. Jackson, Ian Johnston, Dominic Kwiatkowski, Cordelia Langford, John Sillitoe on behalf of the Wellcome Sanger Institute COVID-19 Surveillance Team |
| EPI_ISL_2830132, EPI_ISL_2830133 | Lighthouse Lab in Milton Keynes | Wellcome Sanger Institute for the COVID-19 Genomics UK (COG-UK) Consortium | The Lighthouse Lab in Milton Keynes and Alex Alderton, Roberto Amato, Jeffrey Barrett, Sonia Goncalves, Ewan Harrison, David K. Jackson, Ian Johnston, Dominic Kwiatkowski, Cordelia Langford, John Sillitoe on behalf of the Wellcome Sanger Institute COVID-19 Surveillance Team |
| EPI_ISL_2830134, EPI_ISL_2830135 | Lighthouse Lab in Alderley Park | Wellcome Sanger Institute for the COVID-19 Genomics UK (COG-UK) Consortium | Jacquelyn Wynn, Mairead Hyland, The Lighthouse Lab in Alderley Park and Alex Alderton, Roberto Amato, Jeffrey Barrett, Sonia Goncalves, Ewan Harrison, David K. Jackson, Ian Johnston, Dominic Kwiatkowski, Cordelia Langford, John Sillitoe on behalf of the Wellcome Sanger Institute COVID-19 Surveillance Team |
| EPI_ISL_2830136, EPI_ISL_2830137, EPI_ISL_2830138 | Lighthouse Lab in Milton Keynes | Wellcome Sanger Institute for the COVID-19 Genomics UK (COG-UK) Consortium | The Lighthouse Lab in Milton Keynes and Alex Alderton, Roberto Amato, Jeffrey Barrett, Sonia Goncalves, Ewan Harrison, David K. Jackson, Ian Johnston, Dominic Kwiatkowski, Cordelia Langford, John Sillitoe on behalf of the Wellcome Sanger Institute COVID-19 Surveillance Team |
| EPI_ISL_2830139 | Lighthouse Lab in Glasgow | Wellcome Sanger Institute for the COVID-19 Genomics UK (COG-UK) Consortium | Harper VanSteenhouse, Yumi Kasai, David Gray, Carol Clugston, Anna Dominiczak and Alex Alderton, Roberto Amato, Jeffrey Barrett, Sonia Goncalves, Ewan Harrison, David K. Jackson, Ian Johnston, Dominic Kwiatkowski, Cordelia Langford, John Sillitoe on behalf of the Wellcome Sanger Institute COVID-19 Surveillance Team |
| EPI_ISL_2830140 | Lighthouse Lab in Milton Keynes | Wellcome Sanger Institute for the COVID-19 Genomics UK (COG-UK) Consortium | The Lighthouse Lab in Milton Keynes and Alex Alderton, Roberto Amato, Jeffrey Barrett, Sonia Goncalves, Ewan Harrison, David K. Jackson, Ian Johnston, Dominic Kwiatkowski, Cordelia Langford, John Sillitoe on behalf of the Wellcome Sanger Institute COVID-19 Surveillance Team |
| EPI_ISL_2830141 | Lighthouse Lab in Glasgow | Wellcome Sanger Institute for the COVID-19 Genomics UK (COG-UK) Consortium | Harper VanSteenhouse, Yumi Kasai, David Gray, Carol Clugston, Anna Dominiczak and Alex Alderton, Roberto Amato, Jeffrey Barrett, Sonia Goncalves, Ewan Harrison, David K. Jackson, Ian Johnston, Dominic Kwiatkowski, Cordelia Langford, John Sillitoe on behalf of the Wellcome Sanger Institute COVID-19 Surveillance Team |
| EPI_ISL_2830142 | Lighthouse Lab in Milton Keynes | Wellcome Sanger Institute for the COVID-19 Genomics UK (COG-UK) Consortium | The Lighthouse Lab in Milton Keynes and Alex Alderton, Roberto Amato, Jeffrey Barrett, Sonia Goncalves, Ewan Harrison, David K. Jackson, Ian Johnston, Dominic Kwiatkowski, Cordelia Langford, John Sillitoe on behalf of the Wellcome Sanger Institute COVID-19 Surveillance Team |
| EPI_ISL_2830143, EPI_ISL_2830144 | Lighthouse Lab in Alderley Park | Wellcome Sanger Institute for the COVID-19 Genomics UK (COG-UK) Consortium | Jacquelyn Wynn, Mairead Hyland, The Lighthouse Lab in Alderley Park and Alex Alderton, Roberto Amato, Jeffrey Barrett, Sonia Goncalves, Ewan Harrison, David K. Jackson, Ian Johnston, Dominic Kwiatkowski, Cordelia Langford, John Sillitoe on behalf of the Wellcome Sanger Institute COVID-19 Surveillance Team |
| EPI_ISL_2830145 | Lighthouse Lab in Milton Keynes | Wellcome Sanger Institute for the COVID-19 Genomics UK (COG-UK) Consortium | The Lighthouse Lab in Milton Keynes and Alex Alderton, Roberto Amato, Jeffrey Barrett, Sonia Goncalves, Ewan Harrison, David K. Jackson, Ian Johnston, Dominic Kwiatkowski, Cordelia Langford, John Sillitoe on behalf of the Wellcome Sanger Institute COVID-19 Surveillance Team |
| EPI_ISL_2830146 | Lighthouse Lab in Glasgow | Wellcome Sanger Institute for the COVID-19 Genomics UK (COG-UK) Consortium | Harper VanSteenhouse, Yumi Kasai, David Gray, Carol Clugston, Anna Dominiczak and Alex Alderton, Roberto Amato, Jeffrey Barrett, Sonia Goncalves, Ewan Harrison, David K. Jackson, Ian Johnston, Dominic Kwiatkowski, Cordelia Langford, John Sillitoe on behalf of the Wellcome Sanger Institute COVID-19 Surveillance Team |
| EPI_ISL_2830147, EPI_ISL_2830148, EPI_ISL_2830149 | Lighthouse Lab in Alderley Park | Wellcome Sanger Institute for the COVID-19 Genomics UK (COG-UK) Consortium | Jacquelyn Wynn, Mairead Hyland, The Lighthouse Lab in Alderley Park and Alex Alderton, Roberto Amato, Jeffrey Barrett, Sonia Goncalves, Ewan Harrison, David K. Jackson, Ian Johnston, Dominic Kwiatkowski, Cordelia Langford, John Sillitoe on behalf of the Wellcome Sanger Institute COVID-19 Surveillance Team |
| EPI_ISL_2830150, EPI_ISL_2830151 | Lighthouse Lab in Milton Keynes | Wellcome Sanger Institute for the COVID-19 Genomics UK (COG-UK) Consortium | The Lighthouse Lab in Milton Keynes and Alex Alderton, Roberto Amato, Jeffrey Barrett, Sonia Goncalves, Ewan Harrison, David K. Jackson, Ian Johnston, Dominic Kwiatkowski, Cordelia Langford, John Sillitoe on behalf of the Wellcome Sanger Institute COVID-19 Surveillance Team |
| EPI_ISL_2830152 | Lighthouse Lab in Glasgow | Wellcome Sanger Institute for the COVID-19 Genomics UK (COG-UK) Consortium | Harper VanSteenhouse, Yumi Kasai, David Gray, Carol Clugston, Anna Dominiczak and Alex Alderton, Roberto Amato, Jeffrey Barrett, Sonia Goncalves, Ewan Harrison, David K. Jackson, Ian Johnston, Dominic Kwiatkowski, Cordelia Langford, John Sillitoe on behalf of the Wellcome Sanger Institute COVID-19 Surveillance Team |
| EPI_ISL_2830153 | Lighthouse Lab in Milton Keynes | Wellcome Sanger Institute for the COVID-19 Genomics UK (COG-UK) Consortium | The Lighthouse Lab in Milton Keynes and Alex Alderton, Roberto Amato, Jeffrey Barrett, Sonia Goncalves, Ewan Harrison, David K. Jackson, Ian Johnston, Dominic Kwiatkowski, Cordelia Langford, John Sillitoe on behalf of the Wellcome Sanger Institute COVID-19 Surveillance Team |
| EPI_ISL_2830154 | Lighthouse Lab in Alderley Park | Wellcome Sanger Institute for the COVID-19 Genomics UK (COG-UK) Consortium | Jacquelyn Wynn, Mairead Hyland, The Lighthouse Lab in Alderley Park and Alex Alderton, Roberto Amato, Jeffrey Barrett, Sonia Goncalves, Ewan Harrison, David K. Jackson, Ian Johnston, Dominic Kwiatkowski, Cordelia Langford, John Sillitoe on behalf of the Wellcome Sanger Institute COVID-19 Surveillance Team |
| EPI_ISL_2830155, EPI_ISL_2830156, EPI_ISL_2830157 | Lighthouse Lab in Milton Keynes | Wellcome Sanger Institute for the COVID-19 Genomics UK (COG-UK) Consortium | The Lighthouse Lab in Milton Keynes and Alex Alderton, Roberto Amato, Jeffrey Barrett, Sonia Goncalves, Ewan Harrison, David K. Jackson, Ian Johnston, Dominic Kwiatkowski, Cordelia Langford, John Sillitoe on behalf of the Wellcome Sanger Institute COVID-19 Surveillance Team |
| EPI_ISL_2830158 | Lighthouse Lab in Glasgow | Wellcome Sanger Institute for the COVID-19 Genomics UK (COG-UK) Consortium | Harper VanSteenhouse, Yumi Kasai, David Gray, Carol Clugston, Anna Dominiczak and Alex Alderton, Roberto Amato, Jeffrey Barrett, Sonia Goncalves, Ewan Harrison, David K. Jackson, Ian Johnston, Dominic Kwiatkowski, Cordelia Langford, John Sillitoe on behalf of the Wellcome Sanger Institute COVID-19 Surveillance Team |
| EPI_ISL_2830159 | Lighthouse Lab in Milton Keynes | Wellcome Sanger Institute for the COVID-19 Genomics UK (COG-UK) Consortium | The Lighthouse Lab in Milton Keynes and Alex Alderton, Roberto Amato, Jeffrey Barrett, Sonia Goncalves, Ewan Harrison, David K. Jackson, Ian Johnston, Dominic Kwiatkowski, Cordelia Langford, John Sillitoe on behalf of the Wellcome Sanger Institute COVID-19 Surveillance Team |
| EPI_ISL_2830160, EPI_ISL_2830161 | Lighthouse Lab in Glasgow | Wellcome Sanger Institute for the COVID-19 Genomics UK (COG-UK) Consortium | Harper VanSteenhouse, Yumi Kasai, David Gray, Carol Clugston, Anna Dominiczak and Alex Alderton, Roberto Amato, Jeffrey Barrett, Sonia Goncalves, Ewan Harrison, David K. Jackson, Ian Johnston, Dominic Kwiatkowski, Cordelia Langford, John Sillitoe on behalf of the Wellcome Sanger Institute COVID-19 Surveillance Team |
| EPI_ISL_2830162, EPI_ISL_2830163, EPI_ISL_2830164, EPI_ISL_2830165 | Lighthouse Lab in Alderley Park | Wellcome Sanger Institute for the COVID-19 Genomics UK (COG-UK) Consortium | Jacquelyn Wynn, Mairead Hyland, The Lighthouse Lab in Alderley Park and Alex Alderton, Roberto Amato, Jeffrey Barrett, Sonia Goncalves, Ewan Harrison, David K. Jackson, Ian Johnston, Dominic Kwiatkowski, Cordelia Langford, John Sillitoe on behalf of the Wellcome Sanger Institute COVID-19 Surveillance Team |
| EPI_ISL_2830166 | Lighthouse Lab in Milton Keynes | Wellcome Sanger Institute for the COVID-19 Genomics UK (COG-UK) Consortium | The Lighthouse Lab in Milton Keynes and Alex Alderton, Roberto Amato, Jeffrey Barrett, Sonia Goncalves, Ewan Harrison, David K. Jackson, Ian Johnston, Dominic Kwiatkowski, Cordelia Langford, John Sillitoe on behalf of the Wellcome Sanger Institute COVID-19 Surveillance Team |
| EPI_ISL_2830167 | Lighthouse Lab in Alderley Park | Wellcome Sanger Institute for the COVID-19 Genomics UK (COG-UK) Consortium | Jacquelyn Wynn, Mairead Hyland, The Lighthouse Lab in Alderley Park and Alex Alderton, Roberto Amato, Jeffrey Barrett, Sonia Goncalves, Ewan Harrison, David K. Jackson, Ian Johnston, Dominic Kwiatkowski, Cordelia Langford, John Sillitoe on behalf of the Wellcome Sanger Institute COVID-19 Surveillance Team |
| EPI_ISL_2830168 | Lighthouse Lab in Milton Keynes | Wellcome Sanger Institute for the COVID-19 Genomics UK (COG-UK) Consortium | The Lighthouse Lab in Milton Keynes and Alex Alderton, Roberto Amato, Jeffrey Barrett, Sonia Goncalves, Ewan Harrison, David K. Jackson, Ian Johnston, Dominic Kwiatkowski, Cordelia Langford, John Sillitoe on behalf of the Wellcome Sanger Institute COVID-19 Surveillance Team |
| EPI_ISL_2830169, EPI_ISL_2830170, EPI_ISL_2830171 | Lighthouse Lab in Alderley Park | Wellcome Sanger Institute for the COVID-19 Genomics UK (COG-UK) Consortium | Jacquelyn Wynn, Mairead Hyland, The Lighthouse Lab in Alderley Park and Alex Alderton, Roberto Amato, Jeffrey Barrett, Sonia Goncalves, Ewan Harrison, David K. Jackson, Ian Johnston, Dominic Kwiatkowski, Cordelia Langford, John Sillitoe on behalf of the Wellcome Sanger Institute COVID-19 Surveillance Team |
| EPI_ISL_2830172 | Lighthouse Lab in Milton Keynes | Wellcome Sanger Institute for the COVID-19 Genomics UK (COG-UK) Consortium | The Lighthouse Lab in Milton Keynes and Alex Alderton, Roberto Amato, Jeffrey Barrett, Sonia Goncalves, Ewan Harrison, David K. Jackson, Ian Johnston, Dominic Kwiatkowski, Cordelia Langford, John Sillitoe on behalf of the Wellcome Sanger Institute COVID-19 Surveillance Team |
| EPI_ISL_2830173, EPI_ISL_2830174 | Lighthouse Lab in Alderley Park | Wellcome Sanger Institute for the COVID-19 Genomics UK (COG-UK) Consortium | Jacquelyn Wynn, Mairead Hyland, The Lighthouse Lab in Alderley Park and Alex Alderton, Roberto Amato, Jeffrey Barrett, Sonia Goncalves, Ewan Harrison, David K. Jackson, Ian Johnston, Dominic Kwiatkowski, Cordelia Langford, John Sillitoe on behalf of the Wellcome Sanger Institute COVID-19 Surveillance Team |
| EPI_ISL_2830175 | Lighthouse Lab in Milton Keynes | Wellcome Sanger Institute for the COVID-19 Genomics UK (COG-UK) Consortium | The Lighthouse Lab in Milton Keynes and Alex Alderton, Roberto Amato, Jeffrey Barrett, Sonia Goncalves, Ewan Harrison, David K. Jackson, Ian Johnston, Dominic Kwiatkowski, Cordelia Langford, John Sillitoe on behalf of the Wellcome Sanger Institute COVID-19 Surveillance Team |
| EPI_ISL_2830176 | Lighthouse Lab in Alderley Park | Wellcome Sanger Institute for the COVID-19 Genomics UK (COG-UK) Consortium | Jacquelyn Wynn, Mairead Hyland, The Lighthouse Lab in Alderley Park and Alex Alderton, Roberto Amato, Jeffrey Barrett, Sonia Goncalves, Ewan Harrison, David K. Jackson, Ian Johnston, Dominic Kwiatkowski, Cordelia Langford, John Sillitoe on behalf of the Wellcome Sanger Institute COVID-19 Surveillance Team |
| EPI_ISL_2830177 | Lighthouse Lab in Glasgow | Wellcome Sanger Institute for the COVID-19 Genomics UK (COG-UK) Consortium | Harper VanSteenhouse, Yumi Kasai, David Gray, Carol Clugston, Anna Dominiczak and Alex Alderton, Roberto Amato, Jeffrey Barrett, Sonia Goncalves, Ewan Harrison, David K. Jackson, Ian Johnston, Dominic Kwiatkowski, Cordelia Langford, John Sillitoe on behalf of the Wellcome Sanger Institute COVID-19 Surveillance Team |
| EPI_ISL_2830178, EPI_ISL_2830179 | Lighthouse Lab in Alderley Park | Wellcome Sanger Institute for the COVID-19 Genomics UK (COG-UK) Consortium | Jacquelyn Wynn, Mairead Hyland, The Lighthouse Lab in Alderley Park and Alex Alderton, Roberto Amato, Jeffrey Barrett, Sonia Goncalves, Ewan Harrison, David K. Jackson, Ian Johnston, Dominic Kwiatkowski, Cordelia Langford, John Sillitoe on behalf of the Wellcome Sanger Institute COVID-19 Surveillance Team |
| EPI_ISL_2830180 | Lighthouse Lab in Glasgow | Wellcome Sanger Institute for the COVID-19 Genomics UK (COG-UK) Consortium | Harper VanSteenhouse, Yumi Kasai, David Gray, Carol Clugston, Anna Dominiczak and Alex Alderton, Roberto Amato, Jeffrey Barrett, Sonia Goncalves, Ewan Harrison, David K. Jackson, Ian Johnston, Dominic Kwiatkowski, Cordelia Langford, John Sillitoe on behalf of the Wellcome Sanger Institute COVID-19 Surveillance Team |
| EPI_ISL_2830181, EPI_ISL_2830182 | Lighthouse Lab in Alderley Park | Wellcome Sanger Institute for the COVID-19 Genomics UK (COG-UK) Consortium | Jacquelyn Wynn, Mairead Hyland, The Lighthouse Lab in Alderley Park and Alex Alderton, Roberto Amato, Jeffrey Barrett, Sonia Goncalves, Ewan Harrison, David K. Jackson, Ian Johnston, Dominic Kwiatkowski, Cordelia Langford, John Sillitoe on behalf of the Wellcome Sanger Institute COVID-19 Surveillance Team |
| EPI_ISL_2830183 | Lighthouse Lab in Glasgow | Wellcome Sanger Institute for the COVID-19 Genomics UK (COG-UK) Consortium | Harper VanSteenhouse, Yumi Kasai, David Gray, Carol Clugston, Anna Dominiczak and Alex Alderton, Roberto Amato, Jeffrey Barrett, Sonia Goncalves, Ewan Harrison, David K. Jackson, Ian Johnston, Dominic Kwiatkowski, Cordelia Langford, John Sillitoe on behalf of the Wellcome Sanger Institute COVID-19 Surveillance Team |
| EPI_ISL_2830184, | Lighthouse Lab in Alderley Park | Wellcome Sanger Institute for the COVID-19 Genomics UK (COG-UK) | Jacquelyn Wynn, Mairead Hyland, The Lighthouse Lab in Alderley Park and Alex Alderton, Roberto Amato, Jeffrey Barrett, Sonia Goncalves, Ewan Harrison, David K. Jackson, Ian Johnston, Dominic Kwiatkowski, Cordelia Langford, John Sillitoe on |

| | | | |
|---|---|---|---|
| EPI_ISL_2830185, EPI_ISL_2830186, EPI_ISL_2830187, EPI_ISL_2830188 | | Consortium | behalf of the Wellcome Sanger Institute COVID-19 Surveillance Team |
| EPI_ISL_2830189, EPI_ISL_2830190 | Lighthouse Lab in Glasgow | Wellcome Sanger Institute for the COVID-19 Genomics UK (COG-UK) Consortium | Harper VanSteenhouse, Yumi Kasai, David Gray, Carol Clugston, Anna Dominiczak and Alex Alderton, Roberto Amato, Jeffrey Barrett, Sonia Goncalves, Ewan Harrison, David K. Jackson, Ian Johnston, Dominic Kwiatkowski, Cordelia Langford, John Sillitoe on behalf of the Wellcome Sanger Institute COVID-19 Surveillance Team |
| EPI_ISL_2830191, EPI_ISL_2830192, EPI_ISL_2830193 | Lighthouse Lab in Alderley Park | Wellcome Sanger Institute for the COVID-19 Genomics UK (COG-UK) Consortium | Jacquelyn Wynn, Mairead Hyland, The Lighthouse Lab in Alderley Park and Alex Alderton, Roberto Amato, Jeffrey Barrett, Sonia Goncalves, Ewan Harrison, David K. Jackson, Ian Johnston, Dominic Kwiatkowski, Cordelia Langford, John Sillitoe on behalf of the Wellcome Sanger Institute COVID-19 Surveillance Team |
| EPI_ISL_2830194, EPI_ISL_2830195 | Lighthouse Lab in Glasgow | Wellcome Sanger Institute for the COVID-19 Genomics UK (COG-UK) Consortium | Harper VanSteenhouse, Yumi Kasai, David Gray, Carol Clugston, Anna Dominiczak and Alex Alderton, Roberto Amato, Jeffrey Barrett, Sonia Goncalves, Ewan Harrison, David K. Jackson, Ian Johnston, Dominic Kwiatkowski, Cordelia Langford, John Sillitoe on behalf of the Wellcome Sanger Institute COVID-19 Surveillance Team |
| EPI_ISL_2830196, EPI_ISL_2830198, EPI_ISL_2830199 | Lighthouse Lab in Alderley Park | Wellcome Sanger Institute for the COVID-19 Genomics UK (COG-UK) Consortium | Jacquelyn Wynn, Mairead Hyland, The Lighthouse Lab in Alderley Park and Alex Alderton, Roberto Amato, Jeffrey Barrett, Sonia Goncalves, Ewan Harrison, David K. Jackson, Ian Johnston, Dominic Kwiatkowski, Cordelia Langford, John Sillitoe on behalf of the Wellcome Sanger Institute COVID-19 Surveillance Team |
| EPI_ISL_2830200, EPI_ISL_2830201 | Lighthouse Lab in Glasgow | Wellcome Sanger Institute for the COVID-19 Genomics UK (COG-UK) Consortium | Harper VanSteenhouse, Yumi Kasai, David Gray, Carol Clugston, Anna Dominiczak and Alex Alderton, Roberto Amato, Jeffrey Barrett, Sonia Goncalves, Ewan Harrison, David K. Jackson, Ian Johnston, Dominic Kwiatkowski, Cordelia Langford, John Sillitoe on behalf of the Wellcome Sanger Institute COVID-19 Surveillance Team |
| EPI_ISL_2830202, EPI_ISL_2830203, EPI_ISL_2830204, EPI_ISL_2830205 | Lighthouse Lab in Alderley Park | Wellcome Sanger Institute for the COVID-19 Genomics UK (COG-UK) Consortium | Jacquelyn Wynn, Mairead Hyland, The Lighthouse Lab in Alderley Park and Alex Alderton, Roberto Amato, Jeffrey Barrett, Sonia Goncalves, Ewan Harrison, David K. Jackson, Ian Johnston, Dominic Kwiatkowski, Cordelia Langford, John Sillitoe on behalf of the Wellcome Sanger Institute COVID-19 Surveillance Team |
| EPI_ISL_2830206 | Lighthouse Lab in Glasgow | Wellcome Sanger Institute for the COVID-19 Genomics UK (COG-UK) Consortium | Harper VanSteenhouse, Yumi Kasai, David Gray, Carol Clugston, Anna Dominiczak and Alex Alderton, Roberto Amato, Jeffrey Barrett, Sonia Goncalves, Ewan Harrison, David K. Jackson, Ian Johnston, Dominic Kwiatkowski, Cordelia Langford, John Sillitoe on behalf of the Wellcome Sanger Institute COVID-19 Surveillance Team |
| EPI_ISL_2830207 | Lighthouse Lab in Alderley Park | Wellcome Sanger Institute for the COVID-19 Genomics UK (COG-UK) Consortium | Jacquelyn Wynn, Mairead Hyland, The Lighthouse Lab in Alderley Park and Alex Alderton, Roberto Amato, Jeffrey Barrett, Sonia Goncalves, Ewan Harrison, David K. Jackson, Ian Johnston, Dominic Kwiatkowski, Cordelia Langford, John Sillitoe on behalf of the Wellcome Sanger Institute COVID-19 Surveillance Team |
| EPI_ISL_2830208 | Lighthouse Lab in Glasgow | Wellcome Sanger Institute for the COVID-19 Genomics UK (COG-UK) Consortium | Harper VanSteenhouse, Yumi Kasai, David Gray, Carol Clugston, Anna Dominiczak and Alex Alderton, Roberto Amato, Jeffrey Barrett, Sonia Goncalves, Ewan Harrison, David K. Jackson, Ian Johnston, Dominic Kwiatkowski, Cordelia Langford, John Sillitoe on behalf of the Wellcome Sanger Institute COVID-19 Surveillance Team |
| EPI_ISL_2830209 | Lighthouse Lab in Alderley Park | Wellcome Sanger Institute for the COVID-19 Genomics UK (COG-UK) Consortium | Jacquelyn Wynn, Mairead Hyland, The Lighthouse Lab in Alderley Park and Alex Alderton, Roberto Amato, Jeffrey Barrett, Sonia Goncalves, Ewan Harrison, David K. Jackson, Ian Johnston, Dominic Kwiatkowski, Cordelia Langford, John Sillitoe on behalf of the Wellcome Sanger Institute COVID-19 Surveillance Team |
| EPI_ISL_2830210 | Lighthouse Lab in Glasgow | Wellcome Sanger Institute for the COVID-19 Genomics UK (COG-UK) Consortium | Harper VanSteenhouse, Yumi Kasai, David Gray, Carol Clugston, Anna Dominiczak and Alex Alderton, Roberto Amato, Jeffrey Barrett, Sonia Goncalves, Ewan Harrison, David K. Jackson, Ian Johnston, Dominic Kwiatkowski, Cordelia Langford, John Sillitoe on behalf of the Wellcome Sanger Institute COVID-19 Surveillance Team |
| EPI_ISL_2830211, EPI_ISL_2830212 | Lighthouse Lab in Alderley Park | Wellcome Sanger Institute for the COVID-19 Genomics UK (COG-UK) Consortium | Jacquelyn Wynn, Mairead Hyland, The Lighthouse Lab in Alderley Park and Alex Alderton, Roberto Amato, Jeffrey Barrett, Sonia Goncalves, Ewan Harrison, David K. Jackson, Ian Johnston, Dominic Kwiatkowski, Cordelia Langford, John Sillitoe on behalf of the Wellcome Sanger Institute COVID-19 Surveillance Team |
| EPI_ISL_2830213 | Lighthouse Lab in Glasgow | Wellcome Sanger Institute for the COVID-19 Genomics UK (COG-UK) Consortium | Harper VanSteenhouse, Yumi Kasai, David Gray, Carol Clugston, Anna Dominiczak and Alex Alderton, Roberto Amato, Jeffrey Barrett, Sonia Goncalves, Ewan Harrison, David K. Jackson, Ian Johnston, Dominic Kwiatkowski, Cordelia Langford, John Sillitoe on behalf of the Wellcome Sanger Institute COVID-19 Surveillance Team |
| EPI_ISL_2830214 | Lighthouse Lab in Alderley Park | Wellcome Sanger Institute for the COVID-19 Genomics UK (COG-UK) Consortium | Jacquelyn Wynn, Mairead Hyland, The Lighthouse Lab in Alderley Park and Alex Alderton, Roberto Amato, Jeffrey Barrett, Sonia Goncalves, Ewan Harrison, David K. Jackson, Ian Johnston, Dominic Kwiatkowski, Cordelia Langford, John Sillitoe on behalf of the Wellcome Sanger Institute COVID-19 Surveillance Team |
| EPI_ISL_2830215 | Lighthouse Lab in Glasgow | Wellcome Sanger Institute for the COVID-19 Genomics UK (COG-UK) Consortium | Harper VanSteenhouse, Yumi Kasai, David Gray, Carol Clugston, Anna Dominiczak and Alex Alderton, Roberto Amato, Jeffrey Barrett, Sonia Goncalves, Ewan Harrison, David K. Jackson, Ian Johnston, Dominic Kwiatkowski, Cordelia Langford, John Sillitoe on behalf of the Wellcome Sanger Institute COVID-19 Surveillance Team |
| EPI_ISL_2830216, EPI_ISL_2830217, EPI_ISL_2830218, EPI_ISL_2830219, EPI_ISL_2830220 | Lighthouse Lab in Alderley Park | Wellcome Sanger Institute for the COVID-19 Genomics UK (COG-UK) Consortium | Jacquelyn Wynn, Mairead Hyland, The Lighthouse Lab in Alderley Park and Alex Alderton, Roberto Amato, Jeffrey Barrett, Sonia Goncalves, Ewan Harrison, David K. Jackson, Ian Johnston, Dominic Kwiatkowski, Cordelia Langford, John Sillitoe on behalf of the Wellcome Sanger Institute COVID-19 Surveillance Team |
| EPI_ISL_2830221, EPI_ISL_2830222, EPI_ISL_2830223, EPI_ISL_2830224 | Lighthouse Lab in Glasgow | Wellcome Sanger Institute for the COVID-19 Genomics UK (COG-UK) Consortium | Harper VanSteenhouse, Yumi Kasai, David Gray, Carol Clugston, Anna Dominiczak and Alex Alderton, Roberto Amato, Jeffrey Barrett, Sonia Goncalves, Ewan Harrison, David K. Jackson, Ian Johnston, Dominic Kwiatkowski, Cordelia Langford, John Sillitoe on behalf of the Wellcome Sanger Institute COVID-19 Surveillance Team |
| EPI_ISL_2830225, EPI_ISL_2830226 | Lighthouse Lab in Alderley Park | Wellcome Sanger Institute for the COVID-19 Genomics UK (COG-UK) Consortium | Jacquelyn Wynn, Mairead Hyland, The Lighthouse Lab in Alderley Park and Alex Alderton, Roberto Amato, Jeffrey Barrett, Sonia Goncalves, Ewan Harrison, David K. Jackson, Ian Johnston, Dominic Kwiatkowski, Cordelia Langford, John Sillitoe on behalf of the Wellcome Sanger Institute COVID-19 Surveillance Team |
| EPI_ISL_2830227, EPI_ISL_2830228, EPI_ISL_2830229 | Lighthouse Lab in Glasgow | Wellcome Sanger Institute for the COVID-19 Genomics UK (COG-UK) Consortium | Harper VanSteenhouse, Yumi Kasai, David Gray, Carol Clugston, Anna Dominiczak and Alex Alderton, Roberto Amato, Jeffrey Barrett, Sonia Goncalves, Ewan Harrison, David K. Jackson, Ian Johnston, Dominic Kwiatkowski, Cordelia Langford, John Sillitoe on behalf of the Wellcome Sanger Institute COVID-19 Surveillance Team |
| EPI_ISL_2830230, EPI_ISL_2830231 | Lighthouse Lab in Alderley Park | Wellcome Sanger Institute for the COVID-19 Genomics UK (COG-UK) Consortium | Jacquelyn Wynn, Mairead Hyland, The Lighthouse Lab in Alderley Park and Alex Alderton, Roberto Amato, Jeffrey Barrett, Sonia Goncalves, Ewan Harrison, David K. Jackson, Ian Johnston, Dominic Kwiatkowski, Cordelia Langford, John Sillitoe on behalf of the Wellcome Sanger Institute COVID-19 Surveillance Team |
| EPI_ISL_2830232 | Lighthouse Lab in Glasgow | Wellcome Sanger Institute for the COVID-19 Genomics UK (COG-UK) Consortium | Harper VanSteenhouse, Yumi Kasai, David Gray, Carol Clugston, Anna Dominiczak and Alex Alderton, Roberto Amato, Jeffrey Barrett, Sonia Goncalves, Ewan Harrison, David K. Jackson, Ian Johnston, Dominic Kwiatkowski, Cordelia Langford, John Sillitoe on behalf of the Wellcome Sanger Institute COVID-19 Surveillance Team |
| EPI_ISL_2830233 | Lighthouse Lab in Alderley Park | Wellcome Sanger Institute for the COVID-19 Genomics UK (COG-UK) Consortium | Jacquelyn Wynn, Mairead Hyland, The Lighthouse Lab in Alderley Park and Alex Alderton, Roberto Amato, Jeffrey Barrett, Sonia Goncalves, Ewan Harrison, David K. Jackson, Ian Johnston, Dominic Kwiatkowski, Cordelia Langford, John Sillitoe on behalf of the Wellcome Sanger Institute COVID-19 Surveillance Team |
| EPI_ISL_2830234, EPI_ISL_2830235, EPI_ISL_2830236, EPI_ISL_2830237, EPI_ISL_2830238 | Lighthouse Lab in Glasgow | Wellcome Sanger Institute for the COVID-19 Genomics UK (COG-UK) Consortium | Harper VanSteenhouse, Yumi Kasai, David Gray, Carol Clugston, Anna Dominiczak and Alex Alderton, Roberto Amato, Jeffrey Barrett, Sonia Goncalves, Ewan Harrison, David K. Jackson, Ian Johnston, Dominic Kwiatkowski, Cordelia Langford, John Sillitoe on behalf of the Wellcome Sanger Institute COVID-19 Surveillance Team |
| EPI_ISL_2830239 | Lighthouse Lab in Alderley Park | Wellcome Sanger Institute for the COVID-19 Genomics UK (COG-UK) Consortium | Jacquelyn Wynn, Mairead Hyland, The Lighthouse Lab in Alderley Park and Alex Alderton, Roberto Amato, Jeffrey Barrett, Sonia Goncalves, Ewan Harrison, David K. Jackson, Ian Johnston, Dominic Kwiatkowski, Cordelia Langford, John Sillitoe on behalf of the Wellcome Sanger Institute COVID-19 Surveillance Team |
| EPI_ISL_2830240, EPI_ISL_2830241 | Lighthouse Lab in Glasgow | Wellcome Sanger Institute for the COVID-19 Genomics UK (COG-UK) Consortium | Harper VanSteenhouse, Yumi Kasai, David Gray, Carol Clugston, Anna Dominiczak and Alex Alderton, Roberto Amato, Jeffrey Barrett, Sonia Goncalves, Ewan Harrison, David K. Jackson, Ian Johnston, Dominic Kwiatkowski, Cordelia Langford, John Sillitoe on behalf of the Wellcome Sanger Institute COVID-19 Surveillance Team |
| EPI_ISL_2830242, EPI_ISL_2830243 | Lighthouse Lab in Alderley Park | Wellcome Sanger Institute for the COVID-19 Genomics UK (COG-UK) Consortium | Jacquelyn Wynn, Mairead Hyland, The Lighthouse Lab in Alderley Park and Alex Alderton, Roberto Amato, Jeffrey Barrett, Sonia Goncalves, Ewan Harrison, David K. Jackson, Ian Johnston, Dominic Kwiatkowski, Cordelia Langford, John Sillitoe on behalf of the Wellcome Sanger Institute COVID-19 Surveillance Team |
| EPI_ISL_2830244 | Lighthouse Lab in Glasgow | Wellcome Sanger Institute for the COVID-19 Genomics UK (COG-UK) Consortium | Harper VanSteenhouse, Yumi Kasai, David Gray, Carol Clugston, Anna Dominiczak and Alex Alderton, Roberto Amato, Jeffrey Barrett, Sonia Goncalves, Ewan Harrison, David K. Jackson, Ian Johnston, Dominic Kwiatkowski, Cordelia Langford, John Sillitoe on behalf of the Wellcome Sanger Institute COVID-19 Surveillance Team |
| EPI_ISL_2830245, EPI_ISL_2830246 | Lighthouse Lab in Alderley Park | Wellcome Sanger Institute for the COVID-19 Genomics UK (COG-UK) Consortium | Jacquelyn Wynn, Mairead Hyland, The Lighthouse Lab in Alderley Park and Alex Alderton, Roberto Amato, Jeffrey Barrett, Sonia Goncalves, Ewan Harrison, David K. Jackson, Ian Johnston, Dominic Kwiatkowski, Cordelia Langford, John Sillitoe on behalf of the Wellcome Sanger Institute COVID-19 Surveillance Team |
| EPI_ISL_2830247 | Lighthouse Lab in Glasgow | Wellcome Sanger Institute for the COVID-19 Genomics UK (COG-UK) Consortium | Harper VanSteenhouse, Yumi Kasai, David Gray, Carol Clugston, Anna Dominiczak and Alex Alderton, Roberto Amato, Jeffrey Barrett, Sonia Goncalves, Ewan Harrison, David K. Jackson, Ian Johnston, Dominic Kwiatkowski, Cordelia Langford, John Sillitoe on behalf of the Wellcome Sanger Institute COVID-19 Surveillance Team |
| EPI_ISL_2830248 | Lighthouse Lab in Alderley Park | Wellcome Sanger Institute for the COVID-19 Genomics UK (COG-UK) Consortium | Jacquelyn Wynn, Mairead Hyland, The Lighthouse Lab in Alderley Park and Alex Alderton, Roberto Amato, Jeffrey Barrett, Sonia Goncalves, Ewan Harrison, David K. Jackson, Ian Johnston, Dominic Kwiatkowski, Cordelia Langford, John Sillitoe on behalf of the Wellcome Sanger Institute COVID-19 Surveillance Team |
| EPI_ISL_2830249, EPI_ISL_2830250, EPI_ISL_2830251 | Lighthouse Lab in Glasgow | Wellcome Sanger Institute for the COVID-19 Genomics UK (COG-UK) Consortium | Harper VanSteenhouse, Yumi Kasai, David Gray, Carol Clugston, Anna Dominiczak and Alex Alderton, Roberto Amato, Jeffrey Barrett, Sonia Goncalves, Ewan Harrison, David K. Jackson, Ian Johnston, Dominic Kwiatkowski, Cordelia Langford, John Sillitoe on behalf of the Wellcome Sanger Institute COVID-19 Surveillance Team |
| EPI_ISL_2830252 | Lighthouse Lab in Alderley Park | Wellcome Sanger Institute for the COVID-19 Genomics UK (COG-UK) Consortium | Jacquelyn Wynn, Mairead Hyland, The Lighthouse Lab in Alderley Park and Alex Alderton, Roberto Amato, Jeffrey Barrett, Sonia Goncalves, Ewan Harrison, David K. Jackson, Ian Johnston, Dominic Kwiatkowski, Cordelia Langford, John Sillitoe on behalf of the Wellcome Sanger Institute COVID-19 Surveillance Team |
| EPI_ISL_2830253, EPI_ISL_2830254, EPI_ISL_2830255 | Lighthouse Lab in Glasgow | Wellcome Sanger Institute for the COVID-19 Genomics UK (COG-UK) Consortium | Harper VanSteenhouse, Yumi Kasai, David Gray, Carol Clugston, Anna Dominiczak and Alex Alderton, Roberto Amato, Jeffrey Barrett, Sonia Goncalves, Ewan Harrison, David K. Jackson, Ian Johnston, Dominic Kwiatkowski, Cordelia Langford, John Sillitoe on behalf of the Wellcome Sanger Institute COVID-19 Surveillance Team |
| EPI_ISL_2830256 | Lighthouse Lab in Alderley Park | Wellcome Sanger Institute for the COVID-19 Genomics UK (COG-UK) Consortium | Jacquelyn Wynn, Mairead Hyland, The Lighthouse Lab in Alderley Park and Alex Alderton, Roberto Amato, Jeffrey Barrett, Sonia Goncalves, Ewan Harrison, David K. Jackson, Ian Johnston, Dominic Kwiatkowski, Cordelia Langford, John Sillitoe on behalf of the Wellcome Sanger Institute COVID-19 Surveillance Team |
| EPI_ISL_2830257, EPI_ISL_2830258, EPI_ISL_2830259 | Lighthouse Lab in Glasgow | Wellcome Sanger Institute for the COVID-19 Genomics UK (COG-UK) Consortium | Harper VanSteenhouse, Yumi Kasai, David Gray, Carol Clugston, Anna Dominiczak and Alex Alderton, Roberto Amato, Jeffrey Barrett, Sonia Goncalves, Ewan Harrison, David K. Jackson, Ian Johnston, Dominic Kwiatkowski, Cordelia Langford, John Sillitoe on behalf of the Wellcome Sanger Institute COVID-19 Surveillance Team |
| EPI_ISL_2830260 | Lighthouse Lab in Alderley Park | Wellcome Sanger Institute for the COVID-19 Genomics UK (COG-UK) Consortium | Jacquelyn Wynn, Mairead Hyland, The Lighthouse Lab in Alderley Park and Alex Alderton, Roberto Amato, Jeffrey Barrett, Sonia Goncalves, Ewan Harrison, David K. Jackson, Ian Johnston, Dominic Kwiatkowski, Cordelia Langford, John Sillitoe on behalf of the Wellcome Sanger Institute COVID-19 Surveillance Team |
| EPI_ISL_2830261 | Lighthouse Lab in Glasgow | Wellcome Sanger Institute for the COVID-19 Genomics UK (COG-UK) Consortium | Harper VanSteenhouse, Yumi Kasai, David Gray, Carol Clugston, Anna Dominiczak and Alex Alderton, Roberto Amato, Jeffrey Barrett, Sonia Goncalves, Ewan Harrison, David K. Jackson, Ian Johnston, Dominic Kwiatkowski, Cordelia Langford, John Sillitoe on behalf of the Wellcome Sanger Institute COVID-19 Surveillance Team |

| | | | |
|---|---|---|---|
| EPI_ISL_2830262, EPI_ISL_2830263, EPI_ISL_2830264 | Lighthouse Lab in Alderley Park | Wellcome Sanger Institute for the COVID-19 Genomics UK (COG-UK) Consortium | Jacquelyn Wynn, Mairead Hyland, The Lighthouse Lab in Alderley Park and Alex Alderton, Roberto Amato, Jeffrey Barrett, Sonia Goncalves, Ewan Harrison, David K. Jackson, Ian Johnston, Dominic Kwiatkowski, Cordelia Langford, John Sillitoe on behalf of the Wellcome Sanger Institute COVID-19 Surveillance Team |
| EPI_ISL_2830265 | Lighthouse Lab in Glasgow | Wellcome Sanger Institute for the COVID-19 Genomics UK (COG-UK) Consortium | Harper VanSteenhouse, Yumi Kasai, David Gray, Carol Clugston, Anna Dominiczak and Alex Alderton, Roberto Amato, Jeffrey Barrett, Sonia Goncalves, Ewan Harrison, David K. Jackson, Ian Johnston, Dominic Kwiatkowski, Cordelia Langford, John Sillitoe on behalf of the Wellcome Sanger Institute COVID-19 Surveillance Team |
| EPI_ISL_2830266, EPI_ISL_2830267 | Lighthouse Lab in Alderley Park | Wellcome Sanger Institute for the COVID-19 Genomics UK (COG-UK) Consortium | Jacquelyn Wynn, Mairead Hyland, The Lighthouse Lab in Alderley Park and Alex Alderton, Roberto Amato, Jeffrey Barrett, Sonia Goncalves, Ewan Harrison, David K. Jackson, Ian Johnston, Dominic Kwiatkowski, Cordelia Langford, John Sillitoe on behalf of the Wellcome Sanger Institute COVID-19 Surveillance Team |
| EPI_ISL_2830268, EPI_ISL_2830269 | Lighthouse Lab in Glasgow | Wellcome Sanger Institute for the COVID-19 Genomics UK (COG-UK) Consortium | Harper VanSteenhouse, Yumi Kasai, David Gray, Carol Clugston, Anna Dominiczak and Alex Alderton, Roberto Amato, Jeffrey Barrett, Sonia Goncalves, Ewan Harrison, David K. Jackson, Ian Johnston, Dominic Kwiatkowski, Cordelia Langford, John Sillitoe on behalf of the Wellcome Sanger Institute COVID-19 Surveillance Team |
| EPI_ISL_2830270 | Lighthouse Lab in Alderley Park | Wellcome Sanger Institute for the COVID-19 Genomics UK (COG-UK) Consortium | Jacquelyn Wynn, Mairead Hyland, The Lighthouse Lab in Alderley Park and Alex Alderton, Roberto Amato, Jeffrey Barrett, Sonia Goncalves, Ewan Harrison, David K. Jackson, Ian Johnston, Dominic Kwiatkowski, Cordelia Langford, John Sillitoe on behalf of the Wellcome Sanger Institute COVID-19 Surveillance Team |
| EPI_ISL_2830271 | Lighthouse Lab in Glasgow | Wellcome Sanger Institute for the COVID-19 Genomics UK (COG-UK) Consortium | Harper VanSteenhouse, Yumi Kasai, David Gray, Carol Clugston, Anna Dominiczak and Alex Alderton, Roberto Amato, Jeffrey Barrett, Sonia Goncalves, Ewan Harrison, David K. Jackson, Ian Johnston, Dominic Kwiatkowski, Cordelia Langford, John Sillitoe on behalf of the Wellcome Sanger Institute COVID-19 Surveillance Team |
| EPI_ISL_2830272, EPI_ISL_2830274 | Lighthouse Lab in Alderley Park | Wellcome Sanger Institute for the COVID-19 Genomics UK (COG-UK) Consortium | Jacquelyn Wynn, Mairead Hyland, The Lighthouse Lab in Alderley Park and Alex Alderton, Roberto Amato, Jeffrey Barrett, Sonia Goncalves, Ewan Harrison, David K. Jackson, Ian Johnston, Dominic Kwiatkowski, Cordelia Langford, John Sillitoe on behalf of the Wellcome Sanger Institute COVID-19 Surveillance Team |
| EPI_ISL_2830275 | Lighthouse Lab in Glasgow | Wellcome Sanger Institute for the COVID-19 Genomics UK (COG-UK) Consortium | Harper VanSteenhouse, Yumi Kasai, David Gray, Carol Clugston, Anna Dominiczak and Alex Alderton, Roberto Amato, Jeffrey Barrett, Sonia Goncalves, Ewan Harrison, David K. Jackson, Ian Johnston, Dominic Kwiatkowski, Cordelia Langford, John Sillitoe on behalf of the Wellcome Sanger Institute COVID-19 Surveillance Team |
| EPI_ISL_2830276, EPI_ISL_2830277 | Lighthouse Lab in Alderley Park | Wellcome Sanger Institute for the COVID-19 Genomics UK (COG-UK) Consortium | Jacquelyn Wynn, Mairead Hyland, The Lighthouse Lab in Alderley Park and Alex Alderton, Roberto Amato, Jeffrey Barrett, Sonia Goncalves, Ewan Harrison, David K. Jackson, Ian Johnston, Dominic Kwiatkowski, Cordelia Langford, John Sillitoe on behalf of the Wellcome Sanger Institute COVID-19 Surveillance Team |
| EPI_ISL_2830278 | Lighthouse Lab in Glasgow | Wellcome Sanger Institute for the COVID-19 Genomics UK (COG-UK) Consortium | Harper VanSteenhouse, Yumi Kasai, David Gray, Carol Clugston, Anna Dominiczak and Alex Alderton, Roberto Amato, Jeffrey Barrett, Sonia Goncalves, Ewan Harrison, David K. Jackson, Ian Johnston, Dominic Kwiatkowski, Cordelia Langford, John Sillitoe on behalf of the Wellcome Sanger Institute COVID-19 Surveillance Team |
| EPI_ISL_2830279 | Lighthouse Lab in Alderley Park | Wellcome Sanger Institute for the COVID-19 Genomics UK (COG-UK) Consortium | Jacquelyn Wynn, Mairead Hyland, The Lighthouse Lab in Alderley Park and Alex Alderton, Roberto Amato, Jeffrey Barrett, Sonia Goncalves, Ewan Harrison, David K. Jackson, Ian Johnston, Dominic Kwiatkowski, Cordelia Langford, John Sillitoe on behalf of the Wellcome Sanger Institute COVID-19 Surveillance Team |
| EPI_ISL_2830280 | Lighthouse Lab in Glasgow | Wellcome Sanger Institute for the COVID-19 Genomics UK (COG-UK) Consortium | Harper VanSteenhouse, Yumi Kasai, David Gray, Carol Clugston, Anna Dominiczak and Alex Alderton, Roberto Amato, Jeffrey Barrett, Sonia Goncalves, Ewan Harrison, David K. Jackson, Ian Johnston, Dominic Kwiatkowski, Cordelia Langford, John Sillitoe on behalf of the Wellcome Sanger Institute COVID-19 Surveillance Team |
| EPI_ISL_2830281, EPI_ISL_2830282 | Lighthouse Lab in Alderley Park | Wellcome Sanger Institute for the COVID-19 Genomics UK (COG-UK) Consortium | Jacquelyn Wynn, Mairead Hyland, The Lighthouse Lab in Alderley Park and Alex Alderton, Roberto Amato, Jeffrey Barrett, Sonia Goncalves, Ewan Harrison, David K. Jackson, Ian Johnston, Dominic Kwiatkowski, Cordelia Langford, John Sillitoe on behalf of the Wellcome Sanger Institute COVID-19 Surveillance Team |
| EPI_ISL_2830283 | Lighthouse Lab in Glasgow | Wellcome Sanger Institute for the COVID-19 Genomics UK (COG-UK) Consortium | Harper VanSteenhouse, Yumi Kasai, David Gray, Carol Clugston, Anna Dominiczak and Alex Alderton, Roberto Amato, Jeffrey Barrett, Sonia Goncalves, Ewan Harrison, David K. Jackson, Ian Johnston, Dominic Kwiatkowski, Cordelia Langford, John Sillitoe on behalf of the Wellcome Sanger Institute COVID-19 Surveillance Team |
| EPI_ISL_2830284 | Lighthouse Lab in Alderley Park | Wellcome Sanger Institute for the COVID-19 Genomics UK (COG-UK) Consortium | Jacquelyn Wynn, Mairead Hyland, The Lighthouse Lab in Alderley Park and Alex Alderton, Roberto Amato, Jeffrey Barrett, Sonia Goncalves, Ewan Harrison, David K. Jackson, Ian Johnston, Dominic Kwiatkowski, Cordelia Langford, John Sillitoe on behalf of the Wellcome Sanger Institute COVID-19 Surveillance Team |
| EPI_ISL_2830285 | Lighthouse Lab in Glasgow | Wellcome Sanger Institute for the COVID-19 Genomics UK (COG-UK) Consortium | Harper VanSteenhouse, Yumi Kasai, David Gray, Carol Clugston, Anna Dominiczak and Alex Alderton, Roberto Amato, Jeffrey Barrett, Sonia Goncalves, Ewan Harrison, David K. Jackson, Ian Johnston, Dominic Kwiatkowski, Cordelia Langford, John Sillitoe on behalf of the Wellcome Sanger Institute COVID-19 Surveillance Team |
| EPI_ISL_2830286, EPI_ISL_2830287 | Lighthouse Lab in Alderley Park | Wellcome Sanger Institute for the COVID-19 Genomics UK (COG-UK) Consortium | Jacquelyn Wynn, Mairead Hyland, The Lighthouse Lab in Alderley Park and Alex Alderton, Roberto Amato, Jeffrey Barrett, Sonia Goncalves, Ewan Harrison, David K. Jackson, Ian Johnston, Dominic Kwiatkowski, Cordelia Langford, John Sillitoe on behalf of the Wellcome Sanger Institute COVID-19 Surveillance Team |
| EPI_ISL_2830288 | Lighthouse Lab in Glasgow | Wellcome Sanger Institute for the COVID-19 Genomics UK (COG-UK) Consortium | Harper VanSteenhouse, Yumi Kasai, David Gray, Carol Clugston, Anna Dominiczak and Alex Alderton, Roberto Amato, Jeffrey Barrett, Sonia Goncalves, Ewan Harrison, David K. Jackson, Ian Johnston, Dominic Kwiatkowski, Cordelia Langford, John Sillitoe on behalf of the Wellcome Sanger Institute COVID-19 Surveillance Team |
| EPI_ISL_2830289 | Lighthouse Lab in Alderley Park | Wellcome Sanger Institute for the COVID-19 Genomics UK (COG-UK) Consortium | Jacquelyn Wynn, Mairead Hyland, The Lighthouse Lab in Alderley Park and Alex Alderton, Roberto Amato, Jeffrey Barrett, Sonia Goncalves, Ewan Harrison, David K. Jackson, Ian Johnston, Dominic Kwiatkowski, Cordelia Langford, John Sillitoe on behalf of the Wellcome Sanger Institute COVID-19 Surveillance Team |
| EPI_ISL_2830290 | Lighthouse Lab in Glasgow | Wellcome Sanger Institute for the COVID-19 Genomics UK (COG-UK) Consortium | Harper VanSteenhouse, Yumi Kasai, David Gray, Carol Clugston, Anna Dominiczak and Alex Alderton, Roberto Amato, Jeffrey Barrett, Sonia Goncalves, Ewan Harrison, David K. Jackson, Ian Johnston, Dominic Kwiatkowski, Cordelia Langford, John Sillitoe on behalf of the Wellcome Sanger Institute COVID-19 Surveillance Team |
| EPI_ISL_2830291, EPI_ISL_2830292, EPI_ISL_2830293 | Lighthouse Lab in Alderley Park | Wellcome Sanger Institute for the COVID-19 Genomics UK (COG-UK) Consortium | Jacquelyn Wynn, Mairead Hyland, The Lighthouse Lab in Alderley Park and Alex Alderton, Roberto Amato, Jeffrey Barrett, Sonia Goncalves, Ewan Harrison, David K. Jackson, Ian Johnston, Dominic Kwiatkowski, Cordelia Langford, John Sillitoe on behalf of the Wellcome Sanger Institute COVID-19 Surveillance Team |
| EPI_ISL_2830294, EPI_ISL_2830295 | Lighthouse Lab in Glasgow | Wellcome Sanger Institute for the COVID-19 Genomics UK (COG-UK) Consortium | Harper VanSteenhouse, Yumi Kasai, David Gray, Carol Clugston, Anna Dominiczak and Alex Alderton, Roberto Amato, Jeffrey Barrett, Sonia Goncalves, Ewan Harrison, David K. Jackson, Ian Johnston, Dominic Kwiatkowski, Cordelia Langford, John Sillitoe on behalf of the Wellcome Sanger Institute COVID-19 Surveillance Team |
| EPI_ISL_2830296, EPI_ISL_2830297 | Lighthouse Lab in Alderley Park | Wellcome Sanger Institute for the COVID-19 Genomics UK (COG-UK) Consortium | Jacquelyn Wynn, Mairead Hyland, The Lighthouse Lab in Alderley Park and Alex Alderton, Roberto Amato, Jeffrey Barrett, Sonia Goncalves, Ewan Harrison, David K. Jackson, Ian Johnston, Dominic Kwiatkowski, Cordelia Langford, John Sillitoe on behalf of the Wellcome Sanger Institute COVID-19 Surveillance Team |
| EPI_ISL_2830298 | Lighthouse Lab in Glasgow | Wellcome Sanger Institute for the COVID-19 Genomics UK (COG-UK) Consortium | Harper VanSteenhouse, Yumi Kasai, David Gray, Carol Clugston, Anna Dominiczak and Alex Alderton, Roberto Amato, Jeffrey Barrett, Sonia Goncalves, Ewan Harrison, David K. Jackson, Ian Johnston, Dominic Kwiatkowski, Cordelia Langford, John Sillitoe on behalf of the Wellcome Sanger Institute COVID-19 Surveillance Team |
| EPI_ISL_2830300, EPI_ISL_2830301 | Lighthouse Lab in Alderley Park | Wellcome Sanger Institute for the COVID-19 Genomics UK (COG-UK) Consortium | Jacquelyn Wynn, Mairead Hyland, The Lighthouse Lab in Alderley Park and Alex Alderton, Roberto Amato, Jeffrey Barrett, Sonia Goncalves, Ewan Harrison, David K. Jackson, Ian Johnston, Dominic Kwiatkowski, Cordelia Langford, John Sillitoe on behalf of the Wellcome Sanger Institute COVID-19 Surveillance Team |
| EPI_ISL_2830302, EPI_ISL_2830303, EPI_ISL_2830304, EPI_ISL_2830305 | Lighthouse Lab in Glasgow | Wellcome Sanger Institute for the COVID-19 Genomics UK (COG-UK) Consortium | Harper VanSteenhouse, Yumi Kasai, David Gray, Carol Clugston, Anna Dominiczak and Alex Alderton, Roberto Amato, Jeffrey Barrett, Sonia Goncalves, Ewan Harrison, David K. Jackson, Ian Johnston, Dominic Kwiatkowski, Cordelia Langford, John Sillitoe on behalf of the Wellcome Sanger Institute COVID-19 Surveillance Team |
| EPI_ISL_2830306 | Lighthouse Lab in Alderley Park | Wellcome Sanger Institute for the COVID-19 Genomics UK (COG-UK) Consortium | Jacquelyn Wynn, Mairead Hyland, The Lighthouse Lab in Alderley Park and Alex Alderton, Roberto Amato, Jeffrey Barrett, Sonia Goncalves, Ewan Harrison, David K. Jackson, Ian Johnston, Dominic Kwiatkowski, Cordelia Langford, John Sillitoe on behalf of the Wellcome Sanger Institute COVID-19 Surveillance Team |
| EPI_ISL_2830307 | Lighthouse Lab in Glasgow | Wellcome Sanger Institute for the COVID-19 Genomics UK (COG-UK) Consortium | Harper VanSteenhouse, Yumi Kasai, David Gray, Carol Clugston, Anna Dominiczak and Alex Alderton, Roberto Amato, Jeffrey Barrett, Sonia Goncalves, Ewan Harrison, David K. Jackson, Ian Johnston, Dominic Kwiatkowski, Cordelia Langford, John Sillitoe on behalf of the Wellcome Sanger Institute COVID-19 Surveillance Team |
| EPI_ISL_2830308 | Lighthouse Lab in Alderley Park | Wellcome Sanger Institute for the COVID-19 Genomics UK (COG-UK) Consortium | Jacquelyn Wynn, Mairead Hyland, The Lighthouse Lab in Alderley Park and Alex Alderton, Roberto Amato, Jeffrey Barrett, Sonia Goncalves, Ewan Harrison, David K. Jackson, Ian Johnston, Dominic Kwiatkowski, Cordelia Langford, John Sillitoe on behalf of the Wellcome Sanger Institute COVID-19 Surveillance Team |
| EPI_ISL_2830309, EPI_ISL_2830310 | Lighthouse Lab in Glasgow | Wellcome Sanger Institute for the COVID-19 Genomics UK (COG-UK) Consortium | Harper VanSteenhouse, Yumi Kasai, David Gray, Carol Clugston, Anna Dominiczak and Alex Alderton, Roberto Amato, Jeffrey Barrett, Sonia Goncalves, Ewan Harrison, David K. Jackson, Ian Johnston, Dominic Kwiatkowski, Cordelia Langford, John Sillitoe on behalf of the Wellcome Sanger Institute COVID-19 Surveillance Team |
| EPI_ISL_2830311, EPI_ISL_2830312 | Lighthouse Lab in Alderley Park | Wellcome Sanger Institute for the COVID-19 Genomics UK (COG-UK) Consortium | Jacquelyn Wynn, Mairead Hyland, The Lighthouse Lab in Alderley Park and Alex Alderton, Roberto Amato, Jeffrey Barrett, Sonia Goncalves, Ewan Harrison, David K. Jackson, Ian Johnston, Dominic Kwiatkowski, Cordelia Langford, John Sillitoe on behalf of the Wellcome Sanger Institute COVID-19 Surveillance Team |
| EPI_ISL_2830313, EPI_ISL_2830314 | Lighthouse Lab in Glasgow | Wellcome Sanger Institute for the COVID-19 Genomics UK (COG-UK) Consortium | Harper VanSteenhouse, Yumi Kasai, David Gray, Carol Clugston, Anna Dominiczak and Alex Alderton, Roberto Amato, Jeffrey Barrett, Sonia Goncalves, Ewan Harrison, David K. Jackson, Ian Johnston, Dominic Kwiatkowski, Cordelia Langford, John Sillitoe on behalf of the Wellcome Sanger Institute COVID-19 Surveillance Team |
| EPI_ISL_2830315 | Lighthouse Lab in Alderley Park | Wellcome Sanger Institute for the COVID-19 Genomics UK (COG-UK) Consortium | Jacquelyn Wynn, Mairead Hyland, The Lighthouse Lab in Alderley Park and Alex Alderton, Roberto Amato, Jeffrey Barrett, Sonia Goncalves, Ewan Harrison, David K. Jackson, Ian Johnston, Dominic Kwiatkowski, Cordelia Langford, John Sillitoe on behalf of the Wellcome Sanger Institute COVID-19 Surveillance Team |
| EPI_ISL_2830316 | Lighthouse Lab in Glasgow | Wellcome Sanger Institute for the COVID-19 Genomics UK (COG-UK) Consortium | Harper VanSteenhouse, Yumi Kasai, David Gray, Carol Clugston, Anna Dominiczak and Alex Alderton, Roberto Amato, Jeffrey Barrett, Sonia Goncalves, Ewan Harrison, David K. Jackson, Ian Johnston, Dominic Kwiatkowski, Cordelia Langford, John Sillitoe on behalf of the Wellcome Sanger Institute COVID-19 Surveillance Team |
| EPI_ISL_2830317 | Lighthouse Lab in Alderley Park | Wellcome Sanger Institute for the COVID-19 Genomics UK (COG-UK) Consortium | Jacquelyn Wynn, Mairead Hyland, The Lighthouse Lab in Alderley Park and Alex Alderton, Roberto Amato, Jeffrey Barrett, Sonia Goncalves, Ewan Harrison, David K. Jackson, Ian Johnston, Dominic Kwiatkowski, Cordelia Langford, John Sillitoe on behalf of the Wellcome Sanger Institute COVID-19 Surveillance Team |
| EPI_ISL_2830318 | Lighthouse Lab in Glasgow | Wellcome Sanger Institute for the COVID-19 Genomics UK (COG-UK) Consortium | Harper VanSteenhouse, Yumi Kasai, David Gray, Carol Clugston, Anna Dominiczak and Alex Alderton, Roberto Amato, Jeffrey Barrett, Sonia Goncalves, Ewan Harrison, David K. Jackson, Ian Johnston, Dominic Kwiatkowski, Cordelia Langford, John Sillitoe on behalf of the Wellcome Sanger Institute COVID-19 Surveillance Team |
| EPI_ISL_2830319, EPI_ISL_2830320, EPI_ISL_2830321 | Lighthouse Lab in Alderley Park | Wellcome Sanger Institute for the COVID-19 Genomics UK (COG-UK) Consortium | Jacquelyn Wynn, Mairead Hyland, The Lighthouse Lab in Alderley Park and Alex Alderton, Roberto Amato, Jeffrey Barrett, Sonia Goncalves, Ewan Harrison, David K. Jackson, Ian Johnston, Dominic Kwiatkowski, Cordelia Langford, John Sillitoe on behalf of the Wellcome Sanger Institute COVID-19 Surveillance Team |
| EPI_ISL_2830322, EPI_ISL_2830323, EPI_ISL_2830324, EPI_ISL_2830325, EPI_ISL_2830326, EPI_ISL_2830327 | Lighthouse Lab in Glasgow | Wellcome Sanger Institute for the COVID-19 Genomics UK (COG-UK) Consortium | Harper VanSteenhouse, Yumi Kasai, David Gray, Carol Clugston, Anna Dominiczak and Alex Alderton, Roberto Amato, Jeffrey Barrett, Sonia Goncalves, Ewan Harrison, David K. Jackson, Ian Johnston, Dominic Kwiatkowski, Cordelia Langford, John Sillitoe on behalf of the Wellcome Sanger Institute COVID-19 Surveillance Team |
| EPI_ISL_2830328, EPI_ISL_2830329, EPI_ISL_2830330 | Lighthouse Lab in Alderley Park | Wellcome Sanger Institute for the COVID-19 Genomics UK (COG-UK) Consortium | Jacquelyn Wynn, Mairead Hyland, The Lighthouse Lab in Alderley Park and Alex Alderton, Roberto Amato, Jeffrey Barrett, Sonia Goncalves, Ewan Harrison, David K. Jackson, Ian Johnston, Dominic Kwiatkowski, Cordelia Langford, John Sillitoe on behalf of the Wellcome Sanger Institute COVID-19 Surveillance Team |
| EPI_ISL_2830331 | Lighthouse Lab in Glasgow | Wellcome Sanger Institute for the COVID-19 Genomics UK (COG-UK) Consortium | Harper VanSteenhouse, Yumi Kasai, David Gray, Carol Clugston, Anna Dominiczak and Alex Alderton, Roberto Amato, Jeffrey Barrett, Sonia Goncalves, Ewan Harrison, David K. Jackson, Ian Johnston, Dominic Kwiatkowski, Cordelia Langford, John Sillitoe on behalf of the Wellcome Sanger Institute COVID-19 Surveillance Team |
| EPI_ISL_2830332, EPI_ISL_2830333, | Lighthouse Lab in Milton Keynes | Wellcome Sanger Institute for the COVID-19 Genomics UK (COG-UK) Consortium | The Lighthouse Lab in Milton Keynes and Alex Alderton, Roberto Amato, Jeffrey Barrett, Sonia Goncalves, Ewan Harrison, David K. Jackson, Ian Johnston, Dominic Kwiatkowski, Cordelia Langford, John Sillitoe on behalf of the Wellcome Sanger Institute COVID-19 Surveillance Team |

| | | | |
|---|---|---|---|
| EPI_ISL_2830334, EPI_ISL_2830335 | | | |
| EPI_ISL_2830336 | Lighthouse Lab in Alderley Park | Wellcome Sanger Institute for the COVID-19 Genomics UK (COG-UK) Consortium | Jacquelyn Wynn, Mairead Hyland, The Lighthouse Lab in Alderley Park and Alex Alderton, Roberto Amato, Jeffrey Barrett, Sonia Goncalves, Ewan Harrison, David K. Jackson, Ian Johnston, Dominic Kwiatkowski, Cordelia Langford, John Sillitoe on behalf of the Wellcome Sanger Institute COVID-19 Surveillance Team |
| EPI_ISL_2830337 | Lighthouse Lab in Milton Keynes | Wellcome Sanger Institute for the COVID-19 Genomics UK (COG-UK) Consortium | The Lighthouse Lab in Milton Keynes and Alex Alderton, Roberto Amato, Jeffrey Barrett, Sonia Goncalves, Ewan Harrison, David K. Jackson, Ian Johnston, Dominic Kwiatkowski, Cordelia Langford, John Sillitoe on behalf of the Wellcome Sanger Institute COVID-19 Surveillance Team |
| EPI_ISL_2830338 | Lighthouse Lab in Alderley Park | Wellcome Sanger Institute for the COVID-19 Genomics UK (COG-UK) Consortium | Jacquelyn Wynn, Mairead Hyland, The Lighthouse Lab in Alderley Park and Alex Alderton, Roberto Amato, Jeffrey Barrett, Sonia Goncalves, Ewan Harrison, David K. Jackson, Ian Johnston, Dominic Kwiatkowski, Cordelia Langford, John Sillitoe on behalf of the Wellcome Sanger Institute COVID-19 Surveillance Team |
| EPI_ISL_2830339 | Lighthouse Lab in Milton Keynes | Wellcome Sanger Institute for the COVID-19 Genomics UK (COG-UK) Consortium | The Lighthouse Lab in Milton Keynes and Alex Alderton, Roberto Amato, Jeffrey Barrett, Sonia Goncalves, Ewan Harrison, David K. Jackson, Ian Johnston, Dominic Kwiatkowski, Cordelia Langford, John Sillitoe on behalf of the Wellcome Sanger Institute COVID-19 Surveillance Team |
| EPI_ISL_2830340 | Lighthouse Lab in Alderley Park | Wellcome Sanger Institute for the COVID-19 Genomics UK (COG-UK) Consortium | Jacquelyn Wynn, Mairead Hyland, The Lighthouse Lab in Alderley Park and Alex Alderton, Roberto Amato, Jeffrey Barrett, Sonia Goncalves, Ewan Harrison, David K. Jackson, Ian Johnston, Dominic Kwiatkowski, Cordelia Langford, John Sillitoe on behalf of the Wellcome Sanger Institute COVID-19 Surveillance Team |
| EPI_ISL_2830341, EPI_ISL_2830342, EPI_ISL_2830343 | Lighthouse Lab in Milton Keynes | Wellcome Sanger Institute for the COVID-19 Genomics UK (COG-UK) Consortium | The Lighthouse Lab in Milton Keynes and Alex Alderton, Roberto Amato, Jeffrey Barrett, Sonia Goncalves, Ewan Harrison, David K. Jackson, Ian Johnston, Dominic Kwiatkowski, Cordelia Langford, John Sillitoe on behalf of the Wellcome Sanger Institute COVID-19 Surveillance Team |
| EPI_ISL_2830344 | Lighthouse Lab in Alderley Park | Wellcome Sanger Institute for the COVID-19 Genomics UK (COG-UK) Consortium | Jacquelyn Wynn, Mairead Hyland, The Lighthouse Lab in Alderley Park and Alex Alderton, Roberto Amato, Jeffrey Barrett, Sonia Goncalves, Ewan Harrison, David K. Jackson, Ian Johnston, Dominic Kwiatkowski, Cordelia Langford, John Sillitoe on behalf of the Wellcome Sanger Institute COVID-19 Surveillance Team |
| EPI_ISL_2830345, EPI_ISL_2830346, EPI_ISL_2830347, EPI_ISL_2830348 | Lighthouse Lab in Milton Keynes | Wellcome Sanger Institute for the COVID-19 Genomics UK (COG-UK) Consortium | The Lighthouse Lab in Milton Keynes and Alex Alderton, Roberto Amato, Jeffrey Barrett, Sonia Goncalves, Ewan Harrison, David K. Jackson, Ian Johnston, Dominic Kwiatkowski, Cordelia Langford, John Sillitoe on behalf of the Wellcome Sanger Institute COVID-19 Surveillance Team |
| EPI_ISL_2830349 | Lighthouse Lab in Alderley Park | Wellcome Sanger Institute for the COVID-19 Genomics UK (COG-UK) Consortium | Jacquelyn Wynn, Mairead Hyland, The Lighthouse Lab in Alderley Park and Alex Alderton, Roberto Amato, Jeffrey Barrett, Sonia Goncalves, Ewan Harrison, David K. Jackson, Ian Johnston, Dominic Kwiatkowski, Cordelia Langford, John Sillitoe on behalf of the Wellcome Sanger Institute COVID-19 Surveillance Team |
| EPI_ISL_2830350, EPI_ISL_2830351, EPI_ISL_2830352 | Lighthouse Lab in Milton Keynes | Wellcome Sanger Institute for the COVID-19 Genomics UK (COG-UK) Consortium | The Lighthouse Lab in Milton Keynes and Alex Alderton, Roberto Amato, Jeffrey Barrett, Sonia Goncalves, Ewan Harrison, David K. Jackson, Ian Johnston, Dominic Kwiatkowski, Cordelia Langford, John Sillitoe on behalf of the Wellcome Sanger Institute COVID-19 Surveillance Team |
| EPI_ISL_2830353, EPI_ISL_2830354 | Lighthouse Lab in Alderley Park | Wellcome Sanger Institute for the COVID-19 Genomics UK (COG-UK) Consortium | Jacquelyn Wynn, Mairead Hyland, The Lighthouse Lab in Alderley Park and Alex Alderton, Roberto Amato, Jeffrey Barrett, Sonia Goncalves, Ewan Harrison, David K. Jackson, Ian Johnston, Dominic Kwiatkowski, Cordelia Langford, John Sillitoe on behalf of the Wellcome Sanger Institute COVID-19 Surveillance Team |
| EPI_ISL_2830355 | Lighthouse Lab in Milton Keynes | Wellcome Sanger Institute for the COVID-19 Genomics UK (COG-UK) Consortium | The Lighthouse Lab in Milton Keynes and Alex Alderton, Roberto Amato, Jeffrey Barrett, Sonia Goncalves, Ewan Harrison, David K. Jackson, Ian Johnston, Dominic Kwiatkowski, Cordelia Langford, John Sillitoe on behalf of the Wellcome Sanger Institute COVID-19 Surveillance Team |
| EPI_ISL_2830356 | Lighthouse Lab in Alderley Park | Wellcome Sanger Institute for the COVID-19 Genomics UK (COG-UK) Consortium | Jacquelyn Wynn, Mairead Hyland, The Lighthouse Lab in Alderley Park and Alex Alderton, Roberto Amato, Jeffrey Barrett, Sonia Goncalves, Ewan Harrison, David K. Jackson, Ian Johnston, Dominic Kwiatkowski, Cordelia Langford, John Sillitoe on behalf of the Wellcome Sanger Institute COVID-19 Surveillance Team |
| EPI_ISL_2830357, EPI_ISL_2830358 | Lighthouse Lab in Milton Keynes | Wellcome Sanger Institute for the COVID-19 Genomics UK (COG-UK) Consortium | The Lighthouse Lab in Milton Keynes and Alex Alderton, Roberto Amato, Jeffrey Barrett, Sonia Goncalves, Ewan Harrison, David K. Jackson, Ian Johnston, Dominic Kwiatkowski, Cordelia Langford, John Sillitoe on behalf of the Wellcome Sanger Institute COVID-19 Surveillance Team |
| EPI_ISL_2830359, EPI_ISL_2830360 | Lighthouse Lab in Alderley Park | Wellcome Sanger Institute for the COVID-19 Genomics UK (COG-UK) Consortium | Jacquelyn Wynn, Mairead Hyland, The Lighthouse Lab in Alderley Park and Alex Alderton, Roberto Amato, Jeffrey Barrett, Sonia Goncalves, Ewan Harrison, David K. Jackson, Ian Johnston, Dominic Kwiatkowski, Cordelia Langford, John Sillitoe on behalf of the Wellcome Sanger Institute COVID-19 Surveillance Team |
| EPI_ISL_2830361, EPI_ISL_2830362, EPI_ISL_2830363 | Lighthouse Lab in Milton Keynes | Wellcome Sanger Institute for the COVID-19 Genomics UK (COG-UK) Consortium | The Lighthouse Lab in Milton Keynes and Alex Alderton, Roberto Amato, Jeffrey Barrett, Sonia Goncalves, Ewan Harrison, David K. Jackson, Ian Johnston, Dominic Kwiatkowski, Cordelia Langford, John Sillitoe on behalf of the Wellcome Sanger Institute COVID-19 Surveillance Team |
| EPI_ISL_2830364, EPI_ISL_2830365 | Lighthouse Lab in Alderley Park | Wellcome Sanger Institute for the COVID-19 Genomics UK (COG-UK) Consortium | Jacquelyn Wynn, Mairead Hyland, The Lighthouse Lab in Alderley Park and Alex Alderton, Roberto Amato, Jeffrey Barrett, Sonia Goncalves, Ewan Harrison, David K. Jackson, Ian Johnston, Dominic Kwiatkowski, Cordelia Langford, John Sillitoe on behalf of the Wellcome Sanger Institute COVID-19 Surveillance Team |
| EPI_ISL_2830366 | Lighthouse Lab in Milton Keynes | Wellcome Sanger Institute for the COVID-19 Genomics UK (COG-UK) Consortium | The Lighthouse Lab in Milton Keynes and Alex Alderton, Roberto Amato, Jeffrey Barrett, Sonia Goncalves, Ewan Harrison, David K. Jackson, Ian Johnston, Dominic Kwiatkowski, Cordelia Langford, John Sillitoe on behalf of the Wellcome Sanger Institute COVID-19 Surveillance Team |
| EPI_ISL_2830367 | Lighthouse Lab in Alderley Park | Wellcome Sanger Institute for the COVID-19 Genomics UK (COG-UK) Consortium | Jacquelyn Wynn, Mairead Hyland, The Lighthouse Lab in Alderley Park and Alex Alderton, Roberto Amato, Jeffrey Barrett, Sonia Goncalves, Ewan Harrison, David K. Jackson, Ian Johnston, Dominic Kwiatkowski, Cordelia Langford, John Sillitoe on behalf of the Wellcome Sanger Institute COVID-19 Surveillance Team |
| EPI_ISL_2830368, EPI_ISL_2830369, EPI_ISL_2830370, EPI_ISL_2830371 | Lighthouse Lab in Milton Keynes | Wellcome Sanger Institute for the COVID-19 Genomics UK (COG-UK) Consortium | The Lighthouse Lab in Milton Keynes and Alex Alderton, Roberto Amato, Jeffrey Barrett, Sonia Goncalves, Ewan Harrison, David K. Jackson, Ian Johnston, Dominic Kwiatkowski, Cordelia Langford, John Sillitoe on behalf of the Wellcome Sanger Institute COVID-19 Surveillance Team |
| EPI_ISL_2830372 | Lighthouse Lab in Alderley Park | Wellcome Sanger Institute for the COVID-19 Genomics UK (COG-UK) Consortium | Jacquelyn Wynn, Mairead Hyland, The Lighthouse Lab in Alderley Park and Alex Alderton, Roberto Amato, Jeffrey Barrett, Sonia Goncalves, Ewan Harrison, David K. Jackson, Ian Johnston, Dominic Kwiatkowski, Cordelia Langford, John Sillitoe on behalf of the Wellcome Sanger Institute COVID-19 Surveillance Team |
| EPI_ISL_2830373 | Lighthouse Lab in Milton Keynes | Wellcome Sanger Institute for the COVID-19 Genomics UK (COG-UK) Consortium | The Lighthouse Lab in Milton Keynes and Alex Alderton, Roberto Amato, Jeffrey Barrett, Sonia Goncalves, Ewan Harrison, David K. Jackson, Ian Johnston, Dominic Kwiatkowski, Cordelia Langford, John Sillitoe on behalf of the Wellcome Sanger Institute COVID-19 Surveillance Team |
| EPI_ISL_2830374 | Lighthouse Lab in Alderley Park | Wellcome Sanger Institute for the COVID-19 Genomics UK (COG-UK) Consortium | Jacquelyn Wynn, Mairead Hyland, The Lighthouse Lab in Alderley Park and Alex Alderton, Roberto Amato, Jeffrey Barrett, Sonia Goncalves, Ewan Harrison, David K. Jackson, Ian Johnston, Dominic Kwiatkowski, Cordelia Langford, John Sillitoe on behalf of the Wellcome Sanger Institute COVID-19 Surveillance Team |
| EPI_ISL_2830375, EPI_ISL_2830376, EPI_ISL_2830377 | Lighthouse Lab in Milton Keynes | Wellcome Sanger Institute for the COVID-19 Genomics UK (COG-UK) Consortium | The Lighthouse Lab in Milton Keynes and Alex Alderton, Roberto Amato, Jeffrey Barrett, Sonia Goncalves, Ewan Harrison, David K. Jackson, Ian Johnston, Dominic Kwiatkowski, Cordelia Langford, John Sillitoe on behalf of the Wellcome Sanger Institute COVID-19 Surveillance Team |
| EPI_ISL_2830378 | Lighthouse Lab in Alderley Park | Wellcome Sanger Institute for the COVID-19 Genomics UK (COG-UK) Consortium | Jacquelyn Wynn, Mairead Hyland, The Lighthouse Lab in Alderley Park and Alex Alderton, Roberto Amato, Jeffrey Barrett, Sonia Goncalves, Ewan Harrison, David K. Jackson, Ian Johnston, Dominic Kwiatkowski, Cordelia Langford, John Sillitoe on behalf of the Wellcome Sanger Institute COVID-19 Surveillance Team |
| EPI_ISL_2830379 | Lighthouse Lab in Glasgow | Wellcome Sanger Institute for the COVID-19 Genomics UK (COG-UK) Consortium | Harper VanSteenhouse, Yumi Kasai, David Gray, Carol Clugston, Anna Dominiczak and Alex Alderton, Roberto Amato, Jeffrey Barrett, Sonia Goncalves, Ewan Harrison, David K. Jackson, Ian Johnston, Dominic Kwiatkowski, Cordelia Langford, John Sillitoe on behalf of the Wellcome Sanger Institute COVID-19 Surveillance Team |
| EPI_ISL_2830380 | Lighthouse Lab in Alderley Park | Wellcome Sanger Institute for the COVID-19 Genomics UK (COG-UK) Consortium | Jacquelyn Wynn, Mairead Hyland, The Lighthouse Lab in Alderley Park and Alex Alderton, Roberto Amato, Jeffrey Barrett, Sonia Goncalves, Ewan Harrison, David K. Jackson, Ian Johnston, Dominic Kwiatkowski, Cordelia Langford, John Sillitoe on behalf of the Wellcome Sanger Institute COVID-19 Surveillance Team |
| EPI_ISL_2830381 | Lighthouse Lab in Milton Keynes | Wellcome Sanger Institute for the COVID-19 Genomics UK (COG-UK) Consortium | The Lighthouse Lab in Milton Keynes and Alex Alderton, Roberto Amato, Jeffrey Barrett, Sonia Goncalves, Ewan Harrison, David K. Jackson, Ian Johnston, Dominic Kwiatkowski, Cordelia Langford, John Sillitoe on behalf of the Wellcome Sanger Institute COVID-19 Surveillance Team |
| EPI_ISL_2830382, EPI_ISL_2830383 | Lighthouse Lab in Alderley Park | Wellcome Sanger Institute for the COVID-19 Genomics UK (COG-UK) Consortium | Jacquelyn Wynn, Mairead Hyland, The Lighthouse Lab in Alderley Park and Alex Alderton, Roberto Amato, Jeffrey Barrett, Sonia Goncalves, Ewan Harrison, David K. Jackson, Ian Johnston, Dominic Kwiatkowski, Cordelia Langford, John Sillitoe on behalf of the Wellcome Sanger Institute COVID-19 Surveillance Team |
| EPI_ISL_2830384, EPI_ISL_2830385 | Lighthouse Lab in Milton Keynes | Wellcome Sanger Institute for the COVID-19 Genomics UK (COG-UK) Consortium | The Lighthouse Lab in Milton Keynes and Alex Alderton, Roberto Amato, Jeffrey Barrett, Sonia Goncalves, Ewan Harrison, David K. Jackson, Ian Johnston, Dominic Kwiatkowski, Cordelia Langford, John Sillitoe on behalf of the Wellcome Sanger Institute COVID-19 Surveillance Team |
| EPI_ISL_2830386 | Lighthouse Lab in Alderley Park | Wellcome Sanger Institute for the COVID-19 Genomics UK (COG-UK) Consortium | Jacquelyn Wynn, Mairead Hyland, The Lighthouse Lab in Alderley Park and Alex Alderton, Roberto Amato, Jeffrey Barrett, Sonia Goncalves, Ewan Harrison, David K. Jackson, Ian Johnston, Dominic Kwiatkowski, Cordelia Langford, John Sillitoe on behalf of the Wellcome Sanger Institute COVID-19 Surveillance Team |
| EPI_ISL_2830387, EPI_ISL_2830388, EPI_ISL_2830389 | Lighthouse Lab in Milton Keynes | Wellcome Sanger Institute for the COVID-19 Genomics UK (COG-UK) Consortium | The Lighthouse Lab in Milton Keynes and Alex Alderton, Roberto Amato, Jeffrey Barrett, Sonia Goncalves, Ewan Harrison, David K. Jackson, Ian Johnston, Dominic Kwiatkowski, Cordelia Langford, John Sillitoe on behalf of the Wellcome Sanger Institute COVID-19 Surveillance Team |
| EPI_ISL_2830390 | Lighthouse Lab in Alderley Park | Wellcome Sanger Institute for the COVID-19 Genomics UK (COG-UK) Consortium | Jacquelyn Wynn, Mairead Hyland, The Lighthouse Lab in Alderley Park and Alex Alderton, Roberto Amato, Jeffrey Barrett, Sonia Goncalves, Ewan Harrison, David K. Jackson, Ian Johnston, Dominic Kwiatkowski, Cordelia Langford, John Sillitoe on behalf of the Wellcome Sanger Institute COVID-19 Surveillance Team |
| EPI_ISL_2830391 | Lighthouse Lab in Milton Keynes | Wellcome Sanger Institute for the COVID-19 Genomics UK (COG-UK) Consortium | The Lighthouse Lab in Milton Keynes and Alex Alderton, Roberto Amato, Jeffrey Barrett, Sonia Goncalves, Ewan Harrison, David K. Jackson, Ian Johnston, Dominic Kwiatkowski, Cordelia Langford, John Sillitoe on behalf of the Wellcome Sanger Institute COVID-19 Surveillance Team |
| EPI_ISL_2830392, EPI_ISL_2830393 | Lighthouse Lab in Alderley Park | Wellcome Sanger Institute for the COVID-19 Genomics UK (COG-UK) Consortium | Jacquelyn Wynn, Mairead Hyland, The Lighthouse Lab in Alderley Park and Alex Alderton, Roberto Amato, Jeffrey Barrett, Sonia Goncalves, Ewan Harrison, David K. Jackson, Ian Johnston, Dominic Kwiatkowski, Cordelia Langford, John Sillitoe on behalf of the Wellcome Sanger Institute COVID-19 Surveillance Team |
| EPI_ISL_2830394, EPI_ISL_2830395, EPI_ISL_2830396, EPI_ISL_2830397, EPI_ISL_2830398, EPI_ISL_2830399 | Lighthouse Lab in Milton Keynes | Wellcome Sanger Institute for the COVID-19 Genomics UK (COG-UK) Consortium | The Lighthouse Lab in Milton Keynes and Alex Alderton, Roberto Amato, Jeffrey Barrett, Sonia Goncalves, Ewan Harrison, David K. Jackson, Ian Johnston, Dominic Kwiatkowski, Cordelia Langford, John Sillitoe on behalf of the Wellcome Sanger Institute COVID-19 Surveillance Team |
| EPI_ISL_2830400 | Lighthouse Lab in Alderley Park | Wellcome Sanger Institute for the COVID-19 Genomics UK (COG-UK) Consortium | Jacquelyn Wynn, Mairead Hyland, The Lighthouse Lab in Alderley Park and Alex Alderton, Roberto Amato, Jeffrey Barrett, Sonia Goncalves, Ewan Harrison, David K. Jackson, Ian Johnston, Dominic Kwiatkowski, Cordelia Langford, John Sillitoe on behalf of the Wellcome Sanger Institute COVID-19 Surveillance Team |
| EPI_ISL_2830401, EPI_ISL_2830402, EPI_ISL_2830403 | Lighthouse Lab in Milton Keynes | Wellcome Sanger Institute for the COVID-19 Genomics UK (COG-UK) Consortium | The Lighthouse Lab in Milton Keynes and Alex Alderton, Roberto Amato, Jeffrey Barrett, Sonia Goncalves, Ewan Harrison, David K. Jackson, Ian Johnston, Dominic Kwiatkowski, Cordelia Langford, John Sillitoe on behalf of the Wellcome Sanger Institute COVID-19 Surveillance Team |
| EPI_ISL_2830404, | Lighthouse Lab in Alderley Park | Wellcome Sanger Institute for the COVID-19 Genomics UK (COG-UK) | Jacquelyn Wynn, Mairead Hyland, The Lighthouse Lab in Alderley Park and Alex Alderton, Roberto Amato, Jeffrey Barrett, Sonia Goncalves, Ewan Harrison, David K. Jackson, Ian Johnston, Dominic Kwiatkowski, Cordelia Langford, John Sillitoe on |

| | | | |
|---|---|---|---|
| EPI_ISL_2830406 | | Consortium | behalf of the Wellcome Sanger Institute COVID-19 Surveillance Team |
| EPI_ISL_2830407, EPI_ISL_2830408 | Lighthouse Lab in Milton Keynes | Wellcome Sanger Institute for the COVID-19 Genomics UK (COG-UK) Consortium | The Lighthouse Lab in Milton Keynes and Alex Alderton, Roberto Amato, Jeffrey Barrett, Sonia Goncalves, Ewan Harrison, David K. Jackson, Ian Johnston, Dominic Kwiatkowski, Cordelia Langford, John Sillitoe on behalf of the Wellcome Sanger Institute COVID-19 Surveillance Team |
| EPI_ISL_2830409, EPI_ISL_2830410 | Lighthouse Lab in Alderley Park | Wellcome Sanger Institute for the COVID-19 Genomics UK (COG-UK) Consortium | Jacquelyn Wynn, Mairead Hyland, The Lighthouse Lab in Alderley Park and Alex Alderton, Roberto Amato, Jeffrey Barrett, Sonia Goncalves, Ewan Harrison, David K. Jackson, Ian Johnston, Dominic Kwiatkowski, Cordelia Langford, John Sillitoe on behalf of the Wellcome Sanger Institute COVID-19 Surveillance Team |
| EPI_ISL_2830411, EPI_ISL_2830412, EPI_ISL_2830413, EPI_ISL_2830414, EPI_ISL_2830415, EPI_ISL_2830416, EPI_ISL_2830417 | Lighthouse Lab in Milton Keynes | Wellcome Sanger Institute for the COVID-19 Genomics UK (COG-UK) Consortium | The Lighthouse Lab in Milton Keynes and Alex Alderton, Roberto Amato, Jeffrey Barrett, Sonia Goncalves, Ewan Harrison, David K. Jackson, Ian Johnston, Dominic Kwiatkowski, Cordelia Langford, John Sillitoe on behalf of the Wellcome Sanger Institute COVID-19 Surveillance Team |
| EPI_ISL_2830418 | Lighthouse Lab in Alderley Park | Wellcome Sanger Institute for the COVID-19 Genomics UK (COG-UK) Consortium | Jacquelyn Wynn, Mairead Hyland, The Lighthouse Lab in Alderley Park and Alex Alderton, Roberto Amato, Jeffrey Barrett, Sonia Goncalves, Ewan Harrison, David K. Jackson, Ian Johnston, Dominic Kwiatkowski, Cordelia Langford, John Sillitoe on behalf of the Wellcome Sanger Institute COVID-19 Surveillance Team |
| EPI_ISL_2830419, EPI_ISL_2830420 | Lighthouse Lab in Milton Keynes | Wellcome Sanger Institute for the COVID-19 Genomics UK (COG-UK) Consortium | The Lighthouse Lab in Milton Keynes and Alex Alderton, Roberto Amato, Jeffrey Barrett, Sonia Goncalves, Ewan Harrison, David K. Jackson, Ian Johnston, Dominic Kwiatkowski, Cordelia Langford, John Sillitoe on behalf of the Wellcome Sanger Institute COVID-19 Surveillance Team |
| EPI_ISL_2830421, EPI_ISL_2830422, EPI_ISL_2830423, EPI_ISL_2830424 | Lighthouse Lab in Alderley Park | Wellcome Sanger Institute for the COVID-19 Genomics UK (COG-UK) Consortium | Jacquelyn Wynn, Mairead Hyland, The Lighthouse Lab in Alderley Park and Alex Alderton, Roberto Amato, Jeffrey Barrett, Sonia Goncalves, Ewan Harrison, David K. Jackson, Ian Johnston, Dominic Kwiatkowski, Cordelia Langford, John Sillitoe on behalf of the Wellcome Sanger Institute COVID-19 Surveillance Team |
| EPI_ISL_2830425, EPI_ISL_2830426, EPI_ISL_2830427 | Lighthouse Lab in Milton Keynes | Wellcome Sanger Institute for the COVID-19 Genomics UK (COG-UK) Consortium | The Lighthouse Lab in Milton Keynes and Alex Alderton, Roberto Amato, Jeffrey Barrett, Sonia Goncalves, Ewan Harrison, David K. Jackson, Ian Johnston, Dominic Kwiatkowski, Cordelia Langford, John Sillitoe on behalf of the Wellcome Sanger Institute COVID-19 Surveillance Team |
| EPI_ISL_2830428 | Lighthouse Lab in Alderley Park | Wellcome Sanger Institute for the COVID-19 Genomics UK (COG-UK) Consortium | Jacquelyn Wynn, Mairead Hyland, The Lighthouse Lab in Alderley Park and Alex Alderton, Roberto Amato, Jeffrey Barrett, Sonia Goncalves, Ewan Harrison, David K. Jackson, Ian Johnston, Dominic Kwiatkowski, Cordelia Langford, John Sillitoe on behalf of the Wellcome Sanger Institute COVID-19 Surveillance Team |
| EPI_ISL_2830429, EPI_ISL_2830430 | Lighthouse Lab in Milton Keynes | Wellcome Sanger Institute for the COVID-19 Genomics UK (COG-UK) Consortium | The Lighthouse Lab in Milton Keynes and Alex Alderton, Roberto Amato, Jeffrey Barrett, Sonia Goncalves, Ewan Harrison, David K. Jackson, Ian Johnston, Dominic Kwiatkowski, Cordelia Langford, John Sillitoe on behalf of the Wellcome Sanger Institute COVID-19 Surveillance Team |
| EPI_ISL_2830431 | Lighthouse Lab in Alderley Park | Wellcome Sanger Institute for the COVID-19 Genomics UK (COG-UK) Consortium | Jacquelyn Wynn, Mairead Hyland, The Lighthouse Lab in Alderley Park and Alex Alderton, Roberto Amato, Jeffrey Barrett, Sonia Goncalves, Ewan Harrison, David K. Jackson, Ian Johnston, Dominic Kwiatkowski, Cordelia Langford, John Sillitoe on behalf of the Wellcome Sanger Institute COVID-19 Surveillance Team |
| EPI_ISL_2830432, EPI_ISL_2830433 | Lighthouse Lab in Milton Keynes | Wellcome Sanger Institute for the COVID-19 Genomics UK (COG-UK) Consortium | The Lighthouse Lab in Milton Keynes and Alex Alderton, Roberto Amato, Jeffrey Barrett, Sonia Goncalves, Ewan Harrison, David K. Jackson, Ian Johnston, Dominic Kwiatkowski, Cordelia Langford, John Sillitoe on behalf of the Wellcome Sanger Institute COVID-19 Surveillance Team |
| EPI_ISL_2830434 | Lighthouse Lab in Alderley Park | Wellcome Sanger Institute for the COVID-19 Genomics UK (COG-UK) Consortium | Jacquelyn Wynn, Mairead Hyland, The Lighthouse Lab in Alderley Park and Alex Alderton, Roberto Amato, Jeffrey Barrett, Sonia Goncalves, Ewan Harrison, David K. Jackson, Ian Johnston, Dominic Kwiatkowski, Cordelia Langford, John Sillitoe on behalf of the Wellcome Sanger Institute COVID-19 Surveillance Team |
| EPI_ISL_2830435, EPI_ISL_2830436 | Lighthouse Lab in Milton Keynes | Wellcome Sanger Institute for the COVID-19 Genomics UK (COG-UK) Consortium | The Lighthouse Lab in Milton Keynes and Alex Alderton, Roberto Amato, Jeffrey Barrett, Sonia Goncalves, Ewan Harrison, David K. Jackson, Ian Johnston, Dominic Kwiatkowski, Cordelia Langford, John Sillitoe on behalf of the Wellcome Sanger Institute COVID-19 Surveillance Team |
| EPI_ISL_2830437 | Lighthouse Lab in Alderley Park | Wellcome Sanger Institute for the COVID-19 Genomics UK (COG-UK) Consortium | Jacquelyn Wynn, Mairead Hyland, The Lighthouse Lab in Alderley Park and Alex Alderton, Roberto Amato, Jeffrey Barrett, Sonia Goncalves, Ewan Harrison, David K. Jackson, Ian Johnston, Dominic Kwiatkowski, Cordelia Langford, John Sillitoe on behalf of the Wellcome Sanger Institute COVID-19 Surveillance Team |
| EPI_ISL_2830438, EPI_ISL_2830439, EPI_ISL_2830440, EPI_ISL_2830441, EPI_ISL_2830442, EPI_ISL_2830443, EPI_ISL_2830444, EPI_ISL_2830445 | Lighthouse Lab in Milton Keynes | Wellcome Sanger Institute for the COVID-19 Genomics UK (COG-UK) Consortium | The Lighthouse Lab in Milton Keynes and Alex Alderton, Roberto Amato, Jeffrey Barrett, Sonia Goncalves, Ewan Harrison, David K. Jackson, Ian Johnston, Dominic Kwiatkowski, Cordelia Langford, John Sillitoe on behalf of the Wellcome Sanger Institute COVID-19 Surveillance Team |
| EPI_ISL_2830446, EPI_ISL_2830447 | Lighthouse Lab in Alderley Park | Wellcome Sanger Institute for the COVID-19 Genomics UK (COG-UK) Consortium | Jacquelyn Wynn, Mairead Hyland, The Lighthouse Lab in Alderley Park and Alex Alderton, Roberto Amato, Jeffrey Barrett, Sonia Goncalves, Ewan Harrison, David K. Jackson, Ian Johnston, Dominic Kwiatkowski, Cordelia Langford, John Sillitoe on behalf of the Wellcome Sanger Institute COVID-19 Surveillance Team |
| EPI_ISL_2830448, EPI_ISL_2830449 | Lighthouse Lab in Milton Keynes | Wellcome Sanger Institute for the COVID-19 Genomics UK (COG-UK) Consortium | The Lighthouse Lab in Milton Keynes and Alex Alderton, Roberto Amato, Jeffrey Barrett, Sonia Goncalves, Ewan Harrison, David K. Jackson, Ian Johnston, Dominic Kwiatkowski, Cordelia Langford, John Sillitoe on behalf of the Wellcome Sanger Institute COVID-19 Surveillance Team |
| EPI_ISL_2830450 | Lighthouse Lab in Alderley Park | Wellcome Sanger Institute for the COVID-19 Genomics UK (COG-UK) Consortium | Jacquelyn Wynn, Mairead Hyland, The Lighthouse Lab in Alderley Park and Alex Alderton, Roberto Amato, Jeffrey Barrett, Sonia Goncalves, Ewan Harrison, David K. Jackson, Ian Johnston, Dominic Kwiatkowski, Cordelia Langford, John Sillitoe on behalf of the Wellcome Sanger Institute COVID-19 Surveillance Team |
| EPI_ISL_2830451 | Lighthouse Lab in Milton Keynes | Wellcome Sanger Institute for the COVID-19 Genomics UK (COG-UK) Consortium | The Lighthouse Lab in Milton Keynes and Alex Alderton, Roberto Amato, Jeffrey Barrett, Sonia Goncalves, Ewan Harrison, David K. Jackson, Ian Johnston, Dominic Kwiatkowski, Cordelia Langford, John Sillitoe on behalf of the Wellcome Sanger Institute COVID-19 Surveillance Team |
| EPI_ISL_2830452, EPI_ISL_2830454 | Lighthouse Lab in Alderley Park | Wellcome Sanger Institute for the COVID-19 Genomics UK (COG-UK) Consortium | Jacquelyn Wynn, Mairead Hyland, The Lighthouse Lab in Alderley Park and Alex Alderton, Roberto Amato, Jeffrey Barrett, Sonia Goncalves, Ewan Harrison, David K. Jackson, Ian Johnston, Dominic Kwiatkowski, Cordelia Langford, John Sillitoe on behalf of the Wellcome Sanger Institute COVID-19 Surveillance Team |
| EPI_ISL_2830455 | Lighthouse Lab in Milton Keynes | Wellcome Sanger Institute for the COVID-19 Genomics UK (COG-UK) Consortium | The Lighthouse Lab in Milton Keynes and Alex Alderton, Roberto Amato, Jeffrey Barrett, Sonia Goncalves, Ewan Harrison, David K. Jackson, Ian Johnston, Dominic Kwiatkowski, Cordelia Langford, John Sillitoe on behalf of the Wellcome Sanger Institute COVID-19 Surveillance Team |
| EPI_ISL_2830456, EPI_ISL_2830457 | Lighthouse Lab in Alderley Park | Wellcome Sanger Institute for the COVID-19 Genomics UK (COG-UK) Consortium | Jacquelyn Wynn, Mairead Hyland, The Lighthouse Lab in Alderley Park and Alex Alderton, Roberto Amato, Jeffrey Barrett, Sonia Goncalves, Ewan Harrison, David K. Jackson, Ian Johnston, Dominic Kwiatkowski, Cordelia Langford, John Sillitoe on behalf of the Wellcome Sanger Institute COVID-19 Surveillance Team |
| EPI_ISL_2830458 | Lighthouse Lab in Milton Keynes | Wellcome Sanger Institute for the COVID-19 Genomics UK (COG-UK) Consortium | The Lighthouse Lab in Milton Keynes and Alex Alderton, Roberto Amato, Jeffrey Barrett, Sonia Goncalves, Ewan Harrison, David K. Jackson, Ian Johnston, Dominic Kwiatkowski, Cordelia Langford, John Sillitoe on behalf of the Wellcome Sanger Institute COVID-19 Surveillance Team |
| EPI_ISL_2830459 | Lighthouse Lab in Alderley Park | Wellcome Sanger Institute for the COVID-19 Genomics UK (COG-UK) Consortium | Jacquelyn Wynn, Mairead Hyland, The Lighthouse Lab in Alderley Park and Alex Alderton, Roberto Amato, Jeffrey Barrett, Sonia Goncalves, Ewan Harrison, David K. Jackson, Ian Johnston, Dominic Kwiatkowski, Cordelia Langford, John Sillitoe on behalf of the Wellcome Sanger Institute COVID-19 Surveillance Team |
| EPI_ISL_2830460 | Lighthouse Lab in Milton Keynes | Wellcome Sanger Institute for the COVID-19 Genomics UK (COG-UK) Consortium | The Lighthouse Lab in Milton Keynes and Alex Alderton, Roberto Amato, Jeffrey Barrett, Sonia Goncalves, Ewan Harrison, David K. Jackson, Ian Johnston, Dominic Kwiatkowski, Cordelia Langford, John Sillitoe on behalf of the Wellcome Sanger Institute COVID-19 Surveillance Team |
| EPI_ISL_2830461 | Lighthouse Lab in Alderley Park | Wellcome Sanger Institute for the COVID-19 Genomics UK (COG-UK) Consortium | Jacquelyn Wynn, Mairead Hyland, The Lighthouse Lab in Alderley Park and Alex Alderton, Roberto Amato, Jeffrey Barrett, Sonia Goncalves, Ewan Harrison, David K. Jackson, Ian Johnston, Dominic Kwiatkowski, Cordelia Langford, John Sillitoe on behalf of the Wellcome Sanger Institute COVID-19 Surveillance Team |
| EPI_ISL_2830462, EPI_ISL_2830463, EPI_ISL_2830464, EPI_ISL_2830465 | Lighthouse Lab in Milton Keynes | Wellcome Sanger Institute for the COVID-19 Genomics UK (COG-UK) Consortium | The Lighthouse Lab in Milton Keynes and Alex Alderton, Roberto Amato, Jeffrey Barrett, Sonia Goncalves, Ewan Harrison, David K. Jackson, Ian Johnston, Dominic Kwiatkowski, Cordelia Langford, John Sillitoe on behalf of the Wellcome Sanger Institute COVID-19 Surveillance Team |
| EPI_ISL_2830466, EPI_ISL_2830467 | Lighthouse Lab in Alderley Park | Wellcome Sanger Institute for the COVID-19 Genomics UK (COG-UK) Consortium | Jacquelyn Wynn, Mairead Hyland, The Lighthouse Lab in Alderley Park and Alex Alderton, Roberto Amato, Jeffrey Barrett, Sonia Goncalves, Ewan Harrison, David K. Jackson, Ian Johnston, Dominic Kwiatkowski, Cordelia Langford, John Sillitoe on behalf of the Wellcome Sanger Institute COVID-19 Surveillance Team |
| EPI_ISL_2830468, EPI_ISL_2830469 | Lighthouse Lab in Milton Keynes | Wellcome Sanger Institute for the COVID-19 Genomics UK (COG-UK) Consortium | The Lighthouse Lab in Milton Keynes and Alex Alderton, Roberto Amato, Jeffrey Barrett, Sonia Goncalves, Ewan Harrison, David K. Jackson, Ian Johnston, Dominic Kwiatkowski, Cordelia Langford, John Sillitoe on behalf of the Wellcome Sanger Institute COVID-19 Surveillance Team |
| EPI_ISL_2830470, EPI_ISL_2830471 | Lighthouse Lab in Alderley Park | Wellcome Sanger Institute for the COVID-19 Genomics UK (COG-UK) Consortium | Jacquelyn Wynn, Mairead Hyland, The Lighthouse Lab in Alderley Park and Alex Alderton, Roberto Amato, Jeffrey Barrett, Sonia Goncalves, Ewan Harrison, David K. Jackson, Ian Johnston, Dominic Kwiatkowski, Cordelia Langford, John Sillitoe on behalf of the Wellcome Sanger Institute COVID-19 Surveillance Team |
| EPI_ISL_2830472, EPI_ISL_2830473, EPI_ISL_2830474, EPI_ISL_2830475, EPI_ISL_2830476 | Lighthouse Lab in Milton Keynes | Wellcome Sanger Institute for the COVID-19 Genomics UK (COG-UK) Consortium | The Lighthouse Lab in Milton Keynes and Alex Alderton, Roberto Amato, Jeffrey Barrett, Sonia Goncalves, Ewan Harrison, David K. Jackson, Ian Johnston, Dominic Kwiatkowski, Cordelia Langford, John Sillitoe on behalf of the Wellcome Sanger Institute COVID-19 Surveillance Team |
| EPI_ISL_2830477, EPI_ISL_2830478 | Lighthouse Lab in Alderley Park | Wellcome Sanger Institute for the COVID-19 Genomics UK (COG-UK) Consortium | Jacquelyn Wynn, Mairead Hyland, The Lighthouse Lab in Alderley Park and Alex Alderton, Roberto Amato, Jeffrey Barrett, Sonia Goncalves, Ewan Harrison, David K. Jackson, Ian Johnston, Dominic Kwiatkowski, Cordelia Langford, John Sillitoe on behalf of the Wellcome Sanger Institute COVID-19 Surveillance Team |
| EPI_ISL_2830479 | Lighthouse Lab in Milton Keynes | Wellcome Sanger Institute for the COVID-19 Genomics UK (COG-UK) Consortium | The Lighthouse Lab in Milton Keynes and Alex Alderton, Roberto Amato, Jeffrey Barrett, Sonia Goncalves, Ewan Harrison, David K. Jackson, Ian Johnston, Dominic Kwiatkowski, Cordelia Langford, John Sillitoe on behalf of the Wellcome Sanger Institute COVID-19 Surveillance Team |
| EPI_ISL_2830480, EPI_ISL_2830481 | Lighthouse Lab in Alderley Park | Wellcome Sanger Institute for the COVID-19 Genomics UK (COG-UK) Consortium | Jacquelyn Wynn, Mairead Hyland, The Lighthouse Lab in Alderley Park and Alex Alderton, Roberto Amato, Jeffrey Barrett, Sonia Goncalves, Ewan Harrison, David K. Jackson, Ian Johnston, Dominic Kwiatkowski, Cordelia Langford, John Sillitoe on behalf of the Wellcome Sanger Institute COVID-19 Surveillance Team |
| EPI_ISL_2830482, EPI_ISL_2830483, EPI_ISL_2830484, EPI_ISL_2830485, EPI_ISL_2830486 | Lighthouse Lab in Milton Keynes | Wellcome Sanger Institute for the COVID-19 Genomics UK (COG-UK) Consortium | The Lighthouse Lab in Milton Keynes and Alex Alderton, Roberto Amato, Jeffrey Barrett, Sonia Goncalves, Ewan Harrison, David K. Jackson, Ian Johnston, Dominic Kwiatkowski, Cordelia Langford, John Sillitoe on behalf of the Wellcome Sanger Institute COVID-19 Surveillance Team |
| EPI_ISL_2830487 | Lighthouse Lab in Glasgow | Wellcome Sanger Institute for the COVID-19 Genomics UK (COG-UK) | Harper VanSteenhouse, Yumi Kasai, David Gray, Carol Clugston, Anna Dominiczak and Alex Alderton, Roberto Amato, Jeffrey Barrett, Sonia Goncalves, Ewan Harrison, David K. Jackson, Ian Johnston, Dominic Kwiatkowski, Cordelia Langford, |

| | | | |
|---|---|---|---|
| | | Consortium | John Sillitoe on behalf of the Wellcome Sanger Institute COVID-19 Surveillance Team |
| EPI_ISL_2830488 | Lighthouse Lab in Alderley Park | Wellcome Sanger Institute for the COVID-19 Genomics UK (COG-UK) Consortium | Jacquelyn Wynn, Mairead Hyland, The Lighthouse Lab in Alderley Park and Alex Alderton, Roberto Amato, Jeffrey Barrett, Sonia Goncalves, Ewan Harrison, David K. Jackson, Ian Johnston, Dominic Kwiatkowski, Cordelia Langford, John Sillitoe on behalf of the Wellcome Sanger Institute COVID-19 Surveillance Team |
| EPI_ISL_2830489, EPI_ISL_2830490, EPI_ISL_2830491 | Lighthouse Lab in Milton Keynes | Wellcome Sanger Institute for the COVID-19 Genomics UK (COG-UK) Consortium | The Lighthouse Lab in Milton Keynes and Alex Alderton, Roberto Amato, Jeffrey Barrett, Sonia Goncalves, Ewan Harrison, David K. Jackson, Ian Johnston, Dominic Kwiatkowski, Cordelia Langford, John Sillitoe on behalf of the Wellcome Sanger Institute COVID-19 Surveillance Team |
| EPI_ISL_2830492, EPI_ISL_2830493 | Lighthouse Lab in Alderley Park | Wellcome Sanger Institute for the COVID-19 Genomics UK (COG-UK) Consortium | Jacquelyn Wynn, Mairead Hyland, The Lighthouse Lab in Alderley Park and Alex Alderton, Roberto Amato, Jeffrey Barrett, Sonia Goncalves, Ewan Harrison, David K. Jackson, Ian Johnston, Dominic Kwiatkowski, Cordelia Langford, John Sillitoe on behalf of the Wellcome Sanger Institute COVID-19 Surveillance Team |
| EPI_ISL_2830494, EPI_ISL_2830495, EPI_ISL_2830496, EPI_ISL_2830497, EPI_ISL_2830498, EPI_ISL_2830499, EPI_ISL_2830500, EPI_ISL_2830501 | Lighthouse Lab in Milton Keynes | Wellcome Sanger Institute for the COVID-19 Genomics UK (COG-UK) Consortium | The Lighthouse Lab in Milton Keynes and Alex Alderton, Roberto Amato, Jeffrey Barrett, Sonia Goncalves, Ewan Harrison, David K. Jackson, Ian Johnston, Dominic Kwiatkowski, Cordelia Langford, John Sillitoe on behalf of the Wellcome Sanger Institute COVID-19 Surveillance Team |
| EPI_ISL_2830502, EPI_ISL_2830503, EPI_ISL_2830504, EPI_ISL_2830506, EPI_ISL_2830507 | Lighthouse Lab in Alderley Park | Wellcome Sanger Institute for the COVID-19 Genomics UK (COG-UK) Consortium | Jacquelyn Wynn, Mairead Hyland, The Lighthouse Lab in Alderley Park and Alex Alderton, Roberto Amato, Jeffrey Barrett, Sonia Goncalves, Ewan Harrison, David K. Jackson, Ian Johnston, Dominic Kwiatkowski, Cordelia Langford, John Sillitoe on behalf of the Wellcome Sanger Institute COVID-19 Surveillance Team |
| EPI_ISL_2830508 | Lighthouse Lab in Milton Keynes | Wellcome Sanger Institute for the COVID-19 Genomics UK (COG-UK) Consortium | The Lighthouse Lab in Milton Keynes and Alex Alderton, Roberto Amato, Jeffrey Barrett, Sonia Goncalves, Ewan Harrison, David K. Jackson, Ian Johnston, Dominic Kwiatkowski, Cordelia Langford, John Sillitoe on behalf of the Wellcome Sanger Institute COVID-19 Surveillance Team |
| EPI_ISL_2830509, EPI_ISL_2830510 | Lighthouse Lab in Alderley Park | Wellcome Sanger Institute for the COVID-19 Genomics UK (COG-UK) Consortium | Jacquelyn Wynn, Mairead Hyland, The Lighthouse Lab in Alderley Park and Alex Alderton, Roberto Amato, Jeffrey Barrett, Sonia Goncalves, Ewan Harrison, David K. Jackson, Ian Johnston, Dominic Kwiatkowski, Cordelia Langford, John Sillitoe on behalf of the Wellcome Sanger Institute COVID-19 Surveillance Team |
| EPI_ISL_2830511, EPI_ISL_2830512 | Lighthouse Lab in Milton Keynes | Wellcome Sanger Institute for the COVID-19 Genomics UK (COG-UK) Consortium | The Lighthouse Lab in Milton Keynes and Alex Alderton, Roberto Amato, Jeffrey Barrett, Sonia Goncalves, Ewan Harrison, David K. Jackson, Ian Johnston, Dominic Kwiatkowski, Cordelia Langford, John Sillitoe on behalf of the Wellcome Sanger Institute COVID-19 Surveillance Team |
| EPI_ISL_2830513 | Lighthouse Lab in Alderley Park | Wellcome Sanger Institute for the COVID-19 Genomics UK (COG-UK) Consortium | Jacquelyn Wynn, Mairead Hyland, The Lighthouse Lab in Alderley Park and Alex Alderton, Roberto Amato, Jeffrey Barrett, Sonia Goncalves, Ewan Harrison, David K. Jackson, Ian Johnston, Dominic Kwiatkowski, Cordelia Langford, John Sillitoe on behalf of the Wellcome Sanger Institute COVID-19 Surveillance Team |
| EPI_ISL_2830514 | Lighthouse Lab in Milton Keynes | Wellcome Sanger Institute for the COVID-19 Genomics UK (COG-UK) Consortium | The Lighthouse Lab in Milton Keynes and Alex Alderton, Roberto Amato, Jeffrey Barrett, Sonia Goncalves, Ewan Harrison, David K. Jackson, Ian Johnston, Dominic Kwiatkowski, Cordelia Langford, John Sillitoe on behalf of the Wellcome Sanger Institute COVID-19 Surveillance Team |
| EPI_ISL_2830515, EPI_ISL_2830516 | Lighthouse Lab in Alderley Park | Wellcome Sanger Institute for the COVID-19 Genomics UK (COG-UK) Consortium | Jacquelyn Wynn, Mairead Hyland, The Lighthouse Lab in Alderley Park and Alex Alderton, Roberto Amato, Jeffrey Barrett, Sonia Goncalves, Ewan Harrison, David K. Jackson, Ian Johnston, Dominic Kwiatkowski, Cordelia Langford, John Sillitoe on behalf of the Wellcome Sanger Institute COVID-19 Surveillance Team |
| EPI_ISL_2830517, EPI_ISL_2830518, EPI_ISL_2830519 | Lighthouse Lab in Milton Keynes | Wellcome Sanger Institute for the COVID-19 Genomics UK (COG-UK) Consortium | The Lighthouse Lab in Milton Keynes and Alex Alderton, Roberto Amato, Jeffrey Barrett, Sonia Goncalves, Ewan Harrison, David K. Jackson, Ian Johnston, Dominic Kwiatkowski, Cordelia Langford, John Sillitoe on behalf of the Wellcome Sanger Institute COVID-19 Surveillance Team |
| EPI_ISL_2830520 | Lighthouse Lab in Glasgow | Wellcome Sanger Institute for the COVID-19 Genomics UK (COG-UK) Consortium | Harper VanSteenhouse, Yumi Kasai, David Gray, Carol Clugston, Anna Dominiczak and Alex Alderton, Roberto Amato, Jeffrey Barrett, Sonia Goncalves, Ewan Harrison, David K. Jackson, Ian Johnston, Dominic Kwiatkowski, Cordelia Langford, John Sillitoe on behalf of the Wellcome Sanger Institute COVID-19 Surveillance Team |
| EPI_ISL_2830521, EPI_ISL_2830522, EPI_ISL_2830523, EPI_ISL_2830524 | Lighthouse Lab in Alderley Park | Wellcome Sanger Institute for the COVID-19 Genomics UK (COG-UK) Consortium | Jacquelyn Wynn, Mairead Hyland, The Lighthouse Lab in Alderley Park and Alex Alderton, Roberto Amato, Jeffrey Barrett, Sonia Goncalves, Ewan Harrison, David K. Jackson, Ian Johnston, Dominic Kwiatkowski, Cordelia Langford, John Sillitoe on behalf of the Wellcome Sanger Institute COVID-19 Surveillance Team |
| EPI_ISL_2830525, EPI_ISL_2830526, EPI_ISL_2830527, EPI_ISL_2830528 | Lighthouse Lab in Milton Keynes | Wellcome Sanger Institute for the COVID-19 Genomics UK (COG-UK) Consortium | The Lighthouse Lab in Milton Keynes and Alex Alderton, Roberto Amato, Jeffrey Barrett, Sonia Goncalves, Ewan Harrison, David K. Jackson, Ian Johnston, Dominic Kwiatkowski, Cordelia Langford, John Sillitoe on behalf of the Wellcome Sanger Institute COVID-19 Surveillance Team |
| EPI_ISL_2830529, EPI_ISL_2830530 | Lighthouse Lab in Alderley Park | Wellcome Sanger Institute for the COVID-19 Genomics UK (COG-UK) Consortium | Jacquelyn Wynn, Mairead Hyland, The Lighthouse Lab in Alderley Park and Alex Alderton, Roberto Amato, Jeffrey Barrett, Sonia Goncalves, Ewan Harrison, David K. Jackson, Ian Johnston, Dominic Kwiatkowski, Cordelia Langford, John Sillitoe on behalf of the Wellcome Sanger Institute COVID-19 Surveillance Team |
| EPI_ISL_2830531, EPI_ISL_2830532, EPI_ISL_2830533, EPI_ISL_2830534, EPI_ISL_2830535 | Lighthouse Lab in Milton Keynes | Wellcome Sanger Institute for the COVID-19 Genomics UK (COG-UK) Consortium | The Lighthouse Lab in Milton Keynes and Alex Alderton, Roberto Amato, Jeffrey Barrett, Sonia Goncalves, Ewan Harrison, David K. Jackson, Ian Johnston, Dominic Kwiatkowski, Cordelia Langford, John Sillitoe on behalf of the Wellcome Sanger Institute COVID-19 Surveillance Team |
| EPI_ISL_2830536 | Lighthouse Lab in Alderley Park | Wellcome Sanger Institute for the COVID-19 Genomics UK (COG-UK) Consortium | Jacquelyn Wynn, Mairead Hyland, The Lighthouse Lab in Alderley Park and Alex Alderton, Roberto Amato, Jeffrey Barrett, Sonia Goncalves, Ewan Harrison, David K. Jackson, Ian Johnston, Dominic Kwiatkowski, Cordelia Langford, John Sillitoe on behalf of the Wellcome Sanger Institute COVID-19 Surveillance Team |
| EPI_ISL_2830537 | Lighthouse Lab in Milton Keynes | Wellcome Sanger Institute for the COVID-19 Genomics UK (COG-UK) Consortium | The Lighthouse Lab in Milton Keynes and Alex Alderton, Roberto Amato, Jeffrey Barrett, Sonia Goncalves, Ewan Harrison, David K. Jackson, Ian Johnston, Dominic Kwiatkowski, Cordelia Langford, John Sillitoe on behalf of the Wellcome Sanger Institute COVID-19 Surveillance Team |
| EPI_ISL_2830538, EPI_ISL_2830539 | Lighthouse Lab in Alderley Park | Wellcome Sanger Institute for the COVID-19 Genomics UK (COG-UK) Consortium | Jacquelyn Wynn, Mairead Hyland, The Lighthouse Lab in Alderley Park and Alex Alderton, Roberto Amato, Jeffrey Barrett, Sonia Goncalves, Ewan Harrison, David K. Jackson, Ian Johnston, Dominic Kwiatkowski, Cordelia Langford, John Sillitoe on behalf of the Wellcome Sanger Institute COVID-19 Surveillance Team |
| EPI_ISL_2830540, EPI_ISL_2830541, EPI_ISL_2830542, EPI_ISL_2830543 | Lighthouse Lab in Milton Keynes | Wellcome Sanger Institute for the COVID-19 Genomics UK (COG-UK) Consortium | The Lighthouse Lab in Milton Keynes and Alex Alderton, Roberto Amato, Jeffrey Barrett, Sonia Goncalves, Ewan Harrison, David K. Jackson, Ian Johnston, Dominic Kwiatkowski, Cordelia Langford, John Sillitoe on behalf of the Wellcome Sanger Institute COVID-19 Surveillance Team |
| EPI_ISL_2830544 | Lighthouse Lab in Glasgow | Wellcome Sanger Institute for the COVID-19 Genomics UK (COG-UK) Consortium | Harper VanSteenhouse, Yumi Kasai, David Gray, Carol Clugston, Anna Dominiczak and Alex Alderton, Roberto Amato, Jeffrey Barrett, Sonia Goncalves, Ewan Harrison, David K. Jackson, Ian Johnston, Dominic Kwiatkowski, Cordelia Langford, John Sillitoe on behalf of the Wellcome Sanger Institute COVID-19 Surveillance Team |
| EPI_ISL_2830545 | Lighthouse Lab in Alderley Park | Wellcome Sanger Institute for the COVID-19 Genomics UK (COG-UK) Consortium | Jacquelyn Wynn, Mairead Hyland, The Lighthouse Lab in Alderley Park and Alex Alderton, Roberto Amato, Jeffrey Barrett, Sonia Goncalves, Ewan Harrison, David K. Jackson, Ian Johnston, Dominic Kwiatkowski, Cordelia Langford, John Sillitoe on behalf of the Wellcome Sanger Institute COVID-19 Surveillance Team |
| EPI_ISL_2830546 | Lighthouse Lab in Glasgow | Wellcome Sanger Institute for the COVID-19 Genomics UK (COG-UK) Consortium | Harper VanSteenhouse, Yumi Kasai, David Gray, Carol Clugston, Anna Dominiczak and Alex Alderton, Roberto Amato, Jeffrey Barrett, Sonia Goncalves, Ewan Harrison, David K. Jackson, Ian Johnston, Dominic Kwiatkowski, Cordelia Langford, John Sillitoe on behalf of the Wellcome Sanger Institute COVID-19 Surveillance Team |
| EPI_ISL_2830547, EPI_ISL_2830548 | Lighthouse Lab in Milton Keynes | Wellcome Sanger Institute for the COVID-19 Genomics UK (COG-UK) Consortium | The Lighthouse Lab in Milton Keynes and Alex Alderton, Roberto Amato, Jeffrey Barrett, Sonia Goncalves, Ewan Harrison, David K. Jackson, Ian Johnston, Dominic Kwiatkowski, Cordelia Langford, John Sillitoe on behalf of the Wellcome Sanger Institute COVID-19 Surveillance Team |
| EPI_ISL_2830549 | Lighthouse Lab in Alderley Park | Wellcome Sanger Institute for the COVID-19 Genomics UK (COG-UK) Consortium | Jacquelyn Wynn, Mairead Hyland, The Lighthouse Lab in Alderley Park and Alex Alderton, Roberto Amato, Jeffrey Barrett, Sonia Goncalves, Ewan Harrison, David K. Jackson, Ian Johnston, Dominic Kwiatkowski, Cordelia Langford, John Sillitoe on behalf of the Wellcome Sanger Institute COVID-19 Surveillance Team |
| EPI_ISL_2830550, EPI_ISL_2830551 | Lighthouse Lab in Milton Keynes | Wellcome Sanger Institute for the COVID-19 Genomics UK (COG-UK) Consortium | The Lighthouse Lab in Milton Keynes and Alex Alderton, Roberto Amato, Jeffrey Barrett, Sonia Goncalves, Ewan Harrison, David K. Jackson, Ian Johnston, Dominic Kwiatkowski, Cordelia Langford, John Sillitoe on behalf of the Wellcome Sanger Institute COVID-19 Surveillance Team |
| EPI_ISL_2830552 | Lighthouse Lab in Alderley Park | Wellcome Sanger Institute for the COVID-19 Genomics UK (COG-UK) Consortium | Jacquelyn Wynn, Mairead Hyland, The Lighthouse Lab in Alderley Park and Alex Alderton, Roberto Amato, Jeffrey Barrett, Sonia Goncalves, Ewan Harrison, David K. Jackson, Ian Johnston, Dominic Kwiatkowski, Cordelia Langford, John Sillitoe on behalf of the Wellcome Sanger Institute COVID-19 Surveillance Team |
| EPI_ISL_2830553, EPI_ISL_2830554 | Lighthouse Lab in Milton Keynes | Wellcome Sanger Institute for the COVID-19 Genomics UK (COG-UK) Consortium | The Lighthouse Lab in Milton Keynes and Alex Alderton, Roberto Amato, Jeffrey Barrett, Sonia Goncalves, Ewan Harrison, David K. Jackson, Ian Johnston, Dominic Kwiatkowski, Cordelia Langford, John Sillitoe on behalf of the Wellcome Sanger Institute COVID-19 Surveillance Team |
| EPI_ISL_2830555 | Lighthouse Lab in Alderley Park | Wellcome Sanger Institute for the COVID-19 Genomics UK (COG-UK) Consortium | Jacquelyn Wynn, Mairead Hyland, The Lighthouse Lab in Alderley Park and Alex Alderton, Roberto Amato, Jeffrey Barrett, Sonia Goncalves, Ewan Harrison, David K. Jackson, Ian Johnston, Dominic Kwiatkowski, Cordelia Langford, John Sillitoe on behalf of the Wellcome Sanger Institute COVID-19 Surveillance Team |
| EPI_ISL_2830556, EPI_ISL_2830557, EPI_ISL_2830558, EPI_ISL_2830559, EPI_ISL_2830560 | Lighthouse Lab in Milton Keynes | Wellcome Sanger Institute for the COVID-19 Genomics UK (COG-UK) Consortium | The Lighthouse Lab in Milton Keynes and Alex Alderton, Roberto Amato, Jeffrey Barrett, Sonia Goncalves, Ewan Harrison, David K. Jackson, Ian Johnston, Dominic Kwiatkowski, Cordelia Langford, John Sillitoe on behalf of the Wellcome Sanger Institute COVID-19 Surveillance Team |
| EPI_ISL_2830561 | Lighthouse Lab in Alderley Park | Wellcome Sanger Institute for the COVID-19 Genomics UK (COG-UK) Consortium | Jacquelyn Wynn, Mairead Hyland, The Lighthouse Lab in Alderley Park and Alex Alderton, Roberto Amato, Jeffrey Barrett, Sonia Goncalves, Ewan Harrison, David K. Jackson, Ian Johnston, Dominic Kwiatkowski, Cordelia Langford, John Sillitoe on behalf of the Wellcome Sanger Institute COVID-19 Surveillance Team |
| EPI_ISL_2830562, EPI_ISL_2830563 | Lighthouse Lab in Milton Keynes | Wellcome Sanger Institute for the COVID-19 Genomics UK (COG-UK) Consortium | The Lighthouse Lab in Milton Keynes and Alex Alderton, Roberto Amato, Jeffrey Barrett, Sonia Goncalves, Ewan Harrison, David K. Jackson, Ian Johnston, Dominic Kwiatkowski, Cordelia Langford, John Sillitoe on behalf of the Wellcome Sanger Institute COVID-19 Surveillance Team |
| EPI_ISL_2830564 | Lighthouse Lab in Alderley Park | Wellcome Sanger Institute for the COVID-19 Genomics UK (COG-UK) Consortium | Jacquelyn Wynn, Mairead Hyland, The Lighthouse Lab in Alderley Park and Alex Alderton, Roberto Amato, Jeffrey Barrett, Sonia Goncalves, Ewan Harrison, David K. Jackson, Ian Johnston, Dominic Kwiatkowski, Cordelia Langford, John Sillitoe on behalf of the Wellcome Sanger Institute COVID-19 Surveillance Team |
| EPI_ISL_2830565, EPI_ISL_2830566, | Lighthouse Lab in Milton Keynes | Wellcome Sanger Institute for the COVID-19 Genomics UK (COG-UK) Consortium | The Lighthouse Lab in Milton Keynes and Alex Alderton, Roberto Amato, Jeffrey Barrett, Sonia Goncalves, Ewan Harrison, David K. Jackson, Ian Johnston, Dominic Kwiatkowski, Cordelia Langford, John Sillitoe on behalf of the Wellcome Sanger Institute COVID-19 Surveillance Team |

| | | | |
|---|---|---|---|
| EPI_ISL_2830567 | | | |
| EPI_ISL_2830568 | Lighthouse Lab in Alderley Park | Wellcome Sanger Institute for the COVID-19 Genomics UK (COG-UK) Consortium | Jacquelyn Wynn, Mairead Hyland, The Lighthouse Lab in Alderley Park and Alex Alderton, Roberto Amato, Jeffrey Barrett, Sonia Goncalves, Ewan Harrison, David K. Jackson, Ian Johnston, Dominic Kwiatkowski, Cordelia Langford, John Sillitoe on behalf of the Wellcome Sanger Institute COVID-19 Surveillance Team |
| EPI_ISL_2830569, EPI_ISL_2830570, EPI_ISL_2830571 | Lighthouse Lab in Milton Keynes | Wellcome Sanger Institute for the COVID-19 Genomics UK (COG-UK) Consortium | The Lighthouse Lab in Milton Keynes and Alex Alderton, Roberto Amato, Jeffrey Barrett, Sonia Goncalves, Ewan Harrison, David K. Jackson, Ian Johnston, Dominic Kwiatkowski, Cordelia Langford, John Sillitoe on behalf of the Wellcome Sanger Institute COVID-19 Surveillance Team |
| EPI_ISL_2830572 | Lighthouse Lab in Alderley Park | Wellcome Sanger Institute for the COVID-19 Genomics UK (COG-UK) Consortium | Jacquelyn Wynn, Mairead Hyland, The Lighthouse Lab in Alderley Park and Alex Alderton, Roberto Amato, Jeffrey Barrett, Sonia Goncalves, Ewan Harrison, David K. Jackson, Ian Johnston, Dominic Kwiatkowski, Cordelia Langford, John Sillitoe on behalf of the Wellcome Sanger Institute COVID-19 Surveillance Team |
| EPI_ISL_2830573, EPI_ISL_2830574 | Lighthouse Lab in Milton Keynes | Wellcome Sanger Institute for the COVID-19 Genomics UK (COG-UK) Consortium | The Lighthouse Lab in Milton Keynes and Alex Alderton, Roberto Amato, Jeffrey Barrett, Sonia Goncalves, Ewan Harrison, David K. Jackson, Ian Johnston, Dominic Kwiatkowski, Cordelia Langford, John Sillitoe on behalf of the Wellcome Sanger Institute COVID-19 Surveillance Team |
| EPI_ISL_2830575 | Lighthouse Lab in Alderley Park | Wellcome Sanger Institute for the COVID-19 Genomics UK (COG-UK) Consortium | Jacquelyn Wynn, Mairead Hyland, The Lighthouse Lab in Alderley Park and Alex Alderton, Roberto Amato, Jeffrey Barrett, Sonia Goncalves, Ewan Harrison, David K. Jackson, Ian Johnston, Dominic Kwiatkowski, Cordelia Langford, John Sillitoe on behalf of the Wellcome Sanger Institute COVID-19 Surveillance Team |
| EPI_ISL_2830576 | Lighthouse Lab in Milton Keynes | Wellcome Sanger Institute for the COVID-19 Genomics UK (COG-UK) Consortium | The Lighthouse Lab in Milton Keynes and Alex Alderton, Roberto Amato, Jeffrey Barrett, Sonia Goncalves, Ewan Harrison, David K. Jackson, Ian Johnston, Dominic Kwiatkowski, Cordelia Langford, John Sillitoe on behalf of the Wellcome Sanger Institute COVID-19 Surveillance Team |
| EPI_ISL_2830577 | Lighthouse Lab in Glasgow | Wellcome Sanger Institute for the COVID-19 Genomics UK (COG-UK) Consortium | Harper VanSteenhouse, Yumi Kasai, David Gray, Carol Clugston, Anna Dominiczak and Alex Alderton, Roberto Amato, Jeffrey Barrett, Sonia Goncalves, Ewan Harrison, David K. Jackson, Ian Johnston, Dominic Kwiatkowski, Cordelia Langford, John Sillitoe on behalf of the Wellcome Sanger Institute COVID-19 Surveillance Team |
| EPI_ISL_2830578, EPI_ISL_2830579, EPI_ISL_2830580, EPI_ISL_2830581, EPI_ISL_2830582 | Lighthouse Lab in Milton Keynes | Wellcome Sanger Institute for the COVID-19 Genomics UK (COG-UK) Consortium | The Lighthouse Lab in Milton Keynes and Alex Alderton, Roberto Amato, Jeffrey Barrett, Sonia Goncalves, Ewan Harrison, David K. Jackson, Ian Johnston, Dominic Kwiatkowski, Cordelia Langford, John Sillitoe on behalf of the Wellcome Sanger Institute COVID-19 Surveillance Team |
| EPI_ISL_2830583 | Lighthouse Lab in Glasgow | Wellcome Sanger Institute for the COVID-19 Genomics UK (COG-UK) Consortium | Harper VanSteenhouse, Yumi Kasai, David Gray, Carol Clugston, Anna Dominiczak and Alex Alderton, Roberto Amato, Jeffrey Barrett, Sonia Goncalves, Ewan Harrison, David K. Jackson, Ian Johnston, Dominic Kwiatkowski, Cordelia Langford, John Sillitoe on behalf of the Wellcome Sanger Institute COVID-19 Surveillance Team |
| EPI_ISL_2830584, EPI_ISL_2830585, EPI_ISL_2830586, EPI_ISL_2830587, EPI_ISL_2830588 | Lighthouse Lab in Milton Keynes | Wellcome Sanger Institute for the COVID-19 Genomics UK (COG-UK) Consortium | The Lighthouse Lab in Milton Keynes and Alex Alderton, Roberto Amato, Jeffrey Barrett, Sonia Goncalves, Ewan Harrison, David K. Jackson, Ian Johnston, Dominic Kwiatkowski, Cordelia Langford, John Sillitoe on behalf of the Wellcome Sanger Institute COVID-19 Surveillance Team |
| EPI_ISL_2830589 | Lighthouse Lab in Alderley Park | Wellcome Sanger Institute for the COVID-19 Genomics UK (COG-UK) Consortium | Jacquelyn Wynn, Mairead Hyland, The Lighthouse Lab in Alderley Park and Alex Alderton, Roberto Amato, Jeffrey Barrett, Sonia Goncalves, Ewan Harrison, David K. Jackson, Ian Johnston, Dominic Kwiatkowski, Cordelia Langford, John Sillitoe on behalf of the Wellcome Sanger Institute COVID-19 Surveillance Team |
| EPI_ISL_2830590, EPI_ISL_2830591, EPI_ISL_2830592, EPI_ISL_2830593, EPI_ISL_2830594 | Lighthouse Lab in Milton Keynes | Wellcome Sanger Institute for the COVID-19 Genomics UK (COG-UK) Consortium | The Lighthouse Lab in Milton Keynes and Alex Alderton, Roberto Amato, Jeffrey Barrett, Sonia Goncalves, Ewan Harrison, David K. Jackson, Ian Johnston, Dominic Kwiatkowski, Cordelia Langford, John Sillitoe on behalf of the Wellcome Sanger Institute COVID-19 Surveillance Team |
| EPI_ISL_2830595 | Lighthouse Lab in Alderley Park | Wellcome Sanger Institute for the COVID-19 Genomics UK (COG-UK) Consortium | Jacquelyn Wynn, Mairead Hyland, The Lighthouse Lab in Alderley Park and Alex Alderton, Roberto Amato, Jeffrey Barrett, Sonia Goncalves, Ewan Harrison, David K. Jackson, Ian Johnston, Dominic Kwiatkowski, Cordelia Langford, John Sillitoe on behalf of the Wellcome Sanger Institute COVID-19 Surveillance Team |
| EPI_ISL_2830596, EPI_ISL_2830597, EPI_ISL_2830598, EPI_ISL_2830599 | Lighthouse Lab in Milton Keynes | Wellcome Sanger Institute for the COVID-19 Genomics UK (COG-UK) Consortium | The Lighthouse Lab in Milton Keynes and Alex Alderton, Roberto Amato, Jeffrey Barrett, Sonia Goncalves, Ewan Harrison, David K. Jackson, Ian Johnston, Dominic Kwiatkowski, Cordelia Langford, John Sillitoe on behalf of the Wellcome Sanger Institute COVID-19 Surveillance Team |
| EPI_ISL_2830600 | Lighthouse Lab in Alderley Park | Wellcome Sanger Institute for the COVID-19 Genomics UK (COG-UK) Consortium | Jacquelyn Wynn, Mairead Hyland, The Lighthouse Lab in Alderley Park and Alex Alderton, Roberto Amato, Jeffrey Barrett, Sonia Goncalves, Ewan Harrison, David K. Jackson, Ian Johnston, Dominic Kwiatkowski, Cordelia Langford, John Sillitoe on behalf of the Wellcome Sanger Institute COVID-19 Surveillance Team |
| EPI_ISL_2830601, EPI_ISL_2830602 | Lighthouse Lab in Milton Keynes | Wellcome Sanger Institute for the COVID-19 Genomics UK (COG-UK) Consortium | The Lighthouse Lab in Milton Keynes and Alex Alderton, Roberto Amato, Jeffrey Barrett, Sonia Goncalves, Ewan Harrison, David K. Jackson, Ian Johnston, Dominic Kwiatkowski, Cordelia Langford, John Sillitoe on behalf of the Wellcome Sanger Institute COVID-19 Surveillance Team |
| EPI_ISL_2830603 | Lighthouse Lab in Alderley Park | Wellcome Sanger Institute for the COVID-19 Genomics UK (COG-UK) Consortium | Jacquelyn Wynn, Mairead Hyland, The Lighthouse Lab in Alderley Park and Alex Alderton, Roberto Amato, Jeffrey Barrett, Sonia Goncalves, Ewan Harrison, David K. Jackson, Ian Johnston, Dominic Kwiatkowski, Cordelia Langford, John Sillitoe on behalf of the Wellcome Sanger Institute COVID-19 Surveillance Team |
| EPI_ISL_2830604, EPI_ISL_2830605, EPI_ISL_2830606 | Lighthouse Lab in Milton Keynes | Wellcome Sanger Institute for the COVID-19 Genomics UK (COG-UK) Consortium | The Lighthouse Lab in Milton Keynes and Alex Alderton, Roberto Amato, Jeffrey Barrett, Sonia Goncalves, Ewan Harrison, David K. Jackson, Ian Johnston, Dominic Kwiatkowski, Cordelia Langford, John Sillitoe on behalf of the Wellcome Sanger Institute COVID-19 Surveillance Team |
| EPI_ISL_2830607, EPI_ISL_2830608 | Lighthouse Laboratory Plymouth | Wellcome Sanger Institute for the COVID-19 Genomics UK (COG-UK) Consortium | Lighthouse Laboratory Plymouth and Alex Alderton, Roberto Amato, Jeffrey Barrett, Sonia Goncalves, Ewan Harrison, David K. Jackson, Ian Johnston, Dominic Kwiatkowski, Cordelia Langford, John Sillitoe on behalf of the Wellcome Sanger Institute COVID-19 Surveillance Team |
| EPI_ISL_2830609 | Lighthouse Lab in Milton Keynes | Wellcome Sanger Institute for the COVID-19 Genomics UK (COG-UK) Consortium | The Lighthouse Lab in Milton Keynes and Alex Alderton, Roberto Amato, Jeffrey Barrett, Sonia Goncalves, Ewan Harrison, David K. Jackson, Ian Johnston, Dominic Kwiatkowski, Cordelia Langford, John Sillitoe on behalf of the Wellcome Sanger Institute COVID-19 Surveillance Team |
| EPI_ISL_2830610 | Lighthouse Laboratory Plymouth | Wellcome Sanger Institute for the COVID-19 Genomics UK (COG-UK) Consortium | Lighthouse Laboratory Plymouth and Alex Alderton, Roberto Amato, Jeffrey Barrett, Sonia Goncalves, Ewan Harrison, David K. Jackson, Ian Johnston, Dominic Kwiatkowski, Cordelia Langford, John Sillitoe on behalf of the Wellcome Sanger Institute COVID-19 Surveillance Team |
| EPI_ISL_2830611 | Lighthouse Lab in Glasgow | Wellcome Sanger Institute for the COVID-19 Genomics UK (COG-UK) Consortium | Harper VanSteenhouse, Yumi Kasai, David Gray, Carol Clugston, Anna Dominiczak and Alex Alderton, Roberto Amato, Jeffrey Barrett, Sonia Goncalves, Ewan Harrison, David K. Jackson, Ian Johnston, Dominic Kwiatkowski, Cordelia Langford, John Sillitoe on behalf of the Wellcome Sanger Institute COVID-19 Surveillance Team |
| EPI_ISL_2830612 | Lighthouse Lab in Milton Keynes | Wellcome Sanger Institute for the COVID-19 Genomics UK (COG-UK) Consortium | The Lighthouse Lab in Milton Keynes and Alex Alderton, Roberto Amato, Jeffrey Barrett, Sonia Goncalves, Ewan Harrison, David K. Jackson, Ian Johnston, Dominic Kwiatkowski, Cordelia Langford, John Sillitoe on behalf of the Wellcome Sanger Institute COVID-19 Surveillance Team |
| EPI_ISL_2830613 | Lighthouse Lab in Glasgow | Wellcome Sanger Institute for the COVID-19 Genomics UK (COG-UK) Consortium | Harper VanSteenhouse, Yumi Kasai, David Gray, Carol Clugston, Anna Dominiczak and Alex Alderton, Roberto Amato, Jeffrey Barrett, Sonia Goncalves, Ewan Harrison, David K. Jackson, Ian Johnston, Dominic Kwiatkowski, Cordelia Langford, John Sillitoe on behalf of the Wellcome Sanger Institute COVID-19 Surveillance Team |
| EPI_ISL_2830614 | Lighthouse Laboratory Plymouth | Wellcome Sanger Institute for the COVID-19 Genomics UK (COG-UK) Consortium | Lighthouse Laboratory Plymouth and Alex Alderton, Roberto Amato, Jeffrey Barrett, Sonia Goncalves, Ewan Harrison, David K. Jackson, Ian Johnston, Dominic Kwiatkowski, Cordelia Langford, John Sillitoe on behalf of the Wellcome Sanger Institute COVID-19 Surveillance Team |
| EPI_ISL_2830615 | Lighthouse Lab in Alderley Park | Wellcome Sanger Institute for the COVID-19 Genomics UK (COG-UK) Consortium | Jacquelyn Wynn, Mairead Hyland, The Lighthouse Lab in Alderley Park and Alex Alderton, Roberto Amato, Jeffrey Barrett, Sonia Goncalves, Ewan Harrison, David K. Jackson, Ian Johnston, Dominic Kwiatkowski, Cordelia Langford, John Sillitoe on behalf of the Wellcome Sanger Institute COVID-19 Surveillance Team |
| EPI_ISL_2830616, EPI_ISL_2830617 | Lighthouse Lab in Milton Keynes | Wellcome Sanger Institute for the COVID-19 Genomics UK (COG-UK) Consortium | The Lighthouse Lab in Milton Keynes and Alex Alderton, Roberto Amato, Jeffrey Barrett, Sonia Goncalves, Ewan Harrison, David K. Jackson, Ian Johnston, Dominic Kwiatkowski, Cordelia Langford, John Sillitoe on behalf of the Wellcome Sanger Institute COVID-19 Surveillance Team |
| EPI_ISL_2830618 | Lighthouse Lab in Glasgow | Wellcome Sanger Institute for the COVID-19 Genomics UK (COG-UK) Consortium | Harper VanSteenhouse, Yumi Kasai, David Gray, Carol Clugston, Anna Dominiczak and Alex Alderton, Roberto Amato, Jeffrey Barrett, Sonia Goncalves, Ewan Harrison, David K. Jackson, Ian Johnston, Dominic Kwiatkowski, Cordelia Langford, John Sillitoe on behalf of the Wellcome Sanger Institute COVID-19 Surveillance Team |
| EPI_ISL_2830619, EPI_ISL_2830620 | Lighthouse Lab in Milton Keynes | Wellcome Sanger Institute for the COVID-19 Genomics UK (COG-UK) Consortium | The Lighthouse Lab in Milton Keynes and Alex Alderton, Roberto Amato, Jeffrey Barrett, Sonia Goncalves, Ewan Harrison, David K. Jackson, Ian Johnston, Dominic Kwiatkowski, Cordelia Langford, John Sillitoe on behalf of the Wellcome Sanger Institute COVID-19 Surveillance Team |
| EPI_ISL_2830621, EPI_ISL_2830622, EPI_ISL_2830623, EPI_ISL_2830624 | Lighthouse Lab in Glasgow | Wellcome Sanger Institute for the COVID-19 Genomics UK (COG-UK) Consortium | Harper VanSteenhouse, Yumi Kasai, David Gray, Carol Clugston, Anna Dominiczak and Alex Alderton, Roberto Amato, Jeffrey Barrett, Sonia Goncalves, Ewan Harrison, David K. Jackson, Ian Johnston, Dominic Kwiatkowski, Cordelia Langford, John Sillitoe on behalf of the Wellcome Sanger Institute COVID-19 Surveillance Team |
| EPI_ISL_2830625, EPI_ISL_2830626 | Lighthouse Lab in Milton Keynes | Wellcome Sanger Institute for the COVID-19 Genomics UK (COG-UK) Consortium | The Lighthouse Lab in Milton Keynes and Alex Alderton, Roberto Amato, Jeffrey Barrett, Sonia Goncalves, Ewan Harrison, David K. Jackson, Ian Johnston, Dominic Kwiatkowski, Cordelia Langford, John Sillitoe on behalf of the Wellcome Sanger Institute COVID-19 Surveillance Team |
| EPI_ISL_2830627, EPI_ISL_2830628 | Lighthouse Laboratory Plymouth | Wellcome Sanger Institute for the COVID-19 Genomics UK (COG-UK) Consortium | Lighthouse Laboratory Plymouth and Alex Alderton, Roberto Amato, Jeffrey Barrett, Sonia Goncalves, Ewan Harrison, David K. Jackson, Ian Johnston, Dominic Kwiatkowski, Cordelia Langford, John Sillitoe on behalf of the Wellcome Sanger Institute COVID-19 Surveillance Team |
| EPI_ISL_2830629 | Lighthouse Lab in Glasgow | Wellcome Sanger Institute for the COVID-19 Genomics UK (COG-UK) Consortium | Harper VanSteenhouse, Yumi Kasai, David Gray, Carol Clugston, Anna Dominiczak and Alex Alderton, Roberto Amato, Jeffrey Barrett, Sonia Goncalves, Ewan Harrison, David K. Jackson, Ian Johnston, Dominic Kwiatkowski, Cordelia Langford, John Sillitoe on behalf of the Wellcome Sanger Institute COVID-19 Surveillance Team |
| EPI_ISL_2830630 | Lighthouse Laboratory Plymouth | Wellcome Sanger Institute for the COVID-19 Genomics UK (COG-UK) Consortium | Lighthouse Laboratory Plymouth and Alex Alderton, Roberto Amato, Jeffrey Barrett, Sonia Goncalves, Ewan Harrison, David K. Jackson, Ian Johnston, Dominic Kwiatkowski, Cordelia Langford, John Sillitoe on behalf of the Wellcome Sanger Institute COVID-19 Surveillance Team |
| EPI_ISL_2830631 | Lighthouse Lab in Alderley Park | Wellcome Sanger Institute for the COVID-19 Genomics UK (COG-UK) Consortium | Jacquelyn Wynn, Mairead Hyland, The Lighthouse Lab in Alderley Park and Alex Alderton, Roberto Amato, Jeffrey Barrett, Sonia Goncalves, Ewan Harrison, David K. Jackson, Ian Johnston, Dominic Kwiatkowski, Cordelia Langford, John Sillitoe on behalf of the Wellcome Sanger Institute COVID-19 Surveillance Team |
| EPI_ISL_2830632 | Lighthouse Laboratory Plymouth | Wellcome Sanger Institute for the COVID-19 Genomics UK (COG-UK) Consortium | Lighthouse Laboratory Plymouth and Alex Alderton, Roberto Amato, Jeffrey Barrett, Sonia Goncalves, Ewan Harrison, David K. Jackson, Ian Johnston, Dominic Kwiatkowski, Cordelia Langford, John Sillitoe on behalf of the Wellcome Sanger Institute COVID-19 Surveillance Team |
| EPI_ISL_2830633 | Lighthouse Lab in Milton Keynes | Wellcome Sanger Institute for the COVID-19 Genomics UK (COG-UK) Consortium | The Lighthouse Lab in Milton Keynes and Alex Alderton, Roberto Amato, Jeffrey Barrett, Sonia Goncalves, Ewan Harrison, David K. Jackson, Ian Johnston, Dominic Kwiatkowski, Cordelia Langford, John Sillitoe on behalf of the Wellcome Sanger Institute COVID-19 Surveillance Team |
| EPI_ISL_2830634, EPI_ISL_2830635 | Lighthouse Lab in Alderley Park | Wellcome Sanger Institute for the COVID-19 Genomics UK (COG-UK) Consortium | Jacquelyn Wynn, Mairead Hyland, The Lighthouse Lab in Alderley Park and Alex Alderton, Roberto Amato, Jeffrey Barrett, Sonia Goncalves, Ewan Harrison, David K. Jackson, Ian Johnston, Dominic Kwiatkowski, Cordelia Langford, John Sillitoe on behalf of the Wellcome Sanger Institute COVID-19 Surveillance Team |
| EPI_ISL_2830636 | Lighthouse Lab in Milton Keynes | Wellcome Sanger Institute for the COVID-19 Genomics UK (COG-UK) | The Lighthouse Lab in Milton Keynes and Alex Alderton, Roberto Amato, Jeffrey Barrett, Sonia Goncalves, Ewan Harrison, David K. Jackson, Ian Johnston, Dominic Kwiatkowski, Cordelia Langford, John Sillitoe on behalf of the Wellcome Sanger |

| | | | |
|---|---|---|---|
| | | Consortium | Institute COVID-19 Surveillance Team |
| EPI_ISL_2830637 | Lighthouse Lab in Glasgow | Wellcome Sanger Institute for the COVID-19 Genomics UK (COG-UK) Consortium | Harper VanSteenhouse, Yumi Kasai, David Gray, Carol Clugston, Anna Dominiczak and Alex Alderton, Roberto Amato, Jeffrey Barrett, Sonia Goncalves, Ewan Harrison, David K. Jackson, Ian Johnston, Dominic Kwiatkowski, Cordelia Langford, John Sillitoe on behalf of the Wellcome Sanger Institute COVID-19 Surveillance Team |
| EPI_ISL_2830638 | Lighthouse Laboratory Plymouth | Wellcome Sanger Institute for the COVID-19 Genomics UK (COG-UK) Consortium | Lighthouse Laboratory Plymouth and Alex Alderton, Roberto Amato, Jeffrey Barrett, Sonia Goncalves, Ewan Harrison, David K. Jackson, Ian Johnston, Dominic Kwiatkowski, Cordelia Langford, John Sillitoe on behalf of the Wellcome Sanger Institute COVID-19 Surveillance Team |
| EPI_ISL_2830639, EPI_ISL_2830640 | Lighthouse Lab in Alderley Park | Wellcome Sanger Institute for the COVID-19 Genomics UK (COG-UK) Consortium | Jacquelyn Wynn, Mairead Hyland, The Lighthouse Lab in Alderley Park and Alex Alderton, Roberto Amato, Jeffrey Barrett, Sonia Goncalves, Ewan Harrison, David K. Jackson, Ian Johnston, Dominic Kwiatkowski, Cordelia Langford, John Sillitoe on behalf of the Wellcome Sanger Institute COVID-19 Surveillance Team |
| EPI_ISL_2830641, EPI_ISL_2830642 | Lighthouse Laboratory Plymouth | Wellcome Sanger Institute for the COVID-19 Genomics UK (COG-UK) Consortium | Lighthouse Laboratory Plymouth and Alex Alderton, Roberto Amato, Jeffrey Barrett, Sonia Goncalves, Ewan Harrison, David K. Jackson, Ian Johnston, Dominic Kwiatkowski, Cordelia Langford, John Sillitoe on behalf of the Wellcome Sanger Institute COVID-19 Surveillance Team |
| EPI_ISL_2830643 | Lighthouse Lab in Glasgow | Wellcome Sanger Institute for the COVID-19 Genomics UK (COG-UK) Consortium | Harper VanSteenhouse, Yumi Kasai, David Gray, Carol Clugston, Anna Dominiczak and Alex Alderton, Roberto Amato, Jeffrey Barrett, Sonia Goncalves, Ewan Harrison, David K. Jackson, Ian Johnston, Dominic Kwiatkowski, Cordelia Langford, John Sillitoe on behalf of the Wellcome Sanger Institute COVID-19 Surveillance Team |
| EPI_ISL_2830644 | Lighthouse Lab in Alderley Park | Wellcome Sanger Institute for the COVID-19 Genomics UK (COG-UK) Consortium | Jacquelyn Wynn, Mairead Hyland, The Lighthouse Lab in Alderley Park and Alex Alderton, Roberto Amato, Jeffrey Barrett, Sonia Goncalves, Ewan Harrison, David K. Jackson, Ian Johnston, Dominic Kwiatkowski, Cordelia Langford, John Sillitoe on behalf of the Wellcome Sanger Institute COVID-19 Surveillance Team |
| EPI_ISL_2830645, EPI_ISL_2830646, EPI_ISL_2830647 | Lighthouse Lab in Glasgow | Wellcome Sanger Institute for the COVID-19 Genomics UK (COG-UK) Consortium | Harper VanSteenhouse, Yumi Kasai, David Gray, Carol Clugston, Anna Dominiczak and Alex Alderton, Roberto Amato, Jeffrey Barrett, Sonia Goncalves, Ewan Harrison, David K. Jackson, Ian Johnston, Dominic Kwiatkowski, Cordelia Langford, John Sillitoe on behalf of the Wellcome Sanger Institute COVID-19 Surveillance Team |
| EPI_ISL_2830648 | Lighthouse Laboratory Plymouth | Wellcome Sanger Institute for the COVID-19 Genomics UK (COG-UK) Consortium | Lighthouse Laboratory Plymouth and Alex Alderton, Roberto Amato, Jeffrey Barrett, Sonia Goncalves, Ewan Harrison, David K. Jackson, Ian Johnston, Dominic Kwiatkowski, Cordelia Langford, John Sillitoe on behalf of the Wellcome Sanger Institute COVID-19 Surveillance Team |
| EPI_ISL_2830649 | Lighthouse Lab in Glasgow | Wellcome Sanger Institute for the COVID-19 Genomics UK (COG-UK) Consortium | Harper VanSteenhouse, Yumi Kasai, David Gray, Carol Clugston, Anna Dominiczak and Alex Alderton, Roberto Amato, Jeffrey Barrett, Sonia Goncalves, Ewan Harrison, David K. Jackson, Ian Johnston, Dominic Kwiatkowski, Cordelia Langford, John Sillitoe on behalf of the Wellcome Sanger Institute COVID-19 Surveillance Team |
| EPI_ISL_2830650 | Lighthouse Lab in Milton Keynes | Wellcome Sanger Institute for the COVID-19 Genomics UK (COG-UK) Consortium | The Lighthouse Lab in Milton Keynes and Alex Alderton, Roberto Amato, Jeffrey Barrett, Sonia Goncalves, Ewan Harrison, David K. Jackson, Ian Johnston, Dominic Kwiatkowski, Cordelia Langford, John Sillitoe on behalf of the Wellcome Sanger Institute COVID-19 Surveillance Team |
| EPI_ISL_2830651, EPI_ISL_2830652 | Lighthouse Lab in Glasgow | Wellcome Sanger Institute for the COVID-19 Genomics UK (COG-UK) Consortium | Harper VanSteenhouse, Yumi Kasai, David Gray, Carol Clugston, Anna Dominiczak and Alex Alderton, Roberto Amato, Jeffrey Barrett, Sonia Goncalves, Ewan Harrison, David K. Jackson, Ian Johnston, Dominic Kwiatkowski, Cordelia Langford, John Sillitoe on behalf of the Wellcome Sanger Institute COVID-19 Surveillance Team |
| EPI_ISL_2830653, EPI_ISL_2830654, EPI_ISL_2830655, EPI_ISL_2830656 | Lighthouse Lab in Milton Keynes | Wellcome Sanger Institute for the COVID-19 Genomics UK (COG-UK) Consortium | The Lighthouse Lab in Milton Keynes and Alex Alderton, Roberto Amato, Jeffrey Barrett, Sonia Goncalves, Ewan Harrison, David K. Jackson, Ian Johnston, Dominic Kwiatkowski, Cordelia Langford, John Sillitoe on behalf of the Wellcome Sanger Institute COVID-19 Surveillance Team |
| EPI_ISL_2830657, EPI_ISL_2830658 | Lighthouse Lab in Glasgow | Wellcome Sanger Institute for the COVID-19 Genomics UK (COG-UK) Consortium | Harper VanSteenhouse, Yumi Kasai, David Gray, Carol Clugston, Anna Dominiczak and Alex Alderton, Roberto Amato, Jeffrey Barrett, Sonia Goncalves, Ewan Harrison, David K. Jackson, Ian Johnston, Dominic Kwiatkowski, Cordelia Langford, John Sillitoe on behalf of the Wellcome Sanger Institute COVID-19 Surveillance Team |
| EPI_ISL_2830659 | Lighthouse Laboratory Plymouth | Wellcome Sanger Institute for the COVID-19 Genomics UK (COG-UK) Consortium | Lighthouse Laboratory Plymouth and Alex Alderton, Roberto Amato, Jeffrey Barrett, Sonia Goncalves, Ewan Harrison, David K. Jackson, Ian Johnston, Dominic Kwiatkowski, Cordelia Langford, John Sillitoe on behalf of the Wellcome Sanger Institute COVID-19 Surveillance Team |
| EPI_ISL_2830660 | Lighthouse Lab in Glasgow | Wellcome Sanger Institute for the COVID-19 Genomics UK (COG-UK) Consortium | Harper VanSteenhouse, Yumi Kasai, David Gray, Carol Clugston, Anna Dominiczak and Alex Alderton, Roberto Amato, Jeffrey Barrett, Sonia Goncalves, Ewan Harrison, David K. Jackson, Ian Johnston, Dominic Kwiatkowski, Cordelia Langford, John Sillitoe on behalf of the Wellcome Sanger Institute COVID-19 Surveillance Team |
| EPI_ISL_2830661 | Lighthouse Lab in Alderley Park | Wellcome Sanger Institute for the COVID-19 Genomics UK (COG-UK) Consortium | Jacquelyn Wynn, Mairead Hyland, The Lighthouse Lab in Alderley Park and Alex Alderton, Roberto Amato, Jeffrey Barrett, Sonia Goncalves, Ewan Harrison, David K. Jackson, Ian Johnston, Dominic Kwiatkowski, Cordelia Langford, John Sillitoe on behalf of the Wellcome Sanger Institute COVID-19 Surveillance Team |
| EPI_ISL_2830662 | Lighthouse Lab in Milton Keynes | Wellcome Sanger Institute for the COVID-19 Genomics UK (COG-UK) Consortium | The Lighthouse Lab in Milton Keynes and Alex Alderton, Roberto Amato, Jeffrey Barrett, Sonia Goncalves, Ewan Harrison, David K. Jackson, Ian Johnston, Dominic Kwiatkowski, Cordelia Langford, John Sillitoe on behalf of the Wellcome Sanger Institute COVID-19 Surveillance Team |
| EPI_ISL_2830663 | Lighthouse Laboratory Plymouth | Wellcome Sanger Institute for the COVID-19 Genomics UK (COG-UK) Consortium | Lighthouse Laboratory Plymouth and Alex Alderton, Roberto Amato, Jeffrey Barrett, Sonia Goncalves, Ewan Harrison, David K. Jackson, Ian Johnston, Dominic Kwiatkowski, Cordelia Langford, John Sillitoe on behalf of the Wellcome Sanger Institute COVID-19 Surveillance Team |
| EPI_ISL_2830664 | Lighthouse Lab in Glasgow | Wellcome Sanger Institute for the COVID-19 Genomics UK (COG-UK) Consortium | Harper VanSteenhouse, Yumi Kasai, David Gray, Carol Clugston, Anna Dominiczak and Alex Alderton, Roberto Amato, Jeffrey Barrett, Sonia Goncalves, Ewan Harrison, David K. Jackson, Ian Johnston, Dominic Kwiatkowski, Cordelia Langford, John Sillitoe on behalf of the Wellcome Sanger Institute COVID-19 Surveillance Team |
| EPI_ISL_2830665, EPI_ISL_2830666 | Lighthouse Lab in Milton Keynes | Wellcome Sanger Institute for the COVID-19 Genomics UK (COG-UK) Consortium | The Lighthouse Lab in Milton Keynes and Alex Alderton, Roberto Amato, Jeffrey Barrett, Sonia Goncalves, Ewan Harrison, David K. Jackson, Ian Johnston, Dominic Kwiatkowski, Cordelia Langford, John Sillitoe on behalf of the Wellcome Sanger Institute COVID-19 Surveillance Team |
| EPI_ISL_2830667 | Lighthouse Lab in Glasgow | Wellcome Sanger Institute for the COVID-19 Genomics UK (COG-UK) Consortium | Harper VanSteenhouse, Yumi Kasai, David Gray, Carol Clugston, Anna Dominiczak and Alex Alderton, Roberto Amato, Jeffrey Barrett, Sonia Goncalves, Ewan Harrison, David K. Jackson, Ian Johnston, Dominic Kwiatkowski, Cordelia Langford, John Sillitoe on behalf of the Wellcome Sanger Institute COVID-19 Surveillance Team |
| EPI_ISL_2830668 | Lighthouse Laboratory Plymouth | Wellcome Sanger Institute for the COVID-19 Genomics UK (COG-UK) Consortium | Lighthouse Laboratory Plymouth and Alex Alderton, Roberto Amato, Jeffrey Barrett, Sonia Goncalves, Ewan Harrison, David K. Jackson, Ian Johnston, Dominic Kwiatkowski, Cordelia Langford, John Sillitoe on behalf of the Wellcome Sanger Institute COVID-19 Surveillance Team |
| EPI_ISL_2830669 | Lighthouse Lab in Glasgow | Wellcome Sanger Institute for the COVID-19 Genomics UK (COG-UK) Consortium | Harper VanSteenhouse, Yumi Kasai, David Gray, Carol Clugston, Anna Dominiczak and Alex Alderton, Roberto Amato, Jeffrey Barrett, Sonia Goncalves, Ewan Harrison, David K. Jackson, Ian Johnston, Dominic Kwiatkowski, Cordelia Langford, John Sillitoe on behalf of the Wellcome Sanger Institute COVID-19 Surveillance Team |
| EPI_ISL_2830670 | Lighthouse Lab in Milton Keynes | Wellcome Sanger Institute for the COVID-19 Genomics UK (COG-UK) Consortium | The Lighthouse Lab in Milton Keynes and Alex Alderton, Roberto Amato, Jeffrey Barrett, Sonia Goncalves, Ewan Harrison, David K. Jackson, Ian Johnston, Dominic Kwiatkowski, Cordelia Langford, John Sillitoe on behalf of the Wellcome Sanger Institute COVID-19 Surveillance Team |
| EPI_ISL_2830671 | Lighthouse Lab in Glasgow | Wellcome Sanger Institute for the COVID-19 Genomics UK (COG-UK) Consortium | Harper VanSteenhouse, Yumi Kasai, David Gray, Carol Clugston, Anna Dominiczak and Alex Alderton, Roberto Amato, Jeffrey Barrett, Sonia Goncalves, Ewan Harrison, David K. Jackson, Ian Johnston, Dominic Kwiatkowski, Cordelia Langford, John Sillitoe on behalf of the Wellcome Sanger Institute COVID-19 Surveillance Team |
| EPI_ISL_2830672, EPI_ISL_2830673 | Lighthouse Laboratory Plymouth | Wellcome Sanger Institute for the COVID-19 Genomics UK (COG-UK) Consortium | Lighthouse Laboratory Plymouth and Alex Alderton, Roberto Amato, Jeffrey Barrett, Sonia Goncalves, Ewan Harrison, David K. Jackson, Ian Johnston, Dominic Kwiatkowski, Cordelia Langford, John Sillitoe on behalf of the Wellcome Sanger Institute COVID-19 Surveillance Team |
| EPI_ISL_2830674 | Lighthouse Lab in Alderley Park | Wellcome Sanger Institute for the COVID-19 Genomics UK (COG-UK) Consortium | Jacquelyn Wynn, Mairead Hyland, The Lighthouse Lab in Alderley Park and Alex Alderton, Roberto Amato, Jeffrey Barrett, Sonia Goncalves, Ewan Harrison, David K. Jackson, Ian Johnston, Dominic Kwiatkowski, Cordelia Langford, John Sillitoe on behalf of the Wellcome Sanger Institute COVID-19 Surveillance Team |
| EPI_ISL_2830675, EPI_ISL_2830676, EPI_ISL_2830677 | Lighthouse Lab in Glasgow | Wellcome Sanger Institute for the COVID-19 Genomics UK (COG-UK) Consortium | Harper VanSteenhouse, Yumi Kasai, David Gray, Carol Clugston, Anna Dominiczak and Alex Alderton, Roberto Amato, Jeffrey Barrett, Sonia Goncalves, Ewan Harrison, David K. Jackson, Ian Johnston, Dominic Kwiatkowski, Cordelia Langford, John Sillitoe on behalf of the Wellcome Sanger Institute COVID-19 Surveillance Team |
| EPI_ISL_2830678 | Lighthouse Laboratory Plymouth | Wellcome Sanger Institute for the COVID-19 Genomics UK (COG-UK) Consortium | Lighthouse Laboratory Plymouth and Alex Alderton, Roberto Amato, Jeffrey Barrett, Sonia Goncalves, Ewan Harrison, David K. Jackson, Ian Johnston, Dominic Kwiatkowski, Cordelia Langford, John Sillitoe on behalf of the Wellcome Sanger Institute COVID-19 Surveillance Team |
| EPI_ISL_2830679 | Lighthouse Lab in Glasgow | Wellcome Sanger Institute for the COVID-19 Genomics UK (COG-UK) Consortium | Harper VanSteenhouse, Yumi Kasai, David Gray, Carol Clugston, Anna Dominiczak and Alex Alderton, Roberto Amato, Jeffrey Barrett, Sonia Goncalves, Ewan Harrison, David K. Jackson, Ian Johnston, Dominic Kwiatkowski, Cordelia Langford, John Sillitoe on behalf of the Wellcome Sanger Institute COVID-19 Surveillance Team |
| EPI_ISL_2830680, EPI_ISL_2830681 | Lighthouse Laboratory Plymouth | Wellcome Sanger Institute for the COVID-19 Genomics UK (COG-UK) Consortium | Lighthouse Laboratory Plymouth and Alex Alderton, Roberto Amato, Jeffrey Barrett, Sonia Goncalves, Ewan Harrison, David K. Jackson, Ian Johnston, Dominic Kwiatkowski, Cordelia Langford, John Sillitoe on behalf of the Wellcome Sanger Institute COVID-19 Surveillance Team |
| EPI_ISL_2830682, EPI_ISL_2830683, EPI_ISL_2830684, EPI_ISL_2830685, EPI_ISL_2830686, EPI_ISL_2830687, EPI_ISL_2830688, EPI_ISL_2830689, EPI_ISL_2830690 | Lighthouse Lab in Milton Keynes | Wellcome Sanger Institute for the COVID-19 Genomics UK (COG-UK) Consortium | The Lighthouse Lab in Milton Keynes and Alex Alderton, Roberto Amato, Jeffrey Barrett, Sonia Goncalves, Ewan Harrison, David K. Jackson, Ian Johnston, Dominic Kwiatkowski, Cordelia Langford, John Sillitoe on behalf of the Wellcome Sanger Institute COVID-19 Surveillance Team |
| EPI_ISL_2830691 | Lighthouse Lab in Glasgow | Wellcome Sanger Institute for the COVID-19 Genomics UK (COG-UK) Consortium | Harper VanSteenhouse, Yumi Kasai, David Gray, Carol Clugston, Anna Dominiczak and Alex Alderton, Roberto Amato, Jeffrey Barrett, Sonia Goncalves, Ewan Harrison, David K. Jackson, Ian Johnston, Dominic Kwiatkowski, Cordelia Langford, John Sillitoe on behalf of the Wellcome Sanger Institute COVID-19 Surveillance Team |
| EPI_ISL_2830692 | Lighthouse Lab in Alderley Park | Wellcome Sanger Institute for the COVID-19 Genomics UK (COG-UK) Consortium | Jacquelyn Wynn, Mairead Hyland, The Lighthouse Lab in Alderley Park and Alex Alderton, Roberto Amato, Jeffrey Barrett, Sonia Goncalves, Ewan Harrison, David K. Jackson, Ian Johnston, Dominic Kwiatkowski, Cordelia Langford, John Sillitoe on behalf of the Wellcome Sanger Institute COVID-19 Surveillance Team |
| EPI_ISL_2830693, EPI_ISL_2830694 | Lighthouse Lab in Glasgow | Wellcome Sanger Institute for the COVID-19 Genomics UK (COG-UK) Consortium | Harper VanSteenhouse, Yumi Kasai, David Gray, Carol Clugston, Anna Dominiczak and Alex Alderton, Roberto Amato, Jeffrey Barrett, Sonia Goncalves, Ewan Harrison, David K. Jackson, Ian Johnston, Dominic Kwiatkowski, Cordelia Langford, John Sillitoe on behalf of the Wellcome Sanger Institute COVID-19 Surveillance Team |
| EPI_ISL_2830695 | Lighthouse Lab in Milton Keynes | Wellcome Sanger Institute for the COVID-19 Genomics UK (COG-UK) Consortium | The Lighthouse Lab in Milton Keynes and Alex Alderton, Roberto Amato, Jeffrey Barrett, Sonia Goncalves, Ewan Harrison, David K. Jackson, Ian Johnston, Dominic Kwiatkowski, Cordelia Langford, John Sillitoe on behalf of the Wellcome Sanger Institute COVID-19 Surveillance Team |
| EPI_ISL_2830696 | Lighthouse Laboratory Plymouth | Wellcome Sanger Institute for the COVID-19 Genomics UK (COG-UK) Consortium | Lighthouse Laboratory Plymouth and Alex Alderton, Roberto Amato, Jeffrey Barrett, Sonia Goncalves, Ewan Harrison, David K. Jackson, Ian Johnston, Dominic Kwiatkowski, Cordelia Langford, John Sillitoe on behalf of the Wellcome Sanger Institute COVID-19 Surveillance Team |
| EPI_ISL_2830697 | Lighthouse Lab in Alderley Park | Wellcome Sanger Institute for the COVID-19 Genomics UK (COG-UK) Consortium | Jacquelyn Wynn, Mairead Hyland, The Lighthouse Lab in Alderley Park and Alex Alderton, Roberto Amato, Jeffrey Barrett, Sonia Goncalves, Ewan Harrison, David K. Jackson, Ian Johnston, Dominic Kwiatkowski, Cordelia Langford, John Sillitoe on behalf of the Wellcome Sanger Institute COVID-19 Surveillance Team |
| EPI_ISL_2830698 | Lighthouse Lab in Milton Keynes | Wellcome Sanger Institute for the COVID-19 Genomics UK (COG-UK) Consortium | The Lighthouse Lab in Milton Keynes and Alex Alderton, Roberto Amato, Jeffrey Barrett, Sonia Goncalves, Ewan Harrison, David K. Jackson, Ian Johnston, Dominic Kwiatkowski, Cordelia Langford, John Sillitoe on behalf of the Wellcome Sanger Institute COVID-19 Surveillance Team |
| EPI_ISL_2830699 | Lighthouse Lab in Glasgow | Wellcome Sanger Institute for the COVID-19 Genomics UK (COG-UK) Consortium | Harper VanSteenhouse, Yumi Kasai, David Gray, Carol Clugston, Anna Dominiczak and Alex Alderton, Roberto Amato, Jeffrey Barrett, Sonia Goncalves, Ewan Harrison, David K. Jackson, Ian Johnston, Dominic Kwiatkowski, Cordelia Langford, John Sillitoe on behalf of the Wellcome Sanger Institute COVID-19 Surveillance Team |

| | | | |
|---|---|---|---|
| EPI_ISL_2830700, EPI_ISL_2830701, EPI_ISL_2830702, EPI_ISL_2830703, EPI_ISL_2830704, EPI_ISL_2830705, EPI_ISL_2830706 | Lighthouse Lab in Milton Keynes | Wellcome Sanger Institute for the COVID-19 Genomics UK (COG-UK) Consortium | The Lighthouse Lab in Milton Keynes and Alex Alderton, Roberto Amato, Jeffrey Barrett, Sonia Goncalves, Ewan Harrison, David K. Jackson, Ian Johnston, Dominic Kwiatkowski, Cordelia Langford, John Sillitoe on behalf of the Wellcome Sanger Institute COVID-19 Surveillance Team |
| EPI_ISL_2830707, EPI_ISL_2830708 | Lighthouse Lab in Glasgow | Wellcome Sanger Institute for the COVID-19 Genomics UK (COG-UK) Consortium | Harper VanSteenhouse, Yumi Kasai, David Gray, Carol Clugston, Anna Dominiczak and Alex Alderton, Roberto Amato, Jeffrey Barrett, Sonia Goncalves, Ewan Harrison, David K. Jackson, Ian Johnston, Dominic Kwiatkowski, Cordelia Langford, John Sillitoe on behalf of the Wellcome Sanger Institute COVID-19 Surveillance Team |
| EPI_ISL_2830709 | Lighthouse Lab in Milton Keynes | Wellcome Sanger Institute for the COVID-19 Genomics UK (COG-UK) Consortium | The Lighthouse Lab in Milton Keynes and Alex Alderton, Roberto Amato, Jeffrey Barrett, Sonia Goncalves, Ewan Harrison, David K. Jackson, Ian Johnston, Dominic Kwiatkowski, Cordelia Langford, John Sillitoe on behalf of the Wellcome Sanger Institute COVID-19 Surveillance Team |
| EPI_ISL_2830710 | Lighthouse Lab in Alderley Park | Wellcome Sanger Institute for the COVID-19 Genomics UK (COG-UK) Consortium | Jacquelyn Wynn, Mairead Hyland, The Lighthouse Lab in Alderley Park and Alex Alderton, Roberto Amato, Jeffrey Barrett, Sonia Goncalves, Ewan Harrison, David K. Jackson, Ian Johnston, Dominic Kwiatkowski, Cordelia Langford, John Sillitoe on behalf of the Wellcome Sanger Institute COVID-19 Surveillance Team |
| EPI_ISL_2830711 | Lighthouse Lab in Milton Keynes | Wellcome Sanger Institute for the COVID-19 Genomics UK (COG-UK) Consortium | The Lighthouse Lab in Milton Keynes and Alex Alderton, Roberto Amato, Jeffrey Barrett, Sonia Goncalves, Ewan Harrison, David K. Jackson, Ian Johnston, Dominic Kwiatkowski, Cordelia Langford, John Sillitoe on behalf of the Wellcome Sanger Institute COVID-19 Surveillance Team |
| EPI_ISL_2830712 | Lighthouse Lab in Glasgow | Wellcome Sanger Institute for the COVID-19 Genomics UK (COG-UK) Consortium | Harper VanSteenhouse, Yumi Kasai, David Gray, Carol Clugston, Anna Dominiczak and Alex Alderton, Roberto Amato, Jeffrey Barrett, Sonia Goncalves, Ewan Harrison, David K. Jackson, Ian Johnston, Dominic Kwiatkowski, Cordelia Langford, John Sillitoe on behalf of the Wellcome Sanger Institute COVID-19 Surveillance Team |
| EPI_ISL_2830713, EPI_ISL_2830714 | Lighthouse Lab in Milton Keynes | Wellcome Sanger Institute for the COVID-19 Genomics UK (COG-UK) Consortium | The Lighthouse Lab in Milton Keynes and Alex Alderton, Roberto Amato, Jeffrey Barrett, Sonia Goncalves, Ewan Harrison, David K. Jackson, Ian Johnston, Dominic Kwiatkowski, Cordelia Langford, John Sillitoe on behalf of the Wellcome Sanger Institute COVID-19 Surveillance Team |
| EPI_ISL_2830715 | Lighthouse Lab in Glasgow | Wellcome Sanger Institute for the COVID-19 Genomics UK (COG-UK) Consortium | Harper VanSteenhouse, Yumi Kasai, David Gray, Carol Clugston, Anna Dominiczak and Alex Alderton, Roberto Amato, Jeffrey Barrett, Sonia Goncalves, Ewan Harrison, David K. Jackson, Ian Johnston, Dominic Kwiatkowski, Cordelia Langford, John Sillitoe on behalf of the Wellcome Sanger Institute COVID-19 Surveillance Team |
| EPI_ISL_2830716 | Lighthouse Lab in Alderley Park | Wellcome Sanger Institute for the COVID-19 Genomics UK (COG-UK) Consortium | Jacquelyn Wynn, Mairead Hyland, The Lighthouse Lab in Alderley Park and Alex Alderton, Roberto Amato, Jeffrey Barrett, Sonia Goncalves, Ewan Harrison, David K. Jackson, Ian Johnston, Dominic Kwiatkowski, Cordelia Langford, John Sillitoe on behalf of the Wellcome Sanger Institute COVID-19 Surveillance Team |
| EPI_ISL_2830717 | Lighthouse Lab in Glasgow | Wellcome Sanger Institute for the COVID-19 Genomics UK (COG-UK) Consortium | Harper VanSteenhouse, Yumi Kasai, David Gray, Carol Clugston, Anna Dominiczak and Alex Alderton, Roberto Amato, Jeffrey Barrett, Sonia Goncalves, Ewan Harrison, David K. Jackson, Ian Johnston, Dominic Kwiatkowski, Cordelia Langford, John Sillitoe on behalf of the Wellcome Sanger Institute COVID-19 Surveillance Team |
| EPI_ISL_2830718 | Lighthouse Laboratory Plymouth | Wellcome Sanger Institute for the COVID-19 Genomics UK (COG-UK) Consortium | Lighthouse Laboratory Plymouth and Alex Alderton, Roberto Amato, Jeffrey Barrett, Sonia Goncalves, Ewan Harrison, David K. Jackson, Ian Johnston, Dominic Kwiatkowski, Cordelia Langford, John Sillitoe on behalf of the Wellcome Sanger Institute COVID-19 Surveillance Team |
| EPI_ISL_2830719 | Lighthouse Lab in Glasgow | Wellcome Sanger Institute for the COVID-19 Genomics UK (COG-UK) Consortium | Harper VanSteenhouse, Yumi Kasai, David Gray, Carol Clugston, Anna Dominiczak and Alex Alderton, Roberto Amato, Jeffrey Barrett, Sonia Goncalves, Ewan Harrison, David K. Jackson, Ian Johnston, Dominic Kwiatkowski, Cordelia Langford, John Sillitoe on behalf of the Wellcome Sanger Institute COVID-19 Surveillance Team |
| EPI_ISL_2830720 | Lighthouse Laboratory Plymouth | Wellcome Sanger Institute for the COVID-19 Genomics UK (COG-UK) Consortium | Lighthouse Laboratory Plymouth and Alex Alderton, Roberto Amato, Jeffrey Barrett, Sonia Goncalves, Ewan Harrison, David K. Jackson, Ian Johnston, Dominic Kwiatkowski, Cordelia Langford, John Sillitoe on behalf of the Wellcome Sanger Institute COVID-19 Surveillance Team |
| EPI_ISL_2830721 | Lighthouse Lab in Glasgow | Wellcome Sanger Institute for the COVID-19 Genomics UK (COG-UK) Consortium | Harper VanSteenhouse, Yumi Kasai, David Gray, Carol Clugston, Anna Dominiczak and Alex Alderton, Roberto Amato, Jeffrey Barrett, Sonia Goncalves, Ewan Harrison, David K. Jackson, Ian Johnston, Dominic Kwiatkowski, Cordelia Langford, John Sillitoe on behalf of the Wellcome Sanger Institute COVID-19 Surveillance Team |
| EPI_ISL_2830722 | Lighthouse Laboratory Plymouth | Wellcome Sanger Institute for the COVID-19 Genomics UK (COG-UK) Consortium | Lighthouse Laboratory Plymouth and Alex Alderton, Roberto Amato, Jeffrey Barrett, Sonia Goncalves, Ewan Harrison, David K. Jackson, Ian Johnston, Dominic Kwiatkowski, Cordelia Langford, John Sillitoe on behalf of the Wellcome Sanger Institute COVID-19 Surveillance Team |
| EPI_ISL_2830723 | Lighthouse Lab in Alderley Park | Wellcome Sanger Institute for the COVID-19 Genomics UK (COG-UK) Consortium | Jacquelyn Wynn, Mairead Hyland, The Lighthouse Lab in Alderley Park and Alex Alderton, Roberto Amato, Jeffrey Barrett, Sonia Goncalves, Ewan Harrison, David K. Jackson, Ian Johnston, Dominic Kwiatkowski, Cordelia Langford, John Sillitoe on behalf of the Wellcome Sanger Institute COVID-19 Surveillance Team |
| EPI_ISL_2830724, EPI_ISL_2830725, EPI_ISL_2830726 | Lighthouse Lab in Milton Keynes | Wellcome Sanger Institute for the COVID-19 Genomics UK (COG-UK) Consortium | The Lighthouse Lab in Milton Keynes and Alex Alderton, Roberto Amato, Jeffrey Barrett, Sonia Goncalves, Ewan Harrison, David K. Jackson, Ian Johnston, Dominic Kwiatkowski, Cordelia Langford, John Sillitoe on behalf of the Wellcome Sanger Institute COVID-19 Surveillance Team |
| EPI_ISL_2830727 | Lighthouse Lab in Alderley Park | Wellcome Sanger Institute for the COVID-19 Genomics UK (COG-UK) Consortium | Jacquelyn Wynn, Mairead Hyland, The Lighthouse Lab in Alderley Park and Alex Alderton, Roberto Amato, Jeffrey Barrett, Sonia Goncalves, Ewan Harrison, David K. Jackson, Ian Johnston, Dominic Kwiatkowski, Cordelia Langford, John Sillitoe on behalf of the Wellcome Sanger Institute COVID-19 Surveillance Team |
| EPI_ISL_2830728 | Lighthouse Laboratory Plymouth | Wellcome Sanger Institute for the COVID-19 Genomics UK (COG-UK) Consortium | Lighthouse Laboratory Plymouth and Alex Alderton, Roberto Amato, Jeffrey Barrett, Sonia Goncalves, Ewan Harrison, David K. Jackson, Ian Johnston, Dominic Kwiatkowski, Cordelia Langford, John Sillitoe on behalf of the Wellcome Sanger Institute COVID-19 Surveillance Team |
| EPI_ISL_2830729 | Lighthouse Lab in Glasgow | Wellcome Sanger Institute for the COVID-19 Genomics UK (COG-UK) Consortium | Harper VanSteenhouse, Yumi Kasai, David Gray, Carol Clugston, Anna Dominiczak and Alex Alderton, Roberto Amato, Jeffrey Barrett, Sonia Goncalves, Ewan Harrison, David K. Jackson, Ian Johnston, Dominic Kwiatkowski, Cordelia Langford, John Sillitoe on behalf of the Wellcome Sanger Institute COVID-19 Surveillance Team |
| EPI_ISL_2830730 | Lighthouse Lab in Milton Keynes | Wellcome Sanger Institute for the COVID-19 Genomics UK (COG-UK) Consortium | The Lighthouse Lab in Milton Keynes and Alex Alderton, Roberto Amato, Jeffrey Barrett, Sonia Goncalves, Ewan Harrison, David K. Jackson, Ian Johnston, Dominic Kwiatkowski, Cordelia Langford, John Sillitoe on behalf of the Wellcome Sanger Institute COVID-19 Surveillance Team |
| EPI_ISL_2830731 | Lighthouse Laboratory Plymouth | Wellcome Sanger Institute for the COVID-19 Genomics UK (COG-UK) Consortium | Lighthouse Laboratory Plymouth and Alex Alderton, Roberto Amato, Jeffrey Barrett, Sonia Goncalves, Ewan Harrison, David K. Jackson, Ian Johnston, Dominic Kwiatkowski, Cordelia Langford, John Sillitoe on behalf of the Wellcome Sanger Institute COVID-19 Surveillance Team |
| EPI_ISL_2830732 | Lighthouse Lab in Milton Keynes | Wellcome Sanger Institute for the COVID-19 Genomics UK (COG-UK) Consortium | The Lighthouse Lab in Milton Keynes and Alex Alderton, Roberto Amato, Jeffrey Barrett, Sonia Goncalves, Ewan Harrison, David K. Jackson, Ian Johnston, Dominic Kwiatkowski, Cordelia Langford, John Sillitoe on behalf of the Wellcome Sanger Institute COVID-19 Surveillance Team |
| EPI_ISL_2830733 | Lighthouse Lab in Glasgow | Wellcome Sanger Institute for the COVID-19 Genomics UK (COG-UK) Consortium | Harper VanSteenhouse, Yumi Kasai, David Gray, Carol Clugston, Anna Dominiczak and Alex Alderton, Roberto Amato, Jeffrey Barrett, Sonia Goncalves, Ewan Harrison, David K. Jackson, Ian Johnston, Dominic Kwiatkowski, Cordelia Langford, John Sillitoe on behalf of the Wellcome Sanger Institute COVID-19 Surveillance Team |
| EPI_ISL_2830734 | Lighthouse Lab in Alderley Park | Wellcome Sanger Institute for the COVID-19 Genomics UK (COG-UK) Consortium | Jacquelyn Wynn, Mairead Hyland, The Lighthouse Lab in Alderley Park and Alex Alderton, Roberto Amato, Jeffrey Barrett, Sonia Goncalves, Ewan Harrison, David K. Jackson, Ian Johnston, Dominic Kwiatkowski, Cordelia Langford, John Sillitoe on behalf of the Wellcome Sanger Institute COVID-19 Surveillance Team |
| EPI_ISL_2830735 | Lighthouse Lab in Milton Keynes | Wellcome Sanger Institute for the COVID-19 Genomics UK (COG-UK) Consortium | The Lighthouse Lab in Milton Keynes and Alex Alderton, Roberto Amato, Jeffrey Barrett, Sonia Goncalves, Ewan Harrison, David K. Jackson, Ian Johnston, Dominic Kwiatkowski, Cordelia Langford, John Sillitoe on behalf of the Wellcome Sanger Institute COVID-19 Surveillance Team |
| EPI_ISL_2830736 | Lighthouse Laboratory Plymouth | Wellcome Sanger Institute for the COVID-19 Genomics UK (COG-UK) Consortium | Lighthouse Laboratory Plymouth and Alex Alderton, Roberto Amato, Jeffrey Barrett, Sonia Goncalves, Ewan Harrison, David K. Jackson, Ian Johnston, Dominic Kwiatkowski, Cordelia Langford, John Sillitoe on behalf of the Wellcome Sanger Institute COVID-19 Surveillance Team |
| EPI_ISL_2830737, EPI_ISL_2830738 | Lighthouse Lab in Milton Keynes | Wellcome Sanger Institute for the COVID-19 Genomics UK (COG-UK) Consortium | The Lighthouse Lab in Milton Keynes and Alex Alderton, Roberto Amato, Jeffrey Barrett, Sonia Goncalves, Ewan Harrison, David K. Jackson, Ian Johnston, Dominic Kwiatkowski, Cordelia Langford, John Sillitoe on behalf of the Wellcome Sanger Institute COVID-19 Surveillance Team |
| EPI_ISL_2830739, EPI_ISL_2830740 | Lighthouse Lab in Glasgow | Wellcome Sanger Institute for the COVID-19 Genomics UK (COG-UK) Consortium | Harper VanSteenhouse, Yumi Kasai, David Gray, Carol Clugston, Anna Dominiczak and Alex Alderton, Roberto Amato, Jeffrey Barrett, Sonia Goncalves, Ewan Harrison, David K. Jackson, Ian Johnston, Dominic Kwiatkowski, Cordelia Langford, John Sillitoe on behalf of the Wellcome Sanger Institute COVID-19 Surveillance Team |
| EPI_ISL_2830741 | Lighthouse Lab in Milton Keynes | Wellcome Sanger Institute for the COVID-19 Genomics UK (COG-UK) Consortium | The Lighthouse Lab in Milton Keynes and Alex Alderton, Roberto Amato, Jeffrey Barrett, Sonia Goncalves, Ewan Harrison, David K. Jackson, Ian Johnston, Dominic Kwiatkowski, Cordelia Langford, John Sillitoe on behalf of the Wellcome Sanger Institute COVID-19 Surveillance Team |
| EPI_ISL_2830742 | Lighthouse Lab in Glasgow | Wellcome Sanger Institute for the COVID-19 Genomics UK (COG-UK) Consortium | Harper VanSteenhouse, Yumi Kasai, David Gray, Carol Clugston, Anna Dominiczak and Alex Alderton, Roberto Amato, Jeffrey Barrett, Sonia Goncalves, Ewan Harrison, David K. Jackson, Ian Johnston, Dominic Kwiatkowski, Cordelia Langford, John Sillitoe on behalf of the Wellcome Sanger Institute COVID-19 Surveillance Team |
| EPI_ISL_2830743, EPI_ISL_2830744 | Lighthouse Laboratory Plymouth | Wellcome Sanger Institute for the COVID-19 Genomics UK (COG-UK) Consortium | Lighthouse Laboratory Plymouth and Alex Alderton, Roberto Amato, Jeffrey Barrett, Sonia Goncalves, Ewan Harrison, David K. Jackson, Ian Johnston, Dominic Kwiatkowski, Cordelia Langford, John Sillitoe on behalf of the Wellcome Sanger Institute COVID-19 Surveillance Team |
| EPI_ISL_2830745 | Lighthouse Lab in Milton Keynes | Wellcome Sanger Institute for the COVID-19 Genomics UK (COG-UK) Consortium | The Lighthouse Lab in Milton Keynes and Alex Alderton, Roberto Amato, Jeffrey Barrett, Sonia Goncalves, Ewan Harrison, David K. Jackson, Ian Johnston, Dominic Kwiatkowski, Cordelia Langford, John Sillitoe on behalf of the Wellcome Sanger Institute COVID-19 Surveillance Team |
| EPI_ISL_2830746 | Lighthouse Lab in Glasgow | Wellcome Sanger Institute for the COVID-19 Genomics UK (COG-UK) Consortium | Harper VanSteenhouse, Yumi Kasai, David Gray, Carol Clugston, Anna Dominiczak and Alex Alderton, Roberto Amato, Jeffrey Barrett, Sonia Goncalves, Ewan Harrison, David K. Jackson, Ian Johnston, Dominic Kwiatkowski, Cordelia Langford, John Sillitoe on behalf of the Wellcome Sanger Institute COVID-19 Surveillance Team |
| EPI_ISL_2830747 | Lighthouse Lab in Milton Keynes | Wellcome Sanger Institute for the COVID-19 Genomics UK (COG-UK) Consortium | The Lighthouse Lab in Milton Keynes and Alex Alderton, Roberto Amato, Jeffrey Barrett, Sonia Goncalves, Ewan Harrison, David K. Jackson, Ian Johnston, Dominic Kwiatkowski, Cordelia Langford, John Sillitoe on behalf of the Wellcome Sanger Institute COVID-19 Surveillance Team |
| EPI_ISL_2830748 | Lighthouse Lab in Glasgow | Wellcome Sanger Institute for the COVID-19 Genomics UK (COG-UK) Consortium | Harper VanSteenhouse, Yumi Kasai, David Gray, Carol Clugston, Anna Dominiczak and Alex Alderton, Roberto Amato, Jeffrey Barrett, Sonia Goncalves, Ewan Harrison, David K. Jackson, Ian Johnston, Dominic Kwiatkowski, Cordelia Langford, John Sillitoe on behalf of the Wellcome Sanger Institute COVID-19 Surveillance Team |
| EPI_ISL_2830749 | Lighthouse Laboratory Plymouth | Wellcome Sanger Institute for the COVID-19 Genomics UK (COG-UK) Consortium | Lighthouse Laboratory Plymouth and Alex Alderton, Roberto Amato, Jeffrey Barrett, Sonia Goncalves, Ewan Harrison, David K. Jackson, Ian Johnston, Dominic Kwiatkowski, Cordelia Langford, John Sillitoe on behalf of the Wellcome Sanger Institute COVID-19 Surveillance Team |
| EPI_ISL_2830750 | Lighthouse Lab in Milton Keynes | Wellcome Sanger Institute for the COVID-19 Genomics UK (COG-UK) Consortium | The Lighthouse Lab in Milton Keynes and Alex Alderton, Roberto Amato, Jeffrey Barrett, Sonia Goncalves, Ewan Harrison, David K. Jackson, Ian Johnston, Dominic Kwiatkowski, Cordelia Langford, John Sillitoe on behalf of the Wellcome Sanger Institute COVID-19 Surveillance Team |
| EPI_ISL_2830751 | Lighthouse Lab in Alderley Park | Wellcome Sanger Institute for the COVID-19 Genomics UK (COG-UK) Consortium | Jacquelyn Wynn, Mairead Hyland, The Lighthouse Lab in Alderley Park and Alex Alderton, Roberto Amato, Jeffrey Barrett, Sonia Goncalves, Ewan Harrison, David K. Jackson, Ian Johnston, Dominic Kwiatkowski, Cordelia Langford, John Sillitoe on behalf of the Wellcome Sanger Institute COVID-19 Surveillance Team |
| EPI_ISL_2830752 | Lighthouse Lab in Glasgow | Wellcome Sanger Institute for the COVID-19 Genomics UK (COG-UK) Consortium | Harper VanSteenhouse, Yumi Kasai, David Gray, Carol Clugston, Anna Dominiczak and Alex Alderton, Roberto Amato, Jeffrey Barrett, Sonia Goncalves, Ewan Harrison, David K. Jackson, Ian Johnston, Dominic Kwiatkowski, Cordelia Langford, John Sillitoe on behalf of the Wellcome Sanger Institute COVID-19 Surveillance Team |
| EPI_ISL_2830753, EPI_ISL_2830754 | Lighthouse Laboratory Plymouth | Wellcome Sanger Institute for the COVID-19 Genomics UK (COG-UK) Consortium | Lighthouse Laboratory Plymouth and Alex Alderton, Roberto Amato, Jeffrey Barrett, Sonia Goncalves, Ewan Harrison, David K. Jackson, Ian Johnston, Dominic Kwiatkowski, Cordelia Langford, John Sillitoe on behalf of the Wellcome Sanger Institute COVID-19 Surveillance Team |
| EPI_ISL_2830755 | Lighthouse Lab in Milton Keynes | Wellcome Sanger Institute for the COVID-19 Genomics UK (COG-UK) Consortium | The Lighthouse Lab in Milton Keynes and Alex Alderton, Roberto Amato, Jeffrey Barrett, Sonia Goncalves, Ewan Harrison, David K. Jackson, Ian Johnston, Dominic Kwiatkowski, Cordelia Langford, John Sillitoe on behalf of the Wellcome Sanger Institute COVID-19 Surveillance Team |
| EPI_ISL_2830756 | Lighthouse Lab in Glasgow | Wellcome Sanger Institute for the COVID-19 Genomics UK (COG-UK) | Harper VanSteenhouse, Yumi Kasai, David Gray, Carol Clugston, Anna Dominiczak and Alex Alderton, Roberto Amato, Jeffrey Barrett, Sonia Goncalves, Ewan Harrison, David K. Jackson, Ian Johnston, Dominic Kwiatkowski, Cordelia Langford, |

| | | | |
|---|---|---|---|
| | | Consortium | John Sillitoe on behalf of the Wellcome Sanger Institute COVID-19 Surveillance Team |
| EPI_ISL_2830757, EPI_ISL_2830758, EPI_ISL_2830759, EPI_ISL_2830760 | Lighthouse Lab in Milton Keynes | Wellcome Sanger Institute for the COVID-19 Genomics UK (COG-UK) Consortium | The Lighthouse Lab in Milton Keynes and Alex Alderton, Roberto Amato, Jeffrey Barrett, Sonia Goncalves, Ewan Harrison, David K. Jackson, Ian Johnston, Dominic Kwiatkowski, Cordelia Langford, John Sillitoe on behalf of the Wellcome Sanger Institute COVID-19 Surveillance Team |
| EPI_ISL_2830761 | Lighthouse Lab in Alderley Park | Wellcome Sanger Institute for the COVID-19 Genomics UK (COG-UK) Consortium | Jacquelyn Wynn, Mairead Hyland, The Lighthouse Lab in Alderley Park and Alex Alderton, Roberto Amato, Jeffrey Barrett, Sonia Goncalves, Ewan Harrison, David K. Jackson, Ian Johnston, Dominic Kwiatkowski, Cordelia Langford, John Sillitoe on behalf of the Wellcome Sanger Institute COVID-19 Surveillance Team |
| EPI_ISL_2830762, EPI_ISL_2830763 | Lighthouse Lab in Milton Keynes | Wellcome Sanger Institute for the COVID-19 Genomics UK (COG-UK) Consortium | The Lighthouse Lab in Milton Keynes and Alex Alderton, Roberto Amato, Jeffrey Barrett, Sonia Goncalves, Ewan Harrison, David K. Jackson, Ian Johnston, Dominic Kwiatkowski, Cordelia Langford, John Sillitoe on behalf of the Wellcome Sanger Institute COVID-19 Surveillance Team |
| EPI_ISL_2830764, EPI_ISL_2830765 | Lighthouse Lab in Glasgow | Wellcome Sanger Institute for the COVID-19 Genomics UK (COG-UK) Consortium | Harper VanSteenhouse, Yumi Kasai, David Gray, Carol Clugston, Anna Dominiczak and Alex Alderton, Roberto Amato, Jeffrey Barrett, Sonia Goncalves, Ewan Harrison, David K. Jackson, Ian Johnston, Dominic Kwiatkowski, Cordelia Langford, John Sillitoe on behalf of the Wellcome Sanger Institute COVID-19 Surveillance Team |
| EPI_ISL_2830766 | Lighthouse Laboratory Plymouth | Wellcome Sanger Institute for the COVID-19 Genomics UK (COG-UK) Consortium | Lighthouse Laboratory Plymouth and Alex Alderton, Roberto Amato, Jeffrey Barrett, Sonia Goncalves, Ewan Harrison, David K. Jackson, Ian Johnston, Dominic Kwiatkowski, Cordelia Langford, John Sillitoe on behalf of the Wellcome Sanger Institute COVID-19 Surveillance Team |
| EPI_ISL_2830767, EPI_ISL_2830768 | Lighthouse Lab in Glasgow | Wellcome Sanger Institute for the COVID-19 Genomics UK (COG-UK) Consortium | Harper VanSteenhouse, Yumi Kasai, David Gray, Carol Clugston, Anna Dominiczak and Alex Alderton, Roberto Amato, Jeffrey Barrett, Sonia Goncalves, Ewan Harrison, David K. Jackson, Ian Johnston, Dominic Kwiatkowski, Cordelia Langford, John Sillitoe on behalf of the Wellcome Sanger Institute COVID-19 Surveillance Team |
| EPI_ISL_2830769 | Lighthouse Laboratory Plymouth | Wellcome Sanger Institute for the COVID-19 Genomics UK (COG-UK) Consortium | Lighthouse Laboratory Plymouth and Alex Alderton, Roberto Amato, Jeffrey Barrett, Sonia Goncalves, Ewan Harrison, David K. Jackson, Ian Johnston, Dominic Kwiatkowski, Cordelia Langford, John Sillitoe on behalf of the Wellcome Sanger Institute COVID-19 Surveillance Team |
| EPI_ISL_2830770 | Lighthouse Lab in Milton Keynes | Wellcome Sanger Institute for the COVID-19 Genomics UK (COG-UK) Consortium | The Lighthouse Lab in Milton Keynes and Alex Alderton, Roberto Amato, Jeffrey Barrett, Sonia Goncalves, Ewan Harrison, David K. Jackson, Ian Johnston, Dominic Kwiatkowski, Cordelia Langford, John Sillitoe on behalf of the Wellcome Sanger Institute COVID-19 Surveillance Team |
| EPI_ISL_2830771, EPI_ISL_2830772 | Lighthouse Lab in Glasgow | Wellcome Sanger Institute for the COVID-19 Genomics UK (COG-UK) Consortium | Harper VanSteenhouse, Yumi Kasai, David Gray, Carol Clugston, Anna Dominiczak and Alex Alderton, Roberto Amato, Jeffrey Barrett, Sonia Goncalves, Ewan Harrison, David K. Jackson, Ian Johnston, Dominic Kwiatkowski, Cordelia Langford, John Sillitoe on behalf of the Wellcome Sanger Institute COVID-19 Surveillance Team |
| EPI_ISL_2830773 | Lighthouse Lab in Milton Keynes | Wellcome Sanger Institute for the COVID-19 Genomics UK (COG-UK) Consortium | The Lighthouse Lab in Milton Keynes and Alex Alderton, Roberto Amato, Jeffrey Barrett, Sonia Goncalves, Ewan Harrison, David K. Jackson, Ian Johnston, Dominic Kwiatkowski, Cordelia Langford, John Sillitoe on behalf of the Wellcome Sanger Institute COVID-19 Surveillance Team |
| EPI_ISL_2830774, EPI_ISL_2830775 | Lighthouse Laboratory Plymouth | Wellcome Sanger Institute for the COVID-19 Genomics UK (COG-UK) Consortium | Lighthouse Laboratory Plymouth and Alex Alderton, Roberto Amato, Jeffrey Barrett, Sonia Goncalves, Ewan Harrison, David K. Jackson, Ian Johnston, Dominic Kwiatkowski, Cordelia Langford, John Sillitoe on behalf of the Wellcome Sanger Institute COVID-19 Surveillance Team |
| EPI_ISL_2830776 | Lighthouse Lab in Milton Keynes | Wellcome Sanger Institute for the COVID-19 Genomics UK (COG-UK) Consortium | The Lighthouse Lab in Milton Keynes and Alex Alderton, Roberto Amato, Jeffrey Barrett, Sonia Goncalves, Ewan Harrison, David K. Jackson, Ian Johnston, Dominic Kwiatkowski, Cordelia Langford, John Sillitoe on behalf of the Wellcome Sanger Institute COVID-19 Surveillance Team |
| EPI_ISL_2830777 | Lighthouse Laboratory Plymouth | Wellcome Sanger Institute for the COVID-19 Genomics UK (COG-UK) Consortium | Lighthouse Laboratory Plymouth and Alex Alderton, Roberto Amato, Jeffrey Barrett, Sonia Goncalves, Ewan Harrison, David K. Jackson, Ian Johnston, Dominic Kwiatkowski, Cordelia Langford, John Sillitoe on behalf of the Wellcome Sanger Institute COVID-19 Surveillance Team |
| EPI_ISL_2830778, EPI_ISL_2830779, EPI_ISL_2830780 | Lighthouse Lab in Milton Keynes | Wellcome Sanger Institute for the COVID-19 Genomics UK (COG-UK) Consortium | The Lighthouse Lab in Milton Keynes and Alex Alderton, Roberto Amato, Jeffrey Barrett, Sonia Goncalves, Ewan Harrison, David K. Jackson, Ian Johnston, Dominic Kwiatkowski, Cordelia Langford, John Sillitoe on behalf of the Wellcome Sanger Institute COVID-19 Surveillance Team |
| EPI_ISL_2830781 | Lighthouse Laboratory Plymouth | Wellcome Sanger Institute for the COVID-19 Genomics UK (COG-UK) Consortium | Lighthouse Laboratory Plymouth and Alex Alderton, Roberto Amato, Jeffrey Barrett, Sonia Goncalves, Ewan Harrison, David K. Jackson, Ian Johnston, Dominic Kwiatkowski, Cordelia Langford, John Sillitoe on behalf of the Wellcome Sanger Institute COVID-19 Surveillance Team |
| EPI_ISL_2830782 | Lighthouse Lab in Glasgow | Wellcome Sanger Institute for the COVID-19 Genomics UK (COG-UK) Consortium | Harper VanSteenhouse, Yumi Kasai, David Gray, Carol Clugston, Anna Dominiczak and Alex Alderton, Roberto Amato, Jeffrey Barrett, Sonia Goncalves, Ewan Harrison, David K. Jackson, Ian Johnston, Dominic Kwiatkowski, Cordelia Langford, John Sillitoe on behalf of the Wellcome Sanger Institute COVID-19 Surveillance Team |
| EPI_ISL_2830783 | Lighthouse Laboratory Plymouth | Wellcome Sanger Institute for the COVID-19 Genomics UK (COG-UK) Consortium | Lighthouse Laboratory Plymouth and Alex Alderton, Roberto Amato, Jeffrey Barrett, Sonia Goncalves, Ewan Harrison, David K. Jackson, Ian Johnston, Dominic Kwiatkowski, Cordelia Langford, John Sillitoe on behalf of the Wellcome Sanger Institute COVID-19 Surveillance Team |
| EPI_ISL_2830784 | Lighthouse Lab in Milton Keynes | Wellcome Sanger Institute for the COVID-19 Genomics UK (COG-UK) Consortium | The Lighthouse Lab in Milton Keynes and Alex Alderton, Roberto Amato, Jeffrey Barrett, Sonia Goncalves, Ewan Harrison, David K. Jackson, Ian Johnston, Dominic Kwiatkowski, Cordelia Langford, John Sillitoe on behalf of the Wellcome Sanger Institute COVID-19 Surveillance Team |
| EPI_ISL_2830785, EPI_ISL_2830786 | Lighthouse Lab in Glasgow | Wellcome Sanger Institute for the COVID-19 Genomics UK (COG-UK) Consortium | Harper VanSteenhouse, Yumi Kasai, David Gray, Carol Clugston, Anna Dominiczak and Alex Alderton, Roberto Amato, Jeffrey Barrett, Sonia Goncalves, Ewan Harrison, David K. Jackson, Ian Johnston, Dominic Kwiatkowski, Cordelia Langford, John Sillitoe on behalf of the Wellcome Sanger Institute COVID-19 Surveillance Team |
| EPI_ISL_2830787 | Lighthouse Lab in Milton Keynes | Wellcome Sanger Institute for the COVID-19 Genomics UK (COG-UK) Consortium | The Lighthouse Lab in Milton Keynes and Alex Alderton, Roberto Amato, Jeffrey Barrett, Sonia Goncalves, Ewan Harrison, David K. Jackson, Ian Johnston, Dominic Kwiatkowski, Cordelia Langford, John Sillitoe on behalf of the Wellcome Sanger Institute COVID-19 Surveillance Team |
| EPI_ISL_2830788, EPI_ISL_2830789 | Lighthouse Lab in Glasgow | Wellcome Sanger Institute for the COVID-19 Genomics UK (COG-UK) Consortium | Harper VanSteenhouse, Yumi Kasai, David Gray, Carol Clugston, Anna Dominiczak and Alex Alderton, Roberto Amato, Jeffrey Barrett, Sonia Goncalves, Ewan Harrison, David K. Jackson, Ian Johnston, Dominic Kwiatkowski, Cordelia Langford, John Sillitoe on behalf of the Wellcome Sanger Institute COVID-19 Surveillance Team |
| EPI_ISL_2830790 | Lighthouse Lab in Milton Keynes | Wellcome Sanger Institute for the COVID-19 Genomics UK (COG-UK) Consortium | The Lighthouse Lab in Milton Keynes and Alex Alderton, Roberto Amato, Jeffrey Barrett, Sonia Goncalves, Ewan Harrison, David K. Jackson, Ian Johnston, Dominic Kwiatkowski, Cordelia Langford, John Sillitoe on behalf of the Wellcome Sanger Institute COVID-19 Surveillance Team |
| EPI_ISL_2830791, EPI_ISL_2830792, EPI_ISL_2830793, EPI_ISL_2830794 | Lighthouse Lab in Glasgow | Wellcome Sanger Institute for the COVID-19 Genomics UK (COG-UK) Consortium | Harper VanSteenhouse, Yumi Kasai, David Gray, Carol Clugston, Anna Dominiczak and Alex Alderton, Roberto Amato, Jeffrey Barrett, Sonia Goncalves, Ewan Harrison, David K. Jackson, Ian Johnston, Dominic Kwiatkowski, Cordelia Langford, John Sillitoe on behalf of the Wellcome Sanger Institute COVID-19 Surveillance Team |
| EPI_ISL_2830795, EPI_ISL_2830796 | Lighthouse Lab in Milton Keynes | Wellcome Sanger Institute for the COVID-19 Genomics UK (COG-UK) Consortium | The Lighthouse Lab in Milton Keynes and Alex Alderton, Roberto Amato, Jeffrey Barrett, Sonia Goncalves, Ewan Harrison, David K. Jackson, Ian Johnston, Dominic Kwiatkowski, Cordelia Langford, John Sillitoe on behalf of the Wellcome Sanger Institute COVID-19 Surveillance Team |
| EPI_ISL_2830797 | Lighthouse Lab in Glasgow | Wellcome Sanger Institute for the COVID-19 Genomics UK (COG-UK) Consortium | Harper VanSteenhouse, Yumi Kasai, David Gray, Carol Clugston, Anna Dominiczak and Alex Alderton, Roberto Amato, Jeffrey Barrett, Sonia Goncalves, Ewan Harrison, David K. Jackson, Ian Johnston, Dominic Kwiatkowski, Cordelia Langford, John Sillitoe on behalf of the Wellcome Sanger Institute COVID-19 Surveillance Team |
| EPI_ISL_2830798 | Lighthouse Lab in Milton Keynes | Wellcome Sanger Institute for the COVID-19 Genomics UK (COG-UK) Consortium | The Lighthouse Lab in Milton Keynes and Alex Alderton, Roberto Amato, Jeffrey Barrett, Sonia Goncalves, Ewan Harrison, David K. Jackson, Ian Johnston, Dominic Kwiatkowski, Cordelia Langford, John Sillitoe on behalf of the Wellcome Sanger Institute COVID-19 Surveillance Team |
| EPI_ISL_2830799 | Lighthouse Lab in Alderley Park | Wellcome Sanger Institute for the COVID-19 Genomics UK (COG-UK) Consortium | Jacquelyn Wynn, Mairead Hyland, The Lighthouse Lab in Alderley Park and Alex Alderton, Roberto Amato, Jeffrey Barrett, Sonia Goncalves, Ewan Harrison, David K. Jackson, Ian Johnston, Dominic Kwiatkowski, Cordelia Langford, John Sillitoe on behalf of the Wellcome Sanger Institute COVID-19 Surveillance Team |
| EPI_ISL_2830800 | Lighthouse Laboratory Plymouth | Wellcome Sanger Institute for the COVID-19 Genomics UK (COG-UK) Consortium | Lighthouse Laboratory Plymouth and Alex Alderton, Roberto Amato, Jeffrey Barrett, Sonia Goncalves, Ewan Harrison, David K. Jackson, Ian Johnston, Dominic Kwiatkowski, Cordelia Langford, John Sillitoe on behalf of the Wellcome Sanger Institute COVID-19 Surveillance Team |
| EPI_ISL_2830801 | Lighthouse Lab in Alderley Park | Wellcome Sanger Institute for the COVID-19 Genomics UK (COG-UK) Consortium | Jacquelyn Wynn, Mairead Hyland, The Lighthouse Lab in Alderley Park and Alex Alderton, Roberto Amato, Jeffrey Barrett, Sonia Goncalves, Ewan Harrison, David K. Jackson, Ian Johnston, Dominic Kwiatkowski, Cordelia Langford, John Sillitoe on behalf of the Wellcome Sanger Institute COVID-19 Surveillance Team |
| EPI_ISL_2830802, EPI_ISL_2830803 | Lighthouse Lab in Milton Keynes | Wellcome Sanger Institute for the COVID-19 Genomics UK (COG-UK) Consortium | The Lighthouse Lab in Milton Keynes and Alex Alderton, Roberto Amato, Jeffrey Barrett, Sonia Goncalves, Ewan Harrison, David K. Jackson, Ian Johnston, Dominic Kwiatkowski, Cordelia Langford, John Sillitoe on behalf of the Wellcome Sanger Institute COVID-19 Surveillance Team |
| EPI_ISL_2830804 | Lighthouse Lab in Glasgow | Wellcome Sanger Institute for the COVID-19 Genomics UK (COG-UK) Consortium | Harper VanSteenhouse, Yumi Kasai, David Gray, Carol Clugston, Anna Dominiczak and Alex Alderton, Roberto Amato, Jeffrey Barrett, Sonia Goncalves, Ewan Harrison, David K. Jackson, Ian Johnston, Dominic Kwiatkowski, Cordelia Langford, John Sillitoe on behalf of the Wellcome Sanger Institute COVID-19 Surveillance Team |
| EPI_ISL_2830805, EPI_ISL_2830806 | Lighthouse Lab in Milton Keynes | Wellcome Sanger Institute for the COVID-19 Genomics UK (COG-UK) Consortium | The Lighthouse Lab in Milton Keynes and Alex Alderton, Roberto Amato, Jeffrey Barrett, Sonia Goncalves, Ewan Harrison, David K. Jackson, Ian Johnston, Dominic Kwiatkowski, Cordelia Langford, John Sillitoe on behalf of the Wellcome Sanger Institute COVID-19 Surveillance Team |
| EPI_ISL_2830807 | Lighthouse Lab in Glasgow | Wellcome Sanger Institute for the COVID-19 Genomics UK (COG-UK) Consortium | Harper VanSteenhouse, Yumi Kasai, David Gray, Carol Clugston, Anna Dominiczak and Alex Alderton, Roberto Amato, Jeffrey Barrett, Sonia Goncalves, Ewan Harrison, David K. Jackson, Ian Johnston, Dominic Kwiatkowski, Cordelia Langford, John Sillitoe on behalf of the Wellcome Sanger Institute COVID-19 Surveillance Team |
| EPI_ISL_2830808 | Lighthouse Lab in Milton Keynes | Wellcome Sanger Institute for the COVID-19 Genomics UK (COG-UK) Consortium | The Lighthouse Lab in Milton Keynes and Alex Alderton, Roberto Amato, Jeffrey Barrett, Sonia Goncalves, Ewan Harrison, David K. Jackson, Ian Johnston, Dominic Kwiatkowski, Cordelia Langford, John Sillitoe on behalf of the Wellcome Sanger Institute COVID-19 Surveillance Team |
| EPI_ISL_2830809 | Lighthouse Lab in Glasgow | Wellcome Sanger Institute for the COVID-19 Genomics UK (COG-UK) Consortium | Harper VanSteenhouse, Yumi Kasai, David Gray, Carol Clugston, Anna Dominiczak and Alex Alderton, Roberto Amato, Jeffrey Barrett, Sonia Goncalves, Ewan Harrison, David K. Jackson, Ian Johnston, Dominic Kwiatkowski, Cordelia Langford, John Sillitoe on behalf of the Wellcome Sanger Institute COVID-19 Surveillance Team |
| EPI_ISL_2830810 | Lighthouse Lab in Milton Keynes | Wellcome Sanger Institute for the COVID-19 Genomics UK (COG-UK) Consortium | The Lighthouse Lab in Milton Keynes and Alex Alderton, Roberto Amato, Jeffrey Barrett, Sonia Goncalves, Ewan Harrison, David K. Jackson, Ian Johnston, Dominic Kwiatkowski, Cordelia Langford, John Sillitoe on behalf of the Wellcome Sanger Institute COVID-19 Surveillance Team |
| EPI_ISL_2830811 | Lighthouse Lab in Alderley Park | Wellcome Sanger Institute for the COVID-19 Genomics UK (COG-UK) Consortium | Jacquelyn Wynn, Mairead Hyland, The Lighthouse Lab in Alderley Park and Alex Alderton, Roberto Amato, Jeffrey Barrett, Sonia Goncalves, Ewan Harrison, David K. Jackson, Ian Johnston, Dominic Kwiatkowski, Cordelia Langford, John Sillitoe on behalf of the Wellcome Sanger Institute COVID-19 Surveillance Team |
| EPI_ISL_2830812 | Lighthouse Laboratory Plymouth | Wellcome Sanger Institute for the COVID-19 Genomics UK (COG-UK) Consortium | Lighthouse Laboratory Plymouth and Alex Alderton, Roberto Amato, Jeffrey Barrett, Sonia Goncalves, Ewan Harrison, David K. Jackson, Ian Johnston, Dominic Kwiatkowski, Cordelia Langford, John Sillitoe on behalf of the Wellcome Sanger Institute COVID-19 Surveillance Team |
| EPI_ISL_2830813, EPI_ISL_2830814 | Lighthouse Lab in Alderley Park | Wellcome Sanger Institute for the COVID-19 Genomics UK (COG-UK) Consortium | Jacquelyn Wynn, Mairead Hyland, The Lighthouse Lab in Alderley Park and Alex Alderton, Roberto Amato, Jeffrey Barrett, Sonia Goncalves, Ewan Harrison, David K. Jackson, Ian Johnston, Dominic Kwiatkowski, Cordelia Langford, John Sillitoe on behalf of the Wellcome Sanger Institute COVID-19 Surveillance Team |
| EPI_ISL_2830815 | Lighthouse Lab in Glasgow | Wellcome Sanger Institute for the COVID-19 Genomics UK (COG-UK) Consortium | Harper VanSteenhouse, Yumi Kasai, David Gray, Carol Clugston, Anna Dominiczak and Alex Alderton, Roberto Amato, Jeffrey Barrett, Sonia Goncalves, Ewan Harrison, David K. Jackson, Ian Johnston, Dominic Kwiatkowski, Cordelia Langford, John Sillitoe on behalf of the Wellcome Sanger Institute COVID-19 Surveillance Team |
| EPI_ISL_2830816 | Lighthouse Laboratory Plymouth | Wellcome Sanger Institute for the COVID-19 Genomics UK (COG-UK) Consortium | Lighthouse Laboratory Plymouth and Alex Alderton, Roberto Amato, Jeffrey Barrett, Sonia Goncalves, Ewan Harrison, David K. Jackson, Ian Johnston, Dominic Kwiatkowski, Cordelia Langford, John Sillitoe on behalf of the Wellcome Sanger Institute COVID-19 Surveillance Team |
| EPI_ISL_2830817 | Lighthouse Lab in Milton Keynes | Wellcome Sanger Institute for the COVID-19 Genomics UK (COG-UK) Consortium | The Lighthouse Lab in Milton Keynes and Alex Alderton, Roberto Amato, Jeffrey Barrett, Sonia Goncalves, Ewan Harrison, David K. Jackson, Ian Johnston, Dominic Kwiatkowski, Cordelia Langford, John Sillitoe on behalf of the Wellcome Sanger Institute COVID-19 Surveillance Team |

| | | | |
|---|---|---|---|
| EPI_ISL_2830818 | Lighthouse Lab in Alderley Park | Wellcome Sanger Institute for the COVID-19 Genomics UK (COG-UK) Consortium | Jacquelyn Wynn, Mairead Hyland, The Lighthouse Lab in Alderley Park and Alex Alderton, Roberto Amato, Jeffrey Barrett, Sonia Goncalves, Ewan Harrison, David K. Jackson, Ian Johnston, Dominic Kwiatkowski, Cordelia Langford, John Sillitoe on behalf of the Wellcome Sanger Institute COVID-19 Surveillance Team |
| EPI_ISL_2830819, EPI_ISL_2830820, EPI_ISL_2830821 | Lighthouse Lab in Glasgow | Wellcome Sanger Institute for the COVID-19 Genomics UK (COG-UK) Consortium | Harper VanSteenhouse, Yumi Kasai, David Gray, Carol Clugston, Anna Dominiczak and Alex Alderton, Roberto Amato, Jeffrey Barrett, Sonia Goncalves, Ewan Harrison, David K. Jackson, Ian Johnston, Dominic Kwiatkowski, Cordelia Langford, John Sillitoe on behalf of the Wellcome Sanger Institute COVID-19 Surveillance Team |
| EPI_ISL_2830822, EPI_ISL_2830823 | Lighthouse Lab in Alderley Park | Wellcome Sanger Institute for the COVID-19 Genomics UK (COG-UK) Consortium | Jacquelyn Wynn, Mairead Hyland, The Lighthouse Lab in Alderley Park and Alex Alderton, Roberto Amato, Jeffrey Barrett, Sonia Goncalves, Ewan Harrison, David K. Jackson, Ian Johnston, Dominic Kwiatkowski, Cordelia Langford, John Sillitoe on behalf of the Wellcome Sanger Institute COVID-19 Surveillance Team |
| EPI_ISL_2830824, EPI_ISL_2830825 | Lighthouse Lab in Glasgow | Wellcome Sanger Institute for the COVID-19 Genomics UK (COG-UK) Consortium | Harper VanSteenhouse, Yumi Kasai, David Gray, Carol Clugston, Anna Dominiczak and Alex Alderton, Roberto Amato, Jeffrey Barrett, Sonia Goncalves, Ewan Harrison, David K. Jackson, Ian Johnston, Dominic Kwiatkowski, Cordelia Langford, John Sillitoe on behalf of the Wellcome Sanger Institute COVID-19 Surveillance Team |
| EPI_ISL_2830826 | Lighthouse Lab in Alderley Park | Wellcome Sanger Institute for the COVID-19 Genomics UK (COG-UK) Consortium | Jacquelyn Wynn, Mairead Hyland, The Lighthouse Lab in Alderley Park and Alex Alderton, Roberto Amato, Jeffrey Barrett, Sonia Goncalves, Ewan Harrison, David K. Jackson, Ian Johnston, Dominic Kwiatkowski, Cordelia Langford, John Sillitoe on behalf of the Wellcome Sanger Institute COVID-19 Surveillance Team |
| EPI_ISL_2830828 | Lighthouse Lab in Glasgow | Wellcome Sanger Institute for the COVID-19 Genomics UK (COG-UK) Consortium | Harper VanSteenhouse, Yumi Kasai, David Gray, Carol Clugston, Anna Dominiczak and Alex Alderton, Roberto Amato, Jeffrey Barrett, Sonia Goncalves, Ewan Harrison, David K. Jackson, Ian Johnston, Dominic Kwiatkowski, Cordelia Langford, John Sillitoe on behalf of the Wellcome Sanger Institute COVID-19 Surveillance Team |
| EPI_ISL_2830829, EPI_ISL_2830830 | Lighthouse Lab in Alderley Park | Wellcome Sanger Institute for the COVID-19 Genomics UK (COG-UK) Consortium | Jacquelyn Wynn, Mairead Hyland, The Lighthouse Lab in Alderley Park and Alex Alderton, Roberto Amato, Jeffrey Barrett, Sonia Goncalves, Ewan Harrison, David K. Jackson, Ian Johnston, Dominic Kwiatkowski, Cordelia Langford, John Sillitoe on behalf of the Wellcome Sanger Institute COVID-19 Surveillance Team |
| EPI_ISL_2830831, EPI_ISL_2830832 | Lighthouse Lab in Milton Keynes | Wellcome Sanger Institute for the COVID-19 Genomics UK (COG-UK) Consortium | The Lighthouse Lab in Milton Keynes and Alex Alderton, Roberto Amato, Jeffrey Barrett, Sonia Goncalves, Ewan Harrison, David K. Jackson, Ian Johnston, Dominic Kwiatkowski, Cordelia Langford, John Sillitoe on behalf of the Wellcome Sanger Institute COVID-19 Surveillance Team |
| EPI_ISL_2830833, EPI_ISL_2830834, EPI_ISL_2830835 | Lighthouse Laboratory Plymouth | Wellcome Sanger Institute for the COVID-19 Genomics UK (COG-UK) Consortium | Lighthouse Laboratory Plymouth and Alex Alderton, Roberto Amato, Jeffrey Barrett, Sonia Goncalves, Ewan Harrison, David K. Jackson, Ian Johnston, Dominic Kwiatkowski, Cordelia Langford, John Sillitoe on behalf of the Wellcome Sanger Institute COVID-19 Surveillance Team |
| EPI_ISL_2830836 | Lighthouse Lab in Glasgow | Wellcome Sanger Institute for the COVID-19 Genomics UK (COG-UK) Consortium | Harper VanSteenhouse, Yumi Kasai, David Gray, Carol Clugston, Anna Dominiczak and Alex Alderton, Roberto Amato, Jeffrey Barrett, Sonia Goncalves, Ewan Harrison, David K. Jackson, Ian Johnston, Dominic Kwiatkowski, Cordelia Langford, John Sillitoe on behalf of the Wellcome Sanger Institute COVID-19 Surveillance Team |
| EPI_ISL_2830837 | Lighthouse Lab in Alderley Park | Wellcome Sanger Institute for the COVID-19 Genomics UK (COG-UK) Consortium | Jacquelyn Wynn, Mairead Hyland, The Lighthouse Lab in Alderley Park and Alex Alderton, Roberto Amato, Jeffrey Barrett, Sonia Goncalves, Ewan Harrison, David K. Jackson, Ian Johnston, Dominic Kwiatkowski, Cordelia Langford, John Sillitoe on behalf of the Wellcome Sanger Institute COVID-19 Surveillance Team |
| EPI_ISL_2830838 | Lighthouse Lab in Milton Keynes | Wellcome Sanger Institute for the COVID-19 Genomics UK (COG-UK) Consortium | The Lighthouse Lab in Milton Keynes and Alex Alderton, Roberto Amato, Jeffrey Barrett, Sonia Goncalves, Ewan Harrison, David K. Jackson, Ian Johnston, Dominic Kwiatkowski, Cordelia Langford, John Sillitoe on behalf of the Wellcome Sanger Institute COVID-19 Surveillance Team |
| EPI_ISL_2830839 | Lighthouse Laboratory Plymouth | Wellcome Sanger Institute for the COVID-19 Genomics UK (COG-UK) Consortium | Lighthouse Laboratory Plymouth and Alex Alderton, Roberto Amato, Jeffrey Barrett, Sonia Goncalves, Ewan Harrison, David K. Jackson, Ian Johnston, Dominic Kwiatkowski, Cordelia Langford, John Sillitoe on behalf of the Wellcome Sanger Institute COVID-19 Surveillance Team |
| EPI_ISL_2830840 | Lighthouse Lab in Milton Keynes | Wellcome Sanger Institute for the COVID-19 Genomics UK (COG-UK) Consortium | The Lighthouse Lab in Milton Keynes and Alex Alderton, Roberto Amato, Jeffrey Barrett, Sonia Goncalves, Ewan Harrison, David K. Jackson, Ian Johnston, Dominic Kwiatkowski, Cordelia Langford, John Sillitoe on behalf of the Wellcome Sanger Institute COVID-19 Surveillance Team |
| EPI_ISL_2830841 | Lighthouse Laboratory Plymouth | Wellcome Sanger Institute for the COVID-19 Genomics UK (COG-UK) Consortium | Lighthouse Laboratory Plymouth and Alex Alderton, Roberto Amato, Jeffrey Barrett, Sonia Goncalves, Ewan Harrison, David K. Jackson, Ian Johnston, Dominic Kwiatkowski, Cordelia Langford, John Sillitoe on behalf of the Wellcome Sanger Institute COVID-19 Surveillance Team |
| EPI_ISL_2830842 | Lighthouse Lab in Alderley Park | Wellcome Sanger Institute for the COVID-19 Genomics UK (COG-UK) Consortium | Jacquelyn Wynn, Mairead Hyland, The Lighthouse Lab in Alderley Park and Alex Alderton, Roberto Amato, Jeffrey Barrett, Sonia Goncalves, Ewan Harrison, David K. Jackson, Ian Johnston, Dominic Kwiatkowski, Cordelia Langford, John Sillitoe on behalf of the Wellcome Sanger Institute COVID-19 Surveillance Team |
| EPI_ISL_2830843 | Lighthouse Lab in Milton Keynes | Wellcome Sanger Institute for the COVID-19 Genomics UK (COG-UK) Consortium | The Lighthouse Lab in Milton Keynes and Alex Alderton, Roberto Amato, Jeffrey Barrett, Sonia Goncalves, Ewan Harrison, David K. Jackson, Ian Johnston, Dominic Kwiatkowski, Cordelia Langford, John Sillitoe on behalf of the Wellcome Sanger Institute COVID-19 Surveillance Team |
| EPI_ISL_2830844 | Lighthouse Laboratory Plymouth | Wellcome Sanger Institute for the COVID-19 Genomics UK (COG-UK) Consortium | Lighthouse Laboratory Plymouth and Alex Alderton, Roberto Amato, Jeffrey Barrett, Sonia Goncalves, Ewan Harrison, David K. Jackson, Ian Johnston, Dominic Kwiatkowski, Cordelia Langford, John Sillitoe on behalf of the Wellcome Sanger Institute COVID-19 Surveillance Team |
| EPI_ISL_2830845, EPI_ISL_2830846, EPI_ISL_2830847 | Lighthouse Lab in Glasgow | Wellcome Sanger Institute for the COVID-19 Genomics UK (COG-UK) Consortium | Harper VanSteenhouse, Yumi Kasai, David Gray, Carol Clugston, Anna Dominiczak and Alex Alderton, Roberto Amato, Jeffrey Barrett, Sonia Goncalves, Ewan Harrison, David K. Jackson, Ian Johnston, Dominic Kwiatkowski, Cordelia Langford, John Sillitoe on behalf of the Wellcome Sanger Institute COVID-19 Surveillance Team |
| EPI_ISL_2830848 | Lighthouse Laboratory Plymouth | Wellcome Sanger Institute for the COVID-19 Genomics UK (COG-UK) Consortium | Lighthouse Laboratory Plymouth and Alex Alderton, Roberto Amato, Jeffrey Barrett, Sonia Goncalves, Ewan Harrison, David K. Jackson, Ian Johnston, Dominic Kwiatkowski, Cordelia Langford, John Sillitoe on behalf of the Wellcome Sanger Institute COVID-19 Surveillance Team |
| EPI_ISL_2830849 | Lighthouse Lab in Milton Keynes | Wellcome Sanger Institute for the COVID-19 Genomics UK (COG-UK) Consortium | The Lighthouse Lab in Milton Keynes and Alex Alderton, Roberto Amato, Jeffrey Barrett, Sonia Goncalves, Ewan Harrison, David K. Jackson, Ian Johnston, Dominic Kwiatkowski, Cordelia Langford, John Sillitoe on behalf of the Wellcome Sanger Institute COVID-19 Surveillance Team |
| EPI_ISL_2830851 | Lighthouse Lab in Glasgow | Wellcome Sanger Institute for the COVID-19 Genomics UK (COG-UK) Consortium | Harper VanSteenhouse, Yumi Kasai, David Gray, Carol Clugston, Anna Dominiczak and Alex Alderton, Roberto Amato, Jeffrey Barrett, Sonia Goncalves, Ewan Harrison, David K. Jackson, Ian Johnston, Dominic Kwiatkowski, Cordelia Langford, John Sillitoe on behalf of the Wellcome Sanger Institute COVID-19 Surveillance Team |
| EPI_ISL_2830852 | Lighthouse Laboratory Plymouth | Wellcome Sanger Institute for the COVID-19 Genomics UK (COG-UK) Consortium | Lighthouse Laboratory Plymouth and Alex Alderton, Roberto Amato, Jeffrey Barrett, Sonia Goncalves, Ewan Harrison, David K. Jackson, Ian Johnston, Dominic Kwiatkowski, Cordelia Langford, John Sillitoe on behalf of the Wellcome Sanger Institute COVID-19 Surveillance Team |
| EPI_ISL_2830853, EPI_ISL_2830854, EPI_ISL_2830855, EPI_ISL_2830856 | Lighthouse Lab in Alderley Park | Wellcome Sanger Institute for the COVID-19 Genomics UK (COG-UK) Consortium | Jacquelyn Wynn, Mairead Hyland, The Lighthouse Lab in Alderley Park and Alex Alderton, Roberto Amato, Jeffrey Barrett, Sonia Goncalves, Ewan Harrison, David K. Jackson, Ian Johnston, Dominic Kwiatkowski, Cordelia Langford, John Sillitoe on behalf of the Wellcome Sanger Institute COVID-19 Surveillance Team |
| EPI_ISL_2830857, EPI_ISL_2830858, EPI_ISL_2830859 | Lighthouse Lab in Glasgow | Wellcome Sanger Institute for the COVID-19 Genomics UK (COG-UK) Consortium | Harper VanSteenhouse, Yumi Kasai, David Gray, Carol Clugston, Anna Dominiczak and Alex Alderton, Roberto Amato, Jeffrey Barrett, Sonia Goncalves, Ewan Harrison, David K. Jackson, Ian Johnston, Dominic Kwiatkowski, Cordelia Langford, John Sillitoe on behalf of the Wellcome Sanger Institute COVID-19 Surveillance Team |
| EPI_ISL_2830860 | Lighthouse Lab in Alderley Park | Wellcome Sanger Institute for the COVID-19 Genomics UK (COG-UK) Consortium | Jacquelyn Wynn, Mairead Hyland, The Lighthouse Lab in Alderley Park and Alex Alderton, Roberto Amato, Jeffrey Barrett, Sonia Goncalves, Ewan Harrison, David K. Jackson, Ian Johnston, Dominic Kwiatkowski, Cordelia Langford, John Sillitoe on behalf of the Wellcome Sanger Institute COVID-19 Surveillance Team |
| EPI_ISL_2830861, EPI_ISL_2830862, EPI_ISL_2830863 | Lighthouse Lab in Glasgow | Wellcome Sanger Institute for the COVID-19 Genomics UK (COG-UK) Consortium | Harper VanSteenhouse, Yumi Kasai, David Gray, Carol Clugston, Anna Dominiczak and Alex Alderton, Roberto Amato, Jeffrey Barrett, Sonia Goncalves, Ewan Harrison, David K. Jackson, Ian Johnston, Dominic Kwiatkowski, Cordelia Langford, John Sillitoe on behalf of the Wellcome Sanger Institute COVID-19 Surveillance Team |
| EPI_ISL_2830864 | Lighthouse Lab in Alderley Park | Wellcome Sanger Institute for the COVID-19 Genomics UK (COG-UK) Consortium | Jacquelyn Wynn, Mairead Hyland, The Lighthouse Lab in Alderley Park and Alex Alderton, Roberto Amato, Jeffrey Barrett, Sonia Goncalves, Ewan Harrison, David K. Jackson, Ian Johnston, Dominic Kwiatkowski, Cordelia Langford, John Sillitoe on behalf of the Wellcome Sanger Institute COVID-19 Surveillance Team |
| EPI_ISL_2830865 | Lighthouse Lab in Glasgow | Wellcome Sanger Institute for the COVID-19 Genomics UK (COG-UK) Consortium | Harper VanSteenhouse, Yumi Kasai, David Gray, Carol Clugston, Anna Dominiczak and Alex Alderton, Roberto Amato, Jeffrey Barrett, Sonia Goncalves, Ewan Harrison, David K. Jackson, Ian Johnston, Dominic Kwiatkowski, Cordelia Langford, John Sillitoe on behalf of the Wellcome Sanger Institute COVID-19 Surveillance Team |
| EPI_ISL_2830866, EPI_ISL_2830867 | Lighthouse Lab in Alderley Park | Wellcome Sanger Institute for the COVID-19 Genomics UK (COG-UK) Consortium | Jacquelyn Wynn, Mairead Hyland, The Lighthouse Lab in Alderley Park and Alex Alderton, Roberto Amato, Jeffrey Barrett, Sonia Goncalves, Ewan Harrison, David K. Jackson, Ian Johnston, Dominic Kwiatkowski, Cordelia Langford, John Sillitoe on behalf of the Wellcome Sanger Institute COVID-19 Surveillance Team |
| EPI_ISL_2830868 | Lighthouse Lab in Glasgow | Wellcome Sanger Institute for the COVID-19 Genomics UK (COG-UK) Consortium | Harper VanSteenhouse, Yumi Kasai, David Gray, Carol Clugston, Anna Dominiczak and Alex Alderton, Roberto Amato, Jeffrey Barrett, Sonia Goncalves, Ewan Harrison, David K. Jackson, Ian Johnston, Dominic Kwiatkowski, Cordelia Langford, John Sillitoe on behalf of the Wellcome Sanger Institute COVID-19 Surveillance Team |
| EPI_ISL_2830869, EPI_ISL_2830870, EPI_ISL_2830871, EPI_ISL_2830872 | Lighthouse Lab in Alderley Park | Wellcome Sanger Institute for the COVID-19 Genomics UK (COG-UK) Consortium | Jacquelyn Wynn, Mairead Hyland, The Lighthouse Lab in Alderley Park and Alex Alderton, Roberto Amato, Jeffrey Barrett, Sonia Goncalves, Ewan Harrison, David K. Jackson, Ian Johnston, Dominic Kwiatkowski, Cordelia Langford, John Sillitoe on behalf of the Wellcome Sanger Institute COVID-19 Surveillance Team |
| EPI_ISL_2830873 | Lighthouse Lab in Glasgow | Wellcome Sanger Institute for the COVID-19 Genomics UK (COG-UK) Consortium | Harper VanSteenhouse, Yumi Kasai, David Gray, Carol Clugston, Anna Dominiczak and Alex Alderton, Roberto Amato, Jeffrey Barrett, Sonia Goncalves, Ewan Harrison, David K. Jackson, Ian Johnston, Dominic Kwiatkowski, Cordelia Langford, John Sillitoe on behalf of the Wellcome Sanger Institute COVID-19 Surveillance Team |
| EPI_ISL_2830874 | Lighthouse Lab in Alderley Park | Wellcome Sanger Institute for the COVID-19 Genomics UK (COG-UK) Consortium | Jacquelyn Wynn, Mairead Hyland, The Lighthouse Lab in Alderley Park and Alex Alderton, Roberto Amato, Jeffrey Barrett, Sonia Goncalves, Ewan Harrison, David K. Jackson, Ian Johnston, Dominic Kwiatkowski, Cordelia Langford, John Sillitoe on behalf of the Wellcome Sanger Institute COVID-19 Surveillance Team |
| EPI_ISL_2830875 | Lighthouse Lab in Glasgow | Wellcome Sanger Institute for the COVID-19 Genomics UK (COG-UK) Consortium | Harper VanSteenhouse, Yumi Kasai, David Gray, Carol Clugston, Anna Dominiczak and Alex Alderton, Roberto Amato, Jeffrey Barrett, Sonia Goncalves, Ewan Harrison, David K. Jackson, Ian Johnston, Dominic Kwiatkowski, Cordelia Langford, John Sillitoe on behalf of the Wellcome Sanger Institute COVID-19 Surveillance Team |
| EPI_ISL_2830876, EPI_ISL_2830877 | Lighthouse Lab in Alderley Park | Wellcome Sanger Institute for the COVID-19 Genomics UK (COG-UK) Consortium | Jacquelyn Wynn, Mairead Hyland, The Lighthouse Lab in Alderley Park and Alex Alderton, Roberto Amato, Jeffrey Barrett, Sonia Goncalves, Ewan Harrison, David K. Jackson, Ian Johnston, Dominic Kwiatkowski, Cordelia Langford, John Sillitoe on behalf of the Wellcome Sanger Institute COVID-19 Surveillance Team |
| EPI_ISL_2830878, EPI_ISL_2830879 | Lighthouse Lab in Glasgow | Wellcome Sanger Institute for the COVID-19 Genomics UK (COG-UK) Consortium | Harper VanSteenhouse, Yumi Kasai, David Gray, Carol Clugston, Anna Dominiczak and Alex Alderton, Roberto Amato, Jeffrey Barrett, Sonia Goncalves, Ewan Harrison, David K. Jackson, Ian Johnston, Dominic Kwiatkowski, Cordelia Langford, John Sillitoe on behalf of the Wellcome Sanger Institute COVID-19 Surveillance Team |
| EPI_ISL_2830880 | Lighthouse Lab in Alderley Park | Wellcome Sanger Institute for the COVID-19 Genomics UK (COG-UK) Consortium | Jacquelyn Wynn, Mairead Hyland, The Lighthouse Lab in Alderley Park and Alex Alderton, Roberto Amato, Jeffrey Barrett, Sonia Goncalves, Ewan Harrison, David K. Jackson, Ian Johnston, Dominic Kwiatkowski, Cordelia Langford, John Sillitoe on behalf of the Wellcome Sanger Institute COVID-19 Surveillance Team |
| EPI_ISL_2830881, EPI_ISL_2830882 | Lighthouse Lab in Glasgow | Wellcome Sanger Institute for the COVID-19 Genomics UK (COG-UK) Consortium | Harper VanSteenhouse, Yumi Kasai, David Gray, Carol Clugston, Anna Dominiczak and Alex Alderton, Roberto Amato, Jeffrey Barrett, Sonia Goncalves, Ewan Harrison, David K. Jackson, Ian Johnston, Dominic Kwiatkowski, Cordelia Langford, John Sillitoe on behalf of the Wellcome Sanger Institute COVID-19 Surveillance Team |
| EPI_ISL_2830883, EPI_ISL_2830884, EPI_ISL_2830885, EPI_ISL_2830886, EPI_ISL_2830887 | Lighthouse Lab in Alderley Park | Wellcome Sanger Institute for the COVID-19 Genomics UK (COG-UK) Consortium | Jacquelyn Wynn, Mairead Hyland, The Lighthouse Lab in Alderley Park and Alex Alderton, Roberto Amato, Jeffrey Barrett, Sonia Goncalves, Ewan Harrison, David K. Jackson, Ian Johnston, Dominic Kwiatkowski, Cordelia Langford, John Sillitoe on behalf of the Wellcome Sanger Institute COVID-19 Surveillance Team |
| EPI_ISL_2830888, | Lighthouse Lab in Glasgow | Wellcome Sanger Institute for the COVID-19 Genomics UK (COG-UK) | Harper VanSteenhouse, Yumi Kasai, David Gray, Carol Clugston, Anna Dominiczak and Alex Alderton, Roberto Amato, Jeffrey Barrett, Sonia Goncalves, Ewan Harrison, David K. Jackson, Ian Johnston, Dominic Kwiatkowski, Cordelia Langford, |

| | | | |
|---|---|---|---|
| EPI_ISL_2830889 | | Consortium | John Sillitoe on behalf of the Wellcome Sanger Institute COVID-19 Surveillance Team |
| EPI_ISL_2830890, EPI_ISL_2830891 | Lighthouse Lab in Alderley Park | Wellcome Sanger Institute for the COVID-19 Genomics UK (COG-UK) Consortium | Jacquelyn Wynn, Mairead Hyland, The Lighthouse Lab in Alderley Park and Alex Alderton, Roberto Amato, Jeffrey Barrett, Sonia Goncalves, Ewan Harrison, David K. Jackson, Ian Johnston, Dominic Kwiatkowski, Cordelia Langford, John Sillitoe on behalf of the Wellcome Sanger Institute COVID-19 Surveillance Team |
| EPI_ISL_2830892, EPI_ISL_2830893, EPI_ISL_2830894, EPI_ISL_2830895, EPI_ISL_2830896 | Lighthouse Lab in Glasgow | Wellcome Sanger Institute for the COVID-19 Genomics UK (COG-UK) Consortium | Harper VanSteenhouse, Yumi Kasai, David Gray, Carol Clugston, Anna Dominiczak and Alex Alderton, Roberto Amato, Jeffrey Barrett, Sonia Goncalves, Ewan Harrison, David K. Jackson, Ian Johnston, Dominic Kwiatkowski, Cordelia Langford, John Sillitoe on behalf of the Wellcome Sanger Institute COVID-19 Surveillance Team |
| EPI_ISL_2830897, EPI_ISL_2830898 | Lighthouse Lab in Alderley Park | Wellcome Sanger Institute for the COVID-19 Genomics UK (COG-UK) Consortium | Jacquelyn Wynn, Mairead Hyland, The Lighthouse Lab in Alderley Park and Alex Alderton, Roberto Amato, Jeffrey Barrett, Sonia Goncalves, Ewan Harrison, David K. Jackson, Ian Johnston, Dominic Kwiatkowski, Cordelia Langford, John Sillitoe on behalf of the Wellcome Sanger Institute COVID-19 Surveillance Team |
| EPI_ISL_2830899 | Lighthouse Lab in Glasgow | Wellcome Sanger Institute for the COVID-19 Genomics UK (COG-UK) Consortium | Harper VanSteenhouse, Yumi Kasai, David Gray, Carol Clugston, Anna Dominiczak and Alex Alderton, Roberto Amato, Jeffrey Barrett, Sonia Goncalves, Ewan Harrison, David K. Jackson, Ian Johnston, Dominic Kwiatkowski, Cordelia Langford, John Sillitoe on behalf of the Wellcome Sanger Institute COVID-19 Surveillance Team |
| EPI_ISL_2830900 | Lighthouse Lab in Alderley Park | Wellcome Sanger Institute for the COVID-19 Genomics UK (COG-UK) Consortium | Jacquelyn Wynn, Mairead Hyland, The Lighthouse Lab in Alderley Park and Alex Alderton, Roberto Amato, Jeffrey Barrett, Sonia Goncalves, Ewan Harrison, David K. Jackson, Ian Johnston, Dominic Kwiatkowski, Cordelia Langford, John Sillitoe on behalf of the Wellcome Sanger Institute COVID-19 Surveillance Team |
| EPI_ISL_2830901 | Lighthouse Lab in Glasgow | Wellcome Sanger Institute for the COVID-19 Genomics UK (COG-UK) Consortium | Harper VanSteenhouse, Yumi Kasai, David Gray, Carol Clugston, Anna Dominiczak and Alex Alderton, Roberto Amato, Jeffrey Barrett, Sonia Goncalves, Ewan Harrison, David K. Jackson, Ian Johnston, Dominic Kwiatkowski, Cordelia Langford, John Sillitoe on behalf of the Wellcome Sanger Institute COVID-19 Surveillance Team |
| EPI_ISL_2830902 | Lighthouse Lab in Alderley Park | Wellcome Sanger Institute for the COVID-19 Genomics UK (COG-UK) Consortium | Jacquelyn Wynn, Mairead Hyland, The Lighthouse Lab in Alderley Park and Alex Alderton, Roberto Amato, Jeffrey Barrett, Sonia Goncalves, Ewan Harrison, David K. Jackson, Ian Johnston, Dominic Kwiatkowski, Cordelia Langford, John Sillitoe on behalf of the Wellcome Sanger Institute COVID-19 Surveillance Team |
| EPI_ISL_2830903 | Lighthouse Lab in Glasgow | Wellcome Sanger Institute for the COVID-19 Genomics UK (COG-UK) Consortium | Harper VanSteenhouse, Yumi Kasai, David Gray, Carol Clugston, Anna Dominiczak and Alex Alderton, Roberto Amato, Jeffrey Barrett, Sonia Goncalves, Ewan Harrison, David K. Jackson, Ian Johnston, Dominic Kwiatkowski, Cordelia Langford, John Sillitoe on behalf of the Wellcome Sanger Institute COVID-19 Surveillance Team |
| EPI_ISL_2830904 | Lighthouse Lab in Alderley Park | Wellcome Sanger Institute for the COVID-19 Genomics UK (COG-UK) Consortium | Jacquelyn Wynn, Mairead Hyland, The Lighthouse Lab in Alderley Park and Alex Alderton, Roberto Amato, Jeffrey Barrett, Sonia Goncalves, Ewan Harrison, David K. Jackson, Ian Johnston, Dominic Kwiatkowski, Cordelia Langford, John Sillitoe on behalf of the Wellcome Sanger Institute COVID-19 Surveillance Team |
| EPI_ISL_2830905, EPI_ISL_2830906, EPI_ISL_2830907, EPI_ISL_2830908 | Lighthouse Lab in Glasgow | Wellcome Sanger Institute for the COVID-19 Genomics UK (COG-UK) Consortium | Harper VanSteenhouse, Yumi Kasai, David Gray, Carol Clugston, Anna Dominiczak and Alex Alderton, Roberto Amato, Jeffrey Barrett, Sonia Goncalves, Ewan Harrison, David K. Jackson, Ian Johnston, Dominic Kwiatkowski, Cordelia Langford, John Sillitoe on behalf of the Wellcome Sanger Institute COVID-19 Surveillance Team |
| EPI_ISL_2830909 | Lighthouse Lab in Alderley Park | Wellcome Sanger Institute for the COVID-19 Genomics UK (COG-UK) Consortium | Jacquelyn Wynn, Mairead Hyland, The Lighthouse Lab in Alderley Park and Alex Alderton, Roberto Amato, Jeffrey Barrett, Sonia Goncalves, Ewan Harrison, David K. Jackson, Ian Johnston, Dominic Kwiatkowski, Cordelia Langford, John Sillitoe on behalf of the Wellcome Sanger Institute COVID-19 Surveillance Team |
| EPI_ISL_2830910, EPI_ISL_2830911, EPI_ISL_2830912, EPI_ISL_2830913 | Lighthouse Lab in Glasgow | Wellcome Sanger Institute for the COVID-19 Genomics UK (COG-UK) Consortium | Harper VanSteenhouse, Yumi Kasai, David Gray, Carol Clugston, Anna Dominiczak and Alex Alderton, Roberto Amato, Jeffrey Barrett, Sonia Goncalves, Ewan Harrison, David K. Jackson, Ian Johnston, Dominic Kwiatkowski, Cordelia Langford, John Sillitoe on behalf of the Wellcome Sanger Institute COVID-19 Surveillance Team |
| EPI_ISL_2830914, EPI_ISL_2830915 | Lighthouse Lab in Alderley Park | Wellcome Sanger Institute for the COVID-19 Genomics UK (COG-UK) Consortium | Jacquelyn Wynn, Mairead Hyland, The Lighthouse Lab in Alderley Park and Alex Alderton, Roberto Amato, Jeffrey Barrett, Sonia Goncalves, Ewan Harrison, David K. Jackson, Ian Johnston, Dominic Kwiatkowski, Cordelia Langford, John Sillitoe on behalf of the Wellcome Sanger Institute COVID-19 Surveillance Team |
| EPI_ISL_2830916, EPI_ISL_2830917 | Lighthouse Lab in Glasgow | Wellcome Sanger Institute for the COVID-19 Genomics UK (COG-UK) Consortium | Harper VanSteenhouse, Yumi Kasai, David Gray, Carol Clugston, Anna Dominiczak and Alex Alderton, Roberto Amato, Jeffrey Barrett, Sonia Goncalves, Ewan Harrison, David K. Jackson, Ian Johnston, Dominic Kwiatkowski, Cordelia Langford, John Sillitoe on behalf of the Wellcome Sanger Institute COVID-19 Surveillance Team |
| EPI_ISL_2830918, EPI_ISL_2830919 | Lighthouse Lab in Alderley Park | Wellcome Sanger Institute for the COVID-19 Genomics UK (COG-UK) Consortium | Jacquelyn Wynn, Mairead Hyland, The Lighthouse Lab in Alderley Park and Alex Alderton, Roberto Amato, Jeffrey Barrett, Sonia Goncalves, Ewan Harrison, David K. Jackson, Ian Johnston, Dominic Kwiatkowski, Cordelia Langford, John Sillitoe on behalf of the Wellcome Sanger Institute COVID-19 Surveillance Team |
| EPI_ISL_2830920, EPI_ISL_2830921, EPI_ISL_2830922, EPI_ISL_2830923, EPI_ISL_2830924 | Lighthouse Lab in Glasgow | Wellcome Sanger Institute for the COVID-19 Genomics UK (COG-UK) Consortium | Harper VanSteenhouse, Yumi Kasai, David Gray, Carol Clugston, Anna Dominiczak and Alex Alderton, Roberto Amato, Jeffrey Barrett, Sonia Goncalves, Ewan Harrison, David K. Jackson, Ian Johnston, Dominic Kwiatkowski, Cordelia Langford, John Sillitoe on behalf of the Wellcome Sanger Institute COVID-19 Surveillance Team |
| EPI_ISL_2830925 | Lighthouse Lab in Alderley Park | Wellcome Sanger Institute for the COVID-19 Genomics UK (COG-UK) Consortium | Jacquelyn Wynn, Mairead Hyland, The Lighthouse Lab in Alderley Park and Alex Alderton, Roberto Amato, Jeffrey Barrett, Sonia Goncalves, Ewan Harrison, David K. Jackson, Ian Johnston, Dominic Kwiatkowski, Cordelia Langford, John Sillitoe on behalf of the Wellcome Sanger Institute COVID-19 Surveillance Team |
| EPI_ISL_2830926, EPI_ISL_2830927 | Lighthouse Lab in Glasgow | Wellcome Sanger Institute for the COVID-19 Genomics UK (COG-UK) Consortium | Harper VanSteenhouse, Yumi Kasai, David Gray, Carol Clugston, Anna Dominiczak and Alex Alderton, Roberto Amato, Jeffrey Barrett, Sonia Goncalves, Ewan Harrison, David K. Jackson, Ian Johnston, Dominic Kwiatkowski, Cordelia Langford, John Sillitoe on behalf of the Wellcome Sanger Institute COVID-19 Surveillance Team |
| EPI_ISL_2830928, EPI_ISL_2830929 | Lighthouse Lab in Alderley Park | Wellcome Sanger Institute for the COVID-19 Genomics UK (COG-UK) Consortium | Jacquelyn Wynn, Mairead Hyland, The Lighthouse Lab in Alderley Park and Alex Alderton, Roberto Amato, Jeffrey Barrett, Sonia Goncalves, Ewan Harrison, David K. Jackson, Ian Johnston, Dominic Kwiatkowski, Cordelia Langford, John Sillitoe on behalf of the Wellcome Sanger Institute COVID-19 Surveillance Team |
| EPI_ISL_2830930 | Lighthouse Lab in Glasgow | Wellcome Sanger Institute for the COVID-19 Genomics UK (COG-UK) Consortium | Harper VanSteenhouse, Yumi Kasai, David Gray, Carol Clugston, Anna Dominiczak and Alex Alderton, Roberto Amato, Jeffrey Barrett, Sonia Goncalves, Ewan Harrison, David K. Jackson, Ian Johnston, Dominic Kwiatkowski, Cordelia Langford, John Sillitoe on behalf of the Wellcome Sanger Institute COVID-19 Surveillance Team |
| EPI_ISL_2830931, EPI_ISL_2830932, EPI_ISL_2830933, EPI_ISL_2830934 | Lighthouse Lab in Alderley Park | Wellcome Sanger Institute for the COVID-19 Genomics UK (COG-UK) Consortium | Jacquelyn Wynn, Mairead Hyland, The Lighthouse Lab in Alderley Park and Alex Alderton, Roberto Amato, Jeffrey Barrett, Sonia Goncalves, Ewan Harrison, David K. Jackson, Ian Johnston, Dominic Kwiatkowski, Cordelia Langford, John Sillitoe on behalf of the Wellcome Sanger Institute COVID-19 Surveillance Team |
| EPI_ISL_2830937, EPI_ISL_2830938, EPI_ISL_2830939 | Lighthouse Lab in Glasgow | Wellcome Sanger Institute for the COVID-19 Genomics UK (COG-UK) Consortium | Harper VanSteenhouse, Yumi Kasai, David Gray, Carol Clugston, Anna Dominiczak and Alex Alderton, Roberto Amato, Jeffrey Barrett, Sonia Goncalves, Ewan Harrison, David K. Jackson, Ian Johnston, Dominic Kwiatkowski, Cordelia Langford, John Sillitoe on behalf of the Wellcome Sanger Institute COVID-19 Surveillance Team |
| EPI_ISL_2830940, EPI_ISL_2830941, EPI_ISL_2830942 | Lighthouse Lab in Alderley Park | Wellcome Sanger Institute for the COVID-19 Genomics UK (COG-UK) Consortium | Jacquelyn Wynn, Mairead Hyland, The Lighthouse Lab in Alderley Park and Alex Alderton, Roberto Amato, Jeffrey Barrett, Sonia Goncalves, Ewan Harrison, David K. Jackson, Ian Johnston, Dominic Kwiatkowski, Cordelia Langford, John Sillitoe on behalf of the Wellcome Sanger Institute COVID-19 Surveillance Team |
| EPI_ISL_2830943 | Lighthouse Lab in Glasgow | Wellcome Sanger Institute for the COVID-19 Genomics UK (COG-UK) Consortium | Harper VanSteenhouse, Yumi Kasai, David Gray, Carol Clugston, Anna Dominiczak and Alex Alderton, Roberto Amato, Jeffrey Barrett, Sonia Goncalves, Ewan Harrison, David K. Jackson, Ian Johnston, Dominic Kwiatkowski, Cordelia Langford, John Sillitoe on behalf of the Wellcome Sanger Institute COVID-19 Surveillance Team |
| EPI_ISL_2830944 | Lighthouse Lab in Alderley Park | Wellcome Sanger Institute for the COVID-19 Genomics UK (COG-UK) Consortium | Jacquelyn Wynn, Mairead Hyland, The Lighthouse Lab in Alderley Park and Alex Alderton, Roberto Amato, Jeffrey Barrett, Sonia Goncalves, Ewan Harrison, David K. Jackson, Ian Johnston, Dominic Kwiatkowski, Cordelia Langford, John Sillitoe on behalf of the Wellcome Sanger Institute COVID-19 Surveillance Team |
| EPI_ISL_2830945 | Lighthouse Lab in Glasgow | Wellcome Sanger Institute for the COVID-19 Genomics UK (COG-UK) Consortium | Harper VanSteenhouse, Yumi Kasai, David Gray, Carol Clugston, Anna Dominiczak and Alex Alderton, Roberto Amato, Jeffrey Barrett, Sonia Goncalves, Ewan Harrison, David K. Jackson, Ian Johnston, Dominic Kwiatkowski, Cordelia Langford, John Sillitoe on behalf of the Wellcome Sanger Institute COVID-19 Surveillance Team |
| EPI_ISL_2830946 | Lighthouse Lab in Alderley Park | Wellcome Sanger Institute for the COVID-19 Genomics UK (COG-UK) Consortium | Jacquelyn Wynn, Mairead Hyland, The Lighthouse Lab in Alderley Park and Alex Alderton, Roberto Amato, Jeffrey Barrett, Sonia Goncalves, Ewan Harrison, David K. Jackson, Ian Johnston, Dominic Kwiatkowski, Cordelia Langford, John Sillitoe on behalf of the Wellcome Sanger Institute COVID-19 Surveillance Team |
| EPI_ISL_2830947, EPI_ISL_2830948 | Lighthouse Lab in Glasgow | Wellcome Sanger Institute for the COVID-19 Genomics UK (COG-UK) Consortium | Harper VanSteenhouse, Yumi Kasai, David Gray, Carol Clugston, Anna Dominiczak and Alex Alderton, Roberto Amato, Jeffrey Barrett, Sonia Goncalves, Ewan Harrison, David K. Jackson, Ian Johnston, Dominic Kwiatkowski, Cordelia Langford, John Sillitoe on behalf of the Wellcome Sanger Institute COVID-19 Surveillance Team |
| EPI_ISL_2830949 | Lighthouse Lab in Alderley Park | Wellcome Sanger Institute for the COVID-19 Genomics UK (COG-UK) Consortium | Jacquelyn Wynn, Mairead Hyland, The Lighthouse Lab in Alderley Park and Alex Alderton, Roberto Amato, Jeffrey Barrett, Sonia Goncalves, Ewan Harrison, David K. Jackson, Ian Johnston, Dominic Kwiatkowski, Cordelia Langford, John Sillitoe on behalf of the Wellcome Sanger Institute COVID-19 Surveillance Team |
| EPI_ISL_2830950 | Lighthouse Lab in Glasgow | Wellcome Sanger Institute for the COVID-19 Genomics UK (COG-UK) Consortium | Harper VanSteenhouse, Yumi Kasai, David Gray, Carol Clugston, Anna Dominiczak and Alex Alderton, Roberto Amato, Jeffrey Barrett, Sonia Goncalves, Ewan Harrison, David K. Jackson, Ian Johnston, Dominic Kwiatkowski, Cordelia Langford, John Sillitoe on behalf of the Wellcome Sanger Institute COVID-19 Surveillance Team |
| EPI_ISL_2830951 | Lighthouse Lab in Alderley Park | Wellcome Sanger Institute for the COVID-19 Genomics UK (COG-UK) Consortium | Jacquelyn Wynn, Mairead Hyland, The Lighthouse Lab in Alderley Park and Alex Alderton, Roberto Amato, Jeffrey Barrett, Sonia Goncalves, Ewan Harrison, David K. Jackson, Ian Johnston, Dominic Kwiatkowski, Cordelia Langford, John Sillitoe on behalf of the Wellcome Sanger Institute COVID-19 Surveillance Team |
| EPI_ISL_2830952, EPI_ISL_2830953, EPI_ISL_2830954, EPI_ISL_2830955 | Lighthouse Lab in Glasgow | Wellcome Sanger Institute for the COVID-19 Genomics UK (COG-UK) Consortium | Harper VanSteenhouse, Yumi Kasai, David Gray, Carol Clugston, Anna Dominiczak and Alex Alderton, Roberto Amato, Jeffrey Barrett, Sonia Goncalves, Ewan Harrison, David K. Jackson, Ian Johnston, Dominic Kwiatkowski, Cordelia Langford, John Sillitoe on behalf of the Wellcome Sanger Institute COVID-19 Surveillance Team |
| EPI_ISL_2830956, EPI_ISL_2830957 | Lighthouse Lab in Alderley Park | Wellcome Sanger Institute for the COVID-19 Genomics UK (COG-UK) Consortium | Jacquelyn Wynn, Mairead Hyland, The Lighthouse Lab in Alderley Park and Alex Alderton, Roberto Amato, Jeffrey Barrett, Sonia Goncalves, Ewan Harrison, David K. Jackson, Ian Johnston, Dominic Kwiatkowski, Cordelia Langford, John Sillitoe on behalf of the Wellcome Sanger Institute COVID-19 Surveillance Team |
| EPI_ISL_2830958 | Lighthouse Lab in Glasgow | Wellcome Sanger Institute for the COVID-19 Genomics UK (COG-UK) Consortium | Harper VanSteenhouse, Yumi Kasai, David Gray, Carol Clugston, Anna Dominiczak and Alex Alderton, Roberto Amato, Jeffrey Barrett, Sonia Goncalves, Ewan Harrison, David K. Jackson, Ian Johnston, Dominic Kwiatkowski, Cordelia Langford, John Sillitoe on behalf of the Wellcome Sanger Institute COVID-19 Surveillance Team |
| EPI_ISL_2830959, EPI_ISL_2830960, EPI_ISL_2830961, EPI_ISL_2830962 | Lighthouse Lab in Alderley Park | Wellcome Sanger Institute for the COVID-19 Genomics UK (COG-UK) Consortium | Jacquelyn Wynn, Mairead Hyland, The Lighthouse Lab in Alderley Park and Alex Alderton, Roberto Amato, Jeffrey Barrett, Sonia Goncalves, Ewan Harrison, David K. Jackson, Ian Johnston, Dominic Kwiatkowski, Cordelia Langford, John Sillitoe on behalf of the Wellcome Sanger Institute COVID-19 Surveillance Team |
| EPI_ISL_2830963, EPI_ISL_2830964, EPI_ISL_2830965, EPI_ISL_2830966, EPI_ISL_2830967 | Lighthouse Lab in Glasgow | Wellcome Sanger Institute for the COVID-19 Genomics UK (COG-UK) Consortium | Harper VanSteenhouse, Yumi Kasai, David Gray, Carol Clugston, Anna Dominiczak and Alex Alderton, Roberto Amato, Jeffrey Barrett, Sonia Goncalves, Ewan Harrison, David K. Jackson, Ian Johnston, Dominic Kwiatkowski, Cordelia Langford, John Sillitoe on behalf of the Wellcome Sanger Institute COVID-19 Surveillance Team |

| | | | |
|---|---|---|---|
| EPI_ISL_2830968 | Lighthouse Lab in Alderley Park | Wellcome Sanger Institute for the COVID-19 Genomics UK (COG-UK) Consortium | Jacquelyn Wynn, Mairead Hyland, The Lighthouse Lab in Alderley Park and Alex Alderton, Roberto Amato, Jeffrey Barrett, Sonia Goncalves, Ewan Harrison, David K. Jackson, Ian Johnston, Dominic Kwiatkowski, Cordelia Langford, John Sillitoe on behalf of the Wellcome Sanger Institute COVID-19 Surveillance Team |
| EPI_ISL_2830969, EPI_ISL_2830970, EPI_ISL_2830971, EPI_ISL_2830972 | Lighthouse Lab in Glasgow | Wellcome Sanger Institute for the COVID-19 Genomics UK (COG-UK) Consortium | Harper VanSteenhouse, Yumi Kasai, David Gray, Carol Clugston, Anna Dominiczak and Alex Alderton, Roberto Amato, Jeffrey Barrett, Sonia Goncalves, Ewan Harrison, David K. Jackson, Ian Johnston, Dominic Kwiatkowski, Cordelia Langford, John Sillitoe on behalf of the Wellcome Sanger Institute COVID-19 Surveillance Team |
| EPI_ISL_2830973 | Lighthouse Lab in Alderley Park | Wellcome Sanger Institute for the COVID-19 Genomics UK (COG-UK) Consortium | Jacquelyn Wynn, Mairead Hyland, The Lighthouse Lab in Alderley Park and Alex Alderton, Roberto Amato, Jeffrey Barrett, Sonia Goncalves, Ewan Harrison, David K. Jackson, Ian Johnston, Dominic Kwiatkowski, Cordelia Langford, John Sillitoe on behalf of the Wellcome Sanger Institute COVID-19 Surveillance Team |
| EPI_ISL_2830974, EPI_ISL_2830975, EPI_ISL_2830976 | Lighthouse Lab in Glasgow | Wellcome Sanger Institute for the COVID-19 Genomics UK (COG-UK) Consortium | Harper VanSteenhouse, Yumi Kasai, David Gray, Carol Clugston, Anna Dominiczak and Alex Alderton, Roberto Amato, Jeffrey Barrett, Sonia Goncalves, Ewan Harrison, David K. Jackson, Ian Johnston, Dominic Kwiatkowski, Cordelia Langford, John Sillitoe on behalf of the Wellcome Sanger Institute COVID-19 Surveillance Team |
| EPI_ISL_2830977 | Lighthouse Lab in Alderley Park | Wellcome Sanger Institute for the COVID-19 Genomics UK (COG-UK) Consortium | Jacquelyn Wynn, Mairead Hyland, The Lighthouse Lab in Alderley Park and Alex Alderton, Roberto Amato, Jeffrey Barrett, Sonia Goncalves, Ewan Harrison, David K. Jackson, Ian Johnston, Dominic Kwiatkowski, Cordelia Langford, John Sillitoe on behalf of the Wellcome Sanger Institute COVID-19 Surveillance Team |
| EPI_ISL_2830978 | Lighthouse Lab in Glasgow | Wellcome Sanger Institute for the COVID-19 Genomics UK (COG-UK) Consortium | Harper VanSteenhouse, Yumi Kasai, David Gray, Carol Clugston, Anna Dominiczak and Alex Alderton, Roberto Amato, Jeffrey Barrett, Sonia Goncalves, Ewan Harrison, David K. Jackson, Ian Johnston, Dominic Kwiatkowski, Cordelia Langford, John Sillitoe on behalf of the Wellcome Sanger Institute COVID-19 Surveillance Team |
| EPI_ISL_2830979 | Lighthouse Lab in Alderley Park | Wellcome Sanger Institute for the COVID-19 Genomics UK (COG-UK) Consortium | Jacquelyn Wynn, Mairead Hyland, The Lighthouse Lab in Alderley Park and Alex Alderton, Roberto Amato, Jeffrey Barrett, Sonia Goncalves, Ewan Harrison, David K. Jackson, Ian Johnston, Dominic Kwiatkowski, Cordelia Langford, John Sillitoe on behalf of the Wellcome Sanger Institute COVID-19 Surveillance Team |
| EPI_ISL_2830980, EPI_ISL_2830981 | Lighthouse Lab in Glasgow | Wellcome Sanger Institute for the COVID-19 Genomics UK (COG-UK) Consortium | Harper VanSteenhouse, Yumi Kasai, David Gray, Carol Clugston, Anna Dominiczak and Alex Alderton, Roberto Amato, Jeffrey Barrett, Sonia Goncalves, Ewan Harrison, David K. Jackson, Ian Johnston, Dominic Kwiatkowski, Cordelia Langford, John Sillitoe on behalf of the Wellcome Sanger Institute COVID-19 Surveillance Team |
| EPI_ISL_2830982 | Lighthouse Lab in Alderley Park | Wellcome Sanger Institute for the COVID-19 Genomics UK (COG-UK) Consortium | Jacquelyn Wynn, Mairead Hyland, The Lighthouse Lab in Alderley Park and Alex Alderton, Roberto Amato, Jeffrey Barrett, Sonia Goncalves, Ewan Harrison, David K. Jackson, Ian Johnston, Dominic Kwiatkowski, Cordelia Langford, John Sillitoe on behalf of the Wellcome Sanger Institute COVID-19 Surveillance Team |
| EPI_ISL_2830983, EPI_ISL_2830984 | Lighthouse Lab in Glasgow | Wellcome Sanger Institute for the COVID-19 Genomics UK (COG-UK) Consortium | Harper VanSteenhouse, Yumi Kasai, David Gray, Carol Clugston, Anna Dominiczak and Alex Alderton, Roberto Amato, Jeffrey Barrett, Sonia Goncalves, Ewan Harrison, David K. Jackson, Ian Johnston, Dominic Kwiatkowski, Cordelia Langford, John Sillitoe on behalf of the Wellcome Sanger Institute COVID-19 Surveillance Team |
| EPI_ISL_2830985 | Lighthouse Lab in Alderley Park | Wellcome Sanger Institute for the COVID-19 Genomics UK (COG-UK) Consortium | Jacquelyn Wynn, Mairead Hyland, The Lighthouse Lab in Alderley Park and Alex Alderton, Roberto Amato, Jeffrey Barrett, Sonia Goncalves, Ewan Harrison, David K. Jackson, Ian Johnston, Dominic Kwiatkowski, Cordelia Langford, John Sillitoe on behalf of the Wellcome Sanger Institute COVID-19 Surveillance Team |
| EPI_ISL_2830986, EPI_ISL_2830987 | Lighthouse Lab in Glasgow | Wellcome Sanger Institute for the COVID-19 Genomics UK (COG-UK) Consortium | Harper VanSteenhouse, Yumi Kasai, David Gray, Carol Clugston, Anna Dominiczak and Alex Alderton, Roberto Amato, Jeffrey Barrett, Sonia Goncalves, Ewan Harrison, David K. Jackson, Ian Johnston, Dominic Kwiatkowski, Cordelia Langford, John Sillitoe on behalf of the Wellcome Sanger Institute COVID-19 Surveillance Team |
| EPI_ISL_2830988 | Lighthouse Lab in Alderley Park | Wellcome Sanger Institute for the COVID-19 Genomics UK (COG-UK) Consortium | Jacquelyn Wynn, Mairead Hyland, The Lighthouse Lab in Alderley Park and Alex Alderton, Roberto Amato, Jeffrey Barrett, Sonia Goncalves, Ewan Harrison, David K. Jackson, Ian Johnston, Dominic Kwiatkowski, Cordelia Langford, John Sillitoe on behalf of the Wellcome Sanger Institute COVID-19 Surveillance Team |
| EPI_ISL_2830989, EPI_ISL_2830990, EPI_ISL_2830991 | Lighthouse Lab in Glasgow | Wellcome Sanger Institute for the COVID-19 Genomics UK (COG-UK) Consortium | Harper VanSteenhouse, Yumi Kasai, David Gray, Carol Clugston, Anna Dominiczak and Alex Alderton, Roberto Amato, Jeffrey Barrett, Sonia Goncalves, Ewan Harrison, David K. Jackson, Ian Johnston, Dominic Kwiatkowski, Cordelia Langford, John Sillitoe on behalf of the Wellcome Sanger Institute COVID-19 Surveillance Team |
| EPI_ISL_2830992, EPI_ISL_2830993 | Lighthouse Lab in Alderley Park | Wellcome Sanger Institute for the COVID-19 Genomics UK (COG-UK) Consortium | Jacquelyn Wynn, Mairead Hyland, The Lighthouse Lab in Alderley Park and Alex Alderton, Roberto Amato, Jeffrey Barrett, Sonia Goncalves, Ewan Harrison, David K. Jackson, Ian Johnston, Dominic Kwiatkowski, Cordelia Langford, John Sillitoe on behalf of the Wellcome Sanger Institute COVID-19 Surveillance Team |
| EPI_ISL_2830994, EPI_ISL_2830995, EPI_ISL_2830996 | Lighthouse Lab in Glasgow | Wellcome Sanger Institute for the COVID-19 Genomics UK (COG-UK) Consortium | Harper VanSteenhouse, Yumi Kasai, David Gray, Carol Clugston, Anna Dominiczak and Alex Alderton, Roberto Amato, Jeffrey Barrett, Sonia Goncalves, Ewan Harrison, David K. Jackson, Ian Johnston, Dominic Kwiatkowski, Cordelia Langford, John Sillitoe on behalf of the Wellcome Sanger Institute COVID-19 Surveillance Team |
| EPI_ISL_2830997, EPI_ISL_2830998 | Lighthouse Lab in Alderley Park | Wellcome Sanger Institute for the COVID-19 Genomics UK (COG-UK) Consortium | Jacquelyn Wynn, Mairead Hyland, The Lighthouse Lab in Alderley Park and Alex Alderton, Roberto Amato, Jeffrey Barrett, Sonia Goncalves, Ewan Harrison, David K. Jackson, Ian Johnston, Dominic Kwiatkowski, Cordelia Langford, John Sillitoe on behalf of the Wellcome Sanger Institute COVID-19 Surveillance Team |
| EPI_ISL_2830999, EPI_ISL_2831000 | Lighthouse Lab in Glasgow | Wellcome Sanger Institute for the COVID-19 Genomics UK (COG-UK) Consortium | Harper VanSteenhouse, Yumi Kasai, David Gray, Carol Clugston, Anna Dominiczak and Alex Alderton, Roberto Amato, Jeffrey Barrett, Sonia Goncalves, Ewan Harrison, David K. Jackson, Ian Johnston, Dominic Kwiatkowski, Cordelia Langford, John Sillitoe on behalf of the Wellcome Sanger Institute COVID-19 Surveillance Team |
| EPI_ISL_2831001, EPI_ISL_2831002, EPI_ISL_2831003 | Lighthouse Lab in Alderley Park | Wellcome Sanger Institute for the COVID-19 Genomics UK (COG-UK) Consortium | Jacquelyn Wynn, Mairead Hyland, The Lighthouse Lab in Alderley Park and Alex Alderton, Roberto Amato, Jeffrey Barrett, Sonia Goncalves, Ewan Harrison, David K. Jackson, Ian Johnston, Dominic Kwiatkowski, Cordelia Langford, John Sillitoe on behalf of the Wellcome Sanger Institute COVID-19 Surveillance Team |
| EPI_ISL_2831004 | Lighthouse Lab in Glasgow | Wellcome Sanger Institute for the COVID-19 Genomics UK (COG-UK) Consortium | Harper VanSteenhouse, Yumi Kasai, David Gray, Carol Clugston, Anna Dominiczak and Alex Alderton, Roberto Amato, Jeffrey Barrett, Sonia Goncalves, Ewan Harrison, David K. Jackson, Ian Johnston, Dominic Kwiatkowski, Cordelia Langford, John Sillitoe on behalf of the Wellcome Sanger Institute COVID-19 Surveillance Team |
| EPI_ISL_2831005, EPI_ISL_2831006 | Lighthouse Lab in Alderley Park | Wellcome Sanger Institute for the COVID-19 Genomics UK (COG-UK) Consortium | Jacquelyn Wynn, Mairead Hyland, The Lighthouse Lab in Alderley Park and Alex Alderton, Roberto Amato, Jeffrey Barrett, Sonia Goncalves, Ewan Harrison, David K. Jackson, Ian Johnston, Dominic Kwiatkowski, Cordelia Langford, John Sillitoe on behalf of the Wellcome Sanger Institute COVID-19 Surveillance Team |
| EPI_ISL_2831007 | Lighthouse Lab in Glasgow | Wellcome Sanger Institute for the COVID-19 Genomics UK (COG-UK) Consortium | Harper VanSteenhouse, Yumi Kasai, David Gray, Carol Clugston, Anna Dominiczak and Alex Alderton, Roberto Amato, Jeffrey Barrett, Sonia Goncalves, Ewan Harrison, David K. Jackson, Ian Johnston, Dominic Kwiatkowski, Cordelia Langford, John Sillitoe on behalf of the Wellcome Sanger Institute COVID-19 Surveillance Team |
| EPI_ISL_2831008, EPI_ISL_2831009, EPI_ISL_2831010, EPI_ISL_2831011, EPI_ISL_2831012 | Lighthouse Lab in Alderley Park | Wellcome Sanger Institute for the COVID-19 Genomics UK (COG-UK) Consortium | Jacquelyn Wynn, Mairead Hyland, The Lighthouse Lab in Alderley Park and Alex Alderton, Roberto Amato, Jeffrey Barrett, Sonia Goncalves, Ewan Harrison, David K. Jackson, Ian Johnston, Dominic Kwiatkowski, Cordelia Langford, John Sillitoe on behalf of the Wellcome Sanger Institute COVID-19 Surveillance Team |
| EPI_ISL_2831013 | Lighthouse Lab in Glasgow | Wellcome Sanger Institute for the COVID-19 Genomics UK (COG-UK) Consortium | Harper VanSteenhouse, Yumi Kasai, David Gray, Carol Clugston, Anna Dominiczak and Alex Alderton, Roberto Amato, Jeffrey Barrett, Sonia Goncalves, Ewan Harrison, David K. Jackson, Ian Johnston, Dominic Kwiatkowski, Cordelia Langford, John Sillitoe on behalf of the Wellcome Sanger Institute COVID-19 Surveillance Team |
| EPI_ISL_2831014, EPI_ISL_2831015 | Lighthouse Lab in Alderley Park | Wellcome Sanger Institute for the COVID-19 Genomics UK (COG-UK) Consortium | Jacquelyn Wynn, Mairead Hyland, The Lighthouse Lab in Alderley Park and Alex Alderton, Roberto Amato, Jeffrey Barrett, Sonia Goncalves, Ewan Harrison, David K. Jackson, Ian Johnston, Dominic Kwiatkowski, Cordelia Langford, John Sillitoe on behalf of the Wellcome Sanger Institute COVID-19 Surveillance Team |
| EPI_ISL_2831016, EPI_ISL_2831017, EPI_ISL_2831018, EPI_ISL_2831019, EPI_ISL_2831020 | Lighthouse Lab in Glasgow | Wellcome Sanger Institute for the COVID-19 Genomics UK (COG-UK) Consortium | Harper VanSteenhouse, Yumi Kasai, David Gray, Carol Clugston, Anna Dominiczak and Alex Alderton, Roberto Amato, Jeffrey Barrett, Sonia Goncalves, Ewan Harrison, David K. Jackson, Ian Johnston, Dominic Kwiatkowski, Cordelia Langford, John Sillitoe on behalf of the Wellcome Sanger Institute COVID-19 Surveillance Team |
| EPI_ISL_2831021 | Lighthouse Lab in Alderley Park | Wellcome Sanger Institute for the COVID-19 Genomics UK (COG-UK) Consortium | Jacquelyn Wynn, Mairead Hyland, The Lighthouse Lab in Alderley Park and Alex Alderton, Roberto Amato, Jeffrey Barrett, Sonia Goncalves, Ewan Harrison, David K. Jackson, Ian Johnston, Dominic Kwiatkowski, Cordelia Langford, John Sillitoe on behalf of the Wellcome Sanger Institute COVID-19 Surveillance Team |
| EPI_ISL_2831022, EPI_ISL_2831023 | Lighthouse Lab in Glasgow | Wellcome Sanger Institute for the COVID-19 Genomics UK (COG-UK) Consortium | Harper VanSteenhouse, Yumi Kasai, David Gray, Carol Clugston, Anna Dominiczak and Alex Alderton, Roberto Amato, Jeffrey Barrett, Sonia Goncalves, Ewan Harrison, David K. Jackson, Ian Johnston, Dominic Kwiatkowski, Cordelia Langford, John Sillitoe on behalf of the Wellcome Sanger Institute COVID-19 Surveillance Team |
| EPI_ISL_2831024, EPI_ISL_2831025, EPI_ISL_2831026 | Lighthouse Lab in Alderley Park | Wellcome Sanger Institute for the COVID-19 Genomics UK (COG-UK) Consortium | Jacquelyn Wynn, Mairead Hyland, The Lighthouse Lab in Alderley Park and Alex Alderton, Roberto Amato, Jeffrey Barrett, Sonia Goncalves, Ewan Harrison, David K. Jackson, Ian Johnston, Dominic Kwiatkowski, Cordelia Langford, John Sillitoe on behalf of the Wellcome Sanger Institute COVID-19 Surveillance Team |
| EPI_ISL_2831027 | Lighthouse Lab in Glasgow | Wellcome Sanger Institute for the COVID-19 Genomics UK (COG-UK) Consortium | Harper VanSteenhouse, Yumi Kasai, David Gray, Carol Clugston, Anna Dominiczak and Alex Alderton, Roberto Amato, Jeffrey Barrett, Sonia Goncalves, Ewan Harrison, David K. Jackson, Ian Johnston, Dominic Kwiatkowski, Cordelia Langford, John Sillitoe on behalf of the Wellcome Sanger Institute COVID-19 Surveillance Team |
| EPI_ISL_2831028 | Lighthouse Lab in Alderley Park | Wellcome Sanger Institute for the COVID-19 Genomics UK (COG-UK) Consortium | Jacquelyn Wynn, Mairead Hyland, The Lighthouse Lab in Alderley Park and Alex Alderton, Roberto Amato, Jeffrey Barrett, Sonia Goncalves, Ewan Harrison, David K. Jackson, Ian Johnston, Dominic Kwiatkowski, Cordelia Langford, John Sillitoe on behalf of the Wellcome Sanger Institute COVID-19 Surveillance Team |
| EPI_ISL_2831029, EPI_ISL_2831030 | Lighthouse Lab in Glasgow | Wellcome Sanger Institute for the COVID-19 Genomics UK (COG-UK) Consortium | Harper VanSteenhouse, Yumi Kasai, David Gray, Carol Clugston, Anna Dominiczak and Alex Alderton, Roberto Amato, Jeffrey Barrett, Sonia Goncalves, Ewan Harrison, David K. Jackson, Ian Johnston, Dominic Kwiatkowski, Cordelia Langford, John Sillitoe on behalf of the Wellcome Sanger Institute COVID-19 Surveillance Team |
| EPI_ISL_2831031 | Lighthouse Lab in Alderley Park | Wellcome Sanger Institute for the COVID-19 Genomics UK (COG-UK) Consortium | Jacquelyn Wynn, Mairead Hyland, The Lighthouse Lab in Alderley Park and Alex Alderton, Roberto Amato, Jeffrey Barrett, Sonia Goncalves, Ewan Harrison, David K. Jackson, Ian Johnston, Dominic Kwiatkowski, Cordelia Langford, John Sillitoe on behalf of the Wellcome Sanger Institute COVID-19 Surveillance Team |
| EPI_ISL_2831032 | Lighthouse Lab in Glasgow | Wellcome Sanger Institute for the COVID-19 Genomics UK (COG-UK) Consortium | Harper VanSteenhouse, Yumi Kasai, David Gray, Carol Clugston, Anna Dominiczak and Alex Alderton, Roberto Amato, Jeffrey Barrett, Sonia Goncalves, Ewan Harrison, David K. Jackson, Ian Johnston, Dominic Kwiatkowski, Cordelia Langford, John Sillitoe on behalf of the Wellcome Sanger Institute COVID-19 Surveillance Team |
| EPI_ISL_2831033, EPI_ISL_2831034, EPI_ISL_2831035 | Lighthouse Lab in Alderley Park | Wellcome Sanger Institute for the COVID-19 Genomics UK (COG-UK) Consortium | Jacquelyn Wynn, Mairead Hyland, The Lighthouse Lab in Alderley Park and Alex Alderton, Roberto Amato, Jeffrey Barrett, Sonia Goncalves, Ewan Harrison, David K. Jackson, Ian Johnston, Dominic Kwiatkowski, Cordelia Langford, John Sillitoe on behalf of the Wellcome Sanger Institute COVID-19 Surveillance Team |
| EPI_ISL_2831036 | Lighthouse Lab in Glasgow | Wellcome Sanger Institute for the COVID-19 Genomics UK (COG-UK) Consortium | Harper VanSteenhouse, Yumi Kasai, David Gray, Carol Clugston, Anna Dominiczak and Alex Alderton, Roberto Amato, Jeffrey Barrett, Sonia Goncalves, Ewan Harrison, David K. Jackson, Ian Johnston, Dominic Kwiatkowski, Cordelia Langford, John Sillitoe on behalf of the Wellcome Sanger Institute COVID-19 Surveillance Team |
| EPI_ISL_2831037, EPI_ISL_2831038, EPI_ISL_2831039, EPI_ISL_2831040 | Lighthouse Lab in Alderley Park | Wellcome Sanger Institute for the COVID-19 Genomics UK (COG-UK) Consortium | Jacquelyn Wynn, Mairead Hyland, The Lighthouse Lab in Alderley Park and Alex Alderton, Roberto Amato, Jeffrey Barrett, Sonia Goncalves, Ewan Harrison, David K. Jackson, Ian Johnston, Dominic Kwiatkowski, Cordelia Langford, John Sillitoe on behalf of the Wellcome Sanger Institute COVID-19 Surveillance Team |
| EPI_ISL_2831041 | Lighthouse Lab in Glasgow | Wellcome Sanger Institute for the COVID-19 Genomics UK (COG-UK) Consortium | Harper VanSteenhouse, Yumi Kasai, David Gray, Carol Clugston, Anna Dominiczak and Alex Alderton, Roberto Amato, Jeffrey Barrett, Sonia Goncalves, Ewan Harrison, David K. Jackson, Ian Johnston, Dominic Kwiatkowski, Cordelia Langford, John Sillitoe on behalf of the Wellcome Sanger Institute COVID-19 Surveillance Team |
| EPI_ISL_2831042 | Lighthouse Lab in Alderley Park | Wellcome Sanger Institute for the COVID-19 Genomics UK (COG-UK) | Jacquelyn Wynn, Mairead Hyland, The Lighthouse Lab in Alderley Park and Alex Alderton, Roberto Amato, Jeffrey Barrett, Sonia Goncalves, Ewan Harrison, David K. Jackson, Ian Johnston, Dominic Kwiatkowski, Cordelia Langford, John Sillitoe on |

| | | | |
|---|---|---|---|
| | | Consortium | behalf of the Wellcome Sanger Institute COVID-19 Surveillance Team |
| EPI_ISL_2831043, EPI_ISL_2831044 | Lighthouse Lab in Glasgow | Wellcome Sanger Institute for the COVID-19 Genomics UK (COG-UK) Consortium | Harper VanSteenhouse, Yumi Kasai, David Gray, Carol Clugston, Anna Dominiczak and Alex Alderton, Roberto Amato, Jeffrey Barrett, Sonia Goncalves, Ewan Harrison, David K. Jackson, Ian Johnston, Dominic Kwiatkowski, Cordelia Langford, John Sillitoe on behalf of the Wellcome Sanger Institute COVID-19 Surveillance Team |
| EPI_ISL_2831045 | Lighthouse Lab in Alderley Park | Wellcome Sanger Institute for the COVID-19 Genomics UK (COG-UK) Consortium | Jacquelyn Wynn, Mairead Hyland, The Lighthouse Lab in Alderley Park and Alex Alderton, Roberto Amato, Jeffrey Barrett, Sonia Goncalves, Ewan Harrison, David K. Jackson, Ian Johnston, Dominic Kwiatkowski, Cordelia Langford, John Sillitoe on behalf of the Wellcome Sanger Institute COVID-19 Surveillance Team |
| EPI_ISL_2831046 | Lighthouse Lab in Milton Keynes | Wellcome Sanger Institute for the COVID-19 Genomics UK (COG-UK) Consortium | The Lighthouse Lab in Milton Keynes and Alex Alderton, Roberto Amato, Jeffrey Barrett, Sonia Goncalves, Ewan Harrison, David K. Jackson, Ian Johnston, Dominic Kwiatkowski, Cordelia Langford, John Sillitoe on behalf of the Wellcome Sanger Institute COVID-19 Surveillance Team |
| EPI_ISL_2831047 | Lighthouse Lab in Alderley Park | Wellcome Sanger Institute for the COVID-19 Genomics UK (COG-UK) Consortium | Jacquelyn Wynn, Mairead Hyland, The Lighthouse Lab in Alderley Park and Alex Alderton, Roberto Amato, Jeffrey Barrett, Sonia Goncalves, Ewan Harrison, David K. Jackson, Ian Johnston, Dominic Kwiatkowski, Cordelia Langford, John Sillitoe on behalf of the Wellcome Sanger Institute COVID-19 Surveillance Team |
| EPI_ISL_2831048, EPI_ISL_2831049, EPI_ISL_2831050 | Lighthouse Laboratory Plymouth | Wellcome Sanger Institute for the COVID-19 Genomics UK (COG-UK) Consortium | Lighthouse Laboratory Plymouth and Alex Alderton, Roberto Amato, Jeffrey Barrett, Sonia Goncalves, Ewan Harrison, David K. Jackson, Ian Johnston, Dominic Kwiatkowski, Cordelia Langford, John Sillitoe on behalf of the Wellcome Sanger Institute COVID-19 Surveillance Team |
| EPI_ISL_2831051, EPI_ISL_2831052 | Lighthouse Lab in Glasgow | Wellcome Sanger Institute for the COVID-19 Genomics UK (COG-UK) Consortium | Harper VanSteenhouse, Yumi Kasai, David Gray, Carol Clugston, Anna Dominiczak and Alex Alderton, Roberto Amato, Jeffrey Barrett, Sonia Goncalves, Ewan Harrison, David K. Jackson, Ian Johnston, Dominic Kwiatkowski, Cordelia Langford, John Sillitoe on behalf of the Wellcome Sanger Institute COVID-19 Surveillance Team |
| EPI_ISL_2831053 | Lighthouse Lab in Milton Keynes | Wellcome Sanger Institute for the COVID-19 Genomics UK (COG-UK) Consortium | The Lighthouse Lab in Milton Keynes and Alex Alderton, Roberto Amato, Jeffrey Barrett, Sonia Goncalves, Ewan Harrison, David K. Jackson, Ian Johnston, Dominic Kwiatkowski, Cordelia Langford, John Sillitoe on behalf of the Wellcome Sanger Institute COVID-19 Surveillance Team |
| EPI_ISL_2831054 | Lighthouse Lab in Glasgow | Wellcome Sanger Institute for the COVID-19 Genomics UK (COG-UK) Consortium | Harper VanSteenhouse, Yumi Kasai, David Gray, Carol Clugston, Anna Dominiczak and Alex Alderton, Roberto Amato, Jeffrey Barrett, Sonia Goncalves, Ewan Harrison, David K. Jackson, Ian Johnston, Dominic Kwiatkowski, Cordelia Langford, John Sillitoe on behalf of the Wellcome Sanger Institute COVID-19 Surveillance Team |
| EPI_ISL_2831055 | Lighthouse Lab in Milton Keynes | Wellcome Sanger Institute for the COVID-19 Genomics UK (COG-UK) Consortium | The Lighthouse Lab in Milton Keynes and Alex Alderton, Roberto Amato, Jeffrey Barrett, Sonia Goncalves, Ewan Harrison, David K. Jackson, Ian Johnston, Dominic Kwiatkowski, Cordelia Langford, John Sillitoe on behalf of the Wellcome Sanger Institute COVID-19 Surveillance Team |
| EPI_ISL_2831056 | Lighthouse Lab in Alderley Park | Wellcome Sanger Institute for the COVID-19 Genomics UK (COG-UK) Consortium | Jacquelyn Wynn, Mairead Hyland, The Lighthouse Lab in Alderley Park and Alex Alderton, Roberto Amato, Jeffrey Barrett, Sonia Goncalves, Ewan Harrison, David K. Jackson, Ian Johnston, Dominic Kwiatkowski, Cordelia Langford, John Sillitoe on behalf of the Wellcome Sanger Institute COVID-19 Surveillance Team |
| EPI_ISL_2831057 | Lighthouse Laboratory Plymouth | Wellcome Sanger Institute for the COVID-19 Genomics UK (COG-UK) Consortium | Lighthouse Laboratory Plymouth and Alex Alderton, Roberto Amato, Jeffrey Barrett, Sonia Goncalves, Ewan Harrison, David K. Jackson, Ian Johnston, Dominic Kwiatkowski, Cordelia Langford, John Sillitoe on behalf of the Wellcome Sanger Institute COVID-19 Surveillance Team |
| EPI_ISL_2831058 | Lighthouse Lab in Alderley Park | Wellcome Sanger Institute for the COVID-19 Genomics UK (COG-UK) Consortium | Jacquelyn Wynn, Mairead Hyland, The Lighthouse Lab in Alderley Park and Alex Alderton, Roberto Amato, Jeffrey Barrett, Sonia Goncalves, Ewan Harrison, David K. Jackson, Ian Johnston, Dominic Kwiatkowski, Cordelia Langford, John Sillitoe on behalf of the Wellcome Sanger Institute COVID-19 Surveillance Team |
| EPI_ISL_2831059 | Lighthouse Lab in Milton Keynes | Wellcome Sanger Institute for the COVID-19 Genomics UK (COG-UK) Consortium | The Lighthouse Lab in Milton Keynes and Alex Alderton, Roberto Amato, Jeffrey Barrett, Sonia Goncalves, Ewan Harrison, David K. Jackson, Ian Johnston, Dominic Kwiatkowski, Cordelia Langford, John Sillitoe on behalf of the Wellcome Sanger Institute COVID-19 Surveillance Team |
| EPI_ISL_2831060 | Lighthouse Lab in Alderley Park | Wellcome Sanger Institute for the COVID-19 Genomics UK (COG-UK) Consortium | Jacquelyn Wynn, Mairead Hyland, The Lighthouse Lab in Alderley Park and Alex Alderton, Roberto Amato, Jeffrey Barrett, Sonia Goncalves, Ewan Harrison, David K. Jackson, Ian Johnston, Dominic Kwiatkowski, Cordelia Langford, John Sillitoe on behalf of the Wellcome Sanger Institute COVID-19 Surveillance Team |
| EPI_ISL_2831061 | Lighthouse Lab in Milton Keynes | Wellcome Sanger Institute for the COVID-19 Genomics UK (COG-UK) Consortium | The Lighthouse Lab in Milton Keynes and Alex Alderton, Roberto Amato, Jeffrey Barrett, Sonia Goncalves, Ewan Harrison, David K. Jackson, Ian Johnston, Dominic Kwiatkowski, Cordelia Langford, John Sillitoe on behalf of the Wellcome Sanger Institute COVID-19 Surveillance Team |
| EPI_ISL_2831062 | Lighthouse Lab in Alderley Park | Wellcome Sanger Institute for the COVID-19 Genomics UK (COG-UK) Consortium | Jacquelyn Wynn, Mairead Hyland, The Lighthouse Lab in Alderley Park and Alex Alderton, Roberto Amato, Jeffrey Barrett, Sonia Goncalves, Ewan Harrison, David K. Jackson, Ian Johnston, Dominic Kwiatkowski, Cordelia Langford, John Sillitoe on behalf of the Wellcome Sanger Institute COVID-19 Surveillance Team |
| EPI_ISL_2831063 | Lighthouse Laboratory Plymouth | Wellcome Sanger Institute for the COVID-19 Genomics UK (COG-UK) Consortium | Lighthouse Laboratory Plymouth and Alex Alderton, Roberto Amato, Jeffrey Barrett, Sonia Goncalves, Ewan Harrison, David K. Jackson, Ian Johnston, Dominic Kwiatkowski, Cordelia Langford, John Sillitoe on behalf of the Wellcome Sanger Institute COVID-19 Surveillance Team |
| EPI_ISL_2831064 | Lighthouse Lab in Milton Keynes | Wellcome Sanger Institute for the COVID-19 Genomics UK (COG-UK) Consortium | The Lighthouse Lab in Milton Keynes and Alex Alderton, Roberto Amato, Jeffrey Barrett, Sonia Goncalves, Ewan Harrison, David K. Jackson, Ian Johnston, Dominic Kwiatkowski, Cordelia Langford, John Sillitoe on behalf of the Wellcome Sanger Institute COVID-19 Surveillance Team |
| EPI_ISL_2831065 | Lighthouse Laboratory Plymouth | Wellcome Sanger Institute for the COVID-19 Genomics UK (COG-UK) Consortium | Lighthouse Laboratory Plymouth and Alex Alderton, Roberto Amato, Jeffrey Barrett, Sonia Goncalves, Ewan Harrison, David K. Jackson, Ian Johnston, Dominic Kwiatkowski, Cordelia Langford, John Sillitoe on behalf of the Wellcome Sanger Institute COVID-19 Surveillance Team |
| EPI_ISL_2831066 | Lighthouse Lab in Milton Keynes | Wellcome Sanger Institute for the COVID-19 Genomics UK (COG-UK) Consortium | The Lighthouse Lab in Milton Keynes and Alex Alderton, Roberto Amato, Jeffrey Barrett, Sonia Goncalves, Ewan Harrison, David K. Jackson, Ian Johnston, Dominic Kwiatkowski, Cordelia Langford, John Sillitoe on behalf of the Wellcome Sanger Institute COVID-19 Surveillance Team |
| EPI_ISL_2831067 | Lighthouse Lab in Glasgow | Wellcome Sanger Institute for the COVID-19 Genomics UK (COG-UK) Consortium | Harper VanSteenhouse, Yumi Kasai, David Gray, Carol Clugston, Anna Dominiczak and Alex Alderton, Roberto Amato, Jeffrey Barrett, Sonia Goncalves, Ewan Harrison, David K. Jackson, Ian Johnston, Dominic Kwiatkowski, Cordelia Langford, John Sillitoe on behalf of the Wellcome Sanger Institute COVID-19 Surveillance Team |
| EPI_ISL_2831068 | Lighthouse Lab in Milton Keynes | Wellcome Sanger Institute for the COVID-19 Genomics UK (COG-UK) Consortium | The Lighthouse Lab in Milton Keynes and Alex Alderton, Roberto Amato, Jeffrey Barrett, Sonia Goncalves, Ewan Harrison, David K. Jackson, Ian Johnston, Dominic Kwiatkowski, Cordelia Langford, John Sillitoe on behalf of the Wellcome Sanger Institute COVID-19 Surveillance Team |
| EPI_ISL_2831069 | Lighthouse Lab in Glasgow | Wellcome Sanger Institute for the COVID-19 Genomics UK (COG-UK) Consortium | Harper VanSteenhouse, Yumi Kasai, David Gray, Carol Clugston, Anna Dominiczak and Alex Alderton, Roberto Amato, Jeffrey Barrett, Sonia Goncalves, Ewan Harrison, David K. Jackson, Ian Johnston, Dominic Kwiatkowski, Cordelia Langford, John Sillitoe on behalf of the Wellcome Sanger Institute COVID-19 Surveillance Team |
| EPI_ISL_2831070 | Lighthouse Lab in Alderley Park | Wellcome Sanger Institute for the COVID-19 Genomics UK (COG-UK) Consortium | Jacquelyn Wynn, Mairead Hyland, The Lighthouse Lab in Alderley Park and Alex Alderton, Roberto Amato, Jeffrey Barrett, Sonia Goncalves, Ewan Harrison, David K. Jackson, Ian Johnston, Dominic Kwiatkowski, Cordelia Langford, John Sillitoe on behalf of the Wellcome Sanger Institute COVID-19 Surveillance Team |
| EPI_ISL_2831071, EPI_ISL_2831072 | Lighthouse Laboratory Plymouth | Wellcome Sanger Institute for the COVID-19 Genomics UK (COG-UK) Consortium | Lighthouse Laboratory Plymouth and Alex Alderton, Roberto Amato, Jeffrey Barrett, Sonia Goncalves, Ewan Harrison, David K. Jackson, Ian Johnston, Dominic Kwiatkowski, Cordelia Langford, John Sillitoe on behalf of the Wellcome Sanger Institute COVID-19 Surveillance Team |
| EPI_ISL_2831073 | Lighthouse Lab in Milton Keynes | Wellcome Sanger Institute for the COVID-19 Genomics UK (COG-UK) Consortium | The Lighthouse Lab in Milton Keynes and Alex Alderton, Roberto Amato, Jeffrey Barrett, Sonia Goncalves, Ewan Harrison, David K. Jackson, Ian Johnston, Dominic Kwiatkowski, Cordelia Langford, John Sillitoe on behalf of the Wellcome Sanger Institute COVID-19 Surveillance Team |
| EPI_ISL_2831074, EPI_ISL_2831075 | Lighthouse Lab in Alderley Park | Wellcome Sanger Institute for the COVID-19 Genomics UK (COG-UK) Consortium | Jacquelyn Wynn, Mairead Hyland, The Lighthouse Lab in Alderley Park and Alex Alderton, Roberto Amato, Jeffrey Barrett, Sonia Goncalves, Ewan Harrison, David K. Jackson, Ian Johnston, Dominic Kwiatkowski, Cordelia Langford, John Sillitoe on behalf of the Wellcome Sanger Institute COVID-19 Surveillance Team |
| EPI_ISL_2831076 | Lighthouse Lab in Glasgow | Wellcome Sanger Institute for the COVID-19 Genomics UK (COG-UK) Consortium | Harper VanSteenhouse, Yumi Kasai, David Gray, Carol Clugston, Anna Dominiczak and Alex Alderton, Roberto Amato, Jeffrey Barrett, Sonia Goncalves, Ewan Harrison, David K. Jackson, Ian Johnston, Dominic Kwiatkowski, Cordelia Langford, John Sillitoe on behalf of the Wellcome Sanger Institute COVID-19 Surveillance Team |
| EPI_ISL_2831077 | Lighthouse Lab in Milton Keynes | Wellcome Sanger Institute for the COVID-19 Genomics UK (COG-UK) Consortium | The Lighthouse Lab in Milton Keynes and Alex Alderton, Roberto Amato, Jeffrey Barrett, Sonia Goncalves, Ewan Harrison, David K. Jackson, Ian Johnston, Dominic Kwiatkowski, Cordelia Langford, John Sillitoe on behalf of the Wellcome Sanger Institute COVID-19 Surveillance Team |
| EPI_ISL_2831078, EPI_ISL_2831079 | Lighthouse Lab in Alderley Park | Wellcome Sanger Institute for the COVID-19 Genomics UK (COG-UK) Consortium | Jacquelyn Wynn, Mairead Hyland, The Lighthouse Lab in Alderley Park and Alex Alderton, Roberto Amato, Jeffrey Barrett, Sonia Goncalves, Ewan Harrison, David K. Jackson, Ian Johnston, Dominic Kwiatkowski, Cordelia Langford, John Sillitoe on behalf of the Wellcome Sanger Institute COVID-19 Surveillance Team |
| EPI_ISL_2831080 | Lighthouse Lab in Glasgow | Wellcome Sanger Institute for the COVID-19 Genomics UK (COG-UK) Consortium | Harper VanSteenhouse, Yumi Kasai, David Gray, Carol Clugston, Anna Dominiczak and Alex Alderton, Roberto Amato, Jeffrey Barrett, Sonia Goncalves, Ewan Harrison, David K. Jackson, Ian Johnston, Dominic Kwiatkowski, Cordelia Langford, John Sillitoe on behalf of the Wellcome Sanger Institute COVID-19 Surveillance Team |
| EPI_ISL_2831081 | Lighthouse Lab in Milton Keynes | Wellcome Sanger Institute for the COVID-19 Genomics UK (COG-UK) Consortium | The Lighthouse Lab in Milton Keynes and Alex Alderton, Roberto Amato, Jeffrey Barrett, Sonia Goncalves, Ewan Harrison, David K. Jackson, Ian Johnston, Dominic Kwiatkowski, Cordelia Langford, John Sillitoe on behalf of the Wellcome Sanger Institute COVID-19 Surveillance Team |
| EPI_ISL_2831082 | Lighthouse Lab in Alderley Park | Wellcome Sanger Institute for the COVID-19 Genomics UK (COG-UK) Consortium | Jacquelyn Wynn, Mairead Hyland, The Lighthouse Lab in Alderley Park and Alex Alderton, Roberto Amato, Jeffrey Barrett, Sonia Goncalves, Ewan Harrison, David K. Jackson, Ian Johnston, Dominic Kwiatkowski, Cordelia Langford, John Sillitoe on behalf of the Wellcome Sanger Institute COVID-19 Surveillance Team |
| EPI_ISL_2831084 | Lighthouse Laboratory Plymouth | Wellcome Sanger Institute for the COVID-19 Genomics UK (COG-UK) Consortium | Lighthouse Laboratory Plymouth and Alex Alderton, Roberto Amato, Jeffrey Barrett, Sonia Goncalves, Ewan Harrison, David K. Jackson, Ian Johnston, Dominic Kwiatkowski, Cordelia Langford, John Sillitoe on behalf of the Wellcome Sanger Institute COVID-19 Surveillance Team |
| EPI_ISL_2831085 | Lighthouse Lab in Alderley Park | Wellcome Sanger Institute for the COVID-19 Genomics UK (COG-UK) Consortium | Jacquelyn Wynn, Mairead Hyland, The Lighthouse Lab in Alderley Park and Alex Alderton, Roberto Amato, Jeffrey Barrett, Sonia Goncalves, Ewan Harrison, David K. Jackson, Ian Johnston, Dominic Kwiatkowski, Cordelia Langford, John Sillitoe on behalf of the Wellcome Sanger Institute COVID-19 Surveillance Team |
| EPI_ISL_2831086 | Lighthouse Lab in Milton Keynes | Wellcome Sanger Institute for the COVID-19 Genomics UK (COG-UK) Consortium | The Lighthouse Lab in Milton Keynes and Alex Alderton, Roberto Amato, Jeffrey Barrett, Sonia Goncalves, Ewan Harrison, David K. Jackson, Ian Johnston, Dominic Kwiatkowski, Cordelia Langford, John Sillitoe on behalf of the Wellcome Sanger Institute COVID-19 Surveillance Team |
| EPI_ISL_2831087 | Lighthouse Lab in Alderley Park | Wellcome Sanger Institute for the COVID-19 Genomics UK (COG-UK) Consortium | Jacquelyn Wynn, Mairead Hyland, The Lighthouse Lab in Alderley Park and Alex Alderton, Roberto Amato, Jeffrey Barrett, Sonia Goncalves, Ewan Harrison, David K. Jackson, Ian Johnston, Dominic Kwiatkowski, Cordelia Langford, John Sillitoe on behalf of the Wellcome Sanger Institute COVID-19 Surveillance Team |
| EPI_ISL_2831088 | Lighthouse Lab in Glasgow | Wellcome Sanger Institute for the COVID-19 Genomics UK (COG-UK) Consortium | Harper VanSteenhouse, Yumi Kasai, David Gray, Carol Clugston, Anna Dominiczak and Alex Alderton, Roberto Amato, Jeffrey Barrett, Sonia Goncalves, Ewan Harrison, David K. Jackson, Ian Johnston, Dominic Kwiatkowski, Cordelia Langford, John Sillitoe on behalf of the Wellcome Sanger Institute COVID-19 Surveillance Team |
| EPI_ISL_2831089 | Lighthouse Lab in Milton Keynes | Wellcome Sanger Institute for the COVID-19 Genomics UK (COG-UK) Consortium | The Lighthouse Lab in Milton Keynes and Alex Alderton, Roberto Amato, Jeffrey Barrett, Sonia Goncalves, Ewan Harrison, David K. Jackson, Ian Johnston, Dominic Kwiatkowski, Cordelia Langford, John Sillitoe on behalf of the Wellcome Sanger Institute COVID-19 Surveillance Team |
| EPI_ISL_2831090 | Lighthouse Laboratory Plymouth | Wellcome Sanger Institute for the COVID-19 Genomics UK (COG-UK) Consortium | Lighthouse Laboratory Plymouth and Alex Alderton, Roberto Amato, Jeffrey Barrett, Sonia Goncalves, Ewan Harrison, David K. Jackson, Ian Johnston, Dominic Kwiatkowski, Cordelia Langford, John Sillitoe on behalf of the Wellcome Sanger Institute COVID-19 Surveillance Team |
| EPI_ISL_2831091 | Lighthouse Lab in Glasgow | Wellcome Sanger Institute for the COVID-19 Genomics UK (COG-UK) Consortium | Harper VanSteenhouse, Yumi Kasai, David Gray, Carol Clugston, Anna Dominiczak and Alex Alderton, Roberto Amato, Jeffrey Barrett, Sonia Goncalves, Ewan Harrison, David K. Jackson, Ian Johnston, Dominic Kwiatkowski, Cordelia Langford, John Sillitoe on behalf of the Wellcome Sanger Institute COVID-19 Surveillance Team |
| EPI_ISL_2831092 | Lighthouse Lab in Alderley Park | Wellcome Sanger Institute for the COVID-19 Genomics UK (COG-UK) Consortium | Jacquelyn Wynn, Mairead Hyland, The Lighthouse Lab in Alderley Park and Alex Alderton, Roberto Amato, Jeffrey Barrett, Sonia Goncalves, Ewan Harrison, David K. Jackson, Ian Johnston, Dominic Kwiatkowski, Cordelia Langford, John Sillitoe on behalf of the Wellcome Sanger Institute COVID-19 Surveillance Team |
| EPI_ISL_2831093 | Lighthouse Lab in Milton Keynes | Wellcome Sanger Institute for the COVID-19 Genomics UK (COG-UK) Consortium | The Lighthouse Lab in Milton Keynes and Alex Alderton, Roberto Amato, Jeffrey Barrett, Sonia Goncalves, Ewan Harrison, David K. Jackson, Ian Johnston, Dominic Kwiatkowski, Cordelia Langford, John Sillitoe on behalf of the Wellcome Sanger Institute COVID-19 Surveillance Team |
| EPI_ISL_2831094, EPI_ISL_2831095, | Lighthouse Lab in Alderley Park | Wellcome Sanger Institute for the COVID-19 Genomics UK (COG-UK) Consortium | Jacquelyn Wynn, Mairead Hyland, The Lighthouse Lab in Alderley Park and Alex Alderton, Roberto Amato, Jeffrey Barrett, Sonia Goncalves, Ewan Harrison, David K. Jackson, Ian Johnston, Dominic Kwiatkowski, Cordelia Langford, John Sillitoe on behalf of the Wellcome Sanger Institute COVID-19 Surveillance Team |

| | | | |
|---|---|---|---|
| EPI_ISL_2831096 | | | |
| EPI_ISL_2831097 | Lighthouse Lab in Milton Keynes | Wellcome Sanger Institute for the COVID-19 Genomics UK (COG-UK) Consortium | The Lighthouse Lab in Milton Keynes and Alex Alderton, Roberto Amato, Jeffrey Barrett, Sonia Goncalves, Ewan Harrison, David K. Jackson, Ian Johnston, Dominic Kwiatkowski, Cordelia Langford, John Sillitoe on behalf of the Wellcome Sanger Institute COVID-19 Surveillance Team |
| EPI_ISL_2831098, EPI_ISL_2831099, EPI_ISL_2831100, EPI_ISL_2831101 | Lighthouse Laboratory Plymouth | Wellcome Sanger Institute for the COVID-19 Genomics UK (COG-UK) Consortium | Lighthouse Laboratory Plymouth and Alex Alderton, Roberto Amato, Jeffrey Barrett, Sonia Goncalves, Ewan Harrison, David K. Jackson, Ian Johnston, Dominic Kwiatkowski, Cordelia Langford, John Sillitoe on behalf of the Wellcome Sanger Institute COVID-19 Surveillance Team |
| EPI_ISL_2831102 | Lighthouse Lab in Alderley Park | Wellcome Sanger Institute for the COVID-19 Genomics UK (COG-UK) Consortium | Jacquelyn Wynn, Mairead Hyland, The Lighthouse Lab in Alderley Park and Alex Alderton, Roberto Amato, Jeffrey Barrett, Sonia Goncalves, Ewan Harrison, David K. Jackson, Ian Johnston, Dominic Kwiatkowski, Cordelia Langford, John Sillitoe on behalf of the Wellcome Sanger Institute COVID-19 Surveillance Team |
| EPI_ISL_2831103 | Lighthouse Lab in Milton Keynes | Wellcome Sanger Institute for the COVID-19 Genomics UK (COG-UK) Consortium | The Lighthouse Lab in Milton Keynes and Alex Alderton, Roberto Amato, Jeffrey Barrett, Sonia Goncalves, Ewan Harrison, David K. Jackson, Ian Johnston, Dominic Kwiatkowski, Cordelia Langford, John Sillitoe on behalf of the Wellcome Sanger Institute COVID-19 Surveillance Team |
| EPI_ISL_2831104 | Lighthouse Lab in Glasgow | Wellcome Sanger Institute for the COVID-19 Genomics UK (COG-UK) Consortium | Harper VanSteenhouse, Yumi Kasai, David Gray, Carol Clugston, Anna Dominiczak and Alex Alderton, Roberto Amato, Jeffrey Barrett, Sonia Goncalves, Ewan Harrison, David K. Jackson, Ian Johnston, Dominic Kwiatkowski, Cordelia Langford, John Sillitoe on behalf of the Wellcome Sanger Institute COVID-19 Surveillance Team |
| EPI_ISL_2831105 | Lighthouse Lab in Milton Keynes | Wellcome Sanger Institute for the COVID-19 Genomics UK (COG-UK) Consortium | The Lighthouse Lab in Milton Keynes and Alex Alderton, Roberto Amato, Jeffrey Barrett, Sonia Goncalves, Ewan Harrison, David K. Jackson, Ian Johnston, Dominic Kwiatkowski, Cordelia Langford, John Sillitoe on behalf of the Wellcome Sanger Institute COVID-19 Surveillance Team |
| EPI_ISL_2831106, EPI_ISL_2831107 | Lighthouse Laboratory Plymouth | Wellcome Sanger Institute for the COVID-19 Genomics UK (COG-UK) Consortium | Lighthouse Laboratory Plymouth and Alex Alderton, Roberto Amato, Jeffrey Barrett, Sonia Goncalves, Ewan Harrison, David K. Jackson, Ian Johnston, Dominic Kwiatkowski, Cordelia Langford, John Sillitoe on behalf of the Wellcome Sanger Institute COVID-19 Surveillance Team |
| EPI_ISL_2831108 | Lighthouse Lab in Glasgow | Wellcome Sanger Institute for the COVID-19 Genomics UK (COG-UK) Consortium | Harper VanSteenhouse, Yumi Kasai, David Gray, Carol Clugston, Anna Dominiczak and Alex Alderton, Roberto Amato, Jeffrey Barrett, Sonia Goncalves, Ewan Harrison, David K. Jackson, Ian Johnston, Dominic Kwiatkowski, Cordelia Langford, John Sillitoe on behalf of the Wellcome Sanger Institute COVID-19 Surveillance Team |
| EPI_ISL_2831109, EPI_ISL_2831110 | Lighthouse Lab in Milton Keynes | Wellcome Sanger Institute for the COVID-19 Genomics UK (COG-UK) Consortium | The Lighthouse Lab in Milton Keynes and Alex Alderton, Roberto Amato, Jeffrey Barrett, Sonia Goncalves, Ewan Harrison, David K. Jackson, Ian Johnston, Dominic Kwiatkowski, Cordelia Langford, John Sillitoe on behalf of the Wellcome Sanger Institute COVID-19 Surveillance Team |
| EPI_ISL_2831111 | Lighthouse Laboratory Plymouth | Wellcome Sanger Institute for the COVID-19 Genomics UK (COG-UK) Consortium | Lighthouse Laboratory Plymouth and Alex Alderton, Roberto Amato, Jeffrey Barrett, Sonia Goncalves, Ewan Harrison, David K. Jackson, Ian Johnston, Dominic Kwiatkowski, Cordelia Langford, John Sillitoe on behalf of the Wellcome Sanger Institute COVID-19 Surveillance Team |
| EPI_ISL_2831112 | Lighthouse Lab in Milton Keynes | Wellcome Sanger Institute for the COVID-19 Genomics UK (COG-UK) Consortium | The Lighthouse Lab in Milton Keynes and Alex Alderton, Roberto Amato, Jeffrey Barrett, Sonia Goncalves, Ewan Harrison, David K. Jackson, Ian Johnston, Dominic Kwiatkowski, Cordelia Langford, John Sillitoe on behalf of the Wellcome Sanger Institute COVID-19 Surveillance Team |
| EPI_ISL_2831113, EPI_ISL_2831114 | Lighthouse Lab in Glasgow | Wellcome Sanger Institute for the COVID-19 Genomics UK (COG-UK) Consortium | Harper VanSteenhouse, Yumi Kasai, David Gray, Carol Clugston, Anna Dominiczak and Alex Alderton, Roberto Amato, Jeffrey Barrett, Sonia Goncalves, Ewan Harrison, David K. Jackson, Ian Johnston, Dominic Kwiatkowski, Cordelia Langford, John Sillitoe on behalf of the Wellcome Sanger Institute COVID-19 Surveillance Team |
| EPI_ISL_2831115 | Lighthouse Lab in Milton Keynes | Wellcome Sanger Institute for the COVID-19 Genomics UK (COG-UK) Consortium | The Lighthouse Lab in Milton Keynes and Alex Alderton, Roberto Amato, Jeffrey Barrett, Sonia Goncalves, Ewan Harrison, David K. Jackson, Ian Johnston, Dominic Kwiatkowski, Cordelia Langford, John Sillitoe on behalf of the Wellcome Sanger Institute COVID-19 Surveillance Team |
| EPI_ISL_2831116, EPI_ISL_2831117, EPI_ISL_2831118 | Lighthouse Laboratory Plymouth | Wellcome Sanger Institute for the COVID-19 Genomics UK (COG-UK) Consortium | Lighthouse Laboratory Plymouth and Alex Alderton, Roberto Amato, Jeffrey Barrett, Sonia Goncalves, Ewan Harrison, David K. Jackson, Ian Johnston, Dominic Kwiatkowski, Cordelia Langford, John Sillitoe on behalf of the Wellcome Sanger Institute COVID-19 Surveillance Team |
| EPI_ISL_2831119 | Lighthouse Lab in Alderley Park | Wellcome Sanger Institute for the COVID-19 Genomics UK (COG-UK) Consortium | Jacquelyn Wynn, Mairead Hyland, The Lighthouse Lab in Alderley Park and Alex Alderton, Roberto Amato, Jeffrey Barrett, Sonia Goncalves, Ewan Harrison, David K. Jackson, Ian Johnston, Dominic Kwiatkowski, Cordelia Langford, John Sillitoe on behalf of the Wellcome Sanger Institute COVID-19 Surveillance Team |
| EPI_ISL_2831120 | Lighthouse Lab in Milton Keynes | Wellcome Sanger Institute for the COVID-19 Genomics UK (COG-UK) Consortium | The Lighthouse Lab in Milton Keynes and Alex Alderton, Roberto Amato, Jeffrey Barrett, Sonia Goncalves, Ewan Harrison, David K. Jackson, Ian Johnston, Dominic Kwiatkowski, Cordelia Langford, John Sillitoe on behalf of the Wellcome Sanger Institute COVID-19 Surveillance Team |
| EPI_ISL_2831121, EPI_ISL_2831122 | Lighthouse Lab in Glasgow | Wellcome Sanger Institute for the COVID-19 Genomics UK (COG-UK) Consortium | Harper VanSteenhouse, Yumi Kasai, David Gray, Carol Clugston, Anna Dominiczak and Alex Alderton, Roberto Amato, Jeffrey Barrett, Sonia Goncalves, Ewan Harrison, David K. Jackson, Ian Johnston, Dominic Kwiatkowski, Cordelia Langford, John Sillitoe on behalf of the Wellcome Sanger Institute COVID-19 Surveillance Team |
| EPI_ISL_2831123 | Lighthouse Lab in Milton Keynes | Wellcome Sanger Institute for the COVID-19 Genomics UK (COG-UK) Consortium | The Lighthouse Lab in Milton Keynes and Alex Alderton, Roberto Amato, Jeffrey Barrett, Sonia Goncalves, Ewan Harrison, David K. Jackson, Ian Johnston, Dominic Kwiatkowski, Cordelia Langford, John Sillitoe on behalf of the Wellcome Sanger Institute COVID-19 Surveillance Team |
| EPI_ISL_2831124, EPI_ISL_2831125 | Lighthouse Laboratory Plymouth | Wellcome Sanger Institute for the COVID-19 Genomics UK (COG-UK) Consortium | Lighthouse Laboratory Plymouth and Alex Alderton, Roberto Amato, Jeffrey Barrett, Sonia Goncalves, Ewan Harrison, David K. Jackson, Ian Johnston, Dominic Kwiatkowski, Cordelia Langford, John Sillitoe on behalf of the Wellcome Sanger Institute COVID-19 Surveillance Team |
| EPI_ISL_2831126 | Lighthouse Lab in Milton Keynes | Wellcome Sanger Institute for the COVID-19 Genomics UK (COG-UK) Consortium | The Lighthouse Lab in Milton Keynes and Alex Alderton, Roberto Amato, Jeffrey Barrett, Sonia Goncalves, Ewan Harrison, David K. Jackson, Ian Johnston, Dominic Kwiatkowski, Cordelia Langford, John Sillitoe on behalf of the Wellcome Sanger Institute COVID-19 Surveillance Team |
| EPI_ISL_2831127 | Lighthouse Laboratory Plymouth | Wellcome Sanger Institute for the COVID-19 Genomics UK (COG-UK) Consortium | Lighthouse Laboratory Plymouth and Alex Alderton, Roberto Amato, Jeffrey Barrett, Sonia Goncalves, Ewan Harrison, David K. Jackson, Ian Johnston, Dominic Kwiatkowski, Cordelia Langford, John Sillitoe on behalf of the Wellcome Sanger Institute COVID-19 Surveillance Team |
| EPI_ISL_2831128 | Lighthouse Lab in Glasgow | Wellcome Sanger Institute for the COVID-19 Genomics UK (COG-UK) Consortium | Harper VanSteenhouse, Yumi Kasai, David Gray, Carol Clugston, Anna Dominiczak and Alex Alderton, Roberto Amato, Jeffrey Barrett, Sonia Goncalves, Ewan Harrison, David K. Jackson, Ian Johnston, Dominic Kwiatkowski, Cordelia Langford, John Sillitoe on behalf of the Wellcome Sanger Institute COVID-19 Surveillance Team |
| EPI_ISL_2831129 | Lighthouse Lab in Milton Keynes | Wellcome Sanger Institute for the COVID-19 Genomics UK (COG-UK) Consortium | The Lighthouse Lab in Milton Keynes and Alex Alderton, Roberto Amato, Jeffrey Barrett, Sonia Goncalves, Ewan Harrison, David K. Jackson, Ian Johnston, Dominic Kwiatkowski, Cordelia Langford, John Sillitoe on behalf of the Wellcome Sanger Institute COVID-19 Surveillance Team |
| EPI_ISL_2831130 | Lighthouse Lab in Alderley Park | Wellcome Sanger Institute for the COVID-19 Genomics UK (COG-UK) Consortium | Jacquelyn Wynn, Mairead Hyland, The Lighthouse Lab in Alderley Park and Alex Alderton, Roberto Amato, Jeffrey Barrett, Sonia Goncalves, Ewan Harrison, David K. Jackson, Ian Johnston, Dominic Kwiatkowski, Cordelia Langford, John Sillitoe on behalf of the Wellcome Sanger Institute COVID-19 Surveillance Team |
| EPI_ISL_2831131 | Lighthouse Laboratory Plymouth | Wellcome Sanger Institute for the COVID-19 Genomics UK (COG-UK) Consortium | Lighthouse Laboratory Plymouth and Alex Alderton, Roberto Amato, Jeffrey Barrett, Sonia Goncalves, Ewan Harrison, David K. Jackson, Ian Johnston, Dominic Kwiatkowski, Cordelia Langford, John Sillitoe on behalf of the Wellcome Sanger Institute COVID-19 Surveillance Team |
| EPI_ISL_2831132 | Lighthouse Lab in Milton Keynes | Wellcome Sanger Institute for the COVID-19 Genomics UK (COG-UK) Consortium | The Lighthouse Lab in Milton Keynes and Alex Alderton, Roberto Amato, Jeffrey Barrett, Sonia Goncalves, Ewan Harrison, David K. Jackson, Ian Johnston, Dominic Kwiatkowski, Cordelia Langford, John Sillitoe on behalf of the Wellcome Sanger Institute COVID-19 Surveillance Team |
| EPI_ISL_2831133 | Lighthouse Laboratory Plymouth | Wellcome Sanger Institute for the COVID-19 Genomics UK (COG-UK) Consortium | Lighthouse Laboratory Plymouth and Alex Alderton, Roberto Amato, Jeffrey Barrett, Sonia Goncalves, Ewan Harrison, David K. Jackson, Ian Johnston, Dominic Kwiatkowski, Cordelia Langford, John Sillitoe on behalf of the Wellcome Sanger Institute COVID-19 Surveillance Team |
| EPI_ISL_2831134 | Lighthouse Lab in Alderley Park | Wellcome Sanger Institute for the COVID-19 Genomics UK (COG-UK) Consortium | Jacquelyn Wynn, Mairead Hyland, The Lighthouse Lab in Alderley Park and Alex Alderton, Roberto Amato, Jeffrey Barrett, Sonia Goncalves, Ewan Harrison, David K. Jackson, Ian Johnston, Dominic Kwiatkowski, Cordelia Langford, John Sillitoe on behalf of the Wellcome Sanger Institute COVID-19 Surveillance Team |
| EPI_ISL_2831135, EPI_ISL_2831136 | Lighthouse Laboratory Plymouth | Wellcome Sanger Institute for the COVID-19 Genomics UK (COG-UK) Consortium | Lighthouse Laboratory Plymouth and Alex Alderton, Roberto Amato, Jeffrey Barrett, Sonia Goncalves, Ewan Harrison, David K. Jackson, Ian Johnston, Dominic Kwiatkowski, Cordelia Langford, John Sillitoe on behalf of the Wellcome Sanger Institute COVID-19 Surveillance Team |
| EPI_ISL_2831137 | Lighthouse Lab in Glasgow | Wellcome Sanger Institute for the COVID-19 Genomics UK (COG-UK) Consortium | Harper VanSteenhouse, Yumi Kasai, David Gray, Carol Clugston, Anna Dominiczak and Alex Alderton, Roberto Amato, Jeffrey Barrett, Sonia Goncalves, Ewan Harrison, David K. Jackson, Ian Johnston, Dominic Kwiatkowski, Cordelia Langford, John Sillitoe on behalf of the Wellcome Sanger Institute COVID-19 Surveillance Team |
| EPI_ISL_2831138 | Lighthouse Lab in Alderley Park | Wellcome Sanger Institute for the COVID-19 Genomics UK (COG-UK) Consortium | Jacquelyn Wynn, Mairead Hyland, The Lighthouse Lab in Alderley Park and Alex Alderton, Roberto Amato, Jeffrey Barrett, Sonia Goncalves, Ewan Harrison, David K. Jackson, Ian Johnston, Dominic Kwiatkowski, Cordelia Langford, John Sillitoe on behalf of the Wellcome Sanger Institute COVID-19 Surveillance Team |
| EPI_ISL_2831139 | Lighthouse Laboratory Plymouth | Wellcome Sanger Institute for the COVID-19 Genomics UK (COG-UK) Consortium | Lighthouse Laboratory Plymouth and Alex Alderton, Roberto Amato, Jeffrey Barrett, Sonia Goncalves, Ewan Harrison, David K. Jackson, Ian Johnston, Dominic Kwiatkowski, Cordelia Langford, John Sillitoe on behalf of the Wellcome Sanger Institute COVID-19 Surveillance Team |
| EPI_ISL_2831140 | Lighthouse Lab in Milton Keynes | Wellcome Sanger Institute for the COVID-19 Genomics UK (COG-UK) Consortium | The Lighthouse Lab in Milton Keynes and Alex Alderton, Roberto Amato, Jeffrey Barrett, Sonia Goncalves, Ewan Harrison, David K. Jackson, Ian Johnston, Dominic Kwiatkowski, Cordelia Langford, John Sillitoe on behalf of the Wellcome Sanger Institute COVID-19 Surveillance Team |
| EPI_ISL_2831141 | Lighthouse Laboratory Plymouth | Wellcome Sanger Institute for the COVID-19 Genomics UK (COG-UK) Consortium | Lighthouse Laboratory Plymouth and Alex Alderton, Roberto Amato, Jeffrey Barrett, Sonia Goncalves, Ewan Harrison, David K. Jackson, Ian Johnston, Dominic Kwiatkowski, Cordelia Langford, John Sillitoe on behalf of the Wellcome Sanger Institute COVID-19 Surveillance Team |
| EPI_ISL_2831142 | Lighthouse Lab in Alderley Park | Wellcome Sanger Institute for the COVID-19 Genomics UK (COG-UK) Consortium | Jacquelyn Wynn, Mairead Hyland, The Lighthouse Lab in Alderley Park and Alex Alderton, Roberto Amato, Jeffrey Barrett, Sonia Goncalves, Ewan Harrison, David K. Jackson, Ian Johnston, Dominic Kwiatkowski, Cordelia Langford, John Sillitoe on behalf of the Wellcome Sanger Institute COVID-19 Surveillance Team |
| EPI_ISL_2831143 | Lighthouse Lab in Milton Keynes | Wellcome Sanger Institute for the COVID-19 Genomics UK (COG-UK) Consortium | The Lighthouse Lab in Milton Keynes and Alex Alderton, Roberto Amato, Jeffrey Barrett, Sonia Goncalves, Ewan Harrison, David K. Jackson, Ian Johnston, Dominic Kwiatkowski, Cordelia Langford, John Sillitoe on behalf of the Wellcome Sanger Institute COVID-19 Surveillance Team |
| EPI_ISL_2831144, EPI_ISL_2831145 | Lighthouse Laboratory Plymouth | Wellcome Sanger Institute for the COVID-19 Genomics UK (COG-UK) Consortium | Lighthouse Laboratory Plymouth and Alex Alderton, Roberto Amato, Jeffrey Barrett, Sonia Goncalves, Ewan Harrison, David K. Jackson, Ian Johnston, Dominic Kwiatkowski, Cordelia Langford, John Sillitoe on behalf of the Wellcome Sanger Institute COVID-19 Surveillance Team |
| EPI_ISL_2831146, EPI_ISL_2831147 | Lighthouse Lab in Milton Keynes | Wellcome Sanger Institute for the COVID-19 Genomics UK (COG-UK) Consortium | The Lighthouse Lab in Milton Keynes and Alex Alderton, Roberto Amato, Jeffrey Barrett, Sonia Goncalves, Ewan Harrison, David K. Jackson, Ian Johnston, Dominic Kwiatkowski, Cordelia Langford, John Sillitoe on behalf of the Wellcome Sanger Institute COVID-19 Surveillance Team |
| EPI_ISL_2831148 | Lighthouse Lab in Alderley Park | Wellcome Sanger Institute for the COVID-19 Genomics UK (COG-UK) Consortium | Jacquelyn Wynn, Mairead Hyland, The Lighthouse Lab in Alderley Park and Alex Alderton, Roberto Amato, Jeffrey Barrett, Sonia Goncalves, Ewan Harrison, David K. Jackson, Ian Johnston, Dominic Kwiatkowski, Cordelia Langford, John Sillitoe on behalf of the Wellcome Sanger Institute COVID-19 Surveillance Team |
| EPI_ISL_2831149 | Lighthouse Laboratory Plymouth | Wellcome Sanger Institute for the COVID-19 Genomics UK (COG-UK) Consortium | Lighthouse Laboratory Plymouth and Alex Alderton, Roberto Amato, Jeffrey Barrett, Sonia Goncalves, Ewan Harrison, David K. Jackson, Ian Johnston, Dominic Kwiatkowski, Cordelia Langford, John Sillitoe on behalf of the Wellcome Sanger Institute COVID-19 Surveillance Team |
| EPI_ISL_2831150, EPI_ISL_2831151 | Lighthouse Lab in Alderley Park | Wellcome Sanger Institute for the COVID-19 Genomics UK (COG-UK) Consortium | Jacquelyn Wynn, Mairead Hyland, The Lighthouse Lab in Alderley Park and Alex Alderton, Roberto Amato, Jeffrey Barrett, Sonia Goncalves, Ewan Harrison, David K. Jackson, Ian Johnston, Dominic Kwiatkowski, Cordelia Langford, John Sillitoe on behalf of the Wellcome Sanger Institute COVID-19 Surveillance Team |
| EPI_ISL_2831152, EPI_ISL_2831153 | Lighthouse Laboratory Plymouth | Wellcome Sanger Institute for the COVID-19 Genomics UK (COG-UK) Consortium | Lighthouse Laboratory Plymouth and Alex Alderton, Roberto Amato, Jeffrey Barrett, Sonia Goncalves, Ewan Harrison, David K. Jackson, Ian Johnston, Dominic Kwiatkowski, Cordelia Langford, John Sillitoe on behalf of the Wellcome Sanger Institute COVID-19 Surveillance Team |
| EPI_ISL_2831154 | Lighthouse Lab in Alderley Park | Wellcome Sanger Institute for the COVID-19 Genomics UK (COG-UK) Consortium | Jacquelyn Wynn, Mairead Hyland, The Lighthouse Lab in Alderley Park and Alex Alderton, Roberto Amato, Jeffrey Barrett, Sonia Goncalves, Ewan Harrison, David K. Jackson, Ian Johnston, Dominic Kwiatkowski, Cordelia Langford, John Sillitoe on behalf of the Wellcome Sanger Institute COVID-19 Surveillance Team |

| | | | |
|---|---|---|---|
| EPI_ISL_2831155, EPI_ISL_2831156 | Lighthouse Lab in Milton Keynes | Wellcome Sanger Institute for the COVID-19 Genomics UK (COG-UK) Consortium | The Lighthouse Lab in Milton Keynes and Alex Alderton, Roberto Amato, Jeffrey Barrett, Sonia Goncalves, Ewan Harrison, David K. Jackson, Ian Johnston, Dominic Kwiatkowski, Cordelia Langford, John Sillitoe on behalf of the Wellcome Sanger Institute COVID-19 Surveillance Team |
| EPI_ISL_2831157 | Lighthouse Lab in Alderley Park | Wellcome Sanger Institute for the COVID-19 Genomics UK (COG-UK) Consortium | Jacquelyn Wynn, Mairead Hyland, The Lighthouse Lab in Alderley Park and Alex Alderton, Roberto Amato, Jeffrey Barrett, Sonia Goncalves, Ewan Harrison, David K. Jackson, Ian Johnston, Dominic Kwiatkowski, Cordelia Langford, John Sillitoe on behalf of the Wellcome Sanger Institute COVID-19 Surveillance Team |
| EPI_ISL_2831158 | Lighthouse Lab in Milton Keynes | Wellcome Sanger Institute for the COVID-19 Genomics UK (COG-UK) Consortium | The Lighthouse Lab in Milton Keynes and Alex Alderton, Roberto Amato, Jeffrey Barrett, Sonia Goncalves, Ewan Harrison, David K. Jackson, Ian Johnston, Dominic Kwiatkowski, Cordelia Langford, John Sillitoe on behalf of the Wellcome Sanger Institute COVID-19 Surveillance Team |
| EPI_ISL_2831159, EPI_ISL_2831160 | Lighthouse Lab in Glasgow | Wellcome Sanger Institute for the COVID-19 Genomics UK (COG-UK) Consortium | Harper VanSteenhouse, Yumi Kasai, David Gray, Carol Clugston, Anna Dominiczak and Alex Alderton, Roberto Amato, Jeffrey Barrett, Sonia Goncalves, Ewan Harrison, David K. Jackson, Ian Johnston, Dominic Kwiatkowski, Cordelia Langford, John Sillitoe on behalf of the Wellcome Sanger Institute COVID-19 Surveillance Team |
| EPI_ISL_2831161 | Lighthouse Lab in Alderley Park | Wellcome Sanger Institute for the COVID-19 Genomics UK (COG-UK) Consortium | Jacquelyn Wynn, Mairead Hyland, The Lighthouse Lab in Alderley Park and Alex Alderton, Roberto Amato, Jeffrey Barrett, Sonia Goncalves, Ewan Harrison, David K. Jackson, Ian Johnston, Dominic Kwiatkowski, Cordelia Langford, John Sillitoe on behalf of the Wellcome Sanger Institute COVID-19 Surveillance Team |
| EPI_ISL_2831162 | Lighthouse Lab in Milton Keynes | Wellcome Sanger Institute for the COVID-19 Genomics UK (COG-UK) Consortium | The Lighthouse Lab in Milton Keynes and Alex Alderton, Roberto Amato, Jeffrey Barrett, Sonia Goncalves, Ewan Harrison, David K. Jackson, Ian Johnston, Dominic Kwiatkowski, Cordelia Langford, John Sillitoe on behalf of the Wellcome Sanger Institute COVID-19 Surveillance Team |
| EPI_ISL_2831163 | Lighthouse Lab in Alderley Park | Wellcome Sanger Institute for the COVID-19 Genomics UK (COG-UK) Consortium | Jacquelyn Wynn, Mairead Hyland, The Lighthouse Lab in Alderley Park and Alex Alderton, Roberto Amato, Jeffrey Barrett, Sonia Goncalves, Ewan Harrison, David K. Jackson, Ian Johnston, Dominic Kwiatkowski, Cordelia Langford, John Sillitoe on behalf of the Wellcome Sanger Institute COVID-19 Surveillance Team |
| EPI_ISL_2831164 | Lighthouse Lab in Milton Keynes | Wellcome Sanger Institute for the COVID-19 Genomics UK (COG-UK) Consortium | The Lighthouse Lab in Milton Keynes and Alex Alderton, Roberto Amato, Jeffrey Barrett, Sonia Goncalves, Ewan Harrison, David K. Jackson, Ian Johnston, Dominic Kwiatkowski, Cordelia Langford, John Sillitoe on behalf of the Wellcome Sanger Institute COVID-19 Surveillance Team |
| EPI_ISL_2831165 | Lighthouse Lab in Glasgow | Wellcome Sanger Institute for the COVID-19 Genomics UK (COG-UK) Consortium | Harper VanSteenhouse, Yumi Kasai, David Gray, Carol Clugston, Anna Dominiczak and Alex Alderton, Roberto Amato, Jeffrey Barrett, Sonia Goncalves, Ewan Harrison, David K. Jackson, Ian Johnston, Dominic Kwiatkowski, Cordelia Langford, John Sillitoe on behalf of the Wellcome Sanger Institute COVID-19 Surveillance Team |
| EPI_ISL_2831166 | Lighthouse Lab in Alderley Park | Wellcome Sanger Institute for the COVID-19 Genomics UK (COG-UK) Consortium | Jacquelyn Wynn, Mairead Hyland, The Lighthouse Lab in Alderley Park and Alex Alderton, Roberto Amato, Jeffrey Barrett, Sonia Goncalves, Ewan Harrison, David K. Jackson, Ian Johnston, Dominic Kwiatkowski, Cordelia Langford, John Sillitoe on behalf of the Wellcome Sanger Institute COVID-19 Surveillance Team |
| EPI_ISL_2831167 | Lighthouse Laboratory Plymouth | Wellcome Sanger Institute for the COVID-19 Genomics UK (COG-UK) Consortium | Lighthouse Laboratory Plymouth and Alex Alderton, Roberto Amato, Jeffrey Barrett, Sonia Goncalves, Ewan Harrison, David K. Jackson, Ian Johnston, Dominic Kwiatkowski, Cordelia Langford, John Sillitoe on behalf of the Wellcome Sanger Institute COVID-19 Surveillance Team |
| EPI_ISL_2831168 | Lighthouse Lab in Alderley Park | Wellcome Sanger Institute for the COVID-19 Genomics UK (COG-UK) Consortium | Jacquelyn Wynn, Mairead Hyland, The Lighthouse Lab in Alderley Park and Alex Alderton, Roberto Amato, Jeffrey Barrett, Sonia Goncalves, Ewan Harrison, David K. Jackson, Ian Johnston, Dominic Kwiatkowski, Cordelia Langford, John Sillitoe on behalf of the Wellcome Sanger Institute COVID-19 Surveillance Team |
| EPI_ISL_2831169 | Lighthouse Laboratory Plymouth | Wellcome Sanger Institute for the COVID-19 Genomics UK (COG-UK) Consortium | Lighthouse Laboratory Plymouth and Alex Alderton, Roberto Amato, Jeffrey Barrett, Sonia Goncalves, Ewan Harrison, David K. Jackson, Ian Johnston, Dominic Kwiatkowski, Cordelia Langford, John Sillitoe on behalf of the Wellcome Sanger Institute COVID-19 Surveillance Team |
| EPI_ISL_2831170 | Lighthouse Lab in Glasgow | Wellcome Sanger Institute for the COVID-19 Genomics UK (COG-UK) Consortium | Harper VanSteenhouse, Yumi Kasai, David Gray, Carol Clugston, Anna Dominiczak and Alex Alderton, Roberto Amato, Jeffrey Barrett, Sonia Goncalves, Ewan Harrison, David K. Jackson, Ian Johnston, Dominic Kwiatkowski, Cordelia Langford, John Sillitoe on behalf of the Wellcome Sanger Institute COVID-19 Surveillance Team |
| EPI_ISL_2831171, EPI_ISL_2831172, EPI_ISL_2831173, EPI_ISL_2831174 | Lighthouse Laboratory Plymouth | Wellcome Sanger Institute for the COVID-19 Genomics UK (COG-UK) Consortium | Lighthouse Laboratory Plymouth and Alex Alderton, Roberto Amato, Jeffrey Barrett, Sonia Goncalves, Ewan Harrison, David K. Jackson, Ian Johnston, Dominic Kwiatkowski, Cordelia Langford, John Sillitoe on behalf of the Wellcome Sanger Institute COVID-19 Surveillance Team |
| EPI_ISL_2831175 | Lighthouse Lab in Milton Keynes | Wellcome Sanger Institute for the COVID-19 Genomics UK (COG-UK) Consortium | The Lighthouse Lab in Milton Keynes and Alex Alderton, Roberto Amato, Jeffrey Barrett, Sonia Goncalves, Ewan Harrison, David K. Jackson, Ian Johnston, Dominic Kwiatkowski, Cordelia Langford, John Sillitoe on behalf of the Wellcome Sanger Institute COVID-19 Surveillance Team |
| EPI_ISL_2831176 | Lighthouse Lab in Alderley Park | Wellcome Sanger Institute for the COVID-19 Genomics UK (COG-UK) Consortium | Jacquelyn Wynn, Mairead Hyland, The Lighthouse Lab in Alderley Park and Alex Alderton, Roberto Amato, Jeffrey Barrett, Sonia Goncalves, Ewan Harrison, David K. Jackson, Ian Johnston, Dominic Kwiatkowski, Cordelia Langford, John Sillitoe on behalf of the Wellcome Sanger Institute COVID-19 Surveillance Team |
| EPI_ISL_2831177 | Lighthouse Laboratory Plymouth | Wellcome Sanger Institute for the COVID-19 Genomics UK (COG-UK) Consortium | Lighthouse Laboratory Plymouth and Alex Alderton, Roberto Amato, Jeffrey Barrett, Sonia Goncalves, Ewan Harrison, David K. Jackson, Ian Johnston, Dominic Kwiatkowski, Cordelia Langford, John Sillitoe on behalf of the Wellcome Sanger Institute COVID-19 Surveillance Team |
| EPI_ISL_2831178 | Lighthouse Lab in Milton Keynes | Wellcome Sanger Institute for the COVID-19 Genomics UK (COG-UK) Consortium | The Lighthouse Lab in Milton Keynes and Alex Alderton, Roberto Amato, Jeffrey Barrett, Sonia Goncalves, Ewan Harrison, David K. Jackson, Ian Johnston, Dominic Kwiatkowski, Cordelia Langford, John Sillitoe on behalf of the Wellcome Sanger Institute COVID-19 Surveillance Team |
| EPI_ISL_2831179, EPI_ISL_2831180, EPI_ISL_2831181 | Lighthouse Laboratory Plymouth | Wellcome Sanger Institute for the COVID-19 Genomics UK (COG-UK) Consortium | Lighthouse Laboratory Plymouth and Alex Alderton, Roberto Amato, Jeffrey Barrett, Sonia Goncalves, Ewan Harrison, David K. Jackson, Ian Johnston, Dominic Kwiatkowski, Cordelia Langford, John Sillitoe on behalf of the Wellcome Sanger Institute COVID-19 Surveillance Team |
| EPI_ISL_2831182 | Lighthouse Lab in Glasgow | Wellcome Sanger Institute for the COVID-19 Genomics UK (COG-UK) Consortium | Harper VanSteenhouse, Yumi Kasai, David Gray, Carol Clugston, Anna Dominiczak and Alex Alderton, Roberto Amato, Jeffrey Barrett, Sonia Goncalves, Ewan Harrison, David K. Jackson, Ian Johnston, Dominic Kwiatkowski, Cordelia Langford, John Sillitoe on behalf of the Wellcome Sanger Institute COVID-19 Surveillance Team |
| EPI_ISL_2831183, EPI_ISL_2831184, EPI_ISL_2831185, EPI_ISL_2831186 | Lighthouse Lab in Milton Keynes | Wellcome Sanger Institute for the COVID-19 Genomics UK (COG-UK) Consortium | The Lighthouse Lab in Milton Keynes and Alex Alderton, Roberto Amato, Jeffrey Barrett, Sonia Goncalves, Ewan Harrison, David K. Jackson, Ian Johnston, Dominic Kwiatkowski, Cordelia Langford, John Sillitoe on behalf of the Wellcome Sanger Institute COVID-19 Surveillance Team |
| EPI_ISL_2831187 | Lighthouse Laboratory Plymouth | Wellcome Sanger Institute for the COVID-19 Genomics UK (COG-UK) Consortium | Lighthouse Laboratory Plymouth and Alex Alderton, Roberto Amato, Jeffrey Barrett, Sonia Goncalves, Ewan Harrison, David K. Jackson, Ian Johnston, Dominic Kwiatkowski, Cordelia Langford, John Sillitoe on behalf of the Wellcome Sanger Institute COVID-19 Surveillance Team |
| EPI_ISL_2831188 | Lighthouse Lab in Alderley Park | Wellcome Sanger Institute for the COVID-19 Genomics UK (COG-UK) Consortium | Jacquelyn Wynn, Mairead Hyland, The Lighthouse Lab in Alderley Park and Alex Alderton, Roberto Amato, Jeffrey Barrett, Sonia Goncalves, Ewan Harrison, David K. Jackson, Ian Johnston, Dominic Kwiatkowski, Cordelia Langford, John Sillitoe on behalf of the Wellcome Sanger Institute COVID-19 Surveillance Team |
| EPI_ISL_2831189, EPI_ISL_2831190 | Lighthouse Laboratory Plymouth | Wellcome Sanger Institute for the COVID-19 Genomics UK (COG-UK) Consortium | Lighthouse Laboratory Plymouth and Alex Alderton, Roberto Amato, Jeffrey Barrett, Sonia Goncalves, Ewan Harrison, David K. Jackson, Ian Johnston, Dominic Kwiatkowski, Cordelia Langford, John Sillitoe on behalf of the Wellcome Sanger Institute COVID-19 Surveillance Team |
| EPI_ISL_2831191, EPI_ISL_2831192 | Lighthouse Lab in Milton Keynes | Wellcome Sanger Institute for the COVID-19 Genomics UK (COG-UK) Consortium | The Lighthouse Lab in Milton Keynes and Alex Alderton, Roberto Amato, Jeffrey Barrett, Sonia Goncalves, Ewan Harrison, David K. Jackson, Ian Johnston, Dominic Kwiatkowski, Cordelia Langford, John Sillitoe on behalf of the Wellcome Sanger Institute COVID-19 Surveillance Team |
| EPI_ISL_2831193 | Lighthouse Lab in Glasgow | Wellcome Sanger Institute for the COVID-19 Genomics UK (COG-UK) Consortium | Harper VanSteenhouse, Yumi Kasai, David Gray, Carol Clugston, Anna Dominiczak and Alex Alderton, Roberto Amato, Jeffrey Barrett, Sonia Goncalves, Ewan Harrison, David K. Jackson, Ian Johnston, Dominic Kwiatkowski, Cordelia Langford, John Sillitoe on behalf of the Wellcome Sanger Institute COVID-19 Surveillance Team |
| EPI_ISL_2831194, EPI_ISL_2831195 | Lighthouse Lab in Milton Keynes | Wellcome Sanger Institute for the COVID-19 Genomics UK (COG-UK) Consortium | The Lighthouse Lab in Milton Keynes and Alex Alderton, Roberto Amato, Jeffrey Barrett, Sonia Goncalves, Ewan Harrison, David K. Jackson, Ian Johnston, Dominic Kwiatkowski, Cordelia Langford, John Sillitoe on behalf of the Wellcome Sanger Institute COVID-19 Surveillance Team |
| EPI_ISL_2831196 | Lighthouse Lab in Alderley Park | Wellcome Sanger Institute for the COVID-19 Genomics UK (COG-UK) Consortium | Jacquelyn Wynn, Mairead Hyland, The Lighthouse Lab in Alderley Park and Alex Alderton, Roberto Amato, Jeffrey Barrett, Sonia Goncalves, Ewan Harrison, David K. Jackson, Ian Johnston, Dominic Kwiatkowski, Cordelia Langford, John Sillitoe on behalf of the Wellcome Sanger Institute COVID-19 Surveillance Team |
| EPI_ISL_2831197, EPI_ISL_2831198 | Lighthouse Lab in Milton Keynes | Wellcome Sanger Institute for the COVID-19 Genomics UK (COG-UK) Consortium | The Lighthouse Lab in Milton Keynes and Alex Alderton, Roberto Amato, Jeffrey Barrett, Sonia Goncalves, Ewan Harrison, David K. Jackson, Ian Johnston, Dominic Kwiatkowski, Cordelia Langford, John Sillitoe on behalf of the Wellcome Sanger Institute COVID-19 Surveillance Team |
| EPI_ISL_2831199 | Lighthouse Lab in Glasgow | Wellcome Sanger Institute for the COVID-19 Genomics UK (COG-UK) Consortium | Harper VanSteenhouse, Yumi Kasai, David Gray, Carol Clugston, Anna Dominiczak and Alex Alderton, Roberto Amato, Jeffrey Barrett, Sonia Goncalves, Ewan Harrison, David K. Jackson, Ian Johnston, Dominic Kwiatkowski, Cordelia Langford, John Sillitoe on behalf of the Wellcome Sanger Institute COVID-19 Surveillance Team |
| EPI_ISL_2831200 | Lighthouse Laboratory Plymouth | Wellcome Sanger Institute for the COVID-19 Genomics UK (COG-UK) Consortium | Lighthouse Laboratory Plymouth and Alex Alderton, Roberto Amato, Jeffrey Barrett, Sonia Goncalves, Ewan Harrison, David K. Jackson, Ian Johnston, Dominic Kwiatkowski, Cordelia Langford, John Sillitoe on behalf of the Wellcome Sanger Institute COVID-19 Surveillance Team |
| EPI_ISL_2831201 | Lighthouse Lab in Glasgow | Wellcome Sanger Institute for the COVID-19 Genomics UK (COG-UK) Consortium | Harper VanSteenhouse, Yumi Kasai, David Gray, Carol Clugston, Anna Dominiczak and Alex Alderton, Roberto Amato, Jeffrey Barrett, Sonia Goncalves, Ewan Harrison, David K. Jackson, Ian Johnston, Dominic Kwiatkowski, Cordelia Langford, John Sillitoe on behalf of the Wellcome Sanger Institute COVID-19 Surveillance Team |
| EPI_ISL_2831202 | Lighthouse Lab in Milton Keynes | Wellcome Sanger Institute for the COVID-19 Genomics UK (COG-UK) Consortium | The Lighthouse Lab in Milton Keynes and Alex Alderton, Roberto Amato, Jeffrey Barrett, Sonia Goncalves, Ewan Harrison, David K. Jackson, Ian Johnston, Dominic Kwiatkowski, Cordelia Langford, John Sillitoe on behalf of the Wellcome Sanger Institute COVID-19 Surveillance Team |
| EPI_ISL_2831203 | Lighthouse Lab in Alderley Park | Wellcome Sanger Institute for the COVID-19 Genomics UK (COG-UK) Consortium | Jacquelyn Wynn, Mairead Hyland, The Lighthouse Lab in Alderley Park and Alex Alderton, Roberto Amato, Jeffrey Barrett, Sonia Goncalves, Ewan Harrison, David K. Jackson, Ian Johnston, Dominic Kwiatkowski, Cordelia Langford, John Sillitoe on behalf of the Wellcome Sanger Institute COVID-19 Surveillance Team |
| EPI_ISL_2831204 | Lighthouse Lab in Glasgow | Wellcome Sanger Institute for the COVID-19 Genomics UK (COG-UK) Consortium | Harper VanSteenhouse, Yumi Kasai, David Gray, Carol Clugston, Anna Dominiczak and Alex Alderton, Roberto Amato, Jeffrey Barrett, Sonia Goncalves, Ewan Harrison, David K. Jackson, Ian Johnston, Dominic Kwiatkowski, Cordelia Langford, John Sillitoe on behalf of the Wellcome Sanger Institute COVID-19 Surveillance Team |
| EPI_ISL_2831205 | Lighthouse Lab in Alderley Park | Wellcome Sanger Institute for the COVID-19 Genomics UK (COG-UK) Consortium | Jacquelyn Wynn, Mairead Hyland, The Lighthouse Lab in Alderley Park and Alex Alderton, Roberto Amato, Jeffrey Barrett, Sonia Goncalves, Ewan Harrison, David K. Jackson, Ian Johnston, Dominic Kwiatkowski, Cordelia Langford, John Sillitoe on behalf of the Wellcome Sanger Institute COVID-19 Surveillance Team |
| EPI_ISL_2831206 | Lighthouse Lab in Glasgow | Wellcome Sanger Institute for the COVID-19 Genomics UK (COG-UK) Consortium | Harper VanSteenhouse, Yumi Kasai, David Gray, Carol Clugston, Anna Dominiczak and Alex Alderton, Roberto Amato, Jeffrey Barrett, Sonia Goncalves, Ewan Harrison, David K. Jackson, Ian Johnston, Dominic Kwiatkowski, Cordelia Langford, John Sillitoe on behalf of the Wellcome Sanger Institute COVID-19 Surveillance Team |
| EPI_ISL_2831207 | Lighthouse Lab in Milton Keynes | Wellcome Sanger Institute for the COVID-19 Genomics UK (COG-UK) Consortium | The Lighthouse Lab in Milton Keynes and Alex Alderton, Roberto Amato, Jeffrey Barrett, Sonia Goncalves, Ewan Harrison, David K. Jackson, Ian Johnston, Dominic Kwiatkowski, Cordelia Langford, John Sillitoe on behalf of the Wellcome Sanger Institute COVID-19 Surveillance Team |
| EPI_ISL_2831208 | Lighthouse Lab in Glasgow | Wellcome Sanger Institute for the COVID-19 Genomics UK (COG-UK) Consortium | Harper VanSteenhouse, Yumi Kasai, David Gray, Carol Clugston, Anna Dominiczak and Alex Alderton, Roberto Amato, Jeffrey Barrett, Sonia Goncalves, Ewan Harrison, David K. Jackson, Ian Johnston, Dominic Kwiatkowski, Cordelia Langford, John Sillitoe on behalf of the Wellcome Sanger Institute COVID-19 Surveillance Team |
| EPI_ISL_2831209, EPI_ISL_2831210 | Lighthouse Laboratory Plymouth | Wellcome Sanger Institute for the COVID-19 Genomics UK (COG-UK) Consortium | Lighthouse Laboratory Plymouth and Alex Alderton, Roberto Amato, Jeffrey Barrett, Sonia Goncalves, Ewan Harrison, David K. Jackson, Ian Johnston, Dominic Kwiatkowski, Cordelia Langford, John Sillitoe on behalf of the Wellcome Sanger Institute COVID-19 Surveillance Team |
| EPI_ISL_2831211 | Lighthouse Lab in Alderley Park | Wellcome Sanger Institute for the COVID-19 Genomics UK (COG-UK) Consortium | Jacquelyn Wynn, Mairead Hyland, The Lighthouse Lab in Alderley Park and Alex Alderton, Roberto Amato, Jeffrey Barrett, Sonia Goncalves, Ewan Harrison, David K. Jackson, Ian Johnston, Dominic Kwiatkowski, Cordelia Langford, John Sillitoe on behalf of the Wellcome Sanger Institute COVID-19 Surveillance Team |
| EPI_ISL_2831212 | Lighthouse Lab in Glasgow | Wellcome Sanger Institute for the COVID-19 Genomics UK (COG-UK) Consortium | Harper VanSteenhouse, Yumi Kasai, David Gray, Carol Clugston, Anna Dominiczak and Alex Alderton, Roberto Amato, Jeffrey Barrett, Sonia Goncalves, Ewan Harrison, David K. Jackson, Ian Johnston, Dominic Kwiatkowski, Cordelia Langford, John Sillitoe on behalf of the Wellcome Sanger Institute COVID-19 Surveillance Team |

| | | | |
|---|---|---|---|
| EPI_ISL_2831213, EPI_ISL_2831214, EPI_ISL_2831215 | Lighthouse Laboratory Plymouth | Wellcome Sanger Institute for the COVID-19 Genomics UK (COG-UK) Consortium | Lighthouse Laboratory Plymouth and Alex Alderton, Roberto Amato, Jeffrey Barrett, Sonia Goncalves, Ewan Harrison, David K. Jackson, Ian Johnston, Dominic Kwiatkowski, Cordelia Langford, John Sillitoe on behalf of the Wellcome Sanger Institute COVID-19 Surveillance Team |
| EPI_ISL_2831216, EPI_ISL_2831217 | Lighthouse Lab in Milton Keynes | Wellcome Sanger Institute for the COVID-19 Genomics UK (COG-UK) Consortium | The Lighthouse Lab in Milton Keynes and Alex Alderton, Roberto Amato, Jeffrey Barrett, Sonia Goncalves, Ewan Harrison, David K. Jackson, Ian Johnston, Dominic Kwiatkowski, Cordelia Langford, John Sillitoe on behalf of the Wellcome Sanger Institute COVID-19 Surveillance Team |
| EPI_ISL_2831218 | Lighthouse Laboratory Plymouth | Wellcome Sanger Institute for the COVID-19 Genomics UK (COG-UK) Consortium | Lighthouse Laboratory Plymouth and Alex Alderton, Roberto Amato, Jeffrey Barrett, Sonia Goncalves, Ewan Harrison, David K. Jackson, Ian Johnston, Dominic Kwiatkowski, Cordelia Langford, John Sillitoe on behalf of the Wellcome Sanger Institute COVID-19 Surveillance Team |
| EPI_ISL_2831219 | Lighthouse Lab in Milton Keynes | Wellcome Sanger Institute for the COVID-19 Genomics UK (COG-UK) Consortium | The Lighthouse Lab in Milton Keynes and Alex Alderton, Roberto Amato, Jeffrey Barrett, Sonia Goncalves, Ewan Harrison, David K. Jackson, Ian Johnston, Dominic Kwiatkowski, Cordelia Langford, John Sillitoe on behalf of the Wellcome Sanger Institute COVID-19 Surveillance Team |
| EPI_ISL_2831220 | Lighthouse Lab in Glasgow | Wellcome Sanger Institute for the COVID-19 Genomics UK (COG-UK) Consortium | Harper VanSteenhouse, Yumi Kasai, David Gray, Carol Clugston, Anna Dominiczak and Alex Alderton, Roberto Amato, Jeffrey Barrett, Sonia Goncalves, Ewan Harrison, David K. Jackson, Ian Johnston, Dominic Kwiatkowski, Cordelia Langford, John Sillitoe on behalf of the Wellcome Sanger Institute COVID-19 Surveillance Team |
| EPI_ISL_2831221 | Lighthouse Laboratory Plymouth | Wellcome Sanger Institute for the COVID-19 Genomics UK (COG-UK) Consortium | Lighthouse Laboratory Plymouth and Alex Alderton, Roberto Amato, Jeffrey Barrett, Sonia Goncalves, Ewan Harrison, David K. Jackson, Ian Johnston, Dominic Kwiatkowski, Cordelia Langford, John Sillitoe on behalf of the Wellcome Sanger Institute COVID-19 Surveillance Team |
| EPI_ISL_2831222, EPI_ISL_2831223 | Lighthouse Lab in Alderley Park | Wellcome Sanger Institute for the COVID-19 Genomics UK (COG-UK) Consortium | Jacquelyn Wynn, Mairead Hyland, The Lighthouse Lab in Alderley Park and Alex Alderton, Roberto Amato, Jeffrey Barrett, Sonia Goncalves, Ewan Harrison, David K. Jackson, Ian Johnston, Dominic Kwiatkowski, Cordelia Langford, John Sillitoe on behalf of the Wellcome Sanger Institute COVID-19 Surveillance Team |
| EPI_ISL_2831224 | Lighthouse Lab in Milton Keynes | Wellcome Sanger Institute for the COVID-19 Genomics UK (COG-UK) Consortium | The Lighthouse Lab in Milton Keynes and Alex Alderton, Roberto Amato, Jeffrey Barrett, Sonia Goncalves, Ewan Harrison, David K. Jackson, Ian Johnston, Dominic Kwiatkowski, Cordelia Langford, John Sillitoe on behalf of the Wellcome Sanger Institute COVID-19 Surveillance Team |
| EPI_ISL_2831225 | Lighthouse Lab in Glasgow | Wellcome Sanger Institute for the COVID-19 Genomics UK (COG-UK) Consortium | Harper VanSteenhouse, Yumi Kasai, David Gray, Carol Clugston, Anna Dominiczak and Alex Alderton, Roberto Amato, Jeffrey Barrett, Sonia Goncalves, Ewan Harrison, David K. Jackson, Ian Johnston, Dominic Kwiatkowski, Cordelia Langford, John Sillitoe on behalf of the Wellcome Sanger Institute COVID-19 Surveillance Team |
| EPI_ISL_2831226 | Lighthouse Lab in Milton Keynes | Wellcome Sanger Institute for the COVID-19 Genomics UK (COG-UK) Consortium | The Lighthouse Lab in Milton Keynes and Alex Alderton, Roberto Amato, Jeffrey Barrett, Sonia Goncalves, Ewan Harrison, David K. Jackson, Ian Johnston, Dominic Kwiatkowski, Cordelia Langford, John Sillitoe on behalf of the Wellcome Sanger Institute COVID-19 Surveillance Team |
| EPI_ISL_2831227 | Lighthouse Lab in Glasgow | Wellcome Sanger Institute for the COVID-19 Genomics UK (COG-UK) Consortium | Harper VanSteenhouse, Yumi Kasai, David Gray, Carol Clugston, Anna Dominiczak and Alex Alderton, Roberto Amato, Jeffrey Barrett, Sonia Goncalves, Ewan Harrison, David K. Jackson, Ian Johnston, Dominic Kwiatkowski, Cordelia Langford, John Sillitoe on behalf of the Wellcome Sanger Institute COVID-19 Surveillance Team |
| EPI_ISL_2831228 | Lighthouse Laboratory Plymouth | Wellcome Sanger Institute for the COVID-19 Genomics UK (COG-UK) Consortium | Lighthouse Laboratory Plymouth and Alex Alderton, Roberto Amato, Jeffrey Barrett, Sonia Goncalves, Ewan Harrison, David K. Jackson, Ian Johnston, Dominic Kwiatkowski, Cordelia Langford, John Sillitoe on behalf of the Wellcome Sanger Institute COVID-19 Surveillance Team |
| EPI_ISL_2831229 | Lighthouse Lab in Glasgow | Wellcome Sanger Institute for the COVID-19 Genomics UK (COG-UK) Consortium | Harper VanSteenhouse, Yumi Kasai, David Gray, Carol Clugston, Anna Dominiczak and Alex Alderton, Roberto Amato, Jeffrey Barrett, Sonia Goncalves, Ewan Harrison, David K. Jackson, Ian Johnston, Dominic Kwiatkowski, Cordelia Langford, John Sillitoe on behalf of the Wellcome Sanger Institute COVID-19 Surveillance Team |
| EPI_ISL_2831230 | Lighthouse Laboratory Plymouth | Wellcome Sanger Institute for the COVID-19 Genomics UK (COG-UK) Consortium | Lighthouse Laboratory Plymouth and Alex Alderton, Roberto Amato, Jeffrey Barrett, Sonia Goncalves, Ewan Harrison, David K. Jackson, Ian Johnston, Dominic Kwiatkowski, Cordelia Langford, John Sillitoe on behalf of the Wellcome Sanger Institute COVID-19 Surveillance Team |
| EPI_ISL_2831231 | Lighthouse Lab in Alderley Park | Wellcome Sanger Institute for the COVID-19 Genomics UK (COG-UK) Consortium | Jacquelyn Wynn, Mairead Hyland, The Lighthouse Lab in Alderley Park and Alex Alderton, Roberto Amato, Jeffrey Barrett, Sonia Goncalves, Ewan Harrison, David K. Jackson, Ian Johnston, Dominic Kwiatkowski, Cordelia Langford, John Sillitoe on behalf of the Wellcome Sanger Institute COVID-19 Surveillance Team |
| EPI_ISL_2831232 | Lighthouse Lab in Milton Keynes | Wellcome Sanger Institute for the COVID-19 Genomics UK (COG-UK) Consortium | The Lighthouse Lab in Milton Keynes and Alex Alderton, Roberto Amato, Jeffrey Barrett, Sonia Goncalves, Ewan Harrison, David K. Jackson, Ian Johnston, Dominic Kwiatkowski, Cordelia Langford, John Sillitoe on behalf of the Wellcome Sanger Institute COVID-19 Surveillance Team |
| EPI_ISL_2831233, EPI_ISL_2831234 | Lighthouse Lab in Glasgow | Wellcome Sanger Institute for the COVID-19 Genomics UK (COG-UK) Consortium | Harper VanSteenhouse, Yumi Kasai, David Gray, Carol Clugston, Anna Dominiczak and Alex Alderton, Roberto Amato, Jeffrey Barrett, Sonia Goncalves, Ewan Harrison, David K. Jackson, Ian Johnston, Dominic Kwiatkowski, Cordelia Langford, John Sillitoe on behalf of the Wellcome Sanger Institute COVID-19 Surveillance Team |
| EPI_ISL_2831235, EPI_ISL_2831236 | Lighthouse Lab in Milton Keynes | Wellcome Sanger Institute for the COVID-19 Genomics UK (COG-UK) Consortium | The Lighthouse Lab in Milton Keynes and Alex Alderton, Roberto Amato, Jeffrey Barrett, Sonia Goncalves, Ewan Harrison, David K. Jackson, Ian Johnston, Dominic Kwiatkowski, Cordelia Langford, John Sillitoe on behalf of the Wellcome Sanger Institute COVID-19 Surveillance Team |
| EPI_ISL_2831237 | Lighthouse Lab in Glasgow | Wellcome Sanger Institute for the COVID-19 Genomics UK (COG-UK) Consortium | Harper VanSteenhouse, Yumi Kasai, David Gray, Carol Clugston, Anna Dominiczak and Alex Alderton, Roberto Amato, Jeffrey Barrett, Sonia Goncalves, Ewan Harrison, David K. Jackson, Ian Johnston, Dominic Kwiatkowski, Cordelia Langford, John Sillitoe on behalf of the Wellcome Sanger Institute COVID-19 Surveillance Team |
| EPI_ISL_2831238 | Lighthouse Lab in Milton Keynes | Wellcome Sanger Institute for the COVID-19 Genomics UK (COG-UK) Consortium | The Lighthouse Lab in Milton Keynes and Alex Alderton, Roberto Amato, Jeffrey Barrett, Sonia Goncalves, Ewan Harrison, David K. Jackson, Ian Johnston, Dominic Kwiatkowski, Cordelia Langford, John Sillitoe on behalf of the Wellcome Sanger Institute COVID-19 Surveillance Team |
| EPI_ISL_2831239, EPI_ISL_2831240 | Lighthouse Lab in Alderley Park | Wellcome Sanger Institute for the COVID-19 Genomics UK (COG-UK) Consortium | Jacquelyn Wynn, Mairead Hyland, The Lighthouse Lab in Alderley Park and Alex Alderton, Roberto Amato, Jeffrey Barrett, Sonia Goncalves, Ewan Harrison, David K. Jackson, Ian Johnston, Dominic Kwiatkowski, Cordelia Langford, John Sillitoe on behalf of the Wellcome Sanger Institute COVID-19 Surveillance Team |
| EPI_ISL_2831241 | Lighthouse Lab in Glasgow | Wellcome Sanger Institute for the COVID-19 Genomics UK (COG-UK) Consortium | Harper VanSteenhouse, Yumi Kasai, David Gray, Carol Clugston, Anna Dominiczak and Alex Alderton, Roberto Amato, Jeffrey Barrett, Sonia Goncalves, Ewan Harrison, David K. Jackson, Ian Johnston, Dominic Kwiatkowski, Cordelia Langford, John Sillitoe on behalf of the Wellcome Sanger Institute COVID-19 Surveillance Team |
| EPI_ISL_2831242 | Lighthouse Lab in Alderley Park | Wellcome Sanger Institute for the COVID-19 Genomics UK (COG-UK) Consortium | Jacquelyn Wynn, Mairead Hyland, The Lighthouse Lab in Alderley Park and Alex Alderton, Roberto Amato, Jeffrey Barrett, Sonia Goncalves, Ewan Harrison, David K. Jackson, Ian Johnston, Dominic Kwiatkowski, Cordelia Langford, John Sillitoe on behalf of the Wellcome Sanger Institute COVID-19 Surveillance Team |
| EPI_ISL_2831243 | Lighthouse Laboratory Plymouth | Wellcome Sanger Institute for the COVID-19 Genomics UK (COG-UK) Consortium | Lighthouse Laboratory Plymouth and Alex Alderton, Roberto Amato, Jeffrey Barrett, Sonia Goncalves, Ewan Harrison, David K. Jackson, Ian Johnston, Dominic Kwiatkowski, Cordelia Langford, John Sillitoe on behalf of the Wellcome Sanger Institute COVID-19 Surveillance Team |
| EPI_ISL_2831244 | Lighthouse Lab in Milton Keynes | Wellcome Sanger Institute for the COVID-19 Genomics UK (COG-UK) Consortium | The Lighthouse Lab in Milton Keynes and Alex Alderton, Roberto Amato, Jeffrey Barrett, Sonia Goncalves, Ewan Harrison, David K. Jackson, Ian Johnston, Dominic Kwiatkowski, Cordelia Langford, John Sillitoe on behalf of the Wellcome Sanger Institute COVID-19 Surveillance Team |
| EPI_ISL_2831245 | Lighthouse Lab in Glasgow | Wellcome Sanger Institute for the COVID-19 Genomics UK (COG-UK) Consortium | Harper VanSteenhouse, Yumi Kasai, David Gray, Carol Clugston, Anna Dominiczak and Alex Alderton, Roberto Amato, Jeffrey Barrett, Sonia Goncalves, Ewan Harrison, David K. Jackson, Ian Johnston, Dominic Kwiatkowski, Cordelia Langford, John Sillitoe on behalf of the Wellcome Sanger Institute COVID-19 Surveillance Team |
| EPI_ISL_2831246 | Lighthouse Lab in Alderley Park | Wellcome Sanger Institute for the COVID-19 Genomics UK (COG-UK) Consortium | Jacquelyn Wynn, Mairead Hyland, The Lighthouse Lab in Alderley Park and Alex Alderton, Roberto Amato, Jeffrey Barrett, Sonia Goncalves, Ewan Harrison, David K. Jackson, Ian Johnston, Dominic Kwiatkowski, Cordelia Langford, John Sillitoe on behalf of the Wellcome Sanger Institute COVID-19 Surveillance Team |
| EPI_ISL_2831247 | Lighthouse Lab in Milton Keynes | Wellcome Sanger Institute for the COVID-19 Genomics UK (COG-UK) Consortium | The Lighthouse Lab in Milton Keynes and Alex Alderton, Roberto Amato, Jeffrey Barrett, Sonia Goncalves, Ewan Harrison, David K. Jackson, Ian Johnston, Dominic Kwiatkowski, Cordelia Langford, John Sillitoe on behalf of the Wellcome Sanger Institute COVID-19 Surveillance Team |
| EPI_ISL_2831248 | Lighthouse Lab in Alderley Park | Wellcome Sanger Institute for the COVID-19 Genomics UK (COG-UK) Consortium | Jacquelyn Wynn, Mairead Hyland, The Lighthouse Lab in Alderley Park and Alex Alderton, Roberto Amato, Jeffrey Barrett, Sonia Goncalves, Ewan Harrison, David K. Jackson, Ian Johnston, Dominic Kwiatkowski, Cordelia Langford, John Sillitoe on behalf of the Wellcome Sanger Institute COVID-19 Surveillance Team |
| EPI_ISL_2831249 | Lighthouse Lab in Milton Keynes | Wellcome Sanger Institute for the COVID-19 Genomics UK (COG-UK) Consortium | The Lighthouse Lab in Milton Keynes and Alex Alderton, Roberto Amato, Jeffrey Barrett, Sonia Goncalves, Ewan Harrison, David K. Jackson, Ian Johnston, Dominic Kwiatkowski, Cordelia Langford, John Sillitoe on behalf of the Wellcome Sanger Institute COVID-19 Surveillance Team |
| EPI_ISL_2831250 | Lighthouse Laboratory Plymouth | Wellcome Sanger Institute for the COVID-19 Genomics UK (COG-UK) Consortium | Lighthouse Laboratory Plymouth and Alex Alderton, Roberto Amato, Jeffrey Barrett, Sonia Goncalves, Ewan Harrison, David K. Jackson, Ian Johnston, Dominic Kwiatkowski, Cordelia Langford, John Sillitoe on behalf of the Wellcome Sanger Institute COVID-19 Surveillance Team |
| EPI_ISL_2831251, EPI_ISL_2831252 | Lighthouse Lab in Glasgow | Wellcome Sanger Institute for the COVID-19 Genomics UK (COG-UK) Consortium | Harper VanSteenhouse, Yumi Kasai, David Gray, Carol Clugston, Anna Dominiczak and Alex Alderton, Roberto Amato, Jeffrey Barrett, Sonia Goncalves, Ewan Harrison, David K. Jackson, Ian Johnston, Dominic Kwiatkowski, Cordelia Langford, John Sillitoe on behalf of the Wellcome Sanger Institute COVID-19 Surveillance Team |
| EPI_ISL_2831253 | Lighthouse Laboratory Plymouth | Wellcome Sanger Institute for the COVID-19 Genomics UK (COG-UK) Consortium | Lighthouse Laboratory Plymouth and Alex Alderton, Roberto Amato, Jeffrey Barrett, Sonia Goncalves, Ewan Harrison, David K. Jackson, Ian Johnston, Dominic Kwiatkowski, Cordelia Langford, John Sillitoe on behalf of the Wellcome Sanger Institute COVID-19 Surveillance Team |
| EPI_ISL_2831254 | Lighthouse Lab in Milton Keynes | Wellcome Sanger Institute for the COVID-19 Genomics UK (COG-UK) Consortium | The Lighthouse Lab in Milton Keynes and Alex Alderton, Roberto Amato, Jeffrey Barrett, Sonia Goncalves, Ewan Harrison, David K. Jackson, Ian Johnston, Dominic Kwiatkowski, Cordelia Langford, John Sillitoe on behalf of the Wellcome Sanger Institute COVID-19 Surveillance Team |
| EPI_ISL_2831255 | Lighthouse Lab in Glasgow | Wellcome Sanger Institute for the COVID-19 Genomics UK (COG-UK) Consortium | Harper VanSteenhouse, Yumi Kasai, David Gray, Carol Clugston, Anna Dominiczak and Alex Alderton, Roberto Amato, Jeffrey Barrett, Sonia Goncalves, Ewan Harrison, David K. Jackson, Ian Johnston, Dominic Kwiatkowski, Cordelia Langford, John Sillitoe on behalf of the Wellcome Sanger Institute COVID-19 Surveillance Team |
| EPI_ISL_2831256 | Lighthouse Lab in Alderley Park | Wellcome Sanger Institute for the COVID-19 Genomics UK (COG-UK) Consortium | Jacquelyn Wynn, Mairead Hyland, The Lighthouse Lab in Alderley Park and Alex Alderton, Roberto Amato, Jeffrey Barrett, Sonia Goncalves, Ewan Harrison, David K. Jackson, Ian Johnston, Dominic Kwiatkowski, Cordelia Langford, John Sillitoe on behalf of the Wellcome Sanger Institute COVID-19 Surveillance Team |
| EPI_ISL_2831257 | Lighthouse Lab in Milton Keynes | Wellcome Sanger Institute for the COVID-19 Genomics UK (COG-UK) Consortium | The Lighthouse Lab in Milton Keynes and Alex Alderton, Roberto Amato, Jeffrey Barrett, Sonia Goncalves, Ewan Harrison, David K. Jackson, Ian Johnston, Dominic Kwiatkowski, Cordelia Langford, John Sillitoe on behalf of the Wellcome Sanger Institute COVID-19 Surveillance Team |
| EPI_ISL_2831258 | Lighthouse Lab in Alderley Park | Wellcome Sanger Institute for the COVID-19 Genomics UK (COG-UK) Consortium | Jacquelyn Wynn, Mairead Hyland, The Lighthouse Lab in Alderley Park and Alex Alderton, Roberto Amato, Jeffrey Barrett, Sonia Goncalves, Ewan Harrison, David K. Jackson, Ian Johnston, Dominic Kwiatkowski, Cordelia Langford, John Sillitoe on behalf of the Wellcome Sanger Institute COVID-19 Surveillance Team |
| EPI_ISL_2831259 | Lighthouse Laboratory Plymouth | Wellcome Sanger Institute for the COVID-19 Genomics UK (COG-UK) Consortium | Lighthouse Laboratory Plymouth and Alex Alderton, Roberto Amato, Jeffrey Barrett, Sonia Goncalves, Ewan Harrison, David K. Jackson, Ian Johnston, Dominic Kwiatkowski, Cordelia Langford, John Sillitoe on behalf of the Wellcome Sanger Institute COVID-19 Surveillance Team |
| EPI_ISL_2831260 | Lighthouse Lab in Alderley Park | Wellcome Sanger Institute for the COVID-19 Genomics UK (COG-UK) Consortium | Jacquelyn Wynn, Mairead Hyland, The Lighthouse Lab in Alderley Park and Alex Alderton, Roberto Amato, Jeffrey Barrett, Sonia Goncalves, Ewan Harrison, David K. Jackson, Ian Johnston, Dominic Kwiatkowski, Cordelia Langford, John Sillitoe on behalf of the Wellcome Sanger Institute COVID-19 Surveillance Team |
| EPI_ISL_2831261, EPI_ISL_2831262 | Lighthouse Laboratory Plymouth | Wellcome Sanger Institute for the COVID-19 Genomics UK (COG-UK) Consortium | Lighthouse Laboratory Plymouth and Alex Alderton, Roberto Amato, Jeffrey Barrett, Sonia Goncalves, Ewan Harrison, David K. Jackson, Ian Johnston, Dominic Kwiatkowski, Cordelia Langford, John Sillitoe on behalf of the Wellcome Sanger Institute COVID-19 Surveillance Team |
| EPI_ISL_2831263 | Lighthouse Lab in Milton Keynes | Wellcome Sanger Institute for the COVID-19 Genomics UK (COG-UK) Consortium | The Lighthouse Lab in Milton Keynes and Alex Alderton, Roberto Amato, Jeffrey Barrett, Sonia Goncalves, Ewan Harrison, David K. Jackson, Ian Johnston, Dominic Kwiatkowski, Cordelia Langford, John Sillitoe on behalf of the Wellcome Sanger Institute COVID-19 Surveillance Team |
| EPI_ISL_2831264 | Lighthouse Laboratory Plymouth | Wellcome Sanger Institute for the COVID-19 Genomics UK (COG-UK) Consortium | Lighthouse Laboratory Plymouth and Alex Alderton, Roberto Amato, Jeffrey Barrett, Sonia Goncalves, Ewan Harrison, David K. Jackson, Ian Johnston, Dominic Kwiatkowski, Cordelia Langford, John Sillitoe on behalf of the Wellcome Sanger Institute COVID-19 Surveillance Team |
| EPI_ISL_2831265 | Lighthouse Lab in Glasgow | Wellcome Sanger Institute for the COVID-19 Genomics UK (COG-UK) Consortium | Harper VanSteenhouse, Yumi Kasai, David Gray, Carol Clugston, Anna Dominiczak and Alex Alderton, Roberto Amato, Jeffrey Barrett, Sonia Goncalves, Ewan Harrison, David K. Jackson, Ian Johnston, Dominic Kwiatkowski, Cordelia Langford, John Sillitoe on behalf of the Wellcome Sanger Institute COVID-19 Surveillance Team |
| EPI_ISL_2831266 | Lighthouse Lab in Milton Keynes | Wellcome Sanger Institute for the COVID-19 Genomics UK (COG-UK) Consortium | The Lighthouse Lab in Milton Keynes and Alex Alderton, Roberto Amato, Jeffrey Barrett, Sonia Goncalves, Ewan Harrison, David K. Jackson, Ian Johnston, Dominic Kwiatkowski, Cordelia Langford, John Sillitoe on behalf of the Wellcome Sanger Institute COVID-19 Surveillance Team |

| | | | |
|---|---|---|---|
| EPI_ISL_2831267 | Lighthouse Lab in Glasgow | Wellcome Sanger Institute for the COVID-19 Genomics UK (COG-UK) Consortium | Harper VanSteenhouse, Yumi Kasai, David Gray, Carol Clugston, Anna Dominiczak and Alex Alderton, Roberto Amato, Jeffrey Barrett, Sonia Goncalves, Ewan Harrison, David K. Jackson, Ian Johnston, Dominic Kwiatkowski, Cordelia Langford, John Sillitoe on behalf of the Wellcome Sanger Institute COVID-19 Surveillance Team |
| EPI_ISL_2831268, EPI_ISL_2831269 | Lighthouse Laboratory Plymouth | Wellcome Sanger Institute for the COVID-19 Genomics UK (COG-UK) Consortium | Lighthouse Laboratory Plymouth and Alex Alderton, Roberto Amato, Jeffrey Barrett, Sonia Goncalves, Ewan Harrison, David K. Jackson, Ian Johnston, Dominic Kwiatkowski, Cordelia Langford, John Sillitoe on behalf of the Wellcome Sanger Institute COVID-19 Surveillance Team |
| EPI_ISL_2831270 | Lighthouse Lab in Glasgow | Wellcome Sanger Institute for the COVID-19 Genomics UK (COG-UK) Consortium | Harper VanSteenhouse, Yumi Kasai, David Gray, Carol Clugston, Anna Dominiczak and Alex Alderton, Roberto Amato, Jeffrey Barrett, Sonia Goncalves, Ewan Harrison, David K. Jackson, Ian Johnston, Dominic Kwiatkowski, Cordelia Langford, John Sillitoe on behalf of the Wellcome Sanger Institute COVID-19 Surveillance Team |
| EPI_ISL_2831271 | Lighthouse Laboratory Plymouth | Wellcome Sanger Institute for the COVID-19 Genomics UK (COG-UK) Consortium | Lighthouse Laboratory Plymouth and Alex Alderton, Roberto Amato, Jeffrey Barrett, Sonia Goncalves, Ewan Harrison, David K. Jackson, Ian Johnston, Dominic Kwiatkowski, Cordelia Langford, John Sillitoe on behalf of the Wellcome Sanger Institute COVID-19 Surveillance Team |
| EPI_ISL_2831272 | Lighthouse Lab in Milton Keynes | Wellcome Sanger Institute for the COVID-19 Genomics UK (COG-UK) Consortium | The Lighthouse Lab in Milton Keynes and Alex Alderton, Roberto Amato, Jeffrey Barrett, Sonia Goncalves, Ewan Harrison, David K. Jackson, Ian Johnston, Dominic Kwiatkowski, Cordelia Langford, John Sillitoe on behalf of the Wellcome Sanger Institute COVID-19 Surveillance Team |
| EPI_ISL_2831273 | Lighthouse Laboratory Plymouth | Wellcome Sanger Institute for the COVID-19 Genomics UK (COG-UK) Consortium | Lighthouse Laboratory Plymouth and Alex Alderton, Roberto Amato, Jeffrey Barrett, Sonia Goncalves, Ewan Harrison, David K. Jackson, Ian Johnston, Dominic Kwiatkowski, Cordelia Langford, John Sillitoe on behalf of the Wellcome Sanger Institute COVID-19 Surveillance Team |
| EPI_ISL_2831274 | Lighthouse Lab in Milton Keynes | Wellcome Sanger Institute for the COVID-19 Genomics UK (COG-UK) Consortium | The Lighthouse Lab in Milton Keynes and Alex Alderton, Roberto Amato, Jeffrey Barrett, Sonia Goncalves, Ewan Harrison, David K. Jackson, Ian Johnston, Dominic Kwiatkowski, Cordelia Langford, John Sillitoe on behalf of the Wellcome Sanger Institute COVID-19 Surveillance Team |
| EPI_ISL_2831275 | Lighthouse Lab in Alderley Park | Wellcome Sanger Institute for the COVID-19 Genomics UK (COG-UK) Consortium | Jacquelyn Wynn, Mairead Hyland, The Lighthouse Lab in Alderley Park and Alex Alderton, Roberto Amato, Jeffrey Barrett, Sonia Goncalves, Ewan Harrison, David K. Jackson, Ian Johnston, Dominic Kwiatkowski, Cordelia Langford, John Sillitoe on behalf of the Wellcome Sanger Institute COVID-19 Surveillance Team |
| EPI_ISL_2831276, EPI_ISL_2831277 | Lighthouse Laboratory Plymouth | Wellcome Sanger Institute for the COVID-19 Genomics UK (COG-UK) Consortium | Lighthouse Laboratory Plymouth and Alex Alderton, Roberto Amato, Jeffrey Barrett, Sonia Goncalves, Ewan Harrison, David K. Jackson, Ian Johnston, Dominic Kwiatkowski, Cordelia Langford, John Sillitoe on behalf of the Wellcome Sanger Institute COVID-19 Surveillance Team |
| EPI_ISL_2831278 | Lighthouse Lab in Glasgow | Wellcome Sanger Institute for the COVID-19 Genomics UK (COG-UK) Consortium | Harper VanSteenhouse, Yumi Kasai, David Gray, Carol Clugston, Anna Dominiczak and Alex Alderton, Roberto Amato, Jeffrey Barrett, Sonia Goncalves, Ewan Harrison, David K. Jackson, Ian Johnston, Dominic Kwiatkowski, Cordelia Langford, John Sillitoe on behalf of the Wellcome Sanger Institute COVID-19 Surveillance Team |
| EPI_ISL_2831279, EPI_ISL_2831280 | Lighthouse Lab in Milton Keynes | Wellcome Sanger Institute for the COVID-19 Genomics UK (COG-UK) Consortium | The Lighthouse Lab in Milton Keynes and Alex Alderton, Roberto Amato, Jeffrey Barrett, Sonia Goncalves, Ewan Harrison, David K. Jackson, Ian Johnston, Dominic Kwiatkowski, Cordelia Langford, John Sillitoe on behalf of the Wellcome Sanger Institute COVID-19 Surveillance Team |
| EPI_ISL_2831281 | Lighthouse Laboratory Plymouth | Wellcome Sanger Institute for the COVID-19 Genomics UK (COG-UK) Consortium | Lighthouse Laboratory Plymouth and Alex Alderton, Roberto Amato, Jeffrey Barrett, Sonia Goncalves, Ewan Harrison, David K. Jackson, Ian Johnston, Dominic Kwiatkowski, Cordelia Langford, John Sillitoe on behalf of the Wellcome Sanger Institute COVID-19 Surveillance Team |
| EPI_ISL_2831282 | Lighthouse Lab in Alderley Park | Wellcome Sanger Institute for the COVID-19 Genomics UK (COG-UK) Consortium | Jacquelyn Wynn, Mairead Hyland, The Lighthouse Lab in Alderley Park and Alex Alderton, Roberto Amato, Jeffrey Barrett, Sonia Goncalves, Ewan Harrison, David K. Jackson, Ian Johnston, Dominic Kwiatkowski, Cordelia Langford, John Sillitoe on behalf of the Wellcome Sanger Institute COVID-19 Surveillance Team |
| EPI_ISL_2831283, EPI_ISL_2831284, EPI_ISL_2831285 | Lighthouse Laboratory Plymouth | Wellcome Sanger Institute for the COVID-19 Genomics UK (COG-UK) Consortium | Lighthouse Laboratory Plymouth and Alex Alderton, Roberto Amato, Jeffrey Barrett, Sonia Goncalves, Ewan Harrison, David K. Jackson, Ian Johnston, Dominic Kwiatkowski, Cordelia Langford, John Sillitoe on behalf of the Wellcome Sanger Institute COVID-19 Surveillance Team |
| EPI_ISL_2831286 | Lighthouse Lab in Glasgow | Wellcome Sanger Institute for the COVID-19 Genomics UK (COG-UK) Consortium | Harper VanSteenhouse, Yumi Kasai, David Gray, Carol Clugston, Anna Dominiczak and Alex Alderton, Roberto Amato, Jeffrey Barrett, Sonia Goncalves, Ewan Harrison, David K. Jackson, Ian Johnston, Dominic Kwiatkowski, Cordelia Langford, John Sillitoe on behalf of the Wellcome Sanger Institute COVID-19 Surveillance Team |
| EPI_ISL_2831287 | Lighthouse Lab in Milton Keynes | Wellcome Sanger Institute for the COVID-19 Genomics UK (COG-UK) Consortium | The Lighthouse Lab in Milton Keynes and Alex Alderton, Roberto Amato, Jeffrey Barrett, Sonia Goncalves, Ewan Harrison, David K. Jackson, Ian Johnston, Dominic Kwiatkowski, Cordelia Langford, John Sillitoe on behalf of the Wellcome Sanger Institute COVID-19 Surveillance Team |
| EPI_ISL_2831288 | Lighthouse Lab in Alderley Park | Wellcome Sanger Institute for the COVID-19 Genomics UK (COG-UK) Consortium | Jacquelyn Wynn, Mairead Hyland, The Lighthouse Lab in Alderley Park and Alex Alderton, Roberto Amato, Jeffrey Barrett, Sonia Goncalves, Ewan Harrison, David K. Jackson, Ian Johnston, Dominic Kwiatkowski, Cordelia Langford, John Sillitoe on behalf of the Wellcome Sanger Institute COVID-19 Surveillance Team |
| EPI_ISL_2831289 | Lighthouse Laboratory Plymouth | Wellcome Sanger Institute for the COVID-19 Genomics UK (COG-UK) Consortium | Lighthouse Laboratory Plymouth and Alex Alderton, Roberto Amato, Jeffrey Barrett, Sonia Goncalves, Ewan Harrison, David K. Jackson, Ian Johnston, Dominic Kwiatkowski, Cordelia Langford, John Sillitoe on behalf of the Wellcome Sanger Institute COVID-19 Surveillance Team |
| EPI_ISL_2831290 | Lighthouse Lab in Glasgow | Wellcome Sanger Institute for the COVID-19 Genomics UK (COG-UK) Consortium | Harper VanSteenhouse, Yumi Kasai, David Gray, Carol Clugston, Anna Dominiczak and Alex Alderton, Roberto Amato, Jeffrey Barrett, Sonia Goncalves, Ewan Harrison, David K. Jackson, Ian Johnston, Dominic Kwiatkowski, Cordelia Langford, John Sillitoe on behalf of the Wellcome Sanger Institute COVID-19 Surveillance Team |
| EPI_ISL_2831291, EPI_ISL_2831292, EPI_ISL_2831293 | Lighthouse Laboratory Plymouth | Wellcome Sanger Institute for the COVID-19 Genomics UK (COG-UK) Consortium | Lighthouse Laboratory Plymouth and Alex Alderton, Roberto Amato, Jeffrey Barrett, Sonia Goncalves, Ewan Harrison, David K. Jackson, Ian Johnston, Dominic Kwiatkowski, Cordelia Langford, John Sillitoe on behalf of the Wellcome Sanger Institute COVID-19 Surveillance Team |
| EPI_ISL_2831294 | Lighthouse Lab in Glasgow | Wellcome Sanger Institute for the COVID-19 Genomics UK (COG-UK) Consortium | Harper VanSteenhouse, Yumi Kasai, David Gray, Carol Clugston, Anna Dominiczak and Alex Alderton, Roberto Amato, Jeffrey Barrett, Sonia Goncalves, Ewan Harrison, David K. Jackson, Ian Johnston, Dominic Kwiatkowski, Cordelia Langford, John Sillitoe on behalf of the Wellcome Sanger Institute COVID-19 Surveillance Team |
| EPI_ISL_2831295 | Lighthouse Lab in Alderley Park | Wellcome Sanger Institute for the COVID-19 Genomics UK (COG-UK) Consortium | Jacquelyn Wynn, Mairead Hyland, The Lighthouse Lab in Alderley Park and Alex Alderton, Roberto Amato, Jeffrey Barrett, Sonia Goncalves, Ewan Harrison, David K. Jackson, Ian Johnston, Dominic Kwiatkowski, Cordelia Langford, John Sillitoe on behalf of the Wellcome Sanger Institute COVID-19 Surveillance Team |
| EPI_ISL_2831296 | Lighthouse Lab in Glasgow | Wellcome Sanger Institute for the COVID-19 Genomics UK (COG-UK) Consortium | Harper VanSteenhouse, Yumi Kasai, David Gray, Carol Clugston, Anna Dominiczak and Alex Alderton, Roberto Amato, Jeffrey Barrett, Sonia Goncalves, Ewan Harrison, David K. Jackson, Ian Johnston, Dominic Kwiatkowski, Cordelia Langford, John Sillitoe on behalf of the Wellcome Sanger Institute COVID-19 Surveillance Team |
| EPI_ISL_2831297, EPI_ISL_2831298 | Lighthouse Lab in Alderley Park | Wellcome Sanger Institute for the COVID-19 Genomics UK (COG-UK) Consortium | Jacquelyn Wynn, Mairead Hyland, The Lighthouse Lab in Alderley Park and Alex Alderton, Roberto Amato, Jeffrey Barrett, Sonia Goncalves, Ewan Harrison, David K. Jackson, Ian Johnston, Dominic Kwiatkowski, Cordelia Langford, John Sillitoe on behalf of the Wellcome Sanger Institute COVID-19 Surveillance Team |
| EPI_ISL_2831299 | Lighthouse Lab in Glasgow | Wellcome Sanger Institute for the COVID-19 Genomics UK (COG-UK) Consortium | Harper VanSteenhouse, Yumi Kasai, David Gray, Carol Clugston, Anna Dominiczak and Alex Alderton, Roberto Amato, Jeffrey Barrett, Sonia Goncalves, Ewan Harrison, David K. Jackson, Ian Johnston, Dominic Kwiatkowski, Cordelia Langford, John Sillitoe on behalf of the Wellcome Sanger Institute COVID-19 Surveillance Team |
| EPI_ISL_2831300, EPI_ISL_2831301, EPI_ISL_2831302 | Lighthouse Lab in Milton Keynes | Wellcome Sanger Institute for the COVID-19 Genomics UK (COG-UK) Consortium | The Lighthouse Lab in Milton Keynes and Alex Alderton, Roberto Amato, Jeffrey Barrett, Sonia Goncalves, Ewan Harrison, David K. Jackson, Ian Johnston, Dominic Kwiatkowski, Cordelia Langford, John Sillitoe on behalf of the Wellcome Sanger Institute COVID-19 Surveillance Team |
| EPI_ISL_2831303 | Lighthouse Lab in Glasgow | Wellcome Sanger Institute for the COVID-19 Genomics UK (COG-UK) Consortium | Harper VanSteenhouse, Yumi Kasai, David Gray, Carol Clugston, Anna Dominiczak and Alex Alderton, Roberto Amato, Jeffrey Barrett, Sonia Goncalves, Ewan Harrison, David K. Jackson, Ian Johnston, Dominic Kwiatkowski, Cordelia Langford, John Sillitoe on behalf of the Wellcome Sanger Institute COVID-19 Surveillance Team |
| EPI_ISL_2831304, EPI_ISL_2831305 | Lighthouse Laboratory Plymouth | Wellcome Sanger Institute for the COVID-19 Genomics UK (COG-UK) Consortium | Lighthouse Laboratory Plymouth and Alex Alderton, Roberto Amato, Jeffrey Barrett, Sonia Goncalves, Ewan Harrison, David K. Jackson, Ian Johnston, Dominic Kwiatkowski, Cordelia Langford, John Sillitoe on behalf of the Wellcome Sanger Institute COVID-19 Surveillance Team |
| EPI_ISL_2831306 | Lighthouse Lab in Milton Keynes | Wellcome Sanger Institute for the COVID-19 Genomics UK (COG-UK) Consortium | The Lighthouse Lab in Milton Keynes and Alex Alderton, Roberto Amato, Jeffrey Barrett, Sonia Goncalves, Ewan Harrison, David K. Jackson, Ian Johnston, Dominic Kwiatkowski, Cordelia Langford, John Sillitoe on behalf of the Wellcome Sanger Institute COVID-19 Surveillance Team |
| EPI_ISL_2831307 | Lighthouse Laboratory Plymouth | Wellcome Sanger Institute for the COVID-19 Genomics UK (COG-UK) Consortium | Lighthouse Laboratory Plymouth and Alex Alderton, Roberto Amato, Jeffrey Barrett, Sonia Goncalves, Ewan Harrison, David K. Jackson, Ian Johnston, Dominic Kwiatkowski, Cordelia Langford, John Sillitoe on behalf of the Wellcome Sanger Institute COVID-19 Surveillance Team |
| EPI_ISL_2831308 | Lighthouse Lab in Glasgow | Wellcome Sanger Institute for the COVID-19 Genomics UK (COG-UK) Consortium | Harper VanSteenhouse, Yumi Kasai, David Gray, Carol Clugston, Anna Dominiczak and Alex Alderton, Roberto Amato, Jeffrey Barrett, Sonia Goncalves, Ewan Harrison, David K. Jackson, Ian Johnston, Dominic Kwiatkowski, Cordelia Langford, John Sillitoe on behalf of the Wellcome Sanger Institute COVID-19 Surveillance Team |
| EPI_ISL_2831309, EPI_ISL_2831310 | Lighthouse Lab in Milton Keynes | Wellcome Sanger Institute for the COVID-19 Genomics UK (COG-UK) Consortium | The Lighthouse Lab in Milton Keynes and Alex Alderton, Roberto Amato, Jeffrey Barrett, Sonia Goncalves, Ewan Harrison, David K. Jackson, Ian Johnston, Dominic Kwiatkowski, Cordelia Langford, John Sillitoe on behalf of the Wellcome Sanger Institute COVID-19 Surveillance Team |
| EPI_ISL_2831311 | Lighthouse Laboratory Plymouth | Wellcome Sanger Institute for the COVID-19 Genomics UK (COG-UK) Consortium | Lighthouse Laboratory Plymouth and Alex Alderton, Roberto Amato, Jeffrey Barrett, Sonia Goncalves, Ewan Harrison, David K. Jackson, Ian Johnston, Dominic Kwiatkowski, Cordelia Langford, John Sillitoe on behalf of the Wellcome Sanger Institute COVID-19 Surveillance Team |
| EPI_ISL_2831312 | Lighthouse Lab in Alderley Park | Wellcome Sanger Institute for the COVID-19 Genomics UK (COG-UK) Consortium | Jacquelyn Wynn, Mairead Hyland, The Lighthouse Lab in Alderley Park and Alex Alderton, Roberto Amato, Jeffrey Barrett, Sonia Goncalves, Ewan Harrison, David K. Jackson, Ian Johnston, Dominic Kwiatkowski, Cordelia Langford, John Sillitoe on behalf of the Wellcome Sanger Institute COVID-19 Surveillance Team |
| EPI_ISL_2831313, EPI_ISL_2831314, EPI_ISL_2831315 | Lighthouse Lab in Milton Keynes | Wellcome Sanger Institute for the COVID-19 Genomics UK (COG-UK) Consortium | The Lighthouse Lab in Milton Keynes and Alex Alderton, Roberto Amato, Jeffrey Barrett, Sonia Goncalves, Ewan Harrison, David K. Jackson, Ian Johnston, Dominic Kwiatkowski, Cordelia Langford, John Sillitoe on behalf of the Wellcome Sanger Institute COVID-19 Surveillance Team |
| EPI_ISL_2831316 | Lighthouse Lab in Alderley Park | Wellcome Sanger Institute for the COVID-19 Genomics UK (COG-UK) Consortium | Jacquelyn Wynn, Mairead Hyland, The Lighthouse Lab in Alderley Park and Alex Alderton, Roberto Amato, Jeffrey Barrett, Sonia Goncalves, Ewan Harrison, David K. Jackson, Ian Johnston, Dominic Kwiatkowski, Cordelia Langford, John Sillitoe on behalf of the Wellcome Sanger Institute COVID-19 Surveillance Team |
| EPI_ISL_2831317, EPI_ISL_2831318, EPI_ISL_2831319 | Lighthouse Laboratory Plymouth | Wellcome Sanger Institute for the COVID-19 Genomics UK (COG-UK) Consortium | Lighthouse Laboratory Plymouth and Alex Alderton, Roberto Amato, Jeffrey Barrett, Sonia Goncalves, Ewan Harrison, David K. Jackson, Ian Johnston, Dominic Kwiatkowski, Cordelia Langford, John Sillitoe on behalf of the Wellcome Sanger Institute COVID-19 Surveillance Team |
| EPI_ISL_2831320 | Lighthouse Lab in Milton Keynes | Wellcome Sanger Institute for the COVID-19 Genomics UK (COG-UK) Consortium | The Lighthouse Lab in Milton Keynes and Alex Alderton, Roberto Amato, Jeffrey Barrett, Sonia Goncalves, Ewan Harrison, David K. Jackson, Ian Johnston, Dominic Kwiatkowski, Cordelia Langford, John Sillitoe on behalf of the Wellcome Sanger Institute COVID-19 Surveillance Team |
| EPI_ISL_2831321 | Lighthouse Lab in Alderley Park | Wellcome Sanger Institute for the COVID-19 Genomics UK (COG-UK) Consortium | Jacquelyn Wynn, Mairead Hyland, The Lighthouse Lab in Alderley Park and Alex Alderton, Roberto Amato, Jeffrey Barrett, Sonia Goncalves, Ewan Harrison, David K. Jackson, Ian Johnston, Dominic Kwiatkowski, Cordelia Langford, John Sillitoe on behalf of the Wellcome Sanger Institute COVID-19 Surveillance Team |
| EPI_ISL_2831322 | Lighthouse Lab in Glasgow | Wellcome Sanger Institute for the COVID-19 Genomics UK (COG-UK) Consortium | Harper VanSteenhouse, Yumi Kasai, David Gray, Carol Clugston, Anna Dominiczak and Alex Alderton, Roberto Amato, Jeffrey Barrett, Sonia Goncalves, Ewan Harrison, David K. Jackson, Ian Johnston, Dominic Kwiatkowski, Cordelia Langford, John Sillitoe on behalf of the Wellcome Sanger Institute COVID-19 Surveillance Team |
| EPI_ISL_2831323, EPI_ISL_2831324 | Lighthouse Lab in Alderley Park | Wellcome Sanger Institute for the COVID-19 Genomics UK (COG-UK) Consortium | Jacquelyn Wynn, Mairead Hyland, The Lighthouse Lab in Alderley Park and Alex Alderton, Roberto Amato, Jeffrey Barrett, Sonia Goncalves, Ewan Harrison, David K. Jackson, Ian Johnston, Dominic Kwiatkowski, Cordelia Langford, John Sillitoe on behalf of the Wellcome Sanger Institute COVID-19 Surveillance Team |
| EPI_ISL_2831325 | Lighthouse Laboratory Plymouth | Wellcome Sanger Institute for the COVID-19 Genomics UK (COG-UK) Consortium | Lighthouse Laboratory Plymouth and Alex Alderton, Roberto Amato, Jeffrey Barrett, Sonia Goncalves, Ewan Harrison, David K. Jackson, Ian Johnston, Dominic Kwiatkowski, Cordelia Langford, John Sillitoe on behalf of the Wellcome Sanger Institute COVID-19 Surveillance Team |
| EPI_ISL_2831326 | Lighthouse Lab in Alderley Park | Wellcome Sanger Institute for the COVID-19 Genomics UK (COG-UK) Consortium | Jacquelyn Wynn, Mairead Hyland, The Lighthouse Lab in Alderley Park and Alex Alderton, Roberto Amato, Jeffrey Barrett, Sonia Goncalves, Ewan Harrison, David K. Jackson, Ian Johnston, Dominic Kwiatkowski, Cordelia Langford, John Sillitoe on behalf of the Wellcome Sanger Institute COVID-19 Surveillance Team |

| | | | |
|---|---|---|---|
| EPI_ISL_2831327, EPI_ISL_2831328, EPI_ISL_2831329 | Lighthouse Laboratory Plymouth | Wellcome Sanger Institute for the COVID-19 Genomics UK (COG-UK) Consortium | Lighthouse Laboratory Plymouth and Alex Alderton, Roberto Amato, Jeffrey Barrett, Sonia Goncalves, Ewan Harrison, David K. Jackson, Ian Johnston, Dominic Kwiatkowski, Cordelia Langford, John Sillitoe on behalf of the Wellcome Sanger Institute COVID-19 Surveillance Team |
| EPI_ISL_2831330, EPI_ISL_2831331, EPI_ISL_2831332 | Lighthouse Lab in Alderley Park | Wellcome Sanger Institute for the COVID-19 Genomics UK (COG-UK) Consortium | Jacquelyn Wynn, Mairead Hyland, The Lighthouse Lab in Alderley Park and Alex Alderton, Roberto Amato, Jeffrey Barrett, Sonia Goncalves, Ewan Harrison, David K. Jackson, Ian Johnston, Dominic Kwiatkowski, Cordelia Langford, John Sillitoe on behalf of the Wellcome Sanger Institute COVID-19 Surveillance Team |
| EPI_ISL_2831333 | Lighthouse Laboratory Plymouth | Wellcome Sanger Institute for the COVID-19 Genomics UK (COG-UK) Consortium | Lighthouse Laboratory Plymouth and Alex Alderton, Roberto Amato, Jeffrey Barrett, Sonia Goncalves, Ewan Harrison, David K. Jackson, Ian Johnston, Dominic Kwiatkowski, Cordelia Langford, John Sillitoe on behalf of the Wellcome Sanger Institute COVID-19 Surveillance Team |
| EPI_ISL_2831334, EPI_ISL_2831335, EPI_ISL_2831336 | Lighthouse Lab in Alderley Park | Wellcome Sanger Institute for the COVID-19 Genomics UK (COG-UK) Consortium | Jacquelyn Wynn, Mairead Hyland, The Lighthouse Lab in Alderley Park and Alex Alderton, Roberto Amato, Jeffrey Barrett, Sonia Goncalves, Ewan Harrison, David K. Jackson, Ian Johnston, Dominic Kwiatkowski, Cordelia Langford, John Sillitoe on behalf of the Wellcome Sanger Institute COVID-19 Surveillance Team |
| EPI_ISL_2831337, EPI_ISL_2831338, EPI_ISL_2831339, EPI_ISL_2831340 | Lighthouse Laboratory Plymouth | Wellcome Sanger Institute for the COVID-19 Genomics UK (COG-UK) Consortium | Lighthouse Laboratory Plymouth and Alex Alderton, Roberto Amato, Jeffrey Barrett, Sonia Goncalves, Ewan Harrison, David K. Jackson, Ian Johnston, Dominic Kwiatkowski, Cordelia Langford, John Sillitoe on behalf of the Wellcome Sanger Institute COVID-19 Surveillance Team |
| EPI_ISL_2831341, EPI_ISL_2831342 | Lighthouse Lab in Alderley Park | Wellcome Sanger Institute for the COVID-19 Genomics UK (COG-UK) Consortium | Jacquelyn Wynn, Mairead Hyland, The Lighthouse Lab in Alderley Park and Alex Alderton, Roberto Amato, Jeffrey Barrett, Sonia Goncalves, Ewan Harrison, David K. Jackson, Ian Johnston, Dominic Kwiatkowski, Cordelia Langford, John Sillitoe on behalf of the Wellcome Sanger Institute COVID-19 Surveillance Team |
| EPI_ISL_2831343 | Lighthouse Laboratory Plymouth | Wellcome Sanger Institute for the COVID-19 Genomics UK (COG-UK) Consortium | Lighthouse Laboratory Plymouth and Alex Alderton, Roberto Amato, Jeffrey Barrett, Sonia Goncalves, Ewan Harrison, David K. Jackson, Ian Johnston, Dominic Kwiatkowski, Cordelia Langford, John Sillitoe on behalf of the Wellcome Sanger Institute COVID-19 Surveillance Team |
| EPI_ISL_2831344, EPI_ISL_2831345, EPI_ISL_2831346, EPI_ISL_2831347, EPI_ISL_2831348, EPI_ISL_2831349, EPI_ISL_2831350 | Lighthouse Lab in Alderley Park | Wellcome Sanger Institute for the COVID-19 Genomics UK (COG-UK) Consortium | Jacquelyn Wynn, Mairead Hyland, The Lighthouse Lab in Alderley Park and Alex Alderton, Roberto Amato, Jeffrey Barrett, Sonia Goncalves, Ewan Harrison, David K. Jackson, Ian Johnston, Dominic Kwiatkowski, Cordelia Langford, John Sillitoe on behalf of the Wellcome Sanger Institute COVID-19 Surveillance Team |
| EPI_ISL_2831351 | Lighthouse Laboratory Plymouth | Wellcome Sanger Institute for the COVID-19 Genomics UK (COG-UK) Consortium | Lighthouse Laboratory Plymouth and Alex Alderton, Roberto Amato, Jeffrey Barrett, Sonia Goncalves, Ewan Harrison, David K. Jackson, Ian Johnston, Dominic Kwiatkowski, Cordelia Langford, John Sillitoe on behalf of the Wellcome Sanger Institute COVID-19 Surveillance Team |
| EPI_ISL_2831352 | Lighthouse Lab in Alderley Park | Wellcome Sanger Institute for the COVID-19 Genomics UK (COG-UK) Consortium | Jacquelyn Wynn, Mairead Hyland, The Lighthouse Lab in Alderley Park and Alex Alderton, Roberto Amato, Jeffrey Barrett, Sonia Goncalves, Ewan Harrison, David K. Jackson, Ian Johnston, Dominic Kwiatkowski, Cordelia Langford, John Sillitoe on behalf of the Wellcome Sanger Institute COVID-19 Surveillance Team |
| EPI_ISL_2831353 | Lighthouse Laboratory Plymouth | Wellcome Sanger Institute for the COVID-19 Genomics UK (COG-UK) Consortium | Lighthouse Laboratory Plymouth and Alex Alderton, Roberto Amato, Jeffrey Barrett, Sonia Goncalves, Ewan Harrison, David K. Jackson, Ian Johnston, Dominic Kwiatkowski, Cordelia Langford, John Sillitoe on behalf of the Wellcome Sanger Institute COVID-19 Surveillance Team |
| EPI_ISL_2831354, EPI_ISL_2831355 | Lighthouse Lab in Alderley Park | Wellcome Sanger Institute for the COVID-19 Genomics UK (COG-UK) Consortium | Jacquelyn Wynn, Mairead Hyland, The Lighthouse Lab in Alderley Park and Alex Alderton, Roberto Amato, Jeffrey Barrett, Sonia Goncalves, Ewan Harrison, David K. Jackson, Ian Johnston, Dominic Kwiatkowski, Cordelia Langford, John Sillitoe on behalf of the Wellcome Sanger Institute COVID-19 Surveillance Team |
| EPI_ISL_2831356 | Lighthouse Laboratory Plymouth | Wellcome Sanger Institute for the COVID-19 Genomics UK (COG-UK) Consortium | Lighthouse Laboratory Plymouth and Alex Alderton, Roberto Amato, Jeffrey Barrett, Sonia Goncalves, Ewan Harrison, David K. Jackson, Ian Johnston, Dominic Kwiatkowski, Cordelia Langford, John Sillitoe on behalf of the Wellcome Sanger Institute COVID-19 Surveillance Team |
| EPI_ISL_2831357, EPI_ISL_2831358 | Lighthouse Lab in Alderley Park | Wellcome Sanger Institute for the COVID-19 Genomics UK (COG-UK) Consortium | Jacquelyn Wynn, Mairead Hyland, The Lighthouse Lab in Alderley Park and Alex Alderton, Roberto Amato, Jeffrey Barrett, Sonia Goncalves, Ewan Harrison, David K. Jackson, Ian Johnston, Dominic Kwiatkowski, Cordelia Langford, John Sillitoe on behalf of the Wellcome Sanger Institute COVID-19 Surveillance Team |
| EPI_ISL_2831359 | Lighthouse Laboratory Plymouth | Wellcome Sanger Institute for the COVID-19 Genomics UK (COG-UK) Consortium | Lighthouse Laboratory Plymouth and Alex Alderton, Roberto Amato, Jeffrey Barrett, Sonia Goncalves, Ewan Harrison, David K. Jackson, Ian Johnston, Dominic Kwiatkowski, Cordelia Langford, John Sillitoe on behalf of the Wellcome Sanger Institute COVID-19 Surveillance Team |
| EPI_ISL_2831360, EPI_ISL_2831361, EPI_ISL_2831362 | Lighthouse Lab in Alderley Park | Wellcome Sanger Institute for the COVID-19 Genomics UK (COG-UK) Consortium | Jacquelyn Wynn, Mairead Hyland, The Lighthouse Lab in Alderley Park and Alex Alderton, Roberto Amato, Jeffrey Barrett, Sonia Goncalves, Ewan Harrison, David K. Jackson, Ian Johnston, Dominic Kwiatkowski, Cordelia Langford, John Sillitoe on behalf of the Wellcome Sanger Institute COVID-19 Surveillance Team |
| EPI_ISL_2831363, EPI_ISL_2831364, EPI_ISL_2831365, EPI_ISL_2831366, EPI_ISL_2831367 | Lighthouse Laboratory Plymouth | Wellcome Sanger Institute for the COVID-19 Genomics UK (COG-UK) Consortium | Lighthouse Laboratory Plymouth and Alex Alderton, Roberto Amato, Jeffrey Barrett, Sonia Goncalves, Ewan Harrison, David K. Jackson, Ian Johnston, Dominic Kwiatkowski, Cordelia Langford, John Sillitoe on behalf of the Wellcome Sanger Institute COVID-19 Surveillance Team |
| EPI_ISL_2831368, EPI_ISL_2831369 | Lighthouse Lab in Alderley Park | Wellcome Sanger Institute for the COVID-19 Genomics UK (COG-UK) Consortium | Jacquelyn Wynn, Mairead Hyland, The Lighthouse Lab in Alderley Park and Alex Alderton, Roberto Amato, Jeffrey Barrett, Sonia Goncalves, Ewan Harrison, David K. Jackson, Ian Johnston, Dominic Kwiatkowski, Cordelia Langford, John Sillitoe on behalf of the Wellcome Sanger Institute COVID-19 Surveillance Team |
| EPI_ISL_2831370, EPI_ISL_2831371 | Lighthouse Laboratory Plymouth | Wellcome Sanger Institute for the COVID-19 Genomics UK (COG-UK) Consortium | Lighthouse Laboratory Plymouth and Alex Alderton, Roberto Amato, Jeffrey Barrett, Sonia Goncalves, Ewan Harrison, David K. Jackson, Ian Johnston, Dominic Kwiatkowski, Cordelia Langford, John Sillitoe on behalf of the Wellcome Sanger Institute COVID-19 Surveillance Team |
| EPI_ISL_2831372, EPI_ISL_2831373, EPI_ISL_2831374, EPI_ISL_2831375 | Lighthouse Lab in Alderley Park | Wellcome Sanger Institute for the COVID-19 Genomics UK (COG-UK) Consortium | Jacquelyn Wynn, Mairead Hyland, The Lighthouse Lab in Alderley Park and Alex Alderton, Roberto Amato, Jeffrey Barrett, Sonia Goncalves, Ewan Harrison, David K. Jackson, Ian Johnston, Dominic Kwiatkowski, Cordelia Langford, John Sillitoe on behalf of the Wellcome Sanger Institute COVID-19 Surveillance Team |
| EPI_ISL_2831376 | Lighthouse Laboratory Plymouth | Wellcome Sanger Institute for the COVID-19 Genomics UK (COG-UK) Consortium | Lighthouse Laboratory Plymouth and Alex Alderton, Roberto Amato, Jeffrey Barrett, Sonia Goncalves, Ewan Harrison, David K. Jackson, Ian Johnston, Dominic Kwiatkowski, Cordelia Langford, John Sillitoe on behalf of the Wellcome Sanger Institute COVID-19 Surveillance Team |
| EPI_ISL_2831377 | Lighthouse Lab in Alderley Park | Wellcome Sanger Institute for the COVID-19 Genomics UK (COG-UK) Consortium | Jacquelyn Wynn, Mairead Hyland, The Lighthouse Lab in Alderley Park and Alex Alderton, Roberto Amato, Jeffrey Barrett, Sonia Goncalves, Ewan Harrison, David K. Jackson, Ian Johnston, Dominic Kwiatkowski, Cordelia Langford, John Sillitoe on behalf of the Wellcome Sanger Institute COVID-19 Surveillance Team |
| EPI_ISL_2831378, EPI_ISL_2831379 | Lighthouse Laboratory Plymouth | Wellcome Sanger Institute for the COVID-19 Genomics UK (COG-UK) Consortium | Lighthouse Laboratory Plymouth and Alex Alderton, Roberto Amato, Jeffrey Barrett, Sonia Goncalves, Ewan Harrison, David K. Jackson, Ian Johnston, Dominic Kwiatkowski, Cordelia Langford, John Sillitoe on behalf of the Wellcome Sanger Institute COVID-19 Surveillance Team |
| EPI_ISL_2831380 | Lighthouse Lab in Alderley Park | Wellcome Sanger Institute for the COVID-19 Genomics UK (COG-UK) Consortium | Jacquelyn Wynn, Mairead Hyland, The Lighthouse Lab in Alderley Park and Alex Alderton, Roberto Amato, Jeffrey Barrett, Sonia Goncalves, Ewan Harrison, David K. Jackson, Ian Johnston, Dominic Kwiatkowski, Cordelia Langford, John Sillitoe on behalf of the Wellcome Sanger Institute COVID-19 Surveillance Team |
| EPI_ISL_2831381 | Lighthouse Laboratory Plymouth | Wellcome Sanger Institute for the COVID-19 Genomics UK (COG-UK) Consortium | Lighthouse Laboratory Plymouth and Alex Alderton, Roberto Amato, Jeffrey Barrett, Sonia Goncalves, Ewan Harrison, David K. Jackson, Ian Johnston, Dominic Kwiatkowski, Cordelia Langford, John Sillitoe on behalf of the Wellcome Sanger Institute COVID-19 Surveillance Team |
| EPI_ISL_2831382, EPI_ISL_2831383, EPI_ISL_2831384, EPI_ISL_2831385 | Lighthouse Lab in Alderley Park | Wellcome Sanger Institute for the COVID-19 Genomics UK (COG-UK) Consortium | Jacquelyn Wynn, Mairead Hyland, The Lighthouse Lab in Alderley Park and Alex Alderton, Roberto Amato, Jeffrey Barrett, Sonia Goncalves, Ewan Harrison, David K. Jackson, Ian Johnston, Dominic Kwiatkowski, Cordelia Langford, John Sillitoe on behalf of the Wellcome Sanger Institute COVID-19 Surveillance Team |
| EPI_ISL_2831386, EPI_ISL_2831387 | Lighthouse Laboratory Plymouth | Wellcome Sanger Institute for the COVID-19 Genomics UK (COG-UK) Consortium | Lighthouse Laboratory Plymouth and Alex Alderton, Roberto Amato, Jeffrey Barrett, Sonia Goncalves, Ewan Harrison, David K. Jackson, Ian Johnston, Dominic Kwiatkowski, Cordelia Langford, John Sillitoe on behalf of the Wellcome Sanger Institute COVID-19 Surveillance Team |
| EPI_ISL_2831388 | Lighthouse Lab in Alderley Park | Wellcome Sanger Institute for the COVID-19 Genomics UK (COG-UK) Consortium | Jacquelyn Wynn, Mairead Hyland, The Lighthouse Lab in Alderley Park and Alex Alderton, Roberto Amato, Jeffrey Barrett, Sonia Goncalves, Ewan Harrison, David K. Jackson, Ian Johnston, Dominic Kwiatkowski, Cordelia Langford, John Sillitoe on behalf of the Wellcome Sanger Institute COVID-19 Surveillance Team |
| EPI_ISL_2831389 | Lighthouse Laboratory Plymouth | Wellcome Sanger Institute for the COVID-19 Genomics UK (COG-UK) Consortium | Lighthouse Laboratory Plymouth and Alex Alderton, Roberto Amato, Jeffrey Barrett, Sonia Goncalves, Ewan Harrison, David K. Jackson, Ian Johnston, Dominic Kwiatkowski, Cordelia Langford, John Sillitoe on behalf of the Wellcome Sanger Institute COVID-19 Surveillance Team |
| EPI_ISL_2831390, EPI_ISL_2831391, EPI_ISL_2831392, EPI_ISL_2831393, EPI_ISL_2831394, EPI_ISL_2831395, EPI_ISL_2831396, EPI_ISL_2831397, EPI_ISL_2831398, EPI_ISL_2831399, EPI_ISL_2831400, EPI_ISL_2831401, EPI_ISL_2831402, EPI_ISL_2831403, EPI_ISL_2831404, EPI_ISL_2831405, EPI_ISL_2831406, EPI_ISL_2831407 | | | |
| see above | Lighthouse Lab in Alderley Park | Wellcome Sanger Institute for the COVID-19 Genomics UK (COG-UK) Consortium | Jacquelyn Wynn, Mairead Hyland, The Lighthouse Lab in Alderley Park and Alex Alderton, Roberto Amato, Jeffrey Barrett, Sonia Goncalves, Ewan Harrison, David K. Jackson, Ian Johnston, Dominic Kwiatkowski, Cordelia Langford, John Sillitoe on behalf of the Wellcome Sanger Institute COVID-19 Surveillance Team |
| EPI_ISL_2831408 | Lighthouse Laboratory Plymouth | Wellcome Sanger Institute for the COVID-19 Genomics UK (COG-UK) Consortium | Lighthouse Laboratory Plymouth and Alex Alderton, Roberto Amato, Jeffrey Barrett, Sonia Goncalves, Ewan Harrison, David K. Jackson, Ian Johnston, Dominic Kwiatkowski, Cordelia Langford, John Sillitoe on behalf of the Wellcome Sanger Institute COVID-19 Surveillance Team |
| EPI_ISL_2831409, EPI_ISL_2831410, EPI_ISL_2831411, EPI_ISL_2831412, EPI_ISL_2831413, EPI_ISL_2831414, EPI_ISL_2831415, EPI_ISL_2831416, EPI_ISL_2831417, EPI_ISL_2831418, EPI_ISL_2831419, EPI_ISL_2831420 | | | |
| see above | Lighthouse Lab in Alderley Park | Wellcome Sanger Institute for the COVID-19 Genomics UK (COG-UK) Consortium | Jacquelyn Wynn, Mairead Hyland, The Lighthouse Lab in Alderley Park and Alex Alderton, Roberto Amato, Jeffrey Barrett, Sonia Goncalves, Ewan Harrison, David K. Jackson, Ian Johnston, Dominic Kwiatkowski, Cordelia Langford, John Sillitoe on behalf of the Wellcome Sanger Institute COVID-19 Surveillance Team |
| EPI_ISL_2831421 | Lighthouse Laboratory Plymouth | Wellcome Sanger Institute for the COVID-19 Genomics UK (COG-UK) Consortium | Lighthouse Laboratory Plymouth and Alex Alderton, Roberto Amato, Jeffrey Barrett, Sonia Goncalves, Ewan Harrison, David K. Jackson, Ian Johnston, Dominic Kwiatkowski, Cordelia Langford, John Sillitoe on behalf of the Wellcome Sanger Institute COVID-19 Surveillance Team |
| EPI_ISL_2831422, EPI_ISL_2831423 | Lighthouse Lab in Alderley Park | Wellcome Sanger Institute for the COVID-19 Genomics UK (COG-UK) Consortium | Jacquelyn Wynn, Mairead Hyland, The Lighthouse Lab in Alderley Park and Alex Alderton, Roberto Amato, Jeffrey Barrett, Sonia Goncalves, Ewan Harrison, David K. Jackson, Ian Johnston, Dominic Kwiatkowski, Cordelia Langford, John Sillitoe on behalf of the Wellcome Sanger Institute COVID-19 Surveillance Team |
| EPI_ISL_2831424 | Lighthouse Laboratory Plymouth | Wellcome Sanger Institute for the COVID-19 Genomics UK (COG-UK) Consortium | Lighthouse Laboratory Plymouth and Alex Alderton, Roberto Amato, Jeffrey Barrett, Sonia Goncalves, Ewan Harrison, David K. Jackson, Ian Johnston, Dominic Kwiatkowski, Cordelia Langford, John Sillitoe on behalf of the Wellcome Sanger Institute COVID-19 Surveillance Team |
| EPI_ISL_2831425, EPI_ISL_2831426, EPI_ISL_2831427 | Lighthouse Lab in Alderley Park | Wellcome Sanger Institute for the COVID-19 Genomics UK (COG-UK) Consortium | Jacquelyn Wynn, Mairead Hyland, The Lighthouse Lab in Alderley Park and Alex Alderton, Roberto Amato, Jeffrey Barrett, Sonia Goncalves, Ewan Harrison, David K. Jackson, Ian Johnston, Dominic Kwiatkowski, Cordelia Langford, John Sillitoe on behalf of the Wellcome Sanger Institute COVID-19 Surveillance Team |

| | | | |
|---|---|---|---|
| EPI_ISL_2831428 | Lighthouse Laboratory Plymouth | Wellcome Sanger Institute for the COVID-19 Genomics UK (COG-UK) Consortium | Lighthouse Laboratory Plymouth and Alex Alderton, Roberto Amato, Jeffrey Barrett, Sonia Goncalves, Ewan Harrison, David K. Jackson, Ian Johnston, Dominic Kwiatkowski, Cordelia Langford, John Sillitoe on behalf of the Wellcome Sanger Institute COVID-19 Surveillance Team |
| EPI_ISL_2831429, EPI_ISL_2831430 | Lighthouse Lab in Alderley Park | Wellcome Sanger Institute for the COVID-19 Genomics UK (COG-UK) Consortium | Jacquelyn Wynn, Mairead Hyland, The Lighthouse Lab in Alderley Park and Alex Alderton, Roberto Amato, Jeffrey Barrett, Sonia Goncalves, Ewan Harrison, David K. Jackson, Ian Johnston, Dominic Kwiatkowski, Cordelia Langford, John Sillitoe on behalf of the Wellcome Sanger Institute COVID-19 Surveillance Team |
| EPI_ISL_2831431 | Lighthouse Laboratory Plymouth | Wellcome Sanger Institute for the COVID-19 Genomics UK (COG-UK) Consortium | Lighthouse Laboratory Plymouth and Alex Alderton, Roberto Amato, Jeffrey Barrett, Sonia Goncalves, Ewan Harrison, David K. Jackson, Ian Johnston, Dominic Kwiatkowski, Cordelia Langford, John Sillitoe on behalf of the Wellcome Sanger Institute COVID-19 Surveillance Team |
| EPI_ISL_2831432, EPI_ISL_2831433 | Lighthouse Lab in Alderley Park | Wellcome Sanger Institute for the COVID-19 Genomics UK (COG-UK) Consortium | Jacquelyn Wynn, Mairead Hyland, The Lighthouse Lab in Alderley Park and Alex Alderton, Roberto Amato, Jeffrey Barrett, Sonia Goncalves, Ewan Harrison, David K. Jackson, Ian Johnston, Dominic Kwiatkowski, Cordelia Langford, John Sillitoe on behalf of the Wellcome Sanger Institute COVID-19 Surveillance Team |
| EPI_ISL_2831434 | Lighthouse Laboratory Plymouth | Wellcome Sanger Institute for the COVID-19 Genomics UK (COG-UK) Consortium | Lighthouse Laboratory Plymouth and Alex Alderton, Roberto Amato, Jeffrey Barrett, Sonia Goncalves, Ewan Harrison, David K. Jackson, Ian Johnston, Dominic Kwiatkowski, Cordelia Langford, John Sillitoe on behalf of the Wellcome Sanger Institute COVID-19 Surveillance Team |
| EPI_ISL_2831435, EPI_ISL_2831436, EPI_ISL_2831437 | Lighthouse Lab in Alderley Park | Wellcome Sanger Institute for the COVID-19 Genomics UK (COG-UK) Consortium | Jacquelyn Wynn, Mairead Hyland, The Lighthouse Lab in Alderley Park and Alex Alderton, Roberto Amato, Jeffrey Barrett, Sonia Goncalves, Ewan Harrison, David K. Jackson, Ian Johnston, Dominic Kwiatkowski, Cordelia Langford, John Sillitoe on behalf of the Wellcome Sanger Institute COVID-19 Surveillance Team |
| EPI_ISL_2831438, EPI_ISL_2831439 | Lighthouse Laboratory Plymouth | Wellcome Sanger Institute for the COVID-19 Genomics UK (COG-UK) Consortium | Lighthouse Laboratory Plymouth and Alex Alderton, Roberto Amato, Jeffrey Barrett, Sonia Goncalves, Ewan Harrison, David K. Jackson, Ian Johnston, Dominic Kwiatkowski, Cordelia Langford, John Sillitoe on behalf of the Wellcome Sanger Institute COVID-19 Surveillance Team |
| EPI_ISL_2831440, EPI_ISL_2831441, EPI_ISL_2831442, EPI_ISL_2831443, EPI_ISL_2831444 | Lighthouse Lab in Alderley Park | Wellcome Sanger Institute for the COVID-19 Genomics UK (COG-UK) Consortium | Jacquelyn Wynn, Mairead Hyland, The Lighthouse Lab in Alderley Park and Alex Alderton, Roberto Amato, Jeffrey Barrett, Sonia Goncalves, Ewan Harrison, David K. Jackson, Ian Johnston, Dominic Kwiatkowski, Cordelia Langford, John Sillitoe on behalf of the Wellcome Sanger Institute COVID-19 Surveillance Team |
| EPI_ISL_2831445 | Lighthouse Laboratory Plymouth | Wellcome Sanger Institute for the COVID-19 Genomics UK (COG-UK) Consortium | Lighthouse Laboratory Plymouth and Alex Alderton, Roberto Amato, Jeffrey Barrett, Sonia Goncalves, Ewan Harrison, David K. Jackson, Ian Johnston, Dominic Kwiatkowski, Cordelia Langford, John Sillitoe on behalf of the Wellcome Sanger Institute COVID-19 Surveillance Team |
| EPI_ISL_2831446, EPI_ISL_2831447 | Lighthouse Lab in Alderley Park | Wellcome Sanger Institute for the COVID-19 Genomics UK (COG-UK) Consortium | Jacquelyn Wynn, Mairead Hyland, The Lighthouse Lab in Alderley Park and Alex Alderton, Roberto Amato, Jeffrey Barrett, Sonia Goncalves, Ewan Harrison, David K. Jackson, Ian Johnston, Dominic Kwiatkowski, Cordelia Langford, John Sillitoe on behalf of the Wellcome Sanger Institute COVID-19 Surveillance Team |
| EPI_ISL_2831448 | Lighthouse Laboratory Plymouth | Wellcome Sanger Institute for the COVID-19 Genomics UK (COG-UK) Consortium | Lighthouse Laboratory Plymouth and Alex Alderton, Roberto Amato, Jeffrey Barrett, Sonia Goncalves, Ewan Harrison, David K. Jackson, Ian Johnston, Dominic Kwiatkowski, Cordelia Langford, John Sillitoe on behalf of the Wellcome Sanger Institute COVID-19 Surveillance Team |
| EPI_ISL_2831449, EPI_ISL_2831450 | Lighthouse Lab in Alderley Park | Wellcome Sanger Institute for the COVID-19 Genomics UK (COG-UK) Consortium | Jacquelyn Wynn, Mairead Hyland, The Lighthouse Lab in Alderley Park and Alex Alderton, Roberto Amato, Jeffrey Barrett, Sonia Goncalves, Ewan Harrison, David K. Jackson, Ian Johnston, Dominic Kwiatkowski, Cordelia Langford, John Sillitoe on behalf of the Wellcome Sanger Institute COVID-19 Surveillance Team |
| EPI_ISL_2831451 | Lighthouse Laboratory Plymouth | Wellcome Sanger Institute for the COVID-19 Genomics UK (COG-UK) Consortium | Lighthouse Laboratory Plymouth and Alex Alderton, Roberto Amato, Jeffrey Barrett, Sonia Goncalves, Ewan Harrison, David K. Jackson, Ian Johnston, Dominic Kwiatkowski, Cordelia Langford, John Sillitoe on behalf of the Wellcome Sanger Institute COVID-19 Surveillance Team |
| EPI_ISL_2831452, EPI_ISL_2831453 | Lighthouse Lab in Alderley Park | Wellcome Sanger Institute for the COVID-19 Genomics UK (COG-UK) Consortium | Jacquelyn Wynn, Mairead Hyland, The Lighthouse Lab in Alderley Park and Alex Alderton, Roberto Amato, Jeffrey Barrett, Sonia Goncalves, Ewan Harrison, David K. Jackson, Ian Johnston, Dominic Kwiatkowski, Cordelia Langford, John Sillitoe on behalf of the Wellcome Sanger Institute COVID-19 Surveillance Team |
| EPI_ISL_2831454 | Lighthouse Laboratory Plymouth | Wellcome Sanger Institute for the COVID-19 Genomics UK (COG-UK) Consortium | Lighthouse Laboratory Plymouth and Alex Alderton, Roberto Amato, Jeffrey Barrett, Sonia Goncalves, Ewan Harrison, David K. Jackson, Ian Johnston, Dominic Kwiatkowski, Cordelia Langford, John Sillitoe on behalf of the Wellcome Sanger Institute COVID-19 Surveillance Team |
| EPI_ISL_2831455, EPI_ISL_2831456, EPI_ISL_2831457, EPI_ISL_2831458 | Lighthouse Lab in Alderley Park | Wellcome Sanger Institute for the COVID-19 Genomics UK (COG-UK) Consortium | Jacquelyn Wynn, Mairead Hyland, The Lighthouse Lab in Alderley Park and Alex Alderton, Roberto Amato, Jeffrey Barrett, Sonia Goncalves, Ewan Harrison, David K. Jackson, Ian Johnston, Dominic Kwiatkowski, Cordelia Langford, John Sillitoe on behalf of the Wellcome Sanger Institute COVID-19 Surveillance Team |
| EPI_ISL_2831459, EPI_ISL_2831460, EPI_ISL_2831461 | Lighthouse Laboratory Plymouth | Wellcome Sanger Institute for the COVID-19 Genomics UK (COG-UK) Consortium | Lighthouse Laboratory Plymouth and Alex Alderton, Roberto Amato, Jeffrey Barrett, Sonia Goncalves, Ewan Harrison, David K. Jackson, Ian Johnston, Dominic Kwiatkowski, Cordelia Langford, John Sillitoe on behalf of the Wellcome Sanger Institute COVID-19 Surveillance Team |
| EPI_ISL_2831462, EPI_ISL_2831463, EPI_ISL_2831464 | Lighthouse Lab in Alderley Park | Wellcome Sanger Institute for the COVID-19 Genomics UK (COG-UK) Consortium | Jacquelyn Wynn, Mairead Hyland, The Lighthouse Lab in Alderley Park and Alex Alderton, Roberto Amato, Jeffrey Barrett, Sonia Goncalves, Ewan Harrison, David K. Jackson, Ian Johnston, Dominic Kwiatkowski, Cordelia Langford, John Sillitoe on behalf of the Wellcome Sanger Institute COVID-19 Surveillance Team |
| EPI_ISL_2831465 | Lighthouse Laboratory Plymouth | Wellcome Sanger Institute for the COVID-19 Genomics UK (COG-UK) Consortium | Lighthouse Laboratory Plymouth and Alex Alderton, Roberto Amato, Jeffrey Barrett, Sonia Goncalves, Ewan Harrison, David K. Jackson, Ian Johnston, Dominic Kwiatkowski, Cordelia Langford, John Sillitoe on behalf of the Wellcome Sanger Institute COVID-19 Surveillance Team |
| EPI_ISL_2831466, EPI_ISL_2831467, EPI_ISL_2831468, EPI_ISL_2831469 | Lighthouse Lab in Alderley Park | Wellcome Sanger Institute for the COVID-19 Genomics UK (COG-UK) Consortium | Jacquelyn Wynn, Mairead Hyland, The Lighthouse Lab in Alderley Park and Alex Alderton, Roberto Amato, Jeffrey Barrett, Sonia Goncalves, Ewan Harrison, David K. Jackson, Ian Johnston, Dominic Kwiatkowski, Cordelia Langford, John Sillitoe on behalf of the Wellcome Sanger Institute COVID-19 Surveillance Team |
| EPI_ISL_2831470 | Lighthouse Laboratory Plymouth | Wellcome Sanger Institute for the COVID-19 Genomics UK (COG-UK) Consortium | Lighthouse Laboratory Plymouth and Alex Alderton, Roberto Amato, Jeffrey Barrett, Sonia Goncalves, Ewan Harrison, David K. Jackson, Ian Johnston, Dominic Kwiatkowski, Cordelia Langford, John Sillitoe on behalf of the Wellcome Sanger Institute COVID-19 Surveillance Team |
| EPI_ISL_2831471, EPI_ISL_2831472, EPI_ISL_2831473, EPI_ISL_2831474, EPI_ISL_2831475, EPI_ISL_2831476, EPI_ISL_2831477 | Lighthouse Lab in Alderley Park | Wellcome Sanger Institute for the COVID-19 Genomics UK (COG-UK) Consortium | Jacquelyn Wynn, Mairead Hyland, The Lighthouse Lab in Alderley Park and Alex Alderton, Roberto Amato, Jeffrey Barrett, Sonia Goncalves, Ewan Harrison, David K. Jackson, Ian Johnston, Dominic Kwiatkowski, Cordelia Langford, John Sillitoe on behalf of the Wellcome Sanger Institute COVID-19 Surveillance Team |
| EPI_ISL_2831478, EPI_ISL_2831479 | Lighthouse Laboratory Plymouth | Wellcome Sanger Institute for the COVID-19 Genomics UK (COG-UK) Consortium | Lighthouse Laboratory Plymouth and Alex Alderton, Roberto Amato, Jeffrey Barrett, Sonia Goncalves, Ewan Harrison, David K. Jackson, Ian Johnston, Dominic Kwiatkowski, Cordelia Langford, John Sillitoe on behalf of the Wellcome Sanger Institute COVID-19 Surveillance Team |
| EPI_ISL_2831480, EPI_ISL_2831481 | Lighthouse Lab in Alderley Park | Wellcome Sanger Institute for the COVID-19 Genomics UK (COG-UK) Consortium | Jacquelyn Wynn, Mairead Hyland, The Lighthouse Lab in Alderley Park and Alex Alderton, Roberto Amato, Jeffrey Barrett, Sonia Goncalves, Ewan Harrison, David K. Jackson, Ian Johnston, Dominic Kwiatkowski, Cordelia Langford, John Sillitoe on behalf of the Wellcome Sanger Institute COVID-19 Surveillance Team |
| EPI_ISL_2831482 | Lighthouse Laboratory Plymouth | Wellcome Sanger Institute for the COVID-19 Genomics UK (COG-UK) Consortium | Lighthouse Laboratory Plymouth and Alex Alderton, Roberto Amato, Jeffrey Barrett, Sonia Goncalves, Ewan Harrison, David K. Jackson, Ian Johnston, Dominic Kwiatkowski, Cordelia Langford, John Sillitoe on behalf of the Wellcome Sanger Institute COVID-19 Surveillance Team |
| EPI_ISL_2831483 | Lighthouse Lab in Alderley Park | Wellcome Sanger Institute for the COVID-19 Genomics UK (COG-UK) Consortium | Jacquelyn Wynn, Mairead Hyland, The Lighthouse Lab in Alderley Park and Alex Alderton, Roberto Amato, Jeffrey Barrett, Sonia Goncalves, Ewan Harrison, David K. Jackson, Ian Johnston, Dominic Kwiatkowski, Cordelia Langford, John Sillitoe on behalf of the Wellcome Sanger Institute COVID-19 Surveillance Team |
| EPI_ISL_2831484 | Lighthouse Laboratory Plymouth | Wellcome Sanger Institute for the COVID-19 Genomics UK (COG-UK) Consortium | Lighthouse Laboratory Plymouth and Alex Alderton, Roberto Amato, Jeffrey Barrett, Sonia Goncalves, Ewan Harrison, David K. Jackson, Ian Johnston, Dominic Kwiatkowski, Cordelia Langford, John Sillitoe on behalf of the Wellcome Sanger Institute COVID-19 Surveillance Team |
| EPI_ISL_2831485 | Lighthouse Lab in Alderley Park | Wellcome Sanger Institute for the COVID-19 Genomics UK (COG-UK) Consortium | Jacquelyn Wynn, Mairead Hyland, The Lighthouse Lab in Alderley Park and Alex Alderton, Roberto Amato, Jeffrey Barrett, Sonia Goncalves, Ewan Harrison, David K. Jackson, Ian Johnston, Dominic Kwiatkowski, Cordelia Langford, John Sillitoe on behalf of the Wellcome Sanger Institute COVID-19 Surveillance Team |
| EPI_ISL_2831486 | Lighthouse Laboratory Plymouth | Wellcome Sanger Institute for the COVID-19 Genomics UK (COG-UK) Consortium | Lighthouse Laboratory Plymouth and Alex Alderton, Roberto Amato, Jeffrey Barrett, Sonia Goncalves, Ewan Harrison, David K. Jackson, Ian Johnston, Dominic Kwiatkowski, Cordelia Langford, John Sillitoe on behalf of the Wellcome Sanger Institute COVID-19 Surveillance Team |
| EPI_ISL_2831487 | Lighthouse Lab in Alderley Park | Wellcome Sanger Institute for the COVID-19 Genomics UK (COG-UK) Consortium | Jacquelyn Wynn, Mairead Hyland, The Lighthouse Lab in Alderley Park and Alex Alderton, Roberto Amato, Jeffrey Barrett, Sonia Goncalves, Ewan Harrison, David K. Jackson, Ian Johnston, Dominic Kwiatkowski, Cordelia Langford, John Sillitoe on behalf of the Wellcome Sanger Institute COVID-19 Surveillance Team |
| EPI_ISL_2831488, EPI_ISL_2831489, EPI_ISL_2831490, EPI_ISL_2831491, EPI_ISL_2831492 | Lighthouse Laboratory Plymouth | Wellcome Sanger Institute for the COVID-19 Genomics UK (COG-UK) Consortium | Lighthouse Laboratory Plymouth and Alex Alderton, Roberto Amato, Jeffrey Barrett, Sonia Goncalves, Ewan Harrison, David K. Jackson, Ian Johnston, Dominic Kwiatkowski, Cordelia Langford, John Sillitoe on behalf of the Wellcome Sanger Institute COVID-19 Surveillance Team |
| EPI_ISL_2831493 | Lighthouse Lab in Alderley Park | Wellcome Sanger Institute for the COVID-19 Genomics UK (COG-UK) Consortium | Jacquelyn Wynn, Mairead Hyland, The Lighthouse Lab in Alderley Park and Alex Alderton, Roberto Amato, Jeffrey Barrett, Sonia Goncalves, Ewan Harrison, David K. Jackson, Ian Johnston, Dominic Kwiatkowski, Cordelia Langford, John Sillitoe on behalf of the Wellcome Sanger Institute COVID-19 Surveillance Team |
| EPI_ISL_2831494 | Lighthouse Laboratory Plymouth | Wellcome Sanger Institute for the COVID-19 Genomics UK (COG-UK) Consortium | Lighthouse Laboratory Plymouth and Alex Alderton, Roberto Amato, Jeffrey Barrett, Sonia Goncalves, Ewan Harrison, David K. Jackson, Ian Johnston, Dominic Kwiatkowski, Cordelia Langford, John Sillitoe on behalf of the Wellcome Sanger Institute COVID-19 Surveillance Team |
| EPI_ISL_2831495, EPI_ISL_2831496, EPI_ISL_2831497 | Lighthouse Lab in Alderley Park | Wellcome Sanger Institute for the COVID-19 Genomics UK (COG-UK) Consortium | Jacquelyn Wynn, Mairead Hyland, The Lighthouse Lab in Alderley Park and Alex Alderton, Roberto Amato, Jeffrey Barrett, Sonia Goncalves, Ewan Harrison, David K. Jackson, Ian Johnston, Dominic Kwiatkowski, Cordelia Langford, John Sillitoe on behalf of the Wellcome Sanger Institute COVID-19 Surveillance Team |
| EPI_ISL_2831498 | Lighthouse Laboratory Plymouth | Wellcome Sanger Institute for the COVID-19 Genomics UK (COG-UK) Consortium | Lighthouse Laboratory Plymouth and Alex Alderton, Roberto Amato, Jeffrey Barrett, Sonia Goncalves, Ewan Harrison, David K. Jackson, Ian Johnston, Dominic Kwiatkowski, Cordelia Langford, John Sillitoe on behalf of the Wellcome Sanger Institute COVID-19 Surveillance Team |
| EPI_ISL_2831499, EPI_ISL_2831500, EPI_ISL_2831501 | Lighthouse Lab in Alderley Park | Wellcome Sanger Institute for the COVID-19 Genomics UK (COG-UK) Consortium | Jacquelyn Wynn, Mairead Hyland, The Lighthouse Lab in Alderley Park and Alex Alderton, Roberto Amato, Jeffrey Barrett, Sonia Goncalves, Ewan Harrison, David K. Jackson, Ian Johnston, Dominic Kwiatkowski, Cordelia Langford, John Sillitoe on behalf of the Wellcome Sanger Institute COVID-19 Surveillance Team |
| EPI_ISL_2831502 | Lighthouse Laboratory Plymouth | Wellcome Sanger Institute for the COVID-19 Genomics UK (COG-UK) Consortium | Lighthouse Laboratory Plymouth and Alex Alderton, Roberto Amato, Jeffrey Barrett, Sonia Goncalves, Ewan Harrison, David K. Jackson, Ian Johnston, Dominic Kwiatkowski, Cordelia Langford, John Sillitoe on behalf of the Wellcome Sanger Institute COVID-19 Surveillance Team |

| | | | |
|---|---|---|---|
| EPI_ISL_2831503, EPI_ISL_2831504, EPI_ISL_2831505, EPI_ISL_2831506 | Lighthouse Lab in Alderley Park | Wellcome Sanger Institute for the COVID-19 Genomics UK (COG-UK) Consortium | Jacquelyn Wynn, Mairead Hyland, The Lighthouse Lab in Alderley Park and Alex Alderton, Roberto Amato, Jeffrey Barrett, Sonia Goncalves, Ewan Harrison, David K. Jackson, Ian Johnston, Dominic Kwiatkowski, Cordelia Langford, John Sillitoe on behalf of the Wellcome Sanger Institute COVID-19 Surveillance Team |
| EPI_ISL_2831507 | Lighthouse Laboratory Plymouth | Wellcome Sanger Institute for the COVID-19 Genomics UK (COG-UK) Consortium | Lighthouse Laboratory Plymouth and Alex Alderton, Roberto Amato, Jeffrey Barrett, Sonia Goncalves, Ewan Harrison, David K. Jackson, Ian Johnston, Dominic Kwiatkowski, Cordelia Langford, John Sillitoe on behalf of the Wellcome Sanger Institute COVID-19 Surveillance Team |
| EPI_ISL_2831508, EPI_ISL_2831509 | Lighthouse Lab in Alderley Park | Wellcome Sanger Institute for the COVID-19 Genomics UK (COG-UK) Consortium | Jacquelyn Wynn, Mairead Hyland, The Lighthouse Lab in Alderley Park and Alex Alderton, Roberto Amato, Jeffrey Barrett, Sonia Goncalves, Ewan Harrison, David K. Jackson, Ian Johnston, Dominic Kwiatkowski, Cordelia Langford, John Sillitoe on behalf of the Wellcome Sanger Institute COVID-19 Surveillance Team |
| EPI_ISL_2831510 | Lighthouse Laboratory Plymouth | Wellcome Sanger Institute for the COVID-19 Genomics UK (COG-UK) Consortium | Lighthouse Laboratory Plymouth and Alex Alderton, Roberto Amato, Jeffrey Barrett, Sonia Goncalves, Ewan Harrison, David K. Jackson, Ian Johnston, Dominic Kwiatkowski, Cordelia Langford, John Sillitoe on behalf of the Wellcome Sanger Institute COVID-19 Surveillance Team |
| EPI_ISL_2831511 | Lighthouse Lab in Alderley Park | Wellcome Sanger Institute for the COVID-19 Genomics UK (COG-UK) Consortium | Jacquelyn Wynn, Mairead Hyland, The Lighthouse Lab in Alderley Park and Alex Alderton, Roberto Amato, Jeffrey Barrett, Sonia Goncalves, Ewan Harrison, David K. Jackson, Ian Johnston, Dominic Kwiatkowski, Cordelia Langford, John Sillitoe on behalf of the Wellcome Sanger Institute COVID-19 Surveillance Team |
| EPI_ISL_2831512 | Lighthouse Laboratory Plymouth | Wellcome Sanger Institute for the COVID-19 Genomics UK (COG-UK) Consortium | Lighthouse Laboratory Plymouth and Alex Alderton, Roberto Amato, Jeffrey Barrett, Sonia Goncalves, Ewan Harrison, David K. Jackson, Ian Johnston, Dominic Kwiatkowski, Cordelia Langford, John Sillitoe on behalf of the Wellcome Sanger Institute COVID-19 Surveillance Team |
| EPI_ISL_2831513, EPI_ISL_2831514, EPI_ISL_2831515, EPI_ISL_2831516 | Lighthouse Lab in Alderley Park | Wellcome Sanger Institute for the COVID-19 Genomics UK (COG-UK) Consortium | Jacquelyn Wynn, Mairead Hyland, The Lighthouse Lab in Alderley Park and Alex Alderton, Roberto Amato, Jeffrey Barrett, Sonia Goncalves, Ewan Harrison, David K. Jackson, Ian Johnston, Dominic Kwiatkowski, Cordelia Langford, John Sillitoe on behalf of the Wellcome Sanger Institute COVID-19 Surveillance Team |
| EPI_ISL_2831517 | Lighthouse Laboratory Plymouth | Wellcome Sanger Institute for the COVID-19 Genomics UK (COG-UK) Consortium | Lighthouse Laboratory Plymouth and Alex Alderton, Roberto Amato, Jeffrey Barrett, Sonia Goncalves, Ewan Harrison, David K. Jackson, Ian Johnston, Dominic Kwiatkowski, Cordelia Langford, John Sillitoe on behalf of the Wellcome Sanger Institute COVID-19 Surveillance Team |
| EPI_ISL_2831518, EPI_ISL_2831519 | Lighthouse Lab in Alderley Park | Wellcome Sanger Institute for the COVID-19 Genomics UK (COG-UK) Consortium | Jacquelyn Wynn, Mairead Hyland, The Lighthouse Lab in Alderley Park and Alex Alderton, Roberto Amato, Jeffrey Barrett, Sonia Goncalves, Ewan Harrison, David K. Jackson, Ian Johnston, Dominic Kwiatkowski, Cordelia Langford, John Sillitoe on behalf of the Wellcome Sanger Institute COVID-19 Surveillance Team |
| EPI_ISL_2831520 | Lighthouse Laboratory Plymouth | Wellcome Sanger Institute for the COVID-19 Genomics UK (COG-UK) Consortium | Lighthouse Laboratory Plymouth and Alex Alderton, Roberto Amato, Jeffrey Barrett, Sonia Goncalves, Ewan Harrison, David K. Jackson, Ian Johnston, Dominic Kwiatkowski, Cordelia Langford, John Sillitoe on behalf of the Wellcome Sanger Institute COVID-19 Surveillance Team |
| EPI_ISL_2831521, EPI_ISL_2831522, EPI_ISL_2831523, EPI_ISL_2831524, EPI_ISL_2831525 | Lighthouse Lab in Alderley Park | Wellcome Sanger Institute for the COVID-19 Genomics UK (COG-UK) Consortium | Jacquelyn Wynn, Mairead Hyland, The Lighthouse Lab in Alderley Park and Alex Alderton, Roberto Amato, Jeffrey Barrett, Sonia Goncalves, Ewan Harrison, David K. Jackson, Ian Johnston, Dominic Kwiatkowski, Cordelia Langford, John Sillitoe on behalf of the Wellcome Sanger Institute COVID-19 Surveillance Team |
| EPI_ISL_2831526, EPI_ISL_2831527, EPI_ISL_2831528, EPI_ISL_2831529, EPI_ISL_2831530, EPI_ISL_2831531 | Lighthouse Laboratory Plymouth | Wellcome Sanger Institute for the COVID-19 Genomics UK (COG-UK) Consortium | Lighthouse Laboratory Plymouth and Alex Alderton, Roberto Amato, Jeffrey Barrett, Sonia Goncalves, Ewan Harrison, David K. Jackson, Ian Johnston, Dominic Kwiatkowski, Cordelia Langford, John Sillitoe on behalf of the Wellcome Sanger Institute COVID-19 Surveillance Team |
| EPI_ISL_2831532, EPI_ISL_2831533, EPI_ISL_2831534 | Lighthouse Lab in Alderley Park | Wellcome Sanger Institute for the COVID-19 Genomics UK (COG-UK) Consortium | Jacquelyn Wynn, Mairead Hyland, The Lighthouse Lab in Alderley Park and Alex Alderton, Roberto Amato, Jeffrey Barrett, Sonia Goncalves, Ewan Harrison, David K. Jackson, Ian Johnston, Dominic Kwiatkowski, Cordelia Langford, John Sillitoe on behalf of the Wellcome Sanger Institute COVID-19 Surveillance Team |
| EPI_ISL_2831535 | Lighthouse Laboratory Plymouth | Wellcome Sanger Institute for the COVID-19 Genomics UK (COG-UK) Consortium | Lighthouse Laboratory Plymouth and Alex Alderton, Roberto Amato, Jeffrey Barrett, Sonia Goncalves, Ewan Harrison, David K. Jackson, Ian Johnston, Dominic Kwiatkowski, Cordelia Langford, John Sillitoe on behalf of the Wellcome Sanger Institute COVID-19 Surveillance Team |
| EPI_ISL_2831536, EPI_ISL_2831537, EPI_ISL_2831538 | Lighthouse Lab in Alderley Park | Wellcome Sanger Institute for the COVID-19 Genomics UK (COG-UK) Consortium | Jacquelyn Wynn, Mairead Hyland, The Lighthouse Lab in Alderley Park and Alex Alderton, Roberto Amato, Jeffrey Barrett, Sonia Goncalves, Ewan Harrison, David K. Jackson, Ian Johnston, Dominic Kwiatkowski, Cordelia Langford, John Sillitoe on behalf of the Wellcome Sanger Institute COVID-19 Surveillance Team |
| EPI_ISL_2831539, EPI_ISL_2831540, EPI_ISL_2831541 | Lighthouse Laboratory Plymouth | Wellcome Sanger Institute for the COVID-19 Genomics UK (COG-UK) Consortium | Lighthouse Laboratory Plymouth and Alex Alderton, Roberto Amato, Jeffrey Barrett, Sonia Goncalves, Ewan Harrison, David K. Jackson, Ian Johnston, Dominic Kwiatkowski, Cordelia Langford, John Sillitoe on behalf of the Wellcome Sanger Institute COVID-19 Surveillance Team |
| EPI_ISL_2831542, EPI_ISL_2831543 | Lighthouse Lab in Alderley Park | Wellcome Sanger Institute for the COVID-19 Genomics UK (COG-UK) Consortium | Jacquelyn Wynn, Mairead Hyland, The Lighthouse Lab in Alderley Park and Alex Alderton, Roberto Amato, Jeffrey Barrett, Sonia Goncalves, Ewan Harrison, David K. Jackson, Ian Johnston, Dominic Kwiatkowski, Cordelia Langford, John Sillitoe on behalf of the Wellcome Sanger Institute COVID-19 Surveillance Team |
| EPI_ISL_2831544 | Lighthouse Laboratory Plymouth | Wellcome Sanger Institute for the COVID-19 Genomics UK (COG-UK) Consortium | Lighthouse Laboratory Plymouth and Alex Alderton, Roberto Amato, Jeffrey Barrett, Sonia Goncalves, Ewan Harrison, David K. Jackson, Ian Johnston, Dominic Kwiatkowski, Cordelia Langford, John Sillitoe on behalf of the Wellcome Sanger Institute COVID-19 Surveillance Team |
| EPI_ISL_2831545, EPI_ISL_2831546, EPI_ISL_2831547, EPI_ISL_2831548, EPI_ISL_2831549, EPI_ISL_2831550 | Lighthouse Lab in Alderley Park | Wellcome Sanger Institute for the COVID-19 Genomics UK (COG-UK) Consortium | Jacquelyn Wynn, Mairead Hyland, The Lighthouse Lab in Alderley Park and Alex Alderton, Roberto Amato, Jeffrey Barrett, Sonia Goncalves, Ewan Harrison, David K. Jackson, Ian Johnston, Dominic Kwiatkowski, Cordelia Langford, John Sillitoe on behalf of the Wellcome Sanger Institute COVID-19 Surveillance Team |
| EPI_ISL_2831551, EPI_ISL_2831552 | Lighthouse Laboratory Plymouth | Wellcome Sanger Institute for the COVID-19 Genomics UK (COG-UK) Consortium | Lighthouse Laboratory Plymouth and Alex Alderton, Roberto Amato, Jeffrey Barrett, Sonia Goncalves, Ewan Harrison, David K. Jackson, Ian Johnston, Dominic Kwiatkowski, Cordelia Langford, John Sillitoe on behalf of the Wellcome Sanger Institute COVID-19 Surveillance Team |
| EPI_ISL_2831553, EPI_ISL_2831554, EPI_ISL_2831555, EPI_ISL_2831556, EPI_ISL_2831557 | Lighthouse Lab in Alderley Park | Wellcome Sanger Institute for the COVID-19 Genomics UK (COG-UK) Consortium | Jacquelyn Wynn, Mairead Hyland, The Lighthouse Lab in Alderley Park and Alex Alderton, Roberto Amato, Jeffrey Barrett, Sonia Goncalves, Ewan Harrison, David K. Jackson, Ian Johnston, Dominic Kwiatkowski, Cordelia Langford, John Sillitoe on behalf of the Wellcome Sanger Institute COVID-19 Surveillance Team |
| EPI_ISL_2831558, EPI_ISL_2831559 | Lighthouse Laboratory Plymouth | Wellcome Sanger Institute for the COVID-19 Genomics UK (COG-UK) Consortium | Lighthouse Laboratory Plymouth and Alex Alderton, Roberto Amato, Jeffrey Barrett, Sonia Goncalves, Ewan Harrison, David K. Jackson, Ian Johnston, Dominic Kwiatkowski, Cordelia Langford, John Sillitoe on behalf of the Wellcome Sanger Institute COVID-19 Surveillance Team |
| EPI_ISL_2831560, EPI_ISL_2831561, EPI_ISL_2831562, EPI_ISL_2831563 | Lighthouse Lab in Alderley Park | Wellcome Sanger Institute for the COVID-19 Genomics UK (COG-UK) Consortium | Jacquelyn Wynn, Mairead Hyland, The Lighthouse Lab in Alderley Park and Alex Alderton, Roberto Amato, Jeffrey Barrett, Sonia Goncalves, Ewan Harrison, David K. Jackson, Ian Johnston, Dominic Kwiatkowski, Cordelia Langford, John Sillitoe on behalf of the Wellcome Sanger Institute COVID-19 Surveillance Team |
| EPI_ISL_2831564 | Lighthouse Laboratory Plymouth | Wellcome Sanger Institute for the COVID-19 Genomics UK (COG-UK) Consortium | Lighthouse Laboratory Plymouth and Alex Alderton, Roberto Amato, Jeffrey Barrett, Sonia Goncalves, Ewan Harrison, David K. Jackson, Ian Johnston, Dominic Kwiatkowski, Cordelia Langford, John Sillitoe on behalf of the Wellcome Sanger Institute COVID-19 Surveillance Team |
| EPI_ISL_2831565, EPI_ISL_2831566, EPI_ISL_2831567, EPI_ISL_2831568, EPI_ISL_2831569 | Lighthouse Lab in Alderley Park | Wellcome Sanger Institute for the COVID-19 Genomics UK (COG-UK) Consortium | Jacquelyn Wynn, Mairead Hyland, The Lighthouse Lab in Alderley Park and Alex Alderton, Roberto Amato, Jeffrey Barrett, Sonia Goncalves, Ewan Harrison, David K. Jackson, Ian Johnston, Dominic Kwiatkowski, Cordelia Langford, John Sillitoe on behalf of the Wellcome Sanger Institute COVID-19 Surveillance Team |
| EPI_ISL_2831570 | Lighthouse Laboratory Plymouth | Wellcome Sanger Institute for the COVID-19 Genomics UK (COG-UK) Consortium | Lighthouse Laboratory Plymouth and Alex Alderton, Roberto Amato, Jeffrey Barrett, Sonia Goncalves, Ewan Harrison, David K. Jackson, Ian Johnston, Dominic Kwiatkowski, Cordelia Langford, John Sillitoe on behalf of the Wellcome Sanger Institute COVID-19 Surveillance Team |
| EPI_ISL_2831571 | Lighthouse Lab in Alderley Park | Wellcome Sanger Institute for the COVID-19 Genomics UK (COG-UK) Consortium | Jacquelyn Wynn, Mairead Hyland, The Lighthouse Lab in Alderley Park and Alex Alderton, Roberto Amato, Jeffrey Barrett, Sonia Goncalves, Ewan Harrison, David K. Jackson, Ian Johnston, Dominic Kwiatkowski, Cordelia Langford, John Sillitoe on behalf of the Wellcome Sanger Institute COVID-19 Surveillance Team |
| EPI_ISL_2831572 | Lighthouse Laboratory Plymouth | Wellcome Sanger Institute for the COVID-19 Genomics UK (COG-UK) Consortium | Lighthouse Laboratory Plymouth and Alex Alderton, Roberto Amato, Jeffrey Barrett, Sonia Goncalves, Ewan Harrison, David K. Jackson, Ian Johnston, Dominic Kwiatkowski, Cordelia Langford, John Sillitoe on behalf of the Wellcome Sanger Institute COVID-19 Surveillance Team |
| EPI_ISL_2831573 | Lighthouse Lab in Alderley Park | Wellcome Sanger Institute for the COVID-19 Genomics UK (COG-UK) Consortium | Jacquelyn Wynn, Mairead Hyland, The Lighthouse Lab in Alderley Park and Alex Alderton, Roberto Amato, Jeffrey Barrett, Sonia Goncalves, Ewan Harrison, David K. Jackson, Ian Johnston, Dominic Kwiatkowski, Cordelia Langford, John Sillitoe on behalf of the Wellcome Sanger Institute COVID-19 Surveillance Team |
| EPI_ISL_2831574, EPI_ISL_2831575 | Lighthouse Laboratory Plymouth | Wellcome Sanger Institute for the COVID-19 Genomics UK (COG-UK) Consortium | Lighthouse Laboratory Plymouth and Alex Alderton, Roberto Amato, Jeffrey Barrett, Sonia Goncalves, Ewan Harrison, David K. Jackson, Ian Johnston, Dominic Kwiatkowski, Cordelia Langford, John Sillitoe on behalf of the Wellcome Sanger Institute COVID-19 Surveillance Team |
| EPI_ISL_2831576, EPI_ISL_2831577, EPI_ISL_2831578, EPI_ISL_2831579 | Lighthouse Lab in Alderley Park | Wellcome Sanger Institute for the COVID-19 Genomics UK (COG-UK) Consortium | Jacquelyn Wynn, Mairead Hyland, The Lighthouse Lab in Alderley Park and Alex Alderton, Roberto Amato, Jeffrey Barrett, Sonia Goncalves, Ewan Harrison, David K. Jackson, Ian Johnston, Dominic Kwiatkowski, Cordelia Langford, John Sillitoe on behalf of the Wellcome Sanger Institute COVID-19 Surveillance Team |
| EPI_ISL_2831580, EPI_ISL_2831581 | Lighthouse Laboratory Plymouth | Wellcome Sanger Institute for the COVID-19 Genomics UK (COG-UK) Consortium | Lighthouse Laboratory Plymouth and Alex Alderton, Roberto Amato, Jeffrey Barrett, Sonia Goncalves, Ewan Harrison, David K. Jackson, Ian Johnston, Dominic Kwiatkowski, Cordelia Langford, John Sillitoe on behalf of the Wellcome Sanger Institute COVID-19 Surveillance Team |
| EPI_ISL_2831582, EPI_ISL_2831583, EPI_ISL_2831584, EPI_ISL_2831585, | Lighthouse Lab in Alderley Park | Wellcome Sanger Institute for the COVID-19 Genomics UK (COG-UK) Consortium | Jacquelyn Wynn, Mairead Hyland, The Lighthouse Lab in Alderley Park and Alex Alderton, Roberto Amato, Jeffrey Barrett, Sonia Goncalves, Ewan Harrison, David K. Jackson, Ian Johnston, Dominic Kwiatkowski, Cordelia Langford, John Sillitoe on behalf of the Wellcome Sanger Institute COVID-19 Surveillance Team |

| | | | |
|---|---|---|---|
| EPI_ISL_2831586, EPI_ISL_2831587, EPI_ISL_2831588 | | | |
| EPI_ISL_2831589, EPI_ISL_2831590, EPI_ISL_2831591 | Lighthouse Lab in Glasgow | Wellcome Sanger Institute for the COVID-19 Genomics UK (COG-UK) Consortium | Harper VanSteenhouse, Yumi Kasai, David Gray, Carol Clugston, Anna Dominiczak and Alex Alderton, Roberto Amato, Jeffrey Barrett, Sonia Goncalves, Ewan Harrison, David K. Jackson, Ian Johnston, Dominic Kwiatkowski, Cordelia Langford, John Sillitoe on behalf of the Wellcome Sanger Institute COVID-19 Surveillance Team |
| EPI_ISL_2831592, EPI_ISL_2831593 | Lighthouse Lab in Alderley Park | Wellcome Sanger Institute for the COVID-19 Genomics UK (COG-UK) Consortium | Jacquelyn Wynn, Mairead Hyland, The Lighthouse Lab in Alderley Park and Alex Alderton, Roberto Amato, Jeffrey Barrett, Sonia Goncalves, Ewan Harrison, David K. Jackson, Ian Johnston, Dominic Kwiatkowski, Cordelia Langford, John Sillitoe on behalf of the Wellcome Sanger Institute COVID-19 Surveillance Team |
| EPI_ISL_2831594, EPI_ISL_2831595 | Lighthouse Lab in Glasgow | Wellcome Sanger Institute for the COVID-19 Genomics UK (COG-UK) Consortium | Harper VanSteenhouse, Yumi Kasai, David Gray, Carol Clugston, Anna Dominiczak and Alex Alderton, Roberto Amato, Jeffrey Barrett, Sonia Goncalves, Ewan Harrison, David K. Jackson, Ian Johnston, Dominic Kwiatkowski, Cordelia Langford, John Sillitoe on behalf of the Wellcome Sanger Institute COVID-19 Surveillance Team |
| EPI_ISL_2831596, EPI_ISL_2831597 | Lighthouse Lab in Alderley Park | Wellcome Sanger Institute for the COVID-19 Genomics UK (COG-UK) Consortium | Jacquelyn Wynn, Mairead Hyland, The Lighthouse Lab in Alderley Park and Alex Alderton, Roberto Amato, Jeffrey Barrett, Sonia Goncalves, Ewan Harrison, David K. Jackson, Ian Johnston, Dominic Kwiatkowski, Cordelia Langford, John Sillitoe on behalf of the Wellcome Sanger Institute COVID-19 Surveillance Team |
| EPI_ISL_2831598, EPI_ISL_2831599, EPI_ISL_2831600, EPI_ISL_2831601 | Lighthouse Lab in Glasgow | Wellcome Sanger Institute for the COVID-19 Genomics UK (COG-UK) Consortium | Harper VanSteenhouse, Yumi Kasai, David Gray, Carol Clugston, Anna Dominiczak and Alex Alderton, Roberto Amato, Jeffrey Barrett, Sonia Goncalves, Ewan Harrison, David K. Jackson, Ian Johnston, Dominic Kwiatkowski, Cordelia Langford, John Sillitoe on behalf of the Wellcome Sanger Institute COVID-19 Surveillance Team |
| EPI_ISL_2831602, EPI_ISL_2831603, EPI_ISL_2831604, EPI_ISL_2831605, EPI_ISL_2831606, EPI_ISL_2831608, EPI_ISL_2831609 | Lighthouse Lab in Alderley Park | Wellcome Sanger Institute for the COVID-19 Genomics UK (COG-UK) Consortium | Jacquelyn Wynn, Mairead Hyland, The Lighthouse Lab in Alderley Park and Alex Alderton, Roberto Amato, Jeffrey Barrett, Sonia Goncalves, Ewan Harrison, David K. Jackson, Ian Johnston, Dominic Kwiatkowski, Cordelia Langford, John Sillitoe on behalf of the Wellcome Sanger Institute COVID-19 Surveillance Team |
| EPI_ISL_2831610 | Lighthouse Lab in Glasgow | Wellcome Sanger Institute for the COVID-19 Genomics UK (COG-UK) Consortium | Harper VanSteenhouse, Yumi Kasai, David Gray, Carol Clugston, Anna Dominiczak and Alex Alderton, Roberto Amato, Jeffrey Barrett, Sonia Goncalves, Ewan Harrison, David K. Jackson, Ian Johnston, Dominic Kwiatkowski, Cordelia Langford, John Sillitoe on behalf of the Wellcome Sanger Institute COVID-19 Surveillance Team |
| EPI_ISL_2831611 | Lighthouse Lab in Alderley Park | Wellcome Sanger Institute for the COVID-19 Genomics UK (COG-UK) Consortium | Jacquelyn Wynn, Mairead Hyland, The Lighthouse Lab in Alderley Park and Alex Alderton, Roberto Amato, Jeffrey Barrett, Sonia Goncalves, Ewan Harrison, David K. Jackson, Ian Johnston, Dominic Kwiatkowski, Cordelia Langford, John Sillitoe on behalf of the Wellcome Sanger Institute COVID-19 Surveillance Team |
| EPI_ISL_2831612, EPI_ISL_2831613 | Lighthouse Lab in Glasgow | Wellcome Sanger Institute for the COVID-19 Genomics UK (COG-UK) Consortium | Harper VanSteenhouse, Yumi Kasai, David Gray, Carol Clugston, Anna Dominiczak and Alex Alderton, Roberto Amato, Jeffrey Barrett, Sonia Goncalves, Ewan Harrison, David K. Jackson, Ian Johnston, Dominic Kwiatkowski, Cordelia Langford, John Sillitoe on behalf of the Wellcome Sanger Institute COVID-19 Surveillance Team |
| EPI_ISL_2831615 | Lighthouse Lab in Alderley Park | Wellcome Sanger Institute for the COVID-19 Genomics UK (COG-UK) Consortium | Jacquelyn Wynn, Mairead Hyland, The Lighthouse Lab in Alderley Park and Alex Alderton, Roberto Amato, Jeffrey Barrett, Sonia Goncalves, Ewan Harrison, David K. Jackson, Ian Johnston, Dominic Kwiatkowski, Cordelia Langford, John Sillitoe on behalf of the Wellcome Sanger Institute COVID-19 Surveillance Team |
| EPI_ISL_2831616 | Lighthouse Lab in Glasgow | Wellcome Sanger Institute for the COVID-19 Genomics UK (COG-UK) Consortium | Harper VanSteenhouse, Yumi Kasai, David Gray, Carol Clugston, Anna Dominiczak and Alex Alderton, Roberto Amato, Jeffrey Barrett, Sonia Goncalves, Ewan Harrison, David K. Jackson, Ian Johnston, Dominic Kwiatkowski, Cordelia Langford, John Sillitoe on behalf of the Wellcome Sanger Institute COVID-19 Surveillance Team |
| EPI_ISL_2831617, EPI_ISL_2831618 | Lighthouse Lab in Alderley Park | Wellcome Sanger Institute for the COVID-19 Genomics UK (COG-UK) Consortium | Jacquelyn Wynn, Mairead Hyland, The Lighthouse Lab in Alderley Park and Alex Alderton, Roberto Amato, Jeffrey Barrett, Sonia Goncalves, Ewan Harrison, David K. Jackson, Ian Johnston, Dominic Kwiatkowski, Cordelia Langford, John Sillitoe on behalf of the Wellcome Sanger Institute COVID-19 Surveillance Team |
| EPI_ISL_2831619, EPI_ISL_2831620 | Lighthouse Lab in Glasgow | Wellcome Sanger Institute for the COVID-19 Genomics UK (COG-UK) Consortium | Harper VanSteenhouse, Yumi Kasai, David Gray, Carol Clugston, Anna Dominiczak and Alex Alderton, Roberto Amato, Jeffrey Barrett, Sonia Goncalves, Ewan Harrison, David K. Jackson, Ian Johnston, Dominic Kwiatkowski, Cordelia Langford, John Sillitoe on behalf of the Wellcome Sanger Institute COVID-19 Surveillance Team |
| EPI_ISL_2831621 | Lighthouse Lab in Alderley Park | Wellcome Sanger Institute for the COVID-19 Genomics UK (COG-UK) Consortium | Jacquelyn Wynn, Mairead Hyland, The Lighthouse Lab in Alderley Park and Alex Alderton, Roberto Amato, Jeffrey Barrett, Sonia Goncalves, Ewan Harrison, David K. Jackson, Ian Johnston, Dominic Kwiatkowski, Cordelia Langford, John Sillitoe on behalf of the Wellcome Sanger Institute COVID-19 Surveillance Team |
| EPI_ISL_2831622, EPI_ISL_2831623 | Lighthouse Lab in Glasgow | Wellcome Sanger Institute for the COVID-19 Genomics UK (COG-UK) Consortium | Harper VanSteenhouse, Yumi Kasai, David Gray, Carol Clugston, Anna Dominiczak and Alex Alderton, Roberto Amato, Jeffrey Barrett, Sonia Goncalves, Ewan Harrison, David K. Jackson, Ian Johnston, Dominic Kwiatkowski, Cordelia Langford, John Sillitoe on behalf of the Wellcome Sanger Institute COVID-19 Surveillance Team |
| EPI_ISL_2831624, EPI_ISL_2831625 | Lighthouse Lab in Alderley Park | Wellcome Sanger Institute for the COVID-19 Genomics UK (COG-UK) Consortium | Jacquelyn Wynn, Mairead Hyland, The Lighthouse Lab in Alderley Park and Alex Alderton, Roberto Amato, Jeffrey Barrett, Sonia Goncalves, Ewan Harrison, David K. Jackson, Ian Johnston, Dominic Kwiatkowski, Cordelia Langford, John Sillitoe on behalf of the Wellcome Sanger Institute COVID-19 Surveillance Team |
| EPI_ISL_2831626, EPI_ISL_2831627, EPI_ISL_2831628, EPI_ISL_2831629 | Lighthouse Lab in Glasgow | Wellcome Sanger Institute for the COVID-19 Genomics UK (COG-UK) Consortium | Harper VanSteenhouse, Yumi Kasai, David Gray, Carol Clugston, Anna Dominiczak and Alex Alderton, Roberto Amato, Jeffrey Barrett, Sonia Goncalves, Ewan Harrison, David K. Jackson, Ian Johnston, Dominic Kwiatkowski, Cordelia Langford, John Sillitoe on behalf of the Wellcome Sanger Institute COVID-19 Surveillance Team |
| EPI_ISL_2831630 | Lighthouse Lab in Alderley Park | Wellcome Sanger Institute for the COVID-19 Genomics UK (COG-UK) Consortium | Jacquelyn Wynn, Mairead Hyland, The Lighthouse Lab in Alderley Park and Alex Alderton, Roberto Amato, Jeffrey Barrett, Sonia Goncalves, Ewan Harrison, David K. Jackson, Ian Johnston, Dominic Kwiatkowski, Cordelia Langford, John Sillitoe on behalf of the Wellcome Sanger Institute COVID-19 Surveillance Team |
| EPI_ISL_2831631 | Lighthouse Lab in Glasgow | Wellcome Sanger Institute for the COVID-19 Genomics UK (COG-UK) Consortium | Harper VanSteenhouse, Yumi Kasai, David Gray, Carol Clugston, Anna Dominiczak and Alex Alderton, Roberto Amato, Jeffrey Barrett, Sonia Goncalves, Ewan Harrison, David K. Jackson, Ian Johnston, Dominic Kwiatkowski, Cordelia Langford, John Sillitoe on behalf of the Wellcome Sanger Institute COVID-19 Surveillance Team |
| EPI_ISL_2831632 | Lighthouse Lab in Alderley Park | Wellcome Sanger Institute for the COVID-19 Genomics UK (COG-UK) Consortium | Jacquelyn Wynn, Mairead Hyland, The Lighthouse Lab in Alderley Park and Alex Alderton, Roberto Amato, Jeffrey Barrett, Sonia Goncalves, Ewan Harrison, David K. Jackson, Ian Johnston, Dominic Kwiatkowski, Cordelia Langford, John Sillitoe on behalf of the Wellcome Sanger Institute COVID-19 Surveillance Team |
| EPI_ISL_2831633, EPI_ISL_2831634, EPI_ISL_2831635, EPI_ISL_2831636 | Lighthouse Lab in Glasgow | Wellcome Sanger Institute for the COVID-19 Genomics UK (COG-UK) Consortium | Harper VanSteenhouse, Yumi Kasai, David Gray, Carol Clugston, Anna Dominiczak and Alex Alderton, Roberto Amato, Jeffrey Barrett, Sonia Goncalves, Ewan Harrison, David K. Jackson, Ian Johnston, Dominic Kwiatkowski, Cordelia Langford, John Sillitoe on behalf of the Wellcome Sanger Institute COVID-19 Surveillance Team |
| EPI_ISL_2831637, EPI_ISL_2831638 | Lighthouse Lab in Alderley Park | Wellcome Sanger Institute for the COVID-19 Genomics UK (COG-UK) Consortium | Jacquelyn Wynn, Mairead Hyland, The Lighthouse Lab in Alderley Park and Alex Alderton, Roberto Amato, Jeffrey Barrett, Sonia Goncalves, Ewan Harrison, David K. Jackson, Ian Johnston, Dominic Kwiatkowski, Cordelia Langford, John Sillitoe on behalf of the Wellcome Sanger Institute COVID-19 Surveillance Team |
| EPI_ISL_2831639 | Lighthouse Lab in Glasgow | Wellcome Sanger Institute for the COVID-19 Genomics UK (COG-UK) Consortium | Harper VanSteenhouse, Yumi Kasai, David Gray, Carol Clugston, Anna Dominiczak and Alex Alderton, Roberto Amato, Jeffrey Barrett, Sonia Goncalves, Ewan Harrison, David K. Jackson, Ian Johnston, Dominic Kwiatkowski, Cordelia Langford, John Sillitoe on behalf of the Wellcome Sanger Institute COVID-19 Surveillance Team |
| EPI_ISL_2831640 | Lighthouse Lab in Alderley Park | Wellcome Sanger Institute for the COVID-19 Genomics UK (COG-UK) Consortium | Jacquelyn Wynn, Mairead Hyland, The Lighthouse Lab in Alderley Park and Alex Alderton, Roberto Amato, Jeffrey Barrett, Sonia Goncalves, Ewan Harrison, David K. Jackson, Ian Johnston, Dominic Kwiatkowski, Cordelia Langford, John Sillitoe on behalf of the Wellcome Sanger Institute COVID-19 Surveillance Team |
| EPI_ISL_2831641, EPI_ISL_2831642 | Lighthouse Lab in Glasgow | Wellcome Sanger Institute for the COVID-19 Genomics UK (COG-UK) Consortium | Harper VanSteenhouse, Yumi Kasai, David Gray, Carol Clugston, Anna Dominiczak and Alex Alderton, Roberto Amato, Jeffrey Barrett, Sonia Goncalves, Ewan Harrison, David K. Jackson, Ian Johnston, Dominic Kwiatkowski, Cordelia Langford, John Sillitoe on behalf of the Wellcome Sanger Institute COVID-19 Surveillance Team |
| EPI_ISL_2831643, EPI_ISL_2831644, EPI_ISL_2831645 | Lighthouse Lab in Alderley Park | Wellcome Sanger Institute for the COVID-19 Genomics UK (COG-UK) Consortium | Jacquelyn Wynn, Mairead Hyland, The Lighthouse Lab in Alderley Park and Alex Alderton, Roberto Amato, Jeffrey Barrett, Sonia Goncalves, Ewan Harrison, David K. Jackson, Ian Johnston, Dominic Kwiatkowski, Cordelia Langford, John Sillitoe on behalf of the Wellcome Sanger Institute COVID-19 Surveillance Team |
| EPI_ISL_2831646, EPI_ISL_2831647 | Lighthouse Lab in Glasgow | Wellcome Sanger Institute for the COVID-19 Genomics UK (COG-UK) Consortium | Harper VanSteenhouse, Yumi Kasai, David Gray, Carol Clugston, Anna Dominiczak and Alex Alderton, Roberto Amato, Jeffrey Barrett, Sonia Goncalves, Ewan Harrison, David K. Jackson, Ian Johnston, Dominic Kwiatkowski, Cordelia Langford, John Sillitoe on behalf of the Wellcome Sanger Institute COVID-19 Surveillance Team |
| EPI_ISL_2831648 | Lighthouse Lab in Alderley Park | Wellcome Sanger Institute for the COVID-19 Genomics UK (COG-UK) Consortium | Jacquelyn Wynn, Mairead Hyland, The Lighthouse Lab in Alderley Park and Alex Alderton, Roberto Amato, Jeffrey Barrett, Sonia Goncalves, Ewan Harrison, David K. Jackson, Ian Johnston, Dominic Kwiatkowski, Cordelia Langford, John Sillitoe on behalf of the Wellcome Sanger Institute COVID-19 Surveillance Team |
| EPI_ISL_2831649, EPI_ISL_2831650 | Lighthouse Lab in Glasgow | Wellcome Sanger Institute for the COVID-19 Genomics UK (COG-UK) Consortium | Harper VanSteenhouse, Yumi Kasai, David Gray, Carol Clugston, Anna Dominiczak and Alex Alderton, Roberto Amato, Jeffrey Barrett, Sonia Goncalves, Ewan Harrison, David K. Jackson, Ian Johnston, Dominic Kwiatkowski, Cordelia Langford, John Sillitoe on behalf of the Wellcome Sanger Institute COVID-19 Surveillance Team |
| EPI_ISL_2831651 | Lighthouse Lab in Alderley Park | Wellcome Sanger Institute for the COVID-19 Genomics UK (COG-UK) Consortium | Jacquelyn Wynn, Mairead Hyland, The Lighthouse Lab in Alderley Park and Alex Alderton, Roberto Amato, Jeffrey Barrett, Sonia Goncalves, Ewan Harrison, David K. Jackson, Ian Johnston, Dominic Kwiatkowski, Cordelia Langford, John Sillitoe on behalf of the Wellcome Sanger Institute COVID-19 Surveillance Team |
| EPI_ISL_2831652, EPI_ISL_2831653 | Lighthouse Lab in Glasgow | Wellcome Sanger Institute for the COVID-19 Genomics UK (COG-UK) Consortium | Harper VanSteenhouse, Yumi Kasai, David Gray, Carol Clugston, Anna Dominiczak and Alex Alderton, Roberto Amato, Jeffrey Barrett, Sonia Goncalves, Ewan Harrison, David K. Jackson, Ian Johnston, Dominic Kwiatkowski, Cordelia Langford, John Sillitoe on behalf of the Wellcome Sanger Institute COVID-19 Surveillance Team |
| EPI_ISL_2831654, EPI_ISL_2831655 | Lighthouse Lab in Alderley Park | Wellcome Sanger Institute for the COVID-19 Genomics UK (COG-UK) Consortium | Jacquelyn Wynn, Mairead Hyland, The Lighthouse Lab in Alderley Park and Alex Alderton, Roberto Amato, Jeffrey Barrett, Sonia Goncalves, Ewan Harrison, David K. Jackson, Ian Johnston, Dominic Kwiatkowski, Cordelia Langford, John Sillitoe on behalf of the Wellcome Sanger Institute COVID-19 Surveillance Team |
| EPI_ISL_2831656 | Lighthouse Lab in Glasgow | Wellcome Sanger Institute for the COVID-19 Genomics UK (COG-UK) Consortium | Harper VanSteenhouse, Yumi Kasai, David Gray, Carol Clugston, Anna Dominiczak and Alex Alderton, Roberto Amato, Jeffrey Barrett, Sonia Goncalves, Ewan Harrison, David K. Jackson, Ian Johnston, Dominic Kwiatkowski, Cordelia Langford, John Sillitoe on behalf of the Wellcome Sanger Institute COVID-19 Surveillance Team |
| EPI_ISL_2831657, EPI_ISL_2831658 | Lighthouse Lab in Alderley Park | Wellcome Sanger Institute for the COVID-19 Genomics UK (COG-UK) Consortium | Jacquelyn Wynn, Mairead Hyland, The Lighthouse Lab in Alderley Park and Alex Alderton, Roberto Amato, Jeffrey Barrett, Sonia Goncalves, Ewan Harrison, David K. Jackson, Ian Johnston, Dominic Kwiatkowski, Cordelia Langford, John Sillitoe on behalf of the Wellcome Sanger Institute COVID-19 Surveillance Team |
| EPI_ISL_2831659 | Lighthouse Lab in Glasgow | Wellcome Sanger Institute for the COVID-19 Genomics UK (COG-UK) Consortium | Harper VanSteenhouse, Yumi Kasai, David Gray, Carol Clugston, Anna Dominiczak and Alex Alderton, Roberto Amato, Jeffrey Barrett, Sonia Goncalves, Ewan Harrison, David K. Jackson, Ian Johnston, Dominic Kwiatkowski, Cordelia Langford, John Sillitoe on behalf of the Wellcome Sanger Institute COVID-19 Surveillance Team |
| EPI_ISL_2831660, EPI_ISL_2831661, EPI_ISL_2831662 | Lighthouse Lab in Alderley Park | Wellcome Sanger Institute for the COVID-19 Genomics UK (COG-UK) Consortium | Jacquelyn Wynn, Mairead Hyland, The Lighthouse Lab in Alderley Park and Alex Alderton, Roberto Amato, Jeffrey Barrett, Sonia Goncalves, Ewan Harrison, David K. Jackson, Ian Johnston, Dominic Kwiatkowski, Cordelia Langford, John Sillitoe on behalf of the Wellcome Sanger Institute COVID-19 Surveillance Team |
| EPI_ISL_2831663 | Lighthouse Lab in Glasgow | Wellcome Sanger Institute for the COVID-19 Genomics UK (COG-UK) Consortium | Harper VanSteenhouse, Yumi Kasai, David Gray, Carol Clugston, Anna Dominiczak and Alex Alderton, Roberto Amato, Jeffrey Barrett, Sonia Goncalves, Ewan Harrison, David K. Jackson, Ian Johnston, Dominic Kwiatkowski, Cordelia Langford, John Sillitoe on behalf of the Wellcome Sanger Institute COVID-19 Surveillance Team |
| EPI_ISL_2831664 | Lighthouse Lab in Alderley Park | Wellcome Sanger Institute for the COVID-19 Genomics UK (COG-UK) Consortium | Jacquelyn Wynn, Mairead Hyland, The Lighthouse Lab in Alderley Park and Alex Alderton, Roberto Amato, Jeffrey Barrett, Sonia Goncalves, Ewan Harrison, David K. Jackson, Ian Johnston, Dominic Kwiatkowski, Cordelia Langford, John Sillitoe on behalf of the Wellcome Sanger Institute COVID-19 Surveillance Team |

| | | | |
|---|---|---|---|
| EPI_ISL_2831665, EPI_ISL_2831666 | Lighthouse Lab in Glasgow | Wellcome Sanger Institute for the COVID-19 Genomics UK (COG-UK) Consortium | Harper VanSteenhouse, Yumi Kasai, David Gray, Carol Clugston, Anna Dominiczak and Alex Alderton, Roberto Amato, Jeffrey Barrett, Sonia Goncalves, Ewan Harrison, David K. Jackson, Ian Johnston, Dominic Kwiatkowski, Cordelia Langford, John Sillitoe on behalf of the Wellcome Sanger Institute COVID-19 Surveillance Team |
| EPI_ISL_2831667 | Lighthouse Lab in Alderley Park | Wellcome Sanger Institute for the COVID-19 Genomics UK (COG-UK) Consortium | Jacquelyn Wynn, Mairead Hyland, The Lighthouse Lab in Alderley Park and Alex Alderton, Roberto Amato, Jeffrey Barrett, Sonia Goncalves, Ewan Harrison, David K. Jackson, Ian Johnston, Dominic Kwiatkowski, Cordelia Langford, John Sillitoe on behalf of the Wellcome Sanger Institute COVID-19 Surveillance Team |
| EPI_ISL_2831668 | Lighthouse Lab in Glasgow | Wellcome Sanger Institute for the COVID-19 Genomics UK (COG-UK) Consortium | Harper VanSteenhouse, Yumi Kasai, David Gray, Carol Clugston, Anna Dominiczak and Alex Alderton, Roberto Amato, Jeffrey Barrett, Sonia Goncalves, Ewan Harrison, David K. Jackson, Ian Johnston, Dominic Kwiatkowski, Cordelia Langford, John Sillitoe on behalf of the Wellcome Sanger Institute COVID-19 Surveillance Team |
| EPI_ISL_2831669 | Lighthouse Lab in Alderley Park | Wellcome Sanger Institute for the COVID-19 Genomics UK (COG-UK) Consortium | Jacquelyn Wynn, Mairead Hyland, The Lighthouse Lab in Alderley Park and Alex Alderton, Roberto Amato, Jeffrey Barrett, Sonia Goncalves, Ewan Harrison, David K. Jackson, Ian Johnston, Dominic Kwiatkowski, Cordelia Langford, John Sillitoe on behalf of the Wellcome Sanger Institute COVID-19 Surveillance Team |
| EPI_ISL_2831670, EPI_ISL_2831671, EPI_ISL_2831672, EPI_ISL_2831673 | Lighthouse Lab in Glasgow | Wellcome Sanger Institute for the COVID-19 Genomics UK (COG-UK) Consortium | Harper VanSteenhouse, Yumi Kasai, David Gray, Carol Clugston, Anna Dominiczak and Alex Alderton, Roberto Amato, Jeffrey Barrett, Sonia Goncalves, Ewan Harrison, David K. Jackson, Ian Johnston, Dominic Kwiatkowski, Cordelia Langford, John Sillitoe on behalf of the Wellcome Sanger Institute COVID-19 Surveillance Team |
| EPI_ISL_2831674 | Lighthouse Lab in Alderley Park | Wellcome Sanger Institute for the COVID-19 Genomics UK (COG-UK) Consortium | Jacquelyn Wynn, Mairead Hyland, The Lighthouse Lab in Alderley Park and Alex Alderton, Roberto Amato, Jeffrey Barrett, Sonia Goncalves, Ewan Harrison, David K. Jackson, Ian Johnston, Dominic Kwiatkowski, Cordelia Langford, John Sillitoe on behalf of the Wellcome Sanger Institute COVID-19 Surveillance Team |
| EPI_ISL_2831675 | Lighthouse Lab in Glasgow | Wellcome Sanger Institute for the COVID-19 Genomics UK (COG-UK) Consortium | Harper VanSteenhouse, Yumi Kasai, David Gray, Carol Clugston, Anna Dominiczak and Alex Alderton, Roberto Amato, Jeffrey Barrett, Sonia Goncalves, Ewan Harrison, David K. Jackson, Ian Johnston, Dominic Kwiatkowski, Cordelia Langford, John Sillitoe on behalf of the Wellcome Sanger Institute COVID-19 Surveillance Team |
| EPI_ISL_2831676 | Lighthouse Lab in Alderley Park | Wellcome Sanger Institute for the COVID-19 Genomics UK (COG-UK) Consortium | Jacquelyn Wynn, Mairead Hyland, The Lighthouse Lab in Alderley Park and Alex Alderton, Roberto Amato, Jeffrey Barrett, Sonia Goncalves, Ewan Harrison, David K. Jackson, Ian Johnston, Dominic Kwiatkowski, Cordelia Langford, John Sillitoe on behalf of the Wellcome Sanger Institute COVID-19 Surveillance Team |
| EPI_ISL_2831677, EPI_ISL_2831678, EPI_ISL_2831679, EPI_ISL_2831680 | Lighthouse Lab in Glasgow | Wellcome Sanger Institute for the COVID-19 Genomics UK (COG-UK) Consortium | Harper VanSteenhouse, Yumi Kasai, David Gray, Carol Clugston, Anna Dominiczak and Alex Alderton, Roberto Amato, Jeffrey Barrett, Sonia Goncalves, Ewan Harrison, David K. Jackson, Ian Johnston, Dominic Kwiatkowski, Cordelia Langford, John Sillitoe on behalf of the Wellcome Sanger Institute COVID-19 Surveillance Team |
| EPI_ISL_2831681, EPI_ISL_2831682, EPI_ISL_2831683 | Lighthouse Lab in Alderley Park | Wellcome Sanger Institute for the COVID-19 Genomics UK (COG-UK) Consortium | Jacquelyn Wynn, Mairead Hyland, The Lighthouse Lab in Alderley Park and Alex Alderton, Roberto Amato, Jeffrey Barrett, Sonia Goncalves, Ewan Harrison, David K. Jackson, Ian Johnston, Dominic Kwiatkowski, Cordelia Langford, John Sillitoe on behalf of the Wellcome Sanger Institute COVID-19 Surveillance Team |
| EPI_ISL_2831684 | Lighthouse Lab in Glasgow | Wellcome Sanger Institute for the COVID-19 Genomics UK (COG-UK) Consortium | Harper VanSteenhouse, Yumi Kasai, David Gray, Carol Clugston, Anna Dominiczak and Alex Alderton, Roberto Amato, Jeffrey Barrett, Sonia Goncalves, Ewan Harrison, David K. Jackson, Ian Johnston, Dominic Kwiatkowski, Cordelia Langford, John Sillitoe on behalf of the Wellcome Sanger Institute COVID-19 Surveillance Team |
| EPI_ISL_2831685, EPI_ISL_2831686 | Lighthouse Lab in Alderley Park | Wellcome Sanger Institute for the COVID-19 Genomics UK (COG-UK) Consortium | Jacquelyn Wynn, Mairead Hyland, The Lighthouse Lab in Alderley Park and Alex Alderton, Roberto Amato, Jeffrey Barrett, Sonia Goncalves, Ewan Harrison, David K. Jackson, Ian Johnston, Dominic Kwiatkowski, Cordelia Langford, John Sillitoe on behalf of the Wellcome Sanger Institute COVID-19 Surveillance Team |
| EPI_ISL_2831687, EPI_ISL_2831688, EPI_ISL_2831689, EPI_ISL_2831690 | Lighthouse Lab in Glasgow | Wellcome Sanger Institute for the COVID-19 Genomics UK (COG-UK) Consortium | Harper VanSteenhouse, Yumi Kasai, David Gray, Carol Clugston, Anna Dominiczak and Alex Alderton, Roberto Amato, Jeffrey Barrett, Sonia Goncalves, Ewan Harrison, David K. Jackson, Ian Johnston, Dominic Kwiatkowski, Cordelia Langford, John Sillitoe on behalf of the Wellcome Sanger Institute COVID-19 Surveillance Team |
| EPI_ISL_2831691, EPI_ISL_2831692 | Lighthouse Lab in Alderley Park | Wellcome Sanger Institute for the COVID-19 Genomics UK (COG-UK) Consortium | Jacquelyn Wynn, Mairead Hyland, The Lighthouse Lab in Alderley Park and Alex Alderton, Roberto Amato, Jeffrey Barrett, Sonia Goncalves, Ewan Harrison, David K. Jackson, Ian Johnston, Dominic Kwiatkowski, Cordelia Langford, John Sillitoe on behalf of the Wellcome Sanger Institute COVID-19 Surveillance Team |
| EPI_ISL_2831693 | Lighthouse Lab in Glasgow | Wellcome Sanger Institute for the COVID-19 Genomics UK (COG-UK) Consortium | Harper VanSteenhouse, Yumi Kasai, David Gray, Carol Clugston, Anna Dominiczak and Alex Alderton, Roberto Amato, Jeffrey Barrett, Sonia Goncalves, Ewan Harrison, David K. Jackson, Ian Johnston, Dominic Kwiatkowski, Cordelia Langford, John Sillitoe on behalf of the Wellcome Sanger Institute COVID-19 Surveillance Team |
| EPI_ISL_2831694 | Lighthouse Lab in Alderley Park | Wellcome Sanger Institute for the COVID-19 Genomics UK (COG-UK) Consortium | Jacquelyn Wynn, Mairead Hyland, The Lighthouse Lab in Alderley Park and Alex Alderton, Roberto Amato, Jeffrey Barrett, Sonia Goncalves, Ewan Harrison, David K. Jackson, Ian Johnston, Dominic Kwiatkowski, Cordelia Langford, John Sillitoe on behalf of the Wellcome Sanger Institute COVID-19 Surveillance Team |
| EPI_ISL_2831695, EPI_ISL_2831696 | Lighthouse Lab in Glasgow | Wellcome Sanger Institute for the COVID-19 Genomics UK (COG-UK) Consortium | Harper VanSteenhouse, Yumi Kasai, David Gray, Carol Clugston, Anna Dominiczak and Alex Alderton, Roberto Amato, Jeffrey Barrett, Sonia Goncalves, Ewan Harrison, David K. Jackson, Ian Johnston, Dominic Kwiatkowski, Cordelia Langford, John Sillitoe on behalf of the Wellcome Sanger Institute COVID-19 Surveillance Team |
| EPI_ISL_2831697, EPI_ISL_2831698, EPI_ISL_2831699, EPI_ISL_2831700, EPI_ISL_2831701 | Lighthouse Lab in Alderley Park | Wellcome Sanger Institute for the COVID-19 Genomics UK (COG-UK) Consortium | Jacquelyn Wynn, Mairead Hyland, The Lighthouse Lab in Alderley Park and Alex Alderton, Roberto Amato, Jeffrey Barrett, Sonia Goncalves, Ewan Harrison, David K. Jackson, Ian Johnston, Dominic Kwiatkowski, Cordelia Langford, John Sillitoe on behalf of the Wellcome Sanger Institute COVID-19 Surveillance Team |
| EPI_ISL_2831702 | Lighthouse Lab in Glasgow | Wellcome Sanger Institute for the COVID-19 Genomics UK (COG-UK) Consortium | Harper VanSteenhouse, Yumi Kasai, David Gray, Carol Clugston, Anna Dominiczak and Alex Alderton, Roberto Amato, Jeffrey Barrett, Sonia Goncalves, Ewan Harrison, David K. Jackson, Ian Johnston, Dominic Kwiatkowski, Cordelia Langford, John Sillitoe on behalf of the Wellcome Sanger Institute COVID-19 Surveillance Team |
| EPI_ISL_2831703 | Lighthouse Lab in Alderley Park | Wellcome Sanger Institute for the COVID-19 Genomics UK (COG-UK) Consortium | Jacquelyn Wynn, Mairead Hyland, The Lighthouse Lab in Alderley Park and Alex Alderton, Roberto Amato, Jeffrey Barrett, Sonia Goncalves, Ewan Harrison, David K. Jackson, Ian Johnston, Dominic Kwiatkowski, Cordelia Langford, John Sillitoe on behalf of the Wellcome Sanger Institute COVID-19 Surveillance Team |
| EPI_ISL_2831704 | Lighthouse Lab in Glasgow | Wellcome Sanger Institute for the COVID-19 Genomics UK (COG-UK) Consortium | Harper VanSteenhouse, Yumi Kasai, David Gray, Carol Clugston, Anna Dominiczak and Alex Alderton, Roberto Amato, Jeffrey Barrett, Sonia Goncalves, Ewan Harrison, David K. Jackson, Ian Johnston, Dominic Kwiatkowski, Cordelia Langford, John Sillitoe on behalf of the Wellcome Sanger Institute COVID-19 Surveillance Team |
| EPI_ISL_2831705 | Lighthouse Lab in Alderley Park | Wellcome Sanger Institute for the COVID-19 Genomics UK (COG-UK) Consortium | Jacquelyn Wynn, Mairead Hyland, The Lighthouse Lab in Alderley Park and Alex Alderton, Roberto Amato, Jeffrey Barrett, Sonia Goncalves, Ewan Harrison, David K. Jackson, Ian Johnston, Dominic Kwiatkowski, Cordelia Langford, John Sillitoe on behalf of the Wellcome Sanger Institute COVID-19 Surveillance Team |
| EPI_ISL_2831706, EPI_ISL_2831707, EPI_ISL_2831708, EPI_ISL_2831709, EPI_ISL_2831710 | Lighthouse Lab in Glasgow | Wellcome Sanger Institute for the COVID-19 Genomics UK (COG-UK) Consortium | Harper VanSteenhouse, Yumi Kasai, David Gray, Carol Clugston, Anna Dominiczak and Alex Alderton, Roberto Amato, Jeffrey Barrett, Sonia Goncalves, Ewan Harrison, David K. Jackson, Ian Johnston, Dominic Kwiatkowski, Cordelia Langford, John Sillitoe on behalf of the Wellcome Sanger Institute COVID-19 Surveillance Team |
| EPI_ISL_2831711 | Lighthouse Lab in Alderley Park | Wellcome Sanger Institute for the COVID-19 Genomics UK (COG-UK) Consortium | Jacquelyn Wynn, Mairead Hyland, The Lighthouse Lab in Alderley Park and Alex Alderton, Roberto Amato, Jeffrey Barrett, Sonia Goncalves, Ewan Harrison, David K. Jackson, Ian Johnston, Dominic Kwiatkowski, Cordelia Langford, John Sillitoe on behalf of the Wellcome Sanger Institute COVID-19 Surveillance Team |
| EPI_ISL_2831712, EPI_ISL_2831713, EPI_ISL_2831714 | Lighthouse Lab in Glasgow | Wellcome Sanger Institute for the COVID-19 Genomics UK (COG-UK) Consortium | Harper VanSteenhouse, Yumi Kasai, David Gray, Carol Clugston, Anna Dominiczak and Alex Alderton, Roberto Amato, Jeffrey Barrett, Sonia Goncalves, Ewan Harrison, David K. Jackson, Ian Johnston, Dominic Kwiatkowski, Cordelia Langford, John Sillitoe on behalf of the Wellcome Sanger Institute COVID-19 Surveillance Team |
| EPI_ISL_2831715 | Lighthouse Lab in Alderley Park | Wellcome Sanger Institute for the COVID-19 Genomics UK (COG-UK) Consortium | Jacquelyn Wynn, Mairead Hyland, The Lighthouse Lab in Alderley Park and Alex Alderton, Roberto Amato, Jeffrey Barrett, Sonia Goncalves, Ewan Harrison, David K. Jackson, Ian Johnston, Dominic Kwiatkowski, Cordelia Langford, John Sillitoe on behalf of the Wellcome Sanger Institute COVID-19 Surveillance Team |
| EPI_ISL_2831716, EPI_ISL_2831717 | Lighthouse Lab in Glasgow | Wellcome Sanger Institute for the COVID-19 Genomics UK (COG-UK) Consortium | Harper VanSteenhouse, Yumi Kasai, David Gray, Carol Clugston, Anna Dominiczak and Alex Alderton, Roberto Amato, Jeffrey Barrett, Sonia Goncalves, Ewan Harrison, David K. Jackson, Ian Johnston, Dominic Kwiatkowski, Cordelia Langford, John Sillitoe on behalf of the Wellcome Sanger Institute COVID-19 Surveillance Team |
| EPI_ISL_2831718 | Lighthouse Lab in Alderley Park | Wellcome Sanger Institute for the COVID-19 Genomics UK (COG-UK) Consortium | Jacquelyn Wynn, Mairead Hyland, The Lighthouse Lab in Alderley Park and Alex Alderton, Roberto Amato, Jeffrey Barrett, Sonia Goncalves, Ewan Harrison, David K. Jackson, Ian Johnston, Dominic Kwiatkowski, Cordelia Langford, John Sillitoe on behalf of the Wellcome Sanger Institute COVID-19 Surveillance Team |
| EPI_ISL_2831719 | Lighthouse Lab in Glasgow | Wellcome Sanger Institute for the COVID-19 Genomics UK (COG-UK) Consortium | Harper VanSteenhouse, Yumi Kasai, David Gray, Carol Clugston, Anna Dominiczak and Alex Alderton, Roberto Amato, Jeffrey Barrett, Sonia Goncalves, Ewan Harrison, David K. Jackson, Ian Johnston, Dominic Kwiatkowski, Cordelia Langford, John Sillitoe on behalf of the Wellcome Sanger Institute COVID-19 Surveillance Team |
| EPI_ISL_2831720, EPI_ISL_2831721 | Lighthouse Lab in Alderley Park | Wellcome Sanger Institute for the COVID-19 Genomics UK (COG-UK) Consortium | Jacquelyn Wynn, Mairead Hyland, The Lighthouse Lab in Alderley Park and Alex Alderton, Roberto Amato, Jeffrey Barrett, Sonia Goncalves, Ewan Harrison, David K. Jackson, Ian Johnston, Dominic Kwiatkowski, Cordelia Langford, John Sillitoe on behalf of the Wellcome Sanger Institute COVID-19 Surveillance Team |
| EPI_ISL_2831722, EPI_ISL_2831723, EPI_ISL_2831724 | Lighthouse Lab in Glasgow | Wellcome Sanger Institute for the COVID-19 Genomics UK (COG-UK) Consortium | Harper VanSteenhouse, Yumi Kasai, David Gray, Carol Clugston, Anna Dominiczak and Alex Alderton, Roberto Amato, Jeffrey Barrett, Sonia Goncalves, Ewan Harrison, David K. Jackson, Ian Johnston, Dominic Kwiatkowski, Cordelia Langford, John Sillitoe on behalf of the Wellcome Sanger Institute COVID-19 Surveillance Team |
| EPI_ISL_2831725, EPI_ISL_2831726, EPI_ISL_2831727, EPI_ISL_2831728, EPI_ISL_2831729 | Lighthouse Lab in Alderley Park | Wellcome Sanger Institute for the COVID-19 Genomics UK (COG-UK) Consortium | Jacquelyn Wynn, Mairead Hyland, The Lighthouse Lab in Alderley Park and Alex Alderton, Roberto Amato, Jeffrey Barrett, Sonia Goncalves, Ewan Harrison, David K. Jackson, Ian Johnston, Dominic Kwiatkowski, Cordelia Langford, John Sillitoe on behalf of the Wellcome Sanger Institute COVID-19 Surveillance Team |
| EPI_ISL_2831730, EPI_ISL_2831731, EPI_ISL_2831732 | Lighthouse Lab in Glasgow | Wellcome Sanger Institute for the COVID-19 Genomics UK (COG-UK) Consortium | Harper VanSteenhouse, Yumi Kasai, David Gray, Carol Clugston, Anna Dominiczak and Alex Alderton, Roberto Amato, Jeffrey Barrett, Sonia Goncalves, Ewan Harrison, David K. Jackson, Ian Johnston, Dominic Kwiatkowski, Cordelia Langford, John Sillitoe on behalf of the Wellcome Sanger Institute COVID-19 Surveillance Team |
| EPI_ISL_2831733, EPI_ISL_2831734 | Lighthouse Lab in Alderley Park | Wellcome Sanger Institute for the COVID-19 Genomics UK (COG-UK) Consortium | Jacquelyn Wynn, Mairead Hyland, The Lighthouse Lab in Alderley Park and Alex Alderton, Roberto Amato, Jeffrey Barrett, Sonia Goncalves, Ewan Harrison, David K. Jackson, Ian Johnston, Dominic Kwiatkowski, Cordelia Langford, John Sillitoe on behalf of the Wellcome Sanger Institute COVID-19 Surveillance Team |
| EPI_ISL_2831735 | Lighthouse Lab in Glasgow | Wellcome Sanger Institute for the COVID-19 Genomics UK (COG-UK) Consortium | Harper VanSteenhouse, Yumi Kasai, David Gray, Carol Clugston, Anna Dominiczak and Alex Alderton, Roberto Amato, Jeffrey Barrett, Sonia Goncalves, Ewan Harrison, David K. Jackson, Ian Johnston, Dominic Kwiatkowski, Cordelia Langford, John Sillitoe on behalf of the Wellcome Sanger Institute COVID-19 Surveillance Team |
| EPI_ISL_2831736 | Lighthouse Lab in Alderley Park | Wellcome Sanger Institute for the COVID-19 Genomics UK (COG-UK) Consortium | Jacquelyn Wynn, Mairead Hyland, The Lighthouse Lab in Alderley Park and Alex Alderton, Roberto Amato, Jeffrey Barrett, Sonia Goncalves, Ewan Harrison, David K. Jackson, Ian Johnston, Dominic Kwiatkowski, Cordelia Langford, John Sillitoe on behalf of the Wellcome Sanger Institute COVID-19 Surveillance Team |
| EPI_ISL_2831737, EPI_ISL_2831738, EPI_ISL_2831739 | Lighthouse Lab in Glasgow | Wellcome Sanger Institute for the COVID-19 Genomics UK (COG-UK) Consortium | Harper VanSteenhouse, Yumi Kasai, David Gray, Carol Clugston, Anna Dominiczak and Alex Alderton, Roberto Amato, Jeffrey Barrett, Sonia Goncalves, Ewan Harrison, David K. Jackson, Ian Johnston, Dominic Kwiatkowski, Cordelia Langford, John Sillitoe on behalf of the Wellcome Sanger Institute COVID-19 Surveillance Team |

| | | | |
|---|---|---|---|
| EPI_ISL_2831740 | Lighthouse Lab in Alderley Park | Wellcome Sanger Institute for the COVID-19 Genomics UK (COG-UK) Consortium | Jacquelyn Wynn, Mairead Hyland, The Lighthouse Lab in Alderley Park and Alex Alderton, Roberto Amato, Jeffrey Barrett, Sonia Goncalves, Ewan Harrison, David K. Jackson, Ian Johnston, Dominic Kwiatkowski, Cordelia Langford, John Sillitoe on behalf of the Wellcome Sanger Institute COVID-19 Surveillance Team |
| EPI_ISL_2831741 | Lighthouse Lab in Glasgow | Wellcome Sanger Institute for the COVID-19 Genomics UK (COG-UK) Consortium | Harper VanSteenhouse, Yumi Kasai, David Gray, Carol Clugston, Anna Dominiczak and Alex Alderton, Roberto Amato, Jeffrey Barrett, Sonia Goncalves, Ewan Harrison, David K. Jackson, Ian Johnston, Dominic Kwiatkowski, Cordelia Langford, John Sillitoe on behalf of the Wellcome Sanger Institute COVID-19 Surveillance Team |
| EPI_ISL_2831742, EPI_ISL_2831743 | Lighthouse Lab in Alderley Park | Wellcome Sanger Institute for the COVID-19 Genomics UK (COG-UK) Consortium | Jacquelyn Wynn, Mairead Hyland, The Lighthouse Lab in Alderley Park and Alex Alderton, Roberto Amato, Jeffrey Barrett, Sonia Goncalves, Ewan Harrison, David K. Jackson, Ian Johnston, Dominic Kwiatkowski, Cordelia Langford, John Sillitoe on behalf of the Wellcome Sanger Institute COVID-19 Surveillance Team |
| EPI_ISL_2831744, EPI_ISL_2831745 | Lighthouse Lab in Glasgow | Wellcome Sanger Institute for the COVID-19 Genomics UK (COG-UK) Consortium | Harper VanSteenhouse, Yumi Kasai, David Gray, Carol Clugston, Anna Dominiczak and Alex Alderton, Roberto Amato, Jeffrey Barrett, Sonia Goncalves, Ewan Harrison, David K. Jackson, Ian Johnston, Dominic Kwiatkowski, Cordelia Langford, John Sillitoe on behalf of the Wellcome Sanger Institute COVID-19 Surveillance Team |
| EPI_ISL_2831746 | Lighthouse Lab in Alderley Park | Wellcome Sanger Institute for the COVID-19 Genomics UK (COG-UK) Consortium | Jacquelyn Wynn, Mairead Hyland, The Lighthouse Lab in Alderley Park and Alex Alderton, Roberto Amato, Jeffrey Barrett, Sonia Goncalves, Ewan Harrison, David K. Jackson, Ian Johnston, Dominic Kwiatkowski, Cordelia Langford, John Sillitoe on behalf of the Wellcome Sanger Institute COVID-19 Surveillance Team |
| EPI_ISL_2831747 | Lighthouse Lab in Glasgow | Wellcome Sanger Institute for the COVID-19 Genomics UK (COG-UK) Consortium | Harper VanSteenhouse, Yumi Kasai, David Gray, Carol Clugston, Anna Dominiczak and Alex Alderton, Roberto Amato, Jeffrey Barrett, Sonia Goncalves, Ewan Harrison, David K. Jackson, Ian Johnston, Dominic Kwiatkowski, Cordelia Langford, John Sillitoe on behalf of the Wellcome Sanger Institute COVID-19 Surveillance Team |
| EPI_ISL_2831748 | Lighthouse Lab in Alderley Park | Wellcome Sanger Institute for the COVID-19 Genomics UK (COG-UK) Consortium | Jacquelyn Wynn, Mairead Hyland, The Lighthouse Lab in Alderley Park and Alex Alderton, Roberto Amato, Jeffrey Barrett, Sonia Goncalves, Ewan Harrison, David K. Jackson, Ian Johnston, Dominic Kwiatkowski, Cordelia Langford, John Sillitoe on behalf of the Wellcome Sanger Institute COVID-19 Surveillance Team |
| EPI_ISL_2831749 | Lighthouse Lab in Glasgow | Wellcome Sanger Institute for the COVID-19 Genomics UK (COG-UK) Consortium | Harper VanSteenhouse, Yumi Kasai, David Gray, Carol Clugston, Anna Dominiczak and Alex Alderton, Roberto Amato, Jeffrey Barrett, Sonia Goncalves, Ewan Harrison, David K. Jackson, Ian Johnston, Dominic Kwiatkowski, Cordelia Langford, John Sillitoe on behalf of the Wellcome Sanger Institute COVID-19 Surveillance Team |
| EPI_ISL_2831750, EPI_ISL_2831751, EPI_ISL_2831752, EPI_ISL_2831753, EPI_ISL_2831754 | Lighthouse Lab in Alderley Park | Wellcome Sanger Institute for the COVID-19 Genomics UK (COG-UK) Consortium | Jacquelyn Wynn, Mairead Hyland, The Lighthouse Lab in Alderley Park and Alex Alderton, Roberto Amato, Jeffrey Barrett, Sonia Goncalves, Ewan Harrison, David K. Jackson, Ian Johnston, Dominic Kwiatkowski, Cordelia Langford, John Sillitoe on behalf of the Wellcome Sanger Institute COVID-19 Surveillance Team |
| EPI_ISL_2831755, EPI_ISL_2831756, EPI_ISL_2831757 | Lighthouse Lab in Glasgow | Wellcome Sanger Institute for the COVID-19 Genomics UK (COG-UK) Consortium | Harper VanSteenhouse, Yumi Kasai, David Gray, Carol Clugston, Anna Dominiczak and Alex Alderton, Roberto Amato, Jeffrey Barrett, Sonia Goncalves, Ewan Harrison, David K. Jackson, Ian Johnston, Dominic Kwiatkowski, Cordelia Langford, John Sillitoe on behalf of the Wellcome Sanger Institute COVID-19 Surveillance Team |
| EPI_ISL_2831758, EPI_ISL_2831759, EPI_ISL_2831760, EPI_ISL_2831761, EPI_ISL_2831762 | Lighthouse Lab in Alderley Park | Wellcome Sanger Institute for the COVID-19 Genomics UK (COG-UK) Consortium | Jacquelyn Wynn, Mairead Hyland, The Lighthouse Lab in Alderley Park and Alex Alderton, Roberto Amato, Jeffrey Barrett, Sonia Goncalves, Ewan Harrison, David K. Jackson, Ian Johnston, Dominic Kwiatkowski, Cordelia Langford, John Sillitoe on behalf of the Wellcome Sanger Institute COVID-19 Surveillance Team |
| EPI_ISL_2831763, EPI_ISL_2831764 | Lighthouse Lab in Milton Keynes | Wellcome Sanger Institute for the COVID-19 Genomics UK (COG-UK) Consortium | The Lighthouse Lab in Milton Keynes and Alex Alderton, Roberto Amato, Jeffrey Barrett, Sonia Goncalves, Ewan Harrison, David K. Jackson, Ian Johnston, Dominic Kwiatkowski, Cordelia Langford, John Sillitoe on behalf of the Wellcome Sanger Institute COVID-19 Surveillance Team |
| EPI_ISL_2831765 | Lighthouse Lab in Glasgow | Wellcome Sanger Institute for the COVID-19 Genomics UK (COG-UK) Consortium | Harper VanSteenhouse, Yumi Kasai, David Gray, Carol Clugston, Anna Dominiczak and Alex Alderton, Roberto Amato, Jeffrey Barrett, Sonia Goncalves, Ewan Harrison, David K. Jackson, Ian Johnston, Dominic Kwiatkowski, Cordelia Langford, John Sillitoe on behalf of the Wellcome Sanger Institute COVID-19 Surveillance Team |
| EPI_ISL_2831766 | Lighthouse Lab in Milton Keynes | Wellcome Sanger Institute for the COVID-19 Genomics UK (COG-UK) Consortium | The Lighthouse Lab in Milton Keynes and Alex Alderton, Roberto Amato, Jeffrey Barrett, Sonia Goncalves, Ewan Harrison, David K. Jackson, Ian Johnston, Dominic Kwiatkowski, Cordelia Langford, John Sillitoe on behalf of the Wellcome Sanger Institute COVID-19 Surveillance Team |
| EPI_ISL_2831767 | Lighthouse Laboratory Plymouth | Wellcome Sanger Institute for the COVID-19 Genomics UK (COG-UK) Consortium | Lighthouse Laboratory Plymouth and Alex Alderton, Roberto Amato, Jeffrey Barrett, Sonia Goncalves, Ewan Harrison, David K. Jackson, Ian Johnston, Dominic Kwiatkowski, Cordelia Langford, John Sillitoe on behalf of the Wellcome Sanger Institute COVID-19 Surveillance Team |
| EPI_ISL_2831768, EPI_ISL_2831769, EPI_ISL_2831770 | Lighthouse Lab in Milton Keynes | Wellcome Sanger Institute for the COVID-19 Genomics UK (COG-UK) Consortium | The Lighthouse Lab in Milton Keynes and Alex Alderton, Roberto Amato, Jeffrey Barrett, Sonia Goncalves, Ewan Harrison, David K. Jackson, Ian Johnston, Dominic Kwiatkowski, Cordelia Langford, John Sillitoe on behalf of the Wellcome Sanger Institute COVID-19 Surveillance Team |
| EPI_ISL_2831771 | Lighthouse Laboratory Plymouth | Wellcome Sanger Institute for the COVID-19 Genomics UK (COG-UK) Consortium | Lighthouse Laboratory Plymouth and Alex Alderton, Roberto Amato, Jeffrey Barrett, Sonia Goncalves, Ewan Harrison, David K. Jackson, Ian Johnston, Dominic Kwiatkowski, Cordelia Langford, John Sillitoe on behalf of the Wellcome Sanger Institute COVID-19 Surveillance Team |
| EPI_ISL_2831772, EPI_ISL_2831774 | Lighthouse Lab in Milton Keynes | Wellcome Sanger Institute for the COVID-19 Genomics UK (COG-UK) Consortium | The Lighthouse Lab in Milton Keynes and Alex Alderton, Roberto Amato, Jeffrey Barrett, Sonia Goncalves, Ewan Harrison, David K. Jackson, Ian Johnston, Dominic Kwiatkowski, Cordelia Langford, John Sillitoe on behalf of the Wellcome Sanger Institute COVID-19 Surveillance Team |
| EPI_ISL_2831775 | Lighthouse Laboratory Plymouth | Wellcome Sanger Institute for the COVID-19 Genomics UK (COG-UK) Consortium | Lighthouse Laboratory Plymouth and Alex Alderton, Roberto Amato, Jeffrey Barrett, Sonia Goncalves, Ewan Harrison, David K. Jackson, Ian Johnston, Dominic Kwiatkowski, Cordelia Langford, John Sillitoe on behalf of the Wellcome Sanger Institute COVID-19 Surveillance Team |
| EPI_ISL_2831776, EPI_ISL_2831777 | Lighthouse Lab in Milton Keynes | Wellcome Sanger Institute for the COVID-19 Genomics UK (COG-UK) Consortium | The Lighthouse Lab in Milton Keynes and Alex Alderton, Roberto Amato, Jeffrey Barrett, Sonia Goncalves, Ewan Harrison, David K. Jackson, Ian Johnston, Dominic Kwiatkowski, Cordelia Langford, John Sillitoe on behalf of the Wellcome Sanger Institute COVID-19 Surveillance Team |
| EPI_ISL_2831778 | Lighthouse Laboratory Plymouth | Wellcome Sanger Institute for the COVID-19 Genomics UK (COG-UK) Consortium | Lighthouse Laboratory Plymouth and Alex Alderton, Roberto Amato, Jeffrey Barrett, Sonia Goncalves, Ewan Harrison, David K. Jackson, Ian Johnston, Dominic Kwiatkowski, Cordelia Langford, John Sillitoe on behalf of the Wellcome Sanger Institute COVID-19 Surveillance Team |
| EPI_ISL_2831779 | Lighthouse Lab in Milton Keynes | Wellcome Sanger Institute for the COVID-19 Genomics UK (COG-UK) Consortium | The Lighthouse Lab in Milton Keynes and Alex Alderton, Roberto Amato, Jeffrey Barrett, Sonia Goncalves, Ewan Harrison, David K. Jackson, Ian Johnston, Dominic Kwiatkowski, Cordelia Langford, John Sillitoe on behalf of the Wellcome Sanger Institute COVID-19 Surveillance Team |
| EPI_ISL_2831780 | Lighthouse Laboratory Plymouth | Wellcome Sanger Institute for the COVID-19 Genomics UK (COG-UK) Consortium | Lighthouse Laboratory Plymouth and Alex Alderton, Roberto Amato, Jeffrey Barrett, Sonia Goncalves, Ewan Harrison, David K. Jackson, Ian Johnston, Dominic Kwiatkowski, Cordelia Langford, John Sillitoe on behalf of the Wellcome Sanger Institute COVID-19 Surveillance Team |
| EPI_ISL_2831781, EPI_ISL_2831782, EPI_ISL_2831783, EPI_ISL_2831784, EPI_ISL_2831785 | Lighthouse Lab in Milton Keynes | Wellcome Sanger Institute for the COVID-19 Genomics UK (COG-UK) Consortium | The Lighthouse Lab in Milton Keynes and Alex Alderton, Roberto Amato, Jeffrey Barrett, Sonia Goncalves, Ewan Harrison, David K. Jackson, Ian Johnston, Dominic Kwiatkowski, Cordelia Langford, John Sillitoe on behalf of the Wellcome Sanger Institute COVID-19 Surveillance Team |
| EPI_ISL_2831786 | Lighthouse Laboratory Plymouth | Wellcome Sanger Institute for the COVID-19 Genomics UK (COG-UK) Consortium | Lighthouse Laboratory Plymouth and Alex Alderton, Roberto Amato, Jeffrey Barrett, Sonia Goncalves, Ewan Harrison, David K. Jackson, Ian Johnston, Dominic Kwiatkowski, Cordelia Langford, John Sillitoe on behalf of the Wellcome Sanger Institute COVID-19 Surveillance Team |
| EPI_ISL_2831787 | Lighthouse Lab in Milton Keynes | Wellcome Sanger Institute for the COVID-19 Genomics UK (COG-UK) Consortium | The Lighthouse Lab in Milton Keynes and Alex Alderton, Roberto Amato, Jeffrey Barrett, Sonia Goncalves, Ewan Harrison, David K. Jackson, Ian Johnston, Dominic Kwiatkowski, Cordelia Langford, John Sillitoe on behalf of the Wellcome Sanger Institute COVID-19 Surveillance Team |
| EPI_ISL_2831788, EPI_ISL_2831789 | Lighthouse Laboratory Plymouth | Wellcome Sanger Institute for the COVID-19 Genomics UK (COG-UK) Consortium | Lighthouse Laboratory Plymouth and Alex Alderton, Roberto Amato, Jeffrey Barrett, Sonia Goncalves, Ewan Harrison, David K. Jackson, Ian Johnston, Dominic Kwiatkowski, Cordelia Langford, John Sillitoe on behalf of the Wellcome Sanger Institute COVID-19 Surveillance Team |
| EPI_ISL_2831790 | Lighthouse Lab in Glasgow | Wellcome Sanger Institute for the COVID-19 Genomics UK (COG-UK) Consortium | Harper VanSteenhouse, Yumi Kasai, David Gray, Carol Clugston, Anna Dominiczak and Alex Alderton, Roberto Amato, Jeffrey Barrett, Sonia Goncalves, Ewan Harrison, David K. Jackson, Ian Johnston, Dominic Kwiatkowski, Cordelia Langford, John Sillitoe on behalf of the Wellcome Sanger Institute COVID-19 Surveillance Team |
| EPI_ISL_2831791, EPI_ISL_2831792 | Lighthouse Lab in Milton Keynes | Wellcome Sanger Institute for the COVID-19 Genomics UK (COG-UK) Consortium | The Lighthouse Lab in Milton Keynes and Alex Alderton, Roberto Amato, Jeffrey Barrett, Sonia Goncalves, Ewan Harrison, David K. Jackson, Ian Johnston, Dominic Kwiatkowski, Cordelia Langford, John Sillitoe on behalf of the Wellcome Sanger Institute COVID-19 Surveillance Team |
| EPI_ISL_2831793 | Lighthouse Lab in Glasgow | Wellcome Sanger Institute for the COVID-19 Genomics UK (COG-UK) Consortium | Harper VanSteenhouse, Yumi Kasai, David Gray, Carol Clugston, Anna Dominiczak and Alex Alderton, Roberto Amato, Jeffrey Barrett, Sonia Goncalves, Ewan Harrison, David K. Jackson, Ian Johnston, Dominic Kwiatkowski, Cordelia Langford, John Sillitoe on behalf of the Wellcome Sanger Institute COVID-19 Surveillance Team |
| EPI_ISL_2831794 | Lighthouse Lab in Milton Keynes | Wellcome Sanger Institute for the COVID-19 Genomics UK (COG-UK) Consortium | The Lighthouse Lab in Milton Keynes and Alex Alderton, Roberto Amato, Jeffrey Barrett, Sonia Goncalves, Ewan Harrison, David K. Jackson, Ian Johnston, Dominic Kwiatkowski, Cordelia Langford, John Sillitoe on behalf of the Wellcome Sanger Institute COVID-19 Surveillance Team |
| EPI_ISL_2831795 | Lighthouse Laboratory Plymouth | Wellcome Sanger Institute for the COVID-19 Genomics UK (COG-UK) Consortium | Lighthouse Laboratory Plymouth and Alex Alderton, Roberto Amato, Jeffrey Barrett, Sonia Goncalves, Ewan Harrison, David K. Jackson, Ian Johnston, Dominic Kwiatkowski, Cordelia Langford, John Sillitoe on behalf of the Wellcome Sanger Institute COVID-19 Surveillance Team |
| EPI_ISL_2831796 | Lighthouse Lab in Milton Keynes | Wellcome Sanger Institute for the COVID-19 Genomics UK (COG-UK) Consortium | The Lighthouse Lab in Milton Keynes and Alex Alderton, Roberto Amato, Jeffrey Barrett, Sonia Goncalves, Ewan Harrison, David K. Jackson, Ian Johnston, Dominic Kwiatkowski, Cordelia Langford, John Sillitoe on behalf of the Wellcome Sanger Institute COVID-19 Surveillance Team |
| EPI_ISL_2831797 | Lighthouse Laboratory Plymouth | Wellcome Sanger Institute for the COVID-19 Genomics UK (COG-UK) Consortium | Lighthouse Laboratory Plymouth and Alex Alderton, Roberto Amato, Jeffrey Barrett, Sonia Goncalves, Ewan Harrison, David K. Jackson, Ian Johnston, Dominic Kwiatkowski, Cordelia Langford, John Sillitoe on behalf of the Wellcome Sanger Institute COVID-19 Surveillance Team |
| EPI_ISL_2831798, EPI_ISL_2831799 | Lighthouse Lab in Milton Keynes | Wellcome Sanger Institute for the COVID-19 Genomics UK (COG-UK) Consortium | The Lighthouse Lab in Milton Keynes and Alex Alderton, Roberto Amato, Jeffrey Barrett, Sonia Goncalves, Ewan Harrison, David K. Jackson, Ian Johnston, Dominic Kwiatkowski, Cordelia Langford, John Sillitoe on behalf of the Wellcome Sanger Institute COVID-19 Surveillance Team |
| EPI_ISL_2831800 | Lighthouse Laboratory Plymouth | Wellcome Sanger Institute for the COVID-19 Genomics UK (COG-UK) Consortium | Lighthouse Laboratory Plymouth and Alex Alderton, Roberto Amato, Jeffrey Barrett, Sonia Goncalves, Ewan Harrison, David K. Jackson, Ian Johnston, Dominic Kwiatkowski, Cordelia Langford, John Sillitoe on behalf of the Wellcome Sanger Institute COVID-19 Surveillance Team |
| EPI_ISL_2831801, EPI_ISL_2831802 | Lighthouse Lab in Milton Keynes | Wellcome Sanger Institute for the COVID-19 Genomics UK (COG-UK) Consortium | The Lighthouse Lab in Milton Keynes and Alex Alderton, Roberto Amato, Jeffrey Barrett, Sonia Goncalves, Ewan Harrison, David K. Jackson, Ian Johnston, Dominic Kwiatkowski, Cordelia Langford, John Sillitoe on behalf of the Wellcome Sanger Institute COVID-19 Surveillance Team |
| EPI_ISL_2831803 | Lighthouse Laboratory Plymouth | Wellcome Sanger Institute for the COVID-19 Genomics UK (COG-UK) Consortium | Lighthouse Laboratory Plymouth and Alex Alderton, Roberto Amato, Jeffrey Barrett, Sonia Goncalves, Ewan Harrison, David K. Jackson, Ian Johnston, Dominic Kwiatkowski, Cordelia Langford, John Sillitoe on behalf of the Wellcome Sanger Institute COVID-19 Surveillance Team |
| EPI_ISL_2831804 | Lighthouse Lab in Milton Keynes | Wellcome Sanger Institute for the COVID-19 Genomics UK (COG-UK) Consortium | The Lighthouse Lab in Milton Keynes and Alex Alderton, Roberto Amato, Jeffrey Barrett, Sonia Goncalves, Ewan Harrison, David K. Jackson, Ian Johnston, Dominic Kwiatkowski, Cordelia Langford, John Sillitoe on behalf of the Wellcome Sanger Institute COVID-19 Surveillance Team |
| EPI_ISL_2831805 | Lighthouse Laboratory Plymouth | Wellcome Sanger Institute for the COVID-19 Genomics UK (COG-UK) Consortium | Lighthouse Laboratory Plymouth and Alex Alderton, Roberto Amato, Jeffrey Barrett, Sonia Goncalves, Ewan Harrison, David K. Jackson, Ian Johnston, Dominic Kwiatkowski, Cordelia Langford, John Sillitoe on behalf of the Wellcome Sanger Institute COVID-19 Surveillance Team |
| EPI_ISL_2831806, | Lighthouse Lab in Milton Keynes | Wellcome Sanger Institute for the COVID-19 Genomics UK (COG-UK) | The Lighthouse Lab in Milton Keynes and Alex Alderton, Roberto Amato, Jeffrey Barrett, Sonia Goncalves, Ewan Harrison, David K. Jackson, Ian Johnston, Dominic Kwiatkowski, Cordelia Langford, John Sillitoe on behalf of the Wellcome Sanger |

| | | | |
|---|---|---|---|
| EPI_ISL_2831807 | | Consortium | Institute COVID-19 Surveillance Team |
| EPI_ISL_2831808 | Lighthouse Laboratory Plymouth | Wellcome Sanger Institute for the COVID-19 Genomics UK (COG-UK) Consortium | Lighthouse Laboratory Plymouth and Alex Alderton, Roberto Amato, Jeffrey Barrett, Sonia Goncalves, Ewan Harrison, David K. Jackson, Ian Johnston, Dominic Kwiatkowski, Cordelia Langford, John Sillitoe on behalf of the Wellcome Sanger Institute COVID-19 Surveillance Team |
| EPI_ISL_2831809, EPI_ISL_2831810 | Lighthouse Lab in Milton Keynes | Wellcome Sanger Institute for the COVID-19 Genomics UK (COG-UK) Consortium | The Lighthouse Lab in Milton Keynes and Alex Alderton, Roberto Amato, Jeffrey Barrett, Sonia Goncalves, Ewan Harrison, David K. Jackson, Ian Johnston, Dominic Kwiatkowski, Cordelia Langford, John Sillitoe on behalf of the Wellcome Sanger Institute COVID-19 Surveillance Team |
| EPI_ISL_2831811 | Lighthouse Lab in Glasgow | Wellcome Sanger Institute for the COVID-19 Genomics UK (COG-UK) Consortium | Harper VanSteenhouse, Yumi Kasai, David Gray, Carol Clugston, Anna Dominiczak and Alex Alderton, Roberto Amato, Jeffrey Barrett, Sonia Goncalves, Ewan Harrison, David K. Jackson, Ian Johnston, Dominic Kwiatkowski, Cordelia Langford, John Sillitoe on behalf of the Wellcome Sanger Institute COVID-19 Surveillance Team |
| EPI_ISL_2831812 | Lighthouse Laboratory Plymouth | Wellcome Sanger Institute for the COVID-19 Genomics UK (COG-UK) Consortium | Lighthouse Laboratory Plymouth and Alex Alderton, Roberto Amato, Jeffrey Barrett, Sonia Goncalves, Ewan Harrison, David K. Jackson, Ian Johnston, Dominic Kwiatkowski, Cordelia Langford, John Sillitoe on behalf of the Wellcome Sanger Institute COVID-19 Surveillance Team |
| EPI_ISL_2831813, EPI_ISL_2831814, EPI_ISL_2831815 | Lighthouse Lab in Milton Keynes | Wellcome Sanger Institute for the COVID-19 Genomics UK (COG-UK) Consortium | The Lighthouse Lab in Milton Keynes and Alex Alderton, Roberto Amato, Jeffrey Barrett, Sonia Goncalves, Ewan Harrison, David K. Jackson, Ian Johnston, Dominic Kwiatkowski, Cordelia Langford, John Sillitoe on behalf of the Wellcome Sanger Institute COVID-19 Surveillance Team |
| EPI_ISL_2831816 | Lighthouse Lab in Glasgow | Wellcome Sanger Institute for the COVID-19 Genomics UK (COG-UK) Consortium | Harper VanSteenhouse, Yumi Kasai, David Gray, Carol Clugston, Anna Dominiczak and Alex Alderton, Roberto Amato, Jeffrey Barrett, Sonia Goncalves, Ewan Harrison, David K. Jackson, Ian Johnston, Dominic Kwiatkowski, Cordelia Langford, John Sillitoe on behalf of the Wellcome Sanger Institute COVID-19 Surveillance Team |
| EPI_ISL_2831817, EPI_ISL_2831818 | Lighthouse Lab in Milton Keynes | Wellcome Sanger Institute for the COVID-19 Genomics UK (COG-UK) Consortium | The Lighthouse Lab in Milton Keynes and Alex Alderton, Roberto Amato, Jeffrey Barrett, Sonia Goncalves, Ewan Harrison, David K. Jackson, Ian Johnston, Dominic Kwiatkowski, Cordelia Langford, John Sillitoe on behalf of the Wellcome Sanger Institute COVID-19 Surveillance Team |
| EPI_ISL_2831819, EPI_ISL_2831820 | Lighthouse Laboratory Plymouth | Wellcome Sanger Institute for the COVID-19 Genomics UK (COG-UK) Consortium | Lighthouse Laboratory Plymouth and Alex Alderton, Roberto Amato, Jeffrey Barrett, Sonia Goncalves, Ewan Harrison, David K. Jackson, Ian Johnston, Dominic Kwiatkowski, Cordelia Langford, John Sillitoe on behalf of the Wellcome Sanger Institute COVID-19 Surveillance Team |
| EPI_ISL_2831821 | Lighthouse Lab in Milton Keynes | Wellcome Sanger Institute for the COVID-19 Genomics UK (COG-UK) Consortium | The Lighthouse Lab in Milton Keynes and Alex Alderton, Roberto Amato, Jeffrey Barrett, Sonia Goncalves, Ewan Harrison, David K. Jackson, Ian Johnston, Dominic Kwiatkowski, Cordelia Langford, John Sillitoe on behalf of the Wellcome Sanger Institute COVID-19 Surveillance Team |
| EPI_ISL_2831822 | Lighthouse Laboratory Plymouth | Wellcome Sanger Institute for the COVID-19 Genomics UK (COG-UK) Consortium | Lighthouse Laboratory Plymouth and Alex Alderton, Roberto Amato, Jeffrey Barrett, Sonia Goncalves, Ewan Harrison, David K. Jackson, Ian Johnston, Dominic Kwiatkowski, Cordelia Langford, John Sillitoe on behalf of the Wellcome Sanger Institute COVID-19 Surveillance Team |
| EPI_ISL_2831823 | Lighthouse Lab in Glasgow | Wellcome Sanger Institute for the COVID-19 Genomics UK (COG-UK) Consortium | Harper VanSteenhouse, Yumi Kasai, David Gray, Carol Clugston, Anna Dominiczak and Alex Alderton, Roberto Amato, Jeffrey Barrett, Sonia Goncalves, Ewan Harrison, David K. Jackson, Ian Johnston, Dominic Kwiatkowski, Cordelia Langford, John Sillitoe on behalf of the Wellcome Sanger Institute COVID-19 Surveillance Team |
| EPI_ISL_2831824, EPI_ISL_2831825, EPI_ISL_2831826, EPI_ISL_2831827, EPI_ISL_2831828 | Lighthouse Lab in Milton Keynes | Wellcome Sanger Institute for the COVID-19 Genomics UK (COG-UK) Consortium | The Lighthouse Lab in Milton Keynes and Alex Alderton, Roberto Amato, Jeffrey Barrett, Sonia Goncalves, Ewan Harrison, David K. Jackson, Ian Johnston, Dominic Kwiatkowski, Cordelia Langford, John Sillitoe on behalf of the Wellcome Sanger Institute COVID-19 Surveillance Team |
| EPI_ISL_2831829, EPI_ISL_2831830 | Lighthouse Laboratory Plymouth | Wellcome Sanger Institute for the COVID-19 Genomics UK (COG-UK) Consortium | Lighthouse Laboratory Plymouth and Alex Alderton, Roberto Amato, Jeffrey Barrett, Sonia Goncalves, Ewan Harrison, David K. Jackson, Ian Johnston, Dominic Kwiatkowski, Cordelia Langford, John Sillitoe on behalf of the Wellcome Sanger Institute COVID-19 Surveillance Team |
| EPI_ISL_2831831 | Lighthouse Lab in Glasgow | Wellcome Sanger Institute for the COVID-19 Genomics UK (COG-UK) Consortium | Harper VanSteenhouse, Yumi Kasai, David Gray, Carol Clugston, Anna Dominiczak and Alex Alderton, Roberto Amato, Jeffrey Barrett, Sonia Goncalves, Ewan Harrison, David K. Jackson, Ian Johnston, Dominic Kwiatkowski, Cordelia Langford, John Sillitoe on behalf of the Wellcome Sanger Institute COVID-19 Surveillance Team |
| EPI_ISL_2831832, EPI_ISL_2831833, EPI_ISL_2831834 | Lighthouse Lab in Milton Keynes | Wellcome Sanger Institute for the COVID-19 Genomics UK (COG-UK) Consortium | The Lighthouse Lab in Milton Keynes and Alex Alderton, Roberto Amato, Jeffrey Barrett, Sonia Goncalves, Ewan Harrison, David K. Jackson, Ian Johnston, Dominic Kwiatkowski, Cordelia Langford, John Sillitoe on behalf of the Wellcome Sanger Institute COVID-19 Surveillance Team |
| EPI_ISL_2831835 | Lighthouse Laboratory Plymouth | Wellcome Sanger Institute for the COVID-19 Genomics UK (COG-UK) Consortium | Lighthouse Laboratory Plymouth and Alex Alderton, Roberto Amato, Jeffrey Barrett, Sonia Goncalves, Ewan Harrison, David K. Jackson, Ian Johnston, Dominic Kwiatkowski, Cordelia Langford, John Sillitoe on behalf of the Wellcome Sanger Institute COVID-19 Surveillance Team |
| EPI_ISL_2831836, EPI_ISL_2831838 | Lighthouse Lab in Milton Keynes | Wellcome Sanger Institute for the COVID-19 Genomics UK (COG-UK) Consortium | The Lighthouse Lab in Milton Keynes and Alex Alderton, Roberto Amato, Jeffrey Barrett, Sonia Goncalves, Ewan Harrison, David K. Jackson, Ian Johnston, Dominic Kwiatkowski, Cordelia Langford, John Sillitoe on behalf of the Wellcome Sanger Institute COVID-19 Surveillance Team |
| EPI_ISL_2831839 | Lighthouse Laboratory Plymouth | Wellcome Sanger Institute for the COVID-19 Genomics UK (COG-UK) Consortium | Lighthouse Laboratory Plymouth and Alex Alderton, Roberto Amato, Jeffrey Barrett, Sonia Goncalves, Ewan Harrison, David K. Jackson, Ian Johnston, Dominic Kwiatkowski, Cordelia Langford, John Sillitoe on behalf of the Wellcome Sanger Institute COVID-19 Surveillance Team |
| EPI_ISL_2831840 | Lighthouse Lab in Glasgow | Wellcome Sanger Institute for the COVID-19 Genomics UK (COG-UK) Consortium | Harper VanSteenhouse, Yumi Kasai, David Gray, Carol Clugston, Anna Dominiczak and Alex Alderton, Roberto Amato, Jeffrey Barrett, Sonia Goncalves, Ewan Harrison, David K. Jackson, Ian Johnston, Dominic Kwiatkowski, Cordelia Langford, John Sillitoe on behalf of the Wellcome Sanger Institute COVID-19 Surveillance Team |
| EPI_ISL_2831841 | Lighthouse Laboratory Plymouth | Wellcome Sanger Institute for the COVID-19 Genomics UK (COG-UK) Consortium | Lighthouse Laboratory Plymouth and Alex Alderton, Roberto Amato, Jeffrey Barrett, Sonia Goncalves, Ewan Harrison, David K. Jackson, Ian Johnston, Dominic Kwiatkowski, Cordelia Langford, John Sillitoe on behalf of the Wellcome Sanger Institute COVID-19 Surveillance Team |
| EPI_ISL_2831842, EPI_ISL_2831843 | Lighthouse Lab in Milton Keynes | Wellcome Sanger Institute for the COVID-19 Genomics UK (COG-UK) Consortium | The Lighthouse Lab in Milton Keynes and Alex Alderton, Roberto Amato, Jeffrey Barrett, Sonia Goncalves, Ewan Harrison, David K. Jackson, Ian Johnston, Dominic Kwiatkowski, Cordelia Langford, John Sillitoe on behalf of the Wellcome Sanger Institute COVID-19 Surveillance Team |
| EPI_ISL_2831844 | Lighthouse Lab in Glasgow | Wellcome Sanger Institute for the COVID-19 Genomics UK (COG-UK) Consortium | Harper VanSteenhouse, Yumi Kasai, David Gray, Carol Clugston, Anna Dominiczak and Alex Alderton, Roberto Amato, Jeffrey Barrett, Sonia Goncalves, Ewan Harrison, David K. Jackson, Ian Johnston, Dominic Kwiatkowski, Cordelia Langford, John Sillitoe on behalf of the Wellcome Sanger Institute COVID-19 Surveillance Team |
| EPI_ISL_2831845 | Lighthouse Lab in Milton Keynes | Wellcome Sanger Institute for the COVID-19 Genomics UK (COG-UK) Consortium | The Lighthouse Lab in Milton Keynes and Alex Alderton, Roberto Amato, Jeffrey Barrett, Sonia Goncalves, Ewan Harrison, David K. Jackson, Ian Johnston, Dominic Kwiatkowski, Cordelia Langford, John Sillitoe on behalf of the Wellcome Sanger Institute COVID-19 Surveillance Team |
| EPI_ISL_2831846 | Lighthouse Lab in Glasgow | Wellcome Sanger Institute for the COVID-19 Genomics UK (COG-UK) Consortium | Harper VanSteenhouse, Yumi Kasai, David Gray, Carol Clugston, Anna Dominiczak and Alex Alderton, Roberto Amato, Jeffrey Barrett, Sonia Goncalves, Ewan Harrison, David K. Jackson, Ian Johnston, Dominic Kwiatkowski, Cordelia Langford, John Sillitoe on behalf of the Wellcome Sanger Institute COVID-19 Surveillance Team |
| EPI_ISL_2831847, EPI_ISL_2831848, EPI_ISL_2831849 | Lighthouse Lab in Milton Keynes | Wellcome Sanger Institute for the COVID-19 Genomics UK (COG-UK) Consortium | The Lighthouse Lab in Milton Keynes and Alex Alderton, Roberto Amato, Jeffrey Barrett, Sonia Goncalves, Ewan Harrison, David K. Jackson, Ian Johnston, Dominic Kwiatkowski, Cordelia Langford, John Sillitoe on behalf of the Wellcome Sanger Institute COVID-19 Surveillance Team |
| EPI_ISL_2831850 | Lighthouse Laboratory Plymouth | Wellcome Sanger Institute for the COVID-19 Genomics UK (COG-UK) Consortium | Lighthouse Laboratory Plymouth and Alex Alderton, Roberto Amato, Jeffrey Barrett, Sonia Goncalves, Ewan Harrison, David K. Jackson, Ian Johnston, Dominic Kwiatkowski, Cordelia Langford, John Sillitoe on behalf of the Wellcome Sanger Institute COVID-19 Surveillance Team |
| EPI_ISL_2831851, EPI_ISL_2831852, EPI_ISL_2831853 | Lighthouse Lab in Milton Keynes | Wellcome Sanger Institute for the COVID-19 Genomics UK (COG-UK) Consortium | The Lighthouse Lab in Milton Keynes and Alex Alderton, Roberto Amato, Jeffrey Barrett, Sonia Goncalves, Ewan Harrison, David K. Jackson, Ian Johnston, Dominic Kwiatkowski, Cordelia Langford, John Sillitoe on behalf of the Wellcome Sanger Institute COVID-19 Surveillance Team |
| EPI_ISL_2831854, EPI_ISL_2831855 | Lighthouse Lab in Glasgow | Wellcome Sanger Institute for the COVID-19 Genomics UK (COG-UK) Consortium | Harper VanSteenhouse, Yumi Kasai, David Gray, Carol Clugston, Anna Dominiczak and Alex Alderton, Roberto Amato, Jeffrey Barrett, Sonia Goncalves, Ewan Harrison, David K. Jackson, Ian Johnston, Dominic Kwiatkowski, Cordelia Langford, John Sillitoe on behalf of the Wellcome Sanger Institute COVID-19 Surveillance Team |
| EPI_ISL_2831856, EPI_ISL_2831857 | Lighthouse Laboratory Plymouth | Wellcome Sanger Institute for the COVID-19 Genomics UK (COG-UK) Consortium | Lighthouse Laboratory Plymouth and Alex Alderton, Roberto Amato, Jeffrey Barrett, Sonia Goncalves, Ewan Harrison, David K. Jackson, Ian Johnston, Dominic Kwiatkowski, Cordelia Langford, John Sillitoe on behalf of the Wellcome Sanger Institute COVID-19 Surveillance Team |
| EPI_ISL_2831858 | Lighthouse Lab in Milton Keynes | Wellcome Sanger Institute for the COVID-19 Genomics UK (COG-UK) Consortium | The Lighthouse Lab in Milton Keynes and Alex Alderton, Roberto Amato, Jeffrey Barrett, Sonia Goncalves, Ewan Harrison, David K. Jackson, Ian Johnston, Dominic Kwiatkowski, Cordelia Langford, John Sillitoe on behalf of the Wellcome Sanger Institute COVID-19 Surveillance Team |
| EPI_ISL_2831859, EPI_ISL_2831860 | Lighthouse Laboratory Plymouth | Wellcome Sanger Institute for the COVID-19 Genomics UK (COG-UK) Consortium | Lighthouse Laboratory Plymouth and Alex Alderton, Roberto Amato, Jeffrey Barrett, Sonia Goncalves, Ewan Harrison, David K. Jackson, Ian Johnston, Dominic Kwiatkowski, Cordelia Langford, John Sillitoe on behalf of the Wellcome Sanger Institute COVID-19 Surveillance Team |
| EPI_ISL_2831861 | Lighthouse Lab in Milton Keynes | Wellcome Sanger Institute for the COVID-19 Genomics UK (COG-UK) Consortium | The Lighthouse Lab in Milton Keynes and Alex Alderton, Roberto Amato, Jeffrey Barrett, Sonia Goncalves, Ewan Harrison, David K. Jackson, Ian Johnston, Dominic Kwiatkowski, Cordelia Langford, John Sillitoe on behalf of the Wellcome Sanger Institute COVID-19 Surveillance Team |
| EPI_ISL_2831862 | Lighthouse Laboratory Plymouth | Wellcome Sanger Institute for the COVID-19 Genomics UK (COG-UK) Consortium | Lighthouse Laboratory Plymouth and Alex Alderton, Roberto Amato, Jeffrey Barrett, Sonia Goncalves, Ewan Harrison, David K. Jackson, Ian Johnston, Dominic Kwiatkowski, Cordelia Langford, John Sillitoe on behalf of the Wellcome Sanger Institute COVID-19 Surveillance Team |
| EPI_ISL_2831863, EPI_ISL_2831864 | Lighthouse Lab in Milton Keynes | Wellcome Sanger Institute for the COVID-19 Genomics UK (COG-UK) Consortium | The Lighthouse Lab in Milton Keynes and Alex Alderton, Roberto Amato, Jeffrey Barrett, Sonia Goncalves, Ewan Harrison, David K. Jackson, Ian Johnston, Dominic Kwiatkowski, Cordelia Langford, John Sillitoe on behalf of the Wellcome Sanger Institute COVID-19 Surveillance Team |
| EPI_ISL_2831865, EPI_ISL_2831866 | Lighthouse Laboratory Plymouth | Wellcome Sanger Institute for the COVID-19 Genomics UK (COG-UK) Consortium | Lighthouse Laboratory Plymouth and Alex Alderton, Roberto Amato, Jeffrey Barrett, Sonia Goncalves, Ewan Harrison, David K. Jackson, Ian Johnston, Dominic Kwiatkowski, Cordelia Langford, John Sillitoe on behalf of the Wellcome Sanger Institute COVID-19 Surveillance Team |
| EPI_ISL_2831867, EPI_ISL_2831868, EPI_ISL_2831869, EPI_ISL_2831870, EPI_ISL_2831871, EPI_ISL_2831872, EPI_ISL_2831873, EPI_ISL_2831874, EPI_ISL_2831875 | Lighthouse Lab in Milton Keynes | Wellcome Sanger Institute for the COVID-19 Genomics UK (COG-UK) Consortium | The Lighthouse Lab in Milton Keynes and Alex Alderton, Roberto Amato, Jeffrey Barrett, Sonia Goncalves, Ewan Harrison, David K. Jackson, Ian Johnston, Dominic Kwiatkowski, Cordelia Langford, John Sillitoe on behalf of the Wellcome Sanger Institute COVID-19 Surveillance Team |
| EPI_ISL_2831876 | Lighthouse Lab in Glasgow | Wellcome Sanger Institute for the COVID-19 Genomics UK (COG-UK) Consortium | Harper VanSteenhouse, Yumi Kasai, David Gray, Carol Clugston, Anna Dominiczak and Alex Alderton, Roberto Amato, Jeffrey Barrett, Sonia Goncalves, Ewan Harrison, David K. Jackson, Ian Johnston, Dominic Kwiatkowski, Cordelia Langford, John Sillitoe on behalf of the Wellcome Sanger Institute COVID-19 Surveillance Team |
| EPI_ISL_2831877 | Lighthouse Lab in Milton Keynes | Wellcome Sanger Institute for the COVID-19 Genomics UK (COG-UK) Consortium | The Lighthouse Lab in Milton Keynes and Alex Alderton, Roberto Amato, Jeffrey Barrett, Sonia Goncalves, Ewan Harrison, David K. Jackson, Ian Johnston, Dominic Kwiatkowski, Cordelia Langford, John Sillitoe on behalf of the Wellcome Sanger Institute COVID-19 Surveillance Team |
| EPI_ISL_2831878, | Lighthouse Laboratory Plymouth | Wellcome Sanger Institute for the COVID-19 Genomics UK (COG-UK) | Lighthouse Laboratory Plymouth and Alex Alderton, Roberto Amato, Jeffrey Barrett, Sonia Goncalves, Ewan Harrison, David K. Jackson, Ian Johnston, Dominic Kwiatkowski, Cordelia Langford, John Sillitoe on behalf of the Wellcome Sanger |

| | | | |
|---|---|---|---|
| EPI_ISL_2831879 | | Consortium | Institute COVID-19 Surveillance Team |
| EPI_ISL_2831880 | Lighthouse Lab in Milton Keynes | Wellcome Sanger Institute for the COVID-19 Genomics UK (COG-UK) Consortium | The Lighthouse Lab in Milton Keynes and Alex Alderton, Roberto Amato, Jeffrey Barrett, Sonia Goncalves, Ewan Harrison, David K. Jackson, Ian Johnston, Dominic Kwiatkowski, Cordelia Langford, John Sillitoe on behalf of the Wellcome Sanger Institute COVID-19 Surveillance Team |
| EPI_ISL_2831881 | Lighthouse Laboratory Plymouth | Wellcome Sanger Institute for the COVID-19 Genomics UK (COG-UK) Consortium | Lighthouse Laboratory Plymouth and Alex Alderton, Roberto Amato, Jeffrey Barrett, Sonia Goncalves, Ewan Harrison, David K. Jackson, Ian Johnston, Dominic Kwiatkowski, Cordelia Langford, John Sillitoe on behalf of the Wellcome Sanger Institute COVID-19 Surveillance Team |
| EPI_ISL_2831882 | Lighthouse Lab in Glasgow | Wellcome Sanger Institute for the COVID-19 Genomics UK (COG-UK) Consortium | Harper VanSteenhouse, Yumi Kasai, David Gray, Carol Clugston, Anna Dominiczak and Alex Alderton, Roberto Amato, Jeffrey Barrett, Sonia Goncalves, Ewan Harrison, David K. Jackson, Ian Johnston, Dominic Kwiatkowski, Cordelia Langford, John Sillitoe on behalf of the Wellcome Sanger Institute COVID-19 Surveillance Team |
| EPI_ISL_2831883 | Lighthouse Laboratory Plymouth | Wellcome Sanger Institute for the COVID-19 Genomics UK (COG-UK) Consortium | Lighthouse Laboratory Plymouth and Alex Alderton, Roberto Amato, Jeffrey Barrett, Sonia Goncalves, Ewan Harrison, David K. Jackson, Ian Johnston, Dominic Kwiatkowski, Cordelia Langford, John Sillitoe on behalf of the Wellcome Sanger Institute COVID-19 Surveillance Team |
| EPI_ISL_2831884 | Lighthouse Lab in Milton Keynes | Wellcome Sanger Institute for the COVID-19 Genomics UK (COG-UK) Consortium | The Lighthouse Lab in Milton Keynes and Alex Alderton, Roberto Amato, Jeffrey Barrett, Sonia Goncalves, Ewan Harrison, David K. Jackson, Ian Johnston, Dominic Kwiatkowski, Cordelia Langford, John Sillitoe on behalf of the Wellcome Sanger Institute COVID-19 Surveillance Team |
| EPI_ISL_2831885 | Lighthouse Laboratory Plymouth | Wellcome Sanger Institute for the COVID-19 Genomics UK (COG-UK) Consortium | Lighthouse Laboratory Plymouth and Alex Alderton, Roberto Amato, Jeffrey Barrett, Sonia Goncalves, Ewan Harrison, David K. Jackson, Ian Johnston, Dominic Kwiatkowski, Cordelia Langford, John Sillitoe on behalf of the Wellcome Sanger Institute COVID-19 Surveillance Team |
| EPI_ISL_2831886 | Lighthouse Lab in Glasgow | Wellcome Sanger Institute for the COVID-19 Genomics UK (COG-UK) Consortium | Harper VanSteenhouse, Yumi Kasai, David Gray, Carol Clugston, Anna Dominiczak and Alex Alderton, Roberto Amato, Jeffrey Barrett, Sonia Goncalves, Ewan Harrison, David K. Jackson, Ian Johnston, Dominic Kwiatkowski, Cordelia Langford, John Sillitoe on behalf of the Wellcome Sanger Institute COVID-19 Surveillance Team |
| EPI_ISL_2831887, EPI_ISL_2831888, EPI_ISL_2831889 | Lighthouse Lab in Milton Keynes | Wellcome Sanger Institute for the COVID-19 Genomics UK (COG-UK) Consortium | The Lighthouse Lab in Milton Keynes and Alex Alderton, Roberto Amato, Jeffrey Barrett, Sonia Goncalves, Ewan Harrison, David K. Jackson, Ian Johnston, Dominic Kwiatkowski, Cordelia Langford, John Sillitoe on behalf of the Wellcome Sanger Institute COVID-19 Surveillance Team |
| EPI_ISL_2831890 | Lighthouse Laboratory Plymouth | Wellcome Sanger Institute for the COVID-19 Genomics UK (COG-UK) Consortium | Lighthouse Laboratory Plymouth and Alex Alderton, Roberto Amato, Jeffrey Barrett, Sonia Goncalves, Ewan Harrison, David K. Jackson, Ian Johnston, Dominic Kwiatkowski, Cordelia Langford, John Sillitoe on behalf of the Wellcome Sanger Institute COVID-19 Surveillance Team |
| EPI_ISL_2831891 | Lighthouse Lab in Milton Keynes | Wellcome Sanger Institute for the COVID-19 Genomics UK (COG-UK) Consortium | The Lighthouse Lab in Milton Keynes and Alex Alderton, Roberto Amato, Jeffrey Barrett, Sonia Goncalves, Ewan Harrison, David K. Jackson, Ian Johnston, Dominic Kwiatkowski, Cordelia Langford, John Sillitoe on behalf of the Wellcome Sanger Institute COVID-19 Surveillance Team |
| EPI_ISL_2831892, EPI_ISL_2831893, EPI_ISL_2831894 | Lighthouse Laboratory Plymouth | Wellcome Sanger Institute for the COVID-19 Genomics UK (COG-UK) Consortium | Lighthouse Laboratory Plymouth and Alex Alderton, Roberto Amato, Jeffrey Barrett, Sonia Goncalves, Ewan Harrison, David K. Jackson, Ian Johnston, Dominic Kwiatkowski, Cordelia Langford, John Sillitoe on behalf of the Wellcome Sanger Institute COVID-19 Surveillance Team |
| EPI_ISL_2831895, EPI_ISL_2831896, EPI_ISL_2831897 | Lighthouse Lab in Milton Keynes | Wellcome Sanger Institute for the COVID-19 Genomics UK (COG-UK) Consortium | The Lighthouse Lab in Milton Keynes and Alex Alderton, Roberto Amato, Jeffrey Barrett, Sonia Goncalves, Ewan Harrison, David K. Jackson, Ian Johnston, Dominic Kwiatkowski, Cordelia Langford, John Sillitoe on behalf of the Wellcome Sanger Institute COVID-19 Surveillance Team |
| EPI_ISL_2831898 | Lighthouse Lab in Glasgow | Wellcome Sanger Institute for the COVID-19 Genomics UK (COG-UK) Consortium | Harper VanSteenhouse, Yumi Kasai, David Gray, Carol Clugston, Anna Dominiczak and Alex Alderton, Roberto Amato, Jeffrey Barrett, Sonia Goncalves, Ewan Harrison, David K. Jackson, Ian Johnston, Dominic Kwiatkowski, Cordelia Langford, John Sillitoe on behalf of the Wellcome Sanger Institute COVID-19 Surveillance Team |
| EPI_ISL_2831900, EPI_ISL_2831901, EPI_ISL_2831902, EPI_ISL_2831903 | Lighthouse Lab in Milton Keynes | Wellcome Sanger Institute for the COVID-19 Genomics UK (COG-UK) Consortium | The Lighthouse Lab in Milton Keynes and Alex Alderton, Roberto Amato, Jeffrey Barrett, Sonia Goncalves, Ewan Harrison, David K. Jackson, Ian Johnston, Dominic Kwiatkowski, Cordelia Langford, John Sillitoe on behalf of the Wellcome Sanger Institute COVID-19 Surveillance Team |
| EPI_ISL_2831904, EPI_ISL_2831905, EPI_ISL_2831906 | Lighthouse Laboratory Plymouth | Wellcome Sanger Institute for the COVID-19 Genomics UK (COG-UK) Consortium | Lighthouse Laboratory Plymouth and Alex Alderton, Roberto Amato, Jeffrey Barrett, Sonia Goncalves, Ewan Harrison, David K. Jackson, Ian Johnston, Dominic Kwiatkowski, Cordelia Langford, John Sillitoe on behalf of the Wellcome Sanger Institute COVID-19 Surveillance Team |
| EPI_ISL_2831907 | Lighthouse Lab in Glasgow | Wellcome Sanger Institute for the COVID-19 Genomics UK (COG-UK) Consortium | Harper VanSteenhouse, Yumi Kasai, David Gray, Carol Clugston, Anna Dominiczak and Alex Alderton, Roberto Amato, Jeffrey Barrett, Sonia Goncalves, Ewan Harrison, David K. Jackson, Ian Johnston, Dominic Kwiatkowski, Cordelia Langford, John Sillitoe on behalf of the Wellcome Sanger Institute COVID-19 Surveillance Team |
| EPI_ISL_2831908, EPI_ISL_2831909 | Lighthouse Laboratory Plymouth | Wellcome Sanger Institute for the COVID-19 Genomics UK (COG-UK) Consortium | Lighthouse Laboratory Plymouth and Alex Alderton, Roberto Amato, Jeffrey Barrett, Sonia Goncalves, Ewan Harrison, David K. Jackson, Ian Johnston, Dominic Kwiatkowski, Cordelia Langford, John Sillitoe on behalf of the Wellcome Sanger Institute COVID-19 Surveillance Team |
| EPI_ISL_2831910 | Lighthouse Lab in Milton Keynes | Wellcome Sanger Institute for the COVID-19 Genomics UK (COG-UK) Consortium | The Lighthouse Lab in Milton Keynes and Alex Alderton, Roberto Amato, Jeffrey Barrett, Sonia Goncalves, Ewan Harrison, David K. Jackson, Ian Johnston, Dominic Kwiatkowski, Cordelia Langford, John Sillitoe on behalf of the Wellcome Sanger Institute COVID-19 Surveillance Team |
| EPI_ISL_2831911, EPI_ISL_2831912 | Lighthouse Laboratory Plymouth | Wellcome Sanger Institute for the COVID-19 Genomics UK (COG-UK) Consortium | Lighthouse Laboratory Plymouth and Alex Alderton, Roberto Amato, Jeffrey Barrett, Sonia Goncalves, Ewan Harrison, David K. Jackson, Ian Johnston, Dominic Kwiatkowski, Cordelia Langford, John Sillitoe on behalf of the Wellcome Sanger Institute COVID-19 Surveillance Team |
| EPI_ISL_2831913 | Lighthouse Lab in Milton Keynes | Wellcome Sanger Institute for the COVID-19 Genomics UK (COG-UK) Consortium | The Lighthouse Lab in Milton Keynes and Alex Alderton, Roberto Amato, Jeffrey Barrett, Sonia Goncalves, Ewan Harrison, David K. Jackson, Ian Johnston, Dominic Kwiatkowski, Cordelia Langford, John Sillitoe on behalf of the Wellcome Sanger Institute COVID-19 Surveillance Team |
| EPI_ISL_2831914 | Lighthouse Laboratory Plymouth | Wellcome Sanger Institute for the COVID-19 Genomics UK (COG-UK) Consortium | Lighthouse Laboratory Plymouth and Alex Alderton, Roberto Amato, Jeffrey Barrett, Sonia Goncalves, Ewan Harrison, David K. Jackson, Ian Johnston, Dominic Kwiatkowski, Cordelia Langford, John Sillitoe on behalf of the Wellcome Sanger Institute COVID-19 Surveillance Team |
| EPI_ISL_2831915 | Lighthouse Lab in Milton Keynes | Wellcome Sanger Institute for the COVID-19 Genomics UK (COG-UK) Consortium | The Lighthouse Lab in Milton Keynes and Alex Alderton, Roberto Amato, Jeffrey Barrett, Sonia Goncalves, Ewan Harrison, David K. Jackson, Ian Johnston, Dominic Kwiatkowski, Cordelia Langford, John Sillitoe on behalf of the Wellcome Sanger Institute COVID-19 Surveillance Team |
| EPI_ISL_2831916 | Lighthouse Lab in Glasgow | Wellcome Sanger Institute for the COVID-19 Genomics UK (COG-UK) Consortium | Harper VanSteenhouse, Yumi Kasai, David Gray, Carol Clugston, Anna Dominiczak and Alex Alderton, Roberto Amato, Jeffrey Barrett, Sonia Goncalves, Ewan Harrison, David K. Jackson, Ian Johnston, Dominic Kwiatkowski, Cordelia Langford, John Sillitoe on behalf of the Wellcome Sanger Institute COVID-19 Surveillance Team |
| EPI_ISL_2831917, EPI_ISL_2831918 | Lighthouse Lab in Milton Keynes | Wellcome Sanger Institute for the COVID-19 Genomics UK (COG-UK) Consortium | The Lighthouse Lab in Milton Keynes and Alex Alderton, Roberto Amato, Jeffrey Barrett, Sonia Goncalves, Ewan Harrison, David K. Jackson, Ian Johnston, Dominic Kwiatkowski, Cordelia Langford, John Sillitoe on behalf of the Wellcome Sanger Institute COVID-19 Surveillance Team |
| EPI_ISL_2831919 | Lighthouse Lab in Glasgow | Wellcome Sanger Institute for the COVID-19 Genomics UK (COG-UK) Consortium | Harper VanSteenhouse, Yumi Kasai, David Gray, Carol Clugston, Anna Dominiczak and Alex Alderton, Roberto Amato, Jeffrey Barrett, Sonia Goncalves, Ewan Harrison, David K. Jackson, Ian Johnston, Dominic Kwiatkowski, Cordelia Langford, John Sillitoe on behalf of the Wellcome Sanger Institute COVID-19 Surveillance Team |
| EPI_ISL_2831920 | Lighthouse Laboratory Plymouth | Wellcome Sanger Institute for the COVID-19 Genomics UK (COG-UK) Consortium | Lighthouse Laboratory Plymouth and Alex Alderton, Roberto Amato, Jeffrey Barrett, Sonia Goncalves, Ewan Harrison, David K. Jackson, Ian Johnston, Dominic Kwiatkowski, Cordelia Langford, John Sillitoe on behalf of the Wellcome Sanger Institute COVID-19 Surveillance Team |
| EPI_ISL_2831921, EPI_ISL_2831922, EPI_ISL_2831923, EPI_ISL_2831924 | Lighthouse Lab in Milton Keynes | Wellcome Sanger Institute for the COVID-19 Genomics UK (COG-UK) Consortium | The Lighthouse Lab in Milton Keynes and Alex Alderton, Roberto Amato, Jeffrey Barrett, Sonia Goncalves, Ewan Harrison, David K. Jackson, Ian Johnston, Dominic Kwiatkowski, Cordelia Langford, John Sillitoe on behalf of the Wellcome Sanger Institute COVID-19 Surveillance Team |
| EPI_ISL_2831925, EPI_ISL_2831926 | Lighthouse Laboratory Plymouth | Wellcome Sanger Institute for the COVID-19 Genomics UK (COG-UK) Consortium | Lighthouse Laboratory Plymouth and Alex Alderton, Roberto Amato, Jeffrey Barrett, Sonia Goncalves, Ewan Harrison, David K. Jackson, Ian Johnston, Dominic Kwiatkowski, Cordelia Langford, John Sillitoe on behalf of the Wellcome Sanger Institute COVID-19 Surveillance Team |
| EPI_ISL_2831927, EPI_ISL_2831928, EPI_ISL_2831929, EPI_ISL_2831930 | Lighthouse Lab in Milton Keynes | Wellcome Sanger Institute for the COVID-19 Genomics UK (COG-UK) Consortium | The Lighthouse Lab in Milton Keynes and Alex Alderton, Roberto Amato, Jeffrey Barrett, Sonia Goncalves, Ewan Harrison, David K. Jackson, Ian Johnston, Dominic Kwiatkowski, Cordelia Langford, John Sillitoe on behalf of the Wellcome Sanger Institute COVID-19 Surveillance Team |
| EPI_ISL_2831931, EPI_ISL_2831932, EPI_ISL_2831933, EPI_ISL_2831934 | Lighthouse Laboratory Plymouth | Wellcome Sanger Institute for the COVID-19 Genomics UK (COG-UK) Consortium | Lighthouse Laboratory Plymouth and Alex Alderton, Roberto Amato, Jeffrey Barrett, Sonia Goncalves, Ewan Harrison, David K. Jackson, Ian Johnston, Dominic Kwiatkowski, Cordelia Langford, John Sillitoe on behalf of the Wellcome Sanger Institute COVID-19 Surveillance Team |
| EPI_ISL_2831935, EPI_ISL_2831936 | Lighthouse Lab in Glasgow | Wellcome Sanger Institute for the COVID-19 Genomics UK (COG-UK) Consortium | Harper VanSteenhouse, Yumi Kasai, David Gray, Carol Clugston, Anna Dominiczak and Alex Alderton, Roberto Amato, Jeffrey Barrett, Sonia Goncalves, Ewan Harrison, David K. Jackson, Ian Johnston, Dominic Kwiatkowski, Cordelia Langford, John Sillitoe on behalf of the Wellcome Sanger Institute COVID-19 Surveillance Team |
| EPI_ISL_2831937, EPI_ISL_2831938, EPI_ISL_2831939, EPI_ISL_2831940, EPI_ISL_2831941, EPI_ISL_2831942 | Lighthouse Lab in Milton Keynes | Wellcome Sanger Institute for the COVID-19 Genomics UK (COG-UK) Consortium | The Lighthouse Lab in Milton Keynes and Alex Alderton, Roberto Amato, Jeffrey Barrett, Sonia Goncalves, Ewan Harrison, David K. Jackson, Ian Johnston, Dominic Kwiatkowski, Cordelia Langford, John Sillitoe on behalf of the Wellcome Sanger Institute COVID-19 Surveillance Team |
| EPI_ISL_2831943 | Lighthouse Laboratory Plymouth | Wellcome Sanger Institute for the COVID-19 Genomics UK (COG-UK) Consortium | Lighthouse Laboratory Plymouth and Alex Alderton, Roberto Amato, Jeffrey Barrett, Sonia Goncalves, Ewan Harrison, David K. Jackson, Ian Johnston, Dominic Kwiatkowski, Cordelia Langford, John Sillitoe on behalf of the Wellcome Sanger Institute COVID-19 Surveillance Team |
| EPI_ISL_2831944 | Lighthouse Lab in Milton Keynes | Wellcome Sanger Institute for the COVID-19 Genomics UK (COG-UK) Consortium | The Lighthouse Lab in Milton Keynes and Alex Alderton, Roberto Amato, Jeffrey Barrett, Sonia Goncalves, Ewan Harrison, David K. Jackson, Ian Johnston, Dominic Kwiatkowski, Cordelia Langford, John Sillitoe on behalf of the Wellcome Sanger Institute COVID-19 Surveillance Team |
| EPI_ISL_2831945 | Lighthouse Lab in Glasgow | Wellcome Sanger Institute for the COVID-19 Genomics UK (COG-UK) Consortium | Harper VanSteenhouse, Yumi Kasai, David Gray, Carol Clugston, Anna Dominiczak and Alex Alderton, Roberto Amato, Jeffrey Barrett, Sonia Goncalves, Ewan Harrison, David K. Jackson, Ian Johnston, Dominic Kwiatkowski, Cordelia Langford, John Sillitoe on behalf of the Wellcome Sanger Institute COVID-19 Surveillance Team |
| EPI_ISL_2831946 | Lighthouse Laboratory Plymouth | Wellcome Sanger Institute for the COVID-19 Genomics UK (COG-UK) Consortium | Lighthouse Laboratory Plymouth and Alex Alderton, Roberto Amato, Jeffrey Barrett, Sonia Goncalves, Ewan Harrison, David K. Jackson, Ian Johnston, Dominic Kwiatkowski, Cordelia Langford, John Sillitoe on behalf of the Wellcome Sanger Institute COVID-19 Surveillance Team |
| EPI_ISL_2831947 | Lighthouse Lab in Milton Keynes | Wellcome Sanger Institute for the COVID-19 Genomics UK (COG-UK) Consortium | The Lighthouse Lab in Milton Keynes and Alex Alderton, Roberto Amato, Jeffrey Barrett, Sonia Goncalves, Ewan Harrison, David K. Jackson, Ian Johnston, Dominic Kwiatkowski, Cordelia Langford, John Sillitoe on behalf of the Wellcome Sanger Institute COVID-19 Surveillance Team |
| EPI_ISL_2831948 | Lighthouse Laboratory Plymouth | Wellcome Sanger Institute for the COVID-19 Genomics UK (COG-UK) Consortium | Lighthouse Laboratory Plymouth and Alex Alderton, Roberto Amato, Jeffrey Barrett, Sonia Goncalves, Ewan Harrison, David K. Jackson, Ian Johnston, Dominic Kwiatkowski, Cordelia Langford, John Sillitoe on behalf of the Wellcome Sanger Institute COVID-19 Surveillance Team |

| | | | |
|---|---|---|---|
| EPI_ISL_2831949 | Lighthouse Lab in Milton Keynes | Wellcome Sanger Institute for the COVID-19 Genomics UK (COG-UK) Consortium | The Lighthouse Lab in Milton Keynes and Alex Alderton, Roberto Amato, Jeffrey Barrett, Sonia Goncalves, Ewan Harrison, David K. Jackson, Ian Johnston, Dominic Kwiatkowski, Cordelia Langford, John Sillitoe on behalf of the Wellcome Sanger Institute COVID-19 Surveillance Team |
| EPI_ISL_2831950 | Lighthouse Lab in Glasgow | Wellcome Sanger Institute for the COVID-19 Genomics UK (COG-UK) Consortium | Harper VanSteenhouse, Yumi Kasai, David Gray, Carol Clugston, Anna Dominiczak and Alex Alderton, Roberto Amato, Jeffrey Barrett, Sonia Goncalves, Ewan Harrison, David K. Jackson, Ian Johnston, Dominic Kwiatkowski, Cordelia Langford, John Sillitoe on behalf of the Wellcome Sanger Institute COVID-19 Surveillance Team |
| EPI_ISL_2831951 | Lighthouse Laboratory Plymouth | Wellcome Sanger Institute for the COVID-19 Genomics UK (COG-UK) Consortium | Lighthouse Laboratory Plymouth and Alex Alderton, Roberto Amato, Jeffrey Barrett, Sonia Goncalves, Ewan Harrison, David K. Jackson, Ian Johnston, Dominic Kwiatkowski, Cordelia Langford, John Sillitoe on behalf of the Wellcome Sanger Institute COVID-19 Surveillance Team |
| EPI_ISL_2831952 | Lighthouse Lab in Glasgow | Wellcome Sanger Institute for the COVID-19 Genomics UK (COG-UK) Consortium | Harper VanSteenhouse, Yumi Kasai, David Gray, Carol Clugston, Anna Dominiczak and Alex Alderton, Roberto Amato, Jeffrey Barrett, Sonia Goncalves, Ewan Harrison, David K. Jackson, Ian Johnston, Dominic Kwiatkowski, Cordelia Langford, John Sillitoe on behalf of the Wellcome Sanger Institute COVID-19 Surveillance Team |
| EPI_ISL_2831954 | Lighthouse Lab in Milton Keynes | Wellcome Sanger Institute for the COVID-19 Genomics UK (COG-UK) Consortium | The Lighthouse Lab in Milton Keynes and Alex Alderton, Roberto Amato, Jeffrey Barrett, Sonia Goncalves, Ewan Harrison, David K. Jackson, Ian Johnston, Dominic Kwiatkowski, Cordelia Langford, John Sillitoe on behalf of the Wellcome Sanger Institute COVID-19 Surveillance Team |
| EPI_ISL_2831955 | Lighthouse Laboratory Plymouth | Wellcome Sanger Institute for the COVID-19 Genomics UK (COG-UK) Consortium | Lighthouse Laboratory Plymouth and Alex Alderton, Roberto Amato, Jeffrey Barrett, Sonia Goncalves, Ewan Harrison, David K. Jackson, Ian Johnston, Dominic Kwiatkowski, Cordelia Langford, John Sillitoe on behalf of the Wellcome Sanger Institute COVID-19 Surveillance Team |
| EPI_ISL_2831956, EPI_ISL_2831957 | Lighthouse Lab in Milton Keynes | Wellcome Sanger Institute for the COVID-19 Genomics UK (COG-UK) Consortium | The Lighthouse Lab in Milton Keynes and Alex Alderton, Roberto Amato, Jeffrey Barrett, Sonia Goncalves, Ewan Harrison, David K. Jackson, Ian Johnston, Dominic Kwiatkowski, Cordelia Langford, John Sillitoe on behalf of the Wellcome Sanger Institute COVID-19 Surveillance Team |
| EPI_ISL_2831958 | Lighthouse Laboratory Plymouth | Wellcome Sanger Institute for the COVID-19 Genomics UK (COG-UK) Consortium | Lighthouse Laboratory Plymouth and Alex Alderton, Roberto Amato, Jeffrey Barrett, Sonia Goncalves, Ewan Harrison, David K. Jackson, Ian Johnston, Dominic Kwiatkowski, Cordelia Langford, John Sillitoe on behalf of the Wellcome Sanger Institute COVID-19 Surveillance Team |
| EPI_ISL_2831959, EPI_ISL_2831960 | Lighthouse Lab in Milton Keynes | Wellcome Sanger Institute for the COVID-19 Genomics UK (COG-UK) Consortium | The Lighthouse Lab in Milton Keynes and Alex Alderton, Roberto Amato, Jeffrey Barrett, Sonia Goncalves, Ewan Harrison, David K. Jackson, Ian Johnston, Dominic Kwiatkowski, Cordelia Langford, John Sillitoe on behalf of the Wellcome Sanger Institute COVID-19 Surveillance Team |
| EPI_ISL_2831961, EPI_ISL_2831962 | Lighthouse Laboratory Plymouth | Wellcome Sanger Institute for the COVID-19 Genomics UK (COG-UK) Consortium | Lighthouse Laboratory Plymouth and Alex Alderton, Roberto Amato, Jeffrey Barrett, Sonia Goncalves, Ewan Harrison, David K. Jackson, Ian Johnston, Dominic Kwiatkowski, Cordelia Langford, John Sillitoe on behalf of the Wellcome Sanger Institute COVID-19 Surveillance Team |
| EPI_ISL_2831963 | Lighthouse Lab in Milton Keynes | Wellcome Sanger Institute for the COVID-19 Genomics UK (COG-UK) Consortium | The Lighthouse Lab in Milton Keynes and Alex Alderton, Roberto Amato, Jeffrey Barrett, Sonia Goncalves, Ewan Harrison, David K. Jackson, Ian Johnston, Dominic Kwiatkowski, Cordelia Langford, John Sillitoe on behalf of the Wellcome Sanger Institute COVID-19 Surveillance Team |
| EPI_ISL_2831964, EPI_ISL_2831965 | Lighthouse Laboratory Plymouth | Wellcome Sanger Institute for the COVID-19 Genomics UK (COG-UK) Consortium | Lighthouse Laboratory Plymouth and Alex Alderton, Roberto Amato, Jeffrey Barrett, Sonia Goncalves, Ewan Harrison, David K. Jackson, Ian Johnston, Dominic Kwiatkowski, Cordelia Langford, John Sillitoe on behalf of the Wellcome Sanger Institute COVID-19 Surveillance Team |
| EPI_ISL_2831966, EPI_ISL_2831967 | Lighthouse Lab in Milton Keynes | Wellcome Sanger Institute for the COVID-19 Genomics UK (COG-UK) Consortium | The Lighthouse Lab in Milton Keynes and Alex Alderton, Roberto Amato, Jeffrey Barrett, Sonia Goncalves, Ewan Harrison, David K. Jackson, Ian Johnston, Dominic Kwiatkowski, Cordelia Langford, John Sillitoe on behalf of the Wellcome Sanger Institute COVID-19 Surveillance Team |
| EPI_ISL_2831968 | Lighthouse Lab in Glasgow | Wellcome Sanger Institute for the COVID-19 Genomics UK (COG-UK) Consortium | Harper VanSteenhouse, Yumi Kasai, David Gray, Carol Clugston, Anna Dominiczak and Alex Alderton, Roberto Amato, Jeffrey Barrett, Sonia Goncalves, Ewan Harrison, David K. Jackson, Ian Johnston, Dominic Kwiatkowski, Cordelia Langford, John Sillitoe on behalf of the Wellcome Sanger Institute COVID-19 Surveillance Team |
| EPI_ISL_2831969 | Lighthouse Laboratory Plymouth | Wellcome Sanger Institute for the COVID-19 Genomics UK (COG-UK) Consortium | Lighthouse Laboratory Plymouth and Alex Alderton, Roberto Amato, Jeffrey Barrett, Sonia Goncalves, Ewan Harrison, David K. Jackson, Ian Johnston, Dominic Kwiatkowski, Cordelia Langford, John Sillitoe on behalf of the Wellcome Sanger Institute COVID-19 Surveillance Team |
| EPI_ISL_2831970, EPI_ISL_2831972, EPI_ISL_2831973, EPI_ISL_2831974, EPI_ISL_2831975, EPI_ISL_2831976, EPI_ISL_2831977, EPI_ISL_2831978 | Lighthouse Lab in Milton Keynes | Wellcome Sanger Institute for the COVID-19 Genomics UK (COG-UK) Consortium | The Lighthouse Lab in Milton Keynes and Alex Alderton, Roberto Amato, Jeffrey Barrett, Sonia Goncalves, Ewan Harrison, David K. Jackson, Ian Johnston, Dominic Kwiatkowski, Cordelia Langford, John Sillitoe on behalf of the Wellcome Sanger Institute COVID-19 Surveillance Team |
| EPI_ISL_2831979, EPI_ISL_2831980 | Lighthouse Laboratory Plymouth | Wellcome Sanger Institute for the COVID-19 Genomics UK (COG-UK) Consortium | Lighthouse Laboratory Plymouth and Alex Alderton, Roberto Amato, Jeffrey Barrett, Sonia Goncalves, Ewan Harrison, David K. Jackson, Ian Johnston, Dominic Kwiatkowski, Cordelia Langford, John Sillitoe on behalf of the Wellcome Sanger Institute COVID-19 Surveillance Team |
| EPI_ISL_2831981, EPI_ISL_2831982 | Lighthouse Lab in Milton Keynes | Wellcome Sanger Institute for the COVID-19 Genomics UK (COG-UK) Consortium | The Lighthouse Lab in Milton Keynes and Alex Alderton, Roberto Amato, Jeffrey Barrett, Sonia Goncalves, Ewan Harrison, David K. Jackson, Ian Johnston, Dominic Kwiatkowski, Cordelia Langford, John Sillitoe on behalf of the Wellcome Sanger Institute COVID-19 Surveillance Team |
| EPI_ISL_2831983 | Lighthouse Lab in Glasgow | Wellcome Sanger Institute for the COVID-19 Genomics UK (COG-UK) Consortium | Harper VanSteenhouse, Yumi Kasai, David Gray, Carol Clugston, Anna Dominiczak and Alex Alderton, Roberto Amato, Jeffrey Barrett, Sonia Goncalves, Ewan Harrison, David K. Jackson, Ian Johnston, Dominic Kwiatkowski, Cordelia Langford, John Sillitoe on behalf of the Wellcome Sanger Institute COVID-19 Surveillance Team |
| EPI_ISL_2831984, EPI_ISL_2831985 | Lighthouse Lab in Milton Keynes | Wellcome Sanger Institute for the COVID-19 Genomics UK (COG-UK) Consortium | The Lighthouse Lab in Milton Keynes and Alex Alderton, Roberto Amato, Jeffrey Barrett, Sonia Goncalves, Ewan Harrison, David K. Jackson, Ian Johnston, Dominic Kwiatkowski, Cordelia Langford, John Sillitoe on behalf of the Wellcome Sanger Institute COVID-19 Surveillance Team |
| EPI_ISL_2831986 | Lighthouse Lab in Glasgow | Wellcome Sanger Institute for the COVID-19 Genomics UK (COG-UK) Consortium | Harper VanSteenhouse, Yumi Kasai, David Gray, Carol Clugston, Anna Dominiczak and Alex Alderton, Roberto Amato, Jeffrey Barrett, Sonia Goncalves, Ewan Harrison, David K. Jackson, Ian Johnston, Dominic Kwiatkowski, Cordelia Langford, John Sillitoe on behalf of the Wellcome Sanger Institute COVID-19 Surveillance Team |
| EPI_ISL_2831987, EPI_ISL_2831988, EPI_ISL_2831989, EPI_ISL_2831991, EPI_ISL_2831992 | Lighthouse Lab in Milton Keynes | Wellcome Sanger Institute for the COVID-19 Genomics UK (COG-UK) Consortium | The Lighthouse Lab in Milton Keynes and Alex Alderton, Roberto Amato, Jeffrey Barrett, Sonia Goncalves, Ewan Harrison, David K. Jackson, Ian Johnston, Dominic Kwiatkowski, Cordelia Langford, John Sillitoe on behalf of the Wellcome Sanger Institute COVID-19 Surveillance Team |
| EPI_ISL_2831993, EPI_ISL_2831994 | Lighthouse Laboratory Plymouth | Wellcome Sanger Institute for the COVID-19 Genomics UK (COG-UK) Consortium | Lighthouse Laboratory Plymouth and Alex Alderton, Roberto Amato, Jeffrey Barrett, Sonia Goncalves, Ewan Harrison, David K. Jackson, Ian Johnston, Dominic Kwiatkowski, Cordelia Langford, John Sillitoe on behalf of the Wellcome Sanger Institute COVID-19 Surveillance Team |
| EPI_ISL_2831995, EPI_ISL_2831996 | Lighthouse Lab in Milton Keynes | Wellcome Sanger Institute for the COVID-19 Genomics UK (COG-UK) Consortium | The Lighthouse Lab in Milton Keynes and Alex Alderton, Roberto Amato, Jeffrey Barrett, Sonia Goncalves, Ewan Harrison, David K. Jackson, Ian Johnston, Dominic Kwiatkowski, Cordelia Langford, John Sillitoe on behalf of the Wellcome Sanger Institute COVID-19 Surveillance Team |
| EPI_ISL_2831997, EPI_ISL_2831998 | Lighthouse Laboratory Plymouth | Wellcome Sanger Institute for the COVID-19 Genomics UK (COG-UK) Consortium | Lighthouse Laboratory Plymouth and Alex Alderton, Roberto Amato, Jeffrey Barrett, Sonia Goncalves, Ewan Harrison, David K. Jackson, Ian Johnston, Dominic Kwiatkowski, Cordelia Langford, John Sillitoe on behalf of the Wellcome Sanger Institute COVID-19 Surveillance Team |
| EPI_ISL_2831999, EPI_ISL_2832000, EPI_ISL_2832001 | Lighthouse Lab in Milton Keynes | Wellcome Sanger Institute for the COVID-19 Genomics UK (COG-UK) Consortium | The Lighthouse Lab in Milton Keynes and Alex Alderton, Roberto Amato, Jeffrey Barrett, Sonia Goncalves, Ewan Harrison, David K. Jackson, Ian Johnston, Dominic Kwiatkowski, Cordelia Langford, John Sillitoe on behalf of the Wellcome Sanger Institute COVID-19 Surveillance Team |
| EPI_ISL_2832002 | Lighthouse Laboratory Plymouth | Wellcome Sanger Institute for the COVID-19 Genomics UK (COG-UK) Consortium | Lighthouse Laboratory Plymouth and Alex Alderton, Roberto Amato, Jeffrey Barrett, Sonia Goncalves, Ewan Harrison, David K. Jackson, Ian Johnston, Dominic Kwiatkowski, Cordelia Langford, John Sillitoe on behalf of the Wellcome Sanger Institute COVID-19 Surveillance Team |
| EPI_ISL_2832003, EPI_ISL_2832004, EPI_ISL_2832005, EPI_ISL_2832006, EPI_ISL_2832007, EPI_ISL_2832008, EPI_ISL_2832009 | Lighthouse Lab in Milton Keynes | Wellcome Sanger Institute for the COVID-19 Genomics UK (COG-UK) Consortium | The Lighthouse Lab in Milton Keynes and Alex Alderton, Roberto Amato, Jeffrey Barrett, Sonia Goncalves, Ewan Harrison, David K. Jackson, Ian Johnston, Dominic Kwiatkowski, Cordelia Langford, John Sillitoe on behalf of the Wellcome Sanger Institute COVID-19 Surveillance Team |
| EPI_ISL_2832010 | Lighthouse Laboratory Plymouth | Wellcome Sanger Institute for the COVID-19 Genomics UK (COG-UK) Consortium | Lighthouse Laboratory Plymouth and Alex Alderton, Roberto Amato, Jeffrey Barrett, Sonia Goncalves, Ewan Harrison, David K. Jackson, Ian Johnston, Dominic Kwiatkowski, Cordelia Langford, John Sillitoe on behalf of the Wellcome Sanger Institute COVID-19 Surveillance Team |
| EPI_ISL_2832011, EPI_ISL_2832012, EPI_ISL_2832013 | Lighthouse Lab in Milton Keynes | Wellcome Sanger Institute for the COVID-19 Genomics UK (COG-UK) Consortium | The Lighthouse Lab in Milton Keynes and Alex Alderton, Roberto Amato, Jeffrey Barrett, Sonia Goncalves, Ewan Harrison, David K. Jackson, Ian Johnston, Dominic Kwiatkowski, Cordelia Langford, John Sillitoe on behalf of the Wellcome Sanger Institute COVID-19 Surveillance Team |
| EPI_ISL_2832014, EPI_ISL_2832015 | Lighthouse Laboratory Plymouth | Wellcome Sanger Institute for the COVID-19 Genomics UK (COG-UK) Consortium | Lighthouse Laboratory Plymouth and Alex Alderton, Roberto Amato, Jeffrey Barrett, Sonia Goncalves, Ewan Harrison, David K. Jackson, Ian Johnston, Dominic Kwiatkowski, Cordelia Langford, John Sillitoe on behalf of the Wellcome Sanger Institute COVID-19 Surveillance Team |
| EPI_ISL_2832016, EPI_ISL_2832017 | Lighthouse Lab in Glasgow | Wellcome Sanger Institute for the COVID-19 Genomics UK (COG-UK) Consortium | Harper VanSteenhouse, Yumi Kasai, David Gray, Carol Clugston, Anna Dominiczak and Alex Alderton, Roberto Amato, Jeffrey Barrett, Sonia Goncalves, Ewan Harrison, David K. Jackson, Ian Johnston, Dominic Kwiatkowski, Cordelia Langford, John Sillitoe on behalf of the Wellcome Sanger Institute COVID-19 Surveillance Team |
| EPI_ISL_2832018 | Lighthouse Laboratory Plymouth | Wellcome Sanger Institute for the COVID-19 Genomics UK (COG-UK) Consortium | Lighthouse Laboratory Plymouth and Alex Alderton, Roberto Amato, Jeffrey Barrett, Sonia Goncalves, Ewan Harrison, David K. Jackson, Ian Johnston, Dominic Kwiatkowski, Cordelia Langford, John Sillitoe on behalf of the Wellcome Sanger Institute COVID-19 Surveillance Team |
| EPI_ISL_2832019 | Lighthouse Lab in Milton Keynes | Wellcome Sanger Institute for the COVID-19 Genomics UK (COG-UK) Consortium | The Lighthouse Lab in Milton Keynes and Alex Alderton, Roberto Amato, Jeffrey Barrett, Sonia Goncalves, Ewan Harrison, David K. Jackson, Ian Johnston, Dominic Kwiatkowski, Cordelia Langford, John Sillitoe on behalf of the Wellcome Sanger Institute COVID-19 Surveillance Team |
| EPI_ISL_2832020, EPI_ISL_2832021 | Lighthouse Laboratory Plymouth | Wellcome Sanger Institute for the COVID-19 Genomics UK (COG-UK) Consortium | Lighthouse Laboratory Plymouth and Alex Alderton, Roberto Amato, Jeffrey Barrett, Sonia Goncalves, Ewan Harrison, David K. Jackson, Ian Johnston, Dominic Kwiatkowski, Cordelia Langford, John Sillitoe on behalf of the Wellcome Sanger Institute COVID-19 Surveillance Team |
| EPI_ISL_2832022 | Lighthouse Lab in Glasgow | Wellcome Sanger Institute for the COVID-19 Genomics UK (COG-UK) Consortium | Harper VanSteenhouse, Yumi Kasai, David Gray, Carol Clugston, Anna Dominiczak and Alex Alderton, Roberto Amato, Jeffrey Barrett, Sonia Goncalves, Ewan Harrison, David K. Jackson, Ian Johnston, Dominic Kwiatkowski, Cordelia Langford, John Sillitoe on behalf of the Wellcome Sanger Institute COVID-19 Surveillance Team |
| EPI_ISL_2832023 | Lighthouse Lab in Milton Keynes | Wellcome Sanger Institute for the COVID-19 Genomics UK (COG-UK) Consortium | The Lighthouse Lab in Milton Keynes and Alex Alderton, Roberto Amato, Jeffrey Barrett, Sonia Goncalves, Ewan Harrison, David K. Jackson, Ian Johnston, Dominic Kwiatkowski, Cordelia Langford, John Sillitoe on behalf of the Wellcome Sanger Institute COVID-19 Surveillance Team |
| EPI_ISL_2832024 | Lighthouse Lab in Glasgow | Wellcome Sanger Institute for the COVID-19 Genomics UK (COG-UK) Consortium | Harper VanSteenhouse, Yumi Kasai, David Gray, Carol Clugston, Anna Dominiczak and Alex Alderton, Roberto Amato, Jeffrey Barrett, Sonia Goncalves, Ewan Harrison, David K. Jackson, Ian Johnston, Dominic Kwiatkowski, Cordelia Langford, John Sillitoe on behalf of the Wellcome Sanger Institute COVID-19 Surveillance Team |
| EPI_ISL_2832025 | Lighthouse Lab in Milton Keynes | Wellcome Sanger Institute for the COVID-19 Genomics UK (COG-UK) | The Lighthouse Lab in Milton Keynes and Alex Alderton, Roberto Amato, Jeffrey Barrett, Sonia Goncalves, Ewan Harrison, David K. Jackson, Ian Johnston, Dominic Kwiatkowski, Cordelia Langford, John Sillitoe on behalf of the Wellcome Sanger |

| | | | |
|---|---|---|---|
| | | Consortium | Institute COVID-19 Surveillance Team |
| EPI_ISL_2832026 | Lighthouse Laboratory Plymouth | Wellcome Sanger Institute for the COVID-19 Genomics UK (COG-UK) Consortium | Lighthouse Laboratory Plymouth and Alex Alderton, Roberto Amato, Jeffrey Barrett, Sonia Goncalves, Ewan Harrison, David K. Jackson, Ian Johnston, Dominic Kwiatkowski, Cordelia Langford, John Sillitoe on behalf of the Wellcome Sanger Institute COVID-19 Surveillance Team |
| EPI_ISL_2832027, EPI_ISL_2832028 | Lighthouse Lab in Milton Keynes | Wellcome Sanger Institute for the COVID-19 Genomics UK (COG-UK) Consortium | The Lighthouse Lab in Milton Keynes and Alex Alderton, Roberto Amato, Jeffrey Barrett, Sonia Goncalves, Ewan Harrison, David K. Jackson, Ian Johnston, Dominic Kwiatkowski, Cordelia Langford, John Sillitoe on behalf of the Wellcome Sanger Institute COVID-19 Surveillance Team |
| EPI_ISL_2832029, EPI_ISL_2832030 | Lighthouse Lab in Glasgow | Wellcome Sanger Institute for the COVID-19 Genomics UK (COG-UK) Consortium | Harper VanSteenhouse, Yumi Kasai, David Gray, Carol Clugston, Anna Dominiczak and Alex Alderton, Roberto Amato, Jeffrey Barrett, Sonia Goncalves, Ewan Harrison, David K. Jackson, Ian Johnston, Dominic Kwiatkowski, Cordelia Langford, John Sillitoe on behalf of the Wellcome Sanger Institute COVID-19 Surveillance Team |
| EPI_ISL_2832031, EPI_ISL_2832032, EPI_ISL_2832033, EPI_ISL_2832034, EPI_ISL_2832035, EPI_ISL_2832036, EPI_ISL_2832037 | Lighthouse Lab in Milton Keynes | Wellcome Sanger Institute for the COVID-19 Genomics UK (COG-UK) Consortium | The Lighthouse Lab in Milton Keynes and Alex Alderton, Roberto Amato, Jeffrey Barrett, Sonia Goncalves, Ewan Harrison, David K. Jackson, Ian Johnston, Dominic Kwiatkowski, Cordelia Langford, John Sillitoe on behalf of the Wellcome Sanger Institute COVID-19 Surveillance Team |
| EPI_ISL_2832038 | Lighthouse Laboratory Plymouth | Wellcome Sanger Institute for the COVID-19 Genomics UK (COG-UK) Consortium | Lighthouse Laboratory Plymouth and Alex Alderton, Roberto Amato, Jeffrey Barrett, Sonia Goncalves, Ewan Harrison, David K. Jackson, Ian Johnston, Dominic Kwiatkowski, Cordelia Langford, John Sillitoe on behalf of the Wellcome Sanger Institute COVID-19 Surveillance Team |
| EPI_ISL_2832039 | Lighthouse Lab in Glasgow | Wellcome Sanger Institute for the COVID-19 Genomics UK (COG-UK) Consortium | Harper VanSteenhouse, Yumi Kasai, David Gray, Carol Clugston, Anna Dominiczak and Alex Alderton, Roberto Amato, Jeffrey Barrett, Sonia Goncalves, Ewan Harrison, David K. Jackson, Ian Johnston, Dominic Kwiatkowski, Cordelia Langford, John Sillitoe on behalf of the Wellcome Sanger Institute COVID-19 Surveillance Team |
| EPI_ISL_2832040 | Lighthouse Lab in Milton Keynes | Wellcome Sanger Institute for the COVID-19 Genomics UK (COG-UK) Consortium | The Lighthouse Lab in Milton Keynes and Alex Alderton, Roberto Amato, Jeffrey Barrett, Sonia Goncalves, Ewan Harrison, David K. Jackson, Ian Johnston, Dominic Kwiatkowski, Cordelia Langford, John Sillitoe on behalf of the Wellcome Sanger Institute COVID-19 Surveillance Team |
| EPI_ISL_2832041, EPI_ISL_2832042 | Lighthouse Laboratory Plymouth | Wellcome Sanger Institute for the COVID-19 Genomics UK (COG-UK) Consortium | Lighthouse Laboratory Plymouth and Alex Alderton, Roberto Amato, Jeffrey Barrett, Sonia Goncalves, Ewan Harrison, David K. Jackson, Ian Johnston, Dominic Kwiatkowski, Cordelia Langford, John Sillitoe on behalf of the Wellcome Sanger Institute COVID-19 Surveillance Team |
| EPI_ISL_2832043, EPI_ISL_2832044, EPI_ISL_2832045 | Lighthouse Lab in Milton Keynes | Wellcome Sanger Institute for the COVID-19 Genomics UK (COG-UK) Consortium | The Lighthouse Lab in Milton Keynes and Alex Alderton, Roberto Amato, Jeffrey Barrett, Sonia Goncalves, Ewan Harrison, David K. Jackson, Ian Johnston, Dominic Kwiatkowski, Cordelia Langford, John Sillitoe on behalf of the Wellcome Sanger Institute COVID-19 Surveillance Team |
| EPI_ISL_2832046 | Lighthouse Lab in Glasgow | Wellcome Sanger Institute for the COVID-19 Genomics UK (COG-UK) Consortium | Harper VanSteenhouse, Yumi Kasai, David Gray, Carol Clugston, Anna Dominiczak and Alex Alderton, Roberto Amato, Jeffrey Barrett, Sonia Goncalves, Ewan Harrison, David K. Jackson, Ian Johnston, Dominic Kwiatkowski, Cordelia Langford, John Sillitoe on behalf of the Wellcome Sanger Institute COVID-19 Surveillance Team |
| EPI_ISL_2832047, EPI_ISL_2832048, EPI_ISL_2832049, EPI_ISL_2832050, EPI_ISL_2832051, EPI_ISL_2832052 | Lighthouse Lab in Milton Keynes | Wellcome Sanger Institute for the COVID-19 Genomics UK (COG-UK) Consortium | The Lighthouse Lab in Milton Keynes and Alex Alderton, Roberto Amato, Jeffrey Barrett, Sonia Goncalves, Ewan Harrison, David K. Jackson, Ian Johnston, Dominic Kwiatkowski, Cordelia Langford, John Sillitoe on behalf of the Wellcome Sanger Institute COVID-19 Surveillance Team |
| EPI_ISL_2832053 | Lighthouse Lab in Glasgow | Wellcome Sanger Institute for the COVID-19 Genomics UK (COG-UK) Consortium | Harper VanSteenhouse, Yumi Kasai, David Gray, Carol Clugston, Anna Dominiczak and Alex Alderton, Roberto Amato, Jeffrey Barrett, Sonia Goncalves, Ewan Harrison, David K. Jackson, Ian Johnston, Dominic Kwiatkowski, Cordelia Langford, John Sillitoe on behalf of the Wellcome Sanger Institute COVID-19 Surveillance Team |
| EPI_ISL_2832054 | Lighthouse Lab in Milton Keynes | Wellcome Sanger Institute for the COVID-19 Genomics UK (COG-UK) Consortium | The Lighthouse Lab in Milton Keynes and Alex Alderton, Roberto Amato, Jeffrey Barrett, Sonia Goncalves, Ewan Harrison, David K. Jackson, Ian Johnston, Dominic Kwiatkowski, Cordelia Langford, John Sillitoe on behalf of the Wellcome Sanger Institute COVID-19 Surveillance Team |
| EPI_ISL_2832055 | Lighthouse Laboratory Plymouth | Wellcome Sanger Institute for the COVID-19 Genomics UK (COG-UK) Consortium | Lighthouse Laboratory Plymouth and Alex Alderton, Roberto Amato, Jeffrey Barrett, Sonia Goncalves, Ewan Harrison, David K. Jackson, Ian Johnston, Dominic Kwiatkowski, Cordelia Langford, John Sillitoe on behalf of the Wellcome Sanger Institute COVID-19 Surveillance Team |
| EPI_ISL_2832056, EPI_ISL_2832057, EPI_ISL_2832058, EPI_ISL_2832059, EPI_ISL_2832060 | Lighthouse Lab in Milton Keynes | Wellcome Sanger Institute for the COVID-19 Genomics UK (COG-UK) Consortium | The Lighthouse Lab in Milton Keynes and Alex Alderton, Roberto Amato, Jeffrey Barrett, Sonia Goncalves, Ewan Harrison, David K. Jackson, Ian Johnston, Dominic Kwiatkowski, Cordelia Langford, John Sillitoe on behalf of the Wellcome Sanger Institute COVID-19 Surveillance Team |
| EPI_ISL_2832061 | Lighthouse Laboratory Plymouth | Wellcome Sanger Institute for the COVID-19 Genomics UK (COG-UK) Consortium | Lighthouse Laboratory Plymouth and Alex Alderton, Roberto Amato, Jeffrey Barrett, Sonia Goncalves, Ewan Harrison, David K. Jackson, Ian Johnston, Dominic Kwiatkowski, Cordelia Langford, John Sillitoe on behalf of the Wellcome Sanger Institute COVID-19 Surveillance Team |
| EPI_ISL_2832062 | Lighthouse Lab in Milton Keynes | Wellcome Sanger Institute for the COVID-19 Genomics UK (COG-UK) Consortium | The Lighthouse Lab in Milton Keynes and Alex Alderton, Roberto Amato, Jeffrey Barrett, Sonia Goncalves, Ewan Harrison, David K. Jackson, Ian Johnston, Dominic Kwiatkowski, Cordelia Langford, John Sillitoe on behalf of the Wellcome Sanger Institute COVID-19 Surveillance Team |
| EPI_ISL_2832063 | Lighthouse Laboratory Plymouth | Wellcome Sanger Institute for the COVID-19 Genomics UK (COG-UK) Consortium | Lighthouse Laboratory Plymouth and Alex Alderton, Roberto Amato, Jeffrey Barrett, Sonia Goncalves, Ewan Harrison, David K. Jackson, Ian Johnston, Dominic Kwiatkowski, Cordelia Langford, John Sillitoe on behalf of the Wellcome Sanger Institute COVID-19 Surveillance Team |
| EPI_ISL_2832065 | Lighthouse Lab in Alderley Park | Wellcome Sanger Institute for the COVID-19 Genomics UK (COG-UK) Consortium | Jacquelyn Wynn, Mairead Hyland, The Lighthouse Lab in Alderley Park and Alex Alderton, Roberto Amato, Jeffrey Barrett, Sonia Goncalves, Ewan Harrison, David K. Jackson, Ian Johnston, Dominic Kwiatkowski, Cordelia Langford, John Sillitoe on behalf of the Wellcome Sanger Institute COVID-19 Surveillance Team |
| EPI_ISL_2832066, EPI_ISL_2832067, EPI_ISL_2832068 | Lighthouse Laboratory Plymouth | Wellcome Sanger Institute for the COVID-19 Genomics UK (COG-UK) Consortium | Lighthouse Laboratory Plymouth and Alex Alderton, Roberto Amato, Jeffrey Barrett, Sonia Goncalves, Ewan Harrison, David K. Jackson, Ian Johnston, Dominic Kwiatkowski, Cordelia Langford, John Sillitoe on behalf of the Wellcome Sanger Institute COVID-19 Surveillance Team |
| EPI_ISL_2832069, EPI_ISL_2832070, EPI_ISL_2832071 | Lighthouse Lab in Alderley Park | Wellcome Sanger Institute for the COVID-19 Genomics UK (COG-UK) Consortium | Jacquelyn Wynn, Mairead Hyland, The Lighthouse Lab in Alderley Park and Alex Alderton, Roberto Amato, Jeffrey Barrett, Sonia Goncalves, Ewan Harrison, David K. Jackson, Ian Johnston, Dominic Kwiatkowski, Cordelia Langford, John Sillitoe on behalf of the Wellcome Sanger Institute COVID-19 Surveillance Team |
| EPI_ISL_2832072 | Lighthouse Laboratory Plymouth | Wellcome Sanger Institute for the COVID-19 Genomics UK (COG-UK) Consortium | Lighthouse Laboratory Plymouth and Alex Alderton, Roberto Amato, Jeffrey Barrett, Sonia Goncalves, Ewan Harrison, David K. Jackson, Ian Johnston, Dominic Kwiatkowski, Cordelia Langford, John Sillitoe on behalf of the Wellcome Sanger Institute COVID-19 Surveillance Team |
| EPI_ISL_2832073, EPI_ISL_2832074, EPI_ISL_2832075 | Lighthouse Lab in Alderley Park | Wellcome Sanger Institute for the COVID-19 Genomics UK (COG-UK) Consortium | Jacquelyn Wynn, Mairead Hyland, The Lighthouse Lab in Alderley Park and Alex Alderton, Roberto Amato, Jeffrey Barrett, Sonia Goncalves, Ewan Harrison, David K. Jackson, Ian Johnston, Dominic Kwiatkowski, Cordelia Langford, John Sillitoe on behalf of the Wellcome Sanger Institute COVID-19 Surveillance Team |
| EPI_ISL_2832076 | Lighthouse Lab in Glasgow | Wellcome Sanger Institute for the COVID-19 Genomics UK (COG-UK) Consortium | Harper VanSteenhouse, Yumi Kasai, David Gray, Carol Clugston, Anna Dominiczak and Alex Alderton, Roberto Amato, Jeffrey Barrett, Sonia Goncalves, Ewan Harrison, David K. Jackson, Ian Johnston, Dominic Kwiatkowski, Cordelia Langford, John Sillitoe on behalf of the Wellcome Sanger Institute COVID-19 Surveillance Team |
| EPI_ISL_2832077, EPI_ISL_2832078, EPI_ISL_2832079, EPI_ISL_2832080, EPI_ISL_2832081, EPI_ISL_2832082 | Lighthouse Lab in Alderley Park | Wellcome Sanger Institute for the COVID-19 Genomics UK (COG-UK) Consortium | Jacquelyn Wynn, Mairead Hyland, The Lighthouse Lab in Alderley Park and Alex Alderton, Roberto Amato, Jeffrey Barrett, Sonia Goncalves, Ewan Harrison, David K. Jackson, Ian Johnston, Dominic Kwiatkowski, Cordelia Langford, John Sillitoe on behalf of the Wellcome Sanger Institute COVID-19 Surveillance Team |
| EPI_ISL_2832083 | Lighthouse Lab in Milton Keynes | Wellcome Sanger Institute for the COVID-19 Genomics UK (COG-UK) Consortium | The Lighthouse Lab in Milton Keynes and Alex Alderton, Roberto Amato, Jeffrey Barrett, Sonia Goncalves, Ewan Harrison, David K. Jackson, Ian Johnston, Dominic Kwiatkowski, Cordelia Langford, John Sillitoe on behalf of the Wellcome Sanger Institute COVID-19 Surveillance Team |
| EPI_ISL_2832084, EPI_ISL_2832085 | Lighthouse Laboratory Plymouth | Wellcome Sanger Institute for the COVID-19 Genomics UK (COG-UK) Consortium | Lighthouse Laboratory Plymouth and Alex Alderton, Roberto Amato, Jeffrey Barrett, Sonia Goncalves, Ewan Harrison, David K. Jackson, Ian Johnston, Dominic Kwiatkowski, Cordelia Langford, John Sillitoe on behalf of the Wellcome Sanger Institute COVID-19 Surveillance Team |
| EPI_ISL_2832086, EPI_ISL_2832087 | Lighthouse Lab in Alderley Park | Wellcome Sanger Institute for the COVID-19 Genomics UK (COG-UK) Consortium | Jacquelyn Wynn, Mairead Hyland, The Lighthouse Lab in Alderley Park and Alex Alderton, Roberto Amato, Jeffrey Barrett, Sonia Goncalves, Ewan Harrison, David K. Jackson, Ian Johnston, Dominic Kwiatkowski, Cordelia Langford, John Sillitoe on behalf of the Wellcome Sanger Institute COVID-19 Surveillance Team |
| EPI_ISL_2832088 | Lighthouse Lab in Milton Keynes | Wellcome Sanger Institute for the COVID-19 Genomics UK (COG-UK) Consortium | The Lighthouse Lab in Milton Keynes and Alex Alderton, Roberto Amato, Jeffrey Barrett, Sonia Goncalves, Ewan Harrison, David K. Jackson, Ian Johnston, Dominic Kwiatkowski, Cordelia Langford, John Sillitoe on behalf of the Wellcome Sanger Institute COVID-19 Surveillance Team |
| EPI_ISL_2832089 | Health Services Laboratories | Wellcome Sanger Institute for the COVID-19 Genomics UK (COG-UK) Consortium | Health Services Laboratories and Alex Alderton, Roberto Amato, Jeffrey Barrett, Sonia Goncalves, Ewan Harrison, David K. Jackson, Ian Johnston, Dominic Kwiatkowski, Cordelia Langford, John Sillitoe on behalf of the Wellcome Sanger Institute COVID-19 Surveillance Team |
| EPI_ISL_2832090 | Lighthouse Lab in Milton Keynes | Wellcome Sanger Institute for the COVID-19 Genomics UK (COG-UK) Consortium | The Lighthouse Lab in Milton Keynes and Alex Alderton, Roberto Amato, Jeffrey Barrett, Sonia Goncalves, Ewan Harrison, David K. Jackson, Ian Johnston, Dominic Kwiatkowski, Cordelia Langford, John Sillitoe on behalf of the Wellcome Sanger Institute COVID-19 Surveillance Team |
| EPI_ISL_2832091 | Lighthouse Lab in Glasgow | Wellcome Sanger Institute for the COVID-19 Genomics UK (COG-UK) Consortium | Harper VanSteenhouse, Yumi Kasai, David Gray, Carol Clugston, Anna Dominiczak and Alex Alderton, Roberto Amato, Jeffrey Barrett, Sonia Goncalves, Ewan Harrison, David K. Jackson, Ian Johnston, Dominic Kwiatkowski, Cordelia Langford, John Sillitoe on behalf of the Wellcome Sanger Institute COVID-19 Surveillance Team |
| EPI_ISL_2832092 | Lighthouse Lab in Milton Keynes | Wellcome Sanger Institute for the COVID-19 Genomics UK (COG-UK) Consortium | The Lighthouse Lab in Milton Keynes and Alex Alderton, Roberto Amato, Jeffrey Barrett, Sonia Goncalves, Ewan Harrison, David K. Jackson, Ian Johnston, Dominic Kwiatkowski, Cordelia Langford, John Sillitoe on behalf of the Wellcome Sanger Institute COVID-19 Surveillance Team |
| EPI_ISL_2832093 | Lighthouse Laboratory Plymouth | Wellcome Sanger Institute for the COVID-19 Genomics UK (COG-UK) Consortium | Lighthouse Laboratory Plymouth and Alex Alderton, Roberto Amato, Jeffrey Barrett, Sonia Goncalves, Ewan Harrison, David K. Jackson, Ian Johnston, Dominic Kwiatkowski, Cordelia Langford, John Sillitoe on behalf of the Wellcome Sanger Institute COVID-19 Surveillance Team |
| EPI_ISL_2832094 | Lighthouse Lab in Milton Keynes | Wellcome Sanger Institute for the COVID-19 Genomics UK (COG-UK) Consortium | The Lighthouse Lab in Milton Keynes and Alex Alderton, Roberto Amato, Jeffrey Barrett, Sonia Goncalves, Ewan Harrison, David K. Jackson, Ian Johnston, Dominic Kwiatkowski, Cordelia Langford, John Sillitoe on behalf of the Wellcome Sanger Institute COVID-19 Surveillance Team |
| EPI_ISL_2832095 | Lighthouse Laboratory Plymouth | Wellcome Sanger Institute for the COVID-19 Genomics UK (COG-UK) Consortium | Lighthouse Laboratory Plymouth and Alex Alderton, Roberto Amato, Jeffrey Barrett, Sonia Goncalves, Ewan Harrison, David K. Jackson, Ian Johnston, Dominic Kwiatkowski, Cordelia Langford, John Sillitoe on behalf of the Wellcome Sanger Institute COVID-19 Surveillance Team |

| | | | |
|---|---|---|---|
| EPI_ISL_2832096, EPI_ISL_2832097, EPI_ISL_2832098 | Lighthouse Lab in Alderley Park | Wellcome Sanger Institute for the COVID-19 Genomics UK (COG-UK) Consortium | Jacquelyn Wynn, Mairead Hyland, The Lighthouse Lab in Alderley Park and Alex Alderton, Roberto Amato, Jeffrey Barrett, Sonia Goncalves, Ewan Harrison, David K. Jackson, Ian Johnston, Dominic Kwiatkowski, Cordelia Langford, John Sillitoe on behalf of the Wellcome Sanger Institute COVID-19 Surveillance Team |
| EPI_ISL_2832099 | Lighthouse Lab in Glasgow | Wellcome Sanger Institute for the COVID-19 Genomics UK (COG-UK) Consortium | Harper VanSteenhouse, Yumi Kasai, David Gray, Carol Clugston, Anna Dominiczak and Alex Alderton, Roberto Amato, Jeffrey Barrett, Sonia Goncalves, Ewan Harrison, David K. Jackson, Ian Johnston, Dominic Kwiatkowski, Cordelia Langford, John Sillitoe on behalf of the Wellcome Sanger Institute COVID-19 Surveillance Team |
| EPI_ISL_2832101 | Lighthouse Lab in Alderley Park | Wellcome Sanger Institute for the COVID-19 Genomics UK (COG-UK) Consortium | Jacquelyn Wynn, Mairead Hyland, The Lighthouse Lab in Alderley Park and Alex Alderton, Roberto Amato, Jeffrey Barrett, Sonia Goncalves, Ewan Harrison, David K. Jackson, Ian Johnston, Dominic Kwiatkowski, Cordelia Langford, John Sillitoe on behalf of the Wellcome Sanger Institute COVID-19 Surveillance Team |
| EPI_ISL_2832105, EPI_ISL_2832106 | Laboratory of Genetics and Personalized Medicine, Zan Mitrev Clinic | Laboratory of Genetics and Personalized Medicine, Zan Mitrev Clinic | Kungulovski G. et al. |
| EPI_ISL_2832108 | Hospital Universitari Vall d'Hebron - Vall d'Hebron Institut de Recerca | Hospital Universitari Vall d'Hebron - Vall d'Hebron Institut de Recerca | Cristina Andrés, Maria Piñana, Alejandra González-Sánchez, Damir Garcia-Cehic, Ariadna Rando, Juliana Esperalba, Maria Gema Codina, Carla Castillo, Maria Carmen Martin, Tomàs Pumarola, Josep Quer, Andrés Antón |
| EPI_ISL_2832112 | Microbiologia CATLAB | Can Ruti SARS-CoV-2 Sequencing Hub (HUGTiP/IrsiCaixa/IGTP) | Marc Noguera-Julian, Pilar Armengol, Ignacio Blanco, Antoni E Bordoy, Francesc Catala-Moll, Pere-Joan Cardona, Maria Casadellà, Cristina Casañ, Gemma Clara, Bonaventura Clotet, Montserrat Giménez, Mercedes Guerrero, Anna Not, David Panisello, Roger Paredes, Mariona Parera, Alexia París, Verónica Saludes, Alba Sánchez, Laia Soler, Lauro Sumoy, and Elisa Martró on behalf of the Can Ruti SARS-CoV-2 Sequencing Hub |
| EPI_ISL_2832114 | Parc Taulí Hospital Universitari | Can Ruti SARS-CoV-2 Sequencing Hub (HUGTiP/IrsiCaixa/IGTP) | Marc Noguera-Julian, Pilar Armengol, Ignacio Blanco, Antoni E Bordoy, Francesc Catala-Moll, Pere-Joan Cardona, Maria Casadellà, Cristina Casañ, Gemma Clara, Bonaventura Clotet, Montserrat Giménez, Mercedes Guerrero, Anna Not, David Panisello, Roger Paredes, Mariona Parera, Alexia París, Verónica Saludes, Alba Sánchez, Laia Soler, Lauro Sumoy, and Elisa Martró on behalf of the Can Ruti SARS-CoV-2 Sequencing Hub |
| EPI_ISL_2832118 | Microbiologia CATLAB | Can Ruti SARS-CoV-2 Sequencing Hub (HUGTiP/IrsiCaixa/IGTP) | Marc Noguera-Julian, Pilar Armengol, Ignacio Blanco, Antoni E Bordoy, Francesc Catala-Moll, Pere-Joan Cardona, Maria Casadellà, Cristina Casañ, Gemma Clara, Bonaventura Clotet, Montserrat Giménez, Mercedes Guerrero, Anna Not, David Panisello, Roger Paredes, Mariona Parera, Alexia París, Verónica Saludes, Alba Sánchez, Laia Soler, Lauro Sumoy, and Elisa Martró on behalf of the Can Ruti SARS-CoV-2 Sequencing Hub |
| EPI_ISL_2832120 | Parc Taulí Hospital Universitari | Can Ruti SARS-CoV-2 Sequencing Hub (HUGTiP/IrsiCaixa/IGTP) | Marc Noguera-Julian, Pilar Armengol, Ignacio Blanco, Antoni E Bordoy, Francesc Catala-Moll, Pere-Joan Cardona, Maria Casadellà, Cristina Casañ, Gemma Clara, Bonaventura Clotet, Montserrat Giménez, Mercedes Guerrero, Anna Not, David Panisello, Roger Paredes, Mariona Parera, Alexia París, Verónica Saludes, Alba Sánchez, Laia Soler, Lauro Sumoy, and Elisa Martró on behalf of the Can Ruti SARS-CoV-2 Sequencing Hub |
| EPI_ISL_2832121 | Ospedale San Filippo Neri | INMI Lazzaro Spallanzani IRCCS | F Messina, M Rueca, G Bonfiglio, O Butera, CEM Gruber, F Santini, B Bartolini, E Giombini, M Meledandri, ML Schiavone, A Tamburro, A Di Caro, MR Capobianchi |
| EPI_ISL_2832123, EPI_ISL_2832125 | Microbiologia CATLAB | Can Ruti SARS-CoV-2 Sequencing Hub (HUGTiP/IrsiCaixa/IGTP) | Marc Noguera-Julian, Pilar Armengol, Ignacio Blanco, Antoni E Bordoy, Francesc Catala-Moll, Pere-Joan Cardona, Maria Casadellà, Cristina Casañ, Gemma Clara, Bonaventura Clotet, Montserrat Giménez, Mercedes Guerrero, Anna Not, David Panisello, Roger Paredes, Mariona Parera, Alexia París, Verónica Saludes, Alba Sánchez, Laia Soler, Lauro Sumoy, and Elisa Martró on behalf of the Can Ruti SARS-CoV-2 Sequencing Hub |
| EPI_ISL_2832128 | Parc Taulí Hospital Universitari | Can Ruti SARS-CoV-2 Sequencing Hub (HUGTiP/IrsiCaixa/IGTP) | Marc Noguera-Julian, Pilar Armengol, Ignacio Blanco, Antoni E Bordoy, Francesc Catala-Moll, Pere-Joan Cardona, Maria Casadellà, Cristina Casañ, Gemma Clara, Bonaventura Clotet, Montserrat Giménez, Mercedes Guerrero, Anna Not, David Panisello, Roger Paredes, Mariona Parera, Alexia París, Verónica Saludes, Alba Sánchez, Laia Soler, Lauro Sumoy, and Elisa Martró on behalf of the Can Ruti SARS-CoV-2 Sequencing Hub |
| EPI_ISL_2832134, EPI_ISL_2832135, EPI_ISL_2832136 | Microbiology Department, Laboratori Clínic Metropolitana Nord. Hospital Universitari Germans Trias i Pujol | Can Ruti SARS-CoV-2 Sequencing Hub (HUGTiP/IrsiCaixa/IGTP) | Marc Noguera-Julian, Pilar Armengol, Ignacio Blanco, Antoni E Bordoy, Francesc Catala-Moll, Pere-Joan Cardona, Maria Casadellà, Cristina Casañ, Gemma Clara, Bonaventura Clotet, Montserrat Giménez, Mercedes Guerrero, Anna Not, David Panisello, Roger Paredes, Mariona Parera, Alexia París, Verónica Saludes, Alba Sánchez, Laia Soler, Lauro Sumoy, and Elisa Martró on behalf of the Can Ruti SARS-CoV-2 Sequencing Hub |
| EPI_ISL_2832137 | Microbiologia CATLAB | Can Ruti SARS-CoV-2 Sequencing Hub (HUGTiP/IrsiCaixa/IGTP) | Marc Noguera-Julian, Pilar Armengol, Ignacio Blanco, Antoni E Bordoy, Francesc Catala-Moll, Pere-Joan Cardona, Maria Casadellà, Cristina Casañ, Gemma Clara, Bonaventura Clotet, Montserrat Giménez, Mercedes Guerrero, Anna Not, David Panisello, Roger Paredes, Mariona Parera, Alexia París, Verónica Saludes, Alba Sánchez, Laia Soler, Lauro Sumoy, and Elisa Martró on behalf of the Can Ruti SARS-CoV-2 Sequencing Hub |
| EPI_ISL_2832138, EPI_ISL_2832139, EPI_ISL_2832141, EPI_ISL_2832142, EPI_ISL_2832143, EPI_ISL_2832144, EPI_ISL_2832145, EPI_ISL_2832147, EPI_ISL_2832150, EPI_ISL_2832151, EPI_ISL_2832152, EPI_ISL_2832153, EPI_ISL_2832154, EPI_ISL_2832155, EPI_ISL_2832156, EPI_ISL_2832159, EPI_ISL_2832160, EPI_ISL_2832162, EPI_ISL_2832163, EPI_ISL_2832164, EPI_ISL_2832167, EPI_ISL_2832169, EPI_ISL_2832171, EPI_ISL_2832174, EPI_ISL_2832176, EPI_ISL_2832180, EPI_ISL_2832182, EPI_ISL_2832183 | | | |
| see above | Microbiology Department, Laboratori Clínic Metropolitana Nord. Hospital Universitari Germans Trias i Pujol | Can Ruti SARS-CoV-2 Sequencing Hub (HUGTiP/IrsiCaixa/IGTP) | Marc Noguera-Julian, Pilar Armengol, Ignacio Blanco, Antoni E Bordoy, Francesc Catala-Moll, Pere-Joan Cardona, Maria Casadellà, Cristina Casañ, Gemma Clara, Bonaventura Clotet, Montserrat Giménez, Mercedes Guerrero, Anna Not, David Panisello, Roger Paredes, Mariona Parera, Alexia París, Verónica Saludes, Alba Sánchez, Laia Soler, Lauro Sumoy, and Elisa Martró on behalf of the Can Ruti SARS-CoV-2 Sequencing Hub |
| EPI_ISL_2832184, EPI_ISL_2832185 | Microbiologia CATLAB | Can Ruti SARS-CoV-2 Sequencing Hub (HUGTiP/IrsiCaixa/IGTP) | Marc Noguera-Julian, Pilar Armengol, Ignacio Blanco, Antoni E Bordoy, Francesc Catala-Moll, Pere-Joan Cardona, Maria Casadellà, Cristina Casañ, Gemma Clara, Bonaventura Clotet, Montserrat Giménez, Mercedes Guerrero, Anna Not, David Panisello, Roger Paredes, Mariona Parera, Alexia París, Verónica Saludes, Alba Sánchez, Laia Soler, Lauro Sumoy, and Elisa Martró on behalf of the Can Ruti SARS-CoV-2 Sequencing Hub |
| EPI_ISL_2832186, EPI_ISL_2832187, EPI_ISL_2832191, EPI_ISL_2832192, EPI_ISL_2832194, EPI_ISL_2832195 | Microbiology Department, Laboratori Clínic Metropolitana Nord. Hospital Universitari Germans Trias i Pujol | Can Ruti SARS-CoV-2 Sequencing Hub (HUGTiP/IrsiCaixa/IGTP) | Marc Noguera-Julian, Pilar Armengol, Ignacio Blanco, Antoni E Bordoy, Francesc Catala-Moll, Pere-Joan Cardona, Maria Casadellà, Cristina Casañ, Gemma Clara, Bonaventura Clotet, Montserrat Giménez, Mercedes Guerrero, Anna Not, David Panisello, Roger Paredes, Mariona Parera, Alexia París, Verónica Saludes, Alba Sánchez, Laia Soler, Lauro Sumoy, and Elisa Martró on behalf of the Can Ruti SARS-CoV-2 Sequencing Hub |
| EPI_ISL_2832196 | Microbiologia CATLAB | Can Ruti SARS-CoV-2 Sequencing Hub (HUGTiP/IrsiCaixa/IGTP) | Marc Noguera-Julian, Pilar Armengol, Ignacio Blanco, Antoni E Bordoy, Francesc Catala-Moll, Pere-Joan Cardona, Maria Casadellà, Cristina Casañ, Gemma Clara, Bonaventura Clotet, Montserrat Giménez, Mercedes Guerrero, Anna Not, David Panisello, Roger Paredes, Mariona Parera, Alexia París, Verónica Saludes, Alba Sánchez, Laia Soler, Lauro Sumoy, and Elisa Martró on behalf of the Can Ruti SARS-CoV-2 Sequencing Hub |
| EPI_ISL_2832198, EPI_ISL_2832199, EPI_ISL_2832200, EPI_ISL_2832202, EPI_ISL_2832203, EPI_ISL_2832205, EPI_ISL_2832206 | Microbiology Department, Laboratori Clínic Metropolitana Nord. Hospital Universitari Germans Trias i Pujol | Can Ruti SARS-CoV-2 Sequencing Hub (HUGTiP/IrsiCaixa/IGTP) | Marc Noguera-Julian, Pilar Armengol, Ignacio Blanco, Antoni E Bordoy, Francesc Catala-Moll, Pere-Joan Cardona, Maria Casadellà, Cristina Casañ, Gemma Clara, Bonaventura Clotet, Montserrat Giménez, Mercedes Guerrero, Anna Not, David Panisello, Roger Paredes, Mariona Parera, Alexia París, Verónica Saludes, Alba Sánchez, Laia Soler, Lauro Sumoy, and Elisa Martró on behalf of the Can Ruti SARS-CoV-2 Sequencing Hub |
| EPI_ISL_2832208 | Microbiologia CATLAB | Can Ruti SARS-CoV-2 Sequencing Hub (HUGTiP/IrsiCaixa/IGTP) | Marc Noguera-Julian, Pilar Armengol, Ignacio Blanco, Antoni E Bordoy, Francesc Catala-Moll, Pere-Joan Cardona, Maria Casadellà, Cristina Casañ, Gemma Clara, Bonaventura Clotet, Montserrat Giménez, Mercedes Guerrero, Anna Not, David Panisello, Roger Paredes, Mariona Parera, Alexia París, Verónica Saludes, Alba Sánchez, Laia Soler, Lauro Sumoy, and Elisa Martró on behalf of the Can Ruti SARS-CoV-2 Sequencing Hub |
| EPI_ISL_2832209, EPI_ISL_2832210, EPI_ISL_2832211, EPI_ISL_2832212, EPI_ISL_2832214, EPI_ISL_2832215, EPI_ISL_2832216, EPI_ISL_2832217 | Microbiology Department, Laboratori Clínic Metropolitana Nord. Hospital Universitari Germans Trias i Pujol | Can Ruti SARS-CoV-2 Sequencing Hub (HUGTiP/IrsiCaixa/IGTP) | Marc Noguera-Julian, Pilar Armengol, Ignacio Blanco, Antoni E Bordoy, Francesc Catala-Moll, Pere-Joan Cardona, Maria Casadellà, Cristina Casañ, Gemma Clara, Bonaventura Clotet, Montserrat Giménez, Mercedes Guerrero, Anna Not, David Panisello, Roger Paredes, Mariona Parera, Alexia París, Verónica Saludes, Alba Sánchez, Laia Soler, Lauro Sumoy, and Elisa Martró on behalf of the Can Ruti SARS-CoV-2 Sequencing Hub |
| EPI_ISL_2832219, EPI_ISL_2832220 | Microbiologia CATLAB | Can Ruti SARS-CoV-2 Sequencing Hub (HUGTiP/IrsiCaixa/IGTP) | Marc Noguera-Julian, Pilar Armengol, Ignacio Blanco, Antoni E Bordoy, Francesc Catala-Moll, Pere-Joan Cardona, Maria Casadellà, Cristina Casañ, Gemma Clara, Bonaventura Clotet, Montserrat Giménez, Mercedes Guerrero, Anna Not, David Panisello, Roger Paredes, Mariona Parera, Alexia París, Verónica Saludes, Alba Sánchez, Laia Soler, Lauro Sumoy, and Elisa Martró on behalf of the Can Ruti SARS-CoV-2 Sequencing Hub |
| EPI_ISL_2832221, EPI_ISL_2832222, EPI_ISL_2832223, EPI_ISL_2832224, EPI_ISL_2832225, EPI_ISL_2832226, EPI_ISL_2832227, EPI_ISL_2832228 | Microbiology Department, Laboratori Clínic Metropolitana Nord. Hospital Universitari Germans Trias i Pujol | Can Ruti SARS-CoV-2 Sequencing Hub (HUGTiP/IrsiCaixa/IGTP) | Marc Noguera-Julian, Pilar Armengol, Ignacio Blanco, Antoni E Bordoy, Francesc Catala-Moll, Pere-Joan Cardona, Maria Casadellà, Cristina Casañ, Gemma Clara, Bonaventura Clotet, Montserrat Giménez, Mercedes Guerrero, Anna Not, David Panisello, Roger Paredes, Mariona Parera, Alexia París, Verónica Saludes, Alba Sánchez, Laia Soler, Lauro Sumoy, and Elisa Martró on behalf of the Can Ruti SARS-CoV-2 Sequencing Hub |
| EPI_ISL_2832229, EPI_ISL_2832230, EPI_ISL_2832231, EPI_ISL_2832237, EPI_ISL_2832240, EPI_ISL_2832241, EPI_ISL_2832242 | Plateforme de testing Namuroise | Plateforme de testing Namuroise | Nicolas GILLIARD, Lesly NYINKEU K.; Demars Aurore ; Maschietto Céline ; Nobis Chloé ; Otto Gaetan ; Denis Olivier ; Degosserie Jonathan ; Mullier François |
| EPI_ISL_2832259, EPI_ISL_2832274 | KU Leuven, Rega Institute, Clinical and Epidemiological Virology | KU Leuven, Rega Institute, Clinical and Epidemiological Virology | Tony Wawina-Bokalanga, Bert Vanmechelen, Joan Marti-Carerras, Piet Maes |
| EPI_ISL_2832356 | Department of Virology, Istituto Zooprofilattico Sperimentale del Lazio e della Toscana (IZSLT) | Department of General Diagnostics; Department of Virology; Istituto Zooprofilattico Sperimentale del Lazio e della Toscana (IZSLT) | Patricia Alba, Giuseppe Manna, Elena L. Diaconu, Fabiola Feltrin, Raffaela Conti, Teresa Scicluna, Virginia Carfora, Antonella Cersini, Alessia Franco, Antonio Battisti. |
| EPI_ISL_2832437, EPI_ISL_2832438, EPI_ISL_2832441, EPI_ISL_2832444, EPI_ISL_2832445 | KU Leuven, Rega Institute, Clinical and Epidemiological Virology | KU Leuven, Rega Institute, Clinical and Epidemiological Virology | Tony Wawina-Bokalanga, Bert Vanmechelen, Joan Marti-Carerras, Piet Maes |
| EPI_ISL_2832487 | Curative Labs | New Mexico Department of Health Scientific Laboratory | Ellie Johnson, D'eldra Malone, Jennifer Benoit, Ratheesh Rajan, Linda Salazar, Keila Gutierrez, Anastacia Griego-Fisher |
| EPI_ISL_2832494, EPI_ISL_2832500, EPI_ISL_2832502 | New Mexico Department of Health Scientific Laboratory | New Mexico Department of Health Scientific Laboratory | Ellie Johnson, D'eldra Malone, Jennifer Benoit, Ratheesh Rajan, Linda Salazar, Keila Gutierrez, Anastacia Griego-Fisher |
| EPI_ISL_2832513 | Infinity Biologix | Centers for Disease Control and Prevention Division of Viral Diseases, Pathogen Discovery | Dakota Howard, Dhwani Batra, Peter W. Cook, Kara Moser, Adrian Paskey, Jason Caravas, Benjamin Rambo-Martin, Shatavia Morrison, Christopher Gulvick, Scott Sammons, Yvette Unoarumhi, Darlene Wagner, Matthew Schmerer, Christian Bixby, Yihe Wang, Jonathan Schultz, Chirayu Goswami, Russ Hager, Robin Grimwood, Clinton R. Paden, Duncan MacCannell |
| EPI_ISL_2832518, EPI_ISL_2832519, EPI_ISL_2832524, EPI_ISL_2832528, EPI_ISL_2832530, EPI_ISL_2832531, EPI_ISL_2832533, EPI_ISL_2832537, EPI_ISL_2832539, EPI_ISL_2832540, EPI_ISL_2832541, EPI_ISL_2832542, EPI_ISL_2832545, EPI_ISL_2832549, EPI_ISL_2832552, EPI_ISL_2832553, EPI_ISL_2832554, EPI_ISL_2832555, EPI_ISL_2832557, EPI_ISL_2832564, EPI_ISL_2832565, EPI_ISL_2832570, EPI_ISL_2832572, EPI_ISL_2832575 | | | |
| see above | Aegis Sciences Corporation | Centers for Disease Control and Prevention Division of Viral | Dakota Howard, Dhwani Batra, Peter W. Cook, Kara Moser, Adrian Paskey, Jason Caravas, Benjamin Rambo-Martin, Shatavia Morrison, Christopher Gulvick, Scott Sammons, Yvette Unoarumhi, Darlene Wagner, Matthew Schmerer, Cyndi Clark, |

| | | | |
|---|---|---|---|
| | | Diseases, Pathogen Discovery | Patrick Campbell, Rob Case, Vikramsinha Ghorpade, Holly Houdeshell, Ola Kvalvaag, Dillon Nall, Ethan Sanders, Alec Vest, Shaun Westlund, Matthew Hardison, Clinton R. Paden, Duncan MacCannell |
| EPI_ISL_2832603, EPI_ISL_2832606, EPI_ISL_2832607, EPI_ISL_2832609, EPI_ISL_2832610, EPI_ISL_2832611, EPI_ISL_2832613, EPI_ISL_2832614, EPI_ISL_2832616, EPI_ISL_2832617, EPI_ISL_2832619, EPI_ISL_2832620, EPI_ISL_2832621, EPI_ISL_2832622, EPI_ISL_2832629, EPI_ISL_2832630, EPI_ISL_2832634 | | | |
| see above | KU Leuven, Rega Institute, Clinical and Epidemiological Virology | KU Leuven, Rega Institute, Clinical and Epidemiological Virology | Tony Wawina-Bokalanga, Bert Vanmechelen, Joan Marti-Carerras, Piet Maes |
| EPI_ISL_2832726, EPI_ISL_2832727 | Genomik Solidaritas Indonesia Laboratorium | Genomik Solidaritas Indonesia Laboratorium | Reinhart Gabriel, Anna Christina Brazia, Vania Gavrila Wikasa, Carissa Sintca Wijaya, Dhahlia Agustina Cahyono, Alfin Mohammad Abdillah, Normastuti Adhini Tantyo, Annisa Muthiah Sukirman, Gracia Felias Enos Korompis, Anuraj Shankar, Meutia Ayuputeri Kumaheri, Ariel Pradipta |
| EPI_ISL_2832728, EPI_ISL_2832729 | Genomik Solidaritas Indonesia Laboratorium | Genomik Solidaritas Indonesia Laboratorium | Anna Christina Brazia, Reinhart Gabriel, Vania Gavrila Wikasa, Carissa Sintca Wijaya, Dhahlia Agustina Cahyono, Alfin Mohammad Abdillah, Normastuti Adhini Tantyo, Annisa Muthiah Sukirman, Gracia Felias Enos Korompis, Anuraj Shankar, Meutia Ayuputeri Kumaheri, Ariel Pradipta |
| EPI_ISL_2832730, EPI_ISL_2832731 | Genomik Solidaritas Indonesia Laboratorium | Genomik Solidaritas Indonesia Laboratorium | Vania Gavrila Wikasa, Carissa Sintca Wijaya, Anna Christina Brazia, Reinhart Gabriel, Dhahlia Agustina Cahyono, Alfin Mohammad Abdillah, Normastuti Adhini Tantyo, Annisa Muthiah Sukirman, Gracia Felias Enos Korompis, Anuraj Shankar, Meutia Ayuputeri Kumaheri, Ariel Pradipta |
| EPI_ISL_2832732, EPI_ISL_2832733 | Genomik Solidaritas Indonesia Laboratorium | Genomik Solidaritas Indonesia Laboratorium | Carissa Sintca Wijaya, Vania Gavrila Wikasa, Anna Christina Brazia, Reinhart Gabriel, Dhahlia Agustina Cahyono, Alfin Mohammad Abdillah, Normastuti Adhini Tantyo, Annisa Muthiah Sukirman, Gracia Felias Enos Korompis, Anuraj Shankar, Meutia Ayuputeri Kumaheri, Ariel Pradipta |
| EPI_ISL_2832734, EPI_ISL_2832735, EPI_ISL_2832736 | Genomik Solidaritas Indonesia Laboratorium | Genomik Solidaritas Indonesia Laboratorium | Reinhart Gabriel, Anna Christina Brazia, Vania Gavrila Wikasa, Carissa Sintca Wijaya, Dhahlia Agustina Cahyono, Alfin Mohammad Abdillah, Normastuti Adhini Tantyo, Annisa Muthiah Sukirman, Gracia Felias Enos Korompis, Anuraj Shankar, Meutia Ayuputeri Kumaheri, Ariel Pradipta |
| EPI_ISL_2832737, EPI_ISL_2832738, EPI_ISL_2832739 | Genomik Solidaritas Indonesia Laboratorium | Genomik Solidaritas Indonesia Laboratorium | Anna Christina Brazia, Reinhart Gabriel, Vania Gavrila Wikasa, Carissa Sintca Wijaya, Dhahlia Agustina Cahyono, Alfin Mohammad Abdillah, Normastuti Adhini Tantyo, Annisa Muthiah Sukirman, Gracia Felias Enos Korompis, Anuraj Shankar, Meutia Ayuputeri Kumaheri, Ariel Pradipta |
| EPI_ISL_2832740, EPI_ISL_2832741, EPI_ISL_2832742 | Genomik Solidaritas Indonesia Laboratorium | Genomik Solidaritas Indonesia Laboratorium | Vania Gavrila Wikasa, Carissa Sintca Wijaya, Anna Christina Brazia, Reinhart Gabriel, Dhahlia Agustina Cahyono, Alfin Mohammad Abdillah, Normastuti Adhini Tantyo, Annisa Muthiah Sukirman, Gracia Felias Enos Korompis, Anuraj Shankar, Meutia Ayuputeri Kumaheri, Ariel Pradipta |
| EPI_ISL_2832743, EPI_ISL_2832744, EPI_ISL_2832745 | Genomik Solidaritas Indonesia Laboratorium | Genomik Solidaritas Indonesia Laboratorium | Carissa Sintca Wijaya, Vania Gavrila Wikasa, Anna Christina Brazia, Reinhart Gabriel, Dhahlia Agustina Cahyono, Alfin Mohammad Abdillah, Normastuti Adhini Tantyo, Annisa Muthiah Sukirman, Gracia Felias Enos Korompis, Anuraj Shankar, Meutia Ayuputeri Kumaheri, Ariel Pradipta |
| EPI_ISL_2832755, EPI_ISL_2832757, EPI_ISL_2832764, EPI_ISL_2832769 | Karolinska University Hospital Huddinge | Karolinska University Hospital | Jan Albert ,Tobias Allander ,Annelie Bjerkner ,Sandra Broddesson ,Robert Dyrdak ,Martin Ekman ,Lynda Eneh ,Lina Guerra Blomqvist ,Karolina Ininbergs ,Tanja Normark ,Isak Sylvin ,Zhibing Yun ,Martina Wahlund ,Valtteri Wirta |
| EPI_ISL_2832775, EPI_ISL_2832779 | Unilabs | Karolinska University Hospital | Jan Albert ,Tobias Allander ,Annelie Bjerkner ,Sandra Broddesson ,Robert Dyrdak ,Martin Ekman ,Lynda Eneh ,Lina Guerra Blomqvist ,Karolina Ininbergs ,Tanja Normark ,Isak Sylvin ,Zhibing Yun ,Martina Wahlund ,Valtteri Wirta |
| EPI_ISL_2832780, EPI_ISL_2832781, EPI_ISL_2832782 | Karolinska University Hospital Solna | Karolinska University Hospital | Jan Albert ,Tobias Allander ,Annelie Bjerkner ,Sandra Broddesson ,Robert Dyrdak ,Martin Ekman ,Lynda Eneh ,Lina Guerra Blomqvist ,Karolina Ininbergs ,Tanja Normark ,Isak Sylvin ,Zhibing Yun ,Martina Wahlund ,Valtteri Wirta |
| EPI_ISL_2832783 | Unilabs | Karolinska University Hospital | Jan Albert ,Tobias Allander ,Annelie Bjerkner ,Sandra Broddesson ,Robert Dyrdak ,Martin Ekman ,Lynda Eneh ,Lina Guerra Blomqvist ,Karolina Ininbergs ,Tanja Normark ,Isak Sylvin ,Zhibing Yun ,Martina Wahlund ,Valtteri Wirta |
| EPI_ISL_2832784 | Karolinska University Hospital Solna | Karolinska University Hospital | Jan Albert ,Tobias Allander ,Annelie Bjerkner ,Sandra Broddesson ,Robert Dyrdak ,Martin Ekman ,Lynda Eneh ,Lina Guerra Blomqvist ,Karolina Ininbergs ,Tanja Normark ,Isak Sylvin ,Zhibing Yun ,Martina Wahlund ,Valtteri Wirta |
| EPI_ISL_2832787 | Unilabs | Karolinska University Hospital | Jan Albert ,Tobias Allander ,Annelie Bjerkner ,Sandra Broddesson ,Robert Dyrdak ,Martin Ekman ,Lynda Eneh ,Lina Guerra Blomqvist ,Karolina Ininbergs ,Tanja Normark ,Isak Sylvin ,Zhibing Yun ,Martina Wahlund ,Valtteri Wirta |
| EPI_ISL_2832792, EPI_ISL_2832793 | Karolinska University Hospital Solna | Karolinska University Hospital | Jan Albert ,Tobias Allander ,Annelie Bjerkner ,Sandra Broddesson ,Robert Dyrdak ,Martin Ekman ,Lynda Eneh ,Lina Guerra Blomqvist ,Karolina Ininbergs ,Tanja Normark ,Isak Sylvin ,Zhibing Yun ,Martina Wahlund ,Valtteri Wirta |
| EPI_ISL_2832795 | Unilabs | Karolinska University Hospital | Jan Albert ,Tobias Allander ,Annelie Bjerkner ,Sandra Broddesson ,Robert Dyrdak ,Martin Ekman ,Lynda Eneh ,Lina Guerra Blomqvist ,Karolina Ininbergs ,Tanja Normark ,Isak Sylvin ,Zhibing Yun ,Martina Wahlund ,Valtteri Wirta |
| EPI_ISL_2832796 | Karolinska University Hospital Solna | Karolinska University Hospital | Jan Albert ,Tobias Allander ,Annelie Bjerkner ,Sandra Broddesson ,Robert Dyrdak ,Martin Ekman ,Lynda Eneh ,Lina Guerra Blomqvist ,Karolina Ininbergs ,Tanja Normark ,Isak Sylvin ,Zhibing Yun ,Martina Wahlund ,Valtteri Wirta |
| EPI_ISL_2832797, EPI_ISL_2832798, EPI_ISL_2832804 | Unilabs | Karolinska University Hospital | Jan Albert ,Tobias Allander ,Annelie Bjerkner ,Sandra Broddesson ,Robert Dyrdak ,Martin Ekman ,Lynda Eneh ,Lina Guerra Blomqvist ,Karolina Ininbergs ,Tanja Normark ,Isak Sylvin ,Zhibing Yun ,Martina Wahlund ,Valtteri Wirta |
| EPI_ISL_2832805, EPI_ISL_2832806 | Karolinska University Hospital Solna | Karolinska University Hospital | Jan Albert ,Tobias Allander ,Annelie Bjerkner ,Sandra Broddesson ,Robert Dyrdak ,Martin Ekman ,Lynda Eneh ,Lina Guerra Blomqvist ,Karolina Ininbergs ,Tanja Normark ,Isak Sylvin ,Zhibing Yun ,Martina Wahlund ,Valtteri Wirta |
| EPI_ISL_2832807, EPI_ISL_2832811 | Unilabs | Karolinska University Hospital | Jan Albert ,Tobias Allander ,Annelie Bjerkner ,Sandra Broddesson ,Robert Dyrdak ,Martin Ekman ,Lynda Eneh ,Lina Guerra Blomqvist ,Karolina Ininbergs ,Tanja Normark ,Isak Sylvin ,Zhibing Yun ,Martina Wahlund ,Valtteri Wirta |
| EPI_ISL_2832812, EPI_ISL_2832813, EPI_ISL_2832814 | Karolinska University Hospital Solna | Karolinska University Hospital | Jan Albert ,Tobias Allander ,Annelie Bjerkner ,Sandra Broddesson ,Robert Dyrdak ,Martin Ekman ,Lynda Eneh ,Lina Guerra Blomqvist ,Karolina Ininbergs ,Tanja Normark ,Isak Sylvin ,Zhibing Yun ,Martina Wahlund ,Valtteri Wirta |
| EPI_ISL_2832815 | Unilabs | Karolinska University Hospital | Jan Albert ,Tobias Allander ,Annelie Bjerkner ,Sandra Broddesson ,Robert Dyrdak ,Martin Ekman ,Lynda Eneh ,Lina Guerra Blomqvist ,Karolina Ininbergs ,Tanja Normark ,Isak Sylvin ,Zhibing Yun ,Martina Wahlund ,Valtteri Wirta |
| EPI_ISL_2832816 | Karolinska University Hospital Solna | Karolinska University Hospital | Jan Albert ,Tobias Allander ,Annelie Bjerkner ,Sandra Broddesson ,Robert Dyrdak ,Martin Ekman ,Lynda Eneh ,Lina Guerra Blomqvist ,Karolina Ininbergs ,Tanja Normark ,Isak Sylvin ,Zhibing Yun ,Martina Wahlund ,Valtteri Wirta |
| EPI_ISL_2832817, EPI_ISL_2832818, EPI_ISL_2832822 | Unilabs | Karolinska University Hospital | Jan Albert ,Tobias Allander ,Annelie Bjerkner ,Sandra Broddesson ,Robert Dyrdak ,Martin Ekman ,Lynda Eneh ,Lina Guerra Blomqvist ,Karolina Ininbergs ,Tanja Normark ,Isak Sylvin ,Zhibing Yun ,Martina Wahlund ,Valtteri Wirta |
| EPI_ISL_2832823 | Karolinska University Hospital Solna | Karolinska University Hospital | Jan Albert ,Tobias Allander ,Annelie Bjerkner ,Sandra Broddesson ,Robert Dyrdak ,Martin Ekman ,Lynda Eneh ,Lina Guerra Blomqvist ,Karolina Ininbergs ,Tanja Normark ,Isak Sylvin ,Zhibing Yun ,Martina Wahlund ,Valtteri Wirta |
| EPI_ISL_2832825 | Unilabs | Karolinska University Hospital | Jan Albert ,Tobias Allander ,Annelie Bjerkner ,Sandra Broddesson ,Robert Dyrdak ,Martin Ekman ,Lynda Eneh ,Lina Guerra Blomqvist ,Karolina Ininbergs ,Tanja Normark ,Isak Sylvin ,Zhibing Yun ,Martina Wahlund ,Valtteri Wirta |
| EPI_ISL_2833103, EPI_ISL_2833105, EPI_ISL_2833107, EPI_ISL_2833109, EPI_ISL_2833111, EPI_ISL_2833113, EPI_ISL_2833115, EPI_ISL_2833121, EPI_ISL_2833123, EPI_ISL_2833125, EPI_ISL_2833127, EPI_ISL_2833131, EPI_ISL_2833133, EPI_ISL_2833135 | | | |
| see above | San Gallicano Dermatologic InstituteIRCCS | National Institute for Infectious Diseases (INMI) L. Spallanzani I.R.C.C.S | E Giombini, B Bartolini, M Rueca, G Bonfiglio, O Butera, CEM Gruber, F Santini, F Messina, A Di Caro, MR Capobianchi |
| EPI_ISL_2833202 | KU Leuven, Rega Institute, Clinical and Epidemiological Virology | KU Leuven, Rega Institute, Clinical and Epidemiological Virology | Tony Wawina-Bokalanga, Bert Vanmechelen, Joan Marti-Carerras, Piet Maes |
| EPI_ISL_2833213 | Jessa | Jessa | Berden et al. on behalf of the Jessa_cmdLab |
| EPI_ISL_2833214, EPI_ISL_2833217 | ZOL | Jessa | Berden et al. on behalf of the Jessa_cmdLab |
| EPI_ISL_2833219, EPI_ISL_2833220, EPI_ISL_2833221, EPI_ISL_2833222, EPI_ISL_2833223, EPI_ISL_2833224 | Jessa | Jessa | Berden et al. on behalf of the Jessa_cmdLab |
| EPI_ISL_2833228, EPI_ISL_2833229, EPI_ISL_2833300, EPI_ISL_2833327, EPI_ISL_2833350, EPI_ISL_2833353, EPI_ISL_2833354, EPI_ISL_2833355, EPI_ISL_2833413, EPI_ISL_2833417, EPI_ISL_2833431, EPI_ISL_2833456, EPI_ISL_2833495, EPI_ISL_2833496 | | | |
| see above | KU Leuven, Rega Institute, Clinical and Epidemiological Virology | KU Leuven, Rega Institute, Clinical and Epidemiological Virology | Tony Wawina-Bokalanga, Bert Vanmechelen, Joan Marti-Carerras, Piet Maes |
| EPI_ISL_2833500, EPI_ISL_2833501, EPI_ISL_2833502, EPI_ISL_2833503, EPI_ISL_2833504, EPI_ISL_2833505, EPI_ISL_2833506, EPI_ISL_2833509, EPI_ISL_2833510, EPI_ISL_2833511, EPI_ISL_2833512, EPI_ISL_2833513, EPI_ISL_2833514 | | | |
| see above | A. Krumbholz, Labor Dr. Krause und Kollegen MVZ GmbH, Kiel | Charité Universitätsmedizin Berlin, Institut für Virologie | Victor M Corman, Jörn Beheim-Schwarzbach, Tobias Bleicker, Julia Tesch, Barbara Muehlemann, Andi Krumbholz, Talitha Veith, Julia Schneider, Terry Jones, Christian Drosten |
| EPI_ISL_2833515, EPI_ISL_2833516, EPI_ISL_2833517, EPI_ISL_2833518, EPI_ISL_2833519, EPI_ISL_2833520, EPI_ISL_2833522 | A. Krumbholz, Labor Dr. Krause und Kollegen MVZ GmbH, Kiel | Charité Universitätsmedizin Berlin, Institut für Virologie/Labor Berlin | Peter Menzel, Christine Stephan, Rolf Schwarzer, Victor M Corman, Barbara Mühlemann, Terry Jones, Christian Drosten |
| EPI_ISL_2833524 | Labor Berlin Charité Vivantes GmbH / Institut für Virologie | Charité Universitätsmedizin Berlin, Institut für Virologie/Labor Berlin | Peter Menzel, Christine Stephan, Rolf Schwarzer, Victor M Corman, Barbara Mühlemann, Terry Jones, Christian Drosten |
| EPI_ISL_2833525 | A. Krumbholz, Labor Dr. Krause und Kollegen MVZ GmbH, Kiel | Charité Universitätsmedizin Berlin, Institut für Virologie/Labor Berlin | Peter Menzel, Christine Stephan, Rolf Schwarzer, Victor M Corman, Barbara Mühlemann, Terry Jones, Christian Drosten |
| EPI_ISL_2833527 | Labor Berlin Charité Vivantes GmbH / Institut für Virologie | Charité Universitätsmedizin Berlin, Institut für Virologie/Labor Berlin | Peter Menzel, Christine Stephan, Rolf Schwarzer, Victor M Corman, Barbara Mühlemann, Terry Jones, Christian Drosten |
| EPI_ISL_2833529 | A. Krumbholz, Labor Dr. Krause und Kollegen MVZ GmbH, Kiel | Charité Universitätsmedizin Berlin, Institut für Virologie/Labor Berlin | Peter Menzel, Christine Stephan, Rolf Schwarzer, Victor M Corman, Barbara Mühlemann, Terry Jones, Christian Drosten |
| EPI_ISL_2833531, EPI_ISL_2833534 | Labor Berlin Charité Vivantes GmbH / Institut für Virologie | Charité Universitätsmedizin Berlin, Institut für Virologie/Labor Berlin | Peter Menzel, Christine Stephan, Rolf Schwarzer, Victor M Corman, Barbara Mühlemann, Terry Jones, Christian Drosten |
| EPI_ISL_2833535, EPI_ISL_2833537 | A. Krumbholz, Labor Dr. Krause und Kollegen MVZ GmbH, Kiel | Charité Universitätsmedizin Berlin, Institut für Virologie/Labor Berlin | Peter Menzel, Christine Stephan, Rolf Schwarzer, Victor M Corman, Barbara Mühlemann, Terry Jones, Christian Drosten |
| EPI_ISL_2833538 | Labor Berlin Charité Vivantes GmbH / Institut für Virologie | Charité Universitätsmedizin Berlin, Institut für Virologie/Labor Berlin | Peter Menzel, Christine Stephan, Rolf Schwarzer, Victor M Corman, Barbara Mühlemann, Terry Jones, Christian Drosten |

| | | | |
|---|---|---|---|
| EPI_ISL_2833539, EPI_ISL_2833540 | A. Krumbholz, Labor Dr. Krause und Kollegen MVZ GmbH, Kiel | Charité Universitätsmedizin Berlin, Institut für Virologie/Labor Berlin | Peter Menzel, Christine Stephan, Rolf Schwarzer, Victor M Corman, Barbara Mühlemann, Terry Jones, Christian Drosten |
| EPI_ISL_2833551, EPI_ISL_2833553 | Unilabs | Karolinska University Hospital | Jan Albert ,Tobias Allander ,Annelie Bjerkner ,Sandra Broddesson ,Robert Dyrdak ,Martin Ekman ,Lynda Eneh ,Lina Guerra Blomqvist ,Karolina Ininbergs ,Tanja Normark ,Isak Sylvin ,Zhibing Yun ,Martina Wahlund ,Valtteri Wirta |
| EPI_ISL_2833559 | Karolinska University Hospital Solna | Karolinska University Hospital | Jan Albert ,Tobias Allander ,Annelie Bjerkner ,Sandra Broddesson ,Robert Dyrdak ,Martin Ekman ,Lynda Eneh ,Lina Guerra Blomqvist ,Karolina Ininbergs ,Tanja Normark ,Isak Sylvin ,Zhibing Yun ,Martina Wahlund ,Valtteri Wirta |
| EPI_ISL_2833560 | Unilabs | Karolinska University Hospital | Jan Albert ,Tobias Allander ,Annelie Bjerkner ,Sandra Broddesson ,Robert Dyrdak ,Martin Ekman ,Lynda Eneh ,Lina Guerra Blomqvist ,Karolina Ininbergs ,Tanja Normark ,Isak Sylvin ,Zhibing Yun ,Martina Wahlund ,Valtteri Wirta |
| EPI_ISL_2833562 | Karolinska University Hospital Solna | Karolinska University Hospital | Jan Albert ,Tobias Allander ,Annelie Bjerkner ,Sandra Broddesson ,Robert Dyrdak ,Martin Ekman ,Lynda Eneh ,Lina Guerra Blomqvist ,Karolina Ininbergs ,Tanja Normark ,Isak Sylvin ,Zhibing Yun ,Martina Wahlund ,Valtteri Wirta |
| EPI_ISL_2833564 | Synlab Medilab | Karolinska University Hospital | Jan Albert ,Tobias Allander ,Annelie Bjerkner ,Sandra Broddesson ,Robert Dyrdak ,Martin Ekman ,Lynda Eneh ,Lina Guerra Blomqvist ,Karolina Ininbergs ,Tanja Normark ,Isak Sylvin ,Zhibing Yun ,Martina Wahlund ,Valtteri Wirta |
| EPI_ISL_2833566 | Karolinska University Hospital Solna | Karolinska University Hospital | Jan Albert ,Tobias Allander ,Annelie Bjerkner ,Sandra Broddesson ,Robert Dyrdak ,Martin Ekman ,Lynda Eneh ,Lina Guerra Blomqvist ,Karolina Ininbergs ,Tanja Normark ,Isak Sylvin ,Zhibing Yun ,Martina Wahlund ,Valtteri Wirta |
| EPI_ISL_2833569, EPI_ISL_2833570, EPI_ISL_2833571 | Unilabs | Karolinska University Hospital | Jan Albert ,Tobias Allander ,Annelie Bjerkner ,Sandra Broddesson ,Robert Dyrdak ,Martin Ekman ,Lynda Eneh ,Lina Guerra Blomqvist ,Karolina Ininbergs ,Tanja Normark ,Isak Sylvin ,Zhibing Yun ,Martina Wahlund ,Valtteri Wirta |
| EPI_ISL_2833572 | Karolinska University Hospital Solna | Karolinska University Hospital | Jan Albert ,Tobias Allander ,Annelie Bjerkner ,Sandra Broddesson ,Robert Dyrdak ,Martin Ekman ,Lynda Eneh ,Lina Guerra Blomqvist ,Karolina Ininbergs ,Tanja Normark ,Isak Sylvin ,Zhibing Yun ,Martina Wahlund ,Valtteri Wirta |
| EPI_ISL_2833573, EPI_ISL_2833574, EPI_ISL_2833576, EPI_ISL_2833577 | Unilabs | Karolinska University Hospital | Jan Albert ,Tobias Allander ,Annelie Bjerkner ,Sandra Broddesson ,Robert Dyrdak ,Martin Ekman ,Lynda Eneh ,Lina Guerra Blomqvist ,Karolina Ininbergs ,Tanja Normark ,Isak Sylvin ,Zhibing Yun ,Martina Wahlund ,Valtteri Wirta |
| EPI_ISL_2833580 | Synlab Medilab | Karolinska University Hospital | Jan Albert ,Tobias Allander ,Annelie Bjerkner ,Sandra Broddesson ,Robert Dyrdak ,Martin Ekman ,Lynda Eneh ,Lina Guerra Blomqvist ,Karolina Ininbergs ,Tanja Normark ,Isak Sylvin ,Zhibing Yun ,Martina Wahlund ,Valtteri Wirta |
| EPI_ISL_2833582, EPI_ISL_2833584, EPI_ISL_2833587, EPI_ISL_2833588, EPI_ISL_2833590, EPI_ISL_2833591, EPI_ISL_2833593, EPI_ISL_2833594, EPI_ISL_2833595, EPI_ISL_2833596, EPI_ISL_2833597, EPI_ISL_2833598 | | | |
| see above | Unilabs | Karolinska University Hospital | Jan Albert ,Tobias Allander ,Annelie Bjerkner ,Sandra Broddesson ,Robert Dyrdak ,Martin Ekman ,Lynda Eneh ,Lina Guerra Blomqvist ,Karolina Ininbergs ,Tanja Normark ,Isak Sylvin ,Zhibing Yun ,Martina Wahlund ,Valtteri Wirta |
| EPI_ISL_2833599 | Synlab Medilab | Karolinska University Hospital | Jan Albert ,Tobias Allander ,Annelie Bjerkner ,Sandra Broddesson ,Robert Dyrdak ,Martin Ekman ,Lynda Eneh ,Lina Guerra Blomqvist ,Karolina Ininbergs ,Tanja Normark ,Isak Sylvin ,Zhibing Yun ,Martina Wahlund ,Valtteri Wirta |
| EPI_ISL_2833646, EPI_ISL_2833665 | Department of Microbiology, National Institute for Public Health of Kosova | Charité Universitätsmedizin Berlin, Institut für Virologie | Victor M Corman, Barbara Mühlemann, Jörn Beheim-Schwarzbach, Talitha Veith, Julia Schneider, Zana Deva, Donjeta Hajdari, Aferdita Hyseni, Pranvera Abazi, Nazmi Mehmeti, Blendi Jerliu, Xhevat Jakupi, Terry Jones, Christian Drosten |
| EPI_ISL_2833669, EPI_ISL_2833670 | Felix-Platter Spital | Clinical Bacteriology | Tim Roloff, Fanny Wegner, Helena MB Seth-Smith, Alfredo Mari, Karoline Leuzinger, Julia Bielicki, Manuel Battegay, Hans Hirsch, Adrian Egli |
| EPI_ISL_2833673 | Bürgerspital Solothurn | Institute of Medical Virology | Verena Kufner, Gabriela Ziltener, Maryam Zaheri, Stefan Schmutz, Annette Audigé, Maria Grünberg, Kevin Steiner, Jon Huder, Cyril Shah, Riccarda Capaul, Guido Bloemberg, Jürg Böni, Michael Huber, Alexandra Trkola |
| EPI_ISL_2833680 | Spital Männedorf AG | Institute of Medical Virology | Verena Kufner, Gabriela Ziltener, Maryam Zaheri, Stefan Schmutz, Annette Audigé, Maria Grünberg, Kevin Steiner, Jon Huder, Cyril Shah, Riccarda Capaul, Guido Bloemberg, Jürg Böni, Michael Huber, Alexandra Trkola |
| EPI_ISL_2833681 | Spital Limmattal | Institute of Medical Virology | Verena Kufner, Gabriela Ziltener, Maryam Zaheri, Stefan Schmutz, Annette Audigé, Maria Grünberg, Kevin Steiner, Jon Huder, Cyril Shah, Riccarda Capaul, Guido Bloemberg, Jürg Böni, Michael Huber, Alexandra Trkola |
| EPI_ISL_2833682 | UniversitätsSpital Zürich | Institute of Medical Virology | Verena Kufner, Gabriela Ziltener, Maryam Zaheri, Stefan Schmutz, Annette Audigé, Maria Grünberg, Kevin Steiner, Jon Huder, Cyril Shah, Riccarda Capaul, Guido Bloemberg, Jürg Böni, Michael Huber, Alexandra Trkola |
| EPI_ISL_2833683, EPI_ISL_2833684 | Spital Männedorf AG | Institute of Medical Virology | Verena Kufner, Gabriela Ziltener, Maryam Zaheri, Stefan Schmutz, Annette Audigé, Maria Grünberg, Kevin Steiner, Jon Huder, Cyril Shah, Riccarda Capaul, Guido Bloemberg, Jürg Böni, Michael Huber, Alexandra Trkola |
| EPI_ISL_2833689, EPI_ISL_2833690 | Stadtspital Triemli | Institute of Medical Virology | Verena Kufner, Gabriela Ziltener, Maryam Zaheri, Stefan Schmutz, Annette Audigé, Maria Grünberg, Kevin Steiner, Jon Huder, Cyril Shah, Riccarda Capaul, Guido Bloemberg, Jürg Böni, Michael Huber, Alexandra Trkola |
| EPI_ISL_2833691, EPI_ISL_2833692, EPI_ISL_2833693 | Bioanalytica AG | Institute of Medical Virology | Verena Kufner, Gabriela Ziltener, Maryam Zaheri, Stefan Schmutz, Annette Audigé, Maria Grünberg, Kevin Steiner, Jon Huder, Cyril Shah, Riccarda Capaul, Guido Bloemberg, Jürg Böni, Michael Huber, Alexandra Trkola |
| EPI_ISL_2833694 | Stadtspital Triemli | Institute of Medical Virology | Verena Kufner, Gabriela Ziltener, Maryam Zaheri, Stefan Schmutz, Annette Audigé, Maria Grünberg, Kevin Steiner, Jon Huder, Cyril Shah, Riccarda Capaul, Guido Bloemberg, Jürg Böni, Michael Huber, Alexandra Trkola |
| EPI_ISL_2833713, EPI_ISL_2833724 | KU Leuven, Rega Institute, Clinical and Epidemiological Virology | KU Leuven, Rega Institute, Clinical and Epidemiological Virology | Tony Wawina-Bokalanga, Bert Vanmechelen, Joan Marti-Carerras, Piet Maes |
| EPI_ISL_2833731, EPI_ISL_2833733, EPI_ISL_2833734, EPI_ISL_2833738, EPI_ISL_2833739 | MOH - Jaber Al-Ahmad Hospital (Innovation Research Laboratory) | MOH - Jaber Al-Ahmad Hospital (Innovation Research Laboratory) | Salman Al-Sabah , Mohammad Alghounaim |
| EPI_ISL_2833742, EPI_ISL_2833744, EPI_ISL_2833746, EPI_ISL_2833747, EPI_ISL_2833748, EPI_ISL_2833749, EPI_ISL_2833750, EPI_ISL_2833753 | Adan Hospital | Kuwait Cancer Control Center | Shakir Bahzad, Mona Alateeqi |
| EPI_ISL_2833760, EPI_ISL_2833761, EPI_ISL_2833762, EPI_ISL_2833763, EPI_ISL_2833764, EPI_ISL_2833765, EPI_ISL_2833766, EPI_ISL_2833770, EPI_ISL_2833771, EPI_ISL_2833772, EPI_ISL_2833773, EPI_ISL_2833774, EPI_ISL_2833775 | | | |
| see above | Wyoming Public Health Laboratory | Wyoming Public Health Laboratory | Jim Mildenberger, Taylor Fearing, Channing Weber, Ashley Norberg, Chayse Rowley, Marley Goetz, Brian Dominguez, Elliot Thomasson, Sam Britz, Cari Sloma, Robert Petit, and Rob Christensen |
| EPI_ISL_2833779, EPI_ISL_2833780, EPI_ISL_2833781, EPI_ISL_2833783 | Adan Hospital | Kuwait Cancer Control Center | Shakir Bahzad, Mona Alateeqi |
| EPI_ISL_2833784 | Adan Hospital | Kuwait Cancer control Center | Shakir Bahzad, Mona Alateeqi |
| EPI_ISL_2833791 | Adan Hospital | Kuwait Cancer Control Center | Shakir Bahzad, Mona Alateeqi |
| EPI_ISL_2833815 | Tampa General Hospital Esoteric Lab | Tampa General Hospital Esoteric Research & Development Lab | Jon Faughn, Deanna Becker, Dominic Uy, Jorge Mecias-Frias, Elaine Vendrone, Vicki Healer, Amorce Lima, Suzane Silbert |
| EPI_ISL_2833823 | Adan Hospital | Kuwait cancer Control Center | Shakir Bahzad, Mona Alateeqi |
| EPI_ISL_2833850, EPI_ISL_2833851 | Adan Hospital | Kuwait Cancer Control Center | Shakir Bahzad, Mona Alateeqi |
| EPI_ISL_2833860, EPI_ISL_2833862, EPI_ISL_2833863, EPI_ISL_2833864, EPI_ISL_2833865, EPI_ISL_2833866 | Kansas Health and Environmental Lab | Kansas Health and Environmental Lab | Katherine Wiggins, Mike Grose, Jonathan Barnell, Ben Olsen, and Phil Adam |
| EPI_ISL_2833867 | Adan Hospital | Kuwait Cancer Control Center | Shakir Bahzad, Mona Alateeqi |
| EPI_ISL_2833868, EPI_ISL_2833869, EPI_ISL_2833870, EPI_ISL_2833871, EPI_ISL_2833872, EPI_ISL_2833873, EPI_ISL_2833874, EPI_ISL_2833875, EPI_ISL_2833876, EPI_ISL_2833877, EPI_ISL_2833878, EPI_ISL_2833879, EPI_ISL_2833880, EPI_ISL_2833881, EPI_ISL_2833882, EPI_ISL_2833883, EPI_ISL_2833884, EPI_ISL_2833886, EPI_ISL_2833887 | | | |
| see above | Kansas Health and Environmental Lab | Kansas Health and Environmental Lab | Katherine Wiggins, Mike Grose, Jonathan Barnell, Ben Olsen, and Phil Adam |
| EPI_ISL_2833903 | Adan Hospital | Kuwait Cancer Control Center | Shakir Bahzad, Mona Alateeqi |
| EPI_ISL_2833905, EPI_ISL_2833907, EPI_ISL_2833910 | University of Wisconsin-Madison AIDS Vaccine Research Laboratories | University of Wisconsin-Madison AIDS Vaccine Research Laboratories | Gage Moreno, Katarina Braun, et al. AIDS Vaccine Research Laboratories |
| EPI_ISL_2833913 | Tampa General Hospital Esoteric Lab | Tampa General Hospital Esoteric Research & Development Lab | Jon Faughn, Deanna Becker, Dominic Uy, Jorge Mecias-Frias, Elaine Vendrone, Vicki Healer, Amorce Lima, Suzane Silbert |
| EPI_ISL_2833915, EPI_ISL_2833916 | Adan Hospital | Kuwait Cancer Control Center | Shakir Bahzad, Mona Alateeqi |
| EPI_ISL_2833923, EPI_ISL_2833924, EPI_ISL_2833926, EPI_ISL_2833927, EPI_ISL_2833929, EPI_ISL_2833930, EPI_ISL_2833931, EPI_ISL_2833932, EPI_ISL_2833933, EPI_ISL_2833934, EPI_ISL_2833935, EPI_ISL_2833936, EPI_ISL_2833937 | | | |
| see above | Kansas Health and Environmental Lab | Kansas Health and Environmental Lab | Katherine Wiggins, Mike Grose, Jonathan Barnell, Ben Olsen, and Phil Adam |
| EPI_ISL_2833938, EPI_ISL_2833939, EPI_ISL_2833940, EPI_ISL_2833941, EPI_ISL_2833942, EPI_ISL_2833943, EPI_ISL_2833945, EPI_ISL_2833946, | Wichita State University- Molecular Diagnostic Lab | Kansas Health and Environmental Lab | Katherine Wiggins, Mike Grose, Jonathan Barnell, Ben Olsen, and Phil Adam |

| | | | |
|---|---|---|---|
| EPI_ISL_2833948 | | | |
| EPI_ISL_2833949 | Adan Hospital | Cancer Control Center Center | Shakir Bahzad, Mona Alateeqi |
| EPI_ISL_2833992, EPI_ISL_2833995, EPI_ISL_2833999, EPI_ISL_2834000 | University of Wisconsin-Madison AIDS Vaccine Research Laboratories | University of Wisconsin-Madison AIDS Vaccine Research Laboratories | Gage Moreno, Katarina Braun, et al. AIDS Vaccine Research Laboratories |
| EPI_ISL_2834002, EPI_ISL_2834003, EPI_ISL_2834004 | Department of Acute Infectious Diseases Control and Prevention, Yunnan Provincial Center for Disease Control and Prevention | Department of Acute Infectious Diseases Control and Prevention, Yunnan Provincial Center for Disease Control and Prevention | Meiling Zhang, Jienan Zhou, Zhaosheng Liu, Xiaoqing Fu |
| EPI_ISL_2834009, EPI_ISL_2834018, EPI_ISL_2834021, EPI_ISL_2834023, EPI_ISL_2834025, EPI_ISL_2834027, EPI_ISL_2834028, EPI_ISL_2834029, EPI_ISL_2834031, EPI_ISL_2834032, EPI_ISL_2834033, EPI_ISL_2834034, EPI_ISL_2834035, EPI_ISL_2834036, EPI_ISL_2834039, EPI_ISL_2834041, EPI_ISL_2834042, EPI_ISL_2834043, EPI_ISL_2834045, EPI_ISL_2834048, EPI_ISL_2834049, EPI_ISL_2834050, EPI_ISL_2834051, EPI_ISL_2834054, EPI_ISL_2834057, EPI_ISL_2834060, EPI_ISL_2834062, EPI_ISL_2834063, EPI_ISL_2834064, EPI_ISL_2834065, EPI_ISL_2834066, EPI_ISL_2834067, EPI_ISL_2834069, EPI_ISL_2834070, EPI_ISL_2834071, EPI_ISL_2834072, EPI_ISL_2834073, EPI_ISL_2834074, EPI_ISL_2834075, EPI_ISL_2834076, EPI_ISL_2834077, EPI_ISL_2834078, EPI_ISL_2834079, EPI_ISL_2834080, EPI_ISL_2834081, EPI_ISL_2834082, EPI_ISL_2834083, EPI_ISL_2834084, EPI_ISL_2834085, EPI_ISL_2834086, EPI_ISL_2834088, EPI_ISL_2834089, EPI_ISL_2834090, EPI_ISL_2834091 | | | |
| see above | Servicio de Microbiología. Consorcio Hospital General Universitario de Valencia | SeqCOVID-SPAIN consortium/IBV(CSIC) | María Dolores Ocete, Begoña Fuster Escrivá, Carme Salvador García, Rafael Medina González, Concepción Gimeno Cardona and SeqCOVID-SPAIN consortium |
| EPI_ISL_2834141 | St James's Hospital, Dublin 8, Ireland. | ELDA biotech | Deirdre M. Broderick, Peter R. Flanagan, Brendan Crowley |
| EPI_ISL_2834159, EPI_ISL_2834161, EPI_ISL_2834164, EPI_ISL_2834165, EPI_ISL_2834166, EPI_ISL_2834170, EPI_ISL_2834171, EPI_ISL_2834172, EPI_ISL_2834173, EPI_ISL_2834174, EPI_ISL_2834175, EPI_ISL_2834176, EPI_ISL_2834178, EPI_ISL_2834182, EPI_ISL_2834183, EPI_ISL_2834184, EPI_ISL_2834185, EPI_ISL_2834186, EPI_ISL_2834190, EPI_ISL_2834191, EPI_ISL_2834192, EPI_ISL_2834193, EPI_ISL_2834194, EPI_ISL_2834197, EPI_ISL_2834198, EPI_ISL_2834199, EPI_ISL_2834201, EPI_ISL_2834202, EPI_ISL_2834203, EPI_ISL_2834204, EPI_ISL_2834205, EPI_ISL_2834206 | | | |
| see above | Baylor Scott & White-Temple | Baylor Scott & White-Temple | Ari Rao, Linden Morales, Kimberly Walker, Marcus Volz, Shelby Johnson |
| EPI_ISL_2834245, EPI_ISL_2834246 | The Ohio State University Applied Microbiology Services Laboratory | The Ohio State University Applied Microbiology Services Laboratory | Seth A. Faith PhD |
| EPI_ISL_2834268, EPI_ISL_2834269, EPI_ISL_2834270, EPI_ISL_2834271, EPI_ISL_2834272, EPI_ISL_2834273, EPI_ISL_2834274, EPI_ISL_2834276, EPI_ISL_2834277, EPI_ISL_2834278, EPI_ISL_2834280, EPI_ISL_2834281, EPI_ISL_2834282, EPI_ISL_2834283, EPI_ISL_2834284, EPI_ISL_2834285, EPI_ISL_2834286, EPI_ISL_2834287, EPI_ISL_2834288, EPI_ISL_2834289, EPI_ISL_2834290, EPI_ISL_2834291, EPI_ISL_2834292, EPI_ISL_2834293, EPI_ISL_2834294, EPI_ISL_2834295, EPI_ISL_2834296, EPI_ISL_2834297, EPI_ISL_2834299, EPI_ISL_2834300, EPI_ISL_2834301, EPI_ISL_2834302, EPI_ISL_2834303, EPI_ISL_2834304, EPI_ISL_2834305, EPI_ISL_2834307, EPI_ISL_2834308, EPI_ISL_2834309, EPI_ISL_2834310, EPI_ISL_2834311, EPI_ISL_2834312, EPI_ISL_2834313, EPI_ISL_2834314, EPI_ISL_2834315, EPI_ISL_2834316, EPI_ISL_2834317, EPI_ISL_2834318, EPI_ISL_2834319, EPI_ISL_2834320, EPI_ISL_2834321, EPI_ISL_2834322, EPI_ISL_2834323, EPI_ISL_2834324, EPI_ISL_2834325, EPI_ISL_2834326, EPI_ISL_2834328, EPI_ISL_2834329, EPI_ISL_2834330, EPI_ISL_2834331, EPI_ISL_2834332, EPI_ISL_2834333, EPI_ISL_2834334, EPI_ISL_2834335, EPI_ISL_2834336, EPI_ISL_2834337, EPI_ISL_2834338, EPI_ISL_2834339, EPI_ISL_2834340, EPI_ISL_2834341, EPI_ISL_2834342, EPI_ISL_2834343, EPI_ISL_2834344, EPI_ISL_2834345, EPI_ISL_2834346, EPI_ISL_2834347, EPI_ISL_2834348, EPI_ISL_2834349, EPI_ISL_2834350, EPI_ISL_2834351, EPI_ISL_2834352, EPI_ISL_2834353, EPI_ISL_2834354, EPI_ISL_2834356, EPI_ISL_2834357, EPI_ISL_2834359, EPI_ISL_2834360, EPI_ISL_2834361, EPI_ISL_2834362, EPI_ISL_2834363, EPI_ISL_2834364, EPI_ISL_2834365, EPI_ISL_2834366, EPI_ISL_2834367, EPI_ISL_2834368, EPI_ISL_2834369, EPI_ISL_2834371, EPI_ISL_2834372, EPI_ISL_2834374, EPI_ISL_2834376, EPI_ISL_2834377, EPI_ISL_2834378, EPI_ISL_2834379, EPI_ISL_2834380, EPI_ISL_2834384, EPI_ISL_2834385, EPI_ISL_2834386, EPI_ISL_2834387, EPI_ISL_2834389, EPI_ISL_2834390, EPI_ISL_2834391, EPI_ISL_2834392, EPI_ISL_2834393, EPI_ISL_2834394, EPI_ISL_2834395, EPI_ISL_2834396, EPI_ISL_2834397, EPI_ISL_2834398, EPI_ISL_2834399, EPI_ISL_2834400, EPI_ISL_2834401, EPI_ISL_2834402, EPI_ISL_2834403, EPI_ISL_2834404, EPI_ISL_2834405, EPI_ISL_2834406, EPI_ISL_2834407, EPI_ISL_2834408, EPI_ISL_2834409, EPI_ISL_2834410, EPI_ISL_2834411, EPI_ISL_2834412, EPI_ISL_2834413, EPI_ISL_2834414, EPI_ISL_2834415, EPI_ISL_2834416, EPI_ISL_2834417, EPI_ISL_2834418, EPI_ISL_2834419, EPI_ISL_2834420, EPI_ISL_2834421, EPI_ISL_2834422, EPI_ISL_2834423, EPI_ISL_2834424, EPI_ISL_2834425, EPI_ISL_2834426, EPI_ISL_2834427, EPI_ISL_2834428, EPI_ISL_2834430, EPI_ISL_2834431, EPI_ISL_2834432, EPI_ISL_2834433, EPI_ISL_2834434, EPI_ISL_2834435, EPI_ISL_2834436, EPI_ISL_2834437, EPI_ISL_2834438, EPI_ISL_2834439, EPI_ISL_2834440, EPI_ISL_2834441, EPI_ISL_2834442, EPI_ISL_2834443, EPI_ISL_2834444, EPI_ISL_2834445, EPI_ISL_2834446, EPI_ISL_2834447, EPI_ISL_2834448, EPI_ISL_2834449, EPI_ISL_2834450, EPI_ISL_2834451, EPI_ISL_2834452, EPI_ISL_2834455, EPI_ISL_2834456, EPI_ISL_2834457, EPI_ISL_2834458, EPI_ISL_2834459, EPI_ISL_2834460, EPI_ISL_2834461, EPI_ISL_2834462, EPI_ISL_2834463, EPI_ISL_2834464, EPI_ISL_2834465, EPI_ISL_2834466, EPI_ISL_2834467, EPI_ISL_2834469, EPI_ISL_2834470, EPI_ISL_2834471, EPI_ISL_2834472, EPI_ISL_2834473, EPI_ISL_2834474, EPI_ISL_2834476, EPI_ISL_2834477, EPI_ISL_2834478, EPI_ISL_2834479, EPI_ISL_2834480, EPI_ISL_2834481, EPI_ISL_2834482, EPI_ISL_2834483, EPI_ISL_2834485, EPI_ISL_2834486, EPI_ISL_2834487, EPI_ISL_2834488, EPI_ISL_2834489, EPI_ISL_2834490, EPI_ISL_2834491, EPI_ISL_2834492, EPI_ISL_2834493, EPI_ISL_2834494, EPI_ISL_2834495, EPI_ISL_2834496, EPI_ISL_2834497, EPI_ISL_2834498, EPI_ISL_2834499, EPI_ISL_2834501, EPI_ISL_2834502, EPI_ISL_2834503, EPI_ISL_2834504, EPI_ISL_2834506, EPI_ISL_2834507, EPI_ISL_2834509, EPI_ISL_2834510, EPI_ISL_2834511, EPI_ISL_2834513, EPI_ISL_2834514, EPI_ISL_2834517, EPI_ISL_2834518, EPI_ISL_2834519, EPI_ISL_2834520, EPI_ISL_2834522, EPI_ISL_2834523, EPI_ISL_2834524, EPI_ISL_2834525, EPI_ISL_2834526, EPI_ISL_2834528, EPI_ISL_2834529, EPI_ISL_2834530 | | | |
| see above | Shamir Medical Center (Asaf Harofe) | Shamir Medical Center (Asaf Harofe) | Nir Rainy,Chen Weiner,Reut Sorek Abramovich,Yevgeni Yegorov,Anna Vishnevsky,Patricia Benveniste-Lekovitz,Abu Hamad Ramzia,Adina Bar Chaim |
| EPI_ISL_2834560 | Wyoming Public Health Laboratory | Wyoming Public Health Laboratory | Jim Mildenberger, Taylor Fearing, Channing Weber, Ashley Norberg, Chayse Rowley, Marley Goetz, Brian Dominguez, Elliot Thomasson, Sam Britz, Cari Sloma, Robert Petit, and Rob Christensen |
| EPI_ISL_2834563 | KU Leuven, Rega Institute, Clinical and Epidemiological Virology | KU Leuven, Rega Institute, Clinical and Epidemiological Virology | Tony Wawina-Bokalanga, Bert Vanmechelen, Joan Marti-Carerras, Piet Maes |
| EPI_ISL_2834571, EPI_ISL_2834572, EPI_ISL_2834573, EPI_ISL_2834574, EPI_ISL_2834577, EPI_ISL_2834578, EPI_ISL_2834582, EPI_ISL_2834583, EPI_ISL_2834584, EPI_ISL_2834585, EPI_ISL_2834586, EPI_ISL_2834587, EPI_ISL_2834588, EPI_ISL_2834589, EPI_ISL_2834590 | | | |
| see above | Arizona State Public Health Laboratory | Arizona State Public Health Laboratory | Trung Huynh, Jessica Escobar, Katherine Fullerton, Nobuko Fukushima, Matthew Contursi, Stacy White, Linda Getsinger, Victor Waddell |
| EPI_ISL_2834703, EPI_ISL_2834709, EPI_ISL_2834710, EPI_ISL_2834712, EPI_ISL_2834713, EPI_ISL_2834714, EPI_ISL_2834716, EPI_ISL_2834718, EPI_ISL_2834722 | Washington State Department of Health Public Health Laboratories | Washington State Department of Health Public Health Laboratories | Drew MacKellar, Philip Dykema, Denny Russell, Joenice Gonzalez, Hannah Gray, Geoff Melly, Vanessa De Los Santos, Darren Lucas, JohnAric Peterson, Avi Singh, Rebecca Cao |
| EPI_ISL_2834733, EPI_ISL_2834734, EPI_ISL_2834742, EPI_ISL_2834743 | Ochsner Health | Bioinfoexperts, LLC | Judy Crabtree, Amy Feehan, Rebecca Rose, David J. Nolan, Tessa LaFleur, Julia-Garcia-Diaz, Lucio Miele, Susanna L. Lamers |
| EPI_ISL_2834748, EPI_ISL_2834751, EPI_ISL_2834755, EPI_ISL_2834772, EPI_ISL_2834778, EPI_ISL_2834782, EPI_ISL_2834789, EPI_ISL_2834808, EPI_ISL_2834812 | KU Leuven, Rega Institute, Clinical and Epidemiological Virology | KU Leuven, Rega Institute, Clinical and Epidemiological Virology | Tony Wawina-Bokalanga, Bert Vanmechelen, Joan Marti-Carerras, Piet Maes |
| EPI_ISL_2834925, EPI_ISL_2834927, EPI_ISL_2834928 | Central Health Laboratory/Airport Health Laboratory | Central Health Laboratory ,Victoria Hospital, Candos,Ministry of Health and Wellness, Mauritius | Manraj SS, Sonoo J, Patoo M, Bahadoor BS, Mathur H, Sujeewon C, Jannoo N, Ramuth M |
| EPI_ISL_2834958 | North Memorial | Minnesota Department of Health, Public Health Laboratory | Alexandra Lorentz, Jacob Garfin, Matt Plumb, and Xiong Wang |
| EPI_ISL_2834966 | HealthPartners Central Lab | Minnesota Department of Health, Public Health Laboratory | Alexandra Lorentz, Jacob Garfin, Matt Plumb, and Xiong Wang |
| EPI_ISL_2834967, EPI_ISL_2834968 | Centracare Laboratory Services | Minnesota Department of Health, Public Health Laboratory | Alexandra Lorentz, Jacob Garfin, Matt Plumb, and Xiong Wang |
| EPI_ISL_2834984 | Wyoming Public Health Laboratory | Wyoming Public Health Laboratory | Jim Mildenberger, Taylor Fearing, Channing Weber, Ashley Norberg, Chayse Rowley, Marley Goetz, Brian Dominguez, Elliot Thomasson, Sam Britz, Cari Sloma, Robert Petit, and Rob Christensen |
| EPI_ISL_2835005, EPI_ISL_2835006 | LESP Ciudad de Mexico | Instituto de Diagnostico y Referencia Epidemiologicos (INDRE) | Claudia Wong-Arambula, Abril Rodriguez-Maldonado, Vanessa Rivero-Arredondo, Ariadna Medina-Benitez, Joaquin Quiroz-Mercado, Sergio Rangel-Guerrero, Natividad Cruz-Ortiz, Tatiana Nunez-Garcia, Gisela Barrera-Badillo, Lucia Hernandez-Rivas, Irma Lopez-Martinez, Ernesto Ramirez-Gonzalez. |
| EPI_ISL_2835007 | LESP Veracruz | Instituto de Diagnostico y Referencia Epidemiologicos (INDRE) | Claudia Wong-Arambula, Abril Rodriguez-Maldonado, Vanessa Rivero-Arredondo, Ariadna Medina-Benitez, Joaquin Quiroz-Mercado, Sergio Rangel-Guerrero, Natividad Cruz-Ortiz, Tatiana Nunez-Garcia, Gisela Barrera-Badillo, Lucia Hernandez-Rivas, Irma Lopez-Martinez, Ernesto Ramirez-Gonzalez. |
| EPI_ISL_2835008 | LESP Morelos | Instituto de Diagnostico y Referencia Epidemiologicos (INDRE) | Claudia Wong-Arambula, Abril Rodriguez-Maldonado, Vanessa Rivero-Arredondo, Ariadna Medina-Benitez, Joaquin Quiroz-Mercado, Sergio Rangel-Guerrero, Natividad Cruz-Ortiz, Tatiana Nunez-Garcia, Gisela Barrera-Badillo, Lucia Hernandez-Rivas, Irma Lopez-Martinez, Ernesto Ramirez-Gonzalez. |
| EPI_ISL_2835009, EPI_ISL_2835010, EPI_ISL_2835011, EPI_ISL_2835012, EPI_ISL_2835013 | LESP Ciudad de Mexico | Instituto de Diagnostico y Referencia Epidemiologicos (INDRE) | Claudia Wong-Arambula, Abril Rodriguez-Maldonado, Vanessa Rivero-Arredondo, Ariadna Medina-Benitez, Joaquin Quiroz-Mercado, Sergio Rangel-Guerrero, Natividad Cruz-Ortiz, Tatiana Nunez-Garcia, Gisela Barrera-Badillo, Lucia Hernandez-Rivas, Irma Lopez-Martinez, Ernesto Ramirez-Gonzalez. |
| EPI_ISL_2835014 | LESP Queretaro | Instituto de Diagnostico y Referencia Epidemiologicos (INDRE) | Claudia Wong-Arambula, Abril Rodriguez-Maldonado, Vanessa Rivero-Arredondo, Ariadna Medina-Benitez, Joaquin Quiroz-Mercado, Sergio Rangel-Guerrero, Natividad Cruz-Ortiz, Tatiana Nunez-Garcia, Gisela Barrera-Badillo, Lucia Hernandez-Rivas, Irma Lopez-Martinez, Ernesto Ramirez-Gonzalez. |
| EPI_ISL_2835015 | LESP Tabasco | Instituto de Diagnostico y Referencia Epidemiologicos (INDRE) | Claudia Wong-Arambula, Abril Rodriguez-Maldonado, Vanessa Rivero-Arredondo, Ariadna Medina-Benitez, Joaquin Quiroz-Mercado, Sergio Rangel-Guerrero, Natividad Cruz-Ortiz, Tatiana Nunez-Garcia, Gisela Barrera-Badillo, Lucia Hernandez-Rivas, Irma Lopez-Martinez, Ernesto Ramirez-Gonzalez. |
| EPI_ISL_2835016 | LESP Jalisco | Instituto de Diagnostico y Referencia Epidemiologicos (INDRE) | Claudia Wong-Arambula, Abril Rodriguez-Maldonado, Vanessa Rivero-Arredondo, Ariadna Medina-Benitez, Joaquin Quiroz-Mercado, Sergio Rangel-Guerrero, Natividad Cruz-Ortiz, Tatiana Nunez-Garcia, Gisela Barrera-Badillo, Lucia Hernandez-Rivas, Irma Lopez-Martinez, Ernesto Ramirez-Gonzalez. |
| EPI_ISL_2835017, EPI_ISL_2835018, EPI_ISL_2835019, EPI_ISL_2835020, EPI_ISL_2835021 | LESP Sinaloa | Instituto de Diagnostico y Referencia Epidemiologicos (INDRE) | Claudia Wong-Arambula, Abril Rodriguez-Maldonado, Vanessa Rivero-Arredondo, Ariadna Medina-Benitez, Joaquin Quiroz-Mercado, Sergio Rangel-Guerrero, Natividad Cruz-Ortiz, Tatiana Nunez-Garcia, Gisela Barrera-Badillo, Lucia Hernandez-Rivas, Irma Lopez-Martinez, Ernesto Ramirez-Gonzalez. |
| EPI_ISL_2835022 | LESP Aguascalientes | Instituto de Diagnostico y Referencia Epidemiologicos (INDRE) | Claudia Wong-Arambula, Abril Rodriguez-Maldonado, Vanessa Rivero-Arredondo, Ariadna Medina-Benitez, Joaquin Quiroz-Mercado, Sergio Rangel-Guerrero, Natividad Cruz-Ortiz, Tatiana Nunez-Garcia, Gisela Barrera-Badillo, Lucia Hernandez-Rivas, Irma Lopez-Martinez, Ernesto Ramirez-Gonzalez. |
| EPI_ISL_2835023 | LESP Baja California | Instituto de Diagnostico y Referencia Epidemiologicos (INDRE) | Claudia Wong-Arambula, Abril Rodriguez-Maldonado, Vanessa Rivero-Arredondo, Ariadna Medina-Benitez, Joaquin Quiroz-Mercado, Sergio Rangel-Guerrero, Natividad Cruz-Ortiz, Tatiana Nunez-Garcia, Gisela Barrera-Badillo, Lucia Hernandez-Rivas, Irma Lopez-Martinez, Ernesto Ramirez-Gonzalez. |
| EPI_ISL_2835024 | LESP Campeche | Instituto de Diagnostico y Referencia Epidemiologicos (INDRE) | Claudia Wong-Arambula, Abril Rodriguez-Maldonado, Vanessa Rivero-Arredondo, Ariadna Medina-Benitez, Joaquin Quiroz-Mercado, Sergio Rangel-Guerrero, Natividad Cruz-Ortiz, Tatiana Nunez-Garcia, Gisela Barrera-Badillo, Lucia Hernandez-Rivas, Irma Lopez-Martinez, Ernesto Ramirez-Gonzalez. |

| | | | |
|---|---|---|---|
| EPI_ISL_2835025 | LESP Sonora | Instituto de Diagnostico y Referencia Epidemiologicos (INDRE) | Claudia Wong-Arambula, Abril Rodriguez-Maldonado, Vanessa Rivero-Arredondo, Ariadna Medina-Benitez, Joaquin Quiroz-Mercado, Sergio Rangel-Guerrero, Maribel Gonzalez-Villa, Natividad Cruz-Ortiz, Tatiana Nunez-Garcia, Gisela Barrera-Badillo, Lucia Hernandez-Rivas, Irma Lopez-Martinez, Ernesto Ramirez-Gonzalez. |
| EPI_ISL_2835026, EPI_ISL_2835027, EPI_ISL_2835028, EPI_ISL_2835029, EPI_ISL_2835030, EPI_ISL_2835031, EPI_ISL_2835032, EPI_ISL_2835033, EPI_ISL_2835034, EPI_ISL_2835035, EPI_ISL_2835036, EPI_ISL_2835037, EPI_ISL_2835038, EPI_ISL_2835039 | | | |
| see above | LESP Ciudad de Mexico | Instituto de Diagnostico y Referencia Epidemiologicos (INDRE) | Claudia Wong-Arambula, Abril Rodriguez-Maldonado, Vanessa Rivero-Arredondo, Ariadna Medina-Benitez, Joaquin Quiroz-Mercado, Sergio Rangel-Guerrero, Maribel Gonzalez-Villa, Natividad Cruz-Ortiz, Tatiana Nunez-Garcia, Gisela Barrera-Badillo, Lucia Hernandez-Rivas, Irma Lopez-Martinez, Ernesto Ramirez-Gonzalez. |
| EPI_ISL_2835040 | LESP Estado de Mexico | Instituto de Diagnostico y Referencia Epidemiologicos (INDRE) | Claudia Wong-Arambula, Abril Rodriguez-Maldonado, Vanessa Rivero-Arredondo, Ariadna Medina-Benitez, Joaquin Quiroz-Mercado, Sergio Rangel-Guerrero, Maribel Gonzalez-Villa, Natividad Cruz-Ortiz, Tatiana Nunez-Garcia, Gisela Barrera-Badillo, Lucia Hernandez-Rivas, Irma Lopez-Martinez, Ernesto Ramirez-Gonzalez. |
| EPI_ISL_2835041, EPI_ISL_2835042 | LESP Ciudad de Mexico | Instituto de Diagnostico y Referencia Epidemiologicos (INDRE) | Claudia Wong-Arambula, Abril Rodriguez-Maldonado, Vanessa Rivero-Arredondo, Ariadna Medina-Benitez, Joaquin Quiroz-Mercado, Sergio Rangel-Guerrero, Maribel Gonzalez-Villa, Natividad Cruz-Ortiz, Tatiana Nunez-Garcia, Gisela Barrera-Badillo, Lucia Hernandez-Rivas, Irma Lopez-Martinez, Ernesto Ramirez-Gonzalez. |
| EPI_ISL_2835260, EPI_ISL_2835262, EPI_ISL_2835272 | Rhode Island State Health Laboratory | Rhode Island State Health Laboratory | Ewa King, Richard C. Huard, Kristin Carpenter-Azevedo |
| EPI_ISL_2835552, EPI_ISL_2835553, EPI_ISL_2835554, EPI_ISL_2835555, EPI_ISL_2835559, EPI_ISL_2835561, EPI_ISL_2835563 | Sharp HealthCare Laboratory | Andersen lab at Scripps Research | SEARCH Alliance San Diego with Aaron Harding, Jacquelyn Berumen, Cathy Woerle, Liam McGinnis, Art Mendoza, Omid Bakhtar |
| EPI_ISL_2835600 | NS-QEII Health Sciences Centre | National Microbiology Laboratory (NML) | Anna Majer, Shari Tyson, Grace Seo, Philip Mabon, Elsie Grudeski, Rhiannon Huzarewich, Russell Mandes, Anneliese Landgraff, Jennifer Tanner, Natalie Knox, Morag Graham, Gary Van Domselaar, Todd Hatchette, Jason LeBlanc, Janice Pettipas, Dan Gaston, Nathalie Bastien, Yan Li, Timothy Booth, Darian Hole, Madison Chapel, Kirsten Biggar, CanCOGeN's metadata curation team, Public Health Agency of Canada CanCOGeN team |
| EPI_ISL_2835609 | Clinical Microbiology Laboratory, Faculty of Medicine, Universitas Indonesia | Faculty of Medicine, Universitas Indonesia | Fera Ibrahim, Fadilah, Andi Yasmon, Yulia Rosa, Rafika Indah Paramita, Khaerunissa Anbar Istiadi, Linda Erlina, Hana Apsari Pawestri, Vivi Setiawaty, Fithriyah, Beti Ernawati, Pratiwi Sudarmono, Budi Wiweko, Badriul Hegar, Anis Karuniawati, Ari Fahrial Syam |
| EPI_ISL_2835610 | Clinical Microbiology Laboratory, Faculty of Medicine, Universitas Indonesia | Faculty of Medicine, Universitas Indonesia | Fera Ibrahim, Fadilah, Andi Yasmon, Yulia Rosa, Rafika Indah Paramita, Khaerunissa Anbar Istiadi, Linda Erlina, Hana Apsari Pawestri, Vivi Setiawaty, Fithriyah, Beti Ernawati, Pratiwi Sudarmono, Budi Wiweko, Badriul Hegar, Anis Karuniawati, Ari Fahrial Syam |
| EPI_ISL_2835641 | Ministry of Health-Jaber Al-Ahmad Hospital | Virology Unit, Department of Microbiology, Faculty of Medicine, Kuwait University | Nada Madi, Hussain Safar, Ebaa Al-awadhi, Anfal Al-Adwani |
| EPI_ISL_2835642 | Ministry of Health-Jaber Al-Ahmad Hospital | Virology Unit, Department of Microbiology, Faculty of Medicine, Kuwait University | Nada Madi, Hussain Safar, Ebaa Al-Awadhi, Anfal Al-Adwani |
| EPI_ISL_2835643, EPI_ISL_2835644, EPI_ISL_2835645, EPI_ISL_2835646, EPI_ISL_2835647, EPI_ISL_2835648, EPI_ISL_2835649, EPI_ISL_2835650, EPI_ISL_2835651, EPI_ISL_2835652, EPI_ISL_2835653, EPI_ISL_2835654, EPI_ISL_2835655, EPI_ISL_2835656, EPI_ISL_2835684, EPI_ISL_2835685 | | | |
| see above | Clinical Microbiology Laboratory, Faculty of Medicine, Universitas Indonesia | Faculty of Medicine, Universitas Indonesia | Fera Ibrahim, Fadilah, Andi Yasmon, Yulia Rosa, Rafika Indah Paramita, Khaerunissa Anbar Istiadi, Linda Erlina, Hana Apsari Pawestri, Vivi Setiawaty, Fithriyah, Beti Ernawati, Pratiwi Sudarmono, Budi Wiweko, Badriul Hegar, Anis Karuniawati, Ari Fahrial Syam |
| EPI_ISL_2835686, EPI_ISL_2835687, EPI_ISL_2835688 | Ministry of Health, Jaber Al-Ahmad Hospital | Virology Unit, Department of Microbiology, Faculty of Medicine, Kuwait University | Nada Madi, Hussain Safar, Ebaa Al-Awadhi, Anfal Al-Adwani |
| EPI_ISL_2835689 | Ministry of Health-Jaber Al-Ahmad Hospital | Virology Unit, Department of Microbiology, Faculty of Medicine, Kuwait University | Nada Madi, Hussain Safar, Ebaa Al-Awadi, Anfal Al-Adwani |
| EPI_ISL_2835690, EPI_ISL_2835695, EPI_ISL_2835696, EPI_ISL_2835703 | Wojewodzka Stacja Sanitarno-Epidemiologiczna w Olsztynie, Laboratorium Badan Epidemiologiczno-Klinicznych | Wojewodzka Stacja Sanitarno-Epidemiologiczna w Olsztynie, Laboratorium Badan Epidemiologiczno-Klinicznych | Sylwia Krzetowska, Monika Czerminska, Ewa Liszewska, Tomasz Jakubczak, Marta Lukian, Patryk Bielecki, Aleksandra Kobiatko, Emilia Tarabasz, Paulina Rozycka, Barbara Dolinska |
| EPI_ISL_2835704, EPI_ISL_2835705, EPI_ISL_2835708, EPI_ISL_2835709, EPI_ISL_2835710, EPI_ISL_2835711, EPI_ISL_2835712, EPI_ISL_2835713, EPI_ISL_2835714, EPI_ISL_2835715, EPI_ISL_2835716, EPI_ISL_2835717, EPI_ISL_2835718, EPI_ISL_2835719, EPI_ISL_2835720, EPI_ISL_2835721, EPI_ISL_2835722, EPI_ISL_2835723, EPI_ISL_2835724, EPI_ISL_2835725, EPI_ISL_2835726, EPI_ISL_2835727, EPI_ISL_2835728, EPI_ISL_2835730, EPI_ISL_2835731, EPI_ISL_2835732, EPI_ISL_2835733, EPI_ISL_2835734, EPI_ISL_2835735, EPI_ISL_2835736 | | | |
| see above | Labo Analyses Med | National Reference Center for Viruses of Respiratory Infections, Institut Pasteur, Paris | Marion Barbet, Sylvie Behillil, Méline Bizard, Angela Brisebarre, Camille Capel, Vincent Enouf, Louise Lefrançois, Frédéric Lemoine, Christophe Malabat, Corinne Maufrais, Etienne Simon-Lorière, Maud Vanpeene, Sylvie Van der Werf ,BéNéDicte Roquebert |
| EPI_ISL_2835738 | Hospital | National Reference Center for Viruses of Respiratory Infections, Institut Pasteur, Paris | Marion Barbet, Sylvie Behillil, Méline Bizard, Angela Brisebarre, Camille Capel, Vincent Enouf, Louise Lefrançois, Frédéric Lemoine, Christophe Malabat, Corinne Maufrais, Etienne Simon-Lorière, Maud Vanpeene, Sylvie Van der Werf ,Laurent Bret |
| EPI_ISL_2835743 | Labo Analyses Med | National Reference Center for Viruses of Respiratory Infections, Institut Pasteur, Paris | Marion Barbet, Sylvie Behillil, Méline Bizard, Angela Brisebarre, Camille Capel, Vincent Enouf, Louise Lefrançois, Frédéric Lemoine, Christophe Malabat, Corinne Maufrais, Etienne Simon-Lorière, Maud Vanpeene, Sylvie Van der Werf ,Eve Haguenoer |
| EPI_ISL_2835748 | Labo Analyses Med | National Reference Center for Viruses of Respiratory Infections, Institut Pasteur, Paris | Marion Barbet, Sylvie Behillil, Méline Bizard, Angela Brisebarre, Camille Capel, Vincent Enouf, Louise Lefrançois, Frédéric Lemoine, Christophe Malabat, Corinne Maufrais, Etienne Simon-Lorière, Maud Vanpeene, Sylvie Van der Werf ,Sabine Trombert |
| EPI_ISL_2835749, EPI_ISL_2835756 | Labo Analyses Med | National Reference Center for Viruses of Respiratory Infections, Institut Pasteur, Paris | Marion Barbet, Sylvie Behillil, Méline Bizard, Angela Brisebarre, Camille Capel, Vincent Enouf, Louise Lefrançois, Frédéric Lemoine, Christophe Malabat, Corinne Maufrais, Etienne Simon-Lorière, Maud Vanpeene, Sylvie Van der Werf ,BéNéDicte Roquebert |
| EPI_ISL_2835762, EPI_ISL_2835763 | Labo Analyses Med | National Reference Center for Viruses of Respiratory Infections, Institut Pasteur, Paris | Marion Barbet, Sylvie Behillil, Méline Bizard, Angela Brisebarre, Camille Capel, Vincent Enouf, Louise Lefrançois, Frédéric Lemoine, Christophe Malabat, Corinne Maufrais, Etienne Simon-Lorière, Maud Vanpeene, Sylvie Van der Werf ,Caroline Devaux |
| EPI_ISL_2835777, EPI_ISL_2835783 | Labo Analyses Med | National Reference Center for Viruses of Respiratory Infections, Institut Pasteur, Paris | Marion Barbet, Sylvie Behillil, Méline Bizard, Angela Brisebarre, Camille Capel, Vincent Enouf, Louise Lefrançois, Frédéric Lemoine, Christophe Malabat, Corinne Maufrais, Etienne Simon-Lorière, Maud Vanpeene, Sylvie Van der Werf ,Jean-FrançOis Comes |
| EPI_ISL_2835790, EPI_ISL_2835791, EPI_ISL_2835792, EPI_ISL_2835793, EPI_ISL_2835794, EPI_ISL_2835795, EPI_ISL_2835799, EPI_ISL_2835800, EPI_ISL_2835802, EPI_ISL_2835803, EPI_ISL_2835805, EPI_ISL_2835806 | | | |
| see above | Labo Analyses Med | National Reference Center for Viruses of Respiratory Infections, Institut Pasteur, Paris | Marion Barbet, Sylvie Behillil, Méline Bizard, Angela Brisebarre, Camille Capel, Vincent Enouf, Louise Lefrançois, Frédéric Lemoine, Christophe Malabat, Corinne Maufrais, Etienne Simon-Lorière, Maud Vanpeene, Sylvie Van der Werf ,Anne Holstein |
| EPI_ISL_2835809, EPI_ISL_2835813, EPI_ISL_2835820, EPI_ISL_2835822, EPI_ISL_2835826, EPI_ISL_2835833 | Labo Analyses Med | National Reference Center for Viruses of Respiratory Infections, Institut Pasteur, Paris | Marion Barbet, Sylvie Behillil, Méline Bizard, Angela Brisebarre, Camille Capel, Vincent Enouf, Louise Lefrançois, Frédéric Lemoine, Christophe Malabat, Corinne Maufrais, Etienne Simon-Lorière, Maud Vanpeene, Sylvie Van der Werf ,Vincent Vieillefond |
| EPI_ISL_2835838, EPI_ISL_2835839 | Labo Analyses Med | National Reference Center for Viruses of Respiratory Infections, Institut Pasteur, Paris | Marion Barbet, Sylvie Behillil, Méline Bizard, Angela Brisebarre, Camille Capel, Vincent Enouf, Louise Lefrançois, Frédéric Lemoine, Christophe Malabat, Corinne Maufrais, Etienne Simon-Lorière, Maud Vanpeene, Sylvie Van der Werf ,Eric Grandsire |
| EPI_ISL_2835846, EPI_ISL_2835847 | Hospital | National Reference Center for Viruses of Respiratory Infections, Institut Pasteur, Paris | Marion Barbet, Sylvie Behillil, Méline Bizard, Angela Brisebarre, Camille Capel, Vincent Enouf, Louise Lefrançois, Frédéric Lemoine, Christophe Malabat, Corinne Maufrais, Etienne Simon-Lorière, Maud Vanpeene, Sylvie Van der Werf ,Evelyne Schvoerer |
| EPI_ISL_2835849, EPI_ISL_2835858, EPI_ISL_2835870, EPI_ISL_2835873, EPI_ISL_2835876 | Labo Analyses Med | National Reference Center for Viruses of Respiratory Infections, Institut Pasteur, Paris | Marion Barbet, Sylvie Behillil, Méline Bizard, Angela Brisebarre, Camille Capel, Vincent Enouf, Louise Lefrançois, Frédéric Lemoine, Christophe Malabat, Corinne Maufrais, Etienne Simon-Lorière, Maud Vanpeene, Sylvie Van der Werf ,Sabine Trombert |
| EPI_ISL_2835879 | Hospital | National Reference Center for Viruses of Respiratory Infections, Institut Pasteur, Paris | Marion Barbet, Sylvie Behillil, Méline Bizard, Angela Brisebarre, Camille Capel, Vincent Enouf, Louise Lefrançois, Frédéric Lemoine, Christophe Malabat, Corinne Maufrais, Etienne Simon-Lorière, Maud Vanpeene, Sylvie Van der Werf ,Sylvaine Bastian |
| EPI_ISL_2835892, EPI_ISL_2835893 | Labo Analyses Med | National Reference Center for Viruses of Respiratory Infections, Institut Pasteur, Paris | Marion Barbet, Sylvie Behillil, Méline Bizard, Angela Brisebarre, Camille Capel, Vincent Enouf, Louise Lefrançois, Frédéric Lemoine, Christophe Malabat, Corinne Maufrais, Etienne Simon-Lorière, Maud Vanpeene, Sylvie Van der Werf ,Pierre-Yves Leonard |
| EPI_ISL_2835898 | Hospital | National Reference Center for Viruses of Respiratory Infections, Institut Pasteur, Paris | Marion Barbet, Sylvie Behillil, Méline Bizard, Angela Brisebarre, Camille Capel, Vincent Enouf, Louise Lefrançois, Frédéric Lemoine, Christophe Malabat, Corinne Maufrais, Etienne Simon-Lorière, Maud Vanpeene, Sylvie Van der Werf ,Laurence Louvet |
| EPI_ISL_2835899, EPI_ISL_2835900, EPI_ISL_2835901, EPI_ISL_2835902, EPI_ISL_2835903, EPI_ISL_2835904 | Hospital | National Reference Center for Viruses of Respiratory Infections, Institut Pasteur, Paris | Marion Barbet, Sylvie Behillil, Méline Bizard, Angela Brisebarre, Camille Capel, Vincent Enouf, Louise Lefrançois, Frédéric Lemoine, Christophe Malabat, Corinne Maufrais, Etienne Simon-Lorière, Maud Vanpeene, Sylvie Van der Werf ,Sandrine Castelain |
| EPI_ISL_2835905 | Hospital | National Reference Center for Viruses of Respiratory Infections, Institut Pasteur, Paris | Marion Barbet, Sylvie Behillil, Méline Bizard, Angela Brisebarre, Camille Capel, Vincent Enouf, Louise Lefrançois, Frédéric Lemoine, Christophe Malabat, Corinne Maufrais, Etienne Simon-Lorière, Maud Vanpeene, Sylvie Van der Werf ,Hammas |
| EPI_ISL_2835992 | Eurofins Montjoly | Institut Pasteur de la Guyane | Anne Lavergne, Dominique Rousset, Antoine Enfissi, Arielle Salmier |
| EPI_ISL_2836010 | Epidemiology of Microbial Diseases, Yale School of Public Health | Epidemiology of Microbial Diseases, Yale School of Public Health | Vogels,C.B., Lucas,C., Fauver,J.R., Breban,M.I., Ott,I.M., Alpert,T., Petrone,M.E., Brito,A.F., Watkins,A.E., Kalinich,C.C., Rothman,J.E., Iwasaki,A., Landry,M.L., Grubaugh,N.D. |
| EPI_ISL_2836030, EPI_ISL_2836077, EPI_ISL_2836089, EPI_ISL_2836102, EPI_ISL_2836176, EPI_ISL_2836254, EPI_ISL_2836281, EPI_ISL_2836293, EPI_ISL_2836420, EPI_ISL_2836425, EPI_ISL_2836500, EPI_ISL_2836518, EPI_ISL_2836519, EPI_ISL_2836527, EPI_ISL_2836585, EPI_ISL_2836586, EPI_ISL_2836637, EPI_ISL_2836639, EPI_ISL_2836640, EPI_ISL_2836642, EPI_ISL_2836644, EPI_ISL_2836648, EPI_ISL_2836698, EPI_ISL_2836719, EPI_ISL_2836726, EPI_ISL_2836730, EPI_ISL_2836731, EPI_ISL_2836737, EPI_ISL_2836758, EPI_ISL_2836775, EPI_ISL_2836781, EPI_ISL_2836782, EPI_ISL_2836785, EPI_ISL_2836787, EPI_ISL_2836789, EPI_ISL_2836790, EPI_ISL_2836802, EPI_ISL_2836804, EPI_ISL_2836832, EPI_ISL_2836836, EPI_ISL_2836837, EPI_ISL_2836840, EPI_ISL_2836842, EPI_ISL_2836843 | | | |

| | | | |
|---|---|---|---|
| see above | Florida Bureau of Public Health Laboratories | Florida Bureau of Public Health Laboratories | Sarah Schmedes, Jason Blanton |
| EPI_ISL_2836918, EPI_ISL_2836932, EPI_ISL_2836938, EPI_ISL_2836944, EPI_ISL_2836945, EPI_ISL_2836955, EPI_ISL_2836957 | WACCBIP, University of Ghana, Accra, Ghana | WACCBIP, University of Ghana, Volta Road, Legon-Accra, Ghana | Collins M. Morang'a, Peter K Quashie, Joyce M. Ngoi, Dominic S.Y. Amuzu, Vincent Appiah, Evelyn B. Quansah, Philip M. Soglo, Violette V. M'cormack, Samirah Saiid, Ivy A. Asante, Joseph H.K. Bonney, Evelyn Y. Bonney, John K. Odoom, Nicaise T. Ndam, Frederick Tei-Maya, Mildred Adusei-Poku, Lawrence Ofori-Boadu, Joe K. Mutungi, William K. Ampofo, Yaw Bediako, Lucas N. Amenga-Etego and Gordon A. Awandare |
| EPI_ISL_2837266, EPI_ISL_2837299 | Unidade de apoio ao diagnostico da COVID - UNADIG | Bioinformatics Laboratory / LNCC | Luiz G P de Almeida, Alessandra P Lamarca, Ronaldo da Silva F Jr, Liliane Cavalcante, Alexandra L Gerber, Ana Paula de C Guimaraes, Douglas Terra Machado, Cassia Alves, Diana Mariani, Cintia Policarpo, Gleidson da Silva de Oliveira, Mario Sergio Ribeiro, Silvia Carvalho, Flavio Dias da Silva, Marcio Henrique de Oliveira Garcia, Leandro Magalhaes de Souza, Cristiane Gomes da Silva, Caio Luiz Pereira Ribeiro, Andrea Cony Cavalcanti, Claudia Maria Braga de Mello, Amilcar Tanuri, Ana Tereza R Vasconcelos |
| EPI_ISL_2837693, EPI_ISL_2837899, EPI_ISL_2837907, EPI_ISL_2837919, EPI_ISL_2837920, EPI_ISL_2837921, EPI_ISL_2838017 | California Department of Public Health Valencia Branch Laboratory (CDPH VBL) | California Department of Public Health | CDPH-COVIDNet |
| EPI_ISL_2838048, EPI_ISL_2838049 | Ministry of Health, Jaber Al-Ahmad Hospital | Virology Unit, Department of Microbiology, Faculty of Medicine, Kuwait University | Nada Madi, Hussain Safar, Ebaa Al-Awadhi, Anfal Al-Adwani |
| EPI_ISL_2838050 | UMR190-Unité des virus émergents | UMR190-Unité des virus émergents | C.Baronti |
| EPI_ISL_2838057, EPI_ISL_2838058, EPI_ISL_2838061, EPI_ISL_2838062, EPI_ISL_2838063, EPI_ISL_2838065, EPI_ISL_2838067, EPI_ISL_2838068, EPI_ISL_2838069, EPI_ISL_2838070, EPI_ISL_2838072, EPI_ISL_2838075, EPI_ISL_2838076, EPI_ISL_2838077, EPI_ISL_2838078, EPI_ISL_2838079, EPI_ISL_2838080, EPI_ISL_2838082, EPI_ISL_2838083, EPI_ISL_2838084, EPI_ISL_2838086, EPI_ISL_2838087, EPI_ISL_2838088, EPI_ISL_2838089, EPI_ISL_2838090, EPI_ISL_2838094, EPI_ISL_2838096, EPI_ISL_2838097, EPI_ISL_2838099, EPI_ISL_2838100, EPI_ISL_2838101, EPI_ISL_2838102, EPI_ISL_2838104, EPI_ISL_2838106, EPI_ISL_2838107, EPI_ISL_2838109, EPI_ISL_2838111, EPI_ISL_2838112, EPI_ISL_2838113, EPI_ISL_2838114, EPI_ISL_2838117, EPI_ISL_2838118, EPI_ISL_2838119, EPI_ISL_2838121, EPI_ISL_2838122, EPI_ISL_2838123, EPI_ISL_2838124, EPI_ISL_2838125, EPI_ISL_2838127, EPI_ISL_2838128, EPI_ISL_2838129, EPI_ISL_2838130, EPI_ISL_2838131, EPI_ISL_2838132, EPI_ISL_2838133, EPI_ISL_2838135, EPI_ISL_2838136, EPI_ISL_2838137, EPI_ISL_2838139, EPI_ISL_2838142, EPI_ISL_2838143, EPI_ISL_2838144, EPI_ISL_2838145, EPI_ISL_2838146, EPI_ISL_2838147, EPI_ISL_2838149, EPI_ISL_2838151, EPI_ISL_2838152, EPI_ISL_2838153, EPI_ISL_2838156, EPI_ISL_2838157, EPI_ISL_2838158, EPI_ISL_2838160, EPI_ISL_2838161, EPI_ISL_2838162, EPI_ISL_2838163, EPI_ISL_2838164, EPI_ISL_2838165, EPI_ISL_2838166, EPI_ISL_2838168, EPI_ISL_2838170, EPI_ISL_2838171, EPI_ISL_2838173, EPI_ISL_2838176, EPI_ISL_2838178, EPI_ISL_2838179, EPI_ISL_2838181, EPI_ISL_2838184, EPI_ISL_2838185, EPI_ISL_2838186, EPI_ISL_2838188, EPI_ISL_2838190, EPI_ISL_2838193, EPI_ISL_2838194, EPI_ISL_2838196, EPI_ISL_2838198, EPI_ISL_2838202, EPI_ISL_2838203, EPI_ISL_2838204, EPI_ISL_2838205, EPI_ISL_2838206, EPI_ISL_2838208, EPI_ISL_2838209, EPI_ISL_2838211, EPI_ISL_2838212, EPI_ISL_2838214, EPI_ISL_2838216 | | | |
| see above | Department of Bacteria, Parasites and Fungi, Statens Serum Institut, Copenhagen, Denmark | Statens Serum Institut Bioinformatics and Microbial Genomics | Danish Covid-19 Genome Consortium |
| EPI_ISL_2838217 | Molekylær Medicinsk Afdeling, Aarhus University Hospital, Aarhus, Denmark | Statens Serum Institut Bioinformatics and Microbial Genomics | Danish Covid-19 Genome Consortium |
| EPI_ISL_2838219, EPI_ISL_2838220, EPI_ISL_2838221, EPI_ISL_2838222, EPI_ISL_2838223, EPI_ISL_2838226, EPI_ISL_2838227, EPI_ISL_2838228, EPI_ISL_2838229, EPI_ISL_2838232, EPI_ISL_2838234, EPI_ISL_2838235, EPI_ISL_2838236, EPI_ISL_2838237, EPI_ISL_2838238, EPI_ISL_2838240, EPI_ISL_2838241, EPI_ISL_2838244, EPI_ISL_2838245, EPI_ISL_2838246, EPI_ISL_2838247, EPI_ISL_2838249, EPI_ISL_2838252, EPI_ISL_2838256, EPI_ISL_2838257, EPI_ISL_2838258, EPI_ISL_2838259, EPI_ISL_2838260, EPI_ISL_2838262, EPI_ISL_2838264, EPI_ISL_2838265, EPI_ISL_2838267, EPI_ISL_2838268 | | | |
| see above | Department of Bacteria, Parasites and Fungi, Statens Serum Institut, Copenhagen, Denmark | Statens Serum Institut Bioinformatics and Microbial Genomics | Danish Covid-19 Genome Consortium |
| EPI_ISL_2838269 | Molekylær Medicinsk Afdeling, Aarhus University Hospital, Aarhus, Denmark | Statens Serum Institut Bioinformatics and Microbial Genomics | Danish Covid-19 Genome Consortium |
| EPI_ISL_2838270, EPI_ISL_2838272, EPI_ISL_2838274, EPI_ISL_2838275, EPI_ISL_2838278, EPI_ISL_2838279, EPI_ISL_2838280, EPI_ISL_2838281, EPI_ISL_2838282, EPI_ISL_2838284, EPI_ISL_2838285, EPI_ISL_2838286, EPI_ISL_2838288, EPI_ISL_2838291, EPI_ISL_2838293, EPI_ISL_2838294 | | | |
| see above | Department of Bacteria, Parasites and Fungi, Statens Serum Institut, Copenhagen, Denmark | Statens Serum Institut Bioinformatics and Microbial Genomics | Danish Covid-19 Genome Consortium |
| EPI_ISL_2838295 | Molekylær Medicinsk Afdeling, Aarhus University Hospital, Aarhus, Denmark | Statens Serum Institut Bioinformatics and Microbial Genomics | Danish Covid-19 Genome Consortium |
| EPI_ISL_2838296, EPI_ISL_2838297, EPI_ISL_2838301, EPI_ISL_2838303, EPI_ISL_2838304, EPI_ISL_2838305, EPI_ISL_2838306, EPI_ISL_2838308, EPI_ISL_2838309, EPI_ISL_2838310, EPI_ISL_2838313, EPI_ISL_2838314, EPI_ISL_2838315, EPI_ISL_2838316, EPI_ISL_2838318, EPI_ISL_2838319, EPI_ISL_2838322, EPI_ISL_2838326, EPI_ISL_2838327, EPI_ISL_2838328, EPI_ISL_2838329, EPI_ISL_2838330, EPI_ISL_2838332, EPI_ISL_2838333, EPI_ISL_2838334, EPI_ISL_2838335, EPI_ISL_2838337, EPI_ISL_2838340, EPI_ISL_2838341, EPI_ISL_2838342, EPI_ISL_2838343, EPI_ISL_2838344, EPI_ISL_2838345, EPI_ISL_2838346, EPI_ISL_2838349, EPI_ISL_2838350, EPI_ISL_2838351, EPI_ISL_2838352 | | | |
| see above | Department of Bacteria, Parasites and Fungi, Statens Serum Institut, Copenhagen, Denmark | Statens Serum Institut Bioinformatics and Microbial Genomics | Danish Covid-19 Genome Consortium |
| EPI_ISL_2838886 | OHSU Lab Services Molecular Microbiology Lab | Oregon SARS-CoV-2 Genome Sequencing Center | VLOOKUP(C394,'[UsersoroakBox SyncPHI_SARS-COV-2_Sample_Sheet_infoGISAID_Submission_Infometa_master_2021.xlsx]release20_final'!D:AC,26,FALSE) |
| EPI_ISL_2838887 | OHSU Lab Services Molecular Microbiology Lab | Oregon SARS-CoV-2 Genome Sequencing Center | VLOOKUP(C514,'[UsersoroakBox SyncPHI_SARS-COV-2_Sample_Sheet_infoGISAID_Submission_Infometa_master_2021.xlsx]release20_final'!D:AC,26,FALSE) |
| EPI_ISL_2839094, EPI_ISL_2839095, EPI_ISL_2839096, EPI_ISL_2839097, EPI_ISL_2839098, EPI_ISL_2839099, EPI_ISL_2839113, EPI_ISL_2839114, EPI_ISL_2839119, EPI_ISL_2839124 | Arizona State University | Arizona State University | Peter T. Skidmore, LaRinda A. Holland, Matthew F. Smith, Rabia Maqsood, Nicholas J. Mellor, Joy M. Blain, Valerie Harris, Joshua LaBaer, Vel Murugan, Efrem S. Lim |
| EPI_ISL_2839138, EPI_ISL_2839139, EPI_ISL_2839140 | Oregon State Public Health Laboratory | Oregon State Public Health Laboratory | Rafia Razzaque, Eugene Yeboah, Vanda Makris, Laura Tsaknaridis, John Fontana and Shane Sevey |
| EPI_ISL_2839461, EPI_ISL_2839465 | KU Leuven, Rega Institute, Clinical and Epidemiological Virology | KU Leuven, Rega Institute, Clinical and Epidemiological Virology | Tony Wawina-Bokalanga, Bert Vanmechelen, Joan Marti-Carerras, Piet Maes |
| EPI_ISL_2839562, EPI_ISL_2839563, EPI_ISL_2839564, EPI_ISL_2839565, EPI_ISL_2839566, EPI_ISL_2839567, EPI_ISL_2839568 | Royal Darwin Hospital Pathology | MDU-PHL | Meumann, E., Caly L., Seemann T., Sait, M.L., Druce J., Sherry, N.L. |
| EPI_ISL_2839569, EPI_ISL_2839570 | Microbiological Diagnostic Unit - Public Health Laboratory (MDU-PHL) | MDU-PHL | Seemann T., Sait, M.L., Sherry, N.L. |
| EPI_ISL_2839678, EPI_ISL_2839683, EPI_ISL_2839685, EPI_ISL_2839687, EPI_ISL_2839688, EPI_ISL_2839690, EPI_ISL_2839693, EPI_ISL_2839695, EPI_ISL_2839699, EPI_ISL_2839700, EPI_ISL_2839701, EPI_ISL_2839707, EPI_ISL_2839708, EPI_ISL_2839712, EPI_ISL_2839713, EPI_ISL_2839715, EPI_ISL_2839717, EPI_ISL_2839718, EPI_ISL_2839720, EPI_ISL_2839724, EPI_ISL_2839727, EPI_ISL_2839731, EPI_ISL_2839735, EPI_ISL_2839738, EPI_ISL_2839739, EPI_ISL_2839744, EPI_ISL_2839746, EPI_ISL_2839749, EPI_ISL_2839751, EPI_ISL_2839752, EPI_ISL_2839754, EPI_ISL_2839755, EPI_ISL_2839758, EPI_ISL_2839760, EPI_ISL_2839761, EPI_ISL_2839763, EPI_ISL_2839764, EPI_ISL_2839766, EPI_ISL_2839772, EPI_ISL_2839773, EPI_ISL_2839777, EPI_ISL_2839783, EPI_ISL_2839789, EPI_ISL_2839790, EPI_ISL_2839791, EPI_ISL_2839792, EPI_ISL_2839793, EPI_ISL_2839796, EPI_ISL_2839799, EPI_ISL_2839800, EPI_ISL_2839807, EPI_ISL_2839811, EPI_ISL_2839813, EPI_ISL_2839817, EPI_ISL_2839820, EPI_ISL_2839824, EPI_ISL_2839828, EPI_ISL_2839831, EPI_ISL_2839832, EPI_ISL_2839836, EPI_ISL_2839839, EPI_ISL_2839843, EPI_ISL_2839849, EPI_ISL_2839851, EPI_ISL_2839853, EPI_ISL_2839856, EPI_ISL_2839858, EPI_ISL_2839862, EPI_ISL_2839864, EPI_ISL_2839869, EPI_ISL_2839874, EPI_ISL_2839875, EPI_ISL_2839876, EPI_ISL_2839877, EPI_ISL_2839878, EPI_ISL_2839881, EPI_ISL_2839884, EPI_ISL_2839886, EPI_ISL_2839887, EPI_ISL_2839891, EPI_ISL_2839892, EPI_ISL_2839896, EPI_ISL_2839903, EPI_ISL_2839909, EPI_ISL_2839911, EPI_ISL_2839916, EPI_ISL_2839924, EPI_ISL_2839925, EPI_ISL_2839927, EPI_ISL_2839929, EPI_ISL_2839933, EPI_ISL_2839934, EPI_ISL_2839935, EPI_ISL_2839936, EPI_ISL_2839938, EPI_ISL_2839943, EPI_ISL_2839944, EPI_ISL_2839951, EPI_ISL_2839955, EPI_ISL_2839956, EPI_ISL_2839957, EPI_ISL_2839958, EPI_ISL_2839959, EPI_ISL_2839962, EPI_ISL_2839963, EPI_ISL_2839965, EPI_ISL_2839966, EPI_ISL_2839967, EPI_ISL_2839970, EPI_ISL_2839972, EPI_ISL_2839975, EPI_ISL_2839976, EPI_ISL_2839978 | | | |
| see above | Curative Labs | Curative Labs | Mikhail Hanewich-Hollatz, J.Cesar Ignacio-Espinoza, Elias L. Salfati, Nabjot Sandhu, George Way, Zhiyi Xie, Eugenia Khorosheva, Sophia Quasem, Janet Chen, Vladimir Slepnev |
| EPI_ISL_2839979, EPI_ISL_2839980, EPI_ISL_2839982, EPI_ISL_2839983, EPI_ISL_2839984, EPI_ISL_2839985, EPI_ISL_2839986, EPI_ISL_2839987, EPI_ISL_2839988, EPI_ISL_2839989, EPI_ISL_2839990, EPI_ISL_2839991, EPI_ISL_2839992, EPI_ISL_2839993, EPI_ISL_2839994, EPI_ISL_2839995, EPI_ISL_2839996, EPI_ISL_2839997, EPI_ISL_2839998 | | | |
| see above | SA Pathology | SA Pathology | Caitlin Selway, Lex Leong, Chuan Kok Lim, Mark Turra, Ivan Bastian, Geoff Higgins |
| EPI_ISL_2840060 | Unilabs | Karolinska University Hospital | Jan Albert ,Tobias Allander ,Annelie Bjerkner ,Sandra Broddesson ,Robert Dyrdak ,Martin Ekman ,Lynda Eneh ,Lina Guerra Blomqvist ,Karolina Ininbergs ,Tanja Normark ,Isak Sylvin ,Zhibing Yun ,Martina Wahlund ,Valtteri Wirta |
| EPI_ISL_2840062, EPI_ISL_2840063, EPI_ISL_2840067 | Karolinska University Hospital Solna | Karolinska University Hospital | Jan Albert ,Tobias Allander ,Annelie Bjerkner ,Sandra Broddesson ,Robert Dyrdak ,Martin Ekman ,Lynda Eneh ,Lina Guerra Blomqvist ,Karolina Ininbergs ,Tanja Normark ,Isak Sylvin ,Zhibing Yun ,Martina Wahlund ,Valtteri Wirta |
| EPI_ISL_2840070, EPI_ISL_2840073 | Unilabs | Karolinska University Hospital | Jan Albert ,Tobias Allander ,Annelie Bjerkner ,Sandra Broddesson ,Robert Dyrdak ,Martin Ekman ,Lynda Eneh ,Lina Guerra Blomqvist ,Karolina Ininbergs ,Tanja Normark ,Isak Sylvin ,Zhibing Yun ,Martina Wahlund ,Valtteri Wirta |
| EPI_ISL_2840079, EPI_ISL_2840080 | Karolinska University Hospital Solna | Karolinska University Hospital | Jan Albert ,Tobias Allander ,Annelie Bjerkner ,Sandra Broddesson ,Robert Dyrdak ,Martin Ekman ,Lynda Eneh ,Lina Guerra Blomqvist ,Karolina Ininbergs ,Tanja Normark ,Isak Sylvin ,Zhibing Yun ,Martina Wahlund ,Valtteri Wirta |
| EPI_ISL_2840087, EPI_ISL_2840089, EPI_ISL_2840092, EPI_ISL_2840111, EPI_ISL_2840128, EPI_ISL_2840130, EPI_ISL_2840134, EPI_ISL_2840135 | Unilabs | Karolinska University Hospital | Jan Albert ,Tobias Allander ,Annelie Bjerkner ,Sandra Broddesson ,Robert Dyrdak ,Martin Ekman ,Lynda Eneh ,Lina Guerra Blomqvist ,Karolina Ininbergs ,Tanja Normark ,Isak Sylvin ,Zhibing Yun ,Martina Wahlund ,Valtteri Wirta |

| | | | |
|---|---|---|---|
| EPI_ISL_2840141 | Karolinska University Hospital Solna | Karolinska University Hospital | Jan Albert ,Tobias Allander ,Annelie Bjerkner ,Sandra Broddesson ,Robert Dyrdak ,Martin Ekman ,Lynda Eneh ,Lina Guerra Blomqvist ,Karolina Ininbergs ,Tanja Normark ,Isak Sylvin ,Zhibing Yun ,Martina Wahlund ,Valtteri Wirta |
| EPI_ISL_2840142 | Unilabs | Karolinska University Hospital | Jan Albert ,Tobias Allander ,Annelie Bjerkner ,Sandra Broddesson ,Robert Dyrdak ,Martin Ekman ,Lynda Eneh ,Lina Guerra Blomqvist ,Karolina Ininbergs ,Tanja Normark ,Isak Sylvin ,Zhibing Yun ,Martina Wahlund ,Valtteri Wirta |
| EPI_ISL_2840150 | Karolinska University Hospital Solna | Karolinska University Hospital | Jan Albert ,Tobias Allander ,Annelie Bjerkner ,Sandra Broddesson ,Robert Dyrdak ,Martin Ekman ,Lynda Eneh ,Lina Guerra Blomqvist ,Karolina Ininbergs ,Tanja Normark ,Isak Sylvin ,Zhibing Yun ,Martina Wahlund ,Valtteri Wirta |
| EPI_ISL_2840154, EPI_ISL_2840160 | Synlab Medilab | Karolinska University Hospital | Jan Albert ,Tobias Allander ,Annelie Bjerkner ,Sandra Broddesson ,Robert Dyrdak ,Martin Ekman ,Lynda Eneh ,Lina Guerra Blomqvist ,Karolina Ininbergs ,Tanja Normark ,Isak Sylvin ,Zhibing Yun ,Martina Wahlund ,Valtteri Wirta |
| EPI_ISL_2840161, EPI_ISL_2840164, EPI_ISL_2840167, EPI_ISL_2840168, EPI_ISL_2840169, EPI_ISL_2840170, EPI_ISL_2840172, EPI_ISL_2840173 | Unilabs | Karolinska University Hospital | Jan Albert ,Tobias Allander ,Annelie Bjerkner ,Sandra Broddesson ,Robert Dyrdak ,Martin Ekman ,Lynda Eneh ,Lina Guerra Blomqvist ,Karolina Ininbergs ,Tanja Normark ,Isak Sylvin ,Zhibing Yun ,Martina Wahlund ,Valtteri Wirta |
| EPI_ISL_2840174 | Synlab Medilab | Karolinska University Hospital | Jan Albert ,Tobias Allander ,Annelie Bjerkner ,Sandra Broddesson ,Robert Dyrdak ,Martin Ekman ,Lynda Eneh ,Lina Guerra Blomqvist ,Karolina Ininbergs ,Tanja Normark ,Isak Sylvin ,Zhibing Yun ,Martina Wahlund ,Valtteri Wirta |
| EPI_ISL_2840176, EPI_ISL_2840178, EPI_ISL_2840179 | Unilabs | Karolinska University Hospital | Jan Albert ,Tobias Allander ,Annelie Bjerkner ,Sandra Broddesson ,Robert Dyrdak ,Martin Ekman ,Lynda Eneh ,Lina Guerra Blomqvist ,Karolina Ininbergs ,Tanja Normark ,Isak Sylvin ,Zhibing Yun ,Martina Wahlund ,Valtteri Wirta |
| EPI_ISL_2840259 | Willis-Knighton Medical Center Hospital Laboratory | LSUHS Emerging Viral Threat Laboratory | Lorie M. Atkins, April N. Johnson, Gregory L. Ware, Alexander Mijalis, Jeremy P. Kamil, Jennifer L. Carroll, Maarten Van Diest, Rona S. Scott, Andrew D. Yurochko, Christopher G. Kevil, John A. Vanchiere, Joseph A. Bocchini |
| EPI_ISL_2840269 | "AZ SPHL, Arizona Department of Health Services" | Centers for Disease Control and Prevention Division of Viral Diseases, Pathogen Discovery | Mili Sheth, Sarah Nobles, Jasmine Padilla, Mark Burroughs, Shoshona Le, Katie Dillon, Peter Cook, Clinton R. Paden, Dhwani Batra, Krista Queen, Kristen Knipe, Dakota Howard, Yvette Unoarumhi, Darlene Wagner, Matthew Schmerer, Ben L. Rambo-Martin, Kristine Lacek, Sam Shepard, Alison Laufer Halpin, Dave Wentworth, Vivien Dugan, Suxiang Tong, Justin Lee |
| EPI_ISL_2840270, EPI_ISL_2840272, EPI_ISL_2840273, EPI_ISL_2840275, EPI_ISL_2840276, EPI_ISL_2840277, EPI_ISL_2840278 | WY Public Health Laboratory | Centers for Disease Control and Prevention Division of Viral Diseases, Pathogen Discovery | Mili Sheth, Sarah Nobles, Jasmine Padilla, Mark Burroughs, Shoshona Le, Katie Dillon, Peter Cook, Clinton R. Paden, Dhwani Batra, Krista Queen, Kristen Knipe, Dakota Howard, Yvette Unoarumhi, Darlene Wagner, Matthew Schmerer, Ben L. Rambo-Martin, Kristine Lacek, Sam Shepard, Alison Laufer Halpin, Dave Wentworth, Vivien Dugan, Suxiang Tong, Justin Lee |
| EPI_ISL_2840281 | FL Bureau of Public Health Laboratories-Tampa | Centers for Disease Control and Prevention Division of Viral Diseases, Pathogen Discovery | Mili Sheth, Sarah Nobles, Jasmine Padilla, Mark Burroughs, Shoshona Le, Katie Dillon, Peter Cook, Clinton R. Paden, Dhwani Batra, Krista Queen, Kristen Knipe, Dakota Howard, Yvette Unoarumhi, Darlene Wagner, Matthew Schmerer, Ben L. Rambo-Martin, Kristine Lacek, Sam Shepard, Alison Laufer Halpin, Dave Wentworth, Vivien Dugan, Suxiang Tong, Justin Lee |
| EPI_ISL_2840289 | IL Dept. of Public Health Springfield Laboratory | Centers for Disease Control and Prevention Division of Viral Diseases, Pathogen Discovery | Mili Sheth, Sarah Nobles, Jasmine Padilla, Mark Burroughs, Shoshona Le, Katie Dillon, Peter Cook, Clinton R. Paden, Dhwani Batra, Krista Queen, Kristen Knipe, Dakota Howard, Yvette Unoarumhi, Darlene Wagner, Matthew Schmerer, Ben L. Rambo-Martin, Kristine Lacek, Sam Shepard, Alison Laufer Halpin, Dave Wentworth, Vivien Dugan, Suxiang Tong, Justin Lee |
| EPI_ISL_2840294, EPI_ISL_2840296, EPI_ISL_2840298, EPI_ISL_2840299 | FL Bur. of Public Health Laboratories-Jacksonville | Centers for Disease Control and Prevention Division of Viral Diseases, Pathogen Discovery | Mili Sheth, Sarah Nobles, Jasmine Padilla, Mark Burroughs, Shoshona Le, Katie Dillon, Peter Cook, Clinton R. Paden, Dhwani Batra, Krista Queen, Kristen Knipe, Dakota Howard, Yvette Unoarumhi, Darlene Wagner, Matthew Schmerer, Ben L. Rambo-Martin, Kristine Lacek, Sam Shepard, Alison Laufer Halpin, Dave Wentworth, Vivien Dugan, Suxiang Tong, Justin Lee |
| EPI_ISL_2840325, EPI_ISL_2840326, EPI_ISL_2840328, EPI_ISL_2840329 | MO State Public Health Laboratory | Centers for Disease Control and Prevention Division of Viral Diseases, Pathogen Discovery | Mili Sheth, Sarah Nobles, Jasmine Padilla, Mark Burroughs, Shoshona Le, Katie Dillon, Peter Cook, Clinton R. Paden, Dhwani Batra, Krista Queen, Kristen Knipe, Dakota Howard, Yvette Unoarumhi, Darlene Wagner, Matthew Schmerer, Ben L. Rambo-Martin, Kristine Lacek, Sam Shepard, Alison Laufer Halpin, Dave Wentworth, Vivien Dugan, Suxiang Tong, Justin Lee |
| EPI_ISL_2840348, EPI_ISL_2840349, EPI_ISL_2840350, EPI_ISL_2840352, EPI_ISL_2840353, EPI_ISL_2840355 | NE Public Health Laboratory | Centers for Disease Control and Prevention Division of Viral Diseases, Pathogen Discovery | Mili Sheth, Sarah Nobles, Jasmine Padilla, Mark Burroughs, Shoshona Le, Katie Dillon, Peter Cook, Clinton R. Paden, Dhwani Batra, Krista Queen, Kristen Knipe, Dakota Howard, Yvette Unoarumhi, Darlene Wagner, Matthew Schmerer, Ben L. Rambo-Martin, Kristine Lacek, Sam Shepard, Alison Laufer Halpin, Dave Wentworth, Vivien Dugan, Suxiang Tong, Justin Lee |
| EPI_ISL_2840359, EPI_ISL_2840360, EPI_ISL_2840365, EPI_ISL_2840366, EPI_ISL_2840367, EPI_ISL_2840368 | IA State Hygienic Laboratory | Centers for Disease Control and Prevention Division of Viral Diseases, Pathogen Discovery | Mili Sheth, Sarah Nobles, Jasmine Padilla, Mark Burroughs, Shoshona Le, Katie Dillon, Peter Cook, Clinton R. Paden, Dhwani Batra, Krista Queen, Kristen Knipe, Dakota Howard, Yvette Unoarumhi, Darlene Wagner, Matthew Schmerer, Ben L. Rambo-Martin, Kristine Lacek, Sam Shepard, Alison Laufer Halpin, Dave Wentworth, Vivien Dugan, Suxiang Tong, Justin Lee |
| EPI_ISL_2840383, EPI_ISL_2840385, EPI_ISL_2840386, EPI_ISL_2840387, EPI_ISL_2840388, EPI_ISL_2840391, EPI_ISL_2840392, EPI_ISL_2840393 | "AR Dept. of Health-PHL, Molecular Diagnostics" | Centers for Disease Control and Prevention Division of Viral Diseases, Pathogen Discovery | Mili Sheth, Sarah Nobles, Jasmine Padilla, Mark Burroughs, Shoshona Le, Katie Dillon, Peter Cook, Clinton R. Paden, Dhwani Batra, Krista Queen, Kristen Knipe, Dakota Howard, Yvette Unoarumhi, Darlene Wagner, Matthew Schmerer, Ben L. Rambo-Martin, Kristine Lacek, Sam Shepard, Alison Laufer Halpin, Dave Wentworth, Vivien Dugan, Suxiang Tong, Justin Lee |
| EPI_ISL_2840396, EPI_ISL_2840397, EPI_ISL_2840398 | KS Health and Environmental Laboratories | Centers for Disease Control and Prevention Division of Viral Diseases, Pathogen Discovery | Mili Sheth, Sarah Nobles, Jasmine Padilla, Mark Burroughs, Shoshona Le, Katie Dillon, Peter Cook, Clinton R. Paden, Dhwani Batra, Krista Queen, Kristen Knipe, Dakota Howard, Yvette Unoarumhi, Darlene Wagner, Matthew Schmerer, Ben L. Rambo-Martin, Kristine Lacek, Sam Shepard, Alison Laufer Halpin, Dave Wentworth, Vivien Dugan, Suxiang Tong, Justin Lee |
| EPI_ISL_2840399, EPI_ISL_2840400, EPI_ISL_2840401, EPI_ISL_2840402 | "OK Public Health Laboratory, Oklahoma State DOH" | Centers for Disease Control and Prevention Division of Viral Diseases, Pathogen Discovery | Mili Sheth, Sarah Nobles, Jasmine Padilla, Mark Burroughs, Shoshona Le, Katie Dillon, Peter Cook, Clinton R. Paden, Dhwani Batra, Krista Queen, Kristen Knipe, Dakota Howard, Yvette Unoarumhi, Darlene Wagner, Matthew Schmerer, Ben L. Rambo-Martin, Kristine Lacek, Sam Shepard, Alison Laufer Halpin, Dave Wentworth, Vivien Dugan, Suxiang Tong, Justin Lee |
| EPI_ISL_2840428 | FL Bureau of Public Health Laboratories-Miami | Centers for Disease Control and Prevention Division of Viral Diseases, Pathogen Discovery | Mili Sheth, Sarah Nobles, Jasmine Padilla, Mark Burroughs, Shoshona Le, Katie Dillon, Peter Cook, Clinton R. Paden, Dhwani Batra, Krista Queen, Kristen Knipe, Dakota Howard, Yvette Unoarumhi, Darlene Wagner, Matthew Schmerer, Ben L. Rambo-Martin, Kristine Lacek, Sam Shepard, Alison Laufer Halpin, Dave Wentworth, Vivien Dugan, Suxiang Tong, Justin Lee |
| EPI_ISL_2840429 | CT-Dr. Katherine A. Kelley State Public Health Lab | Centers for Disease Control and Prevention Division of Viral Diseases, Pathogen Discovery | Mili Sheth, Sarah Nobles, Jasmine Padilla, Mark Burroughs, Shoshona Le, Katie Dillon, Peter Cook, Clinton R. Paden, Dhwani Batra, Krista Queen, Kristen Knipe, Dakota Howard, Yvette Unoarumhi, Darlene Wagner, Matthew Schmerer, Ben L. Rambo-Martin, Kristine Lacek, Sam Shepard, Alison Laufer Halpin, Dave Wentworth, Vivien Dugan, Suxiang Tong, Justin Lee |
| EPI_ISL_2840481, EPI_ISL_2840482, EPI_ISL_2840483, EPI_ISL_2840484, EPI_ISL_2840485, EPI_ISL_2840486, EPI_ISL_2840488, EPI_ISL_2840489, EPI_ISL_2840490, EPI_ISL_2840491 | Institute for Infectious Diseases | Institute for Infectious Diseases | Stefan Neuenschwander, Christian Baumann, Miguel A Terrazos Miani, Cora SÃ¤gesser, Pascal Bittel, Peter Keller, Franziska Suter-Riniker, Stephen L Leib, Alban Ramette |
| EPI_ISL_2840534 | KU Leuven, Rega Institute, Clinical and Epidemiological Virology | KU Leuven, Rega Institute, Clinical and Epidemiological Virology | Tony Wawina-Bokalanga, Bert Vanmechelen, Joan Marti-Carerras, Piet Maes |
| EPI_ISL_2840598, EPI_ISL_2840599, EPI_ISL_2840600 | Chulabhorn Hospital | National Institute of Health, Department of Medical Sciences, Ministry of Public Health, Thailand | Pilailuk Okada; Siripaporn Phuygun; Nuttida Thongpramul; Sittiporn Parnmen; Ratana Tacharoenmuang; Pakorn Piromtong; Natchaya Khiadsang; Thanutsapa Thanadachakul; Warawan Wongboot; sirikanda wimol; Sunthareeya Waicharoen; |
| EPI_ISL_2840601, EPI_ISL_2840602, EPI_ISL_2840603, EPI_ISL_2840604 | Urban Institute for Disease Prevention and Control | National Institute of Health, Department of Medical Sciences, Ministry of Public Health, Thailand | Pilailuk Okada; Siripaporn Phuygun; Nuttida Thongpramul; Sittiporn Parnmen; Ratana Tacharoenmuang; Pakorn Piromtong; Natchaya Khiadsang; Thanutsapa Thanadachakul; Warawan Wongboot; sirikanda wimol; Sunthareeya Waicharoen; |
| EPI_ISL_2840605 | Kasemrad international hospital ratthanatibeth | National Institute of Health, Department of Medical Sciences, Ministry of Public Health, Thailand | Pilailuk Okada; Siripaporn Phuygun; Nuttida Thongpramul; Sittiporn Parnmen; Ratana Tacharoenmuang; Pakorn Piromtong; Natchaya Khiadsang; Thanutsapa Thanadachakul; Warawan Wongboot; sirikanda wimol; Sunthareeya Waicharoen; |
| EPI_ISL_2840612, EPI_ISL_2840613, EPI_ISL_2840618, EPI_ISL_2840619, EPI_ISL_2840620 | S.C. Laboratorio Analisi, ASL 3 Liguria | U.O. Igiene, Ospedale Policlinico San Martino | Bruzzone Bianca, Caligiuri Patrizia, De Pace Vanessa, Domnich Alexander, Orsi Andrea, Ricucci Valentina, Spitaleri Antonino, Icardi Giancarlo on behalf of SARS-CoV-2 ITALIAN RESEARCH ENTERPRISE-(SCIRE) Collaborative Group |
| EPI_ISL_2840631, EPI_ISL_2840632, EPI_ISL_2840633 | S.S.D. Microbiologia, Ospedale Santa Corona di Pietra Ligure, ASL 2 Liguria | U.O. Igiene, Ospedale Policlinico San Martino | Bruzzone Bianca, Caligiuri Patrizia, De Pace Vanessa, Domnich Alexander, Orsi Andrea, Ricucci Valentina, Valle Caterina, Icardi Giancarlo on behalf of SARS-CoV-2 ITALIAN RESEARCH ENTERPRISE-(SCIRE) Collaborative Group |

| | | | |
|---|---|---|---|
| EPI_ISL_2840634, EPI_ISL_2840635 | S.C. Patologia Clinica, Ospedale Sant'Andrea, ASL 5 | U.O. Igiene, Ospedale Policlinico San Martino | Bruzzone Bianca, Battolla Enrico, Caligiuri Patrizia, De Pace Vanessa, Domnich Alexander, Orsi Andrea, Ricucci Valentina, Icardi Giancarlo on behalf of SARS-CoV-2 ITALIAN RESEARCH ENTERPRISE-(SCIRE) Collaborative Group |
| EPI_ISL_2840652, EPI_ISL_2840653 | Università degli Studi di Perugia | Istituto Zooprofilattico Sperimentale dell'Abruzzo e Molise "G. Caporale" | Mencacci A, Camilloni B, Biagetti M, Giammarioli M, Lorusso A, Marcacci M, Di Domenico M, Ancora M, Curini V, Mangone I, Rinaldi A, Delli Compagni E, Scialabba S, Di Pasquale A, Cammà C, Puglia I, Calistri P, Savini G |
| EPI_ISL_2840678 | OSPEDALE CIVILE TERAMO - CENTRO TRASFUSIONALE | Istituto Zooprofilattico Sperimentale dell'Abruzzo e Molise "G. Caporale" | Lorusso A, Marcacci M, Di Domenico M, Ancora M, Curini V, Di Lollo Valeria, Mangone I, Rinaldi A, Delli Compagni E, Scialabba S, Caporale M, Di Pasquale A, Cammà C, Puglia I, Calistri P, Savini G |
| EPI_ISL_2840680, EPI_ISL_2840681, EPI_ISL_2840682 | SIESP CHIETI - DRIVE IN LANCIANO | Istituto Zooprofilattico Sperimentale dell'Abruzzo e Molise "G. Caporale" | Lorusso A, Marcacci M, Di Domenico M, Ancora M, Curini V, Di Lollo Valeria, Mangone I, Rinaldi A, Delli Compagni E, Scialabba S, Caporale M, Di Pasquale A, Cammà C, Puglia I, Calistri P, Savini G |
| EPI_ISL_2840683, EPI_ISL_2840684 | DIP. PREV. AVEZZANO SERVIZIO DI IGIENE EPIDEMIOLOGIA E SANITA' PUBBLICA | Istituto Zooprofilattico Sperimentale dell'Abruzzo e Molise "G. Caporale" | Lorusso A, Marcacci M, Di Domenico M, Ancora M, Curini V, Di Lollo Valeria, Mangone I, Rinaldi A, Delli Compagni E, Scialabba S, Caporale M, Di Pasquale A, Cammà C, Puglia I, Calistri P, Savini G |
| EPI_ISL_2840687, EPI_ISL_2840688, EPI_ISL_2840689, EPI_ISL_2840690, EPI_ISL_2840691, EPI_ISL_2840692, EPI_ISL_2840693, EPI_ISL_2840694, EPI_ISL_2840695, EPI_ISL_2840696, EPI_ISL_2840697, EPI_ISL_2840698, EPI_ISL_2840699, EPI_ISL_2840700, EPI_ISL_2840701, EPI_ISL_2840702, EPI_ISL_2840703, EPI_ISL_2840709, EPI_ISL_2840710, EPI_ISL_2840711, EPI_ISL_2840712, EPI_ISL_2840713, EPI_ISL_2840714, EPI_ISL_2840715, EPI_ISL_2840718, EPI_ISL_2840719, EPI_ISL_2840720, EPI_ISL_2840721, EPI_ISL_2840722, EPI_ISL_2840723, EPI_ISL_2840724, EPI_ISL_2840725, EPI_ISL_2840726, EPI_ISL_2840727, EPI_ISL_2840728, EPI_ISL_2840729, EPI_ISL_2840730, EPI_ISL_2840731, EPI_ISL_2840732, EPI_ISL_2840733, EPI_ISL_2840734, EPI_ISL_2840735, EPI_ISL_2840736, EPI_ISL_2840737, EPI_ISL_2840738, EPI_ISL_2840739, EPI_ISL_2840740, EPI_ISL_2840741, EPI_ISL_2840742, EPI_ISL_2840743 | | | |
| see above | OSPEDALE CIVILE TERAMO - CENTRO TRASFUSIONALE | Istituto Zooprofilattico Sperimentale dell'Abruzzo e Molise "G. Caporale" | Lorusso A, Marcacci M, Di Domenico M, Ancora M, Curini V, Di Lollo Valeria, Mangone I, Rinaldi A, Delli Compagni E, Scialabba S, Caporale M, Di Pasquale A, Cammà C, Puglia I, Calistri P, Savini G |
| EPI_ISL_2840853 | Department of Acute Infectious Diseases Control and Prevention, Yunnan Provincial Center for Disease Control and Prevention | Department of Acute Infectious Diseases Control and Prevention, Yunnan Provincial Center for Disease Control and Prevention | Meiling Zhang, Jienan Zhou, Zhaosheng Liu, Xiaoqing Fu |
| EPI_ISL_2840854, EPI_ISL_2840855, EPI_ISL_2840856, EPI_ISL_2840857, EPI_ISL_2840858, EPI_ISL_2840859, EPI_ISL_2840860 | Department of Acute Infectious Diseases Control and Prevention, Yunnan Provincial Center for Disease Control and Prevention | Department of Acute Infectious Diseases Control and Prevention, Yunnan Provincial Center for Disease Control and Prevention | Meiling Zhang, Jienan Zhou, Senquan Jia, Zhaosheng Liu, Xiaoqing Fu |
| EPI_ISL_2840910 | LBM ALPHABIO, Marseille | LBM ALPHABIO, Marseille | Vincent GARCIA |
| EPI_ISL_2840975, EPI_ISL_2840986, EPI_ISL_2841038, EPI_ISL_2841108, EPI_ISL_2841110, EPI_ISL_2841111, EPI_ISL_2841140, EPI_ISL_2841396, EPI_ISL_2841397, EPI_ISL_2841398, EPI_ISL_2841399, EPI_ISL_2841400, EPI_ISL_2841401, EPI_ISL_2841402 | | | |
| see above | National Virus Reference Laboratory | National Virus Reference Laboratory | Zoe Yandle, Charlene Bennett, Gabriel Gonzalez, Michael Carr, Jonathan Dean, Cillian F De Gascun |
| EPI_ISL_2841537, EPI_ISL_2841538 | San Diego County Public Health Laboratory | San Diego County Public Health Laboratory | Ashleigh Murphy Schafer, Brett Austin |
| EPI_ISL_2841544, EPI_ISL_2841545, EPI_ISL_2841546, EPI_ISL_2841548, EPI_ISL_2841549, EPI_ISL_2841550, EPI_ISL_2841551 | Nebraska Public Health Laboratory | NPHL COVID-19 Response Team | NPHL COVID-19 Response Team |
| EPI_ISL_2841613, EPI_ISL_2841615 | San Diego County Public Health Laboratory | San Diego County Public Health Laboratory | Ashleigh Murphy Schafer, Brett Austin |
| EPI_ISL_2841649 | NHLS Charlotte Maxeke Johannesburg Academic Hospital and the University of the Witwatersrand | KRISP, KZn Research Innovation and Sequencing Platform | Florette Treurnicht, Bulelani Manene, Kathleen Subramoney, Giandhari Jennifer, Pillay Sureshnee, Yajna Ramphal, Naidoo Yeshnee, Tshabuila Derek,Tegally Houriiyah, San James, Wilkinson Eduan, de Oliveira Tulio |
| EPI_ISL_2841650 | Netcare The Bay Hospital | KRISP, KZn Research Innovation and Sequencing Platform | Giandhari Jennifer, Pillay Sureshnee, Yajna Ramphal, Naidoo Yeshnee, Tshabuila Derek,Tegally Houriiyah, San James, Wilkinson Eduan, de Oliveira Tulio |
| EPI_ISL_2841651 | Netcare Kingsway Hospital | KRISP, KZn Research Innovation and Sequencing Platform | Giandhari Jennifer, Pillay Sureshnee, Yajna Ramphal, Naidoo Yeshnee, Tshabuila Derek,Tegally Houriiyah, San James, Wilkinson Eduan, de Oliveira Tulio |
| EPI_ISL_2841652, EPI_ISL_2841653, EPI_ISL_2841654 | Instituto Nacional de Investigação em Saúde | CERI, Centre for Epidemic Response and Innvoation, Stellenbosch University and KRISP, KZN Research Innovation and Sequencing Platform, UKZN. | Morais J, Neto Z, Afonso P, Miranda J, David K, Inglês L, Pereira A, Paulo A Carralero RR Paixão JP, Freitas RH, Mufinda M, Lutucuta S, Giandhari J, Pillay S, Naidoo Y, Emmanuel SJ, Tegally H, Wilkinson E, de Oliveira T |
| EPI_ISL_2841660, EPI_ISL_2841661, EPI_ISL_2841662, EPI_ISL_2841663, EPI_ISL_2841664, EPI_ISL_2841665, EPI_ISL_2841666, EPI_ISL_2841667, EPI_ISL_2841669, EPI_ISL_2841670, EPI_ISL_2841671, EPI_ISL_2841672, EPI_ISL_2841673, EPI_ISL_2841674, EPI_ISL_2841675, EPI_ISL_2841676, EPI_ISL_2841678, EPI_ISL_2841679, EPI_ISL_2841680, EPI_ISL_2841681, EPI_ISL_2841682, EPI_ISL_2841683, EPI_ISL_2841684, EPI_ISL_2841685, EPI_ISL_2841686, EPI_ISL_2841687, EPI_ISL_2841688, EPI_ISL_2841689, EPI_ISL_2841690, EPI_ISL_2841691, EPI_ISL_2841692, EPI_ISL_2841693, EPI_ISL_2841695, EPI_ISL_2841696, EPI_ISL_2841697, EPI_ISL_2841698, EPI_ISL_2841699, EPI_ISL_2841700 | | | |
| see above | Wits Vaccines & Infectious Diseases Analytics (VIDA) Research Unit | KRISP, KZn Research Innovation and Sequencing Platform | Baillie Vicky, du Plessis Jeanine, Giandhari Jennifer, Pillay Sureshnee, Naidoo Yeshnee, Tegally Houriiyah, San James, Wilkinson Eduan, de Oliveira Tulio, Madhi Shabir |
| EPI_ISL_2841707 | University Hospital of Northern Norway, Department for Microbiology and Infectious Disease Control | Norwegian Institute of Public Health, Department of Virology | Kathrine Stene-Johansen, Kamilla Heddeland Instefjord, Hilde Elshaug, Garcia Llorente Ignacio, Jon Bråte, Engebretsen Serina Beate,Pedersen Benedikte Nevjen, Line Victoria Moen, Debech Nadia, Atiya R Ali,Marie Paulsen Madsen, Rasmus Riis Kopperud, Hilde Vollan, Karoline Bragstad, Olav Hungnes |
| EPI_ISL_2841718, EPI_ISL_2841726, EPI_ISL_2841728, EPI_ISL_2841730, EPI_ISL_2841732, EPI_ISL_2841733, EPI_ISL_2841734, EPI_ISL_2841735, EPI_ISL_2841737, EPI_ISL_2841738, EPI_ISL_2841751, EPI_ISL_2841753, EPI_ISL_2841756, EPI_ISL_2841757, EPI_ISL_2841758, EPI_ISL_2841760, EPI_ISL_2841761, EPI_ISL_2841762, EPI_ISL_2841763, EPI_ISL_2841764 | | | |
| see above | Nebraska Public Health Laboratory | NPHL COVID-19 Response Team | NPHL COVID-19 Response Team |
| EPI_ISL_2841765, EPI_ISL_2841766, EPI_ISL_2841767, EPI_ISL_2841769, EPI_ISL_2841771, EPI_ISL_2841772, EPI_ISL_2841773, EPI_ISL_2841774, EPI_ISL_2841776, EPI_ISL_2841777, EPI_ISL_2841786 | | | |
| see above | LBM ALPHABIO, Marseille | LBM ALPHABIO, Marseille | Vincent GARCIA |
| EPI_ISL_2841828, EPI_ISL_2841829, EPI_ISL_2841830, EPI_ISL_2841831 | PathWest Laboratory Medicine WA | PathWest Laboratory Medicine WA Microbial Surveillance Unit | PathWest Laboratory Medicine WA Microbial Surveillance Unit |
| EPI_ISL_2842008 | CAP TARADELL | Banc de Sang i Teixits | Francisco Vidal, Irene Corrales, Nina Borràs, Carlos Hobeich, Natália Comes, Lorena Ramírez, Noemí Gonzalez, Silvia Sauleda, Maria Glòria Soria |
| EPI_ISL_2842009, EPI_ISL_2842014, EPI_ISL_2842016 | CAP TONA | Banc de Sang i Teixits | Francisco Vidal, Irene Corrales, Nina Borràs, Carlos Hobeich, Natália Comes, Lorena Ramírez, Noemí Gonzalez, Silvia Sauleda, Maria Glòria Soria |
| EPI_ISL_2842017 | CAP SARRIÀ | Banc de Sang i Teixits | Francisco Vidal, Irene Corrales, Nina Borràs, Carlos Hobeich, Natália Comes, Lorena Ramírez, Noemí Gonzalez, Silvia Sauleda, Maria Glòria Soria |
| EPI_ISL_2842019 | CAP VIC NORD | Banc de Sang i Teixits | Francisco Vidal, Irene Corrales, Nina Borràs, Carlos Hobeich, Natália Comes, Lorena Ramírez, Noemí Gonzalez, Silvia Sauleda, Maria Glòria Soria |
| EPI_ISL_2842025, EPI_ISL_2842026 | CAP SANT BOI DE LLUÇANÈS | Banc de Sang i Teixits | Francisco Vidal, Irene Corrales, Nina Borràs, Carlos Hobeich, Natália Comes, Lorena Ramírez, Noemí Gonzalez, Silvia Sauleda, Maria Glòria Soria |
| EPI_ISL_2842029 | CAP OLOST DE LLUÇANÈS | Banc de Sang i Teixits | Francisco Vidal, Irene Corrales, Nina Borràs, Carlos Hobeich, Natália Comes, Lorena Ramírez, Noemí Gonzalez, Silvia Sauleda, Maria Glòria Soria |
| EPI_ISL_2842030, EPI_ISL_2842037 | CAP TONA | Banc de Sang i Teixits | Francisco Vidal, Irene Corrales, Nina Borràs, Carlos Hobeich, Natália Comes, Lorena Ramírez, Noemí Gonzalez, Silvia Sauleda, Maria Glòria Soria |
| EPI_ISL_2842039 | FUNDACIO ALTHAIA-MANRESA | Banc de Sang i Teixits | Francisco Vidal, Irene Corrales, Nina Borràs, Carlos Hobeich, Natália Comes, Lorena Ramírez, Noemí Gonzalez, Silvia Sauleda, Maria Glòria Soria |
| EPI_ISL_2842044 | CAP SANT QUIRZE DE BESORA | Banc de Sang i Teixits | Francisco Vidal, Irene Corrales, Nina Borràs, Carlos Hobeich, Natália Comes, Lorena Ramírez, Noemí Gonzalez, Silvia Sauleda, Maria Glòria Soria |
| EPI_ISL_2842045 | CAP TONA | Banc de Sang i Teixits | Francisco Vidal, Irene Corrales, Nina Borràs, Carlos Hobeich, Natália Comes, Lorena Ramírez, Noemí Gonzalez, Silvia Sauleda, Maria Glòria Soria |
| EPI_ISL_2842046, EPI_ISL_2842047, EPI_ISL_2842048, EPI_ISL_2842049, EPI_ISL_2842050, EPI_ISL_2842051, EPI_ISL_2842053, EPI_ISL_2842054, EPI_ISL_2842055 | Wichita State University- Molecular Diagnostic Lab | Kansas Health and Environmental Lab | Mike Grose, Katherine Wiggins, Jonathan Barnell, Ben Olsen, and Phil Adam |
| EPI_ISL_2842057, EPI_ISL_2842058, EPI_ISL_2842059, EPI_ISL_2842060, EPI_ISL_2842062, EPI_ISL_2842064, EPI_ISL_2842065, EPI_ISL_2842066, EPI_ISL_2842067, EPI_ISL_2842068, EPI_ISL_2842069, EPI_ISL_2842070, EPI_ISL_2842071 | | | |
| see above | Clinical Reference Laboratory | Kansas Health and Environmental Lab | Mike Grose, Katherine Wiggins, Jonathan Barnell, Ben Olsen, and Phil Adam |
| EPI_ISL_2842073, EPI_ISL_2842075, EPI_ISL_2842076, EPI_ISL_2842077, EPI_ISL_2842079, EPI_ISL_2842080, EPI_ISL_2842081, EPI_ISL_2842082, EPI_ISL_2842084, EPI_ISL_2842085, EPI_ISL_2842086, EPI_ISL_2842087, EPI_ISL_2842088, EPI_ISL_2842089, EPI_ISL_2842090, EPI_ISL_2842091, EPI_ISL_2842092, EPI_ISL_2842093, EPI_ISL_2842094, EPI_ISL_2842095, EPI_ISL_2842098, EPI_ISL_2842099, EPI_ISL_2842100 | | | |
| see above | Kansas Health and Environmental Lab | Kansas Health and Environmental Lab | Mike Grose, Katherine Wiggins, Jonathan Barnell, Ben Olsen, and Phil Adam |
| EPI_ISL_2842101 | NIMHANS | INSACOG-KA, NIMHANS | Chitra Pattabiraman, Anson Kunjumon George,Darshan Sreenivas, Ellango Ramasamy, Sony Sharma, Mahesh Kumar.C.S, Ananthapadmanabha Kotambail, Chetan G K, Gautham Arunachal Udupi, Anita S Desai, V Ravi |

| | | | |
|---|---|---|---|
| EPI_ISL_2842102 | RAILWAY HOSPITAL | INSACOG-KA, NIMHANS | Chitra Pattabiraman, Anson Kunjumon George,Darshan Sreenivas, Ellango Ramasamy, Sony Sharma, Mahesh Kumar.C.S, Ananthapadmanabha Kotambail, Chetan G K, Gautham Arunachal Udupi, Anita S Desai, V Ravi |
| EPI_ISL_2842103 | BBMP Urban | INSACOG-KA, NIMHANS | Chitra Pattabiraman, Anson Kunjumon George,Darshan Sreenivas, Ellango Ramasamy, Sony Sharma, Mahesh Kumar.C.S, Ananthapadmanabha Kotambail, Chetan G K, Gautham Arunachal Udupi, Anita S Desai, V Ravi |
| EPI_ISL_2842104 | DSU KOLAR | INSACOG-KA, NIMHANS | Chitra Pattabiraman, Anson Kunjumon George,Darshan Sreenivas, Ellango Ramasamy, Sony Sharma, Mahesh Kumar.C.S, Ananthapadmanabha Kotambail, Chetan G K, Gautham Arunachal Udupi, Anita S Desai, V Ravi |
| EPI_ISL_2842105, EPI_ISL_2842107, EPI_ISL_2842108, EPI_ISL_2842110, EPI_ISL_2842111, EPI_ISL_2842112 | RAILWAY HOSPITAL | INSACOG-KA, NIMHANS | Chitra Pattabiraman, Anson Kunjumon George,Darshan Sreenivas, Ellango Ramasamy, Sony Sharma, Mahesh Kumar.C.S, Ananthapadmanabha Kotambail, Chetan G K, Gautham Arunachal Udupi, Anita S Desai, V Ravi |
| EPI_ISL_2842113 | SRI JAYADEVA INSTITUTE OF CARDIOLOGY | INSACOG-KA, NIMHANS | Chitra Pattabiraman, Anson Kunjumon George,Darshan Sreenivas, Ellango Ramasamy, Sony Sharma, Mahesh Kumar.C.S, Ananthapadmanabha Kotambail, Chetan G K, Gautham Arunachal Udupi, Anita S Desai, V Ravi |
| EPI_ISL_2842114, EPI_ISL_2842115, EPI_ISL_2842116, EPI_ISL_2842117, EPI_ISL_2842118 | RAILWAY HOSPITAL | INSACOG-KA, NIMHANS | Chitra Pattabiraman, Anson Kunjumon George,Darshan Sreenivas, Ellango Ramasamy, Sony Sharma, Mahesh Kumar.C.S, Ananthapadmanabha Kotambail, Chetan G K, Gautham Arunachal Udupi, Anita S Desai, V Ravi |
| EPI_ISL_2842119 | BBMP Urban | INSACOG-KA, NIMHANS | Chitra Pattabiraman, Anson Kunjumon George,Darshan Sreenivas, Ellango Ramasamy, Sony Sharma, Mahesh Kumar.C.S, Ananthapadmanabha Kotambail, Chetan G K, Gautham Arunachal Udupi, Anita S Desai, V Ravi |
| EPI_ISL_2842120 | RAILWAY HOSPITAL | INSACOG-KA, NIMHANS | Chitra Pattabiraman, Anson Kunjumon George,Darshan Sreenivas, Ellango Ramasamy, Sony Sharma, Mahesh Kumar.C.S, Ananthapadmanabha Kotambail, Chetan G K, Gautham Arunachal Udupi, Anita S Desai, V Ravi |
| EPI_ISL_2842121, EPI_ISL_2842122, EPI_ISL_2842123, EPI_ISL_2842124, EPI_ISL_2842125, EPI_ISL_2842126 | BBMP Urban | INSACOG-KA, NIMHANS | Chitra Pattabiraman, Anson Kunjumon George,Darshan Sreenivas, Ellango Ramasamy, Sony Sharma, Mahesh Kumar.C.S, Ananthapadmanabha Kotambail, Chetan G K, Gautham Arunachal Udupi, Anita S Desai, V Ravi |
| EPI_ISL_2842127 | RAILWAY HOSPITAL | INSACOG-KA, NIMHANS | Chitra Pattabiraman, Anson Kunjumon George,Darshan Sreenivas, Ellango Ramasamy, Sony Sharma, Mahesh Kumar.C.S, Ananthapadmanabha Kotambail, Chetan G K, Gautham Arunachal Udupi, Anita S Desai, V Ravi |
| EPI_ISL_2842128, EPI_ISL_2842129 | NIMHANS | INSACOG-KA, NIMHANS | Chitra Pattabiraman, Anson Kunjumon George,Darshan Sreenivas, Ellango Ramasamy, Sony Sharma, Mahesh Kumar.C.S, Ananthapadmanabha Kotambail, Chetan G K, Gautham Arunachal Udupi, Anita S Desai, V Ravi |
| EPI_ISL_2842130, EPI_ISL_2842131 | BBMP Urban | INSACOG-KA, NIMHANS | Chitra Pattabiraman, Anson Kunjumon George,Darshan Sreenivas, Ellango Ramasamy, Sony Sharma, Mahesh Kumar.C.S, Ananthapadmanabha Kotambail, Chetan G K, Gautham Arunachal Udupi, Anita S Desai, V Ravi |
| EPI_ISL_2842132, EPI_ISL_2842133 | NIMHANS | INSACOG-KA, NIMHANS | Chitra Pattabiraman, Anson Kunjumon George,Darshan Sreenivas, Ellango Ramasamy, Sony Sharma, Mahesh Kumar.C.S, Ananthapadmanabha Kotambail, Chetan G K, Gautham Arunachal Udupi, Anita S Desai, V Ravi |
| EPI_ISL_2842134, EPI_ISL_2842135, EPI_ISL_2842136, EPI_ISL_2842137, EPI_ISL_2842138, EPI_ISL_2842139, EPI_ISL_2842140, EPI_ISL_2842141, EPI_ISL_2842142, EPI_ISL_2842143, EPI_ISL_2842144 | | | |
| see above | VRDL VIJAYAPURA | INSACOG-KA, NIMHANS | Chitra Pattabiraman, Anson Kunjumon George,Darshan Sreenivas, Ellango Ramasamy, Sony Sharma, Mahesh Kumar.C.S, Ananthapadmanabha Kotambail, Chetan G K, Gautham Arunachal Udupi, Anita S Desai, V Ravi |
| EPI_ISL_2842145, EPI_ISL_2842146, EPI_ISL_2842147, EPI_ISL_2842148 | BBMP Urban | INSACOG-KA, NIMHANS | Chitra Pattabiraman, Anson Kunjumon George,Darshan Sreenivas, Ellango Ramasamy, Sony Sharma, Mahesh Kumar.C.S, Ananthapadmanabha Kotambail, Chetan G K, Gautham Arunachal Udupi, Anita S Desai, V Ravi |
| EPI_ISL_2842149 | NIMHANS | INSACOG-KA, NIMHANS | Chitra Pattabiraman, Anson Kunjumon George,Darshan Sreenivas, Ellango Ramasamy, Sony Sharma, Mahesh Kumar.C.S, Ananthapadmanabha Kotambail, Chetan G K, Gautham Arunachal Udupi, Anita S Desai, V Ravi |
| EPI_ISL_2842150, EPI_ISL_2842151, EPI_ISL_2842152, EPI_ISL_2842158, EPI_ISL_2842159, EPI_ISL_2842160, EPI_ISL_2842161, EPI_ISL_2842162, EPI_ISL_2842163, EPI_ISL_2842164 | BBMP Urban | INSACOG-KA, NIMHANS | Chitra Pattabiraman, Anson Kunjumon George,Darshan Sreenivas, Ellango Ramasamy, Sony Sharma, Mahesh Kumar.C.S, Ananthapadmanabha Kotambail, Chetan G K, Gautham Arunachal Udupi, Anita S Desai, V Ravi |
| EPI_ISL_2842165, EPI_ISL_2842166 | NIMHANS | INSACOG-KA, NIMHANS | Chitra Pattabiraman, Anson Kunjumon George,Darshan Sreenivas, Ellango Ramasamy, Sony Sharma, Mahesh Kumar.C.S, Ananthapadmanabha Kotambail, Chetan G K, Gautham Arunachal Udupi, Anita S Desai, V Ravi |
| EPI_ISL_2842167, EPI_ISL_2842168 | RAILWAY HOSPITAL | INSACOG-KA, NIMHANS | Chitra Pattabiraman, Anson Kunjumon George,Darshan Sreenivas, Ellango Ramasamy, Sony Sharma, Mahesh Kumar.C.S, Ananthapadmanabha Kotambail, Chetan G K, Gautham Arunachal Udupi, Anita S Desai, V Ravi |
| EPI_ISL_2842169, EPI_ISL_2842170 | JAYADEVA H | INSACOG-KA, NIMHANS | Chitra Pattabiraman, Anson Kunjumon George,Darshan Sreenivas, Ellango Ramasamy, Sony Sharma, Mahesh Kumar.C.S, Ananthapadmanabha Kotambail, Chetan G K, Gautham Arunachal Udupi, Anita S Desai, V Ravi |
| EPI_ISL_2842171 | NIMHANS | INSACOG-KA, NIMHANS | Chitra Pattabiraman, Anson Kunjumon George,Darshan Sreenivas, Ellango Ramasamy, Sony Sharma, Mahesh Kumar.C.S, Ananthapadmanabha Kotambail, Chetan G K, Gautham Arunachal Udupi, Anita S Desai, V Ravi |
| EPI_ISL_2842172, EPI_ISL_2842173, EPI_ISL_2842174, EPI_ISL_2842175, EPI_ISL_2842176 | BBMP Urban | INSACOG-KA, NIMHANS | Chitra Pattabiraman, Anson Kunjumon George,Darshan Sreenivas, Ellango Ramasamy, Sony Sharma, Mahesh Kumar.C.S, Ananthapadmanabha Kotambail, Chetan G K, Gautham Arunachal Udupi, Anita S Desai, V Ravi |
| EPI_ISL_2842177 | NIMHANS | INSACOG-KA, NIMHANS | Chitra Pattabiraman, Anson Kunjumon George,Darshan Sreenivas, Ellango Ramasamy, Sony Sharma, Mahesh Kumar.C.S, Ananthapadmanabha Kotambail, Chetan G K, Gautham Arunachal Udupi, Anita S Desai, V Ravi |
| EPI_ISL_2842178 | RAILWAY HOSPITAL | INSACOG-KA, NIMHANS | Chitra Pattabiraman, Anson Kunjumon George,Darshan Sreenivas, Ellango Ramasamy, Sony Sharma, Mahesh Kumar.C.S, Ananthapadmanabha Kotambail, Chetan G K, Gautham Arunachal Udupi, Anita S Desai, V Ravi |
| EPI_ISL_2842181, EPI_ISL_2842182 | NIMHANS | INSACOG-KA, NIMHANS | Chitra Pattabiraman, Anson Kunjumon George,Darshan Sreenivas, Ellango Ramasamy, Sony Sharma, Mahesh Kumar.C.S, Ananthapadmanabha Kotambail, Chetan G K, Gautham Arunachal Udupi, Anita S Desai, V Ravi |
| EPI_ISL_2842183, EPI_ISL_2842184, EPI_ISL_2842185, EPI_ISL_2842198, EPI_ISL_2842199, EPI_ISL_2842202 | BBMP Urban | INSACOG-KA, NIMHANS | Chitra Pattabiraman, Anson Kunjumon George,Darshan Sreenivas, Ellango Ramasamy, Sony Sharma, Mahesh Kumar.C.S, Ananthapadmanabha Kotambail, Chetan G K, Gautham Arunachal Udupi, Anita S Desai, V Ravi |
| EPI_ISL_2842206, EPI_ISL_2842209, EPI_ISL_2842210, EPI_ISL_2842211, EPI_ISL_2842212, EPI_ISL_2842213 | NIMHANS | INSACOG-KA, NIMHANS | Chitra Pattabiraman, Anson Kunjumon George,Darshan Sreenivas, Ellango Ramasamy, Sony Sharma, Mahesh Kumar.C.S, Ananthapadmanabha Kotambail, Chetan G K, Gautham Arunachal Udupi, Anita S Desai, V Ravi |
| EPI_ISL_2842216, EPI_ISL_2842218, EPI_ISL_2842219 | KEMPEGOWDA INTERNATIONAL AIRPORT | INSACOG-KA, NIMHANS | Chitra Pattabiraman, Anson Kunjumon George,Darshan Sreenivas, Ellango Ramasamy, Sony Sharma, Mahesh Kumar.C.S, Ananthapadmanabha Kotambail, Chetan G K, Gautham Arunachal Udupi, Anita S Desai, V Ravi |
| EPI_ISL_2842224, EPI_ISL_2842225, EPI_ISL_2842227, EPI_ISL_2842228, EPI_ISL_2842232, EPI_ISL_2842236, EPI_ISL_2842237, EPI_ISL_2842238 | BBMP Urban | INSACOG-KA, NIMHANS | Chitra Pattabiraman, Anson Kunjumon George,Darshan Sreenivas, Ellango Ramasamy, Sony Sharma, Mahesh Kumar.C.S, Ananthapadmanabha Kotambail, Chetan G K, Gautham Arunachal Udupi, Anita S Desai, V Ravi |
| EPI_ISL_2842242, EPI_ISL_2842246, EPI_ISL_2842247 | NIMHANS | INSACOG-KA, NIMHANS | Chitra Pattabiraman, Anson Kunjumon George,Darshan Sreenivas, Ellango Ramasamy, Sony Sharma, Mahesh Kumar.C.S, Ananthapadmanabha Kotambail, Chetan G K, Gautham Arunachal Udupi, Anita S Desai, V Ravi |
| EPI_ISL_2842248, EPI_ISL_2842249, EPI_ISL_2842250, EPI_ISL_2842253, EPI_ISL_2842254 | RAILWAY HOSPITAL | INSACOG-KA, NIMHANS | Chitra Pattabiraman, Anson Kunjumon George,Darshan Sreenivas, Ellango Ramasamy, Sony Sharma, Mahesh Kumar.C.S, Ananthapadmanabha Kotambail, Chetan G K, Gautham Arunachal Udupi, Anita S Desai, V Ravi |
| EPI_ISL_2842257, EPI_ISL_2842258, EPI_ISL_2842259, EPI_ISL_2842261, EPI_ISL_2842264, EPI_ISL_2842265, EPI_ISL_2842266, EPI_ISL_2842267, EPI_ISL_2842271, EPI_ISL_2842272, EPI_ISL_2842273, EPI_ISL_2842275, EPI_ISL_2842276, EPI_ISL_2842279, EPI_ISL_2842280 | | | |
| see above | BBMP Urban | INSACOG-KA, NIMHANS | Chitra Pattabiraman, Anson Kunjumon George,Darshan Sreenivas, Ellango Ramasamy, Sony Sharma, Mahesh Kumar.C.S, Ananthapadmanabha Kotambail, Chetan G K, Gautham Arunachal Udupi, Anita S Desai, V Ravi |

| | | | |
|---|---|---|---|
| EPI_ISL_2842285, EPI_ISL_2842286, EPI_ISL_2842287, EPI_ISL_2842289 | RAILWAY HOSPITAL | INSACOG-KA, NIMHANS | Chitra Pattabiraman, Anson Kunjumon George,Darshan Sreenivas, Ellango Ramasamy, Sony Sharma, Mahesh Kumar.C.S, Ananthapadmanabha Kotambail, Chetan G K, Gautham Arunachal Udupi, Anita S Desai, V Ravi |
| EPI_ISL_2842290 | NIMHANS | INSACOG-KA, NIMHANS | Chitra Pattabiraman, Anson Kunjumon George,Darshan Sreenivas, Ellango Ramasamy, Sony Sharma, Mahesh Kumar.C.S, Ananthapadmanabha Kotambail, Chetan G K, Gautham Arunachal Udupi, Anita S Desai, V Ravi |
| EPI_ISL_2842291, EPI_ISL_2842292, EPI_ISL_2842293 | BBMP Urban | INSACOG-KA, NIMHANS | Chitra Pattabiraman, Anson Kunjumon George,Darshan Sreenivas, Ellango Ramasamy, Sony Sharma, Mahesh Kumar.C.S, Ananthapadmanabha Kotambail, Chetan G K, Gautham Arunachal Udupi, Anita S Desai, V Ravi |
| EPI_ISL_2842294, EPI_ISL_2842295, EPI_ISL_2842296, EPI_ISL_2842297 | RAILWAY HOSPITAL | INSACOG-KA, NIMHANS | Chitra Pattabiraman, Anson Kunjumon George,Darshan Sreenivas, Ellango Ramasamy, Sony Sharma, Mahesh Kumar.C.S, Ananthapadmanabha Kotambail, Chetan G K, Gautham Arunachal Udupi, Anita S Desai, V Ravi |
| EPI_ISL_2842298, EPI_ISL_2842299, EPI_ISL_2842300, EPI_ISL_2842301, EPI_ISL_2842302, EPI_ISL_2842303 | BBMP Urban | INSACOG-KA, NIMHANS | Chitra Pattabiraman, Anson Kunjumon George,Darshan Sreenivas, Ellango Ramasamy, Sony Sharma, Mahesh Kumar.C.S, Ananthapadmanabha Kotambail, Chetan G K, Gautham Arunachal Udupi, Anita S Desai, V Ravi |
| EPI_ISL_2842304 | RAILWAY HOSPITAL | INSACOG-KA, NIMHANS | Chitra Pattabiraman, Anson Kunjumon George,Darshan Sreenivas, Ellango Ramasamy, Sony Sharma, Mahesh Kumar.C.S, Ananthapadmanabha Kotambail, Chetan G K, Gautham Arunachal Udupi, Anita S Desai, V Ravi |
| EPI_ISL_2842306 | JAYADEVA H | INSACOG-KA, NIMHANS | Chitra Pattabiraman, Anson Kunjumon George,Darshan Sreenivas, Ellango Ramasamy, Sony Sharma, Mahesh Kumar.C.S, Ananthapadmanabha Kotambail, Chetan G K, Gautham Arunachal Udupi, Anita S Desai, V Ravi |
| EPI_ISL_2842307, EPI_ISL_2842309, EPI_ISL_2842310, EPI_ISL_2842311, EPI_ISL_2842312 | BBMP Urban | INSACOG-KA, NIMHANS | Chitra Pattabiraman, Anson Kunjumon George,Darshan Sreenivas, Ellango Ramasamy, Sony Sharma, Mahesh Kumar.C.S, Ananthapadmanabha Kotambail, Chetan G K, Gautham Arunachal Udupi, Anita S Desai, V Ravi |
| EPI_ISL_2842314, EPI_ISL_2842315 | RAILWAY HOSPITAL | INSACOG-KA, NIMHANS | Chitra Pattabiraman, Anson Kunjumon George,Darshan Sreenivas, Ellango Ramasamy, Sony Sharma, Mahesh Kumar.C.S, Ananthapadmanabha Kotambail, Chetan G K, Gautham Arunachal Udupi, Anita S Desai, V Ravi |
| EPI_ISL_2842316 | NIMHANS | INSACOG-KA, NIMHANS | Chitra Pattabiraman, Anson Kunjumon George,Darshan Sreenivas, Ellango Ramasamy, Sony Sharma, Mahesh Kumar.C.S, Ananthapadmanabha Kotambail, Chetan G K, Gautham Arunachal Udupi, Anita S Desai, V Ravi |
| EPI_ISL_2842317, EPI_ISL_2842319, EPI_ISL_2842320, EPI_ISL_2842321, EPI_ISL_2842322, EPI_ISL_2842323, EPI_ISL_2842324, EPI_ISL_2842325, EPI_ISL_2842326, EPI_ISL_2842327 | BBMP Urban | INSACOG-KA, NIMHANS | Chitra Pattabiraman, Anson Kunjumon George,Darshan Sreenivas, Ellango Ramasamy, Sony Sharma, Mahesh Kumar.C.S, Ananthapadmanabha Kotambail, Chetan G K, Gautham Arunachal Udupi, Anita S Desai, V Ravi |
| EPI_ISL_2842328 | NIMHANS | INSACOG-KA, NIMHANS | Chitra Pattabiraman, Anson Kunjumon George,Darshan Sreenivas, Ellango Ramasamy, Sony Sharma, Mahesh Kumar.C.S, Ananthapadmanabha Kotambail, Chetan G K, Gautham Arunachal Udupi, Anita S Desai, V Ravi |
| EPI_ISL_2842329, EPI_ISL_2842330, EPI_ISL_2842331, EPI_ISL_2842332, EPI_ISL_2842333, EPI_ISL_2842334, EPI_ISL_2842335, EPI_ISL_2842336, EPI_ISL_2842338, EPI_ISL_2842339, EPI_ISL_2842340, EPI_ISL_2842341, EPI_ISL_2842342, EPI_ISL_2842343, EPI_ISL_2842344, EPI_ISL_2842345, EPI_ISL_2842346 | | | |
| see above | BBMP Urban | INSACOG-KA, NIMHANS | Chitra Pattabiraman, Anson Kunjumon George,Darshan Sreenivas, Ellango Ramasamy, Sony Sharma, Mahesh Kumar.C.S, Ananthapadmanabha Kotambail, Chetan G K, Gautham Arunachal Udupi, Anita S Desai, V Ravi |
| EPI_ISL_2842347 | NIMHANS | INSACOG-KA, NIMHANS | Chitra Pattabiraman, Anson Kunjumon George,Darshan Sreenivas, Ellango Ramasamy, Sony Sharma, Mahesh Kumar.C.S, Ananthapadmanabha Kotambail, Chetan G K, Gautham Arunachal Udupi, Anita S Desai, V Ravi |
| EPI_ISL_2842348, EPI_ISL_2842349, EPI_ISL_2842350, EPI_ISL_2842351, EPI_ISL_2842352, EPI_ISL_2842354 | BBMP Urban | INSACOG-KA, NIMHANS | Chitra Pattabiraman, Anson Kunjumon George,Darshan Sreenivas, Ellango Ramasamy, Sony Sharma, Mahesh Kumar.C.S, Ananthapadmanabha Kotambail, Chetan G K, Gautham Arunachal Udupi, Anita S Desai, V Ravi |
| EPI_ISL_2842355 | NIMHANS | INSACOG-KA, NIMHANS | Chitra Pattabiraman, Anson Kunjumon George,Darshan Sreenivas, Ellango Ramasamy, Sony Sharma, Mahesh Kumar.C.S, Ananthapadmanabha Kotambail, Chetan G K, Gautham Arunachal Udupi, Anita S Desai, V Ravi |
| EPI_ISL_2842356, EPI_ISL_2842357, EPI_ISL_2842358, EPI_ISL_2842359, EPI_ISL_2842360, EPI_ISL_2842361, EPI_ISL_2842362, EPI_ISL_2842363, EPI_ISL_2842364, EPI_ISL_2842365, EPI_ISL_2842367, EPI_ISL_2842368, EPI_ISL_2842369, EPI_ISL_2842370, EPI_ISL_2842371, EPI_ISL_2842372 | | | |
| see above | BBMP Urban | INSACOG-KA, NIMHANS | Chitra Pattabiraman, Anson Kunjumon George,Darshan Sreenivas, Ellango Ramasamy, Sony Sharma, Mahesh Kumar.C.S, Ananthapadmanabha Kotambail, Chetan G K, Gautham Arunachal Udupi, Anita S Desai, V Ravi |
| EPI_ISL_2842373, EPI_ISL_2842374 | RAILWAY HOSPITAL | INSACOG-KA, NIMHANS | Chitra Pattabiraman, Anson Kunjumon George,Darshan Sreenivas, Ellango Ramasamy, Sony Sharma, Mahesh Kumar.C.S, Ananthapadmanabha Kotambail, Chetan G K, Gautham Arunachal Udupi, Anita S Desai, V Ravi |
| EPI_ISL_2842375, EPI_ISL_2842376, EPI_ISL_2842377, EPI_ISL_2842378 | BBMP Urban | INSACOG-KA, NIMHANS | Chitra Pattabiraman, Anson Kunjumon George,Darshan Sreenivas, Ellango Ramasamy, Sony Sharma, Mahesh Kumar.C.S, Ananthapadmanabha Kotambail, Chetan G K, Gautham Arunachal Udupi, Anita S Desai, V Ravi |
| EPI_ISL_2842379 | JAYADEVA H | INSACOG-KA, NIMHANS | Chitra Pattabiraman, Anson Kunjumon George,Darshan Sreenivas, Ellango Ramasamy, Sony Sharma, Mahesh Kumar.C.S, Ananthapadmanabha Kotambail, Chetan G K, Gautham Arunachal Udupi, Anita S Desai, V Ravi |
| EPI_ISL_2842381, EPI_ISL_2842382, EPI_ISL_2842383, EPI_ISL_2842384, EPI_ISL_2842385, EPI_ISL_2842386, EPI_ISL_2842387, EPI_ISL_2842388, EPI_ISL_2842389, EPI_ISL_2842390, EPI_ISL_2842391 | | | |
| see above | BBMP Urban | INSACOG-KA, NIMHANS | Chitra Pattabiraman, Anson Kunjumon George,Darshan Sreenivas, Ellango Ramasamy, Sony Sharma, Mahesh Kumar.C.S, Ananthapadmanabha Kotambail, Chetan G K, Gautham Arunachal Udupi, Anita S Desai, V Ravi |
| EPI_ISL_2842392 | SRI JAYADEVA INSTITUTE OF CARDIOLOGY | INSACOG-KA, NIMHANS | Chitra Pattabiraman, Anson Kunjumon George,Darshan Sreenivas, Ellango Ramasamy, Sony Sharma, Mahesh Kumar.C.S, Ananthapadmanabha Kotambail, Chetan G K, Gautham Arunachal Udupi, Anita S Desai, V Ravi |
| EPI_ISL_2842393, EPI_ISL_2842394, EPI_ISL_2842395 | BBMP Urban | INSACOG-KA, NIMHANS | Chitra Pattabiraman, Anson Kunjumon George,Darshan Sreenivas, Ellango Ramasamy, Sony Sharma, Mahesh Kumar.C.S, Ananthapadmanabha Kotambail, Chetan G K, Gautham Arunachal Udupi, Anita S Desai, V Ravi |
| EPI_ISL_2842396 | NIMHANS | INSACOG-KA, NIMHANS | Chitra Pattabiraman, Anson Kunjumon George,Darshan Sreenivas, Ellango Ramasamy, Sony Sharma, Mahesh Kumar.C.S, Ananthapadmanabha Kotambail, Chetan G K, Gautham Arunachal Udupi, Anita S Desai, V Ravi |
| EPI_ISL_2842397 | RAILWAY HOSPITAL | INSACOG-KA, NIMHANS | Chitra Pattabiraman, Anson Kunjumon George,Darshan Sreenivas, Ellango Ramasamy, Sony Sharma, Mahesh Kumar.C.S, Ananthapadmanabha Kotambail, Chetan G K, Gautham Arunachal Udupi, Anita S Desai, V Ravi |
| EPI_ISL_2842398, EPI_ISL_2842399, EPI_ISL_2842400, EPI_ISL_2842409, EPI_ISL_2842410, EPI_ISL_2842411 | BBMP Urban | INSACOG-KA, NIMHANS | Chitra Pattabiraman, Anson Kunjumon George,Darshan Sreenivas, Ellango Ramasamy, Sony Sharma, Mahesh Kumar.C.S, Ananthapadmanabha Kotambail, Chetan G K, Gautham Arunachal Udupi, Anita S Desai, V Ravi |
| EPI_ISL_2842430, EPI_ISL_2842431, EPI_ISL_2842433, EPI_ISL_2842434, EPI_ISL_2842435, EPI_ISL_2842436 | DSU VIJAYAPURA | INSACOG-KA, NIMHANS | Chitra Pattabiraman, Anson Kunjumon George,Darshan Sreenivas, Ellango Ramasamy, Sony Sharma, Mahesh Kumar.C.S, Ananthapadmanabha Kotambail, Chetan G K, Gautham Arunachal Udupi, Anita S Desai, V Ravi |
| EPI_ISL_2842437 | RAILWAY HOSPITAL | INSACOG-KA, NIMHANS | Chitra Pattabiraman, Anson Kunjumon George,Darshan Sreenivas, Ellango Ramasamy, Sony Sharma, Mahesh Kumar.C.S, Ananthapadmanabha Kotambail, Chetan G K, Gautham Arunachal Udupi, Anita S Desai, V Ravi |
| EPI_ISL_2842438, EPI_ISL_2842439 | BBMP Urban | INSACOG-KA, NIMHANS | Chitra Pattabiraman, Anson Kunjumon George,Darshan Sreenivas, Ellango Ramasamy, Sony Sharma, Mahesh Kumar.C.S, Ananthapadmanabha Kotambail, Chetan G K, Gautham Arunachal Udupi, Anita S Desai, V Ravi |
| EPI_ISL_2842440, EPI_ISL_2842441 | RAILWAY HOSPITAL | INSACOG-KA, NIMHANS | Chitra Pattabiraman, Anson Kunjumon George,Darshan Sreenivas, Ellango Ramasamy, Sony Sharma, Mahesh Kumar.C.S, Ananthapadmanabha Kotambail, Chetan G K, Gautham Arunachal Udupi, Anita S Desai, V Ravi |
| EPI_ISL_2842442 | BBMP Urban | INSACOG-KA, NIMHANS | Chitra Pattabiraman, Anson Kunjumon George,Darshan Sreenivas, Ellango Ramasamy, Sony Sharma, Mahesh Kumar.C.S, Ananthapadmanabha Kotambail, Chetan G K, Gautham Arunachal Udupi, Anita S Desai, V Ravi |
| EPI_ISL_2842443 | NIMHANS | INSACOG-KA, NIMHANS | Chitra Pattabiraman, Anson Kunjumon George,Darshan Sreenivas, Ellango Ramasamy, Sony Sharma, Mahesh Kumar.C.S, Ananthapadmanabha Kotambail, Chetan G K, Gautham Arunachal Udupi, Anita S Desai, V Ravi |
| EPI_ISL_2842444 | RAILWAY HOSPITAL | INSACOG-KA, NIMHANS | Chitra Pattabiraman, Anson Kunjumon George,Darshan Sreenivas, Ellango Ramasamy, Sony Sharma, Mahesh Kumar.C.S, Ananthapadmanabha Kotambail, Chetan G K, Gautham Arunachal Udupi, Anita S Desai, V Ravi |
| EPI_ISL_2842445, EPI_ISL_2842446, EPI_ISL_2842447 | BBMP Urban | INSACOG-KA, NIMHANS | Chitra Pattabiraman, Anson Kunjumon George,Darshan Sreenivas, Ellango Ramasamy, Sony Sharma, Mahesh Kumar.C.S, Ananthapadmanabha Kotambail, Chetan G K, Gautham Arunachal Udupi, Anita S Desai, V Ravi |
| EPI_ISL_2842448 | RAILWAY HOSPITAL | INSACOG-KA, NIMHANS | Chitra Pattabiraman, Anson Kunjumon George,Darshan Sreenivas, Ellango Ramasamy, Sony Sharma, Mahesh Kumar.C.S, Ananthapadmanabha Kotambail, Chetan G K, Gautham Arunachal Udupi, Anita S Desai, V Ravi |
| EPI_ISL_2842449, EPI_ISL_2842450, | BBMP Urban | INSACOG-KA, NIMHANS | Chitra Pattabiraman, Anson Kunjumon George,Darshan Sreenivas, Ellango Ramasamy, Sony Sharma, Mahesh Kumar.C.S, Ananthapadmanabha Kotambail, Chetan G K, Gautham Arunachal Udupi, Anita S Desai, V Ravi |

| | | | |
|---|---|---|---|
| EPI_ISL_2842451 | | | |
| EPI_ISL_2842452 | RAILWAY HOSPITAL | INSACOG-KA, NIMHANS | Chitra Pattabiraman, Anson Kunjumon George,Darshan Sreenivas, Ellango Ramasamy, Sony Sharma, Mahesh Kumar.C.S, Ananthapadmanabha Kotambail, Chetan G K, Gautham Arunachal Udupi, Anita S Desai, V Ravi |
| EPI_ISL_2842453, EPI_ISL_2842454 | BBMP Urban | INSACOG-KA, NIMHANS | Chitra Pattabiraman, Anson Kunjumon George,Darshan Sreenivas, Ellango Ramasamy, Sony Sharma, Mahesh Kumar.C.S, Ananthapadmanabha Kotambail, Chetan G K, Gautham Arunachal Udupi, Anita S Desai, V Ravi |
| EPI_ISL_2842455 | RAILWAY HOSPITAL | INSACOG-KA, NIMHANS | Chitra Pattabiraman, Anson Kunjumon George,Darshan Sreenivas, Ellango Ramasamy, Sony Sharma, Mahesh Kumar.C.S, Ananthapadmanabha Kotambail, Chetan G K, Gautham Arunachal Udupi, Anita S Desai, V Ravi |
| EPI_ISL_2842456, EPI_ISL_2842457 | BBMP Urban | INSACOG-KA, NIMHANS | Chitra Pattabiraman, Anson Kunjumon George,Darshan Sreenivas, Ellango Ramasamy, Sony Sharma, Mahesh Kumar.C.S, Ananthapadmanabha Kotambail, Chetan G K, Gautham Arunachal Udupi, Anita S Desai, V Ravi |
| EPI_ISL_2842458, EPI_ISL_2842460, EPI_ISL_2842463, EPI_ISL_2842464, EPI_ISL_2842465, EPI_ISL_2842466 | VICTORIA HOSPITAL | INSACOG-KA, NIMHANS | Chitra Pattabiraman, Anson Kunjumon George,Darshan Sreenivas, Ellango Ramasamy, Sony Sharma, Mahesh Kumar.C.S, Ananthapadmanabha Kotambail, Chetan G K, Gautham Arunachal Udupi, Anita S Desai, V Ravi |
| EPI_ISL_2842467, EPI_ISL_2842468, EPI_ISL_2842469, EPI_ISL_2842470, EPI_ISL_2842471, EPI_ISL_2842472 | DSU MYSURU | INSACOG-KA, NIMHANS | Chitra Pattabiraman, Anson Kunjumon George,Darshan Sreenivas, Ellango Ramasamy, Sony Sharma, Mahesh Kumar.C.S, Ananthapadmanabha Kotambail, Chetan G K, Gautham Arunachal Udupi, Anita S Desai, V Ravi |
| EPI_ISL_2842473, EPI_ISL_2842474 | DSU MANDYA | INSACOG-KA, NIMHANS | Chitra Pattabiraman, Anson Kunjumon George,Darshan Sreenivas, Ellango Ramasamy, Sony Sharma, Mahesh Kumar.C.S, Ananthapadmanabha Kotambail, Chetan G K, Gautham Arunachal Udupi, Anita S Desai, V Ravi |
| EPI_ISL_2842475, EPI_ISL_2842476, EPI_ISL_2842477 | DSU CHAMRAJANAGARA | INSACOG-KA, NIMHANS | Chitra Pattabiraman, Anson Kunjumon George,Darshan Sreenivas, Ellango Ramasamy, Sony Sharma, Mahesh Kumar.C.S, Ananthapadmanabha Kotambail, Chetan G K, Gautham Arunachal Udupi, Anita S Desai, V Ravi |
| EPI_ISL_2842478, EPI_ISL_2842479, EPI_ISL_2842480 | DSU KODAGU | INSACOG-KA, NIMHANS | Chitra Pattabiraman, Anson Kunjumon George,Darshan Sreenivas, Ellango Ramasamy, Sony Sharma, Mahesh Kumar.C.S, Ananthapadmanabha Kotambail, Chetan G K, Gautham Arunachal Udupi, Anita S Desai, V Ravi |
| EPI_ISL_2842481, EPI_ISL_2842482, EPI_ISL_2842483, EPI_ISL_2842484, EPI_ISL_2842485, EPI_ISL_2842486, EPI_ISL_2842487, EPI_ISL_2842489, EPI_ISL_2842490 | DSU KALABURAGI | INSACOG-KA, NIMHANS | Chitra Pattabiraman, Anson Kunjumon George,Darshan Sreenivas, Ellango Ramasamy, Sony Sharma, Mahesh Kumar.C.S, Ananthapadmanabha Kotambail, Chetan G K, Gautham Arunachal Udupi, Anita S Desai, V Ravi |
| EPI_ISL_2842491, EPI_ISL_2842492, EPI_ISL_2842493, EPI_ISL_2842494, EPI_ISL_2842495, EPI_ISL_2842496, EPI_ISL_2842497, EPI_ISL_2842498, EPI_ISL_2842499 | DSU HASSAN | INSACOG-KA, NIMHANS | Chitra Pattabiraman, Anson Kunjumon George,Darshan Sreenivas, Ellango Ramasamy, Sony Sharma, Mahesh Kumar.C.S, Ananthapadmanabha Kotambail, Chetan G K, Gautham Arunachal Udupi, Anita S Desai, V Ravi |
| EPI_ISL_2842501, EPI_ISL_2842502 | DSU KOLAR | INSACOG-KA, NIMHANS | Chitra Pattabiraman, Anson Kunjumon George,Darshan Sreenivas, Ellango Ramasamy, Sony Sharma, Mahesh Kumar.C.S, Ananthapadmanabha Kotambail, Chetan G K, Gautham Arunachal Udupi, Anita S Desai, V Ravi |
| EPI_ISL_2842514 | Wyoming Public Health Laboratory | Wyoming Public Health Laboratory | Jim Mildenberger, Taylor Fearing, Channing Weber, Ashley Norberg, Chayse Rowley, Marley Goetz, Brian Dominguez, Elliot Thomasson, Sam Britz, Cari Sloma, Robert Petit, and Rob Christensen |
| EPI_ISL_2842619 | Northwestern Memorial Hospital | Northwestern University - Ozer Lab | Ramon Lorenzo-Redondo, Lacy M. Simons, Taylor J. Dean, Chad J. Achenbach, Lawrence J. Jennings, Chao Qi, Michael G. Ison, Judd F. Hultquist, Egon A. Ozer |
| EPI_ISL_2842645, EPI_ISL_2842646, EPI_ISL_2842647, EPI_ISL_2842648, EPI_ISL_2842650, EPI_ISL_2842651, EPI_ISL_2842653, EPI_ISL_2842656, EPI_ISL_2842657, EPI_ISL_2842658, EPI_ISL_2842659, EPI_ISL_2842663, EPI_ISL_2842665, EPI_ISL_2842666, EPI_ISL_2842667, EPI_ISL_2842668, EPI_ISL_2842680, EPI_ISL_2842685, EPI_ISL_2842686, EPI_ISL_2842687, EPI_ISL_2842688, EPI_ISL_2842689, EPI_ISL_2842690 | | | |
| see above | NJDOH, Public Health and Environmental Laboratories | NJ_PHEL | Lindsey Bodnar, Shiv K. Verma, Ryan Pachucki, Chelsea San Filippo, Jacquelyn Deverell, Dana Woell, Allison Roder, Byeong Jeong |
| EPI_ISL_2842735 | Wyoming Public Health Laboratory | Wyoming Public Health Laboratory | Jim Mildenberger, Taylor Fearing, Channing Weber, Ashley Norberg, Chayse Rowley, Marley Goetz, Brian Dominguez, Elliot Thomasson, Sam Britz, Cari Sloma, Robert Petit, and Rob Christensen |
| EPI_ISL_2842784 | Centro Medico ABC | Instituto Nacional de Medicina Genomica | Cedro-Tanda A, Hidalgo-Miranda A, Mendoza-Vargas A, Reyes-Grajeda JP, Roxana Trejo-González, Cesar Lara, Jacomine Reyes-Carrasco, Daniel Aguirre, Gonzalez-Barrera D, Rangel-DeLeon D, Munguia-Garza P, Ramirez-Vega O, Escobar-Arrazola, M, Herrera-Montalvo LA. |
| EPI_ISL_2842808, EPI_ISL_2842812, EPI_ISL_2842813, EPI_ISL_2842814, EPI_ISL_2842815, EPI_ISL_2842816, EPI_ISL_2842818 | California Department of Public Health | California Department of Public Health | CDPH IDLB COVIDNet et al |
| EPI_ISL_2842839, EPI_ISL_2842840, EPI_ISL_2842845 | Queensland Health Forensic and Scientific Services | Queensland Health Forensic and Scientific Services | Son Nguyen |
| EPI_ISL_2842881, EPI_ISL_2842935, EPI_ISL_2842936, EPI_ISL_2842938, EPI_ISL_2842939, EPI_ISL_2842943, EPI_ISL_2842946, EPI_ISL_2842951, EPI_ISL_2842953, EPI_ISL_2842954, EPI_ISL_2842955, EPI_ISL_2842957, EPI_ISL_2842959, EPI_ISL_2842963, EPI_ISL_2842966, EPI_ISL_2842967, EPI_ISL_2842970, EPI_ISL_2842971, EPI_ISL_2842974, EPI_ISL_2842975, EPI_ISL_2842978, EPI_ISL_2842981, EPI_ISL_2842985, EPI_ISL_2842991, EPI_ISL_2842992, EPI_ISL_2842994, EPI_ISL_2842996, EPI_ISL_2842997, EPI_ISL_2842998, EPI_ISL_2843000, EPI_ISL_2843006, EPI_ISL_2843009, EPI_ISL_2843019, EPI_ISL_2843021, EPI_ISL_2843023, EPI_ISL_2843024, EPI_ISL_2843028, EPI_ISL_2843036, EPI_ISL_2843041, EPI_ISL_2843042, EPI_ISL_2843043, EPI_ISL_2843058, EPI_ISL_2843072, EPI_ISL_2843074, EPI_ISL_2843079, EPI_ISL_2843085, EPI_ISL_2843089, EPI_ISL_2843091, EPI_ISL_2843092, EPI_ISL_2843095, EPI_ISL_2843096, EPI_ISL_2843103, EPI_ISL_2843106, EPI_ISL_2843108, EPI_ISL_2843111, EPI_ISL_2843112, EPI_ISL_2843114, EPI_ISL_2843119, EPI_ISL_2843122, EPI_ISL_2843125, EPI_ISL_2843126, EPI_ISL_2843128, EPI_ISL_2843129, EPI_ISL_2843131, EPI_ISL_2843141, EPI_ISL_2843143, EPI_ISL_2843145, EPI_ISL_2843148, EPI_ISL_2843149, EPI_ISL_2843150, EPI_ISL_2843151, EPI_ISL_2843154, EPI_ISL_2843161, EPI_ISL_2843163, EPI_ISL_2843165, EPI_ISL_2843170, EPI_ISL_2843171, EPI_ISL_2843172, EPI_ISL_2843173, EPI_ISL_2843174, EPI_ISL_2843177, EPI_ISL_2843178, EPI_ISL_2843189, EPI_ISL_2843193, EPI_ISL_2843195, EPI_ISL_2843199, EPI_ISL_2843204, EPI_ISL_2843207, EPI_ISL_2843209, EPI_ISL_2843210, EPI_ISL_2843211, EPI_ISL_2843213, EPI_ISL_2843214, EPI_ISL_2843216, EPI_ISL_2843217, EPI_ISL_2843218, EPI_ISL_2843219, EPI_ISL_2843222, EPI_ISL_2843223, EPI_ISL_2843228, EPI_ISL_2843229, EPI_ISL_2843231, EPI_ISL_2843232, EPI_ISL_2843233, EPI_ISL_2843234, EPI_ISL_2843237, EPI_ISL_2843238, EPI_ISL_2843240, EPI_ISL_2843241, EPI_ISL_2843242, EPI_ISL_2843243, EPI_ISL_2843244, EPI_ISL_2843245, EPI_ISL_2843246, EPI_ISL_2843248, EPI_ISL_2843249, EPI_ISL_2843250, EPI_ISL_2843251, EPI_ISL_2843253, EPI_ISL_2843254, EPI_ISL_2843255, EPI_ISL_2843256, EPI_ISL_2843257, EPI_ISL_2843258, EPI_ISL_2843261, EPI_ISL_2843262, EPI_ISL_2843263, EPI_ISL_2843264, EPI_ISL_2843265, EPI_ISL_2843266, EPI_ISL_2843267, EPI_ISL_2843268, EPI_ISL_2843269, EPI_ISL_2843270, EPI_ISL_2843271, EPI_ISL_2843272, EPI_ISL_2843273, EPI_ISL_2843274, EPI_ISL_2843275, EPI_ISL_2843276, EPI_ISL_2843278, EPI_ISL_2843279, EPI_ISL_2843280, EPI_ISL_2843281, EPI_ISL_2843282, EPI_ISL_2843284, EPI_ISL_2843285, EPI_ISL_2843287, EPI_ISL_2843289, EPI_ISL_2843301, EPI_ISL_2843304 | | | |
| see above | Ministry of Public Health / Hamad Medical Corporation | Biomedical Research Center (BRC), Qatar University / Qatar Genome Project (QGP) | BRC: Fatiha M. Benslimane, Heba A. Al-Khatib, Oal Al-Jamal, Dana Al-Batesh, Hadi M. Yassine, Asmaa A. Al-Thani. MOPH and HMC: Abdullatif Al-Khal, Muna A. S. Al-Maslamani, Mashael A. Al-Bader, Hamda Alromaihi, Roberto Bertollini, Peter V. Coyle, Einas A. E. Al-Kuwari, Hamad E. Al-Romaihi, Salih Al-Marri, Mohammed Al-Thani, Reham A. El-Kahlout. QBB: Tasneem Al-Hamad, Dina Elgakhlab QGP: Fatima H. Al-Kuwari, Chadi Saad |
| EPI_ISL_2843670, EPI_ISL_2843671, EPI_ISL_2843672, EPI_ISL_2843673, EPI_ISL_2843677, EPI_ISL_2843678 | MEPHI, Aix Marseille University | MEPHI, Aix Marseille University | Anthony LEVASSEUR |
| EPI_ISL_2844209, EPI_ISL_2844241, EPI_ISL_2844266, EPI_ISL_2844267, EPI_ISL_2844269, EPI_ISL_2844276, EPI_ISL_2844280 | Europe/Sweden/Vastragotaland/Unilabs | Unilabs/Eskilstuna/Sweden | Emma Arvidsson |
| EPI_ISL_2844331, EPI_ISL_2844332, EPI_ISL_2844333, EPI_ISL_2844334, EPI_ISL_2844335, EPI_ISL_2844340, EPI_ISL_2844342, EPI_ISL_2844343, EPI_ISL_2844345, EPI_ISL_2844347, EPI_ISL_2844348, EPI_ISL_2844350, EPI_ISL_2844351, EPI_ISL_2844352, EPI_ISL_2844353, EPI_ISL_2844355, EPI_ISL_2844356, EPI_ISL_2844357, EPI_ISL_2844358, EPI_ISL_2844359, EPI_ISL_2844360, EPI_ISL_2844361, EPI_ISL_2844362, EPI_ISL_2844363, EPI_ISL_2844364, EPI_ISL_2844365, EPI_ISL_2844382, EPI_ISL_2844383, EPI_ISL_2844384, EPI_ISL_2844385, EPI_ISL_2844392, EPI_ISL_2844398, EPI_ISL_2844402, EPI_ISL_2844403, EPI_ISL_2844404, EPI_ISL_2844411, EPI_ISL_2844433, EPI_ISL_2844439, EPI_ISL_2844481, EPI_ISL_2844482, EPI_ISL_2844483 | | | |
| see above | Istituto Zooprofilattico Sperimentale del Mezzogiorno | TIGEM | Antonio Grimaldi Patrizia Annunziata Francesco Panariello Biancamaria Pierri Claudia Tiberio Teresa Giuliano Valentina Bouche Chiara Colantuono Maria Concetta Cuomo Denise Di Concilio Lucio Di Filippo Anna Manfredi Marcello Salvi Antonio Limone Luigi Atripaldi Pellegrino Cerino Andrea Ballabio Davide Cacchiarelli |
| EPI_ISL_2844488, EPI_ISL_2844491, EPI_ISL_2844493, EPI_ISL_2844495, EPI_ISL_2844496, EPI_ISL_2844498, EPI_ISL_2844500, EPI_ISL_2844501, EPI_ISL_2844504, EPI_ISL_2844505, EPI_ISL_2844506, EPI_ISL_2844507, EPI_ISL_2844508, EPI_ISL_2844509, EPI_ISL_2844511, EPI_ISL_2844512, EPI_ISL_2844513, EPI_ISL_2844514, EPI_ISL_2844516, EPI_ISL_2844518, EPI_ISL_2844519, EPI_ISL_2844520, EPI_ISL_2844521, EPI_ISL_2844522, EPI_ISL_2844523, EPI_ISL_2844524 | | | |
| see above | Lab. Microbiologia e Virologia Cotugno A.O. dei Colli - Istituto Zooprofilattico Sperimentale del Mezzogiorno | TIGEM | Antonio Grimaldi Patrizia Annunziata Francesco Panariello Biancamaria Pierri Claudia Tiberio Teresa Giuliano Valentina Bouche Chiara Colantuono Maria Concetta Cuomo Denise Di Concilio Lucio Di Filippo Anna Manfredi Marcello Salvi Antonio Limone Luigi Atripaldi Pellegrino Cerino Andrea Ballabio Davide Cacchiarelli |
| EPI_ISL_2844844 | Jaber Quarantine | Virology Lab, Jaber Al Ahmad Al Sabah Hospital | Dr. Zahrah Buhamad, Dr. Ebaa Al-Awadhi, Adil Afridi, Haroon Masih |
| EPI_ISL_2844846, | Chemische und Veterinäruntersuchungsamt Münsterland- | Robert Koch Institute | unknown |

| | | | |
|---|---|---|---|
| EPI_ISL_2844847, EPI_ISL_2844848, EPI_ISL_2844849 | Emscher-Lippe (CVUA-MEL) | | |
| EPI_ISL_2844850, EPI_ISL_2844851, EPI_ISL_2844852, EPI_ISL_2844853, EPI_ISL_2844854 | Labormedizin Darmstadt | Robert Koch Institute | unknown |
| EPI_ISL_2844855 | IMD Labor Frankfurt | Robert Koch Institute | unknown |
| EPI_ISL_2844858, EPI_ISL_2844859, EPI_ISL_2844860 | Labormedizin Darmstadt | Robert Koch Institute | unknown |
| EPI_ISL_2844861 | IMD Labor Frankfurt | Robert Koch Institute | unknown |
| EPI_ISL_2844862, EPI_ISL_2844863, EPI_ISL_2844864, EPI_ISL_2844865, EPI_ISL_2844866 | Labormedizin Darmstadt | Robert Koch Institute | unknown |
| EPI_ISL_2844867 | IMD Labor Frankfurt | Robert Koch Institute | unknown |
| EPI_ISL_2844868, EPI_ISL_2844869, EPI_ISL_2844870, EPI_ISL_2844872, EPI_ISL_2844873, EPI_ISL_2844874 | Labormedizin Darmstadt | Robert Koch Institute | unknown |
| EPI_ISL_2844875 | IMD Labor Frankfurt | Robert Koch Institute | unknown |
| EPI_ISL_2844876, EPI_ISL_2844878, EPI_ISL_2844879 | Labormedizin Darmstadt | Robert Koch Institute | unknown |
| EPI_ISL_2844882 | Institut für Medizinische Mikrobiologie - Universitätsmedizin Göttingen | Robert Koch Institute | unknown |
| EPI_ISL_2844883 | SYNLAB MVZ Weiden | Robert Koch Institute | unknown |
| EPI_ISL_2844885 | SYNLAB MVZ Leinfelden-Echterdingen | Robert Koch Institute | unknown |
| EPI_ISL_2844886 | SYNLAB MVZ Leverkusen | Robert Koch Institute | unknown |
| EPI_ISL_2844887 | Synlab MVZ Augsburg | Robert Koch Institute | unknown |
| EPI_ISL_2844888, EPI_ISL_2844889, EPI_ISL_2844891, EPI_ISL_2844894 | SYNLAB MVZ Weiden | Robert Koch Institute | unknown |
| EPI_ISL_2844895 | SYNLAB MVZ Leinfelden-Echterdingen | Robert Koch Institute | unknown |
| EPI_ISL_2844896 | Synlab MVZ Augsburg | Robert Koch Institute | unknown |
| EPI_ISL_2844899 | SYNLAB MVZ Leinfelden-Echterdingen | Robert Koch Institute | unknown |
| EPI_ISL_2844901, EPI_ISL_2844902 | SYNLAB MVZ Leverkusen | Robert Koch Institute | unknown |
| EPI_ISL_2844903 | Synlab MVZ Augsburg | Robert Koch Institute | unknown |
| EPI_ISL_2844904 | SYNLAB MVZ Leverkusen | Robert Koch Institute | unknown |
| EPI_ISL_2844908 | SYNLAB MVZ Leinfelden-Echterdingen | Robert Koch Institute | unknown |
| EPI_ISL_2844909 | SYNLAB MVZ Trier | Robert Koch Institute | unknown |
| EPI_ISL_2844911 | SYNLAB MVZ Weiden | Robert Koch Institute | unknown |
| EPI_ISL_2844912 | SYNLAB MVZ Leverkusen | Robert Koch Institute | unknown |
| EPI_ISL_2844913 | Synlab MVZ Augsburg | Robert Koch Institute | unknown |
| EPI_ISL_2844914 | SYNLAB MVZ Leverkusen | Robert Koch Institute | unknown |
| EPI_ISL_2844918 | SYNLAB MVZ Weiden | Robert Koch Institute | unknown |
| EPI_ISL_2844920 | SYNLAB MVZ Neustadt a. d. Weinstraße | Robert Koch Institute | unknown |
| EPI_ISL_2844921 | SYNLAB MVZ Heidelberg | Robert Koch Institute | unknown |
| EPI_ISL_2844922, EPI_ISL_2844924 | SYNLAB MVZ Leinfelden-Echterdingen | Robert Koch Institute | unknown |
| EPI_ISL_2844928, EPI_ISL_2844930, EPI_ISL_2844932 | SYNLAB MVZ Weiden | Robert Koch Institute | unknown |
| EPI_ISL_2844934 | SYNLAB MVZ Trier | Robert Koch Institute | unknown |
| EPI_ISL_2844935 | SYNLAB MVZ Leverkusen | Robert Koch Institute | unknown |
| EPI_ISL_2844936 | SYNLAB MVZ Leinfelden-Echterdingen | Robert Koch Institute | unknown |
| EPI_ISL_2844940, EPI_ISL_2844943, EPI_ISL_2844944 | SYNLAB MVZ Leverkusen | Robert Koch Institute | unknown |
| EPI_ISL_2844945 | SYNLAB MVZ Heidelberg | Robert Koch Institute | unknown |
| EPI_ISL_2844946 | SYNLAB MVZ Weiden | Robert Koch Institute | unknown |
| EPI_ISL_2844948 | SYNLAB MVZ Trier | Robert Koch Institute | unknown |
| EPI_ISL_2844949 | SYNLAB MVZ Leinfelden-Echterdingen | Robert Koch Institute | unknown |
| EPI_ISL_2844951 | Synlab MVZ Augsburg | Robert Koch Institute | unknown |
| EPI_ISL_2844952 | SYNLAB MVZ Weiden | Robert Koch Institute | unknown |
| EPI_ISL_2844953 | SYNLAB MVZ Leinfelden-Echterdingen | Robert Koch Institute | unknown |
| EPI_ISL_2844954 | SYNLAB MVZ Leverkusen | Robert Koch Institute | unknown |
| EPI_ISL_2844955, EPI_ISL_2844957 | SYNLAB MVZ Leinfelden-Echterdingen | Robert Koch Institute | unknown |
| EPI_ISL_2844958, EPI_ISL_2844960, EPI_ISL_2844961 | SYNLAB MVZ Leverkusen | Robert Koch Institute | unknown |
| EPI_ISL_2844962 | SYNLAB MVZ Leinfelden-Echterdingen | Robert Koch Institute | unknown |
| EPI_ISL_2844963 | SYNLAB MVZ Leverkusen | Robert Koch Institute | unknown |
| EPI_ISL_2844966 | SYNLAB MVZ Leinfelden-Echterdingen | Robert Koch Institute | unknown |
| EPI_ISL_2844967, EPI_ISL_2844969 | SYNLAB MVZ Weiden | Robert Koch Institute | unknown |
| EPI_ISL_2844970 | SYNLAB MVZ Leinfelden-Echterdingen | Robert Koch Institute | unknown |
| EPI_ISL_2844972 | SYNLAB MVZ Leverkusen | Robert Koch Institute | unknown |
| EPI_ISL_2844973 | SYNLAB MVZ Leinfelden-Echterdingen | Robert Koch Institute | unknown |

| | | | |
|---|---|---|---|
| EPI_ISL_2844976 | SYNLAB MVZ Weiden | Robert Koch Institute | unknown |
| EPI_ISL_2844977 | SYNLAB MVZ Leinfelden-Echterdingen | Robert Koch Institute | unknown |
| EPI_ISL_2844978 | SYNLAB MVZ Berlin | Robert Koch Institute | unknown |
| EPI_ISL_2844980 | SYNLAB MVZ Leverkusen | Robert Koch Institute | unknown |
| EPI_ISL_2844981, EPI_ISL_2844983 | SYNLAB MVZ Weiden | Robert Koch Institute | unknown |
| EPI_ISL_2844984, EPI_ISL_2844985 | SYNLAB MVZ Leverkusen | Robert Koch Institute | unknown |
| EPI_ISL_2844986 | SYNLAB MVZ Heidelberg | Robert Koch Institute | unknown |
| EPI_ISL_2844987 | SYNLAB MVZ Leinfelden-Echterdingen | Robert Koch Institute | unknown |
| EPI_ISL_2844988, EPI_ISL_2844992 | SYNLAB MVZ Weiden | Robert Koch Institute | unknown |
| EPI_ISL_2844993 | Synlab MVZ Augsburg | Robert Koch Institute | unknown |
| EPI_ISL_2844994 | SYNLAB MVZ Weiden | Robert Koch Institute | unknown |
| EPI_ISL_2844996 | SYNLAB MVZ Leverkusen | Robert Koch Institute | unknown |
| EPI_ISL_2844997 | SYNLAB MVZ Weiden | Robert Koch Institute | unknown |
| EPI_ISL_2844999 | SYNLAB MVZ Leverkusen | Robert Koch Institute | unknown |
| EPI_ISL_2845002, EPI_ISL_2845005 | SYNLAB MVZ Weiden | Robert Koch Institute | unknown |
| EPI_ISL_2845008 | SYNLAB MVZ Leverkusen | Robert Koch Institute | unknown |
| EPI_ISL_2845009, EPI_ISL_2845010 | SYNLAB MVZ Weiden | Robert Koch Institute | unknown |
| EPI_ISL_2845011 | SYNLAB Jena Oncoscreen | Robert Koch Institute | unknown |
| EPI_ISL_2845012, EPI_ISL_2845013 | SYNLAB MVZ Leinfelden-Echterdingen | Robert Koch Institute | unknown |
| EPI_ISL_2845015, EPI_ISL_2845018, EPI_ISL_2845034, EPI_ISL_2845039, EPI_ISL_2845040, EPI_ISL_2845045, EPI_ISL_2845047, EPI_ISL_2845055 | Limbach - MVZ Humangenetik Ulm | Robert Koch Institute | unknown |
| EPI_ISL_2845061, EPI_ISL_2845064, EPI_ISL_2845075, EPI_ISL_2845076 | Medizinische Laboratorien Düsseldorf | Robert Koch Institute | unknown |
| EPI_ISL_2845079 | amedes Medlab Arnold Analytik MVZ GmbH | Robert Koch Institute | unknown |
| EPI_ISL_2845080 | nordlab - Partnerschaftspraxis für Laboratoriumsmedizin | Robert Koch Institute | unknown |
| EPI_ISL_2845083 | amedes MVZ DIAMEDIS Sennestadt | Robert Koch Institute | unknown |
| EPI_ISL_2845084 | amedes MVZ Hannover | Robert Koch Institute | unknown |
| EPI_ISL_2845085 | amedes MVZ für Laboratoriumsdiagnostik Raubling GmbH | Robert Koch Institute | unknown |
| EPI_ISL_2845086 | amedes MVZ Halle/Leipzig | Robert Koch Institute | unknown |
| EPI_ISL_2845087 | nordlab - Partnerschaftspraxis für Laboratoriumsmedizin | Robert Koch Institute | unknown |
| EPI_ISL_2845088 | amedes Medlab Arnold Analytik MVZ GmbH | Robert Koch Institute | unknown |
| EPI_ISL_2845089 | nordlab - Partnerschaftspraxis für Laboratoriumsmedizin | Robert Koch Institute | unknown |
| EPI_ISL_2845090 | amedes MVZ DIAMEDIS Sennestadt | Robert Koch Institute | unknown |
| EPI_ISL_2845091 | amedes MVZ für Laboratoriumsdiagnostik Raubling GmbH | Robert Koch Institute | unknown |
| EPI_ISL_2845093, EPI_ISL_2845095 | MVZ Labor Dr. Limbach & Kollegen GbR | Robert Koch Institute | unknown |
| EPI_ISL_2845096 | MVZ Labor Dr. Reising-Ackermann und Kollegen | Robert Koch Institute | unknown |
| EPI_ISL_2845098 | MVZ Labor Dr. Limbach & Kollegen GbR | Robert Koch Institute | unknown |
| EPI_ISL_2845099 | MVZ Labor Dr. Reising-Ackermann und Kollegen | Robert Koch Institute | unknown |
| EPI_ISL_2845103 | Labor Lübeck bzw. Laborärztliche Gemeinschaftspraxis Lübeck | Robert Koch Institute | unknown |
| EPI_ISL_2845105, EPI_ISL_2845107, EPI_ISL_2845108 | MVZ Dr. Eberhard & Partner Dortmund | Robert Koch Institute | unknown |
| EPI_ISL_2845110 | MVZ für Laboratoriumsmedizin und Mikrobiologie Würzburg | Robert Koch Institute | unknown |
| EPI_ISL_2845111 | MVZ Dr. Eberhard & Partner Dortmund | Robert Koch Institute | unknown |
| EPI_ISL_2845112 | Procomcure Biotech Germany GmbH | Robert Koch Institute | unknown |
| EPI_ISL_2845113 | MVZ für Laboratoriumsmedizin und Mikrobiologie Würzburg | Robert Koch Institute | unknown |
| EPI_ISL_2845114 | MVZ Dr. Eberhard & Partner Dortmund | Robert Koch Institute | unknown |
| EPI_ISL_2845115 | Institut für Klinische Chemie Transfusions- und Laboratoriumsmedizin Berufsgenossenschaftliches Universitätsklinikum Bergmannsheil | Robert Koch Institute | unknown |
| EPI_ISL_2845117 | MVZ für Laboratoriumsmedizin und Mikrobiologie Würzburg | Robert Koch Institute | unknown |
| EPI_ISL_2845118 | Labor Mönchengladbach MVZ Dr. Stein + Kollegen GbR | Robert Koch Institute | unknown |
| EPI_ISL_2845119, EPI_ISL_2845126, EPI_ISL_2845127 | MVZ Labor Dr. Limbach & Kollegen GbR | Robert Koch Institute | unknown |
| EPI_ISL_2845130, EPI_ISL_2845134, EPI_ISL_2845137 | Labor Blackholm MVZ | Robert Koch Institute | unknown |
| EPI_ISL_2845140, EPI_ISL_2845142 | MVZ Labor Dr. Quade & Kollegen GmbH | Robert Koch Institute | unknown |
| EPI_ISL_2845143, EPI_ISL_2845144 | Labor Blackholm MVZ | Robert Koch Institute | unknown |
| EPI_ISL_2845146, EPI_ISL_2845147, EPI_ISL_2845148, EPI_ISL_2845149, EPI_ISL_2845150 | Labor Mönchengladbach MVZ Dr. Stein + Kollegen GbR | Robert Koch Institute | unknown |
| EPI_ISL_2845151 | MVZ Medizinisches Labor Hannover GmbH | Robert Koch Institute | unknown |
| EPI_ISL_2845152 | MVZ für Laboratoriumsmedizin und Mikrobiologie Koblenz-Mittelrhein (Labor Koblenz) | Robert Koch Institute | unknown |
| EPI_ISL_2845153, | MVZ Medizinisches Labor Hannover GmbH | Robert Koch Institute | unknown |

| | | | |
|---|---|---|---|
| EPI_ISL_2845154 | | | |
| EPI_ISL_2845155 | MDI Limbach Berlin GmbH; MVZ Labor Berlin | Robert Koch Institute | unknown |
| EPI_ISL_2845156 | MVZ für Laboratoriumsmedizin und Mikrobiologie Koblenz-Mittelrhein (Labor Koblenz) | Robert Koch Institute | unknown |
| EPI_ISL_2845157 | MDI Limbach Berlin GmbH; MVZ Labor Berlin | Robert Koch Institute | unknown |
| EPI_ISL_2845158 | MVZ für Laboratoriumsmedizin und Mikrobiologie Koblenz-Mittelrhein (Labor Koblenz) | Robert Koch Institute | unknown |
| EPI_ISL_2845159 | MDI Limbach Berlin GmbH; MVZ Labor Berlin | Robert Koch Institute | unknown |
| EPI_ISL_2845160, EPI_ISL_2845161 | MVZ für Laboratoriumsmedizin und Mikrobiologie Koblenz-Mittelrhein (Labor Koblenz) | Robert Koch Institute | unknown |
| EPI_ISL_2845162 | Labor Mönchengladbach MVZ Dr. Stein + Kollegen GbR | Robert Koch Institute | unknown |
| EPI_ISL_2845163 | MVZ für Laboratoriumsmedizin und Mikrobiologie Koblenz-Mittelrhein (Labor Koblenz) | Robert Koch Institute | unknown |
| EPI_ISL_2845164 | Labor Mönchengladbach MVZ Dr. Stein + Kollegen GbR | Robert Koch Institute | unknown |
| EPI_ISL_2845165 | MVZ für Laboratoriumsmedizin und Mikrobiologie Koblenz-Mittelrhein (Labor Koblenz) | Robert Koch Institute | unknown |
| EPI_ISL_2845166 | Labor Mönchengladbach MVZ Dr. Stein + Kollegen GbR | Robert Koch Institute | unknown |
| EPI_ISL_2845167 | MDI Limbach Berlin GmbH; MVZ Labor Berlin | Robert Koch Institute | unknown |
| EPI_ISL_2845168 | Labor Mönchengladbach MVZ Dr. Stein + Kollegen GbR | Robert Koch Institute | unknown |
| EPI_ISL_2845169, EPI_ISL_2845171 | MVZ für Laboratoriumsmedizin und Mikrobiologie Koblenz-Mittelrhein (Labor Koblenz) | Robert Koch Institute | unknown |
| EPI_ISL_2845174 | Labor Mönchengladbach MVZ Dr. Stein + Kollegen GbR | Robert Koch Institute | unknown |
| EPI_ISL_2845175 | MVZ für Laboratoriumsmedizin und Mikrobiologie Koblenz-Mittelrhein (Labor Koblenz) | Robert Koch Institute | unknown |
| EPI_ISL_2845176 | MDI Limbach Berlin GmbH; MVZ Labor Berlin | Robert Koch Institute | unknown |
| EPI_ISL_2845178 | Kerckhoff-Klinik GmbH | Robert Koch Institute | unknown |
| EPI_ISL_2845179 | MVZ für Laboratoriumsmedizin und Mikrobiologie Koblenz-Mittelrhein (Labor Koblenz) | Robert Koch Institute | unknown |
| EPI_ISL_2845180 | Labor Mönchengladbach MVZ Dr. Stein + Kollegen GbR | Robert Koch Institute | unknown |
| EPI_ISL_2845184, EPI_ISL_2845185, EPI_ISL_2845186 | Niedersächsisches Landesgesundheitsamt (NLGA) | Robert Koch Institute | unknown |
| EPI_ISL_2845187, EPI_ISL_2845190 | Labor Mönchengladbach MVZ Dr. Stein + Kollegen GbR | Robert Koch Institute | unknown |
| EPI_ISL_2845197 | labopart - Medizinische Laboratorien Dresden | Robert Koch Institute | unknown |
| EPI_ISL_2845198, EPI_ISL_2845200, EPI_ISL_2845201, EPI_ISL_2845202 | MDI Limbach Berlin GmbH; MVZ Labor Berlin | Robert Koch Institute | unknown |
| EPI_ISL_2845203 | MVZ Labor Dr. Limbach & Kollegen GbR | Robert Koch Institute | unknown |
| EPI_ISL_2845204 | MDI Limbach Berlin GmbH; MVZ Labor Berlin | Robert Koch Institute | unknown |
| EPI_ISL_2845205 | MVZ Labor Dr. Limbach & Kollegen GbR | Robert Koch Institute | unknown |
| EPI_ISL_2845206 | MDI Limbach Berlin GmbH; MVZ Labor Berlin | Robert Koch Institute | unknown |
| EPI_ISL_2845208 | MVZ Labor Dr. Limbach & Kollegen GbR | Robert Koch Institute | unknown |
| EPI_ISL_2845209 | MDI Limbach Berlin GmbH; MVZ Labor Berlin | Robert Koch Institute | unknown |
| EPI_ISL_2845210 | MVZ Labor Dr. Limbach & Kollegen GbR | Robert Koch Institute | unknown |
| EPI_ISL_2845211, EPI_ISL_2845212, EPI_ISL_2845214 | MDI Limbach Berlin GmbH; MVZ Labor Berlin | Robert Koch Institute | unknown |
| EPI_ISL_2845215 | MVZ Labor Dr. Limbach & Kollegen GbR | Robert Koch Institute | unknown |
| EPI_ISL_2845218, EPI_ISL_2845220, EPI_ISL_2845221, EPI_ISL_2845222, EPI_ISL_2845223, EPI_ISL_2845224, EPI_ISL_2845225, EPI_ISL_2845226, EPI_ISL_2845227, EPI_ISL_2845228, EPI_ISL_2845229, EPI_ISL_2845230, EPI_ISL_2845231, EPI_ISL_2845232, EPI_ISL_2845234, EPI_ISL_2845235, EPI_ISL_2845236 | | | |
| see above | Labor Dr. Heidrich & Kollegen MVZ GmbH Hamburg | Robert Koch Institute | unknown |
| EPI_ISL_2845237, EPI_ISL_2845239 | MVZ Dr. Eberhard & Partner Dortmund | Robert Koch Institute | unknown |
| EPI_ISL_2845242 | MVZ für Laboratoriumsmedizin und Mikrobiologie Würzburg | Robert Koch Institute | unknown |
| EPI_ISL_2845243 | Landesamt für Verbraucherschutz des Landes Sachsen-Anhalt | Robert Koch Institute | unknown |
| EPI_ISL_2845244, EPI_ISL_2845245, EPI_ISL_2845246, EPI_ISL_2845247, EPI_ISL_2845248, EPI_ISL_2845249, EPI_ISL_2845250, EPI_ISL_2845251, EPI_ISL_2845252, EPI_ISL_2845253, EPI_ISL_2845254, EPI_ISL_2845255, EPI_ISL_2845256, EPI_ISL_2845257, EPI_ISL_2845258, EPI_ISL_2845259, EPI_ISL_2845260, EPI_ISL_2845262, EPI_ISL_2845263, EPI_ISL_2845264, EPI_ISL_2845265, EPI_ISL_2845266, EPI_ISL_2845267, EPI_ISL_2845268, EPI_ISL_2845269, EPI_ISL_2845270, EPI_ISL_2845271, EPI_ISL_2845272, EPI_ISL_2845273, EPI_ISL_2845274, EPI_ISL_2845275, EPI_ISL_2845276, EPI_ISL_2845277, EPI_ISL_2845278, EPI_ISL_2845279, EPI_ISL_2845280, EPI_ISL_2845281, EPI_ISL_2845283, EPI_ISL_2845284, EPI_ISL_2845286, EPI_ISL_2845287, EPI_ISL_2845288, EPI_ISL_2845289, EPI_ISL_2845290, EPI_ISL_2845291 | | | |
| see above | Labor Dr. Wisplinghoff - Köln | Robert Koch Institute | unknown |
| EPI_ISL_2845292 | MVZ Labor Dr. Limbach & Kollegen GbR | Robert Koch Institute | unknown |
| EPI_ISL_2845294, EPI_ISL_2845295, EPI_ISL_2845296, EPI_ISL_2845298, EPI_ISL_2845299, EPI_ISL_2845300, EPI_ISL_2845301, EPI_ISL_2845305, EPI_ISL_2845308, EPI_ISL_2845310, EPI_ISL_2845311, EPI_ISL_2845312, EPI_ISL_2845313, EPI_ISL_2845314 | | | |
| see above | MVZ Labor Krone GbR | Robert Koch Institute | unknown |
| EPI_ISL_2845317 | Bioscientia MVZ Labor Karlsruhe GmbH | Robert Koch Institute | unknown |
| EPI_ISL_2845318 | Sonic - Labor Dr. von Froreich GmbH | Robert Koch Institute | unknown |
| EPI_ISL_2845320 | Bioscientia Labor Wermsdorf | Robert Koch Institute | unknown |
| EPI_ISL_2845321 | Sonic - Labor Dr. von Froreich GmbH | Robert Koch Institute | unknown |
| EPI_ISL_2845322 | Sonic - MVZ Medizinisches Labor Bremen GmbH | Robert Koch Institute | unknown |
| EPI_ISL_2845323, EPI_ISL_2845325 | Bioscientia Labor Wermsdorf | Robert Koch Institute | unknown |
| EPI_ISL_2845329 | Sonic - MVZ Medizinisches Labor Bremen GmbH | Robert Koch Institute | unknown |
| EPI_ISL_2845330, EPI_ISL_2845334, EPI_ISL_2845335, EPI_ISL_2845336, EPI_ISL_2845338 | Bioscientia Labor Wermsdorf | Robert Koch Institute | unknown |
| EPI_ISL_2845339 | Bioscientia MVZ Labor Karlsruhe GmbH | Robert Koch Institute | unknown |
| EPI_ISL_2845340, EPI_ISL_2845342 | Bioscientia Labor Wermsdorf | Robert Koch Institute | unknown |
| EPI_ISL_2845344 | Bioscientia MVZ Labor Karlsruhe GmbH | Robert Koch Institute | unknown |
| EPI_ISL_2845345 | Bioscientia Labor Wermsdorf | Robert Koch Institute | unknown |
| EPI_ISL_2845346 | Sonic - MVZ Medizinisches Labor Bremen GmbH | Robert Koch Institute | unknown |
| EPI_ISL_2845347, EPI_ISL_2845349 | Bioscientia Labor Wermsdorf | Robert Koch Institute | unknown |
| EPI_ISL_2845351 | Sonic - MVZ Medizinisches Labor Bremen GmbH | Robert Koch Institute | unknown |

| | | | |
|---|---|---|---|
| EPI_ISL_2845352, EPI_ISL_2845353, EPI_ISL_2845354 | Bioscientia Labor Wermsdorf | Robert Koch Institute | unknown |
| EPI_ISL_2845355 | Sonic - MVZ Medizinisches Labor Bremen GmbH | Robert Koch Institute | unknown |
| EPI_ISL_2845356 | ALB FILS KLINIKEN GmbH - Institut für Laboratoriumsmedizin Klinik am Eichert Göppingen | Robert Koch Institute | unknown |
| EPI_ISL_2845357 | Sonic - MVZ Medizinisches Labor Bremen GmbH | Robert Koch Institute | unknown |
| EPI_ISL_2845358 | Bioscientia Labor Wermsdorf | Robert Koch Institute | unknown |
| EPI_ISL_2845359 | Sonic - Labor Dr. von Froreich GmbH | Robert Koch Institute | unknown |
| EPI_ISL_2845361 | Bioscientia Labor Wermsdorf | Robert Koch Institute | unknown |
| EPI_ISL_2845362 | Sonic - MVZ Medizinisches Labor Bremen GmbH | Robert Koch Institute | unknown |
| EPI_ISL_2845363 | Sonic - Labor Dr. von Froreich GmbH | Robert Koch Institute | unknown |
| EPI_ISL_2845366 | Bioscientia Labor Wermsdorf | Robert Koch Institute | unknown |
| EPI_ISL_2845367 | Sonic - MVZ Medizinisches Labor Bremen GmbH | Robert Koch Institute | unknown |
| EPI_ISL_2845369, EPI_ISL_2845371 | Bioscientia Labor Wermsdorf | Robert Koch Institute | unknown |
| EPI_ISL_2845372, EPI_ISL_2845373 | Sonic - MVZ Medizinisches Labor Bremen GmbH | Robert Koch Institute | unknown |
| EPI_ISL_2845374 | Bioscientia MVZ Labor Karlsruhe GmbH | Robert Koch Institute | unknown |
| EPI_ISL_2845376 | Sonic - Labor Staber Dresden (Klipphausen) | Robert Koch Institute | unknown |
| EPI_ISL_2845377 | Bioscientia Labor Wermsdorf | Robert Koch Institute | unknown |
| EPI_ISL_2845380 | LabKom - Labor Augsburg MVZ GmbH | Robert Koch Institute | unknown |
| EPI_ISL_2845381, EPI_ISL_2845382, EPI_ISL_2845384 | Bioscientia Labor Wermsdorf | Robert Koch Institute | unknown |
| EPI_ISL_2845385 | Labor 28 MVZ GmbH | Robert Koch Institute | unknown |
| EPI_ISL_2845386 | LabKom - Labor Mainz MVZ GmbH | Robert Koch Institute | unknown |
| EPI_ISL_2845387, EPI_ISL_2845389 | Bioscientia Labor Wermsdorf | Robert Koch Institute | unknown |
| EPI_ISL_2845390 | LabKom - Labor Augsburg MVZ GmbH | Robert Koch Institute | unknown |
| EPI_ISL_2845394 | Labor 28 MVZ GmbH | Robert Koch Institute | unknown |
| EPI_ISL_2845395 | LabKom - Labor Augsburg MVZ GmbH | Robert Koch Institute | unknown |
| EPI_ISL_2845397 | Universitätsklinikum Frankfurt - Institut für Medizinische Virologie | Robert Koch Institute | unknown |
| EPI_ISL_2845399, EPI_ISL_2845400 | Bioscientia Labor Wermsdorf | Robert Koch Institute | unknown |
| EPI_ISL_2845406 | Bioscientia MVZ Labor Karlsruhe GmbH | Robert Koch Institute | unknown |
| EPI_ISL_2845407 | LabKom - Labor Mainz MVZ GmbH | Robert Koch Institute | unknown |
| EPI_ISL_2845410 | Bioscientia Labor Wermsdorf | Robert Koch Institute | unknown |
| EPI_ISL_2845411 | Klinikum Fulda gAG | Robert Koch Institute | unknown |
| EPI_ISL_2845414 | LabKom - Labor an der Salzbrücke MVZ GmbH | Robert Koch Institute | unknown |
| EPI_ISL_2845418, EPI_ISL_2845419, EPI_ISL_2845421, EPI_ISL_2845422 | Bioscientia Labor Wermsdorf | Robert Koch Institute | unknown |
| EPI_ISL_2845423 | Bioscientia MVZ Labor Karlsruhe GmbH | Robert Koch Institute | unknown |
| EPI_ISL_2845424 | LabKom - MVZ Labor Bochum MLB GmbH | Robert Koch Institute | unknown |
| EPI_ISL_2845425 | LabKom - Labor Augsburg MVZ GmbH | Robert Koch Institute | unknown |
| EPI_ISL_2845426 | Bioscientia Labor Wermsdorf | Robert Koch Institute | unknown |
| EPI_ISL_2845428 | Labor 28 MVZ GmbH | Robert Koch Institute | unknown |
| EPI_ISL_2845432, EPI_ISL_2845433 | Bioscientia Labor Wermsdorf | Robert Koch Institute | unknown |
| EPI_ISL_2845438 | Labor 28 MVZ GmbH | Robert Koch Institute | unknown |
| EPI_ISL_2845439, EPI_ISL_2845440, EPI_ISL_2845441 | Bioscientia Labor Wermsdorf | Robert Koch Institute | unknown |
| EPI_ISL_2845442 | Universitätsklinikum Frankfurt - Institut für Medizinische Virologie | Robert Koch Institute | unknown |
| EPI_ISL_2845443, EPI_ISL_2845446, EPI_ISL_2845450, EPI_ISL_2845451, EPI_ISL_2845452, EPI_ISL_2845453, EPI_ISL_2845456, EPI_ISL_2845457, EPI_ISL_2845463 | Eurofins MVZ Labor Gelsenkirchen | Robert Koch Institute | unknown |
| EPI_ISL_2845469, EPI_ISL_2845472, EPI_ISL_2845474, EPI_ISL_2845475, EPI_ISL_2845476, EPI_ISL_2845478, EPI_ISL_2845481, EPI_ISL_2845484, EPI_ISL_2845486, EPI_ISL_2845487, EPI_ISL_2845489, EPI_ISL_2845493, EPI_ISL_2845494, EPI_ISL_2845495, EPI_ISL_2845496, EPI_ISL_2845499, EPI_ISL_2845500, EPI_ISL_2845502, EPI_ISL_2845504, EPI_ISL_2845505, EPI_ISL_2845506, EPI_ISL_2845509, EPI_ISL_2845511, EPI_ISL_2845512, EPI_ISL_2845513, EPI_ISL_2845514, EPI_ISL_2845515, EPI_ISL_2845517, EPI_ISL_2845518, EPI_ISL_2845520, EPI_ISL_2845524, EPI_ISL_2845527, EPI_ISL_2845528, EPI_ISL_2845529, EPI_ISL_2845532, EPI_ISL_2845534, EPI_ISL_2845536, EPI_ISL_2845537, EPI_ISL_2845538, EPI_ISL_2845539, EPI_ISL_2845540, EPI_ISL_2845542 | | | |
| see above | CENTOGENE Frankfurt Laboratory: Niederlassung Industriepark Höchst | Robert Koch Institute | unknown |
| EPI_ISL_2845650, EPI_ISL_2845653 | Universitätsklinikum Köln; Institut für Virologie | Robert Koch Institute | unknown |
| EPI_ISL_2845679 | IMD - MVZ Labor Martinsried | Robert Koch Institute | unknown |
| EPI_ISL_2845680, EPI_ISL_2845681, EPI_ISL_2845682 | IMD - Labor Oderland | Robert Koch Institute | unknown |
| EPI_ISL_2845683 | IMD - MVZ Labor Martinsried | Robert Koch Institute | unknown |
| EPI_ISL_2845685, EPI_ISL_2845686, EPI_ISL_2845688, EPI_ISL_2845690 | IMD - Labor Oderland | Robert Koch Institute | unknown |
| EPI_ISL_2845691 | IMD - MVZ Labor Martinsried | Robert Koch Institute | unknown |
| EPI_ISL_2845692, EPI_ISL_2845693 | IMD - Labor Oderland | Robert Koch Institute | unknown |
| EPI_ISL_2845696, EPI_ISL_2845705, EPI_ISL_2845709, EPI_ISL_2845712, EPI_ISL_2845713, EPI_ISL_2845719, EPI_ISL_2845721, EPI_ISL_2845723, EPI_ISL_2845724, EPI_ISL_2845725, EPI_ISL_2845728, EPI_ISL_2845729, EPI_ISL_2845730, EPI_ISL_2845731, EPI_ISL_2845733, EPI_ISL_2845736, EPI_ISL_2845738, EPI_ISL_2845739, EPI_ISL_2845740, EPI_ISL_2845741 | | | |

| | | | |
|---|---|---|---|
| see above | Labor Becker & Kollegen (Standort München) | Robert Koch Institute | unknown |
| EPI_ISL_2845747 | Landesuntersuchungsanstalt für das Gesundheits- und Veterinärwesen (LUA) Sachsen | Robert Koch Institute | unknown |
| EPI_ISL_2845750, EPI_ISL_2845751, EPI_ISL_2845754, EPI_ISL_2845758 | DKMS Life Science Lab GmbH Dresden | Robert Koch Institute | unknown |
| EPI_ISL_2845759 | Uniklinikum Carl Gustav Carus an der TU Dresden; Institut für Virologie | Robert Koch Institute | unknown |
| EPI_ISL_2845760 | Städtisches Klinikum Dresden Institut für Klinische Chemie und Laboratoriumsmedizin | Robert Koch Institute | unknown |
| EPI_ISL_2845761 | Landesuntersuchungsanstalt für das Gesundheits- und Veterinärwesen (LUA) Sachsen | Robert Koch Institute | unknown |
| EPI_ISL_2845769, EPI_ISL_2845770, EPI_ISL_2845772, EPI_ISL_2845777 | Universitätsklinikum Heidelberg | Robert Koch Institute | unknown |
| EPI_ISL_2845783, EPI_ISL_2845788 | Dianovis GmbH Greiz | Robert Koch Institute | unknown |
| EPI_ISL_2845789, EPI_ISL_2845791, EPI_ISL_2845793, EPI_ISL_2845795, EPI_ISL_2845797, EPI_ISL_2845798, EPI_ISL_2845800, EPI_ISL_2845801, EPI_ISL_2845804, EPI_ISL_2845807, EPI_ISL_2845810 | | | |
| see above | Labor Becker & Kollegen (Standort München) | Robert Koch Institute | unknown |
| EPI_ISL_2845813, EPI_ISL_2845814, EPI_ISL_2845815, EPI_ISL_2845816, EPI_ISL_2845817, EPI_ISL_2845819, EPI_ISL_2845820, EPI_ISL_2845821 | MVZ Labor Dr. Fenner und Kollegen (Standort Hamburg) | Robert Koch Institute | unknown |
| EPI_ISL_2845834, EPI_ISL_2845836, EPI_ISL_2845838, EPI_ISL_2845839, EPI_ISL_2845840, EPI_ISL_2845842, EPI_ISL_2845843, EPI_ISL_2845844, EPI_ISL_2845845, EPI_ISL_2845846, EPI_ISL_2845849, EPI_ISL_2845850, EPI_ISL_2845851, EPI_ISL_2845853, EPI_ISL_2845856, EPI_ISL_2845857, EPI_ISL_2845860, EPI_ISL_2845861, EPI_ISL_2845862, EPI_ISL_2845865, EPI_ISL_2845866, EPI_ISL_2845867, EPI_ISL_2845868, EPI_ISL_2845869, EPI_ISL_2845870, EPI_ISL_2845871, EPI_ISL_2845875, EPI_ISL_2845876, EPI_ISL_2845878, EPI_ISL_2845879, EPI_ISL_2845880, EPI_ISL_2845881, EPI_ISL_2845884 | | | |
| see above | Eurofins LifeCodexx GmbH | Robert Koch Institute | unknown |
| EPI_ISL_2845886, EPI_ISL_2845889, EPI_ISL_2845890, EPI_ISL_2845891, EPI_ISL_2845894, EPI_ISL_2845897, EPI_ISL_2845898, EPI_ISL_2845899, EPI_ISL_2845900, EPI_ISL_2845901, EPI_ISL_2845902, EPI_ISL_2845904, EPI_ISL_2845905, EPI_ISL_2845906, EPI_ISL_2845907, EPI_ISL_2845908, EPI_ISL_2845909, EPI_ISL_2845912, EPI_ISL_2845914, EPI_ISL_2845915, EPI_ISL_2845916, EPI_ISL_2845917, EPI_ISL_2845920, EPI_ISL_2845922, EPI_ISL_2845923, EPI_ISL_2845924, EPI_ISL_2845925, EPI_ISL_2845926, EPI_ISL_2845928, EPI_ISL_2845929, EPI_ISL_2845932, EPI_ISL_2845934 | | | |
| see above | Bayerisches Landesamt für Gesundheit und Lebensmittelsicherheit (LGL) | Robert Koch Institute | unknown |
| EPI_ISL_2845940, EPI_ISL_2845941, EPI_ISL_2845943, EPI_ISL_2845949 | Clemenshospital - Alexianer Clemenshospital Münster | Robert Koch Institute | unknown |
| EPI_ISL_2845953, EPI_ISL_2845954, EPI_ISL_2845955 | MVZ Labor Dr. Limbach & Kollegen GbR | Robert Koch Institute | unknown |
| EPI_ISL_2845956 | MVZ für Laboratoriumsmedizin und Mikrobiologie Koblenz-Mittelrhein (Labor Koblenz) | Robert Koch Institute | unknown |
| EPI_ISL_2845957, EPI_ISL_2845958, EPI_ISL_2845959 | MVZ Labor Dr. Limbach & Kollegen GbR | Robert Koch Institute | unknown |
| EPI_ISL_2845960 | Labor Prof. Dr. G. Enders MVZ GbR | Robert Koch Institute | unknown |
| EPI_ISL_2845963 | Limbach - MVZ Labor Eveld & Kollegen Essen | Robert Koch Institute | unknown |
| EPI_ISL_2845964 | MVZ Labor Dr. Reising-Ackermann und Kollegen | Robert Koch Institute | unknown |
| EPI_ISL_2845965 | MVZ für Laboratoriumsmedizin und Mikrobiologie Koblenz-Mittelrhein (Labor Koblenz) | Robert Koch Institute | unknown |
| EPI_ISL_2845966 | MVZ Medizinisches Labor Hannover GmbH | Robert Koch Institute | unknown |
| EPI_ISL_2845968, EPI_ISL_2845969 | MVZ für Laboratoriumsmedizin und Mikrobiologie Koblenz-Mittelrhein (Labor Koblenz) | Robert Koch Institute | unknown |
| EPI_ISL_2845970 | MDI Limbach Berlin GmbH; MVZ Labor Berlin | Robert Koch Institute | unknown |
| EPI_ISL_2845974, EPI_ISL_2845975, EPI_ISL_2845976 | MVZ für Laboratoriumsmedizin und Mikrobiologie Koblenz-Mittelrhein (Labor Koblenz) | Robert Koch Institute | unknown |
| EPI_ISL_2845977 | MDI Limbach Berlin GmbH; MVZ Labor Berlin | Robert Koch Institute | unknown |
| EPI_ISL_2845978 | MVZ Medizinisches Labor Hannover GmbH | Robert Koch Institute | unknown |
| EPI_ISL_2845979, EPI_ISL_2845980, EPI_ISL_2845981 | MDI Limbach Berlin GmbH; MVZ Labor Berlin | Robert Koch Institute | unknown |
| EPI_ISL_2845982 | MVZ für Laboratoriumsmedizin und Mikrobiologie Koblenz-Mittelrhein (Labor Koblenz) | Robert Koch Institute | unknown |
| EPI_ISL_2845983, EPI_ISL_2845984, EPI_ISL_2845985, EPI_ISL_2845987 | MDI Limbach Berlin GmbH; MVZ Labor Berlin | Robert Koch Institute | unknown |
| EPI_ISL_2845988 | MVZ Medizinisches Labor Hannover GmbH | Robert Koch Institute | unknown |
| EPI_ISL_2845989 | MDI Limbach Berlin GmbH; MVZ Labor Berlin | Robert Koch Institute | unknown |
| EPI_ISL_2845990 | MVZ für Laboratoriumsmedizin und Mikrobiologie Koblenz-Mittelrhein (Labor Koblenz) | Robert Koch Institute | unknown |
| EPI_ISL_2845991 | MVZ Medizinisches Labor Hannover GmbH | Robert Koch Institute | unknown |
| EPI_ISL_2845992, EPI_ISL_2845993, EPI_ISL_2845994, EPI_ISL_2845998, EPI_ISL_2845999, EPI_ISL_2846000, EPI_ISL_2846001, EPI_ISL_2846002 | MDI Limbach Berlin GmbH; MVZ Labor Berlin | Robert Koch Institute | unknown |
| EPI_ISL_2846003 | MVZ Labor Dr. Limbach & Kollegen GbR | Robert Koch Institute | unknown |
| EPI_ISL_2846004 | Klinikum Fulda gAG | Robert Koch Institute | unknown |
| EPI_ISL_2846007 | Dianovis GmbH Greiz | Robert Koch Institute | unknown |
| EPI_ISL_2846008, EPI_ISL_2846009 | Laborarztpraxis Dres. med. Walther Weindel & Kollegen | Robert Koch Institute | unknown |
| EPI_ISL_2846010, EPI_ISL_2846011, EPI_ISL_2846012, EPI_ISL_2846013, EPI_ISL_2846014, EPI_ISL_2846015, EPI_ISL_2846016, EPI_ISL_2846017, EPI_ISL_2846018, EPI_ISL_2846019, EPI_ISL_2846020, EPI_ISL_2846021 | | | |
| see above | AZ Klina | AZ Klina | Carl Vael - Lynsey Berckmans |

| | | | |
|---|---|---|---|
| EPI_ISL_2846022, EPI_ISL_2846023, EPI_ISL_2846024 | Ministry of Health, Jaber Al-Ahmad Hospital | Virology Unit, Department of Microbiology, Faculty of Medicine, Kuwait University | Nada Madi, Hussain Safar, Ebaa Al-Awadhi, Anfal Al-Adwani |
| EPI_ISL_2846025 | Lighthouse Lab in Alderley Park | Wellcome Sanger Institute for the COVID-19 Genomics UK (COG-UK) Consortium | Jacquelyn Wynn, Mairead Hyland, The Lighthouse Lab in Alderley Park and Alex Alderton, Roberto Amato, Jeffrey Barrett, Sonia Goncalves, Ewan Harrison, David K. Jackson, Ian Johnston, Dominic Kwiatkowski, Cordelia Langford, John Sillitoe on behalf of the Wellcome Sanger Institute COVID-19 Surveillance Team |
| EPI_ISL_2846026 | Lighthouse Lab in Milton Keynes | Wellcome Sanger Institute for the COVID-19 Genomics UK (COG-UK) Consortium | The Lighthouse Lab in Milton Keynes and Alex Alderton, Roberto Amato, Jeffrey Barrett, Sonia Goncalves, Ewan Harrison, David K. Jackson, Ian Johnston, Dominic Kwiatkowski, Cordelia Langford, John Sillitoe on behalf of the Wellcome Sanger Institute COVID-19 Surveillance Team |
| EPI_ISL_2846027 | Lighthouse Lab in Alderley Park | Wellcome Sanger Institute for the COVID-19 Genomics UK (COG-UK) Consortium | Jacquelyn Wynn, Mairead Hyland, The Lighthouse Lab in Alderley Park and Alex Alderton, Roberto Amato, Jeffrey Barrett, Sonia Goncalves, Ewan Harrison, David K. Jackson, Ian Johnston, Dominic Kwiatkowski, Cordelia Langford, John Sillitoe on behalf of the Wellcome Sanger Institute COVID-19 Surveillance Team |
| EPI_ISL_2846028 | Lighthouse Lab in Milton Keynes | Wellcome Sanger Institute for the COVID-19 Genomics UK (COG-UK) Consortium | The Lighthouse Lab in Milton Keynes and Alex Alderton, Roberto Amato, Jeffrey Barrett, Sonia Goncalves, Ewan Harrison, David K. Jackson, Ian Johnston, Dominic Kwiatkowski, Cordelia Langford, John Sillitoe on behalf of the Wellcome Sanger Institute COVID-19 Surveillance Team |
| EPI_ISL_2846029 | Lighthouse Lab in Alderley Park | Wellcome Sanger Institute for the COVID-19 Genomics UK (COG-UK) Consortium | Jacquelyn Wynn, Mairead Hyland, The Lighthouse Lab in Alderley Park and Alex Alderton, Roberto Amato, Jeffrey Barrett, Sonia Goncalves, Ewan Harrison, David K. Jackson, Ian Johnston, Dominic Kwiatkowski, Cordelia Langford, John Sillitoe on behalf of the Wellcome Sanger Institute COVID-19 Surveillance Team |
| EPI_ISL_2846030 | Lighthouse Lab in Milton Keynes | Wellcome Sanger Institute for the COVID-19 Genomics UK (COG-UK) Consortium | The Lighthouse Lab in Milton Keynes and Alex Alderton, Roberto Amato, Jeffrey Barrett, Sonia Goncalves, Ewan Harrison, David K. Jackson, Ian Johnston, Dominic Kwiatkowski, Cordelia Langford, John Sillitoe on behalf of the Wellcome Sanger Institute COVID-19 Surveillance Team |
| EPI_ISL_2846031 | Lighthouse Laboratory Plymouth | Wellcome Sanger Institute for the COVID-19 Genomics UK (COG-UK) Consortium | Lighthouse Laboratory Plymouth and Alex Alderton, Roberto Amato, Jeffrey Barrett, Sonia Goncalves, Ewan Harrison, David K. Jackson, Ian Johnston, Dominic Kwiatkowski, Cordelia Langford, John Sillitoe on behalf of the Wellcome Sanger Institute COVID-19 Surveillance Team |
| EPI_ISL_2846032 | Lighthouse Lab in Alderley Park | Wellcome Sanger Institute for the COVID-19 Genomics UK (COG-UK) Consortium | Jacquelyn Wynn, Mairead Hyland, The Lighthouse Lab in Alderley Park and Alex Alderton, Roberto Amato, Jeffrey Barrett, Sonia Goncalves, Ewan Harrison, David K. Jackson, Ian Johnston, Dominic Kwiatkowski, Cordelia Langford, John Sillitoe on behalf of the Wellcome Sanger Institute COVID-19 Surveillance Team |
| EPI_ISL_2846033 | Lighthouse Lab in Milton Keynes | Wellcome Sanger Institute for the COVID-19 Genomics UK (COG-UK) Consortium | The Lighthouse Lab in Milton Keynes and Alex Alderton, Roberto Amato, Jeffrey Barrett, Sonia Goncalves, Ewan Harrison, David K. Jackson, Ian Johnston, Dominic Kwiatkowski, Cordelia Langford, John Sillitoe on behalf of the Wellcome Sanger Institute COVID-19 Surveillance Team |
| EPI_ISL_2846034, EPI_ISL_2846035 | Lighthouse Lab in Alderley Park | Wellcome Sanger Institute for the COVID-19 Genomics UK (COG-UK) Consortium | Jacquelyn Wynn, Mairead Hyland, The Lighthouse Lab in Alderley Park and Alex Alderton, Roberto Amato, Jeffrey Barrett, Sonia Goncalves, Ewan Harrison, David K. Jackson, Ian Johnston, Dominic Kwiatkowski, Cordelia Langford, John Sillitoe on behalf of the Wellcome Sanger Institute COVID-19 Surveillance Team |
| EPI_ISL_2846036 | Lighthouse Laboratory Plymouth | Wellcome Sanger Institute for the COVID-19 Genomics UK (COG-UK) Consortium | Lighthouse Laboratory Plymouth and Alex Alderton, Roberto Amato, Jeffrey Barrett, Sonia Goncalves, Ewan Harrison, David K. Jackson, Ian Johnston, Dominic Kwiatkowski, Cordelia Langford, John Sillitoe on behalf of the Wellcome Sanger Institute COVID-19 Surveillance Team |
| EPI_ISL_2846037, EPI_ISL_2846038 | Lighthouse Lab in Alderley Park | Wellcome Sanger Institute for the COVID-19 Genomics UK (COG-UK) Consortium | Jacquelyn Wynn, Mairead Hyland, The Lighthouse Lab in Alderley Park and Alex Alderton, Roberto Amato, Jeffrey Barrett, Sonia Goncalves, Ewan Harrison, David K. Jackson, Ian Johnston, Dominic Kwiatkowski, Cordelia Langford, John Sillitoe on behalf of the Wellcome Sanger Institute COVID-19 Surveillance Team |
| EPI_ISL_2846039, EPI_ISL_2846040 | Lighthouse Lab in Milton Keynes | Wellcome Sanger Institute for the COVID-19 Genomics UK (COG-UK) Consortium | The Lighthouse Lab in Milton Keynes and Alex Alderton, Roberto Amato, Jeffrey Barrett, Sonia Goncalves, Ewan Harrison, David K. Jackson, Ian Johnston, Dominic Kwiatkowski, Cordelia Langford, John Sillitoe on behalf of the Wellcome Sanger Institute COVID-19 Surveillance Team |
| EPI_ISL_2846041, EPI_ISL_2846042, EPI_ISL_2846043, EPI_ISL_2846044 | Lighthouse Lab in Alderley Park | Wellcome Sanger Institute for the COVID-19 Genomics UK (COG-UK) Consortium | Jacquelyn Wynn, Mairead Hyland, The Lighthouse Lab in Alderley Park and Alex Alderton, Roberto Amato, Jeffrey Barrett, Sonia Goncalves, Ewan Harrison, David K. Jackson, Ian Johnston, Dominic Kwiatkowski, Cordelia Langford, John Sillitoe on behalf of the Wellcome Sanger Institute COVID-19 Surveillance Team |
| EPI_ISL_2846045 | Lighthouse Laboratory Plymouth | Wellcome Sanger Institute for the COVID-19 Genomics UK (COG-UK) Consortium | Lighthouse Laboratory Plymouth and Alex Alderton, Roberto Amato, Jeffrey Barrett, Sonia Goncalves, Ewan Harrison, David K. Jackson, Ian Johnston, Dominic Kwiatkowski, Cordelia Langford, John Sillitoe on behalf of the Wellcome Sanger Institute COVID-19 Surveillance Team |
| EPI_ISL_2846046 | Lighthouse Lab in Alderley Park | Wellcome Sanger Institute for the COVID-19 Genomics UK (COG-UK) Consortium | Jacquelyn Wynn, Mairead Hyland, The Lighthouse Lab in Alderley Park and Alex Alderton, Roberto Amato, Jeffrey Barrett, Sonia Goncalves, Ewan Harrison, David K. Jackson, Ian Johnston, Dominic Kwiatkowski, Cordelia Langford, John Sillitoe on behalf of the Wellcome Sanger Institute COVID-19 Surveillance Team |
| EPI_ISL_2846047 | Lighthouse Lab in Milton Keynes | Wellcome Sanger Institute for the COVID-19 Genomics UK (COG-UK) Consortium | The Lighthouse Lab in Milton Keynes and Alex Alderton, Roberto Amato, Jeffrey Barrett, Sonia Goncalves, Ewan Harrison, David K. Jackson, Ian Johnston, Dominic Kwiatkowski, Cordelia Langford, John Sillitoe on behalf of the Wellcome Sanger Institute COVID-19 Surveillance Team |
| EPI_ISL_2846048 | Lighthouse Laboratory Plymouth | Wellcome Sanger Institute for the COVID-19 Genomics UK (COG-UK) Consortium | Lighthouse Laboratory Plymouth and Alex Alderton, Roberto Amato, Jeffrey Barrett, Sonia Goncalves, Ewan Harrison, David K. Jackson, Ian Johnston, Dominic Kwiatkowski, Cordelia Langford, John Sillitoe on behalf of the Wellcome Sanger Institute COVID-19 Surveillance Team |
| EPI_ISL_2846049 | Lighthouse Lab in Alderley Park | Wellcome Sanger Institute for the COVID-19 Genomics UK (COG-UK) Consortium | Jacquelyn Wynn, Mairead Hyland, The Lighthouse Lab in Alderley Park and Alex Alderton, Roberto Amato, Jeffrey Barrett, Sonia Goncalves, Ewan Harrison, David K. Jackson, Ian Johnston, Dominic Kwiatkowski, Cordelia Langford, John Sillitoe on behalf of the Wellcome Sanger Institute COVID-19 Surveillance Team |
| EPI_ISL_2846050, EPI_ISL_2846051 | Lighthouse Lab in Milton Keynes | Wellcome Sanger Institute for the COVID-19 Genomics UK (COG-UK) Consortium | The Lighthouse Lab in Milton Keynes and Alex Alderton, Roberto Amato, Jeffrey Barrett, Sonia Goncalves, Ewan Harrison, David K. Jackson, Ian Johnston, Dominic Kwiatkowski, Cordelia Langford, John Sillitoe on behalf of the Wellcome Sanger Institute COVID-19 Surveillance Team |
| EPI_ISL_2846052 | Lighthouse Lab in Alderley Park | Wellcome Sanger Institute for the COVID-19 Genomics UK (COG-UK) Consortium | Jacquelyn Wynn, Mairead Hyland, The Lighthouse Lab in Alderley Park and Alex Alderton, Roberto Amato, Jeffrey Barrett, Sonia Goncalves, Ewan Harrison, David K. Jackson, Ian Johnston, Dominic Kwiatkowski, Cordelia Langford, John Sillitoe on behalf of the Wellcome Sanger Institute COVID-19 Surveillance Team |
| EPI_ISL_2846053 | Lighthouse Lab in Milton Keynes | Wellcome Sanger Institute for the COVID-19 Genomics UK (COG-UK) Consortium | The Lighthouse Lab in Milton Keynes and Alex Alderton, Roberto Amato, Jeffrey Barrett, Sonia Goncalves, Ewan Harrison, David K. Jackson, Ian Johnston, Dominic Kwiatkowski, Cordelia Langford, John Sillitoe on behalf of the Wellcome Sanger Institute COVID-19 Surveillance Team |
| EPI_ISL_2846054 | Lighthouse Lab in Alderley Park | Wellcome Sanger Institute for the COVID-19 Genomics UK (COG-UK) Consortium | Jacquelyn Wynn, Mairead Hyland, The Lighthouse Lab in Alderley Park and Alex Alderton, Roberto Amato, Jeffrey Barrett, Sonia Goncalves, Ewan Harrison, David K. Jackson, Ian Johnston, Dominic Kwiatkowski, Cordelia Langford, John Sillitoe on behalf of the Wellcome Sanger Institute COVID-19 Surveillance Team |
| EPI_ISL_2846055, EPI_ISL_2846056 | Lighthouse Lab in Milton Keynes | Wellcome Sanger Institute for the COVID-19 Genomics UK (COG-UK) Consortium | The Lighthouse Lab in Milton Keynes and Alex Alderton, Roberto Amato, Jeffrey Barrett, Sonia Goncalves, Ewan Harrison, David K. Jackson, Ian Johnston, Dominic Kwiatkowski, Cordelia Langford, John Sillitoe on behalf of the Wellcome Sanger Institute COVID-19 Surveillance Team |
| EPI_ISL_2846057, EPI_ISL_2846058, EPI_ISL_2846059, EPI_ISL_2846060, EPI_ISL_2846061, EPI_ISL_2846062, EPI_ISL_2846063 | Lighthouse Lab in Alderley Park | Wellcome Sanger Institute for the COVID-19 Genomics UK (COG-UK) Consortium | Jacquelyn Wynn, Mairead Hyland, The Lighthouse Lab in Alderley Park and Alex Alderton, Roberto Amato, Jeffrey Barrett, Sonia Goncalves, Ewan Harrison, David K. Jackson, Ian Johnston, Dominic Kwiatkowski, Cordelia Langford, John Sillitoe on behalf of the Wellcome Sanger Institute COVID-19 Surveillance Team |
| EPI_ISL_2846064 | Lighthouse Lab in Milton Keynes | Wellcome Sanger Institute for the COVID-19 Genomics UK (COG-UK) Consortium | The Lighthouse Lab in Milton Keynes and Alex Alderton, Roberto Amato, Jeffrey Barrett, Sonia Goncalves, Ewan Harrison, David K. Jackson, Ian Johnston, Dominic Kwiatkowski, Cordelia Langford, John Sillitoe on behalf of the Wellcome Sanger Institute COVID-19 Surveillance Team |
| EPI_ISL_2846065 | Lighthouse Lab in Alderley Park | Wellcome Sanger Institute for the COVID-19 Genomics UK (COG-UK) Consortium | Jacquelyn Wynn, Mairead Hyland, The Lighthouse Lab in Alderley Park and Alex Alderton, Roberto Amato, Jeffrey Barrett, Sonia Goncalves, Ewan Harrison, David K. Jackson, Ian Johnston, Dominic Kwiatkowski, Cordelia Langford, John Sillitoe on behalf of the Wellcome Sanger Institute COVID-19 Surveillance Team |
| EPI_ISL_2846066, EPI_ISL_2846067, EPI_ISL_2846068, EPI_ISL_2846069, EPI_ISL_2846070 | Lighthouse Laboratory Plymouth | Wellcome Sanger Institute for the COVID-19 Genomics UK (COG-UK) Consortium | Lighthouse Laboratory Plymouth and Alex Alderton, Roberto Amato, Jeffrey Barrett, Sonia Goncalves, Ewan Harrison, David K. Jackson, Ian Johnston, Dominic Kwiatkowski, Cordelia Langford, John Sillitoe on behalf of the Wellcome Sanger Institute COVID-19 Surveillance Team |
| EPI_ISL_2846071 | Lighthouse Lab in Milton Keynes | Wellcome Sanger Institute for the COVID-19 Genomics UK (COG-UK) Consortium | The Lighthouse Lab in Milton Keynes and Alex Alderton, Roberto Amato, Jeffrey Barrett, Sonia Goncalves, Ewan Harrison, David K. Jackson, Ian Johnston, Dominic Kwiatkowski, Cordelia Langford, John Sillitoe on behalf of the Wellcome Sanger Institute COVID-19 Surveillance Team |
| EPI_ISL_2846072, EPI_ISL_2846073 | Lighthouse Lab in Alderley Park | Wellcome Sanger Institute for the COVID-19 Genomics UK (COG-UK) Consortium | Jacquelyn Wynn, Mairead Hyland, The Lighthouse Lab in Alderley Park and Alex Alderton, Roberto Amato, Jeffrey Barrett, Sonia Goncalves, Ewan Harrison, David K. Jackson, Ian Johnston, Dominic Kwiatkowski, Cordelia Langford, John Sillitoe on behalf of the Wellcome Sanger Institute COVID-19 Surveillance Team |
| EPI_ISL_2846074 | Lighthouse Laboratory Plymouth | Wellcome Sanger Institute for the COVID-19 Genomics UK (COG-UK) Consortium | Lighthouse Laboratory Plymouth and Alex Alderton, Roberto Amato, Jeffrey Barrett, Sonia Goncalves, Ewan Harrison, David K. Jackson, Ian Johnston, Dominic Kwiatkowski, Cordelia Langford, John Sillitoe on behalf of the Wellcome Sanger Institute COVID-19 Surveillance Team |
| EPI_ISL_2846075, EPI_ISL_2846076 | Lighthouse Lab in Milton Keynes | Wellcome Sanger Institute for the COVID-19 Genomics UK (COG-UK) Consortium | The Lighthouse Lab in Milton Keynes and Alex Alderton, Roberto Amato, Jeffrey Barrett, Sonia Goncalves, Ewan Harrison, David K. Jackson, Ian Johnston, Dominic Kwiatkowski, Cordelia Langford, John Sillitoe on behalf of the Wellcome Sanger Institute COVID-19 Surveillance Team |
| EPI_ISL_2846077 | Lighthouse Lab in Alderley Park | Wellcome Sanger Institute for the COVID-19 Genomics UK (COG-UK) Consortium | Jacquelyn Wynn, Mairead Hyland, The Lighthouse Lab in Alderley Park and Alex Alderton, Roberto Amato, Jeffrey Barrett, Sonia Goncalves, Ewan Harrison, David K. Jackson, Ian Johnston, Dominic Kwiatkowski, Cordelia Langford, John Sillitoe on behalf of the Wellcome Sanger Institute COVID-19 Surveillance Team |
| EPI_ISL_2846078 | Lighthouse Laboratory Plymouth | Wellcome Sanger Institute for the COVID-19 Genomics UK (COG-UK) Consortium | Lighthouse Laboratory Plymouth and Alex Alderton, Roberto Amato, Jeffrey Barrett, Sonia Goncalves, Ewan Harrison, David K. Jackson, Ian Johnston, Dominic Kwiatkowski, Cordelia Langford, John Sillitoe on behalf of the Wellcome Sanger Institute COVID-19 Surveillance Team |
| EPI_ISL_2846079 | Lighthouse Lab in Alderley Park | Wellcome Sanger Institute for the COVID-19 Genomics UK (COG-UK) Consortium | Jacquelyn Wynn, Mairead Hyland, The Lighthouse Lab in Alderley Park and Alex Alderton, Roberto Amato, Jeffrey Barrett, Sonia Goncalves, Ewan Harrison, David K. Jackson, Ian Johnston, Dominic Kwiatkowski, Cordelia Langford, John Sillitoe on behalf of the Wellcome Sanger Institute COVID-19 Surveillance Team |
| EPI_ISL_2846080 | Lighthouse Laboratory Plymouth | Wellcome Sanger Institute for the COVID-19 Genomics UK (COG-UK) Consortium | Lighthouse Laboratory Plymouth and Alex Alderton, Roberto Amato, Jeffrey Barrett, Sonia Goncalves, Ewan Harrison, David K. Jackson, Ian Johnston, Dominic Kwiatkowski, Cordelia Langford, John Sillitoe on behalf of the Wellcome Sanger Institute COVID-19 Surveillance Team |
| EPI_ISL_2846081 | Lighthouse Lab in Milton Keynes | Wellcome Sanger Institute for the COVID-19 Genomics UK (COG-UK) Consortium | The Lighthouse Lab in Milton Keynes and Alex Alderton, Roberto Amato, Jeffrey Barrett, Sonia Goncalves, Ewan Harrison, David K. Jackson, Ian Johnston, Dominic Kwiatkowski, Cordelia Langford, John Sillitoe on behalf of the Wellcome Sanger Institute COVID-19 Surveillance Team |
| EPI_ISL_2846082, EPI_ISL_2846083 | Lighthouse Lab in Alderley Park | Wellcome Sanger Institute for the COVID-19 Genomics UK (COG-UK) Consortium | Jacquelyn Wynn, Mairead Hyland, The Lighthouse Lab in Alderley Park and Alex Alderton, Roberto Amato, Jeffrey Barrett, Sonia Goncalves, Ewan Harrison, David K. Jackson, Ian Johnston, Dominic Kwiatkowski, Cordelia Langford, John Sillitoe on behalf of the Wellcome Sanger Institute COVID-19 Surveillance Team |
| EPI_ISL_2846084 | Lighthouse Lab in Milton Keynes | Wellcome Sanger Institute for the COVID-19 Genomics UK (COG-UK) Consortium | The Lighthouse Lab in Milton Keynes and Alex Alderton, Roberto Amato, Jeffrey Barrett, Sonia Goncalves, Ewan Harrison, David K. Jackson, Ian Johnston, Dominic Kwiatkowski, Cordelia Langford, John Sillitoe on behalf of the Wellcome Sanger Institute COVID-19 Surveillance Team |
| EPI_ISL_2846085 | Lighthouse Lab in Alderley Park | Wellcome Sanger Institute for the COVID-19 Genomics UK (COG-UK) | Jacquelyn Wynn, Mairead Hyland, The Lighthouse Lab in Alderley Park and Alex Alderton, Roberto Amato, Jeffrey Barrett, Sonia Goncalves, Ewan Harrison, David K. Jackson, Ian Johnston, Dominic Kwiatkowski, Cordelia Langford, John Sillitoe on |

| | | | |
|---|---|---|---|
| | | Consortium | behalf of the Wellcome Sanger Institute COVID-19 Surveillance Team |
| EPI_ISL_2846086 | Lighthouse Lab in Milton Keynes | Wellcome Sanger Institute for the COVID-19 Genomics UK (COG-UK) Consortium | The Lighthouse Lab in Milton Keynes and Alex Alderton, Roberto Amato, Jeffrey Barrett, Sonia Goncalves, Ewan Harrison, David K. Jackson, Ian Johnston, Dominic Kwiatkowski, Cordelia Langford, John Sillitoe on behalf of the Wellcome Sanger Institute COVID-19 Surveillance Team |
| EPI_ISL_2846087, EPI_ISL_2846088 | Lighthouse Lab in Alderley Park | Wellcome Sanger Institute for the COVID-19 Genomics UK (COG-UK) Consortium | Jacquelyn Wynn, Mairead Hyland, The Lighthouse Lab in Alderley Park and Alex Alderton, Roberto Amato, Jeffrey Barrett, Sonia Goncalves, Ewan Harrison, David K. Jackson, Ian Johnston, Dominic Kwiatkowski, Cordelia Langford, John Sillitoe on behalf of the Wellcome Sanger Institute COVID-19 Surveillance Team |
| EPI_ISL_2846089 | Lighthouse Laboratory Plymouth | Wellcome Sanger Institute for the COVID-19 Genomics UK (COG-UK) Consortium | Lighthouse Laboratory Plymouth and Alex Alderton, Roberto Amato, Jeffrey Barrett, Sonia Goncalves, Ewan Harrison, David K. Jackson, Ian Johnston, Dominic Kwiatkowski, Cordelia Langford, John Sillitoe on behalf of the Wellcome Sanger Institute COVID-19 Surveillance Team |
| EPI_ISL_2846090, EPI_ISL_2846091, EPI_ISL_2846092, EPI_ISL_2846093, EPI_ISL_2846094, EPI_ISL_2846095 | Lighthouse Lab in Alderley Park | Wellcome Sanger Institute for the COVID-19 Genomics UK (COG-UK) Consortium | Jacquelyn Wynn, Mairead Hyland, The Lighthouse Lab in Alderley Park and Alex Alderton, Roberto Amato, Jeffrey Barrett, Sonia Goncalves, Ewan Harrison, David K. Jackson, Ian Johnston, Dominic Kwiatkowski, Cordelia Langford, John Sillitoe on behalf of the Wellcome Sanger Institute COVID-19 Surveillance Team |
| EPI_ISL_2846096 | Lighthouse Lab in Milton Keynes | Wellcome Sanger Institute for the COVID-19 Genomics UK (COG-UK) Consortium | The Lighthouse Lab in Milton Keynes and Alex Alderton, Roberto Amato, Jeffrey Barrett, Sonia Goncalves, Ewan Harrison, David K. Jackson, Ian Johnston, Dominic Kwiatkowski, Cordelia Langford, John Sillitoe on behalf of the Wellcome Sanger Institute COVID-19 Surveillance Team |
| EPI_ISL_2846097 | Lighthouse Laboratory Plymouth | Wellcome Sanger Institute for the COVID-19 Genomics UK (COG-UK) Consortium | Lighthouse Laboratory Plymouth and Alex Alderton, Roberto Amato, Jeffrey Barrett, Sonia Goncalves, Ewan Harrison, David K. Jackson, Ian Johnston, Dominic Kwiatkowski, Cordelia Langford, John Sillitoe on behalf of the Wellcome Sanger Institute COVID-19 Surveillance Team |
| EPI_ISL_2846098, EPI_ISL_2846099 | Lighthouse Lab in Alderley Park | Wellcome Sanger Institute for the COVID-19 Genomics UK (COG-UK) Consortium | Jacquelyn Wynn, Mairead Hyland, The Lighthouse Lab in Alderley Park and Alex Alderton, Roberto Amato, Jeffrey Barrett, Sonia Goncalves, Ewan Harrison, David K. Jackson, Ian Johnston, Dominic Kwiatkowski, Cordelia Langford, John Sillitoe on behalf of the Wellcome Sanger Institute COVID-19 Surveillance Team |
| EPI_ISL_2846100 | Lighthouse Laboratory Plymouth | Wellcome Sanger Institute for the COVID-19 Genomics UK (COG-UK) Consortium | Lighthouse Laboratory Plymouth and Alex Alderton, Roberto Amato, Jeffrey Barrett, Sonia Goncalves, Ewan Harrison, David K. Jackson, Ian Johnston, Dominic Kwiatkowski, Cordelia Langford, John Sillitoe on behalf of the Wellcome Sanger Institute COVID-19 Surveillance Team |
| EPI_ISL_2846101, EPI_ISL_2846102 | Lighthouse Lab in Alderley Park | Wellcome Sanger Institute for the COVID-19 Genomics UK (COG-UK) Consortium | Jacquelyn Wynn, Mairead Hyland, The Lighthouse Lab in Alderley Park and Alex Alderton, Roberto Amato, Jeffrey Barrett, Sonia Goncalves, Ewan Harrison, David K. Jackson, Ian Johnston, Dominic Kwiatkowski, Cordelia Langford, John Sillitoe on behalf of the Wellcome Sanger Institute COVID-19 Surveillance Team |
| EPI_ISL_2846103 | Lighthouse Laboratory Plymouth | Wellcome Sanger Institute for the COVID-19 Genomics UK (COG-UK) Consortium | Lighthouse Laboratory Plymouth and Alex Alderton, Roberto Amato, Jeffrey Barrett, Sonia Goncalves, Ewan Harrison, David K. Jackson, Ian Johnston, Dominic Kwiatkowski, Cordelia Langford, John Sillitoe on behalf of the Wellcome Sanger Institute COVID-19 Surveillance Team |
| EPI_ISL_2846104 | Lighthouse Lab in Milton Keynes | Wellcome Sanger Institute for the COVID-19 Genomics UK (COG-UK) Consortium | The Lighthouse Lab in Milton Keynes and Alex Alderton, Roberto Amato, Jeffrey Barrett, Sonia Goncalves, Ewan Harrison, David K. Jackson, Ian Johnston, Dominic Kwiatkowski, Cordelia Langford, John Sillitoe on behalf of the Wellcome Sanger Institute COVID-19 Surveillance Team |
| EPI_ISL_2846105, EPI_ISL_2846106 | Lighthouse Laboratory Plymouth | Wellcome Sanger Institute for the COVID-19 Genomics UK (COG-UK) Consortium | Lighthouse Laboratory Plymouth and Alex Alderton, Roberto Amato, Jeffrey Barrett, Sonia Goncalves, Ewan Harrison, David K. Jackson, Ian Johnston, Dominic Kwiatkowski, Cordelia Langford, John Sillitoe on behalf of the Wellcome Sanger Institute COVID-19 Surveillance Team |
| EPI_ISL_2846107 | Lighthouse Lab in Alderley Park | Wellcome Sanger Institute for the COVID-19 Genomics UK (COG-UK) Consortium | Jacquelyn Wynn, Mairead Hyland, The Lighthouse Lab in Alderley Park and Alex Alderton, Roberto Amato, Jeffrey Barrett, Sonia Goncalves, Ewan Harrison, David K. Jackson, Ian Johnston, Dominic Kwiatkowski, Cordelia Langford, John Sillitoe on behalf of the Wellcome Sanger Institute COVID-19 Surveillance Team |
| EPI_ISL_2846108 | Lighthouse Lab in Milton Keynes | Wellcome Sanger Institute for the COVID-19 Genomics UK (COG-UK) Consortium | The Lighthouse Lab in Milton Keynes and Alex Alderton, Roberto Amato, Jeffrey Barrett, Sonia Goncalves, Ewan Harrison, David K. Jackson, Ian Johnston, Dominic Kwiatkowski, Cordelia Langford, John Sillitoe on behalf of the Wellcome Sanger Institute COVID-19 Surveillance Team |
| EPI_ISL_2846109 | Lighthouse Lab in Alderley Park | Wellcome Sanger Institute for the COVID-19 Genomics UK (COG-UK) Consortium | Jacquelyn Wynn, Mairead Hyland, The Lighthouse Lab in Alderley Park and Alex Alderton, Roberto Amato, Jeffrey Barrett, Sonia Goncalves, Ewan Harrison, David K. Jackson, Ian Johnston, Dominic Kwiatkowski, Cordelia Langford, John Sillitoe on behalf of the Wellcome Sanger Institute COVID-19 Surveillance Team |
| EPI_ISL_2846110 | Lighthouse Laboratory Plymouth | Wellcome Sanger Institute for the COVID-19 Genomics UK (COG-UK) Consortium | Lighthouse Laboratory Plymouth and Alex Alderton, Roberto Amato, Jeffrey Barrett, Sonia Goncalves, Ewan Harrison, David K. Jackson, Ian Johnston, Dominic Kwiatkowski, Cordelia Langford, John Sillitoe on behalf of the Wellcome Sanger Institute COVID-19 Surveillance Team |
| EPI_ISL_2846111 | Lighthouse Lab in Alderley Park | Wellcome Sanger Institute for the COVID-19 Genomics UK (COG-UK) Consortium | Jacquelyn Wynn, Mairead Hyland, The Lighthouse Lab in Alderley Park and Alex Alderton, Roberto Amato, Jeffrey Barrett, Sonia Goncalves, Ewan Harrison, David K. Jackson, Ian Johnston, Dominic Kwiatkowski, Cordelia Langford, John Sillitoe on behalf of the Wellcome Sanger Institute COVID-19 Surveillance Team |
| EPI_ISL_2846112 | Lighthouse Laboratory Plymouth | Wellcome Sanger Institute for the COVID-19 Genomics UK (COG-UK) Consortium | Lighthouse Laboratory Plymouth and Alex Alderton, Roberto Amato, Jeffrey Barrett, Sonia Goncalves, Ewan Harrison, David K. Jackson, Ian Johnston, Dominic Kwiatkowski, Cordelia Langford, John Sillitoe on behalf of the Wellcome Sanger Institute COVID-19 Surveillance Team |
| EPI_ISL_2846113, EPI_ISL_2846114 | Lighthouse Lab in Milton Keynes | Wellcome Sanger Institute for the COVID-19 Genomics UK (COG-UK) Consortium | The Lighthouse Lab in Milton Keynes and Alex Alderton, Roberto Amato, Jeffrey Barrett, Sonia Goncalves, Ewan Harrison, David K. Jackson, Ian Johnston, Dominic Kwiatkowski, Cordelia Langford, John Sillitoe on behalf of the Wellcome Sanger Institute COVID-19 Surveillance Team |
| EPI_ISL_2846115 | Lighthouse Laboratory Plymouth | Wellcome Sanger Institute for the COVID-19 Genomics UK (COG-UK) Consortium | Lighthouse Laboratory Plymouth and Alex Alderton, Roberto Amato, Jeffrey Barrett, Sonia Goncalves, Ewan Harrison, David K. Jackson, Ian Johnston, Dominic Kwiatkowski, Cordelia Langford, John Sillitoe on behalf of the Wellcome Sanger Institute COVID-19 Surveillance Team |
| EPI_ISL_2846116, EPI_ISL_2846117 | Lighthouse Lab in Alderley Park | Wellcome Sanger Institute for the COVID-19 Genomics UK (COG-UK) Consortium | Jacquelyn Wynn, Mairead Hyland, The Lighthouse Lab in Alderley Park and Alex Alderton, Roberto Amato, Jeffrey Barrett, Sonia Goncalves, Ewan Harrison, David K. Jackson, Ian Johnston, Dominic Kwiatkowski, Cordelia Langford, John Sillitoe on behalf of the Wellcome Sanger Institute COVID-19 Surveillance Team |
| EPI_ISL_2846118 | Lighthouse Lab in Milton Keynes | Wellcome Sanger Institute for the COVID-19 Genomics UK (COG-UK) Consortium | The Lighthouse Lab in Milton Keynes and Alex Alderton, Roberto Amato, Jeffrey Barrett, Sonia Goncalves, Ewan Harrison, David K. Jackson, Ian Johnston, Dominic Kwiatkowski, Cordelia Langford, John Sillitoe on behalf of the Wellcome Sanger Institute COVID-19 Surveillance Team |
| EPI_ISL_2846119, EPI_ISL_2846120, EPI_ISL_2846121, EPI_ISL_2846122, EPI_ISL_2846123 | Lighthouse Lab in Alderley Park | Wellcome Sanger Institute for the COVID-19 Genomics UK (COG-UK) Consortium | Jacquelyn Wynn, Mairead Hyland, The Lighthouse Lab in Alderley Park and Alex Alderton, Roberto Amato, Jeffrey Barrett, Sonia Goncalves, Ewan Harrison, David K. Jackson, Ian Johnston, Dominic Kwiatkowski, Cordelia Langford, John Sillitoe on behalf of the Wellcome Sanger Institute COVID-19 Surveillance Team |
| EPI_ISL_2846124 | Lighthouse Lab in Milton Keynes | Wellcome Sanger Institute for the COVID-19 Genomics UK (COG-UK) Consortium | The Lighthouse Lab in Milton Keynes and Alex Alderton, Roberto Amato, Jeffrey Barrett, Sonia Goncalves, Ewan Harrison, David K. Jackson, Ian Johnston, Dominic Kwiatkowski, Cordelia Langford, John Sillitoe on behalf of the Wellcome Sanger Institute COVID-19 Surveillance Team |
| EPI_ISL_2846125, EPI_ISL_2846126, EPI_ISL_2846127, EPI_ISL_2846128, EPI_ISL_2846129 | Lighthouse Lab in Alderley Park | Wellcome Sanger Institute for the COVID-19 Genomics UK (COG-UK) Consortium | Jacquelyn Wynn, Mairead Hyland, The Lighthouse Lab in Alderley Park and Alex Alderton, Roberto Amato, Jeffrey Barrett, Sonia Goncalves, Ewan Harrison, David K. Jackson, Ian Johnston, Dominic Kwiatkowski, Cordelia Langford, John Sillitoe on behalf of the Wellcome Sanger Institute COVID-19 Surveillance Team |
| EPI_ISL_2846130 | Lighthouse Lab in Milton Keynes | Wellcome Sanger Institute for the COVID-19 Genomics UK (COG-UK) Consortium | The Lighthouse Lab in Milton Keynes and Alex Alderton, Roberto Amato, Jeffrey Barrett, Sonia Goncalves, Ewan Harrison, David K. Jackson, Ian Johnston, Dominic Kwiatkowski, Cordelia Langford, John Sillitoe on behalf of the Wellcome Sanger Institute COVID-19 Surveillance Team |
| EPI_ISL_2846131, EPI_ISL_2846132, EPI_ISL_2846133 | Lighthouse Lab in Alderley Park | Wellcome Sanger Institute for the COVID-19 Genomics UK (COG-UK) Consortium | Jacquelyn Wynn, Mairead Hyland, The Lighthouse Lab in Alderley Park and Alex Alderton, Roberto Amato, Jeffrey Barrett, Sonia Goncalves, Ewan Harrison, David K. Jackson, Ian Johnston, Dominic Kwiatkowski, Cordelia Langford, John Sillitoe on behalf of the Wellcome Sanger Institute COVID-19 Surveillance Team |
| EPI_ISL_2846134 | Lighthouse Laboratory Plymouth | Wellcome Sanger Institute for the COVID-19 Genomics UK (COG-UK) Consortium | Lighthouse Laboratory Plymouth and Alex Alderton, Roberto Amato, Jeffrey Barrett, Sonia Goncalves, Ewan Harrison, David K. Jackson, Ian Johnston, Dominic Kwiatkowski, Cordelia Langford, John Sillitoe on behalf of the Wellcome Sanger Institute COVID-19 Surveillance Team |
| EPI_ISL_2846135, EPI_ISL_2846136 | Lighthouse Lab in Milton Keynes | Wellcome Sanger Institute for the COVID-19 Genomics UK (COG-UK) Consortium | The Lighthouse Lab in Milton Keynes and Alex Alderton, Roberto Amato, Jeffrey Barrett, Sonia Goncalves, Ewan Harrison, David K. Jackson, Ian Johnston, Dominic Kwiatkowski, Cordelia Langford, John Sillitoe on behalf of the Wellcome Sanger Institute COVID-19 Surveillance Team |
| EPI_ISL_2846137, EPI_ISL_2846138 | Lighthouse Lab in Alderley Park | Wellcome Sanger Institute for the COVID-19 Genomics UK (COG-UK) Consortium | Jacquelyn Wynn, Mairead Hyland, The Lighthouse Lab in Alderley Park and Alex Alderton, Roberto Amato, Jeffrey Barrett, Sonia Goncalves, Ewan Harrison, David K. Jackson, Ian Johnston, Dominic Kwiatkowski, Cordelia Langford, John Sillitoe on behalf of the Wellcome Sanger Institute COVID-19 Surveillance Team |
| EPI_ISL_2846139 | Lighthouse Lab in Milton Keynes | Wellcome Sanger Institute for the COVID-19 Genomics UK (COG-UK) Consortium | The Lighthouse Lab in Milton Keynes and Alex Alderton, Roberto Amato, Jeffrey Barrett, Sonia Goncalves, Ewan Harrison, David K. Jackson, Ian Johnston, Dominic Kwiatkowski, Cordelia Langford, John Sillitoe on behalf of the Wellcome Sanger Institute COVID-19 Surveillance Team |
| EPI_ISL_2846140 | Lighthouse Lab in Alderley Park | Wellcome Sanger Institute for the COVID-19 Genomics UK (COG-UK) Consortium | Jacquelyn Wynn, Mairead Hyland, The Lighthouse Lab in Alderley Park and Alex Alderton, Roberto Amato, Jeffrey Barrett, Sonia Goncalves, Ewan Harrison, David K. Jackson, Ian Johnston, Dominic Kwiatkowski, Cordelia Langford, John Sillitoe on behalf of the Wellcome Sanger Institute COVID-19 Surveillance Team |
| EPI_ISL_2846141 | Lighthouse Laboratory Plymouth | Wellcome Sanger Institute for the COVID-19 Genomics UK (COG-UK) Consortium | Lighthouse Laboratory Plymouth and Alex Alderton, Roberto Amato, Jeffrey Barrett, Sonia Goncalves, Ewan Harrison, David K. Jackson, Ian Johnston, Dominic Kwiatkowski, Cordelia Langford, John Sillitoe on behalf of the Wellcome Sanger Institute COVID-19 Surveillance Team |
| EPI_ISL_2846142, EPI_ISL_2846143 | Lighthouse Lab in Milton Keynes | Wellcome Sanger Institute for the COVID-19 Genomics UK (COG-UK) Consortium | The Lighthouse Lab in Milton Keynes and Alex Alderton, Roberto Amato, Jeffrey Barrett, Sonia Goncalves, Ewan Harrison, David K. Jackson, Ian Johnston, Dominic Kwiatkowski, Cordelia Langford, John Sillitoe on behalf of the Wellcome Sanger Institute COVID-19 Surveillance Team |
| EPI_ISL_2846144 | Lighthouse Lab in Alderley Park | Wellcome Sanger Institute for the COVID-19 Genomics UK (COG-UK) Consortium | Jacquelyn Wynn, Mairead Hyland, The Lighthouse Lab in Alderley Park and Alex Alderton, Roberto Amato, Jeffrey Barrett, Sonia Goncalves, Ewan Harrison, David K. Jackson, Ian Johnston, Dominic Kwiatkowski, Cordelia Langford, John Sillitoe on behalf of the Wellcome Sanger Institute COVID-19 Surveillance Team |
| EPI_ISL_2846145 | Lighthouse Laboratory Plymouth | Wellcome Sanger Institute for the COVID-19 Genomics UK (COG-UK) Consortium | Lighthouse Laboratory Plymouth and Alex Alderton, Roberto Amato, Jeffrey Barrett, Sonia Goncalves, Ewan Harrison, David K. Jackson, Ian Johnston, Dominic Kwiatkowski, Cordelia Langford, John Sillitoe on behalf of the Wellcome Sanger Institute COVID-19 Surveillance Team |
| EPI_ISL_2846146 | Lighthouse Lab in Alderley Park | Wellcome Sanger Institute for the COVID-19 Genomics UK (COG-UK) Consortium | Jacquelyn Wynn, Mairead Hyland, The Lighthouse Lab in Alderley Park and Alex Alderton, Roberto Amato, Jeffrey Barrett, Sonia Goncalves, Ewan Harrison, David K. Jackson, Ian Johnston, Dominic Kwiatkowski, Cordelia Langford, John Sillitoe on behalf of the Wellcome Sanger Institute COVID-19 Surveillance Team |
| EPI_ISL_2846147 | Lighthouse Laboratory Plymouth | Wellcome Sanger Institute for the COVID-19 Genomics UK (COG-UK) Consortium | Lighthouse Laboratory Plymouth and Alex Alderton, Roberto Amato, Jeffrey Barrett, Sonia Goncalves, Ewan Harrison, David K. Jackson, Ian Johnston, Dominic Kwiatkowski, Cordelia Langford, John Sillitoe on behalf of the Wellcome Sanger Institute COVID-19 Surveillance Team |
| EPI_ISL_2846148 | Lighthouse Lab in Milton Keynes | Wellcome Sanger Institute for the COVID-19 Genomics UK (COG-UK) Consortium | The Lighthouse Lab in Milton Keynes and Alex Alderton, Roberto Amato, Jeffrey Barrett, Sonia Goncalves, Ewan Harrison, David K. Jackson, Ian Johnston, Dominic Kwiatkowski, Cordelia Langford, John Sillitoe on behalf of the Wellcome Sanger Institute COVID-19 Surveillance Team |
| EPI_ISL_2846149 | Lighthouse Lab in Alderley Park | Wellcome Sanger Institute for the COVID-19 Genomics UK (COG-UK) Consortium | Jacquelyn Wynn, Mairead Hyland, The Lighthouse Lab in Alderley Park and Alex Alderton, Roberto Amato, Jeffrey Barrett, Sonia Goncalves, Ewan Harrison, David K. Jackson, Ian Johnston, Dominic Kwiatkowski, Cordelia Langford, John Sillitoe on behalf of the Wellcome Sanger Institute COVID-19 Surveillance Team |

| | | | |
|---|---|---|---|
| EPI_ISL_2846150 | Lighthouse Laboratory Plymouth | Wellcome Sanger Institute for the COVID-19 Genomics UK (COG-UK) Consortium | Lighthouse Laboratory Plymouth and Alex Alderton, Roberto Amato, Jeffrey Barrett, Sonia Goncalves, Ewan Harrison, David K. Jackson, Ian Johnston, Dominic Kwiatkowski, Cordelia Langford, John Sillitoe on behalf of the Wellcome Sanger Institute COVID-19 Surveillance Team |
| EPI_ISL_2846151 | Lighthouse Lab in Alderley Park | Wellcome Sanger Institute for the COVID-19 Genomics UK (COG-UK) Consortium | Jacquelyn Wynn, Mairead Hyland, The Lighthouse Lab in Alderley Park and Alex Alderton, Roberto Amato, Jeffrey Barrett, Sonia Goncalves, Ewan Harrison, David K. Jackson, Ian Johnston, Dominic Kwiatkowski, Cordelia Langford, John Sillitoe on behalf of the Wellcome Sanger Institute COVID-19 Surveillance Team |
| EPI_ISL_2846152 | Lighthouse Laboratory Plymouth | Wellcome Sanger Institute for the COVID-19 Genomics UK (COG-UK) Consortium | Lighthouse Laboratory Plymouth and Alex Alderton, Roberto Amato, Jeffrey Barrett, Sonia Goncalves, Ewan Harrison, David K. Jackson, Ian Johnston, Dominic Kwiatkowski, Cordelia Langford, John Sillitoe on behalf of the Wellcome Sanger Institute COVID-19 Surveillance Team |
| EPI_ISL_2846153 | Lighthouse Lab in Alderley Park | Wellcome Sanger Institute for the COVID-19 Genomics UK (COG-UK) Consortium | Jacquelyn Wynn, Mairead Hyland, The Lighthouse Lab in Alderley Park and Alex Alderton, Roberto Amato, Jeffrey Barrett, Sonia Goncalves, Ewan Harrison, David K. Jackson, Ian Johnston, Dominic Kwiatkowski, Cordelia Langford, John Sillitoe on behalf of the Wellcome Sanger Institute COVID-19 Surveillance Team |
| EPI_ISL_2846154 | Lighthouse Laboratory Plymouth | Wellcome Sanger Institute for the COVID-19 Genomics UK (COG-UK) Consortium | Lighthouse Laboratory Plymouth and Alex Alderton, Roberto Amato, Jeffrey Barrett, Sonia Goncalves, Ewan Harrison, David K. Jackson, Ian Johnston, Dominic Kwiatkowski, Cordelia Langford, John Sillitoe on behalf of the Wellcome Sanger Institute COVID-19 Surveillance Team |
| EPI_ISL_2846155, EPI_ISL_2846156, EPI_ISL_2846157 | Lighthouse Lab in Alderley Park | Wellcome Sanger Institute for the COVID-19 Genomics UK (COG-UK) Consortium | Jacquelyn Wynn, Mairead Hyland, The Lighthouse Lab in Alderley Park and Alex Alderton, Roberto Amato, Jeffrey Barrett, Sonia Goncalves, Ewan Harrison, David K. Jackson, Ian Johnston, Dominic Kwiatkowski, Cordelia Langford, John Sillitoe on behalf of the Wellcome Sanger Institute COVID-19 Surveillance Team |
| EPI_ISL_2846158 | Lighthouse Laboratory Plymouth | Wellcome Sanger Institute for the COVID-19 Genomics UK (COG-UK) Consortium | Lighthouse Laboratory Plymouth and Alex Alderton, Roberto Amato, Jeffrey Barrett, Sonia Goncalves, Ewan Harrison, David K. Jackson, Ian Johnston, Dominic Kwiatkowski, Cordelia Langford, John Sillitoe on behalf of the Wellcome Sanger Institute COVID-19 Surveillance Team |
| EPI_ISL_2846159 | Lighthouse Lab in Milton Keynes | Wellcome Sanger Institute for the COVID-19 Genomics UK (COG-UK) Consortium | The Lighthouse Lab in Milton Keynes and Alex Alderton, Roberto Amato, Jeffrey Barrett, Sonia Goncalves, Ewan Harrison, David K. Jackson, Ian Johnston, Dominic Kwiatkowski, Cordelia Langford, John Sillitoe on behalf of the Wellcome Sanger Institute COVID-19 Surveillance Team |
| EPI_ISL_2846160, EPI_ISL_2846161 | Lighthouse Lab in Alderley Park | Wellcome Sanger Institute for the COVID-19 Genomics UK (COG-UK) Consortium | Jacquelyn Wynn, Mairead Hyland, The Lighthouse Lab in Alderley Park and Alex Alderton, Roberto Amato, Jeffrey Barrett, Sonia Goncalves, Ewan Harrison, David K. Jackson, Ian Johnston, Dominic Kwiatkowski, Cordelia Langford, John Sillitoe on behalf of the Wellcome Sanger Institute COVID-19 Surveillance Team |
| EPI_ISL_2846162, EPI_ISL_2846163 | Lighthouse Laboratory Plymouth | Wellcome Sanger Institute for the COVID-19 Genomics UK (COG-UK) Consortium | Lighthouse Laboratory Plymouth and Alex Alderton, Roberto Amato, Jeffrey Barrett, Sonia Goncalves, Ewan Harrison, David K. Jackson, Ian Johnston, Dominic Kwiatkowski, Cordelia Langford, John Sillitoe on behalf of the Wellcome Sanger Institute COVID-19 Surveillance Team |
| EPI_ISL_2846164 | Lighthouse Lab in Milton Keynes | Wellcome Sanger Institute for the COVID-19 Genomics UK (COG-UK) Consortium | The Lighthouse Lab in Milton Keynes and Alex Alderton, Roberto Amato, Jeffrey Barrett, Sonia Goncalves, Ewan Harrison, David K. Jackson, Ian Johnston, Dominic Kwiatkowski, Cordelia Langford, John Sillitoe on behalf of the Wellcome Sanger Institute COVID-19 Surveillance Team |
| EPI_ISL_2846165 | Lighthouse Lab in Alderley Park | Wellcome Sanger Institute for the COVID-19 Genomics UK (COG-UK) Consortium | Jacquelyn Wynn, Mairead Hyland, The Lighthouse Lab in Alderley Park and Alex Alderton, Roberto Amato, Jeffrey Barrett, Sonia Goncalves, Ewan Harrison, David K. Jackson, Ian Johnston, Dominic Kwiatkowski, Cordelia Langford, John Sillitoe on behalf of the Wellcome Sanger Institute COVID-19 Surveillance Team |
| EPI_ISL_2846166 | Lighthouse Lab in Milton Keynes | Wellcome Sanger Institute for the COVID-19 Genomics UK (COG-UK) Consortium | The Lighthouse Lab in Milton Keynes and Alex Alderton, Roberto Amato, Jeffrey Barrett, Sonia Goncalves, Ewan Harrison, David K. Jackson, Ian Johnston, Dominic Kwiatkowski, Cordelia Langford, John Sillitoe on behalf of the Wellcome Sanger Institute COVID-19 Surveillance Team |
| EPI_ISL_2846167 | Lighthouse Laboratory Plymouth | Wellcome Sanger Institute for the COVID-19 Genomics UK (COG-UK) Consortium | Lighthouse Laboratory Plymouth and Alex Alderton, Roberto Amato, Jeffrey Barrett, Sonia Goncalves, Ewan Harrison, David K. Jackson, Ian Johnston, Dominic Kwiatkowski, Cordelia Langford, John Sillitoe on behalf of the Wellcome Sanger Institute COVID-19 Surveillance Team |
| EPI_ISL_2846168, EPI_ISL_2846169 | Lighthouse Lab in Alderley Park | Wellcome Sanger Institute for the COVID-19 Genomics UK (COG-UK) Consortium | Jacquelyn Wynn, Mairead Hyland, The Lighthouse Lab in Alderley Park and Alex Alderton, Roberto Amato, Jeffrey Barrett, Sonia Goncalves, Ewan Harrison, David K. Jackson, Ian Johnston, Dominic Kwiatkowski, Cordelia Langford, John Sillitoe on behalf of the Wellcome Sanger Institute COVID-19 Surveillance Team |
| EPI_ISL_2846170, EPI_ISL_2846171, EPI_ISL_2846172 | Lighthouse Lab in Milton Keynes | Wellcome Sanger Institute for the COVID-19 Genomics UK (COG-UK) Consortium | The Lighthouse Lab in Milton Keynes and Alex Alderton, Roberto Amato, Jeffrey Barrett, Sonia Goncalves, Ewan Harrison, David K. Jackson, Ian Johnston, Dominic Kwiatkowski, Cordelia Langford, John Sillitoe on behalf of the Wellcome Sanger Institute COVID-19 Surveillance Team |
| EPI_ISL_2846173, EPI_ISL_2846174, EPI_ISL_2846175 | Lighthouse Lab in Alderley Park | Wellcome Sanger Institute for the COVID-19 Genomics UK (COG-UK) Consortium | Jacquelyn Wynn, Mairead Hyland, The Lighthouse Lab in Alderley Park and Alex Alderton, Roberto Amato, Jeffrey Barrett, Sonia Goncalves, Ewan Harrison, David K. Jackson, Ian Johnston, Dominic Kwiatkowski, Cordelia Langford, John Sillitoe on behalf of the Wellcome Sanger Institute COVID-19 Surveillance Team |
| EPI_ISL_2846176 | Lighthouse Lab in Milton Keynes | Wellcome Sanger Institute for the COVID-19 Genomics UK (COG-UK) Consortium | The Lighthouse Lab in Milton Keynes and Alex Alderton, Roberto Amato, Jeffrey Barrett, Sonia Goncalves, Ewan Harrison, David K. Jackson, Ian Johnston, Dominic Kwiatkowski, Cordelia Langford, John Sillitoe on behalf of the Wellcome Sanger Institute COVID-19 Surveillance Team |
| EPI_ISL_2846177 | Lighthouse Lab in Alderley Park | Wellcome Sanger Institute for the COVID-19 Genomics UK (COG-UK) Consortium | Jacquelyn Wynn, Mairead Hyland, The Lighthouse Lab in Alderley Park and Alex Alderton, Roberto Amato, Jeffrey Barrett, Sonia Goncalves, Ewan Harrison, David K. Jackson, Ian Johnston, Dominic Kwiatkowski, Cordelia Langford, John Sillitoe on behalf of the Wellcome Sanger Institute COVID-19 Surveillance Team |
| EPI_ISL_2846178, EPI_ISL_2846179 | Lighthouse Laboratory Plymouth | Wellcome Sanger Institute for the COVID-19 Genomics UK (COG-UK) Consortium | Lighthouse Laboratory Plymouth and Alex Alderton, Roberto Amato, Jeffrey Barrett, Sonia Goncalves, Ewan Harrison, David K. Jackson, Ian Johnston, Dominic Kwiatkowski, Cordelia Langford, John Sillitoe on behalf of the Wellcome Sanger Institute COVID-19 Surveillance Team |
| EPI_ISL_2846180 | Lighthouse Lab in Milton Keynes | Wellcome Sanger Institute for the COVID-19 Genomics UK (COG-UK) Consortium | The Lighthouse Lab in Milton Keynes and Alex Alderton, Roberto Amato, Jeffrey Barrett, Sonia Goncalves, Ewan Harrison, David K. Jackson, Ian Johnston, Dominic Kwiatkowski, Cordelia Langford, John Sillitoe on behalf of the Wellcome Sanger Institute COVID-19 Surveillance Team |
| EPI_ISL_2846181, EPI_ISL_2846182 | Lighthouse Lab in Alderley Park | Wellcome Sanger Institute for the COVID-19 Genomics UK (COG-UK) Consortium | Jacquelyn Wynn, Mairead Hyland, The Lighthouse Lab in Alderley Park and Alex Alderton, Roberto Amato, Jeffrey Barrett, Sonia Goncalves, Ewan Harrison, David K. Jackson, Ian Johnston, Dominic Kwiatkowski, Cordelia Langford, John Sillitoe on behalf of the Wellcome Sanger Institute COVID-19 Surveillance Team |
| EPI_ISL_2846183 | Lighthouse Laboratory Plymouth | Wellcome Sanger Institute for the COVID-19 Genomics UK (COG-UK) Consortium | Lighthouse Laboratory Plymouth and Alex Alderton, Roberto Amato, Jeffrey Barrett, Sonia Goncalves, Ewan Harrison, David K. Jackson, Ian Johnston, Dominic Kwiatkowski, Cordelia Langford, John Sillitoe on behalf of the Wellcome Sanger Institute COVID-19 Surveillance Team |
| EPI_ISL_2846184 | Lighthouse Lab in Alderley Park | Wellcome Sanger Institute for the COVID-19 Genomics UK (COG-UK) Consortium | Jacquelyn Wynn, Mairead Hyland, The Lighthouse Lab in Alderley Park and Alex Alderton, Roberto Amato, Jeffrey Barrett, Sonia Goncalves, Ewan Harrison, David K. Jackson, Ian Johnston, Dominic Kwiatkowski, Cordelia Langford, John Sillitoe on behalf of the Wellcome Sanger Institute COVID-19 Surveillance Team |
| EPI_ISL_2846185 | Lighthouse Laboratory Plymouth | Wellcome Sanger Institute for the COVID-19 Genomics UK (COG-UK) Consortium | Lighthouse Laboratory Plymouth and Alex Alderton, Roberto Amato, Jeffrey Barrett, Sonia Goncalves, Ewan Harrison, David K. Jackson, Ian Johnston, Dominic Kwiatkowski, Cordelia Langford, John Sillitoe on behalf of the Wellcome Sanger Institute COVID-19 Surveillance Team |
| EPI_ISL_2846186 | Lighthouse Lab in Milton Keynes | Wellcome Sanger Institute for the COVID-19 Genomics UK (COG-UK) Consortium | The Lighthouse Lab in Milton Keynes and Alex Alderton, Roberto Amato, Jeffrey Barrett, Sonia Goncalves, Ewan Harrison, David K. Jackson, Ian Johnston, Dominic Kwiatkowski, Cordelia Langford, John Sillitoe on behalf of the Wellcome Sanger Institute COVID-19 Surveillance Team |
| EPI_ISL_2846187, EPI_ISL_2846188 | Lighthouse Lab in Alderley Park | Wellcome Sanger Institute for the COVID-19 Genomics UK (COG-UK) Consortium | Jacquelyn Wynn, Mairead Hyland, The Lighthouse Lab in Alderley Park and Alex Alderton, Roberto Amato, Jeffrey Barrett, Sonia Goncalves, Ewan Harrison, David K. Jackson, Ian Johnston, Dominic Kwiatkowski, Cordelia Langford, John Sillitoe on behalf of the Wellcome Sanger Institute COVID-19 Surveillance Team |
| EPI_ISL_2846189 | Lighthouse Lab in Milton Keynes | Wellcome Sanger Institute for the COVID-19 Genomics UK (COG-UK) Consortium | The Lighthouse Lab in Milton Keynes and Alex Alderton, Roberto Amato, Jeffrey Barrett, Sonia Goncalves, Ewan Harrison, David K. Jackson, Ian Johnston, Dominic Kwiatkowski, Cordelia Langford, John Sillitoe on behalf of the Wellcome Sanger Institute COVID-19 Surveillance Team |
| EPI_ISL_2846190, EPI_ISL_2846191 | Lighthouse Laboratory Plymouth | Wellcome Sanger Institute for the COVID-19 Genomics UK (COG-UK) Consortium | Lighthouse Laboratory Plymouth and Alex Alderton, Roberto Amato, Jeffrey Barrett, Sonia Goncalves, Ewan Harrison, David K. Jackson, Ian Johnston, Dominic Kwiatkowski, Cordelia Langford, John Sillitoe on behalf of the Wellcome Sanger Institute COVID-19 Surveillance Team |
| EPI_ISL_2846192 | Lighthouse Lab in Alderley Park | Wellcome Sanger Institute for the COVID-19 Genomics UK (COG-UK) Consortium | Jacquelyn Wynn, Mairead Hyland, The Lighthouse Lab in Alderley Park and Alex Alderton, Roberto Amato, Jeffrey Barrett, Sonia Goncalves, Ewan Harrison, David K. Jackson, Ian Johnston, Dominic Kwiatkowski, Cordelia Langford, John Sillitoe on behalf of the Wellcome Sanger Institute COVID-19 Surveillance Team |
| EPI_ISL_2846193, EPI_ISL_2846194 | Lighthouse Laboratory Plymouth | Wellcome Sanger Institute for the COVID-19 Genomics UK (COG-UK) Consortium | Lighthouse Laboratory Plymouth and Alex Alderton, Roberto Amato, Jeffrey Barrett, Sonia Goncalves, Ewan Harrison, David K. Jackson, Ian Johnston, Dominic Kwiatkowski, Cordelia Langford, John Sillitoe on behalf of the Wellcome Sanger Institute COVID-19 Surveillance Team |
| EPI_ISL_2846195 | Lighthouse Lab in Alderley Park | Wellcome Sanger Institute for the COVID-19 Genomics UK (COG-UK) Consortium | Jacquelyn Wynn, Mairead Hyland, The Lighthouse Lab in Alderley Park and Alex Alderton, Roberto Amato, Jeffrey Barrett, Sonia Goncalves, Ewan Harrison, David K. Jackson, Ian Johnston, Dominic Kwiatkowski, Cordelia Langford, John Sillitoe on behalf of the Wellcome Sanger Institute COVID-19 Surveillance Team |
| EPI_ISL_2846196 | Lighthouse Lab in Milton Keynes | Wellcome Sanger Institute for the COVID-19 Genomics UK (COG-UK) Consortium | The Lighthouse Lab in Milton Keynes and Alex Alderton, Roberto Amato, Jeffrey Barrett, Sonia Goncalves, Ewan Harrison, David K. Jackson, Ian Johnston, Dominic Kwiatkowski, Cordelia Langford, John Sillitoe on behalf of the Wellcome Sanger Institute COVID-19 Surveillance Team |
| EPI_ISL_2846197 | Lighthouse Laboratory Plymouth | Wellcome Sanger Institute for the COVID-19 Genomics UK (COG-UK) Consortium | Lighthouse Laboratory Plymouth and Alex Alderton, Roberto Amato, Jeffrey Barrett, Sonia Goncalves, Ewan Harrison, David K. Jackson, Ian Johnston, Dominic Kwiatkowski, Cordelia Langford, John Sillitoe on behalf of the Wellcome Sanger Institute COVID-19 Surveillance Team |
| EPI_ISL_2846198 | Lighthouse Lab in Alderley Park | Wellcome Sanger Institute for the COVID-19 Genomics UK (COG-UK) Consortium | Jacquelyn Wynn, Mairead Hyland, The Lighthouse Lab in Alderley Park and Alex Alderton, Roberto Amato, Jeffrey Barrett, Sonia Goncalves, Ewan Harrison, David K. Jackson, Ian Johnston, Dominic Kwiatkowski, Cordelia Langford, John Sillitoe on behalf of the Wellcome Sanger Institute COVID-19 Surveillance Team |
| EPI_ISL_2846199 | Lighthouse Laboratory Plymouth | Wellcome Sanger Institute for the COVID-19 Genomics UK (COG-UK) Consortium | Lighthouse Laboratory Plymouth and Alex Alderton, Roberto Amato, Jeffrey Barrett, Sonia Goncalves, Ewan Harrison, David K. Jackson, Ian Johnston, Dominic Kwiatkowski, Cordelia Langford, John Sillitoe on behalf of the Wellcome Sanger Institute COVID-19 Surveillance Team |
| EPI_ISL_2846200, EPI_ISL_2846201, EPI_ISL_2846202 | Lighthouse Lab in Alderley Park | Wellcome Sanger Institute for the COVID-19 Genomics UK (COG-UK) Consortium | Jacquelyn Wynn, Mairead Hyland, The Lighthouse Lab in Alderley Park and Alex Alderton, Roberto Amato, Jeffrey Barrett, Sonia Goncalves, Ewan Harrison, David K. Jackson, Ian Johnston, Dominic Kwiatkowski, Cordelia Langford, John Sillitoe on behalf of the Wellcome Sanger Institute COVID-19 Surveillance Team |
| EPI_ISL_2846203 | Lighthouse Lab in Milton Keynes | Wellcome Sanger Institute for the COVID-19 Genomics UK (COG-UK) Consortium | The Lighthouse Lab in Milton Keynes and Alex Alderton, Roberto Amato, Jeffrey Barrett, Sonia Goncalves, Ewan Harrison, David K. Jackson, Ian Johnston, Dominic Kwiatkowski, Cordelia Langford, John Sillitoe on behalf of the Wellcome Sanger Institute COVID-19 Surveillance Team |
| EPI_ISL_2846204, EPI_ISL_2846205, EPI_ISL_2846206, EPI_ISL_2846207, EPI_ISL_2846208 | Lighthouse Lab in Alderley Park | Wellcome Sanger Institute for the COVID-19 Genomics UK (COG-UK) Consortium | Jacquelyn Wynn, Mairead Hyland, The Lighthouse Lab in Alderley Park and Alex Alderton, Roberto Amato, Jeffrey Barrett, Sonia Goncalves, Ewan Harrison, David K. Jackson, Ian Johnston, Dominic Kwiatkowski, Cordelia Langford, John Sillitoe on behalf of the Wellcome Sanger Institute COVID-19 Surveillance Team |
| EPI_ISL_2846209 | Lighthouse Lab in Milton Keynes | Wellcome Sanger Institute for the COVID-19 Genomics UK (COG-UK) Consortium | The Lighthouse Lab in Milton Keynes and Alex Alderton, Roberto Amato, Jeffrey Barrett, Sonia Goncalves, Ewan Harrison, David K. Jackson, Ian Johnston, Dominic Kwiatkowski, Cordelia Langford, John Sillitoe on behalf of the Wellcome Sanger Institute COVID-19 Surveillance Team |
| EPI_ISL_2846210, EPI_ISL_2846211 | Lighthouse Laboratory Plymouth | Wellcome Sanger Institute for the COVID-19 Genomics UK (COG-UK) Consortium | Lighthouse Laboratory Plymouth and Alex Alderton, Roberto Amato, Jeffrey Barrett, Sonia Goncalves, Ewan Harrison, David K. Jackson, Ian Johnston, Dominic Kwiatkowski, Cordelia Langford, John Sillitoe on behalf of the Wellcome Sanger Institute COVID-19 Surveillance Team |
| EPI_ISL_2846212 | Lighthouse Lab in Alderley Park | Wellcome Sanger Institute for the COVID-19 Genomics UK (COG-UK) Consortium | Jacquelyn Wynn, Mairead Hyland, The Lighthouse Lab in Alderley Park and Alex Alderton, Roberto Amato, Jeffrey Barrett, Sonia Goncalves, Ewan Harrison, David K. Jackson, Ian Johnston, Dominic Kwiatkowski, Cordelia Langford, John Sillitoe on behalf of the Wellcome Sanger Institute COVID-19 Surveillance Team |
| EPI_ISL_2846213, | Lighthouse Lab in Milton Keynes | Wellcome Sanger Institute for the COVID-19 Genomics UK (COG-UK) | The Lighthouse Lab in Milton Keynes and Alex Alderton, Roberto Amato, Jeffrey Barrett, Sonia Goncalves, Ewan Harrison, David K. Jackson, Ian Johnston, Dominic Kwiatkowski, Cordelia Langford, John Sillitoe on behalf of the Wellcome Sanger |

| | | | |
|---|---|---|---|
| EPI_ISL_2846214 | | Consortium | Institute COVID-19 Surveillance Team |
| EPI_ISL_2846215 | Lighthouse Lab in Alderley Park | Wellcome Sanger Institute for the COVID-19 Genomics UK (COG-UK) Consortium | Jacquelyn Wynn, Mairead Hyland, The Lighthouse Lab in Alderley Park and Alex Alderton, Roberto Amato, Jeffrey Barrett, Sonia Goncalves, Ewan Harrison, David K. Jackson, Ian Johnston, Dominic Kwiatkowski, Cordelia Langford, John Sillitoe on behalf of the Wellcome Sanger Institute COVID-19 Surveillance Team |
| EPI_ISL_2846216, EPI_ISL_2846217 | Lighthouse Lab in Milton Keynes | Wellcome Sanger Institute for the COVID-19 Genomics UK (COG-UK) Consortium | The Lighthouse Lab in Milton Keynes and Alex Alderton, Roberto Amato, Jeffrey Barrett, Sonia Goncalves, Ewan Harrison, David K. Jackson, Ian Johnston, Dominic Kwiatkowski, Cordelia Langford, John Sillitoe on behalf of the Wellcome Sanger Institute COVID-19 Surveillance Team |
| EPI_ISL_2846218 | Lighthouse Lab in Alderley Park | Wellcome Sanger Institute for the COVID-19 Genomics UK (COG-UK) Consortium | Jacquelyn Wynn, Mairead Hyland, The Lighthouse Lab in Alderley Park and Alex Alderton, Roberto Amato, Jeffrey Barrett, Sonia Goncalves, Ewan Harrison, David K. Jackson, Ian Johnston, Dominic Kwiatkowski, Cordelia Langford, John Sillitoe on behalf of the Wellcome Sanger Institute COVID-19 Surveillance Team |
| EPI_ISL_2846219, EPI_ISL_2846220, EPI_ISL_2846221 | Lighthouse Lab in Milton Keynes | Wellcome Sanger Institute for the COVID-19 Genomics UK (COG-UK) Consortium | The Lighthouse Lab in Milton Keynes and Alex Alderton, Roberto Amato, Jeffrey Barrett, Sonia Goncalves, Ewan Harrison, David K. Jackson, Ian Johnston, Dominic Kwiatkowski, Cordelia Langford, John Sillitoe on behalf of the Wellcome Sanger Institute COVID-19 Surveillance Team |
| EPI_ISL_2846222 | Lighthouse Lab in Alderley Park | Wellcome Sanger Institute for the COVID-19 Genomics UK (COG-UK) Consortium | Jacquelyn Wynn, Mairead Hyland, The Lighthouse Lab in Alderley Park and Alex Alderton, Roberto Amato, Jeffrey Barrett, Sonia Goncalves, Ewan Harrison, David K. Jackson, Ian Johnston, Dominic Kwiatkowski, Cordelia Langford, John Sillitoe on behalf of the Wellcome Sanger Institute COVID-19 Surveillance Team |
| EPI_ISL_2846223 | Lighthouse Lab in Milton Keynes | Wellcome Sanger Institute for the COVID-19 Genomics UK (COG-UK) Consortium | The Lighthouse Lab in Milton Keynes and Alex Alderton, Roberto Amato, Jeffrey Barrett, Sonia Goncalves, Ewan Harrison, David K. Jackson, Ian Johnston, Dominic Kwiatkowski, Cordelia Langford, John Sillitoe on behalf of the Wellcome Sanger Institute COVID-19 Surveillance Team |
| EPI_ISL_2846224, EPI_ISL_2846225 | Lighthouse Lab in Alderley Park | Wellcome Sanger Institute for the COVID-19 Genomics UK (COG-UK) Consortium | Jacquelyn Wynn, Mairead Hyland, The Lighthouse Lab in Alderley Park and Alex Alderton, Roberto Amato, Jeffrey Barrett, Sonia Goncalves, Ewan Harrison, David K. Jackson, Ian Johnston, Dominic Kwiatkowski, Cordelia Langford, John Sillitoe on behalf of the Wellcome Sanger Institute COVID-19 Surveillance Team |
| EPI_ISL_2846226 | Lighthouse Laboratory Plymouth | Wellcome Sanger Institute for the COVID-19 Genomics UK (COG-UK) Consortium | Lighthouse Laboratory Plymouth and Alex Alderton, Roberto Amato, Jeffrey Barrett, Sonia Goncalves, Ewan Harrison, David K. Jackson, Ian Johnston, Dominic Kwiatkowski, Cordelia Langford, John Sillitoe on behalf of the Wellcome Sanger Institute COVID-19 Surveillance Team |
| EPI_ISL_2846227, EPI_ISL_2846228 | Lighthouse Lab in Milton Keynes | Wellcome Sanger Institute for the COVID-19 Genomics UK (COG-UK) Consortium | The Lighthouse Lab in Milton Keynes and Alex Alderton, Roberto Amato, Jeffrey Barrett, Sonia Goncalves, Ewan Harrison, David K. Jackson, Ian Johnston, Dominic Kwiatkowski, Cordelia Langford, John Sillitoe on behalf of the Wellcome Sanger Institute COVID-19 Surveillance Team |
| EPI_ISL_2846229 | Lighthouse Laboratory Plymouth | Wellcome Sanger Institute for the COVID-19 Genomics UK (COG-UK) Consortium | Lighthouse Laboratory Plymouth and Alex Alderton, Roberto Amato, Jeffrey Barrett, Sonia Goncalves, Ewan Harrison, David K. Jackson, Ian Johnston, Dominic Kwiatkowski, Cordelia Langford, John Sillitoe on behalf of the Wellcome Sanger Institute COVID-19 Surveillance Team |
| EPI_ISL_2846230, EPI_ISL_2846231 | Lighthouse Lab in Alderley Park | Wellcome Sanger Institute for the COVID-19 Genomics UK (COG-UK) Consortium | Jacquelyn Wynn, Mairead Hyland, The Lighthouse Lab in Alderley Park and Alex Alderton, Roberto Amato, Jeffrey Barrett, Sonia Goncalves, Ewan Harrison, David K. Jackson, Ian Johnston, Dominic Kwiatkowski, Cordelia Langford, John Sillitoe on behalf of the Wellcome Sanger Institute COVID-19 Surveillance Team |
| EPI_ISL_2846232 | Lighthouse Lab in Milton Keynes | Wellcome Sanger Institute for the COVID-19 Genomics UK (COG-UK) Consortium | The Lighthouse Lab in Milton Keynes and Alex Alderton, Roberto Amato, Jeffrey Barrett, Sonia Goncalves, Ewan Harrison, David K. Jackson, Ian Johnston, Dominic Kwiatkowski, Cordelia Langford, John Sillitoe on behalf of the Wellcome Sanger Institute COVID-19 Surveillance Team |
| EPI_ISL_2846233, EPI_ISL_2846234, EPI_ISL_2846235 | Lighthouse Lab in Alderley Park | Wellcome Sanger Institute for the COVID-19 Genomics UK (COG-UK) Consortium | Jacquelyn Wynn, Mairead Hyland, The Lighthouse Lab in Alderley Park and Alex Alderton, Roberto Amato, Jeffrey Barrett, Sonia Goncalves, Ewan Harrison, David K. Jackson, Ian Johnston, Dominic Kwiatkowski, Cordelia Langford, John Sillitoe on behalf of the Wellcome Sanger Institute COVID-19 Surveillance Team |
| EPI_ISL_2846236 | Lighthouse Lab in Milton Keynes | Wellcome Sanger Institute for the COVID-19 Genomics UK (COG-UK) Consortium | The Lighthouse Lab in Milton Keynes and Alex Alderton, Roberto Amato, Jeffrey Barrett, Sonia Goncalves, Ewan Harrison, David K. Jackson, Ian Johnston, Dominic Kwiatkowski, Cordelia Langford, John Sillitoe on behalf of the Wellcome Sanger Institute COVID-19 Surveillance Team |
| EPI_ISL_2846237, EPI_ISL_2846238, EPI_ISL_2846239 | Lighthouse Lab in Alderley Park | Wellcome Sanger Institute for the COVID-19 Genomics UK (COG-UK) Consortium | Jacquelyn Wynn, Mairead Hyland, The Lighthouse Lab in Alderley Park and Alex Alderton, Roberto Amato, Jeffrey Barrett, Sonia Goncalves, Ewan Harrison, David K. Jackson, Ian Johnston, Dominic Kwiatkowski, Cordelia Langford, John Sillitoe on behalf of the Wellcome Sanger Institute COVID-19 Surveillance Team |
| EPI_ISL_2846240, EPI_ISL_2846241 | Lighthouse Lab in Milton Keynes | Wellcome Sanger Institute for the COVID-19 Genomics UK (COG-UK) Consortium | The Lighthouse Lab in Milton Keynes and Alex Alderton, Roberto Amato, Jeffrey Barrett, Sonia Goncalves, Ewan Harrison, David K. Jackson, Ian Johnston, Dominic Kwiatkowski, Cordelia Langford, John Sillitoe on behalf of the Wellcome Sanger Institute COVID-19 Surveillance Team |
| EPI_ISL_2846242 | Lighthouse Lab in Alderley Park | Wellcome Sanger Institute for the COVID-19 Genomics UK (COG-UK) Consortium | Jacquelyn Wynn, Mairead Hyland, The Lighthouse Lab in Alderley Park and Alex Alderton, Roberto Amato, Jeffrey Barrett, Sonia Goncalves, Ewan Harrison, David K. Jackson, Ian Johnston, Dominic Kwiatkowski, Cordelia Langford, John Sillitoe on behalf of the Wellcome Sanger Institute COVID-19 Surveillance Team |
| EPI_ISL_2846243 | Lighthouse Laboratory Plymouth | Wellcome Sanger Institute for the COVID-19 Genomics UK (COG-UK) Consortium | Lighthouse Laboratory Plymouth and Alex Alderton, Roberto Amato, Jeffrey Barrett, Sonia Goncalves, Ewan Harrison, David K. Jackson, Ian Johnston, Dominic Kwiatkowski, Cordelia Langford, John Sillitoe on behalf of the Wellcome Sanger Institute COVID-19 Surveillance Team |
| EPI_ISL_2846244, EPI_ISL_2846245 | Lighthouse Lab in Alderley Park | Wellcome Sanger Institute for the COVID-19 Genomics UK (COG-UK) Consortium | Jacquelyn Wynn, Mairead Hyland, The Lighthouse Lab in Alderley Park and Alex Alderton, Roberto Amato, Jeffrey Barrett, Sonia Goncalves, Ewan Harrison, David K. Jackson, Ian Johnston, Dominic Kwiatkowski, Cordelia Langford, John Sillitoe on behalf of the Wellcome Sanger Institute COVID-19 Surveillance Team |
| EPI_ISL_2846246, EPI_ISL_2846247 | Lighthouse Laboratory Plymouth | Wellcome Sanger Institute for the COVID-19 Genomics UK (COG-UK) Consortium | Lighthouse Laboratory Plymouth and Alex Alderton, Roberto Amato, Jeffrey Barrett, Sonia Goncalves, Ewan Harrison, David K. Jackson, Ian Johnston, Dominic Kwiatkowski, Cordelia Langford, John Sillitoe on behalf of the Wellcome Sanger Institute COVID-19 Surveillance Team |
| EPI_ISL_2846248, EPI_ISL_2846249 | Lighthouse Lab in Alderley Park | Wellcome Sanger Institute for the COVID-19 Genomics UK (COG-UK) Consortium | Jacquelyn Wynn, Mairead Hyland, The Lighthouse Lab in Alderley Park and Alex Alderton, Roberto Amato, Jeffrey Barrett, Sonia Goncalves, Ewan Harrison, David K. Jackson, Ian Johnston, Dominic Kwiatkowski, Cordelia Langford, John Sillitoe on behalf of the Wellcome Sanger Institute COVID-19 Surveillance Team |
| EPI_ISL_2846250 | Lighthouse Lab in Milton Keynes | Wellcome Sanger Institute for the COVID-19 Genomics UK (COG-UK) Consortium | The Lighthouse Lab in Milton Keynes and Alex Alderton, Roberto Amato, Jeffrey Barrett, Sonia Goncalves, Ewan Harrison, David K. Jackson, Ian Johnston, Dominic Kwiatkowski, Cordelia Langford, John Sillitoe on behalf of the Wellcome Sanger Institute COVID-19 Surveillance Team |
| EPI_ISL_2846251, EPI_ISL_2846252 | Lighthouse Laboratory Plymouth | Wellcome Sanger Institute for the COVID-19 Genomics UK (COG-UK) Consortium | Lighthouse Laboratory Plymouth and Alex Alderton, Roberto Amato, Jeffrey Barrett, Sonia Goncalves, Ewan Harrison, David K. Jackson, Ian Johnston, Dominic Kwiatkowski, Cordelia Langford, John Sillitoe on behalf of the Wellcome Sanger Institute COVID-19 Surveillance Team |
| EPI_ISL_2846253 | Lighthouse Lab in Milton Keynes | Wellcome Sanger Institute for the COVID-19 Genomics UK (COG-UK) Consortium | The Lighthouse Lab in Milton Keynes and Alex Alderton, Roberto Amato, Jeffrey Barrett, Sonia Goncalves, Ewan Harrison, David K. Jackson, Ian Johnston, Dominic Kwiatkowski, Cordelia Langford, John Sillitoe on behalf of the Wellcome Sanger Institute COVID-19 Surveillance Team |
| EPI_ISL_2846254 | Lighthouse Lab in Alderley Park | Wellcome Sanger Institute for the COVID-19 Genomics UK (COG-UK) Consortium | Jacquelyn Wynn, Mairead Hyland, The Lighthouse Lab in Alderley Park and Alex Alderton, Roberto Amato, Jeffrey Barrett, Sonia Goncalves, Ewan Harrison, David K. Jackson, Ian Johnston, Dominic Kwiatkowski, Cordelia Langford, John Sillitoe on behalf of the Wellcome Sanger Institute COVID-19 Surveillance Team |
| EPI_ISL_2846255 | Lighthouse Lab in Milton Keynes | Wellcome Sanger Institute for the COVID-19 Genomics UK (COG-UK) Consortium | The Lighthouse Lab in Milton Keynes and Alex Alderton, Roberto Amato, Jeffrey Barrett, Sonia Goncalves, Ewan Harrison, David K. Jackson, Ian Johnston, Dominic Kwiatkowski, Cordelia Langford, John Sillitoe on behalf of the Wellcome Sanger Institute COVID-19 Surveillance Team |
| EPI_ISL_2846256, EPI_ISL_2846257, EPI_ISL_2846258 | Lighthouse Lab in Alderley Park | Wellcome Sanger Institute for the COVID-19 Genomics UK (COG-UK) Consortium | Jacquelyn Wynn, Mairead Hyland, The Lighthouse Lab in Alderley Park and Alex Alderton, Roberto Amato, Jeffrey Barrett, Sonia Goncalves, Ewan Harrison, David K. Jackson, Ian Johnston, Dominic Kwiatkowski, Cordelia Langford, John Sillitoe on behalf of the Wellcome Sanger Institute COVID-19 Surveillance Team |
| EPI_ISL_2846259 | Lighthouse Lab in Milton Keynes | Wellcome Sanger Institute for the COVID-19 Genomics UK (COG-UK) Consortium | The Lighthouse Lab in Milton Keynes and Alex Alderton, Roberto Amato, Jeffrey Barrett, Sonia Goncalves, Ewan Harrison, David K. Jackson, Ian Johnston, Dominic Kwiatkowski, Cordelia Langford, John Sillitoe on behalf of the Wellcome Sanger Institute COVID-19 Surveillance Team |
| EPI_ISL_2846260, EPI_ISL_2846261 | Lighthouse Lab in Alderley Park | Wellcome Sanger Institute for the COVID-19 Genomics UK (COG-UK) Consortium | Jacquelyn Wynn, Mairead Hyland, The Lighthouse Lab in Alderley Park and Alex Alderton, Roberto Amato, Jeffrey Barrett, Sonia Goncalves, Ewan Harrison, David K. Jackson, Ian Johnston, Dominic Kwiatkowski, Cordelia Langford, John Sillitoe on behalf of the Wellcome Sanger Institute COVID-19 Surveillance Team |
| EPI_ISL_2846262 | Lighthouse Lab in Milton Keynes | Wellcome Sanger Institute for the COVID-19 Genomics UK (COG-UK) Consortium | The Lighthouse Lab in Milton Keynes and Alex Alderton, Roberto Amato, Jeffrey Barrett, Sonia Goncalves, Ewan Harrison, David K. Jackson, Ian Johnston, Dominic Kwiatkowski, Cordelia Langford, John Sillitoe on behalf of the Wellcome Sanger Institute COVID-19 Surveillance Team |
| EPI_ISL_2846263 | Lighthouse Lab in Alderley Park | Wellcome Sanger Institute for the COVID-19 Genomics UK (COG-UK) Consortium | Jacquelyn Wynn, Mairead Hyland, The Lighthouse Lab in Alderley Park and Alex Alderton, Roberto Amato, Jeffrey Barrett, Sonia Goncalves, Ewan Harrison, David K. Jackson, Ian Johnston, Dominic Kwiatkowski, Cordelia Langford, John Sillitoe on behalf of the Wellcome Sanger Institute COVID-19 Surveillance Team |
| EPI_ISL_2846264 | Lighthouse Lab in Milton Keynes | Wellcome Sanger Institute for the COVID-19 Genomics UK (COG-UK) Consortium | The Lighthouse Lab in Milton Keynes and Alex Alderton, Roberto Amato, Jeffrey Barrett, Sonia Goncalves, Ewan Harrison, David K. Jackson, Ian Johnston, Dominic Kwiatkowski, Cordelia Langford, John Sillitoe on behalf of the Wellcome Sanger Institute COVID-19 Surveillance Team |
| EPI_ISL_2846265 | Lighthouse Lab in Alderley Park | Wellcome Sanger Institute for the COVID-19 Genomics UK (COG-UK) Consortium | Jacquelyn Wynn, Mairead Hyland, The Lighthouse Lab in Alderley Park and Alex Alderton, Roberto Amato, Jeffrey Barrett, Sonia Goncalves, Ewan Harrison, David K. Jackson, Ian Johnston, Dominic Kwiatkowski, Cordelia Langford, John Sillitoe on behalf of the Wellcome Sanger Institute COVID-19 Surveillance Team |
| EPI_ISL_2846266 | Lighthouse Laboratory Plymouth | Wellcome Sanger Institute for the COVID-19 Genomics UK (COG-UK) Consortium | Lighthouse Laboratory Plymouth and Alex Alderton, Roberto Amato, Jeffrey Barrett, Sonia Goncalves, Ewan Harrison, David K. Jackson, Ian Johnston, Dominic Kwiatkowski, Cordelia Langford, John Sillitoe on behalf of the Wellcome Sanger Institute COVID-19 Surveillance Team |
| EPI_ISL_2846267, EPI_ISL_2846268 | Lighthouse Lab in Milton Keynes | Wellcome Sanger Institute for the COVID-19 Genomics UK (COG-UK) Consortium | The Lighthouse Lab in Milton Keynes and Alex Alderton, Roberto Amato, Jeffrey Barrett, Sonia Goncalves, Ewan Harrison, David K. Jackson, Ian Johnston, Dominic Kwiatkowski, Cordelia Langford, John Sillitoe on behalf of the Wellcome Sanger Institute COVID-19 Surveillance Team |
| EPI_ISL_2846269, EPI_ISL_2846270 | Lighthouse Laboratory Plymouth | Wellcome Sanger Institute for the COVID-19 Genomics UK (COG-UK) Consortium | Lighthouse Laboratory Plymouth and Alex Alderton, Roberto Amato, Jeffrey Barrett, Sonia Goncalves, Ewan Harrison, David K. Jackson, Ian Johnston, Dominic Kwiatkowski, Cordelia Langford, John Sillitoe on behalf of the Wellcome Sanger Institute COVID-19 Surveillance Team |
| EPI_ISL_2846271 | Lighthouse Lab in Milton Keynes | Wellcome Sanger Institute for the COVID-19 Genomics UK (COG-UK) Consortium | The Lighthouse Lab in Milton Keynes and Alex Alderton, Roberto Amato, Jeffrey Barrett, Sonia Goncalves, Ewan Harrison, David K. Jackson, Ian Johnston, Dominic Kwiatkowski, Cordelia Langford, John Sillitoe on behalf of the Wellcome Sanger Institute COVID-19 Surveillance Team |
| EPI_ISL_2846272, EPI_ISL_2846273 | Lighthouse Lab in Alderley Park | Wellcome Sanger Institute for the COVID-19 Genomics UK (COG-UK) Consortium | Jacquelyn Wynn, Mairead Hyland, The Lighthouse Lab in Alderley Park and Alex Alderton, Roberto Amato, Jeffrey Barrett, Sonia Goncalves, Ewan Harrison, David K. Jackson, Ian Johnston, Dominic Kwiatkowski, Cordelia Langford, John Sillitoe on behalf of the Wellcome Sanger Institute COVID-19 Surveillance Team |
| EPI_ISL_2846274 | Lighthouse Laboratory Plymouth | Wellcome Sanger Institute for the COVID-19 Genomics UK (COG-UK) Consortium | Lighthouse Laboratory Plymouth and Alex Alderton, Roberto Amato, Jeffrey Barrett, Sonia Goncalves, Ewan Harrison, David K. Jackson, Ian Johnston, Dominic Kwiatkowski, Cordelia Langford, John Sillitoe on behalf of the Wellcome Sanger Institute COVID-19 Surveillance Team |
| EPI_ISL_2846275 | Lighthouse Lab in Alderley Park | Wellcome Sanger Institute for the COVID-19 Genomics UK (COG-UK) Consortium | Jacquelyn Wynn, Mairead Hyland, The Lighthouse Lab in Alderley Park and Alex Alderton, Roberto Amato, Jeffrey Barrett, Sonia Goncalves, Ewan Harrison, David K. Jackson, Ian Johnston, Dominic Kwiatkowski, Cordelia Langford, John Sillitoe on behalf of the Wellcome Sanger Institute COVID-19 Surveillance Team |
| EPI_ISL_2846276, EPI_ISL_2846277 | Lighthouse Lab in Milton Keynes | Wellcome Sanger Institute for the COVID-19 Genomics UK (COG-UK) Consortium | The Lighthouse Lab in Milton Keynes and Alex Alderton, Roberto Amato, Jeffrey Barrett, Sonia Goncalves, Ewan Harrison, David K. Jackson, Ian Johnston, Dominic Kwiatkowski, Cordelia Langford, John Sillitoe on behalf of the Wellcome Sanger Institute COVID-19 Surveillance Team |
| EPI_ISL_2846278 | Lighthouse Laboratory Plymouth | Wellcome Sanger Institute for the COVID-19 Genomics UK (COG-UK) Consortium | Lighthouse Laboratory Plymouth and Alex Alderton, Roberto Amato, Jeffrey Barrett, Sonia Goncalves, Ewan Harrison, David K. Jackson, Ian Johnston, Dominic Kwiatkowski, Cordelia Langford, John Sillitoe on behalf of the Wellcome Sanger Institute COVID-19 Surveillance Team |
| EPI_ISL_2846279 | Lighthouse Lab in Alderley Park | Wellcome Sanger Institute for the COVID-19 Genomics UK (COG-UK) | Jacquelyn Wynn, Mairead Hyland, The Lighthouse Lab in Alderley Park and Alex Alderton, Roberto Amato, Jeffrey Barrett, Sonia Goncalves, Ewan Harrison, David K. Jackson, Ian Johnston, Dominic Kwiatkowski, Cordelia Langford, John Sillitoe on |

| | | | |
|---|---|---|---|
| | | Consortium | behalf of the Wellcome Sanger Institute COVID-19 Surveillance Team |
| EPI_ISL_2846280 | Lighthouse Lab in Milton Keynes | Wellcome Sanger Institute for the COVID-19 Genomics UK (COG-UK) Consortium | The Lighthouse Lab in Milton Keynes and Alex Alderton, Roberto Amato, Jeffrey Barrett, Sonia Goncalves, Ewan Harrison, David K. Jackson, Ian Johnston, Dominic Kwiatkowski, Cordelia Langford, John Sillitoe on behalf of the Wellcome Sanger Institute COVID-19 Surveillance Team |
| EPI_ISL_2846281 | Lighthouse Laboratory Plymouth | Wellcome Sanger Institute for the COVID-19 Genomics UK (COG-UK) Consortium | Lighthouse Laboratory Plymouth and Alex Alderton, Roberto Amato, Jeffrey Barrett, Sonia Goncalves, Ewan Harrison, David K. Jackson, Ian Johnston, Dominic Kwiatkowski, Cordelia Langford, John Sillitoe on behalf of the Wellcome Sanger Institute COVID-19 Surveillance Team |
| EPI_ISL_2846282, EPI_ISL_2846283, EPI_ISL_2846284 | Lighthouse Lab in Alderley Park | Wellcome Sanger Institute for the COVID-19 Genomics UK (COG-UK) Consortium | Jacquelyn Wynn, Mairead Hyland, The Lighthouse Lab in Alderley Park and Alex Alderton, Roberto Amato, Jeffrey Barrett, Sonia Goncalves, Ewan Harrison, David K. Jackson, Ian Johnston, Dominic Kwiatkowski, Cordelia Langford, John Sillitoe on behalf of the Wellcome Sanger Institute COVID-19 Surveillance Team |
| EPI_ISL_2846285, EPI_ISL_2846286 | Lighthouse Laboratory Plymouth | Wellcome Sanger Institute for the COVID-19 Genomics UK (COG-UK) Consortium | Lighthouse Laboratory Plymouth and Alex Alderton, Roberto Amato, Jeffrey Barrett, Sonia Goncalves, Ewan Harrison, David K. Jackson, Ian Johnston, Dominic Kwiatkowski, Cordelia Langford, John Sillitoe on behalf of the Wellcome Sanger Institute COVID-19 Surveillance Team |
| EPI_ISL_2846287, EPI_ISL_2846288, EPI_ISL_2846289, EPI_ISL_2846290 | Lighthouse Lab in Alderley Park | Wellcome Sanger Institute for the COVID-19 Genomics UK (COG-UK) Consortium | Jacquelyn Wynn, Mairead Hyland, The Lighthouse Lab in Alderley Park and Alex Alderton, Roberto Amato, Jeffrey Barrett, Sonia Goncalves, Ewan Harrison, David K. Jackson, Ian Johnston, Dominic Kwiatkowski, Cordelia Langford, John Sillitoe on behalf of the Wellcome Sanger Institute COVID-19 Surveillance Team |
| EPI_ISL_2846291, EPI_ISL_2846292 | Lighthouse Laboratory Plymouth | Wellcome Sanger Institute for the COVID-19 Genomics UK (COG-UK) Consortium | Lighthouse Laboratory Plymouth and Alex Alderton, Roberto Amato, Jeffrey Barrett, Sonia Goncalves, Ewan Harrison, David K. Jackson, Ian Johnston, Dominic Kwiatkowski, Cordelia Langford, John Sillitoe on behalf of the Wellcome Sanger Institute COVID-19 Surveillance Team |
| EPI_ISL_2846293, EPI_ISL_2846294 | Lighthouse Lab in Milton Keynes | Wellcome Sanger Institute for the COVID-19 Genomics UK (COG-UK) Consortium | The Lighthouse Lab in Milton Keynes and Alex Alderton, Roberto Amato, Jeffrey Barrett, Sonia Goncalves, Ewan Harrison, David K. Jackson, Ian Johnston, Dominic Kwiatkowski, Cordelia Langford, John Sillitoe on behalf of the Wellcome Sanger Institute COVID-19 Surveillance Team |
| EPI_ISL_2846296, EPI_ISL_2846297, EPI_ISL_2846298 | Lighthouse Lab in Alderley Park | Wellcome Sanger Institute for the COVID-19 Genomics UK (COG-UK) Consortium | Jacquelyn Wynn, Mairead Hyland, The Lighthouse Lab in Alderley Park and Alex Alderton, Roberto Amato, Jeffrey Barrett, Sonia Goncalves, Ewan Harrison, David K. Jackson, Ian Johnston, Dominic Kwiatkowski, Cordelia Langford, John Sillitoe on behalf of the Wellcome Sanger Institute COVID-19 Surveillance Team |
| EPI_ISL_2846299, EPI_ISL_2846300 | Lighthouse Laboratory Plymouth | Wellcome Sanger Institute for the COVID-19 Genomics UK (COG-UK) Consortium | Lighthouse Laboratory Plymouth and Alex Alderton, Roberto Amato, Jeffrey Barrett, Sonia Goncalves, Ewan Harrison, David K. Jackson, Ian Johnston, Dominic Kwiatkowski, Cordelia Langford, John Sillitoe on behalf of the Wellcome Sanger Institute COVID-19 Surveillance Team |
| EPI_ISL_2846301 | Lighthouse Lab in Milton Keynes | Wellcome Sanger Institute for the COVID-19 Genomics UK (COG-UK) Consortium | The Lighthouse Lab in Milton Keynes and Alex Alderton, Roberto Amato, Jeffrey Barrett, Sonia Goncalves, Ewan Harrison, David K. Jackson, Ian Johnston, Dominic Kwiatkowski, Cordelia Langford, John Sillitoe on behalf of the Wellcome Sanger Institute COVID-19 Surveillance Team |
| EPI_ISL_2846302, EPI_ISL_2846303 | Lighthouse Laboratory Plymouth | Wellcome Sanger Institute for the COVID-19 Genomics UK (COG-UK) Consortium | Lighthouse Laboratory Plymouth and Alex Alderton, Roberto Amato, Jeffrey Barrett, Sonia Goncalves, Ewan Harrison, David K. Jackson, Ian Johnston, Dominic Kwiatkowski, Cordelia Langford, John Sillitoe on behalf of the Wellcome Sanger Institute COVID-19 Surveillance Team |
| EPI_ISL_2846304 | Lighthouse Lab in Milton Keynes | Wellcome Sanger Institute for the COVID-19 Genomics UK (COG-UK) Consortium | The Lighthouse Lab in Milton Keynes and Alex Alderton, Roberto Amato, Jeffrey Barrett, Sonia Goncalves, Ewan Harrison, David K. Jackson, Ian Johnston, Dominic Kwiatkowski, Cordelia Langford, John Sillitoe on behalf of the Wellcome Sanger Institute COVID-19 Surveillance Team |
| EPI_ISL_2846305, EPI_ISL_2846306, EPI_ISL_2846307 | Lighthouse Lab in Alderley Park | Wellcome Sanger Institute for the COVID-19 Genomics UK (COG-UK) Consortium | Jacquelyn Wynn, Mairead Hyland, The Lighthouse Lab in Alderley Park and Alex Alderton, Roberto Amato, Jeffrey Barrett, Sonia Goncalves, Ewan Harrison, David K. Jackson, Ian Johnston, Dominic Kwiatkowski, Cordelia Langford, John Sillitoe on behalf of the Wellcome Sanger Institute COVID-19 Surveillance Team |
| EPI_ISL_2846308 | Lighthouse Laboratory Plymouth | Wellcome Sanger Institute for the COVID-19 Genomics UK (COG-UK) Consortium | Lighthouse Laboratory Plymouth and Alex Alderton, Roberto Amato, Jeffrey Barrett, Sonia Goncalves, Ewan Harrison, David K. Jackson, Ian Johnston, Dominic Kwiatkowski, Cordelia Langford, John Sillitoe on behalf of the Wellcome Sanger Institute COVID-19 Surveillance Team |
| EPI_ISL_2846309 | Lighthouse Lab in Milton Keynes | Wellcome Sanger Institute for the COVID-19 Genomics UK (COG-UK) Consortium | The Lighthouse Lab in Milton Keynes and Alex Alderton, Roberto Amato, Jeffrey Barrett, Sonia Goncalves, Ewan Harrison, David K. Jackson, Ian Johnston, Dominic Kwiatkowski, Cordelia Langford, John Sillitoe on behalf of the Wellcome Sanger Institute COVID-19 Surveillance Team |
| EPI_ISL_2846310, EPI_ISL_2846311 | Lighthouse Lab in Alderley Park | Wellcome Sanger Institute for the COVID-19 Genomics UK (COG-UK) Consortium | Jacquelyn Wynn, Mairead Hyland, The Lighthouse Lab in Alderley Park and Alex Alderton, Roberto Amato, Jeffrey Barrett, Sonia Goncalves, Ewan Harrison, David K. Jackson, Ian Johnston, Dominic Kwiatkowski, Cordelia Langford, John Sillitoe on behalf of the Wellcome Sanger Institute COVID-19 Surveillance Team |
| EPI_ISL_2846312, EPI_ISL_2846313 | Lighthouse Lab in Milton Keynes | Wellcome Sanger Institute for the COVID-19 Genomics UK (COG-UK) Consortium | The Lighthouse Lab in Milton Keynes and Alex Alderton, Roberto Amato, Jeffrey Barrett, Sonia Goncalves, Ewan Harrison, David K. Jackson, Ian Johnston, Dominic Kwiatkowski, Cordelia Langford, John Sillitoe on behalf of the Wellcome Sanger Institute COVID-19 Surveillance Team |
| EPI_ISL_2846314 | Lighthouse Lab in Alderley Park | Wellcome Sanger Institute for the COVID-19 Genomics UK (COG-UK) Consortium | Jacquelyn Wynn, Mairead Hyland, The Lighthouse Lab in Alderley Park and Alex Alderton, Roberto Amato, Jeffrey Barrett, Sonia Goncalves, Ewan Harrison, David K. Jackson, Ian Johnston, Dominic Kwiatkowski, Cordelia Langford, John Sillitoe on behalf of the Wellcome Sanger Institute COVID-19 Surveillance Team |
| EPI_ISL_2846315 | Lighthouse Lab in Milton Keynes | Wellcome Sanger Institute for the COVID-19 Genomics UK (COG-UK) Consortium | The Lighthouse Lab in Milton Keynes and Alex Alderton, Roberto Amato, Jeffrey Barrett, Sonia Goncalves, Ewan Harrison, David K. Jackson, Ian Johnston, Dominic Kwiatkowski, Cordelia Langford, John Sillitoe on behalf of the Wellcome Sanger Institute COVID-19 Surveillance Team |
| EPI_ISL_2846316 | Lighthouse Lab in Alderley Park | Wellcome Sanger Institute for the COVID-19 Genomics UK (COG-UK) Consortium | Jacquelyn Wynn, Mairead Hyland, The Lighthouse Lab in Alderley Park and Alex Alderton, Roberto Amato, Jeffrey Barrett, Sonia Goncalves, Ewan Harrison, David K. Jackson, Ian Johnston, Dominic Kwiatkowski, Cordelia Langford, John Sillitoe on behalf of the Wellcome Sanger Institute COVID-19 Surveillance Team |
| EPI_ISL_2846317 | Lighthouse Lab in Milton Keynes | Wellcome Sanger Institute for the COVID-19 Genomics UK (COG-UK) Consortium | The Lighthouse Lab in Milton Keynes and Alex Alderton, Roberto Amato, Jeffrey Barrett, Sonia Goncalves, Ewan Harrison, David K. Jackson, Ian Johnston, Dominic Kwiatkowski, Cordelia Langford, John Sillitoe on behalf of the Wellcome Sanger Institute COVID-19 Surveillance Team |
| EPI_ISL_2846318 | Lighthouse Laboratory Plymouth | Wellcome Sanger Institute for the COVID-19 Genomics UK (COG-UK) Consortium | Lighthouse Laboratory Plymouth and Alex Alderton, Roberto Amato, Jeffrey Barrett, Sonia Goncalves, Ewan Harrison, David K. Jackson, Ian Johnston, Dominic Kwiatkowski, Cordelia Langford, John Sillitoe on behalf of the Wellcome Sanger Institute COVID-19 Surveillance Team |
| EPI_ISL_2846319 | Lighthouse Lab in Alderley Park | Wellcome Sanger Institute for the COVID-19 Genomics UK (COG-UK) Consortium | Jacquelyn Wynn, Mairead Hyland, The Lighthouse Lab in Alderley Park and Alex Alderton, Roberto Amato, Jeffrey Barrett, Sonia Goncalves, Ewan Harrison, David K. Jackson, Ian Johnston, Dominic Kwiatkowski, Cordelia Langford, John Sillitoe on behalf of the Wellcome Sanger Institute COVID-19 Surveillance Team |
| EPI_ISL_2846320, EPI_ISL_2846321, EPI_ISL_2846322 | Lighthouse Lab in Milton Keynes | Wellcome Sanger Institute for the COVID-19 Genomics UK (COG-UK) Consortium | The Lighthouse Lab in Milton Keynes and Alex Alderton, Roberto Amato, Jeffrey Barrett, Sonia Goncalves, Ewan Harrison, David K. Jackson, Ian Johnston, Dominic Kwiatkowski, Cordelia Langford, John Sillitoe on behalf of the Wellcome Sanger Institute COVID-19 Surveillance Team |
| EPI_ISL_2846323, EPI_ISL_2846324, EPI_ISL_2846325 | Lighthouse Laboratory Plymouth | Wellcome Sanger Institute for the COVID-19 Genomics UK (COG-UK) Consortium | Lighthouse Laboratory Plymouth and Alex Alderton, Roberto Amato, Jeffrey Barrett, Sonia Goncalves, Ewan Harrison, David K. Jackson, Ian Johnston, Dominic Kwiatkowski, Cordelia Langford, John Sillitoe on behalf of the Wellcome Sanger Institute COVID-19 Surveillance Team |
| EPI_ISL_2846326, EPI_ISL_2846327, EPI_ISL_2846328 | Lighthouse Lab in Alderley Park | Wellcome Sanger Institute for the COVID-19 Genomics UK (COG-UK) Consortium | Jacquelyn Wynn, Mairead Hyland, The Lighthouse Lab in Alderley Park and Alex Alderton, Roberto Amato, Jeffrey Barrett, Sonia Goncalves, Ewan Harrison, David K. Jackson, Ian Johnston, Dominic Kwiatkowski, Cordelia Langford, John Sillitoe on behalf of the Wellcome Sanger Institute COVID-19 Surveillance Team |
| EPI_ISL_2846329 | Lighthouse Lab in Milton Keynes | Wellcome Sanger Institute for the COVID-19 Genomics UK (COG-UK) Consortium | The Lighthouse Lab in Milton Keynes and Alex Alderton, Roberto Amato, Jeffrey Barrett, Sonia Goncalves, Ewan Harrison, David K. Jackson, Ian Johnston, Dominic Kwiatkowski, Cordelia Langford, John Sillitoe on behalf of the Wellcome Sanger Institute COVID-19 Surveillance Team |
| EPI_ISL_2846330, EPI_ISL_2846331 | Lighthouse Lab in Alderley Park | Wellcome Sanger Institute for the COVID-19 Genomics UK (COG-UK) Consortium | Jacquelyn Wynn, Mairead Hyland, The Lighthouse Lab in Alderley Park and Alex Alderton, Roberto Amato, Jeffrey Barrett, Sonia Goncalves, Ewan Harrison, David K. Jackson, Ian Johnston, Dominic Kwiatkowski, Cordelia Langford, John Sillitoe on behalf of the Wellcome Sanger Institute COVID-19 Surveillance Team |
| EPI_ISL_2846332 | Lighthouse Laboratory Plymouth | Wellcome Sanger Institute for the COVID-19 Genomics UK (COG-UK) Consortium | Lighthouse Laboratory Plymouth and Alex Alderton, Roberto Amato, Jeffrey Barrett, Sonia Goncalves, Ewan Harrison, David K. Jackson, Ian Johnston, Dominic Kwiatkowski, Cordelia Langford, John Sillitoe on behalf of the Wellcome Sanger Institute COVID-19 Surveillance Team |
| EPI_ISL_2846333, EPI_ISL_2846334 | Lighthouse Lab in Alderley Park | Wellcome Sanger Institute for the COVID-19 Genomics UK (COG-UK) Consortium | Jacquelyn Wynn, Mairead Hyland, The Lighthouse Lab in Alderley Park and Alex Alderton, Roberto Amato, Jeffrey Barrett, Sonia Goncalves, Ewan Harrison, David K. Jackson, Ian Johnston, Dominic Kwiatkowski, Cordelia Langford, John Sillitoe on behalf of the Wellcome Sanger Institute COVID-19 Surveillance Team |
| EPI_ISL_2846335 | Lighthouse Laboratory Plymouth | Wellcome Sanger Institute for the COVID-19 Genomics UK (COG-UK) Consortium | Lighthouse Laboratory Plymouth and Alex Alderton, Roberto Amato, Jeffrey Barrett, Sonia Goncalves, Ewan Harrison, David K. Jackson, Ian Johnston, Dominic Kwiatkowski, Cordelia Langford, John Sillitoe on behalf of the Wellcome Sanger Institute COVID-19 Surveillance Team |
| EPI_ISL_2846336, EPI_ISL_2846337, EPI_ISL_2846338 | Lighthouse Lab in Alderley Park | Wellcome Sanger Institute for the COVID-19 Genomics UK (COG-UK) Consortium | Jacquelyn Wynn, Mairead Hyland, The Lighthouse Lab in Alderley Park and Alex Alderton, Roberto Amato, Jeffrey Barrett, Sonia Goncalves, Ewan Harrison, David K. Jackson, Ian Johnston, Dominic Kwiatkowski, Cordelia Langford, John Sillitoe on behalf of the Wellcome Sanger Institute COVID-19 Surveillance Team |
| EPI_ISL_2846339 | Lighthouse Laboratory Plymouth | Wellcome Sanger Institute for the COVID-19 Genomics UK (COG-UK) Consortium | Lighthouse Laboratory Plymouth and Alex Alderton, Roberto Amato, Jeffrey Barrett, Sonia Goncalves, Ewan Harrison, David K. Jackson, Ian Johnston, Dominic Kwiatkowski, Cordelia Langford, John Sillitoe on behalf of the Wellcome Sanger Institute COVID-19 Surveillance Team |
| EPI_ISL_2846340, EPI_ISL_2846341 | Lighthouse Lab in Alderley Park | Wellcome Sanger Institute for the COVID-19 Genomics UK (COG-UK) Consortium | Jacquelyn Wynn, Mairead Hyland, The Lighthouse Lab in Alderley Park and Alex Alderton, Roberto Amato, Jeffrey Barrett, Sonia Goncalves, Ewan Harrison, David K. Jackson, Ian Johnston, Dominic Kwiatkowski, Cordelia Langford, John Sillitoe on behalf of the Wellcome Sanger Institute COVID-19 Surveillance Team |
| EPI_ISL_2846342 | AZ Klina | AZ Klina | Carl Vael - Lynsey Berckmans |
| EPI_ISL_2846343, EPI_ISL_2846344, EPI_ISL_2846345, EPI_ISL_2846346 | Lighthouse Lab in Alderley Park | Wellcome Sanger Institute for the COVID-19 Genomics UK (COG-UK) Consortium | Jacquelyn Wynn, Mairead Hyland, The Lighthouse Lab in Alderley Park and Alex Alderton, Roberto Amato, Jeffrey Barrett, Sonia Goncalves, Ewan Harrison, David K. Jackson, Ian Johnston, Dominic Kwiatkowski, Cordelia Langford, John Sillitoe on behalf of the Wellcome Sanger Institute COVID-19 Surveillance Team |
| EPI_ISL_2846347 | Lighthouse Lab in Milton Keynes | Wellcome Sanger Institute for the COVID-19 Genomics UK (COG-UK) Consortium | The Lighthouse Lab in Milton Keynes and Alex Alderton, Roberto Amato, Jeffrey Barrett, Sonia Goncalves, Ewan Harrison, David K. Jackson, Ian Johnston, Dominic Kwiatkowski, Cordelia Langford, John Sillitoe on behalf of the Wellcome Sanger Institute COVID-19 Surveillance Team |
| EPI_ISL_2846348 | Lighthouse Laboratory Plymouth | Wellcome Sanger Institute for the COVID-19 Genomics UK (COG-UK) Consortium | Lighthouse Laboratory Plymouth and Alex Alderton, Roberto Amato, Jeffrey Barrett, Sonia Goncalves, Ewan Harrison, David K. Jackson, Ian Johnston, Dominic Kwiatkowski, Cordelia Langford, John Sillitoe on behalf of the Wellcome Sanger Institute COVID-19 Surveillance Team |
| EPI_ISL_2846349 | Lighthouse Lab in Milton Keynes | Wellcome Sanger Institute for the COVID-19 Genomics UK (COG-UK) Consortium | The Lighthouse Lab in Milton Keynes and Alex Alderton, Roberto Amato, Jeffrey Barrett, Sonia Goncalves, Ewan Harrison, David K. Jackson, Ian Johnston, Dominic Kwiatkowski, Cordelia Langford, John Sillitoe on behalf of the Wellcome Sanger Institute COVID-19 Surveillance Team |
| EPI_ISL_2846350 | Lighthouse Laboratory Plymouth | Wellcome Sanger Institute for the COVID-19 Genomics UK (COG-UK) Consortium | Lighthouse Laboratory Plymouth and Alex Alderton, Roberto Amato, Jeffrey Barrett, Sonia Goncalves, Ewan Harrison, David K. Jackson, Ian Johnston, Dominic Kwiatkowski, Cordelia Langford, John Sillitoe on behalf of the Wellcome Sanger Institute COVID-19 Surveillance Team |
| EPI_ISL_2846351 | Lighthouse Lab in Milton Keynes | Wellcome Sanger Institute for the COVID-19 Genomics UK (COG-UK) | The Lighthouse Lab in Milton Keynes and Alex Alderton, Roberto Amato, Jeffrey Barrett, Sonia Goncalves, Ewan Harrison, David K. Jackson, Ian Johnston, Dominic Kwiatkowski, Cordelia Langford, John Sillitoe on behalf of the Wellcome Sanger |

| | | | |
|---|---|---|---|
| | | Consortium | Institute COVID-19 Surveillance Team |
| EPI_ISL_2846352, EPI_ISL_2846353 | Lighthouse Lab in Alderley Park | Wellcome Sanger Institute for the COVID-19 Genomics UK (COG-UK) Consortium | Jacquelyn Wynn, Mairead Hyland, The Lighthouse Lab in Alderley Park and Alex Alderton, Roberto Amato, Jeffrey Barrett, Sonia Goncalves, Ewan Harrison, David K. Jackson, Ian Johnston, Dominic Kwiatkowski, Cordelia Langford, John Sillitoe on behalf of the Wellcome Sanger Institute COVID-19 Surveillance Team |
| EPI_ISL_2846354, EPI_ISL_2846355 | Lighthouse Laboratory Plymouth | Wellcome Sanger Institute for the COVID-19 Genomics UK (COG-UK) Consortium | Lighthouse Laboratory Plymouth and Alex Alderton, Roberto Amato, Jeffrey Barrett, Sonia Goncalves, Ewan Harrison, David K. Jackson, Ian Johnston, Dominic Kwiatkowski, Cordelia Langford, John Sillitoe on behalf of the Wellcome Sanger Institute COVID-19 Surveillance Team |
| EPI_ISL_2846356, EPI_ISL_2846357 | Lighthouse Lab in Alderley Park | Wellcome Sanger Institute for the COVID-19 Genomics UK (COG-UK) Consortium | Jacquelyn Wynn, Mairead Hyland, The Lighthouse Lab in Alderley Park and Alex Alderton, Roberto Amato, Jeffrey Barrett, Sonia Goncalves, Ewan Harrison, David K. Jackson, Ian Johnston, Dominic Kwiatkowski, Cordelia Langford, John Sillitoe on behalf of the Wellcome Sanger Institute COVID-19 Surveillance Team |
| EPI_ISL_2846358 | Lighthouse Lab in Milton Keynes | Wellcome Sanger Institute for the COVID-19 Genomics UK (COG-UK) Consortium | The Lighthouse Lab in Milton Keynes and Alex Alderton, Roberto Amato, Jeffrey Barrett, Sonia Goncalves, Ewan Harrison, David K. Jackson, Ian Johnston, Dominic Kwiatkowski, Cordelia Langford, John Sillitoe on behalf of the Wellcome Sanger Institute COVID-19 Surveillance Team |
| EPI_ISL_2846359 | Lighthouse Lab in Alderley Park | Wellcome Sanger Institute for the COVID-19 Genomics UK (COG-UK) Consortium | Jacquelyn Wynn, Mairead Hyland, The Lighthouse Lab in Alderley Park and Alex Alderton, Roberto Amato, Jeffrey Barrett, Sonia Goncalves, Ewan Harrison, David K. Jackson, Ian Johnston, Dominic Kwiatkowski, Cordelia Langford, John Sillitoe on behalf of the Wellcome Sanger Institute COVID-19 Surveillance Team |
| EPI_ISL_2846360 | Lighthouse Lab in Milton Keynes | Wellcome Sanger Institute for the COVID-19 Genomics UK (COG-UK) Consortium | The Lighthouse Lab in Milton Keynes and Alex Alderton, Roberto Amato, Jeffrey Barrett, Sonia Goncalves, Ewan Harrison, David K. Jackson, Ian Johnston, Dominic Kwiatkowski, Cordelia Langford, John Sillitoe on behalf of the Wellcome Sanger Institute COVID-19 Surveillance Team |
| EPI_ISL_2846361, EPI_ISL_2846362 | Lighthouse Lab in Glasgow | Wellcome Sanger Institute for the COVID-19 Genomics UK (COG-UK) Consortium | Harper VanSteenhouse, Yumi Kasai, David Gray, Carol Clugston, Anna Dominiczak and Alex Alderton, Roberto Amato, Jeffrey Barrett, Sonia Goncalves, Ewan Harrison, David K. Jackson, Ian Johnston, Dominic Kwiatkowski, Cordelia Langford, John Sillitoe on behalf of the Wellcome Sanger Institute COVID-19 Surveillance Team |
| EPI_ISL_2846363, EPI_ISL_2846364 | Berkshire and Surrey Pathology Services Lighthouse Laboratory | Wellcome Sanger Institute for the COVID-19 Genomics UK (COG-UK) Consortium | Berkshire and Surrey Pathology Services Lighthouse Laboratory and Alex Alderton, Roberto Amato, Jeffrey Barrett, Sonia Goncalves, Ewan Harrison, David K. Jackson, Ian Johnston, Dominic Kwiatkowski, Cordelia Langford, John Sillitoe on behalf of the Wellcome Sanger Institute COVID-19 Surveillance Team |
| EPI_ISL_2846365 | Lighthouse Lab in Glasgow | Wellcome Sanger Institute for the COVID-19 Genomics UK (COG-UK) Consortium | Harper VanSteenhouse, Yumi Kasai, David Gray, Carol Clugston, Anna Dominiczak and Alex Alderton, Roberto Amato, Jeffrey Barrett, Sonia Goncalves, Ewan Harrison, David K. Jackson, Ian Johnston, Dominic Kwiatkowski, Cordelia Langford, John Sillitoe on behalf of the Wellcome Sanger Institute COVID-19 Surveillance Team |
| EPI_ISL_2846366 | Lighthouse Lab in Milton Keynes | Wellcome Sanger Institute for the COVID-19 Genomics UK (COG-UK) Consortium | The Lighthouse Lab in Milton Keynes and Alex Alderton, Roberto Amato, Jeffrey Barrett, Sonia Goncalves, Ewan Harrison, David K. Jackson, Ian Johnston, Dominic Kwiatkowski, Cordelia Langford, John Sillitoe on behalf of the Wellcome Sanger Institute COVID-19 Surveillance Team |
| EPI_ISL_2846367 | Lighthouse Laboratory Plymouth | Wellcome Sanger Institute for the COVID-19 Genomics UK (COG-UK) Consortium | Lighthouse Laboratory Plymouth and Alex Alderton, Roberto Amato, Jeffrey Barrett, Sonia Goncalves, Ewan Harrison, David K. Jackson, Ian Johnston, Dominic Kwiatkowski, Cordelia Langford, John Sillitoe on behalf of the Wellcome Sanger Institute COVID-19 Surveillance Team |
| EPI_ISL_2846368 | Lighthouse Lab in Milton Keynes | Wellcome Sanger Institute for the COVID-19 Genomics UK (COG-UK) Consortium | The Lighthouse Lab in Milton Keynes and Alex Alderton, Roberto Amato, Jeffrey Barrett, Sonia Goncalves, Ewan Harrison, David K. Jackson, Ian Johnston, Dominic Kwiatkowski, Cordelia Langford, John Sillitoe on behalf of the Wellcome Sanger Institute COVID-19 Surveillance Team |
| EPI_ISL_2846369, EPI_ISL_2846370 | Berkshire and Surrey Pathology Services Lighthouse Laboratory | Wellcome Sanger Institute for the COVID-19 Genomics UK (COG-UK) Consortium | Berkshire and Surrey Pathology Services Lighthouse Laboratory and Alex Alderton, Roberto Amato, Jeffrey Barrett, Sonia Goncalves, Ewan Harrison, David K. Jackson, Ian Johnston, Dominic Kwiatkowski, Cordelia Langford, John Sillitoe on behalf of the Wellcome Sanger Institute COVID-19 Surveillance Team |
| EPI_ISL_2846371 | Lighthouse Lab in Glasgow | Wellcome Sanger Institute for the COVID-19 Genomics UK (COG-UK) Consortium | Harper VanSteenhouse, Yumi Kasai, David Gray, Carol Clugston, Anna Dominiczak and Alex Alderton, Roberto Amato, Jeffrey Barrett, Sonia Goncalves, Ewan Harrison, David K. Jackson, Ian Johnston, Dominic Kwiatkowski, Cordelia Langford, John Sillitoe on behalf of the Wellcome Sanger Institute COVID-19 Surveillance Team |
| EPI_ISL_2846372 | Berkshire and Surrey Pathology Services Lighthouse Laboratory | Wellcome Sanger Institute for the COVID-19 Genomics UK (COG-UK) Consortium | Berkshire and Surrey Pathology Services Lighthouse Laboratory and Alex Alderton, Roberto Amato, Jeffrey Barrett, Sonia Goncalves, Ewan Harrison, David K. Jackson, Ian Johnston, Dominic Kwiatkowski, Cordelia Langford, John Sillitoe on behalf of the Wellcome Sanger Institute COVID-19 Surveillance Team |
| EPI_ISL_2846373, EPI_ISL_2846374 | Lighthouse Lab in Milton Keynes | Wellcome Sanger Institute for the COVID-19 Genomics UK (COG-UK) Consortium | The Lighthouse Lab in Milton Keynes and Alex Alderton, Roberto Amato, Jeffrey Barrett, Sonia Goncalves, Ewan Harrison, David K. Jackson, Ian Johnston, Dominic Kwiatkowski, Cordelia Langford, John Sillitoe on behalf of the Wellcome Sanger Institute COVID-19 Surveillance Team |
| EPI_ISL_2846375 | Lighthouse Lab in Glasgow | Wellcome Sanger Institute for the COVID-19 Genomics UK (COG-UK) Consortium | Harper VanSteenhouse, Yumi Kasai, David Gray, Carol Clugston, Anna Dominiczak and Alex Alderton, Roberto Amato, Jeffrey Barrett, Sonia Goncalves, Ewan Harrison, David K. Jackson, Ian Johnston, Dominic Kwiatkowski, Cordelia Langford, John Sillitoe on behalf of the Wellcome Sanger Institute COVID-19 Surveillance Team |
| EPI_ISL_2846376 | Lighthouse Lab in Milton Keynes | Wellcome Sanger Institute for the COVID-19 Genomics UK (COG-UK) Consortium | The Lighthouse Lab in Milton Keynes and Alex Alderton, Roberto Amato, Jeffrey Barrett, Sonia Goncalves, Ewan Harrison, David K. Jackson, Ian Johnston, Dominic Kwiatkowski, Cordelia Langford, John Sillitoe on behalf of the Wellcome Sanger Institute COVID-19 Surveillance Team |
| EPI_ISL_2846377 | Berkshire and Surrey Pathology Services Lighthouse Laboratory | Wellcome Sanger Institute for the COVID-19 Genomics UK (COG-UK) Consortium | Berkshire and Surrey Pathology Services Lighthouse Laboratory and Alex Alderton, Roberto Amato, Jeffrey Barrett, Sonia Goncalves, Ewan Harrison, David K. Jackson, Ian Johnston, Dominic Kwiatkowski, Cordelia Langford, John Sillitoe on behalf of the Wellcome Sanger Institute COVID-19 Surveillance Team |
| EPI_ISL_2846378 | Lighthouse Laboratory Plymouth | Wellcome Sanger Institute for the COVID-19 Genomics UK (COG-UK) Consortium | Lighthouse Laboratory Plymouth and Alex Alderton, Roberto Amato, Jeffrey Barrett, Sonia Goncalves, Ewan Harrison, David K. Jackson, Ian Johnston, Dominic Kwiatkowski, Cordelia Langford, John Sillitoe on behalf of the Wellcome Sanger Institute COVID-19 Surveillance Team |
| EPI_ISL_2846379 | Berkshire and Surrey Pathology Services Lighthouse Laboratory | Wellcome Sanger Institute for the COVID-19 Genomics UK (COG-UK) Consortium | Berkshire and Surrey Pathology Services Lighthouse Laboratory and Alex Alderton, Roberto Amato, Jeffrey Barrett, Sonia Goncalves, Ewan Harrison, David K. Jackson, Ian Johnston, Dominic Kwiatkowski, Cordelia Langford, John Sillitoe on behalf of the Wellcome Sanger Institute COVID-19 Surveillance Team |
| EPI_ISL_2846380, EPI_ISL_2846381, EPI_ISL_2846382, EPI_ISL_2846383 | Lighthouse Lab in Milton Keynes | Wellcome Sanger Institute for the COVID-19 Genomics UK (COG-UK) Consortium | The Lighthouse Lab in Milton Keynes and Alex Alderton, Roberto Amato, Jeffrey Barrett, Sonia Goncalves, Ewan Harrison, David K. Jackson, Ian Johnston, Dominic Kwiatkowski, Cordelia Langford, John Sillitoe on behalf of the Wellcome Sanger Institute COVID-19 Surveillance Team |
| EPI_ISL_2846384, EPI_ISL_2846385 | Lighthouse Lab in Glasgow | Wellcome Sanger Institute for the COVID-19 Genomics UK (COG-UK) Consortium | Harper VanSteenhouse, Yumi Kasai, David Gray, Carol Clugston, Anna Dominiczak and Alex Alderton, Roberto Amato, Jeffrey Barrett, Sonia Goncalves, Ewan Harrison, David K. Jackson, Ian Johnston, Dominic Kwiatkowski, Cordelia Langford, John Sillitoe on behalf of the Wellcome Sanger Institute COVID-19 Surveillance Team |
| EPI_ISL_2846386 | Berkshire and Surrey Pathology Services Lighthouse Laboratory | Wellcome Sanger Institute for the COVID-19 Genomics UK (COG-UK) Consortium | Berkshire and Surrey Pathology Services Lighthouse Laboratory and Alex Alderton, Roberto Amato, Jeffrey Barrett, Sonia Goncalves, Ewan Harrison, David K. Jackson, Ian Johnston, Dominic Kwiatkowski, Cordelia Langford, John Sillitoe on behalf of the Wellcome Sanger Institute COVID-19 Surveillance Team |
| EPI_ISL_2846387 | Lighthouse Lab in Glasgow | Wellcome Sanger Institute for the COVID-19 Genomics UK (COG-UK) Consortium | Harper VanSteenhouse, Yumi Kasai, David Gray, Carol Clugston, Anna Dominiczak and Alex Alderton, Roberto Amato, Jeffrey Barrett, Sonia Goncalves, Ewan Harrison, David K. Jackson, Ian Johnston, Dominic Kwiatkowski, Cordelia Langford, John Sillitoe on behalf of the Wellcome Sanger Institute COVID-19 Surveillance Team |
| EPI_ISL_2846388, EPI_ISL_2846389 | Lighthouse Lab in Milton Keynes | Wellcome Sanger Institute for the COVID-19 Genomics UK (COG-UK) Consortium | The Lighthouse Lab in Milton Keynes and Alex Alderton, Roberto Amato, Jeffrey Barrett, Sonia Goncalves, Ewan Harrison, David K. Jackson, Ian Johnston, Dominic Kwiatkowski, Cordelia Langford, John Sillitoe on behalf of the Wellcome Sanger Institute COVID-19 Surveillance Team |
| EPI_ISL_2846390 | Berkshire and Surrey Pathology Services Lighthouse Laboratory | Wellcome Sanger Institute for the COVID-19 Genomics UK (COG-UK) Consortium | Berkshire and Surrey Pathology Services Lighthouse Laboratory and Alex Alderton, Roberto Amato, Jeffrey Barrett, Sonia Goncalves, Ewan Harrison, David K. Jackson, Ian Johnston, Dominic Kwiatkowski, Cordelia Langford, John Sillitoe on behalf of the Wellcome Sanger Institute COVID-19 Surveillance Team |
| EPI_ISL_2846391 | Lighthouse Lab in Milton Keynes | Wellcome Sanger Institute for the COVID-19 Genomics UK (COG-UK) Consortium | The Lighthouse Lab in Milton Keynes and Alex Alderton, Roberto Amato, Jeffrey Barrett, Sonia Goncalves, Ewan Harrison, David K. Jackson, Ian Johnston, Dominic Kwiatkowski, Cordelia Langford, John Sillitoe on behalf of the Wellcome Sanger Institute COVID-19 Surveillance Team |
| EPI_ISL_2846392, EPI_ISL_2846393, EPI_ISL_2846394 | Berkshire and Surrey Pathology Services Lighthouse Laboratory | Wellcome Sanger Institute for the COVID-19 Genomics UK (COG-UK) Consortium | Berkshire and Surrey Pathology Services Lighthouse Laboratory and Alex Alderton, Roberto Amato, Jeffrey Barrett, Sonia Goncalves, Ewan Harrison, David K. Jackson, Ian Johnston, Dominic Kwiatkowski, Cordelia Langford, John Sillitoe on behalf of the Wellcome Sanger Institute COVID-19 Surveillance Team |
| EPI_ISL_2846395, EPI_ISL_2846396 | Lighthouse Lab in Milton Keynes | Wellcome Sanger Institute for the COVID-19 Genomics UK (COG-UK) Consortium | The Lighthouse Lab in Milton Keynes and Alex Alderton, Roberto Amato, Jeffrey Barrett, Sonia Goncalves, Ewan Harrison, David K. Jackson, Ian Johnston, Dominic Kwiatkowski, Cordelia Langford, John Sillitoe on behalf of the Wellcome Sanger Institute COVID-19 Surveillance Team |
| EPI_ISL_2846397 | Lighthouse Lab in Glasgow | Wellcome Sanger Institute for the COVID-19 Genomics UK (COG-UK) Consortium | Harper VanSteenhouse, Yumi Kasai, David Gray, Carol Clugston, Anna Dominiczak and Alex Alderton, Roberto Amato, Jeffrey Barrett, Sonia Goncalves, Ewan Harrison, David K. Jackson, Ian Johnston, Dominic Kwiatkowski, Cordelia Langford, John Sillitoe on behalf of the Wellcome Sanger Institute COVID-19 Surveillance Team |
| EPI_ISL_2846398, EPI_ISL_2846399 | Berkshire and Surrey Pathology Services Lighthouse Laboratory | Wellcome Sanger Institute for the COVID-19 Genomics UK (COG-UK) Consortium | Berkshire and Surrey Pathology Services Lighthouse Laboratory and Alex Alderton, Roberto Amato, Jeffrey Barrett, Sonia Goncalves, Ewan Harrison, David K. Jackson, Ian Johnston, Dominic Kwiatkowski, Cordelia Langford, John Sillitoe on behalf of the Wellcome Sanger Institute COVID-19 Surveillance Team |
| EPI_ISL_2846400 | Lighthouse Lab in Milton Keynes | Wellcome Sanger Institute for the COVID-19 Genomics UK (COG-UK) Consortium | The Lighthouse Lab in Milton Keynes and Alex Alderton, Roberto Amato, Jeffrey Barrett, Sonia Goncalves, Ewan Harrison, David K. Jackson, Ian Johnston, Dominic Kwiatkowski, Cordelia Langford, John Sillitoe on behalf of the Wellcome Sanger Institute COVID-19 Surveillance Team |
| EPI_ISL_2846401, EPI_ISL_2846402 | Berkshire and Surrey Pathology Services Lighthouse Laboratory | Wellcome Sanger Institute for the COVID-19 Genomics UK (COG-UK) Consortium | Berkshire and Surrey Pathology Services Lighthouse Laboratory and Alex Alderton, Roberto Amato, Jeffrey Barrett, Sonia Goncalves, Ewan Harrison, David K. Jackson, Ian Johnston, Dominic Kwiatkowski, Cordelia Langford, John Sillitoe on behalf of the Wellcome Sanger Institute COVID-19 Surveillance Team |
| EPI_ISL_2846403 | Lighthouse Lab in Glasgow | Wellcome Sanger Institute for the COVID-19 Genomics UK (COG-UK) Consortium | Harper VanSteenhouse, Yumi Kasai, David Gray, Carol Clugston, Anna Dominiczak and Alex Alderton, Roberto Amato, Jeffrey Barrett, Sonia Goncalves, Ewan Harrison, David K. Jackson, Ian Johnston, Dominic Kwiatkowski, Cordelia Langford, John Sillitoe on behalf of the Wellcome Sanger Institute COVID-19 Surveillance Team |
| EPI_ISL_2846404 | Berkshire and Surrey Pathology Services Lighthouse Laboratory | Wellcome Sanger Institute for the COVID-19 Genomics UK (COG-UK) Consortium | Berkshire and Surrey Pathology Services Lighthouse Laboratory and Alex Alderton, Roberto Amato, Jeffrey Barrett, Sonia Goncalves, Ewan Harrison, David K. Jackson, Ian Johnston, Dominic Kwiatkowski, Cordelia Langford, John Sillitoe on behalf of the Wellcome Sanger Institute COVID-19 Surveillance Team |
| EPI_ISL_2846405 | Lighthouse Lab in Glasgow | Wellcome Sanger Institute for the COVID-19 Genomics UK (COG-UK) Consortium | Harper VanSteenhouse, Yumi Kasai, David Gray, Carol Clugston, Anna Dominiczak and Alex Alderton, Roberto Amato, Jeffrey Barrett, Sonia Goncalves, Ewan Harrison, David K. Jackson, Ian Johnston, Dominic Kwiatkowski, Cordelia Langford, John Sillitoe on behalf of the Wellcome Sanger Institute COVID-19 Surveillance Team |
| EPI_ISL_2846406, EPI_ISL_2846407 | Lighthouse Lab in Milton Keynes | Wellcome Sanger Institute for the COVID-19 Genomics UK (COG-UK) Consortium | The Lighthouse Lab in Milton Keynes and Alex Alderton, Roberto Amato, Jeffrey Barrett, Sonia Goncalves, Ewan Harrison, David K. Jackson, Ian Johnston, Dominic Kwiatkowski, Cordelia Langford, John Sillitoe on behalf of the Wellcome Sanger Institute COVID-19 Surveillance Team |
| EPI_ISL_2846408 | Berkshire and Surrey Pathology Services Lighthouse Laboratory | Wellcome Sanger Institute for the COVID-19 Genomics UK (COG-UK) Consortium | Berkshire and Surrey Pathology Services Lighthouse Laboratory and Alex Alderton, Roberto Amato, Jeffrey Barrett, Sonia Goncalves, Ewan Harrison, David K. Jackson, Ian Johnston, Dominic Kwiatkowski, Cordelia Langford, John Sillitoe on behalf of the Wellcome Sanger Institute COVID-19 Surveillance Team |
| EPI_ISL_2846409 | Lighthouse Lab in Milton Keynes | Wellcome Sanger Institute for the COVID-19 Genomics UK (COG-UK) Consortium | The Lighthouse Lab in Milton Keynes and Alex Alderton, Roberto Amato, Jeffrey Barrett, Sonia Goncalves, Ewan Harrison, David K. Jackson, Ian Johnston, Dominic Kwiatkowski, Cordelia Langford, John Sillitoe on behalf of the Wellcome Sanger Institute COVID-19 Surveillance Team |
| EPI_ISL_2846410 | Lighthouse Lab in Glasgow | Wellcome Sanger Institute for the COVID-19 Genomics UK (COG-UK) Consortium | Harper VanSteenhouse, Yumi Kasai, David Gray, Carol Clugston, Anna Dominiczak and Alex Alderton, Roberto Amato, Jeffrey Barrett, Sonia Goncalves, Ewan Harrison, David K. Jackson, Ian Johnston, Dominic Kwiatkowski, Cordelia Langford, John Sillitoe on behalf of the Wellcome Sanger Institute COVID-19 Surveillance Team |
| EPI_ISL_2846411 | Berkshire and Surrey Pathology Services Lighthouse Laboratory | Wellcome Sanger Institute for the COVID-19 Genomics UK (COG-UK) Consortium | Berkshire and Surrey Pathology Services Lighthouse Laboratory and Alex Alderton, Roberto Amato, Jeffrey Barrett, Sonia Goncalves, Ewan Harrison, David K. Jackson, Ian Johnston, Dominic Kwiatkowski, Cordelia Langford, John Sillitoe on behalf of the Wellcome Sanger Institute COVID-19 Surveillance Team |
| EPI_ISL_2846412 | Lighthouse Lab in Milton Keynes | Wellcome Sanger Institute for the COVID-19 Genomics UK (COG-UK) Consortium | The Lighthouse Lab in Milton Keynes and Alex Alderton, Roberto Amato, Jeffrey Barrett, Sonia Goncalves, Ewan Harrison, David K. Jackson, Ian Johnston, Dominic Kwiatkowski, Cordelia Langford, John Sillitoe on behalf of the Wellcome Sanger Institute COVID-19 Surveillance Team |

| | | | |
|---|---|---|---|
| EPI_ISL_2846413 | Berkshire and Surrey Pathology Services Lighthouse Laboratory | Wellcome Sanger Institute for the COVID-19 Genomics UK (COG-UK) Consortium | Berkshire and Surrey Pathology Services Lighthouse Laboratory and Alex Alderton, Roberto Amato, Jeffrey Barrett, Sonia Goncalves, Ewan Harrison, David K. Jackson, Ian Johnston, Dominic Kwiatkowski, Cordelia Langford, John Sillitoe on behalf of the Wellcome Sanger Institute COVID-19 Surveillance Team |
| EPI_ISL_2846414 | Lighthouse Lab in Glasgow | Wellcome Sanger Institute for the COVID-19 Genomics UK (COG-UK) Consortium | Harper VanSteenhouse, Yumi Kasai, David Gray, Carol Clugston, Anna Dominiczak and Alex Alderton, Roberto Amato, Jeffrey Barrett, Sonia Goncalves, Ewan Harrison, David K. Jackson, Ian Johnston, Dominic Kwiatkowski, Cordelia Langford, John Sillitoe on behalf of the Wellcome Sanger Institute COVID-19 Surveillance Team |
| EPI_ISL_2846415, EPI_ISL_2846416 | Berkshire and Surrey Pathology Services Lighthouse Laboratory | Wellcome Sanger Institute for the COVID-19 Genomics UK (COG-UK) Consortium | Berkshire and Surrey Pathology Services Lighthouse Laboratory and Alex Alderton, Roberto Amato, Jeffrey Barrett, Sonia Goncalves, Ewan Harrison, David K. Jackson, Ian Johnston, Dominic Kwiatkowski, Cordelia Langford, John Sillitoe on behalf of the Wellcome Sanger Institute COVID-19 Surveillance Team |
| EPI_ISL_2846417 | Lighthouse Lab in Glasgow | Wellcome Sanger Institute for the COVID-19 Genomics UK (COG-UK) Consortium | Harper VanSteenhouse, Yumi Kasai, David Gray, Carol Clugston, Anna Dominiczak and Alex Alderton, Roberto Amato, Jeffrey Barrett, Sonia Goncalves, Ewan Harrison, David K. Jackson, Ian Johnston, Dominic Kwiatkowski, Cordelia Langford, John Sillitoe on behalf of the Wellcome Sanger Institute COVID-19 Surveillance Team |
| EPI_ISL_2846418 | Berkshire and Surrey Pathology Services Lighthouse Laboratory | Wellcome Sanger Institute for the COVID-19 Genomics UK (COG-UK) Consortium | Berkshire and Surrey Pathology Services Lighthouse Laboratory and Alex Alderton, Roberto Amato, Jeffrey Barrett, Sonia Goncalves, Ewan Harrison, David K. Jackson, Ian Johnston, Dominic Kwiatkowski, Cordelia Langford, John Sillitoe on behalf of the Wellcome Sanger Institute COVID-19 Surveillance Team |
| EPI_ISL_2846419 | Lighthouse Lab in Glasgow | Wellcome Sanger Institute for the COVID-19 Genomics UK (COG-UK) Consortium | Harper VanSteenhouse, Yumi Kasai, David Gray, Carol Clugston, Anna Dominiczak and Alex Alderton, Roberto Amato, Jeffrey Barrett, Sonia Goncalves, Ewan Harrison, David K. Jackson, Ian Johnston, Dominic Kwiatkowski, Cordelia Langford, John Sillitoe on behalf of the Wellcome Sanger Institute COVID-19 Surveillance Team |
| EPI_ISL_2846420, EPI_ISL_2846421, EPI_ISL_2846422 | Lighthouse Lab in Milton Keynes | Wellcome Sanger Institute for the COVID-19 Genomics UK (COG-UK) Consortium | The Lighthouse Lab in Milton Keynes and Alex Alderton, Roberto Amato, Jeffrey Barrett, Sonia Goncalves, Ewan Harrison, David K. Jackson, Ian Johnston, Dominic Kwiatkowski, Cordelia Langford, John Sillitoe on behalf of the Wellcome Sanger Institute COVID-19 Surveillance Team |
| EPI_ISL_2846423 | Berkshire and Surrey Pathology Services Lighthouse Laboratory | Wellcome Sanger Institute for the COVID-19 Genomics UK (COG-UK) Consortium | Berkshire and Surrey Pathology Services Lighthouse Laboratory and Alex Alderton, Roberto Amato, Jeffrey Barrett, Sonia Goncalves, Ewan Harrison, David K. Jackson, Ian Johnston, Dominic Kwiatkowski, Cordelia Langford, John Sillitoe on behalf of the Wellcome Sanger Institute COVID-19 Surveillance Team |
| EPI_ISL_2846424, EPI_ISL_2846425 | Lighthouse Lab in Glasgow | Wellcome Sanger Institute for the COVID-19 Genomics UK (COG-UK) Consortium | Harper VanSteenhouse, Yumi Kasai, David Gray, Carol Clugston, Anna Dominiczak and Alex Alderton, Roberto Amato, Jeffrey Barrett, Sonia Goncalves, Ewan Harrison, David K. Jackson, Ian Johnston, Dominic Kwiatkowski, Cordelia Langford, John Sillitoe on behalf of the Wellcome Sanger Institute COVID-19 Surveillance Team |
| EPI_ISL_2846426, EPI_ISL_2846427 | Lighthouse Lab in Milton Keynes | Wellcome Sanger Institute for the COVID-19 Genomics UK (COG-UK) Consortium | The Lighthouse Lab in Milton Keynes and Alex Alderton, Roberto Amato, Jeffrey Barrett, Sonia Goncalves, Ewan Harrison, David K. Jackson, Ian Johnston, Dominic Kwiatkowski, Cordelia Langford, John Sillitoe on behalf of the Wellcome Sanger Institute COVID-19 Surveillance Team |
| EPI_ISL_2846428 | Lighthouse Lab in Glasgow | Wellcome Sanger Institute for the COVID-19 Genomics UK (COG-UK) Consortium | Harper VanSteenhouse, Yumi Kasai, David Gray, Carol Clugston, Anna Dominiczak and Alex Alderton, Roberto Amato, Jeffrey Barrett, Sonia Goncalves, Ewan Harrison, David K. Jackson, Ian Johnston, Dominic Kwiatkowski, Cordelia Langford, John Sillitoe on behalf of the Wellcome Sanger Institute COVID-19 Surveillance Team |
| EPI_ISL_2846429 | Lighthouse Lab in Milton Keynes | Wellcome Sanger Institute for the COVID-19 Genomics UK (COG-UK) Consortium | The Lighthouse Lab in Milton Keynes and Alex Alderton, Roberto Amato, Jeffrey Barrett, Sonia Goncalves, Ewan Harrison, David K. Jackson, Ian Johnston, Dominic Kwiatkowski, Cordelia Langford, John Sillitoe on behalf of the Wellcome Sanger Institute COVID-19 Surveillance Team |
| EPI_ISL_2846430 | Lighthouse Lab in Glasgow | Wellcome Sanger Institute for the COVID-19 Genomics UK (COG-UK) Consortium | Harper VanSteenhouse, Yumi Kasai, David Gray, Carol Clugston, Anna Dominiczak and Alex Alderton, Roberto Amato, Jeffrey Barrett, Sonia Goncalves, Ewan Harrison, David K. Jackson, Ian Johnston, Dominic Kwiatkowski, Cordelia Langford, John Sillitoe on behalf of the Wellcome Sanger Institute COVID-19 Surveillance Team |
| EPI_ISL_2846431, EPI_ISL_2846432 | Berkshire and Surrey Pathology Services Lighthouse Laboratory | Wellcome Sanger Institute for the COVID-19 Genomics UK (COG-UK) Consortium | Berkshire and Surrey Pathology Services Lighthouse Laboratory and Alex Alderton, Roberto Amato, Jeffrey Barrett, Sonia Goncalves, Ewan Harrison, David K. Jackson, Ian Johnston, Dominic Kwiatkowski, Cordelia Langford, John Sillitoe on behalf of the Wellcome Sanger Institute COVID-19 Surveillance Team |
| EPI_ISL_2846433 | Lighthouse Lab in Milton Keynes | Wellcome Sanger Institute for the COVID-19 Genomics UK (COG-UK) Consortium | The Lighthouse Lab in Milton Keynes and Alex Alderton, Roberto Amato, Jeffrey Barrett, Sonia Goncalves, Ewan Harrison, David K. Jackson, Ian Johnston, Dominic Kwiatkowski, Cordelia Langford, John Sillitoe on behalf of the Wellcome Sanger Institute COVID-19 Surveillance Team |
| EPI_ISL_2846434 | Lighthouse Lab in Glasgow | Wellcome Sanger Institute for the COVID-19 Genomics UK (COG-UK) Consortium | Harper VanSteenhouse, Yumi Kasai, David Gray, Carol Clugston, Anna Dominiczak and Alex Alderton, Roberto Amato, Jeffrey Barrett, Sonia Goncalves, Ewan Harrison, David K. Jackson, Ian Johnston, Dominic Kwiatkowski, Cordelia Langford, John Sillitoe on behalf of the Wellcome Sanger Institute COVID-19 Surveillance Team |
| EPI_ISL_2846435 | Lighthouse Lab in Milton Keynes | Wellcome Sanger Institute for the COVID-19 Genomics UK (COG-UK) Consortium | The Lighthouse Lab in Milton Keynes and Alex Alderton, Roberto Amato, Jeffrey Barrett, Sonia Goncalves, Ewan Harrison, David K. Jackson, Ian Johnston, Dominic Kwiatkowski, Cordelia Langford, John Sillitoe on behalf of the Wellcome Sanger Institute COVID-19 Surveillance Team |
| EPI_ISL_2846436, EPI_ISL_2846437, EPI_ISL_2846438, EPI_ISL_2846439 | Berkshire and Surrey Pathology Services Lighthouse Laboratory | Wellcome Sanger Institute for the COVID-19 Genomics UK (COG-UK) Consortium | Berkshire and Surrey Pathology Services Lighthouse Laboratory and Alex Alderton, Roberto Amato, Jeffrey Barrett, Sonia Goncalves, Ewan Harrison, David K. Jackson, Ian Johnston, Dominic Kwiatkowski, Cordelia Langford, John Sillitoe on behalf of the Wellcome Sanger Institute COVID-19 Surveillance Team |
| EPI_ISL_2846440, EPI_ISL_2846441 | Lighthouse Lab in Milton Keynes | Wellcome Sanger Institute for the COVID-19 Genomics UK (COG-UK) Consortium | The Lighthouse Lab in Milton Keynes and Alex Alderton, Roberto Amato, Jeffrey Barrett, Sonia Goncalves, Ewan Harrison, David K. Jackson, Ian Johnston, Dominic Kwiatkowski, Cordelia Langford, John Sillitoe on behalf of the Wellcome Sanger Institute COVID-19 Surveillance Team |
| EPI_ISL_2846442 | Berkshire and Surrey Pathology Services Lighthouse Laboratory | Wellcome Sanger Institute for the COVID-19 Genomics UK (COG-UK) Consortium | Berkshire and Surrey Pathology Services Lighthouse Laboratory and Alex Alderton, Roberto Amato, Jeffrey Barrett, Sonia Goncalves, Ewan Harrison, David K. Jackson, Ian Johnston, Dominic Kwiatkowski, Cordelia Langford, John Sillitoe on behalf of the Wellcome Sanger Institute COVID-19 Surveillance Team |
| EPI_ISL_2846443 | Lighthouse Lab in Glasgow | Wellcome Sanger Institute for the COVID-19 Genomics UK (COG-UK) Consortium | Harper VanSteenhouse, Yumi Kasai, David Gray, Carol Clugston, Anna Dominiczak and Alex Alderton, Roberto Amato, Jeffrey Barrett, Sonia Goncalves, Ewan Harrison, David K. Jackson, Ian Johnston, Dominic Kwiatkowski, Cordelia Langford, John Sillitoe on behalf of the Wellcome Sanger Institute COVID-19 Surveillance Team |
| EPI_ISL_2846444, EPI_ISL_2846445 | Berkshire and Surrey Pathology Services Lighthouse Laboratory | Wellcome Sanger Institute for the COVID-19 Genomics UK (COG-UK) Consortium | Berkshire and Surrey Pathology Services Lighthouse Laboratory and Alex Alderton, Roberto Amato, Jeffrey Barrett, Sonia Goncalves, Ewan Harrison, David K. Jackson, Ian Johnston, Dominic Kwiatkowski, Cordelia Langford, John Sillitoe on behalf of the Wellcome Sanger Institute COVID-19 Surveillance Team |
| EPI_ISL_2846446 | Lighthouse Lab in Glasgow | Wellcome Sanger Institute for the COVID-19 Genomics UK (COG-UK) Consortium | Harper VanSteenhouse, Yumi Kasai, David Gray, Carol Clugston, Anna Dominiczak and Alex Alderton, Roberto Amato, Jeffrey Barrett, Sonia Goncalves, Ewan Harrison, David K. Jackson, Ian Johnston, Dominic Kwiatkowski, Cordelia Langford, John Sillitoe on behalf of the Wellcome Sanger Institute COVID-19 Surveillance Team |
| EPI_ISL_2846447 | Lighthouse Lab in Milton Keynes | Wellcome Sanger Institute for the COVID-19 Genomics UK (COG-UK) Consortium | The Lighthouse Lab in Milton Keynes and Alex Alderton, Roberto Amato, Jeffrey Barrett, Sonia Goncalves, Ewan Harrison, David K. Jackson, Ian Johnston, Dominic Kwiatkowski, Cordelia Langford, John Sillitoe on behalf of the Wellcome Sanger Institute COVID-19 Surveillance Team |
| EPI_ISL_2846448 | Berkshire and Surrey Pathology Services Lighthouse Laboratory | Wellcome Sanger Institute for the COVID-19 Genomics UK (COG-UK) Consortium | Berkshire and Surrey Pathology Services Lighthouse Laboratory and Alex Alderton, Roberto Amato, Jeffrey Barrett, Sonia Goncalves, Ewan Harrison, David K. Jackson, Ian Johnston, Dominic Kwiatkowski, Cordelia Langford, John Sillitoe on behalf of the Wellcome Sanger Institute COVID-19 Surveillance Team |
| EPI_ISL_2846449 | Lighthouse Lab in Milton Keynes | Wellcome Sanger Institute for the COVID-19 Genomics UK (COG-UK) Consortium | The Lighthouse Lab in Milton Keynes and Alex Alderton, Roberto Amato, Jeffrey Barrett, Sonia Goncalves, Ewan Harrison, David K. Jackson, Ian Johnston, Dominic Kwiatkowski, Cordelia Langford, John Sillitoe on behalf of the Wellcome Sanger Institute COVID-19 Surveillance Team |
| EPI_ISL_2846450, EPI_ISL_2846451 | Lighthouse Lab in Glasgow | Wellcome Sanger Institute for the COVID-19 Genomics UK (COG-UK) Consortium | Harper VanSteenhouse, Yumi Kasai, David Gray, Carol Clugston, Anna Dominiczak and Alex Alderton, Roberto Amato, Jeffrey Barrett, Sonia Goncalves, Ewan Harrison, David K. Jackson, Ian Johnston, Dominic Kwiatkowski, Cordelia Langford, John Sillitoe on behalf of the Wellcome Sanger Institute COVID-19 Surveillance Team |
| EPI_ISL_2846452 | Lighthouse Lab in Milton Keynes | Wellcome Sanger Institute for the COVID-19 Genomics UK (COG-UK) Consortium | The Lighthouse Lab in Milton Keynes and Alex Alderton, Roberto Amato, Jeffrey Barrett, Sonia Goncalves, Ewan Harrison, David K. Jackson, Ian Johnston, Dominic Kwiatkowski, Cordelia Langford, John Sillitoe on behalf of the Wellcome Sanger Institute COVID-19 Surveillance Team |
| EPI_ISL_2846453 | Berkshire and Surrey Pathology Services Lighthouse Laboratory | Wellcome Sanger Institute for the COVID-19 Genomics UK (COG-UK) Consortium | Berkshire and Surrey Pathology Services Lighthouse Laboratory and Alex Alderton, Roberto Amato, Jeffrey Barrett, Sonia Goncalves, Ewan Harrison, David K. Jackson, Ian Johnston, Dominic Kwiatkowski, Cordelia Langford, John Sillitoe on behalf of the Wellcome Sanger Institute COVID-19 Surveillance Team |
| EPI_ISL_2846454 | Lighthouse Lab in Glasgow | Wellcome Sanger Institute for the COVID-19 Genomics UK (COG-UK) Consortium | Harper VanSteenhouse, Yumi Kasai, David Gray, Carol Clugston, Anna Dominiczak and Alex Alderton, Roberto Amato, Jeffrey Barrett, Sonia Goncalves, Ewan Harrison, David K. Jackson, Ian Johnston, Dominic Kwiatkowski, Cordelia Langford, John Sillitoe on behalf of the Wellcome Sanger Institute COVID-19 Surveillance Team |
| EPI_ISL_2846455 | Berkshire and Surrey Pathology Services Lighthouse Laboratory | Wellcome Sanger Institute for the COVID-19 Genomics UK (COG-UK) Consortium | Berkshire and Surrey Pathology Services Lighthouse Laboratory and Alex Alderton, Roberto Amato, Jeffrey Barrett, Sonia Goncalves, Ewan Harrison, David K. Jackson, Ian Johnston, Dominic Kwiatkowski, Cordelia Langford, John Sillitoe on behalf of the Wellcome Sanger Institute COVID-19 Surveillance Team |
| EPI_ISL_2846456 | Lighthouse Lab in Glasgow | Wellcome Sanger Institute for the COVID-19 Genomics UK (COG-UK) Consortium | Harper VanSteenhouse, Yumi Kasai, David Gray, Carol Clugston, Anna Dominiczak and Alex Alderton, Roberto Amato, Jeffrey Barrett, Sonia Goncalves, Ewan Harrison, David K. Jackson, Ian Johnston, Dominic Kwiatkowski, Cordelia Langford, John Sillitoe on behalf of the Wellcome Sanger Institute COVID-19 Surveillance Team |
| EPI_ISL_2846457, EPI_ISL_2846458 | Lighthouse Lab in Milton Keynes | Wellcome Sanger Institute for the COVID-19 Genomics UK (COG-UK) Consortium | The Lighthouse Lab in Milton Keynes and Alex Alderton, Roberto Amato, Jeffrey Barrett, Sonia Goncalves, Ewan Harrison, David K. Jackson, Ian Johnston, Dominic Kwiatkowski, Cordelia Langford, John Sillitoe on behalf of the Wellcome Sanger Institute COVID-19 Surveillance Team |
| EPI_ISL_2846459 | Berkshire and Surrey Pathology Services Lighthouse Laboratory | Wellcome Sanger Institute for the COVID-19 Genomics UK (COG-UK) Consortium | Berkshire and Surrey Pathology Services Lighthouse Laboratory and Alex Alderton, Roberto Amato, Jeffrey Barrett, Sonia Goncalves, Ewan Harrison, David K. Jackson, Ian Johnston, Dominic Kwiatkowski, Cordelia Langford, John Sillitoe on behalf of the Wellcome Sanger Institute COVID-19 Surveillance Team |
| EPI_ISL_2846460 | Lighthouse Lab in Glasgow | Wellcome Sanger Institute for the COVID-19 Genomics UK (COG-UK) Consortium | Harper VanSteenhouse, Yumi Kasai, David Gray, Carol Clugston, Anna Dominiczak and Alex Alderton, Roberto Amato, Jeffrey Barrett, Sonia Goncalves, Ewan Harrison, David K. Jackson, Ian Johnston, Dominic Kwiatkowski, Cordelia Langford, John Sillitoe on behalf of the Wellcome Sanger Institute COVID-19 Surveillance Team |
| EPI_ISL_2846461, EPI_ISL_2846462 | Berkshire and Surrey Pathology Services Lighthouse Laboratory | Wellcome Sanger Institute for the COVID-19 Genomics UK (COG-UK) Consortium | Berkshire and Surrey Pathology Services Lighthouse Laboratory and Alex Alderton, Roberto Amato, Jeffrey Barrett, Sonia Goncalves, Ewan Harrison, David K. Jackson, Ian Johnston, Dominic Kwiatkowski, Cordelia Langford, John Sillitoe on behalf of the Wellcome Sanger Institute COVID-19 Surveillance Team |
| EPI_ISL_2846463 | Lighthouse Lab in Glasgow | Wellcome Sanger Institute for the COVID-19 Genomics UK (COG-UK) Consortium | Harper VanSteenhouse, Yumi Kasai, David Gray, Carol Clugston, Anna Dominiczak and Alex Alderton, Roberto Amato, Jeffrey Barrett, Sonia Goncalves, Ewan Harrison, David K. Jackson, Ian Johnston, Dominic Kwiatkowski, Cordelia Langford, John Sillitoe on behalf of the Wellcome Sanger Institute COVID-19 Surveillance Team |
| EPI_ISL_2846464 | Berkshire and Surrey Pathology Services Lighthouse Laboratory | Wellcome Sanger Institute for the COVID-19 Genomics UK (COG-UK) Consortium | Berkshire and Surrey Pathology Services Lighthouse Laboratory and Alex Alderton, Roberto Amato, Jeffrey Barrett, Sonia Goncalves, Ewan Harrison, David K. Jackson, Ian Johnston, Dominic Kwiatkowski, Cordelia Langford, John Sillitoe on behalf of the Wellcome Sanger Institute COVID-19 Surveillance Team |
| EPI_ISL_2846465 | Lighthouse Lab in Glasgow | Wellcome Sanger Institute for the COVID-19 Genomics UK (COG-UK) Consortium | Harper VanSteenhouse, Yumi Kasai, David Gray, Carol Clugston, Anna Dominiczak and Alex Alderton, Roberto Amato, Jeffrey Barrett, Sonia Goncalves, Ewan Harrison, David K. Jackson, Ian Johnston, Dominic Kwiatkowski, Cordelia Langford, John Sillitoe on behalf of the Wellcome Sanger Institute COVID-19 Surveillance Team |
| EPI_ISL_2846466, EPI_ISL_2846467 | Berkshire and Surrey Pathology Services Lighthouse Laboratory | Wellcome Sanger Institute for the COVID-19 Genomics UK (COG-UK) Consortium | Berkshire and Surrey Pathology Services Lighthouse Laboratory and Alex Alderton, Roberto Amato, Jeffrey Barrett, Sonia Goncalves, Ewan Harrison, David K. Jackson, Ian Johnston, Dominic Kwiatkowski, Cordelia Langford, John Sillitoe on behalf of the Wellcome Sanger Institute COVID-19 Surveillance Team |
| EPI_ISL_2846468, EPI_ISL_2846469 | Lighthouse Lab in Milton Keynes | Wellcome Sanger Institute for the COVID-19 Genomics UK (COG-UK) Consortium | The Lighthouse Lab in Milton Keynes and Alex Alderton, Roberto Amato, Jeffrey Barrett, Sonia Goncalves, Ewan Harrison, David K. Jackson, Ian Johnston, Dominic Kwiatkowski, Cordelia Langford, John Sillitoe on behalf of the Wellcome Sanger Institute COVID-19 Surveillance Team |
| EPI_ISL_2846470 | Berkshire and Surrey Pathology Services Lighthouse Laboratory | Wellcome Sanger Institute for the COVID-19 Genomics UK (COG-UK) Consortium | Berkshire and Surrey Pathology Services Lighthouse Laboratory and Alex Alderton, Roberto Amato, Jeffrey Barrett, Sonia Goncalves, Ewan Harrison, David K. Jackson, Ian Johnston, Dominic Kwiatkowski, Cordelia Langford, John Sillitoe on behalf of the Wellcome Sanger Institute COVID-19 Surveillance Team |
| EPI_ISL_2846471, EPI_ISL_2846472 | Lighthouse Lab in Milton Keynes | Wellcome Sanger Institute for the COVID-19 Genomics UK (COG-UK) Consortium | The Lighthouse Lab in Milton Keynes and Alex Alderton, Roberto Amato, Jeffrey Barrett, Sonia Goncalves, Ewan Harrison, David K. Jackson, Ian Johnston, Dominic Kwiatkowski, Cordelia Langford, John Sillitoe on behalf of the Wellcome Sanger Institute COVID-19 Surveillance Team |
| EPI_ISL_2846473 | Lighthouse Lab in Glasgow | Wellcome Sanger Institute for the COVID-19 Genomics UK (COG-UK) Consortium | Harper VanSteenhouse, Yumi Kasai, David Gray, Carol Clugston, Anna Dominiczak and Alex Alderton, Roberto Amato, Jeffrey Barrett, Sonia Goncalves, Ewan Harrison, David K. Jackson, Ian Johnston, Dominic Kwiatkowski, Cordelia Langford, John Sillitoe on behalf of the Wellcome Sanger Institute COVID-19 Surveillance Team |

| | | | |
|---|---|---|---|
| EPI_ISL_2846474 | Lighthouse Lab in Milton Keynes | Wellcome Sanger Institute for the COVID-19 Genomics UK (COG-UK) Consortium | The Lighthouse Lab in Milton Keynes and Alex Alderton, Roberto Amato, Jeffrey Barrett, Sonia Goncalves, Ewan Harrison, David K. Jackson, Ian Johnston, Dominic Kwiatkowski, Cordelia Langford, John Sillitoe on behalf of the Wellcome Sanger Institute COVID-19 Surveillance Team |
| EPI_ISL_2846475, EPI_ISL_2846476, EPI_ISL_2846477, EPI_ISL_2846478 | Berkshire and Surrey Pathology Services Lighthouse Laboratory | Wellcome Sanger Institute for the COVID-19 Genomics UK (COG-UK) Consortium | Berkshire and Surrey Pathology Services Lighthouse Laboratory and Alex Alderton, Roberto Amato, Jeffrey Barrett, Sonia Goncalves, Ewan Harrison, David K. Jackson, Ian Johnston, Dominic Kwiatkowski, Cordelia Langford, John Sillitoe on behalf of the Wellcome Sanger Institute COVID-19 Surveillance Team |
| EPI_ISL_2846479 | Lighthouse Laboratory Plymouth | Wellcome Sanger Institute for the COVID-19 Genomics UK (COG-UK) Consortium | Lighthouse Laboratory Plymouth and Alex Alderton, Roberto Amato, Jeffrey Barrett, Sonia Goncalves, Ewan Harrison, David K. Jackson, Ian Johnston, Dominic Kwiatkowski, Cordelia Langford, John Sillitoe on behalf of the Wellcome Sanger Institute COVID-19 Surveillance Team |
| EPI_ISL_2846480, EPI_ISL_2846481 | Lighthouse Lab in Milton Keynes | Wellcome Sanger Institute for the COVID-19 Genomics UK (COG-UK) Consortium | The Lighthouse Lab in Milton Keynes and Alex Alderton, Roberto Amato, Jeffrey Barrett, Sonia Goncalves, Ewan Harrison, David K. Jackson, Ian Johnston, Dominic Kwiatkowski, Cordelia Langford, John Sillitoe on behalf of the Wellcome Sanger Institute COVID-19 Surveillance Team |
| EPI_ISL_2846482 | Berkshire and Surrey Pathology Services Lighthouse Laboratory | Wellcome Sanger Institute for the COVID-19 Genomics UK (COG-UK) Consortium | Berkshire and Surrey Pathology Services Lighthouse Laboratory and Alex Alderton, Roberto Amato, Jeffrey Barrett, Sonia Goncalves, Ewan Harrison, David K. Jackson, Ian Johnston, Dominic Kwiatkowski, Cordelia Langford, John Sillitoe on behalf of the Wellcome Sanger Institute COVID-19 Surveillance Team |
| EPI_ISL_2846483 | Lighthouse Lab in Milton Keynes | Wellcome Sanger Institute for the COVID-19 Genomics UK (COG-UK) Consortium | The Lighthouse Lab in Milton Keynes and Alex Alderton, Roberto Amato, Jeffrey Barrett, Sonia Goncalves, Ewan Harrison, David K. Jackson, Ian Johnston, Dominic Kwiatkowski, Cordelia Langford, John Sillitoe on behalf of the Wellcome Sanger Institute COVID-19 Surveillance Team |
| EPI_ISL_2846484 | Lighthouse Laboratory Plymouth | Wellcome Sanger Institute for the COVID-19 Genomics UK (COG-UK) Consortium | Lighthouse Laboratory Plymouth and Alex Alderton, Roberto Amato, Jeffrey Barrett, Sonia Goncalves, Ewan Harrison, David K. Jackson, Ian Johnston, Dominic Kwiatkowski, Cordelia Langford, John Sillitoe on behalf of the Wellcome Sanger Institute COVID-19 Surveillance Team |
| EPI_ISL_2846485 | Lighthouse Lab in Glasgow | Wellcome Sanger Institute for the COVID-19 Genomics UK (COG-UK) Consortium | Harper VanSteenhouse, Yumi Kasai, David Gray, Carol Clugston, Anna Dominiczak and Alex Alderton, Roberto Amato, Jeffrey Barrett, Sonia Goncalves, Ewan Harrison, David K. Jackson, Ian Johnston, Dominic Kwiatkowski, Cordelia Langford, John Sillitoe on behalf of the Wellcome Sanger Institute COVID-19 Surveillance Team |
| EPI_ISL_2846486 | Lighthouse Lab in Milton Keynes | Wellcome Sanger Institute for the COVID-19 Genomics UK (COG-UK) Consortium | The Lighthouse Lab in Milton Keynes and Alex Alderton, Roberto Amato, Jeffrey Barrett, Sonia Goncalves, Ewan Harrison, David K. Jackson, Ian Johnston, Dominic Kwiatkowski, Cordelia Langford, John Sillitoe on behalf of the Wellcome Sanger Institute COVID-19 Surveillance Team |
| EPI_ISL_2846487 | Berkshire and Surrey Pathology Services Lighthouse Laboratory | Wellcome Sanger Institute for the COVID-19 Genomics UK (COG-UK) Consortium | Berkshire and Surrey Pathology Services Lighthouse Laboratory and Alex Alderton, Roberto Amato, Jeffrey Barrett, Sonia Goncalves, Ewan Harrison, David K. Jackson, Ian Johnston, Dominic Kwiatkowski, Cordelia Langford, John Sillitoe on behalf of the Wellcome Sanger Institute COVID-19 Surveillance Team |
| EPI_ISL_2846488, EPI_ISL_2846489, EPI_ISL_2846490 | Lighthouse Lab in Milton Keynes | Wellcome Sanger Institute for the COVID-19 Genomics UK (COG-UK) Consortium | The Lighthouse Lab in Milton Keynes and Alex Alderton, Roberto Amato, Jeffrey Barrett, Sonia Goncalves, Ewan Harrison, David K. Jackson, Ian Johnston, Dominic Kwiatkowski, Cordelia Langford, John Sillitoe on behalf of the Wellcome Sanger Institute COVID-19 Surveillance Team |
| EPI_ISL_2846491 | Lighthouse Lab in Glasgow | Wellcome Sanger Institute for the COVID-19 Genomics UK (COG-UK) Consortium | Harper VanSteenhouse, Yumi Kasai, David Gray, Carol Clugston, Anna Dominiczak and Alex Alderton, Roberto Amato, Jeffrey Barrett, Sonia Goncalves, Ewan Harrison, David K. Jackson, Ian Johnston, Dominic Kwiatkowski, Cordelia Langford, John Sillitoe on behalf of the Wellcome Sanger Institute COVID-19 Surveillance Team |
| EPI_ISL_2846492 | Lighthouse Lab in Milton Keynes | Wellcome Sanger Institute for the COVID-19 Genomics UK (COG-UK) Consortium | The Lighthouse Lab in Milton Keynes and Alex Alderton, Roberto Amato, Jeffrey Barrett, Sonia Goncalves, Ewan Harrison, David K. Jackson, Ian Johnston, Dominic Kwiatkowski, Cordelia Langford, John Sillitoe on behalf of the Wellcome Sanger Institute COVID-19 Surveillance Team |
| EPI_ISL_2846493 | Berkshire and Surrey Pathology Services Lighthouse Laboratory | Wellcome Sanger Institute for the COVID-19 Genomics UK (COG-UK) Consortium | Berkshire and Surrey Pathology Services Lighthouse Laboratory and Alex Alderton, Roberto Amato, Jeffrey Barrett, Sonia Goncalves, Ewan Harrison, David K. Jackson, Ian Johnston, Dominic Kwiatkowski, Cordelia Langford, John Sillitoe on behalf of the Wellcome Sanger Institute COVID-19 Surveillance Team |
| EPI_ISL_2846494 | Lighthouse Lab in Glasgow | Wellcome Sanger Institute for the COVID-19 Genomics UK (COG-UK) Consortium | Harper VanSteenhouse, Yumi Kasai, David Gray, Carol Clugston, Anna Dominiczak and Alex Alderton, Roberto Amato, Jeffrey Barrett, Sonia Goncalves, Ewan Harrison, David K. Jackson, Ian Johnston, Dominic Kwiatkowski, Cordelia Langford, John Sillitoe on behalf of the Wellcome Sanger Institute COVID-19 Surveillance Team |
| EPI_ISL_2846495, EPI_ISL_2846496, EPI_ISL_2846497 | Lighthouse Lab in Milton Keynes | Wellcome Sanger Institute for the COVID-19 Genomics UK (COG-UK) Consortium | The Lighthouse Lab in Milton Keynes and Alex Alderton, Roberto Amato, Jeffrey Barrett, Sonia Goncalves, Ewan Harrison, David K. Jackson, Ian Johnston, Dominic Kwiatkowski, Cordelia Langford, John Sillitoe on behalf of the Wellcome Sanger Institute COVID-19 Surveillance Team |
| EPI_ISL_2846498 | Lighthouse Lab in Glasgow | Wellcome Sanger Institute for the COVID-19 Genomics UK (COG-UK) Consortium | Harper VanSteenhouse, Yumi Kasai, David Gray, Carol Clugston, Anna Dominiczak and Alex Alderton, Roberto Amato, Jeffrey Barrett, Sonia Goncalves, Ewan Harrison, David K. Jackson, Ian Johnston, Dominic Kwiatkowski, Cordelia Langford, John Sillitoe on behalf of the Wellcome Sanger Institute COVID-19 Surveillance Team |
| EPI_ISL_2846499 | Lighthouse Laboratory Plymouth | Wellcome Sanger Institute for the COVID-19 Genomics UK (COG-UK) Consortium | Lighthouse Laboratory Plymouth and Alex Alderton, Roberto Amato, Jeffrey Barrett, Sonia Goncalves, Ewan Harrison, David K. Jackson, Ian Johnston, Dominic Kwiatkowski, Cordelia Langford, John Sillitoe on behalf of the Wellcome Sanger Institute COVID-19 Surveillance Team |
| EPI_ISL_2846500 | Lighthouse Lab in Glasgow | Wellcome Sanger Institute for the COVID-19 Genomics UK (COG-UK) Consortium | Harper VanSteenhouse, Yumi Kasai, David Gray, Carol Clugston, Anna Dominiczak and Alex Alderton, Roberto Amato, Jeffrey Barrett, Sonia Goncalves, Ewan Harrison, David K. Jackson, Ian Johnston, Dominic Kwiatkowski, Cordelia Langford, John Sillitoe on behalf of the Wellcome Sanger Institute COVID-19 Surveillance Team |
| EPI_ISL_2846501 | Lighthouse Laboratory Plymouth | Wellcome Sanger Institute for the COVID-19 Genomics UK (COG-UK) Consortium | Lighthouse Laboratory Plymouth and Alex Alderton, Roberto Amato, Jeffrey Barrett, Sonia Goncalves, Ewan Harrison, David K. Jackson, Ian Johnston, Dominic Kwiatkowski, Cordelia Langford, John Sillitoe on behalf of the Wellcome Sanger Institute COVID-19 Surveillance Team |
| EPI_ISL_2846502 | Lighthouse Lab in Milton Keynes | Wellcome Sanger Institute for the COVID-19 Genomics UK (COG-UK) Consortium | The Lighthouse Lab in Milton Keynes and Alex Alderton, Roberto Amato, Jeffrey Barrett, Sonia Goncalves, Ewan Harrison, David K. Jackson, Ian Johnston, Dominic Kwiatkowski, Cordelia Langford, John Sillitoe on behalf of the Wellcome Sanger Institute COVID-19 Surveillance Team |
| EPI_ISL_2846503 | Lighthouse Lab in Glasgow | Wellcome Sanger Institute for the COVID-19 Genomics UK (COG-UK) Consortium | Harper VanSteenhouse, Yumi Kasai, David Gray, Carol Clugston, Anna Dominiczak and Alex Alderton, Roberto Amato, Jeffrey Barrett, Sonia Goncalves, Ewan Harrison, David K. Jackson, Ian Johnston, Dominic Kwiatkowski, Cordelia Langford, John Sillitoe on behalf of the Wellcome Sanger Institute COVID-19 Surveillance Team |
| EPI_ISL_2846504 | Berkshire and Surrey Pathology Services Lighthouse Laboratory | Wellcome Sanger Institute for the COVID-19 Genomics UK (COG-UK) Consortium | Berkshire and Surrey Pathology Services Lighthouse Laboratory and Alex Alderton, Roberto Amato, Jeffrey Barrett, Sonia Goncalves, Ewan Harrison, David K. Jackson, Ian Johnston, Dominic Kwiatkowski, Cordelia Langford, John Sillitoe on behalf of the Wellcome Sanger Institute COVID-19 Surveillance Team |
| EPI_ISL_2846505 | Lighthouse Lab in Milton Keynes | Wellcome Sanger Institute for the COVID-19 Genomics UK (COG-UK) Consortium | The Lighthouse Lab in Milton Keynes and Alex Alderton, Roberto Amato, Jeffrey Barrett, Sonia Goncalves, Ewan Harrison, David K. Jackson, Ian Johnston, Dominic Kwiatkowski, Cordelia Langford, John Sillitoe on behalf of the Wellcome Sanger Institute COVID-19 Surveillance Team |
| EPI_ISL_2846506 | Berkshire and Surrey Pathology Services Lighthouse Laboratory | Wellcome Sanger Institute for the COVID-19 Genomics UK (COG-UK) Consortium | Berkshire and Surrey Pathology Services Lighthouse Laboratory and Alex Alderton, Roberto Amato, Jeffrey Barrett, Sonia Goncalves, Ewan Harrison, David K. Jackson, Ian Johnston, Dominic Kwiatkowski, Cordelia Langford, John Sillitoe on behalf of the Wellcome Sanger Institute COVID-19 Surveillance Team |
| EPI_ISL_2846507 | Lighthouse Lab in Milton Keynes | Wellcome Sanger Institute for the COVID-19 Genomics UK (COG-UK) Consortium | The Lighthouse Lab in Milton Keynes and Alex Alderton, Roberto Amato, Jeffrey Barrett, Sonia Goncalves, Ewan Harrison, David K. Jackson, Ian Johnston, Dominic Kwiatkowski, Cordelia Langford, John Sillitoe on behalf of the Wellcome Sanger Institute COVID-19 Surveillance Team |
| EPI_ISL_2846508 | Berkshire and Surrey Pathology Services Lighthouse Laboratory | Wellcome Sanger Institute for the COVID-19 Genomics UK (COG-UK) Consortium | Berkshire and Surrey Pathology Services Lighthouse Laboratory and Alex Alderton, Roberto Amato, Jeffrey Barrett, Sonia Goncalves, Ewan Harrison, David K. Jackson, Ian Johnston, Dominic Kwiatkowski, Cordelia Langford, John Sillitoe on behalf of the Wellcome Sanger Institute COVID-19 Surveillance Team |
| EPI_ISL_2846509 | Lighthouse Lab in Glasgow | Wellcome Sanger Institute for the COVID-19 Genomics UK (COG-UK) Consortium | Harper VanSteenhouse, Yumi Kasai, David Gray, Carol Clugston, Anna Dominiczak and Alex Alderton, Roberto Amato, Jeffrey Barrett, Sonia Goncalves, Ewan Harrison, David K. Jackson, Ian Johnston, Dominic Kwiatkowski, Cordelia Langford, John Sillitoe on behalf of the Wellcome Sanger Institute COVID-19 Surveillance Team |
| EPI_ISL_2846510 | Berkshire and Surrey Pathology Services Lighthouse Laboratory | Wellcome Sanger Institute for the COVID-19 Genomics UK (COG-UK) Consortium | Berkshire and Surrey Pathology Services Lighthouse Laboratory and Alex Alderton, Roberto Amato, Jeffrey Barrett, Sonia Goncalves, Ewan Harrison, David K. Jackson, Ian Johnston, Dominic Kwiatkowski, Cordelia Langford, John Sillitoe on behalf of the Wellcome Sanger Institute COVID-19 Surveillance Team |
| EPI_ISL_2846511, EPI_ISL_2846512, EPI_ISL_2846513, EPI_ISL_2846514 | Lighthouse Lab in Milton Keynes | Wellcome Sanger Institute for the COVID-19 Genomics UK (COG-UK) Consortium | The Lighthouse Lab in Milton Keynes and Alex Alderton, Roberto Amato, Jeffrey Barrett, Sonia Goncalves, Ewan Harrison, David K. Jackson, Ian Johnston, Dominic Kwiatkowski, Cordelia Langford, John Sillitoe on behalf of the Wellcome Sanger Institute COVID-19 Surveillance Team |
| EPI_ISL_2846515 | Lighthouse Lab in Glasgow | Wellcome Sanger Institute for the COVID-19 Genomics UK (COG-UK) Consortium | Harper VanSteenhouse, Yumi Kasai, David Gray, Carol Clugston, Anna Dominiczak and Alex Alderton, Roberto Amato, Jeffrey Barrett, Sonia Goncalves, Ewan Harrison, David K. Jackson, Ian Johnston, Dominic Kwiatkowski, Cordelia Langford, John Sillitoe on behalf of the Wellcome Sanger Institute COVID-19 Surveillance Team |
| EPI_ISL_2846516 | Lighthouse Lab in Milton Keynes | Wellcome Sanger Institute for the COVID-19 Genomics UK (COG-UK) Consortium | The Lighthouse Lab in Milton Keynes and Alex Alderton, Roberto Amato, Jeffrey Barrett, Sonia Goncalves, Ewan Harrison, David K. Jackson, Ian Johnston, Dominic Kwiatkowski, Cordelia Langford, John Sillitoe on behalf of the Wellcome Sanger Institute COVID-19 Surveillance Team |
| EPI_ISL_2846517 | Berkshire and Surrey Pathology Services Lighthouse Laboratory | Wellcome Sanger Institute for the COVID-19 Genomics UK (COG-UK) Consortium | Berkshire and Surrey Pathology Services Lighthouse Laboratory and Alex Alderton, Roberto Amato, Jeffrey Barrett, Sonia Goncalves, Ewan Harrison, David K. Jackson, Ian Johnston, Dominic Kwiatkowski, Cordelia Langford, John Sillitoe on behalf of the Wellcome Sanger Institute COVID-19 Surveillance Team |
| EPI_ISL_2846518 | Lighthouse Laboratory Plymouth | Wellcome Sanger Institute for the COVID-19 Genomics UK (COG-UK) Consortium | Lighthouse Laboratory Plymouth and Alex Alderton, Roberto Amato, Jeffrey Barrett, Sonia Goncalves, Ewan Harrison, David K. Jackson, Ian Johnston, Dominic Kwiatkowski, Cordelia Langford, John Sillitoe on behalf of the Wellcome Sanger Institute COVID-19 Surveillance Team |
| EPI_ISL_2846519, EPI_ISL_2846520 | Berkshire and Surrey Pathology Services Lighthouse Laboratory | Wellcome Sanger Institute for the COVID-19 Genomics UK (COG-UK) Consortium | Berkshire and Surrey Pathology Services Lighthouse Laboratory and Alex Alderton, Roberto Amato, Jeffrey Barrett, Sonia Goncalves, Ewan Harrison, David K. Jackson, Ian Johnston, Dominic Kwiatkowski, Cordelia Langford, John Sillitoe on behalf of the Wellcome Sanger Institute COVID-19 Surveillance Team |
| EPI_ISL_2846521 | Lighthouse Lab in Glasgow | Wellcome Sanger Institute for the COVID-19 Genomics UK (COG-UK) Consortium | Harper VanSteenhouse, Yumi Kasai, David Gray, Carol Clugston, Anna Dominiczak and Alex Alderton, Roberto Amato, Jeffrey Barrett, Sonia Goncalves, Ewan Harrison, David K. Jackson, Ian Johnston, Dominic Kwiatkowski, Cordelia Langford, John Sillitoe on behalf of the Wellcome Sanger Institute COVID-19 Surveillance Team |
| EPI_ISL_2846522 | Lighthouse Lab in Milton Keynes | Wellcome Sanger Institute for the COVID-19 Genomics UK (COG-UK) Consortium | The Lighthouse Lab in Milton Keynes and Alex Alderton, Roberto Amato, Jeffrey Barrett, Sonia Goncalves, Ewan Harrison, David K. Jackson, Ian Johnston, Dominic Kwiatkowski, Cordelia Langford, John Sillitoe on behalf of the Wellcome Sanger Institute COVID-19 Surveillance Team |
| EPI_ISL_2846523 | Berkshire and Surrey Pathology Services Lighthouse Laboratory | Wellcome Sanger Institute for the COVID-19 Genomics UK (COG-UK) Consortium | Berkshire and Surrey Pathology Services Lighthouse Laboratory and Alex Alderton, Roberto Amato, Jeffrey Barrett, Sonia Goncalves, Ewan Harrison, David K. Jackson, Ian Johnston, Dominic Kwiatkowski, Cordelia Langford, John Sillitoe on behalf of the Wellcome Sanger Institute COVID-19 Surveillance Team |
| EPI_ISL_2846524 | Lighthouse Lab in Glasgow | Wellcome Sanger Institute for the COVID-19 Genomics UK (COG-UK) Consortium | Harper VanSteenhouse, Yumi Kasai, David Gray, Carol Clugston, Anna Dominiczak and Alex Alderton, Roberto Amato, Jeffrey Barrett, Sonia Goncalves, Ewan Harrison, David K. Jackson, Ian Johnston, Dominic Kwiatkowski, Cordelia Langford, John Sillitoe on behalf of the Wellcome Sanger Institute COVID-19 Surveillance Team |
| EPI_ISL_2846525 | Lighthouse Lab in Milton Keynes | Wellcome Sanger Institute for the COVID-19 Genomics UK (COG-UK) Consortium | The Lighthouse Lab in Milton Keynes and Alex Alderton, Roberto Amato, Jeffrey Barrett, Sonia Goncalves, Ewan Harrison, David K. Jackson, Ian Johnston, Dominic Kwiatkowski, Cordelia Langford, John Sillitoe on behalf of the Wellcome Sanger Institute COVID-19 Surveillance Team |
| EPI_ISL_2846526, EPI_ISL_2846527 | Berkshire and Surrey Pathology Services Lighthouse Laboratory | Wellcome Sanger Institute for the COVID-19 Genomics UK (COG-UK) Consortium | Berkshire and Surrey Pathology Services Lighthouse Laboratory and Alex Alderton, Roberto Amato, Jeffrey Barrett, Sonia Goncalves, Ewan Harrison, David K. Jackson, Ian Johnston, Dominic Kwiatkowski, Cordelia Langford, John Sillitoe on behalf of the Wellcome Sanger Institute COVID-19 Surveillance Team |
| EPI_ISL_2846528 | Lighthouse Lab in Milton Keynes | Wellcome Sanger Institute for the COVID-19 Genomics UK (COG-UK) Consortium | The Lighthouse Lab in Milton Keynes and Alex Alderton, Roberto Amato, Jeffrey Barrett, Sonia Goncalves, Ewan Harrison, David K. Jackson, Ian Johnston, Dominic Kwiatkowski, Cordelia Langford, John Sillitoe on behalf of the Wellcome Sanger Institute COVID-19 Surveillance Team |
| EPI_ISL_2846529, | Berkshire and Surrey Pathology Services Lighthouse Laboratory | Wellcome Sanger Institute for the COVID-19 Genomics UK (COG-UK) | Berkshire and Surrey Pathology Services Lighthouse Laboratory and Alex Alderton, Roberto Amato, Jeffrey Barrett, Sonia Goncalves, Ewan Harrison, David K. Jackson, Ian Johnston, Dominic Kwiatkowski, Cordelia Langford, John Sillitoe on behalf |

| | | | |
|---|---|---|---|
| EPI_ISL_2846530 | | Consortium | of the Wellcome Sanger Institute COVID-19 Surveillance Team |
| EPI_ISL_2846531, EPI_ISL_2846532 | Lighthouse Lab in Milton Keynes | Wellcome Sanger Institute for the COVID-19 Genomics UK (COG-UK) Consortium | The Lighthouse Lab in Milton Keynes and Alex Alderton, Roberto Amato, Jeffrey Barrett, Sonia Goncalves, Ewan Harrison, David K. Jackson, Ian Johnston, Dominic Kwiatkowski, Cordelia Langford, John Sillitoe on behalf of the Wellcome Sanger Institute COVID-19 Surveillance Team |
| EPI_ISL_2846533, EPI_ISL_2846534 | Lighthouse Lab in Glasgow | Wellcome Sanger Institute for the COVID-19 Genomics UK (COG-UK) Consortium | Harper VanSteenhouse, Yumi Kasai, David Gray, Carol Clugston, Anna Dominiczak and Alex Alderton, Roberto Amato, Jeffrey Barrett, Sonia Goncalves, Ewan Harrison, David K. Jackson, Ian Johnston, Dominic Kwiatkowski, Cordelia Langford, John Sillitoe on behalf of the Wellcome Sanger Institute COVID-19 Surveillance Team |
| EPI_ISL_2846535, EPI_ISL_2846536, EPI_ISL_2846537 | Lighthouse Lab in Milton Keynes | Wellcome Sanger Institute for the COVID-19 Genomics UK (COG-UK) Consortium | The Lighthouse Lab in Milton Keynes and Alex Alderton, Roberto Amato, Jeffrey Barrett, Sonia Goncalves, Ewan Harrison, David K. Jackson, Ian Johnston, Dominic Kwiatkowski, Cordelia Langford, John Sillitoe on behalf of the Wellcome Sanger Institute COVID-19 Surveillance Team |
| EPI_ISL_2846538, EPI_ISL_2846539 | Lighthouse Lab in Glasgow | Wellcome Sanger Institute for the COVID-19 Genomics UK (COG-UK) Consortium | Harper VanSteenhouse, Yumi Kasai, David Gray, Carol Clugston, Anna Dominiczak and Alex Alderton, Roberto Amato, Jeffrey Barrett, Sonia Goncalves, Ewan Harrison, David K. Jackson, Ian Johnston, Dominic Kwiatkowski, Cordelia Langford, John Sillitoe on behalf of the Wellcome Sanger Institute COVID-19 Surveillance Team |
| EPI_ISL_2846540 | Berkshire and Surrey Pathology Services Lighthouse Laboratory | Wellcome Sanger Institute for the COVID-19 Genomics UK (COG-UK) Consortium | Berkshire and Surrey Pathology Services Lighthouse Laboratory and Alex Alderton, Roberto Amato, Jeffrey Barrett, Sonia Goncalves, Ewan Harrison, David K. Jackson, Ian Johnston, Dominic Kwiatkowski, Cordelia Langford, John Sillitoe on behalf of the Wellcome Sanger Institute COVID-19 Surveillance Team |
| EPI_ISL_2846541, EPI_ISL_2846542 | Lighthouse Lab in Milton Keynes | Wellcome Sanger Institute for the COVID-19 Genomics UK (COG-UK) Consortium | The Lighthouse Lab in Milton Keynes and Alex Alderton, Roberto Amato, Jeffrey Barrett, Sonia Goncalves, Ewan Harrison, David K. Jackson, Ian Johnston, Dominic Kwiatkowski, Cordelia Langford, John Sillitoe on behalf of the Wellcome Sanger Institute COVID-19 Surveillance Team |
| EPI_ISL_2846543 | Lighthouse Lab in Glasgow | Wellcome Sanger Institute for the COVID-19 Genomics UK (COG-UK) Consortium | Harper VanSteenhouse, Yumi Kasai, David Gray, Carol Clugston, Anna Dominiczak and Alex Alderton, Roberto Amato, Jeffrey Barrett, Sonia Goncalves, Ewan Harrison, David K. Jackson, Ian Johnston, Dominic Kwiatkowski, Cordelia Langford, John Sillitoe on behalf of the Wellcome Sanger Institute COVID-19 Surveillance Team |
| EPI_ISL_2846544, EPI_ISL_2846545 | Lighthouse Lab in Milton Keynes | Wellcome Sanger Institute for the COVID-19 Genomics UK (COG-UK) Consortium | The Lighthouse Lab in Milton Keynes and Alex Alderton, Roberto Amato, Jeffrey Barrett, Sonia Goncalves, Ewan Harrison, David K. Jackson, Ian Johnston, Dominic Kwiatkowski, Cordelia Langford, John Sillitoe on behalf of the Wellcome Sanger Institute COVID-19 Surveillance Team |
| EPI_ISL_2846546, EPI_ISL_2846547 | Berkshire and Surrey Pathology Services Lighthouse Laboratory | Wellcome Sanger Institute for the COVID-19 Genomics UK (COG-UK) Consortium | Berkshire and Surrey Pathology Services Lighthouse Laboratory and Alex Alderton, Roberto Amato, Jeffrey Barrett, Sonia Goncalves, Ewan Harrison, David K. Jackson, Ian Johnston, Dominic Kwiatkowski, Cordelia Langford, John Sillitoe on behalf of the Wellcome Sanger Institute COVID-19 Surveillance Team |
| EPI_ISL_2846548, EPI_ISL_2846549 | Lighthouse Lab in Milton Keynes | Wellcome Sanger Institute for the COVID-19 Genomics UK (COG-UK) Consortium | The Lighthouse Lab in Milton Keynes and Alex Alderton, Roberto Amato, Jeffrey Barrett, Sonia Goncalves, Ewan Harrison, David K. Jackson, Ian Johnston, Dominic Kwiatkowski, Cordelia Langford, John Sillitoe on behalf of the Wellcome Sanger Institute COVID-19 Surveillance Team |
| EPI_ISL_2846550 | Berkshire and Surrey Pathology Services Lighthouse Laboratory | Wellcome Sanger Institute for the COVID-19 Genomics UK (COG-UK) Consortium | Berkshire and Surrey Pathology Services Lighthouse Laboratory and Alex Alderton, Roberto Amato, Jeffrey Barrett, Sonia Goncalves, Ewan Harrison, David K. Jackson, Ian Johnston, Dominic Kwiatkowski, Cordelia Langford, John Sillitoe on behalf of the Wellcome Sanger Institute COVID-19 Surveillance Team |
| EPI_ISL_2846551 | Lighthouse Lab in Milton Keynes | Wellcome Sanger Institute for the COVID-19 Genomics UK (COG-UK) Consortium | The Lighthouse Lab in Milton Keynes and Alex Alderton, Roberto Amato, Jeffrey Barrett, Sonia Goncalves, Ewan Harrison, David K. Jackson, Ian Johnston, Dominic Kwiatkowski, Cordelia Langford, John Sillitoe on behalf of the Wellcome Sanger Institute COVID-19 Surveillance Team |
| EPI_ISL_2846552, EPI_ISL_2846553 | Berkshire and Surrey Pathology Services Lighthouse Laboratory | Wellcome Sanger Institute for the COVID-19 Genomics UK (COG-UK) Consortium | Berkshire and Surrey Pathology Services Lighthouse Laboratory and Alex Alderton, Roberto Amato, Jeffrey Barrett, Sonia Goncalves, Ewan Harrison, David K. Jackson, Ian Johnston, Dominic Kwiatkowski, Cordelia Langford, John Sillitoe on behalf of the Wellcome Sanger Institute COVID-19 Surveillance Team |
| EPI_ISL_2846554, EPI_ISL_2846555 | Lighthouse Lab in Milton Keynes | Wellcome Sanger Institute for the COVID-19 Genomics UK (COG-UK) Consortium | The Lighthouse Lab in Milton Keynes and Alex Alderton, Roberto Amato, Jeffrey Barrett, Sonia Goncalves, Ewan Harrison, David K. Jackson, Ian Johnston, Dominic Kwiatkowski, Cordelia Langford, John Sillitoe on behalf of the Wellcome Sanger Institute COVID-19 Surveillance Team |
| EPI_ISL_2846556 | Lighthouse Lab in Glasgow | Wellcome Sanger Institute for the COVID-19 Genomics UK (COG-UK) Consortium | Harper VanSteenhouse, Yumi Kasai, David Gray, Carol Clugston, Anna Dominiczak and Alex Alderton, Roberto Amato, Jeffrey Barrett, Sonia Goncalves, Ewan Harrison, David K. Jackson, Ian Johnston, Dominic Kwiatkowski, Cordelia Langford, John Sillitoe on behalf of the Wellcome Sanger Institute COVID-19 Surveillance Team |
| EPI_ISL_2846557 | Lighthouse Lab in Milton Keynes | Wellcome Sanger Institute for the COVID-19 Genomics UK (COG-UK) Consortium | The Lighthouse Lab in Milton Keynes and Alex Alderton, Roberto Amato, Jeffrey Barrett, Sonia Goncalves, Ewan Harrison, David K. Jackson, Ian Johnston, Dominic Kwiatkowski, Cordelia Langford, John Sillitoe on behalf of the Wellcome Sanger Institute COVID-19 Surveillance Team |
| EPI_ISL_2846558 | Berkshire and Surrey Pathology Services Lighthouse Laboratory | Wellcome Sanger Institute for the COVID-19 Genomics UK (COG-UK) Consortium | Berkshire and Surrey Pathology Services Lighthouse Laboratory and Alex Alderton, Roberto Amato, Jeffrey Barrett, Sonia Goncalves, Ewan Harrison, David K. Jackson, Ian Johnston, Dominic Kwiatkowski, Cordelia Langford, John Sillitoe on behalf of the Wellcome Sanger Institute COVID-19 Surveillance Team |
| EPI_ISL_2846559, EPI_ISL_2846560 | Lighthouse Lab in Milton Keynes | Wellcome Sanger Institute for the COVID-19 Genomics UK (COG-UK) Consortium | The Lighthouse Lab in Milton Keynes and Alex Alderton, Roberto Amato, Jeffrey Barrett, Sonia Goncalves, Ewan Harrison, David K. Jackson, Ian Johnston, Dominic Kwiatkowski, Cordelia Langford, John Sillitoe on behalf of the Wellcome Sanger Institute COVID-19 Surveillance Team |
| EPI_ISL_2846561, EPI_ISL_2846562, EPI_ISL_2846563 | Berkshire and Surrey Pathology Services Lighthouse Laboratory | Wellcome Sanger Institute for the COVID-19 Genomics UK (COG-UK) Consortium | Berkshire and Surrey Pathology Services Lighthouse Laboratory and Alex Alderton, Roberto Amato, Jeffrey Barrett, Sonia Goncalves, Ewan Harrison, David K. Jackson, Ian Johnston, Dominic Kwiatkowski, Cordelia Langford, John Sillitoe on behalf of the Wellcome Sanger Institute COVID-19 Surveillance Team |
| EPI_ISL_2846564 | Lighthouse Lab in Milton Keynes | Wellcome Sanger Institute for the COVID-19 Genomics UK (COG-UK) Consortium | The Lighthouse Lab in Milton Keynes and Alex Alderton, Roberto Amato, Jeffrey Barrett, Sonia Goncalves, Ewan Harrison, David K. Jackson, Ian Johnston, Dominic Kwiatkowski, Cordelia Langford, John Sillitoe on behalf of the Wellcome Sanger Institute COVID-19 Surveillance Team |
| EPI_ISL_2846565 | Berkshire and Surrey Pathology Services Lighthouse Laboratory | Wellcome Sanger Institute for the COVID-19 Genomics UK (COG-UK) Consortium | Berkshire and Surrey Pathology Services Lighthouse Laboratory and Alex Alderton, Roberto Amato, Jeffrey Barrett, Sonia Goncalves, Ewan Harrison, David K. Jackson, Ian Johnston, Dominic Kwiatkowski, Cordelia Langford, John Sillitoe on behalf of the Wellcome Sanger Institute COVID-19 Surveillance Team |
| EPI_ISL_2846566 | Lighthouse Lab in Milton Keynes | Wellcome Sanger Institute for the COVID-19 Genomics UK (COG-UK) Consortium | The Lighthouse Lab in Milton Keynes and Alex Alderton, Roberto Amato, Jeffrey Barrett, Sonia Goncalves, Ewan Harrison, David K. Jackson, Ian Johnston, Dominic Kwiatkowski, Cordelia Langford, John Sillitoe on behalf of the Wellcome Sanger Institute COVID-19 Surveillance Team |
| EPI_ISL_2846567 | Berkshire and Surrey Pathology Services Lighthouse Laboratory | Wellcome Sanger Institute for the COVID-19 Genomics UK (COG-UK) Consortium | Berkshire and Surrey Pathology Services Lighthouse Laboratory and Alex Alderton, Roberto Amato, Jeffrey Barrett, Sonia Goncalves, Ewan Harrison, David K. Jackson, Ian Johnston, Dominic Kwiatkowski, Cordelia Langford, John Sillitoe on behalf of the Wellcome Sanger Institute COVID-19 Surveillance Team |
| EPI_ISL_2846568 | Lighthouse Lab in Milton Keynes | Wellcome Sanger Institute for the COVID-19 Genomics UK (COG-UK) Consortium | The Lighthouse Lab in Milton Keynes and Alex Alderton, Roberto Amato, Jeffrey Barrett, Sonia Goncalves, Ewan Harrison, David K. Jackson, Ian Johnston, Dominic Kwiatkowski, Cordelia Langford, John Sillitoe on behalf of the Wellcome Sanger Institute COVID-19 Surveillance Team |
| EPI_ISL_2846569 | Lighthouse Lab in Glasgow | Wellcome Sanger Institute for the COVID-19 Genomics UK (COG-UK) Consortium | Harper VanSteenhouse, Yumi Kasai, David Gray, Carol Clugston, Anna Dominiczak and Alex Alderton, Roberto Amato, Jeffrey Barrett, Sonia Goncalves, Ewan Harrison, David K. Jackson, Ian Johnston, Dominic Kwiatkowski, Cordelia Langford, John Sillitoe on behalf of the Wellcome Sanger Institute COVID-19 Surveillance Team |
| EPI_ISL_2846570 | Lighthouse Lab in Milton Keynes | Wellcome Sanger Institute for the COVID-19 Genomics UK (COG-UK) Consortium | The Lighthouse Lab in Milton Keynes and Alex Alderton, Roberto Amato, Jeffrey Barrett, Sonia Goncalves, Ewan Harrison, David K. Jackson, Ian Johnston, Dominic Kwiatkowski, Cordelia Langford, John Sillitoe on behalf of the Wellcome Sanger Institute COVID-19 Surveillance Team |
| EPI_ISL_2846571 | Berkshire and Surrey Pathology Services Lighthouse Laboratory | Wellcome Sanger Institute for the COVID-19 Genomics UK (COG-UK) Consortium | Berkshire and Surrey Pathology Services Lighthouse Laboratory and Alex Alderton, Roberto Amato, Jeffrey Barrett, Sonia Goncalves, Ewan Harrison, David K. Jackson, Ian Johnston, Dominic Kwiatkowski, Cordelia Langford, John Sillitoe on behalf of the Wellcome Sanger Institute COVID-19 Surveillance Team |
| EPI_ISL_2846572, EPI_ISL_2846573 | Lighthouse Lab in Milton Keynes | Wellcome Sanger Institute for the COVID-19 Genomics UK (COG-UK) Consortium | The Lighthouse Lab in Milton Keynes and Alex Alderton, Roberto Amato, Jeffrey Barrett, Sonia Goncalves, Ewan Harrison, David K. Jackson, Ian Johnston, Dominic Kwiatkowski, Cordelia Langford, John Sillitoe on behalf of the Wellcome Sanger Institute COVID-19 Surveillance Team |
| EPI_ISL_2846574, EPI_ISL_2846575, EPI_ISL_2846576 | Lighthouse Lab in Glasgow | Wellcome Sanger Institute for the COVID-19 Genomics UK (COG-UK) Consortium | Harper VanSteenhouse, Yumi Kasai, David Gray, Carol Clugston, Anna Dominiczak and Alex Alderton, Roberto Amato, Jeffrey Barrett, Sonia Goncalves, Ewan Harrison, David K. Jackson, Ian Johnston, Dominic Kwiatkowski, Cordelia Langford, John Sillitoe on behalf of the Wellcome Sanger Institute COVID-19 Surveillance Team |
| EPI_ISL_2846577 | Berkshire and Surrey Pathology Services Lighthouse Laboratory | Wellcome Sanger Institute for the COVID-19 Genomics UK (COG-UK) Consortium | Berkshire and Surrey Pathology Services Lighthouse Laboratory and Alex Alderton, Roberto Amato, Jeffrey Barrett, Sonia Goncalves, Ewan Harrison, David K. Jackson, Ian Johnston, Dominic Kwiatkowski, Cordelia Langford, John Sillitoe on behalf of the Wellcome Sanger Institute COVID-19 Surveillance Team |
| EPI_ISL_2846578 | Lighthouse Lab in Glasgow | Wellcome Sanger Institute for the COVID-19 Genomics UK (COG-UK) Consortium | Harper VanSteenhouse, Yumi Kasai, David Gray, Carol Clugston, Anna Dominiczak and Alex Alderton, Roberto Amato, Jeffrey Barrett, Sonia Goncalves, Ewan Harrison, David K. Jackson, Ian Johnston, Dominic Kwiatkowski, Cordelia Langford, John Sillitoe on behalf of the Wellcome Sanger Institute COVID-19 Surveillance Team |
| EPI_ISL_2846579 | Lighthouse Lab in Milton Keynes | Wellcome Sanger Institute for the COVID-19 Genomics UK (COG-UK) Consortium | The Lighthouse Lab in Milton Keynes and Alex Alderton, Roberto Amato, Jeffrey Barrett, Sonia Goncalves, Ewan Harrison, David K. Jackson, Ian Johnston, Dominic Kwiatkowski, Cordelia Langford, John Sillitoe on behalf of the Wellcome Sanger Institute COVID-19 Surveillance Team |
| EPI_ISL_2846580 | Lighthouse Lab in Glasgow | Wellcome Sanger Institute for the COVID-19 Genomics UK (COG-UK) Consortium | Harper VanSteenhouse, Yumi Kasai, David Gray, Carol Clugston, Anna Dominiczak and Alex Alderton, Roberto Amato, Jeffrey Barrett, Sonia Goncalves, Ewan Harrison, David K. Jackson, Ian Johnston, Dominic Kwiatkowski, Cordelia Langford, John Sillitoe on behalf of the Wellcome Sanger Institute COVID-19 Surveillance Team |
| EPI_ISL_2846581 | Lighthouse Lab in Alderley Park | Wellcome Sanger Institute for the COVID-19 Genomics UK (COG-UK) Consortium | Jacquelyn Wynn, Mairead Hyland, The Lighthouse Lab in Alderley Park and Alex Alderton, Roberto Amato, Jeffrey Barrett, Sonia Goncalves, Ewan Harrison, David K. Jackson, Ian Johnston, Dominic Kwiatkowski, Cordelia Langford, John Sillitoe on behalf of the Wellcome Sanger Institute COVID-19 Surveillance Team |
| EPI_ISL_2846582 | Lighthouse Lab in Milton Keynes | Wellcome Sanger Institute for the COVID-19 Genomics UK (COG-UK) Consortium | The Lighthouse Lab in Milton Keynes and Alex Alderton, Roberto Amato, Jeffrey Barrett, Sonia Goncalves, Ewan Harrison, David K. Jackson, Ian Johnston, Dominic Kwiatkowski, Cordelia Langford, John Sillitoe on behalf of the Wellcome Sanger Institute COVID-19 Surveillance Team |
| EPI_ISL_2846583 | Health Services Laboratories | Wellcome Sanger Institute for the COVID-19 Genomics UK (COG-UK) Consortium | Health Services Laboratories and Alex Alderton, Roberto Amato, Jeffrey Barrett, Sonia Goncalves, Ewan Harrison, David K. Jackson, Ian Johnston, Dominic Kwiatkowski, Cordelia Langford, John Sillitoe on behalf of the Wellcome Sanger Institute COVID-19 Surveillance Team |
| EPI_ISL_2846584 | Lighthouse Laboratory Plymouth | Wellcome Sanger Institute for the COVID-19 Genomics UK (COG-UK) Consortium | Lighthouse Laboratory Plymouth and Alex Alderton, Roberto Amato, Jeffrey Barrett, Sonia Goncalves, Ewan Harrison, David K. Jackson, Ian Johnston, Dominic Kwiatkowski, Cordelia Langford, John Sillitoe on behalf of the Wellcome Sanger Institute COVID-19 Surveillance Team |
| EPI_ISL_2846585, EPI_ISL_2846586, EPI_ISL_2846587 | Lighthouse Lab in Milton Keynes | Wellcome Sanger Institute for the COVID-19 Genomics UK (COG-UK) Consortium | The Lighthouse Lab in Milton Keynes and Alex Alderton, Roberto Amato, Jeffrey Barrett, Sonia Goncalves, Ewan Harrison, David K. Jackson, Ian Johnston, Dominic Kwiatkowski, Cordelia Langford, John Sillitoe on behalf of the Wellcome Sanger Institute COVID-19 Surveillance Team |
| EPI_ISL_2846588, EPI_ISL_2846589 | Lighthouse Lab in Alderley Park | Wellcome Sanger Institute for the COVID-19 Genomics UK (COG-UK) Consortium | Jacquelyn Wynn, Mairead Hyland, The Lighthouse Lab in Alderley Park and Alex Alderton, Roberto Amato, Jeffrey Barrett, Sonia Goncalves, Ewan Harrison, David K. Jackson, Ian Johnston, Dominic Kwiatkowski, Cordelia Langford, John Sillitoe on behalf of the Wellcome Sanger Institute COVID-19 Surveillance Team |
| EPI_ISL_2846590 | Lighthouse Lab in Milton Keynes | Wellcome Sanger Institute for the COVID-19 Genomics UK (COG-UK) Consortium | The Lighthouse Lab in Milton Keynes and Alex Alderton, Roberto Amato, Jeffrey Barrett, Sonia Goncalves, Ewan Harrison, David K. Jackson, Ian Johnston, Dominic Kwiatkowski, Cordelia Langford, John Sillitoe on behalf of the Wellcome Sanger Institute COVID-19 Surveillance Team |
| EPI_ISL_2846591, EPI_ISL_2846592 | Lighthouse Laboratory Plymouth | Wellcome Sanger Institute for the COVID-19 Genomics UK (COG-UK) Consortium | Lighthouse Laboratory Plymouth and Alex Alderton, Roberto Amato, Jeffrey Barrett, Sonia Goncalves, Ewan Harrison, David K. Jackson, Ian Johnston, Dominic Kwiatkowski, Cordelia Langford, John Sillitoe on behalf of the Wellcome Sanger Institute COVID-19 Surveillance Team |
| EPI_ISL_2846593 | Health Services Laboratories | Wellcome Sanger Institute for the COVID-19 Genomics UK (COG-UK) Consortium | Health Services Laboratories and Alex Alderton, Roberto Amato, Jeffrey Barrett, Sonia Goncalves, Ewan Harrison, David K. Jackson, Ian Johnston, Dominic Kwiatkowski, Cordelia Langford, John Sillitoe on behalf of the Wellcome Sanger Institute COVID-19 Surveillance Team |
| EPI_ISL_2846594 | Lighthouse Laboratory Plymouth | Wellcome Sanger Institute for the COVID-19 Genomics UK (COG-UK) | Lighthouse Laboratory Plymouth and Alex Alderton, Roberto Amato, Jeffrey Barrett, Sonia Goncalves, Ewan Harrison, David K. Jackson, Ian Johnston, Dominic Kwiatkowski, Cordelia Langford, John Sillitoe on behalf of the Wellcome Sanger |

| | | | |
|---|---|---|---|
| | | Consortium | Institute COVID-19 Surveillance Team |
| EPI_ISL_2846595 | Lighthouse Lab in Milton Keynes | Wellcome Sanger Institute for the COVID-19 Genomics UK (COG-UK) Consortium | The Lighthouse Lab in Milton Keynes and Alex Alderton, Roberto Amato, Jeffrey Barrett, Sonia Goncalves, Ewan Harrison, David K. Jackson, Ian Johnston, Dominic Kwiatkowski, Cordelia Langford, John Sillitoe on behalf of the Wellcome Sanger Institute COVID-19 Surveillance Team |
| EPI_ISL_2846596, EPI_ISL_2846597 | Lighthouse Lab in Alderley Park | Wellcome Sanger Institute for the COVID-19 Genomics UK (COG-UK) Consortium | Jacquelyn Wynn, Mairead Hyland, The Lighthouse Lab in Alderley Park and Alex Alderton, Roberto Amato, Jeffrey Barrett, Sonia Goncalves, Ewan Harrison, David K. Jackson, Ian Johnston, Dominic Kwiatkowski, Cordelia Langford, John Sillitoe on behalf of the Wellcome Sanger Institute COVID-19 Surveillance Team |
| EPI_ISL_2846598 | Lighthouse Lab in Milton Keynes | Wellcome Sanger Institute for the COVID-19 Genomics UK (COG-UK) Consortium | The Lighthouse Lab in Milton Keynes and Alex Alderton, Roberto Amato, Jeffrey Barrett, Sonia Goncalves, Ewan Harrison, David K. Jackson, Ian Johnston, Dominic Kwiatkowski, Cordelia Langford, John Sillitoe on behalf of the Wellcome Sanger Institute COVID-19 Surveillance Team |
| EPI_ISL_2846599 | Lighthouse Lab in Alderley Park | Wellcome Sanger Institute for the COVID-19 Genomics UK (COG-UK) Consortium | Jacquelyn Wynn, Mairead Hyland, The Lighthouse Lab in Alderley Park and Alex Alderton, Roberto Amato, Jeffrey Barrett, Sonia Goncalves, Ewan Harrison, David K. Jackson, Ian Johnston, Dominic Kwiatkowski, Cordelia Langford, John Sillitoe on behalf of the Wellcome Sanger Institute COVID-19 Surveillance Team |
| EPI_ISL_2846600 | Health Services Laboratories | Wellcome Sanger Institute for the COVID-19 Genomics UK (COG-UK) Consortium | Health Services Laboratories and Alex Alderton, Roberto Amato, Jeffrey Barrett, Sonia Goncalves, Ewan Harrison, David K. Jackson, Ian Johnston, Dominic Kwiatkowski, Cordelia Langford, John Sillitoe on behalf of the Wellcome Sanger Institute COVID-19 Surveillance Team |
| EPI_ISL_2846601 | Lighthouse Lab in Alderley Park | Wellcome Sanger Institute for the COVID-19 Genomics UK (COG-UK) Consortium | Jacquelyn Wynn, Mairead Hyland, The Lighthouse Lab in Alderley Park and Alex Alderton, Roberto Amato, Jeffrey Barrett, Sonia Goncalves, Ewan Harrison, David K. Jackson, Ian Johnston, Dominic Kwiatkowski, Cordelia Langford, John Sillitoe on behalf of the Wellcome Sanger Institute COVID-19 Surveillance Team |
| EPI_ISL_2846602 | Lighthouse Laboratory Plymouth | Wellcome Sanger Institute for the COVID-19 Genomics UK (COG-UK) Consortium | Lighthouse Laboratory Plymouth and Alex Alderton, Roberto Amato, Jeffrey Barrett, Sonia Goncalves, Ewan Harrison, David K. Jackson, Ian Johnston, Dominic Kwiatkowski, Cordelia Langford, John Sillitoe on behalf of the Wellcome Sanger Institute COVID-19 Surveillance Team |
| EPI_ISL_2846603, EPI_ISL_2846604, EPI_ISL_2846605, EPI_ISL_2846606, EPI_ISL_2846607 | Health Services Laboratories | Wellcome Sanger Institute for the COVID-19 Genomics UK (COG-UK) Consortium | Health Services Laboratories and Alex Alderton, Roberto Amato, Jeffrey Barrett, Sonia Goncalves, Ewan Harrison, David K. Jackson, Ian Johnston, Dominic Kwiatkowski, Cordelia Langford, John Sillitoe on behalf of the Wellcome Sanger Institute COVID-19 Surveillance Team |
| EPI_ISL_2846608, EPI_ISL_2846609 | Lighthouse Laboratory Plymouth | Wellcome Sanger Institute for the COVID-19 Genomics UK (COG-UK) Consortium | Lighthouse Laboratory Plymouth and Alex Alderton, Roberto Amato, Jeffrey Barrett, Sonia Goncalves, Ewan Harrison, David K. Jackson, Ian Johnston, Dominic Kwiatkowski, Cordelia Langford, John Sillitoe on behalf of the Wellcome Sanger Institute COVID-19 Surveillance Team |
| EPI_ISL_2846610 | Lighthouse Lab in Milton Keynes | Wellcome Sanger Institute for the COVID-19 Genomics UK (COG-UK) Consortium | The Lighthouse Lab in Milton Keynes and Alex Alderton, Roberto Amato, Jeffrey Barrett, Sonia Goncalves, Ewan Harrison, David K. Jackson, Ian Johnston, Dominic Kwiatkowski, Cordelia Langford, John Sillitoe on behalf of the Wellcome Sanger Institute COVID-19 Surveillance Team |
| EPI_ISL_2846611 | Lighthouse Lab in Alderley Park | Wellcome Sanger Institute for the COVID-19 Genomics UK (COG-UK) Consortium | Jacquelyn Wynn, Mairead Hyland, The Lighthouse Lab in Alderley Park and Alex Alderton, Roberto Amato, Jeffrey Barrett, Sonia Goncalves, Ewan Harrison, David K. Jackson, Ian Johnston, Dominic Kwiatkowski, Cordelia Langford, John Sillitoe on behalf of the Wellcome Sanger Institute COVID-19 Surveillance Team |
| EPI_ISL_2846612 | Lighthouse Lab in Milton Keynes | Wellcome Sanger Institute for the COVID-19 Genomics UK (COG-UK) Consortium | The Lighthouse Lab in Milton Keynes and Alex Alderton, Roberto Amato, Jeffrey Barrett, Sonia Goncalves, Ewan Harrison, David K. Jackson, Ian Johnston, Dominic Kwiatkowski, Cordelia Langford, John Sillitoe on behalf of the Wellcome Sanger Institute COVID-19 Surveillance Team |
| EPI_ISL_2846613 | Lighthouse Lab in Alderley Park | Wellcome Sanger Institute for the COVID-19 Genomics UK (COG-UK) Consortium | Jacquelyn Wynn, Mairead Hyland, The Lighthouse Lab in Alderley Park and Alex Alderton, Roberto Amato, Jeffrey Barrett, Sonia Goncalves, Ewan Harrison, David K. Jackson, Ian Johnston, Dominic Kwiatkowski, Cordelia Langford, John Sillitoe on behalf of the Wellcome Sanger Institute COVID-19 Surveillance Team |
| EPI_ISL_2846614 | Lighthouse Lab in Milton Keynes | Wellcome Sanger Institute for the COVID-19 Genomics UK (COG-UK) Consortium | The Lighthouse Lab in Milton Keynes and Alex Alderton, Roberto Amato, Jeffrey Barrett, Sonia Goncalves, Ewan Harrison, David K. Jackson, Ian Johnston, Dominic Kwiatkowski, Cordelia Langford, John Sillitoe on behalf of the Wellcome Sanger Institute COVID-19 Surveillance Team |
| EPI_ISL_2846615 | Health Services Laboratories | Wellcome Sanger Institute for the COVID-19 Genomics UK (COG-UK) Consortium | Health Services Laboratories and Alex Alderton, Roberto Amato, Jeffrey Barrett, Sonia Goncalves, Ewan Harrison, David K. Jackson, Ian Johnston, Dominic Kwiatkowski, Cordelia Langford, John Sillitoe on behalf of the Wellcome Sanger Institute COVID-19 Surveillance Team |
| EPI_ISL_2846616 | Lighthouse Lab in Milton Keynes | Wellcome Sanger Institute for the COVID-19 Genomics UK (COG-UK) Consortium | The Lighthouse Lab in Milton Keynes and Alex Alderton, Roberto Amato, Jeffrey Barrett, Sonia Goncalves, Ewan Harrison, David K. Jackson, Ian Johnston, Dominic Kwiatkowski, Cordelia Langford, John Sillitoe on behalf of the Wellcome Sanger Institute COVID-19 Surveillance Team |
| EPI_ISL_2846617 | Health Services Laboratories | Wellcome Sanger Institute for the COVID-19 Genomics UK (COG-UK) Consortium | Health Services Laboratories and Alex Alderton, Roberto Amato, Jeffrey Barrett, Sonia Goncalves, Ewan Harrison, David K. Jackson, Ian Johnston, Dominic Kwiatkowski, Cordelia Langford, John Sillitoe on behalf of the Wellcome Sanger Institute COVID-19 Surveillance Team |
| EPI_ISL_2846618, EPI_ISL_2846619 | Lighthouse Lab in Alderley Park | Wellcome Sanger Institute for the COVID-19 Genomics UK (COG-UK) Consortium | Jacquelyn Wynn, Mairead Hyland, The Lighthouse Lab in Alderley Park and Alex Alderton, Roberto Amato, Jeffrey Barrett, Sonia Goncalves, Ewan Harrison, David K. Jackson, Ian Johnston, Dominic Kwiatkowski, Cordelia Langford, John Sillitoe on behalf of the Wellcome Sanger Institute COVID-19 Surveillance Team |
| EPI_ISL_2846620 | Health Services Laboratories | Wellcome Sanger Institute for the COVID-19 Genomics UK (COG-UK) Consortium | Health Services Laboratories and Alex Alderton, Roberto Amato, Jeffrey Barrett, Sonia Goncalves, Ewan Harrison, David K. Jackson, Ian Johnston, Dominic Kwiatkowski, Cordelia Langford, John Sillitoe on behalf of the Wellcome Sanger Institute COVID-19 Surveillance Team |
| EPI_ISL_2846621 | Lighthouse Lab in Milton Keynes | Wellcome Sanger Institute for the COVID-19 Genomics UK (COG-UK) Consortium | The Lighthouse Lab in Milton Keynes and Alex Alderton, Roberto Amato, Jeffrey Barrett, Sonia Goncalves, Ewan Harrison, David K. Jackson, Ian Johnston, Dominic Kwiatkowski, Cordelia Langford, John Sillitoe on behalf of the Wellcome Sanger Institute COVID-19 Surveillance Team |
| EPI_ISL_2846622, EPI_ISL_2846623 | Health Services Laboratories | Wellcome Sanger Institute for the COVID-19 Genomics UK (COG-UK) Consortium | Health Services Laboratories and Alex Alderton, Roberto Amato, Jeffrey Barrett, Sonia Goncalves, Ewan Harrison, David K. Jackson, Ian Johnston, Dominic Kwiatkowski, Cordelia Langford, John Sillitoe on behalf of the Wellcome Sanger Institute COVID-19 Surveillance Team |
| EPI_ISL_2846624 | Lighthouse Laboratory Plymouth | Wellcome Sanger Institute for the COVID-19 Genomics UK (COG-UK) Consortium | Lighthouse Laboratory Plymouth and Alex Alderton, Roberto Amato, Jeffrey Barrett, Sonia Goncalves, Ewan Harrison, David K. Jackson, Ian Johnston, Dominic Kwiatkowski, Cordelia Langford, John Sillitoe on behalf of the Wellcome Sanger Institute COVID-19 Surveillance Team |
| EPI_ISL_2846625, EPI_ISL_2846626 | Lighthouse Lab in Alderley Park | Wellcome Sanger Institute for the COVID-19 Genomics UK (COG-UK) Consortium | Jacquelyn Wynn, Mairead Hyland, The Lighthouse Lab in Alderley Park and Alex Alderton, Roberto Amato, Jeffrey Barrett, Sonia Goncalves, Ewan Harrison, David K. Jackson, Ian Johnston, Dominic Kwiatkowski, Cordelia Langford, John Sillitoe on behalf of the Wellcome Sanger Institute COVID-19 Surveillance Team |
| EPI_ISL_2846627 | Health Services Laboratories | Wellcome Sanger Institute for the COVID-19 Genomics UK (COG-UK) Consortium | Health Services Laboratories and Alex Alderton, Roberto Amato, Jeffrey Barrett, Sonia Goncalves, Ewan Harrison, David K. Jackson, Ian Johnston, Dominic Kwiatkowski, Cordelia Langford, John Sillitoe on behalf of the Wellcome Sanger Institute COVID-19 Surveillance Team |
| EPI_ISL_2846628 | Lighthouse Lab in Milton Keynes | Wellcome Sanger Institute for the COVID-19 Genomics UK (COG-UK) Consortium | The Lighthouse Lab in Milton Keynes and Alex Alderton, Roberto Amato, Jeffrey Barrett, Sonia Goncalves, Ewan Harrison, David K. Jackson, Ian Johnston, Dominic Kwiatkowski, Cordelia Langford, John Sillitoe on behalf of the Wellcome Sanger Institute COVID-19 Surveillance Team |
| EPI_ISL_2846629, EPI_ISL_2846630, EPI_ISL_2846631 | Lighthouse Laboratory Plymouth | Wellcome Sanger Institute for the COVID-19 Genomics UK (COG-UK) Consortium | Lighthouse Laboratory Plymouth and Alex Alderton, Roberto Amato, Jeffrey Barrett, Sonia Goncalves, Ewan Harrison, David K. Jackson, Ian Johnston, Dominic Kwiatkowski, Cordelia Langford, John Sillitoe on behalf of the Wellcome Sanger Institute COVID-19 Surveillance Team |
| EPI_ISL_2846632 | Lighthouse Lab in Milton Keynes | Wellcome Sanger Institute for the COVID-19 Genomics UK (COG-UK) Consortium | The Lighthouse Lab in Milton Keynes and Alex Alderton, Roberto Amato, Jeffrey Barrett, Sonia Goncalves, Ewan Harrison, David K. Jackson, Ian Johnston, Dominic Kwiatkowski, Cordelia Langford, John Sillitoe on behalf of the Wellcome Sanger Institute COVID-19 Surveillance Team |
| EPI_ISL_2846633, EPI_ISL_2846634 | Lighthouse Lab in Alderley Park | Wellcome Sanger Institute for the COVID-19 Genomics UK (COG-UK) Consortium | Jacquelyn Wynn, Mairead Hyland, The Lighthouse Lab in Alderley Park and Alex Alderton, Roberto Amato, Jeffrey Barrett, Sonia Goncalves, Ewan Harrison, David K. Jackson, Ian Johnston, Dominic Kwiatkowski, Cordelia Langford, John Sillitoe on behalf of the Wellcome Sanger Institute COVID-19 Surveillance Team |
| EPI_ISL_2846635 | Health Services Laboratories | Wellcome Sanger Institute for the COVID-19 Genomics UK (COG-UK) Consortium | Health Services Laboratories and Alex Alderton, Roberto Amato, Jeffrey Barrett, Sonia Goncalves, Ewan Harrison, David K. Jackson, Ian Johnston, Dominic Kwiatkowski, Cordelia Langford, John Sillitoe on behalf of the Wellcome Sanger Institute COVID-19 Surveillance Team |
| EPI_ISL_2846636 | Lighthouse Laboratory Plymouth | Wellcome Sanger Institute for the COVID-19 Genomics UK (COG-UK) Consortium | Lighthouse Laboratory Plymouth and Alex Alderton, Roberto Amato, Jeffrey Barrett, Sonia Goncalves, Ewan Harrison, David K. Jackson, Ian Johnston, Dominic Kwiatkowski, Cordelia Langford, John Sillitoe on behalf of the Wellcome Sanger Institute COVID-19 Surveillance Team |
| EPI_ISL_2846637 | Lighthouse Lab in Alderley Park | Wellcome Sanger Institute for the COVID-19 Genomics UK (COG-UK) Consortium | Jacquelyn Wynn, Mairead Hyland, The Lighthouse Lab in Alderley Park and Alex Alderton, Roberto Amato, Jeffrey Barrett, Sonia Goncalves, Ewan Harrison, David K. Jackson, Ian Johnston, Dominic Kwiatkowski, Cordelia Langford, John Sillitoe on behalf of the Wellcome Sanger Institute COVID-19 Surveillance Team |
| EPI_ISL_2846638 | Lighthouse Lab in Milton Keynes | Wellcome Sanger Institute for the COVID-19 Genomics UK (COG-UK) Consortium | The Lighthouse Lab in Milton Keynes and Alex Alderton, Roberto Amato, Jeffrey Barrett, Sonia Goncalves, Ewan Harrison, David K. Jackson, Ian Johnston, Dominic Kwiatkowski, Cordelia Langford, John Sillitoe on behalf of the Wellcome Sanger Institute COVID-19 Surveillance Team |
| EPI_ISL_2846639 | Lighthouse Lab in Alderley Park | Wellcome Sanger Institute for the COVID-19 Genomics UK (COG-UK) Consortium | Jacquelyn Wynn, Mairead Hyland, The Lighthouse Lab in Alderley Park and Alex Alderton, Roberto Amato, Jeffrey Barrett, Sonia Goncalves, Ewan Harrison, David K. Jackson, Ian Johnston, Dominic Kwiatkowski, Cordelia Langford, John Sillitoe on behalf of the Wellcome Sanger Institute COVID-19 Surveillance Team |
| EPI_ISL_2846640 | Lighthouse Lab in Milton Keynes | Wellcome Sanger Institute for the COVID-19 Genomics UK (COG-UK) Consortium | The Lighthouse Lab in Milton Keynes and Alex Alderton, Roberto Amato, Jeffrey Barrett, Sonia Goncalves, Ewan Harrison, David K. Jackson, Ian Johnston, Dominic Kwiatkowski, Cordelia Langford, John Sillitoe on behalf of the Wellcome Sanger Institute COVID-19 Surveillance Team |
| EPI_ISL_2846641 | Health Services Laboratories | Wellcome Sanger Institute for the COVID-19 Genomics UK (COG-UK) Consortium | Health Services Laboratories and Alex Alderton, Roberto Amato, Jeffrey Barrett, Sonia Goncalves, Ewan Harrison, David K. Jackson, Ian Johnston, Dominic Kwiatkowski, Cordelia Langford, John Sillitoe on behalf of the Wellcome Sanger Institute COVID-19 Surveillance Team |
| EPI_ISL_2846642 | Lighthouse Laboratory Plymouth | Wellcome Sanger Institute for the COVID-19 Genomics UK (COG-UK) Consortium | Lighthouse Laboratory Plymouth and Alex Alderton, Roberto Amato, Jeffrey Barrett, Sonia Goncalves, Ewan Harrison, David K. Jackson, Ian Johnston, Dominic Kwiatkowski, Cordelia Langford, John Sillitoe on behalf of the Wellcome Sanger Institute COVID-19 Surveillance Team |
| EPI_ISL_2846643 | Lighthouse Lab in Alderley Park | Wellcome Sanger Institute for the COVID-19 Genomics UK (COG-UK) Consortium | Jacquelyn Wynn, Mairead Hyland, The Lighthouse Lab in Alderley Park and Alex Alderton, Roberto Amato, Jeffrey Barrett, Sonia Goncalves, Ewan Harrison, David K. Jackson, Ian Johnston, Dominic Kwiatkowski, Cordelia Langford, John Sillitoe on behalf of the Wellcome Sanger Institute COVID-19 Surveillance Team |
| EPI_ISL_2846644 | Lighthouse Laboratory Plymouth | Wellcome Sanger Institute for the COVID-19 Genomics UK (COG-UK) Consortium | Lighthouse Laboratory Plymouth and Alex Alderton, Roberto Amato, Jeffrey Barrett, Sonia Goncalves, Ewan Harrison, David K. Jackson, Ian Johnston, Dominic Kwiatkowski, Cordelia Langford, John Sillitoe on behalf of the Wellcome Sanger Institute COVID-19 Surveillance Team |
| EPI_ISL_2846645 | Lighthouse Lab in Alderley Park | Wellcome Sanger Institute for the COVID-19 Genomics UK (COG-UK) Consortium | Jacquelyn Wynn, Mairead Hyland, The Lighthouse Lab in Alderley Park and Alex Alderton, Roberto Amato, Jeffrey Barrett, Sonia Goncalves, Ewan Harrison, David K. Jackson, Ian Johnston, Dominic Kwiatkowski, Cordelia Langford, John Sillitoe on behalf of the Wellcome Sanger Institute COVID-19 Surveillance Team |
| EPI_ISL_2846646, EPI_ISL_2846647, EPI_ISL_2846648 | Lighthouse Lab in Milton Keynes | Wellcome Sanger Institute for the COVID-19 Genomics UK (COG-UK) Consortium | The Lighthouse Lab in Milton Keynes and Alex Alderton, Roberto Amato, Jeffrey Barrett, Sonia Goncalves, Ewan Harrison, David K. Jackson, Ian Johnston, Dominic Kwiatkowski, Cordelia Langford, John Sillitoe on behalf of the Wellcome Sanger Institute COVID-19 Surveillance Team |
| EPI_ISL_2846649, EPI_ISL_2846650 | Lighthouse Lab in Alderley Park | Wellcome Sanger Institute for the COVID-19 Genomics UK (COG-UK) Consortium | Jacquelyn Wynn, Mairead Hyland, The Lighthouse Lab in Alderley Park and Alex Alderton, Roberto Amato, Jeffrey Barrett, Sonia Goncalves, Ewan Harrison, David K. Jackson, Ian Johnston, Dominic Kwiatkowski, Cordelia Langford, John Sillitoe on behalf of the Wellcome Sanger Institute COVID-19 Surveillance Team |
| EPI_ISL_2846651, EPI_ISL_2846652, EPI_ISL_2846653 | Lighthouse Laboratory Plymouth | Wellcome Sanger Institute for the COVID-19 Genomics UK (COG-UK) Consortium | Lighthouse Laboratory Plymouth and Alex Alderton, Roberto Amato, Jeffrey Barrett, Sonia Goncalves, Ewan Harrison, David K. Jackson, Ian Johnston, Dominic Kwiatkowski, Cordelia Langford, John Sillitoe on behalf of the Wellcome Sanger Institute COVID-19 Surveillance Team |

| | | | |
|---|---|---|---|
| EPI_ISL_2846654 | Lighthouse Lab in Alderley Park | Wellcome Sanger Institute for the COVID-19 Genomics UK (COG-UK) Consortium | Jacquelyn Wynn, Mairead Hyland, The Lighthouse Lab in Alderley Park and Alex Alderton, Roberto Amato, Jeffrey Barrett, Sonia Goncalves, Ewan Harrison, David K. Jackson, Ian Johnston, Dominic Kwiatkowski, Cordelia Langford, John Sillitoe on behalf of the Wellcome Sanger Institute COVID-19 Surveillance Team |
| EPI_ISL_2846655 | Health Services Laboratories | Wellcome Sanger Institute for the COVID-19 Genomics UK (COG-UK) Consortium | Health Services Laboratories and Alex Alderton, Roberto Amato, Jeffrey Barrett, Sonia Goncalves, Ewan Harrison, David K. Jackson, Ian Johnston, Dominic Kwiatkowski, Cordelia Langford, John Sillitoe on behalf of the Wellcome Sanger Institute COVID-19 Surveillance Team |
| EPI_ISL_2846656, EPI_ISL_2846657 | Lighthouse Laboratory Plymouth | Wellcome Sanger Institute for the COVID-19 Genomics UK (COG-UK) Consortium | Lighthouse Laboratory Plymouth and Alex Alderton, Roberto Amato, Jeffrey Barrett, Sonia Goncalves, Ewan Harrison, David K. Jackson, Ian Johnston, Dominic Kwiatkowski, Cordelia Langford, John Sillitoe on behalf of the Wellcome Sanger Institute COVID-19 Surveillance Team |
| EPI_ISL_2846658 | Lighthouse Lab in Alderley Park | Wellcome Sanger Institute for the COVID-19 Genomics UK (COG-UK) Consortium | Jacquelyn Wynn, Mairead Hyland, The Lighthouse Lab in Alderley Park and Alex Alderton, Roberto Amato, Jeffrey Barrett, Sonia Goncalves, Ewan Harrison, David K. Jackson, Ian Johnston, Dominic Kwiatkowski, Cordelia Langford, John Sillitoe on behalf of the Wellcome Sanger Institute COVID-19 Surveillance Team |
| EPI_ISL_2846659 | Lighthouse Lab in Milton Keynes | Wellcome Sanger Institute for the COVID-19 Genomics UK (COG-UK) Consortium | The Lighthouse Lab in Milton Keynes and Alex Alderton, Roberto Amato, Jeffrey Barrett, Sonia Goncalves, Ewan Harrison, David K. Jackson, Ian Johnston, Dominic Kwiatkowski, Cordelia Langford, John Sillitoe on behalf of the Wellcome Sanger Institute COVID-19 Surveillance Team |
| EPI_ISL_2846660 | Health Services Laboratories | Wellcome Sanger Institute for the COVID-19 Genomics UK (COG-UK) Consortium | Health Services Laboratories and Alex Alderton, Roberto Amato, Jeffrey Barrett, Sonia Goncalves, Ewan Harrison, David K. Jackson, Ian Johnston, Dominic Kwiatkowski, Cordelia Langford, John Sillitoe on behalf of the Wellcome Sanger Institute COVID-19 Surveillance Team |
| EPI_ISL_2846661 | Lighthouse Lab in Milton Keynes | Wellcome Sanger Institute for the COVID-19 Genomics UK (COG-UK) Consortium | The Lighthouse Lab in Milton Keynes and Alex Alderton, Roberto Amato, Jeffrey Barrett, Sonia Goncalves, Ewan Harrison, David K. Jackson, Ian Johnston, Dominic Kwiatkowski, Cordelia Langford, John Sillitoe on behalf of the Wellcome Sanger Institute COVID-19 Surveillance Team |
| EPI_ISL_2846662 | Lighthouse Laboratory Plymouth | Wellcome Sanger Institute for the COVID-19 Genomics UK (COG-UK) Consortium | Lighthouse Laboratory Plymouth and Alex Alderton, Roberto Amato, Jeffrey Barrett, Sonia Goncalves, Ewan Harrison, David K. Jackson, Ian Johnston, Dominic Kwiatkowski, Cordelia Langford, John Sillitoe on behalf of the Wellcome Sanger Institute COVID-19 Surveillance Team |
| EPI_ISL_2846663 | Lighthouse Lab in Alderley Park | Wellcome Sanger Institute for the COVID-19 Genomics UK (COG-UK) Consortium | Jacquelyn Wynn, Mairead Hyland, The Lighthouse Lab in Alderley Park and Alex Alderton, Roberto Amato, Jeffrey Barrett, Sonia Goncalves, Ewan Harrison, David K. Jackson, Ian Johnston, Dominic Kwiatkowski, Cordelia Langford, John Sillitoe on behalf of the Wellcome Sanger Institute COVID-19 Surveillance Team |
| EPI_ISL_2846665 | Lighthouse Lab in Milton Keynes | Wellcome Sanger Institute for the COVID-19 Genomics UK (COG-UK) Consortium | The Lighthouse Lab in Milton Keynes and Alex Alderton, Roberto Amato, Jeffrey Barrett, Sonia Goncalves, Ewan Harrison, David K. Jackson, Ian Johnston, Dominic Kwiatkowski, Cordelia Langford, John Sillitoe on behalf of the Wellcome Sanger Institute COVID-19 Surveillance Team |
| EPI_ISL_2846666, EPI_ISL_2846667 | Health Services Laboratories | Wellcome Sanger Institute for the COVID-19 Genomics UK (COG-UK) Consortium | Health Services Laboratories and Alex Alderton, Roberto Amato, Jeffrey Barrett, Sonia Goncalves, Ewan Harrison, David K. Jackson, Ian Johnston, Dominic Kwiatkowski, Cordelia Langford, John Sillitoe on behalf of the Wellcome Sanger Institute COVID-19 Surveillance Team |
| EPI_ISL_2846668, EPI_ISL_2846669 | Lighthouse Lab in Alderley Park | Wellcome Sanger Institute for the COVID-19 Genomics UK (COG-UK) Consortium | Jacquelyn Wynn, Mairead Hyland, The Lighthouse Lab in Alderley Park and Alex Alderton, Roberto Amato, Jeffrey Barrett, Sonia Goncalves, Ewan Harrison, David K. Jackson, Ian Johnston, Dominic Kwiatkowski, Cordelia Langford, John Sillitoe on behalf of the Wellcome Sanger Institute COVID-19 Surveillance Team |
| EPI_ISL_2846670 | Lighthouse Lab in Milton Keynes | Wellcome Sanger Institute for the COVID-19 Genomics UK (COG-UK) Consortium | The Lighthouse Lab in Milton Keynes and Alex Alderton, Roberto Amato, Jeffrey Barrett, Sonia Goncalves, Ewan Harrison, David K. Jackson, Ian Johnston, Dominic Kwiatkowski, Cordelia Langford, John Sillitoe on behalf of the Wellcome Sanger Institute COVID-19 Surveillance Team |
| EPI_ISL_2846671 | Lighthouse Laboratory Plymouth | Wellcome Sanger Institute for the COVID-19 Genomics UK (COG-UK) Consortium | Lighthouse Laboratory Plymouth and Alex Alderton, Roberto Amato, Jeffrey Barrett, Sonia Goncalves, Ewan Harrison, David K. Jackson, Ian Johnston, Dominic Kwiatkowski, Cordelia Langford, John Sillitoe on behalf of the Wellcome Sanger Institute COVID-19 Surveillance Team |
| EPI_ISL_2846672 | Lighthouse Lab in Milton Keynes | Wellcome Sanger Institute for the COVID-19 Genomics UK (COG-UK) Consortium | The Lighthouse Lab in Milton Keynes and Alex Alderton, Roberto Amato, Jeffrey Barrett, Sonia Goncalves, Ewan Harrison, David K. Jackson, Ian Johnston, Dominic Kwiatkowski, Cordelia Langford, John Sillitoe on behalf of the Wellcome Sanger Institute COVID-19 Surveillance Team |
| EPI_ISL_2846673 | Lighthouse Lab in Alderley Park | Wellcome Sanger Institute for the COVID-19 Genomics UK (COG-UK) Consortium | Jacquelyn Wynn, Mairead Hyland, The Lighthouse Lab in Alderley Park and Alex Alderton, Roberto Amato, Jeffrey Barrett, Sonia Goncalves, Ewan Harrison, David K. Jackson, Ian Johnston, Dominic Kwiatkowski, Cordelia Langford, John Sillitoe on behalf of the Wellcome Sanger Institute COVID-19 Surveillance Team |
| EPI_ISL_2846674 | Lighthouse Lab in Milton Keynes | Wellcome Sanger Institute for the COVID-19 Genomics UK (COG-UK) Consortium | The Lighthouse Lab in Milton Keynes and Alex Alderton, Roberto Amato, Jeffrey Barrett, Sonia Goncalves, Ewan Harrison, David K. Jackson, Ian Johnston, Dominic Kwiatkowski, Cordelia Langford, John Sillitoe on behalf of the Wellcome Sanger Institute COVID-19 Surveillance Team |
| EPI_ISL_2846675 | Health Services Laboratories | Wellcome Sanger Institute for the COVID-19 Genomics UK (COG-UK) Consortium | Health Services Laboratories and Alex Alderton, Roberto Amato, Jeffrey Barrett, Sonia Goncalves, Ewan Harrison, David K. Jackson, Ian Johnston, Dominic Kwiatkowski, Cordelia Langford, John Sillitoe on behalf of the Wellcome Sanger Institute COVID-19 Surveillance Team |
| EPI_ISL_2846676, EPI_ISL_2846677 | Lighthouse Lab in Alderley Park | Wellcome Sanger Institute for the COVID-19 Genomics UK (COG-UK) Consortium | Jacquelyn Wynn, Mairead Hyland, The Lighthouse Lab in Alderley Park and Alex Alderton, Roberto Amato, Jeffrey Barrett, Sonia Goncalves, Ewan Harrison, David K. Jackson, Ian Johnston, Dominic Kwiatkowski, Cordelia Langford, John Sillitoe on behalf of the Wellcome Sanger Institute COVID-19 Surveillance Team |
| EPI_ISL_2846678 | Lighthouse Lab in Milton Keynes | Wellcome Sanger Institute for the COVID-19 Genomics UK (COG-UK) Consortium | The Lighthouse Lab in Milton Keynes and Alex Alderton, Roberto Amato, Jeffrey Barrett, Sonia Goncalves, Ewan Harrison, David K. Jackson, Ian Johnston, Dominic Kwiatkowski, Cordelia Langford, John Sillitoe on behalf of the Wellcome Sanger Institute COVID-19 Surveillance Team |
| EPI_ISL_2846679 | Health Services Laboratories | Wellcome Sanger Institute for the COVID-19 Genomics UK (COG-UK) Consortium | Health Services Laboratories and Alex Alderton, Roberto Amato, Jeffrey Barrett, Sonia Goncalves, Ewan Harrison, David K. Jackson, Ian Johnston, Dominic Kwiatkowski, Cordelia Langford, John Sillitoe on behalf of the Wellcome Sanger Institute COVID-19 Surveillance Team |
| EPI_ISL_2846680 | Lighthouse Lab in Alderley Park | Wellcome Sanger Institute for the COVID-19 Genomics UK (COG-UK) Consortium | Jacquelyn Wynn, Mairead Hyland, The Lighthouse Lab in Alderley Park and Alex Alderton, Roberto Amato, Jeffrey Barrett, Sonia Goncalves, Ewan Harrison, David K. Jackson, Ian Johnston, Dominic Kwiatkowski, Cordelia Langford, John Sillitoe on behalf of the Wellcome Sanger Institute COVID-19 Surveillance Team |
| EPI_ISL_2846681 | Lighthouse Laboratory Plymouth | Wellcome Sanger Institute for the COVID-19 Genomics UK (COG-UK) Consortium | Lighthouse Laboratory Plymouth and Alex Alderton, Roberto Amato, Jeffrey Barrett, Sonia Goncalves, Ewan Harrison, David K. Jackson, Ian Johnston, Dominic Kwiatkowski, Cordelia Langford, John Sillitoe on behalf of the Wellcome Sanger Institute COVID-19 Surveillance Team |
| EPI_ISL_2846682, EPI_ISL_2846683 | Lighthouse Lab in Alderley Park | Wellcome Sanger Institute for the COVID-19 Genomics UK (COG-UK) Consortium | Jacquelyn Wynn, Mairead Hyland, The Lighthouse Lab in Alderley Park and Alex Alderton, Roberto Amato, Jeffrey Barrett, Sonia Goncalves, Ewan Harrison, David K. Jackson, Ian Johnston, Dominic Kwiatkowski, Cordelia Langford, John Sillitoe on behalf of the Wellcome Sanger Institute COVID-19 Surveillance Team |
| EPI_ISL_2846684, EPI_ISL_2846685 | Health Services Laboratories | Wellcome Sanger Institute for the COVID-19 Genomics UK (COG-UK) Consortium | Health Services Laboratories and Alex Alderton, Roberto Amato, Jeffrey Barrett, Sonia Goncalves, Ewan Harrison, David K. Jackson, Ian Johnston, Dominic Kwiatkowski, Cordelia Langford, John Sillitoe on behalf of the Wellcome Sanger Institute COVID-19 Surveillance Team |
| EPI_ISL_2846686 | Lighthouse Lab in Milton Keynes | Wellcome Sanger Institute for the COVID-19 Genomics UK (COG-UK) Consortium | The Lighthouse Lab in Milton Keynes and Alex Alderton, Roberto Amato, Jeffrey Barrett, Sonia Goncalves, Ewan Harrison, David K. Jackson, Ian Johnston, Dominic Kwiatkowski, Cordelia Langford, John Sillitoe on behalf of the Wellcome Sanger Institute COVID-19 Surveillance Team |
| EPI_ISL_2846687 | Health Services Laboratories | Wellcome Sanger Institute for the COVID-19 Genomics UK (COG-UK) Consortium | Health Services Laboratories and Alex Alderton, Roberto Amato, Jeffrey Barrett, Sonia Goncalves, Ewan Harrison, David K. Jackson, Ian Johnston, Dominic Kwiatkowski, Cordelia Langford, John Sillitoe on behalf of the Wellcome Sanger Institute COVID-19 Surveillance Team |
| EPI_ISL_2846688 | Lighthouse Lab in Alderley Park | Wellcome Sanger Institute for the COVID-19 Genomics UK (COG-UK) Consortium | Jacquelyn Wynn, Mairead Hyland, The Lighthouse Lab in Alderley Park and Alex Alderton, Roberto Amato, Jeffrey Barrett, Sonia Goncalves, Ewan Harrison, David K. Jackson, Ian Johnston, Dominic Kwiatkowski, Cordelia Langford, John Sillitoe on behalf of the Wellcome Sanger Institute COVID-19 Surveillance Team |
| EPI_ISL_2846689 | Lighthouse Laboratory Plymouth | Wellcome Sanger Institute for the COVID-19 Genomics UK (COG-UK) Consortium | Lighthouse Laboratory Plymouth and Alex Alderton, Roberto Amato, Jeffrey Barrett, Sonia Goncalves, Ewan Harrison, David K. Jackson, Ian Johnston, Dominic Kwiatkowski, Cordelia Langford, John Sillitoe on behalf of the Wellcome Sanger Institute COVID-19 Surveillance Team |
| EPI_ISL_2846690 | Health Services Laboratories | Wellcome Sanger Institute for the COVID-19 Genomics UK (COG-UK) Consortium | Health Services Laboratories and Alex Alderton, Roberto Amato, Jeffrey Barrett, Sonia Goncalves, Ewan Harrison, David K. Jackson, Ian Johnston, Dominic Kwiatkowski, Cordelia Langford, John Sillitoe on behalf of the Wellcome Sanger Institute COVID-19 Surveillance Team |
| EPI_ISL_2846691 | Lighthouse Lab in Milton Keynes | Wellcome Sanger Institute for the COVID-19 Genomics UK (COG-UK) Consortium | The Lighthouse Lab in Milton Keynes and Alex Alderton, Roberto Amato, Jeffrey Barrett, Sonia Goncalves, Ewan Harrison, David K. Jackson, Ian Johnston, Dominic Kwiatkowski, Cordelia Langford, John Sillitoe on behalf of the Wellcome Sanger Institute COVID-19 Surveillance Team |
| EPI_ISL_2846692 | Lighthouse Laboratory Plymouth | Wellcome Sanger Institute for the COVID-19 Genomics UK (COG-UK) Consortium | Lighthouse Laboratory Plymouth and Alex Alderton, Roberto Amato, Jeffrey Barrett, Sonia Goncalves, Ewan Harrison, David K. Jackson, Ian Johnston, Dominic Kwiatkowski, Cordelia Langford, John Sillitoe on behalf of the Wellcome Sanger Institute COVID-19 Surveillance Team |
| EPI_ISL_2846693 | Lighthouse Lab in Milton Keynes | Wellcome Sanger Institute for the COVID-19 Genomics UK (COG-UK) Consortium | The Lighthouse Lab in Milton Keynes and Alex Alderton, Roberto Amato, Jeffrey Barrett, Sonia Goncalves, Ewan Harrison, David K. Jackson, Ian Johnston, Dominic Kwiatkowski, Cordelia Langford, John Sillitoe on behalf of the Wellcome Sanger Institute COVID-19 Surveillance Team |
| EPI_ISL_2846694 | Lighthouse Laboratory Plymouth | Wellcome Sanger Institute for the COVID-19 Genomics UK (COG-UK) Consortium | Lighthouse Laboratory Plymouth and Alex Alderton, Roberto Amato, Jeffrey Barrett, Sonia Goncalves, Ewan Harrison, David K. Jackson, Ian Johnston, Dominic Kwiatkowski, Cordelia Langford, John Sillitoe on behalf of the Wellcome Sanger Institute COVID-19 Surveillance Team |
| EPI_ISL_2846695 | Lighthouse Lab in Milton Keynes | Wellcome Sanger Institute for the COVID-19 Genomics UK (COG-UK) Consortium | The Lighthouse Lab in Milton Keynes and Alex Alderton, Roberto Amato, Jeffrey Barrett, Sonia Goncalves, Ewan Harrison, David K. Jackson, Ian Johnston, Dominic Kwiatkowski, Cordelia Langford, John Sillitoe on behalf of the Wellcome Sanger Institute COVID-19 Surveillance Team |
| EPI_ISL_2846696 | Lighthouse Laboratory Plymouth | Wellcome Sanger Institute for the COVID-19 Genomics UK (COG-UK) Consortium | Lighthouse Laboratory Plymouth and Alex Alderton, Roberto Amato, Jeffrey Barrett, Sonia Goncalves, Ewan Harrison, David K. Jackson, Ian Johnston, Dominic Kwiatkowski, Cordelia Langford, John Sillitoe on behalf of the Wellcome Sanger Institute COVID-19 Surveillance Team |
| EPI_ISL_2846697 | Health Services Laboratories | Wellcome Sanger Institute for the COVID-19 Genomics UK (COG-UK) Consortium | Health Services Laboratories and Alex Alderton, Roberto Amato, Jeffrey Barrett, Sonia Goncalves, Ewan Harrison, David K. Jackson, Ian Johnston, Dominic Kwiatkowski, Cordelia Langford, John Sillitoe on behalf of the Wellcome Sanger Institute COVID-19 Surveillance Team |
| EPI_ISL_2846698 | Lighthouse Lab in Alderley Park | Wellcome Sanger Institute for the COVID-19 Genomics UK (COG-UK) Consortium | Jacquelyn Wynn, Mairead Hyland, The Lighthouse Lab in Alderley Park and Alex Alderton, Roberto Amato, Jeffrey Barrett, Sonia Goncalves, Ewan Harrison, David K. Jackson, Ian Johnston, Dominic Kwiatkowski, Cordelia Langford, John Sillitoe on behalf of the Wellcome Sanger Institute COVID-19 Surveillance Team |
| EPI_ISL_2846699 | Lighthouse Laboratory Plymouth | Wellcome Sanger Institute for the COVID-19 Genomics UK (COG-UK) Consortium | Lighthouse Laboratory Plymouth and Alex Alderton, Roberto Amato, Jeffrey Barrett, Sonia Goncalves, Ewan Harrison, David K. Jackson, Ian Johnston, Dominic Kwiatkowski, Cordelia Langford, John Sillitoe on behalf of the Wellcome Sanger Institute COVID-19 Surveillance Team |
| EPI_ISL_2846700, EPI_ISL_2846701 | Lighthouse Lab in Alderley Park | Wellcome Sanger Institute for the COVID-19 Genomics UK (COG-UK) Consortium | Jacquelyn Wynn, Mairead Hyland, The Lighthouse Lab in Alderley Park and Alex Alderton, Roberto Amato, Jeffrey Barrett, Sonia Goncalves, Ewan Harrison, David K. Jackson, Ian Johnston, Dominic Kwiatkowski, Cordelia Langford, John Sillitoe on behalf of the Wellcome Sanger Institute COVID-19 Surveillance Team |
| EPI_ISL_2846702 | Lighthouse Lab in Milton Keynes | Wellcome Sanger Institute for the COVID-19 Genomics UK (COG-UK) Consortium | The Lighthouse Lab in Milton Keynes and Alex Alderton, Roberto Amato, Jeffrey Barrett, Sonia Goncalves, Ewan Harrison, David K. Jackson, Ian Johnston, Dominic Kwiatkowski, Cordelia Langford, John Sillitoe on behalf of the Wellcome Sanger Institute COVID-19 Surveillance Team |
| EPI_ISL_2846703, EPI_ISL_2846704 | Lighthouse Lab in Alderley Park | Wellcome Sanger Institute for the COVID-19 Genomics UK (COG-UK) Consortium | Jacquelyn Wynn, Mairead Hyland, The Lighthouse Lab in Alderley Park and Alex Alderton, Roberto Amato, Jeffrey Barrett, Sonia Goncalves, Ewan Harrison, David K. Jackson, Ian Johnston, Dominic Kwiatkowski, Cordelia Langford, John Sillitoe on behalf of the Wellcome Sanger Institute COVID-19 Surveillance Team |
| EPI_ISL_2846705 | Lighthouse Laboratory Plymouth | Wellcome Sanger Institute for the COVID-19 Genomics UK (COG-UK) Consortium | Lighthouse Laboratory Plymouth and Alex Alderton, Roberto Amato, Jeffrey Barrett, Sonia Goncalves, Ewan Harrison, David K. Jackson, Ian Johnston, Dominic Kwiatkowski, Cordelia Langford, John Sillitoe on behalf of the Wellcome Sanger Institute COVID-19 Surveillance Team |
| EPI_ISL_2846706 | Lighthouse Lab in Milton Keynes | Wellcome Sanger Institute for the COVID-19 Genomics UK (COG-UK) Consortium | The Lighthouse Lab in Milton Keynes and Alex Alderton, Roberto Amato, Jeffrey Barrett, Sonia Goncalves, Ewan Harrison, David K. Jackson, Ian Johnston, Dominic Kwiatkowski, Cordelia Langford, John Sillitoe on behalf of the Wellcome Sanger Institute COVID-19 Surveillance Team |
| EPI_ISL_2846707 | Health Services Laboratories | Wellcome Sanger Institute for the COVID-19 Genomics UK (COG-UK) Consortium | Health Services Laboratories and Alex Alderton, Roberto Amato, Jeffrey Barrett, Sonia Goncalves, Ewan Harrison, David K. Jackson, Ian Johnston, Dominic Kwiatkowski, Cordelia Langford, John Sillitoe on behalf of the Wellcome Sanger Institute COVID-19 Surveillance Team |

| | | | |
|---|---|---|---|
| EPI_ISL_2846708 | Lighthouse Lab in Alderley Park | Wellcome Sanger Institute for the COVID-19 Genomics UK (COG-UK) Consortium | Jacquelyn Wynn, Mairead Hyland, The Lighthouse Lab in Alderley Park and Alex Alderton, Roberto Amato, Jeffrey Barrett, Sonia Goncalves, Ewan Harrison, David K. Jackson, Ian Johnston, Dominic Kwiatkowski, Cordelia Langford, John Sillitoe on behalf of the Wellcome Sanger Institute COVID-19 Surveillance Team |
| EPI_ISL_2846709, EPI_ISL_2846710 | Health Services Laboratories | Wellcome Sanger Institute for the COVID-19 Genomics UK (COG-UK) Consortium | Health Services Laboratories and Alex Alderton, Roberto Amato, Jeffrey Barrett, Sonia Goncalves, Ewan Harrison, David K. Jackson, Ian Johnston, Dominic Kwiatkowski, Cordelia Langford, John Sillitoe on behalf of the Wellcome Sanger Institute COVID-19 Surveillance Team |
| EPI_ISL_2846711 | Lighthouse Lab in Milton Keynes | Wellcome Sanger Institute for the COVID-19 Genomics UK (COG-UK) Consortium | The Lighthouse Lab in Milton Keynes and Alex Alderton, Roberto Amato, Jeffrey Barrett, Sonia Goncalves, Ewan Harrison, David K. Jackson, Ian Johnston, Dominic Kwiatkowski, Cordelia Langford, John Sillitoe on behalf of the Wellcome Sanger Institute COVID-19 Surveillance Team |
| EPI_ISL_2846712 | Lighthouse Laboratory Plymouth | Wellcome Sanger Institute for the COVID-19 Genomics UK (COG-UK) Consortium | Lighthouse Laboratory Plymouth and Alex Alderton, Roberto Amato, Jeffrey Barrett, Sonia Goncalves, Ewan Harrison, David K. Jackson, Ian Johnston, Dominic Kwiatkowski, Cordelia Langford, John Sillitoe on behalf of the Wellcome Sanger Institute COVID-19 Surveillance Team |
| EPI_ISL_2846713 | Lighthouse Lab in Alderley Park | Wellcome Sanger Institute for the COVID-19 Genomics UK (COG-UK) Consortium | Jacquelyn Wynn, Mairead Hyland, The Lighthouse Lab in Alderley Park and Alex Alderton, Roberto Amato, Jeffrey Barrett, Sonia Goncalves, Ewan Harrison, David K. Jackson, Ian Johnston, Dominic Kwiatkowski, Cordelia Langford, John Sillitoe on behalf of the Wellcome Sanger Institute COVID-19 Surveillance Team |
| EPI_ISL_2846714 | Lighthouse Lab in Milton Keynes | Wellcome Sanger Institute for the COVID-19 Genomics UK (COG-UK) Consortium | The Lighthouse Lab in Milton Keynes and Alex Alderton, Roberto Amato, Jeffrey Barrett, Sonia Goncalves, Ewan Harrison, David K. Jackson, Ian Johnston, Dominic Kwiatkowski, Cordelia Langford, John Sillitoe on behalf of the Wellcome Sanger Institute COVID-19 Surveillance Team |
| EPI_ISL_2846715 | Lighthouse Lab in Alderley Park | Wellcome Sanger Institute for the COVID-19 Genomics UK (COG-UK) Consortium | Jacquelyn Wynn, Mairead Hyland, The Lighthouse Lab in Alderley Park and Alex Alderton, Roberto Amato, Jeffrey Barrett, Sonia Goncalves, Ewan Harrison, David K. Jackson, Ian Johnston, Dominic Kwiatkowski, Cordelia Langford, John Sillitoe on behalf of the Wellcome Sanger Institute COVID-19 Surveillance Team |
| EPI_ISL_2846716 | Health Services Laboratories | Wellcome Sanger Institute for the COVID-19 Genomics UK (COG-UK) Consortium | Health Services Laboratories and Alex Alderton, Roberto Amato, Jeffrey Barrett, Sonia Goncalves, Ewan Harrison, David K. Jackson, Ian Johnston, Dominic Kwiatkowski, Cordelia Langford, John Sillitoe on behalf of the Wellcome Sanger Institute COVID-19 Surveillance Team |
| EPI_ISL_2846717, EPI_ISL_2846718 | Lighthouse Lab in Alderley Park | Wellcome Sanger Institute for the COVID-19 Genomics UK (COG-UK) Consortium | Jacquelyn Wynn, Mairead Hyland, The Lighthouse Lab in Alderley Park and Alex Alderton, Roberto Amato, Jeffrey Barrett, Sonia Goncalves, Ewan Harrison, David K. Jackson, Ian Johnston, Dominic Kwiatkowski, Cordelia Langford, John Sillitoe on behalf of the Wellcome Sanger Institute COVID-19 Surveillance Team |
| EPI_ISL_2846719 | Lighthouse Laboratory Plymouth | Wellcome Sanger Institute for the COVID-19 Genomics UK (COG-UK) Consortium | Lighthouse Laboratory Plymouth and Alex Alderton, Roberto Amato, Jeffrey Barrett, Sonia Goncalves, Ewan Harrison, David K. Jackson, Ian Johnston, Dominic Kwiatkowski, Cordelia Langford, John Sillitoe on behalf of the Wellcome Sanger Institute COVID-19 Surveillance Team |
| EPI_ISL_2846720, EPI_ISL_2846721, EPI_ISL_2846722 | Lighthouse Lab in Alderley Park | Wellcome Sanger Institute for the COVID-19 Genomics UK (COG-UK) Consortium | Jacquelyn Wynn, Mairead Hyland, The Lighthouse Lab in Alderley Park and Alex Alderton, Roberto Amato, Jeffrey Barrett, Sonia Goncalves, Ewan Harrison, David K. Jackson, Ian Johnston, Dominic Kwiatkowski, Cordelia Langford, John Sillitoe on behalf of the Wellcome Sanger Institute COVID-19 Surveillance Team |
| EPI_ISL_2846723, EPI_ISL_2846724 | Health Services Laboratories | Wellcome Sanger Institute for the COVID-19 Genomics UK (COG-UK) Consortium | Health Services Laboratories and Alex Alderton, Roberto Amato, Jeffrey Barrett, Sonia Goncalves, Ewan Harrison, David K. Jackson, Ian Johnston, Dominic Kwiatkowski, Cordelia Langford, John Sillitoe on behalf of the Wellcome Sanger Institute COVID-19 Surveillance Team |
| EPI_ISL_2846725 | Lighthouse Laboratory Plymouth | Wellcome Sanger Institute for the COVID-19 Genomics UK (COG-UK) Consortium | Lighthouse Laboratory Plymouth and Alex Alderton, Roberto Amato, Jeffrey Barrett, Sonia Goncalves, Ewan Harrison, David K. Jackson, Ian Johnston, Dominic Kwiatkowski, Cordelia Langford, John Sillitoe on behalf of the Wellcome Sanger Institute COVID-19 Surveillance Team |
| EPI_ISL_2846726 | Health Services Laboratories | Wellcome Sanger Institute for the COVID-19 Genomics UK (COG-UK) Consortium | Health Services Laboratories and Alex Alderton, Roberto Amato, Jeffrey Barrett, Sonia Goncalves, Ewan Harrison, David K. Jackson, Ian Johnston, Dominic Kwiatkowski, Cordelia Langford, John Sillitoe on behalf of the Wellcome Sanger Institute COVID-19 Surveillance Team |
| EPI_ISL_2846727 | Lighthouse Lab in Milton Keynes | Wellcome Sanger Institute for the COVID-19 Genomics UK (COG-UK) Consortium | The Lighthouse Lab in Milton Keynes and Alex Alderton, Roberto Amato, Jeffrey Barrett, Sonia Goncalves, Ewan Harrison, David K. Jackson, Ian Johnston, Dominic Kwiatkowski, Cordelia Langford, John Sillitoe on behalf of the Wellcome Sanger Institute COVID-19 Surveillance Team |
| EPI_ISL_2846728 | Lighthouse Lab in Alderley Park | Wellcome Sanger Institute for the COVID-19 Genomics UK (COG-UK) Consortium | Jacquelyn Wynn, Mairead Hyland, The Lighthouse Lab in Alderley Park and Alex Alderton, Roberto Amato, Jeffrey Barrett, Sonia Goncalves, Ewan Harrison, David K. Jackson, Ian Johnston, Dominic Kwiatkowski, Cordelia Langford, John Sillitoe on behalf of the Wellcome Sanger Institute COVID-19 Surveillance Team |
| EPI_ISL_2846729, EPI_ISL_2846730, EPI_ISL_2846731 | Health Services Laboratories | Wellcome Sanger Institute for the COVID-19 Genomics UK (COG-UK) Consortium | Health Services Laboratories and Alex Alderton, Roberto Amato, Jeffrey Barrett, Sonia Goncalves, Ewan Harrison, David K. Jackson, Ian Johnston, Dominic Kwiatkowski, Cordelia Langford, John Sillitoe on behalf of the Wellcome Sanger Institute COVID-19 Surveillance Team |
| EPI_ISL_2846732, EPI_ISL_2846733 | Lighthouse Lab in Milton Keynes | Wellcome Sanger Institute for the COVID-19 Genomics UK (COG-UK) Consortium | The Lighthouse Lab in Milton Keynes and Alex Alderton, Roberto Amato, Jeffrey Barrett, Sonia Goncalves, Ewan Harrison, David K. Jackson, Ian Johnston, Dominic Kwiatkowski, Cordelia Langford, John Sillitoe on behalf of the Wellcome Sanger Institute COVID-19 Surveillance Team |
| EPI_ISL_2846734, EPI_ISL_2846735 | Lighthouse Laboratory Plymouth | Wellcome Sanger Institute for the COVID-19 Genomics UK (COG-UK) Consortium | Lighthouse Laboratory Plymouth and Alex Alderton, Roberto Amato, Jeffrey Barrett, Sonia Goncalves, Ewan Harrison, David K. Jackson, Ian Johnston, Dominic Kwiatkowski, Cordelia Langford, John Sillitoe on behalf of the Wellcome Sanger Institute COVID-19 Surveillance Team |
| EPI_ISL_2846736 | Health Services Laboratories | Wellcome Sanger Institute for the COVID-19 Genomics UK (COG-UK) Consortium | Health Services Laboratories and Alex Alderton, Roberto Amato, Jeffrey Barrett, Sonia Goncalves, Ewan Harrison, David K. Jackson, Ian Johnston, Dominic Kwiatkowski, Cordelia Langford, John Sillitoe on behalf of the Wellcome Sanger Institute COVID-19 Surveillance Team |
| EPI_ISL_2846737 | Lighthouse Lab in Milton Keynes | Wellcome Sanger Institute for the COVID-19 Genomics UK (COG-UK) Consortium | The Lighthouse Lab in Milton Keynes and Alex Alderton, Roberto Amato, Jeffrey Barrett, Sonia Goncalves, Ewan Harrison, David K. Jackson, Ian Johnston, Dominic Kwiatkowski, Cordelia Langford, John Sillitoe on behalf of the Wellcome Sanger Institute COVID-19 Surveillance Team |
| EPI_ISL_2846738, EPI_ISL_2846739, EPI_ISL_2846740, EPI_ISL_2846741, EPI_ISL_2846742 | Health Services Laboratories | Wellcome Sanger Institute for the COVID-19 Genomics UK (COG-UK) Consortium | Health Services Laboratories and Alex Alderton, Roberto Amato, Jeffrey Barrett, Sonia Goncalves, Ewan Harrison, David K. Jackson, Ian Johnston, Dominic Kwiatkowski, Cordelia Langford, John Sillitoe on behalf of the Wellcome Sanger Institute COVID-19 Surveillance Team |
| EPI_ISL_2846743 | Lighthouse Laboratory Plymouth | Wellcome Sanger Institute for the COVID-19 Genomics UK (COG-UK) Consortium | Lighthouse Laboratory Plymouth and Alex Alderton, Roberto Amato, Jeffrey Barrett, Sonia Goncalves, Ewan Harrison, David K. Jackson, Ian Johnston, Dominic Kwiatkowski, Cordelia Langford, John Sillitoe on behalf of the Wellcome Sanger Institute COVID-19 Surveillance Team |
| EPI_ISL_2846744, EPI_ISL_2846745, EPI_ISL_2846746 | Health Services Laboratories | Wellcome Sanger Institute for the COVID-19 Genomics UK (COG-UK) Consortium | Health Services Laboratories and Alex Alderton, Roberto Amato, Jeffrey Barrett, Sonia Goncalves, Ewan Harrison, David K. Jackson, Ian Johnston, Dominic Kwiatkowski, Cordelia Langford, John Sillitoe on behalf of the Wellcome Sanger Institute COVID-19 Surveillance Team |
| EPI_ISL_2846747 | Lighthouse Laboratory Plymouth | Wellcome Sanger Institute for the COVID-19 Genomics UK (COG-UK) Consortium | Lighthouse Laboratory Plymouth and Alex Alderton, Roberto Amato, Jeffrey Barrett, Sonia Goncalves, Ewan Harrison, David K. Jackson, Ian Johnston, Dominic Kwiatkowski, Cordelia Langford, John Sillitoe on behalf of the Wellcome Sanger Institute COVID-19 Surveillance Team |
| EPI_ISL_2846748 | Lighthouse Lab in Alderley Park | Wellcome Sanger Institute for the COVID-19 Genomics UK (COG-UK) Consortium | Jacquelyn Wynn, Mairead Hyland, The Lighthouse Lab in Alderley Park and Alex Alderton, Roberto Amato, Jeffrey Barrett, Sonia Goncalves, Ewan Harrison, David K. Jackson, Ian Johnston, Dominic Kwiatkowski, Cordelia Langford, John Sillitoe on behalf of the Wellcome Sanger Institute COVID-19 Surveillance Team |
| EPI_ISL_2846749 | Health Services Laboratories | Wellcome Sanger Institute for the COVID-19 Genomics UK (COG-UK) Consortium | Health Services Laboratories and Alex Alderton, Roberto Amato, Jeffrey Barrett, Sonia Goncalves, Ewan Harrison, David K. Jackson, Ian Johnston, Dominic Kwiatkowski, Cordelia Langford, John Sillitoe on behalf of the Wellcome Sanger Institute COVID-19 Surveillance Team |
| EPI_ISL_2846750, EPI_ISL_2846751, EPI_ISL_2846752 | Lighthouse Lab in Milton Keynes | Wellcome Sanger Institute for the COVID-19 Genomics UK (COG-UK) Consortium | The Lighthouse Lab in Milton Keynes and Alex Alderton, Roberto Amato, Jeffrey Barrett, Sonia Goncalves, Ewan Harrison, David K. Jackson, Ian Johnston, Dominic Kwiatkowski, Cordelia Langford, John Sillitoe on behalf of the Wellcome Sanger Institute COVID-19 Surveillance Team |
| EPI_ISL_2846753 | Lighthouse Lab in Alderley Park | Wellcome Sanger Institute for the COVID-19 Genomics UK (COG-UK) Consortium | Jacquelyn Wynn, Mairead Hyland, The Lighthouse Lab in Alderley Park and Alex Alderton, Roberto Amato, Jeffrey Barrett, Sonia Goncalves, Ewan Harrison, David K. Jackson, Ian Johnston, Dominic Kwiatkowski, Cordelia Langford, John Sillitoe on behalf of the Wellcome Sanger Institute COVID-19 Surveillance Team |
| EPI_ISL_2846754, EPI_ISL_2846755 | Health Services Laboratories | Wellcome Sanger Institute for the COVID-19 Genomics UK (COG-UK) Consortium | Health Services Laboratories and Alex Alderton, Roberto Amato, Jeffrey Barrett, Sonia Goncalves, Ewan Harrison, David K. Jackson, Ian Johnston, Dominic Kwiatkowski, Cordelia Langford, John Sillitoe on behalf of the Wellcome Sanger Institute COVID-19 Surveillance Team |
| EPI_ISL_2846756 | Lighthouse Lab in Milton Keynes | Wellcome Sanger Institute for the COVID-19 Genomics UK (COG-UK) Consortium | The Lighthouse Lab in Milton Keynes and Alex Alderton, Roberto Amato, Jeffrey Barrett, Sonia Goncalves, Ewan Harrison, David K. Jackson, Ian Johnston, Dominic Kwiatkowski, Cordelia Langford, John Sillitoe on behalf of the Wellcome Sanger Institute COVID-19 Surveillance Team |
| EPI_ISL_2846757 | Lighthouse Laboratory Plymouth | Wellcome Sanger Institute for the COVID-19 Genomics UK (COG-UK) Consortium | Lighthouse Laboratory Plymouth and Alex Alderton, Roberto Amato, Jeffrey Barrett, Sonia Goncalves, Ewan Harrison, David K. Jackson, Ian Johnston, Dominic Kwiatkowski, Cordelia Langford, John Sillitoe on behalf of the Wellcome Sanger Institute COVID-19 Surveillance Team |
| EPI_ISL_2846758 | Lighthouse Lab in Milton Keynes | Wellcome Sanger Institute for the COVID-19 Genomics UK (COG-UK) Consortium | The Lighthouse Lab in Milton Keynes and Alex Alderton, Roberto Amato, Jeffrey Barrett, Sonia Goncalves, Ewan Harrison, David K. Jackson, Ian Johnston, Dominic Kwiatkowski, Cordelia Langford, John Sillitoe on behalf of the Wellcome Sanger Institute COVID-19 Surveillance Team |
| EPI_ISL_2846759, EPI_ISL_2846760 | Lighthouse Laboratory Plymouth | Wellcome Sanger Institute for the COVID-19 Genomics UK (COG-UK) Consortium | Lighthouse Laboratory Plymouth and Alex Alderton, Roberto Amato, Jeffrey Barrett, Sonia Goncalves, Ewan Harrison, David K. Jackson, Ian Johnston, Dominic Kwiatkowski, Cordelia Langford, John Sillitoe on behalf of the Wellcome Sanger Institute COVID-19 Surveillance Team |
| EPI_ISL_2846761 | Lighthouse Lab in Alderley Park | Wellcome Sanger Institute for the COVID-19 Genomics UK (COG-UK) Consortium | Jacquelyn Wynn, Mairead Hyland, The Lighthouse Lab in Alderley Park and Alex Alderton, Roberto Amato, Jeffrey Barrett, Sonia Goncalves, Ewan Harrison, David K. Jackson, Ian Johnston, Dominic Kwiatkowski, Cordelia Langford, John Sillitoe on behalf of the Wellcome Sanger Institute COVID-19 Surveillance Team |
| EPI_ISL_2846762 | Lighthouse Lab in Milton Keynes | Wellcome Sanger Institute for the COVID-19 Genomics UK (COG-UK) Consortium | The Lighthouse Lab in Milton Keynes and Alex Alderton, Roberto Amato, Jeffrey Barrett, Sonia Goncalves, Ewan Harrison, David K. Jackson, Ian Johnston, Dominic Kwiatkowski, Cordelia Langford, John Sillitoe on behalf of the Wellcome Sanger Institute COVID-19 Surveillance Team |
| EPI_ISL_2846763 | Lighthouse Lab in Alderley Park | Wellcome Sanger Institute for the COVID-19 Genomics UK (COG-UK) Consortium | Jacquelyn Wynn, Mairead Hyland, The Lighthouse Lab in Alderley Park and Alex Alderton, Roberto Amato, Jeffrey Barrett, Sonia Goncalves, Ewan Harrison, David K. Jackson, Ian Johnston, Dominic Kwiatkowski, Cordelia Langford, John Sillitoe on behalf of the Wellcome Sanger Institute COVID-19 Surveillance Team |
| EPI_ISL_2846764, EPI_ISL_2846765 | Health Services Laboratories | Wellcome Sanger Institute for the COVID-19 Genomics UK (COG-UK) Consortium | Health Services Laboratories and Alex Alderton, Roberto Amato, Jeffrey Barrett, Sonia Goncalves, Ewan Harrison, David K. Jackson, Ian Johnston, Dominic Kwiatkowski, Cordelia Langford, John Sillitoe on behalf of the Wellcome Sanger Institute COVID-19 Surveillance Team |
| EPI_ISL_2846766 | Lighthouse Lab in Alderley Park | Wellcome Sanger Institute for the COVID-19 Genomics UK (COG-UK) Consortium | Jacquelyn Wynn, Mairead Hyland, The Lighthouse Lab in Alderley Park and Alex Alderton, Roberto Amato, Jeffrey Barrett, Sonia Goncalves, Ewan Harrison, David K. Jackson, Ian Johnston, Dominic Kwiatkowski, Cordelia Langford, John Sillitoe on behalf of the Wellcome Sanger Institute COVID-19 Surveillance Team |
| EPI_ISL_2846767 | Lighthouse Lab in Milton Keynes | Wellcome Sanger Institute for the COVID-19 Genomics UK (COG-UK) Consortium | The Lighthouse Lab in Milton Keynes and Alex Alderton, Roberto Amato, Jeffrey Barrett, Sonia Goncalves, Ewan Harrison, David K. Jackson, Ian Johnston, Dominic Kwiatkowski, Cordelia Langford, John Sillitoe on behalf of the Wellcome Sanger Institute COVID-19 Surveillance Team |
| EPI_ISL_2846768, EPI_ISL_2846769 | Lighthouse Laboratory Plymouth | Wellcome Sanger Institute for the COVID-19 Genomics UK (COG-UK) Consortium | Lighthouse Laboratory Plymouth and Alex Alderton, Roberto Amato, Jeffrey Barrett, Sonia Goncalves, Ewan Harrison, David K. Jackson, Ian Johnston, Dominic Kwiatkowski, Cordelia Langford, John Sillitoe on behalf of the Wellcome Sanger Institute COVID-19 Surveillance Team |
| EPI_ISL_2846770 | Health Services Laboratories | Wellcome Sanger Institute for the COVID-19 Genomics UK (COG-UK) Consortium | Health Services Laboratories and Alex Alderton, Roberto Amato, Jeffrey Barrett, Sonia Goncalves, Ewan Harrison, David K. Jackson, Ian Johnston, Dominic Kwiatkowski, Cordelia Langford, John Sillitoe on behalf of the Wellcome Sanger Institute COVID-19 Surveillance Team |
| EPI_ISL_2846771, | Lighthouse Lab in Alderley Park | Wellcome Sanger Institute for the COVID-19 Genomics UK (COG-UK) | Jacquelyn Wynn, Mairead Hyland, The Lighthouse Lab in Alderley Park and Alex Alderton, Roberto Amato, Jeffrey Barrett, Sonia Goncalves, Ewan Harrison, David K. Jackson, Ian Johnston, Dominic Kwiatkowski, Cordelia Langford, John Sillitoe on |

| | | | |
|---|---|---|---|
| EPI_ISL_2846772 | | Consortium | behalf of the Wellcome Sanger Institute COVID-19 Surveillance Team |
| EPI_ISL_2846773, EPI_ISL_2846774, EPI_ISL_2846775 | Lighthouse Lab in Milton Keynes | Wellcome Sanger Institute for the COVID-19 Genomics UK (COG-UK) Consortium | The Lighthouse Lab in Milton Keynes and Alex Alderton, Roberto Amato, Jeffrey Barrett, Sonia Goncalves, Ewan Harrison, David K. Jackson, Ian Johnston, Dominic Kwiatkowski, Cordelia Langford, John Sillitoe on behalf of the Wellcome Sanger Institute COVID-19 Surveillance Team |
| EPI_ISL_2846776 | Health Services Laboratories | Wellcome Sanger Institute for the COVID-19 Genomics UK (COG-UK) Consortium | Health Services Laboratories and Alex Alderton, Roberto Amato, Jeffrey Barrett, Sonia Goncalves, Ewan Harrison, David K. Jackson, Ian Johnston, Dominic Kwiatkowski, Cordelia Langford, John Sillitoe on behalf of the Wellcome Sanger Institute COVID-19 Surveillance Team |
| EPI_ISL_2846777, EPI_ISL_2846778 | Lighthouse Laboratory Plymouth | Wellcome Sanger Institute for the COVID-19 Genomics UK (COG-UK) Consortium | Lighthouse Laboratory Plymouth and Alex Alderton, Roberto Amato, Jeffrey Barrett, Sonia Goncalves, Ewan Harrison, David K. Jackson, Ian Johnston, Dominic Kwiatkowski, Cordelia Langford, John Sillitoe on behalf of the Wellcome Sanger Institute COVID-19 Surveillance Team |
| EPI_ISL_2846779 | Health Services Laboratories | Wellcome Sanger Institute for the COVID-19 Genomics UK (COG-UK) Consortium | Health Services Laboratories and Alex Alderton, Roberto Amato, Jeffrey Barrett, Sonia Goncalves, Ewan Harrison, David K. Jackson, Ian Johnston, Dominic Kwiatkowski, Cordelia Langford, John Sillitoe on behalf of the Wellcome Sanger Institute COVID-19 Surveillance Team |
| EPI_ISL_2846780 | Lighthouse Lab in Alderley Park | Wellcome Sanger Institute for the COVID-19 Genomics UK (COG-UK) Consortium | Jacquelyn Wynn, Mairead Hyland, The Lighthouse Lab in Alderley Park and Alex Alderton, Roberto Amato, Jeffrey Barrett, Sonia Goncalves, Ewan Harrison, David K. Jackson, Ian Johnston, Dominic Kwiatkowski, Cordelia Langford, John Sillitoe on behalf of the Wellcome Sanger Institute COVID-19 Surveillance Team |
| EPI_ISL_2846781 | Lighthouse Laboratory Plymouth | Wellcome Sanger Institute for the COVID-19 Genomics UK (COG-UK) Consortium | Lighthouse Laboratory Plymouth and Alex Alderton, Roberto Amato, Jeffrey Barrett, Sonia Goncalves, Ewan Harrison, David K. Jackson, Ian Johnston, Dominic Kwiatkowski, Cordelia Langford, John Sillitoe on behalf of the Wellcome Sanger Institute COVID-19 Surveillance Team |
| EPI_ISL_2846782 | Lighthouse Lab in Alderley Park | Wellcome Sanger Institute for the COVID-19 Genomics UK (COG-UK) Consortium | Jacquelyn Wynn, Mairead Hyland, The Lighthouse Lab in Alderley Park and Alex Alderton, Roberto Amato, Jeffrey Barrett, Sonia Goncalves, Ewan Harrison, David K. Jackson, Ian Johnston, Dominic Kwiatkowski, Cordelia Langford, John Sillitoe on behalf of the Wellcome Sanger Institute COVID-19 Surveillance Team |
| EPI_ISL_2846783, EPI_ISL_2846784, EPI_ISL_2846785 | Lighthouse Laboratory Plymouth | Wellcome Sanger Institute for the COVID-19 Genomics UK (COG-UK) Consortium | Lighthouse Laboratory Plymouth and Alex Alderton, Roberto Amato, Jeffrey Barrett, Sonia Goncalves, Ewan Harrison, David K. Jackson, Ian Johnston, Dominic Kwiatkowski, Cordelia Langford, John Sillitoe on behalf of the Wellcome Sanger Institute COVID-19 Surveillance Team |
| EPI_ISL_2846786 | Lighthouse Lab in Milton Keynes | Wellcome Sanger Institute for the COVID-19 Genomics UK (COG-UK) Consortium | The Lighthouse Lab in Milton Keynes and Alex Alderton, Roberto Amato, Jeffrey Barrett, Sonia Goncalves, Ewan Harrison, David K. Jackson, Ian Johnston, Dominic Kwiatkowski, Cordelia Langford, John Sillitoe on behalf of the Wellcome Sanger Institute COVID-19 Surveillance Team |
| EPI_ISL_2846787 | Lighthouse Lab in Alderley Park | Wellcome Sanger Institute for the COVID-19 Genomics UK (COG-UK) Consortium | Jacquelyn Wynn, Mairead Hyland, The Lighthouse Lab in Alderley Park and Alex Alderton, Roberto Amato, Jeffrey Barrett, Sonia Goncalves, Ewan Harrison, David K. Jackson, Ian Johnston, Dominic Kwiatkowski, Cordelia Langford, John Sillitoe on behalf of the Wellcome Sanger Institute COVID-19 Surveillance Team |
| EPI_ISL_2846788 | Health Services Laboratories | Wellcome Sanger Institute for the COVID-19 Genomics UK (COG-UK) Consortium | Health Services Laboratories and Alex Alderton, Roberto Amato, Jeffrey Barrett, Sonia Goncalves, Ewan Harrison, David K. Jackson, Ian Johnston, Dominic Kwiatkowski, Cordelia Langford, John Sillitoe on behalf of the Wellcome Sanger Institute COVID-19 Surveillance Team |
| EPI_ISL_2846789 | Lighthouse Laboratory Plymouth | Wellcome Sanger Institute for the COVID-19 Genomics UK (COG-UK) Consortium | Lighthouse Laboratory Plymouth and Alex Alderton, Roberto Amato, Jeffrey Barrett, Sonia Goncalves, Ewan Harrison, David K. Jackson, Ian Johnston, Dominic Kwiatkowski, Cordelia Langford, John Sillitoe on behalf of the Wellcome Sanger Institute COVID-19 Surveillance Team |
| EPI_ISL_2846790 | Lighthouse Lab in Alderley Park | Wellcome Sanger Institute for the COVID-19 Genomics UK (COG-UK) Consortium | Jacquelyn Wynn, Mairead Hyland, The Lighthouse Lab in Alderley Park and Alex Alderton, Roberto Amato, Jeffrey Barrett, Sonia Goncalves, Ewan Harrison, David K. Jackson, Ian Johnston, Dominic Kwiatkowski, Cordelia Langford, John Sillitoe on behalf of the Wellcome Sanger Institute COVID-19 Surveillance Team |
| EPI_ISL_2846791 | Lighthouse Laboratory Plymouth | Wellcome Sanger Institute for the COVID-19 Genomics UK (COG-UK) Consortium | Lighthouse Laboratory Plymouth and Alex Alderton, Roberto Amato, Jeffrey Barrett, Sonia Goncalves, Ewan Harrison, David K. Jackson, Ian Johnston, Dominic Kwiatkowski, Cordelia Langford, John Sillitoe on behalf of the Wellcome Sanger Institute COVID-19 Surveillance Team |
| EPI_ISL_2846792 | Health Services Laboratories | Wellcome Sanger Institute for the COVID-19 Genomics UK (COG-UK) Consortium | Health Services Laboratories and Alex Alderton, Roberto Amato, Jeffrey Barrett, Sonia Goncalves, Ewan Harrison, David K. Jackson, Ian Johnston, Dominic Kwiatkowski, Cordelia Langford, John Sillitoe on behalf of the Wellcome Sanger Institute COVID-19 Surveillance Team |
| EPI_ISL_2846793 | Lighthouse Laboratory Plymouth | Wellcome Sanger Institute for the COVID-19 Genomics UK (COG-UK) Consortium | Lighthouse Laboratory Plymouth and Alex Alderton, Roberto Amato, Jeffrey Barrett, Sonia Goncalves, Ewan Harrison, David K. Jackson, Ian Johnston, Dominic Kwiatkowski, Cordelia Langford, John Sillitoe on behalf of the Wellcome Sanger Institute COVID-19 Surveillance Team |
| EPI_ISL_2846794 | Lighthouse Lab in Milton Keynes | Wellcome Sanger Institute for the COVID-19 Genomics UK (COG-UK) Consortium | The Lighthouse Lab in Milton Keynes and Alex Alderton, Roberto Amato, Jeffrey Barrett, Sonia Goncalves, Ewan Harrison, David K. Jackson, Ian Johnston, Dominic Kwiatkowski, Cordelia Langford, John Sillitoe on behalf of the Wellcome Sanger Institute COVID-19 Surveillance Team |
| EPI_ISL_2846795 | Health Services Laboratories | Wellcome Sanger Institute for the COVID-19 Genomics UK (COG-UK) Consortium | Health Services Laboratories and Alex Alderton, Roberto Amato, Jeffrey Barrett, Sonia Goncalves, Ewan Harrison, David K. Jackson, Ian Johnston, Dominic Kwiatkowski, Cordelia Langford, John Sillitoe on behalf of the Wellcome Sanger Institute COVID-19 Surveillance Team |
| EPI_ISL_2846796 | Lighthouse Lab in Milton Keynes | Wellcome Sanger Institute for the COVID-19 Genomics UK (COG-UK) Consortium | The Lighthouse Lab in Milton Keynes and Alex Alderton, Roberto Amato, Jeffrey Barrett, Sonia Goncalves, Ewan Harrison, David K. Jackson, Ian Johnston, Dominic Kwiatkowski, Cordelia Langford, John Sillitoe on behalf of the Wellcome Sanger Institute COVID-19 Surveillance Team |
| EPI_ISL_2846797 | Lighthouse Laboratory Plymouth | Wellcome Sanger Institute for the COVID-19 Genomics UK (COG-UK) Consortium | Lighthouse Laboratory Plymouth and Alex Alderton, Roberto Amato, Jeffrey Barrett, Sonia Goncalves, Ewan Harrison, David K. Jackson, Ian Johnston, Dominic Kwiatkowski, Cordelia Langford, John Sillitoe on behalf of the Wellcome Sanger Institute COVID-19 Surveillance Team |
| EPI_ISL_2846798 | Health Services Laboratories | Wellcome Sanger Institute for the COVID-19 Genomics UK (COG-UK) Consortium | Health Services Laboratories and Alex Alderton, Roberto Amato, Jeffrey Barrett, Sonia Goncalves, Ewan Harrison, David K. Jackson, Ian Johnston, Dominic Kwiatkowski, Cordelia Langford, John Sillitoe on behalf of the Wellcome Sanger Institute COVID-19 Surveillance Team |
| EPI_ISL_2846799 | Lighthouse Laboratory Plymouth | Wellcome Sanger Institute for the COVID-19 Genomics UK (COG-UK) Consortium | Lighthouse Laboratory Plymouth and Alex Alderton, Roberto Amato, Jeffrey Barrett, Sonia Goncalves, Ewan Harrison, David K. Jackson, Ian Johnston, Dominic Kwiatkowski, Cordelia Langford, John Sillitoe on behalf of the Wellcome Sanger Institute COVID-19 Surveillance Team |
| EPI_ISL_2846800 | Lighthouse Lab in Alderley Park | Wellcome Sanger Institute for the COVID-19 Genomics UK (COG-UK) Consortium | Jacquelyn Wynn, Mairead Hyland, The Lighthouse Lab in Alderley Park and Alex Alderton, Roberto Amato, Jeffrey Barrett, Sonia Goncalves, Ewan Harrison, David K. Jackson, Ian Johnston, Dominic Kwiatkowski, Cordelia Langford, John Sillitoe on behalf of the Wellcome Sanger Institute COVID-19 Surveillance Team |
| EPI_ISL_2846801 | Lighthouse Lab in Milton Keynes | Wellcome Sanger Institute for the COVID-19 Genomics UK (COG-UK) Consortium | The Lighthouse Lab in Milton Keynes and Alex Alderton, Roberto Amato, Jeffrey Barrett, Sonia Goncalves, Ewan Harrison, David K. Jackson, Ian Johnston, Dominic Kwiatkowski, Cordelia Langford, John Sillitoe on behalf of the Wellcome Sanger Institute COVID-19 Surveillance Team |
| EPI_ISL_2846802, EPI_ISL_2846803 | Lighthouse Lab in Alderley Park | Wellcome Sanger Institute for the COVID-19 Genomics UK (COG-UK) Consortium | Jacquelyn Wynn, Mairead Hyland, The Lighthouse Lab in Alderley Park and Alex Alderton, Roberto Amato, Jeffrey Barrett, Sonia Goncalves, Ewan Harrison, David K. Jackson, Ian Johnston, Dominic Kwiatkowski, Cordelia Langford, John Sillitoe on behalf of the Wellcome Sanger Institute COVID-19 Surveillance Team |
| EPI_ISL_2846804, EPI_ISL_2846805, EPI_ISL_2846806 | Lighthouse Lab in Milton Keynes | Wellcome Sanger Institute for the COVID-19 Genomics UK (COG-UK) Consortium | The Lighthouse Lab in Milton Keynes and Alex Alderton, Roberto Amato, Jeffrey Barrett, Sonia Goncalves, Ewan Harrison, David K. Jackson, Ian Johnston, Dominic Kwiatkowski, Cordelia Langford, John Sillitoe on behalf of the Wellcome Sanger Institute COVID-19 Surveillance Team |
| EPI_ISL_2846807 | Lighthouse Laboratory Plymouth | Wellcome Sanger Institute for the COVID-19 Genomics UK (COG-UK) Consortium | Lighthouse Laboratory Plymouth and Alex Alderton, Roberto Amato, Jeffrey Barrett, Sonia Goncalves, Ewan Harrison, David K. Jackson, Ian Johnston, Dominic Kwiatkowski, Cordelia Langford, John Sillitoe on behalf of the Wellcome Sanger Institute COVID-19 Surveillance Team |
| EPI_ISL_2846808 | Lighthouse Lab in Alderley Park | Wellcome Sanger Institute for the COVID-19 Genomics UK (COG-UK) Consortium | Jacquelyn Wynn, Mairead Hyland, The Lighthouse Lab in Alderley Park and Alex Alderton, Roberto Amato, Jeffrey Barrett, Sonia Goncalves, Ewan Harrison, David K. Jackson, Ian Johnston, Dominic Kwiatkowski, Cordelia Langford, John Sillitoe on behalf of the Wellcome Sanger Institute COVID-19 Surveillance Team |
| EPI_ISL_2846809 | Health Services Laboratories | Wellcome Sanger Institute for the COVID-19 Genomics UK (COG-UK) Consortium | Health Services Laboratories and Alex Alderton, Roberto Amato, Jeffrey Barrett, Sonia Goncalves, Ewan Harrison, David K. Jackson, Ian Johnston, Dominic Kwiatkowski, Cordelia Langford, John Sillitoe on behalf of the Wellcome Sanger Institute COVID-19 Surveillance Team |
| EPI_ISL_2846810 | Lighthouse Laboratory Plymouth | Wellcome Sanger Institute for the COVID-19 Genomics UK (COG-UK) Consortium | Lighthouse Laboratory Plymouth and Alex Alderton, Roberto Amato, Jeffrey Barrett, Sonia Goncalves, Ewan Harrison, David K. Jackson, Ian Johnston, Dominic Kwiatkowski, Cordelia Langford, John Sillitoe on behalf of the Wellcome Sanger Institute COVID-19 Surveillance Team |
| EPI_ISL_2846811, EPI_ISL_2846812 | Health Services Laboratories | Wellcome Sanger Institute for the COVID-19 Genomics UK (COG-UK) Consortium | Health Services Laboratories and Alex Alderton, Roberto Amato, Jeffrey Barrett, Sonia Goncalves, Ewan Harrison, David K. Jackson, Ian Johnston, Dominic Kwiatkowski, Cordelia Langford, John Sillitoe on behalf of the Wellcome Sanger Institute COVID-19 Surveillance Team |
| EPI_ISL_2846813 | Lighthouse Laboratory Plymouth | Wellcome Sanger Institute for the COVID-19 Genomics UK (COG-UK) Consortium | Lighthouse Laboratory Plymouth and Alex Alderton, Roberto Amato, Jeffrey Barrett, Sonia Goncalves, Ewan Harrison, David K. Jackson, Ian Johnston, Dominic Kwiatkowski, Cordelia Langford, John Sillitoe on behalf of the Wellcome Sanger Institute COVID-19 Surveillance Team |
| EPI_ISL_2846814, EPI_ISL_2846815, EPI_ISL_2846816 | Health Services Laboratories | Wellcome Sanger Institute for the COVID-19 Genomics UK (COG-UK) Consortium | Health Services Laboratories and Alex Alderton, Roberto Amato, Jeffrey Barrett, Sonia Goncalves, Ewan Harrison, David K. Jackson, Ian Johnston, Dominic Kwiatkowski, Cordelia Langford, John Sillitoe on behalf of the Wellcome Sanger Institute COVID-19 Surveillance Team |
| EPI_ISL_2846817 | Lighthouse Lab in Milton Keynes | Wellcome Sanger Institute for the COVID-19 Genomics UK (COG-UK) Consortium | The Lighthouse Lab in Milton Keynes and Alex Alderton, Roberto Amato, Jeffrey Barrett, Sonia Goncalves, Ewan Harrison, David K. Jackson, Ian Johnston, Dominic Kwiatkowski, Cordelia Langford, John Sillitoe on behalf of the Wellcome Sanger Institute COVID-19 Surveillance Team |
| EPI_ISL_2846818 | Health Services Laboratories | Wellcome Sanger Institute for the COVID-19 Genomics UK (COG-UK) Consortium | Health Services Laboratories and Alex Alderton, Roberto Amato, Jeffrey Barrett, Sonia Goncalves, Ewan Harrison, David K. Jackson, Ian Johnston, Dominic Kwiatkowski, Cordelia Langford, John Sillitoe on behalf of the Wellcome Sanger Institute COVID-19 Surveillance Team |
| EPI_ISL_2846819 | Lighthouse Laboratory Plymouth | Wellcome Sanger Institute for the COVID-19 Genomics UK (COG-UK) Consortium | Lighthouse Laboratory Plymouth and Alex Alderton, Roberto Amato, Jeffrey Barrett, Sonia Goncalves, Ewan Harrison, David K. Jackson, Ian Johnston, Dominic Kwiatkowski, Cordelia Langford, John Sillitoe on behalf of the Wellcome Sanger Institute COVID-19 Surveillance Team |
| EPI_ISL_2846820 | Health Services Laboratories | Wellcome Sanger Institute for the COVID-19 Genomics UK (COG-UK) Consortium | Health Services Laboratories and Alex Alderton, Roberto Amato, Jeffrey Barrett, Sonia Goncalves, Ewan Harrison, David K. Jackson, Ian Johnston, Dominic Kwiatkowski, Cordelia Langford, John Sillitoe on behalf of the Wellcome Sanger Institute COVID-19 Surveillance Team |
| EPI_ISL_2846821 | Lighthouse Lab in Milton Keynes | Wellcome Sanger Institute for the COVID-19 Genomics UK (COG-UK) Consortium | The Lighthouse Lab in Milton Keynes and Alex Alderton, Roberto Amato, Jeffrey Barrett, Sonia Goncalves, Ewan Harrison, David K. Jackson, Ian Johnston, Dominic Kwiatkowski, Cordelia Langford, John Sillitoe on behalf of the Wellcome Sanger Institute COVID-19 Surveillance Team |
| EPI_ISL_2846822, EPI_ISL_2846823 | Lighthouse Laboratory Plymouth | Wellcome Sanger Institute for the COVID-19 Genomics UK (COG-UK) Consortium | Lighthouse Laboratory Plymouth and Alex Alderton, Roberto Amato, Jeffrey Barrett, Sonia Goncalves, Ewan Harrison, David K. Jackson, Ian Johnston, Dominic Kwiatkowski, Cordelia Langford, John Sillitoe on behalf of the Wellcome Sanger Institute COVID-19 Surveillance Team |
| EPI_ISL_2846824 | Lighthouse Lab in Milton Keynes | Wellcome Sanger Institute for the COVID-19 Genomics UK (COG-UK) Consortium | The Lighthouse Lab in Milton Keynes and Alex Alderton, Roberto Amato, Jeffrey Barrett, Sonia Goncalves, Ewan Harrison, David K. Jackson, Ian Johnston, Dominic Kwiatkowski, Cordelia Langford, John Sillitoe on behalf of the Wellcome Sanger Institute COVID-19 Surveillance Team |
| EPI_ISL_2846825 | Health Services Laboratories | Wellcome Sanger Institute for the COVID-19 Genomics UK (COG-UK) Consortium | Health Services Laboratories and Alex Alderton, Roberto Amato, Jeffrey Barrett, Sonia Goncalves, Ewan Harrison, David K. Jackson, Ian Johnston, Dominic Kwiatkowski, Cordelia Langford, John Sillitoe on behalf of the Wellcome Sanger Institute COVID-19 Surveillance Team |
| EPI_ISL_2846826 | Lighthouse Lab in Milton Keynes | Wellcome Sanger Institute for the COVID-19 Genomics UK (COG-UK) Consortium | The Lighthouse Lab in Milton Keynes and Alex Alderton, Roberto Amato, Jeffrey Barrett, Sonia Goncalves, Ewan Harrison, David K. Jackson, Ian Johnston, Dominic Kwiatkowski, Cordelia Langford, John Sillitoe on behalf of the Wellcome Sanger Institute COVID-19 Surveillance Team |
| EPI_ISL_2846827 | Health Services Laboratories | Wellcome Sanger Institute for the COVID-19 Genomics UK (COG-UK) Consortium | Health Services Laboratories and Alex Alderton, Roberto Amato, Jeffrey Barrett, Sonia Goncalves, Ewan Harrison, David K. Jackson, Ian Johnston, Dominic Kwiatkowski, Cordelia Langford, John Sillitoe on behalf of the Wellcome Sanger Institute COVID-19 Surveillance Team |

| | | | |
|---|---|---|---|
| EPI_ISL_2846828 | Lighthouse Lab in Alderley Park | Wellcome Sanger Institute for the COVID-19 Genomics UK (COG-UK) Consortium | Jacquelyn Wynn, Mairead Hyland, The Lighthouse Lab in Alderley Park and Alex Alderton, Roberto Amato, Jeffrey Barrett, Sonia Goncalves, Ewan Harrison, David K. Jackson, Ian Johnston, Dominic Kwiatkowski, Cordelia Langford, John Sillitoe on behalf of the Wellcome Sanger Institute COVID-19 Surveillance Team |
| EPI_ISL_2846829 | Lighthouse Lab in Milton Keynes | Wellcome Sanger Institute for the COVID-19 Genomics UK (COG-UK) Consortium | The Lighthouse Lab in Milton Keynes and Alex Alderton, Roberto Amato, Jeffrey Barrett, Sonia Goncalves, Ewan Harrison, David K. Jackson, Ian Johnston, Dominic Kwiatkowski, Cordelia Langford, John Sillitoe on behalf of the Wellcome Sanger Institute COVID-19 Surveillance Team |
| EPI_ISL_2846830 | Lighthouse Laboratory Plymouth | Wellcome Sanger Institute for the COVID-19 Genomics UK (COG-UK) Consortium | Lighthouse Laboratory Plymouth and Alex Alderton, Roberto Amato, Jeffrey Barrett, Sonia Goncalves, Ewan Harrison, David K. Jackson, Ian Johnston, Dominic Kwiatkowski, Cordelia Langford, John Sillitoe on behalf of the Wellcome Sanger Institute COVID-19 Surveillance Team |
| EPI_ISL_2846831 | Lighthouse Lab in Alderley Park | Wellcome Sanger Institute for the COVID-19 Genomics UK (COG-UK) Consortium | Jacquelyn Wynn, Mairead Hyland, The Lighthouse Lab in Alderley Park and Alex Alderton, Roberto Amato, Jeffrey Barrett, Sonia Goncalves, Ewan Harrison, David K. Jackson, Ian Johnston, Dominic Kwiatkowski, Cordelia Langford, John Sillitoe on behalf of the Wellcome Sanger Institute COVID-19 Surveillance Team |
| EPI_ISL_2846832 | Health Services Laboratories | Wellcome Sanger Institute for the COVID-19 Genomics UK (COG-UK) Consortium | Health Services Laboratories and Alex Alderton, Roberto Amato, Jeffrey Barrett, Sonia Goncalves, Ewan Harrison, David K. Jackson, Ian Johnston, Dominic Kwiatkowski, Cordelia Langford, John Sillitoe on behalf of the Wellcome Sanger Institute COVID-19 Surveillance Team |
| EPI_ISL_2846833 | Lighthouse Lab in Alderley Park | Wellcome Sanger Institute for the COVID-19 Genomics UK (COG-UK) Consortium | Jacquelyn Wynn, Mairead Hyland, The Lighthouse Lab in Alderley Park and Alex Alderton, Roberto Amato, Jeffrey Barrett, Sonia Goncalves, Ewan Harrison, David K. Jackson, Ian Johnston, Dominic Kwiatkowski, Cordelia Langford, John Sillitoe on behalf of the Wellcome Sanger Institute COVID-19 Surveillance Team |
| EPI_ISL_2846834, EPI_ISL_2846835 | Lighthouse Lab in Milton Keynes | Wellcome Sanger Institute for the COVID-19 Genomics UK (COG-UK) Consortium | The Lighthouse Lab in Milton Keynes and Alex Alderton, Roberto Amato, Jeffrey Barrett, Sonia Goncalves, Ewan Harrison, David K. Jackson, Ian Johnston, Dominic Kwiatkowski, Cordelia Langford, John Sillitoe on behalf of the Wellcome Sanger Institute COVID-19 Surveillance Team |
| EPI_ISL_2846836 | Lighthouse Laboratory Plymouth | Wellcome Sanger Institute for the COVID-19 Genomics UK (COG-UK) Consortium | Lighthouse Laboratory Plymouth and Alex Alderton, Roberto Amato, Jeffrey Barrett, Sonia Goncalves, Ewan Harrison, David K. Jackson, Ian Johnston, Dominic Kwiatkowski, Cordelia Langford, John Sillitoe on behalf of the Wellcome Sanger Institute COVID-19 Surveillance Team |
| EPI_ISL_2846837 | Health Services Laboratories | Wellcome Sanger Institute for the COVID-19 Genomics UK (COG-UK) Consortium | Health Services Laboratories and Alex Alderton, Roberto Amato, Jeffrey Barrett, Sonia Goncalves, Ewan Harrison, David K. Jackson, Ian Johnston, Dominic Kwiatkowski, Cordelia Langford, John Sillitoe on behalf of the Wellcome Sanger Institute COVID-19 Surveillance Team |
| EPI_ISL_2846838 | Lighthouse Laboratory Plymouth | Wellcome Sanger Institute for the COVID-19 Genomics UK (COG-UK) Consortium | Lighthouse Laboratory Plymouth and Alex Alderton, Roberto Amato, Jeffrey Barrett, Sonia Goncalves, Ewan Harrison, David K. Jackson, Ian Johnston, Dominic Kwiatkowski, Cordelia Langford, John Sillitoe on behalf of the Wellcome Sanger Institute COVID-19 Surveillance Team |
| EPI_ISL_2846839, EPI_ISL_2846840 | Lighthouse Lab in Milton Keynes | Wellcome Sanger Institute for the COVID-19 Genomics UK (COG-UK) Consortium | The Lighthouse Lab in Milton Keynes and Alex Alderton, Roberto Amato, Jeffrey Barrett, Sonia Goncalves, Ewan Harrison, David K. Jackson, Ian Johnston, Dominic Kwiatkowski, Cordelia Langford, John Sillitoe on behalf of the Wellcome Sanger Institute COVID-19 Surveillance Team |
| EPI_ISL_2846841 | Lighthouse Lab in Alderley Park | Wellcome Sanger Institute for the COVID-19 Genomics UK (COG-UK) Consortium | Jacquelyn Wynn, Mairead Hyland, The Lighthouse Lab in Alderley Park and Alex Alderton, Roberto Amato, Jeffrey Barrett, Sonia Goncalves, Ewan Harrison, David K. Jackson, Ian Johnston, Dominic Kwiatkowski, Cordelia Langford, John Sillitoe on behalf of the Wellcome Sanger Institute COVID-19 Surveillance Team |
| EPI_ISL_2846842 | Lighthouse Laboratory Plymouth | Wellcome Sanger Institute for the COVID-19 Genomics UK (COG-UK) Consortium | Lighthouse Laboratory Plymouth and Alex Alderton, Roberto Amato, Jeffrey Barrett, Sonia Goncalves, Ewan Harrison, David K. Jackson, Ian Johnston, Dominic Kwiatkowski, Cordelia Langford, John Sillitoe on behalf of the Wellcome Sanger Institute COVID-19 Surveillance Team |
| EPI_ISL_2846843, EPI_ISL_2846844 | Health Services Laboratories | Wellcome Sanger Institute for the COVID-19 Genomics UK (COG-UK) Consortium | Health Services Laboratories and Alex Alderton, Roberto Amato, Jeffrey Barrett, Sonia Goncalves, Ewan Harrison, David K. Jackson, Ian Johnston, Dominic Kwiatkowski, Cordelia Langford, John Sillitoe on behalf of the Wellcome Sanger Institute COVID-19 Surveillance Team |
| EPI_ISL_2846845, EPI_ISL_2846846, EPI_ISL_2846847 | Lighthouse Lab in Alderley Park | Wellcome Sanger Institute for the COVID-19 Genomics UK (COG-UK) Consortium | Jacquelyn Wynn, Mairead Hyland, The Lighthouse Lab in Alderley Park and Alex Alderton, Roberto Amato, Jeffrey Barrett, Sonia Goncalves, Ewan Harrison, David K. Jackson, Ian Johnston, Dominic Kwiatkowski, Cordelia Langford, John Sillitoe on behalf of the Wellcome Sanger Institute COVID-19 Surveillance Team |
| EPI_ISL_2846848 | Lighthouse Laboratory Plymouth | Wellcome Sanger Institute for the COVID-19 Genomics UK (COG-UK) Consortium | Lighthouse Laboratory Plymouth and Alex Alderton, Roberto Amato, Jeffrey Barrett, Sonia Goncalves, Ewan Harrison, David K. Jackson, Ian Johnston, Dominic Kwiatkowski, Cordelia Langford, John Sillitoe on behalf of the Wellcome Sanger Institute COVID-19 Surveillance Team |
| EPI_ISL_2846849 | Health Services Laboratories | Wellcome Sanger Institute for the COVID-19 Genomics UK (COG-UK) Consortium | Health Services Laboratories and Alex Alderton, Roberto Amato, Jeffrey Barrett, Sonia Goncalves, Ewan Harrison, David K. Jackson, Ian Johnston, Dominic Kwiatkowski, Cordelia Langford, John Sillitoe on behalf of the Wellcome Sanger Institute COVID-19 Surveillance Team |
| EPI_ISL_2846850, EPI_ISL_2846851 | Lighthouse Laboratory Plymouth | Wellcome Sanger Institute for the COVID-19 Genomics UK (COG-UK) Consortium | Lighthouse Laboratory Plymouth and Alex Alderton, Roberto Amato, Jeffrey Barrett, Sonia Goncalves, Ewan Harrison, David K. Jackson, Ian Johnston, Dominic Kwiatkowski, Cordelia Langford, John Sillitoe on behalf of the Wellcome Sanger Institute COVID-19 Surveillance Team |
| EPI_ISL_2846852 | Lighthouse Lab in Milton Keynes | Wellcome Sanger Institute for the COVID-19 Genomics UK (COG-UK) Consortium | The Lighthouse Lab in Milton Keynes and Alex Alderton, Roberto Amato, Jeffrey Barrett, Sonia Goncalves, Ewan Harrison, David K. Jackson, Ian Johnston, Dominic Kwiatkowski, Cordelia Langford, John Sillitoe on behalf of the Wellcome Sanger Institute COVID-19 Surveillance Team |
| EPI_ISL_2846853 | Lighthouse Lab in Alderley Park | Wellcome Sanger Institute for the COVID-19 Genomics UK (COG-UK) Consortium | Jacquelyn Wynn, Mairead Hyland, The Lighthouse Lab in Alderley Park and Alex Alderton, Roberto Amato, Jeffrey Barrett, Sonia Goncalves, Ewan Harrison, David K. Jackson, Ian Johnston, Dominic Kwiatkowski, Cordelia Langford, John Sillitoe on behalf of the Wellcome Sanger Institute COVID-19 Surveillance Team |
| EPI_ISL_2846854 | Health Services Laboratories | Wellcome Sanger Institute for the COVID-19 Genomics UK (COG-UK) Consortium | Health Services Laboratories and Alex Alderton, Roberto Amato, Jeffrey Barrett, Sonia Goncalves, Ewan Harrison, David K. Jackson, Ian Johnston, Dominic Kwiatkowski, Cordelia Langford, John Sillitoe on behalf of the Wellcome Sanger Institute COVID-19 Surveillance Team |
| EPI_ISL_2846855 | Lighthouse Laboratory Plymouth | Wellcome Sanger Institute for the COVID-19 Genomics UK (COG-UK) Consortium | Lighthouse Laboratory Plymouth and Alex Alderton, Roberto Amato, Jeffrey Barrett, Sonia Goncalves, Ewan Harrison, David K. Jackson, Ian Johnston, Dominic Kwiatkowski, Cordelia Langford, John Sillitoe on behalf of the Wellcome Sanger Institute COVID-19 Surveillance Team |
| EPI_ISL_2846856, EPI_ISL_2846857, EPI_ISL_2846858 | Lighthouse Lab in Alderley Park | Wellcome Sanger Institute for the COVID-19 Genomics UK (COG-UK) Consortium | Jacquelyn Wynn, Mairead Hyland, The Lighthouse Lab in Alderley Park and Alex Alderton, Roberto Amato, Jeffrey Barrett, Sonia Goncalves, Ewan Harrison, David K. Jackson, Ian Johnston, Dominic Kwiatkowski, Cordelia Langford, John Sillitoe on behalf of the Wellcome Sanger Institute COVID-19 Surveillance Team |
| EPI_ISL_2846859 | Health Services Laboratories | Wellcome Sanger Institute for the COVID-19 Genomics UK (COG-UK) Consortium | Health Services Laboratories and Alex Alderton, Roberto Amato, Jeffrey Barrett, Sonia Goncalves, Ewan Harrison, David K. Jackson, Ian Johnston, Dominic Kwiatkowski, Cordelia Langford, John Sillitoe on behalf of the Wellcome Sanger Institute COVID-19 Surveillance Team |
| EPI_ISL_2846860 | Lighthouse Lab in Alderley Park | Wellcome Sanger Institute for the COVID-19 Genomics UK (COG-UK) Consortium | Jacquelyn Wynn, Mairead Hyland, The Lighthouse Lab in Alderley Park and Alex Alderton, Roberto Amato, Jeffrey Barrett, Sonia Goncalves, Ewan Harrison, David K. Jackson, Ian Johnston, Dominic Kwiatkowski, Cordelia Langford, John Sillitoe on behalf of the Wellcome Sanger Institute COVID-19 Surveillance Team |
| EPI_ISL_2846861 | Lighthouse Lab in Milton Keynes | Wellcome Sanger Institute for the COVID-19 Genomics UK (COG-UK) Consortium | The Lighthouse Lab in Milton Keynes and Alex Alderton, Roberto Amato, Jeffrey Barrett, Sonia Goncalves, Ewan Harrison, David K. Jackson, Ian Johnston, Dominic Kwiatkowski, Cordelia Langford, John Sillitoe on behalf of the Wellcome Sanger Institute COVID-19 Surveillance Team |
| EPI_ISL_2846862 | Lighthouse Laboratory Plymouth | Wellcome Sanger Institute for the COVID-19 Genomics UK (COG-UK) Consortium | Lighthouse Laboratory Plymouth and Alex Alderton, Roberto Amato, Jeffrey Barrett, Sonia Goncalves, Ewan Harrison, David K. Jackson, Ian Johnston, Dominic Kwiatkowski, Cordelia Langford, John Sillitoe on behalf of the Wellcome Sanger Institute COVID-19 Surveillance Team |
| EPI_ISL_2846863 | Lighthouse Lab in Milton Keynes | Wellcome Sanger Institute for the COVID-19 Genomics UK (COG-UK) Consortium | The Lighthouse Lab in Milton Keynes and Alex Alderton, Roberto Amato, Jeffrey Barrett, Sonia Goncalves, Ewan Harrison, David K. Jackson, Ian Johnston, Dominic Kwiatkowski, Cordelia Langford, John Sillitoe on behalf of the Wellcome Sanger Institute COVID-19 Surveillance Team |
| EPI_ISL_2846864 | Lighthouse Lab in Alderley Park | Wellcome Sanger Institute for the COVID-19 Genomics UK (COG-UK) Consortium | Jacquelyn Wynn, Mairead Hyland, The Lighthouse Lab in Alderley Park and Alex Alderton, Roberto Amato, Jeffrey Barrett, Sonia Goncalves, Ewan Harrison, David K. Jackson, Ian Johnston, Dominic Kwiatkowski, Cordelia Langford, John Sillitoe on behalf of the Wellcome Sanger Institute COVID-19 Surveillance Team |
| EPI_ISL_2846865, EPI_ISL_2846866 | Health Services Laboratories | Wellcome Sanger Institute for the COVID-19 Genomics UK (COG-UK) Consortium | Health Services Laboratories and Alex Alderton, Roberto Amato, Jeffrey Barrett, Sonia Goncalves, Ewan Harrison, David K. Jackson, Ian Johnston, Dominic Kwiatkowski, Cordelia Langford, John Sillitoe on behalf of the Wellcome Sanger Institute COVID-19 Surveillance Team |
| EPI_ISL_2846867 | Lighthouse Lab in Milton Keynes | Wellcome Sanger Institute for the COVID-19 Genomics UK (COG-UK) Consortium | The Lighthouse Lab in Milton Keynes and Alex Alderton, Roberto Amato, Jeffrey Barrett, Sonia Goncalves, Ewan Harrison, David K. Jackson, Ian Johnston, Dominic Kwiatkowski, Cordelia Langford, John Sillitoe on behalf of the Wellcome Sanger Institute COVID-19 Surveillance Team |
| EPI_ISL_2846868 | Lighthouse Lab in Alderley Park | Wellcome Sanger Institute for the COVID-19 Genomics UK (COG-UK) Consortium | Jacquelyn Wynn, Mairead Hyland, The Lighthouse Lab in Alderley Park and Alex Alderton, Roberto Amato, Jeffrey Barrett, Sonia Goncalves, Ewan Harrison, David K. Jackson, Ian Johnston, Dominic Kwiatkowski, Cordelia Langford, John Sillitoe on behalf of the Wellcome Sanger Institute COVID-19 Surveillance Team |
| EPI_ISL_2846869 | Lighthouse Laboratory Plymouth | Wellcome Sanger Institute for the COVID-19 Genomics UK (COG-UK) Consortium | Lighthouse Laboratory Plymouth and Alex Alderton, Roberto Amato, Jeffrey Barrett, Sonia Goncalves, Ewan Harrison, David K. Jackson, Ian Johnston, Dominic Kwiatkowski, Cordelia Langford, John Sillitoe on behalf of the Wellcome Sanger Institute COVID-19 Surveillance Team |
| EPI_ISL_2846870 | Lighthouse Lab in Alderley Park | Wellcome Sanger Institute for the COVID-19 Genomics UK (COG-UK) Consortium | Jacquelyn Wynn, Mairead Hyland, The Lighthouse Lab in Alderley Park and Alex Alderton, Roberto Amato, Jeffrey Barrett, Sonia Goncalves, Ewan Harrison, David K. Jackson, Ian Johnston, Dominic Kwiatkowski, Cordelia Langford, John Sillitoe on behalf of the Wellcome Sanger Institute COVID-19 Surveillance Team |
| EPI_ISL_2846871 | Lighthouse Lab in Milton Keynes | Wellcome Sanger Institute for the COVID-19 Genomics UK (COG-UK) Consortium | The Lighthouse Lab in Milton Keynes and Alex Alderton, Roberto Amato, Jeffrey Barrett, Sonia Goncalves, Ewan Harrison, David K. Jackson, Ian Johnston, Dominic Kwiatkowski, Cordelia Langford, John Sillitoe on behalf of the Wellcome Sanger Institute COVID-19 Surveillance Team |
| EPI_ISL_2846872 | Health Services Laboratories | Wellcome Sanger Institute for the COVID-19 Genomics UK (COG-UK) Consortium | Health Services Laboratories and Alex Alderton, Roberto Amato, Jeffrey Barrett, Sonia Goncalves, Ewan Harrison, David K. Jackson, Ian Johnston, Dominic Kwiatkowski, Cordelia Langford, John Sillitoe on behalf of the Wellcome Sanger Institute COVID-19 Surveillance Team |
| EPI_ISL_2846873 | Lighthouse Lab in Milton Keynes | Wellcome Sanger Institute for the COVID-19 Genomics UK (COG-UK) Consortium | The Lighthouse Lab in Milton Keynes and Alex Alderton, Roberto Amato, Jeffrey Barrett, Sonia Goncalves, Ewan Harrison, David K. Jackson, Ian Johnston, Dominic Kwiatkowski, Cordelia Langford, John Sillitoe on behalf of the Wellcome Sanger Institute COVID-19 Surveillance Team |
| EPI_ISL_2846875, EPI_ISL_2846876, EPI_ISL_2846877 | Lighthouse Laboratory Plymouth | Wellcome Sanger Institute for the COVID-19 Genomics UK (COG-UK) Consortium | Lighthouse Laboratory Plymouth and Alex Alderton, Roberto Amato, Jeffrey Barrett, Sonia Goncalves, Ewan Harrison, David K. Jackson, Ian Johnston, Dominic Kwiatkowski, Cordelia Langford, John Sillitoe on behalf of the Wellcome Sanger Institute COVID-19 Surveillance Team |
| EPI_ISL_2846878 | Lighthouse Lab in Alderley Park | Wellcome Sanger Institute for the COVID-19 Genomics UK (COG-UK) Consortium | Jacquelyn Wynn, Mairead Hyland, The Lighthouse Lab in Alderley Park and Alex Alderton, Roberto Amato, Jeffrey Barrett, Sonia Goncalves, Ewan Harrison, David K. Jackson, Ian Johnston, Dominic Kwiatkowski, Cordelia Langford, John Sillitoe on behalf of the Wellcome Sanger Institute COVID-19 Surveillance Team |
| EPI_ISL_2846879 | Health Services Laboratories | Wellcome Sanger Institute for the COVID-19 Genomics UK (COG-UK) Consortium | Health Services Laboratories and Alex Alderton, Roberto Amato, Jeffrey Barrett, Sonia Goncalves, Ewan Harrison, David K. Jackson, Ian Johnston, Dominic Kwiatkowski, Cordelia Langford, John Sillitoe on behalf of the Wellcome Sanger Institute COVID-19 Surveillance Team |
| EPI_ISL_2846880 | Lighthouse Laboratory Plymouth | Wellcome Sanger Institute for the COVID-19 Genomics UK (COG-UK) Consortium | Lighthouse Laboratory Plymouth and Alex Alderton, Roberto Amato, Jeffrey Barrett, Sonia Goncalves, Ewan Harrison, David K. Jackson, Ian Johnston, Dominic Kwiatkowski, Cordelia Langford, John Sillitoe on behalf of the Wellcome Sanger Institute COVID-19 Surveillance Team |
| EPI_ISL_2846881 | Health Services Laboratories | Wellcome Sanger Institute for the COVID-19 Genomics UK (COG-UK) Consortium | Health Services Laboratories and Alex Alderton, Roberto Amato, Jeffrey Barrett, Sonia Goncalves, Ewan Harrison, David K. Jackson, Ian Johnston, Dominic Kwiatkowski, Cordelia Langford, John Sillitoe on behalf of the Wellcome Sanger Institute COVID-19 Surveillance Team |
| EPI_ISL_2846882 | Lighthouse Lab in Milton Keynes | Wellcome Sanger Institute for the COVID-19 Genomics UK (COG-UK) Consortium | The Lighthouse Lab in Milton Keynes and Alex Alderton, Roberto Amato, Jeffrey Barrett, Sonia Goncalves, Ewan Harrison, David K. Jackson, Ian Johnston, Dominic Kwiatkowski, Cordelia Langford, John Sillitoe on behalf of the Wellcome Sanger Institute COVID-19 Surveillance Team |
| EPI_ISL_2846883 | Lighthouse Lab in Alderley Park | Wellcome Sanger Institute for the COVID-19 Genomics UK (COG-UK) Consortium | Jacquelyn Wynn, Mairead Hyland, The Lighthouse Lab in Alderley Park and Alex Alderton, Roberto Amato, Jeffrey Barrett, Sonia Goncalves, Ewan Harrison, David K. Jackson, Ian Johnston, Dominic Kwiatkowski, Cordelia Langford, John Sillitoe on behalf of the Wellcome Sanger Institute COVID-19 Surveillance Team |

| | | | |
|---|---|---|---|
| EPI_ISL_2846884 | Lighthouse Laboratory Plymouth | Wellcome Sanger Institute for the COVID-19 Genomics UK (COG-UK) Consortium | Lighthouse Laboratory Plymouth and Alex Alderton, Roberto Amato, Jeffrey Barrett, Sonia Goncalves, Ewan Harrison, David K. Jackson, Ian Johnston, Dominic Kwiatkowski, Cordelia Langford, John Sillitoe on behalf of the Wellcome Sanger Institute COVID-19 Surveillance Team |
| EPI_ISL_2846885 | Health Services Laboratories | Wellcome Sanger Institute for the COVID-19 Genomics UK (COG-UK) Consortium | Health Services Laboratories and Alex Alderton, Roberto Amato, Jeffrey Barrett, Sonia Goncalves, Ewan Harrison, David K. Jackson, Ian Johnston, Dominic Kwiatkowski, Cordelia Langford, John Sillitoe on behalf of the Wellcome Sanger Institute COVID-19 Surveillance Team |
| EPI_ISL_2846886 | Lighthouse Lab in Milton Keynes | Wellcome Sanger Institute for the COVID-19 Genomics UK (COG-UK) Consortium | The Lighthouse Lab in Milton Keynes and Alex Alderton, Roberto Amato, Jeffrey Barrett, Sonia Goncalves, Ewan Harrison, David K. Jackson, Ian Johnston, Dominic Kwiatkowski, Cordelia Langford, John Sillitoe on behalf of the Wellcome Sanger Institute COVID-19 Surveillance Team |
| EPI_ISL_2846887 | Lighthouse Lab in Alderley Park | Wellcome Sanger Institute for the COVID-19 Genomics UK (COG-UK) Consortium | Jacquelyn Wynn, Mairead Hyland, The Lighthouse Lab in Alderley Park and Alex Alderton, Roberto Amato, Jeffrey Barrett, Sonia Goncalves, Ewan Harrison, David K. Jackson, Ian Johnston, Dominic Kwiatkowski, Cordelia Langford, John Sillitoe on behalf of the Wellcome Sanger Institute COVID-19 Surveillance Team |
| EPI_ISL_2846888 | Lighthouse Laboratory Plymouth | Wellcome Sanger Institute for the COVID-19 Genomics UK (COG-UK) Consortium | Lighthouse Laboratory Plymouth and Alex Alderton, Roberto Amato, Jeffrey Barrett, Sonia Goncalves, Ewan Harrison, David K. Jackson, Ian Johnston, Dominic Kwiatkowski, Cordelia Langford, John Sillitoe on behalf of the Wellcome Sanger Institute COVID-19 Surveillance Team |
| EPI_ISL_2846889 | Lighthouse Lab in Alderley Park | Wellcome Sanger Institute for the COVID-19 Genomics UK (COG-UK) Consortium | Jacquelyn Wynn, Mairead Hyland, The Lighthouse Lab in Alderley Park and Alex Alderton, Roberto Amato, Jeffrey Barrett, Sonia Goncalves, Ewan Harrison, David K. Jackson, Ian Johnston, Dominic Kwiatkowski, Cordelia Langford, John Sillitoe on behalf of the Wellcome Sanger Institute COVID-19 Surveillance Team |
| EPI_ISL_2846890, EPI_ISL_2846891 | Lighthouse Laboratory Plymouth | Wellcome Sanger Institute for the COVID-19 Genomics UK (COG-UK) Consortium | Lighthouse Laboratory Plymouth and Alex Alderton, Roberto Amato, Jeffrey Barrett, Sonia Goncalves, Ewan Harrison, David K. Jackson, Ian Johnston, Dominic Kwiatkowski, Cordelia Langford, John Sillitoe on behalf of the Wellcome Sanger Institute COVID-19 Surveillance Team |
| EPI_ISL_2846892 | Health Services Laboratories | Wellcome Sanger Institute for the COVID-19 Genomics UK (COG-UK) Consortium | Health Services Laboratories and Alex Alderton, Roberto Amato, Jeffrey Barrett, Sonia Goncalves, Ewan Harrison, David K. Jackson, Ian Johnston, Dominic Kwiatkowski, Cordelia Langford, John Sillitoe on behalf of the Wellcome Sanger Institute COVID-19 Surveillance Team |
| EPI_ISL_2846893 | Lighthouse Lab in Milton Keynes | Wellcome Sanger Institute for the COVID-19 Genomics UK (COG-UK) Consortium | The Lighthouse Lab in Milton Keynes and Alex Alderton, Roberto Amato, Jeffrey Barrett, Sonia Goncalves, Ewan Harrison, David K. Jackson, Ian Johnston, Dominic Kwiatkowski, Cordelia Langford, John Sillitoe on behalf of the Wellcome Sanger Institute COVID-19 Surveillance Team |
| EPI_ISL_2846894 | Lighthouse Laboratory Plymouth | Wellcome Sanger Institute for the COVID-19 Genomics UK (COG-UK) Consortium | Lighthouse Laboratory Plymouth and Alex Alderton, Roberto Amato, Jeffrey Barrett, Sonia Goncalves, Ewan Harrison, David K. Jackson, Ian Johnston, Dominic Kwiatkowski, Cordelia Langford, John Sillitoe on behalf of the Wellcome Sanger Institute COVID-19 Surveillance Team |
| EPI_ISL_2846895 | Health Services Laboratories | Wellcome Sanger Institute for the COVID-19 Genomics UK (COG-UK) Consortium | Health Services Laboratories and Alex Alderton, Roberto Amato, Jeffrey Barrett, Sonia Goncalves, Ewan Harrison, David K. Jackson, Ian Johnston, Dominic Kwiatkowski, Cordelia Langford, John Sillitoe on behalf of the Wellcome Sanger Institute COVID-19 Surveillance Team |
| EPI_ISL_2846896 | Lighthouse Lab in Alderley Park | Wellcome Sanger Institute for the COVID-19 Genomics UK (COG-UK) Consortium | Jacquelyn Wynn, Mairead Hyland, The Lighthouse Lab in Alderley Park and Alex Alderton, Roberto Amato, Jeffrey Barrett, Sonia Goncalves, Ewan Harrison, David K. Jackson, Ian Johnston, Dominic Kwiatkowski, Cordelia Langford, John Sillitoe on behalf of the Wellcome Sanger Institute COVID-19 Surveillance Team |
| EPI_ISL_2846897 | Health Services Laboratories | Wellcome Sanger Institute for the COVID-19 Genomics UK (COG-UK) Consortium | Health Services Laboratories and Alex Alderton, Roberto Amato, Jeffrey Barrett, Sonia Goncalves, Ewan Harrison, David K. Jackson, Ian Johnston, Dominic Kwiatkowski, Cordelia Langford, John Sillitoe on behalf of the Wellcome Sanger Institute COVID-19 Surveillance Team |
| EPI_ISL_2846898 | Lighthouse Lab in Milton Keynes | Wellcome Sanger Institute for the COVID-19 Genomics UK (COG-UK) Consortium | The Lighthouse Lab in Milton Keynes and Alex Alderton, Roberto Amato, Jeffrey Barrett, Sonia Goncalves, Ewan Harrison, David K. Jackson, Ian Johnston, Dominic Kwiatkowski, Cordelia Langford, John Sillitoe on behalf of the Wellcome Sanger Institute COVID-19 Surveillance Team |
| EPI_ISL_2846899 | Health Services Laboratories | Wellcome Sanger Institute for the COVID-19 Genomics UK (COG-UK) Consortium | Health Services Laboratories and Alex Alderton, Roberto Amato, Jeffrey Barrett, Sonia Goncalves, Ewan Harrison, David K. Jackson, Ian Johnston, Dominic Kwiatkowski, Cordelia Langford, John Sillitoe on behalf of the Wellcome Sanger Institute COVID-19 Surveillance Team |
| EPI_ISL_2846900 | Lighthouse Laboratory Plymouth | Wellcome Sanger Institute for the COVID-19 Genomics UK (COG-UK) Consortium | Lighthouse Laboratory Plymouth and Alex Alderton, Roberto Amato, Jeffrey Barrett, Sonia Goncalves, Ewan Harrison, David K. Jackson, Ian Johnston, Dominic Kwiatkowski, Cordelia Langford, John Sillitoe on behalf of the Wellcome Sanger Institute COVID-19 Surveillance Team |
| EPI_ISL_2846901 | Lighthouse Lab in Alderley Park | Wellcome Sanger Institute for the COVID-19 Genomics UK (COG-UK) Consortium | Jacquelyn Wynn, Mairead Hyland, The Lighthouse Lab in Alderley Park and Alex Alderton, Roberto Amato, Jeffrey Barrett, Sonia Goncalves, Ewan Harrison, David K. Jackson, Ian Johnston, Dominic Kwiatkowski, Cordelia Langford, John Sillitoe on behalf of the Wellcome Sanger Institute COVID-19 Surveillance Team |
| EPI_ISL_2846902 | Lighthouse Lab in Milton Keynes | Wellcome Sanger Institute for the COVID-19 Genomics UK (COG-UK) Consortium | The Lighthouse Lab in Milton Keynes and Alex Alderton, Roberto Amato, Jeffrey Barrett, Sonia Goncalves, Ewan Harrison, David K. Jackson, Ian Johnston, Dominic Kwiatkowski, Cordelia Langford, John Sillitoe on behalf of the Wellcome Sanger Institute COVID-19 Surveillance Team |
| EPI_ISL_2846903 | Lighthouse Lab in Alderley Park | Wellcome Sanger Institute for the COVID-19 Genomics UK (COG-UK) Consortium | Jacquelyn Wynn, Mairead Hyland, The Lighthouse Lab in Alderley Park and Alex Alderton, Roberto Amato, Jeffrey Barrett, Sonia Goncalves, Ewan Harrison, David K. Jackson, Ian Johnston, Dominic Kwiatkowski, Cordelia Langford, John Sillitoe on behalf of the Wellcome Sanger Institute COVID-19 Surveillance Team |
| EPI_ISL_2846904, EPI_ISL_2846905, EPI_ISL_2846906, EPI_ISL_2846907 | Health Services Laboratories | Wellcome Sanger Institute for the COVID-19 Genomics UK (COG-UK) Consortium | Health Services Laboratories and Alex Alderton, Roberto Amato, Jeffrey Barrett, Sonia Goncalves, Ewan Harrison, David K. Jackson, Ian Johnston, Dominic Kwiatkowski, Cordelia Langford, John Sillitoe on behalf of the Wellcome Sanger Institute COVID-19 Surveillance Team |
| EPI_ISL_2846908 | Lighthouse Lab in Milton Keynes | Wellcome Sanger Institute for the COVID-19 Genomics UK (COG-UK) Consortium | The Lighthouse Lab in Milton Keynes and Alex Alderton, Roberto Amato, Jeffrey Barrett, Sonia Goncalves, Ewan Harrison, David K. Jackson, Ian Johnston, Dominic Kwiatkowski, Cordelia Langford, John Sillitoe on behalf of the Wellcome Sanger Institute COVID-19 Surveillance Team |
| EPI_ISL_2846909 | Lighthouse Lab in Alderley Park | Wellcome Sanger Institute for the COVID-19 Genomics UK (COG-UK) Consortium | Jacquelyn Wynn, Mairead Hyland, The Lighthouse Lab in Alderley Park and Alex Alderton, Roberto Amato, Jeffrey Barrett, Sonia Goncalves, Ewan Harrison, David K. Jackson, Ian Johnston, Dominic Kwiatkowski, Cordelia Langford, John Sillitoe on behalf of the Wellcome Sanger Institute COVID-19 Surveillance Team |
| EPI_ISL_2846910 | Lighthouse Laboratory Plymouth | Wellcome Sanger Institute for the COVID-19 Genomics UK (COG-UK) Consortium | Lighthouse Laboratory Plymouth and Alex Alderton, Roberto Amato, Jeffrey Barrett, Sonia Goncalves, Ewan Harrison, David K. Jackson, Ian Johnston, Dominic Kwiatkowski, Cordelia Langford, John Sillitoe on behalf of the Wellcome Sanger Institute COVID-19 Surveillance Team |
| EPI_ISL_2846911 | Lighthouse Lab in Alderley Park | Wellcome Sanger Institute for the COVID-19 Genomics UK (COG-UK) Consortium | Jacquelyn Wynn, Mairead Hyland, The Lighthouse Lab in Alderley Park and Alex Alderton, Roberto Amato, Jeffrey Barrett, Sonia Goncalves, Ewan Harrison, David K. Jackson, Ian Johnston, Dominic Kwiatkowski, Cordelia Langford, John Sillitoe on behalf of the Wellcome Sanger Institute COVID-19 Surveillance Team |
| EPI_ISL_2846912 | Lighthouse Laboratory Plymouth | Wellcome Sanger Institute for the COVID-19 Genomics UK (COG-UK) Consortium | Lighthouse Laboratory Plymouth and Alex Alderton, Roberto Amato, Jeffrey Barrett, Sonia Goncalves, Ewan Harrison, David K. Jackson, Ian Johnston, Dominic Kwiatkowski, Cordelia Langford, John Sillitoe on behalf of the Wellcome Sanger Institute COVID-19 Surveillance Team |
| EPI_ISL_2846913 | Lighthouse Lab in Milton Keynes | Wellcome Sanger Institute for the COVID-19 Genomics UK (COG-UK) Consortium | The Lighthouse Lab in Milton Keynes and Alex Alderton, Roberto Amato, Jeffrey Barrett, Sonia Goncalves, Ewan Harrison, David K. Jackson, Ian Johnston, Dominic Kwiatkowski, Cordelia Langford, John Sillitoe on behalf of the Wellcome Sanger Institute COVID-19 Surveillance Team |
| EPI_ISL_2846914 | Health Services Laboratories | Wellcome Sanger Institute for the COVID-19 Genomics UK (COG-UK) Consortium | Health Services Laboratories and Alex Alderton, Roberto Amato, Jeffrey Barrett, Sonia Goncalves, Ewan Harrison, David K. Jackson, Ian Johnston, Dominic Kwiatkowski, Cordelia Langford, John Sillitoe on behalf of the Wellcome Sanger Institute COVID-19 Surveillance Team |
| EPI_ISL_2846915 | Lighthouse Laboratory Plymouth | Wellcome Sanger Institute for the COVID-19 Genomics UK (COG-UK) Consortium | Lighthouse Laboratory Plymouth and Alex Alderton, Roberto Amato, Jeffrey Barrett, Sonia Goncalves, Ewan Harrison, David K. Jackson, Ian Johnston, Dominic Kwiatkowski, Cordelia Langford, John Sillitoe on behalf of the Wellcome Sanger Institute COVID-19 Surveillance Team |
| EPI_ISL_2846916 | Health Services Laboratories | Wellcome Sanger Institute for the COVID-19 Genomics UK (COG-UK) Consortium | Health Services Laboratories and Alex Alderton, Roberto Amato, Jeffrey Barrett, Sonia Goncalves, Ewan Harrison, David K. Jackson, Ian Johnston, Dominic Kwiatkowski, Cordelia Langford, John Sillitoe on behalf of the Wellcome Sanger Institute COVID-19 Surveillance Team |
| EPI_ISL_2846917 | Lighthouse Lab in Alderley Park | Wellcome Sanger Institute for the COVID-19 Genomics UK (COG-UK) Consortium | Jacquelyn Wynn, Mairead Hyland, The Lighthouse Lab in Alderley Park and Alex Alderton, Roberto Amato, Jeffrey Barrett, Sonia Goncalves, Ewan Harrison, David K. Jackson, Ian Johnston, Dominic Kwiatkowski, Cordelia Langford, John Sillitoe on behalf of the Wellcome Sanger Institute COVID-19 Surveillance Team |
| EPI_ISL_2846918 | Lighthouse Laboratory Plymouth | Wellcome Sanger Institute for the COVID-19 Genomics UK (COG-UK) Consortium | Lighthouse Laboratory Plymouth and Alex Alderton, Roberto Amato, Jeffrey Barrett, Sonia Goncalves, Ewan Harrison, David K. Jackson, Ian Johnston, Dominic Kwiatkowski, Cordelia Langford, John Sillitoe on behalf of the Wellcome Sanger Institute COVID-19 Surveillance Team |
| EPI_ISL_2846919 | Lighthouse Lab in Milton Keynes | Wellcome Sanger Institute for the COVID-19 Genomics UK (COG-UK) Consortium | The Lighthouse Lab in Milton Keynes and Alex Alderton, Roberto Amato, Jeffrey Barrett, Sonia Goncalves, Ewan Harrison, David K. Jackson, Ian Johnston, Dominic Kwiatkowski, Cordelia Langford, John Sillitoe on behalf of the Wellcome Sanger Institute COVID-19 Surveillance Team |
| EPI_ISL_2846920 | Lighthouse Lab in Alderley Park | Wellcome Sanger Institute for the COVID-19 Genomics UK (COG-UK) Consortium | Jacquelyn Wynn, Mairead Hyland, The Lighthouse Lab in Alderley Park and Alex Alderton, Roberto Amato, Jeffrey Barrett, Sonia Goncalves, Ewan Harrison, David K. Jackson, Ian Johnston, Dominic Kwiatkowski, Cordelia Langford, John Sillitoe on behalf of the Wellcome Sanger Institute COVID-19 Surveillance Team |
| EPI_ISL_2846921 | Health Services Laboratories | Wellcome Sanger Institute for the COVID-19 Genomics UK (COG-UK) Consortium | Health Services Laboratories and Alex Alderton, Roberto Amato, Jeffrey Barrett, Sonia Goncalves, Ewan Harrison, David K. Jackson, Ian Johnston, Dominic Kwiatkowski, Cordelia Langford, John Sillitoe on behalf of the Wellcome Sanger Institute COVID-19 Surveillance Team |
| EPI_ISL_2846922 | Lighthouse Lab in Milton Keynes | Wellcome Sanger Institute for the COVID-19 Genomics UK (COG-UK) Consortium | The Lighthouse Lab in Milton Keynes and Alex Alderton, Roberto Amato, Jeffrey Barrett, Sonia Goncalves, Ewan Harrison, David K. Jackson, Ian Johnston, Dominic Kwiatkowski, Cordelia Langford, John Sillitoe on behalf of the Wellcome Sanger Institute COVID-19 Surveillance Team |
| EPI_ISL_2846923, EPI_ISL_2846924 | Lighthouse Laboratory Plymouth | Wellcome Sanger Institute for the COVID-19 Genomics UK (COG-UK) Consortium | Lighthouse Laboratory Plymouth and Alex Alderton, Roberto Amato, Jeffrey Barrett, Sonia Goncalves, Ewan Harrison, David K. Jackson, Ian Johnston, Dominic Kwiatkowski, Cordelia Langford, John Sillitoe on behalf of the Wellcome Sanger Institute COVID-19 Surveillance Team |
| EPI_ISL_2846925, EPI_ISL_2846926 | Lighthouse Lab in Milton Keynes | Wellcome Sanger Institute for the COVID-19 Genomics UK (COG-UK) Consortium | The Lighthouse Lab in Milton Keynes and Alex Alderton, Roberto Amato, Jeffrey Barrett, Sonia Goncalves, Ewan Harrison, David K. Jackson, Ian Johnston, Dominic Kwiatkowski, Cordelia Langford, John Sillitoe on behalf of the Wellcome Sanger Institute COVID-19 Surveillance Team |
| EPI_ISL_2846927 | Lighthouse Laboratory Plymouth | Wellcome Sanger Institute for the COVID-19 Genomics UK (COG-UK) Consortium | Lighthouse Laboratory Plymouth and Alex Alderton, Roberto Amato, Jeffrey Barrett, Sonia Goncalves, Ewan Harrison, David K. Jackson, Ian Johnston, Dominic Kwiatkowski, Cordelia Langford, John Sillitoe on behalf of the Wellcome Sanger Institute COVID-19 Surveillance Team |
| EPI_ISL_2846928, EPI_ISL_2846929 | Lighthouse Lab in Milton Keynes | Wellcome Sanger Institute for the COVID-19 Genomics UK (COG-UK) Consortium | The Lighthouse Lab in Milton Keynes and Alex Alderton, Roberto Amato, Jeffrey Barrett, Sonia Goncalves, Ewan Harrison, David K. Jackson, Ian Johnston, Dominic Kwiatkowski, Cordelia Langford, John Sillitoe on behalf of the Wellcome Sanger Institute COVID-19 Surveillance Team |
| EPI_ISL_2846930 | Lighthouse Lab in Alderley Park | Wellcome Sanger Institute for the COVID-19 Genomics UK (COG-UK) Consortium | Jacquelyn Wynn, Mairead Hyland, The Lighthouse Lab in Alderley Park and Alex Alderton, Roberto Amato, Jeffrey Barrett, Sonia Goncalves, Ewan Harrison, David K. Jackson, Ian Johnston, Dominic Kwiatkowski, Cordelia Langford, John Sillitoe on behalf of the Wellcome Sanger Institute COVID-19 Surveillance Team |
| EPI_ISL_2846931, EPI_ISL_2846932 | Lighthouse Lab in Milton Keynes | Wellcome Sanger Institute for the COVID-19 Genomics UK (COG-UK) Consortium | The Lighthouse Lab in Milton Keynes and Alex Alderton, Roberto Amato, Jeffrey Barrett, Sonia Goncalves, Ewan Harrison, David K. Jackson, Ian Johnston, Dominic Kwiatkowski, Cordelia Langford, John Sillitoe on behalf of the Wellcome Sanger Institute COVID-19 Surveillance Team |
| EPI_ISL_2846933 | Lighthouse Lab in Alderley Park | Wellcome Sanger Institute for the COVID-19 Genomics UK (COG-UK) Consortium | Jacquelyn Wynn, Mairead Hyland, The Lighthouse Lab in Alderley Park and Alex Alderton, Roberto Amato, Jeffrey Barrett, Sonia Goncalves, Ewan Harrison, David K. Jackson, Ian Johnston, Dominic Kwiatkowski, Cordelia Langford, John Sillitoe on behalf of the Wellcome Sanger Institute COVID-19 Surveillance Team |
| EPI_ISL_2846934, EPI_ISL_2846935, EPI_ISL_2846936 | Health Services Laboratories | Wellcome Sanger Institute for the COVID-19 Genomics UK (COG-UK) Consortium | Health Services Laboratories and Alex Alderton, Roberto Amato, Jeffrey Barrett, Sonia Goncalves, Ewan Harrison, David K. Jackson, Ian Johnston, Dominic Kwiatkowski, Cordelia Langford, John Sillitoe on behalf of the Wellcome Sanger Institute COVID-19 Surveillance Team |
| EPI_ISL_2846937 | Lighthouse Lab in Glasgow | Wellcome Sanger Institute for the COVID-19 Genomics UK (COG-UK) Consortium | Harper VanSteenhouse, Yumi Kasai, David Gray, Carol Clugston, Anna Dominiczak and Alex Alderton, Roberto Amato, Jeffrey Barrett, Sonia Goncalves, Ewan Harrison, David K. Jackson, Ian Johnston, Dominic Kwiatkowski, Cordelia Langford, John Sillitoe on behalf of the Wellcome Sanger Institute COVID-19 Surveillance Team |

| | | | |
|---|---|---|---|
| EPI_ISL_2846938 | Health Services Laboratories | Wellcome Sanger Institute for the COVID-19 Genomics UK (COG-UK) Consortium | Health Services Laboratories and Alex Alderton, Roberto Amato, Jeffrey Barrett, Sonia Goncalves, Ewan Harrison, David K. Jackson, Ian Johnston, Dominic Kwiatkowski, Cordelia Langford, John Sillitoe on behalf of the Wellcome Sanger Institute COVID-19 Surveillance Team |
| EPI_ISL_2846939, EPI_ISL_2846940, EPI_ISL_2846941 | Lighthouse Lab in Alderley Park | Wellcome Sanger Institute for the COVID-19 Genomics UK (COG-UK) Consortium | Jacquelyn Wynn, Mairead Hyland, The Lighthouse Lab in Alderley Park and Alex Alderton, Roberto Amato, Jeffrey Barrett, Sonia Goncalves, Ewan Harrison, David K. Jackson, Ian Johnston, Dominic Kwiatkowski, Cordelia Langford, John Sillitoe on behalf of the Wellcome Sanger Institute COVID-19 Surveillance Team |
| EPI_ISL_2846942 | Health Services Laboratories | Wellcome Sanger Institute for the COVID-19 Genomics UK (COG-UK) Consortium | Health Services Laboratories and Alex Alderton, Roberto Amato, Jeffrey Barrett, Sonia Goncalves, Ewan Harrison, David K. Jackson, Ian Johnston, Dominic Kwiatkowski, Cordelia Langford, John Sillitoe on behalf of the Wellcome Sanger Institute COVID-19 Surveillance Team |
| EPI_ISL_2846943, EPI_ISL_2846944 | Lighthouse Lab in Milton Keynes | Wellcome Sanger Institute for the COVID-19 Genomics UK (COG-UK) Consortium | The Lighthouse Lab in Milton Keynes and Alex Alderton, Roberto Amato, Jeffrey Barrett, Sonia Goncalves, Ewan Harrison, David K. Jackson, Ian Johnston, Dominic Kwiatkowski, Cordelia Langford, John Sillitoe on behalf of the Wellcome Sanger Institute COVID-19 Surveillance Team |
| EPI_ISL_2846945 | Lighthouse Lab in Glasgow | Wellcome Sanger Institute for the COVID-19 Genomics UK (COG-UK) Consortium | Harper VanSteenhouse, Yumi Kasai, David Gray, Carol Clugston, Anna Dominiczak and Alex Alderton, Roberto Amato, Jeffrey Barrett, Sonia Goncalves, Ewan Harrison, David K. Jackson, Ian Johnston, Dominic Kwiatkowski, Cordelia Langford, John Sillitoe on behalf of the Wellcome Sanger Institute COVID-19 Surveillance Team |
| EPI_ISL_2846946 | Health Services Laboratories | Wellcome Sanger Institute for the COVID-19 Genomics UK (COG-UK) Consortium | Health Services Laboratories and Alex Alderton, Roberto Amato, Jeffrey Barrett, Sonia Goncalves, Ewan Harrison, David K. Jackson, Ian Johnston, Dominic Kwiatkowski, Cordelia Langford, John Sillitoe on behalf of the Wellcome Sanger Institute COVID-19 Surveillance Team |
| EPI_ISL_2846947 | Lighthouse Lab in Milton Keynes | Wellcome Sanger Institute for the COVID-19 Genomics UK (COG-UK) Consortium | The Lighthouse Lab in Milton Keynes and Alex Alderton, Roberto Amato, Jeffrey Barrett, Sonia Goncalves, Ewan Harrison, David K. Jackson, Ian Johnston, Dominic Kwiatkowski, Cordelia Langford, John Sillitoe on behalf of the Wellcome Sanger Institute COVID-19 Surveillance Team |
| EPI_ISL_2846948 | Lighthouse Lab in Alderley Park | Wellcome Sanger Institute for the COVID-19 Genomics UK (COG-UK) Consortium | Jacquelyn Wynn, Mairead Hyland, The Lighthouse Lab in Alderley Park and Alex Alderton, Roberto Amato, Jeffrey Barrett, Sonia Goncalves, Ewan Harrison, David K. Jackson, Ian Johnston, Dominic Kwiatkowski, Cordelia Langford, John Sillitoe on behalf of the Wellcome Sanger Institute COVID-19 Surveillance Team |
| EPI_ISL_2846949, EPI_ISL_2846950, EPI_ISL_2846951, EPI_ISL_2846952 | Health Services Laboratories | Wellcome Sanger Institute for the COVID-19 Genomics UK (COG-UK) Consortium | Health Services Laboratories and Alex Alderton, Roberto Amato, Jeffrey Barrett, Sonia Goncalves, Ewan Harrison, David K. Jackson, Ian Johnston, Dominic Kwiatkowski, Cordelia Langford, John Sillitoe on behalf of the Wellcome Sanger Institute COVID-19 Surveillance Team |
| EPI_ISL_2846953, EPI_ISL_2846954, EPI_ISL_2846955 | Lighthouse Lab in Milton Keynes | Wellcome Sanger Institute for the COVID-19 Genomics UK (COG-UK) Consortium | The Lighthouse Lab in Milton Keynes and Alex Alderton, Roberto Amato, Jeffrey Barrett, Sonia Goncalves, Ewan Harrison, David K. Jackson, Ian Johnston, Dominic Kwiatkowski, Cordelia Langford, John Sillitoe on behalf of the Wellcome Sanger Institute COVID-19 Surveillance Team |
| EPI_ISL_2846956 | Lighthouse Lab in Glasgow | Wellcome Sanger Institute for the COVID-19 Genomics UK (COG-UK) Consortium | Harper VanSteenhouse, Yumi Kasai, David Gray, Carol Clugston, Anna Dominiczak and Alex Alderton, Roberto Amato, Jeffrey Barrett, Sonia Goncalves, Ewan Harrison, David K. Jackson, Ian Johnston, Dominic Kwiatkowski, Cordelia Langford, John Sillitoe on behalf of the Wellcome Sanger Institute COVID-19 Surveillance Team |
| EPI_ISL_2846957 | Lighthouse Lab in Alderley Park | Wellcome Sanger Institute for the COVID-19 Genomics UK (COG-UK) Consortium | Jacquelyn Wynn, Mairead Hyland, The Lighthouse Lab in Alderley Park and Alex Alderton, Roberto Amato, Jeffrey Barrett, Sonia Goncalves, Ewan Harrison, David K. Jackson, Ian Johnston, Dominic Kwiatkowski, Cordelia Langford, John Sillitoe on behalf of the Wellcome Sanger Institute COVID-19 Surveillance Team |
| EPI_ISL_2846958 | Lighthouse Lab in Glasgow | Wellcome Sanger Institute for the COVID-19 Genomics UK (COG-UK) Consortium | Harper VanSteenhouse, Yumi Kasai, David Gray, Carol Clugston, Anna Dominiczak and Alex Alderton, Roberto Amato, Jeffrey Barrett, Sonia Goncalves, Ewan Harrison, David K. Jackson, Ian Johnston, Dominic Kwiatkowski, Cordelia Langford, John Sillitoe on behalf of the Wellcome Sanger Institute COVID-19 Surveillance Team |
| EPI_ISL_2846959, EPI_ISL_2846960, EPI_ISL_2846961 | Lighthouse Lab in Milton Keynes | Wellcome Sanger Institute for the COVID-19 Genomics UK (COG-UK) Consortium | The Lighthouse Lab in Milton Keynes and Alex Alderton, Roberto Amato, Jeffrey Barrett, Sonia Goncalves, Ewan Harrison, David K. Jackson, Ian Johnston, Dominic Kwiatkowski, Cordelia Langford, John Sillitoe on behalf of the Wellcome Sanger Institute COVID-19 Surveillance Team |
| EPI_ISL_2846962, EPI_ISL_2846963 | Lighthouse Lab in Alderley Park | Wellcome Sanger Institute for the COVID-19 Genomics UK (COG-UK) Consortium | Jacquelyn Wynn, Mairead Hyland, The Lighthouse Lab in Alderley Park and Alex Alderton, Roberto Amato, Jeffrey Barrett, Sonia Goncalves, Ewan Harrison, David K. Jackson, Ian Johnston, Dominic Kwiatkowski, Cordelia Langford, John Sillitoe on behalf of the Wellcome Sanger Institute COVID-19 Surveillance Team |
| EPI_ISL_2846964, EPI_ISL_2846965 | Lighthouse Lab in Glasgow | Wellcome Sanger Institute for the COVID-19 Genomics UK (COG-UK) Consortium | Harper VanSteenhouse, Yumi Kasai, David Gray, Carol Clugston, Anna Dominiczak and Alex Alderton, Roberto Amato, Jeffrey Barrett, Sonia Goncalves, Ewan Harrison, David K. Jackson, Ian Johnston, Dominic Kwiatkowski, Cordelia Langford, John Sillitoe on behalf of the Wellcome Sanger Institute COVID-19 Surveillance Team |
| EPI_ISL_2846966 | Health Services Laboratories | Wellcome Sanger Institute for the COVID-19 Genomics UK (COG-UK) Consortium | Health Services Laboratories and Alex Alderton, Roberto Amato, Jeffrey Barrett, Sonia Goncalves, Ewan Harrison, David K. Jackson, Ian Johnston, Dominic Kwiatkowski, Cordelia Langford, John Sillitoe on behalf of the Wellcome Sanger Institute COVID-19 Surveillance Team |
| EPI_ISL_2846967 | Lighthouse Lab in Milton Keynes | Wellcome Sanger Institute for the COVID-19 Genomics UK (COG-UK) Consortium | The Lighthouse Lab in Milton Keynes and Alex Alderton, Roberto Amato, Jeffrey Barrett, Sonia Goncalves, Ewan Harrison, David K. Jackson, Ian Johnston, Dominic Kwiatkowski, Cordelia Langford, John Sillitoe on behalf of the Wellcome Sanger Institute COVID-19 Surveillance Team |
| EPI_ISL_2846968 | Lighthouse Lab in Alderley Park | Wellcome Sanger Institute for the COVID-19 Genomics UK (COG-UK) Consortium | Jacquelyn Wynn, Mairead Hyland, The Lighthouse Lab in Alderley Park and Alex Alderton, Roberto Amato, Jeffrey Barrett, Sonia Goncalves, Ewan Harrison, David K. Jackson, Ian Johnston, Dominic Kwiatkowski, Cordelia Langford, John Sillitoe on behalf of the Wellcome Sanger Institute COVID-19 Surveillance Team |
| EPI_ISL_2846969 | Lighthouse Lab in Glasgow | Wellcome Sanger Institute for the COVID-19 Genomics UK (COG-UK) Consortium | Harper VanSteenhouse, Yumi Kasai, David Gray, Carol Clugston, Anna Dominiczak and Alex Alderton, Roberto Amato, Jeffrey Barrett, Sonia Goncalves, Ewan Harrison, David K. Jackson, Ian Johnston, Dominic Kwiatkowski, Cordelia Langford, John Sillitoe on behalf of the Wellcome Sanger Institute COVID-19 Surveillance Team |
| EPI_ISL_2846970, EPI_ISL_2846971 | Lighthouse Lab in Milton Keynes | Wellcome Sanger Institute for the COVID-19 Genomics UK (COG-UK) Consortium | The Lighthouse Lab in Milton Keynes and Alex Alderton, Roberto Amato, Jeffrey Barrett, Sonia Goncalves, Ewan Harrison, David K. Jackson, Ian Johnston, Dominic Kwiatkowski, Cordelia Langford, John Sillitoe on behalf of the Wellcome Sanger Institute COVID-19 Surveillance Team |
| EPI_ISL_2846972, EPI_ISL_2846973 | Lighthouse Lab in Glasgow | Wellcome Sanger Institute for the COVID-19 Genomics UK (COG-UK) Consortium | Harper VanSteenhouse, Yumi Kasai, David Gray, Carol Clugston, Anna Dominiczak and Alex Alderton, Roberto Amato, Jeffrey Barrett, Sonia Goncalves, Ewan Harrison, David K. Jackson, Ian Johnston, Dominic Kwiatkowski, Cordelia Langford, John Sillitoe on behalf of the Wellcome Sanger Institute COVID-19 Surveillance Team |
| EPI_ISL_2846974 | Lighthouse Lab in Milton Keynes | Wellcome Sanger Institute for the COVID-19 Genomics UK (COG-UK) Consortium | The Lighthouse Lab in Milton Keynes and Alex Alderton, Roberto Amato, Jeffrey Barrett, Sonia Goncalves, Ewan Harrison, David K. Jackson, Ian Johnston, Dominic Kwiatkowski, Cordelia Langford, John Sillitoe on behalf of the Wellcome Sanger Institute COVID-19 Surveillance Team |
| EPI_ISL_2846975 | Lighthouse Lab in Glasgow | Wellcome Sanger Institute for the COVID-19 Genomics UK (COG-UK) Consortium | Harper VanSteenhouse, Yumi Kasai, David Gray, Carol Clugston, Anna Dominiczak and Alex Alderton, Roberto Amato, Jeffrey Barrett, Sonia Goncalves, Ewan Harrison, David K. Jackson, Ian Johnston, Dominic Kwiatkowski, Cordelia Langford, John Sillitoe on behalf of the Wellcome Sanger Institute COVID-19 Surveillance Team |
| EPI_ISL_2846976, EPI_ISL_2846977 | Lighthouse Lab in Alderley Park | Wellcome Sanger Institute for the COVID-19 Genomics UK (COG-UK) Consortium | Jacquelyn Wynn, Mairead Hyland, The Lighthouse Lab in Alderley Park and Alex Alderton, Roberto Amato, Jeffrey Barrett, Sonia Goncalves, Ewan Harrison, David K. Jackson, Ian Johnston, Dominic Kwiatkowski, Cordelia Langford, John Sillitoe on behalf of the Wellcome Sanger Institute COVID-19 Surveillance Team |
| EPI_ISL_2846978 | Lighthouse Lab in Milton Keynes | Wellcome Sanger Institute for the COVID-19 Genomics UK (COG-UK) Consortium | The Lighthouse Lab in Milton Keynes and Alex Alderton, Roberto Amato, Jeffrey Barrett, Sonia Goncalves, Ewan Harrison, David K. Jackson, Ian Johnston, Dominic Kwiatkowski, Cordelia Langford, John Sillitoe on behalf of the Wellcome Sanger Institute COVID-19 Surveillance Team |
| EPI_ISL_2846979 | Lighthouse Lab in Alderley Park | Wellcome Sanger Institute for the COVID-19 Genomics UK (COG-UK) Consortium | Jacquelyn Wynn, Mairead Hyland, The Lighthouse Lab in Alderley Park and Alex Alderton, Roberto Amato, Jeffrey Barrett, Sonia Goncalves, Ewan Harrison, David K. Jackson, Ian Johnston, Dominic Kwiatkowski, Cordelia Langford, John Sillitoe on behalf of the Wellcome Sanger Institute COVID-19 Surveillance Team |
| EPI_ISL_2846980, EPI_ISL_2846981 | Lighthouse Lab in Milton Keynes | Wellcome Sanger Institute for the COVID-19 Genomics UK (COG-UK) Consortium | The Lighthouse Lab in Milton Keynes and Alex Alderton, Roberto Amato, Jeffrey Barrett, Sonia Goncalves, Ewan Harrison, David K. Jackson, Ian Johnston, Dominic Kwiatkowski, Cordelia Langford, John Sillitoe on behalf of the Wellcome Sanger Institute COVID-19 Surveillance Team |
| EPI_ISL_2846982 | Lighthouse Lab in Alderley Park | Wellcome Sanger Institute for the COVID-19 Genomics UK (COG-UK) Consortium | Jacquelyn Wynn, Mairead Hyland, The Lighthouse Lab in Alderley Park and Alex Alderton, Roberto Amato, Jeffrey Barrett, Sonia Goncalves, Ewan Harrison, David K. Jackson, Ian Johnston, Dominic Kwiatkowski, Cordelia Langford, John Sillitoe on behalf of the Wellcome Sanger Institute COVID-19 Surveillance Team |
| EPI_ISL_2846983 | Lighthouse Lab in Milton Keynes | Wellcome Sanger Institute for the COVID-19 Genomics UK (COG-UK) Consortium | The Lighthouse Lab in Milton Keynes and Alex Alderton, Roberto Amato, Jeffrey Barrett, Sonia Goncalves, Ewan Harrison, David K. Jackson, Ian Johnston, Dominic Kwiatkowski, Cordelia Langford, John Sillitoe on behalf of the Wellcome Sanger Institute COVID-19 Surveillance Team |
| EPI_ISL_2846984 | Health Services Laboratories | Wellcome Sanger Institute for the COVID-19 Genomics UK (COG-UK) Consortium | Health Services Laboratories and Alex Alderton, Roberto Amato, Jeffrey Barrett, Sonia Goncalves, Ewan Harrison, David K. Jackson, Ian Johnston, Dominic Kwiatkowski, Cordelia Langford, John Sillitoe on behalf of the Wellcome Sanger Institute COVID-19 Surveillance Team |
| EPI_ISL_2846985, EPI_ISL_2846986 | Lighthouse Lab in Alderley Park | Wellcome Sanger Institute for the COVID-19 Genomics UK (COG-UK) Consortium | Jacquelyn Wynn, Mairead Hyland, The Lighthouse Lab in Alderley Park and Alex Alderton, Roberto Amato, Jeffrey Barrett, Sonia Goncalves, Ewan Harrison, David K. Jackson, Ian Johnston, Dominic Kwiatkowski, Cordelia Langford, John Sillitoe on behalf of the Wellcome Sanger Institute COVID-19 Surveillance Team |
| EPI_ISL_2846987, EPI_ISL_2846988 | Health Services Laboratories | Wellcome Sanger Institute for the COVID-19 Genomics UK (COG-UK) Consortium | Health Services Laboratories and Alex Alderton, Roberto Amato, Jeffrey Barrett, Sonia Goncalves, Ewan Harrison, David K. Jackson, Ian Johnston, Dominic Kwiatkowski, Cordelia Langford, John Sillitoe on behalf of the Wellcome Sanger Institute COVID-19 Surveillance Team |
| EPI_ISL_2846989 | Lighthouse Lab in Milton Keynes | Wellcome Sanger Institute for the COVID-19 Genomics UK (COG-UK) Consortium | The Lighthouse Lab in Milton Keynes and Alex Alderton, Roberto Amato, Jeffrey Barrett, Sonia Goncalves, Ewan Harrison, David K. Jackson, Ian Johnston, Dominic Kwiatkowski, Cordelia Langford, John Sillitoe on behalf of the Wellcome Sanger Institute COVID-19 Surveillance Team |
| EPI_ISL_2846990 | Health Services Laboratories | Wellcome Sanger Institute for the COVID-19 Genomics UK (COG-UK) Consortium | Health Services Laboratories and Alex Alderton, Roberto Amato, Jeffrey Barrett, Sonia Goncalves, Ewan Harrison, David K. Jackson, Ian Johnston, Dominic Kwiatkowski, Cordelia Langford, John Sillitoe on behalf of the Wellcome Sanger Institute COVID-19 Surveillance Team |
| EPI_ISL_2846991, EPI_ISL_2846992 | Lighthouse Lab in Glasgow | Wellcome Sanger Institute for the COVID-19 Genomics UK (COG-UK) Consortium | Harper VanSteenhouse, Yumi Kasai, David Gray, Carol Clugston, Anna Dominiczak and Alex Alderton, Roberto Amato, Jeffrey Barrett, Sonia Goncalves, Ewan Harrison, David K. Jackson, Ian Johnston, Dominic Kwiatkowski, Cordelia Langford, John Sillitoe on behalf of the Wellcome Sanger Institute COVID-19 Surveillance Team |
| EPI_ISL_2846993, EPI_ISL_2846994, EPI_ISL_2846995 | Health Services Laboratories | Wellcome Sanger Institute for the COVID-19 Genomics UK (COG-UK) Consortium | Health Services Laboratories and Alex Alderton, Roberto Amato, Jeffrey Barrett, Sonia Goncalves, Ewan Harrison, David K. Jackson, Ian Johnston, Dominic Kwiatkowski, Cordelia Langford, John Sillitoe on behalf of the Wellcome Sanger Institute COVID-19 Surveillance Team |
| EPI_ISL_2846996 | Lighthouse Lab in Alderley Park | Wellcome Sanger Institute for the COVID-19 Genomics UK (COG-UK) Consortium | Jacquelyn Wynn, Mairead Hyland, The Lighthouse Lab in Alderley Park and Alex Alderton, Roberto Amato, Jeffrey Barrett, Sonia Goncalves, Ewan Harrison, David K. Jackson, Ian Johnston, Dominic Kwiatkowski, Cordelia Langford, John Sillitoe on behalf of the Wellcome Sanger Institute COVID-19 Surveillance Team |
| EPI_ISL_2846997 | Health Services Laboratories | Wellcome Sanger Institute for the COVID-19 Genomics UK (COG-UK) Consortium | Health Services Laboratories and Alex Alderton, Roberto Amato, Jeffrey Barrett, Sonia Goncalves, Ewan Harrison, David K. Jackson, Ian Johnston, Dominic Kwiatkowski, Cordelia Langford, John Sillitoe on behalf of the Wellcome Sanger Institute COVID-19 Surveillance Team |
| EPI_ISL_2846998, EPI_ISL_2846999 | Lighthouse Lab in Glasgow | Wellcome Sanger Institute for the COVID-19 Genomics UK (COG-UK) Consortium | Harper VanSteenhouse, Yumi Kasai, David Gray, Carol Clugston, Anna Dominiczak and Alex Alderton, Roberto Amato, Jeffrey Barrett, Sonia Goncalves, Ewan Harrison, David K. Jackson, Ian Johnston, Dominic Kwiatkowski, Cordelia Langford, John Sillitoe on behalf of the Wellcome Sanger Institute COVID-19 Surveillance Team |
| EPI_ISL_2847000 | Lighthouse Lab in Alderley Park | Wellcome Sanger Institute for the COVID-19 Genomics UK (COG-UK) Consortium | Jacquelyn Wynn, Mairead Hyland, The Lighthouse Lab in Alderley Park and Alex Alderton, Roberto Amato, Jeffrey Barrett, Sonia Goncalves, Ewan Harrison, David K. Jackson, Ian Johnston, Dominic Kwiatkowski, Cordelia Langford, John Sillitoe on behalf of the Wellcome Sanger Institute COVID-19 Surveillance Team |
| EPI_ISL_2847001 | Lighthouse Lab in Milton Keynes | Wellcome Sanger Institute for the COVID-19 Genomics UK (COG-UK) Consortium | The Lighthouse Lab in Milton Keynes and Alex Alderton, Roberto Amato, Jeffrey Barrett, Sonia Goncalves, Ewan Harrison, David K. Jackson, Ian Johnston, Dominic Kwiatkowski, Cordelia Langford, John Sillitoe on behalf of the Wellcome Sanger Institute COVID-19 Surveillance Team |
| EPI_ISL_2847002 | Lighthouse Lab in Alderley Park | Wellcome Sanger Institute for the COVID-19 Genomics UK (COG-UK) | Jacquelyn Wynn, Mairead Hyland, The Lighthouse Lab in Alderley Park and Alex Alderton, Roberto Amato, Jeffrey Barrett, Sonia Goncalves, Ewan Harrison, David K. Jackson, Ian Johnston, Dominic Kwiatkowski, Cordelia Langford, John Sillitoe on |

| | | | |
|---|---|---|---|
| | | Consortium | behalf of the Wellcome Sanger Institute COVID-19 Surveillance Team |
| EPI_ISL_2847003 | Health Services Laboratories | Wellcome Sanger Institute for the COVID-19 Genomics UK (COG-UK) Consortium | Health Services Laboratories and Alex Alderton, Roberto Amato, Jeffrey Barrett, Sonia Goncalves, Ewan Harrison, David K. Jackson, Ian Johnston, Dominic Kwiatkowski, Cordelia Langford, John Sillitoe on behalf of the Wellcome Sanger Institute COVID-19 Surveillance Team |
| EPI_ISL_2847004 | Lighthouse Lab in Glasgow | Wellcome Sanger Institute for the COVID-19 Genomics UK (COG-UK) Consortium | Harper VanSteenhouse, Yumi Kasai, David Gray, Carol Clugston, Anna Dominiczak and Alex Alderton, Roberto Amato, Jeffrey Barrett, Sonia Goncalves, Ewan Harrison, David K. Jackson, Ian Johnston, Dominic Kwiatkowski, Cordelia Langford, John Sillitoe on behalf of the Wellcome Sanger Institute COVID-19 Surveillance Team |
| EPI_ISL_2847005 | Lighthouse Lab in Milton Keynes | Wellcome Sanger Institute for the COVID-19 Genomics UK (COG-UK) Consortium | The Lighthouse Lab in Milton Keynes and Alex Alderton, Roberto Amato, Jeffrey Barrett, Sonia Goncalves, Ewan Harrison, David K. Jackson, Ian Johnston, Dominic Kwiatkowski, Cordelia Langford, John Sillitoe on behalf of the Wellcome Sanger Institute COVID-19 Surveillance Team |
| EPI_ISL_2847006 | Lighthouse Lab in Alderley Park | Wellcome Sanger Institute for the COVID-19 Genomics UK (COG-UK) Consortium | Jacquelyn Wynn, Mairead Hyland, The Lighthouse Lab in Alderley Park and Alex Alderton, Roberto Amato, Jeffrey Barrett, Sonia Goncalves, Ewan Harrison, David K. Jackson, Ian Johnston, Dominic Kwiatkowski, Cordelia Langford, John Sillitoe on behalf of the Wellcome Sanger Institute COVID-19 Surveillance Team |
| EPI_ISL_2847007 | Lighthouse Lab in Glasgow | Wellcome Sanger Institute for the COVID-19 Genomics UK (COG-UK) Consortium | Harper VanSteenhouse, Yumi Kasai, David Gray, Carol Clugston, Anna Dominiczak and Alex Alderton, Roberto Amato, Jeffrey Barrett, Sonia Goncalves, Ewan Harrison, David K. Jackson, Ian Johnston, Dominic Kwiatkowski, Cordelia Langford, John Sillitoe on behalf of the Wellcome Sanger Institute COVID-19 Surveillance Team |
| EPI_ISL_2847008, EPI_ISL_2847009 | Lighthouse Lab in Milton Keynes | Wellcome Sanger Institute for the COVID-19 Genomics UK (COG-UK) Consortium | The Lighthouse Lab in Milton Keynes and Alex Alderton, Roberto Amato, Jeffrey Barrett, Sonia Goncalves, Ewan Harrison, David K. Jackson, Ian Johnston, Dominic Kwiatkowski, Cordelia Langford, John Sillitoe on behalf of the Wellcome Sanger Institute COVID-19 Surveillance Team |
| EPI_ISL_2847010, EPI_ISL_2847011 | Health Services Laboratories | Wellcome Sanger Institute for the COVID-19 Genomics UK (COG-UK) Consortium | Health Services Laboratories and Alex Alderton, Roberto Amato, Jeffrey Barrett, Sonia Goncalves, Ewan Harrison, David K. Jackson, Ian Johnston, Dominic Kwiatkowski, Cordelia Langford, John Sillitoe on behalf of the Wellcome Sanger Institute COVID-19 Surveillance Team |
| EPI_ISL_2847012 | Lighthouse Lab in Glasgow | Wellcome Sanger Institute for the COVID-19 Genomics UK (COG-UK) Consortium | Harper VanSteenhouse, Yumi Kasai, David Gray, Carol Clugston, Anna Dominiczak and Alex Alderton, Roberto Amato, Jeffrey Barrett, Sonia Goncalves, Ewan Harrison, David K. Jackson, Ian Johnston, Dominic Kwiatkowski, Cordelia Langford, John Sillitoe on behalf of the Wellcome Sanger Institute COVID-19 Surveillance Team |
| EPI_ISL_2847013 | Lighthouse Lab in Alderley Park | Wellcome Sanger Institute for the COVID-19 Genomics UK (COG-UK) Consortium | Jacquelyn Wynn, Mairead Hyland, The Lighthouse Lab in Alderley Park and Alex Alderton, Roberto Amato, Jeffrey Barrett, Sonia Goncalves, Ewan Harrison, David K. Jackson, Ian Johnston, Dominic Kwiatkowski, Cordelia Langford, John Sillitoe on behalf of the Wellcome Sanger Institute COVID-19 Surveillance Team |
| EPI_ISL_2847014 | Health Services Laboratories | Wellcome Sanger Institute for the COVID-19 Genomics UK (COG-UK) Consortium | Health Services Laboratories and Alex Alderton, Roberto Amato, Jeffrey Barrett, Sonia Goncalves, Ewan Harrison, David K. Jackson, Ian Johnston, Dominic Kwiatkowski, Cordelia Langford, John Sillitoe on behalf of the Wellcome Sanger Institute COVID-19 Surveillance Team |
| EPI_ISL_2847015 | Lighthouse Lab in Alderley Park | Wellcome Sanger Institute for the COVID-19 Genomics UK (COG-UK) Consortium | Jacquelyn Wynn, Mairead Hyland, The Lighthouse Lab in Alderley Park and Alex Alderton, Roberto Amato, Jeffrey Barrett, Sonia Goncalves, Ewan Harrison, David K. Jackson, Ian Johnston, Dominic Kwiatkowski, Cordelia Langford, John Sillitoe on behalf of the Wellcome Sanger Institute COVID-19 Surveillance Team |
| EPI_ISL_2847016 | Health Services Laboratories | Wellcome Sanger Institute for the COVID-19 Genomics UK (COG-UK) Consortium | Health Services Laboratories and Alex Alderton, Roberto Amato, Jeffrey Barrett, Sonia Goncalves, Ewan Harrison, David K. Jackson, Ian Johnston, Dominic Kwiatkowski, Cordelia Langford, John Sillitoe on behalf of the Wellcome Sanger Institute COVID-19 Surveillance Team |
| EPI_ISL_2847017 | Lighthouse Lab in Alderley Park | Wellcome Sanger Institute for the COVID-19 Genomics UK (COG-UK) Consortium | Jacquelyn Wynn, Mairead Hyland, The Lighthouse Lab in Alderley Park and Alex Alderton, Roberto Amato, Jeffrey Barrett, Sonia Goncalves, Ewan Harrison, David K. Jackson, Ian Johnston, Dominic Kwiatkowski, Cordelia Langford, John Sillitoe on behalf of the Wellcome Sanger Institute COVID-19 Surveillance Team |
| EPI_ISL_2847018 | Lighthouse Lab in Milton Keynes | Wellcome Sanger Institute for the COVID-19 Genomics UK (COG-UK) Consortium | The Lighthouse Lab in Milton Keynes and Alex Alderton, Roberto Amato, Jeffrey Barrett, Sonia Goncalves, Ewan Harrison, David K. Jackson, Ian Johnston, Dominic Kwiatkowski, Cordelia Langford, John Sillitoe on behalf of the Wellcome Sanger Institute COVID-19 Surveillance Team |
| EPI_ISL_2847019 | Health Services Laboratories | Wellcome Sanger Institute for the COVID-19 Genomics UK (COG-UK) Consortium | Health Services Laboratories and Alex Alderton, Roberto Amato, Jeffrey Barrett, Sonia Goncalves, Ewan Harrison, David K. Jackson, Ian Johnston, Dominic Kwiatkowski, Cordelia Langford, John Sillitoe on behalf of the Wellcome Sanger Institute COVID-19 Surveillance Team |
| EPI_ISL_2847020 | Lighthouse Lab in Milton Keynes | Wellcome Sanger Institute for the COVID-19 Genomics UK (COG-UK) Consortium | The Lighthouse Lab in Milton Keynes and Alex Alderton, Roberto Amato, Jeffrey Barrett, Sonia Goncalves, Ewan Harrison, David K. Jackson, Ian Johnston, Dominic Kwiatkowski, Cordelia Langford, John Sillitoe on behalf of the Wellcome Sanger Institute COVID-19 Surveillance Team |
| EPI_ISL_2847021 | Lighthouse Lab in Alderley Park | Wellcome Sanger Institute for the COVID-19 Genomics UK (COG-UK) Consortium | Jacquelyn Wynn, Mairead Hyland, The Lighthouse Lab in Alderley Park and Alex Alderton, Roberto Amato, Jeffrey Barrett, Sonia Goncalves, Ewan Harrison, David K. Jackson, Ian Johnston, Dominic Kwiatkowski, Cordelia Langford, John Sillitoe on behalf of the Wellcome Sanger Institute COVID-19 Surveillance Team |
| EPI_ISL_2847022 | Lighthouse Lab in Glasgow | Wellcome Sanger Institute for the COVID-19 Genomics UK (COG-UK) Consortium | Harper VanSteenhouse, Yumi Kasai, David Gray, Carol Clugston, Anna Dominiczak and Alex Alderton, Roberto Amato, Jeffrey Barrett, Sonia Goncalves, Ewan Harrison, David K. Jackson, Ian Johnston, Dominic Kwiatkowski, Cordelia Langford, John Sillitoe on behalf of the Wellcome Sanger Institute COVID-19 Surveillance Team |
| EPI_ISL_2847023 | Lighthouse Lab in Milton Keynes | Wellcome Sanger Institute for the COVID-19 Genomics UK (COG-UK) Consortium | The Lighthouse Lab in Milton Keynes and Alex Alderton, Roberto Amato, Jeffrey Barrett, Sonia Goncalves, Ewan Harrison, David K. Jackson, Ian Johnston, Dominic Kwiatkowski, Cordelia Langford, John Sillitoe on behalf of the Wellcome Sanger Institute COVID-19 Surveillance Team |
| EPI_ISL_2847024, EPI_ISL_2847025 | Lighthouse Lab in Glasgow | Wellcome Sanger Institute for the COVID-19 Genomics UK (COG-UK) Consortium | Harper VanSteenhouse, Yumi Kasai, David Gray, Carol Clugston, Anna Dominiczak and Alex Alderton, Roberto Amato, Jeffrey Barrett, Sonia Goncalves, Ewan Harrison, David K. Jackson, Ian Johnston, Dominic Kwiatkowski, Cordelia Langford, John Sillitoe on behalf of the Wellcome Sanger Institute COVID-19 Surveillance Team |
| EPI_ISL_2847026 | Lighthouse Lab in Alderley Park | Wellcome Sanger Institute for the COVID-19 Genomics UK (COG-UK) Consortium | Jacquelyn Wynn, Mairead Hyland, The Lighthouse Lab in Alderley Park and Alex Alderton, Roberto Amato, Jeffrey Barrett, Sonia Goncalves, Ewan Harrison, David K. Jackson, Ian Johnston, Dominic Kwiatkowski, Cordelia Langford, John Sillitoe on behalf of the Wellcome Sanger Institute COVID-19 Surveillance Team |
| EPI_ISL_2847027 | Lighthouse Lab in Milton Keynes | Wellcome Sanger Institute for the COVID-19 Genomics UK (COG-UK) Consortium | The Lighthouse Lab in Milton Keynes and Alex Alderton, Roberto Amato, Jeffrey Barrett, Sonia Goncalves, Ewan Harrison, David K. Jackson, Ian Johnston, Dominic Kwiatkowski, Cordelia Langford, John Sillitoe on behalf of the Wellcome Sanger Institute COVID-19 Surveillance Team |
| EPI_ISL_2847028 | Health Services Laboratories | Wellcome Sanger Institute for the COVID-19 Genomics UK (COG-UK) Consortium | Health Services Laboratories and Alex Alderton, Roberto Amato, Jeffrey Barrett, Sonia Goncalves, Ewan Harrison, David K. Jackson, Ian Johnston, Dominic Kwiatkowski, Cordelia Langford, John Sillitoe on behalf of the Wellcome Sanger Institute COVID-19 Surveillance Team |
| EPI_ISL_2847029 | Lighthouse Lab in Milton Keynes | Wellcome Sanger Institute for the COVID-19 Genomics UK (COG-UK) Consortium | The Lighthouse Lab in Milton Keynes and Alex Alderton, Roberto Amato, Jeffrey Barrett, Sonia Goncalves, Ewan Harrison, David K. Jackson, Ian Johnston, Dominic Kwiatkowski, Cordelia Langford, John Sillitoe on behalf of the Wellcome Sanger Institute COVID-19 Surveillance Team |
| EPI_ISL_2847030 | Lighthouse Lab in Glasgow | Wellcome Sanger Institute for the COVID-19 Genomics UK (COG-UK) Consortium | Harper VanSteenhouse, Yumi Kasai, David Gray, Carol Clugston, Anna Dominiczak and Alex Alderton, Roberto Amato, Jeffrey Barrett, Sonia Goncalves, Ewan Harrison, David K. Jackson, Ian Johnston, Dominic Kwiatkowski, Cordelia Langford, John Sillitoe on behalf of the Wellcome Sanger Institute COVID-19 Surveillance Team |
| EPI_ISL_2847031 | Lighthouse Lab in Milton Keynes | Wellcome Sanger Institute for the COVID-19 Genomics UK (COG-UK) Consortium | The Lighthouse Lab in Milton Keynes and Alex Alderton, Roberto Amato, Jeffrey Barrett, Sonia Goncalves, Ewan Harrison, David K. Jackson, Ian Johnston, Dominic Kwiatkowski, Cordelia Langford, John Sillitoe on behalf of the Wellcome Sanger Institute COVID-19 Surveillance Team |
| EPI_ISL_2847032 | Lighthouse Lab in Glasgow | Wellcome Sanger Institute for the COVID-19 Genomics UK (COG-UK) Consortium | Harper VanSteenhouse, Yumi Kasai, David Gray, Carol Clugston, Anna Dominiczak and Alex Alderton, Roberto Amato, Jeffrey Barrett, Sonia Goncalves, Ewan Harrison, David K. Jackson, Ian Johnston, Dominic Kwiatkowski, Cordelia Langford, John Sillitoe on behalf of the Wellcome Sanger Institute COVID-19 Surveillance Team |
| EPI_ISL_2847033 | Lighthouse Lab in Milton Keynes | Wellcome Sanger Institute for the COVID-19 Genomics UK (COG-UK) Consortium | The Lighthouse Lab in Milton Keynes and Alex Alderton, Roberto Amato, Jeffrey Barrett, Sonia Goncalves, Ewan Harrison, David K. Jackson, Ian Johnston, Dominic Kwiatkowski, Cordelia Langford, John Sillitoe on behalf of the Wellcome Sanger Institute COVID-19 Surveillance Team |
| EPI_ISL_2847034 | Lighthouse Lab in Alderley Park | Wellcome Sanger Institute for the COVID-19 Genomics UK (COG-UK) Consortium | Jacquelyn Wynn, Mairead Hyland, The Lighthouse Lab in Alderley Park and Alex Alderton, Roberto Amato, Jeffrey Barrett, Sonia Goncalves, Ewan Harrison, David K. Jackson, Ian Johnston, Dominic Kwiatkowski, Cordelia Langford, John Sillitoe on behalf of the Wellcome Sanger Institute COVID-19 Surveillance Team |
| EPI_ISL_2847035 | Health Services Laboratories | Wellcome Sanger Institute for the COVID-19 Genomics UK (COG-UK) Consortium | Health Services Laboratories and Alex Alderton, Roberto Amato, Jeffrey Barrett, Sonia Goncalves, Ewan Harrison, David K. Jackson, Ian Johnston, Dominic Kwiatkowski, Cordelia Langford, John Sillitoe on behalf of the Wellcome Sanger Institute COVID-19 Surveillance Team |
| EPI_ISL_2847036 | Lighthouse Lab in Glasgow | Wellcome Sanger Institute for the COVID-19 Genomics UK (COG-UK) Consortium | Harper VanSteenhouse, Yumi Kasai, David Gray, Carol Clugston, Anna Dominiczak and Alex Alderton, Roberto Amato, Jeffrey Barrett, Sonia Goncalves, Ewan Harrison, David K. Jackson, Ian Johnston, Dominic Kwiatkowski, Cordelia Langford, John Sillitoe on behalf of the Wellcome Sanger Institute COVID-19 Surveillance Team |
| EPI_ISL_2847037 | Lighthouse Lab in Milton Keynes | Wellcome Sanger Institute for the COVID-19 Genomics UK (COG-UK) Consortium | The Lighthouse Lab in Milton Keynes and Alex Alderton, Roberto Amato, Jeffrey Barrett, Sonia Goncalves, Ewan Harrison, David K. Jackson, Ian Johnston, Dominic Kwiatkowski, Cordelia Langford, John Sillitoe on behalf of the Wellcome Sanger Institute COVID-19 Surveillance Team |
| EPI_ISL_2847038 | Health Services Laboratories | Wellcome Sanger Institute for the COVID-19 Genomics UK (COG-UK) Consortium | Health Services Laboratories and Alex Alderton, Roberto Amato, Jeffrey Barrett, Sonia Goncalves, Ewan Harrison, David K. Jackson, Ian Johnston, Dominic Kwiatkowski, Cordelia Langford, John Sillitoe on behalf of the Wellcome Sanger Institute COVID-19 Surveillance Team |
| EPI_ISL_2847039 | Lighthouse Lab in Milton Keynes | Wellcome Sanger Institute for the COVID-19 Genomics UK (COG-UK) Consortium | The Lighthouse Lab in Milton Keynes and Alex Alderton, Roberto Amato, Jeffrey Barrett, Sonia Goncalves, Ewan Harrison, David K. Jackson, Ian Johnston, Dominic Kwiatkowski, Cordelia Langford, John Sillitoe on behalf of the Wellcome Sanger Institute COVID-19 Surveillance Team |
| EPI_ISL_2847040, EPI_ISL_2847041 | Health Services Laboratories | Wellcome Sanger Institute for the COVID-19 Genomics UK (COG-UK) Consortium | Health Services Laboratories and Alex Alderton, Roberto Amato, Jeffrey Barrett, Sonia Goncalves, Ewan Harrison, David K. Jackson, Ian Johnston, Dominic Kwiatkowski, Cordelia Langford, John Sillitoe on behalf of the Wellcome Sanger Institute COVID-19 Surveillance Team |
| EPI_ISL_2847042 | Lighthouse Lab in Alderley Park | Wellcome Sanger Institute for the COVID-19 Genomics UK (COG-UK) Consortium | Jacquelyn Wynn, Mairead Hyland, The Lighthouse Lab in Alderley Park and Alex Alderton, Roberto Amato, Jeffrey Barrett, Sonia Goncalves, Ewan Harrison, David K. Jackson, Ian Johnston, Dominic Kwiatkowski, Cordelia Langford, John Sillitoe on behalf of the Wellcome Sanger Institute COVID-19 Surveillance Team |
| EPI_ISL_2847043, EPI_ISL_2847044 | Lighthouse Lab in Glasgow | Wellcome Sanger Institute for the COVID-19 Genomics UK (COG-UK) Consortium | Harper VanSteenhouse, Yumi Kasai, David Gray, Carol Clugston, Anna Dominiczak and Alex Alderton, Roberto Amato, Jeffrey Barrett, Sonia Goncalves, Ewan Harrison, David K. Jackson, Ian Johnston, Dominic Kwiatkowski, Cordelia Langford, John Sillitoe on behalf of the Wellcome Sanger Institute COVID-19 Surveillance Team |
| EPI_ISL_2847045 | Lighthouse Lab in Alderley Park | Wellcome Sanger Institute for the COVID-19 Genomics UK (COG-UK) Consortium | Jacquelyn Wynn, Mairead Hyland, The Lighthouse Lab in Alderley Park and Alex Alderton, Roberto Amato, Jeffrey Barrett, Sonia Goncalves, Ewan Harrison, David K. Jackson, Ian Johnston, Dominic Kwiatkowski, Cordelia Langford, John Sillitoe on behalf of the Wellcome Sanger Institute COVID-19 Surveillance Team |
| EPI_ISL_2847046 | Lighthouse Lab in Milton Keynes | Wellcome Sanger Institute for the COVID-19 Genomics UK (COG-UK) Consortium | The Lighthouse Lab in Milton Keynes and Alex Alderton, Roberto Amato, Jeffrey Barrett, Sonia Goncalves, Ewan Harrison, David K. Jackson, Ian Johnston, Dominic Kwiatkowski, Cordelia Langford, John Sillitoe on behalf of the Wellcome Sanger Institute COVID-19 Surveillance Team |
| EPI_ISL_2847047 | Lighthouse Lab in Glasgow | Wellcome Sanger Institute for the COVID-19 Genomics UK (COG-UK) Consortium | Harper VanSteenhouse, Yumi Kasai, David Gray, Carol Clugston, Anna Dominiczak and Alex Alderton, Roberto Amato, Jeffrey Barrett, Sonia Goncalves, Ewan Harrison, David K. Jackson, Ian Johnston, Dominic Kwiatkowski, Cordelia Langford, John Sillitoe on behalf of the Wellcome Sanger Institute COVID-19 Surveillance Team |
| EPI_ISL_2847048 | Lighthouse Lab in Milton Keynes | Wellcome Sanger Institute for the COVID-19 Genomics UK (COG-UK) Consortium | The Lighthouse Lab in Milton Keynes and Alex Alderton, Roberto Amato, Jeffrey Barrett, Sonia Goncalves, Ewan Harrison, David K. Jackson, Ian Johnston, Dominic Kwiatkowski, Cordelia Langford, John Sillitoe on behalf of the Wellcome Sanger Institute COVID-19 Surveillance Team |
| EPI_ISL_2847049 | Lighthouse Lab in Glasgow | Wellcome Sanger Institute for the COVID-19 Genomics UK (COG-UK) Consortium | Harper VanSteenhouse, Yumi Kasai, David Gray, Carol Clugston, Anna Dominiczak and Alex Alderton, Roberto Amato, Jeffrey Barrett, Sonia Goncalves, Ewan Harrison, David K. Jackson, Ian Johnston, Dominic Kwiatkowski, Cordelia Langford, John Sillitoe on behalf of the Wellcome Sanger Institute COVID-19 Surveillance Team |
| EPI_ISL_2847050 | Health Services Laboratories | Wellcome Sanger Institute for the COVID-19 Genomics UK (COG-UK) Consortium | Health Services Laboratories and Alex Alderton, Roberto Amato, Jeffrey Barrett, Sonia Goncalves, Ewan Harrison, David K. Jackson, Ian Johnston, Dominic Kwiatkowski, Cordelia Langford, John Sillitoe on behalf of the Wellcome Sanger Institute COVID-19 Surveillance Team |
| EPI_ISL_2847051 | Lighthouse Lab in Alderley Park | Wellcome Sanger Institute for the COVID-19 Genomics UK (COG-UK) Consortium | Jacquelyn Wynn, Mairead Hyland, The Lighthouse Lab in Alderley Park and Alex Alderton, Roberto Amato, Jeffrey Barrett, Sonia Goncalves, Ewan Harrison, David K. Jackson, Ian Johnston, Dominic Kwiatkowski, Cordelia Langford, John Sillitoe on behalf of the Wellcome Sanger Institute COVID-19 Surveillance Team |
| EPI_ISL_2847052 | Lighthouse Lab in Glasgow | Wellcome Sanger Institute for the COVID-19 Genomics UK (COG-UK) | Harper VanSteenhouse, Yumi Kasai, David Gray, Carol Clugston, Anna Dominiczak and Alex Alderton, Roberto Amato, Jeffrey Barrett, Sonia Goncalves, Ewan Harrison, David K. Jackson, Ian Johnston, Dominic Kwiatkowski, Cordelia Langford, |

| | | | |
|---|---|---|---|
| | | Consortium | John Sillitoe on behalf of the Wellcome Sanger Institute COVID-19 Surveillance Team |
| EPI_ISL_2847053 | Health Services Laboratories | Wellcome Sanger Institute for the COVID-19 Genomics UK (COG-UK) Consortium | Health Services Laboratories and Alex Alderton, Roberto Amato, Jeffrey Barrett, Sonia Goncalves, Ewan Harrison, David K. Jackson, Ian Johnston, Dominic Kwiatkowski, Cordelia Langford, John Sillitoe on behalf of the Wellcome Sanger Institute COVID-19 Surveillance Team |
| EPI_ISL_2847054 | Lighthouse Lab in Glasgow | Wellcome Sanger Institute for the COVID-19 Genomics UK (COG-UK) Consortium | Harper VanSteenhouse, Yumi Kasai, David Gray, Carol Clugston, Anna Dominiczak and Alex Alderton, Roberto Amato, Jeffrey Barrett, Sonia Goncalves, Ewan Harrison, David K. Jackson, Ian Johnston, Dominic Kwiatkowski, Cordelia Langford, John Sillitoe on behalf of the Wellcome Sanger Institute COVID-19 Surveillance Team |
| EPI_ISL_2847055 | Health Services Laboratories | Wellcome Sanger Institute for the COVID-19 Genomics UK (COG-UK) Consortium | Health Services Laboratories and Alex Alderton, Roberto Amato, Jeffrey Barrett, Sonia Goncalves, Ewan Harrison, David K. Jackson, Ian Johnston, Dominic Kwiatkowski, Cordelia Langford, John Sillitoe on behalf of the Wellcome Sanger Institute COVID-19 Surveillance Team |
| EPI_ISL_2847056 | Lighthouse Lab in Glasgow | Wellcome Sanger Institute for the COVID-19 Genomics UK (COG-UK) Consortium | Harper VanSteenhouse, Yumi Kasai, David Gray, Carol Clugston, Anna Dominiczak and Alex Alderton, Roberto Amato, Jeffrey Barrett, Sonia Goncalves, Ewan Harrison, David K. Jackson, Ian Johnston, Dominic Kwiatkowski, Cordelia Langford, John Sillitoe on behalf of the Wellcome Sanger Institute COVID-19 Surveillance Team |
| EPI_ISL_2847057, EPI_ISL_2847058 | Lighthouse Lab in Alderley Park | Wellcome Sanger Institute for the COVID-19 Genomics UK (COG-UK) Consortium | Jacquelyn Wynn, Mairead Hyland, The Lighthouse Lab in Alderley Park and Alex Alderton, Roberto Amato, Jeffrey Barrett, Sonia Goncalves, Ewan Harrison, David K. Jackson, Ian Johnston, Dominic Kwiatkowski, Cordelia Langford, John Sillitoe on behalf of the Wellcome Sanger Institute COVID-19 Surveillance Team |
| EPI_ISL_2847059 | Lighthouse Lab in Glasgow | Wellcome Sanger Institute for the COVID-19 Genomics UK (COG-UK) Consortium | Harper VanSteenhouse, Yumi Kasai, David Gray, Carol Clugston, Anna Dominiczak and Alex Alderton, Roberto Amato, Jeffrey Barrett, Sonia Goncalves, Ewan Harrison, David K. Jackson, Ian Johnston, Dominic Kwiatkowski, Cordelia Langford, John Sillitoe on behalf of the Wellcome Sanger Institute COVID-19 Surveillance Team |
| EPI_ISL_2847060 | Lighthouse Lab in Milton Keynes | Wellcome Sanger Institute for the COVID-19 Genomics UK (COG-UK) Consortium | The Lighthouse Lab in Milton Keynes and Alex Alderton, Roberto Amato, Jeffrey Barrett, Sonia Goncalves, Ewan Harrison, David K. Jackson, Ian Johnston, Dominic Kwiatkowski, Cordelia Langford, John Sillitoe on behalf of the Wellcome Sanger Institute COVID-19 Surveillance Team |
| EPI_ISL_2847061 | Lighthouse Lab in Alderley Park | Wellcome Sanger Institute for the COVID-19 Genomics UK (COG-UK) Consortium | Jacquelyn Wynn, Mairead Hyland, The Lighthouse Lab in Alderley Park and Alex Alderton, Roberto Amato, Jeffrey Barrett, Sonia Goncalves, Ewan Harrison, David K. Jackson, Ian Johnston, Dominic Kwiatkowski, Cordelia Langford, John Sillitoe on behalf of the Wellcome Sanger Institute COVID-19 Surveillance Team |
| EPI_ISL_2847062 | Lighthouse Lab in Glasgow | Wellcome Sanger Institute for the COVID-19 Genomics UK (COG-UK) Consortium | Harper VanSteenhouse, Yumi Kasai, David Gray, Carol Clugston, Anna Dominiczak and Alex Alderton, Roberto Amato, Jeffrey Barrett, Sonia Goncalves, Ewan Harrison, David K. Jackson, Ian Johnston, Dominic Kwiatkowski, Cordelia Langford, John Sillitoe on behalf of the Wellcome Sanger Institute COVID-19 Surveillance Team |
| EPI_ISL_2847063, EPI_ISL_2847064, EPI_ISL_2847065 | Lighthouse Lab in Alderley Park | Wellcome Sanger Institute for the COVID-19 Genomics UK (COG-UK) Consortium | Jacquelyn Wynn, Mairead Hyland, The Lighthouse Lab in Alderley Park and Alex Alderton, Roberto Amato, Jeffrey Barrett, Sonia Goncalves, Ewan Harrison, David K. Jackson, Ian Johnston, Dominic Kwiatkowski, Cordelia Langford, John Sillitoe on behalf of the Wellcome Sanger Institute COVID-19 Surveillance Team |
| EPI_ISL_2847066 | Lighthouse Lab in Milton Keynes | Wellcome Sanger Institute for the COVID-19 Genomics UK (COG-UK) Consortium | The Lighthouse Lab in Milton Keynes and Alex Alderton, Roberto Amato, Jeffrey Barrett, Sonia Goncalves, Ewan Harrison, David K. Jackson, Ian Johnston, Dominic Kwiatkowski, Cordelia Langford, John Sillitoe on behalf of the Wellcome Sanger Institute COVID-19 Surveillance Team |
| EPI_ISL_2847067, EPI_ISL_2847068 | Lighthouse Lab in Glasgow | Wellcome Sanger Institute for the COVID-19 Genomics UK (COG-UK) Consortium | Harper VanSteenhouse, Yumi Kasai, David Gray, Carol Clugston, Anna Dominiczak and Alex Alderton, Roberto Amato, Jeffrey Barrett, Sonia Goncalves, Ewan Harrison, David K. Jackson, Ian Johnston, Dominic Kwiatkowski, Cordelia Langford, John Sillitoe on behalf of the Wellcome Sanger Institute COVID-19 Surveillance Team |
| EPI_ISL_2847069 | Lighthouse Lab in Alderley Park | Wellcome Sanger Institute for the COVID-19 Genomics UK (COG-UK) Consortium | Jacquelyn Wynn, Mairead Hyland, The Lighthouse Lab in Alderley Park and Alex Alderton, Roberto Amato, Jeffrey Barrett, Sonia Goncalves, Ewan Harrison, David K. Jackson, Ian Johnston, Dominic Kwiatkowski, Cordelia Langford, John Sillitoe on behalf of the Wellcome Sanger Institute COVID-19 Surveillance Team |
| EPI_ISL_2847070 | Lighthouse Lab in Milton Keynes | Wellcome Sanger Institute for the COVID-19 Genomics UK (COG-UK) Consortium | The Lighthouse Lab in Milton Keynes and Alex Alderton, Roberto Amato, Jeffrey Barrett, Sonia Goncalves, Ewan Harrison, David K. Jackson, Ian Johnston, Dominic Kwiatkowski, Cordelia Langford, John Sillitoe on behalf of the Wellcome Sanger Institute COVID-19 Surveillance Team |
| EPI_ISL_2847071 | Health Services Laboratories | Wellcome Sanger Institute for the COVID-19 Genomics UK (COG-UK) Consortium | Health Services Laboratories and Alex Alderton, Roberto Amato, Jeffrey Barrett, Sonia Goncalves, Ewan Harrison, David K. Jackson, Ian Johnston, Dominic Kwiatkowski, Cordelia Langford, John Sillitoe on behalf of the Wellcome Sanger Institute COVID-19 Surveillance Team |
| EPI_ISL_2847072 | Lighthouse Lab in Milton Keynes | Wellcome Sanger Institute for the COVID-19 Genomics UK (COG-UK) Consortium | The Lighthouse Lab in Milton Keynes and Alex Alderton, Roberto Amato, Jeffrey Barrett, Sonia Goncalves, Ewan Harrison, David K. Jackson, Ian Johnston, Dominic Kwiatkowski, Cordelia Langford, John Sillitoe on behalf of the Wellcome Sanger Institute COVID-19 Surveillance Team |
| EPI_ISL_2847073 | Lighthouse Lab in Alderley Park | Wellcome Sanger Institute for the COVID-19 Genomics UK (COG-UK) Consortium | Jacquelyn Wynn, Mairead Hyland, The Lighthouse Lab in Alderley Park and Alex Alderton, Roberto Amato, Jeffrey Barrett, Sonia Goncalves, Ewan Harrison, David K. Jackson, Ian Johnston, Dominic Kwiatkowski, Cordelia Langford, John Sillitoe on behalf of the Wellcome Sanger Institute COVID-19 Surveillance Team |
| EPI_ISL_2847074 | Health Services Laboratories | Wellcome Sanger Institute for the COVID-19 Genomics UK (COG-UK) Consortium | Health Services Laboratories and Alex Alderton, Roberto Amato, Jeffrey Barrett, Sonia Goncalves, Ewan Harrison, David K. Jackson, Ian Johnston, Dominic Kwiatkowski, Cordelia Langford, John Sillitoe on behalf of the Wellcome Sanger Institute COVID-19 Surveillance Team |
| EPI_ISL_2847075 | Lighthouse Lab in Glasgow | Wellcome Sanger Institute for the COVID-19 Genomics UK (COG-UK) Consortium | Harper VanSteenhouse, Yumi Kasai, David Gray, Carol Clugston, Anna Dominiczak and Alex Alderton, Roberto Amato, Jeffrey Barrett, Sonia Goncalves, Ewan Harrison, David K. Jackson, Ian Johnston, Dominic Kwiatkowski, Cordelia Langford, John Sillitoe on behalf of the Wellcome Sanger Institute COVID-19 Surveillance Team |
| EPI_ISL_2847076 | Lighthouse Lab in Alderley Park | Wellcome Sanger Institute for the COVID-19 Genomics UK (COG-UK) Consortium | Jacquelyn Wynn, Mairead Hyland, The Lighthouse Lab in Alderley Park and Alex Alderton, Roberto Amato, Jeffrey Barrett, Sonia Goncalves, Ewan Harrison, David K. Jackson, Ian Johnston, Dominic Kwiatkowski, Cordelia Langford, John Sillitoe on behalf of the Wellcome Sanger Institute COVID-19 Surveillance Team |
| EPI_ISL_2847077, EPI_ISL_2847078 | Health Services Laboratories | Wellcome Sanger Institute for the COVID-19 Genomics UK (COG-UK) Consortium | Health Services Laboratories and Alex Alderton, Roberto Amato, Jeffrey Barrett, Sonia Goncalves, Ewan Harrison, David K. Jackson, Ian Johnston, Dominic Kwiatkowski, Cordelia Langford, John Sillitoe on behalf of the Wellcome Sanger Institute COVID-19 Surveillance Team |
| EPI_ISL_2847079 | Lighthouse Lab in Glasgow | Wellcome Sanger Institute for the COVID-19 Genomics UK (COG-UK) Consortium | Harper VanSteenhouse, Yumi Kasai, David Gray, Carol Clugston, Anna Dominiczak and Alex Alderton, Roberto Amato, Jeffrey Barrett, Sonia Goncalves, Ewan Harrison, David K. Jackson, Ian Johnston, Dominic Kwiatkowski, Cordelia Langford, John Sillitoe on behalf of the Wellcome Sanger Institute COVID-19 Surveillance Team |
| EPI_ISL_2847080, EPI_ISL_2847081 | Health Services Laboratories | Wellcome Sanger Institute for the COVID-19 Genomics UK (COG-UK) Consortium | Health Services Laboratories and Alex Alderton, Roberto Amato, Jeffrey Barrett, Sonia Goncalves, Ewan Harrison, David K. Jackson, Ian Johnston, Dominic Kwiatkowski, Cordelia Langford, John Sillitoe on behalf of the Wellcome Sanger Institute COVID-19 Surveillance Team |
| EPI_ISL_2847082, EPI_ISL_2847083 | Lighthouse Lab in Glasgow | Wellcome Sanger Institute for the COVID-19 Genomics UK (COG-UK) Consortium | Harper VanSteenhouse, Yumi Kasai, David Gray, Carol Clugston, Anna Dominiczak and Alex Alderton, Roberto Amato, Jeffrey Barrett, Sonia Goncalves, Ewan Harrison, David K. Jackson, Ian Johnston, Dominic Kwiatkowski, Cordelia Langford, John Sillitoe on behalf of the Wellcome Sanger Institute COVID-19 Surveillance Team |
| EPI_ISL_2847084 | Health Services Laboratories | Wellcome Sanger Institute for the COVID-19 Genomics UK (COG-UK) Consortium | Health Services Laboratories and Alex Alderton, Roberto Amato, Jeffrey Barrett, Sonia Goncalves, Ewan Harrison, David K. Jackson, Ian Johnston, Dominic Kwiatkowski, Cordelia Langford, John Sillitoe on behalf of the Wellcome Sanger Institute COVID-19 Surveillance Team |
| EPI_ISL_2847085, EPI_ISL_2847086, EPI_ISL_2847087 | Lighthouse Lab in Glasgow | Wellcome Sanger Institute for the COVID-19 Genomics UK (COG-UK) Consortium | Harper VanSteenhouse, Yumi Kasai, David Gray, Carol Clugston, Anna Dominiczak and Alex Alderton, Roberto Amato, Jeffrey Barrett, Sonia Goncalves, Ewan Harrison, David K. Jackson, Ian Johnston, Dominic Kwiatkowski, Cordelia Langford, John Sillitoe on behalf of the Wellcome Sanger Institute COVID-19 Surveillance Team |
| EPI_ISL_2847088 | Lighthouse Lab in Milton Keynes | Wellcome Sanger Institute for the COVID-19 Genomics UK (COG-UK) Consortium | The Lighthouse Lab in Milton Keynes and Alex Alderton, Roberto Amato, Jeffrey Barrett, Sonia Goncalves, Ewan Harrison, David K. Jackson, Ian Johnston, Dominic Kwiatkowski, Cordelia Langford, John Sillitoe on behalf of the Wellcome Sanger Institute COVID-19 Surveillance Team |
| EPI_ISL_2847089 | Lighthouse Lab in Glasgow | Wellcome Sanger Institute for the COVID-19 Genomics UK (COG-UK) Consortium | Harper VanSteenhouse, Yumi Kasai, David Gray, Carol Clugston, Anna Dominiczak and Alex Alderton, Roberto Amato, Jeffrey Barrett, Sonia Goncalves, Ewan Harrison, David K. Jackson, Ian Johnston, Dominic Kwiatkowski, Cordelia Langford, John Sillitoe on behalf of the Wellcome Sanger Institute COVID-19 Surveillance Team |
| EPI_ISL_2847090 | Health Services Laboratories | Wellcome Sanger Institute for the COVID-19 Genomics UK (COG-UK) Consortium | Health Services Laboratories and Alex Alderton, Roberto Amato, Jeffrey Barrett, Sonia Goncalves, Ewan Harrison, David K. Jackson, Ian Johnston, Dominic Kwiatkowski, Cordelia Langford, John Sillitoe on behalf of the Wellcome Sanger Institute COVID-19 Surveillance Team |
| EPI_ISL_2847091 | Lighthouse Lab in Glasgow | Wellcome Sanger Institute for the COVID-19 Genomics UK (COG-UK) Consortium | Harper VanSteenhouse, Yumi Kasai, David Gray, Carol Clugston, Anna Dominiczak and Alex Alderton, Roberto Amato, Jeffrey Barrett, Sonia Goncalves, Ewan Harrison, David K. Jackson, Ian Johnston, Dominic Kwiatkowski, Cordelia Langford, John Sillitoe on behalf of the Wellcome Sanger Institute COVID-19 Surveillance Team |
| EPI_ISL_2847092, EPI_ISL_2847093, EPI_ISL_2847094 | Lighthouse Lab in Alderley Park | Wellcome Sanger Institute for the COVID-19 Genomics UK (COG-UK) Consortium | Jacquelyn Wynn, Mairead Hyland, The Lighthouse Lab in Alderley Park and Alex Alderton, Roberto Amato, Jeffrey Barrett, Sonia Goncalves, Ewan Harrison, David K. Jackson, Ian Johnston, Dominic Kwiatkowski, Cordelia Langford, John Sillitoe on behalf of the Wellcome Sanger Institute COVID-19 Surveillance Team |
| EPI_ISL_2847095 | Lighthouse Lab in Glasgow | Wellcome Sanger Institute for the COVID-19 Genomics UK (COG-UK) Consortium | Harper VanSteenhouse, Yumi Kasai, David Gray, Carol Clugston, Anna Dominiczak and Alex Alderton, Roberto Amato, Jeffrey Barrett, Sonia Goncalves, Ewan Harrison, David K. Jackson, Ian Johnston, Dominic Kwiatkowski, Cordelia Langford, John Sillitoe on behalf of the Wellcome Sanger Institute COVID-19 Surveillance Team |
| EPI_ISL_2847096 | Lighthouse Lab in Alderley Park | Wellcome Sanger Institute for the COVID-19 Genomics UK (COG-UK) Consortium | Jacquelyn Wynn, Mairead Hyland, The Lighthouse Lab in Alderley Park and Alex Alderton, Roberto Amato, Jeffrey Barrett, Sonia Goncalves, Ewan Harrison, David K. Jackson, Ian Johnston, Dominic Kwiatkowski, Cordelia Langford, John Sillitoe on behalf of the Wellcome Sanger Institute COVID-19 Surveillance Team |
| EPI_ISL_2847097, EPI_ISL_2847098 | Health Services Laboratories | Wellcome Sanger Institute for the COVID-19 Genomics UK (COG-UK) Consortium | Health Services Laboratories and Alex Alderton, Roberto Amato, Jeffrey Barrett, Sonia Goncalves, Ewan Harrison, David K. Jackson, Ian Johnston, Dominic Kwiatkowski, Cordelia Langford, John Sillitoe on behalf of the Wellcome Sanger Institute COVID-19 Surveillance Team |
| EPI_ISL_2847099 | Lighthouse Lab in Alderley Park | Wellcome Sanger Institute for the COVID-19 Genomics UK (COG-UK) Consortium | Jacquelyn Wynn, Mairead Hyland, The Lighthouse Lab in Alderley Park and Alex Alderton, Roberto Amato, Jeffrey Barrett, Sonia Goncalves, Ewan Harrison, David K. Jackson, Ian Johnston, Dominic Kwiatkowski, Cordelia Langford, John Sillitoe on behalf of the Wellcome Sanger Institute COVID-19 Surveillance Team |
| EPI_ISL_2847100 | Lighthouse Lab in Glasgow | Wellcome Sanger Institute for the COVID-19 Genomics UK (COG-UK) Consortium | Harper VanSteenhouse, Yumi Kasai, David Gray, Carol Clugston, Anna Dominiczak and Alex Alderton, Roberto Amato, Jeffrey Barrett, Sonia Goncalves, Ewan Harrison, David K. Jackson, Ian Johnston, Dominic Kwiatkowski, Cordelia Langford, John Sillitoe on behalf of the Wellcome Sanger Institute COVID-19 Surveillance Team |
| EPI_ISL_2847101 | Lighthouse Lab in Alderley Park | Wellcome Sanger Institute for the COVID-19 Genomics UK (COG-UK) Consortium | Jacquelyn Wynn, Mairead Hyland, The Lighthouse Lab in Alderley Park and Alex Alderton, Roberto Amato, Jeffrey Barrett, Sonia Goncalves, Ewan Harrison, David K. Jackson, Ian Johnston, Dominic Kwiatkowski, Cordelia Langford, John Sillitoe on behalf of the Wellcome Sanger Institute COVID-19 Surveillance Team |
| EPI_ISL_2847102 | Health Services Laboratories | Wellcome Sanger Institute for the COVID-19 Genomics UK (COG-UK) Consortium | Health Services Laboratories and Alex Alderton, Roberto Amato, Jeffrey Barrett, Sonia Goncalves, Ewan Harrison, David K. Jackson, Ian Johnston, Dominic Kwiatkowski, Cordelia Langford, John Sillitoe on behalf of the Wellcome Sanger Institute COVID-19 Surveillance Team |
| EPI_ISL_2847103 | Lighthouse Lab in Glasgow | Wellcome Sanger Institute for the COVID-19 Genomics UK (COG-UK) Consortium | Harper VanSteenhouse, Yumi Kasai, David Gray, Carol Clugston, Anna Dominiczak and Alex Alderton, Roberto Amato, Jeffrey Barrett, Sonia Goncalves, Ewan Harrison, David K. Jackson, Ian Johnston, Dominic Kwiatkowski, Cordelia Langford, John Sillitoe on behalf of the Wellcome Sanger Institute COVID-19 Surveillance Team |
| EPI_ISL_2847104 | Lighthouse Lab in Milton Keynes | Wellcome Sanger Institute for the COVID-19 Genomics UK (COG-UK) Consortium | The Lighthouse Lab in Milton Keynes and Alex Alderton, Roberto Amato, Jeffrey Barrett, Sonia Goncalves, Ewan Harrison, David K. Jackson, Ian Johnston, Dominic Kwiatkowski, Cordelia Langford, John Sillitoe on behalf of the Wellcome Sanger Institute COVID-19 Surveillance Team |
| EPI_ISL_2847105, EPI_ISL_2847106 | Lighthouse Lab in Alderley Park | Wellcome Sanger Institute for the COVID-19 Genomics UK (COG-UK) Consortium | Jacquelyn Wynn, Mairead Hyland, The Lighthouse Lab in Alderley Park and Alex Alderton, Roberto Amato, Jeffrey Barrett, Sonia Goncalves, Ewan Harrison, David K. Jackson, Ian Johnston, Dominic Kwiatkowski, Cordelia Langford, John Sillitoe on behalf of the Wellcome Sanger Institute COVID-19 Surveillance Team |
| EPI_ISL_2847107 | Health Services Laboratories | Wellcome Sanger Institute for the COVID-19 Genomics UK (COG-UK) Consortium | Health Services Laboratories and Alex Alderton, Roberto Amato, Jeffrey Barrett, Sonia Goncalves, Ewan Harrison, David K. Jackson, Ian Johnston, Dominic Kwiatkowski, Cordelia Langford, John Sillitoe on behalf of the Wellcome Sanger Institute COVID-19 Surveillance Team |
| EPI_ISL_2847108, EPI_ISL_2847109 | Lighthouse Lab in Alderley Park | Wellcome Sanger Institute for the COVID-19 Genomics UK (COG-UK) Consortium | Jacquelyn Wynn, Mairead Hyland, The Lighthouse Lab in Alderley Park and Alex Alderton, Roberto Amato, Jeffrey Barrett, Sonia Goncalves, Ewan Harrison, David K. Jackson, Ian Johnston, Dominic Kwiatkowski, Cordelia Langford, John Sillitoe on behalf of the Wellcome Sanger Institute COVID-19 Surveillance Team |
| EPI_ISL_2847110, | Lighthouse Lab in Glasgow | Wellcome Sanger Institute for the COVID-19 Genomics UK (COG-UK) | Harper VanSteenhouse, Yumi Kasai, David Gray, Carol Clugston, Anna Dominiczak and Alex Alderton, Roberto Amato, Jeffrey Barrett, Sonia Goncalves, Ewan Harrison, David K. Jackson, Ian Johnston, Dominic Kwiatkowski, Cordelia Langford, |

| | | | |
|---|---|---|---|
| EPI_ISL_2847111 | | Consortium | John Sillitoe on behalf of the Wellcome Sanger Institute COVID-19 Surveillance Team |
| EPI_ISL_2847112 | Lighthouse Lab in Milton Keynes | Wellcome Sanger Institute for the COVID-19 Genomics UK (COG-UK) Consortium | The Lighthouse Lab in Milton Keynes and Alex Alderton, Roberto Amato, Jeffrey Barrett, Sonia Goncalves, Ewan Harrison, David K. Jackson, Ian Johnston, Dominic Kwiatkowski, Cordelia Langford, John Sillitoe on behalf of the Wellcome Sanger Institute COVID-19 Surveillance Team |
| EPI_ISL_2847113 | Lighthouse Lab in Glasgow | Wellcome Sanger Institute for the COVID-19 Genomics UK (COG-UK) Consortium | Harper VanSteenhouse, Yumi Kasai, David Gray, Carol Clugston, Anna Dominiczak and Alex Alderton, Roberto Amato, Jeffrey Barrett, Sonia Goncalves, Ewan Harrison, David K. Jackson, Ian Johnston, Dominic Kwiatkowski, Cordelia Langford, John Sillitoe on behalf of the Wellcome Sanger Institute COVID-19 Surveillance Team |
| EPI_ISL_2847114 | Lighthouse Lab in Milton Keynes | Wellcome Sanger Institute for the COVID-19 Genomics UK (COG-UK) Consortium | The Lighthouse Lab in Milton Keynes and Alex Alderton, Roberto Amato, Jeffrey Barrett, Sonia Goncalves, Ewan Harrison, David K. Jackson, Ian Johnston, Dominic Kwiatkowski, Cordelia Langford, John Sillitoe on behalf of the Wellcome Sanger Institute COVID-19 Surveillance Team |
| EPI_ISL_2847115 | AZ Klina | AZ Klina | Carl Vael - Lynsey Berckmans |
| EPI_ISL_2847116 | Lighthouse Lab in Milton Keynes | Wellcome Sanger Institute for the COVID-19 Genomics UK (COG-UK) Consortium | The Lighthouse Lab in Milton Keynes and Alex Alderton, Roberto Amato, Jeffrey Barrett, Sonia Goncalves, Ewan Harrison, David K. Jackson, Ian Johnston, Dominic Kwiatkowski, Cordelia Langford, John Sillitoe on behalf of the Wellcome Sanger Institute COVID-19 Surveillance Team |
| EPI_ISL_2847117 | Health Services Laboratories | Wellcome Sanger Institute for the COVID-19 Genomics UK (COG-UK) Consortium | Health Services Laboratories and Alex Alderton, Roberto Amato, Jeffrey Barrett, Sonia Goncalves, Ewan Harrison, David K. Jackson, Ian Johnston, Dominic Kwiatkowski, Cordelia Langford, John Sillitoe on behalf of the Wellcome Sanger Institute COVID-19 Surveillance Team |
| EPI_ISL_2847118, EPI_ISL_2847119 | Lighthouse Lab in Glasgow | Wellcome Sanger Institute for the COVID-19 Genomics UK (COG-UK) Consortium | Harper VanSteenhouse, Yumi Kasai, David Gray, Carol Clugston, Anna Dominiczak and Alex Alderton, Roberto Amato, Jeffrey Barrett, Sonia Goncalves, Ewan Harrison, David K. Jackson, Ian Johnston, Dominic Kwiatkowski, Cordelia Langford, John Sillitoe on behalf of the Wellcome Sanger Institute COVID-19 Surveillance Team |
| EPI_ISL_2847120 | Lighthouse Lab in Alderley Park | Wellcome Sanger Institute for the COVID-19 Genomics UK (COG-UK) Consortium | Jacquelyn Wynn, Mairead Hyland, The Lighthouse Lab in Alderley Park and Alex Alderton, Roberto Amato, Jeffrey Barrett, Sonia Goncalves, Ewan Harrison, David K. Jackson, Ian Johnston, Dominic Kwiatkowski, Cordelia Langford, John Sillitoe on behalf of the Wellcome Sanger Institute COVID-19 Surveillance Team |
| EPI_ISL_2847121, EPI_ISL_2847122 | Lighthouse Lab in Milton Keynes | Wellcome Sanger Institute for the COVID-19 Genomics UK (COG-UK) Consortium | The Lighthouse Lab in Milton Keynes and Alex Alderton, Roberto Amato, Jeffrey Barrett, Sonia Goncalves, Ewan Harrison, David K. Jackson, Ian Johnston, Dominic Kwiatkowski, Cordelia Langford, John Sillitoe on behalf of the Wellcome Sanger Institute COVID-19 Surveillance Team |
| EPI_ISL_2847123 | Lighthouse Lab in Glasgow | Wellcome Sanger Institute for the COVID-19 Genomics UK (COG-UK) Consortium | Harper VanSteenhouse, Yumi Kasai, David Gray, Carol Clugston, Anna Dominiczak and Alex Alderton, Roberto Amato, Jeffrey Barrett, Sonia Goncalves, Ewan Harrison, David K. Jackson, Ian Johnston, Dominic Kwiatkowski, Cordelia Langford, John Sillitoe on behalf of the Wellcome Sanger Institute COVID-19 Surveillance Team |
| EPI_ISL_2847124, EPI_ISL_2847125 | Health Services Laboratories | Wellcome Sanger Institute for the COVID-19 Genomics UK (COG-UK) Consortium | Health Services Laboratories and Alex Alderton, Roberto Amato, Jeffrey Barrett, Sonia Goncalves, Ewan Harrison, David K. Jackson, Ian Johnston, Dominic Kwiatkowski, Cordelia Langford, John Sillitoe on behalf of the Wellcome Sanger Institute COVID-19 Surveillance Team |
| EPI_ISL_2847126 | Lighthouse Lab in Milton Keynes | Wellcome Sanger Institute for the COVID-19 Genomics UK (COG-UK) Consortium | The Lighthouse Lab in Milton Keynes and Alex Alderton, Roberto Amato, Jeffrey Barrett, Sonia Goncalves, Ewan Harrison, David K. Jackson, Ian Johnston, Dominic Kwiatkowski, Cordelia Langford, John Sillitoe on behalf of the Wellcome Sanger Institute COVID-19 Surveillance Team |
| EPI_ISL_2847127 | Lighthouse Lab in Alderley Park | Wellcome Sanger Institute for the COVID-19 Genomics UK (COG-UK) Consortium | Jacquelyn Wynn, Mairead Hyland, The Lighthouse Lab in Alderley Park and Alex Alderton, Roberto Amato, Jeffrey Barrett, Sonia Goncalves, Ewan Harrison, David K. Jackson, Ian Johnston, Dominic Kwiatkowski, Cordelia Langford, John Sillitoe on behalf of the Wellcome Sanger Institute COVID-19 Surveillance Team |
| EPI_ISL_2847128 | Lighthouse Lab in Milton Keynes | Wellcome Sanger Institute for the COVID-19 Genomics UK (COG-UK) Consortium | The Lighthouse Lab in Milton Keynes and Alex Alderton, Roberto Amato, Jeffrey Barrett, Sonia Goncalves, Ewan Harrison, David K. Jackson, Ian Johnston, Dominic Kwiatkowski, Cordelia Langford, John Sillitoe on behalf of the Wellcome Sanger Institute COVID-19 Surveillance Team |
| EPI_ISL_2847129 | Health Services Laboratories | Wellcome Sanger Institute for the COVID-19 Genomics UK (COG-UK) Consortium | Health Services Laboratories and Alex Alderton, Roberto Amato, Jeffrey Barrett, Sonia Goncalves, Ewan Harrison, David K. Jackson, Ian Johnston, Dominic Kwiatkowski, Cordelia Langford, John Sillitoe on behalf of the Wellcome Sanger Institute COVID-19 Surveillance Team |
| EPI_ISL_2847130, EPI_ISL_2847131 | Lighthouse Lab in Alderley Park | Wellcome Sanger Institute for the COVID-19 Genomics UK (COG-UK) Consortium | Jacquelyn Wynn, Mairead Hyland, The Lighthouse Lab in Alderley Park and Alex Alderton, Roberto Amato, Jeffrey Barrett, Sonia Goncalves, Ewan Harrison, David K. Jackson, Ian Johnston, Dominic Kwiatkowski, Cordelia Langford, John Sillitoe on behalf of the Wellcome Sanger Institute COVID-19 Surveillance Team |
| EPI_ISL_2847132 | Health Services Laboratories | Wellcome Sanger Institute for the COVID-19 Genomics UK (COG-UK) Consortium | Health Services Laboratories and Alex Alderton, Roberto Amato, Jeffrey Barrett, Sonia Goncalves, Ewan Harrison, David K. Jackson, Ian Johnston, Dominic Kwiatkowski, Cordelia Langford, John Sillitoe on behalf of the Wellcome Sanger Institute COVID-19 Surveillance Team |
| EPI_ISL_2847133 | Lighthouse Lab in Glasgow | Wellcome Sanger Institute for the COVID-19 Genomics UK (COG-UK) Consortium | Harper VanSteenhouse, Yumi Kasai, David Gray, Carol Clugston, Anna Dominiczak and Alex Alderton, Roberto Amato, Jeffrey Barrett, Sonia Goncalves, Ewan Harrison, David K. Jackson, Ian Johnston, Dominic Kwiatkowski, Cordelia Langford, John Sillitoe on behalf of the Wellcome Sanger Institute COVID-19 Surveillance Team |
| EPI_ISL_2847134 | Lighthouse Lab in Alderley Park | Wellcome Sanger Institute for the COVID-19 Genomics UK (COG-UK) Consortium | Jacquelyn Wynn, Mairead Hyland, The Lighthouse Lab in Alderley Park and Alex Alderton, Roberto Amato, Jeffrey Barrett, Sonia Goncalves, Ewan Harrison, David K. Jackson, Ian Johnston, Dominic Kwiatkowski, Cordelia Langford, John Sillitoe on behalf of the Wellcome Sanger Institute COVID-19 Surveillance Team |
| EPI_ISL_2847135 | Health Services Laboratories | Wellcome Sanger Institute for the COVID-19 Genomics UK (COG-UK) Consortium | Health Services Laboratories and Alex Alderton, Roberto Amato, Jeffrey Barrett, Sonia Goncalves, Ewan Harrison, David K. Jackson, Ian Johnston, Dominic Kwiatkowski, Cordelia Langford, John Sillitoe on behalf of the Wellcome Sanger Institute COVID-19 Surveillance Team |
| EPI_ISL_2847136 | Lighthouse Lab in Milton Keynes | Wellcome Sanger Institute for the COVID-19 Genomics UK (COG-UK) Consortium | The Lighthouse Lab in Milton Keynes and Alex Alderton, Roberto Amato, Jeffrey Barrett, Sonia Goncalves, Ewan Harrison, David K. Jackson, Ian Johnston, Dominic Kwiatkowski, Cordelia Langford, John Sillitoe on behalf of the Wellcome Sanger Institute COVID-19 Surveillance Team |
| EPI_ISL_2847137, EPI_ISL_2847138 | Health Services Laboratories | Wellcome Sanger Institute for the COVID-19 Genomics UK (COG-UK) Consortium | Health Services Laboratories and Alex Alderton, Roberto Amato, Jeffrey Barrett, Sonia Goncalves, Ewan Harrison, David K. Jackson, Ian Johnston, Dominic Kwiatkowski, Cordelia Langford, John Sillitoe on behalf of the Wellcome Sanger Institute COVID-19 Surveillance Team |
| EPI_ISL_2847139 | Lighthouse Lab in Milton Keynes | Wellcome Sanger Institute for the COVID-19 Genomics UK (COG-UK) Consortium | The Lighthouse Lab in Milton Keynes and Alex Alderton, Roberto Amato, Jeffrey Barrett, Sonia Goncalves, Ewan Harrison, David K. Jackson, Ian Johnston, Dominic Kwiatkowski, Cordelia Langford, John Sillitoe on behalf of the Wellcome Sanger Institute COVID-19 Surveillance Team |
| EPI_ISL_2847140 | Lighthouse Lab in Alderley Park | Wellcome Sanger Institute for the COVID-19 Genomics UK (COG-UK) Consortium | Jacquelyn Wynn, Mairead Hyland, The Lighthouse Lab in Alderley Park and Alex Alderton, Roberto Amato, Jeffrey Barrett, Sonia Goncalves, Ewan Harrison, David K. Jackson, Ian Johnston, Dominic Kwiatkowski, Cordelia Langford, John Sillitoe on behalf of the Wellcome Sanger Institute COVID-19 Surveillance Team |
| EPI_ISL_2847141 | Lighthouse Lab in Milton Keynes | Wellcome Sanger Institute for the COVID-19 Genomics UK (COG-UK) Consortium | The Lighthouse Lab in Milton Keynes and Alex Alderton, Roberto Amato, Jeffrey Barrett, Sonia Goncalves, Ewan Harrison, David K. Jackson, Ian Johnston, Dominic Kwiatkowski, Cordelia Langford, John Sillitoe on behalf of the Wellcome Sanger Institute COVID-19 Surveillance Team |
| EPI_ISL_2847142 | Lighthouse Lab in Alderley Park | Wellcome Sanger Institute for the COVID-19 Genomics UK (COG-UK) Consortium | Jacquelyn Wynn, Mairead Hyland, The Lighthouse Lab in Alderley Park and Alex Alderton, Roberto Amato, Jeffrey Barrett, Sonia Goncalves, Ewan Harrison, David K. Jackson, Ian Johnston, Dominic Kwiatkowski, Cordelia Langford, John Sillitoe on behalf of the Wellcome Sanger Institute COVID-19 Surveillance Team |
| EPI_ISL_2847143, EPI_ISL_2847144 | Lighthouse Lab in Milton Keynes | Wellcome Sanger Institute for the COVID-19 Genomics UK (COG-UK) Consortium | The Lighthouse Lab in Milton Keynes and Alex Alderton, Roberto Amato, Jeffrey Barrett, Sonia Goncalves, Ewan Harrison, David K. Jackson, Ian Johnston, Dominic Kwiatkowski, Cordelia Langford, John Sillitoe on behalf of the Wellcome Sanger Institute COVID-19 Surveillance Team |
| EPI_ISL_2847145 | Lighthouse Lab in Alderley Park | Wellcome Sanger Institute for the COVID-19 Genomics UK (COG-UK) Consortium | Jacquelyn Wynn, Mairead Hyland, The Lighthouse Lab in Alderley Park and Alex Alderton, Roberto Amato, Jeffrey Barrett, Sonia Goncalves, Ewan Harrison, David K. Jackson, Ian Johnston, Dominic Kwiatkowski, Cordelia Langford, John Sillitoe on behalf of the Wellcome Sanger Institute COVID-19 Surveillance Team |
| EPI_ISL_2847146 | Lighthouse Lab in Glasgow | Wellcome Sanger Institute for the COVID-19 Genomics UK (COG-UK) Consortium | Harper VanSteenhouse, Yumi Kasai, David Gray, Carol Clugston, Anna Dominiczak and Alex Alderton, Roberto Amato, Jeffrey Barrett, Sonia Goncalves, Ewan Harrison, David K. Jackson, Ian Johnston, Dominic Kwiatkowski, Cordelia Langford, John Sillitoe on behalf of the Wellcome Sanger Institute COVID-19 Surveillance Team |
| EPI_ISL_2847147 | Lighthouse Lab in Milton Keynes | Wellcome Sanger Institute for the COVID-19 Genomics UK (COG-UK) Consortium | The Lighthouse Lab in Milton Keynes and Alex Alderton, Roberto Amato, Jeffrey Barrett, Sonia Goncalves, Ewan Harrison, David K. Jackson, Ian Johnston, Dominic Kwiatkowski, Cordelia Langford, John Sillitoe on behalf of the Wellcome Sanger Institute COVID-19 Surveillance Team |
| EPI_ISL_2847148 | Lighthouse Lab in Alderley Park | Wellcome Sanger Institute for the COVID-19 Genomics UK (COG-UK) Consortium | Jacquelyn Wynn, Mairead Hyland, The Lighthouse Lab in Alderley Park and Alex Alderton, Roberto Amato, Jeffrey Barrett, Sonia Goncalves, Ewan Harrison, David K. Jackson, Ian Johnston, Dominic Kwiatkowski, Cordelia Langford, John Sillitoe on behalf of the Wellcome Sanger Institute COVID-19 Surveillance Team |
| EPI_ISL_2847149 | Lighthouse Lab in Glasgow | Wellcome Sanger Institute for the COVID-19 Genomics UK (COG-UK) Consortium | Harper VanSteenhouse, Yumi Kasai, David Gray, Carol Clugston, Anna Dominiczak and Alex Alderton, Roberto Amato, Jeffrey Barrett, Sonia Goncalves, Ewan Harrison, David K. Jackson, Ian Johnston, Dominic Kwiatkowski, Cordelia Langford, John Sillitoe on behalf of the Wellcome Sanger Institute COVID-19 Surveillance Team |
| EPI_ISL_2847150 | Lighthouse Lab in Alderley Park | Wellcome Sanger Institute for the COVID-19 Genomics UK (COG-UK) Consortium | Jacquelyn Wynn, Mairead Hyland, The Lighthouse Lab in Alderley Park and Alex Alderton, Roberto Amato, Jeffrey Barrett, Sonia Goncalves, Ewan Harrison, David K. Jackson, Ian Johnston, Dominic Kwiatkowski, Cordelia Langford, John Sillitoe on behalf of the Wellcome Sanger Institute COVID-19 Surveillance Team |
| EPI_ISL_2847151 | Lighthouse Lab in Milton Keynes | Wellcome Sanger Institute for the COVID-19 Genomics UK (COG-UK) Consortium | The Lighthouse Lab in Milton Keynes and Alex Alderton, Roberto Amato, Jeffrey Barrett, Sonia Goncalves, Ewan Harrison, David K. Jackson, Ian Johnston, Dominic Kwiatkowski, Cordelia Langford, John Sillitoe on behalf of the Wellcome Sanger Institute COVID-19 Surveillance Team |
| EPI_ISL_2847152 | Lighthouse Lab in Glasgow | Wellcome Sanger Institute for the COVID-19 Genomics UK (COG-UK) Consortium | Harper VanSteenhouse, Yumi Kasai, David Gray, Carol Clugston, Anna Dominiczak and Alex Alderton, Roberto Amato, Jeffrey Barrett, Sonia Goncalves, Ewan Harrison, David K. Jackson, Ian Johnston, Dominic Kwiatkowski, Cordelia Langford, John Sillitoe on behalf of the Wellcome Sanger Institute COVID-19 Surveillance Team |
| EPI_ISL_2847153, EPI_ISL_2847154 | Health Services Laboratories | Wellcome Sanger Institute for the COVID-19 Genomics UK (COG-UK) Consortium | Health Services Laboratories and Alex Alderton, Roberto Amato, Jeffrey Barrett, Sonia Goncalves, Ewan Harrison, David K. Jackson, Ian Johnston, Dominic Kwiatkowski, Cordelia Langford, John Sillitoe on behalf of the Wellcome Sanger Institute COVID-19 Surveillance Team |
| EPI_ISL_2847155 | Lighthouse Lab in Milton Keynes | Wellcome Sanger Institute for the COVID-19 Genomics UK (COG-UK) Consortium | The Lighthouse Lab in Milton Keynes and Alex Alderton, Roberto Amato, Jeffrey Barrett, Sonia Goncalves, Ewan Harrison, David K. Jackson, Ian Johnston, Dominic Kwiatkowski, Cordelia Langford, John Sillitoe on behalf of the Wellcome Sanger Institute COVID-19 Surveillance Team |
| EPI_ISL_2847156 | Health Services Laboratories | Wellcome Sanger Institute for the COVID-19 Genomics UK (COG-UK) Consortium | Health Services Laboratories and Alex Alderton, Roberto Amato, Jeffrey Barrett, Sonia Goncalves, Ewan Harrison, David K. Jackson, Ian Johnston, Dominic Kwiatkowski, Cordelia Langford, John Sillitoe on behalf of the Wellcome Sanger Institute COVID-19 Surveillance Team |
| EPI_ISL_2847157 | Lighthouse Lab in Glasgow | Wellcome Sanger Institute for the COVID-19 Genomics UK (COG-UK) Consortium | Harper VanSteenhouse, Yumi Kasai, David Gray, Carol Clugston, Anna Dominiczak and Alex Alderton, Roberto Amato, Jeffrey Barrett, Sonia Goncalves, Ewan Harrison, David K. Jackson, Ian Johnston, Dominic Kwiatkowski, Cordelia Langford, John Sillitoe on behalf of the Wellcome Sanger Institute COVID-19 Surveillance Team |
| EPI_ISL_2847158 | Lighthouse Lab in Alderley Park | Wellcome Sanger Institute for the COVID-19 Genomics UK (COG-UK) Consortium | Jacquelyn Wynn, Mairead Hyland, The Lighthouse Lab in Alderley Park and Alex Alderton, Roberto Amato, Jeffrey Barrett, Sonia Goncalves, Ewan Harrison, David K. Jackson, Ian Johnston, Dominic Kwiatkowski, Cordelia Langford, John Sillitoe on behalf of the Wellcome Sanger Institute COVID-19 Surveillance Team |
| EPI_ISL_2847159 | Lighthouse Lab in Glasgow | Wellcome Sanger Institute for the COVID-19 Genomics UK (COG-UK) Consortium | Harper VanSteenhouse, Yumi Kasai, David Gray, Carol Clugston, Anna Dominiczak and Alex Alderton, Roberto Amato, Jeffrey Barrett, Sonia Goncalves, Ewan Harrison, David K. Jackson, Ian Johnston, Dominic Kwiatkowski, Cordelia Langford, John Sillitoe on behalf of the Wellcome Sanger Institute COVID-19 Surveillance Team |
| EPI_ISL_2847160 | Lighthouse Lab in Milton Keynes | Wellcome Sanger Institute for the COVID-19 Genomics UK (COG-UK) Consortium | The Lighthouse Lab in Milton Keynes and Alex Alderton, Roberto Amato, Jeffrey Barrett, Sonia Goncalves, Ewan Harrison, David K. Jackson, Ian Johnston, Dominic Kwiatkowski, Cordelia Langford, John Sillitoe on behalf of the Wellcome Sanger Institute COVID-19 Surveillance Team |
| EPI_ISL_2847161, EPI_ISL_2847162, EPI_ISL_2847163 | Lighthouse Lab in Alderley Park | Wellcome Sanger Institute for the COVID-19 Genomics UK (COG-UK) Consortium | Jacquelyn Wynn, Mairead Hyland, The Lighthouse Lab in Alderley Park and Alex Alderton, Roberto Amato, Jeffrey Barrett, Sonia Goncalves, Ewan Harrison, David K. Jackson, Ian Johnston, Dominic Kwiatkowski, Cordelia Langford, John Sillitoe on behalf of the Wellcome Sanger Institute COVID-19 Surveillance Team |
| EPI_ISL_2847164, EPI_ISL_2847165 | Lighthouse Lab in Milton Keynes | Wellcome Sanger Institute for the COVID-19 Genomics UK (COG-UK) Consortium | The Lighthouse Lab in Milton Keynes and Alex Alderton, Roberto Amato, Jeffrey Barrett, Sonia Goncalves, Ewan Harrison, David K. Jackson, Ian Johnston, Dominic Kwiatkowski, Cordelia Langford, John Sillitoe on behalf of the Wellcome Sanger Institute COVID-19 Surveillance Team |

| | | | |
|---|---|---|---|
| EPI_ISL_2847166 | Health Services Laboratories | Wellcome Sanger Institute for the COVID-19 Genomics UK (COG-UK) Consortium | Health Services Laboratories and Alex Alderton, Roberto Amato, Jeffrey Barrett, Sonia Goncalves, Ewan Harrison, David K. Jackson, Ian Johnston, Dominic Kwiatkowski, Cordelia Langford, John Sillitoe on behalf of the Wellcome Sanger Institute COVID-19 Surveillance Team |
| EPI_ISL_2847167 | Lighthouse Lab in Milton Keynes | Wellcome Sanger Institute for the COVID-19 Genomics UK (COG-UK) Consortium | The Lighthouse Lab in Milton Keynes and Alex Alderton, Roberto Amato, Jeffrey Barrett, Sonia Goncalves, Ewan Harrison, David K. Jackson, Ian Johnston, Dominic Kwiatkowski, Cordelia Langford, John Sillitoe on behalf of the Wellcome Sanger Institute COVID-19 Surveillance Team |
| EPI_ISL_2847168 | Lighthouse Lab in Glasgow | Wellcome Sanger Institute for the COVID-19 Genomics UK (COG-UK) Consortium | Harper VanSteenhouse, Yumi Kasai, David Gray, Carol Clugston, Anna Dominiczak and Alex Alderton, Roberto Amato, Jeffrey Barrett, Sonia Goncalves, Ewan Harrison, David K. Jackson, Ian Johnston, Dominic Kwiatkowski, Cordelia Langford, John Sillitoe on behalf of the Wellcome Sanger Institute COVID-19 Surveillance Team |
| EPI_ISL_2847170 | Lighthouse Lab in Alderley Park | Wellcome Sanger Institute for the COVID-19 Genomics UK (COG-UK) Consortium | Jacquelyn Wynn, Mairead Hyland, The Lighthouse Lab in Alderley Park and Alex Alderton, Roberto Amato, Jeffrey Barrett, Sonia Goncalves, Ewan Harrison, David K. Jackson, Ian Johnston, Dominic Kwiatkowski, Cordelia Langford, John Sillitoe on behalf of the Wellcome Sanger Institute COVID-19 Surveillance Team |
| EPI_ISL_2847171 | Lighthouse Lab in Glasgow | Wellcome Sanger Institute for the COVID-19 Genomics UK (COG-UK) Consortium | Harper VanSteenhouse, Yumi Kasai, David Gray, Carol Clugston, Anna Dominiczak and Alex Alderton, Roberto Amato, Jeffrey Barrett, Sonia Goncalves, Ewan Harrison, David K. Jackson, Ian Johnston, Dominic Kwiatkowski, Cordelia Langford, John Sillitoe on behalf of the Wellcome Sanger Institute COVID-19 Surveillance Team |
| EPI_ISL_2847172, EPI_ISL_2847173, EPI_ISL_2847174 | Health Services Laboratories | Wellcome Sanger Institute for the COVID-19 Genomics UK (COG-UK) Consortium | Health Services Laboratories and Alex Alderton, Roberto Amato, Jeffrey Barrett, Sonia Goncalves, Ewan Harrison, David K. Jackson, Ian Johnston, Dominic Kwiatkowski, Cordelia Langford, John Sillitoe on behalf of the Wellcome Sanger Institute COVID-19 Surveillance Team |
| EPI_ISL_2847175 | Lighthouse Lab in Milton Keynes | Wellcome Sanger Institute for the COVID-19 Genomics UK (COG-UK) Consortium | The Lighthouse Lab in Milton Keynes and Alex Alderton, Roberto Amato, Jeffrey Barrett, Sonia Goncalves, Ewan Harrison, David K. Jackson, Ian Johnston, Dominic Kwiatkowski, Cordelia Langford, John Sillitoe on behalf of the Wellcome Sanger Institute COVID-19 Surveillance Team |
| EPI_ISL_2847176, EPI_ISL_2847177 | Lighthouse Lab in Glasgow | Wellcome Sanger Institute for the COVID-19 Genomics UK (COG-UK) Consortium | Harper VanSteenhouse, Yumi Kasai, David Gray, Carol Clugston, Anna Dominiczak and Alex Alderton, Roberto Amato, Jeffrey Barrett, Sonia Goncalves, Ewan Harrison, David K. Jackson, Ian Johnston, Dominic Kwiatkowski, Cordelia Langford, John Sillitoe on behalf of the Wellcome Sanger Institute COVID-19 Surveillance Team |
| EPI_ISL_2847178, EPI_ISL_2847179 | Health Services Laboratories | Wellcome Sanger Institute for the COVID-19 Genomics UK (COG-UK) Consortium | Health Services Laboratories and Alex Alderton, Roberto Amato, Jeffrey Barrett, Sonia Goncalves, Ewan Harrison, David K. Jackson, Ian Johnston, Dominic Kwiatkowski, Cordelia Langford, John Sillitoe on behalf of the Wellcome Sanger Institute COVID-19 Surveillance Team |
| EPI_ISL_2847180 | Lighthouse Lab in Glasgow | Wellcome Sanger Institute for the COVID-19 Genomics UK (COG-UK) Consortium | Harper VanSteenhouse, Yumi Kasai, David Gray, Carol Clugston, Anna Dominiczak and Alex Alderton, Roberto Amato, Jeffrey Barrett, Sonia Goncalves, Ewan Harrison, David K. Jackson, Ian Johnston, Dominic Kwiatkowski, Cordelia Langford, John Sillitoe on behalf of the Wellcome Sanger Institute COVID-19 Surveillance Team |
| EPI_ISL_2847181, EPI_ISL_2847182 | Health Services Laboratories | Wellcome Sanger Institute for the COVID-19 Genomics UK (COG-UK) Consortium | Health Services Laboratories and Alex Alderton, Roberto Amato, Jeffrey Barrett, Sonia Goncalves, Ewan Harrison, David K. Jackson, Ian Johnston, Dominic Kwiatkowski, Cordelia Langford, John Sillitoe on behalf of the Wellcome Sanger Institute COVID-19 Surveillance Team |
| EPI_ISL_2847183 | Lighthouse Lab in Alderley Park | Wellcome Sanger Institute for the COVID-19 Genomics UK (COG-UK) Consortium | Jacquelyn Wynn, Mairead Hyland, The Lighthouse Lab in Alderley Park and Alex Alderton, Roberto Amato, Jeffrey Barrett, Sonia Goncalves, Ewan Harrison, David K. Jackson, Ian Johnston, Dominic Kwiatkowski, Cordelia Langford, John Sillitoe on behalf of the Wellcome Sanger Institute COVID-19 Surveillance Team |
| EPI_ISL_2847184 | Lighthouse Lab in Milton Keynes | Wellcome Sanger Institute for the COVID-19 Genomics UK (COG-UK) Consortium | The Lighthouse Lab in Milton Keynes and Alex Alderton, Roberto Amato, Jeffrey Barrett, Sonia Goncalves, Ewan Harrison, David K. Jackson, Ian Johnston, Dominic Kwiatkowski, Cordelia Langford, John Sillitoe on behalf of the Wellcome Sanger Institute COVID-19 Surveillance Team |
| EPI_ISL_2847185 | Lighthouse Lab in Glasgow | Wellcome Sanger Institute for the COVID-19 Genomics UK (COG-UK) Consortium | Harper VanSteenhouse, Yumi Kasai, David Gray, Carol Clugston, Anna Dominiczak and Alex Alderton, Roberto Amato, Jeffrey Barrett, Sonia Goncalves, Ewan Harrison, David K. Jackson, Ian Johnston, Dominic Kwiatkowski, Cordelia Langford, John Sillitoe on behalf of the Wellcome Sanger Institute COVID-19 Surveillance Team |
| EPI_ISL_2847186 | Lighthouse Lab in Milton Keynes | Wellcome Sanger Institute for the COVID-19 Genomics UK (COG-UK) Consortium | The Lighthouse Lab in Milton Keynes and Alex Alderton, Roberto Amato, Jeffrey Barrett, Sonia Goncalves, Ewan Harrison, David K. Jackson, Ian Johnston, Dominic Kwiatkowski, Cordelia Langford, John Sillitoe on behalf of the Wellcome Sanger Institute COVID-19 Surveillance Team |
| EPI_ISL_2847187, EPI_ISL_2847188 | Health Services Laboratories | Wellcome Sanger Institute for the COVID-19 Genomics UK (COG-UK) Consortium | Health Services Laboratories and Alex Alderton, Roberto Amato, Jeffrey Barrett, Sonia Goncalves, Ewan Harrison, David K. Jackson, Ian Johnston, Dominic Kwiatkowski, Cordelia Langford, John Sillitoe on behalf of the Wellcome Sanger Institute COVID-19 Surveillance Team |
| EPI_ISL_2847189 | Lighthouse Lab in Milton Keynes | Wellcome Sanger Institute for the COVID-19 Genomics UK (COG-UK) Consortium | The Lighthouse Lab in Milton Keynes and Alex Alderton, Roberto Amato, Jeffrey Barrett, Sonia Goncalves, Ewan Harrison, David K. Jackson, Ian Johnston, Dominic Kwiatkowski, Cordelia Langford, John Sillitoe on behalf of the Wellcome Sanger Institute COVID-19 Surveillance Team |
| EPI_ISL_2847190 | Lighthouse Lab in Alderley Park | Wellcome Sanger Institute for the COVID-19 Genomics UK (COG-UK) Consortium | Jacquelyn Wynn, Mairead Hyland, The Lighthouse Lab in Alderley Park and Alex Alderton, Roberto Amato, Jeffrey Barrett, Sonia Goncalves, Ewan Harrison, David K. Jackson, Ian Johnston, Dominic Kwiatkowski, Cordelia Langford, John Sillitoe on behalf of the Wellcome Sanger Institute COVID-19 Surveillance Team |
| EPI_ISL_2847191 | Lighthouse Lab in Glasgow | Wellcome Sanger Institute for the COVID-19 Genomics UK (COG-UK) Consortium | Harper VanSteenhouse, Yumi Kasai, David Gray, Carol Clugston, Anna Dominiczak and Alex Alderton, Roberto Amato, Jeffrey Barrett, Sonia Goncalves, Ewan Harrison, David K. Jackson, Ian Johnston, Dominic Kwiatkowski, Cordelia Langford, John Sillitoe on behalf of the Wellcome Sanger Institute COVID-19 Surveillance Team |
| EPI_ISL_2847192 | Lighthouse Lab in Alderley Park | Wellcome Sanger Institute for the COVID-19 Genomics UK (COG-UK) Consortium | Jacquelyn Wynn, Mairead Hyland, The Lighthouse Lab in Alderley Park and Alex Alderton, Roberto Amato, Jeffrey Barrett, Sonia Goncalves, Ewan Harrison, David K. Jackson, Ian Johnston, Dominic Kwiatkowski, Cordelia Langford, John Sillitoe on behalf of the Wellcome Sanger Institute COVID-19 Surveillance Team |
| EPI_ISL_2847193 | Lighthouse Lab in Glasgow | Wellcome Sanger Institute for the COVID-19 Genomics UK (COG-UK) Consortium | Harper VanSteenhouse, Yumi Kasai, David Gray, Carol Clugston, Anna Dominiczak and Alex Alderton, Roberto Amato, Jeffrey Barrett, Sonia Goncalves, Ewan Harrison, David K. Jackson, Ian Johnston, Dominic Kwiatkowski, Cordelia Langford, John Sillitoe on behalf of the Wellcome Sanger Institute COVID-19 Surveillance Team |
| EPI_ISL_2847194 | Health Services Laboratories | Wellcome Sanger Institute for the COVID-19 Genomics UK (COG-UK) Consortium | Health Services Laboratories and Alex Alderton, Roberto Amato, Jeffrey Barrett, Sonia Goncalves, Ewan Harrison, David K. Jackson, Ian Johnston, Dominic Kwiatkowski, Cordelia Langford, John Sillitoe on behalf of the Wellcome Sanger Institute COVID-19 Surveillance Team |
| EPI_ISL_2847195, EPI_ISL_2847196, EPI_ISL_2847197 | Lighthouse Lab in Alderley Park | Wellcome Sanger Institute for the COVID-19 Genomics UK (COG-UK) Consortium | Jacquelyn Wynn, Mairead Hyland, The Lighthouse Lab in Alderley Park and Alex Alderton, Roberto Amato, Jeffrey Barrett, Sonia Goncalves, Ewan Harrison, David K. Jackson, Ian Johnston, Dominic Kwiatkowski, Cordelia Langford, John Sillitoe on behalf of the Wellcome Sanger Institute COVID-19 Surveillance Team |
| EPI_ISL_2847198 | Lighthouse Lab in Milton Keynes | Wellcome Sanger Institute for the COVID-19 Genomics UK (COG-UK) Consortium | The Lighthouse Lab in Milton Keynes and Alex Alderton, Roberto Amato, Jeffrey Barrett, Sonia Goncalves, Ewan Harrison, David K. Jackson, Ian Johnston, Dominic Kwiatkowski, Cordelia Langford, John Sillitoe on behalf of the Wellcome Sanger Institute COVID-19 Surveillance Team |
| EPI_ISL_2847199 | Lighthouse Lab in Alderley Park | Wellcome Sanger Institute for the COVID-19 Genomics UK (COG-UK) Consortium | Jacquelyn Wynn, Mairead Hyland, The Lighthouse Lab in Alderley Park and Alex Alderton, Roberto Amato, Jeffrey Barrett, Sonia Goncalves, Ewan Harrison, David K. Jackson, Ian Johnston, Dominic Kwiatkowski, Cordelia Langford, John Sillitoe on behalf of the Wellcome Sanger Institute COVID-19 Surveillance Team |
| EPI_ISL_2847200 | Lighthouse Lab in Glasgow | Wellcome Sanger Institute for the COVID-19 Genomics UK (COG-UK) Consortium | Harper VanSteenhouse, Yumi Kasai, David Gray, Carol Clugston, Anna Dominiczak and Alex Alderton, Roberto Amato, Jeffrey Barrett, Sonia Goncalves, Ewan Harrison, David K. Jackson, Ian Johnston, Dominic Kwiatkowski, Cordelia Langford, John Sillitoe on behalf of the Wellcome Sanger Institute COVID-19 Surveillance Team |
| EPI_ISL_2847201 | Health Services Laboratories | Wellcome Sanger Institute for the COVID-19 Genomics UK (COG-UK) Consortium | Health Services Laboratories and Alex Alderton, Roberto Amato, Jeffrey Barrett, Sonia Goncalves, Ewan Harrison, David K. Jackson, Ian Johnston, Dominic Kwiatkowski, Cordelia Langford, John Sillitoe on behalf of the Wellcome Sanger Institute COVID-19 Surveillance Team |
| EPI_ISL_2847202 | Lighthouse Lab in Glasgow | Wellcome Sanger Institute for the COVID-19 Genomics UK (COG-UK) Consortium | Harper VanSteenhouse, Yumi Kasai, David Gray, Carol Clugston, Anna Dominiczak and Alex Alderton, Roberto Amato, Jeffrey Barrett, Sonia Goncalves, Ewan Harrison, David K. Jackson, Ian Johnston, Dominic Kwiatkowski, Cordelia Langford, John Sillitoe on behalf of the Wellcome Sanger Institute COVID-19 Surveillance Team |
| EPI_ISL_2847203 | Lighthouse Lab in Milton Keynes | Wellcome Sanger Institute for the COVID-19 Genomics UK (COG-UK) Consortium | The Lighthouse Lab in Milton Keynes and Alex Alderton, Roberto Amato, Jeffrey Barrett, Sonia Goncalves, Ewan Harrison, David K. Jackson, Ian Johnston, Dominic Kwiatkowski, Cordelia Langford, John Sillitoe on behalf of the Wellcome Sanger Institute COVID-19 Surveillance Team |
| EPI_ISL_2847204 | Health Services Laboratories | Wellcome Sanger Institute for the COVID-19 Genomics UK (COG-UK) Consortium | Health Services Laboratories and Alex Alderton, Roberto Amato, Jeffrey Barrett, Sonia Goncalves, Ewan Harrison, David K. Jackson, Ian Johnston, Dominic Kwiatkowski, Cordelia Langford, John Sillitoe on behalf of the Wellcome Sanger Institute COVID-19 Surveillance Team |
| EPI_ISL_2847205, EPI_ISL_2847206, EPI_ISL_2847207 | Lighthouse Lab in Glasgow | Wellcome Sanger Institute for the COVID-19 Genomics UK (COG-UK) Consortium | Harper VanSteenhouse, Yumi Kasai, David Gray, Carol Clugston, Anna Dominiczak and Alex Alderton, Roberto Amato, Jeffrey Barrett, Sonia Goncalves, Ewan Harrison, David K. Jackson, Ian Johnston, Dominic Kwiatkowski, Cordelia Langford, John Sillitoe on behalf of the Wellcome Sanger Institute COVID-19 Surveillance Team |
| EPI_ISL_2847208 | Lighthouse Lab in Alderley Park | Wellcome Sanger Institute for the COVID-19 Genomics UK (COG-UK) Consortium | Jacquelyn Wynn, Mairead Hyland, The Lighthouse Lab in Alderley Park and Alex Alderton, Roberto Amato, Jeffrey Barrett, Sonia Goncalves, Ewan Harrison, David K. Jackson, Ian Johnston, Dominic Kwiatkowski, Cordelia Langford, John Sillitoe on behalf of the Wellcome Sanger Institute COVID-19 Surveillance Team |
| EPI_ISL_2847209, EPI_ISL_2847210 | Lighthouse Lab in Milton Keynes | Wellcome Sanger Institute for the COVID-19 Genomics UK (COG-UK) Consortium | The Lighthouse Lab in Milton Keynes and Alex Alderton, Roberto Amato, Jeffrey Barrett, Sonia Goncalves, Ewan Harrison, David K. Jackson, Ian Johnston, Dominic Kwiatkowski, Cordelia Langford, John Sillitoe on behalf of the Wellcome Sanger Institute COVID-19 Surveillance Team |
| EPI_ISL_2847211 | Lighthouse Lab in Glasgow | Wellcome Sanger Institute for the COVID-19 Genomics UK (COG-UK) Consortium | Harper VanSteenhouse, Yumi Kasai, David Gray, Carol Clugston, Anna Dominiczak and Alex Alderton, Roberto Amato, Jeffrey Barrett, Sonia Goncalves, Ewan Harrison, David K. Jackson, Ian Johnston, Dominic Kwiatkowski, Cordelia Langford, John Sillitoe on behalf of the Wellcome Sanger Institute COVID-19 Surveillance Team |
| EPI_ISL_2847212 | Lighthouse Lab in Alderley Park | Wellcome Sanger Institute for the COVID-19 Genomics UK (COG-UK) Consortium | Jacquelyn Wynn, Mairead Hyland, The Lighthouse Lab in Alderley Park and Alex Alderton, Roberto Amato, Jeffrey Barrett, Sonia Goncalves, Ewan Harrison, David K. Jackson, Ian Johnston, Dominic Kwiatkowski, Cordelia Langford, John Sillitoe on behalf of the Wellcome Sanger Institute COVID-19 Surveillance Team |
| EPI_ISL_2847213 | Health Services Laboratories | Wellcome Sanger Institute for the COVID-19 Genomics UK (COG-UK) Consortium | Health Services Laboratories and Alex Alderton, Roberto Amato, Jeffrey Barrett, Sonia Goncalves, Ewan Harrison, David K. Jackson, Ian Johnston, Dominic Kwiatkowski, Cordelia Langford, John Sillitoe on behalf of the Wellcome Sanger Institute COVID-19 Surveillance Team |
| EPI_ISL_2847214, EPI_ISL_2847215 | Lighthouse Lab in Glasgow | Wellcome Sanger Institute for the COVID-19 Genomics UK (COG-UK) Consortium | Harper VanSteenhouse, Yumi Kasai, David Gray, Carol Clugston, Anna Dominiczak and Alex Alderton, Roberto Amato, Jeffrey Barrett, Sonia Goncalves, Ewan Harrison, David K. Jackson, Ian Johnston, Dominic Kwiatkowski, Cordelia Langford, John Sillitoe on behalf of the Wellcome Sanger Institute COVID-19 Surveillance Team |
| EPI_ISL_2847216 | Health Services Laboratories | Wellcome Sanger Institute for the COVID-19 Genomics UK (COG-UK) Consortium | Health Services Laboratories and Alex Alderton, Roberto Amato, Jeffrey Barrett, Sonia Goncalves, Ewan Harrison, David K. Jackson, Ian Johnston, Dominic Kwiatkowski, Cordelia Langford, John Sillitoe on behalf of the Wellcome Sanger Institute COVID-19 Surveillance Team |
| EPI_ISL_2847217 | Lighthouse Lab in Alderley Park | Wellcome Sanger Institute for the COVID-19 Genomics UK (COG-UK) Consortium | Jacquelyn Wynn, Mairead Hyland, The Lighthouse Lab in Alderley Park and Alex Alderton, Roberto Amato, Jeffrey Barrett, Sonia Goncalves, Ewan Harrison, David K. Jackson, Ian Johnston, Dominic Kwiatkowski, Cordelia Langford, John Sillitoe on behalf of the Wellcome Sanger Institute COVID-19 Surveillance Team |
| EPI_ISL_2847219, EPI_ISL_2847220 | Lighthouse Lab in Milton Keynes | Wellcome Sanger Institute for the COVID-19 Genomics UK (COG-UK) Consortium | The Lighthouse Lab in Milton Keynes and Alex Alderton, Roberto Amato, Jeffrey Barrett, Sonia Goncalves, Ewan Harrison, David K. Jackson, Ian Johnston, Dominic Kwiatkowski, Cordelia Langford, John Sillitoe on behalf of the Wellcome Sanger Institute COVID-19 Surveillance Team |
| EPI_ISL_2847221, EPI_ISL_2847222 | Lighthouse Lab in Alderley Park | Wellcome Sanger Institute for the COVID-19 Genomics UK (COG-UK) Consortium | Jacquelyn Wynn, Mairead Hyland, The Lighthouse Lab in Alderley Park and Alex Alderton, Roberto Amato, Jeffrey Barrett, Sonia Goncalves, Ewan Harrison, David K. Jackson, Ian Johnston, Dominic Kwiatkowski, Cordelia Langford, John Sillitoe on behalf of the Wellcome Sanger Institute COVID-19 Surveillance Team |
| EPI_ISL_2847223 | Lighthouse Lab in Glasgow | Wellcome Sanger Institute for the COVID-19 Genomics UK (COG-UK) Consortium | Harper VanSteenhouse, Yumi Kasai, David Gray, Carol Clugston, Anna Dominiczak and Alex Alderton, Roberto Amato, Jeffrey Barrett, Sonia Goncalves, Ewan Harrison, David K. Jackson, Ian Johnston, Dominic Kwiatkowski, Cordelia Langford, John Sillitoe on behalf of the Wellcome Sanger Institute COVID-19 Surveillance Team |
| EPI_ISL_2847224 | Lighthouse Lab in Alderley Park | Wellcome Sanger Institute for the COVID-19 Genomics UK (COG-UK) Consortium | Jacquelyn Wynn, Mairead Hyland, The Lighthouse Lab in Alderley Park and Alex Alderton, Roberto Amato, Jeffrey Barrett, Sonia Goncalves, Ewan Harrison, David K. Jackson, Ian Johnston, Dominic Kwiatkowski, Cordelia Langford, John Sillitoe on behalf of the Wellcome Sanger Institute COVID-19 Surveillance Team |
| EPI_ISL_2847225, EPI_ISL_2847226 | Lighthouse Lab in Glasgow | Wellcome Sanger Institute for the COVID-19 Genomics UK (COG-UK) Consortium | Harper VanSteenhouse, Yumi Kasai, David Gray, Carol Clugston, Anna Dominiczak and Alex Alderton, Roberto Amato, Jeffrey Barrett, Sonia Goncalves, Ewan Harrison, David K. Jackson, Ian Johnston, Dominic Kwiatkowski, Cordelia Langford, John Sillitoe on behalf of the Wellcome Sanger Institute COVID-19 Surveillance Team |

| | | | |
|---|---|---|---|
| EPI_ISL_2847227 | Lighthouse Lab in Alderley Park | Wellcome Sanger Institute for the COVID-19 Genomics UK (COG-UK) Consortium | Jacquelyn Wynn, Mairead Hyland, The Lighthouse Lab in Alderley Park and Alex Alderton, Roberto Amato, Jeffrey Barrett, Sonia Goncalves, Ewan Harrison, David K. Jackson, Ian Johnston, Dominic Kwiatkowski, Cordelia Langford, John Sillitoe on behalf of the Wellcome Sanger Institute COVID-19 Surveillance Team |
| EPI_ISL_2847228, EPI_ISL_2847229 | Health Services Laboratories | Wellcome Sanger Institute for the COVID-19 Genomics UK (COG-UK) Consortium | Health Services Laboratories and Alex Alderton, Roberto Amato, Jeffrey Barrett, Sonia Goncalves, Ewan Harrison, David K. Jackson, Ian Johnston, Dominic Kwiatkowski, Cordelia Langford, John Sillitoe on behalf of the Wellcome Sanger Institute COVID-19 Surveillance Team |
| EPI_ISL_2847230 | Lighthouse Lab in Milton Keynes | Wellcome Sanger Institute for the COVID-19 Genomics UK (COG-UK) Consortium | The Lighthouse Lab in Milton Keynes and Alex Alderton, Roberto Amato, Jeffrey Barrett, Sonia Goncalves, Ewan Harrison, David K. Jackson, Ian Johnston, Dominic Kwiatkowski, Cordelia Langford, John Sillitoe on behalf of the Wellcome Sanger Institute COVID-19 Surveillance Team |
| EPI_ISL_2847231, EPI_ISL_2847232 | Lighthouse Lab in Glasgow | Wellcome Sanger Institute for the COVID-19 Genomics UK (COG-UK) Consortium | Harper VanSteenhouse, Yumi Kasai, David Gray, Carol Clugston, Anna Dominiczak and Alex Alderton, Roberto Amato, Jeffrey Barrett, Sonia Goncalves, Ewan Harrison, David K. Jackson, Ian Johnston, Dominic Kwiatkowski, Cordelia Langford, John Sillitoe on behalf of the Wellcome Sanger Institute COVID-19 Surveillance Team |
| EPI_ISL_2847233 | Lighthouse Lab in Alderley Park | Wellcome Sanger Institute for the COVID-19 Genomics UK (COG-UK) Consortium | Jacquelyn Wynn, Mairead Hyland, The Lighthouse Lab in Alderley Park and Alex Alderton, Roberto Amato, Jeffrey Barrett, Sonia Goncalves, Ewan Harrison, David K. Jackson, Ian Johnston, Dominic Kwiatkowski, Cordelia Langford, John Sillitoe on behalf of the Wellcome Sanger Institute COVID-19 Surveillance Team |
| EPI_ISL_2847234 | Lighthouse Lab in Milton Keynes | Wellcome Sanger Institute for the COVID-19 Genomics UK (COG-UK) Consortium | The Lighthouse Lab in Milton Keynes and Alex Alderton, Roberto Amato, Jeffrey Barrett, Sonia Goncalves, Ewan Harrison, David K. Jackson, Ian Johnston, Dominic Kwiatkowski, Cordelia Langford, John Sillitoe on behalf of the Wellcome Sanger Institute COVID-19 Surveillance Team |
| EPI_ISL_2847235 | Lighthouse Lab in Alderley Park | Wellcome Sanger Institute for the COVID-19 Genomics UK (COG-UK) Consortium | Jacquelyn Wynn, Mairead Hyland, The Lighthouse Lab in Alderley Park and Alex Alderton, Roberto Amato, Jeffrey Barrett, Sonia Goncalves, Ewan Harrison, David K. Jackson, Ian Johnston, Dominic Kwiatkowski, Cordelia Langford, John Sillitoe on behalf of the Wellcome Sanger Institute COVID-19 Surveillance Team |
| EPI_ISL_2847236 | Lighthouse Lab in Milton Keynes | Wellcome Sanger Institute for the COVID-19 Genomics UK (COG-UK) Consortium | The Lighthouse Lab in Milton Keynes and Alex Alderton, Roberto Amato, Jeffrey Barrett, Sonia Goncalves, Ewan Harrison, David K. Jackson, Ian Johnston, Dominic Kwiatkowski, Cordelia Langford, John Sillitoe on behalf of the Wellcome Sanger Institute COVID-19 Surveillance Team |
| EPI_ISL_2847237, EPI_ISL_2847238, EPI_ISL_2847239 | Health Services Laboratories | Wellcome Sanger Institute for the COVID-19 Genomics UK (COG-UK) Consortium | Health Services Laboratories and Alex Alderton, Roberto Amato, Jeffrey Barrett, Sonia Goncalves, Ewan Harrison, David K. Jackson, Ian Johnston, Dominic Kwiatkowski, Cordelia Langford, John Sillitoe on behalf of the Wellcome Sanger Institute COVID-19 Surveillance Team |
| EPI_ISL_2847240 | Lighthouse Lab in Alderley Park | Wellcome Sanger Institute for the COVID-19 Genomics UK (COG-UK) Consortium | Jacquelyn Wynn, Mairead Hyland, The Lighthouse Lab in Alderley Park and Alex Alderton, Roberto Amato, Jeffrey Barrett, Sonia Goncalves, Ewan Harrison, David K. Jackson, Ian Johnston, Dominic Kwiatkowski, Cordelia Langford, John Sillitoe on behalf of the Wellcome Sanger Institute COVID-19 Surveillance Team |
| EPI_ISL_2847241, EPI_ISL_2847242, EPI_ISL_2847243 | Lighthouse Lab in Milton Keynes | Wellcome Sanger Institute for the COVID-19 Genomics UK (COG-UK) Consortium | The Lighthouse Lab in Milton Keynes and Alex Alderton, Roberto Amato, Jeffrey Barrett, Sonia Goncalves, Ewan Harrison, David K. Jackson, Ian Johnston, Dominic Kwiatkowski, Cordelia Langford, John Sillitoe on behalf of the Wellcome Sanger Institute COVID-19 Surveillance Team |
| EPI_ISL_2847244, EPI_ISL_2847245 | Lighthouse Lab in Glasgow | Wellcome Sanger Institute for the COVID-19 Genomics UK (COG-UK) Consortium | Harper VanSteenhouse, Yumi Kasai, David Gray, Carol Clugston, Anna Dominiczak and Alex Alderton, Roberto Amato, Jeffrey Barrett, Sonia Goncalves, Ewan Harrison, David K. Jackson, Ian Johnston, Dominic Kwiatkowski, Cordelia Langford, John Sillitoe on behalf of the Wellcome Sanger Institute COVID-19 Surveillance Team |
| EPI_ISL_2847246 | Health Services Laboratories | Wellcome Sanger Institute for the COVID-19 Genomics UK (COG-UK) Consortium | Health Services Laboratories and Alex Alderton, Roberto Amato, Jeffrey Barrett, Sonia Goncalves, Ewan Harrison, David K. Jackson, Ian Johnston, Dominic Kwiatkowski, Cordelia Langford, John Sillitoe on behalf of the Wellcome Sanger Institute COVID-19 Surveillance Team |
| EPI_ISL_2847247 | Lighthouse Lab in Alderley Park | Wellcome Sanger Institute for the COVID-19 Genomics UK (COG-UK) Consortium | Jacquelyn Wynn, Mairead Hyland, The Lighthouse Lab in Alderley Park and Alex Alderton, Roberto Amato, Jeffrey Barrett, Sonia Goncalves, Ewan Harrison, David K. Jackson, Ian Johnston, Dominic Kwiatkowski, Cordelia Langford, John Sillitoe on behalf of the Wellcome Sanger Institute COVID-19 Surveillance Team |
| EPI_ISL_2847248 | Health Services Laboratories | Wellcome Sanger Institute for the COVID-19 Genomics UK (COG-UK) Consortium | Health Services Laboratories and Alex Alderton, Roberto Amato, Jeffrey Barrett, Sonia Goncalves, Ewan Harrison, David K. Jackson, Ian Johnston, Dominic Kwiatkowski, Cordelia Langford, John Sillitoe on behalf of the Wellcome Sanger Institute COVID-19 Surveillance Team |
| EPI_ISL_2847249 | Lighthouse Lab in Alderley Park | Wellcome Sanger Institute for the COVID-19 Genomics UK (COG-UK) Consortium | Jacquelyn Wynn, Mairead Hyland, The Lighthouse Lab in Alderley Park and Alex Alderton, Roberto Amato, Jeffrey Barrett, Sonia Goncalves, Ewan Harrison, David K. Jackson, Ian Johnston, Dominic Kwiatkowski, Cordelia Langford, John Sillitoe on behalf of the Wellcome Sanger Institute COVID-19 Surveillance Team |
| EPI_ISL_2847250 | Health Services Laboratories | Wellcome Sanger Institute for the COVID-19 Genomics UK (COG-UK) Consortium | Health Services Laboratories and Alex Alderton, Roberto Amato, Jeffrey Barrett, Sonia Goncalves, Ewan Harrison, David K. Jackson, Ian Johnston, Dominic Kwiatkowski, Cordelia Langford, John Sillitoe on behalf of the Wellcome Sanger Institute COVID-19 Surveillance Team |
| EPI_ISL_2847251 | Lighthouse Lab in Alderley Park | Wellcome Sanger Institute for the COVID-19 Genomics UK (COG-UK) Consortium | Jacquelyn Wynn, Mairead Hyland, The Lighthouse Lab in Alderley Park and Alex Alderton, Roberto Amato, Jeffrey Barrett, Sonia Goncalves, Ewan Harrison, David K. Jackson, Ian Johnston, Dominic Kwiatkowski, Cordelia Langford, John Sillitoe on behalf of the Wellcome Sanger Institute COVID-19 Surveillance Team |
| EPI_ISL_2847252 | Lighthouse Lab in Glasgow | Wellcome Sanger Institute for the COVID-19 Genomics UK (COG-UK) Consortium | Harper VanSteenhouse, Yumi Kasai, David Gray, Carol Clugston, Anna Dominiczak and Alex Alderton, Roberto Amato, Jeffrey Barrett, Sonia Goncalves, Ewan Harrison, David K. Jackson, Ian Johnston, Dominic Kwiatkowski, Cordelia Langford, John Sillitoe on behalf of the Wellcome Sanger Institute COVID-19 Surveillance Team |
| EPI_ISL_2847253 | Lighthouse Lab in Alderley Park | Wellcome Sanger Institute for the COVID-19 Genomics UK (COG-UK) Consortium | Jacquelyn Wynn, Mairead Hyland, The Lighthouse Lab in Alderley Park and Alex Alderton, Roberto Amato, Jeffrey Barrett, Sonia Goncalves, Ewan Harrison, David K. Jackson, Ian Johnston, Dominic Kwiatkowski, Cordelia Langford, John Sillitoe on behalf of the Wellcome Sanger Institute COVID-19 Surveillance Team |
| EPI_ISL_2847254 | Lighthouse Lab in Milton Keynes | Wellcome Sanger Institute for the COVID-19 Genomics UK (COG-UK) Consortium | The Lighthouse Lab in Milton Keynes and Alex Alderton, Roberto Amato, Jeffrey Barrett, Sonia Goncalves, Ewan Harrison, David K. Jackson, Ian Johnston, Dominic Kwiatkowski, Cordelia Langford, John Sillitoe on behalf of the Wellcome Sanger Institute COVID-19 Surveillance Team |
| EPI_ISL_2847255 | Lighthouse Lab in Glasgow | Wellcome Sanger Institute for the COVID-19 Genomics UK (COG-UK) Consortium | Harper VanSteenhouse, Yumi Kasai, David Gray, Carol Clugston, Anna Dominiczak and Alex Alderton, Roberto Amato, Jeffrey Barrett, Sonia Goncalves, Ewan Harrison, David K. Jackson, Ian Johnston, Dominic Kwiatkowski, Cordelia Langford, John Sillitoe on behalf of the Wellcome Sanger Institute COVID-19 Surveillance Team |
| EPI_ISL_2847256, EPI_ISL_2847257, EPI_ISL_2847258 | Lighthouse Lab in Alderley Park | Wellcome Sanger Institute for the COVID-19 Genomics UK (COG-UK) Consortium | Jacquelyn Wynn, Mairead Hyland, The Lighthouse Lab in Alderley Park and Alex Alderton, Roberto Amato, Jeffrey Barrett, Sonia Goncalves, Ewan Harrison, David K. Jackson, Ian Johnston, Dominic Kwiatkowski, Cordelia Langford, John Sillitoe on behalf of the Wellcome Sanger Institute COVID-19 Surveillance Team |
| EPI_ISL_2847259 | Lighthouse Lab in Milton Keynes | Wellcome Sanger Institute for the COVID-19 Genomics UK (COG-UK) Consortium | The Lighthouse Lab in Milton Keynes and Alex Alderton, Roberto Amato, Jeffrey Barrett, Sonia Goncalves, Ewan Harrison, David K. Jackson, Ian Johnston, Dominic Kwiatkowski, Cordelia Langford, John Sillitoe on behalf of the Wellcome Sanger Institute COVID-19 Surveillance Team |
| EPI_ISL_2847260 | Lighthouse Lab in Alderley Park | Wellcome Sanger Institute for the COVID-19 Genomics UK (COG-UK) Consortium | Jacquelyn Wynn, Mairead Hyland, The Lighthouse Lab in Alderley Park and Alex Alderton, Roberto Amato, Jeffrey Barrett, Sonia Goncalves, Ewan Harrison, David K. Jackson, Ian Johnston, Dominic Kwiatkowski, Cordelia Langford, John Sillitoe on behalf of the Wellcome Sanger Institute COVID-19 Surveillance Team |
| EPI_ISL_2847261 | Health Services Laboratories | Wellcome Sanger Institute for the COVID-19 Genomics UK (COG-UK) Consortium | Health Services Laboratories and Alex Alderton, Roberto Amato, Jeffrey Barrett, Sonia Goncalves, Ewan Harrison, David K. Jackson, Ian Johnston, Dominic Kwiatkowski, Cordelia Langford, John Sillitoe on behalf of the Wellcome Sanger Institute COVID-19 Surveillance Team |
| EPI_ISL_2847262, EPI_ISL_2847263 | Lighthouse Lab in Glasgow | Wellcome Sanger Institute for the COVID-19 Genomics UK (COG-UK) Consortium | Harper VanSteenhouse, Yumi Kasai, David Gray, Carol Clugston, Anna Dominiczak and Alex Alderton, Roberto Amato, Jeffrey Barrett, Sonia Goncalves, Ewan Harrison, David K. Jackson, Ian Johnston, Dominic Kwiatkowski, Cordelia Langford, John Sillitoe on behalf of the Wellcome Sanger Institute COVID-19 Surveillance Team |
| EPI_ISL_2847264 | Lighthouse Lab in Milton Keynes | Wellcome Sanger Institute for the COVID-19 Genomics UK (COG-UK) Consortium | The Lighthouse Lab in Milton Keynes and Alex Alderton, Roberto Amato, Jeffrey Barrett, Sonia Goncalves, Ewan Harrison, David K. Jackson, Ian Johnston, Dominic Kwiatkowski, Cordelia Langford, John Sillitoe on behalf of the Wellcome Sanger Institute COVID-19 Surveillance Team |
| EPI_ISL_2847265 | Health Services Laboratories | Wellcome Sanger Institute for the COVID-19 Genomics UK (COG-UK) Consortium | Health Services Laboratories and Alex Alderton, Roberto Amato, Jeffrey Barrett, Sonia Goncalves, Ewan Harrison, David K. Jackson, Ian Johnston, Dominic Kwiatkowski, Cordelia Langford, John Sillitoe on behalf of the Wellcome Sanger Institute COVID-19 Surveillance Team |
| EPI_ISL_2847266, EPI_ISL_2847267 | Lighthouse Lab in Glasgow | Wellcome Sanger Institute for the COVID-19 Genomics UK (COG-UK) Consortium | Harper VanSteenhouse, Yumi Kasai, David Gray, Carol Clugston, Anna Dominiczak and Alex Alderton, Roberto Amato, Jeffrey Barrett, Sonia Goncalves, Ewan Harrison, David K. Jackson, Ian Johnston, Dominic Kwiatkowski, Cordelia Langford, John Sillitoe on behalf of the Wellcome Sanger Institute COVID-19 Surveillance Team |
| EPI_ISL_2847268, EPI_ISL_2847269 | Lighthouse Lab in Milton Keynes | Wellcome Sanger Institute for the COVID-19 Genomics UK (COG-UK) Consortium | The Lighthouse Lab in Milton Keynes and Alex Alderton, Roberto Amato, Jeffrey Barrett, Sonia Goncalves, Ewan Harrison, David K. Jackson, Ian Johnston, Dominic Kwiatkowski, Cordelia Langford, John Sillitoe on behalf of the Wellcome Sanger Institute COVID-19 Surveillance Team |
| EPI_ISL_2847270 | Health Services Laboratories | Wellcome Sanger Institute for the COVID-19 Genomics UK (COG-UK) Consortium | Health Services Laboratories and Alex Alderton, Roberto Amato, Jeffrey Barrett, Sonia Goncalves, Ewan Harrison, David K. Jackson, Ian Johnston, Dominic Kwiatkowski, Cordelia Langford, John Sillitoe on behalf of the Wellcome Sanger Institute COVID-19 Surveillance Team |
| EPI_ISL_2847271 | Lighthouse Lab in Glasgow | Wellcome Sanger Institute for the COVID-19 Genomics UK (COG-UK) Consortium | Harper VanSteenhouse, Yumi Kasai, David Gray, Carol Clugston, Anna Dominiczak and Alex Alderton, Roberto Amato, Jeffrey Barrett, Sonia Goncalves, Ewan Harrison, David K. Jackson, Ian Johnston, Dominic Kwiatkowski, Cordelia Langford, John Sillitoe on behalf of the Wellcome Sanger Institute COVID-19 Surveillance Team |
| EPI_ISL_2847272 | Lighthouse Lab in Alderley Park | Wellcome Sanger Institute for the COVID-19 Genomics UK (COG-UK) Consortium | Jacquelyn Wynn, Mairead Hyland, The Lighthouse Lab in Alderley Park and Alex Alderton, Roberto Amato, Jeffrey Barrett, Sonia Goncalves, Ewan Harrison, David K. Jackson, Ian Johnston, Dominic Kwiatkowski, Cordelia Langford, John Sillitoe on behalf of the Wellcome Sanger Institute COVID-19 Surveillance Team |
| EPI_ISL_2847273 | Health Services Laboratories | Wellcome Sanger Institute for the COVID-19 Genomics UK (COG-UK) Consortium | Health Services Laboratories and Alex Alderton, Roberto Amato, Jeffrey Barrett, Sonia Goncalves, Ewan Harrison, David K. Jackson, Ian Johnston, Dominic Kwiatkowski, Cordelia Langford, John Sillitoe on behalf of the Wellcome Sanger Institute COVID-19 Surveillance Team |
| EPI_ISL_2847274 | Lighthouse Lab in Milton Keynes | Wellcome Sanger Institute for the COVID-19 Genomics UK (COG-UK) Consortium | The Lighthouse Lab in Milton Keynes and Alex Alderton, Roberto Amato, Jeffrey Barrett, Sonia Goncalves, Ewan Harrison, David K. Jackson, Ian Johnston, Dominic Kwiatkowski, Cordelia Langford, John Sillitoe on behalf of the Wellcome Sanger Institute COVID-19 Surveillance Team |
| EPI_ISL_2847275 | Health Services Laboratories | Wellcome Sanger Institute for the COVID-19 Genomics UK (COG-UK) Consortium | Health Services Laboratories and Alex Alderton, Roberto Amato, Jeffrey Barrett, Sonia Goncalves, Ewan Harrison, David K. Jackson, Ian Johnston, Dominic Kwiatkowski, Cordelia Langford, John Sillitoe on behalf of the Wellcome Sanger Institute COVID-19 Surveillance Team |
| EPI_ISL_2847276 | Lighthouse Lab in Milton Keynes | Wellcome Sanger Institute for the COVID-19 Genomics UK (COG-UK) Consortium | The Lighthouse Lab in Milton Keynes and Alex Alderton, Roberto Amato, Jeffrey Barrett, Sonia Goncalves, Ewan Harrison, David K. Jackson, Ian Johnston, Dominic Kwiatkowski, Cordelia Langford, John Sillitoe on behalf of the Wellcome Sanger Institute COVID-19 Surveillance Team |
| EPI_ISL_2847277 | Lighthouse Lab in Glasgow | Wellcome Sanger Institute for the COVID-19 Genomics UK (COG-UK) Consortium | Harper VanSteenhouse, Yumi Kasai, David Gray, Carol Clugston, Anna Dominiczak and Alex Alderton, Roberto Amato, Jeffrey Barrett, Sonia Goncalves, Ewan Harrison, David K. Jackson, Ian Johnston, Dominic Kwiatkowski, Cordelia Langford, John Sillitoe on behalf of the Wellcome Sanger Institute COVID-19 Surveillance Team |
| EPI_ISL_2847278 | Lighthouse Lab in Alderley Park | Wellcome Sanger Institute for the COVID-19 Genomics UK (COG-UK) Consortium | Jacquelyn Wynn, Mairead Hyland, The Lighthouse Lab in Alderley Park and Alex Alderton, Roberto Amato, Jeffrey Barrett, Sonia Goncalves, Ewan Harrison, David K. Jackson, Ian Johnston, Dominic Kwiatkowski, Cordelia Langford, John Sillitoe on behalf of the Wellcome Sanger Institute COVID-19 Surveillance Team |
| EPI_ISL_2847279 | Lighthouse Lab in Milton Keynes | Wellcome Sanger Institute for the COVID-19 Genomics UK (COG-UK) Consortium | The Lighthouse Lab in Milton Keynes and Alex Alderton, Roberto Amato, Jeffrey Barrett, Sonia Goncalves, Ewan Harrison, David K. Jackson, Ian Johnston, Dominic Kwiatkowski, Cordelia Langford, John Sillitoe on behalf of the Wellcome Sanger Institute COVID-19 Surveillance Team |
| EPI_ISL_2847280 | Lighthouse Lab in Alderley Park | Wellcome Sanger Institute for the COVID-19 Genomics UK (COG-UK) Consortium | Jacquelyn Wynn, Mairead Hyland, The Lighthouse Lab in Alderley Park and Alex Alderton, Roberto Amato, Jeffrey Barrett, Sonia Goncalves, Ewan Harrison, David K. Jackson, Ian Johnston, Dominic Kwiatkowski, Cordelia Langford, John Sillitoe on behalf of the Wellcome Sanger Institute COVID-19 Surveillance Team |
| EPI_ISL_2847281 | Health Services Laboratories | Wellcome Sanger Institute for the COVID-19 Genomics UK (COG-UK) Consortium | Health Services Laboratories and Alex Alderton, Roberto Amato, Jeffrey Barrett, Sonia Goncalves, Ewan Harrison, David K. Jackson, Ian Johnston, Dominic Kwiatkowski, Cordelia Langford, John Sillitoe on behalf of the Wellcome Sanger Institute COVID-19 Surveillance Team |
| EPI_ISL_2847282 | Lighthouse Lab in Glasgow | Wellcome Sanger Institute for the COVID-19 Genomics UK (COG-UK) Consortium | Harper VanSteenhouse, Yumi Kasai, David Gray, Carol Clugston, Anna Dominiczak and Alex Alderton, Roberto Amato, Jeffrey Barrett, Sonia Goncalves, Ewan Harrison, David K. Jackson, Ian Johnston, Dominic Kwiatkowski, Cordelia Langford, John Sillitoe on behalf of the Wellcome Sanger Institute COVID-19 Surveillance Team |

| | | | |
|---|---|---|---|
| EPI_ISL_2847283 | Health Services Laboratories | Wellcome Sanger Institute for the COVID-19 Genomics UK (COG-UK) Consortium | Health Services Laboratories and Alex Alderton, Roberto Amato, Jeffrey Barrett, Sonia Goncalves, Ewan Harrison, David K. Jackson, Ian Johnston, Dominic Kwiatkowski, Cordelia Langford, John Sillitoe on behalf of the Wellcome Sanger Institute COVID-19 Surveillance Team |
| EPI_ISL_2847284 | Lighthouse Lab in Glasgow | Wellcome Sanger Institute for the COVID-19 Genomics UK (COG-UK) Consortium | Harper VanSteenhouse, Yumi Kasai, David Gray, Carol Clugston, Anna Dominiczak and Alex Alderton, Roberto Amato, Jeffrey Barrett, Sonia Goncalves, Ewan Harrison, David K. Jackson, Ian Johnston, Dominic Kwiatkowski, Cordelia Langford, John Sillitoe on behalf of the Wellcome Sanger Institute COVID-19 Surveillance Team |
| EPI_ISL_2847285 | Lighthouse Laboratory Plymouth | Wellcome Sanger Institute for the COVID-19 Genomics UK (COG-UK) Consortium | Lighthouse Laboratory Plymouth and Alex Alderton, Roberto Amato, Jeffrey Barrett, Sonia Goncalves, Ewan Harrison, David K. Jackson, Ian Johnston, Dominic Kwiatkowski, Cordelia Langford, John Sillitoe on behalf of the Wellcome Sanger Institute COVID-19 Surveillance Team |
| EPI_ISL_2847286, EPI_ISL_2847287 | Lighthouse Lab in Milton Keynes | Wellcome Sanger Institute for the COVID-19 Genomics UK (COG-UK) Consortium | The Lighthouse Lab in Milton Keynes and Alex Alderton, Roberto Amato, Jeffrey Barrett, Sonia Goncalves, Ewan Harrison, David K. Jackson, Ian Johnston, Dominic Kwiatkowski, Cordelia Langford, John Sillitoe on behalf of the Wellcome Sanger Institute COVID-19 Surveillance Team |
| EPI_ISL_2847288 | Lighthouse Laboratory Plymouth | Wellcome Sanger Institute for the COVID-19 Genomics UK (COG-UK) Consortium | Lighthouse Laboratory Plymouth and Alex Alderton, Roberto Amato, Jeffrey Barrett, Sonia Goncalves, Ewan Harrison, David K. Jackson, Ian Johnston, Dominic Kwiatkowski, Cordelia Langford, John Sillitoe on behalf of the Wellcome Sanger Institute COVID-19 Surveillance Team |
| EPI_ISL_2847289 | Lighthouse Lab in Glasgow | Wellcome Sanger Institute for the COVID-19 Genomics UK (COG-UK) Consortium | Harper VanSteenhouse, Yumi Kasai, David Gray, Carol Clugston, Anna Dominiczak and Alex Alderton, Roberto Amato, Jeffrey Barrett, Sonia Goncalves, Ewan Harrison, David K. Jackson, Ian Johnston, Dominic Kwiatkowski, Cordelia Langford, John Sillitoe on behalf of the Wellcome Sanger Institute COVID-19 Surveillance Team |
| EPI_ISL_2847290 | Lighthouse Lab in Milton Keynes | Wellcome Sanger Institute for the COVID-19 Genomics UK (COG-UK) Consortium | The Lighthouse Lab in Milton Keynes and Alex Alderton, Roberto Amato, Jeffrey Barrett, Sonia Goncalves, Ewan Harrison, David K. Jackson, Ian Johnston, Dominic Kwiatkowski, Cordelia Langford, John Sillitoe on behalf of the Wellcome Sanger Institute COVID-19 Surveillance Team |
| EPI_ISL_2847291 | Lighthouse Lab in Alderley Park | Wellcome Sanger Institute for the COVID-19 Genomics UK (COG-UK) Consortium | Jacquelyn Wynn, Mairead Hyland, The Lighthouse Lab in Alderley Park and Alex Alderton, Roberto Amato, Jeffrey Barrett, Sonia Goncalves, Ewan Harrison, David K. Jackson, Ian Johnston, Dominic Kwiatkowski, Cordelia Langford, John Sillitoe on behalf of the Wellcome Sanger Institute COVID-19 Surveillance Team |
| EPI_ISL_2847292, EPI_ISL_2847293, EPI_ISL_2847294 | Lighthouse Lab in Milton Keynes | Wellcome Sanger Institute for the COVID-19 Genomics UK (COG-UK) Consortium | The Lighthouse Lab in Milton Keynes and Alex Alderton, Roberto Amato, Jeffrey Barrett, Sonia Goncalves, Ewan Harrison, David K. Jackson, Ian Johnston, Dominic Kwiatkowski, Cordelia Langford, John Sillitoe on behalf of the Wellcome Sanger Institute COVID-19 Surveillance Team |
| EPI_ISL_2847295 | Lighthouse Lab in Glasgow | Wellcome Sanger Institute for the COVID-19 Genomics UK (COG-UK) Consortium | Harper VanSteenhouse, Yumi Kasai, David Gray, Carol Clugston, Anna Dominiczak and Alex Alderton, Roberto Amato, Jeffrey Barrett, Sonia Goncalves, Ewan Harrison, David K. Jackson, Ian Johnston, Dominic Kwiatkowski, Cordelia Langford, John Sillitoe on behalf of the Wellcome Sanger Institute COVID-19 Surveillance Team |
| EPI_ISL_2847297 | Lighthouse Laboratory Plymouth | Wellcome Sanger Institute for the COVID-19 Genomics UK (COG-UK) Consortium | Lighthouse Laboratory Plymouth and Alex Alderton, Roberto Amato, Jeffrey Barrett, Sonia Goncalves, Ewan Harrison, David K. Jackson, Ian Johnston, Dominic Kwiatkowski, Cordelia Langford, John Sillitoe on behalf of the Wellcome Sanger Institute COVID-19 Surveillance Team |
| EPI_ISL_2847298 | Lighthouse Lab in Milton Keynes | Wellcome Sanger Institute for the COVID-19 Genomics UK (COG-UK) Consortium | The Lighthouse Lab in Milton Keynes and Alex Alderton, Roberto Amato, Jeffrey Barrett, Sonia Goncalves, Ewan Harrison, David K. Jackson, Ian Johnston, Dominic Kwiatkowski, Cordelia Langford, John Sillitoe on behalf of the Wellcome Sanger Institute COVID-19 Surveillance Team |
| EPI_ISL_2847299 | Lighthouse Lab in Glasgow | Wellcome Sanger Institute for the COVID-19 Genomics UK (COG-UK) Consortium | Harper VanSteenhouse, Yumi Kasai, David Gray, Carol Clugston, Anna Dominiczak and Alex Alderton, Roberto Amato, Jeffrey Barrett, Sonia Goncalves, Ewan Harrison, David K. Jackson, Ian Johnston, Dominic Kwiatkowski, Cordelia Langford, John Sillitoe on behalf of the Wellcome Sanger Institute COVID-19 Surveillance Team |
| EPI_ISL_2847300, EPI_ISL_2847301 | Lighthouse Lab in Milton Keynes | Wellcome Sanger Institute for the COVID-19 Genomics UK (COG-UK) Consortium | The Lighthouse Lab in Milton Keynes and Alex Alderton, Roberto Amato, Jeffrey Barrett, Sonia Goncalves, Ewan Harrison, David K. Jackson, Ian Johnston, Dominic Kwiatkowski, Cordelia Langford, John Sillitoe on behalf of the Wellcome Sanger Institute COVID-19 Surveillance Team |
| EPI_ISL_2847302, EPI_ISL_2847303 | Lighthouse Lab in Glasgow | Wellcome Sanger Institute for the COVID-19 Genomics UK (COG-UK) Consortium | Harper VanSteenhouse, Yumi Kasai, David Gray, Carol Clugston, Anna Dominiczak and Alex Alderton, Roberto Amato, Jeffrey Barrett, Sonia Goncalves, Ewan Harrison, David K. Jackson, Ian Johnston, Dominic Kwiatkowski, Cordelia Langford, John Sillitoe on behalf of the Wellcome Sanger Institute COVID-19 Surveillance Team |
| EPI_ISL_2847304 | Lighthouse Lab in Milton Keynes | Wellcome Sanger Institute for the COVID-19 Genomics UK (COG-UK) Consortium | The Lighthouse Lab in Milton Keynes and Alex Alderton, Roberto Amato, Jeffrey Barrett, Sonia Goncalves, Ewan Harrison, David K. Jackson, Ian Johnston, Dominic Kwiatkowski, Cordelia Langford, John Sillitoe on behalf of the Wellcome Sanger Institute COVID-19 Surveillance Team |
| EPI_ISL_2847305 | Lighthouse Laboratory Plymouth | Wellcome Sanger Institute for the COVID-19 Genomics UK (COG-UK) Consortium | Lighthouse Laboratory Plymouth and Alex Alderton, Roberto Amato, Jeffrey Barrett, Sonia Goncalves, Ewan Harrison, David K. Jackson, Ian Johnston, Dominic Kwiatkowski, Cordelia Langford, John Sillitoe on behalf of the Wellcome Sanger Institute COVID-19 Surveillance Team |
| EPI_ISL_2847306 | Lighthouse Lab in Alderley Park | Wellcome Sanger Institute for the COVID-19 Genomics UK (COG-UK) Consortium | Jacquelyn Wynn, Mairead Hyland, The Lighthouse Lab in Alderley Park and Alex Alderton, Roberto Amato, Jeffrey Barrett, Sonia Goncalves, Ewan Harrison, David K. Jackson, Ian Johnston, Dominic Kwiatkowski, Cordelia Langford, John Sillitoe on behalf of the Wellcome Sanger Institute COVID-19 Surveillance Team |
| EPI_ISL_2847307 | Lighthouse Lab in Glasgow | Wellcome Sanger Institute for the COVID-19 Genomics UK (COG-UK) Consortium | Harper VanSteenhouse, Yumi Kasai, David Gray, Carol Clugston, Anna Dominiczak and Alex Alderton, Roberto Amato, Jeffrey Barrett, Sonia Goncalves, Ewan Harrison, David K. Jackson, Ian Johnston, Dominic Kwiatkowski, Cordelia Langford, John Sillitoe on behalf of the Wellcome Sanger Institute COVID-19 Surveillance Team |
| EPI_ISL_2847308 | Lighthouse Lab in Alderley Park | Wellcome Sanger Institute for the COVID-19 Genomics UK (COG-UK) Consortium | Jacquelyn Wynn, Mairead Hyland, The Lighthouse Lab in Alderley Park and Alex Alderton, Roberto Amato, Jeffrey Barrett, Sonia Goncalves, Ewan Harrison, David K. Jackson, Ian Johnston, Dominic Kwiatkowski, Cordelia Langford, John Sillitoe on behalf of the Wellcome Sanger Institute COVID-19 Surveillance Team |
| EPI_ISL_2847309, EPI_ISL_2847310 | Lighthouse Lab in Glasgow | Wellcome Sanger Institute for the COVID-19 Genomics UK (COG-UK) Consortium | Harper VanSteenhouse, Yumi Kasai, David Gray, Carol Clugston, Anna Dominiczak and Alex Alderton, Roberto Amato, Jeffrey Barrett, Sonia Goncalves, Ewan Harrison, David K. Jackson, Ian Johnston, Dominic Kwiatkowski, Cordelia Langford, John Sillitoe on behalf of the Wellcome Sanger Institute COVID-19 Surveillance Team |
| EPI_ISL_2847311 | Lighthouse Lab in Alderley Park | Wellcome Sanger Institute for the COVID-19 Genomics UK (COG-UK) Consortium | Jacquelyn Wynn, Mairead Hyland, The Lighthouse Lab in Alderley Park and Alex Alderton, Roberto Amato, Jeffrey Barrett, Sonia Goncalves, Ewan Harrison, David K. Jackson, Ian Johnston, Dominic Kwiatkowski, Cordelia Langford, John Sillitoe on behalf of the Wellcome Sanger Institute COVID-19 Surveillance Team |
| EPI_ISL_2847312 | Lighthouse Lab in Glasgow | Wellcome Sanger Institute for the COVID-19 Genomics UK (COG-UK) Consortium | Harper VanSteenhouse, Yumi Kasai, David Gray, Carol Clugston, Anna Dominiczak and Alex Alderton, Roberto Amato, Jeffrey Barrett, Sonia Goncalves, Ewan Harrison, David K. Jackson, Ian Johnston, Dominic Kwiatkowski, Cordelia Langford, John Sillitoe on behalf of the Wellcome Sanger Institute COVID-19 Surveillance Team |
| EPI_ISL_2847313 | Lighthouse Lab in Milton Keynes | Wellcome Sanger Institute for the COVID-19 Genomics UK (COG-UK) Consortium | The Lighthouse Lab in Milton Keynes and Alex Alderton, Roberto Amato, Jeffrey Barrett, Sonia Goncalves, Ewan Harrison, David K. Jackson, Ian Johnston, Dominic Kwiatkowski, Cordelia Langford, John Sillitoe on behalf of the Wellcome Sanger Institute COVID-19 Surveillance Team |
| EPI_ISL_2847314 | Lighthouse Lab in Glasgow | Wellcome Sanger Institute for the COVID-19 Genomics UK (COG-UK) Consortium | Harper VanSteenhouse, Yumi Kasai, David Gray, Carol Clugston, Anna Dominiczak and Alex Alderton, Roberto Amato, Jeffrey Barrett, Sonia Goncalves, Ewan Harrison, David K. Jackson, Ian Johnston, Dominic Kwiatkowski, Cordelia Langford, John Sillitoe on behalf of the Wellcome Sanger Institute COVID-19 Surveillance Team |
| EPI_ISL_2847315 | Lighthouse Lab in Milton Keynes | Wellcome Sanger Institute for the COVID-19 Genomics UK (COG-UK) Consortium | The Lighthouse Lab in Milton Keynes and Alex Alderton, Roberto Amato, Jeffrey Barrett, Sonia Goncalves, Ewan Harrison, David K. Jackson, Ian Johnston, Dominic Kwiatkowski, Cordelia Langford, John Sillitoe on behalf of the Wellcome Sanger Institute COVID-19 Surveillance Team |
| EPI_ISL_2847316 | Lighthouse Laboratory Plymouth | Wellcome Sanger Institute for the COVID-19 Genomics UK (COG-UK) Consortium | Lighthouse Laboratory Plymouth and Alex Alderton, Roberto Amato, Jeffrey Barrett, Sonia Goncalves, Ewan Harrison, David K. Jackson, Ian Johnston, Dominic Kwiatkowski, Cordelia Langford, John Sillitoe on behalf of the Wellcome Sanger Institute COVID-19 Surveillance Team |
| EPI_ISL_2847317, EPI_ISL_2847318 | Lighthouse Lab in Glasgow | Wellcome Sanger Institute for the COVID-19 Genomics UK (COG-UK) Consortium | Harper VanSteenhouse, Yumi Kasai, David Gray, Carol Clugston, Anna Dominiczak and Alex Alderton, Roberto Amato, Jeffrey Barrett, Sonia Goncalves, Ewan Harrison, David K. Jackson, Ian Johnston, Dominic Kwiatkowski, Cordelia Langford, John Sillitoe on behalf of the Wellcome Sanger Institute COVID-19 Surveillance Team |
| EPI_ISL_2847319, EPI_ISL_2847320 | Lighthouse Laboratory Plymouth | Wellcome Sanger Institute for the COVID-19 Genomics UK (COG-UK) Consortium | Lighthouse Laboratory Plymouth and Alex Alderton, Roberto Amato, Jeffrey Barrett, Sonia Goncalves, Ewan Harrison, David K. Jackson, Ian Johnston, Dominic Kwiatkowski, Cordelia Langford, John Sillitoe on behalf of the Wellcome Sanger Institute COVID-19 Surveillance Team |
| EPI_ISL_2847321, EPI_ISL_2847322 | Lighthouse Lab in Milton Keynes | Wellcome Sanger Institute for the COVID-19 Genomics UK (COG-UK) Consortium | The Lighthouse Lab in Milton Keynes and Alex Alderton, Roberto Amato, Jeffrey Barrett, Sonia Goncalves, Ewan Harrison, David K. Jackson, Ian Johnston, Dominic Kwiatkowski, Cordelia Langford, John Sillitoe on behalf of the Wellcome Sanger Institute COVID-19 Surveillance Team |
| EPI_ISL_2847323 | Lighthouse Laboratory Plymouth | Wellcome Sanger Institute for the COVID-19 Genomics UK (COG-UK) Consortium | Lighthouse Laboratory Plymouth and Alex Alderton, Roberto Amato, Jeffrey Barrett, Sonia Goncalves, Ewan Harrison, David K. Jackson, Ian Johnston, Dominic Kwiatkowski, Cordelia Langford, John Sillitoe on behalf of the Wellcome Sanger Institute COVID-19 Surveillance Team |
| EPI_ISL_2847324 | Lighthouse Lab in Alderley Park | Wellcome Sanger Institute for the COVID-19 Genomics UK (COG-UK) Consortium | Jacquelyn Wynn, Mairead Hyland, The Lighthouse Lab in Alderley Park and Alex Alderton, Roberto Amato, Jeffrey Barrett, Sonia Goncalves, Ewan Harrison, David K. Jackson, Ian Johnston, Dominic Kwiatkowski, Cordelia Langford, John Sillitoe on behalf of the Wellcome Sanger Institute COVID-19 Surveillance Team |
| EPI_ISL_2847325, EPI_ISL_2847326 | Lighthouse Laboratory Plymouth | Wellcome Sanger Institute for the COVID-19 Genomics UK (COG-UK) Consortium | Lighthouse Laboratory Plymouth and Alex Alderton, Roberto Amato, Jeffrey Barrett, Sonia Goncalves, Ewan Harrison, David K. Jackson, Ian Johnston, Dominic Kwiatkowski, Cordelia Langford, John Sillitoe on behalf of the Wellcome Sanger Institute COVID-19 Surveillance Team |
| EPI_ISL_2847327 | Lighthouse Lab in Glasgow | Wellcome Sanger Institute for the COVID-19 Genomics UK (COG-UK) Consortium | Harper VanSteenhouse, Yumi Kasai, David Gray, Carol Clugston, Anna Dominiczak and Alex Alderton, Roberto Amato, Jeffrey Barrett, Sonia Goncalves, Ewan Harrison, David K. Jackson, Ian Johnston, Dominic Kwiatkowski, Cordelia Langford, John Sillitoe on behalf of the Wellcome Sanger Institute COVID-19 Surveillance Team |
| EPI_ISL_2847328 | Lighthouse Laboratory Plymouth | Wellcome Sanger Institute for the COVID-19 Genomics UK (COG-UK) Consortium | Lighthouse Laboratory Plymouth and Alex Alderton, Roberto Amato, Jeffrey Barrett, Sonia Goncalves, Ewan Harrison, David K. Jackson, Ian Johnston, Dominic Kwiatkowski, Cordelia Langford, John Sillitoe on behalf of the Wellcome Sanger Institute COVID-19 Surveillance Team |
| EPI_ISL_2847329, EPI_ISL_2847330 | Lighthouse Lab in Alderley Park | Wellcome Sanger Institute for the COVID-19 Genomics UK (COG-UK) Consortium | Jacquelyn Wynn, Mairead Hyland, The Lighthouse Lab in Alderley Park and Alex Alderton, Roberto Amato, Jeffrey Barrett, Sonia Goncalves, Ewan Harrison, David K. Jackson, Ian Johnston, Dominic Kwiatkowski, Cordelia Langford, John Sillitoe on behalf of the Wellcome Sanger Institute COVID-19 Surveillance Team |
| EPI_ISL_2847331 | Lighthouse Lab in Glasgow | Wellcome Sanger Institute for the COVID-19 Genomics UK (COG-UK) Consortium | Harper VanSteenhouse, Yumi Kasai, David Gray, Carol Clugston, Anna Dominiczak and Alex Alderton, Roberto Amato, Jeffrey Barrett, Sonia Goncalves, Ewan Harrison, David K. Jackson, Ian Johnston, Dominic Kwiatkowski, Cordelia Langford, John Sillitoe on behalf of the Wellcome Sanger Institute COVID-19 Surveillance Team |
| EPI_ISL_2847332 | Lighthouse Lab in Milton Keynes | Wellcome Sanger Institute for the COVID-19 Genomics UK (COG-UK) Consortium | The Lighthouse Lab in Milton Keynes and Alex Alderton, Roberto Amato, Jeffrey Barrett, Sonia Goncalves, Ewan Harrison, David K. Jackson, Ian Johnston, Dominic Kwiatkowski, Cordelia Langford, John Sillitoe on behalf of the Wellcome Sanger Institute COVID-19 Surveillance Team |
| EPI_ISL_2847333 | Lighthouse Lab in Glasgow | Wellcome Sanger Institute for the COVID-19 Genomics UK (COG-UK) Consortium | Harper VanSteenhouse, Yumi Kasai, David Gray, Carol Clugston, Anna Dominiczak and Alex Alderton, Roberto Amato, Jeffrey Barrett, Sonia Goncalves, Ewan Harrison, David K. Jackson, Ian Johnston, Dominic Kwiatkowski, Cordelia Langford, John Sillitoe on behalf of the Wellcome Sanger Institute COVID-19 Surveillance Team |
| EPI_ISL_2847334, EPI_ISL_2847335 | Lighthouse Lab in Alderley Park | Wellcome Sanger Institute for the COVID-19 Genomics UK (COG-UK) Consortium | Jacquelyn Wynn, Mairead Hyland, The Lighthouse Lab in Alderley Park and Alex Alderton, Roberto Amato, Jeffrey Barrett, Sonia Goncalves, Ewan Harrison, David K. Jackson, Ian Johnston, Dominic Kwiatkowski, Cordelia Langford, John Sillitoe on behalf of the Wellcome Sanger Institute COVID-19 Surveillance Team |
| EPI_ISL_2847336 | Lighthouse Lab in Milton Keynes | Wellcome Sanger Institute for the COVID-19 Genomics UK (COG-UK) Consortium | The Lighthouse Lab in Milton Keynes and Alex Alderton, Roberto Amato, Jeffrey Barrett, Sonia Goncalves, Ewan Harrison, David K. Jackson, Ian Johnston, Dominic Kwiatkowski, Cordelia Langford, John Sillitoe on behalf of the Wellcome Sanger Institute COVID-19 Surveillance Team |
| EPI_ISL_2847337 | Lighthouse Lab in Alderley Park | Wellcome Sanger Institute for the COVID-19 Genomics UK (COG-UK) Consortium | Jacquelyn Wynn, Mairead Hyland, The Lighthouse Lab in Alderley Park and Alex Alderton, Roberto Amato, Jeffrey Barrett, Sonia Goncalves, Ewan Harrison, David K. Jackson, Ian Johnston, Dominic Kwiatkowski, Cordelia Langford, John Sillitoe on behalf of the Wellcome Sanger Institute COVID-19 Surveillance Team |
| EPI_ISL_2847338 | Lighthouse Lab in Glasgow | Wellcome Sanger Institute for the COVID-19 Genomics UK (COG-UK) Consortium | Harper VanSteenhouse, Yumi Kasai, David Gray, Carol Clugston, Anna Dominiczak and Alex Alderton, Roberto Amato, Jeffrey Barrett, Sonia Goncalves, Ewan Harrison, David K. Jackson, Ian Johnston, Dominic Kwiatkowski, Cordelia Langford, John Sillitoe on behalf of the Wellcome Sanger Institute COVID-19 Surveillance Team |
| EPI_ISL_2847339 | Lighthouse Laboratory Plymouth | Wellcome Sanger Institute for the COVID-19 Genomics UK (COG-UK) Consortium | Lighthouse Laboratory Plymouth and Alex Alderton, Roberto Amato, Jeffrey Barrett, Sonia Goncalves, Ewan Harrison, David K. Jackson, Ian Johnston, Dominic Kwiatkowski, Cordelia Langford, John Sillitoe on behalf of the Wellcome Sanger Institute COVID-19 Surveillance Team |
| EPI_ISL_2847340, EPI_ISL_2847341 | Lighthouse Lab in Alderley Park | Wellcome Sanger Institute for the COVID-19 Genomics UK (COG-UK) Consortium | Jacquelyn Wynn, Mairead Hyland, The Lighthouse Lab in Alderley Park and Alex Alderton, Roberto Amato, Jeffrey Barrett, Sonia Goncalves, Ewan Harrison, David K. Jackson, Ian Johnston, Dominic Kwiatkowski, Cordelia Langford, John Sillitoe on behalf of the Wellcome Sanger Institute COVID-19 Surveillance Team |

| | | | |
|---|---|---|---|
| EPI_ISL_2847342 | Lighthouse Lab in Glasgow | Wellcome Sanger Institute for the COVID-19 Genomics UK (COG-UK) Consortium | Harper VanSteenhouse, Yumi Kasai, David Gray, Carol Clugston, Anna Dominiczak and Alex Alderton, Roberto Amato, Jeffrey Barrett, Sonia Goncalves, Ewan Harrison, David K. Jackson, Ian Johnston, Dominic Kwiatkowski, Cordelia Langford, John Sillitoe on behalf of the Wellcome Sanger Institute COVID-19 Surveillance Team |
| EPI_ISL_2847343 | Lighthouse Lab in Alderley Park | Wellcome Sanger Institute for the COVID-19 Genomics UK (COG-UK) Consortium | Jacquelyn Wynn, Mairead Hyland, The Lighthouse Lab in Alderley Park and Alex Alderton, Roberto Amato, Jeffrey Barrett, Sonia Goncalves, Ewan Harrison, David K. Jackson, Ian Johnston, Dominic Kwiatkowski, Cordelia Langford, John Sillitoe on behalf of the Wellcome Sanger Institute COVID-19 Surveillance Team |
| EPI_ISL_2847344 | Lighthouse Lab in Milton Keynes | Wellcome Sanger Institute for the COVID-19 Genomics UK (COG-UK) Consortium | The Lighthouse Lab in Milton Keynes and Alex Alderton, Roberto Amato, Jeffrey Barrett, Sonia Goncalves, Ewan Harrison, David K. Jackson, Ian Johnston, Dominic Kwiatkowski, Cordelia Langford, John Sillitoe on behalf of the Wellcome Sanger Institute COVID-19 Surveillance Team |
| EPI_ISL_2847345, EPI_ISL_2847346 | Lighthouse Laboratory Plymouth | Wellcome Sanger Institute for the COVID-19 Genomics UK (COG-UK) Consortium | Lighthouse Laboratory Plymouth and Alex Alderton, Roberto Amato, Jeffrey Barrett, Sonia Goncalves, Ewan Harrison, David K. Jackson, Ian Johnston, Dominic Kwiatkowski, Cordelia Langford, John Sillitoe on behalf of the Wellcome Sanger Institute COVID-19 Surveillance Team |
| EPI_ISL_2847347, EPI_ISL_2847348 | Lighthouse Lab in Milton Keynes | Wellcome Sanger Institute for the COVID-19 Genomics UK (COG-UK) Consortium | The Lighthouse Lab in Milton Keynes and Alex Alderton, Roberto Amato, Jeffrey Barrett, Sonia Goncalves, Ewan Harrison, David K. Jackson, Ian Johnston, Dominic Kwiatkowski, Cordelia Langford, John Sillitoe on behalf of the Wellcome Sanger Institute COVID-19 Surveillance Team |
| EPI_ISL_2847349 | Lighthouse Lab in Glasgow | Wellcome Sanger Institute for the COVID-19 Genomics UK (COG-UK) Consortium | Harper VanSteenhouse, Yumi Kasai, David Gray, Carol Clugston, Anna Dominiczak and Alex Alderton, Roberto Amato, Jeffrey Barrett, Sonia Goncalves, Ewan Harrison, David K. Jackson, Ian Johnston, Dominic Kwiatkowski, Cordelia Langford, John Sillitoe on behalf of the Wellcome Sanger Institute COVID-19 Surveillance Team |
| EPI_ISL_2847350 | Lighthouse Lab in Milton Keynes | Wellcome Sanger Institute for the COVID-19 Genomics UK (COG-UK) Consortium | The Lighthouse Lab in Milton Keynes and Alex Alderton, Roberto Amato, Jeffrey Barrett, Sonia Goncalves, Ewan Harrison, David K. Jackson, Ian Johnston, Dominic Kwiatkowski, Cordelia Langford, John Sillitoe on behalf of the Wellcome Sanger Institute COVID-19 Surveillance Team |
| EPI_ISL_2847351 | Lighthouse Lab in Alderley Park | Wellcome Sanger Institute for the COVID-19 Genomics UK (COG-UK) Consortium | Jacquelyn Wynn, Mairead Hyland, The Lighthouse Lab in Alderley Park and Alex Alderton, Roberto Amato, Jeffrey Barrett, Sonia Goncalves, Ewan Harrison, David K. Jackson, Ian Johnston, Dominic Kwiatkowski, Cordelia Langford, John Sillitoe on behalf of the Wellcome Sanger Institute COVID-19 Surveillance Team |
| EPI_ISL_2847352 | Lighthouse Lab in Milton Keynes | Wellcome Sanger Institute for the COVID-19 Genomics UK (COG-UK) Consortium | The Lighthouse Lab in Milton Keynes and Alex Alderton, Roberto Amato, Jeffrey Barrett, Sonia Goncalves, Ewan Harrison, David K. Jackson, Ian Johnston, Dominic Kwiatkowski, Cordelia Langford, John Sillitoe on behalf of the Wellcome Sanger Institute COVID-19 Surveillance Team |
| EPI_ISL_2847353 | Lighthouse Laboratory Plymouth | Wellcome Sanger Institute for the COVID-19 Genomics UK (COG-UK) Consortium | Lighthouse Laboratory Plymouth and Alex Alderton, Roberto Amato, Jeffrey Barrett, Sonia Goncalves, Ewan Harrison, David K. Jackson, Ian Johnston, Dominic Kwiatkowski, Cordelia Langford, John Sillitoe on behalf of the Wellcome Sanger Institute COVID-19 Surveillance Team |
| EPI_ISL_2847354 | Lighthouse Lab in Milton Keynes | Wellcome Sanger Institute for the COVID-19 Genomics UK (COG-UK) Consortium | The Lighthouse Lab in Milton Keynes and Alex Alderton, Roberto Amato, Jeffrey Barrett, Sonia Goncalves, Ewan Harrison, David K. Jackson, Ian Johnston, Dominic Kwiatkowski, Cordelia Langford, John Sillitoe on behalf of the Wellcome Sanger Institute COVID-19 Surveillance Team |
| EPI_ISL_2847355 | Lighthouse Laboratory Plymouth | Wellcome Sanger Institute for the COVID-19 Genomics UK (COG-UK) Consortium | Lighthouse Laboratory Plymouth and Alex Alderton, Roberto Amato, Jeffrey Barrett, Sonia Goncalves, Ewan Harrison, David K. Jackson, Ian Johnston, Dominic Kwiatkowski, Cordelia Langford, John Sillitoe on behalf of the Wellcome Sanger Institute COVID-19 Surveillance Team |
| EPI_ISL_2847356 | Lighthouse Lab in Glasgow | Wellcome Sanger Institute for the COVID-19 Genomics UK (COG-UK) Consortium | Harper VanSteenhouse, Yumi Kasai, David Gray, Carol Clugston, Anna Dominiczak and Alex Alderton, Roberto Amato, Jeffrey Barrett, Sonia Goncalves, Ewan Harrison, David K. Jackson, Ian Johnston, Dominic Kwiatkowski, Cordelia Langford, John Sillitoe on behalf of the Wellcome Sanger Institute COVID-19 Surveillance Team |
| EPI_ISL_2847357 | Lighthouse Lab in Alderley Park | Wellcome Sanger Institute for the COVID-19 Genomics UK (COG-UK) Consortium | Jacquelyn Wynn, Mairead Hyland, The Lighthouse Lab in Alderley Park and Alex Alderton, Roberto Amato, Jeffrey Barrett, Sonia Goncalves, Ewan Harrison, David K. Jackson, Ian Johnston, Dominic Kwiatkowski, Cordelia Langford, John Sillitoe on behalf of the Wellcome Sanger Institute COVID-19 Surveillance Team |
| EPI_ISL_2847358 | Lighthouse Lab in Milton Keynes | Wellcome Sanger Institute for the COVID-19 Genomics UK (COG-UK) Consortium | The Lighthouse Lab in Milton Keynes and Alex Alderton, Roberto Amato, Jeffrey Barrett, Sonia Goncalves, Ewan Harrison, David K. Jackson, Ian Johnston, Dominic Kwiatkowski, Cordelia Langford, John Sillitoe on behalf of the Wellcome Sanger Institute COVID-19 Surveillance Team |
| EPI_ISL_2847359, EPI_ISL_2847360 | Lighthouse Lab in Alderley Park | Wellcome Sanger Institute for the COVID-19 Genomics UK (COG-UK) Consortium | Jacquelyn Wynn, Mairead Hyland, The Lighthouse Lab in Alderley Park and Alex Alderton, Roberto Amato, Jeffrey Barrett, Sonia Goncalves, Ewan Harrison, David K. Jackson, Ian Johnston, Dominic Kwiatkowski, Cordelia Langford, John Sillitoe on behalf of the Wellcome Sanger Institute COVID-19 Surveillance Team |
| EPI_ISL_2847361 | Lighthouse Lab in Milton Keynes | Wellcome Sanger Institute for the COVID-19 Genomics UK (COG-UK) Consortium | The Lighthouse Lab in Milton Keynes and Alex Alderton, Roberto Amato, Jeffrey Barrett, Sonia Goncalves, Ewan Harrison, David K. Jackson, Ian Johnston, Dominic Kwiatkowski, Cordelia Langford, John Sillitoe on behalf of the Wellcome Sanger Institute COVID-19 Surveillance Team |
| EPI_ISL_2847362 | Lighthouse Lab in Alderley Park | Wellcome Sanger Institute for the COVID-19 Genomics UK (COG-UK) Consortium | Jacquelyn Wynn, Mairead Hyland, The Lighthouse Lab in Alderley Park and Alex Alderton, Roberto Amato, Jeffrey Barrett, Sonia Goncalves, Ewan Harrison, David K. Jackson, Ian Johnston, Dominic Kwiatkowski, Cordelia Langford, John Sillitoe on behalf of the Wellcome Sanger Institute COVID-19 Surveillance Team |
| EPI_ISL_2847363 | Lighthouse Laboratory Plymouth | Wellcome Sanger Institute for the COVID-19 Genomics UK (COG-UK) Consortium | Lighthouse Laboratory Plymouth and Alex Alderton, Roberto Amato, Jeffrey Barrett, Sonia Goncalves, Ewan Harrison, David K. Jackson, Ian Johnston, Dominic Kwiatkowski, Cordelia Langford, John Sillitoe on behalf of the Wellcome Sanger Institute COVID-19 Surveillance Team |
| EPI_ISL_2847364, EPI_ISL_2847365, EPI_ISL_2847366 | Lighthouse Lab in Glasgow | Wellcome Sanger Institute for the COVID-19 Genomics UK (COG-UK) Consortium | Harper VanSteenhouse, Yumi Kasai, David Gray, Carol Clugston, Anna Dominiczak and Alex Alderton, Roberto Amato, Jeffrey Barrett, Sonia Goncalves, Ewan Harrison, David K. Jackson, Ian Johnston, Dominic Kwiatkowski, Cordelia Langford, John Sillitoe on behalf of the Wellcome Sanger Institute COVID-19 Surveillance Team |
| EPI_ISL_2847367 | Lighthouse Lab in Milton Keynes | Wellcome Sanger Institute for the COVID-19 Genomics UK (COG-UK) Consortium | The Lighthouse Lab in Milton Keynes and Alex Alderton, Roberto Amato, Jeffrey Barrett, Sonia Goncalves, Ewan Harrison, David K. Jackson, Ian Johnston, Dominic Kwiatkowski, Cordelia Langford, John Sillitoe on behalf of the Wellcome Sanger Institute COVID-19 Surveillance Team |
| EPI_ISL_2847368, EPI_ISL_2847369 | Lighthouse Laboratory Plymouth | Wellcome Sanger Institute for the COVID-19 Genomics UK (COG-UK) Consortium | Lighthouse Laboratory Plymouth and Alex Alderton, Roberto Amato, Jeffrey Barrett, Sonia Goncalves, Ewan Harrison, David K. Jackson, Ian Johnston, Dominic Kwiatkowski, Cordelia Langford, John Sillitoe on behalf of the Wellcome Sanger Institute COVID-19 Surveillance Team |
| EPI_ISL_2847370, EPI_ISL_2847371 | Lighthouse Lab in Alderley Park | Wellcome Sanger Institute for the COVID-19 Genomics UK (COG-UK) Consortium | Jacquelyn Wynn, Mairead Hyland, The Lighthouse Lab in Alderley Park and Alex Alderton, Roberto Amato, Jeffrey Barrett, Sonia Goncalves, Ewan Harrison, David K. Jackson, Ian Johnston, Dominic Kwiatkowski, Cordelia Langford, John Sillitoe on behalf of the Wellcome Sanger Institute COVID-19 Surveillance Team |
| EPI_ISL_2847372 | Lighthouse Lab in Glasgow | Wellcome Sanger Institute for the COVID-19 Genomics UK (COG-UK) Consortium | Harper VanSteenhouse, Yumi Kasai, David Gray, Carol Clugston, Anna Dominiczak and Alex Alderton, Roberto Amato, Jeffrey Barrett, Sonia Goncalves, Ewan Harrison, David K. Jackson, Ian Johnston, Dominic Kwiatkowski, Cordelia Langford, John Sillitoe on behalf of the Wellcome Sanger Institute COVID-19 Surveillance Team |
| EPI_ISL_2847373 | Lighthouse Laboratory Plymouth | Wellcome Sanger Institute for the COVID-19 Genomics UK (COG-UK) Consortium | Lighthouse Laboratory Plymouth and Alex Alderton, Roberto Amato, Jeffrey Barrett, Sonia Goncalves, Ewan Harrison, David K. Jackson, Ian Johnston, Dominic Kwiatkowski, Cordelia Langford, John Sillitoe on behalf of the Wellcome Sanger Institute COVID-19 Surveillance Team |
| EPI_ISL_2847374 | Lighthouse Lab in Alderley Park | Wellcome Sanger Institute for the COVID-19 Genomics UK (COG-UK) Consortium | Jacquelyn Wynn, Mairead Hyland, The Lighthouse Lab in Alderley Park and Alex Alderton, Roberto Amato, Jeffrey Barrett, Sonia Goncalves, Ewan Harrison, David K. Jackson, Ian Johnston, Dominic Kwiatkowski, Cordelia Langford, John Sillitoe on behalf of the Wellcome Sanger Institute COVID-19 Surveillance Team |
| EPI_ISL_2847375 | Lighthouse Laboratory Plymouth | Wellcome Sanger Institute for the COVID-19 Genomics UK (COG-UK) Consortium | Lighthouse Laboratory Plymouth and Alex Alderton, Roberto Amato, Jeffrey Barrett, Sonia Goncalves, Ewan Harrison, David K. Jackson, Ian Johnston, Dominic Kwiatkowski, Cordelia Langford, John Sillitoe on behalf of the Wellcome Sanger Institute COVID-19 Surveillance Team |
| EPI_ISL_2847376 | Lighthouse Lab in Milton Keynes | Wellcome Sanger Institute for the COVID-19 Genomics UK (COG-UK) Consortium | The Lighthouse Lab in Milton Keynes and Alex Alderton, Roberto Amato, Jeffrey Barrett, Sonia Goncalves, Ewan Harrison, David K. Jackson, Ian Johnston, Dominic Kwiatkowski, Cordelia Langford, John Sillitoe on behalf of the Wellcome Sanger Institute COVID-19 Surveillance Team |
| EPI_ISL_2847377 | Lighthouse Lab in Alderley Park | Wellcome Sanger Institute for the COVID-19 Genomics UK (COG-UK) Consortium | Jacquelyn Wynn, Mairead Hyland, The Lighthouse Lab in Alderley Park and Alex Alderton, Roberto Amato, Jeffrey Barrett, Sonia Goncalves, Ewan Harrison, David K. Jackson, Ian Johnston, Dominic Kwiatkowski, Cordelia Langford, John Sillitoe on behalf of the Wellcome Sanger Institute COVID-19 Surveillance Team |
| EPI_ISL_2847378 | Lighthouse Lab in Milton Keynes | Wellcome Sanger Institute for the COVID-19 Genomics UK (COG-UK) Consortium | The Lighthouse Lab in Milton Keynes and Alex Alderton, Roberto Amato, Jeffrey Barrett, Sonia Goncalves, Ewan Harrison, David K. Jackson, Ian Johnston, Dominic Kwiatkowski, Cordelia Langford, John Sillitoe on behalf of the Wellcome Sanger Institute COVID-19 Surveillance Team |
| EPI_ISL_2847379 | Lighthouse Lab in Alderley Park | Wellcome Sanger Institute for the COVID-19 Genomics UK (COG-UK) Consortium | Jacquelyn Wynn, Mairead Hyland, The Lighthouse Lab in Alderley Park and Alex Alderton, Roberto Amato, Jeffrey Barrett, Sonia Goncalves, Ewan Harrison, David K. Jackson, Ian Johnston, Dominic Kwiatkowski, Cordelia Langford, John Sillitoe on behalf of the Wellcome Sanger Institute COVID-19 Surveillance Team |
| EPI_ISL_2847380, EPI_ISL_2847381 | Lighthouse Lab in Glasgow | Wellcome Sanger Institute for the COVID-19 Genomics UK (COG-UK) Consortium | Harper VanSteenhouse, Yumi Kasai, David Gray, Carol Clugston, Anna Dominiczak and Alex Alderton, Roberto Amato, Jeffrey Barrett, Sonia Goncalves, Ewan Harrison, David K. Jackson, Ian Johnston, Dominic Kwiatkowski, Cordelia Langford, John Sillitoe on behalf of the Wellcome Sanger Institute COVID-19 Surveillance Team |
| EPI_ISL_2847382, EPI_ISL_2847383 | Lighthouse Laboratory Plymouth | Wellcome Sanger Institute for the COVID-19 Genomics UK (COG-UK) Consortium | Lighthouse Laboratory Plymouth and Alex Alderton, Roberto Amato, Jeffrey Barrett, Sonia Goncalves, Ewan Harrison, David K. Jackson, Ian Johnston, Dominic Kwiatkowski, Cordelia Langford, John Sillitoe on behalf of the Wellcome Sanger Institute COVID-19 Surveillance Team |
| EPI_ISL_2847384 | Lighthouse Lab in Alderley Park | Wellcome Sanger Institute for the COVID-19 Genomics UK (COG-UK) Consortium | Jacquelyn Wynn, Mairead Hyland, The Lighthouse Lab in Alderley Park and Alex Alderton, Roberto Amato, Jeffrey Barrett, Sonia Goncalves, Ewan Harrison, David K. Jackson, Ian Johnston, Dominic Kwiatkowski, Cordelia Langford, John Sillitoe on behalf of the Wellcome Sanger Institute COVID-19 Surveillance Team |
| EPI_ISL_2847385 | Lighthouse Lab in Glasgow | Wellcome Sanger Institute for the COVID-19 Genomics UK (COG-UK) Consortium | Harper VanSteenhouse, Yumi Kasai, David Gray, Carol Clugston, Anna Dominiczak and Alex Alderton, Roberto Amato, Jeffrey Barrett, Sonia Goncalves, Ewan Harrison, David K. Jackson, Ian Johnston, Dominic Kwiatkowski, Cordelia Langford, John Sillitoe on behalf of the Wellcome Sanger Institute COVID-19 Surveillance Team |
| EPI_ISL_2847386 | Lighthouse Lab in Alderley Park | Wellcome Sanger Institute for the COVID-19 Genomics UK (COG-UK) Consortium | Jacquelyn Wynn, Mairead Hyland, The Lighthouse Lab in Alderley Park and Alex Alderton, Roberto Amato, Jeffrey Barrett, Sonia Goncalves, Ewan Harrison, David K. Jackson, Ian Johnston, Dominic Kwiatkowski, Cordelia Langford, John Sillitoe on behalf of the Wellcome Sanger Institute COVID-19 Surveillance Team |
| EPI_ISL_2847387 | Lighthouse Lab in Glasgow | Wellcome Sanger Institute for the COVID-19 Genomics UK (COG-UK) Consortium | Harper VanSteenhouse, Yumi Kasai, David Gray, Carol Clugston, Anna Dominiczak and Alex Alderton, Roberto Amato, Jeffrey Barrett, Sonia Goncalves, Ewan Harrison, David K. Jackson, Ian Johnston, Dominic Kwiatkowski, Cordelia Langford, John Sillitoe on behalf of the Wellcome Sanger Institute COVID-19 Surveillance Team |
| EPI_ISL_2847388 | Lighthouse Lab in Milton Keynes | Wellcome Sanger Institute for the COVID-19 Genomics UK (COG-UK) Consortium | The Lighthouse Lab in Milton Keynes and Alex Alderton, Roberto Amato, Jeffrey Barrett, Sonia Goncalves, Ewan Harrison, David K. Jackson, Ian Johnston, Dominic Kwiatkowski, Cordelia Langford, John Sillitoe on behalf of the Wellcome Sanger Institute COVID-19 Surveillance Team |
| EPI_ISL_2847389 | Lighthouse Lab in Alderley Park | Wellcome Sanger Institute for the COVID-19 Genomics UK (COG-UK) Consortium | Jacquelyn Wynn, Mairead Hyland, The Lighthouse Lab in Alderley Park and Alex Alderton, Roberto Amato, Jeffrey Barrett, Sonia Goncalves, Ewan Harrison, David K. Jackson, Ian Johnston, Dominic Kwiatkowski, Cordelia Langford, John Sillitoe on behalf of the Wellcome Sanger Institute COVID-19 Surveillance Team |
| EPI_ISL_2847390 | Lighthouse Lab in Milton Keynes | Wellcome Sanger Institute for the COVID-19 Genomics UK (COG-UK) Consortium | The Lighthouse Lab in Milton Keynes and Alex Alderton, Roberto Amato, Jeffrey Barrett, Sonia Goncalves, Ewan Harrison, David K. Jackson, Ian Johnston, Dominic Kwiatkowski, Cordelia Langford, John Sillitoe on behalf of the Wellcome Sanger Institute COVID-19 Surveillance Team |
| EPI_ISL_2847391 | Lighthouse Laboratory Plymouth | Wellcome Sanger Institute for the COVID-19 Genomics UK (COG-UK) Consortium | Lighthouse Laboratory Plymouth and Alex Alderton, Roberto Amato, Jeffrey Barrett, Sonia Goncalves, Ewan Harrison, David K. Jackson, Ian Johnston, Dominic Kwiatkowski, Cordelia Langford, John Sillitoe on behalf of the Wellcome Sanger Institute COVID-19 Surveillance Team |
| EPI_ISL_2847392 | Lighthouse Lab in Milton Keynes | Wellcome Sanger Institute for the COVID-19 Genomics UK (COG-UK) Consortium | The Lighthouse Lab in Milton Keynes and Alex Alderton, Roberto Amato, Jeffrey Barrett, Sonia Goncalves, Ewan Harrison, David K. Jackson, Ian Johnston, Dominic Kwiatkowski, Cordelia Langford, John Sillitoe on behalf of the Wellcome Sanger Institute COVID-19 Surveillance Team |
| EPI_ISL_2847393 | Lighthouse Laboratory Plymouth | Wellcome Sanger Institute for the COVID-19 Genomics UK (COG-UK) Consortium | Lighthouse Laboratory Plymouth and Alex Alderton, Roberto Amato, Jeffrey Barrett, Sonia Goncalves, Ewan Harrison, David K. Jackson, Ian Johnston, Dominic Kwiatkowski, Cordelia Langford, John Sillitoe on behalf of the Wellcome Sanger Institute COVID-19 Surveillance Team |
| EPI_ISL_2847394, EPI_ISL_2847395 | Lighthouse Lab in Alderley Park | Wellcome Sanger Institute for the COVID-19 Genomics UK (COG-UK) Consortium | Jacquelyn Wynn, Mairead Hyland, The Lighthouse Lab in Alderley Park and Alex Alderton, Roberto Amato, Jeffrey Barrett, Sonia Goncalves, Ewan Harrison, David K. Jackson, Ian Johnston, Dominic Kwiatkowski, Cordelia Langford, John Sillitoe on behalf of the Wellcome Sanger Institute COVID-19 Surveillance Team |

| | | | |
|---|---|---|---|
| EPI_ISL_2847396, EPI_ISL_2847397 | Lighthouse Laboratory Plymouth | Wellcome Sanger Institute for the COVID-19 Genomics UK (COG-UK) Consortium | Lighthouse Laboratory Plymouth and Alex Alderton, Roberto Amato, Jeffrey Barrett, Sonia Goncalves, Ewan Harrison, David K. Jackson, Ian Johnston, Dominic Kwiatkowski, Cordelia Langford, John Sillitoe on behalf of the Wellcome Sanger Institute COVID-19 Surveillance Team |
| EPI_ISL_2847398, EPI_ISL_2847399 | Lighthouse Lab in Alderley Park | Wellcome Sanger Institute for the COVID-19 Genomics UK (COG-UK) Consortium | Jacquelyn Wynn, Mairead Hyland, The Lighthouse Lab in Alderley Park and Alex Alderton, Roberto Amato, Jeffrey Barrett, Sonia Goncalves, Ewan Harrison, David K. Jackson, Ian Johnston, Dominic Kwiatkowski, Cordelia Langford, John Sillitoe on behalf of the Wellcome Sanger Institute COVID-19 Surveillance Team |
| EPI_ISL_2847400 | Lighthouse Lab in Glasgow | Wellcome Sanger Institute for the COVID-19 Genomics UK (COG-UK) Consortium | Harper VanSteenhouse, Yumi Kasai, David Gray, Carol Clugston, Anna Dominiczak and Alex Alderton, Roberto Amato, Jeffrey Barrett, Sonia Goncalves, Ewan Harrison, David K. Jackson, Ian Johnston, Dominic Kwiatkowski, Cordelia Langford, John Sillitoe on behalf of the Wellcome Sanger Institute COVID-19 Surveillance Team |
| EPI_ISL_2847401 | Lighthouse Laboratory Plymouth | Wellcome Sanger Institute for the COVID-19 Genomics UK (COG-UK) Consortium | Lighthouse Laboratory Plymouth and Alex Alderton, Roberto Amato, Jeffrey Barrett, Sonia Goncalves, Ewan Harrison, David K. Jackson, Ian Johnston, Dominic Kwiatkowski, Cordelia Langford, John Sillitoe on behalf of the Wellcome Sanger Institute COVID-19 Surveillance Team |
| EPI_ISL_2847402, EPI_ISL_2847403 | Lighthouse Lab in Milton Keynes | Wellcome Sanger Institute for the COVID-19 Genomics UK (COG-UK) Consortium | The Lighthouse Lab in Milton Keynes and Alex Alderton, Roberto Amato, Jeffrey Barrett, Sonia Goncalves, Ewan Harrison, David K. Jackson, Ian Johnston, Dominic Kwiatkowski, Cordelia Langford, John Sillitoe on behalf of the Wellcome Sanger Institute COVID-19 Surveillance Team |
| EPI_ISL_2847404 | Lighthouse Lab in Alderley Park | Wellcome Sanger Institute for the COVID-19 Genomics UK (COG-UK) Consortium | Jacquelyn Wynn, Mairead Hyland, The Lighthouse Lab in Alderley Park and Alex Alderton, Roberto Amato, Jeffrey Barrett, Sonia Goncalves, Ewan Harrison, David K. Jackson, Ian Johnston, Dominic Kwiatkowski, Cordelia Langford, John Sillitoe on behalf of the Wellcome Sanger Institute COVID-19 Surveillance Team |
| EPI_ISL_2847405 | Lighthouse Lab in Milton Keynes | Wellcome Sanger Institute for the COVID-19 Genomics UK (COG-UK) Consortium | The Lighthouse Lab in Milton Keynes and Alex Alderton, Roberto Amato, Jeffrey Barrett, Sonia Goncalves, Ewan Harrison, David K. Jackson, Ian Johnston, Dominic Kwiatkowski, Cordelia Langford, John Sillitoe on behalf of the Wellcome Sanger Institute COVID-19 Surveillance Team |
| EPI_ISL_2847406 | Lighthouse Lab in Alderley Park | Wellcome Sanger Institute for the COVID-19 Genomics UK (COG-UK) Consortium | Jacquelyn Wynn, Mairead Hyland, The Lighthouse Lab in Alderley Park and Alex Alderton, Roberto Amato, Jeffrey Barrett, Sonia Goncalves, Ewan Harrison, David K. Jackson, Ian Johnston, Dominic Kwiatkowski, Cordelia Langford, John Sillitoe on behalf of the Wellcome Sanger Institute COVID-19 Surveillance Team |
| EPI_ISL_2847407 | Lighthouse Laboratory Plymouth | Wellcome Sanger Institute for the COVID-19 Genomics UK (COG-UK) Consortium | Lighthouse Laboratory Plymouth and Alex Alderton, Roberto Amato, Jeffrey Barrett, Sonia Goncalves, Ewan Harrison, David K. Jackson, Ian Johnston, Dominic Kwiatkowski, Cordelia Langford, John Sillitoe on behalf of the Wellcome Sanger Institute COVID-19 Surveillance Team |
| EPI_ISL_2847408, EPI_ISL_2847409 | Lighthouse Lab in Alderley Park | Wellcome Sanger Institute for the COVID-19 Genomics UK (COG-UK) Consortium | Jacquelyn Wynn, Mairead Hyland, The Lighthouse Lab in Alderley Park and Alex Alderton, Roberto Amato, Jeffrey Barrett, Sonia Goncalves, Ewan Harrison, David K. Jackson, Ian Johnston, Dominic Kwiatkowski, Cordelia Langford, John Sillitoe on behalf of the Wellcome Sanger Institute COVID-19 Surveillance Team |
| EPI_ISL_2847410 | Lighthouse Lab in Glasgow | Wellcome Sanger Institute for the COVID-19 Genomics UK (COG-UK) Consortium | Harper VanSteenhouse, Yumi Kasai, David Gray, Carol Clugston, Anna Dominiczak and Alex Alderton, Roberto Amato, Jeffrey Barrett, Sonia Goncalves, Ewan Harrison, David K. Jackson, Ian Johnston, Dominic Kwiatkowski, Cordelia Langford, John Sillitoe on behalf of the Wellcome Sanger Institute COVID-19 Surveillance Team |
| EPI_ISL_2847411 | Lighthouse Lab in Alderley Park | Wellcome Sanger Institute for the COVID-19 Genomics UK (COG-UK) Consortium | Jacquelyn Wynn, Mairead Hyland, The Lighthouse Lab in Alderley Park and Alex Alderton, Roberto Amato, Jeffrey Barrett, Sonia Goncalves, Ewan Harrison, David K. Jackson, Ian Johnston, Dominic Kwiatkowski, Cordelia Langford, John Sillitoe on behalf of the Wellcome Sanger Institute COVID-19 Surveillance Team |
| EPI_ISL_2847412, EPI_ISL_2847413 | Lighthouse Lab in Milton Keynes | Wellcome Sanger Institute for the COVID-19 Genomics UK (COG-UK) Consortium | The Lighthouse Lab in Milton Keynes and Alex Alderton, Roberto Amato, Jeffrey Barrett, Sonia Goncalves, Ewan Harrison, David K. Jackson, Ian Johnston, Dominic Kwiatkowski, Cordelia Langford, John Sillitoe on behalf of the Wellcome Sanger Institute COVID-19 Surveillance Team |
| EPI_ISL_2847414 | Lighthouse Lab in Alderley Park | Wellcome Sanger Institute for the COVID-19 Genomics UK (COG-UK) Consortium | Jacquelyn Wynn, Mairead Hyland, The Lighthouse Lab in Alderley Park and Alex Alderton, Roberto Amato, Jeffrey Barrett, Sonia Goncalves, Ewan Harrison, David K. Jackson, Ian Johnston, Dominic Kwiatkowski, Cordelia Langford, John Sillitoe on behalf of the Wellcome Sanger Institute COVID-19 Surveillance Team |
| EPI_ISL_2847415 | Lighthouse Lab in Glasgow | Wellcome Sanger Institute for the COVID-19 Genomics UK (COG-UK) Consortium | Harper VanSteenhouse, Yumi Kasai, David Gray, Carol Clugston, Anna Dominiczak and Alex Alderton, Roberto Amato, Jeffrey Barrett, Sonia Goncalves, Ewan Harrison, David K. Jackson, Ian Johnston, Dominic Kwiatkowski, Cordelia Langford, John Sillitoe on behalf of the Wellcome Sanger Institute COVID-19 Surveillance Team |
| EPI_ISL_2847416, EPI_ISL_2847417 | Lighthouse Lab in Alderley Park | Wellcome Sanger Institute for the COVID-19 Genomics UK (COG-UK) Consortium | Jacquelyn Wynn, Mairead Hyland, The Lighthouse Lab in Alderley Park and Alex Alderton, Roberto Amato, Jeffrey Barrett, Sonia Goncalves, Ewan Harrison, David K. Jackson, Ian Johnston, Dominic Kwiatkowski, Cordelia Langford, John Sillitoe on behalf of the Wellcome Sanger Institute COVID-19 Surveillance Team |
| EPI_ISL_2847418 | Lighthouse Lab in Milton Keynes | Wellcome Sanger Institute for the COVID-19 Genomics UK (COG-UK) Consortium | The Lighthouse Lab in Milton Keynes and Alex Alderton, Roberto Amato, Jeffrey Barrett, Sonia Goncalves, Ewan Harrison, David K. Jackson, Ian Johnston, Dominic Kwiatkowski, Cordelia Langford, John Sillitoe on behalf of the Wellcome Sanger Institute COVID-19 Surveillance Team |
| EPI_ISL_2847419 | Lighthouse Laboratory Plymouth | Wellcome Sanger Institute for the COVID-19 Genomics UK (COG-UK) Consortium | Lighthouse Laboratory Plymouth and Alex Alderton, Roberto Amato, Jeffrey Barrett, Sonia Goncalves, Ewan Harrison, David K. Jackson, Ian Johnston, Dominic Kwiatkowski, Cordelia Langford, John Sillitoe on behalf of the Wellcome Sanger Institute COVID-19 Surveillance Team |
| EPI_ISL_2847420 | Lighthouse Lab in Glasgow | Wellcome Sanger Institute for the COVID-19 Genomics UK (COG-UK) Consortium | Harper VanSteenhouse, Yumi Kasai, David Gray, Carol Clugston, Anna Dominiczak and Alex Alderton, Roberto Amato, Jeffrey Barrett, Sonia Goncalves, Ewan Harrison, David K. Jackson, Ian Johnston, Dominic Kwiatkowski, Cordelia Langford, John Sillitoe on behalf of the Wellcome Sanger Institute COVID-19 Surveillance Team |
| EPI_ISL_2847421 | Lighthouse Lab in Alderley Park | Wellcome Sanger Institute for the COVID-19 Genomics UK (COG-UK) Consortium | Jacquelyn Wynn, Mairead Hyland, The Lighthouse Lab in Alderley Park and Alex Alderton, Roberto Amato, Jeffrey Barrett, Sonia Goncalves, Ewan Harrison, David K. Jackson, Ian Johnston, Dominic Kwiatkowski, Cordelia Langford, John Sillitoe on behalf of the Wellcome Sanger Institute COVID-19 Surveillance Team |
| EPI_ISL_2847422 | Lighthouse Lab in Glasgow | Wellcome Sanger Institute for the COVID-19 Genomics UK (COG-UK) Consortium | Harper VanSteenhouse, Yumi Kasai, David Gray, Carol Clugston, Anna Dominiczak and Alex Alderton, Roberto Amato, Jeffrey Barrett, Sonia Goncalves, Ewan Harrison, David K. Jackson, Ian Johnston, Dominic Kwiatkowski, Cordelia Langford, John Sillitoe on behalf of the Wellcome Sanger Institute COVID-19 Surveillance Team |
| EPI_ISL_2847423 | Lighthouse Lab in Alderley Park | Wellcome Sanger Institute for the COVID-19 Genomics UK (COG-UK) Consortium | Jacquelyn Wynn, Mairead Hyland, The Lighthouse Lab in Alderley Park and Alex Alderton, Roberto Amato, Jeffrey Barrett, Sonia Goncalves, Ewan Harrison, David K. Jackson, Ian Johnston, Dominic Kwiatkowski, Cordelia Langford, John Sillitoe on behalf of the Wellcome Sanger Institute COVID-19 Surveillance Team |
| EPI_ISL_2847424 | Lighthouse Laboratory Plymouth | Wellcome Sanger Institute for the COVID-19 Genomics UK (COG-UK) Consortium | Lighthouse Laboratory Plymouth and Alex Alderton, Roberto Amato, Jeffrey Barrett, Sonia Goncalves, Ewan Harrison, David K. Jackson, Ian Johnston, Dominic Kwiatkowski, Cordelia Langford, John Sillitoe on behalf of the Wellcome Sanger Institute COVID-19 Surveillance Team |
| EPI_ISL_2847425 | Lighthouse Lab in Milton Keynes | Wellcome Sanger Institute for the COVID-19 Genomics UK (COG-UK) Consortium | The Lighthouse Lab in Milton Keynes and Alex Alderton, Roberto Amato, Jeffrey Barrett, Sonia Goncalves, Ewan Harrison, David K. Jackson, Ian Johnston, Dominic Kwiatkowski, Cordelia Langford, John Sillitoe on behalf of the Wellcome Sanger Institute COVID-19 Surveillance Team |
| EPI_ISL_2847426 | Lighthouse Lab in Alderley Park | Wellcome Sanger Institute for the COVID-19 Genomics UK (COG-UK) Consortium | Jacquelyn Wynn, Mairead Hyland, The Lighthouse Lab in Alderley Park and Alex Alderton, Roberto Amato, Jeffrey Barrett, Sonia Goncalves, Ewan Harrison, David K. Jackson, Ian Johnston, Dominic Kwiatkowski, Cordelia Langford, John Sillitoe on behalf of the Wellcome Sanger Institute COVID-19 Surveillance Team |
| EPI_ISL_2847427 | Lighthouse Lab in Milton Keynes | Wellcome Sanger Institute for the COVID-19 Genomics UK (COG-UK) Consortium | The Lighthouse Lab in Milton Keynes and Alex Alderton, Roberto Amato, Jeffrey Barrett, Sonia Goncalves, Ewan Harrison, David K. Jackson, Ian Johnston, Dominic Kwiatkowski, Cordelia Langford, John Sillitoe on behalf of the Wellcome Sanger Institute COVID-19 Surveillance Team |
| EPI_ISL_2847428 | Lighthouse Lab in Alderley Park | Wellcome Sanger Institute for the COVID-19 Genomics UK (COG-UK) Consortium | Jacquelyn Wynn, Mairead Hyland, The Lighthouse Lab in Alderley Park and Alex Alderton, Roberto Amato, Jeffrey Barrett, Sonia Goncalves, Ewan Harrison, David K. Jackson, Ian Johnston, Dominic Kwiatkowski, Cordelia Langford, John Sillitoe on behalf of the Wellcome Sanger Institute COVID-19 Surveillance Team |
| EPI_ISL_2847429 | Lighthouse Laboratory Plymouth | Wellcome Sanger Institute for the COVID-19 Genomics UK (COG-UK) Consortium | Lighthouse Laboratory Plymouth and Alex Alderton, Roberto Amato, Jeffrey Barrett, Sonia Goncalves, Ewan Harrison, David K. Jackson, Ian Johnston, Dominic Kwiatkowski, Cordelia Langford, John Sillitoe on behalf of the Wellcome Sanger Institute COVID-19 Surveillance Team |
| EPI_ISL_2847430 | Lighthouse Lab in Milton Keynes | Wellcome Sanger Institute for the COVID-19 Genomics UK (COG-UK) Consortium | The Lighthouse Lab in Milton Keynes and Alex Alderton, Roberto Amato, Jeffrey Barrett, Sonia Goncalves, Ewan Harrison, David K. Jackson, Ian Johnston, Dominic Kwiatkowski, Cordelia Langford, John Sillitoe on behalf of the Wellcome Sanger Institute COVID-19 Surveillance Team |
| EPI_ISL_2847431 | Lighthouse Lab in Alderley Park | Wellcome Sanger Institute for the COVID-19 Genomics UK (COG-UK) Consortium | Jacquelyn Wynn, Mairead Hyland, The Lighthouse Lab in Alderley Park and Alex Alderton, Roberto Amato, Jeffrey Barrett, Sonia Goncalves, Ewan Harrison, David K. Jackson, Ian Johnston, Dominic Kwiatkowski, Cordelia Langford, John Sillitoe on behalf of the Wellcome Sanger Institute COVID-19 Surveillance Team |
| EPI_ISL_2847432 | Lighthouse Laboratory Plymouth | Wellcome Sanger Institute for the COVID-19 Genomics UK (COG-UK) Consortium | Lighthouse Laboratory Plymouth and Alex Alderton, Roberto Amato, Jeffrey Barrett, Sonia Goncalves, Ewan Harrison, David K. Jackson, Ian Johnston, Dominic Kwiatkowski, Cordelia Langford, John Sillitoe on behalf of the Wellcome Sanger Institute COVID-19 Surveillance Team |
| EPI_ISL_2847433 | Lighthouse Lab in Alderley Park | Wellcome Sanger Institute for the COVID-19 Genomics UK (COG-UK) Consortium | Jacquelyn Wynn, Mairead Hyland, The Lighthouse Lab in Alderley Park and Alex Alderton, Roberto Amato, Jeffrey Barrett, Sonia Goncalves, Ewan Harrison, David K. Jackson, Ian Johnston, Dominic Kwiatkowski, Cordelia Langford, John Sillitoe on behalf of the Wellcome Sanger Institute COVID-19 Surveillance Team |
| EPI_ISL_2847434 | Lighthouse Laboratory Plymouth | Wellcome Sanger Institute for the COVID-19 Genomics UK (COG-UK) Consortium | Lighthouse Laboratory Plymouth and Alex Alderton, Roberto Amato, Jeffrey Barrett, Sonia Goncalves, Ewan Harrison, David K. Jackson, Ian Johnston, Dominic Kwiatkowski, Cordelia Langford, John Sillitoe on behalf of the Wellcome Sanger Institute COVID-19 Surveillance Team |
| EPI_ISL_2847435, EPI_ISL_2847436 | Lighthouse Lab in Alderley Park | Wellcome Sanger Institute for the COVID-19 Genomics UK (COG-UK) Consortium | Jacquelyn Wynn, Mairead Hyland, The Lighthouse Lab in Alderley Park and Alex Alderton, Roberto Amato, Jeffrey Barrett, Sonia Goncalves, Ewan Harrison, David K. Jackson, Ian Johnston, Dominic Kwiatkowski, Cordelia Langford, John Sillitoe on behalf of the Wellcome Sanger Institute COVID-19 Surveillance Team |
| EPI_ISL_2847437 | Lighthouse Lab in Milton Keynes | Wellcome Sanger Institute for the COVID-19 Genomics UK (COG-UK) Consortium | The Lighthouse Lab in Milton Keynes and Alex Alderton, Roberto Amato, Jeffrey Barrett, Sonia Goncalves, Ewan Harrison, David K. Jackson, Ian Johnston, Dominic Kwiatkowski, Cordelia Langford, John Sillitoe on behalf of the Wellcome Sanger Institute COVID-19 Surveillance Team |
| EPI_ISL_2847438 | Lighthouse Laboratory Plymouth | Wellcome Sanger Institute for the COVID-19 Genomics UK (COG-UK) Consortium | Lighthouse Laboratory Plymouth and Alex Alderton, Roberto Amato, Jeffrey Barrett, Sonia Goncalves, Ewan Harrison, David K. Jackson, Ian Johnston, Dominic Kwiatkowski, Cordelia Langford, John Sillitoe on behalf of the Wellcome Sanger Institute COVID-19 Surveillance Team |
| EPI_ISL_2847439, EPI_ISL_2847440 | Lighthouse Lab in Glasgow | Wellcome Sanger Institute for the COVID-19 Genomics UK (COG-UK) Consortium | Harper VanSteenhouse, Yumi Kasai, David Gray, Carol Clugston, Anna Dominiczak and Alex Alderton, Roberto Amato, Jeffrey Barrett, Sonia Goncalves, Ewan Harrison, David K. Jackson, Ian Johnston, Dominic Kwiatkowski, Cordelia Langford, John Sillitoe on behalf of the Wellcome Sanger Institute COVID-19 Surveillance Team |
| EPI_ISL_2847441 | Lighthouse Laboratory Plymouth | Wellcome Sanger Institute for the COVID-19 Genomics UK (COG-UK) Consortium | Lighthouse Laboratory Plymouth and Alex Alderton, Roberto Amato, Jeffrey Barrett, Sonia Goncalves, Ewan Harrison, David K. Jackson, Ian Johnston, Dominic Kwiatkowski, Cordelia Langford, John Sillitoe on behalf of the Wellcome Sanger Institute COVID-19 Surveillance Team |
| EPI_ISL_2847442, EPI_ISL_2847443 | Lighthouse Lab in Glasgow | Wellcome Sanger Institute for the COVID-19 Genomics UK (COG-UK) Consortium | Harper VanSteenhouse, Yumi Kasai, David Gray, Carol Clugston, Anna Dominiczak and Alex Alderton, Roberto Amato, Jeffrey Barrett, Sonia Goncalves, Ewan Harrison, David K. Jackson, Ian Johnston, Dominic Kwiatkowski, Cordelia Langford, John Sillitoe on behalf of the Wellcome Sanger Institute COVID-19 Surveillance Team |
| EPI_ISL_2847444, EPI_ISL_2847445 | Lighthouse Lab in Alderley Park | Wellcome Sanger Institute for the COVID-19 Genomics UK (COG-UK) Consortium | Jacquelyn Wynn, Mairead Hyland, The Lighthouse Lab in Alderley Park and Alex Alderton, Roberto Amato, Jeffrey Barrett, Sonia Goncalves, Ewan Harrison, David K. Jackson, Ian Johnston, Dominic Kwiatkowski, Cordelia Langford, John Sillitoe on behalf of the Wellcome Sanger Institute COVID-19 Surveillance Team |
| EPI_ISL_2847446 | Lighthouse Lab in Milton Keynes | Wellcome Sanger Institute for the COVID-19 Genomics UK (COG-UK) Consortium | The Lighthouse Lab in Milton Keynes and Alex Alderton, Roberto Amato, Jeffrey Barrett, Sonia Goncalves, Ewan Harrison, David K. Jackson, Ian Johnston, Dominic Kwiatkowski, Cordelia Langford, John Sillitoe on behalf of the Wellcome Sanger Institute COVID-19 Surveillance Team |
| EPI_ISL_2847447 | Lighthouse Laboratory Plymouth | Wellcome Sanger Institute for the COVID-19 Genomics UK (COG-UK) Consortium | Lighthouse Laboratory Plymouth and Alex Alderton, Roberto Amato, Jeffrey Barrett, Sonia Goncalves, Ewan Harrison, David K. Jackson, Ian Johnston, Dominic Kwiatkowski, Cordelia Langford, John Sillitoe on behalf of the Wellcome Sanger Institute COVID-19 Surveillance Team |
| EPI_ISL_2847448 | Lighthouse Lab in Milton Keynes | Wellcome Sanger Institute for the COVID-19 Genomics UK (COG-UK) Consortium | The Lighthouse Lab in Milton Keynes and Alex Alderton, Roberto Amato, Jeffrey Barrett, Sonia Goncalves, Ewan Harrison, David K. Jackson, Ian Johnston, Dominic Kwiatkowski, Cordelia Langford, John Sillitoe on behalf of the Wellcome Sanger Institute COVID-19 Surveillance Team |
| EPI_ISL_2847449 | Lighthouse Lab in Alderley Park | Wellcome Sanger Institute for the COVID-19 Genomics UK (COG-UK) Consortium | Jacquelyn Wynn, Mairead Hyland, The Lighthouse Lab in Alderley Park and Alex Alderton, Roberto Amato, Jeffrey Barrett, Sonia Goncalves, Ewan Harrison, David K. Jackson, Ian Johnston, Dominic Kwiatkowski, Cordelia Langford, John Sillitoe on behalf of the Wellcome Sanger Institute COVID-19 Surveillance Team |
| EPI_ISL_2847450 | Lighthouse Lab in Milton Keynes | Wellcome Sanger Institute for the COVID-19 Genomics UK (COG-UK) Consortium | The Lighthouse Lab in Milton Keynes and Alex Alderton, Roberto Amato, Jeffrey Barrett, Sonia Goncalves, Ewan Harrison, David K. Jackson, Ian Johnston, Dominic Kwiatkowski, Cordelia Langford, John Sillitoe on behalf of the Wellcome Sanger Institute COVID-19 Surveillance Team |

| | | | |
|---|---|---|---|
| EPI_ISL_2847451 | Lighthouse Lab in Alderley Park | Wellcome Sanger Institute for the COVID-19 Genomics UK (COG-UK) Consortium | Jacquelyn Wynn, Mairead Hyland, The Lighthouse Lab in Alderley Park and Alex Alderton, Roberto Amato, Jeffrey Barrett, Sonia Goncalves, Ewan Harrison, David K. Jackson, Ian Johnston, Dominic Kwiatkowski, Cordelia Langford, John Sillitoe on behalf of the Wellcome Sanger Institute COVID-19 Surveillance Team |
| EPI_ISL_2847452 | Lighthouse Lab in Glasgow | Wellcome Sanger Institute for the COVID-19 Genomics UK (COG-UK) Consortium | Harper VanSteenhouse, Yumi Kasai, David Gray, Carol Clugston, Anna Dominiczak and Alex Alderton, Roberto Amato, Jeffrey Barrett, Sonia Goncalves, Ewan Harrison, David K. Jackson, Ian Johnston, Dominic Kwiatkowski, Cordelia Langford, John Sillitoe on behalf of the Wellcome Sanger Institute COVID-19 Surveillance Team |
| EPI_ISL_2847453 | Lighthouse Lab in Alderley Park | Wellcome Sanger Institute for the COVID-19 Genomics UK (COG-UK) Consortium | Jacquelyn Wynn, Mairead Hyland, The Lighthouse Lab in Alderley Park and Alex Alderton, Roberto Amato, Jeffrey Barrett, Sonia Goncalves, Ewan Harrison, David K. Jackson, Ian Johnston, Dominic Kwiatkowski, Cordelia Langford, John Sillitoe on behalf of the Wellcome Sanger Institute COVID-19 Surveillance Team |
| EPI_ISL_2847454 | Lighthouse Lab in Milton Keynes | Wellcome Sanger Institute for the COVID-19 Genomics UK (COG-UK) Consortium | The Lighthouse Lab in Milton Keynes and Alex Alderton, Roberto Amato, Jeffrey Barrett, Sonia Goncalves, Ewan Harrison, David K. Jackson, Ian Johnston, Dominic Kwiatkowski, Cordelia Langford, John Sillitoe on behalf of the Wellcome Sanger Institute COVID-19 Surveillance Team |
| EPI_ISL_2847455, EPI_ISL_2847456 | Lighthouse Laboratory Plymouth | Wellcome Sanger Institute for the COVID-19 Genomics UK (COG-UK) Consortium | Lighthouse Laboratory Plymouth and Alex Alderton, Roberto Amato, Jeffrey Barrett, Sonia Goncalves, Ewan Harrison, David K. Jackson, Ian Johnston, Dominic Kwiatkowski, Cordelia Langford, John Sillitoe on behalf of the Wellcome Sanger Institute COVID-19 Surveillance Team |
| EPI_ISL_2847457 | Lighthouse Lab in Milton Keynes | Wellcome Sanger Institute for the COVID-19 Genomics UK (COG-UK) Consortium | The Lighthouse Lab in Milton Keynes and Alex Alderton, Roberto Amato, Jeffrey Barrett, Sonia Goncalves, Ewan Harrison, David K. Jackson, Ian Johnston, Dominic Kwiatkowski, Cordelia Langford, John Sillitoe on behalf of the Wellcome Sanger Institute COVID-19 Surveillance Team |
| EPI_ISL_2847458 | Lighthouse Laboratory Plymouth | Wellcome Sanger Institute for the COVID-19 Genomics UK (COG-UK) Consortium | Lighthouse Laboratory Plymouth and Alex Alderton, Roberto Amato, Jeffrey Barrett, Sonia Goncalves, Ewan Harrison, David K. Jackson, Ian Johnston, Dominic Kwiatkowski, Cordelia Langford, John Sillitoe on behalf of the Wellcome Sanger Institute COVID-19 Surveillance Team |
| EPI_ISL_2847459 | Lighthouse Lab in Alderley Park | Wellcome Sanger Institute for the COVID-19 Genomics UK (COG-UK) Consortium | Jacquelyn Wynn, Mairead Hyland, The Lighthouse Lab in Alderley Park and Alex Alderton, Roberto Amato, Jeffrey Barrett, Sonia Goncalves, Ewan Harrison, David K. Jackson, Ian Johnston, Dominic Kwiatkowski, Cordelia Langford, John Sillitoe on behalf of the Wellcome Sanger Institute COVID-19 Surveillance Team |
| EPI_ISL_2847460 | Lighthouse Lab in Milton Keynes | Wellcome Sanger Institute for the COVID-19 Genomics UK (COG-UK) Consortium | The Lighthouse Lab in Milton Keynes and Alex Alderton, Roberto Amato, Jeffrey Barrett, Sonia Goncalves, Ewan Harrison, David K. Jackson, Ian Johnston, Dominic Kwiatkowski, Cordelia Langford, John Sillitoe on behalf of the Wellcome Sanger Institute COVID-19 Surveillance Team |
| EPI_ISL_2847461 | Lighthouse Lab in Glasgow | Wellcome Sanger Institute for the COVID-19 Genomics UK (COG-UK) Consortium | Harper VanSteenhouse, Yumi Kasai, David Gray, Carol Clugston, Anna Dominiczak and Alex Alderton, Roberto Amato, Jeffrey Barrett, Sonia Goncalves, Ewan Harrison, David K. Jackson, Ian Johnston, Dominic Kwiatkowski, Cordelia Langford, John Sillitoe on behalf of the Wellcome Sanger Institute COVID-19 Surveillance Team |
| EPI_ISL_2847462 | Lighthouse Lab in Alderley Park | Wellcome Sanger Institute for the COVID-19 Genomics UK (COG-UK) Consortium | Jacquelyn Wynn, Mairead Hyland, The Lighthouse Lab in Alderley Park and Alex Alderton, Roberto Amato, Jeffrey Barrett, Sonia Goncalves, Ewan Harrison, David K. Jackson, Ian Johnston, Dominic Kwiatkowski, Cordelia Langford, John Sillitoe on behalf of the Wellcome Sanger Institute COVID-19 Surveillance Team |
| EPI_ISL_2847463 | Lighthouse Lab in Milton Keynes | Wellcome Sanger Institute for the COVID-19 Genomics UK (COG-UK) Consortium | The Lighthouse Lab in Milton Keynes and Alex Alderton, Roberto Amato, Jeffrey Barrett, Sonia Goncalves, Ewan Harrison, David K. Jackson, Ian Johnston, Dominic Kwiatkowski, Cordelia Langford, John Sillitoe on behalf of the Wellcome Sanger Institute COVID-19 Surveillance Team |
| EPI_ISL_2847464, EPI_ISL_2847465, EPI_ISL_2847467 | Lighthouse Lab in Alderley Park | Wellcome Sanger Institute for the COVID-19 Genomics UK (COG-UK) Consortium | Jacquelyn Wynn, Mairead Hyland, The Lighthouse Lab in Alderley Park and Alex Alderton, Roberto Amato, Jeffrey Barrett, Sonia Goncalves, Ewan Harrison, David K. Jackson, Ian Johnston, Dominic Kwiatkowski, Cordelia Langford, John Sillitoe on behalf of the Wellcome Sanger Institute COVID-19 Surveillance Team |
| EPI_ISL_2847468 | Lighthouse Laboratory Plymouth | Wellcome Sanger Institute for the COVID-19 Genomics UK (COG-UK) Consortium | Lighthouse Laboratory Plymouth and Alex Alderton, Roberto Amato, Jeffrey Barrett, Sonia Goncalves, Ewan Harrison, David K. Jackson, Ian Johnston, Dominic Kwiatkowski, Cordelia Langford, John Sillitoe on behalf of the Wellcome Sanger Institute COVID-19 Surveillance Team |
| EPI_ISL_2847469, EPI_ISL_2847470, EPI_ISL_2847471 | Lighthouse Lab in Glasgow | Wellcome Sanger Institute for the COVID-19 Genomics UK (COG-UK) Consortium | Harper VanSteenhouse, Yumi Kasai, David Gray, Carol Clugston, Anna Dominiczak and Alex Alderton, Roberto Amato, Jeffrey Barrett, Sonia Goncalves, Ewan Harrison, David K. Jackson, Ian Johnston, Dominic Kwiatkowski, Cordelia Langford, John Sillitoe on behalf of the Wellcome Sanger Institute COVID-19 Surveillance Team |
| EPI_ISL_2847472 | Lighthouse Lab in Milton Keynes | Wellcome Sanger Institute for the COVID-19 Genomics UK (COG-UK) Consortium | The Lighthouse Lab in Milton Keynes and Alex Alderton, Roberto Amato, Jeffrey Barrett, Sonia Goncalves, Ewan Harrison, David K. Jackson, Ian Johnston, Dominic Kwiatkowski, Cordelia Langford, John Sillitoe on behalf of the Wellcome Sanger Institute COVID-19 Surveillance Team |
| EPI_ISL_2847473 | Lighthouse Lab in Glasgow | Wellcome Sanger Institute for the COVID-19 Genomics UK (COG-UK) Consortium | Harper VanSteenhouse, Yumi Kasai, David Gray, Carol Clugston, Anna Dominiczak and Alex Alderton, Roberto Amato, Jeffrey Barrett, Sonia Goncalves, Ewan Harrison, David K. Jackson, Ian Johnston, Dominic Kwiatkowski, Cordelia Langford, John Sillitoe on behalf of the Wellcome Sanger Institute COVID-19 Surveillance Team |
| EPI_ISL_2847474 | Lighthouse Laboratory Plymouth | Wellcome Sanger Institute for the COVID-19 Genomics UK (COG-UK) Consortium | Lighthouse Laboratory Plymouth and Alex Alderton, Roberto Amato, Jeffrey Barrett, Sonia Goncalves, Ewan Harrison, David K. Jackson, Ian Johnston, Dominic Kwiatkowski, Cordelia Langford, John Sillitoe on behalf of the Wellcome Sanger Institute COVID-19 Surveillance Team |
| EPI_ISL_2847475 | Lighthouse Lab in Alderley Park | Wellcome Sanger Institute for the COVID-19 Genomics UK (COG-UK) Consortium | Jacquelyn Wynn, Mairead Hyland, The Lighthouse Lab in Alderley Park and Alex Alderton, Roberto Amato, Jeffrey Barrett, Sonia Goncalves, Ewan Harrison, David K. Jackson, Ian Johnston, Dominic Kwiatkowski, Cordelia Langford, John Sillitoe on behalf of the Wellcome Sanger Institute COVID-19 Surveillance Team |
| EPI_ISL_2847477, EPI_ISL_2847478 | Lighthouse Laboratory Plymouth | Wellcome Sanger Institute for the COVID-19 Genomics UK (COG-UK) Consortium | Lighthouse Laboratory Plymouth and Alex Alderton, Roberto Amato, Jeffrey Barrett, Sonia Goncalves, Ewan Harrison, David K. Jackson, Ian Johnston, Dominic Kwiatkowski, Cordelia Langford, John Sillitoe on behalf of the Wellcome Sanger Institute COVID-19 Surveillance Team |
| EPI_ISL_2847479 | Lighthouse Lab in Alderley Park | Wellcome Sanger Institute for the COVID-19 Genomics UK (COG-UK) Consortium | Jacquelyn Wynn, Mairead Hyland, The Lighthouse Lab in Alderley Park and Alex Alderton, Roberto Amato, Jeffrey Barrett, Sonia Goncalves, Ewan Harrison, David K. Jackson, Ian Johnston, Dominic Kwiatkowski, Cordelia Langford, John Sillitoe on behalf of the Wellcome Sanger Institute COVID-19 Surveillance Team |
| EPI_ISL_2847480, EPI_ISL_2847481 | Lighthouse Lab in Milton Keynes | Wellcome Sanger Institute for the COVID-19 Genomics UK (COG-UK) Consortium | The Lighthouse Lab in Milton Keynes and Alex Alderton, Roberto Amato, Jeffrey Barrett, Sonia Goncalves, Ewan Harrison, David K. Jackson, Ian Johnston, Dominic Kwiatkowski, Cordelia Langford, John Sillitoe on behalf of the Wellcome Sanger Institute COVID-19 Surveillance Team |
| EPI_ISL_2847482 | Lighthouse Lab in Glasgow | Wellcome Sanger Institute for the COVID-19 Genomics UK (COG-UK) Consortium | Harper VanSteenhouse, Yumi Kasai, David Gray, Carol Clugston, Anna Dominiczak and Alex Alderton, Roberto Amato, Jeffrey Barrett, Sonia Goncalves, Ewan Harrison, David K. Jackson, Ian Johnston, Dominic Kwiatkowski, Cordelia Langford, John Sillitoe on behalf of the Wellcome Sanger Institute COVID-19 Surveillance Team |
| EPI_ISL_2847483 | Lighthouse Laboratory Plymouth | Wellcome Sanger Institute for the COVID-19 Genomics UK (COG-UK) Consortium | Lighthouse Laboratory Plymouth and Alex Alderton, Roberto Amato, Jeffrey Barrett, Sonia Goncalves, Ewan Harrison, David K. Jackson, Ian Johnston, Dominic Kwiatkowski, Cordelia Langford, John Sillitoe on behalf of the Wellcome Sanger Institute COVID-19 Surveillance Team |
| EPI_ISL_2847484 | Lighthouse Lab in Glasgow | Wellcome Sanger Institute for the COVID-19 Genomics UK (COG-UK) Consortium | Harper VanSteenhouse, Yumi Kasai, David Gray, Carol Clugston, Anna Dominiczak and Alex Alderton, Roberto Amato, Jeffrey Barrett, Sonia Goncalves, Ewan Harrison, David K. Jackson, Ian Johnston, Dominic Kwiatkowski, Cordelia Langford, John Sillitoe on behalf of the Wellcome Sanger Institute COVID-19 Surveillance Team |
| EPI_ISL_2847485 | Lighthouse Laboratory Plymouth | Wellcome Sanger Institute for the COVID-19 Genomics UK (COG-UK) Consortium | Lighthouse Laboratory Plymouth and Alex Alderton, Roberto Amato, Jeffrey Barrett, Sonia Goncalves, Ewan Harrison, David K. Jackson, Ian Johnston, Dominic Kwiatkowski, Cordelia Langford, John Sillitoe on behalf of the Wellcome Sanger Institute COVID-19 Surveillance Team |
| EPI_ISL_2847486, EPI_ISL_2847487 | Lighthouse Lab in Alderley Park | Wellcome Sanger Institute for the COVID-19 Genomics UK (COG-UK) Consortium | Jacquelyn Wynn, Mairead Hyland, The Lighthouse Lab in Alderley Park and Alex Alderton, Roberto Amato, Jeffrey Barrett, Sonia Goncalves, Ewan Harrison, David K. Jackson, Ian Johnston, Dominic Kwiatkowski, Cordelia Langford, John Sillitoe on behalf of the Wellcome Sanger Institute COVID-19 Surveillance Team |
| EPI_ISL_2847488 | Lighthouse Lab in Glasgow | Wellcome Sanger Institute for the COVID-19 Genomics UK (COG-UK) Consortium | Harper VanSteenhouse, Yumi Kasai, David Gray, Carol Clugston, Anna Dominiczak and Alex Alderton, Roberto Amato, Jeffrey Barrett, Sonia Goncalves, Ewan Harrison, David K. Jackson, Ian Johnston, Dominic Kwiatkowski, Cordelia Langford, John Sillitoe on behalf of the Wellcome Sanger Institute COVID-19 Surveillance Team |
| EPI_ISL_2847489 | Lighthouse Laboratory Plymouth | Wellcome Sanger Institute for the COVID-19 Genomics UK (COG-UK) Consortium | Lighthouse Laboratory Plymouth and Alex Alderton, Roberto Amato, Jeffrey Barrett, Sonia Goncalves, Ewan Harrison, David K. Jackson, Ian Johnston, Dominic Kwiatkowski, Cordelia Langford, John Sillitoe on behalf of the Wellcome Sanger Institute COVID-19 Surveillance Team |
| EPI_ISL_2847490 | Lighthouse Lab in Alderley Park | Wellcome Sanger Institute for the COVID-19 Genomics UK (COG-UK) Consortium | Jacquelyn Wynn, Mairead Hyland, The Lighthouse Lab in Alderley Park and Alex Alderton, Roberto Amato, Jeffrey Barrett, Sonia Goncalves, Ewan Harrison, David K. Jackson, Ian Johnston, Dominic Kwiatkowski, Cordelia Langford, John Sillitoe on behalf of the Wellcome Sanger Institute COVID-19 Surveillance Team |
| EPI_ISL_2847491 | Lighthouse Lab in Glasgow | Wellcome Sanger Institute for the COVID-19 Genomics UK (COG-UK) Consortium | Harper VanSteenhouse, Yumi Kasai, David Gray, Carol Clugston, Anna Dominiczak and Alex Alderton, Roberto Amato, Jeffrey Barrett, Sonia Goncalves, Ewan Harrison, David K. Jackson, Ian Johnston, Dominic Kwiatkowski, Cordelia Langford, John Sillitoe on behalf of the Wellcome Sanger Institute COVID-19 Surveillance Team |
| EPI_ISL_2847492, EPI_ISL_2847493 | Lighthouse Lab in Milton Keynes | Wellcome Sanger Institute for the COVID-19 Genomics UK (COG-UK) Consortium | The Lighthouse Lab in Milton Keynes and Alex Alderton, Roberto Amato, Jeffrey Barrett, Sonia Goncalves, Ewan Harrison, David K. Jackson, Ian Johnston, Dominic Kwiatkowski, Cordelia Langford, John Sillitoe on behalf of the Wellcome Sanger Institute COVID-19 Surveillance Team |
| EPI_ISL_2847494 | Lighthouse Laboratory Plymouth | Wellcome Sanger Institute for the COVID-19 Genomics UK (COG-UK) Consortium | Lighthouse Laboratory Plymouth and Alex Alderton, Roberto Amato, Jeffrey Barrett, Sonia Goncalves, Ewan Harrison, David K. Jackson, Ian Johnston, Dominic Kwiatkowski, Cordelia Langford, John Sillitoe on behalf of the Wellcome Sanger Institute COVID-19 Surveillance Team |
| EPI_ISL_2847495 | Lighthouse Lab in Milton Keynes | Wellcome Sanger Institute for the COVID-19 Genomics UK (COG-UK) Consortium | The Lighthouse Lab in Milton Keynes and Alex Alderton, Roberto Amato, Jeffrey Barrett, Sonia Goncalves, Ewan Harrison, David K. Jackson, Ian Johnston, Dominic Kwiatkowski, Cordelia Langford, John Sillitoe on behalf of the Wellcome Sanger Institute COVID-19 Surveillance Team |
| EPI_ISL_2847496 | Lighthouse Laboratory Plymouth | Wellcome Sanger Institute for the COVID-19 Genomics UK (COG-UK) Consortium | Lighthouse Laboratory Plymouth and Alex Alderton, Roberto Amato, Jeffrey Barrett, Sonia Goncalves, Ewan Harrison, David K. Jackson, Ian Johnston, Dominic Kwiatkowski, Cordelia Langford, John Sillitoe on behalf of the Wellcome Sanger Institute COVID-19 Surveillance Team |
| EPI_ISL_2847497 | Lighthouse Lab in Milton Keynes | Wellcome Sanger Institute for the COVID-19 Genomics UK (COG-UK) Consortium | The Lighthouse Lab in Milton Keynes and Alex Alderton, Roberto Amato, Jeffrey Barrett, Sonia Goncalves, Ewan Harrison, David K. Jackson, Ian Johnston, Dominic Kwiatkowski, Cordelia Langford, John Sillitoe on behalf of the Wellcome Sanger Institute COVID-19 Surveillance Team |
| EPI_ISL_2847498, EPI_ISL_2847499 | Lighthouse Laboratory Plymouth | Wellcome Sanger Institute for the COVID-19 Genomics UK (COG-UK) Consortium | Lighthouse Laboratory Plymouth and Alex Alderton, Roberto Amato, Jeffrey Barrett, Sonia Goncalves, Ewan Harrison, David K. Jackson, Ian Johnston, Dominic Kwiatkowski, Cordelia Langford, John Sillitoe on behalf of the Wellcome Sanger Institute COVID-19 Surveillance Team |
| EPI_ISL_2847500 | Lighthouse Lab in Milton Keynes | Wellcome Sanger Institute for the COVID-19 Genomics UK (COG-UK) Consortium | The Lighthouse Lab in Milton Keynes and Alex Alderton, Roberto Amato, Jeffrey Barrett, Sonia Goncalves, Ewan Harrison, David K. Jackson, Ian Johnston, Dominic Kwiatkowski, Cordelia Langford, John Sillitoe on behalf of the Wellcome Sanger Institute COVID-19 Surveillance Team |
| EPI_ISL_2847501, EPI_ISL_2847502 | Lighthouse Lab in Alderley Park | Wellcome Sanger Institute for the COVID-19 Genomics UK (COG-UK) Consortium | Jacquelyn Wynn, Mairead Hyland, The Lighthouse Lab in Alderley Park and Alex Alderton, Roberto Amato, Jeffrey Barrett, Sonia Goncalves, Ewan Harrison, David K. Jackson, Ian Johnston, Dominic Kwiatkowski, Cordelia Langford, John Sillitoe on behalf of the Wellcome Sanger Institute COVID-19 Surveillance Team |
| EPI_ISL_2847503 | Lighthouse Lab in Milton Keynes | Wellcome Sanger Institute for the COVID-19 Genomics UK (COG-UK) Consortium | The Lighthouse Lab in Milton Keynes and Alex Alderton, Roberto Amato, Jeffrey Barrett, Sonia Goncalves, Ewan Harrison, David K. Jackson, Ian Johnston, Dominic Kwiatkowski, Cordelia Langford, John Sillitoe on behalf of the Wellcome Sanger Institute COVID-19 Surveillance Team |
| EPI_ISL_2847504 | Lighthouse Laboratory Plymouth | Wellcome Sanger Institute for the COVID-19 Genomics UK (COG-UK) Consortium | Lighthouse Laboratory Plymouth and Alex Alderton, Roberto Amato, Jeffrey Barrett, Sonia Goncalves, Ewan Harrison, David K. Jackson, Ian Johnston, Dominic Kwiatkowski, Cordelia Langford, John Sillitoe on behalf of the Wellcome Sanger Institute COVID-19 Surveillance Team |
| EPI_ISL_2847505 | Lighthouse Lab in Glasgow | Wellcome Sanger Institute for the COVID-19 Genomics UK (COG-UK) Consortium | Harper VanSteenhouse, Yumi Kasai, David Gray, Carol Clugston, Anna Dominiczak and Alex Alderton, Roberto Amato, Jeffrey Barrett, Sonia Goncalves, Ewan Harrison, David K. Jackson, Ian Johnston, Dominic Kwiatkowski, Cordelia Langford, John Sillitoe on behalf of the Wellcome Sanger Institute COVID-19 Surveillance Team |
| EPI_ISL_2847506 | Lighthouse Laboratory Plymouth | Wellcome Sanger Institute for the COVID-19 Genomics UK (COG-UK) Consortium | Lighthouse Laboratory Plymouth and Alex Alderton, Roberto Amato, Jeffrey Barrett, Sonia Goncalves, Ewan Harrison, David K. Jackson, Ian Johnston, Dominic Kwiatkowski, Cordelia Langford, John Sillitoe on behalf of the Wellcome Sanger Institute COVID-19 Surveillance Team |
| EPI_ISL_2847507 | Lighthouse Lab in Glasgow | Wellcome Sanger Institute for the COVID-19 Genomics UK (COG-UK) Consortium | Harper VanSteenhouse, Yumi Kasai, David Gray, Carol Clugston, Anna Dominiczak and Alex Alderton, Roberto Amato, Jeffrey Barrett, Sonia Goncalves, Ewan Harrison, David K. Jackson, Ian Johnston, Dominic Kwiatkowski, Cordelia Langford, John Sillitoe on behalf of the Wellcome Sanger Institute COVID-19 Surveillance Team |

| | | | |
|---|---|---|---|
| EPI_ISL_2847508 | Lighthouse Lab in Alderley Park | Wellcome Sanger Institute for the COVID-19 Genomics UK (COG-UK) Consortium | Jacquelyn Wynn, Mairead Hyland, The Lighthouse Lab in Alderley Park and Alex Alderton, Roberto Amato, Jeffrey Barrett, Sonia Goncalves, Ewan Harrison, David K. Jackson, Ian Johnston, Dominic Kwiatkowski, Cordelia Langford, John Sillitoe on behalf of the Wellcome Sanger Institute COVID-19 Surveillance Team |
| EPI_ISL_2847509 | Lighthouse Lab in Milton Keynes | Wellcome Sanger Institute for the COVID-19 Genomics UK (COG-UK) Consortium | The Lighthouse Lab in Milton Keynes and Alex Alderton, Roberto Amato, Jeffrey Barrett, Sonia Goncalves, Ewan Harrison, David K. Jackson, Ian Johnston, Dominic Kwiatkowski, Cordelia Langford, John Sillitoe on behalf of the Wellcome Sanger Institute COVID-19 Surveillance Team |
| EPI_ISL_2847510 | Lighthouse Lab in Glasgow | Wellcome Sanger Institute for the COVID-19 Genomics UK (COG-UK) Consortium | Harper VanSteenhouse, Yumi Kasai, David Gray, Carol Clugston, Anna Dominiczak and Alex Alderton, Roberto Amato, Jeffrey Barrett, Sonia Goncalves, Ewan Harrison, David K. Jackson, Ian Johnston, Dominic Kwiatkowski, Cordelia Langford, John Sillitoe on behalf of the Wellcome Sanger Institute COVID-19 Surveillance Team |
| EPI_ISL_2847511, EPI_ISL_2847512, EPI_ISL_2847513, EPI_ISL_2847514 | Lighthouse Lab in Milton Keynes | Wellcome Sanger Institute for the COVID-19 Genomics UK (COG-UK) Consortium | The Lighthouse Lab in Milton Keynes and Alex Alderton, Roberto Amato, Jeffrey Barrett, Sonia Goncalves, Ewan Harrison, David K. Jackson, Ian Johnston, Dominic Kwiatkowski, Cordelia Langford, John Sillitoe on behalf of the Wellcome Sanger Institute COVID-19 Surveillance Team |
| EPI_ISL_2847515 | Lighthouse Laboratory Plymouth | Wellcome Sanger Institute for the COVID-19 Genomics UK (COG-UK) Consortium | Lighthouse Laboratory Plymouth and Alex Alderton, Roberto Amato, Jeffrey Barrett, Sonia Goncalves, Ewan Harrison, David K. Jackson, Ian Johnston, Dominic Kwiatkowski, Cordelia Langford, John Sillitoe on behalf of the Wellcome Sanger Institute COVID-19 Surveillance Team |
| EPI_ISL_2847516 | Lighthouse Lab in Milton Keynes | Wellcome Sanger Institute for the COVID-19 Genomics UK (COG-UK) Consortium | The Lighthouse Lab in Milton Keynes and Alex Alderton, Roberto Amato, Jeffrey Barrett, Sonia Goncalves, Ewan Harrison, David K. Jackson, Ian Johnston, Dominic Kwiatkowski, Cordelia Langford, John Sillitoe on behalf of the Wellcome Sanger Institute COVID-19 Surveillance Team |
| EPI_ISL_2847517 | Lighthouse Laboratory Plymouth | Wellcome Sanger Institute for the COVID-19 Genomics UK (COG-UK) Consortium | Lighthouse Laboratory Plymouth and Alex Alderton, Roberto Amato, Jeffrey Barrett, Sonia Goncalves, Ewan Harrison, David K. Jackson, Ian Johnston, Dominic Kwiatkowski, Cordelia Langford, John Sillitoe on behalf of the Wellcome Sanger Institute COVID-19 Surveillance Team |
| EPI_ISL_2847518 | Lighthouse Lab in Alderley Park | Wellcome Sanger Institute for the COVID-19 Genomics UK (COG-UK) Consortium | Jacquelyn Wynn, Mairead Hyland, The Lighthouse Lab in Alderley Park and Alex Alderton, Roberto Amato, Jeffrey Barrett, Sonia Goncalves, Ewan Harrison, David K. Jackson, Ian Johnston, Dominic Kwiatkowski, Cordelia Langford, John Sillitoe on behalf of the Wellcome Sanger Institute COVID-19 Surveillance Team |
| EPI_ISL_2847519 | Lighthouse Lab in Milton Keynes | Wellcome Sanger Institute for the COVID-19 Genomics UK (COG-UK) Consortium | The Lighthouse Lab in Milton Keynes and Alex Alderton, Roberto Amato, Jeffrey Barrett, Sonia Goncalves, Ewan Harrison, David K. Jackson, Ian Johnston, Dominic Kwiatkowski, Cordelia Langford, John Sillitoe on behalf of the Wellcome Sanger Institute COVID-19 Surveillance Team |
| EPI_ISL_2847520 | Lighthouse Laboratory Plymouth | Wellcome Sanger Institute for the COVID-19 Genomics UK (COG-UK) Consortium | Lighthouse Laboratory Plymouth and Alex Alderton, Roberto Amato, Jeffrey Barrett, Sonia Goncalves, Ewan Harrison, David K. Jackson, Ian Johnston, Dominic Kwiatkowski, Cordelia Langford, John Sillitoe on behalf of the Wellcome Sanger Institute COVID-19 Surveillance Team |
| EPI_ISL_2847521, EPI_ISL_2847522, EPI_ISL_2847523, EPI_ISL_2847524 | Lighthouse Lab in Alderley Park | Wellcome Sanger Institute for the COVID-19 Genomics UK (COG-UK) Consortium | Jacquelyn Wynn, Mairead Hyland, The Lighthouse Lab in Alderley Park and Alex Alderton, Roberto Amato, Jeffrey Barrett, Sonia Goncalves, Ewan Harrison, David K. Jackson, Ian Johnston, Dominic Kwiatkowski, Cordelia Langford, John Sillitoe on behalf of the Wellcome Sanger Institute COVID-19 Surveillance Team |
| EPI_ISL_2847525, EPI_ISL_2847526, EPI_ISL_2847527 | Lighthouse Lab in Milton Keynes | Wellcome Sanger Institute for the COVID-19 Genomics UK (COG-UK) Consortium | The Lighthouse Lab in Milton Keynes and Alex Alderton, Roberto Amato, Jeffrey Barrett, Sonia Goncalves, Ewan Harrison, David K. Jackson, Ian Johnston, Dominic Kwiatkowski, Cordelia Langford, John Sillitoe on behalf of the Wellcome Sanger Institute COVID-19 Surveillance Team |
| EPI_ISL_2847528 | Lighthouse Lab in Alderley Park | Wellcome Sanger Institute for the COVID-19 Genomics UK (COG-UK) Consortium | Jacquelyn Wynn, Mairead Hyland, The Lighthouse Lab in Alderley Park and Alex Alderton, Roberto Amato, Jeffrey Barrett, Sonia Goncalves, Ewan Harrison, David K. Jackson, Ian Johnston, Dominic Kwiatkowski, Cordelia Langford, John Sillitoe on behalf of the Wellcome Sanger Institute COVID-19 Surveillance Team |
| EPI_ISL_2847529 | Lighthouse Laboratory Plymouth | Wellcome Sanger Institute for the COVID-19 Genomics UK (COG-UK) Consortium | Lighthouse Laboratory Plymouth and Alex Alderton, Roberto Amato, Jeffrey Barrett, Sonia Goncalves, Ewan Harrison, David K. Jackson, Ian Johnston, Dominic Kwiatkowski, Cordelia Langford, John Sillitoe on behalf of the Wellcome Sanger Institute COVID-19 Surveillance Team |
| EPI_ISL_2847530 | Lighthouse Lab in Alderley Park | Wellcome Sanger Institute for the COVID-19 Genomics UK (COG-UK) Consortium | Jacquelyn Wynn, Mairead Hyland, The Lighthouse Lab in Alderley Park and Alex Alderton, Roberto Amato, Jeffrey Barrett, Sonia Goncalves, Ewan Harrison, David K. Jackson, Ian Johnston, Dominic Kwiatkowski, Cordelia Langford, John Sillitoe on behalf of the Wellcome Sanger Institute COVID-19 Surveillance Team |
| EPI_ISL_2847531 | Lighthouse Lab in Milton Keynes | Wellcome Sanger Institute for the COVID-19 Genomics UK (COG-UK) Consortium | The Lighthouse Lab in Milton Keynes and Alex Alderton, Roberto Amato, Jeffrey Barrett, Sonia Goncalves, Ewan Harrison, David K. Jackson, Ian Johnston, Dominic Kwiatkowski, Cordelia Langford, John Sillitoe on behalf of the Wellcome Sanger Institute COVID-19 Surveillance Team |
| EPI_ISL_2847532 | Lighthouse Lab in Glasgow | Wellcome Sanger Institute for the COVID-19 Genomics UK (COG-UK) Consortium | Harper VanSteenhouse, Yumi Kasai, David Gray, Carol Clugston, Anna Dominiczak and Alex Alderton, Roberto Amato, Jeffrey Barrett, Sonia Goncalves, Ewan Harrison, David K. Jackson, Ian Johnston, Dominic Kwiatkowski, Cordelia Langford, John Sillitoe on behalf of the Wellcome Sanger Institute COVID-19 Surveillance Team |
| EPI_ISL_2847533 | Lighthouse Laboratory Plymouth | Wellcome Sanger Institute for the COVID-19 Genomics UK (COG-UK) Consortium | Lighthouse Laboratory Plymouth and Alex Alderton, Roberto Amato, Jeffrey Barrett, Sonia Goncalves, Ewan Harrison, David K. Jackson, Ian Johnston, Dominic Kwiatkowski, Cordelia Langford, John Sillitoe on behalf of the Wellcome Sanger Institute COVID-19 Surveillance Team |
| EPI_ISL_2847534, EPI_ISL_2847535 | Lighthouse Lab in Glasgow | Wellcome Sanger Institute for the COVID-19 Genomics UK (COG-UK) Consortium | Harper VanSteenhouse, Yumi Kasai, David Gray, Carol Clugston, Anna Dominiczak and Alex Alderton, Roberto Amato, Jeffrey Barrett, Sonia Goncalves, Ewan Harrison, David K. Jackson, Ian Johnston, Dominic Kwiatkowski, Cordelia Langford, John Sillitoe on behalf of the Wellcome Sanger Institute COVID-19 Surveillance Team |
| EPI_ISL_2847536, EPI_ISL_2847537 | Lighthouse Lab in Alderley Park | Wellcome Sanger Institute for the COVID-19 Genomics UK (COG-UK) Consortium | Jacquelyn Wynn, Mairead Hyland, The Lighthouse Lab in Alderley Park and Alex Alderton, Roberto Amato, Jeffrey Barrett, Sonia Goncalves, Ewan Harrison, David K. Jackson, Ian Johnston, Dominic Kwiatkowski, Cordelia Langford, John Sillitoe on behalf of the Wellcome Sanger Institute COVID-19 Surveillance Team |
| EPI_ISL_2847538 | Lighthouse Lab in Milton Keynes | Wellcome Sanger Institute for the COVID-19 Genomics UK (COG-UK) Consortium | The Lighthouse Lab in Milton Keynes and Alex Alderton, Roberto Amato, Jeffrey Barrett, Sonia Goncalves, Ewan Harrison, David K. Jackson, Ian Johnston, Dominic Kwiatkowski, Cordelia Langford, John Sillitoe on behalf of the Wellcome Sanger Institute COVID-19 Surveillance Team |
| EPI_ISL_2847539, EPI_ISL_2847540 | Lighthouse Laboratory Plymouth | Wellcome Sanger Institute for the COVID-19 Genomics UK (COG-UK) Consortium | Lighthouse Laboratory Plymouth and Alex Alderton, Roberto Amato, Jeffrey Barrett, Sonia Goncalves, Ewan Harrison, David K. Jackson, Ian Johnston, Dominic Kwiatkowski, Cordelia Langford, John Sillitoe on behalf of the Wellcome Sanger Institute COVID-19 Surveillance Team |
| EPI_ISL_2847541 | Lighthouse Lab in Milton Keynes | Wellcome Sanger Institute for the COVID-19 Genomics UK (COG-UK) Consortium | The Lighthouse Lab in Milton Keynes and Alex Alderton, Roberto Amato, Jeffrey Barrett, Sonia Goncalves, Ewan Harrison, David K. Jackson, Ian Johnston, Dominic Kwiatkowski, Cordelia Langford, John Sillitoe on behalf of the Wellcome Sanger Institute COVID-19 Surveillance Team |
| EPI_ISL_2847542 | Lighthouse Lab in Alderley Park | Wellcome Sanger Institute for the COVID-19 Genomics UK (COG-UK) Consortium | Jacquelyn Wynn, Mairead Hyland, The Lighthouse Lab in Alderley Park and Alex Alderton, Roberto Amato, Jeffrey Barrett, Sonia Goncalves, Ewan Harrison, David K. Jackson, Ian Johnston, Dominic Kwiatkowski, Cordelia Langford, John Sillitoe on behalf of the Wellcome Sanger Institute COVID-19 Surveillance Team |
| EPI_ISL_2847543 | Lighthouse Lab in Glasgow | Wellcome Sanger Institute for the COVID-19 Genomics UK (COG-UK) Consortium | Harper VanSteenhouse, Yumi Kasai, David Gray, Carol Clugston, Anna Dominiczak and Alex Alderton, Roberto Amato, Jeffrey Barrett, Sonia Goncalves, Ewan Harrison, David K. Jackson, Ian Johnston, Dominic Kwiatkowski, Cordelia Langford, John Sillitoe on behalf of the Wellcome Sanger Institute COVID-19 Surveillance Team |
| EPI_ISL_2847544, EPI_ISL_2847545, EPI_ISL_2847546 | Lighthouse Laboratory Plymouth | Wellcome Sanger Institute for the COVID-19 Genomics UK (COG-UK) Consortium | Lighthouse Laboratory Plymouth and Alex Alderton, Roberto Amato, Jeffrey Barrett, Sonia Goncalves, Ewan Harrison, David K. Jackson, Ian Johnston, Dominic Kwiatkowski, Cordelia Langford, John Sillitoe on behalf of the Wellcome Sanger Institute COVID-19 Surveillance Team |
| EPI_ISL_2847547 | Lighthouse Lab in Alderley Park | Wellcome Sanger Institute for the COVID-19 Genomics UK (COG-UK) Consortium | Jacquelyn Wynn, Mairead Hyland, The Lighthouse Lab in Alderley Park and Alex Alderton, Roberto Amato, Jeffrey Barrett, Sonia Goncalves, Ewan Harrison, David K. Jackson, Ian Johnston, Dominic Kwiatkowski, Cordelia Langford, John Sillitoe on behalf of the Wellcome Sanger Institute COVID-19 Surveillance Team |
| EPI_ISL_2847548 | Lighthouse Lab in Glasgow | Wellcome Sanger Institute for the COVID-19 Genomics UK (COG-UK) Consortium | Harper VanSteenhouse, Yumi Kasai, David Gray, Carol Clugston, Anna Dominiczak and Alex Alderton, Roberto Amato, Jeffrey Barrett, Sonia Goncalves, Ewan Harrison, David K. Jackson, Ian Johnston, Dominic Kwiatkowski, Cordelia Langford, John Sillitoe on behalf of the Wellcome Sanger Institute COVID-19 Surveillance Team |
| EPI_ISL_2847549 | Lighthouse Lab in Alderley Park | Wellcome Sanger Institute for the COVID-19 Genomics UK (COG-UK) Consortium | Jacquelyn Wynn, Mairead Hyland, The Lighthouse Lab in Alderley Park and Alex Alderton, Roberto Amato, Jeffrey Barrett, Sonia Goncalves, Ewan Harrison, David K. Jackson, Ian Johnston, Dominic Kwiatkowski, Cordelia Langford, John Sillitoe on behalf of the Wellcome Sanger Institute COVID-19 Surveillance Team |
| EPI_ISL_2847550 | Lighthouse Lab in Milton Keynes | Wellcome Sanger Institute for the COVID-19 Genomics UK (COG-UK) Consortium | The Lighthouse Lab in Milton Keynes and Alex Alderton, Roberto Amato, Jeffrey Barrett, Sonia Goncalves, Ewan Harrison, David K. Jackson, Ian Johnston, Dominic Kwiatkowski, Cordelia Langford, John Sillitoe on behalf of the Wellcome Sanger Institute COVID-19 Surveillance Team |
| EPI_ISL_2847551, EPI_ISL_2847552 | Lighthouse Laboratory Plymouth | Wellcome Sanger Institute for the COVID-19 Genomics UK (COG-UK) Consortium | Lighthouse Laboratory Plymouth and Alex Alderton, Roberto Amato, Jeffrey Barrett, Sonia Goncalves, Ewan Harrison, David K. Jackson, Ian Johnston, Dominic Kwiatkowski, Cordelia Langford, John Sillitoe on behalf of the Wellcome Sanger Institute COVID-19 Surveillance Team |
| EPI_ISL_2847553 | Lighthouse Lab in Alderley Park | Wellcome Sanger Institute for the COVID-19 Genomics UK (COG-UK) Consortium | Jacquelyn Wynn, Mairead Hyland, The Lighthouse Lab in Alderley Park and Alex Alderton, Roberto Amato, Jeffrey Barrett, Sonia Goncalves, Ewan Harrison, David K. Jackson, Ian Johnston, Dominic Kwiatkowski, Cordelia Langford, John Sillitoe on behalf of the Wellcome Sanger Institute COVID-19 Surveillance Team |
| EPI_ISL_2847554 | Lighthouse Lab in Glasgow | Wellcome Sanger Institute for the COVID-19 Genomics UK (COG-UK) Consortium | Harper VanSteenhouse, Yumi Kasai, David Gray, Carol Clugston, Anna Dominiczak and Alex Alderton, Roberto Amato, Jeffrey Barrett, Sonia Goncalves, Ewan Harrison, David K. Jackson, Ian Johnston, Dominic Kwiatkowski, Cordelia Langford, John Sillitoe on behalf of the Wellcome Sanger Institute COVID-19 Surveillance Team |
| EPI_ISL_2847555 | Lighthouse Lab in Alderley Park | Wellcome Sanger Institute for the COVID-19 Genomics UK (COG-UK) Consortium | Jacquelyn Wynn, Mairead Hyland, The Lighthouse Lab in Alderley Park and Alex Alderton, Roberto Amato, Jeffrey Barrett, Sonia Goncalves, Ewan Harrison, David K. Jackson, Ian Johnston, Dominic Kwiatkowski, Cordelia Langford, John Sillitoe on behalf of the Wellcome Sanger Institute COVID-19 Surveillance Team |
| EPI_ISL_2847556 | Lighthouse Lab in Milton Keynes | Wellcome Sanger Institute for the COVID-19 Genomics UK (COG-UK) Consortium | The Lighthouse Lab in Milton Keynes and Alex Alderton, Roberto Amato, Jeffrey Barrett, Sonia Goncalves, Ewan Harrison, David K. Jackson, Ian Johnston, Dominic Kwiatkowski, Cordelia Langford, John Sillitoe on behalf of the Wellcome Sanger Institute COVID-19 Surveillance Team |
| EPI_ISL_2847557 | Lighthouse Laboratory Plymouth | Wellcome Sanger Institute for the COVID-19 Genomics UK (COG-UK) Consortium | Lighthouse Laboratory Plymouth and Alex Alderton, Roberto Amato, Jeffrey Barrett, Sonia Goncalves, Ewan Harrison, David K. Jackson, Ian Johnston, Dominic Kwiatkowski, Cordelia Langford, John Sillitoe on behalf of the Wellcome Sanger Institute COVID-19 Surveillance Team |
| EPI_ISL_2847558 | Lighthouse Lab in Alderley Park | Wellcome Sanger Institute for the COVID-19 Genomics UK (COG-UK) Consortium | Jacquelyn Wynn, Mairead Hyland, The Lighthouse Lab in Alderley Park and Alex Alderton, Roberto Amato, Jeffrey Barrett, Sonia Goncalves, Ewan Harrison, David K. Jackson, Ian Johnston, Dominic Kwiatkowski, Cordelia Langford, John Sillitoe on behalf of the Wellcome Sanger Institute COVID-19 Surveillance Team |
| EPI_ISL_2847559 | Lighthouse Lab in Milton Keynes | Wellcome Sanger Institute for the COVID-19 Genomics UK (COG-UK) Consortium | The Lighthouse Lab in Milton Keynes and Alex Alderton, Roberto Amato, Jeffrey Barrett, Sonia Goncalves, Ewan Harrison, David K. Jackson, Ian Johnston, Dominic Kwiatkowski, Cordelia Langford, John Sillitoe on behalf of the Wellcome Sanger Institute COVID-19 Surveillance Team |
| EPI_ISL_2847560 | Lighthouse Laboratory Plymouth | Wellcome Sanger Institute for the COVID-19 Genomics UK (COG-UK) Consortium | Lighthouse Laboratory Plymouth and Alex Alderton, Roberto Amato, Jeffrey Barrett, Sonia Goncalves, Ewan Harrison, David K. Jackson, Ian Johnston, Dominic Kwiatkowski, Cordelia Langford, John Sillitoe on behalf of the Wellcome Sanger Institute COVID-19 Surveillance Team |
| EPI_ISL_2847561 | Lighthouse Lab in Alderley Park | Wellcome Sanger Institute for the COVID-19 Genomics UK (COG-UK) Consortium | Jacquelyn Wynn, Mairead Hyland, The Lighthouse Lab in Alderley Park and Alex Alderton, Roberto Amato, Jeffrey Barrett, Sonia Goncalves, Ewan Harrison, David K. Jackson, Ian Johnston, Dominic Kwiatkowski, Cordelia Langford, John Sillitoe on behalf of the Wellcome Sanger Institute COVID-19 Surveillance Team |
| EPI_ISL_2847562 | Lighthouse Lab in Milton Keynes | Wellcome Sanger Institute for the COVID-19 Genomics UK (COG-UK) Consortium | The Lighthouse Lab in Milton Keynes and Alex Alderton, Roberto Amato, Jeffrey Barrett, Sonia Goncalves, Ewan Harrison, David K. Jackson, Ian Johnston, Dominic Kwiatkowski, Cordelia Langford, John Sillitoe on behalf of the Wellcome Sanger Institute COVID-19 Surveillance Team |
| EPI_ISL_2847563 | Lighthouse Laboratory Plymouth | Wellcome Sanger Institute for the COVID-19 Genomics UK (COG-UK) Consortium | Lighthouse Laboratory Plymouth and Alex Alderton, Roberto Amato, Jeffrey Barrett, Sonia Goncalves, Ewan Harrison, David K. Jackson, Ian Johnston, Dominic Kwiatkowski, Cordelia Langford, John Sillitoe on behalf of the Wellcome Sanger Institute COVID-19 Surveillance Team |
| EPI_ISL_2847564 | Lighthouse Lab in Milton Keynes | Wellcome Sanger Institute for the COVID-19 Genomics UK (COG-UK) | The Lighthouse Lab in Milton Keynes and Alex Alderton, Roberto Amato, Jeffrey Barrett, Sonia Goncalves, Ewan Harrison, David K. Jackson, Ian Johnston, Dominic Kwiatkowski, Cordelia Langford, John Sillitoe on behalf of the Wellcome Sanger |

| | | | |
|---|---|---|---|
| | | Consortium | Institute COVID-19 Surveillance Team |
| EPI_ISL_2847565, EPI_ISL_2847566, EPI_ISL_2847567 | Lighthouse Laboratory Plymouth | Wellcome Sanger Institute for the COVID-19 Genomics UK (COG-UK) Consortium | Lighthouse Laboratory Plymouth and Alex Alderton, Roberto Amato, Jeffrey Barrett, Sonia Goncalves, Ewan Harrison, David K. Jackson, Ian Johnston, Dominic Kwiatkowski, Cordelia Langford, John Sillitoe on behalf of the Wellcome Sanger Institute COVID-19 Surveillance Team |
| EPI_ISL_2847568 | Lighthouse Lab in Glasgow | Wellcome Sanger Institute for the COVID-19 Genomics UK (COG-UK) Consortium | Harper VanSteenhouse, Yumi Kasai, David Gray, Carol Clugston, Anna Dominiczak and Alex Alderton, Roberto Amato, Jeffrey Barrett, Sonia Goncalves, Ewan Harrison, David K. Jackson, Ian Johnston, Dominic Kwiatkowski, Cordelia Langford, John Sillitoe on behalf of the Wellcome Sanger Institute COVID-19 Surveillance Team |
| EPI_ISL_2847569, EPI_ISL_2847570 | Lighthouse Laboratory Plymouth | Wellcome Sanger Institute for the COVID-19 Genomics UK (COG-UK) Consortium | Lighthouse Laboratory Plymouth and Alex Alderton, Roberto Amato, Jeffrey Barrett, Sonia Goncalves, Ewan Harrison, David K. Jackson, Ian Johnston, Dominic Kwiatkowski, Cordelia Langford, John Sillitoe on behalf of the Wellcome Sanger Institute COVID-19 Surveillance Team |
| EPI_ISL_2847571 | Lighthouse Lab in Milton Keynes | Wellcome Sanger Institute for the COVID-19 Genomics UK (COG-UK) Consortium | The Lighthouse Lab in Milton Keynes and Alex Alderton, Roberto Amato, Jeffrey Barrett, Sonia Goncalves, Ewan Harrison, David K. Jackson, Ian Johnston, Dominic Kwiatkowski, Cordelia Langford, John Sillitoe on behalf of the Wellcome Sanger Institute COVID-19 Surveillance Team |
| EPI_ISL_2847572 | Lighthouse Laboratory Plymouth | Wellcome Sanger Institute for the COVID-19 Genomics UK (COG-UK) Consortium | Lighthouse Laboratory Plymouth and Alex Alderton, Roberto Amato, Jeffrey Barrett, Sonia Goncalves, Ewan Harrison, David K. Jackson, Ian Johnston, Dominic Kwiatkowski, Cordelia Langford, John Sillitoe on behalf of the Wellcome Sanger Institute COVID-19 Surveillance Team |
| EPI_ISL_2847573 | Lighthouse Lab in Alderley Park | Wellcome Sanger Institute for the COVID-19 Genomics UK (COG-UK) Consortium | Jacquelyn Wynn, Mairead Hyland, The Lighthouse Lab in Alderley Park and Alex Alderton, Roberto Amato, Jeffrey Barrett, Sonia Goncalves, Ewan Harrison, David K. Jackson, Ian Johnston, Dominic Kwiatkowski, Cordelia Langford, John Sillitoe on behalf of the Wellcome Sanger Institute COVID-19 Surveillance Team |
| EPI_ISL_2847574 | Lighthouse Lab in Milton Keynes | Wellcome Sanger Institute for the COVID-19 Genomics UK (COG-UK) Consortium | The Lighthouse Lab in Milton Keynes and Alex Alderton, Roberto Amato, Jeffrey Barrett, Sonia Goncalves, Ewan Harrison, David K. Jackson, Ian Johnston, Dominic Kwiatkowski, Cordelia Langford, John Sillitoe on behalf of the Wellcome Sanger Institute COVID-19 Surveillance Team |
| EPI_ISL_2847575 | Lighthouse Lab in Alderley Park | Wellcome Sanger Institute for the COVID-19 Genomics UK (COG-UK) Consortium | Jacquelyn Wynn, Mairead Hyland, The Lighthouse Lab in Alderley Park and Alex Alderton, Roberto Amato, Jeffrey Barrett, Sonia Goncalves, Ewan Harrison, David K. Jackson, Ian Johnston, Dominic Kwiatkowski, Cordelia Langford, John Sillitoe on behalf of the Wellcome Sanger Institute COVID-19 Surveillance Team |
| EPI_ISL_2847576 | Lighthouse Lab in Glasgow | Wellcome Sanger Institute for the COVID-19 Genomics UK (COG-UK) Consortium | Harper VanSteenhouse, Yumi Kasai, David Gray, Carol Clugston, Anna Dominiczak and Alex Alderton, Roberto Amato, Jeffrey Barrett, Sonia Goncalves, Ewan Harrison, David K. Jackson, Ian Johnston, Dominic Kwiatkowski, Cordelia Langford, John Sillitoe on behalf of the Wellcome Sanger Institute COVID-19 Surveillance Team |
| EPI_ISL_2847577, EPI_ISL_2847578 | Lighthouse Laboratory Plymouth | Wellcome Sanger Institute for the COVID-19 Genomics UK (COG-UK) Consortium | Lighthouse Laboratory Plymouth and Alex Alderton, Roberto Amato, Jeffrey Barrett, Sonia Goncalves, Ewan Harrison, David K. Jackson, Ian Johnston, Dominic Kwiatkowski, Cordelia Langford, John Sillitoe on behalf of the Wellcome Sanger Institute COVID-19 Surveillance Team |
| EPI_ISL_2847579 | Lighthouse Lab in Milton Keynes | Wellcome Sanger Institute for the COVID-19 Genomics UK (COG-UK) Consortium | The Lighthouse Lab in Milton Keynes and Alex Alderton, Roberto Amato, Jeffrey Barrett, Sonia Goncalves, Ewan Harrison, David K. Jackson, Ian Johnston, Dominic Kwiatkowski, Cordelia Langford, John Sillitoe on behalf of the Wellcome Sanger Institute COVID-19 Surveillance Team |
| EPI_ISL_2847580, EPI_ISL_2847581 | Lighthouse Lab in Alderley Park | Wellcome Sanger Institute for the COVID-19 Genomics UK (COG-UK) Consortium | Jacquelyn Wynn, Mairead Hyland, The Lighthouse Lab in Alderley Park and Alex Alderton, Roberto Amato, Jeffrey Barrett, Sonia Goncalves, Ewan Harrison, David K. Jackson, Ian Johnston, Dominic Kwiatkowski, Cordelia Langford, John Sillitoe on behalf of the Wellcome Sanger Institute COVID-19 Surveillance Team |
| EPI_ISL_2847582 | Lighthouse Lab in Glasgow | Wellcome Sanger Institute for the COVID-19 Genomics UK (COG-UK) Consortium | Harper VanSteenhouse, Yumi Kasai, David Gray, Carol Clugston, Anna Dominiczak and Alex Alderton, Roberto Amato, Jeffrey Barrett, Sonia Goncalves, Ewan Harrison, David K. Jackson, Ian Johnston, Dominic Kwiatkowski, Cordelia Langford, John Sillitoe on behalf of the Wellcome Sanger Institute COVID-19 Surveillance Team |
| EPI_ISL_2847583 | Lighthouse Laboratory Plymouth | Wellcome Sanger Institute for the COVID-19 Genomics UK (COG-UK) Consortium | Lighthouse Laboratory Plymouth and Alex Alderton, Roberto Amato, Jeffrey Barrett, Sonia Goncalves, Ewan Harrison, David K. Jackson, Ian Johnston, Dominic Kwiatkowski, Cordelia Langford, John Sillitoe on behalf of the Wellcome Sanger Institute COVID-19 Surveillance Team |
| EPI_ISL_2847584 | Lighthouse Lab in Glasgow | Wellcome Sanger Institute for the COVID-19 Genomics UK (COG-UK) Consortium | Harper VanSteenhouse, Yumi Kasai, David Gray, Carol Clugston, Anna Dominiczak and Alex Alderton, Roberto Amato, Jeffrey Barrett, Sonia Goncalves, Ewan Harrison, David K. Jackson, Ian Johnston, Dominic Kwiatkowski, Cordelia Langford, John Sillitoe on behalf of the Wellcome Sanger Institute COVID-19 Surveillance Team |
| EPI_ISL_2847585 | Lighthouse Lab in Milton Keynes | Wellcome Sanger Institute for the COVID-19 Genomics UK (COG-UK) Consortium | The Lighthouse Lab in Milton Keynes and Alex Alderton, Roberto Amato, Jeffrey Barrett, Sonia Goncalves, Ewan Harrison, David K. Jackson, Ian Johnston, Dominic Kwiatkowski, Cordelia Langford, John Sillitoe on behalf of the Wellcome Sanger Institute COVID-19 Surveillance Team |
| EPI_ISL_2847586 | Lighthouse Laboratory Plymouth | Wellcome Sanger Institute for the COVID-19 Genomics UK (COG-UK) Consortium | Lighthouse Laboratory Plymouth and Alex Alderton, Roberto Amato, Jeffrey Barrett, Sonia Goncalves, Ewan Harrison, David K. Jackson, Ian Johnston, Dominic Kwiatkowski, Cordelia Langford, John Sillitoe on behalf of the Wellcome Sanger Institute COVID-19 Surveillance Team |
| EPI_ISL_2847587 | Lighthouse Lab in Glasgow | Wellcome Sanger Institute for the COVID-19 Genomics UK (COG-UK) Consortium | Harper VanSteenhouse, Yumi Kasai, David Gray, Carol Clugston, Anna Dominiczak and Alex Alderton, Roberto Amato, Jeffrey Barrett, Sonia Goncalves, Ewan Harrison, David K. Jackson, Ian Johnston, Dominic Kwiatkowski, Cordelia Langford, John Sillitoe on behalf of the Wellcome Sanger Institute COVID-19 Surveillance Team |
| EPI_ISL_2847588, EPI_ISL_2847589, EPI_ISL_2847590, EPI_ISL_2847591 | Lighthouse Laboratory Plymouth | Wellcome Sanger Institute for the COVID-19 Genomics UK (COG-UK) Consortium | Lighthouse Laboratory Plymouth and Alex Alderton, Roberto Amato, Jeffrey Barrett, Sonia Goncalves, Ewan Harrison, David K. Jackson, Ian Johnston, Dominic Kwiatkowski, Cordelia Langford, John Sillitoe on behalf of the Wellcome Sanger Institute COVID-19 Surveillance Team |
| EPI_ISL_2847592 | Lighthouse Lab in Alderley Park | Wellcome Sanger Institute for the COVID-19 Genomics UK (COG-UK) Consortium | Jacquelyn Wynn, Mairead Hyland, The Lighthouse Lab in Alderley Park and Alex Alderton, Roberto Amato, Jeffrey Barrett, Sonia Goncalves, Ewan Harrison, David K. Jackson, Ian Johnston, Dominic Kwiatkowski, Cordelia Langford, John Sillitoe on behalf of the Wellcome Sanger Institute COVID-19 Surveillance Team |
| EPI_ISL_2847593 | Lighthouse Lab in Milton Keynes | Wellcome Sanger Institute for the COVID-19 Genomics UK (COG-UK) Consortium | The Lighthouse Lab in Milton Keynes and Alex Alderton, Roberto Amato, Jeffrey Barrett, Sonia Goncalves, Ewan Harrison, David K. Jackson, Ian Johnston, Dominic Kwiatkowski, Cordelia Langford, John Sillitoe on behalf of the Wellcome Sanger Institute COVID-19 Surveillance Team |
| EPI_ISL_2847594 | Lighthouse Lab in Glasgow | Wellcome Sanger Institute for the COVID-19 Genomics UK (COG-UK) Consortium | Harper VanSteenhouse, Yumi Kasai, David Gray, Carol Clugston, Anna Dominiczak and Alex Alderton, Roberto Amato, Jeffrey Barrett, Sonia Goncalves, Ewan Harrison, David K. Jackson, Ian Johnston, Dominic Kwiatkowski, Cordelia Langford, John Sillitoe on behalf of the Wellcome Sanger Institute COVID-19 Surveillance Team |
| EPI_ISL_2847595, EPI_ISL_2847596, EPI_ISL_2847597 | Lighthouse Lab in Milton Keynes | Wellcome Sanger Institute for the COVID-19 Genomics UK (COG-UK) Consortium | The Lighthouse Lab in Milton Keynes and Alex Alderton, Roberto Amato, Jeffrey Barrett, Sonia Goncalves, Ewan Harrison, David K. Jackson, Ian Johnston, Dominic Kwiatkowski, Cordelia Langford, John Sillitoe on behalf of the Wellcome Sanger Institute COVID-19 Surveillance Team |
| EPI_ISL_2847598 | Lighthouse Laboratory Plymouth | Wellcome Sanger Institute for the COVID-19 Genomics UK (COG-UK) Consortium | Lighthouse Laboratory Plymouth and Alex Alderton, Roberto Amato, Jeffrey Barrett, Sonia Goncalves, Ewan Harrison, David K. Jackson, Ian Johnston, Dominic Kwiatkowski, Cordelia Langford, John Sillitoe on behalf of the Wellcome Sanger Institute COVID-19 Surveillance Team |
| EPI_ISL_2847599 | Lighthouse Lab in Glasgow | Wellcome Sanger Institute for the COVID-19 Genomics UK (COG-UK) Consortium | Harper VanSteenhouse, Yumi Kasai, David Gray, Carol Clugston, Anna Dominiczak and Alex Alderton, Roberto Amato, Jeffrey Barrett, Sonia Goncalves, Ewan Harrison, David K. Jackson, Ian Johnston, Dominic Kwiatkowski, Cordelia Langford, John Sillitoe on behalf of the Wellcome Sanger Institute COVID-19 Surveillance Team |
| EPI_ISL_2847600 | Lighthouse Lab in Alderley Park | Wellcome Sanger Institute for the COVID-19 Genomics UK (COG-UK) Consortium | Jacquelyn Wynn, Mairead Hyland, The Lighthouse Lab in Alderley Park and Alex Alderton, Roberto Amato, Jeffrey Barrett, Sonia Goncalves, Ewan Harrison, David K. Jackson, Ian Johnston, Dominic Kwiatkowski, Cordelia Langford, John Sillitoe on behalf of the Wellcome Sanger Institute COVID-19 Surveillance Team |
| EPI_ISL_2847601 | Lighthouse Lab in Glasgow | Wellcome Sanger Institute for the COVID-19 Genomics UK (COG-UK) Consortium | Harper VanSteenhouse, Yumi Kasai, David Gray, Carol Clugston, Anna Dominiczak and Alex Alderton, Roberto Amato, Jeffrey Barrett, Sonia Goncalves, Ewan Harrison, David K. Jackson, Ian Johnston, Dominic Kwiatkowski, Cordelia Langford, John Sillitoe on behalf of the Wellcome Sanger Institute COVID-19 Surveillance Team |
| EPI_ISL_2847602, EPI_ISL_2847603 | Lighthouse Lab in Milton Keynes | Wellcome Sanger Institute for the COVID-19 Genomics UK (COG-UK) Consortium | The Lighthouse Lab in Milton Keynes and Alex Alderton, Roberto Amato, Jeffrey Barrett, Sonia Goncalves, Ewan Harrison, David K. Jackson, Ian Johnston, Dominic Kwiatkowski, Cordelia Langford, John Sillitoe on behalf of the Wellcome Sanger Institute COVID-19 Surveillance Team |
| EPI_ISL_2847604 | Lighthouse Lab in Alderley Park | Wellcome Sanger Institute for the COVID-19 Genomics UK (COG-UK) Consortium | Jacquelyn Wynn, Mairead Hyland, The Lighthouse Lab in Alderley Park and Alex Alderton, Roberto Amato, Jeffrey Barrett, Sonia Goncalves, Ewan Harrison, David K. Jackson, Ian Johnston, Dominic Kwiatkowski, Cordelia Langford, John Sillitoe on behalf of the Wellcome Sanger Institute COVID-19 Surveillance Team |
| EPI_ISL_2847605, EPI_ISL_2847606 | Lighthouse Lab in Milton Keynes | Wellcome Sanger Institute for the COVID-19 Genomics UK (COG-UK) Consortium | The Lighthouse Lab in Milton Keynes and Alex Alderton, Roberto Amato, Jeffrey Barrett, Sonia Goncalves, Ewan Harrison, David K. Jackson, Ian Johnston, Dominic Kwiatkowski, Cordelia Langford, John Sillitoe on behalf of the Wellcome Sanger Institute COVID-19 Surveillance Team |
| EPI_ISL_2847607 | Lighthouse Lab in Glasgow | Wellcome Sanger Institute for the COVID-19 Genomics UK (COG-UK) Consortium | Harper VanSteenhouse, Yumi Kasai, David Gray, Carol Clugston, Anna Dominiczak and Alex Alderton, Roberto Amato, Jeffrey Barrett, Sonia Goncalves, Ewan Harrison, David K. Jackson, Ian Johnston, Dominic Kwiatkowski, Cordelia Langford, John Sillitoe on behalf of the Wellcome Sanger Institute COVID-19 Surveillance Team |
| EPI_ISL_2847608, EPI_ISL_2847609, EPI_ISL_2847610 | Lighthouse Laboratory Plymouth | Wellcome Sanger Institute for the COVID-19 Genomics UK (COG-UK) Consortium | Lighthouse Laboratory Plymouth and Alex Alderton, Roberto Amato, Jeffrey Barrett, Sonia Goncalves, Ewan Harrison, David K. Jackson, Ian Johnston, Dominic Kwiatkowski, Cordelia Langford, John Sillitoe on behalf of the Wellcome Sanger Institute COVID-19 Surveillance Team |
| EPI_ISL_2847611 | Lighthouse Lab in Milton Keynes | Wellcome Sanger Institute for the COVID-19 Genomics UK (COG-UK) Consortium | The Lighthouse Lab in Milton Keynes and Alex Alderton, Roberto Amato, Jeffrey Barrett, Sonia Goncalves, Ewan Harrison, David K. Jackson, Ian Johnston, Dominic Kwiatkowski, Cordelia Langford, John Sillitoe on behalf of the Wellcome Sanger Institute COVID-19 Surveillance Team |
| EPI_ISL_2847612, EPI_ISL_2847613, EPI_ISL_2847614, EPI_ISL_2847615 | Lighthouse Lab in Alderley Park | Wellcome Sanger Institute for the COVID-19 Genomics UK (COG-UK) Consortium | Jacquelyn Wynn, Mairead Hyland, The Lighthouse Lab in Alderley Park and Alex Alderton, Roberto Amato, Jeffrey Barrett, Sonia Goncalves, Ewan Harrison, David K. Jackson, Ian Johnston, Dominic Kwiatkowski, Cordelia Langford, John Sillitoe on behalf of the Wellcome Sanger Institute COVID-19 Surveillance Team |
| EPI_ISL_2847616 | Lighthouse Laboratory Plymouth | Wellcome Sanger Institute for the COVID-19 Genomics UK (COG-UK) Consortium | Lighthouse Laboratory Plymouth and Alex Alderton, Roberto Amato, Jeffrey Barrett, Sonia Goncalves, Ewan Harrison, David K. Jackson, Ian Johnston, Dominic Kwiatkowski, Cordelia Langford, John Sillitoe on behalf of the Wellcome Sanger Institute COVID-19 Surveillance Team |
| EPI_ISL_2847617 | Lighthouse Lab in Glasgow | Wellcome Sanger Institute for the COVID-19 Genomics UK (COG-UK) Consortium | Harper VanSteenhouse, Yumi Kasai, David Gray, Carol Clugston, Anna Dominiczak and Alex Alderton, Roberto Amato, Jeffrey Barrett, Sonia Goncalves, Ewan Harrison, David K. Jackson, Ian Johnston, Dominic Kwiatkowski, Cordelia Langford, John Sillitoe on behalf of the Wellcome Sanger Institute COVID-19 Surveillance Team |
| EPI_ISL_2847618 | Lighthouse Lab in Milton Keynes | Wellcome Sanger Institute for the COVID-19 Genomics UK (COG-UK) Consortium | The Lighthouse Lab in Milton Keynes and Alex Alderton, Roberto Amato, Jeffrey Barrett, Sonia Goncalves, Ewan Harrison, David K. Jackson, Ian Johnston, Dominic Kwiatkowski, Cordelia Langford, John Sillitoe on behalf of the Wellcome Sanger Institute COVID-19 Surveillance Team |
| EPI_ISL_2847619, EPI_ISL_2847620, EPI_ISL_2847622, EPI_ISL_2847623, EPI_ISL_2847624, EPI_ISL_2847625 | Lighthouse Laboratory Plymouth | Wellcome Sanger Institute for the COVID-19 Genomics UK (COG-UK) Consortium | Lighthouse Laboratory Plymouth and Alex Alderton, Roberto Amato, Jeffrey Barrett, Sonia Goncalves, Ewan Harrison, David K. Jackson, Ian Johnston, Dominic Kwiatkowski, Cordelia Langford, John Sillitoe on behalf of the Wellcome Sanger Institute COVID-19 Surveillance Team |
| EPI_ISL_2847626 | Lighthouse Lab in Milton Keynes | Wellcome Sanger Institute for the COVID-19 Genomics UK (COG-UK) Consortium | The Lighthouse Lab in Milton Keynes and Alex Alderton, Roberto Amato, Jeffrey Barrett, Sonia Goncalves, Ewan Harrison, David K. Jackson, Ian Johnston, Dominic Kwiatkowski, Cordelia Langford, John Sillitoe on behalf of the Wellcome Sanger Institute COVID-19 Surveillance Team |
| EPI_ISL_2847627 | Lighthouse Laboratory Plymouth | Wellcome Sanger Institute for the COVID-19 Genomics UK (COG-UK) Consortium | Lighthouse Laboratory Plymouth and Alex Alderton, Roberto Amato, Jeffrey Barrett, Sonia Goncalves, Ewan Harrison, David K. Jackson, Ian Johnston, Dominic Kwiatkowski, Cordelia Langford, John Sillitoe on behalf of the Wellcome Sanger Institute COVID-19 Surveillance Team |

| | | | |
|---|---|---|---|
| EPI_ISL_2847628, EPI_ISL_2847629 | Lighthouse Lab in Milton Keynes | Wellcome Sanger Institute for the COVID-19 Genomics UK (COG-UK) Consortium | The Lighthouse Lab in Milton Keynes and Alex Alderton, Roberto Amato, Jeffrey Barrett, Sonia Goncalves, Ewan Harrison, David K. Jackson, Ian Johnston, Dominic Kwiatkowski, Cordelia Langford, John Sillitoe on behalf of the Wellcome Sanger Institute COVID-19 Surveillance Team |
| EPI_ISL_2847630, EPI_ISL_2847631, EPI_ISL_2847632, EPI_ISL_2847633, EPI_ISL_2847634 | Lighthouse Laboratory Plymouth | Wellcome Sanger Institute for the COVID-19 Genomics UK (COG-UK) Consortium | Lighthouse Laboratory Plymouth and Alex Alderton, Roberto Amato, Jeffrey Barrett, Sonia Goncalves, Ewan Harrison, David K. Jackson, Ian Johnston, Dominic Kwiatkowski, Cordelia Langford, John Sillitoe on behalf of the Wellcome Sanger Institute COVID-19 Surveillance Team |
| EPI_ISL_2847635, EPI_ISL_2847636 | Lighthouse Lab in Milton Keynes | Wellcome Sanger Institute for the COVID-19 Genomics UK (COG-UK) Consortium | The Lighthouse Lab in Milton Keynes and Alex Alderton, Roberto Amato, Jeffrey Barrett, Sonia Goncalves, Ewan Harrison, David K. Jackson, Ian Johnston, Dominic Kwiatkowski, Cordelia Langford, John Sillitoe on behalf of the Wellcome Sanger Institute COVID-19 Surveillance Team |
| EPI_ISL_2847637, EPI_ISL_2847638, EPI_ISL_2847639, EPI_ISL_2847640, EPI_ISL_2847641, EPI_ISL_2847642, EPI_ISL_2847643, EPI_ISL_2847644, EPI_ISL_2847645 | Lighthouse Laboratory Plymouth | Wellcome Sanger Institute for the COVID-19 Genomics UK (COG-UK) Consortium | Lighthouse Laboratory Plymouth and Alex Alderton, Roberto Amato, Jeffrey Barrett, Sonia Goncalves, Ewan Harrison, David K. Jackson, Ian Johnston, Dominic Kwiatkowski, Cordelia Langford, John Sillitoe on behalf of the Wellcome Sanger Institute COVID-19 Surveillance Team |
| EPI_ISL_2847647, EPI_ISL_2847648, EPI_ISL_2847649 | Lighthouse Lab in Milton Keynes | Wellcome Sanger Institute for the COVID-19 Genomics UK (COG-UK) Consortium | The Lighthouse Lab in Milton Keynes and Alex Alderton, Roberto Amato, Jeffrey Barrett, Sonia Goncalves, Ewan Harrison, David K. Jackson, Ian Johnston, Dominic Kwiatkowski, Cordelia Langford, John Sillitoe on behalf of the Wellcome Sanger Institute COVID-19 Surveillance Team |
| EPI_ISL_2847650, EPI_ISL_2847651, EPI_ISL_2847652 | Lighthouse Laboratory Plymouth | Wellcome Sanger Institute for the COVID-19 Genomics UK (COG-UK) Consortium | Lighthouse Laboratory Plymouth and Alex Alderton, Roberto Amato, Jeffrey Barrett, Sonia Goncalves, Ewan Harrison, David K. Jackson, Ian Johnston, Dominic Kwiatkowski, Cordelia Langford, John Sillitoe on behalf of the Wellcome Sanger Institute COVID-19 Surveillance Team |
| EPI_ISL_2847653 | Lighthouse Lab in Milton Keynes | Wellcome Sanger Institute for the COVID-19 Genomics UK (COG-UK) Consortium | The Lighthouse Lab in Milton Keynes and Alex Alderton, Roberto Amato, Jeffrey Barrett, Sonia Goncalves, Ewan Harrison, David K. Jackson, Ian Johnston, Dominic Kwiatkowski, Cordelia Langford, John Sillitoe on behalf of the Wellcome Sanger Institute COVID-19 Surveillance Team |
| EPI_ISL_2847654, EPI_ISL_2847655, EPI_ISL_2847656, EPI_ISL_2847657, EPI_ISL_2847658, EPI_ISL_2847659, EPI_ISL_2847660 | Lighthouse Laboratory Plymouth | Wellcome Sanger Institute for the COVID-19 Genomics UK (COG-UK) Consortium | Lighthouse Laboratory Plymouth and Alex Alderton, Roberto Amato, Jeffrey Barrett, Sonia Goncalves, Ewan Harrison, David K. Jackson, Ian Johnston, Dominic Kwiatkowski, Cordelia Langford, John Sillitoe on behalf of the Wellcome Sanger Institute COVID-19 Surveillance Team |
| EPI_ISL_2847661 | Lighthouse Lab in Milton Keynes | Wellcome Sanger Institute for the COVID-19 Genomics UK (COG-UK) Consortium | The Lighthouse Lab in Milton Keynes and Alex Alderton, Roberto Amato, Jeffrey Barrett, Sonia Goncalves, Ewan Harrison, David K. Jackson, Ian Johnston, Dominic Kwiatkowski, Cordelia Langford, John Sillitoe on behalf of the Wellcome Sanger Institute COVID-19 Surveillance Team |
| EPI_ISL_2847662, EPI_ISL_2847663, EPI_ISL_2847664, EPI_ISL_2847665, EPI_ISL_2847666, EPI_ISL_2847667, EPI_ISL_2847668, EPI_ISL_2847669, EPI_ISL_2847670 | Lighthouse Laboratory Plymouth | Wellcome Sanger Institute for the COVID-19 Genomics UK (COG-UK) Consortium | Lighthouse Laboratory Plymouth and Alex Alderton, Roberto Amato, Jeffrey Barrett, Sonia Goncalves, Ewan Harrison, David K. Jackson, Ian Johnston, Dominic Kwiatkowski, Cordelia Langford, John Sillitoe on behalf of the Wellcome Sanger Institute COVID-19 Surveillance Team |
| EPI_ISL_2847671 | Lighthouse Lab in Milton Keynes | Wellcome Sanger Institute for the COVID-19 Genomics UK (COG-UK) Consortium | The Lighthouse Lab in Milton Keynes and Alex Alderton, Roberto Amato, Jeffrey Barrett, Sonia Goncalves, Ewan Harrison, David K. Jackson, Ian Johnston, Dominic Kwiatkowski, Cordelia Langford, John Sillitoe on behalf of the Wellcome Sanger Institute COVID-19 Surveillance Team |
| EPI_ISL_2847672, EPI_ISL_2847673, EPI_ISL_2847674 | Lighthouse Laboratory Plymouth | Wellcome Sanger Institute for the COVID-19 Genomics UK (COG-UK) Consortium | Lighthouse Laboratory Plymouth and Alex Alderton, Roberto Amato, Jeffrey Barrett, Sonia Goncalves, Ewan Harrison, David K. Jackson, Ian Johnston, Dominic Kwiatkowski, Cordelia Langford, John Sillitoe on behalf of the Wellcome Sanger Institute COVID-19 Surveillance Team |
| EPI_ISL_2847675 | Lighthouse Lab in Milton Keynes | Wellcome Sanger Institute for the COVID-19 Genomics UK (COG-UK) Consortium | The Lighthouse Lab in Milton Keynes and Alex Alderton, Roberto Amato, Jeffrey Barrett, Sonia Goncalves, Ewan Harrison, David K. Jackson, Ian Johnston, Dominic Kwiatkowski, Cordelia Langford, John Sillitoe on behalf of the Wellcome Sanger Institute COVID-19 Surveillance Team |
| EPI_ISL_2847676 | Lighthouse Laboratory Plymouth | Wellcome Sanger Institute for the COVID-19 Genomics UK (COG-UK) Consortium | Lighthouse Laboratory Plymouth and Alex Alderton, Roberto Amato, Jeffrey Barrett, Sonia Goncalves, Ewan Harrison, David K. Jackson, Ian Johnston, Dominic Kwiatkowski, Cordelia Langford, John Sillitoe on behalf of the Wellcome Sanger Institute COVID-19 Surveillance Team |
| EPI_ISL_2847677 | Lighthouse Lab in Milton Keynes | Wellcome Sanger Institute for the COVID-19 Genomics UK (COG-UK) Consortium | The Lighthouse Lab in Milton Keynes and Alex Alderton, Roberto Amato, Jeffrey Barrett, Sonia Goncalves, Ewan Harrison, David K. Jackson, Ian Johnston, Dominic Kwiatkowski, Cordelia Langford, John Sillitoe on behalf of the Wellcome Sanger Institute COVID-19 Surveillance Team |
| EPI_ISL_2847678, EPI_ISL_2847679, EPI_ISL_2847680, EPI_ISL_2847681, EPI_ISL_2847682, EPI_ISL_2847683 | Lighthouse Laboratory Plymouth | Wellcome Sanger Institute for the COVID-19 Genomics UK (COG-UK) Consortium | Lighthouse Laboratory Plymouth and Alex Alderton, Roberto Amato, Jeffrey Barrett, Sonia Goncalves, Ewan Harrison, David K. Jackson, Ian Johnston, Dominic Kwiatkowski, Cordelia Langford, John Sillitoe on behalf of the Wellcome Sanger Institute COVID-19 Surveillance Team |
| EPI_ISL_2847684 | Lighthouse Lab in Milton Keynes | Wellcome Sanger Institute for the COVID-19 Genomics UK (COG-UK) Consortium | The Lighthouse Lab in Milton Keynes and Alex Alderton, Roberto Amato, Jeffrey Barrett, Sonia Goncalves, Ewan Harrison, David K. Jackson, Ian Johnston, Dominic Kwiatkowski, Cordelia Langford, John Sillitoe on behalf of the Wellcome Sanger Institute COVID-19 Surveillance Team |
| EPI_ISL_2847685 | Lighthouse Laboratory Plymouth | Wellcome Sanger Institute for the COVID-19 Genomics UK (COG-UK) Consortium | Lighthouse Laboratory Plymouth and Alex Alderton, Roberto Amato, Jeffrey Barrett, Sonia Goncalves, Ewan Harrison, David K. Jackson, Ian Johnston, Dominic Kwiatkowski, Cordelia Langford, John Sillitoe on behalf of the Wellcome Sanger Institute COVID-19 Surveillance Team |
| EPI_ISL_2847686, EPI_ISL_2847687, EPI_ISL_2847688 | Lighthouse Lab in Milton Keynes | Wellcome Sanger Institute for the COVID-19 Genomics UK (COG-UK) Consortium | The Lighthouse Lab in Milton Keynes and Alex Alderton, Roberto Amato, Jeffrey Barrett, Sonia Goncalves, Ewan Harrison, David K. Jackson, Ian Johnston, Dominic Kwiatkowski, Cordelia Langford, John Sillitoe on behalf of the Wellcome Sanger Institute COVID-19 Surveillance Team |
| EPI_ISL_2847689, EPI_ISL_2847690, EPI_ISL_2847691 | Lighthouse Laboratory Plymouth | Wellcome Sanger Institute for the COVID-19 Genomics UK (COG-UK) Consortium | Lighthouse Laboratory Plymouth and Alex Alderton, Roberto Amato, Jeffrey Barrett, Sonia Goncalves, Ewan Harrison, David K. Jackson, Ian Johnston, Dominic Kwiatkowski, Cordelia Langford, John Sillitoe on behalf of the Wellcome Sanger Institute COVID-19 Surveillance Team |
| EPI_ISL_2847692, EPI_ISL_2847693 | Lighthouse Lab in Milton Keynes | Wellcome Sanger Institute for the COVID-19 Genomics UK (COG-UK) Consortium | The Lighthouse Lab in Milton Keynes and Alex Alderton, Roberto Amato, Jeffrey Barrett, Sonia Goncalves, Ewan Harrison, David K. Jackson, Ian Johnston, Dominic Kwiatkowski, Cordelia Langford, John Sillitoe on behalf of the Wellcome Sanger Institute COVID-19 Surveillance Team |
| EPI_ISL_2847694, EPI_ISL_2847695, EPI_ISL_2847696, EPI_ISL_2847697, EPI_ISL_2847698, EPI_ISL_2847699, EPI_ISL_2847700, EPI_ISL_2847701 | Lighthouse Laboratory Plymouth | Wellcome Sanger Institute for the COVID-19 Genomics UK (COG-UK) Consortium | Lighthouse Laboratory Plymouth and Alex Alderton, Roberto Amato, Jeffrey Barrett, Sonia Goncalves, Ewan Harrison, David K. Jackson, Ian Johnston, Dominic Kwiatkowski, Cordelia Langford, John Sillitoe on behalf of the Wellcome Sanger Institute COVID-19 Surveillance Team |
| EPI_ISL_2847702, EPI_ISL_2847703, EPI_ISL_2847704 | Lighthouse Lab in Milton Keynes | Wellcome Sanger Institute for the COVID-19 Genomics UK (COG-UK) Consortium | The Lighthouse Lab in Milton Keynes and Alex Alderton, Roberto Amato, Jeffrey Barrett, Sonia Goncalves, Ewan Harrison, David K. Jackson, Ian Johnston, Dominic Kwiatkowski, Cordelia Langford, John Sillitoe on behalf of the Wellcome Sanger Institute COVID-19 Surveillance Team |
| EPI_ISL_2847705 | Lighthouse Laboratory Plymouth | Wellcome Sanger Institute for the COVID-19 Genomics UK (COG-UK) Consortium | Lighthouse Laboratory Plymouth and Alex Alderton, Roberto Amato, Jeffrey Barrett, Sonia Goncalves, Ewan Harrison, David K. Jackson, Ian Johnston, Dominic Kwiatkowski, Cordelia Langford, John Sillitoe on behalf of the Wellcome Sanger Institute COVID-19 Surveillance Team |
| EPI_ISL_2847706 | Lighthouse Lab in Milton Keynes | Wellcome Sanger Institute for the COVID-19 Genomics UK (COG-UK) Consortium | The Lighthouse Lab in Milton Keynes and Alex Alderton, Roberto Amato, Jeffrey Barrett, Sonia Goncalves, Ewan Harrison, David K. Jackson, Ian Johnston, Dominic Kwiatkowski, Cordelia Langford, John Sillitoe on behalf of the Wellcome Sanger Institute COVID-19 Surveillance Team |
| EPI_ISL_2847707, EPI_ISL_2847708, EPI_ISL_2847709 | Lighthouse Laboratory Plymouth | Wellcome Sanger Institute for the COVID-19 Genomics UK (COG-UK) Consortium | Lighthouse Laboratory Plymouth and Alex Alderton, Roberto Amato, Jeffrey Barrett, Sonia Goncalves, Ewan Harrison, David K. Jackson, Ian Johnston, Dominic Kwiatkowski, Cordelia Langford, John Sillitoe on behalf of the Wellcome Sanger Institute COVID-19 Surveillance Team |
| EPI_ISL_2847711 | Lighthouse Lab in Milton Keynes | Wellcome Sanger Institute for the COVID-19 Genomics UK (COG-UK) Consortium | The Lighthouse Lab in Milton Keynes and Alex Alderton, Roberto Amato, Jeffrey Barrett, Sonia Goncalves, Ewan Harrison, David K. Jackson, Ian Johnston, Dominic Kwiatkowski, Cordelia Langford, John Sillitoe on behalf of the Wellcome Sanger Institute COVID-19 Surveillance Team |
| EPI_ISL_2847712, EPI_ISL_2847713, EPI_ISL_2847714 | Lighthouse Laboratory Plymouth | Wellcome Sanger Institute for the COVID-19 Genomics UK (COG-UK) Consortium | Lighthouse Laboratory Plymouth and Alex Alderton, Roberto Amato, Jeffrey Barrett, Sonia Goncalves, Ewan Harrison, David K. Jackson, Ian Johnston, Dominic Kwiatkowski, Cordelia Langford, John Sillitoe on behalf of the Wellcome Sanger Institute COVID-19 Surveillance Team |
| EPI_ISL_2847715, EPI_ISL_2847717, | Lighthouse Lab in Milton Keynes | Wellcome Sanger Institute for the COVID-19 Genomics UK (COG-UK) Consortium | The Lighthouse Lab in Milton Keynes and Alex Alderton, Roberto Amato, Jeffrey Barrett, Sonia Goncalves, Ewan Harrison, David K. Jackson, Ian Johnston, Dominic Kwiatkowski, Cordelia Langford, John Sillitoe on behalf of the Wellcome Sanger Institute COVID-19 Surveillance Team |

| | | | |
|---|---|---|---|
| EPI_ISL_2847718 | | | |
| EPI_ISL_2847719, EPI_ISL_2847720 | Lighthouse Laboratory Plymouth | Wellcome Sanger Institute for the COVID-19 Genomics UK (COG-UK) Consortium | Lighthouse Laboratory Plymouth and Alex Alderton, Roberto Amato, Jeffrey Barrett, Sonia Goncalves, Ewan Harrison, David K. Jackson, Ian Johnston, Dominic Kwiatkowski, Cordelia Langford, John Sillitoe on behalf of the Wellcome Sanger Institute COVID-19 Surveillance Team |
| EPI_ISL_2847722 | Lighthouse Lab in Milton Keynes | Wellcome Sanger Institute for the COVID-19 Genomics UK (COG-UK) Consortium | The Lighthouse Lab in Milton Keynes and Alex Alderton, Roberto Amato, Jeffrey Barrett, Sonia Goncalves, Ewan Harrison, David K. Jackson, Ian Johnston, Dominic Kwiatkowski, Cordelia Langford, John Sillitoe on behalf of the Wellcome Sanger Institute COVID-19 Surveillance Team |
| EPI_ISL_2847723, EPI_ISL_2847724, EPI_ISL_2847725, EPI_ISL_2847726, EPI_ISL_2847727, EPI_ISL_2847728, EPI_ISL_2847729, EPI_ISL_2847730, EPI_ISL_2847731 | Lighthouse Laboratory Plymouth | Wellcome Sanger Institute for the COVID-19 Genomics UK (COG-UK) Consortium | Lighthouse Laboratory Plymouth and Alex Alderton, Roberto Amato, Jeffrey Barrett, Sonia Goncalves, Ewan Harrison, David K. Jackson, Ian Johnston, Dominic Kwiatkowski, Cordelia Langford, John Sillitoe on behalf of the Wellcome Sanger Institute COVID-19 Surveillance Team |
| EPI_ISL_2847732, EPI_ISL_2847733 | Lighthouse Lab in Milton Keynes | Wellcome Sanger Institute for the COVID-19 Genomics UK (COG-UK) Consortium | The Lighthouse Lab in Milton Keynes and Alex Alderton, Roberto Amato, Jeffrey Barrett, Sonia Goncalves, Ewan Harrison, David K. Jackson, Ian Johnston, Dominic Kwiatkowski, Cordelia Langford, John Sillitoe on behalf of the Wellcome Sanger Institute COVID-19 Surveillance Team |
| EPI_ISL_2847734 | Lighthouse Laboratory Plymouth | Wellcome Sanger Institute for the COVID-19 Genomics UK (COG-UK) Consortium | Lighthouse Laboratory Plymouth and Alex Alderton, Roberto Amato, Jeffrey Barrett, Sonia Goncalves, Ewan Harrison, David K. Jackson, Ian Johnston, Dominic Kwiatkowski, Cordelia Langford, John Sillitoe on behalf of the Wellcome Sanger Institute COVID-19 Surveillance Team |
| EPI_ISL_2847735, EPI_ISL_2847736 | Lighthouse Lab in Milton Keynes | Wellcome Sanger Institute for the COVID-19 Genomics UK (COG-UK) Consortium | The Lighthouse Lab in Milton Keynes and Alex Alderton, Roberto Amato, Jeffrey Barrett, Sonia Goncalves, Ewan Harrison, David K. Jackson, Ian Johnston, Dominic Kwiatkowski, Cordelia Langford, John Sillitoe on behalf of the Wellcome Sanger Institute COVID-19 Surveillance Team |
| EPI_ISL_2847737 | Lighthouse Laboratory Plymouth | Wellcome Sanger Institute for the COVID-19 Genomics UK (COG-UK) Consortium | Lighthouse Laboratory Plymouth and Alex Alderton, Roberto Amato, Jeffrey Barrett, Sonia Goncalves, Ewan Harrison, David K. Jackson, Ian Johnston, Dominic Kwiatkowski, Cordelia Langford, John Sillitoe on behalf of the Wellcome Sanger Institute COVID-19 Surveillance Team |
| EPI_ISL_2847738 | Lighthouse Lab in Milton Keynes | Wellcome Sanger Institute for the COVID-19 Genomics UK (COG-UK) Consortium | The Lighthouse Lab in Milton Keynes and Alex Alderton, Roberto Amato, Jeffrey Barrett, Sonia Goncalves, Ewan Harrison, David K. Jackson, Ian Johnston, Dominic Kwiatkowski, Cordelia Langford, John Sillitoe on behalf of the Wellcome Sanger Institute COVID-19 Surveillance Team |
| EPI_ISL_2847739, EPI_ISL_2847740, EPI_ISL_2847741 | Lighthouse Laboratory Plymouth | Wellcome Sanger Institute for the COVID-19 Genomics UK (COG-UK) Consortium | Lighthouse Laboratory Plymouth and Alex Alderton, Roberto Amato, Jeffrey Barrett, Sonia Goncalves, Ewan Harrison, David K. Jackson, Ian Johnston, Dominic Kwiatkowski, Cordelia Langford, John Sillitoe on behalf of the Wellcome Sanger Institute COVID-19 Surveillance Team |
| EPI_ISL_2847742, EPI_ISL_2847743 | Lighthouse Lab in Milton Keynes | Wellcome Sanger Institute for the COVID-19 Genomics UK (COG-UK) Consortium | The Lighthouse Lab in Milton Keynes and Alex Alderton, Roberto Amato, Jeffrey Barrett, Sonia Goncalves, Ewan Harrison, David K. Jackson, Ian Johnston, Dominic Kwiatkowski, Cordelia Langford, John Sillitoe on behalf of the Wellcome Sanger Institute COVID-19 Surveillance Team |
| EPI_ISL_2847744, EPI_ISL_2847745, EPI_ISL_2847746, EPI_ISL_2847747, EPI_ISL_2847748, EPI_ISL_2847749, EPI_ISL_2847750 | Lighthouse Laboratory Plymouth | Wellcome Sanger Institute for the COVID-19 Genomics UK (COG-UK) Consortium | Lighthouse Laboratory Plymouth and Alex Alderton, Roberto Amato, Jeffrey Barrett, Sonia Goncalves, Ewan Harrison, David K. Jackson, Ian Johnston, Dominic Kwiatkowski, Cordelia Langford, John Sillitoe on behalf of the Wellcome Sanger Institute COVID-19 Surveillance Team |
| EPI_ISL_2847751 | Lighthouse Lab in Milton Keynes | Wellcome Sanger Institute for the COVID-19 Genomics UK (COG-UK) Consortium | The Lighthouse Lab in Milton Keynes and Alex Alderton, Roberto Amato, Jeffrey Barrett, Sonia Goncalves, Ewan Harrison, David K. Jackson, Ian Johnston, Dominic Kwiatkowski, Cordelia Langford, John Sillitoe on behalf of the Wellcome Sanger Institute COVID-19 Surveillance Team |
| EPI_ISL_2847752, EPI_ISL_2847753, EPI_ISL_2847754, EPI_ISL_2847755, EPI_ISL_2847756, EPI_ISL_2847757 | Lighthouse Laboratory Plymouth | Wellcome Sanger Institute for the COVID-19 Genomics UK (COG-UK) Consortium | Lighthouse Laboratory Plymouth and Alex Alderton, Roberto Amato, Jeffrey Barrett, Sonia Goncalves, Ewan Harrison, David K. Jackson, Ian Johnston, Dominic Kwiatkowski, Cordelia Langford, John Sillitoe on behalf of the Wellcome Sanger Institute COVID-19 Surveillance Team |
| EPI_ISL_2847758 | Lighthouse Lab in Milton Keynes | Wellcome Sanger Institute for the COVID-19 Genomics UK (COG-UK) Consortium | The Lighthouse Lab in Milton Keynes and Alex Alderton, Roberto Amato, Jeffrey Barrett, Sonia Goncalves, Ewan Harrison, David K. Jackson, Ian Johnston, Dominic Kwiatkowski, Cordelia Langford, John Sillitoe on behalf of the Wellcome Sanger Institute COVID-19 Surveillance Team |
| EPI_ISL_2847759, EPI_ISL_2847760, EPI_ISL_2847761, EPI_ISL_2847762, EPI_ISL_2847763, EPI_ISL_2847764 | Lighthouse Laboratory Plymouth | Wellcome Sanger Institute for the COVID-19 Genomics UK (COG-UK) Consortium | Lighthouse Laboratory Plymouth and Alex Alderton, Roberto Amato, Jeffrey Barrett, Sonia Goncalves, Ewan Harrison, David K. Jackson, Ian Johnston, Dominic Kwiatkowski, Cordelia Langford, John Sillitoe on behalf of the Wellcome Sanger Institute COVID-19 Surveillance Team |
| EPI_ISL_2847765 | Lighthouse Lab in Milton Keynes | Wellcome Sanger Institute for the COVID-19 Genomics UK (COG-UK) Consortium | The Lighthouse Lab in Milton Keynes and Alex Alderton, Roberto Amato, Jeffrey Barrett, Sonia Goncalves, Ewan Harrison, David K. Jackson, Ian Johnston, Dominic Kwiatkowski, Cordelia Langford, John Sillitoe on behalf of the Wellcome Sanger Institute COVID-19 Surveillance Team |
| EPI_ISL_2847766, EPI_ISL_2847767, EPI_ISL_2847768, EPI_ISL_2847769, EPI_ISL_2847770 | Lighthouse Laboratory Plymouth | Wellcome Sanger Institute for the COVID-19 Genomics UK (COG-UK) Consortium | Lighthouse Laboratory Plymouth and Alex Alderton, Roberto Amato, Jeffrey Barrett, Sonia Goncalves, Ewan Harrison, David K. Jackson, Ian Johnston, Dominic Kwiatkowski, Cordelia Langford, John Sillitoe on behalf of the Wellcome Sanger Institute COVID-19 Surveillance Team |
| EPI_ISL_2847771 | Lighthouse Lab in Milton Keynes | Wellcome Sanger Institute for the COVID-19 Genomics UK (COG-UK) Consortium | The Lighthouse Lab in Milton Keynes and Alex Alderton, Roberto Amato, Jeffrey Barrett, Sonia Goncalves, Ewan Harrison, David K. Jackson, Ian Johnston, Dominic Kwiatkowski, Cordelia Langford, John Sillitoe on behalf of the Wellcome Sanger Institute COVID-19 Surveillance Team |
| EPI_ISL_2847772, EPI_ISL_2847773 | Lighthouse Laboratory Plymouth | Wellcome Sanger Institute for the COVID-19 Genomics UK (COG-UK) Consortium | Lighthouse Laboratory Plymouth and Alex Alderton, Roberto Amato, Jeffrey Barrett, Sonia Goncalves, Ewan Harrison, David K. Jackson, Ian Johnston, Dominic Kwiatkowski, Cordelia Langford, John Sillitoe on behalf of the Wellcome Sanger Institute COVID-19 Surveillance Team |
| EPI_ISL_2847774 | Lighthouse Lab in Milton Keynes | Wellcome Sanger Institute for the COVID-19 Genomics UK (COG-UK) Consortium | The Lighthouse Lab in Milton Keynes and Alex Alderton, Roberto Amato, Jeffrey Barrett, Sonia Goncalves, Ewan Harrison, David K. Jackson, Ian Johnston, Dominic Kwiatkowski, Cordelia Langford, John Sillitoe on behalf of the Wellcome Sanger Institute COVID-19 Surveillance Team |
| EPI_ISL_2847775, EPI_ISL_2847776, EPI_ISL_2847777, EPI_ISL_2847778, EPI_ISL_2847779, EPI_ISL_2847780, EPI_ISL_2847781 | Lighthouse Laboratory Plymouth | Wellcome Sanger Institute for the COVID-19 Genomics UK (COG-UK) Consortium | Lighthouse Laboratory Plymouth and Alex Alderton, Roberto Amato, Jeffrey Barrett, Sonia Goncalves, Ewan Harrison, David K. Jackson, Ian Johnston, Dominic Kwiatkowski, Cordelia Langford, John Sillitoe on behalf of the Wellcome Sanger Institute COVID-19 Surveillance Team |
| EPI_ISL_2847782 | Lighthouse Lab in Milton Keynes | Wellcome Sanger Institute for the COVID-19 Genomics UK (COG-UK) Consortium | The Lighthouse Lab in Milton Keynes and Alex Alderton, Roberto Amato, Jeffrey Barrett, Sonia Goncalves, Ewan Harrison, David K. Jackson, Ian Johnston, Dominic Kwiatkowski, Cordelia Langford, John Sillitoe on behalf of the Wellcome Sanger Institute COVID-19 Surveillance Team |
| EPI_ISL_2847783, EPI_ISL_2847784, EPI_ISL_2847785, EPI_ISL_2847786 | Lighthouse Laboratory Plymouth | Wellcome Sanger Institute for the COVID-19 Genomics UK (COG-UK) Consortium | Lighthouse Laboratory Plymouth and Alex Alderton, Roberto Amato, Jeffrey Barrett, Sonia Goncalves, Ewan Harrison, David K. Jackson, Ian Johnston, Dominic Kwiatkowski, Cordelia Langford, John Sillitoe on behalf of the Wellcome Sanger Institute COVID-19 Surveillance Team |
| EPI_ISL_2847787, EPI_ISL_2847788, EPI_ISL_2847789 | Lighthouse Lab in Milton Keynes | Wellcome Sanger Institute for the COVID-19 Genomics UK (COG-UK) Consortium | The Lighthouse Lab in Milton Keynes and Alex Alderton, Roberto Amato, Jeffrey Barrett, Sonia Goncalves, Ewan Harrison, David K. Jackson, Ian Johnston, Dominic Kwiatkowski, Cordelia Langford, John Sillitoe on behalf of the Wellcome Sanger Institute COVID-19 Surveillance Team |
| EPI_ISL_2847790, EPI_ISL_2847791, EPI_ISL_2847792, EPI_ISL_2847793, EPI_ISL_2847794, EPI_ISL_2847795 | Lighthouse Laboratory Plymouth | Wellcome Sanger Institute for the COVID-19 Genomics UK (COG-UK) Consortium | Lighthouse Laboratory Plymouth and Alex Alderton, Roberto Amato, Jeffrey Barrett, Sonia Goncalves, Ewan Harrison, David K. Jackson, Ian Johnston, Dominic Kwiatkowski, Cordelia Langford, John Sillitoe on behalf of the Wellcome Sanger Institute COVID-19 Surveillance Team |
| EPI_ISL_2847796 | Lighthouse Lab in Milton Keynes | Wellcome Sanger Institute for the COVID-19 Genomics UK (COG-UK) Consortium | The Lighthouse Lab in Milton Keynes and Alex Alderton, Roberto Amato, Jeffrey Barrett, Sonia Goncalves, Ewan Harrison, David K. Jackson, Ian Johnston, Dominic Kwiatkowski, Cordelia Langford, John Sillitoe on behalf of the Wellcome Sanger Institute COVID-19 Surveillance Team |
| EPI_ISL_2847797, EPI_ISL_2847798 | Lighthouse Laboratory Plymouth | Wellcome Sanger Institute for the COVID-19 Genomics UK (COG-UK) Consortium | Lighthouse Laboratory Plymouth and Alex Alderton, Roberto Amato, Jeffrey Barrett, Sonia Goncalves, Ewan Harrison, David K. Jackson, Ian Johnston, Dominic Kwiatkowski, Cordelia Langford, John Sillitoe on behalf of the Wellcome Sanger Institute COVID-19 Surveillance Team |
| EPI_ISL_2847799 | Lighthouse Lab in Milton Keynes | Wellcome Sanger Institute for the COVID-19 Genomics UK (COG-UK) Consortium | The Lighthouse Lab in Milton Keynes and Alex Alderton, Roberto Amato, Jeffrey Barrett, Sonia Goncalves, Ewan Harrison, David K. Jackson, Ian Johnston, Dominic Kwiatkowski, Cordelia Langford, John Sillitoe on behalf of the Wellcome Sanger Institute COVID-19 Surveillance Team |
| EPI_ISL_2847800, EPI_ISL_2847801 | Lighthouse Laboratory Plymouth | Wellcome Sanger Institute for the COVID-19 Genomics UK (COG-UK) Consortium | Lighthouse Laboratory Plymouth and Alex Alderton, Roberto Amato, Jeffrey Barrett, Sonia Goncalves, Ewan Harrison, David K. Jackson, Ian Johnston, Dominic Kwiatkowski, Cordelia Langford, John Sillitoe on behalf of the Wellcome Sanger Institute COVID-19 Surveillance Team |
| EPI_ISL_2847802 | Lighthouse Lab in Milton Keynes | Wellcome Sanger Institute for the COVID-19 Genomics UK (COG-UK) | The Lighthouse Lab in Milton Keynes and Alex Alderton, Roberto Amato, Jeffrey Barrett, Sonia Goncalves, Ewan Harrison, David K. Jackson, Ian Johnston, Dominic Kwiatkowski, Cordelia Langford, John Sillitoe on behalf of the Wellcome Sanger |

| | | | |
|---|---|---|---|
| | | Consortium | Institute COVID-19 Surveillance Team |
| EPI_ISL_2847803, EPI_ISL_2847804, EPI_ISL_2847805 | Lighthouse Laboratory Plymouth | Wellcome Sanger Institute for the COVID-19 Genomics UK (COG-UK) Consortium | Lighthouse Laboratory Plymouth and Alex Alderton, Roberto Amato, Jeffrey Barrett, Sonia Goncalves, Ewan Harrison, David K. Jackson, Ian Johnston, Dominic Kwiatkowski, Cordelia Langford, John Sillitoe on behalf of the Wellcome Sanger Institute COVID-19 Surveillance Team |
| EPI_ISL_2847806 | Lighthouse Lab in Milton Keynes | Wellcome Sanger Institute for the COVID-19 Genomics UK (COG-UK) Consortium | The Lighthouse Lab in Milton Keynes and Alex Alderton, Roberto Amato, Jeffrey Barrett, Sonia Goncalves, Ewan Harrison, David K. Jackson, Ian Johnston, Dominic Kwiatkowski, Cordelia Langford, John Sillitoe on behalf of the Wellcome Sanger Institute COVID-19 Surveillance Team |
| EPI_ISL_2847807 | Lighthouse Laboratory Plymouth | Wellcome Sanger Institute for the COVID-19 Genomics UK (COG-UK) Consortium | Lighthouse Laboratory Plymouth and Alex Alderton, Roberto Amato, Jeffrey Barrett, Sonia Goncalves, Ewan Harrison, David K. Jackson, Ian Johnston, Dominic Kwiatkowski, Cordelia Langford, John Sillitoe on behalf of the Wellcome Sanger Institute COVID-19 Surveillance Team |
| EPI_ISL_2847808 | Lighthouse Lab in Milton Keynes | Wellcome Sanger Institute for the COVID-19 Genomics UK (COG-UK) Consortium | The Lighthouse Lab in Milton Keynes and Alex Alderton, Roberto Amato, Jeffrey Barrett, Sonia Goncalves, Ewan Harrison, David K. Jackson, Ian Johnston, Dominic Kwiatkowski, Cordelia Langford, John Sillitoe on behalf of the Wellcome Sanger Institute COVID-19 Surveillance Team |
| EPI_ISL_2847809 | Lighthouse Laboratory Plymouth | Wellcome Sanger Institute for the COVID-19 Genomics UK (COG-UK) Consortium | Lighthouse Laboratory Plymouth and Alex Alderton, Roberto Amato, Jeffrey Barrett, Sonia Goncalves, Ewan Harrison, David K. Jackson, Ian Johnston, Dominic Kwiatkowski, Cordelia Langford, John Sillitoe on behalf of the Wellcome Sanger Institute COVID-19 Surveillance Team |
| EPI_ISL_2847810, EPI_ISL_2847811 | Lighthouse Lab in Milton Keynes | Wellcome Sanger Institute for the COVID-19 Genomics UK (COG-UK) Consortium | The Lighthouse Lab in Milton Keynes and Alex Alderton, Roberto Amato, Jeffrey Barrett, Sonia Goncalves, Ewan Harrison, David K. Jackson, Ian Johnston, Dominic Kwiatkowski, Cordelia Langford, John Sillitoe on behalf of the Wellcome Sanger Institute COVID-19 Surveillance Team |
| EPI_ISL_2847812, EPI_ISL_2847813, EPI_ISL_2847814, EPI_ISL_2847815, EPI_ISL_2847816, EPI_ISL_2847817 | Lighthouse Laboratory Plymouth | Wellcome Sanger Institute for the COVID-19 Genomics UK (COG-UK) Consortium | Lighthouse Laboratory Plymouth and Alex Alderton, Roberto Amato, Jeffrey Barrett, Sonia Goncalves, Ewan Harrison, David K. Jackson, Ian Johnston, Dominic Kwiatkowski, Cordelia Langford, John Sillitoe on behalf of the Wellcome Sanger Institute COVID-19 Surveillance Team |
| EPI_ISL_2847818 | Lighthouse Lab in Milton Keynes | Wellcome Sanger Institute for the COVID-19 Genomics UK (COG-UK) Consortium | The Lighthouse Lab in Milton Keynes and Alex Alderton, Roberto Amato, Jeffrey Barrett, Sonia Goncalves, Ewan Harrison, David K. Jackson, Ian Johnston, Dominic Kwiatkowski, Cordelia Langford, John Sillitoe on behalf of the Wellcome Sanger Institute COVID-19 Surveillance Team |
| EPI_ISL_2847819, EPI_ISL_2847820, EPI_ISL_2847821, EPI_ISL_2847822, EPI_ISL_2847823 | Lighthouse Laboratory Plymouth | Wellcome Sanger Institute for the COVID-19 Genomics UK (COG-UK) Consortium | Lighthouse Laboratory Plymouth and Alex Alderton, Roberto Amato, Jeffrey Barrett, Sonia Goncalves, Ewan Harrison, David K. Jackson, Ian Johnston, Dominic Kwiatkowski, Cordelia Langford, John Sillitoe on behalf of the Wellcome Sanger Institute COVID-19 Surveillance Team |
| EPI_ISL_2847824 | Lighthouse Lab in Milton Keynes | Wellcome Sanger Institute for the COVID-19 Genomics UK (COG-UK) Consortium | The Lighthouse Lab in Milton Keynes and Alex Alderton, Roberto Amato, Jeffrey Barrett, Sonia Goncalves, Ewan Harrison, David K. Jackson, Ian Johnston, Dominic Kwiatkowski, Cordelia Langford, John Sillitoe on behalf of the Wellcome Sanger Institute COVID-19 Surveillance Team |
| EPI_ISL_2847825, EPI_ISL_2847826, EPI_ISL_2847827, EPI_ISL_2847828, EPI_ISL_2847829, EPI_ISL_2847830, EPI_ISL_2847831 | Lighthouse Laboratory Plymouth | Wellcome Sanger Institute for the COVID-19 Genomics UK (COG-UK) Consortium | Lighthouse Laboratory Plymouth and Alex Alderton, Roberto Amato, Jeffrey Barrett, Sonia Goncalves, Ewan Harrison, David K. Jackson, Ian Johnston, Dominic Kwiatkowski, Cordelia Langford, John Sillitoe on behalf of the Wellcome Sanger Institute COVID-19 Surveillance Team |
| EPI_ISL_2847832 | Lighthouse Lab in Milton Keynes | Wellcome Sanger Institute for the COVID-19 Genomics UK (COG-UK) Consortium | The Lighthouse Lab in Milton Keynes and Alex Alderton, Roberto Amato, Jeffrey Barrett, Sonia Goncalves, Ewan Harrison, David K. Jackson, Ian Johnston, Dominic Kwiatkowski, Cordelia Langford, John Sillitoe on behalf of the Wellcome Sanger Institute COVID-19 Surveillance Team |
| EPI_ISL_2847833, EPI_ISL_2847834 | Lighthouse Laboratory Plymouth | Wellcome Sanger Institute for the COVID-19 Genomics UK (COG-UK) Consortium | Lighthouse Laboratory Plymouth and Alex Alderton, Roberto Amato, Jeffrey Barrett, Sonia Goncalves, Ewan Harrison, David K. Jackson, Ian Johnston, Dominic Kwiatkowski, Cordelia Langford, John Sillitoe on behalf of the Wellcome Sanger Institute COVID-19 Surveillance Team |
| EPI_ISL_2847835 | Lighthouse Lab in Milton Keynes | Wellcome Sanger Institute for the COVID-19 Genomics UK (COG-UK) Consortium | The Lighthouse Lab in Milton Keynes and Alex Alderton, Roberto Amato, Jeffrey Barrett, Sonia Goncalves, Ewan Harrison, David K. Jackson, Ian Johnston, Dominic Kwiatkowski, Cordelia Langford, John Sillitoe on behalf of the Wellcome Sanger Institute COVID-19 Surveillance Team |
| EPI_ISL_2847836 | Lighthouse Laboratory Plymouth | Wellcome Sanger Institute for the COVID-19 Genomics UK (COG-UK) Consortium | Lighthouse Laboratory Plymouth and Alex Alderton, Roberto Amato, Jeffrey Barrett, Sonia Goncalves, Ewan Harrison, David K. Jackson, Ian Johnston, Dominic Kwiatkowski, Cordelia Langford, John Sillitoe on behalf of the Wellcome Sanger Institute COVID-19 Surveillance Team |
| EPI_ISL_2847837, EPI_ISL_2847838 | Lighthouse Lab in Milton Keynes | Wellcome Sanger Institute for the COVID-19 Genomics UK (COG-UK) Consortium | The Lighthouse Lab in Milton Keynes and Alex Alderton, Roberto Amato, Jeffrey Barrett, Sonia Goncalves, Ewan Harrison, David K. Jackson, Ian Johnston, Dominic Kwiatkowski, Cordelia Langford, John Sillitoe on behalf of the Wellcome Sanger Institute COVID-19 Surveillance Team |
| EPI_ISL_2847839 | Lighthouse Laboratory Plymouth | Wellcome Sanger Institute for the COVID-19 Genomics UK (COG-UK) Consortium | Lighthouse Laboratory Plymouth and Alex Alderton, Roberto Amato, Jeffrey Barrett, Sonia Goncalves, Ewan Harrison, David K. Jackson, Ian Johnston, Dominic Kwiatkowski, Cordelia Langford, John Sillitoe on behalf of the Wellcome Sanger Institute COVID-19 Surveillance Team |
| EPI_ISL_2847840 | Lighthouse Lab in Milton Keynes | Wellcome Sanger Institute for the COVID-19 Genomics UK (COG-UK) Consortium | The Lighthouse Lab in Milton Keynes and Alex Alderton, Roberto Amato, Jeffrey Barrett, Sonia Goncalves, Ewan Harrison, David K. Jackson, Ian Johnston, Dominic Kwiatkowski, Cordelia Langford, John Sillitoe on behalf of the Wellcome Sanger Institute COVID-19 Surveillance Team |
| EPI_ISL_2847842, EPI_ISL_2847843 | Lighthouse Laboratory Plymouth | Wellcome Sanger Institute for the COVID-19 Genomics UK (COG-UK) Consortium | Lighthouse Laboratory Plymouth and Alex Alderton, Roberto Amato, Jeffrey Barrett, Sonia Goncalves, Ewan Harrison, David K. Jackson, Ian Johnston, Dominic Kwiatkowski, Cordelia Langford, John Sillitoe on behalf of the Wellcome Sanger Institute COVID-19 Surveillance Team |
| EPI_ISL_2847844, EPI_ISL_2847845 | Lighthouse Lab in Milton Keynes | Wellcome Sanger Institute for the COVID-19 Genomics UK (COG-UK) Consortium | The Lighthouse Lab in Milton Keynes and Alex Alderton, Roberto Amato, Jeffrey Barrett, Sonia Goncalves, Ewan Harrison, David K. Jackson, Ian Johnston, Dominic Kwiatkowski, Cordelia Langford, John Sillitoe on behalf of the Wellcome Sanger Institute COVID-19 Surveillance Team |
| EPI_ISL_2847846 | Lighthouse Laboratory Plymouth | Wellcome Sanger Institute for the COVID-19 Genomics UK (COG-UK) Consortium | Lighthouse Laboratory Plymouth and Alex Alderton, Roberto Amato, Jeffrey Barrett, Sonia Goncalves, Ewan Harrison, David K. Jackson, Ian Johnston, Dominic Kwiatkowski, Cordelia Langford, John Sillitoe on behalf of the Wellcome Sanger Institute COVID-19 Surveillance Team |
| EPI_ISL_2847847 | Lighthouse Lab in Milton Keynes | Wellcome Sanger Institute for the COVID-19 Genomics UK (COG-UK) Consortium | The Lighthouse Lab in Milton Keynes and Alex Alderton, Roberto Amato, Jeffrey Barrett, Sonia Goncalves, Ewan Harrison, David K. Jackson, Ian Johnston, Dominic Kwiatkowski, Cordelia Langford, John Sillitoe on behalf of the Wellcome Sanger Institute COVID-19 Surveillance Team |
| EPI_ISL_2847848 | Lighthouse Laboratory Plymouth | Wellcome Sanger Institute for the COVID-19 Genomics UK (COG-UK) Consortium | Lighthouse Laboratory Plymouth and Alex Alderton, Roberto Amato, Jeffrey Barrett, Sonia Goncalves, Ewan Harrison, David K. Jackson, Ian Johnston, Dominic Kwiatkowski, Cordelia Langford, John Sillitoe on behalf of the Wellcome Sanger Institute COVID-19 Surveillance Team |
| EPI_ISL_2847849, EPI_ISL_2847850 | Lighthouse Lab in Milton Keynes | Wellcome Sanger Institute for the COVID-19 Genomics UK (COG-UK) Consortium | The Lighthouse Lab in Milton Keynes and Alex Alderton, Roberto Amato, Jeffrey Barrett, Sonia Goncalves, Ewan Harrison, David K. Jackson, Ian Johnston, Dominic Kwiatkowski, Cordelia Langford, John Sillitoe on behalf of the Wellcome Sanger Institute COVID-19 Surveillance Team |
| EPI_ISL_2847851, EPI_ISL_2847852, EPI_ISL_2847853 | Lighthouse Laboratory Plymouth | Wellcome Sanger Institute for the COVID-19 Genomics UK (COG-UK) Consortium | Lighthouse Laboratory Plymouth and Alex Alderton, Roberto Amato, Jeffrey Barrett, Sonia Goncalves, Ewan Harrison, David K. Jackson, Ian Johnston, Dominic Kwiatkowski, Cordelia Langford, John Sillitoe on behalf of the Wellcome Sanger Institute COVID-19 Surveillance Team |
| EPI_ISL_2847854 | Lighthouse Lab in Milton Keynes | Wellcome Sanger Institute for the COVID-19 Genomics UK (COG-UK) Consortium | The Lighthouse Lab in Milton Keynes and Alex Alderton, Roberto Amato, Jeffrey Barrett, Sonia Goncalves, Ewan Harrison, David K. Jackson, Ian Johnston, Dominic Kwiatkowski, Cordelia Langford, John Sillitoe on behalf of the Wellcome Sanger Institute COVID-19 Surveillance Team |
| EPI_ISL_2847855, EPI_ISL_2847856, EPI_ISL_2847857 | Lighthouse Laboratory Plymouth | Wellcome Sanger Institute for the COVID-19 Genomics UK (COG-UK) Consortium | Lighthouse Laboratory Plymouth and Alex Alderton, Roberto Amato, Jeffrey Barrett, Sonia Goncalves, Ewan Harrison, David K. Jackson, Ian Johnston, Dominic Kwiatkowski, Cordelia Langford, John Sillitoe on behalf of the Wellcome Sanger Institute COVID-19 Surveillance Team |
| EPI_ISL_2847858, EPI_ISL_2847859 | Lighthouse Lab in Milton Keynes | Wellcome Sanger Institute for the COVID-19 Genomics UK (COG-UK) Consortium | The Lighthouse Lab in Milton Keynes and Alex Alderton, Roberto Amato, Jeffrey Barrett, Sonia Goncalves, Ewan Harrison, David K. Jackson, Ian Johnston, Dominic Kwiatkowski, Cordelia Langford, John Sillitoe on behalf of the Wellcome Sanger Institute COVID-19 Surveillance Team |
| EPI_ISL_2847860, EPI_ISL_2847861, EPI_ISL_2847862, EPI_ISL_2847863 | Lighthouse Laboratory Plymouth | Wellcome Sanger Institute for the COVID-19 Genomics UK (COG-UK) Consortium | Lighthouse Laboratory Plymouth and Alex Alderton, Roberto Amato, Jeffrey Barrett, Sonia Goncalves, Ewan Harrison, David K. Jackson, Ian Johnston, Dominic Kwiatkowski, Cordelia Langford, John Sillitoe on behalf of the Wellcome Sanger Institute COVID-19 Surveillance Team |
| EPI_ISL_2847864 | Lighthouse Lab in Milton Keynes | Wellcome Sanger Institute for the COVID-19 Genomics UK (COG-UK) Consortium | The Lighthouse Lab in Milton Keynes and Alex Alderton, Roberto Amato, Jeffrey Barrett, Sonia Goncalves, Ewan Harrison, David K. Jackson, Ian Johnston, Dominic Kwiatkowski, Cordelia Langford, John Sillitoe on behalf of the Wellcome Sanger Institute COVID-19 Surveillance Team |
| EPI_ISL_2847865, EPI_ISL_2847866 | Lighthouse Laboratory Plymouth | Wellcome Sanger Institute for the COVID-19 Genomics UK (COG-UK) Consortium | Lighthouse Laboratory Plymouth and Alex Alderton, Roberto Amato, Jeffrey Barrett, Sonia Goncalves, Ewan Harrison, David K. Jackson, Ian Johnston, Dominic Kwiatkowski, Cordelia Langford, John Sillitoe on behalf of the Wellcome Sanger Institute COVID-19 Surveillance Team |
| EPI_ISL_2847867, EPI_ISL_2847869 | Lighthouse Lab in Milton Keynes | Wellcome Sanger Institute for the COVID-19 Genomics UK (COG-UK) Consortium | The Lighthouse Lab in Milton Keynes and Alex Alderton, Roberto Amato, Jeffrey Barrett, Sonia Goncalves, Ewan Harrison, David K. Jackson, Ian Johnston, Dominic Kwiatkowski, Cordelia Langford, John Sillitoe on behalf of the Wellcome Sanger Institute COVID-19 Surveillance Team |
| EPI_ISL_2847870 | Lighthouse Laboratory Plymouth | Wellcome Sanger Institute for the COVID-19 Genomics UK (COG-UK) Consortium | Lighthouse Laboratory Plymouth and Alex Alderton, Roberto Amato, Jeffrey Barrett, Sonia Goncalves, Ewan Harrison, David K. Jackson, Ian Johnston, Dominic Kwiatkowski, Cordelia Langford, John Sillitoe on behalf of the Wellcome Sanger Institute COVID-19 Surveillance Team |
| EPI_ISL_2847871 | Lighthouse Lab in Milton Keynes | Wellcome Sanger Institute for the COVID-19 Genomics UK (COG-UK) Consortium | The Lighthouse Lab in Milton Keynes and Alex Alderton, Roberto Amato, Jeffrey Barrett, Sonia Goncalves, Ewan Harrison, David K. Jackson, Ian Johnston, Dominic Kwiatkowski, Cordelia Langford, John Sillitoe on behalf of the Wellcome Sanger Institute COVID-19 Surveillance Team |
| EPI_ISL_2847872, EPI_ISL_2847873 | Lighthouse Laboratory Plymouth | Wellcome Sanger Institute for the COVID-19 Genomics UK (COG-UK) Consortium | Lighthouse Laboratory Plymouth and Alex Alderton, Roberto Amato, Jeffrey Barrett, Sonia Goncalves, Ewan Harrison, David K. Jackson, Ian Johnston, Dominic Kwiatkowski, Cordelia Langford, John Sillitoe on behalf of the Wellcome Sanger Institute COVID-19 Surveillance Team |
| EPI_ISL_2847874 | Lighthouse Lab in Milton Keynes | Wellcome Sanger Institute for the COVID-19 Genomics UK (COG-UK) Consortium | The Lighthouse Lab in Milton Keynes and Alex Alderton, Roberto Amato, Jeffrey Barrett, Sonia Goncalves, Ewan Harrison, David K. Jackson, Ian Johnston, Dominic Kwiatkowski, Cordelia Langford, John Sillitoe on behalf of the Wellcome Sanger Institute COVID-19 Surveillance Team |
| EPI_ISL_2847875, EPI_ISL_2847876, EPI_ISL_2847877 | Lighthouse Laboratory Plymouth | Wellcome Sanger Institute for the COVID-19 Genomics UK (COG-UK) Consortium | Lighthouse Laboratory Plymouth and Alex Alderton, Roberto Amato, Jeffrey Barrett, Sonia Goncalves, Ewan Harrison, David K. Jackson, Ian Johnston, Dominic Kwiatkowski, Cordelia Langford, John Sillitoe on behalf of the Wellcome Sanger Institute COVID-19 Surveillance Team |

| | | | |
|---|---|---|---|
| EPI_ISL_2847878, EPI_ISL_2847879 | Lighthouse Lab in Milton Keynes | Wellcome Sanger Institute for the COVID-19 Genomics UK (COG-UK) Consortium | The Lighthouse Lab in Milton Keynes and Alex Alderton, Roberto Amato, Jeffrey Barrett, Sonia Goncalves, Ewan Harrison, David K. Jackson, Ian Johnston, Dominic Kwiatkowski, Cordelia Langford, John Sillitoe on behalf of the Wellcome Sanger Institute COVID-19 Surveillance Team |
| EPI_ISL_2847880, EPI_ISL_2847881, EPI_ISL_2847882, EPI_ISL_2847883, EPI_ISL_2847884 | Lighthouse Laboratory Plymouth | Wellcome Sanger Institute for the COVID-19 Genomics UK (COG-UK) Consortium | Lighthouse Laboratory Plymouth and Alex Alderton, Roberto Amato, Jeffrey Barrett, Sonia Goncalves, Ewan Harrison, David K. Jackson, Ian Johnston, Dominic Kwiatkowski, Cordelia Langford, John Sillitoe on behalf of the Wellcome Sanger Institute COVID-19 Surveillance Team |
| EPI_ISL_2847885 | Lighthouse Lab in Milton Keynes | Wellcome Sanger Institute for the COVID-19 Genomics UK (COG-UK) Consortium | The Lighthouse Lab in Milton Keynes and Alex Alderton, Roberto Amato, Jeffrey Barrett, Sonia Goncalves, Ewan Harrison, David K. Jackson, Ian Johnston, Dominic Kwiatkowski, Cordelia Langford, John Sillitoe on behalf of the Wellcome Sanger Institute COVID-19 Surveillance Team |
| EPI_ISL_2847886, EPI_ISL_2847887, EPI_ISL_2847888, EPI_ISL_2847889, EPI_ISL_2847890 | Lighthouse Laboratory Plymouth | Wellcome Sanger Institute for the COVID-19 Genomics UK (COG-UK) Consortium | Lighthouse Laboratory Plymouth and Alex Alderton, Roberto Amato, Jeffrey Barrett, Sonia Goncalves, Ewan Harrison, David K. Jackson, Ian Johnston, Dominic Kwiatkowski, Cordelia Langford, John Sillitoe on behalf of the Wellcome Sanger Institute COVID-19 Surveillance Team |
| EPI_ISL_2847891 | Lighthouse Lab in Milton Keynes | Wellcome Sanger Institute for the COVID-19 Genomics UK (COG-UK) Consortium | The Lighthouse Lab in Milton Keynes and Alex Alderton, Roberto Amato, Jeffrey Barrett, Sonia Goncalves, Ewan Harrison, David K. Jackson, Ian Johnston, Dominic Kwiatkowski, Cordelia Langford, John Sillitoe on behalf of the Wellcome Sanger Institute COVID-19 Surveillance Team |
| EPI_ISL_2847892, EPI_ISL_2847893, EPI_ISL_2847894, EPI_ISL_2847895, EPI_ISL_2847896, EPI_ISL_2847897 | Lighthouse Laboratory Plymouth | Wellcome Sanger Institute for the COVID-19 Genomics UK (COG-UK) Consortium | Lighthouse Laboratory Plymouth and Alex Alderton, Roberto Amato, Jeffrey Barrett, Sonia Goncalves, Ewan Harrison, David K. Jackson, Ian Johnston, Dominic Kwiatkowski, Cordelia Langford, John Sillitoe on behalf of the Wellcome Sanger Institute COVID-19 Surveillance Team |
| EPI_ISL_2847898 | Lighthouse Lab in Milton Keynes | Wellcome Sanger Institute for the COVID-19 Genomics UK (COG-UK) Consortium | The Lighthouse Lab in Milton Keynes and Alex Alderton, Roberto Amato, Jeffrey Barrett, Sonia Goncalves, Ewan Harrison, David K. Jackson, Ian Johnston, Dominic Kwiatkowski, Cordelia Langford, John Sillitoe on behalf of the Wellcome Sanger Institute COVID-19 Surveillance Team |
| EPI_ISL_2847899 | Lighthouse Laboratory Plymouth | Wellcome Sanger Institute for the COVID-19 Genomics UK (COG-UK) Consortium | Lighthouse Laboratory Plymouth and Alex Alderton, Roberto Amato, Jeffrey Barrett, Sonia Goncalves, Ewan Harrison, David K. Jackson, Ian Johnston, Dominic Kwiatkowski, Cordelia Langford, John Sillitoe on behalf of the Wellcome Sanger Institute COVID-19 Surveillance Team |
| EPI_ISL_2847900 | Lighthouse Lab in Milton Keynes | Wellcome Sanger Institute for the COVID-19 Genomics UK (COG-UK) Consortium | The Lighthouse Lab in Milton Keynes and Alex Alderton, Roberto Amato, Jeffrey Barrett, Sonia Goncalves, Ewan Harrison, David K. Jackson, Ian Johnston, Dominic Kwiatkowski, Cordelia Langford, John Sillitoe on behalf of the Wellcome Sanger Institute COVID-19 Surveillance Team |
| EPI_ISL_2847901, EPI_ISL_2847902, EPI_ISL_2847903 | Lighthouse Laboratory Plymouth | Wellcome Sanger Institute for the COVID-19 Genomics UK (COG-UK) Consortium | Lighthouse Laboratory Plymouth and Alex Alderton, Roberto Amato, Jeffrey Barrett, Sonia Goncalves, Ewan Harrison, David K. Jackson, Ian Johnston, Dominic Kwiatkowski, Cordelia Langford, John Sillitoe on behalf of the Wellcome Sanger Institute COVID-19 Surveillance Team |
| EPI_ISL_2847904 | Lighthouse Lab in Milton Keynes | Wellcome Sanger Institute for the COVID-19 Genomics UK (COG-UK) Consortium | The Lighthouse Lab in Milton Keynes and Alex Alderton, Roberto Amato, Jeffrey Barrett, Sonia Goncalves, Ewan Harrison, David K. Jackson, Ian Johnston, Dominic Kwiatkowski, Cordelia Langford, John Sillitoe on behalf of the Wellcome Sanger Institute COVID-19 Surveillance Team |
| EPI_ISL_2847905, EPI_ISL_2847906, EPI_ISL_2847907, EPI_ISL_2847908, EPI_ISL_2847909, EPI_ISL_2847910, EPI_ISL_2847911, EPI_ISL_2847912, EPI_ISL_2847913 | Lighthouse Laboratory Plymouth | Wellcome Sanger Institute for the COVID-19 Genomics UK (COG-UK) Consortium | Lighthouse Laboratory Plymouth and Alex Alderton, Roberto Amato, Jeffrey Barrett, Sonia Goncalves, Ewan Harrison, David K. Jackson, Ian Johnston, Dominic Kwiatkowski, Cordelia Langford, John Sillitoe on behalf of the Wellcome Sanger Institute COVID-19 Surveillance Team |
| EPI_ISL_2847914 | Lighthouse Lab in Milton Keynes | Wellcome Sanger Institute for the COVID-19 Genomics UK (COG-UK) Consortium | The Lighthouse Lab in Milton Keynes and Alex Alderton, Roberto Amato, Jeffrey Barrett, Sonia Goncalves, Ewan Harrison, David K. Jackson, Ian Johnston, Dominic Kwiatkowski, Cordelia Langford, John Sillitoe on behalf of the Wellcome Sanger Institute COVID-19 Surveillance Team |
| EPI_ISL_2847915, EPI_ISL_2847916, EPI_ISL_2847917, EPI_ISL_2847918, EPI_ISL_2847919, EPI_ISL_2847920, EPI_ISL_2847921 | Lighthouse Laboratory Plymouth | Wellcome Sanger Institute for the COVID-19 Genomics UK (COG-UK) Consortium | Lighthouse Laboratory Plymouth and Alex Alderton, Roberto Amato, Jeffrey Barrett, Sonia Goncalves, Ewan Harrison, David K. Jackson, Ian Johnston, Dominic Kwiatkowski, Cordelia Langford, John Sillitoe on behalf of the Wellcome Sanger Institute COVID-19 Surveillance Team |
| EPI_ISL_2847922 | Lighthouse Lab in Milton Keynes | Wellcome Sanger Institute for the COVID-19 Genomics UK (COG-UK) Consortium | The Lighthouse Lab in Milton Keynes and Alex Alderton, Roberto Amato, Jeffrey Barrett, Sonia Goncalves, Ewan Harrison, David K. Jackson, Ian Johnston, Dominic Kwiatkowski, Cordelia Langford, John Sillitoe on behalf of the Wellcome Sanger Institute COVID-19 Surveillance Team |
| EPI_ISL_2847923, EPI_ISL_2847924, EPI_ISL_2847925 | Lighthouse Laboratory Plymouth | Wellcome Sanger Institute for the COVID-19 Genomics UK (COG-UK) Consortium | Lighthouse Laboratory Plymouth and Alex Alderton, Roberto Amato, Jeffrey Barrett, Sonia Goncalves, Ewan Harrison, David K. Jackson, Ian Johnston, Dominic Kwiatkowski, Cordelia Langford, John Sillitoe on behalf of the Wellcome Sanger Institute COVID-19 Surveillance Team |
| EPI_ISL_2847926 | Lighthouse Lab in Milton Keynes | Wellcome Sanger Institute for the COVID-19 Genomics UK (COG-UK) Consortium | The Lighthouse Lab in Milton Keynes and Alex Alderton, Roberto Amato, Jeffrey Barrett, Sonia Goncalves, Ewan Harrison, David K. Jackson, Ian Johnston, Dominic Kwiatkowski, Cordelia Langford, John Sillitoe on behalf of the Wellcome Sanger Institute COVID-19 Surveillance Team |
| EPI_ISL_2847927 | Lighthouse Laboratory Plymouth | Wellcome Sanger Institute for the COVID-19 Genomics UK (COG-UK) Consortium | Lighthouse Laboratory Plymouth and Alex Alderton, Roberto Amato, Jeffrey Barrett, Sonia Goncalves, Ewan Harrison, David K. Jackson, Ian Johnston, Dominic Kwiatkowski, Cordelia Langford, John Sillitoe on behalf of the Wellcome Sanger Institute COVID-19 Surveillance Team |
| EPI_ISL_2847928 | Lighthouse Lab in Milton Keynes | Wellcome Sanger Institute for the COVID-19 Genomics UK (COG-UK) Consortium | The Lighthouse Lab in Milton Keynes and Alex Alderton, Roberto Amato, Jeffrey Barrett, Sonia Goncalves, Ewan Harrison, David K. Jackson, Ian Johnston, Dominic Kwiatkowski, Cordelia Langford, John Sillitoe on behalf of the Wellcome Sanger Institute COVID-19 Surveillance Team |
| EPI_ISL_2847929, EPI_ISL_2847930, EPI_ISL_2847931, EPI_ISL_2847932, EPI_ISL_2847933, EPI_ISL_2847934, EPI_ISL_2847935, EPI_ISL_2847936, EPI_ISL_2847937, EPI_ISL_2847938, EPI_ISL_2847939 | | | |
| see above | Lighthouse Laboratory Plymouth | Wellcome Sanger Institute for the COVID-19 Genomics UK (COG-UK) Consortium | Lighthouse Laboratory Plymouth and Alex Alderton, Roberto Amato, Jeffrey Barrett, Sonia Goncalves, Ewan Harrison, David K. Jackson, Ian Johnston, Dominic Kwiatkowski, Cordelia Langford, John Sillitoe on behalf of the Wellcome Sanger Institute COVID-19 Surveillance Team |
| EPI_ISL_2847940, EPI_ISL_2847941 | Lighthouse Lab in Milton Keynes | Wellcome Sanger Institute for the COVID-19 Genomics UK (COG-UK) Consortium | The Lighthouse Lab in Milton Keynes and Alex Alderton, Roberto Amato, Jeffrey Barrett, Sonia Goncalves, Ewan Harrison, David K. Jackson, Ian Johnston, Dominic Kwiatkowski, Cordelia Langford, John Sillitoe on behalf of the Wellcome Sanger Institute COVID-19 Surveillance Team |
| EPI_ISL_2847942 | Lighthouse Laboratory Plymouth | Wellcome Sanger Institute for the COVID-19 Genomics UK (COG-UK) Consortium | Lighthouse Laboratory Plymouth and Alex Alderton, Roberto Amato, Jeffrey Barrett, Sonia Goncalves, Ewan Harrison, David K. Jackson, Ian Johnston, Dominic Kwiatkowski, Cordelia Langford, John Sillitoe on behalf of the Wellcome Sanger Institute COVID-19 Surveillance Team |
| EPI_ISL_2847943 | Lighthouse Lab in Milton Keynes | Wellcome Sanger Institute for the COVID-19 Genomics UK (COG-UK) Consortium | The Lighthouse Lab in Milton Keynes and Alex Alderton, Roberto Amato, Jeffrey Barrett, Sonia Goncalves, Ewan Harrison, David K. Jackson, Ian Johnston, Dominic Kwiatkowski, Cordelia Langford, John Sillitoe on behalf of the Wellcome Sanger Institute COVID-19 Surveillance Team |
| EPI_ISL_2847944, EPI_ISL_2847945, EPI_ISL_2847946, EPI_ISL_2847947, EPI_ISL_2847948, EPI_ISL_2847949 | Lighthouse Laboratory Plymouth | Wellcome Sanger Institute for the COVID-19 Genomics UK (COG-UK) Consortium | Lighthouse Laboratory Plymouth and Alex Alderton, Roberto Amato, Jeffrey Barrett, Sonia Goncalves, Ewan Harrison, David K. Jackson, Ian Johnston, Dominic Kwiatkowski, Cordelia Langford, John Sillitoe on behalf of the Wellcome Sanger Institute COVID-19 Surveillance Team |
| EPI_ISL_2847950, EPI_ISL_2847951 | Lighthouse Lab in Milton Keynes | Wellcome Sanger Institute for the COVID-19 Genomics UK (COG-UK) Consortium | The Lighthouse Lab in Milton Keynes and Alex Alderton, Roberto Amato, Jeffrey Barrett, Sonia Goncalves, Ewan Harrison, David K. Jackson, Ian Johnston, Dominic Kwiatkowski, Cordelia Langford, John Sillitoe on behalf of the Wellcome Sanger Institute COVID-19 Surveillance Team |
| EPI_ISL_2847952, EPI_ISL_2847953, EPI_ISL_2847954, EPI_ISL_2847955, EPI_ISL_2847956 | Lighthouse Laboratory Plymouth | Wellcome Sanger Institute for the COVID-19 Genomics UK (COG-UK) Consortium | Lighthouse Laboratory Plymouth and Alex Alderton, Roberto Amato, Jeffrey Barrett, Sonia Goncalves, Ewan Harrison, David K. Jackson, Ian Johnston, Dominic Kwiatkowski, Cordelia Langford, John Sillitoe on behalf of the Wellcome Sanger Institute COVID-19 Surveillance Team |
| EPI_ISL_2847957, EPI_ISL_2847958 | Lighthouse Lab in Alderley Park | Wellcome Sanger Institute for the COVID-19 Genomics UK (COG-UK) Consortium | Jacquelyn Wynn, Mairead Hyland, The Lighthouse Lab in Alderley Park and Alex Alderton, Roberto Amato, Jeffrey Barrett, Sonia Goncalves, Ewan Harrison, David K. Jackson, Ian Johnston, Dominic Kwiatkowski, Cordelia Langford, John Sillitoe on behalf of the Wellcome Sanger Institute COVID-19 Surveillance Team |
| EPI_ISL_2847959, EPI_ISL_2847960 | Lighthouse Laboratory Plymouth | Wellcome Sanger Institute for the COVID-19 Genomics UK (COG-UK) Consortium | Lighthouse Laboratory Plymouth and Alex Alderton, Roberto Amato, Jeffrey Barrett, Sonia Goncalves, Ewan Harrison, David K. Jackson, Ian Johnston, Dominic Kwiatkowski, Cordelia Langford, John Sillitoe on behalf of the Wellcome Sanger Institute COVID-19 Surveillance Team |
| EPI_ISL_2847961 | Lighthouse Lab in Alderley Park | Wellcome Sanger Institute for the COVID-19 Genomics UK (COG-UK) Consortium | Jacquelyn Wynn, Mairead Hyland, The Lighthouse Lab in Alderley Park and Alex Alderton, Roberto Amato, Jeffrey Barrett, Sonia Goncalves, Ewan Harrison, David K. Jackson, Ian Johnston, Dominic Kwiatkowski, Cordelia Langford, John Sillitoe on behalf of the Wellcome Sanger Institute COVID-19 Surveillance Team |
| EPI_ISL_2847962 | Lighthouse Lab in Milton Keynes | Wellcome Sanger Institute for the COVID-19 Genomics UK (COG-UK) Consortium | The Lighthouse Lab in Milton Keynes and Alex Alderton, Roberto Amato, Jeffrey Barrett, Sonia Goncalves, Ewan Harrison, David K. Jackson, Ian Johnston, Dominic Kwiatkowski, Cordelia Langford, John Sillitoe on behalf of the Wellcome Sanger Institute COVID-19 Surveillance Team |
| EPI_ISL_2847963, EPI_ISL_2847964, EPI_ISL_2847965 | Lighthouse Laboratory Plymouth | Wellcome Sanger Institute for the COVID-19 Genomics UK (COG-UK) Consortium | Lighthouse Laboratory Plymouth and Alex Alderton, Roberto Amato, Jeffrey Barrett, Sonia Goncalves, Ewan Harrison, David K. Jackson, Ian Johnston, Dominic Kwiatkowski, Cordelia Langford, John Sillitoe on behalf of the Wellcome Sanger Institute COVID-19 Surveillance Team |
| EPI_ISL_2847966 | Lighthouse Lab in Alderley Park | Wellcome Sanger Institute for the COVID-19 Genomics UK (COG-UK) | Jacquelyn Wynn, Mairead Hyland, The Lighthouse Lab in Alderley Park and Alex Alderton, Roberto Amato, Jeffrey Barrett, Sonia Goncalves, Ewan Harrison, David K. Jackson, Ian Johnston, Dominic Kwiatkowski, Cordelia Langford, John Sillitoe on |

| | | | |
|---|---|---|---|
| | | Consortium | behalf of the Wellcome Sanger Institute COVID-19 Surveillance Team |
| EPI_ISL_2847967 | Lighthouse Lab in Milton Keynes | Wellcome Sanger Institute for the COVID-19 Genomics UK (COG-UK) Consortium | The Lighthouse Lab in Milton Keynes and Alex Alderton, Roberto Amato, Jeffrey Barrett, Sonia Goncalves, Ewan Harrison, David K. Jackson, Ian Johnston, Dominic Kwiatkowski, Cordelia Langford, John Sillitoe on behalf of the Wellcome Sanger Institute COVID-19 Surveillance Team |
| EPI_ISL_2847968, EPI_ISL_2847969, EPI_ISL_2847970 | Lighthouse Laboratory Plymouth | Wellcome Sanger Institute for the COVID-19 Genomics UK (COG-UK) Consortium | Lighthouse Laboratory Plymouth and Alex Alderton, Roberto Amato, Jeffrey Barrett, Sonia Goncalves, Ewan Harrison, David K. Jackson, Ian Johnston, Dominic Kwiatkowski, Cordelia Langford, John Sillitoe on behalf of the Wellcome Sanger Institute COVID-19 Surveillance Team |
| EPI_ISL_2847971 | Lighthouse Lab in Milton Keynes | Wellcome Sanger Institute for the COVID-19 Genomics UK (COG-UK) Consortium | The Lighthouse Lab in Milton Keynes and Alex Alderton, Roberto Amato, Jeffrey Barrett, Sonia Goncalves, Ewan Harrison, David K. Jackson, Ian Johnston, Dominic Kwiatkowski, Cordelia Langford, John Sillitoe on behalf of the Wellcome Sanger Institute COVID-19 Surveillance Team |
| EPI_ISL_2847972, EPI_ISL_2847973 | Lighthouse Laboratory Plymouth | Wellcome Sanger Institute for the COVID-19 Genomics UK (COG-UK) Consortium | Lighthouse Laboratory Plymouth and Alex Alderton, Roberto Amato, Jeffrey Barrett, Sonia Goncalves, Ewan Harrison, David K. Jackson, Ian Johnston, Dominic Kwiatkowski, Cordelia Langford, John Sillitoe on behalf of the Wellcome Sanger Institute COVID-19 Surveillance Team |
| EPI_ISL_2847974, EPI_ISL_2847975 | Lighthouse Lab in Alderley Park | Wellcome Sanger Institute for the COVID-19 Genomics UK (COG-UK) Consortium | Jacquelyn Wynn, Mairead Hyland, The Lighthouse Lab in Alderley Park and Alex Alderton, Roberto Amato, Jeffrey Barrett, Sonia Goncalves, Ewan Harrison, David K. Jackson, Ian Johnston, Dominic Kwiatkowski, Cordelia Langford, John Sillitoe on behalf of the Wellcome Sanger Institute COVID-19 Surveillance Team |
| EPI_ISL_2847976, EPI_ISL_2847977, EPI_ISL_2847978 | Lighthouse Laboratory Plymouth | Wellcome Sanger Institute for the COVID-19 Genomics UK (COG-UK) Consortium | Lighthouse Laboratory Plymouth and Alex Alderton, Roberto Amato, Jeffrey Barrett, Sonia Goncalves, Ewan Harrison, David K. Jackson, Ian Johnston, Dominic Kwiatkowski, Cordelia Langford, John Sillitoe on behalf of the Wellcome Sanger Institute COVID-19 Surveillance Team |
| EPI_ISL_2847979 | Lighthouse Lab in Milton Keynes | Wellcome Sanger Institute for the COVID-19 Genomics UK (COG-UK) Consortium | The Lighthouse Lab in Milton Keynes and Alex Alderton, Roberto Amato, Jeffrey Barrett, Sonia Goncalves, Ewan Harrison, David K. Jackson, Ian Johnston, Dominic Kwiatkowski, Cordelia Langford, John Sillitoe on behalf of the Wellcome Sanger Institute COVID-19 Surveillance Team |
| EPI_ISL_2847980 | Lighthouse Laboratory Plymouth | Wellcome Sanger Institute for the COVID-19 Genomics UK (COG-UK) Consortium | Lighthouse Laboratory Plymouth and Alex Alderton, Roberto Amato, Jeffrey Barrett, Sonia Goncalves, Ewan Harrison, David K. Jackson, Ian Johnston, Dominic Kwiatkowski, Cordelia Langford, John Sillitoe on behalf of the Wellcome Sanger Institute COVID-19 Surveillance Team |
| EPI_ISL_2847981 | Lighthouse Lab in Alderley Park | Wellcome Sanger Institute for the COVID-19 Genomics UK (COG-UK) Consortium | Jacquelyn Wynn, Mairead Hyland, The Lighthouse Lab in Alderley Park and Alex Alderton, Roberto Amato, Jeffrey Barrett, Sonia Goncalves, Ewan Harrison, David K. Jackson, Ian Johnston, Dominic Kwiatkowski, Cordelia Langford, John Sillitoe on behalf of the Wellcome Sanger Institute COVID-19 Surveillance Team |
| EPI_ISL_2847982 | Lighthouse Laboratory Plymouth | Wellcome Sanger Institute for the COVID-19 Genomics UK (COG-UK) Consortium | Lighthouse Laboratory Plymouth and Alex Alderton, Roberto Amato, Jeffrey Barrett, Sonia Goncalves, Ewan Harrison, David K. Jackson, Ian Johnston, Dominic Kwiatkowski, Cordelia Langford, John Sillitoe on behalf of the Wellcome Sanger Institute COVID-19 Surveillance Team |
| EPI_ISL_2847983 | Lighthouse Lab in Alderley Park | Wellcome Sanger Institute for the COVID-19 Genomics UK (COG-UK) Consortium | Jacquelyn Wynn, Mairead Hyland, The Lighthouse Lab in Alderley Park and Alex Alderton, Roberto Amato, Jeffrey Barrett, Sonia Goncalves, Ewan Harrison, David K. Jackson, Ian Johnston, Dominic Kwiatkowski, Cordelia Langford, John Sillitoe on behalf of the Wellcome Sanger Institute COVID-19 Surveillance Team |
| EPI_ISL_2847984, EPI_ISL_2847985 | Lighthouse Laboratory Plymouth | Wellcome Sanger Institute for the COVID-19 Genomics UK (COG-UK) Consortium | Lighthouse Laboratory Plymouth and Alex Alderton, Roberto Amato, Jeffrey Barrett, Sonia Goncalves, Ewan Harrison, David K. Jackson, Ian Johnston, Dominic Kwiatkowski, Cordelia Langford, John Sillitoe on behalf of the Wellcome Sanger Institute COVID-19 Surveillance Team |
| EPI_ISL_2847986 | Lighthouse Lab in Alderley Park | Wellcome Sanger Institute for the COVID-19 Genomics UK (COG-UK) Consortium | Jacquelyn Wynn, Mairead Hyland, The Lighthouse Lab in Alderley Park and Alex Alderton, Roberto Amato, Jeffrey Barrett, Sonia Goncalves, Ewan Harrison, David K. Jackson, Ian Johnston, Dominic Kwiatkowski, Cordelia Langford, John Sillitoe on behalf of the Wellcome Sanger Institute COVID-19 Surveillance Team |
| EPI_ISL_2847987 | Lighthouse Laboratory Plymouth | Wellcome Sanger Institute for the COVID-19 Genomics UK (COG-UK) Consortium | Lighthouse Laboratory Plymouth and Alex Alderton, Roberto Amato, Jeffrey Barrett, Sonia Goncalves, Ewan Harrison, David K. Jackson, Ian Johnston, Dominic Kwiatkowski, Cordelia Langford, John Sillitoe on behalf of the Wellcome Sanger Institute COVID-19 Surveillance Team |
| EPI_ISL_2847988, EPI_ISL_2847989, EPI_ISL_2847990, EPI_ISL_2847991, EPI_ISL_2847992 | Lighthouse Lab in Milton Keynes | Wellcome Sanger Institute for the COVID-19 Genomics UK (COG-UK) Consortium | The Lighthouse Lab in Milton Keynes and Alex Alderton, Roberto Amato, Jeffrey Barrett, Sonia Goncalves, Ewan Harrison, David K. Jackson, Ian Johnston, Dominic Kwiatkowski, Cordelia Langford, John Sillitoe on behalf of the Wellcome Sanger Institute COVID-19 Surveillance Team |
| EPI_ISL_2847993, EPI_ISL_2847994 | Lighthouse Laboratory Plymouth | Wellcome Sanger Institute for the COVID-19 Genomics UK (COG-UK) Consortium | Lighthouse Laboratory Plymouth and Alex Alderton, Roberto Amato, Jeffrey Barrett, Sonia Goncalves, Ewan Harrison, David K. Jackson, Ian Johnston, Dominic Kwiatkowski, Cordelia Langford, John Sillitoe on behalf of the Wellcome Sanger Institute COVID-19 Surveillance Team |
| EPI_ISL_2847995 | Lighthouse Lab in Milton Keynes | Wellcome Sanger Institute for the COVID-19 Genomics UK (COG-UK) Consortium | The Lighthouse Lab in Milton Keynes and Alex Alderton, Roberto Amato, Jeffrey Barrett, Sonia Goncalves, Ewan Harrison, David K. Jackson, Ian Johnston, Dominic Kwiatkowski, Cordelia Langford, John Sillitoe on behalf of the Wellcome Sanger Institute COVID-19 Surveillance Team |
| EPI_ISL_2847996 | Lighthouse Lab in Alderley Park | Wellcome Sanger Institute for the COVID-19 Genomics UK (COG-UK) Consortium | Jacquelyn Wynn, Mairead Hyland, The Lighthouse Lab in Alderley Park and Alex Alderton, Roberto Amato, Jeffrey Barrett, Sonia Goncalves, Ewan Harrison, David K. Jackson, Ian Johnston, Dominic Kwiatkowski, Cordelia Langford, John Sillitoe on behalf of the Wellcome Sanger Institute COVID-19 Surveillance Team |
| EPI_ISL_2847997 | Lighthouse Lab in Milton Keynes | Wellcome Sanger Institute for the COVID-19 Genomics UK (COG-UK) Consortium | The Lighthouse Lab in Milton Keynes and Alex Alderton, Roberto Amato, Jeffrey Barrett, Sonia Goncalves, Ewan Harrison, David K. Jackson, Ian Johnston, Dominic Kwiatkowski, Cordelia Langford, John Sillitoe on behalf of the Wellcome Sanger Institute COVID-19 Surveillance Team |
| EPI_ISL_2847998 | Lighthouse Lab in Alderley Park | Wellcome Sanger Institute for the COVID-19 Genomics UK (COG-UK) Consortium | Jacquelyn Wynn, Mairead Hyland, The Lighthouse Lab in Alderley Park and Alex Alderton, Roberto Amato, Jeffrey Barrett, Sonia Goncalves, Ewan Harrison, David K. Jackson, Ian Johnston, Dominic Kwiatkowski, Cordelia Langford, John Sillitoe on behalf of the Wellcome Sanger Institute COVID-19 Surveillance Team |
| EPI_ISL_2847999 | Lighthouse Lab in Milton Keynes | Wellcome Sanger Institute for the COVID-19 Genomics UK (COG-UK) Consortium | The Lighthouse Lab in Milton Keynes and Alex Alderton, Roberto Amato, Jeffrey Barrett, Sonia Goncalves, Ewan Harrison, David K. Jackson, Ian Johnston, Dominic Kwiatkowski, Cordelia Langford, John Sillitoe on behalf of the Wellcome Sanger Institute COVID-19 Surveillance Team |
| EPI_ISL_2848000, EPI_ISL_2848001 | Lighthouse Laboratory Plymouth | Wellcome Sanger Institute for the COVID-19 Genomics UK (COG-UK) Consortium | Lighthouse Laboratory Plymouth and Alex Alderton, Roberto Amato, Jeffrey Barrett, Sonia Goncalves, Ewan Harrison, David K. Jackson, Ian Johnston, Dominic Kwiatkowski, Cordelia Langford, John Sillitoe on behalf of the Wellcome Sanger Institute COVID-19 Surveillance Team |
| EPI_ISL_2848002 | Lighthouse Lab in Alderley Park | Wellcome Sanger Institute for the COVID-19 Genomics UK (COG-UK) Consortium | Jacquelyn Wynn, Mairead Hyland, The Lighthouse Lab in Alderley Park and Alex Alderton, Roberto Amato, Jeffrey Barrett, Sonia Goncalves, Ewan Harrison, David K. Jackson, Ian Johnston, Dominic Kwiatkowski, Cordelia Langford, John Sillitoe on behalf of the Wellcome Sanger Institute COVID-19 Surveillance Team |
| EPI_ISL_2848003 | Lighthouse Laboratory Plymouth | Wellcome Sanger Institute for the COVID-19 Genomics UK (COG-UK) Consortium | Lighthouse Laboratory Plymouth and Alex Alderton, Roberto Amato, Jeffrey Barrett, Sonia Goncalves, Ewan Harrison, David K. Jackson, Ian Johnston, Dominic Kwiatkowski, Cordelia Langford, John Sillitoe on behalf of the Wellcome Sanger Institute COVID-19 Surveillance Team |
| EPI_ISL_2848004 | Lighthouse Lab in Milton Keynes | Wellcome Sanger Institute for the COVID-19 Genomics UK (COG-UK) Consortium | The Lighthouse Lab in Milton Keynes and Alex Alderton, Roberto Amato, Jeffrey Barrett, Sonia Goncalves, Ewan Harrison, David K. Jackson, Ian Johnston, Dominic Kwiatkowski, Cordelia Langford, John Sillitoe on behalf of the Wellcome Sanger Institute COVID-19 Surveillance Team |
| EPI_ISL_2848005, EPI_ISL_2848006 | Lighthouse Lab in Alderley Park | Wellcome Sanger Institute for the COVID-19 Genomics UK (COG-UK) Consortium | Jacquelyn Wynn, Mairead Hyland, The Lighthouse Lab in Alderley Park and Alex Alderton, Roberto Amato, Jeffrey Barrett, Sonia Goncalves, Ewan Harrison, David K. Jackson, Ian Johnston, Dominic Kwiatkowski, Cordelia Langford, John Sillitoe on behalf of the Wellcome Sanger Institute COVID-19 Surveillance Team |
| EPI_ISL_2848007, EPI_ISL_2848008, EPI_ISL_2848009 | Lighthouse Laboratory Plymouth | Wellcome Sanger Institute for the COVID-19 Genomics UK (COG-UK) Consortium | Lighthouse Laboratory Plymouth and Alex Alderton, Roberto Amato, Jeffrey Barrett, Sonia Goncalves, Ewan Harrison, David K. Jackson, Ian Johnston, Dominic Kwiatkowski, Cordelia Langford, John Sillitoe on behalf of the Wellcome Sanger Institute COVID-19 Surveillance Team |
| EPI_ISL_2848010 | Lighthouse Lab in Alderley Park | Wellcome Sanger Institute for the COVID-19 Genomics UK (COG-UK) Consortium | Jacquelyn Wynn, Mairead Hyland, The Lighthouse Lab in Alderley Park and Alex Alderton, Roberto Amato, Jeffrey Barrett, Sonia Goncalves, Ewan Harrison, David K. Jackson, Ian Johnston, Dominic Kwiatkowski, Cordelia Langford, John Sillitoe on behalf of the Wellcome Sanger Institute COVID-19 Surveillance Team |
| EPI_ISL_2848011, EPI_ISL_2848012 | Lighthouse Lab in Milton Keynes | Wellcome Sanger Institute for the COVID-19 Genomics UK (COG-UK) Consortium | The Lighthouse Lab in Milton Keynes and Alex Alderton, Roberto Amato, Jeffrey Barrett, Sonia Goncalves, Ewan Harrison, David K. Jackson, Ian Johnston, Dominic Kwiatkowski, Cordelia Langford, John Sillitoe on behalf of the Wellcome Sanger Institute COVID-19 Surveillance Team |
| EPI_ISL_2848013 | Lighthouse Laboratory Plymouth | Wellcome Sanger Institute for the COVID-19 Genomics UK (COG-UK) Consortium | Lighthouse Laboratory Plymouth and Alex Alderton, Roberto Amato, Jeffrey Barrett, Sonia Goncalves, Ewan Harrison, David K. Jackson, Ian Johnston, Dominic Kwiatkowski, Cordelia Langford, John Sillitoe on behalf of the Wellcome Sanger Institute COVID-19 Surveillance Team |
| EPI_ISL_2848014 | Lighthouse Lab in Alderley Park | Wellcome Sanger Institute for the COVID-19 Genomics UK (COG-UK) Consortium | Jacquelyn Wynn, Mairead Hyland, The Lighthouse Lab in Alderley Park and Alex Alderton, Roberto Amato, Jeffrey Barrett, Sonia Goncalves, Ewan Harrison, David K. Jackson, Ian Johnston, Dominic Kwiatkowski, Cordelia Langford, John Sillitoe on behalf of the Wellcome Sanger Institute COVID-19 Surveillance Team |
| EPI_ISL_2848015 | Lighthouse Laboratory Plymouth | Wellcome Sanger Institute for the COVID-19 Genomics UK (COG-UK) Consortium | Lighthouse Laboratory Plymouth and Alex Alderton, Roberto Amato, Jeffrey Barrett, Sonia Goncalves, Ewan Harrison, David K. Jackson, Ian Johnston, Dominic Kwiatkowski, Cordelia Langford, John Sillitoe on behalf of the Wellcome Sanger Institute COVID-19 Surveillance Team |
| EPI_ISL_2848016 | Lighthouse Lab in Milton Keynes | Wellcome Sanger Institute for the COVID-19 Genomics UK (COG-UK) Consortium | The Lighthouse Lab in Milton Keynes and Alex Alderton, Roberto Amato, Jeffrey Barrett, Sonia Goncalves, Ewan Harrison, David K. Jackson, Ian Johnston, Dominic Kwiatkowski, Cordelia Langford, John Sillitoe on behalf of the Wellcome Sanger Institute COVID-19 Surveillance Team |
| EPI_ISL_2848017 | Lighthouse Laboratory Plymouth | Wellcome Sanger Institute for the COVID-19 Genomics UK (COG-UK) Consortium | Lighthouse Laboratory Plymouth and Alex Alderton, Roberto Amato, Jeffrey Barrett, Sonia Goncalves, Ewan Harrison, David K. Jackson, Ian Johnston, Dominic Kwiatkowski, Cordelia Langford, John Sillitoe on behalf of the Wellcome Sanger Institute COVID-19 Surveillance Team |
| EPI_ISL_2848018, EPI_ISL_2848019 | Lighthouse Lab in Milton Keynes | Wellcome Sanger Institute for the COVID-19 Genomics UK (COG-UK) Consortium | The Lighthouse Lab in Milton Keynes and Alex Alderton, Roberto Amato, Jeffrey Barrett, Sonia Goncalves, Ewan Harrison, David K. Jackson, Ian Johnston, Dominic Kwiatkowski, Cordelia Langford, John Sillitoe on behalf of the Wellcome Sanger Institute COVID-19 Surveillance Team |
| EPI_ISL_2848020, EPI_ISL_2848021 | Lighthouse Laboratory Plymouth | Wellcome Sanger Institute for the COVID-19 Genomics UK (COG-UK) Consortium | Lighthouse Laboratory Plymouth and Alex Alderton, Roberto Amato, Jeffrey Barrett, Sonia Goncalves, Ewan Harrison, David K. Jackson, Ian Johnston, Dominic Kwiatkowski, Cordelia Langford, John Sillitoe on behalf of the Wellcome Sanger Institute COVID-19 Surveillance Team |
| EPI_ISL_2848022 | Lighthouse Lab in Alderley Park | Wellcome Sanger Institute for the COVID-19 Genomics UK (COG-UK) Consortium | Jacquelyn Wynn, Mairead Hyland, The Lighthouse Lab in Alderley Park and Alex Alderton, Roberto Amato, Jeffrey Barrett, Sonia Goncalves, Ewan Harrison, David K. Jackson, Ian Johnston, Dominic Kwiatkowski, Cordelia Langford, John Sillitoe on behalf of the Wellcome Sanger Institute COVID-19 Surveillance Team |
| EPI_ISL_2848023 | Lighthouse Laboratory Plymouth | Wellcome Sanger Institute for the COVID-19 Genomics UK (COG-UK) Consortium | Lighthouse Laboratory Plymouth and Alex Alderton, Roberto Amato, Jeffrey Barrett, Sonia Goncalves, Ewan Harrison, David K. Jackson, Ian Johnston, Dominic Kwiatkowski, Cordelia Langford, John Sillitoe on behalf of the Wellcome Sanger Institute COVID-19 Surveillance Team |
| EPI_ISL_2848024, EPI_ISL_2848025, EPI_ISL_2848026 | Lighthouse Lab in Milton Keynes | Wellcome Sanger Institute for the COVID-19 Genomics UK (COG-UK) Consortium | The Lighthouse Lab in Milton Keynes and Alex Alderton, Roberto Amato, Jeffrey Barrett, Sonia Goncalves, Ewan Harrison, David K. Jackson, Ian Johnston, Dominic Kwiatkowski, Cordelia Langford, John Sillitoe on behalf of the Wellcome Sanger Institute COVID-19 Surveillance Team |
| EPI_ISL_2848027 | Lighthouse Laboratory Plymouth | Wellcome Sanger Institute for the COVID-19 Genomics UK (COG-UK) Consortium | Lighthouse Laboratory Plymouth and Alex Alderton, Roberto Amato, Jeffrey Barrett, Sonia Goncalves, Ewan Harrison, David K. Jackson, Ian Johnston, Dominic Kwiatkowski, Cordelia Langford, John Sillitoe on behalf of the Wellcome Sanger Institute COVID-19 Surveillance Team |
| EPI_ISL_2848028 | Lighthouse Lab in Milton Keynes | Wellcome Sanger Institute for the COVID-19 Genomics UK (COG-UK) Consortium | The Lighthouse Lab in Milton Keynes and Alex Alderton, Roberto Amato, Jeffrey Barrett, Sonia Goncalves, Ewan Harrison, David K. Jackson, Ian Johnston, Dominic Kwiatkowski, Cordelia Langford, John Sillitoe on behalf of the Wellcome Sanger Institute COVID-19 Surveillance Team |
| EPI_ISL_2848029 | Lighthouse Lab in Alderley Park | Wellcome Sanger Institute for the COVID-19 Genomics UK (COG-UK) Consortium | Jacquelyn Wynn, Mairead Hyland, The Lighthouse Lab in Alderley Park and Alex Alderton, Roberto Amato, Jeffrey Barrett, Sonia Goncalves, Ewan Harrison, David K. Jackson, Ian Johnston, Dominic Kwiatkowski, Cordelia Langford, John Sillitoe on behalf of the Wellcome Sanger Institute COVID-19 Surveillance Team |

| | | | |
|---|---|---|---|
| EPI_ISL_2848030, EPI_ISL_2848032 | Lighthouse Laboratory Plymouth | Wellcome Sanger Institute for the COVID-19 Genomics UK (COG-UK) Consortium | Lighthouse Laboratory Plymouth and Alex Alderton, Roberto Amato, Jeffrey Barrett, Sonia Goncalves, Ewan Harrison, David K. Jackson, Ian Johnston, Dominic Kwiatkowski, Cordelia Langford, John Sillitoe on behalf of the Wellcome Sanger Institute COVID-19 Surveillance Team |
| EPI_ISL_2848033 | Lighthouse Lab in Milton Keynes | Wellcome Sanger Institute for the COVID-19 Genomics UK (COG-UK) Consortium | The Lighthouse Lab in Milton Keynes and Alex Alderton, Roberto Amato, Jeffrey Barrett, Sonia Goncalves, Ewan Harrison, David K. Jackson, Ian Johnston, Dominic Kwiatkowski, Cordelia Langford, John Sillitoe on behalf of the Wellcome Sanger Institute COVID-19 Surveillance Team |
| EPI_ISL_2848034, EPI_ISL_2848035 | Lighthouse Laboratory Plymouth | Wellcome Sanger Institute for the COVID-19 Genomics UK (COG-UK) Consortium | Lighthouse Laboratory Plymouth and Alex Alderton, Roberto Amato, Jeffrey Barrett, Sonia Goncalves, Ewan Harrison, David K. Jackson, Ian Johnston, Dominic Kwiatkowski, Cordelia Langford, John Sillitoe on behalf of the Wellcome Sanger Institute COVID-19 Surveillance Team |
| EPI_ISL_2848036 | Lighthouse Lab in Alderley Park | Wellcome Sanger Institute for the COVID-19 Genomics UK (COG-UK) Consortium | Jacquelyn Wynn, Mairead Hyland, The Lighthouse Lab in Alderley Park and Alex Alderton, Roberto Amato, Jeffrey Barrett, Sonia Goncalves, Ewan Harrison, David K. Jackson, Ian Johnston, Dominic Kwiatkowski, Cordelia Langford, John Sillitoe on behalf of the Wellcome Sanger Institute COVID-19 Surveillance Team |
| EPI_ISL_2848037 | Lighthouse Laboratory Plymouth | Wellcome Sanger Institute for the COVID-19 Genomics UK (COG-UK) Consortium | Lighthouse Laboratory Plymouth and Alex Alderton, Roberto Amato, Jeffrey Barrett, Sonia Goncalves, Ewan Harrison, David K. Jackson, Ian Johnston, Dominic Kwiatkowski, Cordelia Langford, John Sillitoe on behalf of the Wellcome Sanger Institute COVID-19 Surveillance Team |
| EPI_ISL_2848038, EPI_ISL_2848039 | Lighthouse Lab in Alderley Park | Wellcome Sanger Institute for the COVID-19 Genomics UK (COG-UK) Consortium | Jacquelyn Wynn, Mairead Hyland, The Lighthouse Lab in Alderley Park and Alex Alderton, Roberto Amato, Jeffrey Barrett, Sonia Goncalves, Ewan Harrison, David K. Jackson, Ian Johnston, Dominic Kwiatkowski, Cordelia Langford, John Sillitoe on behalf of the Wellcome Sanger Institute COVID-19 Surveillance Team |
| EPI_ISL_2848040, EPI_ISL_2848041 | Lighthouse Laboratory Plymouth | Wellcome Sanger Institute for the COVID-19 Genomics UK (COG-UK) Consortium | Lighthouse Laboratory Plymouth and Alex Alderton, Roberto Amato, Jeffrey Barrett, Sonia Goncalves, Ewan Harrison, David K. Jackson, Ian Johnston, Dominic Kwiatkowski, Cordelia Langford, John Sillitoe on behalf of the Wellcome Sanger Institute COVID-19 Surveillance Team |
| EPI_ISL_2848042 | Lighthouse Lab in Alderley Park | Wellcome Sanger Institute for the COVID-19 Genomics UK (COG-UK) Consortium | Jacquelyn Wynn, Mairead Hyland, The Lighthouse Lab in Alderley Park and Alex Alderton, Roberto Amato, Jeffrey Barrett, Sonia Goncalves, Ewan Harrison, David K. Jackson, Ian Johnston, Dominic Kwiatkowski, Cordelia Langford, John Sillitoe on behalf of the Wellcome Sanger Institute COVID-19 Surveillance Team |
| EPI_ISL_2848043, EPI_ISL_2848044, EPI_ISL_2848045, EPI_ISL_2848046, EPI_ISL_2848047, EPI_ISL_2848048, EPI_ISL_2848049, EPI_ISL_2848050 | Lighthouse Laboratory Plymouth | Wellcome Sanger Institute for the COVID-19 Genomics UK (COG-UK) Consortium | Lighthouse Laboratory Plymouth and Alex Alderton, Roberto Amato, Jeffrey Barrett, Sonia Goncalves, Ewan Harrison, David K. Jackson, Ian Johnston, Dominic Kwiatkowski, Cordelia Langford, John Sillitoe on behalf of the Wellcome Sanger Institute COVID-19 Surveillance Team |
| EPI_ISL_2848051 | Lighthouse Lab in Alderley Park | Wellcome Sanger Institute for the COVID-19 Genomics UK (COG-UK) Consortium | Jacquelyn Wynn, Mairead Hyland, The Lighthouse Lab in Alderley Park and Alex Alderton, Roberto Amato, Jeffrey Barrett, Sonia Goncalves, Ewan Harrison, David K. Jackson, Ian Johnston, Dominic Kwiatkowski, Cordelia Langford, John Sillitoe on behalf of the Wellcome Sanger Institute COVID-19 Surveillance Team |
| EPI_ISL_2848052, EPI_ISL_2848053, EPI_ISL_2848054 | Lighthouse Laboratory Plymouth | Wellcome Sanger Institute for the COVID-19 Genomics UK (COG-UK) Consortium | Lighthouse Laboratory Plymouth and Alex Alderton, Roberto Amato, Jeffrey Barrett, Sonia Goncalves, Ewan Harrison, David K. Jackson, Ian Johnston, Dominic Kwiatkowski, Cordelia Langford, John Sillitoe on behalf of the Wellcome Sanger Institute COVID-19 Surveillance Team |
| EPI_ISL_2848055 | Lighthouse Lab in Alderley Park | Wellcome Sanger Institute for the COVID-19 Genomics UK (COG-UK) Consortium | Jacquelyn Wynn, Mairead Hyland, The Lighthouse Lab in Alderley Park and Alex Alderton, Roberto Amato, Jeffrey Barrett, Sonia Goncalves, Ewan Harrison, David K. Jackson, Ian Johnston, Dominic Kwiatkowski, Cordelia Langford, John Sillitoe on behalf of the Wellcome Sanger Institute COVID-19 Surveillance Team |
| EPI_ISL_2848056, EPI_ISL_2848057, EPI_ISL_2848058, EPI_ISL_2848059 | Lighthouse Laboratory Plymouth | Wellcome Sanger Institute for the COVID-19 Genomics UK (COG-UK) Consortium | Lighthouse Laboratory Plymouth and Alex Alderton, Roberto Amato, Jeffrey Barrett, Sonia Goncalves, Ewan Harrison, David K. Jackson, Ian Johnston, Dominic Kwiatkowski, Cordelia Langford, John Sillitoe on behalf of the Wellcome Sanger Institute COVID-19 Surveillance Team |
| EPI_ISL_2848060, EPI_ISL_2848061 | Lighthouse Lab in Milton Keynes | Wellcome Sanger Institute for the COVID-19 Genomics UK (COG-UK) Consortium | The Lighthouse Lab in Milton Keynes and Alex Alderton, Roberto Amato, Jeffrey Barrett, Sonia Goncalves, Ewan Harrison, David K. Jackson, Ian Johnston, Dominic Kwiatkowski, Cordelia Langford, John Sillitoe on behalf of the Wellcome Sanger Institute COVID-19 Surveillance Team |
| EPI_ISL_2848062, EPI_ISL_2848063, EPI_ISL_2848064 | Lighthouse Lab in Alderley Park | Wellcome Sanger Institute for the COVID-19 Genomics UK (COG-UK) Consortium | Jacquelyn Wynn, Mairead Hyland, The Lighthouse Lab in Alderley Park and Alex Alderton, Roberto Amato, Jeffrey Barrett, Sonia Goncalves, Ewan Harrison, David K. Jackson, Ian Johnston, Dominic Kwiatkowski, Cordelia Langford, John Sillitoe on behalf of the Wellcome Sanger Institute COVID-19 Surveillance Team |
| EPI_ISL_2848065 | Lighthouse Laboratory Plymouth | Wellcome Sanger Institute for the COVID-19 Genomics UK (COG-UK) Consortium | Lighthouse Laboratory Plymouth and Alex Alderton, Roberto Amato, Jeffrey Barrett, Sonia Goncalves, Ewan Harrison, David K. Jackson, Ian Johnston, Dominic Kwiatkowski, Cordelia Langford, John Sillitoe on behalf of the Wellcome Sanger Institute COVID-19 Surveillance Team |
| EPI_ISL_2848066 | Lighthouse Lab in Milton Keynes | Wellcome Sanger Institute for the COVID-19 Genomics UK (COG-UK) Consortium | The Lighthouse Lab in Milton Keynes and Alex Alderton, Roberto Amato, Jeffrey Barrett, Sonia Goncalves, Ewan Harrison, David K. Jackson, Ian Johnston, Dominic Kwiatkowski, Cordelia Langford, John Sillitoe on behalf of the Wellcome Sanger Institute COVID-19 Surveillance Team |
| EPI_ISL_2848067, EPI_ISL_2848068 | Lighthouse Laboratory Plymouth | Wellcome Sanger Institute for the COVID-19 Genomics UK (COG-UK) Consortium | Lighthouse Laboratory Plymouth and Alex Alderton, Roberto Amato, Jeffrey Barrett, Sonia Goncalves, Ewan Harrison, David K. Jackson, Ian Johnston, Dominic Kwiatkowski, Cordelia Langford, John Sillitoe on behalf of the Wellcome Sanger Institute COVID-19 Surveillance Team |
| EPI_ISL_2848069 | Lighthouse Lab in Milton Keynes | Wellcome Sanger Institute for the COVID-19 Genomics UK (COG-UK) Consortium | The Lighthouse Lab in Milton Keynes and Alex Alderton, Roberto Amato, Jeffrey Barrett, Sonia Goncalves, Ewan Harrison, David K. Jackson, Ian Johnston, Dominic Kwiatkowski, Cordelia Langford, John Sillitoe on behalf of the Wellcome Sanger Institute COVID-19 Surveillance Team |
| EPI_ISL_2848070 | Lighthouse Lab in Alderley Park | Wellcome Sanger Institute for the COVID-19 Genomics UK (COG-UK) Consortium | Jacquelyn Wynn, Mairead Hyland, The Lighthouse Lab in Alderley Park and Alex Alderton, Roberto Amato, Jeffrey Barrett, Sonia Goncalves, Ewan Harrison, David K. Jackson, Ian Johnston, Dominic Kwiatkowski, Cordelia Langford, John Sillitoe on behalf of the Wellcome Sanger Institute COVID-19 Surveillance Team |
| EPI_ISL_2848071, EPI_ISL_2848072 | Lighthouse Laboratory Plymouth | Wellcome Sanger Institute for the COVID-19 Genomics UK (COG-UK) Consortium | Lighthouse Laboratory Plymouth and Alex Alderton, Roberto Amato, Jeffrey Barrett, Sonia Goncalves, Ewan Harrison, David K. Jackson, Ian Johnston, Dominic Kwiatkowski, Cordelia Langford, John Sillitoe on behalf of the Wellcome Sanger Institute COVID-19 Surveillance Team |
| EPI_ISL_2848073, EPI_ISL_2848074 | Lighthouse Lab in Alderley Park | Wellcome Sanger Institute for the COVID-19 Genomics UK (COG-UK) Consortium | Jacquelyn Wynn, Mairead Hyland, The Lighthouse Lab in Alderley Park and Alex Alderton, Roberto Amato, Jeffrey Barrett, Sonia Goncalves, Ewan Harrison, David K. Jackson, Ian Johnston, Dominic Kwiatkowski, Cordelia Langford, John Sillitoe on behalf of the Wellcome Sanger Institute COVID-19 Surveillance Team |
| EPI_ISL_2848075, EPI_ISL_2848076 | Lighthouse Laboratory Plymouth | Wellcome Sanger Institute for the COVID-19 Genomics UK (COG-UK) Consortium | Lighthouse Laboratory Plymouth and Alex Alderton, Roberto Amato, Jeffrey Barrett, Sonia Goncalves, Ewan Harrison, David K. Jackson, Ian Johnston, Dominic Kwiatkowski, Cordelia Langford, John Sillitoe on behalf of the Wellcome Sanger Institute COVID-19 Surveillance Team |
| EPI_ISL_2848077 | Lighthouse Lab in Alderley Park | Wellcome Sanger Institute for the COVID-19 Genomics UK (COG-UK) Consortium | Jacquelyn Wynn, Mairead Hyland, The Lighthouse Lab in Alderley Park and Alex Alderton, Roberto Amato, Jeffrey Barrett, Sonia Goncalves, Ewan Harrison, David K. Jackson, Ian Johnston, Dominic Kwiatkowski, Cordelia Langford, John Sillitoe on behalf of the Wellcome Sanger Institute COVID-19 Surveillance Team |
| EPI_ISL_2848078, EPI_ISL_2848079, EPI_ISL_2848080 | Lighthouse Laboratory Plymouth | Wellcome Sanger Institute for the COVID-19 Genomics UK (COG-UK) Consortium | Lighthouse Laboratory Plymouth and Alex Alderton, Roberto Amato, Jeffrey Barrett, Sonia Goncalves, Ewan Harrison, David K. Jackson, Ian Johnston, Dominic Kwiatkowski, Cordelia Langford, John Sillitoe on behalf of the Wellcome Sanger Institute COVID-19 Surveillance Team |
| EPI_ISL_2848081 | Lighthouse Lab in Alderley Park | Wellcome Sanger Institute for the COVID-19 Genomics UK (COG-UK) Consortium | Jacquelyn Wynn, Mairead Hyland, The Lighthouse Lab in Alderley Park and Alex Alderton, Roberto Amato, Jeffrey Barrett, Sonia Goncalves, Ewan Harrison, David K. Jackson, Ian Johnston, Dominic Kwiatkowski, Cordelia Langford, John Sillitoe on behalf of the Wellcome Sanger Institute COVID-19 Surveillance Team |
| EPI_ISL_2848082, EPI_ISL_2848083 | Lighthouse Laboratory Plymouth | Wellcome Sanger Institute for the COVID-19 Genomics UK (COG-UK) Consortium | Lighthouse Laboratory Plymouth and Alex Alderton, Roberto Amato, Jeffrey Barrett, Sonia Goncalves, Ewan Harrison, David K. Jackson, Ian Johnston, Dominic Kwiatkowski, Cordelia Langford, John Sillitoe on behalf of the Wellcome Sanger Institute COVID-19 Surveillance Team |
| EPI_ISL_2848084 | Lighthouse Lab in Alderley Park | Wellcome Sanger Institute for the COVID-19 Genomics UK (COG-UK) Consortium | Jacquelyn Wynn, Mairead Hyland, The Lighthouse Lab in Alderley Park and Alex Alderton, Roberto Amato, Jeffrey Barrett, Sonia Goncalves, Ewan Harrison, David K. Jackson, Ian Johnston, Dominic Kwiatkowski, Cordelia Langford, John Sillitoe on behalf of the Wellcome Sanger Institute COVID-19 Surveillance Team |
| EPI_ISL_2848085, EPI_ISL_2848086, EPI_ISL_2848087 | Lighthouse Lab in Milton Keynes | Wellcome Sanger Institute for the COVID-19 Genomics UK (COG-UK) Consortium | The Lighthouse Lab in Milton Keynes and Alex Alderton, Roberto Amato, Jeffrey Barrett, Sonia Goncalves, Ewan Harrison, David K. Jackson, Ian Johnston, Dominic Kwiatkowski, Cordelia Langford, John Sillitoe on behalf of the Wellcome Sanger Institute COVID-19 Surveillance Team |
| EPI_ISL_2848088, EPI_ISL_2848089 | Lighthouse Lab in Alderley Park | Wellcome Sanger Institute for the COVID-19 Genomics UK (COG-UK) Consortium | Jacquelyn Wynn, Mairead Hyland, The Lighthouse Lab in Alderley Park and Alex Alderton, Roberto Amato, Jeffrey Barrett, Sonia Goncalves, Ewan Harrison, David K. Jackson, Ian Johnston, Dominic Kwiatkowski, Cordelia Langford, John Sillitoe on behalf of the Wellcome Sanger Institute COVID-19 Surveillance Team |
| EPI_ISL_2848090 | Lighthouse Lab in Milton Keynes | Wellcome Sanger Institute for the COVID-19 Genomics UK (COG-UK) Consortium | The Lighthouse Lab in Milton Keynes and Alex Alderton, Roberto Amato, Jeffrey Barrett, Sonia Goncalves, Ewan Harrison, David K. Jackson, Ian Johnston, Dominic Kwiatkowski, Cordelia Langford, John Sillitoe on behalf of the Wellcome Sanger Institute COVID-19 Surveillance Team |
| EPI_ISL_2848091, EPI_ISL_2848092, EPI_ISL_2848093, EPI_ISL_2848094, EPI_ISL_2848095, EPI_ISL_2848096, EPI_ISL_2848097, EPI_ISL_2848098, EPI_ISL_2848099 | Lighthouse Laboratory Plymouth | Wellcome Sanger Institute for the COVID-19 Genomics UK (COG-UK) Consortium | Lighthouse Laboratory Plymouth and Alex Alderton, Roberto Amato, Jeffrey Barrett, Sonia Goncalves, Ewan Harrison, David K. Jackson, Ian Johnston, Dominic Kwiatkowski, Cordelia Langford, John Sillitoe on behalf of the Wellcome Sanger Institute COVID-19 Surveillance Team |
| EPI_ISL_2848100 | Lighthouse Lab in Alderley Park | Wellcome Sanger Institute for the COVID-19 Genomics UK (COG-UK) Consortium | Jacquelyn Wynn, Mairead Hyland, The Lighthouse Lab in Alderley Park and Alex Alderton, Roberto Amato, Jeffrey Barrett, Sonia Goncalves, Ewan Harrison, David K. Jackson, Ian Johnston, Dominic Kwiatkowski, Cordelia Langford, John Sillitoe on behalf of the Wellcome Sanger Institute COVID-19 Surveillance Team |
| EPI_ISL_2848101, EPI_ISL_2848102, EPI_ISL_2848103 | Lighthouse Laboratory Plymouth | Wellcome Sanger Institute for the COVID-19 Genomics UK (COG-UK) Consortium | Lighthouse Laboratory Plymouth and Alex Alderton, Roberto Amato, Jeffrey Barrett, Sonia Goncalves, Ewan Harrison, David K. Jackson, Ian Johnston, Dominic Kwiatkowski, Cordelia Langford, John Sillitoe on behalf of the Wellcome Sanger Institute COVID-19 Surveillance Team |
| EPI_ISL_2848104 | Lighthouse Lab in Alderley Park | Wellcome Sanger Institute for the COVID-19 Genomics UK (COG-UK) Consortium | Jacquelyn Wynn, Mairead Hyland, The Lighthouse Lab in Alderley Park and Alex Alderton, Roberto Amato, Jeffrey Barrett, Sonia Goncalves, Ewan Harrison, David K. Jackson, Ian Johnston, Dominic Kwiatkowski, Cordelia Langford, John Sillitoe on behalf of the Wellcome Sanger Institute COVID-19 Surveillance Team |
| EPI_ISL_2848105 | Lighthouse Laboratory Plymouth | Wellcome Sanger Institute for the COVID-19 Genomics UK (COG-UK) Consortium | Lighthouse Laboratory Plymouth and Alex Alderton, Roberto Amato, Jeffrey Barrett, Sonia Goncalves, Ewan Harrison, David K. Jackson, Ian Johnston, Dominic Kwiatkowski, Cordelia Langford, John Sillitoe on behalf of the Wellcome Sanger Institute COVID-19 Surveillance Team |
| EPI_ISL_2848106, EPI_ISL_2848107 | Lighthouse Lab in Alderley Park | Wellcome Sanger Institute for the COVID-19 Genomics UK (COG-UK) Consortium | Jacquelyn Wynn, Mairead Hyland, The Lighthouse Lab in Alderley Park and Alex Alderton, Roberto Amato, Jeffrey Barrett, Sonia Goncalves, Ewan Harrison, David K. Jackson, Ian Johnston, Dominic Kwiatkowski, Cordelia Langford, John Sillitoe on behalf of the Wellcome Sanger Institute COVID-19 Surveillance Team |

| | | | |
|---|---|---|---|
| EPI_ISL_2848108, EPI_ISL_2848109, EPI_ISL_2848110 | Lighthouse Laboratory Plymouth | Wellcome Sanger Institute for the COVID-19 Genomics UK (COG-UK) Consortium | Lighthouse Laboratory Plymouth and Alex Alderton, Roberto Amato, Jeffrey Barrett, Sonia Goncalves, Ewan Harrison, David K. Jackson, Ian Johnston, Dominic Kwiatkowski, Cordelia Langford, John Sillitoe on behalf of the Wellcome Sanger Institute COVID-19 Surveillance Team |
| EPI_ISL_2848111 | Lighthouse Lab in Alderley Park | Wellcome Sanger Institute for the COVID-19 Genomics UK (COG-UK) Consortium | Jacquelyn Wynn, Mairead Hyland, The Lighthouse Lab in Alderley Park and Alex Alderton, Roberto Amato, Jeffrey Barrett, Sonia Goncalves, Ewan Harrison, David K. Jackson, Ian Johnston, Dominic Kwiatkowski, Cordelia Langford, John Sillitoe on behalf of the Wellcome Sanger Institute COVID-19 Surveillance Team |
| EPI_ISL_2848112 | Lighthouse Lab in Milton Keynes | Wellcome Sanger Institute for the COVID-19 Genomics UK (COG-UK) Consortium | The Lighthouse Lab in Milton Keynes and Alex Alderton, Roberto Amato, Jeffrey Barrett, Sonia Goncalves, Ewan Harrison, David K. Jackson, Ian Johnston, Dominic Kwiatkowski, Cordelia Langford, John Sillitoe on behalf of the Wellcome Sanger Institute COVID-19 Surveillance Team |
| EPI_ISL_2848113 | Lighthouse Lab in Alderley Park | Wellcome Sanger Institute for the COVID-19 Genomics UK (COG-UK) Consortium | Jacquelyn Wynn, Mairead Hyland, The Lighthouse Lab in Alderley Park and Alex Alderton, Roberto Amato, Jeffrey Barrett, Sonia Goncalves, Ewan Harrison, David K. Jackson, Ian Johnston, Dominic Kwiatkowski, Cordelia Langford, John Sillitoe on behalf of the Wellcome Sanger Institute COVID-19 Surveillance Team |
| EPI_ISL_2848114 | Lighthouse Lab in Milton Keynes | Wellcome Sanger Institute for the COVID-19 Genomics UK (COG-UK) Consortium | The Lighthouse Lab in Milton Keynes and Alex Alderton, Roberto Amato, Jeffrey Barrett, Sonia Goncalves, Ewan Harrison, David K. Jackson, Ian Johnston, Dominic Kwiatkowski, Cordelia Langford, John Sillitoe on behalf of the Wellcome Sanger Institute COVID-19 Surveillance Team |
| EPI_ISL_2848115 | Lighthouse Lab in Alderley Park | Wellcome Sanger Institute for the COVID-19 Genomics UK (COG-UK) Consortium | Jacquelyn Wynn, Mairead Hyland, The Lighthouse Lab in Alderley Park and Alex Alderton, Roberto Amato, Jeffrey Barrett, Sonia Goncalves, Ewan Harrison, David K. Jackson, Ian Johnston, Dominic Kwiatkowski, Cordelia Langford, John Sillitoe on behalf of the Wellcome Sanger Institute COVID-19 Surveillance Team |
| EPI_ISL_2848116 | Lighthouse Lab in Milton Keynes | Wellcome Sanger Institute for the COVID-19 Genomics UK (COG-UK) Consortium | The Lighthouse Lab in Milton Keynes and Alex Alderton, Roberto Amato, Jeffrey Barrett, Sonia Goncalves, Ewan Harrison, David K. Jackson, Ian Johnston, Dominic Kwiatkowski, Cordelia Langford, John Sillitoe on behalf of the Wellcome Sanger Institute COVID-19 Surveillance Team |
| EPI_ISL_2848117, EPI_ISL_2848118 | Lighthouse Laboratory Plymouth | Wellcome Sanger Institute for the COVID-19 Genomics UK (COG-UK) Consortium | Lighthouse Laboratory Plymouth and Alex Alderton, Roberto Amato, Jeffrey Barrett, Sonia Goncalves, Ewan Harrison, David K. Jackson, Ian Johnston, Dominic Kwiatkowski, Cordelia Langford, John Sillitoe on behalf of the Wellcome Sanger Institute COVID-19 Surveillance Team |
| EPI_ISL_2848119 | Lighthouse Lab in Alderley Park | Wellcome Sanger Institute for the COVID-19 Genomics UK (COG-UK) Consortium | Jacquelyn Wynn, Mairead Hyland, The Lighthouse Lab in Alderley Park and Alex Alderton, Roberto Amato, Jeffrey Barrett, Sonia Goncalves, Ewan Harrison, David K. Jackson, Ian Johnston, Dominic Kwiatkowski, Cordelia Langford, John Sillitoe on behalf of the Wellcome Sanger Institute COVID-19 Surveillance Team |
| EPI_ISL_2848120, EPI_ISL_2848121, EPI_ISL_2848122, EPI_ISL_2848123 | Lighthouse Laboratory Plymouth | Wellcome Sanger Institute for the COVID-19 Genomics UK (COG-UK) Consortium | Lighthouse Laboratory Plymouth and Alex Alderton, Roberto Amato, Jeffrey Barrett, Sonia Goncalves, Ewan Harrison, David K. Jackson, Ian Johnston, Dominic Kwiatkowski, Cordelia Langford, John Sillitoe on behalf of the Wellcome Sanger Institute COVID-19 Surveillance Team |
| EPI_ISL_2848124 | Lighthouse Lab in Milton Keynes | Wellcome Sanger Institute for the COVID-19 Genomics UK (COG-UK) Consortium | The Lighthouse Lab in Milton Keynes and Alex Alderton, Roberto Amato, Jeffrey Barrett, Sonia Goncalves, Ewan Harrison, David K. Jackson, Ian Johnston, Dominic Kwiatkowski, Cordelia Langford, John Sillitoe on behalf of the Wellcome Sanger Institute COVID-19 Surveillance Team |
| EPI_ISL_2848125, EPI_ISL_2848126, EPI_ISL_2848127, EPI_ISL_2848128, EPI_ISL_2848129, EPI_ISL_2848130, EPI_ISL_2848131, EPI_ISL_2848132, EPI_ISL_2848133, EPI_ISL_2848134, EPI_ISL_2848135, EPI_ISL_2848136 | | | |
| see above | Lighthouse Laboratory Plymouth | Wellcome Sanger Institute for the COVID-19 Genomics UK (COG-UK) Consortium | Lighthouse Laboratory Plymouth and Alex Alderton, Roberto Amato, Jeffrey Barrett, Sonia Goncalves, Ewan Harrison, David K. Jackson, Ian Johnston, Dominic Kwiatkowski, Cordelia Langford, John Sillitoe on behalf of the Wellcome Sanger Institute COVID-19 Surveillance Team |
| EPI_ISL_2848137 | Lighthouse Lab in Alderley Park | Wellcome Sanger Institute for the COVID-19 Genomics UK (COG-UK) Consortium | Jacquelyn Wynn, Mairead Hyland, The Lighthouse Lab in Alderley Park and Alex Alderton, Roberto Amato, Jeffrey Barrett, Sonia Goncalves, Ewan Harrison, David K. Jackson, Ian Johnston, Dominic Kwiatkowski, Cordelia Langford, John Sillitoe on behalf of the Wellcome Sanger Institute COVID-19 Surveillance Team |
| EPI_ISL_2848138, EPI_ISL_2848139 | Lighthouse Laboratory Plymouth | Wellcome Sanger Institute for the COVID-19 Genomics UK (COG-UK) Consortium | Lighthouse Laboratory Plymouth and Alex Alderton, Roberto Amato, Jeffrey Barrett, Sonia Goncalves, Ewan Harrison, David K. Jackson, Ian Johnston, Dominic Kwiatkowski, Cordelia Langford, John Sillitoe on behalf of the Wellcome Sanger Institute COVID-19 Surveillance Team |
| EPI_ISL_2848140 | Lighthouse Lab in Milton Keynes | Wellcome Sanger Institute for the COVID-19 Genomics UK (COG-UK) Consortium | The Lighthouse Lab in Milton Keynes and Alex Alderton, Roberto Amato, Jeffrey Barrett, Sonia Goncalves, Ewan Harrison, David K. Jackson, Ian Johnston, Dominic Kwiatkowski, Cordelia Langford, John Sillitoe on behalf of the Wellcome Sanger Institute COVID-19 Surveillance Team |
| EPI_ISL_2848141, EPI_ISL_2848142, EPI_ISL_2848143, EPI_ISL_2848144 | Lighthouse Laboratory Plymouth | Wellcome Sanger Institute for the COVID-19 Genomics UK (COG-UK) Consortium | Lighthouse Laboratory Plymouth and Alex Alderton, Roberto Amato, Jeffrey Barrett, Sonia Goncalves, Ewan Harrison, David K. Jackson, Ian Johnston, Dominic Kwiatkowski, Cordelia Langford, John Sillitoe on behalf of the Wellcome Sanger Institute COVID-19 Surveillance Team |
| EPI_ISL_2848145 | Lighthouse Lab in Milton Keynes | Wellcome Sanger Institute for the COVID-19 Genomics UK (COG-UK) Consortium | The Lighthouse Lab in Milton Keynes and Alex Alderton, Roberto Amato, Jeffrey Barrett, Sonia Goncalves, Ewan Harrison, David K. Jackson, Ian Johnston, Dominic Kwiatkowski, Cordelia Langford, John Sillitoe on behalf of the Wellcome Sanger Institute COVID-19 Surveillance Team |
| EPI_ISL_2848146 | Lighthouse Laboratory Plymouth | Wellcome Sanger Institute for the COVID-19 Genomics UK (COG-UK) Consortium | Lighthouse Laboratory Plymouth and Alex Alderton, Roberto Amato, Jeffrey Barrett, Sonia Goncalves, Ewan Harrison, David K. Jackson, Ian Johnston, Dominic Kwiatkowski, Cordelia Langford, John Sillitoe on behalf of the Wellcome Sanger Institute COVID-19 Surveillance Team |
| EPI_ISL_2848147 | Lighthouse Lab in Alderley Park | Wellcome Sanger Institute for the COVID-19 Genomics UK (COG-UK) Consortium | Jacquelyn Wynn, Mairead Hyland, The Lighthouse Lab in Alderley Park and Alex Alderton, Roberto Amato, Jeffrey Barrett, Sonia Goncalves, Ewan Harrison, David K. Jackson, Ian Johnston, Dominic Kwiatkowski, Cordelia Langford, John Sillitoe on behalf of the Wellcome Sanger Institute COVID-19 Surveillance Team |
| EPI_ISL_2848148 | Lighthouse Lab in Milton Keynes | Wellcome Sanger Institute for the COVID-19 Genomics UK (COG-UK) Consortium | The Lighthouse Lab in Milton Keynes and Alex Alderton, Roberto Amato, Jeffrey Barrett, Sonia Goncalves, Ewan Harrison, David K. Jackson, Ian Johnston, Dominic Kwiatkowski, Cordelia Langford, John Sillitoe on behalf of the Wellcome Sanger Institute COVID-19 Surveillance Team |
| EPI_ISL_2848149, EPI_ISL_2848150, EPI_ISL_2848151 | Lighthouse Laboratory Plymouth | Wellcome Sanger Institute for the COVID-19 Genomics UK (COG-UK) Consortium | Lighthouse Laboratory Plymouth and Alex Alderton, Roberto Amato, Jeffrey Barrett, Sonia Goncalves, Ewan Harrison, David K. Jackson, Ian Johnston, Dominic Kwiatkowski, Cordelia Langford, John Sillitoe on behalf of the Wellcome Sanger Institute COVID-19 Surveillance Team |
| EPI_ISL_2848152 | Lighthouse Lab in Milton Keynes | Wellcome Sanger Institute for the COVID-19 Genomics UK (COG-UK) Consortium | The Lighthouse Lab in Milton Keynes and Alex Alderton, Roberto Amato, Jeffrey Barrett, Sonia Goncalves, Ewan Harrison, David K. Jackson, Ian Johnston, Dominic Kwiatkowski, Cordelia Langford, John Sillitoe on behalf of the Wellcome Sanger Institute COVID-19 Surveillance Team |
| EPI_ISL_2848153 | Lighthouse Lab in Alderley Park | Wellcome Sanger Institute for the COVID-19 Genomics UK (COG-UK) Consortium | Jacquelyn Wynn, Mairead Hyland, The Lighthouse Lab in Alderley Park and Alex Alderton, Roberto Amato, Jeffrey Barrett, Sonia Goncalves, Ewan Harrison, David K. Jackson, Ian Johnston, Dominic Kwiatkowski, Cordelia Langford, John Sillitoe on behalf of the Wellcome Sanger Institute COVID-19 Surveillance Team |
| EPI_ISL_2848154, EPI_ISL_2848155 | Lighthouse Laboratory Plymouth | Wellcome Sanger Institute for the COVID-19 Genomics UK (COG-UK) Consortium | Lighthouse Laboratory Plymouth and Alex Alderton, Roberto Amato, Jeffrey Barrett, Sonia Goncalves, Ewan Harrison, David K. Jackson, Ian Johnston, Dominic Kwiatkowski, Cordelia Langford, John Sillitoe on behalf of the Wellcome Sanger Institute COVID-19 Surveillance Team |
| EPI_ISL_2848156 | Lighthouse Lab in Milton Keynes | Wellcome Sanger Institute for the COVID-19 Genomics UK (COG-UK) Consortium | The Lighthouse Lab in Milton Keynes and Alex Alderton, Roberto Amato, Jeffrey Barrett, Sonia Goncalves, Ewan Harrison, David K. Jackson, Ian Johnston, Dominic Kwiatkowski, Cordelia Langford, John Sillitoe on behalf of the Wellcome Sanger Institute COVID-19 Surveillance Team |
| EPI_ISL_2848157 | Lighthouse Laboratory Plymouth | Wellcome Sanger Institute for the COVID-19 Genomics UK (COG-UK) Consortium | Lighthouse Laboratory Plymouth and Alex Alderton, Roberto Amato, Jeffrey Barrett, Sonia Goncalves, Ewan Harrison, David K. Jackson, Ian Johnston, Dominic Kwiatkowski, Cordelia Langford, John Sillitoe on behalf of the Wellcome Sanger Institute COVID-19 Surveillance Team |
| EPI_ISL_2848158, EPI_ISL_2848159 | Lighthouse Lab in Alderley Park | Wellcome Sanger Institute for the COVID-19 Genomics UK (COG-UK) Consortium | Jacquelyn Wynn, Mairead Hyland, The Lighthouse Lab in Alderley Park and Alex Alderton, Roberto Amato, Jeffrey Barrett, Sonia Goncalves, Ewan Harrison, David K. Jackson, Ian Johnston, Dominic Kwiatkowski, Cordelia Langford, John Sillitoe on behalf of the Wellcome Sanger Institute COVID-19 Surveillance Team |
| EPI_ISL_2848160 | Lighthouse Lab in Milton Keynes | Wellcome Sanger Institute for the COVID-19 Genomics UK (COG-UK) Consortium | The Lighthouse Lab in Milton Keynes and Alex Alderton, Roberto Amato, Jeffrey Barrett, Sonia Goncalves, Ewan Harrison, David K. Jackson, Ian Johnston, Dominic Kwiatkowski, Cordelia Langford, John Sillitoe on behalf of the Wellcome Sanger Institute COVID-19 Surveillance Team |
| EPI_ISL_2848161 | Lighthouse Laboratory Plymouth | Wellcome Sanger Institute for the COVID-19 Genomics UK (COG-UK) Consortium | Lighthouse Laboratory Plymouth and Alex Alderton, Roberto Amato, Jeffrey Barrett, Sonia Goncalves, Ewan Harrison, David K. Jackson, Ian Johnston, Dominic Kwiatkowski, Cordelia Langford, John Sillitoe on behalf of the Wellcome Sanger Institute COVID-19 Surveillance Team |
| EPI_ISL_2848162 | Lighthouse Lab in Alderley Park | Wellcome Sanger Institute for the COVID-19 Genomics UK (COG-UK) Consortium | Jacquelyn Wynn, Mairead Hyland, The Lighthouse Lab in Alderley Park and Alex Alderton, Roberto Amato, Jeffrey Barrett, Sonia Goncalves, Ewan Harrison, David K. Jackson, Ian Johnston, Dominic Kwiatkowski, Cordelia Langford, John Sillitoe on behalf of the Wellcome Sanger Institute COVID-19 Surveillance Team |
| EPI_ISL_2848163 | Lighthouse Laboratory Plymouth | Wellcome Sanger Institute for the COVID-19 Genomics UK (COG-UK) Consortium | Lighthouse Laboratory Plymouth and Alex Alderton, Roberto Amato, Jeffrey Barrett, Sonia Goncalves, Ewan Harrison, David K. Jackson, Ian Johnston, Dominic Kwiatkowski, Cordelia Langford, John Sillitoe on behalf of the Wellcome Sanger Institute COVID-19 Surveillance Team |
| EPI_ISL_2848164 | Lighthouse Lab in Alderley Park | Wellcome Sanger Institute for the COVID-19 Genomics UK (COG-UK) Consortium | Jacquelyn Wynn, Mairead Hyland, The Lighthouse Lab in Alderley Park and Alex Alderton, Roberto Amato, Jeffrey Barrett, Sonia Goncalves, Ewan Harrison, David K. Jackson, Ian Johnston, Dominic Kwiatkowski, Cordelia Langford, John Sillitoe on behalf of the Wellcome Sanger Institute COVID-19 Surveillance Team |
| EPI_ISL_2848165 | Lighthouse Laboratory Plymouth | Wellcome Sanger Institute for the COVID-19 Genomics UK (COG-UK) Consortium | Lighthouse Laboratory Plymouth and Alex Alderton, Roberto Amato, Jeffrey Barrett, Sonia Goncalves, Ewan Harrison, David K. Jackson, Ian Johnston, Dominic Kwiatkowski, Cordelia Langford, John Sillitoe on behalf of the Wellcome Sanger Institute COVID-19 Surveillance Team |
| EPI_ISL_2848166 | Lighthouse Lab in Alderley Park | Wellcome Sanger Institute for the COVID-19 Genomics UK (COG-UK) Consortium | Jacquelyn Wynn, Mairead Hyland, The Lighthouse Lab in Alderley Park and Alex Alderton, Roberto Amato, Jeffrey Barrett, Sonia Goncalves, Ewan Harrison, David K. Jackson, Ian Johnston, Dominic Kwiatkowski, Cordelia Langford, John Sillitoe on behalf of the Wellcome Sanger Institute COVID-19 Surveillance Team |
| EPI_ISL_2848167 | Lighthouse Laboratory Plymouth | Wellcome Sanger Institute for the COVID-19 Genomics UK (COG-UK) Consortium | Lighthouse Laboratory Plymouth and Alex Alderton, Roberto Amato, Jeffrey Barrett, Sonia Goncalves, Ewan Harrison, David K. Jackson, Ian Johnston, Dominic Kwiatkowski, Cordelia Langford, John Sillitoe on behalf of the Wellcome Sanger Institute COVID-19 Surveillance Team |
| EPI_ISL_2848168, EPI_ISL_2848169, EPI_ISL_2848170 | Lighthouse Lab in Milton Keynes | Wellcome Sanger Institute for the COVID-19 Genomics UK (COG-UK) Consortium | The Lighthouse Lab in Milton Keynes and Alex Alderton, Roberto Amato, Jeffrey Barrett, Sonia Goncalves, Ewan Harrison, David K. Jackson, Ian Johnston, Dominic Kwiatkowski, Cordelia Langford, John Sillitoe on behalf of the Wellcome Sanger Institute COVID-19 Surveillance Team |
| EPI_ISL_2848171 | Lighthouse Lab in Alderley Park | Wellcome Sanger Institute for the COVID-19 Genomics UK (COG-UK) Consortium | Jacquelyn Wynn, Mairead Hyland, The Lighthouse Lab in Alderley Park and Alex Alderton, Roberto Amato, Jeffrey Barrett, Sonia Goncalves, Ewan Harrison, David K. Jackson, Ian Johnston, Dominic Kwiatkowski, Cordelia Langford, John Sillitoe on behalf of the Wellcome Sanger Institute COVID-19 Surveillance Team |
| EPI_ISL_2848172 | Lighthouse Lab in Milton Keynes | Wellcome Sanger Institute for the COVID-19 Genomics UK (COG-UK) Consortium | The Lighthouse Lab in Milton Keynes and Alex Alderton, Roberto Amato, Jeffrey Barrett, Sonia Goncalves, Ewan Harrison, David K. Jackson, Ian Johnston, Dominic Kwiatkowski, Cordelia Langford, John Sillitoe on behalf of the Wellcome Sanger Institute COVID-19 Surveillance Team |
| EPI_ISL_2848173, EPI_ISL_2848174 | Lighthouse Lab in Alderley Park | Wellcome Sanger Institute for the COVID-19 Genomics UK (COG-UK) Consortium | Jacquelyn Wynn, Mairead Hyland, The Lighthouse Lab in Alderley Park and Alex Alderton, Roberto Amato, Jeffrey Barrett, Sonia Goncalves, Ewan Harrison, David K. Jackson, Ian Johnston, Dominic Kwiatkowski, Cordelia Langford, John Sillitoe on behalf of the Wellcome Sanger Institute COVID-19 Surveillance Team |
| EPI_ISL_2848175 | Lighthouse Laboratory Plymouth | Wellcome Sanger Institute for the COVID-19 Genomics UK (COG-UK) Consortium | Lighthouse Laboratory Plymouth and Alex Alderton, Roberto Amato, Jeffrey Barrett, Sonia Goncalves, Ewan Harrison, David K. Jackson, Ian Johnston, Dominic Kwiatkowski, Cordelia Langford, John Sillitoe on behalf of the Wellcome Sanger Institute COVID-19 Surveillance Team |
| EPI_ISL_2848176 | Lighthouse Lab in Milton Keynes | Wellcome Sanger Institute for the COVID-19 Genomics UK (COG-UK) Consortium | The Lighthouse Lab in Milton Keynes and Alex Alderton, Roberto Amato, Jeffrey Barrett, Sonia Goncalves, Ewan Harrison, David K. Jackson, Ian Johnston, Dominic Kwiatkowski, Cordelia Langford, John Sillitoe on behalf of the Wellcome Sanger Institute COVID-19 Surveillance Team |
| EPI_ISL_2848177 | Lighthouse Laboratory Plymouth | Wellcome Sanger Institute for the COVID-19 Genomics UK (COG-UK) Consortium | Lighthouse Laboratory Plymouth and Alex Alderton, Roberto Amato, Jeffrey Barrett, Sonia Goncalves, Ewan Harrison, David K. Jackson, Ian Johnston, Dominic Kwiatkowski, Cordelia Langford, John Sillitoe on behalf of the Wellcome Sanger Institute COVID-19 Surveillance Team |

| | | | |
|---|---|---|---|
| EPI_ISL_2848178, EPI_ISL_2848179 | Lighthouse Lab in Alderley Park | Wellcome Sanger Institute for the COVID-19 Genomics UK (COG-UK) Consortium | Jacquelyn Wynn, Mairead Hyland, The Lighthouse Lab in Alderley Park and Alex Alderton, Roberto Amato, Jeffrey Barrett, Sonia Goncalves, Ewan Harrison, David K. Jackson, Ian Johnston, Dominic Kwiatkowski, Cordelia Langford, John Sillitoe on behalf of the Wellcome Sanger Institute COVID-19 Surveillance Team |
| EPI_ISL_2848180 | Lighthouse Lab in Milton Keynes | Wellcome Sanger Institute for the COVID-19 Genomics UK (COG-UK) Consortium | The Lighthouse Lab in Milton Keynes and Alex Alderton, Roberto Amato, Jeffrey Barrett, Sonia Goncalves, Ewan Harrison, David K. Jackson, Ian Johnston, Dominic Kwiatkowski, Cordelia Langford, John Sillitoe on behalf of the Wellcome Sanger Institute COVID-19 Surveillance Team |
| EPI_ISL_2848181 | Lighthouse Lab in Alderley Park | Wellcome Sanger Institute for the COVID-19 Genomics UK (COG-UK) Consortium | Jacquelyn Wynn, Mairead Hyland, The Lighthouse Lab in Alderley Park and Alex Alderton, Roberto Amato, Jeffrey Barrett, Sonia Goncalves, Ewan Harrison, David K. Jackson, Ian Johnston, Dominic Kwiatkowski, Cordelia Langford, John Sillitoe on behalf of the Wellcome Sanger Institute COVID-19 Surveillance Team |
| EPI_ISL_2848182 | Lighthouse Laboratory Plymouth | Wellcome Sanger Institute for the COVID-19 Genomics UK (COG-UK) Consortium | Lighthouse Laboratory Plymouth and Alex Alderton, Roberto Amato, Jeffrey Barrett, Sonia Goncalves, Ewan Harrison, David K. Jackson, Ian Johnston, Dominic Kwiatkowski, Cordelia Langford, John Sillitoe on behalf of the Wellcome Sanger Institute COVID-19 Surveillance Team |
| EPI_ISL_2848183 | Lighthouse Lab in Alderley Park | Wellcome Sanger Institute for the COVID-19 Genomics UK (COG-UK) Consortium | Jacquelyn Wynn, Mairead Hyland, The Lighthouse Lab in Alderley Park and Alex Alderton, Roberto Amato, Jeffrey Barrett, Sonia Goncalves, Ewan Harrison, David K. Jackson, Ian Johnston, Dominic Kwiatkowski, Cordelia Langford, John Sillitoe on behalf of the Wellcome Sanger Institute COVID-19 Surveillance Team |
| EPI_ISL_2848184 | Lighthouse Lab in Milton Keynes | Wellcome Sanger Institute for the COVID-19 Genomics UK (COG-UK) Consortium | The Lighthouse Lab in Milton Keynes and Alex Alderton, Roberto Amato, Jeffrey Barrett, Sonia Goncalves, Ewan Harrison, David K. Jackson, Ian Johnston, Dominic Kwiatkowski, Cordelia Langford, John Sillitoe on behalf of the Wellcome Sanger Institute COVID-19 Surveillance Team |
| EPI_ISL_2848185 | Lighthouse Laboratory Plymouth | Wellcome Sanger Institute for the COVID-19 Genomics UK (COG-UK) Consortium | Lighthouse Laboratory Plymouth and Alex Alderton, Roberto Amato, Jeffrey Barrett, Sonia Goncalves, Ewan Harrison, David K. Jackson, Ian Johnston, Dominic Kwiatkowski, Cordelia Langford, John Sillitoe on behalf of the Wellcome Sanger Institute COVID-19 Surveillance Team |
| EPI_ISL_2848186 | Lighthouse Lab in Milton Keynes | Wellcome Sanger Institute for the COVID-19 Genomics UK (COG-UK) Consortium | The Lighthouse Lab in Milton Keynes and Alex Alderton, Roberto Amato, Jeffrey Barrett, Sonia Goncalves, Ewan Harrison, David K. Jackson, Ian Johnston, Dominic Kwiatkowski, Cordelia Langford, John Sillitoe on behalf of the Wellcome Sanger Institute COVID-19 Surveillance Team |
| EPI_ISL_2848187, EPI_ISL_2848188 | Lighthouse Lab in Alderley Park | Wellcome Sanger Institute for the COVID-19 Genomics UK (COG-UK) Consortium | Jacquelyn Wynn, Mairead Hyland, The Lighthouse Lab in Alderley Park and Alex Alderton, Roberto Amato, Jeffrey Barrett, Sonia Goncalves, Ewan Harrison, David K. Jackson, Ian Johnston, Dominic Kwiatkowski, Cordelia Langford, John Sillitoe on behalf of the Wellcome Sanger Institute COVID-19 Surveillance Team |
| EPI_ISL_2848189 | Lighthouse Lab in Milton Keynes | Wellcome Sanger Institute for the COVID-19 Genomics UK (COG-UK) Consortium | The Lighthouse Lab in Milton Keynes and Alex Alderton, Roberto Amato, Jeffrey Barrett, Sonia Goncalves, Ewan Harrison, David K. Jackson, Ian Johnston, Dominic Kwiatkowski, Cordelia Langford, John Sillitoe on behalf of the Wellcome Sanger Institute COVID-19 Surveillance Team |
| EPI_ISL_2848190 | Lighthouse Lab in Alderley Park | Wellcome Sanger Institute for the COVID-19 Genomics UK (COG-UK) Consortium | Jacquelyn Wynn, Mairead Hyland, The Lighthouse Lab in Alderley Park and Alex Alderton, Roberto Amato, Jeffrey Barrett, Sonia Goncalves, Ewan Harrison, David K. Jackson, Ian Johnston, Dominic Kwiatkowski, Cordelia Langford, John Sillitoe on behalf of the Wellcome Sanger Institute COVID-19 Surveillance Team |
| EPI_ISL_2848191 | Lighthouse Laboratory Plymouth | Wellcome Sanger Institute for the COVID-19 Genomics UK (COG-UK) Consortium | Lighthouse Laboratory Plymouth and Alex Alderton, Roberto Amato, Jeffrey Barrett, Sonia Goncalves, Ewan Harrison, David K. Jackson, Ian Johnston, Dominic Kwiatkowski, Cordelia Langford, John Sillitoe on behalf of the Wellcome Sanger Institute COVID-19 Surveillance Team |
| EPI_ISL_2848192 | Lighthouse Lab in Milton Keynes | Wellcome Sanger Institute for the COVID-19 Genomics UK (COG-UK) Consortium | The Lighthouse Lab in Milton Keynes and Alex Alderton, Roberto Amato, Jeffrey Barrett, Sonia Goncalves, Ewan Harrison, David K. Jackson, Ian Johnston, Dominic Kwiatkowski, Cordelia Langford, John Sillitoe on behalf of the Wellcome Sanger Institute COVID-19 Surveillance Team |
| EPI_ISL_2848193 | Lighthouse Lab in Alderley Park | Wellcome Sanger Institute for the COVID-19 Genomics UK (COG-UK) Consortium | Jacquelyn Wynn, Mairead Hyland, The Lighthouse Lab in Alderley Park and Alex Alderton, Roberto Amato, Jeffrey Barrett, Sonia Goncalves, Ewan Harrison, David K. Jackson, Ian Johnston, Dominic Kwiatkowski, Cordelia Langford, John Sillitoe on behalf of the Wellcome Sanger Institute COVID-19 Surveillance Team |
| EPI_ISL_2848194 | Lighthouse Laboratory Plymouth | Wellcome Sanger Institute for the COVID-19 Genomics UK (COG-UK) Consortium | Lighthouse Laboratory Plymouth and Alex Alderton, Roberto Amato, Jeffrey Barrett, Sonia Goncalves, Ewan Harrison, David K. Jackson, Ian Johnston, Dominic Kwiatkowski, Cordelia Langford, John Sillitoe on behalf of the Wellcome Sanger Institute COVID-19 Surveillance Team |
| EPI_ISL_2848195 | Lighthouse Lab in Milton Keynes | Wellcome Sanger Institute for the COVID-19 Genomics UK (COG-UK) Consortium | The Lighthouse Lab in Milton Keynes and Alex Alderton, Roberto Amato, Jeffrey Barrett, Sonia Goncalves, Ewan Harrison, David K. Jackson, Ian Johnston, Dominic Kwiatkowski, Cordelia Langford, John Sillitoe on behalf of the Wellcome Sanger Institute COVID-19 Surveillance Team |
| EPI_ISL_2848196 | Lighthouse Laboratory Plymouth | Wellcome Sanger Institute for the COVID-19 Genomics UK (COG-UK) Consortium | Lighthouse Laboratory Plymouth and Alex Alderton, Roberto Amato, Jeffrey Barrett, Sonia Goncalves, Ewan Harrison, David K. Jackson, Ian Johnston, Dominic Kwiatkowski, Cordelia Langford, John Sillitoe on behalf of the Wellcome Sanger Institute COVID-19 Surveillance Team |
| EPI_ISL_2848197 | Lighthouse Lab in Alderley Park | Wellcome Sanger Institute for the COVID-19 Genomics UK (COG-UK) Consortium | Jacquelyn Wynn, Mairead Hyland, The Lighthouse Lab in Alderley Park and Alex Alderton, Roberto Amato, Jeffrey Barrett, Sonia Goncalves, Ewan Harrison, David K. Jackson, Ian Johnston, Dominic Kwiatkowski, Cordelia Langford, John Sillitoe on behalf of the Wellcome Sanger Institute COVID-19 Surveillance Team |
| EPI_ISL_2848198, EPI_ISL_2848199 | Lighthouse Lab in Milton Keynes | Wellcome Sanger Institute for the COVID-19 Genomics UK (COG-UK) Consortium | The Lighthouse Lab in Milton Keynes and Alex Alderton, Roberto Amato, Jeffrey Barrett, Sonia Goncalves, Ewan Harrison, David K. Jackson, Ian Johnston, Dominic Kwiatkowski, Cordelia Langford, John Sillitoe on behalf of the Wellcome Sanger Institute COVID-19 Surveillance Team |
| EPI_ISL_2848200 | Lighthouse Laboratory Plymouth | Wellcome Sanger Institute for the COVID-19 Genomics UK (COG-UK) Consortium | Lighthouse Laboratory Plymouth and Alex Alderton, Roberto Amato, Jeffrey Barrett, Sonia Goncalves, Ewan Harrison, David K. Jackson, Ian Johnston, Dominic Kwiatkowski, Cordelia Langford, John Sillitoe on behalf of the Wellcome Sanger Institute COVID-19 Surveillance Team |
| EPI_ISL_2848201, EPI_ISL_2848202 | Lighthouse Lab in Milton Keynes | Wellcome Sanger Institute for the COVID-19 Genomics UK (COG-UK) Consortium | The Lighthouse Lab in Milton Keynes and Alex Alderton, Roberto Amato, Jeffrey Barrett, Sonia Goncalves, Ewan Harrison, David K. Jackson, Ian Johnston, Dominic Kwiatkowski, Cordelia Langford, John Sillitoe on behalf of the Wellcome Sanger Institute COVID-19 Surveillance Team |
| EPI_ISL_2848203 | AZ Klina | AZ Klina | Carl Vael - Lynsey Berckmans |
| EPI_ISL_2848204 | Lighthouse Laboratory Plymouth | Wellcome Sanger Institute for the COVID-19 Genomics UK (COG-UK) Consortium | Lighthouse Laboratory Plymouth and Alex Alderton, Roberto Amato, Jeffrey Barrett, Sonia Goncalves, Ewan Harrison, David K. Jackson, Ian Johnston, Dominic Kwiatkowski, Cordelia Langford, John Sillitoe on behalf of the Wellcome Sanger Institute COVID-19 Surveillance Team |
| EPI_ISL_2848205 | Lighthouse Lab in Alderley Park | Wellcome Sanger Institute for the COVID-19 Genomics UK (COG-UK) Consortium | Jacquelyn Wynn, Mairead Hyland, The Lighthouse Lab in Alderley Park and Alex Alderton, Roberto Amato, Jeffrey Barrett, Sonia Goncalves, Ewan Harrison, David K. Jackson, Ian Johnston, Dominic Kwiatkowski, Cordelia Langford, John Sillitoe on behalf of the Wellcome Sanger Institute COVID-19 Surveillance Team |
| EPI_ISL_2848206, EPI_ISL_2848207 | Lighthouse Laboratory Plymouth | Wellcome Sanger Institute for the COVID-19 Genomics UK (COG-UK) Consortium | Lighthouse Laboratory Plymouth and Alex Alderton, Roberto Amato, Jeffrey Barrett, Sonia Goncalves, Ewan Harrison, David K. Jackson, Ian Johnston, Dominic Kwiatkowski, Cordelia Langford, John Sillitoe on behalf of the Wellcome Sanger Institute COVID-19 Surveillance Team |
| EPI_ISL_2848208, EPI_ISL_2848209, EPI_ISL_2848210 | Lighthouse Lab in Milton Keynes | Wellcome Sanger Institute for the COVID-19 Genomics UK (COG-UK) Consortium | The Lighthouse Lab in Milton Keynes and Alex Alderton, Roberto Amato, Jeffrey Barrett, Sonia Goncalves, Ewan Harrison, David K. Jackson, Ian Johnston, Dominic Kwiatkowski, Cordelia Langford, John Sillitoe on behalf of the Wellcome Sanger Institute COVID-19 Surveillance Team |
| EPI_ISL_2848211 | Lighthouse Lab in Alderley Park | Wellcome Sanger Institute for the COVID-19 Genomics UK (COG-UK) Consortium | Jacquelyn Wynn, Mairead Hyland, The Lighthouse Lab in Alderley Park and Alex Alderton, Roberto Amato, Jeffrey Barrett, Sonia Goncalves, Ewan Harrison, David K. Jackson, Ian Johnston, Dominic Kwiatkowski, Cordelia Langford, John Sillitoe on behalf of the Wellcome Sanger Institute COVID-19 Surveillance Team |
| EPI_ISL_2848212 | Lighthouse Laboratory Plymouth | Wellcome Sanger Institute for the COVID-19 Genomics UK (COG-UK) Consortium | Lighthouse Laboratory Plymouth and Alex Alderton, Roberto Amato, Jeffrey Barrett, Sonia Goncalves, Ewan Harrison, David K. Jackson, Ian Johnston, Dominic Kwiatkowski, Cordelia Langford, John Sillitoe on behalf of the Wellcome Sanger Institute COVID-19 Surveillance Team |
| EPI_ISL_2848213 | Lighthouse Lab in Milton Keynes | Wellcome Sanger Institute for the COVID-19 Genomics UK (COG-UK) Consortium | The Lighthouse Lab in Milton Keynes and Alex Alderton, Roberto Amato, Jeffrey Barrett, Sonia Goncalves, Ewan Harrison, David K. Jackson, Ian Johnston, Dominic Kwiatkowski, Cordelia Langford, John Sillitoe on behalf of the Wellcome Sanger Institute COVID-19 Surveillance Team |
| EPI_ISL_2848214 | Lighthouse Laboratory Plymouth | Wellcome Sanger Institute for the COVID-19 Genomics UK (COG-UK) Consortium | Lighthouse Laboratory Plymouth and Alex Alderton, Roberto Amato, Jeffrey Barrett, Sonia Goncalves, Ewan Harrison, David K. Jackson, Ian Johnston, Dominic Kwiatkowski, Cordelia Langford, John Sillitoe on behalf of the Wellcome Sanger Institute COVID-19 Surveillance Team |
| EPI_ISL_2848215 | Lighthouse Lab in Alderley Park | Wellcome Sanger Institute for the COVID-19 Genomics UK (COG-UK) Consortium | Jacquelyn Wynn, Mairead Hyland, The Lighthouse Lab in Alderley Park and Alex Alderton, Roberto Amato, Jeffrey Barrett, Sonia Goncalves, Ewan Harrison, David K. Jackson, Ian Johnston, Dominic Kwiatkowski, Cordelia Langford, John Sillitoe on behalf of the Wellcome Sanger Institute COVID-19 Surveillance Team |
| EPI_ISL_2848216 | Lighthouse Laboratory Plymouth | Wellcome Sanger Institute for the COVID-19 Genomics UK (COG-UK) Consortium | Lighthouse Laboratory Plymouth and Alex Alderton, Roberto Amato, Jeffrey Barrett, Sonia Goncalves, Ewan Harrison, David K. Jackson, Ian Johnston, Dominic Kwiatkowski, Cordelia Langford, John Sillitoe on behalf of the Wellcome Sanger Institute COVID-19 Surveillance Team |
| EPI_ISL_2848217, EPI_ISL_2848218, EPI_ISL_2848219, EPI_ISL_2848220 | Lighthouse Lab in Milton Keynes | Wellcome Sanger Institute for the COVID-19 Genomics UK (COG-UK) Consortium | The Lighthouse Lab in Milton Keynes and Alex Alderton, Roberto Amato, Jeffrey Barrett, Sonia Goncalves, Ewan Harrison, David K. Jackson, Ian Johnston, Dominic Kwiatkowski, Cordelia Langford, John Sillitoe on behalf of the Wellcome Sanger Institute COVID-19 Surveillance Team |
| EPI_ISL_2848221 | Lighthouse Laboratory Plymouth | Wellcome Sanger Institute for the COVID-19 Genomics UK (COG-UK) Consortium | Lighthouse Laboratory Plymouth and Alex Alderton, Roberto Amato, Jeffrey Barrett, Sonia Goncalves, Ewan Harrison, David K. Jackson, Ian Johnston, Dominic Kwiatkowski, Cordelia Langford, John Sillitoe on behalf of the Wellcome Sanger Institute COVID-19 Surveillance Team |
| EPI_ISL_2848222, EPI_ISL_2848223 | Lighthouse Lab in Alderley Park | Wellcome Sanger Institute for the COVID-19 Genomics UK (COG-UK) Consortium | Jacquelyn Wynn, Mairead Hyland, The Lighthouse Lab in Alderley Park and Alex Alderton, Roberto Amato, Jeffrey Barrett, Sonia Goncalves, Ewan Harrison, David K. Jackson, Ian Johnston, Dominic Kwiatkowski, Cordelia Langford, John Sillitoe on behalf of the Wellcome Sanger Institute COVID-19 Surveillance Team |
| EPI_ISL_2848224 | Lighthouse Laboratory Plymouth | Wellcome Sanger Institute for the COVID-19 Genomics UK (COG-UK) Consortium | Lighthouse Laboratory Plymouth and Alex Alderton, Roberto Amato, Jeffrey Barrett, Sonia Goncalves, Ewan Harrison, David K. Jackson, Ian Johnston, Dominic Kwiatkowski, Cordelia Langford, John Sillitoe on behalf of the Wellcome Sanger Institute COVID-19 Surveillance Team |
| EPI_ISL_2848225 | Lighthouse Lab in Milton Keynes | Wellcome Sanger Institute for the COVID-19 Genomics UK (COG-UK) Consortium | The Lighthouse Lab in Milton Keynes and Alex Alderton, Roberto Amato, Jeffrey Barrett, Sonia Goncalves, Ewan Harrison, David K. Jackson, Ian Johnston, Dominic Kwiatkowski, Cordelia Langford, John Sillitoe on behalf of the Wellcome Sanger Institute COVID-19 Surveillance Team |
| EPI_ISL_2848226, EPI_ISL_2848227, EPI_ISL_2848228 | Lighthouse Lab in Alderley Park | Wellcome Sanger Institute for the COVID-19 Genomics UK (COG-UK) Consortium | Jacquelyn Wynn, Mairead Hyland, The Lighthouse Lab in Alderley Park and Alex Alderton, Roberto Amato, Jeffrey Barrett, Sonia Goncalves, Ewan Harrison, David K. Jackson, Ian Johnston, Dominic Kwiatkowski, Cordelia Langford, John Sillitoe on behalf of the Wellcome Sanger Institute COVID-19 Surveillance Team |
| EPI_ISL_2848229, EPI_ISL_2848230 | Lighthouse Laboratory Plymouth | Wellcome Sanger Institute for the COVID-19 Genomics UK (COG-UK) Consortium | Lighthouse Laboratory Plymouth and Alex Alderton, Roberto Amato, Jeffrey Barrett, Sonia Goncalves, Ewan Harrison, David K. Jackson, Ian Johnston, Dominic Kwiatkowski, Cordelia Langford, John Sillitoe on behalf of the Wellcome Sanger Institute COVID-19 Surveillance Team |
| EPI_ISL_2848231, EPI_ISL_2848232 | Lighthouse Lab in Alderley Park | Wellcome Sanger Institute for the COVID-19 Genomics UK (COG-UK) Consortium | Jacquelyn Wynn, Mairead Hyland, The Lighthouse Lab in Alderley Park and Alex Alderton, Roberto Amato, Jeffrey Barrett, Sonia Goncalves, Ewan Harrison, David K. Jackson, Ian Johnston, Dominic Kwiatkowski, Cordelia Langford, John Sillitoe on behalf of the Wellcome Sanger Institute COVID-19 Surveillance Team |
| EPI_ISL_2848233 | Lighthouse Lab in Milton Keynes | Wellcome Sanger Institute for the COVID-19 Genomics UK (COG-UK) Consortium | The Lighthouse Lab in Milton Keynes and Alex Alderton, Roberto Amato, Jeffrey Barrett, Sonia Goncalves, Ewan Harrison, David K. Jackson, Ian Johnston, Dominic Kwiatkowski, Cordelia Langford, John Sillitoe on behalf of the Wellcome Sanger Institute COVID-19 Surveillance Team |
| EPI_ISL_2848234 | Lighthouse Laboratory Plymouth | Wellcome Sanger Institute for the COVID-19 Genomics UK (COG-UK) Consortium | Lighthouse Laboratory Plymouth and Alex Alderton, Roberto Amato, Jeffrey Barrett, Sonia Goncalves, Ewan Harrison, David K. Jackson, Ian Johnston, Dominic Kwiatkowski, Cordelia Langford, John Sillitoe on behalf of the Wellcome Sanger Institute COVID-19 Surveillance Team |
| EPI_ISL_2848235 | Lighthouse Lab in Milton Keynes | Wellcome Sanger Institute for the COVID-19 Genomics UK (COG-UK) Consortium | The Lighthouse Lab in Milton Keynes and Alex Alderton, Roberto Amato, Jeffrey Barrett, Sonia Goncalves, Ewan Harrison, David K. Jackson, Ian Johnston, Dominic Kwiatkowski, Cordelia Langford, John Sillitoe on behalf of the Wellcome Sanger Institute COVID-19 Surveillance Team |
| EPI_ISL_2848236 | Lighthouse Lab in Alderley Park | Wellcome Sanger Institute for the COVID-19 Genomics UK (COG-UK) Consortium | Jacquelyn Wynn, Mairead Hyland, The Lighthouse Lab in Alderley Park and Alex Alderton, Roberto Amato, Jeffrey Barrett, Sonia Goncalves, Ewan Harrison, David K. Jackson, Ian Johnston, Dominic Kwiatkowski, Cordelia Langford, John Sillitoe on behalf of the Wellcome Sanger Institute COVID-19 Surveillance Team |

| | | | |
|---|---|---|---|
| EPI_ISL_2848237 | Lighthouse Laboratory Plymouth | Wellcome Sanger Institute for the COVID-19 Genomics UK (COG-UK) Consortium | Lighthouse Laboratory Plymouth and Alex Alderton, Roberto Amato, Jeffrey Barrett, Sonia Goncalves, Ewan Harrison, David K. Jackson, Ian Johnston, Dominic Kwiatkowski, Cordelia Langford, John Sillitoe on behalf of the Wellcome Sanger Institute COVID-19 Surveillance Team |
| EPI_ISL_2848238 | Lighthouse Lab in Milton Keynes | Wellcome Sanger Institute for the COVID-19 Genomics UK (COG-UK) Consortium | The Lighthouse Lab in Milton Keynes and Alex Alderton, Roberto Amato, Jeffrey Barrett, Sonia Goncalves, Ewan Harrison, David K. Jackson, Ian Johnston, Dominic Kwiatkowski, Cordelia Langford, John Sillitoe on behalf of the Wellcome Sanger Institute COVID-19 Surveillance Team |
| EPI_ISL_2848239 | Lighthouse Lab in Alderley Park | Wellcome Sanger Institute for the COVID-19 Genomics UK (COG-UK) Consortium | Jacquelyn Wynn, Mairead Hyland, The Lighthouse Lab in Alderley Park and Alex Alderton, Roberto Amato, Jeffrey Barrett, Sonia Goncalves, Ewan Harrison, David K. Jackson, Ian Johnston, Dominic Kwiatkowski, Cordelia Langford, John Sillitoe on behalf of the Wellcome Sanger Institute COVID-19 Surveillance Team |
| EPI_ISL_2848240 | Lighthouse Lab in Milton Keynes | Wellcome Sanger Institute for the COVID-19 Genomics UK (COG-UK) Consortium | The Lighthouse Lab in Milton Keynes and Alex Alderton, Roberto Amato, Jeffrey Barrett, Sonia Goncalves, Ewan Harrison, David K. Jackson, Ian Johnston, Dominic Kwiatkowski, Cordelia Langford, John Sillitoe on behalf of the Wellcome Sanger Institute COVID-19 Surveillance Team |
| EPI_ISL_2848241 | Lighthouse Laboratory Plymouth | Wellcome Sanger Institute for the COVID-19 Genomics UK (COG-UK) Consortium | Lighthouse Laboratory Plymouth and Alex Alderton, Roberto Amato, Jeffrey Barrett, Sonia Goncalves, Ewan Harrison, David K. Jackson, Ian Johnston, Dominic Kwiatkowski, Cordelia Langford, John Sillitoe on behalf of the Wellcome Sanger Institute COVID-19 Surveillance Team |
| EPI_ISL_2848242, EPI_ISL_2848243, EPI_ISL_2848244 | Lighthouse Lab in Milton Keynes | Wellcome Sanger Institute for the COVID-19 Genomics UK (COG-UK) Consortium | The Lighthouse Lab in Milton Keynes and Alex Alderton, Roberto Amato, Jeffrey Barrett, Sonia Goncalves, Ewan Harrison, David K. Jackson, Ian Johnston, Dominic Kwiatkowski, Cordelia Langford, John Sillitoe on behalf of the Wellcome Sanger Institute COVID-19 Surveillance Team |
| EPI_ISL_2848245, EPI_ISL_2848246, EPI_ISL_2848247 | Lighthouse Laboratory Plymouth | Wellcome Sanger Institute for the COVID-19 Genomics UK (COG-UK) Consortium | Lighthouse Laboratory Plymouth and Alex Alderton, Roberto Amato, Jeffrey Barrett, Sonia Goncalves, Ewan Harrison, David K. Jackson, Ian Johnston, Dominic Kwiatkowski, Cordelia Langford, John Sillitoe on behalf of the Wellcome Sanger Institute COVID-19 Surveillance Team |
| EPI_ISL_2848248, EPI_ISL_2848249 | Lighthouse Lab in Alderley Park | Wellcome Sanger Institute for the COVID-19 Genomics UK (COG-UK) Consortium | Jacquelyn Wynn, Mairead Hyland, The Lighthouse Lab in Alderley Park and Alex Alderton, Roberto Amato, Jeffrey Barrett, Sonia Goncalves, Ewan Harrison, David K. Jackson, Ian Johnston, Dominic Kwiatkowski, Cordelia Langford, John Sillitoe on behalf of the Wellcome Sanger Institute COVID-19 Surveillance Team |
| EPI_ISL_2848250, EPI_ISL_2848251, EPI_ISL_2848252 | Lighthouse Laboratory Plymouth | Wellcome Sanger Institute for the COVID-19 Genomics UK (COG-UK) Consortium | Lighthouse Laboratory Plymouth and Alex Alderton, Roberto Amato, Jeffrey Barrett, Sonia Goncalves, Ewan Harrison, David K. Jackson, Ian Johnston, Dominic Kwiatkowski, Cordelia Langford, John Sillitoe on behalf of the Wellcome Sanger Institute COVID-19 Surveillance Team |
| EPI_ISL_2848253 | Lighthouse Lab in Alderley Park | Wellcome Sanger Institute for the COVID-19 Genomics UK (COG-UK) Consortium | Jacquelyn Wynn, Mairead Hyland, The Lighthouse Lab in Alderley Park and Alex Alderton, Roberto Amato, Jeffrey Barrett, Sonia Goncalves, Ewan Harrison, David K. Jackson, Ian Johnston, Dominic Kwiatkowski, Cordelia Langford, John Sillitoe on behalf of the Wellcome Sanger Institute COVID-19 Surveillance Team |
| EPI_ISL_2848254, EPI_ISL_2848255, EPI_ISL_2848256 | Lighthouse Laboratory Plymouth | Wellcome Sanger Institute for the COVID-19 Genomics UK (COG-UK) Consortium | Lighthouse Laboratory Plymouth and Alex Alderton, Roberto Amato, Jeffrey Barrett, Sonia Goncalves, Ewan Harrison, David K. Jackson, Ian Johnston, Dominic Kwiatkowski, Cordelia Langford, John Sillitoe on behalf of the Wellcome Sanger Institute COVID-19 Surveillance Team |
| EPI_ISL_2848257, EPI_ISL_2848258 | Lighthouse Lab in Milton Keynes | Wellcome Sanger Institute for the COVID-19 Genomics UK (COG-UK) Consortium | The Lighthouse Lab in Milton Keynes and Alex Alderton, Roberto Amato, Jeffrey Barrett, Sonia Goncalves, Ewan Harrison, David K. Jackson, Ian Johnston, Dominic Kwiatkowski, Cordelia Langford, John Sillitoe on behalf of the Wellcome Sanger Institute COVID-19 Surveillance Team |
| EPI_ISL_2848259 | Lighthouse Laboratory Plymouth | Wellcome Sanger Institute for the COVID-19 Genomics UK (COG-UK) Consortium | Lighthouse Laboratory Plymouth and Alex Alderton, Roberto Amato, Jeffrey Barrett, Sonia Goncalves, Ewan Harrison, David K. Jackson, Ian Johnston, Dominic Kwiatkowski, Cordelia Langford, John Sillitoe on behalf of the Wellcome Sanger Institute COVID-19 Surveillance Team |
| EPI_ISL_2848260 | Lighthouse Lab in Milton Keynes | Wellcome Sanger Institute for the COVID-19 Genomics UK (COG-UK) Consortium | The Lighthouse Lab in Milton Keynes and Alex Alderton, Roberto Amato, Jeffrey Barrett, Sonia Goncalves, Ewan Harrison, David K. Jackson, Ian Johnston, Dominic Kwiatkowski, Cordelia Langford, John Sillitoe on behalf of the Wellcome Sanger Institute COVID-19 Surveillance Team |
| EPI_ISL_2848261 | Lighthouse Laboratory Plymouth | Wellcome Sanger Institute for the COVID-19 Genomics UK (COG-UK) Consortium | Lighthouse Laboratory Plymouth and Alex Alderton, Roberto Amato, Jeffrey Barrett, Sonia Goncalves, Ewan Harrison, David K. Jackson, Ian Johnston, Dominic Kwiatkowski, Cordelia Langford, John Sillitoe on behalf of the Wellcome Sanger Institute COVID-19 Surveillance Team |
| EPI_ISL_2848262, EPI_ISL_2848263 | Lighthouse Lab in Milton Keynes | Wellcome Sanger Institute for the COVID-19 Genomics UK (COG-UK) Consortium | The Lighthouse Lab in Milton Keynes and Alex Alderton, Roberto Amato, Jeffrey Barrett, Sonia Goncalves, Ewan Harrison, David K. Jackson, Ian Johnston, Dominic Kwiatkowski, Cordelia Langford, John Sillitoe on behalf of the Wellcome Sanger Institute COVID-19 Surveillance Team |
| EPI_ISL_2848264, EPI_ISL_2848265, EPI_ISL_2848266 | Lighthouse Laboratory Plymouth | Wellcome Sanger Institute for the COVID-19 Genomics UK (COG-UK) Consortium | Lighthouse Laboratory Plymouth and Alex Alderton, Roberto Amato, Jeffrey Barrett, Sonia Goncalves, Ewan Harrison, David K. Jackson, Ian Johnston, Dominic Kwiatkowski, Cordelia Langford, John Sillitoe on behalf of the Wellcome Sanger Institute COVID-19 Surveillance Team |
| EPI_ISL_2848267 | Lighthouse Lab in Milton Keynes | Wellcome Sanger Institute for the COVID-19 Genomics UK (COG-UK) Consortium | The Lighthouse Lab in Milton Keynes and Alex Alderton, Roberto Amato, Jeffrey Barrett, Sonia Goncalves, Ewan Harrison, David K. Jackson, Ian Johnston, Dominic Kwiatkowski, Cordelia Langford, John Sillitoe on behalf of the Wellcome Sanger Institute COVID-19 Surveillance Team |
| EPI_ISL_2848268, EPI_ISL_2848269 | Lighthouse Laboratory Plymouth | Wellcome Sanger Institute for the COVID-19 Genomics UK (COG-UK) Consortium | Lighthouse Laboratory Plymouth and Alex Alderton, Roberto Amato, Jeffrey Barrett, Sonia Goncalves, Ewan Harrison, David K. Jackson, Ian Johnston, Dominic Kwiatkowski, Cordelia Langford, John Sillitoe on behalf of the Wellcome Sanger Institute COVID-19 Surveillance Team |
| EPI_ISL_2848270 | Lighthouse Lab in Milton Keynes | Wellcome Sanger Institute for the COVID-19 Genomics UK (COG-UK) Consortium | The Lighthouse Lab in Milton Keynes and Alex Alderton, Roberto Amato, Jeffrey Barrett, Sonia Goncalves, Ewan Harrison, David K. Jackson, Ian Johnston, Dominic Kwiatkowski, Cordelia Langford, John Sillitoe on behalf of the Wellcome Sanger Institute COVID-19 Surveillance Team |
| EPI_ISL_2848271, EPI_ISL_2848272, EPI_ISL_2848273, EPI_ISL_2848274, EPI_ISL_2848275 | Lighthouse Laboratory Plymouth | Wellcome Sanger Institute for the COVID-19 Genomics UK (COG-UK) Consortium | Lighthouse Laboratory Plymouth and Alex Alderton, Roberto Amato, Jeffrey Barrett, Sonia Goncalves, Ewan Harrison, David K. Jackson, Ian Johnston, Dominic Kwiatkowski, Cordelia Langford, John Sillitoe on behalf of the Wellcome Sanger Institute COVID-19 Surveillance Team |
| EPI_ISL_2848276 | Lighthouse Lab in Alderley Park | Wellcome Sanger Institute for the COVID-19 Genomics UK (COG-UK) Consortium | Jacquelyn Wynn, Mairead Hyland, The Lighthouse Lab in Alderley Park and Alex Alderton, Roberto Amato, Jeffrey Barrett, Sonia Goncalves, Ewan Harrison, David K. Jackson, Ian Johnston, Dominic Kwiatkowski, Cordelia Langford, John Sillitoe on behalf of the Wellcome Sanger Institute COVID-19 Surveillance Team |
| EPI_ISL_2848277 | Lighthouse Lab in Milton Keynes | Wellcome Sanger Institute for the COVID-19 Genomics UK (COG-UK) Consortium | The Lighthouse Lab in Milton Keynes and Alex Alderton, Roberto Amato, Jeffrey Barrett, Sonia Goncalves, Ewan Harrison, David K. Jackson, Ian Johnston, Dominic Kwiatkowski, Cordelia Langford, John Sillitoe on behalf of the Wellcome Sanger Institute COVID-19 Surveillance Team |
| EPI_ISL_2848278 | Lighthouse Laboratory Plymouth | Wellcome Sanger Institute for the COVID-19 Genomics UK (COG-UK) Consortium | Lighthouse Laboratory Plymouth and Alex Alderton, Roberto Amato, Jeffrey Barrett, Sonia Goncalves, Ewan Harrison, David K. Jackson, Ian Johnston, Dominic Kwiatkowski, Cordelia Langford, John Sillitoe on behalf of the Wellcome Sanger Institute COVID-19 Surveillance Team |
| EPI_ISL_2848279 | Lighthouse Lab in Milton Keynes | Wellcome Sanger Institute for the COVID-19 Genomics UK (COG-UK) Consortium | The Lighthouse Lab in Milton Keynes and Alex Alderton, Roberto Amato, Jeffrey Barrett, Sonia Goncalves, Ewan Harrison, David K. Jackson, Ian Johnston, Dominic Kwiatkowski, Cordelia Langford, John Sillitoe on behalf of the Wellcome Sanger Institute COVID-19 Surveillance Team |
| EPI_ISL_2848280, EPI_ISL_2848281 | Lighthouse Lab in Alderley Park | Wellcome Sanger Institute for the COVID-19 Genomics UK (COG-UK) Consortium | Jacquelyn Wynn, Mairead Hyland, The Lighthouse Lab in Alderley Park and Alex Alderton, Roberto Amato, Jeffrey Barrett, Sonia Goncalves, Ewan Harrison, David K. Jackson, Ian Johnston, Dominic Kwiatkowski, Cordelia Langford, John Sillitoe on behalf of the Wellcome Sanger Institute COVID-19 Surveillance Team |
| EPI_ISL_2848282 | Lighthouse Laboratory Plymouth | Wellcome Sanger Institute for the COVID-19 Genomics UK (COG-UK) Consortium | Lighthouse Laboratory Plymouth and Alex Alderton, Roberto Amato, Jeffrey Barrett, Sonia Goncalves, Ewan Harrison, David K. Jackson, Ian Johnston, Dominic Kwiatkowski, Cordelia Langford, John Sillitoe on behalf of the Wellcome Sanger Institute COVID-19 Surveillance Team |
| EPI_ISL_2848283 | Lighthouse Lab in Alderley Park | Wellcome Sanger Institute for the COVID-19 Genomics UK (COG-UK) Consortium | Jacquelyn Wynn, Mairead Hyland, The Lighthouse Lab in Alderley Park and Alex Alderton, Roberto Amato, Jeffrey Barrett, Sonia Goncalves, Ewan Harrison, David K. Jackson, Ian Johnston, Dominic Kwiatkowski, Cordelia Langford, John Sillitoe on behalf of the Wellcome Sanger Institute COVID-19 Surveillance Team |
| EPI_ISL_2848284, EPI_ISL_2848285, EPI_ISL_2848286 | Lighthouse Laboratory Plymouth | Wellcome Sanger Institute for the COVID-19 Genomics UK (COG-UK) Consortium | Lighthouse Laboratory Plymouth and Alex Alderton, Roberto Amato, Jeffrey Barrett, Sonia Goncalves, Ewan Harrison, David K. Jackson, Ian Johnston, Dominic Kwiatkowski, Cordelia Langford, John Sillitoe on behalf of the Wellcome Sanger Institute COVID-19 Surveillance Team |
| EPI_ISL_2848287 | Lighthouse Lab in Alderley Park | Wellcome Sanger Institute for the COVID-19 Genomics UK (COG-UK) Consortium | Jacquelyn Wynn, Mairead Hyland, The Lighthouse Lab in Alderley Park and Alex Alderton, Roberto Amato, Jeffrey Barrett, Sonia Goncalves, Ewan Harrison, David K. Jackson, Ian Johnston, Dominic Kwiatkowski, Cordelia Langford, John Sillitoe on behalf of the Wellcome Sanger Institute COVID-19 Surveillance Team |
| EPI_ISL_2848288, EPI_ISL_2848289, EPI_ISL_2848290, EPI_ISL_2848291 | Lighthouse Lab in Milton Keynes | Wellcome Sanger Institute for the COVID-19 Genomics UK (COG-UK) Consortium | The Lighthouse Lab in Milton Keynes and Alex Alderton, Roberto Amato, Jeffrey Barrett, Sonia Goncalves, Ewan Harrison, David K. Jackson, Ian Johnston, Dominic Kwiatkowski, Cordelia Langford, John Sillitoe on behalf of the Wellcome Sanger Institute COVID-19 Surveillance Team |
| EPI_ISL_2848292 | Lighthouse Laboratory Plymouth | Wellcome Sanger Institute for the COVID-19 Genomics UK (COG-UK) Consortium | Lighthouse Laboratory Plymouth and Alex Alderton, Roberto Amato, Jeffrey Barrett, Sonia Goncalves, Ewan Harrison, David K. Jackson, Ian Johnston, Dominic Kwiatkowski, Cordelia Langford, John Sillitoe on behalf of the Wellcome Sanger Institute COVID-19 Surveillance Team |
| EPI_ISL_2848293 | Lighthouse Lab in Alderley Park | Wellcome Sanger Institute for the COVID-19 Genomics UK (COG-UK) Consortium | Jacquelyn Wynn, Mairead Hyland, The Lighthouse Lab in Alderley Park and Alex Alderton, Roberto Amato, Jeffrey Barrett, Sonia Goncalves, Ewan Harrison, David K. Jackson, Ian Johnston, Dominic Kwiatkowski, Cordelia Langford, John Sillitoe on behalf of the Wellcome Sanger Institute COVID-19 Surveillance Team |
| EPI_ISL_2848294 | Lighthouse Lab in Milton Keynes | Wellcome Sanger Institute for the COVID-19 Genomics UK (COG-UK) Consortium | The Lighthouse Lab in Milton Keynes and Alex Alderton, Roberto Amato, Jeffrey Barrett, Sonia Goncalves, Ewan Harrison, David K. Jackson, Ian Johnston, Dominic Kwiatkowski, Cordelia Langford, John Sillitoe on behalf of the Wellcome Sanger Institute COVID-19 Surveillance Team |
| EPI_ISL_2848295 | Lighthouse Laboratory Plymouth | Wellcome Sanger Institute for the COVID-19 Genomics UK (COG-UK) Consortium | Lighthouse Laboratory Plymouth and Alex Alderton, Roberto Amato, Jeffrey Barrett, Sonia Goncalves, Ewan Harrison, David K. Jackson, Ian Johnston, Dominic Kwiatkowski, Cordelia Langford, John Sillitoe on behalf of the Wellcome Sanger Institute COVID-19 Surveillance Team |
| EPI_ISL_2848296 | Lighthouse Lab in Milton Keynes | Wellcome Sanger Institute for the COVID-19 Genomics UK (COG-UK) Consortium | The Lighthouse Lab in Milton Keynes and Alex Alderton, Roberto Amato, Jeffrey Barrett, Sonia Goncalves, Ewan Harrison, David K. Jackson, Ian Johnston, Dominic Kwiatkowski, Cordelia Langford, John Sillitoe on behalf of the Wellcome Sanger Institute COVID-19 Surveillance Team |
| EPI_ISL_2848297, EPI_ISL_2848298, EPI_ISL_2848299 | Lighthouse Lab in Alderley Park | Wellcome Sanger Institute for the COVID-19 Genomics UK (COG-UK) Consortium | Jacquelyn Wynn, Mairead Hyland, The Lighthouse Lab in Alderley Park and Alex Alderton, Roberto Amato, Jeffrey Barrett, Sonia Goncalves, Ewan Harrison, David K. Jackson, Ian Johnston, Dominic Kwiatkowski, Cordelia Langford, John Sillitoe on behalf of the Wellcome Sanger Institute COVID-19 Surveillance Team |
| EPI_ISL_2848300 | Lighthouse Laboratory Plymouth | Wellcome Sanger Institute for the COVID-19 Genomics UK (COG-UK) Consortium | Lighthouse Laboratory Plymouth and Alex Alderton, Roberto Amato, Jeffrey Barrett, Sonia Goncalves, Ewan Harrison, David K. Jackson, Ian Johnston, Dominic Kwiatkowski, Cordelia Langford, John Sillitoe on behalf of the Wellcome Sanger Institute COVID-19 Surveillance Team |

EPI_ISL_2848301, EPI_ISL_2848302, EPI_ISL_2848303, EPI_ISL_2848304, EPI_ISL_2848305, EPI_ISL_2848307, EPI_ISL_2848308, EPI_ISL_2848309, EPI_ISL_2848310, EPI_ISL_2848311, EPI_ISL_2848312, EPI_ISL_2848313, EPI_ISL_2848314, EPI_ISL_2848315, EPI_ISL_2848316, EPI_ISL_2848317, EPI_ISL_2848318, EPI_ISL_2848319, EPI_ISL_2848320, EPI_ISL_2848321, EPI_ISL_2848322, EPI_ISL_2848323, EPI_ISL_2848324, EPI_ISL_2848325, EPI_ISL_2848326, EPI_ISL_2848327, EPI_ISL_2848328, EPI_ISL_2848329, EPI_ISL_2848330, EPI_ISL_2848331, EPI_ISL_2848332, EPI_ISL_2848333, EPI_ISL_2848334, EPI_ISL_2848335, EPI_ISL_2848336, EPI_ISL_2848337, EPI_ISL_2848338, EPI_ISL_2848339, EPI_ISL_2848340, EPI_ISL_2848342, EPI_ISL_2848343, EPI_ISL_2848344, EPI_ISL_2848345, EPI_ISL_2848346, EPI_ISL_2848347, EPI_ISL_2848348, EPI_ISL_2848349, EPI_ISL_2848350, EPI_ISL_2848351, EPI_ISL_2848352, EPI_ISL_2848353, EPI_ISL_2848354, EPI_ISL_2848355, EPI_ISL_2848356, EPI_ISL_2848357, EPI_ISL_2848358, EPI_ISL_2848359, EPI_ISL_2848360, EPI_ISL_2848361, EPI_ISL_2848362, EPI_ISL_2848363, EPI_ISL_2848364, EPI_ISL_2848365, EPI_ISL_2848366, EPI_ISL_2848367, EPI_ISL_2848368, EPI_ISL_2848369, EPI_ISL_2848370, EPI_ISL_2848371, EPI_ISL_2848372, EPI_ISL_2848373, EPI_ISL_2848374, EPI_ISL_2848375, EPI_ISL_2848376, EPI_ISL_2848377, EPI_ISL_2848378, EPI_ISL_2848379, EPI_ISL_2848380, EPI_ISL_2848381, EPI_ISL_2848382, EPI_ISL_2848383, EPI_ISL_2848384, EPI_ISL_2848385, EPI_ISL_2848386, EPI_ISL_2848387, EPI_ISL_2848388, EPI_ISL_2848389, EPI_ISL_2848390,

EPI_ISL_2848391, EPI_ISL_2848392, EPI_ISL_2848393, EPI_ISL_2848394, EPI_ISL_2848395, EPI_ISL_2848396, EPI_ISL_2848397, EPI_ISL_2848398, EPI_ISL_2848399, EPI_ISL_2848400, EPI_ISL_2848401, EPI_ISL_2848402, EPI_ISL_2848403, EPI_ISL_2848404, EPI_ISL_2848405, EPI_ISL_2848406, EPI_ISL_2848407, EPI_ISL_2848408, EPI_ISL_2848409, EPI_ISL_2848410, EPI_ISL_2848411, EPI_ISL_2848412, EPI_ISL_2848413, EPI_ISL_2848414, EPI_ISL_2848415, EPI_ISL_2848416, EPI_ISL_2848417, EPI_ISL_2848418, EPI_ISL_2848419, EPI_ISL_2848420, EPI_ISL_2848421, EPI_ISL_2848422, EPI_ISL_2848423, EPI_ISL_2848424, EPI_ISL_2848425, EPI_ISL_2848426, EPI_ISL_2848427, EPI_ISL_2848428, EPI_ISL_2848429, EPI_ISL_2848430, EPI_ISL_2848431, EPI_ISL_2848432, EPI_ISL_2848433, EPI_ISL_2848434, EPI_ISL_2848435, EPI_ISL_2848436, EPI_ISL_2848437, EPI_ISL_2848438, EPI_ISL_2848439, EPI_ISL_2848440, EPI_ISL_2848441, EPI_ISL_2848442, EPI_ISL_2848443, EPI_ISL_2848444, EPI_ISL_2848445, EPI_ISL_2848446, EPI_ISL_2848447, EPI_ISL_2848448, EPI_ISL_2848449, EPI_ISL_2848450, EPI_ISL_2848451, EPI_ISL_2848452, EPI_ISL_2848453, EPI_ISL_2848454, EPI_ISL_2848455, EPI_ISL_2848456, EPI_ISL_2848457, EPI_ISL_2848458, EPI_ISL_2848459, EPI_ISL_2848460, EPI_ISL_2848461, EPI_ISL_2848462, EPI_ISL_2848463, EPI_ISL_2848464, EPI_ISL_2848465, EPI_ISL_2848466, EPI_ISL_2848467, EPI_ISL_2848468, EPI_ISL_2848469, EPI_ISL_2848470, EPI_ISL_2848471, EPI_ISL_2848472, EPI_ISL_2848473, EPI_ISL_2848474, EPI_ISL_2848475, EPI_ISL_2848476, EPI_ISL_2848478, EPI_ISL_2848479, EPI_ISL_2848480, EPI_ISL_2848481, EPI_ISL_2848482, EPI_ISL_2848483, EPI_ISL_2848484, EPI_ISL_2848485, EPI_ISL_2848486, EPI_ISL_2848487, EPI_ISL_2848488, EPI_ISL_2848489, EPI_ISL_2848490, EPI_ISL_2848491, EPI_ISL_2848492, EPI_ISL_2848493, EPI_ISL_2848494, EPI_ISL_2848495, EPI_ISL_2848496, EPI_ISL_2848497, EPI_ISL_2848498, EPI_ISL_2848499, EPI_ISL_2848500, EPI_ISL_2848501, EPI_ISL_2848502, EPI_ISL_2848503, EPI_ISL_2848504, EPI_ISL_2848505, EPI_ISL_2848506, EPI_ISL_2848507, EPI_ISL_2848508, EPI_ISL_2848509, EPI_ISL_2848510, EPI_ISL_2848511, EPI_ISL_2848512, EPI_ISL_2848513, EPI_ISL_2848514, EPI_ISL_2848515, EPI_ISL_2848517, EPI_ISL_2848518, EPI_ISL_2848519, EPI_ISL_2848520, EPI_ISL_2848521, EPI_ISL_2848522, EPI_ISL_2848523, EPI_ISL_2848524, EPI_ISL_2848525, EPI_ISL_2848526, EPI_ISL_2848527, EPI_ISL_2848528, EPI_ISL_2848529, EPI_ISL_2848530, EPI_ISL_2848531, EPI_ISL_2848532, EPI_ISL_2848533, EPI_ISL_2848534, EPI_ISL_2848535, EPI_ISL_2848536, EPI_ISL_2848537, EPI_ISL_2848538, EPI_ISL_2848539, EPI_ISL_2848540, EPI_ISL_2848541, EPI_ISL_2848542, EPI_ISL_2848543, EPI_ISL_2848544, EPI_ISL_2848545, EPI_ISL_2848546, EPI_ISL_2848547, EPI_ISL_2848548, EPI_ISL_2848549, EPI_ISL_2848550, EPI_ISL_2848551, EPI_ISL_2848552, EPI_ISL_2848553, EPI_ISL_2848554, EPI_ISL_2848555, EPI_ISL_2848556, EPI_ISL_2848557, EPI_ISL_2848558, EPI_ISL_2848559, EPI_ISL_2848560, EPI_ISL_2848561, EPI_ISL_2848562, EPI_ISL_2848563, EPI_ISL_2848564, EPI_ISL_2848565, EPI_ISL_2848566, EPI_ISL_2848567, EPI_ISL_2848568, EPI_ISL_2848569, EPI_ISL_2848570, EPI_ISL_2848571, EPI_ISL_2848572, EPI_ISL_2848573, EPI_ISL_2848574, EPI_ISL_2848575, EPI_ISL_2848576, EPI_ISL_2848577, EPI_ISL_2848578, EPI_ISL_2848579, EPI_ISL_2848580, EPI_ISL_2848581, EPI_ISL_2848582, EPI_ISL_2848583, EPI_ISL_2848584, EPI_ISL_2848585, EPI_ISL_2848586, EPI_ISL_2848587, EPI_ISL_2848588, EPI_ISL_2848589, EPI_ISL_2848590, EPI_ISL_2848591, EPI_ISL_2848592, EPI_ISL_2848593, EPI_ISL_2848594, EPI_ISL_2848595, EPI_ISL_2848596, EPI_ISL_2848597, EPI_ISL_2848598, EPI_ISL_2848600, EPI_ISL_2848602, EPI_ISL_2848603, EPI_ISL_2848604, EPI_ISL_2848605, EPI_ISL_2848606, EPI_ISL_2848607, EPI_ISL_2848608, EPI_ISL_2848609, EPI_ISL_2848610, EPI_ISL_2848611, EPI_ISL_2848612, EPI_ISL_2848613, EPI_ISL_2848614, EPI_ISL_2848615, EPI_ISL_2848616, EPI_ISL_2848617, EPI_ISL_2848618, EPI_ISL_2848619, EPI_ISL_2848620, EPI_ISL_2848621, EPI_ISL_2848622, EPI_ISL_2848623, EPI_ISL_2848624, EPI_ISL_2848625, EPI_ISL_2848626, EPI_ISL_2848627, EPI_ISL_2848628, EPI_ISL_2848629, EPI_ISL_2848630, EPI_ISL_2848631, EPI_ISL_2848632, EPI_ISL_2848633, EPI_ISL_2848634, EPI_ISL_2848635, EPI_ISL_2848636, EPI_ISL_2848637, EPI_ISL_2848638, EPI_ISL_2848639, EPI_ISL_2848640, EPI_ISL_2848641, EPI_ISL_2848642, EPI_ISL_2848643, EPI_ISL_2848644, EPI_ISL_2848645, EPI_ISL_2848646, EPI_ISL_2848647, EPI_ISL_2848648, EPI_ISL_2848649, EPI_ISL_2848650, EPI_ISL_2848651, EPI_ISL_2848652, EPI_ISL_2848653, EPI_ISL_2848654, EPI_ISL_2848655, EPI_ISL_2848656

| | | | |
|---|---|---|---|
| see above | Lighthouse Lab in Milton Keynes | Wellcome Sanger Institute for the COVID-19 Genomics UK (COG-UK) Consortium | The Lighthouse Lab in Milton Keynes and Alex Alderton, Roberto Amato, Jeffrey Barrett, Sonia Goncalves, Ewan Harrison, David K. Jackson, Ian Johnston, Dominic Kwiatkowski, Cordelia Langford, John Sillitoe on behalf of the Wellcome Sanger Institute COVID-19 Surveillance Team |
| EPI_ISL_2848657 | Lighthouse Lab in Glasgow | Wellcome Sanger Institute for the COVID-19 Genomics UK (COG-UK) Consortium | Harper VanSteenhouse, Yumi Kasai, David Gray, Carol Clugston, Anna Dominiczak and Alex Alderton, Roberto Amato, Jeffrey Barrett, Sonia Goncalves, Ewan Harrison, David K. Jackson, Ian Johnston, Dominic Kwiatkowski, Cordelia Langford, John Sillitoe on behalf of the Wellcome Sanger Institute COVID-19 Surveillance Team |
| EPI_ISL_2848658, EPI_ISL_2848659, EPI_ISL_2848660, EPI_ISL_2848661, EPI_ISL_2848662 | Lighthouse Lab in Milton Keynes | Wellcome Sanger Institute for the COVID-19 Genomics UK (COG-UK) Consortium | The Lighthouse Lab in Milton Keynes and Alex Alderton, Roberto Amato, Jeffrey Barrett, Sonia Goncalves, Ewan Harrison, David K. Jackson, Ian Johnston, Dominic Kwiatkowski, Cordelia Langford, John Sillitoe on behalf of the Wellcome Sanger Institute COVID-19 Surveillance Team |
| EPI_ISL_2848663 | Lighthouse Lab in Glasgow | Wellcome Sanger Institute for the COVID-19 Genomics UK (COG-UK) Consortium | Harper VanSteenhouse, Yumi Kasai, David Gray, Carol Clugston, Anna Dominiczak and Alex Alderton, Roberto Amato, Jeffrey Barrett, Sonia Goncalves, Ewan Harrison, David K. Jackson, Ian Johnston, Dominic Kwiatkowski, Cordelia Langford, John Sillitoe on behalf of the Wellcome Sanger Institute COVID-19 Surveillance Team |
| EPI_ISL_2848664, EPI_ISL_2848665 | Lighthouse Lab in Milton Keynes | Wellcome Sanger Institute for the COVID-19 Genomics UK (COG-UK) Consortium | The Lighthouse Lab in Milton Keynes and Alex Alderton, Roberto Amato, Jeffrey Barrett, Sonia Goncalves, Ewan Harrison, David K. Jackson, Ian Johnston, Dominic Kwiatkowski, Cordelia Langford, John Sillitoe on behalf of the Wellcome Sanger Institute COVID-19 Surveillance Team |
| EPI_ISL_2848666, EPI_ISL_2848667 | Lighthouse Lab in Glasgow | Wellcome Sanger Institute for the COVID-19 Genomics UK (COG-UK) Consortium | Harper VanSteenhouse, Yumi Kasai, David Gray, Carol Clugston, Anna Dominiczak and Alex Alderton, Roberto Amato, Jeffrey Barrett, Sonia Goncalves, Ewan Harrison, David K. Jackson, Ian Johnston, Dominic Kwiatkowski, Cordelia Langford, John Sillitoe on behalf of the Wellcome Sanger Institute COVID-19 Surveillance Team |
| EPI_ISL_2848668, EPI_ISL_2848669 | Lighthouse Lab in Milton Keynes | Wellcome Sanger Institute for the COVID-19 Genomics UK (COG-UK) Consortium | The Lighthouse Lab in Milton Keynes and Alex Alderton, Roberto Amato, Jeffrey Barrett, Sonia Goncalves, Ewan Harrison, David K. Jackson, Ian Johnston, Dominic Kwiatkowski, Cordelia Langford, John Sillitoe on behalf of the Wellcome Sanger Institute COVID-19 Surveillance Team |
| EPI_ISL_2848670 | Lighthouse Lab in Glasgow | Wellcome Sanger Institute for the COVID-19 Genomics UK (COG-UK) Consortium | Harper VanSteenhouse, Yumi Kasai, David Gray, Carol Clugston, Anna Dominiczak and Alex Alderton, Roberto Amato, Jeffrey Barrett, Sonia Goncalves, Ewan Harrison, David K. Jackson, Ian Johnston, Dominic Kwiatkowski, Cordelia Langford, John Sillitoe on behalf of the Wellcome Sanger Institute COVID-19 Surveillance Team |
| EPI_ISL_2848671, EPI_ISL_2848672, EPI_ISL_2848673, EPI_ISL_2848674, EPI_ISL_2848675, EPI_ISL_2848676, EPI_ISL_2848677, EPI_ISL_2848678 | Lighthouse Lab in Milton Keynes | Wellcome Sanger Institute for the COVID-19 Genomics UK (COG-UK) Consortium | The Lighthouse Lab in Milton Keynes and Alex Alderton, Roberto Amato, Jeffrey Barrett, Sonia Goncalves, Ewan Harrison, David K. Jackson, Ian Johnston, Dominic Kwiatkowski, Cordelia Langford, John Sillitoe on behalf of the Wellcome Sanger Institute COVID-19 Surveillance Team |
| EPI_ISL_2848679 | Lighthouse Lab in Glasgow | Wellcome Sanger Institute for the COVID-19 Genomics UK (COG-UK) Consortium | Harper VanSteenhouse, Yumi Kasai, David Gray, Carol Clugston, Anna Dominiczak and Alex Alderton, Roberto Amato, Jeffrey Barrett, Sonia Goncalves, Ewan Harrison, David K. Jackson, Ian Johnston, Dominic Kwiatkowski, Cordelia Langford, John Sillitoe on behalf of the Wellcome Sanger Institute COVID-19 Surveillance Team |
| EPI_ISL_2848680, EPI_ISL_2848681 | Lighthouse Lab in Milton Keynes | Wellcome Sanger Institute for the COVID-19 Genomics UK (COG-UK) Consortium | The Lighthouse Lab in Milton Keynes and Alex Alderton, Roberto Amato, Jeffrey Barrett, Sonia Goncalves, Ewan Harrison, David K. Jackson, Ian Johnston, Dominic Kwiatkowski, Cordelia Langford, John Sillitoe on behalf of the Wellcome Sanger Institute COVID-19 Surveillance Team |
| EPI_ISL_2848682 | Lighthouse Lab in Glasgow | Wellcome Sanger Institute for the COVID-19 Genomics UK (COG-UK) Consortium | Harper VanSteenhouse, Yumi Kasai, David Gray, Carol Clugston, Anna Dominiczak and Alex Alderton, Roberto Amato, Jeffrey Barrett, Sonia Goncalves, Ewan Harrison, David K. Jackson, Ian Johnston, Dominic Kwiatkowski, Cordelia Langford, John Sillitoe on behalf of the Wellcome Sanger Institute COVID-19 Surveillance Team |
| EPI_ISL_2848683, EPI_ISL_2848684, EPI_ISL_2848685 | Lighthouse Lab in Milton Keynes | Wellcome Sanger Institute for the COVID-19 Genomics UK (COG-UK) Consortium | The Lighthouse Lab in Milton Keynes and Alex Alderton, Roberto Amato, Jeffrey Barrett, Sonia Goncalves, Ewan Harrison, David K. Jackson, Ian Johnston, Dominic Kwiatkowski, Cordelia Langford, John Sillitoe on behalf of the Wellcome Sanger Institute COVID-19 Surveillance Team |
| EPI_ISL_2848686 | Lighthouse Lab in Glasgow | Wellcome Sanger Institute for the COVID-19 Genomics UK (COG-UK) Consortium | Harper VanSteenhouse, Yumi Kasai, David Gray, Carol Clugston, Anna Dominiczak and Alex Alderton, Roberto Amato, Jeffrey Barrett, Sonia Goncalves, Ewan Harrison, David K. Jackson, Ian Johnston, Dominic Kwiatkowski, Cordelia Langford, John Sillitoe on behalf of the Wellcome Sanger Institute COVID-19 Surveillance Team |
| EPI_ISL_2848687, EPI_ISL_2848688, EPI_ISL_2848689 | Lighthouse Lab in Milton Keynes | Wellcome Sanger Institute for the COVID-19 Genomics UK (COG-UK) Consortium | The Lighthouse Lab in Milton Keynes and Alex Alderton, Roberto Amato, Jeffrey Barrett, Sonia Goncalves, Ewan Harrison, David K. Jackson, Ian Johnston, Dominic Kwiatkowski, Cordelia Langford, John Sillitoe on behalf of the Wellcome Sanger Institute COVID-19 Surveillance Team |
| EPI_ISL_2848690, EPI_ISL_2848691, EPI_ISL_2848692 | Lighthouse Lab in Glasgow | Wellcome Sanger Institute for the COVID-19 Genomics UK (COG-UK) Consortium | Harper VanSteenhouse, Yumi Kasai, David Gray, Carol Clugston, Anna Dominiczak and Alex Alderton, Roberto Amato, Jeffrey Barrett, Sonia Goncalves, Ewan Harrison, David K. Jackson, Ian Johnston, Dominic Kwiatkowski, Cordelia Langford, John Sillitoe on behalf of the Wellcome Sanger Institute COVID-19 Surveillance Team |
| EPI_ISL_2848693 | Lighthouse Lab in Milton Keynes | Wellcome Sanger Institute for the COVID-19 Genomics UK (COG-UK) Consortium | The Lighthouse Lab in Milton Keynes and Alex Alderton, Roberto Amato, Jeffrey Barrett, Sonia Goncalves, Ewan Harrison, David K. Jackson, Ian Johnston, Dominic Kwiatkowski, Cordelia Langford, John Sillitoe on behalf of the Wellcome Sanger Institute COVID-19 Surveillance Team |
| EPI_ISL_2848694 | Lighthouse Lab in Glasgow | Wellcome Sanger Institute for the COVID-19 Genomics UK (COG-UK) Consortium | Harper VanSteenhouse, Yumi Kasai, David Gray, Carol Clugston, Anna Dominiczak and Alex Alderton, Roberto Amato, Jeffrey Barrett, Sonia Goncalves, Ewan Harrison, David K. Jackson, Ian Johnston, Dominic Kwiatkowski, Cordelia Langford, John Sillitoe on behalf of the Wellcome Sanger Institute COVID-19 Surveillance Team |
| EPI_ISL_2848695, EPI_ISL_2848696, EPI_ISL_2848697, EPI_ISL_2848698 | Lighthouse Lab in Milton Keynes | Wellcome Sanger Institute for the COVID-19 Genomics UK (COG-UK) Consortium | The Lighthouse Lab in Milton Keynes and Alex Alderton, Roberto Amato, Jeffrey Barrett, Sonia Goncalves, Ewan Harrison, David K. Jackson, Ian Johnston, Dominic Kwiatkowski, Cordelia Langford, John Sillitoe on behalf of the Wellcome Sanger Institute COVID-19 Surveillance Team |
| EPI_ISL_2848699 | Lighthouse Lab in Glasgow | Wellcome Sanger Institute for the COVID-19 Genomics UK (COG-UK) Consortium | Harper VanSteenhouse, Yumi Kasai, David Gray, Carol Clugston, Anna Dominiczak and Alex Alderton, Roberto Amato, Jeffrey Barrett, Sonia Goncalves, Ewan Harrison, David K. Jackson, Ian Johnston, Dominic Kwiatkowski, Cordelia Langford, John Sillitoe on behalf of the Wellcome Sanger Institute COVID-19 Surveillance Team |
| EPI_ISL_2848700, EPI_ISL_2848701, EPI_ISL_2848702 | Lighthouse Lab in Milton Keynes | Wellcome Sanger Institute for the COVID-19 Genomics UK (COG-UK) Consortium | The Lighthouse Lab in Milton Keynes and Alex Alderton, Roberto Amato, Jeffrey Barrett, Sonia Goncalves, Ewan Harrison, David K. Jackson, Ian Johnston, Dominic Kwiatkowski, Cordelia Langford, John Sillitoe on behalf of the Wellcome Sanger Institute COVID-19 Surveillance Team |
| EPI_ISL_2848703 | Lighthouse Lab in Glasgow | Wellcome Sanger Institute for the COVID-19 Genomics UK (COG-UK) Consortium | Harper VanSteenhouse, Yumi Kasai, David Gray, Carol Clugston, Anna Dominiczak and Alex Alderton, Roberto Amato, Jeffrey Barrett, Sonia Goncalves, Ewan Harrison, David K. Jackson, Ian Johnston, Dominic Kwiatkowski, Cordelia Langford, John Sillitoe on behalf of the Wellcome Sanger Institute COVID-19 Surveillance Team |
| EPI_ISL_2848704, EPI_ISL_2848705 | Lighthouse Lab in Milton Keynes | Wellcome Sanger Institute for the COVID-19 Genomics UK (COG-UK) Consortium | The Lighthouse Lab in Milton Keynes and Alex Alderton, Roberto Amato, Jeffrey Barrett, Sonia Goncalves, Ewan Harrison, David K. Jackson, Ian Johnston, Dominic Kwiatkowski, Cordelia Langford, John Sillitoe on behalf of the Wellcome Sanger Institute COVID-19 Surveillance Team |
| EPI_ISL_2848706 | Lighthouse Lab in Glasgow | Wellcome Sanger Institute for the COVID-19 Genomics UK (COG-UK) Consortium | Harper VanSteenhouse, Yumi Kasai, David Gray, Carol Clugston, Anna Dominiczak and Alex Alderton, Roberto Amato, Jeffrey Barrett, Sonia Goncalves, Ewan Harrison, David K. Jackson, Ian Johnston, Dominic Kwiatkowski, Cordelia Langford, John Sillitoe on behalf of the Wellcome Sanger Institute COVID-19 Surveillance Team |
| EPI_ISL_2848707, EPI_ISL_2848708 | Lighthouse Lab in Milton Keynes | Wellcome Sanger Institute for the COVID-19 Genomics UK (COG-UK) Consortium | The Lighthouse Lab in Milton Keynes and Alex Alderton, Roberto Amato, Jeffrey Barrett, Sonia Goncalves, Ewan Harrison, David K. Jackson, Ian Johnston, Dominic Kwiatkowski, Cordelia Langford, John Sillitoe on behalf of the Wellcome Sanger Institute COVID-19 Surveillance Team |
| EPI_ISL_2848709 | AZ Klina | AZ Klina | Carl Vael - Lynsey Berckmans |
| EPI_ISL_2848710, EPI_ISL_2848711, EPI_ISL_2848712, EPI_ISL_2848713 | Lighthouse Lab in Milton Keynes | Wellcome Sanger Institute for the COVID-19 Genomics UK (COG-UK) Consortium | The Lighthouse Lab in Milton Keynes and Alex Alderton, Roberto Amato, Jeffrey Barrett, Sonia Goncalves, Ewan Harrison, David K. Jackson, Ian Johnston, Dominic Kwiatkowski, Cordelia Langford, John Sillitoe on behalf of the Wellcome Sanger Institute COVID-19 Surveillance Team |
| EPI_ISL_2848714, EPI_ISL_2848715 | Lighthouse Lab in Glasgow | Wellcome Sanger Institute for the COVID-19 Genomics UK (COG-UK) Consortium | Harper VanSteenhouse, Yumi Kasai, David Gray, Carol Clugston, Anna Dominiczak and Alex Alderton, Roberto Amato, Jeffrey Barrett, Sonia Goncalves, Ewan Harrison, David K. Jackson, Ian Johnston, Dominic Kwiatkowski, Cordelia Langford, John Sillitoe on behalf of the Wellcome Sanger Institute COVID-19 Surveillance Team |
| EPI_ISL_2848716 | Lighthouse Lab in Milton Keynes | Wellcome Sanger Institute for the COVID-19 Genomics UK (COG-UK) Consortium | The Lighthouse Lab in Milton Keynes and Alex Alderton, Roberto Amato, Jeffrey Barrett, Sonia Goncalves, Ewan Harrison, David K. Jackson, Ian Johnston, Dominic Kwiatkowski, Cordelia Langford, John Sillitoe on behalf of the Wellcome Sanger Institute COVID-19 Surveillance Team |
| EPI_ISL_2848717, EPI_ISL_2848718 | Lighthouse Lab in Glasgow | Wellcome Sanger Institute for the COVID-19 Genomics UK (COG-UK) Consortium | Harper VanSteenhouse, Yumi Kasai, David Gray, Carol Clugston, Anna Dominiczak and Alex Alderton, Roberto Amato, Jeffrey Barrett, Sonia Goncalves, Ewan Harrison, David K. Jackson, Ian Johnston, Dominic Kwiatkowski, Cordelia Langford, John Sillitoe on behalf of the Wellcome Sanger Institute COVID-19 Surveillance Team |
| EPI_ISL_2848719, EPI_ISL_2848720, EPI_ISL_2848721, EPI_ISL_2848722, EPI_ISL_2848723 | Lighthouse Lab in Milton Keynes | Wellcome Sanger Institute for the COVID-19 Genomics UK (COG-UK) Consortium | The Lighthouse Lab in Milton Keynes and Alex Alderton, Roberto Amato, Jeffrey Barrett, Sonia Goncalves, Ewan Harrison, David K. Jackson, Ian Johnston, Dominic Kwiatkowski, Cordelia Langford, John Sillitoe on behalf of the Wellcome Sanger Institute COVID-19 Surveillance Team |
| EPI_ISL_2848724 | Lighthouse Lab in Glasgow | Wellcome Sanger Institute for the COVID-19 Genomics UK (COG-UK) Consortium | Harper VanSteenhouse, Yumi Kasai, David Gray, Carol Clugston, Anna Dominiczak and Alex Alderton, Roberto Amato, Jeffrey Barrett, Sonia Goncalves, Ewan Harrison, David K. Jackson, Ian Johnston, Dominic Kwiatkowski, Cordelia Langford, John Sillitoe on behalf of the Wellcome Sanger Institute COVID-19 Surveillance Team |
| EPI_ISL_2848725 | Lighthouse Lab in Milton Keynes | Wellcome Sanger Institute for the COVID-19 Genomics UK (COG-UK) | The Lighthouse Lab in Milton Keynes and Alex Alderton, Roberto Amato, Jeffrey Barrett, Sonia Goncalves, Ewan Harrison, David K. Jackson, Ian Johnston, Dominic Kwiatkowski, Cordelia Langford, John Sillitoe on behalf of the Wellcome Sanger |

| | | | |
|---|---|---|---|
| | | Consortium | Institute COVID-19 Surveillance Team |
| EPI_ISL_2848726, EPI_ISL_2848727, EPI_ISL_2848728, EPI_ISL_2848729, EPI_ISL_2848730 | Lighthouse Lab in Glasgow | Wellcome Sanger Institute for the COVID-19 Genomics UK (COG-UK) Consortium | Harper VanSteenhouse, Yumi Kasai, David Gray, Carol Clugston, Anna Dominiczak and Alex Alderton, Roberto Amato, Jeffrey Barrett, Sonia Goncalves, Ewan Harrison, David K. Jackson, Ian Johnston, Dominic Kwiatkowski, Cordelia Langford, John Sillitoe on behalf of the Wellcome Sanger Institute COVID-19 Surveillance Team |
| EPI_ISL_2848731 | Lighthouse Lab in Milton Keynes | Wellcome Sanger Institute for the COVID-19 Genomics UK (COG-UK) Consortium | The Lighthouse Lab in Milton Keynes and Alex Alderton, Roberto Amato, Jeffrey Barrett, Sonia Goncalves, Ewan Harrison, David K. Jackson, Ian Johnston, Dominic Kwiatkowski, Cordelia Langford, John Sillitoe on behalf of the Wellcome Sanger Institute COVID-19 Surveillance Team |
| EPI_ISL_2848732 | Lighthouse Lab in Glasgow | Wellcome Sanger Institute for the COVID-19 Genomics UK (COG-UK) Consortium | Harper VanSteenhouse, Yumi Kasai, David Gray, Carol Clugston, Anna Dominiczak and Alex Alderton, Roberto Amato, Jeffrey Barrett, Sonia Goncalves, Ewan Harrison, David K. Jackson, Ian Johnston, Dominic Kwiatkowski, Cordelia Langford, John Sillitoe on behalf of the Wellcome Sanger Institute COVID-19 Surveillance Team |
| EPI_ISL_2848733, EPI_ISL_2848734 | Lighthouse Lab in Milton Keynes | Wellcome Sanger Institute for the COVID-19 Genomics UK (COG-UK) Consortium | The Lighthouse Lab in Milton Keynes and Alex Alderton, Roberto Amato, Jeffrey Barrett, Sonia Goncalves, Ewan Harrison, David K. Jackson, Ian Johnston, Dominic Kwiatkowski, Cordelia Langford, John Sillitoe on behalf of the Wellcome Sanger Institute COVID-19 Surveillance Team |
| EPI_ISL_2848735, EPI_ISL_2848736 | Lighthouse Lab in Glasgow | Wellcome Sanger Institute for the COVID-19 Genomics UK (COG-UK) Consortium | Harper VanSteenhouse, Yumi Kasai, David Gray, Carol Clugston, Anna Dominiczak and Alex Alderton, Roberto Amato, Jeffrey Barrett, Sonia Goncalves, Ewan Harrison, David K. Jackson, Ian Johnston, Dominic Kwiatkowski, Cordelia Langford, John Sillitoe on behalf of the Wellcome Sanger Institute COVID-19 Surveillance Team |
| EPI_ISL_2848737, EPI_ISL_2848738 | Lighthouse Lab in Milton Keynes | Wellcome Sanger Institute for the COVID-19 Genomics UK (COG-UK) Consortium | The Lighthouse Lab in Milton Keynes and Alex Alderton, Roberto Amato, Jeffrey Barrett, Sonia Goncalves, Ewan Harrison, David K. Jackson, Ian Johnston, Dominic Kwiatkowski, Cordelia Langford, John Sillitoe on behalf of the Wellcome Sanger Institute COVID-19 Surveillance Team |
| EPI_ISL_2848739 | Lighthouse Lab in Glasgow | Wellcome Sanger Institute for the COVID-19 Genomics UK (COG-UK) Consortium | Harper VanSteenhouse, Yumi Kasai, David Gray, Carol Clugston, Anna Dominiczak and Alex Alderton, Roberto Amato, Jeffrey Barrett, Sonia Goncalves, Ewan Harrison, David K. Jackson, Ian Johnston, Dominic Kwiatkowski, Cordelia Langford, John Sillitoe on behalf of the Wellcome Sanger Institute COVID-19 Surveillance Team |
| EPI_ISL_2848740, EPI_ISL_2848741, EPI_ISL_2848742, EPI_ISL_2848743, EPI_ISL_2848744, EPI_ISL_2848745, EPI_ISL_2848746, EPI_ISL_2848747, EPI_ISL_2848748, EPI_ISL_2848749, EPI_ISL_2848750, EPI_ISL_2848751, EPI_ISL_2848752, EPI_ISL_2848753 | | | |
| see above | Lighthouse Lab in Milton Keynes | Wellcome Sanger Institute for the COVID-19 Genomics UK (COG-UK) Consortium | The Lighthouse Lab in Milton Keynes and Alex Alderton, Roberto Amato, Jeffrey Barrett, Sonia Goncalves, Ewan Harrison, David K. Jackson, Ian Johnston, Dominic Kwiatkowski, Cordelia Langford, John Sillitoe on behalf of the Wellcome Sanger Institute COVID-19 Surveillance Team |
| EPI_ISL_2848754 | Lighthouse Lab in Glasgow | Wellcome Sanger Institute for the COVID-19 Genomics UK (COG-UK) Consortium | Harper VanSteenhouse, Yumi Kasai, David Gray, Carol Clugston, Anna Dominiczak and Alex Alderton, Roberto Amato, Jeffrey Barrett, Sonia Goncalves, Ewan Harrison, David K. Jackson, Ian Johnston, Dominic Kwiatkowski, Cordelia Langford, John Sillitoe on behalf of the Wellcome Sanger Institute COVID-19 Surveillance Team |
| EPI_ISL_2848755, EPI_ISL_2848756, EPI_ISL_2848757, EPI_ISL_2848758, EPI_ISL_2848759 | Lighthouse Lab in Milton Keynes | Wellcome Sanger Institute for the COVID-19 Genomics UK (COG-UK) Consortium | The Lighthouse Lab in Milton Keynes and Alex Alderton, Roberto Amato, Jeffrey Barrett, Sonia Goncalves, Ewan Harrison, David K. Jackson, Ian Johnston, Dominic Kwiatkowski, Cordelia Langford, John Sillitoe on behalf of the Wellcome Sanger Institute COVID-19 Surveillance Team |
| EPI_ISL_2848760, EPI_ISL_2848761 | Lighthouse Lab in Glasgow | Wellcome Sanger Institute for the COVID-19 Genomics UK (COG-UK) Consortium | Harper VanSteenhouse, Yumi Kasai, David Gray, Carol Clugston, Anna Dominiczak and Alex Alderton, Roberto Amato, Jeffrey Barrett, Sonia Goncalves, Ewan Harrison, David K. Jackson, Ian Johnston, Dominic Kwiatkowski, Cordelia Langford, John Sillitoe on behalf of the Wellcome Sanger Institute COVID-19 Surveillance Team |
| EPI_ISL_2848762 | Lighthouse Lab in Milton Keynes | Wellcome Sanger Institute for the COVID-19 Genomics UK (COG-UK) Consortium | The Lighthouse Lab in Milton Keynes and Alex Alderton, Roberto Amato, Jeffrey Barrett, Sonia Goncalves, Ewan Harrison, David K. Jackson, Ian Johnston, Dominic Kwiatkowski, Cordelia Langford, John Sillitoe on behalf of the Wellcome Sanger Institute COVID-19 Surveillance Team |
| EPI_ISL_2848763 | Lighthouse Lab in Glasgow | Wellcome Sanger Institute for the COVID-19 Genomics UK (COG-UK) Consortium | Harper VanSteenhouse, Yumi Kasai, David Gray, Carol Clugston, Anna Dominiczak and Alex Alderton, Roberto Amato, Jeffrey Barrett, Sonia Goncalves, Ewan Harrison, David K. Jackson, Ian Johnston, Dominic Kwiatkowski, Cordelia Langford, John Sillitoe on behalf of the Wellcome Sanger Institute COVID-19 Surveillance Team |
| EPI_ISL_2848764, EPI_ISL_2848765 | Lighthouse Lab in Milton Keynes | Wellcome Sanger Institute for the COVID-19 Genomics UK (COG-UK) Consortium | The Lighthouse Lab in Milton Keynes and Alex Alderton, Roberto Amato, Jeffrey Barrett, Sonia Goncalves, Ewan Harrison, David K. Jackson, Ian Johnston, Dominic Kwiatkowski, Cordelia Langford, John Sillitoe on behalf of the Wellcome Sanger Institute COVID-19 Surveillance Team |
| EPI_ISL_2848766, EPI_ISL_2848767 | Lighthouse Lab in Glasgow | Wellcome Sanger Institute for the COVID-19 Genomics UK (COG-UK) Consortium | Harper VanSteenhouse, Yumi Kasai, David Gray, Carol Clugston, Anna Dominiczak and Alex Alderton, Roberto Amato, Jeffrey Barrett, Sonia Goncalves, Ewan Harrison, David K. Jackson, Ian Johnston, Dominic Kwiatkowski, Cordelia Langford, John Sillitoe on behalf of the Wellcome Sanger Institute COVID-19 Surveillance Team |
| EPI_ISL_2848768, EPI_ISL_2848769, EPI_ISL_2848770, EPI_ISL_2848771 | Lighthouse Lab in Milton Keynes | Wellcome Sanger Institute for the COVID-19 Genomics UK (COG-UK) Consortium | The Lighthouse Lab in Milton Keynes and Alex Alderton, Roberto Amato, Jeffrey Barrett, Sonia Goncalves, Ewan Harrison, David K. Jackson, Ian Johnston, Dominic Kwiatkowski, Cordelia Langford, John Sillitoe on behalf of the Wellcome Sanger Institute COVID-19 Surveillance Team |
| EPI_ISL_2848772 | Lighthouse Lab in Glasgow | Wellcome Sanger Institute for the COVID-19 Genomics UK (COG-UK) Consortium | Harper VanSteenhouse, Yumi Kasai, David Gray, Carol Clugston, Anna Dominiczak and Alex Alderton, Roberto Amato, Jeffrey Barrett, Sonia Goncalves, Ewan Harrison, David K. Jackson, Ian Johnston, Dominic Kwiatkowski, Cordelia Langford, John Sillitoe on behalf of the Wellcome Sanger Institute COVID-19 Surveillance Team |
| EPI_ISL_2848773, EPI_ISL_2848774, EPI_ISL_2848775, EPI_ISL_2848776, EPI_ISL_2848777, EPI_ISL_2848778, EPI_ISL_2848779, EPI_ISL_2848780, EPI_ISL_2848781, EPI_ISL_2848782 | Lighthouse Lab in Milton Keynes | Wellcome Sanger Institute for the COVID-19 Genomics UK (COG-UK) Consortium | The Lighthouse Lab in Milton Keynes and Alex Alderton, Roberto Amato, Jeffrey Barrett, Sonia Goncalves, Ewan Harrison, David K. Jackson, Ian Johnston, Dominic Kwiatkowski, Cordelia Langford, John Sillitoe on behalf of the Wellcome Sanger Institute COVID-19 Surveillance Team |
| EPI_ISL_2848783 | Lighthouse Lab in Glasgow | Wellcome Sanger Institute for the COVID-19 Genomics UK (COG-UK) Consortium | Harper VanSteenhouse, Yumi Kasai, David Gray, Carol Clugston, Anna Dominiczak and Alex Alderton, Roberto Amato, Jeffrey Barrett, Sonia Goncalves, Ewan Harrison, David K. Jackson, Ian Johnston, Dominic Kwiatkowski, Cordelia Langford, John Sillitoe on behalf of the Wellcome Sanger Institute COVID-19 Surveillance Team |
| EPI_ISL_2848784, EPI_ISL_2848785, EPI_ISL_2848786, EPI_ISL_2848787, EPI_ISL_2848788, EPI_ISL_2848789, EPI_ISL_2848790, EPI_ISL_2848791, EPI_ISL_2848792, EPI_ISL_2848793, EPI_ISL_2848794, EPI_ISL_2848795 | | | |
| see above | Lighthouse Lab in Milton Keynes | Wellcome Sanger Institute for the COVID-19 Genomics UK (COG-UK) Consortium | The Lighthouse Lab in Milton Keynes and Alex Alderton, Roberto Amato, Jeffrey Barrett, Sonia Goncalves, Ewan Harrison, David K. Jackson, Ian Johnston, Dominic Kwiatkowski, Cordelia Langford, John Sillitoe on behalf of the Wellcome Sanger Institute COVID-19 Surveillance Team |
| EPI_ISL_2848796 | Lighthouse Lab in Glasgow | Wellcome Sanger Institute for the COVID-19 Genomics UK (COG-UK) Consortium | Harper VanSteenhouse, Yumi Kasai, David Gray, Carol Clugston, Anna Dominiczak and Alex Alderton, Roberto Amato, Jeffrey Barrett, Sonia Goncalves, Ewan Harrison, David K. Jackson, Ian Johnston, Dominic Kwiatkowski, Cordelia Langford, John Sillitoe on behalf of the Wellcome Sanger Institute COVID-19 Surveillance Team |
| EPI_ISL_2848797, EPI_ISL_2848798 | Lighthouse Lab in Milton Keynes | Wellcome Sanger Institute for the COVID-19 Genomics UK (COG-UK) Consortium | The Lighthouse Lab in Milton Keynes and Alex Alderton, Roberto Amato, Jeffrey Barrett, Sonia Goncalves, Ewan Harrison, David K. Jackson, Ian Johnston, Dominic Kwiatkowski, Cordelia Langford, John Sillitoe on behalf of the Wellcome Sanger Institute COVID-19 Surveillance Team |
| EPI_ISL_2848799 | Lighthouse Lab in Glasgow | Wellcome Sanger Institute for the COVID-19 Genomics UK (COG-UK) Consortium | Harper VanSteenhouse, Yumi Kasai, David Gray, Carol Clugston, Anna Dominiczak and Alex Alderton, Roberto Amato, Jeffrey Barrett, Sonia Goncalves, Ewan Harrison, David K. Jackson, Ian Johnston, Dominic Kwiatkowski, Cordelia Langford, John Sillitoe on behalf of the Wellcome Sanger Institute COVID-19 Surveillance Team |
| EPI_ISL_2848800, EPI_ISL_2848801 | Lighthouse Lab in Milton Keynes | Wellcome Sanger Institute for the COVID-19 Genomics UK (COG-UK) Consortium | The Lighthouse Lab in Milton Keynes and Alex Alderton, Roberto Amato, Jeffrey Barrett, Sonia Goncalves, Ewan Harrison, David K. Jackson, Ian Johnston, Dominic Kwiatkowski, Cordelia Langford, John Sillitoe on behalf of the Wellcome Sanger Institute COVID-19 Surveillance Team |
| EPI_ISL_2848802 | Lighthouse Lab in Glasgow | Wellcome Sanger Institute for the COVID-19 Genomics UK (COG-UK) Consortium | Harper VanSteenhouse, Yumi Kasai, David Gray, Carol Clugston, Anna Dominiczak and Alex Alderton, Roberto Amato, Jeffrey Barrett, Sonia Goncalves, Ewan Harrison, David K. Jackson, Ian Johnston, Dominic Kwiatkowski, Cordelia Langford, John Sillitoe on behalf of the Wellcome Sanger Institute COVID-19 Surveillance Team |
| EPI_ISL_2848803, EPI_ISL_2848804 | Lighthouse Lab in Milton Keynes | Wellcome Sanger Institute for the COVID-19 Genomics UK (COG-UK) Consortium | The Lighthouse Lab in Milton Keynes and Alex Alderton, Roberto Amato, Jeffrey Barrett, Sonia Goncalves, Ewan Harrison, David K. Jackson, Ian Johnston, Dominic Kwiatkowski, Cordelia Langford, John Sillitoe on behalf of the Wellcome Sanger Institute COVID-19 Surveillance Team |
| EPI_ISL_2848805 | Lighthouse Lab in Glasgow | Wellcome Sanger Institute for the COVID-19 Genomics UK (COG-UK) Consortium | Harper VanSteenhouse, Yumi Kasai, David Gray, Carol Clugston, Anna Dominiczak and Alex Alderton, Roberto Amato, Jeffrey Barrett, Sonia Goncalves, Ewan Harrison, David K. Jackson, Ian Johnston, Dominic Kwiatkowski, Cordelia Langford, John Sillitoe on behalf of the Wellcome Sanger Institute COVID-19 Surveillance Team |
| EPI_ISL_2848806, EPI_ISL_2848807 | Lighthouse Lab in Milton Keynes | Wellcome Sanger Institute for the COVID-19 Genomics UK (COG-UK) Consortium | The Lighthouse Lab in Milton Keynes and Alex Alderton, Roberto Amato, Jeffrey Barrett, Sonia Goncalves, Ewan Harrison, David K. Jackson, Ian Johnston, Dominic Kwiatkowski, Cordelia Langford, John Sillitoe on behalf of the Wellcome Sanger Institute COVID-19 Surveillance Team |
| EPI_ISL_2848808, EPI_ISL_2848809 | Lighthouse Lab in Glasgow | Wellcome Sanger Institute for the COVID-19 Genomics UK (COG-UK) Consortium | Harper VanSteenhouse, Yumi Kasai, David Gray, Carol Clugston, Anna Dominiczak and Alex Alderton, Roberto Amato, Jeffrey Barrett, Sonia Goncalves, Ewan Harrison, David K. Jackson, Ian Johnston, Dominic Kwiatkowski, Cordelia Langford, John Sillitoe on behalf of the Wellcome Sanger Institute COVID-19 Surveillance Team |
| EPI_ISL_2848810 | Lighthouse Lab in Milton Keynes | Wellcome Sanger Institute for the COVID-19 Genomics UK (COG-UK) Consortium | The Lighthouse Lab in Milton Keynes and Alex Alderton, Roberto Amato, Jeffrey Barrett, Sonia Goncalves, Ewan Harrison, David K. Jackson, Ian Johnston, Dominic Kwiatkowski, Cordelia Langford, John Sillitoe on behalf of the Wellcome Sanger Institute COVID-19 Surveillance Team |
| EPI_ISL_2848811 | Lighthouse Lab in Glasgow | Wellcome Sanger Institute for the COVID-19 Genomics UK (COG-UK) Consortium | Harper VanSteenhouse, Yumi Kasai, David Gray, Carol Clugston, Anna Dominiczak and Alex Alderton, Roberto Amato, Jeffrey Barrett, Sonia Goncalves, Ewan Harrison, David K. Jackson, Ian Johnston, Dominic Kwiatkowski, Cordelia Langford, John Sillitoe on behalf of the Wellcome Sanger Institute COVID-19 Surveillance Team |
| EPI_ISL_2848812, EPI_ISL_2848813, EPI_ISL_2848814, EPI_ISL_2848815 | Lighthouse Lab in Milton Keynes | Wellcome Sanger Institute for the COVID-19 Genomics UK (COG-UK) Consortium | The Lighthouse Lab in Milton Keynes and Alex Alderton, Roberto Amato, Jeffrey Barrett, Sonia Goncalves, Ewan Harrison, David K. Jackson, Ian Johnston, Dominic Kwiatkowski, Cordelia Langford, John Sillitoe on behalf of the Wellcome Sanger Institute COVID-19 Surveillance Team |
| EPI_ISL_2848816 | Lighthouse Lab in Glasgow | Wellcome Sanger Institute for the COVID-19 Genomics UK (COG-UK) Consortium | Harper VanSteenhouse, Yumi Kasai, David Gray, Carol Clugston, Anna Dominiczak and Alex Alderton, Roberto Amato, Jeffrey Barrett, Sonia Goncalves, Ewan Harrison, David K. Jackson, Ian Johnston, Dominic Kwiatkowski, Cordelia Langford, John Sillitoe on behalf of the Wellcome Sanger Institute COVID-19 Surveillance Team |
| EPI_ISL_2848817, EPI_ISL_2848818, EPI_ISL_2848819, EPI_ISL_2848820 | Lighthouse Lab in Milton Keynes | Wellcome Sanger Institute for the COVID-19 Genomics UK (COG-UK) Consortium | The Lighthouse Lab in Milton Keynes and Alex Alderton, Roberto Amato, Jeffrey Barrett, Sonia Goncalves, Ewan Harrison, David K. Jackson, Ian Johnston, Dominic Kwiatkowski, Cordelia Langford, John Sillitoe on behalf of the Wellcome Sanger Institute COVID-19 Surveillance Team |
| EPI_ISL_2848821 | Lighthouse Lab in Glasgow | Wellcome Sanger Institute for the COVID-19 Genomics UK (COG-UK) | Harper VanSteenhouse, Yumi Kasai, David Gray, Carol Clugston, Anna Dominiczak and Alex Alderton, Roberto Amato, Jeffrey Barrett, Sonia Goncalves, Ewan Harrison, David K. Jackson, Ian Johnston, Dominic Kwiatkowski, Cordelia Langford, |

| | | | |
|---|---|---|---|
| | | Consortium | John Sillitoe on behalf of the Wellcome Sanger Institute COVID-19 Surveillance Team |
| EPI_ISL_2848822, EPI_ISL_2848823, EPI_ISL_2848824, EPI_ISL_2848825, EPI_ISL_2848826, EPI_ISL_2848827 | Lighthouse Lab in Milton Keynes | Wellcome Sanger Institute for the COVID-19 Genomics UK (COG-UK) Consortium | The Lighthouse Lab in Milton Keynes and Alex Alderton, Roberto Amato, Jeffrey Barrett, Sonia Goncalves, Ewan Harrison, David K. Jackson, Ian Johnston, Dominic Kwiatkowski, Cordelia Langford, John Sillitoe on behalf of the Wellcome Sanger Institute COVID-19 Surveillance Team |
| EPI_ISL_2848828 | Lighthouse Lab in Glasgow | Wellcome Sanger Institute for the COVID-19 Genomics UK (COG-UK) Consortium | Harper VanSteenhouse, Yumi Kasai, David Gray, Carol Clugston, Anna Dominiczak and Alex Alderton, Roberto Amato, Jeffrey Barrett, Sonia Goncalves, Ewan Harrison, David K. Jackson, Ian Johnston, Dominic Kwiatkowski, Cordelia Langford, John Sillitoe on behalf of the Wellcome Sanger Institute COVID-19 Surveillance Team |
| EPI_ISL_2848829, EPI_ISL_2848830, EPI_ISL_2848831, EPI_ISL_2848832 | Lighthouse Lab in Milton Keynes | Wellcome Sanger Institute for the COVID-19 Genomics UK (COG-UK) Consortium | The Lighthouse Lab in Milton Keynes and Alex Alderton, Roberto Amato, Jeffrey Barrett, Sonia Goncalves, Ewan Harrison, David K. Jackson, Ian Johnston, Dominic Kwiatkowski, Cordelia Langford, John Sillitoe on behalf of the Wellcome Sanger Institute COVID-19 Surveillance Team |
| EPI_ISL_2848833 | Lighthouse Lab in Glasgow | Wellcome Sanger Institute for the COVID-19 Genomics UK (COG-UK) Consortium | Harper VanSteenhouse, Yumi Kasai, David Gray, Carol Clugston, Anna Dominiczak and Alex Alderton, Roberto Amato, Jeffrey Barrett, Sonia Goncalves, Ewan Harrison, David K. Jackson, Ian Johnston, Dominic Kwiatkowski, Cordelia Langford, John Sillitoe on behalf of the Wellcome Sanger Institute COVID-19 Surveillance Team |
| EPI_ISL_2848834 | Lighthouse Lab in Milton Keynes | Wellcome Sanger Institute for the COVID-19 Genomics UK (COG-UK) Consortium | The Lighthouse Lab in Milton Keynes and Alex Alderton, Roberto Amato, Jeffrey Barrett, Sonia Goncalves, Ewan Harrison, David K. Jackson, Ian Johnston, Dominic Kwiatkowski, Cordelia Langford, John Sillitoe on behalf of the Wellcome Sanger Institute COVID-19 Surveillance Team |
| EPI_ISL_2848835 | Lighthouse Lab in Glasgow | Wellcome Sanger Institute for the COVID-19 Genomics UK (COG-UK) Consortium | Harper VanSteenhouse, Yumi Kasai, David Gray, Carol Clugston, Anna Dominiczak and Alex Alderton, Roberto Amato, Jeffrey Barrett, Sonia Goncalves, Ewan Harrison, David K. Jackson, Ian Johnston, Dominic Kwiatkowski, Cordelia Langford, John Sillitoe on behalf of the Wellcome Sanger Institute COVID-19 Surveillance Team |
| EPI_ISL_2848836, EPI_ISL_2848837 | Lighthouse Lab in Milton Keynes | Wellcome Sanger Institute for the COVID-19 Genomics UK (COG-UK) Consortium | The Lighthouse Lab in Milton Keynes and Alex Alderton, Roberto Amato, Jeffrey Barrett, Sonia Goncalves, Ewan Harrison, David K. Jackson, Ian Johnston, Dominic Kwiatkowski, Cordelia Langford, John Sillitoe on behalf of the Wellcome Sanger Institute COVID-19 Surveillance Team |
| EPI_ISL_2848838, EPI_ISL_2848839 | Lighthouse Lab in Glasgow | Wellcome Sanger Institute for the COVID-19 Genomics UK (COG-UK) Consortium | Harper VanSteenhouse, Yumi Kasai, David Gray, Carol Clugston, Anna Dominiczak and Alex Alderton, Roberto Amato, Jeffrey Barrett, Sonia Goncalves, Ewan Harrison, David K. Jackson, Ian Johnston, Dominic Kwiatkowski, Cordelia Langford, John Sillitoe on behalf of the Wellcome Sanger Institute COVID-19 Surveillance Team |
| EPI_ISL_2848840, EPI_ISL_2848841 | Lighthouse Lab in Milton Keynes | Wellcome Sanger Institute for the COVID-19 Genomics UK (COG-UK) Consortium | The Lighthouse Lab in Milton Keynes and Alex Alderton, Roberto Amato, Jeffrey Barrett, Sonia Goncalves, Ewan Harrison, David K. Jackson, Ian Johnston, Dominic Kwiatkowski, Cordelia Langford, John Sillitoe on behalf of the Wellcome Sanger Institute COVID-19 Surveillance Team |
| EPI_ISL_2848842 | Lighthouse Lab in Glasgow | Wellcome Sanger Institute for the COVID-19 Genomics UK (COG-UK) Consortium | Harper VanSteenhouse, Yumi Kasai, David Gray, Carol Clugston, Anna Dominiczak and Alex Alderton, Roberto Amato, Jeffrey Barrett, Sonia Goncalves, Ewan Harrison, David K. Jackson, Ian Johnston, Dominic Kwiatkowski, Cordelia Langford, John Sillitoe on behalf of the Wellcome Sanger Institute COVID-19 Surveillance Team |
| EPI_ISL_2848843, EPI_ISL_2848844 | Lighthouse Lab in Milton Keynes | Wellcome Sanger Institute for the COVID-19 Genomics UK (COG-UK) Consortium | The Lighthouse Lab in Milton Keynes and Alex Alderton, Roberto Amato, Jeffrey Barrett, Sonia Goncalves, Ewan Harrison, David K. Jackson, Ian Johnston, Dominic Kwiatkowski, Cordelia Langford, John Sillitoe on behalf of the Wellcome Sanger Institute COVID-19 Surveillance Team |
| EPI_ISL_2848845, EPI_ISL_2848846 | Lighthouse Lab in Glasgow | Wellcome Sanger Institute for the COVID-19 Genomics UK (COG-UK) Consortium | Harper VanSteenhouse, Yumi Kasai, David Gray, Carol Clugston, Anna Dominiczak and Alex Alderton, Roberto Amato, Jeffrey Barrett, Sonia Goncalves, Ewan Harrison, David K. Jackson, Ian Johnston, Dominic Kwiatkowski, Cordelia Langford, John Sillitoe on behalf of the Wellcome Sanger Institute COVID-19 Surveillance Team |
| EPI_ISL_2848847, EPI_ISL_2848848, EPI_ISL_2848849, EPI_ISL_2848851, EPI_ISL_2848852 | Lighthouse Lab in Milton Keynes | Wellcome Sanger Institute for the COVID-19 Genomics UK (COG-UK) Consortium | The Lighthouse Lab in Milton Keynes and Alex Alderton, Roberto Amato, Jeffrey Barrett, Sonia Goncalves, Ewan Harrison, David K. Jackson, Ian Johnston, Dominic Kwiatkowski, Cordelia Langford, John Sillitoe on behalf of the Wellcome Sanger Institute COVID-19 Surveillance Team |
| EPI_ISL_2848853 | Lighthouse Lab in Glasgow | Wellcome Sanger Institute for the COVID-19 Genomics UK (COG-UK) Consortium | Harper VanSteenhouse, Yumi Kasai, David Gray, Carol Clugston, Anna Dominiczak and Alex Alderton, Roberto Amato, Jeffrey Barrett, Sonia Goncalves, Ewan Harrison, David K. Jackson, Ian Johnston, Dominic Kwiatkowski, Cordelia Langford, John Sillitoe on behalf of the Wellcome Sanger Institute COVID-19 Surveillance Team |
| EPI_ISL_2848854 | Lighthouse Lab in Milton Keynes | Wellcome Sanger Institute for the COVID-19 Genomics UK (COG-UK) Consortium | The Lighthouse Lab in Milton Keynes and Alex Alderton, Roberto Amato, Jeffrey Barrett, Sonia Goncalves, Ewan Harrison, David K. Jackson, Ian Johnston, Dominic Kwiatkowski, Cordelia Langford, John Sillitoe on behalf of the Wellcome Sanger Institute COVID-19 Surveillance Team |
| EPI_ISL_2848855 | Lighthouse Lab in Glasgow | Wellcome Sanger Institute for the COVID-19 Genomics UK (COG-UK) Consortium | Harper VanSteenhouse, Yumi Kasai, David Gray, Carol Clugston, Anna Dominiczak and Alex Alderton, Roberto Amato, Jeffrey Barrett, Sonia Goncalves, Ewan Harrison, David K. Jackson, Ian Johnston, Dominic Kwiatkowski, Cordelia Langford, John Sillitoe on behalf of the Wellcome Sanger Institute COVID-19 Surveillance Team |
| EPI_ISL_2848856, EPI_ISL_2848857, EPI_ISL_2848858, EPI_ISL_2848859 | Lighthouse Lab in Milton Keynes | Wellcome Sanger Institute for the COVID-19 Genomics UK (COG-UK) Consortium | The Lighthouse Lab in Milton Keynes and Alex Alderton, Roberto Amato, Jeffrey Barrett, Sonia Goncalves, Ewan Harrison, David K. Jackson, Ian Johnston, Dominic Kwiatkowski, Cordelia Langford, John Sillitoe on behalf of the Wellcome Sanger Institute COVID-19 Surveillance Team |
| EPI_ISL_2848860 | Lighthouse Lab in Glasgow | Wellcome Sanger Institute for the COVID-19 Genomics UK (COG-UK) Consortium | Harper VanSteenhouse, Yumi Kasai, David Gray, Carol Clugston, Anna Dominiczak and Alex Alderton, Roberto Amato, Jeffrey Barrett, Sonia Goncalves, Ewan Harrison, David K. Jackson, Ian Johnston, Dominic Kwiatkowski, Cordelia Langford, John Sillitoe on behalf of the Wellcome Sanger Institute COVID-19 Surveillance Team |
| EPI_ISL_2848861, EPI_ISL_2848862, EPI_ISL_2848863, EPI_ISL_2848864 | Lighthouse Lab in Milton Keynes | Wellcome Sanger Institute for the COVID-19 Genomics UK (COG-UK) Consortium | The Lighthouse Lab in Milton Keynes and Alex Alderton, Roberto Amato, Jeffrey Barrett, Sonia Goncalves, Ewan Harrison, David K. Jackson, Ian Johnston, Dominic Kwiatkowski, Cordelia Langford, John Sillitoe on behalf of the Wellcome Sanger Institute COVID-19 Surveillance Team |
| EPI_ISL_2848865 | Lighthouse Lab in Glasgow | Wellcome Sanger Institute for the COVID-19 Genomics UK (COG-UK) Consortium | Harper VanSteenhouse, Yumi Kasai, David Gray, Carol Clugston, Anna Dominiczak and Alex Alderton, Roberto Amato, Jeffrey Barrett, Sonia Goncalves, Ewan Harrison, David K. Jackson, Ian Johnston, Dominic Kwiatkowski, Cordelia Langford, John Sillitoe on behalf of the Wellcome Sanger Institute COVID-19 Surveillance Team |
| EPI_ISL_2848866, EPI_ISL_2848867, EPI_ISL_2848868 | Lighthouse Lab in Milton Keynes | Wellcome Sanger Institute for the COVID-19 Genomics UK (COG-UK) Consortium | The Lighthouse Lab in Milton Keynes and Alex Alderton, Roberto Amato, Jeffrey Barrett, Sonia Goncalves, Ewan Harrison, David K. Jackson, Ian Johnston, Dominic Kwiatkowski, Cordelia Langford, John Sillitoe on behalf of the Wellcome Sanger Institute COVID-19 Surveillance Team |
| EPI_ISL_2848869, EPI_ISL_2848870 | Lighthouse Lab in Glasgow | Wellcome Sanger Institute for the COVID-19 Genomics UK (COG-UK) Consortium | Harper VanSteenhouse, Yumi Kasai, David Gray, Carol Clugston, Anna Dominiczak and Alex Alderton, Roberto Amato, Jeffrey Barrett, Sonia Goncalves, Ewan Harrison, David K. Jackson, Ian Johnston, Dominic Kwiatkowski, Cordelia Langford, John Sillitoe on behalf of the Wellcome Sanger Institute COVID-19 Surveillance Team |
| EPI_ISL_2848871, EPI_ISL_2848872, EPI_ISL_2848873, EPI_ISL_2848874, EPI_ISL_2848875, EPI_ISL_2848876 | Lighthouse Lab in Milton Keynes | Wellcome Sanger Institute for the COVID-19 Genomics UK (COG-UK) Consortium | The Lighthouse Lab in Milton Keynes and Alex Alderton, Roberto Amato, Jeffrey Barrett, Sonia Goncalves, Ewan Harrison, David K. Jackson, Ian Johnston, Dominic Kwiatkowski, Cordelia Langford, John Sillitoe on behalf of the Wellcome Sanger Institute COVID-19 Surveillance Team |
| EPI_ISL_2848877 | Lighthouse Lab in Glasgow | Wellcome Sanger Institute for the COVID-19 Genomics UK (COG-UK) Consortium | Harper VanSteenhouse, Yumi Kasai, David Gray, Carol Clugston, Anna Dominiczak and Alex Alderton, Roberto Amato, Jeffrey Barrett, Sonia Goncalves, Ewan Harrison, David K. Jackson, Ian Johnston, Dominic Kwiatkowski, Cordelia Langford, John Sillitoe on behalf of the Wellcome Sanger Institute COVID-19 Surveillance Team |
| EPI_ISL_2848878, EPI_ISL_2848879, EPI_ISL_2848880, EPI_ISL_2848881, EPI_ISL_2848882, EPI_ISL_2848883 | Lighthouse Lab in Milton Keynes | Wellcome Sanger Institute for the COVID-19 Genomics UK (COG-UK) Consortium | The Lighthouse Lab in Milton Keynes and Alex Alderton, Roberto Amato, Jeffrey Barrett, Sonia Goncalves, Ewan Harrison, David K. Jackson, Ian Johnston, Dominic Kwiatkowski, Cordelia Langford, John Sillitoe on behalf of the Wellcome Sanger Institute COVID-19 Surveillance Team |
| EPI_ISL_2848884 | Lighthouse Lab in Glasgow | Wellcome Sanger Institute for the COVID-19 Genomics UK (COG-UK) Consortium | Harper VanSteenhouse, Yumi Kasai, David Gray, Carol Clugston, Anna Dominiczak and Alex Alderton, Roberto Amato, Jeffrey Barrett, Sonia Goncalves, Ewan Harrison, David K. Jackson, Ian Johnston, Dominic Kwiatkowski, Cordelia Langford, John Sillitoe on behalf of the Wellcome Sanger Institute COVID-19 Surveillance Team |
| EPI_ISL_2848885 | Lighthouse Lab in Milton Keynes | Wellcome Sanger Institute for the COVID-19 Genomics UK (COG-UK) Consortium | The Lighthouse Lab in Milton Keynes and Alex Alderton, Roberto Amato, Jeffrey Barrett, Sonia Goncalves, Ewan Harrison, David K. Jackson, Ian Johnston, Dominic Kwiatkowski, Cordelia Langford, John Sillitoe on behalf of the Wellcome Sanger Institute COVID-19 Surveillance Team |
| EPI_ISL_2848886 | Lighthouse Lab in Glasgow | Wellcome Sanger Institute for the COVID-19 Genomics UK (COG-UK) Consortium | Harper VanSteenhouse, Yumi Kasai, David Gray, Carol Clugston, Anna Dominiczak and Alex Alderton, Roberto Amato, Jeffrey Barrett, Sonia Goncalves, Ewan Harrison, David K. Jackson, Ian Johnston, Dominic Kwiatkowski, Cordelia Langford, John Sillitoe on behalf of the Wellcome Sanger Institute COVID-19 Surveillance Team |
| EPI_ISL_2848887 | Lighthouse Lab in Milton Keynes | Wellcome Sanger Institute for the COVID-19 Genomics UK (COG-UK) Consortium | The Lighthouse Lab in Milton Keynes and Alex Alderton, Roberto Amato, Jeffrey Barrett, Sonia Goncalves, Ewan Harrison, David K. Jackson, Ian Johnston, Dominic Kwiatkowski, Cordelia Langford, John Sillitoe on behalf of the Wellcome Sanger Institute COVID-19 Surveillance Team |
| EPI_ISL_2848888 | Lighthouse Lab in Glasgow | Wellcome Sanger Institute for the COVID-19 Genomics UK (COG-UK) Consortium | Harper VanSteenhouse, Yumi Kasai, David Gray, Carol Clugston, Anna Dominiczak and Alex Alderton, Roberto Amato, Jeffrey Barrett, Sonia Goncalves, Ewan Harrison, David K. Jackson, Ian Johnston, Dominic Kwiatkowski, Cordelia Langford, John Sillitoe on behalf of the Wellcome Sanger Institute COVID-19 Surveillance Team |
| EPI_ISL_2848889, EPI_ISL_2848890 | Lighthouse Lab in Milton Keynes | Wellcome Sanger Institute for the COVID-19 Genomics UK (COG-UK) Consortium | The Lighthouse Lab in Milton Keynes and Alex Alderton, Roberto Amato, Jeffrey Barrett, Sonia Goncalves, Ewan Harrison, David K. Jackson, Ian Johnston, Dominic Kwiatkowski, Cordelia Langford, John Sillitoe on behalf of the Wellcome Sanger Institute COVID-19 Surveillance Team |
| EPI_ISL_2848891, EPI_ISL_2848892 | Lighthouse Lab in Glasgow | Wellcome Sanger Institute for the COVID-19 Genomics UK (COG-UK) Consortium | Harper VanSteenhouse, Yumi Kasai, David Gray, Carol Clugston, Anna Dominiczak and Alex Alderton, Roberto Amato, Jeffrey Barrett, Sonia Goncalves, Ewan Harrison, David K. Jackson, Ian Johnston, Dominic Kwiatkowski, Cordelia Langford, John Sillitoe on behalf of the Wellcome Sanger Institute COVID-19 Surveillance Team |
| EPI_ISL_2848893, EPI_ISL_2848894, EPI_ISL_2848895 | Lighthouse Lab in Milton Keynes | Wellcome Sanger Institute for the COVID-19 Genomics UK (COG-UK) Consortium | The Lighthouse Lab in Milton Keynes and Alex Alderton, Roberto Amato, Jeffrey Barrett, Sonia Goncalves, Ewan Harrison, David K. Jackson, Ian Johnston, Dominic Kwiatkowski, Cordelia Langford, John Sillitoe on behalf of the Wellcome Sanger Institute COVID-19 Surveillance Team |
| EPI_ISL_2848896 | Lighthouse Lab in Glasgow | Wellcome Sanger Institute for the COVID-19 Genomics UK (COG-UK) Consortium | Harper VanSteenhouse, Yumi Kasai, David Gray, Carol Clugston, Anna Dominiczak and Alex Alderton, Roberto Amato, Jeffrey Barrett, Sonia Goncalves, Ewan Harrison, David K. Jackson, Ian Johnston, Dominic Kwiatkowski, Cordelia Langford, John Sillitoe on behalf of the Wellcome Sanger Institute COVID-19 Surveillance Team |
| EPI_ISL_2848897, EPI_ISL_2848898, EPI_ISL_2848899, EPI_ISL_2848900, EPI_ISL_2848901, EPI_ISL_2848902, EPI_ISL_2848903, EPI_ISL_2848904, EPI_ISL_2848905, EPI_ISL_2848906, EPI_ISL_2848907, EPI_ISL_2848908, EPI_ISL_2848909, EPI_ISL_2848910, EPI_ISL_2848911, EPI_ISL_2848912, EPI_ISL_2848913, EPI_ISL_2848914 | | | |
| see above | Lighthouse Lab in Milton Keynes | Wellcome Sanger Institute for the COVID-19 Genomics UK (COG-UK) | The Lighthouse Lab in Milton Keynes and Alex Alderton, Roberto Amato, Jeffrey Barrett, Sonia Goncalves, Ewan Harrison, David K. Jackson, Ian Johnston, Dominic Kwiatkowski, Cordelia Langford, John Sillitoe on behalf of the Wellcome Sanger |

| | | | |
|---|---|---|---|
| | | Consortium | Institute COVID-19 Surveillance Team |
| EPI_ISL_2848915 | Lighthouse Lab in Glasgow | Wellcome Sanger Institute for the COVID-19 Genomics UK (COG-UK) Consortium | Harper VanSteenhouse, Yumi Kasai, David Gray, Carol Clugston, Anna Dominiczak and Alex Alderton, Roberto Amato, Jeffrey Barrett, Sonia Goncalves, Ewan Harrison, David K. Jackson, Ian Johnston, Dominic Kwiatkowski, Cordelia Langford, John Sillitoe on behalf of the Wellcome Sanger Institute COVID-19 Surveillance Team |
| EPI_ISL_2848916 | Lighthouse Lab in Milton Keynes | Wellcome Sanger Institute for the COVID-19 Genomics UK (COG-UK) Consortium | The Lighthouse Lab in Milton Keynes and Alex Alderton, Roberto Amato, Jeffrey Barrett, Sonia Goncalves, Ewan Harrison, David K. Jackson, Ian Johnston, Dominic Kwiatkowski, Cordelia Langford, John Sillitoe on behalf of the Wellcome Sanger Institute COVID-19 Surveillance Team |
| EPI_ISL_2848917, EPI_ISL_2848918 | Lighthouse Lab in Glasgow | Wellcome Sanger Institute for the COVID-19 Genomics UK (COG-UK) Consortium | Harper VanSteenhouse, Yumi Kasai, David Gray, Carol Clugston, Anna Dominiczak and Alex Alderton, Roberto Amato, Jeffrey Barrett, Sonia Goncalves, Ewan Harrison, David K. Jackson, Ian Johnston, Dominic Kwiatkowski, Cordelia Langford, John Sillitoe on behalf of the Wellcome Sanger Institute COVID-19 Surveillance Team |
| EPI_ISL_2848919, EPI_ISL_2848920, EPI_ISL_2848921 | Lighthouse Lab in Milton Keynes | Wellcome Sanger Institute for the COVID-19 Genomics UK (COG-UK) Consortium | The Lighthouse Lab in Milton Keynes and Alex Alderton, Roberto Amato, Jeffrey Barrett, Sonia Goncalves, Ewan Harrison, David K. Jackson, Ian Johnston, Dominic Kwiatkowski, Cordelia Langford, John Sillitoe on behalf of the Wellcome Sanger Institute COVID-19 Surveillance Team |
| EPI_ISL_2848922, EPI_ISL_2848923, EPI_ISL_2848924 | Lighthouse Lab in Glasgow | Wellcome Sanger Institute for the COVID-19 Genomics UK (COG-UK) Consortium | Harper VanSteenhouse, Yumi Kasai, David Gray, Carol Clugston, Anna Dominiczak and Alex Alderton, Roberto Amato, Jeffrey Barrett, Sonia Goncalves, Ewan Harrison, David K. Jackson, Ian Johnston, Dominic Kwiatkowski, Cordelia Langford, John Sillitoe on behalf of the Wellcome Sanger Institute COVID-19 Surveillance Team |
| EPI_ISL_2848925, EPI_ISL_2848926, EPI_ISL_2848927 | Lighthouse Lab in Milton Keynes | Wellcome Sanger Institute for the COVID-19 Genomics UK (COG-UK) Consortium | The Lighthouse Lab in Milton Keynes and Alex Alderton, Roberto Amato, Jeffrey Barrett, Sonia Goncalves, Ewan Harrison, David K. Jackson, Ian Johnston, Dominic Kwiatkowski, Cordelia Langford, John Sillitoe on behalf of the Wellcome Sanger Institute COVID-19 Surveillance Team |
| EPI_ISL_2848928 | Lighthouse Lab in Glasgow | Wellcome Sanger Institute for the COVID-19 Genomics UK (COG-UK) Consortium | Harper VanSteenhouse, Yumi Kasai, David Gray, Carol Clugston, Anna Dominiczak and Alex Alderton, Roberto Amato, Jeffrey Barrett, Sonia Goncalves, Ewan Harrison, David K. Jackson, Ian Johnston, Dominic Kwiatkowski, Cordelia Langford, John Sillitoe on behalf of the Wellcome Sanger Institute COVID-19 Surveillance Team |
| EPI_ISL_2848929 | Lighthouse Lab in Milton Keynes | Wellcome Sanger Institute for the COVID-19 Genomics UK (COG-UK) Consortium | The Lighthouse Lab in Milton Keynes and Alex Alderton, Roberto Amato, Jeffrey Barrett, Sonia Goncalves, Ewan Harrison, David K. Jackson, Ian Johnston, Dominic Kwiatkowski, Cordelia Langford, John Sillitoe on behalf of the Wellcome Sanger Institute COVID-19 Surveillance Team |
| EPI_ISL_2848930 | Lighthouse Lab in Glasgow | Wellcome Sanger Institute for the COVID-19 Genomics UK (COG-UK) Consortium | Harper VanSteenhouse, Yumi Kasai, David Gray, Carol Clugston, Anna Dominiczak and Alex Alderton, Roberto Amato, Jeffrey Barrett, Sonia Goncalves, Ewan Harrison, David K. Jackson, Ian Johnston, Dominic Kwiatkowski, Cordelia Langford, John Sillitoe on behalf of the Wellcome Sanger Institute COVID-19 Surveillance Team |
| EPI_ISL_2848931, EPI_ISL_2848932 | Lighthouse Lab in Milton Keynes | Wellcome Sanger Institute for the COVID-19 Genomics UK (COG-UK) Consortium | The Lighthouse Lab in Milton Keynes and Alex Alderton, Roberto Amato, Jeffrey Barrett, Sonia Goncalves, Ewan Harrison, David K. Jackson, Ian Johnston, Dominic Kwiatkowski, Cordelia Langford, John Sillitoe on behalf of the Wellcome Sanger Institute COVID-19 Surveillance Team |
| EPI_ISL_2848933 | Lighthouse Lab in Glasgow | Wellcome Sanger Institute for the COVID-19 Genomics UK (COG-UK) Consortium | Harper VanSteenhouse, Yumi Kasai, David Gray, Carol Clugston, Anna Dominiczak and Alex Alderton, Roberto Amato, Jeffrey Barrett, Sonia Goncalves, Ewan Harrison, David K. Jackson, Ian Johnston, Dominic Kwiatkowski, Cordelia Langford, John Sillitoe on behalf of the Wellcome Sanger Institute COVID-19 Surveillance Team |
| EPI_ISL_2848934, EPI_ISL_2848935, EPI_ISL_2848936, EPI_ISL_2848937, EPI_ISL_2848938, EPI_ISL_2848939, EPI_ISL_2848940, EPI_ISL_2848941, EPI_ISL_2848942, EPI_ISL_2848943, EPI_ISL_2848944, EPI_ISL_2848945, EPI_ISL_2848946, EPI_ISL_2848947, EPI_ISL_2848948 | | | |
| see above | Lighthouse Lab in Milton Keynes | Wellcome Sanger Institute for the COVID-19 Genomics UK (COG-UK) Consortium | The Lighthouse Lab in Milton Keynes and Alex Alderton, Roberto Amato, Jeffrey Barrett, Sonia Goncalves, Ewan Harrison, David K. Jackson, Ian Johnston, Dominic Kwiatkowski, Cordelia Langford, John Sillitoe on behalf of the Wellcome Sanger Institute COVID-19 Surveillance Team |
| EPI_ISL_2848949 | Lighthouse Lab in Glasgow | Wellcome Sanger Institute for the COVID-19 Genomics UK (COG-UK) Consortium | Harper VanSteenhouse, Yumi Kasai, David Gray, Carol Clugston, Anna Dominiczak and Alex Alderton, Roberto Amato, Jeffrey Barrett, Sonia Goncalves, Ewan Harrison, David K. Jackson, Ian Johnston, Dominic Kwiatkowski, Cordelia Langford, John Sillitoe on behalf of the Wellcome Sanger Institute COVID-19 Surveillance Team |
| EPI_ISL_2848950, EPI_ISL_2848951, EPI_ISL_2848952, EPI_ISL_2848953, EPI_ISL_2848954 | Lighthouse Lab in Milton Keynes | Wellcome Sanger Institute for the COVID-19 Genomics UK (COG-UK) Consortium | The Lighthouse Lab in Milton Keynes and Alex Alderton, Roberto Amato, Jeffrey Barrett, Sonia Goncalves, Ewan Harrison, David K. Jackson, Ian Johnston, Dominic Kwiatkowski, Cordelia Langford, John Sillitoe on behalf of the Wellcome Sanger Institute COVID-19 Surveillance Team |
| EPI_ISL_2848955 | Lighthouse Lab in Glasgow | Wellcome Sanger Institute for the COVID-19 Genomics UK (COG-UK) Consortium | Harper VanSteenhouse, Yumi Kasai, David Gray, Carol Clugston, Anna Dominiczak and Alex Alderton, Roberto Amato, Jeffrey Barrett, Sonia Goncalves, Ewan Harrison, David K. Jackson, Ian Johnston, Dominic Kwiatkowski, Cordelia Langford, John Sillitoe on behalf of the Wellcome Sanger Institute COVID-19 Surveillance Team |
| EPI_ISL_2848956 | Lighthouse Lab in Milton Keynes | Wellcome Sanger Institute for the COVID-19 Genomics UK (COG-UK) Consortium | The Lighthouse Lab in Milton Keynes and Alex Alderton, Roberto Amato, Jeffrey Barrett, Sonia Goncalves, Ewan Harrison, David K. Jackson, Ian Johnston, Dominic Kwiatkowski, Cordelia Langford, John Sillitoe on behalf of the Wellcome Sanger Institute COVID-19 Surveillance Team |
| EPI_ISL_2848957 | Lighthouse Lab in Glasgow | Wellcome Sanger Institute for the COVID-19 Genomics UK (COG-UK) Consortium | Harper VanSteenhouse, Yumi Kasai, David Gray, Carol Clugston, Anna Dominiczak and Alex Alderton, Roberto Amato, Jeffrey Barrett, Sonia Goncalves, Ewan Harrison, David K. Jackson, Ian Johnston, Dominic Kwiatkowski, Cordelia Langford, John Sillitoe on behalf of the Wellcome Sanger Institute COVID-19 Surveillance Team |
| EPI_ISL_2848958, EPI_ISL_2848959, EPI_ISL_2848960, EPI_ISL_2848961, EPI_ISL_2848962, EPI_ISL_2848963, EPI_ISL_2848964 | Lighthouse Lab in Milton Keynes | Wellcome Sanger Institute for the COVID-19 Genomics UK (COG-UK) Consortium | The Lighthouse Lab in Milton Keynes and Alex Alderton, Roberto Amato, Jeffrey Barrett, Sonia Goncalves, Ewan Harrison, David K. Jackson, Ian Johnston, Dominic Kwiatkowski, Cordelia Langford, John Sillitoe on behalf of the Wellcome Sanger Institute COVID-19 Surveillance Team |
| EPI_ISL_2848965, EPI_ISL_2848966 | Lighthouse Lab in Glasgow | Wellcome Sanger Institute for the COVID-19 Genomics UK (COG-UK) Consortium | Harper VanSteenhouse, Yumi Kasai, David Gray, Carol Clugston, Anna Dominiczak and Alex Alderton, Roberto Amato, Jeffrey Barrett, Sonia Goncalves, Ewan Harrison, David K. Jackson, Ian Johnston, Dominic Kwiatkowski, Cordelia Langford, John Sillitoe on behalf of the Wellcome Sanger Institute COVID-19 Surveillance Team |
| EPI_ISL_2848967, EPI_ISL_2848968 | Lighthouse Lab in Milton Keynes | Wellcome Sanger Institute for the COVID-19 Genomics UK (COG-UK) Consortium | The Lighthouse Lab in Milton Keynes and Alex Alderton, Roberto Amato, Jeffrey Barrett, Sonia Goncalves, Ewan Harrison, David K. Jackson, Ian Johnston, Dominic Kwiatkowski, Cordelia Langford, John Sillitoe on behalf of the Wellcome Sanger Institute COVID-19 Surveillance Team |
| EPI_ISL_2848969 | Lighthouse Lab in Glasgow | Wellcome Sanger Institute for the COVID-19 Genomics UK (COG-UK) Consortium | Harper VanSteenhouse, Yumi Kasai, David Gray, Carol Clugston, Anna Dominiczak and Alex Alderton, Roberto Amato, Jeffrey Barrett, Sonia Goncalves, Ewan Harrison, David K. Jackson, Ian Johnston, Dominic Kwiatkowski, Cordelia Langford, John Sillitoe on behalf of the Wellcome Sanger Institute COVID-19 Surveillance Team |
| EPI_ISL_2848970, EPI_ISL_2848971, EPI_ISL_2848972, EPI_ISL_2848973, EPI_ISL_2848974, EPI_ISL_2848975, EPI_ISL_2848976, EPI_ISL_2848977, EPI_ISL_2848978, EPI_ISL_2848979, EPI_ISL_2848980, EPI_ISL_2848981, EPI_ISL_2848982, EPI_ISL_2848983 | | | |
| see above | Lighthouse Lab in Milton Keynes | Wellcome Sanger Institute for the COVID-19 Genomics UK (COG-UK) Consortium | The Lighthouse Lab in Milton Keynes and Alex Alderton, Roberto Amato, Jeffrey Barrett, Sonia Goncalves, Ewan Harrison, David K. Jackson, Ian Johnston, Dominic Kwiatkowski, Cordelia Langford, John Sillitoe on behalf of the Wellcome Sanger Institute COVID-19 Surveillance Team |
| EPI_ISL_2848984 | Lighthouse Lab in Glasgow | Wellcome Sanger Institute for the COVID-19 Genomics UK (COG-UK) Consortium | Harper VanSteenhouse, Yumi Kasai, David Gray, Carol Clugston, Anna Dominiczak and Alex Alderton, Roberto Amato, Jeffrey Barrett, Sonia Goncalves, Ewan Harrison, David K. Jackson, Ian Johnston, Dominic Kwiatkowski, Cordelia Langford, John Sillitoe on behalf of the Wellcome Sanger Institute COVID-19 Surveillance Team |
| EPI_ISL_2848985, EPI_ISL_2848986, EPI_ISL_2848987, EPI_ISL_2848988, EPI_ISL_2848989, EPI_ISL_2848990, EPI_ISL_2848991, EPI_ISL_2848992 | Lighthouse Lab in Milton Keynes | Wellcome Sanger Institute for the COVID-19 Genomics UK (COG-UK) Consortium | The Lighthouse Lab in Milton Keynes and Alex Alderton, Roberto Amato, Jeffrey Barrett, Sonia Goncalves, Ewan Harrison, David K. Jackson, Ian Johnston, Dominic Kwiatkowski, Cordelia Langford, John Sillitoe on behalf of the Wellcome Sanger Institute COVID-19 Surveillance Team |
| EPI_ISL_2848993 | Lighthouse Lab in Glasgow | Wellcome Sanger Institute for the COVID-19 Genomics UK (COG-UK) Consortium | Harper VanSteenhouse, Yumi Kasai, David Gray, Carol Clugston, Anna Dominiczak and Alex Alderton, Roberto Amato, Jeffrey Barrett, Sonia Goncalves, Ewan Harrison, David K. Jackson, Ian Johnston, Dominic Kwiatkowski, Cordelia Langford, John Sillitoe on behalf of the Wellcome Sanger Institute COVID-19 Surveillance Team |
| EPI_ISL_2848994, EPI_ISL_2848995, EPI_ISL_2848996, EPI_ISL_2848997, EPI_ISL_2848998, EPI_ISL_2848999, EPI_ISL_2849000, EPI_ISL_2849001, EPI_ISL_2849002 | Lighthouse Lab in Milton Keynes | Wellcome Sanger Institute for the COVID-19 Genomics UK (COG-UK) Consortium | The Lighthouse Lab in Milton Keynes and Alex Alderton, Roberto Amato, Jeffrey Barrett, Sonia Goncalves, Ewan Harrison, David K. Jackson, Ian Johnston, Dominic Kwiatkowski, Cordelia Langford, John Sillitoe on behalf of the Wellcome Sanger Institute COVID-19 Surveillance Team |
| EPI_ISL_2849003 | Lighthouse Lab in Glasgow | Wellcome Sanger Institute for the COVID-19 Genomics UK (COG-UK) Consortium | Harper VanSteenhouse, Yumi Kasai, David Gray, Carol Clugston, Anna Dominiczak and Alex Alderton, Roberto Amato, Jeffrey Barrett, Sonia Goncalves, Ewan Harrison, David K. Jackson, Ian Johnston, Dominic Kwiatkowski, Cordelia Langford, John Sillitoe on behalf of the Wellcome Sanger Institute COVID-19 Surveillance Team |
| EPI_ISL_2849004, EPI_ISL_2849005, EPI_ISL_2849006, EPI_ISL_2849007, EPI_ISL_2849008 | Lighthouse Lab in Milton Keynes | Wellcome Sanger Institute for the COVID-19 Genomics UK (COG-UK) Consortium | The Lighthouse Lab in Milton Keynes and Alex Alderton, Roberto Amato, Jeffrey Barrett, Sonia Goncalves, Ewan Harrison, David K. Jackson, Ian Johnston, Dominic Kwiatkowski, Cordelia Langford, John Sillitoe on behalf of the Wellcome Sanger Institute COVID-19 Surveillance Team |
| EPI_ISL_2849009 | Lighthouse Lab in Alderley Park | Wellcome Sanger Institute for the COVID-19 Genomics UK (COG-UK) Consortium | Jacquelyn Wynn, Mairead Hyland, The Lighthouse Lab in Alderley Park and Alex Alderton, Roberto Amato, Jeffrey Barrett, Sonia Goncalves, Ewan Harrison, David K. Jackson, Ian Johnston, Dominic Kwiatkowski, Cordelia Langford, John Sillitoe on behalf of the Wellcome Sanger Institute COVID-19 Surveillance Team |
| EPI_ISL_2849010 | Lighthouse Laboratory Plymouth | Wellcome Sanger Institute for the COVID-19 Genomics UK (COG-UK) Consortium | Lighthouse Laboratory Plymouth and Alex Alderton, Roberto Amato, Jeffrey Barrett, Sonia Goncalves, Ewan Harrison, David K. Jackson, Ian Johnston, Dominic Kwiatkowski, Cordelia Langford, John Sillitoe on behalf of the Wellcome Sanger Institute COVID-19 Surveillance Team |
| EPI_ISL_2849011, EPI_ISL_2849012, EPI_ISL_2849013 | Lighthouse Lab in Alderley Park | Wellcome Sanger Institute for the COVID-19 Genomics UK (COG-UK) Consortium | Jacquelyn Wynn, Mairead Hyland, The Lighthouse Lab in Alderley Park and Alex Alderton, Roberto Amato, Jeffrey Barrett, Sonia Goncalves, Ewan Harrison, David K. Jackson, Ian Johnston, Dominic Kwiatkowski, Cordelia Langford, John Sillitoe on behalf of the Wellcome Sanger Institute COVID-19 Surveillance Team |
| EPI_ISL_2849014, EPI_ISL_2849015 | Lighthouse Laboratory Plymouth | Wellcome Sanger Institute for the COVID-19 Genomics UK (COG-UK) Consortium | Lighthouse Laboratory Plymouth and Alex Alderton, Roberto Amato, Jeffrey Barrett, Sonia Goncalves, Ewan Harrison, David K. Jackson, Ian Johnston, Dominic Kwiatkowski, Cordelia Langford, John Sillitoe on behalf of the Wellcome Sanger Institute COVID-19 Surveillance Team |

| | | | |
|---|---|---|---|
| EPI_ISL_2849016, EPI_ISL_2849017 | Lighthouse Lab in Alderley Park | Wellcome Sanger Institute for the COVID-19 Genomics UK (COG-UK) Consortium | Jacquelyn Wynn, Mairead Hyland, The Lighthouse Lab in Alderley Park and Alex Alderton, Roberto Amato, Jeffrey Barrett, Sonia Goncalves, Ewan Harrison, David K. Jackson, Ian Johnston, Dominic Kwiatkowski, Cordelia Langford, John Sillitoe on behalf of the Wellcome Sanger Institute COVID-19 Surveillance Team |
| EPI_ISL_2849018, EPI_ISL_2849019 | Lighthouse Laboratory Plymouth | Wellcome Sanger Institute for the COVID-19 Genomics UK (COG-UK) Consortium | Lighthouse Laboratory Plymouth and Alex Alderton, Roberto Amato, Jeffrey Barrett, Sonia Goncalves, Ewan Harrison, David K. Jackson, Ian Johnston, Dominic Kwiatkowski, Cordelia Langford, John Sillitoe on behalf of the Wellcome Sanger Institute COVID-19 Surveillance Team |
| EPI_ISL_2849020, EPI_ISL_2849021, EPI_ISL_2849022, EPI_ISL_2849023 | Lighthouse Lab in Alderley Park | Wellcome Sanger Institute for the COVID-19 Genomics UK (COG-UK) Consortium | Jacquelyn Wynn, Mairead Hyland, The Lighthouse Lab in Alderley Park and Alex Alderton, Roberto Amato, Jeffrey Barrett, Sonia Goncalves, Ewan Harrison, David K. Jackson, Ian Johnston, Dominic Kwiatkowski, Cordelia Langford, John Sillitoe on behalf of the Wellcome Sanger Institute COVID-19 Surveillance Team |
| EPI_ISL_2849024 | Lighthouse Laboratory Plymouth | Wellcome Sanger Institute for the COVID-19 Genomics UK (COG-UK) Consortium | Lighthouse Laboratory Plymouth and Alex Alderton, Roberto Amato, Jeffrey Barrett, Sonia Goncalves, Ewan Harrison, David K. Jackson, Ian Johnston, Dominic Kwiatkowski, Cordelia Langford, John Sillitoe on behalf of the Wellcome Sanger Institute COVID-19 Surveillance Team |
| EPI_ISL_2849025 | Lighthouse Lab in Alderley Park | Wellcome Sanger Institute for the COVID-19 Genomics UK (COG-UK) Consortium | Jacquelyn Wynn, Mairead Hyland, The Lighthouse Lab in Alderley Park and Alex Alderton, Roberto Amato, Jeffrey Barrett, Sonia Goncalves, Ewan Harrison, David K. Jackson, Ian Johnston, Dominic Kwiatkowski, Cordelia Langford, John Sillitoe on behalf of the Wellcome Sanger Institute COVID-19 Surveillance Team |
| EPI_ISL_2849026 | Lighthouse Laboratory Plymouth | Wellcome Sanger Institute for the COVID-19 Genomics UK (COG-UK) Consortium | Lighthouse Laboratory Plymouth and Alex Alderton, Roberto Amato, Jeffrey Barrett, Sonia Goncalves, Ewan Harrison, David K. Jackson, Ian Johnston, Dominic Kwiatkowski, Cordelia Langford, John Sillitoe on behalf of the Wellcome Sanger Institute COVID-19 Surveillance Team |
| EPI_ISL_2849027, EPI_ISL_2849028, EPI_ISL_2849029 | Lighthouse Lab in Alderley Park | Wellcome Sanger Institute for the COVID-19 Genomics UK (COG-UK) Consortium | Jacquelyn Wynn, Mairead Hyland, The Lighthouse Lab in Alderley Park and Alex Alderton, Roberto Amato, Jeffrey Barrett, Sonia Goncalves, Ewan Harrison, David K. Jackson, Ian Johnston, Dominic Kwiatkowski, Cordelia Langford, John Sillitoe on behalf of the Wellcome Sanger Institute COVID-19 Surveillance Team |
| EPI_ISL_2849030, EPI_ISL_2849031 | Lighthouse Laboratory Plymouth | Wellcome Sanger Institute for the COVID-19 Genomics UK (COG-UK) Consortium | Lighthouse Laboratory Plymouth and Alex Alderton, Roberto Amato, Jeffrey Barrett, Sonia Goncalves, Ewan Harrison, David K. Jackson, Ian Johnston, Dominic Kwiatkowski, Cordelia Langford, John Sillitoe on behalf of the Wellcome Sanger Institute COVID-19 Surveillance Team |
| EPI_ISL_2849032 | Lighthouse Lab in Alderley Park | Wellcome Sanger Institute for the COVID-19 Genomics UK (COG-UK) Consortium | Jacquelyn Wynn, Mairead Hyland, The Lighthouse Lab in Alderley Park and Alex Alderton, Roberto Amato, Jeffrey Barrett, Sonia Goncalves, Ewan Harrison, David K. Jackson, Ian Johnston, Dominic Kwiatkowski, Cordelia Langford, John Sillitoe on behalf of the Wellcome Sanger Institute COVID-19 Surveillance Team |
| EPI_ISL_2849033, EPI_ISL_2849034 | Lighthouse Laboratory Plymouth | Wellcome Sanger Institute for the COVID-19 Genomics UK (COG-UK) Consortium | Lighthouse Laboratory Plymouth and Alex Alderton, Roberto Amato, Jeffrey Barrett, Sonia Goncalves, Ewan Harrison, David K. Jackson, Ian Johnston, Dominic Kwiatkowski, Cordelia Langford, John Sillitoe on behalf of the Wellcome Sanger Institute COVID-19 Surveillance Team |
| EPI_ISL_2849035 | Lighthouse Lab in Alderley Park | Wellcome Sanger Institute for the COVID-19 Genomics UK (COG-UK) Consortium | Jacquelyn Wynn, Mairead Hyland, The Lighthouse Lab in Alderley Park and Alex Alderton, Roberto Amato, Jeffrey Barrett, Sonia Goncalves, Ewan Harrison, David K. Jackson, Ian Johnston, Dominic Kwiatkowski, Cordelia Langford, John Sillitoe on behalf of the Wellcome Sanger Institute COVID-19 Surveillance Team |
| EPI_ISL_2849036, EPI_ISL_2849037, EPI_ISL_2849038 | Lighthouse Laboratory Plymouth | Wellcome Sanger Institute for the COVID-19 Genomics UK (COG-UK) Consortium | Lighthouse Laboratory Plymouth and Alex Alderton, Roberto Amato, Jeffrey Barrett, Sonia Goncalves, Ewan Harrison, David K. Jackson, Ian Johnston, Dominic Kwiatkowski, Cordelia Langford, John Sillitoe on behalf of the Wellcome Sanger Institute COVID-19 Surveillance Team |
| EPI_ISL_2849039, EPI_ISL_2849040 | Lighthouse Lab in Alderley Park | Wellcome Sanger Institute for the COVID-19 Genomics UK (COG-UK) Consortium | Jacquelyn Wynn, Mairead Hyland, The Lighthouse Lab in Alderley Park and Alex Alderton, Roberto Amato, Jeffrey Barrett, Sonia Goncalves, Ewan Harrison, David K. Jackson, Ian Johnston, Dominic Kwiatkowski, Cordelia Langford, John Sillitoe on behalf of the Wellcome Sanger Institute COVID-19 Surveillance Team |
| EPI_ISL_2849041 | Lighthouse Laboratory Plymouth | Wellcome Sanger Institute for the COVID-19 Genomics UK (COG-UK) Consortium | Lighthouse Laboratory Plymouth and Alex Alderton, Roberto Amato, Jeffrey Barrett, Sonia Goncalves, Ewan Harrison, David K. Jackson, Ian Johnston, Dominic Kwiatkowski, Cordelia Langford, John Sillitoe on behalf of the Wellcome Sanger Institute COVID-19 Surveillance Team |
| EPI_ISL_2849042, EPI_ISL_2849043, EPI_ISL_2849044, EPI_ISL_2849045 | Lighthouse Lab in Alderley Park | Wellcome Sanger Institute for the COVID-19 Genomics UK (COG-UK) Consortium | Jacquelyn Wynn, Mairead Hyland, The Lighthouse Lab in Alderley Park and Alex Alderton, Roberto Amato, Jeffrey Barrett, Sonia Goncalves, Ewan Harrison, David K. Jackson, Ian Johnston, Dominic Kwiatkowski, Cordelia Langford, John Sillitoe on behalf of the Wellcome Sanger Institute COVID-19 Surveillance Team |
| EPI_ISL_2849046, EPI_ISL_2849047 | Lighthouse Laboratory Plymouth | Wellcome Sanger Institute for the COVID-19 Genomics UK (COG-UK) Consortium | Lighthouse Laboratory Plymouth and Alex Alderton, Roberto Amato, Jeffrey Barrett, Sonia Goncalves, Ewan Harrison, David K. Jackson, Ian Johnston, Dominic Kwiatkowski, Cordelia Langford, John Sillitoe on behalf of the Wellcome Sanger Institute COVID-19 Surveillance Team |
| EPI_ISL_2849048, EPI_ISL_2849049, EPI_ISL_2849050 | Lighthouse Lab in Alderley Park | Wellcome Sanger Institute for the COVID-19 Genomics UK (COG-UK) Consortium | Jacquelyn Wynn, Mairead Hyland, The Lighthouse Lab in Alderley Park and Alex Alderton, Roberto Amato, Jeffrey Barrett, Sonia Goncalves, Ewan Harrison, David K. Jackson, Ian Johnston, Dominic Kwiatkowski, Cordelia Langford, John Sillitoe on behalf of the Wellcome Sanger Institute COVID-19 Surveillance Team |
| EPI_ISL_2849051 | Lighthouse Laboratory Plymouth | Wellcome Sanger Institute for the COVID-19 Genomics UK (COG-UK) Consortium | Lighthouse Laboratory Plymouth and Alex Alderton, Roberto Amato, Jeffrey Barrett, Sonia Goncalves, Ewan Harrison, David K. Jackson, Ian Johnston, Dominic Kwiatkowski, Cordelia Langford, John Sillitoe on behalf of the Wellcome Sanger Institute COVID-19 Surveillance Team |
| EPI_ISL_2849052 | Lighthouse Lab in Alderley Park | Wellcome Sanger Institute for the COVID-19 Genomics UK (COG-UK) Consortium | Jacquelyn Wynn, Mairead Hyland, The Lighthouse Lab in Alderley Park and Alex Alderton, Roberto Amato, Jeffrey Barrett, Sonia Goncalves, Ewan Harrison, David K. Jackson, Ian Johnston, Dominic Kwiatkowski, Cordelia Langford, John Sillitoe on behalf of the Wellcome Sanger Institute COVID-19 Surveillance Team |
| EPI_ISL_2849053 | Lighthouse Laboratory Plymouth | Wellcome Sanger Institute for the COVID-19 Genomics UK (COG-UK) Consortium | Lighthouse Laboratory Plymouth and Alex Alderton, Roberto Amato, Jeffrey Barrett, Sonia Goncalves, Ewan Harrison, David K. Jackson, Ian Johnston, Dominic Kwiatkowski, Cordelia Langford, John Sillitoe on behalf of the Wellcome Sanger Institute COVID-19 Surveillance Team |
| EPI_ISL_2849054 | Lighthouse Lab in Alderley Park | Wellcome Sanger Institute for the COVID-19 Genomics UK (COG-UK) Consortium | Jacquelyn Wynn, Mairead Hyland, The Lighthouse Lab in Alderley Park and Alex Alderton, Roberto Amato, Jeffrey Barrett, Sonia Goncalves, Ewan Harrison, David K. Jackson, Ian Johnston, Dominic Kwiatkowski, Cordelia Langford, John Sillitoe on behalf of the Wellcome Sanger Institute COVID-19 Surveillance Team |
| EPI_ISL_2849055 | Lighthouse Laboratory Plymouth | Wellcome Sanger Institute for the COVID-19 Genomics UK (COG-UK) Consortium | Lighthouse Laboratory Plymouth and Alex Alderton, Roberto Amato, Jeffrey Barrett, Sonia Goncalves, Ewan Harrison, David K. Jackson, Ian Johnston, Dominic Kwiatkowski, Cordelia Langford, John Sillitoe on behalf of the Wellcome Sanger Institute COVID-19 Surveillance Team |
| EPI_ISL_2849056, EPI_ISL_2849057 | Lighthouse Lab in Alderley Park | Wellcome Sanger Institute for the COVID-19 Genomics UK (COG-UK) Consortium | Jacquelyn Wynn, Mairead Hyland, The Lighthouse Lab in Alderley Park and Alex Alderton, Roberto Amato, Jeffrey Barrett, Sonia Goncalves, Ewan Harrison, David K. Jackson, Ian Johnston, Dominic Kwiatkowski, Cordelia Langford, John Sillitoe on behalf of the Wellcome Sanger Institute COVID-19 Surveillance Team |
| EPI_ISL_2849058, EPI_ISL_2849059 | Lighthouse Laboratory Plymouth | Wellcome Sanger Institute for the COVID-19 Genomics UK (COG-UK) Consortium | Lighthouse Laboratory Plymouth and Alex Alderton, Roberto Amato, Jeffrey Barrett, Sonia Goncalves, Ewan Harrison, David K. Jackson, Ian Johnston, Dominic Kwiatkowski, Cordelia Langford, John Sillitoe on behalf of the Wellcome Sanger Institute COVID-19 Surveillance Team |
| EPI_ISL_2849060 | Lighthouse Lab in Alderley Park | Wellcome Sanger Institute for the COVID-19 Genomics UK (COG-UK) Consortium | Jacquelyn Wynn, Mairead Hyland, The Lighthouse Lab in Alderley Park and Alex Alderton, Roberto Amato, Jeffrey Barrett, Sonia Goncalves, Ewan Harrison, David K. Jackson, Ian Johnston, Dominic Kwiatkowski, Cordelia Langford, John Sillitoe on behalf of the Wellcome Sanger Institute COVID-19 Surveillance Team |
| EPI_ISL_2849061 | Lighthouse Laboratory Plymouth | Wellcome Sanger Institute for the COVID-19 Genomics UK (COG-UK) Consortium | Lighthouse Laboratory Plymouth and Alex Alderton, Roberto Amato, Jeffrey Barrett, Sonia Goncalves, Ewan Harrison, David K. Jackson, Ian Johnston, Dominic Kwiatkowski, Cordelia Langford, John Sillitoe on behalf of the Wellcome Sanger Institute COVID-19 Surveillance Team |
| EPI_ISL_2849062 | Lighthouse Lab in Alderley Park | Wellcome Sanger Institute for the COVID-19 Genomics UK (COG-UK) Consortium | Jacquelyn Wynn, Mairead Hyland, The Lighthouse Lab in Alderley Park and Alex Alderton, Roberto Amato, Jeffrey Barrett, Sonia Goncalves, Ewan Harrison, David K. Jackson, Ian Johnston, Dominic Kwiatkowski, Cordelia Langford, John Sillitoe on behalf of the Wellcome Sanger Institute COVID-19 Surveillance Team |
| EPI_ISL_2849063, EPI_ISL_2849064 | Lighthouse Laboratory Plymouth | Wellcome Sanger Institute for the COVID-19 Genomics UK (COG-UK) Consortium | Lighthouse Laboratory Plymouth and Alex Alderton, Roberto Amato, Jeffrey Barrett, Sonia Goncalves, Ewan Harrison, David K. Jackson, Ian Johnston, Dominic Kwiatkowski, Cordelia Langford, John Sillitoe on behalf of the Wellcome Sanger Institute COVID-19 Surveillance Team |
| EPI_ISL_2849065 | Lighthouse Lab in Alderley Park | Wellcome Sanger Institute for the COVID-19 Genomics UK (COG-UK) Consortium | Jacquelyn Wynn, Mairead Hyland, The Lighthouse Lab in Alderley Park and Alex Alderton, Roberto Amato, Jeffrey Barrett, Sonia Goncalves, Ewan Harrison, David K. Jackson, Ian Johnston, Dominic Kwiatkowski, Cordelia Langford, John Sillitoe on behalf of the Wellcome Sanger Institute COVID-19 Surveillance Team |
| EPI_ISL_2849066 | Lighthouse Laboratory Plymouth | Wellcome Sanger Institute for the COVID-19 Genomics UK (COG-UK) Consortium | Lighthouse Laboratory Plymouth and Alex Alderton, Roberto Amato, Jeffrey Barrett, Sonia Goncalves, Ewan Harrison, David K. Jackson, Ian Johnston, Dominic Kwiatkowski, Cordelia Langford, John Sillitoe on behalf of the Wellcome Sanger Institute COVID-19 Surveillance Team |
| EPI_ISL_2849067, EPI_ISL_2849068, EPI_ISL_2849069, EPI_ISL_2849070, EPI_ISL_2849071 | Lighthouse Lab in Alderley Park | Wellcome Sanger Institute for the COVID-19 Genomics UK (COG-UK) Consortium | Jacquelyn Wynn, Mairead Hyland, The Lighthouse Lab in Alderley Park and Alex Alderton, Roberto Amato, Jeffrey Barrett, Sonia Goncalves, Ewan Harrison, David K. Jackson, Ian Johnston, Dominic Kwiatkowski, Cordelia Langford, John Sillitoe on behalf of the Wellcome Sanger Institute COVID-19 Surveillance Team |
| EPI_ISL_2849072, EPI_ISL_2849073, EPI_ISL_2849074, EPI_ISL_2849075, EPI_ISL_2849076, EPI_ISL_2849077 | Lighthouse Laboratory Plymouth | Wellcome Sanger Institute for the COVID-19 Genomics UK (COG-UK) Consortium | Lighthouse Laboratory Plymouth and Alex Alderton, Roberto Amato, Jeffrey Barrett, Sonia Goncalves, Ewan Harrison, David K. Jackson, Ian Johnston, Dominic Kwiatkowski, Cordelia Langford, John Sillitoe on behalf of the Wellcome Sanger Institute COVID-19 Surveillance Team |
| EPI_ISL_2849078, EPI_ISL_2849079 | Lighthouse Lab in Alderley Park | Wellcome Sanger Institute for the COVID-19 Genomics UK (COG-UK) Consortium | Jacquelyn Wynn, Mairead Hyland, The Lighthouse Lab in Alderley Park and Alex Alderton, Roberto Amato, Jeffrey Barrett, Sonia Goncalves, Ewan Harrison, David K. Jackson, Ian Johnston, Dominic Kwiatkowski, Cordelia Langford, John Sillitoe on behalf of the Wellcome Sanger Institute COVID-19 Surveillance Team |
| EPI_ISL_2849080, EPI_ISL_2849081, EPI_ISL_2849082 | Lighthouse Laboratory Plymouth | Wellcome Sanger Institute for the COVID-19 Genomics UK (COG-UK) Consortium | Lighthouse Laboratory Plymouth and Alex Alderton, Roberto Amato, Jeffrey Barrett, Sonia Goncalves, Ewan Harrison, David K. Jackson, Ian Johnston, Dominic Kwiatkowski, Cordelia Langford, John Sillitoe on behalf of the Wellcome Sanger Institute COVID-19 Surveillance Team |
| EPI_ISL_2849083, EPI_ISL_2849084 | Lighthouse Lab in Alderley Park | Wellcome Sanger Institute for the COVID-19 Genomics UK (COG-UK) Consortium | Jacquelyn Wynn, Mairead Hyland, The Lighthouse Lab in Alderley Park and Alex Alderton, Roberto Amato, Jeffrey Barrett, Sonia Goncalves, Ewan Harrison, David K. Jackson, Ian Johnston, Dominic Kwiatkowski, Cordelia Langford, John Sillitoe on behalf of the Wellcome Sanger Institute COVID-19 Surveillance Team |
| EPI_ISL_2849085 | Lighthouse Laboratory Plymouth | Wellcome Sanger Institute for the COVID-19 Genomics UK (COG-UK) Consortium | Lighthouse Laboratory Plymouth and Alex Alderton, Roberto Amato, Jeffrey Barrett, Sonia Goncalves, Ewan Harrison, David K. Jackson, Ian Johnston, Dominic Kwiatkowski, Cordelia Langford, John Sillitoe on behalf of the Wellcome Sanger Institute COVID-19 Surveillance Team |
| EPI_ISL_2849086 | Lighthouse Lab in Alderley Park | Wellcome Sanger Institute for the COVID-19 Genomics UK (COG-UK) Consortium | Jacquelyn Wynn, Mairead Hyland, The Lighthouse Lab in Alderley Park and Alex Alderton, Roberto Amato, Jeffrey Barrett, Sonia Goncalves, Ewan Harrison, David K. Jackson, Ian Johnston, Dominic Kwiatkowski, Cordelia Langford, John Sillitoe on behalf of the Wellcome Sanger Institute COVID-19 Surveillance Team |
| EPI_ISL_2849087, EPI_ISL_2849088, EPI_ISL_2849089 | Lighthouse Laboratory Plymouth | Wellcome Sanger Institute for the COVID-19 Genomics UK (COG-UK) Consortium | Lighthouse Laboratory Plymouth and Alex Alderton, Roberto Amato, Jeffrey Barrett, Sonia Goncalves, Ewan Harrison, David K. Jackson, Ian Johnston, Dominic Kwiatkowski, Cordelia Langford, John Sillitoe on behalf of the Wellcome Sanger Institute COVID-19 Surveillance Team |
| EPI_ISL_2849090, | Lighthouse Lab in Alderley Park | Wellcome Sanger Institute for the COVID-19 Genomics UK (COG-UK) | Jacquelyn Wynn, Mairead Hyland, The Lighthouse Lab in Alderley Park and Alex Alderton, Roberto Amato, Jeffrey Barrett, Sonia Goncalves, Ewan Harrison, David K. Jackson, Ian Johnston, Dominic Kwiatkowski, Cordelia Langford, John Sillitoe on |

| | | | |
|---|---|---|---|
| EPI_ISL_2849091, EPI_ISL_2849092 | | Consortium | behalf of the Wellcome Sanger Institute COVID-19 Surveillance Team |
| EPI_ISL_2849093 | Lighthouse Laboratory Plymouth | Wellcome Sanger Institute for the COVID-19 Genomics UK (COG-UK) Consortium | Lighthouse Laboratory Plymouth and Alex Alderton, Roberto Amato, Jeffrey Barrett, Sonia Goncalves, Ewan Harrison, David K. Jackson, Ian Johnston, Dominic Kwiatkowski, Cordelia Langford, John Sillitoe on behalf of the Wellcome Sanger Institute COVID-19 Surveillance Team |
| EPI_ISL_2849094 | Lighthouse Lab in Alderley Park | Wellcome Sanger Institute for the COVID-19 Genomics UK (COG-UK) Consortium | Jacquelyn Wynn, Mairead Hyland, The Lighthouse Lab in Alderley Park and Alex Alderton, Roberto Amato, Jeffrey Barrett, Sonia Goncalves, Ewan Harrison, David K. Jackson, Ian Johnston, Dominic Kwiatkowski, Cordelia Langford, John Sillitoe on behalf of the Wellcome Sanger Institute COVID-19 Surveillance Team |
| EPI_ISL_2849095, EPI_ISL_2849096 | Lighthouse Laboratory Plymouth | Wellcome Sanger Institute for the COVID-19 Genomics UK (COG-UK) Consortium | Lighthouse Laboratory Plymouth and Alex Alderton, Roberto Amato, Jeffrey Barrett, Sonia Goncalves, Ewan Harrison, David K. Jackson, Ian Johnston, Dominic Kwiatkowski, Cordelia Langford, John Sillitoe on behalf of the Wellcome Sanger Institute COVID-19 Surveillance Team |
| EPI_ISL_2849097 | Lighthouse Lab in Alderley Park | Wellcome Sanger Institute for the COVID-19 Genomics UK (COG-UK) Consortium | Jacquelyn Wynn, Mairead Hyland, The Lighthouse Lab in Alderley Park and Alex Alderton, Roberto Amato, Jeffrey Barrett, Sonia Goncalves, Ewan Harrison, David K. Jackson, Ian Johnston, Dominic Kwiatkowski, Cordelia Langford, John Sillitoe on behalf of the Wellcome Sanger Institute COVID-19 Surveillance Team |
| EPI_ISL_2849098, EPI_ISL_2849099, EPI_ISL_2849100 | Lighthouse Laboratory Plymouth | Wellcome Sanger Institute for the COVID-19 Genomics UK (COG-UK) Consortium | Lighthouse Laboratory Plymouth and Alex Alderton, Roberto Amato, Jeffrey Barrett, Sonia Goncalves, Ewan Harrison, David K. Jackson, Ian Johnston, Dominic Kwiatkowski, Cordelia Langford, John Sillitoe on behalf of the Wellcome Sanger Institute COVID-19 Surveillance Team |
| EPI_ISL_2849101 | Lighthouse Lab in Alderley Park | Wellcome Sanger Institute for the COVID-19 Genomics UK (COG-UK) Consortium | Jacquelyn Wynn, Mairead Hyland, The Lighthouse Lab in Alderley Park and Alex Alderton, Roberto Amato, Jeffrey Barrett, Sonia Goncalves, Ewan Harrison, David K. Jackson, Ian Johnston, Dominic Kwiatkowski, Cordelia Langford, John Sillitoe on behalf of the Wellcome Sanger Institute COVID-19 Surveillance Team |
| EPI_ISL_2849102 | Lighthouse Laboratory Plymouth | Wellcome Sanger Institute for the COVID-19 Genomics UK (COG-UK) Consortium | Lighthouse Laboratory Plymouth and Alex Alderton, Roberto Amato, Jeffrey Barrett, Sonia Goncalves, Ewan Harrison, David K. Jackson, Ian Johnston, Dominic Kwiatkowski, Cordelia Langford, John Sillitoe on behalf of the Wellcome Sanger Institute COVID-19 Surveillance Team |
| EPI_ISL_2849103, EPI_ISL_2849104, EPI_ISL_2849105, EPI_ISL_2849106, EPI_ISL_2849107, EPI_ISL_2849108 | Lighthouse Lab in Alderley Park | Wellcome Sanger Institute for the COVID-19 Genomics UK (COG-UK) Consortium | Jacquelyn Wynn, Mairead Hyland, The Lighthouse Lab in Alderley Park and Alex Alderton, Roberto Amato, Jeffrey Barrett, Sonia Goncalves, Ewan Harrison, David K. Jackson, Ian Johnston, Dominic Kwiatkowski, Cordelia Langford, John Sillitoe on behalf of the Wellcome Sanger Institute COVID-19 Surveillance Team |
| EPI_ISL_2849109 | Lighthouse Laboratory Plymouth | Wellcome Sanger Institute for the COVID-19 Genomics UK (COG-UK) Consortium | Lighthouse Laboratory Plymouth and Alex Alderton, Roberto Amato, Jeffrey Barrett, Sonia Goncalves, Ewan Harrison, David K. Jackson, Ian Johnston, Dominic Kwiatkowski, Cordelia Langford, John Sillitoe on behalf of the Wellcome Sanger Institute COVID-19 Surveillance Team |
| EPI_ISL_2849110 | Lighthouse Lab in Alderley Park | Wellcome Sanger Institute for the COVID-19 Genomics UK (COG-UK) Consortium | Jacquelyn Wynn, Mairead Hyland, The Lighthouse Lab in Alderley Park and Alex Alderton, Roberto Amato, Jeffrey Barrett, Sonia Goncalves, Ewan Harrison, David K. Jackson, Ian Johnston, Dominic Kwiatkowski, Cordelia Langford, John Sillitoe on behalf of the Wellcome Sanger Institute COVID-19 Surveillance Team |
| EPI_ISL_2849111, EPI_ISL_2849112, EPI_ISL_2849113 | Lighthouse Laboratory Plymouth | Wellcome Sanger Institute for the COVID-19 Genomics UK (COG-UK) Consortium | Lighthouse Laboratory Plymouth and Alex Alderton, Roberto Amato, Jeffrey Barrett, Sonia Goncalves, Ewan Harrison, David K. Jackson, Ian Johnston, Dominic Kwiatkowski, Cordelia Langford, John Sillitoe on behalf of the Wellcome Sanger Institute COVID-19 Surveillance Team |
| EPI_ISL_2849114 | Lighthouse Lab in Alderley Park | Wellcome Sanger Institute for the COVID-19 Genomics UK (COG-UK) Consortium | Jacquelyn Wynn, Mairead Hyland, The Lighthouse Lab in Alderley Park and Alex Alderton, Roberto Amato, Jeffrey Barrett, Sonia Goncalves, Ewan Harrison, David K. Jackson, Ian Johnston, Dominic Kwiatkowski, Cordelia Langford, John Sillitoe on behalf of the Wellcome Sanger Institute COVID-19 Surveillance Team |
| EPI_ISL_2849115, EPI_ISL_2849116, EPI_ISL_2849117, EPI_ISL_2849118 | Lighthouse Laboratory Plymouth | Wellcome Sanger Institute for the COVID-19 Genomics UK (COG-UK) Consortium | Lighthouse Laboratory Plymouth and Alex Alderton, Roberto Amato, Jeffrey Barrett, Sonia Goncalves, Ewan Harrison, David K. Jackson, Ian Johnston, Dominic Kwiatkowski, Cordelia Langford, John Sillitoe on behalf of the Wellcome Sanger Institute COVID-19 Surveillance Team |
| EPI_ISL_2849119, EPI_ISL_2849120, EPI_ISL_2849121 | Lighthouse Lab in Alderley Park | Wellcome Sanger Institute for the COVID-19 Genomics UK (COG-UK) Consortium | Jacquelyn Wynn, Mairead Hyland, The Lighthouse Lab in Alderley Park and Alex Alderton, Roberto Amato, Jeffrey Barrett, Sonia Goncalves, Ewan Harrison, David K. Jackson, Ian Johnston, Dominic Kwiatkowski, Cordelia Langford, John Sillitoe on behalf of the Wellcome Sanger Institute COVID-19 Surveillance Team |
| EPI_ISL_2849122, EPI_ISL_2849123 | Lighthouse Laboratory Plymouth | Wellcome Sanger Institute for the COVID-19 Genomics UK (COG-UK) Consortium | Lighthouse Laboratory Plymouth and Alex Alderton, Roberto Amato, Jeffrey Barrett, Sonia Goncalves, Ewan Harrison, David K. Jackson, Ian Johnston, Dominic Kwiatkowski, Cordelia Langford, John Sillitoe on behalf of the Wellcome Sanger Institute COVID-19 Surveillance Team |
| EPI_ISL_2849124 | Lighthouse Lab in Alderley Park | Wellcome Sanger Institute for the COVID-19 Genomics UK (COG-UK) Consortium | Jacquelyn Wynn, Mairead Hyland, The Lighthouse Lab in Alderley Park and Alex Alderton, Roberto Amato, Jeffrey Barrett, Sonia Goncalves, Ewan Harrison, David K. Jackson, Ian Johnston, Dominic Kwiatkowski, Cordelia Langford, John Sillitoe on behalf of the Wellcome Sanger Institute COVID-19 Surveillance Team |
| EPI_ISL_2849125 | Lighthouse Laboratory Plymouth | Wellcome Sanger Institute for the COVID-19 Genomics UK (COG-UK) Consortium | Lighthouse Laboratory Plymouth and Alex Alderton, Roberto Amato, Jeffrey Barrett, Sonia Goncalves, Ewan Harrison, David K. Jackson, Ian Johnston, Dominic Kwiatkowski, Cordelia Langford, John Sillitoe on behalf of the Wellcome Sanger Institute COVID-19 Surveillance Team |
| EPI_ISL_2849126, EPI_ISL_2849127, EPI_ISL_2849128 | Lighthouse Lab in Alderley Park | Wellcome Sanger Institute for the COVID-19 Genomics UK (COG-UK) Consortium | Jacquelyn Wynn, Mairead Hyland, The Lighthouse Lab in Alderley Park and Alex Alderton, Roberto Amato, Jeffrey Barrett, Sonia Goncalves, Ewan Harrison, David K. Jackson, Ian Johnston, Dominic Kwiatkowski, Cordelia Langford, John Sillitoe on behalf of the Wellcome Sanger Institute COVID-19 Surveillance Team |
| EPI_ISL_2849129 | Lighthouse Laboratory Plymouth | Wellcome Sanger Institute for the COVID-19 Genomics UK (COG-UK) Consortium | Lighthouse Laboratory Plymouth and Alex Alderton, Roberto Amato, Jeffrey Barrett, Sonia Goncalves, Ewan Harrison, David K. Jackson, Ian Johnston, Dominic Kwiatkowski, Cordelia Langford, John Sillitoe on behalf of the Wellcome Sanger Institute COVID-19 Surveillance Team |
| EPI_ISL_2849130 | Lighthouse Lab in Alderley Park | Wellcome Sanger Institute for the COVID-19 Genomics UK (COG-UK) Consortium | Jacquelyn Wynn, Mairead Hyland, The Lighthouse Lab in Alderley Park and Alex Alderton, Roberto Amato, Jeffrey Barrett, Sonia Goncalves, Ewan Harrison, David K. Jackson, Ian Johnston, Dominic Kwiatkowski, Cordelia Langford, John Sillitoe on behalf of the Wellcome Sanger Institute COVID-19 Surveillance Team |
| EPI_ISL_2849131 | Lighthouse Laboratory Plymouth | Wellcome Sanger Institute for the COVID-19 Genomics UK (COG-UK) Consortium | Lighthouse Laboratory Plymouth and Alex Alderton, Roberto Amato, Jeffrey Barrett, Sonia Goncalves, Ewan Harrison, David K. Jackson, Ian Johnston, Dominic Kwiatkowski, Cordelia Langford, John Sillitoe on behalf of the Wellcome Sanger Institute COVID-19 Surveillance Team |
| EPI_ISL_2849132 | Lighthouse Lab in Alderley Park | Wellcome Sanger Institute for the COVID-19 Genomics UK (COG-UK) Consortium | Jacquelyn Wynn, Mairead Hyland, The Lighthouse Lab in Alderley Park and Alex Alderton, Roberto Amato, Jeffrey Barrett, Sonia Goncalves, Ewan Harrison, David K. Jackson, Ian Johnston, Dominic Kwiatkowski, Cordelia Langford, John Sillitoe on behalf of the Wellcome Sanger Institute COVID-19 Surveillance Team |
| EPI_ISL_2849134, EPI_ISL_2849135, EPI_ISL_2849136, EPI_ISL_2849137 | Lighthouse Laboratory Plymouth | Wellcome Sanger Institute for the COVID-19 Genomics UK (COG-UK) Consortium | Lighthouse Laboratory Plymouth and Alex Alderton, Roberto Amato, Jeffrey Barrett, Sonia Goncalves, Ewan Harrison, David K. Jackson, Ian Johnston, Dominic Kwiatkowski, Cordelia Langford, John Sillitoe on behalf of the Wellcome Sanger Institute COVID-19 Surveillance Team |
| EPI_ISL_2849138, EPI_ISL_2849139, EPI_ISL_2849140, EPI_ISL_2849141 | Lighthouse Lab in Alderley Park | Wellcome Sanger Institute for the COVID-19 Genomics UK (COG-UK) Consortium | Jacquelyn Wynn, Mairead Hyland, The Lighthouse Lab in Alderley Park and Alex Alderton, Roberto Amato, Jeffrey Barrett, Sonia Goncalves, Ewan Harrison, David K. Jackson, Ian Johnston, Dominic Kwiatkowski, Cordelia Langford, John Sillitoe on behalf of the Wellcome Sanger Institute COVID-19 Surveillance Team |
| EPI_ISL_2849142 | Lighthouse Laboratory Plymouth | Wellcome Sanger Institute for the COVID-19 Genomics UK (COG-UK) Consortium | Lighthouse Laboratory Plymouth and Alex Alderton, Roberto Amato, Jeffrey Barrett, Sonia Goncalves, Ewan Harrison, David K. Jackson, Ian Johnston, Dominic Kwiatkowski, Cordelia Langford, John Sillitoe on behalf of the Wellcome Sanger Institute COVID-19 Surveillance Team |
| EPI_ISL_2849143, EPI_ISL_2849144 | Lighthouse Lab in Alderley Park | Wellcome Sanger Institute for the COVID-19 Genomics UK (COG-UK) Consortium | Jacquelyn Wynn, Mairead Hyland, The Lighthouse Lab in Alderley Park and Alex Alderton, Roberto Amato, Jeffrey Barrett, Sonia Goncalves, Ewan Harrison, David K. Jackson, Ian Johnston, Dominic Kwiatkowski, Cordelia Langford, John Sillitoe on behalf of the Wellcome Sanger Institute COVID-19 Surveillance Team |
| EPI_ISL_2849145 | Lighthouse Laboratory Plymouth | Wellcome Sanger Institute for the COVID-19 Genomics UK (COG-UK) Consortium | Lighthouse Laboratory Plymouth and Alex Alderton, Roberto Amato, Jeffrey Barrett, Sonia Goncalves, Ewan Harrison, David K. Jackson, Ian Johnston, Dominic Kwiatkowski, Cordelia Langford, John Sillitoe on behalf of the Wellcome Sanger Institute COVID-19 Surveillance Team |
| EPI_ISL_2849146, EPI_ISL_2849147, EPI_ISL_2849148, EPI_ISL_2849149 | Lighthouse Lab in Alderley Park | Wellcome Sanger Institute for the COVID-19 Genomics UK (COG-UK) Consortium | Jacquelyn Wynn, Mairead Hyland, The Lighthouse Lab in Alderley Park and Alex Alderton, Roberto Amato, Jeffrey Barrett, Sonia Goncalves, Ewan Harrison, David K. Jackson, Ian Johnston, Dominic Kwiatkowski, Cordelia Langford, John Sillitoe on behalf of the Wellcome Sanger Institute COVID-19 Surveillance Team |
| EPI_ISL_2849150, EPI_ISL_2849151 | Lighthouse Laboratory Plymouth | Wellcome Sanger Institute for the COVID-19 Genomics UK (COG-UK) Consortium | Lighthouse Laboratory Plymouth and Alex Alderton, Roberto Amato, Jeffrey Barrett, Sonia Goncalves, Ewan Harrison, David K. Jackson, Ian Johnston, Dominic Kwiatkowski, Cordelia Langford, John Sillitoe on behalf of the Wellcome Sanger Institute COVID-19 Surveillance Team |
| EPI_ISL_2849152 | Lighthouse Lab in Alderley Park | Wellcome Sanger Institute for the COVID-19 Genomics UK (COG-UK) Consortium | Jacquelyn Wynn, Mairead Hyland, The Lighthouse Lab in Alderley Park and Alex Alderton, Roberto Amato, Jeffrey Barrett, Sonia Goncalves, Ewan Harrison, David K. Jackson, Ian Johnston, Dominic Kwiatkowski, Cordelia Langford, John Sillitoe on behalf of the Wellcome Sanger Institute COVID-19 Surveillance Team |
| EPI_ISL_2849153 | Lighthouse Laboratory Plymouth | Wellcome Sanger Institute for the COVID-19 Genomics UK (COG-UK) Consortium | Lighthouse Laboratory Plymouth and Alex Alderton, Roberto Amato, Jeffrey Barrett, Sonia Goncalves, Ewan Harrison, David K. Jackson, Ian Johnston, Dominic Kwiatkowski, Cordelia Langford, John Sillitoe on behalf of the Wellcome Sanger Institute COVID-19 Surveillance Team |
| EPI_ISL_2849154, EPI_ISL_2849156, EPI_ISL_2849157 | Lighthouse Lab in Alderley Park | Wellcome Sanger Institute for the COVID-19 Genomics UK (COG-UK) Consortium | Jacquelyn Wynn, Mairead Hyland, The Lighthouse Lab in Alderley Park and Alex Alderton, Roberto Amato, Jeffrey Barrett, Sonia Goncalves, Ewan Harrison, David K. Jackson, Ian Johnston, Dominic Kwiatkowski, Cordelia Langford, John Sillitoe on behalf of the Wellcome Sanger Institute COVID-19 Surveillance Team |
| EPI_ISL_2849158 | Lighthouse Laboratory Plymouth | Wellcome Sanger Institute for the COVID-19 Genomics UK (COG-UK) Consortium | Lighthouse Laboratory Plymouth and Alex Alderton, Roberto Amato, Jeffrey Barrett, Sonia Goncalves, Ewan Harrison, David K. Jackson, Ian Johnston, Dominic Kwiatkowski, Cordelia Langford, John Sillitoe on behalf of the Wellcome Sanger Institute COVID-19 Surveillance Team |
| EPI_ISL_2849159 | Lighthouse Lab in Alderley Park | Wellcome Sanger Institute for the COVID-19 Genomics UK (COG-UK) Consortium | Jacquelyn Wynn, Mairead Hyland, The Lighthouse Lab in Alderley Park and Alex Alderton, Roberto Amato, Jeffrey Barrett, Sonia Goncalves, Ewan Harrison, David K. Jackson, Ian Johnston, Dominic Kwiatkowski, Cordelia Langford, John Sillitoe on behalf of the Wellcome Sanger Institute COVID-19 Surveillance Team |
| EPI_ISL_2849160, EPI_ISL_2849161 | Lighthouse Laboratory Plymouth | Wellcome Sanger Institute for the COVID-19 Genomics UK (COG-UK) Consortium | Lighthouse Laboratory Plymouth and Alex Alderton, Roberto Amato, Jeffrey Barrett, Sonia Goncalves, Ewan Harrison, David K. Jackson, Ian Johnston, Dominic Kwiatkowski, Cordelia Langford, John Sillitoe on behalf of the Wellcome Sanger Institute COVID-19 Surveillance Team |
| EPI_ISL_2849162, EPI_ISL_2849163 | Lighthouse Lab in Alderley Park | Wellcome Sanger Institute for the COVID-19 Genomics UK (COG-UK) Consortium | Jacquelyn Wynn, Mairead Hyland, The Lighthouse Lab in Alderley Park and Alex Alderton, Roberto Amato, Jeffrey Barrett, Sonia Goncalves, Ewan Harrison, David K. Jackson, Ian Johnston, Dominic Kwiatkowski, Cordelia Langford, John Sillitoe on behalf of the Wellcome Sanger Institute COVID-19 Surveillance Team |
| EPI_ISL_2849164, EPI_ISL_2849165 | Lighthouse Laboratory Plymouth | Wellcome Sanger Institute for the COVID-19 Genomics UK (COG-UK) Consortium | Lighthouse Laboratory Plymouth and Alex Alderton, Roberto Amato, Jeffrey Barrett, Sonia Goncalves, Ewan Harrison, David K. Jackson, Ian Johnston, Dominic Kwiatkowski, Cordelia Langford, John Sillitoe on behalf of the Wellcome Sanger Institute COVID-19 Surveillance Team |

| | | | |
|---|---|---|---|
| EPI_ISL_2849166 | Lighthouse Lab in Alderley Park | Wellcome Sanger Institute for the COVID-19 Genomics UK (COG-UK) Consortium | Jacquelyn Wynn, Mairead Hyland, The Lighthouse Lab in Alderley Park and Alex Alderton, Roberto Amato, Jeffrey Barrett, Sonia Goncalves, Ewan Harrison, David K. Jackson, Ian Johnston, Dominic Kwiatkowski, Cordelia Langford, John Sillitoe on behalf of the Wellcome Sanger Institute COVID-19 Surveillance Team |
| EPI_ISL_2849167 | Lighthouse Laboratory Plymouth | Wellcome Sanger Institute for the COVID-19 Genomics UK (COG-UK) Consortium | Lighthouse Laboratory Plymouth and Alex Alderton, Roberto Amato, Jeffrey Barrett, Sonia Goncalves, Ewan Harrison, David K. Jackson, Ian Johnston, Dominic Kwiatkowski, Cordelia Langford, John Sillitoe on behalf of the Wellcome Sanger Institute COVID-19 Surveillance Team |
| EPI_ISL_2849168, EPI_ISL_2849169, EPI_ISL_2849170 | Lighthouse Lab in Alderley Park | Wellcome Sanger Institute for the COVID-19 Genomics UK (COG-UK) Consortium | Jacquelyn Wynn, Mairead Hyland, The Lighthouse Lab in Alderley Park and Alex Alderton, Roberto Amato, Jeffrey Barrett, Sonia Goncalves, Ewan Harrison, David K. Jackson, Ian Johnston, Dominic Kwiatkowski, Cordelia Langford, John Sillitoe on behalf of the Wellcome Sanger Institute COVID-19 Surveillance Team |
| EPI_ISL_2849171, EPI_ISL_2849172, EPI_ISL_2849173, EPI_ISL_2849174 | Lighthouse Laboratory Plymouth | Wellcome Sanger Institute for the COVID-19 Genomics UK (COG-UK) Consortium | Lighthouse Laboratory Plymouth and Alex Alderton, Roberto Amato, Jeffrey Barrett, Sonia Goncalves, Ewan Harrison, David K. Jackson, Ian Johnston, Dominic Kwiatkowski, Cordelia Langford, John Sillitoe on behalf of the Wellcome Sanger Institute COVID-19 Surveillance Team |
| EPI_ISL_2849175 | Lighthouse Lab in Alderley Park | Wellcome Sanger Institute for the COVID-19 Genomics UK (COG-UK) Consortium | Jacquelyn Wynn, Mairead Hyland, The Lighthouse Lab in Alderley Park and Alex Alderton, Roberto Amato, Jeffrey Barrett, Sonia Goncalves, Ewan Harrison, David K. Jackson, Ian Johnston, Dominic Kwiatkowski, Cordelia Langford, John Sillitoe on behalf of the Wellcome Sanger Institute COVID-19 Surveillance Team |
| EPI_ISL_2849176, EPI_ISL_2849177, EPI_ISL_2849178, EPI_ISL_2849179 | Lighthouse Laboratory Plymouth | Wellcome Sanger Institute for the COVID-19 Genomics UK (COG-UK) Consortium | Lighthouse Laboratory Plymouth and Alex Alderton, Roberto Amato, Jeffrey Barrett, Sonia Goncalves, Ewan Harrison, David K. Jackson, Ian Johnston, Dominic Kwiatkowski, Cordelia Langford, John Sillitoe on behalf of the Wellcome Sanger Institute COVID-19 Surveillance Team |
| EPI_ISL_2849180 | Lighthouse Lab in Alderley Park | Wellcome Sanger Institute for the COVID-19 Genomics UK (COG-UK) Consortium | Jacquelyn Wynn, Mairead Hyland, The Lighthouse Lab in Alderley Park and Alex Alderton, Roberto Amato, Jeffrey Barrett, Sonia Goncalves, Ewan Harrison, David K. Jackson, Ian Johnston, Dominic Kwiatkowski, Cordelia Langford, John Sillitoe on behalf of the Wellcome Sanger Institute COVID-19 Surveillance Team |
| EPI_ISL_2849181, EPI_ISL_2849182 | Lighthouse Laboratory Plymouth | Wellcome Sanger Institute for the COVID-19 Genomics UK (COG-UK) Consortium | Lighthouse Laboratory Plymouth and Alex Alderton, Roberto Amato, Jeffrey Barrett, Sonia Goncalves, Ewan Harrison, David K. Jackson, Ian Johnston, Dominic Kwiatkowski, Cordelia Langford, John Sillitoe on behalf of the Wellcome Sanger Institute COVID-19 Surveillance Team |
| EPI_ISL_2849183, EPI_ISL_2849184 | Lighthouse Lab in Alderley Park | Wellcome Sanger Institute for the COVID-19 Genomics UK (COG-UK) Consortium | Jacquelyn Wynn, Mairead Hyland, The Lighthouse Lab in Alderley Park and Alex Alderton, Roberto Amato, Jeffrey Barrett, Sonia Goncalves, Ewan Harrison, David K. Jackson, Ian Johnston, Dominic Kwiatkowski, Cordelia Langford, John Sillitoe on behalf of the Wellcome Sanger Institute COVID-19 Surveillance Team |
| EPI_ISL_2849185, EPI_ISL_2849186 | Lighthouse Laboratory Plymouth | Wellcome Sanger Institute for the COVID-19 Genomics UK (COG-UK) Consortium | Lighthouse Laboratory Plymouth and Alex Alderton, Roberto Amato, Jeffrey Barrett, Sonia Goncalves, Ewan Harrison, David K. Jackson, Ian Johnston, Dominic Kwiatkowski, Cordelia Langford, John Sillitoe on behalf of the Wellcome Sanger Institute COVID-19 Surveillance Team |
| EPI_ISL_2849187, EPI_ISL_2849188 | Lighthouse Lab in Alderley Park | Wellcome Sanger Institute for the COVID-19 Genomics UK (COG-UK) Consortium | Jacquelyn Wynn, Mairead Hyland, The Lighthouse Lab in Alderley Park and Alex Alderton, Roberto Amato, Jeffrey Barrett, Sonia Goncalves, Ewan Harrison, David K. Jackson, Ian Johnston, Dominic Kwiatkowski, Cordelia Langford, John Sillitoe on behalf of the Wellcome Sanger Institute COVID-19 Surveillance Team |
| EPI_ISL_2849189 | Lighthouse Laboratory Plymouth | Wellcome Sanger Institute for the COVID-19 Genomics UK (COG-UK) Consortium | Lighthouse Laboratory Plymouth and Alex Alderton, Roberto Amato, Jeffrey Barrett, Sonia Goncalves, Ewan Harrison, David K. Jackson, Ian Johnston, Dominic Kwiatkowski, Cordelia Langford, John Sillitoe on behalf of the Wellcome Sanger Institute COVID-19 Surveillance Team |
| EPI_ISL_2849190, EPI_ISL_2849191 | Lighthouse Lab in Alderley Park | Wellcome Sanger Institute for the COVID-19 Genomics UK (COG-UK) Consortium | Jacquelyn Wynn, Mairead Hyland, The Lighthouse Lab in Alderley Park and Alex Alderton, Roberto Amato, Jeffrey Barrett, Sonia Goncalves, Ewan Harrison, David K. Jackson, Ian Johnston, Dominic Kwiatkowski, Cordelia Langford, John Sillitoe on behalf of the Wellcome Sanger Institute COVID-19 Surveillance Team |
| EPI_ISL_2849192 | Lighthouse Laboratory Plymouth | Wellcome Sanger Institute for the COVID-19 Genomics UK (COG-UK) Consortium | Lighthouse Laboratory Plymouth and Alex Alderton, Roberto Amato, Jeffrey Barrett, Sonia Goncalves, Ewan Harrison, David K. Jackson, Ian Johnston, Dominic Kwiatkowski, Cordelia Langford, John Sillitoe on behalf of the Wellcome Sanger Institute COVID-19 Surveillance Team |
| EPI_ISL_2849193, EPI_ISL_2849194, EPI_ISL_2849195, EPI_ISL_2849196 | Lighthouse Lab in Alderley Park | Wellcome Sanger Institute for the COVID-19 Genomics UK (COG-UK) Consortium | Jacquelyn Wynn, Mairead Hyland, The Lighthouse Lab in Alderley Park and Alex Alderton, Roberto Amato, Jeffrey Barrett, Sonia Goncalves, Ewan Harrison, David K. Jackson, Ian Johnston, Dominic Kwiatkowski, Cordelia Langford, John Sillitoe on behalf of the Wellcome Sanger Institute COVID-19 Surveillance Team |
| EPI_ISL_2849197, EPI_ISL_2849198 | Lighthouse Laboratory Plymouth | Wellcome Sanger Institute for the COVID-19 Genomics UK (COG-UK) Consortium | Lighthouse Laboratory Plymouth and Alex Alderton, Roberto Amato, Jeffrey Barrett, Sonia Goncalves, Ewan Harrison, David K. Jackson, Ian Johnston, Dominic Kwiatkowski, Cordelia Langford, John Sillitoe on behalf of the Wellcome Sanger Institute COVID-19 Surveillance Team |
| EPI_ISL_2849199 | Lighthouse Lab in Alderley Park | Wellcome Sanger Institute for the COVID-19 Genomics UK (COG-UK) Consortium | Jacquelyn Wynn, Mairead Hyland, The Lighthouse Lab in Alderley Park and Alex Alderton, Roberto Amato, Jeffrey Barrett, Sonia Goncalves, Ewan Harrison, David K. Jackson, Ian Johnston, Dominic Kwiatkowski, Cordelia Langford, John Sillitoe on behalf of the Wellcome Sanger Institute COVID-19 Surveillance Team |
| EPI_ISL_2849200 | Lighthouse Laboratory Plymouth | Wellcome Sanger Institute for the COVID-19 Genomics UK (COG-UK) Consortium | Lighthouse Laboratory Plymouth and Alex Alderton, Roberto Amato, Jeffrey Barrett, Sonia Goncalves, Ewan Harrison, David K. Jackson, Ian Johnston, Dominic Kwiatkowski, Cordelia Langford, John Sillitoe on behalf of the Wellcome Sanger Institute COVID-19 Surveillance Team |
| EPI_ISL_2849201, EPI_ISL_2849202, EPI_ISL_2849203, EPI_ISL_2849204 | Lighthouse Lab in Alderley Park | Wellcome Sanger Institute for the COVID-19 Genomics UK (COG-UK) Consortium | Jacquelyn Wynn, Mairead Hyland, The Lighthouse Lab in Alderley Park and Alex Alderton, Roberto Amato, Jeffrey Barrett, Sonia Goncalves, Ewan Harrison, David K. Jackson, Ian Johnston, Dominic Kwiatkowski, Cordelia Langford, John Sillitoe on behalf of the Wellcome Sanger Institute COVID-19 Surveillance Team |
| EPI_ISL_2849205, EPI_ISL_2849206 | Lighthouse Laboratory Plymouth | Wellcome Sanger Institute for the COVID-19 Genomics UK (COG-UK) Consortium | Lighthouse Laboratory Plymouth and Alex Alderton, Roberto Amato, Jeffrey Barrett, Sonia Goncalves, Ewan Harrison, David K. Jackson, Ian Johnston, Dominic Kwiatkowski, Cordelia Langford, John Sillitoe on behalf of the Wellcome Sanger Institute COVID-19 Surveillance Team |
| EPI_ISL_2849207, EPI_ISL_2849208, EPI_ISL_2849209, EPI_ISL_2849210, EPI_ISL_2849211 | Lighthouse Lab in Alderley Park | Wellcome Sanger Institute for the COVID-19 Genomics UK (COG-UK) Consortium | Jacquelyn Wynn, Mairead Hyland, The Lighthouse Lab in Alderley Park and Alex Alderton, Roberto Amato, Jeffrey Barrett, Sonia Goncalves, Ewan Harrison, David K. Jackson, Ian Johnston, Dominic Kwiatkowski, Cordelia Langford, John Sillitoe on behalf of the Wellcome Sanger Institute COVID-19 Surveillance Team |
| EPI_ISL_2849212, EPI_ISL_2849213 | Lighthouse Laboratory Plymouth | Wellcome Sanger Institute for the COVID-19 Genomics UK (COG-UK) Consortium | Lighthouse Laboratory Plymouth and Alex Alderton, Roberto Amato, Jeffrey Barrett, Sonia Goncalves, Ewan Harrison, David K. Jackson, Ian Johnston, Dominic Kwiatkowski, Cordelia Langford, John Sillitoe on behalf of the Wellcome Sanger Institute COVID-19 Surveillance Team |
| EPI_ISL_2849214 | Lighthouse Lab in Alderley Park | Wellcome Sanger Institute for the COVID-19 Genomics UK (COG-UK) Consortium | Jacquelyn Wynn, Mairead Hyland, The Lighthouse Lab in Alderley Park and Alex Alderton, Roberto Amato, Jeffrey Barrett, Sonia Goncalves, Ewan Harrison, David K. Jackson, Ian Johnston, Dominic Kwiatkowski, Cordelia Langford, John Sillitoe on behalf of the Wellcome Sanger Institute COVID-19 Surveillance Team |
| EPI_ISL_2849215, EPI_ISL_2849216 | Lighthouse Laboratory Plymouth | Wellcome Sanger Institute for the COVID-19 Genomics UK (COG-UK) Consortium | Lighthouse Laboratory Plymouth and Alex Alderton, Roberto Amato, Jeffrey Barrett, Sonia Goncalves, Ewan Harrison, David K. Jackson, Ian Johnston, Dominic Kwiatkowski, Cordelia Langford, John Sillitoe on behalf of the Wellcome Sanger Institute COVID-19 Surveillance Team |
| EPI_ISL_2849217, EPI_ISL_2849218, EPI_ISL_2849219, EPI_ISL_2849220, EPI_ISL_2849221 | Lighthouse Lab in Alderley Park | Wellcome Sanger Institute for the COVID-19 Genomics UK (COG-UK) Consortium | Jacquelyn Wynn, Mairead Hyland, The Lighthouse Lab in Alderley Park and Alex Alderton, Roberto Amato, Jeffrey Barrett, Sonia Goncalves, Ewan Harrison, David K. Jackson, Ian Johnston, Dominic Kwiatkowski, Cordelia Langford, John Sillitoe on behalf of the Wellcome Sanger Institute COVID-19 Surveillance Team |
| EPI_ISL_2849222 | Lighthouse Laboratory Plymouth | Wellcome Sanger Institute for the COVID-19 Genomics UK (COG-UK) Consortium | Lighthouse Laboratory Plymouth and Alex Alderton, Roberto Amato, Jeffrey Barrett, Sonia Goncalves, Ewan Harrison, David K. Jackson, Ian Johnston, Dominic Kwiatkowski, Cordelia Langford, John Sillitoe on behalf of the Wellcome Sanger Institute COVID-19 Surveillance Team |
| EPI_ISL_2849223, EPI_ISL_2849224, EPI_ISL_2849225 | Lighthouse Lab in Alderley Park | Wellcome Sanger Institute for the COVID-19 Genomics UK (COG-UK) Consortium | Jacquelyn Wynn, Mairead Hyland, The Lighthouse Lab in Alderley Park and Alex Alderton, Roberto Amato, Jeffrey Barrett, Sonia Goncalves, Ewan Harrison, David K. Jackson, Ian Johnston, Dominic Kwiatkowski, Cordelia Langford, John Sillitoe on behalf of the Wellcome Sanger Institute COVID-19 Surveillance Team |
| EPI_ISL_2849226 | Lighthouse Laboratory Plymouth | Wellcome Sanger Institute for the COVID-19 Genomics UK (COG-UK) Consortium | Lighthouse Laboratory Plymouth and Alex Alderton, Roberto Amato, Jeffrey Barrett, Sonia Goncalves, Ewan Harrison, David K. Jackson, Ian Johnston, Dominic Kwiatkowski, Cordelia Langford, John Sillitoe on behalf of the Wellcome Sanger Institute COVID-19 Surveillance Team |
| EPI_ISL_2849227 | Lighthouse Lab in Alderley Park | Wellcome Sanger Institute for the COVID-19 Genomics UK (COG-UK) Consortium | Jacquelyn Wynn, Mairead Hyland, The Lighthouse Lab in Alderley Park and Alex Alderton, Roberto Amato, Jeffrey Barrett, Sonia Goncalves, Ewan Harrison, David K. Jackson, Ian Johnston, Dominic Kwiatkowski, Cordelia Langford, John Sillitoe on behalf of the Wellcome Sanger Institute COVID-19 Surveillance Team |
| EPI_ISL_2849228, EPI_ISL_2849229, EPI_ISL_2849230, EPI_ISL_2849231 | Lighthouse Laboratory Plymouth | Wellcome Sanger Institute for the COVID-19 Genomics UK (COG-UK) Consortium | Lighthouse Laboratory Plymouth and Alex Alderton, Roberto Amato, Jeffrey Barrett, Sonia Goncalves, Ewan Harrison, David K. Jackson, Ian Johnston, Dominic Kwiatkowski, Cordelia Langford, John Sillitoe on behalf of the Wellcome Sanger Institute COVID-19 Surveillance Team |
| EPI_ISL_2849232 | Lighthouse Lab in Alderley Park | Wellcome Sanger Institute for the COVID-19 Genomics UK (COG-UK) Consortium | Jacquelyn Wynn, Mairead Hyland, The Lighthouse Lab in Alderley Park and Alex Alderton, Roberto Amato, Jeffrey Barrett, Sonia Goncalves, Ewan Harrison, David K. Jackson, Ian Johnston, Dominic Kwiatkowski, Cordelia Langford, John Sillitoe on behalf of the Wellcome Sanger Institute COVID-19 Surveillance Team |
| EPI_ISL_2849233 | Lighthouse Laboratory Plymouth | Wellcome Sanger Institute for the COVID-19 Genomics UK (COG-UK) Consortium | Lighthouse Laboratory Plymouth and Alex Alderton, Roberto Amato, Jeffrey Barrett, Sonia Goncalves, Ewan Harrison, David K. Jackson, Ian Johnston, Dominic Kwiatkowski, Cordelia Langford, John Sillitoe on behalf of the Wellcome Sanger Institute COVID-19 Surveillance Team |
| EPI_ISL_2849234 | Lighthouse Lab in Alderley Park | Wellcome Sanger Institute for the COVID-19 Genomics UK (COG-UK) Consortium | Jacquelyn Wynn, Mairead Hyland, The Lighthouse Lab in Alderley Park and Alex Alderton, Roberto Amato, Jeffrey Barrett, Sonia Goncalves, Ewan Harrison, David K. Jackson, Ian Johnston, Dominic Kwiatkowski, Cordelia Langford, John Sillitoe on behalf of the Wellcome Sanger Institute COVID-19 Surveillance Team |
| EPI_ISL_2849235, EPI_ISL_2849236, EPI_ISL_2849237 | Lighthouse Laboratory Plymouth | Wellcome Sanger Institute for the COVID-19 Genomics UK (COG-UK) Consortium | Lighthouse Laboratory Plymouth and Alex Alderton, Roberto Amato, Jeffrey Barrett, Sonia Goncalves, Ewan Harrison, David K. Jackson, Ian Johnston, Dominic Kwiatkowski, Cordelia Langford, John Sillitoe on behalf of the Wellcome Sanger Institute COVID-19 Surveillance Team |
| EPI_ISL_2849238, EPI_ISL_2849239, EPI_ISL_2849240, EPI_ISL_2849241, EPI_ISL_2849242 | Lighthouse Lab in Alderley Park | Wellcome Sanger Institute for the COVID-19 Genomics UK (COG-UK) Consortium | Jacquelyn Wynn, Mairead Hyland, The Lighthouse Lab in Alderley Park and Alex Alderton, Roberto Amato, Jeffrey Barrett, Sonia Goncalves, Ewan Harrison, David K. Jackson, Ian Johnston, Dominic Kwiatkowski, Cordelia Langford, John Sillitoe on behalf of the Wellcome Sanger Institute COVID-19 Surveillance Team |
| EPI_ISL_2849243, | Lighthouse Laboratory Plymouth | Wellcome Sanger Institute for the COVID-19 Genomics UK (COG-UK) | Lighthouse Laboratory Plymouth and Alex Alderton, Roberto Amato, Jeffrey Barrett, Sonia Goncalves, Ewan Harrison, David K. Jackson, Ian Johnston, Dominic Kwiatkowski, Cordelia Langford, John Sillitoe on behalf of the Wellcome Sanger |

| | | | |
|---|---|---|---|
| EPI_ISL_2849244, EPI_ISL_2849245, EPI_ISL_2849246, EPI_ISL_2849247, EPI_ISL_2849248 | | Consortium | Institute COVID-19 Surveillance Team |
| EPI_ISL_2849249 | Lighthouse Lab in Alderley Park | Wellcome Sanger Institute for the COVID-19 Genomics UK (COG-UK) Consortium | Jacquelyn Wynn, Mairead Hyland, The Lighthouse Lab in Alderley Park and Alex Alderton, Roberto Amato, Jeffrey Barrett, Sonia Goncalves, Ewan Harrison, David K. Jackson, Ian Johnston, Dominic Kwiatkowski, Cordelia Langford, John Sillitoe on behalf of the Wellcome Sanger Institute COVID-19 Surveillance Team |
| EPI_ISL_2849250 | Lighthouse Laboratory Plymouth | Wellcome Sanger Institute for the COVID-19 Genomics UK (COG-UK) Consortium | Lighthouse Laboratory Plymouth and Alex Alderton, Roberto Amato, Jeffrey Barrett, Sonia Goncalves, Ewan Harrison, David K. Jackson, Ian Johnston, Dominic Kwiatkowski, Cordelia Langford, John Sillitoe on behalf of the Wellcome Sanger Institute COVID-19 Surveillance Team |
| EPI_ISL_2849251 | Lighthouse Lab in Alderley Park | Wellcome Sanger Institute for the COVID-19 Genomics UK (COG-UK) Consortium | Jacquelyn Wynn, Mairead Hyland, The Lighthouse Lab in Alderley Park and Alex Alderton, Roberto Amato, Jeffrey Barrett, Sonia Goncalves, Ewan Harrison, David K. Jackson, Ian Johnston, Dominic Kwiatkowski, Cordelia Langford, John Sillitoe on behalf of the Wellcome Sanger Institute COVID-19 Surveillance Team |
| EPI_ISL_2849252 | Lighthouse Laboratory Plymouth | Wellcome Sanger Institute for the COVID-19 Genomics UK (COG-UK) Consortium | Lighthouse Laboratory Plymouth and Alex Alderton, Roberto Amato, Jeffrey Barrett, Sonia Goncalves, Ewan Harrison, David K. Jackson, Ian Johnston, Dominic Kwiatkowski, Cordelia Langford, John Sillitoe on behalf of the Wellcome Sanger Institute COVID-19 Surveillance Team |
| EPI_ISL_2849253 | Lighthouse Lab in Alderley Park | Wellcome Sanger Institute for the COVID-19 Genomics UK (COG-UK) Consortium | Jacquelyn Wynn, Mairead Hyland, The Lighthouse Lab in Alderley Park and Alex Alderton, Roberto Amato, Jeffrey Barrett, Sonia Goncalves, Ewan Harrison, David K. Jackson, Ian Johnston, Dominic Kwiatkowski, Cordelia Langford, John Sillitoe on behalf of the Wellcome Sanger Institute COVID-19 Surveillance Team |
| EPI_ISL_2849254, EPI_ISL_2849255, EPI_ISL_2849256, EPI_ISL_2849257, EPI_ISL_2849258 | Lighthouse Laboratory Plymouth | Wellcome Sanger Institute for the COVID-19 Genomics UK (COG-UK) Consortium | Lighthouse Laboratory Plymouth and Alex Alderton, Roberto Amato, Jeffrey Barrett, Sonia Goncalves, Ewan Harrison, David K. Jackson, Ian Johnston, Dominic Kwiatkowski, Cordelia Langford, John Sillitoe on behalf of the Wellcome Sanger Institute COVID-19 Surveillance Team |
| EPI_ISL_2849259 | Lighthouse Lab in Alderley Park | Wellcome Sanger Institute for the COVID-19 Genomics UK (COG-UK) Consortium | Jacquelyn Wynn, Mairead Hyland, The Lighthouse Lab in Alderley Park and Alex Alderton, Roberto Amato, Jeffrey Barrett, Sonia Goncalves, Ewan Harrison, David K. Jackson, Ian Johnston, Dominic Kwiatkowski, Cordelia Langford, John Sillitoe on behalf of the Wellcome Sanger Institute COVID-19 Surveillance Team |
| EPI_ISL_2849260 | Lighthouse Laboratory Plymouth | Wellcome Sanger Institute for the COVID-19 Genomics UK (COG-UK) Consortium | Lighthouse Laboratory Plymouth and Alex Alderton, Roberto Amato, Jeffrey Barrett, Sonia Goncalves, Ewan Harrison, David K. Jackson, Ian Johnston, Dominic Kwiatkowski, Cordelia Langford, John Sillitoe on behalf of the Wellcome Sanger Institute COVID-19 Surveillance Team |
| EPI_ISL_2849261 | Lighthouse Lab in Alderley Park | Wellcome Sanger Institute for the COVID-19 Genomics UK (COG-UK) Consortium | Jacquelyn Wynn, Mairead Hyland, The Lighthouse Lab in Alderley Park and Alex Alderton, Roberto Amato, Jeffrey Barrett, Sonia Goncalves, Ewan Harrison, David K. Jackson, Ian Johnston, Dominic Kwiatkowski, Cordelia Langford, John Sillitoe on behalf of the Wellcome Sanger Institute COVID-19 Surveillance Team |
| EPI_ISL_2849262 | Lighthouse Laboratory Plymouth | Wellcome Sanger Institute for the COVID-19 Genomics UK (COG-UK) Consortium | Lighthouse Laboratory Plymouth and Alex Alderton, Roberto Amato, Jeffrey Barrett, Sonia Goncalves, Ewan Harrison, David K. Jackson, Ian Johnston, Dominic Kwiatkowski, Cordelia Langford, John Sillitoe on behalf of the Wellcome Sanger Institute COVID-19 Surveillance Team |
| EPI_ISL_2849263 | Lighthouse Lab in Alderley Park | Wellcome Sanger Institute for the COVID-19 Genomics UK (COG-UK) Consortium | Jacquelyn Wynn, Mairead Hyland, The Lighthouse Lab in Alderley Park and Alex Alderton, Roberto Amato, Jeffrey Barrett, Sonia Goncalves, Ewan Harrison, David K. Jackson, Ian Johnston, Dominic Kwiatkowski, Cordelia Langford, John Sillitoe on behalf of the Wellcome Sanger Institute COVID-19 Surveillance Team |
| EPI_ISL_2849264, EPI_ISL_2849265, EPI_ISL_2849266 | Lighthouse Laboratory Plymouth | Wellcome Sanger Institute for the COVID-19 Genomics UK (COG-UK) Consortium | Lighthouse Laboratory Plymouth and Alex Alderton, Roberto Amato, Jeffrey Barrett, Sonia Goncalves, Ewan Harrison, David K. Jackson, Ian Johnston, Dominic Kwiatkowski, Cordelia Langford, John Sillitoe on behalf of the Wellcome Sanger Institute COVID-19 Surveillance Team |
| EPI_ISL_2849267 | Lighthouse Lab in Alderley Park | Wellcome Sanger Institute for the COVID-19 Genomics UK (COG-UK) Consortium | Jacquelyn Wynn, Mairead Hyland, The Lighthouse Lab in Alderley Park and Alex Alderton, Roberto Amato, Jeffrey Barrett, Sonia Goncalves, Ewan Harrison, David K. Jackson, Ian Johnston, Dominic Kwiatkowski, Cordelia Langford, John Sillitoe on behalf of the Wellcome Sanger Institute COVID-19 Surveillance Team |
| EPI_ISL_2849268, EPI_ISL_2849269, EPI_ISL_2849270, EPI_ISL_2849271 | Lighthouse Laboratory Plymouth | Wellcome Sanger Institute for the COVID-19 Genomics UK (COG-UK) Consortium | Lighthouse Laboratory Plymouth and Alex Alderton, Roberto Amato, Jeffrey Barrett, Sonia Goncalves, Ewan Harrison, David K. Jackson, Ian Johnston, Dominic Kwiatkowski, Cordelia Langford, John Sillitoe on behalf of the Wellcome Sanger Institute COVID-19 Surveillance Team |
| EPI_ISL_2849272 | Lighthouse Lab in Alderley Park | Wellcome Sanger Institute for the COVID-19 Genomics UK (COG-UK) Consortium | Jacquelyn Wynn, Mairead Hyland, The Lighthouse Lab in Alderley Park and Alex Alderton, Roberto Amato, Jeffrey Barrett, Sonia Goncalves, Ewan Harrison, David K. Jackson, Ian Johnston, Dominic Kwiatkowski, Cordelia Langford, John Sillitoe on behalf of the Wellcome Sanger Institute COVID-19 Surveillance Team |
| EPI_ISL_2849273, EPI_ISL_2849274, EPI_ISL_2849275, EPI_ISL_2849276, EPI_ISL_2849277, EPI_ISL_2849278 | Lighthouse Laboratory Plymouth | Wellcome Sanger Institute for the COVID-19 Genomics UK (COG-UK) Consortium | Lighthouse Laboratory Plymouth and Alex Alderton, Roberto Amato, Jeffrey Barrett, Sonia Goncalves, Ewan Harrison, David K. Jackson, Ian Johnston, Dominic Kwiatkowski, Cordelia Langford, John Sillitoe on behalf of the Wellcome Sanger Institute COVID-19 Surveillance Team |
| EPI_ISL_2849279, EPI_ISL_2849280, EPI_ISL_2849281 | Lighthouse Lab in Alderley Park | Wellcome Sanger Institute for the COVID-19 Genomics UK (COG-UK) Consortium | Jacquelyn Wynn, Mairead Hyland, The Lighthouse Lab in Alderley Park and Alex Alderton, Roberto Amato, Jeffrey Barrett, Sonia Goncalves, Ewan Harrison, David K. Jackson, Ian Johnston, Dominic Kwiatkowski, Cordelia Langford, John Sillitoe on behalf of the Wellcome Sanger Institute COVID-19 Surveillance Team |
| EPI_ISL_2849282 | Lighthouse Laboratory Plymouth | Wellcome Sanger Institute for the COVID-19 Genomics UK (COG-UK) Consortium | Lighthouse Laboratory Plymouth and Alex Alderton, Roberto Amato, Jeffrey Barrett, Sonia Goncalves, Ewan Harrison, David K. Jackson, Ian Johnston, Dominic Kwiatkowski, Cordelia Langford, John Sillitoe on behalf of the Wellcome Sanger Institute COVID-19 Surveillance Team |
| EPI_ISL_2849283, EPI_ISL_2849284, EPI_ISL_2849285 | Lighthouse Lab in Alderley Park | Wellcome Sanger Institute for the COVID-19 Genomics UK (COG-UK) Consortium | Jacquelyn Wynn, Mairead Hyland, The Lighthouse Lab in Alderley Park and Alex Alderton, Roberto Amato, Jeffrey Barrett, Sonia Goncalves, Ewan Harrison, David K. Jackson, Ian Johnston, Dominic Kwiatkowski, Cordelia Langford, John Sillitoe on behalf of the Wellcome Sanger Institute COVID-19 Surveillance Team |
| EPI_ISL_2849286 | Lighthouse Laboratory Plymouth | Wellcome Sanger Institute for the COVID-19 Genomics UK (COG-UK) Consortium | Lighthouse Laboratory Plymouth and Alex Alderton, Roberto Amato, Jeffrey Barrett, Sonia Goncalves, Ewan Harrison, David K. Jackson, Ian Johnston, Dominic Kwiatkowski, Cordelia Langford, John Sillitoe on behalf of the Wellcome Sanger Institute COVID-19 Surveillance Team |
| EPI_ISL_2849287 | Lighthouse Lab in Alderley Park | Wellcome Sanger Institute for the COVID-19 Genomics UK (COG-UK) Consortium | Jacquelyn Wynn, Mairead Hyland, The Lighthouse Lab in Alderley Park and Alex Alderton, Roberto Amato, Jeffrey Barrett, Sonia Goncalves, Ewan Harrison, David K. Jackson, Ian Johnston, Dominic Kwiatkowski, Cordelia Langford, John Sillitoe on behalf of the Wellcome Sanger Institute COVID-19 Surveillance Team |
| EPI_ISL_2849288, EPI_ISL_2849289, EPI_ISL_2849290, EPI_ISL_2849291, EPI_ISL_2849292, EPI_ISL_2849293 | Lighthouse Laboratory Plymouth | Wellcome Sanger Institute for the COVID-19 Genomics UK (COG-UK) Consortium | Lighthouse Laboratory Plymouth and Alex Alderton, Roberto Amato, Jeffrey Barrett, Sonia Goncalves, Ewan Harrison, David K. Jackson, Ian Johnston, Dominic Kwiatkowski, Cordelia Langford, John Sillitoe on behalf of the Wellcome Sanger Institute COVID-19 Surveillance Team |
| EPI_ISL_2849294, EPI_ISL_2849295, EPI_ISL_2849296 | Lighthouse Lab in Alderley Park | Wellcome Sanger Institute for the COVID-19 Genomics UK (COG-UK) Consortium | Jacquelyn Wynn, Mairead Hyland, The Lighthouse Lab in Alderley Park and Alex Alderton, Roberto Amato, Jeffrey Barrett, Sonia Goncalves, Ewan Harrison, David K. Jackson, Ian Johnston, Dominic Kwiatkowski, Cordelia Langford, John Sillitoe on behalf of the Wellcome Sanger Institute COVID-19 Surveillance Team |
| EPI_ISL_2849297, EPI_ISL_2849298, EPI_ISL_2849299 | Lighthouse Laboratory Plymouth | Wellcome Sanger Institute for the COVID-19 Genomics UK (COG-UK) Consortium | Lighthouse Laboratory Plymouth and Alex Alderton, Roberto Amato, Jeffrey Barrett, Sonia Goncalves, Ewan Harrison, David K. Jackson, Ian Johnston, Dominic Kwiatkowski, Cordelia Langford, John Sillitoe on behalf of the Wellcome Sanger Institute COVID-19 Surveillance Team |
| EPI_ISL_2849300 | Lighthouse Lab in Alderley Park | Wellcome Sanger Institute for the COVID-19 Genomics UK (COG-UK) Consortium | Jacquelyn Wynn, Mairead Hyland, The Lighthouse Lab in Alderley Park and Alex Alderton, Roberto Amato, Jeffrey Barrett, Sonia Goncalves, Ewan Harrison, David K. Jackson, Ian Johnston, Dominic Kwiatkowski, Cordelia Langford, John Sillitoe on behalf of the Wellcome Sanger Institute COVID-19 Surveillance Team |
| EPI_ISL_2849301, EPI_ISL_2849302 | Lighthouse Laboratory Plymouth | Wellcome Sanger Institute for the COVID-19 Genomics UK (COG-UK) Consortium | Lighthouse Laboratory Plymouth and Alex Alderton, Roberto Amato, Jeffrey Barrett, Sonia Goncalves, Ewan Harrison, David K. Jackson, Ian Johnston, Dominic Kwiatkowski, Cordelia Langford, John Sillitoe on behalf of the Wellcome Sanger Institute COVID-19 Surveillance Team |
| EPI_ISL_2849303 | Lighthouse Lab in Alderley Park | Wellcome Sanger Institute for the COVID-19 Genomics UK (COG-UK) Consortium | Jacquelyn Wynn, Mairead Hyland, The Lighthouse Lab in Alderley Park and Alex Alderton, Roberto Amato, Jeffrey Barrett, Sonia Goncalves, Ewan Harrison, David K. Jackson, Ian Johnston, Dominic Kwiatkowski, Cordelia Langford, John Sillitoe on behalf of the Wellcome Sanger Institute COVID-19 Surveillance Team |
| EPI_ISL_2849304 | Lighthouse Laboratory Plymouth | Wellcome Sanger Institute for the COVID-19 Genomics UK (COG-UK) Consortium | Lighthouse Laboratory Plymouth and Alex Alderton, Roberto Amato, Jeffrey Barrett, Sonia Goncalves, Ewan Harrison, David K. Jackson, Ian Johnston, Dominic Kwiatkowski, Cordelia Langford, John Sillitoe on behalf of the Wellcome Sanger Institute COVID-19 Surveillance Team |
| EPI_ISL_2849305 | Lighthouse Lab in Alderley Park | Wellcome Sanger Institute for the COVID-19 Genomics UK (COG-UK) Consortium | Jacquelyn Wynn, Mairead Hyland, The Lighthouse Lab in Alderley Park and Alex Alderton, Roberto Amato, Jeffrey Barrett, Sonia Goncalves, Ewan Harrison, David K. Jackson, Ian Johnston, Dominic Kwiatkowski, Cordelia Langford, John Sillitoe on behalf of the Wellcome Sanger Institute COVID-19 Surveillance Team |
| EPI_ISL_2849306, EPI_ISL_2849307 | Lighthouse Laboratory Plymouth | Wellcome Sanger Institute for the COVID-19 Genomics UK (COG-UK) Consortium | Lighthouse Laboratory Plymouth and Alex Alderton, Roberto Amato, Jeffrey Barrett, Sonia Goncalves, Ewan Harrison, David K. Jackson, Ian Johnston, Dominic Kwiatkowski, Cordelia Langford, John Sillitoe on behalf of the Wellcome Sanger Institute COVID-19 Surveillance Team |
| EPI_ISL_2849308, EPI_ISL_2849309 | Lighthouse Lab in Alderley Park | Wellcome Sanger Institute for the COVID-19 Genomics UK (COG-UK) Consortium | Jacquelyn Wynn, Mairead Hyland, The Lighthouse Lab in Alderley Park and Alex Alderton, Roberto Amato, Jeffrey Barrett, Sonia Goncalves, Ewan Harrison, David K. Jackson, Ian Johnston, Dominic Kwiatkowski, Cordelia Langford, John Sillitoe on behalf of the Wellcome Sanger Institute COVID-19 Surveillance Team |
| EPI_ISL_2849310, EPI_ISL_2849311 | Lighthouse Laboratory Plymouth | Wellcome Sanger Institute for the COVID-19 Genomics UK (COG-UK) Consortium | Lighthouse Laboratory Plymouth and Alex Alderton, Roberto Amato, Jeffrey Barrett, Sonia Goncalves, Ewan Harrison, David K. Jackson, Ian Johnston, Dominic Kwiatkowski, Cordelia Langford, John Sillitoe on behalf of the Wellcome Sanger Institute COVID-19 Surveillance Team |
| EPI_ISL_2849312, EPI_ISL_2849313 | Lighthouse Lab in Alderley Park | Wellcome Sanger Institute for the COVID-19 Genomics UK (COG-UK) Consortium | Jacquelyn Wynn, Mairead Hyland, The Lighthouse Lab in Alderley Park and Alex Alderton, Roberto Amato, Jeffrey Barrett, Sonia Goncalves, Ewan Harrison, David K. Jackson, Ian Johnston, Dominic Kwiatkowski, Cordelia Langford, John Sillitoe on behalf of the Wellcome Sanger Institute COVID-19 Surveillance Team |
| EPI_ISL_2849314 | Lighthouse Laboratory Plymouth | Wellcome Sanger Institute for the COVID-19 Genomics UK (COG-UK) Consortium | Lighthouse Laboratory Plymouth and Alex Alderton, Roberto Amato, Jeffrey Barrett, Sonia Goncalves, Ewan Harrison, David K. Jackson, Ian Johnston, Dominic Kwiatkowski, Cordelia Langford, John Sillitoe on behalf of the Wellcome Sanger Institute COVID-19 Surveillance Team |
| EPI_ISL_2849315 | Lighthouse Lab in Alderley Park | Wellcome Sanger Institute for the COVID-19 Genomics UK (COG-UK) Consortium | Jacquelyn Wynn, Mairead Hyland, The Lighthouse Lab in Alderley Park and Alex Alderton, Roberto Amato, Jeffrey Barrett, Sonia Goncalves, Ewan Harrison, David K. Jackson, Ian Johnston, Dominic Kwiatkowski, Cordelia Langford, John Sillitoe on behalf of the Wellcome Sanger Institute COVID-19 Surveillance Team |
| EPI_ISL_2849316 | Lighthouse Laboratory Plymouth | Wellcome Sanger Institute for the COVID-19 Genomics UK (COG-UK) | Lighthouse Laboratory Plymouth and Alex Alderton, Roberto Amato, Jeffrey Barrett, Sonia Goncalves, Ewan Harrison, David K. Jackson, Ian Johnston, Dominic Kwiatkowski, Cordelia Langford, John Sillitoe on behalf of the Wellcome Sanger |

| | | | |
|---|---|---|---|
| | | Consortium | Institute COVID-19 Surveillance Team |
| EPI_ISL_2849317 | Lighthouse Lab in Alderley Park | Wellcome Sanger Institute for the COVID-19 Genomics UK (COG-UK) Consortium | Jacquelyn Wynn, Mairead Hyland, The Lighthouse Lab in Alderley Park and Alex Alderton, Roberto Amato, Jeffrey Barrett, Sonia Goncalves, Ewan Harrison, David K. Jackson, Ian Johnston, Dominic Kwiatkowski, Cordelia Langford, John Sillitoe on behalf of the Wellcome Sanger Institute COVID-19 Surveillance Team |
| EPI_ISL_2849318 | Lighthouse Laboratory Plymouth | Wellcome Sanger Institute for the COVID-19 Genomics UK (COG-UK) Consortium | Lighthouse Laboratory Plymouth and Alex Alderton, Roberto Amato, Jeffrey Barrett, Sonia Goncalves, Ewan Harrison, David K. Jackson, Ian Johnston, Dominic Kwiatkowski, Cordelia Langford, John Sillitoe on behalf of the Wellcome Sanger Institute COVID-19 Surveillance Team |
| EPI_ISL_2849319, EPI_ISL_2849320, EPI_ISL_2849321 | Lighthouse Lab in Alderley Park | Wellcome Sanger Institute for the COVID-19 Genomics UK (COG-UK) Consortium | Jacquelyn Wynn, Mairead Hyland, The Lighthouse Lab in Alderley Park and Alex Alderton, Roberto Amato, Jeffrey Barrett, Sonia Goncalves, Ewan Harrison, David K. Jackson, Ian Johnston, Dominic Kwiatkowski, Cordelia Langford, John Sillitoe on behalf of the Wellcome Sanger Institute COVID-19 Surveillance Team |
| EPI_ISL_2849322 | Lighthouse Laboratory Plymouth | Wellcome Sanger Institute for the COVID-19 Genomics UK (COG-UK) Consortium | Lighthouse Laboratory Plymouth and Alex Alderton, Roberto Amato, Jeffrey Barrett, Sonia Goncalves, Ewan Harrison, David K. Jackson, Ian Johnston, Dominic Kwiatkowski, Cordelia Langford, John Sillitoe on behalf of the Wellcome Sanger Institute COVID-19 Surveillance Team |
| EPI_ISL_2849323, EPI_ISL_2849324, EPI_ISL_2849325, EPI_ISL_2849326, EPI_ISL_2849327 | Lighthouse Lab in Alderley Park | Wellcome Sanger Institute for the COVID-19 Genomics UK (COG-UK) Consortium | Jacquelyn Wynn, Mairead Hyland, The Lighthouse Lab in Alderley Park and Alex Alderton, Roberto Amato, Jeffrey Barrett, Sonia Goncalves, Ewan Harrison, David K. Jackson, Ian Johnston, Dominic Kwiatkowski, Cordelia Langford, John Sillitoe on behalf of the Wellcome Sanger Institute COVID-19 Surveillance Team |
| EPI_ISL_2849328, EPI_ISL_2849329 | Lighthouse Laboratory Plymouth | Wellcome Sanger Institute for the COVID-19 Genomics UK (COG-UK) Consortium | Lighthouse Laboratory Plymouth and Alex Alderton, Roberto Amato, Jeffrey Barrett, Sonia Goncalves, Ewan Harrison, David K. Jackson, Ian Johnston, Dominic Kwiatkowski, Cordelia Langford, John Sillitoe on behalf of the Wellcome Sanger Institute COVID-19 Surveillance Team |
| EPI_ISL_2849330, EPI_ISL_2849331, EPI_ISL_2849332, EPI_ISL_2849333 | Lighthouse Lab in Alderley Park | Wellcome Sanger Institute for the COVID-19 Genomics UK (COG-UK) Consortium | Jacquelyn Wynn, Mairead Hyland, The Lighthouse Lab in Alderley Park and Alex Alderton, Roberto Amato, Jeffrey Barrett, Sonia Goncalves, Ewan Harrison, David K. Jackson, Ian Johnston, Dominic Kwiatkowski, Cordelia Langford, John Sillitoe on behalf of the Wellcome Sanger Institute COVID-19 Surveillance Team |
| EPI_ISL_2849334, EPI_ISL_2849335, EPI_ISL_2849336 | Lighthouse Laboratory Plymouth | Wellcome Sanger Institute for the COVID-19 Genomics UK (COG-UK) Consortium | Lighthouse Laboratory Plymouth and Alex Alderton, Roberto Amato, Jeffrey Barrett, Sonia Goncalves, Ewan Harrison, David K. Jackson, Ian Johnston, Dominic Kwiatkowski, Cordelia Langford, John Sillitoe on behalf of the Wellcome Sanger Institute COVID-19 Surveillance Team |
| EPI_ISL_2849337 | Lighthouse Lab in Alderley Park | Wellcome Sanger Institute for the COVID-19 Genomics UK (COG-UK) Consortium | Jacquelyn Wynn, Mairead Hyland, The Lighthouse Lab in Alderley Park and Alex Alderton, Roberto Amato, Jeffrey Barrett, Sonia Goncalves, Ewan Harrison, David K. Jackson, Ian Johnston, Dominic Kwiatkowski, Cordelia Langford, John Sillitoe on behalf of the Wellcome Sanger Institute COVID-19 Surveillance Team |
| EPI_ISL_2849338 | Lighthouse Laboratory Plymouth | Wellcome Sanger Institute for the COVID-19 Genomics UK (COG-UK) Consortium | Lighthouse Laboratory Plymouth and Alex Alderton, Roberto Amato, Jeffrey Barrett, Sonia Goncalves, Ewan Harrison, David K. Jackson, Ian Johnston, Dominic Kwiatkowski, Cordelia Langford, John Sillitoe on behalf of the Wellcome Sanger Institute COVID-19 Surveillance Team |
| EPI_ISL_2849339, EPI_ISL_2849340, EPI_ISL_2849341 | Lighthouse Lab in Alderley Park | Wellcome Sanger Institute for the COVID-19 Genomics UK (COG-UK) Consortium | Jacquelyn Wynn, Mairead Hyland, The Lighthouse Lab in Alderley Park and Alex Alderton, Roberto Amato, Jeffrey Barrett, Sonia Goncalves, Ewan Harrison, David K. Jackson, Ian Johnston, Dominic Kwiatkowski, Cordelia Langford, John Sillitoe on behalf of the Wellcome Sanger Institute COVID-19 Surveillance Team |
| EPI_ISL_2849342 | Lighthouse Laboratory Plymouth | Wellcome Sanger Institute for the COVID-19 Genomics UK (COG-UK) Consortium | Lighthouse Laboratory Plymouth and Alex Alderton, Roberto Amato, Jeffrey Barrett, Sonia Goncalves, Ewan Harrison, David K. Jackson, Ian Johnston, Dominic Kwiatkowski, Cordelia Langford, John Sillitoe on behalf of the Wellcome Sanger Institute COVID-19 Surveillance Team |
| EPI_ISL_2849343 | Lighthouse Lab in Alderley Park | Wellcome Sanger Institute for the COVID-19 Genomics UK (COG-UK) Consortium | Jacquelyn Wynn, Mairead Hyland, The Lighthouse Lab in Alderley Park and Alex Alderton, Roberto Amato, Jeffrey Barrett, Sonia Goncalves, Ewan Harrison, David K. Jackson, Ian Johnston, Dominic Kwiatkowski, Cordelia Langford, John Sillitoe on behalf of the Wellcome Sanger Institute COVID-19 Surveillance Team |
| EPI_ISL_2849344, EPI_ISL_2849345, EPI_ISL_2849346 | Lighthouse Laboratory Plymouth | Wellcome Sanger Institute for the COVID-19 Genomics UK (COG-UK) Consortium | Lighthouse Laboratory Plymouth and Alex Alderton, Roberto Amato, Jeffrey Barrett, Sonia Goncalves, Ewan Harrison, David K. Jackson, Ian Johnston, Dominic Kwiatkowski, Cordelia Langford, John Sillitoe on behalf of the Wellcome Sanger Institute COVID-19 Surveillance Team |
| EPI_ISL_2849347 | Lighthouse Lab in Alderley Park | Wellcome Sanger Institute for the COVID-19 Genomics UK (COG-UK) Consortium | Jacquelyn Wynn, Mairead Hyland, The Lighthouse Lab in Alderley Park and Alex Alderton, Roberto Amato, Jeffrey Barrett, Sonia Goncalves, Ewan Harrison, David K. Jackson, Ian Johnston, Dominic Kwiatkowski, Cordelia Langford, John Sillitoe on behalf of the Wellcome Sanger Institute COVID-19 Surveillance Team |
| EPI_ISL_2849348 | Lighthouse Laboratory Plymouth | Wellcome Sanger Institute for the COVID-19 Genomics UK (COG-UK) Consortium | Lighthouse Laboratory Plymouth and Alex Alderton, Roberto Amato, Jeffrey Barrett, Sonia Goncalves, Ewan Harrison, David K. Jackson, Ian Johnston, Dominic Kwiatkowski, Cordelia Langford, John Sillitoe on behalf of the Wellcome Sanger Institute COVID-19 Surveillance Team |
| EPI_ISL_2849349, EPI_ISL_2849350, EPI_ISL_2849351 | Lighthouse Lab in Alderley Park | Wellcome Sanger Institute for the COVID-19 Genomics UK (COG-UK) Consortium | Jacquelyn Wynn, Mairead Hyland, The Lighthouse Lab in Alderley Park and Alex Alderton, Roberto Amato, Jeffrey Barrett, Sonia Goncalves, Ewan Harrison, David K. Jackson, Ian Johnston, Dominic Kwiatkowski, Cordelia Langford, John Sillitoe on behalf of the Wellcome Sanger Institute COVID-19 Surveillance Team |
| EPI_ISL_2849352, EPI_ISL_2849353 | Lighthouse Laboratory Plymouth | Wellcome Sanger Institute for the COVID-19 Genomics UK (COG-UK) Consortium | Lighthouse Laboratory Plymouth and Alex Alderton, Roberto Amato, Jeffrey Barrett, Sonia Goncalves, Ewan Harrison, David K. Jackson, Ian Johnston, Dominic Kwiatkowski, Cordelia Langford, John Sillitoe on behalf of the Wellcome Sanger Institute COVID-19 Surveillance Team |
| EPI_ISL_2849354 | Lighthouse Lab in Alderley Park | Wellcome Sanger Institute for the COVID-19 Genomics UK (COG-UK) Consortium | Jacquelyn Wynn, Mairead Hyland, The Lighthouse Lab in Alderley Park and Alex Alderton, Roberto Amato, Jeffrey Barrett, Sonia Goncalves, Ewan Harrison, David K. Jackson, Ian Johnston, Dominic Kwiatkowski, Cordelia Langford, John Sillitoe on behalf of the Wellcome Sanger Institute COVID-19 Surveillance Team |
| EPI_ISL_2849355 | Lighthouse Laboratory Plymouth | Wellcome Sanger Institute for the COVID-19 Genomics UK (COG-UK) Consortium | Lighthouse Laboratory Plymouth and Alex Alderton, Roberto Amato, Jeffrey Barrett, Sonia Goncalves, Ewan Harrison, David K. Jackson, Ian Johnston, Dominic Kwiatkowski, Cordelia Langford, John Sillitoe on behalf of the Wellcome Sanger Institute COVID-19 Surveillance Team |
| EPI_ISL_2849356, EPI_ISL_2849357 | Lighthouse Lab in Alderley Park | Wellcome Sanger Institute for the COVID-19 Genomics UK (COG-UK) Consortium | Jacquelyn Wynn, Mairead Hyland, The Lighthouse Lab in Alderley Park and Alex Alderton, Roberto Amato, Jeffrey Barrett, Sonia Goncalves, Ewan Harrison, David K. Jackson, Ian Johnston, Dominic Kwiatkowski, Cordelia Langford, John Sillitoe on behalf of the Wellcome Sanger Institute COVID-19 Surveillance Team |
| EPI_ISL_2849358 | Lighthouse Laboratory Plymouth | Wellcome Sanger Institute for the COVID-19 Genomics UK (COG-UK) Consortium | Lighthouse Laboratory Plymouth and Alex Alderton, Roberto Amato, Jeffrey Barrett, Sonia Goncalves, Ewan Harrison, David K. Jackson, Ian Johnston, Dominic Kwiatkowski, Cordelia Langford, John Sillitoe on behalf of the Wellcome Sanger Institute COVID-19 Surveillance Team |
| EPI_ISL_2849359, EPI_ISL_2849360 | Lighthouse Lab in Alderley Park | Wellcome Sanger Institute for the COVID-19 Genomics UK (COG-UK) Consortium | Jacquelyn Wynn, Mairead Hyland, The Lighthouse Lab in Alderley Park and Alex Alderton, Roberto Amato, Jeffrey Barrett, Sonia Goncalves, Ewan Harrison, David K. Jackson, Ian Johnston, Dominic Kwiatkowski, Cordelia Langford, John Sillitoe on behalf of the Wellcome Sanger Institute COVID-19 Surveillance Team |
| EPI_ISL_2849361, EPI_ISL_2849362, EPI_ISL_2849363, EPI_ISL_2849364, EPI_ISL_2849365 | Lighthouse Laboratory Plymouth | Wellcome Sanger Institute for the COVID-19 Genomics UK (COG-UK) Consortium | Lighthouse Laboratory Plymouth and Alex Alderton, Roberto Amato, Jeffrey Barrett, Sonia Goncalves, Ewan Harrison, David K. Jackson, Ian Johnston, Dominic Kwiatkowski, Cordelia Langford, John Sillitoe on behalf of the Wellcome Sanger Institute COVID-19 Surveillance Team |
| EPI_ISL_2849366, EPI_ISL_2849367 | Lighthouse Lab in Milton Keynes | Wellcome Sanger Institute for the COVID-19 Genomics UK (COG-UK) Consortium | The Lighthouse Lab in Milton Keynes and Alex Alderton, Roberto Amato, Jeffrey Barrett, Sonia Goncalves, Ewan Harrison, David K. Jackson, Ian Johnston, Dominic Kwiatkowski, Cordelia Langford, John Sillitoe on behalf of the Wellcome Sanger Institute COVID-19 Surveillance Team |
| EPI_ISL_2849368 | Lighthouse Lab in Alderley Park | Wellcome Sanger Institute for the COVID-19 Genomics UK (COG-UK) Consortium | Jacquelyn Wynn, Mairead Hyland, The Lighthouse Lab in Alderley Park and Alex Alderton, Roberto Amato, Jeffrey Barrett, Sonia Goncalves, Ewan Harrison, David K. Jackson, Ian Johnston, Dominic Kwiatkowski, Cordelia Langford, John Sillitoe on behalf of the Wellcome Sanger Institute COVID-19 Surveillance Team |
| EPI_ISL_2849369 | Lighthouse Laboratory Plymouth | Wellcome Sanger Institute for the COVID-19 Genomics UK (COG-UK) Consortium | Lighthouse Laboratory Plymouth and Alex Alderton, Roberto Amato, Jeffrey Barrett, Sonia Goncalves, Ewan Harrison, David K. Jackson, Ian Johnston, Dominic Kwiatkowski, Cordelia Langford, John Sillitoe on behalf of the Wellcome Sanger Institute COVID-19 Surveillance Team |
| EPI_ISL_2849370 | Lighthouse Lab in Milton Keynes | Wellcome Sanger Institute for the COVID-19 Genomics UK (COG-UK) Consortium | The Lighthouse Lab in Milton Keynes and Alex Alderton, Roberto Amato, Jeffrey Barrett, Sonia Goncalves, Ewan Harrison, David K. Jackson, Ian Johnston, Dominic Kwiatkowski, Cordelia Langford, John Sillitoe on behalf of the Wellcome Sanger Institute COVID-19 Surveillance Team |
| EPI_ISL_2849371 | Lighthouse Laboratory Plymouth | Wellcome Sanger Institute for the COVID-19 Genomics UK (COG-UK) Consortium | Lighthouse Laboratory Plymouth and Alex Alderton, Roberto Amato, Jeffrey Barrett, Sonia Goncalves, Ewan Harrison, David K. Jackson, Ian Johnston, Dominic Kwiatkowski, Cordelia Langford, John Sillitoe on behalf of the Wellcome Sanger Institute COVID-19 Surveillance Team |
| EPI_ISL_2849372 | Lighthouse Lab in Milton Keynes | Wellcome Sanger Institute for the COVID-19 Genomics UK (COG-UK) Consortium | The Lighthouse Lab in Milton Keynes and Alex Alderton, Roberto Amato, Jeffrey Barrett, Sonia Goncalves, Ewan Harrison, David K. Jackson, Ian Johnston, Dominic Kwiatkowski, Cordelia Langford, John Sillitoe on behalf of the Wellcome Sanger Institute COVID-19 Surveillance Team |
| EPI_ISL_2849373, EPI_ISL_2849374 | Lighthouse Laboratory Plymouth | Wellcome Sanger Institute for the COVID-19 Genomics UK (COG-UK) Consortium | Lighthouse Laboratory Plymouth and Alex Alderton, Roberto Amato, Jeffrey Barrett, Sonia Goncalves, Ewan Harrison, David K. Jackson, Ian Johnston, Dominic Kwiatkowski, Cordelia Langford, John Sillitoe on behalf of the Wellcome Sanger Institute COVID-19 Surveillance Team |
| EPI_ISL_2849375 | Lighthouse Lab in Alderley Park | Wellcome Sanger Institute for the COVID-19 Genomics UK (COG-UK) Consortium | Jacquelyn Wynn, Mairead Hyland, The Lighthouse Lab in Alderley Park and Alex Alderton, Roberto Amato, Jeffrey Barrett, Sonia Goncalves, Ewan Harrison, David K. Jackson, Ian Johnston, Dominic Kwiatkowski, Cordelia Langford, John Sillitoe on behalf of the Wellcome Sanger Institute COVID-19 Surveillance Team |
| EPI_ISL_2849376, EPI_ISL_2849377, EPI_ISL_2849378 | Lighthouse Lab in Milton Keynes | Wellcome Sanger Institute for the COVID-19 Genomics UK (COG-UK) Consortium | The Lighthouse Lab in Milton Keynes and Alex Alderton, Roberto Amato, Jeffrey Barrett, Sonia Goncalves, Ewan Harrison, David K. Jackson, Ian Johnston, Dominic Kwiatkowski, Cordelia Langford, John Sillitoe on behalf of the Wellcome Sanger Institute COVID-19 Surveillance Team |
| EPI_ISL_2849379 | Lighthouse Laboratory Plymouth | Wellcome Sanger Institute for the COVID-19 Genomics UK (COG-UK) Consortium | Lighthouse Laboratory Plymouth and Alex Alderton, Roberto Amato, Jeffrey Barrett, Sonia Goncalves, Ewan Harrison, David K. Jackson, Ian Johnston, Dominic Kwiatkowski, Cordelia Langford, John Sillitoe on behalf of the Wellcome Sanger Institute COVID-19 Surveillance Team |
| EPI_ISL_2849380 | Lighthouse Lab in Milton Keynes | Wellcome Sanger Institute for the COVID-19 Genomics UK (COG-UK) Consortium | The Lighthouse Lab in Milton Keynes and Alex Alderton, Roberto Amato, Jeffrey Barrett, Sonia Goncalves, Ewan Harrison, David K. Jackson, Ian Johnston, Dominic Kwiatkowski, Cordelia Langford, John Sillitoe on behalf of the Wellcome Sanger Institute COVID-19 Surveillance Team |
| EPI_ISL_2849381 | Lighthouse Lab in Alderley Park | Wellcome Sanger Institute for the COVID-19 Genomics UK (COG-UK) Consortium | Jacquelyn Wynn, Mairead Hyland, The Lighthouse Lab in Alderley Park and Alex Alderton, Roberto Amato, Jeffrey Barrett, Sonia Goncalves, Ewan Harrison, David K. Jackson, Ian Johnston, Dominic Kwiatkowski, Cordelia Langford, John Sillitoe on behalf of the Wellcome Sanger Institute COVID-19 Surveillance Team |
| EPI_ISL_2849382 | Lighthouse Lab in Milton Keynes | Wellcome Sanger Institute for the COVID-19 Genomics UK (COG-UK) Consortium | The Lighthouse Lab in Milton Keynes and Alex Alderton, Roberto Amato, Jeffrey Barrett, Sonia Goncalves, Ewan Harrison, David K. Jackson, Ian Johnston, Dominic Kwiatkowski, Cordelia Langford, John Sillitoe on behalf of the Wellcome Sanger Institute COVID-19 Surveillance Team |
| EPI_ISL_2849383 | Lighthouse Laboratory Plymouth | Wellcome Sanger Institute for the COVID-19 Genomics UK (COG-UK) Consortium | Lighthouse Laboratory Plymouth and Alex Alderton, Roberto Amato, Jeffrey Barrett, Sonia Goncalves, Ewan Harrison, David K. Jackson, Ian Johnston, Dominic Kwiatkowski, Cordelia Langford, John Sillitoe on behalf of the Wellcome Sanger Institute COVID-19 Surveillance Team |
| EPI_ISL_2849384, EPI_ISL_2849385 | Lighthouse Lab in Milton Keynes | Wellcome Sanger Institute for the COVID-19 Genomics UK (COG-UK) Consortium | The Lighthouse Lab in Milton Keynes and Alex Alderton, Roberto Amato, Jeffrey Barrett, Sonia Goncalves, Ewan Harrison, David K. Jackson, Ian Johnston, Dominic Kwiatkowski, Cordelia Langford, John Sillitoe on behalf of the Wellcome Sanger Institute COVID-19 Surveillance Team |

| | | | |
|---|---|---|---|
| EPI_ISL_2849386 | Lighthouse Laboratory Plymouth | Wellcome Sanger Institute for the COVID-19 Genomics UK (COG-UK) Consortium | Lighthouse Laboratory Plymouth and Alex Alderton, Roberto Amato, Jeffrey Barrett, Sonia Goncalves, Ewan Harrison, David K. Jackson, Ian Johnston, Dominic Kwiatkowski, Cordelia Langford, John Sillitoe on behalf of the Wellcome Sanger Institute COVID-19 Surveillance Team |
| EPI_ISL_2849387 | Lighthouse Lab in Alderley Park | Wellcome Sanger Institute for the COVID-19 Genomics UK (COG-UK) Consortium | Jacquelyn Wynn, Mairead Hyland, The Lighthouse Lab in Alderley Park and Alex Alderton, Roberto Amato, Jeffrey Barrett, Sonia Goncalves, Ewan Harrison, David K. Jackson, Ian Johnston, Dominic Kwiatkowski, Cordelia Langford, John Sillitoe on behalf of the Wellcome Sanger Institute COVID-19 Surveillance Team |
| EPI_ISL_2849388, EPI_ISL_2849389, EPI_ISL_2849390, EPI_ISL_2849391 | Lighthouse Laboratory Plymouth | Wellcome Sanger Institute for the COVID-19 Genomics UK (COG-UK) Consortium | Lighthouse Laboratory Plymouth and Alex Alderton, Roberto Amato, Jeffrey Barrett, Sonia Goncalves, Ewan Harrison, David K. Jackson, Ian Johnston, Dominic Kwiatkowski, Cordelia Langford, John Sillitoe on behalf of the Wellcome Sanger Institute COVID-19 Surveillance Team |
| EPI_ISL_2849392 | Lighthouse Lab in Milton Keynes | Wellcome Sanger Institute for the COVID-19 Genomics UK (COG-UK) Consortium | The Lighthouse Lab in Milton Keynes and Alex Alderton, Roberto Amato, Jeffrey Barrett, Sonia Goncalves, Ewan Harrison, David K. Jackson, Ian Johnston, Dominic Kwiatkowski, Cordelia Langford, John Sillitoe on behalf of the Wellcome Sanger Institute COVID-19 Surveillance Team |
| EPI_ISL_2849393 | Lighthouse Laboratory Plymouth | Wellcome Sanger Institute for the COVID-19 Genomics UK (COG-UK) Consortium | Lighthouse Laboratory Plymouth and Alex Alderton, Roberto Amato, Jeffrey Barrett, Sonia Goncalves, Ewan Harrison, David K. Jackson, Ian Johnston, Dominic Kwiatkowski, Cordelia Langford, John Sillitoe on behalf of the Wellcome Sanger Institute COVID-19 Surveillance Team |
| EPI_ISL_2849394 | Lighthouse Lab in Alderley Park | Wellcome Sanger Institute for the COVID-19 Genomics UK (COG-UK) Consortium | Jacquelyn Wynn, Mairead Hyland, The Lighthouse Lab in Alderley Park and Alex Alderton, Roberto Amato, Jeffrey Barrett, Sonia Goncalves, Ewan Harrison, David K. Jackson, Ian Johnston, Dominic Kwiatkowski, Cordelia Langford, John Sillitoe on behalf of the Wellcome Sanger Institute COVID-19 Surveillance Team |
| EPI_ISL_2849395 | Lighthouse Laboratory Plymouth | Wellcome Sanger Institute for the COVID-19 Genomics UK (COG-UK) Consortium | Lighthouse Laboratory Plymouth and Alex Alderton, Roberto Amato, Jeffrey Barrett, Sonia Goncalves, Ewan Harrison, David K. Jackson, Ian Johnston, Dominic Kwiatkowski, Cordelia Langford, John Sillitoe on behalf of the Wellcome Sanger Institute COVID-19 Surveillance Team |
| EPI_ISL_2849396, EPI_ISL_2849397 | Lighthouse Lab in Milton Keynes | Wellcome Sanger Institute for the COVID-19 Genomics UK (COG-UK) Consortium | The Lighthouse Lab in Milton Keynes and Alex Alderton, Roberto Amato, Jeffrey Barrett, Sonia Goncalves, Ewan Harrison, David K. Jackson, Ian Johnston, Dominic Kwiatkowski, Cordelia Langford, John Sillitoe on behalf of the Wellcome Sanger Institute COVID-19 Surveillance Team |
| EPI_ISL_2849398 | Lighthouse Laboratory Plymouth | Wellcome Sanger Institute for the COVID-19 Genomics UK (COG-UK) Consortium | Lighthouse Laboratory Plymouth and Alex Alderton, Roberto Amato, Jeffrey Barrett, Sonia Goncalves, Ewan Harrison, David K. Jackson, Ian Johnston, Dominic Kwiatkowski, Cordelia Langford, John Sillitoe on behalf of the Wellcome Sanger Institute COVID-19 Surveillance Team |
| EPI_ISL_2849399 | Lighthouse Lab in Alderley Park | Wellcome Sanger Institute for the COVID-19 Genomics UK (COG-UK) Consortium | Jacquelyn Wynn, Mairead Hyland, The Lighthouse Lab in Alderley Park and Alex Alderton, Roberto Amato, Jeffrey Barrett, Sonia Goncalves, Ewan Harrison, David K. Jackson, Ian Johnston, Dominic Kwiatkowski, Cordelia Langford, John Sillitoe on behalf of the Wellcome Sanger Institute COVID-19 Surveillance Team |
| EPI_ISL_2849400 | Lighthouse Laboratory Plymouth | Wellcome Sanger Institute for the COVID-19 Genomics UK (COG-UK) Consortium | Lighthouse Laboratory Plymouth and Alex Alderton, Roberto Amato, Jeffrey Barrett, Sonia Goncalves, Ewan Harrison, David K. Jackson, Ian Johnston, Dominic Kwiatkowski, Cordelia Langford, John Sillitoe on behalf of the Wellcome Sanger Institute COVID-19 Surveillance Team |
| EPI_ISL_2849401 | Lighthouse Lab in Alderley Park | Wellcome Sanger Institute for the COVID-19 Genomics UK (COG-UK) Consortium | Jacquelyn Wynn, Mairead Hyland, The Lighthouse Lab in Alderley Park and Alex Alderton, Roberto Amato, Jeffrey Barrett, Sonia Goncalves, Ewan Harrison, David K. Jackson, Ian Johnston, Dominic Kwiatkowski, Cordelia Langford, John Sillitoe on behalf of the Wellcome Sanger Institute COVID-19 Surveillance Team |
| EPI_ISL_2849402 | Lighthouse Lab in Milton Keynes | Wellcome Sanger Institute for the COVID-19 Genomics UK (COG-UK) Consortium | The Lighthouse Lab in Milton Keynes and Alex Alderton, Roberto Amato, Jeffrey Barrett, Sonia Goncalves, Ewan Harrison, David K. Jackson, Ian Johnston, Dominic Kwiatkowski, Cordelia Langford, John Sillitoe on behalf of the Wellcome Sanger Institute COVID-19 Surveillance Team |
| EPI_ISL_2849403 | Lighthouse Laboratory Plymouth | Wellcome Sanger Institute for the COVID-19 Genomics UK (COG-UK) Consortium | Lighthouse Laboratory Plymouth and Alex Alderton, Roberto Amato, Jeffrey Barrett, Sonia Goncalves, Ewan Harrison, David K. Jackson, Ian Johnston, Dominic Kwiatkowski, Cordelia Langford, John Sillitoe on behalf of the Wellcome Sanger Institute COVID-19 Surveillance Team |
| EPI_ISL_2849404 | Lighthouse Lab in Alderley Park | Wellcome Sanger Institute for the COVID-19 Genomics UK (COG-UK) Consortium | Jacquelyn Wynn, Mairead Hyland, The Lighthouse Lab in Alderley Park and Alex Alderton, Roberto Amato, Jeffrey Barrett, Sonia Goncalves, Ewan Harrison, David K. Jackson, Ian Johnston, Dominic Kwiatkowski, Cordelia Langford, John Sillitoe on behalf of the Wellcome Sanger Institute COVID-19 Surveillance Team |
| EPI_ISL_2849405 | Lighthouse Laboratory Plymouth | Wellcome Sanger Institute for the COVID-19 Genomics UK (COG-UK) Consortium | Lighthouse Laboratory Plymouth and Alex Alderton, Roberto Amato, Jeffrey Barrett, Sonia Goncalves, Ewan Harrison, David K. Jackson, Ian Johnston, Dominic Kwiatkowski, Cordelia Langford, John Sillitoe on behalf of the Wellcome Sanger Institute COVID-19 Surveillance Team |
| EPI_ISL_2849406 | Lighthouse Lab in Alderley Park | Wellcome Sanger Institute for the COVID-19 Genomics UK (COG-UK) Consortium | Jacquelyn Wynn, Mairead Hyland, The Lighthouse Lab in Alderley Park and Alex Alderton, Roberto Amato, Jeffrey Barrett, Sonia Goncalves, Ewan Harrison, David K. Jackson, Ian Johnston, Dominic Kwiatkowski, Cordelia Langford, John Sillitoe on behalf of the Wellcome Sanger Institute COVID-19 Surveillance Team |
| EPI_ISL_2849407 | Lighthouse Laboratory Plymouth | Wellcome Sanger Institute for the COVID-19 Genomics UK (COG-UK) Consortium | Lighthouse Laboratory Plymouth and Alex Alderton, Roberto Amato, Jeffrey Barrett, Sonia Goncalves, Ewan Harrison, David K. Jackson, Ian Johnston, Dominic Kwiatkowski, Cordelia Langford, John Sillitoe on behalf of the Wellcome Sanger Institute COVID-19 Surveillance Team |
| EPI_ISL_2849408, EPI_ISL_2849409 | Lighthouse Lab in Alderley Park | Wellcome Sanger Institute for the COVID-19 Genomics UK (COG-UK) Consortium | Jacquelyn Wynn, Mairead Hyland, The Lighthouse Lab in Alderley Park and Alex Alderton, Roberto Amato, Jeffrey Barrett, Sonia Goncalves, Ewan Harrison, David K. Jackson, Ian Johnston, Dominic Kwiatkowski, Cordelia Langford, John Sillitoe on behalf of the Wellcome Sanger Institute COVID-19 Surveillance Team |
| EPI_ISL_2849410 | Lighthouse Lab in Milton Keynes | Wellcome Sanger Institute for the COVID-19 Genomics UK (COG-UK) Consortium | The Lighthouse Lab in Milton Keynes and Alex Alderton, Roberto Amato, Jeffrey Barrett, Sonia Goncalves, Ewan Harrison, David K. Jackson, Ian Johnston, Dominic Kwiatkowski, Cordelia Langford, John Sillitoe on behalf of the Wellcome Sanger Institute COVID-19 Surveillance Team |
| EPI_ISL_2849411 | Lighthouse Lab in Alderley Park | Wellcome Sanger Institute for the COVID-19 Genomics UK (COG-UK) Consortium | Jacquelyn Wynn, Mairead Hyland, The Lighthouse Lab in Alderley Park and Alex Alderton, Roberto Amato, Jeffrey Barrett, Sonia Goncalves, Ewan Harrison, David K. Jackson, Ian Johnston, Dominic Kwiatkowski, Cordelia Langford, John Sillitoe on behalf of the Wellcome Sanger Institute COVID-19 Surveillance Team |
| EPI_ISL_2849412 | Lighthouse Lab in Milton Keynes | Wellcome Sanger Institute for the COVID-19 Genomics UK (COG-UK) Consortium | The Lighthouse Lab in Milton Keynes and Alex Alderton, Roberto Amato, Jeffrey Barrett, Sonia Goncalves, Ewan Harrison, David K. Jackson, Ian Johnston, Dominic Kwiatkowski, Cordelia Langford, John Sillitoe on behalf of the Wellcome Sanger Institute COVID-19 Surveillance Team |
| EPI_ISL_2849413 | Lighthouse Laboratory Plymouth | Wellcome Sanger Institute for the COVID-19 Genomics UK (COG-UK) Consortium | Lighthouse Laboratory Plymouth and Alex Alderton, Roberto Amato, Jeffrey Barrett, Sonia Goncalves, Ewan Harrison, David K. Jackson, Ian Johnston, Dominic Kwiatkowski, Cordelia Langford, John Sillitoe on behalf of the Wellcome Sanger Institute COVID-19 Surveillance Team |
| EPI_ISL_2849414, EPI_ISL_2849415, EPI_ISL_2849416, EPI_ISL_2849417 | Lighthouse Lab in Alderley Park | Wellcome Sanger Institute for the COVID-19 Genomics UK (COG-UK) Consortium | Jacquelyn Wynn, Mairead Hyland, The Lighthouse Lab in Alderley Park and Alex Alderton, Roberto Amato, Jeffrey Barrett, Sonia Goncalves, Ewan Harrison, David K. Jackson, Ian Johnston, Dominic Kwiatkowski, Cordelia Langford, John Sillitoe on behalf of the Wellcome Sanger Institute COVID-19 Surveillance Team |
| EPI_ISL_2849418, EPI_ISL_2849419 | Lighthouse Lab in Milton Keynes | Wellcome Sanger Institute for the COVID-19 Genomics UK (COG-UK) Consortium | The Lighthouse Lab in Milton Keynes and Alex Alderton, Roberto Amato, Jeffrey Barrett, Sonia Goncalves, Ewan Harrison, David K. Jackson, Ian Johnston, Dominic Kwiatkowski, Cordelia Langford, John Sillitoe on behalf of the Wellcome Sanger Institute COVID-19 Surveillance Team |
| EPI_ISL_2849420 | Lighthouse Laboratory Plymouth | Wellcome Sanger Institute for the COVID-19 Genomics UK (COG-UK) Consortium | Lighthouse Laboratory Plymouth and Alex Alderton, Roberto Amato, Jeffrey Barrett, Sonia Goncalves, Ewan Harrison, David K. Jackson, Ian Johnston, Dominic Kwiatkowski, Cordelia Langford, John Sillitoe on behalf of the Wellcome Sanger Institute COVID-19 Surveillance Team |
| EPI_ISL_2849421 | Lighthouse Lab in Milton Keynes | Wellcome Sanger Institute for the COVID-19 Genomics UK (COG-UK) Consortium | The Lighthouse Lab in Milton Keynes and Alex Alderton, Roberto Amato, Jeffrey Barrett, Sonia Goncalves, Ewan Harrison, David K. Jackson, Ian Johnston, Dominic Kwiatkowski, Cordelia Langford, John Sillitoe on behalf of the Wellcome Sanger Institute COVID-19 Surveillance Team |
| EPI_ISL_2849422 | Lighthouse Laboratory Plymouth | Wellcome Sanger Institute for the COVID-19 Genomics UK (COG-UK) Consortium | Lighthouse Laboratory Plymouth and Alex Alderton, Roberto Amato, Jeffrey Barrett, Sonia Goncalves, Ewan Harrison, David K. Jackson, Ian Johnston, Dominic Kwiatkowski, Cordelia Langford, John Sillitoe on behalf of the Wellcome Sanger Institute COVID-19 Surveillance Team |
| EPI_ISL_2849423 | Lighthouse Lab in Milton Keynes | Wellcome Sanger Institute for the COVID-19 Genomics UK (COG-UK) Consortium | The Lighthouse Lab in Milton Keynes and Alex Alderton, Roberto Amato, Jeffrey Barrett, Sonia Goncalves, Ewan Harrison, David K. Jackson, Ian Johnston, Dominic Kwiatkowski, Cordelia Langford, John Sillitoe on behalf of the Wellcome Sanger Institute COVID-19 Surveillance Team |
| EPI_ISL_2849424, EPI_ISL_2849425, EPI_ISL_2849426, EPI_ISL_2849427 | Lighthouse Laboratory Plymouth | Wellcome Sanger Institute for the COVID-19 Genomics UK (COG-UK) Consortium | Lighthouse Laboratory Plymouth and Alex Alderton, Roberto Amato, Jeffrey Barrett, Sonia Goncalves, Ewan Harrison, David K. Jackson, Ian Johnston, Dominic Kwiatkowski, Cordelia Langford, John Sillitoe on behalf of the Wellcome Sanger Institute COVID-19 Surveillance Team |
| EPI_ISL_2849428 | Lighthouse Lab in Milton Keynes | Wellcome Sanger Institute for the COVID-19 Genomics UK (COG-UK) Consortium | The Lighthouse Lab in Milton Keynes and Alex Alderton, Roberto Amato, Jeffrey Barrett, Sonia Goncalves, Ewan Harrison, David K. Jackson, Ian Johnston, Dominic Kwiatkowski, Cordelia Langford, John Sillitoe on behalf of the Wellcome Sanger Institute COVID-19 Surveillance Team |
| EPI_ISL_2849429, EPI_ISL_2849430, EPI_ISL_2849431 | Lighthouse Laboratory Plymouth | Wellcome Sanger Institute for the COVID-19 Genomics UK (COG-UK) Consortium | Lighthouse Laboratory Plymouth and Alex Alderton, Roberto Amato, Jeffrey Barrett, Sonia Goncalves, Ewan Harrison, David K. Jackson, Ian Johnston, Dominic Kwiatkowski, Cordelia Langford, John Sillitoe on behalf of the Wellcome Sanger Institute COVID-19 Surveillance Team |
| EPI_ISL_2849432 | Lighthouse Lab in Milton Keynes | Wellcome Sanger Institute for the COVID-19 Genomics UK (COG-UK) Consortium | The Lighthouse Lab in Milton Keynes and Alex Alderton, Roberto Amato, Jeffrey Barrett, Sonia Goncalves, Ewan Harrison, David K. Jackson, Ian Johnston, Dominic Kwiatkowski, Cordelia Langford, John Sillitoe on behalf of the Wellcome Sanger Institute COVID-19 Surveillance Team |
| EPI_ISL_2849433 | Lighthouse Laboratory Plymouth | Wellcome Sanger Institute for the COVID-19 Genomics UK (COG-UK) Consortium | Lighthouse Laboratory Plymouth and Alex Alderton, Roberto Amato, Jeffrey Barrett, Sonia Goncalves, Ewan Harrison, David K. Jackson, Ian Johnston, Dominic Kwiatkowski, Cordelia Langford, John Sillitoe on behalf of the Wellcome Sanger Institute COVID-19 Surveillance Team |
| EPI_ISL_2849434 | Lighthouse Lab in Milton Keynes | Wellcome Sanger Institute for the COVID-19 Genomics UK (COG-UK) Consortium | The Lighthouse Lab in Milton Keynes and Alex Alderton, Roberto Amato, Jeffrey Barrett, Sonia Goncalves, Ewan Harrison, David K. Jackson, Ian Johnston, Dominic Kwiatkowski, Cordelia Langford, John Sillitoe on behalf of the Wellcome Sanger Institute COVID-19 Surveillance Team |
| EPI_ISL_2849435, EPI_ISL_2849436 | Lighthouse Lab in Alderley Park | Wellcome Sanger Institute for the COVID-19 Genomics UK (COG-UK) Consortium | Jacquelyn Wynn, Mairead Hyland, The Lighthouse Lab in Alderley Park and Alex Alderton, Roberto Amato, Jeffrey Barrett, Sonia Goncalves, Ewan Harrison, David K. Jackson, Ian Johnston, Dominic Kwiatkowski, Cordelia Langford, John Sillitoe on behalf of the Wellcome Sanger Institute COVID-19 Surveillance Team |
| EPI_ISL_2849437, EPI_ISL_2849438, EPI_ISL_2849439, EPI_ISL_2849440 | Lighthouse Laboratory Plymouth | Wellcome Sanger Institute for the COVID-19 Genomics UK (COG-UK) Consortium | Lighthouse Laboratory Plymouth and Alex Alderton, Roberto Amato, Jeffrey Barrett, Sonia Goncalves, Ewan Harrison, David K. Jackson, Ian Johnston, Dominic Kwiatkowski, Cordelia Langford, John Sillitoe on behalf of the Wellcome Sanger Institute COVID-19 Surveillance Team |
| EPI_ISL_2849441 | Lighthouse Lab in Milton Keynes | Wellcome Sanger Institute for the COVID-19 Genomics UK (COG-UK) Consortium | The Lighthouse Lab in Milton Keynes and Alex Alderton, Roberto Amato, Jeffrey Barrett, Sonia Goncalves, Ewan Harrison, David K. Jackson, Ian Johnston, Dominic Kwiatkowski, Cordelia Langford, John Sillitoe on behalf of the Wellcome Sanger Institute COVID-19 Surveillance Team |
| EPI_ISL_2849442, EPI_ISL_2849443, EPI_ISL_2849444, EPI_ISL_2849445 | Lighthouse Laboratory Plymouth | Wellcome Sanger Institute for the COVID-19 Genomics UK (COG-UK) Consortium | Lighthouse Laboratory Plymouth and Alex Alderton, Roberto Amato, Jeffrey Barrett, Sonia Goncalves, Ewan Harrison, David K. Jackson, Ian Johnston, Dominic Kwiatkowski, Cordelia Langford, John Sillitoe on behalf of the Wellcome Sanger Institute COVID-19 Surveillance Team |
| EPI_ISL_2849446, EPI_ISL_2849447 | Lighthouse Lab in Alderley Park | Wellcome Sanger Institute for the COVID-19 Genomics UK (COG-UK) Consortium | Jacquelyn Wynn, Mairead Hyland, The Lighthouse Lab in Alderley Park and Alex Alderton, Roberto Amato, Jeffrey Barrett, Sonia Goncalves, Ewan Harrison, David K. Jackson, Ian Johnston, Dominic Kwiatkowski, Cordelia Langford, John Sillitoe on behalf of the Wellcome Sanger Institute COVID-19 Surveillance Team |
| EPI_ISL_2849448 | Lighthouse Lab in Milton Keynes | Wellcome Sanger Institute for the COVID-19 Genomics UK (COG-UK) | The Lighthouse Lab in Milton Keynes and Alex Alderton, Roberto Amato, Jeffrey Barrett, Sonia Goncalves, Ewan Harrison, David K. Jackson, Ian Johnston, Dominic Kwiatkowski, Cordelia Langford, John Sillitoe on behalf of the Wellcome Sanger |

| | | | |
|---|---|---|---|
| | | Consortium | Institute COVID-19 Surveillance Team |
| EPI_ISL_2849449 | Lighthouse Laboratory Plymouth | Wellcome Sanger Institute for the COVID-19 Genomics UK (COG-UK) Consortium | Lighthouse Laboratory Plymouth and Alex Alderton, Roberto Amato, Jeffrey Barrett, Sonia Goncalves, Ewan Harrison, David K. Jackson, Ian Johnston, Dominic Kwiatkowski, Cordelia Langford, John Sillitoe on behalf of the Wellcome Sanger Institute COVID-19 Surveillance Team |
| EPI_ISL_2849450 | Lighthouse Lab in Milton Keynes | Wellcome Sanger Institute for the COVID-19 Genomics UK (COG-UK) Consortium | The Lighthouse Lab in Milton Keynes and Alex Alderton, Roberto Amato, Jeffrey Barrett, Sonia Goncalves, Ewan Harrison, David K. Jackson, Ian Johnston, Dominic Kwiatkowski, Cordelia Langford, John Sillitoe on behalf of the Wellcome Sanger Institute COVID-19 Surveillance Team |
| EPI_ISL_2849451, EPI_ISL_2849452, EPI_ISL_2849453, EPI_ISL_2849454, EPI_ISL_2849455 | Lighthouse Lab in Alderley Park | Wellcome Sanger Institute for the COVID-19 Genomics UK (COG-UK) Consortium | Jacquelyn Wynn, Mairead Hyland, The Lighthouse Lab in Alderley Park and Alex Alderton, Roberto Amato, Jeffrey Barrett, Sonia Goncalves, Ewan Harrison, David K. Jackson, Ian Johnston, Dominic Kwiatkowski, Cordelia Langford, John Sillitoe on behalf of the Wellcome Sanger Institute COVID-19 Surveillance Team |
| EPI_ISL_2849456, EPI_ISL_2849457 | Lighthouse Laboratory Plymouth | Wellcome Sanger Institute for the COVID-19 Genomics UK (COG-UK) Consortium | Lighthouse Laboratory Plymouth and Alex Alderton, Roberto Amato, Jeffrey Barrett, Sonia Goncalves, Ewan Harrison, David K. Jackson, Ian Johnston, Dominic Kwiatkowski, Cordelia Langford, John Sillitoe on behalf of the Wellcome Sanger Institute COVID-19 Surveillance Team |
| EPI_ISL_2849459, EPI_ISL_2849460, EPI_ISL_2849461, EPI_ISL_2849462 | Lighthouse Lab in Alderley Park | Wellcome Sanger Institute for the COVID-19 Genomics UK (COG-UK) Consortium | Jacquelyn Wynn, Mairead Hyland, The Lighthouse Lab in Alderley Park and Alex Alderton, Roberto Amato, Jeffrey Barrett, Sonia Goncalves, Ewan Harrison, David K. Jackson, Ian Johnston, Dominic Kwiatkowski, Cordelia Langford, John Sillitoe on behalf of the Wellcome Sanger Institute COVID-19 Surveillance Team |
| EPI_ISL_2849463 | Lighthouse Laboratory Plymouth | Wellcome Sanger Institute for the COVID-19 Genomics UK (COG-UK) Consortium | Lighthouse Laboratory Plymouth and Alex Alderton, Roberto Amato, Jeffrey Barrett, Sonia Goncalves, Ewan Harrison, David K. Jackson, Ian Johnston, Dominic Kwiatkowski, Cordelia Langford, John Sillitoe on behalf of the Wellcome Sanger Institute COVID-19 Surveillance Team |
| EPI_ISL_2849464 | Lighthouse Lab in Milton Keynes | Wellcome Sanger Institute for the COVID-19 Genomics UK (COG-UK) Consortium | The Lighthouse Lab in Milton Keynes and Alex Alderton, Roberto Amato, Jeffrey Barrett, Sonia Goncalves, Ewan Harrison, David K. Jackson, Ian Johnston, Dominic Kwiatkowski, Cordelia Langford, John Sillitoe on behalf of the Wellcome Sanger Institute COVID-19 Surveillance Team |
| EPI_ISL_2849465 | Lighthouse Laboratory Plymouth | Wellcome Sanger Institute for the COVID-19 Genomics UK (COG-UK) Consortium | Lighthouse Laboratory Plymouth and Alex Alderton, Roberto Amato, Jeffrey Barrett, Sonia Goncalves, Ewan Harrison, David K. Jackson, Ian Johnston, Dominic Kwiatkowski, Cordelia Langford, John Sillitoe on behalf of the Wellcome Sanger Institute COVID-19 Surveillance Team |
| EPI_ISL_2849466, EPI_ISL_2849467 | Lighthouse Lab in Milton Keynes | Wellcome Sanger Institute for the COVID-19 Genomics UK (COG-UK) Consortium | The Lighthouse Lab in Milton Keynes and Alex Alderton, Roberto Amato, Jeffrey Barrett, Sonia Goncalves, Ewan Harrison, David K. Jackson, Ian Johnston, Dominic Kwiatkowski, Cordelia Langford, John Sillitoe on behalf of the Wellcome Sanger Institute COVID-19 Surveillance Team |
| EPI_ISL_2849468 | Lighthouse Lab in Alderley Park | Wellcome Sanger Institute for the COVID-19 Genomics UK (COG-UK) Consortium | Jacquelyn Wynn, Mairead Hyland, The Lighthouse Lab in Alderley Park and Alex Alderton, Roberto Amato, Jeffrey Barrett, Sonia Goncalves, Ewan Harrison, David K. Jackson, Ian Johnston, Dominic Kwiatkowski, Cordelia Langford, John Sillitoe on behalf of the Wellcome Sanger Institute COVID-19 Surveillance Team |
| EPI_ISL_2849469 | Lighthouse Lab in Milton Keynes | Wellcome Sanger Institute for the COVID-19 Genomics UK (COG-UK) Consortium | The Lighthouse Lab in Milton Keynes and Alex Alderton, Roberto Amato, Jeffrey Barrett, Sonia Goncalves, Ewan Harrison, David K. Jackson, Ian Johnston, Dominic Kwiatkowski, Cordelia Langford, John Sillitoe on behalf of the Wellcome Sanger Institute COVID-19 Surveillance Team |
| EPI_ISL_2849470 | Lighthouse Lab in Alderley Park | Wellcome Sanger Institute for the COVID-19 Genomics UK (COG-UK) Consortium | Jacquelyn Wynn, Mairead Hyland, The Lighthouse Lab in Alderley Park and Alex Alderton, Roberto Amato, Jeffrey Barrett, Sonia Goncalves, Ewan Harrison, David K. Jackson, Ian Johnston, Dominic Kwiatkowski, Cordelia Langford, John Sillitoe on behalf of the Wellcome Sanger Institute COVID-19 Surveillance Team |
| EPI_ISL_2849471 | Lighthouse Lab in Milton Keynes | Wellcome Sanger Institute for the COVID-19 Genomics UK (COG-UK) Consortium | The Lighthouse Lab in Milton Keynes and Alex Alderton, Roberto Amato, Jeffrey Barrett, Sonia Goncalves, Ewan Harrison, David K. Jackson, Ian Johnston, Dominic Kwiatkowski, Cordelia Langford, John Sillitoe on behalf of the Wellcome Sanger Institute COVID-19 Surveillance Team |
| EPI_ISL_2849472 | Lighthouse Lab in Alderley Park | Wellcome Sanger Institute for the COVID-19 Genomics UK (COG-UK) Consortium | Jacquelyn Wynn, Mairead Hyland, The Lighthouse Lab in Alderley Park and Alex Alderton, Roberto Amato, Jeffrey Barrett, Sonia Goncalves, Ewan Harrison, David K. Jackson, Ian Johnston, Dominic Kwiatkowski, Cordelia Langford, John Sillitoe on behalf of the Wellcome Sanger Institute COVID-19 Surveillance Team |
| EPI_ISL_2849473, EPI_ISL_2849474 | Lighthouse Laboratory Plymouth | Wellcome Sanger Institute for the COVID-19 Genomics UK (COG-UK) Consortium | Lighthouse Laboratory Plymouth and Alex Alderton, Roberto Amato, Jeffrey Barrett, Sonia Goncalves, Ewan Harrison, David K. Jackson, Ian Johnston, Dominic Kwiatkowski, Cordelia Langford, John Sillitoe on behalf of the Wellcome Sanger Institute COVID-19 Surveillance Team |
| EPI_ISL_2849475, EPI_ISL_2849476 | Lighthouse Lab in Milton Keynes | Wellcome Sanger Institute for the COVID-19 Genomics UK (COG-UK) Consortium | The Lighthouse Lab in Milton Keynes and Alex Alderton, Roberto Amato, Jeffrey Barrett, Sonia Goncalves, Ewan Harrison, David K. Jackson, Ian Johnston, Dominic Kwiatkowski, Cordelia Langford, John Sillitoe on behalf of the Wellcome Sanger Institute COVID-19 Surveillance Team |
| EPI_ISL_2849477 | Lighthouse Laboratory Plymouth | Wellcome Sanger Institute for the COVID-19 Genomics UK (COG-UK) Consortium | Lighthouse Laboratory Plymouth and Alex Alderton, Roberto Amato, Jeffrey Barrett, Sonia Goncalves, Ewan Harrison, David K. Jackson, Ian Johnston, Dominic Kwiatkowski, Cordelia Langford, John Sillitoe on behalf of the Wellcome Sanger Institute COVID-19 Surveillance Team |
| EPI_ISL_2849478 | Lighthouse Lab in Alderley Park | Wellcome Sanger Institute for the COVID-19 Genomics UK (COG-UK) Consortium | Jacquelyn Wynn, Mairead Hyland, The Lighthouse Lab in Alderley Park and Alex Alderton, Roberto Amato, Jeffrey Barrett, Sonia Goncalves, Ewan Harrison, David K. Jackson, Ian Johnston, Dominic Kwiatkowski, Cordelia Langford, John Sillitoe on behalf of the Wellcome Sanger Institute COVID-19 Surveillance Team |
| EPI_ISL_2849479 | Lighthouse Lab in Milton Keynes | Wellcome Sanger Institute for the COVID-19 Genomics UK (COG-UK) Consortium | The Lighthouse Lab in Milton Keynes and Alex Alderton, Roberto Amato, Jeffrey Barrett, Sonia Goncalves, Ewan Harrison, David K. Jackson, Ian Johnston, Dominic Kwiatkowski, Cordelia Langford, John Sillitoe on behalf of the Wellcome Sanger Institute COVID-19 Surveillance Team |
| EPI_ISL_2849480 | Lighthouse Laboratory Plymouth | Wellcome Sanger Institute for the COVID-19 Genomics UK (COG-UK) Consortium | Lighthouse Laboratory Plymouth and Alex Alderton, Roberto Amato, Jeffrey Barrett, Sonia Goncalves, Ewan Harrison, David K. Jackson, Ian Johnston, Dominic Kwiatkowski, Cordelia Langford, John Sillitoe on behalf of the Wellcome Sanger Institute COVID-19 Surveillance Team |
| EPI_ISL_2849481 | Lighthouse Lab in Alderley Park | Wellcome Sanger Institute for the COVID-19 Genomics UK (COG-UK) Consortium | Jacquelyn Wynn, Mairead Hyland, The Lighthouse Lab in Alderley Park and Alex Alderton, Roberto Amato, Jeffrey Barrett, Sonia Goncalves, Ewan Harrison, David K. Jackson, Ian Johnston, Dominic Kwiatkowski, Cordelia Langford, John Sillitoe on behalf of the Wellcome Sanger Institute COVID-19 Surveillance Team |
| EPI_ISL_2849482 | Lighthouse Lab in Milton Keynes | Wellcome Sanger Institute for the COVID-19 Genomics UK (COG-UK) Consortium | The Lighthouse Lab in Milton Keynes and Alex Alderton, Roberto Amato, Jeffrey Barrett, Sonia Goncalves, Ewan Harrison, David K. Jackson, Ian Johnston, Dominic Kwiatkowski, Cordelia Langford, John Sillitoe on behalf of the Wellcome Sanger Institute COVID-19 Surveillance Team |
| EPI_ISL_2849483 | Lighthouse Lab in Alderley Park | Wellcome Sanger Institute for the COVID-19 Genomics UK (COG-UK) Consortium | Jacquelyn Wynn, Mairead Hyland, The Lighthouse Lab in Alderley Park and Alex Alderton, Roberto Amato, Jeffrey Barrett, Sonia Goncalves, Ewan Harrison, David K. Jackson, Ian Johnston, Dominic Kwiatkowski, Cordelia Langford, John Sillitoe on behalf of the Wellcome Sanger Institute COVID-19 Surveillance Team |
| EPI_ISL_2849484 | Lighthouse Laboratory Plymouth | Wellcome Sanger Institute for the COVID-19 Genomics UK (COG-UK) Consortium | Lighthouse Laboratory Plymouth and Alex Alderton, Roberto Amato, Jeffrey Barrett, Sonia Goncalves, Ewan Harrison, David K. Jackson, Ian Johnston, Dominic Kwiatkowski, Cordelia Langford, John Sillitoe on behalf of the Wellcome Sanger Institute COVID-19 Surveillance Team |
| EPI_ISL_2849485, EPI_ISL_2849486, EPI_ISL_2849487 | Lighthouse Lab in Milton Keynes | Wellcome Sanger Institute for the COVID-19 Genomics UK (COG-UK) Consortium | The Lighthouse Lab in Milton Keynes and Alex Alderton, Roberto Amato, Jeffrey Barrett, Sonia Goncalves, Ewan Harrison, David K. Jackson, Ian Johnston, Dominic Kwiatkowski, Cordelia Langford, John Sillitoe on behalf of the Wellcome Sanger Institute COVID-19 Surveillance Team |
| EPI_ISL_2849488 | Lighthouse Lab in Alderley Park | Wellcome Sanger Institute for the COVID-19 Genomics UK (COG-UK) Consortium | Jacquelyn Wynn, Mairead Hyland, The Lighthouse Lab in Alderley Park and Alex Alderton, Roberto Amato, Jeffrey Barrett, Sonia Goncalves, Ewan Harrison, David K. Jackson, Ian Johnston, Dominic Kwiatkowski, Cordelia Langford, John Sillitoe on behalf of the Wellcome Sanger Institute COVID-19 Surveillance Team |
| EPI_ISL_2849489, EPI_ISL_2849490, EPI_ISL_2849492, EPI_ISL_2849493 | Lighthouse Laboratory Plymouth | Wellcome Sanger Institute for the COVID-19 Genomics UK (COG-UK) Consortium | Lighthouse Laboratory Plymouth and Alex Alderton, Roberto Amato, Jeffrey Barrett, Sonia Goncalves, Ewan Harrison, David K. Jackson, Ian Johnston, Dominic Kwiatkowski, Cordelia Langford, John Sillitoe on behalf of the Wellcome Sanger Institute COVID-19 Surveillance Team |
| EPI_ISL_2849494, EPI_ISL_2849495 | Lighthouse Lab in Milton Keynes | Wellcome Sanger Institute for the COVID-19 Genomics UK (COG-UK) Consortium | The Lighthouse Lab in Milton Keynes and Alex Alderton, Roberto Amato, Jeffrey Barrett, Sonia Goncalves, Ewan Harrison, David K. Jackson, Ian Johnston, Dominic Kwiatkowski, Cordelia Langford, John Sillitoe on behalf of the Wellcome Sanger Institute COVID-19 Surveillance Team |
| EPI_ISL_2849496, EPI_ISL_2849497, EPI_ISL_2849498 | Lighthouse Laboratory Plymouth | Wellcome Sanger Institute for the COVID-19 Genomics UK (COG-UK) Consortium | Lighthouse Laboratory Plymouth and Alex Alderton, Roberto Amato, Jeffrey Barrett, Sonia Goncalves, Ewan Harrison, David K. Jackson, Ian Johnston, Dominic Kwiatkowski, Cordelia Langford, John Sillitoe on behalf of the Wellcome Sanger Institute COVID-19 Surveillance Team |
| EPI_ISL_2849499 | Lighthouse Lab in Milton Keynes | Wellcome Sanger Institute for the COVID-19 Genomics UK (COG-UK) Consortium | The Lighthouse Lab in Milton Keynes and Alex Alderton, Roberto Amato, Jeffrey Barrett, Sonia Goncalves, Ewan Harrison, David K. Jackson, Ian Johnston, Dominic Kwiatkowski, Cordelia Langford, John Sillitoe on behalf of the Wellcome Sanger Institute COVID-19 Surveillance Team |
| EPI_ISL_2849500, EPI_ISL_2849501 | Lighthouse Laboratory Plymouth | Wellcome Sanger Institute for the COVID-19 Genomics UK (COG-UK) Consortium | Lighthouse Laboratory Plymouth and Alex Alderton, Roberto Amato, Jeffrey Barrett, Sonia Goncalves, Ewan Harrison, David K. Jackson, Ian Johnston, Dominic Kwiatkowski, Cordelia Langford, John Sillitoe on behalf of the Wellcome Sanger Institute COVID-19 Surveillance Team |
| EPI_ISL_2849502 | Lighthouse Lab in Alderley Park | Wellcome Sanger Institute for the COVID-19 Genomics UK (COG-UK) Consortium | Jacquelyn Wynn, Mairead Hyland, The Lighthouse Lab in Alderley Park and Alex Alderton, Roberto Amato, Jeffrey Barrett, Sonia Goncalves, Ewan Harrison, David K. Jackson, Ian Johnston, Dominic Kwiatkowski, Cordelia Langford, John Sillitoe on behalf of the Wellcome Sanger Institute COVID-19 Surveillance Team |
| EPI_ISL_2849503 | Lighthouse Laboratory Plymouth | Wellcome Sanger Institute for the COVID-19 Genomics UK (COG-UK) Consortium | Lighthouse Laboratory Plymouth and Alex Alderton, Roberto Amato, Jeffrey Barrett, Sonia Goncalves, Ewan Harrison, David K. Jackson, Ian Johnston, Dominic Kwiatkowski, Cordelia Langford, John Sillitoe on behalf of the Wellcome Sanger Institute COVID-19 Surveillance Team |
| EPI_ISL_2849504 | Lighthouse Lab in Milton Keynes | Wellcome Sanger Institute for the COVID-19 Genomics UK (COG-UK) Consortium | The Lighthouse Lab in Milton Keynes and Alex Alderton, Roberto Amato, Jeffrey Barrett, Sonia Goncalves, Ewan Harrison, David K. Jackson, Ian Johnston, Dominic Kwiatkowski, Cordelia Langford, John Sillitoe on behalf of the Wellcome Sanger Institute COVID-19 Surveillance Team |
| EPI_ISL_2849505 | Lighthouse Lab in Alderley Park | Wellcome Sanger Institute for the COVID-19 Genomics UK (COG-UK) Consortium | Jacquelyn Wynn, Mairead Hyland, The Lighthouse Lab in Alderley Park and Alex Alderton, Roberto Amato, Jeffrey Barrett, Sonia Goncalves, Ewan Harrison, David K. Jackson, Ian Johnston, Dominic Kwiatkowski, Cordelia Langford, John Sillitoe on behalf of the Wellcome Sanger Institute COVID-19 Surveillance Team |
| EPI_ISL_2849506, EPI_ISL_2849507 | Lighthouse Laboratory Plymouth | Wellcome Sanger Institute for the COVID-19 Genomics UK (COG-UK) Consortium | Lighthouse Laboratory Plymouth and Alex Alderton, Roberto Amato, Jeffrey Barrett, Sonia Goncalves, Ewan Harrison, David K. Jackson, Ian Johnston, Dominic Kwiatkowski, Cordelia Langford, John Sillitoe on behalf of the Wellcome Sanger Institute COVID-19 Surveillance Team |
| EPI_ISL_2849508, EPI_ISL_2849509 | Lighthouse Lab in Alderley Park | Wellcome Sanger Institute for the COVID-19 Genomics UK (COG-UK) Consortium | Jacquelyn Wynn, Mairead Hyland, The Lighthouse Lab in Alderley Park and Alex Alderton, Roberto Amato, Jeffrey Barrett, Sonia Goncalves, Ewan Harrison, David K. Jackson, Ian Johnston, Dominic Kwiatkowski, Cordelia Langford, John Sillitoe on behalf of the Wellcome Sanger Institute COVID-19 Surveillance Team |
| EPI_ISL_2849510 | Lighthouse Lab in Milton Keynes | Wellcome Sanger Institute for the COVID-19 Genomics UK (COG-UK) Consortium | The Lighthouse Lab in Milton Keynes and Alex Alderton, Roberto Amato, Jeffrey Barrett, Sonia Goncalves, Ewan Harrison, David K. Jackson, Ian Johnston, Dominic Kwiatkowski, Cordelia Langford, John Sillitoe on behalf of the Wellcome Sanger Institute COVID-19 Surveillance Team |
| EPI_ISL_2849511 | Lighthouse Lab in Alderley Park | Wellcome Sanger Institute for the COVID-19 Genomics UK (COG-UK) Consortium | Jacquelyn Wynn, Mairead Hyland, The Lighthouse Lab in Alderley Park and Alex Alderton, Roberto Amato, Jeffrey Barrett, Sonia Goncalves, Ewan Harrison, David K. Jackson, Ian Johnston, Dominic Kwiatkowski, Cordelia Langford, John Sillitoe on behalf of the Wellcome Sanger Institute COVID-19 Surveillance Team |
| EPI_ISL_2849512, EPI_ISL_2849515, EPI_ISL_2849516 | Lighthouse Laboratory Plymouth | Wellcome Sanger Institute for the COVID-19 Genomics UK (COG-UK) Consortium | Lighthouse Laboratory Plymouth and Alex Alderton, Roberto Amato, Jeffrey Barrett, Sonia Goncalves, Ewan Harrison, David K. Jackson, Ian Johnston, Dominic Kwiatkowski, Cordelia Langford, John Sillitoe on behalf of the Wellcome Sanger Institute COVID-19 Surveillance Team |

| | | | |
|---|---|---|---|
| EPI_ISL_2849517 | Lighthouse Lab in Alderley Park | Wellcome Sanger Institute for the COVID-19 Genomics UK (COG-UK) Consortium | Jacquelyn Wynn, Mairead Hyland, The Lighthouse Lab in Alderley Park and Alex Alderton, Roberto Amato, Jeffrey Barrett, Sonia Goncalves, Ewan Harrison, David K. Jackson, Ian Johnston, Dominic Kwiatkowski, Cordelia Langford, John Sillitoe on behalf of the Wellcome Sanger Institute COVID-19 Surveillance Team |
| EPI_ISL_2849518 | Lighthouse Lab in Milton Keynes | Wellcome Sanger Institute for the COVID-19 Genomics UK (COG-UK) Consortium | The Lighthouse Lab in Milton Keynes and Alex Alderton, Roberto Amato, Jeffrey Barrett, Sonia Goncalves, Ewan Harrison, David K. Jackson, Ian Johnston, Dominic Kwiatkowski, Cordelia Langford, John Sillitoe on behalf of the Wellcome Sanger Institute COVID-19 Surveillance Team |
| EPI_ISL_2849519 | Lighthouse Laboratory Plymouth | Wellcome Sanger Institute for the COVID-19 Genomics UK (COG-UK) Consortium | Lighthouse Laboratory Plymouth and Alex Alderton, Roberto Amato, Jeffrey Barrett, Sonia Goncalves, Ewan Harrison, David K. Jackson, Ian Johnston, Dominic Kwiatkowski, Cordelia Langford, John Sillitoe on behalf of the Wellcome Sanger Institute COVID-19 Surveillance Team |
| EPI_ISL_2849520 | Lighthouse Lab in Milton Keynes | Wellcome Sanger Institute for the COVID-19 Genomics UK (COG-UK) Consortium | The Lighthouse Lab in Milton Keynes and Alex Alderton, Roberto Amato, Jeffrey Barrett, Sonia Goncalves, Ewan Harrison, David K. Jackson, Ian Johnston, Dominic Kwiatkowski, Cordelia Langford, John Sillitoe on behalf of the Wellcome Sanger Institute COVID-19 Surveillance Team |
| EPI_ISL_2849521, EPI_ISL_2849522, EPI_ISL_2849523 | Lighthouse Laboratory Plymouth | Wellcome Sanger Institute for the COVID-19 Genomics UK (COG-UK) Consortium | Lighthouse Laboratory Plymouth and Alex Alderton, Roberto Amato, Jeffrey Barrett, Sonia Goncalves, Ewan Harrison, David K. Jackson, Ian Johnston, Dominic Kwiatkowski, Cordelia Langford, John Sillitoe on behalf of the Wellcome Sanger Institute COVID-19 Surveillance Team |
| EPI_ISL_2849524, EPI_ISL_2849525, EPI_ISL_2849526 | Lighthouse Lab in Milton Keynes | Wellcome Sanger Institute for the COVID-19 Genomics UK (COG-UK) Consortium | The Lighthouse Lab in Milton Keynes and Alex Alderton, Roberto Amato, Jeffrey Barrett, Sonia Goncalves, Ewan Harrison, David K. Jackson, Ian Johnston, Dominic Kwiatkowski, Cordelia Langford, John Sillitoe on behalf of the Wellcome Sanger Institute COVID-19 Surveillance Team |
| EPI_ISL_2849527 | Lighthouse Lab in Alderley Park | Wellcome Sanger Institute for the COVID-19 Genomics UK (COG-UK) Consortium | Jacquelyn Wynn, Mairead Hyland, The Lighthouse Lab in Alderley Park and Alex Alderton, Roberto Amato, Jeffrey Barrett, Sonia Goncalves, Ewan Harrison, David K. Jackson, Ian Johnston, Dominic Kwiatkowski, Cordelia Langford, John Sillitoe on behalf of the Wellcome Sanger Institute COVID-19 Surveillance Team |
| EPI_ISL_2849528 | Lighthouse Lab in Milton Keynes | Wellcome Sanger Institute for the COVID-19 Genomics UK (COG-UK) Consortium | The Lighthouse Lab in Milton Keynes and Alex Alderton, Roberto Amato, Jeffrey Barrett, Sonia Goncalves, Ewan Harrison, David K. Jackson, Ian Johnston, Dominic Kwiatkowski, Cordelia Langford, John Sillitoe on behalf of the Wellcome Sanger Institute COVID-19 Surveillance Team |
| EPI_ISL_2849529, EPI_ISL_2849530 | Lighthouse Laboratory Plymouth | Wellcome Sanger Institute for the COVID-19 Genomics UK (COG-UK) Consortium | Lighthouse Laboratory Plymouth and Alex Alderton, Roberto Amato, Jeffrey Barrett, Sonia Goncalves, Ewan Harrison, David K. Jackson, Ian Johnston, Dominic Kwiatkowski, Cordelia Langford, John Sillitoe on behalf of the Wellcome Sanger Institute COVID-19 Surveillance Team |
| EPI_ISL_2849531 | Lighthouse Lab in Milton Keynes | Wellcome Sanger Institute for the COVID-19 Genomics UK (COG-UK) Consortium | The Lighthouse Lab in Milton Keynes and Alex Alderton, Roberto Amato, Jeffrey Barrett, Sonia Goncalves, Ewan Harrison, David K. Jackson, Ian Johnston, Dominic Kwiatkowski, Cordelia Langford, John Sillitoe on behalf of the Wellcome Sanger Institute COVID-19 Surveillance Team |
| EPI_ISL_2849532, EPI_ISL_2849533 | Lighthouse Lab in Alderley Park | Wellcome Sanger Institute for the COVID-19 Genomics UK (COG-UK) Consortium | Jacquelyn Wynn, Mairead Hyland, The Lighthouse Lab in Alderley Park and Alex Alderton, Roberto Amato, Jeffrey Barrett, Sonia Goncalves, Ewan Harrison, David K. Jackson, Ian Johnston, Dominic Kwiatkowski, Cordelia Langford, John Sillitoe on behalf of the Wellcome Sanger Institute COVID-19 Surveillance Team |
| EPI_ISL_2849534 | Lighthouse Lab in Milton Keynes | Wellcome Sanger Institute for the COVID-19 Genomics UK (COG-UK) Consortium | The Lighthouse Lab in Milton Keynes and Alex Alderton, Roberto Amato, Jeffrey Barrett, Sonia Goncalves, Ewan Harrison, David K. Jackson, Ian Johnston, Dominic Kwiatkowski, Cordelia Langford, John Sillitoe on behalf of the Wellcome Sanger Institute COVID-19 Surveillance Team |
| EPI_ISL_2849535 | Lighthouse Lab in Alderley Park | Wellcome Sanger Institute for the COVID-19 Genomics UK (COG-UK) Consortium | Jacquelyn Wynn, Mairead Hyland, The Lighthouse Lab in Alderley Park and Alex Alderton, Roberto Amato, Jeffrey Barrett, Sonia Goncalves, Ewan Harrison, David K. Jackson, Ian Johnston, Dominic Kwiatkowski, Cordelia Langford, John Sillitoe on behalf of the Wellcome Sanger Institute COVID-19 Surveillance Team |
| EPI_ISL_2849536 | Lighthouse Laboratory Plymouth | Wellcome Sanger Institute for the COVID-19 Genomics UK (COG-UK) Consortium | Lighthouse Laboratory Plymouth and Alex Alderton, Roberto Amato, Jeffrey Barrett, Sonia Goncalves, Ewan Harrison, David K. Jackson, Ian Johnston, Dominic Kwiatkowski, Cordelia Langford, John Sillitoe on behalf of the Wellcome Sanger Institute COVID-19 Surveillance Team |
| EPI_ISL_2849537, EPI_ISL_2849538 | Lighthouse Lab in Alderley Park | Wellcome Sanger Institute for the COVID-19 Genomics UK (COG-UK) Consortium | Jacquelyn Wynn, Mairead Hyland, The Lighthouse Lab in Alderley Park and Alex Alderton, Roberto Amato, Jeffrey Barrett, Sonia Goncalves, Ewan Harrison, David K. Jackson, Ian Johnston, Dominic Kwiatkowski, Cordelia Langford, John Sillitoe on behalf of the Wellcome Sanger Institute COVID-19 Surveillance Team |
| EPI_ISL_2849539, EPI_ISL_2849540, EPI_ISL_2849541, EPI_ISL_2849542, EPI_ISL_2849543, EPI_ISL_2849544, EPI_ISL_2849545 | Lighthouse Laboratory Plymouth | Wellcome Sanger Institute for the COVID-19 Genomics UK (COG-UK) Consortium | Lighthouse Laboratory Plymouth and Alex Alderton, Roberto Amato, Jeffrey Barrett, Sonia Goncalves, Ewan Harrison, David K. Jackson, Ian Johnston, Dominic Kwiatkowski, Cordelia Langford, John Sillitoe on behalf of the Wellcome Sanger Institute COVID-19 Surveillance Team |
| EPI_ISL_2849546 | Lighthouse Lab in Alderley Park | Wellcome Sanger Institute for the COVID-19 Genomics UK (COG-UK) Consortium | Jacquelyn Wynn, Mairead Hyland, The Lighthouse Lab in Alderley Park and Alex Alderton, Roberto Amato, Jeffrey Barrett, Sonia Goncalves, Ewan Harrison, David K. Jackson, Ian Johnston, Dominic Kwiatkowski, Cordelia Langford, John Sillitoe on behalf of the Wellcome Sanger Institute COVID-19 Surveillance Team |
| EPI_ISL_2849547, EPI_ISL_2849548, EPI_ISL_2849549, EPI_ISL_2849550 | Lighthouse Laboratory Plymouth | Wellcome Sanger Institute for the COVID-19 Genomics UK (COG-UK) Consortium | Lighthouse Laboratory Plymouth and Alex Alderton, Roberto Amato, Jeffrey Barrett, Sonia Goncalves, Ewan Harrison, David K. Jackson, Ian Johnston, Dominic Kwiatkowski, Cordelia Langford, John Sillitoe on behalf of the Wellcome Sanger Institute COVID-19 Surveillance Team |
| EPI_ISL_2849551 | Lighthouse Lab in Milton Keynes | Wellcome Sanger Institute for the COVID-19 Genomics UK (COG-UK) Consortium | The Lighthouse Lab in Milton Keynes and Alex Alderton, Roberto Amato, Jeffrey Barrett, Sonia Goncalves, Ewan Harrison, David K. Jackson, Ian Johnston, Dominic Kwiatkowski, Cordelia Langford, John Sillitoe on behalf of the Wellcome Sanger Institute COVID-19 Surveillance Team |
| EPI_ISL_2849552, EPI_ISL_2849553 | Lighthouse Laboratory Plymouth | Wellcome Sanger Institute for the COVID-19 Genomics UK (COG-UK) Consortium | Lighthouse Laboratory Plymouth and Alex Alderton, Roberto Amato, Jeffrey Barrett, Sonia Goncalves, Ewan Harrison, David K. Jackson, Ian Johnston, Dominic Kwiatkowski, Cordelia Langford, John Sillitoe on behalf of the Wellcome Sanger Institute COVID-19 Surveillance Team |
| EPI_ISL_2849554 | Lighthouse Lab in Alderley Park | Wellcome Sanger Institute for the COVID-19 Genomics UK (COG-UK) Consortium | Jacquelyn Wynn, Mairead Hyland, The Lighthouse Lab in Alderley Park and Alex Alderton, Roberto Amato, Jeffrey Barrett, Sonia Goncalves, Ewan Harrison, David K. Jackson, Ian Johnston, Dominic Kwiatkowski, Cordelia Langford, John Sillitoe on behalf of the Wellcome Sanger Institute COVID-19 Surveillance Team |
| EPI_ISL_2849555 | Lighthouse Laboratory Plymouth | Wellcome Sanger Institute for the COVID-19 Genomics UK (COG-UK) Consortium | Lighthouse Laboratory Plymouth and Alex Alderton, Roberto Amato, Jeffrey Barrett, Sonia Goncalves, Ewan Harrison, David K. Jackson, Ian Johnston, Dominic Kwiatkowski, Cordelia Langford, John Sillitoe on behalf of the Wellcome Sanger Institute COVID-19 Surveillance Team |
| EPI_ISL_2849556 | Lighthouse Lab in Alderley Park | Wellcome Sanger Institute for the COVID-19 Genomics UK (COG-UK) Consortium | Jacquelyn Wynn, Mairead Hyland, The Lighthouse Lab in Alderley Park and Alex Alderton, Roberto Amato, Jeffrey Barrett, Sonia Goncalves, Ewan Harrison, David K. Jackson, Ian Johnston, Dominic Kwiatkowski, Cordelia Langford, John Sillitoe on behalf of the Wellcome Sanger Institute COVID-19 Surveillance Team |
| EPI_ISL_2849557 | Lighthouse Laboratory Plymouth | Wellcome Sanger Institute for the COVID-19 Genomics UK (COG-UK) Consortium | Lighthouse Laboratory Plymouth and Alex Alderton, Roberto Amato, Jeffrey Barrett, Sonia Goncalves, Ewan Harrison, David K. Jackson, Ian Johnston, Dominic Kwiatkowski, Cordelia Langford, John Sillitoe on behalf of the Wellcome Sanger Institute COVID-19 Surveillance Team |
| EPI_ISL_2849558 | Lighthouse Lab in Alderley Park | Wellcome Sanger Institute for the COVID-19 Genomics UK (COG-UK) Consortium | Jacquelyn Wynn, Mairead Hyland, The Lighthouse Lab in Alderley Park and Alex Alderton, Roberto Amato, Jeffrey Barrett, Sonia Goncalves, Ewan Harrison, David K. Jackson, Ian Johnston, Dominic Kwiatkowski, Cordelia Langford, John Sillitoe on behalf of the Wellcome Sanger Institute COVID-19 Surveillance Team |
| EPI_ISL_2849559 | Lighthouse Laboratory Plymouth | Wellcome Sanger Institute for the COVID-19 Genomics UK (COG-UK) Consortium | Lighthouse Laboratory Plymouth and Alex Alderton, Roberto Amato, Jeffrey Barrett, Sonia Goncalves, Ewan Harrison, David K. Jackson, Ian Johnston, Dominic Kwiatkowski, Cordelia Langford, John Sillitoe on behalf of the Wellcome Sanger Institute COVID-19 Surveillance Team |
| EPI_ISL_2849560 | Lighthouse Lab in Alderley Park | Wellcome Sanger Institute for the COVID-19 Genomics UK (COG-UK) Consortium | Jacquelyn Wynn, Mairead Hyland, The Lighthouse Lab in Alderley Park and Alex Alderton, Roberto Amato, Jeffrey Barrett, Sonia Goncalves, Ewan Harrison, David K. Jackson, Ian Johnston, Dominic Kwiatkowski, Cordelia Langford, John Sillitoe on behalf of the Wellcome Sanger Institute COVID-19 Surveillance Team |
| EPI_ISL_2849561 | Lighthouse Laboratory Plymouth | Wellcome Sanger Institute for the COVID-19 Genomics UK (COG-UK) Consortium | Lighthouse Laboratory Plymouth and Alex Alderton, Roberto Amato, Jeffrey Barrett, Sonia Goncalves, Ewan Harrison, David K. Jackson, Ian Johnston, Dominic Kwiatkowski, Cordelia Langford, John Sillitoe on behalf of the Wellcome Sanger Institute COVID-19 Surveillance Team |
| EPI_ISL_2849562 | Lighthouse Lab in Milton Keynes | Wellcome Sanger Institute for the COVID-19 Genomics UK (COG-UK) Consortium | The Lighthouse Lab in Milton Keynes and Alex Alderton, Roberto Amato, Jeffrey Barrett, Sonia Goncalves, Ewan Harrison, David K. Jackson, Ian Johnston, Dominic Kwiatkowski, Cordelia Langford, John Sillitoe on behalf of the Wellcome Sanger Institute COVID-19 Surveillance Team |
| EPI_ISL_2849563 | Lighthouse Lab in Alderley Park | Wellcome Sanger Institute for the COVID-19 Genomics UK (COG-UK) Consortium | Jacquelyn Wynn, Mairead Hyland, The Lighthouse Lab in Alderley Park and Alex Alderton, Roberto Amato, Jeffrey Barrett, Sonia Goncalves, Ewan Harrison, David K. Jackson, Ian Johnston, Dominic Kwiatkowski, Cordelia Langford, John Sillitoe on behalf of the Wellcome Sanger Institute COVID-19 Surveillance Team |
| EPI_ISL_2849564, EPI_ISL_2849565 | Lighthouse Laboratory Plymouth | Wellcome Sanger Institute for the COVID-19 Genomics UK (COG-UK) Consortium | Lighthouse Laboratory Plymouth and Alex Alderton, Roberto Amato, Jeffrey Barrett, Sonia Goncalves, Ewan Harrison, David K. Jackson, Ian Johnston, Dominic Kwiatkowski, Cordelia Langford, John Sillitoe on behalf of the Wellcome Sanger Institute COVID-19 Surveillance Team |
| EPI_ISL_2849566 | Lighthouse Lab in Alderley Park | Wellcome Sanger Institute for the COVID-19 Genomics UK (COG-UK) Consortium | Jacquelyn Wynn, Mairead Hyland, The Lighthouse Lab in Alderley Park and Alex Alderton, Roberto Amato, Jeffrey Barrett, Sonia Goncalves, Ewan Harrison, David K. Jackson, Ian Johnston, Dominic Kwiatkowski, Cordelia Langford, John Sillitoe on behalf of the Wellcome Sanger Institute COVID-19 Surveillance Team |
| EPI_ISL_2849567 | Lighthouse Laboratory Plymouth | Wellcome Sanger Institute for the COVID-19 Genomics UK (COG-UK) Consortium | Lighthouse Laboratory Plymouth and Alex Alderton, Roberto Amato, Jeffrey Barrett, Sonia Goncalves, Ewan Harrison, David K. Jackson, Ian Johnston, Dominic Kwiatkowski, Cordelia Langford, John Sillitoe on behalf of the Wellcome Sanger Institute COVID-19 Surveillance Team |
| EPI_ISL_2849568, EPI_ISL_2849569 | Lighthouse Lab in Alderley Park | Wellcome Sanger Institute for the COVID-19 Genomics UK (COG-UK) Consortium | Jacquelyn Wynn, Mairead Hyland, The Lighthouse Lab in Alderley Park and Alex Alderton, Roberto Amato, Jeffrey Barrett, Sonia Goncalves, Ewan Harrison, David K. Jackson, Ian Johnston, Dominic Kwiatkowski, Cordelia Langford, John Sillitoe on behalf of the Wellcome Sanger Institute COVID-19 Surveillance Team |
| EPI_ISL_2849570 | Lighthouse Lab in Milton Keynes | Wellcome Sanger Institute for the COVID-19 Genomics UK (COG-UK) Consortium | The Lighthouse Lab in Milton Keynes and Alex Alderton, Roberto Amato, Jeffrey Barrett, Sonia Goncalves, Ewan Harrison, David K. Jackson, Ian Johnston, Dominic Kwiatkowski, Cordelia Langford, John Sillitoe on behalf of the Wellcome Sanger Institute COVID-19 Surveillance Team |
| EPI_ISL_2849571 | Lighthouse Laboratory Plymouth | Wellcome Sanger Institute for the COVID-19 Genomics UK (COG-UK) Consortium | Lighthouse Laboratory Plymouth and Alex Alderton, Roberto Amato, Jeffrey Barrett, Sonia Goncalves, Ewan Harrison, David K. Jackson, Ian Johnston, Dominic Kwiatkowski, Cordelia Langford, John Sillitoe on behalf of the Wellcome Sanger Institute COVID-19 Surveillance Team |
| EPI_ISL_2849572 | Lighthouse Lab in Alderley Park | Wellcome Sanger Institute for the COVID-19 Genomics UK (COG-UK) Consortium | Jacquelyn Wynn, Mairead Hyland, The Lighthouse Lab in Alderley Park and Alex Alderton, Roberto Amato, Jeffrey Barrett, Sonia Goncalves, Ewan Harrison, David K. Jackson, Ian Johnston, Dominic Kwiatkowski, Cordelia Langford, John Sillitoe on behalf of the Wellcome Sanger Institute COVID-19 Surveillance Team |
| EPI_ISL_2849573 | Lighthouse Lab in Milton Keynes | Wellcome Sanger Institute for the COVID-19 Genomics UK (COG-UK) Consortium | The Lighthouse Lab in Milton Keynes and Alex Alderton, Roberto Amato, Jeffrey Barrett, Sonia Goncalves, Ewan Harrison, David K. Jackson, Ian Johnston, Dominic Kwiatkowski, Cordelia Langford, John Sillitoe on behalf of the Wellcome Sanger Institute COVID-19 Surveillance Team |
| EPI_ISL_2849574, EPI_ISL_2849575 | Lighthouse Laboratory Plymouth | Wellcome Sanger Institute for the COVID-19 Genomics UK (COG-UK) Consortium | Lighthouse Laboratory Plymouth and Alex Alderton, Roberto Amato, Jeffrey Barrett, Sonia Goncalves, Ewan Harrison, David K. Jackson, Ian Johnston, Dominic Kwiatkowski, Cordelia Langford, John Sillitoe on behalf of the Wellcome Sanger Institute COVID-19 Surveillance Team |
| EPI_ISL_2849576 | Lighthouse Lab in Milton Keynes | Wellcome Sanger Institute for the COVID-19 Genomics UK (COG-UK) Consortium | The Lighthouse Lab in Milton Keynes and Alex Alderton, Roberto Amato, Jeffrey Barrett, Sonia Goncalves, Ewan Harrison, David K. Jackson, Ian Johnston, Dominic Kwiatkowski, Cordelia Langford, John Sillitoe on behalf of the Wellcome Sanger Institute COVID-19 Surveillance Team |
| EPI_ISL_2849577, EPI_ISL_2849578 | Lighthouse Laboratory Plymouth | Wellcome Sanger Institute for the COVID-19 Genomics UK (COG-UK) Consortium | Lighthouse Laboratory Plymouth and Alex Alderton, Roberto Amato, Jeffrey Barrett, Sonia Goncalves, Ewan Harrison, David K. Jackson, Ian Johnston, Dominic Kwiatkowski, Cordelia Langford, John Sillitoe on behalf of the Wellcome Sanger Institute COVID-19 Surveillance Team |

| | | | |
|---|---|---|---|
| EPI_ISL_2849579 | Lighthouse Lab in Alderley Park | Wellcome Sanger Institute for the COVID-19 Genomics UK (COG-UK) Consortium | Jacquelyn Wynn, Mairead Hyland, The Lighthouse Lab in Alderley Park and Alex Alderton, Roberto Amato, Jeffrey Barrett, Sonia Goncalves, Ewan Harrison, David K. Jackson, Ian Johnston, Dominic Kwiatkowski, Cordelia Langford, John Sillitoe on behalf of the Wellcome Sanger Institute COVID-19 Surveillance Team |
| EPI_ISL_2849580, EPI_ISL_2849581, EPI_ISL_2849582 | Lighthouse Laboratory Plymouth | Wellcome Sanger Institute for the COVID-19 Genomics UK (COG-UK) Consortium | Lighthouse Laboratory Plymouth and Alex Alderton, Roberto Amato, Jeffrey Barrett, Sonia Goncalves, Ewan Harrison, David K. Jackson, Ian Johnston, Dominic Kwiatkowski, Cordelia Langford, John Sillitoe on behalf of the Wellcome Sanger Institute COVID-19 Surveillance Team |
| EPI_ISL_2849583, EPI_ISL_2849584 | Lighthouse Lab in Alderley Park | Wellcome Sanger Institute for the COVID-19 Genomics UK (COG-UK) Consortium | Jacquelyn Wynn, Mairead Hyland, The Lighthouse Lab in Alderley Park and Alex Alderton, Roberto Amato, Jeffrey Barrett, Sonia Goncalves, Ewan Harrison, David K. Jackson, Ian Johnston, Dominic Kwiatkowski, Cordelia Langford, John Sillitoe on behalf of the Wellcome Sanger Institute COVID-19 Surveillance Team |
| EPI_ISL_2849585 | Lighthouse Laboratory Plymouth | Wellcome Sanger Institute for the COVID-19 Genomics UK (COG-UK) Consortium | Lighthouse Laboratory Plymouth and Alex Alderton, Roberto Amato, Jeffrey Barrett, Sonia Goncalves, Ewan Harrison, David K. Jackson, Ian Johnston, Dominic Kwiatkowski, Cordelia Langford, John Sillitoe on behalf of the Wellcome Sanger Institute COVID-19 Surveillance Team |
| EPI_ISL_2849586 | Lighthouse Lab in Milton Keynes | Wellcome Sanger Institute for the COVID-19 Genomics UK (COG-UK) Consortium | The Lighthouse Lab in Milton Keynes and Alex Alderton, Roberto Amato, Jeffrey Barrett, Sonia Goncalves, Ewan Harrison, David K. Jackson, Ian Johnston, Dominic Kwiatkowski, Cordelia Langford, John Sillitoe on behalf of the Wellcome Sanger Institute COVID-19 Surveillance Team |
| EPI_ISL_2849587 | Lighthouse Lab in Alderley Park | Wellcome Sanger Institute for the COVID-19 Genomics UK (COG-UK) Consortium | Jacquelyn Wynn, Mairead Hyland, The Lighthouse Lab in Alderley Park and Alex Alderton, Roberto Amato, Jeffrey Barrett, Sonia Goncalves, Ewan Harrison, David K. Jackson, Ian Johnston, Dominic Kwiatkowski, Cordelia Langford, John Sillitoe on behalf of the Wellcome Sanger Institute COVID-19 Surveillance Team |
| EPI_ISL_2849588 | Lighthouse Laboratory Plymouth | Wellcome Sanger Institute for the COVID-19 Genomics UK (COG-UK) Consortium | Lighthouse Laboratory Plymouth and Alex Alderton, Roberto Amato, Jeffrey Barrett, Sonia Goncalves, Ewan Harrison, David K. Jackson, Ian Johnston, Dominic Kwiatkowski, Cordelia Langford, John Sillitoe on behalf of the Wellcome Sanger Institute COVID-19 Surveillance Team |
| EPI_ISL_2849589 | Lighthouse Lab in Alderley Park | Wellcome Sanger Institute for the COVID-19 Genomics UK (COG-UK) Consortium | Jacquelyn Wynn, Mairead Hyland, The Lighthouse Lab in Alderley Park and Alex Alderton, Roberto Amato, Jeffrey Barrett, Sonia Goncalves, Ewan Harrison, David K. Jackson, Ian Johnston, Dominic Kwiatkowski, Cordelia Langford, John Sillitoe on behalf of the Wellcome Sanger Institute COVID-19 Surveillance Team |
| EPI_ISL_2849590, EPI_ISL_2849591 | Lighthouse Lab in Milton Keynes | Wellcome Sanger Institute for the COVID-19 Genomics UK (COG-UK) Consortium | The Lighthouse Lab in Milton Keynes and Alex Alderton, Roberto Amato, Jeffrey Barrett, Sonia Goncalves, Ewan Harrison, David K. Jackson, Ian Johnston, Dominic Kwiatkowski, Cordelia Langford, John Sillitoe on behalf of the Wellcome Sanger Institute COVID-19 Surveillance Team |
| EPI_ISL_2849592, EPI_ISL_2849593 | Lighthouse Laboratory Plymouth | Wellcome Sanger Institute for the COVID-19 Genomics UK (COG-UK) Consortium | Lighthouse Laboratory Plymouth and Alex Alderton, Roberto Amato, Jeffrey Barrett, Sonia Goncalves, Ewan Harrison, David K. Jackson, Ian Johnston, Dominic Kwiatkowski, Cordelia Langford, John Sillitoe on behalf of the Wellcome Sanger Institute COVID-19 Surveillance Team |
| EPI_ISL_2849594 | Lighthouse Lab in Alderley Park | Wellcome Sanger Institute for the COVID-19 Genomics UK (COG-UK) Consortium | Jacquelyn Wynn, Mairead Hyland, The Lighthouse Lab in Alderley Park and Alex Alderton, Roberto Amato, Jeffrey Barrett, Sonia Goncalves, Ewan Harrison, David K. Jackson, Ian Johnston, Dominic Kwiatkowski, Cordelia Langford, John Sillitoe on behalf of the Wellcome Sanger Institute COVID-19 Surveillance Team |
| EPI_ISL_2849595, EPI_ISL_2849596, EPI_ISL_2849597 | Lighthouse Laboratory Plymouth | Wellcome Sanger Institute for the COVID-19 Genomics UK (COG-UK) Consortium | Lighthouse Laboratory Plymouth and Alex Alderton, Roberto Amato, Jeffrey Barrett, Sonia Goncalves, Ewan Harrison, David K. Jackson, Ian Johnston, Dominic Kwiatkowski, Cordelia Langford, John Sillitoe on behalf of the Wellcome Sanger Institute COVID-19 Surveillance Team |
| EPI_ISL_2849598 | Lighthouse Lab in Milton Keynes | Wellcome Sanger Institute for the COVID-19 Genomics UK (COG-UK) Consortium | The Lighthouse Lab in Milton Keynes and Alex Alderton, Roberto Amato, Jeffrey Barrett, Sonia Goncalves, Ewan Harrison, David K. Jackson, Ian Johnston, Dominic Kwiatkowski, Cordelia Langford, John Sillitoe on behalf of the Wellcome Sanger Institute COVID-19 Surveillance Team |
| EPI_ISL_2849599, EPI_ISL_2849600 | Lighthouse Laboratory Plymouth | Wellcome Sanger Institute for the COVID-19 Genomics UK (COG-UK) Consortium | Lighthouse Laboratory Plymouth and Alex Alderton, Roberto Amato, Jeffrey Barrett, Sonia Goncalves, Ewan Harrison, David K. Jackson, Ian Johnston, Dominic Kwiatkowski, Cordelia Langford, John Sillitoe on behalf of the Wellcome Sanger Institute COVID-19 Surveillance Team |
| EPI_ISL_2849601, EPI_ISL_2849602 | Lighthouse Lab in Milton Keynes | Wellcome Sanger Institute for the COVID-19 Genomics UK (COG-UK) Consortium | The Lighthouse Lab in Milton Keynes and Alex Alderton, Roberto Amato, Jeffrey Barrett, Sonia Goncalves, Ewan Harrison, David K. Jackson, Ian Johnston, Dominic Kwiatkowski, Cordelia Langford, John Sillitoe on behalf of the Wellcome Sanger Institute COVID-19 Surveillance Team |
| EPI_ISL_2849603, EPI_ISL_2849604, EPI_ISL_2849605, EPI_ISL_2849606 | Lighthouse Laboratory Plymouth | Wellcome Sanger Institute for the COVID-19 Genomics UK (COG-UK) Consortium | Lighthouse Laboratory Plymouth and Alex Alderton, Roberto Amato, Jeffrey Barrett, Sonia Goncalves, Ewan Harrison, David K. Jackson, Ian Johnston, Dominic Kwiatkowski, Cordelia Langford, John Sillitoe on behalf of the Wellcome Sanger Institute COVID-19 Surveillance Team |
| EPI_ISL_2849607 | Lighthouse Lab in Alderley Park | Wellcome Sanger Institute for the COVID-19 Genomics UK (COG-UK) Consortium | Jacquelyn Wynn, Mairead Hyland, The Lighthouse Lab in Alderley Park and Alex Alderton, Roberto Amato, Jeffrey Barrett, Sonia Goncalves, Ewan Harrison, David K. Jackson, Ian Johnston, Dominic Kwiatkowski, Cordelia Langford, John Sillitoe on behalf of the Wellcome Sanger Institute COVID-19 Surveillance Team |
| EPI_ISL_2849608 | Lighthouse Lab in Milton Keynes | Wellcome Sanger Institute for the COVID-19 Genomics UK (COG-UK) Consortium | The Lighthouse Lab in Milton Keynes and Alex Alderton, Roberto Amato, Jeffrey Barrett, Sonia Goncalves, Ewan Harrison, David K. Jackson, Ian Johnston, Dominic Kwiatkowski, Cordelia Langford, John Sillitoe on behalf of the Wellcome Sanger Institute COVID-19 Surveillance Team |
| EPI_ISL_2849609, EPI_ISL_2849610 | Lighthouse Laboratory Plymouth | Wellcome Sanger Institute for the COVID-19 Genomics UK (COG-UK) Consortium | Lighthouse Laboratory Plymouth and Alex Alderton, Roberto Amato, Jeffrey Barrett, Sonia Goncalves, Ewan Harrison, David K. Jackson, Ian Johnston, Dominic Kwiatkowski, Cordelia Langford, John Sillitoe on behalf of the Wellcome Sanger Institute COVID-19 Surveillance Team |
| EPI_ISL_2849611 | Lighthouse Lab in Milton Keynes | Wellcome Sanger Institute for the COVID-19 Genomics UK (COG-UK) Consortium | The Lighthouse Lab in Milton Keynes and Alex Alderton, Roberto Amato, Jeffrey Barrett, Sonia Goncalves, Ewan Harrison, David K. Jackson, Ian Johnston, Dominic Kwiatkowski, Cordelia Langford, John Sillitoe on behalf of the Wellcome Sanger Institute COVID-19 Surveillance Team |
| EPI_ISL_2849612, EPI_ISL_2849613 | Lighthouse Laboratory Plymouth | Wellcome Sanger Institute for the COVID-19 Genomics UK (COG-UK) Consortium | Lighthouse Laboratory Plymouth and Alex Alderton, Roberto Amato, Jeffrey Barrett, Sonia Goncalves, Ewan Harrison, David K. Jackson, Ian Johnston, Dominic Kwiatkowski, Cordelia Langford, John Sillitoe on behalf of the Wellcome Sanger Institute COVID-19 Surveillance Team |
| EPI_ISL_2849614 | Lighthouse Lab in Milton Keynes | Wellcome Sanger Institute for the COVID-19 Genomics UK (COG-UK) Consortium | The Lighthouse Lab in Milton Keynes and Alex Alderton, Roberto Amato, Jeffrey Barrett, Sonia Goncalves, Ewan Harrison, David K. Jackson, Ian Johnston, Dominic Kwiatkowski, Cordelia Langford, John Sillitoe on behalf of the Wellcome Sanger Institute COVID-19 Surveillance Team |
| EPI_ISL_2849615 | Lighthouse Laboratory Plymouth | Wellcome Sanger Institute for the COVID-19 Genomics UK (COG-UK) Consortium | Lighthouse Laboratory Plymouth and Alex Alderton, Roberto Amato, Jeffrey Barrett, Sonia Goncalves, Ewan Harrison, David K. Jackson, Ian Johnston, Dominic Kwiatkowski, Cordelia Langford, John Sillitoe on behalf of the Wellcome Sanger Institute COVID-19 Surveillance Team |
| EPI_ISL_2849616 | Lighthouse Lab in Alderley Park | Wellcome Sanger Institute for the COVID-19 Genomics UK (COG-UK) Consortium | Jacquelyn Wynn, Mairead Hyland, The Lighthouse Lab in Alderley Park and Alex Alderton, Roberto Amato, Jeffrey Barrett, Sonia Goncalves, Ewan Harrison, David K. Jackson, Ian Johnston, Dominic Kwiatkowski, Cordelia Langford, John Sillitoe on behalf of the Wellcome Sanger Institute COVID-19 Surveillance Team |
| EPI_ISL_2849617, EPI_ISL_2849618 | Lighthouse Laboratory Plymouth | Wellcome Sanger Institute for the COVID-19 Genomics UK (COG-UK) Consortium | Lighthouse Laboratory Plymouth and Alex Alderton, Roberto Amato, Jeffrey Barrett, Sonia Goncalves, Ewan Harrison, David K. Jackson, Ian Johnston, Dominic Kwiatkowski, Cordelia Langford, John Sillitoe on behalf of the Wellcome Sanger Institute COVID-19 Surveillance Team |
| EPI_ISL_2849619, EPI_ISL_2849620 | Lighthouse Lab in Milton Keynes | Wellcome Sanger Institute for the COVID-19 Genomics UK (COG-UK) Consortium | The Lighthouse Lab in Milton Keynes and Alex Alderton, Roberto Amato, Jeffrey Barrett, Sonia Goncalves, Ewan Harrison, David K. Jackson, Ian Johnston, Dominic Kwiatkowski, Cordelia Langford, John Sillitoe on behalf of the Wellcome Sanger Institute COVID-19 Surveillance Team |
| EPI_ISL_2849621 | Lighthouse Laboratory Plymouth | Wellcome Sanger Institute for the COVID-19 Genomics UK (COG-UK) Consortium | Lighthouse Laboratory Plymouth and Alex Alderton, Roberto Amato, Jeffrey Barrett, Sonia Goncalves, Ewan Harrison, David K. Jackson, Ian Johnston, Dominic Kwiatkowski, Cordelia Langford, John Sillitoe on behalf of the Wellcome Sanger Institute COVID-19 Surveillance Team |
| EPI_ISL_2849622 | Lighthouse Lab in Milton Keynes | Wellcome Sanger Institute for the COVID-19 Genomics UK (COG-UK) Consortium | The Lighthouse Lab in Milton Keynes and Alex Alderton, Roberto Amato, Jeffrey Barrett, Sonia Goncalves, Ewan Harrison, David K. Jackson, Ian Johnston, Dominic Kwiatkowski, Cordelia Langford, John Sillitoe on behalf of the Wellcome Sanger Institute COVID-19 Surveillance Team |
| EPI_ISL_2849623 | Lighthouse Lab in Alderley Park | Wellcome Sanger Institute for the COVID-19 Genomics UK (COG-UK) Consortium | Jacquelyn Wynn, Mairead Hyland, The Lighthouse Lab in Alderley Park and Alex Alderton, Roberto Amato, Jeffrey Barrett, Sonia Goncalves, Ewan Harrison, David K. Jackson, Ian Johnston, Dominic Kwiatkowski, Cordelia Langford, John Sillitoe on behalf of the Wellcome Sanger Institute COVID-19 Surveillance Team |
| EPI_ISL_2849624 | Lighthouse Laboratory Plymouth | Wellcome Sanger Institute for the COVID-19 Genomics UK (COG-UK) Consortium | Lighthouse Laboratory Plymouth and Alex Alderton, Roberto Amato, Jeffrey Barrett, Sonia Goncalves, Ewan Harrison, David K. Jackson, Ian Johnston, Dominic Kwiatkowski, Cordelia Langford, John Sillitoe on behalf of the Wellcome Sanger Institute COVID-19 Surveillance Team |
| EPI_ISL_2849625 | Lighthouse Lab in Milton Keynes | Wellcome Sanger Institute for the COVID-19 Genomics UK (COG-UK) Consortium | The Lighthouse Lab in Milton Keynes and Alex Alderton, Roberto Amato, Jeffrey Barrett, Sonia Goncalves, Ewan Harrison, David K. Jackson, Ian Johnston, Dominic Kwiatkowski, Cordelia Langford, John Sillitoe on behalf of the Wellcome Sanger Institute COVID-19 Surveillance Team |
| EPI_ISL_2849626 | Lighthouse Lab in Alderley Park | Wellcome Sanger Institute for the COVID-19 Genomics UK (COG-UK) Consortium | Jacquelyn Wynn, Mairead Hyland, The Lighthouse Lab in Alderley Park and Alex Alderton, Roberto Amato, Jeffrey Barrett, Sonia Goncalves, Ewan Harrison, David K. Jackson, Ian Johnston, Dominic Kwiatkowski, Cordelia Langford, John Sillitoe on behalf of the Wellcome Sanger Institute COVID-19 Surveillance Team |
| EPI_ISL_2849627 | Lighthouse Laboratory Plymouth | Wellcome Sanger Institute for the COVID-19 Genomics UK (COG-UK) Consortium | Lighthouse Laboratory Plymouth and Alex Alderton, Roberto Amato, Jeffrey Barrett, Sonia Goncalves, Ewan Harrison, David K. Jackson, Ian Johnston, Dominic Kwiatkowski, Cordelia Langford, John Sillitoe on behalf of the Wellcome Sanger Institute COVID-19 Surveillance Team |
| EPI_ISL_2849628 | Lighthouse Lab in Milton Keynes | Wellcome Sanger Institute for the COVID-19 Genomics UK (COG-UK) Consortium | The Lighthouse Lab in Milton Keynes and Alex Alderton, Roberto Amato, Jeffrey Barrett, Sonia Goncalves, Ewan Harrison, David K. Jackson, Ian Johnston, Dominic Kwiatkowski, Cordelia Langford, John Sillitoe on behalf of the Wellcome Sanger Institute COVID-19 Surveillance Team |
| EPI_ISL_2849629 | Lighthouse Laboratory Plymouth | Wellcome Sanger Institute for the COVID-19 Genomics UK (COG-UK) Consortium | Lighthouse Laboratory Plymouth and Alex Alderton, Roberto Amato, Jeffrey Barrett, Sonia Goncalves, Ewan Harrison, David K. Jackson, Ian Johnston, Dominic Kwiatkowski, Cordelia Langford, John Sillitoe on behalf of the Wellcome Sanger Institute COVID-19 Surveillance Team |
| EPI_ISL_2849630, EPI_ISL_2849631 | Lighthouse Lab in Alderley Park | Wellcome Sanger Institute for the COVID-19 Genomics UK (COG-UK) Consortium | Jacquelyn Wynn, Mairead Hyland, The Lighthouse Lab in Alderley Park and Alex Alderton, Roberto Amato, Jeffrey Barrett, Sonia Goncalves, Ewan Harrison, David K. Jackson, Ian Johnston, Dominic Kwiatkowski, Cordelia Langford, John Sillitoe on behalf of the Wellcome Sanger Institute COVID-19 Surveillance Team |
| EPI_ISL_2849632 | Lighthouse Laboratory Plymouth | Wellcome Sanger Institute for the COVID-19 Genomics UK (COG-UK) Consortium | Lighthouse Laboratory Plymouth and Alex Alderton, Roberto Amato, Jeffrey Barrett, Sonia Goncalves, Ewan Harrison, David K. Jackson, Ian Johnston, Dominic Kwiatkowski, Cordelia Langford, John Sillitoe on behalf of the Wellcome Sanger Institute COVID-19 Surveillance Team |
| EPI_ISL_2849633 | Lighthouse Lab in Alderley Park | Wellcome Sanger Institute for the COVID-19 Genomics UK (COG-UK) Consortium | Jacquelyn Wynn, Mairead Hyland, The Lighthouse Lab in Alderley Park and Alex Alderton, Roberto Amato, Jeffrey Barrett, Sonia Goncalves, Ewan Harrison, David K. Jackson, Ian Johnston, Dominic Kwiatkowski, Cordelia Langford, John Sillitoe on behalf of the Wellcome Sanger Institute COVID-19 Surveillance Team |
| EPI_ISL_2849634 | Lighthouse Laboratory Plymouth | Wellcome Sanger Institute for the COVID-19 Genomics UK (COG-UK) Consortium | Lighthouse Laboratory Plymouth and Alex Alderton, Roberto Amato, Jeffrey Barrett, Sonia Goncalves, Ewan Harrison, David K. Jackson, Ian Johnston, Dominic Kwiatkowski, Cordelia Langford, John Sillitoe on behalf of the Wellcome Sanger Institute COVID-19 Surveillance Team |
| EPI_ISL_2849635, EPI_ISL_2849636 | Lighthouse Lab in Milton Keynes | Wellcome Sanger Institute for the COVID-19 Genomics UK (COG-UK) Consortium | The Lighthouse Lab in Milton Keynes and Alex Alderton, Roberto Amato, Jeffrey Barrett, Sonia Goncalves, Ewan Harrison, David K. Jackson, Ian Johnston, Dominic Kwiatkowski, Cordelia Langford, John Sillitoe on behalf of the Wellcome Sanger Institute COVID-19 Surveillance Team |
| EPI_ISL_2849637 | Lighthouse Laboratory Plymouth | Wellcome Sanger Institute for the COVID-19 Genomics UK (COG-UK) Consortium | Lighthouse Laboratory Plymouth and Alex Alderton, Roberto Amato, Jeffrey Barrett, Sonia Goncalves, Ewan Harrison, David K. Jackson, Ian Johnston, Dominic Kwiatkowski, Cordelia Langford, John Sillitoe on behalf of the Wellcome Sanger Institute COVID-19 Surveillance Team |
| EPI_ISL_2849638, EPI_ISL_2849639 | Lighthouse Lab in Alderley Park | Wellcome Sanger Institute for the COVID-19 Genomics UK (COG-UK) Consortium | Jacquelyn Wynn, Mairead Hyland, The Lighthouse Lab in Alderley Park and Alex Alderton, Roberto Amato, Jeffrey Barrett, Sonia Goncalves, Ewan Harrison, David K. Jackson, Ian Johnston, Dominic Kwiatkowski, Cordelia Langford, John Sillitoe on behalf of the Wellcome Sanger Institute COVID-19 Surveillance Team |
| EPI_ISL_2849640 | Lighthouse Laboratory Plymouth | Wellcome Sanger Institute for the COVID-19 Genomics UK (COG-UK) Consortium | Lighthouse Laboratory Plymouth and Alex Alderton, Roberto Amato, Jeffrey Barrett, Sonia Goncalves, Ewan Harrison, David K. Jackson, Ian Johnston, Dominic Kwiatkowski, Cordelia Langford, John Sillitoe on behalf of the Wellcome Sanger Institute COVID-19 Surveillance Team |
| EPI_ISL_2849641, | Lighthouse Lab in Alderley Park | Wellcome Sanger Institute for the COVID-19 Genomics UK (COG-UK) | Jacquelyn Wynn, Mairead Hyland, The Lighthouse Lab in Alderley Park and Alex Alderton, Roberto Amato, Jeffrey Barrett, Sonia Goncalves, Ewan Harrison, David K. Jackson, Ian Johnston, Dominic Kwiatkowski, Cordelia Langford, John Sillitoe on |

| | | | |
|---|---|---|---|
| EPI_ISL_2849642 | | Consortium | behalf of the Wellcome Sanger Institute COVID-19 Surveillance Team |
| EPI_ISL_2849643 | Lighthouse Lab in Milton Keynes | Wellcome Sanger Institute for the COVID-19 Genomics UK (COG-UK) Consortium | The Lighthouse Lab in Milton Keynes and Alex Alderton, Roberto Amato, Jeffrey Barrett, Sonia Goncalves, Ewan Harrison, David K. Jackson, Ian Johnston, Dominic Kwiatkowski, Cordelia Langford, John Sillitoe on behalf of the Wellcome Sanger Institute COVID-19 Surveillance Team |
| EPI_ISL_2849644 | Lighthouse Lab in Alderley Park | Wellcome Sanger Institute for the COVID-19 Genomics UK (COG-UK) Consortium | Jacquelyn Wynn, Mairead Hyland, The Lighthouse Lab in Alderley Park and Alex Alderton, Roberto Amato, Jeffrey Barrett, Sonia Goncalves, Ewan Harrison, David K. Jackson, Ian Johnston, Dominic Kwiatkowski, Cordelia Langford, John Sillitoe on behalf of the Wellcome Sanger Institute COVID-19 Surveillance Team |
| EPI_ISL_2849645, EPI_ISL_2849646 | Lighthouse Laboratory Plymouth | Wellcome Sanger Institute for the COVID-19 Genomics UK (COG-UK) Consortium | Lighthouse Laboratory Plymouth and Alex Alderton, Roberto Amato, Jeffrey Barrett, Sonia Goncalves, Ewan Harrison, David K. Jackson, Ian Johnston, Dominic Kwiatkowski, Cordelia Langford, John Sillitoe on behalf of the Wellcome Sanger Institute COVID-19 Surveillance Team |
| EPI_ISL_2849647 | Lighthouse Lab in Alderley Park | Wellcome Sanger Institute for the COVID-19 Genomics UK (COG-UK) Consortium | Jacquelyn Wynn, Mairead Hyland, The Lighthouse Lab in Alderley Park and Alex Alderton, Roberto Amato, Jeffrey Barrett, Sonia Goncalves, Ewan Harrison, David K. Jackson, Ian Johnston, Dominic Kwiatkowski, Cordelia Langford, John Sillitoe on behalf of the Wellcome Sanger Institute COVID-19 Surveillance Team |
| EPI_ISL_2849648, EPI_ISL_2849649, EPI_ISL_2849650 | Lighthouse Laboratory Plymouth | Wellcome Sanger Institute for the COVID-19 Genomics UK (COG-UK) Consortium | Lighthouse Laboratory Plymouth and Alex Alderton, Roberto Amato, Jeffrey Barrett, Sonia Goncalves, Ewan Harrison, David K. Jackson, Ian Johnston, Dominic Kwiatkowski, Cordelia Langford, John Sillitoe on behalf of the Wellcome Sanger Institute COVID-19 Surveillance Team |
| EPI_ISL_2849651 | Lighthouse Lab in Milton Keynes | Wellcome Sanger Institute for the COVID-19 Genomics UK (COG-UK) Consortium | The Lighthouse Lab in Milton Keynes and Alex Alderton, Roberto Amato, Jeffrey Barrett, Sonia Goncalves, Ewan Harrison, David K. Jackson, Ian Johnston, Dominic Kwiatkowski, Cordelia Langford, John Sillitoe on behalf of the Wellcome Sanger Institute COVID-19 Surveillance Team |
| EPI_ISL_2849652, EPI_ISL_2849653, EPI_ISL_2849654 | Lighthouse Laboratory Plymouth | Wellcome Sanger Institute for the COVID-19 Genomics UK (COG-UK) Consortium | Lighthouse Laboratory Plymouth and Alex Alderton, Roberto Amato, Jeffrey Barrett, Sonia Goncalves, Ewan Harrison, David K. Jackson, Ian Johnston, Dominic Kwiatkowski, Cordelia Langford, John Sillitoe on behalf of the Wellcome Sanger Institute COVID-19 Surveillance Team |
| EPI_ISL_2849655 | Lighthouse Lab in Alderley Park | Wellcome Sanger Institute for the COVID-19 Genomics UK (COG-UK) Consortium | Jacquelyn Wynn, Mairead Hyland, The Lighthouse Lab in Alderley Park and Alex Alderton, Roberto Amato, Jeffrey Barrett, Sonia Goncalves, Ewan Harrison, David K. Jackson, Ian Johnston, Dominic Kwiatkowski, Cordelia Langford, John Sillitoe on behalf of the Wellcome Sanger Institute COVID-19 Surveillance Team |
| EPI_ISL_2849656 | Lighthouse Lab in Milton Keynes | Wellcome Sanger Institute for the COVID-19 Genomics UK (COG-UK) Consortium | The Lighthouse Lab in Milton Keynes and Alex Alderton, Roberto Amato, Jeffrey Barrett, Sonia Goncalves, Ewan Harrison, David K. Jackson, Ian Johnston, Dominic Kwiatkowski, Cordelia Langford, John Sillitoe on behalf of the Wellcome Sanger Institute COVID-19 Surveillance Team |
| EPI_ISL_2849657 | Lighthouse Laboratory Plymouth | Wellcome Sanger Institute for the COVID-19 Genomics UK (COG-UK) Consortium | Lighthouse Laboratory Plymouth and Alex Alderton, Roberto Amato, Jeffrey Barrett, Sonia Goncalves, Ewan Harrison, David K. Jackson, Ian Johnston, Dominic Kwiatkowski, Cordelia Langford, John Sillitoe on behalf of the Wellcome Sanger Institute COVID-19 Surveillance Team |
| EPI_ISL_2849658 | Lighthouse Lab in Milton Keynes | Wellcome Sanger Institute for the COVID-19 Genomics UK (COG-UK) Consortium | The Lighthouse Lab in Milton Keynes and Alex Alderton, Roberto Amato, Jeffrey Barrett, Sonia Goncalves, Ewan Harrison, David K. Jackson, Ian Johnston, Dominic Kwiatkowski, Cordelia Langford, John Sillitoe on behalf of the Wellcome Sanger Institute COVID-19 Surveillance Team |
| EPI_ISL_2849659, EPI_ISL_2849660 | Lighthouse Laboratory Plymouth | Wellcome Sanger Institute for the COVID-19 Genomics UK (COG-UK) Consortium | Lighthouse Laboratory Plymouth and Alex Alderton, Roberto Amato, Jeffrey Barrett, Sonia Goncalves, Ewan Harrison, David K. Jackson, Ian Johnston, Dominic Kwiatkowski, Cordelia Langford, John Sillitoe on behalf of the Wellcome Sanger Institute COVID-19 Surveillance Team |
| EPI_ISL_2849661 | Lighthouse Lab in Milton Keynes | Wellcome Sanger Institute for the COVID-19 Genomics UK (COG-UK) Consortium | The Lighthouse Lab in Milton Keynes and Alex Alderton, Roberto Amato, Jeffrey Barrett, Sonia Goncalves, Ewan Harrison, David K. Jackson, Ian Johnston, Dominic Kwiatkowski, Cordelia Langford, John Sillitoe on behalf of the Wellcome Sanger Institute COVID-19 Surveillance Team |
| EPI_ISL_2849662, EPI_ISL_2849663 | Lighthouse Laboratory Plymouth | Wellcome Sanger Institute for the COVID-19 Genomics UK (COG-UK) Consortium | Lighthouse Laboratory Plymouth and Alex Alderton, Roberto Amato, Jeffrey Barrett, Sonia Goncalves, Ewan Harrison, David K. Jackson, Ian Johnston, Dominic Kwiatkowski, Cordelia Langford, John Sillitoe on behalf of the Wellcome Sanger Institute COVID-19 Surveillance Team |
| EPI_ISL_2849664 | Lighthouse Lab in Alderley Park | Wellcome Sanger Institute for the COVID-19 Genomics UK (COG-UK) Consortium | Jacquelyn Wynn, Mairead Hyland, The Lighthouse Lab in Alderley Park and Alex Alderton, Roberto Amato, Jeffrey Barrett, Sonia Goncalves, Ewan Harrison, David K. Jackson, Ian Johnston, Dominic Kwiatkowski, Cordelia Langford, John Sillitoe on behalf of the Wellcome Sanger Institute COVID-19 Surveillance Team |
| EPI_ISL_2849665 | Lighthouse Lab in Milton Keynes | Wellcome Sanger Institute for the COVID-19 Genomics UK (COG-UK) Consortium | The Lighthouse Lab in Milton Keynes and Alex Alderton, Roberto Amato, Jeffrey Barrett, Sonia Goncalves, Ewan Harrison, David K. Jackson, Ian Johnston, Dominic Kwiatkowski, Cordelia Langford, John Sillitoe on behalf of the Wellcome Sanger Institute COVID-19 Surveillance Team |
| EPI_ISL_2849666 | Lighthouse Lab in Alderley Park | Wellcome Sanger Institute for the COVID-19 Genomics UK (COG-UK) Consortium | Jacquelyn Wynn, Mairead Hyland, The Lighthouse Lab in Alderley Park and Alex Alderton, Roberto Amato, Jeffrey Barrett, Sonia Goncalves, Ewan Harrison, David K. Jackson, Ian Johnston, Dominic Kwiatkowski, Cordelia Langford, John Sillitoe on behalf of the Wellcome Sanger Institute COVID-19 Surveillance Team |
| EPI_ISL_2849667 | Lighthouse Lab in Milton Keynes | Wellcome Sanger Institute for the COVID-19 Genomics UK (COG-UK) Consortium | The Lighthouse Lab in Milton Keynes and Alex Alderton, Roberto Amato, Jeffrey Barrett, Sonia Goncalves, Ewan Harrison, David K. Jackson, Ian Johnston, Dominic Kwiatkowski, Cordelia Langford, John Sillitoe on behalf of the Wellcome Sanger Institute COVID-19 Surveillance Team |
| EPI_ISL_2849668 | Lighthouse Laboratory Plymouth | Wellcome Sanger Institute for the COVID-19 Genomics UK (COG-UK) Consortium | Lighthouse Laboratory Plymouth and Alex Alderton, Roberto Amato, Jeffrey Barrett, Sonia Goncalves, Ewan Harrison, David K. Jackson, Ian Johnston, Dominic Kwiatkowski, Cordelia Langford, John Sillitoe on behalf of the Wellcome Sanger Institute COVID-19 Surveillance Team |
| EPI_ISL_2849669 | Lighthouse Lab in Alderley Park | Wellcome Sanger Institute for the COVID-19 Genomics UK (COG-UK) Consortium | Jacquelyn Wynn, Mairead Hyland, The Lighthouse Lab in Alderley Park and Alex Alderton, Roberto Amato, Jeffrey Barrett, Sonia Goncalves, Ewan Harrison, David K. Jackson, Ian Johnston, Dominic Kwiatkowski, Cordelia Langford, John Sillitoe on behalf of the Wellcome Sanger Institute COVID-19 Surveillance Team |
| EPI_ISL_2849670 | Lighthouse Lab in Milton Keynes | Wellcome Sanger Institute for the COVID-19 Genomics UK (COG-UK) Consortium | The Lighthouse Lab in Milton Keynes and Alex Alderton, Roberto Amato, Jeffrey Barrett, Sonia Goncalves, Ewan Harrison, David K. Jackson, Ian Johnston, Dominic Kwiatkowski, Cordelia Langford, John Sillitoe on behalf of the Wellcome Sanger Institute COVID-19 Surveillance Team |
| EPI_ISL_2849671, EPI_ISL_2849672 | Lighthouse Laboratory Plymouth | Wellcome Sanger Institute for the COVID-19 Genomics UK (COG-UK) Consortium | Lighthouse Laboratory Plymouth and Alex Alderton, Roberto Amato, Jeffrey Barrett, Sonia Goncalves, Ewan Harrison, David K. Jackson, Ian Johnston, Dominic Kwiatkowski, Cordelia Langford, John Sillitoe on behalf of the Wellcome Sanger Institute COVID-19 Surveillance Team |
| EPI_ISL_2849673 | Lighthouse Lab in Alderley Park | Wellcome Sanger Institute for the COVID-19 Genomics UK (COG-UK) Consortium | Jacquelyn Wynn, Mairead Hyland, The Lighthouse Lab in Alderley Park and Alex Alderton, Roberto Amato, Jeffrey Barrett, Sonia Goncalves, Ewan Harrison, David K. Jackson, Ian Johnston, Dominic Kwiatkowski, Cordelia Langford, John Sillitoe on behalf of the Wellcome Sanger Institute COVID-19 Surveillance Team |
| EPI_ISL_2849674 | Lighthouse Laboratory Plymouth | Wellcome Sanger Institute for the COVID-19 Genomics UK (COG-UK) Consortium | Lighthouse Laboratory Plymouth and Alex Alderton, Roberto Amato, Jeffrey Barrett, Sonia Goncalves, Ewan Harrison, David K. Jackson, Ian Johnston, Dominic Kwiatkowski, Cordelia Langford, John Sillitoe on behalf of the Wellcome Sanger Institute COVID-19 Surveillance Team |
| EPI_ISL_2849675, EPI_ISL_2849676 | Lighthouse Lab in Alderley Park | Wellcome Sanger Institute for the COVID-19 Genomics UK (COG-UK) Consortium | Jacquelyn Wynn, Mairead Hyland, The Lighthouse Lab in Alderley Park and Alex Alderton, Roberto Amato, Jeffrey Barrett, Sonia Goncalves, Ewan Harrison, David K. Jackson, Ian Johnston, Dominic Kwiatkowski, Cordelia Langford, John Sillitoe on behalf of the Wellcome Sanger Institute COVID-19 Surveillance Team |
| EPI_ISL_2849677 | AZ Klina | AZ Klina | Carl Vael - Lynsey Berckmans |
| EPI_ISL_2849678, EPI_ISL_2849679, EPI_ISL_2849680 | Lighthouse Laboratory Plymouth | Wellcome Sanger Institute for the COVID-19 Genomics UK (COG-UK) Consortium | Lighthouse Laboratory Plymouth and Alex Alderton, Roberto Amato, Jeffrey Barrett, Sonia Goncalves, Ewan Harrison, David K. Jackson, Ian Johnston, Dominic Kwiatkowski, Cordelia Langford, John Sillitoe on behalf of the Wellcome Sanger Institute COVID-19 Surveillance Team |
| EPI_ISL_2849681 | Lighthouse Lab in Alderley Park | Wellcome Sanger Institute for the COVID-19 Genomics UK (COG-UK) Consortium | Jacquelyn Wynn, Mairead Hyland, The Lighthouse Lab in Alderley Park and Alex Alderton, Roberto Amato, Jeffrey Barrett, Sonia Goncalves, Ewan Harrison, David K. Jackson, Ian Johnston, Dominic Kwiatkowski, Cordelia Langford, John Sillitoe on behalf of the Wellcome Sanger Institute COVID-19 Surveillance Team |
| EPI_ISL_2849682 | Lighthouse Laboratory Plymouth | Wellcome Sanger Institute for the COVID-19 Genomics UK (COG-UK) Consortium | Lighthouse Laboratory Plymouth and Alex Alderton, Roberto Amato, Jeffrey Barrett, Sonia Goncalves, Ewan Harrison, David K. Jackson, Ian Johnston, Dominic Kwiatkowski, Cordelia Langford, John Sillitoe on behalf of the Wellcome Sanger Institute COVID-19 Surveillance Team |
| EPI_ISL_2849683, EPI_ISL_2849684 | Lighthouse Lab in Milton Keynes | Wellcome Sanger Institute for the COVID-19 Genomics UK (COG-UK) Consortium | The Lighthouse Lab in Milton Keynes and Alex Alderton, Roberto Amato, Jeffrey Barrett, Sonia Goncalves, Ewan Harrison, David K. Jackson, Ian Johnston, Dominic Kwiatkowski, Cordelia Langford, John Sillitoe on behalf of the Wellcome Sanger Institute COVID-19 Surveillance Team |
| EPI_ISL_2849685, EPI_ISL_2849686 | Lighthouse Laboratory Plymouth | Wellcome Sanger Institute for the COVID-19 Genomics UK (COG-UK) Consortium | Lighthouse Laboratory Plymouth and Alex Alderton, Roberto Amato, Jeffrey Barrett, Sonia Goncalves, Ewan Harrison, David K. Jackson, Ian Johnston, Dominic Kwiatkowski, Cordelia Langford, John Sillitoe on behalf of the Wellcome Sanger Institute COVID-19 Surveillance Team |
| EPI_ISL_2849687 | Lighthouse Lab in Milton Keynes | Wellcome Sanger Institute for the COVID-19 Genomics UK (COG-UK) Consortium | The Lighthouse Lab in Milton Keynes and Alex Alderton, Roberto Amato, Jeffrey Barrett, Sonia Goncalves, Ewan Harrison, David K. Jackson, Ian Johnston, Dominic Kwiatkowski, Cordelia Langford, John Sillitoe on behalf of the Wellcome Sanger Institute COVID-19 Surveillance Team |
| EPI_ISL_2849688 | Lighthouse Laboratory Plymouth | Wellcome Sanger Institute for the COVID-19 Genomics UK (COG-UK) Consortium | Lighthouse Laboratory Plymouth and Alex Alderton, Roberto Amato, Jeffrey Barrett, Sonia Goncalves, Ewan Harrison, David K. Jackson, Ian Johnston, Dominic Kwiatkowski, Cordelia Langford, John Sillitoe on behalf of the Wellcome Sanger Institute COVID-19 Surveillance Team |
| EPI_ISL_2849689, EPI_ISL_2849690 | Lighthouse Lab in Alderley Park | Wellcome Sanger Institute for the COVID-19 Genomics UK (COG-UK) Consortium | Jacquelyn Wynn, Mairead Hyland, The Lighthouse Lab in Alderley Park and Alex Alderton, Roberto Amato, Jeffrey Barrett, Sonia Goncalves, Ewan Harrison, David K. Jackson, Ian Johnston, Dominic Kwiatkowski, Cordelia Langford, John Sillitoe on behalf of the Wellcome Sanger Institute COVID-19 Surveillance Team |
| EPI_ISL_2849691 | Lighthouse Lab in Milton Keynes | Wellcome Sanger Institute for the COVID-19 Genomics UK (COG-UK) Consortium | The Lighthouse Lab in Milton Keynes and Alex Alderton, Roberto Amato, Jeffrey Barrett, Sonia Goncalves, Ewan Harrison, David K. Jackson, Ian Johnston, Dominic Kwiatkowski, Cordelia Langford, John Sillitoe on behalf of the Wellcome Sanger Institute COVID-19 Surveillance Team |
| EPI_ISL_2849692 | Lighthouse Laboratory Plymouth | Wellcome Sanger Institute for the COVID-19 Genomics UK (COG-UK) Consortium | Lighthouse Laboratory Plymouth and Alex Alderton, Roberto Amato, Jeffrey Barrett, Sonia Goncalves, Ewan Harrison, David K. Jackson, Ian Johnston, Dominic Kwiatkowski, Cordelia Langford, John Sillitoe on behalf of the Wellcome Sanger Institute COVID-19 Surveillance Team |
| EPI_ISL_2849693 | Lighthouse Lab in Alderley Park | Wellcome Sanger Institute for the COVID-19 Genomics UK (COG-UK) Consortium | Jacquelyn Wynn, Mairead Hyland, The Lighthouse Lab in Alderley Park and Alex Alderton, Roberto Amato, Jeffrey Barrett, Sonia Goncalves, Ewan Harrison, David K. Jackson, Ian Johnston, Dominic Kwiatkowski, Cordelia Langford, John Sillitoe on behalf of the Wellcome Sanger Institute COVID-19 Surveillance Team |
| EPI_ISL_2849694 | Lighthouse Laboratory Plymouth | Wellcome Sanger Institute for the COVID-19 Genomics UK (COG-UK) Consortium | Lighthouse Laboratory Plymouth and Alex Alderton, Roberto Amato, Jeffrey Barrett, Sonia Goncalves, Ewan Harrison, David K. Jackson, Ian Johnston, Dominic Kwiatkowski, Cordelia Langford, John Sillitoe on behalf of the Wellcome Sanger Institute COVID-19 Surveillance Team |
| EPI_ISL_2849695, EPI_ISL_2849696 | Lighthouse Lab in Alderley Park | Wellcome Sanger Institute for the COVID-19 Genomics UK (COG-UK) Consortium | Jacquelyn Wynn, Mairead Hyland, The Lighthouse Lab in Alderley Park and Alex Alderton, Roberto Amato, Jeffrey Barrett, Sonia Goncalves, Ewan Harrison, David K. Jackson, Ian Johnston, Dominic Kwiatkowski, Cordelia Langford, John Sillitoe on behalf of the Wellcome Sanger Institute COVID-19 Surveillance Team |
| EPI_ISL_2849697 | Lighthouse Laboratory Plymouth | Wellcome Sanger Institute for the COVID-19 Genomics UK (COG-UK) Consortium | Lighthouse Laboratory Plymouth and Alex Alderton, Roberto Amato, Jeffrey Barrett, Sonia Goncalves, Ewan Harrison, David K. Jackson, Ian Johnston, Dominic Kwiatkowski, Cordelia Langford, John Sillitoe on behalf of the Wellcome Sanger Institute COVID-19 Surveillance Team |
| EPI_ISL_2849698, EPI_ISL_2849699, EPI_ISL_2849700, EPI_ISL_2849701, EPI_ISL_2849702 | Lighthouse Lab in Alderley Park | Wellcome Sanger Institute for the COVID-19 Genomics UK (COG-UK) Consortium | Jacquelyn Wynn, Mairead Hyland, The Lighthouse Lab in Alderley Park and Alex Alderton, Roberto Amato, Jeffrey Barrett, Sonia Goncalves, Ewan Harrison, David K. Jackson, Ian Johnston, Dominic Kwiatkowski, Cordelia Langford, John Sillitoe on behalf of the Wellcome Sanger Institute COVID-19 Surveillance Team |
| EPI_ISL_2849703 | Lighthouse Laboratory Plymouth | Wellcome Sanger Institute for the COVID-19 Genomics UK (COG-UK) Consortium | Lighthouse Laboratory Plymouth and Alex Alderton, Roberto Amato, Jeffrey Barrett, Sonia Goncalves, Ewan Harrison, David K. Jackson, Ian Johnston, Dominic Kwiatkowski, Cordelia Langford, John Sillitoe on behalf of the Wellcome Sanger Institute COVID-19 Surveillance Team |
| EPI_ISL_2849704 | Lighthouse Lab in Alderley Park | Wellcome Sanger Institute for the COVID-19 Genomics UK (COG-UK) | Jacquelyn Wynn, Mairead Hyland, The Lighthouse Lab in Alderley Park and Alex Alderton, Roberto Amato, Jeffrey Barrett, Sonia Goncalves, Ewan Harrison, David K. Jackson, Ian Johnston, Dominic Kwiatkowski, Cordelia Langford, John Sillitoe on |

| | | | |
|---|---|---|---|
| | | Consortium | behalf of the Wellcome Sanger Institute COVID-19 Surveillance Team |
| EPI_ISL_2849705 | Lighthouse Laboratory Plymouth | Wellcome Sanger Institute for the COVID-19 Genomics UK (COG-UK) Consortium | Lighthouse Laboratory Plymouth and Alex Alderton, Roberto Amato, Jeffrey Barrett, Sonia Goncalves, Ewan Harrison, David K. Jackson, Ian Johnston, Dominic Kwiatkowski, Cordelia Langford, John Sillitoe on behalf of the Wellcome Sanger Institute COVID-19 Surveillance Team |
| EPI_ISL_2849706, EPI_ISL_2849707, EPI_ISL_2849708 | Lighthouse Lab in Alderley Park | Wellcome Sanger Institute for the COVID-19 Genomics UK (COG-UK) Consortium | Jacquelyn Wynn, Mairead Hyland, The Lighthouse Lab in Alderley Park and Alex Alderton, Roberto Amato, Jeffrey Barrett, Sonia Goncalves, Ewan Harrison, David K. Jackson, Ian Johnston, Dominic Kwiatkowski, Cordelia Langford, John Sillitoe on behalf of the Wellcome Sanger Institute COVID-19 Surveillance Team |
| EPI_ISL_2849709 | Lighthouse Laboratory Plymouth | Wellcome Sanger Institute for the COVID-19 Genomics UK (COG-UK) Consortium | Lighthouse Laboratory Plymouth and Alex Alderton, Roberto Amato, Jeffrey Barrett, Sonia Goncalves, Ewan Harrison, David K. Jackson, Ian Johnston, Dominic Kwiatkowski, Cordelia Langford, John Sillitoe on behalf of the Wellcome Sanger Institute COVID-19 Surveillance Team |
| EPI_ISL_2849710, EPI_ISL_2849711 | Lighthouse Lab in Alderley Park | Wellcome Sanger Institute for the COVID-19 Genomics UK (COG-UK) Consortium | Jacquelyn Wynn, Mairead Hyland, The Lighthouse Lab in Alderley Park and Alex Alderton, Roberto Amato, Jeffrey Barrett, Sonia Goncalves, Ewan Harrison, David K. Jackson, Ian Johnston, Dominic Kwiatkowski, Cordelia Langford, John Sillitoe on behalf of the Wellcome Sanger Institute COVID-19 Surveillance Team |
| EPI_ISL_2849712, EPI_ISL_2849713, EPI_ISL_2849714 | Lighthouse Laboratory Plymouth | Wellcome Sanger Institute for the COVID-19 Genomics UK (COG-UK) Consortium | Lighthouse Laboratory Plymouth and Alex Alderton, Roberto Amato, Jeffrey Barrett, Sonia Goncalves, Ewan Harrison, David K. Jackson, Ian Johnston, Dominic Kwiatkowski, Cordelia Langford, John Sillitoe on behalf of the Wellcome Sanger Institute COVID-19 Surveillance Team |
| EPI_ISL_2849715 | Lighthouse Lab in Alderley Park | Wellcome Sanger Institute for the COVID-19 Genomics UK (COG-UK) Consortium | Jacquelyn Wynn, Mairead Hyland, The Lighthouse Lab in Alderley Park and Alex Alderton, Roberto Amato, Jeffrey Barrett, Sonia Goncalves, Ewan Harrison, David K. Jackson, Ian Johnston, Dominic Kwiatkowski, Cordelia Langford, John Sillitoe on behalf of the Wellcome Sanger Institute COVID-19 Surveillance Team |
| EPI_ISL_2849716 | Lighthouse Laboratory Plymouth | Wellcome Sanger Institute for the COVID-19 Genomics UK (COG-UK) Consortium | Lighthouse Laboratory Plymouth and Alex Alderton, Roberto Amato, Jeffrey Barrett, Sonia Goncalves, Ewan Harrison, David K. Jackson, Ian Johnston, Dominic Kwiatkowski, Cordelia Langford, John Sillitoe on behalf of the Wellcome Sanger Institute COVID-19 Surveillance Team |
| EPI_ISL_2849717, EPI_ISL_2849718, EPI_ISL_2849719, EPI_ISL_2849720, EPI_ISL_2849721 | Lighthouse Lab in Alderley Park | Wellcome Sanger Institute for the COVID-19 Genomics UK (COG-UK) Consortium | Jacquelyn Wynn, Mairead Hyland, The Lighthouse Lab in Alderley Park and Alex Alderton, Roberto Amato, Jeffrey Barrett, Sonia Goncalves, Ewan Harrison, David K. Jackson, Ian Johnston, Dominic Kwiatkowski, Cordelia Langford, John Sillitoe on behalf of the Wellcome Sanger Institute COVID-19 Surveillance Team |
| EPI_ISL_2849722, EPI_ISL_2849723, EPI_ISL_2849724 | Lighthouse Laboratory Plymouth | Wellcome Sanger Institute for the COVID-19 Genomics UK (COG-UK) Consortium | Lighthouse Laboratory Plymouth and Alex Alderton, Roberto Amato, Jeffrey Barrett, Sonia Goncalves, Ewan Harrison, David K. Jackson, Ian Johnston, Dominic Kwiatkowski, Cordelia Langford, John Sillitoe on behalf of the Wellcome Sanger Institute COVID-19 Surveillance Team |
| EPI_ISL_2849725, EPI_ISL_2849726 | Lighthouse Lab in Alderley Park | Wellcome Sanger Institute for the COVID-19 Genomics UK (COG-UK) Consortium | Jacquelyn Wynn, Mairead Hyland, The Lighthouse Lab in Alderley Park and Alex Alderton, Roberto Amato, Jeffrey Barrett, Sonia Goncalves, Ewan Harrison, David K. Jackson, Ian Johnston, Dominic Kwiatkowski, Cordelia Langford, John Sillitoe on behalf of the Wellcome Sanger Institute COVID-19 Surveillance Team |
| EPI_ISL_2849727 | Lighthouse Laboratory Plymouth | Wellcome Sanger Institute for the COVID-19 Genomics UK (COG-UK) Consortium | Lighthouse Laboratory Plymouth and Alex Alderton, Roberto Amato, Jeffrey Barrett, Sonia Goncalves, Ewan Harrison, David K. Jackson, Ian Johnston, Dominic Kwiatkowski, Cordelia Langford, John Sillitoe on behalf of the Wellcome Sanger Institute COVID-19 Surveillance Team |
| EPI_ISL_2849728, EPI_ISL_2849729, EPI_ISL_2849730, EPI_ISL_2849731, EPI_ISL_2849732 | Lighthouse Lab in Alderley Park | Wellcome Sanger Institute for the COVID-19 Genomics UK (COG-UK) Consortium | Jacquelyn Wynn, Mairead Hyland, The Lighthouse Lab in Alderley Park and Alex Alderton, Roberto Amato, Jeffrey Barrett, Sonia Goncalves, Ewan Harrison, David K. Jackson, Ian Johnston, Dominic Kwiatkowski, Cordelia Langford, John Sillitoe on behalf of the Wellcome Sanger Institute COVID-19 Surveillance Team |
| EPI_ISL_2849733, EPI_ISL_2849734 | Lighthouse Laboratory Plymouth | Wellcome Sanger Institute for the COVID-19 Genomics UK (COG-UK) Consortium | Lighthouse Laboratory Plymouth and Alex Alderton, Roberto Amato, Jeffrey Barrett, Sonia Goncalves, Ewan Harrison, David K. Jackson, Ian Johnston, Dominic Kwiatkowski, Cordelia Langford, John Sillitoe on behalf of the Wellcome Sanger Institute COVID-19 Surveillance Team |
| EPI_ISL_2849735, EPI_ISL_2849736 | Lighthouse Lab in Alderley Park | Wellcome Sanger Institute for the COVID-19 Genomics UK (COG-UK) Consortium | Jacquelyn Wynn, Mairead Hyland, The Lighthouse Lab in Alderley Park and Alex Alderton, Roberto Amato, Jeffrey Barrett, Sonia Goncalves, Ewan Harrison, David K. Jackson, Ian Johnston, Dominic Kwiatkowski, Cordelia Langford, John Sillitoe on behalf of the Wellcome Sanger Institute COVID-19 Surveillance Team |
| EPI_ISL_2849737 | Lighthouse Laboratory Plymouth | Wellcome Sanger Institute for the COVID-19 Genomics UK (COG-UK) Consortium | Lighthouse Laboratory Plymouth and Alex Alderton, Roberto Amato, Jeffrey Barrett, Sonia Goncalves, Ewan Harrison, David K. Jackson, Ian Johnston, Dominic Kwiatkowski, Cordelia Langford, John Sillitoe on behalf of the Wellcome Sanger Institute COVID-19 Surveillance Team |
| EPI_ISL_2849738 | Lighthouse Lab in Alderley Park | Wellcome Sanger Institute for the COVID-19 Genomics UK (COG-UK) Consortium | Jacquelyn Wynn, Mairead Hyland, The Lighthouse Lab in Alderley Park and Alex Alderton, Roberto Amato, Jeffrey Barrett, Sonia Goncalves, Ewan Harrison, David K. Jackson, Ian Johnston, Dominic Kwiatkowski, Cordelia Langford, John Sillitoe on behalf of the Wellcome Sanger Institute COVID-19 Surveillance Team |
| EPI_ISL_2849739 | Lighthouse Laboratory Plymouth | Wellcome Sanger Institute for the COVID-19 Genomics UK (COG-UK) Consortium | Lighthouse Laboratory Plymouth and Alex Alderton, Roberto Amato, Jeffrey Barrett, Sonia Goncalves, Ewan Harrison, David K. Jackson, Ian Johnston, Dominic Kwiatkowski, Cordelia Langford, John Sillitoe on behalf of the Wellcome Sanger Institute COVID-19 Surveillance Team |
| EPI_ISL_2849740 | Lighthouse Lab in Alderley Park | Wellcome Sanger Institute for the COVID-19 Genomics UK (COG-UK) Consortium | Jacquelyn Wynn, Mairead Hyland, The Lighthouse Lab in Alderley Park and Alex Alderton, Roberto Amato, Jeffrey Barrett, Sonia Goncalves, Ewan Harrison, David K. Jackson, Ian Johnston, Dominic Kwiatkowski, Cordelia Langford, John Sillitoe on behalf of the Wellcome Sanger Institute COVID-19 Surveillance Team |
| EPI_ISL_2849741, EPI_ISL_2849742 | Lighthouse Laboratory Plymouth | Wellcome Sanger Institute for the COVID-19 Genomics UK (COG-UK) Consortium | Lighthouse Laboratory Plymouth and Alex Alderton, Roberto Amato, Jeffrey Barrett, Sonia Goncalves, Ewan Harrison, David K. Jackson, Ian Johnston, Dominic Kwiatkowski, Cordelia Langford, John Sillitoe on behalf of the Wellcome Sanger Institute COVID-19 Surveillance Team |
| EPI_ISL_2849743 | Lighthouse Lab in Alderley Park | Wellcome Sanger Institute for the COVID-19 Genomics UK (COG-UK) Consortium | Jacquelyn Wynn, Mairead Hyland, The Lighthouse Lab in Alderley Park and Alex Alderton, Roberto Amato, Jeffrey Barrett, Sonia Goncalves, Ewan Harrison, David K. Jackson, Ian Johnston, Dominic Kwiatkowski, Cordelia Langford, John Sillitoe on behalf of the Wellcome Sanger Institute COVID-19 Surveillance Team |
| EPI_ISL_2849744 | Lighthouse Laboratory Plymouth | Wellcome Sanger Institute for the COVID-19 Genomics UK (COG-UK) Consortium | Lighthouse Laboratory Plymouth and Alex Alderton, Roberto Amato, Jeffrey Barrett, Sonia Goncalves, Ewan Harrison, David K. Jackson, Ian Johnston, Dominic Kwiatkowski, Cordelia Langford, John Sillitoe on behalf of the Wellcome Sanger Institute COVID-19 Surveillance Team |
| EPI_ISL_2849745, EPI_ISL_2849746, EPI_ISL_2849747, EPI_ISL_2849748, EPI_ISL_2849749, EPI_ISL_2849750, EPI_ISL_2849751, EPI_ISL_2849752, EPI_ISL_2849753, EPI_ISL_2849754 | Lighthouse Lab in Alderley Park | Wellcome Sanger Institute for the COVID-19 Genomics UK (COG-UK) Consortium | Jacquelyn Wynn, Mairead Hyland, The Lighthouse Lab in Alderley Park and Alex Alderton, Roberto Amato, Jeffrey Barrett, Sonia Goncalves, Ewan Harrison, David K. Jackson, Ian Johnston, Dominic Kwiatkowski, Cordelia Langford, John Sillitoe on behalf of the Wellcome Sanger Institute COVID-19 Surveillance Team |
| EPI_ISL_2849755, EPI_ISL_2849756 | Lighthouse Laboratory Plymouth | Wellcome Sanger Institute for the COVID-19 Genomics UK (COG-UK) Consortium | Lighthouse Laboratory Plymouth and Alex Alderton, Roberto Amato, Jeffrey Barrett, Sonia Goncalves, Ewan Harrison, David K. Jackson, Ian Johnston, Dominic Kwiatkowski, Cordelia Langford, John Sillitoe on behalf of the Wellcome Sanger Institute COVID-19 Surveillance Team |
| EPI_ISL_2849757, EPI_ISL_2849758 | Lighthouse Lab in Alderley Park | Wellcome Sanger Institute for the COVID-19 Genomics UK (COG-UK) Consortium | Jacquelyn Wynn, Mairead Hyland, The Lighthouse Lab in Alderley Park and Alex Alderton, Roberto Amato, Jeffrey Barrett, Sonia Goncalves, Ewan Harrison, David K. Jackson, Ian Johnston, Dominic Kwiatkowski, Cordelia Langford, John Sillitoe on behalf of the Wellcome Sanger Institute COVID-19 Surveillance Team |
| EPI_ISL_2849759 | Lighthouse Laboratory Plymouth | Wellcome Sanger Institute for the COVID-19 Genomics UK (COG-UK) Consortium | Lighthouse Laboratory Plymouth and Alex Alderton, Roberto Amato, Jeffrey Barrett, Sonia Goncalves, Ewan Harrison, David K. Jackson, Ian Johnston, Dominic Kwiatkowski, Cordelia Langford, John Sillitoe on behalf of the Wellcome Sanger Institute COVID-19 Surveillance Team |
| EPI_ISL_2849760, EPI_ISL_2849761 | Lighthouse Lab in Alderley Park | Wellcome Sanger Institute for the COVID-19 Genomics UK (COG-UK) Consortium | Jacquelyn Wynn, Mairead Hyland, The Lighthouse Lab in Alderley Park and Alex Alderton, Roberto Amato, Jeffrey Barrett, Sonia Goncalves, Ewan Harrison, David K. Jackson, Ian Johnston, Dominic Kwiatkowski, Cordelia Langford, John Sillitoe on behalf of the Wellcome Sanger Institute COVID-19 Surveillance Team |
| EPI_ISL_2849762 | Lighthouse Laboratory Plymouth | Wellcome Sanger Institute for the COVID-19 Genomics UK (COG-UK) Consortium | Lighthouse Laboratory Plymouth and Alex Alderton, Roberto Amato, Jeffrey Barrett, Sonia Goncalves, Ewan Harrison, David K. Jackson, Ian Johnston, Dominic Kwiatkowski, Cordelia Langford, John Sillitoe on behalf of the Wellcome Sanger Institute COVID-19 Surveillance Team |
| EPI_ISL_2849763 | Artic Network workflow-EPI2ME labs | King Khalid Bin Abdul Aziz street, South Surra, Kuwait City | Nada Madi, Hussain Safar, Ebaa Al-Awadhi, Anfal Al-Adwani |
| EPI_ISL_2849764 | Lighthouse Laboratory Plymouth | Wellcome Sanger Institute for the COVID-19 Genomics UK (COG-UK) Consortium | Lighthouse Laboratory Plymouth and Alex Alderton, Roberto Amato, Jeffrey Barrett, Sonia Goncalves, Ewan Harrison, David K. Jackson, Ian Johnston, Dominic Kwiatkowski, Cordelia Langford, John Sillitoe on behalf of the Wellcome Sanger Institute COVID-19 Surveillance Team |
| EPI_ISL_2849765 | Lighthouse Lab in Alderley Park | Wellcome Sanger Institute for the COVID-19 Genomics UK (COG-UK) Consortium | Jacquelyn Wynn, Mairead Hyland, The Lighthouse Lab in Alderley Park and Alex Alderton, Roberto Amato, Jeffrey Barrett, Sonia Goncalves, Ewan Harrison, David K. Jackson, Ian Johnston, Dominic Kwiatkowski, Cordelia Langford, John Sillitoe on behalf of the Wellcome Sanger Institute COVID-19 Surveillance Team |
| EPI_ISL_2849766 | Lighthouse Laboratory Plymouth | Wellcome Sanger Institute for the COVID-19 Genomics UK (COG-UK) Consortium | Lighthouse Laboratory Plymouth and Alex Alderton, Roberto Amato, Jeffrey Barrett, Sonia Goncalves, Ewan Harrison, David K. Jackson, Ian Johnston, Dominic Kwiatkowski, Cordelia Langford, John Sillitoe on behalf of the Wellcome Sanger Institute COVID-19 Surveillance Team |
| EPI_ISL_2849767, EPI_ISL_2849768 | Lighthouse Lab in Alderley Park | Wellcome Sanger Institute for the COVID-19 Genomics UK (COG-UK) Consortium | Jacquelyn Wynn, Mairead Hyland, The Lighthouse Lab in Alderley Park and Alex Alderton, Roberto Amato, Jeffrey Barrett, Sonia Goncalves, Ewan Harrison, David K. Jackson, Ian Johnston, Dominic Kwiatkowski, Cordelia Langford, John Sillitoe on behalf of the Wellcome Sanger Institute COVID-19 Surveillance Team |
| EPI_ISL_2849769 | Lighthouse Laboratory Plymouth | Wellcome Sanger Institute for the COVID-19 Genomics UK (COG-UK) Consortium | Lighthouse Laboratory Plymouth and Alex Alderton, Roberto Amato, Jeffrey Barrett, Sonia Goncalves, Ewan Harrison, David K. Jackson, Ian Johnston, Dominic Kwiatkowski, Cordelia Langford, John Sillitoe on behalf of the Wellcome Sanger Institute COVID-19 Surveillance Team |
| EPI_ISL_2849770, EPI_ISL_2849771, EPI_ISL_2849772, EPI_ISL_2849773, EPI_ISL_2849774 | Lighthouse Lab in Alderley Park | Wellcome Sanger Institute for the COVID-19 Genomics UK (COG-UK) Consortium | Jacquelyn Wynn, Mairead Hyland, The Lighthouse Lab in Alderley Park and Alex Alderton, Roberto Amato, Jeffrey Barrett, Sonia Goncalves, Ewan Harrison, David K. Jackson, Ian Johnston, Dominic Kwiatkowski, Cordelia Langford, John Sillitoe on behalf of the Wellcome Sanger Institute COVID-19 Surveillance Team |
| EPI_ISL_2849775 | Lighthouse Laboratory Plymouth | Wellcome Sanger Institute for the COVID-19 Genomics UK (COG-UK) Consortium | Lighthouse Laboratory Plymouth and Alex Alderton, Roberto Amato, Jeffrey Barrett, Sonia Goncalves, Ewan Harrison, David K. Jackson, Ian Johnston, Dominic Kwiatkowski, Cordelia Langford, John Sillitoe on behalf of the Wellcome Sanger Institute COVID-19 Surveillance Team |
| EPI_ISL_2849776, EPI_ISL_2849777 | Lighthouse Lab in Alderley Park | Wellcome Sanger Institute for the COVID-19 Genomics UK (COG-UK) Consortium | Jacquelyn Wynn, Mairead Hyland, The Lighthouse Lab in Alderley Park and Alex Alderton, Roberto Amato, Jeffrey Barrett, Sonia Goncalves, Ewan Harrison, David K. Jackson, Ian Johnston, Dominic Kwiatkowski, Cordelia Langford, John Sillitoe on behalf of the Wellcome Sanger Institute COVID-19 Surveillance Team |
| EPI_ISL_2849778 | Lighthouse Laboratory Plymouth | Wellcome Sanger Institute for the COVID-19 Genomics UK (COG-UK) Consortium | Lighthouse Laboratory Plymouth and Alex Alderton, Roberto Amato, Jeffrey Barrett, Sonia Goncalves, Ewan Harrison, David K. Jackson, Ian Johnston, Dominic Kwiatkowski, Cordelia Langford, John Sillitoe on behalf of the Wellcome Sanger Institute COVID-19 Surveillance Team |

| | | | |
|---|---|---|---|
| EPI_ISL_2849779, EPI_ISL_2849780 | Lighthouse Lab in Alderley Park | Wellcome Sanger Institute for the COVID-19 Genomics UK (COG-UK) Consortium | Jacquelyn Wynn, Mairead Hyland, The Lighthouse Lab in Alderley Park and Alex Alderton, Roberto Amato, Jeffrey Barrett, Sonia Goncalves, Ewan Harrison, David K. Jackson, Ian Johnston, Dominic Kwiatkowski, Cordelia Langford, John Sillitoe on behalf of the Wellcome Sanger Institute COVID-19 Surveillance Team |
| EPI_ISL_2849781, EPI_ISL_2849782 | Lighthouse Laboratory Plymouth | Wellcome Sanger Institute for the COVID-19 Genomics UK (COG-UK) Consortium | Lighthouse Laboratory Plymouth and Alex Alderton, Roberto Amato, Jeffrey Barrett, Sonia Goncalves, Ewan Harrison, David K. Jackson, Ian Johnston, Dominic Kwiatkowski, Cordelia Langford, John Sillitoe on behalf of the Wellcome Sanger Institute COVID-19 Surveillance Team |
| EPI_ISL_2849783, EPI_ISL_2849784 | Lighthouse Lab in Alderley Park | Wellcome Sanger Institute for the COVID-19 Genomics UK (COG-UK) Consortium | Jacquelyn Wynn, Mairead Hyland, The Lighthouse Lab in Alderley Park and Alex Alderton, Roberto Amato, Jeffrey Barrett, Sonia Goncalves, Ewan Harrison, David K. Jackson, Ian Johnston, Dominic Kwiatkowski, Cordelia Langford, John Sillitoe on behalf of the Wellcome Sanger Institute COVID-19 Surveillance Team |
| EPI_ISL_2849785, EPI_ISL_2849786 | Lighthouse Laboratory Plymouth | Wellcome Sanger Institute for the COVID-19 Genomics UK (COG-UK) Consortium | Lighthouse Laboratory Plymouth and Alex Alderton, Roberto Amato, Jeffrey Barrett, Sonia Goncalves, Ewan Harrison, David K. Jackson, Ian Johnston, Dominic Kwiatkowski, Cordelia Langford, John Sillitoe on behalf of the Wellcome Sanger Institute COVID-19 Surveillance Team |
| EPI_ISL_2849787 | Lighthouse Lab in Alderley Park | Wellcome Sanger Institute for the COVID-19 Genomics UK (COG-UK) Consortium | Jacquelyn Wynn, Mairead Hyland, The Lighthouse Lab in Alderley Park and Alex Alderton, Roberto Amato, Jeffrey Barrett, Sonia Goncalves, Ewan Harrison, David K. Jackson, Ian Johnston, Dominic Kwiatkowski, Cordelia Langford, John Sillitoe on behalf of the Wellcome Sanger Institute COVID-19 Surveillance Team |
| EPI_ISL_2849788, EPI_ISL_2849789 | Lighthouse Laboratory Plymouth | Wellcome Sanger Institute for the COVID-19 Genomics UK (COG-UK) Consortium | Lighthouse Laboratory Plymouth and Alex Alderton, Roberto Amato, Jeffrey Barrett, Sonia Goncalves, Ewan Harrison, David K. Jackson, Ian Johnston, Dominic Kwiatkowski, Cordelia Langford, John Sillitoe on behalf of the Wellcome Sanger Institute COVID-19 Surveillance Team |
| EPI_ISL_2849790, EPI_ISL_2849791 | Lighthouse Lab in Alderley Park | Wellcome Sanger Institute for the COVID-19 Genomics UK (COG-UK) Consortium | Jacquelyn Wynn, Mairead Hyland, The Lighthouse Lab in Alderley Park and Alex Alderton, Roberto Amato, Jeffrey Barrett, Sonia Goncalves, Ewan Harrison, David K. Jackson, Ian Johnston, Dominic Kwiatkowski, Cordelia Langford, John Sillitoe on behalf of the Wellcome Sanger Institute COVID-19 Surveillance Team |
| EPI_ISL_2849792 | Lighthouse Laboratory Plymouth | Wellcome Sanger Institute for the COVID-19 Genomics UK (COG-UK) Consortium | Lighthouse Laboratory Plymouth and Alex Alderton, Roberto Amato, Jeffrey Barrett, Sonia Goncalves, Ewan Harrison, David K. Jackson, Ian Johnston, Dominic Kwiatkowski, Cordelia Langford, John Sillitoe on behalf of the Wellcome Sanger Institute COVID-19 Surveillance Team |
| EPI_ISL_2849793 | Lighthouse Lab in Alderley Park | Wellcome Sanger Institute for the COVID-19 Genomics UK (COG-UK) Consortium | Jacquelyn Wynn, Mairead Hyland, The Lighthouse Lab in Alderley Park and Alex Alderton, Roberto Amato, Jeffrey Barrett, Sonia Goncalves, Ewan Harrison, David K. Jackson, Ian Johnston, Dominic Kwiatkowski, Cordelia Langford, John Sillitoe on behalf of the Wellcome Sanger Institute COVID-19 Surveillance Team |
| EPI_ISL_2849794 | Lighthouse Laboratory Plymouth | Wellcome Sanger Institute for the COVID-19 Genomics UK (COG-UK) Consortium | Lighthouse Laboratory Plymouth and Alex Alderton, Roberto Amato, Jeffrey Barrett, Sonia Goncalves, Ewan Harrison, David K. Jackson, Ian Johnston, Dominic Kwiatkowski, Cordelia Langford, John Sillitoe on behalf of the Wellcome Sanger Institute COVID-19 Surveillance Team |
| EPI_ISL_2849795, EPI_ISL_2849796, EPI_ISL_2849797 | Lighthouse Lab in Alderley Park | Wellcome Sanger Institute for the COVID-19 Genomics UK (COG-UK) Consortium | Jacquelyn Wynn, Mairead Hyland, The Lighthouse Lab in Alderley Park and Alex Alderton, Roberto Amato, Jeffrey Barrett, Sonia Goncalves, Ewan Harrison, David K. Jackson, Ian Johnston, Dominic Kwiatkowski, Cordelia Langford, John Sillitoe on behalf of the Wellcome Sanger Institute COVID-19 Surveillance Team |
| EPI_ISL_2849798, EPI_ISL_2849799, EPI_ISL_2849800 | Lighthouse Laboratory Plymouth | Wellcome Sanger Institute for the COVID-19 Genomics UK (COG-UK) Consortium | Lighthouse Laboratory Plymouth and Alex Alderton, Roberto Amato, Jeffrey Barrett, Sonia Goncalves, Ewan Harrison, David K. Jackson, Ian Johnston, Dominic Kwiatkowski, Cordelia Langford, John Sillitoe on behalf of the Wellcome Sanger Institute COVID-19 Surveillance Team |
| EPI_ISL_2849801 | Lighthouse Lab in Alderley Park | Wellcome Sanger Institute for the COVID-19 Genomics UK (COG-UK) Consortium | Jacquelyn Wynn, Mairead Hyland, The Lighthouse Lab in Alderley Park and Alex Alderton, Roberto Amato, Jeffrey Barrett, Sonia Goncalves, Ewan Harrison, David K. Jackson, Ian Johnston, Dominic Kwiatkowski, Cordelia Langford, John Sillitoe on behalf of the Wellcome Sanger Institute COVID-19 Surveillance Team |
| EPI_ISL_2849802, EPI_ISL_2849803, EPI_ISL_2849804 | Lighthouse Laboratory Plymouth | Wellcome Sanger Institute for the COVID-19 Genomics UK (COG-UK) Consortium | Lighthouse Laboratory Plymouth and Alex Alderton, Roberto Amato, Jeffrey Barrett, Sonia Goncalves, Ewan Harrison, David K. Jackson, Ian Johnston, Dominic Kwiatkowski, Cordelia Langford, John Sillitoe on behalf of the Wellcome Sanger Institute COVID-19 Surveillance Team |
| EPI_ISL_2849805, EPI_ISL_2849806 | Lighthouse Lab in Alderley Park | Wellcome Sanger Institute for the COVID-19 Genomics UK (COG-UK) Consortium | Jacquelyn Wynn, Mairead Hyland, The Lighthouse Lab in Alderley Park and Alex Alderton, Roberto Amato, Jeffrey Barrett, Sonia Goncalves, Ewan Harrison, David K. Jackson, Ian Johnston, Dominic Kwiatkowski, Cordelia Langford, John Sillitoe on behalf of the Wellcome Sanger Institute COVID-19 Surveillance Team |
| EPI_ISL_2849807 | Lighthouse Laboratory Plymouth | Wellcome Sanger Institute for the COVID-19 Genomics UK (COG-UK) Consortium | Lighthouse Laboratory Plymouth and Alex Alderton, Roberto Amato, Jeffrey Barrett, Sonia Goncalves, Ewan Harrison, David K. Jackson, Ian Johnston, Dominic Kwiatkowski, Cordelia Langford, John Sillitoe on behalf of the Wellcome Sanger Institute COVID-19 Surveillance Team |
| EPI_ISL_2849808 | Lighthouse Lab in Alderley Park | Wellcome Sanger Institute for the COVID-19 Genomics UK (COG-UK) Consortium | Jacquelyn Wynn, Mairead Hyland, The Lighthouse Lab in Alderley Park and Alex Alderton, Roberto Amato, Jeffrey Barrett, Sonia Goncalves, Ewan Harrison, David K. Jackson, Ian Johnston, Dominic Kwiatkowski, Cordelia Langford, John Sillitoe on behalf of the Wellcome Sanger Institute COVID-19 Surveillance Team |
| EPI_ISL_2849809 | Lighthouse Laboratory Plymouth | Wellcome Sanger Institute for the COVID-19 Genomics UK (COG-UK) Consortium | Lighthouse Laboratory Plymouth and Alex Alderton, Roberto Amato, Jeffrey Barrett, Sonia Goncalves, Ewan Harrison, David K. Jackson, Ian Johnston, Dominic Kwiatkowski, Cordelia Langford, John Sillitoe on behalf of the Wellcome Sanger Institute COVID-19 Surveillance Team |
| EPI_ISL_2849810 | Lighthouse Lab in Alderley Park | Wellcome Sanger Institute for the COVID-19 Genomics UK (COG-UK) Consortium | Jacquelyn Wynn, Mairead Hyland, The Lighthouse Lab in Alderley Park and Alex Alderton, Roberto Amato, Jeffrey Barrett, Sonia Goncalves, Ewan Harrison, David K. Jackson, Ian Johnston, Dominic Kwiatkowski, Cordelia Langford, John Sillitoe on behalf of the Wellcome Sanger Institute COVID-19 Surveillance Team |
| EPI_ISL_2849811 | Lighthouse Laboratory Plymouth | Wellcome Sanger Institute for the COVID-19 Genomics UK (COG-UK) Consortium | Lighthouse Laboratory Plymouth and Alex Alderton, Roberto Amato, Jeffrey Barrett, Sonia Goncalves, Ewan Harrison, David K. Jackson, Ian Johnston, Dominic Kwiatkowski, Cordelia Langford, John Sillitoe on behalf of the Wellcome Sanger Institute COVID-19 Surveillance Team |
| EPI_ISL_2849812, EPI_ISL_2849813, EPI_ISL_2849814, EPI_ISL_2849815 | Lighthouse Lab in Alderley Park | Wellcome Sanger Institute for the COVID-19 Genomics UK (COG-UK) Consortium | Jacquelyn Wynn, Mairead Hyland, The Lighthouse Lab in Alderley Park and Alex Alderton, Roberto Amato, Jeffrey Barrett, Sonia Goncalves, Ewan Harrison, David K. Jackson, Ian Johnston, Dominic Kwiatkowski, Cordelia Langford, John Sillitoe on behalf of the Wellcome Sanger Institute COVID-19 Surveillance Team |
| EPI_ISL_2849816, EPI_ISL_2849817 | Lighthouse Laboratory Plymouth | Wellcome Sanger Institute for the COVID-19 Genomics UK (COG-UK) Consortium | Lighthouse Laboratory Plymouth and Alex Alderton, Roberto Amato, Jeffrey Barrett, Sonia Goncalves, Ewan Harrison, David K. Jackson, Ian Johnston, Dominic Kwiatkowski, Cordelia Langford, John Sillitoe on behalf of the Wellcome Sanger Institute COVID-19 Surveillance Team |
| EPI_ISL_2849818, EPI_ISL_2849819 | Lighthouse Lab in Alderley Park | Wellcome Sanger Institute for the COVID-19 Genomics UK (COG-UK) Consortium | Jacquelyn Wynn, Mairead Hyland, The Lighthouse Lab in Alderley Park and Alex Alderton, Roberto Amato, Jeffrey Barrett, Sonia Goncalves, Ewan Harrison, David K. Jackson, Ian Johnston, Dominic Kwiatkowski, Cordelia Langford, John Sillitoe on behalf of the Wellcome Sanger Institute COVID-19 Surveillance Team |
| EPI_ISL_2849820 | Lighthouse Laboratory Plymouth | Wellcome Sanger Institute for the COVID-19 Genomics UK (COG-UK) Consortium | Lighthouse Laboratory Plymouth and Alex Alderton, Roberto Amato, Jeffrey Barrett, Sonia Goncalves, Ewan Harrison, David K. Jackson, Ian Johnston, Dominic Kwiatkowski, Cordelia Langford, John Sillitoe on behalf of the Wellcome Sanger Institute COVID-19 Surveillance Team |
| EPI_ISL_2849821, EPI_ISL_2849822 | Lighthouse Lab in Alderley Park | Wellcome Sanger Institute for the COVID-19 Genomics UK (COG-UK) Consortium | Jacquelyn Wynn, Mairead Hyland, The Lighthouse Lab in Alderley Park and Alex Alderton, Roberto Amato, Jeffrey Barrett, Sonia Goncalves, Ewan Harrison, David K. Jackson, Ian Johnston, Dominic Kwiatkowski, Cordelia Langford, John Sillitoe on behalf of the Wellcome Sanger Institute COVID-19 Surveillance Team |
| EPI_ISL_2849823 | Lighthouse Laboratory Plymouth | Wellcome Sanger Institute for the COVID-19 Genomics UK (COG-UK) Consortium | Lighthouse Laboratory Plymouth and Alex Alderton, Roberto Amato, Jeffrey Barrett, Sonia Goncalves, Ewan Harrison, David K. Jackson, Ian Johnston, Dominic Kwiatkowski, Cordelia Langford, John Sillitoe on behalf of the Wellcome Sanger Institute COVID-19 Surveillance Team |
| EPI_ISL_2849824 | Lighthouse Lab in Alderley Park | Wellcome Sanger Institute for the COVID-19 Genomics UK (COG-UK) Consortium | Jacquelyn Wynn, Mairead Hyland, The Lighthouse Lab in Alderley Park and Alex Alderton, Roberto Amato, Jeffrey Barrett, Sonia Goncalves, Ewan Harrison, David K. Jackson, Ian Johnston, Dominic Kwiatkowski, Cordelia Langford, John Sillitoe on behalf of the Wellcome Sanger Institute COVID-19 Surveillance Team |
| EPI_ISL_2849825 | Lighthouse Laboratory Plymouth | Wellcome Sanger Institute for the COVID-19 Genomics UK (COG-UK) Consortium | Lighthouse Laboratory Plymouth and Alex Alderton, Roberto Amato, Jeffrey Barrett, Sonia Goncalves, Ewan Harrison, David K. Jackson, Ian Johnston, Dominic Kwiatkowski, Cordelia Langford, John Sillitoe on behalf of the Wellcome Sanger Institute COVID-19 Surveillance Team |
| EPI_ISL_2849826, EPI_ISL_2849827 | Lighthouse Lab in Alderley Park | Wellcome Sanger Institute for the COVID-19 Genomics UK (COG-UK) Consortium | Jacquelyn Wynn, Mairead Hyland, The Lighthouse Lab in Alderley Park and Alex Alderton, Roberto Amato, Jeffrey Barrett, Sonia Goncalves, Ewan Harrison, David K. Jackson, Ian Johnston, Dominic Kwiatkowski, Cordelia Langford, John Sillitoe on behalf of the Wellcome Sanger Institute COVID-19 Surveillance Team |
| EPI_ISL_2849828, EPI_ISL_2849829, EPI_ISL_2849830, EPI_ISL_2849831, EPI_ISL_2849832, EPI_ISL_2849833 | Lighthouse Laboratory Plymouth | Wellcome Sanger Institute for the COVID-19 Genomics UK (COG-UK) Consortium | Lighthouse Laboratory Plymouth and Alex Alderton, Roberto Amato, Jeffrey Barrett, Sonia Goncalves, Ewan Harrison, David K. Jackson, Ian Johnston, Dominic Kwiatkowski, Cordelia Langford, John Sillitoe on behalf of the Wellcome Sanger Institute COVID-19 Surveillance Team |
| EPI_ISL_2849834, EPI_ISL_2849835 | Lighthouse Lab in Alderley Park | Wellcome Sanger Institute for the COVID-19 Genomics UK (COG-UK) Consortium | Jacquelyn Wynn, Mairead Hyland, The Lighthouse Lab in Alderley Park and Alex Alderton, Roberto Amato, Jeffrey Barrett, Sonia Goncalves, Ewan Harrison, David K. Jackson, Ian Johnston, Dominic Kwiatkowski, Cordelia Langford, John Sillitoe on behalf of the Wellcome Sanger Institute COVID-19 Surveillance Team |
| EPI_ISL_2849836 | Lighthouse Laboratory Plymouth | Wellcome Sanger Institute for the COVID-19 Genomics UK (COG-UK) Consortium | Lighthouse Laboratory Plymouth and Alex Alderton, Roberto Amato, Jeffrey Barrett, Sonia Goncalves, Ewan Harrison, David K. Jackson, Ian Johnston, Dominic Kwiatkowski, Cordelia Langford, John Sillitoe on behalf of the Wellcome Sanger Institute COVID-19 Surveillance Team |
| EPI_ISL_2849838, EPI_ISL_2849839 | Lighthouse Lab in Alderley Park | Wellcome Sanger Institute for the COVID-19 Genomics UK (COG-UK) Consortium | Jacquelyn Wynn, Mairead Hyland, The Lighthouse Lab in Alderley Park and Alex Alderton, Roberto Amato, Jeffrey Barrett, Sonia Goncalves, Ewan Harrison, David K. Jackson, Ian Johnston, Dominic Kwiatkowski, Cordelia Langford, John Sillitoe on behalf of the Wellcome Sanger Institute COVID-19 Surveillance Team |
| EPI_ISL_2849840, EPI_ISL_2849841 | Lighthouse Laboratory Plymouth | Wellcome Sanger Institute for the COVID-19 Genomics UK (COG-UK) Consortium | Lighthouse Laboratory Plymouth and Alex Alderton, Roberto Amato, Jeffrey Barrett, Sonia Goncalves, Ewan Harrison, David K. Jackson, Ian Johnston, Dominic Kwiatkowski, Cordelia Langford, John Sillitoe on behalf of the Wellcome Sanger Institute COVID-19 Surveillance Team |
| EPI_ISL_2849842 | Lighthouse Lab in Alderley Park | Wellcome Sanger Institute for the COVID-19 Genomics UK (COG-UK) Consortium | Jacquelyn Wynn, Mairead Hyland, The Lighthouse Lab in Alderley Park and Alex Alderton, Roberto Amato, Jeffrey Barrett, Sonia Goncalves, Ewan Harrison, David K. Jackson, Ian Johnston, Dominic Kwiatkowski, Cordelia Langford, John Sillitoe on behalf of the Wellcome Sanger Institute COVID-19 Surveillance Team |
| EPI_ISL_2849843, EPI_ISL_2849844 | Lighthouse Laboratory Plymouth | Wellcome Sanger Institute for the COVID-19 Genomics UK (COG-UK) Consortium | Lighthouse Laboratory Plymouth and Alex Alderton, Roberto Amato, Jeffrey Barrett, Sonia Goncalves, Ewan Harrison, David K. Jackson, Ian Johnston, Dominic Kwiatkowski, Cordelia Langford, John Sillitoe on behalf of the Wellcome Sanger Institute COVID-19 Surveillance Team |
| EPI_ISL_2849845 | Lighthouse Lab in Alderley Park | Wellcome Sanger Institute for the COVID-19 Genomics UK (COG-UK) Consortium | Jacquelyn Wynn, Mairead Hyland, The Lighthouse Lab in Alderley Park and Alex Alderton, Roberto Amato, Jeffrey Barrett, Sonia Goncalves, Ewan Harrison, David K. Jackson, Ian Johnston, Dominic Kwiatkowski, Cordelia Langford, John Sillitoe on behalf of the Wellcome Sanger Institute COVID-19 Surveillance Team |
| EPI_ISL_2849846 | Lighthouse Laboratory Plymouth | Wellcome Sanger Institute for the COVID-19 Genomics UK (COG-UK) Consortium | Lighthouse Laboratory Plymouth and Alex Alderton, Roberto Amato, Jeffrey Barrett, Sonia Goncalves, Ewan Harrison, David K. Jackson, Ian Johnston, Dominic Kwiatkowski, Cordelia Langford, John Sillitoe on behalf of the Wellcome Sanger Institute COVID-19 Surveillance Team |
| EPI_ISL_2849847 | Lighthouse Lab in Alderley Park | Wellcome Sanger Institute for the COVID-19 Genomics UK (COG-UK) Consortium | Jacquelyn Wynn, Mairead Hyland, The Lighthouse Lab in Alderley Park and Alex Alderton, Roberto Amato, Jeffrey Barrett, Sonia Goncalves, Ewan Harrison, David K. Jackson, Ian Johnston, Dominic Kwiatkowski, Cordelia Langford, John Sillitoe on behalf of the Wellcome Sanger Institute COVID-19 Surveillance Team |
| EPI_ISL_2849848, EPI_ISL_2849849, EPI_ISL_2849850 | Lighthouse Laboratory Plymouth | Wellcome Sanger Institute for the COVID-19 Genomics UK (COG-UK) Consortium | Lighthouse Laboratory Plymouth and Alex Alderton, Roberto Amato, Jeffrey Barrett, Sonia Goncalves, Ewan Harrison, David K. Jackson, Ian Johnston, Dominic Kwiatkowski, Cordelia Langford, John Sillitoe on behalf of the Wellcome Sanger Institute COVID-19 Surveillance Team |
| EPI_ISL_2849851 | Lighthouse Lab in Alderley Park | Wellcome Sanger Institute for the COVID-19 Genomics UK (COG-UK) Consortium | Jacquelyn Wynn, Mairead Hyland, The Lighthouse Lab in Alderley Park and Alex Alderton, Roberto Amato, Jeffrey Barrett, Sonia Goncalves, Ewan Harrison, David K. Jackson, Ian Johnston, Dominic Kwiatkowski, Cordelia Langford, John Sillitoe on behalf of the Wellcome Sanger Institute COVID-19 Surveillance Team |

| | | | |
|---|---|---|---|
| EPI_ISL_2849852 | Lighthouse Laboratory Plymouth | Wellcome Sanger Institute for the COVID-19 Genomics UK (COG-UK) Consortium | Lighthouse Laboratory Plymouth and Alex Alderton, Roberto Amato, Jeffrey Barrett, Sonia Goncalves, Ewan Harrison, David K. Jackson, Ian Johnston, Dominic Kwiatkowski, Cordelia Langford, John Sillitoe on behalf of the Wellcome Sanger Institute COVID-19 Surveillance Team |
| EPI_ISL_2849853, EPI_ISL_2849854, EPI_ISL_2849855 | Lighthouse Lab in Alderley Park | Wellcome Sanger Institute for the COVID-19 Genomics UK (COG-UK) Consortium | Jacquelyn Wynn, Mairead Hyland, The Lighthouse Lab in Alderley Park and Alex Alderton, Roberto Amato, Jeffrey Barrett, Sonia Goncalves, Ewan Harrison, David K. Jackson, Ian Johnston, Dominic Kwiatkowski, Cordelia Langford, John Sillitoe on behalf of the Wellcome Sanger Institute COVID-19 Surveillance Team |
| EPI_ISL_2849856, EPI_ISL_2849857 | Lighthouse Laboratory Plymouth | Wellcome Sanger Institute for the COVID-19 Genomics UK (COG-UK) Consortium | Lighthouse Laboratory Plymouth and Alex Alderton, Roberto Amato, Jeffrey Barrett, Sonia Goncalves, Ewan Harrison, David K. Jackson, Ian Johnston, Dominic Kwiatkowski, Cordelia Langford, John Sillitoe on behalf of the Wellcome Sanger Institute COVID-19 Surveillance Team |
| EPI_ISL_2849858 | Lighthouse Lab in Alderley Park | Wellcome Sanger Institute for the COVID-19 Genomics UK (COG-UK) Consortium | Jacquelyn Wynn, Mairead Hyland, The Lighthouse Lab in Alderley Park and Alex Alderton, Roberto Amato, Jeffrey Barrett, Sonia Goncalves, Ewan Harrison, David K. Jackson, Ian Johnston, Dominic Kwiatkowski, Cordelia Langford, John Sillitoe on behalf of the Wellcome Sanger Institute COVID-19 Surveillance Team |
| EPI_ISL_2849859 | Lighthouse Laboratory Plymouth | Wellcome Sanger Institute for the COVID-19 Genomics UK (COG-UK) Consortium | Lighthouse Laboratory Plymouth and Alex Alderton, Roberto Amato, Jeffrey Barrett, Sonia Goncalves, Ewan Harrison, David K. Jackson, Ian Johnston, Dominic Kwiatkowski, Cordelia Langford, John Sillitoe on behalf of the Wellcome Sanger Institute COVID-19 Surveillance Team |
| EPI_ISL_2849860 | Lighthouse Lab in Alderley Park | Wellcome Sanger Institute for the COVID-19 Genomics UK (COG-UK) Consortium | Jacquelyn Wynn, Mairead Hyland, The Lighthouse Lab in Alderley Park and Alex Alderton, Roberto Amato, Jeffrey Barrett, Sonia Goncalves, Ewan Harrison, David K. Jackson, Ian Johnston, Dominic Kwiatkowski, Cordelia Langford, John Sillitoe on behalf of the Wellcome Sanger Institute COVID-19 Surveillance Team |
| EPI_ISL_2849861 | Lighthouse Laboratory Plymouth | Wellcome Sanger Institute for the COVID-19 Genomics UK (COG-UK) Consortium | Lighthouse Laboratory Plymouth and Alex Alderton, Roberto Amato, Jeffrey Barrett, Sonia Goncalves, Ewan Harrison, David K. Jackson, Ian Johnston, Dominic Kwiatkowski, Cordelia Langford, John Sillitoe on behalf of the Wellcome Sanger Institute COVID-19 Surveillance Team |
| EPI_ISL_2849862, EPI_ISL_2849863 | Lighthouse Lab in Alderley Park | Wellcome Sanger Institute for the COVID-19 Genomics UK (COG-UK) Consortium | Jacquelyn Wynn, Mairead Hyland, The Lighthouse Lab in Alderley Park and Alex Alderton, Roberto Amato, Jeffrey Barrett, Sonia Goncalves, Ewan Harrison, David K. Jackson, Ian Johnston, Dominic Kwiatkowski, Cordelia Langford, John Sillitoe on behalf of the Wellcome Sanger Institute COVID-19 Surveillance Team |
| EPI_ISL_2849864, EPI_ISL_2849865, EPI_ISL_2849866 | Lighthouse Laboratory Plymouth | Wellcome Sanger Institute for the COVID-19 Genomics UK (COG-UK) Consortium | Lighthouse Laboratory Plymouth and Alex Alderton, Roberto Amato, Jeffrey Barrett, Sonia Goncalves, Ewan Harrison, David K. Jackson, Ian Johnston, Dominic Kwiatkowski, Cordelia Langford, John Sillitoe on behalf of the Wellcome Sanger Institute COVID-19 Surveillance Team |
| EPI_ISL_2849867, EPI_ISL_2849868, EPI_ISL_2849869 | Lighthouse Lab in Alderley Park | Wellcome Sanger Institute for the COVID-19 Genomics UK (COG-UK) Consortium | Jacquelyn Wynn, Mairead Hyland, The Lighthouse Lab in Alderley Park and Alex Alderton, Roberto Amato, Jeffrey Barrett, Sonia Goncalves, Ewan Harrison, David K. Jackson, Ian Johnston, Dominic Kwiatkowski, Cordelia Langford, John Sillitoe on behalf of the Wellcome Sanger Institute COVID-19 Surveillance Team |
| EPI_ISL_2849870 | Lighthouse Laboratory Plymouth | Wellcome Sanger Institute for the COVID-19 Genomics UK (COG-UK) Consortium | Lighthouse Laboratory Plymouth and Alex Alderton, Roberto Amato, Jeffrey Barrett, Sonia Goncalves, Ewan Harrison, David K. Jackson, Ian Johnston, Dominic Kwiatkowski, Cordelia Langford, John Sillitoe on behalf of the Wellcome Sanger Institute COVID-19 Surveillance Team |
| EPI_ISL_2849871 | Lighthouse Lab in Alderley Park | Wellcome Sanger Institute for the COVID-19 Genomics UK (COG-UK) Consortium | Jacquelyn Wynn, Mairead Hyland, The Lighthouse Lab in Alderley Park and Alex Alderton, Roberto Amato, Jeffrey Barrett, Sonia Goncalves, Ewan Harrison, David K. Jackson, Ian Johnston, Dominic Kwiatkowski, Cordelia Langford, John Sillitoe on behalf of the Wellcome Sanger Institute COVID-19 Surveillance Team |
| EPI_ISL_2849872, EPI_ISL_2849873, EPI_ISL_2849874, EPI_ISL_2849875, EPI_ISL_2849876, EPI_ISL_2849877 | Lighthouse Laboratory Plymouth | Wellcome Sanger Institute for the COVID-19 Genomics UK (COG-UK) Consortium | Lighthouse Laboratory Plymouth and Alex Alderton, Roberto Amato, Jeffrey Barrett, Sonia Goncalves, Ewan Harrison, David K. Jackson, Ian Johnston, Dominic Kwiatkowski, Cordelia Langford, John Sillitoe on behalf of the Wellcome Sanger Institute COVID-19 Surveillance Team |
| EPI_ISL_2849878, EPI_ISL_2849879 | Lighthouse Lab in Alderley Park | Wellcome Sanger Institute for the COVID-19 Genomics UK (COG-UK) Consortium | Jacquelyn Wynn, Mairead Hyland, The Lighthouse Lab in Alderley Park and Alex Alderton, Roberto Amato, Jeffrey Barrett, Sonia Goncalves, Ewan Harrison, David K. Jackson, Ian Johnston, Dominic Kwiatkowski, Cordelia Langford, John Sillitoe on behalf of the Wellcome Sanger Institute COVID-19 Surveillance Team |
| EPI_ISL_2849880 | Lighthouse Laboratory Plymouth | Wellcome Sanger Institute for the COVID-19 Genomics UK (COG-UK) Consortium | Lighthouse Laboratory Plymouth and Alex Alderton, Roberto Amato, Jeffrey Barrett, Sonia Goncalves, Ewan Harrison, David K. Jackson, Ian Johnston, Dominic Kwiatkowski, Cordelia Langford, John Sillitoe on behalf of the Wellcome Sanger Institute COVID-19 Surveillance Team |
| EPI_ISL_2849881 | Lighthouse Lab in Alderley Park | Wellcome Sanger Institute for the COVID-19 Genomics UK (COG-UK) Consortium | Jacquelyn Wynn, Mairead Hyland, The Lighthouse Lab in Alderley Park and Alex Alderton, Roberto Amato, Jeffrey Barrett, Sonia Goncalves, Ewan Harrison, David K. Jackson, Ian Johnston, Dominic Kwiatkowski, Cordelia Langford, John Sillitoe on behalf of the Wellcome Sanger Institute COVID-19 Surveillance Team |
| EPI_ISL_2849882, EPI_ISL_2849883, EPI_ISL_2849884 | Lighthouse Laboratory Plymouth | Wellcome Sanger Institute for the COVID-19 Genomics UK (COG-UK) Consortium | Lighthouse Laboratory Plymouth and Alex Alderton, Roberto Amato, Jeffrey Barrett, Sonia Goncalves, Ewan Harrison, David K. Jackson, Ian Johnston, Dominic Kwiatkowski, Cordelia Langford, John Sillitoe on behalf of the Wellcome Sanger Institute COVID-19 Surveillance Team |
| EPI_ISL_2849885, EPI_ISL_2849886 | Lighthouse Lab in Alderley Park | Wellcome Sanger Institute for the COVID-19 Genomics UK (COG-UK) Consortium | Jacquelyn Wynn, Mairead Hyland, The Lighthouse Lab in Alderley Park and Alex Alderton, Roberto Amato, Jeffrey Barrett, Sonia Goncalves, Ewan Harrison, David K. Jackson, Ian Johnston, Dominic Kwiatkowski, Cordelia Langford, John Sillitoe on behalf of the Wellcome Sanger Institute COVID-19 Surveillance Team |
| EPI_ISL_2849887, EPI_ISL_2849888, EPI_ISL_2849889, EPI_ISL_2849890, EPI_ISL_2849891 | Lighthouse Laboratory Plymouth | Wellcome Sanger Institute for the COVID-19 Genomics UK (COG-UK) Consortium | Lighthouse Laboratory Plymouth and Alex Alderton, Roberto Amato, Jeffrey Barrett, Sonia Goncalves, Ewan Harrison, David K. Jackson, Ian Johnston, Dominic Kwiatkowski, Cordelia Langford, John Sillitoe on behalf of the Wellcome Sanger Institute COVID-19 Surveillance Team |
| EPI_ISL_2849892, EPI_ISL_2849893, EPI_ISL_2849894 | Lighthouse Lab in Alderley Park | Wellcome Sanger Institute for the COVID-19 Genomics UK (COG-UK) Consortium | Jacquelyn Wynn, Mairead Hyland, The Lighthouse Lab in Alderley Park and Alex Alderton, Roberto Amato, Jeffrey Barrett, Sonia Goncalves, Ewan Harrison, David K. Jackson, Ian Johnston, Dominic Kwiatkowski, Cordelia Langford, John Sillitoe on behalf of the Wellcome Sanger Institute COVID-19 Surveillance Team |
| EPI_ISL_2849895, EPI_ISL_2849896 | Lighthouse Laboratory Plymouth | Wellcome Sanger Institute for the COVID-19 Genomics UK (COG-UK) Consortium | Lighthouse Laboratory Plymouth and Alex Alderton, Roberto Amato, Jeffrey Barrett, Sonia Goncalves, Ewan Harrison, David K. Jackson, Ian Johnston, Dominic Kwiatkowski, Cordelia Langford, John Sillitoe on behalf of the Wellcome Sanger Institute COVID-19 Surveillance Team |
| EPI_ISL_2849897 | Lighthouse Lab in Alderley Park | Wellcome Sanger Institute for the COVID-19 Genomics UK (COG-UK) Consortium | Jacquelyn Wynn, Mairead Hyland, The Lighthouse Lab in Alderley Park and Alex Alderton, Roberto Amato, Jeffrey Barrett, Sonia Goncalves, Ewan Harrison, David K. Jackson, Ian Johnston, Dominic Kwiatkowski, Cordelia Langford, John Sillitoe on behalf of the Wellcome Sanger Institute COVID-19 Surveillance Team |
| EPI_ISL_2849898, EPI_ISL_2849899, EPI_ISL_2849900 | Lighthouse Laboratory Plymouth | Wellcome Sanger Institute for the COVID-19 Genomics UK (COG-UK) Consortium | Lighthouse Laboratory Plymouth and Alex Alderton, Roberto Amato, Jeffrey Barrett, Sonia Goncalves, Ewan Harrison, David K. Jackson, Ian Johnston, Dominic Kwiatkowski, Cordelia Langford, John Sillitoe on behalf of the Wellcome Sanger Institute COVID-19 Surveillance Team |
| EPI_ISL_2849901, EPI_ISL_2849902 | Lighthouse Lab in Alderley Park | Wellcome Sanger Institute for the COVID-19 Genomics UK (COG-UK) Consortium | Jacquelyn Wynn, Mairead Hyland, The Lighthouse Lab in Alderley Park and Alex Alderton, Roberto Amato, Jeffrey Barrett, Sonia Goncalves, Ewan Harrison, David K. Jackson, Ian Johnston, Dominic Kwiatkowski, Cordelia Langford, John Sillitoe on behalf of the Wellcome Sanger Institute COVID-19 Surveillance Team |
| EPI_ISL_2849903 | Lighthouse Laboratory Plymouth | Wellcome Sanger Institute for the COVID-19 Genomics UK (COG-UK) Consortium | Lighthouse Laboratory Plymouth and Alex Alderton, Roberto Amato, Jeffrey Barrett, Sonia Goncalves, Ewan Harrison, David K. Jackson, Ian Johnston, Dominic Kwiatkowski, Cordelia Langford, John Sillitoe on behalf of the Wellcome Sanger Institute COVID-19 Surveillance Team |
| EPI_ISL_2849904, EPI_ISL_2849905 | Lighthouse Lab in Alderley Park | Wellcome Sanger Institute for the COVID-19 Genomics UK (COG-UK) Consortium | Jacquelyn Wynn, Mairead Hyland, The Lighthouse Lab in Alderley Park and Alex Alderton, Roberto Amato, Jeffrey Barrett, Sonia Goncalves, Ewan Harrison, David K. Jackson, Ian Johnston, Dominic Kwiatkowski, Cordelia Langford, John Sillitoe on behalf of the Wellcome Sanger Institute COVID-19 Surveillance Team |
| EPI_ISL_2849906, EPI_ISL_2849907 | Lighthouse Laboratory Plymouth | Wellcome Sanger Institute for the COVID-19 Genomics UK (COG-UK) Consortium | Lighthouse Laboratory Plymouth and Alex Alderton, Roberto Amato, Jeffrey Barrett, Sonia Goncalves, Ewan Harrison, David K. Jackson, Ian Johnston, Dominic Kwiatkowski, Cordelia Langford, John Sillitoe on behalf of the Wellcome Sanger Institute COVID-19 Surveillance Team |
| EPI_ISL_2849908, EPI_ISL_2849909, EPI_ISL_2849910 | Lighthouse Lab in Alderley Park | Wellcome Sanger Institute for the COVID-19 Genomics UK (COG-UK) Consortium | Jacquelyn Wynn, Mairead Hyland, The Lighthouse Lab in Alderley Park and Alex Alderton, Roberto Amato, Jeffrey Barrett, Sonia Goncalves, Ewan Harrison, David K. Jackson, Ian Johnston, Dominic Kwiatkowski, Cordelia Langford, John Sillitoe on behalf of the Wellcome Sanger Institute COVID-19 Surveillance Team |
| EPI_ISL_2849911, EPI_ISL_2849912, EPI_ISL_2849913, EPI_ISL_2849914 | Lighthouse Laboratory Plymouth | Wellcome Sanger Institute for the COVID-19 Genomics UK (COG-UK) Consortium | Lighthouse Laboratory Plymouth and Alex Alderton, Roberto Amato, Jeffrey Barrett, Sonia Goncalves, Ewan Harrison, David K. Jackson, Ian Johnston, Dominic Kwiatkowski, Cordelia Langford, John Sillitoe on behalf of the Wellcome Sanger Institute COVID-19 Surveillance Team |
| EPI_ISL_2849915 | Lighthouse Lab in Alderley Park | Wellcome Sanger Institute for the COVID-19 Genomics UK (COG-UK) Consortium | Jacquelyn Wynn, Mairead Hyland, The Lighthouse Lab in Alderley Park and Alex Alderton, Roberto Amato, Jeffrey Barrett, Sonia Goncalves, Ewan Harrison, David K. Jackson, Ian Johnston, Dominic Kwiatkowski, Cordelia Langford, John Sillitoe on behalf of the Wellcome Sanger Institute COVID-19 Surveillance Team |
| EPI_ISL_2849916 | Lighthouse Laboratory Plymouth | Wellcome Sanger Institute for the COVID-19 Genomics UK (COG-UK) Consortium | Lighthouse Laboratory Plymouth and Alex Alderton, Roberto Amato, Jeffrey Barrett, Sonia Goncalves, Ewan Harrison, David K. Jackson, Ian Johnston, Dominic Kwiatkowski, Cordelia Langford, John Sillitoe on behalf of the Wellcome Sanger Institute COVID-19 Surveillance Team |
| EPI_ISL_2849917 | Lighthouse Lab in Alderley Park | Wellcome Sanger Institute for the COVID-19 Genomics UK (COG-UK) Consortium | Jacquelyn Wynn, Mairead Hyland, The Lighthouse Lab in Alderley Park and Alex Alderton, Roberto Amato, Jeffrey Barrett, Sonia Goncalves, Ewan Harrison, David K. Jackson, Ian Johnston, Dominic Kwiatkowski, Cordelia Langford, John Sillitoe on behalf of the Wellcome Sanger Institute COVID-19 Surveillance Team |
| EPI_ISL_2849918 | Lighthouse Laboratory Plymouth | Wellcome Sanger Institute for the COVID-19 Genomics UK (COG-UK) Consortium | Lighthouse Laboratory Plymouth and Alex Alderton, Roberto Amato, Jeffrey Barrett, Sonia Goncalves, Ewan Harrison, David K. Jackson, Ian Johnston, Dominic Kwiatkowski, Cordelia Langford, John Sillitoe on behalf of the Wellcome Sanger Institute COVID-19 Surveillance Team |
| EPI_ISL_2849919, EPI_ISL_2849920 | Lighthouse Lab in Alderley Park | Wellcome Sanger Institute for the COVID-19 Genomics UK (COG-UK) Consortium | Jacquelyn Wynn, Mairead Hyland, The Lighthouse Lab in Alderley Park and Alex Alderton, Roberto Amato, Jeffrey Barrett, Sonia Goncalves, Ewan Harrison, David K. Jackson, Ian Johnston, Dominic Kwiatkowski, Cordelia Langford, John Sillitoe on behalf of the Wellcome Sanger Institute COVID-19 Surveillance Team |
| EPI_ISL_2849921 | Lighthouse Laboratory Plymouth | Wellcome Sanger Institute for the COVID-19 Genomics UK (COG-UK) Consortium | Lighthouse Laboratory Plymouth and Alex Alderton, Roberto Amato, Jeffrey Barrett, Sonia Goncalves, Ewan Harrison, David K. Jackson, Ian Johnston, Dominic Kwiatkowski, Cordelia Langford, John Sillitoe on behalf of the Wellcome Sanger Institute COVID-19 Surveillance Team |
| EPI_ISL_2849922, EPI_ISL_2849923 | Lighthouse Lab in Alderley Park | Wellcome Sanger Institute for the COVID-19 Genomics UK (COG-UK) Consortium | Jacquelyn Wynn, Mairead Hyland, The Lighthouse Lab in Alderley Park and Alex Alderton, Roberto Amato, Jeffrey Barrett, Sonia Goncalves, Ewan Harrison, David K. Jackson, Ian Johnston, Dominic Kwiatkowski, Cordelia Langford, John Sillitoe on behalf of the Wellcome Sanger Institute COVID-19 Surveillance Team |
| EPI_ISL_2849924, EPI_ISL_2849925 | Lighthouse Laboratory Plymouth | Wellcome Sanger Institute for the COVID-19 Genomics UK (COG-UK) Consortium | Lighthouse Laboratory Plymouth and Alex Alderton, Roberto Amato, Jeffrey Barrett, Sonia Goncalves, Ewan Harrison, David K. Jackson, Ian Johnston, Dominic Kwiatkowski, Cordelia Langford, John Sillitoe on behalf of the Wellcome Sanger Institute COVID-19 Surveillance Team |
| EPI_ISL_2849926, EPI_ISL_2849927, EPI_ISL_2849928 | Lighthouse Lab in Alderley Park | Wellcome Sanger Institute for the COVID-19 Genomics UK (COG-UK) Consortium | Jacquelyn Wynn, Mairead Hyland, The Lighthouse Lab in Alderley Park and Alex Alderton, Roberto Amato, Jeffrey Barrett, Sonia Goncalves, Ewan Harrison, David K. Jackson, Ian Johnston, Dominic Kwiatkowski, Cordelia Langford, John Sillitoe on behalf of the Wellcome Sanger Institute COVID-19 Surveillance Team |

| | | | |
|---|---|---|---|
| EPI_ISL_2849929, EPI_ISL_2849930, EPI_ISL_2849931 | Lighthouse Laboratory Plymouth | Wellcome Sanger Institute for the COVID-19 Genomics UK (COG-UK) Consortium | Lighthouse Laboratory Plymouth and Alex Alderton, Roberto Amato, Jeffrey Barrett, Sonia Goncalves, Ewan Harrison, David K. Jackson, Ian Johnston, Dominic Kwiatkowski, Cordelia Langford, John Sillitoe on behalf of the Wellcome Sanger Institute COVID-19 Surveillance Team |
| EPI_ISL_2849932, EPI_ISL_2849933, EPI_ISL_2849934 | Lighthouse Lab in Alderley Park | Wellcome Sanger Institute for the COVID-19 Genomics UK (COG-UK) Consortium | Jacquelyn Wynn, Mairead Hyland, The Lighthouse Lab in Alderley Park and Alex Alderton, Roberto Amato, Jeffrey Barrett, Sonia Goncalves, Ewan Harrison, David K. Jackson, Ian Johnston, Dominic Kwiatkowski, Cordelia Langford, John Sillitoe on behalf of the Wellcome Sanger Institute COVID-19 Surveillance Team |
| EPI_ISL_2849935, EPI_ISL_2849936, EPI_ISL_2849937, EPI_ISL_2849938 | Lighthouse Laboratory Plymouth | Wellcome Sanger Institute for the COVID-19 Genomics UK (COG-UK) Consortium | Lighthouse Laboratory Plymouth and Alex Alderton, Roberto Amato, Jeffrey Barrett, Sonia Goncalves, Ewan Harrison, David K. Jackson, Ian Johnston, Dominic Kwiatkowski, Cordelia Langford, John Sillitoe on behalf of the Wellcome Sanger Institute COVID-19 Surveillance Team |
| EPI_ISL_2849939, EPI_ISL_2849940, EPI_ISL_2849941, EPI_ISL_2849942 | Lighthouse Lab in Alderley Park | Wellcome Sanger Institute for the COVID-19 Genomics UK (COG-UK) Consortium | Jacquelyn Wynn, Mairead Hyland, The Lighthouse Lab in Alderley Park and Alex Alderton, Roberto Amato, Jeffrey Barrett, Sonia Goncalves, Ewan Harrison, David K. Jackson, Ian Johnston, Dominic Kwiatkowski, Cordelia Langford, John Sillitoe on behalf of the Wellcome Sanger Institute COVID-19 Surveillance Team |
| EPI_ISL_2849943, EPI_ISL_2849944 | Lighthouse Laboratory Plymouth | Wellcome Sanger Institute for the COVID-19 Genomics UK (COG-UK) Consortium | Lighthouse Laboratory Plymouth and Alex Alderton, Roberto Amato, Jeffrey Barrett, Sonia Goncalves, Ewan Harrison, David K. Jackson, Ian Johnston, Dominic Kwiatkowski, Cordelia Langford, John Sillitoe on behalf of the Wellcome Sanger Institute COVID-19 Surveillance Team |
| EPI_ISL_2849945 | Lighthouse Lab in Alderley Park | Wellcome Sanger Institute for the COVID-19 Genomics UK (COG-UK) Consortium | Jacquelyn Wynn, Mairead Hyland, The Lighthouse Lab in Alderley Park and Alex Alderton, Roberto Amato, Jeffrey Barrett, Sonia Goncalves, Ewan Harrison, David K. Jackson, Ian Johnston, Dominic Kwiatkowski, Cordelia Langford, John Sillitoe on behalf of the Wellcome Sanger Institute COVID-19 Surveillance Team |
| EPI_ISL_2849946 | Lighthouse Laboratory Plymouth | Wellcome Sanger Institute for the COVID-19 Genomics UK (COG-UK) Consortium | Lighthouse Laboratory Plymouth and Alex Alderton, Roberto Amato, Jeffrey Barrett, Sonia Goncalves, Ewan Harrison, David K. Jackson, Ian Johnston, Dominic Kwiatkowski, Cordelia Langford, John Sillitoe on behalf of the Wellcome Sanger Institute COVID-19 Surveillance Team |
| EPI_ISL_2849947, EPI_ISL_2849948 | Lighthouse Lab in Alderley Park | Wellcome Sanger Institute for the COVID-19 Genomics UK (COG-UK) Consortium | Jacquelyn Wynn, Mairead Hyland, The Lighthouse Lab in Alderley Park and Alex Alderton, Roberto Amato, Jeffrey Barrett, Sonia Goncalves, Ewan Harrison, David K. Jackson, Ian Johnston, Dominic Kwiatkowski, Cordelia Langford, John Sillitoe on behalf of the Wellcome Sanger Institute COVID-19 Surveillance Team |
| EPI_ISL_2849949, EPI_ISL_2849950 | Lighthouse Laboratory Plymouth | Wellcome Sanger Institute for the COVID-19 Genomics UK (COG-UK) Consortium | Lighthouse Laboratory Plymouth and Alex Alderton, Roberto Amato, Jeffrey Barrett, Sonia Goncalves, Ewan Harrison, David K. Jackson, Ian Johnston, Dominic Kwiatkowski, Cordelia Langford, John Sillitoe on behalf of the Wellcome Sanger Institute COVID-19 Surveillance Team |
| EPI_ISL_2849951, EPI_ISL_2849952 | Lighthouse Lab in Alderley Park | Wellcome Sanger Institute for the COVID-19 Genomics UK (COG-UK) Consortium | Jacquelyn Wynn, Mairead Hyland, The Lighthouse Lab in Alderley Park and Alex Alderton, Roberto Amato, Jeffrey Barrett, Sonia Goncalves, Ewan Harrison, David K. Jackson, Ian Johnston, Dominic Kwiatkowski, Cordelia Langford, John Sillitoe on behalf of the Wellcome Sanger Institute COVID-19 Surveillance Team |
| EPI_ISL_2849953, EPI_ISL_2849954, EPI_ISL_2849955 | Lighthouse Laboratory Plymouth | Wellcome Sanger Institute for the COVID-19 Genomics UK (COG-UK) Consortium | Lighthouse Laboratory Plymouth and Alex Alderton, Roberto Amato, Jeffrey Barrett, Sonia Goncalves, Ewan Harrison, David K. Jackson, Ian Johnston, Dominic Kwiatkowski, Cordelia Langford, John Sillitoe on behalf of the Wellcome Sanger Institute COVID-19 Surveillance Team |
| EPI_ISL_2849956, EPI_ISL_2849957, EPI_ISL_2849958, EPI_ISL_2849959, EPI_ISL_2849960, EPI_ISL_2849961 | Lighthouse Lab in Alderley Park | Wellcome Sanger Institute for the COVID-19 Genomics UK (COG-UK) Consortium | Jacquelyn Wynn, Mairead Hyland, The Lighthouse Lab in Alderley Park and Alex Alderton, Roberto Amato, Jeffrey Barrett, Sonia Goncalves, Ewan Harrison, David K. Jackson, Ian Johnston, Dominic Kwiatkowski, Cordelia Langford, John Sillitoe on behalf of the Wellcome Sanger Institute COVID-19 Surveillance Team |
| EPI_ISL_2849962 | Lighthouse Laboratory Plymouth | Wellcome Sanger Institute for the COVID-19 Genomics UK (COG-UK) Consortium | Lighthouse Laboratory Plymouth and Alex Alderton, Roberto Amato, Jeffrey Barrett, Sonia Goncalves, Ewan Harrison, David K. Jackson, Ian Johnston, Dominic Kwiatkowski, Cordelia Langford, John Sillitoe on behalf of the Wellcome Sanger Institute COVID-19 Surveillance Team |
| EPI_ISL_2849963, EPI_ISL_2849964, EPI_ISL_2849965 | Lighthouse Lab in Alderley Park | Wellcome Sanger Institute for the COVID-19 Genomics UK (COG-UK) Consortium | Jacquelyn Wynn, Mairead Hyland, The Lighthouse Lab in Alderley Park and Alex Alderton, Roberto Amato, Jeffrey Barrett, Sonia Goncalves, Ewan Harrison, David K. Jackson, Ian Johnston, Dominic Kwiatkowski, Cordelia Langford, John Sillitoe on behalf of the Wellcome Sanger Institute COVID-19 Surveillance Team |
| EPI_ISL_2849966, EPI_ISL_2849967 | Lighthouse Laboratory Plymouth | Wellcome Sanger Institute for the COVID-19 Genomics UK (COG-UK) Consortium | Lighthouse Laboratory Plymouth and Alex Alderton, Roberto Amato, Jeffrey Barrett, Sonia Goncalves, Ewan Harrison, David K. Jackson, Ian Johnston, Dominic Kwiatkowski, Cordelia Langford, John Sillitoe on behalf of the Wellcome Sanger Institute COVID-19 Surveillance Team |
| EPI_ISL_2849968, EPI_ISL_2849969 | Lighthouse Lab in Alderley Park | Wellcome Sanger Institute for the COVID-19 Genomics UK (COG-UK) Consortium | Jacquelyn Wynn, Mairead Hyland, The Lighthouse Lab in Alderley Park and Alex Alderton, Roberto Amato, Jeffrey Barrett, Sonia Goncalves, Ewan Harrison, David K. Jackson, Ian Johnston, Dominic Kwiatkowski, Cordelia Langford, John Sillitoe on behalf of the Wellcome Sanger Institute COVID-19 Surveillance Team |
| EPI_ISL_2849970, EPI_ISL_2849971 | Lighthouse Laboratory Plymouth | Wellcome Sanger Institute for the COVID-19 Genomics UK (COG-UK) Consortium | Lighthouse Laboratory Plymouth and Alex Alderton, Roberto Amato, Jeffrey Barrett, Sonia Goncalves, Ewan Harrison, David K. Jackson, Ian Johnston, Dominic Kwiatkowski, Cordelia Langford, John Sillitoe on behalf of the Wellcome Sanger Institute COVID-19 Surveillance Team |
| EPI_ISL_2849972, EPI_ISL_2849973 | Lighthouse Lab in Alderley Park | Wellcome Sanger Institute for the COVID-19 Genomics UK (COG-UK) Consortium | Jacquelyn Wynn, Mairead Hyland, The Lighthouse Lab in Alderley Park and Alex Alderton, Roberto Amato, Jeffrey Barrett, Sonia Goncalves, Ewan Harrison, David K. Jackson, Ian Johnston, Dominic Kwiatkowski, Cordelia Langford, John Sillitoe on behalf of the Wellcome Sanger Institute COVID-19 Surveillance Team |
| EPI_ISL_2849974, EPI_ISL_2849975, EPI_ISL_2849976 | Lighthouse Laboratory Plymouth | Wellcome Sanger Institute for the COVID-19 Genomics UK (COG-UK) Consortium | Lighthouse Laboratory Plymouth and Alex Alderton, Roberto Amato, Jeffrey Barrett, Sonia Goncalves, Ewan Harrison, David K. Jackson, Ian Johnston, Dominic Kwiatkowski, Cordelia Langford, John Sillitoe on behalf of the Wellcome Sanger Institute COVID-19 Surveillance Team |
| EPI_ISL_2849977 | Lighthouse Lab in Alderley Park | Wellcome Sanger Institute for the COVID-19 Genomics UK (COG-UK) Consortium | Jacquelyn Wynn, Mairead Hyland, The Lighthouse Lab in Alderley Park and Alex Alderton, Roberto Amato, Jeffrey Barrett, Sonia Goncalves, Ewan Harrison, David K. Jackson, Ian Johnston, Dominic Kwiatkowski, Cordelia Langford, John Sillitoe on behalf of the Wellcome Sanger Institute COVID-19 Surveillance Team |
| EPI_ISL_2849978, EPI_ISL_2849979, EPI_ISL_2849980, EPI_ISL_2849981, EPI_ISL_2849982, EPI_ISL_2849983, EPI_ISL_2849984, EPI_ISL_2849985 | Lighthouse Laboratory Plymouth | Wellcome Sanger Institute for the COVID-19 Genomics UK (COG-UK) Consortium | Lighthouse Laboratory Plymouth and Alex Alderton, Roberto Amato, Jeffrey Barrett, Sonia Goncalves, Ewan Harrison, David K. Jackson, Ian Johnston, Dominic Kwiatkowski, Cordelia Langford, John Sillitoe on behalf of the Wellcome Sanger Institute COVID-19 Surveillance Team |
| EPI_ISL_2849986, EPI_ISL_2849987, EPI_ISL_2849988 | Lighthouse Lab in Alderley Park | Wellcome Sanger Institute for the COVID-19 Genomics UK (COG-UK) Consortium | Jacquelyn Wynn, Mairead Hyland, The Lighthouse Lab in Alderley Park and Alex Alderton, Roberto Amato, Jeffrey Barrett, Sonia Goncalves, Ewan Harrison, David K. Jackson, Ian Johnston, Dominic Kwiatkowski, Cordelia Langford, John Sillitoe on behalf of the Wellcome Sanger Institute COVID-19 Surveillance Team |
| EPI_ISL_2849989 | Lighthouse Laboratory Plymouth | Wellcome Sanger Institute for the COVID-19 Genomics UK (COG-UK) Consortium | Lighthouse Laboratory Plymouth and Alex Alderton, Roberto Amato, Jeffrey Barrett, Sonia Goncalves, Ewan Harrison, David K. Jackson, Ian Johnston, Dominic Kwiatkowski, Cordelia Langford, John Sillitoe on behalf of the Wellcome Sanger Institute COVID-19 Surveillance Team |
| EPI_ISL_2849990 | Lighthouse Lab in Alderley Park | Wellcome Sanger Institute for the COVID-19 Genomics UK (COG-UK) Consortium | Jacquelyn Wynn, Mairead Hyland, The Lighthouse Lab in Alderley Park and Alex Alderton, Roberto Amato, Jeffrey Barrett, Sonia Goncalves, Ewan Harrison, David K. Jackson, Ian Johnston, Dominic Kwiatkowski, Cordelia Langford, John Sillitoe on behalf of the Wellcome Sanger Institute COVID-19 Surveillance Team |
| EPI_ISL_2849991 | Lighthouse Laboratory Plymouth | Wellcome Sanger Institute for the COVID-19 Genomics UK (COG-UK) Consortium | Lighthouse Laboratory Plymouth and Alex Alderton, Roberto Amato, Jeffrey Barrett, Sonia Goncalves, Ewan Harrison, David K. Jackson, Ian Johnston, Dominic Kwiatkowski, Cordelia Langford, John Sillitoe on behalf of the Wellcome Sanger Institute COVID-19 Surveillance Team |
| EPI_ISL_2849992 | Lighthouse Lab in Alderley Park | Wellcome Sanger Institute for the COVID-19 Genomics UK (COG-UK) Consortium | Jacquelyn Wynn, Mairead Hyland, The Lighthouse Lab in Alderley Park and Alex Alderton, Roberto Amato, Jeffrey Barrett, Sonia Goncalves, Ewan Harrison, David K. Jackson, Ian Johnston, Dominic Kwiatkowski, Cordelia Langford, John Sillitoe on behalf of the Wellcome Sanger Institute COVID-19 Surveillance Team |
| EPI_ISL_2849993 | Lighthouse Laboratory Plymouth | Wellcome Sanger Institute for the COVID-19 Genomics UK (COG-UK) Consortium | Lighthouse Laboratory Plymouth and Alex Alderton, Roberto Amato, Jeffrey Barrett, Sonia Goncalves, Ewan Harrison, David K. Jackson, Ian Johnston, Dominic Kwiatkowski, Cordelia Langford, John Sillitoe on behalf of the Wellcome Sanger Institute COVID-19 Surveillance Team |
| EPI_ISL_2849994, EPI_ISL_2849995 | Lighthouse Lab in Alderley Park | Wellcome Sanger Institute for the COVID-19 Genomics UK (COG-UK) Consortium | Jacquelyn Wynn, Mairead Hyland, The Lighthouse Lab in Alderley Park and Alex Alderton, Roberto Amato, Jeffrey Barrett, Sonia Goncalves, Ewan Harrison, David K. Jackson, Ian Johnston, Dominic Kwiatkowski, Cordelia Langford, John Sillitoe on behalf of the Wellcome Sanger Institute COVID-19 Surveillance Team |
| EPI_ISL_2849996 | Lighthouse Laboratory Plymouth | Wellcome Sanger Institute for the COVID-19 Genomics UK (COG-UK) Consortium | Lighthouse Laboratory Plymouth and Alex Alderton, Roberto Amato, Jeffrey Barrett, Sonia Goncalves, Ewan Harrison, David K. Jackson, Ian Johnston, Dominic Kwiatkowski, Cordelia Langford, John Sillitoe on behalf of the Wellcome Sanger Institute COVID-19 Surveillance Team |
| EPI_ISL_2849997, EPI_ISL_2849998 | Lighthouse Lab in Alderley Park | Wellcome Sanger Institute for the COVID-19 Genomics UK (COG-UK) Consortium | Jacquelyn Wynn, Mairead Hyland, The Lighthouse Lab in Alderley Park and Alex Alderton, Roberto Amato, Jeffrey Barrett, Sonia Goncalves, Ewan Harrison, David K. Jackson, Ian Johnston, Dominic Kwiatkowski, Cordelia Langford, John Sillitoe on behalf of the Wellcome Sanger Institute COVID-19 Surveillance Team |
| EPI_ISL_2849999, EPI_ISL_2850000, EPI_ISL_2850001, EPI_ISL_2850002, EPI_ISL_2850003 | Lighthouse Laboratory Plymouth | Wellcome Sanger Institute for the COVID-19 Genomics UK (COG-UK) Consortium | Lighthouse Laboratory Plymouth and Alex Alderton, Roberto Amato, Jeffrey Barrett, Sonia Goncalves, Ewan Harrison, David K. Jackson, Ian Johnston, Dominic Kwiatkowski, Cordelia Langford, John Sillitoe on behalf of the Wellcome Sanger Institute COVID-19 Surveillance Team |
| EPI_ISL_2850004, EPI_ISL_2850005, EPI_ISL_2850006 | Lighthouse Lab in Alderley Park | Wellcome Sanger Institute for the COVID-19 Genomics UK (COG-UK) Consortium | Jacquelyn Wynn, Mairead Hyland, The Lighthouse Lab in Alderley Park and Alex Alderton, Roberto Amato, Jeffrey Barrett, Sonia Goncalves, Ewan Harrison, David K. Jackson, Ian Johnston, Dominic Kwiatkowski, Cordelia Langford, John Sillitoe on behalf of the Wellcome Sanger Institute COVID-19 Surveillance Team |
| EPI_ISL_2850007 | Lighthouse Laboratory Plymouth | Wellcome Sanger Institute for the COVID-19 Genomics UK (COG-UK) Consortium | Lighthouse Laboratory Plymouth and Alex Alderton, Roberto Amato, Jeffrey Barrett, Sonia Goncalves, Ewan Harrison, David K. Jackson, Ian Johnston, Dominic Kwiatkowski, Cordelia Langford, John Sillitoe on behalf of the Wellcome Sanger Institute COVID-19 Surveillance Team |
| EPI_ISL_2850008 | Lighthouse Lab in Alderley Park | Wellcome Sanger Institute for the COVID-19 Genomics UK (COG-UK) Consortium | Jacquelyn Wynn, Mairead Hyland, The Lighthouse Lab in Alderley Park and Alex Alderton, Roberto Amato, Jeffrey Barrett, Sonia Goncalves, Ewan Harrison, David K. Jackson, Ian Johnston, Dominic Kwiatkowski, Cordelia Langford, John Sillitoe on behalf of the Wellcome Sanger Institute COVID-19 Surveillance Team |
| EPI_ISL_2850009 | Lighthouse Laboratory Plymouth | Wellcome Sanger Institute for the COVID-19 Genomics UK (COG-UK) Consortium | Lighthouse Laboratory Plymouth and Alex Alderton, Roberto Amato, Jeffrey Barrett, Sonia Goncalves, Ewan Harrison, David K. Jackson, Ian Johnston, Dominic Kwiatkowski, Cordelia Langford, John Sillitoe on behalf of the Wellcome Sanger Institute COVID-19 Surveillance Team |
| EPI_ISL_2850010, | Lighthouse Lab in Alderley Park | Wellcome Sanger Institute for the COVID-19 Genomics UK (COG-UK) | Jacquelyn Wynn, Mairead Hyland, The Lighthouse Lab in Alderley Park and Alex Alderton, Roberto Amato, Jeffrey Barrett, Sonia Goncalves, Ewan Harrison, David K. Jackson, Ian Johnston, Dominic Kwiatkowski, Cordelia Langford, John Sillitoe on |

| | | | |
|---|---|---|---|
| EPI_ISL_2850011 | | Consortium | behalf of the Wellcome Sanger Institute COVID-19 Surveillance Team |
| EPI_ISL_2850012 | Lighthouse Laboratory Plymouth | Wellcome Sanger Institute for the COVID-19 Genomics UK (COG-UK) Consortium | Lighthouse Laboratory Plymouth and Alex Alderton, Roberto Amato, Jeffrey Barrett, Sonia Goncalves, Ewan Harrison, David K. Jackson, Ian Johnston, Dominic Kwiatkowski, Cordelia Langford, John Sillitoe on behalf of the Wellcome Sanger Institute COVID-19 Surveillance Team |
| EPI_ISL_2850013 | Lighthouse Lab in Alderley Park | Wellcome Sanger Institute for the COVID-19 Genomics UK (COG-UK) Consortium | Jacquelyn Wynn, Mairead Hyland, The Lighthouse Lab in Alderley Park and Alex Alderton, Roberto Amato, Jeffrey Barrett, Sonia Goncalves, Ewan Harrison, David K. Jackson, Ian Johnston, Dominic Kwiatkowski, Cordelia Langford, John Sillitoe on behalf of the Wellcome Sanger Institute COVID-19 Surveillance Team |
| EPI_ISL_2850014, EPI_ISL_2850015, EPI_ISL_2850016, EPI_ISL_2850017, EPI_ISL_2850018, EPI_ISL_2850019 | Lighthouse Laboratory Plymouth | Wellcome Sanger Institute for the COVID-19 Genomics UK (COG-UK) Consortium | Lighthouse Laboratory Plymouth and Alex Alderton, Roberto Amato, Jeffrey Barrett, Sonia Goncalves, Ewan Harrison, David K. Jackson, Ian Johnston, Dominic Kwiatkowski, Cordelia Langford, John Sillitoe on behalf of the Wellcome Sanger Institute COVID-19 Surveillance Team |
| EPI_ISL_2850020, EPI_ISL_2850021 | Lighthouse Lab in Alderley Park | Wellcome Sanger Institute for the COVID-19 Genomics UK (COG-UK) Consortium | Jacquelyn Wynn, Mairead Hyland, The Lighthouse Lab in Alderley Park and Alex Alderton, Roberto Amato, Jeffrey Barrett, Sonia Goncalves, Ewan Harrison, David K. Jackson, Ian Johnston, Dominic Kwiatkowski, Cordelia Langford, John Sillitoe on behalf of the Wellcome Sanger Institute COVID-19 Surveillance Team |
| EPI_ISL_2850022, EPI_ISL_2850023, EPI_ISL_2850024, EPI_ISL_2850025, EPI_ISL_2850026 | Lighthouse Laboratory Plymouth | Wellcome Sanger Institute for the COVID-19 Genomics UK (COG-UK) Consortium | Lighthouse Laboratory Plymouth and Alex Alderton, Roberto Amato, Jeffrey Barrett, Sonia Goncalves, Ewan Harrison, David K. Jackson, Ian Johnston, Dominic Kwiatkowski, Cordelia Langford, John Sillitoe on behalf of the Wellcome Sanger Institute COVID-19 Surveillance Team |
| EPI_ISL_2850027, EPI_ISL_2850028, EPI_ISL_2850029 | Lighthouse Lab in Alderley Park | Wellcome Sanger Institute for the COVID-19 Genomics UK (COG-UK) Consortium | Jacquelyn Wynn, Mairead Hyland, The Lighthouse Lab in Alderley Park and Alex Alderton, Roberto Amato, Jeffrey Barrett, Sonia Goncalves, Ewan Harrison, David K. Jackson, Ian Johnston, Dominic Kwiatkowski, Cordelia Langford, John Sillitoe on behalf of the Wellcome Sanger Institute COVID-19 Surveillance Team |
| EPI_ISL_2850030 | Lighthouse Laboratory Plymouth | Wellcome Sanger Institute for the COVID-19 Genomics UK (COG-UK) Consortium | Lighthouse Laboratory Plymouth and Alex Alderton, Roberto Amato, Jeffrey Barrett, Sonia Goncalves, Ewan Harrison, David K. Jackson, Ian Johnston, Dominic Kwiatkowski, Cordelia Langford, John Sillitoe on behalf of the Wellcome Sanger Institute COVID-19 Surveillance Team |
| EPI_ISL_2850031, EPI_ISL_2850033, EPI_ISL_2850034, EPI_ISL_2850035, EPI_ISL_2850036 | Lighthouse Lab in Alderley Park | Wellcome Sanger Institute for the COVID-19 Genomics UK (COG-UK) Consortium | Jacquelyn Wynn, Mairead Hyland, The Lighthouse Lab in Alderley Park and Alex Alderton, Roberto Amato, Jeffrey Barrett, Sonia Goncalves, Ewan Harrison, David K. Jackson, Ian Johnston, Dominic Kwiatkowski, Cordelia Langford, John Sillitoe on behalf of the Wellcome Sanger Institute COVID-19 Surveillance Team |
| EPI_ISL_2850037, EPI_ISL_2850038 | Lighthouse Lab in Glasgow | Wellcome Sanger Institute for the COVID-19 Genomics UK (COG-UK) Consortium | Harper VanSteenhouse, Yumi Kasai, David Gray, Carol Clugston, Anna Dominiczak and Alex Alderton, Roberto Amato, Jeffrey Barrett, Sonia Goncalves, Ewan Harrison, David K. Jackson, Ian Johnston, Dominic Kwiatkowski, Cordelia Langford, John Sillitoe on behalf of the Wellcome Sanger Institute COVID-19 Surveillance Team |
| EPI_ISL_2850039 | Lighthouse Laboratory Plymouth | Wellcome Sanger Institute for the COVID-19 Genomics UK (COG-UK) Consortium | Lighthouse Laboratory Plymouth and Alex Alderton, Roberto Amato, Jeffrey Barrett, Sonia Goncalves, Ewan Harrison, David K. Jackson, Ian Johnston, Dominic Kwiatkowski, Cordelia Langford, John Sillitoe on behalf of the Wellcome Sanger Institute COVID-19 Surveillance Team |
| EPI_ISL_2850040, EPI_ISL_2850041, EPI_ISL_2850042 | Lighthouse Lab in Glasgow | Wellcome Sanger Institute for the COVID-19 Genomics UK (COG-UK) Consortium | Harper VanSteenhouse, Yumi Kasai, David Gray, Carol Clugston, Anna Dominiczak and Alex Alderton, Roberto Amato, Jeffrey Barrett, Sonia Goncalves, Ewan Harrison, David K. Jackson, Ian Johnston, Dominic Kwiatkowski, Cordelia Langford, John Sillitoe on behalf of the Wellcome Sanger Institute COVID-19 Surveillance Team |
| EPI_ISL_2850043 | Lighthouse Laboratory Plymouth | Wellcome Sanger Institute for the COVID-19 Genomics UK (COG-UK) Consortium | Lighthouse Laboratory Plymouth and Alex Alderton, Roberto Amato, Jeffrey Barrett, Sonia Goncalves, Ewan Harrison, David K. Jackson, Ian Johnston, Dominic Kwiatkowski, Cordelia Langford, John Sillitoe on behalf of the Wellcome Sanger Institute COVID-19 Surveillance Team |
| EPI_ISL_2850044, EPI_ISL_2850045, EPI_ISL_2850046, EPI_ISL_2850047, EPI_ISL_2850048, EPI_ISL_2850049 | Lighthouse Lab in Glasgow | Wellcome Sanger Institute for the COVID-19 Genomics UK (COG-UK) Consortium | Harper VanSteenhouse, Yumi Kasai, David Gray, Carol Clugston, Anna Dominiczak and Alex Alderton, Roberto Amato, Jeffrey Barrett, Sonia Goncalves, Ewan Harrison, David K. Jackson, Ian Johnston, Dominic Kwiatkowski, Cordelia Langford, John Sillitoe on behalf of the Wellcome Sanger Institute COVID-19 Surveillance Team |
| EPI_ISL_2850050 | Lighthouse Laboratory Plymouth | Wellcome Sanger Institute for the COVID-19 Genomics UK (COG-UK) Consortium | Lighthouse Laboratory Plymouth and Alex Alderton, Roberto Amato, Jeffrey Barrett, Sonia Goncalves, Ewan Harrison, David K. Jackson, Ian Johnston, Dominic Kwiatkowski, Cordelia Langford, John Sillitoe on behalf of the Wellcome Sanger Institute COVID-19 Surveillance Team |
| EPI_ISL_2850051, EPI_ISL_2850052, EPI_ISL_2850053, EPI_ISL_2850054, EPI_ISL_2850055, EPI_ISL_2850056, EPI_ISL_2850057, EPI_ISL_2850058, EPI_ISL_2850059 | Lighthouse Lab in Glasgow | Wellcome Sanger Institute for the COVID-19 Genomics UK (COG-UK) Consortium | Harper VanSteenhouse, Yumi Kasai, David Gray, Carol Clugston, Anna Dominiczak and Alex Alderton, Roberto Amato, Jeffrey Barrett, Sonia Goncalves, Ewan Harrison, David K. Jackson, Ian Johnston, Dominic Kwiatkowski, Cordelia Langford, John Sillitoe on behalf of the Wellcome Sanger Institute COVID-19 Surveillance Team |
| EPI_ISL_2850060 | Lighthouse Laboratory Plymouth | Wellcome Sanger Institute for the COVID-19 Genomics UK (COG-UK) Consortium | Lighthouse Laboratory Plymouth and Alex Alderton, Roberto Amato, Jeffrey Barrett, Sonia Goncalves, Ewan Harrison, David K. Jackson, Ian Johnston, Dominic Kwiatkowski, Cordelia Langford, John Sillitoe on behalf of the Wellcome Sanger Institute COVID-19 Surveillance Team |
| EPI_ISL_2850061, EPI_ISL_2850062 | Lighthouse Lab in Glasgow | Wellcome Sanger Institute for the COVID-19 Genomics UK (COG-UK) Consortium | Harper VanSteenhouse, Yumi Kasai, David Gray, Carol Clugston, Anna Dominiczak and Alex Alderton, Roberto Amato, Jeffrey Barrett, Sonia Goncalves, Ewan Harrison, David K. Jackson, Ian Johnston, Dominic Kwiatkowski, Cordelia Langford, John Sillitoe on behalf of the Wellcome Sanger Institute COVID-19 Surveillance Team |
| EPI_ISL_2850064, EPI_ISL_2850065 | Lighthouse Laboratory Plymouth | Wellcome Sanger Institute for the COVID-19 Genomics UK (COG-UK) Consortium | Lighthouse Laboratory Plymouth and Alex Alderton, Roberto Amato, Jeffrey Barrett, Sonia Goncalves, Ewan Harrison, David K. Jackson, Ian Johnston, Dominic Kwiatkowski, Cordelia Langford, John Sillitoe on behalf of the Wellcome Sanger Institute COVID-19 Surveillance Team |
| EPI_ISL_2850066, EPI_ISL_2850067, EPI_ISL_2850068 | Lighthouse Lab in Glasgow | Wellcome Sanger Institute for the COVID-19 Genomics UK (COG-UK) Consortium | Harper VanSteenhouse, Yumi Kasai, David Gray, Carol Clugston, Anna Dominiczak and Alex Alderton, Roberto Amato, Jeffrey Barrett, Sonia Goncalves, Ewan Harrison, David K. Jackson, Ian Johnston, Dominic Kwiatkowski, Cordelia Langford, John Sillitoe on behalf of the Wellcome Sanger Institute COVID-19 Surveillance Team |
| EPI_ISL_2850069, EPI_ISL_2850070 | Lighthouse Laboratory Plymouth | Wellcome Sanger Institute for the COVID-19 Genomics UK (COG-UK) Consortium | Lighthouse Laboratory Plymouth and Alex Alderton, Roberto Amato, Jeffrey Barrett, Sonia Goncalves, Ewan Harrison, David K. Jackson, Ian Johnston, Dominic Kwiatkowski, Cordelia Langford, John Sillitoe on behalf of the Wellcome Sanger Institute COVID-19 Surveillance Team |
| EPI_ISL_2850071, EPI_ISL_2850072, EPI_ISL_2850073, EPI_ISL_2850074 | Lighthouse Lab in Glasgow | Wellcome Sanger Institute for the COVID-19 Genomics UK (COG-UK) Consortium | Harper VanSteenhouse, Yumi Kasai, David Gray, Carol Clugston, Anna Dominiczak and Alex Alderton, Roberto Amato, Jeffrey Barrett, Sonia Goncalves, Ewan Harrison, David K. Jackson, Ian Johnston, Dominic Kwiatkowski, Cordelia Langford, John Sillitoe on behalf of the Wellcome Sanger Institute COVID-19 Surveillance Team |
| EPI_ISL_2850075 | Lighthouse Laboratory Plymouth | Wellcome Sanger Institute for the COVID-19 Genomics UK (COG-UK) Consortium | Lighthouse Laboratory Plymouth and Alex Alderton, Roberto Amato, Jeffrey Barrett, Sonia Goncalves, Ewan Harrison, David K. Jackson, Ian Johnston, Dominic Kwiatkowski, Cordelia Langford, John Sillitoe on behalf of the Wellcome Sanger Institute COVID-19 Surveillance Team |
| EPI_ISL_2850076 | Lighthouse Lab in Glasgow | Wellcome Sanger Institute for the COVID-19 Genomics UK (COG-UK) Consortium | Harper VanSteenhouse, Yumi Kasai, David Gray, Carol Clugston, Anna Dominiczak and Alex Alderton, Roberto Amato, Jeffrey Barrett, Sonia Goncalves, Ewan Harrison, David K. Jackson, Ian Johnston, Dominic Kwiatkowski, Cordelia Langford, John Sillitoe on behalf of the Wellcome Sanger Institute COVID-19 Surveillance Team |
| EPI_ISL_2850077 | Lighthouse Laboratory Plymouth | Wellcome Sanger Institute for the COVID-19 Genomics UK (COG-UK) Consortium | Lighthouse Laboratory Plymouth and Alex Alderton, Roberto Amato, Jeffrey Barrett, Sonia Goncalves, Ewan Harrison, David K. Jackson, Ian Johnston, Dominic Kwiatkowski, Cordelia Langford, John Sillitoe on behalf of the Wellcome Sanger Institute COVID-19 Surveillance Team |
| EPI_ISL_2850078 | Lighthouse Lab in Glasgow | Wellcome Sanger Institute for the COVID-19 Genomics UK (COG-UK) Consortium | Harper VanSteenhouse, Yumi Kasai, David Gray, Carol Clugston, Anna Dominiczak and Alex Alderton, Roberto Amato, Jeffrey Barrett, Sonia Goncalves, Ewan Harrison, David K. Jackson, Ian Johnston, Dominic Kwiatkowski, Cordelia Langford, John Sillitoe on behalf of the Wellcome Sanger Institute COVID-19 Surveillance Team |
| EPI_ISL_2850079 | Lighthouse Laboratory Plymouth | Wellcome Sanger Institute for the COVID-19 Genomics UK (COG-UK) Consortium | Lighthouse Laboratory Plymouth and Alex Alderton, Roberto Amato, Jeffrey Barrett, Sonia Goncalves, Ewan Harrison, David K. Jackson, Ian Johnston, Dominic Kwiatkowski, Cordelia Langford, John Sillitoe on behalf of the Wellcome Sanger Institute COVID-19 Surveillance Team |
| EPI_ISL_2850080 | Lighthouse Lab in Glasgow | Wellcome Sanger Institute for the COVID-19 Genomics UK (COG-UK) Consortium | Harper VanSteenhouse, Yumi Kasai, David Gray, Carol Clugston, Anna Dominiczak and Alex Alderton, Roberto Amato, Jeffrey Barrett, Sonia Goncalves, Ewan Harrison, David K. Jackson, Ian Johnston, Dominic Kwiatkowski, Cordelia Langford, John Sillitoe on behalf of the Wellcome Sanger Institute COVID-19 Surveillance Team |
| EPI_ISL_2850081 | Lighthouse Laboratory Plymouth | Wellcome Sanger Institute for the COVID-19 Genomics UK (COG-UK) Consortium | Lighthouse Laboratory Plymouth and Alex Alderton, Roberto Amato, Jeffrey Barrett, Sonia Goncalves, Ewan Harrison, David K. Jackson, Ian Johnston, Dominic Kwiatkowski, Cordelia Langford, John Sillitoe on behalf of the Wellcome Sanger Institute COVID-19 Surveillance Team |
| EPI_ISL_2850082, EPI_ISL_2850083, EPI_ISL_2850084, EPI_ISL_2850085, EPI_ISL_2850086, EPI_ISL_2850087, EPI_ISL_2850088 | Lighthouse Lab in Glasgow | Wellcome Sanger Institute for the COVID-19 Genomics UK (COG-UK) Consortium | Harper VanSteenhouse, Yumi Kasai, David Gray, Carol Clugston, Anna Dominiczak and Alex Alderton, Roberto Amato, Jeffrey Barrett, Sonia Goncalves, Ewan Harrison, David K. Jackson, Ian Johnston, Dominic Kwiatkowski, Cordelia Langford, John Sillitoe on behalf of the Wellcome Sanger Institute COVID-19 Surveillance Team |
| EPI_ISL_2850089, EPI_ISL_2850090 | Lighthouse Laboratory Plymouth | Wellcome Sanger Institute for the COVID-19 Genomics UK (COG-UK) Consortium | Lighthouse Laboratory Plymouth and Alex Alderton, Roberto Amato, Jeffrey Barrett, Sonia Goncalves, Ewan Harrison, David K. Jackson, Ian Johnston, Dominic Kwiatkowski, Cordelia Langford, John Sillitoe on behalf of the Wellcome Sanger Institute COVID-19 Surveillance Team |
| EPI_ISL_2850091 | Lighthouse Lab in Glasgow | Wellcome Sanger Institute for the COVID-19 Genomics UK (COG-UK) Consortium | Harper VanSteenhouse, Yumi Kasai, David Gray, Carol Clugston, Anna Dominiczak and Alex Alderton, Roberto Amato, Jeffrey Barrett, Sonia Goncalves, Ewan Harrison, David K. Jackson, Ian Johnston, Dominic Kwiatkowski, Cordelia Langford, John Sillitoe on behalf of the Wellcome Sanger Institute COVID-19 Surveillance Team |
| EPI_ISL_2850092 | Lighthouse Laboratory Plymouth | Wellcome Sanger Institute for the COVID-19 Genomics UK (COG-UK) | Lighthouse Laboratory Plymouth and Alex Alderton, Roberto Amato, Jeffrey Barrett, Sonia Goncalves, Ewan Harrison, David K. Jackson, Ian Johnston, Dominic Kwiatkowski, Cordelia Langford, John Sillitoe on behalf of the Wellcome Sanger |

| | | | |
|---|---|---|---|
| | | Consortium | Institute COVID-19 Surveillance Team |
| EPI_ISL_2850093, EPI_ISL_2850094, EPI_ISL_2850095, EPI_ISL_2850096, EPI_ISL_2850097, EPI_ISL_2850098 | Lighthouse Lab in Glasgow | Wellcome Sanger Institute for the COVID-19 Genomics UK (COG-UK) Consortium | Harper VanSteenhouse, Yumi Kasai, David Gray, Carol Clugston, Anna Dominiczak and Alex Alderton, Roberto Amato, Jeffrey Barrett, Sonia Goncalves, Ewan Harrison, David K. Jackson, Ian Johnston, Dominic Kwiatkowski, Cordelia Langford, John Sillitoe on behalf of the Wellcome Sanger Institute COVID-19 Surveillance Team |
| EPI_ISL_2850099 | Lighthouse Laboratory Plymouth | Wellcome Sanger Institute for the COVID-19 Genomics UK (COG-UK) Consortium | Lighthouse Laboratory Plymouth and Alex Alderton, Roberto Amato, Jeffrey Barrett, Sonia Goncalves, Ewan Harrison, David K. Jackson, Ian Johnston, Dominic Kwiatkowski, Cordelia Langford, John Sillitoe on behalf of the Wellcome Sanger Institute COVID-19 Surveillance Team |
| EPI_ISL_2850100 | Lighthouse Lab in Glasgow | Wellcome Sanger Institute for the COVID-19 Genomics UK (COG-UK) Consortium | Harper VanSteenhouse, Yumi Kasai, David Gray, Carol Clugston, Anna Dominiczak and Alex Alderton, Roberto Amato, Jeffrey Barrett, Sonia Goncalves, Ewan Harrison, David K. Jackson, Ian Johnston, Dominic Kwiatkowski, Cordelia Langford, John Sillitoe on behalf of the Wellcome Sanger Institute COVID-19 Surveillance Team |
| EPI_ISL_2850101, EPI_ISL_2850102 | Lighthouse Laboratory Plymouth | Wellcome Sanger Institute for the COVID-19 Genomics UK (COG-UK) Consortium | Lighthouse Laboratory Plymouth and Alex Alderton, Roberto Amato, Jeffrey Barrett, Sonia Goncalves, Ewan Harrison, David K. Jackson, Ian Johnston, Dominic Kwiatkowski, Cordelia Langford, John Sillitoe on behalf of the Wellcome Sanger Institute COVID-19 Surveillance Team |
| EPI_ISL_2850103 | Lighthouse Lab in Glasgow | Wellcome Sanger Institute for the COVID-19 Genomics UK (COG-UK) Consortium | Harper VanSteenhouse, Yumi Kasai, David Gray, Carol Clugston, Anna Dominiczak and Alex Alderton, Roberto Amato, Jeffrey Barrett, Sonia Goncalves, Ewan Harrison, David K. Jackson, Ian Johnston, Dominic Kwiatkowski, Cordelia Langford, John Sillitoe on behalf of the Wellcome Sanger Institute COVID-19 Surveillance Team |
| EPI_ISL_2850104, EPI_ISL_2850105 | Lighthouse Laboratory Plymouth | Wellcome Sanger Institute for the COVID-19 Genomics UK (COG-UK) Consortium | Lighthouse Laboratory Plymouth and Alex Alderton, Roberto Amato, Jeffrey Barrett, Sonia Goncalves, Ewan Harrison, David K. Jackson, Ian Johnston, Dominic Kwiatkowski, Cordelia Langford, John Sillitoe on behalf of the Wellcome Sanger Institute COVID-19 Surveillance Team |
| EPI_ISL_2850106, EPI_ISL_2850107, EPI_ISL_2850108, EPI_ISL_2850109, EPI_ISL_2850110, EPI_ISL_2850111, EPI_ISL_2850112, EPI_ISL_2850113 | Lighthouse Lab in Glasgow | Wellcome Sanger Institute for the COVID-19 Genomics UK (COG-UK) Consortium | Harper VanSteenhouse, Yumi Kasai, David Gray, Carol Clugston, Anna Dominiczak and Alex Alderton, Roberto Amato, Jeffrey Barrett, Sonia Goncalves, Ewan Harrison, David K. Jackson, Ian Johnston, Dominic Kwiatkowski, Cordelia Langford, John Sillitoe on behalf of the Wellcome Sanger Institute COVID-19 Surveillance Team |
| EPI_ISL_2850114 | Lighthouse Laboratory Plymouth | Wellcome Sanger Institute for the COVID-19 Genomics UK (COG-UK) Consortium | Lighthouse Laboratory Plymouth and Alex Alderton, Roberto Amato, Jeffrey Barrett, Sonia Goncalves, Ewan Harrison, David K. Jackson, Ian Johnston, Dominic Kwiatkowski, Cordelia Langford, John Sillitoe on behalf of the Wellcome Sanger Institute COVID-19 Surveillance Team |
| EPI_ISL_2850115, EPI_ISL_2850116, EPI_ISL_2850117 | Lighthouse Lab in Glasgow | Wellcome Sanger Institute for the COVID-19 Genomics UK (COG-UK) Consortium | Harper VanSteenhouse, Yumi Kasai, David Gray, Carol Clugston, Anna Dominiczak and Alex Alderton, Roberto Amato, Jeffrey Barrett, Sonia Goncalves, Ewan Harrison, David K. Jackson, Ian Johnston, Dominic Kwiatkowski, Cordelia Langford, John Sillitoe on behalf of the Wellcome Sanger Institute COVID-19 Surveillance Team |
| EPI_ISL_2850118 | Lighthouse Laboratory Plymouth | Wellcome Sanger Institute for the COVID-19 Genomics UK (COG-UK) Consortium | Lighthouse Laboratory Plymouth and Alex Alderton, Roberto Amato, Jeffrey Barrett, Sonia Goncalves, Ewan Harrison, David K. Jackson, Ian Johnston, Dominic Kwiatkowski, Cordelia Langford, John Sillitoe on behalf of the Wellcome Sanger Institute COVID-19 Surveillance Team |
| EPI_ISL_2850119 | Lighthouse Lab in Glasgow | Wellcome Sanger Institute for the COVID-19 Genomics UK (COG-UK) Consortium | Harper VanSteenhouse, Yumi Kasai, David Gray, Carol Clugston, Anna Dominiczak and Alex Alderton, Roberto Amato, Jeffrey Barrett, Sonia Goncalves, Ewan Harrison, David K. Jackson, Ian Johnston, Dominic Kwiatkowski, Cordelia Langford, John Sillitoe on behalf of the Wellcome Sanger Institute COVID-19 Surveillance Team |
| EPI_ISL_2850120 | Lighthouse Laboratory Plymouth | Wellcome Sanger Institute for the COVID-19 Genomics UK (COG-UK) Consortium | Lighthouse Laboratory Plymouth and Alex Alderton, Roberto Amato, Jeffrey Barrett, Sonia Goncalves, Ewan Harrison, David K. Jackson, Ian Johnston, Dominic Kwiatkowski, Cordelia Langford, John Sillitoe on behalf of the Wellcome Sanger Institute COVID-19 Surveillance Team |
| EPI_ISL_2850121 | Lighthouse Lab in Glasgow | Wellcome Sanger Institute for the COVID-19 Genomics UK (COG-UK) Consortium | Harper VanSteenhouse, Yumi Kasai, David Gray, Carol Clugston, Anna Dominiczak and Alex Alderton, Roberto Amato, Jeffrey Barrett, Sonia Goncalves, Ewan Harrison, David K. Jackson, Ian Johnston, Dominic Kwiatkowski, Cordelia Langford, John Sillitoe on behalf of the Wellcome Sanger Institute COVID-19 Surveillance Team |
| EPI_ISL_2850122, EPI_ISL_2850123 | Lighthouse Laboratory Plymouth | Wellcome Sanger Institute for the COVID-19 Genomics UK (COG-UK) Consortium | Lighthouse Laboratory Plymouth and Alex Alderton, Roberto Amato, Jeffrey Barrett, Sonia Goncalves, Ewan Harrison, David K. Jackson, Ian Johnston, Dominic Kwiatkowski, Cordelia Langford, John Sillitoe on behalf of the Wellcome Sanger Institute COVID-19 Surveillance Team |
| EPI_ISL_2850124, EPI_ISL_2850125, EPI_ISL_2850126, EPI_ISL_2850127, EPI_ISL_2850128, EPI_ISL_2850129, EPI_ISL_2850130 | Lighthouse Lab in Glasgow | Wellcome Sanger Institute for the COVID-19 Genomics UK (COG-UK) Consortium | Harper VanSteenhouse, Yumi Kasai, David Gray, Carol Clugston, Anna Dominiczak and Alex Alderton, Roberto Amato, Jeffrey Barrett, Sonia Goncalves, Ewan Harrison, David K. Jackson, Ian Johnston, Dominic Kwiatkowski, Cordelia Langford, John Sillitoe on behalf of the Wellcome Sanger Institute COVID-19 Surveillance Team |
| EPI_ISL_2850131 | Lighthouse Laboratory Plymouth | Wellcome Sanger Institute for the COVID-19 Genomics UK (COG-UK) Consortium | Lighthouse Laboratory Plymouth and Alex Alderton, Roberto Amato, Jeffrey Barrett, Sonia Goncalves, Ewan Harrison, David K. Jackson, Ian Johnston, Dominic Kwiatkowski, Cordelia Langford, John Sillitoe on behalf of the Wellcome Sanger Institute COVID-19 Surveillance Team |
| EPI_ISL_2850132, EPI_ISL_2850133, EPI_ISL_2850134 | Lighthouse Lab in Glasgow | Wellcome Sanger Institute for the COVID-19 Genomics UK (COG-UK) Consortium | Harper VanSteenhouse, Yumi Kasai, David Gray, Carol Clugston, Anna Dominiczak and Alex Alderton, Roberto Amato, Jeffrey Barrett, Sonia Goncalves, Ewan Harrison, David K. Jackson, Ian Johnston, Dominic Kwiatkowski, Cordelia Langford, John Sillitoe on behalf of the Wellcome Sanger Institute COVID-19 Surveillance Team |
| EPI_ISL_2850135, EPI_ISL_2850136 | Lighthouse Laboratory Plymouth | Wellcome Sanger Institute for the COVID-19 Genomics UK (COG-UK) Consortium | Lighthouse Laboratory Plymouth and Alex Alderton, Roberto Amato, Jeffrey Barrett, Sonia Goncalves, Ewan Harrison, David K. Jackson, Ian Johnston, Dominic Kwiatkowski, Cordelia Langford, John Sillitoe on behalf of the Wellcome Sanger Institute COVID-19 Surveillance Team |
| EPI_ISL_2850137, EPI_ISL_2850138 | Lighthouse Lab in Glasgow | Wellcome Sanger Institute for the COVID-19 Genomics UK (COG-UK) Consortium | Harper VanSteenhouse, Yumi Kasai, David Gray, Carol Clugston, Anna Dominiczak and Alex Alderton, Roberto Amato, Jeffrey Barrett, Sonia Goncalves, Ewan Harrison, David K. Jackson, Ian Johnston, Dominic Kwiatkowski, Cordelia Langford, John Sillitoe on behalf of the Wellcome Sanger Institute COVID-19 Surveillance Team |
| EPI_ISL_2850139, EPI_ISL_2850140 | Lighthouse Laboratory Plymouth | Wellcome Sanger Institute for the COVID-19 Genomics UK (COG-UK) Consortium | Lighthouse Laboratory Plymouth and Alex Alderton, Roberto Amato, Jeffrey Barrett, Sonia Goncalves, Ewan Harrison, David K. Jackson, Ian Johnston, Dominic Kwiatkowski, Cordelia Langford, John Sillitoe on behalf of the Wellcome Sanger Institute COVID-19 Surveillance Team |
| EPI_ISL_2850141, EPI_ISL_2850142, EPI_ISL_2850143, EPI_ISL_2850144, EPI_ISL_2850145 | Lighthouse Lab in Glasgow | Wellcome Sanger Institute for the COVID-19 Genomics UK (COG-UK) Consortium | Harper VanSteenhouse, Yumi Kasai, David Gray, Carol Clugston, Anna Dominiczak and Alex Alderton, Roberto Amato, Jeffrey Barrett, Sonia Goncalves, Ewan Harrison, David K. Jackson, Ian Johnston, Dominic Kwiatkowski, Cordelia Langford, John Sillitoe on behalf of the Wellcome Sanger Institute COVID-19 Surveillance Team |
| EPI_ISL_2850146 | Lighthouse Laboratory Plymouth | Wellcome Sanger Institute for the COVID-19 Genomics UK (COG-UK) Consortium | Lighthouse Laboratory Plymouth and Alex Alderton, Roberto Amato, Jeffrey Barrett, Sonia Goncalves, Ewan Harrison, David K. Jackson, Ian Johnston, Dominic Kwiatkowski, Cordelia Langford, John Sillitoe on behalf of the Wellcome Sanger Institute COVID-19 Surveillance Team |
| EPI_ISL_2850147, EPI_ISL_2850148, EPI_ISL_2850149 | Lighthouse Lab in Glasgow | Wellcome Sanger Institute for the COVID-19 Genomics UK (COG-UK) Consortium | Harper VanSteenhouse, Yumi Kasai, David Gray, Carol Clugston, Anna Dominiczak and Alex Alderton, Roberto Amato, Jeffrey Barrett, Sonia Goncalves, Ewan Harrison, David K. Jackson, Ian Johnston, Dominic Kwiatkowski, Cordelia Langford, John Sillitoe on behalf of the Wellcome Sanger Institute COVID-19 Surveillance Team |
| EPI_ISL_2850150 | Lighthouse Laboratory Plymouth | Wellcome Sanger Institute for the COVID-19 Genomics UK (COG-UK) Consortium | Lighthouse Laboratory Plymouth and Alex Alderton, Roberto Amato, Jeffrey Barrett, Sonia Goncalves, Ewan Harrison, David K. Jackson, Ian Johnston, Dominic Kwiatkowski, Cordelia Langford, John Sillitoe on behalf of the Wellcome Sanger Institute COVID-19 Surveillance Team |
| EPI_ISL_2850151, EPI_ISL_2850152, EPI_ISL_2850153, EPI_ISL_2850154, EPI_ISL_2850155, EPI_ISL_2850156, EPI_ISL_2850157, EPI_ISL_2850158, EPI_ISL_2850159, EPI_ISL_2850160, EPI_ISL_2850161, EPI_ISL_2850162, EPI_ISL_2850163, EPI_ISL_2850164, EPI_ISL_2850165, EPI_ISL_2850166, EPI_ISL_2850167 | | | |
| see above | Lighthouse Lab in Glasgow | Wellcome Sanger Institute for the COVID-19 Genomics UK (COG-UK) Consortium | Harper VanSteenhouse, Yumi Kasai, David Gray, Carol Clugston, Anna Dominiczak and Alex Alderton, Roberto Amato, Jeffrey Barrett, Sonia Goncalves, Ewan Harrison, David K. Jackson, Ian Johnston, Dominic Kwiatkowski, Cordelia Langford, John Sillitoe on behalf of the Wellcome Sanger Institute COVID-19 Surveillance Team |
| EPI_ISL_2850168 | Lighthouse Laboratory Plymouth | Wellcome Sanger Institute for the COVID-19 Genomics UK (COG-UK) Consortium | Lighthouse Laboratory Plymouth and Alex Alderton, Roberto Amato, Jeffrey Barrett, Sonia Goncalves, Ewan Harrison, David K. Jackson, Ian Johnston, Dominic Kwiatkowski, Cordelia Langford, John Sillitoe on behalf of the Wellcome Sanger Institute COVID-19 Surveillance Team |
| EPI_ISL_2850169, EPI_ISL_2850170, EPI_ISL_2850171, EPI_ISL_2850172, EPI_ISL_2850173, EPI_ISL_2850174, EPI_ISL_2850175 | Lighthouse Lab in Glasgow | Wellcome Sanger Institute for the COVID-19 Genomics UK (COG-UK) Consortium | Harper VanSteenhouse, Yumi Kasai, David Gray, Carol Clugston, Anna Dominiczak and Alex Alderton, Roberto Amato, Jeffrey Barrett, Sonia Goncalves, Ewan Harrison, David K. Jackson, Ian Johnston, Dominic Kwiatkowski, Cordelia Langford, John Sillitoe on behalf of the Wellcome Sanger Institute COVID-19 Surveillance Team |
| EPI_ISL_2850176 | Lighthouse Laboratory Plymouth | Wellcome Sanger Institute for the COVID-19 Genomics UK (COG-UK) Consortium | Lighthouse Laboratory Plymouth and Alex Alderton, Roberto Amato, Jeffrey Barrett, Sonia Goncalves, Ewan Harrison, David K. Jackson, Ian Johnston, Dominic Kwiatkowski, Cordelia Langford, John Sillitoe on behalf of the Wellcome Sanger Institute COVID-19 Surveillance Team |
| EPI_ISL_2850177 | Lighthouse Lab in Glasgow | Wellcome Sanger Institute for the COVID-19 Genomics UK (COG-UK) Consortium | Harper VanSteenhouse, Yumi Kasai, David Gray, Carol Clugston, Anna Dominiczak and Alex Alderton, Roberto Amato, Jeffrey Barrett, Sonia Goncalves, Ewan Harrison, David K. Jackson, Ian Johnston, Dominic Kwiatkowski, Cordelia Langford, John Sillitoe on behalf of the Wellcome Sanger Institute COVID-19 Surveillance Team |
| EPI_ISL_2850178 | Lighthouse Laboratory Plymouth | Wellcome Sanger Institute for the COVID-19 Genomics UK (COG-UK) Consortium | Lighthouse Laboratory Plymouth and Alex Alderton, Roberto Amato, Jeffrey Barrett, Sonia Goncalves, Ewan Harrison, David K. Jackson, Ian Johnston, Dominic Kwiatkowski, Cordelia Langford, John Sillitoe on behalf of the Wellcome Sanger Institute COVID-19 Surveillance Team |
| EPI_ISL_2850179, EPI_ISL_2850180 | Lighthouse Lab in Glasgow | Wellcome Sanger Institute for the COVID-19 Genomics UK (COG-UK) Consortium | Harper VanSteenhouse, Yumi Kasai, David Gray, Carol Clugston, Anna Dominiczak and Alex Alderton, Roberto Amato, Jeffrey Barrett, Sonia Goncalves, Ewan Harrison, David K. Jackson, Ian Johnston, Dominic Kwiatkowski, Cordelia Langford, John Sillitoe on behalf of the Wellcome Sanger Institute COVID-19 Surveillance Team |
| EPI_ISL_2850181, EPI_ISL_2850182 | Lighthouse Laboratory Plymouth | Wellcome Sanger Institute for the COVID-19 Genomics UK (COG-UK) Consortium | Lighthouse Laboratory Plymouth and Alex Alderton, Roberto Amato, Jeffrey Barrett, Sonia Goncalves, Ewan Harrison, David K. Jackson, Ian Johnston, Dominic Kwiatkowski, Cordelia Langford, John Sillitoe on behalf of the Wellcome Sanger Institute COVID-19 Surveillance Team |
| EPI_ISL_2850183, EPI_ISL_2850184, EPI_ISL_2850185, | Lighthouse Lab in Glasgow | Wellcome Sanger Institute for the COVID-19 Genomics UK (COG-UK) Consortium | Harper VanSteenhouse, Yumi Kasai, David Gray, Carol Clugston, Anna Dominiczak and Alex Alderton, Roberto Amato, Jeffrey Barrett, Sonia Goncalves, Ewan Harrison, David K. Jackson, Ian Johnston, Dominic Kwiatkowski, Cordelia Langford, John Sillitoe on behalf of the Wellcome Sanger Institute COVID-19 Surveillance Team |

| | | | |
|---|---|---|---|
| EPI_ISL_2850186 | | | |
| EPI_ISL_2850187 | Lighthouse Laboratory Plymouth | Wellcome Sanger Institute for the COVID-19 Genomics UK (COG-UK) Consortium | Lighthouse Laboratory Plymouth and Alex Alderton, Roberto Amato, Jeffrey Barrett, Sonia Goncalves, Ewan Harrison, David K. Jackson, Ian Johnston, Dominic Kwiatkowski, Cordelia Langford, John Sillitoe on behalf of the Wellcome Sanger Institute COVID-19 Surveillance Team |
| EPI_ISL_2850188, EPI_ISL_2850189, EPI_ISL_2850190, EPI_ISL_2850191, EPI_ISL_2850192 | Lighthouse Lab in Glasgow | Wellcome Sanger Institute for the COVID-19 Genomics UK (COG-UK) Consortium | Harper VanSteenhouse, Yumi Kasai, David Gray, Carol Clugston, Anna Dominiczak and Alex Alderton, Roberto Amato, Jeffrey Barrett, Sonia Goncalves, Ewan Harrison, David K. Jackson, Ian Johnston, Dominic Kwiatkowski, Cordelia Langford, John Sillitoe on behalf of the Wellcome Sanger Institute COVID-19 Surveillance Team |
| EPI_ISL_2850193 | AZ Klina | AZ Klina | Carl Vael - Lynsey Berckmans |
| EPI_ISL_2850194, EPI_ISL_2850195, EPI_ISL_2850196, EPI_ISL_2850197 | Lighthouse Lab in Glasgow | Wellcome Sanger Institute for the COVID-19 Genomics UK (COG-UK) Consortium | Harper VanSteenhouse, Yumi Kasai, David Gray, Carol Clugston, Anna Dominiczak and Alex Alderton, Roberto Amato, Jeffrey Barrett, Sonia Goncalves, Ewan Harrison, David K. Jackson, Ian Johnston, Dominic Kwiatkowski, Cordelia Langford, John Sillitoe on behalf of the Wellcome Sanger Institute COVID-19 Surveillance Team |
| EPI_ISL_2850198 | Lighthouse Laboratory Plymouth | Wellcome Sanger Institute for the COVID-19 Genomics UK (COG-UK) Consortium | Lighthouse Laboratory Plymouth and Alex Alderton, Roberto Amato, Jeffrey Barrett, Sonia Goncalves, Ewan Harrison, David K. Jackson, Ian Johnston, Dominic Kwiatkowski, Cordelia Langford, John Sillitoe on behalf of the Wellcome Sanger Institute COVID-19 Surveillance Team |
| EPI_ISL_2850199, EPI_ISL_2850200, EPI_ISL_2850201, EPI_ISL_2850202, EPI_ISL_2850203, EPI_ISL_2850204, EPI_ISL_2850205, EPI_ISL_2850206, EPI_ISL_2850207, EPI_ISL_2850208, EPI_ISL_2850209, EPI_ISL_2850210, EPI_ISL_2850211, EPI_ISL_2850212 | | | |
| see above | Lighthouse Lab in Glasgow | Wellcome Sanger Institute for the COVID-19 Genomics UK (COG-UK) Consortium | Harper VanSteenhouse, Yumi Kasai, David Gray, Carol Clugston, Anna Dominiczak and Alex Alderton, Roberto Amato, Jeffrey Barrett, Sonia Goncalves, Ewan Harrison, David K. Jackson, Ian Johnston, Dominic Kwiatkowski, Cordelia Langford, John Sillitoe on behalf of the Wellcome Sanger Institute COVID-19 Surveillance Team |
| EPI_ISL_2850213 | Lighthouse Laboratory Plymouth | Wellcome Sanger Institute for the COVID-19 Genomics UK (COG-UK) Consortium | Lighthouse Laboratory Plymouth and Alex Alderton, Roberto Amato, Jeffrey Barrett, Sonia Goncalves, Ewan Harrison, David K. Jackson, Ian Johnston, Dominic Kwiatkowski, Cordelia Langford, John Sillitoe on behalf of the Wellcome Sanger Institute COVID-19 Surveillance Team |
| EPI_ISL_2850214, EPI_ISL_2850215, EPI_ISL_2850216, EPI_ISL_2850217 | Lighthouse Lab in Glasgow | Wellcome Sanger Institute for the COVID-19 Genomics UK (COG-UK) Consortium | Harper VanSteenhouse, Yumi Kasai, David Gray, Carol Clugston, Anna Dominiczak and Alex Alderton, Roberto Amato, Jeffrey Barrett, Sonia Goncalves, Ewan Harrison, David K. Jackson, Ian Johnston, Dominic Kwiatkowski, Cordelia Langford, John Sillitoe on behalf of the Wellcome Sanger Institute COVID-19 Surveillance Team |
| EPI_ISL_2850218 | Lighthouse Laboratory Plymouth | Wellcome Sanger Institute for the COVID-19 Genomics UK (COG-UK) Consortium | Lighthouse Laboratory Plymouth and Alex Alderton, Roberto Amato, Jeffrey Barrett, Sonia Goncalves, Ewan Harrison, David K. Jackson, Ian Johnston, Dominic Kwiatkowski, Cordelia Langford, John Sillitoe on behalf of the Wellcome Sanger Institute COVID-19 Surveillance Team |
| EPI_ISL_2850219, EPI_ISL_2850220 | Lighthouse Lab in Glasgow | Wellcome Sanger Institute for the COVID-19 Genomics UK (COG-UK) Consortium | Harper VanSteenhouse, Yumi Kasai, David Gray, Carol Clugston, Anna Dominiczak and Alex Alderton, Roberto Amato, Jeffrey Barrett, Sonia Goncalves, Ewan Harrison, David K. Jackson, Ian Johnston, Dominic Kwiatkowski, Cordelia Langford, John Sillitoe on behalf of the Wellcome Sanger Institute COVID-19 Surveillance Team |
| EPI_ISL_2850221 | Lighthouse Laboratory Plymouth | Wellcome Sanger Institute for the COVID-19 Genomics UK (COG-UK) Consortium | Lighthouse Laboratory Plymouth and Alex Alderton, Roberto Amato, Jeffrey Barrett, Sonia Goncalves, Ewan Harrison, David K. Jackson, Ian Johnston, Dominic Kwiatkowski, Cordelia Langford, John Sillitoe on behalf of the Wellcome Sanger Institute COVID-19 Surveillance Team |
| EPI_ISL_2850222, EPI_ISL_2850223, EPI_ISL_2850224, EPI_ISL_2850225, EPI_ISL_2850226, EPI_ISL_2850227, EPI_ISL_2850228, EPI_ISL_2850229 | Lighthouse Lab in Glasgow | Wellcome Sanger Institute for the COVID-19 Genomics UK (COG-UK) Consortium | Harper VanSteenhouse, Yumi Kasai, David Gray, Carol Clugston, Anna Dominiczak and Alex Alderton, Roberto Amato, Jeffrey Barrett, Sonia Goncalves, Ewan Harrison, David K. Jackson, Ian Johnston, Dominic Kwiatkowski, Cordelia Langford, John Sillitoe on behalf of the Wellcome Sanger Institute COVID-19 Surveillance Team |
| EPI_ISL_2850230, EPI_ISL_2850231, EPI_ISL_2850232 | Lighthouse Laboratory Plymouth | Wellcome Sanger Institute for the COVID-19 Genomics UK (COG-UK) Consortium | Lighthouse Laboratory Plymouth and Alex Alderton, Roberto Amato, Jeffrey Barrett, Sonia Goncalves, Ewan Harrison, David K. Jackson, Ian Johnston, Dominic Kwiatkowski, Cordelia Langford, John Sillitoe on behalf of the Wellcome Sanger Institute COVID-19 Surveillance Team |
| EPI_ISL_2850233, EPI_ISL_2850234, EPI_ISL_2850235, EPI_ISL_2850236, EPI_ISL_2850237, EPI_ISL_2850238, EPI_ISL_2850239, EPI_ISL_2850240, EPI_ISL_2850241 | Lighthouse Lab in Glasgow | Wellcome Sanger Institute for the COVID-19 Genomics UK (COG-UK) Consortium | Harper VanSteenhouse, Yumi Kasai, David Gray, Carol Clugston, Anna Dominiczak and Alex Alderton, Roberto Amato, Jeffrey Barrett, Sonia Goncalves, Ewan Harrison, David K. Jackson, Ian Johnston, Dominic Kwiatkowski, Cordelia Langford, John Sillitoe on behalf of the Wellcome Sanger Institute COVID-19 Surveillance Team |
| EPI_ISL_2850242 | Lighthouse Laboratory Plymouth | Wellcome Sanger Institute for the COVID-19 Genomics UK (COG-UK) Consortium | Lighthouse Laboratory Plymouth and Alex Alderton, Roberto Amato, Jeffrey Barrett, Sonia Goncalves, Ewan Harrison, David K. Jackson, Ian Johnston, Dominic Kwiatkowski, Cordelia Langford, John Sillitoe on behalf of the Wellcome Sanger Institute COVID-19 Surveillance Team |
| EPI_ISL_2850243 | Lighthouse Lab in Glasgow | Wellcome Sanger Institute for the COVID-19 Genomics UK (COG-UK) Consortium | Harper VanSteenhouse, Yumi Kasai, David Gray, Carol Clugston, Anna Dominiczak and Alex Alderton, Roberto Amato, Jeffrey Barrett, Sonia Goncalves, Ewan Harrison, David K. Jackson, Ian Johnston, Dominic Kwiatkowski, Cordelia Langford, John Sillitoe on behalf of the Wellcome Sanger Institute COVID-19 Surveillance Team |
| EPI_ISL_2850244 | Lighthouse Laboratory Plymouth | Wellcome Sanger Institute for the COVID-19 Genomics UK (COG-UK) Consortium | Lighthouse Laboratory Plymouth and Alex Alderton, Roberto Amato, Jeffrey Barrett, Sonia Goncalves, Ewan Harrison, David K. Jackson, Ian Johnston, Dominic Kwiatkowski, Cordelia Langford, John Sillitoe on behalf of the Wellcome Sanger Institute COVID-19 Surveillance Team |
| EPI_ISL_2850245 | Lighthouse Lab in Glasgow | Wellcome Sanger Institute for the COVID-19 Genomics UK (COG-UK) Consortium | Harper VanSteenhouse, Yumi Kasai, David Gray, Carol Clugston, Anna Dominiczak and Alex Alderton, Roberto Amato, Jeffrey Barrett, Sonia Goncalves, Ewan Harrison, David K. Jackson, Ian Johnston, Dominic Kwiatkowski, Cordelia Langford, John Sillitoe on behalf of the Wellcome Sanger Institute COVID-19 Surveillance Team |
| EPI_ISL_2850246 | Lighthouse Laboratory Plymouth | Wellcome Sanger Institute for the COVID-19 Genomics UK (COG-UK) Consortium | Lighthouse Laboratory Plymouth and Alex Alderton, Roberto Amato, Jeffrey Barrett, Sonia Goncalves, Ewan Harrison, David K. Jackson, Ian Johnston, Dominic Kwiatkowski, Cordelia Langford, John Sillitoe on behalf of the Wellcome Sanger Institute COVID-19 Surveillance Team |
| EPI_ISL_2850247, EPI_ISL_2850248, EPI_ISL_2850249, EPI_ISL_2850250, EPI_ISL_2850251, EPI_ISL_2850252, EPI_ISL_2850253, EPI_ISL_2850254, EPI_ISL_2850255, EPI_ISL_2850256 | Lighthouse Lab in Glasgow | Wellcome Sanger Institute for the COVID-19 Genomics UK (COG-UK) Consortium | Harper VanSteenhouse, Yumi Kasai, David Gray, Carol Clugston, Anna Dominiczak and Alex Alderton, Roberto Amato, Jeffrey Barrett, Sonia Goncalves, Ewan Harrison, David K. Jackson, Ian Johnston, Dominic Kwiatkowski, Cordelia Langford, John Sillitoe on behalf of the Wellcome Sanger Institute COVID-19 Surveillance Team |
| EPI_ISL_2850257 | Lighthouse Laboratory Plymouth | Wellcome Sanger Institute for the COVID-19 Genomics UK (COG-UK) Consortium | Lighthouse Laboratory Plymouth and Alex Alderton, Roberto Amato, Jeffrey Barrett, Sonia Goncalves, Ewan Harrison, David K. Jackson, Ian Johnston, Dominic Kwiatkowski, Cordelia Langford, John Sillitoe on behalf of the Wellcome Sanger Institute COVID-19 Surveillance Team |
| EPI_ISL_2850258, EPI_ISL_2850259, EPI_ISL_2850260, EPI_ISL_2850261 | Lighthouse Lab in Glasgow | Wellcome Sanger Institute for the COVID-19 Genomics UK (COG-UK) Consortium | Harper VanSteenhouse, Yumi Kasai, David Gray, Carol Clugston, Anna Dominiczak and Alex Alderton, Roberto Amato, Jeffrey Barrett, Sonia Goncalves, Ewan Harrison, David K. Jackson, Ian Johnston, Dominic Kwiatkowski, Cordelia Langford, John Sillitoe on behalf of the Wellcome Sanger Institute COVID-19 Surveillance Team |
| EPI_ISL_2850262, EPI_ISL_2850263 | Lighthouse Laboratory Plymouth | Wellcome Sanger Institute for the COVID-19 Genomics UK (COG-UK) Consortium | Lighthouse Laboratory Plymouth and Alex Alderton, Roberto Amato, Jeffrey Barrett, Sonia Goncalves, Ewan Harrison, David K. Jackson, Ian Johnston, Dominic Kwiatkowski, Cordelia Langford, John Sillitoe on behalf of the Wellcome Sanger Institute COVID-19 Surveillance Team |
| EPI_ISL_2850264, EPI_ISL_2850265, EPI_ISL_2850266, EPI_ISL_2850267, EPI_ISL_2850268, EPI_ISL_2850269, EPI_ISL_2850270, EPI_ISL_2850271, EPI_ISL_2850272, EPI_ISL_2850273, EPI_ISL_2850274, EPI_ISL_2850275 | | | |
| see above | Lighthouse Lab in Glasgow | Wellcome Sanger Institute for the COVID-19 Genomics UK (COG-UK) Consortium | Harper VanSteenhouse, Yumi Kasai, David Gray, Carol Clugston, Anna Dominiczak and Alex Alderton, Roberto Amato, Jeffrey Barrett, Sonia Goncalves, Ewan Harrison, David K. Jackson, Ian Johnston, Dominic Kwiatkowski, Cordelia Langford, John Sillitoe on behalf of the Wellcome Sanger Institute COVID-19 Surveillance Team |
| EPI_ISL_2850276, EPI_ISL_2850277 | Lighthouse Laboratory Plymouth | Wellcome Sanger Institute for the COVID-19 Genomics UK (COG-UK) Consortium | Lighthouse Laboratory Plymouth and Alex Alderton, Roberto Amato, Jeffrey Barrett, Sonia Goncalves, Ewan Harrison, David K. Jackson, Ian Johnston, Dominic Kwiatkowski, Cordelia Langford, John Sillitoe on behalf of the Wellcome Sanger Institute COVID-19 Surveillance Team |
| EPI_ISL_2850278, EPI_ISL_2850279, EPI_ISL_2850280, EPI_ISL_2850281, EPI_ISL_2850282, EPI_ISL_2850283, EPI_ISL_2850284, EPI_ISL_2850285, EPI_ISL_2850286, EPI_ISL_2850287, EPI_ISL_2850288 | | | |
| see above | Lighthouse Lab in Glasgow | Wellcome Sanger Institute for the COVID-19 Genomics UK (COG-UK) Consortium | Harper VanSteenhouse, Yumi Kasai, David Gray, Carol Clugston, Anna Dominiczak and Alex Alderton, Roberto Amato, Jeffrey Barrett, Sonia Goncalves, Ewan Harrison, David K. Jackson, Ian Johnston, Dominic Kwiatkowski, Cordelia Langford, John Sillitoe on behalf of the Wellcome Sanger Institute COVID-19 Surveillance Team |
| EPI_ISL_2850289, EPI_ISL_2850290 | Lighthouse Laboratory Plymouth | Wellcome Sanger Institute for the COVID-19 Genomics UK (COG-UK) Consortium | Lighthouse Laboratory Plymouth and Alex Alderton, Roberto Amato, Jeffrey Barrett, Sonia Goncalves, Ewan Harrison, David K. Jackson, Ian Johnston, Dominic Kwiatkowski, Cordelia Langford, John Sillitoe on behalf of the Wellcome Sanger Institute COVID-19 Surveillance Team |
| EPI_ISL_2850291, EPI_ISL_2850292 | Lighthouse Lab in Glasgow | Wellcome Sanger Institute for the COVID-19 Genomics UK (COG-UK) Consortium | Harper VanSteenhouse, Yumi Kasai, David Gray, Carol Clugston, Anna Dominiczak and Alex Alderton, Roberto Amato, Jeffrey Barrett, Sonia Goncalves, Ewan Harrison, David K. Jackson, Ian Johnston, Dominic Kwiatkowski, Cordelia Langford, John Sillitoe on behalf of the Wellcome Sanger Institute COVID-19 Surveillance Team |
| EPI_ISL_2850293 | Lighthouse Laboratory Plymouth | Wellcome Sanger Institute for the COVID-19 Genomics UK (COG-UK) Consortium | Lighthouse Laboratory Plymouth and Alex Alderton, Roberto Amato, Jeffrey Barrett, Sonia Goncalves, Ewan Harrison, David K. Jackson, Ian Johnston, Dominic Kwiatkowski, Cordelia Langford, John Sillitoe on behalf of the Wellcome Sanger Institute COVID-19 Surveillance Team |
| EPI_ISL_2850294, EPI_ISL_2850295, EPI_ISL_2850296, | Lighthouse Lab in Glasgow | Wellcome Sanger Institute for the COVID-19 Genomics UK (COG-UK) Consortium | Harper VanSteenhouse, Yumi Kasai, David Gray, Carol Clugston, Anna Dominiczak and Alex Alderton, Roberto Amato, Jeffrey Barrett, Sonia Goncalves, Ewan Harrison, David K. Jackson, Ian Johnston, Dominic Kwiatkowski, Cordelia Langford, John Sillitoe on behalf of the Wellcome Sanger Institute COVID-19 Surveillance Team |

| | | | |
|---|---|---|---|
| EPI_ISL_2850297, EPI_ISL_2850298, EPI_ISL_2850299 | | | |
| EPI_ISL_2850300, EPI_ISL_2850301, EPI_ISL_2850302 | Lighthouse Laboratory Plymouth | Wellcome Sanger Institute for the COVID-19 Genomics UK (COG-UK) Consortium | Lighthouse Laboratory Plymouth and Alex Alderton, Roberto Amato, Jeffrey Barrett, Sonia Goncalves, Ewan Harrison, David K. Jackson, Ian Johnston, Dominic Kwiatkowski, Cordelia Langford, John Sillitoe on behalf of the Wellcome Sanger Institute COVID-19 Surveillance Team |
| EPI_ISL_2850303, EPI_ISL_2850304, EPI_ISL_2850305 | Lighthouse Lab in Glasgow | Wellcome Sanger Institute for the COVID-19 Genomics UK (COG-UK) Consortium | Harper VanSteenhouse, Yumi Kasai, David Gray, Carol Clugston, Anna Dominiczak and Alex Alderton, Roberto Amato, Jeffrey Barrett, Sonia Goncalves, Ewan Harrison, David K. Jackson, Ian Johnston, Dominic Kwiatkowski, Cordelia Langford, John Sillitoe on behalf of the Wellcome Sanger Institute COVID-19 Surveillance Team |
| EPI_ISL_2850306 | Lighthouse Laboratory Plymouth | Wellcome Sanger Institute for the COVID-19 Genomics UK (COG-UK) Consortium | Lighthouse Laboratory Plymouth and Alex Alderton, Roberto Amato, Jeffrey Barrett, Sonia Goncalves, Ewan Harrison, David K. Jackson, Ian Johnston, Dominic Kwiatkowski, Cordelia Langford, John Sillitoe on behalf of the Wellcome Sanger Institute COVID-19 Surveillance Team |
| EPI_ISL_2850307, EPI_ISL_2850308, EPI_ISL_2850309, EPI_ISL_2850310, EPI_ISL_2850311 | Lighthouse Lab in Glasgow | Wellcome Sanger Institute for the COVID-19 Genomics UK (COG-UK) Consortium | Harper VanSteenhouse, Yumi Kasai, David Gray, Carol Clugston, Anna Dominiczak and Alex Alderton, Roberto Amato, Jeffrey Barrett, Sonia Goncalves, Ewan Harrison, David K. Jackson, Ian Johnston, Dominic Kwiatkowski, Cordelia Langford, John Sillitoe on behalf of the Wellcome Sanger Institute COVID-19 Surveillance Team |
| EPI_ISL_2850312 | Lighthouse Laboratory Plymouth | Wellcome Sanger Institute for the COVID-19 Genomics UK (COG-UK) Consortium | Lighthouse Laboratory Plymouth and Alex Alderton, Roberto Amato, Jeffrey Barrett, Sonia Goncalves, Ewan Harrison, David K. Jackson, Ian Johnston, Dominic Kwiatkowski, Cordelia Langford, John Sillitoe on behalf of the Wellcome Sanger Institute COVID-19 Surveillance Team |
| EPI_ISL_2850313, EPI_ISL_2850314, EPI_ISL_2850315, EPI_ISL_2850316, EPI_ISL_2850317, EPI_ISL_2850318, EPI_ISL_2850319, EPI_ISL_2850320, EPI_ISL_2850321, EPI_ISL_2850322, EPI_ISL_2850323, EPI_ISL_2850324, EPI_ISL_2850325 | | | |
| see above | Lighthouse Lab in Glasgow | Wellcome Sanger Institute for the COVID-19 Genomics UK (COG-UK) Consortium | Harper VanSteenhouse, Yumi Kasai, David Gray, Carol Clugston, Anna Dominiczak and Alex Alderton, Roberto Amato, Jeffrey Barrett, Sonia Goncalves, Ewan Harrison, David K. Jackson, Ian Johnston, Dominic Kwiatkowski, Cordelia Langford, John Sillitoe on behalf of the Wellcome Sanger Institute COVID-19 Surveillance Team |
| EPI_ISL_2850326 | Lighthouse Laboratory Plymouth | Wellcome Sanger Institute for the COVID-19 Genomics UK (COG-UK) Consortium | Lighthouse Laboratory Plymouth and Alex Alderton, Roberto Amato, Jeffrey Barrett, Sonia Goncalves, Ewan Harrison, David K. Jackson, Ian Johnston, Dominic Kwiatkowski, Cordelia Langford, John Sillitoe on behalf of the Wellcome Sanger Institute COVID-19 Surveillance Team |
| EPI_ISL_2850327, EPI_ISL_2850328 | Lighthouse Lab in Glasgow | Wellcome Sanger Institute for the COVID-19 Genomics UK (COG-UK) Consortium | Harper VanSteenhouse, Yumi Kasai, David Gray, Carol Clugston, Anna Dominiczak and Alex Alderton, Roberto Amato, Jeffrey Barrett, Sonia Goncalves, Ewan Harrison, David K. Jackson, Ian Johnston, Dominic Kwiatkowski, Cordelia Langford, John Sillitoe on behalf of the Wellcome Sanger Institute COVID-19 Surveillance Team |
| EPI_ISL_2850329 | Lighthouse Laboratory Plymouth | Wellcome Sanger Institute for the COVID-19 Genomics UK (COG-UK) Consortium | Lighthouse Laboratory Plymouth and Alex Alderton, Roberto Amato, Jeffrey Barrett, Sonia Goncalves, Ewan Harrison, David K. Jackson, Ian Johnston, Dominic Kwiatkowski, Cordelia Langford, John Sillitoe on behalf of the Wellcome Sanger Institute COVID-19 Surveillance Team |
| EPI_ISL_2850330 | Lighthouse Lab in Glasgow | Wellcome Sanger Institute for the COVID-19 Genomics UK (COG-UK) Consortium | Harper VanSteenhouse, Yumi Kasai, David Gray, Carol Clugston, Anna Dominiczak and Alex Alderton, Roberto Amato, Jeffrey Barrett, Sonia Goncalves, Ewan Harrison, David K. Jackson, Ian Johnston, Dominic Kwiatkowski, Cordelia Langford, John Sillitoe on behalf of the Wellcome Sanger Institute COVID-19 Surveillance Team |
| EPI_ISL_2850331 | Lighthouse Laboratory Plymouth | Wellcome Sanger Institute for the COVID-19 Genomics UK (COG-UK) Consortium | Lighthouse Laboratory Plymouth and Alex Alderton, Roberto Amato, Jeffrey Barrett, Sonia Goncalves, Ewan Harrison, David K. Jackson, Ian Johnston, Dominic Kwiatkowski, Cordelia Langford, John Sillitoe on behalf of the Wellcome Sanger Institute COVID-19 Surveillance Team |
| EPI_ISL_2850332 | Lighthouse Lab in Glasgow | Wellcome Sanger Institute for the COVID-19 Genomics UK (COG-UK) Consortium | Harper VanSteenhouse, Yumi Kasai, David Gray, Carol Clugston, Anna Dominiczak and Alex Alderton, Roberto Amato, Jeffrey Barrett, Sonia Goncalves, Ewan Harrison, David K. Jackson, Ian Johnston, Dominic Kwiatkowski, Cordelia Langford, John Sillitoe on behalf of the Wellcome Sanger Institute COVID-19 Surveillance Team |
| EPI_ISL_2850333 | Lighthouse Laboratory Plymouth | Wellcome Sanger Institute for the COVID-19 Genomics UK (COG-UK) Consortium | Lighthouse Laboratory Plymouth and Alex Alderton, Roberto Amato, Jeffrey Barrett, Sonia Goncalves, Ewan Harrison, David K. Jackson, Ian Johnston, Dominic Kwiatkowski, Cordelia Langford, John Sillitoe on behalf of the Wellcome Sanger Institute COVID-19 Surveillance Team |
| EPI_ISL_2850334, EPI_ISL_2850335, EPI_ISL_2850336 | Lighthouse Lab in Glasgow | Wellcome Sanger Institute for the COVID-19 Genomics UK (COG-UK) Consortium | Harper VanSteenhouse, Yumi Kasai, David Gray, Carol Clugston, Anna Dominiczak and Alex Alderton, Roberto Amato, Jeffrey Barrett, Sonia Goncalves, Ewan Harrison, David K. Jackson, Ian Johnston, Dominic Kwiatkowski, Cordelia Langford, John Sillitoe on behalf of the Wellcome Sanger Institute COVID-19 Surveillance Team |
| EPI_ISL_2850337 | Lighthouse Laboratory Plymouth | Wellcome Sanger Institute for the COVID-19 Genomics UK (COG-UK) Consortium | Lighthouse Laboratory Plymouth and Alex Alderton, Roberto Amato, Jeffrey Barrett, Sonia Goncalves, Ewan Harrison, David K. Jackson, Ian Johnston, Dominic Kwiatkowski, Cordelia Langford, John Sillitoe on behalf of the Wellcome Sanger Institute COVID-19 Surveillance Team |
| EPI_ISL_2850338, EPI_ISL_2850339, EPI_ISL_2850340 | Lighthouse Lab in Glasgow | Wellcome Sanger Institute for the COVID-19 Genomics UK (COG-UK) Consortium | Harper VanSteenhouse, Yumi Kasai, David Gray, Carol Clugston, Anna Dominiczak and Alex Alderton, Roberto Amato, Jeffrey Barrett, Sonia Goncalves, Ewan Harrison, David K. Jackson, Ian Johnston, Dominic Kwiatkowski, Cordelia Langford, John Sillitoe on behalf of the Wellcome Sanger Institute COVID-19 Surveillance Team |
| EPI_ISL_2850341 | Lighthouse Laboratory Plymouth | Wellcome Sanger Institute for the COVID-19 Genomics UK (COG-UK) Consortium | Lighthouse Laboratory Plymouth and Alex Alderton, Roberto Amato, Jeffrey Barrett, Sonia Goncalves, Ewan Harrison, David K. Jackson, Ian Johnston, Dominic Kwiatkowski, Cordelia Langford, John Sillitoe on behalf of the Wellcome Sanger Institute COVID-19 Surveillance Team |
| EPI_ISL_2850342, EPI_ISL_2850343 | Lighthouse Lab in Glasgow | Wellcome Sanger Institute for the COVID-19 Genomics UK (COG-UK) Consortium | Harper VanSteenhouse, Yumi Kasai, David Gray, Carol Clugston, Anna Dominiczak and Alex Alderton, Roberto Amato, Jeffrey Barrett, Sonia Goncalves, Ewan Harrison, David K. Jackson, Ian Johnston, Dominic Kwiatkowski, Cordelia Langford, John Sillitoe on behalf of the Wellcome Sanger Institute COVID-19 Surveillance Team |
| EPI_ISL_2850344 | Lighthouse Laboratory Plymouth | Wellcome Sanger Institute for the COVID-19 Genomics UK (COG-UK) Consortium | Lighthouse Laboratory Plymouth and Alex Alderton, Roberto Amato, Jeffrey Barrett, Sonia Goncalves, Ewan Harrison, David K. Jackson, Ian Johnston, Dominic Kwiatkowski, Cordelia Langford, John Sillitoe on behalf of the Wellcome Sanger Institute COVID-19 Surveillance Team |
| EPI_ISL_2850345 | Lighthouse Lab in Glasgow | Wellcome Sanger Institute for the COVID-19 Genomics UK (COG-UK) Consortium | Harper VanSteenhouse, Yumi Kasai, David Gray, Carol Clugston, Anna Dominiczak and Alex Alderton, Roberto Amato, Jeffrey Barrett, Sonia Goncalves, Ewan Harrison, David K. Jackson, Ian Johnston, Dominic Kwiatkowski, Cordelia Langford, John Sillitoe on behalf of the Wellcome Sanger Institute COVID-19 Surveillance Team |
| EPI_ISL_2850346 | Lighthouse Laboratory Plymouth | Wellcome Sanger Institute for the COVID-19 Genomics UK (COG-UK) Consortium | Lighthouse Laboratory Plymouth and Alex Alderton, Roberto Amato, Jeffrey Barrett, Sonia Goncalves, Ewan Harrison, David K. Jackson, Ian Johnston, Dominic Kwiatkowski, Cordelia Langford, John Sillitoe on behalf of the Wellcome Sanger Institute COVID-19 Surveillance Team |
| EPI_ISL_2850347, EPI_ISL_2850348, EPI_ISL_2850349, EPI_ISL_2850350 | Lighthouse Lab in Glasgow | Wellcome Sanger Institute for the COVID-19 Genomics UK (COG-UK) Consortium | Harper VanSteenhouse, Yumi Kasai, David Gray, Carol Clugston, Anna Dominiczak and Alex Alderton, Roberto Amato, Jeffrey Barrett, Sonia Goncalves, Ewan Harrison, David K. Jackson, Ian Johnston, Dominic Kwiatkowski, Cordelia Langford, John Sillitoe on behalf of the Wellcome Sanger Institute COVID-19 Surveillance Team |
| EPI_ISL_2850351 | Lighthouse Laboratory Plymouth | Wellcome Sanger Institute for the COVID-19 Genomics UK (COG-UK) Consortium | Lighthouse Laboratory Plymouth and Alex Alderton, Roberto Amato, Jeffrey Barrett, Sonia Goncalves, Ewan Harrison, David K. Jackson, Ian Johnston, Dominic Kwiatkowski, Cordelia Langford, John Sillitoe on behalf of the Wellcome Sanger Institute COVID-19 Surveillance Team |
| EPI_ISL_2850352, EPI_ISL_2850353, EPI_ISL_2850354, EPI_ISL_2850355, EPI_ISL_2850356, EPI_ISL_2850357 | Lighthouse Lab in Glasgow | Wellcome Sanger Institute for the COVID-19 Genomics UK (COG-UK) Consortium | Harper VanSteenhouse, Yumi Kasai, David Gray, Carol Clugston, Anna Dominiczak and Alex Alderton, Roberto Amato, Jeffrey Barrett, Sonia Goncalves, Ewan Harrison, David K. Jackson, Ian Johnston, Dominic Kwiatkowski, Cordelia Langford, John Sillitoe on behalf of the Wellcome Sanger Institute COVID-19 Surveillance Team |
| EPI_ISL_2850358 | Lighthouse Laboratory Plymouth | Wellcome Sanger Institute for the COVID-19 Genomics UK (COG-UK) Consortium | Lighthouse Laboratory Plymouth and Alex Alderton, Roberto Amato, Jeffrey Barrett, Sonia Goncalves, Ewan Harrison, David K. Jackson, Ian Johnston, Dominic Kwiatkowski, Cordelia Langford, John Sillitoe on behalf of the Wellcome Sanger Institute COVID-19 Surveillance Team |
| EPI_ISL_2850359, EPI_ISL_2850360, EPI_ISL_2850361, EPI_ISL_2850362 | Lighthouse Lab in Glasgow | Wellcome Sanger Institute for the COVID-19 Genomics UK (COG-UK) Consortium | Harper VanSteenhouse, Yumi Kasai, David Gray, Carol Clugston, Anna Dominiczak and Alex Alderton, Roberto Amato, Jeffrey Barrett, Sonia Goncalves, Ewan Harrison, David K. Jackson, Ian Johnston, Dominic Kwiatkowski, Cordelia Langford, John Sillitoe on behalf of the Wellcome Sanger Institute COVID-19 Surveillance Team |
| EPI_ISL_2850363 | Lighthouse Laboratory Plymouth | Wellcome Sanger Institute for the COVID-19 Genomics UK (COG-UK) Consortium | Lighthouse Laboratory Plymouth and Alex Alderton, Roberto Amato, Jeffrey Barrett, Sonia Goncalves, Ewan Harrison, David K. Jackson, Ian Johnston, Dominic Kwiatkowski, Cordelia Langford, John Sillitoe on behalf of the Wellcome Sanger Institute COVID-19 Surveillance Team |
| EPI_ISL_2850364 | Lighthouse Lab in Glasgow | Wellcome Sanger Institute for the COVID-19 Genomics UK (COG-UK) Consortium | Harper VanSteenhouse, Yumi Kasai, David Gray, Carol Clugston, Anna Dominiczak and Alex Alderton, Roberto Amato, Jeffrey Barrett, Sonia Goncalves, Ewan Harrison, David K. Jackson, Ian Johnston, Dominic Kwiatkowski, Cordelia Langford, John Sillitoe on behalf of the Wellcome Sanger Institute COVID-19 Surveillance Team |
| EPI_ISL_2850365 | Lighthouse Laboratory Plymouth | Wellcome Sanger Institute for the COVID-19 Genomics UK (COG-UK) Consortium | Lighthouse Laboratory Plymouth and Alex Alderton, Roberto Amato, Jeffrey Barrett, Sonia Goncalves, Ewan Harrison, David K. Jackson, Ian Johnston, Dominic Kwiatkowski, Cordelia Langford, John Sillitoe on behalf of the Wellcome Sanger Institute COVID-19 Surveillance Team |
| EPI_ISL_2850366, EPI_ISL_2850367 | Lighthouse Lab in Glasgow | Wellcome Sanger Institute for the COVID-19 Genomics UK (COG-UK) Consortium | Harper VanSteenhouse, Yumi Kasai, David Gray, Carol Clugston, Anna Dominiczak and Alex Alderton, Roberto Amato, Jeffrey Barrett, Sonia Goncalves, Ewan Harrison, David K. Jackson, Ian Johnston, Dominic Kwiatkowski, Cordelia Langford, John Sillitoe on behalf of the Wellcome Sanger Institute COVID-19 Surveillance Team |
| EPI_ISL_2850368 | Lighthouse Laboratory Plymouth | Wellcome Sanger Institute for the COVID-19 Genomics UK (COG-UK) Consortium | Lighthouse Laboratory Plymouth and Alex Alderton, Roberto Amato, Jeffrey Barrett, Sonia Goncalves, Ewan Harrison, David K. Jackson, Ian Johnston, Dominic Kwiatkowski, Cordelia Langford, John Sillitoe on behalf of the Wellcome Sanger Institute COVID-19 Surveillance Team |
| EPI_ISL_2850369, EPI_ISL_2850370, EPI_ISL_2850371 | Lighthouse Lab in Glasgow | Wellcome Sanger Institute for the COVID-19 Genomics UK (COG-UK) Consortium | Harper VanSteenhouse, Yumi Kasai, David Gray, Carol Clugston, Anna Dominiczak and Alex Alderton, Roberto Amato, Jeffrey Barrett, Sonia Goncalves, Ewan Harrison, David K. Jackson, Ian Johnston, Dominic Kwiatkowski, Cordelia Langford, John Sillitoe on behalf of the Wellcome Sanger Institute COVID-19 Surveillance Team |
| EPI_ISL_2850372 | Lighthouse Laboratory Plymouth | Wellcome Sanger Institute for the COVID-19 Genomics UK (COG-UK) Consortium | Lighthouse Laboratory Plymouth and Alex Alderton, Roberto Amato, Jeffrey Barrett, Sonia Goncalves, Ewan Harrison, David K. Jackson, Ian Johnston, Dominic Kwiatkowski, Cordelia Langford, John Sillitoe on behalf of the Wellcome Sanger Institute COVID-19 Surveillance Team |
| EPI_ISL_2850373 | Lighthouse Lab in Glasgow | Wellcome Sanger Institute for the COVID-19 Genomics UK (COG-UK) Consortium | Harper VanSteenhouse, Yumi Kasai, David Gray, Carol Clugston, Anna Dominiczak and Alex Alderton, Roberto Amato, Jeffrey Barrett, Sonia Goncalves, Ewan Harrison, David K. Jackson, Ian Johnston, Dominic Kwiatkowski, Cordelia Langford, John Sillitoe on behalf of the Wellcome Sanger Institute COVID-19 Surveillance Team |
| EPI_ISL_2850374 | Lighthouse Lab in Alderley Park | Wellcome Sanger Institute for the COVID-19 Genomics UK (COG-UK) Consortium | Jacquelyn Wynn, Mairead Hyland, The Lighthouse Lab in Alderley Park and Alex Alderton, Roberto Amato, Jeffrey Barrett, Sonia Goncalves, Ewan Harrison, David K. Jackson, Ian Johnston, Dominic Kwiatkowski, Cordelia Langford, John Sillitoe on behalf of the Wellcome Sanger Institute COVID-19 Surveillance Team |
| EPI_ISL_2850375 | Lighthouse Laboratory Plymouth | Wellcome Sanger Institute for the COVID-19 Genomics UK (COG-UK) Consortium | Lighthouse Laboratory Plymouth and Alex Alderton, Roberto Amato, Jeffrey Barrett, Sonia Goncalves, Ewan Harrison, David K. Jackson, Ian Johnston, Dominic Kwiatkowski, Cordelia Langford, John Sillitoe on behalf of the Wellcome Sanger Institute COVID-19 Surveillance Team |
| EPI_ISL_2850376 | Lighthouse Lab in Milton Keynes | Wellcome Sanger Institute for the COVID-19 Genomics UK (COG-UK) | The Lighthouse Lab in Milton Keynes and Alex Alderton, Roberto Amato, Jeffrey Barrett, Sonia Goncalves, Ewan Harrison, David K. Jackson, Ian Johnston, Dominic Kwiatkowski, Cordelia Langford, John Sillitoe on behalf of the Wellcome Sanger |

| | | | |
|---|---|---|---|
| | | Consortium | Institute COVID-19 Surveillance Team |
| EPI_ISL_2850377, EPI_ISL_2850378 | Lighthouse Lab in Alderley Park | Wellcome Sanger Institute for the COVID-19 Genomics UK (COG-UK) Consortium | Jacquelyn Wynn, Mairead Hyland, The Lighthouse Lab in Alderley Park and Alex Alderton, Roberto Amato, Jeffrey Barrett, Sonia Goncalves, Ewan Harrison, David K. Jackson, Ian Johnston, Dominic Kwiatkowski, Cordelia Langford, John Sillitoe on behalf of the Wellcome Sanger Institute COVID-19 Surveillance Team |
| EPI_ISL_2850379 | Lighthouse Lab in Milton Keynes | Wellcome Sanger Institute for the COVID-19 Genomics UK (COG-UK) Consortium | The Lighthouse Lab in Milton Keynes and Alex Alderton, Roberto Amato, Jeffrey Barrett, Sonia Goncalves, Ewan Harrison, David K. Jackson, Ian Johnston, Dominic Kwiatkowski, Cordelia Langford, John Sillitoe on behalf of the Wellcome Sanger Institute COVID-19 Surveillance Team |
| EPI_ISL_2850380, EPI_ISL_2850381 | Lighthouse Laboratory Plymouth | Wellcome Sanger Institute for the COVID-19 Genomics UK (COG-UK) Consortium | Lighthouse Laboratory Plymouth and Alex Alderton, Roberto Amato, Jeffrey Barrett, Sonia Goncalves, Ewan Harrison, David K. Jackson, Ian Johnston, Dominic Kwiatkowski, Cordelia Langford, John Sillitoe on behalf of the Wellcome Sanger Institute COVID-19 Surveillance Team |
| EPI_ISL_2850382, EPI_ISL_2850383, EPI_ISL_2850384, EPI_ISL_2850385 | Lighthouse Lab in Alderley Park | Wellcome Sanger Institute for the COVID-19 Genomics UK (COG-UK) Consortium | Jacquelyn Wynn, Mairead Hyland, The Lighthouse Lab in Alderley Park and Alex Alderton, Roberto Amato, Jeffrey Barrett, Sonia Goncalves, Ewan Harrison, David K. Jackson, Ian Johnston, Dominic Kwiatkowski, Cordelia Langford, John Sillitoe on behalf of the Wellcome Sanger Institute COVID-19 Surveillance Team |
| EPI_ISL_2850386, EPI_ISL_2850387 | Lighthouse Laboratory Plymouth | Wellcome Sanger Institute for the COVID-19 Genomics UK (COG-UK) Consortium | Lighthouse Laboratory Plymouth and Alex Alderton, Roberto Amato, Jeffrey Barrett, Sonia Goncalves, Ewan Harrison, David K. Jackson, Ian Johnston, Dominic Kwiatkowski, Cordelia Langford, John Sillitoe on behalf of the Wellcome Sanger Institute COVID-19 Surveillance Team |
| EPI_ISL_2850388 | Lighthouse Lab in Alderley Park | Wellcome Sanger Institute for the COVID-19 Genomics UK (COG-UK) Consortium | Jacquelyn Wynn, Mairead Hyland, The Lighthouse Lab in Alderley Park and Alex Alderton, Roberto Amato, Jeffrey Barrett, Sonia Goncalves, Ewan Harrison, David K. Jackson, Ian Johnston, Dominic Kwiatkowski, Cordelia Langford, John Sillitoe on behalf of the Wellcome Sanger Institute COVID-19 Surveillance Team |
| EPI_ISL_2850389 | Lighthouse Lab in Milton Keynes | Wellcome Sanger Institute for the COVID-19 Genomics UK (COG-UK) Consortium | The Lighthouse Lab in Milton Keynes and Alex Alderton, Roberto Amato, Jeffrey Barrett, Sonia Goncalves, Ewan Harrison, David K. Jackson, Ian Johnston, Dominic Kwiatkowski, Cordelia Langford, John Sillitoe on behalf of the Wellcome Sanger Institute COVID-19 Surveillance Team |
| EPI_ISL_2850390 | Lighthouse Laboratory Plymouth | Wellcome Sanger Institute for the COVID-19 Genomics UK (COG-UK) Consortium | Lighthouse Laboratory Plymouth and Alex Alderton, Roberto Amato, Jeffrey Barrett, Sonia Goncalves, Ewan Harrison, David K. Jackson, Ian Johnston, Dominic Kwiatkowski, Cordelia Langford, John Sillitoe on behalf of the Wellcome Sanger Institute COVID-19 Surveillance Team |
| EPI_ISL_2850391, EPI_ISL_2850392, EPI_ISL_2850393 | Lighthouse Lab in Alderley Park | Wellcome Sanger Institute for the COVID-19 Genomics UK (COG-UK) Consortium | Jacquelyn Wynn, Mairead Hyland, The Lighthouse Lab in Alderley Park and Alex Alderton, Roberto Amato, Jeffrey Barrett, Sonia Goncalves, Ewan Harrison, David K. Jackson, Ian Johnston, Dominic Kwiatkowski, Cordelia Langford, John Sillitoe on behalf of the Wellcome Sanger Institute COVID-19 Surveillance Team |
| EPI_ISL_2850394 | Lighthouse Laboratory Plymouth | Wellcome Sanger Institute for the COVID-19 Genomics UK (COG-UK) Consortium | Lighthouse Laboratory Plymouth and Alex Alderton, Roberto Amato, Jeffrey Barrett, Sonia Goncalves, Ewan Harrison, David K. Jackson, Ian Johnston, Dominic Kwiatkowski, Cordelia Langford, John Sillitoe on behalf of the Wellcome Sanger Institute COVID-19 Surveillance Team |
| EPI_ISL_2850395 | Lighthouse Lab in Alderley Park | Wellcome Sanger Institute for the COVID-19 Genomics UK (COG-UK) Consortium | Jacquelyn Wynn, Mairead Hyland, The Lighthouse Lab in Alderley Park and Alex Alderton, Roberto Amato, Jeffrey Barrett, Sonia Goncalves, Ewan Harrison, David K. Jackson, Ian Johnston, Dominic Kwiatkowski, Cordelia Langford, John Sillitoe on behalf of the Wellcome Sanger Institute COVID-19 Surveillance Team |
| EPI_ISL_2850396 | Lighthouse Lab in Milton Keynes | Wellcome Sanger Institute for the COVID-19 Genomics UK (COG-UK) Consortium | The Lighthouse Lab in Milton Keynes and Alex Alderton, Roberto Amato, Jeffrey Barrett, Sonia Goncalves, Ewan Harrison, David K. Jackson, Ian Johnston, Dominic Kwiatkowski, Cordelia Langford, John Sillitoe on behalf of the Wellcome Sanger Institute COVID-19 Surveillance Team |
| EPI_ISL_2850397 | Lighthouse Lab in Alderley Park | Wellcome Sanger Institute for the COVID-19 Genomics UK (COG-UK) Consortium | Jacquelyn Wynn, Mairead Hyland, The Lighthouse Lab in Alderley Park and Alex Alderton, Roberto Amato, Jeffrey Barrett, Sonia Goncalves, Ewan Harrison, David K. Jackson, Ian Johnston, Dominic Kwiatkowski, Cordelia Langford, John Sillitoe on behalf of the Wellcome Sanger Institute COVID-19 Surveillance Team |
| EPI_ISL_2850398, EPI_ISL_2850399 | Lighthouse Laboratory Plymouth | Wellcome Sanger Institute for the COVID-19 Genomics UK (COG-UK) Consortium | Lighthouse Laboratory Plymouth and Alex Alderton, Roberto Amato, Jeffrey Barrett, Sonia Goncalves, Ewan Harrison, David K. Jackson, Ian Johnston, Dominic Kwiatkowski, Cordelia Langford, John Sillitoe on behalf of the Wellcome Sanger Institute COVID-19 Surveillance Team |
| EPI_ISL_2850400 | Lighthouse Lab in Alderley Park | Wellcome Sanger Institute for the COVID-19 Genomics UK (COG-UK) Consortium | Jacquelyn Wynn, Mairead Hyland, The Lighthouse Lab in Alderley Park and Alex Alderton, Roberto Amato, Jeffrey Barrett, Sonia Goncalves, Ewan Harrison, David K. Jackson, Ian Johnston, Dominic Kwiatkowski, Cordelia Langford, John Sillitoe on behalf of the Wellcome Sanger Institute COVID-19 Surveillance Team |
| EPI_ISL_2850401 | Lighthouse Laboratory Plymouth | Wellcome Sanger Institute for the COVID-19 Genomics UK (COG-UK) Consortium | Lighthouse Laboratory Plymouth and Alex Alderton, Roberto Amato, Jeffrey Barrett, Sonia Goncalves, Ewan Harrison, David K. Jackson, Ian Johnston, Dominic Kwiatkowski, Cordelia Langford, John Sillitoe on behalf of the Wellcome Sanger Institute COVID-19 Surveillance Team |
| EPI_ISL_2850402 | Lighthouse Lab in Milton Keynes | Wellcome Sanger Institute for the COVID-19 Genomics UK (COG-UK) Consortium | The Lighthouse Lab in Milton Keynes and Alex Alderton, Roberto Amato, Jeffrey Barrett, Sonia Goncalves, Ewan Harrison, David K. Jackson, Ian Johnston, Dominic Kwiatkowski, Cordelia Langford, John Sillitoe on behalf of the Wellcome Sanger Institute COVID-19 Surveillance Team |
| EPI_ISL_2850403 | Lighthouse Lab in Alderley Park | Wellcome Sanger Institute for the COVID-19 Genomics UK (COG-UK) Consortium | Jacquelyn Wynn, Mairead Hyland, The Lighthouse Lab in Alderley Park and Alex Alderton, Roberto Amato, Jeffrey Barrett, Sonia Goncalves, Ewan Harrison, David K. Jackson, Ian Johnston, Dominic Kwiatkowski, Cordelia Langford, John Sillitoe on behalf of the Wellcome Sanger Institute COVID-19 Surveillance Team |
| EPI_ISL_2850404, EPI_ISL_2850405 | Lighthouse Laboratory Plymouth | Wellcome Sanger Institute for the COVID-19 Genomics UK (COG-UK) Consortium | Lighthouse Laboratory Plymouth and Alex Alderton, Roberto Amato, Jeffrey Barrett, Sonia Goncalves, Ewan Harrison, David K. Jackson, Ian Johnston, Dominic Kwiatkowski, Cordelia Langford, John Sillitoe on behalf of the Wellcome Sanger Institute COVID-19 Surveillance Team |
| EPI_ISL_2850406 | Lighthouse Lab in Milton Keynes | Wellcome Sanger Institute for the COVID-19 Genomics UK (COG-UK) Consortium | The Lighthouse Lab in Milton Keynes and Alex Alderton, Roberto Amato, Jeffrey Barrett, Sonia Goncalves, Ewan Harrison, David K. Jackson, Ian Johnston, Dominic Kwiatkowski, Cordelia Langford, John Sillitoe on behalf of the Wellcome Sanger Institute COVID-19 Surveillance Team |
| EPI_ISL_2850407 | Lighthouse Lab in Alderley Park | Wellcome Sanger Institute for the COVID-19 Genomics UK (COG-UK) Consortium | Jacquelyn Wynn, Mairead Hyland, The Lighthouse Lab in Alderley Park and Alex Alderton, Roberto Amato, Jeffrey Barrett, Sonia Goncalves, Ewan Harrison, David K. Jackson, Ian Johnston, Dominic Kwiatkowski, Cordelia Langford, John Sillitoe on behalf of the Wellcome Sanger Institute COVID-19 Surveillance Team |
| EPI_ISL_2850408 | Lighthouse Laboratory Plymouth | Wellcome Sanger Institute for the COVID-19 Genomics UK (COG-UK) Consortium | Lighthouse Laboratory Plymouth and Alex Alderton, Roberto Amato, Jeffrey Barrett, Sonia Goncalves, Ewan Harrison, David K. Jackson, Ian Johnston, Dominic Kwiatkowski, Cordelia Langford, John Sillitoe on behalf of the Wellcome Sanger Institute COVID-19 Surveillance Team |
| EPI_ISL_2850409 | Lighthouse Lab in Milton Keynes | Wellcome Sanger Institute for the COVID-19 Genomics UK (COG-UK) Consortium | The Lighthouse Lab in Milton Keynes and Alex Alderton, Roberto Amato, Jeffrey Barrett, Sonia Goncalves, Ewan Harrison, David K. Jackson, Ian Johnston, Dominic Kwiatkowski, Cordelia Langford, John Sillitoe on behalf of the Wellcome Sanger Institute COVID-19 Surveillance Team |
| EPI_ISL_2850410 | Lighthouse Laboratory Plymouth | Wellcome Sanger Institute for the COVID-19 Genomics UK (COG-UK) Consortium | Lighthouse Laboratory Plymouth and Alex Alderton, Roberto Amato, Jeffrey Barrett, Sonia Goncalves, Ewan Harrison, David K. Jackson, Ian Johnston, Dominic Kwiatkowski, Cordelia Langford, John Sillitoe on behalf of the Wellcome Sanger Institute COVID-19 Surveillance Team |
| EPI_ISL_2850411, EPI_ISL_2850412 | Lighthouse Lab in Milton Keynes | Wellcome Sanger Institute for the COVID-19 Genomics UK (COG-UK) Consortium | The Lighthouse Lab in Milton Keynes and Alex Alderton, Roberto Amato, Jeffrey Barrett, Sonia Goncalves, Ewan Harrison, David K. Jackson, Ian Johnston, Dominic Kwiatkowski, Cordelia Langford, John Sillitoe on behalf of the Wellcome Sanger Institute COVID-19 Surveillance Team |
| EPI_ISL_2850413 | Lighthouse Lab in Alderley Park | Wellcome Sanger Institute for the COVID-19 Genomics UK (COG-UK) Consortium | Jacquelyn Wynn, Mairead Hyland, The Lighthouse Lab in Alderley Park and Alex Alderton, Roberto Amato, Jeffrey Barrett, Sonia Goncalves, Ewan Harrison, David K. Jackson, Ian Johnston, Dominic Kwiatkowski, Cordelia Langford, John Sillitoe on behalf of the Wellcome Sanger Institute COVID-19 Surveillance Team |
| EPI_ISL_2850414 | Lighthouse Laboratory Plymouth | Wellcome Sanger Institute for the COVID-19 Genomics UK (COG-UK) Consortium | Lighthouse Laboratory Plymouth and Alex Alderton, Roberto Amato, Jeffrey Barrett, Sonia Goncalves, Ewan Harrison, David K. Jackson, Ian Johnston, Dominic Kwiatkowski, Cordelia Langford, John Sillitoe on behalf of the Wellcome Sanger Institute COVID-19 Surveillance Team |
| EPI_ISL_2850415 | Lighthouse Lab in Milton Keynes | Wellcome Sanger Institute for the COVID-19 Genomics UK (COG-UK) Consortium | The Lighthouse Lab in Milton Keynes and Alex Alderton, Roberto Amato, Jeffrey Barrett, Sonia Goncalves, Ewan Harrison, David K. Jackson, Ian Johnston, Dominic Kwiatkowski, Cordelia Langford, John Sillitoe on behalf of the Wellcome Sanger Institute COVID-19 Surveillance Team |
| EPI_ISL_2850416 | Lighthouse Laboratory Plymouth | Wellcome Sanger Institute for the COVID-19 Genomics UK (COG-UK) Consortium | Lighthouse Laboratory Plymouth and Alex Alderton, Roberto Amato, Jeffrey Barrett, Sonia Goncalves, Ewan Harrison, David K. Jackson, Ian Johnston, Dominic Kwiatkowski, Cordelia Langford, John Sillitoe on behalf of the Wellcome Sanger Institute COVID-19 Surveillance Team |
| EPI_ISL_2850417, EPI_ISL_2850418, EPI_ISL_2850419, EPI_ISL_2850420, EPI_ISL_2850421 | Lighthouse Lab in Alderley Park | Wellcome Sanger Institute for the COVID-19 Genomics UK (COG-UK) Consortium | Jacquelyn Wynn, Mairead Hyland, The Lighthouse Lab in Alderley Park and Alex Alderton, Roberto Amato, Jeffrey Barrett, Sonia Goncalves, Ewan Harrison, David K. Jackson, Ian Johnston, Dominic Kwiatkowski, Cordelia Langford, John Sillitoe on behalf of the Wellcome Sanger Institute COVID-19 Surveillance Team |
| EPI_ISL_2850422 | Lighthouse Lab in Milton Keynes | Wellcome Sanger Institute for the COVID-19 Genomics UK (COG-UK) Consortium | The Lighthouse Lab in Milton Keynes and Alex Alderton, Roberto Amato, Jeffrey Barrett, Sonia Goncalves, Ewan Harrison, David K. Jackson, Ian Johnston, Dominic Kwiatkowski, Cordelia Langford, John Sillitoe on behalf of the Wellcome Sanger Institute COVID-19 Surveillance Team |
| EPI_ISL_2850423, EPI_ISL_2850424, EPI_ISL_2850425 | Lighthouse Lab in Alderley Park | Wellcome Sanger Institute for the COVID-19 Genomics UK (COG-UK) Consortium | Jacquelyn Wynn, Mairead Hyland, The Lighthouse Lab in Alderley Park and Alex Alderton, Roberto Amato, Jeffrey Barrett, Sonia Goncalves, Ewan Harrison, David K. Jackson, Ian Johnston, Dominic Kwiatkowski, Cordelia Langford, John Sillitoe on behalf of the Wellcome Sanger Institute COVID-19 Surveillance Team |
| EPI_ISL_2850426 | Lighthouse Laboratory Plymouth | Wellcome Sanger Institute for the COVID-19 Genomics UK (COG-UK) Consortium | Lighthouse Laboratory Plymouth and Alex Alderton, Roberto Amato, Jeffrey Barrett, Sonia Goncalves, Ewan Harrison, David K. Jackson, Ian Johnston, Dominic Kwiatkowski, Cordelia Langford, John Sillitoe on behalf of the Wellcome Sanger Institute COVID-19 Surveillance Team |
| EPI_ISL_2850427 | Lighthouse Lab in Milton Keynes | Wellcome Sanger Institute for the COVID-19 Genomics UK (COG-UK) Consortium | The Lighthouse Lab in Milton Keynes and Alex Alderton, Roberto Amato, Jeffrey Barrett, Sonia Goncalves, Ewan Harrison, David K. Jackson, Ian Johnston, Dominic Kwiatkowski, Cordelia Langford, John Sillitoe on behalf of the Wellcome Sanger Institute COVID-19 Surveillance Team |
| EPI_ISL_2850428 | Lighthouse Laboratory Plymouth | Wellcome Sanger Institute for the COVID-19 Genomics UK (COG-UK) Consortium | Lighthouse Laboratory Plymouth and Alex Alderton, Roberto Amato, Jeffrey Barrett, Sonia Goncalves, Ewan Harrison, David K. Jackson, Ian Johnston, Dominic Kwiatkowski, Cordelia Langford, John Sillitoe on behalf of the Wellcome Sanger Institute COVID-19 Surveillance Team |
| EPI_ISL_2850429, EPI_ISL_2850430, EPI_ISL_2850431, EPI_ISL_2850432, EPI_ISL_2850433, EPI_ISL_2850434 | Lighthouse Lab in Alderley Park | Wellcome Sanger Institute for the COVID-19 Genomics UK (COG-UK) Consortium | Jacquelyn Wynn, Mairead Hyland, The Lighthouse Lab in Alderley Park and Alex Alderton, Roberto Amato, Jeffrey Barrett, Sonia Goncalves, Ewan Harrison, David K. Jackson, Ian Johnston, Dominic Kwiatkowski, Cordelia Langford, John Sillitoe on behalf of the Wellcome Sanger Institute COVID-19 Surveillance Team |
| EPI_ISL_2850435 | Lighthouse Laboratory Plymouth | Wellcome Sanger Institute for the COVID-19 Genomics UK (COG-UK) Consortium | Lighthouse Laboratory Plymouth and Alex Alderton, Roberto Amato, Jeffrey Barrett, Sonia Goncalves, Ewan Harrison, David K. Jackson, Ian Johnston, Dominic Kwiatkowski, Cordelia Langford, John Sillitoe on behalf of the Wellcome Sanger Institute COVID-19 Surveillance Team |
| EPI_ISL_2850436 | Lighthouse Lab in Milton Keynes | Wellcome Sanger Institute for the COVID-19 Genomics UK (COG-UK) Consortium | The Lighthouse Lab in Milton Keynes and Alex Alderton, Roberto Amato, Jeffrey Barrett, Sonia Goncalves, Ewan Harrison, David K. Jackson, Ian Johnston, Dominic Kwiatkowski, Cordelia Langford, John Sillitoe on behalf of the Wellcome Sanger Institute COVID-19 Surveillance Team |
| EPI_ISL_2850437, EPI_ISL_2850438 | Lighthouse Lab in Alderley Park | Wellcome Sanger Institute for the COVID-19 Genomics UK (COG-UK) Consortium | Jacquelyn Wynn, Mairead Hyland, The Lighthouse Lab in Alderley Park and Alex Alderton, Roberto Amato, Jeffrey Barrett, Sonia Goncalves, Ewan Harrison, David K. Jackson, Ian Johnston, Dominic Kwiatkowski, Cordelia Langford, John Sillitoe on behalf of the Wellcome Sanger Institute COVID-19 Surveillance Team |
| EPI_ISL_2850439 | Lighthouse Lab in Milton Keynes | Wellcome Sanger Institute for the COVID-19 Genomics UK (COG-UK) Consortium | The Lighthouse Lab in Milton Keynes and Alex Alderton, Roberto Amato, Jeffrey Barrett, Sonia Goncalves, Ewan Harrison, David K. Jackson, Ian Johnston, Dominic Kwiatkowski, Cordelia Langford, John Sillitoe on behalf of the Wellcome Sanger Institute COVID-19 Surveillance Team |

| | | | |
|---|---|---|---|
| EPI_ISL_2850440, EPI_ISL_2850441 | Lighthouse Lab in Alderley Park | Wellcome Sanger Institute for the COVID-19 Genomics UK (COG-UK) Consortium | Jacquelyn Wynn, Mairead Hyland, The Lighthouse Lab in Alderley Park and Alex Alderton, Roberto Amato, Jeffrey Barrett, Sonia Goncalves, Ewan Harrison, David K. Jackson, Ian Johnston, Dominic Kwiatkowski, Cordelia Langford, John Sillitoe on behalf of the Wellcome Sanger Institute COVID-19 Surveillance Team |
| EPI_ISL_2850442 | Lighthouse Lab in Milton Keynes | Wellcome Sanger Institute for the COVID-19 Genomics UK (COG-UK) Consortium | The Lighthouse Lab in Milton Keynes and Alex Alderton, Roberto Amato, Jeffrey Barrett, Sonia Goncalves, Ewan Harrison, David K. Jackson, Ian Johnston, Dominic Kwiatkowski, Cordelia Langford, John Sillitoe on behalf of the Wellcome Sanger Institute COVID-19 Surveillance Team |
| EPI_ISL_2850443, EPI_ISL_2850444 | Lighthouse Laboratory Plymouth | Wellcome Sanger Institute for the COVID-19 Genomics UK (COG-UK) Consortium | Lighthouse Laboratory Plymouth and Alex Alderton, Roberto Amato, Jeffrey Barrett, Sonia Goncalves, Ewan Harrison, David K. Jackson, Ian Johnston, Dominic Kwiatkowski, Cordelia Langford, John Sillitoe on behalf of the Wellcome Sanger Institute COVID-19 Surveillance Team |
| EPI_ISL_2850445 | Lighthouse Lab in Milton Keynes | Wellcome Sanger Institute for the COVID-19 Genomics UK (COG-UK) Consortium | The Lighthouse Lab in Milton Keynes and Alex Alderton, Roberto Amato, Jeffrey Barrett, Sonia Goncalves, Ewan Harrison, David K. Jackson, Ian Johnston, Dominic Kwiatkowski, Cordelia Langford, John Sillitoe on behalf of the Wellcome Sanger Institute COVID-19 Surveillance Team |
| EPI_ISL_2850446, EPI_ISL_2850447, EPI_ISL_2850448 | Lighthouse Lab in Alderley Park | Wellcome Sanger Institute for the COVID-19 Genomics UK (COG-UK) Consortium | Jacquelyn Wynn, Mairead Hyland, The Lighthouse Lab in Alderley Park and Alex Alderton, Roberto Amato, Jeffrey Barrett, Sonia Goncalves, Ewan Harrison, David K. Jackson, Ian Johnston, Dominic Kwiatkowski, Cordelia Langford, John Sillitoe on behalf of the Wellcome Sanger Institute COVID-19 Surveillance Team |
| EPI_ISL_2850449 | Lighthouse Lab in Milton Keynes | Wellcome Sanger Institute for the COVID-19 Genomics UK (COG-UK) Consortium | The Lighthouse Lab in Milton Keynes and Alex Alderton, Roberto Amato, Jeffrey Barrett, Sonia Goncalves, Ewan Harrison, David K. Jackson, Ian Johnston, Dominic Kwiatkowski, Cordelia Langford, John Sillitoe on behalf of the Wellcome Sanger Institute COVID-19 Surveillance Team |
| EPI_ISL_2850450 | Lighthouse Laboratory Plymouth | Wellcome Sanger Institute for the COVID-19 Genomics UK (COG-UK) Consortium | Lighthouse Laboratory Plymouth and Alex Alderton, Roberto Amato, Jeffrey Barrett, Sonia Goncalves, Ewan Harrison, David K. Jackson, Ian Johnston, Dominic Kwiatkowski, Cordelia Langford, John Sillitoe on behalf of the Wellcome Sanger Institute COVID-19 Surveillance Team |
| EPI_ISL_2850451 | Lighthouse Lab in Alderley Park | Wellcome Sanger Institute for the COVID-19 Genomics UK (COG-UK) Consortium | Jacquelyn Wynn, Mairead Hyland, The Lighthouse Lab in Alderley Park and Alex Alderton, Roberto Amato, Jeffrey Barrett, Sonia Goncalves, Ewan Harrison, David K. Jackson, Ian Johnston, Dominic Kwiatkowski, Cordelia Langford, John Sillitoe on behalf of the Wellcome Sanger Institute COVID-19 Surveillance Team |
| EPI_ISL_2850452 | Lighthouse Laboratory Plymouth | Wellcome Sanger Institute for the COVID-19 Genomics UK (COG-UK) Consortium | Lighthouse Laboratory Plymouth and Alex Alderton, Roberto Amato, Jeffrey Barrett, Sonia Goncalves, Ewan Harrison, David K. Jackson, Ian Johnston, Dominic Kwiatkowski, Cordelia Langford, John Sillitoe on behalf of the Wellcome Sanger Institute COVID-19 Surveillance Team |
| EPI_ISL_2850453, EPI_ISL_2850454, EPI_ISL_2850455, EPI_ISL_2850456, EPI_ISL_2850457 | Lighthouse Lab in Alderley Park | Wellcome Sanger Institute for the COVID-19 Genomics UK (COG-UK) Consortium | Jacquelyn Wynn, Mairead Hyland, The Lighthouse Lab in Alderley Park and Alex Alderton, Roberto Amato, Jeffrey Barrett, Sonia Goncalves, Ewan Harrison, David K. Jackson, Ian Johnston, Dominic Kwiatkowski, Cordelia Langford, John Sillitoe on behalf of the Wellcome Sanger Institute COVID-19 Surveillance Team |
| EPI_ISL_2850458 | Lighthouse Lab in Milton Keynes | Wellcome Sanger Institute for the COVID-19 Genomics UK (COG-UK) Consortium | The Lighthouse Lab in Milton Keynes and Alex Alderton, Roberto Amato, Jeffrey Barrett, Sonia Goncalves, Ewan Harrison, David K. Jackson, Ian Johnston, Dominic Kwiatkowski, Cordelia Langford, John Sillitoe on behalf of the Wellcome Sanger Institute COVID-19 Surveillance Team |
| EPI_ISL_2850459 | Lighthouse Lab in Alderley Park | Wellcome Sanger Institute for the COVID-19 Genomics UK (COG-UK) Consortium | Jacquelyn Wynn, Mairead Hyland, The Lighthouse Lab in Alderley Park and Alex Alderton, Roberto Amato, Jeffrey Barrett, Sonia Goncalves, Ewan Harrison, David K. Jackson, Ian Johnston, Dominic Kwiatkowski, Cordelia Langford, John Sillitoe on behalf of the Wellcome Sanger Institute COVID-19 Surveillance Team |
| EPI_ISL_2850460, EPI_ISL_2850461 | Lighthouse Lab in Milton Keynes | Wellcome Sanger Institute for the COVID-19 Genomics UK (COG-UK) Consortium | The Lighthouse Lab in Milton Keynes and Alex Alderton, Roberto Amato, Jeffrey Barrett, Sonia Goncalves, Ewan Harrison, David K. Jackson, Ian Johnston, Dominic Kwiatkowski, Cordelia Langford, John Sillitoe on behalf of the Wellcome Sanger Institute COVID-19 Surveillance Team |
| EPI_ISL_2850462 | Lighthouse Laboratory Plymouth | Wellcome Sanger Institute for the COVID-19 Genomics UK (COG-UK) Consortium | Lighthouse Laboratory Plymouth and Alex Alderton, Roberto Amato, Jeffrey Barrett, Sonia Goncalves, Ewan Harrison, David K. Jackson, Ian Johnston, Dominic Kwiatkowski, Cordelia Langford, John Sillitoe on behalf of the Wellcome Sanger Institute COVID-19 Surveillance Team |
| EPI_ISL_2850463, EPI_ISL_2850464 | Lighthouse Lab in Milton Keynes | Wellcome Sanger Institute for the COVID-19 Genomics UK (COG-UK) Consortium | The Lighthouse Lab in Milton Keynes and Alex Alderton, Roberto Amato, Jeffrey Barrett, Sonia Goncalves, Ewan Harrison, David K. Jackson, Ian Johnston, Dominic Kwiatkowski, Cordelia Langford, John Sillitoe on behalf of the Wellcome Sanger Institute COVID-19 Surveillance Team |
| EPI_ISL_2850465, EPI_ISL_2850466 | Lighthouse Lab in Alderley Park | Wellcome Sanger Institute for the COVID-19 Genomics UK (COG-UK) Consortium | Jacquelyn Wynn, Mairead Hyland, The Lighthouse Lab in Alderley Park and Alex Alderton, Roberto Amato, Jeffrey Barrett, Sonia Goncalves, Ewan Harrison, David K. Jackson, Ian Johnston, Dominic Kwiatkowski, Cordelia Langford, John Sillitoe on behalf of the Wellcome Sanger Institute COVID-19 Surveillance Team |
| EPI_ISL_2850467 | Lighthouse Laboratory Plymouth | Wellcome Sanger Institute for the COVID-19 Genomics UK (COG-UK) Consortium | Lighthouse Laboratory Plymouth and Alex Alderton, Roberto Amato, Jeffrey Barrett, Sonia Goncalves, Ewan Harrison, David K. Jackson, Ian Johnston, Dominic Kwiatkowski, Cordelia Langford, John Sillitoe on behalf of the Wellcome Sanger Institute COVID-19 Surveillance Team |
| EPI_ISL_2850468 | Lighthouse Lab in Alderley Park | Wellcome Sanger Institute for the COVID-19 Genomics UK (COG-UK) Consortium | Jacquelyn Wynn, Mairead Hyland, The Lighthouse Lab in Alderley Park and Alex Alderton, Roberto Amato, Jeffrey Barrett, Sonia Goncalves, Ewan Harrison, David K. Jackson, Ian Johnston, Dominic Kwiatkowski, Cordelia Langford, John Sillitoe on behalf of the Wellcome Sanger Institute COVID-19 Surveillance Team |
| EPI_ISL_2850469 | Lighthouse Lab in Milton Keynes | Wellcome Sanger Institute for the COVID-19 Genomics UK (COG-UK) Consortium | The Lighthouse Lab in Milton Keynes and Alex Alderton, Roberto Amato, Jeffrey Barrett, Sonia Goncalves, Ewan Harrison, David K. Jackson, Ian Johnston, Dominic Kwiatkowski, Cordelia Langford, John Sillitoe on behalf of the Wellcome Sanger Institute COVID-19 Surveillance Team |
| EPI_ISL_2850470 | Lighthouse Laboratory Plymouth | Wellcome Sanger Institute for the COVID-19 Genomics UK (COG-UK) Consortium | Lighthouse Laboratory Plymouth and Alex Alderton, Roberto Amato, Jeffrey Barrett, Sonia Goncalves, Ewan Harrison, David K. Jackson, Ian Johnston, Dominic Kwiatkowski, Cordelia Langford, John Sillitoe on behalf of the Wellcome Sanger Institute COVID-19 Surveillance Team |
| EPI_ISL_2850471, EPI_ISL_2850472, EPI_ISL_2850473 | Lighthouse Lab in Alderley Park | Wellcome Sanger Institute for the COVID-19 Genomics UK (COG-UK) Consortium | Jacquelyn Wynn, Mairead Hyland, The Lighthouse Lab in Alderley Park and Alex Alderton, Roberto Amato, Jeffrey Barrett, Sonia Goncalves, Ewan Harrison, David K. Jackson, Ian Johnston, Dominic Kwiatkowski, Cordelia Langford, John Sillitoe on behalf of the Wellcome Sanger Institute COVID-19 Surveillance Team |
| EPI_ISL_2850474 | Lighthouse Lab in Milton Keynes | Wellcome Sanger Institute for the COVID-19 Genomics UK (COG-UK) Consortium | The Lighthouse Lab in Milton Keynes and Alex Alderton, Roberto Amato, Jeffrey Barrett, Sonia Goncalves, Ewan Harrison, David K. Jackson, Ian Johnston, Dominic Kwiatkowski, Cordelia Langford, John Sillitoe on behalf of the Wellcome Sanger Institute COVID-19 Surveillance Team |
| EPI_ISL_2850475, EPI_ISL_2850476 | Lighthouse Lab in Alderley Park | Wellcome Sanger Institute for the COVID-19 Genomics UK (COG-UK) Consortium | Jacquelyn Wynn, Mairead Hyland, The Lighthouse Lab in Alderley Park and Alex Alderton, Roberto Amato, Jeffrey Barrett, Sonia Goncalves, Ewan Harrison, David K. Jackson, Ian Johnston, Dominic Kwiatkowski, Cordelia Langford, John Sillitoe on behalf of the Wellcome Sanger Institute COVID-19 Surveillance Team |
| EPI_ISL_2850477, EPI_ISL_2850478, EPI_ISL_2850479 | Lighthouse Lab in Milton Keynes | Wellcome Sanger Institute for the COVID-19 Genomics UK (COG-UK) Consortium | The Lighthouse Lab in Milton Keynes and Alex Alderton, Roberto Amato, Jeffrey Barrett, Sonia Goncalves, Ewan Harrison, David K. Jackson, Ian Johnston, Dominic Kwiatkowski, Cordelia Langford, John Sillitoe on behalf of the Wellcome Sanger Institute COVID-19 Surveillance Team |
| EPI_ISL_2850480 | Lighthouse Lab in Alderley Park | Wellcome Sanger Institute for the COVID-19 Genomics UK (COG-UK) Consortium | Jacquelyn Wynn, Mairead Hyland, The Lighthouse Lab in Alderley Park and Alex Alderton, Roberto Amato, Jeffrey Barrett, Sonia Goncalves, Ewan Harrison, David K. Jackson, Ian Johnston, Dominic Kwiatkowski, Cordelia Langford, John Sillitoe on behalf of the Wellcome Sanger Institute COVID-19 Surveillance Team |
| EPI_ISL_2850481 | Lighthouse Lab in Milton Keynes | Wellcome Sanger Institute for the COVID-19 Genomics UK (COG-UK) Consortium | The Lighthouse Lab in Milton Keynes and Alex Alderton, Roberto Amato, Jeffrey Barrett, Sonia Goncalves, Ewan Harrison, David K. Jackson, Ian Johnston, Dominic Kwiatkowski, Cordelia Langford, John Sillitoe on behalf of the Wellcome Sanger Institute COVID-19 Surveillance Team |
| EPI_ISL_2850482, EPI_ISL_2850483, EPI_ISL_2850484 | Lighthouse Lab in Alderley Park | Wellcome Sanger Institute for the COVID-19 Genomics UK (COG-UK) Consortium | Jacquelyn Wynn, Mairead Hyland, The Lighthouse Lab in Alderley Park and Alex Alderton, Roberto Amato, Jeffrey Barrett, Sonia Goncalves, Ewan Harrison, David K. Jackson, Ian Johnston, Dominic Kwiatkowski, Cordelia Langford, John Sillitoe on behalf of the Wellcome Sanger Institute COVID-19 Surveillance Team |
| EPI_ISL_2850485 | Lighthouse Laboratory Plymouth | Wellcome Sanger Institute for the COVID-19 Genomics UK (COG-UK) Consortium | Lighthouse Laboratory Plymouth and Alex Alderton, Roberto Amato, Jeffrey Barrett, Sonia Goncalves, Ewan Harrison, David K. Jackson, Ian Johnston, Dominic Kwiatkowski, Cordelia Langford, John Sillitoe on behalf of the Wellcome Sanger Institute COVID-19 Surveillance Team |
| EPI_ISL_2850486, EPI_ISL_2850487, EPI_ISL_2850488 | Lighthouse Lab in Alderley Park | Wellcome Sanger Institute for the COVID-19 Genomics UK (COG-UK) Consortium | Jacquelyn Wynn, Mairead Hyland, The Lighthouse Lab in Alderley Park and Alex Alderton, Roberto Amato, Jeffrey Barrett, Sonia Goncalves, Ewan Harrison, David K. Jackson, Ian Johnston, Dominic Kwiatkowski, Cordelia Langford, John Sillitoe on behalf of the Wellcome Sanger Institute COVID-19 Surveillance Team |
| EPI_ISL_2850489 | Lighthouse Laboratory Plymouth | Wellcome Sanger Institute for the COVID-19 Genomics UK (COG-UK) Consortium | Lighthouse Laboratory Plymouth and Alex Alderton, Roberto Amato, Jeffrey Barrett, Sonia Goncalves, Ewan Harrison, David K. Jackson, Ian Johnston, Dominic Kwiatkowski, Cordelia Langford, John Sillitoe on behalf of the Wellcome Sanger Institute COVID-19 Surveillance Team |
| EPI_ISL_2850490, EPI_ISL_2850491 | Lighthouse Lab in Alderley Park | Wellcome Sanger Institute for the COVID-19 Genomics UK (COG-UK) Consortium | Jacquelyn Wynn, Mairead Hyland, The Lighthouse Lab in Alderley Park and Alex Alderton, Roberto Amato, Jeffrey Barrett, Sonia Goncalves, Ewan Harrison, David K. Jackson, Ian Johnston, Dominic Kwiatkowski, Cordelia Langford, John Sillitoe on behalf of the Wellcome Sanger Institute COVID-19 Surveillance Team |
| EPI_ISL_2850492 | Lighthouse Lab in Milton Keynes | Wellcome Sanger Institute for the COVID-19 Genomics UK (COG-UK) Consortium | The Lighthouse Lab in Milton Keynes and Alex Alderton, Roberto Amato, Jeffrey Barrett, Sonia Goncalves, Ewan Harrison, David K. Jackson, Ian Johnston, Dominic Kwiatkowski, Cordelia Langford, John Sillitoe on behalf of the Wellcome Sanger Institute COVID-19 Surveillance Team |
| EPI_ISL_2850493 | Lighthouse Laboratory Plymouth | Wellcome Sanger Institute for the COVID-19 Genomics UK (COG-UK) Consortium | Lighthouse Laboratory Plymouth and Alex Alderton, Roberto Amato, Jeffrey Barrett, Sonia Goncalves, Ewan Harrison, David K. Jackson, Ian Johnston, Dominic Kwiatkowski, Cordelia Langford, John Sillitoe on behalf of the Wellcome Sanger Institute COVID-19 Surveillance Team |
| EPI_ISL_2850494 | Lighthouse Lab in Milton Keynes | Wellcome Sanger Institute for the COVID-19 Genomics UK (COG-UK) Consortium | The Lighthouse Lab in Milton Keynes and Alex Alderton, Roberto Amato, Jeffrey Barrett, Sonia Goncalves, Ewan Harrison, David K. Jackson, Ian Johnston, Dominic Kwiatkowski, Cordelia Langford, John Sillitoe on behalf of the Wellcome Sanger Institute COVID-19 Surveillance Team |
| EPI_ISL_2850495 | Lighthouse Lab in Alderley Park | Wellcome Sanger Institute for the COVID-19 Genomics UK (COG-UK) Consortium | Jacquelyn Wynn, Mairead Hyland, The Lighthouse Lab in Alderley Park and Alex Alderton, Roberto Amato, Jeffrey Barrett, Sonia Goncalves, Ewan Harrison, David K. Jackson, Ian Johnston, Dominic Kwiatkowski, Cordelia Langford, John Sillitoe on behalf of the Wellcome Sanger Institute COVID-19 Surveillance Team |
| EPI_ISL_2850496, EPI_ISL_2850497, EPI_ISL_2850498 | Lighthouse Lab in Milton Keynes | Wellcome Sanger Institute for the COVID-19 Genomics UK (COG-UK) Consortium | The Lighthouse Lab in Milton Keynes and Alex Alderton, Roberto Amato, Jeffrey Barrett, Sonia Goncalves, Ewan Harrison, David K. Jackson, Ian Johnston, Dominic Kwiatkowski, Cordelia Langford, John Sillitoe on behalf of the Wellcome Sanger Institute COVID-19 Surveillance Team |
| EPI_ISL_2850499 | Lighthouse Laboratory Plymouth | Wellcome Sanger Institute for the COVID-19 Genomics UK (COG-UK) Consortium | Lighthouse Laboratory Plymouth and Alex Alderton, Roberto Amato, Jeffrey Barrett, Sonia Goncalves, Ewan Harrison, David K. Jackson, Ian Johnston, Dominic Kwiatkowski, Cordelia Langford, John Sillitoe on behalf of the Wellcome Sanger Institute COVID-19 Surveillance Team |
| EPI_ISL_2850500, EPI_ISL_2850501 | Lighthouse Lab in Alderley Park | Wellcome Sanger Institute for the COVID-19 Genomics UK (COG-UK) Consortium | Jacquelyn Wynn, Mairead Hyland, The Lighthouse Lab in Alderley Park and Alex Alderton, Roberto Amato, Jeffrey Barrett, Sonia Goncalves, Ewan Harrison, David K. Jackson, Ian Johnston, Dominic Kwiatkowski, Cordelia Langford, John Sillitoe on behalf of the Wellcome Sanger Institute COVID-19 Surveillance Team |
| EPI_ISL_2850502, EPI_ISL_2850503 | Lighthouse Laboratory Plymouth | Wellcome Sanger Institute for the COVID-19 Genomics UK (COG-UK) Consortium | Lighthouse Laboratory Plymouth and Alex Alderton, Roberto Amato, Jeffrey Barrett, Sonia Goncalves, Ewan Harrison, David K. Jackson, Ian Johnston, Dominic Kwiatkowski, Cordelia Langford, John Sillitoe on behalf of the Wellcome Sanger Institute COVID-19 Surveillance Team |
| EPI_ISL_2850504 | Lighthouse Lab in Milton Keynes | Wellcome Sanger Institute for the COVID-19 Genomics UK (COG-UK) Consortium | The Lighthouse Lab in Milton Keynes and Alex Alderton, Roberto Amato, Jeffrey Barrett, Sonia Goncalves, Ewan Harrison, David K. Jackson, Ian Johnston, Dominic Kwiatkowski, Cordelia Langford, John Sillitoe on behalf of the Wellcome Sanger Institute COVID-19 Surveillance Team |
| EPI_ISL_2850505 | Lighthouse Laboratory Plymouth | Wellcome Sanger Institute for the COVID-19 Genomics UK (COG-UK) Consortium | Lighthouse Laboratory Plymouth and Alex Alderton, Roberto Amato, Jeffrey Barrett, Sonia Goncalves, Ewan Harrison, David K. Jackson, Ian Johnston, Dominic Kwiatkowski, Cordelia Langford, John Sillitoe on behalf of the Wellcome Sanger Institute COVID-19 Surveillance Team |
| EPI_ISL_2850506, | Lighthouse Lab in Alderley Park | Wellcome Sanger Institute for the COVID-19 Genomics UK (COG-UK) | Jacquelyn Wynn, Mairead Hyland, The Lighthouse Lab in Alderley Park and Alex Alderton, Roberto Amato, Jeffrey Barrett, Sonia Goncalves, Ewan Harrison, David K. Jackson, Ian Johnston, Dominic Kwiatkowski, Cordelia Langford, John Sillitoe on |

| | | | |
|---|---|---|---|
| EPI_ISL_2850507, EPI_ISL_2850508 | | Consortium | behalf of the Wellcome Sanger Institute COVID-19 Surveillance Team |
| EPI_ISL_2850509, EPI_ISL_2850510, EPI_ISL_2850511, EPI_ISL_2850513 | Lighthouse Laboratory Plymouth | Wellcome Sanger Institute for the COVID-19 Genomics UK (COG-UK) Consortium | Lighthouse Laboratory Plymouth and Alex Alderton, Roberto Amato, Jeffrey Barrett, Sonia Goncalves, Ewan Harrison, David K. Jackson, Ian Johnston, Dominic Kwiatkowski, Cordelia Langford, John Sillitoe on behalf of the Wellcome Sanger Institute COVID-19 Surveillance Team |
| EPI_ISL_2850514 | Lighthouse Lab in Alderley Park | Wellcome Sanger Institute for the COVID-19 Genomics UK (COG-UK) Consortium | Jacquelyn Wynn, Mairead Hyland, The Lighthouse Lab in Alderley Park and Alex Alderton, Roberto Amato, Jeffrey Barrett, Sonia Goncalves, Ewan Harrison, David K. Jackson, Ian Johnston, Dominic Kwiatkowski, Cordelia Langford, John Sillitoe on behalf of the Wellcome Sanger Institute COVID-19 Surveillance Team |
| EPI_ISL_2850515, EPI_ISL_2850516 | Lighthouse Laboratory Plymouth | Wellcome Sanger Institute for the COVID-19 Genomics UK (COG-UK) Consortium | Lighthouse Laboratory Plymouth and Alex Alderton, Roberto Amato, Jeffrey Barrett, Sonia Goncalves, Ewan Harrison, David K. Jackson, Ian Johnston, Dominic Kwiatkowski, Cordelia Langford, John Sillitoe on behalf of the Wellcome Sanger Institute COVID-19 Surveillance Team |
| EPI_ISL_2850517, EPI_ISL_2850518, EPI_ISL_2850519, EPI_ISL_2850520, EPI_ISL_2850521 | Lighthouse Lab in Alderley Park | Wellcome Sanger Institute for the COVID-19 Genomics UK (COG-UK) Consortium | Jacquelyn Wynn, Mairead Hyland, The Lighthouse Lab in Alderley Park and Alex Alderton, Roberto Amato, Jeffrey Barrett, Sonia Goncalves, Ewan Harrison, David K. Jackson, Ian Johnston, Dominic Kwiatkowski, Cordelia Langford, John Sillitoe on behalf of the Wellcome Sanger Institute COVID-19 Surveillance Team |
| EPI_ISL_2850522 | Lighthouse Lab in Milton Keynes | Wellcome Sanger Institute for the COVID-19 Genomics UK (COG-UK) Consortium | The Lighthouse Lab in Milton Keynes and Alex Alderton, Roberto Amato, Jeffrey Barrett, Sonia Goncalves, Ewan Harrison, David K. Jackson, Ian Johnston, Dominic Kwiatkowski, Cordelia Langford, John Sillitoe on behalf of the Wellcome Sanger Institute COVID-19 Surveillance Team |
| EPI_ISL_2850523 | Lighthouse Lab in Alderley Park | Wellcome Sanger Institute for the COVID-19 Genomics UK (COG-UK) Consortium | Jacquelyn Wynn, Mairead Hyland, The Lighthouse Lab in Alderley Park and Alex Alderton, Roberto Amato, Jeffrey Barrett, Sonia Goncalves, Ewan Harrison, David K. Jackson, Ian Johnston, Dominic Kwiatkowski, Cordelia Langford, John Sillitoe on behalf of the Wellcome Sanger Institute COVID-19 Surveillance Team |
| EPI_ISL_2850524 | Lighthouse Lab in Milton Keynes | Wellcome Sanger Institute for the COVID-19 Genomics UK (COG-UK) Consortium | The Lighthouse Lab in Milton Keynes and Alex Alderton, Roberto Amato, Jeffrey Barrett, Sonia Goncalves, Ewan Harrison, David K. Jackson, Ian Johnston, Dominic Kwiatkowski, Cordelia Langford, John Sillitoe on behalf of the Wellcome Sanger Institute COVID-19 Surveillance Team |
| EPI_ISL_2850525, EPI_ISL_2850526 | Lighthouse Lab in Alderley Park | Wellcome Sanger Institute for the COVID-19 Genomics UK (COG-UK) Consortium | Jacquelyn Wynn, Mairead Hyland, The Lighthouse Lab in Alderley Park and Alex Alderton, Roberto Amato, Jeffrey Barrett, Sonia Goncalves, Ewan Harrison, David K. Jackson, Ian Johnston, Dominic Kwiatkowski, Cordelia Langford, John Sillitoe on behalf of the Wellcome Sanger Institute COVID-19 Surveillance Team |
| EPI_ISL_2850527 | Lighthouse Laboratory Plymouth | Wellcome Sanger Institute for the COVID-19 Genomics UK (COG-UK) Consortium | Lighthouse Laboratory Plymouth and Alex Alderton, Roberto Amato, Jeffrey Barrett, Sonia Goncalves, Ewan Harrison, David K. Jackson, Ian Johnston, Dominic Kwiatkowski, Cordelia Langford, John Sillitoe on behalf of the Wellcome Sanger Institute COVID-19 Surveillance Team |
| EPI_ISL_2850528 | Lighthouse Lab in Milton Keynes | Wellcome Sanger Institute for the COVID-19 Genomics UK (COG-UK) Consortium | The Lighthouse Lab in Milton Keynes and Alex Alderton, Roberto Amato, Jeffrey Barrett, Sonia Goncalves, Ewan Harrison, David K. Jackson, Ian Johnston, Dominic Kwiatkowski, Cordelia Langford, John Sillitoe on behalf of the Wellcome Sanger Institute COVID-19 Surveillance Team |
| EPI_ISL_2850529 | Lighthouse Lab in Alderley Park | Wellcome Sanger Institute for the COVID-19 Genomics UK (COG-UK) Consortium | Jacquelyn Wynn, Mairead Hyland, The Lighthouse Lab in Alderley Park and Alex Alderton, Roberto Amato, Jeffrey Barrett, Sonia Goncalves, Ewan Harrison, David K. Jackson, Ian Johnston, Dominic Kwiatkowski, Cordelia Langford, John Sillitoe on behalf of the Wellcome Sanger Institute COVID-19 Surveillance Team |
| EPI_ISL_2850530 | Lighthouse Laboratory Plymouth | Wellcome Sanger Institute for the COVID-19 Genomics UK (COG-UK) Consortium | Lighthouse Laboratory Plymouth and Alex Alderton, Roberto Amato, Jeffrey Barrett, Sonia Goncalves, Ewan Harrison, David K. Jackson, Ian Johnston, Dominic Kwiatkowski, Cordelia Langford, John Sillitoe on behalf of the Wellcome Sanger Institute COVID-19 Surveillance Team |
| EPI_ISL_2850531 | Lighthouse Lab in Milton Keynes | Wellcome Sanger Institute for the COVID-19 Genomics UK (COG-UK) Consortium | The Lighthouse Lab in Milton Keynes and Alex Alderton, Roberto Amato, Jeffrey Barrett, Sonia Goncalves, Ewan Harrison, David K. Jackson, Ian Johnston, Dominic Kwiatkowski, Cordelia Langford, John Sillitoe on behalf of the Wellcome Sanger Institute COVID-19 Surveillance Team |
| EPI_ISL_2850532 | Lighthouse Laboratory Plymouth | Wellcome Sanger Institute for the COVID-19 Genomics UK (COG-UK) Consortium | Lighthouse Laboratory Plymouth and Alex Alderton, Roberto Amato, Jeffrey Barrett, Sonia Goncalves, Ewan Harrison, David K. Jackson, Ian Johnston, Dominic Kwiatkowski, Cordelia Langford, John Sillitoe on behalf of the Wellcome Sanger Institute COVID-19 Surveillance Team |
| EPI_ISL_2850533, EPI_ISL_2850534 | Lighthouse Lab in Alderley Park | Wellcome Sanger Institute for the COVID-19 Genomics UK (COG-UK) Consortium | Jacquelyn Wynn, Mairead Hyland, The Lighthouse Lab in Alderley Park and Alex Alderton, Roberto Amato, Jeffrey Barrett, Sonia Goncalves, Ewan Harrison, David K. Jackson, Ian Johnston, Dominic Kwiatkowski, Cordelia Langford, John Sillitoe on behalf of the Wellcome Sanger Institute COVID-19 Surveillance Team |
| EPI_ISL_2850535 | Lighthouse Lab in Milton Keynes | Wellcome Sanger Institute for the COVID-19 Genomics UK (COG-UK) Consortium | The Lighthouse Lab in Milton Keynes and Alex Alderton, Roberto Amato, Jeffrey Barrett, Sonia Goncalves, Ewan Harrison, David K. Jackson, Ian Johnston, Dominic Kwiatkowski, Cordelia Langford, John Sillitoe on behalf of the Wellcome Sanger Institute COVID-19 Surveillance Team |
| EPI_ISL_2850536 | Lighthouse Laboratory Plymouth | Wellcome Sanger Institute for the COVID-19 Genomics UK (COG-UK) Consortium | Lighthouse Laboratory Plymouth and Alex Alderton, Roberto Amato, Jeffrey Barrett, Sonia Goncalves, Ewan Harrison, David K. Jackson, Ian Johnston, Dominic Kwiatkowski, Cordelia Langford, John Sillitoe on behalf of the Wellcome Sanger Institute COVID-19 Surveillance Team |
| EPI_ISL_2850537 | Lighthouse Lab in Milton Keynes | Wellcome Sanger Institute for the COVID-19 Genomics UK (COG-UK) Consortium | The Lighthouse Lab in Milton Keynes and Alex Alderton, Roberto Amato, Jeffrey Barrett, Sonia Goncalves, Ewan Harrison, David K. Jackson, Ian Johnston, Dominic Kwiatkowski, Cordelia Langford, John Sillitoe on behalf of the Wellcome Sanger Institute COVID-19 Surveillance Team |
| EPI_ISL_2850538 | Lighthouse Laboratory Plymouth | Wellcome Sanger Institute for the COVID-19 Genomics UK (COG-UK) Consortium | Lighthouse Laboratory Plymouth and Alex Alderton, Roberto Amato, Jeffrey Barrett, Sonia Goncalves, Ewan Harrison, David K. Jackson, Ian Johnston, Dominic Kwiatkowski, Cordelia Langford, John Sillitoe on behalf of the Wellcome Sanger Institute COVID-19 Surveillance Team |
| EPI_ISL_2850539 | Lighthouse Lab in Alderley Park | Wellcome Sanger Institute for the COVID-19 Genomics UK (COG-UK) Consortium | Jacquelyn Wynn, Mairead Hyland, The Lighthouse Lab in Alderley Park and Alex Alderton, Roberto Amato, Jeffrey Barrett, Sonia Goncalves, Ewan Harrison, David K. Jackson, Ian Johnston, Dominic Kwiatkowski, Cordelia Langford, John Sillitoe on behalf of the Wellcome Sanger Institute COVID-19 Surveillance Team |
| EPI_ISL_2850540 | Lighthouse Lab in Milton Keynes | Wellcome Sanger Institute for the COVID-19 Genomics UK (COG-UK) Consortium | The Lighthouse Lab in Milton Keynes and Alex Alderton, Roberto Amato, Jeffrey Barrett, Sonia Goncalves, Ewan Harrison, David K. Jackson, Ian Johnston, Dominic Kwiatkowski, Cordelia Langford, John Sillitoe on behalf of the Wellcome Sanger Institute COVID-19 Surveillance Team |
| EPI_ISL_2850541 | Lighthouse Laboratory Plymouth | Wellcome Sanger Institute for the COVID-19 Genomics UK (COG-UK) Consortium | Lighthouse Laboratory Plymouth and Alex Alderton, Roberto Amato, Jeffrey Barrett, Sonia Goncalves, Ewan Harrison, David K. Jackson, Ian Johnston, Dominic Kwiatkowski, Cordelia Langford, John Sillitoe on behalf of the Wellcome Sanger Institute COVID-19 Surveillance Team |
| EPI_ISL_2850542 | Lighthouse Lab in Alderley Park | Wellcome Sanger Institute for the COVID-19 Genomics UK (COG-UK) Consortium | Jacquelyn Wynn, Mairead Hyland, The Lighthouse Lab in Alderley Park and Alex Alderton, Roberto Amato, Jeffrey Barrett, Sonia Goncalves, Ewan Harrison, David K. Jackson, Ian Johnston, Dominic Kwiatkowski, Cordelia Langford, John Sillitoe on behalf of the Wellcome Sanger Institute COVID-19 Surveillance Team |
| EPI_ISL_2850543 | Lighthouse Laboratory Plymouth | Wellcome Sanger Institute for the COVID-19 Genomics UK (COG-UK) Consortium | Lighthouse Laboratory Plymouth and Alex Alderton, Roberto Amato, Jeffrey Barrett, Sonia Goncalves, Ewan Harrison, David K. Jackson, Ian Johnston, Dominic Kwiatkowski, Cordelia Langford, John Sillitoe on behalf of the Wellcome Sanger Institute COVID-19 Surveillance Team |
| EPI_ISL_2850544, EPI_ISL_2850545 | Lighthouse Lab in Alderley Park | Wellcome Sanger Institute for the COVID-19 Genomics UK (COG-UK) Consortium | Jacquelyn Wynn, Mairead Hyland, The Lighthouse Lab in Alderley Park and Alex Alderton, Roberto Amato, Jeffrey Barrett, Sonia Goncalves, Ewan Harrison, David K. Jackson, Ian Johnston, Dominic Kwiatkowski, Cordelia Langford, John Sillitoe on behalf of the Wellcome Sanger Institute COVID-19 Surveillance Team |
| EPI_ISL_2850546, EPI_ISL_2850547 | Lighthouse Lab in Milton Keynes | Wellcome Sanger Institute for the COVID-19 Genomics UK (COG-UK) Consortium | The Lighthouse Lab in Milton Keynes and Alex Alderton, Roberto Amato, Jeffrey Barrett, Sonia Goncalves, Ewan Harrison, David K. Jackson, Ian Johnston, Dominic Kwiatkowski, Cordelia Langford, John Sillitoe on behalf of the Wellcome Sanger Institute COVID-19 Surveillance Team |
| EPI_ISL_2850548 | Lighthouse Lab in Alderley Park | Wellcome Sanger Institute for the COVID-19 Genomics UK (COG-UK) Consortium | Jacquelyn Wynn, Mairead Hyland, The Lighthouse Lab in Alderley Park and Alex Alderton, Roberto Amato, Jeffrey Barrett, Sonia Goncalves, Ewan Harrison, David K. Jackson, Ian Johnston, Dominic Kwiatkowski, Cordelia Langford, John Sillitoe on behalf of the Wellcome Sanger Institute COVID-19 Surveillance Team |
| EPI_ISL_2850549 | Lighthouse Lab in Milton Keynes | Wellcome Sanger Institute for the COVID-19 Genomics UK (COG-UK) Consortium | The Lighthouse Lab in Milton Keynes and Alex Alderton, Roberto Amato, Jeffrey Barrett, Sonia Goncalves, Ewan Harrison, David K. Jackson, Ian Johnston, Dominic Kwiatkowski, Cordelia Langford, John Sillitoe on behalf of the Wellcome Sanger Institute COVID-19 Surveillance Team |
| EPI_ISL_2850550 | Lighthouse Lab in Alderley Park | Wellcome Sanger Institute for the COVID-19 Genomics UK (COG-UK) Consortium | Jacquelyn Wynn, Mairead Hyland, The Lighthouse Lab in Alderley Park and Alex Alderton, Roberto Amato, Jeffrey Barrett, Sonia Goncalves, Ewan Harrison, David K. Jackson, Ian Johnston, Dominic Kwiatkowski, Cordelia Langford, John Sillitoe on behalf of the Wellcome Sanger Institute COVID-19 Surveillance Team |
| EPI_ISL_2850551 | Lighthouse Lab in Milton Keynes | Wellcome Sanger Institute for the COVID-19 Genomics UK (COG-UK) Consortium | The Lighthouse Lab in Milton Keynes and Alex Alderton, Roberto Amato, Jeffrey Barrett, Sonia Goncalves, Ewan Harrison, David K. Jackson, Ian Johnston, Dominic Kwiatkowski, Cordelia Langford, John Sillitoe on behalf of the Wellcome Sanger Institute COVID-19 Surveillance Team |
| EPI_ISL_2850552 | Lighthouse Laboratory Plymouth | Wellcome Sanger Institute for the COVID-19 Genomics UK (COG-UK) Consortium | Lighthouse Laboratory Plymouth and Alex Alderton, Roberto Amato, Jeffrey Barrett, Sonia Goncalves, Ewan Harrison, David K. Jackson, Ian Johnston, Dominic Kwiatkowski, Cordelia Langford, John Sillitoe on behalf of the Wellcome Sanger Institute COVID-19 Surveillance Team |
| EPI_ISL_2850553 | Lighthouse Lab in Alderley Park | Wellcome Sanger Institute for the COVID-19 Genomics UK (COG-UK) Consortium | Jacquelyn Wynn, Mairead Hyland, The Lighthouse Lab in Alderley Park and Alex Alderton, Roberto Amato, Jeffrey Barrett, Sonia Goncalves, Ewan Harrison, David K. Jackson, Ian Johnston, Dominic Kwiatkowski, Cordelia Langford, John Sillitoe on behalf of the Wellcome Sanger Institute COVID-19 Surveillance Team |
| EPI_ISL_2850554 | Lighthouse Lab in Milton Keynes | Wellcome Sanger Institute for the COVID-19 Genomics UK (COG-UK) Consortium | The Lighthouse Lab in Milton Keynes and Alex Alderton, Roberto Amato, Jeffrey Barrett, Sonia Goncalves, Ewan Harrison, David K. Jackson, Ian Johnston, Dominic Kwiatkowski, Cordelia Langford, John Sillitoe on behalf of the Wellcome Sanger Institute COVID-19 Surveillance Team |
| EPI_ISL_2850555 | Lighthouse Laboratory Plymouth | Wellcome Sanger Institute for the COVID-19 Genomics UK (COG-UK) Consortium | Lighthouse Laboratory Plymouth and Alex Alderton, Roberto Amato, Jeffrey Barrett, Sonia Goncalves, Ewan Harrison, David K. Jackson, Ian Johnston, Dominic Kwiatkowski, Cordelia Langford, John Sillitoe on behalf of the Wellcome Sanger Institute COVID-19 Surveillance Team |
| EPI_ISL_2850556 | Lighthouse Lab in Milton Keynes | Wellcome Sanger Institute for the COVID-19 Genomics UK (COG-UK) Consortium | The Lighthouse Lab in Milton Keynes and Alex Alderton, Roberto Amato, Jeffrey Barrett, Sonia Goncalves, Ewan Harrison, David K. Jackson, Ian Johnston, Dominic Kwiatkowski, Cordelia Langford, John Sillitoe on behalf of the Wellcome Sanger Institute COVID-19 Surveillance Team |
| EPI_ISL_2850557, EPI_ISL_2850558, EPI_ISL_2850559, EPI_ISL_2850560 | Lighthouse Lab in Alderley Park | Wellcome Sanger Institute for the COVID-19 Genomics UK (COG-UK) Consortium | Jacquelyn Wynn, Mairead Hyland, The Lighthouse Lab in Alderley Park and Alex Alderton, Roberto Amato, Jeffrey Barrett, Sonia Goncalves, Ewan Harrison, David K. Jackson, Ian Johnston, Dominic Kwiatkowski, Cordelia Langford, John Sillitoe on behalf of the Wellcome Sanger Institute COVID-19 Surveillance Team |
| EPI_ISL_2850561, EPI_ISL_2850562, EPI_ISL_2850563 | Lighthouse Lab in Milton Keynes | Wellcome Sanger Institute for the COVID-19 Genomics UK (COG-UK) Consortium | The Lighthouse Lab in Milton Keynes and Alex Alderton, Roberto Amato, Jeffrey Barrett, Sonia Goncalves, Ewan Harrison, David K. Jackson, Ian Johnston, Dominic Kwiatkowski, Cordelia Langford, John Sillitoe on behalf of the Wellcome Sanger Institute COVID-19 Surveillance Team |
| EPI_ISL_2850564, EPI_ISL_2850565 | Lighthouse Laboratory Plymouth | Wellcome Sanger Institute for the COVID-19 Genomics UK (COG-UK) Consortium | Lighthouse Laboratory Plymouth and Alex Alderton, Roberto Amato, Jeffrey Barrett, Sonia Goncalves, Ewan Harrison, David K. Jackson, Ian Johnston, Dominic Kwiatkowski, Cordelia Langford, John Sillitoe on behalf of the Wellcome Sanger Institute COVID-19 Surveillance Team |
| EPI_ISL_2850566, EPI_ISL_2850567, EPI_ISL_2850568, EPI_ISL_2850569, EPI_ISL_2850570, EPI_ISL_2850571 | Lighthouse Lab in Alderley Park | Wellcome Sanger Institute for the COVID-19 Genomics UK (COG-UK) Consortium | Jacquelyn Wynn, Mairead Hyland, The Lighthouse Lab in Alderley Park and Alex Alderton, Roberto Amato, Jeffrey Barrett, Sonia Goncalves, Ewan Harrison, David K. Jackson, Ian Johnston, Dominic Kwiatkowski, Cordelia Langford, John Sillitoe on behalf of the Wellcome Sanger Institute COVID-19 Surveillance Team |

| | | | |
|---|---|---|---|
| EPI_ISL_2850572, EPI_ISL_2850573 | Lighthouse Lab in Milton Keynes | Wellcome Sanger Institute for the COVID-19 Genomics UK (COG-UK) Consortium | The Lighthouse Lab in Milton Keynes and Alex Alderton, Roberto Amato, Jeffrey Barrett, Sonia Goncalves, Ewan Harrison, David K. Jackson, Ian Johnston, Dominic Kwiatkowski, Cordelia Langford, John Sillitoe on behalf of the Wellcome Sanger Institute COVID-19 Surveillance Team |
| EPI_ISL_2850574, EPI_ISL_2850575 | Lighthouse Laboratory Plymouth | Wellcome Sanger Institute for the COVID-19 Genomics UK (COG-UK) Consortium | Lighthouse Laboratory Plymouth and Alex Alderton, Roberto Amato, Jeffrey Barrett, Sonia Goncalves, Ewan Harrison, David K. Jackson, Ian Johnston, Dominic Kwiatkowski, Cordelia Langford, John Sillitoe on behalf of the Wellcome Sanger Institute COVID-19 Surveillance Team |
| EPI_ISL_2850576 | Lighthouse Lab in Alderley Park | Wellcome Sanger Institute for the COVID-19 Genomics UK (COG-UK) Consortium | Jacquelyn Wynn, Mairead Hyland, The Lighthouse Lab in Alderley Park and Alex Alderton, Roberto Amato, Jeffrey Barrett, Sonia Goncalves, Ewan Harrison, David K. Jackson, Ian Johnston, Dominic Kwiatkowski, Cordelia Langford, John Sillitoe on behalf of the Wellcome Sanger Institute COVID-19 Surveillance Team |
| EPI_ISL_2850577 | Lighthouse Laboratory Plymouth | Wellcome Sanger Institute for the COVID-19 Genomics UK (COG-UK) Consortium | Lighthouse Laboratory Plymouth and Alex Alderton, Roberto Amato, Jeffrey Barrett, Sonia Goncalves, Ewan Harrison, David K. Jackson, Ian Johnston, Dominic Kwiatkowski, Cordelia Langford, John Sillitoe on behalf of the Wellcome Sanger Institute COVID-19 Surveillance Team |
| EPI_ISL_2850578, EPI_ISL_2850579, EPI_ISL_2850580, EPI_ISL_2850581 | Lighthouse Lab in Milton Keynes | Wellcome Sanger Institute for the COVID-19 Genomics UK (COG-UK) Consortium | The Lighthouse Lab in Milton Keynes and Alex Alderton, Roberto Amato, Jeffrey Barrett, Sonia Goncalves, Ewan Harrison, David K. Jackson, Ian Johnston, Dominic Kwiatkowski, Cordelia Langford, John Sillitoe on behalf of the Wellcome Sanger Institute COVID-19 Surveillance Team |
| EPI_ISL_2850582, EPI_ISL_2850583, EPI_ISL_2850584, EPI_ISL_2850585 | Lighthouse Lab in Alderley Park | Wellcome Sanger Institute for the COVID-19 Genomics UK (COG-UK) Consortium | Jacquelyn Wynn, Mairead Hyland, The Lighthouse Lab in Alderley Park and Alex Alderton, Roberto Amato, Jeffrey Barrett, Sonia Goncalves, Ewan Harrison, David K. Jackson, Ian Johnston, Dominic Kwiatkowski, Cordelia Langford, John Sillitoe on behalf of the Wellcome Sanger Institute COVID-19 Surveillance Team |
| EPI_ISL_2850586 | Lighthouse Lab in Milton Keynes | Wellcome Sanger Institute for the COVID-19 Genomics UK (COG-UK) Consortium | The Lighthouse Lab in Milton Keynes and Alex Alderton, Roberto Amato, Jeffrey Barrett, Sonia Goncalves, Ewan Harrison, David K. Jackson, Ian Johnston, Dominic Kwiatkowski, Cordelia Langford, John Sillitoe on behalf of the Wellcome Sanger Institute COVID-19 Surveillance Team |
| EPI_ISL_2850587 | Lighthouse Lab in Alderley Park | Wellcome Sanger Institute for the COVID-19 Genomics UK (COG-UK) Consortium | Jacquelyn Wynn, Mairead Hyland, The Lighthouse Lab in Alderley Park and Alex Alderton, Roberto Amato, Jeffrey Barrett, Sonia Goncalves, Ewan Harrison, David K. Jackson, Ian Johnston, Dominic Kwiatkowski, Cordelia Langford, John Sillitoe on behalf of the Wellcome Sanger Institute COVID-19 Surveillance Team |
| EPI_ISL_2850588 | Lighthouse Laboratory Plymouth | Wellcome Sanger Institute for the COVID-19 Genomics UK (COG-UK) Consortium | Lighthouse Laboratory Plymouth and Alex Alderton, Roberto Amato, Jeffrey Barrett, Sonia Goncalves, Ewan Harrison, David K. Jackson, Ian Johnston, Dominic Kwiatkowski, Cordelia Langford, John Sillitoe on behalf of the Wellcome Sanger Institute COVID-19 Surveillance Team |
| EPI_ISL_2850589, EPI_ISL_2850590 | Lighthouse Lab in Alderley Park | Wellcome Sanger Institute for the COVID-19 Genomics UK (COG-UK) Consortium | Jacquelyn Wynn, Mairead Hyland, The Lighthouse Lab in Alderley Park and Alex Alderton, Roberto Amato, Jeffrey Barrett, Sonia Goncalves, Ewan Harrison, David K. Jackson, Ian Johnston, Dominic Kwiatkowski, Cordelia Langford, John Sillitoe on behalf of the Wellcome Sanger Institute COVID-19 Surveillance Team |
| EPI_ISL_2850591 | Lighthouse Laboratory Plymouth | Wellcome Sanger Institute for the COVID-19 Genomics UK (COG-UK) Consortium | Lighthouse Laboratory Plymouth and Alex Alderton, Roberto Amato, Jeffrey Barrett, Sonia Goncalves, Ewan Harrison, David K. Jackson, Ian Johnston, Dominic Kwiatkowski, Cordelia Langford, John Sillitoe on behalf of the Wellcome Sanger Institute COVID-19 Surveillance Team |
| EPI_ISL_2850592 | Lighthouse Lab in Alderley Park | Wellcome Sanger Institute for the COVID-19 Genomics UK (COG-UK) Consortium | Jacquelyn Wynn, Mairead Hyland, The Lighthouse Lab in Alderley Park and Alex Alderton, Roberto Amato, Jeffrey Barrett, Sonia Goncalves, Ewan Harrison, David K. Jackson, Ian Johnston, Dominic Kwiatkowski, Cordelia Langford, John Sillitoe on behalf of the Wellcome Sanger Institute COVID-19 Surveillance Team |
| EPI_ISL_2850593 | Lighthouse Laboratory Plymouth | Wellcome Sanger Institute for the COVID-19 Genomics UK (COG-UK) Consortium | Lighthouse Laboratory Plymouth and Alex Alderton, Roberto Amato, Jeffrey Barrett, Sonia Goncalves, Ewan Harrison, David K. Jackson, Ian Johnston, Dominic Kwiatkowski, Cordelia Langford, John Sillitoe on behalf of the Wellcome Sanger Institute COVID-19 Surveillance Team |
| EPI_ISL_2850594 | Lighthouse Lab in Alderley Park | Wellcome Sanger Institute for the COVID-19 Genomics UK (COG-UK) Consortium | Jacquelyn Wynn, Mairead Hyland, The Lighthouse Lab in Alderley Park and Alex Alderton, Roberto Amato, Jeffrey Barrett, Sonia Goncalves, Ewan Harrison, David K. Jackson, Ian Johnston, Dominic Kwiatkowski, Cordelia Langford, John Sillitoe on behalf of the Wellcome Sanger Institute COVID-19 Surveillance Team |
| EPI_ISL_2850595 | Lighthouse Lab in Milton Keynes | Wellcome Sanger Institute for the COVID-19 Genomics UK (COG-UK) Consortium | The Lighthouse Lab in Milton Keynes and Alex Alderton, Roberto Amato, Jeffrey Barrett, Sonia Goncalves, Ewan Harrison, David K. Jackson, Ian Johnston, Dominic Kwiatkowski, Cordelia Langford, John Sillitoe on behalf of the Wellcome Sanger Institute COVID-19 Surveillance Team |
| EPI_ISL_2850596 | Lighthouse Lab in Alderley Park | Wellcome Sanger Institute for the COVID-19 Genomics UK (COG-UK) Consortium | Jacquelyn Wynn, Mairead Hyland, The Lighthouse Lab in Alderley Park and Alex Alderton, Roberto Amato, Jeffrey Barrett, Sonia Goncalves, Ewan Harrison, David K. Jackson, Ian Johnston, Dominic Kwiatkowski, Cordelia Langford, John Sillitoe on behalf of the Wellcome Sanger Institute COVID-19 Surveillance Team |
| EPI_ISL_2850597 | Lighthouse Lab in Milton Keynes | Wellcome Sanger Institute for the COVID-19 Genomics UK (COG-UK) Consortium | The Lighthouse Lab in Milton Keynes and Alex Alderton, Roberto Amato, Jeffrey Barrett, Sonia Goncalves, Ewan Harrison, David K. Jackson, Ian Johnston, Dominic Kwiatkowski, Cordelia Langford, John Sillitoe on behalf of the Wellcome Sanger Institute COVID-19 Surveillance Team |
| EPI_ISL_2850598 | Lighthouse Laboratory Plymouth | Wellcome Sanger Institute for the COVID-19 Genomics UK (COG-UK) Consortium | Lighthouse Laboratory Plymouth and Alex Alderton, Roberto Amato, Jeffrey Barrett, Sonia Goncalves, Ewan Harrison, David K. Jackson, Ian Johnston, Dominic Kwiatkowski, Cordelia Langford, John Sillitoe on behalf of the Wellcome Sanger Institute COVID-19 Surveillance Team |
| EPI_ISL_2850599, EPI_ISL_2850600, EPI_ISL_2850601 | Lighthouse Lab in Alderley Park | Wellcome Sanger Institute for the COVID-19 Genomics UK (COG-UK) Consortium | Jacquelyn Wynn, Mairead Hyland, The Lighthouse Lab in Alderley Park and Alex Alderton, Roberto Amato, Jeffrey Barrett, Sonia Goncalves, Ewan Harrison, David K. Jackson, Ian Johnston, Dominic Kwiatkowski, Cordelia Langford, John Sillitoe on behalf of the Wellcome Sanger Institute COVID-19 Surveillance Team |
| EPI_ISL_2850602 | Lighthouse Laboratory Plymouth | Wellcome Sanger Institute for the COVID-19 Genomics UK (COG-UK) Consortium | Lighthouse Laboratory Plymouth and Alex Alderton, Roberto Amato, Jeffrey Barrett, Sonia Goncalves, Ewan Harrison, David K. Jackson, Ian Johnston, Dominic Kwiatkowski, Cordelia Langford, John Sillitoe on behalf of the Wellcome Sanger Institute COVID-19 Surveillance Team |
| EPI_ISL_2850603 | Lighthouse Lab in Alderley Park | Wellcome Sanger Institute for the COVID-19 Genomics UK (COG-UK) Consortium | Jacquelyn Wynn, Mairead Hyland, The Lighthouse Lab in Alderley Park and Alex Alderton, Roberto Amato, Jeffrey Barrett, Sonia Goncalves, Ewan Harrison, David K. Jackson, Ian Johnston, Dominic Kwiatkowski, Cordelia Langford, John Sillitoe on behalf of the Wellcome Sanger Institute COVID-19 Surveillance Team |
| EPI_ISL_2850604 | Lighthouse Laboratory Plymouth | Wellcome Sanger Institute for the COVID-19 Genomics UK (COG-UK) Consortium | Lighthouse Laboratory Plymouth and Alex Alderton, Roberto Amato, Jeffrey Barrett, Sonia Goncalves, Ewan Harrison, David K. Jackson, Ian Johnston, Dominic Kwiatkowski, Cordelia Langford, John Sillitoe on behalf of the Wellcome Sanger Institute COVID-19 Surveillance Team |
| EPI_ISL_2850605 | Lighthouse Lab in Milton Keynes | Wellcome Sanger Institute for the COVID-19 Genomics UK (COG-UK) Consortium | The Lighthouse Lab in Milton Keynes and Alex Alderton, Roberto Amato, Jeffrey Barrett, Sonia Goncalves, Ewan Harrison, David K. Jackson, Ian Johnston, Dominic Kwiatkowski, Cordelia Langford, John Sillitoe on behalf of the Wellcome Sanger Institute COVID-19 Surveillance Team |
| EPI_ISL_2850606 | Lighthouse Lab in Alderley Park | Wellcome Sanger Institute for the COVID-19 Genomics UK (COG-UK) Consortium | Jacquelyn Wynn, Mairead Hyland, The Lighthouse Lab in Alderley Park and Alex Alderton, Roberto Amato, Jeffrey Barrett, Sonia Goncalves, Ewan Harrison, David K. Jackson, Ian Johnston, Dominic Kwiatkowski, Cordelia Langford, John Sillitoe on behalf of the Wellcome Sanger Institute COVID-19 Surveillance Team |
| EPI_ISL_2850607, EPI_ISL_2850608 | Lighthouse Laboratory Plymouth | Wellcome Sanger Institute for the COVID-19 Genomics UK (COG-UK) Consortium | Lighthouse Laboratory Plymouth and Alex Alderton, Roberto Amato, Jeffrey Barrett, Sonia Goncalves, Ewan Harrison, David K. Jackson, Ian Johnston, Dominic Kwiatkowski, Cordelia Langford, John Sillitoe on behalf of the Wellcome Sanger Institute COVID-19 Surveillance Team |
| EPI_ISL_2850609 | Lighthouse Lab in Milton Keynes | Wellcome Sanger Institute for the COVID-19 Genomics UK (COG-UK) Consortium | The Lighthouse Lab in Milton Keynes and Alex Alderton, Roberto Amato, Jeffrey Barrett, Sonia Goncalves, Ewan Harrison, David K. Jackson, Ian Johnston, Dominic Kwiatkowski, Cordelia Langford, John Sillitoe on behalf of the Wellcome Sanger Institute COVID-19 Surveillance Team |
| EPI_ISL_2850610 | Lighthouse Laboratory Plymouth | Wellcome Sanger Institute for the COVID-19 Genomics UK (COG-UK) Consortium | Lighthouse Laboratory Plymouth and Alex Alderton, Roberto Amato, Jeffrey Barrett, Sonia Goncalves, Ewan Harrison, David K. Jackson, Ian Johnston, Dominic Kwiatkowski, Cordelia Langford, John Sillitoe on behalf of the Wellcome Sanger Institute COVID-19 Surveillance Team |
| EPI_ISL_2850611 | Lighthouse Lab in Alderley Park | Wellcome Sanger Institute for the COVID-19 Genomics UK (COG-UK) Consortium | Jacquelyn Wynn, Mairead Hyland, The Lighthouse Lab in Alderley Park and Alex Alderton, Roberto Amato, Jeffrey Barrett, Sonia Goncalves, Ewan Harrison, David K. Jackson, Ian Johnston, Dominic Kwiatkowski, Cordelia Langford, John Sillitoe on behalf of the Wellcome Sanger Institute COVID-19 Surveillance Team |
| EPI_ISL_2850612, EPI_ISL_2850613 | Lighthouse Lab in Milton Keynes | Wellcome Sanger Institute for the COVID-19 Genomics UK (COG-UK) Consortium | The Lighthouse Lab in Milton Keynes and Alex Alderton, Roberto Amato, Jeffrey Barrett, Sonia Goncalves, Ewan Harrison, David K. Jackson, Ian Johnston, Dominic Kwiatkowski, Cordelia Langford, John Sillitoe on behalf of the Wellcome Sanger Institute COVID-19 Surveillance Team |
| EPI_ISL_2850614 | Lighthouse Laboratory Plymouth | Wellcome Sanger Institute for the COVID-19 Genomics UK (COG-UK) Consortium | Lighthouse Laboratory Plymouth and Alex Alderton, Roberto Amato, Jeffrey Barrett, Sonia Goncalves, Ewan Harrison, David K. Jackson, Ian Johnston, Dominic Kwiatkowski, Cordelia Langford, John Sillitoe on behalf of the Wellcome Sanger Institute COVID-19 Surveillance Team |
| EPI_ISL_2850615 | Lighthouse Lab in Milton Keynes | Wellcome Sanger Institute for the COVID-19 Genomics UK (COG-UK) Consortium | The Lighthouse Lab in Milton Keynes and Alex Alderton, Roberto Amato, Jeffrey Barrett, Sonia Goncalves, Ewan Harrison, David K. Jackson, Ian Johnston, Dominic Kwiatkowski, Cordelia Langford, John Sillitoe on behalf of the Wellcome Sanger Institute COVID-19 Surveillance Team |
| EPI_ISL_2850616, EPI_ISL_2850617, EPI_ISL_2850618, EPI_ISL_2850619 | Lighthouse Lab in Alderley Park | Wellcome Sanger Institute for the COVID-19 Genomics UK (COG-UK) Consortium | Jacquelyn Wynn, Mairead Hyland, The Lighthouse Lab in Alderley Park and Alex Alderton, Roberto Amato, Jeffrey Barrett, Sonia Goncalves, Ewan Harrison, David K. Jackson, Ian Johnston, Dominic Kwiatkowski, Cordelia Langford, John Sillitoe on behalf of the Wellcome Sanger Institute COVID-19 Surveillance Team |
| EPI_ISL_2850620 | Lighthouse Lab in Milton Keynes | Wellcome Sanger Institute for the COVID-19 Genomics UK (COG-UK) Consortium | The Lighthouse Lab in Milton Keynes and Alex Alderton, Roberto Amato, Jeffrey Barrett, Sonia Goncalves, Ewan Harrison, David K. Jackson, Ian Johnston, Dominic Kwiatkowski, Cordelia Langford, John Sillitoe on behalf of the Wellcome Sanger Institute COVID-19 Surveillance Team |
| EPI_ISL_2850621, EPI_ISL_2850622, EPI_ISL_2850623 | Lighthouse Lab in Alderley Park | Wellcome Sanger Institute for the COVID-19 Genomics UK (COG-UK) Consortium | Jacquelyn Wynn, Mairead Hyland, The Lighthouse Lab in Alderley Park and Alex Alderton, Roberto Amato, Jeffrey Barrett, Sonia Goncalves, Ewan Harrison, David K. Jackson, Ian Johnston, Dominic Kwiatkowski, Cordelia Langford, John Sillitoe on behalf of the Wellcome Sanger Institute COVID-19 Surveillance Team |
| EPI_ISL_2850624 | Lighthouse Laboratory Plymouth | Wellcome Sanger Institute for the COVID-19 Genomics UK (COG-UK) Consortium | Lighthouse Laboratory Plymouth and Alex Alderton, Roberto Amato, Jeffrey Barrett, Sonia Goncalves, Ewan Harrison, David K. Jackson, Ian Johnston, Dominic Kwiatkowski, Cordelia Langford, John Sillitoe on behalf of the Wellcome Sanger Institute COVID-19 Surveillance Team |
| EPI_ISL_2850625 | Lighthouse Lab in Milton Keynes | Wellcome Sanger Institute for the COVID-19 Genomics UK (COG-UK) Consortium | The Lighthouse Lab in Milton Keynes and Alex Alderton, Roberto Amato, Jeffrey Barrett, Sonia Goncalves, Ewan Harrison, David K. Jackson, Ian Johnston, Dominic Kwiatkowski, Cordelia Langford, John Sillitoe on behalf of the Wellcome Sanger Institute COVID-19 Surveillance Team |
| EPI_ISL_2850626, EPI_ISL_2850627, EPI_ISL_2850628 | Lighthouse Lab in Alderley Park | Wellcome Sanger Institute for the COVID-19 Genomics UK (COG-UK) Consortium | Jacquelyn Wynn, Mairead Hyland, The Lighthouse Lab in Alderley Park and Alex Alderton, Roberto Amato, Jeffrey Barrett, Sonia Goncalves, Ewan Harrison, David K. Jackson, Ian Johnston, Dominic Kwiatkowski, Cordelia Langford, John Sillitoe on behalf of the Wellcome Sanger Institute COVID-19 Surveillance Team |
| EPI_ISL_2850629 | Lighthouse Lab in Milton Keynes | Wellcome Sanger Institute for the COVID-19 Genomics UK (COG-UK) Consortium | The Lighthouse Lab in Milton Keynes and Alex Alderton, Roberto Amato, Jeffrey Barrett, Sonia Goncalves, Ewan Harrison, David K. Jackson, Ian Johnston, Dominic Kwiatkowski, Cordelia Langford, John Sillitoe on behalf of the Wellcome Sanger Institute COVID-19 Surveillance Team |
| EPI_ISL_2850630, EPI_ISL_2850631 | Lighthouse Lab in Alderley Park | Wellcome Sanger Institute for the COVID-19 Genomics UK (COG-UK) Consortium | Jacquelyn Wynn, Mairead Hyland, The Lighthouse Lab in Alderley Park and Alex Alderton, Roberto Amato, Jeffrey Barrett, Sonia Goncalves, Ewan Harrison, David K. Jackson, Ian Johnston, Dominic Kwiatkowski, Cordelia Langford, John Sillitoe on behalf of the Wellcome Sanger Institute COVID-19 Surveillance Team |
| EPI_ISL_2850632, EPI_ISL_2850633 | Lighthouse Lab in Milton Keynes | Wellcome Sanger Institute for the COVID-19 Genomics UK (COG-UK) Consortium | The Lighthouse Lab in Milton Keynes and Alex Alderton, Roberto Amato, Jeffrey Barrett, Sonia Goncalves, Ewan Harrison, David K. Jackson, Ian Johnston, Dominic Kwiatkowski, Cordelia Langford, John Sillitoe on behalf of the Wellcome Sanger Institute COVID-19 Surveillance Team |
| EPI_ISL_2850634 | Lighthouse Laboratory Plymouth | Wellcome Sanger Institute for the COVID-19 Genomics UK (COG-UK) Consortium | Lighthouse Laboratory Plymouth and Alex Alderton, Roberto Amato, Jeffrey Barrett, Sonia Goncalves, Ewan Harrison, David K. Jackson, Ian Johnston, Dominic Kwiatkowski, Cordelia Langford, John Sillitoe on behalf of the Wellcome Sanger Institute COVID-19 Surveillance Team |
| EPI_ISL_2850635 | Lighthouse Lab in Alderley Park | Wellcome Sanger Institute for the COVID-19 Genomics UK (COG-UK) | Jacquelyn Wynn, Mairead Hyland, The Lighthouse Lab in Alderley Park and Alex Alderton, Roberto Amato, Jeffrey Barrett, Sonia Goncalves, Ewan Harrison, David K. Jackson, Ian Johnston, Dominic Kwiatkowski, Cordelia Langford, John Sillitoe on |

| | | | |
|---|---|---|---|
| | | Consortium | behalf of the Wellcome Sanger Institute COVID-19 Surveillance Team |
| EPI_ISL_2850636 | Lighthouse Lab in Milton Keynes | Wellcome Sanger Institute for the COVID-19 Genomics UK (COG-UK) Consortium | The Lighthouse Lab in Milton Keynes and Alex Alderton, Roberto Amato, Jeffrey Barrett, Sonia Goncalves, Ewan Harrison, David K. Jackson, Ian Johnston, Dominic Kwiatkowski, Cordelia Langford, John Sillitoe on behalf of the Wellcome Sanger Institute COVID-19 Surveillance Team |
| EPI_ISL_2850637 | Lighthouse Lab in Alderley Park | Wellcome Sanger Institute for the COVID-19 Genomics UK (COG-UK) Consortium | Jacquelyn Wynn, Mairead Hyland, The Lighthouse Lab in Alderley Park and Alex Alderton, Roberto Amato, Jeffrey Barrett, Sonia Goncalves, Ewan Harrison, David K. Jackson, Ian Johnston, Dominic Kwiatkowski, Cordelia Langford, John Sillitoe on behalf of the Wellcome Sanger Institute COVID-19 Surveillance Team |
| EPI_ISL_2850638 | Lighthouse Lab in Milton Keynes | Wellcome Sanger Institute for the COVID-19 Genomics UK (COG-UK) Consortium | The Lighthouse Lab in Milton Keynes and Alex Alderton, Roberto Amato, Jeffrey Barrett, Sonia Goncalves, Ewan Harrison, David K. Jackson, Ian Johnston, Dominic Kwiatkowski, Cordelia Langford, John Sillitoe on behalf of the Wellcome Sanger Institute COVID-19 Surveillance Team |
| EPI_ISL_2850639 | Lighthouse Lab in Alderley Park | Wellcome Sanger Institute for the COVID-19 Genomics UK (COG-UK) Consortium | Jacquelyn Wynn, Mairead Hyland, The Lighthouse Lab in Alderley Park and Alex Alderton, Roberto Amato, Jeffrey Barrett, Sonia Goncalves, Ewan Harrison, David K. Jackson, Ian Johnston, Dominic Kwiatkowski, Cordelia Langford, John Sillitoe on behalf of the Wellcome Sanger Institute COVID-19 Surveillance Team |
| EPI_ISL_2850640 | Lighthouse Laboratory Plymouth | Wellcome Sanger Institute for the COVID-19 Genomics UK (COG-UK) Consortium | Lighthouse Laboratory Plymouth and Alex Alderton, Roberto Amato, Jeffrey Barrett, Sonia Goncalves, Ewan Harrison, David K. Jackson, Ian Johnston, Dominic Kwiatkowski, Cordelia Langford, John Sillitoe on behalf of the Wellcome Sanger Institute COVID-19 Surveillance Team |
| EPI_ISL_2850641 | Lighthouse Lab in Milton Keynes | Wellcome Sanger Institute for the COVID-19 Genomics UK (COG-UK) Consortium | The Lighthouse Lab in Milton Keynes and Alex Alderton, Roberto Amato, Jeffrey Barrett, Sonia Goncalves, Ewan Harrison, David K. Jackson, Ian Johnston, Dominic Kwiatkowski, Cordelia Langford, John Sillitoe on behalf of the Wellcome Sanger Institute COVID-19 Surveillance Team |
| EPI_ISL_2850642, EPI_ISL_2850643, EPI_ISL_2850644, EPI_ISL_2850645, EPI_ISL_2850646, EPI_ISL_2850647, EPI_ISL_2850648, EPI_ISL_2850649, EPI_ISL_2850650 | Lighthouse Lab in Alderley Park | Wellcome Sanger Institute for the COVID-19 Genomics UK (COG-UK) Consortium | Jacquelyn Wynn, Mairead Hyland, The Lighthouse Lab in Alderley Park and Alex Alderton, Roberto Amato, Jeffrey Barrett, Sonia Goncalves, Ewan Harrison, David K. Jackson, Ian Johnston, Dominic Kwiatkowski, Cordelia Langford, John Sillitoe on behalf of the Wellcome Sanger Institute COVID-19 Surveillance Team |
| EPI_ISL_2850651 | Lighthouse Laboratory Plymouth | Wellcome Sanger Institute for the COVID-19 Genomics UK (COG-UK) Consortium | Lighthouse Laboratory Plymouth and Alex Alderton, Roberto Amato, Jeffrey Barrett, Sonia Goncalves, Ewan Harrison, David K. Jackson, Ian Johnston, Dominic Kwiatkowski, Cordelia Langford, John Sillitoe on behalf of the Wellcome Sanger Institute COVID-19 Surveillance Team |
| EPI_ISL_2850652, EPI_ISL_2850653 | Lighthouse Lab in Alderley Park | Wellcome Sanger Institute for the COVID-19 Genomics UK (COG-UK) Consortium | Jacquelyn Wynn, Mairead Hyland, The Lighthouse Lab in Alderley Park and Alex Alderton, Roberto Amato, Jeffrey Barrett, Sonia Goncalves, Ewan Harrison, David K. Jackson, Ian Johnston, Dominic Kwiatkowski, Cordelia Langford, John Sillitoe on behalf of the Wellcome Sanger Institute COVID-19 Surveillance Team |
| EPI_ISL_2850654, EPI_ISL_2850655 | Lighthouse Laboratory Plymouth | Wellcome Sanger Institute for the COVID-19 Genomics UK (COG-UK) Consortium | Lighthouse Laboratory Plymouth and Alex Alderton, Roberto Amato, Jeffrey Barrett, Sonia Goncalves, Ewan Harrison, David K. Jackson, Ian Johnston, Dominic Kwiatkowski, Cordelia Langford, John Sillitoe on behalf of the Wellcome Sanger Institute COVID-19 Surveillance Team |
| EPI_ISL_2850656, EPI_ISL_2850657, EPI_ISL_2850658 | Lighthouse Lab in Milton Keynes | Wellcome Sanger Institute for the COVID-19 Genomics UK (COG-UK) Consortium | The Lighthouse Lab in Milton Keynes and Alex Alderton, Roberto Amato, Jeffrey Barrett, Sonia Goncalves, Ewan Harrison, David K. Jackson, Ian Johnston, Dominic Kwiatkowski, Cordelia Langford, John Sillitoe on behalf of the Wellcome Sanger Institute COVID-19 Surveillance Team |
| EPI_ISL_2850659 | Lighthouse Laboratory Plymouth | Wellcome Sanger Institute for the COVID-19 Genomics UK (COG-UK) Consortium | Lighthouse Laboratory Plymouth and Alex Alderton, Roberto Amato, Jeffrey Barrett, Sonia Goncalves, Ewan Harrison, David K. Jackson, Ian Johnston, Dominic Kwiatkowski, Cordelia Langford, John Sillitoe on behalf of the Wellcome Sanger Institute COVID-19 Surveillance Team |
| EPI_ISL_2850660, EPI_ISL_2850661 | Lighthouse Lab in Milton Keynes | Wellcome Sanger Institute for the COVID-19 Genomics UK (COG-UK) Consortium | The Lighthouse Lab in Milton Keynes and Alex Alderton, Roberto Amato, Jeffrey Barrett, Sonia Goncalves, Ewan Harrison, David K. Jackson, Ian Johnston, Dominic Kwiatkowski, Cordelia Langford, John Sillitoe on behalf of the Wellcome Sanger Institute COVID-19 Surveillance Team |
| EPI_ISL_2850662, EPI_ISL_2850663, EPI_ISL_2850664, EPI_ISL_2850665 | Lighthouse Laboratory Plymouth | Wellcome Sanger Institute for the COVID-19 Genomics UK (COG-UK) Consortium | Lighthouse Laboratory Plymouth and Alex Alderton, Roberto Amato, Jeffrey Barrett, Sonia Goncalves, Ewan Harrison, David K. Jackson, Ian Johnston, Dominic Kwiatkowski, Cordelia Langford, John Sillitoe on behalf of the Wellcome Sanger Institute COVID-19 Surveillance Team |
| EPI_ISL_2850666, EPI_ISL_2850667 | Lighthouse Lab in Alderley Park | Wellcome Sanger Institute for the COVID-19 Genomics UK (COG-UK) Consortium | Jacquelyn Wynn, Mairead Hyland, The Lighthouse Lab in Alderley Park and Alex Alderton, Roberto Amato, Jeffrey Barrett, Sonia Goncalves, Ewan Harrison, David K. Jackson, Ian Johnston, Dominic Kwiatkowski, Cordelia Langford, John Sillitoe on behalf of the Wellcome Sanger Institute COVID-19 Surveillance Team |
| EPI_ISL_2850668 | Lighthouse Laboratory Plymouth | Wellcome Sanger Institute for the COVID-19 Genomics UK (COG-UK) Consortium | Lighthouse Laboratory Plymouth and Alex Alderton, Roberto Amato, Jeffrey Barrett, Sonia Goncalves, Ewan Harrison, David K. Jackson, Ian Johnston, Dominic Kwiatkowski, Cordelia Langford, John Sillitoe on behalf of the Wellcome Sanger Institute COVID-19 Surveillance Team |
| EPI_ISL_2850669, EPI_ISL_2850670 | Lighthouse Lab in Alderley Park | Wellcome Sanger Institute for the COVID-19 Genomics UK (COG-UK) Consortium | Jacquelyn Wynn, Mairead Hyland, The Lighthouse Lab in Alderley Park and Alex Alderton, Roberto Amato, Jeffrey Barrett, Sonia Goncalves, Ewan Harrison, David K. Jackson, Ian Johnston, Dominic Kwiatkowski, Cordelia Langford, John Sillitoe on behalf of the Wellcome Sanger Institute COVID-19 Surveillance Team |
| EPI_ISL_2850671 | AZ Klina | AZ Klina | Carl Vael - Lynsey Berckmans |
| EPI_ISL_2850672, EPI_ISL_2850673, EPI_ISL_2850674 | Lighthouse Lab in Alderley Park | Wellcome Sanger Institute for the COVID-19 Genomics UK (COG-UK) Consortium | Jacquelyn Wynn, Mairead Hyland, The Lighthouse Lab in Alderley Park and Alex Alderton, Roberto Amato, Jeffrey Barrett, Sonia Goncalves, Ewan Harrison, David K. Jackson, Ian Johnston, Dominic Kwiatkowski, Cordelia Langford, John Sillitoe on behalf of the Wellcome Sanger Institute COVID-19 Surveillance Team |
| EPI_ISL_2850675, EPI_ISL_2850676 | Lighthouse Laboratory Plymouth | Wellcome Sanger Institute for the COVID-19 Genomics UK (COG-UK) Consortium | Lighthouse Laboratory Plymouth and Alex Alderton, Roberto Amato, Jeffrey Barrett, Sonia Goncalves, Ewan Harrison, David K. Jackson, Ian Johnston, Dominic Kwiatkowski, Cordelia Langford, John Sillitoe on behalf of the Wellcome Sanger Institute COVID-19 Surveillance Team |
| EPI_ISL_2850677 | Lighthouse Lab in Milton Keynes | Wellcome Sanger Institute for the COVID-19 Genomics UK (COG-UK) Consortium | The Lighthouse Lab in Milton Keynes and Alex Alderton, Roberto Amato, Jeffrey Barrett, Sonia Goncalves, Ewan Harrison, David K. Jackson, Ian Johnston, Dominic Kwiatkowski, Cordelia Langford, John Sillitoe on behalf of the Wellcome Sanger Institute COVID-19 Surveillance Team |
| EPI_ISL_2850678, EPI_ISL_2850679 | Lighthouse Lab in Alderley Park | Wellcome Sanger Institute for the COVID-19 Genomics UK (COG-UK) Consortium | Jacquelyn Wynn, Mairead Hyland, The Lighthouse Lab in Alderley Park and Alex Alderton, Roberto Amato, Jeffrey Barrett, Sonia Goncalves, Ewan Harrison, David K. Jackson, Ian Johnston, Dominic Kwiatkowski, Cordelia Langford, John Sillitoe on behalf of the Wellcome Sanger Institute COVID-19 Surveillance Team |
| EPI_ISL_2850680 | Lighthouse Laboratory Plymouth | Wellcome Sanger Institute for the COVID-19 Genomics UK (COG-UK) Consortium | Lighthouse Laboratory Plymouth and Alex Alderton, Roberto Amato, Jeffrey Barrett, Sonia Goncalves, Ewan Harrison, David K. Jackson, Ian Johnston, Dominic Kwiatkowski, Cordelia Langford, John Sillitoe on behalf of the Wellcome Sanger Institute COVID-19 Surveillance Team |
| EPI_ISL_2850681, EPI_ISL_2850682 | Lighthouse Lab in Alderley Park | Wellcome Sanger Institute for the COVID-19 Genomics UK (COG-UK) Consortium | Jacquelyn Wynn, Mairead Hyland, The Lighthouse Lab in Alderley Park and Alex Alderton, Roberto Amato, Jeffrey Barrett, Sonia Goncalves, Ewan Harrison, David K. Jackson, Ian Johnston, Dominic Kwiatkowski, Cordelia Langford, John Sillitoe on behalf of the Wellcome Sanger Institute COVID-19 Surveillance Team |
| EPI_ISL_2850683, EPI_ISL_2850684, EPI_ISL_2850685 | Lighthouse Laboratory Plymouth | Wellcome Sanger Institute for the COVID-19 Genomics UK (COG-UK) Consortium | Lighthouse Laboratory Plymouth and Alex Alderton, Roberto Amato, Jeffrey Barrett, Sonia Goncalves, Ewan Harrison, David K. Jackson, Ian Johnston, Dominic Kwiatkowski, Cordelia Langford, John Sillitoe on behalf of the Wellcome Sanger Institute COVID-19 Surveillance Team |
| EPI_ISL_2850686, EPI_ISL_2850687, EPI_ISL_2850688 | Lighthouse Lab in Alderley Park | Wellcome Sanger Institute for the COVID-19 Genomics UK (COG-UK) Consortium | Jacquelyn Wynn, Mairead Hyland, The Lighthouse Lab in Alderley Park and Alex Alderton, Roberto Amato, Jeffrey Barrett, Sonia Goncalves, Ewan Harrison, David K. Jackson, Ian Johnston, Dominic Kwiatkowski, Cordelia Langford, John Sillitoe on behalf of the Wellcome Sanger Institute COVID-19 Surveillance Team |
| EPI_ISL_2850689 | Lighthouse Laboratory Plymouth | Wellcome Sanger Institute for the COVID-19 Genomics UK (COG-UK) Consortium | Lighthouse Laboratory Plymouth and Alex Alderton, Roberto Amato, Jeffrey Barrett, Sonia Goncalves, Ewan Harrison, David K. Jackson, Ian Johnston, Dominic Kwiatkowski, Cordelia Langford, John Sillitoe on behalf of the Wellcome Sanger Institute COVID-19 Surveillance Team |
| EPI_ISL_2850690 | Lighthouse Lab in Alderley Park | Wellcome Sanger Institute for the COVID-19 Genomics UK (COG-UK) Consortium | Jacquelyn Wynn, Mairead Hyland, The Lighthouse Lab in Alderley Park and Alex Alderton, Roberto Amato, Jeffrey Barrett, Sonia Goncalves, Ewan Harrison, David K. Jackson, Ian Johnston, Dominic Kwiatkowski, Cordelia Langford, John Sillitoe on behalf of the Wellcome Sanger Institute COVID-19 Surveillance Team |
| EPI_ISL_2850691 | Lighthouse Lab in Milton Keynes | Wellcome Sanger Institute for the COVID-19 Genomics UK (COG-UK) Consortium | The Lighthouse Lab in Milton Keynes and Alex Alderton, Roberto Amato, Jeffrey Barrett, Sonia Goncalves, Ewan Harrison, David K. Jackson, Ian Johnston, Dominic Kwiatkowski, Cordelia Langford, John Sillitoe on behalf of the Wellcome Sanger Institute COVID-19 Surveillance Team |
| EPI_ISL_2850692, EPI_ISL_2850693 | Lighthouse Lab in Alderley Park | Wellcome Sanger Institute for the COVID-19 Genomics UK (COG-UK) Consortium | Jacquelyn Wynn, Mairead Hyland, The Lighthouse Lab in Alderley Park and Alex Alderton, Roberto Amato, Jeffrey Barrett, Sonia Goncalves, Ewan Harrison, David K. Jackson, Ian Johnston, Dominic Kwiatkowski, Cordelia Langford, John Sillitoe on behalf of the Wellcome Sanger Institute COVID-19 Surveillance Team |
| EPI_ISL_2850694 | Lighthouse Lab in Milton Keynes | Wellcome Sanger Institute for the COVID-19 Genomics UK (COG-UK) Consortium | The Lighthouse Lab in Milton Keynes and Alex Alderton, Roberto Amato, Jeffrey Barrett, Sonia Goncalves, Ewan Harrison, David K. Jackson, Ian Johnston, Dominic Kwiatkowski, Cordelia Langford, John Sillitoe on behalf of the Wellcome Sanger Institute COVID-19 Surveillance Team |
| EPI_ISL_2850695, EPI_ISL_2850696 | Lighthouse Lab in Alderley Park | Wellcome Sanger Institute for the COVID-19 Genomics UK (COG-UK) Consortium | Jacquelyn Wynn, Mairead Hyland, The Lighthouse Lab in Alderley Park and Alex Alderton, Roberto Amato, Jeffrey Barrett, Sonia Goncalves, Ewan Harrison, David K. Jackson, Ian Johnston, Dominic Kwiatkowski, Cordelia Langford, John Sillitoe on behalf of the Wellcome Sanger Institute COVID-19 Surveillance Team |
| EPI_ISL_2850697 | Lighthouse Laboratory Plymouth | Wellcome Sanger Institute for the COVID-19 Genomics UK (COG-UK) Consortium | Lighthouse Laboratory Plymouth and Alex Alderton, Roberto Amato, Jeffrey Barrett, Sonia Goncalves, Ewan Harrison, David K. Jackson, Ian Johnston, Dominic Kwiatkowski, Cordelia Langford, John Sillitoe on behalf of the Wellcome Sanger Institute COVID-19 Surveillance Team |
| EPI_ISL_2850698, EPI_ISL_2850699 | Lighthouse Lab in Milton Keynes | Wellcome Sanger Institute for the COVID-19 Genomics UK (COG-UK) Consortium | The Lighthouse Lab in Milton Keynes and Alex Alderton, Roberto Amato, Jeffrey Barrett, Sonia Goncalves, Ewan Harrison, David K. Jackson, Ian Johnston, Dominic Kwiatkowski, Cordelia Langford, John Sillitoe on behalf of the Wellcome Sanger Institute COVID-19 Surveillance Team |
| EPI_ISL_2850700 | Lighthouse Lab in Alderley Park | Wellcome Sanger Institute for the COVID-19 Genomics UK (COG-UK) Consortium | Jacquelyn Wynn, Mairead Hyland, The Lighthouse Lab in Alderley Park and Alex Alderton, Roberto Amato, Jeffrey Barrett, Sonia Goncalves, Ewan Harrison, David K. Jackson, Ian Johnston, Dominic Kwiatkowski, Cordelia Langford, John Sillitoe on behalf of the Wellcome Sanger Institute COVID-19 Surveillance Team |
| EPI_ISL_2850701 | Lighthouse Lab in Milton Keynes | Wellcome Sanger Institute for the COVID-19 Genomics UK (COG-UK) Consortium | The Lighthouse Lab in Milton Keynes and Alex Alderton, Roberto Amato, Jeffrey Barrett, Sonia Goncalves, Ewan Harrison, David K. Jackson, Ian Johnston, Dominic Kwiatkowski, Cordelia Langford, John Sillitoe on behalf of the Wellcome Sanger Institute COVID-19 Surveillance Team |
| EPI_ISL_2850702, EPI_ISL_2850703 | Lighthouse Laboratory Plymouth | Wellcome Sanger Institute for the COVID-19 Genomics UK (COG-UK) Consortium | Lighthouse Laboratory Plymouth and Alex Alderton, Roberto Amato, Jeffrey Barrett, Sonia Goncalves, Ewan Harrison, David K. Jackson, Ian Johnston, Dominic Kwiatkowski, Cordelia Langford, John Sillitoe on behalf of the Wellcome Sanger Institute COVID-19 Surveillance Team |
| EPI_ISL_2850704 | Lighthouse Lab in Alderley Park | Wellcome Sanger Institute for the COVID-19 Genomics UK (COG-UK) Consortium | Jacquelyn Wynn, Mairead Hyland, The Lighthouse Lab in Alderley Park and Alex Alderton, Roberto Amato, Jeffrey Barrett, Sonia Goncalves, Ewan Harrison, David K. Jackson, Ian Johnston, Dominic Kwiatkowski, Cordelia Langford, John Sillitoe on behalf of the Wellcome Sanger Institute COVID-19 Surveillance Team |
| EPI_ISL_2850705 | Lighthouse Lab in Milton Keynes | Wellcome Sanger Institute for the COVID-19 Genomics UK (COG-UK) Consortium | The Lighthouse Lab in Milton Keynes and Alex Alderton, Roberto Amato, Jeffrey Barrett, Sonia Goncalves, Ewan Harrison, David K. Jackson, Ian Johnston, Dominic Kwiatkowski, Cordelia Langford, John Sillitoe on behalf of the Wellcome Sanger Institute COVID-19 Surveillance Team |
| EPI_ISL_2850706 | Lighthouse Lab in Alderley Park | Wellcome Sanger Institute for the COVID-19 Genomics UK (COG-UK) | Jacquelyn Wynn, Mairead Hyland, The Lighthouse Lab in Alderley Park and Alex Alderton, Roberto Amato, Jeffrey Barrett, Sonia Goncalves, Ewan Harrison, David K. Jackson, Ian Johnston, Dominic Kwiatkowski, Cordelia Langford, John Sillitoe on |

| | | | |
|---|---|---|---|
| | | Consortium | behalf of the Wellcome Sanger Institute COVID-19 Surveillance Team |
| EPI_ISL_2850707 | Lighthouse Lab in Milton Keynes | Wellcome Sanger Institute for the COVID-19 Genomics UK (COG-UK) Consortium | The Lighthouse Lab in Milton Keynes and Alex Alderton, Roberto Amato, Jeffrey Barrett, Sonia Goncalves, Ewan Harrison, David K. Jackson, Ian Johnston, Dominic Kwiatkowski, Cordelia Langford, John Sillitoe on behalf of the Wellcome Sanger Institute COVID-19 Surveillance Team |
| EPI_ISL_2850708, EPI_ISL_2850709 | Lighthouse Laboratory Plymouth | Wellcome Sanger Institute for the COVID-19 Genomics UK (COG-UK) Consortium | Lighthouse Laboratory Plymouth and Alex Alderton, Roberto Amato, Jeffrey Barrett, Sonia Goncalves, Ewan Harrison, David K. Jackson, Ian Johnston, Dominic Kwiatkowski, Cordelia Langford, John Sillitoe on behalf of the Wellcome Sanger Institute COVID-19 Surveillance Team |
| EPI_ISL_2850710, EPI_ISL_2850711, EPI_ISL_2850712, EPI_ISL_2850713, EPI_ISL_2850714 | Lighthouse Lab in Alderley Park | Wellcome Sanger Institute for the COVID-19 Genomics UK (COG-UK) Consortium | Jacquelyn Wynn, Mairead Hyland, The Lighthouse Lab in Alderley Park and Alex Alderton, Roberto Amato, Jeffrey Barrett, Sonia Goncalves, Ewan Harrison, David K. Jackson, Ian Johnston, Dominic Kwiatkowski, Cordelia Langford, John Sillitoe on behalf of the Wellcome Sanger Institute COVID-19 Surveillance Team |
| EPI_ISL_2850715 | Lighthouse Laboratory Plymouth | Wellcome Sanger Institute for the COVID-19 Genomics UK (COG-UK) Consortium | Lighthouse Laboratory Plymouth and Alex Alderton, Roberto Amato, Jeffrey Barrett, Sonia Goncalves, Ewan Harrison, David K. Jackson, Ian Johnston, Dominic Kwiatkowski, Cordelia Langford, John Sillitoe on behalf of the Wellcome Sanger Institute COVID-19 Surveillance Team |
| EPI_ISL_2850716, EPI_ISL_2850717 | Lighthouse Lab in Alderley Park | Wellcome Sanger Institute for the COVID-19 Genomics UK (COG-UK) Consortium | Jacquelyn Wynn, Mairead Hyland, The Lighthouse Lab in Alderley Park and Alex Alderton, Roberto Amato, Jeffrey Barrett, Sonia Goncalves, Ewan Harrison, David K. Jackson, Ian Johnston, Dominic Kwiatkowski, Cordelia Langford, John Sillitoe on behalf of the Wellcome Sanger Institute COVID-19 Surveillance Team |
| EPI_ISL_2850718 | Lighthouse Laboratory Plymouth | Wellcome Sanger Institute for the COVID-19 Genomics UK (COG-UK) Consortium | Lighthouse Laboratory Plymouth and Alex Alderton, Roberto Amato, Jeffrey Barrett, Sonia Goncalves, Ewan Harrison, David K. Jackson, Ian Johnston, Dominic Kwiatkowski, Cordelia Langford, John Sillitoe on behalf of the Wellcome Sanger Institute COVID-19 Surveillance Team |
| EPI_ISL_2850719 | Lighthouse Lab in Milton Keynes | Wellcome Sanger Institute for the COVID-19 Genomics UK (COG-UK) Consortium | The Lighthouse Lab in Milton Keynes and Alex Alderton, Roberto Amato, Jeffrey Barrett, Sonia Goncalves, Ewan Harrison, David K. Jackson, Ian Johnston, Dominic Kwiatkowski, Cordelia Langford, John Sillitoe on behalf of the Wellcome Sanger Institute COVID-19 Surveillance Team |
| EPI_ISL_2850720 | Lighthouse Lab in Alderley Park | Wellcome Sanger Institute for the COVID-19 Genomics UK (COG-UK) Consortium | Jacquelyn Wynn, Mairead Hyland, The Lighthouse Lab in Alderley Park and Alex Alderton, Roberto Amato, Jeffrey Barrett, Sonia Goncalves, Ewan Harrison, David K. Jackson, Ian Johnston, Dominic Kwiatkowski, Cordelia Langford, John Sillitoe on behalf of the Wellcome Sanger Institute COVID-19 Surveillance Team |
| EPI_ISL_2850721 | Lighthouse Laboratory Plymouth | Wellcome Sanger Institute for the COVID-19 Genomics UK (COG-UK) Consortium | Lighthouse Laboratory Plymouth and Alex Alderton, Roberto Amato, Jeffrey Barrett, Sonia Goncalves, Ewan Harrison, David K. Jackson, Ian Johnston, Dominic Kwiatkowski, Cordelia Langford, John Sillitoe on behalf of the Wellcome Sanger Institute COVID-19 Surveillance Team |
| EPI_ISL_2850722 | Lighthouse Lab in Alderley Park | Wellcome Sanger Institute for the COVID-19 Genomics UK (COG-UK) Consortium | Jacquelyn Wynn, Mairead Hyland, The Lighthouse Lab in Alderley Park and Alex Alderton, Roberto Amato, Jeffrey Barrett, Sonia Goncalves, Ewan Harrison, David K. Jackson, Ian Johnston, Dominic Kwiatkowski, Cordelia Langford, John Sillitoe on behalf of the Wellcome Sanger Institute COVID-19 Surveillance Team |
| EPI_ISL_2850723 | Lighthouse Lab in Milton Keynes | Wellcome Sanger Institute for the COVID-19 Genomics UK (COG-UK) Consortium | The Lighthouse Lab in Milton Keynes and Alex Alderton, Roberto Amato, Jeffrey Barrett, Sonia Goncalves, Ewan Harrison, David K. Jackson, Ian Johnston, Dominic Kwiatkowski, Cordelia Langford, John Sillitoe on behalf of the Wellcome Sanger Institute COVID-19 Surveillance Team |
| EPI_ISL_2850724, EPI_ISL_2850725, EPI_ISL_2850726, EPI_ISL_2850727, EPI_ISL_2850728 | Lighthouse Lab in Alderley Park | Wellcome Sanger Institute for the COVID-19 Genomics UK (COG-UK) Consortium | Jacquelyn Wynn, Mairead Hyland, The Lighthouse Lab in Alderley Park and Alex Alderton, Roberto Amato, Jeffrey Barrett, Sonia Goncalves, Ewan Harrison, David K. Jackson, Ian Johnston, Dominic Kwiatkowski, Cordelia Langford, John Sillitoe on behalf of the Wellcome Sanger Institute COVID-19 Surveillance Team |
| EPI_ISL_2850729 | Lighthouse Lab in Milton Keynes | Wellcome Sanger Institute for the COVID-19 Genomics UK (COG-UK) Consortium | The Lighthouse Lab in Milton Keynes and Alex Alderton, Roberto Amato, Jeffrey Barrett, Sonia Goncalves, Ewan Harrison, David K. Jackson, Ian Johnston, Dominic Kwiatkowski, Cordelia Langford, John Sillitoe on behalf of the Wellcome Sanger Institute COVID-19 Surveillance Team |
| EPI_ISL_2850730, EPI_ISL_2850731, EPI_ISL_2850732 | Lighthouse Lab in Alderley Park | Wellcome Sanger Institute for the COVID-19 Genomics UK (COG-UK) Consortium | Jacquelyn Wynn, Mairead Hyland, The Lighthouse Lab in Alderley Park and Alex Alderton, Roberto Amato, Jeffrey Barrett, Sonia Goncalves, Ewan Harrison, David K. Jackson, Ian Johnston, Dominic Kwiatkowski, Cordelia Langford, John Sillitoe on behalf of the Wellcome Sanger Institute COVID-19 Surveillance Team |
| EPI_ISL_2850733 | Lighthouse Lab in Milton Keynes | Wellcome Sanger Institute for the COVID-19 Genomics UK (COG-UK) Consortium | The Lighthouse Lab in Milton Keynes and Alex Alderton, Roberto Amato, Jeffrey Barrett, Sonia Goncalves, Ewan Harrison, David K. Jackson, Ian Johnston, Dominic Kwiatkowski, Cordelia Langford, John Sillitoe on behalf of the Wellcome Sanger Institute COVID-19 Surveillance Team |
| EPI_ISL_2850734 | Lighthouse Laboratory Plymouth | Wellcome Sanger Institute for the COVID-19 Genomics UK (COG-UK) Consortium | Lighthouse Laboratory Plymouth and Alex Alderton, Roberto Amato, Jeffrey Barrett, Sonia Goncalves, Ewan Harrison, David K. Jackson, Ian Johnston, Dominic Kwiatkowski, Cordelia Langford, John Sillitoe on behalf of the Wellcome Sanger Institute COVID-19 Surveillance Team |
| EPI_ISL_2850735, EPI_ISL_2850736, EPI_ISL_2850737 | Lighthouse Lab in Alderley Park | Wellcome Sanger Institute for the COVID-19 Genomics UK (COG-UK) Consortium | Jacquelyn Wynn, Mairead Hyland, The Lighthouse Lab in Alderley Park and Alex Alderton, Roberto Amato, Jeffrey Barrett, Sonia Goncalves, Ewan Harrison, David K. Jackson, Ian Johnston, Dominic Kwiatkowski, Cordelia Langford, John Sillitoe on behalf of the Wellcome Sanger Institute COVID-19 Surveillance Team |
| EPI_ISL_2850738 | Lighthouse Lab in Milton Keynes | Wellcome Sanger Institute for the COVID-19 Genomics UK (COG-UK) Consortium | The Lighthouse Lab in Milton Keynes and Alex Alderton, Roberto Amato, Jeffrey Barrett, Sonia Goncalves, Ewan Harrison, David K. Jackson, Ian Johnston, Dominic Kwiatkowski, Cordelia Langford, John Sillitoe on behalf of the Wellcome Sanger Institute COVID-19 Surveillance Team |
| EPI_ISL_2850739, EPI_ISL_2850740, EPI_ISL_2850741, EPI_ISL_2850742, EPI_ISL_2850743, EPI_ISL_2850744, EPI_ISL_2850745, EPI_ISL_2850746, EPI_ISL_2850747, EPI_ISL_2850748, EPI_ISL_2850749, EPI_ISL_2850750, EPI_ISL_2850751, EPI_ISL_2850752, EPI_ISL_2850753, EPI_ISL_2850754, EPI_ISL_2850755, EPI_ISL_2850756, EPI_ISL_2850757, EPI_ISL_2850758, EPI_ISL_2850759, EPI_ISL_2850760, EPI_ISL_2850761, EPI_ISL_2850762, EPI_ISL_2850763, EPI_ISL_2850764, EPI_ISL_2850765, EPI_ISL_2850766, EPI_ISL_2850767, EPI_ISL_2850768, EPI_ISL_2850769, EPI_ISL_2850770, EPI_ISL_2850771, EPI_ISL_2850772, EPI_ISL_2850773, EPI_ISL_2850774, EPI_ISL_2850775, EPI_ISL_2850776, EPI_ISL_2850777, EPI_ISL_2850778, EPI_ISL_2850779, EPI_ISL_2850780, EPI_ISL_2850781, EPI_ISL_2850782, EPI_ISL_2850783, EPI_ISL_2850784, EPI_ISL_2850785, EPI_ISL_2850786, EPI_ISL_2850787, EPI_ISL_2850788, EPI_ISL_2850789, EPI_ISL_2850790, EPI_ISL_2850791, EPI_ISL_2850792, EPI_ISL_2850793, EPI_ISL_2850794, EPI_ISL_2850795, EPI_ISL_2850796, EPI_ISL_2850797, EPI_ISL_2850798, EPI_ISL_2850799, EPI_ISL_2850800, EPI_ISL_2850801, EPI_ISL_2850802, EPI_ISL_2850803, EPI_ISL_2850804, EPI_ISL_2850805, EPI_ISL_2850806, EPI_ISL_2850807, EPI_ISL_2850808, EPI_ISL_2850809, EPI_ISL_2850810, EPI_ISL_2850811, EPI_ISL_2850812, EPI_ISL_2850813, EPI_ISL_2850814, EPI_ISL_2850815, EPI_ISL_2850816, EPI_ISL_2850817, EPI_ISL_2850818, EPI_ISL_2850819, EPI_ISL_2850820, EPI_ISL_2850821, EPI_ISL_2850822, EPI_ISL_2850823, EPI_ISL_2850824, EPI_ISL_2850825, EPI_ISL_2850826, EPI_ISL_2850827, EPI_ISL_2850828, EPI_ISL_2850829, EPI_ISL_2850830, EPI_ISL_2850831, EPI_ISL_2850832, EPI_ISL_2850833, EPI_ISL_2850834, EPI_ISL_2850835, EPI_ISL_2850836, EPI_ISL_2850837, EPI_ISL_2850838, EPI_ISL_2850839, EPI_ISL_2850840, EPI_ISL_2850841, EPI_ISL_2850842, EPI_ISL_2850843, EPI_ISL_2850844, EPI_ISL_2850845, EPI_ISL_2850846, EPI_ISL_2850847, EPI_ISL_2850848, EPI_ISL_2850849, EPI_ISL_2850850, EPI_ISL_2850851, EPI_ISL_2850852, EPI_ISL_2850853, EPI_ISL_2850854, EPI_ISL_2850855 | | | |
| see above | Lighthouse Lab in Glasgow | Wellcome Sanger Institute for the COVID-19 Genomics UK (COG-UK) Consortium | Harper VanSteenhouse, Yumi Kasai, David Gray, Carol Clugston, Anna Dominiczak and Alex Alderton, Roberto Amato, Jeffrey Barrett, Sonia Goncalves, Ewan Harrison, David K. Jackson, Ian Johnston, Dominic Kwiatkowski, Cordelia Langford, John Sillitoe on behalf of the Wellcome Sanger Institute COVID-19 Surveillance Team |
| EPI_ISL_2850856 | Lighthouse Lab in Milton Keynes | Wellcome Sanger Institute for the COVID-19 Genomics UK (COG-UK) Consortium | The Lighthouse Lab in Milton Keynes and Alex Alderton, Roberto Amato, Jeffrey Barrett, Sonia Goncalves, Ewan Harrison, David K. Jackson, Ian Johnston, Dominic Kwiatkowski, Cordelia Langford, John Sillitoe on behalf of the Wellcome Sanger Institute COVID-19 Surveillance Team |
| EPI_ISL_2850857, EPI_ISL_2850858, EPI_ISL_2850859, EPI_ISL_2850860, EPI_ISL_2850861, EPI_ISL_2850862, EPI_ISL_2850863, EPI_ISL_2850864, EPI_ISL_2850865, EPI_ISL_2850866, EPI_ISL_2850867, EPI_ISL_2850868, EPI_ISL_2850869, EPI_ISL_2850870, EPI_ISL_2850871, EPI_ISL_2850872, EPI_ISL_2850873, EPI_ISL_2850874, EPI_ISL_2850875, EPI_ISL_2850876, EPI_ISL_2850877, EPI_ISL_2850878, EPI_ISL_2850879, EPI_ISL_2850880, EPI_ISL_2850881, EPI_ISL_2850882, EPI_ISL_2850883, EPI_ISL_2850884, EPI_ISL_2850885, EPI_ISL_2850886, EPI_ISL_2850887, EPI_ISL_2850888, EPI_ISL_2850889, EPI_ISL_2850890, EPI_ISL_2850891, EPI_ISL_2850892, EPI_ISL_2850893, EPI_ISL_2850894, EPI_ISL_2850895, EPI_ISL_2850896, EPI_ISL_2850897, EPI_ISL_2850898, EPI_ISL_2850899, EPI_ISL_2850900, EPI_ISL_2850901, EPI_ISL_2850902, EPI_ISL_2850903, EPI_ISL_2850904, EPI_ISL_2850905, EPI_ISL_2850906, EPI_ISL_2850907, EPI_ISL_2850908, EPI_ISL_2850909, EPI_ISL_2850910, EPI_ISL_2850911, EPI_ISL_2850912, EPI_ISL_2850913, EPI_ISL_2850914, EPI_ISL_2850915, EPI_ISL_2850916, EPI_ISL_2850917, EPI_ISL_2850918, EPI_ISL_2850919, EPI_ISL_2850920, EPI_ISL_2850921 | | | |
| see above | Lighthouse Lab in Glasgow | Wellcome Sanger Institute for the COVID-19 Genomics UK (COG-UK) Consortium | Harper VanSteenhouse, Yumi Kasai, David Gray, Carol Clugston, Anna Dominiczak and Alex Alderton, Roberto Amato, Jeffrey Barrett, Sonia Goncalves, Ewan Harrison, David K. Jackson, Ian Johnston, Dominic Kwiatkowski, Cordelia Langford, John Sillitoe on behalf of the Wellcome Sanger Institute COVID-19 Surveillance Team |
| EPI_ISL_2850922 | Lighthouse Lab in Alderley Park | Wellcome Sanger Institute for the COVID-19 Genomics UK (COG-UK) Consortium | Jacquelyn Wynn, Mairead Hyland, The Lighthouse Lab in Alderley Park and Alex Alderton, Roberto Amato, Jeffrey Barrett, Sonia Goncalves, Ewan Harrison, David K. Jackson, Ian Johnston, Dominic Kwiatkowski, Cordelia Langford, John Sillitoe on behalf of the Wellcome Sanger Institute COVID-19 Surveillance Team |
| EPI_ISL_2850923, EPI_ISL_2850924, EPI_ISL_2850925, EPI_ISL_2850926, EPI_ISL_2850927, EPI_ISL_2850928, EPI_ISL_2850929, EPI_ISL_2850930, EPI_ISL_2850931, EPI_ISL_2850932, EPI_ISL_2850933, EPI_ISL_2850934, EPI_ISL_2850935, EPI_ISL_2850936, EPI_ISL_2850937, EPI_ISL_2850938, EPI_ISL_2850939, EPI_ISL_2850940, EPI_ISL_2850941, EPI_ISL_2850942, EPI_ISL_2850943, EPI_ISL_2850944, EPI_ISL_2850945, EPI_ISL_2850946, EPI_ISL_2850947, EPI_ISL_2850948, EPI_ISL_2850949, EPI_ISL_2850950, EPI_ISL_2850951, EPI_ISL_2850952, EPI_ISL_2850953, EPI_ISL_2850954, EPI_ISL_2850955, EPI_ISL_2850956, EPI_ISL_2850957, EPI_ISL_2850958, EPI_ISL_2850959, EPI_ISL_2850960, EPI_ISL_2850961, EPI_ISL_2850962, EPI_ISL_2850963, EPI_ISL_2850964, EPI_ISL_2850965, EPI_ISL_2850966, EPI_ISL_2850967, EPI_ISL_2850968, EPI_ISL_2850969, EPI_ISL_2850970, EPI_ISL_2850971, EPI_ISL_2850972, EPI_ISL_2850973, EPI_ISL_2850974, EPI_ISL_2850975, EPI_ISL_2850976, EPI_ISL_2850977, EPI_ISL_2850978, EPI_ISL_2850979, EPI_ISL_2850980, EPI_ISL_2850981, EPI_ISL_2850982, EPI_ISL_2850983, EPI_ISL_2850984, EPI_ISL_2850985, EPI_ISL_2850986, EPI_ISL_2850987, EPI_ISL_2850988, EPI_ISL_2850989, EPI_ISL_2850990, EPI_ISL_2850991, EPI_ISL_2850992, EPI_ISL_2850993, EPI_ISL_2850994, EPI_ISL_2850995, EPI_ISL_2850996, EPI_ISL_2850997, EPI_ISL_2850998, EPI_ISL_2850999, EPI_ISL_2851000, EPI_ISL_2851001, EPI_ISL_2851002, EPI_ISL_2851003, EPI_ISL_2851004, EPI_ISL_2851005, EPI_ISL_2851006, EPI_ISL_2851007, EPI_ISL_2851008, EPI_ISL_2851009, EPI_ISL_2851010, EPI_ISL_2851011, EPI_ISL_2851012, EPI_ISL_2851013, EPI_ISL_2851014, EPI_ISL_2851015 | | | |
| see above | Lighthouse Lab in Glasgow | Wellcome Sanger Institute for the COVID-19 Genomics UK (COG-UK) Consortium | Harper VanSteenhouse, Yumi Kasai, David Gray, Carol Clugston, Anna Dominiczak and Alex Alderton, Roberto Amato, Jeffrey Barrett, Sonia Goncalves, Ewan Harrison, David K. Jackson, Ian Johnston, Dominic Kwiatkowski, Cordelia Langford, John Sillitoe on behalf of the Wellcome Sanger Institute COVID-19 Surveillance Team |
| EPI_ISL_2851016 | Lighthouse Lab in Alderley Park | Wellcome Sanger Institute for the COVID-19 Genomics UK (COG-UK) Consortium | Jacquelyn Wynn, Mairead Hyland, The Lighthouse Lab in Alderley Park and Alex Alderton, Roberto Amato, Jeffrey Barrett, Sonia Goncalves, Ewan Harrison, David K. Jackson, Ian Johnston, Dominic Kwiatkowski, Cordelia Langford, John Sillitoe on behalf of the Wellcome Sanger Institute COVID-19 Surveillance Team |
| EPI_ISL_2851017, EPI_ISL_2851018, EPI_ISL_2851019, EPI_ISL_2851020, EPI_ISL_2851021, EPI_ISL_2851022, EPI_ISL_2851023, EPI_ISL_2851024, EPI_ISL_2851025, EPI_ISL_2851026, EPI_ISL_2851027, EPI_ISL_2851028, EPI_ISL_2851029, EPI_ISL_2851030, EPI_ISL_2851031, EPI_ISL_2851032, EPI_ISL_2851033, EPI_ISL_2851034, EPI_ISL_2851035 | | | |
| see above | Lighthouse Lab in Glasgow | Wellcome Sanger Institute for the COVID-19 Genomics UK (COG-UK) Consortium | Harper VanSteenhouse, Yumi Kasai, David Gray, Carol Clugston, Anna Dominiczak and Alex Alderton, Roberto Amato, Jeffrey Barrett, Sonia Goncalves, Ewan Harrison, David K. Jackson, Ian Johnston, Dominic Kwiatkowski, Cordelia Langford, John Sillitoe on behalf of the Wellcome Sanger Institute COVID-19 Surveillance Team |
| EPI_ISL_2851036 | Ministry of Health, Jaber Al-Ahmad Hospital | Virology Unit, Department of Microbiology, Faculty of Medicine, Kuwait University | Nada Madi, Hussain Safar, Ebaa Al-Awadhi, Anfal Al-Adwani |
| EPI_ISL_2851037, EPI_ISL_2851038, EPI_ISL_2851039, EPI_ISL_2851040, EPI_ISL_2851041, EPI_ISL_2851042, EPI_ISL_2851043, EPI_ISL_2851044, EPI_ISL_2851045, EPI_ISL_2851046, EPI_ISL_2851047, EPI_ISL_2851048, EPI_ISL_2851049, EPI_ISL_2851050, EPI_ISL_2851051, EPI_ISL_2851052, EPI_ISL_2851053, EPI_ISL_2851054, EPI_ISL_2851055, EPI_ISL_2851056, EPI_ISL_2851057, EPI_ISL_2851058, EPI_ISL_2851059, EPI_ISL_2851060, EPI_ISL_2851061, EPI_ISL_2851062, EPI_ISL_2851063, EPI_ISL_2851064, EPI_ISL_2851065, EPI_ISL_2851066, EPI_ISL_2851067, EPI_ISL_2851068, EPI_ISL_2851069, EPI_ISL_2851070, EPI_ISL_2851071, EPI_ISL_2851072, EPI_ISL_2851073, EPI_ISL_2851074, EPI_ISL_2851075, EPI_ISL_2851076, EPI_ISL_2851077, EPI_ISL_2851078 | | | |
| see above | Lighthouse Lab in Glasgow | Wellcome Sanger Institute for the COVID-19 Genomics UK (COG-UK) Consortium | Harper VanSteenhouse, Yumi Kasai, David Gray, Carol Clugston, Anna Dominiczak and Alex Alderton, Roberto Amato, Jeffrey Barrett, Sonia Goncalves, Ewan Harrison, David K. Jackson, Ian Johnston, Dominic Kwiatkowski, Cordelia Langford, John Sillitoe on behalf of the Wellcome Sanger Institute COVID-19 Surveillance Team |
| EPI_ISL_2851079 | Lighthouse Lab in Alderley Park | Wellcome Sanger Institute for the COVID-19 Genomics UK (COG-UK) Consortium | Jacquelyn Wynn, Mairead Hyland, The Lighthouse Lab in Alderley Park and Alex Alderton, Roberto Amato, Jeffrey Barrett, Sonia Goncalves, Ewan Harrison, David K. Jackson, Ian Johnston, Dominic Kwiatkowski, Cordelia Langford, John Sillitoe on behalf of the Wellcome Sanger Institute COVID-19 Surveillance Team |
| EPI_ISL_2851080, EPI_ISL_2851081, EPI_ISL_2851082, EPI_ISL_2851083, EPI_ISL_2851084, EPI_ISL_2851085, EPI_ISL_2851086, EPI_ISL_2851087, EPI_ISL_2851088, EPI_ISL_2851089, EPI_ISL_2851090, EPI_ISL_2851091, EPI_ISL_2851092, EPI_ISL_2851093, EPI_ISL_2851094, EPI_ISL_2851095, EPI_ISL_2851096, EPI_ISL_2851097, EPI_ISL_2851098, EPI_ISL_2851099, EPI_ISL_2851100, EPI_ISL_2851101, EPI_ISL_2851102, EPI_ISL_2851103, EPI_ISL_2851104, EPI_ISL_2851105, EPI_ISL_2851106, EPI_ISL_2851107, EPI_ISL_2851108, EPI_ISL_2851109, EPI_ISL_2851110, EPI_ISL_2851111, EPI_ISL_2851112, EPI_ISL_2851113, EPI_ISL_2851114, EPI_ISL_2851115, EPI_ISL_2851116, EPI_ISL_2851117, EPI_ISL_2851118, EPI_ISL_2851119, EPI_ISL_2851120, EPI_ISL_2851121, EPI_ISL_2851122, EPI_ISL_2851123, EPI_ISL_2851124, EPI_ISL_2851125, EPI_ISL_2851126, EPI_ISL_2851127, EPI_ISL_2851128, EPI_ISL_2851129, EPI_ISL_2851130, EPI_ISL_2851131, EPI_ISL_2851132 | | | |
| see above | Lighthouse Lab in Glasgow | Wellcome Sanger Institute for the COVID-19 Genomics UK (COG-UK) Consortium | Harper VanSteenhouse, Yumi Kasai, David Gray, Carol Clugston, Anna Dominiczak and Alex Alderton, Roberto Amato, Jeffrey Barrett, Sonia Goncalves, Ewan Harrison, David K. Jackson, Ian Johnston, Dominic Kwiatkowski, Cordelia Langford, John Sillitoe on behalf of the Wellcome Sanger Institute COVID-19 Surveillance Team |
| EPI_ISL_2851133 | AZ Klina | AZ Klina | Carl Vael - Lynsey Berckmans |
| EPI_ISL_2851134, EPI_ISL_2851135, EPI_ISL_2851136, EPI_ISL_2851137, EPI_ISL_2851138, EPI_ISL_2851139, EPI_ISL_2851140, EPI_ISL_2851141, EPI_ISL_2851142, EPI_ISL_2851143, EPI_ISL_2851144, EPI_ISL_2851145, EPI_ISL_2851146, EPI_ISL_2851147, EPI_ISL_2851148, EPI_ISL_2851149, EPI_ISL_2851150, EPI_ISL_2851151, EPI_ISL_2851152, EPI_ISL_2851153, EPI_ISL_2851154, EPI_ISL_2851155, EPI_ISL_2851156, EPI_ISL_2851157, EPI_ISL_2851158, EPI_ISL_2851159, EPI_ISL_2851160, EPI_ISL_2851161, EPI_ISL_2851162, EPI_ISL_2851163, EPI_ISL_2851164, EPI_ISL_2851165, EPI_ISL_2851166, EPI_ISL_2851167, EPI_ISL_2851168, EPI_ISL_2851169, EPI_ISL_2851170, EPI_ISL_2851171, EPI_ISL_2851172, EPI_ISL_2851173, EPI_ISL_2851174, EPI_ISL_2851175, EPI_ISL_2851176, EPI_ISL_2851177, EPI_ISL_2851178, EPI_ISL_2851179, EPI_ISL_2851180, EPI_ISL_2851181, EPI_ISL_2851182, EPI_ISL_2851183, EPI_ISL_2851184, EPI_ISL_2851185, EPI_ISL_2851186, EPI_ISL_2851187, EPI_ISL_2851188 | | | |
| see above | Lighthouse Lab in Glasgow | Wellcome Sanger Institute for the COVID-19 Genomics UK (COG-UK) Consortium | Harper VanSteenhouse, Yumi Kasai, David Gray, Carol Clugston, Anna Dominiczak and Alex Alderton, Roberto Amato, Jeffrey Barrett, Sonia Goncalves, Ewan Harrison, David K. Jackson, Ian Johnston, Dominic Kwiatkowski, Cordelia Langford, John Sillitoe on behalf of the Wellcome Sanger Institute COVID-19 Surveillance Team |

| | | | |
|---|---|---|---|
| EPI_ISL_2851189 | Lighthouse Lab in Alderley Park | Wellcome Sanger Institute for the COVID-19 Genomics UK (COG-UK) Consortium | Jacquelyn Wynn, Mairead Hyland, The Lighthouse Lab in Alderley Park and Alex Alderton, Roberto Amato, Jeffrey Barrett, Sonia Goncalves, Ewan Harrison, David K. Jackson, Ian Johnston, Dominic Kwiatkowski, Cordelia Langford, John Sillitoe on behalf of the Wellcome Sanger Institute COVID-19 Surveillance Team |
| EPI_ISL_2851190, EPI_ISL_2851191, EPI_ISL_2851192, EPI_ISL_2851193, EPI_ISL_2851194, EPI_ISL_2851195, EPI_ISL_2851196, EPI_ISL_2851197, EPI_ISL_2851198, EPI_ISL_2851199, EPI_ISL_2851200, EPI_ISL_2851201, EPI_ISL_2851202, EPI_ISL_2851203, EPI_ISL_2851204, EPI_ISL_2851205, EPI_ISL_2851206, EPI_ISL_2851207, EPI_ISL_2851208, EPI_ISL_2851209, EPI_ISL_2851210, EPI_ISL_2851211, EPI_ISL_2851212, EPI_ISL_2851213, EPI_ISL_2851214, EPI_ISL_2851215, EPI_ISL_2851216, EPI_ISL_2851217, EPI_ISL_2851218, EPI_ISL_2851219, EPI_ISL_2851220, EPI_ISL_2851221, EPI_ISL_2851222, EPI_ISL_2851223, EPI_ISL_2851224, EPI_ISL_2851225, EPI_ISL_2851226, EPI_ISL_2851227, EPI_ISL_2851228, EPI_ISL_2851229, EPI_ISL_2851230, EPI_ISL_2851231, EPI_ISL_2851232, EPI_ISL_2851233, EPI_ISL_2851234, EPI_ISL_2851235, EPI_ISL_2851236, EPI_ISL_2851237, EPI_ISL_2851238, EPI_ISL_2851239, EPI_ISL_2851240, EPI_ISL_2851241, EPI_ISL_2851242, EPI_ISL_2851243, EPI_ISL_2851244 | | | |
| see above | Lighthouse Lab in Glasgow | Wellcome Sanger Institute for the COVID-19 Genomics UK (COG-UK) Consortium | Harper VanSteenhouse, Yumi Kasai, David Gray, Carol Clugston, Anna Dominiczak and Alex Alderton, Roberto Amato, Jeffrey Barrett, Sonia Goncalves, Ewan Harrison, David K. Jackson, Ian Johnston, Dominic Kwiatkowski, Cordelia Langford, John Sillitoe on behalf of the Wellcome Sanger Institute COVID-19 Surveillance Team |
| EPI_ISL_2851245 | Lighthouse Lab in Alderley Park | Wellcome Sanger Institute for the COVID-19 Genomics UK (COG-UK) Consortium | Jacquelyn Wynn, Mairead Hyland, The Lighthouse Lab in Alderley Park and Alex Alderton, Roberto Amato, Jeffrey Barrett, Sonia Goncalves, Ewan Harrison, David K. Jackson, Ian Johnston, Dominic Kwiatkowski, Cordelia Langford, John Sillitoe on behalf of the Wellcome Sanger Institute COVID-19 Surveillance Team |
| EPI_ISL_2851246, EPI_ISL_2851247, EPI_ISL_2851248, EPI_ISL_2851249, EPI_ISL_2851250, EPI_ISL_2851251, EPI_ISL_2851252, EPI_ISL_2851253, EPI_ISL_2851254, EPI_ISL_2851255, EPI_ISL_2851256, EPI_ISL_2851257, EPI_ISL_2851258, EPI_ISL_2851259, EPI_ISL_2851260, EPI_ISL_2851261, EPI_ISL_2851262, EPI_ISL_2851263, EPI_ISL_2851264, EPI_ISL_2851265, EPI_ISL_2851266, EPI_ISL_2851267, EPI_ISL_2851268, EPI_ISL_2851269, EPI_ISL_2851270, EPI_ISL_2851271, EPI_ISL_2851272, EPI_ISL_2851273, EPI_ISL_2851274, EPI_ISL_2851275, EPI_ISL_2851276, EPI_ISL_2851277, EPI_ISL_2851278, EPI_ISL_2851279, EPI_ISL_2851280, EPI_ISL_2851281, EPI_ISL_2851282, EPI_ISL_2851283, EPI_ISL_2851284, EPI_ISL_2851285, EPI_ISL_2851286, EPI_ISL_2851287, EPI_ISL_2851288, EPI_ISL_2851289, EPI_ISL_2851290, EPI_ISL_2851291, EPI_ISL_2851293, EPI_ISL_2851294, EPI_ISL_2851295, EPI_ISL_2851296, EPI_ISL_2851297, EPI_ISL_2851298, EPI_ISL_2851299, EPI_ISL_2851300, EPI_ISL_2851301, EPI_ISL_2851302, EPI_ISL_2851303, EPI_ISL_2851304, EPI_ISL_2851305, EPI_ISL_2851306, EPI_ISL_2851307, EPI_ISL_2851308, EPI_ISL_2851309, EPI_ISL_2851310, EPI_ISL_2851311, EPI_ISL_2851312, EPI_ISL_2851313, EPI_ISL_2851314, EPI_ISL_2851315, EPI_ISL_2851316, EPI_ISL_2851317, EPI_ISL_2851318, EPI_ISL_2851319, EPI_ISL_2851320, EPI_ISL_2851321, EPI_ISL_2851322, EPI_ISL_2851323, EPI_ISL_2851324, EPI_ISL_2851325, EPI_ISL_2851326, EPI_ISL_2851327, EPI_ISL_2851328, EPI_ISL_2851329, EPI_ISL_2851330, EPI_ISL_2851331, EPI_ISL_2851332, EPI_ISL_2851333, EPI_ISL_2851334, EPI_ISL_2851335, EPI_ISL_2851336, EPI_ISL_2851337, EPI_ISL_2851338, EPI_ISL_2851339, EPI_ISL_2851340, EPI_ISL_2851341, EPI_ISL_2851342, EPI_ISL_2851343, EPI_ISL_2851344, EPI_ISL_2851345, EPI_ISL_2851346, EPI_ISL_2851347, EPI_ISL_2851348, EPI_ISL_2851349, EPI_ISL_2851350, EPI_ISL_2851351, EPI_ISL_2851352, EPI_ISL_2851353, EPI_ISL_2851354, EPI_ISL_2851355, EPI_ISL_2851356, EPI_ISL_2851357, EPI_ISL_2851358, EPI_ISL_2851359, EPI_ISL_2851360, EPI_ISL_2851361, EPI_ISL_2851362, EPI_ISL_2851363, EPI_ISL_2851364, EPI_ISL_2851365, EPI_ISL_2851366, EPI_ISL_2851367, EPI_ISL_2851368, EPI_ISL_2851369, EPI_ISL_2851370, EPI_ISL_2851371, EPI_ISL_2851372, EPI_ISL_2851373, EPI_ISL_2851374, EPI_ISL_2851375, EPI_ISL_2851376 | | | |
| see above | Lighthouse Lab in Glasgow | Wellcome Sanger Institute for the COVID-19 Genomics UK (COG-UK) Consortium | Harper VanSteenhouse, Yumi Kasai, David Gray, Carol Clugston, Anna Dominiczak and Alex Alderton, Roberto Amato, Jeffrey Barrett, Sonia Goncalves, Ewan Harrison, David K. Jackson, Ian Johnston, Dominic Kwiatkowski, Cordelia Langford, John Sillitoe on behalf of the Wellcome Sanger Institute COVID-19 Surveillance Team |
| EPI_ISL_2851377 | Lighthouse Lab in Alderley Park | Wellcome Sanger Institute for the COVID-19 Genomics UK (COG-UK) Consortium | Jacquelyn Wynn, Mairead Hyland, The Lighthouse Lab in Alderley Park and Alex Alderton, Roberto Amato, Jeffrey Barrett, Sonia Goncalves, Ewan Harrison, David K. Jackson, Ian Johnston, Dominic Kwiatkowski, Cordelia Langford, John Sillitoe on behalf of the Wellcome Sanger Institute COVID-19 Surveillance Team |
| EPI_ISL_2851378, EPI_ISL_2851379, EPI_ISL_2851380, EPI_ISL_2851381, EPI_ISL_2851382, EPI_ISL_2851383, EPI_ISL_2851384, EPI_ISL_2851385, EPI_ISL_2851387, EPI_ISL_2851388, EPI_ISL_2851389, EPI_ISL_2851390, EPI_ISL_2851391, EPI_ISL_2851392, EPI_ISL_2851393, EPI_ISL_2851395, EPI_ISL_2851396, EPI_ISL_2851397, EPI_ISL_2851398, EPI_ISL_2851399, EPI_ISL_2851400, EPI_ISL_2851401, EPI_ISL_2851402, EPI_ISL_2851403, EPI_ISL_2851404, EPI_ISL_2851405, EPI_ISL_2851406, EPI_ISL_2851407, EPI_ISL_2851408, EPI_ISL_2851409 | | | |
| see above | Lighthouse Lab in Glasgow | Wellcome Sanger Institute for the COVID-19 Genomics UK (COG-UK) Consortium | Harper VanSteenhouse, Yumi Kasai, David Gray, Carol Clugston, Anna Dominiczak and Alex Alderton, Roberto Amato, Jeffrey Barrett, Sonia Goncalves, Ewan Harrison, David K. Jackson, Ian Johnston, Dominic Kwiatkowski, Cordelia Langford, John Sillitoe on behalf of the Wellcome Sanger Institute COVID-19 Surveillance Team |
| EPI_ISL_2851410 | Lighthouse Lab in Milton Keynes | Wellcome Sanger Institute for the COVID-19 Genomics UK (COG-UK) Consortium | The Lighthouse Lab in Milton Keynes and Alex Alderton, Roberto Amato, Jeffrey Barrett, Sonia Goncalves, Ewan Harrison, David K. Jackson, Ian Johnston, Dominic Kwiatkowski, Cordelia Langford, John Sillitoe on behalf of the Wellcome Sanger Institute COVID-19 Surveillance Team |
| EPI_ISL_2851411 | Lighthouse Lab in Alderley Park | Wellcome Sanger Institute for the COVID-19 Genomics UK (COG-UK) Consortium | Jacquelyn Wynn, Mairead Hyland, The Lighthouse Lab in Alderley Park and Alex Alderton, Roberto Amato, Jeffrey Barrett, Sonia Goncalves, Ewan Harrison, David K. Jackson, Ian Johnston, Dominic Kwiatkowski, Cordelia Langford, John Sillitoe on behalf of the Wellcome Sanger Institute COVID-19 Surveillance Team |
| EPI_ISL_2851412, EPI_ISL_2851413 | Health Services Laboratories | Wellcome Sanger Institute for the COVID-19 Genomics UK (COG-UK) Consortium | Health Services Laboratories and Alex Alderton, Roberto Amato, Jeffrey Barrett, Sonia Goncalves, Ewan Harrison, David K. Jackson, Ian Johnston, Dominic Kwiatkowski, Cordelia Langford, John Sillitoe on behalf of the Wellcome Sanger Institute COVID-19 Surveillance Team |
| EPI_ISL_2851414 | Lighthouse Lab in Alderley Park | Wellcome Sanger Institute for the COVID-19 Genomics UK (COG-UK) Consortium | Jacquelyn Wynn, Mairead Hyland, The Lighthouse Lab in Alderley Park and Alex Alderton, Roberto Amato, Jeffrey Barrett, Sonia Goncalves, Ewan Harrison, David K. Jackson, Ian Johnston, Dominic Kwiatkowski, Cordelia Langford, John Sillitoe on behalf of the Wellcome Sanger Institute COVID-19 Surveillance Team |
| EPI_ISL_2851415, EPI_ISL_2851416, EPI_ISL_2851417, EPI_ISL_2851418, EPI_ISL_2851419, EPI_ISL_2851420, EPI_ISL_2851421 | Health Services Laboratories | Wellcome Sanger Institute for the COVID-19 Genomics UK (COG-UK) Consortium | Health Services Laboratories and Alex Alderton, Roberto Amato, Jeffrey Barrett, Sonia Goncalves, Ewan Harrison, David K. Jackson, Ian Johnston, Dominic Kwiatkowski, Cordelia Langford, John Sillitoe on behalf of the Wellcome Sanger Institute COVID-19 Surveillance Team |
| EPI_ISL_2851422, EPI_ISL_2851423, EPI_ISL_2851424 | Lighthouse Lab in Milton Keynes | Wellcome Sanger Institute for the COVID-19 Genomics UK (COG-UK) Consortium | The Lighthouse Lab in Milton Keynes and Alex Alderton, Roberto Amato, Jeffrey Barrett, Sonia Goncalves, Ewan Harrison, David K. Jackson, Ian Johnston, Dominic Kwiatkowski, Cordelia Langford, John Sillitoe on behalf of the Wellcome Sanger Institute COVID-19 Surveillance Team |
| EPI_ISL_2851425 | Lighthouse Lab in Alderley Park | Wellcome Sanger Institute for the COVID-19 Genomics UK (COG-UK) Consortium | Jacquelyn Wynn, Mairead Hyland, The Lighthouse Lab in Alderley Park and Alex Alderton, Roberto Amato, Jeffrey Barrett, Sonia Goncalves, Ewan Harrison, David K. Jackson, Ian Johnston, Dominic Kwiatkowski, Cordelia Langford, John Sillitoe on behalf of the Wellcome Sanger Institute COVID-19 Surveillance Team |
| EPI_ISL_2851426 | Health Services Laboratories | Wellcome Sanger Institute for the COVID-19 Genomics UK (COG-UK) Consortium | Health Services Laboratories and Alex Alderton, Roberto Amato, Jeffrey Barrett, Sonia Goncalves, Ewan Harrison, David K. Jackson, Ian Johnston, Dominic Kwiatkowski, Cordelia Langford, John Sillitoe on behalf of the Wellcome Sanger Institute COVID-19 Surveillance Team |
| EPI_ISL_2851427 | Lighthouse Lab in Alderley Park | Wellcome Sanger Institute for the COVID-19 Genomics UK (COG-UK) Consortium | Jacquelyn Wynn, Mairead Hyland, The Lighthouse Lab in Alderley Park and Alex Alderton, Roberto Amato, Jeffrey Barrett, Sonia Goncalves, Ewan Harrison, David K. Jackson, Ian Johnston, Dominic Kwiatkowski, Cordelia Langford, John Sillitoe on behalf of the Wellcome Sanger Institute COVID-19 Surveillance Team |
| EPI_ISL_2851428, EPI_ISL_2851429 | Health Services Laboratories | Wellcome Sanger Institute for the COVID-19 Genomics UK (COG-UK) Consortium | Health Services Laboratories and Alex Alderton, Roberto Amato, Jeffrey Barrett, Sonia Goncalves, Ewan Harrison, David K. Jackson, Ian Johnston, Dominic Kwiatkowski, Cordelia Langford, John Sillitoe on behalf of the Wellcome Sanger Institute COVID-19 Surveillance Team |
| EPI_ISL_2851430, EPI_ISL_2851431 | Lighthouse Lab in Alderley Park | Wellcome Sanger Institute for the COVID-19 Genomics UK (COG-UK) Consortium | Jacquelyn Wynn, Mairead Hyland, The Lighthouse Lab in Alderley Park and Alex Alderton, Roberto Amato, Jeffrey Barrett, Sonia Goncalves, Ewan Harrison, David K. Jackson, Ian Johnston, Dominic Kwiatkowski, Cordelia Langford, John Sillitoe on behalf of the Wellcome Sanger Institute COVID-19 Surveillance Team |
| EPI_ISL_2851432, EPI_ISL_2851433 | Health Services Laboratories | Wellcome Sanger Institute for the COVID-19 Genomics UK (COG-UK) Consortium | Health Services Laboratories and Alex Alderton, Roberto Amato, Jeffrey Barrett, Sonia Goncalves, Ewan Harrison, David K. Jackson, Ian Johnston, Dominic Kwiatkowski, Cordelia Langford, John Sillitoe on behalf of the Wellcome Sanger Institute COVID-19 Surveillance Team |
| EPI_ISL_2851434 | Lighthouse Lab in Alderley Park | Wellcome Sanger Institute for the COVID-19 Genomics UK (COG-UK) Consortium | Jacquelyn Wynn, Mairead Hyland, The Lighthouse Lab in Alderley Park and Alex Alderton, Roberto Amato, Jeffrey Barrett, Sonia Goncalves, Ewan Harrison, David K. Jackson, Ian Johnston, Dominic Kwiatkowski, Cordelia Langford, John Sillitoe on behalf of the Wellcome Sanger Institute COVID-19 Surveillance Team |
| EPI_ISL_2851435 | Health Services Laboratories | Wellcome Sanger Institute for the COVID-19 Genomics UK (COG-UK) Consortium | Health Services Laboratories and Alex Alderton, Roberto Amato, Jeffrey Barrett, Sonia Goncalves, Ewan Harrison, David K. Jackson, Ian Johnston, Dominic Kwiatkowski, Cordelia Langford, John Sillitoe on behalf of the Wellcome Sanger Institute COVID-19 Surveillance Team |
| EPI_ISL_2851436 | Lighthouse Lab in Milton Keynes | Wellcome Sanger Institute for the COVID-19 Genomics UK (COG-UK) Consortium | The Lighthouse Lab in Milton Keynes and Alex Alderton, Roberto Amato, Jeffrey Barrett, Sonia Goncalves, Ewan Harrison, David K. Jackson, Ian Johnston, Dominic Kwiatkowski, Cordelia Langford, John Sillitoe on behalf of the Wellcome Sanger Institute COVID-19 Surveillance Team |
| EPI_ISL_2851437 | Lighthouse Lab in Alderley Park | Wellcome Sanger Institute for the COVID-19 Genomics UK (COG-UK) Consortium | Jacquelyn Wynn, Mairead Hyland, The Lighthouse Lab in Alderley Park and Alex Alderton, Roberto Amato, Jeffrey Barrett, Sonia Goncalves, Ewan Harrison, David K. Jackson, Ian Johnston, Dominic Kwiatkowski, Cordelia Langford, John Sillitoe on behalf of the Wellcome Sanger Institute COVID-19 Surveillance Team |
| EPI_ISL_2851438, EPI_ISL_2851439, EPI_ISL_2851440, EPI_ISL_2851441, EPI_ISL_2851442, EPI_ISL_2851443 | Health Services Laboratories | Wellcome Sanger Institute for the COVID-19 Genomics UK (COG-UK) Consortium | Health Services Laboratories and Alex Alderton, Roberto Amato, Jeffrey Barrett, Sonia Goncalves, Ewan Harrison, David K. Jackson, Ian Johnston, Dominic Kwiatkowski, Cordelia Langford, John Sillitoe on behalf of the Wellcome Sanger Institute COVID-19 Surveillance Team |
| EPI_ISL_2851444 | Lighthouse Lab in Alderley Park | Wellcome Sanger Institute for the COVID-19 Genomics UK (COG-UK) Consortium | Jacquelyn Wynn, Mairead Hyland, The Lighthouse Lab in Alderley Park and Alex Alderton, Roberto Amato, Jeffrey Barrett, Sonia Goncalves, Ewan Harrison, David K. Jackson, Ian Johnston, Dominic Kwiatkowski, Cordelia Langford, John Sillitoe on behalf of the Wellcome Sanger Institute COVID-19 Surveillance Team |
| EPI_ISL_2851445 | Health Services Laboratories | Wellcome Sanger Institute for the COVID-19 Genomics UK (COG-UK) Consortium | Health Services Laboratories and Alex Alderton, Roberto Amato, Jeffrey Barrett, Sonia Goncalves, Ewan Harrison, David K. Jackson, Ian Johnston, Dominic Kwiatkowski, Cordelia Langford, John Sillitoe on behalf of the Wellcome Sanger Institute COVID-19 Surveillance Team |
| EPI_ISL_2851446, EPI_ISL_2851447 | Lighthouse Lab in Milton Keynes | Wellcome Sanger Institute for the COVID-19 Genomics UK (COG-UK) Consortium | The Lighthouse Lab in Milton Keynes and Alex Alderton, Roberto Amato, Jeffrey Barrett, Sonia Goncalves, Ewan Harrison, David K. Jackson, Ian Johnston, Dominic Kwiatkowski, Cordelia Langford, John Sillitoe on behalf of the Wellcome Sanger Institute COVID-19 Surveillance Team |
| EPI_ISL_2851448, EPI_ISL_2851449, EPI_ISL_2851450 | Health Services Laboratories | Wellcome Sanger Institute for the COVID-19 Genomics UK (COG-UK) Consortium | Health Services Laboratories and Alex Alderton, Roberto Amato, Jeffrey Barrett, Sonia Goncalves, Ewan Harrison, David K. Jackson, Ian Johnston, Dominic Kwiatkowski, Cordelia Langford, John Sillitoe on behalf of the Wellcome Sanger Institute COVID-19 Surveillance Team |
| EPI_ISL_2851451 | Lighthouse Lab in Alderley Park | Wellcome Sanger Institute for the COVID-19 Genomics UK (COG-UK) Consortium | Jacquelyn Wynn, Mairead Hyland, The Lighthouse Lab in Alderley Park and Alex Alderton, Roberto Amato, Jeffrey Barrett, Sonia Goncalves, Ewan Harrison, David K. Jackson, Ian Johnston, Dominic Kwiatkowski, Cordelia Langford, John Sillitoe on behalf of the Wellcome Sanger Institute COVID-19 Surveillance Team |
| EPI_ISL_2851452, EPI_ISL_2851453, EPI_ISL_2851454 | Lighthouse Lab in Milton Keynes | Wellcome Sanger Institute for the COVID-19 Genomics UK (COG-UK) Consortium | The Lighthouse Lab in Milton Keynes and Alex Alderton, Roberto Amato, Jeffrey Barrett, Sonia Goncalves, Ewan Harrison, David K. Jackson, Ian Johnston, Dominic Kwiatkowski, Cordelia Langford, John Sillitoe on behalf of the Wellcome Sanger Institute COVID-19 Surveillance Team |
| EPI_ISL_2851455, EPI_ISL_2851456 | Lighthouse Lab in Alderley Park | Wellcome Sanger Institute for the COVID-19 Genomics UK (COG-UK) Consortium | Jacquelyn Wynn, Mairead Hyland, The Lighthouse Lab in Alderley Park and Alex Alderton, Roberto Amato, Jeffrey Barrett, Sonia Goncalves, Ewan Harrison, David K. Jackson, Ian Johnston, Dominic Kwiatkowski, Cordelia Langford, John Sillitoe on behalf of the Wellcome Sanger Institute COVID-19 Surveillance Team |
| EPI_ISL_2851457 | Health Services Laboratories | Wellcome Sanger Institute for the COVID-19 Genomics UK (COG-UK) Consortium | Health Services Laboratories and Alex Alderton, Roberto Amato, Jeffrey Barrett, Sonia Goncalves, Ewan Harrison, David K. Jackson, Ian Johnston, Dominic Kwiatkowski, Cordelia Langford, John Sillitoe on behalf of the Wellcome Sanger Institute COVID-19 Surveillance Team |
| EPI_ISL_2851458, EPI_ISL_2851459, EPI_ISL_2851460 | Lighthouse Lab in Alderley Park | Wellcome Sanger Institute for the COVID-19 Genomics UK (COG-UK) Consortium | Jacquelyn Wynn, Mairead Hyland, The Lighthouse Lab in Alderley Park and Alex Alderton, Roberto Amato, Jeffrey Barrett, Sonia Goncalves, Ewan Harrison, David K. Jackson, Ian Johnston, Dominic Kwiatkowski, Cordelia Langford, John Sillitoe on behalf of the Wellcome Sanger Institute COVID-19 Surveillance Team |
| EPI_ISL_2851461, EPI_ISL_2851462, EPI_ISL_2851463 | Health Services Laboratories | Wellcome Sanger Institute for the COVID-19 Genomics UK (COG-UK) Consortium | Health Services Laboratories and Alex Alderton, Roberto Amato, Jeffrey Barrett, Sonia Goncalves, Ewan Harrison, David K. Jackson, Ian Johnston, Dominic Kwiatkowski, Cordelia Langford, John Sillitoe on behalf of the Wellcome Sanger Institute COVID-19 Surveillance Team |
| EPI_ISL_2851464, EPI_ISL_2851465 | Lighthouse Lab in Alderley Park | Wellcome Sanger Institute for the COVID-19 Genomics UK (COG-UK) Consortium | Jacquelyn Wynn, Mairead Hyland, The Lighthouse Lab in Alderley Park and Alex Alderton, Roberto Amato, Jeffrey Barrett, Sonia Goncalves, Ewan Harrison, David K. Jackson, Ian Johnston, Dominic Kwiatkowski, Cordelia Langford, John Sillitoe on behalf of the Wellcome Sanger Institute COVID-19 Surveillance Team |
| EPI_ISL_2851466 | Health Services Laboratories | Wellcome Sanger Institute for the COVID-19 Genomics UK (COG-UK) | Health Services Laboratories and Alex Alderton, Roberto Amato, Jeffrey Barrett, Sonia Goncalves, Ewan Harrison, David K. Jackson, Ian Johnston, Dominic Kwiatkowski, Cordelia Langford, John Sillitoe on behalf of the Wellcome Sanger Institute |

| | | | |
|---|---|---|---|
| | | Consortium | COVID-19 Surveillance Team |
| EPI_ISL_2851467, EPI_ISL_2851468 | Lighthouse Lab in Alderley Park | Wellcome Sanger Institute for the COVID-19 Genomics UK (COG-UK) Consortium | Jacquelyn Wynn, Mairead Hyland, The Lighthouse Lab in Alderley Park and Alex Alderton, Roberto Amato, Jeffrey Barrett, Sonia Goncalves, Ewan Harrison, David K. Jackson, Ian Johnston, Dominic Kwiatkowski, Cordelia Langford, John Sillitoe on behalf of the Wellcome Sanger Institute COVID-19 Surveillance Team |
| EPI_ISL_2851469 | Health Services Laboratories | Wellcome Sanger Institute for the COVID-19 Genomics UK (COG-UK) Consortium | Health Services Laboratories and Alex Alderton, Roberto Amato, Jeffrey Barrett, Sonia Goncalves, Ewan Harrison, David K. Jackson, Ian Johnston, Dominic Kwiatkowski, Cordelia Langford, John Sillitoe on behalf of the Wellcome Sanger Institute COVID-19 Surveillance Team |
| EPI_ISL_2851470 | Lighthouse Lab in Alderley Park | Wellcome Sanger Institute for the COVID-19 Genomics UK (COG-UK) Consortium | Jacquelyn Wynn, Mairead Hyland, The Lighthouse Lab in Alderley Park and Alex Alderton, Roberto Amato, Jeffrey Barrett, Sonia Goncalves, Ewan Harrison, David K. Jackson, Ian Johnston, Dominic Kwiatkowski, Cordelia Langford, John Sillitoe on behalf of the Wellcome Sanger Institute COVID-19 Surveillance Team |
| EPI_ISL_2851471, EPI_ISL_2851472 | Health Services Laboratories | Wellcome Sanger Institute for the COVID-19 Genomics UK (COG-UK) Consortium | Health Services Laboratories and Alex Alderton, Roberto Amato, Jeffrey Barrett, Sonia Goncalves, Ewan Harrison, David K. Jackson, Ian Johnston, Dominic Kwiatkowski, Cordelia Langford, John Sillitoe on behalf of the Wellcome Sanger Institute COVID-19 Surveillance Team |
| EPI_ISL_2851473 | Lighthouse Lab in Milton Keynes | Wellcome Sanger Institute for the COVID-19 Genomics UK (COG-UK) Consortium | The Lighthouse Lab in Milton Keynes and Alex Alderton, Roberto Amato, Jeffrey Barrett, Sonia Goncalves, Ewan Harrison, David K. Jackson, Ian Johnston, Dominic Kwiatkowski, Cordelia Langford, John Sillitoe on behalf of the Wellcome Sanger Institute COVID-19 Surveillance Team |
| EPI_ISL_2851474, EPI_ISL_2851475, EPI_ISL_2851476, EPI_ISL_2851477, EPI_ISL_2851478 | Health Services Laboratories | Wellcome Sanger Institute for the COVID-19 Genomics UK (COG-UK) Consortium | Health Services Laboratories and Alex Alderton, Roberto Amato, Jeffrey Barrett, Sonia Goncalves, Ewan Harrison, David K. Jackson, Ian Johnston, Dominic Kwiatkowski, Cordelia Langford, John Sillitoe on behalf of the Wellcome Sanger Institute COVID-19 Surveillance Team |
| EPI_ISL_2851479 | Lighthouse Lab in Milton Keynes | Wellcome Sanger Institute for the COVID-19 Genomics UK (COG-UK) Consortium | The Lighthouse Lab in Milton Keynes and Alex Alderton, Roberto Amato, Jeffrey Barrett, Sonia Goncalves, Ewan Harrison, David K. Jackson, Ian Johnston, Dominic Kwiatkowski, Cordelia Langford, John Sillitoe on behalf of the Wellcome Sanger Institute COVID-19 Surveillance Team |
| EPI_ISL_2851480, EPI_ISL_2851481, EPI_ISL_2851482 | Health Services Laboratories | Wellcome Sanger Institute for the COVID-19 Genomics UK (COG-UK) Consortium | Health Services Laboratories and Alex Alderton, Roberto Amato, Jeffrey Barrett, Sonia Goncalves, Ewan Harrison, David K. Jackson, Ian Johnston, Dominic Kwiatkowski, Cordelia Langford, John Sillitoe on behalf of the Wellcome Sanger Institute COVID-19 Surveillance Team |
| EPI_ISL_2851483 | Lighthouse Lab in Alderley Park | Wellcome Sanger Institute for the COVID-19 Genomics UK (COG-UK) Consortium | Jacquelyn Wynn, Mairead Hyland, The Lighthouse Lab in Alderley Park and Alex Alderton, Roberto Amato, Jeffrey Barrett, Sonia Goncalves, Ewan Harrison, David K. Jackson, Ian Johnston, Dominic Kwiatkowski, Cordelia Langford, John Sillitoe on behalf of the Wellcome Sanger Institute COVID-19 Surveillance Team |
| EPI_ISL_2851484, EPI_ISL_2851485, EPI_ISL_2851486 | Health Services Laboratories | Wellcome Sanger Institute for the COVID-19 Genomics UK (COG-UK) Consortium | Health Services Laboratories and Alex Alderton, Roberto Amato, Jeffrey Barrett, Sonia Goncalves, Ewan Harrison, David K. Jackson, Ian Johnston, Dominic Kwiatkowski, Cordelia Langford, John Sillitoe on behalf of the Wellcome Sanger Institute COVID-19 Surveillance Team |
| EPI_ISL_2851487 | Lighthouse Lab in Alderley Park | Wellcome Sanger Institute for the COVID-19 Genomics UK (COG-UK) Consortium | Jacquelyn Wynn, Mairead Hyland, The Lighthouse Lab in Alderley Park and Alex Alderton, Roberto Amato, Jeffrey Barrett, Sonia Goncalves, Ewan Harrison, David K. Jackson, Ian Johnston, Dominic Kwiatkowski, Cordelia Langford, John Sillitoe on behalf of the Wellcome Sanger Institute COVID-19 Surveillance Team |
| EPI_ISL_2851488, EPI_ISL_2851489, EPI_ISL_2851490, EPI_ISL_2851491, EPI_ISL_2851492 | Health Services Laboratories | Wellcome Sanger Institute for the COVID-19 Genomics UK (COG-UK) Consortium | Health Services Laboratories and Alex Alderton, Roberto Amato, Jeffrey Barrett, Sonia Goncalves, Ewan Harrison, David K. Jackson, Ian Johnston, Dominic Kwiatkowski, Cordelia Langford, John Sillitoe on behalf of the Wellcome Sanger Institute COVID-19 Surveillance Team |
| EPI_ISL_2851493 | Lighthouse Lab in Alderley Park | Wellcome Sanger Institute for the COVID-19 Genomics UK (COG-UK) Consortium | Jacquelyn Wynn, Mairead Hyland, The Lighthouse Lab in Alderley Park and Alex Alderton, Roberto Amato, Jeffrey Barrett, Sonia Goncalves, Ewan Harrison, David K. Jackson, Ian Johnston, Dominic Kwiatkowski, Cordelia Langford, John Sillitoe on behalf of the Wellcome Sanger Institute COVID-19 Surveillance Team |
| EPI_ISL_2851494, EPI_ISL_2851495 | Health Services Laboratories | Wellcome Sanger Institute for the COVID-19 Genomics UK (COG-UK) Consortium | Health Services Laboratories and Alex Alderton, Roberto Amato, Jeffrey Barrett, Sonia Goncalves, Ewan Harrison, David K. Jackson, Ian Johnston, Dominic Kwiatkowski, Cordelia Langford, John Sillitoe on behalf of the Wellcome Sanger Institute COVID-19 Surveillance Team |
| EPI_ISL_2851496, EPI_ISL_2851497, EPI_ISL_2851498 | Lighthouse Lab in Alderley Park | Wellcome Sanger Institute for the COVID-19 Genomics UK (COG-UK) Consortium | Jacquelyn Wynn, Mairead Hyland, The Lighthouse Lab in Alderley Park and Alex Alderton, Roberto Amato, Jeffrey Barrett, Sonia Goncalves, Ewan Harrison, David K. Jackson, Ian Johnston, Dominic Kwiatkowski, Cordelia Langford, John Sillitoe on behalf of the Wellcome Sanger Institute COVID-19 Surveillance Team |
| EPI_ISL_2851499, EPI_ISL_2851500 | Health Services Laboratories | Wellcome Sanger Institute for the COVID-19 Genomics UK (COG-UK) Consortium | Health Services Laboratories and Alex Alderton, Roberto Amato, Jeffrey Barrett, Sonia Goncalves, Ewan Harrison, David K. Jackson, Ian Johnston, Dominic Kwiatkowski, Cordelia Langford, John Sillitoe on behalf of the Wellcome Sanger Institute COVID-19 Surveillance Team |
| EPI_ISL_2851501, EPI_ISL_2851502, EPI_ISL_2851503 | Lighthouse Lab in Milton Keynes | Wellcome Sanger Institute for the COVID-19 Genomics UK (COG-UK) Consortium | The Lighthouse Lab in Milton Keynes and Alex Alderton, Roberto Amato, Jeffrey Barrett, Sonia Goncalves, Ewan Harrison, David K. Jackson, Ian Johnston, Dominic Kwiatkowski, Cordelia Langford, John Sillitoe on behalf of the Wellcome Sanger Institute COVID-19 Surveillance Team |
| EPI_ISL_2851504 | Lighthouse Lab in Alderley Park | Wellcome Sanger Institute for the COVID-19 Genomics UK (COG-UK) Consortium | Jacquelyn Wynn, Mairead Hyland, The Lighthouse Lab in Alderley Park and Alex Alderton, Roberto Amato, Jeffrey Barrett, Sonia Goncalves, Ewan Harrison, David K. Jackson, Ian Johnston, Dominic Kwiatkowski, Cordelia Langford, John Sillitoe on behalf of the Wellcome Sanger Institute COVID-19 Surveillance Team |
| EPI_ISL_2851505 | Health Services Laboratories | Wellcome Sanger Institute for the COVID-19 Genomics UK (COG-UK) Consortium | Health Services Laboratories and Alex Alderton, Roberto Amato, Jeffrey Barrett, Sonia Goncalves, Ewan Harrison, David K. Jackson, Ian Johnston, Dominic Kwiatkowski, Cordelia Langford, John Sillitoe on behalf of the Wellcome Sanger Institute COVID-19 Surveillance Team |
| EPI_ISL_2851506 | Lighthouse Lab in Alderley Park | Wellcome Sanger Institute for the COVID-19 Genomics UK (COG-UK) Consortium | Jacquelyn Wynn, Mairead Hyland, The Lighthouse Lab in Alderley Park and Alex Alderton, Roberto Amato, Jeffrey Barrett, Sonia Goncalves, Ewan Harrison, David K. Jackson, Ian Johnston, Dominic Kwiatkowski, Cordelia Langford, John Sillitoe on behalf of the Wellcome Sanger Institute COVID-19 Surveillance Team |
| EPI_ISL_2851507, EPI_ISL_2851508 | Health Services Laboratories | Wellcome Sanger Institute for the COVID-19 Genomics UK (COG-UK) Consortium | Health Services Laboratories and Alex Alderton, Roberto Amato, Jeffrey Barrett, Sonia Goncalves, Ewan Harrison, David K. Jackson, Ian Johnston, Dominic Kwiatkowski, Cordelia Langford, John Sillitoe on behalf of the Wellcome Sanger Institute COVID-19 Surveillance Team |
| EPI_ISL_2851509, EPI_ISL_2851510 | Lighthouse Lab in Alderley Park | Wellcome Sanger Institute for the COVID-19 Genomics UK (COG-UK) Consortium | Jacquelyn Wynn, Mairead Hyland, The Lighthouse Lab in Alderley Park and Alex Alderton, Roberto Amato, Jeffrey Barrett, Sonia Goncalves, Ewan Harrison, David K. Jackson, Ian Johnston, Dominic Kwiatkowski, Cordelia Langford, John Sillitoe on behalf of the Wellcome Sanger Institute COVID-19 Surveillance Team |
| EPI_ISL_2851511, EPI_ISL_2851512 | Lighthouse Lab in Milton Keynes | Wellcome Sanger Institute for the COVID-19 Genomics UK (COG-UK) Consortium | The Lighthouse Lab in Milton Keynes and Alex Alderton, Roberto Amato, Jeffrey Barrett, Sonia Goncalves, Ewan Harrison, David K. Jackson, Ian Johnston, Dominic Kwiatkowski, Cordelia Langford, John Sillitoe on behalf of the Wellcome Sanger Institute COVID-19 Surveillance Team |
| EPI_ISL_2851513 | Health Services Laboratories | Wellcome Sanger Institute for the COVID-19 Genomics UK (COG-UK) Consortium | Health Services Laboratories and Alex Alderton, Roberto Amato, Jeffrey Barrett, Sonia Goncalves, Ewan Harrison, David K. Jackson, Ian Johnston, Dominic Kwiatkowski, Cordelia Langford, John Sillitoe on behalf of the Wellcome Sanger Institute COVID-19 Surveillance Team |
| EPI_ISL_2851514 | Lighthouse Lab in Milton Keynes | Wellcome Sanger Institute for the COVID-19 Genomics UK (COG-UK) Consortium | The Lighthouse Lab in Milton Keynes and Alex Alderton, Roberto Amato, Jeffrey Barrett, Sonia Goncalves, Ewan Harrison, David K. Jackson, Ian Johnston, Dominic Kwiatkowski, Cordelia Langford, John Sillitoe on behalf of the Wellcome Sanger Institute COVID-19 Surveillance Team |
| EPI_ISL_2851515 | Lighthouse Lab in Alderley Park | Wellcome Sanger Institute for the COVID-19 Genomics UK (COG-UK) Consortium | Jacquelyn Wynn, Mairead Hyland, The Lighthouse Lab in Alderley Park and Alex Alderton, Roberto Amato, Jeffrey Barrett, Sonia Goncalves, Ewan Harrison, David K. Jackson, Ian Johnston, Dominic Kwiatkowski, Cordelia Langford, John Sillitoe on behalf of the Wellcome Sanger Institute COVID-19 Surveillance Team |
| EPI_ISL_2851516, EPI_ISL_2851517 | Health Services Laboratories | Wellcome Sanger Institute for the COVID-19 Genomics UK (COG-UK) Consortium | Health Services Laboratories and Alex Alderton, Roberto Amato, Jeffrey Barrett, Sonia Goncalves, Ewan Harrison, David K. Jackson, Ian Johnston, Dominic Kwiatkowski, Cordelia Langford, John Sillitoe on behalf of the Wellcome Sanger Institute COVID-19 Surveillance Team |
| EPI_ISL_2851518 | Lighthouse Lab in Alderley Park | Wellcome Sanger Institute for the COVID-19 Genomics UK (COG-UK) Consortium | Jacquelyn Wynn, Mairead Hyland, The Lighthouse Lab in Alderley Park and Alex Alderton, Roberto Amato, Jeffrey Barrett, Sonia Goncalves, Ewan Harrison, David K. Jackson, Ian Johnston, Dominic Kwiatkowski, Cordelia Langford, John Sillitoe on behalf of the Wellcome Sanger Institute COVID-19 Surveillance Team |
| EPI_ISL_2851519 | Lighthouse Lab in Milton Keynes | Wellcome Sanger Institute for the COVID-19 Genomics UK (COG-UK) Consortium | The Lighthouse Lab in Milton Keynes and Alex Alderton, Roberto Amato, Jeffrey Barrett, Sonia Goncalves, Ewan Harrison, David K. Jackson, Ian Johnston, Dominic Kwiatkowski, Cordelia Langford, John Sillitoe on behalf of the Wellcome Sanger Institute COVID-19 Surveillance Team |
| EPI_ISL_2851520 | Health Services Laboratories | Wellcome Sanger Institute for the COVID-19 Genomics UK (COG-UK) Consortium | Health Services Laboratories and Alex Alderton, Roberto Amato, Jeffrey Barrett, Sonia Goncalves, Ewan Harrison, David K. Jackson, Ian Johnston, Dominic Kwiatkowski, Cordelia Langford, John Sillitoe on behalf of the Wellcome Sanger Institute COVID-19 Surveillance Team |
| EPI_ISL_2851521 | Lighthouse Lab in Alderley Park | Wellcome Sanger Institute for the COVID-19 Genomics UK (COG-UK) Consortium | Jacquelyn Wynn, Mairead Hyland, The Lighthouse Lab in Alderley Park and Alex Alderton, Roberto Amato, Jeffrey Barrett, Sonia Goncalves, Ewan Harrison, David K. Jackson, Ian Johnston, Dominic Kwiatkowski, Cordelia Langford, John Sillitoe on behalf of the Wellcome Sanger Institute COVID-19 Surveillance Team |
| EPI_ISL_2851522 | Health Services Laboratories | Wellcome Sanger Institute for the COVID-19 Genomics UK (COG-UK) Consortium | Health Services Laboratories and Alex Alderton, Roberto Amato, Jeffrey Barrett, Sonia Goncalves, Ewan Harrison, David K. Jackson, Ian Johnston, Dominic Kwiatkowski, Cordelia Langford, John Sillitoe on behalf of the Wellcome Sanger Institute COVID-19 Surveillance Team |
| EPI_ISL_2851523, EPI_ISL_2851524 | Lighthouse Lab in Milton Keynes | Wellcome Sanger Institute for the COVID-19 Genomics UK (COG-UK) Consortium | The Lighthouse Lab in Milton Keynes and Alex Alderton, Roberto Amato, Jeffrey Barrett, Sonia Goncalves, Ewan Harrison, David K. Jackson, Ian Johnston, Dominic Kwiatkowski, Cordelia Langford, John Sillitoe on behalf of the Wellcome Sanger Institute COVID-19 Surveillance Team |
| EPI_ISL_2851525 | Health Services Laboratories | Wellcome Sanger Institute for the COVID-19 Genomics UK (COG-UK) Consortium | Health Services Laboratories and Alex Alderton, Roberto Amato, Jeffrey Barrett, Sonia Goncalves, Ewan Harrison, David K. Jackson, Ian Johnston, Dominic Kwiatkowski, Cordelia Langford, John Sillitoe on behalf of the Wellcome Sanger Institute COVID-19 Surveillance Team |
| EPI_ISL_2851526 | Lighthouse Lab in Milton Keynes | Wellcome Sanger Institute for the COVID-19 Genomics UK (COG-UK) Consortium | The Lighthouse Lab in Milton Keynes and Alex Alderton, Roberto Amato, Jeffrey Barrett, Sonia Goncalves, Ewan Harrison, David K. Jackson, Ian Johnston, Dominic Kwiatkowski, Cordelia Langford, John Sillitoe on behalf of the Wellcome Sanger Institute COVID-19 Surveillance Team |
| EPI_ISL_2851527, EPI_ISL_2851528 | Health Services Laboratories | Wellcome Sanger Institute for the COVID-19 Genomics UK (COG-UK) Consortium | Health Services Laboratories and Alex Alderton, Roberto Amato, Jeffrey Barrett, Sonia Goncalves, Ewan Harrison, David K. Jackson, Ian Johnston, Dominic Kwiatkowski, Cordelia Langford, John Sillitoe on behalf of the Wellcome Sanger Institute COVID-19 Surveillance Team |
| EPI_ISL_2851529, EPI_ISL_2851530, EPI_ISL_2851531 | Lighthouse Lab in Milton Keynes | Wellcome Sanger Institute for the COVID-19 Genomics UK (COG-UK) Consortium | The Lighthouse Lab in Milton Keynes and Alex Alderton, Roberto Amato, Jeffrey Barrett, Sonia Goncalves, Ewan Harrison, David K. Jackson, Ian Johnston, Dominic Kwiatkowski, Cordelia Langford, John Sillitoe on behalf of the Wellcome Sanger Institute COVID-19 Surveillance Team |
| EPI_ISL_2851532 | Health Services Laboratories | Wellcome Sanger Institute for the COVID-19 Genomics UK (COG-UK) Consortium | Health Services Laboratories and Alex Alderton, Roberto Amato, Jeffrey Barrett, Sonia Goncalves, Ewan Harrison, David K. Jackson, Ian Johnston, Dominic Kwiatkowski, Cordelia Langford, John Sillitoe on behalf of the Wellcome Sanger Institute COVID-19 Surveillance Team |
| EPI_ISL_2851533 | Lighthouse Lab in Milton Keynes | Wellcome Sanger Institute for the COVID-19 Genomics UK (COG-UK) Consortium | The Lighthouse Lab in Milton Keynes and Alex Alderton, Roberto Amato, Jeffrey Barrett, Sonia Goncalves, Ewan Harrison, David K. Jackson, Ian Johnston, Dominic Kwiatkowski, Cordelia Langford, John Sillitoe on behalf of the Wellcome Sanger Institute COVID-19 Surveillance Team |
| EPI_ISL_2851534 | Lighthouse Lab in Alderley Park | Wellcome Sanger Institute for the COVID-19 Genomics UK (COG-UK) Consortium | Jacquelyn Wynn, Mairead Hyland, The Lighthouse Lab in Alderley Park and Alex Alderton, Roberto Amato, Jeffrey Barrett, Sonia Goncalves, Ewan Harrison, David K. Jackson, Ian Johnston, Dominic Kwiatkowski, Cordelia Langford, John Sillitoe on behalf of the Wellcome Sanger Institute COVID-19 Surveillance Team |

| | | | |
|---|---|---|---|
| EPI_ISL_2851535 | Health Services Laboratories | Wellcome Sanger Institute for the COVID-19 Genomics UK (COG-UK) Consortium | Health Services Laboratories and Alex Alderton, Roberto Amato, Jeffrey Barrett, Sonia Goncalves, Ewan Harrison, David K. Jackson, Ian Johnston, Dominic Kwiatkowski, Cordelia Langford, John Sillitoe on behalf of the Wellcome Sanger Institute COVID-19 Surveillance Team |
| EPI_ISL_2851536 | Lighthouse Lab in Milton Keynes | Wellcome Sanger Institute for the COVID-19 Genomics UK (COG-UK) Consortium | The Lighthouse Lab in Milton Keynes and Alex Alderton, Roberto Amato, Jeffrey Barrett, Sonia Goncalves, Ewan Harrison, David K. Jackson, Ian Johnston, Dominic Kwiatkowski, Cordelia Langford, John Sillitoe on behalf of the Wellcome Sanger Institute COVID-19 Surveillance Team |
| EPI_ISL_2851537 | Health Services Laboratories | Wellcome Sanger Institute for the COVID-19 Genomics UK (COG-UK) Consortium | Health Services Laboratories and Alex Alderton, Roberto Amato, Jeffrey Barrett, Sonia Goncalves, Ewan Harrison, David K. Jackson, Ian Johnston, Dominic Kwiatkowski, Cordelia Langford, John Sillitoe on behalf of the Wellcome Sanger Institute COVID-19 Surveillance Team |
| EPI_ISL_2851538 | Lighthouse Lab in Alderley Park | Wellcome Sanger Institute for the COVID-19 Genomics UK (COG-UK) Consortium | Jacquelyn Wynn, Mairead Hyland, The Lighthouse Lab in Alderley Park and Alex Alderton, Roberto Amato, Jeffrey Barrett, Sonia Goncalves, Ewan Harrison, David K. Jackson, Ian Johnston, Dominic Kwiatkowski, Cordelia Langford, John Sillitoe on behalf of the Wellcome Sanger Institute COVID-19 Surveillance Team |
| EPI_ISL_2851539 | Lighthouse Lab in Milton Keynes | Wellcome Sanger Institute for the COVID-19 Genomics UK (COG-UK) Consortium | The Lighthouse Lab in Milton Keynes and Alex Alderton, Roberto Amato, Jeffrey Barrett, Sonia Goncalves, Ewan Harrison, David K. Jackson, Ian Johnston, Dominic Kwiatkowski, Cordelia Langford, John Sillitoe on behalf of the Wellcome Sanger Institute COVID-19 Surveillance Team |
| EPI_ISL_2851540 | Health Services Laboratories | Wellcome Sanger Institute for the COVID-19 Genomics UK (COG-UK) Consortium | Health Services Laboratories and Alex Alderton, Roberto Amato, Jeffrey Barrett, Sonia Goncalves, Ewan Harrison, David K. Jackson, Ian Johnston, Dominic Kwiatkowski, Cordelia Langford, John Sillitoe on behalf of the Wellcome Sanger Institute COVID-19 Surveillance Team |
| EPI_ISL_2851541 | Lighthouse Lab in Alderley Park | Wellcome Sanger Institute for the COVID-19 Genomics UK (COG-UK) Consortium | Jacquelyn Wynn, Mairead Hyland, The Lighthouse Lab in Alderley Park and Alex Alderton, Roberto Amato, Jeffrey Barrett, Sonia Goncalves, Ewan Harrison, David K. Jackson, Ian Johnston, Dominic Kwiatkowski, Cordelia Langford, John Sillitoe on behalf of the Wellcome Sanger Institute COVID-19 Surveillance Team |
| EPI_ISL_2851542, EPI_ISL_2851543 | Health Services Laboratories | Wellcome Sanger Institute for the COVID-19 Genomics UK (COG-UK) Consortium | Health Services Laboratories and Alex Alderton, Roberto Amato, Jeffrey Barrett, Sonia Goncalves, Ewan Harrison, David K. Jackson, Ian Johnston, Dominic Kwiatkowski, Cordelia Langford, John Sillitoe on behalf of the Wellcome Sanger Institute COVID-19 Surveillance Team |
| EPI_ISL_2851544, EPI_ISL_2851545, EPI_ISL_2851546 | Lighthouse Lab in Alderley Park | Wellcome Sanger Institute for the COVID-19 Genomics UK (COG-UK) Consortium | Jacquelyn Wynn, Mairead Hyland, The Lighthouse Lab in Alderley Park and Alex Alderton, Roberto Amato, Jeffrey Barrett, Sonia Goncalves, Ewan Harrison, David K. Jackson, Ian Johnston, Dominic Kwiatkowski, Cordelia Langford, John Sillitoe on behalf of the Wellcome Sanger Institute COVID-19 Surveillance Team |
| EPI_ISL_2851547, EPI_ISL_2851548 | Health Services Laboratories | Wellcome Sanger Institute for the COVID-19 Genomics UK (COG-UK) Consortium | Health Services Laboratories and Alex Alderton, Roberto Amato, Jeffrey Barrett, Sonia Goncalves, Ewan Harrison, David K. Jackson, Ian Johnston, Dominic Kwiatkowski, Cordelia Langford, John Sillitoe on behalf of the Wellcome Sanger Institute COVID-19 Surveillance Team |
| EPI_ISL_2851549 | Lighthouse Lab in Alderley Park | Wellcome Sanger Institute for the COVID-19 Genomics UK (COG-UK) Consortium | Jacquelyn Wynn, Mairead Hyland, The Lighthouse Lab in Alderley Park and Alex Alderton, Roberto Amato, Jeffrey Barrett, Sonia Goncalves, Ewan Harrison, David K. Jackson, Ian Johnston, Dominic Kwiatkowski, Cordelia Langford, John Sillitoe on behalf of the Wellcome Sanger Institute COVID-19 Surveillance Team |
| EPI_ISL_2851550, EPI_ISL_2851551, EPI_ISL_2851552, EPI_ISL_2851553, EPI_ISL_2851554 | Health Services Laboratories | Wellcome Sanger Institute for the COVID-19 Genomics UK (COG-UK) Consortium | Health Services Laboratories and Alex Alderton, Roberto Amato, Jeffrey Barrett, Sonia Goncalves, Ewan Harrison, David K. Jackson, Ian Johnston, Dominic Kwiatkowski, Cordelia Langford, John Sillitoe on behalf of the Wellcome Sanger Institute COVID-19 Surveillance Team |
| EPI_ISL_2851555 | Lighthouse Lab in Milton Keynes | Wellcome Sanger Institute for the COVID-19 Genomics UK (COG-UK) Consortium | The Lighthouse Lab in Milton Keynes and Alex Alderton, Roberto Amato, Jeffrey Barrett, Sonia Goncalves, Ewan Harrison, David K. Jackson, Ian Johnston, Dominic Kwiatkowski, Cordelia Langford, John Sillitoe on behalf of the Wellcome Sanger Institute COVID-19 Surveillance Team |
| EPI_ISL_2851556 | Lighthouse Lab in Alderley Park | Wellcome Sanger Institute for the COVID-19 Genomics UK (COG-UK) Consortium | Jacquelyn Wynn, Mairead Hyland, The Lighthouse Lab in Alderley Park and Alex Alderton, Roberto Amato, Jeffrey Barrett, Sonia Goncalves, Ewan Harrison, David K. Jackson, Ian Johnston, Dominic Kwiatkowski, Cordelia Langford, John Sillitoe on behalf of the Wellcome Sanger Institute COVID-19 Surveillance Team |
| EPI_ISL_2851557 | Health Services Laboratories | Wellcome Sanger Institute for the COVID-19 Genomics UK (COG-UK) Consortium | Health Services Laboratories and Alex Alderton, Roberto Amato, Jeffrey Barrett, Sonia Goncalves, Ewan Harrison, David K. Jackson, Ian Johnston, Dominic Kwiatkowski, Cordelia Langford, John Sillitoe on behalf of the Wellcome Sanger Institute COVID-19 Surveillance Team |
| EPI_ISL_2851558 | Lighthouse Lab in Milton Keynes | Wellcome Sanger Institute for the COVID-19 Genomics UK (COG-UK) Consortium | The Lighthouse Lab in Milton Keynes and Alex Alderton, Roberto Amato, Jeffrey Barrett, Sonia Goncalves, Ewan Harrison, David K. Jackson, Ian Johnston, Dominic Kwiatkowski, Cordelia Langford, John Sillitoe on behalf of the Wellcome Sanger Institute COVID-19 Surveillance Team |
| EPI_ISL_2851559 | Health Services Laboratories | Wellcome Sanger Institute for the COVID-19 Genomics UK (COG-UK) Consortium | Health Services Laboratories and Alex Alderton, Roberto Amato, Jeffrey Barrett, Sonia Goncalves, Ewan Harrison, David K. Jackson, Ian Johnston, Dominic Kwiatkowski, Cordelia Langford, John Sillitoe on behalf of the Wellcome Sanger Institute COVID-19 Surveillance Team |
| EPI_ISL_2851560 | Lighthouse Lab in Alderley Park | Wellcome Sanger Institute for the COVID-19 Genomics UK (COG-UK) Consortium | Jacquelyn Wynn, Mairead Hyland, The Lighthouse Lab in Alderley Park and Alex Alderton, Roberto Amato, Jeffrey Barrett, Sonia Goncalves, Ewan Harrison, David K. Jackson, Ian Johnston, Dominic Kwiatkowski, Cordelia Langford, John Sillitoe on behalf of the Wellcome Sanger Institute COVID-19 Surveillance Team |
| EPI_ISL_2851561 | Lighthouse Lab in Milton Keynes | Wellcome Sanger Institute for the COVID-19 Genomics UK (COG-UK) Consortium | The Lighthouse Lab in Milton Keynes and Alex Alderton, Roberto Amato, Jeffrey Barrett, Sonia Goncalves, Ewan Harrison, David K. Jackson, Ian Johnston, Dominic Kwiatkowski, Cordelia Langford, John Sillitoe on behalf of the Wellcome Sanger Institute COVID-19 Surveillance Team |
| EPI_ISL_2851562 | Lighthouse Lab in Alderley Park | Wellcome Sanger Institute for the COVID-19 Genomics UK (COG-UK) Consortium | Jacquelyn Wynn, Mairead Hyland, The Lighthouse Lab in Alderley Park and Alex Alderton, Roberto Amato, Jeffrey Barrett, Sonia Goncalves, Ewan Harrison, David K. Jackson, Ian Johnston, Dominic Kwiatkowski, Cordelia Langford, John Sillitoe on behalf of the Wellcome Sanger Institute COVID-19 Surveillance Team |
| EPI_ISL_2851563 | Lighthouse Lab in Milton Keynes | Wellcome Sanger Institute for the COVID-19 Genomics UK (COG-UK) Consortium | The Lighthouse Lab in Milton Keynes and Alex Alderton, Roberto Amato, Jeffrey Barrett, Sonia Goncalves, Ewan Harrison, David K. Jackson, Ian Johnston, Dominic Kwiatkowski, Cordelia Langford, John Sillitoe on behalf of the Wellcome Sanger Institute COVID-19 Surveillance Team |
| EPI_ISL_2851564, EPI_ISL_2851565, EPI_ISL_2851566, EPI_ISL_2851567 | Health Services Laboratories | Wellcome Sanger Institute for the COVID-19 Genomics UK (COG-UK) Consortium | Health Services Laboratories and Alex Alderton, Roberto Amato, Jeffrey Barrett, Sonia Goncalves, Ewan Harrison, David K. Jackson, Ian Johnston, Dominic Kwiatkowski, Cordelia Langford, John Sillitoe on behalf of the Wellcome Sanger Institute COVID-19 Surveillance Team |
| EPI_ISL_2851568 | Lighthouse Lab in Alderley Park | Wellcome Sanger Institute for the COVID-19 Genomics UK (COG-UK) Consortium | Jacquelyn Wynn, Mairead Hyland, The Lighthouse Lab in Alderley Park and Alex Alderton, Roberto Amato, Jeffrey Barrett, Sonia Goncalves, Ewan Harrison, David K. Jackson, Ian Johnston, Dominic Kwiatkowski, Cordelia Langford, John Sillitoe on behalf of the Wellcome Sanger Institute COVID-19 Surveillance Team |
| EPI_ISL_2851569 | Lighthouse Lab in Milton Keynes | Wellcome Sanger Institute for the COVID-19 Genomics UK (COG-UK) Consortium | The Lighthouse Lab in Milton Keynes and Alex Alderton, Roberto Amato, Jeffrey Barrett, Sonia Goncalves, Ewan Harrison, David K. Jackson, Ian Johnston, Dominic Kwiatkowski, Cordelia Langford, John Sillitoe on behalf of the Wellcome Sanger Institute COVID-19 Surveillance Team |
| EPI_ISL_2851570 | Lighthouse Lab in Alderley Park | Wellcome Sanger Institute for the COVID-19 Genomics UK (COG-UK) Consortium | Jacquelyn Wynn, Mairead Hyland, The Lighthouse Lab in Alderley Park and Alex Alderton, Roberto Amato, Jeffrey Barrett, Sonia Goncalves, Ewan Harrison, David K. Jackson, Ian Johnston, Dominic Kwiatkowski, Cordelia Langford, John Sillitoe on behalf of the Wellcome Sanger Institute COVID-19 Surveillance Team |
| EPI_ISL_2851571 | Lighthouse Lab in Milton Keynes | Wellcome Sanger Institute for the COVID-19 Genomics UK (COG-UK) Consortium | The Lighthouse Lab in Milton Keynes and Alex Alderton, Roberto Amato, Jeffrey Barrett, Sonia Goncalves, Ewan Harrison, David K. Jackson, Ian Johnston, Dominic Kwiatkowski, Cordelia Langford, John Sillitoe on behalf of the Wellcome Sanger Institute COVID-19 Surveillance Team |
| EPI_ISL_2851572 | Health Services Laboratories | Wellcome Sanger Institute for the COVID-19 Genomics UK (COG-UK) Consortium | Health Services Laboratories and Alex Alderton, Roberto Amato, Jeffrey Barrett, Sonia Goncalves, Ewan Harrison, David K. Jackson, Ian Johnston, Dominic Kwiatkowski, Cordelia Langford, John Sillitoe on behalf of the Wellcome Sanger Institute COVID-19 Surveillance Team |
| EPI_ISL_2851573, EPI_ISL_2851574 | Lighthouse Lab in Milton Keynes | Wellcome Sanger Institute for the COVID-19 Genomics UK (COG-UK) Consortium | The Lighthouse Lab in Milton Keynes and Alex Alderton, Roberto Amato, Jeffrey Barrett, Sonia Goncalves, Ewan Harrison, David K. Jackson, Ian Johnston, Dominic Kwiatkowski, Cordelia Langford, John Sillitoe on behalf of the Wellcome Sanger Institute COVID-19 Surveillance Team |
| EPI_ISL_2851575 | Lighthouse Lab in Alderley Park | Wellcome Sanger Institute for the COVID-19 Genomics UK (COG-UK) Consortium | Jacquelyn Wynn, Mairead Hyland, The Lighthouse Lab in Alderley Park and Alex Alderton, Roberto Amato, Jeffrey Barrett, Sonia Goncalves, Ewan Harrison, David K. Jackson, Ian Johnston, Dominic Kwiatkowski, Cordelia Langford, John Sillitoe on behalf of the Wellcome Sanger Institute COVID-19 Surveillance Team |
| EPI_ISL_2851576, EPI_ISL_2851577 | Health Services Laboratories | Wellcome Sanger Institute for the COVID-19 Genomics UK (COG-UK) Consortium | Health Services Laboratories and Alex Alderton, Roberto Amato, Jeffrey Barrett, Sonia Goncalves, Ewan Harrison, David K. Jackson, Ian Johnston, Dominic Kwiatkowski, Cordelia Langford, John Sillitoe on behalf of the Wellcome Sanger Institute COVID-19 Surveillance Team |
| EPI_ISL_2851578 | Lighthouse Lab in Alderley Park | Wellcome Sanger Institute for the COVID-19 Genomics UK (COG-UK) Consortium | Jacquelyn Wynn, Mairead Hyland, The Lighthouse Lab in Alderley Park and Alex Alderton, Roberto Amato, Jeffrey Barrett, Sonia Goncalves, Ewan Harrison, David K. Jackson, Ian Johnston, Dominic Kwiatkowski, Cordelia Langford, John Sillitoe on behalf of the Wellcome Sanger Institute COVID-19 Surveillance Team |
| EPI_ISL_2851579 | Lighthouse Lab in Milton Keynes | Wellcome Sanger Institute for the COVID-19 Genomics UK (COG-UK) Consortium | The Lighthouse Lab in Milton Keynes and Alex Alderton, Roberto Amato, Jeffrey Barrett, Sonia Goncalves, Ewan Harrison, David K. Jackson, Ian Johnston, Dominic Kwiatkowski, Cordelia Langford, John Sillitoe on behalf of the Wellcome Sanger Institute COVID-19 Surveillance Team |
| EPI_ISL_2851580 | Health Services Laboratories | Wellcome Sanger Institute for the COVID-19 Genomics UK (COG-UK) Consortium | Health Services Laboratories and Alex Alderton, Roberto Amato, Jeffrey Barrett, Sonia Goncalves, Ewan Harrison, David K. Jackson, Ian Johnston, Dominic Kwiatkowski, Cordelia Langford, John Sillitoe on behalf of the Wellcome Sanger Institute COVID-19 Surveillance Team |
| EPI_ISL_2851581 | Lighthouse Lab in Alderley Park | Wellcome Sanger Institute for the COVID-19 Genomics UK (COG-UK) Consortium | Jacquelyn Wynn, Mairead Hyland, The Lighthouse Lab in Alderley Park and Alex Alderton, Roberto Amato, Jeffrey Barrett, Sonia Goncalves, Ewan Harrison, David K. Jackson, Ian Johnston, Dominic Kwiatkowski, Cordelia Langford, John Sillitoe on behalf of the Wellcome Sanger Institute COVID-19 Surveillance Team |
| EPI_ISL_2851582 | Health Services Laboratories | Wellcome Sanger Institute for the COVID-19 Genomics UK (COG-UK) Consortium | Health Services Laboratories and Alex Alderton, Roberto Amato, Jeffrey Barrett, Sonia Goncalves, Ewan Harrison, David K. Jackson, Ian Johnston, Dominic Kwiatkowski, Cordelia Langford, John Sillitoe on behalf of the Wellcome Sanger Institute COVID-19 Surveillance Team |
| EPI_ISL_2851583, EPI_ISL_2851584 | Lighthouse Lab in Alderley Park | Wellcome Sanger Institute for the COVID-19 Genomics UK (COG-UK) Consortium | Jacquelyn Wynn, Mairead Hyland, The Lighthouse Lab in Alderley Park and Alex Alderton, Roberto Amato, Jeffrey Barrett, Sonia Goncalves, Ewan Harrison, David K. Jackson, Ian Johnston, Dominic Kwiatkowski, Cordelia Langford, John Sillitoe on behalf of the Wellcome Sanger Institute COVID-19 Surveillance Team |
| EPI_ISL_2851585, EPI_ISL_2851586, EPI_ISL_2851587 | Health Services Laboratories | Wellcome Sanger Institute for the COVID-19 Genomics UK (COG-UK) Consortium | Health Services Laboratories and Alex Alderton, Roberto Amato, Jeffrey Barrett, Sonia Goncalves, Ewan Harrison, David K. Jackson, Ian Johnston, Dominic Kwiatkowski, Cordelia Langford, John Sillitoe on behalf of the Wellcome Sanger Institute COVID-19 Surveillance Team |
| EPI_ISL_2851588 | Lighthouse Lab in Milton Keynes | Wellcome Sanger Institute for the COVID-19 Genomics UK (COG-UK) Consortium | The Lighthouse Lab in Milton Keynes and Alex Alderton, Roberto Amato, Jeffrey Barrett, Sonia Goncalves, Ewan Harrison, David K. Jackson, Ian Johnston, Dominic Kwiatkowski, Cordelia Langford, John Sillitoe on behalf of the Wellcome Sanger Institute COVID-19 Surveillance Team |
| EPI_ISL_2851589 | Lighthouse Lab in Alderley Park | Wellcome Sanger Institute for the COVID-19 Genomics UK (COG-UK) Consortium | Jacquelyn Wynn, Mairead Hyland, The Lighthouse Lab in Alderley Park and Alex Alderton, Roberto Amato, Jeffrey Barrett, Sonia Goncalves, Ewan Harrison, David K. Jackson, Ian Johnston, Dominic Kwiatkowski, Cordelia Langford, John Sillitoe on behalf of the Wellcome Sanger Institute COVID-19 Surveillance Team |
| EPI_ISL_2851590 | Health Services Laboratories | Wellcome Sanger Institute for the COVID-19 Genomics UK (COG-UK) Consortium | Health Services Laboratories and Alex Alderton, Roberto Amato, Jeffrey Barrett, Sonia Goncalves, Ewan Harrison, David K. Jackson, Ian Johnston, Dominic Kwiatkowski, Cordelia Langford, John Sillitoe on behalf of the Wellcome Sanger Institute COVID-19 Surveillance Team |
| EPI_ISL_2851591 | Lighthouse Lab in Alderley Park | Wellcome Sanger Institute for the COVID-19 Genomics UK (COG-UK) Consortium | Jacquelyn Wynn, Mairead Hyland, The Lighthouse Lab in Alderley Park and Alex Alderton, Roberto Amato, Jeffrey Barrett, Sonia Goncalves, Ewan Harrison, David K. Jackson, Ian Johnston, Dominic Kwiatkowski, Cordelia Langford, John Sillitoe on behalf of the Wellcome Sanger Institute COVID-19 Surveillance Team |
| EPI_ISL_2851592 | Lighthouse Lab in Milton Keynes | Wellcome Sanger Institute for the COVID-19 Genomics UK (COG-UK) Consortium | The Lighthouse Lab in Milton Keynes and Alex Alderton, Roberto Amato, Jeffrey Barrett, Sonia Goncalves, Ewan Harrison, David K. Jackson, Ian Johnston, Dominic Kwiatkowski, Cordelia Langford, John Sillitoe on behalf of the Wellcome Sanger Institute COVID-19 Surveillance Team |
| EPI_ISL_2851593 | Lighthouse Lab in Alderley Park | Wellcome Sanger Institute for the COVID-19 Genomics UK (COG-UK) | Jacquelyn Wynn, Mairead Hyland, The Lighthouse Lab in Alderley Park and Alex Alderton, Roberto Amato, Jeffrey Barrett, Sonia Goncalves, Ewan Harrison, David K. Jackson, Ian Johnston, Dominic Kwiatkowski, Cordelia Langford, John Sillitoe on |

| | | | |
|---|---|---|---|
| | | Consortium | behalf of the Wellcome Sanger Institute COVID-19 Surveillance Team |
| EPI_ISL_2851594, EPI_ISL_2851595, EPI_ISL_2851596, EPI_ISL_2851597 | Health Services Laboratories | Wellcome Sanger Institute for the COVID-19 Genomics UK (COG-UK) Consortium | Health Services Laboratories and Alex Alderton, Roberto Amato, Jeffrey Barrett, Sonia Goncalves, Ewan Harrison, David K. Jackson, Ian Johnston, Dominic Kwiatkowski, Cordelia Langford, John Sillitoe on behalf of the Wellcome Sanger Institute COVID-19 Surveillance Team |
| EPI_ISL_2851598 | Lighthouse Lab in Alderley Park | Wellcome Sanger Institute for the COVID-19 Genomics UK (COG-UK) Consortium | Jacquelyn Wynn, Mairead Hyland, The Lighthouse Lab in Alderley Park and Alex Alderton, Roberto Amato, Jeffrey Barrett, Sonia Goncalves, Ewan Harrison, David K. Jackson, Ian Johnston, Dominic Kwiatkowski, Cordelia Langford, John Sillitoe on behalf of the Wellcome Sanger Institute COVID-19 Surveillance Team |
| EPI_ISL_2851599 | Lighthouse Lab in Milton Keynes | Wellcome Sanger Institute for the COVID-19 Genomics UK (COG-UK) Consortium | The Lighthouse Lab in Milton Keynes and Alex Alderton, Roberto Amato, Jeffrey Barrett, Sonia Goncalves, Ewan Harrison, David K. Jackson, Ian Johnston, Dominic Kwiatkowski, Cordelia Langford, John Sillitoe on behalf of the Wellcome Sanger Institute COVID-19 Surveillance Team |
| EPI_ISL_2851600 | Lighthouse Lab in Alderley Park | Wellcome Sanger Institute for the COVID-19 Genomics UK (COG-UK) Consortium | Jacquelyn Wynn, Mairead Hyland, The Lighthouse Lab in Alderley Park and Alex Alderton, Roberto Amato, Jeffrey Barrett, Sonia Goncalves, Ewan Harrison, David K. Jackson, Ian Johnston, Dominic Kwiatkowski, Cordelia Langford, John Sillitoe on behalf of the Wellcome Sanger Institute COVID-19 Surveillance Team |
| EPI_ISL_2851601 | Lighthouse Lab in Milton Keynes | Wellcome Sanger Institute for the COVID-19 Genomics UK (COG-UK) Consortium | The Lighthouse Lab in Milton Keynes and Alex Alderton, Roberto Amato, Jeffrey Barrett, Sonia Goncalves, Ewan Harrison, David K. Jackson, Ian Johnston, Dominic Kwiatkowski, Cordelia Langford, John Sillitoe on behalf of the Wellcome Sanger Institute COVID-19 Surveillance Team |
| EPI_ISL_2851602 | Lighthouse Lab in Alderley Park | Wellcome Sanger Institute for the COVID-19 Genomics UK (COG-UK) Consortium | Jacquelyn Wynn, Mairead Hyland, The Lighthouse Lab in Alderley Park and Alex Alderton, Roberto Amato, Jeffrey Barrett, Sonia Goncalves, Ewan Harrison, David K. Jackson, Ian Johnston, Dominic Kwiatkowski, Cordelia Langford, John Sillitoe on behalf of the Wellcome Sanger Institute COVID-19 Surveillance Team |
| EPI_ISL_2851603 | Lighthouse Lab in Milton Keynes | Wellcome Sanger Institute for the COVID-19 Genomics UK (COG-UK) Consortium | The Lighthouse Lab in Milton Keynes and Alex Alderton, Roberto Amato, Jeffrey Barrett, Sonia Goncalves, Ewan Harrison, David K. Jackson, Ian Johnston, Dominic Kwiatkowski, Cordelia Langford, John Sillitoe on behalf of the Wellcome Sanger Institute COVID-19 Surveillance Team |
| EPI_ISL_2851604 | Lighthouse Lab in Alderley Park | Wellcome Sanger Institute for the COVID-19 Genomics UK (COG-UK) Consortium | Jacquelyn Wynn, Mairead Hyland, The Lighthouse Lab in Alderley Park and Alex Alderton, Roberto Amato, Jeffrey Barrett, Sonia Goncalves, Ewan Harrison, David K. Jackson, Ian Johnston, Dominic Kwiatkowski, Cordelia Langford, John Sillitoe on behalf of the Wellcome Sanger Institute COVID-19 Surveillance Team |
| EPI_ISL_2851605 | Lighthouse Lab in Milton Keynes | Wellcome Sanger Institute for the COVID-19 Genomics UK (COG-UK) Consortium | The Lighthouse Lab in Milton Keynes and Alex Alderton, Roberto Amato, Jeffrey Barrett, Sonia Goncalves, Ewan Harrison, David K. Jackson, Ian Johnston, Dominic Kwiatkowski, Cordelia Langford, John Sillitoe on behalf of the Wellcome Sanger Institute COVID-19 Surveillance Team |
| EPI_ISL_2851607 | Health Services Laboratories | Wellcome Sanger Institute for the COVID-19 Genomics UK (COG-UK) Consortium | Health Services Laboratories and Alex Alderton, Roberto Amato, Jeffrey Barrett, Sonia Goncalves, Ewan Harrison, David K. Jackson, Ian Johnston, Dominic Kwiatkowski, Cordelia Langford, John Sillitoe on behalf of the Wellcome Sanger Institute COVID-19 Surveillance Team |
| EPI_ISL_2851608 | Lighthouse Lab in Milton Keynes | Wellcome Sanger Institute for the COVID-19 Genomics UK (COG-UK) Consortium | The Lighthouse Lab in Milton Keynes and Alex Alderton, Roberto Amato, Jeffrey Barrett, Sonia Goncalves, Ewan Harrison, David K. Jackson, Ian Johnston, Dominic Kwiatkowski, Cordelia Langford, John Sillitoe on behalf of the Wellcome Sanger Institute COVID-19 Surveillance Team |
| EPI_ISL_2851609 | Health Services Laboratories | Wellcome Sanger Institute for the COVID-19 Genomics UK (COG-UK) Consortium | Health Services Laboratories and Alex Alderton, Roberto Amato, Jeffrey Barrett, Sonia Goncalves, Ewan Harrison, David K. Jackson, Ian Johnston, Dominic Kwiatkowski, Cordelia Langford, John Sillitoe on behalf of the Wellcome Sanger Institute COVID-19 Surveillance Team |
| EPI_ISL_2851610 | Lighthouse Lab in Alderley Park | Wellcome Sanger Institute for the COVID-19 Genomics UK (COG-UK) Consortium | Jacquelyn Wynn, Mairead Hyland, The Lighthouse Lab in Alderley Park and Alex Alderton, Roberto Amato, Jeffrey Barrett, Sonia Goncalves, Ewan Harrison, David K. Jackson, Ian Johnston, Dominic Kwiatkowski, Cordelia Langford, John Sillitoe on behalf of the Wellcome Sanger Institute COVID-19 Surveillance Team |
| EPI_ISL_2851611 | Lighthouse Lab in Milton Keynes | Wellcome Sanger Institute for the COVID-19 Genomics UK (COG-UK) Consortium | The Lighthouse Lab in Milton Keynes and Alex Alderton, Roberto Amato, Jeffrey Barrett, Sonia Goncalves, Ewan Harrison, David K. Jackson, Ian Johnston, Dominic Kwiatkowski, Cordelia Langford, John Sillitoe on behalf of the Wellcome Sanger Institute COVID-19 Surveillance Team |
| EPI_ISL_2851612 | Health Services Laboratories | Wellcome Sanger Institute for the COVID-19 Genomics UK (COG-UK) Consortium | Health Services Laboratories and Alex Alderton, Roberto Amato, Jeffrey Barrett, Sonia Goncalves, Ewan Harrison, David K. Jackson, Ian Johnston, Dominic Kwiatkowski, Cordelia Langford, John Sillitoe on behalf of the Wellcome Sanger Institute COVID-19 Surveillance Team |
| EPI_ISL_2851613, EPI_ISL_2851614, EPI_ISL_2851615 | Lighthouse Lab in Milton Keynes | Wellcome Sanger Institute for the COVID-19 Genomics UK (COG-UK) Consortium | The Lighthouse Lab in Milton Keynes and Alex Alderton, Roberto Amato, Jeffrey Barrett, Sonia Goncalves, Ewan Harrison, David K. Jackson, Ian Johnston, Dominic Kwiatkowski, Cordelia Langford, John Sillitoe on behalf of the Wellcome Sanger Institute COVID-19 Surveillance Team |
| EPI_ISL_2851616 | AZ Klina | AZ Klina | Carl Vael - Lynsey Berckmans |
| EPI_ISL_2851617 | Health Services Laboratories | Wellcome Sanger Institute for the COVID-19 Genomics UK (COG-UK) Consortium | Health Services Laboratories and Alex Alderton, Roberto Amato, Jeffrey Barrett, Sonia Goncalves, Ewan Harrison, David K. Jackson, Ian Johnston, Dominic Kwiatkowski, Cordelia Langford, John Sillitoe on behalf of the Wellcome Sanger Institute COVID-19 Surveillance Team |
| EPI_ISL_2851618 | Lighthouse Lab in Milton Keynes | Wellcome Sanger Institute for the COVID-19 Genomics UK (COG-UK) Consortium | The Lighthouse Lab in Milton Keynes and Alex Alderton, Roberto Amato, Jeffrey Barrett, Sonia Goncalves, Ewan Harrison, David K. Jackson, Ian Johnston, Dominic Kwiatkowski, Cordelia Langford, John Sillitoe on behalf of the Wellcome Sanger Institute COVID-19 Surveillance Team |
| EPI_ISL_2851620 | Lighthouse Lab in Alderley Park | Wellcome Sanger Institute for the COVID-19 Genomics UK (COG-UK) Consortium | Jacquelyn Wynn, Mairead Hyland, The Lighthouse Lab in Alderley Park and Alex Alderton, Roberto Amato, Jeffrey Barrett, Sonia Goncalves, Ewan Harrison, David K. Jackson, Ian Johnston, Dominic Kwiatkowski, Cordelia Langford, John Sillitoe on behalf of the Wellcome Sanger Institute COVID-19 Surveillance Team |
| EPI_ISL_2851621 | Health Services Laboratories | Wellcome Sanger Institute for the COVID-19 Genomics UK (COG-UK) Consortium | Health Services Laboratories and Alex Alderton, Roberto Amato, Jeffrey Barrett, Sonia Goncalves, Ewan Harrison, David K. Jackson, Ian Johnston, Dominic Kwiatkowski, Cordelia Langford, John Sillitoe on behalf of the Wellcome Sanger Institute COVID-19 Surveillance Team |
| EPI_ISL_2851622 | Lighthouse Lab in Alderley Park | Wellcome Sanger Institute for the COVID-19 Genomics UK (COG-UK) Consortium | Jacquelyn Wynn, Mairead Hyland, The Lighthouse Lab in Alderley Park and Alex Alderton, Roberto Amato, Jeffrey Barrett, Sonia Goncalves, Ewan Harrison, David K. Jackson, Ian Johnston, Dominic Kwiatkowski, Cordelia Langford, John Sillitoe on behalf of the Wellcome Sanger Institute COVID-19 Surveillance Team |
| EPI_ISL_2851623 | Lighthouse Lab in Milton Keynes | Wellcome Sanger Institute for the COVID-19 Genomics UK (COG-UK) Consortium | The Lighthouse Lab in Milton Keynes and Alex Alderton, Roberto Amato, Jeffrey Barrett, Sonia Goncalves, Ewan Harrison, David K. Jackson, Ian Johnston, Dominic Kwiatkowski, Cordelia Langford, John Sillitoe on behalf of the Wellcome Sanger Institute COVID-19 Surveillance Team |
| EPI_ISL_2851624, EPI_ISL_2851625 | Health Services Laboratories | Wellcome Sanger Institute for the COVID-19 Genomics UK (COG-UK) Consortium | Health Services Laboratories and Alex Alderton, Roberto Amato, Jeffrey Barrett, Sonia Goncalves, Ewan Harrison, David K. Jackson, Ian Johnston, Dominic Kwiatkowski, Cordelia Langford, John Sillitoe on behalf of the Wellcome Sanger Institute COVID-19 Surveillance Team |
| EPI_ISL_2851626 | Lighthouse Lab in Milton Keynes | Wellcome Sanger Institute for the COVID-19 Genomics UK (COG-UK) Consortium | The Lighthouse Lab in Milton Keynes and Alex Alderton, Roberto Amato, Jeffrey Barrett, Sonia Goncalves, Ewan Harrison, David K. Jackson, Ian Johnston, Dominic Kwiatkowski, Cordelia Langford, John Sillitoe on behalf of the Wellcome Sanger Institute COVID-19 Surveillance Team |
| EPI_ISL_2851627 | Health Services Laboratories | Wellcome Sanger Institute for the COVID-19 Genomics UK (COG-UK) Consortium | Health Services Laboratories and Alex Alderton, Roberto Amato, Jeffrey Barrett, Sonia Goncalves, Ewan Harrison, David K. Jackson, Ian Johnston, Dominic Kwiatkowski, Cordelia Langford, John Sillitoe on behalf of the Wellcome Sanger Institute COVID-19 Surveillance Team |
| EPI_ISL_2851628, EPI_ISL_2851629 | Lighthouse Lab in Alderley Park | Wellcome Sanger Institute for the COVID-19 Genomics UK (COG-UK) Consortium | Jacquelyn Wynn, Mairead Hyland, The Lighthouse Lab in Alderley Park and Alex Alderton, Roberto Amato, Jeffrey Barrett, Sonia Goncalves, Ewan Harrison, David K. Jackson, Ian Johnston, Dominic Kwiatkowski, Cordelia Langford, John Sillitoe on behalf of the Wellcome Sanger Institute COVID-19 Surveillance Team |
| EPI_ISL_2851630, EPI_ISL_2851631, EPI_ISL_2851632, EPI_ISL_2851633 | Health Services Laboratories | Wellcome Sanger Institute for the COVID-19 Genomics UK (COG-UK) Consortium | Health Services Laboratories and Alex Alderton, Roberto Amato, Jeffrey Barrett, Sonia Goncalves, Ewan Harrison, David K. Jackson, Ian Johnston, Dominic Kwiatkowski, Cordelia Langford, John Sillitoe on behalf of the Wellcome Sanger Institute COVID-19 Surveillance Team |
| EPI_ISL_2851634 | Lighthouse Lab in Milton Keynes | Wellcome Sanger Institute for the COVID-19 Genomics UK (COG-UK) Consortium | The Lighthouse Lab in Milton Keynes and Alex Alderton, Roberto Amato, Jeffrey Barrett, Sonia Goncalves, Ewan Harrison, David K. Jackson, Ian Johnston, Dominic Kwiatkowski, Cordelia Langford, John Sillitoe on behalf of the Wellcome Sanger Institute COVID-19 Surveillance Team |
| EPI_ISL_2851635 | Lighthouse Lab in Alderley Park | Wellcome Sanger Institute for the COVID-19 Genomics UK (COG-UK) Consortium | Jacquelyn Wynn, Mairead Hyland, The Lighthouse Lab in Alderley Park and Alex Alderton, Roberto Amato, Jeffrey Barrett, Sonia Goncalves, Ewan Harrison, David K. Jackson, Ian Johnston, Dominic Kwiatkowski, Cordelia Langford, John Sillitoe on behalf of the Wellcome Sanger Institute COVID-19 Surveillance Team |
| EPI_ISL_2851636 | Lighthouse Lab in Milton Keynes | Wellcome Sanger Institute for the COVID-19 Genomics UK (COG-UK) Consortium | The Lighthouse Lab in Milton Keynes and Alex Alderton, Roberto Amato, Jeffrey Barrett, Sonia Goncalves, Ewan Harrison, David K. Jackson, Ian Johnston, Dominic Kwiatkowski, Cordelia Langford, John Sillitoe on behalf of the Wellcome Sanger Institute COVID-19 Surveillance Team |
| EPI_ISL_2851637, EPI_ISL_2851638 | Health Services Laboratories | Wellcome Sanger Institute for the COVID-19 Genomics UK (COG-UK) Consortium | Health Services Laboratories and Alex Alderton, Roberto Amato, Jeffrey Barrett, Sonia Goncalves, Ewan Harrison, David K. Jackson, Ian Johnston, Dominic Kwiatkowski, Cordelia Langford, John Sillitoe on behalf of the Wellcome Sanger Institute COVID-19 Surveillance Team |
| EPI_ISL_2851639, EPI_ISL_2851640, EPI_ISL_2851641 | Lighthouse Lab in Milton Keynes | Wellcome Sanger Institute for the COVID-19 Genomics UK (COG-UK) Consortium | The Lighthouse Lab in Milton Keynes and Alex Alderton, Roberto Amato, Jeffrey Barrett, Sonia Goncalves, Ewan Harrison, David K. Jackson, Ian Johnston, Dominic Kwiatkowski, Cordelia Langford, John Sillitoe on behalf of the Wellcome Sanger Institute COVID-19 Surveillance Team |
| EPI_ISL_2851642 | Lighthouse Lab in Alderley Park | Wellcome Sanger Institute for the COVID-19 Genomics UK (COG-UK) Consortium | Jacquelyn Wynn, Mairead Hyland, The Lighthouse Lab in Alderley Park and Alex Alderton, Roberto Amato, Jeffrey Barrett, Sonia Goncalves, Ewan Harrison, David K. Jackson, Ian Johnston, Dominic Kwiatkowski, Cordelia Langford, John Sillitoe on behalf of the Wellcome Sanger Institute COVID-19 Surveillance Team |
| EPI_ISL_2851643, EPI_ISL_2851644, EPI_ISL_2851645, EPI_ISL_2851646, EPI_ISL_2851647, EPI_ISL_2851648 | Health Services Laboratories | Wellcome Sanger Institute for the COVID-19 Genomics UK (COG-UK) Consortium | Health Services Laboratories and Alex Alderton, Roberto Amato, Jeffrey Barrett, Sonia Goncalves, Ewan Harrison, David K. Jackson, Ian Johnston, Dominic Kwiatkowski, Cordelia Langford, John Sillitoe on behalf of the Wellcome Sanger Institute COVID-19 Surveillance Team |
| EPI_ISL_2851649 | Lighthouse Lab in Milton Keynes | Wellcome Sanger Institute for the COVID-19 Genomics UK (COG-UK) Consortium | The Lighthouse Lab in Milton Keynes and Alex Alderton, Roberto Amato, Jeffrey Barrett, Sonia Goncalves, Ewan Harrison, David K. Jackson, Ian Johnston, Dominic Kwiatkowski, Cordelia Langford, John Sillitoe on behalf of the Wellcome Sanger Institute COVID-19 Surveillance Team |
| EPI_ISL_2851650, EPI_ISL_2851651, EPI_ISL_2851652 | Health Services Laboratories | Wellcome Sanger Institute for the COVID-19 Genomics UK (COG-UK) Consortium | Health Services Laboratories and Alex Alderton, Roberto Amato, Jeffrey Barrett, Sonia Goncalves, Ewan Harrison, David K. Jackson, Ian Johnston, Dominic Kwiatkowski, Cordelia Langford, John Sillitoe on behalf of the Wellcome Sanger Institute COVID-19 Surveillance Team |
| EPI_ISL_2851653 | Lighthouse Lab in Milton Keynes | Wellcome Sanger Institute for the COVID-19 Genomics UK (COG-UK) Consortium | The Lighthouse Lab in Milton Keynes and Alex Alderton, Roberto Amato, Jeffrey Barrett, Sonia Goncalves, Ewan Harrison, David K. Jackson, Ian Johnston, Dominic Kwiatkowski, Cordelia Langford, John Sillitoe on behalf of the Wellcome Sanger Institute COVID-19 Surveillance Team |
| EPI_ISL_2851654 | Health Services Laboratories | Wellcome Sanger Institute for the COVID-19 Genomics UK (COG-UK) Consortium | Health Services Laboratories and Alex Alderton, Roberto Amato, Jeffrey Barrett, Sonia Goncalves, Ewan Harrison, David K. Jackson, Ian Johnston, Dominic Kwiatkowski, Cordelia Langford, John Sillitoe on behalf of the Wellcome Sanger Institute COVID-19 Surveillance Team |
| EPI_ISL_2851655, EPI_ISL_2851656, EPI_ISL_2851657 | Lighthouse Lab in Milton Keynes | Wellcome Sanger Institute for the COVID-19 Genomics UK (COG-UK) Consortium | The Lighthouse Lab in Milton Keynes and Alex Alderton, Roberto Amato, Jeffrey Barrett, Sonia Goncalves, Ewan Harrison, David K. Jackson, Ian Johnston, Dominic Kwiatkowski, Cordelia Langford, John Sillitoe on behalf of the Wellcome Sanger Institute COVID-19 Surveillance Team |

| | | | |
|---|---|---|---|
| EPI_ISL_2851658, EPI_ISL_2851659 | Health Services Laboratories | Wellcome Sanger Institute for the COVID-19 Genomics UK (COG-UK) Consortium | Health Services Laboratories and Alex Alderton, Roberto Amato, Jeffrey Barrett, Sonia Goncalves, Ewan Harrison, David K. Jackson, Ian Johnston, Dominic Kwiatkowski, Cordelia Langford, John Sillitoe on behalf of the Wellcome Sanger Institute COVID-19 Surveillance Team |
| EPI_ISL_2851660 | Lighthouse Lab in Alderley Park | Wellcome Sanger Institute for the COVID-19 Genomics UK (COG-UK) Consortium | Jacquelyn Wynn, Mairead Hyland, The Lighthouse Lab in Alderley Park and Alex Alderton, Roberto Amato, Jeffrey Barrett, Sonia Goncalves, Ewan Harrison, David K. Jackson, Ian Johnston, Dominic Kwiatkowski, Cordelia Langford, John Sillitoe on behalf of the Wellcome Sanger Institute COVID-19 Surveillance Team |
| EPI_ISL_2851661 | Lighthouse Lab in Milton Keynes | Wellcome Sanger Institute for the COVID-19 Genomics UK (COG-UK) Consortium | The Lighthouse Lab in Milton Keynes and Alex Alderton, Roberto Amato, Jeffrey Barrett, Sonia Goncalves, Ewan Harrison, David K. Jackson, Ian Johnston, Dominic Kwiatkowski, Cordelia Langford, John Sillitoe on behalf of the Wellcome Sanger Institute COVID-19 Surveillance Team |
| EPI_ISL_2851662, EPI_ISL_2851663 | Health Services Laboratories | Wellcome Sanger Institute for the COVID-19 Genomics UK (COG-UK) Consortium | Health Services Laboratories and Alex Alderton, Roberto Amato, Jeffrey Barrett, Sonia Goncalves, Ewan Harrison, David K. Jackson, Ian Johnston, Dominic Kwiatkowski, Cordelia Langford, John Sillitoe on behalf of the Wellcome Sanger Institute COVID-19 Surveillance Team |
| EPI_ISL_2851664 | Lighthouse Lab in Milton Keynes | Wellcome Sanger Institute for the COVID-19 Genomics UK (COG-UK) Consortium | The Lighthouse Lab in Milton Keynes and Alex Alderton, Roberto Amato, Jeffrey Barrett, Sonia Goncalves, Ewan Harrison, David K. Jackson, Ian Johnston, Dominic Kwiatkowski, Cordelia Langford, John Sillitoe on behalf of the Wellcome Sanger Institute COVID-19 Surveillance Team |
| EPI_ISL_2851665 | Health Services Laboratories | Wellcome Sanger Institute for the COVID-19 Genomics UK (COG-UK) Consortium | Health Services Laboratories and Alex Alderton, Roberto Amato, Jeffrey Barrett, Sonia Goncalves, Ewan Harrison, David K. Jackson, Ian Johnston, Dominic Kwiatkowski, Cordelia Langford, John Sillitoe on behalf of the Wellcome Sanger Institute COVID-19 Surveillance Team |
| EPI_ISL_2851666 | Lighthouse Lab in Milton Keynes | Wellcome Sanger Institute for the COVID-19 Genomics UK (COG-UK) Consortium | The Lighthouse Lab in Milton Keynes and Alex Alderton, Roberto Amato, Jeffrey Barrett, Sonia Goncalves, Ewan Harrison, David K. Jackson, Ian Johnston, Dominic Kwiatkowski, Cordelia Langford, John Sillitoe on behalf of the Wellcome Sanger Institute COVID-19 Surveillance Team |
| EPI_ISL_2851667 | Health Services Laboratories | Wellcome Sanger Institute for the COVID-19 Genomics UK (COG-UK) Consortium | Health Services Laboratories and Alex Alderton, Roberto Amato, Jeffrey Barrett, Sonia Goncalves, Ewan Harrison, David K. Jackson, Ian Johnston, Dominic Kwiatkowski, Cordelia Langford, John Sillitoe on behalf of the Wellcome Sanger Institute COVID-19 Surveillance Team |
| EPI_ISL_2851668 | Lighthouse Lab in Alderley Park | Wellcome Sanger Institute for the COVID-19 Genomics UK (COG-UK) Consortium | Jacquelyn Wynn, Mairead Hyland, The Lighthouse Lab in Alderley Park and Alex Alderton, Roberto Amato, Jeffrey Barrett, Sonia Goncalves, Ewan Harrison, David K. Jackson, Ian Johnston, Dominic Kwiatkowski, Cordelia Langford, John Sillitoe on behalf of the Wellcome Sanger Institute COVID-19 Surveillance Team |
| EPI_ISL_2851669, EPI_ISL_2851670 | Lighthouse Lab in Milton Keynes | Wellcome Sanger Institute for the COVID-19 Genomics UK (COG-UK) Consortium | The Lighthouse Lab in Milton Keynes and Alex Alderton, Roberto Amato, Jeffrey Barrett, Sonia Goncalves, Ewan Harrison, David K. Jackson, Ian Johnston, Dominic Kwiatkowski, Cordelia Langford, John Sillitoe on behalf of the Wellcome Sanger Institute COVID-19 Surveillance Team |
| EPI_ISL_2851671 | Health Services Laboratories | Wellcome Sanger Institute for the COVID-19 Genomics UK (COG-UK) Consortium | Health Services Laboratories and Alex Alderton, Roberto Amato, Jeffrey Barrett, Sonia Goncalves, Ewan Harrison, David K. Jackson, Ian Johnston, Dominic Kwiatkowski, Cordelia Langford, John Sillitoe on behalf of the Wellcome Sanger Institute COVID-19 Surveillance Team |
| EPI_ISL_2851672 | Lighthouse Lab in Alderley Park | Wellcome Sanger Institute for the COVID-19 Genomics UK (COG-UK) Consortium | Jacquelyn Wynn, Mairead Hyland, The Lighthouse Lab in Alderley Park and Alex Alderton, Roberto Amato, Jeffrey Barrett, Sonia Goncalves, Ewan Harrison, David K. Jackson, Ian Johnston, Dominic Kwiatkowski, Cordelia Langford, John Sillitoe on behalf of the Wellcome Sanger Institute COVID-19 Surveillance Team |
| EPI_ISL_2851673, EPI_ISL_2851674 | Health Services Laboratories | Wellcome Sanger Institute for the COVID-19 Genomics UK (COG-UK) Consortium | Health Services Laboratories and Alex Alderton, Roberto Amato, Jeffrey Barrett, Sonia Goncalves, Ewan Harrison, David K. Jackson, Ian Johnston, Dominic Kwiatkowski, Cordelia Langford, John Sillitoe on behalf of the Wellcome Sanger Institute COVID-19 Surveillance Team |
| EPI_ISL_2851675 | Lighthouse Lab in Milton Keynes | Wellcome Sanger Institute for the COVID-19 Genomics UK (COG-UK) Consortium | The Lighthouse Lab in Milton Keynes and Alex Alderton, Roberto Amato, Jeffrey Barrett, Sonia Goncalves, Ewan Harrison, David K. Jackson, Ian Johnston, Dominic Kwiatkowski, Cordelia Langford, John Sillitoe on behalf of the Wellcome Sanger Institute COVID-19 Surveillance Team |
| EPI_ISL_2851676, EPI_ISL_2851677 | Health Services Laboratories | Wellcome Sanger Institute for the COVID-19 Genomics UK (COG-UK) Consortium | Health Services Laboratories and Alex Alderton, Roberto Amato, Jeffrey Barrett, Sonia Goncalves, Ewan Harrison, David K. Jackson, Ian Johnston, Dominic Kwiatkowski, Cordelia Langford, John Sillitoe on behalf of the Wellcome Sanger Institute COVID-19 Surveillance Team |
| EPI_ISL_2851678 | Lighthouse Lab in Milton Keynes | Wellcome Sanger Institute for the COVID-19 Genomics UK (COG-UK) Consortium | The Lighthouse Lab in Milton Keynes and Alex Alderton, Roberto Amato, Jeffrey Barrett, Sonia Goncalves, Ewan Harrison, David K. Jackson, Ian Johnston, Dominic Kwiatkowski, Cordelia Langford, John Sillitoe on behalf of the Wellcome Sanger Institute COVID-19 Surveillance Team |
| EPI_ISL_2851679, EPI_ISL_2851680 | Health Services Laboratories | Wellcome Sanger Institute for the COVID-19 Genomics UK (COG-UK) Consortium | Health Services Laboratories and Alex Alderton, Roberto Amato, Jeffrey Barrett, Sonia Goncalves, Ewan Harrison, David K. Jackson, Ian Johnston, Dominic Kwiatkowski, Cordelia Langford, John Sillitoe on behalf of the Wellcome Sanger Institute COVID-19 Surveillance Team |
| EPI_ISL_2851681 | Lighthouse Lab in Alderley Park | Wellcome Sanger Institute for the COVID-19 Genomics UK (COG-UK) Consortium | Jacquelyn Wynn, Mairead Hyland, The Lighthouse Lab in Alderley Park and Alex Alderton, Roberto Amato, Jeffrey Barrett, Sonia Goncalves, Ewan Harrison, David K. Jackson, Ian Johnston, Dominic Kwiatkowski, Cordelia Langford, John Sillitoe on behalf of the Wellcome Sanger Institute COVID-19 Surveillance Team |
| EPI_ISL_2851682, EPI_ISL_2851683 | Health Services Laboratories | Wellcome Sanger Institute for the COVID-19 Genomics UK (COG-UK) Consortium | Health Services Laboratories and Alex Alderton, Roberto Amato, Jeffrey Barrett, Sonia Goncalves, Ewan Harrison, David K. Jackson, Ian Johnston, Dominic Kwiatkowski, Cordelia Langford, John Sillitoe on behalf of the Wellcome Sanger Institute COVID-19 Surveillance Team |
| EPI_ISL_2851684, EPI_ISL_2851685 | Lighthouse Lab in Alderley Park | Wellcome Sanger Institute for the COVID-19 Genomics UK (COG-UK) Consortium | Jacquelyn Wynn, Mairead Hyland, The Lighthouse Lab in Alderley Park and Alex Alderton, Roberto Amato, Jeffrey Barrett, Sonia Goncalves, Ewan Harrison, David K. Jackson, Ian Johnston, Dominic Kwiatkowski, Cordelia Langford, John Sillitoe on behalf of the Wellcome Sanger Institute COVID-19 Surveillance Team |
| EPI_ISL_2851686 | Lighthouse Lab in Milton Keynes | Wellcome Sanger Institute for the COVID-19 Genomics UK (COG-UK) Consortium | The Lighthouse Lab in Milton Keynes and Alex Alderton, Roberto Amato, Jeffrey Barrett, Sonia Goncalves, Ewan Harrison, David K. Jackson, Ian Johnston, Dominic Kwiatkowski, Cordelia Langford, John Sillitoe on behalf of the Wellcome Sanger Institute COVID-19 Surveillance Team |
| EPI_ISL_2851687, EPI_ISL_2851688, EPI_ISL_2851689, EPI_ISL_2851690 | Health Services Laboratories | Wellcome Sanger Institute for the COVID-19 Genomics UK (COG-UK) Consortium | Health Services Laboratories and Alex Alderton, Roberto Amato, Jeffrey Barrett, Sonia Goncalves, Ewan Harrison, David K. Jackson, Ian Johnston, Dominic Kwiatkowski, Cordelia Langford, John Sillitoe on behalf of the Wellcome Sanger Institute COVID-19 Surveillance Team |
| EPI_ISL_2851691 | Lighthouse Lab in Milton Keynes | Wellcome Sanger Institute for the COVID-19 Genomics UK (COG-UK) Consortium | The Lighthouse Lab in Milton Keynes and Alex Alderton, Roberto Amato, Jeffrey Barrett, Sonia Goncalves, Ewan Harrison, David K. Jackson, Ian Johnston, Dominic Kwiatkowski, Cordelia Langford, John Sillitoe on behalf of the Wellcome Sanger Institute COVID-19 Surveillance Team |
| EPI_ISL_2851692 | Health Services Laboratories | Wellcome Sanger Institute for the COVID-19 Genomics UK (COG-UK) Consortium | Health Services Laboratories and Alex Alderton, Roberto Amato, Jeffrey Barrett, Sonia Goncalves, Ewan Harrison, David K. Jackson, Ian Johnston, Dominic Kwiatkowski, Cordelia Langford, John Sillitoe on behalf of the Wellcome Sanger Institute COVID-19 Surveillance Team |
| EPI_ISL_2851693 | Lighthouse Lab in Alderley Park | Wellcome Sanger Institute for the COVID-19 Genomics UK (COG-UK) Consortium | Jacquelyn Wynn, Mairead Hyland, The Lighthouse Lab in Alderley Park and Alex Alderton, Roberto Amato, Jeffrey Barrett, Sonia Goncalves, Ewan Harrison, David K. Jackson, Ian Johnston, Dominic Kwiatkowski, Cordelia Langford, John Sillitoe on behalf of the Wellcome Sanger Institute COVID-19 Surveillance Team |
| EPI_ISL_2851694 | Health Services Laboratories | Wellcome Sanger Institute for the COVID-19 Genomics UK (COG-UK) Consortium | Health Services Laboratories and Alex Alderton, Roberto Amato, Jeffrey Barrett, Sonia Goncalves, Ewan Harrison, David K. Jackson, Ian Johnston, Dominic Kwiatkowski, Cordelia Langford, John Sillitoe on behalf of the Wellcome Sanger Institute COVID-19 Surveillance Team |
| EPI_ISL_2851695 | Lighthouse Lab in Milton Keynes | Wellcome Sanger Institute for the COVID-19 Genomics UK (COG-UK) Consortium | The Lighthouse Lab in Milton Keynes and Alex Alderton, Roberto Amato, Jeffrey Barrett, Sonia Goncalves, Ewan Harrison, David K. Jackson, Ian Johnston, Dominic Kwiatkowski, Cordelia Langford, John Sillitoe on behalf of the Wellcome Sanger Institute COVID-19 Surveillance Team |
| EPI_ISL_2851696, EPI_ISL_2851697, EPI_ISL_2851698 | Health Services Laboratories | Wellcome Sanger Institute for the COVID-19 Genomics UK (COG-UK) Consortium | Health Services Laboratories and Alex Alderton, Roberto Amato, Jeffrey Barrett, Sonia Goncalves, Ewan Harrison, David K. Jackson, Ian Johnston, Dominic Kwiatkowski, Cordelia Langford, John Sillitoe on behalf of the Wellcome Sanger Institute COVID-19 Surveillance Team |
| EPI_ISL_2851699 | Lighthouse Lab in Milton Keynes | Wellcome Sanger Institute for the COVID-19 Genomics UK (COG-UK) Consortium | The Lighthouse Lab in Milton Keynes and Alex Alderton, Roberto Amato, Jeffrey Barrett, Sonia Goncalves, Ewan Harrison, David K. Jackson, Ian Johnston, Dominic Kwiatkowski, Cordelia Langford, John Sillitoe on behalf of the Wellcome Sanger Institute COVID-19 Surveillance Team |
| EPI_ISL_2851700 | Health Services Laboratories | Wellcome Sanger Institute for the COVID-19 Genomics UK (COG-UK) Consortium | Health Services Laboratories and Alex Alderton, Roberto Amato, Jeffrey Barrett, Sonia Goncalves, Ewan Harrison, David K. Jackson, Ian Johnston, Dominic Kwiatkowski, Cordelia Langford, John Sillitoe on behalf of the Wellcome Sanger Institute COVID-19 Surveillance Team |
| EPI_ISL_2851701 | Lighthouse Lab in Milton Keynes | Wellcome Sanger Institute for the COVID-19 Genomics UK (COG-UK) Consortium | The Lighthouse Lab in Milton Keynes and Alex Alderton, Roberto Amato, Jeffrey Barrett, Sonia Goncalves, Ewan Harrison, David K. Jackson, Ian Johnston, Dominic Kwiatkowski, Cordelia Langford, John Sillitoe on behalf of the Wellcome Sanger Institute COVID-19 Surveillance Team |
| EPI_ISL_2851702 | Health Services Laboratories | Wellcome Sanger Institute for the COVID-19 Genomics UK (COG-UK) Consortium | Health Services Laboratories and Alex Alderton, Roberto Amato, Jeffrey Barrett, Sonia Goncalves, Ewan Harrison, David K. Jackson, Ian Johnston, Dominic Kwiatkowski, Cordelia Langford, John Sillitoe on behalf of the Wellcome Sanger Institute COVID-19 Surveillance Team |
| EPI_ISL_2851703 | Lighthouse Lab in Milton Keynes | Wellcome Sanger Institute for the COVID-19 Genomics UK (COG-UK) Consortium | The Lighthouse Lab in Milton Keynes and Alex Alderton, Roberto Amato, Jeffrey Barrett, Sonia Goncalves, Ewan Harrison, David K. Jackson, Ian Johnston, Dominic Kwiatkowski, Cordelia Langford, John Sillitoe on behalf of the Wellcome Sanger Institute COVID-19 Surveillance Team |
| EPI_ISL_2851704 | Lighthouse Lab in Alderley Park | Wellcome Sanger Institute for the COVID-19 Genomics UK (COG-UK) Consortium | Jacquelyn Wynn, Mairead Hyland, The Lighthouse Lab in Alderley Park and Alex Alderton, Roberto Amato, Jeffrey Barrett, Sonia Goncalves, Ewan Harrison, David K. Jackson, Ian Johnston, Dominic Kwiatkowski, Cordelia Langford, John Sillitoe on behalf of the Wellcome Sanger Institute COVID-19 Surveillance Team |
| EPI_ISL_2851705 | Lighthouse Lab in Milton Keynes | Wellcome Sanger Institute for the COVID-19 Genomics UK (COG-UK) Consortium | The Lighthouse Lab in Milton Keynes and Alex Alderton, Roberto Amato, Jeffrey Barrett, Sonia Goncalves, Ewan Harrison, David K. Jackson, Ian Johnston, Dominic Kwiatkowski, Cordelia Langford, John Sillitoe on behalf of the Wellcome Sanger Institute COVID-19 Surveillance Team |
| EPI_ISL_2851706 | Lighthouse Lab in Alderley Park | Wellcome Sanger Institute for the COVID-19 Genomics UK (COG-UK) Consortium | Jacquelyn Wynn, Mairead Hyland, The Lighthouse Lab in Alderley Park and Alex Alderton, Roberto Amato, Jeffrey Barrett, Sonia Goncalves, Ewan Harrison, David K. Jackson, Ian Johnston, Dominic Kwiatkowski, Cordelia Langford, John Sillitoe on behalf of the Wellcome Sanger Institute COVID-19 Surveillance Team |
| EPI_ISL_2851707, EPI_ISL_2851708, EPI_ISL_2851709, EPI_ISL_2851710 | Health Services Laboratories | Wellcome Sanger Institute for the COVID-19 Genomics UK (COG-UK) Consortium | Health Services Laboratories and Alex Alderton, Roberto Amato, Jeffrey Barrett, Sonia Goncalves, Ewan Harrison, David K. Jackson, Ian Johnston, Dominic Kwiatkowski, Cordelia Langford, John Sillitoe on behalf of the Wellcome Sanger Institute COVID-19 Surveillance Team |
| EPI_ISL_2851711, EPI_ISL_2851712 | Lighthouse Lab in Milton Keynes | Wellcome Sanger Institute for the COVID-19 Genomics UK (COG-UK) Consortium | The Lighthouse Lab in Milton Keynes and Alex Alderton, Roberto Amato, Jeffrey Barrett, Sonia Goncalves, Ewan Harrison, David K. Jackson, Ian Johnston, Dominic Kwiatkowski, Cordelia Langford, John Sillitoe on behalf of the Wellcome Sanger Institute COVID-19 Surveillance Team |
| EPI_ISL_2851713 | Lighthouse Lab in Alderley Park | Wellcome Sanger Institute for the COVID-19 Genomics UK (COG-UK) Consortium | Jacquelyn Wynn, Mairead Hyland, The Lighthouse Lab in Alderley Park and Alex Alderton, Roberto Amato, Jeffrey Barrett, Sonia Goncalves, Ewan Harrison, David K. Jackson, Ian Johnston, Dominic Kwiatkowski, Cordelia Langford, John Sillitoe on behalf of the Wellcome Sanger Institute COVID-19 Surveillance Team |
| EPI_ISL_2851714, EPI_ISL_2851715, EPI_ISL_2851716, EPI_ISL_2851717, EPI_ISL_2851718 | Health Services Laboratories | Wellcome Sanger Institute for the COVID-19 Genomics UK (COG-UK) Consortium | Health Services Laboratories and Alex Alderton, Roberto Amato, Jeffrey Barrett, Sonia Goncalves, Ewan Harrison, David K. Jackson, Ian Johnston, Dominic Kwiatkowski, Cordelia Langford, John Sillitoe on behalf of the Wellcome Sanger Institute COVID-19 Surveillance Team |
| EPI_ISL_2851719 | Lighthouse Lab in Alderley Park | Wellcome Sanger Institute for the COVID-19 Genomics UK (COG-UK) Consortium | Jacquelyn Wynn, Mairead Hyland, The Lighthouse Lab in Alderley Park and Alex Alderton, Roberto Amato, Jeffrey Barrett, Sonia Goncalves, Ewan Harrison, David K. Jackson, Ian Johnston, Dominic Kwiatkowski, Cordelia Langford, John Sillitoe on behalf of the Wellcome Sanger Institute COVID-19 Surveillance Team |
| EPI_ISL_2851720 | Health Services Laboratories | Wellcome Sanger Institute for the COVID-19 Genomics UK (COG-UK) Consortium | Health Services Laboratories and Alex Alderton, Roberto Amato, Jeffrey Barrett, Sonia Goncalves, Ewan Harrison, David K. Jackson, Ian Johnston, Dominic Kwiatkowski, Cordelia Langford, John Sillitoe on behalf of the Wellcome Sanger Institute COVID-19 Surveillance Team |

| | | | |
|---|---|---|---|
| EPI_ISL_2851721, EPI_ISL_2851722 | Lighthouse Lab in Alderley Park | Wellcome Sanger Institute for the COVID-19 Genomics UK (COG-UK) Consortium | Jacquelyn Wynn, Mairead Hyland, The Lighthouse Lab in Alderley Park and Alex Alderton, Roberto Amato, Jeffrey Barrett, Sonia Goncalves, Ewan Harrison, David K. Jackson, Ian Johnston, Dominic Kwiatkowski, Cordelia Langford, John Sillitoe on behalf of the Wellcome Sanger Institute COVID-19 Surveillance Team |
| EPI_ISL_2851723, EPI_ISL_2851724, EPI_ISL_2851725 | Health Services Laboratories | Wellcome Sanger Institute for the COVID-19 Genomics UK (COG-UK) Consortium | Health Services Laboratories and Alex Alderton, Roberto Amato, Jeffrey Barrett, Sonia Goncalves, Ewan Harrison, David K. Jackson, Ian Johnston, Dominic Kwiatkowski, Cordelia Langford, John Sillitoe on behalf of the Wellcome Sanger Institute COVID-19 Surveillance Team |
| EPI_ISL_2851726 | Lighthouse Lab in Alderley Park | Wellcome Sanger Institute for the COVID-19 Genomics UK (COG-UK) Consortium | Jacquelyn Wynn, Mairead Hyland, The Lighthouse Lab in Alderley Park and Alex Alderton, Roberto Amato, Jeffrey Barrett, Sonia Goncalves, Ewan Harrison, David K. Jackson, Ian Johnston, Dominic Kwiatkowski, Cordelia Langford, John Sillitoe on behalf of the Wellcome Sanger Institute COVID-19 Surveillance Team |
| EPI_ISL_2851727 | Health Services Laboratories | Wellcome Sanger Institute for the COVID-19 Genomics UK (COG-UK) Consortium | Health Services Laboratories and Alex Alderton, Roberto Amato, Jeffrey Barrett, Sonia Goncalves, Ewan Harrison, David K. Jackson, Ian Johnston, Dominic Kwiatkowski, Cordelia Langford, John Sillitoe on behalf of the Wellcome Sanger Institute COVID-19 Surveillance Team |
| EPI_ISL_2851728 | Lighthouse Lab in Alderley Park | Wellcome Sanger Institute for the COVID-19 Genomics UK (COG-UK) Consortium | Jacquelyn Wynn, Mairead Hyland, The Lighthouse Lab in Alderley Park and Alex Alderton, Roberto Amato, Jeffrey Barrett, Sonia Goncalves, Ewan Harrison, David K. Jackson, Ian Johnston, Dominic Kwiatkowski, Cordelia Langford, John Sillitoe on behalf of the Wellcome Sanger Institute COVID-19 Surveillance Team |
| EPI_ISL_2851729 | Lighthouse Lab in Milton Keynes | Wellcome Sanger Institute for the COVID-19 Genomics UK (COG-UK) Consortium | The Lighthouse Lab in Milton Keynes and Alex Alderton, Roberto Amato, Jeffrey Barrett, Sonia Goncalves, Ewan Harrison, David K. Jackson, Ian Johnston, Dominic Kwiatkowski, Cordelia Langford, John Sillitoe on behalf of the Wellcome Sanger Institute COVID-19 Surveillance Team |
| EPI_ISL_2851730, EPI_ISL_2851731 | Lighthouse Lab in Alderley Park | Wellcome Sanger Institute for the COVID-19 Genomics UK (COG-UK) Consortium | Jacquelyn Wynn, Mairead Hyland, The Lighthouse Lab in Alderley Park and Alex Alderton, Roberto Amato, Jeffrey Barrett, Sonia Goncalves, Ewan Harrison, David K. Jackson, Ian Johnston, Dominic Kwiatkowski, Cordelia Langford, John Sillitoe on behalf of the Wellcome Sanger Institute COVID-19 Surveillance Team |
| EPI_ISL_2851732, EPI_ISL_2851733, EPI_ISL_2851734, EPI_ISL_2851735, EPI_ISL_2851736 | Health Services Laboratories | Wellcome Sanger Institute for the COVID-19 Genomics UK (COG-UK) Consortium | Health Services Laboratories and Alex Alderton, Roberto Amato, Jeffrey Barrett, Sonia Goncalves, Ewan Harrison, David K. Jackson, Ian Johnston, Dominic Kwiatkowski, Cordelia Langford, John Sillitoe on behalf of the Wellcome Sanger Institute COVID-19 Surveillance Team |
| EPI_ISL_2851737, EPI_ISL_2851738 | Lighthouse Lab in Alderley Park | Wellcome Sanger Institute for the COVID-19 Genomics UK (COG-UK) Consortium | Jacquelyn Wynn, Mairead Hyland, The Lighthouse Lab in Alderley Park and Alex Alderton, Roberto Amato, Jeffrey Barrett, Sonia Goncalves, Ewan Harrison, David K. Jackson, Ian Johnston, Dominic Kwiatkowski, Cordelia Langford, John Sillitoe on behalf of the Wellcome Sanger Institute COVID-19 Surveillance Team |
| EPI_ISL_2851739, EPI_ISL_2851740, EPI_ISL_2851741 | Health Services Laboratories | Wellcome Sanger Institute for the COVID-19 Genomics UK (COG-UK) Consortium | Health Services Laboratories and Alex Alderton, Roberto Amato, Jeffrey Barrett, Sonia Goncalves, Ewan Harrison, David K. Jackson, Ian Johnston, Dominic Kwiatkowski, Cordelia Langford, John Sillitoe on behalf of the Wellcome Sanger Institute COVID-19 Surveillance Team |
| EPI_ISL_2851742, EPI_ISL_2851743 | Lighthouse Lab in Milton Keynes | Wellcome Sanger Institute for the COVID-19 Genomics UK (COG-UK) Consortium | The Lighthouse Lab in Milton Keynes and Alex Alderton, Roberto Amato, Jeffrey Barrett, Sonia Goncalves, Ewan Harrison, David K. Jackson, Ian Johnston, Dominic Kwiatkowski, Cordelia Langford, John Sillitoe on behalf of the Wellcome Sanger Institute COVID-19 Surveillance Team |
| EPI_ISL_2851744 | Health Services Laboratories | Wellcome Sanger Institute for the COVID-19 Genomics UK (COG-UK) Consortium | Health Services Laboratories and Alex Alderton, Roberto Amato, Jeffrey Barrett, Sonia Goncalves, Ewan Harrison, David K. Jackson, Ian Johnston, Dominic Kwiatkowski, Cordelia Langford, John Sillitoe on behalf of the Wellcome Sanger Institute COVID-19 Surveillance Team |
| EPI_ISL_2851745 | Lighthouse Lab in Milton Keynes | Wellcome Sanger Institute for the COVID-19 Genomics UK (COG-UK) Consortium | The Lighthouse Lab in Milton Keynes and Alex Alderton, Roberto Amato, Jeffrey Barrett, Sonia Goncalves, Ewan Harrison, David K. Jackson, Ian Johnston, Dominic Kwiatkowski, Cordelia Langford, John Sillitoe on behalf of the Wellcome Sanger Institute COVID-19 Surveillance Team |
| EPI_ISL_2851746, EPI_ISL_2851747, EPI_ISL_2851748, EPI_ISL_2851749, EPI_ISL_2851750 | Health Services Laboratories | Wellcome Sanger Institute for the COVID-19 Genomics UK (COG-UK) Consortium | Health Services Laboratories and Alex Alderton, Roberto Amato, Jeffrey Barrett, Sonia Goncalves, Ewan Harrison, David K. Jackson, Ian Johnston, Dominic Kwiatkowski, Cordelia Langford, John Sillitoe on behalf of the Wellcome Sanger Institute COVID-19 Surveillance Team |
| EPI_ISL_2851751 | Lighthouse Lab in Alderley Park | Wellcome Sanger Institute for the COVID-19 Genomics UK (COG-UK) Consortium | Jacquelyn Wynn, Mairead Hyland, The Lighthouse Lab in Alderley Park and Alex Alderton, Roberto Amato, Jeffrey Barrett, Sonia Goncalves, Ewan Harrison, David K. Jackson, Ian Johnston, Dominic Kwiatkowski, Cordelia Langford, John Sillitoe on behalf of the Wellcome Sanger Institute COVID-19 Surveillance Team |
| EPI_ISL_2851752 | Health Services Laboratories | Wellcome Sanger Institute for the COVID-19 Genomics UK (COG-UK) Consortium | Health Services Laboratories and Alex Alderton, Roberto Amato, Jeffrey Barrett, Sonia Goncalves, Ewan Harrison, David K. Jackson, Ian Johnston, Dominic Kwiatkowski, Cordelia Langford, John Sillitoe on behalf of the Wellcome Sanger Institute COVID-19 Surveillance Team |
| EPI_ISL_2851753, EPI_ISL_2851754 | Lighthouse Lab in Milton Keynes | Wellcome Sanger Institute for the COVID-19 Genomics UK (COG-UK) Consortium | The Lighthouse Lab in Milton Keynes and Alex Alderton, Roberto Amato, Jeffrey Barrett, Sonia Goncalves, Ewan Harrison, David K. Jackson, Ian Johnston, Dominic Kwiatkowski, Cordelia Langford, John Sillitoe on behalf of the Wellcome Sanger Institute COVID-19 Surveillance Team |
| EPI_ISL_2851755 | Health Services Laboratories | Wellcome Sanger Institute for the COVID-19 Genomics UK (COG-UK) Consortium | Health Services Laboratories and Alex Alderton, Roberto Amato, Jeffrey Barrett, Sonia Goncalves, Ewan Harrison, David K. Jackson, Ian Johnston, Dominic Kwiatkowski, Cordelia Langford, John Sillitoe on behalf of the Wellcome Sanger Institute COVID-19 Surveillance Team |
| EPI_ISL_2851756, EPI_ISL_2851757 | Lighthouse Lab in Milton Keynes | Wellcome Sanger Institute for the COVID-19 Genomics UK (COG-UK) Consortium | The Lighthouse Lab in Milton Keynes and Alex Alderton, Roberto Amato, Jeffrey Barrett, Sonia Goncalves, Ewan Harrison, David K. Jackson, Ian Johnston, Dominic Kwiatkowski, Cordelia Langford, John Sillitoe on behalf of the Wellcome Sanger Institute COVID-19 Surveillance Team |
| EPI_ISL_2851758, EPI_ISL_2851759 | Lighthouse Lab in Alderley Park | Wellcome Sanger Institute for the COVID-19 Genomics UK (COG-UK) Consortium | Jacquelyn Wynn, Mairead Hyland, The Lighthouse Lab in Alderley Park and Alex Alderton, Roberto Amato, Jeffrey Barrett, Sonia Goncalves, Ewan Harrison, David K. Jackson, Ian Johnston, Dominic Kwiatkowski, Cordelia Langford, John Sillitoe on behalf of the Wellcome Sanger Institute COVID-19 Surveillance Team |
| EPI_ISL_2851760, EPI_ISL_2851761, EPI_ISL_2851762 | Health Services Laboratories | Wellcome Sanger Institute for the COVID-19 Genomics UK (COG-UK) Consortium | Health Services Laboratories and Alex Alderton, Roberto Amato, Jeffrey Barrett, Sonia Goncalves, Ewan Harrison, David K. Jackson, Ian Johnston, Dominic Kwiatkowski, Cordelia Langford, John Sillitoe on behalf of the Wellcome Sanger Institute COVID-19 Surveillance Team |
| EPI_ISL_2851763 | Lighthouse Lab in Milton Keynes | Wellcome Sanger Institute for the COVID-19 Genomics UK (COG-UK) Consortium | The Lighthouse Lab in Milton Keynes and Alex Alderton, Roberto Amato, Jeffrey Barrett, Sonia Goncalves, Ewan Harrison, David K. Jackson, Ian Johnston, Dominic Kwiatkowski, Cordelia Langford, John Sillitoe on behalf of the Wellcome Sanger Institute COVID-19 Surveillance Team |
| EPI_ISL_2851764, EPI_ISL_2851765 | Health Services Laboratories | Wellcome Sanger Institute for the COVID-19 Genomics UK (COG-UK) Consortium | Health Services Laboratories and Alex Alderton, Roberto Amato, Jeffrey Barrett, Sonia Goncalves, Ewan Harrison, David K. Jackson, Ian Johnston, Dominic Kwiatkowski, Cordelia Langford, John Sillitoe on behalf of the Wellcome Sanger Institute COVID-19 Surveillance Team |
| EPI_ISL_2851766 | Lighthouse Lab in Alderley Park | Wellcome Sanger Institute for the COVID-19 Genomics UK (COG-UK) Consortium | Jacquelyn Wynn, Mairead Hyland, The Lighthouse Lab in Alderley Park and Alex Alderton, Roberto Amato, Jeffrey Barrett, Sonia Goncalves, Ewan Harrison, David K. Jackson, Ian Johnston, Dominic Kwiatkowski, Cordelia Langford, John Sillitoe on behalf of the Wellcome Sanger Institute COVID-19 Surveillance Team |
| EPI_ISL_2851767, EPI_ISL_2851768 | Lighthouse Lab in Milton Keynes | Wellcome Sanger Institute for the COVID-19 Genomics UK (COG-UK) Consortium | The Lighthouse Lab in Milton Keynes and Alex Alderton, Roberto Amato, Jeffrey Barrett, Sonia Goncalves, Ewan Harrison, David K. Jackson, Ian Johnston, Dominic Kwiatkowski, Cordelia Langford, John Sillitoe on behalf of the Wellcome Sanger Institute COVID-19 Surveillance Team |
| EPI_ISL_2851769 | Health Services Laboratories | Wellcome Sanger Institute for the COVID-19 Genomics UK (COG-UK) Consortium | Health Services Laboratories and Alex Alderton, Roberto Amato, Jeffrey Barrett, Sonia Goncalves, Ewan Harrison, David K. Jackson, Ian Johnston, Dominic Kwiatkowski, Cordelia Langford, John Sillitoe on behalf of the Wellcome Sanger Institute COVID-19 Surveillance Team |
| EPI_ISL_2851770 | Lighthouse Lab in Alderley Park | Wellcome Sanger Institute for the COVID-19 Genomics UK (COG-UK) Consortium | Jacquelyn Wynn, Mairead Hyland, The Lighthouse Lab in Alderley Park and Alex Alderton, Roberto Amato, Jeffrey Barrett, Sonia Goncalves, Ewan Harrison, David K. Jackson, Ian Johnston, Dominic Kwiatkowski, Cordelia Langford, John Sillitoe on behalf of the Wellcome Sanger Institute COVID-19 Surveillance Team |
| EPI_ISL_2851771 | Health Services Laboratories | Wellcome Sanger Institute for the COVID-19 Genomics UK (COG-UK) Consortium | Health Services Laboratories and Alex Alderton, Roberto Amato, Jeffrey Barrett, Sonia Goncalves, Ewan Harrison, David K. Jackson, Ian Johnston, Dominic Kwiatkowski, Cordelia Langford, John Sillitoe on behalf of the Wellcome Sanger Institute COVID-19 Surveillance Team |
| EPI_ISL_2851772 | Lighthouse Lab in Milton Keynes | Wellcome Sanger Institute for the COVID-19 Genomics UK (COG-UK) Consortium | The Lighthouse Lab in Milton Keynes and Alex Alderton, Roberto Amato, Jeffrey Barrett, Sonia Goncalves, Ewan Harrison, David K. Jackson, Ian Johnston, Dominic Kwiatkowski, Cordelia Langford, John Sillitoe on behalf of the Wellcome Sanger Institute COVID-19 Surveillance Team |
| EPI_ISL_2851773 | Lighthouse Lab in Alderley Park | Wellcome Sanger Institute for the COVID-19 Genomics UK (COG-UK) Consortium | Jacquelyn Wynn, Mairead Hyland, The Lighthouse Lab in Alderley Park and Alex Alderton, Roberto Amato, Jeffrey Barrett, Sonia Goncalves, Ewan Harrison, David K. Jackson, Ian Johnston, Dominic Kwiatkowski, Cordelia Langford, John Sillitoe on behalf of the Wellcome Sanger Institute COVID-19 Surveillance Team |
| EPI_ISL_2851775, EPI_ISL_2851776 | Lighthouse Lab in Glasgow | Wellcome Sanger Institute for the COVID-19 Genomics UK (COG-UK) Consortium | Harper VanSteenhouse, Yumi Kasai, David Gray, Carol Clugston, Anna Dominiczak and Alex Alderton, Roberto Amato, Jeffrey Barrett, Sonia Goncalves, Ewan Harrison, David K. Jackson, Ian Johnston, Dominic Kwiatkowski, Cordelia Langford, John Sillitoe on behalf of the Wellcome Sanger Institute COVID-19 Surveillance Team |
| EPI_ISL_2851777 | Lighthouse Lab in Milton Keynes | Wellcome Sanger Institute for the COVID-19 Genomics UK (COG-UK) Consortium | The Lighthouse Lab in Milton Keynes and Alex Alderton, Roberto Amato, Jeffrey Barrett, Sonia Goncalves, Ewan Harrison, David K. Jackson, Ian Johnston, Dominic Kwiatkowski, Cordelia Langford, John Sillitoe on behalf of the Wellcome Sanger Institute COVID-19 Surveillance Team |
| EPI_ISL_2851778, EPI_ISL_2851779, EPI_ISL_2851780, EPI_ISL_2851781, EPI_ISL_2851782, EPI_ISL_2851783 | Lighthouse Lab in Glasgow | Wellcome Sanger Institute for the COVID-19 Genomics UK (COG-UK) Consortium | Harper VanSteenhouse, Yumi Kasai, David Gray, Carol Clugston, Anna Dominiczak and Alex Alderton, Roberto Amato, Jeffrey Barrett, Sonia Goncalves, Ewan Harrison, David K. Jackson, Ian Johnston, Dominic Kwiatkowski, Cordelia Langford, John Sillitoe on behalf of the Wellcome Sanger Institute COVID-19 Surveillance Team |
| EPI_ISL_2851784 | Lighthouse Lab in Alderley Park | Wellcome Sanger Institute for the COVID-19 Genomics UK (COG-UK) Consortium | Jacquelyn Wynn, Mairead Hyland, The Lighthouse Lab in Alderley Park and Alex Alderton, Roberto Amato, Jeffrey Barrett, Sonia Goncalves, Ewan Harrison, David K. Jackson, Ian Johnston, Dominic Kwiatkowski, Cordelia Langford, John Sillitoe on behalf of the Wellcome Sanger Institute COVID-19 Surveillance Team |
| EPI_ISL_2851785 | Lighthouse Lab in Glasgow | Wellcome Sanger Institute for the COVID-19 Genomics UK (COG-UK) Consortium | Harper VanSteenhouse, Yumi Kasai, David Gray, Carol Clugston, Anna Dominiczak and Alex Alderton, Roberto Amato, Jeffrey Barrett, Sonia Goncalves, Ewan Harrison, David K. Jackson, Ian Johnston, Dominic Kwiatkowski, Cordelia Langford, John Sillitoe on behalf of the Wellcome Sanger Institute COVID-19 Surveillance Team |
| EPI_ISL_2851786 | Lighthouse Lab in Milton Keynes | Wellcome Sanger Institute for the COVID-19 Genomics UK (COG-UK) Consortium | The Lighthouse Lab in Milton Keynes and Alex Alderton, Roberto Amato, Jeffrey Barrett, Sonia Goncalves, Ewan Harrison, David K. Jackson, Ian Johnston, Dominic Kwiatkowski, Cordelia Langford, John Sillitoe on behalf of the Wellcome Sanger Institute COVID-19 Surveillance Team |
| EPI_ISL_2851787, EPI_ISL_2851788, EPI_ISL_2851789 | Lighthouse Lab in Glasgow | Wellcome Sanger Institute for the COVID-19 Genomics UK (COG-UK) Consortium | Harper VanSteenhouse, Yumi Kasai, David Gray, Carol Clugston, Anna Dominiczak and Alex Alderton, Roberto Amato, Jeffrey Barrett, Sonia Goncalves, Ewan Harrison, David K. Jackson, Ian Johnston, Dominic Kwiatkowski, Cordelia Langford, John Sillitoe on behalf of the Wellcome Sanger Institute COVID-19 Surveillance Team |
| EPI_ISL_2851790, EPI_ISL_2851791 | Lighthouse Lab in Alderley Park | Wellcome Sanger Institute for the COVID-19 Genomics UK (COG-UK) Consortium | Jacquelyn Wynn, Mairead Hyland, The Lighthouse Lab in Alderley Park and Alex Alderton, Roberto Amato, Jeffrey Barrett, Sonia Goncalves, Ewan Harrison, David K. Jackson, Ian Johnston, Dominic Kwiatkowski, Cordelia Langford, John Sillitoe on behalf of the Wellcome Sanger Institute COVID-19 Surveillance Team |
| EPI_ISL_2851792 | Lighthouse Lab in Milton Keynes | Wellcome Sanger Institute for the COVID-19 Genomics UK (COG-UK) Consortium | The Lighthouse Lab in Milton Keynes and Alex Alderton, Roberto Amato, Jeffrey Barrett, Sonia Goncalves, Ewan Harrison, David K. Jackson, Ian Johnston, Dominic Kwiatkowski, Cordelia Langford, John Sillitoe on behalf of the Wellcome Sanger Institute COVID-19 Surveillance Team |

| | | | |
|---|---|---|---|
| EPI_ISL_2851793 | Lighthouse Lab in Alderley Park | Wellcome Sanger Institute for the COVID-19 Genomics UK (COG-UK) Consortium | Jacquelyn Wynn, Mairead Hyland, The Lighthouse Lab in Alderley Park and Alex Alderton, Roberto Amato, Jeffrey Barrett, Sonia Goncalves, Ewan Harrison, David K. Jackson, Ian Johnston, Dominic Kwiatkowski, Cordelia Langford, John Sillitoe on behalf of the Wellcome Sanger Institute COVID-19 Surveillance Team |
| EPI_ISL_2851794, EPI_ISL_2851795 | Lighthouse Lab in Glasgow | Wellcome Sanger Institute for the COVID-19 Genomics UK (COG-UK) Consortium | Harper VanSteenhouse, Yumi Kasai, David Gray, Carol Clugston, Anna Dominiczak and Alex Alderton, Roberto Amato, Jeffrey Barrett, Sonia Goncalves, Ewan Harrison, David K. Jackson, Ian Johnston, Dominic Kwiatkowski, Cordelia Langford, John Sillitoe on behalf of the Wellcome Sanger Institute COVID-19 Surveillance Team |
| EPI_ISL_2851796 | Lighthouse Lab in Alderley Park | Wellcome Sanger Institute for the COVID-19 Genomics UK (COG-UK) Consortium | Jacquelyn Wynn, Mairead Hyland, The Lighthouse Lab in Alderley Park and Alex Alderton, Roberto Amato, Jeffrey Barrett, Sonia Goncalves, Ewan Harrison, David K. Jackson, Ian Johnston, Dominic Kwiatkowski, Cordelia Langford, John Sillitoe on behalf of the Wellcome Sanger Institute COVID-19 Surveillance Team |
| EPI_ISL_2851797 | Lighthouse Lab in Glasgow | Wellcome Sanger Institute for the COVID-19 Genomics UK (COG-UK) Consortium | Harper VanSteenhouse, Yumi Kasai, David Gray, Carol Clugston, Anna Dominiczak and Alex Alderton, Roberto Amato, Jeffrey Barrett, Sonia Goncalves, Ewan Harrison, David K. Jackson, Ian Johnston, Dominic Kwiatkowski, Cordelia Langford, John Sillitoe on behalf of the Wellcome Sanger Institute COVID-19 Surveillance Team |
| EPI_ISL_2851798 | Lighthouse Lab in Milton Keynes | Wellcome Sanger Institute for the COVID-19 Genomics UK (COG-UK) Consortium | The Lighthouse Lab in Milton Keynes and Alex Alderton, Roberto Amato, Jeffrey Barrett, Sonia Goncalves, Ewan Harrison, David K. Jackson, Ian Johnston, Dominic Kwiatkowski, Cordelia Langford, John Sillitoe on behalf of the Wellcome Sanger Institute COVID-19 Surveillance Team |
| EPI_ISL_2851799 | Lighthouse Lab in Glasgow | Wellcome Sanger Institute for the COVID-19 Genomics UK (COG-UK) Consortium | Harper VanSteenhouse, Yumi Kasai, David Gray, Carol Clugston, Anna Dominiczak and Alex Alderton, Roberto Amato, Jeffrey Barrett, Sonia Goncalves, Ewan Harrison, David K. Jackson, Ian Johnston, Dominic Kwiatkowski, Cordelia Langford, John Sillitoe on behalf of the Wellcome Sanger Institute COVID-19 Surveillance Team |
| EPI_ISL_2851800, EPI_ISL_2851801 | Lighthouse Lab in Milton Keynes | Wellcome Sanger Institute for the COVID-19 Genomics UK (COG-UK) Consortium | The Lighthouse Lab in Milton Keynes and Alex Alderton, Roberto Amato, Jeffrey Barrett, Sonia Goncalves, Ewan Harrison, David K. Jackson, Ian Johnston, Dominic Kwiatkowski, Cordelia Langford, John Sillitoe on behalf of the Wellcome Sanger Institute COVID-19 Surveillance Team |
| EPI_ISL_2851802 | Lighthouse Lab in Alderley Park | Wellcome Sanger Institute for the COVID-19 Genomics UK (COG-UK) Consortium | Jacquelyn Wynn, Mairead Hyland, The Lighthouse Lab in Alderley Park and Alex Alderton, Roberto Amato, Jeffrey Barrett, Sonia Goncalves, Ewan Harrison, David K. Jackson, Ian Johnston, Dominic Kwiatkowski, Cordelia Langford, John Sillitoe on behalf of the Wellcome Sanger Institute COVID-19 Surveillance Team |
| EPI_ISL_2851803 | Lighthouse Lab in Milton Keynes | Wellcome Sanger Institute for the COVID-19 Genomics UK (COG-UK) Consortium | The Lighthouse Lab in Milton Keynes and Alex Alderton, Roberto Amato, Jeffrey Barrett, Sonia Goncalves, Ewan Harrison, David K. Jackson, Ian Johnston, Dominic Kwiatkowski, Cordelia Langford, John Sillitoe on behalf of the Wellcome Sanger Institute COVID-19 Surveillance Team |
| EPI_ISL_2851804 | Lighthouse Lab in Alderley Park | Wellcome Sanger Institute for the COVID-19 Genomics UK (COG-UK) Consortium | Jacquelyn Wynn, Mairead Hyland, The Lighthouse Lab in Alderley Park and Alex Alderton, Roberto Amato, Jeffrey Barrett, Sonia Goncalves, Ewan Harrison, David K. Jackson, Ian Johnston, Dominic Kwiatkowski, Cordelia Langford, John Sillitoe on behalf of the Wellcome Sanger Institute COVID-19 Surveillance Team |
| EPI_ISL_2851805, EPI_ISL_2851806 | Lighthouse Lab in Glasgow | Wellcome Sanger Institute for the COVID-19 Genomics UK (COG-UK) Consortium | Harper VanSteenhouse, Yumi Kasai, David Gray, Carol Clugston, Anna Dominiczak and Alex Alderton, Roberto Amato, Jeffrey Barrett, Sonia Goncalves, Ewan Harrison, David K. Jackson, Ian Johnston, Dominic Kwiatkowski, Cordelia Langford, John Sillitoe on behalf of the Wellcome Sanger Institute COVID-19 Surveillance Team |
| EPI_ISL_2851807, EPI_ISL_2851808 | Lighthouse Lab in Milton Keynes | Wellcome Sanger Institute for the COVID-19 Genomics UK (COG-UK) Consortium | The Lighthouse Lab in Milton Keynes and Alex Alderton, Roberto Amato, Jeffrey Barrett, Sonia Goncalves, Ewan Harrison, David K. Jackson, Ian Johnston, Dominic Kwiatkowski, Cordelia Langford, John Sillitoe on behalf of the Wellcome Sanger Institute COVID-19 Surveillance Team |
| EPI_ISL_2851809, EPI_ISL_2851810, EPI_ISL_2851811 | Lighthouse Lab in Glasgow | Wellcome Sanger Institute for the COVID-19 Genomics UK (COG-UK) Consortium | Harper VanSteenhouse, Yumi Kasai, David Gray, Carol Clugston, Anna Dominiczak and Alex Alderton, Roberto Amato, Jeffrey Barrett, Sonia Goncalves, Ewan Harrison, David K. Jackson, Ian Johnston, Dominic Kwiatkowski, Cordelia Langford, John Sillitoe on behalf of the Wellcome Sanger Institute COVID-19 Surveillance Team |
| EPI_ISL_2851812 | Lighthouse Lab in Milton Keynes | Wellcome Sanger Institute for the COVID-19 Genomics UK (COG-UK) Consortium | The Lighthouse Lab in Milton Keynes and Alex Alderton, Roberto Amato, Jeffrey Barrett, Sonia Goncalves, Ewan Harrison, David K. Jackson, Ian Johnston, Dominic Kwiatkowski, Cordelia Langford, John Sillitoe on behalf of the Wellcome Sanger Institute COVID-19 Surveillance Team |
| EPI_ISL_2851813 | Lighthouse Lab in Alderley Park | Wellcome Sanger Institute for the COVID-19 Genomics UK (COG-UK) Consortium | Jacquelyn Wynn, Mairead Hyland, The Lighthouse Lab in Alderley Park and Alex Alderton, Roberto Amato, Jeffrey Barrett, Sonia Goncalves, Ewan Harrison, David K. Jackson, Ian Johnston, Dominic Kwiatkowski, Cordelia Langford, John Sillitoe on behalf of the Wellcome Sanger Institute COVID-19 Surveillance Team |
| EPI_ISL_2851814 | Lighthouse Lab in Milton Keynes | Wellcome Sanger Institute for the COVID-19 Genomics UK (COG-UK) Consortium | The Lighthouse Lab in Milton Keynes and Alex Alderton, Roberto Amato, Jeffrey Barrett, Sonia Goncalves, Ewan Harrison, David K. Jackson, Ian Johnston, Dominic Kwiatkowski, Cordelia Langford, John Sillitoe on behalf of the Wellcome Sanger Institute COVID-19 Surveillance Team |
| EPI_ISL_2851815 | Lighthouse Lab in Glasgow | Wellcome Sanger Institute for the COVID-19 Genomics UK (COG-UK) Consortium | Harper VanSteenhouse, Yumi Kasai, David Gray, Carol Clugston, Anna Dominiczak and Alex Alderton, Roberto Amato, Jeffrey Barrett, Sonia Goncalves, Ewan Harrison, David K. Jackson, Ian Johnston, Dominic Kwiatkowski, Cordelia Langford, John Sillitoe on behalf of the Wellcome Sanger Institute COVID-19 Surveillance Team |
| EPI_ISL_2851816 | Lighthouse Lab in Milton Keynes | Wellcome Sanger Institute for the COVID-19 Genomics UK (COG-UK) Consortium | The Lighthouse Lab in Milton Keynes and Alex Alderton, Roberto Amato, Jeffrey Barrett, Sonia Goncalves, Ewan Harrison, David K. Jackson, Ian Johnston, Dominic Kwiatkowski, Cordelia Langford, John Sillitoe on behalf of the Wellcome Sanger Institute COVID-19 Surveillance Team |
| EPI_ISL_2851817, EPI_ISL_2851818 | Lighthouse Lab in Glasgow | Wellcome Sanger Institute for the COVID-19 Genomics UK (COG-UK) Consortium | Harper VanSteenhouse, Yumi Kasai, David Gray, Carol Clugston, Anna Dominiczak and Alex Alderton, Roberto Amato, Jeffrey Barrett, Sonia Goncalves, Ewan Harrison, David K. Jackson, Ian Johnston, Dominic Kwiatkowski, Cordelia Langford, John Sillitoe on behalf of the Wellcome Sanger Institute COVID-19 Surveillance Team |
| EPI_ISL_2851819, EPI_ISL_2851820 | Lighthouse Lab in Milton Keynes | Wellcome Sanger Institute for the COVID-19 Genomics UK (COG-UK) Consortium | The Lighthouse Lab in Milton Keynes and Alex Alderton, Roberto Amato, Jeffrey Barrett, Sonia Goncalves, Ewan Harrison, David K. Jackson, Ian Johnston, Dominic Kwiatkowski, Cordelia Langford, John Sillitoe on behalf of the Wellcome Sanger Institute COVID-19 Surveillance Team |
| EPI_ISL_2851821 | Lighthouse Lab in Glasgow | Wellcome Sanger Institute for the COVID-19 Genomics UK (COG-UK) Consortium | Harper VanSteenhouse, Yumi Kasai, David Gray, Carol Clugston, Anna Dominiczak and Alex Alderton, Roberto Amato, Jeffrey Barrett, Sonia Goncalves, Ewan Harrison, David K. Jackson, Ian Johnston, Dominic Kwiatkowski, Cordelia Langford, John Sillitoe on behalf of the Wellcome Sanger Institute COVID-19 Surveillance Team |
| EPI_ISL_2851822 | Lighthouse Lab in Milton Keynes | Wellcome Sanger Institute for the COVID-19 Genomics UK (COG-UK) Consortium | The Lighthouse Lab in Milton Keynes and Alex Alderton, Roberto Amato, Jeffrey Barrett, Sonia Goncalves, Ewan Harrison, David K. Jackson, Ian Johnston, Dominic Kwiatkowski, Cordelia Langford, John Sillitoe on behalf of the Wellcome Sanger Institute COVID-19 Surveillance Team |
| EPI_ISL_2851823 | Lighthouse Lab in Glasgow | Wellcome Sanger Institute for the COVID-19 Genomics UK (COG-UK) Consortium | Harper VanSteenhouse, Yumi Kasai, David Gray, Carol Clugston, Anna Dominiczak and Alex Alderton, Roberto Amato, Jeffrey Barrett, Sonia Goncalves, Ewan Harrison, David K. Jackson, Ian Johnston, Dominic Kwiatkowski, Cordelia Langford, John Sillitoe on behalf of the Wellcome Sanger Institute COVID-19 Surveillance Team |
| EPI_ISL_2851824 | Lighthouse Lab in Alderley Park | Wellcome Sanger Institute for the COVID-19 Genomics UK (COG-UK) Consortium | Jacquelyn Wynn, Mairead Hyland, The Lighthouse Lab in Alderley Park and Alex Alderton, Roberto Amato, Jeffrey Barrett, Sonia Goncalves, Ewan Harrison, David K. Jackson, Ian Johnston, Dominic Kwiatkowski, Cordelia Langford, John Sillitoe on behalf of the Wellcome Sanger Institute COVID-19 Surveillance Team |
| EPI_ISL_2851825 | Lighthouse Lab in Milton Keynes | Wellcome Sanger Institute for the COVID-19 Genomics UK (COG-UK) Consortium | The Lighthouse Lab in Milton Keynes and Alex Alderton, Roberto Amato, Jeffrey Barrett, Sonia Goncalves, Ewan Harrison, David K. Jackson, Ian Johnston, Dominic Kwiatkowski, Cordelia Langford, John Sillitoe on behalf of the Wellcome Sanger Institute COVID-19 Surveillance Team |
| EPI_ISL_2851826 | Lighthouse Lab in Alderley Park | Wellcome Sanger Institute for the COVID-19 Genomics UK (COG-UK) Consortium | Jacquelyn Wynn, Mairead Hyland, The Lighthouse Lab in Alderley Park and Alex Alderton, Roberto Amato, Jeffrey Barrett, Sonia Goncalves, Ewan Harrison, David K. Jackson, Ian Johnston, Dominic Kwiatkowski, Cordelia Langford, John Sillitoe on behalf of the Wellcome Sanger Institute COVID-19 Surveillance Team |
| EPI_ISL_2851827 | Lighthouse Lab in Glasgow | Wellcome Sanger Institute for the COVID-19 Genomics UK (COG-UK) Consortium | Harper VanSteenhouse, Yumi Kasai, David Gray, Carol Clugston, Anna Dominiczak and Alex Alderton, Roberto Amato, Jeffrey Barrett, Sonia Goncalves, Ewan Harrison, David K. Jackson, Ian Johnston, Dominic Kwiatkowski, Cordelia Langford, John Sillitoe on behalf of the Wellcome Sanger Institute COVID-19 Surveillance Team |
| EPI_ISL_2851828 | Lighthouse Lab in Milton Keynes | Wellcome Sanger Institute for the COVID-19 Genomics UK (COG-UK) Consortium | The Lighthouse Lab in Milton Keynes and Alex Alderton, Roberto Amato, Jeffrey Barrett, Sonia Goncalves, Ewan Harrison, David K. Jackson, Ian Johnston, Dominic Kwiatkowski, Cordelia Langford, John Sillitoe on behalf of the Wellcome Sanger Institute COVID-19 Surveillance Team |
| EPI_ISL_2851829, EPI_ISL_2851830, EPI_ISL_2851831, EPI_ISL_2851832, EPI_ISL_2851833, EPI_ISL_2851834 | Lighthouse Lab in Glasgow | Wellcome Sanger Institute for the COVID-19 Genomics UK (COG-UK) Consortium | Harper VanSteenhouse, Yumi Kasai, David Gray, Carol Clugston, Anna Dominiczak and Alex Alderton, Roberto Amato, Jeffrey Barrett, Sonia Goncalves, Ewan Harrison, David K. Jackson, Ian Johnston, Dominic Kwiatkowski, Cordelia Langford, John Sillitoe on behalf of the Wellcome Sanger Institute COVID-19 Surveillance Team |
| EPI_ISL_2851835 | Lighthouse Lab in Alderley Park | Wellcome Sanger Institute for the COVID-19 Genomics UK (COG-UK) Consortium | Jacquelyn Wynn, Mairead Hyland, The Lighthouse Lab in Alderley Park and Alex Alderton, Roberto Amato, Jeffrey Barrett, Sonia Goncalves, Ewan Harrison, David K. Jackson, Ian Johnston, Dominic Kwiatkowski, Cordelia Langford, John Sillitoe on behalf of the Wellcome Sanger Institute COVID-19 Surveillance Team |
| EPI_ISL_2851836 | Lighthouse Lab in Glasgow | Wellcome Sanger Institute for the COVID-19 Genomics UK (COG-UK) Consortium | Harper VanSteenhouse, Yumi Kasai, David Gray, Carol Clugston, Anna Dominiczak and Alex Alderton, Roberto Amato, Jeffrey Barrett, Sonia Goncalves, Ewan Harrison, David K. Jackson, Ian Johnston, Dominic Kwiatkowski, Cordelia Langford, John Sillitoe on behalf of the Wellcome Sanger Institute COVID-19 Surveillance Team |
| EPI_ISL_2851837 | Lighthouse Lab in Milton Keynes | Wellcome Sanger Institute for the COVID-19 Genomics UK (COG-UK) Consortium | The Lighthouse Lab in Milton Keynes and Alex Alderton, Roberto Amato, Jeffrey Barrett, Sonia Goncalves, Ewan Harrison, David K. Jackson, Ian Johnston, Dominic Kwiatkowski, Cordelia Langford, John Sillitoe on behalf of the Wellcome Sanger Institute COVID-19 Surveillance Team |
| EPI_ISL_2851838, EPI_ISL_2851839 | Lighthouse Lab in Glasgow | Wellcome Sanger Institute for the COVID-19 Genomics UK (COG-UK) Consortium | Harper VanSteenhouse, Yumi Kasai, David Gray, Carol Clugston, Anna Dominiczak and Alex Alderton, Roberto Amato, Jeffrey Barrett, Sonia Goncalves, Ewan Harrison, David K. Jackson, Ian Johnston, Dominic Kwiatkowski, Cordelia Langford, John Sillitoe on behalf of the Wellcome Sanger Institute COVID-19 Surveillance Team |
| EPI_ISL_2851840 | Lighthouse Lab in Alderley Park | Wellcome Sanger Institute for the COVID-19 Genomics UK (COG-UK) Consortium | Jacquelyn Wynn, Mairead Hyland, The Lighthouse Lab in Alderley Park and Alex Alderton, Roberto Amato, Jeffrey Barrett, Sonia Goncalves, Ewan Harrison, David K. Jackson, Ian Johnston, Dominic Kwiatkowski, Cordelia Langford, John Sillitoe on behalf of the Wellcome Sanger Institute COVID-19 Surveillance Team |
| EPI_ISL_2851841, EPI_ISL_2851842, EPI_ISL_2851843, EPI_ISL_2851844, EPI_ISL_2851845 | Lighthouse Lab in Glasgow | Wellcome Sanger Institute for the COVID-19 Genomics UK (COG-UK) Consortium | Harper VanSteenhouse, Yumi Kasai, David Gray, Carol Clugston, Anna Dominiczak and Alex Alderton, Roberto Amato, Jeffrey Barrett, Sonia Goncalves, Ewan Harrison, David K. Jackson, Ian Johnston, Dominic Kwiatkowski, Cordelia Langford, John Sillitoe on behalf of the Wellcome Sanger Institute COVID-19 Surveillance Team |
| EPI_ISL_2851846 | Lighthouse Lab in Alderley Park | Wellcome Sanger Institute for the COVID-19 Genomics UK (COG-UK) Consortium | Jacquelyn Wynn, Mairead Hyland, The Lighthouse Lab in Alderley Park and Alex Alderton, Roberto Amato, Jeffrey Barrett, Sonia Goncalves, Ewan Harrison, David K. Jackson, Ian Johnston, Dominic Kwiatkowski, Cordelia Langford, John Sillitoe on behalf of the Wellcome Sanger Institute COVID-19 Surveillance Team |
| EPI_ISL_2851847, EPI_ISL_2851848, EPI_ISL_2851849, EPI_ISL_2851850, EPI_ISL_2851851, EPI_ISL_2851852, EPI_ISL_2851853, EPI_ISL_2851854, EPI_ISL_2851855, EPI_ISL_2851856, EPI_ISL_2851857, EPI_ISL_2851858, EPI_ISL_2851859, EPI_ISL_2851860, EPI_ISL_2851861 | | | |
| see above | Lighthouse Lab in Glasgow | Wellcome Sanger Institute for the COVID-19 Genomics UK (COG-UK) Consortium | Harper VanSteenhouse, Yumi Kasai, David Gray, Carol Clugston, Anna Dominiczak and Alex Alderton, Roberto Amato, Jeffrey Barrett, Sonia Goncalves, Ewan Harrison, David K. Jackson, Ian Johnston, Dominic Kwiatkowski, Cordelia Langford, John Sillitoe on behalf of the Wellcome Sanger Institute COVID-19 Surveillance Team |
| EPI_ISL_2851862 | Lighthouse Lab in Milton Keynes | Wellcome Sanger Institute for the COVID-19 Genomics UK (COG-UK) Consortium | The Lighthouse Lab in Milton Keynes and Alex Alderton, Roberto Amato, Jeffrey Barrett, Sonia Goncalves, Ewan Harrison, David K. Jackson, Ian Johnston, Dominic Kwiatkowski, Cordelia Langford, John Sillitoe on behalf of the Wellcome Sanger Institute COVID-19 Surveillance Team |
| EPI_ISL_2851863 | Lighthouse Lab in Alderley Park | Wellcome Sanger Institute for the COVID-19 Genomics UK (COG-UK) Consortium | Jacquelyn Wynn, Mairead Hyland, The Lighthouse Lab in Alderley Park and Alex Alderton, Roberto Amato, Jeffrey Barrett, Sonia Goncalves, Ewan Harrison, David K. Jackson, Ian Johnston, Dominic Kwiatkowski, Cordelia Langford, John Sillitoe on behalf of the Wellcome Sanger Institute COVID-19 Surveillance Team |
| EPI_ISL_2851864, EPI_ISL_2851865, EPI_ISL_2851866 | Lighthouse Lab in Milton Keynes | Wellcome Sanger Institute for the COVID-19 Genomics UK (COG-UK) Consortium | The Lighthouse Lab in Milton Keynes and Alex Alderton, Roberto Amato, Jeffrey Barrett, Sonia Goncalves, Ewan Harrison, David K. Jackson, Ian Johnston, Dominic Kwiatkowski, Cordelia Langford, John Sillitoe on behalf of the Wellcome Sanger Institute COVID-19 Surveillance Team |
| EPI_ISL_2851867, | Lighthouse Lab in Glasgow | Wellcome Sanger Institute for the COVID-19 Genomics UK (COG-UK) | Harper VanSteenhouse, Yumi Kasai, David Gray, Carol Clugston, Anna Dominiczak and Alex Alderton, Roberto Amato, Jeffrey Barrett, Sonia Goncalves, Ewan Harrison, David K. Jackson, Ian Johnston, Dominic Kwiatkowski, Cordelia Langford, |

| | | | |
|---|---|---|---|
| EPI_ISL_2851868 | | Consortium | John Sillitoe on behalf of the Wellcome Sanger Institute COVID-19 Surveillance Team |
| EPI_ISL_2851869 | Lighthouse Lab in Alderley Park | Wellcome Sanger Institute for the COVID-19 Genomics UK (COG-UK) Consortium | Jacquelyn Wynn, Mairead Hyland, The Lighthouse Lab in Alderley Park and Alex Alderton, Roberto Amato, Jeffrey Barrett, Sonia Goncalves, Ewan Harrison, David K. Jackson, Ian Johnston, Dominic Kwiatkowski, Cordelia Langford, John Sillitoe on behalf of the Wellcome Sanger Institute COVID-19 Surveillance Team |
| EPI_ISL_2851870, EPI_ISL_2851871 | Lighthouse Lab in Glasgow | Wellcome Sanger Institute for the COVID-19 Genomics UK (COG-UK) Consortium | Harper VanSteenhouse, Yumi Kasai, David Gray, Carol Clugston, Anna Dominiczak and Alex Alderton, Roberto Amato, Jeffrey Barrett, Sonia Goncalves, Ewan Harrison, David K. Jackson, Ian Johnston, Dominic Kwiatkowski, Cordelia Langford, John Sillitoe on behalf of the Wellcome Sanger Institute COVID-19 Surveillance Team |
| EPI_ISL_2851872 | Lighthouse Lab in Milton Keynes | Wellcome Sanger Institute for the COVID-19 Genomics UK (COG-UK) Consortium | The Lighthouse Lab in Milton Keynes and Alex Alderton, Roberto Amato, Jeffrey Barrett, Sonia Goncalves, Ewan Harrison, David K. Jackson, Ian Johnston, Dominic Kwiatkowski, Cordelia Langford, John Sillitoe on behalf of the Wellcome Sanger Institute COVID-19 Surveillance Team |
| EPI_ISL_2851873, EPI_ISL_2851874 | Lighthouse Lab in Alderley Park | Wellcome Sanger Institute for the COVID-19 Genomics UK (COG-UK) Consortium | Jacquelyn Wynn, Mairead Hyland, The Lighthouse Lab in Alderley Park and Alex Alderton, Roberto Amato, Jeffrey Barrett, Sonia Goncalves, Ewan Harrison, David K. Jackson, Ian Johnston, Dominic Kwiatkowski, Cordelia Langford, John Sillitoe on behalf of the Wellcome Sanger Institute COVID-19 Surveillance Team |
| EPI_ISL_2851875 | Lighthouse Lab in Glasgow | Wellcome Sanger Institute for the COVID-19 Genomics UK (COG-UK) Consortium | Harper VanSteenhouse, Yumi Kasai, David Gray, Carol Clugston, Anna Dominiczak and Alex Alderton, Roberto Amato, Jeffrey Barrett, Sonia Goncalves, Ewan Harrison, David K. Jackson, Ian Johnston, Dominic Kwiatkowski, Cordelia Langford, John Sillitoe on behalf of the Wellcome Sanger Institute COVID-19 Surveillance Team |
| EPI_ISL_2851876 | Lighthouse Lab in Alderley Park | Wellcome Sanger Institute for the COVID-19 Genomics UK (COG-UK) Consortium | Jacquelyn Wynn, Mairead Hyland, The Lighthouse Lab in Alderley Park and Alex Alderton, Roberto Amato, Jeffrey Barrett, Sonia Goncalves, Ewan Harrison, David K. Jackson, Ian Johnston, Dominic Kwiatkowski, Cordelia Langford, John Sillitoe on behalf of the Wellcome Sanger Institute COVID-19 Surveillance Team |
| EPI_ISL_2851877, EPI_ISL_2851878, EPI_ISL_2851879, EPI_ISL_2851880 | Lighthouse Lab in Milton Keynes | Wellcome Sanger Institute for the COVID-19 Genomics UK (COG-UK) Consortium | The Lighthouse Lab in Milton Keynes and Alex Alderton, Roberto Amato, Jeffrey Barrett, Sonia Goncalves, Ewan Harrison, David K. Jackson, Ian Johnston, Dominic Kwiatkowski, Cordelia Langford, John Sillitoe on behalf of the Wellcome Sanger Institute COVID-19 Surveillance Team |
| EPI_ISL_2851881, EPI_ISL_2851882 | Lighthouse Lab in Alderley Park | Wellcome Sanger Institute for the COVID-19 Genomics UK (COG-UK) Consortium | Jacquelyn Wynn, Mairead Hyland, The Lighthouse Lab in Alderley Park and Alex Alderton, Roberto Amato, Jeffrey Barrett, Sonia Goncalves, Ewan Harrison, David K. Jackson, Ian Johnston, Dominic Kwiatkowski, Cordelia Langford, John Sillitoe on behalf of the Wellcome Sanger Institute COVID-19 Surveillance Team |
| EPI_ISL_2851883 | Lighthouse Lab in Glasgow | Wellcome Sanger Institute for the COVID-19 Genomics UK (COG-UK) Consortium | Harper VanSteenhouse, Yumi Kasai, David Gray, Carol Clugston, Anna Dominiczak and Alex Alderton, Roberto Amato, Jeffrey Barrett, Sonia Goncalves, Ewan Harrison, David K. Jackson, Ian Johnston, Dominic Kwiatkowski, Cordelia Langford, John Sillitoe on behalf of the Wellcome Sanger Institute COVID-19 Surveillance Team |
| EPI_ISL_2851884, EPI_ISL_2851885 | Lighthouse Lab in Milton Keynes | Wellcome Sanger Institute for the COVID-19 Genomics UK (COG-UK) Consortium | The Lighthouse Lab in Milton Keynes and Alex Alderton, Roberto Amato, Jeffrey Barrett, Sonia Goncalves, Ewan Harrison, David K. Jackson, Ian Johnston, Dominic Kwiatkowski, Cordelia Langford, John Sillitoe on behalf of the Wellcome Sanger Institute COVID-19 Surveillance Team |
| EPI_ISL_2851886 | Lighthouse Lab in Glasgow | Wellcome Sanger Institute for the COVID-19 Genomics UK (COG-UK) Consortium | Harper VanSteenhouse, Yumi Kasai, David Gray, Carol Clugston, Anna Dominiczak and Alex Alderton, Roberto Amato, Jeffrey Barrett, Sonia Goncalves, Ewan Harrison, David K. Jackson, Ian Johnston, Dominic Kwiatkowski, Cordelia Langford, John Sillitoe on behalf of the Wellcome Sanger Institute COVID-19 Surveillance Team |
| EPI_ISL_2851887 | Lighthouse Lab in Milton Keynes | Wellcome Sanger Institute for the COVID-19 Genomics UK (COG-UK) Consortium | The Lighthouse Lab in Milton Keynes and Alex Alderton, Roberto Amato, Jeffrey Barrett, Sonia Goncalves, Ewan Harrison, David K. Jackson, Ian Johnston, Dominic Kwiatkowski, Cordelia Langford, John Sillitoe on behalf of the Wellcome Sanger Institute COVID-19 Surveillance Team |
| EPI_ISL_2851888, EPI_ISL_2851889, EPI_ISL_2851890, EPI_ISL_2851891 | Lighthouse Lab in Glasgow | Wellcome Sanger Institute for the COVID-19 Genomics UK (COG-UK) Consortium | Harper VanSteenhouse, Yumi Kasai, David Gray, Carol Clugston, Anna Dominiczak and Alex Alderton, Roberto Amato, Jeffrey Barrett, Sonia Goncalves, Ewan Harrison, David K. Jackson, Ian Johnston, Dominic Kwiatkowski, Cordelia Langford, John Sillitoe on behalf of the Wellcome Sanger Institute COVID-19 Surveillance Team |
| EPI_ISL_2851892, EPI_ISL_2851893 | Lighthouse Lab in Alderley Park | Wellcome Sanger Institute for the COVID-19 Genomics UK (COG-UK) Consortium | Jacquelyn Wynn, Mairead Hyland, The Lighthouse Lab in Alderley Park and Alex Alderton, Roberto Amato, Jeffrey Barrett, Sonia Goncalves, Ewan Harrison, David K. Jackson, Ian Johnston, Dominic Kwiatkowski, Cordelia Langford, John Sillitoe on behalf of the Wellcome Sanger Institute COVID-19 Surveillance Team |
| EPI_ISL_2851894, EPI_ISL_2851895 | Lighthouse Lab in Glasgow | Wellcome Sanger Institute for the COVID-19 Genomics UK (COG-UK) Consortium | Harper VanSteenhouse, Yumi Kasai, David Gray, Carol Clugston, Anna Dominiczak and Alex Alderton, Roberto Amato, Jeffrey Barrett, Sonia Goncalves, Ewan Harrison, David K. Jackson, Ian Johnston, Dominic Kwiatkowski, Cordelia Langford, John Sillitoe on behalf of the Wellcome Sanger Institute COVID-19 Surveillance Team |
| EPI_ISL_2851896, EPI_ISL_2851897 | Lighthouse Lab in Alderley Park | Wellcome Sanger Institute for the COVID-19 Genomics UK (COG-UK) Consortium | Jacquelyn Wynn, Mairead Hyland, The Lighthouse Lab in Alderley Park and Alex Alderton, Roberto Amato, Jeffrey Barrett, Sonia Goncalves, Ewan Harrison, David K. Jackson, Ian Johnston, Dominic Kwiatkowski, Cordelia Langford, John Sillitoe on behalf of the Wellcome Sanger Institute COVID-19 Surveillance Team |
| EPI_ISL_2851898, EPI_ISL_2851899, EPI_ISL_2851900 | Lighthouse Lab in Glasgow | Wellcome Sanger Institute for the COVID-19 Genomics UK (COG-UK) Consortium | Harper VanSteenhouse, Yumi Kasai, David Gray, Carol Clugston, Anna Dominiczak and Alex Alderton, Roberto Amato, Jeffrey Barrett, Sonia Goncalves, Ewan Harrison, David K. Jackson, Ian Johnston, Dominic Kwiatkowski, Cordelia Langford, John Sillitoe on behalf of the Wellcome Sanger Institute COVID-19 Surveillance Team |
| EPI_ISL_2851901 | Lighthouse Lab in Alderley Park | Wellcome Sanger Institute for the COVID-19 Genomics UK (COG-UK) Consortium | Jacquelyn Wynn, Mairead Hyland, The Lighthouse Lab in Alderley Park and Alex Alderton, Roberto Amato, Jeffrey Barrett, Sonia Goncalves, Ewan Harrison, David K. Jackson, Ian Johnston, Dominic Kwiatkowski, Cordelia Langford, John Sillitoe on behalf of the Wellcome Sanger Institute COVID-19 Surveillance Team |
| EPI_ISL_2851902, EPI_ISL_2851904, EPI_ISL_2851905 | Lighthouse Lab in Milton Keynes | Wellcome Sanger Institute for the COVID-19 Genomics UK (COG-UK) Consortium | The Lighthouse Lab in Milton Keynes and Alex Alderton, Roberto Amato, Jeffrey Barrett, Sonia Goncalves, Ewan Harrison, David K. Jackson, Ian Johnston, Dominic Kwiatkowski, Cordelia Langford, John Sillitoe on behalf of the Wellcome Sanger Institute COVID-19 Surveillance Team |
| EPI_ISL_2851906 | Lighthouse Lab in Alderley Park | Wellcome Sanger Institute for the COVID-19 Genomics UK (COG-UK) Consortium | Jacquelyn Wynn, Mairead Hyland, The Lighthouse Lab in Alderley Park and Alex Alderton, Roberto Amato, Jeffrey Barrett, Sonia Goncalves, Ewan Harrison, David K. Jackson, Ian Johnston, Dominic Kwiatkowski, Cordelia Langford, John Sillitoe on behalf of the Wellcome Sanger Institute COVID-19 Surveillance Team |
| EPI_ISL_2851907 | Lighthouse Lab in Milton Keynes | Wellcome Sanger Institute for the COVID-19 Genomics UK (COG-UK) Consortium | The Lighthouse Lab in Milton Keynes and Alex Alderton, Roberto Amato, Jeffrey Barrett, Sonia Goncalves, Ewan Harrison, David K. Jackson, Ian Johnston, Dominic Kwiatkowski, Cordelia Langford, John Sillitoe on behalf of the Wellcome Sanger Institute COVID-19 Surveillance Team |
| EPI_ISL_2851908 | Lighthouse Lab in Glasgow | Wellcome Sanger Institute for the COVID-19 Genomics UK (COG-UK) Consortium | Harper VanSteenhouse, Yumi Kasai, David Gray, Carol Clugston, Anna Dominiczak and Alex Alderton, Roberto Amato, Jeffrey Barrett, Sonia Goncalves, Ewan Harrison, David K. Jackson, Ian Johnston, Dominic Kwiatkowski, Cordelia Langford, John Sillitoe on behalf of the Wellcome Sanger Institute COVID-19 Surveillance Team |
| EPI_ISL_2851909 | Lighthouse Lab in Alderley Park | Wellcome Sanger Institute for the COVID-19 Genomics UK (COG-UK) Consortium | Jacquelyn Wynn, Mairead Hyland, The Lighthouse Lab in Alderley Park and Alex Alderton, Roberto Amato, Jeffrey Barrett, Sonia Goncalves, Ewan Harrison, David K. Jackson, Ian Johnston, Dominic Kwiatkowski, Cordelia Langford, John Sillitoe on behalf of the Wellcome Sanger Institute COVID-19 Surveillance Team |
| EPI_ISL_2851910, EPI_ISL_2851911, EPI_ISL_2851912 | Lighthouse Lab in Glasgow | Wellcome Sanger Institute for the COVID-19 Genomics UK (COG-UK) Consortium | Harper VanSteenhouse, Yumi Kasai, David Gray, Carol Clugston, Anna Dominiczak and Alex Alderton, Roberto Amato, Jeffrey Barrett, Sonia Goncalves, Ewan Harrison, David K. Jackson, Ian Johnston, Dominic Kwiatkowski, Cordelia Langford, John Sillitoe on behalf of the Wellcome Sanger Institute COVID-19 Surveillance Team |
| EPI_ISL_2851913, EPI_ISL_2851914 | Lighthouse Lab in Milton Keynes | Wellcome Sanger Institute for the COVID-19 Genomics UK (COG-UK) Consortium | The Lighthouse Lab in Milton Keynes and Alex Alderton, Roberto Amato, Jeffrey Barrett, Sonia Goncalves, Ewan Harrison, David K. Jackson, Ian Johnston, Dominic Kwiatkowski, Cordelia Langford, John Sillitoe on behalf of the Wellcome Sanger Institute COVID-19 Surveillance Team |
| EPI_ISL_2851915, EPI_ISL_2851916, EPI_ISL_2851917 | Lighthouse Lab in Glasgow | Wellcome Sanger Institute for the COVID-19 Genomics UK (COG-UK) Consortium | Harper VanSteenhouse, Yumi Kasai, David Gray, Carol Clugston, Anna Dominiczak and Alex Alderton, Roberto Amato, Jeffrey Barrett, Sonia Goncalves, Ewan Harrison, David K. Jackson, Ian Johnston, Dominic Kwiatkowski, Cordelia Langford, John Sillitoe on behalf of the Wellcome Sanger Institute COVID-19 Surveillance Team |
| EPI_ISL_2851918 | Lighthouse Lab in Milton Keynes | Wellcome Sanger Institute for the COVID-19 Genomics UK (COG-UK) Consortium | The Lighthouse Lab in Milton Keynes and Alex Alderton, Roberto Amato, Jeffrey Barrett, Sonia Goncalves, Ewan Harrison, David K. Jackson, Ian Johnston, Dominic Kwiatkowski, Cordelia Langford, John Sillitoe on behalf of the Wellcome Sanger Institute COVID-19 Surveillance Team |
| EPI_ISL_2851919 | Lighthouse Lab in Glasgow | Wellcome Sanger Institute for the COVID-19 Genomics UK (COG-UK) Consortium | Harper VanSteenhouse, Yumi Kasai, David Gray, Carol Clugston, Anna Dominiczak and Alex Alderton, Roberto Amato, Jeffrey Barrett, Sonia Goncalves, Ewan Harrison, David K. Jackson, Ian Johnston, Dominic Kwiatkowski, Cordelia Langford, John Sillitoe on behalf of the Wellcome Sanger Institute COVID-19 Surveillance Team |
| EPI_ISL_2851920 | Lighthouse Lab in Milton Keynes | Wellcome Sanger Institute for the COVID-19 Genomics UK (COG-UK) Consortium | The Lighthouse Lab in Milton Keynes and Alex Alderton, Roberto Amato, Jeffrey Barrett, Sonia Goncalves, Ewan Harrison, David K. Jackson, Ian Johnston, Dominic Kwiatkowski, Cordelia Langford, John Sillitoe on behalf of the Wellcome Sanger Institute COVID-19 Surveillance Team |
| EPI_ISL_2851921 | Lighthouse Lab in Glasgow | Wellcome Sanger Institute for the COVID-19 Genomics UK (COG-UK) Consortium | Harper VanSteenhouse, Yumi Kasai, David Gray, Carol Clugston, Anna Dominiczak and Alex Alderton, Roberto Amato, Jeffrey Barrett, Sonia Goncalves, Ewan Harrison, David K. Jackson, Ian Johnston, Dominic Kwiatkowski, Cordelia Langford, John Sillitoe on behalf of the Wellcome Sanger Institute COVID-19 Surveillance Team |
| EPI_ISL_2851922 | Lighthouse Lab in Milton Keynes | Wellcome Sanger Institute for the COVID-19 Genomics UK (COG-UK) Consortium | The Lighthouse Lab in Milton Keynes and Alex Alderton, Roberto Amato, Jeffrey Barrett, Sonia Goncalves, Ewan Harrison, David K. Jackson, Ian Johnston, Dominic Kwiatkowski, Cordelia Langford, John Sillitoe on behalf of the Wellcome Sanger Institute COVID-19 Surveillance Team |
| EPI_ISL_2851923, EPI_ISL_2851924, EPI_ISL_2851925, EPI_ISL_2851926, EPI_ISL_2851927, EPI_ISL_2851928 | Lighthouse Lab in Glasgow | Wellcome Sanger Institute for the COVID-19 Genomics UK (COG-UK) Consortium | Harper VanSteenhouse, Yumi Kasai, David Gray, Carol Clugston, Anna Dominiczak and Alex Alderton, Roberto Amato, Jeffrey Barrett, Sonia Goncalves, Ewan Harrison, David K. Jackson, Ian Johnston, Dominic Kwiatkowski, Cordelia Langford, John Sillitoe on behalf of the Wellcome Sanger Institute COVID-19 Surveillance Team |
| EPI_ISL_2851929 | Lighthouse Lab in Milton Keynes | Wellcome Sanger Institute for the COVID-19 Genomics UK (COG-UK) Consortium | The Lighthouse Lab in Milton Keynes and Alex Alderton, Roberto Amato, Jeffrey Barrett, Sonia Goncalves, Ewan Harrison, David K. Jackson, Ian Johnston, Dominic Kwiatkowski, Cordelia Langford, John Sillitoe on behalf of the Wellcome Sanger Institute COVID-19 Surveillance Team |
| EPI_ISL_2851930 | Lighthouse Lab in Glasgow | Wellcome Sanger Institute for the COVID-19 Genomics UK (COG-UK) Consortium | Harper VanSteenhouse, Yumi Kasai, David Gray, Carol Clugston, Anna Dominiczak and Alex Alderton, Roberto Amato, Jeffrey Barrett, Sonia Goncalves, Ewan Harrison, David K. Jackson, Ian Johnston, Dominic Kwiatkowski, Cordelia Langford, John Sillitoe on behalf of the Wellcome Sanger Institute COVID-19 Surveillance Team |
| EPI_ISL_2851931 | Lighthouse Lab in Milton Keynes | Wellcome Sanger Institute for the COVID-19 Genomics UK (COG-UK) Consortium | The Lighthouse Lab in Milton Keynes and Alex Alderton, Roberto Amato, Jeffrey Barrett, Sonia Goncalves, Ewan Harrison, David K. Jackson, Ian Johnston, Dominic Kwiatkowski, Cordelia Langford, John Sillitoe on behalf of the Wellcome Sanger Institute COVID-19 Surveillance Team |
| EPI_ISL_2851932, EPI_ISL_2851933, EPI_ISL_2851934 | Lighthouse Lab in Glasgow | Wellcome Sanger Institute for the COVID-19 Genomics UK (COG-UK) Consortium | Harper VanSteenhouse, Yumi Kasai, David Gray, Carol Clugston, Anna Dominiczak and Alex Alderton, Roberto Amato, Jeffrey Barrett, Sonia Goncalves, Ewan Harrison, David K. Jackson, Ian Johnston, Dominic Kwiatkowski, Cordelia Langford, John Sillitoe on behalf of the Wellcome Sanger Institute COVID-19 Surveillance Team |
| EPI_ISL_2851935 | Lighthouse Lab in Alderley Park | Wellcome Sanger Institute for the COVID-19 Genomics UK (COG-UK) Consortium | Jacquelyn Wynn, Mairead Hyland, The Lighthouse Lab in Alderley Park and Alex Alderton, Roberto Amato, Jeffrey Barrett, Sonia Goncalves, Ewan Harrison, David K. Jackson, Ian Johnston, Dominic Kwiatkowski, Cordelia Langford, John Sillitoe on behalf of the Wellcome Sanger Institute COVID-19 Surveillance Team |
| EPI_ISL_2851936 | Lighthouse Lab in Milton Keynes | Wellcome Sanger Institute for the COVID-19 Genomics UK (COG-UK) Consortium | The Lighthouse Lab in Milton Keynes and Alex Alderton, Roberto Amato, Jeffrey Barrett, Sonia Goncalves, Ewan Harrison, David K. Jackson, Ian Johnston, Dominic Kwiatkowski, Cordelia Langford, John Sillitoe on behalf of the Wellcome Sanger Institute COVID-19 Surveillance Team |
| EPI_ISL_2851937, EPI_ISL_2851938 | Lighthouse Lab in Glasgow | Wellcome Sanger Institute for the COVID-19 Genomics UK (COG-UK) Consortium | Harper VanSteenhouse, Yumi Kasai, David Gray, Carol Clugston, Anna Dominiczak and Alex Alderton, Roberto Amato, Jeffrey Barrett, Sonia Goncalves, Ewan Harrison, David K. Jackson, Ian Johnston, Dominic Kwiatkowski, Cordelia Langford, John Sillitoe on behalf of the Wellcome Sanger Institute COVID-19 Surveillance Team |

| | | | |
|---|---|---|---|
| EPI_ISL_2851939 | Lighthouse Lab in Alderley Park | Wellcome Sanger Institute for the COVID-19 Genomics UK (COG-UK) Consortium | Jacquelyn Wynn, Mairead Hyland, The Lighthouse Lab in Alderley Park and Alex Alderton, Roberto Amato, Jeffrey Barrett, Sonia Goncalves, Ewan Harrison, David K. Jackson, Ian Johnston, Dominic Kwiatkowski, Cordelia Langford, John Sillitoe on behalf of the Wellcome Sanger Institute COVID-19 Surveillance Team |
| EPI_ISL_2851940, EPI_ISL_2851941, EPI_ISL_2851942 | Lighthouse Lab in Glasgow | Wellcome Sanger Institute for the COVID-19 Genomics UK (COG-UK) Consortium | Harper VanSteenhouse, Yumi Kasai, David Gray, Carol Clugston, Anna Dominiczak and Alex Alderton, Roberto Amato, Jeffrey Barrett, Sonia Goncalves, Ewan Harrison, David K. Jackson, Ian Johnston, Dominic Kwiatkowski, Cordelia Langford, John Sillitoe on behalf of the Wellcome Sanger Institute COVID-19 Surveillance Team |
| EPI_ISL_2851943 | Lighthouse Lab in Alderley Park | Wellcome Sanger Institute for the COVID-19 Genomics UK (COG-UK) Consortium | Jacquelyn Wynn, Mairead Hyland, The Lighthouse Lab in Alderley Park and Alex Alderton, Roberto Amato, Jeffrey Barrett, Sonia Goncalves, Ewan Harrison, David K. Jackson, Ian Johnston, Dominic Kwiatkowski, Cordelia Langford, John Sillitoe on behalf of the Wellcome Sanger Institute COVID-19 Surveillance Team |
| EPI_ISL_2851944 | Lighthouse Lab in Glasgow | Wellcome Sanger Institute for the COVID-19 Genomics UK (COG-UK) Consortium | Harper VanSteenhouse, Yumi Kasai, David Gray, Carol Clugston, Anna Dominiczak and Alex Alderton, Roberto Amato, Jeffrey Barrett, Sonia Goncalves, Ewan Harrison, David K. Jackson, Ian Johnston, Dominic Kwiatkowski, Cordelia Langford, John Sillitoe on behalf of the Wellcome Sanger Institute COVID-19 Surveillance Team |
| EPI_ISL_2851945 | Lighthouse Lab in Milton Keynes | Wellcome Sanger Institute for the COVID-19 Genomics UK (COG-UK) Consortium | The Lighthouse Lab in Milton Keynes and Alex Alderton, Roberto Amato, Jeffrey Barrett, Sonia Goncalves, Ewan Harrison, David K. Jackson, Ian Johnston, Dominic Kwiatkowski, Cordelia Langford, John Sillitoe on behalf of the Wellcome Sanger Institute COVID-19 Surveillance Team |
| EPI_ISL_2851946 | Lighthouse Lab in Glasgow | Wellcome Sanger Institute for the COVID-19 Genomics UK (COG-UK) Consortium | Harper VanSteenhouse, Yumi Kasai, David Gray, Carol Clugston, Anna Dominiczak and Alex Alderton, Roberto Amato, Jeffrey Barrett, Sonia Goncalves, Ewan Harrison, David K. Jackson, Ian Johnston, Dominic Kwiatkowski, Cordelia Langford, John Sillitoe on behalf of the Wellcome Sanger Institute COVID-19 Surveillance Team |
| EPI_ISL_2851947 | Lighthouse Lab in Alderley Park | Wellcome Sanger Institute for the COVID-19 Genomics UK (COG-UK) Consortium | Jacquelyn Wynn, Mairead Hyland, The Lighthouse Lab in Alderley Park and Alex Alderton, Roberto Amato, Jeffrey Barrett, Sonia Goncalves, Ewan Harrison, David K. Jackson, Ian Johnston, Dominic Kwiatkowski, Cordelia Langford, John Sillitoe on behalf of the Wellcome Sanger Institute COVID-19 Surveillance Team |
| EPI_ISL_2851948, EPI_ISL_2851949 | Lighthouse Lab in Glasgow | Wellcome Sanger Institute for the COVID-19 Genomics UK (COG-UK) Consortium | Harper VanSteenhouse, Yumi Kasai, David Gray, Carol Clugston, Anna Dominiczak and Alex Alderton, Roberto Amato, Jeffrey Barrett, Sonia Goncalves, Ewan Harrison, David K. Jackson, Ian Johnston, Dominic Kwiatkowski, Cordelia Langford, John Sillitoe on behalf of the Wellcome Sanger Institute COVID-19 Surveillance Team |
| EPI_ISL_2851950 | Lighthouse Lab in Milton Keynes | Wellcome Sanger Institute for the COVID-19 Genomics UK (COG-UK) Consortium | The Lighthouse Lab in Milton Keynes and Alex Alderton, Roberto Amato, Jeffrey Barrett, Sonia Goncalves, Ewan Harrison, David K. Jackson, Ian Johnston, Dominic Kwiatkowski, Cordelia Langford, John Sillitoe on behalf of the Wellcome Sanger Institute COVID-19 Surveillance Team |
| EPI_ISL_2851951, EPI_ISL_2851952 | Lighthouse Lab in Glasgow | Wellcome Sanger Institute for the COVID-19 Genomics UK (COG-UK) Consortium | Harper VanSteenhouse, Yumi Kasai, David Gray, Carol Clugston, Anna Dominiczak and Alex Alderton, Roberto Amato, Jeffrey Barrett, Sonia Goncalves, Ewan Harrison, David K. Jackson, Ian Johnston, Dominic Kwiatkowski, Cordelia Langford, John Sillitoe on behalf of the Wellcome Sanger Institute COVID-19 Surveillance Team |
| EPI_ISL_2851953 | Lighthouse Lab in Milton Keynes | Wellcome Sanger Institute for the COVID-19 Genomics UK (COG-UK) Consortium | The Lighthouse Lab in Milton Keynes and Alex Alderton, Roberto Amato, Jeffrey Barrett, Sonia Goncalves, Ewan Harrison, David K. Jackson, Ian Johnston, Dominic Kwiatkowski, Cordelia Langford, John Sillitoe on behalf of the Wellcome Sanger Institute COVID-19 Surveillance Team |
| EPI_ISL_2851954 | Lighthouse Lab in Glasgow | Wellcome Sanger Institute for the COVID-19 Genomics UK (COG-UK) Consortium | Harper VanSteenhouse, Yumi Kasai, David Gray, Carol Clugston, Anna Dominiczak and Alex Alderton, Roberto Amato, Jeffrey Barrett, Sonia Goncalves, Ewan Harrison, David K. Jackson, Ian Johnston, Dominic Kwiatkowski, Cordelia Langford, John Sillitoe on behalf of the Wellcome Sanger Institute COVID-19 Surveillance Team |
| EPI_ISL_2851955 | Lighthouse Lab in Milton Keynes | Wellcome Sanger Institute for the COVID-19 Genomics UK (COG-UK) Consortium | The Lighthouse Lab in Milton Keynes and Alex Alderton, Roberto Amato, Jeffrey Barrett, Sonia Goncalves, Ewan Harrison, David K. Jackson, Ian Johnston, Dominic Kwiatkowski, Cordelia Langford, John Sillitoe on behalf of the Wellcome Sanger Institute COVID-19 Surveillance Team |
| EPI_ISL_2851956 | Lighthouse Lab in Glasgow | Wellcome Sanger Institute for the COVID-19 Genomics UK (COG-UK) Consortium | Harper VanSteenhouse, Yumi Kasai, David Gray, Carol Clugston, Anna Dominiczak and Alex Alderton, Roberto Amato, Jeffrey Barrett, Sonia Goncalves, Ewan Harrison, David K. Jackson, Ian Johnston, Dominic Kwiatkowski, Cordelia Langford, John Sillitoe on behalf of the Wellcome Sanger Institute COVID-19 Surveillance Team |
| EPI_ISL_2851957 | Lighthouse Lab in Milton Keynes | Wellcome Sanger Institute for the COVID-19 Genomics UK (COG-UK) Consortium | The Lighthouse Lab in Milton Keynes and Alex Alderton, Roberto Amato, Jeffrey Barrett, Sonia Goncalves, Ewan Harrison, David K. Jackson, Ian Johnston, Dominic Kwiatkowski, Cordelia Langford, John Sillitoe on behalf of the Wellcome Sanger Institute COVID-19 Surveillance Team |
| EPI_ISL_2851958 | Lighthouse Lab in Alderley Park | Wellcome Sanger Institute for the COVID-19 Genomics UK (COG-UK) Consortium | Jacquelyn Wynn, Mairead Hyland, The Lighthouse Lab in Alderley Park and Alex Alderton, Roberto Amato, Jeffrey Barrett, Sonia Goncalves, Ewan Harrison, David K. Jackson, Ian Johnston, Dominic Kwiatkowski, Cordelia Langford, John Sillitoe on behalf of the Wellcome Sanger Institute COVID-19 Surveillance Team |
| EPI_ISL_2851959 | Lighthouse Lab in Glasgow | Wellcome Sanger Institute for the COVID-19 Genomics UK (COG-UK) Consortium | Harper VanSteenhouse, Yumi Kasai, David Gray, Carol Clugston, Anna Dominiczak and Alex Alderton, Roberto Amato, Jeffrey Barrett, Sonia Goncalves, Ewan Harrison, David K. Jackson, Ian Johnston, Dominic Kwiatkowski, Cordelia Langford, John Sillitoe on behalf of the Wellcome Sanger Institute COVID-19 Surveillance Team |
| EPI_ISL_2851960, EPI_ISL_2851961, EPI_ISL_2851962 | Lighthouse Lab in Milton Keynes | Wellcome Sanger Institute for the COVID-19 Genomics UK (COG-UK) Consortium | The Lighthouse Lab in Milton Keynes and Alex Alderton, Roberto Amato, Jeffrey Barrett, Sonia Goncalves, Ewan Harrison, David K. Jackson, Ian Johnston, Dominic Kwiatkowski, Cordelia Langford, John Sillitoe on behalf of the Wellcome Sanger Institute COVID-19 Surveillance Team |
| EPI_ISL_2851963, EPI_ISL_2851964 | Lighthouse Lab in Glasgow | Wellcome Sanger Institute for the COVID-19 Genomics UK (COG-UK) Consortium | Harper VanSteenhouse, Yumi Kasai, David Gray, Carol Clugston, Anna Dominiczak and Alex Alderton, Roberto Amato, Jeffrey Barrett, Sonia Goncalves, Ewan Harrison, David K. Jackson, Ian Johnston, Dominic Kwiatkowski, Cordelia Langford, John Sillitoe on behalf of the Wellcome Sanger Institute COVID-19 Surveillance Team |
| EPI_ISL_2851965, EPI_ISL_2851966 | Lighthouse Lab in Alderley Park | Wellcome Sanger Institute for the COVID-19 Genomics UK (COG-UK) Consortium | Jacquelyn Wynn, Mairead Hyland, The Lighthouse Lab in Alderley Park and Alex Alderton, Roberto Amato, Jeffrey Barrett, Sonia Goncalves, Ewan Harrison, David K. Jackson, Ian Johnston, Dominic Kwiatkowski, Cordelia Langford, John Sillitoe on behalf of the Wellcome Sanger Institute COVID-19 Surveillance Team |
| EPI_ISL_2851967 | Lighthouse Lab in Milton Keynes | Wellcome Sanger Institute for the COVID-19 Genomics UK (COG-UK) Consortium | The Lighthouse Lab in Milton Keynes and Alex Alderton, Roberto Amato, Jeffrey Barrett, Sonia Goncalves, Ewan Harrison, David K. Jackson, Ian Johnston, Dominic Kwiatkowski, Cordelia Langford, John Sillitoe on behalf of the Wellcome Sanger Institute COVID-19 Surveillance Team |
| EPI_ISL_2851968 | Lighthouse Lab in Glasgow | Wellcome Sanger Institute for the COVID-19 Genomics UK (COG-UK) Consortium | Harper VanSteenhouse, Yumi Kasai, David Gray, Carol Clugston, Anna Dominiczak and Alex Alderton, Roberto Amato, Jeffrey Barrett, Sonia Goncalves, Ewan Harrison, David K. Jackson, Ian Johnston, Dominic Kwiatkowski, Cordelia Langford, John Sillitoe on behalf of the Wellcome Sanger Institute COVID-19 Surveillance Team |
| EPI_ISL_2851969 | Lighthouse Lab in Milton Keynes | Wellcome Sanger Institute for the COVID-19 Genomics UK (COG-UK) Consortium | The Lighthouse Lab in Milton Keynes and Alex Alderton, Roberto Amato, Jeffrey Barrett, Sonia Goncalves, Ewan Harrison, David K. Jackson, Ian Johnston, Dominic Kwiatkowski, Cordelia Langford, John Sillitoe on behalf of the Wellcome Sanger Institute COVID-19 Surveillance Team |
| EPI_ISL_2851970, EPI_ISL_2851971, EPI_ISL_2851972 | Lighthouse Lab in Glasgow | Wellcome Sanger Institute for the COVID-19 Genomics UK (COG-UK) Consortium | Harper VanSteenhouse, Yumi Kasai, David Gray, Carol Clugston, Anna Dominiczak and Alex Alderton, Roberto Amato, Jeffrey Barrett, Sonia Goncalves, Ewan Harrison, David K. Jackson, Ian Johnston, Dominic Kwiatkowski, Cordelia Langford, John Sillitoe on behalf of the Wellcome Sanger Institute COVID-19 Surveillance Team |
| EPI_ISL_2851973 | Lighthouse Lab in Alderley Park | Wellcome Sanger Institute for the COVID-19 Genomics UK (COG-UK) Consortium | Jacquelyn Wynn, Mairead Hyland, The Lighthouse Lab in Alderley Park and Alex Alderton, Roberto Amato, Jeffrey Barrett, Sonia Goncalves, Ewan Harrison, David K. Jackson, Ian Johnston, Dominic Kwiatkowski, Cordelia Langford, John Sillitoe on behalf of the Wellcome Sanger Institute COVID-19 Surveillance Team |
| EPI_ISL_2851974, EPI_ISL_2851975 | Lighthouse Lab in Glasgow | Wellcome Sanger Institute for the COVID-19 Genomics UK (COG-UK) Consortium | Harper VanSteenhouse, Yumi Kasai, David Gray, Carol Clugston, Anna Dominiczak and Alex Alderton, Roberto Amato, Jeffrey Barrett, Sonia Goncalves, Ewan Harrison, David K. Jackson, Ian Johnston, Dominic Kwiatkowski, Cordelia Langford, John Sillitoe on behalf of the Wellcome Sanger Institute COVID-19 Surveillance Team |
| EPI_ISL_2851976 | Lighthouse Lab in Milton Keynes | Wellcome Sanger Institute for the COVID-19 Genomics UK (COG-UK) Consortium | The Lighthouse Lab in Milton Keynes and Alex Alderton, Roberto Amato, Jeffrey Barrett, Sonia Goncalves, Ewan Harrison, David K. Jackson, Ian Johnston, Dominic Kwiatkowski, Cordelia Langford, John Sillitoe on behalf of the Wellcome Sanger Institute COVID-19 Surveillance Team |
| EPI_ISL_2851977, EPI_ISL_2851978 | Lighthouse Lab in Alderley Park | Wellcome Sanger Institute for the COVID-19 Genomics UK (COG-UK) Consortium | Jacquelyn Wynn, Mairead Hyland, The Lighthouse Lab in Alderley Park and Alex Alderton, Roberto Amato, Jeffrey Barrett, Sonia Goncalves, Ewan Harrison, David K. Jackson, Ian Johnston, Dominic Kwiatkowski, Cordelia Langford, John Sillitoe on behalf of the Wellcome Sanger Institute COVID-19 Surveillance Team |
| EPI_ISL_2851979, EPI_ISL_2851980, EPI_ISL_2851981, EPI_ISL_2851982 | Lighthouse Lab in Glasgow | Wellcome Sanger Institute for the COVID-19 Genomics UK (COG-UK) Consortium | Harper VanSteenhouse, Yumi Kasai, David Gray, Carol Clugston, Anna Dominiczak and Alex Alderton, Roberto Amato, Jeffrey Barrett, Sonia Goncalves, Ewan Harrison, David K. Jackson, Ian Johnston, Dominic Kwiatkowski, Cordelia Langford, John Sillitoe on behalf of the Wellcome Sanger Institute COVID-19 Surveillance Team |
| EPI_ISL_2851983, EPI_ISL_2851984 | Lighthouse Lab in Milton Keynes | Wellcome Sanger Institute for the COVID-19 Genomics UK (COG-UK) Consortium | The Lighthouse Lab in Milton Keynes and Alex Alderton, Roberto Amato, Jeffrey Barrett, Sonia Goncalves, Ewan Harrison, David K. Jackson, Ian Johnston, Dominic Kwiatkowski, Cordelia Langford, John Sillitoe on behalf of the Wellcome Sanger Institute COVID-19 Surveillance Team |
| EPI_ISL_2851985, EPI_ISL_2851986, EPI_ISL_2851987 | Lighthouse Lab in Glasgow | Wellcome Sanger Institute for the COVID-19 Genomics UK (COG-UK) Consortium | Harper VanSteenhouse, Yumi Kasai, David Gray, Carol Clugston, Anna Dominiczak and Alex Alderton, Roberto Amato, Jeffrey Barrett, Sonia Goncalves, Ewan Harrison, David K. Jackson, Ian Johnston, Dominic Kwiatkowski, Cordelia Langford, John Sillitoe on behalf of the Wellcome Sanger Institute COVID-19 Surveillance Team |
| EPI_ISL_2851988 | Lighthouse Lab in Milton Keynes | Wellcome Sanger Institute for the COVID-19 Genomics UK (COG-UK) Consortium | The Lighthouse Lab in Milton Keynes and Alex Alderton, Roberto Amato, Jeffrey Barrett, Sonia Goncalves, Ewan Harrison, David K. Jackson, Ian Johnston, Dominic Kwiatkowski, Cordelia Langford, John Sillitoe on behalf of the Wellcome Sanger Institute COVID-19 Surveillance Team |
| EPI_ISL_2851989 | Lighthouse Lab in Alderley Park | Wellcome Sanger Institute for the COVID-19 Genomics UK (COG-UK) Consortium | Jacquelyn Wynn, Mairead Hyland, The Lighthouse Lab in Alderley Park and Alex Alderton, Roberto Amato, Jeffrey Barrett, Sonia Goncalves, Ewan Harrison, David K. Jackson, Ian Johnston, Dominic Kwiatkowski, Cordelia Langford, John Sillitoe on behalf of the Wellcome Sanger Institute COVID-19 Surveillance Team |
| EPI_ISL_2851990 | Lighthouse Lab in Milton Keynes | Wellcome Sanger Institute for the COVID-19 Genomics UK (COG-UK) Consortium | The Lighthouse Lab in Milton Keynes and Alex Alderton, Roberto Amato, Jeffrey Barrett, Sonia Goncalves, Ewan Harrison, David K. Jackson, Ian Johnston, Dominic Kwiatkowski, Cordelia Langford, John Sillitoe on behalf of the Wellcome Sanger Institute COVID-19 Surveillance Team |
| EPI_ISL_2851991 | Lighthouse Lab in Alderley Park | Wellcome Sanger Institute for the COVID-19 Genomics UK (COG-UK) Consortium | Jacquelyn Wynn, Mairead Hyland, The Lighthouse Lab in Alderley Park and Alex Alderton, Roberto Amato, Jeffrey Barrett, Sonia Goncalves, Ewan Harrison, David K. Jackson, Ian Johnston, Dominic Kwiatkowski, Cordelia Langford, John Sillitoe on behalf of the Wellcome Sanger Institute COVID-19 Surveillance Team |
| EPI_ISL_2851992, EPI_ISL_2851993 | Lighthouse Lab in Glasgow | Wellcome Sanger Institute for the COVID-19 Genomics UK (COG-UK) Consortium | Harper VanSteenhouse, Yumi Kasai, David Gray, Carol Clugston, Anna Dominiczak and Alex Alderton, Roberto Amato, Jeffrey Barrett, Sonia Goncalves, Ewan Harrison, David K. Jackson, Ian Johnston, Dominic Kwiatkowski, Cordelia Langford, John Sillitoe on behalf of the Wellcome Sanger Institute COVID-19 Surveillance Team |
| EPI_ISL_2851994 | Lighthouse Lab in Alderley Park | Wellcome Sanger Institute for the COVID-19 Genomics UK (COG-UK) Consortium | Jacquelyn Wynn, Mairead Hyland, The Lighthouse Lab in Alderley Park and Alex Alderton, Roberto Amato, Jeffrey Barrett, Sonia Goncalves, Ewan Harrison, David K. Jackson, Ian Johnston, Dominic Kwiatkowski, Cordelia Langford, John Sillitoe on behalf of the Wellcome Sanger Institute COVID-19 Surveillance Team |
| EPI_ISL_2851995 | Lighthouse Lab in Glasgow | Wellcome Sanger Institute for the COVID-19 Genomics UK (COG-UK) Consortium | Harper VanSteenhouse, Yumi Kasai, David Gray, Carol Clugston, Anna Dominiczak and Alex Alderton, Roberto Amato, Jeffrey Barrett, Sonia Goncalves, Ewan Harrison, David K. Jackson, Ian Johnston, Dominic Kwiatkowski, Cordelia Langford, John Sillitoe on behalf of the Wellcome Sanger Institute COVID-19 Surveillance Team |
| EPI_ISL_2851996 | Lighthouse Lab in Alderley Park | Wellcome Sanger Institute for the COVID-19 Genomics UK (COG-UK) Consortium | Jacquelyn Wynn, Mairead Hyland, The Lighthouse Lab in Alderley Park and Alex Alderton, Roberto Amato, Jeffrey Barrett, Sonia Goncalves, Ewan Harrison, David K. Jackson, Ian Johnston, Dominic Kwiatkowski, Cordelia Langford, John Sillitoe on behalf of the Wellcome Sanger Institute COVID-19 Surveillance Team |
| EPI_ISL_2851997 | Lighthouse Lab in Glasgow | Wellcome Sanger Institute for the COVID-19 Genomics UK (COG-UK) Consortium | Harper VanSteenhouse, Yumi Kasai, David Gray, Carol Clugston, Anna Dominiczak and Alex Alderton, Roberto Amato, Jeffrey Barrett, Sonia Goncalves, Ewan Harrison, David K. Jackson, Ian Johnston, Dominic Kwiatkowski, Cordelia Langford, John Sillitoe on behalf of the Wellcome Sanger Institute COVID-19 Surveillance Team |
| EPI_ISL_2851998 | Lighthouse Lab in Milton Keynes | Wellcome Sanger Institute for the COVID-19 Genomics UK (COG-UK) Consortium | The Lighthouse Lab in Milton Keynes and Alex Alderton, Roberto Amato, Jeffrey Barrett, Sonia Goncalves, Ewan Harrison, David K. Jackson, Ian Johnston, Dominic Kwiatkowski, Cordelia Langford, John Sillitoe on behalf of the Wellcome Sanger Institute COVID-19 Surveillance Team |
| EPI_ISL_2851999, EPI_ISL_2852000, EPI_ISL_2852001, EPI_ISL_2852002, | Lighthouse Lab in Alderley Park | Wellcome Sanger Institute for the COVID-19 Genomics UK (COG-UK) Consortium | Jacquelyn Wynn, Mairead Hyland, The Lighthouse Lab in Alderley Park and Alex Alderton, Roberto Amato, Jeffrey Barrett, Sonia Goncalves, Ewan Harrison, David K. Jackson, Ian Johnston, Dominic Kwiatkowski, Cordelia Langford, John Sillitoe on behalf of the Wellcome Sanger Institute COVID-19 Surveillance Team |

| | | | |
|---|---|---|---|
| EPI_ISL_2852003, EPI_ISL_2852004, EPI_ISL_2852005 | | | |
| EPI_ISL_2852006 | Lighthouse Lab in Glasgow | Wellcome Sanger Institute for the COVID-19 Genomics UK (COG-UK) Consortium | Harper VanSteenhouse, Yumi Kasai, David Gray, Carol Clugston, Anna Dominiczak and Alex Alderton, Roberto Amato, Jeffrey Barrett, Sonia Goncalves, Ewan Harrison, David K. Jackson, Ian Johnston, Dominic Kwiatkowski, Cordelia Langford, John Sillitoe on behalf of the Wellcome Sanger Institute COVID-19 Surveillance Team |
| EPI_ISL_2852007 | Lighthouse Lab in Milton Keynes | Wellcome Sanger Institute for the COVID-19 Genomics UK (COG-UK) Consortium | The Lighthouse Lab in Milton Keynes and Alex Alderton, Roberto Amato, Jeffrey Barrett, Sonia Goncalves, Ewan Harrison, David K. Jackson, Ian Johnston, Dominic Kwiatkowski, Cordelia Langford, John Sillitoe on behalf of the Wellcome Sanger Institute COVID-19 Surveillance Team |
| EPI_ISL_2852008 | Lighthouse Lab in Alderley Park | Wellcome Sanger Institute for the COVID-19 Genomics UK (COG-UK) Consortium | Jacquelyn Wynn, Mairead Hyland, The Lighthouse Lab in Alderley Park and Alex Alderton, Roberto Amato, Jeffrey Barrett, Sonia Goncalves, Ewan Harrison, David K. Jackson, Ian Johnston, Dominic Kwiatkowski, Cordelia Langford, John Sillitoe on behalf of the Wellcome Sanger Institute COVID-19 Surveillance Team |
| EPI_ISL_2852009, EPI_ISL_2852010 | Lighthouse Lab in Glasgow | Wellcome Sanger Institute for the COVID-19 Genomics UK (COG-UK) Consortium | Harper VanSteenhouse, Yumi Kasai, David Gray, Carol Clugston, Anna Dominiczak and Alex Alderton, Roberto Amato, Jeffrey Barrett, Sonia Goncalves, Ewan Harrison, David K. Jackson, Ian Johnston, Dominic Kwiatkowski, Cordelia Langford, John Sillitoe on behalf of the Wellcome Sanger Institute COVID-19 Surveillance Team |
| EPI_ISL_2852011 | Lighthouse Lab in Alderley Park | Wellcome Sanger Institute for the COVID-19 Genomics UK (COG-UK) Consortium | Jacquelyn Wynn, Mairead Hyland, The Lighthouse Lab in Alderley Park and Alex Alderton, Roberto Amato, Jeffrey Barrett, Sonia Goncalves, Ewan Harrison, David K. Jackson, Ian Johnston, Dominic Kwiatkowski, Cordelia Langford, John Sillitoe on behalf of the Wellcome Sanger Institute COVID-19 Surveillance Team |
| EPI_ISL_2852012 | Lighthouse Lab in Glasgow | Wellcome Sanger Institute for the COVID-19 Genomics UK (COG-UK) Consortium | Harper VanSteenhouse, Yumi Kasai, David Gray, Carol Clugston, Anna Dominiczak and Alex Alderton, Roberto Amato, Jeffrey Barrett, Sonia Goncalves, Ewan Harrison, David K. Jackson, Ian Johnston, Dominic Kwiatkowski, Cordelia Langford, John Sillitoe on behalf of the Wellcome Sanger Institute COVID-19 Surveillance Team |
| EPI_ISL_2852013 | Lighthouse Lab in Milton Keynes | Wellcome Sanger Institute for the COVID-19 Genomics UK (COG-UK) Consortium | The Lighthouse Lab in Milton Keynes and Alex Alderton, Roberto Amato, Jeffrey Barrett, Sonia Goncalves, Ewan Harrison, David K. Jackson, Ian Johnston, Dominic Kwiatkowski, Cordelia Langford, John Sillitoe on behalf of the Wellcome Sanger Institute COVID-19 Surveillance Team |
| EPI_ISL_2852014 | Lighthouse Lab in Alderley Park | Wellcome Sanger Institute for the COVID-19 Genomics UK (COG-UK) Consortium | Jacquelyn Wynn, Mairead Hyland, The Lighthouse Lab in Alderley Park and Alex Alderton, Roberto Amato, Jeffrey Barrett, Sonia Goncalves, Ewan Harrison, David K. Jackson, Ian Johnston, Dominic Kwiatkowski, Cordelia Langford, John Sillitoe on behalf of the Wellcome Sanger Institute COVID-19 Surveillance Team |
| EPI_ISL_2852015, EPI_ISL_2852016, EPI_ISL_2852018, EPI_ISL_2852019 | Lighthouse Lab in Glasgow | Wellcome Sanger Institute for the COVID-19 Genomics UK (COG-UK) Consortium | Harper VanSteenhouse, Yumi Kasai, David Gray, Carol Clugston, Anna Dominiczak and Alex Alderton, Roberto Amato, Jeffrey Barrett, Sonia Goncalves, Ewan Harrison, David K. Jackson, Ian Johnston, Dominic Kwiatkowski, Cordelia Langford, John Sillitoe on behalf of the Wellcome Sanger Institute COVID-19 Surveillance Team |
| EPI_ISL_2852020 | Lighthouse Lab in Alderley Park | Wellcome Sanger Institute for the COVID-19 Genomics UK (COG-UK) Consortium | Jacquelyn Wynn, Mairead Hyland, The Lighthouse Lab in Alderley Park and Alex Alderton, Roberto Amato, Jeffrey Barrett, Sonia Goncalves, Ewan Harrison, David K. Jackson, Ian Johnston, Dominic Kwiatkowski, Cordelia Langford, John Sillitoe on behalf of the Wellcome Sanger Institute COVID-19 Surveillance Team |
| EPI_ISL_2852021 | Lighthouse Lab in Glasgow | Wellcome Sanger Institute for the COVID-19 Genomics UK (COG-UK) Consortium | Harper VanSteenhouse, Yumi Kasai, David Gray, Carol Clugston, Anna Dominiczak and Alex Alderton, Roberto Amato, Jeffrey Barrett, Sonia Goncalves, Ewan Harrison, David K. Jackson, Ian Johnston, Dominic Kwiatkowski, Cordelia Langford, John Sillitoe on behalf of the Wellcome Sanger Institute COVID-19 Surveillance Team |
| EPI_ISL_2852022, EPI_ISL_2852023 | Lighthouse Lab in Alderley Park | Wellcome Sanger Institute for the COVID-19 Genomics UK (COG-UK) Consortium | Jacquelyn Wynn, Mairead Hyland, The Lighthouse Lab in Alderley Park and Alex Alderton, Roberto Amato, Jeffrey Barrett, Sonia Goncalves, Ewan Harrison, David K. Jackson, Ian Johnston, Dominic Kwiatkowski, Cordelia Langford, John Sillitoe on behalf of the Wellcome Sanger Institute COVID-19 Surveillance Team |
| EPI_ISL_2852024 | Lighthouse Lab in Glasgow | Wellcome Sanger Institute for the COVID-19 Genomics UK (COG-UK) Consortium | Harper VanSteenhouse, Yumi Kasai, David Gray, Carol Clugston, Anna Dominiczak and Alex Alderton, Roberto Amato, Jeffrey Barrett, Sonia Goncalves, Ewan Harrison, David K. Jackson, Ian Johnston, Dominic Kwiatkowski, Cordelia Langford, John Sillitoe on behalf of the Wellcome Sanger Institute COVID-19 Surveillance Team |
| EPI_ISL_2852025, EPI_ISL_2852026 | Lighthouse Lab in Alderley Park | Wellcome Sanger Institute for the COVID-19 Genomics UK (COG-UK) Consortium | Jacquelyn Wynn, Mairead Hyland, The Lighthouse Lab in Alderley Park and Alex Alderton, Roberto Amato, Jeffrey Barrett, Sonia Goncalves, Ewan Harrison, David K. Jackson, Ian Johnston, Dominic Kwiatkowski, Cordelia Langford, John Sillitoe on behalf of the Wellcome Sanger Institute COVID-19 Surveillance Team |
| EPI_ISL_2852027 | Lighthouse Lab in Glasgow | Wellcome Sanger Institute for the COVID-19 Genomics UK (COG-UK) Consortium | Harper VanSteenhouse, Yumi Kasai, David Gray, Carol Clugston, Anna Dominiczak and Alex Alderton, Roberto Amato, Jeffrey Barrett, Sonia Goncalves, Ewan Harrison, David K. Jackson, Ian Johnston, Dominic Kwiatkowski, Cordelia Langford, John Sillitoe on behalf of the Wellcome Sanger Institute COVID-19 Surveillance Team |
| EPI_ISL_2852028 | Lighthouse Lab in Alderley Park | Wellcome Sanger Institute for the COVID-19 Genomics UK (COG-UK) Consortium | Jacquelyn Wynn, Mairead Hyland, The Lighthouse Lab in Alderley Park and Alex Alderton, Roberto Amato, Jeffrey Barrett, Sonia Goncalves, Ewan Harrison, David K. Jackson, Ian Johnston, Dominic Kwiatkowski, Cordelia Langford, John Sillitoe on behalf of the Wellcome Sanger Institute COVID-19 Surveillance Team |
| EPI_ISL_2852029 | Lighthouse Lab in Milton Keynes | Wellcome Sanger Institute for the COVID-19 Genomics UK (COG-UK) Consortium | The Lighthouse Lab in Milton Keynes and Alex Alderton, Roberto Amato, Jeffrey Barrett, Sonia Goncalves, Ewan Harrison, David K. Jackson, Ian Johnston, Dominic Kwiatkowski, Cordelia Langford, John Sillitoe on behalf of the Wellcome Sanger Institute COVID-19 Surveillance Team |
| EPI_ISL_2852030, EPI_ISL_2852031 | Lighthouse Lab in Glasgow | Wellcome Sanger Institute for the COVID-19 Genomics UK (COG-UK) Consortium | Harper VanSteenhouse, Yumi Kasai, David Gray, Carol Clugston, Anna Dominiczak and Alex Alderton, Roberto Amato, Jeffrey Barrett, Sonia Goncalves, Ewan Harrison, David K. Jackson, Ian Johnston, Dominic Kwiatkowski, Cordelia Langford, John Sillitoe on behalf of the Wellcome Sanger Institute COVID-19 Surveillance Team |
| EPI_ISL_2852032 | Lighthouse Lab in Milton Keynes | Wellcome Sanger Institute for the COVID-19 Genomics UK (COG-UK) Consortium | The Lighthouse Lab in Milton Keynes and Alex Alderton, Roberto Amato, Jeffrey Barrett, Sonia Goncalves, Ewan Harrison, David K. Jackson, Ian Johnston, Dominic Kwiatkowski, Cordelia Langford, John Sillitoe on behalf of the Wellcome Sanger Institute COVID-19 Surveillance Team |
| EPI_ISL_2852033, EPI_ISL_2852034 | Lighthouse Lab in Glasgow | Wellcome Sanger Institute for the COVID-19 Genomics UK (COG-UK) Consortium | Harper VanSteenhouse, Yumi Kasai, David Gray, Carol Clugston, Anna Dominiczak and Alex Alderton, Roberto Amato, Jeffrey Barrett, Sonia Goncalves, Ewan Harrison, David K. Jackson, Ian Johnston, Dominic Kwiatkowski, Cordelia Langford, John Sillitoe on behalf of the Wellcome Sanger Institute COVID-19 Surveillance Team |
| EPI_ISL_2852035 | Lighthouse Lab in Milton Keynes | Wellcome Sanger Institute for the COVID-19 Genomics UK (COG-UK) Consortium | The Lighthouse Lab in Milton Keynes and Alex Alderton, Roberto Amato, Jeffrey Barrett, Sonia Goncalves, Ewan Harrison, David K. Jackson, Ian Johnston, Dominic Kwiatkowski, Cordelia Langford, John Sillitoe on behalf of the Wellcome Sanger Institute COVID-19 Surveillance Team |
| EPI_ISL_2852036 | Lighthouse Lab in Glasgow | Wellcome Sanger Institute for the COVID-19 Genomics UK (COG-UK) Consortium | Harper VanSteenhouse, Yumi Kasai, David Gray, Carol Clugston, Anna Dominiczak and Alex Alderton, Roberto Amato, Jeffrey Barrett, Sonia Goncalves, Ewan Harrison, David K. Jackson, Ian Johnston, Dominic Kwiatkowski, Cordelia Langford, John Sillitoe on behalf of the Wellcome Sanger Institute COVID-19 Surveillance Team |
| EPI_ISL_2852037 | Lighthouse Lab in Alderley Park | Wellcome Sanger Institute for the COVID-19 Genomics UK (COG-UK) Consortium | Jacquelyn Wynn, Mairead Hyland, The Lighthouse Lab in Alderley Park and Alex Alderton, Roberto Amato, Jeffrey Barrett, Sonia Goncalves, Ewan Harrison, David K. Jackson, Ian Johnston, Dominic Kwiatkowski, Cordelia Langford, John Sillitoe on behalf of the Wellcome Sanger Institute COVID-19 Surveillance Team |
| EPI_ISL_2852038 | Lighthouse Lab in Milton Keynes | Wellcome Sanger Institute for the COVID-19 Genomics UK (COG-UK) Consortium | The Lighthouse Lab in Milton Keynes and Alex Alderton, Roberto Amato, Jeffrey Barrett, Sonia Goncalves, Ewan Harrison, David K. Jackson, Ian Johnston, Dominic Kwiatkowski, Cordelia Langford, John Sillitoe on behalf of the Wellcome Sanger Institute COVID-19 Surveillance Team |
| EPI_ISL_2852039, EPI_ISL_2852040 | Lighthouse Lab in Alderley Park | Wellcome Sanger Institute for the COVID-19 Genomics UK (COG-UK) Consortium | Jacquelyn Wynn, Mairead Hyland, The Lighthouse Lab in Alderley Park and Alex Alderton, Roberto Amato, Jeffrey Barrett, Sonia Goncalves, Ewan Harrison, David K. Jackson, Ian Johnston, Dominic Kwiatkowski, Cordelia Langford, John Sillitoe on behalf of the Wellcome Sanger Institute COVID-19 Surveillance Team |
| EPI_ISL_2852041, EPI_ISL_2852042, EPI_ISL_2852043 | Lighthouse Lab in Glasgow | Wellcome Sanger Institute for the COVID-19 Genomics UK (COG-UK) Consortium | Harper VanSteenhouse, Yumi Kasai, David Gray, Carol Clugston, Anna Dominiczak and Alex Alderton, Roberto Amato, Jeffrey Barrett, Sonia Goncalves, Ewan Harrison, David K. Jackson, Ian Johnston, Dominic Kwiatkowski, Cordelia Langford, John Sillitoe on behalf of the Wellcome Sanger Institute COVID-19 Surveillance Team |
| EPI_ISL_2852044, EPI_ISL_2852045 | Lighthouse Lab in Alderley Park | Wellcome Sanger Institute for the COVID-19 Genomics UK (COG-UK) Consortium | Jacquelyn Wynn, Mairead Hyland, The Lighthouse Lab in Alderley Park and Alex Alderton, Roberto Amato, Jeffrey Barrett, Sonia Goncalves, Ewan Harrison, David K. Jackson, Ian Johnston, Dominic Kwiatkowski, Cordelia Langford, John Sillitoe on behalf of the Wellcome Sanger Institute COVID-19 Surveillance Team |
| EPI_ISL_2852046, EPI_ISL_2852047 | Lighthouse Lab in Glasgow | Wellcome Sanger Institute for the COVID-19 Genomics UK (COG-UK) Consortium | Harper VanSteenhouse, Yumi Kasai, David Gray, Carol Clugston, Anna Dominiczak and Alex Alderton, Roberto Amato, Jeffrey Barrett, Sonia Goncalves, Ewan Harrison, David K. Jackson, Ian Johnston, Dominic Kwiatkowski, Cordelia Langford, John Sillitoe on behalf of the Wellcome Sanger Institute COVID-19 Surveillance Team |
| EPI_ISL_2852048, EPI_ISL_2852049 | Lighthouse Lab in Alderley Park | Wellcome Sanger Institute for the COVID-19 Genomics UK (COG-UK) Consortium | Jacquelyn Wynn, Mairead Hyland, The Lighthouse Lab in Alderley Park and Alex Alderton, Roberto Amato, Jeffrey Barrett, Sonia Goncalves, Ewan Harrison, David K. Jackson, Ian Johnston, Dominic Kwiatkowski, Cordelia Langford, John Sillitoe on behalf of the Wellcome Sanger Institute COVID-19 Surveillance Team |
| EPI_ISL_2852050, EPI_ISL_2852051 | Lighthouse Lab in Glasgow | Wellcome Sanger Institute for the COVID-19 Genomics UK (COG-UK) Consortium | Harper VanSteenhouse, Yumi Kasai, David Gray, Carol Clugston, Anna Dominiczak and Alex Alderton, Roberto Amato, Jeffrey Barrett, Sonia Goncalves, Ewan Harrison, David K. Jackson, Ian Johnston, Dominic Kwiatkowski, Cordelia Langford, John Sillitoe on behalf of the Wellcome Sanger Institute COVID-19 Surveillance Team |
| EPI_ISL_2852052 | Lighthouse Lab in Alderley Park | Wellcome Sanger Institute for the COVID-19 Genomics UK (COG-UK) Consortium | Jacquelyn Wynn, Mairead Hyland, The Lighthouse Lab in Alderley Park and Alex Alderton, Roberto Amato, Jeffrey Barrett, Sonia Goncalves, Ewan Harrison, David K. Jackson, Ian Johnston, Dominic Kwiatkowski, Cordelia Langford, John Sillitoe on behalf of the Wellcome Sanger Institute COVID-19 Surveillance Team |
| EPI_ISL_2852053, EPI_ISL_2852054 | Lighthouse Lab in Milton Keynes | Wellcome Sanger Institute for the COVID-19 Genomics UK (COG-UK) Consortium | The Lighthouse Lab in Milton Keynes and Alex Alderton, Roberto Amato, Jeffrey Barrett, Sonia Goncalves, Ewan Harrison, David K. Jackson, Ian Johnston, Dominic Kwiatkowski, Cordelia Langford, John Sillitoe on behalf of the Wellcome Sanger Institute COVID-19 Surveillance Team |
| EPI_ISL_2852055, EPI_ISL_2852056 | Lighthouse Lab in Alderley Park | Wellcome Sanger Institute for the COVID-19 Genomics UK (COG-UK) Consortium | Jacquelyn Wynn, Mairead Hyland, The Lighthouse Lab in Alderley Park and Alex Alderton, Roberto Amato, Jeffrey Barrett, Sonia Goncalves, Ewan Harrison, David K. Jackson, Ian Johnston, Dominic Kwiatkowski, Cordelia Langford, John Sillitoe on behalf of the Wellcome Sanger Institute COVID-19 Surveillance Team |
| EPI_ISL_2852057 | Lighthouse Lab in Glasgow | Wellcome Sanger Institute for the COVID-19 Genomics UK (COG-UK) Consortium | Harper VanSteenhouse, Yumi Kasai, David Gray, Carol Clugston, Anna Dominiczak and Alex Alderton, Roberto Amato, Jeffrey Barrett, Sonia Goncalves, Ewan Harrison, David K. Jackson, Ian Johnston, Dominic Kwiatkowski, Cordelia Langford, John Sillitoe on behalf of the Wellcome Sanger Institute COVID-19 Surveillance Team |
| EPI_ISL_2852058 | Lighthouse Lab in Milton Keynes | Wellcome Sanger Institute for the COVID-19 Genomics UK (COG-UK) Consortium | The Lighthouse Lab in Milton Keynes and Alex Alderton, Roberto Amato, Jeffrey Barrett, Sonia Goncalves, Ewan Harrison, David K. Jackson, Ian Johnston, Dominic Kwiatkowski, Cordelia Langford, John Sillitoe on behalf of the Wellcome Sanger Institute COVID-19 Surveillance Team |
| EPI_ISL_2852059, EPI_ISL_2852060 | Lighthouse Lab in Glasgow | Wellcome Sanger Institute for the COVID-19 Genomics UK (COG-UK) Consortium | Harper VanSteenhouse, Yumi Kasai, David Gray, Carol Clugston, Anna Dominiczak and Alex Alderton, Roberto Amato, Jeffrey Barrett, Sonia Goncalves, Ewan Harrison, David K. Jackson, Ian Johnston, Dominic Kwiatkowski, Cordelia Langford, John Sillitoe on behalf of the Wellcome Sanger Institute COVID-19 Surveillance Team |
| EPI_ISL_2852061 | Lighthouse Lab in Alderley Park | Wellcome Sanger Institute for the COVID-19 Genomics UK (COG-UK) Consortium | Jacquelyn Wynn, Mairead Hyland, The Lighthouse Lab in Alderley Park and Alex Alderton, Roberto Amato, Jeffrey Barrett, Sonia Goncalves, Ewan Harrison, David K. Jackson, Ian Johnston, Dominic Kwiatkowski, Cordelia Langford, John Sillitoe on behalf of the Wellcome Sanger Institute COVID-19 Surveillance Team |
| EPI_ISL_2852062 | Lighthouse Lab in Glasgow | Wellcome Sanger Institute for the COVID-19 Genomics UK (COG-UK) Consortium | Harper VanSteenhouse, Yumi Kasai, David Gray, Carol Clugston, Anna Dominiczak and Alex Alderton, Roberto Amato, Jeffrey Barrett, Sonia Goncalves, Ewan Harrison, David K. Jackson, Ian Johnston, Dominic Kwiatkowski, Cordelia Langford, John Sillitoe on behalf of the Wellcome Sanger Institute COVID-19 Surveillance Team |
| EPI_ISL_2852063, EPI_ISL_2852064, EPI_ISL_2852065 | Lighthouse Lab in Alderley Park | Wellcome Sanger Institute for the COVID-19 Genomics UK (COG-UK) Consortium | Jacquelyn Wynn, Mairead Hyland, The Lighthouse Lab in Alderley Park and Alex Alderton, Roberto Amato, Jeffrey Barrett, Sonia Goncalves, Ewan Harrison, David K. Jackson, Ian Johnston, Dominic Kwiatkowski, Cordelia Langford, John Sillitoe on behalf of the Wellcome Sanger Institute COVID-19 Surveillance Team |
| EPI_ISL_2852066 | Lighthouse Lab in Glasgow | Wellcome Sanger Institute for the COVID-19 Genomics UK (COG-UK) Consortium | Harper VanSteenhouse, Yumi Kasai, David Gray, Carol Clugston, Anna Dominiczak and Alex Alderton, Roberto Amato, Jeffrey Barrett, Sonia Goncalves, Ewan Harrison, David K. Jackson, Ian Johnston, Dominic Kwiatkowski, Cordelia Langford, John Sillitoe on behalf of the Wellcome Sanger Institute COVID-19 Surveillance Team |
| EPI_ISL_2852067, EPI_ISL_2852068, EPI_ISL_2852069, EPI_ISL_2852070 | Lighthouse Lab in Alderley Park | Wellcome Sanger Institute for the COVID-19 Genomics UK (COG-UK) Consortium | Jacquelyn Wynn, Mairead Hyland, The Lighthouse Lab in Alderley Park and Alex Alderton, Roberto Amato, Jeffrey Barrett, Sonia Goncalves, Ewan Harrison, David K. Jackson, Ian Johnston, Dominic Kwiatkowski, Cordelia Langford, John Sillitoe on behalf of the Wellcome Sanger Institute COVID-19 Surveillance Team |

| | | | |
|---|---|---|---|
| EPI_ISL_2852071, EPI_ISL_2852072, EPI_ISL_2852073 | Lighthouse Lab in Glasgow | Wellcome Sanger Institute for the COVID-19 Genomics UK (COG-UK) Consortium | Harper VanSteenhouse, Yumi Kasai, David Gray, Carol Clugston, Anna Dominiczak and Alex Alderton, Roberto Amato, Jeffrey Barrett, Sonia Goncalves, Ewan Harrison, David K. Jackson, Ian Johnston, Dominic Kwiatkowski, Cordelia Langford, John Sillitoe on behalf of the Wellcome Sanger Institute COVID-19 Surveillance Team |
| EPI_ISL_2852074 | Lighthouse Lab in Milton Keynes | Wellcome Sanger Institute for the COVID-19 Genomics UK (COG-UK) Consortium | The Lighthouse Lab in Milton Keynes and Alex Alderton, Roberto Amato, Jeffrey Barrett, Sonia Goncalves, Ewan Harrison, David K. Jackson, Ian Johnston, Dominic Kwiatkowski, Cordelia Langford, John Sillitoe on behalf of the Wellcome Sanger Institute COVID-19 Surveillance Team |
| EPI_ISL_2852075, EPI_ISL_2852076 | Lighthouse Lab in Alderley Park | Wellcome Sanger Institute for the COVID-19 Genomics UK (COG-UK) Consortium | Jacquelyn Wynn, Mairead Hyland, The Lighthouse Lab in Alderley Park and Alex Alderton, Roberto Amato, Jeffrey Barrett, Sonia Goncalves, Ewan Harrison, David K. Jackson, Ian Johnston, Dominic Kwiatkowski, Cordelia Langford, John Sillitoe on behalf of the Wellcome Sanger Institute COVID-19 Surveillance Team |
| EPI_ISL_2852077 | Lighthouse Lab in Milton Keynes | Wellcome Sanger Institute for the COVID-19 Genomics UK (COG-UK) Consortium | The Lighthouse Lab in Milton Keynes and Alex Alderton, Roberto Amato, Jeffrey Barrett, Sonia Goncalves, Ewan Harrison, David K. Jackson, Ian Johnston, Dominic Kwiatkowski, Cordelia Langford, John Sillitoe on behalf of the Wellcome Sanger Institute COVID-19 Surveillance Team |
| EPI_ISL_2852078 | Lighthouse Lab in Glasgow | Wellcome Sanger Institute for the COVID-19 Genomics UK (COG-UK) Consortium | Harper VanSteenhouse, Yumi Kasai, David Gray, Carol Clugston, Anna Dominiczak and Alex Alderton, Roberto Amato, Jeffrey Barrett, Sonia Goncalves, Ewan Harrison, David K. Jackson, Ian Johnston, Dominic Kwiatkowski, Cordelia Langford, John Sillitoe on behalf of the Wellcome Sanger Institute COVID-19 Surveillance Team |
| EPI_ISL_2852079 | Lighthouse Lab in Milton Keynes | Wellcome Sanger Institute for the COVID-19 Genomics UK (COG-UK) Consortium | The Lighthouse Lab in Milton Keynes and Alex Alderton, Roberto Amato, Jeffrey Barrett, Sonia Goncalves, Ewan Harrison, David K. Jackson, Ian Johnston, Dominic Kwiatkowski, Cordelia Langford, John Sillitoe on behalf of the Wellcome Sanger Institute COVID-19 Surveillance Team |
| EPI_ISL_2852080 | Lighthouse Lab in Alderley Park | Wellcome Sanger Institute for the COVID-19 Genomics UK (COG-UK) Consortium | Jacquelyn Wynn, Mairead Hyland, The Lighthouse Lab in Alderley Park and Alex Alderton, Roberto Amato, Jeffrey Barrett, Sonia Goncalves, Ewan Harrison, David K. Jackson, Ian Johnston, Dominic Kwiatkowski, Cordelia Langford, John Sillitoe on behalf of the Wellcome Sanger Institute COVID-19 Surveillance Team |
| EPI_ISL_2852081, EPI_ISL_2852082 | Lighthouse Lab in Milton Keynes | Wellcome Sanger Institute for the COVID-19 Genomics UK (COG-UK) Consortium | The Lighthouse Lab in Milton Keynes and Alex Alderton, Roberto Amato, Jeffrey Barrett, Sonia Goncalves, Ewan Harrison, David K. Jackson, Ian Johnston, Dominic Kwiatkowski, Cordelia Langford, John Sillitoe on behalf of the Wellcome Sanger Institute COVID-19 Surveillance Team |
| EPI_ISL_2852083, EPI_ISL_2852084 | Lighthouse Lab in Glasgow | Wellcome Sanger Institute for the COVID-19 Genomics UK (COG-UK) Consortium | Harper VanSteenhouse, Yumi Kasai, David Gray, Carol Clugston, Anna Dominiczak and Alex Alderton, Roberto Amato, Jeffrey Barrett, Sonia Goncalves, Ewan Harrison, David K. Jackson, Ian Johnston, Dominic Kwiatkowski, Cordelia Langford, John Sillitoe on behalf of the Wellcome Sanger Institute COVID-19 Surveillance Team |
| EPI_ISL_2852085 | Lighthouse Lab in Milton Keynes | Wellcome Sanger Institute for the COVID-19 Genomics UK (COG-UK) Consortium | The Lighthouse Lab in Milton Keynes and Alex Alderton, Roberto Amato, Jeffrey Barrett, Sonia Goncalves, Ewan Harrison, David K. Jackson, Ian Johnston, Dominic Kwiatkowski, Cordelia Langford, John Sillitoe on behalf of the Wellcome Sanger Institute COVID-19 Surveillance Team |
| EPI_ISL_2852086, EPI_ISL_2852087, EPI_ISL_2852088, EPI_ISL_2852089 | Lighthouse Lab in Glasgow | Wellcome Sanger Institute for the COVID-19 Genomics UK (COG-UK) Consortium | Harper VanSteenhouse, Yumi Kasai, David Gray, Carol Clugston, Anna Dominiczak and Alex Alderton, Roberto Amato, Jeffrey Barrett, Sonia Goncalves, Ewan Harrison, David K. Jackson, Ian Johnston, Dominic Kwiatkowski, Cordelia Langford, John Sillitoe on behalf of the Wellcome Sanger Institute COVID-19 Surveillance Team |
| EPI_ISL_2852090 | Lighthouse Lab in Alderley Park | Wellcome Sanger Institute for the COVID-19 Genomics UK (COG-UK) Consortium | Jacquelyn Wynn, Mairead Hyland, The Lighthouse Lab in Alderley Park and Alex Alderton, Roberto Amato, Jeffrey Barrett, Sonia Goncalves, Ewan Harrison, David K. Jackson, Ian Johnston, Dominic Kwiatkowski, Cordelia Langford, John Sillitoe on behalf of the Wellcome Sanger Institute COVID-19 Surveillance Team |
| EPI_ISL_2852091 | Lighthouse Lab in Glasgow | Wellcome Sanger Institute for the COVID-19 Genomics UK (COG-UK) Consortium | Harper VanSteenhouse, Yumi Kasai, David Gray, Carol Clugston, Anna Dominiczak and Alex Alderton, Roberto Amato, Jeffrey Barrett, Sonia Goncalves, Ewan Harrison, David K. Jackson, Ian Johnston, Dominic Kwiatkowski, Cordelia Langford, John Sillitoe on behalf of the Wellcome Sanger Institute COVID-19 Surveillance Team |
| EPI_ISL_2852092 | Lighthouse Lab in Alderley Park | Wellcome Sanger Institute for the COVID-19 Genomics UK (COG-UK) Consortium | Jacquelyn Wynn, Mairead Hyland, The Lighthouse Lab in Alderley Park and Alex Alderton, Roberto Amato, Jeffrey Barrett, Sonia Goncalves, Ewan Harrison, David K. Jackson, Ian Johnston, Dominic Kwiatkowski, Cordelia Langford, John Sillitoe on behalf of the Wellcome Sanger Institute COVID-19 Surveillance Team |
| EPI_ISL_2852093, EPI_ISL_2852094 | Lighthouse Lab in Glasgow | Wellcome Sanger Institute for the COVID-19 Genomics UK (COG-UK) Consortium | Harper VanSteenhouse, Yumi Kasai, David Gray, Carol Clugston, Anna Dominiczak and Alex Alderton, Roberto Amato, Jeffrey Barrett, Sonia Goncalves, Ewan Harrison, David K. Jackson, Ian Johnston, Dominic Kwiatkowski, Cordelia Langford, John Sillitoe on behalf of the Wellcome Sanger Institute COVID-19 Surveillance Team |
| EPI_ISL_2852095, EPI_ISL_2852096, EPI_ISL_2852097, EPI_ISL_2852098 | Lighthouse Lab in Milton Keynes | Wellcome Sanger Institute for the COVID-19 Genomics UK (COG-UK) Consortium | The Lighthouse Lab in Milton Keynes and Alex Alderton, Roberto Amato, Jeffrey Barrett, Sonia Goncalves, Ewan Harrison, David K. Jackson, Ian Johnston, Dominic Kwiatkowski, Cordelia Langford, John Sillitoe on behalf of the Wellcome Sanger Institute COVID-19 Surveillance Team |
| EPI_ISL_2852099 | Lighthouse Lab in Glasgow | Wellcome Sanger Institute for the COVID-19 Genomics UK (COG-UK) Consortium | Harper VanSteenhouse, Yumi Kasai, David Gray, Carol Clugston, Anna Dominiczak and Alex Alderton, Roberto Amato, Jeffrey Barrett, Sonia Goncalves, Ewan Harrison, David K. Jackson, Ian Johnston, Dominic Kwiatkowski, Cordelia Langford, John Sillitoe on behalf of the Wellcome Sanger Institute COVID-19 Surveillance Team |
| EPI_ISL_2852100 | Lighthouse Lab in Milton Keynes | Wellcome Sanger Institute for the COVID-19 Genomics UK (COG-UK) Consortium | The Lighthouse Lab in Milton Keynes and Alex Alderton, Roberto Amato, Jeffrey Barrett, Sonia Goncalves, Ewan Harrison, David K. Jackson, Ian Johnston, Dominic Kwiatkowski, Cordelia Langford, John Sillitoe on behalf of the Wellcome Sanger Institute COVID-19 Surveillance Team |
| EPI_ISL_2852101, EPI_ISL_2852102, EPI_ISL_2852103 | Lighthouse Lab in Alderley Park | Wellcome Sanger Institute for the COVID-19 Genomics UK (COG-UK) Consortium | Jacquelyn Wynn, Mairead Hyland, The Lighthouse Lab in Alderley Park and Alex Alderton, Roberto Amato, Jeffrey Barrett, Sonia Goncalves, Ewan Harrison, David K. Jackson, Ian Johnston, Dominic Kwiatkowski, Cordelia Langford, John Sillitoe on behalf of the Wellcome Sanger Institute COVID-19 Surveillance Team |
| EPI_ISL_2852104 | Lighthouse Lab in Glasgow | Wellcome Sanger Institute for the COVID-19 Genomics UK (COG-UK) Consortium | Harper VanSteenhouse, Yumi Kasai, David Gray, Carol Clugston, Anna Dominiczak and Alex Alderton, Roberto Amato, Jeffrey Barrett, Sonia Goncalves, Ewan Harrison, David K. Jackson, Ian Johnston, Dominic Kwiatkowski, Cordelia Langford, John Sillitoe on behalf of the Wellcome Sanger Institute COVID-19 Surveillance Team |
| EPI_ISL_2852105 | Lighthouse Lab in Milton Keynes | Wellcome Sanger Institute for the COVID-19 Genomics UK (COG-UK) Consortium | The Lighthouse Lab in Milton Keynes and Alex Alderton, Roberto Amato, Jeffrey Barrett, Sonia Goncalves, Ewan Harrison, David K. Jackson, Ian Johnston, Dominic Kwiatkowski, Cordelia Langford, John Sillitoe on behalf of the Wellcome Sanger Institute COVID-19 Surveillance Team |
| EPI_ISL_2852106 | Lighthouse Lab in Glasgow | Wellcome Sanger Institute for the COVID-19 Genomics UK (COG-UK) Consortium | Harper VanSteenhouse, Yumi Kasai, David Gray, Carol Clugston, Anna Dominiczak and Alex Alderton, Roberto Amato, Jeffrey Barrett, Sonia Goncalves, Ewan Harrison, David K. Jackson, Ian Johnston, Dominic Kwiatkowski, Cordelia Langford, John Sillitoe on behalf of the Wellcome Sanger Institute COVID-19 Surveillance Team |
| EPI_ISL_2852107, EPI_ISL_2852108, EPI_ISL_2852109 | Lighthouse Lab in Milton Keynes | Wellcome Sanger Institute for the COVID-19 Genomics UK (COG-UK) Consortium | The Lighthouse Lab in Milton Keynes and Alex Alderton, Roberto Amato, Jeffrey Barrett, Sonia Goncalves, Ewan Harrison, David K. Jackson, Ian Johnston, Dominic Kwiatkowski, Cordelia Langford, John Sillitoe on behalf of the Wellcome Sanger Institute COVID-19 Surveillance Team |
| EPI_ISL_2852110 | AZ Klina | AZ Klina | Carl Vael - Lynsey Berckmans |
| EPI_ISL_2852111 | Lighthouse Lab in Alderley Park | Wellcome Sanger Institute for the COVID-19 Genomics UK (COG-UK) Consortium | Jacquelyn Wynn, Mairead Hyland, The Lighthouse Lab in Alderley Park and Alex Alderton, Roberto Amato, Jeffrey Barrett, Sonia Goncalves, Ewan Harrison, David K. Jackson, Ian Johnston, Dominic Kwiatkowski, Cordelia Langford, John Sillitoe on behalf of the Wellcome Sanger Institute COVID-19 Surveillance Team |
| EPI_ISL_2852112 | Lighthouse Lab in Glasgow | Wellcome Sanger Institute for the COVID-19 Genomics UK (COG-UK) Consortium | Harper VanSteenhouse, Yumi Kasai, David Gray, Carol Clugston, Anna Dominiczak and Alex Alderton, Roberto Amato, Jeffrey Barrett, Sonia Goncalves, Ewan Harrison, David K. Jackson, Ian Johnston, Dominic Kwiatkowski, Cordelia Langford, John Sillitoe on behalf of the Wellcome Sanger Institute COVID-19 Surveillance Team |
| EPI_ISL_2852113 | Lighthouse Lab in Milton Keynes | Wellcome Sanger Institute for the COVID-19 Genomics UK (COG-UK) Consortium | The Lighthouse Lab in Milton Keynes and Alex Alderton, Roberto Amato, Jeffrey Barrett, Sonia Goncalves, Ewan Harrison, David K. Jackson, Ian Johnston, Dominic Kwiatkowski, Cordelia Langford, John Sillitoe on behalf of the Wellcome Sanger Institute COVID-19 Surveillance Team |
| EPI_ISL_2852114 | Lighthouse Lab in Glasgow | Wellcome Sanger Institute for the COVID-19 Genomics UK (COG-UK) Consortium | Harper VanSteenhouse, Yumi Kasai, David Gray, Carol Clugston, Anna Dominiczak and Alex Alderton, Roberto Amato, Jeffrey Barrett, Sonia Goncalves, Ewan Harrison, David K. Jackson, Ian Johnston, Dominic Kwiatkowski, Cordelia Langford, John Sillitoe on behalf of the Wellcome Sanger Institute COVID-19 Surveillance Team |
| EPI_ISL_2852115 | Lighthouse Lab in Alderley Park | Wellcome Sanger Institute for the COVID-19 Genomics UK (COG-UK) Consortium | Jacquelyn Wynn, Mairead Hyland, The Lighthouse Lab in Alderley Park and Alex Alderton, Roberto Amato, Jeffrey Barrett, Sonia Goncalves, Ewan Harrison, David K. Jackson, Ian Johnston, Dominic Kwiatkowski, Cordelia Langford, John Sillitoe on behalf of the Wellcome Sanger Institute COVID-19 Surveillance Team |
| EPI_ISL_2852116, EPI_ISL_2852117 | Lighthouse Lab in Glasgow | Wellcome Sanger Institute for the COVID-19 Genomics UK (COG-UK) Consortium | Harper VanSteenhouse, Yumi Kasai, David Gray, Carol Clugston, Anna Dominiczak and Alex Alderton, Roberto Amato, Jeffrey Barrett, Sonia Goncalves, Ewan Harrison, David K. Jackson, Ian Johnston, Dominic Kwiatkowski, Cordelia Langford, John Sillitoe on behalf of the Wellcome Sanger Institute COVID-19 Surveillance Team |
| EPI_ISL_2852118, EPI_ISL_2852119 | Lighthouse Lab in Milton Keynes | Wellcome Sanger Institute for the COVID-19 Genomics UK (COG-UK) Consortium | The Lighthouse Lab in Milton Keynes and Alex Alderton, Roberto Amato, Jeffrey Barrett, Sonia Goncalves, Ewan Harrison, David K. Jackson, Ian Johnston, Dominic Kwiatkowski, Cordelia Langford, John Sillitoe on behalf of the Wellcome Sanger Institute COVID-19 Surveillance Team |
| EPI_ISL_2852120 | Lighthouse Lab in Glasgow | Wellcome Sanger Institute for the COVID-19 Genomics UK (COG-UK) Consortium | Harper VanSteenhouse, Yumi Kasai, David Gray, Carol Clugston, Anna Dominiczak and Alex Alderton, Roberto Amato, Jeffrey Barrett, Sonia Goncalves, Ewan Harrison, David K. Jackson, Ian Johnston, Dominic Kwiatkowski, Cordelia Langford, John Sillitoe on behalf of the Wellcome Sanger Institute COVID-19 Surveillance Team |
| EPI_ISL_2852121 | Lighthouse Lab in Alderley Park | Wellcome Sanger Institute for the COVID-19 Genomics UK (COG-UK) Consortium | Jacquelyn Wynn, Mairead Hyland, The Lighthouse Lab in Alderley Park and Alex Alderton, Roberto Amato, Jeffrey Barrett, Sonia Goncalves, Ewan Harrison, David K. Jackson, Ian Johnston, Dominic Kwiatkowski, Cordelia Langford, John Sillitoe on behalf of the Wellcome Sanger Institute COVID-19 Surveillance Team |
| EPI_ISL_2852122 | Lighthouse Lab in Glasgow | Wellcome Sanger Institute for the COVID-19 Genomics UK (COG-UK) Consortium | Harper VanSteenhouse, Yumi Kasai, David Gray, Carol Clugston, Anna Dominiczak and Alex Alderton, Roberto Amato, Jeffrey Barrett, Sonia Goncalves, Ewan Harrison, David K. Jackson, Ian Johnston, Dominic Kwiatkowski, Cordelia Langford, John Sillitoe on behalf of the Wellcome Sanger Institute COVID-19 Surveillance Team |
| EPI_ISL_2852123 | Lighthouse Lab in Alderley Park | Wellcome Sanger Institute for the COVID-19 Genomics UK (COG-UK) Consortium | Jacquelyn Wynn, Mairead Hyland, The Lighthouse Lab in Alderley Park and Alex Alderton, Roberto Amato, Jeffrey Barrett, Sonia Goncalves, Ewan Harrison, David K. Jackson, Ian Johnston, Dominic Kwiatkowski, Cordelia Langford, John Sillitoe on behalf of the Wellcome Sanger Institute COVID-19 Surveillance Team |
| EPI_ISL_2852124, EPI_ISL_2852125 | Lighthouse Lab in Glasgow | Wellcome Sanger Institute for the COVID-19 Genomics UK (COG-UK) Consortium | Harper VanSteenhouse, Yumi Kasai, David Gray, Carol Clugston, Anna Dominiczak and Alex Alderton, Roberto Amato, Jeffrey Barrett, Sonia Goncalves, Ewan Harrison, David K. Jackson, Ian Johnston, Dominic Kwiatkowski, Cordelia Langford, John Sillitoe on behalf of the Wellcome Sanger Institute COVID-19 Surveillance Team |
| EPI_ISL_2852126 | Lighthouse Lab in Milton Keynes | Wellcome Sanger Institute for the COVID-19 Genomics UK (COG-UK) Consortium | The Lighthouse Lab in Milton Keynes and Alex Alderton, Roberto Amato, Jeffrey Barrett, Sonia Goncalves, Ewan Harrison, David K. Jackson, Ian Johnston, Dominic Kwiatkowski, Cordelia Langford, John Sillitoe on behalf of the Wellcome Sanger Institute COVID-19 Surveillance Team |
| EPI_ISL_2852127 | Lighthouse Lab in Alderley Park | Wellcome Sanger Institute for the COVID-19 Genomics UK (COG-UK) Consortium | Jacquelyn Wynn, Mairead Hyland, The Lighthouse Lab in Alderley Park and Alex Alderton, Roberto Amato, Jeffrey Barrett, Sonia Goncalves, Ewan Harrison, David K. Jackson, Ian Johnston, Dominic Kwiatkowski, Cordelia Langford, John Sillitoe on behalf of the Wellcome Sanger Institute COVID-19 Surveillance Team |
| EPI_ISL_2852128 | Lighthouse Lab in Glasgow | Wellcome Sanger Institute for the COVID-19 Genomics UK (COG-UK) Consortium | Harper VanSteenhouse, Yumi Kasai, David Gray, Carol Clugston, Anna Dominiczak and Alex Alderton, Roberto Amato, Jeffrey Barrett, Sonia Goncalves, Ewan Harrison, David K. Jackson, Ian Johnston, Dominic Kwiatkowski, Cordelia Langford, John Sillitoe on behalf of the Wellcome Sanger Institute COVID-19 Surveillance Team |
| EPI_ISL_2852129 | Lighthouse Lab in Alderley Park | Wellcome Sanger Institute for the COVID-19 Genomics UK (COG-UK) Consortium | Jacquelyn Wynn, Mairead Hyland, The Lighthouse Lab in Alderley Park and Alex Alderton, Roberto Amato, Jeffrey Barrett, Sonia Goncalves, Ewan Harrison, David K. Jackson, Ian Johnston, Dominic Kwiatkowski, Cordelia Langford, John Sillitoe on behalf of the Wellcome Sanger Institute COVID-19 Surveillance Team |
| EPI_ISL_2852130, EPI_ISL_2852131, EPI_ISL_2852132 | Lighthouse Lab in Glasgow | Wellcome Sanger Institute for the COVID-19 Genomics UK (COG-UK) Consortium | Harper VanSteenhouse, Yumi Kasai, David Gray, Carol Clugston, Anna Dominiczak and Alex Alderton, Roberto Amato, Jeffrey Barrett, Sonia Goncalves, Ewan Harrison, David K. Jackson, Ian Johnston, Dominic Kwiatkowski, Cordelia Langford, John Sillitoe on behalf of the Wellcome Sanger Institute COVID-19 Surveillance Team |
| EPI_ISL_2852133 | Lighthouse Lab in Milton Keynes | Wellcome Sanger Institute for the COVID-19 Genomics UK (COG-UK) Consortium | The Lighthouse Lab in Milton Keynes and Alex Alderton, Roberto Amato, Jeffrey Barrett, Sonia Goncalves, Ewan Harrison, David K. Jackson, Ian Johnston, Dominic Kwiatkowski, Cordelia Langford, John Sillitoe on behalf of the Wellcome Sanger Institute COVID-19 Surveillance Team |

| | | | |
|---|---|---|---|
| EPI_ISL_2852134, EPI_ISL_2852135 | Lighthouse Lab in Glasgow | Wellcome Sanger Institute for the COVID-19 Genomics UK (COG-UK) Consortium | Harper VanSteenhouse, Yumi Kasai, David Gray, Carol Clugston, Anna Dominiczak and Alex Alderton, Roberto Amato, Jeffrey Barrett, Sonia Goncalves, Ewan Harrison, David K. Jackson, Ian Johnston, Dominic Kwiatkowski, Cordelia Langford, John Sillitoe on behalf of the Wellcome Sanger Institute COVID-19 Surveillance Team |
| EPI_ISL_2852136 | Lighthouse Lab in Alderley Park | Wellcome Sanger Institute for the COVID-19 Genomics UK (COG-UK) Consortium | Jacquelyn Wynn, Mairead Hyland, The Lighthouse Lab in Alderley Park and Alex Alderton, Roberto Amato, Jeffrey Barrett, Sonia Goncalves, Ewan Harrison, David K. Jackson, Ian Johnston, Dominic Kwiatkowski, Cordelia Langford, John Sillitoe on behalf of the Wellcome Sanger Institute COVID-19 Surveillance Team |
| EPI_ISL_2852137, EPI_ISL_2852138 | Lighthouse Lab in Milton Keynes | Wellcome Sanger Institute for the COVID-19 Genomics UK (COG-UK) Consortium | The Lighthouse Lab in Milton Keynes and Alex Alderton, Roberto Amato, Jeffrey Barrett, Sonia Goncalves, Ewan Harrison, David K. Jackson, Ian Johnston, Dominic Kwiatkowski, Cordelia Langford, John Sillitoe on behalf of the Wellcome Sanger Institute COVID-19 Surveillance Team |
| EPI_ISL_2852139 | Lighthouse Lab in Alderley Park | Wellcome Sanger Institute for the COVID-19 Genomics UK (COG-UK) Consortium | Jacquelyn Wynn, Mairead Hyland, The Lighthouse Lab in Alderley Park and Alex Alderton, Roberto Amato, Jeffrey Barrett, Sonia Goncalves, Ewan Harrison, David K. Jackson, Ian Johnston, Dominic Kwiatkowski, Cordelia Langford, John Sillitoe on behalf of the Wellcome Sanger Institute COVID-19 Surveillance Team |
| EPI_ISL_2852140, EPI_ISL_2852141 | Lighthouse Lab in Glasgow | Wellcome Sanger Institute for the COVID-19 Genomics UK (COG-UK) Consortium | Harper VanSteenhouse, Yumi Kasai, David Gray, Carol Clugston, Anna Dominiczak and Alex Alderton, Roberto Amato, Jeffrey Barrett, Sonia Goncalves, Ewan Harrison, David K. Jackson, Ian Johnston, Dominic Kwiatkowski, Cordelia Langford, John Sillitoe on behalf of the Wellcome Sanger Institute COVID-19 Surveillance Team |
| EPI_ISL_2852142 | Lighthouse Laboratory Plymouth | Wellcome Sanger Institute for the COVID-19 Genomics UK (COG-UK) Consortium | Lighthouse Laboratory Plymouth and Alex Alderton, Roberto Amato, Jeffrey Barrett, Sonia Goncalves, Ewan Harrison, David K. Jackson, Ian Johnston, Dominic Kwiatkowski, Cordelia Langford, John Sillitoe on behalf of the Wellcome Sanger Institute COVID-19 Surveillance Team |
| EPI_ISL_2852143, EPI_ISL_2852144, EPI_ISL_2852145, EPI_ISL_2852146, EPI_ISL_2852147, EPI_ISL_2852148, EPI_ISL_2852149, EPI_ISL_2852150, EPI_ISL_2852151 | Lighthouse Lab in Glasgow | Wellcome Sanger Institute for the COVID-19 Genomics UK (COG-UK) Consortium | Harper VanSteenhouse, Yumi Kasai, David Gray, Carol Clugston, Anna Dominiczak and Alex Alderton, Roberto Amato, Jeffrey Barrett, Sonia Goncalves, Ewan Harrison, David K. Jackson, Ian Johnston, Dominic Kwiatkowski, Cordelia Langford, John Sillitoe on behalf of the Wellcome Sanger Institute COVID-19 Surveillance Team |
| EPI_ISL_2852152 | Lighthouse Laboratory Plymouth | Wellcome Sanger Institute for the COVID-19 Genomics UK (COG-UK) Consortium | Lighthouse Laboratory Plymouth and Alex Alderton, Roberto Amato, Jeffrey Barrett, Sonia Goncalves, Ewan Harrison, David K. Jackson, Ian Johnston, Dominic Kwiatkowski, Cordelia Langford, John Sillitoe on behalf of the Wellcome Sanger Institute COVID-19 Surveillance Team |
| EPI_ISL_2852153, EPI_ISL_2852154 | Lighthouse Lab in Glasgow | Wellcome Sanger Institute for the COVID-19 Genomics UK (COG-UK) Consortium | Harper VanSteenhouse, Yumi Kasai, David Gray, Carol Clugston, Anna Dominiczak and Alex Alderton, Roberto Amato, Jeffrey Barrett, Sonia Goncalves, Ewan Harrison, David K. Jackson, Ian Johnston, Dominic Kwiatkowski, Cordelia Langford, John Sillitoe on behalf of the Wellcome Sanger Institute COVID-19 Surveillance Team |
| EPI_ISL_2852155 | Lighthouse Laboratory Plymouth | Wellcome Sanger Institute for the COVID-19 Genomics UK (COG-UK) Consortium | Lighthouse Laboratory Plymouth and Alex Alderton, Roberto Amato, Jeffrey Barrett, Sonia Goncalves, Ewan Harrison, David K. Jackson, Ian Johnston, Dominic Kwiatkowski, Cordelia Langford, John Sillitoe on behalf of the Wellcome Sanger Institute COVID-19 Surveillance Team |
| EPI_ISL_2852156 | Lighthouse Lab in Glasgow | Wellcome Sanger Institute for the COVID-19 Genomics UK (COG-UK) Consortium | Harper VanSteenhouse, Yumi Kasai, David Gray, Carol Clugston, Anna Dominiczak and Alex Alderton, Roberto Amato, Jeffrey Barrett, Sonia Goncalves, Ewan Harrison, David K. Jackson, Ian Johnston, Dominic Kwiatkowski, Cordelia Langford, John Sillitoe on behalf of the Wellcome Sanger Institute COVID-19 Surveillance Team |
| EPI_ISL_2852157, EPI_ISL_2852158, EPI_ISL_2852159 | Lighthouse Laboratory Plymouth | Wellcome Sanger Institute for the COVID-19 Genomics UK (COG-UK) Consortium | Lighthouse Laboratory Plymouth and Alex Alderton, Roberto Amato, Jeffrey Barrett, Sonia Goncalves, Ewan Harrison, David K. Jackson, Ian Johnston, Dominic Kwiatkowski, Cordelia Langford, John Sillitoe on behalf of the Wellcome Sanger Institute COVID-19 Surveillance Team |
| EPI_ISL_2852160, EPI_ISL_2852161, EPI_ISL_2852162, EPI_ISL_2852163, EPI_ISL_2852164, EPI_ISL_2852165, EPI_ISL_2852166 | Lighthouse Lab in Glasgow | Wellcome Sanger Institute for the COVID-19 Genomics UK (COG-UK) Consortium | Harper VanSteenhouse, Yumi Kasai, David Gray, Carol Clugston, Anna Dominiczak and Alex Alderton, Roberto Amato, Jeffrey Barrett, Sonia Goncalves, Ewan Harrison, David K. Jackson, Ian Johnston, Dominic Kwiatkowski, Cordelia Langford, John Sillitoe on behalf of the Wellcome Sanger Institute COVID-19 Surveillance Team |
| EPI_ISL_2852167 | Lighthouse Laboratory Plymouth | Wellcome Sanger Institute for the COVID-19 Genomics UK (COG-UK) Consortium | Lighthouse Laboratory Plymouth and Alex Alderton, Roberto Amato, Jeffrey Barrett, Sonia Goncalves, Ewan Harrison, David K. Jackson, Ian Johnston, Dominic Kwiatkowski, Cordelia Langford, John Sillitoe on behalf of the Wellcome Sanger Institute COVID-19 Surveillance Team |
| EPI_ISL_2852168 | Lighthouse Lab in Glasgow | Wellcome Sanger Institute for the COVID-19 Genomics UK (COG-UK) Consortium | Harper VanSteenhouse, Yumi Kasai, David Gray, Carol Clugston, Anna Dominiczak and Alex Alderton, Roberto Amato, Jeffrey Barrett, Sonia Goncalves, Ewan Harrison, David K. Jackson, Ian Johnston, Dominic Kwiatkowski, Cordelia Langford, John Sillitoe on behalf of the Wellcome Sanger Institute COVID-19 Surveillance Team |
| EPI_ISL_2852169, EPI_ISL_2852170 | Lighthouse Laboratory Plymouth | Wellcome Sanger Institute for the COVID-19 Genomics UK (COG-UK) Consortium | Lighthouse Laboratory Plymouth and Alex Alderton, Roberto Amato, Jeffrey Barrett, Sonia Goncalves, Ewan Harrison, David K. Jackson, Ian Johnston, Dominic Kwiatkowski, Cordelia Langford, John Sillitoe on behalf of the Wellcome Sanger Institute COVID-19 Surveillance Team |
| EPI_ISL_2852171, EPI_ISL_2852172, EPI_ISL_2852173, EPI_ISL_2852174, EPI_ISL_2852175, EPI_ISL_2852176, EPI_ISL_2852177, EPI_ISL_2852178, EPI_ISL_2852179 | Lighthouse Lab in Glasgow | Wellcome Sanger Institute for the COVID-19 Genomics UK (COG-UK) Consortium | Harper VanSteenhouse, Yumi Kasai, David Gray, Carol Clugston, Anna Dominiczak and Alex Alderton, Roberto Amato, Jeffrey Barrett, Sonia Goncalves, Ewan Harrison, David K. Jackson, Ian Johnston, Dominic Kwiatkowski, Cordelia Langford, John Sillitoe on behalf of the Wellcome Sanger Institute COVID-19 Surveillance Team |
| EPI_ISL_2852180 | Lighthouse Laboratory Plymouth | Wellcome Sanger Institute for the COVID-19 Genomics UK (COG-UK) Consortium | Lighthouse Laboratory Plymouth and Alex Alderton, Roberto Amato, Jeffrey Barrett, Sonia Goncalves, Ewan Harrison, David K. Jackson, Ian Johnston, Dominic Kwiatkowski, Cordelia Langford, John Sillitoe on behalf of the Wellcome Sanger Institute COVID-19 Surveillance Team |
| EPI_ISL_2852181, EPI_ISL_2852182 | Lighthouse Lab in Glasgow | Wellcome Sanger Institute for the COVID-19 Genomics UK (COG-UK) Consortium | Harper VanSteenhouse, Yumi Kasai, David Gray, Carol Clugston, Anna Dominiczak and Alex Alderton, Roberto Amato, Jeffrey Barrett, Sonia Goncalves, Ewan Harrison, David K. Jackson, Ian Johnston, Dominic Kwiatkowski, Cordelia Langford, John Sillitoe on behalf of the Wellcome Sanger Institute COVID-19 Surveillance Team |
| EPI_ISL_2852183 | Lighthouse Laboratory Plymouth | Wellcome Sanger Institute for the COVID-19 Genomics UK (COG-UK) Consortium | Lighthouse Laboratory Plymouth and Alex Alderton, Roberto Amato, Jeffrey Barrett, Sonia Goncalves, Ewan Harrison, David K. Jackson, Ian Johnston, Dominic Kwiatkowski, Cordelia Langford, John Sillitoe on behalf of the Wellcome Sanger Institute COVID-19 Surveillance Team |
| EPI_ISL_2852184, EPI_ISL_2852185 | Lighthouse Lab in Glasgow | Wellcome Sanger Institute for the COVID-19 Genomics UK (COG-UK) Consortium | Harper VanSteenhouse, Yumi Kasai, David Gray, Carol Clugston, Anna Dominiczak and Alex Alderton, Roberto Amato, Jeffrey Barrett, Sonia Goncalves, Ewan Harrison, David K. Jackson, Ian Johnston, Dominic Kwiatkowski, Cordelia Langford, John Sillitoe on behalf of the Wellcome Sanger Institute COVID-19 Surveillance Team |
| EPI_ISL_2852186, EPI_ISL_2852187 | Lighthouse Laboratory Plymouth | Wellcome Sanger Institute for the COVID-19 Genomics UK (COG-UK) Consortium | Lighthouse Laboratory Plymouth and Alex Alderton, Roberto Amato, Jeffrey Barrett, Sonia Goncalves, Ewan Harrison, David K. Jackson, Ian Johnston, Dominic Kwiatkowski, Cordelia Langford, John Sillitoe on behalf of the Wellcome Sanger Institute COVID-19 Surveillance Team |
| EPI_ISL_2852188, EPI_ISL_2852189, EPI_ISL_2852190, EPI_ISL_2852191 | Lighthouse Lab in Glasgow | Wellcome Sanger Institute for the COVID-19 Genomics UK (COG-UK) Consortium | Harper VanSteenhouse, Yumi Kasai, David Gray, Carol Clugston, Anna Dominiczak and Alex Alderton, Roberto Amato, Jeffrey Barrett, Sonia Goncalves, Ewan Harrison, David K. Jackson, Ian Johnston, Dominic Kwiatkowski, Cordelia Langford, John Sillitoe on behalf of the Wellcome Sanger Institute COVID-19 Surveillance Team |
| EPI_ISL_2852192, EPI_ISL_2852193 | Lighthouse Laboratory Plymouth | Wellcome Sanger Institute for the COVID-19 Genomics UK (COG-UK) Consortium | Lighthouse Laboratory Plymouth and Alex Alderton, Roberto Amato, Jeffrey Barrett, Sonia Goncalves, Ewan Harrison, David K. Jackson, Ian Johnston, Dominic Kwiatkowski, Cordelia Langford, John Sillitoe on behalf of the Wellcome Sanger Institute COVID-19 Surveillance Team |
| EPI_ISL_2852194 | Lighthouse Lab in Glasgow | Wellcome Sanger Institute for the COVID-19 Genomics UK (COG-UK) Consortium | Harper VanSteenhouse, Yumi Kasai, David Gray, Carol Clugston, Anna Dominiczak and Alex Alderton, Roberto Amato, Jeffrey Barrett, Sonia Goncalves, Ewan Harrison, David K. Jackson, Ian Johnston, Dominic Kwiatkowski, Cordelia Langford, John Sillitoe on behalf of the Wellcome Sanger Institute COVID-19 Surveillance Team |
| EPI_ISL_2852195 | Lighthouse Laboratory Plymouth | Wellcome Sanger Institute for the COVID-19 Genomics UK (COG-UK) Consortium | Lighthouse Laboratory Plymouth and Alex Alderton, Roberto Amato, Jeffrey Barrett, Sonia Goncalves, Ewan Harrison, David K. Jackson, Ian Johnston, Dominic Kwiatkowski, Cordelia Langford, John Sillitoe on behalf of the Wellcome Sanger Institute COVID-19 Surveillance Team |
| EPI_ISL_2852196 | Lighthouse Lab in Glasgow | Wellcome Sanger Institute for the COVID-19 Genomics UK (COG-UK) Consortium | Harper VanSteenhouse, Yumi Kasai, David Gray, Carol Clugston, Anna Dominiczak and Alex Alderton, Roberto Amato, Jeffrey Barrett, Sonia Goncalves, Ewan Harrison, David K. Jackson, Ian Johnston, Dominic Kwiatkowski, Cordelia Langford, John Sillitoe on behalf of the Wellcome Sanger Institute COVID-19 Surveillance Team |
| EPI_ISL_2852197 | Lighthouse Laboratory Plymouth | Wellcome Sanger Institute for the COVID-19 Genomics UK (COG-UK) Consortium | Lighthouse Laboratory Plymouth and Alex Alderton, Roberto Amato, Jeffrey Barrett, Sonia Goncalves, Ewan Harrison, David K. Jackson, Ian Johnston, Dominic Kwiatkowski, Cordelia Langford, John Sillitoe on behalf of the Wellcome Sanger Institute COVID-19 Surveillance Team |
| EPI_ISL_2852198, EPI_ISL_2852199, EPI_ISL_2852200, EPI_ISL_2852201, EPI_ISL_2852202, EPI_ISL_2852203, EPI_ISL_2852204, EPI_ISL_2852205 | Lighthouse Lab in Glasgow | Wellcome Sanger Institute for the COVID-19 Genomics UK (COG-UK) Consortium | Harper VanSteenhouse, Yumi Kasai, David Gray, Carol Clugston, Anna Dominiczak and Alex Alderton, Roberto Amato, Jeffrey Barrett, Sonia Goncalves, Ewan Harrison, David K. Jackson, Ian Johnston, Dominic Kwiatkowski, Cordelia Langford, John Sillitoe on behalf of the Wellcome Sanger Institute COVID-19 Surveillance Team |
| EPI_ISL_2852206 | Lighthouse Laboratory Plymouth | Wellcome Sanger Institute for the COVID-19 Genomics UK (COG-UK) Consortium | Lighthouse Laboratory Plymouth and Alex Alderton, Roberto Amato, Jeffrey Barrett, Sonia Goncalves, Ewan Harrison, David K. Jackson, Ian Johnston, Dominic Kwiatkowski, Cordelia Langford, John Sillitoe on behalf of the Wellcome Sanger Institute COVID-19 Surveillance Team |
| EPI_ISL_2852207, EPI_ISL_2852208, EPI_ISL_2852209, EPI_ISL_2852210 | Lighthouse Lab in Glasgow | Wellcome Sanger Institute for the COVID-19 Genomics UK (COG-UK) Consortium | Harper VanSteenhouse, Yumi Kasai, David Gray, Carol Clugston, Anna Dominiczak and Alex Alderton, Roberto Amato, Jeffrey Barrett, Sonia Goncalves, Ewan Harrison, David K. Jackson, Ian Johnston, Dominic Kwiatkowski, Cordelia Langford, John Sillitoe on behalf of the Wellcome Sanger Institute COVID-19 Surveillance Team |
| EPI_ISL_2852211 | Lighthouse Laboratory Plymouth | Wellcome Sanger Institute for the COVID-19 Genomics UK (COG-UK) Consortium | Lighthouse Laboratory Plymouth and Alex Alderton, Roberto Amato, Jeffrey Barrett, Sonia Goncalves, Ewan Harrison, David K. Jackson, Ian Johnston, Dominic Kwiatkowski, Cordelia Langford, John Sillitoe on behalf of the Wellcome Sanger Institute COVID-19 Surveillance Team |

EPI_ISL_2852212, EPI_ISL_2852213, EPI_ISL_2852214, EPI_ISL_2852215, EPI_ISL_2852216, EPI_ISL_2852217, EPI_ISL_2852218, EPI_ISL_2852219, EPI_ISL_2852220, EPI_ISL_2852221, EPI_ISL_2852222, EPI_ISL_2852223, EPI_ISL_2852224

| | | | |
|---|---|---|---|
| see above | Lighthouse Lab in Glasgow | Wellcome Sanger Institute for the COVID-19 Genomics UK (COG-UK) Consortium | Harper VanSteenhouse, Yumi Kasai, David Gray, Carol Clugston, Anna Dominiczak and Alex Alderton, Roberto Amato, Jeffrey Barrett, Sonia Goncalves, Ewan Harrison, David K. Jackson, Ian Johnston, Dominic Kwiatkowski, Cordelia Langford, John Sillitoe on behalf of the Wellcome Sanger Institute COVID-19 Surveillance Team |
| EPI_ISL_2852225 | Lighthouse Laboratory Plymouth | Wellcome Sanger Institute for the COVID-19 Genomics UK (COG-UK) Consortium | Lighthouse Laboratory Plymouth and Alex Alderton, Roberto Amato, Jeffrey Barrett, Sonia Goncalves, Ewan Harrison, David K. Jackson, Ian Johnston, Dominic Kwiatkowski, Cordelia Langford, John Sillitoe on behalf of the Wellcome Sanger Institute COVID-19 Surveillance Team |
| EPI_ISL_2852226 | Lighthouse Lab in Glasgow | Wellcome Sanger Institute for the COVID-19 Genomics UK (COG-UK) Consortium | Harper VanSteenhouse, Yumi Kasai, David Gray, Carol Clugston, Anna Dominiczak and Alex Alderton, Roberto Amato, Jeffrey Barrett, Sonia Goncalves, Ewan Harrison, David K. Jackson, Ian Johnston, Dominic Kwiatkowski, Cordelia Langford, John Sillitoe on behalf of the Wellcome Sanger Institute COVID-19 Surveillance Team |
| EPI_ISL_2852227 | Lighthouse Laboratory Plymouth | Wellcome Sanger Institute for the COVID-19 Genomics UK (COG-UK) Consortium | Lighthouse Laboratory Plymouth and Alex Alderton, Roberto Amato, Jeffrey Barrett, Sonia Goncalves, Ewan Harrison, David K. Jackson, Ian Johnston, Dominic Kwiatkowski, Cordelia Langford, John Sillitoe on behalf of the Wellcome Sanger Institute COVID-19 Surveillance Team |
| EPI_ISL_2852228, EPI_ISL_2852229, EPI_ISL_2852230 | Lighthouse Lab in Glasgow | Wellcome Sanger Institute for the COVID-19 Genomics UK (COG-UK) Consortium | Harper VanSteenhouse, Yumi Kasai, David Gray, Carol Clugston, Anna Dominiczak and Alex Alderton, Roberto Amato, Jeffrey Barrett, Sonia Goncalves, Ewan Harrison, David K. Jackson, Ian Johnston, Dominic Kwiatkowski, Cordelia Langford, John Sillitoe on behalf of the Wellcome Sanger Institute COVID-19 Surveillance Team |
| EPI_ISL_2852231 | Lighthouse Laboratory Plymouth | Wellcome Sanger Institute for the COVID-19 Genomics UK (COG-UK) Consortium | Lighthouse Laboratory Plymouth and Alex Alderton, Roberto Amato, Jeffrey Barrett, Sonia Goncalves, Ewan Harrison, David K. Jackson, Ian Johnston, Dominic Kwiatkowski, Cordelia Langford, John Sillitoe on behalf of the Wellcome Sanger Institute COVID-19 Surveillance Team |
| EPI_ISL_2852232, EPI_ISL_2852233, EPI_ISL_2852234, EPI_ISL_2852235, EPI_ISL_2852236 | Lighthouse Lab in Glasgow | Wellcome Sanger Institute for the COVID-19 Genomics UK (COG-UK) Consortium | Harper VanSteenhouse, Yumi Kasai, David Gray, Carol Clugston, Anna Dominiczak and Alex Alderton, Roberto Amato, Jeffrey Barrett, Sonia Goncalves, Ewan Harrison, David K. Jackson, Ian Johnston, Dominic Kwiatkowski, Cordelia Langford, John Sillitoe on behalf of the Wellcome Sanger Institute COVID-19 Surveillance Team |
| EPI_ISL_2852237, EPI_ISL_2852238 | Lighthouse Laboratory Plymouth | Wellcome Sanger Institute for the COVID-19 Genomics UK (COG-UK) Consortium | Lighthouse Laboratory Plymouth and Alex Alderton, Roberto Amato, Jeffrey Barrett, Sonia Goncalves, Ewan Harrison, David K. Jackson, Ian Johnston, Dominic Kwiatkowski, Cordelia Langford, John Sillitoe on behalf of the Wellcome Sanger Institute COVID-19 Surveillance Team |
| EPI_ISL_2852239, EPI_ISL_2852240, EPI_ISL_2852241, EPI_ISL_2852242, EPI_ISL_2852243, EPI_ISL_2852244, EPI_ISL_2852245, EPI_ISL_2852246, EPI_ISL_2852247 | Lighthouse Lab in Glasgow | Wellcome Sanger Institute for the COVID-19 Genomics UK (COG-UK) Consortium | Harper VanSteenhouse, Yumi Kasai, David Gray, Carol Clugston, Anna Dominiczak and Alex Alderton, Roberto Amato, Jeffrey Barrett, Sonia Goncalves, Ewan Harrison, David K. Jackson, Ian Johnston, Dominic Kwiatkowski, Cordelia Langford, John Sillitoe on behalf of the Wellcome Sanger Institute COVID-19 Surveillance Team |
| EPI_ISL_2852248, EPI_ISL_2852249, EPI_ISL_2852250, EPI_ISL_2852251 | Lighthouse Laboratory Plymouth | Wellcome Sanger Institute for the COVID-19 Genomics UK (COG-UK) Consortium | Lighthouse Laboratory Plymouth and Alex Alderton, Roberto Amato, Jeffrey Barrett, Sonia Goncalves, Ewan Harrison, David K. Jackson, Ian Johnston, Dominic Kwiatkowski, Cordelia Langford, John Sillitoe on behalf of the Wellcome Sanger Institute COVID-19 Surveillance Team |
| EPI_ISL_2852252, EPI_ISL_2852253, EPI_ISL_2852254 | Lighthouse Lab in Glasgow | Wellcome Sanger Institute for the COVID-19 Genomics UK (COG-UK) Consortium | Harper VanSteenhouse, Yumi Kasai, David Gray, Carol Clugston, Anna Dominiczak and Alex Alderton, Roberto Amato, Jeffrey Barrett, Sonia Goncalves, Ewan Harrison, David K. Jackson, Ian Johnston, Dominic Kwiatkowski, Cordelia Langford, John Sillitoe on behalf of the Wellcome Sanger Institute COVID-19 Surveillance Team |
| EPI_ISL_2852255 | Lighthouse Laboratory Plymouth | Wellcome Sanger Institute for the COVID-19 Genomics UK (COG-UK) Consortium | Lighthouse Laboratory Plymouth and Alex Alderton, Roberto Amato, Jeffrey Barrett, Sonia Goncalves, Ewan Harrison, David K. Jackson, Ian Johnston, Dominic Kwiatkowski, Cordelia Langford, John Sillitoe on behalf of the Wellcome Sanger Institute COVID-19 Surveillance Team |
| EPI_ISL_2852256, EPI_ISL_2852257, EPI_ISL_2852258 | Lighthouse Lab in Glasgow | Wellcome Sanger Institute for the COVID-19 Genomics UK (COG-UK) Consortium | Harper VanSteenhouse, Yumi Kasai, David Gray, Carol Clugston, Anna Dominiczak and Alex Alderton, Roberto Amato, Jeffrey Barrett, Sonia Goncalves, Ewan Harrison, David K. Jackson, Ian Johnston, Dominic Kwiatkowski, Cordelia Langford, John Sillitoe on behalf of the Wellcome Sanger Institute COVID-19 Surveillance Team |
| EPI_ISL_2852259 | Lighthouse Laboratory Plymouth | Wellcome Sanger Institute for the COVID-19 Genomics UK (COG-UK) Consortium | Lighthouse Laboratory Plymouth and Alex Alderton, Roberto Amato, Jeffrey Barrett, Sonia Goncalves, Ewan Harrison, David K. Jackson, Ian Johnston, Dominic Kwiatkowski, Cordelia Langford, John Sillitoe on behalf of the Wellcome Sanger Institute COVID-19 Surveillance Team |
| EPI_ISL_2852260, EPI_ISL_2852261, EPI_ISL_2852262 | Lighthouse Lab in Glasgow | Wellcome Sanger Institute for the COVID-19 Genomics UK (COG-UK) Consortium | Harper VanSteenhouse, Yumi Kasai, David Gray, Carol Clugston, Anna Dominiczak and Alex Alderton, Roberto Amato, Jeffrey Barrett, Sonia Goncalves, Ewan Harrison, David K. Jackson, Ian Johnston, Dominic Kwiatkowski, Cordelia Langford, John Sillitoe on behalf of the Wellcome Sanger Institute COVID-19 Surveillance Team |
| EPI_ISL_2852263 | Lighthouse Laboratory Plymouth | Wellcome Sanger Institute for the COVID-19 Genomics UK (COG-UK) Consortium | Lighthouse Laboratory Plymouth and Alex Alderton, Roberto Amato, Jeffrey Barrett, Sonia Goncalves, Ewan Harrison, David K. Jackson, Ian Johnston, Dominic Kwiatkowski, Cordelia Langford, John Sillitoe on behalf of the Wellcome Sanger Institute COVID-19 Surveillance Team |
| EPI_ISL_2852264, EPI_ISL_2852265, EPI_ISL_2852268, EPI_ISL_2852269, EPI_ISL_2852270, EPI_ISL_2852271, EPI_ISL_2852272, EPI_ISL_2852273, EPI_ISL_2852274, EPI_ISL_2852275, EPI_ISL_2852276, EPI_ISL_2852277, EPI_ISL_2852278, EPI_ISL_2852280, EPI_ISL_2852281 | | | |
| see above | Lighthouse Lab in Glasgow | Wellcome Sanger Institute for the COVID-19 Genomics UK (COG-UK) Consortium | Harper VanSteenhouse, Yumi Kasai, David Gray, Carol Clugston, Anna Dominiczak and Alex Alderton, Roberto Amato, Jeffrey Barrett, Sonia Goncalves, Ewan Harrison, David K. Jackson, Ian Johnston, Dominic Kwiatkowski, Cordelia Langford, John Sillitoe on behalf of the Wellcome Sanger Institute COVID-19 Surveillance Team |
| EPI_ISL_2852282 | Lighthouse Laboratory Plymouth | Wellcome Sanger Institute for the COVID-19 Genomics UK (COG-UK) Consortium | Lighthouse Laboratory Plymouth and Alex Alderton, Roberto Amato, Jeffrey Barrett, Sonia Goncalves, Ewan Harrison, David K. Jackson, Ian Johnston, Dominic Kwiatkowski, Cordelia Langford, John Sillitoe on behalf of the Wellcome Sanger Institute COVID-19 Surveillance Team |
| EPI_ISL_2852283, EPI_ISL_2852284, EPI_ISL_2852285 | Lighthouse Lab in Glasgow | Wellcome Sanger Institute for the COVID-19 Genomics UK (COG-UK) Consortium | Harper VanSteenhouse, Yumi Kasai, David Gray, Carol Clugston, Anna Dominiczak and Alex Alderton, Roberto Amato, Jeffrey Barrett, Sonia Goncalves, Ewan Harrison, David K. Jackson, Ian Johnston, Dominic Kwiatkowski, Cordelia Langford, John Sillitoe on behalf of the Wellcome Sanger Institute COVID-19 Surveillance Team |
| EPI_ISL_2852286 | Lighthouse Laboratory Plymouth | Wellcome Sanger Institute for the COVID-19 Genomics UK (COG-UK) Consortium | Lighthouse Laboratory Plymouth and Alex Alderton, Roberto Amato, Jeffrey Barrett, Sonia Goncalves, Ewan Harrison, David K. Jackson, Ian Johnston, Dominic Kwiatkowski, Cordelia Langford, John Sillitoe on behalf of the Wellcome Sanger Institute COVID-19 Surveillance Team |
| EPI_ISL_2852287, EPI_ISL_2852288, EPI_ISL_2852289, EPI_ISL_2852290, EPI_ISL_2852291, EPI_ISL_2852292, EPI_ISL_2852293, EPI_ISL_2852294, EPI_ISL_2852295, EPI_ISL_2852296, EPI_ISL_2852297, EPI_ISL_2852298, EPI_ISL_2852299, EPI_ISL_2852300, EPI_ISL_2852301 | | | |
| see above | Lighthouse Lab in Glasgow | Wellcome Sanger Institute for the COVID-19 Genomics UK (COG-UK) Consortium | Harper VanSteenhouse, Yumi Kasai, David Gray, Carol Clugston, Anna Dominiczak and Alex Alderton, Roberto Amato, Jeffrey Barrett, Sonia Goncalves, Ewan Harrison, David K. Jackson, Ian Johnston, Dominic Kwiatkowski, Cordelia Langford, John Sillitoe on behalf of the Wellcome Sanger Institute COVID-19 Surveillance Team |
| EPI_ISL_2852302, EPI_ISL_2852303 | Lighthouse Laboratory Plymouth | Wellcome Sanger Institute for the COVID-19 Genomics UK (COG-UK) Consortium | Lighthouse Laboratory Plymouth and Alex Alderton, Roberto Amato, Jeffrey Barrett, Sonia Goncalves, Ewan Harrison, David K. Jackson, Ian Johnston, Dominic Kwiatkowski, Cordelia Langford, John Sillitoe on behalf of the Wellcome Sanger Institute COVID-19 Surveillance Team |
| EPI_ISL_2852304, EPI_ISL_2852305, EPI_ISL_2852306, EPI_ISL_2852307, EPI_ISL_2852308, EPI_ISL_2852309 | Lighthouse Lab in Glasgow | Wellcome Sanger Institute for the COVID-19 Genomics UK (COG-UK) Consortium | Harper VanSteenhouse, Yumi Kasai, David Gray, Carol Clugston, Anna Dominiczak and Alex Alderton, Roberto Amato, Jeffrey Barrett, Sonia Goncalves, Ewan Harrison, David K. Jackson, Ian Johnston, Dominic Kwiatkowski, Cordelia Langford, John Sillitoe on behalf of the Wellcome Sanger Institute COVID-19 Surveillance Team |
| EPI_ISL_2852310 | Lighthouse Laboratory Plymouth | Wellcome Sanger Institute for the COVID-19 Genomics UK (COG-UK) Consortium | Lighthouse Laboratory Plymouth and Alex Alderton, Roberto Amato, Jeffrey Barrett, Sonia Goncalves, Ewan Harrison, David K. Jackson, Ian Johnston, Dominic Kwiatkowski, Cordelia Langford, John Sillitoe on behalf of the Wellcome Sanger Institute COVID-19 Surveillance Team |
| EPI_ISL_2852311, EPI_ISL_2852312 | Lighthouse Lab in Glasgow | Wellcome Sanger Institute for the COVID-19 Genomics UK (COG-UK) Consortium | Harper VanSteenhouse, Yumi Kasai, David Gray, Carol Clugston, Anna Dominiczak and Alex Alderton, Roberto Amato, Jeffrey Barrett, Sonia Goncalves, Ewan Harrison, David K. Jackson, Ian Johnston, Dominic Kwiatkowski, Cordelia Langford, John Sillitoe on behalf of the Wellcome Sanger Institute COVID-19 Surveillance Team |
| EPI_ISL_2852313 | Lighthouse Laboratory Plymouth | Wellcome Sanger Institute for the COVID-19 Genomics UK (COG-UK) Consortium | Lighthouse Laboratory Plymouth and Alex Alderton, Roberto Amato, Jeffrey Barrett, Sonia Goncalves, Ewan Harrison, David K. Jackson, Ian Johnston, Dominic Kwiatkowski, Cordelia Langford, John Sillitoe on behalf of the Wellcome Sanger Institute COVID-19 Surveillance Team |
| EPI_ISL_2852314, EPI_ISL_2852315 | Lighthouse Lab in Glasgow | Wellcome Sanger Institute for the COVID-19 Genomics UK (COG-UK) Consortium | Harper VanSteenhouse, Yumi Kasai, David Gray, Carol Clugston, Anna Dominiczak and Alex Alderton, Roberto Amato, Jeffrey Barrett, Sonia Goncalves, Ewan Harrison, David K. Jackson, Ian Johnston, Dominic Kwiatkowski, Cordelia Langford, John Sillitoe on behalf of the Wellcome Sanger Institute COVID-19 Surveillance Team |
| EPI_ISL_2852316 | Lighthouse Laboratory Plymouth | Wellcome Sanger Institute for the COVID-19 Genomics UK (COG-UK) Consortium | Lighthouse Laboratory Plymouth and Alex Alderton, Roberto Amato, Jeffrey Barrett, Sonia Goncalves, Ewan Harrison, David K. Jackson, Ian Johnston, Dominic Kwiatkowski, Cordelia Langford, John Sillitoe on behalf of the Wellcome Sanger Institute COVID-19 Surveillance Team |
| EPI_ISL_2852317, EPI_ISL_2852318 | Lighthouse Lab in Glasgow | Wellcome Sanger Institute for the COVID-19 Genomics UK (COG-UK) Consortium | Harper VanSteenhouse, Yumi Kasai, David Gray, Carol Clugston, Anna Dominiczak and Alex Alderton, Roberto Amato, Jeffrey Barrett, Sonia Goncalves, Ewan Harrison, David K. Jackson, Ian Johnston, Dominic Kwiatkowski, Cordelia Langford, John Sillitoe on behalf of the Wellcome Sanger Institute COVID-19 Surveillance Team |
| EPI_ISL_2852319, EPI_ISL_2852320, EPI_ISL_2852321 | Lighthouse Laboratory Plymouth | Wellcome Sanger Institute for the COVID-19 Genomics UK (COG-UK) Consortium | Lighthouse Laboratory Plymouth and Alex Alderton, Roberto Amato, Jeffrey Barrett, Sonia Goncalves, Ewan Harrison, David K. Jackson, Ian Johnston, Dominic Kwiatkowski, Cordelia Langford, John Sillitoe on behalf of the Wellcome Sanger Institute COVID-19 Surveillance Team |
| EPI_ISL_2852322 | Lighthouse Lab in Glasgow | Wellcome Sanger Institute for the COVID-19 Genomics UK (COG-UK) Consortium | Harper VanSteenhouse, Yumi Kasai, David Gray, Carol Clugston, Anna Dominiczak and Alex Alderton, Roberto Amato, Jeffrey Barrett, Sonia Goncalves, Ewan Harrison, David K. Jackson, Ian Johnston, Dominic Kwiatkowski, Cordelia Langford, John Sillitoe on behalf of the Wellcome Sanger Institute COVID-19 Surveillance Team |
| EPI_ISL_2852323, EPI_ISL_2852324 | Lighthouse Laboratory Plymouth | Wellcome Sanger Institute for the COVID-19 Genomics UK (COG-UK) Consortium | Lighthouse Laboratory Plymouth and Alex Alderton, Roberto Amato, Jeffrey Barrett, Sonia Goncalves, Ewan Harrison, David K. Jackson, Ian Johnston, Dominic Kwiatkowski, Cordelia Langford, John Sillitoe on behalf of the Wellcome Sanger Institute COVID-19 Surveillance Team |
| EPI_ISL_2852325, EPI_ISL_2852326, EPI_ISL_2852327, EPI_ISL_2852328, EPI_ISL_2852329, EPI_ISL_2852330, | Lighthouse Lab in Glasgow | Wellcome Sanger Institute for the COVID-19 Genomics UK (COG-UK) Consortium | Harper VanSteenhouse, Yumi Kasai, David Gray, Carol Clugston, Anna Dominiczak and Alex Alderton, Roberto Amato, Jeffrey Barrett, Sonia Goncalves, Ewan Harrison, David K. Jackson, Ian Johnston, Dominic Kwiatkowski, Cordelia Langford, John Sillitoe on behalf of the Wellcome Sanger Institute COVID-19 Surveillance Team |

| | | | |
|---|---|---|---|
| EPI_ISL_2852331 | | | |
| EPI_ISL_2852332 | Lighthouse Laboratory Plymouth | Wellcome Sanger Institute for the COVID-19 Genomics UK (COG-UK) Consortium | Lighthouse Laboratory Plymouth and Alex Alderton, Roberto Amato, Jeffrey Barrett, Sonia Goncalves, Ewan Harrison, David K. Jackson, Ian Johnston, Dominic Kwiatkowski, Cordelia Langford, John Sillitoe on behalf of the Wellcome Sanger Institute COVID-19 Surveillance Team |
| EPI_ISL_2852333, EPI_ISL_2852334, EPI_ISL_2852335, EPI_ISL_2852336, EPI_ISL_2852337, EPI_ISL_2852338, EPI_ISL_2852339, EPI_ISL_2852340 | Lighthouse Lab in Glasgow | Wellcome Sanger Institute for the COVID-19 Genomics UK (COG-UK) Consortium | Harper VanSteenhouse, Yumi Kasai, David Gray, Carol Clugston, Anna Dominiczak and Alex Alderton, Roberto Amato, Jeffrey Barrett, Sonia Goncalves, Ewan Harrison, David K. Jackson, Ian Johnston, Dominic Kwiatkowski, Cordelia Langford, John Sillitoe on behalf of the Wellcome Sanger Institute COVID-19 Surveillance Team |
| EPI_ISL_2852341 | Lighthouse Laboratory Plymouth | Wellcome Sanger Institute for the COVID-19 Genomics UK (COG-UK) Consortium | Lighthouse Laboratory Plymouth and Alex Alderton, Roberto Amato, Jeffrey Barrett, Sonia Goncalves, Ewan Harrison, David K. Jackson, Ian Johnston, Dominic Kwiatkowski, Cordelia Langford, John Sillitoe on behalf of the Wellcome Sanger Institute COVID-19 Surveillance Team |
| EPI_ISL_2852342, EPI_ISL_2852343 | Lighthouse Lab in Glasgow | Wellcome Sanger Institute for the COVID-19 Genomics UK (COG-UK) Consortium | Harper VanSteenhouse, Yumi Kasai, David Gray, Carol Clugston, Anna Dominiczak and Alex Alderton, Roberto Amato, Jeffrey Barrett, Sonia Goncalves, Ewan Harrison, David K. Jackson, Ian Johnston, Dominic Kwiatkowski, Cordelia Langford, John Sillitoe on behalf of the Wellcome Sanger Institute COVID-19 Surveillance Team |
| EPI_ISL_2852344, EPI_ISL_2852345 | Lighthouse Laboratory Plymouth | Wellcome Sanger Institute for the COVID-19 Genomics UK (COG-UK) Consortium | Lighthouse Laboratory Plymouth and Alex Alderton, Roberto Amato, Jeffrey Barrett, Sonia Goncalves, Ewan Harrison, David K. Jackson, Ian Johnston, Dominic Kwiatkowski, Cordelia Langford, John Sillitoe on behalf of the Wellcome Sanger Institute COVID-19 Surveillance Team |
| EPI_ISL_2852346, EPI_ISL_2852347, EPI_ISL_2852348 | Lighthouse Lab in Glasgow | Wellcome Sanger Institute for the COVID-19 Genomics UK (COG-UK) Consortium | Harper VanSteenhouse, Yumi Kasai, David Gray, Carol Clugston, Anna Dominiczak and Alex Alderton, Roberto Amato, Jeffrey Barrett, Sonia Goncalves, Ewan Harrison, David K. Jackson, Ian Johnston, Dominic Kwiatkowski, Cordelia Langford, John Sillitoe on behalf of the Wellcome Sanger Institute COVID-19 Surveillance Team |
| EPI_ISL_2852349, EPI_ISL_2852350 | Lighthouse Laboratory Plymouth | Wellcome Sanger Institute for the COVID-19 Genomics UK (COG-UK) Consortium | Lighthouse Laboratory Plymouth and Alex Alderton, Roberto Amato, Jeffrey Barrett, Sonia Goncalves, Ewan Harrison, David K. Jackson, Ian Johnston, Dominic Kwiatkowski, Cordelia Langford, John Sillitoe on behalf of the Wellcome Sanger Institute COVID-19 Surveillance Team |
| EPI_ISL_2852351, EPI_ISL_2852352, EPI_ISL_2852353, EPI_ISL_2852354 | Lighthouse Lab in Glasgow | Wellcome Sanger Institute for the COVID-19 Genomics UK (COG-UK) Consortium | Harper VanSteenhouse, Yumi Kasai, David Gray, Carol Clugston, Anna Dominiczak and Alex Alderton, Roberto Amato, Jeffrey Barrett, Sonia Goncalves, Ewan Harrison, David K. Jackson, Ian Johnston, Dominic Kwiatkowski, Cordelia Langford, John Sillitoe on behalf of the Wellcome Sanger Institute COVID-19 Surveillance Team |
| EPI_ISL_2852355 | Lighthouse Laboratory Plymouth | Wellcome Sanger Institute for the COVID-19 Genomics UK (COG-UK) Consortium | Lighthouse Laboratory Plymouth and Alex Alderton, Roberto Amato, Jeffrey Barrett, Sonia Goncalves, Ewan Harrison, David K. Jackson, Ian Johnston, Dominic Kwiatkowski, Cordelia Langford, John Sillitoe on behalf of the Wellcome Sanger Institute COVID-19 Surveillance Team |
| EPI_ISL_2852356 | Lighthouse Lab in Glasgow | Wellcome Sanger Institute for the COVID-19 Genomics UK (COG-UK) Consortium | Harper VanSteenhouse, Yumi Kasai, David Gray, Carol Clugston, Anna Dominiczak and Alex Alderton, Roberto Amato, Jeffrey Barrett, Sonia Goncalves, Ewan Harrison, David K. Jackson, Ian Johnston, Dominic Kwiatkowski, Cordelia Langford, John Sillitoe on behalf of the Wellcome Sanger Institute COVID-19 Surveillance Team |
| EPI_ISL_2852357, EPI_ISL_2852358 | Lighthouse Laboratory Plymouth | Wellcome Sanger Institute for the COVID-19 Genomics UK (COG-UK) Consortium | Lighthouse Laboratory Plymouth and Alex Alderton, Roberto Amato, Jeffrey Barrett, Sonia Goncalves, Ewan Harrison, David K. Jackson, Ian Johnston, Dominic Kwiatkowski, Cordelia Langford, John Sillitoe on behalf of the Wellcome Sanger Institute COVID-19 Surveillance Team |
| EPI_ISL_2852359, EPI_ISL_2852360 | Lighthouse Lab in Glasgow | Wellcome Sanger Institute for the COVID-19 Genomics UK (COG-UK) Consortium | Harper VanSteenhouse, Yumi Kasai, David Gray, Carol Clugston, Anna Dominiczak and Alex Alderton, Roberto Amato, Jeffrey Barrett, Sonia Goncalves, Ewan Harrison, David K. Jackson, Ian Johnston, Dominic Kwiatkowski, Cordelia Langford, John Sillitoe on behalf of the Wellcome Sanger Institute COVID-19 Surveillance Team |
| EPI_ISL_2852361 | Lighthouse Laboratory Plymouth | Wellcome Sanger Institute for the COVID-19 Genomics UK (COG-UK) Consortium | Lighthouse Laboratory Plymouth and Alex Alderton, Roberto Amato, Jeffrey Barrett, Sonia Goncalves, Ewan Harrison, David K. Jackson, Ian Johnston, Dominic Kwiatkowski, Cordelia Langford, John Sillitoe on behalf of the Wellcome Sanger Institute COVID-19 Surveillance Team |
| EPI_ISL_2852362, EPI_ISL_2852363, EPI_ISL_2852364, EPI_ISL_2852365 | Lighthouse Lab in Glasgow | Wellcome Sanger Institute for the COVID-19 Genomics UK (COG-UK) Consortium | Harper VanSteenhouse, Yumi Kasai, David Gray, Carol Clugston, Anna Dominiczak and Alex Alderton, Roberto Amato, Jeffrey Barrett, Sonia Goncalves, Ewan Harrison, David K. Jackson, Ian Johnston, Dominic Kwiatkowski, Cordelia Langford, John Sillitoe on behalf of the Wellcome Sanger Institute COVID-19 Surveillance Team |
| EPI_ISL_2852366 | Lighthouse Laboratory Plymouth | Wellcome Sanger Institute for the COVID-19 Genomics UK (COG-UK) Consortium | Lighthouse Laboratory Plymouth and Alex Alderton, Roberto Amato, Jeffrey Barrett, Sonia Goncalves, Ewan Harrison, David K. Jackson, Ian Johnston, Dominic Kwiatkowski, Cordelia Langford, John Sillitoe on behalf of the Wellcome Sanger Institute COVID-19 Surveillance Team |
| EPI_ISL_2852367, EPI_ISL_2852368, EPI_ISL_2852369, EPI_ISL_2852370 | Lighthouse Lab in Glasgow | Wellcome Sanger Institute for the COVID-19 Genomics UK (COG-UK) Consortium | Harper VanSteenhouse, Yumi Kasai, David Gray, Carol Clugston, Anna Dominiczak and Alex Alderton, Roberto Amato, Jeffrey Barrett, Sonia Goncalves, Ewan Harrison, David K. Jackson, Ian Johnston, Dominic Kwiatkowski, Cordelia Langford, John Sillitoe on behalf of the Wellcome Sanger Institute COVID-19 Surveillance Team |
| EPI_ISL_2852371 | Lighthouse Laboratory Plymouth | Wellcome Sanger Institute for the COVID-19 Genomics UK (COG-UK) Consortium | Lighthouse Laboratory Plymouth and Alex Alderton, Roberto Amato, Jeffrey Barrett, Sonia Goncalves, Ewan Harrison, David K. Jackson, Ian Johnston, Dominic Kwiatkowski, Cordelia Langford, John Sillitoe on behalf of the Wellcome Sanger Institute COVID-19 Surveillance Team |
| EPI_ISL_2852372, EPI_ISL_2852373, EPI_ISL_2852375, EPI_ISL_2852376, EPI_ISL_2852377, EPI_ISL_2852378, EPI_ISL_2852379 | Lighthouse Lab in Glasgow | Wellcome Sanger Institute for the COVID-19 Genomics UK (COG-UK) Consortium | Harper VanSteenhouse, Yumi Kasai, David Gray, Carol Clugston, Anna Dominiczak and Alex Alderton, Roberto Amato, Jeffrey Barrett, Sonia Goncalves, Ewan Harrison, David K. Jackson, Ian Johnston, Dominic Kwiatkowski, Cordelia Langford, John Sillitoe on behalf of the Wellcome Sanger Institute COVID-19 Surveillance Team |
| EPI_ISL_2852380 | Lighthouse Laboratory Plymouth | Wellcome Sanger Institute for the COVID-19 Genomics UK (COG-UK) Consortium | Lighthouse Laboratory Plymouth and Alex Alderton, Roberto Amato, Jeffrey Barrett, Sonia Goncalves, Ewan Harrison, David K. Jackson, Ian Johnston, Dominic Kwiatkowski, Cordelia Langford, John Sillitoe on behalf of the Wellcome Sanger Institute COVID-19 Surveillance Team |
| EPI_ISL_2852381, EPI_ISL_2852382, EPI_ISL_2852383, EPI_ISL_2852384, EPI_ISL_2852385, EPI_ISL_2852386 | Lighthouse Lab in Glasgow | Wellcome Sanger Institute for the COVID-19 Genomics UK (COG-UK) Consortium | Harper VanSteenhouse, Yumi Kasai, David Gray, Carol Clugston, Anna Dominiczak and Alex Alderton, Roberto Amato, Jeffrey Barrett, Sonia Goncalves, Ewan Harrison, David K. Jackson, Ian Johnston, Dominic Kwiatkowski, Cordelia Langford, John Sillitoe on behalf of the Wellcome Sanger Institute COVID-19 Surveillance Team |
| EPI_ISL_2852387, EPI_ISL_2852388 | Lighthouse Laboratory Plymouth | Wellcome Sanger Institute for the COVID-19 Genomics UK (COG-UK) Consortium | Lighthouse Laboratory Plymouth and Alex Alderton, Roberto Amato, Jeffrey Barrett, Sonia Goncalves, Ewan Harrison, David K. Jackson, Ian Johnston, Dominic Kwiatkowski, Cordelia Langford, John Sillitoe on behalf of the Wellcome Sanger Institute COVID-19 Surveillance Team |
| EPI_ISL_2852389 | Lighthouse Lab in Glasgow | Wellcome Sanger Institute for the COVID-19 Genomics UK (COG-UK) Consortium | Harper VanSteenhouse, Yumi Kasai, David Gray, Carol Clugston, Anna Dominiczak and Alex Alderton, Roberto Amato, Jeffrey Barrett, Sonia Goncalves, Ewan Harrison, David K. Jackson, Ian Johnston, Dominic Kwiatkowski, Cordelia Langford, John Sillitoe on behalf of the Wellcome Sanger Institute COVID-19 Surveillance Team |
| EPI_ISL_2852390 | Lighthouse Laboratory Plymouth | Wellcome Sanger Institute for the COVID-19 Genomics UK (COG-UK) Consortium | Lighthouse Laboratory Plymouth and Alex Alderton, Roberto Amato, Jeffrey Barrett, Sonia Goncalves, Ewan Harrison, David K. Jackson, Ian Johnston, Dominic Kwiatkowski, Cordelia Langford, John Sillitoe on behalf of the Wellcome Sanger Institute COVID-19 Surveillance Team |

EPI_ISL_2852391, EPI_ISL_2852392, EPI_ISL_2852393, EPI_ISL_2852394, EPI_ISL_2852395, EPI_ISL_2852396, EPI_ISL_2852397, EPI_ISL_2852398, EPI_ISL_2852399, EPI_ISL_2852400, EPI_ISL_2852401, EPI_ISL_2852402, EPI_ISL_2852403

| | | | |
|---|---|---|---|
| see above | Lighthouse Lab in Glasgow | Wellcome Sanger Institute for the COVID-19 Genomics UK (COG-UK) Consortium | Harper VanSteenhouse, Yumi Kasai, David Gray, Carol Clugston, Anna Dominiczak and Alex Alderton, Roberto Amato, Jeffrey Barrett, Sonia Goncalves, Ewan Harrison, David K. Jackson, Ian Johnston, Dominic Kwiatkowski, Cordelia Langford, John Sillitoe on behalf of the Wellcome Sanger Institute COVID-19 Surveillance Team |
| EPI_ISL_2852404 | Lighthouse Laboratory Plymouth | Wellcome Sanger Institute for the COVID-19 Genomics UK (COG-UK) Consortium | Lighthouse Laboratory Plymouth and Alex Alderton, Roberto Amato, Jeffrey Barrett, Sonia Goncalves, Ewan Harrison, David K. Jackson, Ian Johnston, Dominic Kwiatkowski, Cordelia Langford, John Sillitoe on behalf of the Wellcome Sanger Institute COVID-19 Surveillance Team |
| EPI_ISL_2852405, EPI_ISL_2852407, EPI_ISL_2852408, EPI_ISL_2852409 | Lighthouse Lab in Glasgow | Wellcome Sanger Institute for the COVID-19 Genomics UK (COG-UK) Consortium | Harper VanSteenhouse, Yumi Kasai, David Gray, Carol Clugston, Anna Dominiczak and Alex Alderton, Roberto Amato, Jeffrey Barrett, Sonia Goncalves, Ewan Harrison, David K. Jackson, Ian Johnston, Dominic Kwiatkowski, Cordelia Langford, John Sillitoe on behalf of the Wellcome Sanger Institute COVID-19 Surveillance Team |
| EPI_ISL_2852410 | Lighthouse Laboratory Plymouth | Wellcome Sanger Institute for the COVID-19 Genomics UK (COG-UK) Consortium | Lighthouse Laboratory Plymouth and Alex Alderton, Roberto Amato, Jeffrey Barrett, Sonia Goncalves, Ewan Harrison, David K. Jackson, Ian Johnston, Dominic Kwiatkowski, Cordelia Langford, John Sillitoe on behalf of the Wellcome Sanger Institute COVID-19 Surveillance Team |

EPI_ISL_2852411, EPI_ISL_2852412, EPI_ISL_2852413, EPI_ISL_2852414, EPI_ISL_2852415, EPI_ISL_2852416, EPI_ISL_2852417, EPI_ISL_2852418, EPI_ISL_2852419, EPI_ISL_2852420, EPI_ISL_2852421, EPI_ISL_2852422, EPI_ISL_2852423

| | | | |
|---|---|---|---|
| see above | Lighthouse Lab in Glasgow | Wellcome Sanger Institute for the COVID-19 Genomics UK (COG-UK) Consortium | Harper VanSteenhouse, Yumi Kasai, David Gray, Carol Clugston, Anna Dominiczak and Alex Alderton, Roberto Amato, Jeffrey Barrett, Sonia Goncalves, Ewan Harrison, David K. Jackson, Ian Johnston, Dominic Kwiatkowski, Cordelia Langford, John Sillitoe on behalf of the Wellcome Sanger Institute COVID-19 Surveillance Team |
| EPI_ISL_2852424 | Lighthouse Laboratory Plymouth | Wellcome Sanger Institute for the COVID-19 Genomics UK (COG-UK) Consortium | Lighthouse Laboratory Plymouth and Alex Alderton, Roberto Amato, Jeffrey Barrett, Sonia Goncalves, Ewan Harrison, David K. Jackson, Ian Johnston, Dominic Kwiatkowski, Cordelia Langford, John Sillitoe on behalf of the Wellcome Sanger Institute COVID-19 Surveillance Team |
| EPI_ISL_2852425, EPI_ISL_2852426, EPI_ISL_2852427 | Lighthouse Lab in Glasgow | Wellcome Sanger Institute for the COVID-19 Genomics UK (COG-UK) Consortium | Harper VanSteenhouse, Yumi Kasai, David Gray, Carol Clugston, Anna Dominiczak and Alex Alderton, Roberto Amato, Jeffrey Barrett, Sonia Goncalves, Ewan Harrison, David K. Jackson, Ian Johnston, Dominic Kwiatkowski, Cordelia Langford, John Sillitoe on behalf of the Wellcome Sanger Institute COVID-19 Surveillance Team |
| EPI_ISL_2852428 | Lighthouse Laboratory Plymouth | Wellcome Sanger Institute for the COVID-19 Genomics UK (COG-UK) Consortium | Lighthouse Laboratory Plymouth and Alex Alderton, Roberto Amato, Jeffrey Barrett, Sonia Goncalves, Ewan Harrison, David K. Jackson, Ian Johnston, Dominic Kwiatkowski, Cordelia Langford, John Sillitoe on behalf of the Wellcome Sanger Institute COVID-19 Surveillance Team |
| EPI_ISL_2852429, EPI_ISL_2852430, EPI_ISL_2852431, EPI_ISL_2852432, EPI_ISL_2852433, | Lighthouse Lab in Glasgow | Wellcome Sanger Institute for the COVID-19 Genomics UK (COG-UK) Consortium | Harper VanSteenhouse, Yumi Kasai, David Gray, Carol Clugston, Anna Dominiczak and Alex Alderton, Roberto Amato, Jeffrey Barrett, Sonia Goncalves, Ewan Harrison, David K. Jackson, Ian Johnston, Dominic Kwiatkowski, Cordelia Langford, John Sillitoe on behalf of the Wellcome Sanger Institute COVID-19 Surveillance Team |

| | | | |
|---|---|---|---|
| EPI_ISL_2852434, EPI_ISL_2852435, EPI_ISL_2852436 | | | |
| EPI_ISL_2852437 | Lighthouse Laboratory Plymouth | Wellcome Sanger Institute for the COVID-19 Genomics UK (COG-UK) Consortium | Lighthouse Laboratory Plymouth and Alex Alderton, Roberto Amato, Jeffrey Barrett, Sonia Goncalves, Ewan Harrison, David K. Jackson, Ian Johnston, Dominic Kwiatkowski, Cordelia Langford, John Sillitoe on behalf of the Wellcome Sanger Institute COVID-19 Surveillance Team |
| EPI_ISL_2852438, EPI_ISL_2852439 | Lighthouse Lab in Glasgow | Wellcome Sanger Institute for the COVID-19 Genomics UK (COG-UK) Consortium | Harper VanSteenhouse, Yumi Kasai, David Gray, Carol Clugston, Anna Dominiczak and Alex Alderton, Roberto Amato, Jeffrey Barrett, Sonia Goncalves, Ewan Harrison, David K. Jackson, Ian Johnston, Dominic Kwiatkowski, Cordelia Langford, John Sillitoe on behalf of the Wellcome Sanger Institute COVID-19 Surveillance Team |
| EPI_ISL_2852440 | Lighthouse Laboratory Plymouth | Wellcome Sanger Institute for the COVID-19 Genomics UK (COG-UK) Consortium | Lighthouse Laboratory Plymouth and Alex Alderton, Roberto Amato, Jeffrey Barrett, Sonia Goncalves, Ewan Harrison, David K. Jackson, Ian Johnston, Dominic Kwiatkowski, Cordelia Langford, John Sillitoe on behalf of the Wellcome Sanger Institute COVID-19 Surveillance Team |
| EPI_ISL_2852441, EPI_ISL_2852442, EPI_ISL_2852443, EPI_ISL_2852444, EPI_ISL_2852445 | Lighthouse Lab in Glasgow | Wellcome Sanger Institute for the COVID-19 Genomics UK (COG-UK) Consortium | Harper VanSteenhouse, Yumi Kasai, David Gray, Carol Clugston, Anna Dominiczak and Alex Alderton, Roberto Amato, Jeffrey Barrett, Sonia Goncalves, Ewan Harrison, David K. Jackson, Ian Johnston, Dominic Kwiatkowski, Cordelia Langford, John Sillitoe on behalf of the Wellcome Sanger Institute COVID-19 Surveillance Team |
| EPI_ISL_2852446 | Lighthouse Laboratory Plymouth | Wellcome Sanger Institute for the COVID-19 Genomics UK (COG-UK) Consortium | Lighthouse Laboratory Plymouth and Alex Alderton, Roberto Amato, Jeffrey Barrett, Sonia Goncalves, Ewan Harrison, David K. Jackson, Ian Johnston, Dominic Kwiatkowski, Cordelia Langford, John Sillitoe on behalf of the Wellcome Sanger Institute COVID-19 Surveillance Team |
| EPI_ISL_2852447 | Lighthouse Lab in Glasgow | Wellcome Sanger Institute for the COVID-19 Genomics UK (COG-UK) Consortium | Harper VanSteenhouse, Yumi Kasai, David Gray, Carol Clugston, Anna Dominiczak and Alex Alderton, Roberto Amato, Jeffrey Barrett, Sonia Goncalves, Ewan Harrison, David K. Jackson, Ian Johnston, Dominic Kwiatkowski, Cordelia Langford, John Sillitoe on behalf of the Wellcome Sanger Institute COVID-19 Surveillance Team |
| EPI_ISL_2852448 | Lighthouse Laboratory Plymouth | Wellcome Sanger Institute for the COVID-19 Genomics UK (COG-UK) Consortium | Lighthouse Laboratory Plymouth and Alex Alderton, Roberto Amato, Jeffrey Barrett, Sonia Goncalves, Ewan Harrison, David K. Jackson, Ian Johnston, Dominic Kwiatkowski, Cordelia Langford, John Sillitoe on behalf of the Wellcome Sanger Institute COVID-19 Surveillance Team |
| EPI_ISL_2852449, EPI_ISL_2852450 | Lighthouse Lab in Glasgow | Wellcome Sanger Institute for the COVID-19 Genomics UK (COG-UK) Consortium | Harper VanSteenhouse, Yumi Kasai, David Gray, Carol Clugston, Anna Dominiczak and Alex Alderton, Roberto Amato, Jeffrey Barrett, Sonia Goncalves, Ewan Harrison, David K. Jackson, Ian Johnston, Dominic Kwiatkowski, Cordelia Langford, John Sillitoe on behalf of the Wellcome Sanger Institute COVID-19 Surveillance Team |
| EPI_ISL_2852451, EPI_ISL_2852452 | Lighthouse Laboratory Plymouth | Wellcome Sanger Institute for the COVID-19 Genomics UK (COG-UK) Consortium | Lighthouse Laboratory Plymouth and Alex Alderton, Roberto Amato, Jeffrey Barrett, Sonia Goncalves, Ewan Harrison, David K. Jackson, Ian Johnston, Dominic Kwiatkowski, Cordelia Langford, John Sillitoe on behalf of the Wellcome Sanger Institute COVID-19 Surveillance Team |
| EPI_ISL_2852453, EPI_ISL_2852454, EPI_ISL_2852455, EPI_ISL_2852456, EPI_ISL_2852457, EPI_ISL_2852458, EPI_ISL_2852459 | Lighthouse Lab in Glasgow | Wellcome Sanger Institute for the COVID-19 Genomics UK (COG-UK) Consortium | Harper VanSteenhouse, Yumi Kasai, David Gray, Carol Clugston, Anna Dominiczak and Alex Alderton, Roberto Amato, Jeffrey Barrett, Sonia Goncalves, Ewan Harrison, David K. Jackson, Ian Johnston, Dominic Kwiatkowski, Cordelia Langford, John Sillitoe on behalf of the Wellcome Sanger Institute COVID-19 Surveillance Team |
| EPI_ISL_2852460 | Lighthouse Laboratory Plymouth | Wellcome Sanger Institute for the COVID-19 Genomics UK (COG-UK) Consortium | Lighthouse Laboratory Plymouth and Alex Alderton, Roberto Amato, Jeffrey Barrett, Sonia Goncalves, Ewan Harrison, David K. Jackson, Ian Johnston, Dominic Kwiatkowski, Cordelia Langford, John Sillitoe on behalf of the Wellcome Sanger Institute COVID-19 Surveillance Team |
| EPI_ISL_2852461, EPI_ISL_2852462 | Lighthouse Lab in Glasgow | Wellcome Sanger Institute for the COVID-19 Genomics UK (COG-UK) Consortium | Harper VanSteenhouse, Yumi Kasai, David Gray, Carol Clugston, Anna Dominiczak and Alex Alderton, Roberto Amato, Jeffrey Barrett, Sonia Goncalves, Ewan Harrison, David K. Jackson, Ian Johnston, Dominic Kwiatkowski, Cordelia Langford, John Sillitoe on behalf of the Wellcome Sanger Institute COVID-19 Surveillance Team |
| EPI_ISL_2852463 | Lighthouse Laboratory Plymouth | Wellcome Sanger Institute for the COVID-19 Genomics UK (COG-UK) Consortium | Lighthouse Laboratory Plymouth and Alex Alderton, Roberto Amato, Jeffrey Barrett, Sonia Goncalves, Ewan Harrison, David K. Jackson, Ian Johnston, Dominic Kwiatkowski, Cordelia Langford, John Sillitoe on behalf of the Wellcome Sanger Institute COVID-19 Surveillance Team |
| EPI_ISL_2852464, EPI_ISL_2852465 | Lighthouse Lab in Glasgow | Wellcome Sanger Institute for the COVID-19 Genomics UK (COG-UK) Consortium | Harper VanSteenhouse, Yumi Kasai, David Gray, Carol Clugston, Anna Dominiczak and Alex Alderton, Roberto Amato, Jeffrey Barrett, Sonia Goncalves, Ewan Harrison, David K. Jackson, Ian Johnston, Dominic Kwiatkowski, Cordelia Langford, John Sillitoe on behalf of the Wellcome Sanger Institute COVID-19 Surveillance Team |
| EPI_ISL_2852466 | Lighthouse Laboratory Plymouth | Wellcome Sanger Institute for the COVID-19 Genomics UK (COG-UK) Consortium | Lighthouse Laboratory Plymouth and Alex Alderton, Roberto Amato, Jeffrey Barrett, Sonia Goncalves, Ewan Harrison, David K. Jackson, Ian Johnston, Dominic Kwiatkowski, Cordelia Langford, John Sillitoe on behalf of the Wellcome Sanger Institute COVID-19 Surveillance Team |
| EPI_ISL_2852467, EPI_ISL_2852468, EPI_ISL_2852469, EPI_ISL_2852470, EPI_ISL_2852471, EPI_ISL_2852472, EPI_ISL_2852473, EPI_ISL_2852474, EPI_ISL_2852475, EPI_ISL_2852476, EPI_ISL_2852477, EPI_ISL_2852478, EPI_ISL_2852479 | | | |
| see above | Lighthouse Lab in Glasgow | Wellcome Sanger Institute for the COVID-19 Genomics UK (COG-UK) Consortium | Harper VanSteenhouse, Yumi Kasai, David Gray, Carol Clugston, Anna Dominiczak and Alex Alderton, Roberto Amato, Jeffrey Barrett, Sonia Goncalves, Ewan Harrison, David K. Jackson, Ian Johnston, Dominic Kwiatkowski, Cordelia Langford, John Sillitoe on behalf of the Wellcome Sanger Institute COVID-19 Surveillance Team |
| EPI_ISL_2852480 | Lighthouse Lab in Milton Keynes | Wellcome Sanger Institute for the COVID-19 Genomics UK (COG-UK) Consortium | The Lighthouse Lab in Milton Keynes and Alex Alderton, Roberto Amato, Jeffrey Barrett, Sonia Goncalves, Ewan Harrison, David K. Jackson, Ian Johnston, Dominic Kwiatkowski, Cordelia Langford, John Sillitoe on behalf of the Wellcome Sanger Institute COVID-19 Surveillance Team |
| EPI_ISL_2852481, EPI_ISL_2852482 | Lighthouse Laboratory Plymouth | Wellcome Sanger Institute for the COVID-19 Genomics UK (COG-UK) Consortium | Lighthouse Laboratory Plymouth and Alex Alderton, Roberto Amato, Jeffrey Barrett, Sonia Goncalves, Ewan Harrison, David K. Jackson, Ian Johnston, Dominic Kwiatkowski, Cordelia Langford, John Sillitoe on behalf of the Wellcome Sanger Institute COVID-19 Surveillance Team |
| EPI_ISL_2852483, EPI_ISL_2852484, EPI_ISL_2852485, EPI_ISL_2852486, EPI_ISL_2852487, EPI_ISL_2852488, EPI_ISL_2852489 | Lighthouse Lab in Glasgow | Wellcome Sanger Institute for the COVID-19 Genomics UK (COG-UK) Consortium | Harper VanSteenhouse, Yumi Kasai, David Gray, Carol Clugston, Anna Dominiczak and Alex Alderton, Roberto Amato, Jeffrey Barrett, Sonia Goncalves, Ewan Harrison, David K. Jackson, Ian Johnston, Dominic Kwiatkowski, Cordelia Langford, John Sillitoe on behalf of the Wellcome Sanger Institute COVID-19 Surveillance Team |
| EPI_ISL_2852490 | Lighthouse Lab in Alderley Park | Wellcome Sanger Institute for the COVID-19 Genomics UK (COG-UK) Consortium | Jacquelyn Wynn, Mairead Hyland, The Lighthouse Lab in Alderley Park and Alex Alderton, Roberto Amato, Jeffrey Barrett, Sonia Goncalves, Ewan Harrison, David K. Jackson, Ian Johnston, Dominic Kwiatkowski, Cordelia Langford, John Sillitoe on behalf of the Wellcome Sanger Institute COVID-19 Surveillance Team |
| EPI_ISL_2852491, EPI_ISL_2852492, EPI_ISL_2852493, EPI_ISL_2852494, EPI_ISL_2852495, EPI_ISL_2852496, EPI_ISL_2852497, EPI_ISL_2852498, EPI_ISL_2852499, EPI_ISL_2852500, EPI_ISL_2852501, EPI_ISL_2852502, EPI_ISL_2852503, EPI_ISL_2852504, EPI_ISL_2852505, EPI_ISL_2852506, EPI_ISL_2852507, EPI_ISL_2852508, EPI_ISL_2852509, EPI_ISL_2852510, EPI_ISL_2852511, EPI_ISL_2852512, EPI_ISL_2852513, EPI_ISL_2852514, EPI_ISL_2852515, EPI_ISL_2852516, EPI_ISL_2852517, EPI_ISL_2852518, EPI_ISL_2852519, EPI_ISL_2852520, EPI_ISL_2852521, EPI_ISL_2852522, EPI_ISL_2852523, EPI_ISL_2852524, EPI_ISL_2852525, EPI_ISL_2852526, EPI_ISL_2852527, EPI_ISL_2852528, EPI_ISL_2852529, EPI_ISL_2852530, EPI_ISL_2852531, EPI_ISL_2852532, EPI_ISL_2852533, EPI_ISL_2852534, EPI_ISL_2852535, EPI_ISL_2852536, EPI_ISL_2852537, EPI_ISL_2852538, EPI_ISL_2852539, EPI_ISL_2852540, EPI_ISL_2852541, EPI_ISL_2852542, EPI_ISL_2852543, EPI_ISL_2852544, EPI_ISL_2852545, EPI_ISL_2852546, EPI_ISL_2852547, EPI_ISL_2852548, EPI_ISL_2852549, EPI_ISL_2852550, EPI_ISL_2852551, EPI_ISL_2852552, EPI_ISL_2852553, EPI_ISL_2852554, EPI_ISL_2852555, EPI_ISL_2852556, EPI_ISL_2852557, EPI_ISL_2852558, EPI_ISL_2852559, EPI_ISL_2852560, EPI_ISL_2852561, EPI_ISL_2852562, EPI_ISL_2852563, EPI_ISL_2852564, EPI_ISL_2852565, EPI_ISL_2852566, EPI_ISL_2852567, EPI_ISL_2852568, EPI_ISL_2852569, EPI_ISL_2852570, EPI_ISL_2852571, EPI_ISL_2852572, EPI_ISL_2852573, EPI_ISL_2852574, EPI_ISL_2852575 | | | |
| see above | Lighthouse Lab in Glasgow | Wellcome Sanger Institute for the COVID-19 Genomics UK (COG-UK) Consortium | Harper VanSteenhouse, Yumi Kasai, David Gray, Carol Clugston, Anna Dominiczak and Alex Alderton, Roberto Amato, Jeffrey Barrett, Sonia Goncalves, Ewan Harrison, David K. Jackson, Ian Johnston, Dominic Kwiatkowski, Cordelia Langford, John Sillitoe on behalf of the Wellcome Sanger Institute COVID-19 Surveillance Team |
| EPI_ISL_2852576 | Lighthouse Lab in Alderley Park | Wellcome Sanger Institute for the COVID-19 Genomics UK (COG-UK) Consortium | Jacquelyn Wynn, Mairead Hyland, The Lighthouse Lab in Alderley Park and Alex Alderton, Roberto Amato, Jeffrey Barrett, Sonia Goncalves, Ewan Harrison, David K. Jackson, Ian Johnston, Dominic Kwiatkowski, Cordelia Langford, John Sillitoe on behalf of the Wellcome Sanger Institute COVID-19 Surveillance Team |
| EPI_ISL_2852577, EPI_ISL_2852579, EPI_ISL_2852580, EPI_ISL_2852581, EPI_ISL_2852582, EPI_ISL_2852583, EPI_ISL_2852584, EPI_ISL_2852585, EPI_ISL_2852586, EPI_ISL_2852587, EPI_ISL_2852588, EPI_ISL_2852589, EPI_ISL_2852590, EPI_ISL_2852592, EPI_ISL_2852593, EPI_ISL_2852594, EPI_ISL_2852595, EPI_ISL_2852596, EPI_ISL_2852597, EPI_ISL_2852598, EPI_ISL_2852599, EPI_ISL_2852600, EPI_ISL_2852601, EPI_ISL_2852602, EPI_ISL_2852603, EPI_ISL_2852604, EPI_ISL_2852605, EPI_ISL_2852606, EPI_ISL_2852607, EPI_ISL_2852608, EPI_ISL_2852609, EPI_ISL_2852610, EPI_ISL_2852611, EPI_ISL_2852612, EPI_ISL_2852613, EPI_ISL_2852614, EPI_ISL_2852615, EPI_ISL_2852616, EPI_ISL_2852617, EPI_ISL_2852618, EPI_ISL_2852619, EPI_ISL_2852620, EPI_ISL_2852621, EPI_ISL_2852622, EPI_ISL_2852623, EPI_ISL_2852624, EPI_ISL_2852625, EPI_ISL_2852626, EPI_ISL_2852627, EPI_ISL_2852628, EPI_ISL_2852629, EPI_ISL_2852630, EPI_ISL_2852631, EPI_ISL_2852632, EPI_ISL_2852633, EPI_ISL_2852634, EPI_ISL_2852635, EPI_ISL_2852636, EPI_ISL_2852637, EPI_ISL_2852638, EPI_ISL_2852639, EPI_ISL_2852640, EPI_ISL_2852641, EPI_ISL_2852642, EPI_ISL_2852643, EPI_ISL_2852644, EPI_ISL_2852645, EPI_ISL_2852646, EPI_ISL_2852647, EPI_ISL_2852648, EPI_ISL_2852649, EPI_ISL_2852650, EPI_ISL_2852651, EPI_ISL_2852652, EPI_ISL_2852653, EPI_ISL_2852654, EPI_ISL_2852655, EPI_ISL_2852656, EPI_ISL_2852657, EPI_ISL_2852658, EPI_ISL_2852659, EPI_ISL_2852660, EPI_ISL_2852661, EPI_ISL_2852662, EPI_ISL_2852663, EPI_ISL_2852664, EPI_ISL_2852665, EPI_ISL_2852666, EPI_ISL_2852667, EPI_ISL_2852668, EPI_ISL_2852669, EPI_ISL_2852670, EPI_ISL_2852671, EPI_ISL_2852672, EPI_ISL_2852673, EPI_ISL_2852674, EPI_ISL_2852675, EPI_ISL_2852676, EPI_ISL_2852677 | | | |
| see above | Lighthouse Lab in Glasgow | Wellcome Sanger Institute for the COVID-19 Genomics UK (COG-UK) Consortium | Harper VanSteenhouse, Yumi Kasai, David Gray, Carol Clugston, Anna Dominiczak and Alex Alderton, Roberto Amato, Jeffrey Barrett, Sonia Goncalves, Ewan Harrison, David K. Jackson, Ian Johnston, Dominic Kwiatkowski, Cordelia Langford, John Sillitoe on behalf of the Wellcome Sanger Institute COVID-19 Surveillance Team |
| EPI_ISL_2852678 | Lighthouse Lab in Alderley Park | Wellcome Sanger Institute for the COVID-19 Genomics UK (COG-UK) Consortium | Jacquelyn Wynn, Mairead Hyland, The Lighthouse Lab in Alderley Park and Alex Alderton, Roberto Amato, Jeffrey Barrett, Sonia Goncalves, Ewan Harrison, David K. Jackson, Ian Johnston, Dominic Kwiatkowski, Cordelia Langford, John Sillitoe on behalf of the Wellcome Sanger Institute COVID-19 Surveillance Team |
| EPI_ISL_2852679, EPI_ISL_2852680, EPI_ISL_2852681, EPI_ISL_2852682, EPI_ISL_2852683, EPI_ISL_2852684, EPI_ISL_2852685, EPI_ISL_2852686, EPI_ISL_2852687, EPI_ISL_2852688, EPI_ISL_2852689, EPI_ISL_2852690, EPI_ISL_2852691, EPI_ISL_2852692, EPI_ISL_2852693, EPI_ISL_2852694, EPI_ISL_2852695, EPI_ISL_2852696, EPI_ISL_2852697 | | | |
| see above | Lighthouse Lab in Glasgow | Wellcome Sanger Institute for the COVID-19 Genomics UK (COG-UK) Consortium | Harper VanSteenhouse, Yumi Kasai, David Gray, Carol Clugston, Anna Dominiczak and Alex Alderton, Roberto Amato, Jeffrey Barrett, Sonia Goncalves, Ewan Harrison, David K. Jackson, Ian Johnston, Dominic Kwiatkowski, Cordelia Langford, John Sillitoe on behalf of the Wellcome Sanger Institute COVID-19 Surveillance Team |
| EPI_ISL_2852698 | Lighthouse Lab in Alderley Park | Wellcome Sanger Institute for the COVID-19 Genomics UK (COG-UK) Consortium | Jacquelyn Wynn, Mairead Hyland, The Lighthouse Lab in Alderley Park and Alex Alderton, Roberto Amato, Jeffrey Barrett, Sonia Goncalves, Ewan Harrison, David K. Jackson, Ian Johnston, Dominic Kwiatkowski, Cordelia Langford, John Sillitoe on behalf of the Wellcome Sanger Institute COVID-19 Surveillance Team |
| EPI_ISL_2852699, EPI_ISL_2852700, EPI_ISL_2852701, EPI_ISL_2852702, EPI_ISL_2852703, EPI_ISL_2852704, EPI_ISL_2852705, EPI_ISL_2852706, EPI_ISL_2852707, EPI_ISL_2852708, EPI_ISL_2852709, EPI_ISL_2852710, EPI_ISL_2852711, EPI_ISL_2852712, EPI_ISL_2852713, EPI_ISL_2852714, EPI_ISL_2852715, EPI_ISL_2852716, EPI_ISL_2852717, EPI_ISL_2852718, EPI_ISL_2852719, EPI_ISL_2852720, EPI_ISL_2852721, EPI_ISL_2852722, EPI_ISL_2852723, EPI_ISL_2852724, EPI_ISL_2852725, EPI_ISL_2852726, EPI_ISL_2852727, EPI_ISL_2852728, EPI_ISL_2852729, EPI_ISL_2852730, EPI_ISL_2852731, EPI_ISL_2852732, EPI_ISL_2852733, EPI_ISL_2852734, EPI_ISL_2852735, EPI_ISL_2852736, EPI_ISL_2852737, EPI_ISL_2852738, EPI_ISL_2852739, EPI_ISL_2852740, EPI_ISL_2852741, EPI_ISL_2852742, EPI_ISL_2852743, EPI_ISL_2852744, EPI_ISL_2852745, EPI_ISL_2852746, EPI_ISL_2852747, EPI_ISL_2852748, EPI_ISL_2852749, EPI_ISL_2852750, EPI_ISL_2852751, EPI_ISL_2852752, EPI_ISL_2852753, EPI_ISL_2852754, EPI_ISL_2852755, EPI_ISL_2852756, EPI_ISL_2852757, EPI_ISL_2852758, EPI_ISL_2852759, EPI_ISL_2852760, EPI_ISL_2852761, EPI_ISL_2852762, EPI_ISL_2852763, EPI_ISL_2852764, EPI_ISL_2852765, EPI_ISL_2852766, EPI_ISL_2852767, EPI_ISL_2852768, EPI_ISL_2852769, EPI_ISL_2852770, EPI_ISL_2852771, EPI_ISL_2852772, EPI_ISL_2852773, EPI_ISL_2852774, EPI_ISL_2852775, EPI_ISL_2852776, EPI_ISL_2852777, EPI_ISL_2852778, EPI_ISL_2852779, EPI_ISL_2852780, EPI_ISL_2852781, EPI_ISL_2852782, EPI_ISL_2852783, EPI_ISL_2852784, EPI_ISL_2852785, EPI_ISL_2852786, EPI_ISL_2852787, EPI_ISL_2852788, EPI_ISL_2852789, EPI_ISL_2852790, EPI_ISL_2852791, EPI_ISL_2852792, EPI_ISL_2852793, EPI_ISL_2852794, EPI_ISL_2852795, EPI_ISL_2852796, EPI_ISL_2852797, EPI_ISL_2852798, EPI_ISL_2852799, EPI_ISL_2852800, EPI_ISL_2852801, EPI_ISL_2852802, EPI_ISL_2852803, EPI_ISL_2852804, EPI_ISL_2852805, EPI_ISL_2852806, EPI_ISL_2852807, EPI_ISL_2852808, EPI_ISL_2852809, EPI_ISL_2852810, EPI_ISL_2852811, EPI_ISL_2852812, EPI_ISL_2852813, EPI_ISL_2852814, EPI_ISL_2852815, EPI_ISL_2852816, EPI_ISL_2852817, EPI_ISL_2852818 | | | |
| see above | Lighthouse Lab in Glasgow | Wellcome Sanger Institute for the COVID-19 Genomics UK (COG-UK) Consortium | Harper VanSteenhouse, Yumi Kasai, David Gray, Carol Clugston, Anna Dominiczak and Alex Alderton, Roberto Amato, Jeffrey Barrett, Sonia Goncalves, Ewan Harrison, David K. Jackson, Ian Johnston, Dominic Kwiatkowski, Cordelia Langford, John Sillitoe on behalf of the Wellcome Sanger Institute COVID-19 Surveillance Team |
| EPI_ISL_2852819 | Lighthouse Laboratory Plymouth | Wellcome Sanger Institute for the COVID-19 Genomics UK (COG-UK) Consortium | Lighthouse Laboratory Plymouth and Alex Alderton, Roberto Amato, Jeffrey Barrett, Sonia Goncalves, Ewan Harrison, David K. Jackson, Ian Johnston, Dominic Kwiatkowski, Cordelia Langford, John Sillitoe on behalf of the Wellcome Sanger Institute COVID-19 Surveillance Team |
| EPI_ISL_2852820, EPI_ISL_2852821, EPI_ISL_2852822, EPI_ISL_2852823, EPI_ISL_2852824, EPI_ISL_2852825, EPI_ISL_2852826, EPI_ISL_2852827, EPI_ISL_2852828, EPI_ISL_2852829, EPI_ISL_2852830, EPI_ISL_2852831, EPI_ISL_2852832, EPI_ISL_2852833, EPI_ISL_2852834, EPI_ISL_2852835, EPI_ISL_2852836, EPI_ISL_2852837, EPI_ISL_2852838, EPI_ISL_2852839, EPI_ISL_2852840, EPI_ISL_2852841, EPI_ISL_2852842, EPI_ISL_2852843, EPI_ISL_2852844, EPI_ISL_2852845, EPI_ISL_2852846, EPI_ISL_2852847, EPI_ISL_2852848, EPI_ISL_2852849, EPI_ISL_2852850, EPI_ISL_2852851 | | | |
| see above | Lighthouse Lab in Glasgow | Wellcome Sanger Institute for the COVID-19 Genomics UK (COG-UK) Consortium | Harper VanSteenhouse, Yumi Kasai, David Gray, Carol Clugston, Anna Dominiczak and Alex Alderton, Roberto Amato, Jeffrey Barrett, Sonia Goncalves, Ewan Harrison, David K. Jackson, Ian Johnston, Dominic Kwiatkowski, Cordelia Langford, John Sillitoe on behalf of the Wellcome Sanger Institute COVID-19 Surveillance Team |
| EPI_ISL_2852852, | Lighthouse Lab in Alderley Park | Wellcome Sanger Institute for the COVID-19 Genomics UK (COG-UK) | Jacquelyn Wynn, Mairead Hyland, The Lighthouse Lab in Alderley Park and Alex Alderton, Roberto Amato, Jeffrey Barrett, Sonia Goncalves, Ewan Harrison, David K. Jackson, Ian Johnston, Dominic Kwiatkowski, Cordelia Langford, John Sillitoe on |

| | | | |
|---|---|---|---|
| EPI_ISL_2852853, EPI_ISL_2852854 | | Consortium | behalf of the Wellcome Sanger Institute COVID-19 Surveillance Team |
| EPI_ISL_2852855 | Lighthouse Lab in Milton Keynes | Wellcome Sanger Institute for the COVID-19 Genomics UK (COG-UK) Consortium | The Lighthouse Lab in Milton Keynes and Alex Alderton, Roberto Amato, Jeffrey Barrett, Sonia Goncalves, Ewan Harrison, David K. Jackson, Ian Johnston, Dominic Kwiatkowski, Cordelia Langford, John Sillitoe on behalf of the Wellcome Sanger Institute COVID-19 Surveillance Team |
| EPI_ISL_2852856, EPI_ISL_2852857, EPI_ISL_2852858, EPI_ISL_2852859, EPI_ISL_2852860, EPI_ISL_2852861, EPI_ISL_2852862, EPI_ISL_2852863, EPI_ISL_2852864, EPI_ISL_2852865, EPI_ISL_2852867, EPI_ISL_2852868, EPI_ISL_2852869, EPI_ISL_2852870, EPI_ISL_2852871, EPI_ISL_2852872, EPI_ISL_2852873, EPI_ISL_2852874, EPI_ISL_2852875, EPI_ISL_2852876, EPI_ISL_2852877, EPI_ISL_2852878, EPI_ISL_2852879, EPI_ISL_2852880, EPI_ISL_2852881, EPI_ISL_2852882, EPI_ISL_2852883, EPI_ISL_2852884, EPI_ISL_2852885, EPI_ISL_2852886, EPI_ISL_2852887, EPI_ISL_2852888, EPI_ISL_2852889, EPI_ISL_2852890, EPI_ISL_2852891, EPI_ISL_2852892, EPI_ISL_2852893, EPI_ISL_2852894, EPI_ISL_2852895, EPI_ISL_2852896, EPI_ISL_2852897, EPI_ISL_2852898, EPI_ISL_2852899, EPI_ISL_2852900, EPI_ISL_2852901, EPI_ISL_2852902, EPI_ISL_2852903, EPI_ISL_2852904, EPI_ISL_2852905, EPI_ISL_2852906, EPI_ISL_2852907, EPI_ISL_2852908, EPI_ISL_2852909, EPI_ISL_2852910, EPI_ISL_2852911, EPI_ISL_2852912, EPI_ISL_2852913, EPI_ISL_2852914, EPI_ISL_2852915, EPI_ISL_2852916, EPI_ISL_2852917, EPI_ISL_2852918, EPI_ISL_2852919, EPI_ISL_2852920, EPI_ISL_2852921, EPI_ISL_2852922, EPI_ISL_2852923, EPI_ISL_2852924, EPI_ISL_2852925, EPI_ISL_2852926, EPI_ISL_2852927, EPI_ISL_2852928, EPI_ISL_2852929, EPI_ISL_2852930, EPI_ISL_2852932, EPI_ISL_2852933, EPI_ISL_2852934, EPI_ISL_2852935, EPI_ISL_2852936, EPI_ISL_2852937, EPI_ISL_2852938, EPI_ISL_2852939, EPI_ISL_2852940, EPI_ISL_2852941, EPI_ISL_2852942, EPI_ISL_2852943, EPI_ISL_2852944, EPI_ISL_2852945, EPI_ISL_2852946, EPI_ISL_2852947, EPI_ISL_2852948, EPI_ISL_2852949, EPI_ISL_2852950, EPI_ISL_2852951, EPI_ISL_2852952, EPI_ISL_2852953, EPI_ISL_2852954, EPI_ISL_2852955, EPI_ISL_2852956, EPI_ISL_2852957, EPI_ISL_2852958, EPI_ISL_2852959, EPI_ISL_2852960, EPI_ISL_2852961, EPI_ISL_2852962, EPI_ISL_2852963, EPI_ISL_2852964, EPI_ISL_2852966, EPI_ISL_2852967, EPI_ISL_2852968, EPI_ISL_2852969, EPI_ISL_2852970, EPI_ISL_2852971, EPI_ISL_2852973, EPI_ISL_2852974, EPI_ISL_2852975, EPI_ISL_2852976, EPI_ISL_2852977, EPI_ISL_2852978, EPI_ISL_2852979, EPI_ISL_2852980, EPI_ISL_2852981, EPI_ISL_2852982, EPI_ISL_2852983, EPI_ISL_2852984, EPI_ISL_2852985, EPI_ISL_2852986, EPI_ISL_2852987, EPI_ISL_2852988, EPI_ISL_2852989, EPI_ISL_2852990, EPI_ISL_2852991, EPI_ISL_2852992, EPI_ISL_2852993, EPI_ISL_2852994, EPI_ISL_2852995, EPI_ISL_2852996, EPI_ISL_2852997, EPI_ISL_2852998, EPI_ISL_2852999, EPI_ISL_2853000, EPI_ISL_2853001, EPI_ISL_2853002, EPI_ISL_2853003, EPI_ISL_2853004, EPI_ISL_2853005, EPI_ISL_2853006, EPI_ISL_2853008, EPI_ISL_2853009, EPI_ISL_2853010, EPI_ISL_2853011, EPI_ISL_2853012, EPI_ISL_2853013 | | | |
| see above | Lighthouse Lab in Glasgow | Wellcome Sanger Institute for the COVID-19 Genomics UK (COG-UK) Consortium | Harper VanSteenhouse, Yumi Kasai, David Gray, Carol Clugston, Anna Dominiczak and Alex Alderton, Roberto Amato, Jeffrey Barrett, Sonia Goncalves, Ewan Harrison, David K. Jackson, Ian Johnston, Dominic Kwiatkowski, Cordelia Langford, John Sillitoe on behalf of the Wellcome Sanger Institute COVID-19 Surveillance Team |
| EPI_ISL_2853014 | Lighthouse Laboratory Plymouth | Wellcome Sanger Institute for the COVID-19 Genomics UK (COG-UK) Consortium | Lighthouse Laboratory Plymouth and Alex Alderton, Roberto Amato, Jeffrey Barrett, Sonia Goncalves, Ewan Harrison, David K. Jackson, Ian Johnston, Dominic Kwiatkowski, Cordelia Langford, John Sillitoe on behalf of the Wellcome Sanger Institute COVID-19 Surveillance Team |
| EPI_ISL_2853015, EPI_ISL_2853016, EPI_ISL_2853017, EPI_ISL_2853018 | Lighthouse Lab in Glasgow | Wellcome Sanger Institute for the COVID-19 Genomics UK (COG-UK) Consortium | Harper VanSteenhouse, Yumi Kasai, David Gray, Carol Clugston, Anna Dominiczak and Alex Alderton, Roberto Amato, Jeffrey Barrett, Sonia Goncalves, Ewan Harrison, David K. Jackson, Ian Johnston, Dominic Kwiatkowski, Cordelia Langford, John Sillitoe on behalf of the Wellcome Sanger Institute COVID-19 Surveillance Team |
| EPI_ISL_2853019 | Lighthouse Lab in Milton Keynes | Wellcome Sanger Institute for the COVID-19 Genomics UK (COG-UK) Consortium | The Lighthouse Lab in Milton Keynes and Alex Alderton, Roberto Amato, Jeffrey Barrett, Sonia Goncalves, Ewan Harrison, David K. Jackson, Ian Johnston, Dominic Kwiatkowski, Cordelia Langford, John Sillitoe on behalf of the Wellcome Sanger Institute COVID-19 Surveillance Team |
| EPI_ISL_2853020, EPI_ISL_2853021, EPI_ISL_2853022, EPI_ISL_2853023, EPI_ISL_2853024, EPI_ISL_2853025, EPI_ISL_2853026, EPI_ISL_2853027, EPI_ISL_2853028, EPI_ISL_2853029, EPI_ISL_2853030, EPI_ISL_2853031, EPI_ISL_2853032, EPI_ISL_2853033, EPI_ISL_2853034, EPI_ISL_2853035, EPI_ISL_2853036, EPI_ISL_2853037, EPI_ISL_2853038, EPI_ISL_2853039, EPI_ISL_2853040, EPI_ISL_2853041, EPI_ISL_2853042, EPI_ISL_2853043, EPI_ISL_2853044, EPI_ISL_2853045, EPI_ISL_2853046, EPI_ISL_2853047, EPI_ISL_2853048, EPI_ISL_2853049, EPI_ISL_2853050, EPI_ISL_2853051, EPI_ISL_2853052, EPI_ISL_2853053, EPI_ISL_2853054, EPI_ISL_2853055, EPI_ISL_2853056, EPI_ISL_2853057, EPI_ISL_2853058, EPI_ISL_2853059, EPI_ISL_2853060, EPI_ISL_2853061, EPI_ISL_2853062 | | | |
| see above | Lighthouse Lab in Glasgow | Wellcome Sanger Institute for the COVID-19 Genomics UK (COG-UK) Consortium | Harper VanSteenhouse, Yumi Kasai, David Gray, Carol Clugston, Anna Dominiczak and Alex Alderton, Roberto Amato, Jeffrey Barrett, Sonia Goncalves, Ewan Harrison, David K. Jackson, Ian Johnston, Dominic Kwiatkowski, Cordelia Langford, John Sillitoe on behalf of the Wellcome Sanger Institute COVID-19 Surveillance Team |
| EPI_ISL_2853063 | Lighthouse Laboratory Plymouth | Wellcome Sanger Institute for the COVID-19 Genomics UK (COG-UK) Consortium | Lighthouse Laboratory Plymouth and Alex Alderton, Roberto Amato, Jeffrey Barrett, Sonia Goncalves, Ewan Harrison, David K. Jackson, Ian Johnston, Dominic Kwiatkowski, Cordelia Langford, John Sillitoe on behalf of the Wellcome Sanger Institute COVID-19 Surveillance Team |
| EPI_ISL_2853064, EPI_ISL_2853065, EPI_ISL_2853066, EPI_ISL_2853068, EPI_ISL_2853069, EPI_ISL_2853070, EPI_ISL_2853071, EPI_ISL_2853072, EPI_ISL_2853073, EPI_ISL_2853074, EPI_ISL_2853075, EPI_ISL_2853076, EPI_ISL_2853077, EPI_ISL_2853078, EPI_ISL_2853079, EPI_ISL_2853080, EPI_ISL_2853081, EPI_ISL_2853082, EPI_ISL_2853083, EPI_ISL_2853084, EPI_ISL_2853085, EPI_ISL_2853086, EPI_ISL_2853087, EPI_ISL_2853088, EPI_ISL_2853089, EPI_ISL_2853090, EPI_ISL_2853091, EPI_ISL_2853092, EPI_ISL_2853093, EPI_ISL_2853094, EPI_ISL_2853096, EPI_ISL_2853097, EPI_ISL_2853098, EPI_ISL_2853099 | | | |
| see above | Lighthouse Lab in Glasgow | Wellcome Sanger Institute for the COVID-19 Genomics UK (COG-UK) Consortium | Harper VanSteenhouse, Yumi Kasai, David Gray, Carol Clugston, Anna Dominiczak and Alex Alderton, Roberto Amato, Jeffrey Barrett, Sonia Goncalves, Ewan Harrison, David K. Jackson, Ian Johnston, Dominic Kwiatkowski, Cordelia Langford, John Sillitoe on behalf of the Wellcome Sanger Institute COVID-19 Surveillance Team |
| EPI_ISL_2853100 | Lighthouse Lab in Alderley Park | Wellcome Sanger Institute for the COVID-19 Genomics UK (COG-UK) Consortium | Jacquelyn Wynn, Mairead Hyland, The Lighthouse Lab in Alderley Park and Alex Alderton, Roberto Amato, Jeffrey Barrett, Sonia Goncalves, Ewan Harrison, David K. Jackson, Ian Johnston, Dominic Kwiatkowski, Cordelia Langford, John Sillitoe on behalf of the Wellcome Sanger Institute COVID-19 Surveillance Team |
| EPI_ISL_2853101, EPI_ISL_2853102, EPI_ISL_2853103, EPI_ISL_2853104, EPI_ISL_2853105, EPI_ISL_2853106, EPI_ISL_2853107, EPI_ISL_2853108, EPI_ISL_2853109, EPI_ISL_2853110, EPI_ISL_2853111, EPI_ISL_2853112, EPI_ISL_2853113, EPI_ISL_2853114, EPI_ISL_2853115, EPI_ISL_2853116, EPI_ISL_2853117, EPI_ISL_2853118, EPI_ISL_2853119, EPI_ISL_2853120, EPI_ISL_2853121, EPI_ISL_2853122, EPI_ISL_2853123, EPI_ISL_2853124, EPI_ISL_2853125, EPI_ISL_2853126, EPI_ISL_2853127, EPI_ISL_2853129, EPI_ISL_2853130, EPI_ISL_2853131, EPI_ISL_2853132, EPI_ISL_2853133, EPI_ISL_2853134, EPI_ISL_2853135, EPI_ISL_2853136, EPI_ISL_2853137, EPI_ISL_2853138, EPI_ISL_2853139, EPI_ISL_2853140, EPI_ISL_2853141, EPI_ISL_2853142, EPI_ISL_2853143, EPI_ISL_2853144, EPI_ISL_2853145, EPI_ISL_2853146, EPI_ISL_2853147, EPI_ISL_2853148, EPI_ISL_2853149, EPI_ISL_2853150, EPI_ISL_2853151, EPI_ISL_2853152, EPI_ISL_2853153, EPI_ISL_2853154, EPI_ISL_2853155, EPI_ISL_2853156, EPI_ISL_2853157, EPI_ISL_2853158, EPI_ISL_2853159, EPI_ISL_2853160, EPI_ISL_2853161, EPI_ISL_2853162, EPI_ISL_2853163, EPI_ISL_2853164, EPI_ISL_2853166, EPI_ISL_2853167, EPI_ISL_2853168, EPI_ISL_2853169, EPI_ISL_2853170, EPI_ISL_2853171, EPI_ISL_2853172, EPI_ISL_2853173, EPI_ISL_2853174, EPI_ISL_2853175, EPI_ISL_2853176, EPI_ISL_2853177, EPI_ISL_2853178, EPI_ISL_2853179, EPI_ISL_2853180, EPI_ISL_2853181, EPI_ISL_2853182, EPI_ISL_2853183, EPI_ISL_2853184, EPI_ISL_2853185, EPI_ISL_2853186, EPI_ISL_2853187, EPI_ISL_2853188, EPI_ISL_2853189, EPI_ISL_2853190, EPI_ISL_2853191, EPI_ISL_2853192, EPI_ISL_2853193, EPI_ISL_2853194, EPI_ISL_2853195, EPI_ISL_2853196, EPI_ISL_2853197, EPI_ISL_2853198, EPI_ISL_2853199, EPI_ISL_2853200, EPI_ISL_2853201, EPI_ISL_2853202, EPI_ISL_2853203, EPI_ISL_2853204, EPI_ISL_2853205, EPI_ISL_2853206, EPI_ISL_2853207, EPI_ISL_2853208, EPI_ISL_2853209, EPI_ISL_2853210, EPI_ISL_2853211, EPI_ISL_2853212, EPI_ISL_2853213, EPI_ISL_2853214, EPI_ISL_2853215, EPI_ISL_2853216, EPI_ISL_2853217, EPI_ISL_2853218, EPI_ISL_2853219, EPI_ISL_2853220, EPI_ISL_2853221, EPI_ISL_2853222, EPI_ISL_2853223, EPI_ISL_2853224, EPI_ISL_2853225, EPI_ISL_2853226, EPI_ISL_2853227, EPI_ISL_2853228, EPI_ISL_2853229, EPI_ISL_2853230, EPI_ISL_2853231, EPI_ISL_2853232, EPI_ISL_2853233, EPI_ISL_2853234, EPI_ISL_2853235, EPI_ISL_2853236, EPI_ISL_2853237, EPI_ISL_2853238, EPI_ISL_2853239, EPI_ISL_2853240, EPI_ISL_2853241, EPI_ISL_2853242, EPI_ISL_2853243, EPI_ISL_2853244, EPI_ISL_2853245, EPI_ISL_2853246, EPI_ISL_2853247, EPI_ISL_2853248, EPI_ISL_2853249, EPI_ISL_2853250, EPI_ISL_2853251, EPI_ISL_2853252, EPI_ISL_2853253, EPI_ISL_2853254, EPI_ISL_2853255, EPI_ISL_2853256, EPI_ISL_2853257, EPI_ISL_2853258 | | | |
| see above | Lighthouse Lab in Glasgow | Wellcome Sanger Institute for the COVID-19 Genomics UK (COG-UK) Consortium | Harper VanSteenhouse, Yumi Kasai, David Gray, Carol Clugston, Anna Dominiczak and Alex Alderton, Roberto Amato, Jeffrey Barrett, Sonia Goncalves, Ewan Harrison, David K. Jackson, Ian Johnston, Dominic Kwiatkowski, Cordelia Langford, John Sillitoe on behalf of the Wellcome Sanger Institute COVID-19 Surveillance Team |
| EPI_ISL_2853259 | Lighthouse Laboratory Plymouth | Wellcome Sanger Institute for the COVID-19 Genomics UK (COG-UK) Consortium | Lighthouse Laboratory Plymouth and Alex Alderton, Roberto Amato, Jeffrey Barrett, Sonia Goncalves, Ewan Harrison, David K. Jackson, Ian Johnston, Dominic Kwiatkowski, Cordelia Langford, John Sillitoe on behalf of the Wellcome Sanger Institute COVID-19 Surveillance Team |
| EPI_ISL_2853260, EPI_ISL_2853261, EPI_ISL_2853262 | Lighthouse Lab in Alderley Park | Wellcome Sanger Institute for the COVID-19 Genomics UK (COG-UK) Consortium | Jacquelyn Wynn, Mairead Hyland, The Lighthouse Lab in Alderley Park and Alex Alderton, Roberto Amato, Jeffrey Barrett, Sonia Goncalves, Ewan Harrison, David K. Jackson, Ian Johnston, Dominic Kwiatkowski, Cordelia Langford, John Sillitoe on behalf of the Wellcome Sanger Institute COVID-19 Surveillance Team |
| EPI_ISL_2853263, EPI_ISL_2853264, EPI_ISL_2853265 | Lighthouse Laboratory Plymouth | Wellcome Sanger Institute for the COVID-19 Genomics UK (COG-UK) Consortium | Lighthouse Laboratory Plymouth and Alex Alderton, Roberto Amato, Jeffrey Barrett, Sonia Goncalves, Ewan Harrison, David K. Jackson, Ian Johnston, Dominic Kwiatkowski, Cordelia Langford, John Sillitoe on behalf of the Wellcome Sanger Institute COVID-19 Surveillance Team |
| EPI_ISL_2853266, EPI_ISL_2853267 | Lighthouse Lab in Alderley Park | Wellcome Sanger Institute for the COVID-19 Genomics UK (COG-UK) Consortium | Jacquelyn Wynn, Mairead Hyland, The Lighthouse Lab in Alderley Park and Alex Alderton, Roberto Amato, Jeffrey Barrett, Sonia Goncalves, Ewan Harrison, David K. Jackson, Ian Johnston, Dominic Kwiatkowski, Cordelia Langford, John Sillitoe on behalf of the Wellcome Sanger Institute COVID-19 Surveillance Team |
| EPI_ISL_2853268 | Lighthouse Laboratory Plymouth | Wellcome Sanger Institute for the COVID-19 Genomics UK (COG-UK) Consortium | Lighthouse Laboratory Plymouth and Alex Alderton, Roberto Amato, Jeffrey Barrett, Sonia Goncalves, Ewan Harrison, David K. Jackson, Ian Johnston, Dominic Kwiatkowski, Cordelia Langford, John Sillitoe on behalf of the Wellcome Sanger Institute COVID-19 Surveillance Team |
| EPI_ISL_2853269, EPI_ISL_2853270, EPI_ISL_2853271, EPI_ISL_2853272, EPI_ISL_2853274 | Lighthouse Lab in Alderley Park | Wellcome Sanger Institute for the COVID-19 Genomics UK (COG-UK) Consortium | Jacquelyn Wynn, Mairead Hyland, The Lighthouse Lab in Alderley Park and Alex Alderton, Roberto Amato, Jeffrey Barrett, Sonia Goncalves, Ewan Harrison, David K. Jackson, Ian Johnston, Dominic Kwiatkowski, Cordelia Langford, John Sillitoe on behalf of the Wellcome Sanger Institute COVID-19 Surveillance Team |
| EPI_ISL_2853275 | Lighthouse Laboratory Plymouth | Wellcome Sanger Institute for the COVID-19 Genomics UK (COG-UK) Consortium | Lighthouse Laboratory Plymouth and Alex Alderton, Roberto Amato, Jeffrey Barrett, Sonia Goncalves, Ewan Harrison, David K. Jackson, Ian Johnston, Dominic Kwiatkowski, Cordelia Langford, John Sillitoe on behalf of the Wellcome Sanger Institute COVID-19 Surveillance Team |
| EPI_ISL_2853276, EPI_ISL_2853277, EPI_ISL_2853278, EPI_ISL_2853279, EPI_ISL_2853280 | Lighthouse Lab in Alderley Park | Wellcome Sanger Institute for the COVID-19 Genomics UK (COG-UK) Consortium | Jacquelyn Wynn, Mairead Hyland, The Lighthouse Lab in Alderley Park and Alex Alderton, Roberto Amato, Jeffrey Barrett, Sonia Goncalves, Ewan Harrison, David K. Jackson, Ian Johnston, Dominic Kwiatkowski, Cordelia Langford, John Sillitoe on behalf of the Wellcome Sanger Institute COVID-19 Surveillance Team |
| EPI_ISL_2853281 | Lighthouse Laboratory Plymouth | Wellcome Sanger Institute for the COVID-19 Genomics UK (COG-UK) Consortium | Lighthouse Laboratory Plymouth and Alex Alderton, Roberto Amato, Jeffrey Barrett, Sonia Goncalves, Ewan Harrison, David K. Jackson, Ian Johnston, Dominic Kwiatkowski, Cordelia Langford, John Sillitoe on behalf of the Wellcome Sanger Institute COVID-19 Surveillance Team |
| EPI_ISL_2853282, EPI_ISL_2853283, EPI_ISL_2853284 | Lighthouse Lab in Alderley Park | Wellcome Sanger Institute for the COVID-19 Genomics UK (COG-UK) Consortium | Jacquelyn Wynn, Mairead Hyland, The Lighthouse Lab in Alderley Park and Alex Alderton, Roberto Amato, Jeffrey Barrett, Sonia Goncalves, Ewan Harrison, David K. Jackson, Ian Johnston, Dominic Kwiatkowski, Cordelia Langford, John Sillitoe on behalf of the Wellcome Sanger Institute COVID-19 Surveillance Team |
| EPI_ISL_2853285 | Lighthouse Laboratory Plymouth | Wellcome Sanger Institute for the COVID-19 Genomics UK (COG-UK) Consortium | Lighthouse Laboratory Plymouth and Alex Alderton, Roberto Amato, Jeffrey Barrett, Sonia Goncalves, Ewan Harrison, David K. Jackson, Ian Johnston, Dominic Kwiatkowski, Cordelia Langford, John Sillitoe on behalf of the Wellcome Sanger Institute COVID-19 Surveillance Team |
| EPI_ISL_2853286, EPI_ISL_2853287 | Lighthouse Lab in Alderley Park | Wellcome Sanger Institute for the COVID-19 Genomics UK (COG-UK) Consortium | Jacquelyn Wynn, Mairead Hyland, The Lighthouse Lab in Alderley Park and Alex Alderton, Roberto Amato, Jeffrey Barrett, Sonia Goncalves, Ewan Harrison, David K. Jackson, Ian Johnston, Dominic Kwiatkowski, Cordelia Langford, John Sillitoe on behalf of the Wellcome Sanger Institute COVID-19 Surveillance Team |
| EPI_ISL_2853288 | Lighthouse Laboratory Plymouth | Wellcome Sanger Institute for the COVID-19 Genomics UK (COG-UK) Consortium | Lighthouse Laboratory Plymouth and Alex Alderton, Roberto Amato, Jeffrey Barrett, Sonia Goncalves, Ewan Harrison, David K. Jackson, Ian Johnston, Dominic Kwiatkowski, Cordelia Langford, John Sillitoe on behalf of the Wellcome Sanger Institute COVID-19 Surveillance Team |
| EPI_ISL_2853289 | Lighthouse Lab in Alderley Park | Wellcome Sanger Institute for the COVID-19 Genomics UK (COG-UK) Consortium | Jacquelyn Wynn, Mairead Hyland, The Lighthouse Lab in Alderley Park and Alex Alderton, Roberto Amato, Jeffrey Barrett, Sonia Goncalves, Ewan Harrison, David K. Jackson, Ian Johnston, Dominic Kwiatkowski, Cordelia Langford, John Sillitoe on behalf of the Wellcome Sanger Institute COVID-19 Surveillance Team |
| EPI_ISL_2853290 | Lighthouse Laboratory Plymouth | Wellcome Sanger Institute for the COVID-19 Genomics UK (COG-UK) Consortium | Lighthouse Laboratory Plymouth and Alex Alderton, Roberto Amato, Jeffrey Barrett, Sonia Goncalves, Ewan Harrison, David K. Jackson, Ian Johnston, Dominic Kwiatkowski, Cordelia Langford, John Sillitoe on behalf of the Wellcome Sanger Institute COVID-19 Surveillance Team |
| EPI_ISL_2853291, EPI_ISL_2853292, EPI_ISL_2853293, EPI_ISL_2853294 | Lighthouse Lab in Alderley Park | Wellcome Sanger Institute for the COVID-19 Genomics UK (COG-UK) Consortium | Jacquelyn Wynn, Mairead Hyland, The Lighthouse Lab in Alderley Park and Alex Alderton, Roberto Amato, Jeffrey Barrett, Sonia Goncalves, Ewan Harrison, David K. Jackson, Ian Johnston, Dominic Kwiatkowski, Cordelia Langford, John Sillitoe on behalf of the Wellcome Sanger Institute COVID-19 Surveillance Team |
| EPI_ISL_2853295, EPI_ISL_2853296, EPI_ISL_2853297 | Lighthouse Laboratory Plymouth | Wellcome Sanger Institute for the COVID-19 Genomics UK (COG-UK) Consortium | Lighthouse Laboratory Plymouth and Alex Alderton, Roberto Amato, Jeffrey Barrett, Sonia Goncalves, Ewan Harrison, David K. Jackson, Ian Johnston, Dominic Kwiatkowski, Cordelia Langford, John Sillitoe on behalf of the Wellcome Sanger Institute COVID-19 Surveillance Team |
| EPI_ISL_2853298 | Lighthouse Lab in Alderley Park | Wellcome Sanger Institute for the COVID-19 Genomics UK (COG-UK) Consortium | Jacquelyn Wynn, Mairead Hyland, The Lighthouse Lab in Alderley Park and Alex Alderton, Roberto Amato, Jeffrey Barrett, Sonia Goncalves, Ewan Harrison, David K. Jackson, Ian Johnston, Dominic Kwiatkowski, Cordelia Langford, John Sillitoe on behalf of the Wellcome Sanger Institute COVID-19 Surveillance Team |
| EPI_ISL_2853299, EPI_ISL_2853300, EPI_ISL_2853301 | Lighthouse Laboratory Plymouth | Wellcome Sanger Institute for the COVID-19 Genomics UK (COG-UK) Consortium | Lighthouse Laboratory Plymouth and Alex Alderton, Roberto Amato, Jeffrey Barrett, Sonia Goncalves, Ewan Harrison, David K. Jackson, Ian Johnston, Dominic Kwiatkowski, Cordelia Langford, John Sillitoe on behalf of the Wellcome Sanger Institute COVID-19 Surveillance Team |
| EPI_ISL_2853302, | Lighthouse Lab in Alderley Park | Wellcome Sanger Institute for the COVID-19 Genomics UK (COG-UK) | Jacquelyn Wynn, Mairead Hyland, The Lighthouse Lab in Alderley Park and Alex Alderton, Roberto Amato, Jeffrey Barrett, Sonia Goncalves, Ewan Harrison, David K. Jackson, Ian Johnston, Dominic Kwiatkowski, Cordelia Langford, John Sillitoe on |

| | | | |
|---|---|---|---|
| EPI_ISL_2853303, EPI_ISL_2853304, EPI_ISL_2853305, EPI_ISL_2853306 | | Consortium | behalf of the Wellcome Sanger Institute COVID-19 Surveillance Team |
| EPI_ISL_2853307 | Lighthouse Laboratory Plymouth | Wellcome Sanger Institute for the COVID-19 Genomics UK (COG-UK) Consortium | Lighthouse Laboratory Plymouth and Alex Alderton, Roberto Amato, Jeffrey Barrett, Sonia Goncalves, Ewan Harrison, David K. Jackson, Ian Johnston, Dominic Kwiatkowski, Cordelia Langford, John Sillitoe on behalf of the Wellcome Sanger Institute COVID-19 Surveillance Team |
| EPI_ISL_2853308, EPI_ISL_2853309, EPI_ISL_2853310 | Lighthouse Lab in Alderley Park | Wellcome Sanger Institute for the COVID-19 Genomics UK (COG-UK) Consortium | Jacquelyn Wynn, Mairead Hyland, The Lighthouse Lab in Alderley Park and Alex Alderton, Roberto Amato, Jeffrey Barrett, Sonia Goncalves, Ewan Harrison, David K. Jackson, Ian Johnston, Dominic Kwiatkowski, Cordelia Langford, John Sillitoe on behalf of the Wellcome Sanger Institute COVID-19 Surveillance Team |
| EPI_ISL_2853311, EPI_ISL_2853312 | Lighthouse Laboratory Plymouth | Wellcome Sanger Institute for the COVID-19 Genomics UK (COG-UK) Consortium | Lighthouse Laboratory Plymouth and Alex Alderton, Roberto Amato, Jeffrey Barrett, Sonia Goncalves, Ewan Harrison, David K. Jackson, Ian Johnston, Dominic Kwiatkowski, Cordelia Langford, John Sillitoe on behalf of the Wellcome Sanger Institute COVID-19 Surveillance Team |
| EPI_ISL_2853313 | Lighthouse Lab in Alderley Park | Wellcome Sanger Institute for the COVID-19 Genomics UK (COG-UK) Consortium | Jacquelyn Wynn, Mairead Hyland, The Lighthouse Lab in Alderley Park and Alex Alderton, Roberto Amato, Jeffrey Barrett, Sonia Goncalves, Ewan Harrison, David K. Jackson, Ian Johnston, Dominic Kwiatkowski, Cordelia Langford, John Sillitoe on behalf of the Wellcome Sanger Institute COVID-19 Surveillance Team |
| EPI_ISL_2853314, EPI_ISL_2853315 | Lighthouse Laboratory Plymouth | Wellcome Sanger Institute for the COVID-19 Genomics UK (COG-UK) Consortium | Lighthouse Laboratory Plymouth and Alex Alderton, Roberto Amato, Jeffrey Barrett, Sonia Goncalves, Ewan Harrison, David K. Jackson, Ian Johnston, Dominic Kwiatkowski, Cordelia Langford, John Sillitoe on behalf of the Wellcome Sanger Institute COVID-19 Surveillance Team |
| EPI_ISL_2853316, EPI_ISL_2853317, EPI_ISL_2853318, EPI_ISL_2853319, EPI_ISL_2853320, EPI_ISL_2853321 | Lighthouse Lab in Alderley Park | Wellcome Sanger Institute for the COVID-19 Genomics UK (COG-UK) Consortium | Jacquelyn Wynn, Mairead Hyland, The Lighthouse Lab in Alderley Park and Alex Alderton, Roberto Amato, Jeffrey Barrett, Sonia Goncalves, Ewan Harrison, David K. Jackson, Ian Johnston, Dominic Kwiatkowski, Cordelia Langford, John Sillitoe on behalf of the Wellcome Sanger Institute COVID-19 Surveillance Team |
| EPI_ISL_2853322 | Lighthouse Laboratory Plymouth | Wellcome Sanger Institute for the COVID-19 Genomics UK (COG-UK) Consortium | Lighthouse Laboratory Plymouth and Alex Alderton, Roberto Amato, Jeffrey Barrett, Sonia Goncalves, Ewan Harrison, David K. Jackson, Ian Johnston, Dominic Kwiatkowski, Cordelia Langford, John Sillitoe on behalf of the Wellcome Sanger Institute COVID-19 Surveillance Team |
| EPI_ISL_2853323, EPI_ISL_2853324, EPI_ISL_2853325, EPI_ISL_2853326 | Lighthouse Lab in Alderley Park | Wellcome Sanger Institute for the COVID-19 Genomics UK (COG-UK) Consortium | Jacquelyn Wynn, Mairead Hyland, The Lighthouse Lab in Alderley Park and Alex Alderton, Roberto Amato, Jeffrey Barrett, Sonia Goncalves, Ewan Harrison, David K. Jackson, Ian Johnston, Dominic Kwiatkowski, Cordelia Langford, John Sillitoe on behalf of the Wellcome Sanger Institute COVID-19 Surveillance Team |
| EPI_ISL_2853327 | Lighthouse Laboratory Plymouth | Wellcome Sanger Institute for the COVID-19 Genomics UK (COG-UK) Consortium | Lighthouse Laboratory Plymouth and Alex Alderton, Roberto Amato, Jeffrey Barrett, Sonia Goncalves, Ewan Harrison, David K. Jackson, Ian Johnston, Dominic Kwiatkowski, Cordelia Langford, John Sillitoe on behalf of the Wellcome Sanger Institute COVID-19 Surveillance Team |
| EPI_ISL_2853328, EPI_ISL_2853329, EPI_ISL_2853330 | Lighthouse Lab in Alderley Park | Wellcome Sanger Institute for the COVID-19 Genomics UK (COG-UK) Consortium | Jacquelyn Wynn, Mairead Hyland, The Lighthouse Lab in Alderley Park and Alex Alderton, Roberto Amato, Jeffrey Barrett, Sonia Goncalves, Ewan Harrison, David K. Jackson, Ian Johnston, Dominic Kwiatkowski, Cordelia Langford, John Sillitoe on behalf of the Wellcome Sanger Institute COVID-19 Surveillance Team |
| EPI_ISL_2853331 | Lighthouse Laboratory Plymouth | Wellcome Sanger Institute for the COVID-19 Genomics UK (COG-UK) Consortium | Lighthouse Laboratory Plymouth and Alex Alderton, Roberto Amato, Jeffrey Barrett, Sonia Goncalves, Ewan Harrison, David K. Jackson, Ian Johnston, Dominic Kwiatkowski, Cordelia Langford, John Sillitoe on behalf of the Wellcome Sanger Institute COVID-19 Surveillance Team |
| EPI_ISL_2853332, EPI_ISL_2853333, EPI_ISL_2853334 | Lighthouse Lab in Alderley Park | Wellcome Sanger Institute for the COVID-19 Genomics UK (COG-UK) Consortium | Jacquelyn Wynn, Mairead Hyland, The Lighthouse Lab in Alderley Park and Alex Alderton, Roberto Amato, Jeffrey Barrett, Sonia Goncalves, Ewan Harrison, David K. Jackson, Ian Johnston, Dominic Kwiatkowski, Cordelia Langford, John Sillitoe on behalf of the Wellcome Sanger Institute COVID-19 Surveillance Team |
| EPI_ISL_2853335, EPI_ISL_2853336 | Lighthouse Laboratory Plymouth | Wellcome Sanger Institute for the COVID-19 Genomics UK (COG-UK) Consortium | Lighthouse Laboratory Plymouth and Alex Alderton, Roberto Amato, Jeffrey Barrett, Sonia Goncalves, Ewan Harrison, David K. Jackson, Ian Johnston, Dominic Kwiatkowski, Cordelia Langford, John Sillitoe on behalf of the Wellcome Sanger Institute COVID-19 Surveillance Team |
| EPI_ISL_2853337, EPI_ISL_2853338, EPI_ISL_2853339, EPI_ISL_2853340, EPI_ISL_2853341, EPI_ISL_2853342, EPI_ISL_2853343, EPI_ISL_2853344, EPI_ISL_2853345 | Lighthouse Lab in Alderley Park | Wellcome Sanger Institute for the COVID-19 Genomics UK (COG-UK) Consortium | Jacquelyn Wynn, Mairead Hyland, The Lighthouse Lab in Alderley Park and Alex Alderton, Roberto Amato, Jeffrey Barrett, Sonia Goncalves, Ewan Harrison, David K. Jackson, Ian Johnston, Dominic Kwiatkowski, Cordelia Langford, John Sillitoe on behalf of the Wellcome Sanger Institute COVID-19 Surveillance Team |
| EPI_ISL_2853346, EPI_ISL_2853347 | Lighthouse Laboratory Plymouth | Wellcome Sanger Institute for the COVID-19 Genomics UK (COG-UK) Consortium | Lighthouse Laboratory Plymouth and Alex Alderton, Roberto Amato, Jeffrey Barrett, Sonia Goncalves, Ewan Harrison, David K. Jackson, Ian Johnston, Dominic Kwiatkowski, Cordelia Langford, John Sillitoe on behalf of the Wellcome Sanger Institute COVID-19 Surveillance Team |
| EPI_ISL_2853349, EPI_ISL_2853350, EPI_ISL_2853351, EPI_ISL_2853352, EPI_ISL_2853353, EPI_ISL_2853354, EPI_ISL_2853355 | Lighthouse Lab in Alderley Park | Wellcome Sanger Institute for the COVID-19 Genomics UK (COG-UK) Consortium | Jacquelyn Wynn, Mairead Hyland, The Lighthouse Lab in Alderley Park and Alex Alderton, Roberto Amato, Jeffrey Barrett, Sonia Goncalves, Ewan Harrison, David K. Jackson, Ian Johnston, Dominic Kwiatkowski, Cordelia Langford, John Sillitoe on behalf of the Wellcome Sanger Institute COVID-19 Surveillance Team |
| EPI_ISL_2853356 | Lighthouse Laboratory Plymouth | Wellcome Sanger Institute for the COVID-19 Genomics UK (COG-UK) Consortium | Lighthouse Laboratory Plymouth and Alex Alderton, Roberto Amato, Jeffrey Barrett, Sonia Goncalves, Ewan Harrison, David K. Jackson, Ian Johnston, Dominic Kwiatkowski, Cordelia Langford, John Sillitoe on behalf of the Wellcome Sanger Institute COVID-19 Surveillance Team |
| EPI_ISL_2853357, EPI_ISL_2853358, EPI_ISL_2853360 | Lighthouse Lab in Alderley Park | Wellcome Sanger Institute for the COVID-19 Genomics UK (COG-UK) Consortium | Jacquelyn Wynn, Mairead Hyland, The Lighthouse Lab in Alderley Park and Alex Alderton, Roberto Amato, Jeffrey Barrett, Sonia Goncalves, Ewan Harrison, David K. Jackson, Ian Johnston, Dominic Kwiatkowski, Cordelia Langford, John Sillitoe on behalf of the Wellcome Sanger Institute COVID-19 Surveillance Team |
| EPI_ISL_2853361 | Lighthouse Laboratory Plymouth | Wellcome Sanger Institute for the COVID-19 Genomics UK (COG-UK) Consortium | Lighthouse Laboratory Plymouth and Alex Alderton, Roberto Amato, Jeffrey Barrett, Sonia Goncalves, Ewan Harrison, David K. Jackson, Ian Johnston, Dominic Kwiatkowski, Cordelia Langford, John Sillitoe on behalf of the Wellcome Sanger Institute COVID-19 Surveillance Team |
| EPI_ISL_2853362 | Lighthouse Lab in Alderley Park | Wellcome Sanger Institute for the COVID-19 Genomics UK (COG-UK) Consortium | Jacquelyn Wynn, Mairead Hyland, The Lighthouse Lab in Alderley Park and Alex Alderton, Roberto Amato, Jeffrey Barrett, Sonia Goncalves, Ewan Harrison, David K. Jackson, Ian Johnston, Dominic Kwiatkowski, Cordelia Langford, John Sillitoe on behalf of the Wellcome Sanger Institute COVID-19 Surveillance Team |
| EPI_ISL_2853363, EPI_ISL_2853364, EPI_ISL_2853365 | Lighthouse Laboratory Plymouth | Wellcome Sanger Institute for the COVID-19 Genomics UK (COG-UK) Consortium | Lighthouse Laboratory Plymouth and Alex Alderton, Roberto Amato, Jeffrey Barrett, Sonia Goncalves, Ewan Harrison, David K. Jackson, Ian Johnston, Dominic Kwiatkowski, Cordelia Langford, John Sillitoe on behalf of the Wellcome Sanger Institute COVID-19 Surveillance Team |
| EPI_ISL_2853366, EPI_ISL_2853367 | Lighthouse Lab in Alderley Park | Wellcome Sanger Institute for the COVID-19 Genomics UK (COG-UK) Consortium | Jacquelyn Wynn, Mairead Hyland, The Lighthouse Lab in Alderley Park and Alex Alderton, Roberto Amato, Jeffrey Barrett, Sonia Goncalves, Ewan Harrison, David K. Jackson, Ian Johnston, Dominic Kwiatkowski, Cordelia Langford, John Sillitoe on behalf of the Wellcome Sanger Institute COVID-19 Surveillance Team |
| EPI_ISL_2853368 | Lighthouse Laboratory Plymouth | Wellcome Sanger Institute for the COVID-19 Genomics UK (COG-UK) Consortium | Lighthouse Laboratory Plymouth and Alex Alderton, Roberto Amato, Jeffrey Barrett, Sonia Goncalves, Ewan Harrison, David K. Jackson, Ian Johnston, Dominic Kwiatkowski, Cordelia Langford, John Sillitoe on behalf of the Wellcome Sanger Institute COVID-19 Surveillance Team |
| EPI_ISL_2853369, EPI_ISL_2853370 | Lighthouse Lab in Alderley Park | Wellcome Sanger Institute for the COVID-19 Genomics UK (COG-UK) Consortium | Jacquelyn Wynn, Mairead Hyland, The Lighthouse Lab in Alderley Park and Alex Alderton, Roberto Amato, Jeffrey Barrett, Sonia Goncalves, Ewan Harrison, David K. Jackson, Ian Johnston, Dominic Kwiatkowski, Cordelia Langford, John Sillitoe on behalf of the Wellcome Sanger Institute COVID-19 Surveillance Team |
| EPI_ISL_2853371, EPI_ISL_2853372 | Lighthouse Laboratory Plymouth | Wellcome Sanger Institute for the COVID-19 Genomics UK (COG-UK) Consortium | Lighthouse Laboratory Plymouth and Alex Alderton, Roberto Amato, Jeffrey Barrett, Sonia Goncalves, Ewan Harrison, David K. Jackson, Ian Johnston, Dominic Kwiatkowski, Cordelia Langford, John Sillitoe on behalf of the Wellcome Sanger Institute COVID-19 Surveillance Team |
| EPI_ISL_2853373 | Lighthouse Lab in Alderley Park | Wellcome Sanger Institute for the COVID-19 Genomics UK (COG-UK) Consortium | Jacquelyn Wynn, Mairead Hyland, The Lighthouse Lab in Alderley Park and Alex Alderton, Roberto Amato, Jeffrey Barrett, Sonia Goncalves, Ewan Harrison, David K. Jackson, Ian Johnston, Dominic Kwiatkowski, Cordelia Langford, John Sillitoe on behalf of the Wellcome Sanger Institute COVID-19 Surveillance Team |
| EPI_ISL_2853374 | Lighthouse Laboratory Plymouth | Wellcome Sanger Institute for the COVID-19 Genomics UK (COG-UK) Consortium | Lighthouse Laboratory Plymouth and Alex Alderton, Roberto Amato, Jeffrey Barrett, Sonia Goncalves, Ewan Harrison, David K. Jackson, Ian Johnston, Dominic Kwiatkowski, Cordelia Langford, John Sillitoe on behalf of the Wellcome Sanger Institute COVID-19 Surveillance Team |
| EPI_ISL_2853375, EPI_ISL_2853376 | Lighthouse Lab in Alderley Park | Wellcome Sanger Institute for the COVID-19 Genomics UK (COG-UK) Consortium | Jacquelyn Wynn, Mairead Hyland, The Lighthouse Lab in Alderley Park and Alex Alderton, Roberto Amato, Jeffrey Barrett, Sonia Goncalves, Ewan Harrison, David K. Jackson, Ian Johnston, Dominic Kwiatkowski, Cordelia Langford, John Sillitoe on behalf of the Wellcome Sanger Institute COVID-19 Surveillance Team |
| EPI_ISL_2853377 | Lighthouse Laboratory Plymouth | Wellcome Sanger Institute for the COVID-19 Genomics UK (COG-UK) Consortium | Lighthouse Laboratory Plymouth and Alex Alderton, Roberto Amato, Jeffrey Barrett, Sonia Goncalves, Ewan Harrison, David K. Jackson, Ian Johnston, Dominic Kwiatkowski, Cordelia Langford, John Sillitoe on behalf of the Wellcome Sanger Institute COVID-19 Surveillance Team |
| EPI_ISL_2853378, EPI_ISL_2853379, EPI_ISL_2853380, EPI_ISL_2853381, EPI_ISL_2853382 | Lighthouse Lab in Alderley Park | Wellcome Sanger Institute for the COVID-19 Genomics UK (COG-UK) Consortium | Jacquelyn Wynn, Mairead Hyland, The Lighthouse Lab in Alderley Park and Alex Alderton, Roberto Amato, Jeffrey Barrett, Sonia Goncalves, Ewan Harrison, David K. Jackson, Ian Johnston, Dominic Kwiatkowski, Cordelia Langford, John Sillitoe on behalf of the Wellcome Sanger Institute COVID-19 Surveillance Team |
| EPI_ISL_2853383 | Lighthouse Laboratory Plymouth | Wellcome Sanger Institute for the COVID-19 Genomics UK (COG-UK) Consortium | Lighthouse Laboratory Plymouth and Alex Alderton, Roberto Amato, Jeffrey Barrett, Sonia Goncalves, Ewan Harrison, David K. Jackson, Ian Johnston, Dominic Kwiatkowski, Cordelia Langford, John Sillitoe on behalf of the Wellcome Sanger Institute COVID-19 Surveillance Team |
| EPI_ISL_2853384, EPI_ISL_2853385, EPI_ISL_2853386, | Lighthouse Lab in Alderley Park | Wellcome Sanger Institute for the COVID-19 Genomics UK (COG-UK) Consortium | Jacquelyn Wynn, Mairead Hyland, The Lighthouse Lab in Alderley Park and Alex Alderton, Roberto Amato, Jeffrey Barrett, Sonia Goncalves, Ewan Harrison, David K. Jackson, Ian Johnston, Dominic Kwiatkowski, Cordelia Langford, John Sillitoe on behalf of the Wellcome Sanger Institute COVID-19 Surveillance Team |

| | | | |
|---|---|---|---|
| EPI_ISL_2853387, EPI_ISL_2853388, EPI_ISL_2853389, EPI_ISL_2853390, EPI_ISL_2853391, EPI_ISL_2853392, EPI_ISL_2853393 | | | |
| EPI_ISL_2853394 | Lighthouse Laboratory Plymouth | Wellcome Sanger Institute for the COVID-19 Genomics UK (COG-UK) Consortium | Lighthouse Laboratory Plymouth and Alex Alderton, Roberto Amato, Jeffrey Barrett, Sonia Goncalves, Ewan Harrison, David K. Jackson, Ian Johnston, Dominic Kwiatkowski, Cordelia Langford, John Sillitoe on behalf of the Wellcome Sanger Institute COVID-19 Surveillance Team |
| EPI_ISL_2853395, EPI_ISL_2853396, EPI_ISL_2853397, EPI_ISL_2853398 | Lighthouse Lab in Alderley Park | Wellcome Sanger Institute for the COVID-19 Genomics UK (COG-UK) Consortium | Jacquelyn Wynn, Mairead Hyland, The Lighthouse Lab in Alderley Park and Alex Alderton, Roberto Amato, Jeffrey Barrett, Sonia Goncalves, Ewan Harrison, David K. Jackson, Ian Johnston, Dominic Kwiatkowski, Cordelia Langford, John Sillitoe on behalf of the Wellcome Sanger Institute COVID-19 Surveillance Team |
| EPI_ISL_2853399, EPI_ISL_2853400, EPI_ISL_2853401 | Lighthouse Laboratory Plymouth | Wellcome Sanger Institute for the COVID-19 Genomics UK (COG-UK) Consortium | Lighthouse Laboratory Plymouth and Alex Alderton, Roberto Amato, Jeffrey Barrett, Sonia Goncalves, Ewan Harrison, David K. Jackson, Ian Johnston, Dominic Kwiatkowski, Cordelia Langford, John Sillitoe on behalf of the Wellcome Sanger Institute COVID-19 Surveillance Team |
| EPI_ISL_2853402 | Lighthouse Lab in Alderley Park | Wellcome Sanger Institute for the COVID-19 Genomics UK (COG-UK) Consortium | Jacquelyn Wynn, Mairead Hyland, The Lighthouse Lab in Alderley Park and Alex Alderton, Roberto Amato, Jeffrey Barrett, Sonia Goncalves, Ewan Harrison, David K. Jackson, Ian Johnston, Dominic Kwiatkowski, Cordelia Langford, John Sillitoe on behalf of the Wellcome Sanger Institute COVID-19 Surveillance Team |
| EPI_ISL_2853403 | Lighthouse Laboratory Plymouth | Wellcome Sanger Institute for the COVID-19 Genomics UK (COG-UK) Consortium | Lighthouse Laboratory Plymouth and Alex Alderton, Roberto Amato, Jeffrey Barrett, Sonia Goncalves, Ewan Harrison, David K. Jackson, Ian Johnston, Dominic Kwiatkowski, Cordelia Langford, John Sillitoe on behalf of the Wellcome Sanger Institute COVID-19 Surveillance Team |
| EPI_ISL_2853404, EPI_ISL_2853405, EPI_ISL_2853406, EPI_ISL_2853407, EPI_ISL_2853408, EPI_ISL_2853409, EPI_ISL_2853410 | Lighthouse Lab in Alderley Park | Wellcome Sanger Institute for the COVID-19 Genomics UK (COG-UK) Consortium | Jacquelyn Wynn, Mairead Hyland, The Lighthouse Lab in Alderley Park and Alex Alderton, Roberto Amato, Jeffrey Barrett, Sonia Goncalves, Ewan Harrison, David K. Jackson, Ian Johnston, Dominic Kwiatkowski, Cordelia Langford, John Sillitoe on behalf of the Wellcome Sanger Institute COVID-19 Surveillance Team |
| EPI_ISL_2853411 | Lighthouse Laboratory Plymouth | Wellcome Sanger Institute for the COVID-19 Genomics UK (COG-UK) Consortium | Lighthouse Laboratory Plymouth and Alex Alderton, Roberto Amato, Jeffrey Barrett, Sonia Goncalves, Ewan Harrison, David K. Jackson, Ian Johnston, Dominic Kwiatkowski, Cordelia Langford, John Sillitoe on behalf of the Wellcome Sanger Institute COVID-19 Surveillance Team |
| EPI_ISL_2853412, EPI_ISL_2853413, EPI_ISL_2853414, EPI_ISL_2853415, EPI_ISL_2853416 | Lighthouse Lab in Alderley Park | Wellcome Sanger Institute for the COVID-19 Genomics UK (COG-UK) Consortium | Jacquelyn Wynn, Mairead Hyland, The Lighthouse Lab in Alderley Park and Alex Alderton, Roberto Amato, Jeffrey Barrett, Sonia Goncalves, Ewan Harrison, David K. Jackson, Ian Johnston, Dominic Kwiatkowski, Cordelia Langford, John Sillitoe on behalf of the Wellcome Sanger Institute COVID-19 Surveillance Team |
| EPI_ISL_2853417 | Lighthouse Laboratory Plymouth | Wellcome Sanger Institute for the COVID-19 Genomics UK (COG-UK) Consortium | Lighthouse Laboratory Plymouth and Alex Alderton, Roberto Amato, Jeffrey Barrett, Sonia Goncalves, Ewan Harrison, David K. Jackson, Ian Johnston, Dominic Kwiatkowski, Cordelia Langford, John Sillitoe on behalf of the Wellcome Sanger Institute COVID-19 Surveillance Team |
| EPI_ISL_2853418, EPI_ISL_2853419 | Lighthouse Lab in Alderley Park | Wellcome Sanger Institute for the COVID-19 Genomics UK (COG-UK) Consortium | Jacquelyn Wynn, Mairead Hyland, The Lighthouse Lab in Alderley Park and Alex Alderton, Roberto Amato, Jeffrey Barrett, Sonia Goncalves, Ewan Harrison, David K. Jackson, Ian Johnston, Dominic Kwiatkowski, Cordelia Langford, John Sillitoe on behalf of the Wellcome Sanger Institute COVID-19 Surveillance Team |
| EPI_ISL_2853420 | Lighthouse Laboratory Plymouth | Wellcome Sanger Institute for the COVID-19 Genomics UK (COG-UK) Consortium | Lighthouse Laboratory Plymouth and Alex Alderton, Roberto Amato, Jeffrey Barrett, Sonia Goncalves, Ewan Harrison, David K. Jackson, Ian Johnston, Dominic Kwiatkowski, Cordelia Langford, John Sillitoe on behalf of the Wellcome Sanger Institute COVID-19 Surveillance Team |
| EPI_ISL_2853421, EPI_ISL_2853423 | Lighthouse Lab in Alderley Park | Wellcome Sanger Institute for the COVID-19 Genomics UK (COG-UK) Consortium | Jacquelyn Wynn, Mairead Hyland, The Lighthouse Lab in Alderley Park and Alex Alderton, Roberto Amato, Jeffrey Barrett, Sonia Goncalves, Ewan Harrison, David K. Jackson, Ian Johnston, Dominic Kwiatkowski, Cordelia Langford, John Sillitoe on behalf of the Wellcome Sanger Institute COVID-19 Surveillance Team |
| EPI_ISL_2853424 | Lighthouse Laboratory Plymouth | Wellcome Sanger Institute for the COVID-19 Genomics UK (COG-UK) Consortium | Lighthouse Laboratory Plymouth and Alex Alderton, Roberto Amato, Jeffrey Barrett, Sonia Goncalves, Ewan Harrison, David K. Jackson, Ian Johnston, Dominic Kwiatkowski, Cordelia Langford, John Sillitoe on behalf of the Wellcome Sanger Institute COVID-19 Surveillance Team |
| EPI_ISL_2853425, EPI_ISL_2853426, EPI_ISL_2853427 | Lighthouse Lab in Alderley Park | Wellcome Sanger Institute for the COVID-19 Genomics UK (COG-UK) Consortium | Jacquelyn Wynn, Mairead Hyland, The Lighthouse Lab in Alderley Park and Alex Alderton, Roberto Amato, Jeffrey Barrett, Sonia Goncalves, Ewan Harrison, David K. Jackson, Ian Johnston, Dominic Kwiatkowski, Cordelia Langford, John Sillitoe on behalf of the Wellcome Sanger Institute COVID-19 Surveillance Team |
| EPI_ISL_2853428, EPI_ISL_2853429 | Lighthouse Laboratory Plymouth | Wellcome Sanger Institute for the COVID-19 Genomics UK (COG-UK) Consortium | Lighthouse Laboratory Plymouth and Alex Alderton, Roberto Amato, Jeffrey Barrett, Sonia Goncalves, Ewan Harrison, David K. Jackson, Ian Johnston, Dominic Kwiatkowski, Cordelia Langford, John Sillitoe on behalf of the Wellcome Sanger Institute COVID-19 Surveillance Team |
| EPI_ISL_2853430, EPI_ISL_2853431, EPI_ISL_2853432 | Lighthouse Lab in Alderley Park | Wellcome Sanger Institute for the COVID-19 Genomics UK (COG-UK) Consortium | Jacquelyn Wynn, Mairead Hyland, The Lighthouse Lab in Alderley Park and Alex Alderton, Roberto Amato, Jeffrey Barrett, Sonia Goncalves, Ewan Harrison, David K. Jackson, Ian Johnston, Dominic Kwiatkowski, Cordelia Langford, John Sillitoe on behalf of the Wellcome Sanger Institute COVID-19 Surveillance Team |
| EPI_ISL_2853433 | Lighthouse Laboratory Plymouth | Wellcome Sanger Institute for the COVID-19 Genomics UK (COG-UK) Consortium | Lighthouse Laboratory Plymouth and Alex Alderton, Roberto Amato, Jeffrey Barrett, Sonia Goncalves, Ewan Harrison, David K. Jackson, Ian Johnston, Dominic Kwiatkowski, Cordelia Langford, John Sillitoe on behalf of the Wellcome Sanger Institute COVID-19 Surveillance Team |
| EPI_ISL_2853434, EPI_ISL_2853435, EPI_ISL_2853436, EPI_ISL_2853437 | Lighthouse Lab in Alderley Park | Wellcome Sanger Institute for the COVID-19 Genomics UK (COG-UK) Consortium | Jacquelyn Wynn, Mairead Hyland, The Lighthouse Lab in Alderley Park and Alex Alderton, Roberto Amato, Jeffrey Barrett, Sonia Goncalves, Ewan Harrison, David K. Jackson, Ian Johnston, Dominic Kwiatkowski, Cordelia Langford, John Sillitoe on behalf of the Wellcome Sanger Institute COVID-19 Surveillance Team |
| EPI_ISL_2853438 | Lighthouse Laboratory Plymouth | Wellcome Sanger Institute for the COVID-19 Genomics UK (COG-UK) Consortium | Lighthouse Laboratory Plymouth and Alex Alderton, Roberto Amato, Jeffrey Barrett, Sonia Goncalves, Ewan Harrison, David K. Jackson, Ian Johnston, Dominic Kwiatkowski, Cordelia Langford, John Sillitoe on behalf of the Wellcome Sanger Institute COVID-19 Surveillance Team |
| EPI_ISL_2853439, EPI_ISL_2853440, EPI_ISL_2853441, EPI_ISL_2853443, EPI_ISL_2853444, EPI_ISL_2853445, EPI_ISL_2853446 | Lighthouse Lab in Alderley Park | Wellcome Sanger Institute for the COVID-19 Genomics UK (COG-UK) Consortium | Jacquelyn Wynn, Mairead Hyland, The Lighthouse Lab in Alderley Park and Alex Alderton, Roberto Amato, Jeffrey Barrett, Sonia Goncalves, Ewan Harrison, David K. Jackson, Ian Johnston, Dominic Kwiatkowski, Cordelia Langford, John Sillitoe on behalf of the Wellcome Sanger Institute COVID-19 Surveillance Team |
| EPI_ISL_2853447 | Lighthouse Laboratory Plymouth | Wellcome Sanger Institute for the COVID-19 Genomics UK (COG-UK) Consortium | Lighthouse Laboratory Plymouth and Alex Alderton, Roberto Amato, Jeffrey Barrett, Sonia Goncalves, Ewan Harrison, David K. Jackson, Ian Johnston, Dominic Kwiatkowski, Cordelia Langford, John Sillitoe on behalf of the Wellcome Sanger Institute COVID-19 Surveillance Team |
| EPI_ISL_2853448, EPI_ISL_2853449, EPI_ISL_2853450, EPI_ISL_2853451, EPI_ISL_2853452, EPI_ISL_2853453 | Lighthouse Lab in Alderley Park | Wellcome Sanger Institute for the COVID-19 Genomics UK (COG-UK) Consortium | Jacquelyn Wynn, Mairead Hyland, The Lighthouse Lab in Alderley Park and Alex Alderton, Roberto Amato, Jeffrey Barrett, Sonia Goncalves, Ewan Harrison, David K. Jackson, Ian Johnston, Dominic Kwiatkowski, Cordelia Langford, John Sillitoe on behalf of the Wellcome Sanger Institute COVID-19 Surveillance Team |
| EPI_ISL_2853454 | Lighthouse Laboratory Plymouth | Wellcome Sanger Institute for the COVID-19 Genomics UK (COG-UK) Consortium | Lighthouse Laboratory Plymouth and Alex Alderton, Roberto Amato, Jeffrey Barrett, Sonia Goncalves, Ewan Harrison, David K. Jackson, Ian Johnston, Dominic Kwiatkowski, Cordelia Langford, John Sillitoe on behalf of the Wellcome Sanger Institute COVID-19 Surveillance Team |
| EPI_ISL_2853455 | Lighthouse Lab in Alderley Park | Wellcome Sanger Institute for the COVID-19 Genomics UK (COG-UK) Consortium | Jacquelyn Wynn, Mairead Hyland, The Lighthouse Lab in Alderley Park and Alex Alderton, Roberto Amato, Jeffrey Barrett, Sonia Goncalves, Ewan Harrison, David K. Jackson, Ian Johnston, Dominic Kwiatkowski, Cordelia Langford, John Sillitoe on behalf of the Wellcome Sanger Institute COVID-19 Surveillance Team |
| EPI_ISL_2853456, EPI_ISL_2853457, EPI_ISL_2853458, EPI_ISL_2853459, EPI_ISL_2853460, EPI_ISL_2853461 | Lighthouse Laboratory Plymouth | Wellcome Sanger Institute for the COVID-19 Genomics UK (COG-UK) Consortium | Lighthouse Laboratory Plymouth and Alex Alderton, Roberto Amato, Jeffrey Barrett, Sonia Goncalves, Ewan Harrison, David K. Jackson, Ian Johnston, Dominic Kwiatkowski, Cordelia Langford, John Sillitoe on behalf of the Wellcome Sanger Institute COVID-19 Surveillance Team |
| EPI_ISL_2853462, EPI_ISL_2853463, EPI_ISL_2853464, EPI_ISL_2853465, EPI_ISL_2853466 | Lighthouse Lab in Alderley Park | Wellcome Sanger Institute for the COVID-19 Genomics UK (COG-UK) Consortium | Jacquelyn Wynn, Mairead Hyland, The Lighthouse Lab in Alderley Park and Alex Alderton, Roberto Amato, Jeffrey Barrett, Sonia Goncalves, Ewan Harrison, David K. Jackson, Ian Johnston, Dominic Kwiatkowski, Cordelia Langford, John Sillitoe on behalf of the Wellcome Sanger Institute COVID-19 Surveillance Team |
| EPI_ISL_2853467 | Lighthouse Laboratory Plymouth | Wellcome Sanger Institute for the COVID-19 Genomics UK (COG-UK) Consortium | Lighthouse Laboratory Plymouth and Alex Alderton, Roberto Amato, Jeffrey Barrett, Sonia Goncalves, Ewan Harrison, David K. Jackson, Ian Johnston, Dominic Kwiatkowski, Cordelia Langford, John Sillitoe on behalf of the Wellcome Sanger Institute COVID-19 Surveillance Team |
| EPI_ISL_2853468, EPI_ISL_2853469, EPI_ISL_2853470, EPI_ISL_2853471, EPI_ISL_2853472 | Lighthouse Lab in Alderley Park | Wellcome Sanger Institute for the COVID-19 Genomics UK (COG-UK) Consortium | Jacquelyn Wynn, Mairead Hyland, The Lighthouse Lab in Alderley Park and Alex Alderton, Roberto Amato, Jeffrey Barrett, Sonia Goncalves, Ewan Harrison, David K. Jackson, Ian Johnston, Dominic Kwiatkowski, Cordelia Langford, John Sillitoe on behalf of the Wellcome Sanger Institute COVID-19 Surveillance Team |

| | | | |
|---|---|---|---|
| EPI_ISL_2853473 | Lighthouse Laboratory Plymouth | Wellcome Sanger Institute for the COVID-19 Genomics UK (COG-UK) Consortium | Lighthouse Laboratory Plymouth and Alex Alderton, Roberto Amato, Jeffrey Barrett, Sonia Goncalves, Ewan Harrison, David K. Jackson, Ian Johnston, Dominic Kwiatkowski, Cordelia Langford, John Sillitoe on behalf of the Wellcome Sanger Institute COVID-19 Surveillance Team |
| EPI_ISL_2853474, EPI_ISL_2853475 | Lighthouse Lab in Alderley Park | Wellcome Sanger Institute for the COVID-19 Genomics UK (COG-UK) Consortium | Jacquelyn Wynn, Mairead Hyland, The Lighthouse Lab in Alderley Park and Alex Alderton, Roberto Amato, Jeffrey Barrett, Sonia Goncalves, Ewan Harrison, David K. Jackson, Ian Johnston, Dominic Kwiatkowski, Cordelia Langford, John Sillitoe on behalf of the Wellcome Sanger Institute COVID-19 Surveillance Team |
| EPI_ISL_2853476 | Lighthouse Laboratory Plymouth | Wellcome Sanger Institute for the COVID-19 Genomics UK (COG-UK) Consortium | Lighthouse Laboratory Plymouth and Alex Alderton, Roberto Amato, Jeffrey Barrett, Sonia Goncalves, Ewan Harrison, David K. Jackson, Ian Johnston, Dominic Kwiatkowski, Cordelia Langford, John Sillitoe on behalf of the Wellcome Sanger Institute COVID-19 Surveillance Team |
| EPI_ISL_2853477, EPI_ISL_2853478 | Lighthouse Lab in Alderley Park | Wellcome Sanger Institute for the COVID-19 Genomics UK (COG-UK) Consortium | Jacquelyn Wynn, Mairead Hyland, The Lighthouse Lab in Alderley Park and Alex Alderton, Roberto Amato, Jeffrey Barrett, Sonia Goncalves, Ewan Harrison, David K. Jackson, Ian Johnston, Dominic Kwiatkowski, Cordelia Langford, John Sillitoe on behalf of the Wellcome Sanger Institute COVID-19 Surveillance Team |
| EPI_ISL_2853479 | Lighthouse Laboratory Plymouth | Wellcome Sanger Institute for the COVID-19 Genomics UK (COG-UK) Consortium | Lighthouse Laboratory Plymouth and Alex Alderton, Roberto Amato, Jeffrey Barrett, Sonia Goncalves, Ewan Harrison, David K. Jackson, Ian Johnston, Dominic Kwiatkowski, Cordelia Langford, John Sillitoe on behalf of the Wellcome Sanger Institute COVID-19 Surveillance Team |
| EPI_ISL_2853480, EPI_ISL_2853481, EPI_ISL_2853482 | Lighthouse Lab in Alderley Park | Wellcome Sanger Institute for the COVID-19 Genomics UK (COG-UK) Consortium | Jacquelyn Wynn, Mairead Hyland, The Lighthouse Lab in Alderley Park and Alex Alderton, Roberto Amato, Jeffrey Barrett, Sonia Goncalves, Ewan Harrison, David K. Jackson, Ian Johnston, Dominic Kwiatkowski, Cordelia Langford, John Sillitoe on behalf of the Wellcome Sanger Institute COVID-19 Surveillance Team |
| EPI_ISL_2853483 | Lighthouse Laboratory Plymouth | Wellcome Sanger Institute for the COVID-19 Genomics UK (COG-UK) Consortium | Lighthouse Laboratory Plymouth and Alex Alderton, Roberto Amato, Jeffrey Barrett, Sonia Goncalves, Ewan Harrison, David K. Jackson, Ian Johnston, Dominic Kwiatkowski, Cordelia Langford, John Sillitoe on behalf of the Wellcome Sanger Institute COVID-19 Surveillance Team |
| EPI_ISL_2853484, EPI_ISL_2853485, EPI_ISL_2853486 | Lighthouse Lab in Alderley Park | Wellcome Sanger Institute for the COVID-19 Genomics UK (COG-UK) Consortium | Jacquelyn Wynn, Mairead Hyland, The Lighthouse Lab in Alderley Park and Alex Alderton, Roberto Amato, Jeffrey Barrett, Sonia Goncalves, Ewan Harrison, David K. Jackson, Ian Johnston, Dominic Kwiatkowski, Cordelia Langford, John Sillitoe on behalf of the Wellcome Sanger Institute COVID-19 Surveillance Team |
| EPI_ISL_2853487 | Lighthouse Laboratory Plymouth | Wellcome Sanger Institute for the COVID-19 Genomics UK (COG-UK) Consortium | Lighthouse Laboratory Plymouth and Alex Alderton, Roberto Amato, Jeffrey Barrett, Sonia Goncalves, Ewan Harrison, David K. Jackson, Ian Johnston, Dominic Kwiatkowski, Cordelia Langford, John Sillitoe on behalf of the Wellcome Sanger Institute COVID-19 Surveillance Team |
| EPI_ISL_2853488, EPI_ISL_2853489 | Lighthouse Lab in Alderley Park | Wellcome Sanger Institute for the COVID-19 Genomics UK (COG-UK) Consortium | Jacquelyn Wynn, Mairead Hyland, The Lighthouse Lab in Alderley Park and Alex Alderton, Roberto Amato, Jeffrey Barrett, Sonia Goncalves, Ewan Harrison, David K. Jackson, Ian Johnston, Dominic Kwiatkowski, Cordelia Langford, John Sillitoe on behalf of the Wellcome Sanger Institute COVID-19 Surveillance Team |
| EPI_ISL_2853490, EPI_ISL_2853491 | Lighthouse Laboratory Plymouth | Wellcome Sanger Institute for the COVID-19 Genomics UK (COG-UK) Consortium | Lighthouse Laboratory Plymouth and Alex Alderton, Roberto Amato, Jeffrey Barrett, Sonia Goncalves, Ewan Harrison, David K. Jackson, Ian Johnston, Dominic Kwiatkowski, Cordelia Langford, John Sillitoe on behalf of the Wellcome Sanger Institute COVID-19 Surveillance Team |
| EPI_ISL_2853492 | Lighthouse Lab in Alderley Park | Wellcome Sanger Institute for the COVID-19 Genomics UK (COG-UK) Consortium | Jacquelyn Wynn, Mairead Hyland, The Lighthouse Lab in Alderley Park and Alex Alderton, Roberto Amato, Jeffrey Barrett, Sonia Goncalves, Ewan Harrison, David K. Jackson, Ian Johnston, Dominic Kwiatkowski, Cordelia Langford, John Sillitoe on behalf of the Wellcome Sanger Institute COVID-19 Surveillance Team |
| EPI_ISL_2853493 | Lighthouse Laboratory Plymouth | Wellcome Sanger Institute for the COVID-19 Genomics UK (COG-UK) Consortium | Lighthouse Laboratory Plymouth and Alex Alderton, Roberto Amato, Jeffrey Barrett, Sonia Goncalves, Ewan Harrison, David K. Jackson, Ian Johnston, Dominic Kwiatkowski, Cordelia Langford, John Sillitoe on behalf of the Wellcome Sanger Institute COVID-19 Surveillance Team |
| EPI_ISL_2853494 | Lighthouse Lab in Alderley Park | Wellcome Sanger Institute for the COVID-19 Genomics UK (COG-UK) Consortium | Jacquelyn Wynn, Mairead Hyland, The Lighthouse Lab in Alderley Park and Alex Alderton, Roberto Amato, Jeffrey Barrett, Sonia Goncalves, Ewan Harrison, David K. Jackson, Ian Johnston, Dominic Kwiatkowski, Cordelia Langford, John Sillitoe on behalf of the Wellcome Sanger Institute COVID-19 Surveillance Team |
| EPI_ISL_2853495 | Lighthouse Laboratory Plymouth | Wellcome Sanger Institute for the COVID-19 Genomics UK (COG-UK) Consortium | Lighthouse Laboratory Plymouth and Alex Alderton, Roberto Amato, Jeffrey Barrett, Sonia Goncalves, Ewan Harrison, David K. Jackson, Ian Johnston, Dominic Kwiatkowski, Cordelia Langford, John Sillitoe on behalf of the Wellcome Sanger Institute COVID-19 Surveillance Team |
| EPI_ISL_2853496, EPI_ISL_2853497, EPI_ISL_2853498, EPI_ISL_2853499, EPI_ISL_2853500 | Lighthouse Lab in Alderley Park | Wellcome Sanger Institute for the COVID-19 Genomics UK (COG-UK) Consortium | Jacquelyn Wynn, Mairead Hyland, The Lighthouse Lab in Alderley Park and Alex Alderton, Roberto Amato, Jeffrey Barrett, Sonia Goncalves, Ewan Harrison, David K. Jackson, Ian Johnston, Dominic Kwiatkowski, Cordelia Langford, John Sillitoe on behalf of the Wellcome Sanger Institute COVID-19 Surveillance Team |
| EPI_ISL_2853501, EPI_ISL_2853502 | Lighthouse Laboratory Plymouth | Wellcome Sanger Institute for the COVID-19 Genomics UK (COG-UK) Consortium | Lighthouse Laboratory Plymouth and Alex Alderton, Roberto Amato, Jeffrey Barrett, Sonia Goncalves, Ewan Harrison, David K. Jackson, Ian Johnston, Dominic Kwiatkowski, Cordelia Langford, John Sillitoe on behalf of the Wellcome Sanger Institute COVID-19 Surveillance Team |
| EPI_ISL_2853503, EPI_ISL_2853504, EPI_ISL_2853505, EPI_ISL_2853506, EPI_ISL_2853507 | Lighthouse Lab in Alderley Park | Wellcome Sanger Institute for the COVID-19 Genomics UK (COG-UK) Consortium | Jacquelyn Wynn, Mairead Hyland, The Lighthouse Lab in Alderley Park and Alex Alderton, Roberto Amato, Jeffrey Barrett, Sonia Goncalves, Ewan Harrison, David K. Jackson, Ian Johnston, Dominic Kwiatkowski, Cordelia Langford, John Sillitoe on behalf of the Wellcome Sanger Institute COVID-19 Surveillance Team |
| EPI_ISL_2853508 | Lighthouse Laboratory Plymouth | Wellcome Sanger Institute for the COVID-19 Genomics UK (COG-UK) Consortium | Lighthouse Laboratory Plymouth and Alex Alderton, Roberto Amato, Jeffrey Barrett, Sonia Goncalves, Ewan Harrison, David K. Jackson, Ian Johnston, Dominic Kwiatkowski, Cordelia Langford, John Sillitoe on behalf of the Wellcome Sanger Institute COVID-19 Surveillance Team |
| EPI_ISL_2853509, EPI_ISL_2853510, EPI_ISL_2853511, EPI_ISL_2853512, EPI_ISL_2853513 | Lighthouse Lab in Alderley Park | Wellcome Sanger Institute for the COVID-19 Genomics UK (COG-UK) Consortium | Jacquelyn Wynn, Mairead Hyland, The Lighthouse Lab in Alderley Park and Alex Alderton, Roberto Amato, Jeffrey Barrett, Sonia Goncalves, Ewan Harrison, David K. Jackson, Ian Johnston, Dominic Kwiatkowski, Cordelia Langford, John Sillitoe on behalf of the Wellcome Sanger Institute COVID-19 Surveillance Team |
| EPI_ISL_2853514 | Lighthouse Laboratory Plymouth | Wellcome Sanger Institute for the COVID-19 Genomics UK (COG-UK) Consortium | Lighthouse Laboratory Plymouth and Alex Alderton, Roberto Amato, Jeffrey Barrett, Sonia Goncalves, Ewan Harrison, David K. Jackson, Ian Johnston, Dominic Kwiatkowski, Cordelia Langford, John Sillitoe on behalf of the Wellcome Sanger Institute COVID-19 Surveillance Team |
| EPI_ISL_2853515 | Lighthouse Lab in Alderley Park | Wellcome Sanger Institute for the COVID-19 Genomics UK (COG-UK) Consortium | Jacquelyn Wynn, Mairead Hyland, The Lighthouse Lab in Alderley Park and Alex Alderton, Roberto Amato, Jeffrey Barrett, Sonia Goncalves, Ewan Harrison, David K. Jackson, Ian Johnston, Dominic Kwiatkowski, Cordelia Langford, John Sillitoe on behalf of the Wellcome Sanger Institute COVID-19 Surveillance Team |
| EPI_ISL_2853516 | Lighthouse Laboratory Plymouth | Wellcome Sanger Institute for the COVID-19 Genomics UK (COG-UK) Consortium | Lighthouse Laboratory Plymouth and Alex Alderton, Roberto Amato, Jeffrey Barrett, Sonia Goncalves, Ewan Harrison, David K. Jackson, Ian Johnston, Dominic Kwiatkowski, Cordelia Langford, John Sillitoe on behalf of the Wellcome Sanger Institute COVID-19 Surveillance Team |
| EPI_ISL_2853517, EPI_ISL_2853518, EPI_ISL_2853519, EPI_ISL_2853520, EPI_ISL_2853521, EPI_ISL_2853522, EPI_ISL_2853523, EPI_ISL_2853524, EPI_ISL_2853525 | Lighthouse Lab in Alderley Park | Wellcome Sanger Institute for the COVID-19 Genomics UK (COG-UK) Consortium | Jacquelyn Wynn, Mairead Hyland, The Lighthouse Lab in Alderley Park and Alex Alderton, Roberto Amato, Jeffrey Barrett, Sonia Goncalves, Ewan Harrison, David K. Jackson, Ian Johnston, Dominic Kwiatkowski, Cordelia Langford, John Sillitoe on behalf of the Wellcome Sanger Institute COVID-19 Surveillance Team |
| EPI_ISL_2853526 | Lighthouse Laboratory Plymouth | Wellcome Sanger Institute for the COVID-19 Genomics UK (COG-UK) Consortium | Lighthouse Laboratory Plymouth and Alex Alderton, Roberto Amato, Jeffrey Barrett, Sonia Goncalves, Ewan Harrison, David K. Jackson, Ian Johnston, Dominic Kwiatkowski, Cordelia Langford, John Sillitoe on behalf of the Wellcome Sanger Institute COVID-19 Surveillance Team |
| EPI_ISL_2853527, EPI_ISL_2853528, EPI_ISL_2853529, EPI_ISL_2853530, EPI_ISL_2853531, EPI_ISL_2853532 | Lighthouse Lab in Alderley Park | Wellcome Sanger Institute for the COVID-19 Genomics UK (COG-UK) Consortium | Jacquelyn Wynn, Mairead Hyland, The Lighthouse Lab in Alderley Park and Alex Alderton, Roberto Amato, Jeffrey Barrett, Sonia Goncalves, Ewan Harrison, David K. Jackson, Ian Johnston, Dominic Kwiatkowski, Cordelia Langford, John Sillitoe on behalf of the Wellcome Sanger Institute COVID-19 Surveillance Team |
| EPI_ISL_2853533, EPI_ISL_2853534 | Lighthouse Laboratory Plymouth | Wellcome Sanger Institute for the COVID-19 Genomics UK (COG-UK) Consortium | Lighthouse Laboratory Plymouth and Alex Alderton, Roberto Amato, Jeffrey Barrett, Sonia Goncalves, Ewan Harrison, David K. Jackson, Ian Johnston, Dominic Kwiatkowski, Cordelia Langford, John Sillitoe on behalf of the Wellcome Sanger Institute COVID-19 Surveillance Team |
| EPI_ISL_2853535 | Lighthouse Lab in Alderley Park | Wellcome Sanger Institute for the COVID-19 Genomics UK (COG-UK) Consortium | Jacquelyn Wynn, Mairead Hyland, The Lighthouse Lab in Alderley Park and Alex Alderton, Roberto Amato, Jeffrey Barrett, Sonia Goncalves, Ewan Harrison, David K. Jackson, Ian Johnston, Dominic Kwiatkowski, Cordelia Langford, John Sillitoe on behalf of the Wellcome Sanger Institute COVID-19 Surveillance Team |
| EPI_ISL_2853536 | Lighthouse Laboratory Plymouth | Wellcome Sanger Institute for the COVID-19 Genomics UK (COG-UK) Consortium | Lighthouse Laboratory Plymouth and Alex Alderton, Roberto Amato, Jeffrey Barrett, Sonia Goncalves, Ewan Harrison, David K. Jackson, Ian Johnston, Dominic Kwiatkowski, Cordelia Langford, John Sillitoe on behalf of the Wellcome Sanger Institute COVID-19 Surveillance Team |
| EPI_ISL_2853537, EPI_ISL_2853538, EPI_ISL_2853539 | Lighthouse Lab in Alderley Park | Wellcome Sanger Institute for the COVID-19 Genomics UK (COG-UK) Consortium | Jacquelyn Wynn, Mairead Hyland, The Lighthouse Lab in Alderley Park and Alex Alderton, Roberto Amato, Jeffrey Barrett, Sonia Goncalves, Ewan Harrison, David K. Jackson, Ian Johnston, Dominic Kwiatkowski, Cordelia Langford, John Sillitoe on behalf of the Wellcome Sanger Institute COVID-19 Surveillance Team |
| EPI_ISL_2853540 | Lighthouse Laboratory Plymouth | Wellcome Sanger Institute for the COVID-19 Genomics UK (COG-UK) Consortium | Lighthouse Laboratory Plymouth and Alex Alderton, Roberto Amato, Jeffrey Barrett, Sonia Goncalves, Ewan Harrison, David K. Jackson, Ian Johnston, Dominic Kwiatkowski, Cordelia Langford, John Sillitoe on behalf of the Wellcome Sanger Institute COVID-19 Surveillance Team |
| EPI_ISL_2853541, EPI_ISL_2853542 | Lighthouse Lab in Alderley Park | Wellcome Sanger Institute for the COVID-19 Genomics UK (COG-UK) Consortium | Jacquelyn Wynn, Mairead Hyland, The Lighthouse Lab in Alderley Park and Alex Alderton, Roberto Amato, Jeffrey Barrett, Sonia Goncalves, Ewan Harrison, David K. Jackson, Ian Johnston, Dominic Kwiatkowski, Cordelia Langford, John Sillitoe on behalf of the Wellcome Sanger Institute COVID-19 Surveillance Team |
| EPI_ISL_2853543, EPI_ISL_2853544, EPI_ISL_2853545 | Lighthouse Laboratory Plymouth | Wellcome Sanger Institute for the COVID-19 Genomics UK (COG-UK) Consortium | Lighthouse Laboratory Plymouth and Alex Alderton, Roberto Amato, Jeffrey Barrett, Sonia Goncalves, Ewan Harrison, David K. Jackson, Ian Johnston, Dominic Kwiatkowski, Cordelia Langford, John Sillitoe on behalf of the Wellcome Sanger Institute COVID-19 Surveillance Team |
| EPI_ISL_2853546 | Lighthouse Lab in Alderley Park | Wellcome Sanger Institute for the COVID-19 Genomics UK (COG-UK) Consortium | Jacquelyn Wynn, Mairead Hyland, The Lighthouse Lab in Alderley Park and Alex Alderton, Roberto Amato, Jeffrey Barrett, Sonia Goncalves, Ewan Harrison, David K. Jackson, Ian Johnston, Dominic Kwiatkowski, Cordelia Langford, John Sillitoe on behalf of the Wellcome Sanger Institute COVID-19 Surveillance Team |
| EPI_ISL_2853547 | Lighthouse Laboratory Plymouth | Wellcome Sanger Institute for the COVID-19 Genomics UK (COG-UK) Consortium | Lighthouse Laboratory Plymouth and Alex Alderton, Roberto Amato, Jeffrey Barrett, Sonia Goncalves, Ewan Harrison, David K. Jackson, Ian Johnston, Dominic Kwiatkowski, Cordelia Langford, John Sillitoe on behalf of the Wellcome Sanger Institute COVID-19 Surveillance Team |
| EPI_ISL_2853548, | Lighthouse Lab in Alderley Park | Wellcome Sanger Institute for the COVID-19 Genomics UK (COG-UK) | Jacquelyn Wynn, Mairead Hyland, The Lighthouse Lab in Alderley Park and Alex Alderton, Roberto Amato, Jeffrey Barrett, Sonia Goncalves, Ewan Harrison, David K. Jackson, Ian Johnston, Dominic Kwiatkowski, Cordelia Langford, John Sillitoe on |

| | | | |
|---|---|---|---|
| EPI_ISL_2853549, EPI_ISL_2853551, EPI_ISL_2853552, EPI_ISL_2853553, EPI_ISL_2853554, EPI_ISL_2853555, EPI_ISL_2853556 | | Consortium | behalf of the Wellcome Sanger Institute COVID-19 Surveillance Team |
| EPI_ISL_2853557 | Lighthouse Laboratory Plymouth | Wellcome Sanger Institute for the COVID-19 Genomics UK (COG-UK) Consortium | Lighthouse Laboratory Plymouth and Alex Alderton, Roberto Amato, Jeffrey Barrett, Sonia Goncalves, Ewan Harrison, David K. Jackson, Ian Johnston, Dominic Kwiatkowski, Cordelia Langford, John Sillitoe on behalf of the Wellcome Sanger Institute COVID-19 Surveillance Team |
| EPI_ISL_2853558, EPI_ISL_2853559, EPI_ISL_2853560, EPI_ISL_2853561, EPI_ISL_2853562, EPI_ISL_2853563, EPI_ISL_2853564, EPI_ISL_2853565, EPI_ISL_2853566, EPI_ISL_2853567, EPI_ISL_2853568, EPI_ISL_2853569, EPI_ISL_2853570, EPI_ISL_2853571, EPI_ISL_2853572, EPI_ISL_2853573 | | | |
| see above | Lighthouse Lab in Alderley Park | Wellcome Sanger Institute for the COVID-19 Genomics UK (COG-UK) Consortium | Jacquelyn Wynn, Mairead Hyland, The Lighthouse Lab in Alderley Park and Alex Alderton, Roberto Amato, Jeffrey Barrett, Sonia Goncalves, Ewan Harrison, David K. Jackson, Ian Johnston, Dominic Kwiatkowski, Cordelia Langford, John Sillitoe on behalf of the Wellcome Sanger Institute COVID-19 Surveillance Team |
| EPI_ISL_2853574 | Lighthouse Laboratory Plymouth | Wellcome Sanger Institute for the COVID-19 Genomics UK (COG-UK) Consortium | Lighthouse Laboratory Plymouth and Alex Alderton, Roberto Amato, Jeffrey Barrett, Sonia Goncalves, Ewan Harrison, David K. Jackson, Ian Johnston, Dominic Kwiatkowski, Cordelia Langford, John Sillitoe on behalf of the Wellcome Sanger Institute COVID-19 Surveillance Team |
| EPI_ISL_2853575, EPI_ISL_2853576, EPI_ISL_2853577, EPI_ISL_2853578, EPI_ISL_2853579, EPI_ISL_2853580, EPI_ISL_2853581 | Lighthouse Lab in Alderley Park | Wellcome Sanger Institute for the COVID-19 Genomics UK (COG-UK) Consortium | Jacquelyn Wynn, Mairead Hyland, The Lighthouse Lab in Alderley Park and Alex Alderton, Roberto Amato, Jeffrey Barrett, Sonia Goncalves, Ewan Harrison, David K. Jackson, Ian Johnston, Dominic Kwiatkowski, Cordelia Langford, John Sillitoe on behalf of the Wellcome Sanger Institute COVID-19 Surveillance Team |
| EPI_ISL_2853582 | Lighthouse Laboratory Plymouth | Wellcome Sanger Institute for the COVID-19 Genomics UK (COG-UK) Consortium | Lighthouse Laboratory Plymouth and Alex Alderton, Roberto Amato, Jeffrey Barrett, Sonia Goncalves, Ewan Harrison, David K. Jackson, Ian Johnston, Dominic Kwiatkowski, Cordelia Langford, John Sillitoe on behalf of the Wellcome Sanger Institute COVID-19 Surveillance Team |
| EPI_ISL_2853583, EPI_ISL_2853584, EPI_ISL_2853585 | Lighthouse Lab in Alderley Park | Wellcome Sanger Institute for the COVID-19 Genomics UK (COG-UK) Consortium | Jacquelyn Wynn, Mairead Hyland, The Lighthouse Lab in Alderley Park and Alex Alderton, Roberto Amato, Jeffrey Barrett, Sonia Goncalves, Ewan Harrison, David K. Jackson, Ian Johnston, Dominic Kwiatkowski, Cordelia Langford, John Sillitoe on behalf of the Wellcome Sanger Institute COVID-19 Surveillance Team |
| EPI_ISL_2853586 | Ministry of Health, Jaber Al-Ahmad Hospital | Virology Unit, Department of Microbiology, Faculty of Medicine, Kuwait University | Nada Madi, Hussain Safar, Ebaa Al-Awadhi, Anfal Al-Adwani |
| EPI_ISL_2854048 | Labuan Hospital | Institute for Medical Research, Infectious Disease Research Centre, National Institutes of Health, Ministry of Health Malaysia | Suppiah J, Kamel K, Mohd Zawawi Z, Azizan MA, Ramly N, Robert F, Anasir MI, Thayan R |
| EPI_ISL_2854052 | Ampang Hospital | Institute for Medical Research, Infectious Disease Research Centre, National Institutes of Health, Ministry of Health Malaysia | Suppiah J, Kamel K, Mohd Zawawi Z, Azizan MA, Ramly N, Robert F, Anasir MI, Thayan R |
| EPI_ISL_2854057, EPI_ISL_2854066 | Microbiology and Virology Laboratory, "Policlinico Riuniti, Azienda Ospedaliero Universitaria, Foggia" | Microbiology and Virology Laboratory, "Policlinico Riuniti, Azienda Ospedaliero Universitaria, Foggia" | Fabio Arena, Settimia Altamura, Donatella Cedola, Valeria Delli Carri, Annamaria D'Aprile, Maria Rosaria Lipsi, Loris Micelli, Maria Pia Patrizio, Daniela Pisanelli, Tiziana Rollo, Maurizio Margaglione and Rosella De Nittis. |
| EPI_ISL_2854187 | Kuala Terengganu District Health Office | Institute for Medical Research, Infectious Disease Research Centre, National Institutes of Health, Ministry of Health Malaysia | Suppiah J, Kamel K, Mohd Zawawi Z, Azizan MA, Ramly N, Robert F, Anasir MI, Thayan R |
| EPI_ISL_2854189, EPI_ISL_2854190, EPI_ISL_2854191, EPI_ISL_2854194, EPI_ISL_2854196, EPI_ISL_2854197, EPI_ISL_2854202, EPI_ISL_2854203, EPI_ISL_2854204, EPI_ISL_2854205, EPI_ISL_2854206, EPI_ISL_2854207 | | | |
| see above | Arizona State Public Health Laboratory | Arizona State Public Health Laboratory | Trung Huynh, Jessica Escobar, Katherine Fullerton, Nobuko Fukushima, Matthew Contursi, Stacy White, Linda Getsinger, Victor Waddell |
| EPI_ISL_2854210 | Genome Information Research Center, Research Institute for Microbial Diseases, Osaka University | Genome Information Research Center, Research Institute for Microbial Diseases, Osaka University | Daisuke Motooka, Kentaro Tanaka, Hiroya Oki, and Shota Nakamura |
| EPI_ISL_2854230, EPI_ISL_2854233, EPI_ISL_2854234, EPI_ISL_2854246, EPI_ISL_2854251, EPI_ISL_2854253, EPI_ISL_2854254, EPI_ISL_2854255, EPI_ISL_2854256 | NCSLPH | NCSLPH | Chase K, Miller MC, Greene S, Glover W |
| EPI_ISL_2854269 | Wojewodzka Stacja Sanitarno-Epidemiologiczna w Lodzi, Oddzial Laboratoryjny Mikrobiologii i Parazytologii | Wojewodzka Stacja Sanitarno-Epidemiologiczna w Lodzi, Oddzial Laboratoryjny Mikrobiologii i Parazytologii | Elian Terpo, Szymon Konkolewski, Marta Gawronska, Martyna Matera, Agnieszka Karnowska, Danuta Bartczak, Katarzyna Kosowska, Lucyna Kochanowska, Aleksandra Krawczyk |
| EPI_ISL_2854272, EPI_ISL_2854273, EPI_ISL_2854275, EPI_ISL_2854276, EPI_ISL_2854277, EPI_ISL_2854280, EPI_ISL_2854282, EPI_ISL_2854284, EPI_ISL_2854285, EPI_ISL_2854287, EPI_ISL_2854288, EPI_ISL_2854289, EPI_ISL_2854290, EPI_ISL_2854291, EPI_ISL_2854292, EPI_ISL_2854293, EPI_ISL_2854294, EPI_ISL_2854296, EPI_ISL_2854297, EPI_ISL_2854299, EPI_ISL_2854300, EPI_ISL_2854301, EPI_ISL_2854302, EPI_ISL_2854303, EPI_ISL_2854306, EPI_ISL_2854307, EPI_ISL_2854308, EPI_ISL_2854309, EPI_ISL_2854310, EPI_ISL_2854311, EPI_ISL_2854312, EPI_ISL_2854313, EPI_ISL_2854315, EPI_ISL_2854316, EPI_ISL_2854317, EPI_ISL_2854318, EPI_ISL_2854320, EPI_ISL_2854321, EPI_ISL_2854322, EPI_ISL_2854323, EPI_ISL_2854324, EPI_ISL_2854325, EPI_ISL_2854326, EPI_ISL_2854327, EPI_ISL_2854328, EPI_ISL_2854329, EPI_ISL_2854330, EPI_ISL_2854331, EPI_ISL_2854333, EPI_ISL_2854334, EPI_ISL_2854337, EPI_ISL_2854338, EPI_ISL_2854339, EPI_ISL_2854340, EPI_ISL_2854341, EPI_ISL_2854342, EPI_ISL_2854343, EPI_ISL_2854344, EPI_ISL_2854345, EPI_ISL_2854346, EPI_ISL_2854349, EPI_ISL_2854350, EPI_ISL_2854351, EPI_ISL_2854352, EPI_ISL_2854353, EPI_ISL_2854354, EPI_ISL_2854355, EPI_ISL_2854356, EPI_ISL_2854358, EPI_ISL_2854359, EPI_ISL_2854360, EPI_ISL_2854364, EPI_ISL_2854365, EPI_ISL_2854366, EPI_ISL_2854367, EPI_ISL_2854368, EPI_ISL_2854369, EPI_ISL_2854370, EPI_ISL_2854371, EPI_ISL_2854372, EPI_ISL_2854373, EPI_ISL_2854374, EPI_ISL_2854376, EPI_ISL_2854377, EPI_ISL_2854378, EPI_ISL_2854380, EPI_ISL_2854381, EPI_ISL_2854382, EPI_ISL_2854384, EPI_ISL_2854385, EPI_ISL_2854386, EPI_ISL_2854387, EPI_ISL_2854388, EPI_ISL_2854389, EPI_ISL_2854390, EPI_ISL_2854391, EPI_ISL_2854392, EPI_ISL_2854393, EPI_ISL_2854394, EPI_ISL_2854395, EPI_ISL_2854396, EPI_ISL_2854397, EPI_ISL_2854398, EPI_ISL_2854400, EPI_ISL_2854402, EPI_ISL_2854403, EPI_ISL_2854405, EPI_ISL_2854406, EPI_ISL_2854407, EPI_ISL_2854408, EPI_ISL_2854409, EPI_ISL_2854411, EPI_ISL_2854412, EPI_ISL_2854413, EPI_ISL_2854414, EPI_ISL_2854415, EPI_ISL_2854416, EPI_ISL_2854417, EPI_ISL_2854418, EPI_ISL_2854419, EPI_ISL_2854420, EPI_ISL_2854421, EPI_ISL_2854422, EPI_ISL_2854423, EPI_ISL_2854424, EPI_ISL_2854425, EPI_ISL_2854426, EPI_ISL_2854427, EPI_ISL_2854428, EPI_ISL_2854429, EPI_ISL_2854430, EPI_ISL_2854431, EPI_ISL_2854432, EPI_ISL_2854433, EPI_ISL_2854434, EPI_ISL_2854435, EPI_ISL_2854436, EPI_ISL_2854437, EPI_ISL_2854438, EPI_ISL_2854440, EPI_ISL_2854441, EPI_ISL_2854443, EPI_ISL_2854445, EPI_ISL_2854447, EPI_ISL_2854448, EPI_ISL_2854449, EPI_ISL_2854450, EPI_ISL_2854451, EPI_ISL_2854452, EPI_ISL_2854453, EPI_ISL_2854454, EPI_ISL_2854455, EPI_ISL_2854456, EPI_ISL_2854457, EPI_ISL_2854458, EPI_ISL_2854459, EPI_ISL_2854460, EPI_ISL_2854461, EPI_ISL_2854462, EPI_ISL_2854463, EPI_ISL_2854464, EPI_ISL_2854465, EPI_ISL_2854466, EPI_ISL_2854467, EPI_ISL_2854468, EPI_ISL_2854469, EPI_ISL_2854470, EPI_ISL_2854471, EPI_ISL_2854472, EPI_ISL_2854473, EPI_ISL_2854474, EPI_ISL_2854475, EPI_ISL_2854478, EPI_ISL_2854479, EPI_ISL_2854480, EPI_ISL_2854482, EPI_ISL_2854483, EPI_ISL_2854484, EPI_ISL_2854485, EPI_ISL_2854487, EPI_ISL_2854488, EPI_ISL_2854489, EPI_ISL_2854490, EPI_ISL_2854491, EPI_ISL_2854492, EPI_ISL_2854493, EPI_ISL_2854494, EPI_ISL_2854497, EPI_ISL_2854498, EPI_ISL_2854499, EPI_ISL_2854500, EPI_ISL_2854501, EPI_ISL_2854502, EPI_ISL_2854503, EPI_ISL_2854505, EPI_ISL_2854506, EPI_ISL_2854507, EPI_ISL_2854508, EPI_ISL_2854509, EPI_ISL_2854510, EPI_ISL_2854511, EPI_ISL_2854512, EPI_ISL_2854513, EPI_ISL_2854514, EPI_ISL_2854516, EPI_ISL_2854517, EPI_ISL_2854518, EPI_ISL_2854520, EPI_ISL_2854521, EPI_ISL_2854522, EPI_ISL_2854524, EPI_ISL_2854525, EPI_ISL_2854526, EPI_ISL_2854527, EPI_ISL_2854528, EPI_ISL_2854529, EPI_ISL_2854530, EPI_ISL_2854531, EPI_ISL_2854532, EPI_ISL_2854533, EPI_ISL_2854534, EPI_ISL_2854535, EPI_ISL_2854536, EPI_ISL_2854537, EPI_ISL_2854538, EPI_ISL_2854539, EPI_ISL_2854540, EPI_ISL_2854541, EPI_ISL_2854542, EPI_ISL_2854543, EPI_ISL_2854544, EPI_ISL_2854545, EPI_ISL_2854546, EPI_ISL_2854547, EPI_ISL_2854548, EPI_ISL_2854549, EPI_ISL_2854550, EPI_ISL_2854551, EPI_ISL_2854553, EPI_ISL_2854554, EPI_ISL_2854556, EPI_ISL_2854557, EPI_ISL_2854558, EPI_ISL_2854559, EPI_ISL_2854560, EPI_ISL_2854562, EPI_ISL_2854563, EPI_ISL_2854564, EPI_ISL_2854565, EPI_ISL_2854566, EPI_ISL_2854569, EPI_ISL_2854570, EPI_ISL_2854572, EPI_ISL_2854574, EPI_ISL_2854575, EPI_ISL_2854577, EPI_ISL_2854579, EPI_ISL_2854580, EPI_ISL_2854581, EPI_ISL_2854582, EPI_ISL_2854583, EPI_ISL_2854584, EPI_ISL_2854586, EPI_ISL_2854587, EPI_ISL_2854588, EPI_ISL_2854589, EPI_ISL_2854590, EPI_ISL_2854591, EPI_ISL_2854593, EPI_ISL_2854594, EPI_ISL_2854596, EPI_ISL_2854598, EPI_ISL_2854600, EPI_ISL_2854601, EPI_ISL_2854603, EPI_ISL_2854604, EPI_ISL_2854605, EPI_ISL_2854606, EPI_ISL_2854608, EPI_ISL_2854609, EPI_ISL_2854611, EPI_ISL_2854612, EPI_ISL_2854613, EPI_ISL_2854614, EPI_ISL_2854615, EPI_ISL_2854617, EPI_ISL_2854618, EPI_ISL_2854619, EPI_ISL_2854620, EPI_ISL_2854621, EPI_ISL_2854622, EPI_ISL_2854623, EPI_ISL_2854624, EPI_ISL_2854625, EPI_ISL_2854626, EPI_ISL_2854628, EPI_ISL_2854629, EPI_ISL_2854630, EPI_ISL_2854631, EPI_ISL_2854632, EPI_ISL_2854633, EPI_ISL_2854635, EPI_ISL_2854637, EPI_ISL_2854638, EPI_ISL_2854639, EPI_ISL_2854640, EPI_ISL_2854641, EPI_ISL_2854642, EPI_ISL_2854645, EPI_ISL_2854646, EPI_ISL_2854648, EPI_ISL_2854649, EPI_ISL_2854650 | | | |
| see above | Department of Bacteria, Parasites and Fungi, Statens Serum Institut, Copenhagen, Denmark | Statens Serum Institut Bioinformatics and Microbial Genomics | Danish Covid-19 Genome Consortium |
| EPI_ISL_2854653, EPI_ISL_2854663, EPI_ISL_2854666 | Molecular Diagnostics Pathology Department Mater Dei Hospital Malta | Molecular Diagnostics Pathology Department Mater Dei Hospital Malta | G Zahra, R Borg, C Cilia, L Grech, M Briffa |
| EPI_ISL_2854707 | UPTD LABKESDA Kota Samarinda | National Institute of Health Research and Development | Vivi Setiawaty, Hana Apsari Pawestri, Subangkit, Kartika Dewi Puspa, Arie Ardiansyah Nugraha, Hartanti Dian Ikawati, Krisna Pangesti, Triyani Soekarso, Nelly Puspandari |
| EPI_ISL_2854724 | RS Hermina Depok | National Institute of Health Research and Development | Subangkit, Hana Apsari Pawestri, , Kartika Dewi Puspa, Arie Ardiansyah Nugraha, Hartanti Dian Ikawati, Krisna Pangesti, Triyani Soekarso, Nelly Puspandari, Vivi Setiawaty |
| EPI_ISL_2854726 | Labkesda Kota Tangerang | National Institute of Health Research and Development | Subangkit, Hana Apsari Pawestri, , Kartika Dewi Puspa, Arie Ardiansyah Nugraha, Hartanti Dian Ikawati, Krisna Pangesti, Triyani Soekarso, Nelly Puspandari, Vivi Setiawaty |
| EPI_ISL_2854729 | LAB RS Mitra Kemayoran | National Institute of Health Research and Development | Subangkit, Hana Apsari Pawestri, , Kartika Dewi Puspa, Arie Ardiansyah Nugraha, Hartanti Dian Ikawati, Krisna Pangesti, Triyani Soekarso, Nelly Puspandari, Vivi Setiawaty |
| EPI_ISL_2854741, EPI_ISL_2854742, EPI_ISL_2854743, EPI_ISL_2854744 | RSU Kartika Pulomas | National Institute of Health Research and Development | Subangkit, Hana Apsari Pawestri, , Kartika Dewi Puspa, Arie Ardiansyah Nugraha, Hartanti Dian Ikawati, Krisna Pangesti, Triyani Soekarso, Nelly Puspandari, Vivi Setiawaty |
| EPI_ISL_2854747 | RSUD Ciawi | National Institute of Health Research and Development | Subangkit, Hana Apsari Pawestri, , Kartika Dewi Puspa, Arie Ardiansyah Nugraha, Hartanti Dian Ikawati, Krisna Pangesti, Triyani Soekarso, Nelly Puspandari, Vivi Setiawaty |
| EPI_ISL_2854748 | RS Ichsan Medical Centre Bintaro Tangsel | National Institute of Health Research and Development | Subangkit, Hana Apsari Pawestri, , Kartika Dewi Puspa, Arie Ardiansyah Nugraha, Hartanti Dian Ikawati, Krisna Pangesti, Triyani Soekarso, Nelly Puspandari, Vivi Setiawaty |
| EPI_ISL_2854749 | RS Kartika Pulomas | National Institute of Health Research and Development | Hana Apsari Pawestri, Subangkit, Kartika Dewi Puspa, Arie Ardiansyah Nugraha, Hartanti Dian Ikawati, Krisna Pangesti, Triyani Soekarso, Nelly Puspandari, Vivi Setiawaty |
| EPI_ISL_2854757, EPI_ISL_2854758, EPI_ISL_2854759 | Kantor Kesehatan Pelabuhan Kelas III Bengkulu | National Institute of Health Research and Development | Hana Apsari Pawestri, Subangkit, Kartika Dewi Puspa, Arie Ardiansyah Nugraha, Hartanti Dian Ikawati, Krisna Pangesti, Triyani Soekarso, Nelly Puspandari, Vivi Setiawaty |
| EPI_ISL_2854776 | RSU St Carolus | National Institute of Health Research and Development | Hana Apsari Pawestri, Subangkit, Kartika Dewi Puspa, Arie Ardiansyah Nugraha, Hartanti Dian Ikawati, Krisna Pangesti, Triyani Soekarso, Nelly Puspandari, Vivi Setiawaty |
| EPI_ISL_2854777 | RSU Permata Depok | National Institute of Health Research and Development | Hana Apsari Pawestri, Subangkit, Kartika Dewi Puspa, Arie Ardiansyah Nugraha, Hartanti Dian Ikawati, Krisna Pangesti, Triyani Soekarso, Nelly Puspandari, Vivi Setiawaty |
| EPI_ISL_2854778 | RS Krakatau Medika, Cilegon | National Institute of Health Research and Development | Hana Apsari Pawestri, Subangkit, Kartika Dewi Puspa, Arie Ardiansyah Nugraha, Hartanti Dian Ikawati, Krisna Pangesti, Triyani Soekarso, Nelly Puspandari, Vivi Setiawaty |
| EPI_ISL_2854779 | KKP Kelas II Cilacap | National Institute of Health Research and Development | Hana Apsari Pawestri, Subangkit, Kartika Dewi Puspa, Arie Ardiansyah Nugraha, Hartanti Dian Ikawati, Krisna Pangesti, Triyani Soekarso, Nelly Puspandari, Vivi Setiawaty |
| EPI_ISL_2854780 | Puskesmas kec Menteng | National Institute of Health Research and Development | Hana Apsari Pawestri, Subangkit, Kartika Dewi Puspa, Arie Ardiansyah Nugraha, Hartanti Dian Ikawati, Krisna Pangesti, Triyani Soekarso, Nelly Puspandari, Vivi Setiawaty |
| EPI_ISL_2854781, EPI_ISL_2854782, EPI_ISL_2854783, EPI_ISL_2854784, EPI_ISL_2854785, EPI_ISL_2854786, EPI_ISL_2854787, EPI_ISL_2854788, EPI_ISL_2854789, EPI_ISL_2854790, EPI_ISL_2854791, EPI_ISL_2854792, EPI_ISL_2854793, EPI_ISL_2854794, EPI_ISL_2854795, EPI_ISL_2854796 | | | |
| see above | Kantor Kesehatan Pelabuhan Kelas I Medan | National Institute of Health Research and Development | Hana Apsari Pawestri, Subangkit, Kartika Dewi Puspa, Arie Ardiansyah Nugraha, Hartanti Dian Ikawati, Krisna Pangesti, Triyani Soekarso, Nelly Puspandari, Vivi Setiawaty |
| EPI_ISL_2854797, | Molecular Diagnostics Pathology Department Mater Dei Hospital | Molecular Diagnostics Pathology Department Mater Dei Hospital | G Zahra, R Borg, C Cilia, L Grech, M Briffa |

| | | | |
|---|---|---|---|
| EPI_ISL_2854798 | Malta | Malta | |
| EPI_ISL_2854799 | Molecular Diagnostics Pathology Department Mater Dei Hospital Malta | Molecular Diagnostics Pathology Department Mater Dei Hospital Malta | G Zahra, R Borg, C Cilia, L Grech, M briffa |
| EPI_ISL_2854800, EPI_ISL_2854802 | Molecular Diagnostics Pathology Department Mater Dei Hospital Malta | Molecular Diagnostics Pathology Department Mater Dei Hospital Malta | G Zahra, R Borg, C Cilia, L Grech, M Briffa |
| EPI_ISL_2854803 | Jaber Quarantine | Virology Lab, Jaber Al Ahmad Hospital | Dr. Zahrah Buhamad, Dr. Ebaa Al-Awadhi, Adil Afridi, Haroon Masih |
| EPI_ISL_2854883, EPI_ISL_2854894, EPI_ISL_2854925, EPI_ISL_2854948, EPI_ISL_2854952 | Gravity Diagnostics, LLC | Gravity Diagnostics, LLC | Gravity Diagnostics |
| EPI_ISL_2854976, EPI_ISL_2855444, EPI_ISL_2855852 | Jaber Quarantine | Virology Lab, Jaber Al Ahmad Hospital | Dr. Zahrah Buhamad, Dr. Ebaa Al-Awadhi, Adil Afridi, Haroon Masih |
| EPI_ISL_2855867, EPI_ISL_2855870, EPI_ISL_2855872, EPI_ISL_2855874, EPI_ISL_2855909, EPI_ISL_2855911, EPI_ISL_2855977, EPI_ISL_2855994, EPI_ISL_2856026, EPI_ISL_2856035, EPI_ISL_2856038, EPI_ISL_2856051, EPI_ISL_2856067, EPI_ISL_2856069, EPI_ISL_2856096, EPI_ISL_2856119, EPI_ISL_2856122, EPI_ISL_2856123 | | | |
| see above | Utah Public Health Laboratory | Utah Public Health Laboratory | Olinto Linares-Perdomo, Erin L. Young, Kelly F. Oakeson, Tara Gallagher |
| EPI_ISL_2856149 | Jaber Quarantine | Virology Lab, Jaber Al Ahmad Hospital | Dr. Zahrah Buhamad, Dr. Ebaa Al-Awadhi, Adil Afridi, Haroon Masih |
| EPI_ISL_2856176, EPI_ISL_2856184, EPI_ISL_2856185, EPI_ISL_2856187, EPI_ISL_2856188, EPI_ISL_2856189, EPI_ISL_2856190, EPI_ISL_2856191, EPI_ISL_2856192, EPI_ISL_2856193, EPI_ISL_2856195, EPI_ISL_2856196, EPI_ISL_2856197, EPI_ISL_2856199, EPI_ISL_2856200, EPI_ISL_2856201, EPI_ISL_2856204, EPI_ISL_2856205, EPI_ISL_2856206, EPI_ISL_2856208, EPI_ISL_2856209, EPI_ISL_2856210, EPI_ISL_2856212, EPI_ISL_2856213, EPI_ISL_2856214, EPI_ISL_2856215, EPI_ISL_2856218, EPI_ISL_2856219, EPI_ISL_2856220, EPI_ISL_2856221, EPI_ISL_2856222, EPI_ISL_2856223, EPI_ISL_2856224, EPI_ISL_2856225, EPI_ISL_2856226, EPI_ISL_2856227, EPI_ISL_2856228, EPI_ISL_2856230, EPI_ISL_2856231, EPI_ISL_2856234, EPI_ISL_2856236, EPI_ISL_2856238, EPI_ISL_2856240, EPI_ISL_2856241, EPI_ISL_2856242, EPI_ISL_2856247, EPI_ISL_2856251, EPI_ISL_2856252, EPI_ISL_2856253, EPI_ISL_2856254, EPI_ISL_2856256, EPI_ISL_2856260, EPI_ISL_2856261, EPI_ISL_2856262, EPI_ISL_2856263, EPI_ISL_2856264, EPI_ISL_2856265, EPI_ISL_2856266, EPI_ISL_2856267, EPI_ISL_2856268, EPI_ISL_2856269, EPI_ISL_2856270, EPI_ISL_2856271, EPI_ISL_2856272, EPI_ISL_2856278, EPI_ISL_2856279, EPI_ISL_2856280, EPI_ISL_2856281, EPI_ISL_2856282, EPI_ISL_2856283, EPI_ISL_2856284, EPI_ISL_2856285, EPI_ISL_2856286, EPI_ISL_2856290, EPI_ISL_2856295, EPI_ISL_2856296, EPI_ISL_2856297, EPI_ISL_2856299, EPI_ISL_2856300, EPI_ISL_2856303, EPI_ISL_2856305, EPI_ISL_2856307, EPI_ISL_2856308, EPI_ISL_2856309, EPI_ISL_2856310, EPI_ISL_2856312, EPI_ISL_2856316, EPI_ISL_2856317, EPI_ISL_2856318, EPI_ISL_2856320, EPI_ISL_2856321, EPI_ISL_2856323, EPI_ISL_2856324, EPI_ISL_2856325, EPI_ISL_2856326, EPI_ISL_2856327, EPI_ISL_2856329, EPI_ISL_2856330, EPI_ISL_2856331, EPI_ISL_2856333, EPI_ISL_2856334, EPI_ISL_2856335, EPI_ISL_2856336, EPI_ISL_2856337, EPI_ISL_2856338, EPI_ISL_2856339, EPI_ISL_2856341, EPI_ISL_2856342, EPI_ISL_2856345, EPI_ISL_2856367, EPI_ISL_2856368, EPI_ISL_2856377, EPI_ISL_2856382, EPI_ISL_2856387, EPI_ISL_2856388, EPI_ISL_2856392, EPI_ISL_2856400, EPI_ISL_2856401, EPI_ISL_2856415, EPI_ISL_2856418, EPI_ISL_2856424, EPI_ISL_2856427, EPI_ISL_2856428, EPI_ISL_2856438, EPI_ISL_2856441, EPI_ISL_2856456, EPI_ISL_2856461, EPI_ISL_2856462, EPI_ISL_2856477, EPI_ISL_2856481, EPI_ISL_2856485, EPI_ISL_2856489, EPI_ISL_2856495, EPI_ISL_2856500, EPI_ISL_2856501, EPI_ISL_2856502, EPI_ISL_2856504, EPI_ISL_2856506, EPI_ISL_2856508, EPI_ISL_2856509, EPI_ISL_2856511, EPI_ISL_2856514, EPI_ISL_2856518, EPI_ISL_2856519, EPI_ISL_2856521, EPI_ISL_2856522, EPI_ISL_2856523, EPI_ISL_2856524, EPI_ISL_2856525, EPI_ISL_2856527, EPI_ISL_2856528, EPI_ISL_2856529, EPI_ISL_2856530, EPI_ISL_2856531, EPI_ISL_2856532, EPI_ISL_2856533, EPI_ISL_2856534, EPI_ISL_2856535, EPI_ISL_2856536, EPI_ISL_2856537, EPI_ISL_2856538, EPI_ISL_2856539, EPI_ISL_2856540, EPI_ISL_2856541, EPI_ISL_2856542, EPI_ISL_2856543, EPI_ISL_2856544, EPI_ISL_2856545, EPI_ISL_2856547, EPI_ISL_2856549, EPI_ISL_2856552, EPI_ISL_2856553, EPI_ISL_2856554, EPI_ISL_2856558, EPI_ISL_2856559, EPI_ISL_2856560, EPI_ISL_2856562, EPI_ISL_2856564, EPI_ISL_2856565, EPI_ISL_2856566, EPI_ISL_2856568, EPI_ISL_2856569, EPI_ISL_2856572, EPI_ISL_2856573, EPI_ISL_2856574, EPI_ISL_2856575, EPI_ISL_2856576, EPI_ISL_2856577, EPI_ISL_2856578, EPI_ISL_2856579, EPI_ISL_2856581, EPI_ISL_2856583, EPI_ISL_2856584, EPI_ISL_2856586, EPI_ISL_2856587, EPI_ISL_2856588, EPI_ISL_2856590, EPI_ISL_2856591, EPI_ISL_2856592, EPI_ISL_2856593, EPI_ISL_2856594, EPI_ISL_2856595, EPI_ISL_2856596, EPI_ISL_2856598, EPI_ISL_2856600, EPI_ISL_2856604, EPI_ISL_2856605, EPI_ISL_2856606, EPI_ISL_2856607, EPI_ISL_2856608, EPI_ISL_2856609, EPI_ISL_2856610, EPI_ISL_2856611, EPI_ISL_2856612, EPI_ISL_2856613, EPI_ISL_2856614, EPI_ISL_2856615, EPI_ISL_2856617, EPI_ISL_2856619, EPI_ISL_2856620, EPI_ISL_2856622, EPI_ISL_2856623, EPI_ISL_2856624, EPI_ISL_2856625, EPI_ISL_2856626, EPI_ISL_2856627, EPI_ISL_2856628, EPI_ISL_2856631, EPI_ISL_2856632, EPI_ISL_2856637, EPI_ISL_2856639, EPI_ISL_2856640, EPI_ISL_2856641, EPI_ISL_2856646, EPI_ISL_2856648, EPI_ISL_2856653, EPI_ISL_2856654, EPI_ISL_2856655, EPI_ISL_2856656, EPI_ISL_2856657, EPI_ISL_2856661, EPI_ISL_2856662, EPI_ISL_2856664, EPI_ISL_2856667, EPI_ISL_2856670, EPI_ISL_2856673, EPI_ISL_2856674, EPI_ISL_2856685, EPI_ISL_2856686, EPI_ISL_2856688, EPI_ISL_2856690, EPI_ISL_2856696, EPI_ISL_2856697, EPI_ISL_2856698, EPI_ISL_2856699, EPI_ISL_2856706, EPI_ISL_2856707, EPI_ISL_2856708, EPI_ISL_2856709, EPI_ISL_2856710, EPI_ISL_2856712, EPI_ISL_2856714, EPI_ISL_2856715, EPI_ISL_2856716, EPI_ISL_2856717, EPI_ISL_2856719, EPI_ISL_2856720, EPI_ISL_2856722, EPI_ISL_2856726, EPI_ISL_2856729, EPI_ISL_2856730, EPI_ISL_2856734, EPI_ISL_2856737, EPI_ISL_2856738, EPI_ISL_2856741, EPI_ISL_2856742, EPI_ISL_2856743, EPI_ISL_2856744, EPI_ISL_2856746, EPI_ISL_2856747, EPI_ISL_2856749, EPI_ISL_2856751, EPI_ISL_2856752, EPI_ISL_2856754, EPI_ISL_2856755, EPI_ISL_2856758, EPI_ISL_2856762, EPI_ISL_2856763, EPI_ISL_2856765, EPI_ISL_2856766, EPI_ISL_2856767, EPI_ISL_2856769, EPI_ISL_2856770, EPI_ISL_2856771, EPI_ISL_2856772, EPI_ISL_2856773, EPI_ISL_2856774, EPI_ISL_2856775, EPI_ISL_2856776, EPI_ISL_2856778, EPI_ISL_2856779, EPI_ISL_2856780, EPI_ISL_2856782, EPI_ISL_2856786, EPI_ISL_2856788, EPI_ISL_2856790, EPI_ISL_2856791, EPI_ISL_2856793, EPI_ISL_2856794, EPI_ISL_2856795, EPI_ISL_2856797, EPI_ISL_2856798, EPI_ISL_2856799, EPI_ISL_2856802, EPI_ISL_2856803, EPI_ISL_2856804, EPI_ISL_2856806, EPI_ISL_2856807, EPI_ISL_2856812, EPI_ISL_2856816, EPI_ISL_2856817, EPI_ISL_2856818, EPI_ISL_2856821, EPI_ISL_2856822, EPI_ISL_2856824, EPI_ISL_2856825, EPI_ISL_2856827, EPI_ISL_2856828, EPI_ISL_2856829, EPI_ISL_2856830, EPI_ISL_2856831, EPI_ISL_2856832, EPI_ISL_2856833, EPI_ISL_2856834, EPI_ISL_2856835, EPI_ISL_2856836, EPI_ISL_2856837, EPI_ISL_2856838, EPI_ISL_2856839, EPI_ISL_2856840, EPI_ISL_2856841, EPI_ISL_2856842, EPI_ISL_2856843, EPI_ISL_2856845, EPI_ISL_2856846, EPI_ISL_2856847, EPI_ISL_2856848, EPI_ISL_2856850, EPI_ISL_2856852, EPI_ISL_2856855, EPI_ISL_2856856, EPI_ISL_2856857, EPI_ISL_2856858, EPI_ISL_2856859, EPI_ISL_2856860, EPI_ISL_2856861, EPI_ISL_2856865, EPI_ISL_2856866, EPI_ISL_2856874, EPI_ISL_2856877, EPI_ISL_2856879, EPI_ISL_2856880, EPI_ISL_2856882, EPI_ISL_2856887, EPI_ISL_2856889, EPI_ISL_2856890, EPI_ISL_2856891, EPI_ISL_2856901, EPI_ISL_2856903, EPI_ISL_2856905, EPI_ISL_2856906, EPI_ISL_2856907, EPI_ISL_2856908, EPI_ISL_2856911, EPI_ISL_2856912, EPI_ISL_2856914, EPI_ISL_2856915, EPI_ISL_2856916, EPI_ISL_2856920, EPI_ISL_2856921, EPI_ISL_2856922, EPI_ISL_2856924, EPI_ISL_2856926, EPI_ISL_2856927, EPI_ISL_2856928, EPI_ISL_2856930, EPI_ISL_2856933, EPI_ISL_2856935, EPI_ISL_2856936, EPI_ISL_2856937, EPI_ISL_2856938, EPI_ISL_2856939, EPI_ISL_2856940, EPI_ISL_2856941, EPI_ISL_2856942, EPI_ISL_2856943, EPI_ISL_2856944, EPI_ISL_2856945, EPI_ISL_2856946, EPI_ISL_2856947, EPI_ISL_2856948, EPI_ISL_2856949, EPI_ISL_2856950, EPI_ISL_2856951, EPI_ISL_2856952, EPI_ISL_2856953, EPI_ISL_2856954, EPI_ISL_2856955, EPI_ISL_2856956, EPI_ISL_2856957, EPI_ISL_2856959, EPI_ISL_2856960, EPI_ISL_2856961, EPI_ISL_2856962, EPI_ISL_2856963, EPI_ISL_2856965, EPI_ISL_2856967, EPI_ISL_2856968, EPI_ISL_2856969, EPI_ISL_2856970, EPI_ISL_2856971, EPI_ISL_2856973, EPI_ISL_2856974, EPI_ISL_2856976, EPI_ISL_2856977, EPI_ISL_2856978, EPI_ISL_2856979, EPI_ISL_2856980, EPI_ISL_2856981, EPI_ISL_2856982, EPI_ISL_2856983, EPI_ISL_2856985, EPI_ISL_2856986, EPI_ISL_2856987, EPI_ISL_2856988, EPI_ISL_2856989, EPI_ISL_2856990, EPI_ISL_2856991, EPI_ISL_2856992, EPI_ISL_2856993, EPI_ISL_2856994, EPI_ISL_2856995, EPI_ISL_2856996, EPI_ISL_2856997, EPI_ISL_2856998, EPI_ISL_2857001, EPI_ISL_2857003, EPI_ISL_2857004, EPI_ISL_2857005, EPI_ISL_2857006, EPI_ISL_2857007, EPI_ISL_2857008, EPI_ISL_2857011, EPI_ISL_2857012, EPI_ISL_2857013, EPI_ISL_2857014, EPI_ISL_2857015, EPI_ISL_2857016, EPI_ISL_2857017, EPI_ISL_2857020, EPI_ISL_2857021, EPI_ISL_2857023, EPI_ISL_2857025, EPI_ISL_2857026, EPI_ISL_2857027, EPI_ISL_2857029, EPI_ISL_2857030, EPI_ISL_2857031, EPI_ISL_2857036, EPI_ISL_2857038, EPI_ISL_2857041, EPI_ISL_2857043, EPI_ISL_2857047, EPI_ISL_2857048, EPI_ISL_2857049, EPI_ISL_2857050, EPI_ISL_2857052, EPI_ISL_2857057, EPI_ISL_2857058, EPI_ISL_2857059, EPI_ISL_2857060, EPI_ISL_2857061, EPI_ISL_2857062, EPI_ISL_2857063, EPI_ISL_2857064, EPI_ISL_2857065, EPI_ISL_2857066, EPI_ISL_2857067, EPI_ISL_2857068, EPI_ISL_2857069, EPI_ISL_2857072, EPI_ISL_2857073, EPI_ISL_2857075, EPI_ISL_2857078, EPI_ISL_2857080, EPI_ISL_2857081, EPI_ISL_2857083, EPI_ISL_2857084, EPI_ISL_2857085, EPI_ISL_2857086, EPI_ISL_2857087, EPI_ISL_2857088, EPI_ISL_2857089, EPI_ISL_2857090, EPI_ISL_2857091, EPI_ISL_2857092, EPI_ISL_2857094, EPI_ISL_2857095, EPI_ISL_2857096, EPI_ISL_2857099, EPI_ISL_2857100, EPI_ISL_2857101, EPI_ISL_2857102, EPI_ISL_2857103, EPI_ISL_2857105, EPI_ISL_2857107, EPI_ISL_2857108, EPI_ISL_2857109, EPI_ISL_2857110, EPI_ISL_2857111, EPI_ISL_2857112, EPI_ISL_2857114, EPI_ISL_2857115, EPI_ISL_2857116, EPI_ISL_2857117, EPI_ISL_2857118, EPI_ISL_2857119, EPI_ISL_2857120, EPI_ISL_2857122, EPI_ISL_2857123, EPI_ISL_2857124, EPI_ISL_2857126, EPI_ISL_2857127, EPI_ISL_2857128, EPI_ISL_2857131, EPI_ISL_2857132, EPI_ISL_2857134, EPI_ISL_2857135, EPI_ISL_2857136, EPI_ISL_2857137, EPI_ISL_2857141, EPI_ISL_2857142, EPI_ISL_2857144, EPI_ISL_2857145, EPI_ISL_2857146, EPI_ISL_2857147, EPI_ISL_2857150, EPI_ISL_2857151, EPI_ISL_2857154, EPI_ISL_2857155, EPI_ISL_2857157, EPI_ISL_2857158, EPI_ISL_2857159, EPI_ISL_2857160, EPI_ISL_2857161, EPI_ISL_2857163, EPI_ISL_2857165, EPI_ISL_2857168, EPI_ISL_2857169, EPI_ISL_2857170, EPI_ISL_2857171, EPI_ISL_2857173, EPI_ISL_2857174, EPI_ISL_2857176, EPI_ISL_2857177, EPI_ISL_2857178, EPI_ISL_2857179, EPI_ISL_2857180, EPI_ISL_2857182, EPI_ISL_2857183, EPI_ISL_2857184, EPI_ISL_2857185, EPI_ISL_2857187, EPI_ISL_2857188, EPI_ISL_2857190, EPI_ISL_2857191, EPI_ISL_2857192, EPI_ISL_2857193, EPI_ISL_2857194, EPI_ISL_2857195, EPI_ISL_2857196, EPI_ISL_2857197, EPI_ISL_2857198, EPI_ISL_2857201, EPI_ISL_2857204, EPI_ISL_2857205, EPI_ISL_2857207, EPI_ISL_2857208, EPI_ISL_2857211, EPI_ISL_2857212, EPI_ISL_2857213, EPI_ISL_2857214, EPI_ISL_2857217, EPI_ISL_2857221, EPI_ISL_2857222, EPI_ISL_2857223, EPI_ISL_2857224, EPI_ISL_2857225, EPI_ISL_2857226, EPI_ISL_2857227, EPI_ISL_2857229, EPI_ISL_2857230, EPI_ISL_2857231, EPI_ISL_2857233, EPI_ISL_2857236, EPI_ISL_2857241, EPI_ISL_2857243, EPI_ISL_2857244, EPI_ISL_2857245, EPI_ISL_2857246, EPI_ISL_2857247, EPI_ISL_2857248, EPI_ISL_2857249, EPI_ISL_2857252, EPI_ISL_2857253, EPI_ISL_2857254, EPI_ISL_2857255, EPI_ISL_2857256, EPI_ISL_2857258, EPI_ISL_2857259, EPI_ISL_2857260, EPI_ISL_2857261, EPI_ISL_2857262, EPI_ISL_2857263, EPI_ISL_2857264, EPI_ISL_2857265, EPI_ISL_2857267, EPI_ISL_2857268, EPI_ISL_2857269, EPI_ISL_2857270, EPI_ISL_2857278, EPI_ISL_2857279, EPI_ISL_2857280, EPI_ISL_2857281, EPI_ISL_2857283, EPI_ISL_2857284, EPI_ISL_2857286, EPI_ISL_2857287, EPI_ISL_2857288, EPI_ISL_2857289, EPI_ISL_2857290, EPI_ISL_2857291, EPI_ISL_2857292, EPI_ISL_2857293, EPI_ISL_2857294, EPI_ISL_2857296, EPI_ISL_2857297, EPI_ISL_2857298, EPI_ISL_2857299, EPI_ISL_2857300, EPI_ISL_2857301, EPI_ISL_2857302, EPI_ISL_2857303, EPI_ISL_2857304, EPI_ISL_2857305, EPI_ISL_2857306, EPI_ISL_2857307, EPI_ISL_2857308, EPI_ISL_2857309, EPI_ISL_2857310, EPI_ISL_2857312, EPI_ISL_2857313, EPI_ISL_2857314, EPI_ISL_2857316, EPI_ISL_2857317, EPI_ISL_2857318, EPI_ISL_2857319, EPI_ISL_2857321, EPI_ISL_2857322, EPI_ISL_2857323, EPI_ISL_2857324, EPI_ISL_2857325, EPI_ISL_2857326, EPI_ISL_2857327, EPI_ISL_2857328, EPI_ISL_2857329, EPI_ISL_2857330, EPI_ISL_2857331, EPI_ISL_2857332, EPI_ISL_2857333, EPI_ISL_2857334, EPI_ISL_2857335, EPI_ISL_2857361, EPI_ISL_2857364, EPI_ISL_2857365, EPI_ISL_2857366, EPI_ISL_2857367, EPI_ISL_2857369, EPI_ISL_2857370, EPI_ISL_2857371, EPI_ISL_2857372, EPI_ISL_2857373, EPI_ISL_2857374, EPI_ISL_2857375, EPI_ISL_2857376, EPI_ISL_2857377, EPI_ISL_2857378, EPI_ISL_2857379, EPI_ISL_2857380, EPI_ISL_2857381, EPI_ISL_2857383, EPI_ISL_2857384, EPI_ISL_2857385, EPI_ISL_2857387, EPI_ISL_2857388, EPI_ISL_2857389, EPI_ISL_2857390, EPI_ISL_2857391, EPI_ISL_2857395, EPI_ISL_2857396, EPI_ISL_2857397, EPI_ISL_2857398, EPI_ISL_2857399, EPI_ISL_2857400, EPI_ISL_2857401, EPI_ISL_2857402, EPI_ISL_2857403, EPI_ISL_2857404, EPI_ISL_2857405, EPI_ISL_2857406, EPI_ISL_2857407, EPI_ISL_2857408, EPI_ISL_2857409, EPI_ISL_2857410, EPI_ISL_2857411, EPI_ISL_2857412, EPI_ISL_2857413, EPI_ISL_2857414, EPI_ISL_2857415, EPI_ISL_2857417, EPI_ISL_2857418, EPI_ISL_2857419, EPI_ISL_2857420, EPI_ISL_2857421, EPI_ISL_2857422, EPI_ISL_2857425, EPI_ISL_2857426, EPI_ISL_2857427, EPI_ISL_2857428, EPI_ISL_2857429, EPI_ISL_2857430, EPI_ISL_2857431, EPI_ISL_2857432, EPI_ISL_2857433, EPI_ISL_2857434, EPI_ISL_2857435, EPI_ISL_2857436, EPI_ISL_2857437, EPI_ISL_2857438, EPI_ISL_2857439, EPI_ISL_2857440, EPI_ISL_2857441, EPI_ISL_2857442, EPI_ISL_2857443, EPI_ISL_2857444, EPI_ISL_2857446, EPI_ISL_2857447, EPI_ISL_2857449, EPI_ISL_2857450, EPI_ISL_2857451, EPI_ISL_2857452, EPI_ISL_2857453, EPI_ISL_2857479, EPI_ISL_2857545, EPI_ISL_2857546, EPI_ISL_2857547, EPI_ISL_2857548, EPI_ISL_2857549, EPI_ISL_2857550, EPI_ISL_2857551, EPI_ISL_2857552, EPI_ISL_2857553, EPI_ISL_2857555, EPI_ISL_2857556, EPI_ISL_2857557, EPI_ISL_2857559, EPI_ISL_2857560, EPI_ISL_2857561, EPI_ISL_2857562, EPI_ISL_2857563, EPI_ISL_2857564, EPI_ISL_2857565, EPI_ISL_2857566, EPI_ISL_2857567, EPI_ISL_2857568, EPI_ISL_2857570, EPI_ISL_2857571, EPI_ISL_2857572, EPI_ISL_2857573, EPI_ISL_2857574, EPI_ISL_2857575, EPI_ISL_2857578, EPI_ISL_2857579, EPI_ISL_2857580, EPI_ISL_2857581, EPI_ISL_2857584, EPI_ISL_2857587, EPI_ISL_2857588, EPI_ISL_2857589, EPI_ISL_2857590, EPI_ISL_2857591, EPI_ISL_2857593, EPI_ISL_2857594, EPI_ISL_2857595, EPI_ISL_2857596, EPI_ISL_2857597, EPI_ISL_2857598, EPI_ISL_2857599, EPI_ISL_2857600, EPI_ISL_2857601, EPI_ISL_2857604, EPI_ISL_2857605, EPI_ISL_2857606, EPI_ISL_2857607, EPI_ISL_2857609, EPI_ISL_2857610, EPI_ISL_2857611, EPI_ISL_2857613, EPI_ISL_2857614 | | | |
| see above | Utah Public Health Laboratory | Utah Public Health Laboratory | Olinto Linares-Perdomo, Erin L. Young, Kelly F. Oakeson, Tara Gallagher |
| EPI_ISL_2857615 | Jaber Quarantine | Virology Lab, Jaber Al Ahmad Hospital | Dr. Zahrah Buhamad, Dr. Ebaa Al-Awadhi, Adil Afridi, Haroon Masih |
| EPI_ISL_2857616, EPI_ISL_2857617, EPI_ISL_2857618, EPI_ISL_2857619, EPI_ISL_2857620, EPI_ISL_2857622, EPI_ISL_2857623, EPI_ISL_2857625, EPI_ISL_2857626, EPI_ISL_2857627, EPI_ISL_2857629, EPI_ISL_2857630, EPI_ISL_2857631, EPI_ISL_2857633, EPI_ISL_2857634, EPI_ISL_2857635, EPI_ISL_2857636, EPI_ISL_2857637, EPI_ISL_2857638, EPI_ISL_2857639, EPI_ISL_2857640, EPI_ISL_2857641, EPI_ISL_2857642, EPI_ISL_2857643, EPI_ISL_2857644, EPI_ISL_2857645, EPI_ISL_2857646, EPI_ISL_2857647, EPI_ISL_2857648, EPI_ISL_2857649, EPI_ISL_2857651, EPI_ISL_2857652, EPI_ISL_2857653, EPI_ISL_2857654, EPI_ISL_2857656, EPI_ISL_2857657, EPI_ISL_2857660, EPI_ISL_2857661, EPI_ISL_2857663, EPI_ISL_2857664, EPI_ISL_2857665, EPI_ISL_2857666, EPI_ISL_2857668, EPI_ISL_2857669, EPI_ISL_2857670, EPI_ISL_2857671, EPI_ISL_2857672, EPI_ISL_2857673, EPI_ISL_2857674, EPI_ISL_2857675, EPI_ISL_2857676, EPI_ISL_2857677, EPI_ISL_2857678, EPI_ISL_2857679, EPI_ISL_2857680, EPI_ISL_2857681, EPI_ISL_2857682, EPI_ISL_2857683, EPI_ISL_2857685, EPI_ISL_2857686, EPI_ISL_2857687, EPI_ISL_2857688, EPI_ISL_2857689, EPI_ISL_2857690, EPI_ISL_2857691, EPI_ISL_2857693, EPI_ISL_2857694, EPI_ISL_2857695, EPI_ISL_2857696, EPI_ISL_2857697, EPI_ISL_2857698, EPI_ISL_2857699, EPI_ISL_2857700, EPI_ISL_2857703, EPI_ISL_2857705, EPI_ISL_2857707, EPI_ISL_2857708, EPI_ISL_2857709, EPI_ISL_2857710, EPI_ISL_2857711, EPI_ISL_2857712, EPI_ISL_2857713, EPI_ISL_2857714, EPI_ISL_2857715, EPI_ISL_2857716, EPI_ISL_2857718, EPI_ISL_2857719, EPI_ISL_2857720, EPI_ISL_2857721, EPI_ISL_2857722, EPI_ISL_2857723, EPI_ISL_2857724, EPI_ISL_2857725, EPI_ISL_2857727, EPI_ISL_2857728, EPI_ISL_2857729, EPI_ISL_2857730, EPI_ISL_2857731, EPI_ISL_2857732, EPI_ISL_2857733, EPI_ISL_2857734, EPI_ISL_2857735, EPI_ISL_2857736, EPI_ISL_2857737, EPI_ISL_2857738, EPI_ISL_2857739, EPI_ISL_2857740, EPI_ISL_2857741, EPI_ISL_2857742, EPI_ISL_2857743, EPI_ISL_2857744, EPI_ISL_2857745, EPI_ISL_2857748, EPI_ISL_2857750, EPI_ISL_2857751, EPI_ISL_2857752, EPI_ISL_2857753, EPI_ISL_2857754, EPI_ISL_2857755, EPI_ISL_2857756, EPI_ISL_2857757, EPI_ISL_2857758, EPI_ISL_2857759, EPI_ISL_2857760, EPI_ISL_2857761, EPI_ISL_2857762, EPI_ISL_2857765, EPI_ISL_2857766, EPI_ISL_2857768, EPI_ISL_2857769, EPI_ISL_2857770, EPI_ISL_2857771, EPI_ISL_2857772, EPI_ISL_2857773, EPI_ISL_2857774, EPI_ISL_2857775, EPI_ISL_2857776, EPI_ISL_2857777, EPI_ISL_2857778, EPI_ISL_2857779, EPI_ISL_2857781, EPI_ISL_2857782, EPI_ISL_2857784, EPI_ISL_2857785, EPI_ISL_2857786, EPI_ISL_2857787, EPI_ISL_2857788, EPI_ISL_2857790, EPI_ISL_2857791, EPI_ISL_2857792, EPI_ISL_2857793, EPI_ISL_2857794, EPI_ISL_2857795, EPI_ISL_2857796, EPI_ISL_2857797, EPI_ISL_2857798, EPI_ISL_2857799, EPI_ISL_2857800, EPI_ISL_2857802, EPI_ISL_2857803, EPI_ISL_2857804, EPI_ISL_2857805, EPI_ISL_2857806, EPI_ISL_2857807, EPI_ISL_2857808, EPI_ISL_2857809, EPI_ISL_2857810, EPI_ISL_2857811, EPI_ISL_2857812, EPI_ISL_2857813, EPI_ISL_2857814, EPI_ISL_2857815, EPI_ISL_2857816, EPI_ISL_2857817, EPI_ISL_2857818, EPI_ISL_2857819, EPI_ISL_2857820, EPI_ISL_2857821, EPI_ISL_2857822, EPI_ISL_2857824, EPI_ISL_2857825, EPI_ISL_2857826, EPI_ISL_2857827, EPI_ISL_2857828, EPI_ISL_2857829, EPI_ISL_2857830, EPI_ISL_2857831, EPI_ISL_2857832, EPI_ISL_2857833, EPI_ISL_2857834, EPI_ISL_2857835, EPI_ISL_2857836, EPI_ISL_2857837, EPI_ISL_2857838, EPI_ISL_2857839, EPI_ISL_2857840, EPI_ISL_2857841, EPI_ISL_2857842, EPI_ISL_2857843, EPI_ISL_2857844, EPI_ISL_2857845, EPI_ISL_2857846, EPI_ISL_2857847, EPI_ISL_2857848, EPI_ISL_2857850, EPI_ISL_2857851, EPI_ISL_2857852, EPI_ISL_2857853, EPI_ISL_2857854, EPI_ISL_2857855, EPI_ISL_2857856, EPI_ISL_2857857, EPI_ISL_2857858, EPI_ISL_2857859, EPI_ISL_2857860, EPI_ISL_2857861, EPI_ISL_2857862, EPI_ISL_2857864, EPI_ISL_2857865, EPI_ISL_2857866, EPI_ISL_2857867, EPI_ISL_2857868, EPI_ISL_2857869, EPI_ISL_2857870, EPI_ISL_2857871, EPI_ISL_2857872, EPI_ISL_2857873, EPI_ISL_2857874, EPI_ISL_2857875, EPI_ISL_2857876, EPI_ISL_2857877, EPI_ISL_2857878, EPI_ISL_2857879, EPI_ISL_2857880, EPI_ISL_2857881, EPI_ISL_2857882, EPI_ISL_2857883, EPI_ISL_2857884, EPI_ISL_2857885, EPI_ISL_2857886, EPI_ISL_2857887, EPI_ISL_2857888, EPI_ISL_2857889, EPI_ISL_2857890, EPI_ISL_2857891, EPI_ISL_2857892, EPI_ISL_2857894, EPI_ISL_2857895, EPI_ISL_2857896, EPI_ISL_2857897, EPI_ISL_2857898, EPI_ISL_2857899, EPI_ISL_2857900, EPI_ISL_2857901, EPI_ISL_2857902, EPI_ISL_2857903, EPI_ISL_2857905, EPI_ISL_2857906, EPI_ISL_2857907, EPI_ISL_2857908, EPI_ISL_2857909, EPI_ISL_2857910, EPI_ISL_2857911, EPI_ISL_2857912, EPI_ISL_2857913, EPI_ISL_2857922, EPI_ISL_2857923, EPI_ISL_2857926, EPI_ISL_2857933, EPI_ISL_2857934, EPI_ISL_2857942, EPI_ISL_2857945, EPI_ISL_2857947, EPI_ISL_2857966, EPI_ISL_2857967, EPI_ISL_2857970, EPI_ISL_2857991, EPI_ISL_2857994, EPI_ISL_2858008 | | | |
| see above | Utah Public Health Laboratory | Utah Public Health Laboratory | Olinto Linares-Perdomo, Erin L. Young, Kelly F. Oakeson, Tara Gallagher |
| EPI_ISL_2858250, EPI_ISL_2858252, EPI_ISL_2858253 | Jaber Quarantine | Virology Lab, Jaber Al Ahmad Hospital | Dr. Zahrah Buhamad, Dr. Ebaa Al-Awadhi, Adil Afridi, Haroon Masih |
| EPI_ISL_2858697 | Spital Männedorf AG | Institute of Medical Virology | Verena Kufner, Gabriela Ziltener, Maryam Zaheri, Stefan Schmutz, Annette Audigé, Maria Grünberg, Kevin Steiner, Jon Huder, Cyril Shah, Riccarda Capaul, Guido Bloemberg, Jürg Böni, Michael Huber, Alexandra Trkola |
| EPI_ISL_2858698 | UniversitätsSpital Zürich | Institute of Medical Virology | Verena Kufner, Gabriela Ziltener, Maryam Zaheri, Stefan Schmutz, Annette Audigé, Maria Grünberg, Kevin Steiner, Jon Huder, Cyril Shah, Riccarda Capaul, Guido Bloemberg, Jürg Böni, Michael Huber, Alexandra Trkola |
| EPI_ISL_2858700 | RSPI PROF DR. SULIANTI SAROSO | Genomik Solidaritas Indonesia Laboratorium | Anna Christina Brazia, Himawan Masyhuri, Normastuti Adhini Tantyo, Alfin Muhammad Abdillah, Dhahlia Agustina Cahyono, Ahmad Zoebad Foeady, Annisa Muthiah Sukirman, Gracia Felias Enos Korompis, Teguh Sarry Hartono, Louisa Markus, Anuraj Shankar, Meutia Ayuputeri Kumaheri, Anindya Pradipta Susanto, Ariel Pradipta |
| EPI_ISL_2858764, EPI_ISL_2858765 | North Dakota Department of Health, Public Health Laboratory | North Dakota Department of Health, Public Health Laboratory | Lisa Wingerter |
| EPI_ISL_2858768, EPI_ISL_2858770, EPI_ISL_2858772, EPI_ISL_2858774, EPI_ISL_2858775, EPI_ISL_2858776, EPI_ISL_2858777, EPI_ISL_2858778, EPI_ISL_2858779, EPI_ISL_2858780, EPI_ISL_2858781, EPI_ISL_2858783, EPI_ISL_2858784, EPI_ISL_2858785, EPI_ISL_2858786, EPI_ISL_2858787, EPI_ISL_2858788 | | | |
| see above | University of Mississippi Medical Center, Department of Pathology | University of Mississippi Medical Center, Molecular and Genomics Core Facility | Ashley C. Johnson, Wenjie Wu, Ithiel J. Frame, Krishna K. Ayyalasomayajula, Michael R. Garrett, D. Ashley Robinson |
| EPI_ISL_2858792, EPI_ISL_2858793, EPI_ISL_2858794, EPI_ISL_2858796, EPI_ISL_2858797, EPI_ISL_2858798, EPI_ISL_2858799, EPI_ISL_2858800, EPI_ISL_2858802, EPI_ISL_2858805, EPI_ISL_2858807, EPI_ISL_2858808, EPI_ISL_2858809, EPI_ISL_2858810 | | | |
| see above | MS Public Health Laboratory | University of Mississippi Medical Center, Molecular and Genomics Core Facility | Ashley C. Johnson, Wenjie Wu, Ithiel J. Frame, Krishna K. Ayyalasomayajula, Michael R. Garrett, D. Ashley Robinson |
| EPI_ISL_2858834, EPI_ISL_2858835, EPI_ISL_2858836, EPI_ISL_2858841, EPI_ISL_2858842, EPI_ISL_2858845, EPI_ISL_2858848, EPI_ISL_2858850, EPI_ISL_2858854, EPI_ISL_2858855, EPI_ISL_2858857, EPI_ISL_2858858, EPI_ISL_2858864, EPI_ISL_2858869, EPI_ISL_2858870 | | | |
| see above | North Dakota Department of Health, Public Health Laboratory | North Dakota Department of Health, Public Health Laboratory | Lisa Wingerter |
| EPI_ISL_2858872 | Genomic Research Laboratory, Bangabandhu Sheikh Mujib Medical University | Genomic Research Laboratory, BSMMU | Laila Anjuman Banu, Md.Saydur Rahman, Sharadindu Kanti Sinha, Arifa Akram, Md.Sharfuddin Ahmed |
| EPI_ISL_2858873, EPI_ISL_2858874, EPI_ISL_2858875, EPI_ISL_2858876, EPI_ISL_2858878, EPI_ISL_2858879, EPI_ISL_2858880, EPI_ISL_2858881, EPI_ISL_2858882, EPI_ISL_2858884, EPI_ISL_2858885, EPI_ISL_2858886, EPI_ISL_2858888, EPI_ISL_2858889, EPI_ISL_2858890, EPI_ISL_2858891 | | | |
| see above | TXDSHS | TXDSHS | Rashmi Tuladhar, Bonnie Oh, Jenny Zhang, Maliha Rahman, Mayela Pedrueza, Anita Pokharel, Karen Bobier, Lorraine Rodriguez, Myong Koag, Chun Wang, Rachel Lee, Grace Kubin |
| EPI_ISL_2858898, EPI_ISL_2858911, EPI_ISL_2858916, EPI_ISL_2858922 | New Mexico Department of Health Scientific Laboratory | New Mexico Department of Health Scientific Laboratory | Ellie Johnson, D'eldra Malone, Jennifer Benoit, Ratheesh Rajan, Linda Salazar, Keila Gutierrez, Anastacia Griego-Fisher |

| | | | |
|---|---|---|---|
| EPI_ISL_2859431, EPI_ISL_2859432, EPI_ISL_2859433, EPI_ISL_2859435, EPI_ISL_2859438, EPI_ISL_2859439, EPI_ISL_2859446, EPI_ISL_2859447, EPI_ISL_2859448, EPI_ISL_2859449 | Washington State Department of Health Public Health Laboratories | Washington State Department of Health Public Health Laboratories | Drew MacKellar, Philip Dykema, Denny Russell, Joenice Gonzalez, Hannah Gray, Geoff Melly, Vanessa De Los Santos, Darren Lucas, JohnAric Peterson, Avi Singh, Rebecca Cao |
| EPI_ISL_2860278, EPI_ISL_2860279, EPI_ISL_2860280 | Badan Besar Rehabilitasi BNN | Biosafety Level-3 Laboratory, Indonesian Institute of Sciences (LIPI) | Indriawati, Masrukhin, Idris, Yuliawati, Listiana Oktavia, Anik B. Dharmayanthi, Andri Wardiana, Ratih A. Ningrum |
| EPI_ISL_2860281 | Klinik Primera Clinica | Biosafety Level-3 Laboratory, Indonesian Institute of Sciences (LIPI) | Indriawati, Masrukhin, Idris, Yuliawati, Listiana Oktavia, Anik B. Dharmayanthi, Andri Wardiana, Ratih A. Ningrum |
| EPI_ISL_2860285 | Klinik Primera Clinica | Biosafety Level-3 Laboratory, Indonesian Institute of Sciences (LIPI) | Syam B. Iryanto, Yuliawati, Masrukhin, Indriawati, Andri Wardiana, Anik B. Dharmayanthi, Sugiyono Saputra |
| EPI_ISL_2860286, EPI_ISL_2860287 | Klinik Primera Clinica | Biosafety Level-3 Laboratory, Indonesian Institute of Sciences (LIPI) | Masrukhin, Idris, Yuliawati, Indriawati, Listiana Oktavia, Anik B. Dharmayanthi, Asep M.Ridwanulloh, Sugiyono Saputra |
| EPI_ISL_2860291, EPI_ISL_2860293 | RS Umum Trimitra | Biosafety Level-3 Laboratory, Indonesian Institute of Sciences (LIPI) | Yuliawati, Indriawati, Listiana Oktavia, Anik B. Dharmayanthi, Asep M.Ridwanulloh, Andri Wardiana, Syam B. Iryanto |
| EPI_ISL_2860294, EPI_ISL_2860295 | Klinik Dr Nurdin | Biosafety Level-3 Laboratory, Indonesian Institute of Sciences (LIPI) | Yuliawati, Indriawati, Listiana Oktavia, Anik B. Dharmayanthi, Asep M.Ridwanulloh, Andri Wardiana, Syam B. Iryanto |
| EPI_ISL_2860298 | Rumah Sakit Umum Medika Dramaga | Biosafety Level-3 Laboratory, Indonesian Institute of Sciences (LIPI) | Listiana Oktavia, Yuliawati, Indriawati, Anik B. Dharmayanthi, Asep M.Ridwanulloh, Andri Wardiana, Syam B. Iryanto |
| EPI_ISL_2860299 | Rumah Sakit Umum Medika Dramaga | Biosafety Level-3 Laboratory, Indonesian Institute of Sciences (LIPI) | Listiana Oktavia, Yuliawati, Indriawati, Andri Wardiana, Syam B. Iryanto, Sugiyono Saputra |
| EPI_ISL_2860305, EPI_ISL_2860307, EPI_ISL_2860308, EPI_ISL_2860311, EPI_ISL_2860313, EPI_ISL_2860314, EPI_ISL_2860315, EPI_ISL_2860317 | Yale Clinical Virology Lab | Grubaugh Lab - Yale School of Public Health | Joseph Fauver, Mallery Breban, Isabel Ott, Tara Alpert, Mary Petrone, Anderson Brito, Chantal Vogels, Annie Watkins, Chaney Kalinich, Jessica Rothman, Marie L. Landry, Nathan Grubaugh |
| EPI_ISL_2860320, EPI_ISL_2860323 | UCC | Grubaugh Lab - Yale School of Public Health | Joseph Fauver, Mallery Breban, Isabel Ott, Tara Alpert, Mary Petrone, Anderson Brito, Chantal Vogels, Annie Watkins, Chaney Kalinich, Jessica Rothman, Kendall Billig, Tobias Koch, Fabiola Cruz-Lopez, David Rivera Aponte, Fabiola Fontanet, Natalie Machargo, Andrea Firpo, Vianca Aponte, Ernesto Vazquez, Patricia Serrano, Tania Mitwalli, Karolane Gonzalez, Nathan Grubaugh |
| EPI_ISL_2860332, EPI_ISL_2860333, EPI_ISL_2860334, EPI_ISL_2860335, EPI_ISL_2860336 | RS Umum Trimitra | Biosafety Level-3 Laboratory, Indonesian Institute of Sciences (LIPI) | Fahrurrozi, Linda Sukmarini, Nova D. Yanthi, Syam B. Iryanto, Michael Saut Parulian, Yuhoniz A. Maryam, Andri Wardiana, Asep M. Ridwanulloh, Sugiyono Saputra |
| EPI_ISL_2860337, EPI_ISL_2860338, EPI_ISL_2860339 | RS Umum Trimitra | Biosafety Level-3 Laboratory, Indonesian Institute of Sciences (LIPI) | Linda Sukmarini, Nova D. Yanthi, Fahrurrozi, Ocha P. Meitha, Willery Yeo, Syam B. Iryanto, Andri Wardiana, Asep M. Ridwanulloh, Ratih A. Ningrum |
| EPI_ISL_2860340, EPI_ISL_2860341, EPI_ISL_2860342 | RS Umum Trimitra | Biosafety Level-3 Laboratory, Indonesian Institute of Sciences (LIPI) | Nova D. Yanthi, Fahrurrozi, Syam B. Iryanto, Andri Wardiana, Annisa D. Septiani, Lusiana, Maya Komalasari, Linda Sukmarini, Asep M. Ridwanulloh, Puspita Lisdiyanti |
| EPI_ISL_2860343, EPI_ISL_2860344, EPI_ISL_2860345, EPI_ISL_2860346 | RS Umum Trimitra | Biosafety Level-3 Laboratory, Indonesian Institute of Sciences (LIPI) | Syam B. Iryanto, Ruby Setiawan, Linda Sukmarini, Nova D. Yanthi, Fahrurrozi, Asep M. Ridwanulloh, Ratih A. Ningrum, Sugiyono Saputra |
| EPI_ISL_2860347 | Klinik Dr Nurdin | Biosafety Level-3 Laboratory, Indonesian Institute of Sciences (LIPI) | Syam B. Iryanto, Ruby Setiawan, Linda Sukmarini, Nova D. Yanthi, Fahrurrozi, Asep M. Ridwanulloh, Anggia Prasetyoputri, Anik B. Dharmayanthi |
| EPI_ISL_2860348, EPI_ISL_2860349 | Klinik Primera Clinica | Biosafety Level-3 Laboratory, Indonesian Institute of Sciences (LIPI) | Syam B. Iryanto, Ruby Setiawan, Linda Sukmarini, Nova D. Yanthi, Fahrurrozi, Asep M. Ridwanulloh, Anggia Prasetyoputri, Anik B. Dharmayanthi |
| EPI_ISL_2860350 | Labkesda Depok | Biosafety Level-3 Laboratory, Indonesian Institute of Sciences (LIPI) | Syaiful Rizal, Sri Swasthikawati, Ruby Setiawan, Masrukhin, Elin Herliana, Kania R. Astuti, Kurniyawati, Harto W. Tamara, Sugiyono Saputra, Herman Irawan, Syam B. Iryanto, Ratih A. Ningrum |
| EPI_ISL_2860351 | Labkesda Depok | Biosafety Level-3 Laboratory, Indonesian Institute of Sciences (LIPI) | Sri Swasthikawati, Ruby Setiawan, Masrukhin, Elin Herliana, Syaiful Rizal, Kania R. Astuti, Kurniyawati, Harto W. Tamara, Herman Irawan, Sugeng Suhadi, Syam B. Iryanto, Ratih A. Ningrum |
| EPI_ISL_2860352 | Labkesda Depok | Biosafety Level-3 Laboratory, Indonesian Institute of Sciences (LIPI) | Sri Swasthikawati, Ruby Setiawan, Masrukhin, Elin Herliana, Syaiful Rizal, Kania R. Astuti, Kurniyawati, Andri Wardiana, Herman Irawan, Sugeng Suhadi, Syam B. Iryanto, Puspita Lisdiyanti |
| EPI_ISL_2860353 | Labkesda Depok | Biosafety Level-3 Laboratory, Indonesian Institute of Sciences (LIPI) | Sri Swasthikawati, Ruby Setiawan, Masrukhin, Ratih A. Ningrum, Syaiful Rizal, Kania R. Astuti, Kurniyawati, Harto W. Tamara, Herman Irawan, Sugeng Suhadi, Syam B. Iryanto, Puspita Lisdiyanti |
| EPI_ISL_2860354 | Labkesda Depok | Biosafety Level-3 Laboratory, Indonesian Institute of Sciences (LIPI) | Ruby Setiawan, Masrukhin, Elin Herliana, Herman Irawan, Syaiful Rizal, Kania R. Astuti, Sri Swasthikawati, Kurniyawati, Harto W. Tamara, Sugeng Suhadi, Sugiyono Saputra, Puspita Lisdiyanti |
| EPI_ISL_2860355 | Labkesda Depok | Biosafety Level-3 Laboratory, Indonesian Institute of Sciences (LIPI) | Ruby Setiawan, Masrukhin, Elin Herliana, Herman Irawan, Syaiful Rizal, Kania R. Astuti, Sri Swasthikawati, Syam B. Iryanto, Harto W. Tamara, Sugeng Suhadi, Anggia Prasetyoputri, Sugiyono Saputra |
| EPI_ISL_2860356 | Labkesda Depok | Biosafety Level-3 Laboratory, Indonesian Institute of Sciences (LIPI) | Ruby Setiawan, Masrukhin, Elin Herliana, Herman Irawan, Syaiful Rizal, Kania R. Astuti, Sri Swasthikawati, Kurniyawati, Harto W. Tamara, Sugeng Suhadi, Anggia Prasetyoputri, Sugiyono Saputra |
| EPI_ISL_2860357 | Labkesda Depok | Biosafety Level-3 Laboratory, Indonesian Institute of Sciences (LIPI) | Masrukhin, Herman Irawan, Elin Herliana, Syaiful Rizal, Ruby Setiawan, Ratih A. Ningrum, Sri Swasthikawati, Syam B. Iryanto, Harto W. Tamara, Sugeng Suhadi, Andri Wardiana, Sugiyono Saputra |
| EPI_ISL_2860358, EPI_ISL_2860359 | Labkesda Depok | Biosafety Level-3 Laboratory, Indonesian Institute of Sciences (LIPI) | Masrukhin, Herman Irawan, Elin Herliana, Syaiful Rizal, Ruby Setiawan, Kania R. Astuti, Sri Swasthikawati, Kurniyawati, Harto W. Tamara, Sugeng Suhadi, Ratih A. Ningrum, Anik B. Dharmayanthi |
| EPI_ISL_2860360, EPI_ISL_2860361 | RS Umum Trimitra | Biosafety Level-3 Laboratory, Indonesian Institute of Sciences (LIPI) | Sri Swasthikawati, Ruby Setiawan, Masrukhin, Michael S. Parulian, Yuhoniz A. Maryam, Syam B. Iryanto, Ratih A. Ningrum, Anggia Prasetyoputri, Sugiyono Saputra |
| EPI_ISL_2860362, EPI_ISL_2860363 | RS Umum Trimitra | Biosafety Level-3 Laboratory, Indonesian Institute of Sciences (LIPI) | Ruby Setiawan, Masrukhin, Sri Swasthikawati, Ocha P. Meitha, Willery Yeo, Syam B. Iryanto, Anggia Prasetyoputri, Anik B. Dharmayanthi, Wien Kusharyoto |
| EPI_ISL_2860364, EPI_ISL_2860365 | RS Umum Trimitra | Biosafety Level-3 Laboratory, Indonesian Institute of Sciences (LIPI) | Masrukhin, Sri Swasthikawati, Ruby Setiawan, Annisa D. Septiani, Lusiana, Maya Komalasari, Syam B. Iryanto, Ade Andriani, Asep M. Ridwanulloh, Puspita Lisdiyanti |
| EPI_ISL_2860366, EPI_ISL_2860367 | RS Umum Trimitra | Biosafety Level-3 Laboratory, Indonesian Institute of Sciences (LIPI) | Syam B. Iryanto, Sri Swasthikawati, Ruby Setiawan, Masrukhin, Annisa D. Septiani, Lusiana, Maya Komalasari, Andri Wardiana, Anik B. Dharmayanthi, Ratih A. Ningrum |
| EPI_ISL_2860368 | Klinik Primera Clinica | Biosafety Level-3 Laboratory, Indonesian Institute of Sciences (LIPI) | Syam B. Iryanto, Sri Swasthikawati, Ruby Setiawan, Masrukhin, Annisa D. Septiani, Lusiana, Maya Komalasari, Andri Wardiana, Anik B. Dharmayanthi, Ratih A. Ningrum |
| EPI_ISL_2860369, EPI_ISL_2860370, EPI_ISL_2860371, EPI_ISL_2860372, EPI_ISL_2860374, EPI_ISL_2860375, EPI_ISL_2860376, EPI_ISL_2860378, EPI_ISL_2860379, EPI_ISL_2860380, EPI_ISL_2860381, EPI_ISL_2860382, EPI_ISL_2860383, EPI_ISL_2860384, EPI_ISL_2860385, EPI_ISL_2860386, EPI_ISL_2860387, EPI_ISL_2860388, EPI_ISL_2860389, EPI_ISL_2860391, EPI_ISL_2860392, EPI_ISL_2860393, EPI_ISL_2860394, EPI_ISL_2860395, EPI_ISL_2860396, EPI_ISL_2860397, EPI_ISL_2860398, EPI_ISL_2860399 | | | |
| see above | IN State Department of Health Laboratory Services | IN State Department of Health Laboratory Services | Cassandra Campion, Jamie Yeadon, Brian Pope, Lixia Liu, Kyle Brownlee, Melissa Hindenlang, Mark Glazier |
| EPI_ISL_2860423, EPI_ISL_2860425, EPI_ISL_2860430, EPI_ISL_2860435, EPI_ISL_2860458, EPI_ISL_2860461, EPI_ISL_2860462, EPI_ISL_2860463, EPI_ISL_2860464, EPI_ISL_2860465, EPI_ISL_2860466, EPI_ISL_2860467, EPI_ISL_2860470, EPI_ISL_2860471, EPI_ISL_2860475, EPI_ISL_2860508, EPI_ISL_2860509, EPI_ISL_2860522, EPI_ISL_2860527, EPI_ISL_2860528, EPI_ISL_2860529, EPI_ISL_2860537, EPI_ISL_2860538, EPI_ISL_2860539, EPI_ISL_2860540, EPI_ISL_2860541, EPI_ISL_2860543, EPI_ISL_2860547, EPI_ISL_2860549, EPI_ISL_2860551, EPI_ISL_2860552, EPI_ISL_2860553, EPI_ISL_2860554, EPI_ISL_2860563, EPI_ISL_2860564, EPI_ISL_2860565, EPI_ISL_2860567, EPI_ISL_2860568, EPI_ISL_2860571, EPI_ISL_2860573, EPI_ISL_2860620, EPI_ISL_2860621, EPI_ISL_2860623, EPI_ISL_2860624, EPI_ISL_2860625, EPI_ISL_2860626, EPI_ISL_2860627, EPI_ISL_2860628, EPI_ISL_2860629, EPI_ISL_2860630, EPI_ISL_2860631, EPI_ISL_2860632, EPI_ISL_2860633, EPI_ISL_2860634 | | | |
| see above | Dutch COVID-19 response team | Erasmus Medical Center | Bas Oude Munnink, Reina Sikkema, David Nieuwenhuijse, Irina Chestakova, Anne van der Linden, Marjan Boter, Emmanuelle Munger, Corine GeurtsvanKessel, Annemiek van der Eijk, Richard Molenkamp, Marion Koopmans, on behalf of the Dutch national COVID-19 respo |
| EPI_ISL_2860648, EPI_ISL_2860656, EPI_ISL_2860657, EPI_ISL_2860668, EPI_ISL_2860669, EPI_ISL_2860763, EPI_ISL_2860764 | Hyogo Prefectural Institute of Public Health science | Pathogen Genomics Center, National Institute of Infectious Diseases | Kenichi Ogita, Tomohiro Oshibe, Miki Ogi, Denshi Takai, Etsuko Saito, Kyoko Suzuki, Masatsugu Chikahira, Yumi Akiyama, Tsuyoshi Sekizuka, Kentaro Itokawa, Rina Tanaka, Masanori Hashino, Makoto Kuroda |
| EPI_ISL_2860930, EPI_ISL_2860931, EPI_ISL_2860932, EPI_ISL_2860933, EPI_ISL_2860934, EPI_ISL_2860935, EPI_ISL_2860936, EPI_ISL_2860937, EPI_ISL_2860938, EPI_ISL_2860939, EPI_ISL_2860940, EPI_ISL_2860941, EPI_ISL_2860942, EPI_ISL_2860943, EPI_ISL_2860944, EPI_ISL_2860945, EPI_ISL_2860946, EPI_ISL_2860947, EPI_ISL_2860949, EPI_ISL_2860950, EPI_ISL_2860951, EPI_ISL_2860952, EPI_ISL_2860953, EPI_ISL_2860954, EPI_ISL_2860955, EPI_ISL_2860956, EPI_ISL_2860958, EPI_ISL_2860959, EPI_ISL_2860961, EPI_ISL_2860963, EPI_ISL_2860964, EPI_ISL_2860965, EPI_ISL_2860967, EPI_ISL_2860969, EPI_ISL_2860970, EPI_ISL_2860971, EPI_ISL_2860972, EPI_ISL_2860975, EPI_ISL_2860976, EPI_ISL_2860978, EPI_ISL_2860979, EPI_ISL_2860980, EPI_ISL_2860981, EPI_ISL_2860982, EPI_ISL_2860983, EPI_ISL_2860984, EPI_ISL_2860985, EPI_ISL_2860986, EPI_ISL_2860987, EPI_ISL_2860988, EPI_ISL_2860989, EPI_ISL_2860990, EPI_ISL_2860991, EPI_ISL_2860992, EPI_ISL_2860993, EPI_ISL_2860994, EPI_ISL_2860995, EPI_ISL_2860996, EPI_ISL_2860997, EPI_ISL_2860998, EPI_ISL_2860999, EPI_ISL_2861000, EPI_ISL_2861001, EPI_ISL_2861002, EPI_ISL_2861003, EPI_ISL_2861004, EPI_ISL_2861005, EPI_ISL_2861006, EPI_ISL_2861007, EPI_ISL_2861008, EPI_ISL_2861009 | | | |
| see above | INSACOG-WB | National Institute of Biomedical Genomics – INSACOG | Arindam Maitra, Bhaswati Bandyopadhyay, Nidhan Kumar Biswas, Tamal Ghosh, Sreedhar Chinnaswamy, Ajay Chakraborti, Saumitra Das |
| EPI_ISL_2861037, EPI_ISL_2861038, EPI_ISL_2861039, EPI_ISL_2861040, EPI_ISL_2861041, EPI_ISL_2861046, EPI_ISL_2861048, EPI_ISL_2861050, EPI_ISL_2861051, EPI_ISL_2861052, EPI_ISL_2861055, EPI_ISL_2861058, EPI_ISL_2861066, EPI_ISL_2861067, EPI_ISL_2861068, EPI_ISL_2861069, EPI_ISL_2861072, EPI_ISL_2861073, EPI_ISL_2861075 | | | |
| see above | Servicio de Microbiologia Hospital Ramon y Cajal | Servicio de Microbiologia Hospital Ramon y Cajal | Ponce M, Galan JC, Martinez L. Abreu M, y Gonzalez-Alba JM |

| | | | |
|---|---|---|---|
| EPI_ISL_2861076, EPI_ISL_2861078, EPI_ISL_2861083, EPI_ISL_2861085, EPI_ISL_2861086, EPI_ISL_2861087, EPI_ISL_2861088, EPI_ISL_2861089, EPI_ISL_2861091, EPI_ISL_2861092, EPI_ISL_2861093, EPI_ISL_2861095, EPI_ISL_2861096, EPI_ISL_2861118, EPI_ISL_2861119, EPI_ISL_2861121, EPI_ISL_2861122 | | | |
| see above | Mobile County Health Department | Synergy Laboratories | Megan Cornwell |
| EPI_ISL_2861127 | Synergy Laboratories | Synergy Laboratories | Megan Cornwell |
| EPI_ISL_2861140, EPI_ISL_2861143, EPI_ISL_2861146 | Mobile County Health Department | Synergy Laboratories | Megan Cornwell |
| EPI_ISL_2861158 | LESP Ciudad de Mexico | Instituto de Diagnostico y Referencia Epidemiologicos (INDRE) | Claudia Wong-Arambula, Abril Rodriguez-Maldonado, Vanessa Rivero-Arredondo, Ariadna Medina-Benitez, Joaquin Quiroz-Mercado, Sergio Rangel-Guerrero, Maribel Gonzalez-Villa, Natividad Cruz-Ortiz, Tatiana Nunez-Garcia, Gisela Barrera-Badillo, Lucia Hernandez-Rivas, Irma Lopez-Martinez, Ernesto Ramirez-Gonzalez. |
| EPI_ISL_2861197, EPI_ISL_2861198, EPI_ISL_2861200, EPI_ISL_2861206, EPI_ISL_2861208, EPI_ISL_2861209 | MD PHL | MD PHL | Maryland Department of Health Laboratories Administration |
| EPI_ISL_2861250 | Liverpool Hospital | NSW Health Pathology - Institute of Clinical Pathology and Medical Research; Westmead Hospital; University of Sydney | CIDM-PH et al. |
| EPI_ISL_2861251, EPI_ISL_2861252 | Southern IML | NSW Health Pathology - Institute of Clinical Pathology and Medical Research; Westmead Hospital; University of Sydney | CIDM-PH et al. |
| EPI_ISL_2861253, EPI_ISL_2861254 | Laverty Pathology | NSW Health Pathology - Institute of Clinical Pathology and Medical Research; Westmead Hospital; University of Sydney | CIDM-PH et al. |
| EPI_ISL_2861255 | St Vincent's Pathology (SydPath) | NSW Health Pathology - Institute of Clinical Pathology and Medical Research; Westmead Hospital; University of Sydney | CIDM-PH et al. |
| EPI_ISL_2861256 | Sydney South West Pathology Service (SSWPS) - Liverpool Hospital - NSW Health Pathology | NSW Health Pathology - Institute of Clinical Pathology and Medical Research; Westmead Hospital; University of Sydney | CIDM-PH et al. |
| EPI_ISL_2861262, EPI_ISL_2861263, EPI_ISL_2861265 | Dr. Yoshitaka Tamura Department of Clinical Laboratory, Osaka Habikino Medical Center | Department of Developmental Medicine, Research Institute, Osaka Women's and Children's Hospital | Itaru Yanagihara, Yukiko Nakura |
| EPI_ISL_2861270 | PathWest Laboratory Medicine WA | PathWest Laboratory Medicine WA Microbial Surveillance Unit | PathWest Laboratory Medicine WA Microbial Surveillance Unit |
| EPI_ISL_2861273, EPI_ISL_2861274, EPI_ISL_2861275 | AZ Zeno | AZ Sint Jan | Marijke Reynders, Patrick Descheemaeker, Laurien Hoornaert, Thomas Van Landschoot, Marie Madeleine Chabert-Consen, Jorn Hellemans |
| EPI_ISL_2861276 | AZ Sint-Jan campus Oostende | AZ Sint Jan | Marijke Reynders, Patrick Descheemaeker, Laurien Hoornaert, Thomas Van Landschoot, Marie Madeleine Chabert-Consen, Jorn Hellemans |
| EPI_ISL_2861277, EPI_ISL_2861278 | AZ Zeno | AZ Sint Jan | Marijke Reynders, Patrick Descheemaeker, Laurien Hoornaert, Thomas Van Landschoot, Marie Madeleine Chabert-Consen, Jorn Hellemans |
| EPI_ISL_2861288, EPI_ISL_2861289, EPI_ISL_2861290, EPI_ISL_2861291, EPI_ISL_2861292, EPI_ISL_2861293, EPI_ISL_2861294, EPI_ISL_2861295, EPI_ISL_2861296, EPI_ISL_2861298, EPI_ISL_2861299, EPI_ISL_2861300, EPI_ISL_2861301, EPI_ISL_2861302, EPI_ISL_2861304, EPI_ISL_2861305, EPI_ISL_2861306, EPI_ISL_2861307, EPI_ISL_2861308, EPI_ISL_2861309, EPI_ISL_2861310, EPI_ISL_2861311, EPI_ISL_2861312, EPI_ISL_2861313, EPI_ISL_2861314, EPI_ISL_2861315, EPI_ISL_2861316 | | | |
| see above | Microbiology Department. Complexo Hospitalario Universitario de Vigo | Microbiology Department. Complexo Hospitalario Universitario de Vigo | Alfaya N, Alonso I, Alvarez M, Cabrera JJ, Carballo R, Cores O, Cortizo S, del-Campo V, Martinez L, Mediero G, Perez S, Davina C, Potel C, Regueiro B, Rey S, Vasallo FJ |
| EPI_ISL_2861320, EPI_ISL_2861321, EPI_ISL_2861322, EPI_ISL_2861323 | Microbiology Department, University Hospital Donostia | Microbiology Department, University Hospital Donostia | Marimon JM, Montes M, Piñeiro L, Martin-Peñaranda T, Gomez M, Sorarrain A, Cilla G. |
| EPI_ISL_2861339 | Istituto Zooprofilattico Sperimentale della Puglia e della Basilicata | Istituto Zooprofilattico Sperimentale della Puglia e della Basilicata | Parisi A., Bianco A., Capozzi L., Del Sambro L., Simone D., Difato L., Santoro A., Ridolfi D., Giannico A., Manzari C., Pesole G., Pace L., Galante D. |
| EPI_ISL_2861344, EPI_ISL_2861350, EPI_ISL_2861357, EPI_ISL_2861358, EPI_ISL_2861363, EPI_ISL_2861369 | AZDelta | AZ Delta Medical Laboratories in Roeselare, Belgium | Geert Martens, Dieter De Smet, Merijn Vanhee, on behalf of AZ Delta COVID-19 Genomics core (member of Genomic surveillance of SARS-CoV-2 in Belgium network) |
| EPI_ISL_2861376 | St James's Hospital, Dublin 8, Ireland. | ELDA biotech | Deirdre M. Broderick, Peter R. Flanagan, Brendan Crowley |
| EPI_ISL_2861381, EPI_ISL_2861385, EPI_ISL_2861399, EPI_ISL_2861400, EPI_ISL_2861402, EPI_ISL_2861408, EPI_ISL_2861420, EPI_ISL_2861424, EPI_ISL_2861425, EPI_ISL_2861426, EPI_ISL_2861431, EPI_ISL_2861441, EPI_ISL_2861449, EPI_ISL_2861451, EPI_ISL_2861452, EPI_ISL_2861457, EPI_ISL_2861458, EPI_ISL_2861460, EPI_ISL_2861462, EPI_ISL_2861463, EPI_ISL_2861467, EPI_ISL_2861468, EPI_ISL_2861469, EPI_ISL_2861470, EPI_ISL_2861474, EPI_ISL_2861475, EPI_ISL_2861476, EPI_ISL_2861477, EPI_ISL_2861478, EPI_ISL_2861481, EPI_ISL_2861482, EPI_ISL_2861484, EPI_ISL_2861486, EPI_ISL_2861487, EPI_ISL_2861488, EPI_ISL_2861490, EPI_ISL_2861492, EPI_ISL_2861493, EPI_ISL_2861494, EPI_ISL_2861495, EPI_ISL_2861496, EPI_ISL_2861497, EPI_ISL_2861498, EPI_ISL_2861500, EPI_ISL_2861501, EPI_ISL_2861502, EPI_ISL_2861505, EPI_ISL_2861507, EPI_ISL_2861508, EPI_ISL_2861509, EPI_ISL_2861510, EPI_ISL_2861511, EPI_ISL_2861512, EPI_ISL_2861513, EPI_ISL_2861514, EPI_ISL_2861515, EPI_ISL_2861516, EPI_ISL_2861517, EPI_ISL_2861518, EPI_ISL_2861519, EPI_ISL_2861520, EPI_ISL_2861521, EPI_ISL_2861522, EPI_ISL_2861523, EPI_ISL_2861524, EPI_ISL_2861525, EPI_ISL_2861526, EPI_ISL_2861527, EPI_ISL_2861528, EPI_ISL_2861529, EPI_ISL_2861530, EPI_ISL_2861531, EPI_ISL_2861532, EPI_ISL_2861533, EPI_ISL_2861535, EPI_ISL_2861536, EPI_ISL_2861537, EPI_ISL_2861539, EPI_ISL_2861540, EPI_ISL_2861541, EPI_ISL_2861543, EPI_ISL_2861544, EPI_ISL_2861545, EPI_ISL_2861546, EPI_ISL_2861547, EPI_ISL_2861548, EPI_ISL_2861549, EPI_ISL_2861550, EPI_ISL_2861551, EPI_ISL_2861552, EPI_ISL_2861553, EPI_ISL_2861554, EPI_ISL_2861555, EPI_ISL_2861556, EPI_ISL_2861557, EPI_ISL_2861558, EPI_ISL_2861559, EPI_ISL_2861560 | | | |
| see above | Lab voor klinische biologie | Lab voor klinische biologie | Marija Janevska, Hannelore Hamerlinck, Bruno Verhasselt |
| EPI_ISL_2861575, EPI_ISL_2861577, EPI_ISL_2861578, EPI_ISL_2861579, EPI_ISL_2861580, EPI_ISL_2861581, EPI_ISL_2861582, EPI_ISL_2861584, EPI_ISL_2861585, EPI_ISL_2861586, EPI_ISL_2861587, EPI_ISL_2861588, EPI_ISL_2861589, EPI_ISL_2861591, EPI_ISL_2861593, EPI_ISL_2861594, EPI_ISL_2861595, EPI_ISL_2861596, EPI_ISL_2861598, EPI_ISL_2861600, EPI_ISL_2861601, EPI_ISL_2861602, EPI_ISL_2861606, EPI_ISL_2861607, EPI_ISL_2861608, EPI_ISL_2861609, EPI_ISL_2861610, EPI_ISL_2861611 | | | |
| see above | Platform BIS UZA/UAntwerpen | Labo Klinische Biologie, UZA | Marie Le Mercier, Jasmine Coppens, Kathleen Holemans, Ines Verbesselt, Basil Britto Xavier, Christine Lammens, Veerle Matheeussen, Herman Goossens |
| EPI_ISL_2861616, EPI_ISL_2861617 | Algemeen Medisch Labo | Labo Klinische Biologie, UZA | Marie Le Mercier, Jasmine Coppens, Kathleen Holemans, Ines Verbesselt, Basil Britto Xavier, Christine Lammens, Veerle Matheeussen, Herman Goossens |
| EPI_ISL_2861637, EPI_ISL_2861643, EPI_ISL_2861647 | Platform BIS UZA/UAntwerpen | Labo Klinische Biologie, UZA | Marie Le Mercier, Jasmine Coppens, Kathleen Holemans, Ines Verbesselt, Basil Britto Xavier, Christine Lammens, Veerle Matheeussen, Herman Goossens |
| EPI_ISL_2861648 | Algemeen Klinisch Labo | Labo Klinische Biologie, UZA | Marie Le Mercier, Jasmine Coppens, Kathleen Holemans, Ines Verbesselt, Basil Britto Xavier, Christine Lammens, Veerle Matheeussen, Herman Goossens |
| EPI_ISL_2861649, EPI_ISL_2861652, EPI_ISL_2861654, EPI_ISL_2861655, EPI_ISL_2861656, EPI_ISL_2861659, EPI_ISL_2861660, EPI_ISL_2861661, EPI_ISL_2861662, EPI_ISL_2861663, EPI_ISL_2861664, EPI_ISL_2861665, EPI_ISL_2861667, EPI_ISL_2861668, EPI_ISL_2861669, EPI_ISL_2861670, EPI_ISL_2861671, EPI_ISL_2861673, EPI_ISL_2861674, EPI_ISL_2861675 | | | |
| see above | Platform BIS UZA/UAntwerpen | Labo Klinische Biologie, UZA | Marie Le Mercier, Jasmine Coppens, Kathleen Holemans, Ines Verbesselt, Basil Britto Xavier, Christine Lammens, Veerle Matheeussen, Herman Goossens |
| EPI_ISL_2861676 | Algemeen Klinisch Labo | Labo Klinische Biologie, UZA | Marie Le Mercier, Jasmine Coppens, Kathleen Holemans, Ines Verbesselt, Basil Britto Xavier, Christine Lammens, Veerle Matheeussen, Herman Goossens |
| EPI_ISL_2861681, EPI_ISL_2861682, EPI_ISL_2861683, EPI_ISL_2861684, EPI_ISL_2861686 | LHUB-ULB | Labo Klinische Biologie, UZA | Marie Le Mercier, Jasmine Coppens, Kathleen Holemans, Ines Verbesselt, Basil Britto Xavier, Christine Lammens, Veerle Matheeussen, Herman Goossens |
| EPI_ISL_2861687, EPI_ISL_2861688, EPI_ISL_2861689, EPI_ISL_2861690, EPI_ISL_2861691, EPI_ISL_2861693, EPI_ISL_2861694, EPI_ISL_2861695, EPI_ISL_2861696, EPI_ISL_2861697, EPI_ISL_2861699, EPI_ISL_2861700, EPI_ISL_2861701, EPI_ISL_2861703, EPI_ISL_2861704, EPI_ISL_2861706, EPI_ISL_2861708, EPI_ISL_2861709, EPI_ISL_2861710, EPI_ISL_2861711, EPI_ISL_2861713 | | | |
| see above | Platform BIS UZA/UAntwerpen | Labo Klinische Biologie, UZA | Marie Le Mercier, Jasmine Coppens, Kathleen Holemans, Ines Verbesselt, Basil Britto Xavier, Christine Lammens, Veerle Matheeussen, Herman Goossens |
| EPI_ISL_2861718 | Genome Information Research Center, Research Institute for Microbial Diseases, Osaka University | Genome Information Research Center, Research Institute for Microbial Diseases, Osaka University | Daisuke Motooka, Kentaro Tanaka, Hiroya Oki, and Shota Nakamura |
| EPI_ISL_2861726, EPI_ISL_2861728, EPI_ISL_2861729, EPI_ISL_2861731, EPI_ISL_2861737 | CLILAB | Microbiology Department | Sara Marti, Aida Gonzalez-Diaz, Laura Calatayud, Jordi Niubó, Miguel Fernandez-Huerta, Carmen Ardanuy, Jordi Camara, M Angeles Domínguez |
| EPI_ISL_2861738, EPI_ISL_2861739, EPI_ISL_2861741, EPI_ISL_2861742, EPI_ISL_2861743, EPI_ISL_2861744, EPI_ISL_2861746, EPI_ISL_2861749, EPI_ISL_2861750, EPI_ISL_2861751, EPI_ISL_2861752, EPI_ISL_2861755, EPI_ISL_2861756, EPI_ISL_2861757, EPI_ISL_2861758, EPI_ISL_2861759, EPI_ISL_2861761, EPI_ISL_2861763, EPI_ISL_2861764, EPI_ISL_2861765, EPI_ISL_2861767, EPI_ISL_2861768, EPI_ISL_2861769, EPI_ISL_2861770, EPI_ISL_2861771, EPI_ISL_2861772, EPI_ISL_2861774, EPI_ISL_2861775, EPI_ISL_2861778, EPI_ISL_2861782, EPI_ISL_2861783, EPI_ISL_2861784, EPI_ISL_2861788, EPI_ISL_2861789, EPI_ISL_2861790, EPI_ISL_2861791, EPI_ISL_2861792, EPI_ISL_2861793, EPI_ISL_2861794, EPI_ISL_2861795, EPI_ISL_2861796, EPI_ISL_2861797 | | | |
| see above | Hospital Universitari Bellvitge | Microbiology Department | Sara Marti, Aida Gonzalez-Diaz, Laura Calatayud, Jordi Niubó, Miguel Fernandez-Huerta, Carmen Ardanuy, Jordi Camara, M Angeles Domínguez |
| EPI_ISL_2861815, EPI_ISL_2861817, EPI_ISL_2861819, EPI_ISL_2861820, EPI_ISL_2861822 | Laboratorio di Riferimento Regionale della Sicilia Occidentale per l'Emergenza COVID-19 | Laboratorio di Riferimento Regionale della Sicilia Occidentale per l'Emergenza COVID-19 | Fabio Tramuto, Carmelo Massimo Maida, Daniela Di Naro, Giulia Randazzo, Walter Mazzucco, Giorgio Graziano, Vincenzo Restivo, Claudio Costantino, Francesco Vitale |
| EPI_ISL_2861854, EPI_ISL_2861861, EPI_ISL_2861862, EPI_ISL_2861867, EPI_ISL_2861868, EPI_ISL_2861869, EPI_ISL_2861870, EPI_ISL_2861871, EPI_ISL_2861873, EPI_ISL_2861874, EPI_ISL_2861876, EPI_ISL_2861878, EPI_ISL_2861879, EPI_ISL_2861880, EPI_ISL_2861886, EPI_ISL_2861887, EPI_ISL_2861889, EPI_ISL_2861893, EPI_ISL_2861895, EPI_ISL_2861899, EPI_ISL_2861900, EPI_ISL_2861901, | | | |

| | | | |
|---|---|---|---|
| EPI_ISL_2861903, EPI_ISL_2861904, EPI_ISL_2861905, EPI_ISL_2861907, EPI_ISL_2861908, EPI_ISL_2861910, EPI_ISL_2861911, EPI_ISL_2861914, EPI_ISL_2861917 | | | |
| see above | ASL Napoli 1 Centro | AMES Centro Polidiagnostico Strumentale S.r.l. | Giovanni Savarese, Raffaella Ruggiero, Eloisa Evangelista, Antonella Di Carlo, Luisa Circelli, Luigi D'Amore, Roberto Sirica,Monica Ianniello, Nadia Petrillo, Maurizio D'Amora, Antonio Fico |
| EPI_ISL_2862010 | Genomic Research Laboratory, Bangabandhu Sheikh Mujib Medical University | Genomic Research Laboratory, BSMMU | Laila Anjuman Banu, Md.Saydur Rahman, Sharadindu Kanti Sinha, Arifa Akram, Md.Sharfuddin Ahmed |
| EPI_ISL_2862015 | Genomic Research Laboratory, Bangabandhu Sheikh Mujib Medical University (BSMMU) Shahbag, Dhaka_1000 | Genomic Research Laboratory, BSMMU | Laila Anjuman Banu, Md.Saydur Rahman, Sharadindu Kanti Sinha, Arifa Akram, Md.Sharfuddin Ahmed |
| EPI_ISL_2862016 | Genomic Research Laboratory, Bangabandhu Sheikh Mujib Medical University | Genomic Research Laboratory, BSMMU | Laila Anjuman Banu, Md.Saydur Rahman, Sharadindu Kanti Sinha, Arifa Akram, Md.Sharfuddin Ahmed |
| EPI_ISL_2862023, EPI_ISL_2862030, EPI_ISL_2862032 | MD PHL | MD PHL | Maryland Department of Health Laboratories Administration |
| EPI_ISL_2862048 | Genomic Research Laboratory, Bangabandhu Sheikh Mujib Medical University | Genomic Research Laboratory, BSMMU | Laila Anjuman Banu, Md.Saydur Rahman, Sharadindu Kanti Sinha, Arifa Akram, Md.Sharfuddin Ahmed |
| EPI_ISL_2862049, EPI_ISL_2862051, EPI_ISL_2862052, EPI_ISL_2862072, EPI_ISL_2862074, EPI_ISL_2862076, EPI_ISL_2862079, EPI_ISL_2862080 | National Virus Reference Laboratory | National Virus Reference Laboratory | Zoe Yandle, Charlene Bennett, Gabriel Gonzalez, Michael Carr, Jonathan Dean, Cillian F De Gascun |
| EPI_ISL_2862085, EPI_ISL_2862086, EPI_ISL_2862087 | National Virus Reference Laboratory | National Virus Reference Laboratory | Fiona Crispie, Calum Walsh, Matthew McCabe, Zoe Yandle, Charlene Bennet, Gabriel Gonzalez, Michael Carr, Jonathan Dean, Paul Cotter, Cillian F De Gascun |
| EPI_ISL_2862094 | National Virus Reference Laboratory | National Virus Reference Laboratory | Zoe Yandle, Charlene Bennett, Gabriel Gonzalez, Michael Carr, Jonathan Dean, Cillian F De Gascun |
| EPI_ISL_2862111, EPI_ISL_2862112, EPI_ISL_2862113, EPI_ISL_2862114, EPI_ISL_2862115, EPI_ISL_2862116, EPI_ISL_2862117, EPI_ISL_2862118, EPI_ISL_2862119, EPI_ISL_2862120 | National Virus Reference Laboratory | National Virus Reference Laboratory | Fiona Crispie, Calum Walsh, Matthew McCabe, Zoe Yandle, Charlene Bennet, Gabriel Gonzalez, Michael Carr, Jonathan Dean, Paul Cotter, Cillian F De Gascun |
| EPI_ISL_2862121, EPI_ISL_2862130, EPI_ISL_2862131, EPI_ISL_2862133, EPI_ISL_2862337, EPI_ISL_2862340, EPI_ISL_2862341, EPI_ISL_2862342, EPI_ISL_2862344 | National Virus Reference Laboratory | National Virus Reference Laboratory | Zoe Yandle, Charlene Bennett, Gabriel Gonzalez, Michael Carr, Jonathan Dean, Cillian F De Gascun |
| EPI_ISL_2862351 | National Virus Reference Laboratory | National Virus Reference Laboratory | Fiona Crispie, Calum Walsh, Matthew McCabe, Zoe Yandle, Charlene Bennet, Gabriel Gonzalez, Michael Carr, Jonathan Dean, Paul Cotter, Cillian F De Gascun |
| EPI_ISL_2862354, EPI_ISL_2862355, EPI_ISL_2862356, EPI_ISL_2862357, EPI_ISL_2862358, EPI_ISL_2862359, EPI_ISL_2862360, EPI_ISL_2862361, EPI_ISL_2862362, EPI_ISL_2862363, EPI_ISL_2862364, EPI_ISL_2862365, EPI_ISL_2862366, EPI_ISL_2862367, EPI_ISL_2862368, EPI_ISL_2862369, EPI_ISL_2862370, EPI_ISL_2862371, EPI_ISL_2862372, EPI_ISL_2862373, EPI_ISL_2862374, EPI_ISL_2862375, EPI_ISL_2862376, EPI_ISL_2862377, EPI_ISL_2862378, EPI_ISL_2862379, EPI_ISL_2862380, EPI_ISL_2862381, EPI_ISL_2862382, EPI_ISL_2862383, EPI_ISL_2862384, EPI_ISL_2862385, EPI_ISL_2862386, EPI_ISL_2862387, EPI_ISL_2862388, EPI_ISL_2862389, EPI_ISL_2862390, EPI_ISL_2862391, EPI_ISL_2862392, EPI_ISL_2862393, EPI_ISL_2862394, EPI_ISL_2862395, EPI_ISL_2862396, EPI_ISL_2862397, EPI_ISL_2862398, EPI_ISL_2862399, EPI_ISL_2862400, EPI_ISL_2862401, EPI_ISL_2862402, EPI_ISL_2862403, EPI_ISL_2862404, EPI_ISL_2862405, EPI_ISL_2862406, EPI_ISL_2862407, EPI_ISL_2862408, EPI_ISL_2862409, EPI_ISL_2862410, EPI_ISL_2862411, EPI_ISL_2862412, EPI_ISL_2862413, EPI_ISL_2862414, EPI_ISL_2862415, EPI_ISL_2862416, EPI_ISL_2862417, EPI_ISL_2862418, EPI_ISL_2862419, EPI_ISL_2862420, EPI_ISL_2862421, EPI_ISL_2862422, EPI_ISL_2862423, EPI_ISL_2862424, EPI_ISL_2862425, EPI_ISL_2862426 | | | |
| see above | National Virus Reference Laboratory | National Virus Reference Laboratory | Zoe Yandle, Charlene Bennett, Gabriel Gonzalez, Michael Carr, Jonathan Dean, Cillian F De Gascun |
| EPI_ISL_2862427, EPI_ISL_2862428 | National Virus Reference Laboratory | National Virus Reference Laboratory | Fiona Crispie, Calum Walsh, Matthew McCabe, Zoe Yandle, Charlene Bennet, Gabriel Gonzalez, Michael Carr, Jonathan Dean, Paul Cotter, Cillian F De Gascun |
| EPI_ISL_2862435, EPI_ISL_2862441 | Dutch COVID-19 response team | National Institute for Public Health and the Environment (RIVM) | Adam Meijer, Harry Vennema, Dirk Eggink, Jeroen Cremer, Sharon van den Brink, Bas van der Veer, AnneMarie van den Brandt, Lisa Wijsman, Kim Freriks, Ryanne Jaarsma, Euníce Then, Lynn Aarts, Sanne Bos, Melissa van Tuil, Florian Zwagemaker, Dennis Schmitz, Annelies Kroneman, Karim Hajji, Chantal Reusken, on behalf of the national COVID-19 response team |
| EPI_ISL_2862662, EPI_ISL_2862664, EPI_ISL_2862666, EPI_ISL_2862668, EPI_ISL_2862670, EPI_ISL_2862672, EPI_ISL_2862674, EPI_ISL_2862676, EPI_ISL_2862678, EPI_ISL_2862680, EPI_ISL_2862682, EPI_ISL_2862684, EPI_ISL_2862686, EPI_ISL_2862688, EPI_ISL_2862690, EPI_ISL_2862692, EPI_ISL_2862694, EPI_ISL_2862697, EPI_ISL_2862699, EPI_ISL_2862701, EPI_ISL_2862703, EPI_ISL_2862705 | | | |
| see above | Department of Bacteria, Parasites and Fungi, Statens Serum Institut, Copenhagen, Denmark | Statens Serum Institut Bioinformatics and Microbial Genomics | Danish Covid-19 Genome Consortium |
| EPI_ISL_2862706, EPI_ISL_2862707 | Dutch COVID-19 response team | National Institute for Public Health and the Environment (RIVM) | Adam Meijer, Harry Vennema, Dirk Eggink, Jeroen Cremer, Sharon van den Brink, Bas van der Veer, AnneMarie van den Brandt, Lisa Wijsman, Kim Freriks, Ryanne Jaarsma, Euníce Then, Lynn Aarts, Sanne Bos, Melissa van Tuil, Florian Zwagemaker, Dennis Schmitz, Annelies Kroneman, Karim Hajji, Chantal Reusken, on behalf of the national COVID-19 response team |
| EPI_ISL_2862708 | Department of Bacteria, Parasites and Fungi, Statens Serum Institut, Copenhagen, Denmark | Statens Serum Institut Bioinformatics and Microbial Genomics | Danish Covid-19 Genome Consortium |
| EPI_ISL_2862709 | Dutch COVID-19 response team | National Institute for Public Health and the Environment (RIVM) | Adam Meijer, Harry Vennema, Dirk Eggink, Jeroen Cremer, Sharon van den Brink, Bas van der Veer, AnneMarie van den Brandt, Lisa Wijsman, Kim Freriks, Ryanne Jaarsma, Euníce Then, Lynn Aarts, Sanne Bos, Melissa van Tuil, Florian Zwagemaker, Dennis Schmitz, Annelies Kroneman, Karim Hajji, Chantal Reusken, on behalf of the national COVID-19 response team |
| EPI_ISL_2862710, EPI_ISL_2862712, EPI_ISL_2862714, EPI_ISL_2862716, EPI_ISL_2862717, EPI_ISL_2862722, EPI_ISL_2862724, EPI_ISL_2862726, EPI_ISL_2862728, EPI_ISL_2862730, EPI_ISL_2862732, EPI_ISL_2862734, EPI_ISL_2862736, EPI_ISL_2862738, EPI_ISL_2862740, EPI_ISL_2862742, EPI_ISL_2862744 | | | |
| see above | Department of Bacteria, Parasites and Fungi, Statens Serum Institut, Copenhagen, Denmark | Statens Serum Institut Bioinformatics and Microbial Genomics | Danish Covid-19 Genome Consortium |
| EPI_ISL_2862746 | Department of Clinical Microbiology, Odense University Hospital, Odense, Denmark | Statens Serum Institut Bioinformatics and Microbial Genomics | Danish Covid-19 Genome Consortium |
| EPI_ISL_2862747, EPI_ISL_2862749, EPI_ISL_2862751 | Department of Bacteria, Parasites and Fungi, Statens Serum Institut, Copenhagen, Denmark | Statens Serum Institut Bioinformatics and Microbial Genomics | Danish Covid-19 Genome Consortium |
| EPI_ISL_2862752 | Dutch COVID-19 response team | National Institute for Public Health and the Environment (RIVM) | Adam Meijer, Harry Vennema, Dirk Eggink, Jeroen Cremer, Sharon van den Brink, Bas van der Veer, AnneMarie van den Brandt, Lisa Wijsman, Kim Freriks, Ryanne Jaarsma, Euníce Then, Lynn Aarts, Sanne Bos, Melissa van Tuil, Florian Zwagemaker, Dennis Schmitz, Annelies Kroneman, Karim Hajji, Chantal Reusken, on behalf of the national COVID-19 response team |
| EPI_ISL_2862753 | Department of Bacteria, Parasites and Fungi, Statens Serum Institut, Copenhagen, Denmark | Statens Serum Institut Bioinformatics and Microbial Genomics | Danish Covid-19 Genome Consortium |
| EPI_ISL_2862754 | Dutch COVID-19 response team | National Institute for Public Health and the Environment (RIVM) | Adam Meijer, Harry Vennema, Dirk Eggink, Jeroen Cremer, Sharon van den Brink, Bas van der Veer, AnneMarie van den Brandt, Lisa Wijsman, Kim Freriks, Ryanne Jaarsma, Euníce Then, Lynn Aarts, Sanne Bos, Melissa van Tuil, Florian Zwagemaker, Dennis Schmitz, Annelies Kroneman, Karim Hajji, Chantal Reusken, on behalf of the national COVID-19 response team |
| EPI_ISL_2862755 | Department of Bacteria, Parasites and Fungi, Statens Serum Institut, Copenhagen, Denmark | Statens Serum Institut Bioinformatics and Microbial Genomics | Danish Covid-19 Genome Consortium |
| EPI_ISL_2862756 | Dutch COVID-19 response team | National Institute for Public Health and the Environment (RIVM) | Adam Meijer, Harry Vennema, Dirk Eggink, Jeroen Cremer, Sharon van den Brink, Bas van der Veer, AnneMarie van den Brandt, Lisa Wijsman, Kim Freriks, Ryanne Jaarsma, Euníce Then, Lynn Aarts, Sanne Bos, Melissa van Tuil, Florian Zwagemaker, Dennis Schmitz, Annelies Kroneman, Karim Hajji, Chantal Reusken, on behalf of the national COVID-19 response team |
| EPI_ISL_2862757 | Department of Bacteria, Parasites and Fungi, Statens Serum Institut, Copenhagen, Denmark | Statens Serum Institut Bioinformatics and Microbial Genomics | Danish Covid-19 Genome Consortium |
| EPI_ISL_2862758 | Dutch COVID-19 response team | National Institute for Public Health and the Environment (RIVM) | Adam Meijer, Harry Vennema, Dirk Eggink, Jeroen Cremer, Sharon van den Brink, Bas van der Veer, AnneMarie van den Brandt, Lisa Wijsman, Kim Freriks, Ryanne Jaarsma, Euníce Then, Lynn Aarts, Sanne Bos, Melissa van Tuil, Florian Zwagemaker, Dennis Schmitz, Annelies Kroneman, Karim Hajji, Chantal Reusken, on behalf of the national COVID-19 response team |
| EPI_ISL_2862760, EPI_ISL_2862762, EPI_ISL_2862764, EPI_ISL_2862766, EPI_ISL_2862768, EPI_ISL_2862770, EPI_ISL_2862772, EPI_ISL_2862774, EPI_ISL_2862776, EPI_ISL_2862778, EPI_ISL_2862780, EPI_ISL_2862782, EPI_ISL_2862784, EPI_ISL_2862786, EPI_ISL_2862788, EPI_ISL_2862791, EPI_ISL_2862793, EPI_ISL_2862795, EPI_ISL_2862797, EPI_ISL_2862799, EPI_ISL_2862801, EPI_ISL_2862803, EPI_ISL_2862805, EPI_ISL_2862807, EPI_ISL_2862809, EPI_ISL_2862812, EPI_ISL_2862814, EPI_ISL_2862816, EPI_ISL_2862818, EPI_ISL_2862820, EPI_ISL_2862822, EPI_ISL_2862825, EPI_ISL_2862826, EPI_ISL_2862829, EPI_ISL_2862831, EPI_ISL_2862833, EPI_ISL_2862835, EPI_ISL_2862837, EPI_ISL_2862839, EPI_ISL_2862841, EPI_ISL_2862843, EPI_ISL_2862845, EPI_ISL_2862848, EPI_ISL_2862850, EPI_ISL_2862852, EPI_ISL_2862854, EPI_ISL_2862856, EPI_ISL_2862858, EPI_ISL_2862860, EPI_ISL_2862862, EPI_ISL_2862864, EPI_ISL_2862866, EPI_ISL_2862868, EPI_ISL_2862870, EPI_ISL_2862872, EPI_ISL_2862874, EPI_ISL_2862877, EPI_ISL_2862879, EPI_ISL_2862881, EPI_ISL_2862883, EPI_ISL_2862885 | | | |
| see above | Department of Bacteria, Parasites and Fungi, Statens Serum Institut, Copenhagen, Denmark | Statens Serum Institut Bioinformatics and Microbial Genomics | Danish Covid-19 Genome Consortium |
| EPI_ISL_2862886 | Dutch COVID-19 response team | National Institute for Public Health and the Environment (RIVM) | Adam Meijer, Harry Vennema, Dirk Eggink, Jeroen Cremer, Sharon van den Brink, Bas van der Veer, AnneMarie van den Brandt, Lisa Wijsman, Kim Freriks, Ryanne Jaarsma, Euníce Then, Lynn Aarts, Sanne Bos, Melissa van Tuil, Florian Zwagemaker, Dennis Schmitz, Annelies Kroneman, Karim Hajji, Chantal Reusken, on behalf of the national COVID-19 response team |
| EPI_ISL_2862887, | Department of Bacteria, Parasites and Fungi, Statens Serum | Statens Serum Institut Bioinformatics and Microbial Genomics | Danish Covid-19 Genome Consortium |

| | | | |
|---|---|---|---|
| EPI_ISL_2862889 | Institut, Copenhagen, Denmark | | |
| EPI_ISL_2862890 | Dutch COVID-19 response team | National Institute for Public Health and the Environment (RIVM) | Adam Meijer, Harry Vennema, Dirk Eggink, Jeroen Cremer, Sharon van den Brink, Bas van der Veer, AnneMarie van den Brandt, Lisa Wijsman, Kim Freriks, Ryanne Jaarsma, Euníce Then, Lynn Aarts, Sanne Bos, Melissa van Tuil, Florian Zwagemaker, Dennis Schmitz, Annelies Kroneman, Karim Hajji, Chantal Reusken, on behalf of the national COVID-19 response team |
| EPI_ISL_2862891 | Department of Bacteria, Parasites and Fungi, Statens Serum Institut, Copenhagen, Denmark | Statens Serum Institut Bioinformatics and Microbial Genomics | Danish Covid-19 Genome Consortium |
| EPI_ISL_2862892, EPI_ISL_2862893 | Dutch COVID-19 response team | National Institute for Public Health and the Environment (RIVM) | Adam Meijer, Harry Vennema, Dirk Eggink, Jeroen Cremer, Sharon van den Brink, Bas van der Veer, AnneMarie van den Brandt, Lisa Wijsman, Kim Freriks, Ryanne Jaarsma, Euníce Then, Lynn Aarts, Sanne Bos, Melissa van Tuil, Florian Zwagemaker, Dennis Schmitz, Annelies Kroneman, Karim Hajji, Chantal Reusken, on behalf of the national COVID-19 response team |
| EPI_ISL_2862894 | Department of Bacteria, Parasites and Fungi, Statens Serum Institut, Copenhagen, Denmark | Statens Serum Institut Bioinformatics and Microbial Genomics | Danish Covid-19 Genome Consortium |
| EPI_ISL_2862895 | Dutch COVID-19 response team | National Institute for Public Health and the Environment (RIVM) | Adam Meijer, Harry Vennema, Dirk Eggink, Jeroen Cremer, Sharon van den Brink, Bas van der Veer, AnneMarie van den Brandt, Lisa Wijsman, Kim Freriks, Ryanne Jaarsma, Euníce Then, Lynn Aarts, Sanne Bos, Melissa van Tuil, Florian Zwagemaker, Dennis Schmitz, Annelies Kroneman, Karim Hajji, Chantal Reusken, on behalf of the national COVID-19 response team |
| EPI_ISL_2862896 | Department of Bacteria, Parasites and Fungi, Statens Serum Institut, Copenhagen, Denmark | Statens Serum Institut Bioinformatics and Microbial Genomics | Danish Covid-19 Genome Consortium |
| EPI_ISL_2862897 | Dutch COVID-19 response team | National Institute for Public Health and the Environment (RIVM) | Adam Meijer, Harry Vennema, Dirk Eggink, Jeroen Cremer, Sharon van den Brink, Bas van der Veer, AnneMarie van den Brandt, Lisa Wijsman, Kim Freriks, Ryanne Jaarsma, Euníce Then, Lynn Aarts, Sanne Bos, Melissa van Tuil, Florian Zwagemaker, Dennis Schmitz, Annelies Kroneman, Karim Hajji, Chantal Reusken, on behalf of the national COVID-19 response team |
| EPI_ISL_2862898 | Department of Bacteria, Parasites and Fungi, Statens Serum Institut, Copenhagen, Denmark | Statens Serum Institut Bioinformatics and Microbial Genomics | Danish Covid-19 Genome Consortium |
| EPI_ISL_2862899 | Dutch COVID-19 response team | National Institute for Public Health and the Environment (RIVM) | Adam Meijer, Harry Vennema, Dirk Eggink, Jeroen Cremer, Sharon van den Brink, Bas van der Veer, AnneMarie van den Brandt, Lisa Wijsman, Kim Freriks, Ryanne Jaarsma, Euníce Then, Lynn Aarts, Sanne Bos, Melissa van Tuil, Florian Zwagemaker, Dennis Schmitz, Annelies Kroneman, Karim Hajji, Chantal Reusken, on behalf of the national COVID-19 response team |
| EPI_ISL_2862900 | Department of Bacteria, Parasites and Fungi, Statens Serum Institut, Copenhagen, Denmark | Statens Serum Institut Bioinformatics and Microbial Genomics | Danish Covid-19 Genome Consortium |
| EPI_ISL_2862901 | Dutch COVID-19 response team | National Institute for Public Health and the Environment (RIVM) | Adam Meijer, Harry Vennema, Dirk Eggink, Jeroen Cremer, Sharon van den Brink, Bas van der Veer, AnneMarie van den Brandt, Lisa Wijsman, Kim Freriks, Ryanne Jaarsma, Euníce Then, Lynn Aarts, Sanne Bos, Melissa van Tuil, Florian Zwagemaker, Dennis Schmitz, Annelies Kroneman, Karim Hajji, Chantal Reusken, on behalf of the national COVID-19 response team |
| EPI_ISL_2862902 | Department of Bacteria, Parasites and Fungi, Statens Serum Institut, Copenhagen, Denmark | Statens Serum Institut Bioinformatics and Microbial Genomics | Danish Covid-19 Genome Consortium |
| EPI_ISL_2862903 | Dutch COVID-19 response team | National Institute for Public Health and the Environment (RIVM) | Adam Meijer, Harry Vennema, Dirk Eggink, Jeroen Cremer, Sharon van den Brink, Bas van der Veer, AnneMarie van den Brandt, Lisa Wijsman, Kim Freriks, Ryanne Jaarsma, Euníce Then, Lynn Aarts, Sanne Bos, Melissa van Tuil, Florian Zwagemaker, Dennis Schmitz, Annelies Kroneman, Karim Hajji, Chantal Reusken, on behalf of the national COVID-19 response team |
| EPI_ISL_2862904 | Department of Bacteria, Parasites and Fungi, Statens Serum Institut, Copenhagen, Denmark | Statens Serum Institut Bioinformatics and Microbial Genomics | Danish Covid-19 Genome Consortium |
| EPI_ISL_2862905 | Dutch COVID-19 response team | National Institute for Public Health and the Environment (RIVM) | Adam Meijer, Harry Vennema, Dirk Eggink, Jeroen Cremer, Sharon van den Brink, Bas van der Veer, AnneMarie van den Brandt, Lisa Wijsman, Kim Freriks, Ryanne Jaarsma, Euníce Then, Lynn Aarts, Sanne Bos, Melissa van Tuil, Florian Zwagemaker, Dennis Schmitz, Annelies Kroneman, Karim Hajji, Chantal Reusken, on behalf of the national COVID-19 response team |
| EPI_ISL_2862906, EPI_ISL_2862908 | Department of Bacteria, Parasites and Fungi, Statens Serum Institut, Copenhagen, Denmark | Statens Serum Institut Bioinformatics and Microbial Genomics | Danish Covid-19 Genome Consortium |
| EPI_ISL_2862909, EPI_ISL_2862910 | Dutch COVID-19 response team | National Institute for Public Health and the Environment (RIVM) | Adam Meijer, Harry Vennema, Dirk Eggink, Jeroen Cremer, Sharon van den Brink, Bas van der Veer, AnneMarie van den Brandt, Lisa Wijsman, Kim Freriks, Ryanne Jaarsma, Euníce Then, Lynn Aarts, Sanne Bos, Melissa van Tuil, Florian Zwagemaker, Dennis Schmitz, Annelies Kroneman, Karim Hajji, Chantal Reusken, on behalf of the national COVID-19 response team |
| EPI_ISL_2862911 | Department of Bacteria, Parasites and Fungi, Statens Serum Institut, Copenhagen, Denmark | Statens Serum Institut Bioinformatics and Microbial Genomics | Danish Covid-19 Genome Consortium |
| EPI_ISL_2862912 | Dutch COVID-19 response team | National Institute for Public Health and the Environment (RIVM) | Adam Meijer, Harry Vennema, Dirk Eggink, Jeroen Cremer, Sharon van den Brink, Bas van der Veer, AnneMarie van den Brandt, Lisa Wijsman, Kim Freriks, Ryanne Jaarsma, Euníce Then, Lynn Aarts, Sanne Bos, Melissa van Tuil, Florian Zwagemaker, Dennis Schmitz, Annelies Kroneman, Karim Hajji, Chantal Reusken, on behalf of the national COVID-19 response team |
| EPI_ISL_2862913 | Department of Bacteria, Parasites and Fungi, Statens Serum Institut, Copenhagen, Denmark | Statens Serum Institut Bioinformatics and Microbial Genomics | Danish Covid-19 Genome Consortium |
| EPI_ISL_2862914 | Dutch COVID-19 response team | National Institute for Public Health and the Environment (RIVM) | Adam Meijer, Harry Vennema, Dirk Eggink, Jeroen Cremer, Sharon van den Brink, Bas van der Veer, AnneMarie van den Brandt, Lisa Wijsman, Kim Freriks, Ryanne Jaarsma, Euníce Then, Lynn Aarts, Sanne Bos, Melissa van Tuil, Florian Zwagemaker, Dennis Schmitz, Annelies Kroneman, Karim Hajji, Chantal Reusken, on behalf of the national COVID-19 response team |
| EPI_ISL_2862915 | Department of Bacteria, Parasites and Fungi, Statens Serum Institut, Copenhagen, Denmark | Statens Serum Institut Bioinformatics and Microbial Genomics | Danish Covid-19 Genome Consortium |
| EPI_ISL_2862916 | Dutch COVID-19 response team | National Institute for Public Health and the Environment (RIVM) | Adam Meijer, Harry Vennema, Dirk Eggink, Jeroen Cremer, Sharon van den Brink, Bas van der Veer, AnneMarie van den Brandt, Lisa Wijsman, Kim Freriks, Ryanne Jaarsma, Euníce Then, Lynn Aarts, Sanne Bos, Melissa van Tuil, Florian Zwagemaker, Dennis Schmitz, Annelies Kroneman, Karim Hajji, Chantal Reusken, on behalf of the national COVID-19 response team |
| EPI_ISL_2862917 | Department of Bacteria, Parasites and Fungi, Statens Serum Institut, Copenhagen, Denmark | Statens Serum Institut Bioinformatics and Microbial Genomics | Danish Covid-19 Genome Consortium |
| EPI_ISL_2862918 | Dutch COVID-19 response team | National Institute for Public Health and the Environment (RIVM) | Adam Meijer, Harry Vennema, Dirk Eggink, Jeroen Cremer, Sharon van den Brink, Bas van der Veer, AnneMarie van den Brandt, Lisa Wijsman, Kim Freriks, Ryanne Jaarsma, Euníce Then, Lynn Aarts, Sanne Bos, Melissa van Tuil, Florian Zwagemaker, Dennis Schmitz, Annelies Kroneman, Karim Hajji, Chantal Reusken, on behalf of the national COVID-19 response team |
| EPI_ISL_2862919 | Department of Bacteria, Parasites and Fungi, Statens Serum Institut, Copenhagen, Denmark | Statens Serum Institut Bioinformatics and Microbial Genomics | Danish Covid-19 Genome Consortium |
| EPI_ISL_2862920 | Dutch COVID-19 response team | National Institute for Public Health and the Environment (RIVM) | Adam Meijer, Harry Vennema, Dirk Eggink, Jeroen Cremer, Sharon van den Brink, Bas van der Veer, AnneMarie van den Brandt, Lisa Wijsman, Kim Freriks, Ryanne Jaarsma, Euníce Then, Lynn Aarts, Sanne Bos, Melissa van Tuil, Florian Zwagemaker, Dennis Schmitz, Annelies Kroneman, Karim Hajji, Chantal Reusken, on behalf of the national COVID-19 response team |
| EPI_ISL_2862921 | Department of Bacteria, Parasites and Fungi, Statens Serum Institut, Copenhagen, Denmark | Statens Serum Institut Bioinformatics and Microbial Genomics | Danish Covid-19 Genome Consortium |
| EPI_ISL_2862922 | Dutch COVID-19 response team | National Institute for Public Health and the Environment (RIVM) | Adam Meijer, Harry Vennema, Dirk Eggink, Jeroen Cremer, Sharon van den Brink, Bas van der Veer, AnneMarie van den Brandt, Lisa Wijsman, Kim Freriks, Ryanne Jaarsma, Euníce Then, Lynn Aarts, Sanne Bos, Melissa van Tuil, Florian Zwagemaker, Dennis Schmitz, Annelies Kroneman, Karim Hajji, Chantal Reusken, on behalf of the national COVID-19 response team |
| EPI_ISL_2862923 | Department of Bacteria, Parasites and Fungi, Statens Serum Institut, Copenhagen, Denmark | Statens Serum Institut Bioinformatics and Microbial Genomics | Danish Covid-19 Genome Consortium |
| EPI_ISL_2862924 | Dutch COVID-19 response team | National Institute for Public Health and the Environment (RIVM) | Adam Meijer, Harry Vennema, Dirk Eggink, Jeroen Cremer, Sharon van den Brink, Bas van der Veer, AnneMarie van den Brandt, Lisa Wijsman, Kim Freriks, Ryanne Jaarsma, Euníce Then, Lynn Aarts, Sanne Bos, Melissa van Tuil, Florian Zwagemaker, Dennis Schmitz, Annelies Kroneman, Karim Hajji, Chantal Reusken, on behalf of the national COVID-19 response team |
| EPI_ISL_2862925 | Department of Bacteria, Parasites and Fungi, Statens Serum Institut, Copenhagen, Denmark | Statens Serum Institut Bioinformatics and Microbial Genomics | Danish Covid-19 Genome Consortium |
| EPI_ISL_2862926 | Dutch COVID-19 response team | National Institute for Public Health and the Environment (RIVM) | Adam Meijer, Harry Vennema, Dirk Eggink, Jeroen Cremer, Sharon van den Brink, Bas van der Veer, AnneMarie van den Brandt, Lisa Wijsman, Kim Freriks, Ryanne Jaarsma, Euníce Then, Lynn Aarts, Sanne Bos, Melissa van Tuil, Florian Zwagemaker, Dennis Schmitz, Annelies Kroneman, Karim Hajji, Chantal Reusken, on behalf of the national COVID-19 response team |
| EPI_ISL_2862927 | Department of Bacteria, Parasites and Fungi, Statens Serum Institut, Copenhagen, Denmark | Statens Serum Institut Bioinformatics and Microbial Genomics | Danish Covid-19 Genome Consortium |
| EPI_ISL_2862928 | Dutch COVID-19 response team | National Institute for Public Health and the Environment (RIVM) | Adam Meijer, Harry Vennema, Dirk Eggink, Jeroen Cremer, Sharon van den Brink, Bas van der Veer, AnneMarie van den Brandt, Lisa Wijsman, Kim Freriks, Ryanne Jaarsma, Euníce Then, Lynn Aarts, Sanne Bos, Melissa van Tuil, Florian Zwagemaker, Dennis Schmitz, Annelies Kroneman, Karim Hajji, Chantal Reusken, on behalf of the national COVID-19 response team |
| EPI_ISL_2862929 | Department of Bacteria, Parasites and Fungi, Statens Serum Institut, Copenhagen, Denmark | Statens Serum Institut Bioinformatics and Microbial Genomics | Danish Covid-19 Genome Consortium |
| EPI_ISL_2862930 | Dutch COVID-19 response team | National Institute for Public Health and the Environment (RIVM) | Adam Meijer, Harry Vennema, Dirk Eggink, Jeroen Cremer, Sharon van den Brink, Bas van der Veer, AnneMarie van den Brandt, Lisa Wijsman, Kim Freriks, Ryanne Jaarsma, Euníce Then, Lynn Aarts, Sanne Bos, Melissa van Tuil, Florian Zwagemaker, Dennis Schmitz, Annelies Kroneman, Karim Hajji, Chantal Reusken, on behalf of the national COVID-19 response team |
| EPI_ISL_2862931 | Department of Bacteria, Parasites and Fungi, Statens Serum Institut, Copenhagen, Denmark | Statens Serum Institut Bioinformatics and Microbial Genomics | Danish Covid-19 Genome Consortium |
| EPI_ISL_2862932 | Dutch COVID-19 response team | National Institute for Public Health and the Environment (RIVM) | Adam Meijer, Harry Vennema, Dirk Eggink, Jeroen Cremer, Sharon van den Brink, Bas van der Veer, AnneMarie van den Brandt, Lisa Wijsman, Kim Freriks, Ryanne Jaarsma, Euníce Then, Lynn Aarts, Sanne Bos, Melissa van Tuil, Florian Zwagemaker, Dennis Schmitz, Annelies Kroneman, Karim Hajji, Chantal Reusken, on behalf of the national COVID-19 response team |
| EPI_ISL_2862933 | Department of Bacteria, Parasites and Fungi, Statens Serum Institut, Copenhagen, Denmark | Statens Serum Institut Bioinformatics and Microbial Genomics | Danish Covid-19 Genome Consortium |
| EPI_ISL_2862935 | Dutch COVID-19 response team | National Institute for Public Health and the Environment (RIVM) | Adam Meijer, Harry Vennema, Dirk Eggink, Jeroen Cremer, Sharon van den Brink, Bas van der Veer, AnneMarie van den Brandt, Lisa Wijsman, Kim Freriks, Ryanne Jaarsma, Euníce Then, Lynn Aarts, Sanne Bos, Melissa van Tuil, Florian Zwagemaker, Dennis Schmitz, Annelies Kroneman, Karim Hajji, Chantal Reusken, on behalf of the national COVID-19 response team |
| EPI_ISL_2862936 | Department of Bacteria, Parasites and Fungi, Statens Serum Institut, Copenhagen, Denmark | Statens Serum Institut Bioinformatics and Microbial Genomics | Danish Covid-19 Genome Consortium |
| EPI_ISL_2862937 | Dutch COVID-19 response team | National Institute for Public Health and the Environment (RIVM) | Adam Meijer, Harry Vennema, Dirk Eggink, Jeroen Cremer, Sharon van den Brink, Bas van der Veer, AnneMarie van den Brandt, Lisa Wijsman, Kim Freriks, Ryanne Jaarsma, Euníce Then, Lynn Aarts, Sanne Bos, Melissa van Tuil, Florian Zwagemaker, Dennis Schmitz, Annelies Kroneman, Karim Hajji, Chantal Reusken, on behalf of the national COVID-19 response team |
| EPI_ISL_2862938, EPI_ISL_2862940 | Department of Bacteria, Parasites and Fungi, Statens Serum Institut, Copenhagen, Denmark | Statens Serum Institut Bioinformatics and Microbial Genomics | Danish Covid-19 Genome Consortium |
| EPI_ISL_2862941 | Dutch COVID-19 response team | National Institute for Public Health and the Environment (RIVM) | Adam Meijer, Harry Vennema, Dirk Eggink, Jeroen Cremer, Sharon van den Brink, Bas van der Veer, AnneMarie van den Brandt, Lisa Wijsman, Kim Freriks, Ryanne Jaarsma, Euníce Then, Lynn Aarts, Sanne Bos, Melissa van Tuil, Florian |

| | | | |
|---|---|---|---|
| | | | Zwagemaker, Dennis Schmitz, Annelies Kroneman, Karim Hajji, Chantal Reusken, on behalf of the national COVID-19 response team |
| EPI_ISL_2862942, EPI_ISL_2862944, EPI_ISL_2862946, EPI_ISL_2862948, EPI_ISL_2862950, EPI_ISL_2862952, EPI_ISL_2862954, EPI_ISL_2862956, EPI_ISL_2862958, EPI_ISL_2862960 | Department of Bacteria, Parasites and Fungi, Statens Serum Institut, Copenhagen, Denmark | Statens Serum Institut Bioinformatics and Microbial Genomics | Danish Covid-19 Genome Consortium |
| EPI_ISL_2862961 | Dutch COVID-19 response team | National Institute for Public Health and the Environment (RIVM) | Adam Meijer, Harry Vennema, Dirk Eggink, Jeroen Cremer, Sharon van den Brink, Bas van der Veer, AnneMarie van den Brandt, Lisa Wijsman, Kim Freriks, Ryanne Jaarsma, Euníce Then, Lynn Aarts, Sanne Bos, Melissa van Tuil, Florian Zwagemaker, Dennis Schmitz, Annelies Kroneman, Karim Hajji, Chantal Reusken, on behalf of the national COVID-19 response team |
| EPI_ISL_2862962 | Department of Bacteria, Parasites and Fungi, Statens Serum Institut, Copenhagen, Denmark | Statens Serum Institut Bioinformatics and Microbial Genomics | Danish Covid-19 Genome Consortium |
| EPI_ISL_2862963 | Dutch COVID-19 response team | National Institute for Public Health and the Environment (RIVM) | Adam Meijer, Harry Vennema, Dirk Eggink, Jeroen Cremer, Sharon van den Brink, Bas van der Veer, AnneMarie van den Brandt, Lisa Wijsman, Kim Freriks, Ryanne Jaarsma, Euníce Then, Lynn Aarts, Sanne Bos, Melissa van Tuil, Florian Zwagemaker, Dennis Schmitz, Annelies Kroneman, Karim Hajji, Chantal Reusken, on behalf of the national COVID-19 response team |
| EPI_ISL_2862964 | Department of Bacteria, Parasites and Fungi, Statens Serum Institut, Copenhagen, Denmark | Statens Serum Institut Bioinformatics and Microbial Genomics | Danish Covid-19 Genome Consortium |
| EPI_ISL_2862965, EPI_ISL_2862966 | Dutch COVID-19 response team | National Institute for Public Health and the Environment (RIVM) | Adam Meijer, Harry Vennema, Dirk Eggink, Jeroen Cremer, Sharon van den Brink, Bas van der Veer, AnneMarie van den Brandt, Lisa Wijsman, Kim Freriks, Ryanne Jaarsma, Euníce Then, Lynn Aarts, Sanne Bos, Melissa van Tuil, Florian Zwagemaker, Dennis Schmitz, Annelies Kroneman, Karim Hajji, Chantal Reusken, on behalf of the national COVID-19 response team |
| EPI_ISL_2862967 | Department of Bacteria, Parasites and Fungi, Statens Serum Institut, Copenhagen, Denmark | Statens Serum Institut Bioinformatics and Microbial Genomics | Danish Covid-19 Genome Consortium |
| EPI_ISL_2862968 | Dutch COVID-19 response team | National Institute for Public Health and the Environment (RIVM) | Adam Meijer, Harry Vennema, Dirk Eggink, Jeroen Cremer, Sharon van den Brink, Bas van der Veer, AnneMarie van den Brandt, Lisa Wijsman, Kim Freriks, Ryanne Jaarsma, Euníce Then, Lynn Aarts, Sanne Bos, Melissa van Tuil, Florian Zwagemaker, Dennis Schmitz, Annelies Kroneman, Karim Hajji, Chantal Reusken, on behalf of the national COVID-19 response team |
| EPI_ISL_2862969 | Department of Bacteria, Parasites and Fungi, Statens Serum Institut, Copenhagen, Denmark | Statens Serum Institut Bioinformatics and Microbial Genomics | Danish Covid-19 Genome Consortium |
| EPI_ISL_2862970 | Dutch COVID-19 response team | National Institute for Public Health and the Environment (RIVM) | Adam Meijer, Harry Vennema, Dirk Eggink, Jeroen Cremer, Sharon van den Brink, Bas van der Veer, AnneMarie van den Brandt, Lisa Wijsman, Kim Freriks, Ryanne Jaarsma, Euníce Then, Lynn Aarts, Sanne Bos, Melissa van Tuil, Florian Zwagemaker, Dennis Schmitz, Annelies Kroneman, Karim Hajji, Chantal Reusken, on behalf of the national COVID-19 response team |
| EPI_ISL_2862971 | Department of Bacteria, Parasites and Fungi, Statens Serum Institut, Copenhagen, Denmark | Statens Serum Institut Bioinformatics and Microbial Genomics | Danish Covid-19 Genome Consortium |
| EPI_ISL_2862972 | Dutch COVID-19 response team | National Institute for Public Health and the Environment (RIVM) | Adam Meijer, Harry Vennema, Dirk Eggink, Jeroen Cremer, Sharon van den Brink, Bas van der Veer, AnneMarie van den Brandt, Lisa Wijsman, Kim Freriks, Ryanne Jaarsma, Euníce Then, Lynn Aarts, Sanne Bos, Melissa van Tuil, Florian Zwagemaker, Dennis Schmitz, Annelies Kroneman, Karim Hajji, Chantal Reusken, on behalf of the national COVID-19 response team |
| EPI_ISL_2862973 | Department of Bacteria, Parasites and Fungi, Statens Serum Institut, Copenhagen, Denmark | Statens Serum Institut Bioinformatics and Microbial Genomics | Danish Covid-19 Genome Consortium |
| EPI_ISL_2862974 | Dutch COVID-19 response team | National Institute for Public Health and the Environment (RIVM) | Adam Meijer, Harry Vennema, Dirk Eggink, Jeroen Cremer, Sharon van den Brink, Bas van der Veer, AnneMarie van den Brandt, Lisa Wijsman, Kim Freriks, Ryanne Jaarsma, Euníce Then, Lynn Aarts, Sanne Bos, Melissa van Tuil, Florian Zwagemaker, Dennis Schmitz, Annelies Kroneman, Karim Hajji, Chantal Reusken, on behalf of the national COVID-19 response team |
| EPI_ISL_2862975 | Department of Bacteria, Parasites and Fungi, Statens Serum Institut, Copenhagen, Denmark | Statens Serum Institut Bioinformatics and Microbial Genomics | Danish Covid-19 Genome Consortium |
| EPI_ISL_2862976 | Dutch COVID-19 response team | National Institute for Public Health and the Environment (RIVM) | Adam Meijer, Harry Vennema, Dirk Eggink, Jeroen Cremer, Sharon van den Brink, Bas van der Veer, AnneMarie van den Brandt, Lisa Wijsman, Kim Freriks, Ryanne Jaarsma, Euníce Then, Lynn Aarts, Sanne Bos, Melissa van Tuil, Florian Zwagemaker, Dennis Schmitz, Annelies Kroneman, Karim Hajji, Chantal Reusken, on behalf of the national COVID-19 response team |
| EPI_ISL_2862977 | Department of Bacteria, Parasites and Fungi, Statens Serum Institut, Copenhagen, Denmark | Statens Serum Institut Bioinformatics and Microbial Genomics | Danish Covid-19 Genome Consortium |
| EPI_ISL_2862978 | Dutch COVID-19 response team | National Institute for Public Health and the Environment (RIVM) | Adam Meijer, Harry Vennema, Dirk Eggink, Jeroen Cremer, Sharon van den Brink, Bas van der Veer, AnneMarie van den Brandt, Lisa Wijsman, Kim Freriks, Ryanne Jaarsma, Euníce Then, Lynn Aarts, Sanne Bos, Melissa van Tuil, Florian Zwagemaker, Dennis Schmitz, Annelies Kroneman, Karim Hajji, Chantal Reusken, on behalf of the national COVID-19 response team |
| EPI_ISL_2862979 | Department of Bacteria, Parasites and Fungi, Statens Serum Institut, Copenhagen, Denmark | Statens Serum Institut Bioinformatics and Microbial Genomics | Danish Covid-19 Genome Consortium |
| EPI_ISL_2862980, EPI_ISL_2862981 | Dutch COVID-19 response team | National Institute for Public Health and the Environment (RIVM) | Adam Meijer, Harry Vennema, Dirk Eggink, Jeroen Cremer, Sharon van den Brink, Bas van der Veer, AnneMarie van den Brandt, Lisa Wijsman, Kim Freriks, Ryanne Jaarsma, Euníce Then, Lynn Aarts, Sanne Bos, Melissa van Tuil, Florian Zwagemaker, Dennis Schmitz, Annelies Kroneman, Karim Hajji, Chantal Reusken, on behalf of the national COVID-19 response team |
| EPI_ISL_2862982, EPI_ISL_2862983 | Department of Bacteria, Parasites and Fungi, Statens Serum Institut, Copenhagen, Denmark | Statens Serum Institut Bioinformatics and Microbial Genomics | Danish Covid-19 Genome Consortium |
| EPI_ISL_2862984, EPI_ISL_2862985 | Dutch COVID-19 response team | National Institute for Public Health and the Environment (RIVM) | Adam Meijer, Harry Vennema, Dirk Eggink, Jeroen Cremer, Sharon van den Brink, Bas van der Veer, AnneMarie van den Brandt, Lisa Wijsman, Kim Freriks, Ryanne Jaarsma, Euníce Then, Lynn Aarts, Sanne Bos, Melissa van Tuil, Florian Zwagemaker, Dennis Schmitz, Annelies Kroneman, Karim Hajji, Chantal Reusken, on behalf of the national COVID-19 response team |
| EPI_ISL_2862986 | Department of Bacteria, Parasites and Fungi, Statens Serum Institut, Copenhagen, Denmark | Statens Serum Institut Bioinformatics and Microbial Genomics | Danish Covid-19 Genome Consortium |
| EPI_ISL_2862987 | Dutch COVID-19 response team | National Institute for Public Health and the Environment (RIVM) | Adam Meijer, Harry Vennema, Dirk Eggink, Jeroen Cremer, Sharon van den Brink, Bas van der Veer, AnneMarie van den Brandt, Lisa Wijsman, Kim Freriks, Ryanne Jaarsma, Euníce Then, Lynn Aarts, Sanne Bos, Melissa van Tuil, Florian Zwagemaker, Dennis Schmitz, Annelies Kroneman, Karim Hajji, Chantal Reusken, on behalf of the national COVID-19 response team |
| EPI_ISL_2862988 | Department of Bacteria, Parasites and Fungi, Statens Serum Institut, Copenhagen, Denmark | Statens Serum Institut Bioinformatics and Microbial Genomics | Danish Covid-19 Genome Consortium |
| EPI_ISL_2862989 | Dutch COVID-19 response team | National Institute for Public Health and the Environment (RIVM) | Adam Meijer, Harry Vennema, Dirk Eggink, Jeroen Cremer, Sharon van den Brink, Bas van der Veer, AnneMarie van den Brandt, Lisa Wijsman, Kim Freriks, Ryanne Jaarsma, Euníce Then, Lynn Aarts, Sanne Bos, Melissa van Tuil, Florian Zwagemaker, Dennis Schmitz, Annelies Kroneman, Karim Hajji, Chantal Reusken, on behalf of the national COVID-19 response team |
| EPI_ISL_2862990 | Department of Bacteria, Parasites and Fungi, Statens Serum Institut, Copenhagen, Denmark | Statens Serum Institut Bioinformatics and Microbial Genomics | Danish Covid-19 Genome Consortium |
| EPI_ISL_2862991 | Dutch COVID-19 response team | National Institute for Public Health and the Environment (RIVM) | Adam Meijer, Harry Vennema, Dirk Eggink, Jeroen Cremer, Sharon van den Brink, Bas van der Veer, AnneMarie van den Brandt, Lisa Wijsman, Kim Freriks, Ryanne Jaarsma, Euníce Then, Lynn Aarts, Sanne Bos, Melissa van Tuil, Florian Zwagemaker, Dennis Schmitz, Annelies Kroneman, Karim Hajji, Chantal Reusken, on behalf of the national COVID-19 response team |
| EPI_ISL_2862992 | Department of Bacteria, Parasites and Fungi, Statens Serum Institut, Copenhagen, Denmark | Statens Serum Institut Bioinformatics and Microbial Genomics | Danish Covid-19 Genome Consortium |
| EPI_ISL_2862993 | Dutch COVID-19 response team | National Institute for Public Health and the Environment (RIVM) | Adam Meijer, Harry Vennema, Dirk Eggink, Jeroen Cremer, Sharon van den Brink, Bas van der Veer, AnneMarie van den Brandt, Lisa Wijsman, Kim Freriks, Ryanne Jaarsma, Euníce Then, Lynn Aarts, Sanne Bos, Melissa van Tuil, Florian Zwagemaker, Dennis Schmitz, Annelies Kroneman, Karim Hajji, Chantal Reusken, on behalf of the national COVID-19 response team |
| EPI_ISL_2862994 | Department of Bacteria, Parasites and Fungi, Statens Serum Institut, Copenhagen, Denmark | Statens Serum Institut Bioinformatics and Microbial Genomics | Danish Covid-19 Genome Consortium |
| EPI_ISL_2862995 | Dutch COVID-19 response team | National Institute for Public Health and the Environment (RIVM) | Adam Meijer, Harry Vennema, Dirk Eggink, Jeroen Cremer, Sharon van den Brink, Bas van der Veer, AnneMarie van den Brandt, Lisa Wijsman, Kim Freriks, Ryanne Jaarsma, Euníce Then, Lynn Aarts, Sanne Bos, Melissa van Tuil, Florian Zwagemaker, Dennis Schmitz, Annelies Kroneman, Karim Hajji, Chantal Reusken, on behalf of the national COVID-19 response team |
| EPI_ISL_2862996 | Department of Bacteria, Parasites and Fungi, Statens Serum Institut, Copenhagen, Denmark | Statens Serum Institut Bioinformatics and Microbial Genomics | Danish Covid-19 Genome Consortium |
| EPI_ISL_2862997 | Dutch COVID-19 response team | National Institute for Public Health and the Environment (RIVM) | Adam Meijer, Harry Vennema, Dirk Eggink, Jeroen Cremer, Sharon van den Brink, Bas van der Veer, AnneMarie van den Brandt, Lisa Wijsman, Kim Freriks, Ryanne Jaarsma, Euníce Then, Lynn Aarts, Sanne Bos, Melissa van Tuil, Florian Zwagemaker, Dennis Schmitz, Annelies Kroneman, Karim Hajji, Chantal Reusken, on behalf of the national COVID-19 response team |
| EPI_ISL_2862998 | Department of Bacteria, Parasites and Fungi, Statens Serum Institut, Copenhagen, Denmark | Statens Serum Institut Bioinformatics and Microbial Genomics | Danish Covid-19 Genome Consortium |
| EPI_ISL_2862999 | Dutch COVID-19 response team | National Institute for Public Health and the Environment (RIVM) | Adam Meijer, Harry Vennema, Dirk Eggink, Jeroen Cremer, Sharon van den Brink, Bas van der Veer, AnneMarie van den Brandt, Lisa Wijsman, Kim Freriks, Ryanne Jaarsma, Euníce Then, Lynn Aarts, Sanne Bos, Melissa van Tuil, Florian Zwagemaker, Dennis Schmitz, Annelies Kroneman, Karim Hajji, Chantal Reusken, on behalf of the national COVID-19 response team |
| EPI_ISL_2863000 | Department of Bacteria, Parasites and Fungi, Statens Serum Institut, Copenhagen, Denmark | Statens Serum Institut Bioinformatics and Microbial Genomics | Danish Covid-19 Genome Consortium |
| EPI_ISL_2863001 | Dutch COVID-19 response team | National Institute for Public Health and the Environment (RIVM) | Adam Meijer, Harry Vennema, Dirk Eggink, Jeroen Cremer, Sharon van den Brink, Bas van der Veer, AnneMarie van den Brandt, Lisa Wijsman, Kim Freriks, Ryanne Jaarsma, Euníce Then, Lynn Aarts, Sanne Bos, Melissa van Tuil, Florian Zwagemaker, Dennis Schmitz, Annelies Kroneman, Karim Hajji, Chantal Reusken, on behalf of the national COVID-19 response team |
| EPI_ISL_2863002 | Department of Bacteria, Parasites and Fungi, Statens Serum Institut, Copenhagen, Denmark | Statens Serum Institut Bioinformatics and Microbial Genomics | Danish Covid-19 Genome Consortium |
| EPI_ISL_2863003 | Dutch COVID-19 response team | National Institute for Public Health and the Environment (RIVM) | Adam Meijer, Harry Vennema, Dirk Eggink, Jeroen Cremer, Sharon van den Brink, Bas van der Veer, AnneMarie van den Brandt, Lisa Wijsman, Kim Freriks, Ryanne Jaarsma, Euníce Then, Lynn Aarts, Sanne Bos, Melissa van Tuil, Florian Zwagemaker, Dennis Schmitz, Annelies Kroneman, Karim Hajji, Chantal Reusken, on behalf of the national COVID-19 response team |
| EPI_ISL_2863004 | Department of Bacteria, Parasites and Fungi, Statens Serum Institut, Copenhagen, Denmark | Statens Serum Institut Bioinformatics and Microbial Genomics | Danish Covid-19 Genome Consortium |

| | | | |
|---|---|---|---|
| EPI_ISL_2863005 | Dutch COVID-19 response team | National Institute for Public Health and the Environment (RIVM) | Adam Meijer, Harry Vennema, Dirk Eggink, Jeroen Cremer, Sharon van den Brink, Bas van der Veer, AnneMarie van den Brandt, Lisa Wijsman, Kim Freriks, Ryanne Jaarsma, Euníce Then, Lynn Aarts, Sanne Bos, Melissa van Tuil, Florian Zwagemaker, Dennis Schmitz, Annelies Kroneman, Karim Hajji, Chantal Reusken, on behalf of the national COVID-19 response team |
| EPI_ISL_2863006 | Department of Bacteria, Parasites and Fungi, Statens Serum Institut, Copenhagen, Denmark | Statens Serum Institut Bioinformatics and Microbial Genomics | Danish Covid-19 Genome Consortium |
| EPI_ISL_2863007 | Dutch COVID-19 response team | National Institute for Public Health and the Environment (RIVM) | Adam Meijer, Harry Vennema, Dirk Eggink, Jeroen Cremer, Sharon van den Brink, Bas van der Veer, AnneMarie van den Brandt, Lisa Wijsman, Kim Freriks, Ryanne Jaarsma, Euníce Then, Lynn Aarts, Sanne Bos, Melissa van Tuil, Florian Zwagemaker, Dennis Schmitz, Annelies Kroneman, Karim Hajji, Chantal Reusken, on behalf of the national COVID-19 response team |
| EPI_ISL_2863008 | Department of Bacteria, Parasites and Fungi, Statens Serum Institut, Copenhagen, Denmark | Statens Serum Institut Bioinformatics and Microbial Genomics | Danish Covid-19 Genome Consortium |
| EPI_ISL_2863009 | Dutch COVID-19 response team | National Institute for Public Health and the Environment (RIVM) | Adam Meijer, Harry Vennema, Dirk Eggink, Jeroen Cremer, Sharon van den Brink, Bas van der Veer, AnneMarie van den Brandt, Lisa Wijsman, Kim Freriks, Ryanne Jaarsma, Euníce Then, Lynn Aarts, Sanne Bos, Melissa van Tuil, Florian Zwagemaker, Dennis Schmitz, Annelies Kroneman, Karim Hajji, Chantal Reusken, on behalf of the national COVID-19 response team |
| EPI_ISL_2863010 | Department of Bacteria, Parasites and Fungi, Statens Serum Institut, Copenhagen, Denmark | Statens Serum Institut Bioinformatics and Microbial Genomics | Danish Covid-19 Genome Consortium |
| EPI_ISL_2863011, EPI_ISL_2863012 | Dutch COVID-19 response team | National Institute for Public Health and the Environment (RIVM) | Adam Meijer, Harry Vennema, Dirk Eggink, Jeroen Cremer, Sharon van den Brink, Bas van der Veer, AnneMarie van den Brandt, Lisa Wijsman, Kim Freriks, Ryanne Jaarsma, Euníce Then, Lynn Aarts, Sanne Bos, Melissa van Tuil, Florian Zwagemaker, Dennis Schmitz, Annelies Kroneman, Karim Hajji, Chantal Reusken, on behalf of the national COVID-19 response team |
| EPI_ISL_2863013 | Department of Bacteria, Parasites and Fungi, Statens Serum Institut, Copenhagen, Denmark | Statens Serum Institut Bioinformatics and Microbial Genomics | Danish Covid-19 Genome Consortium |
| EPI_ISL_2863014 | Dutch COVID-19 response team | National Institute for Public Health and the Environment (RIVM) | Adam Meijer, Harry Vennema, Dirk Eggink, Jeroen Cremer, Sharon van den Brink, Bas van der Veer, AnneMarie van den Brandt, Lisa Wijsman, Kim Freriks, Ryanne Jaarsma, Euníce Then, Lynn Aarts, Sanne Bos, Melissa van Tuil, Florian Zwagemaker, Dennis Schmitz, Annelies Kroneman, Karim Hajji, Chantal Reusken, on behalf of the national COVID-19 response team |
| EPI_ISL_2863015 | Department of Bacteria, Parasites and Fungi, Statens Serum Institut, Copenhagen, Denmark | Statens Serum Institut Bioinformatics and Microbial Genomics | Danish Covid-19 Genome Consortium |
| EPI_ISL_2863016 | Dutch COVID-19 response team | National Institute for Public Health and the Environment (RIVM) | Adam Meijer, Harry Vennema, Dirk Eggink, Jeroen Cremer, Sharon van den Brink, Bas van der Veer, AnneMarie van den Brandt, Lisa Wijsman, Kim Freriks, Ryanne Jaarsma, Euníce Then, Lynn Aarts, Sanne Bos, Melissa van Tuil, Florian Zwagemaker, Dennis Schmitz, Annelies Kroneman, Karim Hajji, Chantal Reusken, on behalf of the national COVID-19 response team |
| EPI_ISL_2863017 | Department of Bacteria, Parasites and Fungi, Statens Serum Institut, Copenhagen, Denmark | Statens Serum Institut Bioinformatics and Microbial Genomics | Danish Covid-19 Genome Consortium |
| EPI_ISL_2863018, EPI_ISL_2863020 | Dutch COVID-19 response team | National Institute for Public Health and the Environment (RIVM) | Adam Meijer, Harry Vennema, Dirk Eggink, Jeroen Cremer, Sharon van den Brink, Bas van der Veer, AnneMarie van den Brandt, Lisa Wijsman, Kim Freriks, Ryanne Jaarsma, Euníce Then, Lynn Aarts, Sanne Bos, Melissa van Tuil, Florian Zwagemaker, Dennis Schmitz, Annelies Kroneman, Karim Hajji, Chantal Reusken, on behalf of the national COVID-19 response team |
| EPI_ISL_2863021 | Department of Bacteria, Parasites and Fungi, Statens Serum Institut, Copenhagen, Denmark | Statens Serum Institut Bioinformatics and Microbial Genomics | Danish Covid-19 Genome Consortium |
| EPI_ISL_2863022 | Dutch COVID-19 response team | National Institute for Public Health and the Environment (RIVM) | Adam Meijer, Harry Vennema, Dirk Eggink, Jeroen Cremer, Sharon van den Brink, Bas van der Veer, AnneMarie van den Brandt, Lisa Wijsman, Kim Freriks, Ryanne Jaarsma, Euníce Then, Lynn Aarts, Sanne Bos, Melissa van Tuil, Florian Zwagemaker, Dennis Schmitz, Annelies Kroneman, Karim Hajji, Chantal Reusken, on behalf of the national COVID-19 response team |
| EPI_ISL_2863023 | Department of Bacteria, Parasites and Fungi, Statens Serum Institut, Copenhagen, Denmark | Statens Serum Institut Bioinformatics and Microbial Genomics | Danish Covid-19 Genome Consortium |
| EPI_ISL_2863024 | Dutch COVID-19 response team | National Institute for Public Health and the Environment (RIVM) | Adam Meijer, Harry Vennema, Dirk Eggink, Jeroen Cremer, Sharon van den Brink, Bas van der Veer, AnneMarie van den Brandt, Lisa Wijsman, Kim Freriks, Ryanne Jaarsma, Euníce Then, Lynn Aarts, Sanne Bos, Melissa van Tuil, Florian Zwagemaker, Dennis Schmitz, Annelies Kroneman, Karim Hajji, Chantal Reusken, on behalf of the national COVID-19 response team |
| EPI_ISL_2863025 | Department of Bacteria, Parasites and Fungi, Statens Serum Institut, Copenhagen, Denmark | Statens Serum Institut Bioinformatics and Microbial Genomics | Danish Covid-19 Genome Consortium |
| EPI_ISL_2863026 | Dutch COVID-19 response team | National Institute for Public Health and the Environment (RIVM) | Adam Meijer, Harry Vennema, Dirk Eggink, Jeroen Cremer, Sharon van den Brink, Bas van der Veer, AnneMarie van den Brandt, Lisa Wijsman, Kim Freriks, Ryanne Jaarsma, Euníce Then, Lynn Aarts, Sanne Bos, Melissa van Tuil, Florian Zwagemaker, Dennis Schmitz, Annelies Kroneman, Karim Hajji, Chantal Reusken, on behalf of the national COVID-19 response team |
| EPI_ISL_2863027 | Department of Bacteria, Parasites and Fungi, Statens Serum Institut, Copenhagen, Denmark | Statens Serum Institut Bioinformatics and Microbial Genomics | Danish Covid-19 Genome Consortium |
| EPI_ISL_2863028 | Dutch COVID-19 response team | National Institute for Public Health and the Environment (RIVM) | Adam Meijer, Harry Vennema, Dirk Eggink, Jeroen Cremer, Sharon van den Brink, Bas van der Veer, AnneMarie van den Brandt, Lisa Wijsman, Kim Freriks, Ryanne Jaarsma, Euníce Then, Lynn Aarts, Sanne Bos, Melissa van Tuil, Florian Zwagemaker, Dennis Schmitz, Annelies Kroneman, Karim Hajji, Chantal Reusken, on behalf of the national COVID-19 response team |
| EPI_ISL_2863029 | Department of Bacteria, Parasites and Fungi, Statens Serum Institut, Copenhagen, Denmark | Statens Serum Institut Bioinformatics and Microbial Genomics | Danish Covid-19 Genome Consortium |
| EPI_ISL_2863030 | Dutch COVID-19 response team | National Institute for Public Health and the Environment (RIVM) | Adam Meijer, Harry Vennema, Dirk Eggink, Jeroen Cremer, Sharon van den Brink, Bas van der Veer, AnneMarie van den Brandt, Lisa Wijsman, Kim Freriks, Ryanne Jaarsma, Euníce Then, Lynn Aarts, Sanne Bos, Melissa van Tuil, Florian Zwagemaker, Dennis Schmitz, Annelies Kroneman, Karim Hajji, Chantal Reusken, on behalf of the national COVID-19 response team |
| EPI_ISL_2863031 | Department of Bacteria, Parasites and Fungi, Statens Serum Institut, Copenhagen, Denmark | Statens Serum Institut Bioinformatics and Microbial Genomics | Danish Covid-19 Genome Consortium |
| EPI_ISL_2863032 | Dutch COVID-19 response team | National Institute for Public Health and the Environment (RIVM) | Adam Meijer, Harry Vennema, Dirk Eggink, Jeroen Cremer, Sharon van den Brink, Bas van der Veer, AnneMarie van den Brandt, Lisa Wijsman, Kim Freriks, Ryanne Jaarsma, Euníce Then, Lynn Aarts, Sanne Bos, Melissa van Tuil, Florian Zwagemaker, Dennis Schmitz, Annelies Kroneman, Karim Hajji, Chantal Reusken, on behalf of the national COVID-19 response team |
| EPI_ISL_2863033 | Department of Bacteria, Parasites and Fungi, Statens Serum Institut, Copenhagen, Denmark | Statens Serum Institut Bioinformatics and Microbial Genomics | Danish Covid-19 Genome Consortium |
| EPI_ISL_2863034 | Dutch COVID-19 response team | National Institute for Public Health and the Environment (RIVM) | Adam Meijer, Harry Vennema, Dirk Eggink, Jeroen Cremer, Sharon van den Brink, Bas van der Veer, AnneMarie van den Brandt, Lisa Wijsman, Kim Freriks, Ryanne Jaarsma, Euníce Then, Lynn Aarts, Sanne Bos, Melissa van Tuil, Florian Zwagemaker, Dennis Schmitz, Annelies Kroneman, Karim Hajji, Chantal Reusken, on behalf of the national COVID-19 response team |
| EPI_ISL_2863035 | Department of Bacteria, Parasites and Fungi, Statens Serum Institut, Copenhagen, Denmark | Statens Serum Institut Bioinformatics and Microbial Genomics | Danish Covid-19 Genome Consortium |
| EPI_ISL_2863036 | Dutch COVID-19 response team | National Institute for Public Health and the Environment (RIVM) | Adam Meijer, Harry Vennema, Dirk Eggink, Jeroen Cremer, Sharon van den Brink, Bas van der Veer, AnneMarie van den Brandt, Lisa Wijsman, Kim Freriks, Ryanne Jaarsma, Euníce Then, Lynn Aarts, Sanne Bos, Melissa van Tuil, Florian Zwagemaker, Dennis Schmitz, Annelies Kroneman, Karim Hajji, Chantal Reusken, on behalf of the national COVID-19 response team |
| EPI_ISL_2863037 | Department of Bacteria, Parasites and Fungi, Statens Serum Institut, Copenhagen, Denmark | Statens Serum Institut Bioinformatics and Microbial Genomics | Danish Covid-19 Genome Consortium |
| EPI_ISL_2863038 | Dutch COVID-19 response team | National Institute for Public Health and the Environment (RIVM) | Adam Meijer, Harry Vennema, Dirk Eggink, Jeroen Cremer, Sharon van den Brink, Bas van der Veer, AnneMarie van den Brandt, Lisa Wijsman, Kim Freriks, Ryanne Jaarsma, Euníce Then, Lynn Aarts, Sanne Bos, Melissa van Tuil, Florian Zwagemaker, Dennis Schmitz, Annelies Kroneman, Karim Hajji, Chantal Reusken, on behalf of the national COVID-19 response team |
| EPI_ISL_2863039 | Department of Bacteria, Parasites and Fungi, Statens Serum Institut, Copenhagen, Denmark | Statens Serum Institut Bioinformatics and Microbial Genomics | Danish Covid-19 Genome Consortium |
| EPI_ISL_2863040 | Dutch COVID-19 response team | National Institute for Public Health and the Environment (RIVM) | Adam Meijer, Harry Vennema, Dirk Eggink, Jeroen Cremer, Sharon van den Brink, Bas van der Veer, AnneMarie van den Brandt, Lisa Wijsman, Kim Freriks, Ryanne Jaarsma, Euníce Then, Lynn Aarts, Sanne Bos, Melissa van Tuil, Florian Zwagemaker, Dennis Schmitz, Annelies Kroneman, Karim Hajji, Chantal Reusken, on behalf of the national COVID-19 response team |
| EPI_ISL_2863041 | Department of Bacteria, Parasites and Fungi, Statens Serum Institut, Copenhagen, Denmark | Statens Serum Institut Bioinformatics and Microbial Genomics | Danish Covid-19 Genome Consortium |
| EPI_ISL_2863042 | Dutch COVID-19 response team | National Institute for Public Health and the Environment (RIVM) | Adam Meijer, Harry Vennema, Dirk Eggink, Jeroen Cremer, Sharon van den Brink, Bas van der Veer, AnneMarie van den Brandt, Lisa Wijsman, Kim Freriks, Ryanne Jaarsma, Euníce Then, Lynn Aarts, Sanne Bos, Melissa van Tuil, Florian Zwagemaker, Dennis Schmitz, Annelies Kroneman, Karim Hajji, Chantal Reusken, on behalf of the national COVID-19 response team |
| EPI_ISL_2863043 | Department of Bacteria, Parasites and Fungi, Statens Serum Institut, Copenhagen, Denmark | Statens Serum Institut Bioinformatics and Microbial Genomics | Danish Covid-19 Genome Consortium |
| EPI_ISL_2863044 | Dutch COVID-19 response team | National Institute for Public Health and the Environment (RIVM) | Adam Meijer, Harry Vennema, Dirk Eggink, Jeroen Cremer, Sharon van den Brink, Bas van der Veer, AnneMarie van den Brandt, Lisa Wijsman, Kim Freriks, Ryanne Jaarsma, Euníce Then, Lynn Aarts, Sanne Bos, Melissa van Tuil, Florian Zwagemaker, Dennis Schmitz, Annelies Kroneman, Karim Hajji, Chantal Reusken, on behalf of the national COVID-19 response team |
| EPI_ISL_2863045 | Department of Bacteria, Parasites and Fungi, Statens Serum Institut, Copenhagen, Denmark | Statens Serum Institut Bioinformatics and Microbial Genomics | Danish Covid-19 Genome Consortium |
| EPI_ISL_2863046 | Dutch COVID-19 response team | National Institute for Public Health and the Environment (RIVM) | Adam Meijer, Harry Vennema, Dirk Eggink, Jeroen Cremer, Sharon van den Brink, Bas van der Veer, AnneMarie van den Brandt, Lisa Wijsman, Kim Freriks, Ryanne Jaarsma, Euníce Then, Lynn Aarts, Sanne Bos, Melissa van Tuil, Florian Zwagemaker, Dennis Schmitz, Annelies Kroneman, Karim Hajji, Chantal Reusken, on behalf of the national COVID-19 response team |
| EPI_ISL_2863047 | Department of Bacteria, Parasites and Fungi, Statens Serum Institut, Copenhagen, Denmark | Statens Serum Institut Bioinformatics and Microbial Genomics | Danish Covid-19 Genome Consortium |
| EPI_ISL_2863048 | Dutch COVID-19 response team | National Institute for Public Health and the Environment (RIVM) | Adam Meijer, Harry Vennema, Dirk Eggink, Jeroen Cremer, Sharon van den Brink, Bas van der Veer, AnneMarie van den Brandt, Lisa Wijsman, Kim Freriks, Ryanne Jaarsma, Euníce Then, Lynn Aarts, Sanne Bos, Melissa van Tuil, Florian Zwagemaker, Dennis Schmitz, Annelies Kroneman, Karim Hajji, Chantal Reusken, on behalf of the national COVID-19 response team |
| EPI_ISL_2863049 | Department of Bacteria, Parasites and Fungi, Statens Serum Institut, Copenhagen, Denmark | Statens Serum Institut Bioinformatics and Microbial Genomics | Danish Covid-19 Genome Consortium |
| EPI_ISL_2863050, EPI_ISL_2863051 | Dutch COVID-19 response team | National Institute for Public Health and the Environment (RIVM) | Adam Meijer, Harry Vennema, Dirk Eggink, Jeroen Cremer, Sharon van den Brink, Bas van der Veer, AnneMarie van den Brandt, Lisa Wijsman, Kim Freriks, Ryanne Jaarsma, Euníce Then, Lynn Aarts, Sanne Bos, Melissa van Tuil, Florian Zwagemaker, Dennis Schmitz, Annelies Kroneman, Karim Hajji, Chantal Reusken, on behalf of the national COVID-19 response team |
| EPI_ISL_2863052, EPI_ISL_2863054 | Department of Bacteria, Parasites and Fungi, Statens Serum Institut, Copenhagen, Denmark | Statens Serum Institut Bioinformatics and Microbial Genomics | Danish Covid-19 Genome Consortium |
| EPI_ISL_2863055 | Dutch COVID-19 response team | National Institute for Public Health and the Environment (RIVM) | Adam Meijer, Harry Vennema, Dirk Eggink, Jeroen Cremer, Sharon van den Brink, Bas van der Veer, AnneMarie van den Brandt, Lisa Wijsman, Kim Freriks, Ryanne Jaarsma, Euníce Then, Lynn Aarts, Sanne Bos, Melissa van Tuil, Florian Zwagemaker, Dennis Schmitz, Annelies Kroneman, Karim Hajji, Chantal Reusken, on behalf of the national COVID-19 response team |

| | | | |
|---|---|---|---|
| EPI_ISL_2863056, EPI_ISL_2863058, EPI_ISL_2863060 | Department of Bacteria, Parasites and Fungi, Statens Serum Institut, Copenhagen, Denmark | Statens Serum Institut Bioinformatics and Microbial Genomics | Danish Covid-19 Genome Consortium |
| EPI_ISL_2863061 | Dutch COVID-19 response team | National Institute for Public Health and the Environment (RIVM) | Adam Meijer, Harry Vennema, Dirk Eggink, Jeroen Cremer, Sharon van den Brink, Bas van der Veer, AnneMarie van den Brandt, Lisa Wijsman, Kim Freriks, Ryanne Jaarsma, Euníce Then, Lynn Aarts, Sanne Bos, Melissa van Tuil, Florian Zwagemaker, Dennis Schmitz, Annelies Kroneman, Karim Hajji, Chantal Reusken, on behalf of the national COVID-19 response team |
| EPI_ISL_2863062 | Department of Bacteria, Parasites and Fungi, Statens Serum Institut, Copenhagen, Denmark | Statens Serum Institut Bioinformatics and Microbial Genomics | Danish Covid-19 Genome Consortium |
| EPI_ISL_2863063 | Dutch COVID-19 response team | National Institute for Public Health and the Environment (RIVM) | Adam Meijer, Harry Vennema, Dirk Eggink, Jeroen Cremer, Sharon van den Brink, Bas van der Veer, AnneMarie van den Brandt, Lisa Wijsman, Kim Freriks, Ryanne Jaarsma, Euníce Then, Lynn Aarts, Sanne Bos, Melissa van Tuil, Florian Zwagemaker, Dennis Schmitz, Annelies Kroneman, Karim Hajji, Chantal Reusken, on behalf of the national COVID-19 response team |
| EPI_ISL_2863064 | Department of Bacteria, Parasites and Fungi, Statens Serum Institut, Copenhagen, Denmark | Statens Serum Institut Bioinformatics and Microbial Genomics | Danish Covid-19 Genome Consortium |
| EPI_ISL_2863065 | Dutch COVID-19 response team | National Institute for Public Health and the Environment (RIVM) | Adam Meijer, Harry Vennema, Dirk Eggink, Jeroen Cremer, Sharon van den Brink, Bas van der Veer, AnneMarie van den Brandt, Lisa Wijsman, Kim Freriks, Ryanne Jaarsma, Euníce Then, Lynn Aarts, Sanne Bos, Melissa van Tuil, Florian Zwagemaker, Dennis Schmitz, Annelies Kroneman, Karim Hajji, Chantal Reusken, on behalf of the national COVID-19 response team |
| EPI_ISL_2863066 | Department of Bacteria, Parasites and Fungi, Statens Serum Institut, Copenhagen, Denmark | Statens Serum Institut Bioinformatics and Microbial Genomics | Danish Covid-19 Genome Consortium |
| EPI_ISL_2863067 | Dutch COVID-19 response team | National Institute for Public Health and the Environment (RIVM) | Adam Meijer, Harry Vennema, Dirk Eggink, Jeroen Cremer, Sharon van den Brink, Bas van der Veer, AnneMarie van den Brandt, Lisa Wijsman, Kim Freriks, Ryanne Jaarsma, Euníce Then, Lynn Aarts, Sanne Bos, Melissa van Tuil, Florian Zwagemaker, Dennis Schmitz, Annelies Kroneman, Karim Hajji, Chantal Reusken, on behalf of the national COVID-19 response team |
| EPI_ISL_2863068 | Department of Bacteria, Parasites and Fungi, Statens Serum Institut, Copenhagen, Denmark | Statens Serum Institut Bioinformatics and Microbial Genomics | Danish Covid-19 Genome Consortium |
| EPI_ISL_2863069 | Dutch COVID-19 response team | National Institute for Public Health and the Environment (RIVM) | Adam Meijer, Harry Vennema, Dirk Eggink, Jeroen Cremer, Sharon van den Brink, Bas van der Veer, AnneMarie van den Brandt, Lisa Wijsman, Kim Freriks, Ryanne Jaarsma, Euníce Then, Lynn Aarts, Sanne Bos, Melissa van Tuil, Florian Zwagemaker, Dennis Schmitz, Annelies Kroneman, Karim Hajji, Chantal Reusken, on behalf of the national COVID-19 response team |
| EPI_ISL_2863070 | Department of Bacteria, Parasites and Fungi, Statens Serum Institut, Copenhagen, Denmark | Statens Serum Institut Bioinformatics and Microbial Genomics | Danish Covid-19 Genome Consortium |
| EPI_ISL_2863071 | Dutch COVID-19 response team | National Institute for Public Health and the Environment (RIVM) | Adam Meijer, Harry Vennema, Dirk Eggink, Jeroen Cremer, Sharon van den Brink, Bas van der Veer, AnneMarie van den Brandt, Lisa Wijsman, Kim Freriks, Ryanne Jaarsma, Euníce Then, Lynn Aarts, Sanne Bos, Melissa van Tuil, Florian Zwagemaker, Dennis Schmitz, Annelies Kroneman, Karim Hajji, Chantal Reusken, on behalf of the national COVID-19 response team |
| EPI_ISL_2863072 | Department of Bacteria, Parasites and Fungi, Statens Serum Institut, Copenhagen, Denmark | Statens Serum Institut Bioinformatics and Microbial Genomics | Danish Covid-19 Genome Consortium |
| EPI_ISL_2863073 | Dutch COVID-19 response team | National Institute for Public Health and the Environment (RIVM) | Adam Meijer, Harry Vennema, Dirk Eggink, Jeroen Cremer, Sharon van den Brink, Bas van der Veer, AnneMarie van den Brandt, Lisa Wijsman, Kim Freriks, Ryanne Jaarsma, Euníce Then, Lynn Aarts, Sanne Bos, Melissa van Tuil, Florian Zwagemaker, Dennis Schmitz, Annelies Kroneman, Karim Hajji, Chantal Reusken, on behalf of the national COVID-19 response team |
| EPI_ISL_2863074 | Department of Bacteria, Parasites and Fungi, Statens Serum Institut, Copenhagen, Denmark | Statens Serum Institut Bioinformatics and Microbial Genomics | Danish Covid-19 Genome Consortium |
| EPI_ISL_2863075 | Dutch COVID-19 response team | National Institute for Public Health and the Environment (RIVM) | Adam Meijer, Harry Vennema, Dirk Eggink, Jeroen Cremer, Sharon van den Brink, Bas van der Veer, AnneMarie van den Brandt, Lisa Wijsman, Kim Freriks, Ryanne Jaarsma, Euníce Then, Lynn Aarts, Sanne Bos, Melissa van Tuil, Florian Zwagemaker, Dennis Schmitz, Annelies Kroneman, Karim Hajji, Chantal Reusken, on behalf of the national COVID-19 response team |
| EPI_ISL_2863076 | Department of Bacteria, Parasites and Fungi, Statens Serum Institut, Copenhagen, Denmark | Statens Serum Institut Bioinformatics and Microbial Genomics | Danish Covid-19 Genome Consortium |
| EPI_ISL_2863077 | Dutch COVID-19 response team | National Institute for Public Health and the Environment (RIVM) | Adam Meijer, Harry Vennema, Dirk Eggink, Jeroen Cremer, Sharon van den Brink, Bas van der Veer, AnneMarie van den Brandt, Lisa Wijsman, Kim Freriks, Ryanne Jaarsma, Euníce Then, Lynn Aarts, Sanne Bos, Melissa van Tuil, Florian Zwagemaker, Dennis Schmitz, Annelies Kroneman, Karim Hajji, Chantal Reusken, on behalf of the national COVID-19 response team |
| EPI_ISL_2863078 | Department of Bacteria, Parasites and Fungi, Statens Serum Institut, Copenhagen, Denmark | Statens Serum Institut Bioinformatics and Microbial Genomics | Danish Covid-19 Genome Consortium |
| EPI_ISL_2863079, EPI_ISL_2863080 | Dutch COVID-19 response team | National Institute for Public Health and the Environment (RIVM) | Adam Meijer, Harry Vennema, Dirk Eggink, Jeroen Cremer, Sharon van den Brink, Bas van der Veer, AnneMarie van den Brandt, Lisa Wijsman, Kim Freriks, Ryanne Jaarsma, Euníce Then, Lynn Aarts, Sanne Bos, Melissa van Tuil, Florian Zwagemaker, Dennis Schmitz, Annelies Kroneman, Karim Hajji, Chantal Reusken, on behalf of the national COVID-19 response team |
| EPI_ISL_2863081 | Department of Bacteria, Parasites and Fungi, Statens Serum Institut, Copenhagen, Denmark | Statens Serum Institut Bioinformatics and Microbial Genomics | Danish Covid-19 Genome Consortium |
| EPI_ISL_2863082 | Dutch COVID-19 response team | National Institute for Public Health and the Environment (RIVM) | Adam Meijer, Harry Vennema, Dirk Eggink, Jeroen Cremer, Sharon van den Brink, Bas van der Veer, AnneMarie van den Brandt, Lisa Wijsman, Kim Freriks, Ryanne Jaarsma, Euníce Then, Lynn Aarts, Sanne Bos, Melissa van Tuil, Florian Zwagemaker, Dennis Schmitz, Annelies Kroneman, Karim Hajji, Chantal Reusken, on behalf of the national COVID-19 response team |
| EPI_ISL_2863083 | Department of Bacteria, Parasites and Fungi, Statens Serum Institut, Copenhagen, Denmark | Statens Serum Institut Bioinformatics and Microbial Genomics | Danish Covid-19 Genome Consortium |
| EPI_ISL_2863084 | Dutch COVID-19 response team | National Institute for Public Health and the Environment (RIVM) | Adam Meijer, Harry Vennema, Dirk Eggink, Jeroen Cremer, Sharon van den Brink, Bas van der Veer, AnneMarie van den Brandt, Lisa Wijsman, Kim Freriks, Ryanne Jaarsma, Euníce Then, Lynn Aarts, Sanne Bos, Melissa van Tuil, Florian Zwagemaker, Dennis Schmitz, Annelies Kroneman, Karim Hajji, Chantal Reusken, on behalf of the national COVID-19 response team |
| EPI_ISL_2863085 | Department of Bacteria, Parasites and Fungi, Statens Serum Institut, Copenhagen, Denmark | Statens Serum Institut Bioinformatics and Microbial Genomics | Danish Covid-19 Genome Consortium |
| EPI_ISL_2863086 | Dutch COVID-19 response team | National Institute for Public Health and the Environment (RIVM) | Adam Meijer, Harry Vennema, Dirk Eggink, Jeroen Cremer, Sharon van den Brink, Bas van der Veer, AnneMarie van den Brandt, Lisa Wijsman, Kim Freriks, Ryanne Jaarsma, Euníce Then, Lynn Aarts, Sanne Bos, Melissa van Tuil, Florian Zwagemaker, Dennis Schmitz, Annelies Kroneman, Karim Hajji, Chantal Reusken, on behalf of the national COVID-19 response team |
| EPI_ISL_2863087 | Department of Bacteria, Parasites and Fungi, Statens Serum Institut, Copenhagen, Denmark | Statens Serum Institut Bioinformatics and Microbial Genomics | Danish Covid-19 Genome Consortium |
| EPI_ISL_2863088 | Dutch COVID-19 response team | National Institute for Public Health and the Environment (RIVM) | Adam Meijer, Harry Vennema, Dirk Eggink, Jeroen Cremer, Sharon van den Brink, Bas van der Veer, AnneMarie van den Brandt, Lisa Wijsman, Kim Freriks, Ryanne Jaarsma, Euníce Then, Lynn Aarts, Sanne Bos, Melissa van Tuil, Florian Zwagemaker, Dennis Schmitz, Annelies Kroneman, Karim Hajji, Chantal Reusken, on behalf of the national COVID-19 response team |
| EPI_ISL_2863089 | Department of Bacteria, Parasites and Fungi, Statens Serum Institut, Copenhagen, Denmark | Statens Serum Institut Bioinformatics and Microbial Genomics | Danish Covid-19 Genome Consortium |
| EPI_ISL_2863090 | Dutch COVID-19 response team | National Institute for Public Health and the Environment (RIVM) | Adam Meijer, Harry Vennema, Dirk Eggink, Jeroen Cremer, Sharon van den Brink, Bas van der Veer, AnneMarie van den Brandt, Lisa Wijsman, Kim Freriks, Ryanne Jaarsma, Euníce Then, Lynn Aarts, Sanne Bos, Melissa van Tuil, Florian Zwagemaker, Dennis Schmitz, Annelies Kroneman, Karim Hajji, Chantal Reusken, on behalf of the national COVID-19 response team |
| EPI_ISL_2863091 | Department of Bacteria, Parasites and Fungi, Statens Serum Institut, Copenhagen, Denmark | Statens Serum Institut Bioinformatics and Microbial Genomics | Danish Covid-19 Genome Consortium |
| EPI_ISL_2863092, EPI_ISL_2863093 | Dutch COVID-19 response team | National Institute for Public Health and the Environment (RIVM) | Adam Meijer, Harry Vennema, Dirk Eggink, Jeroen Cremer, Sharon van den Brink, Bas van der Veer, AnneMarie van den Brandt, Lisa Wijsman, Kim Freriks, Ryanne Jaarsma, Euníce Then, Lynn Aarts, Sanne Bos, Melissa van Tuil, Florian Zwagemaker, Dennis Schmitz, Annelies Kroneman, Karim Hajji, Chantal Reusken, on behalf of the national COVID-19 response team |
| EPI_ISL_2863094 | Department of Bacteria, Parasites and Fungi, Statens Serum Institut, Copenhagen, Denmark | Statens Serum Institut Bioinformatics and Microbial Genomics | Danish Covid-19 Genome Consortium |
| EPI_ISL_2863095 | Dutch COVID-19 response team | National Institute for Public Health and the Environment (RIVM) | Adam Meijer, Harry Vennema, Dirk Eggink, Jeroen Cremer, Sharon van den Brink, Bas van der Veer, AnneMarie van den Brandt, Lisa Wijsman, Kim Freriks, Ryanne Jaarsma, Euníce Then, Lynn Aarts, Sanne Bos, Melissa van Tuil, Florian Zwagemaker, Dennis Schmitz, Annelies Kroneman, Karim Hajji, Chantal Reusken, on behalf of the national COVID-19 response team |
| EPI_ISL_2863096 | Department of Bacteria, Parasites and Fungi, Statens Serum Institut, Copenhagen, Denmark | Statens Serum Institut Bioinformatics and Microbial Genomics | Danish Covid-19 Genome Consortium |
| EPI_ISL_2863097 | Dutch COVID-19 response team | National Institute for Public Health and the Environment (RIVM) | Adam Meijer, Harry Vennema, Dirk Eggink, Jeroen Cremer, Sharon van den Brink, Bas van der Veer, AnneMarie van den Brandt, Lisa Wijsman, Kim Freriks, Ryanne Jaarsma, Euníce Then, Lynn Aarts, Sanne Bos, Melissa van Tuil, Florian Zwagemaker, Dennis Schmitz, Annelies Kroneman, Karim Hajji, Chantal Reusken, on behalf of the national COVID-19 response team |
| EPI_ISL_2863098 | Department of Bacteria, Parasites and Fungi, Statens Serum Institut, Copenhagen, Denmark | Statens Serum Institut Bioinformatics and Microbial Genomics | Danish Covid-19 Genome Consortium |
| EPI_ISL_2863099 | Dutch COVID-19 response team | National Institute for Public Health and the Environment (RIVM) | Adam Meijer, Harry Vennema, Dirk Eggink, Jeroen Cremer, Sharon van den Brink, Bas van der Veer, AnneMarie van den Brandt, Lisa Wijsman, Kim Freriks, Ryanne Jaarsma, Euníce Then, Lynn Aarts, Sanne Bos, Melissa van Tuil, Florian Zwagemaker, Dennis Schmitz, Annelies Kroneman, Karim Hajji, Chantal Reusken, on behalf of the national COVID-19 response team |
| EPI_ISL_2863100 | Department of Bacteria, Parasites and Fungi, Statens Serum Institut, Copenhagen, Denmark | Statens Serum Institut Bioinformatics and Microbial Genomics | Danish Covid-19 Genome Consortium |
| EPI_ISL_2863101 | Dutch COVID-19 response team | National Institute for Public Health and the Environment (RIVM) | Adam Meijer, Harry Vennema, Dirk Eggink, Jeroen Cremer, Sharon van den Brink, Bas van der Veer, AnneMarie van den Brandt, Lisa Wijsman, Kim Freriks, Ryanne Jaarsma, Euníce Then, Lynn Aarts, Sanne Bos, Melissa van Tuil, Florian Zwagemaker, Dennis Schmitz, Annelies Kroneman, Karim Hajji, Chantal Reusken, on behalf of the national COVID-19 response team |
| EPI_ISL_2863102 | Department of Bacteria, Parasites and Fungi, Statens Serum Institut, Copenhagen, Denmark | Statens Serum Institut Bioinformatics and Microbial Genomics | Danish Covid-19 Genome Consortium |
| EPI_ISL_2863103 | Dutch COVID-19 response team | National Institute for Public Health and the Environment (RIVM) | Adam Meijer, Harry Vennema, Dirk Eggink, Jeroen Cremer, Sharon van den Brink, Bas van der Veer, AnneMarie van den Brandt, Lisa Wijsman, Kim Freriks, Ryanne Jaarsma, Euníce Then, Lynn Aarts, Sanne Bos, Melissa van Tuil, Florian Zwagemaker, Dennis Schmitz, Annelies Kroneman, Karim Hajji, Chantal Reusken, on behalf of the national COVID-19 response team |
| EPI_ISL_2863104 | Department of Bacteria, Parasites and Fungi, Statens Serum Institut, Copenhagen, Denmark | Statens Serum Institut Bioinformatics and Microbial Genomics | Danish Covid-19 Genome Consortium |
| EPI_ISL_2863105 | Dutch COVID-19 response team | National Institute for Public Health and the Environment (RIVM) | Adam Meijer, Harry Vennema, Dirk Eggink, Jeroen Cremer, Sharon van den Brink, Bas van der Veer, AnneMarie van den Brandt, Lisa Wijsman, Kim Freriks, Ryanne Jaarsma, Euníce Then, Lynn Aarts, Sanne Bos, Melissa van Tuil, Florian Zwagemaker, Dennis Schmitz, Annelies Kroneman, Karim Hajji, Chantal Reusken, on behalf of the national COVID-19 response team |
| EPI_ISL_2863106 | Department of Bacteria, Parasites and Fungi, Statens Serum Institut, Copenhagen, Denmark | Statens Serum Institut Bioinformatics and Microbial Genomics | Danish Covid-19 Genome Consortium |

| | | | |
|---|---|---|---|
| EPI_ISL_2863107, EPI_ISL_2863108 | Dutch COVID-19 response team | National Institute for Public Health and the Environment (RIVM) | Adam Meijer, Harry Vennema, Dirk Eggink, Jeroen Cremer, Sharon van den Brink, Bas van der Veer, AnneMarie van den Brandt, Lisa Wijsman, Kim Freriks, Ryanne Jaarsma, Euníce Then, Lynn Aarts, Sanne Bos, Melissa van Tuil, Florian Zwagemaker, Dennis Schmitz, Annelies Kroneman, Karim Hajji, Chantal Reusken, on behalf of the national COVID-19 response team |
| EPI_ISL_2863109 | Department of Bacteria, Parasites and Fungi, Statens Serum Institut, Copenhagen, Denmark | Statens Serum Institut Bioinformatics and Microbial Genomics | Danish Covid-19 Genome Consortium |
| EPI_ISL_2863110 | Dutch COVID-19 response team | National Institute for Public Health and the Environment (RIVM) | Adam Meijer, Harry Vennema, Dirk Eggink, Jeroen Cremer, Sharon van den Brink, Bas van der Veer, AnneMarie van den Brandt, Lisa Wijsman, Kim Freriks, Ryanne Jaarsma, Euníce Then, Lynn Aarts, Sanne Bos, Melissa van Tuil, Florian Zwagemaker, Dennis Schmitz, Annelies Kroneman, Karim Hajji, Chantal Reusken, on behalf of the national COVID-19 response team |
| EPI_ISL_2863111 | Department of Bacteria, Parasites and Fungi, Statens Serum Institut, Copenhagen, Denmark | Statens Serum Institut Bioinformatics and Microbial Genomics | Danish Covid-19 Genome Consortium |
| EPI_ISL_2863112 | Dutch COVID-19 response team | National Institute for Public Health and the Environment (RIVM) | Adam Meijer, Harry Vennema, Dirk Eggink, Jeroen Cremer, Sharon van den Brink, Bas van der Veer, AnneMarie van den Brandt, Lisa Wijsman, Kim Freriks, Ryanne Jaarsma, Euníce Then, Lynn Aarts, Sanne Bos, Melissa van Tuil, Florian Zwagemaker, Dennis Schmitz, Annelies Kroneman, Karim Hajji, Chantal Reusken, on behalf of the national COVID-19 response team |
| EPI_ISL_2863113 | Department of Bacteria, Parasites and Fungi, Statens Serum Institut, Copenhagen, Denmark | Statens Serum Institut Bioinformatics and Microbial Genomics | Danish Covid-19 Genome Consortium |
| EPI_ISL_2863114 | Dutch COVID-19 response team | National Institute for Public Health and the Environment (RIVM) | Adam Meijer, Harry Vennema, Dirk Eggink, Jeroen Cremer, Sharon van den Brink, Bas van der Veer, AnneMarie van den Brandt, Lisa Wijsman, Kim Freriks, Ryanne Jaarsma, Euníce Then, Lynn Aarts, Sanne Bos, Melissa van Tuil, Florian Zwagemaker, Dennis Schmitz, Annelies Kroneman, Karim Hajji, Chantal Reusken, on behalf of the national COVID-19 response team |
| EPI_ISL_2863115 | Department of Bacteria, Parasites and Fungi, Statens Serum Institut, Copenhagen, Denmark | Statens Serum Institut Bioinformatics and Microbial Genomics | Danish Covid-19 Genome Consortium |
| EPI_ISL_2863116 | Dutch COVID-19 response team | National Institute for Public Health and the Environment (RIVM) | Adam Meijer, Harry Vennema, Dirk Eggink, Jeroen Cremer, Sharon van den Brink, Bas van der Veer, AnneMarie van den Brandt, Lisa Wijsman, Kim Freriks, Ryanne Jaarsma, Euníce Then, Lynn Aarts, Sanne Bos, Melissa van Tuil, Florian Zwagemaker, Dennis Schmitz, Annelies Kroneman, Karim Hajji, Chantal Reusken, on behalf of the national COVID-19 response team |
| EPI_ISL_2863117 | Department of Bacteria, Parasites and Fungi, Statens Serum Institut, Copenhagen, Denmark | Statens Serum Institut Bioinformatics and Microbial Genomics | Danish Covid-19 Genome Consortium |
| EPI_ISL_2863118 | Dutch COVID-19 response team | National Institute for Public Health and the Environment (RIVM) | Adam Meijer, Harry Vennema, Dirk Eggink, Jeroen Cremer, Sharon van den Brink, Bas van der Veer, AnneMarie van den Brandt, Lisa Wijsman, Kim Freriks, Ryanne Jaarsma, Euníce Then, Lynn Aarts, Sanne Bos, Melissa van Tuil, Florian Zwagemaker, Dennis Schmitz, Annelies Kroneman, Karim Hajji, Chantal Reusken, on behalf of the national COVID-19 response team |
| EPI_ISL_2863119 | Department of Bacteria, Parasites and Fungi, Statens Serum Institut, Copenhagen, Denmark | Statens Serum Institut Bioinformatics and Microbial Genomics | Danish Covid-19 Genome Consortium |
| EPI_ISL_2863120 | Dutch COVID-19 response team | National Institute for Public Health and the Environment (RIVM) | Adam Meijer, Harry Vennema, Dirk Eggink, Jeroen Cremer, Sharon van den Brink, Bas van der Veer, AnneMarie van den Brandt, Lisa Wijsman, Kim Freriks, Ryanne Jaarsma, Euníce Then, Lynn Aarts, Sanne Bos, Melissa van Tuil, Florian Zwagemaker, Dennis Schmitz, Annelies Kroneman, Karim Hajji, Chantal Reusken, on behalf of the national COVID-19 response team |
| EPI_ISL_2863121 | Department of Bacteria, Parasites and Fungi, Statens Serum Institut, Copenhagen, Denmark | Statens Serum Institut Bioinformatics and Microbial Genomics | Danish Covid-19 Genome Consortium |
| EPI_ISL_2863122 | Dutch COVID-19 response team | National Institute for Public Health and the Environment (RIVM) | Adam Meijer, Harry Vennema, Dirk Eggink, Jeroen Cremer, Sharon van den Brink, Bas van der Veer, AnneMarie van den Brandt, Lisa Wijsman, Kim Freriks, Ryanne Jaarsma, Euníce Then, Lynn Aarts, Sanne Bos, Melissa van Tuil, Florian Zwagemaker, Dennis Schmitz, Annelies Kroneman, Karim Hajji, Chantal Reusken, on behalf of the national COVID-19 response team |
| EPI_ISL_2863123 | Department of Bacteria, Parasites and Fungi, Statens Serum Institut, Copenhagen, Denmark | Statens Serum Institut Bioinformatics and Microbial Genomics | Danish Covid-19 Genome Consortium |
| EPI_ISL_2863124 | Dutch COVID-19 response team | National Institute for Public Health and the Environment (RIVM) | Adam Meijer, Harry Vennema, Dirk Eggink, Jeroen Cremer, Sharon van den Brink, Bas van der Veer, AnneMarie van den Brandt, Lisa Wijsman, Kim Freriks, Ryanne Jaarsma, Euníce Then, Lynn Aarts, Sanne Bos, Melissa van Tuil, Florian Zwagemaker, Dennis Schmitz, Annelies Kroneman, Karim Hajji, Chantal Reusken, on behalf of the national COVID-19 response team |
| EPI_ISL_2863125 | Department of Bacteria, Parasites and Fungi, Statens Serum Institut, Copenhagen, Denmark | Statens Serum Institut Bioinformatics and Microbial Genomics | Danish Covid-19 Genome Consortium |
| EPI_ISL_2863126 | Dutch COVID-19 response team | National Institute for Public Health and the Environment (RIVM) | Adam Meijer, Harry Vennema, Dirk Eggink, Jeroen Cremer, Sharon van den Brink, Bas van der Veer, AnneMarie van den Brandt, Lisa Wijsman, Kim Freriks, Ryanne Jaarsma, Euníce Then, Lynn Aarts, Sanne Bos, Melissa van Tuil, Florian Zwagemaker, Dennis Schmitz, Annelies Kroneman, Karim Hajji, Chantal Reusken, on behalf of the national COVID-19 response team |
| EPI_ISL_2863127 | Department of Bacteria, Parasites and Fungi, Statens Serum Institut, Copenhagen, Denmark | Statens Serum Institut Bioinformatics and Microbial Genomics | Danish Covid-19 Genome Consortium |
| EPI_ISL_2863128, EPI_ISL_2863129 | Dutch COVID-19 response team | National Institute for Public Health and the Environment (RIVM) | Adam Meijer, Harry Vennema, Dirk Eggink, Jeroen Cremer, Sharon van den Brink, Bas van der Veer, AnneMarie van den Brandt, Lisa Wijsman, Kim Freriks, Ryanne Jaarsma, Euníce Then, Lynn Aarts, Sanne Bos, Melissa van Tuil, Florian Zwagemaker, Dennis Schmitz, Annelies Kroneman, Karim Hajji, Chantal Reusken, on behalf of the national COVID-19 response team |
| EPI_ISL_2863130 | Department of Bacteria, Parasites and Fungi, Statens Serum Institut, Copenhagen, Denmark | Statens Serum Institut Bioinformatics and Microbial Genomics | Danish Covid-19 Genome Consortium |
| EPI_ISL_2863131 | Dutch COVID-19 response team | National Institute for Public Health and the Environment (RIVM) | Adam Meijer, Harry Vennema, Dirk Eggink, Jeroen Cremer, Sharon van den Brink, Bas van der Veer, AnneMarie van den Brandt, Lisa Wijsman, Kim Freriks, Ryanne Jaarsma, Euníce Then, Lynn Aarts, Sanne Bos, Melissa van Tuil, Florian Zwagemaker, Dennis Schmitz, Annelies Kroneman, Karim Hajji, Chantal Reusken, on behalf of the national COVID-19 response team |
| EPI_ISL_2863132 | Department of Bacteria, Parasites and Fungi, Statens Serum Institut, Copenhagen, Denmark | Statens Serum Institut Bioinformatics and Microbial Genomics | Danish Covid-19 Genome Consortium |
| EPI_ISL_2863133 | Dutch COVID-19 response team | National Institute for Public Health and the Environment (RIVM) | Adam Meijer, Harry Vennema, Dirk Eggink, Jeroen Cremer, Sharon van den Brink, Bas van der Veer, AnneMarie van den Brandt, Lisa Wijsman, Kim Freriks, Ryanne Jaarsma, Euníce Then, Lynn Aarts, Sanne Bos, Melissa van Tuil, Florian Zwagemaker, Dennis Schmitz, Annelies Kroneman, Karim Hajji, Chantal Reusken, on behalf of the national COVID-19 response team |
| EPI_ISL_2863134 | Department of Bacteria, Parasites and Fungi, Statens Serum Institut, Copenhagen, Denmark | Statens Serum Institut Bioinformatics and Microbial Genomics | Danish Covid-19 Genome Consortium |
| EPI_ISL_2863135 | Dutch COVID-19 response team | National Institute for Public Health and the Environment (RIVM) | Adam Meijer, Harry Vennema, Dirk Eggink, Jeroen Cremer, Sharon van den Brink, Bas van der Veer, AnneMarie van den Brandt, Lisa Wijsman, Kim Freriks, Ryanne Jaarsma, Euníce Then, Lynn Aarts, Sanne Bos, Melissa van Tuil, Florian Zwagemaker, Dennis Schmitz, Annelies Kroneman, Karim Hajji, Chantal Reusken, on behalf of the national COVID-19 response team |
| EPI_ISL_2863136 | Department of Bacteria, Parasites and Fungi, Statens Serum Institut, Copenhagen, Denmark | Statens Serum Institut Bioinformatics and Microbial Genomics | Danish Covid-19 Genome Consortium |
| EPI_ISL_2863137 | Dutch COVID-19 response team | National Institute for Public Health and the Environment (RIVM) | Adam Meijer, Harry Vennema, Dirk Eggink, Jeroen Cremer, Sharon van den Brink, Bas van der Veer, AnneMarie van den Brandt, Lisa Wijsman, Kim Freriks, Ryanne Jaarsma, Euníce Then, Lynn Aarts, Sanne Bos, Melissa van Tuil, Florian Zwagemaker, Dennis Schmitz, Annelies Kroneman, Karim Hajji, Chantal Reusken, on behalf of the national COVID-19 response team |
| EPI_ISL_2863138 | Department of Bacteria, Parasites and Fungi, Statens Serum Institut, Copenhagen, Denmark | Statens Serum Institut Bioinformatics and Microbial Genomics | Danish Covid-19 Genome Consortium |
| EPI_ISL_2863139 | Dutch COVID-19 response team | National Institute for Public Health and the Environment (RIVM) | Adam Meijer, Harry Vennema, Dirk Eggink, Jeroen Cremer, Sharon van den Brink, Bas van der Veer, AnneMarie van den Brandt, Lisa Wijsman, Kim Freriks, Ryanne Jaarsma, Euníce Then, Lynn Aarts, Sanne Bos, Melissa van Tuil, Florian Zwagemaker, Dennis Schmitz, Annelies Kroneman, Karim Hajji, Chantal Reusken, on behalf of the national COVID-19 response team |
| EPI_ISL_2863140 | Department of Bacteria, Parasites and Fungi, Statens Serum Institut, Copenhagen, Denmark | Statens Serum Institut Bioinformatics and Microbial Genomics | Danish Covid-19 Genome Consortium |
| EPI_ISL_2863141 | Dutch COVID-19 response team | National Institute for Public Health and the Environment (RIVM) | Adam Meijer, Harry Vennema, Dirk Eggink, Jeroen Cremer, Sharon van den Brink, Bas van der Veer, AnneMarie van den Brandt, Lisa Wijsman, Kim Freriks, Ryanne Jaarsma, Euníce Then, Lynn Aarts, Sanne Bos, Melissa van Tuil, Florian Zwagemaker, Dennis Schmitz, Annelies Kroneman, Karim Hajji, Chantal Reusken, on behalf of the national COVID-19 response team |
| EPI_ISL_2863142 | Department of Bacteria, Parasites and Fungi, Statens Serum Institut, Copenhagen, Denmark | Statens Serum Institut Bioinformatics and Microbial Genomics | Danish Covid-19 Genome Consortium |
| EPI_ISL_2863143 | Dutch COVID-19 response team | National Institute for Public Health and the Environment (RIVM) | Adam Meijer, Harry Vennema, Dirk Eggink, Jeroen Cremer, Sharon van den Brink, Bas van der Veer, AnneMarie van den Brandt, Lisa Wijsman, Kim Freriks, Ryanne Jaarsma, Euníce Then, Lynn Aarts, Sanne Bos, Melissa van Tuil, Florian Zwagemaker, Dennis Schmitz, Annelies Kroneman, Karim Hajji, Chantal Reusken, on behalf of the national COVID-19 response team |
| EPI_ISL_2863144 | Department of Bacteria, Parasites and Fungi, Statens Serum Institut, Copenhagen, Denmark | Statens Serum Institut Bioinformatics and Microbial Genomics | Danish Covid-19 Genome Consortium |
| EPI_ISL_2863145 | Dutch COVID-19 response team | National Institute for Public Health and the Environment (RIVM) | Adam Meijer, Harry Vennema, Dirk Eggink, Jeroen Cremer, Sharon van den Brink, Bas van der Veer, AnneMarie van den Brandt, Lisa Wijsman, Kim Freriks, Ryanne Jaarsma, Euníce Then, Lynn Aarts, Sanne Bos, Melissa van Tuil, Florian Zwagemaker, Dennis Schmitz, Annelies Kroneman, Karim Hajji, Chantal Reusken, on behalf of the national COVID-19 response team |
| EPI_ISL_2863146 | Department of Bacteria, Parasites and Fungi, Statens Serum Institut, Copenhagen, Denmark | Statens Serum Institut Bioinformatics and Microbial Genomics | Danish Covid-19 Genome Consortium |
| EPI_ISL_2863147 | Dutch COVID-19 response team | National Institute for Public Health and the Environment (RIVM) | Adam Meijer, Harry Vennema, Dirk Eggink, Jeroen Cremer, Sharon van den Brink, Bas van der Veer, AnneMarie van den Brandt, Lisa Wijsman, Kim Freriks, Ryanne Jaarsma, Euníce Then, Lynn Aarts, Sanne Bos, Melissa van Tuil, Florian Zwagemaker, Dennis Schmitz, Annelies Kroneman, Karim Hajji, Chantal Reusken, on behalf of the national COVID-19 response team |
| EPI_ISL_2863148 | Department of Bacteria, Parasites and Fungi, Statens Serum Institut, Copenhagen, Denmark | Statens Serum Institut Bioinformatics and Microbial Genomics | Danish Covid-19 Genome Consortium |
| EPI_ISL_2863149 | Dutch COVID-19 response team | National Institute for Public Health and the Environment (RIVM) | Adam Meijer, Harry Vennema, Dirk Eggink, Jeroen Cremer, Sharon van den Brink, Bas van der Veer, AnneMarie van den Brandt, Lisa Wijsman, Kim Freriks, Ryanne Jaarsma, Euníce Then, Lynn Aarts, Sanne Bos, Melissa van Tuil, Florian Zwagemaker, Dennis Schmitz, Annelies Kroneman, Karim Hajji, Chantal Reusken, on behalf of the national COVID-19 response team |
| EPI_ISL_2863150 | Department of Bacteria, Parasites and Fungi, Statens Serum Institut, Copenhagen, Denmark | Statens Serum Institut Bioinformatics and Microbial Genomics | Danish Covid-19 Genome Consortium |
| EPI_ISL_2863151 | Dutch COVID-19 response team | National Institute for Public Health and the Environment (RIVM) | Adam Meijer, Harry Vennema, Dirk Eggink, Jeroen Cremer, Sharon van den Brink, Bas van der Veer, AnneMarie van den Brandt, Lisa Wijsman, Kim Freriks, Ryanne Jaarsma, Euníce Then, Lynn Aarts, Sanne Bos, Melissa van Tuil, Florian Zwagemaker, Dennis Schmitz, Annelies Kroneman, Karim Hajji, Chantal Reusken, on behalf of the national COVID-19 response team |
| EPI_ISL_2863152 | Department of Bacteria, Parasites and Fungi, Statens Serum Institut, Copenhagen, Denmark | Statens Serum Institut Bioinformatics and Microbial Genomics | Danish Covid-19 Genome Consortium |
| EPI_ISL_2863153 | Dutch COVID-19 response team | National Institute for Public Health and the Environment (RIVM) | Adam Meijer, Harry Vennema, Dirk Eggink, Jeroen Cremer, Sharon van den Brink, Bas van der Veer, AnneMarie van den Brandt, Lisa Wijsman, Kim Freriks, Ryanne Jaarsma, Euníce Then, Lynn Aarts, Sanne Bos, Melissa van Tuil, Florian Zwagemaker, Dennis Schmitz, Annelies Kroneman, Karim Hajji, Chantal Reusken, on behalf of the national COVID-19 response team |

| | | | |
|---|---|---|---|
| EPI_ISL_2863154 | Department of Bacteria, Parasites and Fungi, Statens Serum Institut, Copenhagen, Denmark | Statens Serum Institut Bioinformatics and Microbial Genomics | Danish Covid-19 Genome Consortium |
| EPI_ISL_2863155, EPI_ISL_2863156 | Dutch COVID-19 response team | National Institute for Public Health and the Environment (RIVM) | Adam Meijer, Harry Vennema, Dirk Eggink, Jeroen Cremer, Sharon van den Brink, Bas van der Veer, AnneMarie van den Brandt, Lisa Wijsman, Kim Freriks, Ryanne Jaarsma, Euníce Then, Lynn Aarts, Sanne Bos, Melissa van Tuil, Florian Zwagemaker, Dennis Schmitz, Annelies Kroneman, Karim Hajji, Chantal Reusken, on behalf of the national COVID-19 response team |
| EPI_ISL_2863157 | Department of Bacteria, Parasites and Fungi, Statens Serum Institut, Copenhagen, Denmark | Statens Serum Institut Bioinformatics and Microbial Genomics | Danish Covid-19 Genome Consortium |
| EPI_ISL_2863158 | Dutch COVID-19 response team | National Institute for Public Health and the Environment (RIVM) | Adam Meijer, Harry Vennema, Dirk Eggink, Jeroen Cremer, Sharon van den Brink, Bas van der Veer, AnneMarie van den Brandt, Lisa Wijsman, Kim Freriks, Ryanne Jaarsma, Euníce Then, Lynn Aarts, Sanne Bos, Melissa van Tuil, Florian Zwagemaker, Dennis Schmitz, Annelies Kroneman, Karim Hajji, Chantal Reusken, on behalf of the national COVID-19 response team |
| EPI_ISL_2863159 | Department of Bacteria, Parasites and Fungi, Statens Serum Institut, Copenhagen, Denmark | Statens Serum Institut Bioinformatics and Microbial Genomics | Danish Covid-19 Genome Consortium |
| EPI_ISL_2863160, EPI_ISL_2863162 | Dutch COVID-19 response team | National Institute for Public Health and the Environment (RIVM) | Adam Meijer, Harry Vennema, Dirk Eggink, Jeroen Cremer, Sharon van den Brink, Bas van der Veer, AnneMarie van den Brandt, Lisa Wijsman, Kim Freriks, Ryanne Jaarsma, Euníce Then, Lynn Aarts, Sanne Bos, Melissa van Tuil, Florian Zwagemaker, Dennis Schmitz, Annelies Kroneman, Karim Hajji, Chantal Reusken, on behalf of the national COVID-19 response team |
| EPI_ISL_2863163 | Department of Bacteria, Parasites and Fungi, Statens Serum Institut, Copenhagen, Denmark | Statens Serum Institut Bioinformatics and Microbial Genomics | Danish Covid-19 Genome Consortium |
| EPI_ISL_2863164 | Dutch COVID-19 response team | National Institute for Public Health and the Environment (RIVM) | Adam Meijer, Harry Vennema, Dirk Eggink, Jeroen Cremer, Sharon van den Brink, Bas van der Veer, AnneMarie van den Brandt, Lisa Wijsman, Kim Freriks, Ryanne Jaarsma, Euníce Then, Lynn Aarts, Sanne Bos, Melissa van Tuil, Florian Zwagemaker, Dennis Schmitz, Annelies Kroneman, Karim Hajji, Chantal Reusken, on behalf of the national COVID-19 response team |
| EPI_ISL_2863165 | Department of Bacteria, Parasites and Fungi, Statens Serum Institut, Copenhagen, Denmark | Statens Serum Institut Bioinformatics and Microbial Genomics | Danish Covid-19 Genome Consortium |
| EPI_ISL_2863166 | Dutch COVID-19 response team | National Institute for Public Health and the Environment (RIVM) | Adam Meijer, Harry Vennema, Dirk Eggink, Jeroen Cremer, Sharon van den Brink, Bas van der Veer, AnneMarie van den Brandt, Lisa Wijsman, Kim Freriks, Ryanne Jaarsma, Euníce Then, Lynn Aarts, Sanne Bos, Melissa van Tuil, Florian Zwagemaker, Dennis Schmitz, Annelies Kroneman, Karim Hajji, Chantal Reusken, on behalf of the national COVID-19 response team |
| EPI_ISL_2863167 | Department of Bacteria, Parasites and Fungi, Statens Serum Institut, Copenhagen, Denmark | Statens Serum Institut Bioinformatics and Microbial Genomics | Danish Covid-19 Genome Consortium |
| EPI_ISL_2863168 | Dutch COVID-19 response team | National Institute for Public Health and the Environment (RIVM) | Adam Meijer, Harry Vennema, Dirk Eggink, Jeroen Cremer, Sharon van den Brink, Bas van der Veer, AnneMarie van den Brandt, Lisa Wijsman, Kim Freriks, Ryanne Jaarsma, Euníce Then, Lynn Aarts, Sanne Bos, Melissa van Tuil, Florian Zwagemaker, Dennis Schmitz, Annelies Kroneman, Karim Hajji, Chantal Reusken, on behalf of the national COVID-19 response team |
| EPI_ISL_2863169 | Department of Bacteria, Parasites and Fungi, Statens Serum Institut, Copenhagen, Denmark | Statens Serum Institut Bioinformatics and Microbial Genomics | Danish Covid-19 Genome Consortium |
| EPI_ISL_2863170 | Dutch COVID-19 response team | National Institute for Public Health and the Environment (RIVM) | Adam Meijer, Harry Vennema, Dirk Eggink, Jeroen Cremer, Sharon van den Brink, Bas van der Veer, AnneMarie van den Brandt, Lisa Wijsman, Kim Freriks, Ryanne Jaarsma, Euníce Then, Lynn Aarts, Sanne Bos, Melissa van Tuil, Florian Zwagemaker, Dennis Schmitz, Annelies Kroneman, Karim Hajji, Chantal Reusken, on behalf of the national COVID-19 response team |
| EPI_ISL_2863171 | Department of Bacteria, Parasites and Fungi, Statens Serum Institut, Copenhagen, Denmark | Statens Serum Institut Bioinformatics and Microbial Genomics | Danish Covid-19 Genome Consortium |
| EPI_ISL_2863172 | Dutch COVID-19 response team | National Institute for Public Health and the Environment (RIVM) | Adam Meijer, Harry Vennema, Dirk Eggink, Jeroen Cremer, Sharon van den Brink, Bas van der Veer, AnneMarie van den Brandt, Lisa Wijsman, Kim Freriks, Ryanne Jaarsma, Euníce Then, Lynn Aarts, Sanne Bos, Melissa van Tuil, Florian Zwagemaker, Dennis Schmitz, Annelies Kroneman, Karim Hajji, Chantal Reusken, on behalf of the national COVID-19 response team |
| EPI_ISL_2863173 | Department of Bacteria, Parasites and Fungi, Statens Serum Institut, Copenhagen, Denmark | Statens Serum Institut Bioinformatics and Microbial Genomics | Danish Covid-19 Genome Consortium |
| EPI_ISL_2863174 | Dutch COVID-19 response team | National Institute for Public Health and the Environment (RIVM) | Adam Meijer, Harry Vennema, Dirk Eggink, Jeroen Cremer, Sharon van den Brink, Bas van der Veer, AnneMarie van den Brandt, Lisa Wijsman, Kim Freriks, Ryanne Jaarsma, Euníce Then, Lynn Aarts, Sanne Bos, Melissa van Tuil, Florian Zwagemaker, Dennis Schmitz, Annelies Kroneman, Karim Hajji, Chantal Reusken, on behalf of the national COVID-19 response team |
| EPI_ISL_2863175 | Department of Bacteria, Parasites and Fungi, Statens Serum Institut, Copenhagen, Denmark | Statens Serum Institut Bioinformatics and Microbial Genomics | Danish Covid-19 Genome Consortium |
| EPI_ISL_2863176 | Dutch COVID-19 response team | National Institute for Public Health and the Environment (RIVM) | Adam Meijer, Harry Vennema, Dirk Eggink, Jeroen Cremer, Sharon van den Brink, Bas van der Veer, AnneMarie van den Brandt, Lisa Wijsman, Kim Freriks, Ryanne Jaarsma, Euníce Then, Lynn Aarts, Sanne Bos, Melissa van Tuil, Florian Zwagemaker, Dennis Schmitz, Annelies Kroneman, Karim Hajji, Chantal Reusken, on behalf of the national COVID-19 response team |
| EPI_ISL_2863177 | Department of Bacteria, Parasites and Fungi, Statens Serum Institut, Copenhagen, Denmark | Statens Serum Institut Bioinformatics and Microbial Genomics | Danish Covid-19 Genome Consortium |
| EPI_ISL_2863178 | Dutch COVID-19 response team | National Institute for Public Health and the Environment (RIVM) | Adam Meijer, Harry Vennema, Dirk Eggink, Jeroen Cremer, Sharon van den Brink, Bas van der Veer, AnneMarie van den Brandt, Lisa Wijsman, Kim Freriks, Ryanne Jaarsma, Euníce Then, Lynn Aarts, Sanne Bos, Melissa van Tuil, Florian Zwagemaker, Dennis Schmitz, Annelies Kroneman, Karim Hajji, Chantal Reusken, on behalf of the national COVID-19 response team |
| EPI_ISL_2863179 | Department of Bacteria, Parasites and Fungi, Statens Serum Institut, Copenhagen, Denmark | Statens Serum Institut Bioinformatics and Microbial Genomics | Danish Covid-19 Genome Consortium |
| EPI_ISL_2863180, EPI_ISL_2863181 | Dutch COVID-19 response team | National Institute for Public Health and the Environment (RIVM) | Adam Meijer, Harry Vennema, Dirk Eggink, Jeroen Cremer, Sharon van den Brink, Bas van der Veer, AnneMarie van den Brandt, Lisa Wijsman, Kim Freriks, Ryanne Jaarsma, Euníce Then, Lynn Aarts, Sanne Bos, Melissa van Tuil, Florian Zwagemaker, Dennis Schmitz, Annelies Kroneman, Karim Hajji, Chantal Reusken, on behalf of the national COVID-19 response team |
| EPI_ISL_2863182 | Department of Bacteria, Parasites and Fungi, Statens Serum Institut, Copenhagen, Denmark | Statens Serum Institut Bioinformatics and Microbial Genomics | Danish Covid-19 Genome Consortium |
| EPI_ISL_2863183 | Dutch COVID-19 response team | National Institute for Public Health and the Environment (RIVM) | Adam Meijer, Harry Vennema, Dirk Eggink, Jeroen Cremer, Sharon van den Brink, Bas van der Veer, AnneMarie van den Brandt, Lisa Wijsman, Kim Freriks, Ryanne Jaarsma, Euníce Then, Lynn Aarts, Sanne Bos, Melissa van Tuil, Florian Zwagemaker, Dennis Schmitz, Annelies Kroneman, Karim Hajji, Chantal Reusken, on behalf of the national COVID-19 response team |
| EPI_ISL_2863184 | Department of Bacteria, Parasites and Fungi, Statens Serum Institut, Copenhagen, Denmark | Statens Serum Institut Bioinformatics and Microbial Genomics | Danish Covid-19 Genome Consortium |
| EPI_ISL_2863185 | Dutch COVID-19 response team | National Institute for Public Health and the Environment (RIVM) | Adam Meijer, Harry Vennema, Dirk Eggink, Jeroen Cremer, Sharon van den Brink, Bas van der Veer, AnneMarie van den Brandt, Lisa Wijsman, Kim Freriks, Ryanne Jaarsma, Euníce Then, Lynn Aarts, Sanne Bos, Melissa van Tuil, Florian Zwagemaker, Dennis Schmitz, Annelies Kroneman, Karim Hajji, Chantal Reusken, on behalf of the national COVID-19 response team |
| EPI_ISL_2863186 | Department of Bacteria, Parasites and Fungi, Statens Serum Institut, Copenhagen, Denmark | Statens Serum Institut Bioinformatics and Microbial Genomics | Danish Covid-19 Genome Consortium |
| EPI_ISL_2863187 | Dutch COVID-19 response team | National Institute for Public Health and the Environment (RIVM) | Adam Meijer, Harry Vennema, Dirk Eggink, Jeroen Cremer, Sharon van den Brink, Bas van der Veer, AnneMarie van den Brandt, Lisa Wijsman, Kim Freriks, Ryanne Jaarsma, Euníce Then, Lynn Aarts, Sanne Bos, Melissa van Tuil, Florian Zwagemaker, Dennis Schmitz, Annelies Kroneman, Karim Hajji, Chantal Reusken, on behalf of the national COVID-19 response team |
| EPI_ISL_2863188 | Department of Bacteria, Parasites and Fungi, Statens Serum Institut, Copenhagen, Denmark | Statens Serum Institut Bioinformatics and Microbial Genomics | Danish Covid-19 Genome Consortium |
| EPI_ISL_2863189 | Dutch COVID-19 response team | National Institute for Public Health and the Environment (RIVM) | Adam Meijer, Harry Vennema, Dirk Eggink, Jeroen Cremer, Sharon van den Brink, Bas van der Veer, AnneMarie van den Brandt, Lisa Wijsman, Kim Freriks, Ryanne Jaarsma, Euníce Then, Lynn Aarts, Sanne Bos, Melissa van Tuil, Florian Zwagemaker, Dennis Schmitz, Annelies Kroneman, Karim Hajji, Chantal Reusken, on behalf of the national COVID-19 response team |
| EPI_ISL_2863190 | Department of Bacteria, Parasites and Fungi, Statens Serum Institut, Copenhagen, Denmark | Statens Serum Institut Bioinformatics and Microbial Genomics | Danish Covid-19 Genome Consortium |
| EPI_ISL_2863191 | Dutch COVID-19 response team | National Institute for Public Health and the Environment (RIVM) | Adam Meijer, Harry Vennema, Dirk Eggink, Jeroen Cremer, Sharon van den Brink, Bas van der Veer, AnneMarie van den Brandt, Lisa Wijsman, Kim Freriks, Ryanne Jaarsma, Euníce Then, Lynn Aarts, Sanne Bos, Melissa van Tuil, Florian Zwagemaker, Dennis Schmitz, Annelies Kroneman, Karim Hajji, Chantal Reusken, on behalf of the national COVID-19 response team |
| EPI_ISL_2863192 | Department of Bacteria, Parasites and Fungi, Statens Serum Institut, Copenhagen, Denmark | Statens Serum Institut Bioinformatics and Microbial Genomics | Danish Covid-19 Genome Consortium |
| EPI_ISL_2863193 | Dutch COVID-19 response team | National Institute for Public Health and the Environment (RIVM) | Adam Meijer, Harry Vennema, Dirk Eggink, Jeroen Cremer, Sharon van den Brink, Bas van der Veer, AnneMarie van den Brandt, Lisa Wijsman, Kim Freriks, Ryanne Jaarsma, Euníce Then, Lynn Aarts, Sanne Bos, Melissa van Tuil, Florian Zwagemaker, Dennis Schmitz, Annelies Kroneman, Karim Hajji, Chantal Reusken, on behalf of the national COVID-19 response team |
| EPI_ISL_2863194 | Department of Bacteria, Parasites and Fungi, Statens Serum Institut, Copenhagen, Denmark | Statens Serum Institut Bioinformatics and Microbial Genomics | Danish Covid-19 Genome Consortium |
| EPI_ISL_2863195 | Dutch COVID-19 response team | National Institute for Public Health and the Environment (RIVM) | Adam Meijer, Harry Vennema, Dirk Eggink, Jeroen Cremer, Sharon van den Brink, Bas van der Veer, AnneMarie van den Brandt, Lisa Wijsman, Kim Freriks, Ryanne Jaarsma, Euníce Then, Lynn Aarts, Sanne Bos, Melissa van Tuil, Florian Zwagemaker, Dennis Schmitz, Annelies Kroneman, Karim Hajji, Chantal Reusken, on behalf of the national COVID-19 response team |
| EPI_ISL_2863196 | Department of Bacteria, Parasites and Fungi, Statens Serum Institut, Copenhagen, Denmark | Statens Serum Institut Bioinformatics and Microbial Genomics | Danish Covid-19 Genome Consortium |
| EPI_ISL_2863197 | Dutch COVID-19 response team | National Institute for Public Health and the Environment (RIVM) | Adam Meijer, Harry Vennema, Dirk Eggink, Jeroen Cremer, Sharon van den Brink, Bas van der Veer, AnneMarie van den Brandt, Lisa Wijsman, Kim Freriks, Ryanne Jaarsma, Euníce Then, Lynn Aarts, Sanne Bos, Melissa van Tuil, Florian Zwagemaker, Dennis Schmitz, Annelies Kroneman, Karim Hajji, Chantal Reusken, on behalf of the national COVID-19 response team |
| EPI_ISL_2863198 | Department of Bacteria, Parasites and Fungi, Statens Serum Institut, Copenhagen, Denmark | Statens Serum Institut Bioinformatics and Microbial Genomics | Danish Covid-19 Genome Consortium |
| EPI_ISL_2863199 | Dutch COVID-19 response team | National Institute for Public Health and the Environment (RIVM) | Adam Meijer, Harry Vennema, Dirk Eggink, Jeroen Cremer, Sharon van den Brink, Bas van der Veer, AnneMarie van den Brandt, Lisa Wijsman, Kim Freriks, Ryanne Jaarsma, Euníce Then, Lynn Aarts, Sanne Bos, Melissa van Tuil, Florian Zwagemaker, Dennis Schmitz, Annelies Kroneman, Karim Hajji, Chantal Reusken, on behalf of the national COVID-19 response team |
| EPI_ISL_2863200 | Department of Bacteria, Parasites and Fungi, Statens Serum Institut, Copenhagen, Denmark | Statens Serum Institut Bioinformatics and Microbial Genomics | Danish Covid-19 Genome Consortium |
| EPI_ISL_2863201 | Dutch COVID-19 response team | National Institute for Public Health and the Environment (RIVM) | Adam Meijer, Harry Vennema, Dirk Eggink, Jeroen Cremer, Sharon van den Brink, Bas van der Veer, AnneMarie van den Brandt, Lisa Wijsman, Kim Freriks, Ryanne Jaarsma, Euníce Then, Lynn Aarts, Sanne Bos, Melissa van Tuil, Florian Zwagemaker, Dennis Schmitz, Annelies Kroneman, Karim Hajji, Chantal Reusken, on behalf of the national COVID-19 response team |
| EPI_ISL_2863202 | Department of Bacteria, Parasites and Fungi, Statens Serum Institut, Copenhagen, Denmark | Statens Serum Institut Bioinformatics and Microbial Genomics | Danish Covid-19 Genome Consortium |

| | | | |
|---|---|---|---|
| EPI_ISL_2863203 | Dutch COVID-19 response team | National Institute for Public Health and the Environment (RIVM) | Adam Meijer, Harry Vennema, Dirk Eggink, Jeroen Cremer, Sharon van den Brink, Bas van der Veer, AnneMarie van den Brandt, Lisa Wijsman, Kim Freriks, Ryanne Jaarsma, Euníce Then, Lynn Aarts, Sanne Bos, Melissa van Tuil, Florian Zwagemaker, Dennis Schmitz, Annelies Kroneman, Karim Hajji, Chantal Reusken, on behalf of the national COVID-19 response team |
| EPI_ISL_2863204 | Department of Bacteria, Parasites and Fungi, Statens Serum Institut, Copenhagen, Denmark | Statens Serum Institut Bioinformatics and Microbial Genomics | Danish Covid-19 Genome Consortium |
| EPI_ISL_2863205 | Dutch COVID-19 response team | National Institute for Public Health and the Environment (RIVM) | Adam Meijer, Harry Vennema, Dirk Eggink, Jeroen Cremer, Sharon van den Brink, Bas van der Veer, AnneMarie van den Brandt, Lisa Wijsman, Kim Freriks, Ryanne Jaarsma, Euníce Then, Lynn Aarts, Sanne Bos, Melissa van Tuil, Florian Zwagemaker, Dennis Schmitz, Annelies Kroneman, Karim Hajji, Chantal Reusken, on behalf of the national COVID-19 response team |
| EPI_ISL_2863206 | Department of Bacteria, Parasites and Fungi, Statens Serum Institut, Copenhagen, Denmark | Statens Serum Institut Bioinformatics and Microbial Genomics | Danish Covid-19 Genome Consortium |
| EPI_ISL_2863207 | Dutch COVID-19 response team | National Institute for Public Health and the Environment (RIVM) | Adam Meijer, Harry Vennema, Dirk Eggink, Jeroen Cremer, Sharon van den Brink, Bas van der Veer, AnneMarie van den Brandt, Lisa Wijsman, Kim Freriks, Ryanne Jaarsma, Euníce Then, Lynn Aarts, Sanne Bos, Melissa van Tuil, Florian Zwagemaker, Dennis Schmitz, Annelies Kroneman, Karim Hajji, Chantal Reusken, on behalf of the national COVID-19 response team |
| EPI_ISL_2863208 | Department of Bacteria, Parasites and Fungi, Statens Serum Institut, Copenhagen, Denmark | Statens Serum Institut Bioinformatics and Microbial Genomics | Danish Covid-19 Genome Consortium |
| EPI_ISL_2863209, EPI_ISL_2863210 | Dutch COVID-19 response team | National Institute for Public Health and the Environment (RIVM) | Adam Meijer, Harry Vennema, Dirk Eggink, Jeroen Cremer, Sharon van den Brink, Bas van der Veer, AnneMarie van den Brandt, Lisa Wijsman, Kim Freriks, Ryanne Jaarsma, Euníce Then, Lynn Aarts, Sanne Bos, Melissa van Tuil, Florian Zwagemaker, Dennis Schmitz, Annelies Kroneman, Karim Hajji, Chantal Reusken, on behalf of the national COVID-19 response team |
| EPI_ISL_2863211 | Department of Bacteria, Parasites and Fungi, Statens Serum Institut, Copenhagen, Denmark | Statens Serum Institut Bioinformatics and Microbial Genomics | Danish Covid-19 Genome Consortium |
| EPI_ISL_2863212 | Dutch COVID-19 response team | National Institute for Public Health and the Environment (RIVM) | Adam Meijer, Harry Vennema, Dirk Eggink, Jeroen Cremer, Sharon van den Brink, Bas van der Veer, AnneMarie van den Brandt, Lisa Wijsman, Kim Freriks, Ryanne Jaarsma, Euníce Then, Lynn Aarts, Sanne Bos, Melissa van Tuil, Florian Zwagemaker, Dennis Schmitz, Annelies Kroneman, Karim Hajji, Chantal Reusken, on behalf of the national COVID-19 response team |
| EPI_ISL_2863213 | Department of Bacteria, Parasites and Fungi, Statens Serum Institut, Copenhagen, Denmark | Statens Serum Institut Bioinformatics and Microbial Genomics | Danish Covid-19 Genome Consortium |
| EPI_ISL_2863214 | Dutch COVID-19 response team | National Institute for Public Health and the Environment (RIVM) | Adam Meijer, Harry Vennema, Dirk Eggink, Jeroen Cremer, Sharon van den Brink, Bas van der Veer, AnneMarie van den Brandt, Lisa Wijsman, Kim Freriks, Ryanne Jaarsma, Euníce Then, Lynn Aarts, Sanne Bos, Melissa van Tuil, Florian Zwagemaker, Dennis Schmitz, Annelies Kroneman, Karim Hajji, Chantal Reusken, on behalf of the national COVID-19 response team |
| EPI_ISL_2863215 | Department of Bacteria, Parasites and Fungi, Statens Serum Institut, Copenhagen, Denmark | Statens Serum Institut Bioinformatics and Microbial Genomics | Danish Covid-19 Genome Consortium |
| EPI_ISL_2863216 | Dutch COVID-19 response team | National Institute for Public Health and the Environment (RIVM) | Adam Meijer, Harry Vennema, Dirk Eggink, Jeroen Cremer, Sharon van den Brink, Bas van der Veer, AnneMarie van den Brandt, Lisa Wijsman, Kim Freriks, Ryanne Jaarsma, Euníce Then, Lynn Aarts, Sanne Bos, Melissa van Tuil, Florian Zwagemaker, Dennis Schmitz, Annelies Kroneman, Karim Hajji, Chantal Reusken, on behalf of the national COVID-19 response team |
| EPI_ISL_2863217 | Department of Bacteria, Parasites and Fungi, Statens Serum Institut, Copenhagen, Denmark | Statens Serum Institut Bioinformatics and Microbial Genomics | Danish Covid-19 Genome Consortium |
| EPI_ISL_2863218 | Dutch COVID-19 response team | National Institute for Public Health and the Environment (RIVM) | Adam Meijer, Harry Vennema, Dirk Eggink, Jeroen Cremer, Sharon van den Brink, Bas van der Veer, AnneMarie van den Brandt, Lisa Wijsman, Kim Freriks, Ryanne Jaarsma, Euníce Then, Lynn Aarts, Sanne Bos, Melissa van Tuil, Florian Zwagemaker, Dennis Schmitz, Annelies Kroneman, Karim Hajji, Chantal Reusken, on behalf of the national COVID-19 response team |
| EPI_ISL_2863219 | Department of Bacteria, Parasites and Fungi, Statens Serum Institut, Copenhagen, Denmark | Statens Serum Institut Bioinformatics and Microbial Genomics | Danish Covid-19 Genome Consortium |
| EPI_ISL_2863220 | Dutch COVID-19 response team | National Institute for Public Health and the Environment (RIVM) | Adam Meijer, Harry Vennema, Dirk Eggink, Jeroen Cremer, Sharon van den Brink, Bas van der Veer, AnneMarie van den Brandt, Lisa Wijsman, Kim Freriks, Ryanne Jaarsma, Euníce Then, Lynn Aarts, Sanne Bos, Melissa van Tuil, Florian Zwagemaker, Dennis Schmitz, Annelies Kroneman, Karim Hajji, Chantal Reusken, on behalf of the national COVID-19 response team |
| EPI_ISL_2863221 | Department of Bacteria, Parasites and Fungi, Statens Serum Institut, Copenhagen, Denmark | Statens Serum Institut Bioinformatics and Microbial Genomics | Danish Covid-19 Genome Consortium |
| EPI_ISL_2863222 | Dutch COVID-19 response team | National Institute for Public Health and the Environment (RIVM) | Adam Meijer, Harry Vennema, Dirk Eggink, Jeroen Cremer, Sharon van den Brink, Bas van der Veer, AnneMarie van den Brandt, Lisa Wijsman, Kim Freriks, Ryanne Jaarsma, Euníce Then, Lynn Aarts, Sanne Bos, Melissa van Tuil, Florian Zwagemaker, Dennis Schmitz, Annelies Kroneman, Karim Hajji, Chantal Reusken, on behalf of the national COVID-19 response team |
| EPI_ISL_2863223 | Department of Bacteria, Parasites and Fungi, Statens Serum Institut, Copenhagen, Denmark | Statens Serum Institut Bioinformatics and Microbial Genomics | Danish Covid-19 Genome Consortium |
| EPI_ISL_2863224 | Dutch COVID-19 response team | National Institute for Public Health and the Environment (RIVM) | Adam Meijer, Harry Vennema, Dirk Eggink, Jeroen Cremer, Sharon van den Brink, Bas van der Veer, AnneMarie van den Brandt, Lisa Wijsman, Kim Freriks, Ryanne Jaarsma, Euníce Then, Lynn Aarts, Sanne Bos, Melissa van Tuil, Florian Zwagemaker, Dennis Schmitz, Annelies Kroneman, Karim Hajji, Chantal Reusken, on behalf of the national COVID-19 response team |
| EPI_ISL_2863225 | Department of Bacteria, Parasites and Fungi, Statens Serum Institut, Copenhagen, Denmark | Statens Serum Institut Bioinformatics and Microbial Genomics | Danish Covid-19 Genome Consortium |
| EPI_ISL_2863226 | Dutch COVID-19 response team | National Institute for Public Health and the Environment (RIVM) | Adam Meijer, Harry Vennema, Dirk Eggink, Jeroen Cremer, Sharon van den Brink, Bas van der Veer, AnneMarie van den Brandt, Lisa Wijsman, Kim Freriks, Ryanne Jaarsma, Euníce Then, Lynn Aarts, Sanne Bos, Melissa van Tuil, Florian Zwagemaker, Dennis Schmitz, Annelies Kroneman, Karim Hajji, Chantal Reusken, on behalf of the national COVID-19 response team |
| EPI_ISL_2863227 | Department of Bacteria, Parasites and Fungi, Statens Serum Institut, Copenhagen, Denmark | Statens Serum Institut Bioinformatics and Microbial Genomics | Danish Covid-19 Genome Consortium |
| EPI_ISL_2863228, EPI_ISL_2863229 | Dutch COVID-19 response team | National Institute for Public Health and the Environment (RIVM) | Adam Meijer, Harry Vennema, Dirk Eggink, Jeroen Cremer, Sharon van den Brink, Bas van der Veer, AnneMarie van den Brandt, Lisa Wijsman, Kim Freriks, Ryanne Jaarsma, Euníce Then, Lynn Aarts, Sanne Bos, Melissa van Tuil, Florian Zwagemaker, Dennis Schmitz, Annelies Kroneman, Karim Hajji, Chantal Reusken, on behalf of the national COVID-19 response team |
| EPI_ISL_2863230 | Department of Bacteria, Parasites and Fungi, Statens Serum Institut, Copenhagen, Denmark | Statens Serum Institut Bioinformatics and Microbial Genomics | Danish Covid-19 Genome Consortium |
| EPI_ISL_2863231 | Dutch COVID-19 response team | National Institute for Public Health and the Environment (RIVM) | Adam Meijer, Harry Vennema, Dirk Eggink, Jeroen Cremer, Sharon van den Brink, Bas van der Veer, AnneMarie van den Brandt, Lisa Wijsman, Kim Freriks, Ryanne Jaarsma, Euníce Then, Lynn Aarts, Sanne Bos, Melissa van Tuil, Florian Zwagemaker, Dennis Schmitz, Annelies Kroneman, Karim Hajji, Chantal Reusken, on behalf of the national COVID-19 response team |
| EPI_ISL_2863232 | Department of Bacteria, Parasites and Fungi, Statens Serum Institut, Copenhagen, Denmark | Statens Serum Institut Bioinformatics and Microbial Genomics | Danish Covid-19 Genome Consortium |
| EPI_ISL_2863233 | Dutch COVID-19 response team | National Institute for Public Health and the Environment (RIVM) | Adam Meijer, Harry Vennema, Dirk Eggink, Jeroen Cremer, Sharon van den Brink, Bas van der Veer, AnneMarie van den Brandt, Lisa Wijsman, Kim Freriks, Ryanne Jaarsma, Euníce Then, Lynn Aarts, Sanne Bos, Melissa van Tuil, Florian Zwagemaker, Dennis Schmitz, Annelies Kroneman, Karim Hajji, Chantal Reusken, on behalf of the national COVID-19 response team |
| EPI_ISL_2863234 | Department of Bacteria, Parasites and Fungi, Statens Serum Institut, Copenhagen, Denmark | Statens Serum Institut Bioinformatics and Microbial Genomics | Danish Covid-19 Genome Consortium |
| EPI_ISL_2863235 | Dutch COVID-19 response team | National Institute for Public Health and the Environment (RIVM) | Adam Meijer, Harry Vennema, Dirk Eggink, Jeroen Cremer, Sharon van den Brink, Bas van der Veer, AnneMarie van den Brandt, Lisa Wijsman, Kim Freriks, Ryanne Jaarsma, Euníce Then, Lynn Aarts, Sanne Bos, Melissa van Tuil, Florian Zwagemaker, Dennis Schmitz, Annelies Kroneman, Karim Hajji, Chantal Reusken, on behalf of the national COVID-19 response team |
| EPI_ISL_2863236 | Department of Bacteria, Parasites and Fungi, Statens Serum Institut, Copenhagen, Denmark | Statens Serum Institut Bioinformatics and Microbial Genomics | Danish Covid-19 Genome Consortium |
| EPI_ISL_2863237 | Dutch COVID-19 response team | National Institute for Public Health and the Environment (RIVM) | Adam Meijer, Harry Vennema, Dirk Eggink, Jeroen Cremer, Sharon van den Brink, Bas van der Veer, AnneMarie van den Brandt, Lisa Wijsman, Kim Freriks, Ryanne Jaarsma, Euníce Then, Lynn Aarts, Sanne Bos, Melissa van Tuil, Florian Zwagemaker, Dennis Schmitz, Annelies Kroneman, Karim Hajji, Chantal Reusken, on behalf of the national COVID-19 response team |
| EPI_ISL_2863238 | Department of Bacteria, Parasites and Fungi, Statens Serum Institut, Copenhagen, Denmark | Statens Serum Institut Bioinformatics and Microbial Genomics | Danish Covid-19 Genome Consortium |
| EPI_ISL_2863239 | Dutch COVID-19 response team | National Institute for Public Health and the Environment (RIVM) | Adam Meijer, Harry Vennema, Dirk Eggink, Jeroen Cremer, Sharon van den Brink, Bas van der Veer, AnneMarie van den Brandt, Lisa Wijsman, Kim Freriks, Ryanne Jaarsma, Euníce Then, Lynn Aarts, Sanne Bos, Melissa van Tuil, Florian Zwagemaker, Dennis Schmitz, Annelies Kroneman, Karim Hajji, Chantal Reusken, on behalf of the national COVID-19 response team |
| EPI_ISL_2863240 | Department of Bacteria, Parasites and Fungi, Statens Serum Institut, Copenhagen, Denmark | Statens Serum Institut Bioinformatics and Microbial Genomics | Danish Covid-19 Genome Consortium |
| EPI_ISL_2863241, EPI_ISL_2863242 | Dutch COVID-19 response team | National Institute for Public Health and the Environment (RIVM) | Adam Meijer, Harry Vennema, Dirk Eggink, Jeroen Cremer, Sharon van den Brink, Bas van der Veer, AnneMarie van den Brandt, Lisa Wijsman, Kim Freriks, Ryanne Jaarsma, Euníce Then, Lynn Aarts, Sanne Bos, Melissa van Tuil, Florian Zwagemaker, Dennis Schmitz, Annelies Kroneman, Karim Hajji, Chantal Reusken, on behalf of the national COVID-19 response team |
| EPI_ISL_2863243 | Department of Bacteria, Parasites and Fungi, Statens Serum Institut, Copenhagen, Denmark | Statens Serum Institut Bioinformatics and Microbial Genomics | Danish Covid-19 Genome Consortium |
| EPI_ISL_2863244 | Dutch COVID-19 response team | National Institute for Public Health and the Environment (RIVM) | Adam Meijer, Harry Vennema, Dirk Eggink, Jeroen Cremer, Sharon van den Brink, Bas van der Veer, AnneMarie van den Brandt, Lisa Wijsman, Kim Freriks, Ryanne Jaarsma, Euníce Then, Lynn Aarts, Sanne Bos, Melissa van Tuil, Florian Zwagemaker, Dennis Schmitz, Annelies Kroneman, Karim Hajji, Chantal Reusken, on behalf of the national COVID-19 response team |
| EPI_ISL_2863245 | Department of Bacteria, Parasites and Fungi, Statens Serum Institut, Copenhagen, Denmark | Statens Serum Institut Bioinformatics and Microbial Genomics | Danish Covid-19 Genome Consortium |
| EPI_ISL_2863246 | Dutch COVID-19 response team | National Institute for Public Health and the Environment (RIVM) | Adam Meijer, Harry Vennema, Dirk Eggink, Jeroen Cremer, Sharon van den Brink, Bas van der Veer, AnneMarie van den Brandt, Lisa Wijsman, Kim Freriks, Ryanne Jaarsma, Euníce Then, Lynn Aarts, Sanne Bos, Melissa van Tuil, Florian Zwagemaker, Dennis Schmitz, Annelies Kroneman, Karim Hajji, Chantal Reusken, on behalf of the national COVID-19 response team |
| EPI_ISL_2863247 | Department of Bacteria, Parasites and Fungi, Statens Serum Institut, Copenhagen, Denmark | Statens Serum Institut Bioinformatics and Microbial Genomics | Danish Covid-19 Genome Consortium |
| EPI_ISL_2863248 | Dutch COVID-19 response team | National Institute for Public Health and the Environment (RIVM) | Adam Meijer, Harry Vennema, Dirk Eggink, Jeroen Cremer, Sharon van den Brink, Bas van der Veer, AnneMarie van den Brandt, Lisa Wijsman, Kim Freriks, Ryanne Jaarsma, Euníce Then, Lynn Aarts, Sanne Bos, Melissa van Tuil, Florian Zwagemaker, Dennis Schmitz, Annelies Kroneman, Karim Hajji, Chantal Reusken, on behalf of the national COVID-19 response team |
| EPI_ISL_2863249 | Department of Bacteria, Parasites and Fungi, Statens Serum Institut, Copenhagen, Denmark | Statens Serum Institut Bioinformatics and Microbial Genomics | Danish Covid-19 Genome Consortium |
| EPI_ISL_2863250 | Dutch COVID-19 response team | National Institute for Public Health and the Environment (RIVM) | Adam Meijer, Harry Vennema, Dirk Eggink, Jeroen Cremer, Sharon van den Brink, Bas van der Veer, AnneMarie van den Brandt, Lisa Wijsman, Kim Freriks, Ryanne Jaarsma, Euníce Then, Lynn Aarts, Sanne Bos, Melissa van Tuil, Florian Zwagemaker, Dennis Schmitz, Annelies Kroneman, Karim Hajji, Chantal Reusken, on behalf of the national COVID-19 response team |

| | | | |
|---|---|---|---|
| EPI_ISL_2863251 | Department of Bacteria, Parasites and Fungi, Statens Serum Institut, Copenhagen, Denmark | Statens Serum Institut Bioinformatics and Microbial Genomics | Danish Covid-19 Genome Consortium |
| EPI_ISL_2863252 | Dutch COVID-19 response team | National Institute for Public Health and the Environment (RIVM) | Adam Meijer, Harry Vennema, Dirk Eggink, Jeroen Cremer, Sharon van den Brink, Bas van der Veer, AnneMarie van den Brandt, Lisa Wijsman, Kim Freriks, Ryanne Jaarsma, Euníce Then, Lynn Aarts, Sanne Bos, Melissa van Tuil, Florian Zwagemaker, Dennis Schmitz, Annelies Kroneman, Karim Hajji, Chantal Reusken, on behalf of the national COVID-19 response team |
| EPI_ISL_2863253 | Department of Bacteria, Parasites and Fungi, Statens Serum Institut, Copenhagen, Denmark | Statens Serum Institut Bioinformatics and Microbial Genomics | Danish Covid-19 Genome Consortium |
| EPI_ISL_2863254 | Dutch COVID-19 response team | National Institute for Public Health and the Environment (RIVM) | Adam Meijer, Harry Vennema, Dirk Eggink, Jeroen Cremer, Sharon van den Brink, Bas van der Veer, AnneMarie van den Brandt, Lisa Wijsman, Kim Freriks, Ryanne Jaarsma, Euníce Then, Lynn Aarts, Sanne Bos, Melissa van Tuil, Florian Zwagemaker, Dennis Schmitz, Annelies Kroneman, Karim Hajji, Chantal Reusken, on behalf of the national COVID-19 response team |
| EPI_ISL_2863255 | Department of Bacteria, Parasites and Fungi, Statens Serum Institut, Copenhagen, Denmark | Statens Serum Institut Bioinformatics and Microbial Genomics | Danish Covid-19 Genome Consortium |
| EPI_ISL_2863256 | Dutch COVID-19 response team | National Institute for Public Health and the Environment (RIVM) | Adam Meijer, Harry Vennema, Dirk Eggink, Jeroen Cremer, Sharon van den Brink, Bas van der Veer, AnneMarie van den Brandt, Lisa Wijsman, Kim Freriks, Ryanne Jaarsma, Euníce Then, Lynn Aarts, Sanne Bos, Melissa van Tuil, Florian Zwagemaker, Dennis Schmitz, Annelies Kroneman, Karim Hajji, Chantal Reusken, on behalf of the national COVID-19 response team |
| EPI_ISL_2863257 | Department of Bacteria, Parasites and Fungi, Statens Serum Institut, Copenhagen, Denmark | Statens Serum Institut Bioinformatics and Microbial Genomics | Danish Covid-19 Genome Consortium |
| EPI_ISL_2863258 | Dutch COVID-19 response team | National Institute for Public Health and the Environment (RIVM) | Adam Meijer, Harry Vennema, Dirk Eggink, Jeroen Cremer, Sharon van den Brink, Bas van der Veer, AnneMarie van den Brandt, Lisa Wijsman, Kim Freriks, Ryanne Jaarsma, Euníce Then, Lynn Aarts, Sanne Bos, Melissa van Tuil, Florian Zwagemaker, Dennis Schmitz, Annelies Kroneman, Karim Hajji, Chantal Reusken, on behalf of the national COVID-19 response team |
| EPI_ISL_2863259 | Department of Bacteria, Parasites and Fungi, Statens Serum Institut, Copenhagen, Denmark | Statens Serum Institut Bioinformatics and Microbial Genomics | Danish Covid-19 Genome Consortium |
| EPI_ISL_2863260 | Dutch COVID-19 response team | National Institute for Public Health and the Environment (RIVM) | Adam Meijer, Harry Vennema, Dirk Eggink, Jeroen Cremer, Sharon van den Brink, Bas van der Veer, AnneMarie van den Brandt, Lisa Wijsman, Kim Freriks, Ryanne Jaarsma, Euníce Then, Lynn Aarts, Sanne Bos, Melissa van Tuil, Florian Zwagemaker, Dennis Schmitz, Annelies Kroneman, Karim Hajji, Chantal Reusken, on behalf of the national COVID-19 response team |
| EPI_ISL_2863261 | Department of Bacteria, Parasites and Fungi, Statens Serum Institut, Copenhagen, Denmark | Statens Serum Institut Bioinformatics and Microbial Genomics | Danish Covid-19 Genome Consortium |
| EPI_ISL_2863262 | Dutch COVID-19 response team | National Institute for Public Health and the Environment (RIVM) | Adam Meijer, Harry Vennema, Dirk Eggink, Jeroen Cremer, Sharon van den Brink, Bas van der Veer, AnneMarie van den Brandt, Lisa Wijsman, Kim Freriks, Ryanne Jaarsma, Euníce Then, Lynn Aarts, Sanne Bos, Melissa van Tuil, Florian Zwagemaker, Dennis Schmitz, Annelies Kroneman, Karim Hajji, Chantal Reusken, on behalf of the national COVID-19 response team |
| EPI_ISL_2863263 | Department of Bacteria, Parasites and Fungi, Statens Serum Institut, Copenhagen, Denmark | Statens Serum Institut Bioinformatics and Microbial Genomics | Danish Covid-19 Genome Consortium |
| EPI_ISL_2863264, EPI_ISL_2863265 | Dutch COVID-19 response team | National Institute for Public Health and the Environment (RIVM) | Adam Meijer, Harry Vennema, Dirk Eggink, Jeroen Cremer, Sharon van den Brink, Bas van der Veer, AnneMarie van den Brandt, Lisa Wijsman, Kim Freriks, Ryanne Jaarsma, Euníce Then, Lynn Aarts, Sanne Bos, Melissa van Tuil, Florian Zwagemaker, Dennis Schmitz, Annelies Kroneman, Karim Hajji, Chantal Reusken, on behalf of the national COVID-19 response team |
| EPI_ISL_2863266 | Department of Bacteria, Parasites and Fungi, Statens Serum Institut, Copenhagen, Denmark | Statens Serum Institut Bioinformatics and Microbial Genomics | Danish Covid-19 Genome Consortium |
| EPI_ISL_2863267 | Dutch COVID-19 response team | National Institute for Public Health and the Environment (RIVM) | Adam Meijer, Harry Vennema, Dirk Eggink, Jeroen Cremer, Sharon van den Brink, Bas van der Veer, AnneMarie van den Brandt, Lisa Wijsman, Kim Freriks, Ryanne Jaarsma, Euníce Then, Lynn Aarts, Sanne Bos, Melissa van Tuil, Florian Zwagemaker, Dennis Schmitz, Annelies Kroneman, Karim Hajji, Chantal Reusken, on behalf of the national COVID-19 response team |
| EPI_ISL_2863268 | Department of Bacteria, Parasites and Fungi, Statens Serum Institut, Copenhagen, Denmark | Statens Serum Institut Bioinformatics and Microbial Genomics | Danish Covid-19 Genome Consortium |
| EPI_ISL_2863269 | Dutch COVID-19 response team | National Institute for Public Health and the Environment (RIVM) | Adam Meijer, Harry Vennema, Dirk Eggink, Jeroen Cremer, Sharon van den Brink, Bas van der Veer, AnneMarie van den Brandt, Lisa Wijsman, Kim Freriks, Ryanne Jaarsma, Euníce Then, Lynn Aarts, Sanne Bos, Melissa van Tuil, Florian Zwagemaker, Dennis Schmitz, Annelies Kroneman, Karim Hajji, Chantal Reusken, on behalf of the national COVID-19 response team |
| EPI_ISL_2863270 | Department of Bacteria, Parasites and Fungi, Statens Serum Institut, Copenhagen, Denmark | Statens Serum Institut Bioinformatics and Microbial Genomics | Danish Covid-19 Genome Consortium |
| EPI_ISL_2863271 | Dutch COVID-19 response team | National Institute for Public Health and the Environment (RIVM) | Adam Meijer, Harry Vennema, Dirk Eggink, Jeroen Cremer, Sharon van den Brink, Bas van der Veer, AnneMarie van den Brandt, Lisa Wijsman, Kim Freriks, Ryanne Jaarsma, Euníce Then, Lynn Aarts, Sanne Bos, Melissa van Tuil, Florian Zwagemaker, Dennis Schmitz, Annelies Kroneman, Karim Hajji, Chantal Reusken, on behalf of the national COVID-19 response team |
| EPI_ISL_2863272 | Department of Bacteria, Parasites and Fungi, Statens Serum Institut, Copenhagen, Denmark | Statens Serum Institut Bioinformatics and Microbial Genomics | Danish Covid-19 Genome Consortium |
| EPI_ISL_2863273 | Dutch COVID-19 response team | National Institute for Public Health and the Environment (RIVM) | Adam Meijer, Harry Vennema, Dirk Eggink, Jeroen Cremer, Sharon van den Brink, Bas van der Veer, AnneMarie van den Brandt, Lisa Wijsman, Kim Freriks, Ryanne Jaarsma, Euníce Then, Lynn Aarts, Sanne Bos, Melissa van Tuil, Florian Zwagemaker, Dennis Schmitz, Annelies Kroneman, Karim Hajji, Chantal Reusken, on behalf of the national COVID-19 response team |
| EPI_ISL_2863274 | Department of Bacteria, Parasites and Fungi, Statens Serum Institut, Copenhagen, Denmark | Statens Serum Institut Bioinformatics and Microbial Genomics | Danish Covid-19 Genome Consortium |
| EPI_ISL_2863275 | Dutch COVID-19 response team | National Institute for Public Health and the Environment (RIVM) | Adam Meijer, Harry Vennema, Dirk Eggink, Jeroen Cremer, Sharon van den Brink, Bas van der Veer, AnneMarie van den Brandt, Lisa Wijsman, Kim Freriks, Ryanne Jaarsma, Euníce Then, Lynn Aarts, Sanne Bos, Melissa van Tuil, Florian Zwagemaker, Dennis Schmitz, Annelies Kroneman, Karim Hajji, Chantal Reusken, on behalf of the national COVID-19 response team |
| EPI_ISL_2863276 | Department of Bacteria, Parasites and Fungi, Statens Serum Institut, Copenhagen, Denmark | Statens Serum Institut Bioinformatics and Microbial Genomics | Danish Covid-19 Genome Consortium |
| EPI_ISL_2863277 | Dutch COVID-19 response team | National Institute for Public Health and the Environment (RIVM) | Adam Meijer, Harry Vennema, Dirk Eggink, Jeroen Cremer, Sharon van den Brink, Bas van der Veer, AnneMarie van den Brandt, Lisa Wijsman, Kim Freriks, Ryanne Jaarsma, Euníce Then, Lynn Aarts, Sanne Bos, Melissa van Tuil, Florian Zwagemaker, Dennis Schmitz, Annelies Kroneman, Karim Hajji, Chantal Reusken, on behalf of the national COVID-19 response team |
| EPI_ISL_2863278 | Department of Bacteria, Parasites and Fungi, Statens Serum Institut, Copenhagen, Denmark | Statens Serum Institut Bioinformatics and Microbial Genomics | Danish Covid-19 Genome Consortium |
| EPI_ISL_2863279 | Dutch COVID-19 response team | National Institute for Public Health and the Environment (RIVM) | Adam Meijer, Harry Vennema, Dirk Eggink, Jeroen Cremer, Sharon van den Brink, Bas van der Veer, AnneMarie van den Brandt, Lisa Wijsman, Kim Freriks, Ryanne Jaarsma, Euníce Then, Lynn Aarts, Sanne Bos, Melissa van Tuil, Florian Zwagemaker, Dennis Schmitz, Annelies Kroneman, Karim Hajji, Chantal Reusken, on behalf of the national COVID-19 response team |
| EPI_ISL_2863280 | Department of Bacteria, Parasites and Fungi, Statens Serum Institut, Copenhagen, Denmark | Statens Serum Institut Bioinformatics and Microbial Genomics | Danish Covid-19 Genome Consortium |
| EPI_ISL_2863281 | Dutch COVID-19 response team | National Institute for Public Health and the Environment (RIVM) | Adam Meijer, Harry Vennema, Dirk Eggink, Jeroen Cremer, Sharon van den Brink, Bas van der Veer, AnneMarie van den Brandt, Lisa Wijsman, Kim Freriks, Ryanne Jaarsma, Euníce Then, Lynn Aarts, Sanne Bos, Melissa van Tuil, Florian Zwagemaker, Dennis Schmitz, Annelies Kroneman, Karim Hajji, Chantal Reusken, on behalf of the national COVID-19 response team |
| EPI_ISL_2863282 | Department of Bacteria, Parasites and Fungi, Statens Serum Institut, Copenhagen, Denmark | Statens Serum Institut Bioinformatics and Microbial Genomics | Danish Covid-19 Genome Consortium |
| EPI_ISL_2863283 | Dutch COVID-19 response team | National Institute for Public Health and the Environment (RIVM) | Adam Meijer, Harry Vennema, Dirk Eggink, Jeroen Cremer, Sharon van den Brink, Bas van der Veer, AnneMarie van den Brandt, Lisa Wijsman, Kim Freriks, Ryanne Jaarsma, Euníce Then, Lynn Aarts, Sanne Bos, Melissa van Tuil, Florian Zwagemaker, Dennis Schmitz, Annelies Kroneman, Karim Hajji, Chantal Reusken, on behalf of the national COVID-19 response team |
| EPI_ISL_2863284 | Department of Bacteria, Parasites and Fungi, Statens Serum Institut, Copenhagen, Denmark | Statens Serum Institut Bioinformatics and Microbial Genomics | Danish Covid-19 Genome Consortium |
| EPI_ISL_2863285 | Dutch COVID-19 response team | National Institute for Public Health and the Environment (RIVM) | Adam Meijer, Harry Vennema, Dirk Eggink, Jeroen Cremer, Sharon van den Brink, Bas van der Veer, AnneMarie van den Brandt, Lisa Wijsman, Kim Freriks, Ryanne Jaarsma, Euníce Then, Lynn Aarts, Sanne Bos, Melissa van Tuil, Florian Zwagemaker, Dennis Schmitz, Annelies Kroneman, Karim Hajji, Chantal Reusken, on behalf of the national COVID-19 response team |
| EPI_ISL_2863286, EPI_ISL_2863287 | Department of Bacteria, Parasites and Fungi, Statens Serum Institut, Copenhagen, Denmark | Statens Serum Institut Bioinformatics and Microbial Genomics | Danish Covid-19 Genome Consortium |
| EPI_ISL_2863288, EPI_ISL_2863289 | Dutch COVID-19 response team | National Institute for Public Health and the Environment (RIVM) | Adam Meijer, Harry Vennema, Dirk Eggink, Jeroen Cremer, Sharon van den Brink, Bas van der Veer, AnneMarie van den Brandt, Lisa Wijsman, Kim Freriks, Ryanne Jaarsma, Euníce Then, Lynn Aarts, Sanne Bos, Melissa van Tuil, Florian Zwagemaker, Dennis Schmitz, Annelies Kroneman, Karim Hajji, Chantal Reusken, on behalf of the national COVID-19 response team |
| EPI_ISL_2863290 | Department of Bacteria, Parasites and Fungi, Statens Serum Institut, Copenhagen, Denmark | Statens Serum Institut Bioinformatics and Microbial Genomics | Danish Covid-19 Genome Consortium |
| EPI_ISL_2863291 | Dutch COVID-19 response team | National Institute for Public Health and the Environment (RIVM) | Adam Meijer, Harry Vennema, Dirk Eggink, Jeroen Cremer, Sharon van den Brink, Bas van der Veer, AnneMarie van den Brandt, Lisa Wijsman, Kim Freriks, Ryanne Jaarsma, Euníce Then, Lynn Aarts, Sanne Bos, Melissa van Tuil, Florian Zwagemaker, Dennis Schmitz, Annelies Kroneman, Karim Hajji, Chantal Reusken, on behalf of the national COVID-19 response team |
| EPI_ISL_2863292 | Department of Bacteria, Parasites and Fungi, Statens Serum Institut, Copenhagen, Denmark | Statens Serum Institut Bioinformatics and Microbial Genomics | Danish Covid-19 Genome Consortium |
| EPI_ISL_2863293 | Dutch COVID-19 response team | National Institute for Public Health and the Environment (RIVM) | Adam Meijer, Harry Vennema, Dirk Eggink, Jeroen Cremer, Sharon van den Brink, Bas van der Veer, AnneMarie van den Brandt, Lisa Wijsman, Kim Freriks, Ryanne Jaarsma, Euníce Then, Lynn Aarts, Sanne Bos, Melissa van Tuil, Florian Zwagemaker, Dennis Schmitz, Annelies Kroneman, Karim Hajji, Chantal Reusken, on behalf of the national COVID-19 response team |
| EPI_ISL_2863294 | Department of Bacteria, Parasites and Fungi, Statens Serum Institut, Copenhagen, Denmark | Statens Serum Institut Bioinformatics and Microbial Genomics | Danish Covid-19 Genome Consortium |
| EPI_ISL_2863295 | Dutch COVID-19 response team | National Institute for Public Health and the Environment (RIVM) | Adam Meijer, Harry Vennema, Dirk Eggink, Jeroen Cremer, Sharon van den Brink, Bas van der Veer, AnneMarie van den Brandt, Lisa Wijsman, Kim Freriks, Ryanne Jaarsma, Euníce Then, Lynn Aarts, Sanne Bos, Melissa van Tuil, Florian Zwagemaker, Dennis Schmitz, Annelies Kroneman, Karim Hajji, Chantal Reusken, on behalf of the national COVID-19 response team |
| EPI_ISL_2863296 | Department of Bacteria, Parasites and Fungi, Statens Serum Institut, Copenhagen, Denmark | Statens Serum Institut Bioinformatics and Microbial Genomics | Danish Covid-19 Genome Consortium |
| EPI_ISL_2863297 | Dutch COVID-19 response team | National Institute for Public Health and the Environment (RIVM) | Adam Meijer, Harry Vennema, Dirk Eggink, Jeroen Cremer, Sharon van den Brink, Bas van der Veer, AnneMarie van den Brandt, Lisa Wijsman, Kim Freriks, Ryanne Jaarsma, Euníce Then, Lynn Aarts, Sanne Bos, Melissa van Tuil, Florian Zwagemaker, Dennis Schmitz, Annelies Kroneman, Karim Hajji, Chantal Reusken, on behalf of the national COVID-19 response team |
| EPI_ISL_2863298 | Department of Bacteria, Parasites and Fungi, Statens Serum Institut, Copenhagen, Denmark | Statens Serum Institut Bioinformatics and Microbial Genomics | Danish Covid-19 Genome Consortium |

| | | | |
|---|---|---|---|
| EPI_ISL_2863299 | Dutch COVID-19 response team | National Institute for Public Health and the Environment (RIVM) | Adam Meijer, Harry Vennema, Dirk Eggink, Jeroen Cremer, Sharon van den Brink, Bas van der Veer, AnneMarie van den Brandt, Lisa Wijsman, Kim Freriks, Ryanne Jaarsma, Euníce Then, Lynn Aarts, Sanne Bos, Melissa van Tuil, Florian Zwagemaker, Dennis Schmitz, Annelies Kroneman, Karim Hajji, Chantal Reusken, on behalf of the national COVID-19 response team |
| EPI_ISL_2863300 | Department of Bacteria, Parasites and Fungi, Statens Serum Institut, Copenhagen, Denmark | Statens Serum Institut Bioinformatics and Microbial Genomics | Danish Covid-19 Genome Consortium |
| EPI_ISL_2863301 | Dutch COVID-19 response team | National Institute for Public Health and the Environment (RIVM) | Adam Meijer, Harry Vennema, Dirk Eggink, Jeroen Cremer, Sharon van den Brink, Bas van der Veer, AnneMarie van den Brandt, Lisa Wijsman, Kim Freriks, Ryanne Jaarsma, Euníce Then, Lynn Aarts, Sanne Bos, Melissa van Tuil, Florian Zwagemaker, Dennis Schmitz, Annelies Kroneman, Karim Hajji, Chantal Reusken, on behalf of the national COVID-19 response team |
| EPI_ISL_2863302 | Department of Bacteria, Parasites and Fungi, Statens Serum Institut, Copenhagen, Denmark | Statens Serum Institut Bioinformatics and Microbial Genomics | Danish Covid-19 Genome Consortium |
| EPI_ISL_2863303 | Dutch COVID-19 response team | National Institute for Public Health and the Environment (RIVM) | Adam Meijer, Harry Vennema, Dirk Eggink, Jeroen Cremer, Sharon van den Brink, Bas van der Veer, AnneMarie van den Brandt, Lisa Wijsman, Kim Freriks, Ryanne Jaarsma, Euníce Then, Lynn Aarts, Sanne Bos, Melissa van Tuil, Florian Zwagemaker, Dennis Schmitz, Annelies Kroneman, Karim Hajji, Chantal Reusken, on behalf of the national COVID-19 response team |
| EPI_ISL_2863304 | Department of Bacteria, Parasites and Fungi, Statens Serum Institut, Copenhagen, Denmark | Statens Serum Institut Bioinformatics and Microbial Genomics | Danish Covid-19 Genome Consortium |
| EPI_ISL_2863305 | Dutch COVID-19 response team | National Institute for Public Health and the Environment (RIVM) | Adam Meijer, Harry Vennema, Dirk Eggink, Jeroen Cremer, Sharon van den Brink, Bas van der Veer, AnneMarie van den Brandt, Lisa Wijsman, Kim Freriks, Ryanne Jaarsma, Euníce Then, Lynn Aarts, Sanne Bos, Melissa van Tuil, Florian Zwagemaker, Dennis Schmitz, Annelies Kroneman, Karim Hajji, Chantal Reusken, on behalf of the national COVID-19 response team |
| EPI_ISL_2863306 | Department of Bacteria, Parasites and Fungi, Statens Serum Institut, Copenhagen, Denmark | Statens Serum Institut Bioinformatics and Microbial Genomics | Danish Covid-19 Genome Consortium |
| EPI_ISL_2863307 | Dutch COVID-19 response team | National Institute for Public Health and the Environment (RIVM) | Adam Meijer, Harry Vennema, Dirk Eggink, Jeroen Cremer, Sharon van den Brink, Bas van der Veer, AnneMarie van den Brandt, Lisa Wijsman, Kim Freriks, Ryanne Jaarsma, Euníce Then, Lynn Aarts, Sanne Bos, Melissa van Tuil, Florian Zwagemaker, Dennis Schmitz, Annelies Kroneman, Karim Hajji, Chantal Reusken, on behalf of the national COVID-19 response team |
| EPI_ISL_2863308 | Department of Bacteria, Parasites and Fungi, Statens Serum Institut, Copenhagen, Denmark | Statens Serum Institut Bioinformatics and Microbial Genomics | Danish Covid-19 Genome Consortium |
| EPI_ISL_2863309 | Dutch COVID-19 response team | National Institute for Public Health and the Environment (RIVM) | Adam Meijer, Harry Vennema, Dirk Eggink, Jeroen Cremer, Sharon van den Brink, Bas van der Veer, AnneMarie van den Brandt, Lisa Wijsman, Kim Freriks, Ryanne Jaarsma, Euníce Then, Lynn Aarts, Sanne Bos, Melissa van Tuil, Florian Zwagemaker, Dennis Schmitz, Annelies Kroneman, Karim Hajji, Chantal Reusken, on behalf of the national COVID-19 response team |
| EPI_ISL_2863310 | Department of Bacteria, Parasites and Fungi, Statens Serum Institut, Copenhagen, Denmark | Statens Serum Institut Bioinformatics and Microbial Genomics | Danish Covid-19 Genome Consortium |
| EPI_ISL_2863311, EPI_ISL_2863313 | Dutch COVID-19 response team | National Institute for Public Health and the Environment (RIVM) | Adam Meijer, Harry Vennema, Dirk Eggink, Jeroen Cremer, Sharon van den Brink, Bas van der Veer, AnneMarie van den Brandt, Lisa Wijsman, Kim Freriks, Ryanne Jaarsma, Euníce Then, Lynn Aarts, Sanne Bos, Melissa van Tuil, Florian Zwagemaker, Dennis Schmitz, Annelies Kroneman, Karim Hajji, Chantal Reusken, on behalf of the national COVID-19 response team |
| EPI_ISL_2863314 | Department of Bacteria, Parasites and Fungi, Statens Serum Institut, Copenhagen, Denmark | Statens Serum Institut Bioinformatics and Microbial Genomics | Danish Covid-19 Genome Consortium |
| EPI_ISL_2863315 | Dutch COVID-19 response team | National Institute for Public Health and the Environment (RIVM) | Adam Meijer, Harry Vennema, Dirk Eggink, Jeroen Cremer, Sharon van den Brink, Bas van der Veer, AnneMarie van den Brandt, Lisa Wijsman, Kim Freriks, Ryanne Jaarsma, Euníce Then, Lynn Aarts, Sanne Bos, Melissa van Tuil, Florian Zwagemaker, Dennis Schmitz, Annelies Kroneman, Karim Hajji, Chantal Reusken, on behalf of the national COVID-19 response team |
| EPI_ISL_2863316 | Department of Bacteria, Parasites and Fungi, Statens Serum Institut, Copenhagen, Denmark | Statens Serum Institut Bioinformatics and Microbial Genomics | Danish Covid-19 Genome Consortium |
| EPI_ISL_2863317 | Dutch COVID-19 response team | National Institute for Public Health and the Environment (RIVM) | Adam Meijer, Harry Vennema, Dirk Eggink, Jeroen Cremer, Sharon van den Brink, Bas van der Veer, AnneMarie van den Brandt, Lisa Wijsman, Kim Freriks, Ryanne Jaarsma, Euníce Then, Lynn Aarts, Sanne Bos, Melissa van Tuil, Florian Zwagemaker, Dennis Schmitz, Annelies Kroneman, Karim Hajji, Chantal Reusken, on behalf of the national COVID-19 response team |
| EPI_ISL_2863318 | Department of Bacteria, Parasites and Fungi, Statens Serum Institut, Copenhagen, Denmark | Statens Serum Institut Bioinformatics and Microbial Genomics | Danish Covid-19 Genome Consortium |
| EPI_ISL_2863319 | Dutch COVID-19 response team | National Institute for Public Health and the Environment (RIVM) | Adam Meijer, Harry Vennema, Dirk Eggink, Jeroen Cremer, Sharon van den Brink, Bas van der Veer, AnneMarie van den Brandt, Lisa Wijsman, Kim Freriks, Ryanne Jaarsma, Euníce Then, Lynn Aarts, Sanne Bos, Melissa van Tuil, Florian Zwagemaker, Dennis Schmitz, Annelies Kroneman, Karim Hajji, Chantal Reusken, on behalf of the national COVID-19 response team |
| EPI_ISL_2863320 | Department of Bacteria, Parasites and Fungi, Statens Serum Institut, Copenhagen, Denmark | Statens Serum Institut Bioinformatics and Microbial Genomics | Danish Covid-19 Genome Consortium |
| EPI_ISL_2863321 | Dutch COVID-19 response team | National Institute for Public Health and the Environment (RIVM) | Adam Meijer, Harry Vennema, Dirk Eggink, Jeroen Cremer, Sharon van den Brink, Bas van der Veer, AnneMarie van den Brandt, Lisa Wijsman, Kim Freriks, Ryanne Jaarsma, Euníce Then, Lynn Aarts, Sanne Bos, Melissa van Tuil, Florian Zwagemaker, Dennis Schmitz, Annelies Kroneman, Karim Hajji, Chantal Reusken, on behalf of the national COVID-19 response team |
| EPI_ISL_2863322 | Department of Bacteria, Parasites and Fungi, Statens Serum Institut, Copenhagen, Denmark | Statens Serum Institut Bioinformatics and Microbial Genomics | Danish Covid-19 Genome Consortium |
| EPI_ISL_2863323 | Dutch COVID-19 response team | National Institute for Public Health and the Environment (RIVM) | Adam Meijer, Harry Vennema, Dirk Eggink, Jeroen Cremer, Sharon van den Brink, Bas van der Veer, AnneMarie van den Brandt, Lisa Wijsman, Kim Freriks, Ryanne Jaarsma, Euníce Then, Lynn Aarts, Sanne Bos, Melissa van Tuil, Florian Zwagemaker, Dennis Schmitz, Annelies Kroneman, Karim Hajji, Chantal Reusken, on behalf of the national COVID-19 response team |
| EPI_ISL_2863324 | Department of Bacteria, Parasites and Fungi, Statens Serum Institut, Copenhagen, Denmark | Statens Serum Institut Bioinformatics and Microbial Genomics | Danish Covid-19 Genome Consortium |
| EPI_ISL_2863325 | Dutch COVID-19 response team | National Institute for Public Health and the Environment (RIVM) | Adam Meijer, Harry Vennema, Dirk Eggink, Jeroen Cremer, Sharon van den Brink, Bas van der Veer, AnneMarie van den Brandt, Lisa Wijsman, Kim Freriks, Ryanne Jaarsma, Euníce Then, Lynn Aarts, Sanne Bos, Melissa van Tuil, Florian Zwagemaker, Dennis Schmitz, Annelies Kroneman, Karim Hajji, Chantal Reusken, on behalf of the national COVID-19 response team |
| EPI_ISL_2863326 | Department of Bacteria, Parasites and Fungi, Statens Serum Institut, Copenhagen, Denmark | Statens Serum Institut Bioinformatics and Microbial Genomics | Danish Covid-19 Genome Consortium |
| EPI_ISL_2863327 | Dutch COVID-19 response team | National Institute for Public Health and the Environment (RIVM) | Adam Meijer, Harry Vennema, Dirk Eggink, Jeroen Cremer, Sharon van den Brink, Bas van der Veer, AnneMarie van den Brandt, Lisa Wijsman, Kim Freriks, Ryanne Jaarsma, Euníce Then, Lynn Aarts, Sanne Bos, Melissa van Tuil, Florian Zwagemaker, Dennis Schmitz, Annelies Kroneman, Karim Hajji, Chantal Reusken, on behalf of the national COVID-19 response team |
| EPI_ISL_2863328 | Department of Bacteria, Parasites and Fungi, Statens Serum Institut, Copenhagen, Denmark | Statens Serum Institut Bioinformatics and Microbial Genomics | Danish Covid-19 Genome Consortium |
| EPI_ISL_2863329 | Dutch COVID-19 response team | National Institute for Public Health and the Environment (RIVM) | Adam Meijer, Harry Vennema, Dirk Eggink, Jeroen Cremer, Sharon van den Brink, Bas van der Veer, AnneMarie van den Brandt, Lisa Wijsman, Kim Freriks, Ryanne Jaarsma, Euníce Then, Lynn Aarts, Sanne Bos, Melissa van Tuil, Florian Zwagemaker, Dennis Schmitz, Annelies Kroneman, Karim Hajji, Chantal Reusken, on behalf of the national COVID-19 response team |
| EPI_ISL_2863330 | Department of Bacteria, Parasites and Fungi, Statens Serum Institut, Copenhagen, Denmark | Statens Serum Institut Bioinformatics and Microbial Genomics | Danish Covid-19 Genome Consortium |
| EPI_ISL_2863331 | Dutch COVID-19 response team | National Institute for Public Health and the Environment (RIVM) | Adam Meijer, Harry Vennema, Dirk Eggink, Jeroen Cremer, Sharon van den Brink, Bas van der Veer, AnneMarie van den Brandt, Lisa Wijsman, Kim Freriks, Ryanne Jaarsma, Euníce Then, Lynn Aarts, Sanne Bos, Melissa van Tuil, Florian Zwagemaker, Dennis Schmitz, Annelies Kroneman, Karim Hajji, Chantal Reusken, on behalf of the national COVID-19 response team |
| EPI_ISL_2863332 | Department of Bacteria, Parasites and Fungi, Statens Serum Institut, Copenhagen, Denmark | Statens Serum Institut Bioinformatics and Microbial Genomics | Danish Covid-19 Genome Consortium |
| EPI_ISL_2863333 | Dutch COVID-19 response team | National Institute for Public Health and the Environment (RIVM) | Adam Meijer, Harry Vennema, Dirk Eggink, Jeroen Cremer, Sharon van den Brink, Bas van der Veer, AnneMarie van den Brandt, Lisa Wijsman, Kim Freriks, Ryanne Jaarsma, Euníce Then, Lynn Aarts, Sanne Bos, Melissa van Tuil, Florian Zwagemaker, Dennis Schmitz, Annelies Kroneman, Karim Hajji, Chantal Reusken, on behalf of the national COVID-19 response team |
| EPI_ISL_2863334 | Department of Bacteria, Parasites and Fungi, Statens Serum Institut, Copenhagen, Denmark | Statens Serum Institut Bioinformatics and Microbial Genomics | Danish Covid-19 Genome Consortium |
| EPI_ISL_2863335 | Dutch COVID-19 response team | National Institute for Public Health and the Environment (RIVM) | Adam Meijer, Harry Vennema, Dirk Eggink, Jeroen Cremer, Sharon van den Brink, Bas van der Veer, AnneMarie van den Brandt, Lisa Wijsman, Kim Freriks, Ryanne Jaarsma, Euníce Then, Lynn Aarts, Sanne Bos, Melissa van Tuil, Florian Zwagemaker, Dennis Schmitz, Annelies Kroneman, Karim Hajji, Chantal Reusken, on behalf of the national COVID-19 response team |
| EPI_ISL_2863336 | Department of Bacteria, Parasites and Fungi, Statens Serum Institut, Copenhagen, Denmark | Statens Serum Institut Bioinformatics and Microbial Genomics | Danish Covid-19 Genome Consortium |
| EPI_ISL_2863337 | Dutch COVID-19 response team | National Institute for Public Health and the Environment (RIVM) | Adam Meijer, Harry Vennema, Dirk Eggink, Jeroen Cremer, Sharon van den Brink, Bas van der Veer, AnneMarie van den Brandt, Lisa Wijsman, Kim Freriks, Ryanne Jaarsma, Euníce Then, Lynn Aarts, Sanne Bos, Melissa van Tuil, Florian Zwagemaker, Dennis Schmitz, Annelies Kroneman, Karim Hajji, Chantal Reusken, on behalf of the national COVID-19 response team |
| EPI_ISL_2863338 | Department of Bacteria, Parasites and Fungi, Statens Serum Institut, Copenhagen, Denmark | Statens Serum Institut Bioinformatics and Microbial Genomics | Danish Covid-19 Genome Consortium |
| EPI_ISL_2863339, EPI_ISL_2863340 | Dutch COVID-19 response team | National Institute for Public Health and the Environment (RIVM) | Adam Meijer, Harry Vennema, Dirk Eggink, Jeroen Cremer, Sharon van den Brink, Bas van der Veer, AnneMarie van den Brandt, Lisa Wijsman, Kim Freriks, Ryanne Jaarsma, Euníce Then, Lynn Aarts, Sanne Bos, Melissa van Tuil, Florian Zwagemaker, Dennis Schmitz, Annelies Kroneman, Karim Hajji, Chantal Reusken, on behalf of the national COVID-19 response team |
| EPI_ISL_2863341 | Department of Bacteria, Parasites and Fungi, Statens Serum Institut, Copenhagen, Denmark | Statens Serum Institut Bioinformatics and Microbial Genomics | Danish Covid-19 Genome Consortium |
| EPI_ISL_2863342 | Dutch COVID-19 response team | National Institute for Public Health and the Environment (RIVM) | Adam Meijer, Harry Vennema, Dirk Eggink, Jeroen Cremer, Sharon van den Brink, Bas van der Veer, AnneMarie van den Brandt, Lisa Wijsman, Kim Freriks, Ryanne Jaarsma, Euníce Then, Lynn Aarts, Sanne Bos, Melissa van Tuil, Florian Zwagemaker, Dennis Schmitz, Annelies Kroneman, Karim Hajji, Chantal Reusken, on behalf of the national COVID-19 response team |
| EPI_ISL_2863343 | Department of Bacteria, Parasites and Fungi, Statens Serum Institut, Copenhagen, Denmark | Statens Serum Institut Bioinformatics and Microbial Genomics | Danish Covid-19 Genome Consortium |
| EPI_ISL_2863344 | Dutch COVID-19 response team | National Institute for Public Health and the Environment (RIVM) | Adam Meijer, Harry Vennema, Dirk Eggink, Jeroen Cremer, Sharon van den Brink, Bas van der Veer, AnneMarie van den Brandt, Lisa Wijsman, Kim Freriks, Ryanne Jaarsma, Euníce Then, Lynn Aarts, Sanne Bos, Melissa van Tuil, Florian Zwagemaker, Dennis Schmitz, Annelies Kroneman, Karim Hajji, Chantal Reusken, on behalf of the national COVID-19 response team |
| EPI_ISL_2863345 | Department of Bacteria, Parasites and Fungi, Statens Serum Institut, Copenhagen, Denmark | Statens Serum Institut Bioinformatics and Microbial Genomics | Danish Covid-19 Genome Consortium |
| EPI_ISL_2863346 | Dutch COVID-19 response team | National Institute for Public Health and the Environment (RIVM) | Adam Meijer, Harry Vennema, Dirk Eggink, Jeroen Cremer, Sharon van den Brink, Bas van der Veer, AnneMarie van den Brandt, Lisa Wijsman, Kim Freriks, Ryanne Jaarsma, Euníce Then, Lynn Aarts, Sanne Bos, Melissa van Tuil, Florian Zwagemaker, Dennis Schmitz, Annelies Kroneman, Karim Hajji, Chantal Reusken, on behalf of the national COVID-19 response team |

| | | | |
|---|---|---|---|
| EPI_ISL_2863347 | Department of Bacteria, Parasites and Fungi, Statens Serum Institut, Copenhagen, Denmark | Statens Serum Institut Bioinformatics and Microbial Genomics | Danish Covid-19 Genome Consortium |
| EPI_ISL_2863348 | Dutch COVID-19 response team | National Institute for Public Health and the Environment (RIVM) | Adam Meijer, Harry Vennema, Dirk Eggink, Jeroen Cremer, Sharon van den Brink, Bas van der Veer, AnneMarie van den Brandt, Lisa Wijsman, Kim Freriks, Ryanne Jaarsma, Euníce Then, Lynn Aarts, Sanne Bos, Melissa van Tuil, Florian Zwagemaker, Dennis Schmitz, Annelies Kroneman, Karim Hajji, Chantal Reusken, on behalf of the national COVID-19 response team |
| EPI_ISL_2863349 | Department of Bacteria, Parasites and Fungi, Statens Serum Institut, Copenhagen, Denmark | Statens Serum Institut Bioinformatics and Microbial Genomics | Danish Covid-19 Genome Consortium |
| EPI_ISL_2863350 | Dutch COVID-19 response team | National Institute for Public Health and the Environment (RIVM) | Adam Meijer, Harry Vennema, Dirk Eggink, Jeroen Cremer, Sharon van den Brink, Bas van der Veer, AnneMarie van den Brandt, Lisa Wijsman, Kim Freriks, Ryanne Jaarsma, Euníce Then, Lynn Aarts, Sanne Bos, Melissa van Tuil, Florian Zwagemaker, Dennis Schmitz, Annelies Kroneman, Karim Hajji, Chantal Reusken, on behalf of the national COVID-19 response team |
| EPI_ISL_2863351 | Department of Bacteria, Parasites and Fungi, Statens Serum Institut, Copenhagen, Denmark | Statens Serum Institut Bioinformatics and Microbial Genomics | Danish Covid-19 Genome Consortium |
| EPI_ISL_2863352 | Dutch COVID-19 response team | National Institute for Public Health and the Environment (RIVM) | Adam Meijer, Harry Vennema, Dirk Eggink, Jeroen Cremer, Sharon van den Brink, Bas van der Veer, AnneMarie van den Brandt, Lisa Wijsman, Kim Freriks, Ryanne Jaarsma, Euníce Then, Lynn Aarts, Sanne Bos, Melissa van Tuil, Florian Zwagemaker, Dennis Schmitz, Annelies Kroneman, Karim Hajji, Chantal Reusken, on behalf of the national COVID-19 response team |
| EPI_ISL_2863353 | Department of Bacteria, Parasites and Fungi, Statens Serum Institut, Copenhagen, Denmark | Statens Serum Institut Bioinformatics and Microbial Genomics | Danish Covid-19 Genome Consortium |
| EPI_ISL_2863354 | Dutch COVID-19 response team | National Institute for Public Health and the Environment (RIVM) | Adam Meijer, Harry Vennema, Dirk Eggink, Jeroen Cremer, Sharon van den Brink, Bas van der Veer, AnneMarie van den Brandt, Lisa Wijsman, Kim Freriks, Ryanne Jaarsma, Euníce Then, Lynn Aarts, Sanne Bos, Melissa van Tuil, Florian Zwagemaker, Dennis Schmitz, Annelies Kroneman, Karim Hajji, Chantal Reusken, on behalf of the national COVID-19 response team |
| EPI_ISL_2863355 | Department of Clinical Microbiology, Odense University Hospital, Odense, Denmark | Statens Serum Institut Bioinformatics and Microbial Genomics | Danish Covid-19 Genome Consortium |
| EPI_ISL_2863356 | Dutch COVID-19 response team | National Institute for Public Health and the Environment (RIVM) | Adam Meijer, Harry Vennema, Dirk Eggink, Jeroen Cremer, Sharon van den Brink, Bas van der Veer, AnneMarie van den Brandt, Lisa Wijsman, Kim Freriks, Ryanne Jaarsma, Euníce Then, Lynn Aarts, Sanne Bos, Melissa van Tuil, Florian Zwagemaker, Dennis Schmitz, Annelies Kroneman, Karim Hajji, Chantal Reusken, on behalf of the national COVID-19 response team |
| EPI_ISL_2863357 | Department of Bacteria, Parasites and Fungi, Statens Serum Institut, Copenhagen, Denmark | Statens Serum Institut Bioinformatics and Microbial Genomics | Danish Covid-19 Genome Consortium |
| EPI_ISL_2863358 | Dutch COVID-19 response team | National Institute for Public Health and the Environment (RIVM) | Adam Meijer, Harry Vennema, Dirk Eggink, Jeroen Cremer, Sharon van den Brink, Bas van der Veer, AnneMarie van den Brandt, Lisa Wijsman, Kim Freriks, Ryanne Jaarsma, Euníce Then, Lynn Aarts, Sanne Bos, Melissa van Tuil, Florian Zwagemaker, Dennis Schmitz, Annelies Kroneman, Karim Hajji, Chantal Reusken, on behalf of the national COVID-19 response team |
| EPI_ISL_2863359 | Department of Bacteria, Parasites and Fungi, Statens Serum Institut, Copenhagen, Denmark | Statens Serum Institut Bioinformatics and Microbial Genomics | Danish Covid-19 Genome Consortium |
| EPI_ISL_2863360, EPI_ISL_2863361 | Dutch COVID-19 response team | National Institute for Public Health and the Environment (RIVM) | Adam Meijer, Harry Vennema, Dirk Eggink, Jeroen Cremer, Sharon van den Brink, Bas van der Veer, AnneMarie van den Brandt, Lisa Wijsman, Kim Freriks, Ryanne Jaarsma, Euníce Then, Lynn Aarts, Sanne Bos, Melissa van Tuil, Florian Zwagemaker, Dennis Schmitz, Annelies Kroneman, Karim Hajji, Chantal Reusken, on behalf of the national COVID-19 response team |
| EPI_ISL_2863362 | Department of Bacteria, Parasites and Fungi, Statens Serum Institut, Copenhagen, Denmark | Statens Serum Institut Bioinformatics and Microbial Genomics | Danish Covid-19 Genome Consortium |
| EPI_ISL_2863363 | Dutch COVID-19 response team | National Institute for Public Health and the Environment (RIVM) | Adam Meijer, Harry Vennema, Dirk Eggink, Jeroen Cremer, Sharon van den Brink, Bas van der Veer, AnneMarie van den Brandt, Lisa Wijsman, Kim Freriks, Ryanne Jaarsma, Euníce Then, Lynn Aarts, Sanne Bos, Melissa van Tuil, Florian Zwagemaker, Dennis Schmitz, Annelies Kroneman, Karim Hajji, Chantal Reusken, on behalf of the national COVID-19 response team |
| EPI_ISL_2863364 | Department of Bacteria, Parasites and Fungi, Statens Serum Institut, Copenhagen, Denmark | Statens Serum Institut Bioinformatics and Microbial Genomics | Danish Covid-19 Genome Consortium |
| EPI_ISL_2863365 | Dutch COVID-19 response team | National Institute for Public Health and the Environment (RIVM) | Adam Meijer, Harry Vennema, Dirk Eggink, Jeroen Cremer, Sharon van den Brink, Bas van der Veer, AnneMarie van den Brandt, Lisa Wijsman, Kim Freriks, Ryanne Jaarsma, Euníce Then, Lynn Aarts, Sanne Bos, Melissa van Tuil, Florian Zwagemaker, Dennis Schmitz, Annelies Kroneman, Karim Hajji, Chantal Reusken, on behalf of the national COVID-19 response team |
| EPI_ISL_2863366 | Department of Bacteria, Parasites and Fungi, Statens Serum Institut, Copenhagen, Denmark | Statens Serum Institut Bioinformatics and Microbial Genomics | Danish Covid-19 Genome Consortium |
| EPI_ISL_2863367 | Dutch COVID-19 response team | National Institute for Public Health and the Environment (RIVM) | Adam Meijer, Harry Vennema, Dirk Eggink, Jeroen Cremer, Sharon van den Brink, Bas van der Veer, AnneMarie van den Brandt, Lisa Wijsman, Kim Freriks, Ryanne Jaarsma, Euníce Then, Lynn Aarts, Sanne Bos, Melissa van Tuil, Florian Zwagemaker, Dennis Schmitz, Annelies Kroneman, Karim Hajji, Chantal Reusken, on behalf of the national COVID-19 response team |
| EPI_ISL_2863368 | Department of Bacteria, Parasites and Fungi, Statens Serum Institut, Copenhagen, Denmark | Statens Serum Institut Bioinformatics and Microbial Genomics | Danish Covid-19 Genome Consortium |
| EPI_ISL_2863369 | Dutch COVID-19 response team | National Institute for Public Health and the Environment (RIVM) | Adam Meijer, Harry Vennema, Dirk Eggink, Jeroen Cremer, Sharon van den Brink, Bas van der Veer, AnneMarie van den Brandt, Lisa Wijsman, Kim Freriks, Ryanne Jaarsma, Euníce Then, Lynn Aarts, Sanne Bos, Melissa van Tuil, Florian Zwagemaker, Dennis Schmitz, Annelies Kroneman, Karim Hajji, Chantal Reusken, on behalf of the national COVID-19 response team |
| EPI_ISL_2863370 | Department of Bacteria, Parasites and Fungi, Statens Serum Institut, Copenhagen, Denmark | Statens Serum Institut Bioinformatics and Microbial Genomics | Danish Covid-19 Genome Consortium |
| EPI_ISL_2863371 | Dutch COVID-19 response team | National Institute for Public Health and the Environment (RIVM) | Adam Meijer, Harry Vennema, Dirk Eggink, Jeroen Cremer, Sharon van den Brink, Bas van der Veer, AnneMarie van den Brandt, Lisa Wijsman, Kim Freriks, Ryanne Jaarsma, Euníce Then, Lynn Aarts, Sanne Bos, Melissa van Tuil, Florian Zwagemaker, Dennis Schmitz, Annelies Kroneman, Karim Hajji, Chantal Reusken, on behalf of the national COVID-19 response team |
| EPI_ISL_2863372 | Department of Bacteria, Parasites and Fungi, Statens Serum Institut, Copenhagen, Denmark | Statens Serum Institut Bioinformatics and Microbial Genomics | Danish Covid-19 Genome Consortium |
| EPI_ISL_2863373 | Dutch COVID-19 response team | National Institute for Public Health and the Environment (RIVM) | Adam Meijer, Harry Vennema, Dirk Eggink, Jeroen Cremer, Sharon van den Brink, Bas van der Veer, AnneMarie van den Brandt, Lisa Wijsman, Kim Freriks, Ryanne Jaarsma, Euníce Then, Lynn Aarts, Sanne Bos, Melissa van Tuil, Florian Zwagemaker, Dennis Schmitz, Annelies Kroneman, Karim Hajji, Chantal Reusken, on behalf of the national COVID-19 response team |
| EPI_ISL_2863374 | Department of Bacteria, Parasites and Fungi, Statens Serum Institut, Copenhagen, Denmark | Statens Serum Institut Bioinformatics and Microbial Genomics | Danish Covid-19 Genome Consortium |
| EPI_ISL_2863375 | Dutch COVID-19 response team | National Institute for Public Health and the Environment (RIVM) | Adam Meijer, Harry Vennema, Dirk Eggink, Jeroen Cremer, Sharon van den Brink, Bas van der Veer, AnneMarie van den Brandt, Lisa Wijsman, Kim Freriks, Ryanne Jaarsma, Euníce Then, Lynn Aarts, Sanne Bos, Melissa van Tuil, Florian Zwagemaker, Dennis Schmitz, Annelies Kroneman, Karim Hajji, Chantal Reusken, on behalf of the national COVID-19 response team |
| EPI_ISL_2863376 | Department of Bacteria, Parasites and Fungi, Statens Serum Institut, Copenhagen, Denmark | Statens Serum Institut Bioinformatics and Microbial Genomics | Danish Covid-19 Genome Consortium |
| EPI_ISL_2863377 | Dutch COVID-19 response team | National Institute for Public Health and the Environment (RIVM) | Adam Meijer, Harry Vennema, Dirk Eggink, Jeroen Cremer, Sharon van den Brink, Bas van der Veer, AnneMarie van den Brandt, Lisa Wijsman, Kim Freriks, Ryanne Jaarsma, Euníce Then, Lynn Aarts, Sanne Bos, Melissa van Tuil, Florian Zwagemaker, Dennis Schmitz, Annelies Kroneman, Karim Hajji, Chantal Reusken, on behalf of the national COVID-19 response team |
| EPI_ISL_2863378 | Department of Bacteria, Parasites and Fungi, Statens Serum Institut, Copenhagen, Denmark | Statens Serum Institut Bioinformatics and Microbial Genomics | Danish Covid-19 Genome Consortium |
| EPI_ISL_2863379 | Dutch COVID-19 response team | National Institute for Public Health and the Environment (RIVM) | Adam Meijer, Harry Vennema, Dirk Eggink, Jeroen Cremer, Sharon van den Brink, Bas van der Veer, AnneMarie van den Brandt, Lisa Wijsman, Kim Freriks, Ryanne Jaarsma, Euníce Then, Lynn Aarts, Sanne Bos, Melissa van Tuil, Florian Zwagemaker, Dennis Schmitz, Annelies Kroneman, Karim Hajji, Chantal Reusken, on behalf of the national COVID-19 response team |
| EPI_ISL_2863380 | Department of Bacteria, Parasites and Fungi, Statens Serum Institut, Copenhagen, Denmark | Statens Serum Institut Bioinformatics and Microbial Genomics | Danish Covid-19 Genome Consortium |
| EPI_ISL_2863381 | Dutch COVID-19 response team | National Institute for Public Health and the Environment (RIVM) | Adam Meijer, Harry Vennema, Dirk Eggink, Jeroen Cremer, Sharon van den Brink, Bas van der Veer, AnneMarie van den Brandt, Lisa Wijsman, Kim Freriks, Ryanne Jaarsma, Euníce Then, Lynn Aarts, Sanne Bos, Melissa van Tuil, Florian Zwagemaker, Dennis Schmitz, Annelies Kroneman, Karim Hajji, Chantal Reusken, on behalf of the national COVID-19 response team |
| EPI_ISL_2863382 | Department of Bacteria, Parasites and Fungi, Statens Serum Institut, Copenhagen, Denmark | Statens Serum Institut Bioinformatics and Microbial Genomics | Danish Covid-19 Genome Consortium |
| EPI_ISL_2863383 | Dutch COVID-19 response team | National Institute for Public Health and the Environment (RIVM) | Adam Meijer, Harry Vennema, Dirk Eggink, Jeroen Cremer, Sharon van den Brink, Bas van der Veer, AnneMarie van den Brandt, Lisa Wijsman, Kim Freriks, Ryanne Jaarsma, Euníce Then, Lynn Aarts, Sanne Bos, Melissa van Tuil, Florian Zwagemaker, Dennis Schmitz, Annelies Kroneman, Karim Hajji, Chantal Reusken, on behalf of the national COVID-19 response team |
| EPI_ISL_2863384 | Department of Bacteria, Parasites and Fungi, Statens Serum Institut, Copenhagen, Denmark | Statens Serum Institut Bioinformatics and Microbial Genomics | Danish Covid-19 Genome Consortium |
| EPI_ISL_2863385 | Dutch COVID-19 response team | National Institute for Public Health and the Environment (RIVM) | Adam Meijer, Harry Vennema, Dirk Eggink, Jeroen Cremer, Sharon van den Brink, Bas van der Veer, AnneMarie van den Brandt, Lisa Wijsman, Kim Freriks, Ryanne Jaarsma, Euníce Then, Lynn Aarts, Sanne Bos, Melissa van Tuil, Florian Zwagemaker, Dennis Schmitz, Annelies Kroneman, Karim Hajji, Chantal Reusken, on behalf of the national COVID-19 response team |
| EPI_ISL_2863386 | Department of Bacteria, Parasites and Fungi, Statens Serum Institut, Copenhagen, Denmark | Statens Serum Institut Bioinformatics and Microbial Genomics | Danish Covid-19 Genome Consortium |
| EPI_ISL_2863387 | Dutch COVID-19 response team | National Institute for Public Health and the Environment (RIVM) | Adam Meijer, Harry Vennema, Dirk Eggink, Jeroen Cremer, Sharon van den Brink, Bas van der Veer, AnneMarie van den Brandt, Lisa Wijsman, Kim Freriks, Ryanne Jaarsma, Euníce Then, Lynn Aarts, Sanne Bos, Melissa van Tuil, Florian Zwagemaker, Dennis Schmitz, Annelies Kroneman, Karim Hajji, Chantal Reusken, on behalf of the national COVID-19 response team |
| EPI_ISL_2863388 | Department of Bacteria, Parasites and Fungi, Statens Serum Institut, Copenhagen, Denmark | Statens Serum Institut Bioinformatics and Microbial Genomics | Danish Covid-19 Genome Consortium |
| EPI_ISL_2863389 | Dutch COVID-19 response team | National Institute for Public Health and the Environment (RIVM) | Adam Meijer, Harry Vennema, Dirk Eggink, Jeroen Cremer, Sharon van den Brink, Bas van der Veer, AnneMarie van den Brandt, Lisa Wijsman, Kim Freriks, Ryanne Jaarsma, Euníce Then, Lynn Aarts, Sanne Bos, Melissa van Tuil, Florian Zwagemaker, Dennis Schmitz, Annelies Kroneman, Karim Hajji, Chantal Reusken, on behalf of the national COVID-19 response team |
| EPI_ISL_2863390 | Department of Bacteria, Parasites and Fungi, Statens Serum Institut, Copenhagen, Denmark | Statens Serum Institut Bioinformatics and Microbial Genomics | Danish Covid-19 Genome Consortium |
| EPI_ISL_2863391 | Dutch COVID-19 response team | National Institute for Public Health and the Environment (RIVM) | Adam Meijer, Harry Vennema, Dirk Eggink, Jeroen Cremer, Sharon van den Brink, Bas van der Veer, AnneMarie van den Brandt, Lisa Wijsman, Kim Freriks, Ryanne Jaarsma, Euníce Then, Lynn Aarts, Sanne Bos, Melissa van Tuil, Florian Zwagemaker, Dennis Schmitz, Annelies Kroneman, Karim Hajji, Chantal Reusken, on behalf of the national COVID-19 response team |
| EPI_ISL_2863392 | Department of Bacteria, Parasites and Fungi, Statens Serum Institut, Copenhagen, Denmark | Statens Serum Institut Bioinformatics and Microbial Genomics | Danish Covid-19 Genome Consortium |

| | | | |
|---|---|---|---|
| EPI_ISL_2863393 | Dutch COVID-19 response team | National Institute for Public Health and the Environment (RIVM) | Adam Meijer, Harry Vennema, Dirk Eggink, Jeroen Cremer, Sharon van den Brink, Bas van der Veer, AnneMarie van den Brandt, Lisa Wijsman, Kim Freriks, Ryanne Jaarsma, Euníce Then, Lynn Aarts, Sanne Bos, Melissa van Tuil, Florian Zwagemaker, Dennis Schmitz, Annelies Kroneman, Karim Hajji, Chantal Reusken, on behalf of the national COVID-19 response team |
| EPI_ISL_2863394 | Department of Bacteria, Parasites and Fungi, Statens Serum Institut, Copenhagen, Denmark | Statens Serum Institut Bioinformatics and Microbial Genomics | Danish Covid-19 Genome Consortium |
| EPI_ISL_2863395 | Dutch COVID-19 response team | National Institute for Public Health and the Environment (RIVM) | Adam Meijer, Harry Vennema, Dirk Eggink, Jeroen Cremer, Sharon van den Brink, Bas van der Veer, AnneMarie van den Brandt, Lisa Wijsman, Kim Freriks, Ryanne Jaarsma, Euníce Then, Lynn Aarts, Sanne Bos, Melissa van Tuil, Florian Zwagemaker, Dennis Schmitz, Annelies Kroneman, Karim Hajji, Chantal Reusken, on behalf of the national COVID-19 response team |
| EPI_ISL_2863396 | Department of Bacteria, Parasites and Fungi, Statens Serum Institut, Copenhagen, Denmark | Statens Serum Institut Bioinformatics and Microbial Genomics | Danish Covid-19 Genome Consortium |
| EPI_ISL_2863397 | Dutch COVID-19 response team | National Institute for Public Health and the Environment (RIVM) | Adam Meijer, Harry Vennema, Dirk Eggink, Jeroen Cremer, Sharon van den Brink, Bas van der Veer, AnneMarie van den Brandt, Lisa Wijsman, Kim Freriks, Ryanne Jaarsma, Euníce Then, Lynn Aarts, Sanne Bos, Melissa van Tuil, Florian Zwagemaker, Dennis Schmitz, Annelies Kroneman, Karim Hajji, Chantal Reusken, on behalf of the national COVID-19 response team |
| EPI_ISL_2863398 | Department of Bacteria, Parasites and Fungi, Statens Serum Institut, Copenhagen, Denmark | Statens Serum Institut Bioinformatics and Microbial Genomics | Danish Covid-19 Genome Consortium |
| EPI_ISL_2863399, EPI_ISL_2863400 | Dutch COVID-19 response team | National Institute for Public Health and the Environment (RIVM) | Adam Meijer, Harry Vennema, Dirk Eggink, Jeroen Cremer, Sharon van den Brink, Bas van der Veer, AnneMarie van den Brandt, Lisa Wijsman, Kim Freriks, Ryanne Jaarsma, Euníce Then, Lynn Aarts, Sanne Bos, Melissa van Tuil, Florian Zwagemaker, Dennis Schmitz, Annelies Kroneman, Karim Hajji, Chantal Reusken, on behalf of the national COVID-19 response team |
| EPI_ISL_2863401 | Department of Bacteria, Parasites and Fungi, Statens Serum Institut, Copenhagen, Denmark | Statens Serum Institut Bioinformatics and Microbial Genomics | Danish Covid-19 Genome Consortium |
| EPI_ISL_2863402 | Dutch COVID-19 response team | National Institute for Public Health and the Environment (RIVM) | Adam Meijer, Harry Vennema, Dirk Eggink, Jeroen Cremer, Sharon van den Brink, Bas van der Veer, AnneMarie van den Brandt, Lisa Wijsman, Kim Freriks, Ryanne Jaarsma, Euníce Then, Lynn Aarts, Sanne Bos, Melissa van Tuil, Florian Zwagemaker, Dennis Schmitz, Annelies Kroneman, Karim Hajji, Chantal Reusken, on behalf of the national COVID-19 response team |
| EPI_ISL_2863403 | Department of Bacteria, Parasites and Fungi, Statens Serum Institut, Copenhagen, Denmark | Statens Serum Institut Bioinformatics and Microbial Genomics | Danish Covid-19 Genome Consortium |
| EPI_ISL_2863404 | Dutch COVID-19 response team | National Institute for Public Health and the Environment (RIVM) | Adam Meijer, Harry Vennema, Dirk Eggink, Jeroen Cremer, Sharon van den Brink, Bas van der Veer, AnneMarie van den Brandt, Lisa Wijsman, Kim Freriks, Ryanne Jaarsma, Euníce Then, Lynn Aarts, Sanne Bos, Melissa van Tuil, Florian Zwagemaker, Dennis Schmitz, Annelies Kroneman, Karim Hajji, Chantal Reusken, on behalf of the national COVID-19 response team |
| EPI_ISL_2863405 | Department of Bacteria, Parasites and Fungi, Statens Serum Institut, Copenhagen, Denmark | Statens Serum Institut Bioinformatics and Microbial Genomics | Danish Covid-19 Genome Consortium |
| EPI_ISL_2863406 | Dutch COVID-19 response team | National Institute for Public Health and the Environment (RIVM) | Adam Meijer, Harry Vennema, Dirk Eggink, Jeroen Cremer, Sharon van den Brink, Bas van der Veer, AnneMarie van den Brandt, Lisa Wijsman, Kim Freriks, Ryanne Jaarsma, Euníce Then, Lynn Aarts, Sanne Bos, Melissa van Tuil, Florian Zwagemaker, Dennis Schmitz, Annelies Kroneman, Karim Hajji, Chantal Reusken, on behalf of the national COVID-19 response team |
| EPI_ISL_2863407 | Department of Bacteria, Parasites and Fungi, Statens Serum Institut, Copenhagen, Denmark | Statens Serum Institut Bioinformatics and Microbial Genomics | Danish Covid-19 Genome Consortium |
| EPI_ISL_2863408 | Dutch COVID-19 response team | National Institute for Public Health and the Environment (RIVM) | Adam Meijer, Harry Vennema, Dirk Eggink, Jeroen Cremer, Sharon van den Brink, Bas van der Veer, AnneMarie van den Brandt, Lisa Wijsman, Kim Freriks, Ryanne Jaarsma, Euníce Then, Lynn Aarts, Sanne Bos, Melissa van Tuil, Florian Zwagemaker, Dennis Schmitz, Annelies Kroneman, Karim Hajji, Chantal Reusken, on behalf of the national COVID-19 response team |
| EPI_ISL_2863409 | Department of Bacteria, Parasites and Fungi, Statens Serum Institut, Copenhagen, Denmark | Statens Serum Institut Bioinformatics and Microbial Genomics | Danish Covid-19 Genome Consortium |
| EPI_ISL_2863410 | Dutch COVID-19 response team | National Institute for Public Health and the Environment (RIVM) | Adam Meijer, Harry Vennema, Dirk Eggink, Jeroen Cremer, Sharon van den Brink, Bas van der Veer, AnneMarie van den Brandt, Lisa Wijsman, Kim Freriks, Ryanne Jaarsma, Euníce Then, Lynn Aarts, Sanne Bos, Melissa van Tuil, Florian Zwagemaker, Dennis Schmitz, Annelies Kroneman, Karim Hajji, Chantal Reusken, on behalf of the national COVID-19 response team |
| EPI_ISL_2863411 | Department of Bacteria, Parasites and Fungi, Statens Serum Institut, Copenhagen, Denmark | Statens Serum Institut Bioinformatics and Microbial Genomics | Danish Covid-19 Genome Consortium |
| EPI_ISL_2863412 | Dutch COVID-19 response team | National Institute for Public Health and the Environment (RIVM) | Adam Meijer, Harry Vennema, Dirk Eggink, Jeroen Cremer, Sharon van den Brink, Bas van der Veer, AnneMarie van den Brandt, Lisa Wijsman, Kim Freriks, Ryanne Jaarsma, Euníce Then, Lynn Aarts, Sanne Bos, Melissa van Tuil, Florian Zwagemaker, Dennis Schmitz, Annelies Kroneman, Karim Hajji, Chantal Reusken, on behalf of the national COVID-19 response team |
| EPI_ISL_2863413 | Department of Bacteria, Parasites and Fungi, Statens Serum Institut, Copenhagen, Denmark | Statens Serum Institut Bioinformatics and Microbial Genomics | Danish Covid-19 Genome Consortium |
| EPI_ISL_2863414 | Dutch COVID-19 response team | National Institute for Public Health and the Environment (RIVM) | Adam Meijer, Harry Vennema, Dirk Eggink, Jeroen Cremer, Sharon van den Brink, Bas van der Veer, AnneMarie van den Brandt, Lisa Wijsman, Kim Freriks, Ryanne Jaarsma, Euníce Then, Lynn Aarts, Sanne Bos, Melissa van Tuil, Florian Zwagemaker, Dennis Schmitz, Annelies Kroneman, Karim Hajji, Chantal Reusken, on behalf of the national COVID-19 response team |
| EPI_ISL_2863415 | Department of Bacteria, Parasites and Fungi, Statens Serum Institut, Copenhagen, Denmark | Statens Serum Institut Bioinformatics and Microbial Genomics | Danish Covid-19 Genome Consortium |
| EPI_ISL_2863416, EPI_ISL_2863417 | Dutch COVID-19 response team | National Institute for Public Health and the Environment (RIVM) | Adam Meijer, Harry Vennema, Dirk Eggink, Jeroen Cremer, Sharon van den Brink, Bas van der Veer, AnneMarie van den Brandt, Lisa Wijsman, Kim Freriks, Ryanne Jaarsma, Euníce Then, Lynn Aarts, Sanne Bos, Melissa van Tuil, Florian Zwagemaker, Dennis Schmitz, Annelies Kroneman, Karim Hajji, Chantal Reusken, on behalf of the national COVID-19 response team |
| EPI_ISL_2863418 | Department of Bacteria, Parasites and Fungi, Statens Serum Institut, Copenhagen, Denmark | Statens Serum Institut Bioinformatics and Microbial Genomics | Danish Covid-19 Genome Consortium |
| EPI_ISL_2863419 | Dutch COVID-19 response team | National Institute for Public Health and the Environment (RIVM) | Adam Meijer, Harry Vennema, Dirk Eggink, Jeroen Cremer, Sharon van den Brink, Bas van der Veer, AnneMarie van den Brandt, Lisa Wijsman, Kim Freriks, Ryanne Jaarsma, Euníce Then, Lynn Aarts, Sanne Bos, Melissa van Tuil, Florian Zwagemaker, Dennis Schmitz, Annelies Kroneman, Karim Hajji, Chantal Reusken, on behalf of the national COVID-19 response team |
| EPI_ISL_2863420 | Department of Bacteria, Parasites and Fungi, Statens Serum Institut, Copenhagen, Denmark | Statens Serum Institut Bioinformatics and Microbial Genomics | Danish Covid-19 Genome Consortium |
| EPI_ISL_2863421 | Dutch COVID-19 response team | National Institute for Public Health and the Environment (RIVM) | Adam Meijer, Harry Vennema, Dirk Eggink, Jeroen Cremer, Sharon van den Brink, Bas van der Veer, AnneMarie van den Brandt, Lisa Wijsman, Kim Freriks, Ryanne Jaarsma, Euníce Then, Lynn Aarts, Sanne Bos, Melissa van Tuil, Florian Zwagemaker, Dennis Schmitz, Annelies Kroneman, Karim Hajji, Chantal Reusken, on behalf of the national COVID-19 response team |
| EPI_ISL_2863422 | Department of Bacteria, Parasites and Fungi, Statens Serum Institut, Copenhagen, Denmark | Statens Serum Institut Bioinformatics and Microbial Genomics | Danish Covid-19 Genome Consortium |
| EPI_ISL_2863423 | Dutch COVID-19 response team | National Institute for Public Health and the Environment (RIVM) | Adam Meijer, Harry Vennema, Dirk Eggink, Jeroen Cremer, Sharon van den Brink, Bas van der Veer, AnneMarie van den Brandt, Lisa Wijsman, Kim Freriks, Ryanne Jaarsma, Euníce Then, Lynn Aarts, Sanne Bos, Melissa van Tuil, Florian Zwagemaker, Dennis Schmitz, Annelies Kroneman, Karim Hajji, Chantal Reusken, on behalf of the national COVID-19 response team |
| EPI_ISL_2863424 | Department of Bacteria, Parasites and Fungi, Statens Serum Institut, Copenhagen, Denmark | Statens Serum Institut Bioinformatics and Microbial Genomics | Danish Covid-19 Genome Consortium |
| EPI_ISL_2863425 | Dutch COVID-19 response team | National Institute for Public Health and the Environment (RIVM) | Adam Meijer, Harry Vennema, Dirk Eggink, Jeroen Cremer, Sharon van den Brink, Bas van der Veer, AnneMarie van den Brandt, Lisa Wijsman, Kim Freriks, Ryanne Jaarsma, Euníce Then, Lynn Aarts, Sanne Bos, Melissa van Tuil, Florian Zwagemaker, Dennis Schmitz, Annelies Kroneman, Karim Hajji, Chantal Reusken, on behalf of the national COVID-19 response team |
| EPI_ISL_2863426 | Department of Bacteria, Parasites and Fungi, Statens Serum Institut, Copenhagen, Denmark | Statens Serum Institut Bioinformatics and Microbial Genomics | Danish Covid-19 Genome Consortium |
| EPI_ISL_2863427 | Dutch COVID-19 response team | National Institute for Public Health and the Environment (RIVM) | Adam Meijer, Harry Vennema, Dirk Eggink, Jeroen Cremer, Sharon van den Brink, Bas van der Veer, AnneMarie van den Brandt, Lisa Wijsman, Kim Freriks, Ryanne Jaarsma, Euníce Then, Lynn Aarts, Sanne Bos, Melissa van Tuil, Florian Zwagemaker, Dennis Schmitz, Annelies Kroneman, Karim Hajji, Chantal Reusken, on behalf of the national COVID-19 response team |
| EPI_ISL_2863428 | Department of Bacteria, Parasites and Fungi, Statens Serum Institut, Copenhagen, Denmark | Statens Serum Institut Bioinformatics and Microbial Genomics | Danish Covid-19 Genome Consortium |
| EPI_ISL_2863429 | Dutch COVID-19 response team | National Institute for Public Health and the Environment (RIVM) | Adam Meijer, Harry Vennema, Dirk Eggink, Jeroen Cremer, Sharon van den Brink, Bas van der Veer, AnneMarie van den Brandt, Lisa Wijsman, Kim Freriks, Ryanne Jaarsma, Euníce Then, Lynn Aarts, Sanne Bos, Melissa van Tuil, Florian Zwagemaker, Dennis Schmitz, Annelies Kroneman, Karim Hajji, Chantal Reusken, on behalf of the national COVID-19 response team |
| EPI_ISL_2863430 | Department of Bacteria, Parasites and Fungi, Statens Serum Institut, Copenhagen, Denmark | Statens Serum Institut Bioinformatics and Microbial Genomics | Danish Covid-19 Genome Consortium |
| EPI_ISL_2863431 | Dutch COVID-19 response team | National Institute for Public Health and the Environment (RIVM) | Adam Meijer, Harry Vennema, Dirk Eggink, Jeroen Cremer, Sharon van den Brink, Bas van der Veer, AnneMarie van den Brandt, Lisa Wijsman, Kim Freriks, Ryanne Jaarsma, Euníce Then, Lynn Aarts, Sanne Bos, Melissa van Tuil, Florian Zwagemaker, Dennis Schmitz, Annelies Kroneman, Karim Hajji, Chantal Reusken, on behalf of the national COVID-19 response team |
| EPI_ISL_2863432 | Department of Bacteria, Parasites and Fungi, Statens Serum Institut, Copenhagen, Denmark | Statens Serum Institut Bioinformatics and Microbial Genomics | Danish Covid-19 Genome Consortium |
| EPI_ISL_2863433 | Dutch COVID-19 response team | National Institute for Public Health and the Environment (RIVM) | Adam Meijer, Harry Vennema, Dirk Eggink, Jeroen Cremer, Sharon van den Brink, Bas van der Veer, AnneMarie van den Brandt, Lisa Wijsman, Kim Freriks, Ryanne Jaarsma, Euníce Then, Lynn Aarts, Sanne Bos, Melissa van Tuil, Florian Zwagemaker, Dennis Schmitz, Annelies Kroneman, Karim Hajji, Chantal Reusken, on behalf of the national COVID-19 response team |
| EPI_ISL_2863434 | Department of Bacteria, Parasites and Fungi, Statens Serum Institut, Copenhagen, Denmark | Statens Serum Institut Bioinformatics and Microbial Genomics | Danish Covid-19 Genome Consortium |
| EPI_ISL_2863435, EPI_ISL_2863436 | Dutch COVID-19 response team | National Institute for Public Health and the Environment (RIVM) | Adam Meijer, Harry Vennema, Dirk Eggink, Jeroen Cremer, Sharon van den Brink, Bas van der Veer, AnneMarie van den Brandt, Lisa Wijsman, Kim Freriks, Ryanne Jaarsma, Euníce Then, Lynn Aarts, Sanne Bos, Melissa van Tuil, Florian Zwagemaker, Dennis Schmitz, Annelies Kroneman, Karim Hajji, Chantal Reusken, on behalf of the national COVID-19 response team |

EPI_ISL_2863437, EPI_ISL_2863439, EPI_ISL_2863441, EPI_ISL_2863443, EPI_ISL_2863445, EPI_ISL_2863447, EPI_ISL_2863449, EPI_ISL_2863451, EPI_ISL_2863453, EPI_ISL_2863455, EPI_ISL_2863457, EPI_ISL_2863459, EPI_ISL_2863461, EPI_ISL_2863462, EPI_ISL_2863463, EPI_ISL_2863464, EPI_ISL_2863465, EPI_ISL_2863466, EPI_ISL_2863467, EPI_ISL_2863468, EPI_ISL_2863469, EPI_ISL_2863470, EPI_ISL_2863471, EPI_ISL_2863472, EPI_ISL_2863473, EPI_ISL_2863474, EPI_ISL_2863475, EPI_ISL_2863476, EPI_ISL_2863477, EPI_ISL_2863478, EPI_ISL_2863479, EPI_ISL_2863480, EPI_ISL_2863481, EPI_ISL_2863482, EPI_ISL_2863483, EPI_ISL_2863484, EPI_ISL_2863485, EPI_ISL_2863486, EPI_ISL_2863487, EPI_ISL_2863488, EPI_ISL_2863489, EPI_ISL_2863490, EPI_ISL_2863491, EPI_ISL_2863492, EPI_ISL_2863493, EPI_ISL_2863494, EPI_ISL_2863495, EPI_ISL_2863496, EPI_ISL_2863497, EPI_ISL_2863498, EPI_ISL_2863499, EPI_ISL_2863500, EPI_ISL_2863501, EPI_ISL_2863502, EPI_ISL_2863504, EPI_ISL_2863505, EPI_ISL_2863506, EPI_ISL_2863507, EPI_ISL_2863508, EPI_ISL_2863509, EPI_ISL_2863510, EPI_ISL_2863511, EPI_ISL_2863512, EPI_ISL_2863513, EPI_ISL_2863514, EPI_ISL_2863515, EPI_ISL_2863516, EPI_ISL_2863517, EPI_ISL_2863518, EPI_ISL_2863519, EPI_ISL_2863520, EPI_ISL_2863521, EPI_ISL_2863522, EPI_ISL_2863523, EPI_ISL_2863524, EPI_ISL_2863525, EPI_ISL_2863526, EPI_ISL_2863527, EPI_ISL_2863528, EPI_ISL_2863529, EPI_ISL_2863530, EPI_ISL_2863531, EPI_ISL_2863532, EPI_ISL_2863533, EPI_ISL_2863534, EPI_ISL_2863535, EPI_ISL_2863536, EPI_ISL_2863537, EPI_ISL_2863538, EPI_ISL_2863539, EPI_ISL_2863540, EPI_ISL_2863541, EPI_ISL_2863542, EPI_ISL_2863543, EPI_ISL_2863544, EPI_ISL_2863545, EPI_ISL_2863546, EPI_ISL_2863547, EPI_ISL_2863548, EPI_ISL_2863549, EPI_ISL_2863550, EPI_ISL_2863551, EPI_ISL_2863552, EPI_ISL_2863553, EPI_ISL_2863554, EPI_ISL_2863555, EPI_ISL_2863556, EPI_ISL_2863557, EPI_ISL_2863558, EPI_ISL_2863559, EPI_ISL_2863560, EPI_ISL_2863561, EPI_ISL_2863562, EPI_ISL_2863563, EPI_ISL_2863564, EPI_ISL_2863565, EPI_ISL_2863566, EPI_ISL_2863567, EPI_ISL_2863568, EPI_ISL_2863570, EPI_ISL_2863572, EPI_ISL_2863574, EPI_ISL_2863577, EPI_ISL_2863579, EPI_ISL_2863581, EPI_ISL_2863583, EPI_ISL_2863590, EPI_ISL_2863592, EPI_ISL_2863594, EPI_ISL_2863597, EPI_ISL_2863599, EPI_ISL_2863601, EPI_ISL_2863603, EPI_ISL_2863606, EPI_ISL_2863610, EPI_ISL_2863612, EPI_ISL_2863615, EPI_ISL_2863617, EPI_ISL_2863620, EPI_ISL_2863622, EPI_ISL_2863624, EPI_ISL_2863627, EPI_ISL_2863629, EPI_ISL_2863631, EPI_ISL_2863633, EPI_ISL_2863636, EPI_ISL_2863638, EPI_ISL_2863640, EPI_ISL_2863642, EPI_ISL_2863644, EPI_ISL_2863646, EPI_ISL_2863649, EPI_ISL_2863650, EPI_ISL_2863652, EPI_ISL_2863654, EPI_ISL_2863657, EPI_ISL_2863659, EPI_ISL_2863661, EPI_ISL_2863663, EPI_ISL_2863665, EPI_ISL_2863668, EPI_ISL_2863670, EPI_ISL_2863672, EPI_ISL_2863674, EPI_ISL_2863676, EPI_ISL_2863678, EPI_ISL_2863681, EPI_ISL_2863683, EPI_ISL_2863686, EPI_ISL_2863688, EPI_ISL_2863690, EPI_ISL_2863692, EPI_ISL_2863694, EPI_ISL_2863697, EPI_ISL_2863699, EPI_ISL_2863701, EPI_ISL_2863703, EPI_ISL_2863705, EPI_ISL_2863708, EPI_ISL_2863710, EPI_ISL_2863713, EPI_ISL_2863715, EPI_ISL_2863718, EPI_ISL_2863720, EPI_ISL_2863722, EPI_ISL_2863724, EPI_ISL_2863726, EPI_ISL_2863730, EPI_ISL_2863731, EPI_ISL_2863732, EPI_ISL_2863733, EPI_ISL_2863734, EPI_ISL_2863735, EPI_ISL_2863736, EPI_ISL_2863737, EPI_ISL_2863738, EPI_ISL_2863739, EPI_ISL_2863740,

| | | | |
|---|---|---|---|
| EPI_ISL_2863741, EPI_ISL_2863742, EPI_ISL_2863743, EPI_ISL_2863744, EPI_ISL_2863745, EPI_ISL_2863746, EPI_ISL_2863747, EPI_ISL_2863748, EPI_ISL_2863749, EPI_ISL_2863750, EPI_ISL_2863751, EPI_ISL_2863752, EPI_ISL_2863753, EPI_ISL_2863754, EPI_ISL_2863755, EPI_ISL_2863756, EPI_ISL_2863757, EPI_ISL_2863758, EPI_ISL_2863759, EPI_ISL_2863760, EPI_ISL_2863761, EPI_ISL_2863762, EPI_ISL_2863763, EPI_ISL_2863764, EPI_ISL_2863765, EPI_ISL_2863766, EPI_ISL_2863767, EPI_ISL_2863768, EPI_ISL_2863769, EPI_ISL_2863770, EPI_ISL_2863771, EPI_ISL_2863772, EPI_ISL_2863773, EPI_ISL_2863774, EPI_ISL_2863775, EPI_ISL_2863776, EPI_ISL_2863777, EPI_ISL_2863778, EPI_ISL_2863779, EPI_ISL_2863780, EPI_ISL_2863781, EPI_ISL_2863782, EPI_ISL_2863783, EPI_ISL_2863784, EPI_ISL_2863785, EPI_ISL_2863786, EPI_ISL_2863787, EPI_ISL_2863788, EPI_ISL_2863789, EPI_ISL_2863790, EPI_ISL_2863791, EPI_ISL_2863792, EPI_ISL_2863793, EPI_ISL_2863794 | | | |
| see above | Department of Bacteria, Parasites and Fungi, Statens Serum Institut, Copenhagen, Denmark | Statens Serum Institut Bioinformatics and Microbial Genomics | Danish Covid-19 Genome Consortium |
| EPI_ISL_2863898, EPI_ISL_2863899 | Lboratorio Central de Saude Publica do Estado do Parana (LACEN/PR) | Laboratory of Respiratory Viruses and Measles, Oswaldo Cruz Institute, FIOCRUZ | Paola Resende, Luciana Appolinario, Fernando Motta, Anna Carolina Paixao, Ana Carolina Mendonca, Alice Sampaio Rocha, Taina Venas, Elisa Cavalcante Pereira, Renata Serrano Lopes, Agatha Cristinne Prudencio Soares, Ighor Leonardo Arantes Gomes, Irina Riediger, Marilda Siqueira on behalf of the Fiocruz COVID-19 Genomic Surveillance Network |
| EPI_ISL_2863902, EPI_ISL_2863903, EPI_ISL_2863905, EPI_ISL_2863906, EPI_ISL_2863907, EPI_ISL_2863909, EPI_ISL_2863914, EPI_ISL_2863916, EPI_ISL_2863923, EPI_ISL_2863924, EPI_ISL_2863927, EPI_ISL_2863928 | | | |
| see above | Arizona State University | Arizona State University | Peter T. Skidmore, LaRinda A. Holland, Matthew F. Smith, Rabia Maqsood, Nicholas J. Mellor, Joy M. Blain, Valerie Harris, Joshua LaBaer, Vel Murugan, Efrem S. Lim |
| EPI_ISL_2863929, EPI_ISL_2863930, EPI_ISL_2863931, EPI_ISL_2863932, EPI_ISL_2863933, EPI_ISL_2863934, EPI_ISL_2863935 | Department of Acute Infectious Diseases Control and Prevention, Yunnan Provincial Center for Disease Control and Prevention | Department of Acute Infectious Diseases Control and Prevention, Yunnan Provincial Center for Disease Control and Prevention | Meiling Zhang, Jienan Zhou, Zhaosheng Liu, Xiaoqing Fu |
| EPI_ISL_2863936, EPI_ISL_2863937, EPI_ISL_2863938, EPI_ISL_2863939, EPI_ISL_2863942 | Nucleic Acid Testing, National Reference Laboratory | GIGA Medical Genomics | Yvan Butera, Keith Durkin, Maria Artesi, Bouchra Boujemla, Robert Rutayisire, Reuben Sindayiheba, Patrick Tuyisenge, Esperence Umumararungu, Sébastien Bontems, Marie-Pierre Hayette, Nathalie Renotte, Swaibu Gatare, Jacob Souopgui, Sabin Nsanzimana, Vincent Bours, Léon Mutesa |
| EPI_ISL_2863943, EPI_ISL_2863944, EPI_ISL_2863946, EPI_ISL_2863947, EPI_ISL_2863950, EPI_ISL_2863951, EPI_ISL_2863952, EPI_ISL_2863953, EPI_ISL_2863954, EPI_ISL_2863955, EPI_ISL_2863956 | | | |
| see above | Fondazione IRCCS Ca' Granda Ospedale Maggiore Policlinico | Fondazione IRCCS Ca' Granda Ospedale Maggiore Policlinico | Ferruccio Ceriotti, Sara Uceda Renteria |
| EPI_ISL_2864126, EPI_ISL_2864127, EPI_ISL_2864128, EPI_ISL_2864129, EPI_ISL_2864130, EPI_ISL_2864131, EPI_ISL_2864132, EPI_ISL_2864133, EPI_ISL_2864134, EPI_ISL_2864135, EPI_ISL_2864136, EPI_ISL_2864137, EPI_ISL_2864138, EPI_ISL_2864139, EPI_ISL_2864140 | | | |
| see above | Azienda Sanitaria Locale di Piacenza - Presidio Ospedaliero - Laboratorio di Microbiologia | Istituto Zooprofilattico Sperimentale della Lombardia e dell'Emilia Romagna (IZSLER), Risk Analysis and Genomic Epidemiology Unit | Lo Cascio Giuliana, Schiavo Roberta, Peroni Irene, Marina Morganti, Ilaria Menozzi, Erika Scaltriti, Stefano Pongolini |
| EPI_ISL_2864141, EPI_ISL_2864142, EPI_ISL_2864143, EPI_ISL_2864144, EPI_ISL_2864145, EPI_ISL_2864146, EPI_ISL_2864147, EPI_ISL_2864148 | Microbiology Department. Complexo Hospitalario Universitario de Vigo | Microbiology Department. Complexo Hospitalario Universitario de Vigo | Alfaya N, Alonso I, Alvarez M, Cabrera JJ, Carballo R, Cores O, Cortizo S, del-Campo V, Martinez L, Mediero G, Perez S, Davina C, Potel C, Regueiro B, Rey S, Vasallo FJ |
| EPI_ISL_2864207, EPI_ISL_2864210, EPI_ISL_2864238, EPI_ISL_2864254, EPI_ISL_2864261, EPI_ISL_2864304, EPI_ISL_2864305, EPI_ISL_2864306, EPI_ISL_2864307, EPI_ISL_2864310, EPI_ISL_2864311, EPI_ISL_2864340 | | | |
| see above | KU Leuven, Rega Institute, Clinical and Epidemiological Virology | KU Leuven, Rega Institute, Clinical and Epidemiological Virology | Tony Wawina-Bokalanga, Bert Vanmechelen, Joan Marti-Carerras, Piet Maes |
| EPI_ISL_2864380, EPI_ISL_2864416, EPI_ISL_2864431, EPI_ISL_2864432, EPI_ISL_2864433, EPI_ISL_2864434, EPI_ISL_2864435, EPI_ISL_2864436, EPI_ISL_2864437, EPI_ISL_2864438, EPI_ISL_2864439, EPI_ISL_2864440, EPI_ISL_2864441, EPI_ISL_2864442, EPI_ISL_2864443, EPI_ISL_2864444, EPI_ISL_2864445, EPI_ISL_2864446, EPI_ISL_2864447, EPI_ISL_2864448, EPI_ISL_2864449, EPI_ISL_2864450, EPI_ISL_2864451, EPI_ISL_2864452, EPI_ISL_2864453 | | | |
| see above | Arcispedale Santa Maria Nuova, Autoimmunità, Allergologia e Biotecnologie Innovative | Istituto Zooprofilattico Sperimentale della Lombardia e dell'Emilia Romagna (IZSLER), Risk Analysis and Genomic Epidemiology Unit | Alessandro Zerbini, Lucia Belloni, Stefania Croci, Marina Morganti, Ilaria Menozzi, Erika Scaltriti, Stefano Pongolini |
| EPI_ISL_2864459, EPI_ISL_2864473, EPI_ISL_2864474, EPI_ISL_2864478, EPI_ISL_2864483, EPI_ISL_2864485, EPI_ISL_2864489, EPI_ISL_2864493 | KU Leuven, Rega Institute, Clinical and Epidemiological Virology | KU Leuven, Rega Institute, Clinical and Epidemiological Virology | Tony Wawina-Bokalanga, Bert Vanmechelen, Joan Marti-Carerras, Piet Maes |
| EPI_ISL_2864525, EPI_ISL_2864541, EPI_ISL_2864542, EPI_ISL_2864543, EPI_ISL_2864544, EPI_ISL_2864545, EPI_ISL_2864546, EPI_ISL_2864547, EPI_ISL_2864548, EPI_ISL_2864549, EPI_ISL_2864550, EPI_ISL_2864551, EPI_ISL_2864552, EPI_ISL_2864554, EPI_ISL_2864555, EPI_ISL_2864556, EPI_ISL_2864557, EPI_ISL_2864558, EPI_ISL_2864559, EPI_ISL_2864560, EPI_ISL_2864561 | | | |
| see above | UW Virology Lab | UW Virology Lab | Pavitra Roychoudhury, Hong Xie, Lasata Shrestha, Shah Mohamed Bakhash, Tien V. Nguyen, Noah R. Baker, Sean Ellis, Meei-Li Huang, Keith R Jerome, Alexander Greninger |
| EPI_ISL_2864567, EPI_ISL_2864573, EPI_ISL_2864574, EPI_ISL_2864575, EPI_ISL_2864576, EPI_ISL_2864629, EPI_ISL_2864641, EPI_ISL_2864645, EPI_ISL_2864647, EPI_ISL_2864691, EPI_ISL_2864696, EPI_ISL_2864707, EPI_ISL_2864708, EPI_ISL_2864709, EPI_ISL_2864710, EPI_ISL_2864711 | | | |
| see above | KU Leuven, Rega Institute, Clinical and Epidemiological Virology | KU Leuven, Rega Institute, Clinical and Epidemiological Virology | Tony Wawina-Bokalanga, Bert Vanmechelen, Joan Marti-Carerras, Piet Maes |
| EPI_ISL_2864713 | Genomic Research Laboratory, Bangabandhu Sheikh Mujib Medical University | Genomic Research Laboratory, BSMMU | Laila Anjuman Banu, Md.Saydur Rahman, Sharadindu Kanti Sinha, Arifa Akram, Md.Sharfuddin Ahmed |
| EPI_ISL_2864714, EPI_ISL_2864715, EPI_ISL_2864717, EPI_ISL_2864718, EPI_ISL_2864719, EPI_ISL_2864720, EPI_ISL_2864721, EPI_ISL_2864762, EPI_ISL_2864771 | KU Leuven, Rega Institute, Clinical and Epidemiological Virology | KU Leuven, Rega Institute, Clinical and Epidemiological Virology | Tony Wawina-Bokalanga, Bert Vanmechelen, Joan Marti-Carerras, Piet Maes |
| EPI_ISL_2864777 | Lighthouse Laboratory Plymouth | Wellcome Sanger Institute for the COVID-19 Genomics UK (COG-UK) Consortium | Lighthouse Laboratory Plymouth and Alex Alderton, Roberto Amato, Jeffrey Barrett, Sonia Goncalves, Ewan Harrison, David K. Jackson, Ian Johnston, Dominic Kwiatkowski, Cordelia Langford, John Sillitoe on behalf of the Wellcome Sanger Institute COVID-19 Surveillance Team |
| EPI_ISL_2864778 | Berkshire and Surrey Pathology Services Lighthouse Laboratory | Wellcome Sanger Institute for the COVID-19 Genomics UK (COG-UK) Consortium | Berkshire and Surrey Pathology Services Lighthouse Laboratory and Alex Alderton, Roberto Amato, Jeffrey Barrett, Sonia Goncalves, Ewan Harrison, David K. Jackson, Ian Johnston, Dominic Kwiatkowski, Cordelia Langford, John Sillitoe on behalf of the Wellcome Sanger Institute COVID-19 Surveillance Team |
| EPI_ISL_2864779 | Lighthouse Laboratory Plymouth | Wellcome Sanger Institute for the COVID-19 Genomics UK (COG-UK) Consortium | Lighthouse Laboratory Plymouth and Alex Alderton, Roberto Amato, Jeffrey Barrett, Sonia Goncalves, Ewan Harrison, David K. Jackson, Ian Johnston, Dominic Kwiatkowski, Cordelia Langford, John Sillitoe on behalf of the Wellcome Sanger Institute COVID-19 Surveillance Team |
| EPI_ISL_2864780 | Lighthouse Lab in Glasgow | Wellcome Sanger Institute for the COVID-19 Genomics UK (COG-UK) Consortium | Harper VanSteenhouse, Yumi Kasai, David Gray, Carol Clugston, Anna Dominiczak and Alex Alderton, Roberto Amato, Jeffrey Barrett, Sonia Goncalves, Ewan Harrison, David K. Jackson, Ian Johnston, Dominic Kwiatkowski, Cordelia Langford, John Sillitoe on behalf of the Wellcome Sanger Institute COVID-19 Surveillance Team |
| EPI_ISL_2864781, EPI_ISL_2864782 | Lighthouse Laboratory Plymouth | Wellcome Sanger Institute for the COVID-19 Genomics UK (COG-UK) Consortium | Lighthouse Laboratory Plymouth and Alex Alderton, Roberto Amato, Jeffrey Barrett, Sonia Goncalves, Ewan Harrison, David K. Jackson, Ian Johnston, Dominic Kwiatkowski, Cordelia Langford, John Sillitoe on behalf of the Wellcome Sanger Institute COVID-19 Surveillance Team |
| EPI_ISL_2864783 | Berkshire and Surrey Pathology Services Lighthouse Laboratory | Wellcome Sanger Institute for the COVID-19 Genomics UK (COG-UK) Consortium | Berkshire and Surrey Pathology Services Lighthouse Laboratory and Alex Alderton, Roberto Amato, Jeffrey Barrett, Sonia Goncalves, Ewan Harrison, David K. Jackson, Ian Johnston, Dominic Kwiatkowski, Cordelia Langford, John Sillitoe on behalf of the Wellcome Sanger Institute COVID-19 Surveillance Team |
| EPI_ISL_2864784, EPI_ISL_2864785, EPI_ISL_2864786, EPI_ISL_2864787 | Lighthouse Laboratory Plymouth | Wellcome Sanger Institute for the COVID-19 Genomics UK (COG-UK) Consortium | Lighthouse Laboratory Plymouth and Alex Alderton, Roberto Amato, Jeffrey Barrett, Sonia Goncalves, Ewan Harrison, David K. Jackson, Ian Johnston, Dominic Kwiatkowski, Cordelia Langford, John Sillitoe on behalf of the Wellcome Sanger Institute COVID-19 Surveillance Team |
| EPI_ISL_2864788 | Lighthouse Lab in Glasgow | Wellcome Sanger Institute for the COVID-19 Genomics UK (COG-UK) Consortium | Harper VanSteenhouse, Yumi Kasai, David Gray, Carol Clugston, Anna Dominiczak and Alex Alderton, Roberto Amato, Jeffrey Barrett, Sonia Goncalves, Ewan Harrison, David K. Jackson, Ian Johnston, Dominic Kwiatkowski, Cordelia Langford, John Sillitoe on behalf of the Wellcome Sanger Institute COVID-19 Surveillance Team |
| EPI_ISL_2864789 | Lighthouse Laboratory Plymouth | Wellcome Sanger Institute for the COVID-19 Genomics UK (COG-UK) Consortium | Lighthouse Laboratory Plymouth and Alex Alderton, Roberto Amato, Jeffrey Barrett, Sonia Goncalves, Ewan Harrison, David K. Jackson, Ian Johnston, Dominic Kwiatkowski, Cordelia Langford, John Sillitoe on behalf of the Wellcome Sanger Institute COVID-19 Surveillance Team |
| EPI_ISL_2864790, EPI_ISL_2864791 | Berkshire and Surrey Pathology Services Lighthouse Laboratory | Wellcome Sanger Institute for the COVID-19 Genomics UK (COG-UK) Consortium | Berkshire and Surrey Pathology Services Lighthouse Laboratory and Alex Alderton, Roberto Amato, Jeffrey Barrett, Sonia Goncalves, Ewan Harrison, David K. Jackson, Ian Johnston, Dominic Kwiatkowski, Cordelia Langford, John Sillitoe on behalf of the Wellcome Sanger Institute COVID-19 Surveillance Team |
| EPI_ISL_2864792 | Lighthouse Laboratory Plymouth | Wellcome Sanger Institute for the COVID-19 Genomics UK (COG-UK) Consortium | Lighthouse Laboratory Plymouth and Alex Alderton, Roberto Amato, Jeffrey Barrett, Sonia Goncalves, Ewan Harrison, David K. Jackson, Ian Johnston, Dominic Kwiatkowski, Cordelia Langford, John Sillitoe on behalf of the Wellcome Sanger Institute COVID-19 Surveillance Team |
| EPI_ISL_2864793, EPI_ISL_2864794 | Lighthouse Lab in Glasgow | Wellcome Sanger Institute for the COVID-19 Genomics UK (COG-UK) Consortium | Harper VanSteenhouse, Yumi Kasai, David Gray, Carol Clugston, Anna Dominiczak and Alex Alderton, Roberto Amato, Jeffrey Barrett, Sonia Goncalves, Ewan Harrison, David K. Jackson, Ian Johnston, Dominic Kwiatkowski, Cordelia Langford, John Sillitoe on behalf of the Wellcome Sanger Institute COVID-19 Surveillance Team |
| EPI_ISL_2864795, EPI_ISL_2864796 | Berkshire and Surrey Pathology Services Lighthouse Laboratory | Wellcome Sanger Institute for the COVID-19 Genomics UK (COG-UK) Consortium | Berkshire and Surrey Pathology Services Lighthouse Laboratory and Alex Alderton, Roberto Amato, Jeffrey Barrett, Sonia Goncalves, Ewan Harrison, David K. Jackson, Ian Johnston, Dominic Kwiatkowski, Cordelia Langford, John Sillitoe on behalf of the Wellcome Sanger Institute COVID-19 Surveillance Team |
| EPI_ISL_2864797, EPI_ISL_2864798, | Lighthouse Laboratory Plymouth | Wellcome Sanger Institute for the COVID-19 Genomics UK (COG-UK) Consortium | Lighthouse Laboratory Plymouth and Alex Alderton, Roberto Amato, Jeffrey Barrett, Sonia Goncalves, Ewan Harrison, David K. Jackson, Ian Johnston, Dominic Kwiatkowski, Cordelia Langford, John Sillitoe on behalf of the Wellcome Sanger Institute COVID-19 Surveillance Team |

| | | | |
|---|---|---|---|
| EPI_ISL_2864799 | | | |
| EPI_ISL_2864800, EPI_ISL_2864801, EPI_ISL_2864802 | Berkshire and Surrey Pathology Services Lighthouse Laboratory | Wellcome Sanger Institute for the COVID-19 Genomics UK (COG-UK) Consortium | Berkshire and Surrey Pathology Services Lighthouse Laboratory and Alex Alderton, Roberto Amato, Jeffrey Barrett, Sonia Goncalves, Ewan Harrison, David K. Jackson, Ian Johnston, Dominic Kwiatkowski, Cordelia Langford, John Sillitoe on behalf of the Wellcome Sanger Institute COVID-19 Surveillance Team |
| EPI_ISL_2864803, EPI_ISL_2864804, EPI_ISL_2864805, EPI_ISL_2864806, EPI_ISL_2864807 | Lighthouse Laboratory Plymouth | Wellcome Sanger Institute for the COVID-19 Genomics UK (COG-UK) Consortium | Lighthouse Laboratory Plymouth and Alex Alderton, Roberto Amato, Jeffrey Barrett, Sonia Goncalves, Ewan Harrison, David K. Jackson, Ian Johnston, Dominic Kwiatkowski, Cordelia Langford, John Sillitoe on behalf of the Wellcome Sanger Institute COVID-19 Surveillance Team |
| EPI_ISL_2864808 | Berkshire and Surrey Pathology Services Lighthouse Laboratory | Wellcome Sanger Institute for the COVID-19 Genomics UK (COG-UK) Consortium | Berkshire and Surrey Pathology Services Lighthouse Laboratory and Alex Alderton, Roberto Amato, Jeffrey Barrett, Sonia Goncalves, Ewan Harrison, David K. Jackson, Ian Johnston, Dominic Kwiatkowski, Cordelia Langford, John Sillitoe on behalf of the Wellcome Sanger Institute COVID-19 Surveillance Team |
| EPI_ISL_2864809 | Lighthouse Laboratory Plymouth | Wellcome Sanger Institute for the COVID-19 Genomics UK (COG-UK) Consortium | Lighthouse Laboratory Plymouth and Alex Alderton, Roberto Amato, Jeffrey Barrett, Sonia Goncalves, Ewan Harrison, David K. Jackson, Ian Johnston, Dominic Kwiatkowski, Cordelia Langford, John Sillitoe on behalf of the Wellcome Sanger Institute COVID-19 Surveillance Team |
| EPI_ISL_2864810 | Berkshire and Surrey Pathology Services Lighthouse Laboratory | Wellcome Sanger Institute for the COVID-19 Genomics UK (COG-UK) Consortium | Berkshire and Surrey Pathology Services Lighthouse Laboratory and Alex Alderton, Roberto Amato, Jeffrey Barrett, Sonia Goncalves, Ewan Harrison, David K. Jackson, Ian Johnston, Dominic Kwiatkowski, Cordelia Langford, John Sillitoe on behalf of the Wellcome Sanger Institute COVID-19 Surveillance Team |
| EPI_ISL_2864811 | Lighthouse Laboratory Plymouth | Wellcome Sanger Institute for the COVID-19 Genomics UK (COG-UK) Consortium | Lighthouse Laboratory Plymouth and Alex Alderton, Roberto Amato, Jeffrey Barrett, Sonia Goncalves, Ewan Harrison, David K. Jackson, Ian Johnston, Dominic Kwiatkowski, Cordelia Langford, John Sillitoe on behalf of the Wellcome Sanger Institute COVID-19 Surveillance Team |
| EPI_ISL_2864812 | Lighthouse Lab in Glasgow | Wellcome Sanger Institute for the COVID-19 Genomics UK (COG-UK) Consortium | Harper VanSteenhouse, Yumi Kasai, David Gray, Carol Clugston, Anna Dominiczak and Alex Alderton, Roberto Amato, Jeffrey Barrett, Sonia Goncalves, Ewan Harrison, David K. Jackson, Ian Johnston, Dominic Kwiatkowski, Cordelia Langford, John Sillitoe on behalf of the Wellcome Sanger Institute COVID-19 Surveillance Team |
| EPI_ISL_2864813 | Berkshire and Surrey Pathology Services Lighthouse Laboratory | Wellcome Sanger Institute for the COVID-19 Genomics UK (COG-UK) Consortium | Berkshire and Surrey Pathology Services Lighthouse Laboratory and Alex Alderton, Roberto Amato, Jeffrey Barrett, Sonia Goncalves, Ewan Harrison, David K. Jackson, Ian Johnston, Dominic Kwiatkowski, Cordelia Langford, John Sillitoe on behalf of the Wellcome Sanger Institute COVID-19 Surveillance Team |
| EPI_ISL_2864814, EPI_ISL_2864815 | Lighthouse Laboratory Plymouth | Wellcome Sanger Institute for the COVID-19 Genomics UK (COG-UK) Consortium | Lighthouse Laboratory Plymouth and Alex Alderton, Roberto Amato, Jeffrey Barrett, Sonia Goncalves, Ewan Harrison, David K. Jackson, Ian Johnston, Dominic Kwiatkowski, Cordelia Langford, John Sillitoe on behalf of the Wellcome Sanger Institute COVID-19 Surveillance Team |
| EPI_ISL_2864816 | Berkshire and Surrey Pathology Services Lighthouse Laboratory | Wellcome Sanger Institute for the COVID-19 Genomics UK (COG-UK) Consortium | Berkshire and Surrey Pathology Services Lighthouse Laboratory and Alex Alderton, Roberto Amato, Jeffrey Barrett, Sonia Goncalves, Ewan Harrison, David K. Jackson, Ian Johnston, Dominic Kwiatkowski, Cordelia Langford, John Sillitoe on behalf of the Wellcome Sanger Institute COVID-19 Surveillance Team |
| EPI_ISL_2864817, EPI_ISL_2864818 | Lighthouse Laboratory Plymouth | Wellcome Sanger Institute for the COVID-19 Genomics UK (COG-UK) Consortium | Lighthouse Laboratory Plymouth and Alex Alderton, Roberto Amato, Jeffrey Barrett, Sonia Goncalves, Ewan Harrison, David K. Jackson, Ian Johnston, Dominic Kwiatkowski, Cordelia Langford, John Sillitoe on behalf of the Wellcome Sanger Institute COVID-19 Surveillance Team |
| EPI_ISL_2864819 | Lighthouse Lab in Glasgow | Wellcome Sanger Institute for the COVID-19 Genomics UK (COG-UK) Consortium | Harper VanSteenhouse, Yumi Kasai, David Gray, Carol Clugston, Anna Dominiczak and Alex Alderton, Roberto Amato, Jeffrey Barrett, Sonia Goncalves, Ewan Harrison, David K. Jackson, Ian Johnston, Dominic Kwiatkowski, Cordelia Langford, John Sillitoe on behalf of the Wellcome Sanger Institute COVID-19 Surveillance Team |
| EPI_ISL_2864820, EPI_ISL_2864821 | Lighthouse Laboratory Plymouth | Wellcome Sanger Institute for the COVID-19 Genomics UK (COG-UK) Consortium | Lighthouse Laboratory Plymouth and Alex Alderton, Roberto Amato, Jeffrey Barrett, Sonia Goncalves, Ewan Harrison, David K. Jackson, Ian Johnston, Dominic Kwiatkowski, Cordelia Langford, John Sillitoe on behalf of the Wellcome Sanger Institute COVID-19 Surveillance Team |
| EPI_ISL_2864822 | Berkshire and Surrey Pathology Services Lighthouse Laboratory | Wellcome Sanger Institute for the COVID-19 Genomics UK (COG-UK) Consortium | Berkshire and Surrey Pathology Services Lighthouse Laboratory and Alex Alderton, Roberto Amato, Jeffrey Barrett, Sonia Goncalves, Ewan Harrison, David K. Jackson, Ian Johnston, Dominic Kwiatkowski, Cordelia Langford, John Sillitoe on behalf of the Wellcome Sanger Institute COVID-19 Surveillance Team |
| EPI_ISL_2864823, EPI_ISL_2864824, EPI_ISL_2864825, EPI_ISL_2864826 | Lighthouse Laboratory Plymouth | Wellcome Sanger Institute for the COVID-19 Genomics UK (COG-UK) Consortium | Lighthouse Laboratory Plymouth and Alex Alderton, Roberto Amato, Jeffrey Barrett, Sonia Goncalves, Ewan Harrison, David K. Jackson, Ian Johnston, Dominic Kwiatkowski, Cordelia Langford, John Sillitoe on behalf of the Wellcome Sanger Institute COVID-19 Surveillance Team |
| EPI_ISL_2864827 | Berkshire and Surrey Pathology Services Lighthouse Laboratory | Wellcome Sanger Institute for the COVID-19 Genomics UK (COG-UK) Consortium | Berkshire and Surrey Pathology Services Lighthouse Laboratory and Alex Alderton, Roberto Amato, Jeffrey Barrett, Sonia Goncalves, Ewan Harrison, David K. Jackson, Ian Johnston, Dominic Kwiatkowski, Cordelia Langford, John Sillitoe on behalf of the Wellcome Sanger Institute COVID-19 Surveillance Team |
| EPI_ISL_2864828, EPI_ISL_2864829 | Lighthouse Laboratory Plymouth | Wellcome Sanger Institute for the COVID-19 Genomics UK (COG-UK) Consortium | Lighthouse Laboratory Plymouth and Alex Alderton, Roberto Amato, Jeffrey Barrett, Sonia Goncalves, Ewan Harrison, David K. Jackson, Ian Johnston, Dominic Kwiatkowski, Cordelia Langford, John Sillitoe on behalf of the Wellcome Sanger Institute COVID-19 Surveillance Team |
| EPI_ISL_2864830 | Berkshire and Surrey Pathology Services Lighthouse Laboratory | Wellcome Sanger Institute for the COVID-19 Genomics UK (COG-UK) Consortium | Berkshire and Surrey Pathology Services Lighthouse Laboratory and Alex Alderton, Roberto Amato, Jeffrey Barrett, Sonia Goncalves, Ewan Harrison, David K. Jackson, Ian Johnston, Dominic Kwiatkowski, Cordelia Langford, John Sillitoe on behalf of the Wellcome Sanger Institute COVID-19 Surveillance Team |
| EPI_ISL_2864831, EPI_ISL_2864832, EPI_ISL_2864833, EPI_ISL_2864834 | Lighthouse Laboratory Plymouth | Wellcome Sanger Institute for the COVID-19 Genomics UK (COG-UK) Consortium | Lighthouse Laboratory Plymouth and Alex Alderton, Roberto Amato, Jeffrey Barrett, Sonia Goncalves, Ewan Harrison, David K. Jackson, Ian Johnston, Dominic Kwiatkowski, Cordelia Langford, John Sillitoe on behalf of the Wellcome Sanger Institute COVID-19 Surveillance Team |
| EPI_ISL_2864835 | Berkshire and Surrey Pathology Services Lighthouse Laboratory | Wellcome Sanger Institute for the COVID-19 Genomics UK (COG-UK) Consortium | Berkshire and Surrey Pathology Services Lighthouse Laboratory and Alex Alderton, Roberto Amato, Jeffrey Barrett, Sonia Goncalves, Ewan Harrison, David K. Jackson, Ian Johnston, Dominic Kwiatkowski, Cordelia Langford, John Sillitoe on behalf of the Wellcome Sanger Institute COVID-19 Surveillance Team |
| EPI_ISL_2864836, EPI_ISL_2864837 | Lighthouse Laboratory Plymouth | Wellcome Sanger Institute for the COVID-19 Genomics UK (COG-UK) Consortium | Lighthouse Laboratory Plymouth and Alex Alderton, Roberto Amato, Jeffrey Barrett, Sonia Goncalves, Ewan Harrison, David K. Jackson, Ian Johnston, Dominic Kwiatkowski, Cordelia Langford, John Sillitoe on behalf of the Wellcome Sanger Institute COVID-19 Surveillance Team |
| EPI_ISL_2864838, EPI_ISL_2864839 | Berkshire and Surrey Pathology Services Lighthouse Laboratory | Wellcome Sanger Institute for the COVID-19 Genomics UK (COG-UK) Consortium | Berkshire and Surrey Pathology Services Lighthouse Laboratory and Alex Alderton, Roberto Amato, Jeffrey Barrett, Sonia Goncalves, Ewan Harrison, David K. Jackson, Ian Johnston, Dominic Kwiatkowski, Cordelia Langford, John Sillitoe on behalf of the Wellcome Sanger Institute COVID-19 Surveillance Team |
| EPI_ISL_2864840 | Lighthouse Lab in Glasgow | Wellcome Sanger Institute for the COVID-19 Genomics UK (COG-UK) Consortium | Harper VanSteenhouse, Yumi Kasai, David Gray, Carol Clugston, Anna Dominiczak and Alex Alderton, Roberto Amato, Jeffrey Barrett, Sonia Goncalves, Ewan Harrison, David K. Jackson, Ian Johnston, Dominic Kwiatkowski, Cordelia Langford, John Sillitoe on behalf of the Wellcome Sanger Institute COVID-19 Surveillance Team |
| EPI_ISL_2864841 | Berkshire and Surrey Pathology Services Lighthouse Laboratory | Wellcome Sanger Institute for the COVID-19 Genomics UK (COG-UK) Consortium | Berkshire and Surrey Pathology Services Lighthouse Laboratory and Alex Alderton, Roberto Amato, Jeffrey Barrett, Sonia Goncalves, Ewan Harrison, David K. Jackson, Ian Johnston, Dominic Kwiatkowski, Cordelia Langford, John Sillitoe on behalf of the Wellcome Sanger Institute COVID-19 Surveillance Team |
| EPI_ISL_2864842, EPI_ISL_2864843 | Lighthouse Laboratory Plymouth | Wellcome Sanger Institute for the COVID-19 Genomics UK (COG-UK) Consortium | Lighthouse Laboratory Plymouth and Alex Alderton, Roberto Amato, Jeffrey Barrett, Sonia Goncalves, Ewan Harrison, David K. Jackson, Ian Johnston, Dominic Kwiatkowski, Cordelia Langford, John Sillitoe on behalf of the Wellcome Sanger Institute COVID-19 Surveillance Team |
| EPI_ISL_2864844 | Berkshire and Surrey Pathology Services Lighthouse Laboratory | Wellcome Sanger Institute for the COVID-19 Genomics UK (COG-UK) Consortium | Berkshire and Surrey Pathology Services Lighthouse Laboratory and Alex Alderton, Roberto Amato, Jeffrey Barrett, Sonia Goncalves, Ewan Harrison, David K. Jackson, Ian Johnston, Dominic Kwiatkowski, Cordelia Langford, John Sillitoe on behalf of the Wellcome Sanger Institute COVID-19 Surveillance Team |
| EPI_ISL_2864845 | Lighthouse Laboratory Plymouth | Wellcome Sanger Institute for the COVID-19 Genomics UK (COG-UK) Consortium | Lighthouse Laboratory Plymouth and Alex Alderton, Roberto Amato, Jeffrey Barrett, Sonia Goncalves, Ewan Harrison, David K. Jackson, Ian Johnston, Dominic Kwiatkowski, Cordelia Langford, John Sillitoe on behalf of the Wellcome Sanger Institute COVID-19 Surveillance Team |
| EPI_ISL_2864846 | Lighthouse Lab in Glasgow | Wellcome Sanger Institute for the COVID-19 Genomics UK (COG-UK) Consortium | Harper VanSteenhouse, Yumi Kasai, David Gray, Carol Clugston, Anna Dominiczak and Alex Alderton, Roberto Amato, Jeffrey Barrett, Sonia Goncalves, Ewan Harrison, David K. Jackson, Ian Johnston, Dominic Kwiatkowski, Cordelia Langford, John Sillitoe on behalf of the Wellcome Sanger Institute COVID-19 Surveillance Team |
| EPI_ISL_2864847 | Berkshire and Surrey Pathology Services Lighthouse Laboratory | Wellcome Sanger Institute for the COVID-19 Genomics UK (COG-UK) Consortium | Berkshire and Surrey Pathology Services Lighthouse Laboratory and Alex Alderton, Roberto Amato, Jeffrey Barrett, Sonia Goncalves, Ewan Harrison, David K. Jackson, Ian Johnston, Dominic Kwiatkowski, Cordelia Langford, John Sillitoe on behalf of the Wellcome Sanger Institute COVID-19 Surveillance Team |
| EPI_ISL_2864848 | Lighthouse Laboratory Plymouth | Wellcome Sanger Institute for the COVID-19 Genomics UK (COG-UK) Consortium | Lighthouse Laboratory Plymouth and Alex Alderton, Roberto Amato, Jeffrey Barrett, Sonia Goncalves, Ewan Harrison, David K. Jackson, Ian Johnston, Dominic Kwiatkowski, Cordelia Langford, John Sillitoe on behalf of the Wellcome Sanger Institute COVID-19 Surveillance Team |
| EPI_ISL_2864849 | Berkshire and Surrey Pathology Services Lighthouse Laboratory | Wellcome Sanger Institute for the COVID-19 Genomics UK (COG-UK) Consortium | Berkshire and Surrey Pathology Services Lighthouse Laboratory and Alex Alderton, Roberto Amato, Jeffrey Barrett, Sonia Goncalves, Ewan Harrison, David K. Jackson, Ian Johnston, Dominic Kwiatkowski, Cordelia Langford, John Sillitoe on behalf of the Wellcome Sanger Institute COVID-19 Surveillance Team |
| EPI_ISL_2864850, EPI_ISL_2864851, EPI_ISL_2864852 | Lighthouse Laboratory Plymouth | Wellcome Sanger Institute for the COVID-19 Genomics UK (COG-UK) Consortium | Lighthouse Laboratory Plymouth and Alex Alderton, Roberto Amato, Jeffrey Barrett, Sonia Goncalves, Ewan Harrison, David K. Jackson, Ian Johnston, Dominic Kwiatkowski, Cordelia Langford, John Sillitoe on behalf of the Wellcome Sanger Institute COVID-19 Surveillance Team |
| EPI_ISL_2864853 | Berkshire and Surrey Pathology Services Lighthouse Laboratory | Wellcome Sanger Institute for the COVID-19 Genomics UK (COG-UK) Consortium | Berkshire and Surrey Pathology Services Lighthouse Laboratory and Alex Alderton, Roberto Amato, Jeffrey Barrett, Sonia Goncalves, Ewan Harrison, David K. Jackson, Ian Johnston, Dominic Kwiatkowski, Cordelia Langford, John Sillitoe on behalf of the Wellcome Sanger Institute COVID-19 Surveillance Team |
| EPI_ISL_2864854 | Lighthouse Laboratory Plymouth | Wellcome Sanger Institute for the COVID-19 Genomics UK (COG-UK) Consortium | Lighthouse Laboratory Plymouth and Alex Alderton, Roberto Amato, Jeffrey Barrett, Sonia Goncalves, Ewan Harrison, David K. Jackson, Ian Johnston, Dominic Kwiatkowski, Cordelia Langford, John Sillitoe on behalf of the Wellcome Sanger Institute COVID-19 Surveillance Team |
| EPI_ISL_2864855 | Lighthouse Lab in Glasgow | Wellcome Sanger Institute for the COVID-19 Genomics UK (COG-UK) Consortium | Harper VanSteenhouse, Yumi Kasai, David Gray, Carol Clugston, Anna Dominiczak and Alex Alderton, Roberto Amato, Jeffrey Barrett, Sonia Goncalves, Ewan Harrison, David K. Jackson, Ian Johnston, Dominic Kwiatkowski, Cordelia Langford, John Sillitoe on behalf of the Wellcome Sanger Institute COVID-19 Surveillance Team |
| EPI_ISL_2864856 | Lighthouse Laboratory Plymouth | Wellcome Sanger Institute for the COVID-19 Genomics UK (COG-UK) Consortium | Lighthouse Laboratory Plymouth and Alex Alderton, Roberto Amato, Jeffrey Barrett, Sonia Goncalves, Ewan Harrison, David K. Jackson, Ian Johnston, Dominic Kwiatkowski, Cordelia Langford, John Sillitoe on behalf of the Wellcome Sanger Institute COVID-19 Surveillance Team |
| EPI_ISL_2864857 | Berkshire and Surrey Pathology Services Lighthouse Laboratory | Wellcome Sanger Institute for the COVID-19 Genomics UK (COG-UK) Consortium | Berkshire and Surrey Pathology Services Lighthouse Laboratory and Alex Alderton, Roberto Amato, Jeffrey Barrett, Sonia Goncalves, Ewan Harrison, David K. Jackson, Ian Johnston, Dominic Kwiatkowski, Cordelia Langford, John Sillitoe on behalf of the Wellcome Sanger Institute COVID-19 Surveillance Team |
| EPI_ISL_2864858, EPI_ISL_2864859, EPI_ISL_2864860 | Lighthouse Laboratory Plymouth | Wellcome Sanger Institute for the COVID-19 Genomics UK (COG-UK) Consortium | Lighthouse Laboratory Plymouth and Alex Alderton, Roberto Amato, Jeffrey Barrett, Sonia Goncalves, Ewan Harrison, David K. Jackson, Ian Johnston, Dominic Kwiatkowski, Cordelia Langford, John Sillitoe on behalf of the Wellcome Sanger Institute COVID-19 Surveillance Team |
| EPI_ISL_2864861 | Lighthouse Lab in Glasgow | Wellcome Sanger Institute for the COVID-19 Genomics UK (COG-UK) Consortium | Harper VanSteenhouse, Yumi Kasai, David Gray, Carol Clugston, Anna Dominiczak and Alex Alderton, Roberto Amato, Jeffrey Barrett, Sonia Goncalves, Ewan Harrison, David K. Jackson, Ian Johnston, Dominic Kwiatkowski, Cordelia Langford, John Sillitoe on behalf of the Wellcome Sanger Institute COVID-19 Surveillance Team |
| EPI_ISL_2864862 | Lighthouse Laboratory Plymouth | Wellcome Sanger Institute for the COVID-19 Genomics UK (COG-UK) Consortium | Lighthouse Laboratory Plymouth and Alex Alderton, Roberto Amato, Jeffrey Barrett, Sonia Goncalves, Ewan Harrison, David K. Jackson, Ian Johnston, Dominic Kwiatkowski, Cordelia Langford, John Sillitoe on behalf of the Wellcome Sanger Institute COVID-19 Surveillance Team |

| | | | |
|---|---|---|---|
| EPI_ISL_2864863, EPI_ISL_2864864 | Berkshire and Surrey Pathology Services Lighthouse Laboratory | Wellcome Sanger Institute for the COVID-19 Genomics UK (COG-UK) Consortium | Berkshire and Surrey Pathology Services Lighthouse Laboratory and Alex Alderton, Roberto Amato, Jeffrey Barrett, Sonia Goncalves, Ewan Harrison, David K. Jackson, Ian Johnston, Dominic Kwiatkowski, Cordelia Langford, John Sillitoe on behalf of the Wellcome Sanger Institute COVID-19 Surveillance Team |
| EPI_ISL_2864865 | Lighthouse Laboratory Plymouth | Wellcome Sanger Institute for the COVID-19 Genomics UK (COG-UK) Consortium | Lighthouse Laboratory Plymouth and Alex Alderton, Roberto Amato, Jeffrey Barrett, Sonia Goncalves, Ewan Harrison, David K. Jackson, Ian Johnston, Dominic Kwiatkowski, Cordelia Langford, John Sillitoe on behalf of the Wellcome Sanger Institute COVID-19 Surveillance Team |
| EPI_ISL_2864866 | Lighthouse Lab in Glasgow | Wellcome Sanger Institute for the COVID-19 Genomics UK (COG-UK) Consortium | Harper VanSteenhouse, Yumi Kasai, David Gray, Carol Clugston, Anna Dominiczak and Alex Alderton, Roberto Amato, Jeffrey Barrett, Sonia Goncalves, Ewan Harrison, David K. Jackson, Ian Johnston, Dominic Kwiatkowski, Cordelia Langford, John Sillitoe on behalf of the Wellcome Sanger Institute COVID-19 Surveillance Team |
| EPI_ISL_2864867, EPI_ISL_2864868, EPI_ISL_2864869 | Lighthouse Laboratory Plymouth | Wellcome Sanger Institute for the COVID-19 Genomics UK (COG-UK) Consortium | Lighthouse Laboratory Plymouth and Alex Alderton, Roberto Amato, Jeffrey Barrett, Sonia Goncalves, Ewan Harrison, David K. Jackson, Ian Johnston, Dominic Kwiatkowski, Cordelia Langford, John Sillitoe on behalf of the Wellcome Sanger Institute COVID-19 Surveillance Team |
| EPI_ISL_2864870 | Lighthouse Lab in Glasgow | Wellcome Sanger Institute for the COVID-19 Genomics UK (COG-UK) Consortium | Harper VanSteenhouse, Yumi Kasai, David Gray, Carol Clugston, Anna Dominiczak and Alex Alderton, Roberto Amato, Jeffrey Barrett, Sonia Goncalves, Ewan Harrison, David K. Jackson, Ian Johnston, Dominic Kwiatkowski, Cordelia Langford, John Sillitoe on behalf of the Wellcome Sanger Institute COVID-19 Surveillance Team |
| EPI_ISL_2864871, EPI_ISL_2864873 | Berkshire and Surrey Pathology Services Lighthouse Laboratory | Wellcome Sanger Institute for the COVID-19 Genomics UK (COG-UK) Consortium | Berkshire and Surrey Pathology Services Lighthouse Laboratory and Alex Alderton, Roberto Amato, Jeffrey Barrett, Sonia Goncalves, Ewan Harrison, David K. Jackson, Ian Johnston, Dominic Kwiatkowski, Cordelia Langford, John Sillitoe on behalf of the Wellcome Sanger Institute COVID-19 Surveillance Team |
| EPI_ISL_2864874, EPI_ISL_2864875, EPI_ISL_2864876, EPI_ISL_2864877, EPI_ISL_2864878, EPI_ISL_2864879, EPI_ISL_2864880, EPI_ISL_2864881, EPI_ISL_2864882 | Lighthouse Laboratory Plymouth | Wellcome Sanger Institute for the COVID-19 Genomics UK (COG-UK) Consortium | Lighthouse Laboratory Plymouth and Alex Alderton, Roberto Amato, Jeffrey Barrett, Sonia Goncalves, Ewan Harrison, David K. Jackson, Ian Johnston, Dominic Kwiatkowski, Cordelia Langford, John Sillitoe on behalf of the Wellcome Sanger Institute COVID-19 Surveillance Team |
| EPI_ISL_2864883 | Lighthouse Lab in Glasgow | Wellcome Sanger Institute for the COVID-19 Genomics UK (COG-UK) Consortium | Harper VanSteenhouse, Yumi Kasai, David Gray, Carol Clugston, Anna Dominiczak and Alex Alderton, Roberto Amato, Jeffrey Barrett, Sonia Goncalves, Ewan Harrison, David K. Jackson, Ian Johnston, Dominic Kwiatkowski, Cordelia Langford, John Sillitoe on behalf of the Wellcome Sanger Institute COVID-19 Surveillance Team |
| EPI_ISL_2864884 | Lighthouse Laboratory Plymouth | Wellcome Sanger Institute for the COVID-19 Genomics UK (COG-UK) Consortium | Lighthouse Laboratory Plymouth and Alex Alderton, Roberto Amato, Jeffrey Barrett, Sonia Goncalves, Ewan Harrison, David K. Jackson, Ian Johnston, Dominic Kwiatkowski, Cordelia Langford, John Sillitoe on behalf of the Wellcome Sanger Institute COVID-19 Surveillance Team |
| EPI_ISL_2864885, EPI_ISL_2864886 | Berkshire and Surrey Pathology Services Lighthouse Laboratory | Wellcome Sanger Institute for the COVID-19 Genomics UK (COG-UK) Consortium | Berkshire and Surrey Pathology Services Lighthouse Laboratory and Alex Alderton, Roberto Amato, Jeffrey Barrett, Sonia Goncalves, Ewan Harrison, David K. Jackson, Ian Johnston, Dominic Kwiatkowski, Cordelia Langford, John Sillitoe on behalf of the Wellcome Sanger Institute COVID-19 Surveillance Team |
| EPI_ISL_2864887, EPI_ISL_2864888 | Lighthouse Laboratory Plymouth | Wellcome Sanger Institute for the COVID-19 Genomics UK (COG-UK) Consortium | Lighthouse Laboratory Plymouth and Alex Alderton, Roberto Amato, Jeffrey Barrett, Sonia Goncalves, Ewan Harrison, David K. Jackson, Ian Johnston, Dominic Kwiatkowski, Cordelia Langford, John Sillitoe on behalf of the Wellcome Sanger Institute COVID-19 Surveillance Team |
| EPI_ISL_2864889, EPI_ISL_2864890 | Berkshire and Surrey Pathology Services Lighthouse Laboratory | Wellcome Sanger Institute for the COVID-19 Genomics UK (COG-UK) Consortium | Berkshire and Surrey Pathology Services Lighthouse Laboratory and Alex Alderton, Roberto Amato, Jeffrey Barrett, Sonia Goncalves, Ewan Harrison, David K. Jackson, Ian Johnston, Dominic Kwiatkowski, Cordelia Langford, John Sillitoe on behalf of the Wellcome Sanger Institute COVID-19 Surveillance Team |
| EPI_ISL_2864891, EPI_ISL_2864892, EPI_ISL_2864893 | Lighthouse Laboratory Plymouth | Wellcome Sanger Institute for the COVID-19 Genomics UK (COG-UK) Consortium | Lighthouse Laboratory Plymouth and Alex Alderton, Roberto Amato, Jeffrey Barrett, Sonia Goncalves, Ewan Harrison, David K. Jackson, Ian Johnston, Dominic Kwiatkowski, Cordelia Langford, John Sillitoe on behalf of the Wellcome Sanger Institute COVID-19 Surveillance Team |
| EPI_ISL_2864894, EPI_ISL_2864895 | Berkshire and Surrey Pathology Services Lighthouse Laboratory | Wellcome Sanger Institute for the COVID-19 Genomics UK (COG-UK) Consortium | Berkshire and Surrey Pathology Services Lighthouse Laboratory and Alex Alderton, Roberto Amato, Jeffrey Barrett, Sonia Goncalves, Ewan Harrison, David K. Jackson, Ian Johnston, Dominic Kwiatkowski, Cordelia Langford, John Sillitoe on behalf of the Wellcome Sanger Institute COVID-19 Surveillance Team |
| EPI_ISL_2864896, EPI_ISL_2864897 | Lighthouse Laboratory Plymouth | Wellcome Sanger Institute for the COVID-19 Genomics UK (COG-UK) Consortium | Lighthouse Laboratory Plymouth and Alex Alderton, Roberto Amato, Jeffrey Barrett, Sonia Goncalves, Ewan Harrison, David K. Jackson, Ian Johnston, Dominic Kwiatkowski, Cordelia Langford, John Sillitoe on behalf of the Wellcome Sanger Institute COVID-19 Surveillance Team |
| EPI_ISL_2864898 | Lighthouse Lab in Glasgow | Wellcome Sanger Institute for the COVID-19 Genomics UK (COG-UK) Consortium | Harper VanSteenhouse, Yumi Kasai, David Gray, Carol Clugston, Anna Dominiczak and Alex Alderton, Roberto Amato, Jeffrey Barrett, Sonia Goncalves, Ewan Harrison, David K. Jackson, Ian Johnston, Dominic Kwiatkowski, Cordelia Langford, John Sillitoe on behalf of the Wellcome Sanger Institute COVID-19 Surveillance Team |
| EPI_ISL_2864899, EPI_ISL_2864900, EPI_ISL_2864901, EPI_ISL_2864902 | Lighthouse Laboratory Plymouth | Wellcome Sanger Institute for the COVID-19 Genomics UK (COG-UK) Consortium | Lighthouse Laboratory Plymouth and Alex Alderton, Roberto Amato, Jeffrey Barrett, Sonia Goncalves, Ewan Harrison, David K. Jackson, Ian Johnston, Dominic Kwiatkowski, Cordelia Langford, John Sillitoe on behalf of the Wellcome Sanger Institute COVID-19 Surveillance Team |
| EPI_ISL_2864903 | Berkshire and Surrey Pathology Services Lighthouse Laboratory | Wellcome Sanger Institute for the COVID-19 Genomics UK (COG-UK) Consortium | Berkshire and Surrey Pathology Services Lighthouse Laboratory and Alex Alderton, Roberto Amato, Jeffrey Barrett, Sonia Goncalves, Ewan Harrison, David K. Jackson, Ian Johnston, Dominic Kwiatkowski, Cordelia Langford, John Sillitoe on behalf of the Wellcome Sanger Institute COVID-19 Surveillance Team |
| EPI_ISL_2864904, EPI_ISL_2864905, EPI_ISL_2864906, EPI_ISL_2864907, EPI_ISL_2864908, EPI_ISL_2864909, EPI_ISL_2864910 | Lighthouse Laboratory Plymouth | Wellcome Sanger Institute for the COVID-19 Genomics UK (COG-UK) Consortium | Lighthouse Laboratory Plymouth and Alex Alderton, Roberto Amato, Jeffrey Barrett, Sonia Goncalves, Ewan Harrison, David K. Jackson, Ian Johnston, Dominic Kwiatkowski, Cordelia Langford, John Sillitoe on behalf of the Wellcome Sanger Institute COVID-19 Surveillance Team |
| EPI_ISL_2864911 | Berkshire and Surrey Pathology Services Lighthouse Laboratory | Wellcome Sanger Institute for the COVID-19 Genomics UK (COG-UK) Consortium | Berkshire and Surrey Pathology Services Lighthouse Laboratory and Alex Alderton, Roberto Amato, Jeffrey Barrett, Sonia Goncalves, Ewan Harrison, David K. Jackson, Ian Johnston, Dominic Kwiatkowski, Cordelia Langford, John Sillitoe on behalf of the Wellcome Sanger Institute COVID-19 Surveillance Team |
| EPI_ISL_2864912 | Lighthouse Lab in Glasgow | Wellcome Sanger Institute for the COVID-19 Genomics UK (COG-UK) Consortium | Harper VanSteenhouse, Yumi Kasai, David Gray, Carol Clugston, Anna Dominiczak and Alex Alderton, Roberto Amato, Jeffrey Barrett, Sonia Goncalves, Ewan Harrison, David K. Jackson, Ian Johnston, Dominic Kwiatkowski, Cordelia Langford, John Sillitoe on behalf of the Wellcome Sanger Institute COVID-19 Surveillance Team |
| EPI_ISL_2864913, EPI_ISL_2864914, EPI_ISL_2864915, EPI_ISL_2864916, EPI_ISL_2864917 | Lighthouse Laboratory Plymouth | Wellcome Sanger Institute for the COVID-19 Genomics UK (COG-UK) Consortium | Lighthouse Laboratory Plymouth and Alex Alderton, Roberto Amato, Jeffrey Barrett, Sonia Goncalves, Ewan Harrison, David K. Jackson, Ian Johnston, Dominic Kwiatkowski, Cordelia Langford, John Sillitoe on behalf of the Wellcome Sanger Institute COVID-19 Surveillance Team |
| EPI_ISL_2864918 | Lighthouse Lab in Glasgow | Wellcome Sanger Institute for the COVID-19 Genomics UK (COG-UK) Consortium | Harper VanSteenhouse, Yumi Kasai, David Gray, Carol Clugston, Anna Dominiczak and Alex Alderton, Roberto Amato, Jeffrey Barrett, Sonia Goncalves, Ewan Harrison, David K. Jackson, Ian Johnston, Dominic Kwiatkowski, Cordelia Langford, John Sillitoe on behalf of the Wellcome Sanger Institute COVID-19 Surveillance Team |
| EPI_ISL_2864919 | Berkshire and Surrey Pathology Services Lighthouse Laboratory | Wellcome Sanger Institute for the COVID-19 Genomics UK (COG-UK) Consortium | Berkshire and Surrey Pathology Services Lighthouse Laboratory and Alex Alderton, Roberto Amato, Jeffrey Barrett, Sonia Goncalves, Ewan Harrison, David K. Jackson, Ian Johnston, Dominic Kwiatkowski, Cordelia Langford, John Sillitoe on behalf of the Wellcome Sanger Institute COVID-19 Surveillance Team |
| EPI_ISL_2864920 | Lighthouse Laboratory Plymouth | Wellcome Sanger Institute for the COVID-19 Genomics UK (COG-UK) Consortium | Lighthouse Laboratory Plymouth and Alex Alderton, Roberto Amato, Jeffrey Barrett, Sonia Goncalves, Ewan Harrison, David K. Jackson, Ian Johnston, Dominic Kwiatkowski, Cordelia Langford, John Sillitoe on behalf of the Wellcome Sanger Institute COVID-19 Surveillance Team |
| EPI_ISL_2864921, EPI_ISL_2864922, EPI_ISL_2864923 | Berkshire and Surrey Pathology Services Lighthouse Laboratory | Wellcome Sanger Institute for the COVID-19 Genomics UK (COG-UK) Consortium | Berkshire and Surrey Pathology Services Lighthouse Laboratory and Alex Alderton, Roberto Amato, Jeffrey Barrett, Sonia Goncalves, Ewan Harrison, David K. Jackson, Ian Johnston, Dominic Kwiatkowski, Cordelia Langford, John Sillitoe on behalf of the Wellcome Sanger Institute COVID-19 Surveillance Team |
| EPI_ISL_2864924, EPI_ISL_2864925, EPI_ISL_2864926, EPI_ISL_2864927, EPI_ISL_2864928 | Lighthouse Laboratory Plymouth | Wellcome Sanger Institute for the COVID-19 Genomics UK (COG-UK) Consortium | Lighthouse Laboratory Plymouth and Alex Alderton, Roberto Amato, Jeffrey Barrett, Sonia Goncalves, Ewan Harrison, David K. Jackson, Ian Johnston, Dominic Kwiatkowski, Cordelia Langford, John Sillitoe on behalf of the Wellcome Sanger Institute COVID-19 Surveillance Team |
| EPI_ISL_2864929 | Berkshire and Surrey Pathology Services Lighthouse Laboratory | Wellcome Sanger Institute for the COVID-19 Genomics UK (COG-UK) Consortium | Berkshire and Surrey Pathology Services Lighthouse Laboratory and Alex Alderton, Roberto Amato, Jeffrey Barrett, Sonia Goncalves, Ewan Harrison, David K. Jackson, Ian Johnston, Dominic Kwiatkowski, Cordelia Langford, John Sillitoe on behalf of the Wellcome Sanger Institute COVID-19 Surveillance Team |
| EPI_ISL_2864930 | Lighthouse Lab in Glasgow | Wellcome Sanger Institute for the COVID-19 Genomics UK (COG-UK) Consortium | Harper VanSteenhouse, Yumi Kasai, David Gray, Carol Clugston, Anna Dominiczak and Alex Alderton, Roberto Amato, Jeffrey Barrett, Sonia Goncalves, Ewan Harrison, David K. Jackson, Ian Johnston, Dominic Kwiatkowski, Cordelia Langford, John Sillitoe on behalf of the Wellcome Sanger Institute COVID-19 Surveillance Team |
| EPI_ISL_2864931 | Lighthouse Laboratory Plymouth | Wellcome Sanger Institute for the COVID-19 Genomics UK (COG-UK) Consortium | Lighthouse Laboratory Plymouth and Alex Alderton, Roberto Amato, Jeffrey Barrett, Sonia Goncalves, Ewan Harrison, David K. Jackson, Ian Johnston, Dominic Kwiatkowski, Cordelia Langford, John Sillitoe on behalf of the Wellcome Sanger Institute COVID-19 Surveillance Team |
| EPI_ISL_2864932 | Lighthouse Lab in Glasgow | Wellcome Sanger Institute for the COVID-19 Genomics UK (COG-UK) Consortium | Harper VanSteenhouse, Yumi Kasai, David Gray, Carol Clugston, Anna Dominiczak and Alex Alderton, Roberto Amato, Jeffrey Barrett, Sonia Goncalves, Ewan Harrison, David K. Jackson, Ian Johnston, Dominic Kwiatkowski, Cordelia Langford, John Sillitoe on behalf of the Wellcome Sanger Institute COVID-19 Surveillance Team |
| EPI_ISL_2864933, EPI_ISL_2864934 | Lighthouse Laboratory Plymouth | Wellcome Sanger Institute for the COVID-19 Genomics UK (COG-UK) Consortium | Lighthouse Laboratory Plymouth and Alex Alderton, Roberto Amato, Jeffrey Barrett, Sonia Goncalves, Ewan Harrison, David K. Jackson, Ian Johnston, Dominic Kwiatkowski, Cordelia Langford, John Sillitoe on behalf of the Wellcome Sanger Institute COVID-19 Surveillance Team |
| EPI_ISL_2864935, EPI_ISL_2864936 | Berkshire and Surrey Pathology Services Lighthouse Laboratory | Wellcome Sanger Institute for the COVID-19 Genomics UK (COG-UK) Consortium | Berkshire and Surrey Pathology Services Lighthouse Laboratory and Alex Alderton, Roberto Amato, Jeffrey Barrett, Sonia Goncalves, Ewan Harrison, David K. Jackson, Ian Johnston, Dominic Kwiatkowski, Cordelia Langford, John Sillitoe on behalf of the Wellcome Sanger Institute COVID-19 Surveillance Team |
| EPI_ISL_2864937 | Lighthouse Laboratory Plymouth | Wellcome Sanger Institute for the COVID-19 Genomics UK (COG-UK) Consortium | Lighthouse Laboratory Plymouth and Alex Alderton, Roberto Amato, Jeffrey Barrett, Sonia Goncalves, Ewan Harrison, David K. Jackson, Ian Johnston, Dominic Kwiatkowski, Cordelia Langford, John Sillitoe on behalf of the Wellcome Sanger Institute COVID-19 Surveillance Team |
| EPI_ISL_2864938 | Lighthouse Lab in Glasgow | Wellcome Sanger Institute for the COVID-19 Genomics UK (COG-UK) Consortium | Harper VanSteenhouse, Yumi Kasai, David Gray, Carol Clugston, Anna Dominiczak and Alex Alderton, Roberto Amato, Jeffrey Barrett, Sonia Goncalves, Ewan Harrison, David K. Jackson, Ian Johnston, Dominic Kwiatkowski, Cordelia Langford, John Sillitoe on behalf of the Wellcome Sanger Institute COVID-19 Surveillance Team |
| EPI_ISL_2864939, | Berkshire and Surrey Pathology Services Lighthouse Laboratory | Wellcome Sanger Institute for the COVID-19 Genomics UK (COG-UK) | Berkshire and Surrey Pathology Services Lighthouse Laboratory and Alex Alderton, Roberto Amato, Jeffrey Barrett, Sonia Goncalves, Ewan Harrison, David K. Jackson, Ian Johnston, Dominic Kwiatkowski, Cordelia Langford, John Sillitoe on behalf |

| | | | |
|---|---|---|---|
| EPI_ISL_2864940, EPI_ISL_2864941 | | Consortium | of the Wellcome Sanger Institute COVID-19 Surveillance Team |
| EPI_ISL_2864942 | Lighthouse Laboratory Plymouth | Wellcome Sanger Institute for the COVID-19 Genomics UK (COG-UK) Consortium | Lighthouse Laboratory Plymouth and Alex Alderton, Roberto Amato, Jeffrey Barrett, Sonia Goncalves, Ewan Harrison, David K. Jackson, Ian Johnston, Dominic Kwiatkowski, Cordelia Langford, John Sillitoe on behalf of the Wellcome Sanger Institute COVID-19 Surveillance Team |
| EPI_ISL_2864943, EPI_ISL_2864944 | Lighthouse Lab in Glasgow | Wellcome Sanger Institute for the COVID-19 Genomics UK (COG-UK) Consortium | Harper VanSteenhouse, Yumi Kasai, David Gray, Carol Clugston, Anna Dominiczak and Alex Alderton, Roberto Amato, Jeffrey Barrett, Sonia Goncalves, Ewan Harrison, David K. Jackson, Ian Johnston, Dominic Kwiatkowski, Cordelia Langford, John Sillitoe on behalf of the Wellcome Sanger Institute COVID-19 Surveillance Team |
| EPI_ISL_2864945, EPI_ISL_2864946 | Lighthouse Laboratory Plymouth | Wellcome Sanger Institute for the COVID-19 Genomics UK (COG-UK) Consortium | Lighthouse Laboratory Plymouth and Alex Alderton, Roberto Amato, Jeffrey Barrett, Sonia Goncalves, Ewan Harrison, David K. Jackson, Ian Johnston, Dominic Kwiatkowski, Cordelia Langford, John Sillitoe on behalf of the Wellcome Sanger Institute COVID-19 Surveillance Team |
| EPI_ISL_2864947 | Berkshire and Surrey Pathology Services Lighthouse Laboratory | Wellcome Sanger Institute for the COVID-19 Genomics UK (COG-UK) Consortium | Berkshire and Surrey Pathology Services Lighthouse Laboratory and Alex Alderton, Roberto Amato, Jeffrey Barrett, Sonia Goncalves, Ewan Harrison, David K. Jackson, Ian Johnston, Dominic Kwiatkowski, Cordelia Langford, John Sillitoe on behalf of the Wellcome Sanger Institute COVID-19 Surveillance Team |
| EPI_ISL_2864948 | Lighthouse Laboratory Plymouth | Wellcome Sanger Institute for the COVID-19 Genomics UK (COG-UK) Consortium | Lighthouse Laboratory Plymouth and Alex Alderton, Roberto Amato, Jeffrey Barrett, Sonia Goncalves, Ewan Harrison, David K. Jackson, Ian Johnston, Dominic Kwiatkowski, Cordelia Langford, John Sillitoe on behalf of the Wellcome Sanger Institute COVID-19 Surveillance Team |
| EPI_ISL_2864949, EPI_ISL_2864950 | Berkshire and Surrey Pathology Services Lighthouse Laboratory | Wellcome Sanger Institute for the COVID-19 Genomics UK (COG-UK) Consortium | Berkshire and Surrey Pathology Services Lighthouse Laboratory and Alex Alderton, Roberto Amato, Jeffrey Barrett, Sonia Goncalves, Ewan Harrison, David K. Jackson, Ian Johnston, Dominic Kwiatkowski, Cordelia Langford, John Sillitoe on behalf of the Wellcome Sanger Institute COVID-19 Surveillance Team |
| EPI_ISL_2864951 | Lighthouse Lab in Glasgow | Wellcome Sanger Institute for the COVID-19 Genomics UK (COG-UK) Consortium | Harper VanSteenhouse, Yumi Kasai, David Gray, Carol Clugston, Anna Dominiczak and Alex Alderton, Roberto Amato, Jeffrey Barrett, Sonia Goncalves, Ewan Harrison, David K. Jackson, Ian Johnston, Dominic Kwiatkowski, Cordelia Langford, John Sillitoe on behalf of the Wellcome Sanger Institute COVID-19 Surveillance Team |
| EPI_ISL_2864952 | Berkshire and Surrey Pathology Services Lighthouse Laboratory | Wellcome Sanger Institute for the COVID-19 Genomics UK (COG-UK) Consortium | Berkshire and Surrey Pathology Services Lighthouse Laboratory and Alex Alderton, Roberto Amato, Jeffrey Barrett, Sonia Goncalves, Ewan Harrison, David K. Jackson, Ian Johnston, Dominic Kwiatkowski, Cordelia Langford, John Sillitoe on behalf of the Wellcome Sanger Institute COVID-19 Surveillance Team |
| EPI_ISL_2864953 | Lighthouse Laboratory Plymouth | Wellcome Sanger Institute for the COVID-19 Genomics UK (COG-UK) Consortium | Lighthouse Laboratory Plymouth and Alex Alderton, Roberto Amato, Jeffrey Barrett, Sonia Goncalves, Ewan Harrison, David K. Jackson, Ian Johnston, Dominic Kwiatkowski, Cordelia Langford, John Sillitoe on behalf of the Wellcome Sanger Institute COVID-19 Surveillance Team |
| EPI_ISL_2864954 | Lighthouse Lab in Glasgow | Wellcome Sanger Institute for the COVID-19 Genomics UK (COG-UK) Consortium | Harper VanSteenhouse, Yumi Kasai, David Gray, Carol Clugston, Anna Dominiczak and Alex Alderton, Roberto Amato, Jeffrey Barrett, Sonia Goncalves, Ewan Harrison, David K. Jackson, Ian Johnston, Dominic Kwiatkowski, Cordelia Langford, John Sillitoe on behalf of the Wellcome Sanger Institute COVID-19 Surveillance Team |
| EPI_ISL_2864955 | Berkshire and Surrey Pathology Services Lighthouse Laboratory | Wellcome Sanger Institute for the COVID-19 Genomics UK (COG-UK) Consortium | Berkshire and Surrey Pathology Services Lighthouse Laboratory and Alex Alderton, Roberto Amato, Jeffrey Barrett, Sonia Goncalves, Ewan Harrison, David K. Jackson, Ian Johnston, Dominic Kwiatkowski, Cordelia Langford, John Sillitoe on behalf of the Wellcome Sanger Institute COVID-19 Surveillance Team |
| EPI_ISL_2864956 | Lighthouse Laboratory Plymouth | Wellcome Sanger Institute for the COVID-19 Genomics UK (COG-UK) Consortium | Lighthouse Laboratory Plymouth and Alex Alderton, Roberto Amato, Jeffrey Barrett, Sonia Goncalves, Ewan Harrison, David K. Jackson, Ian Johnston, Dominic Kwiatkowski, Cordelia Langford, John Sillitoe on behalf of the Wellcome Sanger Institute COVID-19 Surveillance Team |
| EPI_ISL_2864957, EPI_ISL_2864958 | Berkshire and Surrey Pathology Services Lighthouse Laboratory | Wellcome Sanger Institute for the COVID-19 Genomics UK (COG-UK) Consortium | Berkshire and Surrey Pathology Services Lighthouse Laboratory and Alex Alderton, Roberto Amato, Jeffrey Barrett, Sonia Goncalves, Ewan Harrison, David K. Jackson, Ian Johnston, Dominic Kwiatkowski, Cordelia Langford, John Sillitoe on behalf of the Wellcome Sanger Institute COVID-19 Surveillance Team |
| EPI_ISL_2864959, EPI_ISL_2864960 | Lighthouse Laboratory Plymouth | Wellcome Sanger Institute for the COVID-19 Genomics UK (COG-UK) Consortium | Lighthouse Laboratory Plymouth and Alex Alderton, Roberto Amato, Jeffrey Barrett, Sonia Goncalves, Ewan Harrison, David K. Jackson, Ian Johnston, Dominic Kwiatkowski, Cordelia Langford, John Sillitoe on behalf of the Wellcome Sanger Institute COVID-19 Surveillance Team |
| EPI_ISL_2864961 | Berkshire and Surrey Pathology Services Lighthouse Laboratory | Wellcome Sanger Institute for the COVID-19 Genomics UK (COG-UK) Consortium | Berkshire and Surrey Pathology Services Lighthouse Laboratory and Alex Alderton, Roberto Amato, Jeffrey Barrett, Sonia Goncalves, Ewan Harrison, David K. Jackson, Ian Johnston, Dominic Kwiatkowski, Cordelia Langford, John Sillitoe on behalf of the Wellcome Sanger Institute COVID-19 Surveillance Team |
| EPI_ISL_2864962, EPI_ISL_2864963 | Lighthouse Laboratory Plymouth | Wellcome Sanger Institute for the COVID-19 Genomics UK (COG-UK) Consortium | Lighthouse Laboratory Plymouth and Alex Alderton, Roberto Amato, Jeffrey Barrett, Sonia Goncalves, Ewan Harrison, David K. Jackson, Ian Johnston, Dominic Kwiatkowski, Cordelia Langford, John Sillitoe on behalf of the Wellcome Sanger Institute COVID-19 Surveillance Team |
| EPI_ISL_2864964 | Berkshire and Surrey Pathology Services Lighthouse Laboratory | Wellcome Sanger Institute for the COVID-19 Genomics UK (COG-UK) Consortium | Berkshire and Surrey Pathology Services Lighthouse Laboratory and Alex Alderton, Roberto Amato, Jeffrey Barrett, Sonia Goncalves, Ewan Harrison, David K. Jackson, Ian Johnston, Dominic Kwiatkowski, Cordelia Langford, John Sillitoe on behalf of the Wellcome Sanger Institute COVID-19 Surveillance Team |
| EPI_ISL_2864965 | Lighthouse Laboratory Plymouth | Wellcome Sanger Institute for the COVID-19 Genomics UK (COG-UK) Consortium | Lighthouse Laboratory Plymouth and Alex Alderton, Roberto Amato, Jeffrey Barrett, Sonia Goncalves, Ewan Harrison, David K. Jackson, Ian Johnston, Dominic Kwiatkowski, Cordelia Langford, John Sillitoe on behalf of the Wellcome Sanger Institute COVID-19 Surveillance Team |
| EPI_ISL_2864966, EPI_ISL_2864967 | Berkshire and Surrey Pathology Services Lighthouse Laboratory | Wellcome Sanger Institute for the COVID-19 Genomics UK (COG-UK) Consortium | Berkshire and Surrey Pathology Services Lighthouse Laboratory and Alex Alderton, Roberto Amato, Jeffrey Barrett, Sonia Goncalves, Ewan Harrison, David K. Jackson, Ian Johnston, Dominic Kwiatkowski, Cordelia Langford, John Sillitoe on behalf of the Wellcome Sanger Institute COVID-19 Surveillance Team |
| EPI_ISL_2864968 | Lighthouse Laboratory Plymouth | Wellcome Sanger Institute for the COVID-19 Genomics UK (COG-UK) Consortium | Lighthouse Laboratory Plymouth and Alex Alderton, Roberto Amato, Jeffrey Barrett, Sonia Goncalves, Ewan Harrison, David K. Jackson, Ian Johnston, Dominic Kwiatkowski, Cordelia Langford, John Sillitoe on behalf of the Wellcome Sanger Institute COVID-19 Surveillance Team |
| EPI_ISL_2864969, EPI_ISL_2864970, EPI_ISL_2864971, EPI_ISL_2864972 | Lighthouse Lab in Glasgow | Wellcome Sanger Institute for the COVID-19 Genomics UK (COG-UK) Consortium | Harper VanSteenhouse, Yumi Kasai, David Gray, Carol Clugston, Anna Dominiczak and Alex Alderton, Roberto Amato, Jeffrey Barrett, Sonia Goncalves, Ewan Harrison, David K. Jackson, Ian Johnston, Dominic Kwiatkowski, Cordelia Langford, John Sillitoe on behalf of the Wellcome Sanger Institute COVID-19 Surveillance Team |
| EPI_ISL_2864973, EPI_ISL_2864974, EPI_ISL_2864975 | Lighthouse Laboratory Plymouth | Wellcome Sanger Institute for the COVID-19 Genomics UK (COG-UK) Consortium | Lighthouse Laboratory Plymouth and Alex Alderton, Roberto Amato, Jeffrey Barrett, Sonia Goncalves, Ewan Harrison, David K. Jackson, Ian Johnston, Dominic Kwiatkowski, Cordelia Langford, John Sillitoe on behalf of the Wellcome Sanger Institute COVID-19 Surveillance Team |
| EPI_ISL_2864976 | Lighthouse Lab in Glasgow | Wellcome Sanger Institute for the COVID-19 Genomics UK (COG-UK) Consortium | Harper VanSteenhouse, Yumi Kasai, David Gray, Carol Clugston, Anna Dominiczak and Alex Alderton, Roberto Amato, Jeffrey Barrett, Sonia Goncalves, Ewan Harrison, David K. Jackson, Ian Johnston, Dominic Kwiatkowski, Cordelia Langford, John Sillitoe on behalf of the Wellcome Sanger Institute COVID-19 Surveillance Team |
| EPI_ISL_2864977 | Berkshire and Surrey Pathology Services Lighthouse Laboratory | Wellcome Sanger Institute for the COVID-19 Genomics UK (COG-UK) Consortium | Berkshire and Surrey Pathology Services Lighthouse Laboratory and Alex Alderton, Roberto Amato, Jeffrey Barrett, Sonia Goncalves, Ewan Harrison, David K. Jackson, Ian Johnston, Dominic Kwiatkowski, Cordelia Langford, John Sillitoe on behalf of the Wellcome Sanger Institute COVID-19 Surveillance Team |
| EPI_ISL_2864978, EPI_ISL_2864979, EPI_ISL_2864980, EPI_ISL_2864981, EPI_ISL_2864982, EPI_ISL_2864983, EPI_ISL_2864984, EPI_ISL_2864985, EPI_ISL_2864986 | Lighthouse Laboratory Plymouth | Wellcome Sanger Institute for the COVID-19 Genomics UK (COG-UK) Consortium | Lighthouse Laboratory Plymouth and Alex Alderton, Roberto Amato, Jeffrey Barrett, Sonia Goncalves, Ewan Harrison, David K. Jackson, Ian Johnston, Dominic Kwiatkowski, Cordelia Langford, John Sillitoe on behalf of the Wellcome Sanger Institute COVID-19 Surveillance Team |
| EPI_ISL_2864987 | Berkshire and Surrey Pathology Services Lighthouse Laboratory | Wellcome Sanger Institute for the COVID-19 Genomics UK (COG-UK) Consortium | Berkshire and Surrey Pathology Services Lighthouse Laboratory and Alex Alderton, Roberto Amato, Jeffrey Barrett, Sonia Goncalves, Ewan Harrison, David K. Jackson, Ian Johnston, Dominic Kwiatkowski, Cordelia Langford, John Sillitoe on behalf of the Wellcome Sanger Institute COVID-19 Surveillance Team |
| EPI_ISL_2864988 | Lighthouse Lab in Glasgow | Wellcome Sanger Institute for the COVID-19 Genomics UK (COG-UK) Consortium | Harper VanSteenhouse, Yumi Kasai, David Gray, Carol Clugston, Anna Dominiczak and Alex Alderton, Roberto Amato, Jeffrey Barrett, Sonia Goncalves, Ewan Harrison, David K. Jackson, Ian Johnston, Dominic Kwiatkowski, Cordelia Langford, John Sillitoe on behalf of the Wellcome Sanger Institute COVID-19 Surveillance Team |
| EPI_ISL_2864989, EPI_ISL_2864990 | Berkshire and Surrey Pathology Services Lighthouse Laboratory | Wellcome Sanger Institute for the COVID-19 Genomics UK (COG-UK) Consortium | Berkshire and Surrey Pathology Services Lighthouse Laboratory and Alex Alderton, Roberto Amato, Jeffrey Barrett, Sonia Goncalves, Ewan Harrison, David K. Jackson, Ian Johnston, Dominic Kwiatkowski, Cordelia Langford, John Sillitoe on behalf of the Wellcome Sanger Institute COVID-19 Surveillance Team |
| EPI_ISL_2864991 | Lighthouse Laboratory Plymouth | Wellcome Sanger Institute for the COVID-19 Genomics UK (COG-UK) Consortium | Lighthouse Laboratory Plymouth and Alex Alderton, Roberto Amato, Jeffrey Barrett, Sonia Goncalves, Ewan Harrison, David K. Jackson, Ian Johnston, Dominic Kwiatkowski, Cordelia Langford, John Sillitoe on behalf of the Wellcome Sanger Institute COVID-19 Surveillance Team |
| EPI_ISL_2864992 | Berkshire and Surrey Pathology Services Lighthouse Laboratory | Wellcome Sanger Institute for the COVID-19 Genomics UK (COG-UK) Consortium | Berkshire and Surrey Pathology Services Lighthouse Laboratory and Alex Alderton, Roberto Amato, Jeffrey Barrett, Sonia Goncalves, Ewan Harrison, David K. Jackson, Ian Johnston, Dominic Kwiatkowski, Cordelia Langford, John Sillitoe on behalf of the Wellcome Sanger Institute COVID-19 Surveillance Team |
| EPI_ISL_2864993 | Lighthouse Laboratory Plymouth | Wellcome Sanger Institute for the COVID-19 Genomics UK (COG-UK) Consortium | Lighthouse Laboratory Plymouth and Alex Alderton, Roberto Amato, Jeffrey Barrett, Sonia Goncalves, Ewan Harrison, David K. Jackson, Ian Johnston, Dominic Kwiatkowski, Cordelia Langford, John Sillitoe on behalf of the Wellcome Sanger Institute COVID-19 Surveillance Team |
| EPI_ISL_2864994 | Berkshire and Surrey Pathology Services Lighthouse Laboratory | Wellcome Sanger Institute for the COVID-19 Genomics UK (COG-UK) Consortium | Berkshire and Surrey Pathology Services Lighthouse Laboratory and Alex Alderton, Roberto Amato, Jeffrey Barrett, Sonia Goncalves, Ewan Harrison, David K. Jackson, Ian Johnston, Dominic Kwiatkowski, Cordelia Langford, John Sillitoe on behalf of the Wellcome Sanger Institute COVID-19 Surveillance Team |
| EPI_ISL_2864995, EPI_ISL_2864996, EPI_ISL_2864997, EPI_ISL_2864998, EPI_ISL_2864999, EPI_ISL_2865000, EPI_ISL_2865001, EPI_ISL_2865002 | Lighthouse Laboratory Plymouth | Wellcome Sanger Institute for the COVID-19 Genomics UK (COG-UK) Consortium | Lighthouse Laboratory Plymouth and Alex Alderton, Roberto Amato, Jeffrey Barrett, Sonia Goncalves, Ewan Harrison, David K. Jackson, Ian Johnston, Dominic Kwiatkowski, Cordelia Langford, John Sillitoe on behalf of the Wellcome Sanger Institute COVID-19 Surveillance Team |
| EPI_ISL_2865003 | Berkshire and Surrey Pathology Services Lighthouse Laboratory | Wellcome Sanger Institute for the COVID-19 Genomics UK (COG-UK) Consortium | Berkshire and Surrey Pathology Services Lighthouse Laboratory and Alex Alderton, Roberto Amato, Jeffrey Barrett, Sonia Goncalves, Ewan Harrison, David K. Jackson, Ian Johnston, Dominic Kwiatkowski, Cordelia Langford, John Sillitoe on behalf of the Wellcome Sanger Institute COVID-19 Surveillance Team |
| EPI_ISL_2865004 | Lighthouse Laboratory Plymouth | Wellcome Sanger Institute for the COVID-19 Genomics UK (COG-UK) Consortium | Lighthouse Laboratory Plymouth and Alex Alderton, Roberto Amato, Jeffrey Barrett, Sonia Goncalves, Ewan Harrison, David K. Jackson, Ian Johnston, Dominic Kwiatkowski, Cordelia Langford, John Sillitoe on behalf of the Wellcome Sanger Institute COVID-19 Surveillance Team |
| EPI_ISL_2865005 | Berkshire and Surrey Pathology Services Lighthouse Laboratory | Wellcome Sanger Institute for the COVID-19 Genomics UK (COG-UK) Consortium | Berkshire and Surrey Pathology Services Lighthouse Laboratory and Alex Alderton, Roberto Amato, Jeffrey Barrett, Sonia Goncalves, Ewan Harrison, David K. Jackson, Ian Johnston, Dominic Kwiatkowski, Cordelia Langford, John Sillitoe on behalf of the Wellcome Sanger Institute COVID-19 Surveillance Team |
| EPI_ISL_2865006 | Lighthouse Laboratory Plymouth | Wellcome Sanger Institute for the COVID-19 Genomics UK (COG-UK) Consortium | Lighthouse Laboratory Plymouth and Alex Alderton, Roberto Amato, Jeffrey Barrett, Sonia Goncalves, Ewan Harrison, David K. Jackson, Ian Johnston, Dominic Kwiatkowski, Cordelia Langford, John Sillitoe on behalf of the Wellcome Sanger Institute COVID-19 Surveillance Team |
| EPI_ISL_2865007, | Berkshire and Surrey Pathology Services Lighthouse Laboratory | Wellcome Sanger Institute for the COVID-19 Genomics UK (COG-UK) | Berkshire and Surrey Pathology Services Lighthouse Laboratory and Alex Alderton, Roberto Amato, Jeffrey Barrett, Sonia Goncalves, Ewan Harrison, David K. Jackson, Ian Johnston, Dominic Kwiatkowski, Cordelia Langford, John Sillitoe on behalf |

| | | | |
|---|---|---|---|
| EPI_ISL_2865008, EPI_ISL_2865009 | | Consortium | of the Wellcome Sanger Institute COVID-19 Surveillance Team |
| EPI_ISL_2865010 | Lighthouse Lab in Glasgow | Wellcome Sanger Institute for the COVID-19 Genomics UK (COG-UK) Consortium | Harper VanSteenhouse, Yumi Kasai, David Gray, Carol Clugston, Anna Dominiczak and Alex Alderton, Roberto Amato, Jeffrey Barrett, Sonia Goncalves, Ewan Harrison, David K. Jackson, Ian Johnston, Dominic Kwiatkowski, Cordelia Langford, John Sillitoe on behalf of the Wellcome Sanger Institute COVID-19 Surveillance Team |
| EPI_ISL_2865011, EPI_ISL_2865012 | Lighthouse Laboratory Plymouth | Wellcome Sanger Institute for the COVID-19 Genomics UK (COG-UK) Consortium | Lighthouse Laboratory Plymouth and Alex Alderton, Roberto Amato, Jeffrey Barrett, Sonia Goncalves, Ewan Harrison, David K. Jackson, Ian Johnston, Dominic Kwiatkowski, Cordelia Langford, John Sillitoe on behalf of the Wellcome Sanger Institute COVID-19 Surveillance Team |
| EPI_ISL_2865013, EPI_ISL_2865014, EPI_ISL_2865015 | Lighthouse Lab in Glasgow | Wellcome Sanger Institute for the COVID-19 Genomics UK (COG-UK) Consortium | Harper VanSteenhouse, Yumi Kasai, David Gray, Carol Clugston, Anna Dominiczak and Alex Alderton, Roberto Amato, Jeffrey Barrett, Sonia Goncalves, Ewan Harrison, David K. Jackson, Ian Johnston, Dominic Kwiatkowski, Cordelia Langford, John Sillitoe on behalf of the Wellcome Sanger Institute COVID-19 Surveillance Team |
| EPI_ISL_2865016 | Berkshire and Surrey Pathology Services Lighthouse Laboratory | Wellcome Sanger Institute for the COVID-19 Genomics UK (COG-UK) Consortium | Berkshire and Surrey Pathology Services Lighthouse Laboratory and Alex Alderton, Roberto Amato, Jeffrey Barrett, Sonia Goncalves, Ewan Harrison, David K. Jackson, Ian Johnston, Dominic Kwiatkowski, Cordelia Langford, John Sillitoe on behalf of the Wellcome Sanger Institute COVID-19 Surveillance Team |
| EPI_ISL_2865017 | Lighthouse Lab in Glasgow | Wellcome Sanger Institute for the COVID-19 Genomics UK (COG-UK) Consortium | Harper VanSteenhouse, Yumi Kasai, David Gray, Carol Clugston, Anna Dominiczak and Alex Alderton, Roberto Amato, Jeffrey Barrett, Sonia Goncalves, Ewan Harrison, David K. Jackson, Ian Johnston, Dominic Kwiatkowski, Cordelia Langford, John Sillitoe on behalf of the Wellcome Sanger Institute COVID-19 Surveillance Team |
| EPI_ISL_2865018 | Lighthouse Laboratory Plymouth | Wellcome Sanger Institute for the COVID-19 Genomics UK (COG-UK) Consortium | Lighthouse Laboratory Plymouth and Alex Alderton, Roberto Amato, Jeffrey Barrett, Sonia Goncalves, Ewan Harrison, David K. Jackson, Ian Johnston, Dominic Kwiatkowski, Cordelia Langford, John Sillitoe on behalf of the Wellcome Sanger Institute COVID-19 Surveillance Team |
| EPI_ISL_2865019 | Berkshire and Surrey Pathology Services Lighthouse Laboratory | Wellcome Sanger Institute for the COVID-19 Genomics UK (COG-UK) Consortium | Berkshire and Surrey Pathology Services Lighthouse Laboratory and Alex Alderton, Roberto Amato, Jeffrey Barrett, Sonia Goncalves, Ewan Harrison, David K. Jackson, Ian Johnston, Dominic Kwiatkowski, Cordelia Langford, John Sillitoe on behalf of the Wellcome Sanger Institute COVID-19 Surveillance Team |
| EPI_ISL_2865020, EPI_ISL_2865021, EPI_ISL_2865022 | Lighthouse Laboratory Plymouth | Wellcome Sanger Institute for the COVID-19 Genomics UK (COG-UK) Consortium | Lighthouse Laboratory Plymouth and Alex Alderton, Roberto Amato, Jeffrey Barrett, Sonia Goncalves, Ewan Harrison, David K. Jackson, Ian Johnston, Dominic Kwiatkowski, Cordelia Langford, John Sillitoe on behalf of the Wellcome Sanger Institute COVID-19 Surveillance Team |
| EPI_ISL_2865023 | Berkshire and Surrey Pathology Services Lighthouse Laboratory | Wellcome Sanger Institute for the COVID-19 Genomics UK (COG-UK) Consortium | Berkshire and Surrey Pathology Services Lighthouse Laboratory and Alex Alderton, Roberto Amato, Jeffrey Barrett, Sonia Goncalves, Ewan Harrison, David K. Jackson, Ian Johnston, Dominic Kwiatkowski, Cordelia Langford, John Sillitoe on behalf of the Wellcome Sanger Institute COVID-19 Surveillance Team |
| EPI_ISL_2865024, EPI_ISL_2865025 | Lighthouse Laboratory Plymouth | Wellcome Sanger Institute for the COVID-19 Genomics UK (COG-UK) Consortium | Lighthouse Laboratory Plymouth and Alex Alderton, Roberto Amato, Jeffrey Barrett, Sonia Goncalves, Ewan Harrison, David K. Jackson, Ian Johnston, Dominic Kwiatkowski, Cordelia Langford, John Sillitoe on behalf of the Wellcome Sanger Institute COVID-19 Surveillance Team |
| EPI_ISL_2865026 | Lighthouse Lab in Glasgow | Wellcome Sanger Institute for the COVID-19 Genomics UK (COG-UK) Consortium | Harper VanSteenhouse, Yumi Kasai, David Gray, Carol Clugston, Anna Dominiczak and Alex Alderton, Roberto Amato, Jeffrey Barrett, Sonia Goncalves, Ewan Harrison, David K. Jackson, Ian Johnston, Dominic Kwiatkowski, Cordelia Langford, John Sillitoe on behalf of the Wellcome Sanger Institute COVID-19 Surveillance Team |
| EPI_ISL_2865027, EPI_ISL_2865028, EPI_ISL_2865029 | Berkshire and Surrey Pathology Services Lighthouse Laboratory | Wellcome Sanger Institute for the COVID-19 Genomics UK (COG-UK) Consortium | Berkshire and Surrey Pathology Services Lighthouse Laboratory and Alex Alderton, Roberto Amato, Jeffrey Barrett, Sonia Goncalves, Ewan Harrison, David K. Jackson, Ian Johnston, Dominic Kwiatkowski, Cordelia Langford, John Sillitoe on behalf of the Wellcome Sanger Institute COVID-19 Surveillance Team |
| EPI_ISL_2865030, EPI_ISL_2865031 | Lighthouse Laboratory Plymouth | Wellcome Sanger Institute for the COVID-19 Genomics UK (COG-UK) Consortium | Lighthouse Laboratory Plymouth and Alex Alderton, Roberto Amato, Jeffrey Barrett, Sonia Goncalves, Ewan Harrison, David K. Jackson, Ian Johnston, Dominic Kwiatkowski, Cordelia Langford, John Sillitoe on behalf of the Wellcome Sanger Institute COVID-19 Surveillance Team |
| EPI_ISL_2865032 | Lighthouse Lab in Glasgow | Wellcome Sanger Institute for the COVID-19 Genomics UK (COG-UK) Consortium | Harper VanSteenhouse, Yumi Kasai, David Gray, Carol Clugston, Anna Dominiczak and Alex Alderton, Roberto Amato, Jeffrey Barrett, Sonia Goncalves, Ewan Harrison, David K. Jackson, Ian Johnston, Dominic Kwiatkowski, Cordelia Langford, John Sillitoe on behalf of the Wellcome Sanger Institute COVID-19 Surveillance Team |
| EPI_ISL_2865033, EPI_ISL_2865034, EPI_ISL_2865035 | Lighthouse Laboratory Plymouth | Wellcome Sanger Institute for the COVID-19 Genomics UK (COG-UK) Consortium | Lighthouse Laboratory Plymouth and Alex Alderton, Roberto Amato, Jeffrey Barrett, Sonia Goncalves, Ewan Harrison, David K. Jackson, Ian Johnston, Dominic Kwiatkowski, Cordelia Langford, John Sillitoe on behalf of the Wellcome Sanger Institute COVID-19 Surveillance Team |
| EPI_ISL_2865036 | Berkshire and Surrey Pathology Services Lighthouse Laboratory | Wellcome Sanger Institute for the COVID-19 Genomics UK (COG-UK) Consortium | Berkshire and Surrey Pathology Services Lighthouse Laboratory and Alex Alderton, Roberto Amato, Jeffrey Barrett, Sonia Goncalves, Ewan Harrison, David K. Jackson, Ian Johnston, Dominic Kwiatkowski, Cordelia Langford, John Sillitoe on behalf of the Wellcome Sanger Institute COVID-19 Surveillance Team |
| EPI_ISL_2865037, EPI_ISL_2865038, EPI_ISL_2865039, EPI_ISL_2865040, EPI_ISL_2865041, EPI_ISL_2865042, EPI_ISL_2865043 | Lighthouse Laboratory Plymouth | Wellcome Sanger Institute for the COVID-19 Genomics UK (COG-UK) Consortium | Lighthouse Laboratory Plymouth and Alex Alderton, Roberto Amato, Jeffrey Barrett, Sonia Goncalves, Ewan Harrison, David K. Jackson, Ian Johnston, Dominic Kwiatkowski, Cordelia Langford, John Sillitoe on behalf of the Wellcome Sanger Institute COVID-19 Surveillance Team |
| EPI_ISL_2865044 | Berkshire and Surrey Pathology Services Lighthouse Laboratory | Wellcome Sanger Institute for the COVID-19 Genomics UK (COG-UK) Consortium | Berkshire and Surrey Pathology Services Lighthouse Laboratory and Alex Alderton, Roberto Amato, Jeffrey Barrett, Sonia Goncalves, Ewan Harrison, David K. Jackson, Ian Johnston, Dominic Kwiatkowski, Cordelia Langford, John Sillitoe on behalf of the Wellcome Sanger Institute COVID-19 Surveillance Team |
| EPI_ISL_2865045, EPI_ISL_2865046, EPI_ISL_2865047, EPI_ISL_2865048, EPI_ISL_2865049, EPI_ISL_2865050 | Lighthouse Laboratory Plymouth | Wellcome Sanger Institute for the COVID-19 Genomics UK (COG-UK) Consortium | Lighthouse Laboratory Plymouth and Alex Alderton, Roberto Amato, Jeffrey Barrett, Sonia Goncalves, Ewan Harrison, David K. Jackson, Ian Johnston, Dominic Kwiatkowski, Cordelia Langford, John Sillitoe on behalf of the Wellcome Sanger Institute COVID-19 Surveillance Team |
| EPI_ISL_2865051, EPI_ISL_2865052 | Berkshire and Surrey Pathology Services Lighthouse Laboratory | Wellcome Sanger Institute for the COVID-19 Genomics UK (COG-UK) Consortium | Berkshire and Surrey Pathology Services Lighthouse Laboratory and Alex Alderton, Roberto Amato, Jeffrey Barrett, Sonia Goncalves, Ewan Harrison, David K. Jackson, Ian Johnston, Dominic Kwiatkowski, Cordelia Langford, John Sillitoe on behalf of the Wellcome Sanger Institute COVID-19 Surveillance Team |
| EPI_ISL_2865053, EPI_ISL_2865054, EPI_ISL_2865055 | Lighthouse Laboratory Plymouth | Wellcome Sanger Institute for the COVID-19 Genomics UK (COG-UK) Consortium | Lighthouse Laboratory Plymouth and Alex Alderton, Roberto Amato, Jeffrey Barrett, Sonia Goncalves, Ewan Harrison, David K. Jackson, Ian Johnston, Dominic Kwiatkowski, Cordelia Langford, John Sillitoe on behalf of the Wellcome Sanger Institute COVID-19 Surveillance Team |
| EPI_ISL_2865056 | Berkshire and Surrey Pathology Services Lighthouse Laboratory | Wellcome Sanger Institute for the COVID-19 Genomics UK (COG-UK) Consortium | Berkshire and Surrey Pathology Services Lighthouse Laboratory and Alex Alderton, Roberto Amato, Jeffrey Barrett, Sonia Goncalves, Ewan Harrison, David K. Jackson, Ian Johnston, Dominic Kwiatkowski, Cordelia Langford, John Sillitoe on behalf of the Wellcome Sanger Institute COVID-19 Surveillance Team |
| EPI_ISL_2865057, EPI_ISL_2865058, EPI_ISL_2865059 | Lighthouse Laboratory Plymouth | Wellcome Sanger Institute for the COVID-19 Genomics UK (COG-UK) Consortium | Lighthouse Laboratory Plymouth and Alex Alderton, Roberto Amato, Jeffrey Barrett, Sonia Goncalves, Ewan Harrison, David K. Jackson, Ian Johnston, Dominic Kwiatkowski, Cordelia Langford, John Sillitoe on behalf of the Wellcome Sanger Institute COVID-19 Surveillance Team |
| EPI_ISL_2865060, EPI_ISL_2865061 | Berkshire and Surrey Pathology Services Lighthouse Laboratory | Wellcome Sanger Institute for the COVID-19 Genomics UK (COG-UK) Consortium | Berkshire and Surrey Pathology Services Lighthouse Laboratory and Alex Alderton, Roberto Amato, Jeffrey Barrett, Sonia Goncalves, Ewan Harrison, David K. Jackson, Ian Johnston, Dominic Kwiatkowski, Cordelia Langford, John Sillitoe on behalf of the Wellcome Sanger Institute COVID-19 Surveillance Team |
| EPI_ISL_2865062 | Lighthouse Lab in Milton Keynes | Wellcome Sanger Institute for the COVID-19 Genomics UK (COG-UK) Consortium | The Lighthouse Lab in Milton Keynes and Alex Alderton, Roberto Amato, Jeffrey Barrett, Sonia Goncalves, Ewan Harrison, David K. Jackson, Ian Johnston, Dominic Kwiatkowski, Cordelia Langford, John Sillitoe on behalf of the Wellcome Sanger Institute COVID-19 Surveillance Team |
| EPI_ISL_2865063 | Lighthouse Lab in Alderley Park | Wellcome Sanger Institute for the COVID-19 Genomics UK (COG-UK) Consortium | Jacquelyn Wynn, Mairead Hyland, The Lighthouse Lab in Alderley Park and Alex Alderton, Roberto Amato, Jeffrey Barrett, Sonia Goncalves, Ewan Harrison, David K. Jackson, Ian Johnston, Dominic Kwiatkowski, Cordelia Langford, John Sillitoe on behalf of the Wellcome Sanger Institute COVID-19 Surveillance Team |
| EPI_ISL_2865064 | Berkshire and Surrey Pathology Services Lighthouse Laboratory | Wellcome Sanger Institute for the COVID-19 Genomics UK (COG-UK) Consortium | Berkshire and Surrey Pathology Services Lighthouse Laboratory and Alex Alderton, Roberto Amato, Jeffrey Barrett, Sonia Goncalves, Ewan Harrison, David K. Jackson, Ian Johnston, Dominic Kwiatkowski, Cordelia Langford, John Sillitoe on behalf of the Wellcome Sanger Institute COVID-19 Surveillance Team |
| EPI_ISL_2865065, EPI_ISL_2865066 | Lighthouse Lab in Milton Keynes | Wellcome Sanger Institute for the COVID-19 Genomics UK (COG-UK) Consortium | The Lighthouse Lab in Milton Keynes and Alex Alderton, Roberto Amato, Jeffrey Barrett, Sonia Goncalves, Ewan Harrison, David K. Jackson, Ian Johnston, Dominic Kwiatkowski, Cordelia Langford, John Sillitoe on behalf of the Wellcome Sanger Institute COVID-19 Surveillance Team |
| EPI_ISL_2865067 | Lighthouse Lab in Alderley Park | Wellcome Sanger Institute for the COVID-19 Genomics UK (COG-UK) Consortium | Jacquelyn Wynn, Mairead Hyland, The Lighthouse Lab in Alderley Park and Alex Alderton, Roberto Amato, Jeffrey Barrett, Sonia Goncalves, Ewan Harrison, David K. Jackson, Ian Johnston, Dominic Kwiatkowski, Cordelia Langford, John Sillitoe on behalf of the Wellcome Sanger Institute COVID-19 Surveillance Team |
| EPI_ISL_2865068 | Berkshire and Surrey Pathology Services Lighthouse Laboratory | Wellcome Sanger Institute for the COVID-19 Genomics UK (COG-UK) Consortium | Berkshire and Surrey Pathology Services Lighthouse Laboratory and Alex Alderton, Roberto Amato, Jeffrey Barrett, Sonia Goncalves, Ewan Harrison, David K. Jackson, Ian Johnston, Dominic Kwiatkowski, Cordelia Langford, John Sillitoe on behalf of the Wellcome Sanger Institute COVID-19 Surveillance Team |
| EPI_ISL_2865069 | Lighthouse Lab in Milton Keynes | Wellcome Sanger Institute for the COVID-19 Genomics UK (COG-UK) Consortium | The Lighthouse Lab in Milton Keynes and Alex Alderton, Roberto Amato, Jeffrey Barrett, Sonia Goncalves, Ewan Harrison, David K. Jackson, Ian Johnston, Dominic Kwiatkowski, Cordelia Langford, John Sillitoe on behalf of the Wellcome Sanger Institute COVID-19 Surveillance Team |
| EPI_ISL_2865070 | Lighthouse Lab in Alderley Park | Wellcome Sanger Institute for the COVID-19 Genomics UK (COG-UK) Consortium | Jacquelyn Wynn, Mairead Hyland, The Lighthouse Lab in Alderley Park and Alex Alderton, Roberto Amato, Jeffrey Barrett, Sonia Goncalves, Ewan Harrison, David K. Jackson, Ian Johnston, Dominic Kwiatkowski, Cordelia Langford, John Sillitoe on behalf of the Wellcome Sanger Institute COVID-19 Surveillance Team |
| EPI_ISL_2865071 | Lighthouse Lab in Milton Keynes | Wellcome Sanger Institute for the COVID-19 Genomics UK (COG-UK) Consortium | The Lighthouse Lab in Milton Keynes and Alex Alderton, Roberto Amato, Jeffrey Barrett, Sonia Goncalves, Ewan Harrison, David K. Jackson, Ian Johnston, Dominic Kwiatkowski, Cordelia Langford, John Sillitoe on behalf of the Wellcome Sanger Institute COVID-19 Surveillance Team |
| EPI_ISL_2865072 | Lighthouse Lab in Alderley Park | Wellcome Sanger Institute for the COVID-19 Genomics UK (COG-UK) Consortium | Jacquelyn Wynn, Mairead Hyland, The Lighthouse Lab in Alderley Park and Alex Alderton, Roberto Amato, Jeffrey Barrett, Sonia Goncalves, Ewan Harrison, David K. Jackson, Ian Johnston, Dominic Kwiatkowski, Cordelia Langford, John Sillitoe on behalf of the Wellcome Sanger Institute COVID-19 Surveillance Team |
| EPI_ISL_2865073 | Berkshire and Surrey Pathology Services Lighthouse Laboratory | Wellcome Sanger Institute for the COVID-19 Genomics UK (COG-UK) Consortium | Berkshire and Surrey Pathology Services Lighthouse Laboratory and Alex Alderton, Roberto Amato, Jeffrey Barrett, Sonia Goncalves, Ewan Harrison, David K. Jackson, Ian Johnston, Dominic Kwiatkowski, Cordelia Langford, John Sillitoe on behalf of the Wellcome Sanger Institute COVID-19 Surveillance Team |
| EPI_ISL_2865074, EPI_ISL_2865075 | Lighthouse Lab in Alderley Park | Wellcome Sanger Institute for the COVID-19 Genomics UK (COG-UK) Consortium | Jacquelyn Wynn, Mairead Hyland, The Lighthouse Lab in Alderley Park and Alex Alderton, Roberto Amato, Jeffrey Barrett, Sonia Goncalves, Ewan Harrison, David K. Jackson, Ian Johnston, Dominic Kwiatkowski, Cordelia Langford, John Sillitoe on behalf of the Wellcome Sanger Institute COVID-19 Surveillance Team |
| EPI_ISL_2865076, EPI_ISL_2865077 | Berkshire and Surrey Pathology Services Lighthouse Laboratory | Wellcome Sanger Institute for the COVID-19 Genomics UK (COG-UK) Consortium | Berkshire and Surrey Pathology Services Lighthouse Laboratory and Alex Alderton, Roberto Amato, Jeffrey Barrett, Sonia Goncalves, Ewan Harrison, David K. Jackson, Ian Johnston, Dominic Kwiatkowski, Cordelia Langford, John Sillitoe on behalf of the Wellcome Sanger Institute COVID-19 Surveillance Team |
| EPI_ISL_2865078, EPI_ISL_2865079, | Lighthouse Lab in Alderley Park | Wellcome Sanger Institute for the COVID-19 Genomics UK (COG-UK) Consortium | Jacquelyn Wynn, Mairead Hyland, The Lighthouse Lab in Alderley Park and Alex Alderton, Roberto Amato, Jeffrey Barrett, Sonia Goncalves, Ewan Harrison, David K. Jackson, Ian Johnston, Dominic Kwiatkowski, Cordelia Langford, John Sillitoe on behalf of the Wellcome Sanger Institute COVID-19 Surveillance Team |

| | | | |
|---|---|---|---|
| EPI_ISL_2865080, EPI_ISL_2865081 | | | |
| EPI_ISL_2865082 | Berkshire and Surrey Pathology Services Lighthouse Laboratory | Wellcome Sanger Institute for the COVID-19 Genomics UK (COG-UK) Consortium | Berkshire and Surrey Pathology Services Lighthouse Laboratory and Alex Alderton, Roberto Amato, Jeffrey Barrett, Sonia Goncalves, Ewan Harrison, David K. Jackson, Ian Johnston, Dominic Kwiatkowski, Cordelia Langford, John Sillitoe on behalf of the Wellcome Sanger Institute COVID-19 Surveillance Team |
| EPI_ISL_2865083 | Lighthouse Lab in Alderley Park | Wellcome Sanger Institute for the COVID-19 Genomics UK (COG-UK) Consortium | Jacquelyn Wynn, Mairead Hyland, The Lighthouse Lab in Alderley Park and Alex Alderton, Roberto Amato, Jeffrey Barrett, Sonia Goncalves, Ewan Harrison, David K. Jackson, Ian Johnston, Dominic Kwiatkowski, Cordelia Langford, John Sillitoe on behalf of the Wellcome Sanger Institute COVID-19 Surveillance Team |
| EPI_ISL_2865084 | Lighthouse Lab in Milton Keynes | Wellcome Sanger Institute for the COVID-19 Genomics UK (COG-UK) Consortium | The Lighthouse Lab in Milton Keynes and Alex Alderton, Roberto Amato, Jeffrey Barrett, Sonia Goncalves, Ewan Harrison, David K. Jackson, Ian Johnston, Dominic Kwiatkowski, Cordelia Langford, John Sillitoe on behalf of the Wellcome Sanger Institute COVID-19 Surveillance Team |
| EPI_ISL_2865085 | Berkshire and Surrey Pathology Services Lighthouse Laboratory | Wellcome Sanger Institute for the COVID-19 Genomics UK (COG-UK) Consortium | Berkshire and Surrey Pathology Services Lighthouse Laboratory and Alex Alderton, Roberto Amato, Jeffrey Barrett, Sonia Goncalves, Ewan Harrison, David K. Jackson, Ian Johnston, Dominic Kwiatkowski, Cordelia Langford, John Sillitoe on behalf of the Wellcome Sanger Institute COVID-19 Surveillance Team |
| EPI_ISL_2865086 | Lighthouse Lab in Milton Keynes | Wellcome Sanger Institute for the COVID-19 Genomics UK (COG-UK) Consortium | The Lighthouse Lab in Milton Keynes and Alex Alderton, Roberto Amato, Jeffrey Barrett, Sonia Goncalves, Ewan Harrison, David K. Jackson, Ian Johnston, Dominic Kwiatkowski, Cordelia Langford, John Sillitoe on behalf of the Wellcome Sanger Institute COVID-19 Surveillance Team |
| EPI_ISL_2865087 | Lighthouse Lab in Alderley Park | Wellcome Sanger Institute for the COVID-19 Genomics UK (COG-UK) Consortium | Jacquelyn Wynn, Mairead Hyland, The Lighthouse Lab in Alderley Park and Alex Alderton, Roberto Amato, Jeffrey Barrett, Sonia Goncalves, Ewan Harrison, David K. Jackson, Ian Johnston, Dominic Kwiatkowski, Cordelia Langford, John Sillitoe on behalf of the Wellcome Sanger Institute COVID-19 Surveillance Team |
| EPI_ISL_2865088 | Berkshire and Surrey Pathology Services Lighthouse Laboratory | Wellcome Sanger Institute for the COVID-19 Genomics UK (COG-UK) Consortium | Berkshire and Surrey Pathology Services Lighthouse Laboratory and Alex Alderton, Roberto Amato, Jeffrey Barrett, Sonia Goncalves, Ewan Harrison, David K. Jackson, Ian Johnston, Dominic Kwiatkowski, Cordelia Langford, John Sillitoe on behalf of the Wellcome Sanger Institute COVID-19 Surveillance Team |
| EPI_ISL_2865089 | Lighthouse Lab in Alderley Park | Wellcome Sanger Institute for the COVID-19 Genomics UK (COG-UK) Consortium | Jacquelyn Wynn, Mairead Hyland, The Lighthouse Lab in Alderley Park and Alex Alderton, Roberto Amato, Jeffrey Barrett, Sonia Goncalves, Ewan Harrison, David K. Jackson, Ian Johnston, Dominic Kwiatkowski, Cordelia Langford, John Sillitoe on behalf of the Wellcome Sanger Institute COVID-19 Surveillance Team |
| EPI_ISL_2865090 | Lighthouse Lab in Milton Keynes | Wellcome Sanger Institute for the COVID-19 Genomics UK (COG-UK) Consortium | The Lighthouse Lab in Milton Keynes and Alex Alderton, Roberto Amato, Jeffrey Barrett, Sonia Goncalves, Ewan Harrison, David K. Jackson, Ian Johnston, Dominic Kwiatkowski, Cordelia Langford, John Sillitoe on behalf of the Wellcome Sanger Institute COVID-19 Surveillance Team |
| EPI_ISL_2865091 | Lighthouse Lab in Alderley Park | Wellcome Sanger Institute for the COVID-19 Genomics UK (COG-UK) Consortium | Jacquelyn Wynn, Mairead Hyland, The Lighthouse Lab in Alderley Park and Alex Alderton, Roberto Amato, Jeffrey Barrett, Sonia Goncalves, Ewan Harrison, David K. Jackson, Ian Johnston, Dominic Kwiatkowski, Cordelia Langford, John Sillitoe on behalf of the Wellcome Sanger Institute COVID-19 Surveillance Team |
| EPI_ISL_2865092 | Lighthouse Lab in Milton Keynes | Wellcome Sanger Institute for the COVID-19 Genomics UK (COG-UK) Consortium | The Lighthouse Lab in Milton Keynes and Alex Alderton, Roberto Amato, Jeffrey Barrett, Sonia Goncalves, Ewan Harrison, David K. Jackson, Ian Johnston, Dominic Kwiatkowski, Cordelia Langford, John Sillitoe on behalf of the Wellcome Sanger Institute COVID-19 Surveillance Team |
| EPI_ISL_2865093 | Lighthouse Lab in Alderley Park | Wellcome Sanger Institute for the COVID-19 Genomics UK (COG-UK) Consortium | Jacquelyn Wynn, Mairead Hyland, The Lighthouse Lab in Alderley Park and Alex Alderton, Roberto Amato, Jeffrey Barrett, Sonia Goncalves, Ewan Harrison, David K. Jackson, Ian Johnston, Dominic Kwiatkowski, Cordelia Langford, John Sillitoe on behalf of the Wellcome Sanger Institute COVID-19 Surveillance Team |
| EPI_ISL_2865094 | Lighthouse Lab in Milton Keynes | Wellcome Sanger Institute for the COVID-19 Genomics UK (COG-UK) Consortium | The Lighthouse Lab in Milton Keynes and Alex Alderton, Roberto Amato, Jeffrey Barrett, Sonia Goncalves, Ewan Harrison, David K. Jackson, Ian Johnston, Dominic Kwiatkowski, Cordelia Langford, John Sillitoe on behalf of the Wellcome Sanger Institute COVID-19 Surveillance Team |
| EPI_ISL_2865095 | Lighthouse Lab in Alderley Park | Wellcome Sanger Institute for the COVID-19 Genomics UK (COG-UK) Consortium | Jacquelyn Wynn, Mairead Hyland, The Lighthouse Lab in Alderley Park and Alex Alderton, Roberto Amato, Jeffrey Barrett, Sonia Goncalves, Ewan Harrison, David K. Jackson, Ian Johnston, Dominic Kwiatkowski, Cordelia Langford, John Sillitoe on behalf of the Wellcome Sanger Institute COVID-19 Surveillance Team |
| EPI_ISL_2865096 | Lighthouse Lab in Milton Keynes | Wellcome Sanger Institute for the COVID-19 Genomics UK (COG-UK) Consortium | The Lighthouse Lab in Milton Keynes and Alex Alderton, Roberto Amato, Jeffrey Barrett, Sonia Goncalves, Ewan Harrison, David K. Jackson, Ian Johnston, Dominic Kwiatkowski, Cordelia Langford, John Sillitoe on behalf of the Wellcome Sanger Institute COVID-19 Surveillance Team |
| EPI_ISL_2865097 | Berkshire and Surrey Pathology Services Lighthouse Laboratory | Wellcome Sanger Institute for the COVID-19 Genomics UK (COG-UK) Consortium | Berkshire and Surrey Pathology Services Lighthouse Laboratory and Alex Alderton, Roberto Amato, Jeffrey Barrett, Sonia Goncalves, Ewan Harrison, David K. Jackson, Ian Johnston, Dominic Kwiatkowski, Cordelia Langford, John Sillitoe on behalf of the Wellcome Sanger Institute COVID-19 Surveillance Team |
| EPI_ISL_2865098, EPI_ISL_2865099 | Lighthouse Lab in Alderley Park | Wellcome Sanger Institute for the COVID-19 Genomics UK (COG-UK) Consortium | Jacquelyn Wynn, Mairead Hyland, The Lighthouse Lab in Alderley Park and Alex Alderton, Roberto Amato, Jeffrey Barrett, Sonia Goncalves, Ewan Harrison, David K. Jackson, Ian Johnston, Dominic Kwiatkowski, Cordelia Langford, John Sillitoe on behalf of the Wellcome Sanger Institute COVID-19 Surveillance Team |
| EPI_ISL_2865100 | Berkshire and Surrey Pathology Services Lighthouse Laboratory | Wellcome Sanger Institute for the COVID-19 Genomics UK (COG-UK) Consortium | Berkshire and Surrey Pathology Services Lighthouse Laboratory and Alex Alderton, Roberto Amato, Jeffrey Barrett, Sonia Goncalves, Ewan Harrison, David K. Jackson, Ian Johnston, Dominic Kwiatkowski, Cordelia Langford, John Sillitoe on behalf of the Wellcome Sanger Institute COVID-19 Surveillance Team |
| EPI_ISL_2865101 | Lighthouse Lab in Alderley Park | Wellcome Sanger Institute for the COVID-19 Genomics UK (COG-UK) Consortium | Jacquelyn Wynn, Mairead Hyland, The Lighthouse Lab in Alderley Park and Alex Alderton, Roberto Amato, Jeffrey Barrett, Sonia Goncalves, Ewan Harrison, David K. Jackson, Ian Johnston, Dominic Kwiatkowski, Cordelia Langford, John Sillitoe on behalf of the Wellcome Sanger Institute COVID-19 Surveillance Team |
| EPI_ISL_2865102 | Berkshire and Surrey Pathology Services Lighthouse Laboratory | Wellcome Sanger Institute for the COVID-19 Genomics UK (COG-UK) Consortium | Berkshire and Surrey Pathology Services Lighthouse Laboratory and Alex Alderton, Roberto Amato, Jeffrey Barrett, Sonia Goncalves, Ewan Harrison, David K. Jackson, Ian Johnston, Dominic Kwiatkowski, Cordelia Langford, John Sillitoe on behalf of the Wellcome Sanger Institute COVID-19 Surveillance Team |
| EPI_ISL_2865103, EPI_ISL_2865104 | Lighthouse Lab in Alderley Park | Wellcome Sanger Institute for the COVID-19 Genomics UK (COG-UK) Consortium | Jacquelyn Wynn, Mairead Hyland, The Lighthouse Lab in Alderley Park and Alex Alderton, Roberto Amato, Jeffrey Barrett, Sonia Goncalves, Ewan Harrison, David K. Jackson, Ian Johnston, Dominic Kwiatkowski, Cordelia Langford, John Sillitoe on behalf of the Wellcome Sanger Institute COVID-19 Surveillance Team |
| EPI_ISL_2865105 | Lighthouse Lab in Milton Keynes | Wellcome Sanger Institute for the COVID-19 Genomics UK (COG-UK) Consortium | The Lighthouse Lab in Milton Keynes and Alex Alderton, Roberto Amato, Jeffrey Barrett, Sonia Goncalves, Ewan Harrison, David K. Jackson, Ian Johnston, Dominic Kwiatkowski, Cordelia Langford, John Sillitoe on behalf of the Wellcome Sanger Institute COVID-19 Surveillance Team |
| EPI_ISL_2865106 | Berkshire and Surrey Pathology Services Lighthouse Laboratory | Wellcome Sanger Institute for the COVID-19 Genomics UK (COG-UK) Consortium | Berkshire and Surrey Pathology Services Lighthouse Laboratory and Alex Alderton, Roberto Amato, Jeffrey Barrett, Sonia Goncalves, Ewan Harrison, David K. Jackson, Ian Johnston, Dominic Kwiatkowski, Cordelia Langford, John Sillitoe on behalf of the Wellcome Sanger Institute COVID-19 Surveillance Team |
| EPI_ISL_2865107 | Lighthouse Lab in Alderley Park | Wellcome Sanger Institute for the COVID-19 Genomics UK (COG-UK) Consortium | Jacquelyn Wynn, Mairead Hyland, The Lighthouse Lab in Alderley Park and Alex Alderton, Roberto Amato, Jeffrey Barrett, Sonia Goncalves, Ewan Harrison, David K. Jackson, Ian Johnston, Dominic Kwiatkowski, Cordelia Langford, John Sillitoe on behalf of the Wellcome Sanger Institute COVID-19 Surveillance Team |
| EPI_ISL_2865108 | Lighthouse Lab in Glasgow | Wellcome Sanger Institute for the COVID-19 Genomics UK (COG-UK) Consortium | Harper VanSteenhouse, Yumi Kasai, David Gray, Carol Clugston, Anna Dominiczak and Alex Alderton, Roberto Amato, Jeffrey Barrett, Sonia Goncalves, Ewan Harrison, David K. Jackson, Ian Johnston, Dominic Kwiatkowski, Cordelia Langford, John Sillitoe on behalf of the Wellcome Sanger Institute COVID-19 Surveillance Team |
| EPI_ISL_2865109, EPI_ISL_2865110 | Berkshire and Surrey Pathology Services Lighthouse Laboratory | Wellcome Sanger Institute for the COVID-19 Genomics UK (COG-UK) Consortium | Berkshire and Surrey Pathology Services Lighthouse Laboratory and Alex Alderton, Roberto Amato, Jeffrey Barrett, Sonia Goncalves, Ewan Harrison, David K. Jackson, Ian Johnston, Dominic Kwiatkowski, Cordelia Langford, John Sillitoe on behalf of the Wellcome Sanger Institute COVID-19 Surveillance Team |
| EPI_ISL_2865111 | Lighthouse Lab in Glasgow | Wellcome Sanger Institute for the COVID-19 Genomics UK (COG-UK) Consortium | Harper VanSteenhouse, Yumi Kasai, David Gray, Carol Clugston, Anna Dominiczak and Alex Alderton, Roberto Amato, Jeffrey Barrett, Sonia Goncalves, Ewan Harrison, David K. Jackson, Ian Johnston, Dominic Kwiatkowski, Cordelia Langford, John Sillitoe on behalf of the Wellcome Sanger Institute COVID-19 Surveillance Team |
| EPI_ISL_2865112 | Berkshire and Surrey Pathology Services Lighthouse Laboratory | Wellcome Sanger Institute for the COVID-19 Genomics UK (COG-UK) Consortium | Berkshire and Surrey Pathology Services Lighthouse Laboratory and Alex Alderton, Roberto Amato, Jeffrey Barrett, Sonia Goncalves, Ewan Harrison, David K. Jackson, Ian Johnston, Dominic Kwiatkowski, Cordelia Langford, John Sillitoe on behalf of the Wellcome Sanger Institute COVID-19 Surveillance Team |
| EPI_ISL_2865113 | Lighthouse Lab in Alderley Park | Wellcome Sanger Institute for the COVID-19 Genomics UK (COG-UK) Consortium | Jacquelyn Wynn, Mairead Hyland, The Lighthouse Lab in Alderley Park and Alex Alderton, Roberto Amato, Jeffrey Barrett, Sonia Goncalves, Ewan Harrison, David K. Jackson, Ian Johnston, Dominic Kwiatkowski, Cordelia Langford, John Sillitoe on behalf of the Wellcome Sanger Institute COVID-19 Surveillance Team |
| EPI_ISL_2865114 | Lighthouse Lab in Glasgow | Wellcome Sanger Institute for the COVID-19 Genomics UK (COG-UK) Consortium | Harper VanSteenhouse, Yumi Kasai, David Gray, Carol Clugston, Anna Dominiczak and Alex Alderton, Roberto Amato, Jeffrey Barrett, Sonia Goncalves, Ewan Harrison, David K. Jackson, Ian Johnston, Dominic Kwiatkowski, Cordelia Langford, John Sillitoe on behalf of the Wellcome Sanger Institute COVID-19 Surveillance Team |
| EPI_ISL_2865115 | Lighthouse Lab in Alderley Park | Wellcome Sanger Institute for the COVID-19 Genomics UK (COG-UK) Consortium | Jacquelyn Wynn, Mairead Hyland, The Lighthouse Lab in Alderley Park and Alex Alderton, Roberto Amato, Jeffrey Barrett, Sonia Goncalves, Ewan Harrison, David K. Jackson, Ian Johnston, Dominic Kwiatkowski, Cordelia Langford, John Sillitoe on behalf of the Wellcome Sanger Institute COVID-19 Surveillance Team |
| EPI_ISL_2865116 | Lighthouse Lab in Glasgow | Wellcome Sanger Institute for the COVID-19 Genomics UK (COG-UK) Consortium | Harper VanSteenhouse, Yumi Kasai, David Gray, Carol Clugston, Anna Dominiczak and Alex Alderton, Roberto Amato, Jeffrey Barrett, Sonia Goncalves, Ewan Harrison, David K. Jackson, Ian Johnston, Dominic Kwiatkowski, Cordelia Langford, John Sillitoe on behalf of the Wellcome Sanger Institute COVID-19 Surveillance Team |
| EPI_ISL_2865117 | Lighthouse Lab in Milton Keynes | Wellcome Sanger Institute for the COVID-19 Genomics UK (COG-UK) Consortium | The Lighthouse Lab in Milton Keynes and Alex Alderton, Roberto Amato, Jeffrey Barrett, Sonia Goncalves, Ewan Harrison, David K. Jackson, Ian Johnston, Dominic Kwiatkowski, Cordelia Langford, John Sillitoe on behalf of the Wellcome Sanger Institute COVID-19 Surveillance Team |
| EPI_ISL_2865118 | Lighthouse Lab in Alderley Park | Wellcome Sanger Institute for the COVID-19 Genomics UK (COG-UK) Consortium | Jacquelyn Wynn, Mairead Hyland, The Lighthouse Lab in Alderley Park and Alex Alderton, Roberto Amato, Jeffrey Barrett, Sonia Goncalves, Ewan Harrison, David K. Jackson, Ian Johnston, Dominic Kwiatkowski, Cordelia Langford, John Sillitoe on behalf of the Wellcome Sanger Institute COVID-19 Surveillance Team |
| EPI_ISL_2865119 | Lighthouse Lab in Milton Keynes | Wellcome Sanger Institute for the COVID-19 Genomics UK (COG-UK) Consortium | The Lighthouse Lab in Milton Keynes and Alex Alderton, Roberto Amato, Jeffrey Barrett, Sonia Goncalves, Ewan Harrison, David K. Jackson, Ian Johnston, Dominic Kwiatkowski, Cordelia Langford, John Sillitoe on behalf of the Wellcome Sanger Institute COVID-19 Surveillance Team |
| EPI_ISL_2865120, EPI_ISL_2865121 | Lighthouse Lab in Alderley Park | Wellcome Sanger Institute for the COVID-19 Genomics UK (COG-UK) Consortium | Jacquelyn Wynn, Mairead Hyland, The Lighthouse Lab in Alderley Park and Alex Alderton, Roberto Amato, Jeffrey Barrett, Sonia Goncalves, Ewan Harrison, David K. Jackson, Ian Johnston, Dominic Kwiatkowski, Cordelia Langford, John Sillitoe on behalf of the Wellcome Sanger Institute COVID-19 Surveillance Team |
| EPI_ISL_2865122 | Lighthouse Lab in Milton Keynes | Wellcome Sanger Institute for the COVID-19 Genomics UK (COG-UK) Consortium | The Lighthouse Lab in Milton Keynes and Alex Alderton, Roberto Amato, Jeffrey Barrett, Sonia Goncalves, Ewan Harrison, David K. Jackson, Ian Johnston, Dominic Kwiatkowski, Cordelia Langford, John Sillitoe on behalf of the Wellcome Sanger Institute COVID-19 Surveillance Team |
| EPI_ISL_2865123 | Lighthouse Lab in Glasgow | Wellcome Sanger Institute for the COVID-19 Genomics UK (COG-UK) Consortium | Harper VanSteenhouse, Yumi Kasai, David Gray, Carol Clugston, Anna Dominiczak and Alex Alderton, Roberto Amato, Jeffrey Barrett, Sonia Goncalves, Ewan Harrison, David K. Jackson, Ian Johnston, Dominic Kwiatkowski, Cordelia Langford, John Sillitoe on behalf of the Wellcome Sanger Institute COVID-19 Surveillance Team |
| EPI_ISL_2865124 | Lighthouse Lab in Alderley Park | Wellcome Sanger Institute for the COVID-19 Genomics UK (COG-UK) Consortium | Jacquelyn Wynn, Mairead Hyland, The Lighthouse Lab in Alderley Park and Alex Alderton, Roberto Amato, Jeffrey Barrett, Sonia Goncalves, Ewan Harrison, David K. Jackson, Ian Johnston, Dominic Kwiatkowski, Cordelia Langford, John Sillitoe on behalf of the Wellcome Sanger Institute COVID-19 Surveillance Team |
| EPI_ISL_2865125 | Lighthouse Lab in Glasgow | Wellcome Sanger Institute for the COVID-19 Genomics UK (COG-UK) Consortium | Harper VanSteenhouse, Yumi Kasai, David Gray, Carol Clugston, Anna Dominiczak and Alex Alderton, Roberto Amato, Jeffrey Barrett, Sonia Goncalves, Ewan Harrison, David K. Jackson, Ian Johnston, Dominic Kwiatkowski, Cordelia Langford, John Sillitoe on behalf of the Wellcome Sanger Institute COVID-19 Surveillance Team |
| EPI_ISL_2865126 | Lighthouse Lab in Milton Keynes | Wellcome Sanger Institute for the COVID-19 Genomics UK (COG-UK) Consortium | The Lighthouse Lab in Milton Keynes and Alex Alderton, Roberto Amato, Jeffrey Barrett, Sonia Goncalves, Ewan Harrison, David K. Jackson, Ian Johnston, Dominic Kwiatkowski, Cordelia Langford, John Sillitoe on behalf of the Wellcome Sanger Institute COVID-19 Surveillance Team |
| EPI_ISL_2865127 | Lighthouse Lab in Alderley Park | Wellcome Sanger Institute for the COVID-19 Genomics UK (COG-UK) Consortium | Jacquelyn Wynn, Mairead Hyland, The Lighthouse Lab in Alderley Park and Alex Alderton, Roberto Amato, Jeffrey Barrett, Sonia Goncalves, Ewan Harrison, David K. Jackson, Ian Johnston, Dominic Kwiatkowski, Cordelia Langford, John Sillitoe on behalf of the Wellcome Sanger Institute COVID-19 Surveillance Team |
| EPI_ISL_2865128 | Berkshire and Surrey Pathology Services Lighthouse Laboratory | Wellcome Sanger Institute for the COVID-19 Genomics UK (COG-UK) Consortium | Berkshire and Surrey Pathology Services Lighthouse Laboratory and Alex Alderton, Roberto Amato, Jeffrey Barrett, Sonia Goncalves, Ewan Harrison, David K. Jackson, Ian Johnston, Dominic Kwiatkowski, Cordelia Langford, John Sillitoe on behalf of the Wellcome Sanger Institute COVID-19 Surveillance Team |
| EPI_ISL_2865129 | Lighthouse Lab in Glasgow | Wellcome Sanger Institute for the COVID-19 Genomics UK (COG-UK) Consortium | Harper VanSteenhouse, Yumi Kasai, David Gray, Carol Clugston, Anna Dominiczak and Alex Alderton, Roberto Amato, Jeffrey Barrett, Sonia Goncalves, Ewan Harrison, David K. Jackson, Ian Johnston, Dominic Kwiatkowski, Cordelia Langford, John Sillitoe on behalf of the Wellcome Sanger Institute COVID-19 Surveillance Team |

| | | | |
|---|---|---|---|
| EPI_ISL_2865130 | Berkshire and Surrey Pathology Services Lighthouse Laboratory | Wellcome Sanger Institute for the COVID-19 Genomics UK (COG-UK) Consortium | Berkshire and Surrey Pathology Services Lighthouse Laboratory and Alex Alderton, Roberto Amato, Jeffrey Barrett, Sonia Goncalves, Ewan Harrison, David K. Jackson, Ian Johnston, Dominic Kwiatkowski, Cordelia Langford, John Sillitoe on behalf of the Wellcome Sanger Institute COVID-19 Surveillance Team |
| EPI_ISL_2865131 | Lighthouse Lab in Milton Keynes | Wellcome Sanger Institute for the COVID-19 Genomics UK (COG-UK) Consortium | The Lighthouse Lab in Milton Keynes and Alex Alderton, Roberto Amato, Jeffrey Barrett, Sonia Goncalves, Ewan Harrison, David K. Jackson, Ian Johnston, Dominic Kwiatkowski, Cordelia Langford, John Sillitoe on behalf of the Wellcome Sanger Institute COVID-19 Surveillance Team |
| EPI_ISL_2865132 | Berkshire and Surrey Pathology Services Lighthouse Laboratory | Wellcome Sanger Institute for the COVID-19 Genomics UK (COG-UK) Consortium | Berkshire and Surrey Pathology Services Lighthouse Laboratory and Alex Alderton, Roberto Amato, Jeffrey Barrett, Sonia Goncalves, Ewan Harrison, David K. Jackson, Ian Johnston, Dominic Kwiatkowski, Cordelia Langford, John Sillitoe on behalf of the Wellcome Sanger Institute COVID-19 Surveillance Team |
| EPI_ISL_2865133 | Lighthouse Lab in Alderley Park | Wellcome Sanger Institute for the COVID-19 Genomics UK (COG-UK) Consortium | Jacquelyn Wynn, Mairead Hyland, The Lighthouse Lab in Alderley Park and Alex Alderton, Roberto Amato, Jeffrey Barrett, Sonia Goncalves, Ewan Harrison, David K. Jackson, Ian Johnston, Dominic Kwiatkowski, Cordelia Langford, John Sillitoe on behalf of the Wellcome Sanger Institute COVID-19 Surveillance Team |
| EPI_ISL_2865134 | Berkshire and Surrey Pathology Services Lighthouse Laboratory | Wellcome Sanger Institute for the COVID-19 Genomics UK (COG-UK) Consortium | Berkshire and Surrey Pathology Services Lighthouse Laboratory and Alex Alderton, Roberto Amato, Jeffrey Barrett, Sonia Goncalves, Ewan Harrison, David K. Jackson, Ian Johnston, Dominic Kwiatkowski, Cordelia Langford, John Sillitoe on behalf of the Wellcome Sanger Institute COVID-19 Surveillance Team |
| EPI_ISL_2865135, EPI_ISL_2865136 | Lighthouse Lab in Alderley Park | Wellcome Sanger Institute for the COVID-19 Genomics UK (COG-UK) Consortium | Jacquelyn Wynn, Mairead Hyland, The Lighthouse Lab in Alderley Park and Alex Alderton, Roberto Amato, Jeffrey Barrett, Sonia Goncalves, Ewan Harrison, David K. Jackson, Ian Johnston, Dominic Kwiatkowski, Cordelia Langford, John Sillitoe on behalf of the Wellcome Sanger Institute COVID-19 Surveillance Team |
| EPI_ISL_2865137 | Lighthouse Lab in Milton Keynes | Wellcome Sanger Institute for the COVID-19 Genomics UK (COG-UK) Consortium | The Lighthouse Lab in Milton Keynes and Alex Alderton, Roberto Amato, Jeffrey Barrett, Sonia Goncalves, Ewan Harrison, David K. Jackson, Ian Johnston, Dominic Kwiatkowski, Cordelia Langford, John Sillitoe on behalf of the Wellcome Sanger Institute COVID-19 Surveillance Team |
| EPI_ISL_2865138, EPI_ISL_2865139 | Berkshire and Surrey Pathology Services Lighthouse Laboratory | Wellcome Sanger Institute for the COVID-19 Genomics UK (COG-UK) Consortium | Berkshire and Surrey Pathology Services Lighthouse Laboratory and Alex Alderton, Roberto Amato, Jeffrey Barrett, Sonia Goncalves, Ewan Harrison, David K. Jackson, Ian Johnston, Dominic Kwiatkowski, Cordelia Langford, John Sillitoe on behalf of the Wellcome Sanger Institute COVID-19 Surveillance Team |
| EPI_ISL_2865140, EPI_ISL_2865141 | Lighthouse Lab in Alderley Park | Wellcome Sanger Institute for the COVID-19 Genomics UK (COG-UK) Consortium | Jacquelyn Wynn, Mairead Hyland, The Lighthouse Lab in Alderley Park and Alex Alderton, Roberto Amato, Jeffrey Barrett, Sonia Goncalves, Ewan Harrison, David K. Jackson, Ian Johnston, Dominic Kwiatkowski, Cordelia Langford, John Sillitoe on behalf of the Wellcome Sanger Institute COVID-19 Surveillance Team |
| EPI_ISL_2865142 | Lighthouse Lab in Glasgow | Wellcome Sanger Institute for the COVID-19 Genomics UK (COG-UK) Consortium | Harper VanSteenhouse, Yumi Kasai, David Gray, Carol Clugston, Anna Dominiczak and Alex Alderton, Roberto Amato, Jeffrey Barrett, Sonia Goncalves, Ewan Harrison, David K. Jackson, Ian Johnston, Dominic Kwiatkowski, Cordelia Langford, John Sillitoe on behalf of the Wellcome Sanger Institute COVID-19 Surveillance Team |
| EPI_ISL_2865143 | Berkshire and Surrey Pathology Services Lighthouse Laboratory | Wellcome Sanger Institute for the COVID-19 Genomics UK (COG-UK) Consortium | Berkshire and Surrey Pathology Services Lighthouse Laboratory and Alex Alderton, Roberto Amato, Jeffrey Barrett, Sonia Goncalves, Ewan Harrison, David K. Jackson, Ian Johnston, Dominic Kwiatkowski, Cordelia Langford, John Sillitoe on behalf of the Wellcome Sanger Institute COVID-19 Surveillance Team |
| EPI_ISL_2865144 | Lighthouse Lab in Glasgow | Wellcome Sanger Institute for the COVID-19 Genomics UK (COG-UK) Consortium | Harper VanSteenhouse, Yumi Kasai, David Gray, Carol Clugston, Anna Dominiczak and Alex Alderton, Roberto Amato, Jeffrey Barrett, Sonia Goncalves, Ewan Harrison, David K. Jackson, Ian Johnston, Dominic Kwiatkowski, Cordelia Langford, John Sillitoe on behalf of the Wellcome Sanger Institute COVID-19 Surveillance Team |
| EPI_ISL_2865145 | Berkshire and Surrey Pathology Services Lighthouse Laboratory | Wellcome Sanger Institute for the COVID-19 Genomics UK (COG-UK) Consortium | Berkshire and Surrey Pathology Services Lighthouse Laboratory and Alex Alderton, Roberto Amato, Jeffrey Barrett, Sonia Goncalves, Ewan Harrison, David K. Jackson, Ian Johnston, Dominic Kwiatkowski, Cordelia Langford, John Sillitoe on behalf of the Wellcome Sanger Institute COVID-19 Surveillance Team |
| EPI_ISL_2865146 | Lighthouse Lab in Glasgow | Wellcome Sanger Institute for the COVID-19 Genomics UK (COG-UK) Consortium | Harper VanSteenhouse, Yumi Kasai, David Gray, Carol Clugston, Anna Dominiczak and Alex Alderton, Roberto Amato, Jeffrey Barrett, Sonia Goncalves, Ewan Harrison, David K. Jackson, Ian Johnston, Dominic Kwiatkowski, Cordelia Langford, John Sillitoe on behalf of the Wellcome Sanger Institute COVID-19 Surveillance Team |
| EPI_ISL_2865147 | Berkshire and Surrey Pathology Services Lighthouse Laboratory | Wellcome Sanger Institute for the COVID-19 Genomics UK (COG-UK) Consortium | Berkshire and Surrey Pathology Services Lighthouse Laboratory and Alex Alderton, Roberto Amato, Jeffrey Barrett, Sonia Goncalves, Ewan Harrison, David K. Jackson, Ian Johnston, Dominic Kwiatkowski, Cordelia Langford, John Sillitoe on behalf of the Wellcome Sanger Institute COVID-19 Surveillance Team |
| EPI_ISL_2865148 | Lighthouse Lab in Milton Keynes | Wellcome Sanger Institute for the COVID-19 Genomics UK (COG-UK) Consortium | The Lighthouse Lab in Milton Keynes and Alex Alderton, Roberto Amato, Jeffrey Barrett, Sonia Goncalves, Ewan Harrison, David K. Jackson, Ian Johnston, Dominic Kwiatkowski, Cordelia Langford, John Sillitoe on behalf of the Wellcome Sanger Institute COVID-19 Surveillance Team |
| EPI_ISL_2865149 | Berkshire and Surrey Pathology Services Lighthouse Laboratory | Wellcome Sanger Institute for the COVID-19 Genomics UK (COG-UK) Consortium | Berkshire and Surrey Pathology Services Lighthouse Laboratory and Alex Alderton, Roberto Amato, Jeffrey Barrett, Sonia Goncalves, Ewan Harrison, David K. Jackson, Ian Johnston, Dominic Kwiatkowski, Cordelia Langford, John Sillitoe on behalf of the Wellcome Sanger Institute COVID-19 Surveillance Team |
| EPI_ISL_2865150 | Lighthouse Lab in Milton Keynes | Wellcome Sanger Institute for the COVID-19 Genomics UK (COG-UK) Consortium | The Lighthouse Lab in Milton Keynes and Alex Alderton, Roberto Amato, Jeffrey Barrett, Sonia Goncalves, Ewan Harrison, David K. Jackson, Ian Johnston, Dominic Kwiatkowski, Cordelia Langford, John Sillitoe on behalf of the Wellcome Sanger Institute COVID-19 Surveillance Team |
| EPI_ISL_2865151 | Lighthouse Lab in Glasgow | Wellcome Sanger Institute for the COVID-19 Genomics UK (COG-UK) Consortium | Harper VanSteenhouse, Yumi Kasai, David Gray, Carol Clugston, Anna Dominiczak and Alex Alderton, Roberto Amato, Jeffrey Barrett, Sonia Goncalves, Ewan Harrison, David K. Jackson, Ian Johnston, Dominic Kwiatkowski, Cordelia Langford, John Sillitoe on behalf of the Wellcome Sanger Institute COVID-19 Surveillance Team |
| EPI_ISL_2865152 | Lighthouse Lab in Alderley Park | Wellcome Sanger Institute for the COVID-19 Genomics UK (COG-UK) Consortium | Jacquelyn Wynn, Mairead Hyland, The Lighthouse Lab in Alderley Park and Alex Alderton, Roberto Amato, Jeffrey Barrett, Sonia Goncalves, Ewan Harrison, David K. Jackson, Ian Johnston, Dominic Kwiatkowski, Cordelia Langford, John Sillitoe on behalf of the Wellcome Sanger Institute COVID-19 Surveillance Team |
| EPI_ISL_2865153 | Lighthouse Lab in Milton Keynes | Wellcome Sanger Institute for the COVID-19 Genomics UK (COG-UK) Consortium | The Lighthouse Lab in Milton Keynes and Alex Alderton, Roberto Amato, Jeffrey Barrett, Sonia Goncalves, Ewan Harrison, David K. Jackson, Ian Johnston, Dominic Kwiatkowski, Cordelia Langford, John Sillitoe on behalf of the Wellcome Sanger Institute COVID-19 Surveillance Team |
| EPI_ISL_2865154, EPI_ISL_2865155 | Berkshire and Surrey Pathology Services Lighthouse Laboratory | Wellcome Sanger Institute for the COVID-19 Genomics UK (COG-UK) Consortium | Berkshire and Surrey Pathology Services Lighthouse Laboratory and Alex Alderton, Roberto Amato, Jeffrey Barrett, Sonia Goncalves, Ewan Harrison, David K. Jackson, Ian Johnston, Dominic Kwiatkowski, Cordelia Langford, John Sillitoe on behalf of the Wellcome Sanger Institute COVID-19 Surveillance Team |
| EPI_ISL_2865156 | Lighthouse Lab in Alderley Park | Wellcome Sanger Institute for the COVID-19 Genomics UK (COG-UK) Consortium | Jacquelyn Wynn, Mairead Hyland, The Lighthouse Lab in Alderley Park and Alex Alderton, Roberto Amato, Jeffrey Barrett, Sonia Goncalves, Ewan Harrison, David K. Jackson, Ian Johnston, Dominic Kwiatkowski, Cordelia Langford, John Sillitoe on behalf of the Wellcome Sanger Institute COVID-19 Surveillance Team |
| EPI_ISL_2865157, EPI_ISL_2865158 | Lighthouse Lab in Milton Keynes | Wellcome Sanger Institute for the COVID-19 Genomics UK (COG-UK) Consortium | The Lighthouse Lab in Milton Keynes and Alex Alderton, Roberto Amato, Jeffrey Barrett, Sonia Goncalves, Ewan Harrison, David K. Jackson, Ian Johnston, Dominic Kwiatkowski, Cordelia Langford, John Sillitoe on behalf of the Wellcome Sanger Institute COVID-19 Surveillance Team |
| EPI_ISL_2865159, EPI_ISL_2865160, EPI_ISL_2865161, EPI_ISL_2865162 | Berkshire and Surrey Pathology Services Lighthouse Laboratory | Wellcome Sanger Institute for the COVID-19 Genomics UK (COG-UK) Consortium | Berkshire and Surrey Pathology Services Lighthouse Laboratory and Alex Alderton, Roberto Amato, Jeffrey Barrett, Sonia Goncalves, Ewan Harrison, David K. Jackson, Ian Johnston, Dominic Kwiatkowski, Cordelia Langford, John Sillitoe on behalf of the Wellcome Sanger Institute COVID-19 Surveillance Team |
| EPI_ISL_2865163 | Lighthouse Lab in Milton Keynes | Wellcome Sanger Institute for the COVID-19 Genomics UK (COG-UK) Consortium | The Lighthouse Lab in Milton Keynes and Alex Alderton, Roberto Amato, Jeffrey Barrett, Sonia Goncalves, Ewan Harrison, David K. Jackson, Ian Johnston, Dominic Kwiatkowski, Cordelia Langford, John Sillitoe on behalf of the Wellcome Sanger Institute COVID-19 Surveillance Team |
| EPI_ISL_2865164 | Lighthouse Lab in Glasgow | Wellcome Sanger Institute for the COVID-19 Genomics UK (COG-UK) Consortium | Harper VanSteenhouse, Yumi Kasai, David Gray, Carol Clugston, Anna Dominiczak and Alex Alderton, Roberto Amato, Jeffrey Barrett, Sonia Goncalves, Ewan Harrison, David K. Jackson, Ian Johnston, Dominic Kwiatkowski, Cordelia Langford, John Sillitoe on behalf of the Wellcome Sanger Institute COVID-19 Surveillance Team |
| EPI_ISL_2865165 | Berkshire and Surrey Pathology Services Lighthouse Laboratory | Wellcome Sanger Institute for the COVID-19 Genomics UK (COG-UK) Consortium | Berkshire and Surrey Pathology Services Lighthouse Laboratory and Alex Alderton, Roberto Amato, Jeffrey Barrett, Sonia Goncalves, Ewan Harrison, David K. Jackson, Ian Johnston, Dominic Kwiatkowski, Cordelia Langford, John Sillitoe on behalf of the Wellcome Sanger Institute COVID-19 Surveillance Team |
| EPI_ISL_2865166 | Lighthouse Lab in Milton Keynes | Wellcome Sanger Institute for the COVID-19 Genomics UK (COG-UK) Consortium | The Lighthouse Lab in Milton Keynes and Alex Alderton, Roberto Amato, Jeffrey Barrett, Sonia Goncalves, Ewan Harrison, David K. Jackson, Ian Johnston, Dominic Kwiatkowski, Cordelia Langford, John Sillitoe on behalf of the Wellcome Sanger Institute COVID-19 Surveillance Team |
| EPI_ISL_2865167, EPI_ISL_2865168, EPI_ISL_2865169 | Lighthouse Lab in Alderley Park | Wellcome Sanger Institute for the COVID-19 Genomics UK (COG-UK) Consortium | Jacquelyn Wynn, Mairead Hyland, The Lighthouse Lab in Alderley Park and Alex Alderton, Roberto Amato, Jeffrey Barrett, Sonia Goncalves, Ewan Harrison, David K. Jackson, Ian Johnston, Dominic Kwiatkowski, Cordelia Langford, John Sillitoe on behalf of the Wellcome Sanger Institute COVID-19 Surveillance Team |
| EPI_ISL_2865170 | Berkshire and Surrey Pathology Services Lighthouse Laboratory | Wellcome Sanger Institute for the COVID-19 Genomics UK (COG-UK) Consortium | Berkshire and Surrey Pathology Services Lighthouse Laboratory and Alex Alderton, Roberto Amato, Jeffrey Barrett, Sonia Goncalves, Ewan Harrison, David K. Jackson, Ian Johnston, Dominic Kwiatkowski, Cordelia Langford, John Sillitoe on behalf of the Wellcome Sanger Institute COVID-19 Surveillance Team |
| EPI_ISL_2865171 | Lighthouse Lab in Milton Keynes | Wellcome Sanger Institute for the COVID-19 Genomics UK (COG-UK) Consortium | The Lighthouse Lab in Milton Keynes and Alex Alderton, Roberto Amato, Jeffrey Barrett, Sonia Goncalves, Ewan Harrison, David K. Jackson, Ian Johnston, Dominic Kwiatkowski, Cordelia Langford, John Sillitoe on behalf of the Wellcome Sanger Institute COVID-19 Surveillance Team |
| EPI_ISL_2865172 | Lighthouse Lab in Alderley Park | Wellcome Sanger Institute for the COVID-19 Genomics UK (COG-UK) Consortium | Jacquelyn Wynn, Mairead Hyland, The Lighthouse Lab in Alderley Park and Alex Alderton, Roberto Amato, Jeffrey Barrett, Sonia Goncalves, Ewan Harrison, David K. Jackson, Ian Johnston, Dominic Kwiatkowski, Cordelia Langford, John Sillitoe on behalf of the Wellcome Sanger Institute COVID-19 Surveillance Team |
| EPI_ISL_2865173 | Lighthouse Lab in Milton Keynes | Wellcome Sanger Institute for the COVID-19 Genomics UK (COG-UK) Consortium | The Lighthouse Lab in Milton Keynes and Alex Alderton, Roberto Amato, Jeffrey Barrett, Sonia Goncalves, Ewan Harrison, David K. Jackson, Ian Johnston, Dominic Kwiatkowski, Cordelia Langford, John Sillitoe on behalf of the Wellcome Sanger Institute COVID-19 Surveillance Team |
| EPI_ISL_2865174 | Berkshire and Surrey Pathology Services Lighthouse Laboratory | Wellcome Sanger Institute for the COVID-19 Genomics UK (COG-UK) Consortium | Berkshire and Surrey Pathology Services Lighthouse Laboratory and Alex Alderton, Roberto Amato, Jeffrey Barrett, Sonia Goncalves, Ewan Harrison, David K. Jackson, Ian Johnston, Dominic Kwiatkowski, Cordelia Langford, John Sillitoe on behalf of the Wellcome Sanger Institute COVID-19 Surveillance Team |
| EPI_ISL_2865175 | Lighthouse Lab in Alderley Park | Wellcome Sanger Institute for the COVID-19 Genomics UK (COG-UK) Consortium | Jacquelyn Wynn, Mairead Hyland, The Lighthouse Lab in Alderley Park and Alex Alderton, Roberto Amato, Jeffrey Barrett, Sonia Goncalves, Ewan Harrison, David K. Jackson, Ian Johnston, Dominic Kwiatkowski, Cordelia Langford, John Sillitoe on behalf of the Wellcome Sanger Institute COVID-19 Surveillance Team |
| EPI_ISL_2865176 | Lighthouse Lab in Milton Keynes | Wellcome Sanger Institute for the COVID-19 Genomics UK (COG-UK) Consortium | The Lighthouse Lab in Milton Keynes and Alex Alderton, Roberto Amato, Jeffrey Barrett, Sonia Goncalves, Ewan Harrison, David K. Jackson, Ian Johnston, Dominic Kwiatkowski, Cordelia Langford, John Sillitoe on behalf of the Wellcome Sanger Institute COVID-19 Surveillance Team |
| EPI_ISL_2865177 | Berkshire and Surrey Pathology Services Lighthouse Laboratory | Wellcome Sanger Institute for the COVID-19 Genomics UK (COG-UK) Consortium | Berkshire and Surrey Pathology Services Lighthouse Laboratory and Alex Alderton, Roberto Amato, Jeffrey Barrett, Sonia Goncalves, Ewan Harrison, David K. Jackson, Ian Johnston, Dominic Kwiatkowski, Cordelia Langford, John Sillitoe on behalf of the Wellcome Sanger Institute COVID-19 Surveillance Team |
| EPI_ISL_2865178 | Lighthouse Lab in Glasgow | Wellcome Sanger Institute for the COVID-19 Genomics UK (COG-UK) Consortium | Harper VanSteenhouse, Yumi Kasai, David Gray, Carol Clugston, Anna Dominiczak and Alex Alderton, Roberto Amato, Jeffrey Barrett, Sonia Goncalves, Ewan Harrison, David K. Jackson, Ian Johnston, Dominic Kwiatkowski, Cordelia Langford, John Sillitoe on behalf of the Wellcome Sanger Institute COVID-19 Surveillance Team |
| EPI_ISL_2865179 | Berkshire and Surrey Pathology Services Lighthouse Laboratory | Wellcome Sanger Institute for the COVID-19 Genomics UK (COG-UK) Consortium | Berkshire and Surrey Pathology Services Lighthouse Laboratory and Alex Alderton, Roberto Amato, Jeffrey Barrett, Sonia Goncalves, Ewan Harrison, David K. Jackson, Ian Johnston, Dominic Kwiatkowski, Cordelia Langford, John Sillitoe on behalf of the Wellcome Sanger Institute COVID-19 Surveillance Team |
| EPI_ISL_2865180, EPI_ISL_2865181 | Lighthouse Lab in Milton Keynes | Wellcome Sanger Institute for the COVID-19 Genomics UK (COG-UK) Consortium | The Lighthouse Lab in Milton Keynes and Alex Alderton, Roberto Amato, Jeffrey Barrett, Sonia Goncalves, Ewan Harrison, David K. Jackson, Ian Johnston, Dominic Kwiatkowski, Cordelia Langford, John Sillitoe on behalf of the Wellcome Sanger Institute COVID-19 Surveillance Team |
| EPI_ISL_2865182 | Berkshire and Surrey Pathology Services Lighthouse Laboratory | Wellcome Sanger Institute for the COVID-19 Genomics UK (COG-UK) Consortium | Berkshire and Surrey Pathology Services Lighthouse Laboratory and Alex Alderton, Roberto Amato, Jeffrey Barrett, Sonia Goncalves, Ewan Harrison, David K. Jackson, Ian Johnston, Dominic Kwiatkowski, Cordelia Langford, John Sillitoe on behalf of the Wellcome Sanger Institute COVID-19 Surveillance Team |
| EPI_ISL_2865183 | Lighthouse Lab in Milton Keynes | Wellcome Sanger Institute for the COVID-19 Genomics UK (COG-UK) Consortium | The Lighthouse Lab in Milton Keynes and Alex Alderton, Roberto Amato, Jeffrey Barrett, Sonia Goncalves, Ewan Harrison, David K. Jackson, Ian Johnston, Dominic Kwiatkowski, Cordelia Langford, John Sillitoe on behalf of the Wellcome Sanger Institute COVID-19 Surveillance Team |
| EPI_ISL_2865184 | Berkshire and Surrey Pathology Services Lighthouse Laboratory | Wellcome Sanger Institute for the COVID-19 Genomics UK (COG-UK) Consortium | Berkshire and Surrey Pathology Services Lighthouse Laboratory and Alex Alderton, Roberto Amato, Jeffrey Barrett, Sonia Goncalves, Ewan Harrison, David K. Jackson, Ian Johnston, Dominic Kwiatkowski, Cordelia Langford, John Sillitoe on behalf of the Wellcome Sanger Institute COVID-19 Surveillance Team |

| | | | |
|---|---|---|---|
| EPI_ISL_2865185 | Lighthouse Lab in Milton Keynes | Wellcome Sanger Institute for the COVID-19 Genomics UK (COG-UK) Consortium | The Lighthouse Lab in Milton Keynes and Alex Alderton, Roberto Amato, Jeffrey Barrett, Sonia Goncalves, Ewan Harrison, David K. Jackson, Ian Johnston, Dominic Kwiatkowski, Cordelia Langford, John Sillitoe on behalf of the Wellcome Sanger Institute COVID-19 Surveillance Team |
| EPI_ISL_2865186 | Berkshire and Surrey Pathology Services Lighthouse Laboratory | Wellcome Sanger Institute for the COVID-19 Genomics UK (COG-UK) Consortium | Berkshire and Surrey Pathology Services Lighthouse Laboratory and Alex Alderton, Roberto Amato, Jeffrey Barrett, Sonia Goncalves, Ewan Harrison, David K. Jackson, Ian Johnston, Dominic Kwiatkowski, Cordelia Langford, John Sillitoe on behalf of the Wellcome Sanger Institute COVID-19 Surveillance Team |
| EPI_ISL_2865187 | Lighthouse Lab in Milton Keynes | Wellcome Sanger Institute for the COVID-19 Genomics UK (COG-UK) Consortium | The Lighthouse Lab in Milton Keynes and Alex Alderton, Roberto Amato, Jeffrey Barrett, Sonia Goncalves, Ewan Harrison, David K. Jackson, Ian Johnston, Dominic Kwiatkowski, Cordelia Langford, John Sillitoe on behalf of the Wellcome Sanger Institute COVID-19 Surveillance Team |
| EPI_ISL_2865188 | Lighthouse Lab in Glasgow | Wellcome Sanger Institute for the COVID-19 Genomics UK (COG-UK) Consortium | Harper VanSteenhouse, Yumi Kasai, David Gray, Carol Clugston, Anna Dominiczak and Alex Alderton, Roberto Amato, Jeffrey Barrett, Sonia Goncalves, Ewan Harrison, David K. Jackson, Ian Johnston, Dominic Kwiatkowski, Cordelia Langford, John Sillitoe on behalf of the Wellcome Sanger Institute COVID-19 Surveillance Team |
| EPI_ISL_2865189, EPI_ISL_2865190 | Berkshire and Surrey Pathology Services Lighthouse Laboratory | Wellcome Sanger Institute for the COVID-19 Genomics UK (COG-UK) Consortium | Berkshire and Surrey Pathology Services Lighthouse Laboratory and Alex Alderton, Roberto Amato, Jeffrey Barrett, Sonia Goncalves, Ewan Harrison, David K. Jackson, Ian Johnston, Dominic Kwiatkowski, Cordelia Langford, John Sillitoe on behalf of the Wellcome Sanger Institute COVID-19 Surveillance Team |
| EPI_ISL_2865191 | Lighthouse Lab in Glasgow | Wellcome Sanger Institute for the COVID-19 Genomics UK (COG-UK) Consortium | Harper VanSteenhouse, Yumi Kasai, David Gray, Carol Clugston, Anna Dominiczak and Alex Alderton, Roberto Amato, Jeffrey Barrett, Sonia Goncalves, Ewan Harrison, David K. Jackson, Ian Johnston, Dominic Kwiatkowski, Cordelia Langford, John Sillitoe on behalf of the Wellcome Sanger Institute COVID-19 Surveillance Team |
| EPI_ISL_2865192 | Lighthouse Lab in Milton Keynes | Wellcome Sanger Institute for the COVID-19 Genomics UK (COG-UK) Consortium | The Lighthouse Lab in Milton Keynes and Alex Alderton, Roberto Amato, Jeffrey Barrett, Sonia Goncalves, Ewan Harrison, David K. Jackson, Ian Johnston, Dominic Kwiatkowski, Cordelia Langford, John Sillitoe on behalf of the Wellcome Sanger Institute COVID-19 Surveillance Team |
| EPI_ISL_2865193 | Berkshire and Surrey Pathology Services Lighthouse Laboratory | Wellcome Sanger Institute for the COVID-19 Genomics UK (COG-UK) Consortium | Berkshire and Surrey Pathology Services Lighthouse Laboratory and Alex Alderton, Roberto Amato, Jeffrey Barrett, Sonia Goncalves, Ewan Harrison, David K. Jackson, Ian Johnston, Dominic Kwiatkowski, Cordelia Langford, John Sillitoe on behalf of the Wellcome Sanger Institute COVID-19 Surveillance Team |
| EPI_ISL_2865194, EPI_ISL_2865195 | Lighthouse Lab in Milton Keynes | Wellcome Sanger Institute for the COVID-19 Genomics UK (COG-UK) Consortium | The Lighthouse Lab in Milton Keynes and Alex Alderton, Roberto Amato, Jeffrey Barrett, Sonia Goncalves, Ewan Harrison, David K. Jackson, Ian Johnston, Dominic Kwiatkowski, Cordelia Langford, John Sillitoe on behalf of the Wellcome Sanger Institute COVID-19 Surveillance Team |
| EPI_ISL_2865196 | Lighthouse Lab in Alderley Park | Wellcome Sanger Institute for the COVID-19 Genomics UK (COG-UK) Consortium | Jacquelyn Wynn, Mairead Hyland, The Lighthouse Lab in Alderley Park and Alex Alderton, Roberto Amato, Jeffrey Barrett, Sonia Goncalves, Ewan Harrison, David K. Jackson, Ian Johnston, Dominic Kwiatkowski, Cordelia Langford, John Sillitoe on behalf of the Wellcome Sanger Institute COVID-19 Surveillance Team |
| EPI_ISL_2865197 | Berkshire and Surrey Pathology Services Lighthouse Laboratory | Wellcome Sanger Institute for the COVID-19 Genomics UK (COG-UK) Consortium | Berkshire and Surrey Pathology Services Lighthouse Laboratory and Alex Alderton, Roberto Amato, Jeffrey Barrett, Sonia Goncalves, Ewan Harrison, David K. Jackson, Ian Johnston, Dominic Kwiatkowski, Cordelia Langford, John Sillitoe on behalf of the Wellcome Sanger Institute COVID-19 Surveillance Team |
| EPI_ISL_2865198 | Lighthouse Lab in Milton Keynes | Wellcome Sanger Institute for the COVID-19 Genomics UK (COG-UK) Consortium | The Lighthouse Lab in Milton Keynes and Alex Alderton, Roberto Amato, Jeffrey Barrett, Sonia Goncalves, Ewan Harrison, David K. Jackson, Ian Johnston, Dominic Kwiatkowski, Cordelia Langford, John Sillitoe on behalf of the Wellcome Sanger Institute COVID-19 Surveillance Team |
| EPI_ISL_2865199 | Berkshire and Surrey Pathology Services Lighthouse Laboratory | Wellcome Sanger Institute for the COVID-19 Genomics UK (COG-UK) Consortium | Berkshire and Surrey Pathology Services Lighthouse Laboratory and Alex Alderton, Roberto Amato, Jeffrey Barrett, Sonia Goncalves, Ewan Harrison, David K. Jackson, Ian Johnston, Dominic Kwiatkowski, Cordelia Langford, John Sillitoe on behalf of the Wellcome Sanger Institute COVID-19 Surveillance Team |
| EPI_ISL_2865200 | Lighthouse Lab in Glasgow | Wellcome Sanger Institute for the COVID-19 Genomics UK (COG-UK) Consortium | Harper VanSteenhouse, Yumi Kasai, David Gray, Carol Clugston, Anna Dominiczak and Alex Alderton, Roberto Amato, Jeffrey Barrett, Sonia Goncalves, Ewan Harrison, David K. Jackson, Ian Johnston, Dominic Kwiatkowski, Cordelia Langford, John Sillitoe on behalf of the Wellcome Sanger Institute COVID-19 Surveillance Team |
| EPI_ISL_2865201, EPI_ISL_2865202 | Lighthouse Lab in Alderley Park | Wellcome Sanger Institute for the COVID-19 Genomics UK (COG-UK) Consortium | Jacquelyn Wynn, Mairead Hyland, The Lighthouse Lab in Alderley Park and Alex Alderton, Roberto Amato, Jeffrey Barrett, Sonia Goncalves, Ewan Harrison, David K. Jackson, Ian Johnston, Dominic Kwiatkowski, Cordelia Langford, John Sillitoe on behalf of the Wellcome Sanger Institute COVID-19 Surveillance Team |
| EPI_ISL_2865203 | Berkshire and Surrey Pathology Services Lighthouse Laboratory | Wellcome Sanger Institute for the COVID-19 Genomics UK (COG-UK) Consortium | Berkshire and Surrey Pathology Services Lighthouse Laboratory and Alex Alderton, Roberto Amato, Jeffrey Barrett, Sonia Goncalves, Ewan Harrison, David K. Jackson, Ian Johnston, Dominic Kwiatkowski, Cordelia Langford, John Sillitoe on behalf of the Wellcome Sanger Institute COVID-19 Surveillance Team |
| EPI_ISL_2865204 | Lighthouse Lab in Milton Keynes | Wellcome Sanger Institute for the COVID-19 Genomics UK (COG-UK) Consortium | The Lighthouse Lab in Milton Keynes and Alex Alderton, Roberto Amato, Jeffrey Barrett, Sonia Goncalves, Ewan Harrison, David K. Jackson, Ian Johnston, Dominic Kwiatkowski, Cordelia Langford, John Sillitoe on behalf of the Wellcome Sanger Institute COVID-19 Surveillance Team |
| EPI_ISL_2865205 | Lighthouse Lab in Alderley Park | Wellcome Sanger Institute for the COVID-19 Genomics UK (COG-UK) Consortium | Jacquelyn Wynn, Mairead Hyland, The Lighthouse Lab in Alderley Park and Alex Alderton, Roberto Amato, Jeffrey Barrett, Sonia Goncalves, Ewan Harrison, David K. Jackson, Ian Johnston, Dominic Kwiatkowski, Cordelia Langford, John Sillitoe on behalf of the Wellcome Sanger Institute COVID-19 Surveillance Team |
| EPI_ISL_2865206, EPI_ISL_2865207 | Lighthouse Lab in Milton Keynes | Wellcome Sanger Institute for the COVID-19 Genomics UK (COG-UK) Consortium | The Lighthouse Lab in Milton Keynes and Alex Alderton, Roberto Amato, Jeffrey Barrett, Sonia Goncalves, Ewan Harrison, David K. Jackson, Ian Johnston, Dominic Kwiatkowski, Cordelia Langford, John Sillitoe on behalf of the Wellcome Sanger Institute COVID-19 Surveillance Team |
| EPI_ISL_2865208 | Lighthouse Lab in Glasgow | Wellcome Sanger Institute for the COVID-19 Genomics UK (COG-UK) Consortium | Harper VanSteenhouse, Yumi Kasai, David Gray, Carol Clugston, Anna Dominiczak and Alex Alderton, Roberto Amato, Jeffrey Barrett, Sonia Goncalves, Ewan Harrison, David K. Jackson, Ian Johnston, Dominic Kwiatkowski, Cordelia Langford, John Sillitoe on behalf of the Wellcome Sanger Institute COVID-19 Surveillance Team |
| EPI_ISL_2865209 | Lighthouse Lab in Milton Keynes | Wellcome Sanger Institute for the COVID-19 Genomics UK (COG-UK) Consortium | The Lighthouse Lab in Milton Keynes and Alex Alderton, Roberto Amato, Jeffrey Barrett, Sonia Goncalves, Ewan Harrison, David K. Jackson, Ian Johnston, Dominic Kwiatkowski, Cordelia Langford, John Sillitoe on behalf of the Wellcome Sanger Institute COVID-19 Surveillance Team |
| EPI_ISL_2865210, EPI_ISL_2865211 | Berkshire and Surrey Pathology Services Lighthouse Laboratory | Wellcome Sanger Institute for the COVID-19 Genomics UK (COG-UK) Consortium | Berkshire and Surrey Pathology Services Lighthouse Laboratory and Alex Alderton, Roberto Amato, Jeffrey Barrett, Sonia Goncalves, Ewan Harrison, David K. Jackson, Ian Johnston, Dominic Kwiatkowski, Cordelia Langford, John Sillitoe on behalf of the Wellcome Sanger Institute COVID-19 Surveillance Team |
| EPI_ISL_2865212, EPI_ISL_2865213, EPI_ISL_2865214 | Lighthouse Lab in Alderley Park | Wellcome Sanger Institute for the COVID-19 Genomics UK (COG-UK) Consortium | Jacquelyn Wynn, Mairead Hyland, The Lighthouse Lab in Alderley Park and Alex Alderton, Roberto Amato, Jeffrey Barrett, Sonia Goncalves, Ewan Harrison, David K. Jackson, Ian Johnston, Dominic Kwiatkowski, Cordelia Langford, John Sillitoe on behalf of the Wellcome Sanger Institute COVID-19 Surveillance Team |
| EPI_ISL_2865215, EPI_ISL_2865216 | Berkshire and Surrey Pathology Services Lighthouse Laboratory | Wellcome Sanger Institute for the COVID-19 Genomics UK (COG-UK) Consortium | Berkshire and Surrey Pathology Services Lighthouse Laboratory and Alex Alderton, Roberto Amato, Jeffrey Barrett, Sonia Goncalves, Ewan Harrison, David K. Jackson, Ian Johnston, Dominic Kwiatkowski, Cordelia Langford, John Sillitoe on behalf of the Wellcome Sanger Institute COVID-19 Surveillance Team |
| EPI_ISL_2865217 | Lighthouse Lab in Glasgow | Wellcome Sanger Institute for the COVID-19 Genomics UK (COG-UK) Consortium | Harper VanSteenhouse, Yumi Kasai, David Gray, Carol Clugston, Anna Dominiczak and Alex Alderton, Roberto Amato, Jeffrey Barrett, Sonia Goncalves, Ewan Harrison, David K. Jackson, Ian Johnston, Dominic Kwiatkowski, Cordelia Langford, John Sillitoe on behalf of the Wellcome Sanger Institute COVID-19 Surveillance Team |
| EPI_ISL_2865218 | Lighthouse Lab in Milton Keynes | Wellcome Sanger Institute for the COVID-19 Genomics UK (COG-UK) Consortium | The Lighthouse Lab in Milton Keynes and Alex Alderton, Roberto Amato, Jeffrey Barrett, Sonia Goncalves, Ewan Harrison, David K. Jackson, Ian Johnston, Dominic Kwiatkowski, Cordelia Langford, John Sillitoe on behalf of the Wellcome Sanger Institute COVID-19 Surveillance Team |
| EPI_ISL_2865219, EPI_ISL_2865220 | Berkshire and Surrey Pathology Services Lighthouse Laboratory | Wellcome Sanger Institute for the COVID-19 Genomics UK (COG-UK) Consortium | Berkshire and Surrey Pathology Services Lighthouse Laboratory and Alex Alderton, Roberto Amato, Jeffrey Barrett, Sonia Goncalves, Ewan Harrison, David K. Jackson, Ian Johnston, Dominic Kwiatkowski, Cordelia Langford, John Sillitoe on behalf of the Wellcome Sanger Institute COVID-19 Surveillance Team |
| EPI_ISL_2865221, EPI_ISL_2865222 | Lighthouse Lab in Milton Keynes | Wellcome Sanger Institute for the COVID-19 Genomics UK (COG-UK) Consortium | The Lighthouse Lab in Milton Keynes and Alex Alderton, Roberto Amato, Jeffrey Barrett, Sonia Goncalves, Ewan Harrison, David K. Jackson, Ian Johnston, Dominic Kwiatkowski, Cordelia Langford, John Sillitoe on behalf of the Wellcome Sanger Institute COVID-19 Surveillance Team |
| EPI_ISL_2865223 | Berkshire and Surrey Pathology Services Lighthouse Laboratory | Wellcome Sanger Institute for the COVID-19 Genomics UK (COG-UK) Consortium | Berkshire and Surrey Pathology Services Lighthouse Laboratory and Alex Alderton, Roberto Amato, Jeffrey Barrett, Sonia Goncalves, Ewan Harrison, David K. Jackson, Ian Johnston, Dominic Kwiatkowski, Cordelia Langford, John Sillitoe on behalf of the Wellcome Sanger Institute COVID-19 Surveillance Team |
| EPI_ISL_2865224, EPI_ISL_2865225, EPI_ISL_2865226, EPI_ISL_2865227, EPI_ISL_2865228 | Lighthouse Lab in Alderley Park | Wellcome Sanger Institute for the COVID-19 Genomics UK (COG-UK) Consortium | Jacquelyn Wynn, Mairead Hyland, The Lighthouse Lab in Alderley Park and Alex Alderton, Roberto Amato, Jeffrey Barrett, Sonia Goncalves, Ewan Harrison, David K. Jackson, Ian Johnston, Dominic Kwiatkowski, Cordelia Langford, John Sillitoe on behalf of the Wellcome Sanger Institute COVID-19 Surveillance Team |
| EPI_ISL_2865229 | Berkshire and Surrey Pathology Services Lighthouse Laboratory | Wellcome Sanger Institute for the COVID-19 Genomics UK (COG-UK) Consortium | Berkshire and Surrey Pathology Services Lighthouse Laboratory and Alex Alderton, Roberto Amato, Jeffrey Barrett, Sonia Goncalves, Ewan Harrison, David K. Jackson, Ian Johnston, Dominic Kwiatkowski, Cordelia Langford, John Sillitoe on behalf of the Wellcome Sanger Institute COVID-19 Surveillance Team |
| EPI_ISL_2865230 | Lighthouse Lab in Milton Keynes | Wellcome Sanger Institute for the COVID-19 Genomics UK (COG-UK) Consortium | The Lighthouse Lab in Milton Keynes and Alex Alderton, Roberto Amato, Jeffrey Barrett, Sonia Goncalves, Ewan Harrison, David K. Jackson, Ian Johnston, Dominic Kwiatkowski, Cordelia Langford, John Sillitoe on behalf of the Wellcome Sanger Institute COVID-19 Surveillance Team |
| EPI_ISL_2865231 | Berkshire and Surrey Pathology Services Lighthouse Laboratory | Wellcome Sanger Institute for the COVID-19 Genomics UK (COG-UK) Consortium | Berkshire and Surrey Pathology Services Lighthouse Laboratory and Alex Alderton, Roberto Amato, Jeffrey Barrett, Sonia Goncalves, Ewan Harrison, David K. Jackson, Ian Johnston, Dominic Kwiatkowski, Cordelia Langford, John Sillitoe on behalf of the Wellcome Sanger Institute COVID-19 Surveillance Team |
| EPI_ISL_2865232 | Lighthouse Lab in Milton Keynes | Wellcome Sanger Institute for the COVID-19 Genomics UK (COG-UK) Consortium | The Lighthouse Lab in Milton Keynes and Alex Alderton, Roberto Amato, Jeffrey Barrett, Sonia Goncalves, Ewan Harrison, David K. Jackson, Ian Johnston, Dominic Kwiatkowski, Cordelia Langford, John Sillitoe on behalf of the Wellcome Sanger Institute COVID-19 Surveillance Team |
| EPI_ISL_2865233 | Berkshire and Surrey Pathology Services Lighthouse Laboratory | Wellcome Sanger Institute for the COVID-19 Genomics UK (COG-UK) Consortium | Berkshire and Surrey Pathology Services Lighthouse Laboratory and Alex Alderton, Roberto Amato, Jeffrey Barrett, Sonia Goncalves, Ewan Harrison, David K. Jackson, Ian Johnston, Dominic Kwiatkowski, Cordelia Langford, John Sillitoe on behalf of the Wellcome Sanger Institute COVID-19 Surveillance Team |
| EPI_ISL_2865234 | Lighthouse Lab in Alderley Park | Wellcome Sanger Institute for the COVID-19 Genomics UK (COG-UK) Consortium | Jacquelyn Wynn, Mairead Hyland, The Lighthouse Lab in Alderley Park and Alex Alderton, Roberto Amato, Jeffrey Barrett, Sonia Goncalves, Ewan Harrison, David K. Jackson, Ian Johnston, Dominic Kwiatkowski, Cordelia Langford, John Sillitoe on behalf of the Wellcome Sanger Institute COVID-19 Surveillance Team |
| EPI_ISL_2865235 | Berkshire and Surrey Pathology Services Lighthouse Laboratory | Wellcome Sanger Institute for the COVID-19 Genomics UK (COG-UK) Consortium | Berkshire and Surrey Pathology Services Lighthouse Laboratory and Alex Alderton, Roberto Amato, Jeffrey Barrett, Sonia Goncalves, Ewan Harrison, David K. Jackson, Ian Johnston, Dominic Kwiatkowski, Cordelia Langford, John Sillitoe on behalf of the Wellcome Sanger Institute COVID-19 Surveillance Team |
| EPI_ISL_2865236 | Lighthouse Lab in Milton Keynes | Wellcome Sanger Institute for the COVID-19 Genomics UK (COG-UK) Consortium | The Lighthouse Lab in Milton Keynes and Alex Alderton, Roberto Amato, Jeffrey Barrett, Sonia Goncalves, Ewan Harrison, David K. Jackson, Ian Johnston, Dominic Kwiatkowski, Cordelia Langford, John Sillitoe on behalf of the Wellcome Sanger Institute COVID-19 Surveillance Team |
| EPI_ISL_2865237, EPI_ISL_2865238 | Berkshire and Surrey Pathology Services Lighthouse Laboratory | Wellcome Sanger Institute for the COVID-19 Genomics UK (COG-UK) Consortium | Berkshire and Surrey Pathology Services Lighthouse Laboratory and Alex Alderton, Roberto Amato, Jeffrey Barrett, Sonia Goncalves, Ewan Harrison, David K. Jackson, Ian Johnston, Dominic Kwiatkowski, Cordelia Langford, John Sillitoe on behalf of the Wellcome Sanger Institute COVID-19 Surveillance Team |
| EPI_ISL_2865239 | Lighthouse Lab in Alderley Park | Wellcome Sanger Institute for the COVID-19 Genomics UK (COG-UK) Consortium | Jacquelyn Wynn, Mairead Hyland, The Lighthouse Lab in Alderley Park and Alex Alderton, Roberto Amato, Jeffrey Barrett, Sonia Goncalves, Ewan Harrison, David K. Jackson, Ian Johnston, Dominic Kwiatkowski, Cordelia Langford, John Sillitoe on behalf of the Wellcome Sanger Institute COVID-19 Surveillance Team |
| EPI_ISL_2865240 | Lighthouse Lab in Milton Keynes | Wellcome Sanger Institute for the COVID-19 Genomics UK (COG-UK) Consortium | The Lighthouse Lab in Milton Keynes and Alex Alderton, Roberto Amato, Jeffrey Barrett, Sonia Goncalves, Ewan Harrison, David K. Jackson, Ian Johnston, Dominic Kwiatkowski, Cordelia Langford, John Sillitoe on behalf of the Wellcome Sanger Institute COVID-19 Surveillance Team |
| EPI_ISL_2865241 | Berkshire and Surrey Pathology Services Lighthouse Laboratory | Wellcome Sanger Institute for the COVID-19 Genomics UK (COG-UK) Consortium | Berkshire and Surrey Pathology Services Lighthouse Laboratory and Alex Alderton, Roberto Amato, Jeffrey Barrett, Sonia Goncalves, Ewan Harrison, David K. Jackson, Ian Johnston, Dominic Kwiatkowski, Cordelia Langford, John Sillitoe on behalf of the Wellcome Sanger Institute COVID-19 Surveillance Team |
| EPI_ISL_2865242 | Lighthouse Lab in Milton Keynes | Wellcome Sanger Institute for the COVID-19 Genomics UK (COG-UK) Consortium | The Lighthouse Lab in Milton Keynes and Alex Alderton, Roberto Amato, Jeffrey Barrett, Sonia Goncalves, Ewan Harrison, David K. Jackson, Ian Johnston, Dominic Kwiatkowski, Cordelia Langford, John Sillitoe on behalf of the Wellcome Sanger Institute COVID-19 Surveillance Team |
| EPI_ISL_2865243 | Lighthouse Lab in Alderley Park | Wellcome Sanger Institute for the COVID-19 Genomics UK (COG-UK) | Jacquelyn Wynn, Mairead Hyland, The Lighthouse Lab in Alderley Park and Alex Alderton, Roberto Amato, Jeffrey Barrett, Sonia Goncalves, Ewan Harrison, David K. Jackson, Ian Johnston, Dominic Kwiatkowski, Cordelia Langford, John Sillitoe on |

| | | | |
|---|---|---|---|
| | | Consortium | behalf of the Wellcome Sanger Institute COVID-19 Surveillance Team |
| EPI_ISL_2865244 | Lighthouse Lab in Glasgow | Wellcome Sanger Institute for the COVID-19 Genomics UK (COG-UK) Consortium | Harper VanSteenhouse, Yumi Kasai, David Gray, Carol Clugston, Anna Dominiczak and Alex Alderton, Roberto Amato, Jeffrey Barrett, Sonia Goncalves, Ewan Harrison, David K. Jackson, Ian Johnston, Dominic Kwiatkowski, Cordelia Langford, John Sillitoe on behalf of the Wellcome Sanger Institute COVID-19 Surveillance Team |
| EPI_ISL_2865245, EPI_ISL_2865246 | Lighthouse Lab in Milton Keynes | Wellcome Sanger Institute for the COVID-19 Genomics UK (COG-UK) Consortium | The Lighthouse Lab in Milton Keynes and Alex Alderton, Roberto Amato, Jeffrey Barrett, Sonia Goncalves, Ewan Harrison, David K. Jackson, Ian Johnston, Dominic Kwiatkowski, Cordelia Langford, John Sillitoe on behalf of the Wellcome Sanger Institute COVID-19 Surveillance Team |
| EPI_ISL_2865247 | Lighthouse Lab in Glasgow | Wellcome Sanger Institute for the COVID-19 Genomics UK (COG-UK) Consortium | Harper VanSteenhouse, Yumi Kasai, David Gray, Carol Clugston, Anna Dominiczak and Alex Alderton, Roberto Amato, Jeffrey Barrett, Sonia Goncalves, Ewan Harrison, David K. Jackson, Ian Johnston, Dominic Kwiatkowski, Cordelia Langford, John Sillitoe on behalf of the Wellcome Sanger Institute COVID-19 Surveillance Team |
| EPI_ISL_2865248 | Lighthouse Lab in Milton Keynes | Wellcome Sanger Institute for the COVID-19 Genomics UK (COG-UK) Consortium | The Lighthouse Lab in Milton Keynes and Alex Alderton, Roberto Amato, Jeffrey Barrett, Sonia Goncalves, Ewan Harrison, David K. Jackson, Ian Johnston, Dominic Kwiatkowski, Cordelia Langford, John Sillitoe on behalf of the Wellcome Sanger Institute COVID-19 Surveillance Team |
| EPI_ISL_2865249 | Lighthouse Lab in Alderley Park | Wellcome Sanger Institute for the COVID-19 Genomics UK (COG-UK) Consortium | Jacquelyn Wynn, Mairead Hyland, The Lighthouse Lab in Alderley Park and Alex Alderton, Roberto Amato, Jeffrey Barrett, Sonia Goncalves, Ewan Harrison, David K. Jackson, Ian Johnston, Dominic Kwiatkowski, Cordelia Langford, John Sillitoe on behalf of the Wellcome Sanger Institute COVID-19 Surveillance Team |
| EPI_ISL_2865250, EPI_ISL_2865251 | Lighthouse Lab in Milton Keynes | Wellcome Sanger Institute for the COVID-19 Genomics UK (COG-UK) Consortium | The Lighthouse Lab in Milton Keynes and Alex Alderton, Roberto Amato, Jeffrey Barrett, Sonia Goncalves, Ewan Harrison, David K. Jackson, Ian Johnston, Dominic Kwiatkowski, Cordelia Langford, John Sillitoe on behalf of the Wellcome Sanger Institute COVID-19 Surveillance Team |
| EPI_ISL_2865252 | Lighthouse Lab in Glasgow | Wellcome Sanger Institute for the COVID-19 Genomics UK (COG-UK) Consortium | Harper VanSteenhouse, Yumi Kasai, David Gray, Carol Clugston, Anna Dominiczak and Alex Alderton, Roberto Amato, Jeffrey Barrett, Sonia Goncalves, Ewan Harrison, David K. Jackson, Ian Johnston, Dominic Kwiatkowski, Cordelia Langford, John Sillitoe on behalf of the Wellcome Sanger Institute COVID-19 Surveillance Team |
| EPI_ISL_2865253 | Lighthouse Lab in Alderley Park | Wellcome Sanger Institute for the COVID-19 Genomics UK (COG-UK) Consortium | Jacquelyn Wynn, Mairead Hyland, The Lighthouse Lab in Alderley Park and Alex Alderton, Roberto Amato, Jeffrey Barrett, Sonia Goncalves, Ewan Harrison, David K. Jackson, Ian Johnston, Dominic Kwiatkowski, Cordelia Langford, John Sillitoe on behalf of the Wellcome Sanger Institute COVID-19 Surveillance Team |
| EPI_ISL_2865254, EPI_ISL_2865255 | Lighthouse Lab in Milton Keynes | Wellcome Sanger Institute for the COVID-19 Genomics UK (COG-UK) Consortium | The Lighthouse Lab in Milton Keynes and Alex Alderton, Roberto Amato, Jeffrey Barrett, Sonia Goncalves, Ewan Harrison, David K. Jackson, Ian Johnston, Dominic Kwiatkowski, Cordelia Langford, John Sillitoe on behalf of the Wellcome Sanger Institute COVID-19 Surveillance Team |
| EPI_ISL_2865256 | Berkshire and Surrey Pathology Services Lighthouse Laboratory | Wellcome Sanger Institute for the COVID-19 Genomics UK (COG-UK) Consortium | Berkshire and Surrey Pathology Services Lighthouse Laboratory and Alex Alderton, Roberto Amato, Jeffrey Barrett, Sonia Goncalves, Ewan Harrison, David K. Jackson, Ian Johnston, Dominic Kwiatkowski, Cordelia Langford, John Sillitoe on behalf of the Wellcome Sanger Institute COVID-19 Surveillance Team |
| EPI_ISL_2865257 | Lighthouse Lab in Milton Keynes | Wellcome Sanger Institute for the COVID-19 Genomics UK (COG-UK) Consortium | The Lighthouse Lab in Milton Keynes and Alex Alderton, Roberto Amato, Jeffrey Barrett, Sonia Goncalves, Ewan Harrison, David K. Jackson, Ian Johnston, Dominic Kwiatkowski, Cordelia Langford, John Sillitoe on behalf of the Wellcome Sanger Institute COVID-19 Surveillance Team |
| EPI_ISL_2865258 | Lighthouse Lab in Alderley Park | Wellcome Sanger Institute for the COVID-19 Genomics UK (COG-UK) Consortium | Jacquelyn Wynn, Mairead Hyland, The Lighthouse Lab in Alderley Park and Alex Alderton, Roberto Amato, Jeffrey Barrett, Sonia Goncalves, Ewan Harrison, David K. Jackson, Ian Johnston, Dominic Kwiatkowski, Cordelia Langford, John Sillitoe on behalf of the Wellcome Sanger Institute COVID-19 Surveillance Team |
| EPI_ISL_2865259 | Lighthouse Lab in Glasgow | Wellcome Sanger Institute for the COVID-19 Genomics UK (COG-UK) Consortium | Harper VanSteenhouse, Yumi Kasai, David Gray, Carol Clugston, Anna Dominiczak and Alex Alderton, Roberto Amato, Jeffrey Barrett, Sonia Goncalves, Ewan Harrison, David K. Jackson, Ian Johnston, Dominic Kwiatkowski, Cordelia Langford, John Sillitoe on behalf of the Wellcome Sanger Institute COVID-19 Surveillance Team |
| EPI_ISL_2865260 | Lighthouse Lab in Alderley Park | Wellcome Sanger Institute for the COVID-19 Genomics UK (COG-UK) Consortium | Jacquelyn Wynn, Mairead Hyland, The Lighthouse Lab in Alderley Park and Alex Alderton, Roberto Amato, Jeffrey Barrett, Sonia Goncalves, Ewan Harrison, David K. Jackson, Ian Johnston, Dominic Kwiatkowski, Cordelia Langford, John Sillitoe on behalf of the Wellcome Sanger Institute COVID-19 Surveillance Team |
| EPI_ISL_2865261 | Lighthouse Lab in Glasgow | Wellcome Sanger Institute for the COVID-19 Genomics UK (COG-UK) Consortium | Harper VanSteenhouse, Yumi Kasai, David Gray, Carol Clugston, Anna Dominiczak and Alex Alderton, Roberto Amato, Jeffrey Barrett, Sonia Goncalves, Ewan Harrison, David K. Jackson, Ian Johnston, Dominic Kwiatkowski, Cordelia Langford, John Sillitoe on behalf of the Wellcome Sanger Institute COVID-19 Surveillance Team |
| EPI_ISL_2865262 | Lighthouse Lab in Milton Keynes | Wellcome Sanger Institute for the COVID-19 Genomics UK (COG-UK) Consortium | The Lighthouse Lab in Milton Keynes and Alex Alderton, Roberto Amato, Jeffrey Barrett, Sonia Goncalves, Ewan Harrison, David K. Jackson, Ian Johnston, Dominic Kwiatkowski, Cordelia Langford, John Sillitoe on behalf of the Wellcome Sanger Institute COVID-19 Surveillance Team |
| EPI_ISL_2865263 | Lighthouse Lab in Alderley Park | Wellcome Sanger Institute for the COVID-19 Genomics UK (COG-UK) Consortium | Jacquelyn Wynn, Mairead Hyland, The Lighthouse Lab in Alderley Park and Alex Alderton, Roberto Amato, Jeffrey Barrett, Sonia Goncalves, Ewan Harrison, David K. Jackson, Ian Johnston, Dominic Kwiatkowski, Cordelia Langford, John Sillitoe on behalf of the Wellcome Sanger Institute COVID-19 Surveillance Team |
| EPI_ISL_2865264 | Berkshire and Surrey Pathology Services Lighthouse Laboratory | Wellcome Sanger Institute for the COVID-19 Genomics UK (COG-UK) Consortium | Berkshire and Surrey Pathology Services Lighthouse Laboratory and Alex Alderton, Roberto Amato, Jeffrey Barrett, Sonia Goncalves, Ewan Harrison, David K. Jackson, Ian Johnston, Dominic Kwiatkowski, Cordelia Langford, John Sillitoe on behalf of the Wellcome Sanger Institute COVID-19 Surveillance Team |
| EPI_ISL_2865265 | Lighthouse Lab in Alderley Park | Wellcome Sanger Institute for the COVID-19 Genomics UK (COG-UK) Consortium | Jacquelyn Wynn, Mairead Hyland, The Lighthouse Lab in Alderley Park and Alex Alderton, Roberto Amato, Jeffrey Barrett, Sonia Goncalves, Ewan Harrison, David K. Jackson, Ian Johnston, Dominic Kwiatkowski, Cordelia Langford, John Sillitoe on behalf of the Wellcome Sanger Institute COVID-19 Surveillance Team |
| EPI_ISL_2865266, EPI_ISL_2865267 | Lighthouse Lab in Milton Keynes | Wellcome Sanger Institute for the COVID-19 Genomics UK (COG-UK) Consortium | The Lighthouse Lab in Milton Keynes and Alex Alderton, Roberto Amato, Jeffrey Barrett, Sonia Goncalves, Ewan Harrison, David K. Jackson, Ian Johnston, Dominic Kwiatkowski, Cordelia Langford, John Sillitoe on behalf of the Wellcome Sanger Institute COVID-19 Surveillance Team |
| EPI_ISL_2865268 | Lighthouse Lab in Glasgow | Wellcome Sanger Institute for the COVID-19 Genomics UK (COG-UK) Consortium | Harper VanSteenhouse, Yumi Kasai, David Gray, Carol Clugston, Anna Dominiczak and Alex Alderton, Roberto Amato, Jeffrey Barrett, Sonia Goncalves, Ewan Harrison, David K. Jackson, Ian Johnston, Dominic Kwiatkowski, Cordelia Langford, John Sillitoe on behalf of the Wellcome Sanger Institute COVID-19 Surveillance Team |
| EPI_ISL_2865269 | Berkshire and Surrey Pathology Services Lighthouse Laboratory | Wellcome Sanger Institute for the COVID-19 Genomics UK (COG-UK) Consortium | Berkshire and Surrey Pathology Services Lighthouse Laboratory and Alex Alderton, Roberto Amato, Jeffrey Barrett, Sonia Goncalves, Ewan Harrison, David K. Jackson, Ian Johnston, Dominic Kwiatkowski, Cordelia Langford, John Sillitoe on behalf of the Wellcome Sanger Institute COVID-19 Surveillance Team |
| EPI_ISL_2865270 | Lighthouse Lab in Glasgow | Wellcome Sanger Institute for the COVID-19 Genomics UK (COG-UK) Consortium | Harper VanSteenhouse, Yumi Kasai, David Gray, Carol Clugston, Anna Dominiczak and Alex Alderton, Roberto Amato, Jeffrey Barrett, Sonia Goncalves, Ewan Harrison, David K. Jackson, Ian Johnston, Dominic Kwiatkowski, Cordelia Langford, John Sillitoe on behalf of the Wellcome Sanger Institute COVID-19 Surveillance Team |
| EPI_ISL_2865271 | Lighthouse Lab in Milton Keynes | Wellcome Sanger Institute for the COVID-19 Genomics UK (COG-UK) Consortium | The Lighthouse Lab in Milton Keynes and Alex Alderton, Roberto Amato, Jeffrey Barrett, Sonia Goncalves, Ewan Harrison, David K. Jackson, Ian Johnston, Dominic Kwiatkowski, Cordelia Langford, John Sillitoe on behalf of the Wellcome Sanger Institute COVID-19 Surveillance Team |
| EPI_ISL_2865272 | Berkshire and Surrey Pathology Services Lighthouse Laboratory | Wellcome Sanger Institute for the COVID-19 Genomics UK (COG-UK) Consortium | Berkshire and Surrey Pathology Services Lighthouse Laboratory and Alex Alderton, Roberto Amato, Jeffrey Barrett, Sonia Goncalves, Ewan Harrison, David K. Jackson, Ian Johnston, Dominic Kwiatkowski, Cordelia Langford, John Sillitoe on behalf of the Wellcome Sanger Institute COVID-19 Surveillance Team |
| EPI_ISL_2865273 | Lighthouse Lab in Glasgow | Wellcome Sanger Institute for the COVID-19 Genomics UK (COG-UK) Consortium | Harper VanSteenhouse, Yumi Kasai, David Gray, Carol Clugston, Anna Dominiczak and Alex Alderton, Roberto Amato, Jeffrey Barrett, Sonia Goncalves, Ewan Harrison, David K. Jackson, Ian Johnston, Dominic Kwiatkowski, Cordelia Langford, John Sillitoe on behalf of the Wellcome Sanger Institute COVID-19 Surveillance Team |
| EPI_ISL_2865274 | Lighthouse Lab in Alderley Park | Wellcome Sanger Institute for the COVID-19 Genomics UK (COG-UK) Consortium | Jacquelyn Wynn, Mairead Hyland, The Lighthouse Lab in Alderley Park and Alex Alderton, Roberto Amato, Jeffrey Barrett, Sonia Goncalves, Ewan Harrison, David K. Jackson, Ian Johnston, Dominic Kwiatkowski, Cordelia Langford, John Sillitoe on behalf of the Wellcome Sanger Institute COVID-19 Surveillance Team |
| EPI_ISL_2865275 | Lighthouse Lab in Milton Keynes | Wellcome Sanger Institute for the COVID-19 Genomics UK (COG-UK) Consortium | The Lighthouse Lab in Milton Keynes and Alex Alderton, Roberto Amato, Jeffrey Barrett, Sonia Goncalves, Ewan Harrison, David K. Jackson, Ian Johnston, Dominic Kwiatkowski, Cordelia Langford, John Sillitoe on behalf of the Wellcome Sanger Institute COVID-19 Surveillance Team |
| EPI_ISL_2865276 | Berkshire and Surrey Pathology Services Lighthouse Laboratory | Wellcome Sanger Institute for the COVID-19 Genomics UK (COG-UK) Consortium | Berkshire and Surrey Pathology Services Lighthouse Laboratory and Alex Alderton, Roberto Amato, Jeffrey Barrett, Sonia Goncalves, Ewan Harrison, David K. Jackson, Ian Johnston, Dominic Kwiatkowski, Cordelia Langford, John Sillitoe on behalf of the Wellcome Sanger Institute COVID-19 Surveillance Team |
| EPI_ISL_2865277, EPI_ISL_2865278, EPI_ISL_2865279 | Lighthouse Lab in Milton Keynes | Wellcome Sanger Institute for the COVID-19 Genomics UK (COG-UK) Consortium | The Lighthouse Lab in Milton Keynes and Alex Alderton, Roberto Amato, Jeffrey Barrett, Sonia Goncalves, Ewan Harrison, David K. Jackson, Ian Johnston, Dominic Kwiatkowski, Cordelia Langford, John Sillitoe on behalf of the Wellcome Sanger Institute COVID-19 Surveillance Team |
| EPI_ISL_2865280 | Lighthouse Lab in Alderley Park | Wellcome Sanger Institute for the COVID-19 Genomics UK (COG-UK) Consortium | Jacquelyn Wynn, Mairead Hyland, The Lighthouse Lab in Alderley Park and Alex Alderton, Roberto Amato, Jeffrey Barrett, Sonia Goncalves, Ewan Harrison, David K. Jackson, Ian Johnston, Dominic Kwiatkowski, Cordelia Langford, John Sillitoe on behalf of the Wellcome Sanger Institute COVID-19 Surveillance Team |
| EPI_ISL_2865281 | Berkshire and Surrey Pathology Services Lighthouse Laboratory | Wellcome Sanger Institute for the COVID-19 Genomics UK (COG-UK) Consortium | Berkshire and Surrey Pathology Services Lighthouse Laboratory and Alex Alderton, Roberto Amato, Jeffrey Barrett, Sonia Goncalves, Ewan Harrison, David K. Jackson, Ian Johnston, Dominic Kwiatkowski, Cordelia Langford, John Sillitoe on behalf of the Wellcome Sanger Institute COVID-19 Surveillance Team |
| EPI_ISL_2865282 | Lighthouse Lab in Alderley Park | Wellcome Sanger Institute for the COVID-19 Genomics UK (COG-UK) Consortium | Jacquelyn Wynn, Mairead Hyland, The Lighthouse Lab in Alderley Park and Alex Alderton, Roberto Amato, Jeffrey Barrett, Sonia Goncalves, Ewan Harrison, David K. Jackson, Ian Johnston, Dominic Kwiatkowski, Cordelia Langford, John Sillitoe on behalf of the Wellcome Sanger Institute COVID-19 Surveillance Team |
| EPI_ISL_2865283 | Lighthouse Lab in Milton Keynes | Wellcome Sanger Institute for the COVID-19 Genomics UK (COG-UK) Consortium | The Lighthouse Lab in Milton Keynes and Alex Alderton, Roberto Amato, Jeffrey Barrett, Sonia Goncalves, Ewan Harrison, David K. Jackson, Ian Johnston, Dominic Kwiatkowski, Cordelia Langford, John Sillitoe on behalf of the Wellcome Sanger Institute COVID-19 Surveillance Team |
| EPI_ISL_2865284, EPI_ISL_2865285 | Lighthouse Lab in Alderley Park | Wellcome Sanger Institute for the COVID-19 Genomics UK (COG-UK) Consortium | Jacquelyn Wynn, Mairead Hyland, The Lighthouse Lab in Alderley Park and Alex Alderton, Roberto Amato, Jeffrey Barrett, Sonia Goncalves, Ewan Harrison, David K. Jackson, Ian Johnston, Dominic Kwiatkowski, Cordelia Langford, John Sillitoe on behalf of the Wellcome Sanger Institute COVID-19 Surveillance Team |
| EPI_ISL_2865286, EPI_ISL_2865287 | Berkshire and Surrey Pathology Services Lighthouse Laboratory | Wellcome Sanger Institute for the COVID-19 Genomics UK (COG-UK) Consortium | Berkshire and Surrey Pathology Services Lighthouse Laboratory and Alex Alderton, Roberto Amato, Jeffrey Barrett, Sonia Goncalves, Ewan Harrison, David K. Jackson, Ian Johnston, Dominic Kwiatkowski, Cordelia Langford, John Sillitoe on behalf of the Wellcome Sanger Institute COVID-19 Surveillance Team |
| EPI_ISL_2865288 | Lighthouse Lab in Alderley Park | Wellcome Sanger Institute for the COVID-19 Genomics UK (COG-UK) Consortium | Jacquelyn Wynn, Mairead Hyland, The Lighthouse Lab in Alderley Park and Alex Alderton, Roberto Amato, Jeffrey Barrett, Sonia Goncalves, Ewan Harrison, David K. Jackson, Ian Johnston, Dominic Kwiatkowski, Cordelia Langford, John Sillitoe on behalf of the Wellcome Sanger Institute COVID-19 Surveillance Team |
| EPI_ISL_2865289 | Berkshire and Surrey Pathology Services Lighthouse Laboratory | Wellcome Sanger Institute for the COVID-19 Genomics UK (COG-UK) Consortium | Berkshire and Surrey Pathology Services Lighthouse Laboratory and Alex Alderton, Roberto Amato, Jeffrey Barrett, Sonia Goncalves, Ewan Harrison, David K. Jackson, Ian Johnston, Dominic Kwiatkowski, Cordelia Langford, John Sillitoe on behalf of the Wellcome Sanger Institute COVID-19 Surveillance Team |
| EPI_ISL_2865290, EPI_ISL_2865291, EPI_ISL_2865292 | Lighthouse Lab in Milton Keynes | Wellcome Sanger Institute for the COVID-19 Genomics UK (COG-UK) Consortium | The Lighthouse Lab in Milton Keynes and Alex Alderton, Roberto Amato, Jeffrey Barrett, Sonia Goncalves, Ewan Harrison, David K. Jackson, Ian Johnston, Dominic Kwiatkowski, Cordelia Langford, John Sillitoe on behalf of the Wellcome Sanger Institute COVID-19 Surveillance Team |
| EPI_ISL_2865293 | Lighthouse Lab in Alderley Park | Wellcome Sanger Institute for the COVID-19 Genomics UK (COG-UK) Consortium | Jacquelyn Wynn, Mairead Hyland, The Lighthouse Lab in Alderley Park and Alex Alderton, Roberto Amato, Jeffrey Barrett, Sonia Goncalves, Ewan Harrison, David K. Jackson, Ian Johnston, Dominic Kwiatkowski, Cordelia Langford, John Sillitoe on behalf of the Wellcome Sanger Institute COVID-19 Surveillance Team |
| EPI_ISL_2865294, EPI_ISL_2865295 | Lighthouse Lab in Milton Keynes | Wellcome Sanger Institute for the COVID-19 Genomics UK (COG-UK) Consortium | The Lighthouse Lab in Milton Keynes and Alex Alderton, Roberto Amato, Jeffrey Barrett, Sonia Goncalves, Ewan Harrison, David K. Jackson, Ian Johnston, Dominic Kwiatkowski, Cordelia Langford, John Sillitoe on behalf of the Wellcome Sanger Institute COVID-19 Surveillance Team |
| EPI_ISL_2865296 | Berkshire and Surrey Pathology Services Lighthouse Laboratory | Wellcome Sanger Institute for the COVID-19 Genomics UK (COG-UK) Consortium | Berkshire and Surrey Pathology Services Lighthouse Laboratory and Alex Alderton, Roberto Amato, Jeffrey Barrett, Sonia Goncalves, Ewan Harrison, David K. Jackson, Ian Johnston, Dominic Kwiatkowski, Cordelia Langford, John Sillitoe on behalf of the Wellcome Sanger Institute COVID-19 Surveillance Team |
| EPI_ISL_2865297, EPI_ISL_2865298, EPI_ISL_2865299 | Lighthouse Lab in Alderley Park | Wellcome Sanger Institute for the COVID-19 Genomics UK (COG-UK) Consortium | Jacquelyn Wynn, Mairead Hyland, The Lighthouse Lab in Alderley Park and Alex Alderton, Roberto Amato, Jeffrey Barrett, Sonia Goncalves, Ewan Harrison, David K. Jackson, Ian Johnston, Dominic Kwiatkowski, Cordelia Langford, John Sillitoe on behalf of the Wellcome Sanger Institute COVID-19 Surveillance Team |
| EPI_ISL_2865300 | Berkshire and Surrey Pathology Services Lighthouse Laboratory | Wellcome Sanger Institute for the COVID-19 Genomics UK (COG-UK) | Berkshire and Surrey Pathology Services Lighthouse Laboratory and Alex Alderton, Roberto Amato, Jeffrey Barrett, Sonia Goncalves, Ewan Harrison, David K. Jackson, Ian Johnston, Dominic Kwiatkowski, Cordelia Langford, John Sillitoe on behalf |

| | | | |
|---|---|---|---|
| | | Consortium | of the Wellcome Sanger Institute COVID-19 Surveillance Team |
| EPI_ISL_2865301, EPI_ISL_2865302 | Lighthouse Lab in Glasgow | Wellcome Sanger Institute for the COVID-19 Genomics UK (COG-UK) Consortium | Harper VanSteenhouse, Yumi Kasai, David Gray, Carol Clugston, Anna Dominiczak and Alex Alderton, Roberto Amato, Jeffrey Barrett, Sonia Goncalves, Ewan Harrison, David K. Jackson, Ian Johnston, Dominic Kwiatkowski, Cordelia Langford, John Sillitoe on behalf of the Wellcome Sanger Institute COVID-19 Surveillance Team |
| EPI_ISL_2865303 | Lighthouse Lab in Milton Keynes | Wellcome Sanger Institute for the COVID-19 Genomics UK (COG-UK) Consortium | The Lighthouse Lab in Milton Keynes and Alex Alderton, Roberto Amato, Jeffrey Barrett, Sonia Goncalves, Ewan Harrison, David K. Jackson, Ian Johnston, Dominic Kwiatkowski, Cordelia Langford, John Sillitoe on behalf of the Wellcome Sanger Institute COVID-19 Surveillance Team |
| EPI_ISL_2865304 | Berkshire and Surrey Pathology Services Lighthouse Laboratory | Wellcome Sanger Institute for the COVID-19 Genomics UK (COG-UK) Consortium | Berkshire and Surrey Pathology Services Lighthouse Laboratory and Alex Alderton, Roberto Amato, Jeffrey Barrett, Sonia Goncalves, Ewan Harrison, David K. Jackson, Ian Johnston, Dominic Kwiatkowski, Cordelia Langford, John Sillitoe on behalf of the Wellcome Sanger Institute COVID-19 Surveillance Team |
| EPI_ISL_2865305 | Lighthouse Lab in Milton Keynes | Wellcome Sanger Institute for the COVID-19 Genomics UK (COG-UK) Consortium | The Lighthouse Lab in Milton Keynes and Alex Alderton, Roberto Amato, Jeffrey Barrett, Sonia Goncalves, Ewan Harrison, David K. Jackson, Ian Johnston, Dominic Kwiatkowski, Cordelia Langford, John Sillitoe on behalf of the Wellcome Sanger Institute COVID-19 Surveillance Team |
| EPI_ISL_2865306, EPI_ISL_2865307 | Lighthouse Lab in Alderley Park | Wellcome Sanger Institute for the COVID-19 Genomics UK (COG-UK) Consortium | Jacquelyn Wynn, Mairead Hyland, The Lighthouse Lab in Alderley Park and Alex Alderton, Roberto Amato, Jeffrey Barrett, Sonia Goncalves, Ewan Harrison, David K. Jackson, Ian Johnston, Dominic Kwiatkowski, Cordelia Langford, John Sillitoe on behalf of the Wellcome Sanger Institute COVID-19 Surveillance Team |
| EPI_ISL_2865308 | Berkshire and Surrey Pathology Services Lighthouse Laboratory | Wellcome Sanger Institute for the COVID-19 Genomics UK (COG-UK) Consortium | Berkshire and Surrey Pathology Services Lighthouse Laboratory and Alex Alderton, Roberto Amato, Jeffrey Barrett, Sonia Goncalves, Ewan Harrison, David K. Jackson, Ian Johnston, Dominic Kwiatkowski, Cordelia Langford, John Sillitoe on behalf of the Wellcome Sanger Institute COVID-19 Surveillance Team |
| EPI_ISL_2865309, EPI_ISL_2865310 | Lighthouse Lab in Alderley Park | Wellcome Sanger Institute for the COVID-19 Genomics UK (COG-UK) Consortium | Jacquelyn Wynn, Mairead Hyland, The Lighthouse Lab in Alderley Park and Alex Alderton, Roberto Amato, Jeffrey Barrett, Sonia Goncalves, Ewan Harrison, David K. Jackson, Ian Johnston, Dominic Kwiatkowski, Cordelia Langford, John Sillitoe on behalf of the Wellcome Sanger Institute COVID-19 Surveillance Team |
| EPI_ISL_2865311, EPI_ISL_2865312 | Lighthouse Lab in Glasgow | Wellcome Sanger Institute for the COVID-19 Genomics UK (COG-UK) Consortium | Harper VanSteenhouse, Yumi Kasai, David Gray, Carol Clugston, Anna Dominiczak and Alex Alderton, Roberto Amato, Jeffrey Barrett, Sonia Goncalves, Ewan Harrison, David K. Jackson, Ian Johnston, Dominic Kwiatkowski, Cordelia Langford, John Sillitoe on behalf of the Wellcome Sanger Institute COVID-19 Surveillance Team |
| EPI_ISL_2865313 | Lighthouse Lab in Alderley Park | Wellcome Sanger Institute for the COVID-19 Genomics UK (COG-UK) Consortium | Jacquelyn Wynn, Mairead Hyland, The Lighthouse Lab in Alderley Park and Alex Alderton, Roberto Amato, Jeffrey Barrett, Sonia Goncalves, Ewan Harrison, David K. Jackson, Ian Johnston, Dominic Kwiatkowski, Cordelia Langford, John Sillitoe on behalf of the Wellcome Sanger Institute COVID-19 Surveillance Team |
| EPI_ISL_2865314 | Berkshire and Surrey Pathology Services Lighthouse Laboratory | Wellcome Sanger Institute for the COVID-19 Genomics UK (COG-UK) Consortium | Berkshire and Surrey Pathology Services Lighthouse Laboratory and Alex Alderton, Roberto Amato, Jeffrey Barrett, Sonia Goncalves, Ewan Harrison, David K. Jackson, Ian Johnston, Dominic Kwiatkowski, Cordelia Langford, John Sillitoe on behalf of the Wellcome Sanger Institute COVID-19 Surveillance Team |
| EPI_ISL_2865315, EPI_ISL_2865316 | Lighthouse Lab in Milton Keynes | Wellcome Sanger Institute for the COVID-19 Genomics UK (COG-UK) Consortium | The Lighthouse Lab in Milton Keynes and Alex Alderton, Roberto Amato, Jeffrey Barrett, Sonia Goncalves, Ewan Harrison, David K. Jackson, Ian Johnston, Dominic Kwiatkowski, Cordelia Langford, John Sillitoe on behalf of the Wellcome Sanger Institute COVID-19 Surveillance Team |
| EPI_ISL_2865317 | Lighthouse Lab in Glasgow | Wellcome Sanger Institute for the COVID-19 Genomics UK (COG-UK) Consortium | Harper VanSteenhouse, Yumi Kasai, David Gray, Carol Clugston, Anna Dominiczak and Alex Alderton, Roberto Amato, Jeffrey Barrett, Sonia Goncalves, Ewan Harrison, David K. Jackson, Ian Johnston, Dominic Kwiatkowski, Cordelia Langford, John Sillitoe on behalf of the Wellcome Sanger Institute COVID-19 Surveillance Team |
| EPI_ISL_2865318 | Lighthouse Lab in Alderley Park | Wellcome Sanger Institute for the COVID-19 Genomics UK (COG-UK) Consortium | Jacquelyn Wynn, Mairead Hyland, The Lighthouse Lab in Alderley Park and Alex Alderton, Roberto Amato, Jeffrey Barrett, Sonia Goncalves, Ewan Harrison, David K. Jackson, Ian Johnston, Dominic Kwiatkowski, Cordelia Langford, John Sillitoe on behalf of the Wellcome Sanger Institute COVID-19 Surveillance Team |
| EPI_ISL_2865319 | Berkshire and Surrey Pathology Services Lighthouse Laboratory | Wellcome Sanger Institute for the COVID-19 Genomics UK (COG-UK) Consortium | Berkshire and Surrey Pathology Services Lighthouse Laboratory and Alex Alderton, Roberto Amato, Jeffrey Barrett, Sonia Goncalves, Ewan Harrison, David K. Jackson, Ian Johnston, Dominic Kwiatkowski, Cordelia Langford, John Sillitoe on behalf of the Wellcome Sanger Institute COVID-19 Surveillance Team |
| EPI_ISL_2865320, EPI_ISL_2865321 | Lighthouse Lab in Alderley Park | Wellcome Sanger Institute for the COVID-19 Genomics UK (COG-UK) Consortium | Jacquelyn Wynn, Mairead Hyland, The Lighthouse Lab in Alderley Park and Alex Alderton, Roberto Amato, Jeffrey Barrett, Sonia Goncalves, Ewan Harrison, David K. Jackson, Ian Johnston, Dominic Kwiatkowski, Cordelia Langford, John Sillitoe on behalf of the Wellcome Sanger Institute COVID-19 Surveillance Team |
| EPI_ISL_2865322 | Berkshire and Surrey Pathology Services Lighthouse Laboratory | Wellcome Sanger Institute for the COVID-19 Genomics UK (COG-UK) Consortium | Berkshire and Surrey Pathology Services Lighthouse Laboratory and Alex Alderton, Roberto Amato, Jeffrey Barrett, Sonia Goncalves, Ewan Harrison, David K. Jackson, Ian Johnston, Dominic Kwiatkowski, Cordelia Langford, John Sillitoe on behalf of the Wellcome Sanger Institute COVID-19 Surveillance Team |
| EPI_ISL_2865323 | Lighthouse Lab in Milton Keynes | Wellcome Sanger Institute for the COVID-19 Genomics UK (COG-UK) Consortium | The Lighthouse Lab in Milton Keynes and Alex Alderton, Roberto Amato, Jeffrey Barrett, Sonia Goncalves, Ewan Harrison, David K. Jackson, Ian Johnston, Dominic Kwiatkowski, Cordelia Langford, John Sillitoe on behalf of the Wellcome Sanger Institute COVID-19 Surveillance Team |
| EPI_ISL_2865324, EPI_ISL_2865325 | Lighthouse Lab in Glasgow | Wellcome Sanger Institute for the COVID-19 Genomics UK (COG-UK) Consortium | Harper VanSteenhouse, Yumi Kasai, David Gray, Carol Clugston, Anna Dominiczak and Alex Alderton, Roberto Amato, Jeffrey Barrett, Sonia Goncalves, Ewan Harrison, David K. Jackson, Ian Johnston, Dominic Kwiatkowski, Cordelia Langford, John Sillitoe on behalf of the Wellcome Sanger Institute COVID-19 Surveillance Team |
| EPI_ISL_2865326 | Lighthouse Lab in Alderley Park | Wellcome Sanger Institute for the COVID-19 Genomics UK (COG-UK) Consortium | Jacquelyn Wynn, Mairead Hyland, The Lighthouse Lab in Alderley Park and Alex Alderton, Roberto Amato, Jeffrey Barrett, Sonia Goncalves, Ewan Harrison, David K. Jackson, Ian Johnston, Dominic Kwiatkowski, Cordelia Langford, John Sillitoe on behalf of the Wellcome Sanger Institute COVID-19 Surveillance Team |
| EPI_ISL_2865327, EPI_ISL_2865328 | Lighthouse Lab in Glasgow | Wellcome Sanger Institute for the COVID-19 Genomics UK (COG-UK) Consortium | Harper VanSteenhouse, Yumi Kasai, David Gray, Carol Clugston, Anna Dominiczak and Alex Alderton, Roberto Amato, Jeffrey Barrett, Sonia Goncalves, Ewan Harrison, David K. Jackson, Ian Johnston, Dominic Kwiatkowski, Cordelia Langford, John Sillitoe on behalf of the Wellcome Sanger Institute COVID-19 Surveillance Team |
| EPI_ISL_2865329 | Lighthouse Lab in Milton Keynes | Wellcome Sanger Institute for the COVID-19 Genomics UK (COG-UK) Consortium | The Lighthouse Lab in Milton Keynes and Alex Alderton, Roberto Amato, Jeffrey Barrett, Sonia Goncalves, Ewan Harrison, David K. Jackson, Ian Johnston, Dominic Kwiatkowski, Cordelia Langford, John Sillitoe on behalf of the Wellcome Sanger Institute COVID-19 Surveillance Team |
| EPI_ISL_2865330 | Lighthouse Lab in Alderley Park | Wellcome Sanger Institute for the COVID-19 Genomics UK (COG-UK) Consortium | Jacquelyn Wynn, Mairead Hyland, The Lighthouse Lab in Alderley Park and Alex Alderton, Roberto Amato, Jeffrey Barrett, Sonia Goncalves, Ewan Harrison, David K. Jackson, Ian Johnston, Dominic Kwiatkowski, Cordelia Langford, John Sillitoe on behalf of the Wellcome Sanger Institute COVID-19 Surveillance Team |
| EPI_ISL_2865331 | Lighthouse Lab in Milton Keynes | Wellcome Sanger Institute for the COVID-19 Genomics UK (COG-UK) Consortium | The Lighthouse Lab in Milton Keynes and Alex Alderton, Roberto Amato, Jeffrey Barrett, Sonia Goncalves, Ewan Harrison, David K. Jackson, Ian Johnston, Dominic Kwiatkowski, Cordelia Langford, John Sillitoe on behalf of the Wellcome Sanger Institute COVID-19 Surveillance Team |
| EPI_ISL_2865332 | Lighthouse Lab in Alderley Park | Wellcome Sanger Institute for the COVID-19 Genomics UK (COG-UK) Consortium | Jacquelyn Wynn, Mairead Hyland, The Lighthouse Lab in Alderley Park and Alex Alderton, Roberto Amato, Jeffrey Barrett, Sonia Goncalves, Ewan Harrison, David K. Jackson, Ian Johnston, Dominic Kwiatkowski, Cordelia Langford, John Sillitoe on behalf of the Wellcome Sanger Institute COVID-19 Surveillance Team |
| EPI_ISL_2865333, EPI_ISL_2865334 | Lighthouse Lab in Glasgow | Wellcome Sanger Institute for the COVID-19 Genomics UK (COG-UK) Consortium | Harper VanSteenhouse, Yumi Kasai, David Gray, Carol Clugston, Anna Dominiczak and Alex Alderton, Roberto Amato, Jeffrey Barrett, Sonia Goncalves, Ewan Harrison, David K. Jackson, Ian Johnston, Dominic Kwiatkowski, Cordelia Langford, John Sillitoe on behalf of the Wellcome Sanger Institute COVID-19 Surveillance Team |
| EPI_ISL_2865335 | Lighthouse Lab in Milton Keynes | Wellcome Sanger Institute for the COVID-19 Genomics UK (COG-UK) Consortium | The Lighthouse Lab in Milton Keynes and Alex Alderton, Roberto Amato, Jeffrey Barrett, Sonia Goncalves, Ewan Harrison, David K. Jackson, Ian Johnston, Dominic Kwiatkowski, Cordelia Langford, John Sillitoe on behalf of the Wellcome Sanger Institute COVID-19 Surveillance Team |
| EPI_ISL_2865336 | Lighthouse Lab in Alderley Park | Wellcome Sanger Institute for the COVID-19 Genomics UK (COG-UK) Consortium | Jacquelyn Wynn, Mairead Hyland, The Lighthouse Lab in Alderley Park and Alex Alderton, Roberto Amato, Jeffrey Barrett, Sonia Goncalves, Ewan Harrison, David K. Jackson, Ian Johnston, Dominic Kwiatkowski, Cordelia Langford, John Sillitoe on behalf of the Wellcome Sanger Institute COVID-19 Surveillance Team |
| EPI_ISL_2865337, EPI_ISL_2865338 | Lighthouse Lab in Milton Keynes | Wellcome Sanger Institute for the COVID-19 Genomics UK (COG-UK) Consortium | The Lighthouse Lab in Milton Keynes and Alex Alderton, Roberto Amato, Jeffrey Barrett, Sonia Goncalves, Ewan Harrison, David K. Jackson, Ian Johnston, Dominic Kwiatkowski, Cordelia Langford, John Sillitoe on behalf of the Wellcome Sanger Institute COVID-19 Surveillance Team |
| EPI_ISL_2865339, EPI_ISL_2865340 | Berkshire and Surrey Pathology Services Lighthouse Laboratory | Wellcome Sanger Institute for the COVID-19 Genomics UK (COG-UK) Consortium | Berkshire and Surrey Pathology Services Lighthouse Laboratory and Alex Alderton, Roberto Amato, Jeffrey Barrett, Sonia Goncalves, Ewan Harrison, David K. Jackson, Ian Johnston, Dominic Kwiatkowski, Cordelia Langford, John Sillitoe on behalf of the Wellcome Sanger Institute COVID-19 Surveillance Team |
| EPI_ISL_2865341 | Lighthouse Lab in Alderley Park | Wellcome Sanger Institute for the COVID-19 Genomics UK (COG-UK) Consortium | Jacquelyn Wynn, Mairead Hyland, The Lighthouse Lab in Alderley Park and Alex Alderton, Roberto Amato, Jeffrey Barrett, Sonia Goncalves, Ewan Harrison, David K. Jackson, Ian Johnston, Dominic Kwiatkowski, Cordelia Langford, John Sillitoe on behalf of the Wellcome Sanger Institute COVID-19 Surveillance Team |
| EPI_ISL_2865342, EPI_ISL_2865343, EPI_ISL_2865344 | Lighthouse Lab in Milton Keynes | Wellcome Sanger Institute for the COVID-19 Genomics UK (COG-UK) Consortium | The Lighthouse Lab in Milton Keynes and Alex Alderton, Roberto Amato, Jeffrey Barrett, Sonia Goncalves, Ewan Harrison, David K. Jackson, Ian Johnston, Dominic Kwiatkowski, Cordelia Langford, John Sillitoe on behalf of the Wellcome Sanger Institute COVID-19 Surveillance Team |
| EPI_ISL_2865345 | Berkshire and Surrey Pathology Services Lighthouse Laboratory | Wellcome Sanger Institute for the COVID-19 Genomics UK (COG-UK) Consortium | Berkshire and Surrey Pathology Services Lighthouse Laboratory and Alex Alderton, Roberto Amato, Jeffrey Barrett, Sonia Goncalves, Ewan Harrison, David K. Jackson, Ian Johnston, Dominic Kwiatkowski, Cordelia Langford, John Sillitoe on behalf of the Wellcome Sanger Institute COVID-19 Surveillance Team |
| EPI_ISL_2865346 | Lighthouse Lab in Glasgow | Wellcome Sanger Institute for the COVID-19 Genomics UK (COG-UK) Consortium | Harper VanSteenhouse, Yumi Kasai, David Gray, Carol Clugston, Anna Dominiczak and Alex Alderton, Roberto Amato, Jeffrey Barrett, Sonia Goncalves, Ewan Harrison, David K. Jackson, Ian Johnston, Dominic Kwiatkowski, Cordelia Langford, John Sillitoe on behalf of the Wellcome Sanger Institute COVID-19 Surveillance Team |
| EPI_ISL_2865347 | Lighthouse Lab in Alderley Park | Wellcome Sanger Institute for the COVID-19 Genomics UK (COG-UK) Consortium | Jacquelyn Wynn, Mairead Hyland, The Lighthouse Lab in Alderley Park and Alex Alderton, Roberto Amato, Jeffrey Barrett, Sonia Goncalves, Ewan Harrison, David K. Jackson, Ian Johnston, Dominic Kwiatkowski, Cordelia Langford, John Sillitoe on behalf of the Wellcome Sanger Institute COVID-19 Surveillance Team |
| EPI_ISL_2865348 | Lighthouse Lab in Glasgow | Wellcome Sanger Institute for the COVID-19 Genomics UK (COG-UK) Consortium | Harper VanSteenhouse, Yumi Kasai, David Gray, Carol Clugston, Anna Dominiczak and Alex Alderton, Roberto Amato, Jeffrey Barrett, Sonia Goncalves, Ewan Harrison, David K. Jackson, Ian Johnston, Dominic Kwiatkowski, Cordelia Langford, John Sillitoe on behalf of the Wellcome Sanger Institute COVID-19 Surveillance Team |
| EPI_ISL_2865349, EPI_ISL_2865350 | Lighthouse Lab in Milton Keynes | Wellcome Sanger Institute for the COVID-19 Genomics UK (COG-UK) Consortium | The Lighthouse Lab in Milton Keynes and Alex Alderton, Roberto Amato, Jeffrey Barrett, Sonia Goncalves, Ewan Harrison, David K. Jackson, Ian Johnston, Dominic Kwiatkowski, Cordelia Langford, John Sillitoe on behalf of the Wellcome Sanger Institute COVID-19 Surveillance Team |
| EPI_ISL_2865351 | Berkshire and Surrey Pathology Services Lighthouse Laboratory | Wellcome Sanger Institute for the COVID-19 Genomics UK (COG-UK) Consortium | Berkshire and Surrey Pathology Services Lighthouse Laboratory and Alex Alderton, Roberto Amato, Jeffrey Barrett, Sonia Goncalves, Ewan Harrison, David K. Jackson, Ian Johnston, Dominic Kwiatkowski, Cordelia Langford, John Sillitoe on behalf of the Wellcome Sanger Institute COVID-19 Surveillance Team |
| EPI_ISL_2865352 | Lighthouse Lab in Alderley Park | Wellcome Sanger Institute for the COVID-19 Genomics UK (COG-UK) Consortium | Jacquelyn Wynn, Mairead Hyland, The Lighthouse Lab in Alderley Park and Alex Alderton, Roberto Amato, Jeffrey Barrett, Sonia Goncalves, Ewan Harrison, David K. Jackson, Ian Johnston, Dominic Kwiatkowski, Cordelia Langford, John Sillitoe on behalf of the Wellcome Sanger Institute COVID-19 Surveillance Team |
| EPI_ISL_2865353, EPI_ISL_2865354, EPI_ISL_2865355 | Berkshire and Surrey Pathology Services Lighthouse Laboratory | Wellcome Sanger Institute for the COVID-19 Genomics UK (COG-UK) Consortium | Berkshire and Surrey Pathology Services Lighthouse Laboratory and Alex Alderton, Roberto Amato, Jeffrey Barrett, Sonia Goncalves, Ewan Harrison, David K. Jackson, Ian Johnston, Dominic Kwiatkowski, Cordelia Langford, John Sillitoe on behalf of the Wellcome Sanger Institute COVID-19 Surveillance Team |
| EPI_ISL_2865356 | Lighthouse Lab in Alderley Park | Wellcome Sanger Institute for the COVID-19 Genomics UK (COG-UK) Consortium | Jacquelyn Wynn, Mairead Hyland, The Lighthouse Lab in Alderley Park and Alex Alderton, Roberto Amato, Jeffrey Barrett, Sonia Goncalves, Ewan Harrison, David K. Jackson, Ian Johnston, Dominic Kwiatkowski, Cordelia Langford, John Sillitoe on behalf of the Wellcome Sanger Institute COVID-19 Surveillance Team |
| EPI_ISL_2865357 | Lighthouse Lab in Glasgow | Wellcome Sanger Institute for the COVID-19 Genomics UK (COG-UK) Consortium | Harper VanSteenhouse, Yumi Kasai, David Gray, Carol Clugston, Anna Dominiczak and Alex Alderton, Roberto Amato, Jeffrey Barrett, Sonia Goncalves, Ewan Harrison, David K. Jackson, Ian Johnston, Dominic Kwiatkowski, Cordelia Langford, John Sillitoe on behalf of the Wellcome Sanger Institute COVID-19 Surveillance Team |
| EPI_ISL_2865358 | Lighthouse Lab in Milton Keynes | Wellcome Sanger Institute for the COVID-19 Genomics UK (COG-UK) Consortium | The Lighthouse Lab in Milton Keynes and Alex Alderton, Roberto Amato, Jeffrey Barrett, Sonia Goncalves, Ewan Harrison, David K. Jackson, Ian Johnston, Dominic Kwiatkowski, Cordelia Langford, John Sillitoe on behalf of the Wellcome Sanger Institute COVID-19 Surveillance Team |
| EPI_ISL_2865359 | Berkshire and Surrey Pathology Services Lighthouse Laboratory | Wellcome Sanger Institute for the COVID-19 Genomics UK (COG-UK) Consortium | Berkshire and Surrey Pathology Services Lighthouse Laboratory and Alex Alderton, Roberto Amato, Jeffrey Barrett, Sonia Goncalves, Ewan Harrison, David K. Jackson, Ian Johnston, Dominic Kwiatkowski, Cordelia Langford, John Sillitoe on behalf of the Wellcome Sanger Institute COVID-19 Surveillance Team |
| EPI_ISL_2865360 | Lighthouse Lab in Alderley Park | Wellcome Sanger Institute for the COVID-19 Genomics UK (COG-UK) Consortium | Jacquelyn Wynn, Mairead Hyland, The Lighthouse Lab in Alderley Park and Alex Alderton, Roberto Amato, Jeffrey Barrett, Sonia Goncalves, Ewan Harrison, David K. Jackson, Ian Johnston, Dominic Kwiatkowski, Cordelia Langford, John Sillitoe on behalf of the Wellcome Sanger Institute COVID-19 Surveillance Team |

| | | | |
|---|---|---|---|
| EPI_ISL_2865361 | Berkshire and Surrey Pathology Services Lighthouse Laboratory | Wellcome Sanger Institute for the COVID-19 Genomics UK (COG-UK) Consortium | Berkshire and Surrey Pathology Services Lighthouse Laboratory and Alex Alderton, Roberto Amato, Jeffrey Barrett, Sonia Goncalves, Ewan Harrison, David K. Jackson, Ian Johnston, Dominic Kwiatkowski, Cordelia Langford, John Sillitoe on behalf of the Wellcome Sanger Institute COVID-19 Surveillance Team |
| EPI_ISL_2865362 | Lighthouse Lab in Milton Keynes | Wellcome Sanger Institute for the COVID-19 Genomics UK (COG-UK) Consortium | The Lighthouse Lab in Milton Keynes and Alex Alderton, Roberto Amato, Jeffrey Barrett, Sonia Goncalves, Ewan Harrison, David K. Jackson, Ian Johnston, Dominic Kwiatkowski, Cordelia Langford, John Sillitoe on behalf of the Wellcome Sanger Institute COVID-19 Surveillance Team |
| EPI_ISL_2865363 | Lighthouse Lab in Glasgow | Wellcome Sanger Institute for the COVID-19 Genomics UK (COG-UK) Consortium | Harper VanSteenhouse, Yumi Kasai, David Gray, Carol Clugston, Anna Dominiczak and Alex Alderton, Roberto Amato, Jeffrey Barrett, Sonia Goncalves, Ewan Harrison, David K. Jackson, Ian Johnston, Dominic Kwiatkowski, Cordelia Langford, John Sillitoe on behalf of the Wellcome Sanger Institute COVID-19 Surveillance Team |
| EPI_ISL_2865364, EPI_ISL_2865365 | Lighthouse Lab in Milton Keynes | Wellcome Sanger Institute for the COVID-19 Genomics UK (COG-UK) Consortium | The Lighthouse Lab in Milton Keynes and Alex Alderton, Roberto Amato, Jeffrey Barrett, Sonia Goncalves, Ewan Harrison, David K. Jackson, Ian Johnston, Dominic Kwiatkowski, Cordelia Langford, John Sillitoe on behalf of the Wellcome Sanger Institute COVID-19 Surveillance Team |
| EPI_ISL_2865366 | Lighthouse Lab in Glasgow | Wellcome Sanger Institute for the COVID-19 Genomics UK (COG-UK) Consortium | Harper VanSteenhouse, Yumi Kasai, David Gray, Carol Clugston, Anna Dominiczak and Alex Alderton, Roberto Amato, Jeffrey Barrett, Sonia Goncalves, Ewan Harrison, David K. Jackson, Ian Johnston, Dominic Kwiatkowski, Cordelia Langford, John Sillitoe on behalf of the Wellcome Sanger Institute COVID-19 Surveillance Team |
| EPI_ISL_2865367, EPI_ISL_2865368 | Lighthouse Lab in Milton Keynes | Wellcome Sanger Institute for the COVID-19 Genomics UK (COG-UK) Consortium | The Lighthouse Lab in Milton Keynes and Alex Alderton, Roberto Amato, Jeffrey Barrett, Sonia Goncalves, Ewan Harrison, David K. Jackson, Ian Johnston, Dominic Kwiatkowski, Cordelia Langford, John Sillitoe on behalf of the Wellcome Sanger Institute COVID-19 Surveillance Team |
| EPI_ISL_2865369 | Lighthouse Lab in Glasgow | Wellcome Sanger Institute for the COVID-19 Genomics UK (COG-UK) Consortium | Harper VanSteenhouse, Yumi Kasai, David Gray, Carol Clugston, Anna Dominiczak and Alex Alderton, Roberto Amato, Jeffrey Barrett, Sonia Goncalves, Ewan Harrison, David K. Jackson, Ian Johnston, Dominic Kwiatkowski, Cordelia Langford, John Sillitoe on behalf of the Wellcome Sanger Institute COVID-19 Surveillance Team |
| EPI_ISL_2865370 | Lighthouse Lab in Milton Keynes | Wellcome Sanger Institute for the COVID-19 Genomics UK (COG-UK) Consortium | The Lighthouse Lab in Milton Keynes and Alex Alderton, Roberto Amato, Jeffrey Barrett, Sonia Goncalves, Ewan Harrison, David K. Jackson, Ian Johnston, Dominic Kwiatkowski, Cordelia Langford, John Sillitoe on behalf of the Wellcome Sanger Institute COVID-19 Surveillance Team |
| EPI_ISL_2865371 | Lighthouse Lab in Alderley Park | Wellcome Sanger Institute for the COVID-19 Genomics UK (COG-UK) Consortium | Jacquelyn Wynn, Mairead Hyland, The Lighthouse Lab in Alderley Park and Alex Alderton, Roberto Amato, Jeffrey Barrett, Sonia Goncalves, Ewan Harrison, David K. Jackson, Ian Johnston, Dominic Kwiatkowski, Cordelia Langford, John Sillitoe on behalf of the Wellcome Sanger Institute COVID-19 Surveillance Team |
| EPI_ISL_2865372, EPI_ISL_2865373, EPI_ISL_2865374 | Lighthouse Lab in Glasgow | Wellcome Sanger Institute for the COVID-19 Genomics UK (COG-UK) Consortium | Harper VanSteenhouse, Yumi Kasai, David Gray, Carol Clugston, Anna Dominiczak and Alex Alderton, Roberto Amato, Jeffrey Barrett, Sonia Goncalves, Ewan Harrison, David K. Jackson, Ian Johnston, Dominic Kwiatkowski, Cordelia Langford, John Sillitoe on behalf of the Wellcome Sanger Institute COVID-19 Surveillance Team |
| EPI_ISL_2865375, EPI_ISL_2865376 | Lighthouse Lab in Milton Keynes | Wellcome Sanger Institute for the COVID-19 Genomics UK (COG-UK) Consortium | The Lighthouse Lab in Milton Keynes and Alex Alderton, Roberto Amato, Jeffrey Barrett, Sonia Goncalves, Ewan Harrison, David K. Jackson, Ian Johnston, Dominic Kwiatkowski, Cordelia Langford, John Sillitoe on behalf of the Wellcome Sanger Institute COVID-19 Surveillance Team |
| EPI_ISL_2865377, EPI_ISL_2865378, EPI_ISL_2865379 | Lighthouse Lab in Glasgow | Wellcome Sanger Institute for the COVID-19 Genomics UK (COG-UK) Consortium | Harper VanSteenhouse, Yumi Kasai, David Gray, Carol Clugston, Anna Dominiczak and Alex Alderton, Roberto Amato, Jeffrey Barrett, Sonia Goncalves, Ewan Harrison, David K. Jackson, Ian Johnston, Dominic Kwiatkowski, Cordelia Langford, John Sillitoe on behalf of the Wellcome Sanger Institute COVID-19 Surveillance Team |
| EPI_ISL_2865380 | Lighthouse Lab in Milton Keynes | Wellcome Sanger Institute for the COVID-19 Genomics UK (COG-UK) Consortium | The Lighthouse Lab in Milton Keynes and Alex Alderton, Roberto Amato, Jeffrey Barrett, Sonia Goncalves, Ewan Harrison, David K. Jackson, Ian Johnston, Dominic Kwiatkowski, Cordelia Langford, John Sillitoe on behalf of the Wellcome Sanger Institute COVID-19 Surveillance Team |
| EPI_ISL_2865381 | Lighthouse Lab in Glasgow | Wellcome Sanger Institute for the COVID-19 Genomics UK (COG-UK) Consortium | Harper VanSteenhouse, Yumi Kasai, David Gray, Carol Clugston, Anna Dominiczak and Alex Alderton, Roberto Amato, Jeffrey Barrett, Sonia Goncalves, Ewan Harrison, David K. Jackson, Ian Johnston, Dominic Kwiatkowski, Cordelia Langford, John Sillitoe on behalf of the Wellcome Sanger Institute COVID-19 Surveillance Team |
| EPI_ISL_2865382 | Lighthouse Lab in Milton Keynes | Wellcome Sanger Institute for the COVID-19 Genomics UK (COG-UK) Consortium | The Lighthouse Lab in Milton Keynes and Alex Alderton, Roberto Amato, Jeffrey Barrett, Sonia Goncalves, Ewan Harrison, David K. Jackson, Ian Johnston, Dominic Kwiatkowski, Cordelia Langford, John Sillitoe on behalf of the Wellcome Sanger Institute COVID-19 Surveillance Team |
| EPI_ISL_2865383 | Lighthouse Lab in Glasgow | Wellcome Sanger Institute for the COVID-19 Genomics UK (COG-UK) Consortium | Harper VanSteenhouse, Yumi Kasai, David Gray, Carol Clugston, Anna Dominiczak and Alex Alderton, Roberto Amato, Jeffrey Barrett, Sonia Goncalves, Ewan Harrison, David K. Jackson, Ian Johnston, Dominic Kwiatkowski, Cordelia Langford, John Sillitoe on behalf of the Wellcome Sanger Institute COVID-19 Surveillance Team |
| EPI_ISL_2865384 | Lighthouse Lab in Alderley Park | Wellcome Sanger Institute for the COVID-19 Genomics UK (COG-UK) Consortium | Jacquelyn Wynn, Mairead Hyland, The Lighthouse Lab in Alderley Park and Alex Alderton, Roberto Amato, Jeffrey Barrett, Sonia Goncalves, Ewan Harrison, David K. Jackson, Ian Johnston, Dominic Kwiatkowski, Cordelia Langford, John Sillitoe on behalf of the Wellcome Sanger Institute COVID-19 Surveillance Team |
| EPI_ISL_2865385 | Lighthouse Lab in Milton Keynes | Wellcome Sanger Institute for the COVID-19 Genomics UK (COG-UK) Consortium | The Lighthouse Lab in Milton Keynes and Alex Alderton, Roberto Amato, Jeffrey Barrett, Sonia Goncalves, Ewan Harrison, David K. Jackson, Ian Johnston, Dominic Kwiatkowski, Cordelia Langford, John Sillitoe on behalf of the Wellcome Sanger Institute COVID-19 Surveillance Team |
| EPI_ISL_2865386, EPI_ISL_2865387 | Lighthouse Lab in Glasgow | Wellcome Sanger Institute for the COVID-19 Genomics UK (COG-UK) Consortium | Harper VanSteenhouse, Yumi Kasai, David Gray, Carol Clugston, Anna Dominiczak and Alex Alderton, Roberto Amato, Jeffrey Barrett, Sonia Goncalves, Ewan Harrison, David K. Jackson, Ian Johnston, Dominic Kwiatkowski, Cordelia Langford, John Sillitoe on behalf of the Wellcome Sanger Institute COVID-19 Surveillance Team |
| EPI_ISL_2865388 | Lighthouse Lab in Milton Keynes | Wellcome Sanger Institute for the COVID-19 Genomics UK (COG-UK) Consortium | The Lighthouse Lab in Milton Keynes and Alex Alderton, Roberto Amato, Jeffrey Barrett, Sonia Goncalves, Ewan Harrison, David K. Jackson, Ian Johnston, Dominic Kwiatkowski, Cordelia Langford, John Sillitoe on behalf of the Wellcome Sanger Institute COVID-19 Surveillance Team |
| EPI_ISL_2865389 | Lighthouse Lab in Glasgow | Wellcome Sanger Institute for the COVID-19 Genomics UK (COG-UK) Consortium | Harper VanSteenhouse, Yumi Kasai, David Gray, Carol Clugston, Anna Dominiczak and Alex Alderton, Roberto Amato, Jeffrey Barrett, Sonia Goncalves, Ewan Harrison, David K. Jackson, Ian Johnston, Dominic Kwiatkowski, Cordelia Langford, John Sillitoe on behalf of the Wellcome Sanger Institute COVID-19 Surveillance Team |
| EPI_ISL_2865390 | Lighthouse Lab in Milton Keynes | Wellcome Sanger Institute for the COVID-19 Genomics UK (COG-UK) Consortium | The Lighthouse Lab in Milton Keynes and Alex Alderton, Roberto Amato, Jeffrey Barrett, Sonia Goncalves, Ewan Harrison, David K. Jackson, Ian Johnston, Dominic Kwiatkowski, Cordelia Langford, John Sillitoe on behalf of the Wellcome Sanger Institute COVID-19 Surveillance Team |
| EPI_ISL_2865391, EPI_ISL_2865392 | Lighthouse Lab in Glasgow | Wellcome Sanger Institute for the COVID-19 Genomics UK (COG-UK) Consortium | Harper VanSteenhouse, Yumi Kasai, David Gray, Carol Clugston, Anna Dominiczak and Alex Alderton, Roberto Amato, Jeffrey Barrett, Sonia Goncalves, Ewan Harrison, David K. Jackson, Ian Johnston, Dominic Kwiatkowski, Cordelia Langford, John Sillitoe on behalf of the Wellcome Sanger Institute COVID-19 Surveillance Team |
| EPI_ISL_2865393, EPI_ISL_2865394, EPI_ISL_2865395, EPI_ISL_2865396 | Lighthouse Lab in Milton Keynes | Wellcome Sanger Institute for the COVID-19 Genomics UK (COG-UK) Consortium | The Lighthouse Lab in Milton Keynes and Alex Alderton, Roberto Amato, Jeffrey Barrett, Sonia Goncalves, Ewan Harrison, David K. Jackson, Ian Johnston, Dominic Kwiatkowski, Cordelia Langford, John Sillitoe on behalf of the Wellcome Sanger Institute COVID-19 Surveillance Team |
| EPI_ISL_2865397 | Lighthouse Lab in Glasgow | Wellcome Sanger Institute for the COVID-19 Genomics UK (COG-UK) Consortium | Harper VanSteenhouse, Yumi Kasai, David Gray, Carol Clugston, Anna Dominiczak and Alex Alderton, Roberto Amato, Jeffrey Barrett, Sonia Goncalves, Ewan Harrison, David K. Jackson, Ian Johnston, Dominic Kwiatkowski, Cordelia Langford, John Sillitoe on behalf of the Wellcome Sanger Institute COVID-19 Surveillance Team |
| EPI_ISL_2865398 | Lighthouse Lab in Milton Keynes | Wellcome Sanger Institute for the COVID-19 Genomics UK (COG-UK) Consortium | The Lighthouse Lab in Milton Keynes and Alex Alderton, Roberto Amato, Jeffrey Barrett, Sonia Goncalves, Ewan Harrison, David K. Jackson, Ian Johnston, Dominic Kwiatkowski, Cordelia Langford, John Sillitoe on behalf of the Wellcome Sanger Institute COVID-19 Surveillance Team |
| EPI_ISL_2865399 | Lighthouse Lab in Glasgow | Wellcome Sanger Institute for the COVID-19 Genomics UK (COG-UK) Consortium | Harper VanSteenhouse, Yumi Kasai, David Gray, Carol Clugston, Anna Dominiczak and Alex Alderton, Roberto Amato, Jeffrey Barrett, Sonia Goncalves, Ewan Harrison, David K. Jackson, Ian Johnston, Dominic Kwiatkowski, Cordelia Langford, John Sillitoe on behalf of the Wellcome Sanger Institute COVID-19 Surveillance Team |
| EPI_ISL_2865400 | Lighthouse Lab in Milton Keynes | Wellcome Sanger Institute for the COVID-19 Genomics UK (COG-UK) Consortium | The Lighthouse Lab in Milton Keynes and Alex Alderton, Roberto Amato, Jeffrey Barrett, Sonia Goncalves, Ewan Harrison, David K. Jackson, Ian Johnston, Dominic Kwiatkowski, Cordelia Langford, John Sillitoe on behalf of the Wellcome Sanger Institute COVID-19 Surveillance Team |
| EPI_ISL_2865401 | Lighthouse Lab in Glasgow | Wellcome Sanger Institute for the COVID-19 Genomics UK (COG-UK) Consortium | Harper VanSteenhouse, Yumi Kasai, David Gray, Carol Clugston, Anna Dominiczak and Alex Alderton, Roberto Amato, Jeffrey Barrett, Sonia Goncalves, Ewan Harrison, David K. Jackson, Ian Johnston, Dominic Kwiatkowski, Cordelia Langford, John Sillitoe on behalf of the Wellcome Sanger Institute COVID-19 Surveillance Team |
| EPI_ISL_2865402 | Lighthouse Lab in Milton Keynes | Wellcome Sanger Institute for the COVID-19 Genomics UK (COG-UK) Consortium | The Lighthouse Lab in Milton Keynes and Alex Alderton, Roberto Amato, Jeffrey Barrett, Sonia Goncalves, Ewan Harrison, David K. Jackson, Ian Johnston, Dominic Kwiatkowski, Cordelia Langford, John Sillitoe on behalf of the Wellcome Sanger Institute COVID-19 Surveillance Team |
| EPI_ISL_2865403 | Lighthouse Lab in Glasgow | Wellcome Sanger Institute for the COVID-19 Genomics UK (COG-UK) Consortium | Harper VanSteenhouse, Yumi Kasai, David Gray, Carol Clugston, Anna Dominiczak and Alex Alderton, Roberto Amato, Jeffrey Barrett, Sonia Goncalves, Ewan Harrison, David K. Jackson, Ian Johnston, Dominic Kwiatkowski, Cordelia Langford, John Sillitoe on behalf of the Wellcome Sanger Institute COVID-19 Surveillance Team |
| EPI_ISL_2865404 | Lighthouse Lab in Milton Keynes | Wellcome Sanger Institute for the COVID-19 Genomics UK (COG-UK) Consortium | The Lighthouse Lab in Milton Keynes and Alex Alderton, Roberto Amato, Jeffrey Barrett, Sonia Goncalves, Ewan Harrison, David K. Jackson, Ian Johnston, Dominic Kwiatkowski, Cordelia Langford, John Sillitoe on behalf of the Wellcome Sanger Institute COVID-19 Surveillance Team |
| EPI_ISL_2865405 | Lighthouse Lab in Glasgow | Wellcome Sanger Institute for the COVID-19 Genomics UK (COG-UK) Consortium | Harper VanSteenhouse, Yumi Kasai, David Gray, Carol Clugston, Anna Dominiczak and Alex Alderton, Roberto Amato, Jeffrey Barrett, Sonia Goncalves, Ewan Harrison, David K. Jackson, Ian Johnston, Dominic Kwiatkowski, Cordelia Langford, John Sillitoe on behalf of the Wellcome Sanger Institute COVID-19 Surveillance Team |
| EPI_ISL_2865406, EPI_ISL_2865407, EPI_ISL_2865408, EPI_ISL_2865409, EPI_ISL_2865411, EPI_ISL_2865412, EPI_ISL_2865413, EPI_ISL_2865414 | Lighthouse Lab in Milton Keynes | Wellcome Sanger Institute for the COVID-19 Genomics UK (COG-UK) Consortium | The Lighthouse Lab in Milton Keynes and Alex Alderton, Roberto Amato, Jeffrey Barrett, Sonia Goncalves, Ewan Harrison, David K. Jackson, Ian Johnston, Dominic Kwiatkowski, Cordelia Langford, John Sillitoe on behalf of the Wellcome Sanger Institute COVID-19 Surveillance Team |
| EPI_ISL_2865415 | Lighthouse Lab in Glasgow | Wellcome Sanger Institute for the COVID-19 Genomics UK (COG-UK) Consortium | Harper VanSteenhouse, Yumi Kasai, David Gray, Carol Clugston, Anna Dominiczak and Alex Alderton, Roberto Amato, Jeffrey Barrett, Sonia Goncalves, Ewan Harrison, David K. Jackson, Ian Johnston, Dominic Kwiatkowski, Cordelia Langford, John Sillitoe on behalf of the Wellcome Sanger Institute COVID-19 Surveillance Team |
| EPI_ISL_2865416 | Lighthouse Lab in Milton Keynes | Wellcome Sanger Institute for the COVID-19 Genomics UK (COG-UK) Consortium | The Lighthouse Lab in Milton Keynes and Alex Alderton, Roberto Amato, Jeffrey Barrett, Sonia Goncalves, Ewan Harrison, David K. Jackson, Ian Johnston, Dominic Kwiatkowski, Cordelia Langford, John Sillitoe on behalf of the Wellcome Sanger Institute COVID-19 Surveillance Team |
| EPI_ISL_2865417 | Lighthouse Lab in Glasgow | Wellcome Sanger Institute for the COVID-19 Genomics UK (COG-UK) Consortium | Harper VanSteenhouse, Yumi Kasai, David Gray, Carol Clugston, Anna Dominiczak and Alex Alderton, Roberto Amato, Jeffrey Barrett, Sonia Goncalves, Ewan Harrison, David K. Jackson, Ian Johnston, Dominic Kwiatkowski, Cordelia Langford, John Sillitoe on behalf of the Wellcome Sanger Institute COVID-19 Surveillance Team |
| EPI_ISL_2865418, EPI_ISL_2865419 | Lighthouse Lab in Milton Keynes | Wellcome Sanger Institute for the COVID-19 Genomics UK (COG-UK) Consortium | The Lighthouse Lab in Milton Keynes and Alex Alderton, Roberto Amato, Jeffrey Barrett, Sonia Goncalves, Ewan Harrison, David K. Jackson, Ian Johnston, Dominic Kwiatkowski, Cordelia Langford, John Sillitoe on behalf of the Wellcome Sanger Institute COVID-19 Surveillance Team |
| EPI_ISL_2865420 | Lighthouse Lab in Glasgow | Wellcome Sanger Institute for the COVID-19 Genomics UK (COG-UK) Consortium | Harper VanSteenhouse, Yumi Kasai, David Gray, Carol Clugston, Anna Dominiczak and Alex Alderton, Roberto Amato, Jeffrey Barrett, Sonia Goncalves, Ewan Harrison, David K. Jackson, Ian Johnston, Dominic Kwiatkowski, Cordelia Langford, John Sillitoe on behalf of the Wellcome Sanger Institute COVID-19 Surveillance Team |
| EPI_ISL_2865421, EPI_ISL_2865422 | Lighthouse Lab in Milton Keynes | Wellcome Sanger Institute for the COVID-19 Genomics UK (COG-UK) Consortium | The Lighthouse Lab in Milton Keynes and Alex Alderton, Roberto Amato, Jeffrey Barrett, Sonia Goncalves, Ewan Harrison, David K. Jackson, Ian Johnston, Dominic Kwiatkowski, Cordelia Langford, John Sillitoe on behalf of the Wellcome Sanger Institute COVID-19 Surveillance Team |
| EPI_ISL_2865423 | Lighthouse Lab in Glasgow | Wellcome Sanger Institute for the COVID-19 Genomics UK (COG-UK) Consortium | Harper VanSteenhouse, Yumi Kasai, David Gray, Carol Clugston, Anna Dominiczak and Alex Alderton, Roberto Amato, Jeffrey Barrett, Sonia Goncalves, Ewan Harrison, David K. Jackson, Ian Johnston, Dominic Kwiatkowski, Cordelia Langford, John Sillitoe on behalf of the Wellcome Sanger Institute COVID-19 Surveillance Team |

| | | | |
|---|---|---|---|
| EPI_ISL_2865424, EPI_ISL_2865425 | Lighthouse Lab in Milton Keynes | Wellcome Sanger Institute for the COVID-19 Genomics UK (COG-UK) Consortium | The Lighthouse Lab in Milton Keynes and Alex Alderton, Roberto Amato, Jeffrey Barrett, Sonia Goncalves, Ewan Harrison, David K. Jackson, Ian Johnston, Dominic Kwiatkowski, Cordelia Langford, John Sillitoe on behalf of the Wellcome Sanger Institute COVID-19 Surveillance Team |
| EPI_ISL_2865426, EPI_ISL_2865427 | Lighthouse Lab in Glasgow | Wellcome Sanger Institute for the COVID-19 Genomics UK (COG-UK) Consortium | Harper VanSteenhouse, Yumi Kasai, David Gray, Carol Clugston, Anna Dominiczak and Alex Alderton, Roberto Amato, Jeffrey Barrett, Sonia Goncalves, Ewan Harrison, David K. Jackson, Ian Johnston, Dominic Kwiatkowski, Cordelia Langford, John Sillitoe on behalf of the Wellcome Sanger Institute COVID-19 Surveillance Team |
| EPI_ISL_2865428, EPI_ISL_2865429, EPI_ISL_2865430 | Lighthouse Lab in Milton Keynes | Wellcome Sanger Institute for the COVID-19 Genomics UK (COG-UK) Consortium | The Lighthouse Lab in Milton Keynes and Alex Alderton, Roberto Amato, Jeffrey Barrett, Sonia Goncalves, Ewan Harrison, David K. Jackson, Ian Johnston, Dominic Kwiatkowski, Cordelia Langford, John Sillitoe on behalf of the Wellcome Sanger Institute COVID-19 Surveillance Team |
| EPI_ISL_2865431, EPI_ISL_2865432 | Lighthouse Lab in Glasgow | Wellcome Sanger Institute for the COVID-19 Genomics UK (COG-UK) Consortium | Harper VanSteenhouse, Yumi Kasai, David Gray, Carol Clugston, Anna Dominiczak and Alex Alderton, Roberto Amato, Jeffrey Barrett, Sonia Goncalves, Ewan Harrison, David K. Jackson, Ian Johnston, Dominic Kwiatkowski, Cordelia Langford, John Sillitoe on behalf of the Wellcome Sanger Institute COVID-19 Surveillance Team |
| EPI_ISL_2865433 | Lighthouse Lab in Alderley Park | Wellcome Sanger Institute for the COVID-19 Genomics UK (COG-UK) Consortium | Jacquelyn Wynn, Mairead Hyland, The Lighthouse Lab in Alderley Park and Alex Alderton, Roberto Amato, Jeffrey Barrett, Sonia Goncalves, Ewan Harrison, David K. Jackson, Ian Johnston, Dominic Kwiatkowski, Cordelia Langford, John Sillitoe on behalf of the Wellcome Sanger Institute COVID-19 Surveillance Team |
| EPI_ISL_2865434 | Lighthouse Lab in Milton Keynes | Wellcome Sanger Institute for the COVID-19 Genomics UK (COG-UK) Consortium | The Lighthouse Lab in Milton Keynes and Alex Alderton, Roberto Amato, Jeffrey Barrett, Sonia Goncalves, Ewan Harrison, David K. Jackson, Ian Johnston, Dominic Kwiatkowski, Cordelia Langford, John Sillitoe on behalf of the Wellcome Sanger Institute COVID-19 Surveillance Team |
| EPI_ISL_2865435 | Lighthouse Lab in Glasgow | Wellcome Sanger Institute for the COVID-19 Genomics UK (COG-UK) Consortium | Harper VanSteenhouse, Yumi Kasai, David Gray, Carol Clugston, Anna Dominiczak and Alex Alderton, Roberto Amato, Jeffrey Barrett, Sonia Goncalves, Ewan Harrison, David K. Jackson, Ian Johnston, Dominic Kwiatkowski, Cordelia Langford, John Sillitoe on behalf of the Wellcome Sanger Institute COVID-19 Surveillance Team |
| EPI_ISL_2865436 | Lighthouse Lab in Milton Keynes | Wellcome Sanger Institute for the COVID-19 Genomics UK (COG-UK) Consortium | The Lighthouse Lab in Milton Keynes and Alex Alderton, Roberto Amato, Jeffrey Barrett, Sonia Goncalves, Ewan Harrison, David K. Jackson, Ian Johnston, Dominic Kwiatkowski, Cordelia Langford, John Sillitoe on behalf of the Wellcome Sanger Institute COVID-19 Surveillance Team |
| EPI_ISL_2865437, EPI_ISL_2865438, EPI_ISL_2865439 | Lighthouse Lab in Glasgow | Wellcome Sanger Institute for the COVID-19 Genomics UK (COG-UK) Consortium | Harper VanSteenhouse, Yumi Kasai, David Gray, Carol Clugston, Anna Dominiczak and Alex Alderton, Roberto Amato, Jeffrey Barrett, Sonia Goncalves, Ewan Harrison, David K. Jackson, Ian Johnston, Dominic Kwiatkowski, Cordelia Langford, John Sillitoe on behalf of the Wellcome Sanger Institute COVID-19 Surveillance Team |
| EPI_ISL_2865440, EPI_ISL_2865441, EPI_ISL_2865442, EPI_ISL_2865443 | Lighthouse Lab in Milton Keynes | Wellcome Sanger Institute for the COVID-19 Genomics UK (COG-UK) Consortium | The Lighthouse Lab in Milton Keynes and Alex Alderton, Roberto Amato, Jeffrey Barrett, Sonia Goncalves, Ewan Harrison, David K. Jackson, Ian Johnston, Dominic Kwiatkowski, Cordelia Langford, John Sillitoe on behalf of the Wellcome Sanger Institute COVID-19 Surveillance Team |
| EPI_ISL_2865444 | Lighthouse Lab in Glasgow | Wellcome Sanger Institute for the COVID-19 Genomics UK (COG-UK) Consortium | Harper VanSteenhouse, Yumi Kasai, David Gray, Carol Clugston, Anna Dominiczak and Alex Alderton, Roberto Amato, Jeffrey Barrett, Sonia Goncalves, Ewan Harrison, David K. Jackson, Ian Johnston, Dominic Kwiatkowski, Cordelia Langford, John Sillitoe on behalf of the Wellcome Sanger Institute COVID-19 Surveillance Team |
| EPI_ISL_2865446 | Lighthouse Lab in Milton Keynes | Wellcome Sanger Institute for the COVID-19 Genomics UK (COG-UK) Consortium | The Lighthouse Lab in Milton Keynes and Alex Alderton, Roberto Amato, Jeffrey Barrett, Sonia Goncalves, Ewan Harrison, David K. Jackson, Ian Johnston, Dominic Kwiatkowski, Cordelia Langford, John Sillitoe on behalf of the Wellcome Sanger Institute COVID-19 Surveillance Team |
| EPI_ISL_2865447 | Lighthouse Lab in Glasgow | Wellcome Sanger Institute for the COVID-19 Genomics UK (COG-UK) Consortium | Harper VanSteenhouse, Yumi Kasai, David Gray, Carol Clugston, Anna Dominiczak and Alex Alderton, Roberto Amato, Jeffrey Barrett, Sonia Goncalves, Ewan Harrison, David K. Jackson, Ian Johnston, Dominic Kwiatkowski, Cordelia Langford, John Sillitoe on behalf of the Wellcome Sanger Institute COVID-19 Surveillance Team |
| EPI_ISL_2865448, EPI_ISL_2865449, EPI_ISL_2865450, EPI_ISL_2865451 | Lighthouse Lab in Milton Keynes | Wellcome Sanger Institute for the COVID-19 Genomics UK (COG-UK) Consortium | The Lighthouse Lab in Milton Keynes and Alex Alderton, Roberto Amato, Jeffrey Barrett, Sonia Goncalves, Ewan Harrison, David K. Jackson, Ian Johnston, Dominic Kwiatkowski, Cordelia Langford, John Sillitoe on behalf of the Wellcome Sanger Institute COVID-19 Surveillance Team |
| EPI_ISL_2865452 | Lighthouse Lab in Glasgow | Wellcome Sanger Institute for the COVID-19 Genomics UK (COG-UK) Consortium | Harper VanSteenhouse, Yumi Kasai, David Gray, Carol Clugston, Anna Dominiczak and Alex Alderton, Roberto Amato, Jeffrey Barrett, Sonia Goncalves, Ewan Harrison, David K. Jackson, Ian Johnston, Dominic Kwiatkowski, Cordelia Langford, John Sillitoe on behalf of the Wellcome Sanger Institute COVID-19 Surveillance Team |
| EPI_ISL_2865453 | Lighthouse Lab in Alderley Park | Wellcome Sanger Institute for the COVID-19 Genomics UK (COG-UK) Consortium | Jacquelyn Wynn, Mairead Hyland, The Lighthouse Lab in Alderley Park and Alex Alderton, Roberto Amato, Jeffrey Barrett, Sonia Goncalves, Ewan Harrison, David K. Jackson, Ian Johnston, Dominic Kwiatkowski, Cordelia Langford, John Sillitoe on behalf of the Wellcome Sanger Institute COVID-19 Surveillance Team |
| EPI_ISL_2865454, EPI_ISL_2865455 | Lighthouse Lab in Glasgow | Wellcome Sanger Institute for the COVID-19 Genomics UK (COG-UK) Consortium | Harper VanSteenhouse, Yumi Kasai, David Gray, Carol Clugston, Anna Dominiczak and Alex Alderton, Roberto Amato, Jeffrey Barrett, Sonia Goncalves, Ewan Harrison, David K. Jackson, Ian Johnston, Dominic Kwiatkowski, Cordelia Langford, John Sillitoe on behalf of the Wellcome Sanger Institute COVID-19 Surveillance Team |
| EPI_ISL_2865456, EPI_ISL_2865457, EPI_ISL_2865458, EPI_ISL_2865459 | Lighthouse Lab in Milton Keynes | Wellcome Sanger Institute for the COVID-19 Genomics UK (COG-UK) Consortium | The Lighthouse Lab in Milton Keynes and Alex Alderton, Roberto Amato, Jeffrey Barrett, Sonia Goncalves, Ewan Harrison, David K. Jackson, Ian Johnston, Dominic Kwiatkowski, Cordelia Langford, John Sillitoe on behalf of the Wellcome Sanger Institute COVID-19 Surveillance Team |
| EPI_ISL_2865460 | Lighthouse Lab in Glasgow | Wellcome Sanger Institute for the COVID-19 Genomics UK (COG-UK) Consortium | Harper VanSteenhouse, Yumi Kasai, David Gray, Carol Clugston, Anna Dominiczak and Alex Alderton, Roberto Amato, Jeffrey Barrett, Sonia Goncalves, Ewan Harrison, David K. Jackson, Ian Johnston, Dominic Kwiatkowski, Cordelia Langford, John Sillitoe on behalf of the Wellcome Sanger Institute COVID-19 Surveillance Team |
| EPI_ISL_2865461 | Lighthouse Lab in Alderley Park | Wellcome Sanger Institute for the COVID-19 Genomics UK (COG-UK) Consortium | Jacquelyn Wynn, Mairead Hyland, The Lighthouse Lab in Alderley Park and Alex Alderton, Roberto Amato, Jeffrey Barrett, Sonia Goncalves, Ewan Harrison, David K. Jackson, Ian Johnston, Dominic Kwiatkowski, Cordelia Langford, John Sillitoe on behalf of the Wellcome Sanger Institute COVID-19 Surveillance Team |
| EPI_ISL_2865462, EPI_ISL_2865463, EPI_ISL_2865464, EPI_ISL_2865465 | Lighthouse Lab in Milton Keynes | Wellcome Sanger Institute for the COVID-19 Genomics UK (COG-UK) Consortium | The Lighthouse Lab in Milton Keynes and Alex Alderton, Roberto Amato, Jeffrey Barrett, Sonia Goncalves, Ewan Harrison, David K. Jackson, Ian Johnston, Dominic Kwiatkowski, Cordelia Langford, John Sillitoe on behalf of the Wellcome Sanger Institute COVID-19 Surveillance Team |
| EPI_ISL_2865466, EPI_ISL_2865467 | Lighthouse Lab in Glasgow | Wellcome Sanger Institute for the COVID-19 Genomics UK (COG-UK) Consortium | Harper VanSteenhouse, Yumi Kasai, David Gray, Carol Clugston, Anna Dominiczak and Alex Alderton, Roberto Amato, Jeffrey Barrett, Sonia Goncalves, Ewan Harrison, David K. Jackson, Ian Johnston, Dominic Kwiatkowski, Cordelia Langford, John Sillitoe on behalf of the Wellcome Sanger Institute COVID-19 Surveillance Team |
| EPI_ISL_2865468 | Lighthouse Lab in Milton Keynes | Wellcome Sanger Institute for the COVID-19 Genomics UK (COG-UK) Consortium | The Lighthouse Lab in Milton Keynes and Alex Alderton, Roberto Amato, Jeffrey Barrett, Sonia Goncalves, Ewan Harrison, David K. Jackson, Ian Johnston, Dominic Kwiatkowski, Cordelia Langford, John Sillitoe on behalf of the Wellcome Sanger Institute COVID-19 Surveillance Team |
| EPI_ISL_2865469 | Lighthouse Lab in Glasgow | Wellcome Sanger Institute for the COVID-19 Genomics UK (COG-UK) Consortium | Harper VanSteenhouse, Yumi Kasai, David Gray, Carol Clugston, Anna Dominiczak and Alex Alderton, Roberto Amato, Jeffrey Barrett, Sonia Goncalves, Ewan Harrison, David K. Jackson, Ian Johnston, Dominic Kwiatkowski, Cordelia Langford, John Sillitoe on behalf of the Wellcome Sanger Institute COVID-19 Surveillance Team |
| EPI_ISL_2865470 | Lighthouse Lab in Alderley Park | Wellcome Sanger Institute for the COVID-19 Genomics UK (COG-UK) Consortium | Jacquelyn Wynn, Mairead Hyland, The Lighthouse Lab in Alderley Park and Alex Alderton, Roberto Amato, Jeffrey Barrett, Sonia Goncalves, Ewan Harrison, David K. Jackson, Ian Johnston, Dominic Kwiatkowski, Cordelia Langford, John Sillitoe on behalf of the Wellcome Sanger Institute COVID-19 Surveillance Team |
| EPI_ISL_2865471, EPI_ISL_2865472, EPI_ISL_2865473, EPI_ISL_2865474 | Lighthouse Lab in Glasgow | Wellcome Sanger Institute for the COVID-19 Genomics UK (COG-UK) Consortium | Harper VanSteenhouse, Yumi Kasai, David Gray, Carol Clugston, Anna Dominiczak and Alex Alderton, Roberto Amato, Jeffrey Barrett, Sonia Goncalves, Ewan Harrison, David K. Jackson, Ian Johnston, Dominic Kwiatkowski, Cordelia Langford, John Sillitoe on behalf of the Wellcome Sanger Institute COVID-19 Surveillance Team |
| EPI_ISL_2865475 | Lighthouse Lab in Milton Keynes | Wellcome Sanger Institute for the COVID-19 Genomics UK (COG-UK) Consortium | The Lighthouse Lab in Milton Keynes and Alex Alderton, Roberto Amato, Jeffrey Barrett, Sonia Goncalves, Ewan Harrison, David K. Jackson, Ian Johnston, Dominic Kwiatkowski, Cordelia Langford, John Sillitoe on behalf of the Wellcome Sanger Institute COVID-19 Surveillance Team |
| EPI_ISL_2865476 | Lighthouse Lab in Alderley Park | Wellcome Sanger Institute for the COVID-19 Genomics UK (COG-UK) Consortium | Jacquelyn Wynn, Mairead Hyland, The Lighthouse Lab in Alderley Park and Alex Alderton, Roberto Amato, Jeffrey Barrett, Sonia Goncalves, Ewan Harrison, David K. Jackson, Ian Johnston, Dominic Kwiatkowski, Cordelia Langford, John Sillitoe on behalf of the Wellcome Sanger Institute COVID-19 Surveillance Team |
| EPI_ISL_2865477 | Lighthouse Lab in Glasgow | Wellcome Sanger Institute for the COVID-19 Genomics UK (COG-UK) Consortium | Harper VanSteenhouse, Yumi Kasai, David Gray, Carol Clugston, Anna Dominiczak and Alex Alderton, Roberto Amato, Jeffrey Barrett, Sonia Goncalves, Ewan Harrison, David K. Jackson, Ian Johnston, Dominic Kwiatkowski, Cordelia Langford, John Sillitoe on behalf of the Wellcome Sanger Institute COVID-19 Surveillance Team |
| EPI_ISL_2865478, EPI_ISL_2865479 | Lighthouse Lab in Milton Keynes | Wellcome Sanger Institute for the COVID-19 Genomics UK (COG-UK) Consortium | The Lighthouse Lab in Milton Keynes and Alex Alderton, Roberto Amato, Jeffrey Barrett, Sonia Goncalves, Ewan Harrison, David K. Jackson, Ian Johnston, Dominic Kwiatkowski, Cordelia Langford, John Sillitoe on behalf of the Wellcome Sanger Institute COVID-19 Surveillance Team |
| EPI_ISL_2865480, EPI_ISL_2865481, EPI_ISL_2865482 | Lighthouse Lab in Glasgow | Wellcome Sanger Institute for the COVID-19 Genomics UK (COG-UK) Consortium | Harper VanSteenhouse, Yumi Kasai, David Gray, Carol Clugston, Anna Dominiczak and Alex Alderton, Roberto Amato, Jeffrey Barrett, Sonia Goncalves, Ewan Harrison, David K. Jackson, Ian Johnston, Dominic Kwiatkowski, Cordelia Langford, John Sillitoe on behalf of the Wellcome Sanger Institute COVID-19 Surveillance Team |
| EPI_ISL_2865483 | Lighthouse Lab in Milton Keynes | Wellcome Sanger Institute for the COVID-19 Genomics UK (COG-UK) Consortium | The Lighthouse Lab in Milton Keynes and Alex Alderton, Roberto Amato, Jeffrey Barrett, Sonia Goncalves, Ewan Harrison, David K. Jackson, Ian Johnston, Dominic Kwiatkowski, Cordelia Langford, John Sillitoe on behalf of the Wellcome Sanger Institute COVID-19 Surveillance Team |
| EPI_ISL_2865484 | Lighthouse Lab in Glasgow | Wellcome Sanger Institute for the COVID-19 Genomics UK (COG-UK) Consortium | Harper VanSteenhouse, Yumi Kasai, David Gray, Carol Clugston, Anna Dominiczak and Alex Alderton, Roberto Amato, Jeffrey Barrett, Sonia Goncalves, Ewan Harrison, David K. Jackson, Ian Johnston, Dominic Kwiatkowski, Cordelia Langford, John Sillitoe on behalf of the Wellcome Sanger Institute COVID-19 Surveillance Team |
| EPI_ISL_2865485 | Lighthouse Lab in Milton Keynes | Wellcome Sanger Institute for the COVID-19 Genomics UK (COG-UK) Consortium | The Lighthouse Lab in Milton Keynes and Alex Alderton, Roberto Amato, Jeffrey Barrett, Sonia Goncalves, Ewan Harrison, David K. Jackson, Ian Johnston, Dominic Kwiatkowski, Cordelia Langford, John Sillitoe on behalf of the Wellcome Sanger Institute COVID-19 Surveillance Team |
| EPI_ISL_2865486 | Lighthouse Lab in Glasgow | Wellcome Sanger Institute for the COVID-19 Genomics UK (COG-UK) Consortium | Harper VanSteenhouse, Yumi Kasai, David Gray, Carol Clugston, Anna Dominiczak and Alex Alderton, Roberto Amato, Jeffrey Barrett, Sonia Goncalves, Ewan Harrison, David K. Jackson, Ian Johnston, Dominic Kwiatkowski, Cordelia Langford, John Sillitoe on behalf of the Wellcome Sanger Institute COVID-19 Surveillance Team |
| EPI_ISL_2865487, EPI_ISL_2865488 | Lighthouse Lab in Milton Keynes | Wellcome Sanger Institute for the COVID-19 Genomics UK (COG-UK) Consortium | The Lighthouse Lab in Milton Keynes and Alex Alderton, Roberto Amato, Jeffrey Barrett, Sonia Goncalves, Ewan Harrison, David K. Jackson, Ian Johnston, Dominic Kwiatkowski, Cordelia Langford, John Sillitoe on behalf of the Wellcome Sanger Institute COVID-19 Surveillance Team |
| EPI_ISL_2865489, EPI_ISL_2865490 | Lighthouse Lab in Glasgow | Wellcome Sanger Institute for the COVID-19 Genomics UK (COG-UK) Consortium | Harper VanSteenhouse, Yumi Kasai, David Gray, Carol Clugston, Anna Dominiczak and Alex Alderton, Roberto Amato, Jeffrey Barrett, Sonia Goncalves, Ewan Harrison, David K. Jackson, Ian Johnston, Dominic Kwiatkowski, Cordelia Langford, John Sillitoe on behalf of the Wellcome Sanger Institute COVID-19 Surveillance Team |
| EPI_ISL_2865491, EPI_ISL_2865492 | Lighthouse Lab in Milton Keynes | Wellcome Sanger Institute for the COVID-19 Genomics UK (COG-UK) Consortium | The Lighthouse Lab in Milton Keynes and Alex Alderton, Roberto Amato, Jeffrey Barrett, Sonia Goncalves, Ewan Harrison, David K. Jackson, Ian Johnston, Dominic Kwiatkowski, Cordelia Langford, John Sillitoe on behalf of the Wellcome Sanger Institute COVID-19 Surveillance Team |
| EPI_ISL_2865493 | Lighthouse Lab in Glasgow | Wellcome Sanger Institute for the COVID-19 Genomics UK (COG-UK) Consortium | Harper VanSteenhouse, Yumi Kasai, David Gray, Carol Clugston, Anna Dominiczak and Alex Alderton, Roberto Amato, Jeffrey Barrett, Sonia Goncalves, Ewan Harrison, David K. Jackson, Ian Johnston, Dominic Kwiatkowski, Cordelia Langford, John Sillitoe on behalf of the Wellcome Sanger Institute COVID-19 Surveillance Team |
| EPI_ISL_2865494 | Lighthouse Lab in Milton Keynes | Wellcome Sanger Institute for the COVID-19 Genomics UK (COG-UK) Consortium | The Lighthouse Lab in Milton Keynes and Alex Alderton, Roberto Amato, Jeffrey Barrett, Sonia Goncalves, Ewan Harrison, David K. Jackson, Ian Johnston, Dominic Kwiatkowski, Cordelia Langford, John Sillitoe on behalf of the Wellcome Sanger Institute COVID-19 Surveillance Team |

| | | | |
|---|---|---|---|
| EPI_ISL_2865495 | Lighthouse Lab in Glasgow | Wellcome Sanger Institute for the COVID-19 Genomics UK (COG-UK) Consortium | Harper VanSteenhouse, Yumi Kasai, David Gray, Carol Clugston, Anna Dominiczak and Alex Alderton, Roberto Amato, Jeffrey Barrett, Sonia Goncalves, Ewan Harrison, David K. Jackson, Ian Johnston, Dominic Kwiatkowski, Cordelia Langford, John Sillitoe on behalf of the Wellcome Sanger Institute COVID-19 Surveillance Team |
| EPI_ISL_2865496 | Lighthouse Lab in Alderley Park | Wellcome Sanger Institute for the COVID-19 Genomics UK (COG-UK) Consortium | Jacquelyn Wynn, Mairead Hyland, The Lighthouse Lab in Alderley Park and Alex Alderton, Roberto Amato, Jeffrey Barrett, Sonia Goncalves, Ewan Harrison, David K. Jackson, Ian Johnston, Dominic Kwiatkowski, Cordelia Langford, John Sillitoe on behalf of the Wellcome Sanger Institute COVID-19 Surveillance Team |
| EPI_ISL_2865497 | Lighthouse Lab in Glasgow | Wellcome Sanger Institute for the COVID-19 Genomics UK (COG-UK) Consortium | Harper VanSteenhouse, Yumi Kasai, David Gray, Carol Clugston, Anna Dominiczak and Alex Alderton, Roberto Amato, Jeffrey Barrett, Sonia Goncalves, Ewan Harrison, David K. Jackson, Ian Johnston, Dominic Kwiatkowski, Cordelia Langford, John Sillitoe on behalf of the Wellcome Sanger Institute COVID-19 Surveillance Team |
| EPI_ISL_2865498 | Lighthouse Lab in Milton Keynes | Wellcome Sanger Institute for the COVID-19 Genomics UK (COG-UK) Consortium | The Lighthouse Lab in Milton Keynes and Alex Alderton, Roberto Amato, Jeffrey Barrett, Sonia Goncalves, Ewan Harrison, David K. Jackson, Ian Johnston, Dominic Kwiatkowski, Cordelia Langford, John Sillitoe on behalf of the Wellcome Sanger Institute COVID-19 Surveillance Team |
| EPI_ISL_2865499 | Lighthouse Lab in Alderley Park | Wellcome Sanger Institute for the COVID-19 Genomics UK (COG-UK) Consortium | Jacquelyn Wynn, Mairead Hyland, The Lighthouse Lab in Alderley Park and Alex Alderton, Roberto Amato, Jeffrey Barrett, Sonia Goncalves, Ewan Harrison, David K. Jackson, Ian Johnston, Dominic Kwiatkowski, Cordelia Langford, John Sillitoe on behalf of the Wellcome Sanger Institute COVID-19 Surveillance Team |
| EPI_ISL_2865500, EPI_ISL_2865501, EPI_ISL_2865502, EPI_ISL_2865503 | Lighthouse Lab in Glasgow | Wellcome Sanger Institute for the COVID-19 Genomics UK (COG-UK) Consortium | Harper VanSteenhouse, Yumi Kasai, David Gray, Carol Clugston, Anna Dominiczak and Alex Alderton, Roberto Amato, Jeffrey Barrett, Sonia Goncalves, Ewan Harrison, David K. Jackson, Ian Johnston, Dominic Kwiatkowski, Cordelia Langford, John Sillitoe on behalf of the Wellcome Sanger Institute COVID-19 Surveillance Team |
| EPI_ISL_2865504 | Lighthouse Lab in Milton Keynes | Wellcome Sanger Institute for the COVID-19 Genomics UK (COG-UK) Consortium | The Lighthouse Lab in Milton Keynes and Alex Alderton, Roberto Amato, Jeffrey Barrett, Sonia Goncalves, Ewan Harrison, David K. Jackson, Ian Johnston, Dominic Kwiatkowski, Cordelia Langford, John Sillitoe on behalf of the Wellcome Sanger Institute COVID-19 Surveillance Team |
| EPI_ISL_2865505, EPI_ISL_2865506, EPI_ISL_2865507, EPI_ISL_2865508 | Lighthouse Lab in Glasgow | Wellcome Sanger Institute for the COVID-19 Genomics UK (COG-UK) Consortium | Harper VanSteenhouse, Yumi Kasai, David Gray, Carol Clugston, Anna Dominiczak and Alex Alderton, Roberto Amato, Jeffrey Barrett, Sonia Goncalves, Ewan Harrison, David K. Jackson, Ian Johnston, Dominic Kwiatkowski, Cordelia Langford, John Sillitoe on behalf of the Wellcome Sanger Institute COVID-19 Surveillance Team |
| EPI_ISL_2865509 | Lighthouse Lab in Milton Keynes | Wellcome Sanger Institute for the COVID-19 Genomics UK (COG-UK) Consortium | The Lighthouse Lab in Milton Keynes and Alex Alderton, Roberto Amato, Jeffrey Barrett, Sonia Goncalves, Ewan Harrison, David K. Jackson, Ian Johnston, Dominic Kwiatkowski, Cordelia Langford, John Sillitoe on behalf of the Wellcome Sanger Institute COVID-19 Surveillance Team |
| EPI_ISL_2865510 | Lighthouse Lab in Glasgow | Wellcome Sanger Institute for the COVID-19 Genomics UK (COG-UK) Consortium | Harper VanSteenhouse, Yumi Kasai, David Gray, Carol Clugston, Anna Dominiczak and Alex Alderton, Roberto Amato, Jeffrey Barrett, Sonia Goncalves, Ewan Harrison, David K. Jackson, Ian Johnston, Dominic Kwiatkowski, Cordelia Langford, John Sillitoe on behalf of the Wellcome Sanger Institute COVID-19 Surveillance Team |
| EPI_ISL_2865511 | Lighthouse Lab in Milton Keynes | Wellcome Sanger Institute for the COVID-19 Genomics UK (COG-UK) Consortium | The Lighthouse Lab in Milton Keynes and Alex Alderton, Roberto Amato, Jeffrey Barrett, Sonia Goncalves, Ewan Harrison, David K. Jackson, Ian Johnston, Dominic Kwiatkowski, Cordelia Langford, John Sillitoe on behalf of the Wellcome Sanger Institute COVID-19 Surveillance Team |
| EPI_ISL_2865512 | Lighthouse Lab in Glasgow | Wellcome Sanger Institute for the COVID-19 Genomics UK (COG-UK) Consortium | Harper VanSteenhouse, Yumi Kasai, David Gray, Carol Clugston, Anna Dominiczak and Alex Alderton, Roberto Amato, Jeffrey Barrett, Sonia Goncalves, Ewan Harrison, David K. Jackson, Ian Johnston, Dominic Kwiatkowski, Cordelia Langford, John Sillitoe on behalf of the Wellcome Sanger Institute COVID-19 Surveillance Team |
| EPI_ISL_2865513, EPI_ISL_2865514, EPI_ISL_2865515, EPI_ISL_2865516 | Lighthouse Lab in Milton Keynes | Wellcome Sanger Institute for the COVID-19 Genomics UK (COG-UK) Consortium | The Lighthouse Lab in Milton Keynes and Alex Alderton, Roberto Amato, Jeffrey Barrett, Sonia Goncalves, Ewan Harrison, David K. Jackson, Ian Johnston, Dominic Kwiatkowski, Cordelia Langford, John Sillitoe on behalf of the Wellcome Sanger Institute COVID-19 Surveillance Team |
| EPI_ISL_2865517 | Lighthouse Lab in Glasgow | Wellcome Sanger Institute for the COVID-19 Genomics UK (COG-UK) Consortium | Harper VanSteenhouse, Yumi Kasai, David Gray, Carol Clugston, Anna Dominiczak and Alex Alderton, Roberto Amato, Jeffrey Barrett, Sonia Goncalves, Ewan Harrison, David K. Jackson, Ian Johnston, Dominic Kwiatkowski, Cordelia Langford, John Sillitoe on behalf of the Wellcome Sanger Institute COVID-19 Surveillance Team |
| EPI_ISL_2865518, EPI_ISL_2865519, EPI_ISL_2865520 | Lighthouse Lab in Milton Keynes | Wellcome Sanger Institute for the COVID-19 Genomics UK (COG-UK) Consortium | The Lighthouse Lab in Milton Keynes and Alex Alderton, Roberto Amato, Jeffrey Barrett, Sonia Goncalves, Ewan Harrison, David K. Jackson, Ian Johnston, Dominic Kwiatkowski, Cordelia Langford, John Sillitoe on behalf of the Wellcome Sanger Institute COVID-19 Surveillance Team |
| EPI_ISL_2865521 | Lighthouse Lab in Glasgow | Wellcome Sanger Institute for the COVID-19 Genomics UK (COG-UK) Consortium | Harper VanSteenhouse, Yumi Kasai, David Gray, Carol Clugston, Anna Dominiczak and Alex Alderton, Roberto Amato, Jeffrey Barrett, Sonia Goncalves, Ewan Harrison, David K. Jackson, Ian Johnston, Dominic Kwiatkowski, Cordelia Langford, John Sillitoe on behalf of the Wellcome Sanger Institute COVID-19 Surveillance Team |
| EPI_ISL_2865522, EPI_ISL_2865523 | Lighthouse Lab in Milton Keynes | Wellcome Sanger Institute for the COVID-19 Genomics UK (COG-UK) Consortium | The Lighthouse Lab in Milton Keynes and Alex Alderton, Roberto Amato, Jeffrey Barrett, Sonia Goncalves, Ewan Harrison, David K. Jackson, Ian Johnston, Dominic Kwiatkowski, Cordelia Langford, John Sillitoe on behalf of the Wellcome Sanger Institute COVID-19 Surveillance Team |
| EPI_ISL_2865524, EPI_ISL_2865525 | Lighthouse Lab in Glasgow | Wellcome Sanger Institute for the COVID-19 Genomics UK (COG-UK) Consortium | Harper VanSteenhouse, Yumi Kasai, David Gray, Carol Clugston, Anna Dominiczak and Alex Alderton, Roberto Amato, Jeffrey Barrett, Sonia Goncalves, Ewan Harrison, David K. Jackson, Ian Johnston, Dominic Kwiatkowski, Cordelia Langford, John Sillitoe on behalf of the Wellcome Sanger Institute COVID-19 Surveillance Team |
| EPI_ISL_2865526 | Lighthouse Lab in Milton Keynes | Wellcome Sanger Institute for the COVID-19 Genomics UK (COG-UK) Consortium | The Lighthouse Lab in Milton Keynes and Alex Alderton, Roberto Amato, Jeffrey Barrett, Sonia Goncalves, Ewan Harrison, David K. Jackson, Ian Johnston, Dominic Kwiatkowski, Cordelia Langford, John Sillitoe on behalf of the Wellcome Sanger Institute COVID-19 Surveillance Team |
| EPI_ISL_2865527 | Lighthouse Lab in Alderley Park | Wellcome Sanger Institute for the COVID-19 Genomics UK (COG-UK) Consortium | Jacquelyn Wynn, Mairead Hyland, The Lighthouse Lab in Alderley Park and Alex Alderton, Roberto Amato, Jeffrey Barrett, Sonia Goncalves, Ewan Harrison, David K. Jackson, Ian Johnston, Dominic Kwiatkowski, Cordelia Langford, John Sillitoe on behalf of the Wellcome Sanger Institute COVID-19 Surveillance Team |
| EPI_ISL_2865528 | Lighthouse Lab in Glasgow | Wellcome Sanger Institute for the COVID-19 Genomics UK (COG-UK) Consortium | Harper VanSteenhouse, Yumi Kasai, David Gray, Carol Clugston, Anna Dominiczak and Alex Alderton, Roberto Amato, Jeffrey Barrett, Sonia Goncalves, Ewan Harrison, David K. Jackson, Ian Johnston, Dominic Kwiatkowski, Cordelia Langford, John Sillitoe on behalf of the Wellcome Sanger Institute COVID-19 Surveillance Team |
| EPI_ISL_2865529 | Lighthouse Lab in Milton Keynes | Wellcome Sanger Institute for the COVID-19 Genomics UK (COG-UK) Consortium | The Lighthouse Lab in Milton Keynes and Alex Alderton, Roberto Amato, Jeffrey Barrett, Sonia Goncalves, Ewan Harrison, David K. Jackson, Ian Johnston, Dominic Kwiatkowski, Cordelia Langford, John Sillitoe on behalf of the Wellcome Sanger Institute COVID-19 Surveillance Team |
| EPI_ISL_2865530 | Lighthouse Lab in Glasgow | Wellcome Sanger Institute for the COVID-19 Genomics UK (COG-UK) Consortium | Harper VanSteenhouse, Yumi Kasai, David Gray, Carol Clugston, Anna Dominiczak and Alex Alderton, Roberto Amato, Jeffrey Barrett, Sonia Goncalves, Ewan Harrison, David K. Jackson, Ian Johnston, Dominic Kwiatkowski, Cordelia Langford, John Sillitoe on behalf of the Wellcome Sanger Institute COVID-19 Surveillance Team |
| EPI_ISL_2865531 | Lighthouse Lab in Milton Keynes | Wellcome Sanger Institute for the COVID-19 Genomics UK (COG-UK) Consortium | The Lighthouse Lab in Milton Keynes and Alex Alderton, Roberto Amato, Jeffrey Barrett, Sonia Goncalves, Ewan Harrison, David K. Jackson, Ian Johnston, Dominic Kwiatkowski, Cordelia Langford, John Sillitoe on behalf of the Wellcome Sanger Institute COVID-19 Surveillance Team |
| EPI_ISL_2865532, EPI_ISL_2865533 | Lighthouse Lab in Glasgow | Wellcome Sanger Institute for the COVID-19 Genomics UK (COG-UK) Consortium | Harper VanSteenhouse, Yumi Kasai, David Gray, Carol Clugston, Anna Dominiczak and Alex Alderton, Roberto Amato, Jeffrey Barrett, Sonia Goncalves, Ewan Harrison, David K. Jackson, Ian Johnston, Dominic Kwiatkowski, Cordelia Langford, John Sillitoe on behalf of the Wellcome Sanger Institute COVID-19 Surveillance Team |
| EPI_ISL_2865534 | Lighthouse Lab in Milton Keynes | Wellcome Sanger Institute for the COVID-19 Genomics UK (COG-UK) Consortium | The Lighthouse Lab in Milton Keynes and Alex Alderton, Roberto Amato, Jeffrey Barrett, Sonia Goncalves, Ewan Harrison, David K. Jackson, Ian Johnston, Dominic Kwiatkowski, Cordelia Langford, John Sillitoe on behalf of the Wellcome Sanger Institute COVID-19 Surveillance Team |
| EPI_ISL_2865535 | Lighthouse Lab in Glasgow | Wellcome Sanger Institute for the COVID-19 Genomics UK (COG-UK) Consortium | Harper VanSteenhouse, Yumi Kasai, David Gray, Carol Clugston, Anna Dominiczak and Alex Alderton, Roberto Amato, Jeffrey Barrett, Sonia Goncalves, Ewan Harrison, David K. Jackson, Ian Johnston, Dominic Kwiatkowski, Cordelia Langford, John Sillitoe on behalf of the Wellcome Sanger Institute COVID-19 Surveillance Team |
| EPI_ISL_2865536, EPI_ISL_2865537, EPI_ISL_2865538 | Lighthouse Lab in Milton Keynes | Wellcome Sanger Institute for the COVID-19 Genomics UK (COG-UK) Consortium | The Lighthouse Lab in Milton Keynes and Alex Alderton, Roberto Amato, Jeffrey Barrett, Sonia Goncalves, Ewan Harrison, David K. Jackson, Ian Johnston, Dominic Kwiatkowski, Cordelia Langford, John Sillitoe on behalf of the Wellcome Sanger Institute COVID-19 Surveillance Team |
| EPI_ISL_2865539, EPI_ISL_2865540, EPI_ISL_2865541 | Lighthouse Lab in Glasgow | Wellcome Sanger Institute for the COVID-19 Genomics UK (COG-UK) Consortium | Harper VanSteenhouse, Yumi Kasai, David Gray, Carol Clugston, Anna Dominiczak and Alex Alderton, Roberto Amato, Jeffrey Barrett, Sonia Goncalves, Ewan Harrison, David K. Jackson, Ian Johnston, Dominic Kwiatkowski, Cordelia Langford, John Sillitoe on behalf of the Wellcome Sanger Institute COVID-19 Surveillance Team |
| EPI_ISL_2865542 | Lighthouse Lab in Milton Keynes | Wellcome Sanger Institute for the COVID-19 Genomics UK (COG-UK) Consortium | The Lighthouse Lab in Milton Keynes and Alex Alderton, Roberto Amato, Jeffrey Barrett, Sonia Goncalves, Ewan Harrison, David K. Jackson, Ian Johnston, Dominic Kwiatkowski, Cordelia Langford, John Sillitoe on behalf of the Wellcome Sanger Institute COVID-19 Surveillance Team |
| EPI_ISL_2865543, EPI_ISL_2865544 | Lighthouse Lab in Glasgow | Wellcome Sanger Institute for the COVID-19 Genomics UK (COG-UK) Consortium | Harper VanSteenhouse, Yumi Kasai, David Gray, Carol Clugston, Anna Dominiczak and Alex Alderton, Roberto Amato, Jeffrey Barrett, Sonia Goncalves, Ewan Harrison, David K. Jackson, Ian Johnston, Dominic Kwiatkowski, Cordelia Langford, John Sillitoe on behalf of the Wellcome Sanger Institute COVID-19 Surveillance Team |
| EPI_ISL_2865545 | Lighthouse Lab in Milton Keynes | Wellcome Sanger Institute for the COVID-19 Genomics UK (COG-UK) Consortium | The Lighthouse Lab in Milton Keynes and Alex Alderton, Roberto Amato, Jeffrey Barrett, Sonia Goncalves, Ewan Harrison, David K. Jackson, Ian Johnston, Dominic Kwiatkowski, Cordelia Langford, John Sillitoe on behalf of the Wellcome Sanger Institute COVID-19 Surveillance Team |
| EPI_ISL_2865546 | Lighthouse Lab in Glasgow | Wellcome Sanger Institute for the COVID-19 Genomics UK (COG-UK) Consortium | Harper VanSteenhouse, Yumi Kasai, David Gray, Carol Clugston, Anna Dominiczak and Alex Alderton, Roberto Amato, Jeffrey Barrett, Sonia Goncalves, Ewan Harrison, David K. Jackson, Ian Johnston, Dominic Kwiatkowski, Cordelia Langford, John Sillitoe on behalf of the Wellcome Sanger Institute COVID-19 Surveillance Team |
| EPI_ISL_2865547 | Lighthouse Lab in Alderley Park | Wellcome Sanger Institute for the COVID-19 Genomics UK (COG-UK) Consortium | Jacquelyn Wynn, Mairead Hyland, The Lighthouse Lab in Alderley Park and Alex Alderton, Roberto Amato, Jeffrey Barrett, Sonia Goncalves, Ewan Harrison, David K. Jackson, Ian Johnston, Dominic Kwiatkowski, Cordelia Langford, John Sillitoe on behalf of the Wellcome Sanger Institute COVID-19 Surveillance Team |
| EPI_ISL_2865548 | Lighthouse Lab in Milton Keynes | Wellcome Sanger Institute for the COVID-19 Genomics UK (COG-UK) Consortium | The Lighthouse Lab in Milton Keynes and Alex Alderton, Roberto Amato, Jeffrey Barrett, Sonia Goncalves, Ewan Harrison, David K. Jackson, Ian Johnston, Dominic Kwiatkowski, Cordelia Langford, John Sillitoe on behalf of the Wellcome Sanger Institute COVID-19 Surveillance Team |
| EPI_ISL_2865549 | Lighthouse Lab in Glasgow | Wellcome Sanger Institute for the COVID-19 Genomics UK (COG-UK) Consortium | Harper VanSteenhouse, Yumi Kasai, David Gray, Carol Clugston, Anna Dominiczak and Alex Alderton, Roberto Amato, Jeffrey Barrett, Sonia Goncalves, Ewan Harrison, David K. Jackson, Ian Johnston, Dominic Kwiatkowski, Cordelia Langford, John Sillitoe on behalf of the Wellcome Sanger Institute COVID-19 Surveillance Team |
| EPI_ISL_2865550 | Lighthouse Lab in Milton Keynes | Wellcome Sanger Institute for the COVID-19 Genomics UK (COG-UK) Consortium | The Lighthouse Lab in Milton Keynes and Alex Alderton, Roberto Amato, Jeffrey Barrett, Sonia Goncalves, Ewan Harrison, David K. Jackson, Ian Johnston, Dominic Kwiatkowski, Cordelia Langford, John Sillitoe on behalf of the Wellcome Sanger Institute COVID-19 Surveillance Team |
| EPI_ISL_2865551, EPI_ISL_2865552 | Lighthouse Lab in Glasgow | Wellcome Sanger Institute for the COVID-19 Genomics UK (COG-UK) Consortium | Harper VanSteenhouse, Yumi Kasai, David Gray, Carol Clugston, Anna Dominiczak and Alex Alderton, Roberto Amato, Jeffrey Barrett, Sonia Goncalves, Ewan Harrison, David K. Jackson, Ian Johnston, Dominic Kwiatkowski, Cordelia Langford, John Sillitoe on behalf of the Wellcome Sanger Institute COVID-19 Surveillance Team |
| EPI_ISL_2865553 | Lighthouse Lab in Milton Keynes | Wellcome Sanger Institute for the COVID-19 Genomics UK (COG-UK) Consortium | The Lighthouse Lab in Milton Keynes and Alex Alderton, Roberto Amato, Jeffrey Barrett, Sonia Goncalves, Ewan Harrison, David K. Jackson, Ian Johnston, Dominic Kwiatkowski, Cordelia Langford, John Sillitoe on behalf of the Wellcome Sanger Institute COVID-19 Surveillance Team |
| EPI_ISL_2865554, EPI_ISL_2865555 | Lighthouse Lab in Glasgow | Wellcome Sanger Institute for the COVID-19 Genomics UK (COG-UK) Consortium | Harper VanSteenhouse, Yumi Kasai, David Gray, Carol Clugston, Anna Dominiczak and Alex Alderton, Roberto Amato, Jeffrey Barrett, Sonia Goncalves, Ewan Harrison, David K. Jackson, Ian Johnston, Dominic Kwiatkowski, Cordelia Langford, John Sillitoe on behalf of the Wellcome Sanger Institute COVID-19 Surveillance Team |
| EPI_ISL_2865556 | Lighthouse Lab in Milton Keynes | Wellcome Sanger Institute for the COVID-19 Genomics UK (COG-UK) Consortium | The Lighthouse Lab in Milton Keynes and Alex Alderton, Roberto Amato, Jeffrey Barrett, Sonia Goncalves, Ewan Harrison, David K. Jackson, Ian Johnston, Dominic Kwiatkowski, Cordelia Langford, John Sillitoe on behalf of the Wellcome Sanger Institute COVID-19 Surveillance Team |
| EPI_ISL_2865557 | Lighthouse Lab in Glasgow | Wellcome Sanger Institute for the COVID-19 Genomics UK (COG-UK) | Harper VanSteenhouse, Yumi Kasai, David Gray, Carol Clugston, Anna Dominiczak and Alex Alderton, Roberto Amato, Jeffrey Barrett, Sonia Goncalves, Ewan Harrison, David K. Jackson, Ian Johnston, Dominic Kwiatkowski, Cordelia Langford, |

| | | | |
|---|---|---|---|
| | | Consortium | John Sillitoe on behalf of the Wellcome Sanger Institute COVID-19 Surveillance Team |
| EPI_ISL_2865558 | Lighthouse Lab in Milton Keynes | Wellcome Sanger Institute for the COVID-19 Genomics UK (COG-UK) Consortium | The Lighthouse Lab in Milton Keynes and Alex Alderton, Roberto Amato, Jeffrey Barrett, Sonia Goncalves, Ewan Harrison, David K. Jackson, Ian Johnston, Dominic Kwiatkowski, Cordelia Langford, John Sillitoe on behalf of the Wellcome Sanger Institute COVID-19 Surveillance Team |
| EPI_ISL_2865559 | Lighthouse Lab in Glasgow | Wellcome Sanger Institute for the COVID-19 Genomics UK (COG-UK) Consortium | Harper VanSteenhouse, Yumi Kasai, David Gray, Carol Clugston, Anna Dominiczak and Alex Alderton, Roberto Amato, Jeffrey Barrett, Sonia Goncalves, Ewan Harrison, David K. Jackson, Ian Johnston, Dominic Kwiatkowski, Cordelia Langford, John Sillitoe on behalf of the Wellcome Sanger Institute COVID-19 Surveillance Team |
| EPI_ISL_2865560 | Lighthouse Lab in Milton Keynes | Wellcome Sanger Institute for the COVID-19 Genomics UK (COG-UK) Consortium | The Lighthouse Lab in Milton Keynes and Alex Alderton, Roberto Amato, Jeffrey Barrett, Sonia Goncalves, Ewan Harrison, David K. Jackson, Ian Johnston, Dominic Kwiatkowski, Cordelia Langford, John Sillitoe on behalf of the Wellcome Sanger Institute COVID-19 Surveillance Team |
| EPI_ISL_2865561, EPI_ISL_2865562, EPI_ISL_2865563 | Lighthouse Lab in Glasgow | Wellcome Sanger Institute for the COVID-19 Genomics UK (COG-UK) Consortium | Harper VanSteenhouse, Yumi Kasai, David Gray, Carol Clugston, Anna Dominiczak and Alex Alderton, Roberto Amato, Jeffrey Barrett, Sonia Goncalves, Ewan Harrison, David K. Jackson, Ian Johnston, Dominic Kwiatkowski, Cordelia Langford, John Sillitoe on behalf of the Wellcome Sanger Institute COVID-19 Surveillance Team |
| EPI_ISL_2865564, EPI_ISL_2865565, EPI_ISL_2865566 | Lighthouse Lab in Milton Keynes | Wellcome Sanger Institute for the COVID-19 Genomics UK (COG-UK) Consortium | The Lighthouse Lab in Milton Keynes and Alex Alderton, Roberto Amato, Jeffrey Barrett, Sonia Goncalves, Ewan Harrison, David K. Jackson, Ian Johnston, Dominic Kwiatkowski, Cordelia Langford, John Sillitoe on behalf of the Wellcome Sanger Institute COVID-19 Surveillance Team |
| EPI_ISL_2865567, EPI_ISL_2865568, EPI_ISL_2865569 | Lighthouse Lab in Glasgow | Wellcome Sanger Institute for the COVID-19 Genomics UK (COG-UK) Consortium | Harper VanSteenhouse, Yumi Kasai, David Gray, Carol Clugston, Anna Dominiczak and Alex Alderton, Roberto Amato, Jeffrey Barrett, Sonia Goncalves, Ewan Harrison, David K. Jackson, Ian Johnston, Dominic Kwiatkowski, Cordelia Langford, John Sillitoe on behalf of the Wellcome Sanger Institute COVID-19 Surveillance Team |
| EPI_ISL_2865570 | Lighthouse Lab in Milton Keynes | Wellcome Sanger Institute for the COVID-19 Genomics UK (COG-UK) Consortium | The Lighthouse Lab in Milton Keynes and Alex Alderton, Roberto Amato, Jeffrey Barrett, Sonia Goncalves, Ewan Harrison, David K. Jackson, Ian Johnston, Dominic Kwiatkowski, Cordelia Langford, John Sillitoe on behalf of the Wellcome Sanger Institute COVID-19 Surveillance Team |
| EPI_ISL_2865571, EPI_ISL_2865572 | Lighthouse Lab in Glasgow | Wellcome Sanger Institute for the COVID-19 Genomics UK (COG-UK) Consortium | Harper VanSteenhouse, Yumi Kasai, David Gray, Carol Clugston, Anna Dominiczak and Alex Alderton, Roberto Amato, Jeffrey Barrett, Sonia Goncalves, Ewan Harrison, David K. Jackson, Ian Johnston, Dominic Kwiatkowski, Cordelia Langford, John Sillitoe on behalf of the Wellcome Sanger Institute COVID-19 Surveillance Team |
| EPI_ISL_2865573 | Lighthouse Lab in Milton Keynes | Wellcome Sanger Institute for the COVID-19 Genomics UK (COG-UK) Consortium | The Lighthouse Lab in Milton Keynes and Alex Alderton, Roberto Amato, Jeffrey Barrett, Sonia Goncalves, Ewan Harrison, David K. Jackson, Ian Johnston, Dominic Kwiatkowski, Cordelia Langford, John Sillitoe on behalf of the Wellcome Sanger Institute COVID-19 Surveillance Team |
| EPI_ISL_2865574, EPI_ISL_2865575 | Lighthouse Lab in Glasgow | Wellcome Sanger Institute for the COVID-19 Genomics UK (COG-UK) Consortium | Harper VanSteenhouse, Yumi Kasai, David Gray, Carol Clugston, Anna Dominiczak and Alex Alderton, Roberto Amato, Jeffrey Barrett, Sonia Goncalves, Ewan Harrison, David K. Jackson, Ian Johnston, Dominic Kwiatkowski, Cordelia Langford, John Sillitoe on behalf of the Wellcome Sanger Institute COVID-19 Surveillance Team |
| EPI_ISL_2865576 | Lighthouse Lab in Milton Keynes | Wellcome Sanger Institute for the COVID-19 Genomics UK (COG-UK) Consortium | The Lighthouse Lab in Milton Keynes and Alex Alderton, Roberto Amato, Jeffrey Barrett, Sonia Goncalves, Ewan Harrison, David K. Jackson, Ian Johnston, Dominic Kwiatkowski, Cordelia Langford, John Sillitoe on behalf of the Wellcome Sanger Institute COVID-19 Surveillance Team |
| EPI_ISL_2865577 | Lighthouse Lab in Glasgow | Wellcome Sanger Institute for the COVID-19 Genomics UK (COG-UK) Consortium | Harper VanSteenhouse, Yumi Kasai, David Gray, Carol Clugston, Anna Dominiczak and Alex Alderton, Roberto Amato, Jeffrey Barrett, Sonia Goncalves, Ewan Harrison, David K. Jackson, Ian Johnston, Dominic Kwiatkowski, Cordelia Langford, John Sillitoe on behalf of the Wellcome Sanger Institute COVID-19 Surveillance Team |
| EPI_ISL_2865578 | Lighthouse Lab in Milton Keynes | Wellcome Sanger Institute for the COVID-19 Genomics UK (COG-UK) Consortium | The Lighthouse Lab in Milton Keynes and Alex Alderton, Roberto Amato, Jeffrey Barrett, Sonia Goncalves, Ewan Harrison, David K. Jackson, Ian Johnston, Dominic Kwiatkowski, Cordelia Langford, John Sillitoe on behalf of the Wellcome Sanger Institute COVID-19 Surveillance Team |
| EPI_ISL_2865579, EPI_ISL_2865580, EPI_ISL_2865581, EPI_ISL_2865582 | Lighthouse Lab in Glasgow | Wellcome Sanger Institute for the COVID-19 Genomics UK (COG-UK) Consortium | Harper VanSteenhouse, Yumi Kasai, David Gray, Carol Clugston, Anna Dominiczak and Alex Alderton, Roberto Amato, Jeffrey Barrett, Sonia Goncalves, Ewan Harrison, David K. Jackson, Ian Johnston, Dominic Kwiatkowski, Cordelia Langford, John Sillitoe on behalf of the Wellcome Sanger Institute COVID-19 Surveillance Team |
| EPI_ISL_2865583, EPI_ISL_2865584 | Lighthouse Lab in Milton Keynes | Wellcome Sanger Institute for the COVID-19 Genomics UK (COG-UK) Consortium | The Lighthouse Lab in Milton Keynes and Alex Alderton, Roberto Amato, Jeffrey Barrett, Sonia Goncalves, Ewan Harrison, David K. Jackson, Ian Johnston, Dominic Kwiatkowski, Cordelia Langford, John Sillitoe on behalf of the Wellcome Sanger Institute COVID-19 Surveillance Team |
| EPI_ISL_2865585 | Lighthouse Lab in Glasgow | Wellcome Sanger Institute for the COVID-19 Genomics UK (COG-UK) Consortium | Harper VanSteenhouse, Yumi Kasai, David Gray, Carol Clugston, Anna Dominiczak and Alex Alderton, Roberto Amato, Jeffrey Barrett, Sonia Goncalves, Ewan Harrison, David K. Jackson, Ian Johnston, Dominic Kwiatkowski, Cordelia Langford, John Sillitoe on behalf of the Wellcome Sanger Institute COVID-19 Surveillance Team |
| EPI_ISL_2865586, EPI_ISL_2865587 | Lighthouse Lab in Milton Keynes | Wellcome Sanger Institute for the COVID-19 Genomics UK (COG-UK) Consortium | The Lighthouse Lab in Milton Keynes and Alex Alderton, Roberto Amato, Jeffrey Barrett, Sonia Goncalves, Ewan Harrison, David K. Jackson, Ian Johnston, Dominic Kwiatkowski, Cordelia Langford, John Sillitoe on behalf of the Wellcome Sanger Institute COVID-19 Surveillance Team |
| EPI_ISL_2865588, EPI_ISL_2865589 | Lighthouse Lab in Glasgow | Wellcome Sanger Institute for the COVID-19 Genomics UK (COG-UK) Consortium | Harper VanSteenhouse, Yumi Kasai, David Gray, Carol Clugston, Anna Dominiczak and Alex Alderton, Roberto Amato, Jeffrey Barrett, Sonia Goncalves, Ewan Harrison, David K. Jackson, Ian Johnston, Dominic Kwiatkowski, Cordelia Langford, John Sillitoe on behalf of the Wellcome Sanger Institute COVID-19 Surveillance Team |
| EPI_ISL_2865590 | Lighthouse Lab in Milton Keynes | Wellcome Sanger Institute for the COVID-19 Genomics UK (COG-UK) Consortium | The Lighthouse Lab in Milton Keynes and Alex Alderton, Roberto Amato, Jeffrey Barrett, Sonia Goncalves, Ewan Harrison, David K. Jackson, Ian Johnston, Dominic Kwiatkowski, Cordelia Langford, John Sillitoe on behalf of the Wellcome Sanger Institute COVID-19 Surveillance Team |
| EPI_ISL_2865591 | Lighthouse Lab in Glasgow | Wellcome Sanger Institute for the COVID-19 Genomics UK (COG-UK) Consortium | Harper VanSteenhouse, Yumi Kasai, David Gray, Carol Clugston, Anna Dominiczak and Alex Alderton, Roberto Amato, Jeffrey Barrett, Sonia Goncalves, Ewan Harrison, David K. Jackson, Ian Johnston, Dominic Kwiatkowski, Cordelia Langford, John Sillitoe on behalf of the Wellcome Sanger Institute COVID-19 Surveillance Team |
| EPI_ISL_2865592, EPI_ISL_2865593 | Lighthouse Lab in Milton Keynes | Wellcome Sanger Institute for the COVID-19 Genomics UK (COG-UK) Consortium | The Lighthouse Lab in Milton Keynes and Alex Alderton, Roberto Amato, Jeffrey Barrett, Sonia Goncalves, Ewan Harrison, David K. Jackson, Ian Johnston, Dominic Kwiatkowski, Cordelia Langford, John Sillitoe on behalf of the Wellcome Sanger Institute COVID-19 Surveillance Team |
| EPI_ISL_2865594 | Lighthouse Lab in Glasgow | Wellcome Sanger Institute for the COVID-19 Genomics UK (COG-UK) Consortium | Harper VanSteenhouse, Yumi Kasai, David Gray, Carol Clugston, Anna Dominiczak and Alex Alderton, Roberto Amato, Jeffrey Barrett, Sonia Goncalves, Ewan Harrison, David K. Jackson, Ian Johnston, Dominic Kwiatkowski, Cordelia Langford, John Sillitoe on behalf of the Wellcome Sanger Institute COVID-19 Surveillance Team |
| EPI_ISL_2865595, EPI_ISL_2865596 | Lighthouse Lab in Milton Keynes | Wellcome Sanger Institute for the COVID-19 Genomics UK (COG-UK) Consortium | The Lighthouse Lab in Milton Keynes and Alex Alderton, Roberto Amato, Jeffrey Barrett, Sonia Goncalves, Ewan Harrison, David K. Jackson, Ian Johnston, Dominic Kwiatkowski, Cordelia Langford, John Sillitoe on behalf of the Wellcome Sanger Institute COVID-19 Surveillance Team |
| EPI_ISL_2865597 | Lighthouse Lab in Glasgow | Wellcome Sanger Institute for the COVID-19 Genomics UK (COG-UK) Consortium | Harper VanSteenhouse, Yumi Kasai, David Gray, Carol Clugston, Anna Dominiczak and Alex Alderton, Roberto Amato, Jeffrey Barrett, Sonia Goncalves, Ewan Harrison, David K. Jackson, Ian Johnston, Dominic Kwiatkowski, Cordelia Langford, John Sillitoe on behalf of the Wellcome Sanger Institute COVID-19 Surveillance Team |
| EPI_ISL_2865598, EPI_ISL_2865599, EPI_ISL_2865600, EPI_ISL_2865601 | Lighthouse Lab in Milton Keynes | Wellcome Sanger Institute for the COVID-19 Genomics UK (COG-UK) Consortium | The Lighthouse Lab in Milton Keynes and Alex Alderton, Roberto Amato, Jeffrey Barrett, Sonia Goncalves, Ewan Harrison, David K. Jackson, Ian Johnston, Dominic Kwiatkowski, Cordelia Langford, John Sillitoe on behalf of the Wellcome Sanger Institute COVID-19 Surveillance Team |
| EPI_ISL_2865602, EPI_ISL_2865603, EPI_ISL_2865604, EPI_ISL_2865605 | Lighthouse Lab in Glasgow | Wellcome Sanger Institute for the COVID-19 Genomics UK (COG-UK) Consortium | Harper VanSteenhouse, Yumi Kasai, David Gray, Carol Clugston, Anna Dominiczak and Alex Alderton, Roberto Amato, Jeffrey Barrett, Sonia Goncalves, Ewan Harrison, David K. Jackson, Ian Johnston, Dominic Kwiatkowski, Cordelia Langford, John Sillitoe on behalf of the Wellcome Sanger Institute COVID-19 Surveillance Team |
| EPI_ISL_2865606, EPI_ISL_2865607 | Lighthouse Lab in Milton Keynes | Wellcome Sanger Institute for the COVID-19 Genomics UK (COG-UK) Consortium | The Lighthouse Lab in Milton Keynes and Alex Alderton, Roberto Amato, Jeffrey Barrett, Sonia Goncalves, Ewan Harrison, David K. Jackson, Ian Johnston, Dominic Kwiatkowski, Cordelia Langford, John Sillitoe on behalf of the Wellcome Sanger Institute COVID-19 Surveillance Team |
| EPI_ISL_2865608 | Lighthouse Lab in Alderley Park | Wellcome Sanger Institute for the COVID-19 Genomics UK (COG-UK) Consortium | Jacquelyn Wynn, Mairead Hyland, The Lighthouse Lab in Alderley Park and Alex Alderton, Roberto Amato, Jeffrey Barrett, Sonia Goncalves, Ewan Harrison, David K. Jackson, Ian Johnston, Dominic Kwiatkowski, Cordelia Langford, John Sillitoe on behalf of the Wellcome Sanger Institute COVID-19 Surveillance Team |
| EPI_ISL_2865609, EPI_ISL_2865610 | Lighthouse Lab in Milton Keynes | Wellcome Sanger Institute for the COVID-19 Genomics UK (COG-UK) Consortium | The Lighthouse Lab in Milton Keynes and Alex Alderton, Roberto Amato, Jeffrey Barrett, Sonia Goncalves, Ewan Harrison, David K. Jackson, Ian Johnston, Dominic Kwiatkowski, Cordelia Langford, John Sillitoe on behalf of the Wellcome Sanger Institute COVID-19 Surveillance Team |
| EPI_ISL_2865611, EPI_ISL_2865612 | Lighthouse Lab in Glasgow | Wellcome Sanger Institute for the COVID-19 Genomics UK (COG-UK) Consortium | Harper VanSteenhouse, Yumi Kasai, David Gray, Carol Clugston, Anna Dominiczak and Alex Alderton, Roberto Amato, Jeffrey Barrett, Sonia Goncalves, Ewan Harrison, David K. Jackson, Ian Johnston, Dominic Kwiatkowski, Cordelia Langford, John Sillitoe on behalf of the Wellcome Sanger Institute COVID-19 Surveillance Team |
| EPI_ISL_2865613 | Lighthouse Lab in Milton Keynes | Wellcome Sanger Institute for the COVID-19 Genomics UK (COG-UK) Consortium | The Lighthouse Lab in Milton Keynes and Alex Alderton, Roberto Amato, Jeffrey Barrett, Sonia Goncalves, Ewan Harrison, David K. Jackson, Ian Johnston, Dominic Kwiatkowski, Cordelia Langford, John Sillitoe on behalf of the Wellcome Sanger Institute COVID-19 Surveillance Team |
| EPI_ISL_2865614, EPI_ISL_2865615 | Lighthouse Lab in Glasgow | Wellcome Sanger Institute for the COVID-19 Genomics UK (COG-UK) Consortium | Harper VanSteenhouse, Yumi Kasai, David Gray, Carol Clugston, Anna Dominiczak and Alex Alderton, Roberto Amato, Jeffrey Barrett, Sonia Goncalves, Ewan Harrison, David K. Jackson, Ian Johnston, Dominic Kwiatkowski, Cordelia Langford, John Sillitoe on behalf of the Wellcome Sanger Institute COVID-19 Surveillance Team |
| EPI_ISL_2865616 | Lighthouse Lab in Milton Keynes | Wellcome Sanger Institute for the COVID-19 Genomics UK (COG-UK) Consortium | The Lighthouse Lab in Milton Keynes and Alex Alderton, Roberto Amato, Jeffrey Barrett, Sonia Goncalves, Ewan Harrison, David K. Jackson, Ian Johnston, Dominic Kwiatkowski, Cordelia Langford, John Sillitoe on behalf of the Wellcome Sanger Institute COVID-19 Surveillance Team |
| EPI_ISL_2865617, EPI_ISL_2865618, EPI_ISL_2865619, EPI_ISL_2865620 | Lighthouse Lab in Glasgow | Wellcome Sanger Institute for the COVID-19 Genomics UK (COG-UK) Consortium | Harper VanSteenhouse, Yumi Kasai, David Gray, Carol Clugston, Anna Dominiczak and Alex Alderton, Roberto Amato, Jeffrey Barrett, Sonia Goncalves, Ewan Harrison, David K. Jackson, Ian Johnston, Dominic Kwiatkowski, Cordelia Langford, John Sillitoe on behalf of the Wellcome Sanger Institute COVID-19 Surveillance Team |
| EPI_ISL_2865621, EPI_ISL_2865622, EPI_ISL_2865623, EPI_ISL_2865624 | Lighthouse Lab in Milton Keynes | Wellcome Sanger Institute for the COVID-19 Genomics UK (COG-UK) Consortium | The Lighthouse Lab in Milton Keynes and Alex Alderton, Roberto Amato, Jeffrey Barrett, Sonia Goncalves, Ewan Harrison, David K. Jackson, Ian Johnston, Dominic Kwiatkowski, Cordelia Langford, John Sillitoe on behalf of the Wellcome Sanger Institute COVID-19 Surveillance Team |
| EPI_ISL_2865625 | Lighthouse Lab in Glasgow | Wellcome Sanger Institute for the COVID-19 Genomics UK (COG-UK) Consortium | Harper VanSteenhouse, Yumi Kasai, David Gray, Carol Clugston, Anna Dominiczak and Alex Alderton, Roberto Amato, Jeffrey Barrett, Sonia Goncalves, Ewan Harrison, David K. Jackson, Ian Johnston, Dominic Kwiatkowski, Cordelia Langford, John Sillitoe on behalf of the Wellcome Sanger Institute COVID-19 Surveillance Team |
| EPI_ISL_2865626 | Lighthouse Lab in Milton Keynes | Wellcome Sanger Institute for the COVID-19 Genomics UK (COG-UK) Consortium | The Lighthouse Lab in Milton Keynes and Alex Alderton, Roberto Amato, Jeffrey Barrett, Sonia Goncalves, Ewan Harrison, David K. Jackson, Ian Johnston, Dominic Kwiatkowski, Cordelia Langford, John Sillitoe on behalf of the Wellcome Sanger Institute COVID-19 Surveillance Team |
| EPI_ISL_2865627 | Lighthouse Lab in Glasgow | Wellcome Sanger Institute for the COVID-19 Genomics UK (COG-UK) Consortium | Harper VanSteenhouse, Yumi Kasai, David Gray, Carol Clugston, Anna Dominiczak and Alex Alderton, Roberto Amato, Jeffrey Barrett, Sonia Goncalves, Ewan Harrison, David K. Jackson, Ian Johnston, Dominic Kwiatkowski, Cordelia Langford, John Sillitoe on behalf of the Wellcome Sanger Institute COVID-19 Surveillance Team |
| EPI_ISL_2865628 | Lighthouse Lab in Milton Keynes | Wellcome Sanger Institute for the COVID-19 Genomics UK (COG-UK) Consortium | The Lighthouse Lab in Milton Keynes and Alex Alderton, Roberto Amato, Jeffrey Barrett, Sonia Goncalves, Ewan Harrison, David K. Jackson, Ian Johnston, Dominic Kwiatkowski, Cordelia Langford, John Sillitoe on behalf of the Wellcome Sanger Institute COVID-19 Surveillance Team |

| | | | |
|---|---|---|---|
| EPI_ISL_2865629 | Lighthouse Lab in Glasgow | Wellcome Sanger Institute for the COVID-19 Genomics UK (COG-UK) Consortium | Harper VanSteenhouse, Yumi Kasai, David Gray, Carol Clugston, Anna Dominiczak and Alex Alderton, Roberto Amato, Jeffrey Barrett, Sonia Goncalves, Ewan Harrison, David K. Jackson, Ian Johnston, Dominic Kwiatkowski, Cordelia Langford, John Sillitoe on behalf of the Wellcome Sanger Institute COVID-19 Surveillance Team |
| EPI_ISL_2865630, EPI_ISL_2865631 | Lighthouse Lab in Milton Keynes | Wellcome Sanger Institute for the COVID-19 Genomics UK (COG-UK) Consortium | The Lighthouse Lab in Milton Keynes and Alex Alderton, Roberto Amato, Jeffrey Barrett, Sonia Goncalves, Ewan Harrison, David K. Jackson, Ian Johnston, Dominic Kwiatkowski, Cordelia Langford, John Sillitoe on behalf of the Wellcome Sanger Institute COVID-19 Surveillance Team |
| EPI_ISL_2865632, EPI_ISL_2865633, EPI_ISL_2865634 | Lighthouse Lab in Glasgow | Wellcome Sanger Institute for the COVID-19 Genomics UK (COG-UK) Consortium | Harper VanSteenhouse, Yumi Kasai, David Gray, Carol Clugston, Anna Dominiczak and Alex Alderton, Roberto Amato, Jeffrey Barrett, Sonia Goncalves, Ewan Harrison, David K. Jackson, Ian Johnston, Dominic Kwiatkowski, Cordelia Langford, John Sillitoe on behalf of the Wellcome Sanger Institute COVID-19 Surveillance Team |
| EPI_ISL_2865635 | Lighthouse Lab in Milton Keynes | Wellcome Sanger Institute for the COVID-19 Genomics UK (COG-UK) Consortium | The Lighthouse Lab in Milton Keynes and Alex Alderton, Roberto Amato, Jeffrey Barrett, Sonia Goncalves, Ewan Harrison, David K. Jackson, Ian Johnston, Dominic Kwiatkowski, Cordelia Langford, John Sillitoe on behalf of the Wellcome Sanger Institute COVID-19 Surveillance Team |
| EPI_ISL_2865636 | Lighthouse Lab in Glasgow | Wellcome Sanger Institute for the COVID-19 Genomics UK (COG-UK) Consortium | Harper VanSteenhouse, Yumi Kasai, David Gray, Carol Clugston, Anna Dominiczak and Alex Alderton, Roberto Amato, Jeffrey Barrett, Sonia Goncalves, Ewan Harrison, David K. Jackson, Ian Johnston, Dominic Kwiatkowski, Cordelia Langford, John Sillitoe on behalf of the Wellcome Sanger Institute COVID-19 Surveillance Team |
| EPI_ISL_2865637, EPI_ISL_2865638 | Lighthouse Lab in Milton Keynes | Wellcome Sanger Institute for the COVID-19 Genomics UK (COG-UK) Consortium | The Lighthouse Lab in Milton Keynes and Alex Alderton, Roberto Amato, Jeffrey Barrett, Sonia Goncalves, Ewan Harrison, David K. Jackson, Ian Johnston, Dominic Kwiatkowski, Cordelia Langford, John Sillitoe on behalf of the Wellcome Sanger Institute COVID-19 Surveillance Team |
| EPI_ISL_2865639 | Lighthouse Lab in Glasgow | Wellcome Sanger Institute for the COVID-19 Genomics UK (COG-UK) Consortium | Harper VanSteenhouse, Yumi Kasai, David Gray, Carol Clugston, Anna Dominiczak and Alex Alderton, Roberto Amato, Jeffrey Barrett, Sonia Goncalves, Ewan Harrison, David K. Jackson, Ian Johnston, Dominic Kwiatkowski, Cordelia Langford, John Sillitoe on behalf of the Wellcome Sanger Institute COVID-19 Surveillance Team |
| EPI_ISL_2865640 | Lighthouse Lab in Milton Keynes | Wellcome Sanger Institute for the COVID-19 Genomics UK (COG-UK) Consortium | The Lighthouse Lab in Milton Keynes and Alex Alderton, Roberto Amato, Jeffrey Barrett, Sonia Goncalves, Ewan Harrison, David K. Jackson, Ian Johnston, Dominic Kwiatkowski, Cordelia Langford, John Sillitoe on behalf of the Wellcome Sanger Institute COVID-19 Surveillance Team |
| EPI_ISL_2865641, EPI_ISL_2865642 | Lighthouse Lab in Glasgow | Wellcome Sanger Institute for the COVID-19 Genomics UK (COG-UK) Consortium | Harper VanSteenhouse, Yumi Kasai, David Gray, Carol Clugston, Anna Dominiczak and Alex Alderton, Roberto Amato, Jeffrey Barrett, Sonia Goncalves, Ewan Harrison, David K. Jackson, Ian Johnston, Dominic Kwiatkowski, Cordelia Langford, John Sillitoe on behalf of the Wellcome Sanger Institute COVID-19 Surveillance Team |
| EPI_ISL_2865643, EPI_ISL_2865644 | Lighthouse Lab in Milton Keynes | Wellcome Sanger Institute for the COVID-19 Genomics UK (COG-UK) Consortium | The Lighthouse Lab in Milton Keynes and Alex Alderton, Roberto Amato, Jeffrey Barrett, Sonia Goncalves, Ewan Harrison, David K. Jackson, Ian Johnston, Dominic Kwiatkowski, Cordelia Langford, John Sillitoe on behalf of the Wellcome Sanger Institute COVID-19 Surveillance Team |
| EPI_ISL_2865645 | Lighthouse Lab in Glasgow | Wellcome Sanger Institute for the COVID-19 Genomics UK (COG-UK) Consortium | Harper VanSteenhouse, Yumi Kasai, David Gray, Carol Clugston, Anna Dominiczak and Alex Alderton, Roberto Amato, Jeffrey Barrett, Sonia Goncalves, Ewan Harrison, David K. Jackson, Ian Johnston, Dominic Kwiatkowski, Cordelia Langford, John Sillitoe on behalf of the Wellcome Sanger Institute COVID-19 Surveillance Team |
| EPI_ISL_2865646 | Lighthouse Lab in Milton Keynes | Wellcome Sanger Institute for the COVID-19 Genomics UK (COG-UK) Consortium | The Lighthouse Lab in Milton Keynes and Alex Alderton, Roberto Amato, Jeffrey Barrett, Sonia Goncalves, Ewan Harrison, David K. Jackson, Ian Johnston, Dominic Kwiatkowski, Cordelia Langford, John Sillitoe on behalf of the Wellcome Sanger Institute COVID-19 Surveillance Team |
| EPI_ISL_2865647 | Lighthouse Lab in Alderley Park | Wellcome Sanger Institute for the COVID-19 Genomics UK (COG-UK) Consortium | Jacquelyn Wynn, Mairead Hyland, The Lighthouse Lab in Alderley Park and Alex Alderton, Roberto Amato, Jeffrey Barrett, Sonia Goncalves, Ewan Harrison, David K. Jackson, Ian Johnston, Dominic Kwiatkowski, Cordelia Langford, John Sillitoe on behalf of the Wellcome Sanger Institute COVID-19 Surveillance Team |
| EPI_ISL_2865648 | Lighthouse Lab in Glasgow | Wellcome Sanger Institute for the COVID-19 Genomics UK (COG-UK) Consortium | Harper VanSteenhouse, Yumi Kasai, David Gray, Carol Clugston, Anna Dominiczak and Alex Alderton, Roberto Amato, Jeffrey Barrett, Sonia Goncalves, Ewan Harrison, David K. Jackson, Ian Johnston, Dominic Kwiatkowski, Cordelia Langford, John Sillitoe on behalf of the Wellcome Sanger Institute COVID-19 Surveillance Team |
| EPI_ISL_2865649, EPI_ISL_2865650, EPI_ISL_2865651 | Lighthouse Lab in Milton Keynes | Wellcome Sanger Institute for the COVID-19 Genomics UK (COG-UK) Consortium | The Lighthouse Lab in Milton Keynes and Alex Alderton, Roberto Amato, Jeffrey Barrett, Sonia Goncalves, Ewan Harrison, David K. Jackson, Ian Johnston, Dominic Kwiatkowski, Cordelia Langford, John Sillitoe on behalf of the Wellcome Sanger Institute COVID-19 Surveillance Team |
| EPI_ISL_2865652 | Lighthouse Lab in Glasgow | Wellcome Sanger Institute for the COVID-19 Genomics UK (COG-UK) Consortium | Harper VanSteenhouse, Yumi Kasai, David Gray, Carol Clugston, Anna Dominiczak and Alex Alderton, Roberto Amato, Jeffrey Barrett, Sonia Goncalves, Ewan Harrison, David K. Jackson, Ian Johnston, Dominic Kwiatkowski, Cordelia Langford, John Sillitoe on behalf of the Wellcome Sanger Institute COVID-19 Surveillance Team |
| EPI_ISL_2865653, EPI_ISL_2865654 | Lighthouse Lab in Milton Keynes | Wellcome Sanger Institute for the COVID-19 Genomics UK (COG-UK) Consortium | The Lighthouse Lab in Milton Keynes and Alex Alderton, Roberto Amato, Jeffrey Barrett, Sonia Goncalves, Ewan Harrison, David K. Jackson, Ian Johnston, Dominic Kwiatkowski, Cordelia Langford, John Sillitoe on behalf of the Wellcome Sanger Institute COVID-19 Surveillance Team |
| EPI_ISL_2865655, EPI_ISL_2865656 | Lighthouse Lab in Glasgow | Wellcome Sanger Institute for the COVID-19 Genomics UK (COG-UK) Consortium | Harper VanSteenhouse, Yumi Kasai, David Gray, Carol Clugston, Anna Dominiczak and Alex Alderton, Roberto Amato, Jeffrey Barrett, Sonia Goncalves, Ewan Harrison, David K. Jackson, Ian Johnston, Dominic Kwiatkowski, Cordelia Langford, John Sillitoe on behalf of the Wellcome Sanger Institute COVID-19 Surveillance Team |
| EPI_ISL_2865657, EPI_ISL_2865658 | Lighthouse Lab in Milton Keynes | Wellcome Sanger Institute for the COVID-19 Genomics UK (COG-UK) Consortium | The Lighthouse Lab in Milton Keynes and Alex Alderton, Roberto Amato, Jeffrey Barrett, Sonia Goncalves, Ewan Harrison, David K. Jackson, Ian Johnston, Dominic Kwiatkowski, Cordelia Langford, John Sillitoe on behalf of the Wellcome Sanger Institute COVID-19 Surveillance Team |
| EPI_ISL_2865659 | Lighthouse Lab in Glasgow | Wellcome Sanger Institute for the COVID-19 Genomics UK (COG-UK) Consortium | Harper VanSteenhouse, Yumi Kasai, David Gray, Carol Clugston, Anna Dominiczak and Alex Alderton, Roberto Amato, Jeffrey Barrett, Sonia Goncalves, Ewan Harrison, David K. Jackson, Ian Johnston, Dominic Kwiatkowski, Cordelia Langford, John Sillitoe on behalf of the Wellcome Sanger Institute COVID-19 Surveillance Team |
| EPI_ISL_2865660, EPI_ISL_2865661 | Lighthouse Lab in Milton Keynes | Wellcome Sanger Institute for the COVID-19 Genomics UK (COG-UK) Consortium | The Lighthouse Lab in Milton Keynes and Alex Alderton, Roberto Amato, Jeffrey Barrett, Sonia Goncalves, Ewan Harrison, David K. Jackson, Ian Johnston, Dominic Kwiatkowski, Cordelia Langford, John Sillitoe on behalf of the Wellcome Sanger Institute COVID-19 Surveillance Team |
| EPI_ISL_2865662 | Lighthouse Lab in Alderley Park | Wellcome Sanger Institute for the COVID-19 Genomics UK (COG-UK) Consortium | Jacquelyn Wynn, Mairead Hyland, The Lighthouse Lab in Alderley Park and Alex Alderton, Roberto Amato, Jeffrey Barrett, Sonia Goncalves, Ewan Harrison, David K. Jackson, Ian Johnston, Dominic Kwiatkowski, Cordelia Langford, John Sillitoe on behalf of the Wellcome Sanger Institute COVID-19 Surveillance Team |
| EPI_ISL_2865663, EPI_ISL_2865664, EPI_ISL_2865665 | Lighthouse Lab in Milton Keynes | Wellcome Sanger Institute for the COVID-19 Genomics UK (COG-UK) Consortium | The Lighthouse Lab in Milton Keynes and Alex Alderton, Roberto Amato, Jeffrey Barrett, Sonia Goncalves, Ewan Harrison, David K. Jackson, Ian Johnston, Dominic Kwiatkowski, Cordelia Langford, John Sillitoe on behalf of the Wellcome Sanger Institute COVID-19 Surveillance Team |
| EPI_ISL_2865666, EPI_ISL_2865667, EPI_ISL_2865668, EPI_ISL_2865669, EPI_ISL_2865670, EPI_ISL_2865671 | Lighthouse Lab in Glasgow | Wellcome Sanger Institute for the COVID-19 Genomics UK (COG-UK) Consortium | Harper VanSteenhouse, Yumi Kasai, David Gray, Carol Clugston, Anna Dominiczak and Alex Alderton, Roberto Amato, Jeffrey Barrett, Sonia Goncalves, Ewan Harrison, David K. Jackson, Ian Johnston, Dominic Kwiatkowski, Cordelia Langford, John Sillitoe on behalf of the Wellcome Sanger Institute COVID-19 Surveillance Team |
| EPI_ISL_2865672, EPI_ISL_2865673 | Lighthouse Lab in Milton Keynes | Wellcome Sanger Institute for the COVID-19 Genomics UK (COG-UK) Consortium | The Lighthouse Lab in Milton Keynes and Alex Alderton, Roberto Amato, Jeffrey Barrett, Sonia Goncalves, Ewan Harrison, David K. Jackson, Ian Johnston, Dominic Kwiatkowski, Cordelia Langford, John Sillitoe on behalf of the Wellcome Sanger Institute COVID-19 Surveillance Team |
| EPI_ISL_2865674, EPI_ISL_2865675 | Lighthouse Lab in Glasgow | Wellcome Sanger Institute for the COVID-19 Genomics UK (COG-UK) Consortium | Harper VanSteenhouse, Yumi Kasai, David Gray, Carol Clugston, Anna Dominiczak and Alex Alderton, Roberto Amato, Jeffrey Barrett, Sonia Goncalves, Ewan Harrison, David K. Jackson, Ian Johnston, Dominic Kwiatkowski, Cordelia Langford, John Sillitoe on behalf of the Wellcome Sanger Institute COVID-19 Surveillance Team |
| EPI_ISL_2865676 | Lighthouse Lab in Alderley Park | Wellcome Sanger Institute for the COVID-19 Genomics UK (COG-UK) Consortium | Jacquelyn Wynn, Mairead Hyland, The Lighthouse Lab in Alderley Park and Alex Alderton, Roberto Amato, Jeffrey Barrett, Sonia Goncalves, Ewan Harrison, David K. Jackson, Ian Johnston, Dominic Kwiatkowski, Cordelia Langford, John Sillitoe on behalf of the Wellcome Sanger Institute COVID-19 Surveillance Team |
| EPI_ISL_2865677 | Lighthouse Lab in Milton Keynes | Wellcome Sanger Institute for the COVID-19 Genomics UK (COG-UK) Consortium | The Lighthouse Lab in Milton Keynes and Alex Alderton, Roberto Amato, Jeffrey Barrett, Sonia Goncalves, Ewan Harrison, David K. Jackson, Ian Johnston, Dominic Kwiatkowski, Cordelia Langford, John Sillitoe on behalf of the Wellcome Sanger Institute COVID-19 Surveillance Team |
| EPI_ISL_2865678 | Lighthouse Lab in Glasgow | Wellcome Sanger Institute for the COVID-19 Genomics UK (COG-UK) Consortium | Harper VanSteenhouse, Yumi Kasai, David Gray, Carol Clugston, Anna Dominiczak and Alex Alderton, Roberto Amato, Jeffrey Barrett, Sonia Goncalves, Ewan Harrison, David K. Jackson, Ian Johnston, Dominic Kwiatkowski, Cordelia Langford, John Sillitoe on behalf of the Wellcome Sanger Institute COVID-19 Surveillance Team |
| EPI_ISL_2865679, EPI_ISL_2865680 | Lighthouse Lab in Milton Keynes | Wellcome Sanger Institute for the COVID-19 Genomics UK (COG-UK) Consortium | The Lighthouse Lab in Milton Keynes and Alex Alderton, Roberto Amato, Jeffrey Barrett, Sonia Goncalves, Ewan Harrison, David K. Jackson, Ian Johnston, Dominic Kwiatkowski, Cordelia Langford, John Sillitoe on behalf of the Wellcome Sanger Institute COVID-19 Surveillance Team |
| EPI_ISL_2865681 | Lighthouse Lab in Glasgow | Wellcome Sanger Institute for the COVID-19 Genomics UK (COG-UK) Consortium | Harper VanSteenhouse, Yumi Kasai, David Gray, Carol Clugston, Anna Dominiczak and Alex Alderton, Roberto Amato, Jeffrey Barrett, Sonia Goncalves, Ewan Harrison, David K. Jackson, Ian Johnston, Dominic Kwiatkowski, Cordelia Langford, John Sillitoe on behalf of the Wellcome Sanger Institute COVID-19 Surveillance Team |
| EPI_ISL_2865682 | Lighthouse Lab in Milton Keynes | Wellcome Sanger Institute for the COVID-19 Genomics UK (COG-UK) Consortium | The Lighthouse Lab in Milton Keynes and Alex Alderton, Roberto Amato, Jeffrey Barrett, Sonia Goncalves, Ewan Harrison, David K. Jackson, Ian Johnston, Dominic Kwiatkowski, Cordelia Langford, John Sillitoe on behalf of the Wellcome Sanger Institute COVID-19 Surveillance Team |
| EPI_ISL_2865683, EPI_ISL_2865684, EPI_ISL_2865685, EPI_ISL_2865686 | Lighthouse Lab in Glasgow | Wellcome Sanger Institute for the COVID-19 Genomics UK (COG-UK) Consortium | Harper VanSteenhouse, Yumi Kasai, David Gray, Carol Clugston, Anna Dominiczak and Alex Alderton, Roberto Amato, Jeffrey Barrett, Sonia Goncalves, Ewan Harrison, David K. Jackson, Ian Johnston, Dominic Kwiatkowski, Cordelia Langford, John Sillitoe on behalf of the Wellcome Sanger Institute COVID-19 Surveillance Team |
| EPI_ISL_2865687, EPI_ISL_2865688 | Lighthouse Lab in Milton Keynes | Wellcome Sanger Institute for the COVID-19 Genomics UK (COG-UK) Consortium | The Lighthouse Lab in Milton Keynes and Alex Alderton, Roberto Amato, Jeffrey Barrett, Sonia Goncalves, Ewan Harrison, David K. Jackson, Ian Johnston, Dominic Kwiatkowski, Cordelia Langford, John Sillitoe on behalf of the Wellcome Sanger Institute COVID-19 Surveillance Team |
| EPI_ISL_2865689, EPI_ISL_2865690, EPI_ISL_2865691, EPI_ISL_2865692, EPI_ISL_2865693 | Lighthouse Lab in Glasgow | Wellcome Sanger Institute for the COVID-19 Genomics UK (COG-UK) Consortium | Harper VanSteenhouse, Yumi Kasai, David Gray, Carol Clugston, Anna Dominiczak and Alex Alderton, Roberto Amato, Jeffrey Barrett, Sonia Goncalves, Ewan Harrison, David K. Jackson, Ian Johnston, Dominic Kwiatkowski, Cordelia Langford, John Sillitoe on behalf of the Wellcome Sanger Institute COVID-19 Surveillance Team |
| EPI_ISL_2865694, EPI_ISL_2865695 | Lighthouse Lab in Milton Keynes | Wellcome Sanger Institute for the COVID-19 Genomics UK (COG-UK) Consortium | The Lighthouse Lab in Milton Keynes and Alex Alderton, Roberto Amato, Jeffrey Barrett, Sonia Goncalves, Ewan Harrison, David K. Jackson, Ian Johnston, Dominic Kwiatkowski, Cordelia Langford, John Sillitoe on behalf of the Wellcome Sanger Institute COVID-19 Surveillance Team |
| EPI_ISL_2865696 | Lighthouse Lab in Glasgow | Wellcome Sanger Institute for the COVID-19 Genomics UK (COG-UK) Consortium | Harper VanSteenhouse, Yumi Kasai, David Gray, Carol Clugston, Anna Dominiczak and Alex Alderton, Roberto Amato, Jeffrey Barrett, Sonia Goncalves, Ewan Harrison, David K. Jackson, Ian Johnston, Dominic Kwiatkowski, Cordelia Langford, John Sillitoe on behalf of the Wellcome Sanger Institute COVID-19 Surveillance Team |
| EPI_ISL_2865697 | Lighthouse Lab in Milton Keynes | Wellcome Sanger Institute for the COVID-19 Genomics UK (COG-UK) Consortium | The Lighthouse Lab in Milton Keynes and Alex Alderton, Roberto Amato, Jeffrey Barrett, Sonia Goncalves, Ewan Harrison, David K. Jackson, Ian Johnston, Dominic Kwiatkowski, Cordelia Langford, John Sillitoe on behalf of the Wellcome Sanger Institute COVID-19 Surveillance Team |
| EPI_ISL_2865698, EPI_ISL_2865699 | Lighthouse Lab in Glasgow | Wellcome Sanger Institute for the COVID-19 Genomics UK (COG-UK) Consortium | Harper VanSteenhouse, Yumi Kasai, David Gray, Carol Clugston, Anna Dominiczak and Alex Alderton, Roberto Amato, Jeffrey Barrett, Sonia Goncalves, Ewan Harrison, David K. Jackson, Ian Johnston, Dominic Kwiatkowski, Cordelia Langford, John Sillitoe on behalf of the Wellcome Sanger Institute COVID-19 Surveillance Team |
| EPI_ISL_2865700, EPI_ISL_2865701, | Lighthouse Lab in Milton Keynes | Wellcome Sanger Institute for the COVID-19 Genomics UK (COG-UK) Consortium | The Lighthouse Lab in Milton Keynes and Alex Alderton, Roberto Amato, Jeffrey Barrett, Sonia Goncalves, Ewan Harrison, David K. Jackson, Ian Johnston, Dominic Kwiatkowski, Cordelia Langford, John Sillitoe on behalf of the Wellcome Sanger Institute COVID-19 Surveillance Team |

| | | | |
|---|---|---|---|
| EPI_ISL_2865702, EPI_ISL_2865703 | | | |
| EPI_ISL_2865704 | Lighthouse Lab in Glasgow | Wellcome Sanger Institute for the COVID-19 Genomics UK (COG-UK) Consortium | Harper VanSteenhouse, Yumi Kasai, David Gray, Carol Clugston, Anna Dominiczak and Alex Alderton, Roberto Amato, Jeffrey Barrett, Sonia Goncalves, Ewan Harrison, David K. Jackson, Ian Johnston, Dominic Kwiatkowski, Cordelia Langford, John Sillitoe on behalf of the Wellcome Sanger Institute COVID-19 Surveillance Team |
| EPI_ISL_2865705 | Lighthouse Lab in Milton Keynes | Wellcome Sanger Institute for the COVID-19 Genomics UK (COG-UK) Consortium | The Lighthouse Lab in Milton Keynes and Alex Alderton, Roberto Amato, Jeffrey Barrett, Sonia Goncalves, Ewan Harrison, David K. Jackson, Ian Johnston, Dominic Kwiatkowski, Cordelia Langford, John Sillitoe on behalf of the Wellcome Sanger Institute COVID-19 Surveillance Team |
| EPI_ISL_2865706 | Lighthouse Lab in Glasgow | Wellcome Sanger Institute for the COVID-19 Genomics UK (COG-UK) Consortium | Harper VanSteenhouse, Yumi Kasai, David Gray, Carol Clugston, Anna Dominiczak and Alex Alderton, Roberto Amato, Jeffrey Barrett, Sonia Goncalves, Ewan Harrison, David K. Jackson, Ian Johnston, Dominic Kwiatkowski, Cordelia Langford, John Sillitoe on behalf of the Wellcome Sanger Institute COVID-19 Surveillance Team |
| EPI_ISL_2865707 | Lighthouse Lab in Milton Keynes | Wellcome Sanger Institute for the COVID-19 Genomics UK (COG-UK) Consortium | The Lighthouse Lab in Milton Keynes and Alex Alderton, Roberto Amato, Jeffrey Barrett, Sonia Goncalves, Ewan Harrison, David K. Jackson, Ian Johnston, Dominic Kwiatkowski, Cordelia Langford, John Sillitoe on behalf of the Wellcome Sanger Institute COVID-19 Surveillance Team |
| EPI_ISL_2865708, EPI_ISL_2865709, EPI_ISL_2865710 | Lighthouse Lab in Glasgow | Wellcome Sanger Institute for the COVID-19 Genomics UK (COG-UK) Consortium | Harper VanSteenhouse, Yumi Kasai, David Gray, Carol Clugston, Anna Dominiczak and Alex Alderton, Roberto Amato, Jeffrey Barrett, Sonia Goncalves, Ewan Harrison, David K. Jackson, Ian Johnston, Dominic Kwiatkowski, Cordelia Langford, John Sillitoe on behalf of the Wellcome Sanger Institute COVID-19 Surveillance Team |
| EPI_ISL_2865711 | Lighthouse Lab in Alderley Park | Wellcome Sanger Institute for the COVID-19 Genomics UK (COG-UK) Consortium | Jacquelyn Wynn, Mairead Hyland, The Lighthouse Lab in Alderley Park and Alex Alderton, Roberto Amato, Jeffrey Barrett, Sonia Goncalves, Ewan Harrison, David K. Jackson, Ian Johnston, Dominic Kwiatkowski, Cordelia Langford, John Sillitoe on behalf of the Wellcome Sanger Institute COVID-19 Surveillance Team |
| EPI_ISL_2865712 | Lighthouse Lab in Milton Keynes | Wellcome Sanger Institute for the COVID-19 Genomics UK (COG-UK) Consortium | The Lighthouse Lab in Milton Keynes and Alex Alderton, Roberto Amato, Jeffrey Barrett, Sonia Goncalves, Ewan Harrison, David K. Jackson, Ian Johnston, Dominic Kwiatkowski, Cordelia Langford, John Sillitoe on behalf of the Wellcome Sanger Institute COVID-19 Surveillance Team |
| EPI_ISL_2865713 | Lighthouse Lab in Alderley Park | Wellcome Sanger Institute for the COVID-19 Genomics UK (COG-UK) Consortium | Jacquelyn Wynn, Mairead Hyland, The Lighthouse Lab in Alderley Park and Alex Alderton, Roberto Amato, Jeffrey Barrett, Sonia Goncalves, Ewan Harrison, David K. Jackson, Ian Johnston, Dominic Kwiatkowski, Cordelia Langford, John Sillitoe on behalf of the Wellcome Sanger Institute COVID-19 Surveillance Team |
| EPI_ISL_2865714 | Lighthouse Lab in Milton Keynes | Wellcome Sanger Institute for the COVID-19 Genomics UK (COG-UK) Consortium | The Lighthouse Lab in Milton Keynes and Alex Alderton, Roberto Amato, Jeffrey Barrett, Sonia Goncalves, Ewan Harrison, David K. Jackson, Ian Johnston, Dominic Kwiatkowski, Cordelia Langford, John Sillitoe on behalf of the Wellcome Sanger Institute COVID-19 Surveillance Team |
| EPI_ISL_2865715 | Lighthouse Lab in Alderley Park | Wellcome Sanger Institute for the COVID-19 Genomics UK (COG-UK) Consortium | Jacquelyn Wynn, Mairead Hyland, The Lighthouse Lab in Alderley Park and Alex Alderton, Roberto Amato, Jeffrey Barrett, Sonia Goncalves, Ewan Harrison, David K. Jackson, Ian Johnston, Dominic Kwiatkowski, Cordelia Langford, John Sillitoe on behalf of the Wellcome Sanger Institute COVID-19 Surveillance Team |
| EPI_ISL_2865716 | Lighthouse Lab in Glasgow | Wellcome Sanger Institute for the COVID-19 Genomics UK (COG-UK) Consortium | Harper VanSteenhouse, Yumi Kasai, David Gray, Carol Clugston, Anna Dominiczak and Alex Alderton, Roberto Amato, Jeffrey Barrett, Sonia Goncalves, Ewan Harrison, David K. Jackson, Ian Johnston, Dominic Kwiatkowski, Cordelia Langford, John Sillitoe on behalf of the Wellcome Sanger Institute COVID-19 Surveillance Team |
| EPI_ISL_2865717 | Lighthouse Lab in Alderley Park | Wellcome Sanger Institute for the COVID-19 Genomics UK (COG-UK) Consortium | Jacquelyn Wynn, Mairead Hyland, The Lighthouse Lab in Alderley Park and Alex Alderton, Roberto Amato, Jeffrey Barrett, Sonia Goncalves, Ewan Harrison, David K. Jackson, Ian Johnston, Dominic Kwiatkowski, Cordelia Langford, John Sillitoe on behalf of the Wellcome Sanger Institute COVID-19 Surveillance Team |
| EPI_ISL_2865718 | Lighthouse Lab in Milton Keynes | Wellcome Sanger Institute for the COVID-19 Genomics UK (COG-UK) Consortium | The Lighthouse Lab in Milton Keynes and Alex Alderton, Roberto Amato, Jeffrey Barrett, Sonia Goncalves, Ewan Harrison, David K. Jackson, Ian Johnston, Dominic Kwiatkowski, Cordelia Langford, John Sillitoe on behalf of the Wellcome Sanger Institute COVID-19 Surveillance Team |
| EPI_ISL_2865719, EPI_ISL_2865721 | Lighthouse Lab in Glasgow | Wellcome Sanger Institute for the COVID-19 Genomics UK (COG-UK) Consortium | Harper VanSteenhouse, Yumi Kasai, David Gray, Carol Clugston, Anna Dominiczak and Alex Alderton, Roberto Amato, Jeffrey Barrett, Sonia Goncalves, Ewan Harrison, David K. Jackson, Ian Johnston, Dominic Kwiatkowski, Cordelia Langford, John Sillitoe on behalf of the Wellcome Sanger Institute COVID-19 Surveillance Team |
| EPI_ISL_2865722 | Lighthouse Lab in Milton Keynes | Wellcome Sanger Institute for the COVID-19 Genomics UK (COG-UK) Consortium | The Lighthouse Lab in Milton Keynes and Alex Alderton, Roberto Amato, Jeffrey Barrett, Sonia Goncalves, Ewan Harrison, David K. Jackson, Ian Johnston, Dominic Kwiatkowski, Cordelia Langford, John Sillitoe on behalf of the Wellcome Sanger Institute COVID-19 Surveillance Team |
| EPI_ISL_2865723, EPI_ISL_2865724, EPI_ISL_2865725 | Lighthouse Lab in Alderley Park | Wellcome Sanger Institute for the COVID-19 Genomics UK (COG-UK) Consortium | Jacquelyn Wynn, Mairead Hyland, The Lighthouse Lab in Alderley Park and Alex Alderton, Roberto Amato, Jeffrey Barrett, Sonia Goncalves, Ewan Harrison, David K. Jackson, Ian Johnston, Dominic Kwiatkowski, Cordelia Langford, John Sillitoe on behalf of the Wellcome Sanger Institute COVID-19 Surveillance Team |
| EPI_ISL_2865726 | Lighthouse Lab in Milton Keynes | Wellcome Sanger Institute for the COVID-19 Genomics UK (COG-UK) Consortium | The Lighthouse Lab in Milton Keynes and Alex Alderton, Roberto Amato, Jeffrey Barrett, Sonia Goncalves, Ewan Harrison, David K. Jackson, Ian Johnston, Dominic Kwiatkowski, Cordelia Langford, John Sillitoe on behalf of the Wellcome Sanger Institute COVID-19 Surveillance Team |
| EPI_ISL_2865727, EPI_ISL_2865728 | Lighthouse Lab in Glasgow | Wellcome Sanger Institute for the COVID-19 Genomics UK (COG-UK) Consortium | Harper VanSteenhouse, Yumi Kasai, David Gray, Carol Clugston, Anna Dominiczak and Alex Alderton, Roberto Amato, Jeffrey Barrett, Sonia Goncalves, Ewan Harrison, David K. Jackson, Ian Johnston, Dominic Kwiatkowski, Cordelia Langford, John Sillitoe on behalf of the Wellcome Sanger Institute COVID-19 Surveillance Team |
| EPI_ISL_2865729 | Lighthouse Lab in Milton Keynes | Wellcome Sanger Institute for the COVID-19 Genomics UK (COG-UK) Consortium | The Lighthouse Lab in Milton Keynes and Alex Alderton, Roberto Amato, Jeffrey Barrett, Sonia Goncalves, Ewan Harrison, David K. Jackson, Ian Johnston, Dominic Kwiatkowski, Cordelia Langford, John Sillitoe on behalf of the Wellcome Sanger Institute COVID-19 Surveillance Team |
| EPI_ISL_2865730, EPI_ISL_2865731 | Lighthouse Lab in Alderley Park | Wellcome Sanger Institute for the COVID-19 Genomics UK (COG-UK) Consortium | Jacquelyn Wynn, Mairead Hyland, The Lighthouse Lab in Alderley Park and Alex Alderton, Roberto Amato, Jeffrey Barrett, Sonia Goncalves, Ewan Harrison, David K. Jackson, Ian Johnston, Dominic Kwiatkowski, Cordelia Langford, John Sillitoe on behalf of the Wellcome Sanger Institute COVID-19 Surveillance Team |
| EPI_ISL_2865732 | Lighthouse Lab in Glasgow | Wellcome Sanger Institute for the COVID-19 Genomics UK (COG-UK) Consortium | Harper VanSteenhouse, Yumi Kasai, David Gray, Carol Clugston, Anna Dominiczak and Alex Alderton, Roberto Amato, Jeffrey Barrett, Sonia Goncalves, Ewan Harrison, David K. Jackson, Ian Johnston, Dominic Kwiatkowski, Cordelia Langford, John Sillitoe on behalf of the Wellcome Sanger Institute COVID-19 Surveillance Team |
| EPI_ISL_2865733 | Lighthouse Lab in Alderley Park | Wellcome Sanger Institute for the COVID-19 Genomics UK (COG-UK) Consortium | Jacquelyn Wynn, Mairead Hyland, The Lighthouse Lab in Alderley Park and Alex Alderton, Roberto Amato, Jeffrey Barrett, Sonia Goncalves, Ewan Harrison, David K. Jackson, Ian Johnston, Dominic Kwiatkowski, Cordelia Langford, John Sillitoe on behalf of the Wellcome Sanger Institute COVID-19 Surveillance Team |
| EPI_ISL_2865734 | Lighthouse Lab in Glasgow | Wellcome Sanger Institute for the COVID-19 Genomics UK (COG-UK) Consortium | Harper VanSteenhouse, Yumi Kasai, David Gray, Carol Clugston, Anna Dominiczak and Alex Alderton, Roberto Amato, Jeffrey Barrett, Sonia Goncalves, Ewan Harrison, David K. Jackson, Ian Johnston, Dominic Kwiatkowski, Cordelia Langford, John Sillitoe on behalf of the Wellcome Sanger Institute COVID-19 Surveillance Team |
| EPI_ISL_2865735 | Lighthouse Lab in Milton Keynes | Wellcome Sanger Institute for the COVID-19 Genomics UK (COG-UK) Consortium | The Lighthouse Lab in Milton Keynes and Alex Alderton, Roberto Amato, Jeffrey Barrett, Sonia Goncalves, Ewan Harrison, David K. Jackson, Ian Johnston, Dominic Kwiatkowski, Cordelia Langford, John Sillitoe on behalf of the Wellcome Sanger Institute COVID-19 Surveillance Team |
| EPI_ISL_2865736, EPI_ISL_2865737, EPI_ISL_2865738 | Lighthouse Lab in Glasgow | Wellcome Sanger Institute for the COVID-19 Genomics UK (COG-UK) Consortium | Harper VanSteenhouse, Yumi Kasai, David Gray, Carol Clugston, Anna Dominiczak and Alex Alderton, Roberto Amato, Jeffrey Barrett, Sonia Goncalves, Ewan Harrison, David K. Jackson, Ian Johnston, Dominic Kwiatkowski, Cordelia Langford, John Sillitoe on behalf of the Wellcome Sanger Institute COVID-19 Surveillance Team |
| EPI_ISL_2865739 | Lighthouse Lab in Milton Keynes | Wellcome Sanger Institute for the COVID-19 Genomics UK (COG-UK) Consortium | The Lighthouse Lab in Milton Keynes and Alex Alderton, Roberto Amato, Jeffrey Barrett, Sonia Goncalves, Ewan Harrison, David K. Jackson, Ian Johnston, Dominic Kwiatkowski, Cordelia Langford, John Sillitoe on behalf of the Wellcome Sanger Institute COVID-19 Surveillance Team |
| EPI_ISL_2865740, EPI_ISL_2865741, EPI_ISL_2865742, EPI_ISL_2865743, EPI_ISL_2865744, EPI_ISL_2865745, EPI_ISL_2865746 | Lighthouse Lab in Glasgow | Wellcome Sanger Institute for the COVID-19 Genomics UK (COG-UK) Consortium | Harper VanSteenhouse, Yumi Kasai, David Gray, Carol Clugston, Anna Dominiczak and Alex Alderton, Roberto Amato, Jeffrey Barrett, Sonia Goncalves, Ewan Harrison, David K. Jackson, Ian Johnston, Dominic Kwiatkowski, Cordelia Langford, John Sillitoe on behalf of the Wellcome Sanger Institute COVID-19 Surveillance Team |
| EPI_ISL_2865748 | Lighthouse Laboratory Plymouth | Wellcome Sanger Institute for the COVID-19 Genomics UK (COG-UK) Consortium | Lighthouse Laboratory Plymouth and Alex Alderton, Roberto Amato, Jeffrey Barrett, Sonia Goncalves, Ewan Harrison, David K. Jackson, Ian Johnston, Dominic Kwiatkowski, Cordelia Langford, John Sillitoe on behalf of the Wellcome Sanger Institute COVID-19 Surveillance Team |
| EPI_ISL_2865749 | Lighthouse Lab in Alderley Park | Wellcome Sanger Institute for the COVID-19 Genomics UK (COG-UK) Consortium | Jacquelyn Wynn, Mairead Hyland, The Lighthouse Lab in Alderley Park and Alex Alderton, Roberto Amato, Jeffrey Barrett, Sonia Goncalves, Ewan Harrison, David K. Jackson, Ian Johnston, Dominic Kwiatkowski, Cordelia Langford, John Sillitoe on behalf of the Wellcome Sanger Institute COVID-19 Surveillance Team |
| EPI_ISL_2865750 | Lighthouse Lab in Glasgow | Wellcome Sanger Institute for the COVID-19 Genomics UK (COG-UK) Consortium | Harper VanSteenhouse, Yumi Kasai, David Gray, Carol Clugston, Anna Dominiczak and Alex Alderton, Roberto Amato, Jeffrey Barrett, Sonia Goncalves, Ewan Harrison, David K. Jackson, Ian Johnston, Dominic Kwiatkowski, Cordelia Langford, John Sillitoe on behalf of the Wellcome Sanger Institute COVID-19 Surveillance Team |
| EPI_ISL_2865751, EPI_ISL_2865752 | Lighthouse Lab in Milton Keynes | Wellcome Sanger Institute for the COVID-19 Genomics UK (COG-UK) Consortium | The Lighthouse Lab in Milton Keynes and Alex Alderton, Roberto Amato, Jeffrey Barrett, Sonia Goncalves, Ewan Harrison, David K. Jackson, Ian Johnston, Dominic Kwiatkowski, Cordelia Langford, John Sillitoe on behalf of the Wellcome Sanger Institute COVID-19 Surveillance Team |
| EPI_ISL_2865753 | Lighthouse Lab in Alderley Park | Wellcome Sanger Institute for the COVID-19 Genomics UK (COG-UK) Consortium | Jacquelyn Wynn, Mairead Hyland, The Lighthouse Lab in Alderley Park and Alex Alderton, Roberto Amato, Jeffrey Barrett, Sonia Goncalves, Ewan Harrison, David K. Jackson, Ian Johnston, Dominic Kwiatkowski, Cordelia Langford, John Sillitoe on behalf of the Wellcome Sanger Institute COVID-19 Surveillance Team |
| EPI_ISL_2865754, EPI_ISL_2865755 | Lighthouse Lab in Glasgow | Wellcome Sanger Institute for the COVID-19 Genomics UK (COG-UK) Consortium | Harper VanSteenhouse, Yumi Kasai, David Gray, Carol Clugston, Anna Dominiczak and Alex Alderton, Roberto Amato, Jeffrey Barrett, Sonia Goncalves, Ewan Harrison, David K. Jackson, Ian Johnston, Dominic Kwiatkowski, Cordelia Langford, John Sillitoe on behalf of the Wellcome Sanger Institute COVID-19 Surveillance Team |
| EPI_ISL_2865756, EPI_ISL_2865757 | Lighthouse Lab in Milton Keynes | Wellcome Sanger Institute for the COVID-19 Genomics UK (COG-UK) Consortium | The Lighthouse Lab in Milton Keynes and Alex Alderton, Roberto Amato, Jeffrey Barrett, Sonia Goncalves, Ewan Harrison, David K. Jackson, Ian Johnston, Dominic Kwiatkowski, Cordelia Langford, John Sillitoe on behalf of the Wellcome Sanger Institute COVID-19 Surveillance Team |
| EPI_ISL_2865758, EPI_ISL_2865759 | Lighthouse Lab in Glasgow | Wellcome Sanger Institute for the COVID-19 Genomics UK (COG-UK) Consortium | Harper VanSteenhouse, Yumi Kasai, David Gray, Carol Clugston, Anna Dominiczak and Alex Alderton, Roberto Amato, Jeffrey Barrett, Sonia Goncalves, Ewan Harrison, David K. Jackson, Ian Johnston, Dominic Kwiatkowski, Cordelia Langford, John Sillitoe on behalf of the Wellcome Sanger Institute COVID-19 Surveillance Team |
| EPI_ISL_2865760, EPI_ISL_2865761 | Lighthouse Lab in Alderley Park | Wellcome Sanger Institute for the COVID-19 Genomics UK (COG-UK) Consortium | Jacquelyn Wynn, Mairead Hyland, The Lighthouse Lab in Alderley Park and Alex Alderton, Roberto Amato, Jeffrey Barrett, Sonia Goncalves, Ewan Harrison, David K. Jackson, Ian Johnston, Dominic Kwiatkowski, Cordelia Langford, John Sillitoe on behalf of the Wellcome Sanger Institute COVID-19 Surveillance Team |
| EPI_ISL_2865763, EPI_ISL_2865764 | Lighthouse Lab in Glasgow | Wellcome Sanger Institute for the COVID-19 Genomics UK (COG-UK) Consortium | Harper VanSteenhouse, Yumi Kasai, David Gray, Carol Clugston, Anna Dominiczak and Alex Alderton, Roberto Amato, Jeffrey Barrett, Sonia Goncalves, Ewan Harrison, David K. Jackson, Ian Johnston, Dominic Kwiatkowski, Cordelia Langford, John Sillitoe on behalf of the Wellcome Sanger Institute COVID-19 Surveillance Team |
| EPI_ISL_2865765, EPI_ISL_2865766 | Lighthouse Lab in Milton Keynes | Wellcome Sanger Institute for the COVID-19 Genomics UK (COG-UK) Consortium | The Lighthouse Lab in Milton Keynes and Alex Alderton, Roberto Amato, Jeffrey Barrett, Sonia Goncalves, Ewan Harrison, David K. Jackson, Ian Johnston, Dominic Kwiatkowski, Cordelia Langford, John Sillitoe on behalf of the Wellcome Sanger Institute COVID-19 Surveillance Team |
| EPI_ISL_2865767 | Lighthouse Lab in Glasgow | Wellcome Sanger Institute for the COVID-19 Genomics UK (COG-UK) Consortium | Harper VanSteenhouse, Yumi Kasai, David Gray, Carol Clugston, Anna Dominiczak and Alex Alderton, Roberto Amato, Jeffrey Barrett, Sonia Goncalves, Ewan Harrison, David K. Jackson, Ian Johnston, Dominic Kwiatkowski, Cordelia Langford, John Sillitoe on behalf of the Wellcome Sanger Institute COVID-19 Surveillance Team |
| EPI_ISL_2865768 | Lighthouse Lab in Milton Keynes | Wellcome Sanger Institute for the COVID-19 Genomics UK (COG-UK) Consortium | The Lighthouse Lab in Milton Keynes and Alex Alderton, Roberto Amato, Jeffrey Barrett, Sonia Goncalves, Ewan Harrison, David K. Jackson, Ian Johnston, Dominic Kwiatkowski, Cordelia Langford, John Sillitoe on behalf of the Wellcome Sanger Institute COVID-19 Surveillance Team |
| EPI_ISL_2865769, EPI_ISL_2865770, | Lighthouse Lab in Glasgow | Wellcome Sanger Institute for the COVID-19 Genomics UK (COG-UK) Consortium | Harper VanSteenhouse, Yumi Kasai, David Gray, Carol Clugston, Anna Dominiczak and Alex Alderton, Roberto Amato, Jeffrey Barrett, Sonia Goncalves, Ewan Harrison, David K. Jackson, Ian Johnston, Dominic Kwiatkowski, Cordelia Langford, John Sillitoe on behalf of the Wellcome Sanger Institute COVID-19 Surveillance Team |

| | | | |
|---|---|---|---|
| EPI_ISL_2865771, EPI_ISL_2865772, EPI_ISL_2865773 | | | |
| EPI_ISL_2865774 | Lighthouse Lab in Alderley Park | Wellcome Sanger Institute for the COVID-19 Genomics UK (COG-UK) Consortium | Jacquelyn Wynn, Mairead Hyland, The Lighthouse Lab in Alderley Park and Alex Alderton, Roberto Amato, Jeffrey Barrett, Sonia Goncalves, Ewan Harrison, David K. Jackson, Ian Johnston, Dominic Kwiatkowski, Cordelia Langford, John Sillitoe on behalf of the Wellcome Sanger Institute COVID-19 Surveillance Team |
| EPI_ISL_2865775, EPI_ISL_2865776, EPI_ISL_2865777 | Lighthouse Lab in Glasgow | Wellcome Sanger Institute for the COVID-19 Genomics UK (COG-UK) Consortium | Harper VanSteenhouse, Yumi Kasai, David Gray, Carol Clugston, Anna Dominiczak and Alex Alderton, Roberto Amato, Jeffrey Barrett, Sonia Goncalves, Ewan Harrison, David K. Jackson, Ian Johnston, Dominic Kwiatkowski, Cordelia Langford, John Sillitoe on behalf of the Wellcome Sanger Institute COVID-19 Surveillance Team |
| EPI_ISL_2865778, EPI_ISL_2865779 | Lighthouse Lab in Alderley Park | Wellcome Sanger Institute for the COVID-19 Genomics UK (COG-UK) Consortium | Jacquelyn Wynn, Mairead Hyland, The Lighthouse Lab in Alderley Park and Alex Alderton, Roberto Amato, Jeffrey Barrett, Sonia Goncalves, Ewan Harrison, David K. Jackson, Ian Johnston, Dominic Kwiatkowski, Cordelia Langford, John Sillitoe on behalf of the Wellcome Sanger Institute COVID-19 Surveillance Team |
| EPI_ISL_2865780, EPI_ISL_2865781, EPI_ISL_2865782, EPI_ISL_2865783 | Lighthouse Lab in Glasgow | Wellcome Sanger Institute for the COVID-19 Genomics UK (COG-UK) Consortium | Harper VanSteenhouse, Yumi Kasai, David Gray, Carol Clugston, Anna Dominiczak and Alex Alderton, Roberto Amato, Jeffrey Barrett, Sonia Goncalves, Ewan Harrison, David K. Jackson, Ian Johnston, Dominic Kwiatkowski, Cordelia Langford, John Sillitoe on behalf of the Wellcome Sanger Institute COVID-19 Surveillance Team |
| EPI_ISL_2865784 | Lighthouse Lab in Milton Keynes | Wellcome Sanger Institute for the COVID-19 Genomics UK (COG-UK) Consortium | The Lighthouse Lab in Milton Keynes and Alex Alderton, Roberto Amato, Jeffrey Barrett, Sonia Goncalves, Ewan Harrison, David K. Jackson, Ian Johnston, Dominic Kwiatkowski, Cordelia Langford, John Sillitoe on behalf of the Wellcome Sanger Institute COVID-19 Surveillance Team |
| EPI_ISL_2865785, EPI_ISL_2865786 | Lighthouse Lab in Glasgow | Wellcome Sanger Institute for the COVID-19 Genomics UK (COG-UK) Consortium | Harper VanSteenhouse, Yumi Kasai, David Gray, Carol Clugston, Anna Dominiczak and Alex Alderton, Roberto Amato, Jeffrey Barrett, Sonia Goncalves, Ewan Harrison, David K. Jackson, Ian Johnston, Dominic Kwiatkowski, Cordelia Langford, John Sillitoe on behalf of the Wellcome Sanger Institute COVID-19 Surveillance Team |
| EPI_ISL_2865787 | Lighthouse Lab in Milton Keynes | Wellcome Sanger Institute for the COVID-19 Genomics UK (COG-UK) Consortium | The Lighthouse Lab in Milton Keynes and Alex Alderton, Roberto Amato, Jeffrey Barrett, Sonia Goncalves, Ewan Harrison, David K. Jackson, Ian Johnston, Dominic Kwiatkowski, Cordelia Langford, John Sillitoe on behalf of the Wellcome Sanger Institute COVID-19 Surveillance Team |
| EPI_ISL_2865788, EPI_ISL_2865789 | Lighthouse Lab in Glasgow | Wellcome Sanger Institute for the COVID-19 Genomics UK (COG-UK) Consortium | Harper VanSteenhouse, Yumi Kasai, David Gray, Carol Clugston, Anna Dominiczak and Alex Alderton, Roberto Amato, Jeffrey Barrett, Sonia Goncalves, Ewan Harrison, David K. Jackson, Ian Johnston, Dominic Kwiatkowski, Cordelia Langford, John Sillitoe on behalf of the Wellcome Sanger Institute COVID-19 Surveillance Team |
| EPI_ISL_2865790 | Lighthouse Lab in Milton Keynes | Wellcome Sanger Institute for the COVID-19 Genomics UK (COG-UK) Consortium | The Lighthouse Lab in Milton Keynes and Alex Alderton, Roberto Amato, Jeffrey Barrett, Sonia Goncalves, Ewan Harrison, David K. Jackson, Ian Johnston, Dominic Kwiatkowski, Cordelia Langford, John Sillitoe on behalf of the Wellcome Sanger Institute COVID-19 Surveillance Team |
| EPI_ISL_2865791 | Lighthouse Lab in Alderley Park | Wellcome Sanger Institute for the COVID-19 Genomics UK (COG-UK) Consortium | Jacquelyn Wynn, Mairead Hyland, The Lighthouse Lab in Alderley Park and Alex Alderton, Roberto Amato, Jeffrey Barrett, Sonia Goncalves, Ewan Harrison, David K. Jackson, Ian Johnston, Dominic Kwiatkowski, Cordelia Langford, John Sillitoe on behalf of the Wellcome Sanger Institute COVID-19 Surveillance Team |
| EPI_ISL_2865792 | Lighthouse Lab in Glasgow | Wellcome Sanger Institute for the COVID-19 Genomics UK (COG-UK) Consortium | Harper VanSteenhouse, Yumi Kasai, David Gray, Carol Clugston, Anna Dominiczak and Alex Alderton, Roberto Amato, Jeffrey Barrett, Sonia Goncalves, Ewan Harrison, David K. Jackson, Ian Johnston, Dominic Kwiatkowski, Cordelia Langford, John Sillitoe on behalf of the Wellcome Sanger Institute COVID-19 Surveillance Team |
| EPI_ISL_2865793 | Lighthouse Lab in Alderley Park | Wellcome Sanger Institute for the COVID-19 Genomics UK (COG-UK) Consortium | Jacquelyn Wynn, Mairead Hyland, The Lighthouse Lab in Alderley Park and Alex Alderton, Roberto Amato, Jeffrey Barrett, Sonia Goncalves, Ewan Harrison, David K. Jackson, Ian Johnston, Dominic Kwiatkowski, Cordelia Langford, John Sillitoe on behalf of the Wellcome Sanger Institute COVID-19 Surveillance Team |
| EPI_ISL_2865794 | Lighthouse Lab in Milton Keynes | Wellcome Sanger Institute for the COVID-19 Genomics UK (COG-UK) Consortium | The Lighthouse Lab in Milton Keynes and Alex Alderton, Roberto Amato, Jeffrey Barrett, Sonia Goncalves, Ewan Harrison, David K. Jackson, Ian Johnston, Dominic Kwiatkowski, Cordelia Langford, John Sillitoe on behalf of the Wellcome Sanger Institute COVID-19 Surveillance Team |
| EPI_ISL_2865795 | Lighthouse Lab in Alderley Park | Wellcome Sanger Institute for the COVID-19 Genomics UK (COG-UK) Consortium | Jacquelyn Wynn, Mairead Hyland, The Lighthouse Lab in Alderley Park and Alex Alderton, Roberto Amato, Jeffrey Barrett, Sonia Goncalves, Ewan Harrison, David K. Jackson, Ian Johnston, Dominic Kwiatkowski, Cordelia Langford, John Sillitoe on behalf of the Wellcome Sanger Institute COVID-19 Surveillance Team |
| EPI_ISL_2865796, EPI_ISL_2865797 | Lighthouse Lab in Milton Keynes | Wellcome Sanger Institute for the COVID-19 Genomics UK (COG-UK) Consortium | The Lighthouse Lab in Milton Keynes and Alex Alderton, Roberto Amato, Jeffrey Barrett, Sonia Goncalves, Ewan Harrison, David K. Jackson, Ian Johnston, Dominic Kwiatkowski, Cordelia Langford, John Sillitoe on behalf of the Wellcome Sanger Institute COVID-19 Surveillance Team |
| EPI_ISL_2865798 | Lighthouse Lab in Alderley Park | Wellcome Sanger Institute for the COVID-19 Genomics UK (COG-UK) Consortium | Jacquelyn Wynn, Mairead Hyland, The Lighthouse Lab in Alderley Park and Alex Alderton, Roberto Amato, Jeffrey Barrett, Sonia Goncalves, Ewan Harrison, David K. Jackson, Ian Johnston, Dominic Kwiatkowski, Cordelia Langford, John Sillitoe on behalf of the Wellcome Sanger Institute COVID-19 Surveillance Team |
| EPI_ISL_2865799 | Lighthouse Lab in Milton Keynes | Wellcome Sanger Institute for the COVID-19 Genomics UK (COG-UK) Consortium | The Lighthouse Lab in Milton Keynes and Alex Alderton, Roberto Amato, Jeffrey Barrett, Sonia Goncalves, Ewan Harrison, David K. Jackson, Ian Johnston, Dominic Kwiatkowski, Cordelia Langford, John Sillitoe on behalf of the Wellcome Sanger Institute COVID-19 Surveillance Team |
| EPI_ISL_2865800 | Lighthouse Lab in Alderley Park | Wellcome Sanger Institute for the COVID-19 Genomics UK (COG-UK) Consortium | Jacquelyn Wynn, Mairead Hyland, The Lighthouse Lab in Alderley Park and Alex Alderton, Roberto Amato, Jeffrey Barrett, Sonia Goncalves, Ewan Harrison, David K. Jackson, Ian Johnston, Dominic Kwiatkowski, Cordelia Langford, John Sillitoe on behalf of the Wellcome Sanger Institute COVID-19 Surveillance Team |
| EPI_ISL_2865801 | Lighthouse Lab in Milton Keynes | Wellcome Sanger Institute for the COVID-19 Genomics UK (COG-UK) Consortium | The Lighthouse Lab in Milton Keynes and Alex Alderton, Roberto Amato, Jeffrey Barrett, Sonia Goncalves, Ewan Harrison, David K. Jackson, Ian Johnston, Dominic Kwiatkowski, Cordelia Langford, John Sillitoe on behalf of the Wellcome Sanger Institute COVID-19 Surveillance Team |
| EPI_ISL_2865802, EPI_ISL_2865803 | Lighthouse Lab in Glasgow | Wellcome Sanger Institute for the COVID-19 Genomics UK (COG-UK) Consortium | Harper VanSteenhouse, Yumi Kasai, David Gray, Carol Clugston, Anna Dominiczak and Alex Alderton, Roberto Amato, Jeffrey Barrett, Sonia Goncalves, Ewan Harrison, David K. Jackson, Ian Johnston, Dominic Kwiatkowski, Cordelia Langford, John Sillitoe on behalf of the Wellcome Sanger Institute COVID-19 Surveillance Team |
| EPI_ISL_2865804 | Lighthouse Lab in Milton Keynes | Wellcome Sanger Institute for the COVID-19 Genomics UK (COG-UK) Consortium | The Lighthouse Lab in Milton Keynes and Alex Alderton, Roberto Amato, Jeffrey Barrett, Sonia Goncalves, Ewan Harrison, David K. Jackson, Ian Johnston, Dominic Kwiatkowski, Cordelia Langford, John Sillitoe on behalf of the Wellcome Sanger Institute COVID-19 Surveillance Team |
| EPI_ISL_2865805 | Lighthouse Lab in Alderley Park | Wellcome Sanger Institute for the COVID-19 Genomics UK (COG-UK) Consortium | Jacquelyn Wynn, Mairead Hyland, The Lighthouse Lab in Alderley Park and Alex Alderton, Roberto Amato, Jeffrey Barrett, Sonia Goncalves, Ewan Harrison, David K. Jackson, Ian Johnston, Dominic Kwiatkowski, Cordelia Langford, John Sillitoe on behalf of the Wellcome Sanger Institute COVID-19 Surveillance Team |
| EPI_ISL_2865806 | Lighthouse Lab in Milton Keynes | Wellcome Sanger Institute for the COVID-19 Genomics UK (COG-UK) Consortium | The Lighthouse Lab in Milton Keynes and Alex Alderton, Roberto Amato, Jeffrey Barrett, Sonia Goncalves, Ewan Harrison, David K. Jackson, Ian Johnston, Dominic Kwiatkowski, Cordelia Langford, John Sillitoe on behalf of the Wellcome Sanger Institute COVID-19 Surveillance Team |
| EPI_ISL_2865807 | Lighthouse Laboratory Plymouth | Wellcome Sanger Institute for the COVID-19 Genomics UK (COG-UK) Consortium | Lighthouse Laboratory Plymouth and Alex Alderton, Roberto Amato, Jeffrey Barrett, Sonia Goncalves, Ewan Harrison, David K. Jackson, Ian Johnston, Dominic Kwiatkowski, Cordelia Langford, John Sillitoe on behalf of the Wellcome Sanger Institute COVID-19 Surveillance Team |
| EPI_ISL_2865808, EPI_ISL_2865809, EPI_ISL_2865810 | Lighthouse Lab in Glasgow | Wellcome Sanger Institute for the COVID-19 Genomics UK (COG-UK) Consortium | Harper VanSteenhouse, Yumi Kasai, David Gray, Carol Clugston, Anna Dominiczak and Alex Alderton, Roberto Amato, Jeffrey Barrett, Sonia Goncalves, Ewan Harrison, David K. Jackson, Ian Johnston, Dominic Kwiatkowski, Cordelia Langford, John Sillitoe on behalf of the Wellcome Sanger Institute COVID-19 Surveillance Team |
| EPI_ISL_2865811 | Lighthouse Lab in Alderley Park | Wellcome Sanger Institute for the COVID-19 Genomics UK (COG-UK) Consortium | Jacquelyn Wynn, Mairead Hyland, The Lighthouse Lab in Alderley Park and Alex Alderton, Roberto Amato, Jeffrey Barrett, Sonia Goncalves, Ewan Harrison, David K. Jackson, Ian Johnston, Dominic Kwiatkowski, Cordelia Langford, John Sillitoe on behalf of the Wellcome Sanger Institute COVID-19 Surveillance Team |
| EPI_ISL_2865812 | Lighthouse Lab in Milton Keynes | Wellcome Sanger Institute for the COVID-19 Genomics UK (COG-UK) Consortium | The Lighthouse Lab in Milton Keynes and Alex Alderton, Roberto Amato, Jeffrey Barrett, Sonia Goncalves, Ewan Harrison, David K. Jackson, Ian Johnston, Dominic Kwiatkowski, Cordelia Langford, John Sillitoe on behalf of the Wellcome Sanger Institute COVID-19 Surveillance Team |
| EPI_ISL_2865813 | Lighthouse Lab in Glasgow | Wellcome Sanger Institute for the COVID-19 Genomics UK (COG-UK) Consortium | Harper VanSteenhouse, Yumi Kasai, David Gray, Carol Clugston, Anna Dominiczak and Alex Alderton, Roberto Amato, Jeffrey Barrett, Sonia Goncalves, Ewan Harrison, David K. Jackson, Ian Johnston, Dominic Kwiatkowski, Cordelia Langford, John Sillitoe on behalf of the Wellcome Sanger Institute COVID-19 Surveillance Team |
| EPI_ISL_2865814 | Lighthouse Lab in Milton Keynes | Wellcome Sanger Institute for the COVID-19 Genomics UK (COG-UK) Consortium | The Lighthouse Lab in Milton Keynes and Alex Alderton, Roberto Amato, Jeffrey Barrett, Sonia Goncalves, Ewan Harrison, David K. Jackson, Ian Johnston, Dominic Kwiatkowski, Cordelia Langford, John Sillitoe on behalf of the Wellcome Sanger Institute COVID-19 Surveillance Team |
| EPI_ISL_2865815 | Lighthouse Lab in Alderley Park | Wellcome Sanger Institute for the COVID-19 Genomics UK (COG-UK) Consortium | Jacquelyn Wynn, Mairead Hyland, The Lighthouse Lab in Alderley Park and Alex Alderton, Roberto Amato, Jeffrey Barrett, Sonia Goncalves, Ewan Harrison, David K. Jackson, Ian Johnston, Dominic Kwiatkowski, Cordelia Langford, John Sillitoe on behalf of the Wellcome Sanger Institute COVID-19 Surveillance Team |
| EPI_ISL_2865816, EPI_ISL_2865817 | Lighthouse Lab in Glasgow | Wellcome Sanger Institute for the COVID-19 Genomics UK (COG-UK) Consortium | Harper VanSteenhouse, Yumi Kasai, David Gray, Carol Clugston, Anna Dominiczak and Alex Alderton, Roberto Amato, Jeffrey Barrett, Sonia Goncalves, Ewan Harrison, David K. Jackson, Ian Johnston, Dominic Kwiatkowski, Cordelia Langford, John Sillitoe on behalf of the Wellcome Sanger Institute COVID-19 Surveillance Team |
| EPI_ISL_2865818, EPI_ISL_2865819 | Lighthouse Lab in Alderley Park | Wellcome Sanger Institute for the COVID-19 Genomics UK (COG-UK) Consortium | Jacquelyn Wynn, Mairead Hyland, The Lighthouse Lab in Alderley Park and Alex Alderton, Roberto Amato, Jeffrey Barrett, Sonia Goncalves, Ewan Harrison, David K. Jackson, Ian Johnston, Dominic Kwiatkowski, Cordelia Langford, John Sillitoe on behalf of the Wellcome Sanger Institute COVID-19 Surveillance Team |
| EPI_ISL_2865820 | Lighthouse Lab in Glasgow | Wellcome Sanger Institute for the COVID-19 Genomics UK (COG-UK) Consortium | Harper VanSteenhouse, Yumi Kasai, David Gray, Carol Clugston, Anna Dominiczak and Alex Alderton, Roberto Amato, Jeffrey Barrett, Sonia Goncalves, Ewan Harrison, David K. Jackson, Ian Johnston, Dominic Kwiatkowski, Cordelia Langford, John Sillitoe on behalf of the Wellcome Sanger Institute COVID-19 Surveillance Team |
| EPI_ISL_2865821 | Lighthouse Lab in Milton Keynes | Wellcome Sanger Institute for the COVID-19 Genomics UK (COG-UK) Consortium | The Lighthouse Lab in Milton Keynes and Alex Alderton, Roberto Amato, Jeffrey Barrett, Sonia Goncalves, Ewan Harrison, David K. Jackson, Ian Johnston, Dominic Kwiatkowski, Cordelia Langford, John Sillitoe on behalf of the Wellcome Sanger Institute COVID-19 Surveillance Team |
| EPI_ISL_2865822 | Lighthouse Lab in Glasgow | Wellcome Sanger Institute for the COVID-19 Genomics UK (COG-UK) Consortium | Harper VanSteenhouse, Yumi Kasai, David Gray, Carol Clugston, Anna Dominiczak and Alex Alderton, Roberto Amato, Jeffrey Barrett, Sonia Goncalves, Ewan Harrison, David K. Jackson, Ian Johnston, Dominic Kwiatkowski, Cordelia Langford, John Sillitoe on behalf of the Wellcome Sanger Institute COVID-19 Surveillance Team |
| EPI_ISL_2865823 | Lighthouse Lab in Milton Keynes | Wellcome Sanger Institute for the COVID-19 Genomics UK (COG-UK) Consortium | The Lighthouse Lab in Milton Keynes and Alex Alderton, Roberto Amato, Jeffrey Barrett, Sonia Goncalves, Ewan Harrison, David K. Jackson, Ian Johnston, Dominic Kwiatkowski, Cordelia Langford, John Sillitoe on behalf of the Wellcome Sanger Institute COVID-19 Surveillance Team |
| EPI_ISL_2865824, EPI_ISL_2865825, EPI_ISL_2865826, EPI_ISL_2865827 | Lighthouse Lab in Glasgow | Wellcome Sanger Institute for the COVID-19 Genomics UK (COG-UK) Consortium | Harper VanSteenhouse, Yumi Kasai, David Gray, Carol Clugston, Anna Dominiczak and Alex Alderton, Roberto Amato, Jeffrey Barrett, Sonia Goncalves, Ewan Harrison, David K. Jackson, Ian Johnston, Dominic Kwiatkowski, Cordelia Langford, John Sillitoe on behalf of the Wellcome Sanger Institute COVID-19 Surveillance Team |
| EPI_ISL_2865828 | Lighthouse Lab in Alderley Park | Wellcome Sanger Institute for the COVID-19 Genomics UK (COG-UK) Consortium | Jacquelyn Wynn, Mairead Hyland, The Lighthouse Lab in Alderley Park and Alex Alderton, Roberto Amato, Jeffrey Barrett, Sonia Goncalves, Ewan Harrison, David K. Jackson, Ian Johnston, Dominic Kwiatkowski, Cordelia Langford, John Sillitoe on behalf of the Wellcome Sanger Institute COVID-19 Surveillance Team |
| EPI_ISL_2865829, EPI_ISL_2865830 | Lighthouse Lab in Milton Keynes | Wellcome Sanger Institute for the COVID-19 Genomics UK (COG-UK) Consortium | The Lighthouse Lab in Milton Keynes and Alex Alderton, Roberto Amato, Jeffrey Barrett, Sonia Goncalves, Ewan Harrison, David K. Jackson, Ian Johnston, Dominic Kwiatkowski, Cordelia Langford, John Sillitoe on behalf of the Wellcome Sanger Institute COVID-19 Surveillance Team |
| EPI_ISL_2865831, EPI_ISL_2865832, EPI_ISL_2865833 | Lighthouse Lab in Glasgow | Wellcome Sanger Institute for the COVID-19 Genomics UK (COG-UK) Consortium | Harper VanSteenhouse, Yumi Kasai, David Gray, Carol Clugston, Anna Dominiczak and Alex Alderton, Roberto Amato, Jeffrey Barrett, Sonia Goncalves, Ewan Harrison, David K. Jackson, Ian Johnston, Dominic Kwiatkowski, Cordelia Langford, John Sillitoe on behalf of the Wellcome Sanger Institute COVID-19 Surveillance Team |
| EPI_ISL_2865834 | Lighthouse Lab in Milton Keynes | Wellcome Sanger Institute for the COVID-19 Genomics UK (COG-UK) Consortium | The Lighthouse Lab in Milton Keynes and Alex Alderton, Roberto Amato, Jeffrey Barrett, Sonia Goncalves, Ewan Harrison, David K. Jackson, Ian Johnston, Dominic Kwiatkowski, Cordelia Langford, John Sillitoe on behalf of the Wellcome Sanger Institute COVID-19 Surveillance Team |

| | | | |
|---|---|---|---|
| EPI_ISL_2865835, EPI_ISL_2865836 | Lighthouse Lab in Glasgow | Wellcome Sanger Institute for the COVID-19 Genomics UK (COG-UK) Consortium | Harper VanSteenhouse, Yumi Kasai, David Gray, Carol Clugston, Anna Dominiczak and Alex Alderton, Roberto Amato, Jeffrey Barrett, Sonia Goncalves, Ewan Harrison, David K. Jackson, Ian Johnston, Dominic Kwiatkowski, Cordelia Langford, John Sillitoe on behalf of the Wellcome Sanger Institute COVID-19 Surveillance Team |
| EPI_ISL_2865837 | Lighthouse Laboratory Plymouth | Wellcome Sanger Institute for the COVID-19 Genomics UK (COG-UK) Consortium | Lighthouse Laboratory Plymouth and Alex Alderton, Roberto Amato, Jeffrey Barrett, Sonia Goncalves, Ewan Harrison, David K. Jackson, Ian Johnston, Dominic Kwiatkowski, Cordelia Langford, John Sillitoe on behalf of the Wellcome Sanger Institute COVID-19 Surveillance Team |
| EPI_ISL_2865838 | Lighthouse Lab in Alderley Park | Wellcome Sanger Institute for the COVID-19 Genomics UK (COG-UK) Consortium | Jacquelyn Wynn, Mairead Hyland, The Lighthouse Lab in Alderley Park and Alex Alderton, Roberto Amato, Jeffrey Barrett, Sonia Goncalves, Ewan Harrison, David K. Jackson, Ian Johnston, Dominic Kwiatkowski, Cordelia Langford, John Sillitoe on behalf of the Wellcome Sanger Institute COVID-19 Surveillance Team |
| EPI_ISL_2865839 | Lighthouse Lab in Glasgow | Wellcome Sanger Institute for the COVID-19 Genomics UK (COG-UK) Consortium | Harper VanSteenhouse, Yumi Kasai, David Gray, Carol Clugston, Anna Dominiczak and Alex Alderton, Roberto Amato, Jeffrey Barrett, Sonia Goncalves, Ewan Harrison, David K. Jackson, Ian Johnston, Dominic Kwiatkowski, Cordelia Langford, John Sillitoe on behalf of the Wellcome Sanger Institute COVID-19 Surveillance Team |
| EPI_ISL_2865840 | Lighthouse Lab in Milton Keynes | Wellcome Sanger Institute for the COVID-19 Genomics UK (COG-UK) Consortium | The Lighthouse Lab in Milton Keynes and Alex Alderton, Roberto Amato, Jeffrey Barrett, Sonia Goncalves, Ewan Harrison, David K. Jackson, Ian Johnston, Dominic Kwiatkowski, Cordelia Langford, John Sillitoe on behalf of the Wellcome Sanger Institute COVID-19 Surveillance Team |
| EPI_ISL_2865841 | Lighthouse Lab in Glasgow | Wellcome Sanger Institute for the COVID-19 Genomics UK (COG-UK) Consortium | Harper VanSteenhouse, Yumi Kasai, David Gray, Carol Clugston, Anna Dominiczak and Alex Alderton, Roberto Amato, Jeffrey Barrett, Sonia Goncalves, Ewan Harrison, David K. Jackson, Ian Johnston, Dominic Kwiatkowski, Cordelia Langford, John Sillitoe on behalf of the Wellcome Sanger Institute COVID-19 Surveillance Team |
| EPI_ISL_2865842 | Lighthouse Laboratory Plymouth | Wellcome Sanger Institute for the COVID-19 Genomics UK (COG-UK) Consortium | Lighthouse Laboratory Plymouth and Alex Alderton, Roberto Amato, Jeffrey Barrett, Sonia Goncalves, Ewan Harrison, David K. Jackson, Ian Johnston, Dominic Kwiatkowski, Cordelia Langford, John Sillitoe on behalf of the Wellcome Sanger Institute COVID-19 Surveillance Team |
| EPI_ISL_2865843 | Lighthouse Lab in Milton Keynes | Wellcome Sanger Institute for the COVID-19 Genomics UK (COG-UK) Consortium | The Lighthouse Lab in Milton Keynes and Alex Alderton, Roberto Amato, Jeffrey Barrett, Sonia Goncalves, Ewan Harrison, David K. Jackson, Ian Johnston, Dominic Kwiatkowski, Cordelia Langford, John Sillitoe on behalf of the Wellcome Sanger Institute COVID-19 Surveillance Team |
| EPI_ISL_2865844, EPI_ISL_2865845, EPI_ISL_2865846, EPI_ISL_2865847, EPI_ISL_2865848 | Lighthouse Lab in Glasgow | Wellcome Sanger Institute for the COVID-19 Genomics UK (COG-UK) Consortium | Harper VanSteenhouse, Yumi Kasai, David Gray, Carol Clugston, Anna Dominiczak and Alex Alderton, Roberto Amato, Jeffrey Barrett, Sonia Goncalves, Ewan Harrison, David K. Jackson, Ian Johnston, Dominic Kwiatkowski, Cordelia Langford, John Sillitoe on behalf of the Wellcome Sanger Institute COVID-19 Surveillance Team |
| EPI_ISL_2865849 | Lighthouse Lab in Milton Keynes | Wellcome Sanger Institute for the COVID-19 Genomics UK (COG-UK) Consortium | The Lighthouse Lab in Milton Keynes and Alex Alderton, Roberto Amato, Jeffrey Barrett, Sonia Goncalves, Ewan Harrison, David K. Jackson, Ian Johnston, Dominic Kwiatkowski, Cordelia Langford, John Sillitoe on behalf of the Wellcome Sanger Institute COVID-19 Surveillance Team |
| EPI_ISL_2865850, EPI_ISL_2865851, EPI_ISL_2865852 | Lighthouse Lab in Glasgow | Wellcome Sanger Institute for the COVID-19 Genomics UK (COG-UK) Consortium | Harper VanSteenhouse, Yumi Kasai, David Gray, Carol Clugston, Anna Dominiczak and Alex Alderton, Roberto Amato, Jeffrey Barrett, Sonia Goncalves, Ewan Harrison, David K. Jackson, Ian Johnston, Dominic Kwiatkowski, Cordelia Langford, John Sillitoe on behalf of the Wellcome Sanger Institute COVID-19 Surveillance Team |
| EPI_ISL_2865853 | Lighthouse Lab in Milton Keynes | Wellcome Sanger Institute for the COVID-19 Genomics UK (COG-UK) Consortium | The Lighthouse Lab in Milton Keynes and Alex Alderton, Roberto Amato, Jeffrey Barrett, Sonia Goncalves, Ewan Harrison, David K. Jackson, Ian Johnston, Dominic Kwiatkowski, Cordelia Langford, John Sillitoe on behalf of the Wellcome Sanger Institute COVID-19 Surveillance Team |
| EPI_ISL_2865854 | Lighthouse Lab in Alderley Park | Wellcome Sanger Institute for the COVID-19 Genomics UK (COG-UK) Consortium | Jacquelyn Wynn, Mairead Hyland, The Lighthouse Lab in Alderley Park and Alex Alderton, Roberto Amato, Jeffrey Barrett, Sonia Goncalves, Ewan Harrison, David K. Jackson, Ian Johnston, Dominic Kwiatkowski, Cordelia Langford, John Sillitoe on behalf of the Wellcome Sanger Institute COVID-19 Surveillance Team |
| EPI_ISL_2865855 | Lighthouse Lab in Milton Keynes | Wellcome Sanger Institute for the COVID-19 Genomics UK (COG-UK) Consortium | The Lighthouse Lab in Milton Keynes and Alex Alderton, Roberto Amato, Jeffrey Barrett, Sonia Goncalves, Ewan Harrison, David K. Jackson, Ian Johnston, Dominic Kwiatkowski, Cordelia Langford, John Sillitoe on behalf of the Wellcome Sanger Institute COVID-19 Surveillance Team |
| EPI_ISL_2865856 | Lighthouse Lab in Glasgow | Wellcome Sanger Institute for the COVID-19 Genomics UK (COG-UK) Consortium | Harper VanSteenhouse, Yumi Kasai, David Gray, Carol Clugston, Anna Dominiczak and Alex Alderton, Roberto Amato, Jeffrey Barrett, Sonia Goncalves, Ewan Harrison, David K. Jackson, Ian Johnston, Dominic Kwiatkowski, Cordelia Langford, John Sillitoe on behalf of the Wellcome Sanger Institute COVID-19 Surveillance Team |
| EPI_ISL_2865857 | Lighthouse Lab in Milton Keynes | Wellcome Sanger Institute for the COVID-19 Genomics UK (COG-UK) Consortium | The Lighthouse Lab in Milton Keynes and Alex Alderton, Roberto Amato, Jeffrey Barrett, Sonia Goncalves, Ewan Harrison, David K. Jackson, Ian Johnston, Dominic Kwiatkowski, Cordelia Langford, John Sillitoe on behalf of the Wellcome Sanger Institute COVID-19 Surveillance Team |
| EPI_ISL_2865858 | Lighthouse Lab in Alderley Park | Wellcome Sanger Institute for the COVID-19 Genomics UK (COG-UK) Consortium | Jacquelyn Wynn, Mairead Hyland, The Lighthouse Lab in Alderley Park and Alex Alderton, Roberto Amato, Jeffrey Barrett, Sonia Goncalves, Ewan Harrison, David K. Jackson, Ian Johnston, Dominic Kwiatkowski, Cordelia Langford, John Sillitoe on behalf of the Wellcome Sanger Institute COVID-19 Surveillance Team |
| EPI_ISL_2865859, EPI_ISL_2865860, EPI_ISL_2865861, EPI_ISL_2865862 | Lighthouse Lab in Glasgow | Wellcome Sanger Institute for the COVID-19 Genomics UK (COG-UK) Consortium | Harper VanSteenhouse, Yumi Kasai, David Gray, Carol Clugston, Anna Dominiczak and Alex Alderton, Roberto Amato, Jeffrey Barrett, Sonia Goncalves, Ewan Harrison, David K. Jackson, Ian Johnston, Dominic Kwiatkowski, Cordelia Langford, John Sillitoe on behalf of the Wellcome Sanger Institute COVID-19 Surveillance Team |
| EPI_ISL_2865863, EPI_ISL_2865864, EPI_ISL_2865865 | Lighthouse Lab in Alderley Park | Wellcome Sanger Institute for the COVID-19 Genomics UK (COG-UK) Consortium | Jacquelyn Wynn, Mairead Hyland, The Lighthouse Lab in Alderley Park and Alex Alderton, Roberto Amato, Jeffrey Barrett, Sonia Goncalves, Ewan Harrison, David K. Jackson, Ian Johnston, Dominic Kwiatkowski, Cordelia Langford, John Sillitoe on behalf of the Wellcome Sanger Institute COVID-19 Surveillance Team |
| EPI_ISL_2865866 | Lighthouse Lab in Milton Keynes | Wellcome Sanger Institute for the COVID-19 Genomics UK (COG-UK) Consortium | The Lighthouse Lab in Milton Keynes and Alex Alderton, Roberto Amato, Jeffrey Barrett, Sonia Goncalves, Ewan Harrison, David K. Jackson, Ian Johnston, Dominic Kwiatkowski, Cordelia Langford, John Sillitoe on behalf of the Wellcome Sanger Institute COVID-19 Surveillance Team |
| EPI_ISL_2865867 | Lighthouse Lab in Glasgow | Wellcome Sanger Institute for the COVID-19 Genomics UK (COG-UK) Consortium | Harper VanSteenhouse, Yumi Kasai, David Gray, Carol Clugston, Anna Dominiczak and Alex Alderton, Roberto Amato, Jeffrey Barrett, Sonia Goncalves, Ewan Harrison, David K. Jackson, Ian Johnston, Dominic Kwiatkowski, Cordelia Langford, John Sillitoe on behalf of the Wellcome Sanger Institute COVID-19 Surveillance Team |
| EPI_ISL_2865868 | Lighthouse Lab in Alderley Park | Wellcome Sanger Institute for the COVID-19 Genomics UK (COG-UK) Consortium | Jacquelyn Wynn, Mairead Hyland, The Lighthouse Lab in Alderley Park and Alex Alderton, Roberto Amato, Jeffrey Barrett, Sonia Goncalves, Ewan Harrison, David K. Jackson, Ian Johnston, Dominic Kwiatkowski, Cordelia Langford, John Sillitoe on behalf of the Wellcome Sanger Institute COVID-19 Surveillance Team |
| EPI_ISL_2865869 | Lighthouse Lab in Glasgow | Wellcome Sanger Institute for the COVID-19 Genomics UK (COG-UK) Consortium | Harper VanSteenhouse, Yumi Kasai, David Gray, Carol Clugston, Anna Dominiczak and Alex Alderton, Roberto Amato, Jeffrey Barrett, Sonia Goncalves, Ewan Harrison, David K. Jackson, Ian Johnston, Dominic Kwiatkowski, Cordelia Langford, John Sillitoe on behalf of the Wellcome Sanger Institute COVID-19 Surveillance Team |
| EPI_ISL_2865870 | Lighthouse Laboratory Plymouth | Wellcome Sanger Institute for the COVID-19 Genomics UK (COG-UK) Consortium | Lighthouse Laboratory Plymouth and Alex Alderton, Roberto Amato, Jeffrey Barrett, Sonia Goncalves, Ewan Harrison, David K. Jackson, Ian Johnston, Dominic Kwiatkowski, Cordelia Langford, John Sillitoe on behalf of the Wellcome Sanger Institute COVID-19 Surveillance Team |
| EPI_ISL_2865871 | Lighthouse Lab in Glasgow | Wellcome Sanger Institute for the COVID-19 Genomics UK (COG-UK) Consortium | Harper VanSteenhouse, Yumi Kasai, David Gray, Carol Clugston, Anna Dominiczak and Alex Alderton, Roberto Amato, Jeffrey Barrett, Sonia Goncalves, Ewan Harrison, David K. Jackson, Ian Johnston, Dominic Kwiatkowski, Cordelia Langford, John Sillitoe on behalf of the Wellcome Sanger Institute COVID-19 Surveillance Team |
| EPI_ISL_2865872, EPI_ISL_2865873 | Lighthouse Lab in Milton Keynes | Wellcome Sanger Institute for the COVID-19 Genomics UK (COG-UK) Consortium | The Lighthouse Lab in Milton Keynes and Alex Alderton, Roberto Amato, Jeffrey Barrett, Sonia Goncalves, Ewan Harrison, David K. Jackson, Ian Johnston, Dominic Kwiatkowski, Cordelia Langford, John Sillitoe on behalf of the Wellcome Sanger Institute COVID-19 Surveillance Team |
| EPI_ISL_2865874 | Lighthouse Lab in Glasgow | Wellcome Sanger Institute for the COVID-19 Genomics UK (COG-UK) Consortium | Harper VanSteenhouse, Yumi Kasai, David Gray, Carol Clugston, Anna Dominiczak and Alex Alderton, Roberto Amato, Jeffrey Barrett, Sonia Goncalves, Ewan Harrison, David K. Jackson, Ian Johnston, Dominic Kwiatkowski, Cordelia Langford, John Sillitoe on behalf of the Wellcome Sanger Institute COVID-19 Surveillance Team |
| EPI_ISL_2865875 | Lighthouse Lab in Milton Keynes | Wellcome Sanger Institute for the COVID-19 Genomics UK (COG-UK) Consortium | The Lighthouse Lab in Milton Keynes and Alex Alderton, Roberto Amato, Jeffrey Barrett, Sonia Goncalves, Ewan Harrison, David K. Jackson, Ian Johnston, Dominic Kwiatkowski, Cordelia Langford, John Sillitoe on behalf of the Wellcome Sanger Institute COVID-19 Surveillance Team |
| EPI_ISL_2865876, EPI_ISL_2865877 | Lighthouse Lab in Glasgow | Wellcome Sanger Institute for the COVID-19 Genomics UK (COG-UK) Consortium | Harper VanSteenhouse, Yumi Kasai, David Gray, Carol Clugston, Anna Dominiczak and Alex Alderton, Roberto Amato, Jeffrey Barrett, Sonia Goncalves, Ewan Harrison, David K. Jackson, Ian Johnston, Dominic Kwiatkowski, Cordelia Langford, John Sillitoe on behalf of the Wellcome Sanger Institute COVID-19 Surveillance Team |
| EPI_ISL_2865878, EPI_ISL_2865879 | Lighthouse Lab in Alderley Park | Wellcome Sanger Institute for the COVID-19 Genomics UK (COG-UK) Consortium | Jacquelyn Wynn, Mairead Hyland, The Lighthouse Lab in Alderley Park and Alex Alderton, Roberto Amato, Jeffrey Barrett, Sonia Goncalves, Ewan Harrison, David K. Jackson, Ian Johnston, Dominic Kwiatkowski, Cordelia Langford, John Sillitoe on behalf of the Wellcome Sanger Institute COVID-19 Surveillance Team |
| EPI_ISL_2865880, EPI_ISL_2865881 | Lighthouse Lab in Glasgow | Wellcome Sanger Institute for the COVID-19 Genomics UK (COG-UK) Consortium | Harper VanSteenhouse, Yumi Kasai, David Gray, Carol Clugston, Anna Dominiczak and Alex Alderton, Roberto Amato, Jeffrey Barrett, Sonia Goncalves, Ewan Harrison, David K. Jackson, Ian Johnston, Dominic Kwiatkowski, Cordelia Langford, John Sillitoe on behalf of the Wellcome Sanger Institute COVID-19 Surveillance Team |
| EPI_ISL_2865882, EPI_ISL_2865883, EPI_ISL_2865884 | Lighthouse Lab in Alderley Park | Wellcome Sanger Institute for the COVID-19 Genomics UK (COG-UK) Consortium | Jacquelyn Wynn, Mairead Hyland, The Lighthouse Lab in Alderley Park and Alex Alderton, Roberto Amato, Jeffrey Barrett, Sonia Goncalves, Ewan Harrison, David K. Jackson, Ian Johnston, Dominic Kwiatkowski, Cordelia Langford, John Sillitoe on behalf of the Wellcome Sanger Institute COVID-19 Surveillance Team |
| EPI_ISL_2865885 | Lighthouse Lab in Milton Keynes | Wellcome Sanger Institute for the COVID-19 Genomics UK (COG-UK) Consortium | The Lighthouse Lab in Milton Keynes and Alex Alderton, Roberto Amato, Jeffrey Barrett, Sonia Goncalves, Ewan Harrison, David K. Jackson, Ian Johnston, Dominic Kwiatkowski, Cordelia Langford, John Sillitoe on behalf of the Wellcome Sanger Institute COVID-19 Surveillance Team |
| EPI_ISL_2865886 | Lighthouse Lab in Glasgow | Wellcome Sanger Institute for the COVID-19 Genomics UK (COG-UK) Consortium | Harper VanSteenhouse, Yumi Kasai, David Gray, Carol Clugston, Anna Dominiczak and Alex Alderton, Roberto Amato, Jeffrey Barrett, Sonia Goncalves, Ewan Harrison, David K. Jackson, Ian Johnston, Dominic Kwiatkowski, Cordelia Langford, John Sillitoe on behalf of the Wellcome Sanger Institute COVID-19 Surveillance Team |
| EPI_ISL_2865887 | Lighthouse Lab in Alderley Park | Wellcome Sanger Institute for the COVID-19 Genomics UK (COG-UK) Consortium | Jacquelyn Wynn, Mairead Hyland, The Lighthouse Lab in Alderley Park and Alex Alderton, Roberto Amato, Jeffrey Barrett, Sonia Goncalves, Ewan Harrison, David K. Jackson, Ian Johnston, Dominic Kwiatkowski, Cordelia Langford, John Sillitoe on behalf of the Wellcome Sanger Institute COVID-19 Surveillance Team |
| EPI_ISL_2865888, EPI_ISL_2865889 | Lighthouse Lab in Glasgow | Wellcome Sanger Institute for the COVID-19 Genomics UK (COG-UK) Consortium | Harper VanSteenhouse, Yumi Kasai, David Gray, Carol Clugston, Anna Dominiczak and Alex Alderton, Roberto Amato, Jeffrey Barrett, Sonia Goncalves, Ewan Harrison, David K. Jackson, Ian Johnston, Dominic Kwiatkowski, Cordelia Langford, John Sillitoe on behalf of the Wellcome Sanger Institute COVID-19 Surveillance Team |
| EPI_ISL_2865890 | Lighthouse Lab in Alderley Park | Wellcome Sanger Institute for the COVID-19 Genomics UK (COG-UK) Consortium | Jacquelyn Wynn, Mairead Hyland, The Lighthouse Lab in Alderley Park and Alex Alderton, Roberto Amato, Jeffrey Barrett, Sonia Goncalves, Ewan Harrison, David K. Jackson, Ian Johnston, Dominic Kwiatkowski, Cordelia Langford, John Sillitoe on behalf of the Wellcome Sanger Institute COVID-19 Surveillance Team |
| EPI_ISL_2865891 | Lighthouse Lab in Glasgow | Wellcome Sanger Institute for the COVID-19 Genomics UK (COG-UK) Consortium | Harper VanSteenhouse, Yumi Kasai, David Gray, Carol Clugston, Anna Dominiczak and Alex Alderton, Roberto Amato, Jeffrey Barrett, Sonia Goncalves, Ewan Harrison, David K. Jackson, Ian Johnston, Dominic Kwiatkowski, Cordelia Langford, John Sillitoe on behalf of the Wellcome Sanger Institute COVID-19 Surveillance Team |
| EPI_ISL_2865892 | Lighthouse Lab in Milton Keynes | Wellcome Sanger Institute for the COVID-19 Genomics UK (COG-UK) Consortium | The Lighthouse Lab in Milton Keynes and Alex Alderton, Roberto Amato, Jeffrey Barrett, Sonia Goncalves, Ewan Harrison, David K. Jackson, Ian Johnston, Dominic Kwiatkowski, Cordelia Langford, John Sillitoe on behalf of the Wellcome Sanger Institute COVID-19 Surveillance Team |
| EPI_ISL_2865893, EPI_ISL_2865894 | Lighthouse Lab in Alderley Park | Wellcome Sanger Institute for the COVID-19 Genomics UK (COG-UK) Consortium | Jacquelyn Wynn, Mairead Hyland, The Lighthouse Lab in Alderley Park and Alex Alderton, Roberto Amato, Jeffrey Barrett, Sonia Goncalves, Ewan Harrison, David K. Jackson, Ian Johnston, Dominic Kwiatkowski, Cordelia Langford, John Sillitoe on behalf of the Wellcome Sanger Institute COVID-19 Surveillance Team |
| EPI_ISL_2865895 | Lighthouse Lab in Glasgow | Wellcome Sanger Institute for the COVID-19 Genomics UK (COG-UK) Consortium | Harper VanSteenhouse, Yumi Kasai, David Gray, Carol Clugston, Anna Dominiczak and Alex Alderton, Roberto Amato, Jeffrey Barrett, Sonia Goncalves, Ewan Harrison, David K. Jackson, Ian Johnston, Dominic Kwiatkowski, Cordelia Langford, John Sillitoe on behalf of the Wellcome Sanger Institute COVID-19 Surveillance Team |
| EPI_ISL_2865896 | Lighthouse Lab in Alderley Park | Wellcome Sanger Institute for the COVID-19 Genomics UK (COG-UK) Consortium | Jacquelyn Wynn, Mairead Hyland, The Lighthouse Lab in Alderley Park and Alex Alderton, Roberto Amato, Jeffrey Barrett, Sonia Goncalves, Ewan Harrison, David K. Jackson, Ian Johnston, Dominic Kwiatkowski, Cordelia Langford, John Sillitoe on behalf of the Wellcome Sanger Institute COVID-19 Surveillance Team |

| | | | |
|---|---|---|---|
| EPI_ISL_2865897 | Lighthouse Lab in Milton Keynes | Wellcome Sanger Institute for the COVID-19 Genomics UK (COG-UK) Consortium | The Lighthouse Lab in Milton Keynes and Alex Alderton, Roberto Amato, Jeffrey Barrett, Sonia Goncalves, Ewan Harrison, David K. Jackson, Ian Johnston, Dominic Kwiatkowski, Cordelia Langford, John Sillitoe on behalf of the Wellcome Sanger Institute COVID-19 Surveillance Team |
| EPI_ISL_2865898, EPI_ISL_2865899 | Lighthouse Lab in Glasgow | Wellcome Sanger Institute for the COVID-19 Genomics UK (COG-UK) Consortium | Harper VanSteenhouse, Yumi Kasai, David Gray, Carol Clugston, Anna Dominiczak and Alex Alderton, Roberto Amato, Jeffrey Barrett, Sonia Goncalves, Ewan Harrison, David K. Jackson, Ian Johnston, Dominic Kwiatkowski, Cordelia Langford, John Sillitoe on behalf of the Wellcome Sanger Institute COVID-19 Surveillance Team |
| EPI_ISL_2865900, EPI_ISL_2865901 | Lighthouse Lab in Alderley Park | Wellcome Sanger Institute for the COVID-19 Genomics UK (COG-UK) Consortium | Jacquelyn Wynn, Mairead Hyland, The Lighthouse Lab in Alderley Park and Alex Alderton, Roberto Amato, Jeffrey Barrett, Sonia Goncalves, Ewan Harrison, David K. Jackson, Ian Johnston, Dominic Kwiatkowski, Cordelia Langford, John Sillitoe on behalf of the Wellcome Sanger Institute COVID-19 Surveillance Team |
| EPI_ISL_2865902 | Lighthouse Lab in Glasgow | Wellcome Sanger Institute for the COVID-19 Genomics UK (COG-UK) Consortium | Harper VanSteenhouse, Yumi Kasai, David Gray, Carol Clugston, Anna Dominiczak and Alex Alderton, Roberto Amato, Jeffrey Barrett, Sonia Goncalves, Ewan Harrison, David K. Jackson, Ian Johnston, Dominic Kwiatkowski, Cordelia Langford, John Sillitoe on behalf of the Wellcome Sanger Institute COVID-19 Surveillance Team |
| EPI_ISL_2865903 | Lighthouse Lab in Alderley Park | Wellcome Sanger Institute for the COVID-19 Genomics UK (COG-UK) Consortium | Jacquelyn Wynn, Mairead Hyland, The Lighthouse Lab in Alderley Park and Alex Alderton, Roberto Amato, Jeffrey Barrett, Sonia Goncalves, Ewan Harrison, David K. Jackson, Ian Johnston, Dominic Kwiatkowski, Cordelia Langford, John Sillitoe on behalf of the Wellcome Sanger Institute COVID-19 Surveillance Team |
| EPI_ISL_2865904, EPI_ISL_2865905 | Lighthouse Lab in Glasgow | Wellcome Sanger Institute for the COVID-19 Genomics UK (COG-UK) Consortium | Harper VanSteenhouse, Yumi Kasai, David Gray, Carol Clugston, Anna Dominiczak and Alex Alderton, Roberto Amato, Jeffrey Barrett, Sonia Goncalves, Ewan Harrison, David K. Jackson, Ian Johnston, Dominic Kwiatkowski, Cordelia Langford, John Sillitoe on behalf of the Wellcome Sanger Institute COVID-19 Surveillance Team |
| EPI_ISL_2865906 | Lighthouse Lab in Alderley Park | Wellcome Sanger Institute for the COVID-19 Genomics UK (COG-UK) Consortium | Jacquelyn Wynn, Mairead Hyland, The Lighthouse Lab in Alderley Park and Alex Alderton, Roberto Amato, Jeffrey Barrett, Sonia Goncalves, Ewan Harrison, David K. Jackson, Ian Johnston, Dominic Kwiatkowski, Cordelia Langford, John Sillitoe on behalf of the Wellcome Sanger Institute COVID-19 Surveillance Team |
| EPI_ISL_2865907 | Lighthouse Lab in Glasgow | Wellcome Sanger Institute for the COVID-19 Genomics UK (COG-UK) Consortium | Harper VanSteenhouse, Yumi Kasai, David Gray, Carol Clugston, Anna Dominiczak and Alex Alderton, Roberto Amato, Jeffrey Barrett, Sonia Goncalves, Ewan Harrison, David K. Jackson, Ian Johnston, Dominic Kwiatkowski, Cordelia Langford, John Sillitoe on behalf of the Wellcome Sanger Institute COVID-19 Surveillance Team |
| EPI_ISL_2865908, EPI_ISL_2865909, EPI_ISL_2865910 | Lighthouse Lab in Milton Keynes | Wellcome Sanger Institute for the COVID-19 Genomics UK (COG-UK) Consortium | The Lighthouse Lab in Milton Keynes and Alex Alderton, Roberto Amato, Jeffrey Barrett, Sonia Goncalves, Ewan Harrison, David K. Jackson, Ian Johnston, Dominic Kwiatkowski, Cordelia Langford, John Sillitoe on behalf of the Wellcome Sanger Institute COVID-19 Surveillance Team |
| EPI_ISL_2865911, EPI_ISL_2865912, EPI_ISL_2865913, EPI_ISL_2865914, EPI_ISL_2865915 | Lighthouse Lab in Glasgow | Wellcome Sanger Institute for the COVID-19 Genomics UK (COG-UK) Consortium | Harper VanSteenhouse, Yumi Kasai, David Gray, Carol Clugston, Anna Dominiczak and Alex Alderton, Roberto Amato, Jeffrey Barrett, Sonia Goncalves, Ewan Harrison, David K. Jackson, Ian Johnston, Dominic Kwiatkowski, Cordelia Langford, John Sillitoe on behalf of the Wellcome Sanger Institute COVID-19 Surveillance Team |
| EPI_ISL_2865916 | Lighthouse Lab in Milton Keynes | Wellcome Sanger Institute for the COVID-19 Genomics UK (COG-UK) Consortium | The Lighthouse Lab in Milton Keynes and Alex Alderton, Roberto Amato, Jeffrey Barrett, Sonia Goncalves, Ewan Harrison, David K. Jackson, Ian Johnston, Dominic Kwiatkowski, Cordelia Langford, John Sillitoe on behalf of the Wellcome Sanger Institute COVID-19 Surveillance Team |
| EPI_ISL_2865918 | Lighthouse Lab in Glasgow | Wellcome Sanger Institute for the COVID-19 Genomics UK (COG-UK) Consortium | Harper VanSteenhouse, Yumi Kasai, David Gray, Carol Clugston, Anna Dominiczak and Alex Alderton, Roberto Amato, Jeffrey Barrett, Sonia Goncalves, Ewan Harrison, David K. Jackson, Ian Johnston, Dominic Kwiatkowski, Cordelia Langford, John Sillitoe on behalf of the Wellcome Sanger Institute COVID-19 Surveillance Team |
| EPI_ISL_2865919 | Lighthouse Lab in Milton Keynes | Wellcome Sanger Institute for the COVID-19 Genomics UK (COG-UK) Consortium | The Lighthouse Lab in Milton Keynes and Alex Alderton, Roberto Amato, Jeffrey Barrett, Sonia Goncalves, Ewan Harrison, David K. Jackson, Ian Johnston, Dominic Kwiatkowski, Cordelia Langford, John Sillitoe on behalf of the Wellcome Sanger Institute COVID-19 Surveillance Team |
| EPI_ISL_2865920 | Lighthouse Lab in Alderley Park | Wellcome Sanger Institute for the COVID-19 Genomics UK (COG-UK) Consortium | Jacquelyn Wynn, Mairead Hyland, The Lighthouse Lab in Alderley Park and Alex Alderton, Roberto Amato, Jeffrey Barrett, Sonia Goncalves, Ewan Harrison, David K. Jackson, Ian Johnston, Dominic Kwiatkowski, Cordelia Langford, John Sillitoe on behalf of the Wellcome Sanger Institute COVID-19 Surveillance Team |
| EPI_ISL_2865922 | Lighthouse Lab in Milton Keynes | Wellcome Sanger Institute for the COVID-19 Genomics UK (COG-UK) Consortium | The Lighthouse Lab in Milton Keynes and Alex Alderton, Roberto Amato, Jeffrey Barrett, Sonia Goncalves, Ewan Harrison, David K. Jackson, Ian Johnston, Dominic Kwiatkowski, Cordelia Langford, John Sillitoe on behalf of the Wellcome Sanger Institute COVID-19 Surveillance Team |
| EPI_ISL_2865923, EPI_ISL_2865924 | Lighthouse Lab in Glasgow | Wellcome Sanger Institute for the COVID-19 Genomics UK (COG-UK) Consortium | Harper VanSteenhouse, Yumi Kasai, David Gray, Carol Clugston, Anna Dominiczak and Alex Alderton, Roberto Amato, Jeffrey Barrett, Sonia Goncalves, Ewan Harrison, David K. Jackson, Ian Johnston, Dominic Kwiatkowski, Cordelia Langford, John Sillitoe on behalf of the Wellcome Sanger Institute COVID-19 Surveillance Team |
| EPI_ISL_2865925 | Lighthouse Lab in Milton Keynes | Wellcome Sanger Institute for the COVID-19 Genomics UK (COG-UK) Consortium | The Lighthouse Lab in Milton Keynes and Alex Alderton, Roberto Amato, Jeffrey Barrett, Sonia Goncalves, Ewan Harrison, David K. Jackson, Ian Johnston, Dominic Kwiatkowski, Cordelia Langford, John Sillitoe on behalf of the Wellcome Sanger Institute COVID-19 Surveillance Team |
| EPI_ISL_2865926 | Lighthouse Lab in Alderley Park | Wellcome Sanger Institute for the COVID-19 Genomics UK (COG-UK) Consortium | Jacquelyn Wynn, Mairead Hyland, The Lighthouse Lab in Alderley Park and Alex Alderton, Roberto Amato, Jeffrey Barrett, Sonia Goncalves, Ewan Harrison, David K. Jackson, Ian Johnston, Dominic Kwiatkowski, Cordelia Langford, John Sillitoe on behalf of the Wellcome Sanger Institute COVID-19 Surveillance Team |
| EPI_ISL_2865927 | Lighthouse Lab in Milton Keynes | Wellcome Sanger Institute for the COVID-19 Genomics UK (COG-UK) Consortium | The Lighthouse Lab in Milton Keynes and Alex Alderton, Roberto Amato, Jeffrey Barrett, Sonia Goncalves, Ewan Harrison, David K. Jackson, Ian Johnston, Dominic Kwiatkowski, Cordelia Langford, John Sillitoe on behalf of the Wellcome Sanger Institute COVID-19 Surveillance Team |
| EPI_ISL_2865928 | Lighthouse Lab in Glasgow | Wellcome Sanger Institute for the COVID-19 Genomics UK (COG-UK) Consortium | Harper VanSteenhouse, Yumi Kasai, David Gray, Carol Clugston, Anna Dominiczak and Alex Alderton, Roberto Amato, Jeffrey Barrett, Sonia Goncalves, Ewan Harrison, David K. Jackson, Ian Johnston, Dominic Kwiatkowski, Cordelia Langford, John Sillitoe on behalf of the Wellcome Sanger Institute COVID-19 Surveillance Team |
| EPI_ISL_2865929 | Lighthouse Laboratory Plymouth | Wellcome Sanger Institute for the COVID-19 Genomics UK (COG-UK) Consortium | Lighthouse Laboratory Plymouth and Alex Alderton, Roberto Amato, Jeffrey Barrett, Sonia Goncalves, Ewan Harrison, David K. Jackson, Ian Johnston, Dominic Kwiatkowski, Cordelia Langford, John Sillitoe on behalf of the Wellcome Sanger Institute COVID-19 Surveillance Team |
| EPI_ISL_2865930, EPI_ISL_2865931 | Lighthouse Lab in Milton Keynes | Wellcome Sanger Institute for the COVID-19 Genomics UK (COG-UK) Consortium | The Lighthouse Lab in Milton Keynes and Alex Alderton, Roberto Amato, Jeffrey Barrett, Sonia Goncalves, Ewan Harrison, David K. Jackson, Ian Johnston, Dominic Kwiatkowski, Cordelia Langford, John Sillitoe on behalf of the Wellcome Sanger Institute COVID-19 Surveillance Team |
| EPI_ISL_2865932 | Lighthouse Lab in Glasgow | Wellcome Sanger Institute for the COVID-19 Genomics UK (COG-UK) Consortium | Harper VanSteenhouse, Yumi Kasai, David Gray, Carol Clugston, Anna Dominiczak and Alex Alderton, Roberto Amato, Jeffrey Barrett, Sonia Goncalves, Ewan Harrison, David K. Jackson, Ian Johnston, Dominic Kwiatkowski, Cordelia Langford, John Sillitoe on behalf of the Wellcome Sanger Institute COVID-19 Surveillance Team |
| EPI_ISL_2865933 | Lighthouse Lab in Milton Keynes | Wellcome Sanger Institute for the COVID-19 Genomics UK (COG-UK) Consortium | The Lighthouse Lab in Milton Keynes and Alex Alderton, Roberto Amato, Jeffrey Barrett, Sonia Goncalves, Ewan Harrison, David K. Jackson, Ian Johnston, Dominic Kwiatkowski, Cordelia Langford, John Sillitoe on behalf of the Wellcome Sanger Institute COVID-19 Surveillance Team |
| EPI_ISL_2865934 | Lighthouse Lab in Glasgow | Wellcome Sanger Institute for the COVID-19 Genomics UK (COG-UK) Consortium | Harper VanSteenhouse, Yumi Kasai, David Gray, Carol Clugston, Anna Dominiczak and Alex Alderton, Roberto Amato, Jeffrey Barrett, Sonia Goncalves, Ewan Harrison, David K. Jackson, Ian Johnston, Dominic Kwiatkowski, Cordelia Langford, John Sillitoe on behalf of the Wellcome Sanger Institute COVID-19 Surveillance Team |
| EPI_ISL_2865935 | Lighthouse Lab in Alderley Park | Wellcome Sanger Institute for the COVID-19 Genomics UK (COG-UK) Consortium | Jacquelyn Wynn, Mairead Hyland, The Lighthouse Lab in Alderley Park and Alex Alderton, Roberto Amato, Jeffrey Barrett, Sonia Goncalves, Ewan Harrison, David K. Jackson, Ian Johnston, Dominic Kwiatkowski, Cordelia Langford, John Sillitoe on behalf of the Wellcome Sanger Institute COVID-19 Surveillance Team |
| EPI_ISL_2865936, EPI_ISL_2865937 | Lighthouse Lab in Glasgow | Wellcome Sanger Institute for the COVID-19 Genomics UK (COG-UK) Consortium | Harper VanSteenhouse, Yumi Kasai, David Gray, Carol Clugston, Anna Dominiczak and Alex Alderton, Roberto Amato, Jeffrey Barrett, Sonia Goncalves, Ewan Harrison, David K. Jackson, Ian Johnston, Dominic Kwiatkowski, Cordelia Langford, John Sillitoe on behalf of the Wellcome Sanger Institute COVID-19 Surveillance Team |
| EPI_ISL_2865938 | Lighthouse Lab in Alderley Park | Wellcome Sanger Institute for the COVID-19 Genomics UK (COG-UK) Consortium | Jacquelyn Wynn, Mairead Hyland, The Lighthouse Lab in Alderley Park and Alex Alderton, Roberto Amato, Jeffrey Barrett, Sonia Goncalves, Ewan Harrison, David K. Jackson, Ian Johnston, Dominic Kwiatkowski, Cordelia Langford, John Sillitoe on behalf of the Wellcome Sanger Institute COVID-19 Surveillance Team |
| EPI_ISL_2865939 | Lighthouse Lab in Glasgow | Wellcome Sanger Institute for the COVID-19 Genomics UK (COG-UK) Consortium | Harper VanSteenhouse, Yumi Kasai, David Gray, Carol Clugston, Anna Dominiczak and Alex Alderton, Roberto Amato, Jeffrey Barrett, Sonia Goncalves, Ewan Harrison, David K. Jackson, Ian Johnston, Dominic Kwiatkowski, Cordelia Langford, John Sillitoe on behalf of the Wellcome Sanger Institute COVID-19 Surveillance Team |
| EPI_ISL_2865940, EPI_ISL_2865941 | Lighthouse Lab in Milton Keynes | Wellcome Sanger Institute for the COVID-19 Genomics UK (COG-UK) Consortium | The Lighthouse Lab in Milton Keynes and Alex Alderton, Roberto Amato, Jeffrey Barrett, Sonia Goncalves, Ewan Harrison, David K. Jackson, Ian Johnston, Dominic Kwiatkowski, Cordelia Langford, John Sillitoe on behalf of the Wellcome Sanger Institute COVID-19 Surveillance Team |
| EPI_ISL_2865942 | Lighthouse Laboratory Plymouth | Wellcome Sanger Institute for the COVID-19 Genomics UK (COG-UK) Consortium | Lighthouse Laboratory Plymouth and Alex Alderton, Roberto Amato, Jeffrey Barrett, Sonia Goncalves, Ewan Harrison, David K. Jackson, Ian Johnston, Dominic Kwiatkowski, Cordelia Langford, John Sillitoe on behalf of the Wellcome Sanger Institute COVID-19 Surveillance Team |
| EPI_ISL_2865943 | Lighthouse Lab in Glasgow | Wellcome Sanger Institute for the COVID-19 Genomics UK (COG-UK) Consortium | Harper VanSteenhouse, Yumi Kasai, David Gray, Carol Clugston, Anna Dominiczak and Alex Alderton, Roberto Amato, Jeffrey Barrett, Sonia Goncalves, Ewan Harrison, David K. Jackson, Ian Johnston, Dominic Kwiatkowski, Cordelia Langford, John Sillitoe on behalf of the Wellcome Sanger Institute COVID-19 Surveillance Team |
| EPI_ISL_2865944 | Lighthouse Lab in Milton Keynes | Wellcome Sanger Institute for the COVID-19 Genomics UK (COG-UK) Consortium | The Lighthouse Lab in Milton Keynes and Alex Alderton, Roberto Amato, Jeffrey Barrett, Sonia Goncalves, Ewan Harrison, David K. Jackson, Ian Johnston, Dominic Kwiatkowski, Cordelia Langford, John Sillitoe on behalf of the Wellcome Sanger Institute COVID-19 Surveillance Team |
| EPI_ISL_2865945 | Lighthouse Laboratory Plymouth | Wellcome Sanger Institute for the COVID-19 Genomics UK (COG-UK) Consortium | Lighthouse Laboratory Plymouth and Alex Alderton, Roberto Amato, Jeffrey Barrett, Sonia Goncalves, Ewan Harrison, David K. Jackson, Ian Johnston, Dominic Kwiatkowski, Cordelia Langford, John Sillitoe on behalf of the Wellcome Sanger Institute COVID-19 Surveillance Team |
| EPI_ISL_2865946 | Lighthouse Lab in Milton Keynes | Wellcome Sanger Institute for the COVID-19 Genomics UK (COG-UK) Consortium | The Lighthouse Lab in Milton Keynes and Alex Alderton, Roberto Amato, Jeffrey Barrett, Sonia Goncalves, Ewan Harrison, David K. Jackson, Ian Johnston, Dominic Kwiatkowski, Cordelia Langford, John Sillitoe on behalf of the Wellcome Sanger Institute COVID-19 Surveillance Team |
| EPI_ISL_2865947 | Lighthouse Lab in Glasgow | Wellcome Sanger Institute for the COVID-19 Genomics UK (COG-UK) Consortium | Harper VanSteenhouse, Yumi Kasai, David Gray, Carol Clugston, Anna Dominiczak and Alex Alderton, Roberto Amato, Jeffrey Barrett, Sonia Goncalves, Ewan Harrison, David K. Jackson, Ian Johnston, Dominic Kwiatkowski, Cordelia Langford, John Sillitoe on behalf of the Wellcome Sanger Institute COVID-19 Surveillance Team |
| EPI_ISL_2865948 | Lighthouse Lab in Alderley Park | Wellcome Sanger Institute for the COVID-19 Genomics UK (COG-UK) Consortium | Jacquelyn Wynn, Mairead Hyland, The Lighthouse Lab in Alderley Park and Alex Alderton, Roberto Amato, Jeffrey Barrett, Sonia Goncalves, Ewan Harrison, David K. Jackson, Ian Johnston, Dominic Kwiatkowski, Cordelia Langford, John Sillitoe on behalf of the Wellcome Sanger Institute COVID-19 Surveillance Team |
| EPI_ISL_2865949 | Lighthouse Lab in Milton Keynes | Wellcome Sanger Institute for the COVID-19 Genomics UK (COG-UK) Consortium | The Lighthouse Lab in Milton Keynes and Alex Alderton, Roberto Amato, Jeffrey Barrett, Sonia Goncalves, Ewan Harrison, David K. Jackson, Ian Johnston, Dominic Kwiatkowski, Cordelia Langford, John Sillitoe on behalf of the Wellcome Sanger Institute COVID-19 Surveillance Team |
| EPI_ISL_2865950 | Lighthouse Lab in Glasgow | Wellcome Sanger Institute for the COVID-19 Genomics UK (COG-UK) Consortium | Harper VanSteenhouse, Yumi Kasai, David Gray, Carol Clugston, Anna Dominiczak and Alex Alderton, Roberto Amato, Jeffrey Barrett, Sonia Goncalves, Ewan Harrison, David K. Jackson, Ian Johnston, Dominic Kwiatkowski, Cordelia Langford, John Sillitoe on behalf of the Wellcome Sanger Institute COVID-19 Surveillance Team |
| EPI_ISL_2865951, EPI_ISL_2865952 | Lighthouse Lab in Milton Keynes | Wellcome Sanger Institute for the COVID-19 Genomics UK (COG-UK) Consortium | The Lighthouse Lab in Milton Keynes and Alex Alderton, Roberto Amato, Jeffrey Barrett, Sonia Goncalves, Ewan Harrison, David K. Jackson, Ian Johnston, Dominic Kwiatkowski, Cordelia Langford, John Sillitoe on behalf of the Wellcome Sanger Institute COVID-19 Surveillance Team |
| EPI_ISL_2865953 | Lighthouse Lab in Alderley Park | Wellcome Sanger Institute for the COVID-19 Genomics UK (COG-UK) Consortium | Jacquelyn Wynn, Mairead Hyland, The Lighthouse Lab in Alderley Park and Alex Alderton, Roberto Amato, Jeffrey Barrett, Sonia Goncalves, Ewan Harrison, David K. Jackson, Ian Johnston, Dominic Kwiatkowski, Cordelia Langford, John Sillitoe on behalf of the Wellcome Sanger Institute COVID-19 Surveillance Team |
| EPI_ISL_2865954 | Lighthouse Lab in Glasgow | Wellcome Sanger Institute for the COVID-19 Genomics UK (COG-UK) Consortium | Harper VanSteenhouse, Yumi Kasai, David Gray, Carol Clugston, Anna Dominiczak and Alex Alderton, Roberto Amato, Jeffrey Barrett, Sonia Goncalves, Ewan Harrison, David K. Jackson, Ian Johnston, Dominic Kwiatkowski, Cordelia Langford, John Sillitoe on behalf of the Wellcome Sanger Institute COVID-19 Surveillance Team |
| EPI_ISL_2865955, EPI_ISL_2865956, | Lighthouse Lab in Alderley Park | Wellcome Sanger Institute for the COVID-19 Genomics UK (COG-UK) Consortium | Jacquelyn Wynn, Mairead Hyland, The Lighthouse Lab in Alderley Park and Alex Alderton, Roberto Amato, Jeffrey Barrett, Sonia Goncalves, Ewan Harrison, David K. Jackson, Ian Johnston, Dominic Kwiatkowski, Cordelia Langford, John Sillitoe on behalf of the Wellcome Sanger Institute COVID-19 Surveillance Team |

| | | | |
|---|---|---|---|
| EPI_ISL_2865957 | | | |
| EPI_ISL_2865958 | Lighthouse Lab in Glasgow | Wellcome Sanger Institute for the COVID-19 Genomics UK (COG-UK) Consortium | Harper VanSteenhouse, Yumi Kasai, David Gray, Carol Clugston, Anna Dominiczak and Alex Alderton, Roberto Amato, Jeffrey Barrett, Sonia Goncalves, Ewan Harrison, David K. Jackson, Ian Johnston, Dominic Kwiatkowski, Cordelia Langford, John Sillitoe on behalf of the Wellcome Sanger Institute COVID-19 Surveillance Team |
| EPI_ISL_2865959 | Lighthouse Lab in Milton Keynes | Wellcome Sanger Institute for the COVID-19 Genomics UK (COG-UK) Consortium | The Lighthouse Lab in Milton Keynes and Alex Alderton, Roberto Amato, Jeffrey Barrett, Sonia Goncalves, Ewan Harrison, David K. Jackson, Ian Johnston, Dominic Kwiatkowski, Cordelia Langford, John Sillitoe on behalf of the Wellcome Sanger Institute COVID-19 Surveillance Team |
| EPI_ISL_2865960 | Lighthouse Lab in Alderley Park | Wellcome Sanger Institute for the COVID-19 Genomics UK (COG-UK) Consortium | Jacquelyn Wynn, Mairead Hyland, The Lighthouse Lab in Alderley Park and Alex Alderton, Roberto Amato, Jeffrey Barrett, Sonia Goncalves, Ewan Harrison, David K. Jackson, Ian Johnston, Dominic Kwiatkowski, Cordelia Langford, John Sillitoe on behalf of the Wellcome Sanger Institute COVID-19 Surveillance Team |
| EPI_ISL_2865961 | Lighthouse Lab in Glasgow | Wellcome Sanger Institute for the COVID-19 Genomics UK (COG-UK) Consortium | Harper VanSteenhouse, Yumi Kasai, David Gray, Carol Clugston, Anna Dominiczak and Alex Alderton, Roberto Amato, Jeffrey Barrett, Sonia Goncalves, Ewan Harrison, David K. Jackson, Ian Johnston, Dominic Kwiatkowski, Cordelia Langford, John Sillitoe on behalf of the Wellcome Sanger Institute COVID-19 Surveillance Team |
| EPI_ISL_2865962 | Lighthouse Lab in Milton Keynes | Wellcome Sanger Institute for the COVID-19 Genomics UK (COG-UK) Consortium | The Lighthouse Lab in Milton Keynes and Alex Alderton, Roberto Amato, Jeffrey Barrett, Sonia Goncalves, Ewan Harrison, David K. Jackson, Ian Johnston, Dominic Kwiatkowski, Cordelia Langford, John Sillitoe on behalf of the Wellcome Sanger Institute COVID-19 Surveillance Team |
| EPI_ISL_2865963 | Lighthouse Lab in Alderley Park | Wellcome Sanger Institute for the COVID-19 Genomics UK (COG-UK) Consortium | Jacquelyn Wynn, Mairead Hyland, The Lighthouse Lab in Alderley Park and Alex Alderton, Roberto Amato, Jeffrey Barrett, Sonia Goncalves, Ewan Harrison, David K. Jackson, Ian Johnston, Dominic Kwiatkowski, Cordelia Langford, John Sillitoe on behalf of the Wellcome Sanger Institute COVID-19 Surveillance Team |
| EPI_ISL_2865964 | Lighthouse Lab in Milton Keynes | Wellcome Sanger Institute for the COVID-19 Genomics UK (COG-UK) Consortium | The Lighthouse Lab in Milton Keynes and Alex Alderton, Roberto Amato, Jeffrey Barrett, Sonia Goncalves, Ewan Harrison, David K. Jackson, Ian Johnston, Dominic Kwiatkowski, Cordelia Langford, John Sillitoe on behalf of the Wellcome Sanger Institute COVID-19 Surveillance Team |
| EPI_ISL_2865965, EPI_ISL_2865966 | Lighthouse Lab in Glasgow | Wellcome Sanger Institute for the COVID-19 Genomics UK (COG-UK) Consortium | Harper VanSteenhouse, Yumi Kasai, David Gray, Carol Clugston, Anna Dominiczak and Alex Alderton, Roberto Amato, Jeffrey Barrett, Sonia Goncalves, Ewan Harrison, David K. Jackson, Ian Johnston, Dominic Kwiatkowski, Cordelia Langford, John Sillitoe on behalf of the Wellcome Sanger Institute COVID-19 Surveillance Team |
| EPI_ISL_2865967 | Lighthouse Lab in Milton Keynes | Wellcome Sanger Institute for the COVID-19 Genomics UK (COG-UK) Consortium | The Lighthouse Lab in Milton Keynes and Alex Alderton, Roberto Amato, Jeffrey Barrett, Sonia Goncalves, Ewan Harrison, David K. Jackson, Ian Johnston, Dominic Kwiatkowski, Cordelia Langford, John Sillitoe on behalf of the Wellcome Sanger Institute COVID-19 Surveillance Team |
| EPI_ISL_2865968 | Lighthouse Lab in Alderley Park | Wellcome Sanger Institute for the COVID-19 Genomics UK (COG-UK) Consortium | Jacquelyn Wynn, Mairead Hyland, The Lighthouse Lab in Alderley Park and Alex Alderton, Roberto Amato, Jeffrey Barrett, Sonia Goncalves, Ewan Harrison, David K. Jackson, Ian Johnston, Dominic Kwiatkowski, Cordelia Langford, John Sillitoe on behalf of the Wellcome Sanger Institute COVID-19 Surveillance Team |
| EPI_ISL_2865969, EPI_ISL_2865970 | Lighthouse Lab in Milton Keynes | Wellcome Sanger Institute for the COVID-19 Genomics UK (COG-UK) Consortium | The Lighthouse Lab in Milton Keynes and Alex Alderton, Roberto Amato, Jeffrey Barrett, Sonia Goncalves, Ewan Harrison, David K. Jackson, Ian Johnston, Dominic Kwiatkowski, Cordelia Langford, John Sillitoe on behalf of the Wellcome Sanger Institute COVID-19 Surveillance Team |
| EPI_ISL_2865971, EPI_ISL_2865972, EPI_ISL_2865973, EPI_ISL_2865974, EPI_ISL_2865975 | Lighthouse Lab in Glasgow | Wellcome Sanger Institute for the COVID-19 Genomics UK (COG-UK) Consortium | Harper VanSteenhouse, Yumi Kasai, David Gray, Carol Clugston, Anna Dominiczak and Alex Alderton, Roberto Amato, Jeffrey Barrett, Sonia Goncalves, Ewan Harrison, David K. Jackson, Ian Johnston, Dominic Kwiatkowski, Cordelia Langford, John Sillitoe on behalf of the Wellcome Sanger Institute COVID-19 Surveillance Team |
| EPI_ISL_2865976 | Lighthouse Lab in Milton Keynes | Wellcome Sanger Institute for the COVID-19 Genomics UK (COG-UK) Consortium | The Lighthouse Lab in Milton Keynes and Alex Alderton, Roberto Amato, Jeffrey Barrett, Sonia Goncalves, Ewan Harrison, David K. Jackson, Ian Johnston, Dominic Kwiatkowski, Cordelia Langford, John Sillitoe on behalf of the Wellcome Sanger Institute COVID-19 Surveillance Team |
| EPI_ISL_2865977, EPI_ISL_2865978 | Lighthouse Lab in Glasgow | Wellcome Sanger Institute for the COVID-19 Genomics UK (COG-UK) Consortium | Harper VanSteenhouse, Yumi Kasai, David Gray, Carol Clugston, Anna Dominiczak and Alex Alderton, Roberto Amato, Jeffrey Barrett, Sonia Goncalves, Ewan Harrison, David K. Jackson, Ian Johnston, Dominic Kwiatkowski, Cordelia Langford, John Sillitoe on behalf of the Wellcome Sanger Institute COVID-19 Surveillance Team |
| EPI_ISL_2865979 | Lighthouse Laboratory Plymouth | Wellcome Sanger Institute for the COVID-19 Genomics UK (COG-UK) Consortium | Lighthouse Laboratory Plymouth and Alex Alderton, Roberto Amato, Jeffrey Barrett, Sonia Goncalves, Ewan Harrison, David K. Jackson, Ian Johnston, Dominic Kwiatkowski, Cordelia Langford, John Sillitoe on behalf of the Wellcome Sanger Institute COVID-19 Surveillance Team |
| EPI_ISL_2865980, EPI_ISL_2865981, EPI_ISL_2865982, EPI_ISL_2865983 | Lighthouse Lab in Glasgow | Wellcome Sanger Institute for the COVID-19 Genomics UK (COG-UK) Consortium | Harper VanSteenhouse, Yumi Kasai, David Gray, Carol Clugston, Anna Dominiczak and Alex Alderton, Roberto Amato, Jeffrey Barrett, Sonia Goncalves, Ewan Harrison, David K. Jackson, Ian Johnston, Dominic Kwiatkowski, Cordelia Langford, John Sillitoe on behalf of the Wellcome Sanger Institute COVID-19 Surveillance Team |
| EPI_ISL_2865985 | Lighthouse Lab in Alderley Park | Wellcome Sanger Institute for the COVID-19 Genomics UK (COG-UK) Consortium | Jacquelyn Wynn, Mairead Hyland, The Lighthouse Lab in Alderley Park and Alex Alderton, Roberto Amato, Jeffrey Barrett, Sonia Goncalves, Ewan Harrison, David K. Jackson, Ian Johnston, Dominic Kwiatkowski, Cordelia Langford, John Sillitoe on behalf of the Wellcome Sanger Institute COVID-19 Surveillance Team |
| EPI_ISL_2865986, EPI_ISL_2865987 | Lighthouse Lab in Glasgow | Wellcome Sanger Institute for the COVID-19 Genomics UK (COG-UK) Consortium | Harper VanSteenhouse, Yumi Kasai, David Gray, Carol Clugston, Anna Dominiczak and Alex Alderton, Roberto Amato, Jeffrey Barrett, Sonia Goncalves, Ewan Harrison, David K. Jackson, Ian Johnston, Dominic Kwiatkowski, Cordelia Langford, John Sillitoe on behalf of the Wellcome Sanger Institute COVID-19 Surveillance Team |
| EPI_ISL_2865988 | Lighthouse Lab in Milton Keynes | Wellcome Sanger Institute for the COVID-19 Genomics UK (COG-UK) Consortium | The Lighthouse Lab in Milton Keynes and Alex Alderton, Roberto Amato, Jeffrey Barrett, Sonia Goncalves, Ewan Harrison, David K. Jackson, Ian Johnston, Dominic Kwiatkowski, Cordelia Langford, John Sillitoe on behalf of the Wellcome Sanger Institute COVID-19 Surveillance Team |
| EPI_ISL_2865989, EPI_ISL_2865990, EPI_ISL_2865991, EPI_ISL_2865992 | Lighthouse Lab in Glasgow | Wellcome Sanger Institute for the COVID-19 Genomics UK (COG-UK) Consortium | Harper VanSteenhouse, Yumi Kasai, David Gray, Carol Clugston, Anna Dominiczak and Alex Alderton, Roberto Amato, Jeffrey Barrett, Sonia Goncalves, Ewan Harrison, David K. Jackson, Ian Johnston, Dominic Kwiatkowski, Cordelia Langford, John Sillitoe on behalf of the Wellcome Sanger Institute COVID-19 Surveillance Team |
| EPI_ISL_2865993 | Lighthouse Lab in Milton Keynes | Wellcome Sanger Institute for the COVID-19 Genomics UK (COG-UK) Consortium | The Lighthouse Lab in Milton Keynes and Alex Alderton, Roberto Amato, Jeffrey Barrett, Sonia Goncalves, Ewan Harrison, David K. Jackson, Ian Johnston, Dominic Kwiatkowski, Cordelia Langford, John Sillitoe on behalf of the Wellcome Sanger Institute COVID-19 Surveillance Team |
| EPI_ISL_2865994, EPI_ISL_2865995 | Lighthouse Lab in Glasgow | Wellcome Sanger Institute for the COVID-19 Genomics UK (COG-UK) Consortium | Harper VanSteenhouse, Yumi Kasai, David Gray, Carol Clugston, Anna Dominiczak and Alex Alderton, Roberto Amato, Jeffrey Barrett, Sonia Goncalves, Ewan Harrison, David K. Jackson, Ian Johnston, Dominic Kwiatkowski, Cordelia Langford, John Sillitoe on behalf of the Wellcome Sanger Institute COVID-19 Surveillance Team |
| EPI_ISL_2865996 | Lighthouse Lab in Alderley Park | Wellcome Sanger Institute for the COVID-19 Genomics UK (COG-UK) Consortium | Jacquelyn Wynn, Mairead Hyland, The Lighthouse Lab in Alderley Park and Alex Alderton, Roberto Amato, Jeffrey Barrett, Sonia Goncalves, Ewan Harrison, David K. Jackson, Ian Johnston, Dominic Kwiatkowski, Cordelia Langford, John Sillitoe on behalf of the Wellcome Sanger Institute COVID-19 Surveillance Team |
| EPI_ISL_2865997 | Lighthouse Lab in Milton Keynes | Wellcome Sanger Institute for the COVID-19 Genomics UK (COG-UK) Consortium | The Lighthouse Lab in Milton Keynes and Alex Alderton, Roberto Amato, Jeffrey Barrett, Sonia Goncalves, Ewan Harrison, David K. Jackson, Ian Johnston, Dominic Kwiatkowski, Cordelia Langford, John Sillitoe on behalf of the Wellcome Sanger Institute COVID-19 Surveillance Team |
| EPI_ISL_2865998, EPI_ISL_2865999 | Lighthouse Lab in Alderley Park | Wellcome Sanger Institute for the COVID-19 Genomics UK (COG-UK) Consortium | Jacquelyn Wynn, Mairead Hyland, The Lighthouse Lab in Alderley Park and Alex Alderton, Roberto Amato, Jeffrey Barrett, Sonia Goncalves, Ewan Harrison, David K. Jackson, Ian Johnston, Dominic Kwiatkowski, Cordelia Langford, John Sillitoe on behalf of the Wellcome Sanger Institute COVID-19 Surveillance Team |
| EPI_ISL_2866000 | Lighthouse Lab in Milton Keynes | Wellcome Sanger Institute for the COVID-19 Genomics UK (COG-UK) Consortium | The Lighthouse Lab in Milton Keynes and Alex Alderton, Roberto Amato, Jeffrey Barrett, Sonia Goncalves, Ewan Harrison, David K. Jackson, Ian Johnston, Dominic Kwiatkowski, Cordelia Langford, John Sillitoe on behalf of the Wellcome Sanger Institute COVID-19 Surveillance Team |
| EPI_ISL_2866001, EPI_ISL_2866002, EPI_ISL_2866003, EPI_ISL_2866004 | Lighthouse Lab in Glasgow | Wellcome Sanger Institute for the COVID-19 Genomics UK (COG-UK) Consortium | Harper VanSteenhouse, Yumi Kasai, David Gray, Carol Clugston, Anna Dominiczak and Alex Alderton, Roberto Amato, Jeffrey Barrett, Sonia Goncalves, Ewan Harrison, David K. Jackson, Ian Johnston, Dominic Kwiatkowski, Cordelia Langford, John Sillitoe on behalf of the Wellcome Sanger Institute COVID-19 Surveillance Team |
| EPI_ISL_2866005 | Lighthouse Lab in Milton Keynes | Wellcome Sanger Institute for the COVID-19 Genomics UK (COG-UK) Consortium | The Lighthouse Lab in Milton Keynes and Alex Alderton, Roberto Amato, Jeffrey Barrett, Sonia Goncalves, Ewan Harrison, David K. Jackson, Ian Johnston, Dominic Kwiatkowski, Cordelia Langford, John Sillitoe on behalf of the Wellcome Sanger Institute COVID-19 Surveillance Team |
| EPI_ISL_2866006, EPI_ISL_2866007 | Lighthouse Lab in Glasgow | Wellcome Sanger Institute for the COVID-19 Genomics UK (COG-UK) Consortium | Harper VanSteenhouse, Yumi Kasai, David Gray, Carol Clugston, Anna Dominiczak and Alex Alderton, Roberto Amato, Jeffrey Barrett, Sonia Goncalves, Ewan Harrison, David K. Jackson, Ian Johnston, Dominic Kwiatkowski, Cordelia Langford, John Sillitoe on behalf of the Wellcome Sanger Institute COVID-19 Surveillance Team |
| EPI_ISL_2866008 | Lighthouse Lab in Milton Keynes | Wellcome Sanger Institute for the COVID-19 Genomics UK (COG-UK) Consortium | The Lighthouse Lab in Milton Keynes and Alex Alderton, Roberto Amato, Jeffrey Barrett, Sonia Goncalves, Ewan Harrison, David K. Jackson, Ian Johnston, Dominic Kwiatkowski, Cordelia Langford, John Sillitoe on behalf of the Wellcome Sanger Institute COVID-19 Surveillance Team |
| EPI_ISL_2866009 | Lighthouse Lab in Alderley Park | Wellcome Sanger Institute for the COVID-19 Genomics UK (COG-UK) Consortium | Jacquelyn Wynn, Mairead Hyland, The Lighthouse Lab in Alderley Park and Alex Alderton, Roberto Amato, Jeffrey Barrett, Sonia Goncalves, Ewan Harrison, David K. Jackson, Ian Johnston, Dominic Kwiatkowski, Cordelia Langford, John Sillitoe on behalf of the Wellcome Sanger Institute COVID-19 Surveillance Team |
| EPI_ISL_2866010 | Lighthouse Lab in Glasgow | Wellcome Sanger Institute for the COVID-19 Genomics UK (COG-UK) Consortium | Harper VanSteenhouse, Yumi Kasai, David Gray, Carol Clugston, Anna Dominiczak and Alex Alderton, Roberto Amato, Jeffrey Barrett, Sonia Goncalves, Ewan Harrison, David K. Jackson, Ian Johnston, Dominic Kwiatkowski, Cordelia Langford, John Sillitoe on behalf of the Wellcome Sanger Institute COVID-19 Surveillance Team |
| EPI_ISL_2866011 | Lighthouse Lab in Milton Keynes | Wellcome Sanger Institute for the COVID-19 Genomics UK (COG-UK) Consortium | The Lighthouse Lab in Milton Keynes and Alex Alderton, Roberto Amato, Jeffrey Barrett, Sonia Goncalves, Ewan Harrison, David K. Jackson, Ian Johnston, Dominic Kwiatkowski, Cordelia Langford, John Sillitoe on behalf of the Wellcome Sanger Institute COVID-19 Surveillance Team |
| EPI_ISL_2866012 | Lighthouse Lab in Alderley Park | Wellcome Sanger Institute for the COVID-19 Genomics UK (COG-UK) Consortium | Jacquelyn Wynn, Mairead Hyland, The Lighthouse Lab in Alderley Park and Alex Alderton, Roberto Amato, Jeffrey Barrett, Sonia Goncalves, Ewan Harrison, David K. Jackson, Ian Johnston, Dominic Kwiatkowski, Cordelia Langford, John Sillitoe on behalf of the Wellcome Sanger Institute COVID-19 Surveillance Team |
| EPI_ISL_2866013 | Lighthouse Lab in Glasgow | Wellcome Sanger Institute for the COVID-19 Genomics UK (COG-UK) Consortium | Harper VanSteenhouse, Yumi Kasai, David Gray, Carol Clugston, Anna Dominiczak and Alex Alderton, Roberto Amato, Jeffrey Barrett, Sonia Goncalves, Ewan Harrison, David K. Jackson, Ian Johnston, Dominic Kwiatkowski, Cordelia Langford, John Sillitoe on behalf of the Wellcome Sanger Institute COVID-19 Surveillance Team |
| EPI_ISL_2866014 | Lighthouse Laboratory Plymouth | Wellcome Sanger Institute for the COVID-19 Genomics UK (COG-UK) Consortium | Lighthouse Laboratory Plymouth and Alex Alderton, Roberto Amato, Jeffrey Barrett, Sonia Goncalves, Ewan Harrison, David K. Jackson, Ian Johnston, Dominic Kwiatkowski, Cordelia Langford, John Sillitoe on behalf of the Wellcome Sanger Institute COVID-19 Surveillance Team |
| EPI_ISL_2866015, EPI_ISL_2866016, EPI_ISL_2866017, EPI_ISL_2866018 | Lighthouse Lab in Alderley Park | Wellcome Sanger Institute for the COVID-19 Genomics UK (COG-UK) Consortium | Jacquelyn Wynn, Mairead Hyland, The Lighthouse Lab in Alderley Park and Alex Alderton, Roberto Amato, Jeffrey Barrett, Sonia Goncalves, Ewan Harrison, David K. Jackson, Ian Johnston, Dominic Kwiatkowski, Cordelia Langford, John Sillitoe on behalf of the Wellcome Sanger Institute COVID-19 Surveillance Team |
| EPI_ISL_2866019, EPI_ISL_2866020, EPI_ISL_2866021, EPI_ISL_2866022, EPI_ISL_2866023 | Lighthouse Lab in Glasgow | Wellcome Sanger Institute for the COVID-19 Genomics UK (COG-UK) Consortium | Harper VanSteenhouse, Yumi Kasai, David Gray, Carol Clugston, Anna Dominiczak and Alex Alderton, Roberto Amato, Jeffrey Barrett, Sonia Goncalves, Ewan Harrison, David K. Jackson, Ian Johnston, Dominic Kwiatkowski, Cordelia Langford, John Sillitoe on behalf of the Wellcome Sanger Institute COVID-19 Surveillance Team |
| EPI_ISL_2866024 | Lighthouse Laboratory Plymouth | Wellcome Sanger Institute for the COVID-19 Genomics UK (COG-UK) Consortium | Lighthouse Laboratory Plymouth and Alex Alderton, Roberto Amato, Jeffrey Barrett, Sonia Goncalves, Ewan Harrison, David K. Jackson, Ian Johnston, Dominic Kwiatkowski, Cordelia Langford, John Sillitoe on behalf of the Wellcome Sanger Institute COVID-19 Surveillance Team |

| | | | |
|---|---|---|---|
| EPI_ISL_2866025, EPI_ISL_2866026, EPI_ISL_2866027, EPI_ISL_2866028 | Lighthouse Lab in Glasgow | Wellcome Sanger Institute for the COVID-19 Genomics UK (COG-UK) Consortium | Harper VanSteenhouse, Yumi Kasai, David Gray, Carol Clugston, Anna Dominiczak and Alex Alderton, Roberto Amato, Jeffrey Barrett, Sonia Goncalves, Ewan Harrison, David K. Jackson, Ian Johnston, Dominic Kwiatkowski, Cordelia Langford, John Sillitoe on behalf of the Wellcome Sanger Institute COVID-19 Surveillance Team |
| EPI_ISL_2866029 | Lighthouse Lab in Milton Keynes | Wellcome Sanger Institute for the COVID-19 Genomics UK (COG-UK) Consortium | The Lighthouse Lab in Milton Keynes and Alex Alderton, Roberto Amato, Jeffrey Barrett, Sonia Goncalves, Ewan Harrison, David K. Jackson, Ian Johnston, Dominic Kwiatkowski, Cordelia Langford, John Sillitoe on behalf of the Wellcome Sanger Institute COVID-19 Surveillance Team |
| EPI_ISL_2866030 | Lighthouse Lab in Glasgow | Wellcome Sanger Institute for the COVID-19 Genomics UK (COG-UK) Consortium | Harper VanSteenhouse, Yumi Kasai, David Gray, Carol Clugston, Anna Dominiczak and Alex Alderton, Roberto Amato, Jeffrey Barrett, Sonia Goncalves, Ewan Harrison, David K. Jackson, Ian Johnston, Dominic Kwiatkowski, Cordelia Langford, John Sillitoe on behalf of the Wellcome Sanger Institute COVID-19 Surveillance Team |
| EPI_ISL_2866031 | Lighthouse Lab in Milton Keynes | Wellcome Sanger Institute for the COVID-19 Genomics UK (COG-UK) Consortium | The Lighthouse Lab in Milton Keynes and Alex Alderton, Roberto Amato, Jeffrey Barrett, Sonia Goncalves, Ewan Harrison, David K. Jackson, Ian Johnston, Dominic Kwiatkowski, Cordelia Langford, John Sillitoe on behalf of the Wellcome Sanger Institute COVID-19 Surveillance Team |
| EPI_ISL_2866032 | Lighthouse Laboratory Plymouth | Wellcome Sanger Institute for the COVID-19 Genomics UK (COG-UK) Consortium | Lighthouse Laboratory Plymouth and Alex Alderton, Roberto Amato, Jeffrey Barrett, Sonia Goncalves, Ewan Harrison, David K. Jackson, Ian Johnston, Dominic Kwiatkowski, Cordelia Langford, John Sillitoe on behalf of the Wellcome Sanger Institute COVID-19 Surveillance Team |
| EPI_ISL_2866033 | Lighthouse Lab in Glasgow | Wellcome Sanger Institute for the COVID-19 Genomics UK (COG-UK) Consortium | Harper VanSteenhouse, Yumi Kasai, David Gray, Carol Clugston, Anna Dominiczak and Alex Alderton, Roberto Amato, Jeffrey Barrett, Sonia Goncalves, Ewan Harrison, David K. Jackson, Ian Johnston, Dominic Kwiatkowski, Cordelia Langford, John Sillitoe on behalf of the Wellcome Sanger Institute COVID-19 Surveillance Team |
| EPI_ISL_2866034 | Lighthouse Lab in Milton Keynes | Wellcome Sanger Institute for the COVID-19 Genomics UK (COG-UK) Consortium | The Lighthouse Lab in Milton Keynes and Alex Alderton, Roberto Amato, Jeffrey Barrett, Sonia Goncalves, Ewan Harrison, David K. Jackson, Ian Johnston, Dominic Kwiatkowski, Cordelia Langford, John Sillitoe on behalf of the Wellcome Sanger Institute COVID-19 Surveillance Team |
| EPI_ISL_2866035, EPI_ISL_2866036, EPI_ISL_2866037, EPI_ISL_2866038, EPI_ISL_2866039, EPI_ISL_2866040 | Lighthouse Lab in Glasgow | Wellcome Sanger Institute for the COVID-19 Genomics UK (COG-UK) Consortium | Harper VanSteenhouse, Yumi Kasai, David Gray, Carol Clugston, Anna Dominiczak and Alex Alderton, Roberto Amato, Jeffrey Barrett, Sonia Goncalves, Ewan Harrison, David K. Jackson, Ian Johnston, Dominic Kwiatkowski, Cordelia Langford, John Sillitoe on behalf of the Wellcome Sanger Institute COVID-19 Surveillance Team |
| EPI_ISL_2866041, EPI_ISL_2866042, EPI_ISL_2866043 | Lighthouse Lab in Milton Keynes | Wellcome Sanger Institute for the COVID-19 Genomics UK (COG-UK) Consortium | The Lighthouse Lab in Milton Keynes and Alex Alderton, Roberto Amato, Jeffrey Barrett, Sonia Goncalves, Ewan Harrison, David K. Jackson, Ian Johnston, Dominic Kwiatkowski, Cordelia Langford, John Sillitoe on behalf of the Wellcome Sanger Institute COVID-19 Surveillance Team |
| EPI_ISL_2866044, EPI_ISL_2866045, EPI_ISL_2866046, EPI_ISL_2866047 | Lighthouse Lab in Glasgow | Wellcome Sanger Institute for the COVID-19 Genomics UK (COG-UK) Consortium | Harper VanSteenhouse, Yumi Kasai, David Gray, Carol Clugston, Anna Dominiczak and Alex Alderton, Roberto Amato, Jeffrey Barrett, Sonia Goncalves, Ewan Harrison, David K. Jackson, Ian Johnston, Dominic Kwiatkowski, Cordelia Langford, John Sillitoe on behalf of the Wellcome Sanger Institute COVID-19 Surveillance Team |
| EPI_ISL_2866048 | Lighthouse Lab in Alderley Park | Wellcome Sanger Institute for the COVID-19 Genomics UK (COG-UK) Consortium | Jacquelyn Wynn, Mairead Hyland, The Lighthouse Lab in Alderley Park and Alex Alderton, Roberto Amato, Jeffrey Barrett, Sonia Goncalves, Ewan Harrison, David K. Jackson, Ian Johnston, Dominic Kwiatkowski, Cordelia Langford, John Sillitoe on behalf of the Wellcome Sanger Institute COVID-19 Surveillance Team |
| EPI_ISL_2866049, EPI_ISL_2866050 | Lighthouse Lab in Glasgow | Wellcome Sanger Institute for the COVID-19 Genomics UK (COG-UK) Consortium | Harper VanSteenhouse, Yumi Kasai, David Gray, Carol Clugston, Anna Dominiczak and Alex Alderton, Roberto Amato, Jeffrey Barrett, Sonia Goncalves, Ewan Harrison, David K. Jackson, Ian Johnston, Dominic Kwiatkowski, Cordelia Langford, John Sillitoe on behalf of the Wellcome Sanger Institute COVID-19 Surveillance Team |
| EPI_ISL_2866051 | Lighthouse Lab in Milton Keynes | Wellcome Sanger Institute for the COVID-19 Genomics UK (COG-UK) Consortium | The Lighthouse Lab in Milton Keynes and Alex Alderton, Roberto Amato, Jeffrey Barrett, Sonia Goncalves, Ewan Harrison, David K. Jackson, Ian Johnston, Dominic Kwiatkowski, Cordelia Langford, John Sillitoe on behalf of the Wellcome Sanger Institute COVID-19 Surveillance Team |
| EPI_ISL_2866052, EPI_ISL_2866053, EPI_ISL_2866054, EPI_ISL_2866055, EPI_ISL_2866056 | Lighthouse Lab in Glasgow | Wellcome Sanger Institute for the COVID-19 Genomics UK (COG-UK) Consortium | Harper VanSteenhouse, Yumi Kasai, David Gray, Carol Clugston, Anna Dominiczak and Alex Alderton, Roberto Amato, Jeffrey Barrett, Sonia Goncalves, Ewan Harrison, David K. Jackson, Ian Johnston, Dominic Kwiatkowski, Cordelia Langford, John Sillitoe on behalf of the Wellcome Sanger Institute COVID-19 Surveillance Team |
| EPI_ISL_2866057 | Lighthouse Lab in Milton Keynes | Wellcome Sanger Institute for the COVID-19 Genomics UK (COG-UK) Consortium | The Lighthouse Lab in Milton Keynes and Alex Alderton, Roberto Amato, Jeffrey Barrett, Sonia Goncalves, Ewan Harrison, David K. Jackson, Ian Johnston, Dominic Kwiatkowski, Cordelia Langford, John Sillitoe on behalf of the Wellcome Sanger Institute COVID-19 Surveillance Team |
| EPI_ISL_2866058 | Lighthouse Lab in Alderley Park | Wellcome Sanger Institute for the COVID-19 Genomics UK (COG-UK) Consortium | Jacquelyn Wynn, Mairead Hyland, The Lighthouse Lab in Alderley Park and Alex Alderton, Roberto Amato, Jeffrey Barrett, Sonia Goncalves, Ewan Harrison, David K. Jackson, Ian Johnston, Dominic Kwiatkowski, Cordelia Langford, John Sillitoe on behalf of the Wellcome Sanger Institute COVID-19 Surveillance Team |
| EPI_ISL_2866059 | Lighthouse Lab in Milton Keynes | Wellcome Sanger Institute for the COVID-19 Genomics UK (COG-UK) Consortium | The Lighthouse Lab in Milton Keynes and Alex Alderton, Roberto Amato, Jeffrey Barrett, Sonia Goncalves, Ewan Harrison, David K. Jackson, Ian Johnston, Dominic Kwiatkowski, Cordelia Langford, John Sillitoe on behalf of the Wellcome Sanger Institute COVID-19 Surveillance Team |
| EPI_ISL_2866060, EPI_ISL_2866061 | Lighthouse Lab in Alderley Park | Wellcome Sanger Institute for the COVID-19 Genomics UK (COG-UK) Consortium | Jacquelyn Wynn, Mairead Hyland, The Lighthouse Lab in Alderley Park and Alex Alderton, Roberto Amato, Jeffrey Barrett, Sonia Goncalves, Ewan Harrison, David K. Jackson, Ian Johnston, Dominic Kwiatkowski, Cordelia Langford, John Sillitoe on behalf of the Wellcome Sanger Institute COVID-19 Surveillance Team |
| EPI_ISL_2866062 | Lighthouse Lab in Glasgow | Wellcome Sanger Institute for the COVID-19 Genomics UK (COG-UK) Consortium | Harper VanSteenhouse, Yumi Kasai, David Gray, Carol Clugston, Anna Dominiczak and Alex Alderton, Roberto Amato, Jeffrey Barrett, Sonia Goncalves, Ewan Harrison, David K. Jackson, Ian Johnston, Dominic Kwiatkowski, Cordelia Langford, John Sillitoe on behalf of the Wellcome Sanger Institute COVID-19 Surveillance Team |
| EPI_ISL_2866063 | Lighthouse Lab in Alderley Park | Wellcome Sanger Institute for the COVID-19 Genomics UK (COG-UK) Consortium | Jacquelyn Wynn, Mairead Hyland, The Lighthouse Lab in Alderley Park and Alex Alderton, Roberto Amato, Jeffrey Barrett, Sonia Goncalves, Ewan Harrison, David K. Jackson, Ian Johnston, Dominic Kwiatkowski, Cordelia Langford, John Sillitoe on behalf of the Wellcome Sanger Institute COVID-19 Surveillance Team |
| EPI_ISL_2866064 | Lighthouse Laboratory Plymouth | Wellcome Sanger Institute for the COVID-19 Genomics UK (COG-UK) Consortium | Lighthouse Laboratory Plymouth and Alex Alderton, Roberto Amato, Jeffrey Barrett, Sonia Goncalves, Ewan Harrison, David K. Jackson, Ian Johnston, Dominic Kwiatkowski, Cordelia Langford, John Sillitoe on behalf of the Wellcome Sanger Institute COVID-19 Surveillance Team |
| EPI_ISL_2866065, EPI_ISL_2866066, EPI_ISL_2866067 | Lighthouse Lab in Milton Keynes | Wellcome Sanger Institute for the COVID-19 Genomics UK (COG-UK) Consortium | The Lighthouse Lab in Milton Keynes and Alex Alderton, Roberto Amato, Jeffrey Barrett, Sonia Goncalves, Ewan Harrison, David K. Jackson, Ian Johnston, Dominic Kwiatkowski, Cordelia Langford, John Sillitoe on behalf of the Wellcome Sanger Institute COVID-19 Surveillance Team |
| EPI_ISL_2866068, EPI_ISL_2866069, EPI_ISL_2866070 | Lighthouse Laboratory Plymouth | Wellcome Sanger Institute for the COVID-19 Genomics UK (COG-UK) Consortium | Lighthouse Laboratory Plymouth and Alex Alderton, Roberto Amato, Jeffrey Barrett, Sonia Goncalves, Ewan Harrison, David K. Jackson, Ian Johnston, Dominic Kwiatkowski, Cordelia Langford, John Sillitoe on behalf of the Wellcome Sanger Institute COVID-19 Surveillance Team |
| EPI_ISL_2866071 | Lighthouse Lab in Milton Keynes | Wellcome Sanger Institute for the COVID-19 Genomics UK (COG-UK) Consortium | The Lighthouse Lab in Milton Keynes and Alex Alderton, Roberto Amato, Jeffrey Barrett, Sonia Goncalves, Ewan Harrison, David K. Jackson, Ian Johnston, Dominic Kwiatkowski, Cordelia Langford, John Sillitoe on behalf of the Wellcome Sanger Institute COVID-19 Surveillance Team |
| EPI_ISL_2866072, EPI_ISL_2866073 | Lighthouse Laboratory Plymouth | Wellcome Sanger Institute for the COVID-19 Genomics UK (COG-UK) Consortium | Lighthouse Laboratory Plymouth and Alex Alderton, Roberto Amato, Jeffrey Barrett, Sonia Goncalves, Ewan Harrison, David K. Jackson, Ian Johnston, Dominic Kwiatkowski, Cordelia Langford, John Sillitoe on behalf of the Wellcome Sanger Institute COVID-19 Surveillance Team |
| EPI_ISL_2866074 | Lighthouse Lab in Milton Keynes | Wellcome Sanger Institute for the COVID-19 Genomics UK (COG-UK) Consortium | The Lighthouse Lab in Milton Keynes and Alex Alderton, Roberto Amato, Jeffrey Barrett, Sonia Goncalves, Ewan Harrison, David K. Jackson, Ian Johnston, Dominic Kwiatkowski, Cordelia Langford, John Sillitoe on behalf of the Wellcome Sanger Institute COVID-19 Surveillance Team |
| EPI_ISL_2866075 | Lighthouse Laboratory Plymouth | Wellcome Sanger Institute for the COVID-19 Genomics UK (COG-UK) Consortium | Lighthouse Laboratory Plymouth and Alex Alderton, Roberto Amato, Jeffrey Barrett, Sonia Goncalves, Ewan Harrison, David K. Jackson, Ian Johnston, Dominic Kwiatkowski, Cordelia Langford, John Sillitoe on behalf of the Wellcome Sanger Institute COVID-19 Surveillance Team |
| EPI_ISL_2866076 | Lighthouse Lab in Milton Keynes | Wellcome Sanger Institute for the COVID-19 Genomics UK (COG-UK) Consortium | The Lighthouse Lab in Milton Keynes and Alex Alderton, Roberto Amato, Jeffrey Barrett, Sonia Goncalves, Ewan Harrison, David K. Jackson, Ian Johnston, Dominic Kwiatkowski, Cordelia Langford, John Sillitoe on behalf of the Wellcome Sanger Institute COVID-19 Surveillance Team |
| EPI_ISL_2866077 | Lighthouse Laboratory Plymouth | Wellcome Sanger Institute for the COVID-19 Genomics UK (COG-UK) Consortium | Lighthouse Laboratory Plymouth and Alex Alderton, Roberto Amato, Jeffrey Barrett, Sonia Goncalves, Ewan Harrison, David K. Jackson, Ian Johnston, Dominic Kwiatkowski, Cordelia Langford, John Sillitoe on behalf of the Wellcome Sanger Institute COVID-19 Surveillance Team |
| EPI_ISL_2866078 | Lighthouse Lab in Milton Keynes | Wellcome Sanger Institute for the COVID-19 Genomics UK (COG-UK) Consortium | The Lighthouse Lab in Milton Keynes and Alex Alderton, Roberto Amato, Jeffrey Barrett, Sonia Goncalves, Ewan Harrison, David K. Jackson, Ian Johnston, Dominic Kwiatkowski, Cordelia Langford, John Sillitoe on behalf of the Wellcome Sanger Institute COVID-19 Surveillance Team |
| EPI_ISL_2866079 | Lighthouse Laboratory Plymouth | Wellcome Sanger Institute for the COVID-19 Genomics UK (COG-UK) Consortium | Lighthouse Laboratory Plymouth and Alex Alderton, Roberto Amato, Jeffrey Barrett, Sonia Goncalves, Ewan Harrison, David K. Jackson, Ian Johnston, Dominic Kwiatkowski, Cordelia Langford, John Sillitoe on behalf of the Wellcome Sanger Institute COVID-19 Surveillance Team |
| EPI_ISL_2866080, EPI_ISL_2866081 | Lighthouse Lab in Milton Keynes | Wellcome Sanger Institute for the COVID-19 Genomics UK (COG-UK) Consortium | The Lighthouse Lab in Milton Keynes and Alex Alderton, Roberto Amato, Jeffrey Barrett, Sonia Goncalves, Ewan Harrison, David K. Jackson, Ian Johnston, Dominic Kwiatkowski, Cordelia Langford, John Sillitoe on behalf of the Wellcome Sanger Institute COVID-19 Surveillance Team |
| EPI_ISL_2866082, EPI_ISL_2866083, EPI_ISL_2866084, EPI_ISL_2866085, EPI_ISL_2866086, EPI_ISL_2866087 | Lighthouse Laboratory Plymouth | Wellcome Sanger Institute for the COVID-19 Genomics UK (COG-UK) Consortium | Lighthouse Laboratory Plymouth and Alex Alderton, Roberto Amato, Jeffrey Barrett, Sonia Goncalves, Ewan Harrison, David K. Jackson, Ian Johnston, Dominic Kwiatkowski, Cordelia Langford, John Sillitoe on behalf of the Wellcome Sanger Institute COVID-19 Surveillance Team |
| EPI_ISL_2866088 | Lighthouse Lab in Milton Keynes | Wellcome Sanger Institute for the COVID-19 Genomics UK (COG-UK) Consortium | The Lighthouse Lab in Milton Keynes and Alex Alderton, Roberto Amato, Jeffrey Barrett, Sonia Goncalves, Ewan Harrison, David K. Jackson, Ian Johnston, Dominic Kwiatkowski, Cordelia Langford, John Sillitoe on behalf of the Wellcome Sanger Institute COVID-19 Surveillance Team |
| EPI_ISL_2866089 | Lighthouse Laboratory Plymouth | Wellcome Sanger Institute for the COVID-19 Genomics UK (COG-UK) Consortium | Lighthouse Laboratory Plymouth and Alex Alderton, Roberto Amato, Jeffrey Barrett, Sonia Goncalves, Ewan Harrison, David K. Jackson, Ian Johnston, Dominic Kwiatkowski, Cordelia Langford, John Sillitoe on behalf of the Wellcome Sanger Institute COVID-19 Surveillance Team |
| EPI_ISL_2866090 | Lighthouse Lab in Milton Keynes | Wellcome Sanger Institute for the COVID-19 Genomics UK (COG-UK) Consortium | The Lighthouse Lab in Milton Keynes and Alex Alderton, Roberto Amato, Jeffrey Barrett, Sonia Goncalves, Ewan Harrison, David K. Jackson, Ian Johnston, Dominic Kwiatkowski, Cordelia Langford, John Sillitoe on behalf of the Wellcome Sanger Institute COVID-19 Surveillance Team |
| EPI_ISL_2866091, EPI_ISL_2866092, EPI_ISL_2866093, | Lighthouse Laboratory Plymouth | Wellcome Sanger Institute for the COVID-19 Genomics UK (COG-UK) Consortium | Lighthouse Laboratory Plymouth and Alex Alderton, Roberto Amato, Jeffrey Barrett, Sonia Goncalves, Ewan Harrison, David K. Jackson, Ian Johnston, Dominic Kwiatkowski, Cordelia Langford, John Sillitoe on behalf of the Wellcome Sanger Institute COVID-19 Surveillance Team |

| | | | |
|---|---|---|---|
| EPI_ISL_2866094, EPI_ISL_2866095 | | | |
| EPI_ISL_2866096, EPI_ISL_2866097 | Lighthouse Lab in Milton Keynes | Wellcome Sanger Institute for the COVID-19 Genomics UK (COG-UK) Consortium | The Lighthouse Lab in Milton Keynes and Alex Alderton, Roberto Amato, Jeffrey Barrett, Sonia Goncalves, Ewan Harrison, David K. Jackson, Ian Johnston, Dominic Kwiatkowski, Cordelia Langford, John Sillitoe on behalf of the Wellcome Sanger Institute COVID-19 Surveillance Team |
| EPI_ISL_2866098, EPI_ISL_2866099, EPI_ISL_2866100, EPI_ISL_2866101, EPI_ISL_2866102, EPI_ISL_2866103, EPI_ISL_2866104, EPI_ISL_2866105 | Lighthouse Laboratory Plymouth | Wellcome Sanger Institute for the COVID-19 Genomics UK (COG-UK) Consortium | Lighthouse Laboratory Plymouth and Alex Alderton, Roberto Amato, Jeffrey Barrett, Sonia Goncalves, Ewan Harrison, David K. Jackson, Ian Johnston, Dominic Kwiatkowski, Cordelia Langford, John Sillitoe on behalf of the Wellcome Sanger Institute COVID-19 Surveillance Team |
| EPI_ISL_2866106, EPI_ISL_2866107, EPI_ISL_2866108, EPI_ISL_2866109, EPI_ISL_2866110, EPI_ISL_2866111 | Lighthouse Lab in Milton Keynes | Wellcome Sanger Institute for the COVID-19 Genomics UK (COG-UK) Consortium | The Lighthouse Lab in Milton Keynes and Alex Alderton, Roberto Amato, Jeffrey Barrett, Sonia Goncalves, Ewan Harrison, David K. Jackson, Ian Johnston, Dominic Kwiatkowski, Cordelia Langford, John Sillitoe on behalf of the Wellcome Sanger Institute COVID-19 Surveillance Team |
| EPI_ISL_2866112, EPI_ISL_2866113 | Lighthouse Laboratory Plymouth | Wellcome Sanger Institute for the COVID-19 Genomics UK (COG-UK) Consortium | Lighthouse Laboratory Plymouth and Alex Alderton, Roberto Amato, Jeffrey Barrett, Sonia Goncalves, Ewan Harrison, David K. Jackson, Ian Johnston, Dominic Kwiatkowski, Cordelia Langford, John Sillitoe on behalf of the Wellcome Sanger Institute COVID-19 Surveillance Team |
| EPI_ISL_2866114 | Lighthouse Lab in Milton Keynes | Wellcome Sanger Institute for the COVID-19 Genomics UK (COG-UK) Consortium | The Lighthouse Lab in Milton Keynes and Alex Alderton, Roberto Amato, Jeffrey Barrett, Sonia Goncalves, Ewan Harrison, David K. Jackson, Ian Johnston, Dominic Kwiatkowski, Cordelia Langford, John Sillitoe on behalf of the Wellcome Sanger Institute COVID-19 Surveillance Team |
| EPI_ISL_2866115 | Lighthouse Laboratory Plymouth | Wellcome Sanger Institute for the COVID-19 Genomics UK (COG-UK) Consortium | Lighthouse Laboratory Plymouth and Alex Alderton, Roberto Amato, Jeffrey Barrett, Sonia Goncalves, Ewan Harrison, David K. Jackson, Ian Johnston, Dominic Kwiatkowski, Cordelia Langford, John Sillitoe on behalf of the Wellcome Sanger Institute COVID-19 Surveillance Team |
| EPI_ISL_2866116 | Lighthouse Lab in Milton Keynes | Wellcome Sanger Institute for the COVID-19 Genomics UK (COG-UK) Consortium | The Lighthouse Lab in Milton Keynes and Alex Alderton, Roberto Amato, Jeffrey Barrett, Sonia Goncalves, Ewan Harrison, David K. Jackson, Ian Johnston, Dominic Kwiatkowski, Cordelia Langford, John Sillitoe on behalf of the Wellcome Sanger Institute COVID-19 Surveillance Team |
| EPI_ISL_2866117 | Lighthouse Laboratory Plymouth | Wellcome Sanger Institute for the COVID-19 Genomics UK (COG-UK) Consortium | Lighthouse Laboratory Plymouth and Alex Alderton, Roberto Amato, Jeffrey Barrett, Sonia Goncalves, Ewan Harrison, David K. Jackson, Ian Johnston, Dominic Kwiatkowski, Cordelia Langford, John Sillitoe on behalf of the Wellcome Sanger Institute COVID-19 Surveillance Team |
| EPI_ISL_2866118, EPI_ISL_2866119 | Lighthouse Lab in Milton Keynes | Wellcome Sanger Institute for the COVID-19 Genomics UK (COG-UK) Consortium | The Lighthouse Lab in Milton Keynes and Alex Alderton, Roberto Amato, Jeffrey Barrett, Sonia Goncalves, Ewan Harrison, David K. Jackson, Ian Johnston, Dominic Kwiatkowski, Cordelia Langford, John Sillitoe on behalf of the Wellcome Sanger Institute COVID-19 Surveillance Team |
| EPI_ISL_2866120, EPI_ISL_2866121, EPI_ISL_2866122, EPI_ISL_2866123, EPI_ISL_2866124, EPI_ISL_2866125, EPI_ISL_2866126, EPI_ISL_2866127 | Lighthouse Laboratory Plymouth | Wellcome Sanger Institute for the COVID-19 Genomics UK (COG-UK) Consortium | Lighthouse Laboratory Plymouth and Alex Alderton, Roberto Amato, Jeffrey Barrett, Sonia Goncalves, Ewan Harrison, David K. Jackson, Ian Johnston, Dominic Kwiatkowski, Cordelia Langford, John Sillitoe on behalf of the Wellcome Sanger Institute COVID-19 Surveillance Team |
| EPI_ISL_2866128 | Lighthouse Lab in Milton Keynes | Wellcome Sanger Institute for the COVID-19 Genomics UK (COG-UK) Consortium | The Lighthouse Lab in Milton Keynes and Alex Alderton, Roberto Amato, Jeffrey Barrett, Sonia Goncalves, Ewan Harrison, David K. Jackson, Ian Johnston, Dominic Kwiatkowski, Cordelia Langford, John Sillitoe on behalf of the Wellcome Sanger Institute COVID-19 Surveillance Team |
| EPI_ISL_2866129 | Lighthouse Laboratory Plymouth | Wellcome Sanger Institute for the COVID-19 Genomics UK (COG-UK) Consortium | Lighthouse Laboratory Plymouth and Alex Alderton, Roberto Amato, Jeffrey Barrett, Sonia Goncalves, Ewan Harrison, David K. Jackson, Ian Johnston, Dominic Kwiatkowski, Cordelia Langford, John Sillitoe on behalf of the Wellcome Sanger Institute COVID-19 Surveillance Team |
| EPI_ISL_2866130 | Lighthouse Lab in Milton Keynes | Wellcome Sanger Institute for the COVID-19 Genomics UK (COG-UK) Consortium | The Lighthouse Lab in Milton Keynes and Alex Alderton, Roberto Amato, Jeffrey Barrett, Sonia Goncalves, Ewan Harrison, David K. Jackson, Ian Johnston, Dominic Kwiatkowski, Cordelia Langford, John Sillitoe on behalf of the Wellcome Sanger Institute COVID-19 Surveillance Team |
| EPI_ISL_2866131, EPI_ISL_2866132, EPI_ISL_2866133 | Lighthouse Laboratory Plymouth | Wellcome Sanger Institute for the COVID-19 Genomics UK (COG-UK) Consortium | Lighthouse Laboratory Plymouth and Alex Alderton, Roberto Amato, Jeffrey Barrett, Sonia Goncalves, Ewan Harrison, David K. Jackson, Ian Johnston, Dominic Kwiatkowski, Cordelia Langford, John Sillitoe on behalf of the Wellcome Sanger Institute COVID-19 Surveillance Team |
| EPI_ISL_2866134 | Lighthouse Lab in Milton Keynes | Wellcome Sanger Institute for the COVID-19 Genomics UK (COG-UK) Consortium | The Lighthouse Lab in Milton Keynes and Alex Alderton, Roberto Amato, Jeffrey Barrett, Sonia Goncalves, Ewan Harrison, David K. Jackson, Ian Johnston, Dominic Kwiatkowski, Cordelia Langford, John Sillitoe on behalf of the Wellcome Sanger Institute COVID-19 Surveillance Team |
| EPI_ISL_2866135, EPI_ISL_2866136 | Lighthouse Laboratory Plymouth | Wellcome Sanger Institute for the COVID-19 Genomics UK (COG-UK) Consortium | Lighthouse Laboratory Plymouth and Alex Alderton, Roberto Amato, Jeffrey Barrett, Sonia Goncalves, Ewan Harrison, David K. Jackson, Ian Johnston, Dominic Kwiatkowski, Cordelia Langford, John Sillitoe on behalf of the Wellcome Sanger Institute COVID-19 Surveillance Team |
| EPI_ISL_2866137, EPI_ISL_2866138, EPI_ISL_2866139 | Lighthouse Lab in Milton Keynes | Wellcome Sanger Institute for the COVID-19 Genomics UK (COG-UK) Consortium | The Lighthouse Lab in Milton Keynes and Alex Alderton, Roberto Amato, Jeffrey Barrett, Sonia Goncalves, Ewan Harrison, David K. Jackson, Ian Johnston, Dominic Kwiatkowski, Cordelia Langford, John Sillitoe on behalf of the Wellcome Sanger Institute COVID-19 Surveillance Team |
| EPI_ISL_2866140, EPI_ISL_2866141 | Lighthouse Laboratory Plymouth | Wellcome Sanger Institute for the COVID-19 Genomics UK (COG-UK) Consortium | Lighthouse Laboratory Plymouth and Alex Alderton, Roberto Amato, Jeffrey Barrett, Sonia Goncalves, Ewan Harrison, David K. Jackson, Ian Johnston, Dominic Kwiatkowski, Cordelia Langford, John Sillitoe on behalf of the Wellcome Sanger Institute COVID-19 Surveillance Team |
| EPI_ISL_2866142 | Lighthouse Lab in Milton Keynes | Wellcome Sanger Institute for the COVID-19 Genomics UK (COG-UK) Consortium | The Lighthouse Lab in Milton Keynes and Alex Alderton, Roberto Amato, Jeffrey Barrett, Sonia Goncalves, Ewan Harrison, David K. Jackson, Ian Johnston, Dominic Kwiatkowski, Cordelia Langford, John Sillitoe on behalf of the Wellcome Sanger Institute COVID-19 Surveillance Team |
| EPI_ISL_2866143, EPI_ISL_2866144 | Lighthouse Laboratory Plymouth | Wellcome Sanger Institute for the COVID-19 Genomics UK (COG-UK) Consortium | Lighthouse Laboratory Plymouth and Alex Alderton, Roberto Amato, Jeffrey Barrett, Sonia Goncalves, Ewan Harrison, David K. Jackson, Ian Johnston, Dominic Kwiatkowski, Cordelia Langford, John Sillitoe on behalf of the Wellcome Sanger Institute COVID-19 Surveillance Team |
| EPI_ISL_2866145, EPI_ISL_2866146, EPI_ISL_2866147 | Lighthouse Lab in Milton Keynes | Wellcome Sanger Institute for the COVID-19 Genomics UK (COG-UK) Consortium | The Lighthouse Lab in Milton Keynes and Alex Alderton, Roberto Amato, Jeffrey Barrett, Sonia Goncalves, Ewan Harrison, David K. Jackson, Ian Johnston, Dominic Kwiatkowski, Cordelia Langford, John Sillitoe on behalf of the Wellcome Sanger Institute COVID-19 Surveillance Team |
| EPI_ISL_2866148 | Lighthouse Laboratory Plymouth | Wellcome Sanger Institute for the COVID-19 Genomics UK (COG-UK) Consortium | Lighthouse Laboratory Plymouth and Alex Alderton, Roberto Amato, Jeffrey Barrett, Sonia Goncalves, Ewan Harrison, David K. Jackson, Ian Johnston, Dominic Kwiatkowski, Cordelia Langford, John Sillitoe on behalf of the Wellcome Sanger Institute COVID-19 Surveillance Team |
| EPI_ISL_2866149 | Lighthouse Lab in Milton Keynes | Wellcome Sanger Institute for the COVID-19 Genomics UK (COG-UK) Consortium | The Lighthouse Lab in Milton Keynes and Alex Alderton, Roberto Amato, Jeffrey Barrett, Sonia Goncalves, Ewan Harrison, David K. Jackson, Ian Johnston, Dominic Kwiatkowski, Cordelia Langford, John Sillitoe on behalf of the Wellcome Sanger Institute COVID-19 Surveillance Team |
| EPI_ISL_2866150, EPI_ISL_2866151, EPI_ISL_2866152, EPI_ISL_2866153 | Lighthouse Laboratory Plymouth | Wellcome Sanger Institute for the COVID-19 Genomics UK (COG-UK) Consortium | Lighthouse Laboratory Plymouth and Alex Alderton, Roberto Amato, Jeffrey Barrett, Sonia Goncalves, Ewan Harrison, David K. Jackson, Ian Johnston, Dominic Kwiatkowski, Cordelia Langford, John Sillitoe on behalf of the Wellcome Sanger Institute COVID-19 Surveillance Team |
| EPI_ISL_2866154, EPI_ISL_2866155 | Lighthouse Lab in Milton Keynes | Wellcome Sanger Institute for the COVID-19 Genomics UK (COG-UK) Consortium | The Lighthouse Lab in Milton Keynes and Alex Alderton, Roberto Amato, Jeffrey Barrett, Sonia Goncalves, Ewan Harrison, David K. Jackson, Ian Johnston, Dominic Kwiatkowski, Cordelia Langford, John Sillitoe on behalf of the Wellcome Sanger Institute COVID-19 Surveillance Team |
| EPI_ISL_2866156 | Lighthouse Laboratory Plymouth | Wellcome Sanger Institute for the COVID-19 Genomics UK (COG-UK) Consortium | Lighthouse Laboratory Plymouth and Alex Alderton, Roberto Amato, Jeffrey Barrett, Sonia Goncalves, Ewan Harrison, David K. Jackson, Ian Johnston, Dominic Kwiatkowski, Cordelia Langford, John Sillitoe on behalf of the Wellcome Sanger Institute COVID-19 Surveillance Team |
| EPI_ISL_2866157 | Lighthouse Lab in Milton Keynes | Wellcome Sanger Institute for the COVID-19 Genomics UK (COG-UK) Consortium | The Lighthouse Lab in Milton Keynes and Alex Alderton, Roberto Amato, Jeffrey Barrett, Sonia Goncalves, Ewan Harrison, David K. Jackson, Ian Johnston, Dominic Kwiatkowski, Cordelia Langford, John Sillitoe on behalf of the Wellcome Sanger Institute COVID-19 Surveillance Team |
| EPI_ISL_2866158 | Lighthouse Laboratory Plymouth | Wellcome Sanger Institute for the COVID-19 Genomics UK (COG-UK) Consortium | Lighthouse Laboratory Plymouth and Alex Alderton, Roberto Amato, Jeffrey Barrett, Sonia Goncalves, Ewan Harrison, David K. Jackson, Ian Johnston, Dominic Kwiatkowski, Cordelia Langford, John Sillitoe on behalf of the Wellcome Sanger Institute COVID-19 Surveillance Team |
| EPI_ISL_2866159 | Lighthouse Lab in Milton Keynes | Wellcome Sanger Institute for the COVID-19 Genomics UK (COG-UK) Consortium | The Lighthouse Lab in Milton Keynes and Alex Alderton, Roberto Amato, Jeffrey Barrett, Sonia Goncalves, Ewan Harrison, David K. Jackson, Ian Johnston, Dominic Kwiatkowski, Cordelia Langford, John Sillitoe on behalf of the Wellcome Sanger Institute COVID-19 Surveillance Team |
| EPI_ISL_2866160, EPI_ISL_2866161, EPI_ISL_2866162, EPI_ISL_2866163, EPI_ISL_2866164, EPI_ISL_2866165, EPI_ISL_2866166 | Lighthouse Laboratory Plymouth | Wellcome Sanger Institute for the COVID-19 Genomics UK (COG-UK) Consortium | Lighthouse Laboratory Plymouth and Alex Alderton, Roberto Amato, Jeffrey Barrett, Sonia Goncalves, Ewan Harrison, David K. Jackson, Ian Johnston, Dominic Kwiatkowski, Cordelia Langford, John Sillitoe on behalf of the Wellcome Sanger Institute COVID-19 Surveillance Team |
| EPI_ISL_2866167 | Lighthouse Lab in Milton Keynes | Wellcome Sanger Institute for the COVID-19 Genomics UK (COG-UK) Consortium | The Lighthouse Lab in Milton Keynes and Alex Alderton, Roberto Amato, Jeffrey Barrett, Sonia Goncalves, Ewan Harrison, David K. Jackson, Ian Johnston, Dominic Kwiatkowski, Cordelia Langford, John Sillitoe on behalf of the Wellcome Sanger Institute COVID-19 Surveillance Team |
| EPI_ISL_2866168 | Lighthouse Laboratory Plymouth | Wellcome Sanger Institute for the COVID-19 Genomics UK (COG-UK) Consortium | Lighthouse Laboratory Plymouth and Alex Alderton, Roberto Amato, Jeffrey Barrett, Sonia Goncalves, Ewan Harrison, David K. Jackson, Ian Johnston, Dominic Kwiatkowski, Cordelia Langford, John Sillitoe on behalf of the Wellcome Sanger Institute COVID-19 Surveillance Team |
| EPI_ISL_2866169 | Lighthouse Lab in Milton Keynes | Wellcome Sanger Institute for the COVID-19 Genomics UK (COG-UK) Consortium | The Lighthouse Lab in Milton Keynes and Alex Alderton, Roberto Amato, Jeffrey Barrett, Sonia Goncalves, Ewan Harrison, David K. Jackson, Ian Johnston, Dominic Kwiatkowski, Cordelia Langford, John Sillitoe on behalf of the Wellcome Sanger Institute COVID-19 Surveillance Team |

| | | | |
|---|---|---|---|
| EPI_ISL_2866170 | Lighthouse Laboratory Plymouth | Wellcome Sanger Institute for the COVID-19 Genomics UK (COG-UK) Consortium | Lighthouse Laboratory Plymouth and Alex Alderton, Roberto Amato, Jeffrey Barrett, Sonia Goncalves, Ewan Harrison, David K. Jackson, Ian Johnston, Dominic Kwiatkowski, Cordelia Langford, John Sillitoe on behalf of the Wellcome Sanger Institute COVID-19 Surveillance Team |
| EPI_ISL_2866171 | Lighthouse Lab in Milton Keynes | Wellcome Sanger Institute for the COVID-19 Genomics UK (COG-UK) Consortium | The Lighthouse Lab in Milton Keynes and Alex Alderton, Roberto Amato, Jeffrey Barrett, Sonia Goncalves, Ewan Harrison, David K. Jackson, Ian Johnston, Dominic Kwiatkowski, Cordelia Langford, John Sillitoe on behalf of the Wellcome Sanger Institute COVID-19 Surveillance Team |
| EPI_ISL_2866172 | Lighthouse Laboratory Plymouth | Wellcome Sanger Institute for the COVID-19 Genomics UK (COG-UK) Consortium | Lighthouse Laboratory Plymouth and Alex Alderton, Roberto Amato, Jeffrey Barrett, Sonia Goncalves, Ewan Harrison, David K. Jackson, Ian Johnston, Dominic Kwiatkowski, Cordelia Langford, John Sillitoe on behalf of the Wellcome Sanger Institute COVID-19 Surveillance Team |
| EPI_ISL_2866173 | Lighthouse Lab in Milton Keynes | Wellcome Sanger Institute for the COVID-19 Genomics UK (COG-UK) Consortium | The Lighthouse Lab in Milton Keynes and Alex Alderton, Roberto Amato, Jeffrey Barrett, Sonia Goncalves, Ewan Harrison, David K. Jackson, Ian Johnston, Dominic Kwiatkowski, Cordelia Langford, John Sillitoe on behalf of the Wellcome Sanger Institute COVID-19 Surveillance Team |
| EPI_ISL_2866174, EPI_ISL_2866175, EPI_ISL_2866176, EPI_ISL_2866177, EPI_ISL_2866178, EPI_ISL_2866179, EPI_ISL_2866180 | Lighthouse Laboratory Plymouth | Wellcome Sanger Institute for the COVID-19 Genomics UK (COG-UK) Consortium | Lighthouse Laboratory Plymouth and Alex Alderton, Roberto Amato, Jeffrey Barrett, Sonia Goncalves, Ewan Harrison, David K. Jackson, Ian Johnston, Dominic Kwiatkowski, Cordelia Langford, John Sillitoe on behalf of the Wellcome Sanger Institute COVID-19 Surveillance Team |
| EPI_ISL_2866181 | Lighthouse Lab in Milton Keynes | Wellcome Sanger Institute for the COVID-19 Genomics UK (COG-UK) Consortium | The Lighthouse Lab in Milton Keynes and Alex Alderton, Roberto Amato, Jeffrey Barrett, Sonia Goncalves, Ewan Harrison, David K. Jackson, Ian Johnston, Dominic Kwiatkowski, Cordelia Langford, John Sillitoe on behalf of the Wellcome Sanger Institute COVID-19 Surveillance Team |
| EPI_ISL_2866182, EPI_ISL_2866183, EPI_ISL_2866184, EPI_ISL_2866185, EPI_ISL_2866187, EPI_ISL_2866188 | Lighthouse Laboratory Plymouth | Wellcome Sanger Institute for the COVID-19 Genomics UK (COG-UK) Consortium | Lighthouse Laboratory Plymouth and Alex Alderton, Roberto Amato, Jeffrey Barrett, Sonia Goncalves, Ewan Harrison, David K. Jackson, Ian Johnston, Dominic Kwiatkowski, Cordelia Langford, John Sillitoe on behalf of the Wellcome Sanger Institute COVID-19 Surveillance Team |
| EPI_ISL_2866189, EPI_ISL_2866190 | Lighthouse Lab in Milton Keynes | Wellcome Sanger Institute for the COVID-19 Genomics UK (COG-UK) Consortium | The Lighthouse Lab in Milton Keynes and Alex Alderton, Roberto Amato, Jeffrey Barrett, Sonia Goncalves, Ewan Harrison, David K. Jackson, Ian Johnston, Dominic Kwiatkowski, Cordelia Langford, John Sillitoe on behalf of the Wellcome Sanger Institute COVID-19 Surveillance Team |
| EPI_ISL_2866191 | Lighthouse Laboratory Plymouth | Wellcome Sanger Institute for the COVID-19 Genomics UK (COG-UK) Consortium | Lighthouse Laboratory Plymouth and Alex Alderton, Roberto Amato, Jeffrey Barrett, Sonia Goncalves, Ewan Harrison, David K. Jackson, Ian Johnston, Dominic Kwiatkowski, Cordelia Langford, John Sillitoe on behalf of the Wellcome Sanger Institute COVID-19 Surveillance Team |
| EPI_ISL_2866192 | Lighthouse Lab in Milton Keynes | Wellcome Sanger Institute for the COVID-19 Genomics UK (COG-UK) Consortium | The Lighthouse Lab in Milton Keynes and Alex Alderton, Roberto Amato, Jeffrey Barrett, Sonia Goncalves, Ewan Harrison, David K. Jackson, Ian Johnston, Dominic Kwiatkowski, Cordelia Langford, John Sillitoe on behalf of the Wellcome Sanger Institute COVID-19 Surveillance Team |
| EPI_ISL_2866193, EPI_ISL_2866194, EPI_ISL_2866195, EPI_ISL_2866196 | Lighthouse Laboratory Plymouth | Wellcome Sanger Institute for the COVID-19 Genomics UK (COG-UK) Consortium | Lighthouse Laboratory Plymouth and Alex Alderton, Roberto Amato, Jeffrey Barrett, Sonia Goncalves, Ewan Harrison, David K. Jackson, Ian Johnston, Dominic Kwiatkowski, Cordelia Langford, John Sillitoe on behalf of the Wellcome Sanger Institute COVID-19 Surveillance Team |
| EPI_ISL_2866197 | Lighthouse Lab in Milton Keynes | Wellcome Sanger Institute for the COVID-19 Genomics UK (COG-UK) Consortium | The Lighthouse Lab in Milton Keynes and Alex Alderton, Roberto Amato, Jeffrey Barrett, Sonia Goncalves, Ewan Harrison, David K. Jackson, Ian Johnston, Dominic Kwiatkowski, Cordelia Langford, John Sillitoe on behalf of the Wellcome Sanger Institute COVID-19 Surveillance Team |
| EPI_ISL_2866198 | Lighthouse Laboratory Plymouth | Wellcome Sanger Institute for the COVID-19 Genomics UK (COG-UK) Consortium | Lighthouse Laboratory Plymouth and Alex Alderton, Roberto Amato, Jeffrey Barrett, Sonia Goncalves, Ewan Harrison, David K. Jackson, Ian Johnston, Dominic Kwiatkowski, Cordelia Langford, John Sillitoe on behalf of the Wellcome Sanger Institute COVID-19 Surveillance Team |
| EPI_ISL_2866199, EPI_ISL_2866200 | Lighthouse Lab in Milton Keynes | Wellcome Sanger Institute for the COVID-19 Genomics UK (COG-UK) Consortium | The Lighthouse Lab in Milton Keynes and Alex Alderton, Roberto Amato, Jeffrey Barrett, Sonia Goncalves, Ewan Harrison, David K. Jackson, Ian Johnston, Dominic Kwiatkowski, Cordelia Langford, John Sillitoe on behalf of the Wellcome Sanger Institute COVID-19 Surveillance Team |
| EPI_ISL_2866201, EPI_ISL_2866202 | Lighthouse Laboratory Plymouth | Wellcome Sanger Institute for the COVID-19 Genomics UK (COG-UK) Consortium | Lighthouse Laboratory Plymouth and Alex Alderton, Roberto Amato, Jeffrey Barrett, Sonia Goncalves, Ewan Harrison, David K. Jackson, Ian Johnston, Dominic Kwiatkowski, Cordelia Langford, John Sillitoe on behalf of the Wellcome Sanger Institute COVID-19 Surveillance Team |
| EPI_ISL_2866203 | Lighthouse Lab in Milton Keynes | Wellcome Sanger Institute for the COVID-19 Genomics UK (COG-UK) Consortium | The Lighthouse Lab in Milton Keynes and Alex Alderton, Roberto Amato, Jeffrey Barrett, Sonia Goncalves, Ewan Harrison, David K. Jackson, Ian Johnston, Dominic Kwiatkowski, Cordelia Langford, John Sillitoe on behalf of the Wellcome Sanger Institute COVID-19 Surveillance Team |
| EPI_ISL_2866204 | Lighthouse Laboratory Plymouth | Wellcome Sanger Institute for the COVID-19 Genomics UK (COG-UK) Consortium | Lighthouse Laboratory Plymouth and Alex Alderton, Roberto Amato, Jeffrey Barrett, Sonia Goncalves, Ewan Harrison, David K. Jackson, Ian Johnston, Dominic Kwiatkowski, Cordelia Langford, John Sillitoe on behalf of the Wellcome Sanger Institute COVID-19 Surveillance Team |
| EPI_ISL_2866205, EPI_ISL_2866206, EPI_ISL_2866207 | Lighthouse Lab in Milton Keynes | Wellcome Sanger Institute for the COVID-19 Genomics UK (COG-UK) Consortium | The Lighthouse Lab in Milton Keynes and Alex Alderton, Roberto Amato, Jeffrey Barrett, Sonia Goncalves, Ewan Harrison, David K. Jackson, Ian Johnston, Dominic Kwiatkowski, Cordelia Langford, John Sillitoe on behalf of the Wellcome Sanger Institute COVID-19 Surveillance Team |
| EPI_ISL_2866208, EPI_ISL_2866209 | Lighthouse Laboratory Plymouth | Wellcome Sanger Institute for the COVID-19 Genomics UK (COG-UK) Consortium | Lighthouse Laboratory Plymouth and Alex Alderton, Roberto Amato, Jeffrey Barrett, Sonia Goncalves, Ewan Harrison, David K. Jackson, Ian Johnston, Dominic Kwiatkowski, Cordelia Langford, John Sillitoe on behalf of the Wellcome Sanger Institute COVID-19 Surveillance Team |
| EPI_ISL_2866210, EPI_ISL_2866211, EPI_ISL_2866212 | Lighthouse Lab in Milton Keynes | Wellcome Sanger Institute for the COVID-19 Genomics UK (COG-UK) Consortium | The Lighthouse Lab in Milton Keynes and Alex Alderton, Roberto Amato, Jeffrey Barrett, Sonia Goncalves, Ewan Harrison, David K. Jackson, Ian Johnston, Dominic Kwiatkowski, Cordelia Langford, John Sillitoe on behalf of the Wellcome Sanger Institute COVID-19 Surveillance Team |
| EPI_ISL_2866213 | Lighthouse Laboratory Plymouth | Wellcome Sanger Institute for the COVID-19 Genomics UK (COG-UK) Consortium | Lighthouse Laboratory Plymouth and Alex Alderton, Roberto Amato, Jeffrey Barrett, Sonia Goncalves, Ewan Harrison, David K. Jackson, Ian Johnston, Dominic Kwiatkowski, Cordelia Langford, John Sillitoe on behalf of the Wellcome Sanger Institute COVID-19 Surveillance Team |
| EPI_ISL_2866214 | Lighthouse Lab in Milton Keynes | Wellcome Sanger Institute for the COVID-19 Genomics UK (COG-UK) Consortium | The Lighthouse Lab in Milton Keynes and Alex Alderton, Roberto Amato, Jeffrey Barrett, Sonia Goncalves, Ewan Harrison, David K. Jackson, Ian Johnston, Dominic Kwiatkowski, Cordelia Langford, John Sillitoe on behalf of the Wellcome Sanger Institute COVID-19 Surveillance Team |
| EPI_ISL_2866215, EPI_ISL_2866216, EPI_ISL_2866217 | Lighthouse Laboratory Plymouth | Wellcome Sanger Institute for the COVID-19 Genomics UK (COG-UK) Consortium | Lighthouse Laboratory Plymouth and Alex Alderton, Roberto Amato, Jeffrey Barrett, Sonia Goncalves, Ewan Harrison, David K. Jackson, Ian Johnston, Dominic Kwiatkowski, Cordelia Langford, John Sillitoe on behalf of the Wellcome Sanger Institute COVID-19 Surveillance Team |
| EPI_ISL_2866218 | Lighthouse Lab in Milton Keynes | Wellcome Sanger Institute for the COVID-19 Genomics UK (COG-UK) Consortium | The Lighthouse Lab in Milton Keynes and Alex Alderton, Roberto Amato, Jeffrey Barrett, Sonia Goncalves, Ewan Harrison, David K. Jackson, Ian Johnston, Dominic Kwiatkowski, Cordelia Langford, John Sillitoe on behalf of the Wellcome Sanger Institute COVID-19 Surveillance Team |
| EPI_ISL_2866219, EPI_ISL_2866220, EPI_ISL_2866221, EPI_ISL_2866222 | Lighthouse Laboratory Plymouth | Wellcome Sanger Institute for the COVID-19 Genomics UK (COG-UK) Consortium | Lighthouse Laboratory Plymouth and Alex Alderton, Roberto Amato, Jeffrey Barrett, Sonia Goncalves, Ewan Harrison, David K. Jackson, Ian Johnston, Dominic Kwiatkowski, Cordelia Langford, John Sillitoe on behalf of the Wellcome Sanger Institute COVID-19 Surveillance Team |
| EPI_ISL_2866223, EPI_ISL_2866224 | Lighthouse Lab in Milton Keynes | Wellcome Sanger Institute for the COVID-19 Genomics UK (COG-UK) Consortium | The Lighthouse Lab in Milton Keynes and Alex Alderton, Roberto Amato, Jeffrey Barrett, Sonia Goncalves, Ewan Harrison, David K. Jackson, Ian Johnston, Dominic Kwiatkowski, Cordelia Langford, John Sillitoe on behalf of the Wellcome Sanger Institute COVID-19 Surveillance Team |
| EPI_ISL_2866225 | Lighthouse Laboratory Plymouth | Wellcome Sanger Institute for the COVID-19 Genomics UK (COG-UK) Consortium | Lighthouse Laboratory Plymouth and Alex Alderton, Roberto Amato, Jeffrey Barrett, Sonia Goncalves, Ewan Harrison, David K. Jackson, Ian Johnston, Dominic Kwiatkowski, Cordelia Langford, John Sillitoe on behalf of the Wellcome Sanger Institute COVID-19 Surveillance Team |
| EPI_ISL_2866226 | Lighthouse Lab in Milton Keynes | Wellcome Sanger Institute for the COVID-19 Genomics UK (COG-UK) Consortium | The Lighthouse Lab in Milton Keynes and Alex Alderton, Roberto Amato, Jeffrey Barrett, Sonia Goncalves, Ewan Harrison, David K. Jackson, Ian Johnston, Dominic Kwiatkowski, Cordelia Langford, John Sillitoe on behalf of the Wellcome Sanger Institute COVID-19 Surveillance Team |
| EPI_ISL_2866227 | Lighthouse Laboratory Plymouth | Wellcome Sanger Institute for the COVID-19 Genomics UK (COG-UK) Consortium | Lighthouse Laboratory Plymouth and Alex Alderton, Roberto Amato, Jeffrey Barrett, Sonia Goncalves, Ewan Harrison, David K. Jackson, Ian Johnston, Dominic Kwiatkowski, Cordelia Langford, John Sillitoe on behalf of the Wellcome Sanger Institute COVID-19 Surveillance Team |
| EPI_ISL_2866228 | Lighthouse Lab in Milton Keynes | Wellcome Sanger Institute for the COVID-19 Genomics UK (COG-UK) Consortium | The Lighthouse Lab in Milton Keynes and Alex Alderton, Roberto Amato, Jeffrey Barrett, Sonia Goncalves, Ewan Harrison, David K. Jackson, Ian Johnston, Dominic Kwiatkowski, Cordelia Langford, John Sillitoe on behalf of the Wellcome Sanger Institute COVID-19 Surveillance Team |
| EPI_ISL_2866229 | Lighthouse Laboratory Plymouth | Wellcome Sanger Institute for the COVID-19 Genomics UK (COG-UK) Consortium | Lighthouse Laboratory Plymouth and Alex Alderton, Roberto Amato, Jeffrey Barrett, Sonia Goncalves, Ewan Harrison, David K. Jackson, Ian Johnston, Dominic Kwiatkowski, Cordelia Langford, John Sillitoe on behalf of the Wellcome Sanger Institute COVID-19 Surveillance Team |
| EPI_ISL_2866230, EPI_ISL_2866231, EPI_ISL_2866232 | Lighthouse Lab in Milton Keynes | Wellcome Sanger Institute for the COVID-19 Genomics UK (COG-UK) Consortium | The Lighthouse Lab in Milton Keynes and Alex Alderton, Roberto Amato, Jeffrey Barrett, Sonia Goncalves, Ewan Harrison, David K. Jackson, Ian Johnston, Dominic Kwiatkowski, Cordelia Langford, John Sillitoe on behalf of the Wellcome Sanger Institute COVID-19 Surveillance Team |
| EPI_ISL_2866233 | Lighthouse Laboratory Plymouth | Wellcome Sanger Institute for the COVID-19 Genomics UK (COG-UK) Consortium | Lighthouse Laboratory Plymouth and Alex Alderton, Roberto Amato, Jeffrey Barrett, Sonia Goncalves, Ewan Harrison, David K. Jackson, Ian Johnston, Dominic Kwiatkowski, Cordelia Langford, John Sillitoe on behalf of the Wellcome Sanger Institute COVID-19 Surveillance Team |
| EPI_ISL_2866234 | Lighthouse Lab in Milton Keynes | Wellcome Sanger Institute for the COVID-19 Genomics UK (COG-UK) Consortium | The Lighthouse Lab in Milton Keynes and Alex Alderton, Roberto Amato, Jeffrey Barrett, Sonia Goncalves, Ewan Harrison, David K. Jackson, Ian Johnston, Dominic Kwiatkowski, Cordelia Langford, John Sillitoe on behalf of the Wellcome Sanger Institute COVID-19 Surveillance Team |
| EPI_ISL_2866235, EPI_ISL_2866236, EPI_ISL_2866237 | Lighthouse Laboratory Plymouth | Wellcome Sanger Institute for the COVID-19 Genomics UK (COG-UK) Consortium | Lighthouse Laboratory Plymouth and Alex Alderton, Roberto Amato, Jeffrey Barrett, Sonia Goncalves, Ewan Harrison, David K. Jackson, Ian Johnston, Dominic Kwiatkowski, Cordelia Langford, John Sillitoe on behalf of the Wellcome Sanger Institute COVID-19 Surveillance Team |
| EPI_ISL_2866238 | Lighthouse Lab in Milton Keynes | Wellcome Sanger Institute for the COVID-19 Genomics UK (COG-UK) Consortium | The Lighthouse Lab in Milton Keynes and Alex Alderton, Roberto Amato, Jeffrey Barrett, Sonia Goncalves, Ewan Harrison, David K. Jackson, Ian Johnston, Dominic Kwiatkowski, Cordelia Langford, John Sillitoe on behalf of the Wellcome Sanger Institute COVID-19 Surveillance Team |
| EPI_ISL_2866239, EPI_ISL_2866240 | Lighthouse Laboratory Plymouth | Wellcome Sanger Institute for the COVID-19 Genomics UK (COG-UK) Consortium | Lighthouse Laboratory Plymouth and Alex Alderton, Roberto Amato, Jeffrey Barrett, Sonia Goncalves, Ewan Harrison, David K. Jackson, Ian Johnston, Dominic Kwiatkowski, Cordelia Langford, John Sillitoe on behalf of the Wellcome Sanger Institute COVID-19 Surveillance Team |
| EPI_ISL_2866241 | Lighthouse Lab in Milton Keynes | Wellcome Sanger Institute for the COVID-19 Genomics UK (COG-UK) | The Lighthouse Lab in Milton Keynes and Alex Alderton, Roberto Amato, Jeffrey Barrett, Sonia Goncalves, Ewan Harrison, David K. Jackson, Ian Johnston, Dominic Kwiatkowski, Cordelia Langford, John Sillitoe on behalf of the Wellcome Sanger |

| | | | |
|---|---|---|---|
| | | Consortium | Institute COVID-19 Surveillance Team |
| EPI_ISL_2866242, EPI_ISL_2866243 | Lighthouse Laboratory Plymouth | Wellcome Sanger Institute for the COVID-19 Genomics UK (COG-UK) Consortium | Lighthouse Laboratory Plymouth and Alex Alderton, Roberto Amato, Jeffrey Barrett, Sonia Goncalves, Ewan Harrison, David K. Jackson, Ian Johnston, Dominic Kwiatkowski, Cordelia Langford, John Sillitoe on behalf of the Wellcome Sanger Institute COVID-19 Surveillance Team |
| EPI_ISL_2866244 | Lighthouse Lab in Milton Keynes | Wellcome Sanger Institute for the COVID-19 Genomics UK (COG-UK) Consortium | The Lighthouse Lab in Milton Keynes and Alex Alderton, Roberto Amato, Jeffrey Barrett, Sonia Goncalves, Ewan Harrison, David K. Jackson, Ian Johnston, Dominic Kwiatkowski, Cordelia Langford, John Sillitoe on behalf of the Wellcome Sanger Institute COVID-19 Surveillance Team |
| EPI_ISL_2866245, EPI_ISL_2866246, EPI_ISL_2866247, EPI_ISL_2866248, EPI_ISL_2866249 | Lighthouse Laboratory Plymouth | Wellcome Sanger Institute for the COVID-19 Genomics UK (COG-UK) Consortium | Lighthouse Laboratory Plymouth and Alex Alderton, Roberto Amato, Jeffrey Barrett, Sonia Goncalves, Ewan Harrison, David K. Jackson, Ian Johnston, Dominic Kwiatkowski, Cordelia Langford, John Sillitoe on behalf of the Wellcome Sanger Institute COVID-19 Surveillance Team |
| EPI_ISL_2866250 | Lighthouse Lab in Milton Keynes | Wellcome Sanger Institute for the COVID-19 Genomics UK (COG-UK) Consortium | The Lighthouse Lab in Milton Keynes and Alex Alderton, Roberto Amato, Jeffrey Barrett, Sonia Goncalves, Ewan Harrison, David K. Jackson, Ian Johnston, Dominic Kwiatkowski, Cordelia Langford, John Sillitoe on behalf of the Wellcome Sanger Institute COVID-19 Surveillance Team |
| EPI_ISL_2866251 | Lighthouse Laboratory Plymouth | Wellcome Sanger Institute for the COVID-19 Genomics UK (COG-UK) Consortium | Lighthouse Laboratory Plymouth and Alex Alderton, Roberto Amato, Jeffrey Barrett, Sonia Goncalves, Ewan Harrison, David K. Jackson, Ian Johnston, Dominic Kwiatkowski, Cordelia Langford, John Sillitoe on behalf of the Wellcome Sanger Institute COVID-19 Surveillance Team |
| EPI_ISL_2866252 | Lighthouse Lab in Milton Keynes | Wellcome Sanger Institute for the COVID-19 Genomics UK (COG-UK) Consortium | The Lighthouse Lab in Milton Keynes and Alex Alderton, Roberto Amato, Jeffrey Barrett, Sonia Goncalves, Ewan Harrison, David K. Jackson, Ian Johnston, Dominic Kwiatkowski, Cordelia Langford, John Sillitoe on behalf of the Wellcome Sanger Institute COVID-19 Surveillance Team |
| EPI_ISL_2866253, EPI_ISL_2866254, EPI_ISL_2866255, EPI_ISL_2866256, EPI_ISL_2866257, EPI_ISL_2866258, EPI_ISL_2866259, EPI_ISL_2866260 | Lighthouse Laboratory Plymouth | Wellcome Sanger Institute for the COVID-19 Genomics UK (COG-UK) Consortium | Lighthouse Laboratory Plymouth and Alex Alderton, Roberto Amato, Jeffrey Barrett, Sonia Goncalves, Ewan Harrison, David K. Jackson, Ian Johnston, Dominic Kwiatkowski, Cordelia Langford, John Sillitoe on behalf of the Wellcome Sanger Institute COVID-19 Surveillance Team |
| EPI_ISL_2866261, EPI_ISL_2866262 | Lighthouse Lab in Milton Keynes | Wellcome Sanger Institute for the COVID-19 Genomics UK (COG-UK) Consortium | The Lighthouse Lab in Milton Keynes and Alex Alderton, Roberto Amato, Jeffrey Barrett, Sonia Goncalves, Ewan Harrison, David K. Jackson, Ian Johnston, Dominic Kwiatkowski, Cordelia Langford, John Sillitoe on behalf of the Wellcome Sanger Institute COVID-19 Surveillance Team |
| EPI_ISL_2866263 | Lighthouse Laboratory Plymouth | Wellcome Sanger Institute for the COVID-19 Genomics UK (COG-UK) Consortium | Lighthouse Laboratory Plymouth and Alex Alderton, Roberto Amato, Jeffrey Barrett, Sonia Goncalves, Ewan Harrison, David K. Jackson, Ian Johnston, Dominic Kwiatkowski, Cordelia Langford, John Sillitoe on behalf of the Wellcome Sanger Institute COVID-19 Surveillance Team |
| EPI_ISL_2866264 | Lighthouse Lab in Milton Keynes | Wellcome Sanger Institute for the COVID-19 Genomics UK (COG-UK) Consortium | The Lighthouse Lab in Milton Keynes and Alex Alderton, Roberto Amato, Jeffrey Barrett, Sonia Goncalves, Ewan Harrison, David K. Jackson, Ian Johnston, Dominic Kwiatkowski, Cordelia Langford, John Sillitoe on behalf of the Wellcome Sanger Institute COVID-19 Surveillance Team |
| EPI_ISL_2866265 | Lighthouse Laboratory Plymouth | Wellcome Sanger Institute for the COVID-19 Genomics UK (COG-UK) Consortium | Lighthouse Laboratory Plymouth and Alex Alderton, Roberto Amato, Jeffrey Barrett, Sonia Goncalves, Ewan Harrison, David K. Jackson, Ian Johnston, Dominic Kwiatkowski, Cordelia Langford, John Sillitoe on behalf of the Wellcome Sanger Institute COVID-19 Surveillance Team |
| EPI_ISL_2866266 | Lighthouse Lab in Milton Keynes | Wellcome Sanger Institute for the COVID-19 Genomics UK (COG-UK) Consortium | The Lighthouse Lab in Milton Keynes and Alex Alderton, Roberto Amato, Jeffrey Barrett, Sonia Goncalves, Ewan Harrison, David K. Jackson, Ian Johnston, Dominic Kwiatkowski, Cordelia Langford, John Sillitoe on behalf of the Wellcome Sanger Institute COVID-19 Surveillance Team |
| EPI_ISL_2866267, EPI_ISL_2866268, EPI_ISL_2866269, EPI_ISL_2866270, EPI_ISL_2866271, EPI_ISL_2866272, EPI_ISL_2866273, EPI_ISL_2866274, EPI_ISL_2866275 | Lighthouse Laboratory Plymouth | Wellcome Sanger Institute for the COVID-19 Genomics UK (COG-UK) Consortium | Lighthouse Laboratory Plymouth and Alex Alderton, Roberto Amato, Jeffrey Barrett, Sonia Goncalves, Ewan Harrison, David K. Jackson, Ian Johnston, Dominic Kwiatkowski, Cordelia Langford, John Sillitoe on behalf of the Wellcome Sanger Institute COVID-19 Surveillance Team |
| EPI_ISL_2866276, EPI_ISL_2866277, EPI_ISL_2866278 | Lighthouse Lab in Milton Keynes | Wellcome Sanger Institute for the COVID-19 Genomics UK (COG-UK) Consortium | The Lighthouse Lab in Milton Keynes and Alex Alderton, Roberto Amato, Jeffrey Barrett, Sonia Goncalves, Ewan Harrison, David K. Jackson, Ian Johnston, Dominic Kwiatkowski, Cordelia Langford, John Sillitoe on behalf of the Wellcome Sanger Institute COVID-19 Surveillance Team |
| EPI_ISL_2866279, EPI_ISL_2866280 | Lighthouse Laboratory Plymouth | Wellcome Sanger Institute for the COVID-19 Genomics UK (COG-UK) Consortium | Lighthouse Laboratory Plymouth and Alex Alderton, Roberto Amato, Jeffrey Barrett, Sonia Goncalves, Ewan Harrison, David K. Jackson, Ian Johnston, Dominic Kwiatkowski, Cordelia Langford, John Sillitoe on behalf of the Wellcome Sanger Institute COVID-19 Surveillance Team |
| EPI_ISL_2866281 | Lighthouse Lab in Milton Keynes | Wellcome Sanger Institute for the COVID-19 Genomics UK (COG-UK) Consortium | The Lighthouse Lab in Milton Keynes and Alex Alderton, Roberto Amato, Jeffrey Barrett, Sonia Goncalves, Ewan Harrison, David K. Jackson, Ian Johnston, Dominic Kwiatkowski, Cordelia Langford, John Sillitoe on behalf of the Wellcome Sanger Institute COVID-19 Surveillance Team |
| EPI_ISL_2866282 | Lighthouse Laboratory Plymouth | Wellcome Sanger Institute for the COVID-19 Genomics UK (COG-UK) Consortium | Lighthouse Laboratory Plymouth and Alex Alderton, Roberto Amato, Jeffrey Barrett, Sonia Goncalves, Ewan Harrison, David K. Jackson, Ian Johnston, Dominic Kwiatkowski, Cordelia Langford, John Sillitoe on behalf of the Wellcome Sanger Institute COVID-19 Surveillance Team |
| EPI_ISL_2866283 | Lighthouse Lab in Milton Keynes | Wellcome Sanger Institute for the COVID-19 Genomics UK (COG-UK) Consortium | The Lighthouse Lab in Milton Keynes and Alex Alderton, Roberto Amato, Jeffrey Barrett, Sonia Goncalves, Ewan Harrison, David K. Jackson, Ian Johnston, Dominic Kwiatkowski, Cordelia Langford, John Sillitoe on behalf of the Wellcome Sanger Institute COVID-19 Surveillance Team |
| EPI_ISL_2866284, EPI_ISL_2866285, EPI_ISL_2866286 | Lighthouse Laboratory Plymouth | Wellcome Sanger Institute for the COVID-19 Genomics UK (COG-UK) Consortium | Lighthouse Laboratory Plymouth and Alex Alderton, Roberto Amato, Jeffrey Barrett, Sonia Goncalves, Ewan Harrison, David K. Jackson, Ian Johnston, Dominic Kwiatkowski, Cordelia Langford, John Sillitoe on behalf of the Wellcome Sanger Institute COVID-19 Surveillance Team |
| EPI_ISL_2866287, EPI_ISL_2866288 | Lighthouse Lab in Milton Keynes | Wellcome Sanger Institute for the COVID-19 Genomics UK (COG-UK) Consortium | The Lighthouse Lab in Milton Keynes and Alex Alderton, Roberto Amato, Jeffrey Barrett, Sonia Goncalves, Ewan Harrison, David K. Jackson, Ian Johnston, Dominic Kwiatkowski, Cordelia Langford, John Sillitoe on behalf of the Wellcome Sanger Institute COVID-19 Surveillance Team |
| EPI_ISL_2866289, EPI_ISL_2866290 | Lighthouse Laboratory Plymouth | Wellcome Sanger Institute for the COVID-19 Genomics UK (COG-UK) Consortium | Lighthouse Laboratory Plymouth and Alex Alderton, Roberto Amato, Jeffrey Barrett, Sonia Goncalves, Ewan Harrison, David K. Jackson, Ian Johnston, Dominic Kwiatkowski, Cordelia Langford, John Sillitoe on behalf of the Wellcome Sanger Institute COVID-19 Surveillance Team |
| EPI_ISL_2866291 | Lighthouse Lab in Milton Keynes | Wellcome Sanger Institute for the COVID-19 Genomics UK (COG-UK) Consortium | The Lighthouse Lab in Milton Keynes and Alex Alderton, Roberto Amato, Jeffrey Barrett, Sonia Goncalves, Ewan Harrison, David K. Jackson, Ian Johnston, Dominic Kwiatkowski, Cordelia Langford, John Sillitoe on behalf of the Wellcome Sanger Institute COVID-19 Surveillance Team |
| EPI_ISL_2866292, EPI_ISL_2866293, EPI_ISL_2866294, EPI_ISL_2866295 | Lighthouse Laboratory Plymouth | Wellcome Sanger Institute for the COVID-19 Genomics UK (COG-UK) Consortium | Lighthouse Laboratory Plymouth and Alex Alderton, Roberto Amato, Jeffrey Barrett, Sonia Goncalves, Ewan Harrison, David K. Jackson, Ian Johnston, Dominic Kwiatkowski, Cordelia Langford, John Sillitoe on behalf of the Wellcome Sanger Institute COVID-19 Surveillance Team |
| EPI_ISL_2866296 | Lighthouse Lab in Milton Keynes | Wellcome Sanger Institute for the COVID-19 Genomics UK (COG-UK) Consortium | The Lighthouse Lab in Milton Keynes and Alex Alderton, Roberto Amato, Jeffrey Barrett, Sonia Goncalves, Ewan Harrison, David K. Jackson, Ian Johnston, Dominic Kwiatkowski, Cordelia Langford, John Sillitoe on behalf of the Wellcome Sanger Institute COVID-19 Surveillance Team |
| EPI_ISL_2866297 | Lighthouse Laboratory Plymouth | Wellcome Sanger Institute for the COVID-19 Genomics UK (COG-UK) Consortium | Lighthouse Laboratory Plymouth and Alex Alderton, Roberto Amato, Jeffrey Barrett, Sonia Goncalves, Ewan Harrison, David K. Jackson, Ian Johnston, Dominic Kwiatkowski, Cordelia Langford, John Sillitoe on behalf of the Wellcome Sanger Institute COVID-19 Surveillance Team |
| EPI_ISL_2866298, EPI_ISL_2866299, EPI_ISL_2866300 | Lighthouse Lab in Milton Keynes | Wellcome Sanger Institute for the COVID-19 Genomics UK (COG-UK) Consortium | The Lighthouse Lab in Milton Keynes and Alex Alderton, Roberto Amato, Jeffrey Barrett, Sonia Goncalves, Ewan Harrison, David K. Jackson, Ian Johnston, Dominic Kwiatkowski, Cordelia Langford, John Sillitoe on behalf of the Wellcome Sanger Institute COVID-19 Surveillance Team |
| EPI_ISL_2866301, EPI_ISL_2866302, EPI_ISL_2866303 | Lighthouse Laboratory Plymouth | Wellcome Sanger Institute for the COVID-19 Genomics UK (COG-UK) Consortium | Lighthouse Laboratory Plymouth and Alex Alderton, Roberto Amato, Jeffrey Barrett, Sonia Goncalves, Ewan Harrison, David K. Jackson, Ian Johnston, Dominic Kwiatkowski, Cordelia Langford, John Sillitoe on behalf of the Wellcome Sanger Institute COVID-19 Surveillance Team |
| EPI_ISL_2866304 | Lighthouse Lab in Milton Keynes | Wellcome Sanger Institute for the COVID-19 Genomics UK (COG-UK) Consortium | The Lighthouse Lab in Milton Keynes and Alex Alderton, Roberto Amato, Jeffrey Barrett, Sonia Goncalves, Ewan Harrison, David K. Jackson, Ian Johnston, Dominic Kwiatkowski, Cordelia Langford, John Sillitoe on behalf of the Wellcome Sanger Institute COVID-19 Surveillance Team |
| EPI_ISL_2866305, EPI_ISL_2866306, EPI_ISL_2866307 | Lighthouse Laboratory Plymouth | Wellcome Sanger Institute for the COVID-19 Genomics UK (COG-UK) Consortium | Lighthouse Laboratory Plymouth and Alex Alderton, Roberto Amato, Jeffrey Barrett, Sonia Goncalves, Ewan Harrison, David K. Jackson, Ian Johnston, Dominic Kwiatkowski, Cordelia Langford, John Sillitoe on behalf of the Wellcome Sanger Institute COVID-19 Surveillance Team |
| EPI_ISL_2866308 | Lighthouse Lab in Milton Keynes | Wellcome Sanger Institute for the COVID-19 Genomics UK (COG-UK) Consortium | The Lighthouse Lab in Milton Keynes and Alex Alderton, Roberto Amato, Jeffrey Barrett, Sonia Goncalves, Ewan Harrison, David K. Jackson, Ian Johnston, Dominic Kwiatkowski, Cordelia Langford, John Sillitoe on behalf of the Wellcome Sanger Institute COVID-19 Surveillance Team |
| EPI_ISL_2866309 | Lighthouse Laboratory Plymouth | Wellcome Sanger Institute for the COVID-19 Genomics UK (COG-UK) Consortium | Lighthouse Laboratory Plymouth and Alex Alderton, Roberto Amato, Jeffrey Barrett, Sonia Goncalves, Ewan Harrison, David K. Jackson, Ian Johnston, Dominic Kwiatkowski, Cordelia Langford, John Sillitoe on behalf of the Wellcome Sanger Institute COVID-19 Surveillance Team |
| EPI_ISL_2866310 | Lighthouse Lab in Milton Keynes | Wellcome Sanger Institute for the COVID-19 Genomics UK (COG-UK) Consortium | The Lighthouse Lab in Milton Keynes and Alex Alderton, Roberto Amato, Jeffrey Barrett, Sonia Goncalves, Ewan Harrison, David K. Jackson, Ian Johnston, Dominic Kwiatkowski, Cordelia Langford, John Sillitoe on behalf of the Wellcome Sanger Institute COVID-19 Surveillance Team |
| EPI_ISL_2866311, EPI_ISL_2866312, EPI_ISL_2866313, EPI_ISL_2866314 | Lighthouse Laboratory Plymouth | Wellcome Sanger Institute for the COVID-19 Genomics UK (COG-UK) Consortium | Lighthouse Laboratory Plymouth and Alex Alderton, Roberto Amato, Jeffrey Barrett, Sonia Goncalves, Ewan Harrison, David K. Jackson, Ian Johnston, Dominic Kwiatkowski, Cordelia Langford, John Sillitoe on behalf of the Wellcome Sanger Institute COVID-19 Surveillance Team |
| EPI_ISL_2866316 | Lighthouse Lab in Milton Keynes | Wellcome Sanger Institute for the COVID-19 Genomics UK (COG-UK) Consortium | The Lighthouse Lab in Milton Keynes and Alex Alderton, Roberto Amato, Jeffrey Barrett, Sonia Goncalves, Ewan Harrison, David K. Jackson, Ian Johnston, Dominic Kwiatkowski, Cordelia Langford, John Sillitoe on behalf of the Wellcome Sanger Institute COVID-19 Surveillance Team |
| EPI_ISL_2866317, | Lighthouse Laboratory Plymouth | Wellcome Sanger Institute for the COVID-19 Genomics UK (COG-UK) | Lighthouse Laboratory Plymouth and Alex Alderton, Roberto Amato, Jeffrey Barrett, Sonia Goncalves, Ewan Harrison, David K. Jackson, Ian Johnston, Dominic Kwiatkowski, Cordelia Langford, John Sillitoe on behalf of the Wellcome Sanger |

| | | | |
|---|---|---|---|
| EPI_ISL_2866318, EPI_ISL_2866319, EPI_ISL_2866320 | | Consortium | Institute COVID-19 Surveillance Team |
| EPI_ISL_2866321, EPI_ISL_2866322 | Lighthouse Lab in Milton Keynes | Wellcome Sanger Institute for the COVID-19 Genomics UK (COG-UK) Consortium | The Lighthouse Lab in Milton Keynes and Alex Alderton, Roberto Amato, Jeffrey Barrett, Sonia Goncalves, Ewan Harrison, David K. Jackson, Ian Johnston, Dominic Kwiatkowski, Cordelia Langford, John Sillitoe on behalf of the Wellcome Sanger Institute COVID-19 Surveillance Team |
| EPI_ISL_2866323, EPI_ISL_2866324, EPI_ISL_2866325, EPI_ISL_2866326 | Lighthouse Laboratory Plymouth | Wellcome Sanger Institute for the COVID-19 Genomics UK (COG-UK) Consortium | Lighthouse Laboratory Plymouth and Alex Alderton, Roberto Amato, Jeffrey Barrett, Sonia Goncalves, Ewan Harrison, David K. Jackson, Ian Johnston, Dominic Kwiatkowski, Cordelia Langford, John Sillitoe on behalf of the Wellcome Sanger Institute COVID-19 Surveillance Team |
| EPI_ISL_2866327 | Lighthouse Lab in Milton Keynes | Wellcome Sanger Institute for the COVID-19 Genomics UK (COG-UK) Consortium | The Lighthouse Lab in Milton Keynes and Alex Alderton, Roberto Amato, Jeffrey Barrett, Sonia Goncalves, Ewan Harrison, David K. Jackson, Ian Johnston, Dominic Kwiatkowski, Cordelia Langford, John Sillitoe on behalf of the Wellcome Sanger Institute COVID-19 Surveillance Team |
| EPI_ISL_2866328 | Lighthouse Laboratory Plymouth | Wellcome Sanger Institute for the COVID-19 Genomics UK (COG-UK) Consortium | Lighthouse Laboratory Plymouth and Alex Alderton, Roberto Amato, Jeffrey Barrett, Sonia Goncalves, Ewan Harrison, David K. Jackson, Ian Johnston, Dominic Kwiatkowski, Cordelia Langford, John Sillitoe on behalf of the Wellcome Sanger Institute COVID-19 Surveillance Team |
| EPI_ISL_2866329, EPI_ISL_2866330, EPI_ISL_2866331 | Lighthouse Lab in Milton Keynes | Wellcome Sanger Institute for the COVID-19 Genomics UK (COG-UK) Consortium | The Lighthouse Lab in Milton Keynes and Alex Alderton, Roberto Amato, Jeffrey Barrett, Sonia Goncalves, Ewan Harrison, David K. Jackson, Ian Johnston, Dominic Kwiatkowski, Cordelia Langford, John Sillitoe on behalf of the Wellcome Sanger Institute COVID-19 Surveillance Team |
| EPI_ISL_2866332, EPI_ISL_2866333, EPI_ISL_2866334 | Lighthouse Laboratory Plymouth | Wellcome Sanger Institute for the COVID-19 Genomics UK (COG-UK) Consortium | Lighthouse Laboratory Plymouth and Alex Alderton, Roberto Amato, Jeffrey Barrett, Sonia Goncalves, Ewan Harrison, David K. Jackson, Ian Johnston, Dominic Kwiatkowski, Cordelia Langford, John Sillitoe on behalf of the Wellcome Sanger Institute COVID-19 Surveillance Team |
| EPI_ISL_2866335 | Lighthouse Lab in Milton Keynes | Wellcome Sanger Institute for the COVID-19 Genomics UK (COG-UK) Consortium | The Lighthouse Lab in Milton Keynes and Alex Alderton, Roberto Amato, Jeffrey Barrett, Sonia Goncalves, Ewan Harrison, David K. Jackson, Ian Johnston, Dominic Kwiatkowski, Cordelia Langford, John Sillitoe on behalf of the Wellcome Sanger Institute COVID-19 Surveillance Team |
| EPI_ISL_2866336 | Lighthouse Laboratory Plymouth | Wellcome Sanger Institute for the COVID-19 Genomics UK (COG-UK) Consortium | Lighthouse Laboratory Plymouth and Alex Alderton, Roberto Amato, Jeffrey Barrett, Sonia Goncalves, Ewan Harrison, David K. Jackson, Ian Johnston, Dominic Kwiatkowski, Cordelia Langford, John Sillitoe on behalf of the Wellcome Sanger Institute COVID-19 Surveillance Team |
| EPI_ISL_2866337, EPI_ISL_2866338 | Lighthouse Lab in Milton Keynes | Wellcome Sanger Institute for the COVID-19 Genomics UK (COG-UK) Consortium | The Lighthouse Lab in Milton Keynes and Alex Alderton, Roberto Amato, Jeffrey Barrett, Sonia Goncalves, Ewan Harrison, David K. Jackson, Ian Johnston, Dominic Kwiatkowski, Cordelia Langford, John Sillitoe on behalf of the Wellcome Sanger Institute COVID-19 Surveillance Team |
| EPI_ISL_2866339, EPI_ISL_2866340, EPI_ISL_2866341 | Lighthouse Laboratory Plymouth | Wellcome Sanger Institute for the COVID-19 Genomics UK (COG-UK) Consortium | Lighthouse Laboratory Plymouth and Alex Alderton, Roberto Amato, Jeffrey Barrett, Sonia Goncalves, Ewan Harrison, David K. Jackson, Ian Johnston, Dominic Kwiatkowski, Cordelia Langford, John Sillitoe on behalf of the Wellcome Sanger Institute COVID-19 Surveillance Team |
| EPI_ISL_2866342 | Lighthouse Lab in Milton Keynes | Wellcome Sanger Institute for the COVID-19 Genomics UK (COG-UK) Consortium | The Lighthouse Lab in Milton Keynes and Alex Alderton, Roberto Amato, Jeffrey Barrett, Sonia Goncalves, Ewan Harrison, David K. Jackson, Ian Johnston, Dominic Kwiatkowski, Cordelia Langford, John Sillitoe on behalf of the Wellcome Sanger Institute COVID-19 Surveillance Team |
| EPI_ISL_2866343 | Lighthouse Laboratory Plymouth | Wellcome Sanger Institute for the COVID-19 Genomics UK (COG-UK) Consortium | Lighthouse Laboratory Plymouth and Alex Alderton, Roberto Amato, Jeffrey Barrett, Sonia Goncalves, Ewan Harrison, David K. Jackson, Ian Johnston, Dominic Kwiatkowski, Cordelia Langford, John Sillitoe on behalf of the Wellcome Sanger Institute COVID-19 Surveillance Team |
| EPI_ISL_2866344 | Lighthouse Lab in Milton Keynes | Wellcome Sanger Institute for the COVID-19 Genomics UK (COG-UK) Consortium | The Lighthouse Lab in Milton Keynes and Alex Alderton, Roberto Amato, Jeffrey Barrett, Sonia Goncalves, Ewan Harrison, David K. Jackson, Ian Johnston, Dominic Kwiatkowski, Cordelia Langford, John Sillitoe on behalf of the Wellcome Sanger Institute COVID-19 Surveillance Team |
| EPI_ISL_2866345 | Lighthouse Laboratory Plymouth | Wellcome Sanger Institute for the COVID-19 Genomics UK (COG-UK) Consortium | Lighthouse Laboratory Plymouth and Alex Alderton, Roberto Amato, Jeffrey Barrett, Sonia Goncalves, Ewan Harrison, David K. Jackson, Ian Johnston, Dominic Kwiatkowski, Cordelia Langford, John Sillitoe on behalf of the Wellcome Sanger Institute COVID-19 Surveillance Team |
| EPI_ISL_2866346 | Lighthouse Lab in Milton Keynes | Wellcome Sanger Institute for the COVID-19 Genomics UK (COG-UK) Consortium | The Lighthouse Lab in Milton Keynes and Alex Alderton, Roberto Amato, Jeffrey Barrett, Sonia Goncalves, Ewan Harrison, David K. Jackson, Ian Johnston, Dominic Kwiatkowski, Cordelia Langford, John Sillitoe on behalf of the Wellcome Sanger Institute COVID-19 Surveillance Team |
| EPI_ISL_2866347 | Lighthouse Laboratory Plymouth | Wellcome Sanger Institute for the COVID-19 Genomics UK (COG-UK) Consortium | Lighthouse Laboratory Plymouth and Alex Alderton, Roberto Amato, Jeffrey Barrett, Sonia Goncalves, Ewan Harrison, David K. Jackson, Ian Johnston, Dominic Kwiatkowski, Cordelia Langford, John Sillitoe on behalf of the Wellcome Sanger Institute COVID-19 Surveillance Team |
| EPI_ISL_2866348 | Lighthouse Lab in Milton Keynes | Wellcome Sanger Institute for the COVID-19 Genomics UK (COG-UK) Consortium | The Lighthouse Lab in Milton Keynes and Alex Alderton, Roberto Amato, Jeffrey Barrett, Sonia Goncalves, Ewan Harrison, David K. Jackson, Ian Johnston, Dominic Kwiatkowski, Cordelia Langford, John Sillitoe on behalf of the Wellcome Sanger Institute COVID-19 Surveillance Team |
| EPI_ISL_2866349, EPI_ISL_2866350 | Lighthouse Laboratory Plymouth | Wellcome Sanger Institute for the COVID-19 Genomics UK (COG-UK) Consortium | Lighthouse Laboratory Plymouth and Alex Alderton, Roberto Amato, Jeffrey Barrett, Sonia Goncalves, Ewan Harrison, David K. Jackson, Ian Johnston, Dominic Kwiatkowski, Cordelia Langford, John Sillitoe on behalf of the Wellcome Sanger Institute COVID-19 Surveillance Team |
| EPI_ISL_2866351, EPI_ISL_2866352, EPI_ISL_2866353, EPI_ISL_2866354 | Lighthouse Lab in Milton Keynes | Wellcome Sanger Institute for the COVID-19 Genomics UK (COG-UK) Consortium | The Lighthouse Lab in Milton Keynes and Alex Alderton, Roberto Amato, Jeffrey Barrett, Sonia Goncalves, Ewan Harrison, David K. Jackson, Ian Johnston, Dominic Kwiatkowski, Cordelia Langford, John Sillitoe on behalf of the Wellcome Sanger Institute COVID-19 Surveillance Team |
| EPI_ISL_2866355 | Lighthouse Laboratory Plymouth | Wellcome Sanger Institute for the COVID-19 Genomics UK (COG-UK) Consortium | Lighthouse Laboratory Plymouth and Alex Alderton, Roberto Amato, Jeffrey Barrett, Sonia Goncalves, Ewan Harrison, David K. Jackson, Ian Johnston, Dominic Kwiatkowski, Cordelia Langford, John Sillitoe on behalf of the Wellcome Sanger Institute COVID-19 Surveillance Team |
| EPI_ISL_2866356 | Lighthouse Lab in Milton Keynes | Wellcome Sanger Institute for the COVID-19 Genomics UK (COG-UK) Consortium | The Lighthouse Lab in Milton Keynes and Alex Alderton, Roberto Amato, Jeffrey Barrett, Sonia Goncalves, Ewan Harrison, David K. Jackson, Ian Johnston, Dominic Kwiatkowski, Cordelia Langford, John Sillitoe on behalf of the Wellcome Sanger Institute COVID-19 Surveillance Team |
| EPI_ISL_2866357, EPI_ISL_2866358, EPI_ISL_2866359 | Lighthouse Laboratory Plymouth | Wellcome Sanger Institute for the COVID-19 Genomics UK (COG-UK) Consortium | Lighthouse Laboratory Plymouth and Alex Alderton, Roberto Amato, Jeffrey Barrett, Sonia Goncalves, Ewan Harrison, David K. Jackson, Ian Johnston, Dominic Kwiatkowski, Cordelia Langford, John Sillitoe on behalf of the Wellcome Sanger Institute COVID-19 Surveillance Team |
| EPI_ISL_2866360, EPI_ISL_2866361, EPI_ISL_2866362 | Lighthouse Lab in Milton Keynes | Wellcome Sanger Institute for the COVID-19 Genomics UK (COG-UK) Consortium | The Lighthouse Lab in Milton Keynes and Alex Alderton, Roberto Amato, Jeffrey Barrett, Sonia Goncalves, Ewan Harrison, David K. Jackson, Ian Johnston, Dominic Kwiatkowski, Cordelia Langford, John Sillitoe on behalf of the Wellcome Sanger Institute COVID-19 Surveillance Team |
| EPI_ISL_2866363, EPI_ISL_2866364 | Lighthouse Laboratory Plymouth | Wellcome Sanger Institute for the COVID-19 Genomics UK (COG-UK) Consortium | Lighthouse Laboratory Plymouth and Alex Alderton, Roberto Amato, Jeffrey Barrett, Sonia Goncalves, Ewan Harrison, David K. Jackson, Ian Johnston, Dominic Kwiatkowski, Cordelia Langford, John Sillitoe on behalf of the Wellcome Sanger Institute COVID-19 Surveillance Team |
| EPI_ISL_2866365 | Lighthouse Lab in Milton Keynes | Wellcome Sanger Institute for the COVID-19 Genomics UK (COG-UK) Consortium | The Lighthouse Lab in Milton Keynes and Alex Alderton, Roberto Amato, Jeffrey Barrett, Sonia Goncalves, Ewan Harrison, David K. Jackson, Ian Johnston, Dominic Kwiatkowski, Cordelia Langford, John Sillitoe on behalf of the Wellcome Sanger Institute COVID-19 Surveillance Team |
| EPI_ISL_2866366 | Lighthouse Laboratory Plymouth | Wellcome Sanger Institute for the COVID-19 Genomics UK (COG-UK) Consortium | Lighthouse Laboratory Plymouth and Alex Alderton, Roberto Amato, Jeffrey Barrett, Sonia Goncalves, Ewan Harrison, David K. Jackson, Ian Johnston, Dominic Kwiatkowski, Cordelia Langford, John Sillitoe on behalf of the Wellcome Sanger Institute COVID-19 Surveillance Team |
| EPI_ISL_2866367 | Lighthouse Lab in Milton Keynes | Wellcome Sanger Institute for the COVID-19 Genomics UK (COG-UK) Consortium | The Lighthouse Lab in Milton Keynes and Alex Alderton, Roberto Amato, Jeffrey Barrett, Sonia Goncalves, Ewan Harrison, David K. Jackson, Ian Johnston, Dominic Kwiatkowski, Cordelia Langford, John Sillitoe on behalf of the Wellcome Sanger Institute COVID-19 Surveillance Team |
| EPI_ISL_2866368 | Lighthouse Laboratory Plymouth | Wellcome Sanger Institute for the COVID-19 Genomics UK (COG-UK) Consortium | Lighthouse Laboratory Plymouth and Alex Alderton, Roberto Amato, Jeffrey Barrett, Sonia Goncalves, Ewan Harrison, David K. Jackson, Ian Johnston, Dominic Kwiatkowski, Cordelia Langford, John Sillitoe on behalf of the Wellcome Sanger Institute COVID-19 Surveillance Team |
| EPI_ISL_2866369, EPI_ISL_2866370 | Lighthouse Lab in Milton Keynes | Wellcome Sanger Institute for the COVID-19 Genomics UK (COG-UK) Consortium | The Lighthouse Lab in Milton Keynes and Alex Alderton, Roberto Amato, Jeffrey Barrett, Sonia Goncalves, Ewan Harrison, David K. Jackson, Ian Johnston, Dominic Kwiatkowski, Cordelia Langford, John Sillitoe on behalf of the Wellcome Sanger Institute COVID-19 Surveillance Team |
| EPI_ISL_2866371, EPI_ISL_2866372, EPI_ISL_2866373 | Lighthouse Laboratory Plymouth | Wellcome Sanger Institute for the COVID-19 Genomics UK (COG-UK) Consortium | Lighthouse Laboratory Plymouth and Alex Alderton, Roberto Amato, Jeffrey Barrett, Sonia Goncalves, Ewan Harrison, David K. Jackson, Ian Johnston, Dominic Kwiatkowski, Cordelia Langford, John Sillitoe on behalf of the Wellcome Sanger Institute COVID-19 Surveillance Team |
| EPI_ISL_2866374 | Lighthouse Lab in Milton Keynes | Wellcome Sanger Institute for the COVID-19 Genomics UK (COG-UK) Consortium | The Lighthouse Lab in Milton Keynes and Alex Alderton, Roberto Amato, Jeffrey Barrett, Sonia Goncalves, Ewan Harrison, David K. Jackson, Ian Johnston, Dominic Kwiatkowski, Cordelia Langford, John Sillitoe on behalf of the Wellcome Sanger Institute COVID-19 Surveillance Team |
| EPI_ISL_2866375, EPI_ISL_2866376, EPI_ISL_2866377, EPI_ISL_2866378 | Lighthouse Laboratory Plymouth | Wellcome Sanger Institute for the COVID-19 Genomics UK (COG-UK) Consortium | Lighthouse Laboratory Plymouth and Alex Alderton, Roberto Amato, Jeffrey Barrett, Sonia Goncalves, Ewan Harrison, David K. Jackson, Ian Johnston, Dominic Kwiatkowski, Cordelia Langford, John Sillitoe on behalf of the Wellcome Sanger Institute COVID-19 Surveillance Team |
| EPI_ISL_2866379 | Lighthouse Lab in Milton Keynes | Wellcome Sanger Institute for the COVID-19 Genomics UK (COG-UK) Consortium | The Lighthouse Lab in Milton Keynes and Alex Alderton, Roberto Amato, Jeffrey Barrett, Sonia Goncalves, Ewan Harrison, David K. Jackson, Ian Johnston, Dominic Kwiatkowski, Cordelia Langford, John Sillitoe on behalf of the Wellcome Sanger Institute COVID-19 Surveillance Team |
| EPI_ISL_2866380, EPI_ISL_2866381, EPI_ISL_2866382 | Lighthouse Laboratory Plymouth | Wellcome Sanger Institute for the COVID-19 Genomics UK (COG-UK) Consortium | Lighthouse Laboratory Plymouth and Alex Alderton, Roberto Amato, Jeffrey Barrett, Sonia Goncalves, Ewan Harrison, David K. Jackson, Ian Johnston, Dominic Kwiatkowski, Cordelia Langford, John Sillitoe on behalf of the Wellcome Sanger Institute COVID-19 Surveillance Team |
| EPI_ISL_2866383, EPI_ISL_2866384 | Lighthouse Lab in Milton Keynes | Wellcome Sanger Institute for the COVID-19 Genomics UK (COG-UK) Consortium | The Lighthouse Lab in Milton Keynes and Alex Alderton, Roberto Amato, Jeffrey Barrett, Sonia Goncalves, Ewan Harrison, David K. Jackson, Ian Johnston, Dominic Kwiatkowski, Cordelia Langford, John Sillitoe on behalf of the Wellcome Sanger Institute COVID-19 Surveillance Team |
| EPI_ISL_2866385, EPI_ISL_2866386, EPI_ISL_2866387 | Lighthouse Laboratory Plymouth | Wellcome Sanger Institute for the COVID-19 Genomics UK (COG-UK) Consortium | Lighthouse Laboratory Plymouth and Alex Alderton, Roberto Amato, Jeffrey Barrett, Sonia Goncalves, Ewan Harrison, David K. Jackson, Ian Johnston, Dominic Kwiatkowski, Cordelia Langford, John Sillitoe on behalf of the Wellcome Sanger Institute COVID-19 Surveillance Team |
| EPI_ISL_2866388 | Lighthouse Lab in Milton Keynes | Wellcome Sanger Institute for the COVID-19 Genomics UK (COG-UK) Consortium | The Lighthouse Lab in Milton Keynes and Alex Alderton, Roberto Amato, Jeffrey Barrett, Sonia Goncalves, Ewan Harrison, David K. Jackson, Ian Johnston, Dominic Kwiatkowski, Cordelia Langford, John Sillitoe on behalf of the Wellcome Sanger Institute COVID-19 Surveillance Team |
| EPI_ISL_2866389, EPI_ISL_2866390 | Lighthouse Laboratory Plymouth | Wellcome Sanger Institute for the COVID-19 Genomics UK (COG-UK) Consortium | Lighthouse Laboratory Plymouth and Alex Alderton, Roberto Amato, Jeffrey Barrett, Sonia Goncalves, Ewan Harrison, David K. Jackson, Ian Johnston, Dominic Kwiatkowski, Cordelia Langford, John Sillitoe on behalf of the Wellcome Sanger Institute COVID-19 Surveillance Team |

| | | | |
|---|---|---|---|
| EPI_ISL_2866391 | Lighthouse Lab in Milton Keynes | Wellcome Sanger Institute for the COVID-19 Genomics UK (COG-UK) Consortium | The Lighthouse Lab in Milton Keynes and Alex Alderton, Roberto Amato, Jeffrey Barrett, Sonia Goncalves, Ewan Harrison, David K. Jackson, Ian Johnston, Dominic Kwiatkowski, Cordelia Langford, John Sillitoe on behalf of the Wellcome Sanger Institute COVID-19 Surveillance Team |
| EPI_ISL_2866392 | Lighthouse Laboratory Plymouth | Wellcome Sanger Institute for the COVID-19 Genomics UK (COG-UK) Consortium | Lighthouse Laboratory Plymouth and Alex Alderton, Roberto Amato, Jeffrey Barrett, Sonia Goncalves, Ewan Harrison, David K. Jackson, Ian Johnston, Dominic Kwiatkowski, Cordelia Langford, John Sillitoe on behalf of the Wellcome Sanger Institute COVID-19 Surveillance Team |
| EPI_ISL_2866393, EPI_ISL_2866394, EPI_ISL_2866395 | Lighthouse Lab in Milton Keynes | Wellcome Sanger Institute for the COVID-19 Genomics UK (COG-UK) Consortium | The Lighthouse Lab in Milton Keynes and Alex Alderton, Roberto Amato, Jeffrey Barrett, Sonia Goncalves, Ewan Harrison, David K. Jackson, Ian Johnston, Dominic Kwiatkowski, Cordelia Langford, John Sillitoe on behalf of the Wellcome Sanger Institute COVID-19 Surveillance Team |
| EPI_ISL_2866396 | Lighthouse Laboratory Plymouth | Wellcome Sanger Institute for the COVID-19 Genomics UK (COG-UK) Consortium | Lighthouse Laboratory Plymouth and Alex Alderton, Roberto Amato, Jeffrey Barrett, Sonia Goncalves, Ewan Harrison, David K. Jackson, Ian Johnston, Dominic Kwiatkowski, Cordelia Langford, John Sillitoe on behalf of the Wellcome Sanger Institute COVID-19 Surveillance Team |
| EPI_ISL_2866397, EPI_ISL_2866398 | Lighthouse Lab in Milton Keynes | Wellcome Sanger Institute for the COVID-19 Genomics UK (COG-UK) Consortium | The Lighthouse Lab in Milton Keynes and Alex Alderton, Roberto Amato, Jeffrey Barrett, Sonia Goncalves, Ewan Harrison, David K. Jackson, Ian Johnston, Dominic Kwiatkowski, Cordelia Langford, John Sillitoe on behalf of the Wellcome Sanger Institute COVID-19 Surveillance Team |
| EPI_ISL_2866399 | Lighthouse Laboratory Plymouth | Wellcome Sanger Institute for the COVID-19 Genomics UK (COG-UK) Consortium | Lighthouse Laboratory Plymouth and Alex Alderton, Roberto Amato, Jeffrey Barrett, Sonia Goncalves, Ewan Harrison, David K. Jackson, Ian Johnston, Dominic Kwiatkowski, Cordelia Langford, John Sillitoe on behalf of the Wellcome Sanger Institute COVID-19 Surveillance Team |
| EPI_ISL_2866400, EPI_ISL_2866401, EPI_ISL_2866402 | Lighthouse Lab in Milton Keynes | Wellcome Sanger Institute for the COVID-19 Genomics UK (COG-UK) Consortium | The Lighthouse Lab in Milton Keynes and Alex Alderton, Roberto Amato, Jeffrey Barrett, Sonia Goncalves, Ewan Harrison, David K. Jackson, Ian Johnston, Dominic Kwiatkowski, Cordelia Langford, John Sillitoe on behalf of the Wellcome Sanger Institute COVID-19 Surveillance Team |
| EPI_ISL_2866403, EPI_ISL_2866404, EPI_ISL_2866405 | Lighthouse Laboratory Plymouth | Wellcome Sanger Institute for the COVID-19 Genomics UK (COG-UK) Consortium | Lighthouse Laboratory Plymouth and Alex Alderton, Roberto Amato, Jeffrey Barrett, Sonia Goncalves, Ewan Harrison, David K. Jackson, Ian Johnston, Dominic Kwiatkowski, Cordelia Langford, John Sillitoe on behalf of the Wellcome Sanger Institute COVID-19 Surveillance Team |
| EPI_ISL_2866406 | Lighthouse Lab in Milton Keynes | Wellcome Sanger Institute for the COVID-19 Genomics UK (COG-UK) Consortium | The Lighthouse Lab in Milton Keynes and Alex Alderton, Roberto Amato, Jeffrey Barrett, Sonia Goncalves, Ewan Harrison, David K. Jackson, Ian Johnston, Dominic Kwiatkowski, Cordelia Langford, John Sillitoe on behalf of the Wellcome Sanger Institute COVID-19 Surveillance Team |
| EPI_ISL_2866407, EPI_ISL_2866408, EPI_ISL_2866409, EPI_ISL_2866410, EPI_ISL_2866411 | Lighthouse Laboratory Plymouth | Wellcome Sanger Institute for the COVID-19 Genomics UK (COG-UK) Consortium | Lighthouse Laboratory Plymouth and Alex Alderton, Roberto Amato, Jeffrey Barrett, Sonia Goncalves, Ewan Harrison, David K. Jackson, Ian Johnston, Dominic Kwiatkowski, Cordelia Langford, John Sillitoe on behalf of the Wellcome Sanger Institute COVID-19 Surveillance Team |
| EPI_ISL_2866412 | Lighthouse Lab in Milton Keynes | Wellcome Sanger Institute for the COVID-19 Genomics UK (COG-UK) Consortium | The Lighthouse Lab in Milton Keynes and Alex Alderton, Roberto Amato, Jeffrey Barrett, Sonia Goncalves, Ewan Harrison, David K. Jackson, Ian Johnston, Dominic Kwiatkowski, Cordelia Langford, John Sillitoe on behalf of the Wellcome Sanger Institute COVID-19 Surveillance Team |
| EPI_ISL_2866413, EPI_ISL_2866414, EPI_ISL_2866415 | Lighthouse Laboratory Plymouth | Wellcome Sanger Institute for the COVID-19 Genomics UK (COG-UK) Consortium | Lighthouse Laboratory Plymouth and Alex Alderton, Roberto Amato, Jeffrey Barrett, Sonia Goncalves, Ewan Harrison, David K. Jackson, Ian Johnston, Dominic Kwiatkowski, Cordelia Langford, John Sillitoe on behalf of the Wellcome Sanger Institute COVID-19 Surveillance Team |
| EPI_ISL_2866416, EPI_ISL_2866417 | Lighthouse Lab in Milton Keynes | Wellcome Sanger Institute for the COVID-19 Genomics UK (COG-UK) Consortium | The Lighthouse Lab in Milton Keynes and Alex Alderton, Roberto Amato, Jeffrey Barrett, Sonia Goncalves, Ewan Harrison, David K. Jackson, Ian Johnston, Dominic Kwiatkowski, Cordelia Langford, John Sillitoe on behalf of the Wellcome Sanger Institute COVID-19 Surveillance Team |
| EPI_ISL_2866418 | Lighthouse Laboratory Plymouth | Wellcome Sanger Institute for the COVID-19 Genomics UK (COG-UK) Consortium | Lighthouse Laboratory Plymouth and Alex Alderton, Roberto Amato, Jeffrey Barrett, Sonia Goncalves, Ewan Harrison, David K. Jackson, Ian Johnston, Dominic Kwiatkowski, Cordelia Langford, John Sillitoe on behalf of the Wellcome Sanger Institute COVID-19 Surveillance Team |
| EPI_ISL_2866419 | Lighthouse Lab in Milton Keynes | Wellcome Sanger Institute for the COVID-19 Genomics UK (COG-UK) Consortium | The Lighthouse Lab in Milton Keynes and Alex Alderton, Roberto Amato, Jeffrey Barrett, Sonia Goncalves, Ewan Harrison, David K. Jackson, Ian Johnston, Dominic Kwiatkowski, Cordelia Langford, John Sillitoe on behalf of the Wellcome Sanger Institute COVID-19 Surveillance Team |
| EPI_ISL_2866420, EPI_ISL_2866421, EPI_ISL_2866422, EPI_ISL_2866423, EPI_ISL_2866424 | Lighthouse Laboratory Plymouth | Wellcome Sanger Institute for the COVID-19 Genomics UK (COG-UK) Consortium | Lighthouse Laboratory Plymouth and Alex Alderton, Roberto Amato, Jeffrey Barrett, Sonia Goncalves, Ewan Harrison, David K. Jackson, Ian Johnston, Dominic Kwiatkowski, Cordelia Langford, John Sillitoe on behalf of the Wellcome Sanger Institute COVID-19 Surveillance Team |
| EPI_ISL_2866425 | Lighthouse Lab in Milton Keynes | Wellcome Sanger Institute for the COVID-19 Genomics UK (COG-UK) Consortium | The Lighthouse Lab in Milton Keynes and Alex Alderton, Roberto Amato, Jeffrey Barrett, Sonia Goncalves, Ewan Harrison, David K. Jackson, Ian Johnston, Dominic Kwiatkowski, Cordelia Langford, John Sillitoe on behalf of the Wellcome Sanger Institute COVID-19 Surveillance Team |
| EPI_ISL_2866426 | Lighthouse Laboratory Plymouth | Wellcome Sanger Institute for the COVID-19 Genomics UK (COG-UK) Consortium | Lighthouse Laboratory Plymouth and Alex Alderton, Roberto Amato, Jeffrey Barrett, Sonia Goncalves, Ewan Harrison, David K. Jackson, Ian Johnston, Dominic Kwiatkowski, Cordelia Langford, John Sillitoe on behalf of the Wellcome Sanger Institute COVID-19 Surveillance Team |
| EPI_ISL_2866427, EPI_ISL_2866428, EPI_ISL_2866429, EPI_ISL_2866430, EPI_ISL_2866431, EPI_ISL_2866432 | Lighthouse Lab in Milton Keynes | Wellcome Sanger Institute for the COVID-19 Genomics UK (COG-UK) Consortium | The Lighthouse Lab in Milton Keynes and Alex Alderton, Roberto Amato, Jeffrey Barrett, Sonia Goncalves, Ewan Harrison, David K. Jackson, Ian Johnston, Dominic Kwiatkowski, Cordelia Langford, John Sillitoe on behalf of the Wellcome Sanger Institute COVID-19 Surveillance Team |
| EPI_ISL_2866433, EPI_ISL_2866434 | Lighthouse Laboratory Plymouth | Wellcome Sanger Institute for the COVID-19 Genomics UK (COG-UK) Consortium | Lighthouse Laboratory Plymouth and Alex Alderton, Roberto Amato, Jeffrey Barrett, Sonia Goncalves, Ewan Harrison, David K. Jackson, Ian Johnston, Dominic Kwiatkowski, Cordelia Langford, John Sillitoe on behalf of the Wellcome Sanger Institute COVID-19 Surveillance Team |
| EPI_ISL_2866435, EPI_ISL_2866436 | Lighthouse Lab in Milton Keynes | Wellcome Sanger Institute for the COVID-19 Genomics UK (COG-UK) Consortium | The Lighthouse Lab in Milton Keynes and Alex Alderton, Roberto Amato, Jeffrey Barrett, Sonia Goncalves, Ewan Harrison, David K. Jackson, Ian Johnston, Dominic Kwiatkowski, Cordelia Langford, John Sillitoe on behalf of the Wellcome Sanger Institute COVID-19 Surveillance Team |
| EPI_ISL_2866437 | Lighthouse Laboratory Plymouth | Wellcome Sanger Institute for the COVID-19 Genomics UK (COG-UK) Consortium | Lighthouse Laboratory Plymouth and Alex Alderton, Roberto Amato, Jeffrey Barrett, Sonia Goncalves, Ewan Harrison, David K. Jackson, Ian Johnston, Dominic Kwiatkowski, Cordelia Langford, John Sillitoe on behalf of the Wellcome Sanger Institute COVID-19 Surveillance Team |
| EPI_ISL_2866438 | Lighthouse Lab in Milton Keynes | Wellcome Sanger Institute for the COVID-19 Genomics UK (COG-UK) Consortium | The Lighthouse Lab in Milton Keynes and Alex Alderton, Roberto Amato, Jeffrey Barrett, Sonia Goncalves, Ewan Harrison, David K. Jackson, Ian Johnston, Dominic Kwiatkowski, Cordelia Langford, John Sillitoe on behalf of the Wellcome Sanger Institute COVID-19 Surveillance Team |
| EPI_ISL_2866439 | Lighthouse Laboratory Plymouth | Wellcome Sanger Institute for the COVID-19 Genomics UK (COG-UK) Consortium | Lighthouse Laboratory Plymouth and Alex Alderton, Roberto Amato, Jeffrey Barrett, Sonia Goncalves, Ewan Harrison, David K. Jackson, Ian Johnston, Dominic Kwiatkowski, Cordelia Langford, John Sillitoe on behalf of the Wellcome Sanger Institute COVID-19 Surveillance Team |
| EPI_ISL_2866440 | Lighthouse Lab in Milton Keynes | Wellcome Sanger Institute for the COVID-19 Genomics UK (COG-UK) Consortium | The Lighthouse Lab in Milton Keynes and Alex Alderton, Roberto Amato, Jeffrey Barrett, Sonia Goncalves, Ewan Harrison, David K. Jackson, Ian Johnston, Dominic Kwiatkowski, Cordelia Langford, John Sillitoe on behalf of the Wellcome Sanger Institute COVID-19 Surveillance Team |
| EPI_ISL_2866441 | Lighthouse Laboratory Plymouth | Wellcome Sanger Institute for the COVID-19 Genomics UK (COG-UK) Consortium | Lighthouse Laboratory Plymouth and Alex Alderton, Roberto Amato, Jeffrey Barrett, Sonia Goncalves, Ewan Harrison, David K. Jackson, Ian Johnston, Dominic Kwiatkowski, Cordelia Langford, John Sillitoe on behalf of the Wellcome Sanger Institute COVID-19 Surveillance Team |
| EPI_ISL_2866442, EPI_ISL_2866443 | Lighthouse Lab in Milton Keynes | Wellcome Sanger Institute for the COVID-19 Genomics UK (COG-UK) Consortium | The Lighthouse Lab in Milton Keynes and Alex Alderton, Roberto Amato, Jeffrey Barrett, Sonia Goncalves, Ewan Harrison, David K. Jackson, Ian Johnston, Dominic Kwiatkowski, Cordelia Langford, John Sillitoe on behalf of the Wellcome Sanger Institute COVID-19 Surveillance Team |
| EPI_ISL_2866444, EPI_ISL_2866445 | Lighthouse Laboratory Plymouth | Wellcome Sanger Institute for the COVID-19 Genomics UK (COG-UK) Consortium | Lighthouse Laboratory Plymouth and Alex Alderton, Roberto Amato, Jeffrey Barrett, Sonia Goncalves, Ewan Harrison, David K. Jackson, Ian Johnston, Dominic Kwiatkowski, Cordelia Langford, John Sillitoe on behalf of the Wellcome Sanger Institute COVID-19 Surveillance Team |
| EPI_ISL_2866446, EPI_ISL_2866447, EPI_ISL_2866448, EPI_ISL_2866449 | Lighthouse Lab in Milton Keynes | Wellcome Sanger Institute for the COVID-19 Genomics UK (COG-UK) Consortium | The Lighthouse Lab in Milton Keynes and Alex Alderton, Roberto Amato, Jeffrey Barrett, Sonia Goncalves, Ewan Harrison, David K. Jackson, Ian Johnston, Dominic Kwiatkowski, Cordelia Langford, John Sillitoe on behalf of the Wellcome Sanger Institute COVID-19 Surveillance Team |
| EPI_ISL_2866450, EPI_ISL_2866451, EPI_ISL_2866452 | Lighthouse Laboratory Plymouth | Wellcome Sanger Institute for the COVID-19 Genomics UK (COG-UK) Consortium | Lighthouse Laboratory Plymouth and Alex Alderton, Roberto Amato, Jeffrey Barrett, Sonia Goncalves, Ewan Harrison, David K. Jackson, Ian Johnston, Dominic Kwiatkowski, Cordelia Langford, John Sillitoe on behalf of the Wellcome Sanger Institute COVID-19 Surveillance Team |
| EPI_ISL_2866453 | Lighthouse Lab in Milton Keynes | Wellcome Sanger Institute for the COVID-19 Genomics UK (COG-UK) Consortium | The Lighthouse Lab in Milton Keynes and Alex Alderton, Roberto Amato, Jeffrey Barrett, Sonia Goncalves, Ewan Harrison, David K. Jackson, Ian Johnston, Dominic Kwiatkowski, Cordelia Langford, John Sillitoe on behalf of the Wellcome Sanger Institute COVID-19 Surveillance Team |
| EPI_ISL_2866454, EPI_ISL_2866455 | Lighthouse Laboratory Plymouth | Wellcome Sanger Institute for the COVID-19 Genomics UK (COG-UK) Consortium | Lighthouse Laboratory Plymouth and Alex Alderton, Roberto Amato, Jeffrey Barrett, Sonia Goncalves, Ewan Harrison, David K. Jackson, Ian Johnston, Dominic Kwiatkowski, Cordelia Langford, John Sillitoe on behalf of the Wellcome Sanger Institute COVID-19 Surveillance Team |
| EPI_ISL_2866456 | Lighthouse Lab in Milton Keynes | Wellcome Sanger Institute for the COVID-19 Genomics UK (COG-UK) Consortium | The Lighthouse Lab in Milton Keynes and Alex Alderton, Roberto Amato, Jeffrey Barrett, Sonia Goncalves, Ewan Harrison, David K. Jackson, Ian Johnston, Dominic Kwiatkowski, Cordelia Langford, John Sillitoe on behalf of the Wellcome Sanger Institute COVID-19 Surveillance Team |
| EPI_ISL_2866457 | Lighthouse Laboratory Plymouth | Wellcome Sanger Institute for the COVID-19 Genomics UK (COG-UK) Consortium | Lighthouse Laboratory Plymouth and Alex Alderton, Roberto Amato, Jeffrey Barrett, Sonia Goncalves, Ewan Harrison, David K. Jackson, Ian Johnston, Dominic Kwiatkowski, Cordelia Langford, John Sillitoe on behalf of the Wellcome Sanger Institute COVID-19 Surveillance Team |
| EPI_ISL_2866458 | Lighthouse Lab in Milton Keynes | Wellcome Sanger Institute for the COVID-19 Genomics UK (COG-UK) Consortium | The Lighthouse Lab in Milton Keynes and Alex Alderton, Roberto Amato, Jeffrey Barrett, Sonia Goncalves, Ewan Harrison, David K. Jackson, Ian Johnston, Dominic Kwiatkowski, Cordelia Langford, John Sillitoe on behalf of the Wellcome Sanger Institute COVID-19 Surveillance Team |
| EPI_ISL_2866459 | Lighthouse Laboratory Plymouth | Wellcome Sanger Institute for the COVID-19 Genomics UK (COG-UK) Consortium | Lighthouse Laboratory Plymouth and Alex Alderton, Roberto Amato, Jeffrey Barrett, Sonia Goncalves, Ewan Harrison, David K. Jackson, Ian Johnston, Dominic Kwiatkowski, Cordelia Langford, John Sillitoe on behalf of the Wellcome Sanger Institute COVID-19 Surveillance Team |
| EPI_ISL_2866460, EPI_ISL_2866461 | Lighthouse Lab in Milton Keynes | Wellcome Sanger Institute for the COVID-19 Genomics UK (COG-UK) Consortium | The Lighthouse Lab in Milton Keynes and Alex Alderton, Roberto Amato, Jeffrey Barrett, Sonia Goncalves, Ewan Harrison, David K. Jackson, Ian Johnston, Dominic Kwiatkowski, Cordelia Langford, John Sillitoe on behalf of the Wellcome Sanger Institute COVID-19 Surveillance Team |
| EPI_ISL_2866462 | Lighthouse Laboratory Plymouth | Wellcome Sanger Institute for the COVID-19 Genomics UK (COG-UK) Consortium | Lighthouse Laboratory Plymouth and Alex Alderton, Roberto Amato, Jeffrey Barrett, Sonia Goncalves, Ewan Harrison, David K. Jackson, Ian Johnston, Dominic Kwiatkowski, Cordelia Langford, John Sillitoe on behalf of the Wellcome Sanger Institute COVID-19 Surveillance Team |

| | | | |
|---|---|---|---|
| EPI_ISL_2866463, EPI_ISL_2866464, EPI_ISL_2866465 | Lighthouse Lab in Milton Keynes | Wellcome Sanger Institute for the COVID-19 Genomics UK (COG-UK) Consortium | The Lighthouse Lab in Milton Keynes and Alex Alderton, Roberto Amato, Jeffrey Barrett, Sonia Goncalves, Ewan Harrison, David K. Jackson, Ian Johnston, Dominic Kwiatkowski, Cordelia Langford, John Sillitoe on behalf of the Wellcome Sanger Institute COVID-19 Surveillance Team |
| EPI_ISL_2866466, EPI_ISL_2866467 | Lighthouse Laboratory Plymouth | Wellcome Sanger Institute for the COVID-19 Genomics UK (COG-UK) Consortium | Lighthouse Laboratory Plymouth and Alex Alderton, Roberto Amato, Jeffrey Barrett, Sonia Goncalves, Ewan Harrison, David K. Jackson, Ian Johnston, Dominic Kwiatkowski, Cordelia Langford, John Sillitoe on behalf of the Wellcome Sanger Institute COVID-19 Surveillance Team |
| EPI_ISL_2866468, EPI_ISL_2866469, EPI_ISL_2866470, EPI_ISL_2866471, EPI_ISL_2866472, EPI_ISL_2866473 | Lighthouse Lab in Milton Keynes | Wellcome Sanger Institute for the COVID-19 Genomics UK (COG-UK) Consortium | The Lighthouse Lab in Milton Keynes and Alex Alderton, Roberto Amato, Jeffrey Barrett, Sonia Goncalves, Ewan Harrison, David K. Jackson, Ian Johnston, Dominic Kwiatkowski, Cordelia Langford, John Sillitoe on behalf of the Wellcome Sanger Institute COVID-19 Surveillance Team |
| EPI_ISL_2866474 | Lighthouse Laboratory Plymouth | Wellcome Sanger Institute for the COVID-19 Genomics UK (COG-UK) Consortium | Lighthouse Laboratory Plymouth and Alex Alderton, Roberto Amato, Jeffrey Barrett, Sonia Goncalves, Ewan Harrison, David K. Jackson, Ian Johnston, Dominic Kwiatkowski, Cordelia Langford, John Sillitoe on behalf of the Wellcome Sanger Institute COVID-19 Surveillance Team |
| EPI_ISL_2866475 | Lighthouse Lab in Milton Keynes | Wellcome Sanger Institute for the COVID-19 Genomics UK (COG-UK) Consortium | The Lighthouse Lab in Milton Keynes and Alex Alderton, Roberto Amato, Jeffrey Barrett, Sonia Goncalves, Ewan Harrison, David K. Jackson, Ian Johnston, Dominic Kwiatkowski, Cordelia Langford, John Sillitoe on behalf of the Wellcome Sanger Institute COVID-19 Surveillance Team |
| EPI_ISL_2866476 | Lighthouse Laboratory Plymouth | Wellcome Sanger Institute for the COVID-19 Genomics UK (COG-UK) Consortium | Lighthouse Laboratory Plymouth and Alex Alderton, Roberto Amato, Jeffrey Barrett, Sonia Goncalves, Ewan Harrison, David K. Jackson, Ian Johnston, Dominic Kwiatkowski, Cordelia Langford, John Sillitoe on behalf of the Wellcome Sanger Institute COVID-19 Surveillance Team |
| EPI_ISL_2866477 | Lighthouse Lab in Milton Keynes | Wellcome Sanger Institute for the COVID-19 Genomics UK (COG-UK) Consortium | The Lighthouse Lab in Milton Keynes and Alex Alderton, Roberto Amato, Jeffrey Barrett, Sonia Goncalves, Ewan Harrison, David K. Jackson, Ian Johnston, Dominic Kwiatkowski, Cordelia Langford, John Sillitoe on behalf of the Wellcome Sanger Institute COVID-19 Surveillance Team |
| EPI_ISL_2866478 | Lighthouse Laboratory Plymouth | Wellcome Sanger Institute for the COVID-19 Genomics UK (COG-UK) Consortium | Lighthouse Laboratory Plymouth and Alex Alderton, Roberto Amato, Jeffrey Barrett, Sonia Goncalves, Ewan Harrison, David K. Jackson, Ian Johnston, Dominic Kwiatkowski, Cordelia Langford, John Sillitoe on behalf of the Wellcome Sanger Institute COVID-19 Surveillance Team |
| EPI_ISL_2866479, EPI_ISL_2866480, EPI_ISL_2866481 | Lighthouse Lab in Milton Keynes | Wellcome Sanger Institute for the COVID-19 Genomics UK (COG-UK) Consortium | The Lighthouse Lab in Milton Keynes and Alex Alderton, Roberto Amato, Jeffrey Barrett, Sonia Goncalves, Ewan Harrison, David K. Jackson, Ian Johnston, Dominic Kwiatkowski, Cordelia Langford, John Sillitoe on behalf of the Wellcome Sanger Institute COVID-19 Surveillance Team |
| EPI_ISL_2866482, EPI_ISL_2866483, EPI_ISL_2866484, EPI_ISL_2866485 | Lighthouse Laboratory Plymouth | Wellcome Sanger Institute for the COVID-19 Genomics UK (COG-UK) Consortium | Lighthouse Laboratory Plymouth and Alex Alderton, Roberto Amato, Jeffrey Barrett, Sonia Goncalves, Ewan Harrison, David K. Jackson, Ian Johnston, Dominic Kwiatkowski, Cordelia Langford, John Sillitoe on behalf of the Wellcome Sanger Institute COVID-19 Surveillance Team |
| EPI_ISL_2866486, EPI_ISL_2866487, EPI_ISL_2866488 | Lighthouse Lab in Milton Keynes | Wellcome Sanger Institute for the COVID-19 Genomics UK (COG-UK) Consortium | The Lighthouse Lab in Milton Keynes and Alex Alderton, Roberto Amato, Jeffrey Barrett, Sonia Goncalves, Ewan Harrison, David K. Jackson, Ian Johnston, Dominic Kwiatkowski, Cordelia Langford, John Sillitoe on behalf of the Wellcome Sanger Institute COVID-19 Surveillance Team |
| EPI_ISL_2866489, EPI_ISL_2866490 | Lighthouse Laboratory Plymouth | Wellcome Sanger Institute for the COVID-19 Genomics UK (COG-UK) Consortium | Lighthouse Laboratory Plymouth and Alex Alderton, Roberto Amato, Jeffrey Barrett, Sonia Goncalves, Ewan Harrison, David K. Jackson, Ian Johnston, Dominic Kwiatkowski, Cordelia Langford, John Sillitoe on behalf of the Wellcome Sanger Institute COVID-19 Surveillance Team |
| EPI_ISL_2866491 | Lighthouse Lab in Milton Keynes | Wellcome Sanger Institute for the COVID-19 Genomics UK (COG-UK) Consortium | The Lighthouse Lab in Milton Keynes and Alex Alderton, Roberto Amato, Jeffrey Barrett, Sonia Goncalves, Ewan Harrison, David K. Jackson, Ian Johnston, Dominic Kwiatkowski, Cordelia Langford, John Sillitoe on behalf of the Wellcome Sanger Institute COVID-19 Surveillance Team |
| EPI_ISL_2866492 | Lighthouse Laboratory Plymouth | Wellcome Sanger Institute for the COVID-19 Genomics UK (COG-UK) Consortium | Lighthouse Laboratory Plymouth and Alex Alderton, Roberto Amato, Jeffrey Barrett, Sonia Goncalves, Ewan Harrison, David K. Jackson, Ian Johnston, Dominic Kwiatkowski, Cordelia Langford, John Sillitoe on behalf of the Wellcome Sanger Institute COVID-19 Surveillance Team |
| EPI_ISL_2866493, EPI_ISL_2866494 | Lighthouse Lab in Milton Keynes | Wellcome Sanger Institute for the COVID-19 Genomics UK (COG-UK) Consortium | The Lighthouse Lab in Milton Keynes and Alex Alderton, Roberto Amato, Jeffrey Barrett, Sonia Goncalves, Ewan Harrison, David K. Jackson, Ian Johnston, Dominic Kwiatkowski, Cordelia Langford, John Sillitoe on behalf of the Wellcome Sanger Institute COVID-19 Surveillance Team |
| EPI_ISL_2866495 | Lighthouse Laboratory Plymouth | Wellcome Sanger Institute for the COVID-19 Genomics UK (COG-UK) Consortium | Lighthouse Laboratory Plymouth and Alex Alderton, Roberto Amato, Jeffrey Barrett, Sonia Goncalves, Ewan Harrison, David K. Jackson, Ian Johnston, Dominic Kwiatkowski, Cordelia Langford, John Sillitoe on behalf of the Wellcome Sanger Institute COVID-19 Surveillance Team |
| EPI_ISL_2866496, EPI_ISL_2866497 | Lighthouse Lab in Milton Keynes | Wellcome Sanger Institute for the COVID-19 Genomics UK (COG-UK) Consortium | The Lighthouse Lab in Milton Keynes and Alex Alderton, Roberto Amato, Jeffrey Barrett, Sonia Goncalves, Ewan Harrison, David K. Jackson, Ian Johnston, Dominic Kwiatkowski, Cordelia Langford, John Sillitoe on behalf of the Wellcome Sanger Institute COVID-19 Surveillance Team |
| EPI_ISL_2866498 | Lighthouse Laboratory Plymouth | Wellcome Sanger Institute for the COVID-19 Genomics UK (COG-UK) Consortium | Lighthouse Laboratory Plymouth and Alex Alderton, Roberto Amato, Jeffrey Barrett, Sonia Goncalves, Ewan Harrison, David K. Jackson, Ian Johnston, Dominic Kwiatkowski, Cordelia Langford, John Sillitoe on behalf of the Wellcome Sanger Institute COVID-19 Surveillance Team |
| EPI_ISL_2866499, EPI_ISL_2866500, EPI_ISL_2866501 | Lighthouse Lab in Milton Keynes | Wellcome Sanger Institute for the COVID-19 Genomics UK (COG-UK) Consortium | The Lighthouse Lab in Milton Keynes and Alex Alderton, Roberto Amato, Jeffrey Barrett, Sonia Goncalves, Ewan Harrison, David K. Jackson, Ian Johnston, Dominic Kwiatkowski, Cordelia Langford, John Sillitoe on behalf of the Wellcome Sanger Institute COVID-19 Surveillance Team |
| EPI_ISL_2866502, EPI_ISL_2866503 | Lighthouse Laboratory Plymouth | Wellcome Sanger Institute for the COVID-19 Genomics UK (COG-UK) Consortium | Lighthouse Laboratory Plymouth and Alex Alderton, Roberto Amato, Jeffrey Barrett, Sonia Goncalves, Ewan Harrison, David K. Jackson, Ian Johnston, Dominic Kwiatkowski, Cordelia Langford, John Sillitoe on behalf of the Wellcome Sanger Institute COVID-19 Surveillance Team |
| EPI_ISL_2866504 | Lighthouse Lab in Milton Keynes | Wellcome Sanger Institute for the COVID-19 Genomics UK (COG-UK) Consortium | The Lighthouse Lab in Milton Keynes and Alex Alderton, Roberto Amato, Jeffrey Barrett, Sonia Goncalves, Ewan Harrison, David K. Jackson, Ian Johnston, Dominic Kwiatkowski, Cordelia Langford, John Sillitoe on behalf of the Wellcome Sanger Institute COVID-19 Surveillance Team |
| EPI_ISL_2866505, EPI_ISL_2866506, EPI_ISL_2866507, EPI_ISL_2866508 | Lighthouse Laboratory Plymouth | Wellcome Sanger Institute for the COVID-19 Genomics UK (COG-UK) Consortium | Lighthouse Laboratory Plymouth and Alex Alderton, Roberto Amato, Jeffrey Barrett, Sonia Goncalves, Ewan Harrison, David K. Jackson, Ian Johnston, Dominic Kwiatkowski, Cordelia Langford, John Sillitoe on behalf of the Wellcome Sanger Institute COVID-19 Surveillance Team |
| EPI_ISL_2866509, EPI_ISL_2866510 | Lighthouse Lab in Milton Keynes | Wellcome Sanger Institute for the COVID-19 Genomics UK (COG-UK) Consortium | The Lighthouse Lab in Milton Keynes and Alex Alderton, Roberto Amato, Jeffrey Barrett, Sonia Goncalves, Ewan Harrison, David K. Jackson, Ian Johnston, Dominic Kwiatkowski, Cordelia Langford, John Sillitoe on behalf of the Wellcome Sanger Institute COVID-19 Surveillance Team |
| EPI_ISL_2866511, EPI_ISL_2866512 | Lighthouse Laboratory Plymouth | Wellcome Sanger Institute for the COVID-19 Genomics UK (COG-UK) Consortium | Lighthouse Laboratory Plymouth and Alex Alderton, Roberto Amato, Jeffrey Barrett, Sonia Goncalves, Ewan Harrison, David K. Jackson, Ian Johnston, Dominic Kwiatkowski, Cordelia Langford, John Sillitoe on behalf of the Wellcome Sanger Institute COVID-19 Surveillance Team |
| EPI_ISL_2866513, EPI_ISL_2866514, EPI_ISL_2866515 | Lighthouse Lab in Milton Keynes | Wellcome Sanger Institute for the COVID-19 Genomics UK (COG-UK) Consortium | The Lighthouse Lab in Milton Keynes and Alex Alderton, Roberto Amato, Jeffrey Barrett, Sonia Goncalves, Ewan Harrison, David K. Jackson, Ian Johnston, Dominic Kwiatkowski, Cordelia Langford, John Sillitoe on behalf of the Wellcome Sanger Institute COVID-19 Surveillance Team |
| EPI_ISL_2866516, EPI_ISL_2866517 | Lighthouse Laboratory Plymouth | Wellcome Sanger Institute for the COVID-19 Genomics UK (COG-UK) Consortium | Lighthouse Laboratory Plymouth and Alex Alderton, Roberto Amato, Jeffrey Barrett, Sonia Goncalves, Ewan Harrison, David K. Jackson, Ian Johnston, Dominic Kwiatkowski, Cordelia Langford, John Sillitoe on behalf of the Wellcome Sanger Institute COVID-19 Surveillance Team |
| EPI_ISL_2866518, EPI_ISL_2866519 | Lighthouse Lab in Milton Keynes | Wellcome Sanger Institute for the COVID-19 Genomics UK (COG-UK) Consortium | The Lighthouse Lab in Milton Keynes and Alex Alderton, Roberto Amato, Jeffrey Barrett, Sonia Goncalves, Ewan Harrison, David K. Jackson, Ian Johnston, Dominic Kwiatkowski, Cordelia Langford, John Sillitoe on behalf of the Wellcome Sanger Institute COVID-19 Surveillance Team |
| EPI_ISL_2866520 | Lighthouse Laboratory Plymouth | Wellcome Sanger Institute for the COVID-19 Genomics UK (COG-UK) Consortium | Lighthouse Laboratory Plymouth and Alex Alderton, Roberto Amato, Jeffrey Barrett, Sonia Goncalves, Ewan Harrison, David K. Jackson, Ian Johnston, Dominic Kwiatkowski, Cordelia Langford, John Sillitoe on behalf of the Wellcome Sanger Institute COVID-19 Surveillance Team |
| EPI_ISL_2866521 | Lighthouse Lab in Milton Keynes | Wellcome Sanger Institute for the COVID-19 Genomics UK (COG-UK) Consortium | The Lighthouse Lab in Milton Keynes and Alex Alderton, Roberto Amato, Jeffrey Barrett, Sonia Goncalves, Ewan Harrison, David K. Jackson, Ian Johnston, Dominic Kwiatkowski, Cordelia Langford, John Sillitoe on behalf of the Wellcome Sanger Institute COVID-19 Surveillance Team |
| EPI_ISL_2866522 | Lighthouse Laboratory Plymouth | Wellcome Sanger Institute for the COVID-19 Genomics UK (COG-UK) Consortium | Lighthouse Laboratory Plymouth and Alex Alderton, Roberto Amato, Jeffrey Barrett, Sonia Goncalves, Ewan Harrison, David K. Jackson, Ian Johnston, Dominic Kwiatkowski, Cordelia Langford, John Sillitoe on behalf of the Wellcome Sanger Institute COVID-19 Surveillance Team |
| EPI_ISL_2866523, EPI_ISL_2866524 | Lighthouse Lab in Milton Keynes | Wellcome Sanger Institute for the COVID-19 Genomics UK (COG-UK) Consortium | The Lighthouse Lab in Milton Keynes and Alex Alderton, Roberto Amato, Jeffrey Barrett, Sonia Goncalves, Ewan Harrison, David K. Jackson, Ian Johnston, Dominic Kwiatkowski, Cordelia Langford, John Sillitoe on behalf of the Wellcome Sanger Institute COVID-19 Surveillance Team |
| EPI_ISL_2866525, EPI_ISL_2866526, EPI_ISL_2866527, EPI_ISL_2866528 | Lighthouse Laboratory Plymouth | Wellcome Sanger Institute for the COVID-19 Genomics UK (COG-UK) Consortium | Lighthouse Laboratory Plymouth and Alex Alderton, Roberto Amato, Jeffrey Barrett, Sonia Goncalves, Ewan Harrison, David K. Jackson, Ian Johnston, Dominic Kwiatkowski, Cordelia Langford, John Sillitoe on behalf of the Wellcome Sanger Institute COVID-19 Surveillance Team |
| EPI_ISL_2866529 | Lighthouse Lab in Milton Keynes | Wellcome Sanger Institute for the COVID-19 Genomics UK (COG-UK) Consortium | The Lighthouse Lab in Milton Keynes and Alex Alderton, Roberto Amato, Jeffrey Barrett, Sonia Goncalves, Ewan Harrison, David K. Jackson, Ian Johnston, Dominic Kwiatkowski, Cordelia Langford, John Sillitoe on behalf of the Wellcome Sanger Institute COVID-19 Surveillance Team |
| EPI_ISL_2866530, EPI_ISL_2866531 | Lighthouse Laboratory Plymouth | Wellcome Sanger Institute for the COVID-19 Genomics UK (COG-UK) Consortium | Lighthouse Laboratory Plymouth and Alex Alderton, Roberto Amato, Jeffrey Barrett, Sonia Goncalves, Ewan Harrison, David K. Jackson, Ian Johnston, Dominic Kwiatkowski, Cordelia Langford, John Sillitoe on behalf of the Wellcome Sanger Institute COVID-19 Surveillance Team |
| EPI_ISL_2866532 | Lighthouse Lab in Milton Keynes | Wellcome Sanger Institute for the COVID-19 Genomics UK (COG-UK) Consortium | The Lighthouse Lab in Milton Keynes and Alex Alderton, Roberto Amato, Jeffrey Barrett, Sonia Goncalves, Ewan Harrison, David K. Jackson, Ian Johnston, Dominic Kwiatkowski, Cordelia Langford, John Sillitoe on behalf of the Wellcome Sanger Institute COVID-19 Surveillance Team |
| EPI_ISL_2866533, EPI_ISL_2866534 | Lighthouse Laboratory Plymouth | Wellcome Sanger Institute for the COVID-19 Genomics UK (COG-UK) Consortium | Lighthouse Laboratory Plymouth and Alex Alderton, Roberto Amato, Jeffrey Barrett, Sonia Goncalves, Ewan Harrison, David K. Jackson, Ian Johnston, Dominic Kwiatkowski, Cordelia Langford, John Sillitoe on behalf of the Wellcome Sanger Institute COVID-19 Surveillance Team |
| EPI_ISL_2866535 | Lighthouse Lab in Milton Keynes | Wellcome Sanger Institute for the COVID-19 Genomics UK (COG-UK) Consortium | The Lighthouse Lab in Milton Keynes and Alex Alderton, Roberto Amato, Jeffrey Barrett, Sonia Goncalves, Ewan Harrison, David K. Jackson, Ian Johnston, Dominic Kwiatkowski, Cordelia Langford, John Sillitoe on behalf of the Wellcome Sanger Institute COVID-19 Surveillance Team |
| EPI_ISL_2866536, EPI_ISL_2866537, EPI_ISL_2866538 | Lighthouse Laboratory Plymouth | Wellcome Sanger Institute for the COVID-19 Genomics UK (COG-UK) Consortium | Lighthouse Laboratory Plymouth and Alex Alderton, Roberto Amato, Jeffrey Barrett, Sonia Goncalves, Ewan Harrison, David K. Jackson, Ian Johnston, Dominic Kwiatkowski, Cordelia Langford, John Sillitoe on behalf of the Wellcome Sanger Institute COVID-19 Surveillance Team |
| EPI_ISL_2866539 | Lighthouse Lab in Milton Keynes | Wellcome Sanger Institute for the COVID-19 Genomics UK (COG-UK) | The Lighthouse Lab in Milton Keynes and Alex Alderton, Roberto Amato, Jeffrey Barrett, Sonia Goncalves, Ewan Harrison, David K. Jackson, Ian Johnston, Dominic Kwiatkowski, Cordelia Langford, John Sillitoe on behalf of the Wellcome Sanger |

| | | | |
|---|---|---|---|
| | | Consortium | Institute COVID-19 Surveillance Team |
| EPI_ISL_2866540, EPI_ISL_2866541 | Lighthouse Laboratory Plymouth | Wellcome Sanger Institute for the COVID-19 Genomics UK (COG-UK) Consortium | Lighthouse Laboratory Plymouth and Alex Alderton, Roberto Amato, Jeffrey Barrett, Sonia Goncalves, Ewan Harrison, David K. Jackson, Ian Johnston, Dominic Kwiatkowski, Cordelia Langford, John Sillitoe on behalf of the Wellcome Sanger Institute COVID-19 Surveillance Team |
| EPI_ISL_2866542 | Lighthouse Lab in Milton Keynes | Wellcome Sanger Institute for the COVID-19 Genomics UK (COG-UK) Consortium | The Lighthouse Lab in Milton Keynes and Alex Alderton, Roberto Amato, Jeffrey Barrett, Sonia Goncalves, Ewan Harrison, David K. Jackson, Ian Johnston, Dominic Kwiatkowski, Cordelia Langford, John Sillitoe on behalf of the Wellcome Sanger Institute COVID-19 Surveillance Team |
| EPI_ISL_2866543, EPI_ISL_2866544 | Lighthouse Laboratory Plymouth | Wellcome Sanger Institute for the COVID-19 Genomics UK (COG-UK) Consortium | Lighthouse Laboratory Plymouth and Alex Alderton, Roberto Amato, Jeffrey Barrett, Sonia Goncalves, Ewan Harrison, David K. Jackson, Ian Johnston, Dominic Kwiatkowski, Cordelia Langford, John Sillitoe on behalf of the Wellcome Sanger Institute COVID-19 Surveillance Team |
| EPI_ISL_2866545, EPI_ISL_2866546 | Lighthouse Lab in Milton Keynes | Wellcome Sanger Institute for the COVID-19 Genomics UK (COG-UK) Consortium | The Lighthouse Lab in Milton Keynes and Alex Alderton, Roberto Amato, Jeffrey Barrett, Sonia Goncalves, Ewan Harrison, David K. Jackson, Ian Johnston, Dominic Kwiatkowski, Cordelia Langford, John Sillitoe on behalf of the Wellcome Sanger Institute COVID-19 Surveillance Team |
| EPI_ISL_2866547 | Lighthouse Laboratory Plymouth | Wellcome Sanger Institute for the COVID-19 Genomics UK (COG-UK) Consortium | Lighthouse Laboratory Plymouth and Alex Alderton, Roberto Amato, Jeffrey Barrett, Sonia Goncalves, Ewan Harrison, David K. Jackson, Ian Johnston, Dominic Kwiatkowski, Cordelia Langford, John Sillitoe on behalf of the Wellcome Sanger Institute COVID-19 Surveillance Team |
| EPI_ISL_2866548, EPI_ISL_2866549, EPI_ISL_2866550, EPI_ISL_2866551 | Lighthouse Lab in Milton Keynes | Wellcome Sanger Institute for the COVID-19 Genomics UK (COG-UK) Consortium | The Lighthouse Lab in Milton Keynes and Alex Alderton, Roberto Amato, Jeffrey Barrett, Sonia Goncalves, Ewan Harrison, David K. Jackson, Ian Johnston, Dominic Kwiatkowski, Cordelia Langford, John Sillitoe on behalf of the Wellcome Sanger Institute COVID-19 Surveillance Team |
| EPI_ISL_2866552, EPI_ISL_2866553 | Lighthouse Laboratory Plymouth | Wellcome Sanger Institute for the COVID-19 Genomics UK (COG-UK) Consortium | Lighthouse Laboratory Plymouth and Alex Alderton, Roberto Amato, Jeffrey Barrett, Sonia Goncalves, Ewan Harrison, David K. Jackson, Ian Johnston, Dominic Kwiatkowski, Cordelia Langford, John Sillitoe on behalf of the Wellcome Sanger Institute COVID-19 Surveillance Team |
| EPI_ISL_2866554 | Lighthouse Lab in Milton Keynes | Wellcome Sanger Institute for the COVID-19 Genomics UK (COG-UK) Consortium | The Lighthouse Lab in Milton Keynes and Alex Alderton, Roberto Amato, Jeffrey Barrett, Sonia Goncalves, Ewan Harrison, David K. Jackson, Ian Johnston, Dominic Kwiatkowski, Cordelia Langford, John Sillitoe on behalf of the Wellcome Sanger Institute COVID-19 Surveillance Team |
| EPI_ISL_2866555 | Lighthouse Laboratory Plymouth | Wellcome Sanger Institute for the COVID-19 Genomics UK (COG-UK) Consortium | Lighthouse Laboratory Plymouth and Alex Alderton, Roberto Amato, Jeffrey Barrett, Sonia Goncalves, Ewan Harrison, David K. Jackson, Ian Johnston, Dominic Kwiatkowski, Cordelia Langford, John Sillitoe on behalf of the Wellcome Sanger Institute COVID-19 Surveillance Team |
| EPI_ISL_2866556, EPI_ISL_2866557, EPI_ISL_2866558, EPI_ISL_2866559 | Lighthouse Lab in Milton Keynes | Wellcome Sanger Institute for the COVID-19 Genomics UK (COG-UK) Consortium | The Lighthouse Lab in Milton Keynes and Alex Alderton, Roberto Amato, Jeffrey Barrett, Sonia Goncalves, Ewan Harrison, David K. Jackson, Ian Johnston, Dominic Kwiatkowski, Cordelia Langford, John Sillitoe on behalf of the Wellcome Sanger Institute COVID-19 Surveillance Team |
| EPI_ISL_2866560, EPI_ISL_2866561 | Lighthouse Laboratory Plymouth | Wellcome Sanger Institute for the COVID-19 Genomics UK (COG-UK) Consortium | Lighthouse Laboratory Plymouth and Alex Alderton, Roberto Amato, Jeffrey Barrett, Sonia Goncalves, Ewan Harrison, David K. Jackson, Ian Johnston, Dominic Kwiatkowski, Cordelia Langford, John Sillitoe on behalf of the Wellcome Sanger Institute COVID-19 Surveillance Team |
| EPI_ISL_2866562 | Lighthouse Lab in Milton Keynes | Wellcome Sanger Institute for the COVID-19 Genomics UK (COG-UK) Consortium | The Lighthouse Lab in Milton Keynes and Alex Alderton, Roberto Amato, Jeffrey Barrett, Sonia Goncalves, Ewan Harrison, David K. Jackson, Ian Johnston, Dominic Kwiatkowski, Cordelia Langford, John Sillitoe on behalf of the Wellcome Sanger Institute COVID-19 Surveillance Team |
| EPI_ISL_2866563 | Lighthouse Laboratory Plymouth | Wellcome Sanger Institute for the COVID-19 Genomics UK (COG-UK) Consortium | Lighthouse Laboratory Plymouth and Alex Alderton, Roberto Amato, Jeffrey Barrett, Sonia Goncalves, Ewan Harrison, David K. Jackson, Ian Johnston, Dominic Kwiatkowski, Cordelia Langford, John Sillitoe on behalf of the Wellcome Sanger Institute COVID-19 Surveillance Team |
| EPI_ISL_2866564, EPI_ISL_2866565 | Lighthouse Lab in Milton Keynes | Wellcome Sanger Institute for the COVID-19 Genomics UK (COG-UK) Consortium | The Lighthouse Lab in Milton Keynes and Alex Alderton, Roberto Amato, Jeffrey Barrett, Sonia Goncalves, Ewan Harrison, David K. Jackson, Ian Johnston, Dominic Kwiatkowski, Cordelia Langford, John Sillitoe on behalf of the Wellcome Sanger Institute COVID-19 Surveillance Team |
| EPI_ISL_2866566, EPI_ISL_2866567 | Lighthouse Laboratory Plymouth | Wellcome Sanger Institute for the COVID-19 Genomics UK (COG-UK) Consortium | Lighthouse Laboratory Plymouth and Alex Alderton, Roberto Amato, Jeffrey Barrett, Sonia Goncalves, Ewan Harrison, David K. Jackson, Ian Johnston, Dominic Kwiatkowski, Cordelia Langford, John Sillitoe on behalf of the Wellcome Sanger Institute COVID-19 Surveillance Team |
| EPI_ISL_2866568 | Lighthouse Lab in Milton Keynes | Wellcome Sanger Institute for the COVID-19 Genomics UK (COG-UK) Consortium | The Lighthouse Lab in Milton Keynes and Alex Alderton, Roberto Amato, Jeffrey Barrett, Sonia Goncalves, Ewan Harrison, David K. Jackson, Ian Johnston, Dominic Kwiatkowski, Cordelia Langford, John Sillitoe on behalf of the Wellcome Sanger Institute COVID-19 Surveillance Team |
| EPI_ISL_2866569, EPI_ISL_2866570, EPI_ISL_2866571 | Lighthouse Laboratory Plymouth | Wellcome Sanger Institute for the COVID-19 Genomics UK (COG-UK) Consortium | Lighthouse Laboratory Plymouth and Alex Alderton, Roberto Amato, Jeffrey Barrett, Sonia Goncalves, Ewan Harrison, David K. Jackson, Ian Johnston, Dominic Kwiatkowski, Cordelia Langford, John Sillitoe on behalf of the Wellcome Sanger Institute COVID-19 Surveillance Team |
| EPI_ISL_2866572, EPI_ISL_2866573 | Lighthouse Lab in Milton Keynes | Wellcome Sanger Institute for the COVID-19 Genomics UK (COG-UK) Consortium | The Lighthouse Lab in Milton Keynes and Alex Alderton, Roberto Amato, Jeffrey Barrett, Sonia Goncalves, Ewan Harrison, David K. Jackson, Ian Johnston, Dominic Kwiatkowski, Cordelia Langford, John Sillitoe on behalf of the Wellcome Sanger Institute COVID-19 Surveillance Team |
| EPI_ISL_2866574, EPI_ISL_2866575, EPI_ISL_2866576 | Lighthouse Laboratory Plymouth | Wellcome Sanger Institute for the COVID-19 Genomics UK (COG-UK) Consortium | Lighthouse Laboratory Plymouth and Alex Alderton, Roberto Amato, Jeffrey Barrett, Sonia Goncalves, Ewan Harrison, David K. Jackson, Ian Johnston, Dominic Kwiatkowski, Cordelia Langford, John Sillitoe on behalf of the Wellcome Sanger Institute COVID-19 Surveillance Team |
| EPI_ISL_2866577, EPI_ISL_2866578, EPI_ISL_2866579, EPI_ISL_2866580, EPI_ISL_2866581, EPI_ISL_2866582, EPI_ISL_2866583, EPI_ISL_2866584, EPI_ISL_2866585, EPI_ISL_2866586, EPI_ISL_2866587, EPI_ISL_2866588 | | | |
| see above | Lighthouse Lab in Milton Keynes | Wellcome Sanger Institute for the COVID-19 Genomics UK (COG-UK) Consortium | The Lighthouse Lab in Milton Keynes and Alex Alderton, Roberto Amato, Jeffrey Barrett, Sonia Goncalves, Ewan Harrison, David K. Jackson, Ian Johnston, Dominic Kwiatkowski, Cordelia Langford, John Sillitoe on behalf of the Wellcome Sanger Institute COVID-19 Surveillance Team |
| EPI_ISL_2866589, EPI_ISL_2866590 | Lighthouse Laboratory Plymouth | Wellcome Sanger Institute for the COVID-19 Genomics UK (COG-UK) Consortium | Lighthouse Laboratory Plymouth and Alex Alderton, Roberto Amato, Jeffrey Barrett, Sonia Goncalves, Ewan Harrison, David K. Jackson, Ian Johnston, Dominic Kwiatkowski, Cordelia Langford, John Sillitoe on behalf of the Wellcome Sanger Institute COVID-19 Surveillance Team |
| EPI_ISL_2866591, EPI_ISL_2866592 | Lighthouse Lab in Milton Keynes | Wellcome Sanger Institute for the COVID-19 Genomics UK (COG-UK) Consortium | The Lighthouse Lab in Milton Keynes and Alex Alderton, Roberto Amato, Jeffrey Barrett, Sonia Goncalves, Ewan Harrison, David K. Jackson, Ian Johnston, Dominic Kwiatkowski, Cordelia Langford, John Sillitoe on behalf of the Wellcome Sanger Institute COVID-19 Surveillance Team |
| EPI_ISL_2866593 | Lighthouse Laboratory Plymouth | Wellcome Sanger Institute for the COVID-19 Genomics UK (COG-UK) Consortium | Lighthouse Laboratory Plymouth and Alex Alderton, Roberto Amato, Jeffrey Barrett, Sonia Goncalves, Ewan Harrison, David K. Jackson, Ian Johnston, Dominic Kwiatkowski, Cordelia Langford, John Sillitoe on behalf of the Wellcome Sanger Institute COVID-19 Surveillance Team |
| EPI_ISL_2866594, EPI_ISL_2866595, EPI_ISL_2866596, EPI_ISL_2866597, EPI_ISL_2866598 | Lighthouse Lab in Milton Keynes | Wellcome Sanger Institute for the COVID-19 Genomics UK (COG-UK) Consortium | The Lighthouse Lab in Milton Keynes and Alex Alderton, Roberto Amato, Jeffrey Barrett, Sonia Goncalves, Ewan Harrison, David K. Jackson, Ian Johnston, Dominic Kwiatkowski, Cordelia Langford, John Sillitoe on behalf of the Wellcome Sanger Institute COVID-19 Surveillance Team |
| EPI_ISL_2866599 | Lighthouse Laboratory Plymouth | Wellcome Sanger Institute for the COVID-19 Genomics UK (COG-UK) Consortium | Lighthouse Laboratory Plymouth and Alex Alderton, Roberto Amato, Jeffrey Barrett, Sonia Goncalves, Ewan Harrison, David K. Jackson, Ian Johnston, Dominic Kwiatkowski, Cordelia Langford, John Sillitoe on behalf of the Wellcome Sanger Institute COVID-19 Surveillance Team |
| EPI_ISL_2866600 | Lighthouse Lab in Milton Keynes | Wellcome Sanger Institute for the COVID-19 Genomics UK (COG-UK) Consortium | The Lighthouse Lab in Milton Keynes and Alex Alderton, Roberto Amato, Jeffrey Barrett, Sonia Goncalves, Ewan Harrison, David K. Jackson, Ian Johnston, Dominic Kwiatkowski, Cordelia Langford, John Sillitoe on behalf of the Wellcome Sanger Institute COVID-19 Surveillance Team |
| EPI_ISL_2866601 | Lighthouse Laboratory Plymouth | Wellcome Sanger Institute for the COVID-19 Genomics UK (COG-UK) Consortium | Lighthouse Laboratory Plymouth and Alex Alderton, Roberto Amato, Jeffrey Barrett, Sonia Goncalves, Ewan Harrison, David K. Jackson, Ian Johnston, Dominic Kwiatkowski, Cordelia Langford, John Sillitoe on behalf of the Wellcome Sanger Institute COVID-19 Surveillance Team |
| EPI_ISL_2866602 | Lighthouse Lab in Milton Keynes | Wellcome Sanger Institute for the COVID-19 Genomics UK (COG-UK) Consortium | The Lighthouse Lab in Milton Keynes and Alex Alderton, Roberto Amato, Jeffrey Barrett, Sonia Goncalves, Ewan Harrison, David K. Jackson, Ian Johnston, Dominic Kwiatkowski, Cordelia Langford, John Sillitoe on behalf of the Wellcome Sanger Institute COVID-19 Surveillance Team |
| EPI_ISL_2866603 | Lighthouse Laboratory Plymouth | Wellcome Sanger Institute for the COVID-19 Genomics UK (COG-UK) Consortium | Lighthouse Laboratory Plymouth and Alex Alderton, Roberto Amato, Jeffrey Barrett, Sonia Goncalves, Ewan Harrison, David K. Jackson, Ian Johnston, Dominic Kwiatkowski, Cordelia Langford, John Sillitoe on behalf of the Wellcome Sanger Institute COVID-19 Surveillance Team |
| EPI_ISL_2866604 | Lighthouse Lab in Milton Keynes | Wellcome Sanger Institute for the COVID-19 Genomics UK (COG-UK) Consortium | The Lighthouse Lab in Milton Keynes and Alex Alderton, Roberto Amato, Jeffrey Barrett, Sonia Goncalves, Ewan Harrison, David K. Jackson, Ian Johnston, Dominic Kwiatkowski, Cordelia Langford, John Sillitoe on behalf of the Wellcome Sanger Institute COVID-19 Surveillance Team |
| EPI_ISL_2866605 | Lighthouse Laboratory Plymouth | Wellcome Sanger Institute for the COVID-19 Genomics UK (COG-UK) Consortium | Lighthouse Laboratory Plymouth and Alex Alderton, Roberto Amato, Jeffrey Barrett, Sonia Goncalves, Ewan Harrison, David K. Jackson, Ian Johnston, Dominic Kwiatkowski, Cordelia Langford, John Sillitoe on behalf of the Wellcome Sanger Institute COVID-19 Surveillance Team |
| EPI_ISL_2866606 | Lighthouse Lab in Milton Keynes | Wellcome Sanger Institute for the COVID-19 Genomics UK (COG-UK) Consortium | The Lighthouse Lab in Milton Keynes and Alex Alderton, Roberto Amato, Jeffrey Barrett, Sonia Goncalves, Ewan Harrison, David K. Jackson, Ian Johnston, Dominic Kwiatkowski, Cordelia Langford, John Sillitoe on behalf of the Wellcome Sanger Institute COVID-19 Surveillance Team |
| EPI_ISL_2866607 | Lighthouse Laboratory Plymouth | Wellcome Sanger Institute for the COVID-19 Genomics UK (COG-UK) Consortium | Lighthouse Laboratory Plymouth and Alex Alderton, Roberto Amato, Jeffrey Barrett, Sonia Goncalves, Ewan Harrison, David K. Jackson, Ian Johnston, Dominic Kwiatkowski, Cordelia Langford, John Sillitoe on behalf of the Wellcome Sanger Institute COVID-19 Surveillance Team |
| EPI_ISL_2866608 | Lighthouse Lab in Milton Keynes | Wellcome Sanger Institute for the COVID-19 Genomics UK (COG-UK) Consortium | The Lighthouse Lab in Milton Keynes and Alex Alderton, Roberto Amato, Jeffrey Barrett, Sonia Goncalves, Ewan Harrison, David K. Jackson, Ian Johnston, Dominic Kwiatkowski, Cordelia Langford, John Sillitoe on behalf of the Wellcome Sanger Institute COVID-19 Surveillance Team |
| EPI_ISL_2866609 | Lighthouse Laboratory Plymouth | Wellcome Sanger Institute for the COVID-19 Genomics UK (COG-UK) Consortium | Lighthouse Laboratory Plymouth and Alex Alderton, Roberto Amato, Jeffrey Barrett, Sonia Goncalves, Ewan Harrison, David K. Jackson, Ian Johnston, Dominic Kwiatkowski, Cordelia Langford, John Sillitoe on behalf of the Wellcome Sanger Institute COVID-19 Surveillance Team |
| EPI_ISL_2866610 | Lighthouse Lab in Milton Keynes | Wellcome Sanger Institute for the COVID-19 Genomics UK (COG-UK) Consortium | The Lighthouse Lab in Milton Keynes and Alex Alderton, Roberto Amato, Jeffrey Barrett, Sonia Goncalves, Ewan Harrison, David K. Jackson, Ian Johnston, Dominic Kwiatkowski, Cordelia Langford, John Sillitoe on behalf of the Wellcome Sanger Institute COVID-19 Surveillance Team |
| EPI_ISL_2866611, EPI_ISL_2866612, EPI_ISL_2866613 | Lighthouse Laboratory Plymouth | Wellcome Sanger Institute for the COVID-19 Genomics UK (COG-UK) Consortium | Lighthouse Laboratory Plymouth and Alex Alderton, Roberto Amato, Jeffrey Barrett, Sonia Goncalves, Ewan Harrison, David K. Jackson, Ian Johnston, Dominic Kwiatkowski, Cordelia Langford, John Sillitoe on behalf of the Wellcome Sanger Institute COVID-19 Surveillance Team |
| EPI_ISL_2866614, EPI_ISL_2866615, EPI_ISL_2866616, EPI_ISL_2866617 | Lighthouse Lab in Milton Keynes | Wellcome Sanger Institute for the COVID-19 Genomics UK (COG-UK) Consortium | The Lighthouse Lab in Milton Keynes and Alex Alderton, Roberto Amato, Jeffrey Barrett, Sonia Goncalves, Ewan Harrison, David K. Jackson, Ian Johnston, Dominic Kwiatkowski, Cordelia Langford, John Sillitoe on behalf of the Wellcome Sanger Institute COVID-19 Surveillance Team |
| EPI_ISL_2866618, EPI_ISL_2866619, EPI_ISL_2866620, | Lighthouse Laboratory Plymouth | Wellcome Sanger Institute for the COVID-19 Genomics UK (COG-UK) Consortium | Lighthouse Laboratory Plymouth and Alex Alderton, Roberto Amato, Jeffrey Barrett, Sonia Goncalves, Ewan Harrison, David K. Jackson, Ian Johnston, Dominic Kwiatkowski, Cordelia Langford, John Sillitoe on behalf of the Wellcome Sanger Institute COVID-19 Surveillance Team |

| | | | |
|---|---|---|---|
| EPI_ISL_2866621 | | | |
| EPI_ISL_2866622 | Lighthouse Lab in Milton Keynes | Wellcome Sanger Institute for the COVID-19 Genomics UK (COG-UK) Consortium | The Lighthouse Lab in Milton Keynes and Alex Alderton, Roberto Amato, Jeffrey Barrett, Sonia Goncalves, Ewan Harrison, David K. Jackson, Ian Johnston, Dominic Kwiatkowski, Cordelia Langford, John Sillitoe on behalf of the Wellcome Sanger Institute COVID-19 Surveillance Team |
| EPI_ISL_2866623, EPI_ISL_2866624 | Lighthouse Laboratory Plymouth | Wellcome Sanger Institute for the COVID-19 Genomics UK (COG-UK) Consortium | Lighthouse Laboratory Plymouth and Alex Alderton, Roberto Amato, Jeffrey Barrett, Sonia Goncalves, Ewan Harrison, David K. Jackson, Ian Johnston, Dominic Kwiatkowski, Cordelia Langford, John Sillitoe on behalf of the Wellcome Sanger Institute COVID-19 Surveillance Team |
| EPI_ISL_2866625, EPI_ISL_2866626 | Lighthouse Lab in Milton Keynes | Wellcome Sanger Institute for the COVID-19 Genomics UK (COG-UK) Consortium | The Lighthouse Lab in Milton Keynes and Alex Alderton, Roberto Amato, Jeffrey Barrett, Sonia Goncalves, Ewan Harrison, David K. Jackson, Ian Johnston, Dominic Kwiatkowski, Cordelia Langford, John Sillitoe on behalf of the Wellcome Sanger Institute COVID-19 Surveillance Team |
| EPI_ISL_2866627, EPI_ISL_2866628 | Lighthouse Laboratory Plymouth | Wellcome Sanger Institute for the COVID-19 Genomics UK (COG-UK) Consortium | Lighthouse Laboratory Plymouth and Alex Alderton, Roberto Amato, Jeffrey Barrett, Sonia Goncalves, Ewan Harrison, David K. Jackson, Ian Johnston, Dominic Kwiatkowski, Cordelia Langford, John Sillitoe on behalf of the Wellcome Sanger Institute COVID-19 Surveillance Team |
| EPI_ISL_2866629, EPI_ISL_2866630, EPI_ISL_2866631, EPI_ISL_2866632 | Lighthouse Lab in Milton Keynes | Wellcome Sanger Institute for the COVID-19 Genomics UK (COG-UK) Consortium | The Lighthouse Lab in Milton Keynes and Alex Alderton, Roberto Amato, Jeffrey Barrett, Sonia Goncalves, Ewan Harrison, David K. Jackson, Ian Johnston, Dominic Kwiatkowski, Cordelia Langford, John Sillitoe on behalf of the Wellcome Sanger Institute COVID-19 Surveillance Team |
| EPI_ISL_2866633, EPI_ISL_2866634, EPI_ISL_2866635 | Lighthouse Laboratory Plymouth | Wellcome Sanger Institute for the COVID-19 Genomics UK (COG-UK) Consortium | Lighthouse Laboratory Plymouth and Alex Alderton, Roberto Amato, Jeffrey Barrett, Sonia Goncalves, Ewan Harrison, David K. Jackson, Ian Johnston, Dominic Kwiatkowski, Cordelia Langford, John Sillitoe on behalf of the Wellcome Sanger Institute COVID-19 Surveillance Team |
| EPI_ISL_2866636 | Lighthouse Lab in Milton Keynes | Wellcome Sanger Institute for the COVID-19 Genomics UK (COG-UK) Consortium | The Lighthouse Lab in Milton Keynes and Alex Alderton, Roberto Amato, Jeffrey Barrett, Sonia Goncalves, Ewan Harrison, David K. Jackson, Ian Johnston, Dominic Kwiatkowski, Cordelia Langford, John Sillitoe on behalf of the Wellcome Sanger Institute COVID-19 Surveillance Team |
| EPI_ISL_2866637, EPI_ISL_2866638 | Lighthouse Laboratory Plymouth | Wellcome Sanger Institute for the COVID-19 Genomics UK (COG-UK) Consortium | Lighthouse Laboratory Plymouth and Alex Alderton, Roberto Amato, Jeffrey Barrett, Sonia Goncalves, Ewan Harrison, David K. Jackson, Ian Johnston, Dominic Kwiatkowski, Cordelia Langford, John Sillitoe on behalf of the Wellcome Sanger Institute COVID-19 Surveillance Team |
| EPI_ISL_2866639 | Lighthouse Lab in Milton Keynes | Wellcome Sanger Institute for the COVID-19 Genomics UK (COG-UK) Consortium | The Lighthouse Lab in Milton Keynes and Alex Alderton, Roberto Amato, Jeffrey Barrett, Sonia Goncalves, Ewan Harrison, David K. Jackson, Ian Johnston, Dominic Kwiatkowski, Cordelia Langford, John Sillitoe on behalf of the Wellcome Sanger Institute COVID-19 Surveillance Team |
| EPI_ISL_2866640, EPI_ISL_2866641 | Lighthouse Laboratory Plymouth | Wellcome Sanger Institute for the COVID-19 Genomics UK (COG-UK) Consortium | Lighthouse Laboratory Plymouth and Alex Alderton, Roberto Amato, Jeffrey Barrett, Sonia Goncalves, Ewan Harrison, David K. Jackson, Ian Johnston, Dominic Kwiatkowski, Cordelia Langford, John Sillitoe on behalf of the Wellcome Sanger Institute COVID-19 Surveillance Team |
| EPI_ISL_2866642, EPI_ISL_2866643, EPI_ISL_2866644 | Lighthouse Lab in Milton Keynes | Wellcome Sanger Institute for the COVID-19 Genomics UK (COG-UK) Consortium | The Lighthouse Lab in Milton Keynes and Alex Alderton, Roberto Amato, Jeffrey Barrett, Sonia Goncalves, Ewan Harrison, David K. Jackson, Ian Johnston, Dominic Kwiatkowski, Cordelia Langford, John Sillitoe on behalf of the Wellcome Sanger Institute COVID-19 Surveillance Team |
| EPI_ISL_2866645 | Lighthouse Laboratory Plymouth | Wellcome Sanger Institute for the COVID-19 Genomics UK (COG-UK) Consortium | Lighthouse Laboratory Plymouth and Alex Alderton, Roberto Amato, Jeffrey Barrett, Sonia Goncalves, Ewan Harrison, David K. Jackson, Ian Johnston, Dominic Kwiatkowski, Cordelia Langford, John Sillitoe on behalf of the Wellcome Sanger Institute COVID-19 Surveillance Team |
| EPI_ISL_2866646 | Lighthouse Lab in Milton Keynes | Wellcome Sanger Institute for the COVID-19 Genomics UK (COG-UK) Consortium | The Lighthouse Lab in Milton Keynes and Alex Alderton, Roberto Amato, Jeffrey Barrett, Sonia Goncalves, Ewan Harrison, David K. Jackson, Ian Johnston, Dominic Kwiatkowski, Cordelia Langford, John Sillitoe on behalf of the Wellcome Sanger Institute COVID-19 Surveillance Team |
| EPI_ISL_2866647 | Lighthouse Laboratory Plymouth | Wellcome Sanger Institute for the COVID-19 Genomics UK (COG-UK) Consortium | Lighthouse Laboratory Plymouth and Alex Alderton, Roberto Amato, Jeffrey Barrett, Sonia Goncalves, Ewan Harrison, David K. Jackson, Ian Johnston, Dominic Kwiatkowski, Cordelia Langford, John Sillitoe on behalf of the Wellcome Sanger Institute COVID-19 Surveillance Team |
| EPI_ISL_2866648, EPI_ISL_2866649 | Lighthouse Lab in Milton Keynes | Wellcome Sanger Institute for the COVID-19 Genomics UK (COG-UK) Consortium | The Lighthouse Lab in Milton Keynes and Alex Alderton, Roberto Amato, Jeffrey Barrett, Sonia Goncalves, Ewan Harrison, David K. Jackson, Ian Johnston, Dominic Kwiatkowski, Cordelia Langford, John Sillitoe on behalf of the Wellcome Sanger Institute COVID-19 Surveillance Team |
| EPI_ISL_2866650, EPI_ISL_2866651 | Lighthouse Laboratory Plymouth | Wellcome Sanger Institute for the COVID-19 Genomics UK (COG-UK) Consortium | Lighthouse Laboratory Plymouth and Alex Alderton, Roberto Amato, Jeffrey Barrett, Sonia Goncalves, Ewan Harrison, David K. Jackson, Ian Johnston, Dominic Kwiatkowski, Cordelia Langford, John Sillitoe on behalf of the Wellcome Sanger Institute COVID-19 Surveillance Team |
| EPI_ISL_2866652 | Lighthouse Lab in Milton Keynes | Wellcome Sanger Institute for the COVID-19 Genomics UK (COG-UK) Consortium | The Lighthouse Lab in Milton Keynes and Alex Alderton, Roberto Amato, Jeffrey Barrett, Sonia Goncalves, Ewan Harrison, David K. Jackson, Ian Johnston, Dominic Kwiatkowski, Cordelia Langford, John Sillitoe on behalf of the Wellcome Sanger Institute COVID-19 Surveillance Team |
| EPI_ISL_2866653 | Lighthouse Laboratory Plymouth | Wellcome Sanger Institute for the COVID-19 Genomics UK (COG-UK) Consortium | Lighthouse Laboratory Plymouth and Alex Alderton, Roberto Amato, Jeffrey Barrett, Sonia Goncalves, Ewan Harrison, David K. Jackson, Ian Johnston, Dominic Kwiatkowski, Cordelia Langford, John Sillitoe on behalf of the Wellcome Sanger Institute COVID-19 Surveillance Team |
| EPI_ISL_2866654, EPI_ISL_2866655 | Lighthouse Lab in Milton Keynes | Wellcome Sanger Institute for the COVID-19 Genomics UK (COG-UK) Consortium | The Lighthouse Lab in Milton Keynes and Alex Alderton, Roberto Amato, Jeffrey Barrett, Sonia Goncalves, Ewan Harrison, David K. Jackson, Ian Johnston, Dominic Kwiatkowski, Cordelia Langford, John Sillitoe on behalf of the Wellcome Sanger Institute COVID-19 Surveillance Team |
| EPI_ISL_2866656, EPI_ISL_2866657, EPI_ISL_2866658, EPI_ISL_2866659 | Lighthouse Laboratory Plymouth | Wellcome Sanger Institute for the COVID-19 Genomics UK (COG-UK) Consortium | Lighthouse Laboratory Plymouth and Alex Alderton, Roberto Amato, Jeffrey Barrett, Sonia Goncalves, Ewan Harrison, David K. Jackson, Ian Johnston, Dominic Kwiatkowski, Cordelia Langford, John Sillitoe on behalf of the Wellcome Sanger Institute COVID-19 Surveillance Team |
| EPI_ISL_2866660 | Lighthouse Lab in Milton Keynes | Wellcome Sanger Institute for the COVID-19 Genomics UK (COG-UK) Consortium | The Lighthouse Lab in Milton Keynes and Alex Alderton, Roberto Amato, Jeffrey Barrett, Sonia Goncalves, Ewan Harrison, David K. Jackson, Ian Johnston, Dominic Kwiatkowski, Cordelia Langford, John Sillitoe on behalf of the Wellcome Sanger Institute COVID-19 Surveillance Team |
| EPI_ISL_2866661 | Lighthouse Laboratory Plymouth | Wellcome Sanger Institute for the COVID-19 Genomics UK (COG-UK) Consortium | Lighthouse Laboratory Plymouth and Alex Alderton, Roberto Amato, Jeffrey Barrett, Sonia Goncalves, Ewan Harrison, David K. Jackson, Ian Johnston, Dominic Kwiatkowski, Cordelia Langford, John Sillitoe on behalf of the Wellcome Sanger Institute COVID-19 Surveillance Team |
| EPI_ISL_2866662 | Lighthouse Lab in Milton Keynes | Wellcome Sanger Institute for the COVID-19 Genomics UK (COG-UK) Consortium | The Lighthouse Lab in Milton Keynes and Alex Alderton, Roberto Amato, Jeffrey Barrett, Sonia Goncalves, Ewan Harrison, David K. Jackson, Ian Johnston, Dominic Kwiatkowski, Cordelia Langford, John Sillitoe on behalf of the Wellcome Sanger Institute COVID-19 Surveillance Team |
| EPI_ISL_2866663 | Lighthouse Laboratory Plymouth | Wellcome Sanger Institute for the COVID-19 Genomics UK (COG-UK) Consortium | Lighthouse Laboratory Plymouth and Alex Alderton, Roberto Amato, Jeffrey Barrett, Sonia Goncalves, Ewan Harrison, David K. Jackson, Ian Johnston, Dominic Kwiatkowski, Cordelia Langford, John Sillitoe on behalf of the Wellcome Sanger Institute COVID-19 Surveillance Team |
| EPI_ISL_2866664 | Lighthouse Lab in Milton Keynes | Wellcome Sanger Institute for the COVID-19 Genomics UK (COG-UK) Consortium | The Lighthouse Lab in Milton Keynes and Alex Alderton, Roberto Amato, Jeffrey Barrett, Sonia Goncalves, Ewan Harrison, David K. Jackson, Ian Johnston, Dominic Kwiatkowski, Cordelia Langford, John Sillitoe on behalf of the Wellcome Sanger Institute COVID-19 Surveillance Team |
| EPI_ISL_2866665, EPI_ISL_2866666, EPI_ISL_2866667, EPI_ISL_2866668, EPI_ISL_2866669 | Lighthouse Laboratory Plymouth | Wellcome Sanger Institute for the COVID-19 Genomics UK (COG-UK) Consortium | Lighthouse Laboratory Plymouth and Alex Alderton, Roberto Amato, Jeffrey Barrett, Sonia Goncalves, Ewan Harrison, David K. Jackson, Ian Johnston, Dominic Kwiatkowski, Cordelia Langford, John Sillitoe on behalf of the Wellcome Sanger Institute COVID-19 Surveillance Team |
| EPI_ISL_2866670 | Lighthouse Lab in Milton Keynes | Wellcome Sanger Institute for the COVID-19 Genomics UK (COG-UK) Consortium | The Lighthouse Lab in Milton Keynes and Alex Alderton, Roberto Amato, Jeffrey Barrett, Sonia Goncalves, Ewan Harrison, David K. Jackson, Ian Johnston, Dominic Kwiatkowski, Cordelia Langford, John Sillitoe on behalf of the Wellcome Sanger Institute COVID-19 Surveillance Team |
| EPI_ISL_2866671, EPI_ISL_2866672, EPI_ISL_2866673 | Lighthouse Laboratory Plymouth | Wellcome Sanger Institute for the COVID-19 Genomics UK (COG-UK) Consortium | Lighthouse Laboratory Plymouth and Alex Alderton, Roberto Amato, Jeffrey Barrett, Sonia Goncalves, Ewan Harrison, David K. Jackson, Ian Johnston, Dominic Kwiatkowski, Cordelia Langford, John Sillitoe on behalf of the Wellcome Sanger Institute COVID-19 Surveillance Team |
| EPI_ISL_2866674 | Lighthouse Lab in Glasgow | Wellcome Sanger Institute for the COVID-19 Genomics UK (COG-UK) Consortium | Harper VanSteenhouse, Yumi Kasai, David Gray, Carol Clugston, Anna Dominiczak and Alex Alderton, Roberto Amato, Jeffrey Barrett, Sonia Goncalves, Ewan Harrison, David K. Jackson, Ian Johnston, Dominic Kwiatkowski, Cordelia Langford, John Sillitoe on behalf of the Wellcome Sanger Institute COVID-19 Surveillance Team |
| EPI_ISL_2866675 | Lighthouse Lab in Milton Keynes | Wellcome Sanger Institute for the COVID-19 Genomics UK (COG-UK) Consortium | The Lighthouse Lab in Milton Keynes and Alex Alderton, Roberto Amato, Jeffrey Barrett, Sonia Goncalves, Ewan Harrison, David K. Jackson, Ian Johnston, Dominic Kwiatkowski, Cordelia Langford, John Sillitoe on behalf of the Wellcome Sanger Institute COVID-19 Surveillance Team |
| EPI_ISL_2866676 | Lighthouse Laboratory Plymouth | Wellcome Sanger Institute for the COVID-19 Genomics UK (COG-UK) Consortium | Lighthouse Laboratory Plymouth and Alex Alderton, Roberto Amato, Jeffrey Barrett, Sonia Goncalves, Ewan Harrison, David K. Jackson, Ian Johnston, Dominic Kwiatkowski, Cordelia Langford, John Sillitoe on behalf of the Wellcome Sanger Institute COVID-19 Surveillance Team |
| EPI_ISL_2866677, EPI_ISL_2866678 | Lighthouse Lab in Glasgow | Wellcome Sanger Institute for the COVID-19 Genomics UK (COG-UK) Consortium | Harper VanSteenhouse, Yumi Kasai, David Gray, Carol Clugston, Anna Dominiczak and Alex Alderton, Roberto Amato, Jeffrey Barrett, Sonia Goncalves, Ewan Harrison, David K. Jackson, Ian Johnston, Dominic Kwiatkowski, Cordelia Langford, John Sillitoe on behalf of the Wellcome Sanger Institute COVID-19 Surveillance Team |
| EPI_ISL_2866679 | Lighthouse Laboratory Plymouth | Wellcome Sanger Institute for the COVID-19 Genomics UK (COG-UK) Consortium | Lighthouse Laboratory Plymouth and Alex Alderton, Roberto Amato, Jeffrey Barrett, Sonia Goncalves, Ewan Harrison, David K. Jackson, Ian Johnston, Dominic Kwiatkowski, Cordelia Langford, John Sillitoe on behalf of the Wellcome Sanger Institute COVID-19 Surveillance Team |
| EPI_ISL_2866680 | Lighthouse Lab in Glasgow | Wellcome Sanger Institute for the COVID-19 Genomics UK (COG-UK) Consortium | Harper VanSteenhouse, Yumi Kasai, David Gray, Carol Clugston, Anna Dominiczak and Alex Alderton, Roberto Amato, Jeffrey Barrett, Sonia Goncalves, Ewan Harrison, David K. Jackson, Ian Johnston, Dominic Kwiatkowski, Cordelia Langford, John Sillitoe on behalf of the Wellcome Sanger Institute COVID-19 Surveillance Team |
| EPI_ISL_2866681 | Lighthouse Laboratory Plymouth | Wellcome Sanger Institute for the COVID-19 Genomics UK (COG-UK) Consortium | Lighthouse Laboratory Plymouth and Alex Alderton, Roberto Amato, Jeffrey Barrett, Sonia Goncalves, Ewan Harrison, David K. Jackson, Ian Johnston, Dominic Kwiatkowski, Cordelia Langford, John Sillitoe on behalf of the Wellcome Sanger Institute COVID-19 Surveillance Team |
| EPI_ISL_2866682 | Lighthouse Lab in Glasgow | Wellcome Sanger Institute for the COVID-19 Genomics UK (COG-UK) Consortium | Harper VanSteenhouse, Yumi Kasai, David Gray, Carol Clugston, Anna Dominiczak and Alex Alderton, Roberto Amato, Jeffrey Barrett, Sonia Goncalves, Ewan Harrison, David K. Jackson, Ian Johnston, Dominic Kwiatkowski, Cordelia Langford, John Sillitoe on behalf of the Wellcome Sanger Institute COVID-19 Surveillance Team |
| EPI_ISL_2866683, EPI_ISL_2866684 | Lighthouse Lab in Milton Keynes | Wellcome Sanger Institute for the COVID-19 Genomics UK (COG-UK) Consortium | The Lighthouse Lab in Milton Keynes and Alex Alderton, Roberto Amato, Jeffrey Barrett, Sonia Goncalves, Ewan Harrison, David K. Jackson, Ian Johnston, Dominic Kwiatkowski, Cordelia Langford, John Sillitoe on behalf of the Wellcome Sanger Institute COVID-19 Surveillance Team |
| EPI_ISL_2866685, EPI_ISL_2866686 | Lighthouse Lab in Glasgow | Wellcome Sanger Institute for the COVID-19 Genomics UK (COG-UK) Consortium | Harper VanSteenhouse, Yumi Kasai, David Gray, Carol Clugston, Anna Dominiczak and Alex Alderton, Roberto Amato, Jeffrey Barrett, Sonia Goncalves, Ewan Harrison, David K. Jackson, Ian Johnston, Dominic Kwiatkowski, Cordelia Langford, John Sillitoe on behalf of the Wellcome Sanger Institute COVID-19 Surveillance Team |
| EPI_ISL_2866687, EPI_ISL_2866688 | Lighthouse Laboratory Plymouth | Wellcome Sanger Institute for the COVID-19 Genomics UK (COG-UK) Consortium | Lighthouse Laboratory Plymouth and Alex Alderton, Roberto Amato, Jeffrey Barrett, Sonia Goncalves, Ewan Harrison, David K. Jackson, Ian Johnston, Dominic Kwiatkowski, Cordelia Langford, John Sillitoe on behalf of the Wellcome Sanger Institute COVID-19 Surveillance Team |
| EPI_ISL_2866689, EPI_ISL_2866690, EPI_ISL_2866691, | Lighthouse Lab in Glasgow | Wellcome Sanger Institute for the COVID-19 Genomics UK (COG-UK) Consortium | Harper VanSteenhouse, Yumi Kasai, David Gray, Carol Clugston, Anna Dominiczak and Alex Alderton, Roberto Amato, Jeffrey Barrett, Sonia Goncalves, Ewan Harrison, David K. Jackson, Ian Johnston, Dominic Kwiatkowski, Cordelia Langford, John Sillitoe on behalf of the Wellcome Sanger Institute COVID-19 Surveillance Team |

| | | | |
|---|---|---|---|
| EPI_ISL_2866692, EPI_ISL_2866693 | | | |
| EPI_ISL_2866694, EPI_ISL_2866695, EPI_ISL_2866696 | Lighthouse Laboratory Plymouth | Wellcome Sanger Institute for the COVID-19 Genomics UK (COG-UK) Consortium | Lighthouse Laboratory Plymouth and Alex Alderton, Roberto Amato, Jeffrey Barrett, Sonia Goncalves, Ewan Harrison, David K. Jackson, Ian Johnston, Dominic Kwiatkowski, Cordelia Langford, John Sillitoe on behalf of the Wellcome Sanger Institute COVID-19 Surveillance Team |
| EPI_ISL_2866697, EPI_ISL_2866698, EPI_ISL_2866699, EPI_ISL_2866700 | Lighthouse Lab in Glasgow | Wellcome Sanger Institute for the COVID-19 Genomics UK (COG-UK) Consortium | Harper VanSteenhouse, Yumi Kasai, David Gray, Carol Clugston, Anna Dominiczak and Alex Alderton, Roberto Amato, Jeffrey Barrett, Sonia Goncalves, Ewan Harrison, David K. Jackson, Ian Johnston, Dominic Kwiatkowski, Cordelia Langford, John Sillitoe on behalf of the Wellcome Sanger Institute COVID-19 Surveillance Team |
| EPI_ISL_2866701 | Lighthouse Lab in Milton Keynes | Wellcome Sanger Institute for the COVID-19 Genomics UK (COG-UK) Consortium | The Lighthouse Lab in Milton Keynes and Alex Alderton, Roberto Amato, Jeffrey Barrett, Sonia Goncalves, Ewan Harrison, David K. Jackson, Ian Johnston, Dominic Kwiatkowski, Cordelia Langford, John Sillitoe on behalf of the Wellcome Sanger Institute COVID-19 Surveillance Team |
| EPI_ISL_2866702, EPI_ISL_2866703 | Lighthouse Lab in Glasgow | Wellcome Sanger Institute for the COVID-19 Genomics UK (COG-UK) Consortium | Harper VanSteenhouse, Yumi Kasai, David Gray, Carol Clugston, Anna Dominiczak and Alex Alderton, Roberto Amato, Jeffrey Barrett, Sonia Goncalves, Ewan Harrison, David K. Jackson, Ian Johnston, Dominic Kwiatkowski, Cordelia Langford, John Sillitoe on behalf of the Wellcome Sanger Institute COVID-19 Surveillance Team |
| EPI_ISL_2866704 | Lighthouse Laboratory Plymouth | Wellcome Sanger Institute for the COVID-19 Genomics UK (COG-UK) Consortium | Lighthouse Laboratory Plymouth and Alex Alderton, Roberto Amato, Jeffrey Barrett, Sonia Goncalves, Ewan Harrison, David K. Jackson, Ian Johnston, Dominic Kwiatkowski, Cordelia Langford, John Sillitoe on behalf of the Wellcome Sanger Institute COVID-19 Surveillance Team |
| EPI_ISL_2866705 | Lighthouse Lab in Glasgow | Wellcome Sanger Institute for the COVID-19 Genomics UK (COG-UK) Consortium | Harper VanSteenhouse, Yumi Kasai, David Gray, Carol Clugston, Anna Dominiczak and Alex Alderton, Roberto Amato, Jeffrey Barrett, Sonia Goncalves, Ewan Harrison, David K. Jackson, Ian Johnston, Dominic Kwiatkowski, Cordelia Langford, John Sillitoe on behalf of the Wellcome Sanger Institute COVID-19 Surveillance Team |
| EPI_ISL_2866706 | Lighthouse Lab in Milton Keynes | Wellcome Sanger Institute for the COVID-19 Genomics UK (COG-UK) Consortium | The Lighthouse Lab in Milton Keynes and Alex Alderton, Roberto Amato, Jeffrey Barrett, Sonia Goncalves, Ewan Harrison, David K. Jackson, Ian Johnston, Dominic Kwiatkowski, Cordelia Langford, John Sillitoe on behalf of the Wellcome Sanger Institute COVID-19 Surveillance Team |
| EPI_ISL_2866708 | Lighthouse Lab in Glasgow | Wellcome Sanger Institute for the COVID-19 Genomics UK (COG-UK) Consortium | Harper VanSteenhouse, Yumi Kasai, David Gray, Carol Clugston, Anna Dominiczak and Alex Alderton, Roberto Amato, Jeffrey Barrett, Sonia Goncalves, Ewan Harrison, David K. Jackson, Ian Johnston, Dominic Kwiatkowski, Cordelia Langford, John Sillitoe on behalf of the Wellcome Sanger Institute COVID-19 Surveillance Team |
| EPI_ISL_2866709, EPI_ISL_2866710, EPI_ISL_2866711 | Lighthouse Laboratory Plymouth | Wellcome Sanger Institute for the COVID-19 Genomics UK (COG-UK) Consortium | Lighthouse Laboratory Plymouth and Alex Alderton, Roberto Amato, Jeffrey Barrett, Sonia Goncalves, Ewan Harrison, David K. Jackson, Ian Johnston, Dominic Kwiatkowski, Cordelia Langford, John Sillitoe on behalf of the Wellcome Sanger Institute COVID-19 Surveillance Team |
| EPI_ISL_2866712, EPI_ISL_2866713, EPI_ISL_2866714 | Lighthouse Lab in Glasgow | Wellcome Sanger Institute for the COVID-19 Genomics UK (COG-UK) Consortium | Harper VanSteenhouse, Yumi Kasai, David Gray, Carol Clugston, Anna Dominiczak and Alex Alderton, Roberto Amato, Jeffrey Barrett, Sonia Goncalves, Ewan Harrison, David K. Jackson, Ian Johnston, Dominic Kwiatkowski, Cordelia Langford, John Sillitoe on behalf of the Wellcome Sanger Institute COVID-19 Surveillance Team |
| EPI_ISL_2866715 | Lighthouse Laboratory Plymouth | Wellcome Sanger Institute for the COVID-19 Genomics UK (COG-UK) Consortium | Lighthouse Laboratory Plymouth and Alex Alderton, Roberto Amato, Jeffrey Barrett, Sonia Goncalves, Ewan Harrison, David K. Jackson, Ian Johnston, Dominic Kwiatkowski, Cordelia Langford, John Sillitoe on behalf of the Wellcome Sanger Institute COVID-19 Surveillance Team |
| EPI_ISL_2866716, EPI_ISL_2866717, EPI_ISL_2866718 | Lighthouse Lab in Glasgow | Wellcome Sanger Institute for the COVID-19 Genomics UK (COG-UK) Consortium | Harper VanSteenhouse, Yumi Kasai, David Gray, Carol Clugston, Anna Dominiczak and Alex Alderton, Roberto Amato, Jeffrey Barrett, Sonia Goncalves, Ewan Harrison, David K. Jackson, Ian Johnston, Dominic Kwiatkowski, Cordelia Langford, John Sillitoe on behalf of the Wellcome Sanger Institute COVID-19 Surveillance Team |
| EPI_ISL_2866719, EPI_ISL_2866720 | Lighthouse Laboratory Plymouth | Wellcome Sanger Institute for the COVID-19 Genomics UK (COG-UK) Consortium | Lighthouse Laboratory Plymouth and Alex Alderton, Roberto Amato, Jeffrey Barrett, Sonia Goncalves, Ewan Harrison, David K. Jackson, Ian Johnston, Dominic Kwiatkowski, Cordelia Langford, John Sillitoe on behalf of the Wellcome Sanger Institute COVID-19 Surveillance Team |
| EPI_ISL_2866721, EPI_ISL_2866722 | Lighthouse Lab in Glasgow | Wellcome Sanger Institute for the COVID-19 Genomics UK (COG-UK) Consortium | Harper VanSteenhouse, Yumi Kasai, David Gray, Carol Clugston, Anna Dominiczak and Alex Alderton, Roberto Amato, Jeffrey Barrett, Sonia Goncalves, Ewan Harrison, David K. Jackson, Ian Johnston, Dominic Kwiatkowski, Cordelia Langford, John Sillitoe on behalf of the Wellcome Sanger Institute COVID-19 Surveillance Team |
| EPI_ISL_2866723, EPI_ISL_2866724 | Lighthouse Lab in Milton Keynes | Wellcome Sanger Institute for the COVID-19 Genomics UK (COG-UK) Consortium | The Lighthouse Lab in Milton Keynes and Alex Alderton, Roberto Amato, Jeffrey Barrett, Sonia Goncalves, Ewan Harrison, David K. Jackson, Ian Johnston, Dominic Kwiatkowski, Cordelia Langford, John Sillitoe on behalf of the Wellcome Sanger Institute COVID-19 Surveillance Team |
| EPI_ISL_2866725 | Lighthouse Laboratory Plymouth | Wellcome Sanger Institute for the COVID-19 Genomics UK (COG-UK) Consortium | Lighthouse Laboratory Plymouth and Alex Alderton, Roberto Amato, Jeffrey Barrett, Sonia Goncalves, Ewan Harrison, David K. Jackson, Ian Johnston, Dominic Kwiatkowski, Cordelia Langford, John Sillitoe on behalf of the Wellcome Sanger Institute COVID-19 Surveillance Team |
| EPI_ISL_2866726 | Lighthouse Lab in Glasgow | Wellcome Sanger Institute for the COVID-19 Genomics UK (COG-UK) Consortium | Harper VanSteenhouse, Yumi Kasai, David Gray, Carol Clugston, Anna Dominiczak and Alex Alderton, Roberto Amato, Jeffrey Barrett, Sonia Goncalves, Ewan Harrison, David K. Jackson, Ian Johnston, Dominic Kwiatkowski, Cordelia Langford, John Sillitoe on behalf of the Wellcome Sanger Institute COVID-19 Surveillance Team |
| EPI_ISL_2866727 | Lighthouse Laboratory Plymouth | Wellcome Sanger Institute for the COVID-19 Genomics UK (COG-UK) Consortium | Lighthouse Laboratory Plymouth and Alex Alderton, Roberto Amato, Jeffrey Barrett, Sonia Goncalves, Ewan Harrison, David K. Jackson, Ian Johnston, Dominic Kwiatkowski, Cordelia Langford, John Sillitoe on behalf of the Wellcome Sanger Institute COVID-19 Surveillance Team |
| EPI_ISL_2866728 | Lighthouse Lab in Milton Keynes | Wellcome Sanger Institute for the COVID-19 Genomics UK (COG-UK) Consortium | The Lighthouse Lab in Milton Keynes and Alex Alderton, Roberto Amato, Jeffrey Barrett, Sonia Goncalves, Ewan Harrison, David K. Jackson, Ian Johnston, Dominic Kwiatkowski, Cordelia Langford, John Sillitoe on behalf of the Wellcome Sanger Institute COVID-19 Surveillance Team |
| EPI_ISL_2866729, EPI_ISL_2866730 | Lighthouse Lab in Glasgow | Wellcome Sanger Institute for the COVID-19 Genomics UK (COG-UK) Consortium | Harper VanSteenhouse, Yumi Kasai, David Gray, Carol Clugston, Anna Dominiczak and Alex Alderton, Roberto Amato, Jeffrey Barrett, Sonia Goncalves, Ewan Harrison, David K. Jackson, Ian Johnston, Dominic Kwiatkowski, Cordelia Langford, John Sillitoe on behalf of the Wellcome Sanger Institute COVID-19 Surveillance Team |
| EPI_ISL_2866731 | Lighthouse Lab in Milton Keynes | Wellcome Sanger Institute for the COVID-19 Genomics UK (COG-UK) Consortium | The Lighthouse Lab in Milton Keynes and Alex Alderton, Roberto Amato, Jeffrey Barrett, Sonia Goncalves, Ewan Harrison, David K. Jackson, Ian Johnston, Dominic Kwiatkowski, Cordelia Langford, John Sillitoe on behalf of the Wellcome Sanger Institute COVID-19 Surveillance Team |
| EPI_ISL_2866732 | Lighthouse Lab in Glasgow | Wellcome Sanger Institute for the COVID-19 Genomics UK (COG-UK) Consortium | Harper VanSteenhouse, Yumi Kasai, David Gray, Carol Clugston, Anna Dominiczak and Alex Alderton, Roberto Amato, Jeffrey Barrett, Sonia Goncalves, Ewan Harrison, David K. Jackson, Ian Johnston, Dominic Kwiatkowski, Cordelia Langford, John Sillitoe on behalf of the Wellcome Sanger Institute COVID-19 Surveillance Team |
| EPI_ISL_2866733 | Lighthouse Lab in Milton Keynes | Wellcome Sanger Institute for the COVID-19 Genomics UK (COG-UK) Consortium | The Lighthouse Lab in Milton Keynes and Alex Alderton, Roberto Amato, Jeffrey Barrett, Sonia Goncalves, Ewan Harrison, David K. Jackson, Ian Johnston, Dominic Kwiatkowski, Cordelia Langford, John Sillitoe on behalf of the Wellcome Sanger Institute COVID-19 Surveillance Team |
| EPI_ISL_2866734, EPI_ISL_2866735 | Lighthouse Lab in Glasgow | Wellcome Sanger Institute for the COVID-19 Genomics UK (COG-UK) Consortium | Harper VanSteenhouse, Yumi Kasai, David Gray, Carol Clugston, Anna Dominiczak and Alex Alderton, Roberto Amato, Jeffrey Barrett, Sonia Goncalves, Ewan Harrison, David K. Jackson, Ian Johnston, Dominic Kwiatkowski, Cordelia Langford, John Sillitoe on behalf of the Wellcome Sanger Institute COVID-19 Surveillance Team |
| EPI_ISL_2866736 | Lighthouse Lab in Milton Keynes | Wellcome Sanger Institute for the COVID-19 Genomics UK (COG-UK) Consortium | The Lighthouse Lab in Milton Keynes and Alex Alderton, Roberto Amato, Jeffrey Barrett, Sonia Goncalves, Ewan Harrison, David K. Jackson, Ian Johnston, Dominic Kwiatkowski, Cordelia Langford, John Sillitoe on behalf of the Wellcome Sanger Institute COVID-19 Surveillance Team |
| EPI_ISL_2866737, EPI_ISL_2866738, EPI_ISL_2866739 | Lighthouse Lab in Glasgow | Wellcome Sanger Institute for the COVID-19 Genomics UK (COG-UK) Consortium | Harper VanSteenhouse, Yumi Kasai, David Gray, Carol Clugston, Anna Dominiczak and Alex Alderton, Roberto Amato, Jeffrey Barrett, Sonia Goncalves, Ewan Harrison, David K. Jackson, Ian Johnston, Dominic Kwiatkowski, Cordelia Langford, John Sillitoe on behalf of the Wellcome Sanger Institute COVID-19 Surveillance Team |
| EPI_ISL_2866740 | Lighthouse Laboratory Plymouth | Wellcome Sanger Institute for the COVID-19 Genomics UK (COG-UK) Consortium | Lighthouse Laboratory Plymouth and Alex Alderton, Roberto Amato, Jeffrey Barrett, Sonia Goncalves, Ewan Harrison, David K. Jackson, Ian Johnston, Dominic Kwiatkowski, Cordelia Langford, John Sillitoe on behalf of the Wellcome Sanger Institute COVID-19 Surveillance Team |
| EPI_ISL_2866741, EPI_ISL_2866742 | Lighthouse Lab in Glasgow | Wellcome Sanger Institute for the COVID-19 Genomics UK (COG-UK) Consortium | Harper VanSteenhouse, Yumi Kasai, David Gray, Carol Clugston, Anna Dominiczak and Alex Alderton, Roberto Amato, Jeffrey Barrett, Sonia Goncalves, Ewan Harrison, David K. Jackson, Ian Johnston, Dominic Kwiatkowski, Cordelia Langford, John Sillitoe on behalf of the Wellcome Sanger Institute COVID-19 Surveillance Team |
| EPI_ISL_2866743 | Lighthouse Lab in Milton Keynes | Wellcome Sanger Institute for the COVID-19 Genomics UK (COG-UK) Consortium | The Lighthouse Lab in Milton Keynes and Alex Alderton, Roberto Amato, Jeffrey Barrett, Sonia Goncalves, Ewan Harrison, David K. Jackson, Ian Johnston, Dominic Kwiatkowski, Cordelia Langford, John Sillitoe on behalf of the Wellcome Sanger Institute COVID-19 Surveillance Team |
| EPI_ISL_2866744, EPI_ISL_2866745, EPI_ISL_2866746 | Lighthouse Lab in Glasgow | Wellcome Sanger Institute for the COVID-19 Genomics UK (COG-UK) Consortium | Harper VanSteenhouse, Yumi Kasai, David Gray, Carol Clugston, Anna Dominiczak and Alex Alderton, Roberto Amato, Jeffrey Barrett, Sonia Goncalves, Ewan Harrison, David K. Jackson, Ian Johnston, Dominic Kwiatkowski, Cordelia Langford, John Sillitoe on behalf of the Wellcome Sanger Institute COVID-19 Surveillance Team |
| EPI_ISL_2866747, EPI_ISL_2866748 | Lighthouse Lab in Milton Keynes | Wellcome Sanger Institute for the COVID-19 Genomics UK (COG-UK) Consortium | The Lighthouse Lab in Milton Keynes and Alex Alderton, Roberto Amato, Jeffrey Barrett, Sonia Goncalves, Ewan Harrison, David K. Jackson, Ian Johnston, Dominic Kwiatkowski, Cordelia Langford, John Sillitoe on behalf of the Wellcome Sanger Institute COVID-19 Surveillance Team |
| EPI_ISL_2866749 | Lighthouse Lab in Glasgow | Wellcome Sanger Institute for the COVID-19 Genomics UK (COG-UK) Consortium | Harper VanSteenhouse, Yumi Kasai, David Gray, Carol Clugston, Anna Dominiczak and Alex Alderton, Roberto Amato, Jeffrey Barrett, Sonia Goncalves, Ewan Harrison, David K. Jackson, Ian Johnston, Dominic Kwiatkowski, Cordelia Langford, John Sillitoe on behalf of the Wellcome Sanger Institute COVID-19 Surveillance Team |
| EPI_ISL_2866750 | Lighthouse Lab in Milton Keynes | Wellcome Sanger Institute for the COVID-19 Genomics UK (COG-UK) Consortium | The Lighthouse Lab in Milton Keynes and Alex Alderton, Roberto Amato, Jeffrey Barrett, Sonia Goncalves, Ewan Harrison, David K. Jackson, Ian Johnston, Dominic Kwiatkowski, Cordelia Langford, John Sillitoe on behalf of the Wellcome Sanger Institute COVID-19 Surveillance Team |
| EPI_ISL_2866751, EPI_ISL_2866752, EPI_ISL_2866753 | Lighthouse Lab in Glasgow | Wellcome Sanger Institute for the COVID-19 Genomics UK (COG-UK) Consortium | Harper VanSteenhouse, Yumi Kasai, David Gray, Carol Clugston, Anna Dominiczak and Alex Alderton, Roberto Amato, Jeffrey Barrett, Sonia Goncalves, Ewan Harrison, David K. Jackson, Ian Johnston, Dominic Kwiatkowski, Cordelia Langford, John Sillitoe on behalf of the Wellcome Sanger Institute COVID-19 Surveillance Team |
| EPI_ISL_2866754, EPI_ISL_2866755 | Lighthouse Lab in Milton Keynes | Wellcome Sanger Institute for the COVID-19 Genomics UK (COG-UK) Consortium | The Lighthouse Lab in Milton Keynes and Alex Alderton, Roberto Amato, Jeffrey Barrett, Sonia Goncalves, Ewan Harrison, David K. Jackson, Ian Johnston, Dominic Kwiatkowski, Cordelia Langford, John Sillitoe on behalf of the Wellcome Sanger Institute COVID-19 Surveillance Team |
| EPI_ISL_2866756, EPI_ISL_2866757 | Lighthouse Lab in Glasgow | Wellcome Sanger Institute for the COVID-19 Genomics UK (COG-UK) Consortium | Harper VanSteenhouse, Yumi Kasai, David Gray, Carol Clugston, Anna Dominiczak and Alex Alderton, Roberto Amato, Jeffrey Barrett, Sonia Goncalves, Ewan Harrison, David K. Jackson, Ian Johnston, Dominic Kwiatkowski, Cordelia Langford, John Sillitoe on behalf of the Wellcome Sanger Institute COVID-19 Surveillance Team |
| EPI_ISL_2866758 | Lighthouse Lab in Milton Keynes | Wellcome Sanger Institute for the COVID-19 Genomics UK (COG-UK) Consortium | The Lighthouse Lab in Milton Keynes and Alex Alderton, Roberto Amato, Jeffrey Barrett, Sonia Goncalves, Ewan Harrison, David K. Jackson, Ian Johnston, Dominic Kwiatkowski, Cordelia Langford, John Sillitoe on behalf of the Wellcome Sanger Institute COVID-19 Surveillance Team |
| EPI_ISL_2866759 | Lighthouse Lab in Glasgow | Wellcome Sanger Institute for the COVID-19 Genomics UK (COG-UK) Consortium | Harper VanSteenhouse, Yumi Kasai, David Gray, Carol Clugston, Anna Dominiczak and Alex Alderton, Roberto Amato, Jeffrey Barrett, Sonia Goncalves, Ewan Harrison, David K. Jackson, Ian Johnston, Dominic Kwiatkowski, Cordelia Langford, John Sillitoe on behalf of the Wellcome Sanger Institute COVID-19 Surveillance Team |
| EPI_ISL_2866760 | Lighthouse Laboratory Plymouth | Wellcome Sanger Institute for the COVID-19 Genomics UK (COG-UK) Consortium | Lighthouse Laboratory Plymouth and Alex Alderton, Roberto Amato, Jeffrey Barrett, Sonia Goncalves, Ewan Harrison, David K. Jackson, Ian Johnston, Dominic Kwiatkowski, Cordelia Langford, John Sillitoe on behalf of the Wellcome Sanger Institute COVID-19 Surveillance Team |
| EPI_ISL_2866761, EPI_ISL_2866762, EPI_ISL_2866763, | Lighthouse Lab in Glasgow | Wellcome Sanger Institute for the COVID-19 Genomics UK (COG-UK) Consortium | Harper VanSteenhouse, Yumi Kasai, David Gray, Carol Clugston, Anna Dominiczak and Alex Alderton, Roberto Amato, Jeffrey Barrett, Sonia Goncalves, Ewan Harrison, David K. Jackson, Ian Johnston, Dominic Kwiatkowski, Cordelia Langford, John Sillitoe on behalf of the Wellcome Sanger Institute COVID-19 Surveillance Team |

| | | | |
|---|---|---|---|
| EPI_ISL_2866764 | | | |
| EPI_ISL_2866765 | Lighthouse Laboratory Plymouth | Wellcome Sanger Institute for the COVID-19 Genomics UK (COG-UK) Consortium | Lighthouse Laboratory Plymouth and Alex Alderton, Roberto Amato, Jeffrey Barrett, Sonia Goncalves, Ewan Harrison, David K. Jackson, Ian Johnston, Dominic Kwiatkowski, Cordelia Langford, John Sillitoe on behalf of the Wellcome Sanger Institute COVID-19 Surveillance Team |
| EPI_ISL_2866766, EPI_ISL_2866767 | Lighthouse Lab in Glasgow | Wellcome Sanger Institute for the COVID-19 Genomics UK (COG-UK) Consortium | Harper VanSteenhouse, Yumi Kasai, David Gray, Carol Clugston, Anna Dominiczak and Alex Alderton, Roberto Amato, Jeffrey Barrett, Sonia Goncalves, Ewan Harrison, David K. Jackson, Ian Johnston, Dominic Kwiatkowski, Cordelia Langford, John Sillitoe on behalf of the Wellcome Sanger Institute COVID-19 Surveillance Team |
| EPI_ISL_2866768 | Lighthouse Lab in Milton Keynes | Wellcome Sanger Institute for the COVID-19 Genomics UK (COG-UK) Consortium | The Lighthouse Lab in Milton Keynes and Alex Alderton, Roberto Amato, Jeffrey Barrett, Sonia Goncalves, Ewan Harrison, David K. Jackson, Ian Johnston, Dominic Kwiatkowski, Cordelia Langford, John Sillitoe on behalf of the Wellcome Sanger Institute COVID-19 Surveillance Team |
| EPI_ISL_2866769, EPI_ISL_2866770, EPI_ISL_2866771, EPI_ISL_2866772, EPI_ISL_2866773 | Lighthouse Lab in Glasgow | Wellcome Sanger Institute for the COVID-19 Genomics UK (COG-UK) Consortium | Harper VanSteenhouse, Yumi Kasai, David Gray, Carol Clugston, Anna Dominiczak and Alex Alderton, Roberto Amato, Jeffrey Barrett, Sonia Goncalves, Ewan Harrison, David K. Jackson, Ian Johnston, Dominic Kwiatkowski, Cordelia Langford, John Sillitoe on behalf of the Wellcome Sanger Institute COVID-19 Surveillance Team |
| EPI_ISL_2866774, EPI_ISL_2866775 | Lighthouse Laboratory Plymouth | Wellcome Sanger Institute for the COVID-19 Genomics UK (COG-UK) Consortium | Lighthouse Laboratory Plymouth and Alex Alderton, Roberto Amato, Jeffrey Barrett, Sonia Goncalves, Ewan Harrison, David K. Jackson, Ian Johnston, Dominic Kwiatkowski, Cordelia Langford, John Sillitoe on behalf of the Wellcome Sanger Institute COVID-19 Surveillance Team |
| EPI_ISL_2866776 | Lighthouse Lab in Glasgow | Wellcome Sanger Institute for the COVID-19 Genomics UK (COG-UK) Consortium | Harper VanSteenhouse, Yumi Kasai, David Gray, Carol Clugston, Anna Dominiczak and Alex Alderton, Roberto Amato, Jeffrey Barrett, Sonia Goncalves, Ewan Harrison, David K. Jackson, Ian Johnston, Dominic Kwiatkowski, Cordelia Langford, John Sillitoe on behalf of the Wellcome Sanger Institute COVID-19 Surveillance Team |
| EPI_ISL_2866777 | Lighthouse Laboratory Plymouth | Wellcome Sanger Institute for the COVID-19 Genomics UK (COG-UK) Consortium | Lighthouse Laboratory Plymouth and Alex Alderton, Roberto Amato, Jeffrey Barrett, Sonia Goncalves, Ewan Harrison, David K. Jackson, Ian Johnston, Dominic Kwiatkowski, Cordelia Langford, John Sillitoe on behalf of the Wellcome Sanger Institute COVID-19 Surveillance Team |
| EPI_ISL_2866778 | Lighthouse Lab in Glasgow | Wellcome Sanger Institute for the COVID-19 Genomics UK (COG-UK) Consortium | Harper VanSteenhouse, Yumi Kasai, David Gray, Carol Clugston, Anna Dominiczak and Alex Alderton, Roberto Amato, Jeffrey Barrett, Sonia Goncalves, Ewan Harrison, David K. Jackson, Ian Johnston, Dominic Kwiatkowski, Cordelia Langford, John Sillitoe on behalf of the Wellcome Sanger Institute COVID-19 Surveillance Team |
| EPI_ISL_2866779 | Lighthouse Laboratory Plymouth | Wellcome Sanger Institute for the COVID-19 Genomics UK (COG-UK) Consortium | Lighthouse Laboratory Plymouth and Alex Alderton, Roberto Amato, Jeffrey Barrett, Sonia Goncalves, Ewan Harrison, David K. Jackson, Ian Johnston, Dominic Kwiatkowski, Cordelia Langford, John Sillitoe on behalf of the Wellcome Sanger Institute COVID-19 Surveillance Team |
| EPI_ISL_2866781, EPI_ISL_2866782 | Lighthouse Lab in Glasgow | Wellcome Sanger Institute for the COVID-19 Genomics UK (COG-UK) Consortium | Harper VanSteenhouse, Yumi Kasai, David Gray, Carol Clugston, Anna Dominiczak and Alex Alderton, Roberto Amato, Jeffrey Barrett, Sonia Goncalves, Ewan Harrison, David K. Jackson, Ian Johnston, Dominic Kwiatkowski, Cordelia Langford, John Sillitoe on behalf of the Wellcome Sanger Institute COVID-19 Surveillance Team |
| EPI_ISL_2866783 | Lighthouse Lab in Milton Keynes | Wellcome Sanger Institute for the COVID-19 Genomics UK (COG-UK) Consortium | The Lighthouse Lab in Milton Keynes and Alex Alderton, Roberto Amato, Jeffrey Barrett, Sonia Goncalves, Ewan Harrison, David K. Jackson, Ian Johnston, Dominic Kwiatkowski, Cordelia Langford, John Sillitoe on behalf of the Wellcome Sanger Institute COVID-19 Surveillance Team |
| EPI_ISL_2866784, EPI_ISL_2866785, EPI_ISL_2866786, EPI_ISL_2866787, EPI_ISL_2866788 | Lighthouse Lab in Glasgow | Wellcome Sanger Institute for the COVID-19 Genomics UK (COG-UK) Consortium | Harper VanSteenhouse, Yumi Kasai, David Gray, Carol Clugston, Anna Dominiczak and Alex Alderton, Roberto Amato, Jeffrey Barrett, Sonia Goncalves, Ewan Harrison, David K. Jackson, Ian Johnston, Dominic Kwiatkowski, Cordelia Langford, John Sillitoe on behalf of the Wellcome Sanger Institute COVID-19 Surveillance Team |
| EPI_ISL_2866789 | Lighthouse Lab in Milton Keynes | Wellcome Sanger Institute for the COVID-19 Genomics UK (COG-UK) Consortium | The Lighthouse Lab in Milton Keynes and Alex Alderton, Roberto Amato, Jeffrey Barrett, Sonia Goncalves, Ewan Harrison, David K. Jackson, Ian Johnston, Dominic Kwiatkowski, Cordelia Langford, John Sillitoe on behalf of the Wellcome Sanger Institute COVID-19 Surveillance Team |
| EPI_ISL_2866790 | Lighthouse Laboratory Plymouth | Wellcome Sanger Institute for the COVID-19 Genomics UK (COG-UK) Consortium | Lighthouse Laboratory Plymouth and Alex Alderton, Roberto Amato, Jeffrey Barrett, Sonia Goncalves, Ewan Harrison, David K. Jackson, Ian Johnston, Dominic Kwiatkowski, Cordelia Langford, John Sillitoe on behalf of the Wellcome Sanger Institute COVID-19 Surveillance Team |
| EPI_ISL_2866791, EPI_ISL_2866792 | Lighthouse Lab in Glasgow | Wellcome Sanger Institute for the COVID-19 Genomics UK (COG-UK) Consortium | Harper VanSteenhouse, Yumi Kasai, David Gray, Carol Clugston, Anna Dominiczak and Alex Alderton, Roberto Amato, Jeffrey Barrett, Sonia Goncalves, Ewan Harrison, David K. Jackson, Ian Johnston, Dominic Kwiatkowski, Cordelia Langford, John Sillitoe on behalf of the Wellcome Sanger Institute COVID-19 Surveillance Team |
| EPI_ISL_2866793 | Lighthouse Laboratory Plymouth | Wellcome Sanger Institute for the COVID-19 Genomics UK (COG-UK) Consortium | Lighthouse Laboratory Plymouth and Alex Alderton, Roberto Amato, Jeffrey Barrett, Sonia Goncalves, Ewan Harrison, David K. Jackson, Ian Johnston, Dominic Kwiatkowski, Cordelia Langford, John Sillitoe on behalf of the Wellcome Sanger Institute COVID-19 Surveillance Team |
| EPI_ISL_2866794, EPI_ISL_2866795 | Lighthouse Lab in Glasgow | Wellcome Sanger Institute for the COVID-19 Genomics UK (COG-UK) Consortium | Harper VanSteenhouse, Yumi Kasai, David Gray, Carol Clugston, Anna Dominiczak and Alex Alderton, Roberto Amato, Jeffrey Barrett, Sonia Goncalves, Ewan Harrison, David K. Jackson, Ian Johnston, Dominic Kwiatkowski, Cordelia Langford, John Sillitoe on behalf of the Wellcome Sanger Institute COVID-19 Surveillance Team |
| EPI_ISL_2866796 | Lighthouse Laboratory Plymouth | Wellcome Sanger Institute for the COVID-19 Genomics UK (COG-UK) Consortium | Lighthouse Laboratory Plymouth and Alex Alderton, Roberto Amato, Jeffrey Barrett, Sonia Goncalves, Ewan Harrison, David K. Jackson, Ian Johnston, Dominic Kwiatkowski, Cordelia Langford, John Sillitoe on behalf of the Wellcome Sanger Institute COVID-19 Surveillance Team |
| EPI_ISL_2866797, EPI_ISL_2866798 | Lighthouse Lab in Milton Keynes | Wellcome Sanger Institute for the COVID-19 Genomics UK (COG-UK) Consortium | The Lighthouse Lab in Milton Keynes and Alex Alderton, Roberto Amato, Jeffrey Barrett, Sonia Goncalves, Ewan Harrison, David K. Jackson, Ian Johnston, Dominic Kwiatkowski, Cordelia Langford, John Sillitoe on behalf of the Wellcome Sanger Institute COVID-19 Surveillance Team |
| EPI_ISL_2866800 | Lighthouse Lab in Glasgow | Wellcome Sanger Institute for the COVID-19 Genomics UK (COG-UK) Consortium | Harper VanSteenhouse, Yumi Kasai, David Gray, Carol Clugston, Anna Dominiczak and Alex Alderton, Roberto Amato, Jeffrey Barrett, Sonia Goncalves, Ewan Harrison, David K. Jackson, Ian Johnston, Dominic Kwiatkowski, Cordelia Langford, John Sillitoe on behalf of the Wellcome Sanger Institute COVID-19 Surveillance Team |
| EPI_ISL_2866801 | Lighthouse Lab in Milton Keynes | Wellcome Sanger Institute for the COVID-19 Genomics UK (COG-UK) Consortium | The Lighthouse Lab in Milton Keynes and Alex Alderton, Roberto Amato, Jeffrey Barrett, Sonia Goncalves, Ewan Harrison, David K. Jackson, Ian Johnston, Dominic Kwiatkowski, Cordelia Langford, John Sillitoe on behalf of the Wellcome Sanger Institute COVID-19 Surveillance Team |
| EPI_ISL_2866802, EPI_ISL_2866803 | Lighthouse Lab in Glasgow | Wellcome Sanger Institute for the COVID-19 Genomics UK (COG-UK) Consortium | Harper VanSteenhouse, Yumi Kasai, David Gray, Carol Clugston, Anna Dominiczak and Alex Alderton, Roberto Amato, Jeffrey Barrett, Sonia Goncalves, Ewan Harrison, David K. Jackson, Ian Johnston, Dominic Kwiatkowski, Cordelia Langford, John Sillitoe on behalf of the Wellcome Sanger Institute COVID-19 Surveillance Team |
| EPI_ISL_2866804, EPI_ISL_2866805 | Lighthouse Lab in Milton Keynes | Wellcome Sanger Institute for the COVID-19 Genomics UK (COG-UK) Consortium | The Lighthouse Lab in Milton Keynes and Alex Alderton, Roberto Amato, Jeffrey Barrett, Sonia Goncalves, Ewan Harrison, David K. Jackson, Ian Johnston, Dominic Kwiatkowski, Cordelia Langford, John Sillitoe on behalf of the Wellcome Sanger Institute COVID-19 Surveillance Team |
| EPI_ISL_2866806, EPI_ISL_2866807 | Lighthouse Lab in Glasgow | Wellcome Sanger Institute for the COVID-19 Genomics UK (COG-UK) Consortium | Harper VanSteenhouse, Yumi Kasai, David Gray, Carol Clugston, Anna Dominiczak and Alex Alderton, Roberto Amato, Jeffrey Barrett, Sonia Goncalves, Ewan Harrison, David K. Jackson, Ian Johnston, Dominic Kwiatkowski, Cordelia Langford, John Sillitoe on behalf of the Wellcome Sanger Institute COVID-19 Surveillance Team |
| EPI_ISL_2866808 | Lighthouse Laboratory Plymouth | Wellcome Sanger Institute for the COVID-19 Genomics UK (COG-UK) Consortium | Lighthouse Laboratory Plymouth and Alex Alderton, Roberto Amato, Jeffrey Barrett, Sonia Goncalves, Ewan Harrison, David K. Jackson, Ian Johnston, Dominic Kwiatkowski, Cordelia Langford, John Sillitoe on behalf of the Wellcome Sanger Institute COVID-19 Surveillance Team |
| EPI_ISL_2866809, EPI_ISL_2866810 | Lighthouse Lab in Milton Keynes | Wellcome Sanger Institute for the COVID-19 Genomics UK (COG-UK) Consortium | The Lighthouse Lab in Milton Keynes and Alex Alderton, Roberto Amato, Jeffrey Barrett, Sonia Goncalves, Ewan Harrison, David K. Jackson, Ian Johnston, Dominic Kwiatkowski, Cordelia Langford, John Sillitoe on behalf of the Wellcome Sanger Institute COVID-19 Surveillance Team |
| EPI_ISL_2866811, EPI_ISL_2866812 | Lighthouse Lab in Glasgow | Wellcome Sanger Institute for the COVID-19 Genomics UK (COG-UK) Consortium | Harper VanSteenhouse, Yumi Kasai, David Gray, Carol Clugston, Anna Dominiczak and Alex Alderton, Roberto Amato, Jeffrey Barrett, Sonia Goncalves, Ewan Harrison, David K. Jackson, Ian Johnston, Dominic Kwiatkowski, Cordelia Langford, John Sillitoe on behalf of the Wellcome Sanger Institute COVID-19 Surveillance Team |
| EPI_ISL_2866813 | Lighthouse Laboratory Plymouth | Wellcome Sanger Institute for the COVID-19 Genomics UK (COG-UK) Consortium | Lighthouse Laboratory Plymouth and Alex Alderton, Roberto Amato, Jeffrey Barrett, Sonia Goncalves, Ewan Harrison, David K. Jackson, Ian Johnston, Dominic Kwiatkowski, Cordelia Langford, John Sillitoe on behalf of the Wellcome Sanger Institute COVID-19 Surveillance Team |
| EPI_ISL_2866814, EPI_ISL_2866815 | Lighthouse Lab in Milton Keynes | Wellcome Sanger Institute for the COVID-19 Genomics UK (COG-UK) Consortium | The Lighthouse Lab in Milton Keynes and Alex Alderton, Roberto Amato, Jeffrey Barrett, Sonia Goncalves, Ewan Harrison, David K. Jackson, Ian Johnston, Dominic Kwiatkowski, Cordelia Langford, John Sillitoe on behalf of the Wellcome Sanger Institute COVID-19 Surveillance Team |
| EPI_ISL_2866816 | Lighthouse Laboratory Plymouth | Wellcome Sanger Institute for the COVID-19 Genomics UK (COG-UK) Consortium | Lighthouse Laboratory Plymouth and Alex Alderton, Roberto Amato, Jeffrey Barrett, Sonia Goncalves, Ewan Harrison, David K. Jackson, Ian Johnston, Dominic Kwiatkowski, Cordelia Langford, John Sillitoe on behalf of the Wellcome Sanger Institute COVID-19 Surveillance Team |
| EPI_ISL_2866817 | Lighthouse Lab in Milton Keynes | Wellcome Sanger Institute for the COVID-19 Genomics UK (COG-UK) Consortium | The Lighthouse Lab in Milton Keynes and Alex Alderton, Roberto Amato, Jeffrey Barrett, Sonia Goncalves, Ewan Harrison, David K. Jackson, Ian Johnston, Dominic Kwiatkowski, Cordelia Langford, John Sillitoe on behalf of the Wellcome Sanger Institute COVID-19 Surveillance Team |
| EPI_ISL_2866818, EPI_ISL_2866819, EPI_ISL_2866820, EPI_ISL_2866821, EPI_ISL_2866822, EPI_ISL_2866823 | Lighthouse Lab in Glasgow | Wellcome Sanger Institute for the COVID-19 Genomics UK (COG-UK) Consortium | Harper VanSteenhouse, Yumi Kasai, David Gray, Carol Clugston, Anna Dominiczak and Alex Alderton, Roberto Amato, Jeffrey Barrett, Sonia Goncalves, Ewan Harrison, David K. Jackson, Ian Johnston, Dominic Kwiatkowski, Cordelia Langford, John Sillitoe on behalf of the Wellcome Sanger Institute COVID-19 Surveillance Team |
| EPI_ISL_2866824 | Lighthouse Laboratory Plymouth | Wellcome Sanger Institute for the COVID-19 Genomics UK (COG-UK) Consortium | Lighthouse Laboratory Plymouth and Alex Alderton, Roberto Amato, Jeffrey Barrett, Sonia Goncalves, Ewan Harrison, David K. Jackson, Ian Johnston, Dominic Kwiatkowski, Cordelia Langford, John Sillitoe on behalf of the Wellcome Sanger Institute COVID-19 Surveillance Team |
| EPI_ISL_2866825 | Lighthouse Lab in Milton Keynes | Wellcome Sanger Institute for the COVID-19 Genomics UK (COG-UK) Consortium | The Lighthouse Lab in Milton Keynes and Alex Alderton, Roberto Amato, Jeffrey Barrett, Sonia Goncalves, Ewan Harrison, David K. Jackson, Ian Johnston, Dominic Kwiatkowski, Cordelia Langford, John Sillitoe on behalf of the Wellcome Sanger Institute COVID-19 Surveillance Team |
| EPI_ISL_2866826 | Lighthouse Laboratory Plymouth | Wellcome Sanger Institute for the COVID-19 Genomics UK (COG-UK) Consortium | Lighthouse Laboratory Plymouth and Alex Alderton, Roberto Amato, Jeffrey Barrett, Sonia Goncalves, Ewan Harrison, David K. Jackson, Ian Johnston, Dominic Kwiatkowski, Cordelia Langford, John Sillitoe on behalf of the Wellcome Sanger Institute COVID-19 Surveillance Team |
| EPI_ISL_2866827 | Lighthouse Lab in Glasgow | Wellcome Sanger Institute for the COVID-19 Genomics UK (COG-UK) Consortium | Harper VanSteenhouse, Yumi Kasai, David Gray, Carol Clugston, Anna Dominiczak and Alex Alderton, Roberto Amato, Jeffrey Barrett, Sonia Goncalves, Ewan Harrison, David K. Jackson, Ian Johnston, Dominic Kwiatkowski, Cordelia Langford, John Sillitoe on behalf of the Wellcome Sanger Institute COVID-19 Surveillance Team |
| EPI_ISL_2866828 | Lighthouse Laboratory Plymouth | Wellcome Sanger Institute for the COVID-19 Genomics UK (COG-UK) Consortium | Lighthouse Laboratory Plymouth and Alex Alderton, Roberto Amato, Jeffrey Barrett, Sonia Goncalves, Ewan Harrison, David K. Jackson, Ian Johnston, Dominic Kwiatkowski, Cordelia Langford, John Sillitoe on behalf of the Wellcome Sanger Institute COVID-19 Surveillance Team |
| EPI_ISL_2866829 | Lighthouse Lab in Glasgow | Wellcome Sanger Institute for the COVID-19 Genomics UK (COG-UK) Consortium | Harper VanSteenhouse, Yumi Kasai, David Gray, Carol Clugston, Anna Dominiczak and Alex Alderton, Roberto Amato, Jeffrey Barrett, Sonia Goncalves, Ewan Harrison, David K. Jackson, Ian Johnston, Dominic Kwiatkowski, Cordelia Langford, John Sillitoe on behalf of the Wellcome Sanger Institute COVID-19 Surveillance Team |
| EPI_ISL_2866830 | Lighthouse Laboratory Plymouth | Wellcome Sanger Institute for the COVID-19 Genomics UK (COG-UK) Consortium | Lighthouse Laboratory Plymouth and Alex Alderton, Roberto Amato, Jeffrey Barrett, Sonia Goncalves, Ewan Harrison, David K. Jackson, Ian Johnston, Dominic Kwiatkowski, Cordelia Langford, John Sillitoe on behalf of the Wellcome Sanger Institute COVID-19 Surveillance Team |
| EPI_ISL_2866831 | Lighthouse Lab in Glasgow | Wellcome Sanger Institute for the COVID-19 Genomics UK (COG-UK) Consortium | Harper VanSteenhouse, Yumi Kasai, David Gray, Carol Clugston, Anna Dominiczak and Alex Alderton, Roberto Amato, Jeffrey Barrett, Sonia Goncalves, Ewan Harrison, David K. Jackson, Ian Johnston, Dominic Kwiatkowski, Cordelia Langford, John Sillitoe on behalf of the Wellcome Sanger Institute COVID-19 Surveillance Team |

| | | | |
|---|---|---|---|
| EPI_ISL_2866832, EPI_ISL_2866834 | Lighthouse Laboratory Plymouth | Wellcome Sanger Institute for the COVID-19 Genomics UK (COG-UK) Consortium | Lighthouse Laboratory Plymouth and Alex Alderton, Roberto Amato, Jeffrey Barrett, Sonia Goncalves, Ewan Harrison, David K. Jackson, Ian Johnston, Dominic Kwiatkowski, Cordelia Langford, John Sillitoe on behalf of the Wellcome Sanger Institute COVID-19 Surveillance Team |
| EPI_ISL_2866835, EPI_ISL_2866836, EPI_ISL_2866837 | Lighthouse Lab in Milton Keynes | Wellcome Sanger Institute for the COVID-19 Genomics UK (COG-UK) Consortium | The Lighthouse Lab in Milton Keynes and Alex Alderton, Roberto Amato, Jeffrey Barrett, Sonia Goncalves, Ewan Harrison, David K. Jackson, Ian Johnston, Dominic Kwiatkowski, Cordelia Langford, John Sillitoe on behalf of the Wellcome Sanger Institute COVID-19 Surveillance Team |
| EPI_ISL_2866838 | Lighthouse Lab in Glasgow | Wellcome Sanger Institute for the COVID-19 Genomics UK (COG-UK) Consortium | Harper VanSteenhouse, Yumi Kasai, David Gray, Carol Clugston, Anna Dominiczak and Alex Alderton, Roberto Amato, Jeffrey Barrett, Sonia Goncalves, Ewan Harrison, David K. Jackson, Ian Johnston, Dominic Kwiatkowski, Cordelia Langford, John Sillitoe on behalf of the Wellcome Sanger Institute COVID-19 Surveillance Team |
| EPI_ISL_2866839 | Lighthouse Lab in Milton Keynes | Wellcome Sanger Institute for the COVID-19 Genomics UK (COG-UK) Consortium | The Lighthouse Lab in Milton Keynes and Alex Alderton, Roberto Amato, Jeffrey Barrett, Sonia Goncalves, Ewan Harrison, David K. Jackson, Ian Johnston, Dominic Kwiatkowski, Cordelia Langford, John Sillitoe on behalf of the Wellcome Sanger Institute COVID-19 Surveillance Team |
| EPI_ISL_2866840 | Lighthouse Laboratory Plymouth | Wellcome Sanger Institute for the COVID-19 Genomics UK (COG-UK) Consortium | Lighthouse Laboratory Plymouth and Alex Alderton, Roberto Amato, Jeffrey Barrett, Sonia Goncalves, Ewan Harrison, David K. Jackson, Ian Johnston, Dominic Kwiatkowski, Cordelia Langford, John Sillitoe on behalf of the Wellcome Sanger Institute COVID-19 Surveillance Team |
| EPI_ISL_2866841, EPI_ISL_2866842 | Lighthouse Lab in Glasgow | Wellcome Sanger Institute for the COVID-19 Genomics UK (COG-UK) Consortium | Harper VanSteenhouse, Yumi Kasai, David Gray, Carol Clugston, Anna Dominiczak and Alex Alderton, Roberto Amato, Jeffrey Barrett, Sonia Goncalves, Ewan Harrison, David K. Jackson, Ian Johnston, Dominic Kwiatkowski, Cordelia Langford, John Sillitoe on behalf of the Wellcome Sanger Institute COVID-19 Surveillance Team |
| EPI_ISL_2866843 | Lighthouse Lab in Milton Keynes | Wellcome Sanger Institute for the COVID-19 Genomics UK (COG-UK) Consortium | The Lighthouse Lab in Milton Keynes and Alex Alderton, Roberto Amato, Jeffrey Barrett, Sonia Goncalves, Ewan Harrison, David K. Jackson, Ian Johnston, Dominic Kwiatkowski, Cordelia Langford, John Sillitoe on behalf of the Wellcome Sanger Institute COVID-19 Surveillance Team |
| EPI_ISL_2866844 | Lighthouse Lab in Glasgow | Wellcome Sanger Institute for the COVID-19 Genomics UK (COG-UK) Consortium | Harper VanSteenhouse, Yumi Kasai, David Gray, Carol Clugston, Anna Dominiczak and Alex Alderton, Roberto Amato, Jeffrey Barrett, Sonia Goncalves, Ewan Harrison, David K. Jackson, Ian Johnston, Dominic Kwiatkowski, Cordelia Langford, John Sillitoe on behalf of the Wellcome Sanger Institute COVID-19 Surveillance Team |
| EPI_ISL_2866845 | Lighthouse Lab in Milton Keynes | Wellcome Sanger Institute for the COVID-19 Genomics UK (COG-UK) Consortium | The Lighthouse Lab in Milton Keynes and Alex Alderton, Roberto Amato, Jeffrey Barrett, Sonia Goncalves, Ewan Harrison, David K. Jackson, Ian Johnston, Dominic Kwiatkowski, Cordelia Langford, John Sillitoe on behalf of the Wellcome Sanger Institute COVID-19 Surveillance Team |
| EPI_ISL_2866846, EPI_ISL_2866847, EPI_ISL_2866848 | Lighthouse Lab in Glasgow | Wellcome Sanger Institute for the COVID-19 Genomics UK (COG-UK) Consortium | Harper VanSteenhouse, Yumi Kasai, David Gray, Carol Clugston, Anna Dominiczak and Alex Alderton, Roberto Amato, Jeffrey Barrett, Sonia Goncalves, Ewan Harrison, David K. Jackson, Ian Johnston, Dominic Kwiatkowski, Cordelia Langford, John Sillitoe on behalf of the Wellcome Sanger Institute COVID-19 Surveillance Team |
| EPI_ISL_2866849 | Lighthouse Lab in Milton Keynes | Wellcome Sanger Institute for the COVID-19 Genomics UK (COG-UK) Consortium | The Lighthouse Lab in Milton Keynes and Alex Alderton, Roberto Amato, Jeffrey Barrett, Sonia Goncalves, Ewan Harrison, David K. Jackson, Ian Johnston, Dominic Kwiatkowski, Cordelia Langford, John Sillitoe on behalf of the Wellcome Sanger Institute COVID-19 Surveillance Team |
| EPI_ISL_2866850, EPI_ISL_2866851 | Lighthouse Lab in Glasgow | Wellcome Sanger Institute for the COVID-19 Genomics UK (COG-UK) Consortium | Harper VanSteenhouse, Yumi Kasai, David Gray, Carol Clugston, Anna Dominiczak and Alex Alderton, Roberto Amato, Jeffrey Barrett, Sonia Goncalves, Ewan Harrison, David K. Jackson, Ian Johnston, Dominic Kwiatkowski, Cordelia Langford, John Sillitoe on behalf of the Wellcome Sanger Institute COVID-19 Surveillance Team |
| EPI_ISL_2866852 | Lighthouse Laboratory Plymouth | Wellcome Sanger Institute for the COVID-19 Genomics UK (COG-UK) Consortium | Lighthouse Laboratory Plymouth and Alex Alderton, Roberto Amato, Jeffrey Barrett, Sonia Goncalves, Ewan Harrison, David K. Jackson, Ian Johnston, Dominic Kwiatkowski, Cordelia Langford, John Sillitoe on behalf of the Wellcome Sanger Institute COVID-19 Surveillance Team |
| EPI_ISL_2866853 | Lighthouse Lab in Milton Keynes | Wellcome Sanger Institute for the COVID-19 Genomics UK (COG-UK) Consortium | The Lighthouse Lab in Milton Keynes and Alex Alderton, Roberto Amato, Jeffrey Barrett, Sonia Goncalves, Ewan Harrison, David K. Jackson, Ian Johnston, Dominic Kwiatkowski, Cordelia Langford, John Sillitoe on behalf of the Wellcome Sanger Institute COVID-19 Surveillance Team |
| EPI_ISL_2866854, EPI_ISL_2866856 | Lighthouse Lab in Glasgow | Wellcome Sanger Institute for the COVID-19 Genomics UK (COG-UK) Consortium | Harper VanSteenhouse, Yumi Kasai, David Gray, Carol Clugston, Anna Dominiczak and Alex Alderton, Roberto Amato, Jeffrey Barrett, Sonia Goncalves, Ewan Harrison, David K. Jackson, Ian Johnston, Dominic Kwiatkowski, Cordelia Langford, John Sillitoe on behalf of the Wellcome Sanger Institute COVID-19 Surveillance Team |
| EPI_ISL_2866857 | Lighthouse Laboratory Plymouth | Wellcome Sanger Institute for the COVID-19 Genomics UK (COG-UK) Consortium | Lighthouse Laboratory Plymouth and Alex Alderton, Roberto Amato, Jeffrey Barrett, Sonia Goncalves, Ewan Harrison, David K. Jackson, Ian Johnston, Dominic Kwiatkowski, Cordelia Langford, John Sillitoe on behalf of the Wellcome Sanger Institute COVID-19 Surveillance Team |
| EPI_ISL_2866858, EPI_ISL_2866859, EPI_ISL_2866860, EPI_ISL_2866861, EPI_ISL_2866862 | Lighthouse Lab in Glasgow | Wellcome Sanger Institute for the COVID-19 Genomics UK (COG-UK) Consortium | Harper VanSteenhouse, Yumi Kasai, David Gray, Carol Clugston, Anna Dominiczak and Alex Alderton, Roberto Amato, Jeffrey Barrett, Sonia Goncalves, Ewan Harrison, David K. Jackson, Ian Johnston, Dominic Kwiatkowski, Cordelia Langford, John Sillitoe on behalf of the Wellcome Sanger Institute COVID-19 Surveillance Team |
| EPI_ISL_2866863 | Lighthouse Laboratory Plymouth | Wellcome Sanger Institute for the COVID-19 Genomics UK (COG-UK) Consortium | Lighthouse Laboratory Plymouth and Alex Alderton, Roberto Amato, Jeffrey Barrett, Sonia Goncalves, Ewan Harrison, David K. Jackson, Ian Johnston, Dominic Kwiatkowski, Cordelia Langford, John Sillitoe on behalf of the Wellcome Sanger Institute COVID-19 Surveillance Team |
| EPI_ISL_2866864 | Lighthouse Lab in Glasgow | Wellcome Sanger Institute for the COVID-19 Genomics UK (COG-UK) Consortium | Harper VanSteenhouse, Yumi Kasai, David Gray, Carol Clugston, Anna Dominiczak and Alex Alderton, Roberto Amato, Jeffrey Barrett, Sonia Goncalves, Ewan Harrison, David K. Jackson, Ian Johnston, Dominic Kwiatkowski, Cordelia Langford, John Sillitoe on behalf of the Wellcome Sanger Institute COVID-19 Surveillance Team |
| EPI_ISL_2866865 | Lighthouse Laboratory Plymouth | Wellcome Sanger Institute for the COVID-19 Genomics UK (COG-UK) Consortium | Lighthouse Laboratory Plymouth and Alex Alderton, Roberto Amato, Jeffrey Barrett, Sonia Goncalves, Ewan Harrison, David K. Jackson, Ian Johnston, Dominic Kwiatkowski, Cordelia Langford, John Sillitoe on behalf of the Wellcome Sanger Institute COVID-19 Surveillance Team |
| EPI_ISL_2866866 | Lighthouse Lab in Glasgow | Wellcome Sanger Institute for the COVID-19 Genomics UK (COG-UK) Consortium | Harper VanSteenhouse, Yumi Kasai, David Gray, Carol Clugston, Anna Dominiczak and Alex Alderton, Roberto Amato, Jeffrey Barrett, Sonia Goncalves, Ewan Harrison, David K. Jackson, Ian Johnston, Dominic Kwiatkowski, Cordelia Langford, John Sillitoe on behalf of the Wellcome Sanger Institute COVID-19 Surveillance Team |
| EPI_ISL_2866867 | Lighthouse Lab in Milton Keynes | Wellcome Sanger Institute for the COVID-19 Genomics UK (COG-UK) Consortium | The Lighthouse Lab in Milton Keynes and Alex Alderton, Roberto Amato, Jeffrey Barrett, Sonia Goncalves, Ewan Harrison, David K. Jackson, Ian Johnston, Dominic Kwiatkowski, Cordelia Langford, John Sillitoe on behalf of the Wellcome Sanger Institute COVID-19 Surveillance Team |
| EPI_ISL_2866868 | Lighthouse Laboratory Plymouth | Wellcome Sanger Institute for the COVID-19 Genomics UK (COG-UK) Consortium | Lighthouse Laboratory Plymouth and Alex Alderton, Roberto Amato, Jeffrey Barrett, Sonia Goncalves, Ewan Harrison, David K. Jackson, Ian Johnston, Dominic Kwiatkowski, Cordelia Langford, John Sillitoe on behalf of the Wellcome Sanger Institute COVID-19 Surveillance Team |
| EPI_ISL_2866869, EPI_ISL_2866870 | Lighthouse Lab in Glasgow | Wellcome Sanger Institute for the COVID-19 Genomics UK (COG-UK) Consortium | Harper VanSteenhouse, Yumi Kasai, David Gray, Carol Clugston, Anna Dominiczak and Alex Alderton, Roberto Amato, Jeffrey Barrett, Sonia Goncalves, Ewan Harrison, David K. Jackson, Ian Johnston, Dominic Kwiatkowski, Cordelia Langford, John Sillitoe on behalf of the Wellcome Sanger Institute COVID-19 Surveillance Team |
| EPI_ISL_2866871 | Lighthouse Lab in Milton Keynes | Wellcome Sanger Institute for the COVID-19 Genomics UK (COG-UK) Consortium | The Lighthouse Lab in Milton Keynes and Alex Alderton, Roberto Amato, Jeffrey Barrett, Sonia Goncalves, Ewan Harrison, David K. Jackson, Ian Johnston, Dominic Kwiatkowski, Cordelia Langford, John Sillitoe on behalf of the Wellcome Sanger Institute COVID-19 Surveillance Team |
| EPI_ISL_2866872 | Lighthouse Lab in Glasgow | Wellcome Sanger Institute for the COVID-19 Genomics UK (COG-UK) Consortium | Harper VanSteenhouse, Yumi Kasai, David Gray, Carol Clugston, Anna Dominiczak and Alex Alderton, Roberto Amato, Jeffrey Barrett, Sonia Goncalves, Ewan Harrison, David K. Jackson, Ian Johnston, Dominic Kwiatkowski, Cordelia Langford, John Sillitoe on behalf of the Wellcome Sanger Institute COVID-19 Surveillance Team |
| EPI_ISL_2866873 | Lighthouse Lab in Milton Keynes | Wellcome Sanger Institute for the COVID-19 Genomics UK (COG-UK) Consortium | The Lighthouse Lab in Milton Keynes and Alex Alderton, Roberto Amato, Jeffrey Barrett, Sonia Goncalves, Ewan Harrison, David K. Jackson, Ian Johnston, Dominic Kwiatkowski, Cordelia Langford, John Sillitoe on behalf of the Wellcome Sanger Institute COVID-19 Surveillance Team |
| EPI_ISL_2866874 | Lighthouse Lab in Glasgow | Wellcome Sanger Institute for the COVID-19 Genomics UK (COG-UK) Consortium | Harper VanSteenhouse, Yumi Kasai, David Gray, Carol Clugston, Anna Dominiczak and Alex Alderton, Roberto Amato, Jeffrey Barrett, Sonia Goncalves, Ewan Harrison, David K. Jackson, Ian Johnston, Dominic Kwiatkowski, Cordelia Langford, John Sillitoe on behalf of the Wellcome Sanger Institute COVID-19 Surveillance Team |
| EPI_ISL_2866875, EPI_ISL_2866876, EPI_ISL_2866877 | Lighthouse Laboratory Plymouth | Wellcome Sanger Institute for the COVID-19 Genomics UK (COG-UK) Consortium | Lighthouse Laboratory Plymouth and Alex Alderton, Roberto Amato, Jeffrey Barrett, Sonia Goncalves, Ewan Harrison, David K. Jackson, Ian Johnston, Dominic Kwiatkowski, Cordelia Langford, John Sillitoe on behalf of the Wellcome Sanger Institute COVID-19 Surveillance Team |
| EPI_ISL_2866878 | Lighthouse Lab in Glasgow | Wellcome Sanger Institute for the COVID-19 Genomics UK (COG-UK) Consortium | Harper VanSteenhouse, Yumi Kasai, David Gray, Carol Clugston, Anna Dominiczak and Alex Alderton, Roberto Amato, Jeffrey Barrett, Sonia Goncalves, Ewan Harrison, David K. Jackson, Ian Johnston, Dominic Kwiatkowski, Cordelia Langford, John Sillitoe on behalf of the Wellcome Sanger Institute COVID-19 Surveillance Team |
| EPI_ISL_2866879 | Lighthouse Lab in Milton Keynes | Wellcome Sanger Institute for the COVID-19 Genomics UK (COG-UK) Consortium | The Lighthouse Lab in Milton Keynes and Alex Alderton, Roberto Amato, Jeffrey Barrett, Sonia Goncalves, Ewan Harrison, David K. Jackson, Ian Johnston, Dominic Kwiatkowski, Cordelia Langford, John Sillitoe on behalf of the Wellcome Sanger Institute COVID-19 Surveillance Team |
| EPI_ISL_2866880 | Lighthouse Lab in Glasgow | Wellcome Sanger Institute for the COVID-19 Genomics UK (COG-UK) Consortium | Harper VanSteenhouse, Yumi Kasai, David Gray, Carol Clugston, Anna Dominiczak and Alex Alderton, Roberto Amato, Jeffrey Barrett, Sonia Goncalves, Ewan Harrison, David K. Jackson, Ian Johnston, Dominic Kwiatkowski, Cordelia Langford, John Sillitoe on behalf of the Wellcome Sanger Institute COVID-19 Surveillance Team |
| EPI_ISL_2866881, EPI_ISL_2866882 | Lighthouse Laboratory Plymouth | Wellcome Sanger Institute for the COVID-19 Genomics UK (COG-UK) Consortium | Lighthouse Laboratory Plymouth and Alex Alderton, Roberto Amato, Jeffrey Barrett, Sonia Goncalves, Ewan Harrison, David K. Jackson, Ian Johnston, Dominic Kwiatkowski, Cordelia Langford, John Sillitoe on behalf of the Wellcome Sanger Institute COVID-19 Surveillance Team |
| EPI_ISL_2866883 | Lighthouse Lab in Milton Keynes | Wellcome Sanger Institute for the COVID-19 Genomics UK (COG-UK) Consortium | The Lighthouse Lab in Milton Keynes and Alex Alderton, Roberto Amato, Jeffrey Barrett, Sonia Goncalves, Ewan Harrison, David K. Jackson, Ian Johnston, Dominic Kwiatkowski, Cordelia Langford, John Sillitoe on behalf of the Wellcome Sanger Institute COVID-19 Surveillance Team |
| EPI_ISL_2866884 | Lighthouse Lab in Glasgow | Wellcome Sanger Institute for the COVID-19 Genomics UK (COG-UK) Consortium | Harper VanSteenhouse, Yumi Kasai, David Gray, Carol Clugston, Anna Dominiczak and Alex Alderton, Roberto Amato, Jeffrey Barrett, Sonia Goncalves, Ewan Harrison, David K. Jackson, Ian Johnston, Dominic Kwiatkowski, Cordelia Langford, John Sillitoe on behalf of the Wellcome Sanger Institute COVID-19 Surveillance Team |
| EPI_ISL_2866885 | Lighthouse Lab in Milton Keynes | Wellcome Sanger Institute for the COVID-19 Genomics UK (COG-UK) Consortium | The Lighthouse Lab in Milton Keynes and Alex Alderton, Roberto Amato, Jeffrey Barrett, Sonia Goncalves, Ewan Harrison, David K. Jackson, Ian Johnston, Dominic Kwiatkowski, Cordelia Langford, John Sillitoe on behalf of the Wellcome Sanger Institute COVID-19 Surveillance Team |
| EPI_ISL_2866886, EPI_ISL_2866887 | Lighthouse Lab in Glasgow | Wellcome Sanger Institute for the COVID-19 Genomics UK (COG-UK) Consortium | Harper VanSteenhouse, Yumi Kasai, David Gray, Carol Clugston, Anna Dominiczak and Alex Alderton, Roberto Amato, Jeffrey Barrett, Sonia Goncalves, Ewan Harrison, David K. Jackson, Ian Johnston, Dominic Kwiatkowski, Cordelia Langford, John Sillitoe on behalf of the Wellcome Sanger Institute COVID-19 Surveillance Team |
| EPI_ISL_2866888 | Lighthouse Lab in Milton Keynes | Wellcome Sanger Institute for the COVID-19 Genomics UK (COG-UK) Consortium | The Lighthouse Lab in Milton Keynes and Alex Alderton, Roberto Amato, Jeffrey Barrett, Sonia Goncalves, Ewan Harrison, David K. Jackson, Ian Johnston, Dominic Kwiatkowski, Cordelia Langford, John Sillitoe on behalf of the Wellcome Sanger Institute COVID-19 Surveillance Team |
| EPI_ISL_2866889 | Lighthouse Laboratory Plymouth | Wellcome Sanger Institute for the COVID-19 Genomics UK (COG-UK) Consortium | Lighthouse Laboratory Plymouth and Alex Alderton, Roberto Amato, Jeffrey Barrett, Sonia Goncalves, Ewan Harrison, David K. Jackson, Ian Johnston, Dominic Kwiatkowski, Cordelia Langford, John Sillitoe on behalf of the Wellcome Sanger Institute COVID-19 Surveillance Team |
| EPI_ISL_2866890, EPI_ISL_2866891 | Lighthouse Lab in Milton Keynes | Wellcome Sanger Institute for the COVID-19 Genomics UK (COG-UK) Consortium | The Lighthouse Lab in Milton Keynes and Alex Alderton, Roberto Amato, Jeffrey Barrett, Sonia Goncalves, Ewan Harrison, David K. Jackson, Ian Johnston, Dominic Kwiatkowski, Cordelia Langford, John Sillitoe on behalf of the Wellcome Sanger Institute COVID-19 Surveillance Team |
| EPI_ISL_2866892 | Lighthouse Lab in Glasgow | Wellcome Sanger Institute for the COVID-19 Genomics UK (COG-UK) Consortium | Harper VanSteenhouse, Yumi Kasai, David Gray, Carol Clugston, Anna Dominiczak and Alex Alderton, Roberto Amato, Jeffrey Barrett, Sonia Goncalves, Ewan Harrison, David K. Jackson, Ian Johnston, Dominic Kwiatkowski, Cordelia Langford, John Sillitoe on behalf of the Wellcome Sanger Institute COVID-19 Surveillance Team |
| EPI_ISL_2866893 | Lighthouse Laboratory Plymouth | Wellcome Sanger Institute for the COVID-19 Genomics UK (COG-UK) Consortium | Lighthouse Laboratory Plymouth and Alex Alderton, Roberto Amato, Jeffrey Barrett, Sonia Goncalves, Ewan Harrison, David K. Jackson, Ian Johnston, Dominic Kwiatkowski, Cordelia Langford, John Sillitoe on behalf of the Wellcome Sanger Institute COVID-19 Surveillance Team |

| | | | |
|---|---|---|---|
| EPI_ISL_2866894, EPI_ISL_2866895 | Lighthouse Lab in Glasgow | Wellcome Sanger Institute for the COVID-19 Genomics UK (COG-UK) Consortium | Harper VanSteenhouse, Yumi Kasai, David Gray, Carol Clugston, Anna Dominiczak and Alex Alderton, Roberto Amato, Jeffrey Barrett, Sonia Goncalves, Ewan Harrison, David K. Jackson, Ian Johnston, Dominic Kwiatkowski, Cordelia Langford, John Sillitoe on behalf of the Wellcome Sanger Institute COVID-19 Surveillance Team |
| EPI_ISL_2866896 | Lighthouse Laboratory Plymouth | Wellcome Sanger Institute for the COVID-19 Genomics UK (COG-UK) Consortium | Lighthouse Laboratory Plymouth and Alex Alderton, Roberto Amato, Jeffrey Barrett, Sonia Goncalves, Ewan Harrison, David K. Jackson, Ian Johnston, Dominic Kwiatkowski, Cordelia Langford, John Sillitoe on behalf of the Wellcome Sanger Institute COVID-19 Surveillance Team |
| EPI_ISL_2866897 | Lighthouse Lab in Milton Keynes | Wellcome Sanger Institute for the COVID-19 Genomics UK (COG-UK) Consortium | The Lighthouse Lab in Milton Keynes and Alex Alderton, Roberto Amato, Jeffrey Barrett, Sonia Goncalves, Ewan Harrison, David K. Jackson, Ian Johnston, Dominic Kwiatkowski, Cordelia Langford, John Sillitoe on behalf of the Wellcome Sanger Institute COVID-19 Surveillance Team |
| EPI_ISL_2866898, EPI_ISL_2866899, EPI_ISL_2866900, EPI_ISL_2866901 | Lighthouse Lab in Glasgow | Wellcome Sanger Institute for the COVID-19 Genomics UK (COG-UK) Consortium | Harper VanSteenhouse, Yumi Kasai, David Gray, Carol Clugston, Anna Dominiczak and Alex Alderton, Roberto Amato, Jeffrey Barrett, Sonia Goncalves, Ewan Harrison, David K. Jackson, Ian Johnston, Dominic Kwiatkowski, Cordelia Langford, John Sillitoe on behalf of the Wellcome Sanger Institute COVID-19 Surveillance Team |
| EPI_ISL_2866902 | Lighthouse Laboratory Plymouth | Wellcome Sanger Institute for the COVID-19 Genomics UK (COG-UK) Consortium | Lighthouse Laboratory Plymouth and Alex Alderton, Roberto Amato, Jeffrey Barrett, Sonia Goncalves, Ewan Harrison, David K. Jackson, Ian Johnston, Dominic Kwiatkowski, Cordelia Langford, John Sillitoe on behalf of the Wellcome Sanger Institute COVID-19 Surveillance Team |
| EPI_ISL_2866903, EPI_ISL_2866904 | Lighthouse Lab in Glasgow | Wellcome Sanger Institute for the COVID-19 Genomics UK (COG-UK) Consortium | Harper VanSteenhouse, Yumi Kasai, David Gray, Carol Clugston, Anna Dominiczak and Alex Alderton, Roberto Amato, Jeffrey Barrett, Sonia Goncalves, Ewan Harrison, David K. Jackson, Ian Johnston, Dominic Kwiatkowski, Cordelia Langford, John Sillitoe on behalf of the Wellcome Sanger Institute COVID-19 Surveillance Team |
| EPI_ISL_2866905 | Lighthouse Lab in Milton Keynes | Wellcome Sanger Institute for the COVID-19 Genomics UK (COG-UK) Consortium | The Lighthouse Lab in Milton Keynes and Alex Alderton, Roberto Amato, Jeffrey Barrett, Sonia Goncalves, Ewan Harrison, David K. Jackson, Ian Johnston, Dominic Kwiatkowski, Cordelia Langford, John Sillitoe on behalf of the Wellcome Sanger Institute COVID-19 Surveillance Team |
| EPI_ISL_2866906, EPI_ISL_2866907 | Lighthouse Lab in Glasgow | Wellcome Sanger Institute for the COVID-19 Genomics UK (COG-UK) Consortium | Harper VanSteenhouse, Yumi Kasai, David Gray, Carol Clugston, Anna Dominiczak and Alex Alderton, Roberto Amato, Jeffrey Barrett, Sonia Goncalves, Ewan Harrison, David K. Jackson, Ian Johnston, Dominic Kwiatkowski, Cordelia Langford, John Sillitoe on behalf of the Wellcome Sanger Institute COVID-19 Surveillance Team |
| EPI_ISL_2866908, EPI_ISL_2866909, EPI_ISL_2866910 | Lighthouse Lab in Milton Keynes | Wellcome Sanger Institute for the COVID-19 Genomics UK (COG-UK) Consortium | The Lighthouse Lab in Milton Keynes and Alex Alderton, Roberto Amato, Jeffrey Barrett, Sonia Goncalves, Ewan Harrison, David K. Jackson, Ian Johnston, Dominic Kwiatkowski, Cordelia Langford, John Sillitoe on behalf of the Wellcome Sanger Institute COVID-19 Surveillance Team |
| EPI_ISL_2866911 | Lighthouse Lab in Glasgow | Wellcome Sanger Institute for the COVID-19 Genomics UK (COG-UK) Consortium | Harper VanSteenhouse, Yumi Kasai, David Gray, Carol Clugston, Anna Dominiczak and Alex Alderton, Roberto Amato, Jeffrey Barrett, Sonia Goncalves, Ewan Harrison, David K. Jackson, Ian Johnston, Dominic Kwiatkowski, Cordelia Langford, John Sillitoe on behalf of the Wellcome Sanger Institute COVID-19 Surveillance Team |
| EPI_ISL_2866912 | Lighthouse Lab in Milton Keynes | Wellcome Sanger Institute for the COVID-19 Genomics UK (COG-UK) Consortium | The Lighthouse Lab in Milton Keynes and Alex Alderton, Roberto Amato, Jeffrey Barrett, Sonia Goncalves, Ewan Harrison, David K. Jackson, Ian Johnston, Dominic Kwiatkowski, Cordelia Langford, John Sillitoe on behalf of the Wellcome Sanger Institute COVID-19 Surveillance Team |
| EPI_ISL_2866913 | Lighthouse Lab in Glasgow | Wellcome Sanger Institute for the COVID-19 Genomics UK (COG-UK) Consortium | Harper VanSteenhouse, Yumi Kasai, David Gray, Carol Clugston, Anna Dominiczak and Alex Alderton, Roberto Amato, Jeffrey Barrett, Sonia Goncalves, Ewan Harrison, David K. Jackson, Ian Johnston, Dominic Kwiatkowski, Cordelia Langford, John Sillitoe on behalf of the Wellcome Sanger Institute COVID-19 Surveillance Team |
| EPI_ISL_2866914 | Lighthouse Laboratory Plymouth | Wellcome Sanger Institute for the COVID-19 Genomics UK (COG-UK) Consortium | Lighthouse Laboratory Plymouth and Alex Alderton, Roberto Amato, Jeffrey Barrett, Sonia Goncalves, Ewan Harrison, David K. Jackson, Ian Johnston, Dominic Kwiatkowski, Cordelia Langford, John Sillitoe on behalf of the Wellcome Sanger Institute COVID-19 Surveillance Team |
| EPI_ISL_2866915, EPI_ISL_2866916 | Lighthouse Lab in Glasgow | Wellcome Sanger Institute for the COVID-19 Genomics UK (COG-UK) Consortium | Harper VanSteenhouse, Yumi Kasai, David Gray, Carol Clugston, Anna Dominiczak and Alex Alderton, Roberto Amato, Jeffrey Barrett, Sonia Goncalves, Ewan Harrison, David K. Jackson, Ian Johnston, Dominic Kwiatkowski, Cordelia Langford, John Sillitoe on behalf of the Wellcome Sanger Institute COVID-19 Surveillance Team |
| EPI_ISL_2866917 | Lighthouse Laboratory Plymouth | Wellcome Sanger Institute for the COVID-19 Genomics UK (COG-UK) Consortium | Lighthouse Laboratory Plymouth and Alex Alderton, Roberto Amato, Jeffrey Barrett, Sonia Goncalves, Ewan Harrison, David K. Jackson, Ian Johnston, Dominic Kwiatkowski, Cordelia Langford, John Sillitoe on behalf of the Wellcome Sanger Institute COVID-19 Surveillance Team |
| EPI_ISL_2866918 | Lighthouse Lab in Milton Keynes | Wellcome Sanger Institute for the COVID-19 Genomics UK (COG-UK) Consortium | The Lighthouse Lab in Milton Keynes and Alex Alderton, Roberto Amato, Jeffrey Barrett, Sonia Goncalves, Ewan Harrison, David K. Jackson, Ian Johnston, Dominic Kwiatkowski, Cordelia Langford, John Sillitoe on behalf of the Wellcome Sanger Institute COVID-19 Surveillance Team |
| EPI_ISL_2866919, EPI_ISL_2866920, EPI_ISL_2866921, EPI_ISL_2866922 | Lighthouse Laboratory Plymouth | Wellcome Sanger Institute for the COVID-19 Genomics UK (COG-UK) Consortium | Lighthouse Laboratory Plymouth and Alex Alderton, Roberto Amato, Jeffrey Barrett, Sonia Goncalves, Ewan Harrison, David K. Jackson, Ian Johnston, Dominic Kwiatkowski, Cordelia Langford, John Sillitoe on behalf of the Wellcome Sanger Institute COVID-19 Surveillance Team |
| EPI_ISL_2866923 | Lighthouse Lab in Glasgow | Wellcome Sanger Institute for the COVID-19 Genomics UK (COG-UK) Consortium | Harper VanSteenhouse, Yumi Kasai, David Gray, Carol Clugston, Anna Dominiczak and Alex Alderton, Roberto Amato, Jeffrey Barrett, Sonia Goncalves, Ewan Harrison, David K. Jackson, Ian Johnston, Dominic Kwiatkowski, Cordelia Langford, John Sillitoe on behalf of the Wellcome Sanger Institute COVID-19 Surveillance Team |
| EPI_ISL_2866924 | Lighthouse Lab in Milton Keynes | Wellcome Sanger Institute for the COVID-19 Genomics UK (COG-UK) Consortium | The Lighthouse Lab in Milton Keynes and Alex Alderton, Roberto Amato, Jeffrey Barrett, Sonia Goncalves, Ewan Harrison, David K. Jackson, Ian Johnston, Dominic Kwiatkowski, Cordelia Langford, John Sillitoe on behalf of the Wellcome Sanger Institute COVID-19 Surveillance Team |
| EPI_ISL_2866925 | Lighthouse Lab in Glasgow | Wellcome Sanger Institute for the COVID-19 Genomics UK (COG-UK) Consortium | Harper VanSteenhouse, Yumi Kasai, David Gray, Carol Clugston, Anna Dominiczak and Alex Alderton, Roberto Amato, Jeffrey Barrett, Sonia Goncalves, Ewan Harrison, David K. Jackson, Ian Johnston, Dominic Kwiatkowski, Cordelia Langford, John Sillitoe on behalf of the Wellcome Sanger Institute COVID-19 Surveillance Team |
| EPI_ISL_2866926 | Lighthouse Lab in Milton Keynes | Wellcome Sanger Institute for the COVID-19 Genomics UK (COG-UK) Consortium | The Lighthouse Lab in Milton Keynes and Alex Alderton, Roberto Amato, Jeffrey Barrett, Sonia Goncalves, Ewan Harrison, David K. Jackson, Ian Johnston, Dominic Kwiatkowski, Cordelia Langford, John Sillitoe on behalf of the Wellcome Sanger Institute COVID-19 Surveillance Team |
| EPI_ISL_2866927, EPI_ISL_2866928, EPI_ISL_2866929, EPI_ISL_2866930, EPI_ISL_2866931 | Lighthouse Lab in Glasgow | Wellcome Sanger Institute for the COVID-19 Genomics UK (COG-UK) Consortium | Harper VanSteenhouse, Yumi Kasai, David Gray, Carol Clugston, Anna Dominiczak and Alex Alderton, Roberto Amato, Jeffrey Barrett, Sonia Goncalves, Ewan Harrison, David K. Jackson, Ian Johnston, Dominic Kwiatkowski, Cordelia Langford, John Sillitoe on behalf of the Wellcome Sanger Institute COVID-19 Surveillance Team |
| EPI_ISL_2866932 | Lighthouse Laboratory Plymouth | Wellcome Sanger Institute for the COVID-19 Genomics UK (COG-UK) Consortium | Lighthouse Laboratory Plymouth and Alex Alderton, Roberto Amato, Jeffrey Barrett, Sonia Goncalves, Ewan Harrison, David K. Jackson, Ian Johnston, Dominic Kwiatkowski, Cordelia Langford, John Sillitoe on behalf of the Wellcome Sanger Institute COVID-19 Surveillance Team |
| EPI_ISL_2866933, EPI_ISL_2866934, EPI_ISL_2866935, EPI_ISL_2866936 | Lighthouse Lab in Glasgow | Wellcome Sanger Institute for the COVID-19 Genomics UK (COG-UK) Consortium | Harper VanSteenhouse, Yumi Kasai, David Gray, Carol Clugston, Anna Dominiczak and Alex Alderton, Roberto Amato, Jeffrey Barrett, Sonia Goncalves, Ewan Harrison, David K. Jackson, Ian Johnston, Dominic Kwiatkowski, Cordelia Langford, John Sillitoe on behalf of the Wellcome Sanger Institute COVID-19 Surveillance Team |
| EPI_ISL_2866937 | Lighthouse Lab in Milton Keynes | Wellcome Sanger Institute for the COVID-19 Genomics UK (COG-UK) Consortium | The Lighthouse Lab in Milton Keynes and Alex Alderton, Roberto Amato, Jeffrey Barrett, Sonia Goncalves, Ewan Harrison, David K. Jackson, Ian Johnston, Dominic Kwiatkowski, Cordelia Langford, John Sillitoe on behalf of the Wellcome Sanger Institute COVID-19 Surveillance Team |
| EPI_ISL_2866938, EPI_ISL_2866939 | Lighthouse Laboratory Plymouth | Wellcome Sanger Institute for the COVID-19 Genomics UK (COG-UK) Consortium | Lighthouse Laboratory Plymouth and Alex Alderton, Roberto Amato, Jeffrey Barrett, Sonia Goncalves, Ewan Harrison, David K. Jackson, Ian Johnston, Dominic Kwiatkowski, Cordelia Langford, John Sillitoe on behalf of the Wellcome Sanger Institute COVID-19 Surveillance Team |
| EPI_ISL_2866940 | Lighthouse Lab in Milton Keynes | Wellcome Sanger Institute for the COVID-19 Genomics UK (COG-UK) Consortium | The Lighthouse Lab in Milton Keynes and Alex Alderton, Roberto Amato, Jeffrey Barrett, Sonia Goncalves, Ewan Harrison, David K. Jackson, Ian Johnston, Dominic Kwiatkowski, Cordelia Langford, John Sillitoe on behalf of the Wellcome Sanger Institute COVID-19 Surveillance Team |
| EPI_ISL_2866941 | Lighthouse Lab in Glasgow | Wellcome Sanger Institute for the COVID-19 Genomics UK (COG-UK) Consortium | Harper VanSteenhouse, Yumi Kasai, David Gray, Carol Clugston, Anna Dominiczak and Alex Alderton, Roberto Amato, Jeffrey Barrett, Sonia Goncalves, Ewan Harrison, David K. Jackson, Ian Johnston, Dominic Kwiatkowski, Cordelia Langford, John Sillitoe on behalf of the Wellcome Sanger Institute COVID-19 Surveillance Team |
| EPI_ISL_2866942, EPI_ISL_2866943 | Lighthouse Laboratory Plymouth | Wellcome Sanger Institute for the COVID-19 Genomics UK (COG-UK) Consortium | Lighthouse Laboratory Plymouth and Alex Alderton, Roberto Amato, Jeffrey Barrett, Sonia Goncalves, Ewan Harrison, David K. Jackson, Ian Johnston, Dominic Kwiatkowski, Cordelia Langford, John Sillitoe on behalf of the Wellcome Sanger Institute COVID-19 Surveillance Team |
| EPI_ISL_2866944 | Lighthouse Lab in Milton Keynes | Wellcome Sanger Institute for the COVID-19 Genomics UK (COG-UK) Consortium | The Lighthouse Lab in Milton Keynes and Alex Alderton, Roberto Amato, Jeffrey Barrett, Sonia Goncalves, Ewan Harrison, David K. Jackson, Ian Johnston, Dominic Kwiatkowski, Cordelia Langford, John Sillitoe on behalf of the Wellcome Sanger Institute COVID-19 Surveillance Team |
| EPI_ISL_2866945 | Lighthouse Laboratory Plymouth | Wellcome Sanger Institute for the COVID-19 Genomics UK (COG-UK) Consortium | Lighthouse Laboratory Plymouth and Alex Alderton, Roberto Amato, Jeffrey Barrett, Sonia Goncalves, Ewan Harrison, David K. Jackson, Ian Johnston, Dominic Kwiatkowski, Cordelia Langford, John Sillitoe on behalf of the Wellcome Sanger Institute COVID-19 Surveillance Team |
| EPI_ISL_2866946 | Lighthouse Lab in Milton Keynes | Wellcome Sanger Institute for the COVID-19 Genomics UK (COG-UK) Consortium | The Lighthouse Lab in Milton Keynes and Alex Alderton, Roberto Amato, Jeffrey Barrett, Sonia Goncalves, Ewan Harrison, David K. Jackson, Ian Johnston, Dominic Kwiatkowski, Cordelia Langford, John Sillitoe on behalf of the Wellcome Sanger Institute COVID-19 Surveillance Team |
| EPI_ISL_2866947 | Lighthouse Lab in Glasgow | Wellcome Sanger Institute for the COVID-19 Genomics UK (COG-UK) Consortium | Harper VanSteenhouse, Yumi Kasai, David Gray, Carol Clugston, Anna Dominiczak and Alex Alderton, Roberto Amato, Jeffrey Barrett, Sonia Goncalves, Ewan Harrison, David K. Jackson, Ian Johnston, Dominic Kwiatkowski, Cordelia Langford, John Sillitoe on behalf of the Wellcome Sanger Institute COVID-19 Surveillance Team |
| EPI_ISL_2866948 | Lighthouse Laboratory Plymouth | Wellcome Sanger Institute for the COVID-19 Genomics UK (COG-UK) Consortium | Lighthouse Laboratory Plymouth and Alex Alderton, Roberto Amato, Jeffrey Barrett, Sonia Goncalves, Ewan Harrison, David K. Jackson, Ian Johnston, Dominic Kwiatkowski, Cordelia Langford, John Sillitoe on behalf of the Wellcome Sanger Institute COVID-19 Surveillance Team |
| EPI_ISL_2866949 | Lighthouse Lab in Glasgow | Wellcome Sanger Institute for the COVID-19 Genomics UK (COG-UK) Consortium | Harper VanSteenhouse, Yumi Kasai, David Gray, Carol Clugston, Anna Dominiczak and Alex Alderton, Roberto Amato, Jeffrey Barrett, Sonia Goncalves, Ewan Harrison, David K. Jackson, Ian Johnston, Dominic Kwiatkowski, Cordelia Langford, John Sillitoe on behalf of the Wellcome Sanger Institute COVID-19 Surveillance Team |
| EPI_ISL_2866950 | Lighthouse Laboratory Plymouth | Wellcome Sanger Institute for the COVID-19 Genomics UK (COG-UK) Consortium | Lighthouse Laboratory Plymouth and Alex Alderton, Roberto Amato, Jeffrey Barrett, Sonia Goncalves, Ewan Harrison, David K. Jackson, Ian Johnston, Dominic Kwiatkowski, Cordelia Langford, John Sillitoe on behalf of the Wellcome Sanger Institute COVID-19 Surveillance Team |
| EPI_ISL_2866951 | Lighthouse Lab in Glasgow | Wellcome Sanger Institute for the COVID-19 Genomics UK (COG-UK) Consortium | Harper VanSteenhouse, Yumi Kasai, David Gray, Carol Clugston, Anna Dominiczak and Alex Alderton, Roberto Amato, Jeffrey Barrett, Sonia Goncalves, Ewan Harrison, David K. Jackson, Ian Johnston, Dominic Kwiatkowski, Cordelia Langford, John Sillitoe on behalf of the Wellcome Sanger Institute COVID-19 Surveillance Team |
| EPI_ISL_2866952 | Lighthouse Laboratory Plymouth | Wellcome Sanger Institute for the COVID-19 Genomics UK (COG-UK) Consortium | Lighthouse Laboratory Plymouth and Alex Alderton, Roberto Amato, Jeffrey Barrett, Sonia Goncalves, Ewan Harrison, David K. Jackson, Ian Johnston, Dominic Kwiatkowski, Cordelia Langford, John Sillitoe on behalf of the Wellcome Sanger Institute COVID-19 Surveillance Team |
| EPI_ISL_2866953, EPI_ISL_2866954 | Lighthouse Lab in Milton Keynes | Wellcome Sanger Institute for the COVID-19 Genomics UK (COG-UK) Consortium | The Lighthouse Lab in Milton Keynes and Alex Alderton, Roberto Amato, Jeffrey Barrett, Sonia Goncalves, Ewan Harrison, David K. Jackson, Ian Johnston, Dominic Kwiatkowski, Cordelia Langford, John Sillitoe on behalf of the Wellcome Sanger Institute COVID-19 Surveillance Team |
| EPI_ISL_2866955 | Lighthouse Lab in Glasgow | Wellcome Sanger Institute for the COVID-19 Genomics UK (COG-UK) Consortium | Harper VanSteenhouse, Yumi Kasai, David Gray, Carol Clugston, Anna Dominiczak and Alex Alderton, Roberto Amato, Jeffrey Barrett, Sonia Goncalves, Ewan Harrison, David K. Jackson, Ian Johnston, Dominic Kwiatkowski, Cordelia Langford, John Sillitoe on behalf of the Wellcome Sanger Institute COVID-19 Surveillance Team |
| EPI_ISL_2866956 | Lighthouse Laboratory Plymouth | Wellcome Sanger Institute for the COVID-19 Genomics UK (COG-UK) Consortium | Lighthouse Laboratory Plymouth and Alex Alderton, Roberto Amato, Jeffrey Barrett, Sonia Goncalves, Ewan Harrison, David K. Jackson, Ian Johnston, Dominic Kwiatkowski, Cordelia Langford, John Sillitoe on behalf of the Wellcome Sanger Institute COVID-19 Surveillance Team |

| | | | |
|---|---|---|---|
| EPI_ISL_2866957, EPI_ISL_2866958 | Lighthouse Lab in Glasgow | Wellcome Sanger Institute for the COVID-19 Genomics UK (COG-UK) Consortium | Harper VanSteenhouse, Yumi Kasai, David Gray, Carol Clugston, Anna Dominiczak and Alex Alderton, Roberto Amato, Jeffrey Barrett, Sonia Goncalves, Ewan Harrison, David K. Jackson, Ian Johnston, Dominic Kwiatkowski, Cordelia Langford, John Sillitoe on behalf of the Wellcome Sanger Institute COVID-19 Surveillance Team |
| EPI_ISL_2866959 | Lighthouse Laboratory Plymouth | Wellcome Sanger Institute for the COVID-19 Genomics UK (COG-UK) Consortium | Lighthouse Laboratory Plymouth and Alex Alderton, Roberto Amato, Jeffrey Barrett, Sonia Goncalves, Ewan Harrison, David K. Jackson, Ian Johnston, Dominic Kwiatkowski, Cordelia Langford, John Sillitoe on behalf of the Wellcome Sanger Institute COVID-19 Surveillance Team |
| EPI_ISL_2866960 | Lighthouse Lab in Milton Keynes | Wellcome Sanger Institute for the COVID-19 Genomics UK (COG-UK) Consortium | The Lighthouse Lab in Milton Keynes and Alex Alderton, Roberto Amato, Jeffrey Barrett, Sonia Goncalves, Ewan Harrison, David K. Jackson, Ian Johnston, Dominic Kwiatkowski, Cordelia Langford, John Sillitoe on behalf of the Wellcome Sanger Institute COVID-19 Surveillance Team |
| EPI_ISL_2866961, EPI_ISL_2866962 | Lighthouse Lab in Glasgow | Wellcome Sanger Institute for the COVID-19 Genomics UK (COG-UK) Consortium | Harper VanSteenhouse, Yumi Kasai, David Gray, Carol Clugston, Anna Dominiczak and Alex Alderton, Roberto Amato, Jeffrey Barrett, Sonia Goncalves, Ewan Harrison, David K. Jackson, Ian Johnston, Dominic Kwiatkowski, Cordelia Langford, John Sillitoe on behalf of the Wellcome Sanger Institute COVID-19 Surveillance Team |
| EPI_ISL_2866963 | Lighthouse Laboratory Plymouth | Wellcome Sanger Institute for the COVID-19 Genomics UK (COG-UK) Consortium | Lighthouse Laboratory Plymouth and Alex Alderton, Roberto Amato, Jeffrey Barrett, Sonia Goncalves, Ewan Harrison, David K. Jackson, Ian Johnston, Dominic Kwiatkowski, Cordelia Langford, John Sillitoe on behalf of the Wellcome Sanger Institute COVID-19 Surveillance Team |
| EPI_ISL_2866964, EPI_ISL_2866965, EPI_ISL_2866966 | Lighthouse Lab in Glasgow | Wellcome Sanger Institute for the COVID-19 Genomics UK (COG-UK) Consortium | Harper VanSteenhouse, Yumi Kasai, David Gray, Carol Clugston, Anna Dominiczak and Alex Alderton, Roberto Amato, Jeffrey Barrett, Sonia Goncalves, Ewan Harrison, David K. Jackson, Ian Johnston, Dominic Kwiatkowski, Cordelia Langford, John Sillitoe on behalf of the Wellcome Sanger Institute COVID-19 Surveillance Team |
| EPI_ISL_2866967 | Lighthouse Laboratory Plymouth | Wellcome Sanger Institute for the COVID-19 Genomics UK (COG-UK) Consortium | Lighthouse Laboratory Plymouth and Alex Alderton, Roberto Amato, Jeffrey Barrett, Sonia Goncalves, Ewan Harrison, David K. Jackson, Ian Johnston, Dominic Kwiatkowski, Cordelia Langford, John Sillitoe on behalf of the Wellcome Sanger Institute COVID-19 Surveillance Team |
| EPI_ISL_2866968 | Lighthouse Lab in Glasgow | Wellcome Sanger Institute for the COVID-19 Genomics UK (COG-UK) Consortium | Harper VanSteenhouse, Yumi Kasai, David Gray, Carol Clugston, Anna Dominiczak and Alex Alderton, Roberto Amato, Jeffrey Barrett, Sonia Goncalves, Ewan Harrison, David K. Jackson, Ian Johnston, Dominic Kwiatkowski, Cordelia Langford, John Sillitoe on behalf of the Wellcome Sanger Institute COVID-19 Surveillance Team |
| EPI_ISL_2866969 | Lighthouse Lab in Milton Keynes | Wellcome Sanger Institute for the COVID-19 Genomics UK (COG-UK) Consortium | The Lighthouse Lab in Milton Keynes and Alex Alderton, Roberto Amato, Jeffrey Barrett, Sonia Goncalves, Ewan Harrison, David K. Jackson, Ian Johnston, Dominic Kwiatkowski, Cordelia Langford, John Sillitoe on behalf of the Wellcome Sanger Institute COVID-19 Surveillance Team |
| EPI_ISL_2866970 | Lighthouse Lab in Glasgow | Wellcome Sanger Institute for the COVID-19 Genomics UK (COG-UK) Consortium | Harper VanSteenhouse, Yumi Kasai, David Gray, Carol Clugston, Anna Dominiczak and Alex Alderton, Roberto Amato, Jeffrey Barrett, Sonia Goncalves, Ewan Harrison, David K. Jackson, Ian Johnston, Dominic Kwiatkowski, Cordelia Langford, John Sillitoe on behalf of the Wellcome Sanger Institute COVID-19 Surveillance Team |
| EPI_ISL_2866971 | Lighthouse Lab in Milton Keynes | Wellcome Sanger Institute for the COVID-19 Genomics UK (COG-UK) Consortium | The Lighthouse Lab in Milton Keynes and Alex Alderton, Roberto Amato, Jeffrey Barrett, Sonia Goncalves, Ewan Harrison, David K. Jackson, Ian Johnston, Dominic Kwiatkowski, Cordelia Langford, John Sillitoe on behalf of the Wellcome Sanger Institute COVID-19 Surveillance Team |
| EPI_ISL_2866972 | Lighthouse Lab in Glasgow | Wellcome Sanger Institute for the COVID-19 Genomics UK (COG-UK) Consortium | Harper VanSteenhouse, Yumi Kasai, David Gray, Carol Clugston, Anna Dominiczak and Alex Alderton, Roberto Amato, Jeffrey Barrett, Sonia Goncalves, Ewan Harrison, David K. Jackson, Ian Johnston, Dominic Kwiatkowski, Cordelia Langford, John Sillitoe on behalf of the Wellcome Sanger Institute COVID-19 Surveillance Team |
| EPI_ISL_2866973 | Lighthouse Lab in Milton Keynes | Wellcome Sanger Institute for the COVID-19 Genomics UK (COG-UK) Consortium | The Lighthouse Lab in Milton Keynes and Alex Alderton, Roberto Amato, Jeffrey Barrett, Sonia Goncalves, Ewan Harrison, David K. Jackson, Ian Johnston, Dominic Kwiatkowski, Cordelia Langford, John Sillitoe on behalf of the Wellcome Sanger Institute COVID-19 Surveillance Team |
| EPI_ISL_2866974 | Lighthouse Laboratory Plymouth | Wellcome Sanger Institute for the COVID-19 Genomics UK (COG-UK) Consortium | Lighthouse Laboratory Plymouth and Alex Alderton, Roberto Amato, Jeffrey Barrett, Sonia Goncalves, Ewan Harrison, David K. Jackson, Ian Johnston, Dominic Kwiatkowski, Cordelia Langford, John Sillitoe on behalf of the Wellcome Sanger Institute COVID-19 Surveillance Team |
| EPI_ISL_2866975 | Lighthouse Lab in Milton Keynes | Wellcome Sanger Institute for the COVID-19 Genomics UK (COG-UK) Consortium | The Lighthouse Lab in Milton Keynes and Alex Alderton, Roberto Amato, Jeffrey Barrett, Sonia Goncalves, Ewan Harrison, David K. Jackson, Ian Johnston, Dominic Kwiatkowski, Cordelia Langford, John Sillitoe on behalf of the Wellcome Sanger Institute COVID-19 Surveillance Team |
| EPI_ISL_2866976 | Lighthouse Laboratory Plymouth | Wellcome Sanger Institute for the COVID-19 Genomics UK (COG-UK) Consortium | Lighthouse Laboratory Plymouth and Alex Alderton, Roberto Amato, Jeffrey Barrett, Sonia Goncalves, Ewan Harrison, David K. Jackson, Ian Johnston, Dominic Kwiatkowski, Cordelia Langford, John Sillitoe on behalf of the Wellcome Sanger Institute COVID-19 Surveillance Team |
| EPI_ISL_2866977 | Lighthouse Lab in Glasgow | Wellcome Sanger Institute for the COVID-19 Genomics UK (COG-UK) Consortium | Harper VanSteenhouse, Yumi Kasai, David Gray, Carol Clugston, Anna Dominiczak and Alex Alderton, Roberto Amato, Jeffrey Barrett, Sonia Goncalves, Ewan Harrison, David K. Jackson, Ian Johnston, Dominic Kwiatkowski, Cordelia Langford, John Sillitoe on behalf of the Wellcome Sanger Institute COVID-19 Surveillance Team |
| EPI_ISL_2866978 | Lighthouse Lab in Milton Keynes | Wellcome Sanger Institute for the COVID-19 Genomics UK (COG-UK) Consortium | The Lighthouse Lab in Milton Keynes and Alex Alderton, Roberto Amato, Jeffrey Barrett, Sonia Goncalves, Ewan Harrison, David K. Jackson, Ian Johnston, Dominic Kwiatkowski, Cordelia Langford, John Sillitoe on behalf of the Wellcome Sanger Institute COVID-19 Surveillance Team |
| EPI_ISL_2866979 | Lighthouse Laboratory Plymouth | Wellcome Sanger Institute for the COVID-19 Genomics UK (COG-UK) Consortium | Lighthouse Laboratory Plymouth and Alex Alderton, Roberto Amato, Jeffrey Barrett, Sonia Goncalves, Ewan Harrison, David K. Jackson, Ian Johnston, Dominic Kwiatkowski, Cordelia Langford, John Sillitoe on behalf of the Wellcome Sanger Institute COVID-19 Surveillance Team |
| EPI_ISL_2866980, EPI_ISL_2866981 | Lighthouse Lab in Milton Keynes | Wellcome Sanger Institute for the COVID-19 Genomics UK (COG-UK) Consortium | The Lighthouse Lab in Milton Keynes and Alex Alderton, Roberto Amato, Jeffrey Barrett, Sonia Goncalves, Ewan Harrison, David K. Jackson, Ian Johnston, Dominic Kwiatkowski, Cordelia Langford, John Sillitoe on behalf of the Wellcome Sanger Institute COVID-19 Surveillance Team |
| EPI_ISL_2866982, EPI_ISL_2866983 | Lighthouse Lab in Glasgow | Wellcome Sanger Institute for the COVID-19 Genomics UK (COG-UK) Consortium | Harper VanSteenhouse, Yumi Kasai, David Gray, Carol Clugston, Anna Dominiczak and Alex Alderton, Roberto Amato, Jeffrey Barrett, Sonia Goncalves, Ewan Harrison, David K. Jackson, Ian Johnston, Dominic Kwiatkowski, Cordelia Langford, John Sillitoe on behalf of the Wellcome Sanger Institute COVID-19 Surveillance Team |
| EPI_ISL_2866984 | Lighthouse Lab in Milton Keynes | Wellcome Sanger Institute for the COVID-19 Genomics UK (COG-UK) Consortium | The Lighthouse Lab in Milton Keynes and Alex Alderton, Roberto Amato, Jeffrey Barrett, Sonia Goncalves, Ewan Harrison, David K. Jackson, Ian Johnston, Dominic Kwiatkowski, Cordelia Langford, John Sillitoe on behalf of the Wellcome Sanger Institute COVID-19 Surveillance Team |
| EPI_ISL_2866985 | Lighthouse Lab in Glasgow | Wellcome Sanger Institute for the COVID-19 Genomics UK (COG-UK) Consortium | Harper VanSteenhouse, Yumi Kasai, David Gray, Carol Clugston, Anna Dominiczak and Alex Alderton, Roberto Amato, Jeffrey Barrett, Sonia Goncalves, Ewan Harrison, David K. Jackson, Ian Johnston, Dominic Kwiatkowski, Cordelia Langford, John Sillitoe on behalf of the Wellcome Sanger Institute COVID-19 Surveillance Team |
| EPI_ISL_2866986 | Lighthouse Lab in Milton Keynes | Wellcome Sanger Institute for the COVID-19 Genomics UK (COG-UK) Consortium | The Lighthouse Lab in Milton Keynes and Alex Alderton, Roberto Amato, Jeffrey Barrett, Sonia Goncalves, Ewan Harrison, David K. Jackson, Ian Johnston, Dominic Kwiatkowski, Cordelia Langford, John Sillitoe on behalf of the Wellcome Sanger Institute COVID-19 Surveillance Team |
| EPI_ISL_2866987, EPI_ISL_2866988 | Lighthouse Lab in Glasgow | Wellcome Sanger Institute for the COVID-19 Genomics UK (COG-UK) Consortium | Harper VanSteenhouse, Yumi Kasai, David Gray, Carol Clugston, Anna Dominiczak and Alex Alderton, Roberto Amato, Jeffrey Barrett, Sonia Goncalves, Ewan Harrison, David K. Jackson, Ian Johnston, Dominic Kwiatkowski, Cordelia Langford, John Sillitoe on behalf of the Wellcome Sanger Institute COVID-19 Surveillance Team |
| EPI_ISL_2866989 | Lighthouse Laboratory Plymouth | Wellcome Sanger Institute for the COVID-19 Genomics UK (COG-UK) Consortium | Lighthouse Laboratory Plymouth and Alex Alderton, Roberto Amato, Jeffrey Barrett, Sonia Goncalves, Ewan Harrison, David K. Jackson, Ian Johnston, Dominic Kwiatkowski, Cordelia Langford, John Sillitoe on behalf of the Wellcome Sanger Institute COVID-19 Surveillance Team |
| EPI_ISL_2866990 | Lighthouse Lab in Milton Keynes | Wellcome Sanger Institute for the COVID-19 Genomics UK (COG-UK) Consortium | The Lighthouse Lab in Milton Keynes and Alex Alderton, Roberto Amato, Jeffrey Barrett, Sonia Goncalves, Ewan Harrison, David K. Jackson, Ian Johnston, Dominic Kwiatkowski, Cordelia Langford, John Sillitoe on behalf of the Wellcome Sanger Institute COVID-19 Surveillance Team |
| EPI_ISL_2866991, EPI_ISL_2866992, EPI_ISL_2866993 | Lighthouse Lab in Glasgow | Wellcome Sanger Institute for the COVID-19 Genomics UK (COG-UK) Consortium | Harper VanSteenhouse, Yumi Kasai, David Gray, Carol Clugston, Anna Dominiczak and Alex Alderton, Roberto Amato, Jeffrey Barrett, Sonia Goncalves, Ewan Harrison, David K. Jackson, Ian Johnston, Dominic Kwiatkowski, Cordelia Langford, John Sillitoe on behalf of the Wellcome Sanger Institute COVID-19 Surveillance Team |
| EPI_ISL_2866994 | Lighthouse Laboratory Plymouth | Wellcome Sanger Institute for the COVID-19 Genomics UK (COG-UK) Consortium | Lighthouse Laboratory Plymouth and Alex Alderton, Roberto Amato, Jeffrey Barrett, Sonia Goncalves, Ewan Harrison, David K. Jackson, Ian Johnston, Dominic Kwiatkowski, Cordelia Langford, John Sillitoe on behalf of the Wellcome Sanger Institute COVID-19 Surveillance Team |
| EPI_ISL_2866995, EPI_ISL_2866996, EPI_ISL_2866997 | Lighthouse Lab in Milton Keynes | Wellcome Sanger Institute for the COVID-19 Genomics UK (COG-UK) Consortium | The Lighthouse Lab in Milton Keynes and Alex Alderton, Roberto Amato, Jeffrey Barrett, Sonia Goncalves, Ewan Harrison, David K. Jackson, Ian Johnston, Dominic Kwiatkowski, Cordelia Langford, John Sillitoe on behalf of the Wellcome Sanger Institute COVID-19 Surveillance Team |
| EPI_ISL_2866998 | Lighthouse Lab in Glasgow | Wellcome Sanger Institute for the COVID-19 Genomics UK (COG-UK) Consortium | Harper VanSteenhouse, Yumi Kasai, David Gray, Carol Clugston, Anna Dominiczak and Alex Alderton, Roberto Amato, Jeffrey Barrett, Sonia Goncalves, Ewan Harrison, David K. Jackson, Ian Johnston, Dominic Kwiatkowski, Cordelia Langford, John Sillitoe on behalf of the Wellcome Sanger Institute COVID-19 Surveillance Team |
| EPI_ISL_2866999 | Lighthouse Laboratory Plymouth | Wellcome Sanger Institute for the COVID-19 Genomics UK (COG-UK) Consortium | Lighthouse Laboratory Plymouth and Alex Alderton, Roberto Amato, Jeffrey Barrett, Sonia Goncalves, Ewan Harrison, David K. Jackson, Ian Johnston, Dominic Kwiatkowski, Cordelia Langford, John Sillitoe on behalf of the Wellcome Sanger Institute COVID-19 Surveillance Team |
| EPI_ISL_2867000, EPI_ISL_2867001 | Lighthouse Lab in Milton Keynes | Wellcome Sanger Institute for the COVID-19 Genomics UK (COG-UK) Consortium | The Lighthouse Lab in Milton Keynes and Alex Alderton, Roberto Amato, Jeffrey Barrett, Sonia Goncalves, Ewan Harrison, David K. Jackson, Ian Johnston, Dominic Kwiatkowski, Cordelia Langford, John Sillitoe on behalf of the Wellcome Sanger Institute COVID-19 Surveillance Team |
| EPI_ISL_2867002 | Lighthouse Laboratory Plymouth | Wellcome Sanger Institute for the COVID-19 Genomics UK (COG-UK) Consortium | Lighthouse Laboratory Plymouth and Alex Alderton, Roberto Amato, Jeffrey Barrett, Sonia Goncalves, Ewan Harrison, David K. Jackson, Ian Johnston, Dominic Kwiatkowski, Cordelia Langford, John Sillitoe on behalf of the Wellcome Sanger Institute COVID-19 Surveillance Team |
| EPI_ISL_2867003 | Lighthouse Lab in Milton Keynes | Wellcome Sanger Institute for the COVID-19 Genomics UK (COG-UK) Consortium | The Lighthouse Lab in Milton Keynes and Alex Alderton, Roberto Amato, Jeffrey Barrett, Sonia Goncalves, Ewan Harrison, David K. Jackson, Ian Johnston, Dominic Kwiatkowski, Cordelia Langford, John Sillitoe on behalf of the Wellcome Sanger Institute COVID-19 Surveillance Team |
| EPI_ISL_2867004, EPI_ISL_2867005, EPI_ISL_2867006, EPI_ISL_2867007, EPI_ISL_2867008, EPI_ISL_2867009 | Lighthouse Lab in Glasgow | Wellcome Sanger Institute for the COVID-19 Genomics UK (COG-UK) Consortium | Harper VanSteenhouse, Yumi Kasai, David Gray, Carol Clugston, Anna Dominiczak and Alex Alderton, Roberto Amato, Jeffrey Barrett, Sonia Goncalves, Ewan Harrison, David K. Jackson, Ian Johnston, Dominic Kwiatkowski, Cordelia Langford, John Sillitoe on behalf of the Wellcome Sanger Institute COVID-19 Surveillance Team |
| EPI_ISL_2867010 | Lighthouse Lab in Milton Keynes | Wellcome Sanger Institute for the COVID-19 Genomics UK (COG-UK) Consortium | The Lighthouse Lab in Milton Keynes and Alex Alderton, Roberto Amato, Jeffrey Barrett, Sonia Goncalves, Ewan Harrison, David K. Jackson, Ian Johnston, Dominic Kwiatkowski, Cordelia Langford, John Sillitoe on behalf of the Wellcome Sanger Institute COVID-19 Surveillance Team |
| EPI_ISL_2867011, EPI_ISL_2867012, EPI_ISL_2867013, EPI_ISL_2867014, EPI_ISL_2867015, EPI_ISL_2867016 | Lighthouse Lab in Glasgow | Wellcome Sanger Institute for the COVID-19 Genomics UK (COG-UK) Consortium | Harper VanSteenhouse, Yumi Kasai, David Gray, Carol Clugston, Anna Dominiczak and Alex Alderton, Roberto Amato, Jeffrey Barrett, Sonia Goncalves, Ewan Harrison, David K. Jackson, Ian Johnston, Dominic Kwiatkowski, Cordelia Langford, John Sillitoe on behalf of the Wellcome Sanger Institute COVID-19 Surveillance Team |
| EPI_ISL_2867017 | Lighthouse Laboratory Plymouth | Wellcome Sanger Institute for the COVID-19 Genomics UK (COG-UK) Consortium | Lighthouse Laboratory Plymouth and Alex Alderton, Roberto Amato, Jeffrey Barrett, Sonia Goncalves, Ewan Harrison, David K. Jackson, Ian Johnston, Dominic Kwiatkowski, Cordelia Langford, John Sillitoe on behalf of the Wellcome Sanger Institute COVID-19 Surveillance Team |
| EPI_ISL_2867018 | Lighthouse Lab in Glasgow | Wellcome Sanger Institute for the COVID-19 Genomics UK (COG-UK) Consortium | Harper VanSteenhouse, Yumi Kasai, David Gray, Carol Clugston, Anna Dominiczak and Alex Alderton, Roberto Amato, Jeffrey Barrett, Sonia Goncalves, Ewan Harrison, David K. Jackson, Ian Johnston, Dominic Kwiatkowski, Cordelia Langford, John Sillitoe on behalf of the Wellcome Sanger Institute COVID-19 Surveillance Team |

| | | | |
|---|---|---|---|
| EPI_ISL_2867019, EPI_ISL_2867020 | Lighthouse Laboratory Plymouth | Wellcome Sanger Institute for the COVID-19 Genomics UK (COG-UK) Consortium | Lighthouse Laboratory Plymouth and Alex Alderton, Roberto Amato, Jeffrey Barrett, Sonia Goncalves, Ewan Harrison, David K. Jackson, Ian Johnston, Dominic Kwiatkowski, Cordelia Langford, John Sillitoe on behalf of the Wellcome Sanger Institute COVID-19 Surveillance Team |
| EPI_ISL_2867021, EPI_ISL_2867022 | Lighthouse Lab in Glasgow | Wellcome Sanger Institute for the COVID-19 Genomics UK (COG-UK) Consortium | Harper VanSteenhouse, Yumi Kasai, David Gray, Carol Clugston, Anna Dominiczak and Alex Alderton, Roberto Amato, Jeffrey Barrett, Sonia Goncalves, Ewan Harrison, David K. Jackson, Ian Johnston, Dominic Kwiatkowski, Cordelia Langford, John Sillitoe on behalf of the Wellcome Sanger Institute COVID-19 Surveillance Team |
| EPI_ISL_2867023 | Lighthouse Lab in Milton Keynes | Wellcome Sanger Institute for the COVID-19 Genomics UK (COG-UK) Consortium | The Lighthouse Lab in Milton Keynes and Alex Alderton, Roberto Amato, Jeffrey Barrett, Sonia Goncalves, Ewan Harrison, David K. Jackson, Ian Johnston, Dominic Kwiatkowski, Cordelia Langford, John Sillitoe on behalf of the Wellcome Sanger Institute COVID-19 Surveillance Team |
| EPI_ISL_2867024, EPI_ISL_2867025 | Lighthouse Lab in Glasgow | Wellcome Sanger Institute for the COVID-19 Genomics UK (COG-UK) Consortium | Harper VanSteenhouse, Yumi Kasai, David Gray, Carol Clugston, Anna Dominiczak and Alex Alderton, Roberto Amato, Jeffrey Barrett, Sonia Goncalves, Ewan Harrison, David K. Jackson, Ian Johnston, Dominic Kwiatkowski, Cordelia Langford, John Sillitoe on behalf of the Wellcome Sanger Institute COVID-19 Surveillance Team |
| EPI_ISL_2867026, EPI_ISL_2867027 | Lighthouse Lab in Milton Keynes | Wellcome Sanger Institute for the COVID-19 Genomics UK (COG-UK) Consortium | The Lighthouse Lab in Milton Keynes and Alex Alderton, Roberto Amato, Jeffrey Barrett, Sonia Goncalves, Ewan Harrison, David K. Jackson, Ian Johnston, Dominic Kwiatkowski, Cordelia Langford, John Sillitoe on behalf of the Wellcome Sanger Institute COVID-19 Surveillance Team |
| EPI_ISL_2867028 | Lighthouse Lab in Glasgow | Wellcome Sanger Institute for the COVID-19 Genomics UK (COG-UK) Consortium | Harper VanSteenhouse, Yumi Kasai, David Gray, Carol Clugston, Anna Dominiczak and Alex Alderton, Roberto Amato, Jeffrey Barrett, Sonia Goncalves, Ewan Harrison, David K. Jackson, Ian Johnston, Dominic Kwiatkowski, Cordelia Langford, John Sillitoe on behalf of the Wellcome Sanger Institute COVID-19 Surveillance Team |
| EPI_ISL_2867029 | Lighthouse Laboratory Plymouth | Wellcome Sanger Institute for the COVID-19 Genomics UK (COG-UK) Consortium | Lighthouse Laboratory Plymouth and Alex Alderton, Roberto Amato, Jeffrey Barrett, Sonia Goncalves, Ewan Harrison, David K. Jackson, Ian Johnston, Dominic Kwiatkowski, Cordelia Langford, John Sillitoe on behalf of the Wellcome Sanger Institute COVID-19 Surveillance Team |
| EPI_ISL_2867030 | Berkshire and Surrey Pathology Services Lighthouse Laboratory | Wellcome Sanger Institute for the COVID-19 Genomics UK (COG-UK) Consortium | Berkshire and Surrey Pathology Services Lighthouse Laboratory and Alex Alderton, Roberto Amato, Jeffrey Barrett, Sonia Goncalves, Ewan Harrison, David K. Jackson, Ian Johnston, Dominic Kwiatkowski, Cordelia Langford, John Sillitoe on behalf of the Wellcome Sanger Institute COVID-19 Surveillance Team |
| EPI_ISL_2867031 | Lighthouse Lab in Alderley Park | Wellcome Sanger Institute for the COVID-19 Genomics UK (COG-UK) Consortium | Jacquelyn Wynn, Mairead Hyland, The Lighthouse Lab in Alderley Park and Alex Alderton, Roberto Amato, Jeffrey Barrett, Sonia Goncalves, Ewan Harrison, David K. Jackson, Ian Johnston, Dominic Kwiatkowski, Cordelia Langford, John Sillitoe on behalf of the Wellcome Sanger Institute COVID-19 Surveillance Team |
| EPI_ISL_2867032 | Berkshire and Surrey Pathology Services Lighthouse Laboratory | Wellcome Sanger Institute for the COVID-19 Genomics UK (COG-UK) Consortium | Berkshire and Surrey Pathology Services Lighthouse Laboratory and Alex Alderton, Roberto Amato, Jeffrey Barrett, Sonia Goncalves, Ewan Harrison, David K. Jackson, Ian Johnston, Dominic Kwiatkowski, Cordelia Langford, John Sillitoe on behalf of the Wellcome Sanger Institute COVID-19 Surveillance Team |
| EPI_ISL_2867033 | Lighthouse Lab in Alderley Park | Wellcome Sanger Institute for the COVID-19 Genomics UK (COG-UK) Consortium | Jacquelyn Wynn, Mairead Hyland, The Lighthouse Lab in Alderley Park and Alex Alderton, Roberto Amato, Jeffrey Barrett, Sonia Goncalves, Ewan Harrison, David K. Jackson, Ian Johnston, Dominic Kwiatkowski, Cordelia Langford, John Sillitoe on behalf of the Wellcome Sanger Institute COVID-19 Surveillance Team |
| EPI_ISL_2867034, EPI_ISL_2867035 | Berkshire and Surrey Pathology Services Lighthouse Laboratory | Wellcome Sanger Institute for the COVID-19 Genomics UK (COG-UK) Consortium | Berkshire and Surrey Pathology Services Lighthouse Laboratory and Alex Alderton, Roberto Amato, Jeffrey Barrett, Sonia Goncalves, Ewan Harrison, David K. Jackson, Ian Johnston, Dominic Kwiatkowski, Cordelia Langford, John Sillitoe on behalf of the Wellcome Sanger Institute COVID-19 Surveillance Team |
| EPI_ISL_2867036 | Lighthouse Laboratory Plymouth | Wellcome Sanger Institute for the COVID-19 Genomics UK (COG-UK) Consortium | Lighthouse Laboratory Plymouth and Alex Alderton, Roberto Amato, Jeffrey Barrett, Sonia Goncalves, Ewan Harrison, David K. Jackson, Ian Johnston, Dominic Kwiatkowski, Cordelia Langford, John Sillitoe on behalf of the Wellcome Sanger Institute COVID-19 Surveillance Team |
| EPI_ISL_2867037 | Berkshire and Surrey Pathology Services Lighthouse Laboratory | Wellcome Sanger Institute for the COVID-19 Genomics UK (COG-UK) Consortium | Berkshire and Surrey Pathology Services Lighthouse Laboratory and Alex Alderton, Roberto Amato, Jeffrey Barrett, Sonia Goncalves, Ewan Harrison, David K. Jackson, Ian Johnston, Dominic Kwiatkowski, Cordelia Langford, John Sillitoe on behalf of the Wellcome Sanger Institute COVID-19 Surveillance Team |
| EPI_ISL_2867038 | Lighthouse Lab in Alderley Park | Wellcome Sanger Institute for the COVID-19 Genomics UK (COG-UK) Consortium | Jacquelyn Wynn, Mairead Hyland, The Lighthouse Lab in Alderley Park and Alex Alderton, Roberto Amato, Jeffrey Barrett, Sonia Goncalves, Ewan Harrison, David K. Jackson, Ian Johnston, Dominic Kwiatkowski, Cordelia Langford, John Sillitoe on behalf of the Wellcome Sanger Institute COVID-19 Surveillance Team |
| EPI_ISL_2867039 | Berkshire and Surrey Pathology Services Lighthouse Laboratory | Wellcome Sanger Institute for the COVID-19 Genomics UK (COG-UK) Consortium | Berkshire and Surrey Pathology Services Lighthouse Laboratory and Alex Alderton, Roberto Amato, Jeffrey Barrett, Sonia Goncalves, Ewan Harrison, David K. Jackson, Ian Johnston, Dominic Kwiatkowski, Cordelia Langford, John Sillitoe on behalf of the Wellcome Sanger Institute COVID-19 Surveillance Team |
| EPI_ISL_2867040, EPI_ISL_2867041, EPI_ISL_2867042 | Lighthouse Lab in Alderley Park | Wellcome Sanger Institute for the COVID-19 Genomics UK (COG-UK) Consortium | Jacquelyn Wynn, Mairead Hyland, The Lighthouse Lab in Alderley Park and Alex Alderton, Roberto Amato, Jeffrey Barrett, Sonia Goncalves, Ewan Harrison, David K. Jackson, Ian Johnston, Dominic Kwiatkowski, Cordelia Langford, John Sillitoe on behalf of the Wellcome Sanger Institute COVID-19 Surveillance Team |
| EPI_ISL_2867043 | Berkshire and Surrey Pathology Services Lighthouse Laboratory | Wellcome Sanger Institute for the COVID-19 Genomics UK (COG-UK) Consortium | Berkshire and Surrey Pathology Services Lighthouse Laboratory and Alex Alderton, Roberto Amato, Jeffrey Barrett, Sonia Goncalves, Ewan Harrison, David K. Jackson, Ian Johnston, Dominic Kwiatkowski, Cordelia Langford, John Sillitoe on behalf of the Wellcome Sanger Institute COVID-19 Surveillance Team |
| EPI_ISL_2867044 | Lighthouse Lab in Alderley Park | Wellcome Sanger Institute for the COVID-19 Genomics UK (COG-UK) Consortium | Jacquelyn Wynn, Mairead Hyland, The Lighthouse Lab in Alderley Park and Alex Alderton, Roberto Amato, Jeffrey Barrett, Sonia Goncalves, Ewan Harrison, David K. Jackson, Ian Johnston, Dominic Kwiatkowski, Cordelia Langford, John Sillitoe on behalf of the Wellcome Sanger Institute COVID-19 Surveillance Team |
| EPI_ISL_2867045, EPI_ISL_2867046, EPI_ISL_2867047, EPI_ISL_2867048 | Berkshire and Surrey Pathology Services Lighthouse Laboratory | Wellcome Sanger Institute for the COVID-19 Genomics UK (COG-UK) Consortium | Berkshire and Surrey Pathology Services Lighthouse Laboratory and Alex Alderton, Roberto Amato, Jeffrey Barrett, Sonia Goncalves, Ewan Harrison, David K. Jackson, Ian Johnston, Dominic Kwiatkowski, Cordelia Langford, John Sillitoe on behalf of the Wellcome Sanger Institute COVID-19 Surveillance Team |
| EPI_ISL_2867049 | Lighthouse Lab in Alderley Park | Wellcome Sanger Institute for the COVID-19 Genomics UK (COG-UK) Consortium | Jacquelyn Wynn, Mairead Hyland, The Lighthouse Lab in Alderley Park and Alex Alderton, Roberto Amato, Jeffrey Barrett, Sonia Goncalves, Ewan Harrison, David K. Jackson, Ian Johnston, Dominic Kwiatkowski, Cordelia Langford, John Sillitoe on behalf of the Wellcome Sanger Institute COVID-19 Surveillance Team |
| EPI_ISL_2867050 | Lighthouse Laboratory Plymouth | Wellcome Sanger Institute for the COVID-19 Genomics UK (COG-UK) Consortium | Lighthouse Laboratory Plymouth and Alex Alderton, Roberto Amato, Jeffrey Barrett, Sonia Goncalves, Ewan Harrison, David K. Jackson, Ian Johnston, Dominic Kwiatkowski, Cordelia Langford, John Sillitoe on behalf of the Wellcome Sanger Institute COVID-19 Surveillance Team |
| EPI_ISL_2867051 | Berkshire and Surrey Pathology Services Lighthouse Laboratory | Wellcome Sanger Institute for the COVID-19 Genomics UK (COG-UK) Consortium | Berkshire and Surrey Pathology Services Lighthouse Laboratory and Alex Alderton, Roberto Amato, Jeffrey Barrett, Sonia Goncalves, Ewan Harrison, David K. Jackson, Ian Johnston, Dominic Kwiatkowski, Cordelia Langford, John Sillitoe on behalf of the Wellcome Sanger Institute COVID-19 Surveillance Team |
| EPI_ISL_2867052 | Lighthouse Lab in Alderley Park | Wellcome Sanger Institute for the COVID-19 Genomics UK (COG-UK) Consortium | Jacquelyn Wynn, Mairead Hyland, The Lighthouse Lab in Alderley Park and Alex Alderton, Roberto Amato, Jeffrey Barrett, Sonia Goncalves, Ewan Harrison, David K. Jackson, Ian Johnston, Dominic Kwiatkowski, Cordelia Langford, John Sillitoe on behalf of the Wellcome Sanger Institute COVID-19 Surveillance Team |
| EPI_ISL_2867053 | Lighthouse Laboratory Plymouth | Wellcome Sanger Institute for the COVID-19 Genomics UK (COG-UK) Consortium | Lighthouse Laboratory Plymouth and Alex Alderton, Roberto Amato, Jeffrey Barrett, Sonia Goncalves, Ewan Harrison, David K. Jackson, Ian Johnston, Dominic Kwiatkowski, Cordelia Langford, John Sillitoe on behalf of the Wellcome Sanger Institute COVID-19 Surveillance Team |
| EPI_ISL_2867054 | Lighthouse Lab in Alderley Park | Wellcome Sanger Institute for the COVID-19 Genomics UK (COG-UK) Consortium | Jacquelyn Wynn, Mairead Hyland, The Lighthouse Lab in Alderley Park and Alex Alderton, Roberto Amato, Jeffrey Barrett, Sonia Goncalves, Ewan Harrison, David K. Jackson, Ian Johnston, Dominic Kwiatkowski, Cordelia Langford, John Sillitoe on behalf of the Wellcome Sanger Institute COVID-19 Surveillance Team |
| EPI_ISL_2867055, EPI_ISL_2867056 | Berkshire and Surrey Pathology Services Lighthouse Laboratory | Wellcome Sanger Institute for the COVID-19 Genomics UK (COG-UK) Consortium | Berkshire and Surrey Pathology Services Lighthouse Laboratory and Alex Alderton, Roberto Amato, Jeffrey Barrett, Sonia Goncalves, Ewan Harrison, David K. Jackson, Ian Johnston, Dominic Kwiatkowski, Cordelia Langford, John Sillitoe on behalf of the Wellcome Sanger Institute COVID-19 Surveillance Team |
| EPI_ISL_2867057, EPI_ISL_2867058, EPI_ISL_2867059, EPI_ISL_2867060, EPI_ISL_2867061 | Lighthouse Laboratory Plymouth | Wellcome Sanger Institute for the COVID-19 Genomics UK (COG-UK) Consortium | Lighthouse Laboratory Plymouth and Alex Alderton, Roberto Amato, Jeffrey Barrett, Sonia Goncalves, Ewan Harrison, David K. Jackson, Ian Johnston, Dominic Kwiatkowski, Cordelia Langford, John Sillitoe on behalf of the Wellcome Sanger Institute COVID-19 Surveillance Team |
| EPI_ISL_2867062, EPI_ISL_2867063 | Lighthouse Lab in Alderley Park | Wellcome Sanger Institute for the COVID-19 Genomics UK (COG-UK) Consortium | Jacquelyn Wynn, Mairead Hyland, The Lighthouse Lab in Alderley Park and Alex Alderton, Roberto Amato, Jeffrey Barrett, Sonia Goncalves, Ewan Harrison, David K. Jackson, Ian Johnston, Dominic Kwiatkowski, Cordelia Langford, John Sillitoe on behalf of the Wellcome Sanger Institute COVID-19 Surveillance Team |
| EPI_ISL_2867064, EPI_ISL_2867065, EPI_ISL_2867066, EPI_ISL_2867067 | Berkshire and Surrey Pathology Services Lighthouse Laboratory | Wellcome Sanger Institute for the COVID-19 Genomics UK (COG-UK) Consortium | Berkshire and Surrey Pathology Services Lighthouse Laboratory and Alex Alderton, Roberto Amato, Jeffrey Barrett, Sonia Goncalves, Ewan Harrison, David K. Jackson, Ian Johnston, Dominic Kwiatkowski, Cordelia Langford, John Sillitoe on behalf of the Wellcome Sanger Institute COVID-19 Surveillance Team |
| EPI_ISL_2867068 | Lighthouse Lab in Alderley Park | Wellcome Sanger Institute for the COVID-19 Genomics UK (COG-UK) Consortium | Jacquelyn Wynn, Mairead Hyland, The Lighthouse Lab in Alderley Park and Alex Alderton, Roberto Amato, Jeffrey Barrett, Sonia Goncalves, Ewan Harrison, David K. Jackson, Ian Johnston, Dominic Kwiatkowski, Cordelia Langford, John Sillitoe on behalf of the Wellcome Sanger Institute COVID-19 Surveillance Team |
| EPI_ISL_2867069, EPI_ISL_2867070 | Berkshire and Surrey Pathology Services Lighthouse Laboratory | Wellcome Sanger Institute for the COVID-19 Genomics UK (COG-UK) Consortium | Berkshire and Surrey Pathology Services Lighthouse Laboratory and Alex Alderton, Roberto Amato, Jeffrey Barrett, Sonia Goncalves, Ewan Harrison, David K. Jackson, Ian Johnston, Dominic Kwiatkowski, Cordelia Langford, John Sillitoe on behalf of the Wellcome Sanger Institute COVID-19 Surveillance Team |
| EPI_ISL_2867071 | Lighthouse Laboratory Plymouth | Wellcome Sanger Institute for the COVID-19 Genomics UK (COG-UK) Consortium | Lighthouse Laboratory Plymouth and Alex Alderton, Roberto Amato, Jeffrey Barrett, Sonia Goncalves, Ewan Harrison, David K. Jackson, Ian Johnston, Dominic Kwiatkowski, Cordelia Langford, John Sillitoe on behalf of the Wellcome Sanger Institute COVID-19 Surveillance Team |
| EPI_ISL_2867072 | Berkshire and Surrey Pathology Services Lighthouse Laboratory | Wellcome Sanger Institute for the COVID-19 Genomics UK (COG-UK) Consortium | Berkshire and Surrey Pathology Services Lighthouse Laboratory and Alex Alderton, Roberto Amato, Jeffrey Barrett, Sonia Goncalves, Ewan Harrison, David K. Jackson, Ian Johnston, Dominic Kwiatkowski, Cordelia Langford, John Sillitoe on behalf of the Wellcome Sanger Institute COVID-19 Surveillance Team |
| EPI_ISL_2867073 | Lighthouse Lab in Alderley Park | Wellcome Sanger Institute for the COVID-19 Genomics UK (COG-UK) Consortium | Jacquelyn Wynn, Mairead Hyland, The Lighthouse Lab in Alderley Park and Alex Alderton, Roberto Amato, Jeffrey Barrett, Sonia Goncalves, Ewan Harrison, David K. Jackson, Ian Johnston, Dominic Kwiatkowski, Cordelia Langford, John Sillitoe on behalf of the Wellcome Sanger Institute COVID-19 Surveillance Team |
| EPI_ISL_2867074, EPI_ISL_2867075 | Berkshire and Surrey Pathology Services Lighthouse Laboratory | Wellcome Sanger Institute for the COVID-19 Genomics UK (COG-UK) Consortium | Berkshire and Surrey Pathology Services Lighthouse Laboratory and Alex Alderton, Roberto Amato, Jeffrey Barrett, Sonia Goncalves, Ewan Harrison, David K. Jackson, Ian Johnston, Dominic Kwiatkowski, Cordelia Langford, John Sillitoe on behalf of the Wellcome Sanger Institute COVID-19 Surveillance Team |
| EPI_ISL_2867076, EPI_ISL_2867077 | Lighthouse Lab in Alderley Park | Wellcome Sanger Institute for the COVID-19 Genomics UK (COG-UK) Consortium | Jacquelyn Wynn, Mairead Hyland, The Lighthouse Lab in Alderley Park and Alex Alderton, Roberto Amato, Jeffrey Barrett, Sonia Goncalves, Ewan Harrison, David K. Jackson, Ian Johnston, Dominic Kwiatkowski, Cordelia Langford, John Sillitoe on behalf of the Wellcome Sanger Institute COVID-19 Surveillance Team |
| EPI_ISL_2867078 | Berkshire and Surrey Pathology Services Lighthouse Laboratory | Wellcome Sanger Institute for the COVID-19 Genomics UK (COG-UK) Consortium | Berkshire and Surrey Pathology Services Lighthouse Laboratory and Alex Alderton, Roberto Amato, Jeffrey Barrett, Sonia Goncalves, Ewan Harrison, David K. Jackson, Ian Johnston, Dominic Kwiatkowski, Cordelia Langford, John Sillitoe on behalf of the Wellcome Sanger Institute COVID-19 Surveillance Team |
| EPI_ISL_2867079 | Lighthouse Laboratory Plymouth | Wellcome Sanger Institute for the COVID-19 Genomics UK (COG-UK) Consortium | Lighthouse Laboratory Plymouth and Alex Alderton, Roberto Amato, Jeffrey Barrett, Sonia Goncalves, Ewan Harrison, David K. Jackson, Ian Johnston, Dominic Kwiatkowski, Cordelia Langford, John Sillitoe on behalf of the Wellcome Sanger Institute COVID-19 Surveillance Team |
| EPI_ISL_2867080 | Berkshire and Surrey Pathology Services Lighthouse Laboratory | Wellcome Sanger Institute for the COVID-19 Genomics UK (COG-UK) Consortium | Berkshire and Surrey Pathology Services Lighthouse Laboratory and Alex Alderton, Roberto Amato, Jeffrey Barrett, Sonia Goncalves, Ewan Harrison, David K. Jackson, Ian Johnston, Dominic Kwiatkowski, Cordelia Langford, John Sillitoe on behalf of the Wellcome Sanger Institute COVID-19 Surveillance Team |
| EPI_ISL_2867081, EPI_ISL_2867082 | Lighthouse Lab in Alderley Park | Wellcome Sanger Institute for the COVID-19 Genomics UK (COG-UK) Consortium | Jacquelyn Wynn, Mairead Hyland, The Lighthouse Lab in Alderley Park and Alex Alderton, Roberto Amato, Jeffrey Barrett, Sonia Goncalves, Ewan Harrison, David K. Jackson, Ian Johnston, Dominic Kwiatkowski, Cordelia Langford, John Sillitoe on behalf of the Wellcome Sanger Institute COVID-19 Surveillance Team |
| EPI_ISL_2867083, | Berkshire and Surrey Pathology Services Lighthouse Laboratory | Wellcome Sanger Institute for the COVID-19 Genomics UK (COG-UK) | Berkshire and Surrey Pathology Services Lighthouse Laboratory and Alex Alderton, Roberto Amato, Jeffrey Barrett, Sonia Goncalves, Ewan Harrison, David K. Jackson, Ian Johnston, Dominic Kwiatkowski, Cordelia Langford, John Sillitoe on behalf |

| | | | |
|---|---|---|---|
| EPI_ISL_2867084 | | Consortium | of the Wellcome Sanger Institute COVID-19 Surveillance Team |
| EPI_ISL_2867085 | Lighthouse Laboratory Plymouth | Wellcome Sanger Institute for the COVID-19 Genomics UK (COG-UK) Consortium | Lighthouse Laboratory Plymouth and Alex Alderton, Roberto Amato, Jeffrey Barrett, Sonia Goncalves, Ewan Harrison, David K. Jackson, Ian Johnston, Dominic Kwiatkowski, Cordelia Langford, John Sillitoe on behalf of the Wellcome Sanger Institute COVID-19 Surveillance Team |
| EPI_ISL_2867086 | Lighthouse Lab in Alderley Park | Wellcome Sanger Institute for the COVID-19 Genomics UK (COG-UK) Consortium | Jacquelyn Wynn, Mairead Hyland, The Lighthouse Lab in Alderley Park and Alex Alderton, Roberto Amato, Jeffrey Barrett, Sonia Goncalves, Ewan Harrison, David K. Jackson, Ian Johnston, Dominic Kwiatkowski, Cordelia Langford, John Sillitoe on behalf of the Wellcome Sanger Institute COVID-19 Surveillance Team |
| EPI_ISL_2867087, EPI_ISL_2867088 | Lighthouse Laboratory Plymouth | Wellcome Sanger Institute for the COVID-19 Genomics UK (COG-UK) Consortium | Lighthouse Laboratory Plymouth and Alex Alderton, Roberto Amato, Jeffrey Barrett, Sonia Goncalves, Ewan Harrison, David K. Jackson, Ian Johnston, Dominic Kwiatkowski, Cordelia Langford, John Sillitoe on behalf of the Wellcome Sanger Institute COVID-19 Surveillance Team |
| EPI_ISL_2867089, EPI_ISL_2867090 | Berkshire and Surrey Pathology Services Lighthouse Laboratory | Wellcome Sanger Institute for the COVID-19 Genomics UK (COG-UK) Consortium | Berkshire and Surrey Pathology Services Lighthouse Laboratory and Alex Alderton, Roberto Amato, Jeffrey Barrett, Sonia Goncalves, Ewan Harrison, David K. Jackson, Ian Johnston, Dominic Kwiatkowski, Cordelia Langford, John Sillitoe on behalf of the Wellcome Sanger Institute COVID-19 Surveillance Team |
| EPI_ISL_2867091 | Lighthouse Laboratory Plymouth | Wellcome Sanger Institute for the COVID-19 Genomics UK (COG-UK) Consortium | Lighthouse Laboratory Plymouth and Alex Alderton, Roberto Amato, Jeffrey Barrett, Sonia Goncalves, Ewan Harrison, David K. Jackson, Ian Johnston, Dominic Kwiatkowski, Cordelia Langford, John Sillitoe on behalf of the Wellcome Sanger Institute COVID-19 Surveillance Team |
| EPI_ISL_2867092, EPI_ISL_2867093 | Berkshire and Surrey Pathology Services Lighthouse Laboratory | Wellcome Sanger Institute for the COVID-19 Genomics UK (COG-UK) Consortium | Berkshire and Surrey Pathology Services Lighthouse Laboratory and Alex Alderton, Roberto Amato, Jeffrey Barrett, Sonia Goncalves, Ewan Harrison, David K. Jackson, Ian Johnston, Dominic Kwiatkowski, Cordelia Langford, John Sillitoe on behalf of the Wellcome Sanger Institute COVID-19 Surveillance Team |
| EPI_ISL_2867094 | Lighthouse Laboratory Plymouth | Wellcome Sanger Institute for the COVID-19 Genomics UK (COG-UK) Consortium | Lighthouse Laboratory Plymouth and Alex Alderton, Roberto Amato, Jeffrey Barrett, Sonia Goncalves, Ewan Harrison, David K. Jackson, Ian Johnston, Dominic Kwiatkowski, Cordelia Langford, John Sillitoe on behalf of the Wellcome Sanger Institute COVID-19 Surveillance Team |
| EPI_ISL_2867095, EPI_ISL_2867096 | Berkshire and Surrey Pathology Services Lighthouse Laboratory | Wellcome Sanger Institute for the COVID-19 Genomics UK (COG-UK) Consortium | Berkshire and Surrey Pathology Services Lighthouse Laboratory and Alex Alderton, Roberto Amato, Jeffrey Barrett, Sonia Goncalves, Ewan Harrison, David K. Jackson, Ian Johnston, Dominic Kwiatkowski, Cordelia Langford, John Sillitoe on behalf of the Wellcome Sanger Institute COVID-19 Surveillance Team |
| EPI_ISL_2867097 | Lighthouse Lab in Alderley Park | Wellcome Sanger Institute for the COVID-19 Genomics UK (COG-UK) Consortium | Jacquelyn Wynn, Mairead Hyland, The Lighthouse Lab in Alderley Park and Alex Alderton, Roberto Amato, Jeffrey Barrett, Sonia Goncalves, Ewan Harrison, David K. Jackson, Ian Johnston, Dominic Kwiatkowski, Cordelia Langford, John Sillitoe on behalf of the Wellcome Sanger Institute COVID-19 Surveillance Team |
| EPI_ISL_2867098, EPI_ISL_2867099 | Berkshire and Surrey Pathology Services Lighthouse Laboratory | Wellcome Sanger Institute for the COVID-19 Genomics UK (COG-UK) Consortium | Berkshire and Surrey Pathology Services Lighthouse Laboratory and Alex Alderton, Roberto Amato, Jeffrey Barrett, Sonia Goncalves, Ewan Harrison, David K. Jackson, Ian Johnston, Dominic Kwiatkowski, Cordelia Langford, John Sillitoe on behalf of the Wellcome Sanger Institute COVID-19 Surveillance Team |
| EPI_ISL_2867100 | Lighthouse Lab in Alderley Park | Wellcome Sanger Institute for the COVID-19 Genomics UK (COG-UK) Consortium | Jacquelyn Wynn, Mairead Hyland, The Lighthouse Lab in Alderley Park and Alex Alderton, Roberto Amato, Jeffrey Barrett, Sonia Goncalves, Ewan Harrison, David K. Jackson, Ian Johnston, Dominic Kwiatkowski, Cordelia Langford, John Sillitoe on behalf of the Wellcome Sanger Institute COVID-19 Surveillance Team |
| EPI_ISL_2867101, EPI_ISL_2867102 | Berkshire and Surrey Pathology Services Lighthouse Laboratory | Wellcome Sanger Institute for the COVID-19 Genomics UK (COG-UK) Consortium | Berkshire and Surrey Pathology Services Lighthouse Laboratory and Alex Alderton, Roberto Amato, Jeffrey Barrett, Sonia Goncalves, Ewan Harrison, David K. Jackson, Ian Johnston, Dominic Kwiatkowski, Cordelia Langford, John Sillitoe on behalf of the Wellcome Sanger Institute COVID-19 Surveillance Team |
| EPI_ISL_2867103 | Lighthouse Laboratory Plymouth | Wellcome Sanger Institute for the COVID-19 Genomics UK (COG-UK) Consortium | Lighthouse Laboratory Plymouth and Alex Alderton, Roberto Amato, Jeffrey Barrett, Sonia Goncalves, Ewan Harrison, David K. Jackson, Ian Johnston, Dominic Kwiatkowski, Cordelia Langford, John Sillitoe on behalf of the Wellcome Sanger Institute COVID-19 Surveillance Team |
| EPI_ISL_2867104, EPI_ISL_2867105 | Berkshire and Surrey Pathology Services Lighthouse Laboratory | Wellcome Sanger Institute for the COVID-19 Genomics UK (COG-UK) Consortium | Berkshire and Surrey Pathology Services Lighthouse Laboratory and Alex Alderton, Roberto Amato, Jeffrey Barrett, Sonia Goncalves, Ewan Harrison, David K. Jackson, Ian Johnston, Dominic Kwiatkowski, Cordelia Langford, John Sillitoe on behalf of the Wellcome Sanger Institute COVID-19 Surveillance Team |
| EPI_ISL_2867106 | Lighthouse Lab in Alderley Park | Wellcome Sanger Institute for the COVID-19 Genomics UK (COG-UK) Consortium | Jacquelyn Wynn, Mairead Hyland, The Lighthouse Lab in Alderley Park and Alex Alderton, Roberto Amato, Jeffrey Barrett, Sonia Goncalves, Ewan Harrison, David K. Jackson, Ian Johnston, Dominic Kwiatkowski, Cordelia Langford, John Sillitoe on behalf of the Wellcome Sanger Institute COVID-19 Surveillance Team |
| EPI_ISL_2867107 | Berkshire and Surrey Pathology Services Lighthouse Laboratory | Wellcome Sanger Institute for the COVID-19 Genomics UK (COG-UK) Consortium | Berkshire and Surrey Pathology Services Lighthouse Laboratory and Alex Alderton, Roberto Amato, Jeffrey Barrett, Sonia Goncalves, Ewan Harrison, David K. Jackson, Ian Johnston, Dominic Kwiatkowski, Cordelia Langford, John Sillitoe on behalf of the Wellcome Sanger Institute COVID-19 Surveillance Team |
| EPI_ISL_2867108 | Lighthouse Lab in Alderley Park | Wellcome Sanger Institute for the COVID-19 Genomics UK (COG-UK) Consortium | Jacquelyn Wynn, Mairead Hyland, The Lighthouse Lab in Alderley Park and Alex Alderton, Roberto Amato, Jeffrey Barrett, Sonia Goncalves, Ewan Harrison, David K. Jackson, Ian Johnston, Dominic Kwiatkowski, Cordelia Langford, John Sillitoe on behalf of the Wellcome Sanger Institute COVID-19 Surveillance Team |
| EPI_ISL_2867109 | Berkshire and Surrey Pathology Services Lighthouse Laboratory | Wellcome Sanger Institute for the COVID-19 Genomics UK (COG-UK) Consortium | Berkshire and Surrey Pathology Services Lighthouse Laboratory and Alex Alderton, Roberto Amato, Jeffrey Barrett, Sonia Goncalves, Ewan Harrison, David K. Jackson, Ian Johnston, Dominic Kwiatkowski, Cordelia Langford, John Sillitoe on behalf of the Wellcome Sanger Institute COVID-19 Surveillance Team |
| EPI_ISL_2867110, EPI_ISL_2867111, EPI_ISL_2867112, EPI_ISL_2867113 | Lighthouse Laboratory Plymouth | Wellcome Sanger Institute for the COVID-19 Genomics UK (COG-UK) Consortium | Lighthouse Laboratory Plymouth and Alex Alderton, Roberto Amato, Jeffrey Barrett, Sonia Goncalves, Ewan Harrison, David K. Jackson, Ian Johnston, Dominic Kwiatkowski, Cordelia Langford, John Sillitoe on behalf of the Wellcome Sanger Institute COVID-19 Surveillance Team |
| EPI_ISL_2867114, EPI_ISL_2867115 | Berkshire and Surrey Pathology Services Lighthouse Laboratory | Wellcome Sanger Institute for the COVID-19 Genomics UK (COG-UK) Consortium | Berkshire and Surrey Pathology Services Lighthouse Laboratory and Alex Alderton, Roberto Amato, Jeffrey Barrett, Sonia Goncalves, Ewan Harrison, David K. Jackson, Ian Johnston, Dominic Kwiatkowski, Cordelia Langford, John Sillitoe on behalf of the Wellcome Sanger Institute COVID-19 Surveillance Team |
| EPI_ISL_2867116 | Lighthouse Lab in Alderley Park | Wellcome Sanger Institute for the COVID-19 Genomics UK (COG-UK) Consortium | Jacquelyn Wynn, Mairead Hyland, The Lighthouse Lab in Alderley Park and Alex Alderton, Roberto Amato, Jeffrey Barrett, Sonia Goncalves, Ewan Harrison, David K. Jackson, Ian Johnston, Dominic Kwiatkowski, Cordelia Langford, John Sillitoe on behalf of the Wellcome Sanger Institute COVID-19 Surveillance Team |
| EPI_ISL_2867117, EPI_ISL_2867118, EPI_ISL_2867119 | Berkshire and Surrey Pathology Services Lighthouse Laboratory | Wellcome Sanger Institute for the COVID-19 Genomics UK (COG-UK) Consortium | Berkshire and Surrey Pathology Services Lighthouse Laboratory and Alex Alderton, Roberto Amato, Jeffrey Barrett, Sonia Goncalves, Ewan Harrison, David K. Jackson, Ian Johnston, Dominic Kwiatkowski, Cordelia Langford, John Sillitoe on behalf of the Wellcome Sanger Institute COVID-19 Surveillance Team |
| EPI_ISL_2867120 | Lighthouse Lab in Alderley Park | Wellcome Sanger Institute for the COVID-19 Genomics UK (COG-UK) Consortium | Jacquelyn Wynn, Mairead Hyland, The Lighthouse Lab in Alderley Park and Alex Alderton, Roberto Amato, Jeffrey Barrett, Sonia Goncalves, Ewan Harrison, David K. Jackson, Ian Johnston, Dominic Kwiatkowski, Cordelia Langford, John Sillitoe on behalf of the Wellcome Sanger Institute COVID-19 Surveillance Team |
| EPI_ISL_2867121 | Berkshire and Surrey Pathology Services Lighthouse Laboratory | Wellcome Sanger Institute for the COVID-19 Genomics UK (COG-UK) Consortium | Berkshire and Surrey Pathology Services Lighthouse Laboratory and Alex Alderton, Roberto Amato, Jeffrey Barrett, Sonia Goncalves, Ewan Harrison, David K. Jackson, Ian Johnston, Dominic Kwiatkowski, Cordelia Langford, John Sillitoe on behalf of the Wellcome Sanger Institute COVID-19 Surveillance Team |
| EPI_ISL_2867122 | Lighthouse Laboratory Plymouth | Wellcome Sanger Institute for the COVID-19 Genomics UK (COG-UK) Consortium | Lighthouse Laboratory Plymouth and Alex Alderton, Roberto Amato, Jeffrey Barrett, Sonia Goncalves, Ewan Harrison, David K. Jackson, Ian Johnston, Dominic Kwiatkowski, Cordelia Langford, John Sillitoe on behalf of the Wellcome Sanger Institute COVID-19 Surveillance Team |
| EPI_ISL_2867123 | Berkshire and Surrey Pathology Services Lighthouse Laboratory | Wellcome Sanger Institute for the COVID-19 Genomics UK (COG-UK) Consortium | Berkshire and Surrey Pathology Services Lighthouse Laboratory and Alex Alderton, Roberto Amato, Jeffrey Barrett, Sonia Goncalves, Ewan Harrison, David K. Jackson, Ian Johnston, Dominic Kwiatkowski, Cordelia Langford, John Sillitoe on behalf of the Wellcome Sanger Institute COVID-19 Surveillance Team |
| EPI_ISL_2867124 | Lighthouse Lab in Alderley Park | Wellcome Sanger Institute for the COVID-19 Genomics UK (COG-UK) Consortium | Jacquelyn Wynn, Mairead Hyland, The Lighthouse Lab in Alderley Park and Alex Alderton, Roberto Amato, Jeffrey Barrett, Sonia Goncalves, Ewan Harrison, David K. Jackson, Ian Johnston, Dominic Kwiatkowski, Cordelia Langford, John Sillitoe on behalf of the Wellcome Sanger Institute COVID-19 Surveillance Team |
| EPI_ISL_2867125 | Lighthouse Laboratory Plymouth | Wellcome Sanger Institute for the COVID-19 Genomics UK (COG-UK) Consortium | Lighthouse Laboratory Plymouth and Alex Alderton, Roberto Amato, Jeffrey Barrett, Sonia Goncalves, Ewan Harrison, David K. Jackson, Ian Johnston, Dominic Kwiatkowski, Cordelia Langford, John Sillitoe on behalf of the Wellcome Sanger Institute COVID-19 Surveillance Team |
| EPI_ISL_2867126, EPI_ISL_2867127 | Berkshire and Surrey Pathology Services Lighthouse Laboratory | Wellcome Sanger Institute for the COVID-19 Genomics UK (COG-UK) Consortium | Berkshire and Surrey Pathology Services Lighthouse Laboratory and Alex Alderton, Roberto Amato, Jeffrey Barrett, Sonia Goncalves, Ewan Harrison, David K. Jackson, Ian Johnston, Dominic Kwiatkowski, Cordelia Langford, John Sillitoe on behalf of the Wellcome Sanger Institute COVID-19 Surveillance Team |
| EPI_ISL_2867128, EPI_ISL_2867129 | Lighthouse Laboratory Plymouth | Wellcome Sanger Institute for the COVID-19 Genomics UK (COG-UK) Consortium | Lighthouse Laboratory Plymouth and Alex Alderton, Roberto Amato, Jeffrey Barrett, Sonia Goncalves, Ewan Harrison, David K. Jackson, Ian Johnston, Dominic Kwiatkowski, Cordelia Langford, John Sillitoe on behalf of the Wellcome Sanger Institute COVID-19 Surveillance Team |
| EPI_ISL_2867130 | Berkshire and Surrey Pathology Services Lighthouse Laboratory | Wellcome Sanger Institute for the COVID-19 Genomics UK (COG-UK) Consortium | Berkshire and Surrey Pathology Services Lighthouse Laboratory and Alex Alderton, Roberto Amato, Jeffrey Barrett, Sonia Goncalves, Ewan Harrison, David K. Jackson, Ian Johnston, Dominic Kwiatkowski, Cordelia Langford, John Sillitoe on behalf of the Wellcome Sanger Institute COVID-19 Surveillance Team |
| EPI_ISL_2867131 | Lighthouse Laboratory Plymouth | Wellcome Sanger Institute for the COVID-19 Genomics UK (COG-UK) Consortium | Lighthouse Laboratory Plymouth and Alex Alderton, Roberto Amato, Jeffrey Barrett, Sonia Goncalves, Ewan Harrison, David K. Jackson, Ian Johnston, Dominic Kwiatkowski, Cordelia Langford, John Sillitoe on behalf of the Wellcome Sanger Institute COVID-19 Surveillance Team |
| EPI_ISL_2867132 | Lighthouse Lab in Alderley Park | Wellcome Sanger Institute for the COVID-19 Genomics UK (COG-UK) Consortium | Jacquelyn Wynn, Mairead Hyland, The Lighthouse Lab in Alderley Park and Alex Alderton, Roberto Amato, Jeffrey Barrett, Sonia Goncalves, Ewan Harrison, David K. Jackson, Ian Johnston, Dominic Kwiatkowski, Cordelia Langford, John Sillitoe on behalf of the Wellcome Sanger Institute COVID-19 Surveillance Team |
| EPI_ISL_2867133, EPI_ISL_2867134, EPI_ISL_2867135 | Berkshire and Surrey Pathology Services Lighthouse Laboratory | Wellcome Sanger Institute for the COVID-19 Genomics UK (COG-UK) Consortium | Berkshire and Surrey Pathology Services Lighthouse Laboratory and Alex Alderton, Roberto Amato, Jeffrey Barrett, Sonia Goncalves, Ewan Harrison, David K. Jackson, Ian Johnston, Dominic Kwiatkowski, Cordelia Langford, John Sillitoe on behalf of the Wellcome Sanger Institute COVID-19 Surveillance Team |
| EPI_ISL_2867136, EPI_ISL_2867137 | Lighthouse Laboratory Plymouth | Wellcome Sanger Institute for the COVID-19 Genomics UK (COG-UK) Consortium | Lighthouse Laboratory Plymouth and Alex Alderton, Roberto Amato, Jeffrey Barrett, Sonia Goncalves, Ewan Harrison, David K. Jackson, Ian Johnston, Dominic Kwiatkowski, Cordelia Langford, John Sillitoe on behalf of the Wellcome Sanger Institute COVID-19 Surveillance Team |
| EPI_ISL_2867138 | Berkshire and Surrey Pathology Services Lighthouse Laboratory | Wellcome Sanger Institute for the COVID-19 Genomics UK (COG-UK) Consortium | Berkshire and Surrey Pathology Services Lighthouse Laboratory and Alex Alderton, Roberto Amato, Jeffrey Barrett, Sonia Goncalves, Ewan Harrison, David K. Jackson, Ian Johnston, Dominic Kwiatkowski, Cordelia Langford, John Sillitoe on behalf of the Wellcome Sanger Institute COVID-19 Surveillance Team |
| EPI_ISL_2867139, EPI_ISL_2867140, EPI_ISL_2867141 | Lighthouse Lab in Alderley Park | Wellcome Sanger Institute for the COVID-19 Genomics UK (COG-UK) Consortium | Jacquelyn Wynn, Mairead Hyland, The Lighthouse Lab in Alderley Park and Alex Alderton, Roberto Amato, Jeffrey Barrett, Sonia Goncalves, Ewan Harrison, David K. Jackson, Ian Johnston, Dominic Kwiatkowski, Cordelia Langford, John Sillitoe on behalf of the Wellcome Sanger Institute COVID-19 Surveillance Team |
| EPI_ISL_2867142 | Berkshire and Surrey Pathology Services Lighthouse Laboratory | Wellcome Sanger Institute for the COVID-19 Genomics UK (COG-UK) Consortium | Berkshire and Surrey Pathology Services Lighthouse Laboratory and Alex Alderton, Roberto Amato, Jeffrey Barrett, Sonia Goncalves, Ewan Harrison, David K. Jackson, Ian Johnston, Dominic Kwiatkowski, Cordelia Langford, John Sillitoe on behalf of the Wellcome Sanger Institute COVID-19 Surveillance Team |
| EPI_ISL_2867143 | Lighthouse Lab in Alderley Park | Wellcome Sanger Institute for the COVID-19 Genomics UK (COG-UK) Consortium | Jacquelyn Wynn, Mairead Hyland, The Lighthouse Lab in Alderley Park and Alex Alderton, Roberto Amato, Jeffrey Barrett, Sonia Goncalves, Ewan Harrison, David K. Jackson, Ian Johnston, Dominic Kwiatkowski, Cordelia Langford, John Sillitoe on behalf of the Wellcome Sanger Institute COVID-19 Surveillance Team |
| EPI_ISL_2867144, EPI_ISL_2867145, EPI_ISL_2867146 | Berkshire and Surrey Pathology Services Lighthouse Laboratory | Wellcome Sanger Institute for the COVID-19 Genomics UK (COG-UK) Consortium | Berkshire and Surrey Pathology Services Lighthouse Laboratory and Alex Alderton, Roberto Amato, Jeffrey Barrett, Sonia Goncalves, Ewan Harrison, David K. Jackson, Ian Johnston, Dominic Kwiatkowski, Cordelia Langford, John Sillitoe on behalf of the Wellcome Sanger Institute COVID-19 Surveillance Team |
| EPI_ISL_2867147 | Lighthouse Lab in Alderley Park | Wellcome Sanger Institute for the COVID-19 Genomics UK (COG-UK) Consortium | Jacquelyn Wynn, Mairead Hyland, The Lighthouse Lab in Alderley Park and Alex Alderton, Roberto Amato, Jeffrey Barrett, Sonia Goncalves, Ewan Harrison, David K. Jackson, Ian Johnston, Dominic Kwiatkowski, Cordelia Langford, John Sillitoe on behalf of the Wellcome Sanger Institute COVID-19 Surveillance Team |
| EPI_ISL_2867148 | Berkshire and Surrey Pathology Services Lighthouse Laboratory | Wellcome Sanger Institute for the COVID-19 Genomics UK (COG-UK) Consortium | Berkshire and Surrey Pathology Services Lighthouse Laboratory and Alex Alderton, Roberto Amato, Jeffrey Barrett, Sonia Goncalves, Ewan Harrison, David K. Jackson, Ian Johnston, Dominic Kwiatkowski, Cordelia Langford, John Sillitoe on behalf of the Wellcome Sanger Institute COVID-19 Surveillance Team |

| | | | |
|---|---|---|---|
| EPI_ISL_2867149, EPI_ISL_2867150 | Lighthouse Lab in Alderley Park | Wellcome Sanger Institute for the COVID-19 Genomics UK (COG-UK) Consortium | Jacquelyn Wynn, Mairead Hyland, The Lighthouse Lab in Alderley Park and Alex Alderton, Roberto Amato, Jeffrey Barrett, Sonia Goncalves, Ewan Harrison, David K. Jackson, Ian Johnston, Dominic Kwiatkowski, Cordelia Langford, John Sillitoe on behalf of the Wellcome Sanger Institute COVID-19 Surveillance Team |
| EPI_ISL_2867151, EPI_ISL_2867152 | Lighthouse Laboratory Plymouth | Wellcome Sanger Institute for the COVID-19 Genomics UK (COG-UK) Consortium | Lighthouse Laboratory Plymouth and Alex Alderton, Roberto Amato, Jeffrey Barrett, Sonia Goncalves, Ewan Harrison, David K. Jackson, Ian Johnston, Dominic Kwiatkowski, Cordelia Langford, John Sillitoe on behalf of the Wellcome Sanger Institute COVID-19 Surveillance Team |
| EPI_ISL_2867153, EPI_ISL_2867154 | Berkshire and Surrey Pathology Services Lighthouse Laboratory | Wellcome Sanger Institute for the COVID-19 Genomics UK (COG-UK) Consortium | Berkshire and Surrey Pathology Services Lighthouse Laboratory and Alex Alderton, Roberto Amato, Jeffrey Barrett, Sonia Goncalves, Ewan Harrison, David K. Jackson, Ian Johnston, Dominic Kwiatkowski, Cordelia Langford, John Sillitoe on behalf of the Wellcome Sanger Institute COVID-19 Surveillance Team |
| EPI_ISL_2867155, EPI_ISL_2867156 | Lighthouse Lab in Alderley Park | Wellcome Sanger Institute for the COVID-19 Genomics UK (COG-UK) Consortium | Jacquelyn Wynn, Mairead Hyland, The Lighthouse Lab in Alderley Park and Alex Alderton, Roberto Amato, Jeffrey Barrett, Sonia Goncalves, Ewan Harrison, David K. Jackson, Ian Johnston, Dominic Kwiatkowski, Cordelia Langford, John Sillitoe on behalf of the Wellcome Sanger Institute COVID-19 Surveillance Team |
| EPI_ISL_2867157, EPI_ISL_2867158 | Berkshire and Surrey Pathology Services Lighthouse Laboratory | Wellcome Sanger Institute for the COVID-19 Genomics UK (COG-UK) Consortium | Berkshire and Surrey Pathology Services Lighthouse Laboratory and Alex Alderton, Roberto Amato, Jeffrey Barrett, Sonia Goncalves, Ewan Harrison, David K. Jackson, Ian Johnston, Dominic Kwiatkowski, Cordelia Langford, John Sillitoe on behalf of the Wellcome Sanger Institute COVID-19 Surveillance Team |
| EPI_ISL_2867159 | Lighthouse Laboratory Plymouth | Wellcome Sanger Institute for the COVID-19 Genomics UK (COG-UK) Consortium | Lighthouse Laboratory Plymouth and Alex Alderton, Roberto Amato, Jeffrey Barrett, Sonia Goncalves, Ewan Harrison, David K. Jackson, Ian Johnston, Dominic Kwiatkowski, Cordelia Langford, John Sillitoe on behalf of the Wellcome Sanger Institute COVID-19 Surveillance Team |
| EPI_ISL_2867160 | Lighthouse Lab in Alderley Park | Wellcome Sanger Institute for the COVID-19 Genomics UK (COG-UK) Consortium | Jacquelyn Wynn, Mairead Hyland, The Lighthouse Lab in Alderley Park and Alex Alderton, Roberto Amato, Jeffrey Barrett, Sonia Goncalves, Ewan Harrison, David K. Jackson, Ian Johnston, Dominic Kwiatkowski, Cordelia Langford, John Sillitoe on behalf of the Wellcome Sanger Institute COVID-19 Surveillance Team |
| EPI_ISL_2867161, EPI_ISL_2867162 | Berkshire and Surrey Pathology Services Lighthouse Laboratory | Wellcome Sanger Institute for the COVID-19 Genomics UK (COG-UK) Consortium | Berkshire and Surrey Pathology Services Lighthouse Laboratory and Alex Alderton, Roberto Amato, Jeffrey Barrett, Sonia Goncalves, Ewan Harrison, David K. Jackson, Ian Johnston, Dominic Kwiatkowski, Cordelia Langford, John Sillitoe on behalf of the Wellcome Sanger Institute COVID-19 Surveillance Team |
| EPI_ISL_2867163, EPI_ISL_2867164, EPI_ISL_2867165 | Lighthouse Lab in Alderley Park | Wellcome Sanger Institute for the COVID-19 Genomics UK (COG-UK) Consortium | Jacquelyn Wynn, Mairead Hyland, The Lighthouse Lab in Alderley Park and Alex Alderton, Roberto Amato, Jeffrey Barrett, Sonia Goncalves, Ewan Harrison, David K. Jackson, Ian Johnston, Dominic Kwiatkowski, Cordelia Langford, John Sillitoe on behalf of the Wellcome Sanger Institute COVID-19 Surveillance Team |
| EPI_ISL_2867166 | Lighthouse Laboratory Plymouth | Wellcome Sanger Institute for the COVID-19 Genomics UK (COG-UK) Consortium | Lighthouse Laboratory Plymouth and Alex Alderton, Roberto Amato, Jeffrey Barrett, Sonia Goncalves, Ewan Harrison, David K. Jackson, Ian Johnston, Dominic Kwiatkowski, Cordelia Langford, John Sillitoe on behalf of the Wellcome Sanger Institute COVID-19 Surveillance Team |
| EPI_ISL_2867167 | Berkshire and Surrey Pathology Services Lighthouse Laboratory | Wellcome Sanger Institute for the COVID-19 Genomics UK (COG-UK) Consortium | Berkshire and Surrey Pathology Services Lighthouse Laboratory and Alex Alderton, Roberto Amato, Jeffrey Barrett, Sonia Goncalves, Ewan Harrison, David K. Jackson, Ian Johnston, Dominic Kwiatkowski, Cordelia Langford, John Sillitoe on behalf of the Wellcome Sanger Institute COVID-19 Surveillance Team |
| EPI_ISL_2867168 | Lighthouse Laboratory Plymouth | Wellcome Sanger Institute for the COVID-19 Genomics UK (COG-UK) Consortium | Lighthouse Laboratory Plymouth and Alex Alderton, Roberto Amato, Jeffrey Barrett, Sonia Goncalves, Ewan Harrison, David K. Jackson, Ian Johnston, Dominic Kwiatkowski, Cordelia Langford, John Sillitoe on behalf of the Wellcome Sanger Institute COVID-19 Surveillance Team |
| EPI_ISL_2867169 | Berkshire and Surrey Pathology Services Lighthouse Laboratory | Wellcome Sanger Institute for the COVID-19 Genomics UK (COG-UK) Consortium | Berkshire and Surrey Pathology Services Lighthouse Laboratory and Alex Alderton, Roberto Amato, Jeffrey Barrett, Sonia Goncalves, Ewan Harrison, David K. Jackson, Ian Johnston, Dominic Kwiatkowski, Cordelia Langford, John Sillitoe on behalf of the Wellcome Sanger Institute COVID-19 Surveillance Team |
| EPI_ISL_2867170, EPI_ISL_2867171, EPI_ISL_2867172, EPI_ISL_2867173 | Lighthouse Laboratory Plymouth | Wellcome Sanger Institute for the COVID-19 Genomics UK (COG-UK) Consortium | Lighthouse Laboratory Plymouth and Alex Alderton, Roberto Amato, Jeffrey Barrett, Sonia Goncalves, Ewan Harrison, David K. Jackson, Ian Johnston, Dominic Kwiatkowski, Cordelia Langford, John Sillitoe on behalf of the Wellcome Sanger Institute COVID-19 Surveillance Team |
| EPI_ISL_2867174 | Berkshire and Surrey Pathology Services Lighthouse Laboratory | Wellcome Sanger Institute for the COVID-19 Genomics UK (COG-UK) Consortium | Berkshire and Surrey Pathology Services Lighthouse Laboratory and Alex Alderton, Roberto Amato, Jeffrey Barrett, Sonia Goncalves, Ewan Harrison, David K. Jackson, Ian Johnston, Dominic Kwiatkowski, Cordelia Langford, John Sillitoe on behalf of the Wellcome Sanger Institute COVID-19 Surveillance Team |
| EPI_ISL_2867175 | Lighthouse Laboratory Plymouth | Wellcome Sanger Institute for the COVID-19 Genomics UK (COG-UK) Consortium | Lighthouse Laboratory Plymouth and Alex Alderton, Roberto Amato, Jeffrey Barrett, Sonia Goncalves, Ewan Harrison, David K. Jackson, Ian Johnston, Dominic Kwiatkowski, Cordelia Langford, John Sillitoe on behalf of the Wellcome Sanger Institute COVID-19 Surveillance Team |
| EPI_ISL_2867176, EPI_ISL_2867177 | Lighthouse Lab in Alderley Park | Wellcome Sanger Institute for the COVID-19 Genomics UK (COG-UK) Consortium | Jacquelyn Wynn, Mairead Hyland, The Lighthouse Lab in Alderley Park and Alex Alderton, Roberto Amato, Jeffrey Barrett, Sonia Goncalves, Ewan Harrison, David K. Jackson, Ian Johnston, Dominic Kwiatkowski, Cordelia Langford, John Sillitoe on behalf of the Wellcome Sanger Institute COVID-19 Surveillance Team |
| EPI_ISL_2867178 | Lighthouse Laboratory Plymouth | Wellcome Sanger Institute for the COVID-19 Genomics UK (COG-UK) Consortium | Lighthouse Laboratory Plymouth and Alex Alderton, Roberto Amato, Jeffrey Barrett, Sonia Goncalves, Ewan Harrison, David K. Jackson, Ian Johnston, Dominic Kwiatkowski, Cordelia Langford, John Sillitoe on behalf of the Wellcome Sanger Institute COVID-19 Surveillance Team |
| EPI_ISL_2867179 | Berkshire and Surrey Pathology Services Lighthouse Laboratory | Wellcome Sanger Institute for the COVID-19 Genomics UK (COG-UK) Consortium | Berkshire and Surrey Pathology Services Lighthouse Laboratory and Alex Alderton, Roberto Amato, Jeffrey Barrett, Sonia Goncalves, Ewan Harrison, David K. Jackson, Ian Johnston, Dominic Kwiatkowski, Cordelia Langford, John Sillitoe on behalf of the Wellcome Sanger Institute COVID-19 Surveillance Team |
| EPI_ISL_2867180 | Lighthouse Laboratory Plymouth | Wellcome Sanger Institute for the COVID-19 Genomics UK (COG-UK) Consortium | Lighthouse Laboratory Plymouth and Alex Alderton, Roberto Amato, Jeffrey Barrett, Sonia Goncalves, Ewan Harrison, David K. Jackson, Ian Johnston, Dominic Kwiatkowski, Cordelia Langford, John Sillitoe on behalf of the Wellcome Sanger Institute COVID-19 Surveillance Team |
| EPI_ISL_2867181, EPI_ISL_2867182, EPI_ISL_2867183, EPI_ISL_2867184, EPI_ISL_2867185 | Berkshire and Surrey Pathology Services Lighthouse Laboratory | Wellcome Sanger Institute for the COVID-19 Genomics UK (COG-UK) Consortium | Berkshire and Surrey Pathology Services Lighthouse Laboratory and Alex Alderton, Roberto Amato, Jeffrey Barrett, Sonia Goncalves, Ewan Harrison, David K. Jackson, Ian Johnston, Dominic Kwiatkowski, Cordelia Langford, John Sillitoe on behalf of the Wellcome Sanger Institute COVID-19 Surveillance Team |
| EPI_ISL_2867186 | Lighthouse Laboratory Plymouth | Wellcome Sanger Institute for the COVID-19 Genomics UK (COG-UK) Consortium | Lighthouse Laboratory Plymouth and Alex Alderton, Roberto Amato, Jeffrey Barrett, Sonia Goncalves, Ewan Harrison, David K. Jackson, Ian Johnston, Dominic Kwiatkowski, Cordelia Langford, John Sillitoe on behalf of the Wellcome Sanger Institute COVID-19 Surveillance Team |
| EPI_ISL_2867187 | Lighthouse Lab in Alderley Park | Wellcome Sanger Institute for the COVID-19 Genomics UK (COG-UK) Consortium | Jacquelyn Wynn, Mairead Hyland, The Lighthouse Lab in Alderley Park and Alex Alderton, Roberto Amato, Jeffrey Barrett, Sonia Goncalves, Ewan Harrison, David K. Jackson, Ian Johnston, Dominic Kwiatkowski, Cordelia Langford, John Sillitoe on behalf of the Wellcome Sanger Institute COVID-19 Surveillance Team |
| EPI_ISL_2867188 | Lighthouse Laboratory Plymouth | Wellcome Sanger Institute for the COVID-19 Genomics UK (COG-UK) Consortium | Lighthouse Laboratory Plymouth and Alex Alderton, Roberto Amato, Jeffrey Barrett, Sonia Goncalves, Ewan Harrison, David K. Jackson, Ian Johnston, Dominic Kwiatkowski, Cordelia Langford, John Sillitoe on behalf of the Wellcome Sanger Institute COVID-19 Surveillance Team |
| EPI_ISL_2867189 | Berkshire and Surrey Pathology Services Lighthouse Laboratory | Wellcome Sanger Institute for the COVID-19 Genomics UK (COG-UK) Consortium | Berkshire and Surrey Pathology Services Lighthouse Laboratory and Alex Alderton, Roberto Amato, Jeffrey Barrett, Sonia Goncalves, Ewan Harrison, David K. Jackson, Ian Johnston, Dominic Kwiatkowski, Cordelia Langford, John Sillitoe on behalf of the Wellcome Sanger Institute COVID-19 Surveillance Team |
| EPI_ISL_2867190 | Lighthouse Lab in Alderley Park | Wellcome Sanger Institute for the COVID-19 Genomics UK (COG-UK) Consortium | Jacquelyn Wynn, Mairead Hyland, The Lighthouse Lab in Alderley Park and Alex Alderton, Roberto Amato, Jeffrey Barrett, Sonia Goncalves, Ewan Harrison, David K. Jackson, Ian Johnston, Dominic Kwiatkowski, Cordelia Langford, John Sillitoe on behalf of the Wellcome Sanger Institute COVID-19 Surveillance Team |
| EPI_ISL_2867191, EPI_ISL_2867192, EPI_ISL_2867193, EPI_ISL_2867194 | Berkshire and Surrey Pathology Services Lighthouse Laboratory | Wellcome Sanger Institute for the COVID-19 Genomics UK (COG-UK) Consortium | Berkshire and Surrey Pathology Services Lighthouse Laboratory and Alex Alderton, Roberto Amato, Jeffrey Barrett, Sonia Goncalves, Ewan Harrison, David K. Jackson, Ian Johnston, Dominic Kwiatkowski, Cordelia Langford, John Sillitoe on behalf of the Wellcome Sanger Institute COVID-19 Surveillance Team |
| EPI_ISL_2867195 | Lighthouse Laboratory Plymouth | Wellcome Sanger Institute for the COVID-19 Genomics UK (COG-UK) Consortium | Lighthouse Laboratory Plymouth and Alex Alderton, Roberto Amato, Jeffrey Barrett, Sonia Goncalves, Ewan Harrison, David K. Jackson, Ian Johnston, Dominic Kwiatkowski, Cordelia Langford, John Sillitoe on behalf of the Wellcome Sanger Institute COVID-19 Surveillance Team |
| EPI_ISL_2867196 | Lighthouse Lab in Alderley Park | Wellcome Sanger Institute for the COVID-19 Genomics UK (COG-UK) Consortium | Jacquelyn Wynn, Mairead Hyland, The Lighthouse Lab in Alderley Park and Alex Alderton, Roberto Amato, Jeffrey Barrett, Sonia Goncalves, Ewan Harrison, David K. Jackson, Ian Johnston, Dominic Kwiatkowski, Cordelia Langford, John Sillitoe on behalf of the Wellcome Sanger Institute COVID-19 Surveillance Team |
| EPI_ISL_2867197, EPI_ISL_2867198 | Berkshire and Surrey Pathology Services Lighthouse Laboratory | Wellcome Sanger Institute for the COVID-19 Genomics UK (COG-UK) Consortium | Berkshire and Surrey Pathology Services Lighthouse Laboratory and Alex Alderton, Roberto Amato, Jeffrey Barrett, Sonia Goncalves, Ewan Harrison, David K. Jackson, Ian Johnston, Dominic Kwiatkowski, Cordelia Langford, John Sillitoe on behalf of the Wellcome Sanger Institute COVID-19 Surveillance Team |
| EPI_ISL_2867199 | Lighthouse Lab in Alderley Park | Wellcome Sanger Institute for the COVID-19 Genomics UK (COG-UK) Consortium | Jacquelyn Wynn, Mairead Hyland, The Lighthouse Lab in Alderley Park and Alex Alderton, Roberto Amato, Jeffrey Barrett, Sonia Goncalves, Ewan Harrison, David K. Jackson, Ian Johnston, Dominic Kwiatkowski, Cordelia Langford, John Sillitoe on behalf of the Wellcome Sanger Institute COVID-19 Surveillance Team |
| EPI_ISL_2867200, EPI_ISL_2867201 | Berkshire and Surrey Pathology Services Lighthouse Laboratory | Wellcome Sanger Institute for the COVID-19 Genomics UK (COG-UK) Consortium | Berkshire and Surrey Pathology Services Lighthouse Laboratory and Alex Alderton, Roberto Amato, Jeffrey Barrett, Sonia Goncalves, Ewan Harrison, David K. Jackson, Ian Johnston, Dominic Kwiatkowski, Cordelia Langford, John Sillitoe on behalf of the Wellcome Sanger Institute COVID-19 Surveillance Team |
| EPI_ISL_2867202 | Lighthouse Lab in Alderley Park | Wellcome Sanger Institute for the COVID-19 Genomics UK (COG-UK) Consortium | Jacquelyn Wynn, Mairead Hyland, The Lighthouse Lab in Alderley Park and Alex Alderton, Roberto Amato, Jeffrey Barrett, Sonia Goncalves, Ewan Harrison, David K. Jackson, Ian Johnston, Dominic Kwiatkowski, Cordelia Langford, John Sillitoe on behalf of the Wellcome Sanger Institute COVID-19 Surveillance Team |
| EPI_ISL_2867203 | Berkshire and Surrey Pathology Services Lighthouse Laboratory | Wellcome Sanger Institute for the COVID-19 Genomics UK (COG-UK) Consortium | Berkshire and Surrey Pathology Services Lighthouse Laboratory and Alex Alderton, Roberto Amato, Jeffrey Barrett, Sonia Goncalves, Ewan Harrison, David K. Jackson, Ian Johnston, Dominic Kwiatkowski, Cordelia Langford, John Sillitoe on behalf of the Wellcome Sanger Institute COVID-19 Surveillance Team |
| EPI_ISL_2867204 | Lighthouse Laboratory Plymouth | Wellcome Sanger Institute for the COVID-19 Genomics UK (COG-UK) Consortium | Lighthouse Laboratory Plymouth and Alex Alderton, Roberto Amato, Jeffrey Barrett, Sonia Goncalves, Ewan Harrison, David K. Jackson, Ian Johnston, Dominic Kwiatkowski, Cordelia Langford, John Sillitoe on behalf of the Wellcome Sanger Institute COVID-19 Surveillance Team |
| EPI_ISL_2867205 | Lighthouse Lab in Alderley Park | Wellcome Sanger Institute for the COVID-19 Genomics UK (COG-UK) Consortium | Jacquelyn Wynn, Mairead Hyland, The Lighthouse Lab in Alderley Park and Alex Alderton, Roberto Amato, Jeffrey Barrett, Sonia Goncalves, Ewan Harrison, David K. Jackson, Ian Johnston, Dominic Kwiatkowski, Cordelia Langford, John Sillitoe on behalf of the Wellcome Sanger Institute COVID-19 Surveillance Team |
| EPI_ISL_2867206, EPI_ISL_2867207, EPI_ISL_2867208 | Berkshire and Surrey Pathology Services Lighthouse Laboratory | Wellcome Sanger Institute for the COVID-19 Genomics UK (COG-UK) Consortium | Berkshire and Surrey Pathology Services Lighthouse Laboratory and Alex Alderton, Roberto Amato, Jeffrey Barrett, Sonia Goncalves, Ewan Harrison, David K. Jackson, Ian Johnston, Dominic Kwiatkowski, Cordelia Langford, John Sillitoe on behalf of the Wellcome Sanger Institute COVID-19 Surveillance Team |
| EPI_ISL_2867209 | Lighthouse Laboratory Plymouth | Wellcome Sanger Institute for the COVID-19 Genomics UK (COG-UK) Consortium | Lighthouse Laboratory Plymouth and Alex Alderton, Roberto Amato, Jeffrey Barrett, Sonia Goncalves, Ewan Harrison, David K. Jackson, Ian Johnston, Dominic Kwiatkowski, Cordelia Langford, John Sillitoe on behalf of the Wellcome Sanger Institute COVID-19 Surveillance Team |
| EPI_ISL_2867210, EPI_ISL_2867211 | Berkshire and Surrey Pathology Services Lighthouse Laboratory | Wellcome Sanger Institute for the COVID-19 Genomics UK (COG-UK) Consortium | Berkshire and Surrey Pathology Services Lighthouse Laboratory and Alex Alderton, Roberto Amato, Jeffrey Barrett, Sonia Goncalves, Ewan Harrison, David K. Jackson, Ian Johnston, Dominic Kwiatkowski, Cordelia Langford, John Sillitoe on behalf of the Wellcome Sanger Institute COVID-19 Surveillance Team |
| EPI_ISL_2867212 | Lighthouse Laboratory Plymouth | Wellcome Sanger Institute for the COVID-19 Genomics UK (COG-UK) Consortium | Lighthouse Laboratory Plymouth and Alex Alderton, Roberto Amato, Jeffrey Barrett, Sonia Goncalves, Ewan Harrison, David K. Jackson, Ian Johnston, Dominic Kwiatkowski, Cordelia Langford, John Sillitoe on behalf of the Wellcome Sanger Institute COVID-19 Surveillance Team |
| EPI_ISL_2867213 | Berkshire and Surrey Pathology Services Lighthouse Laboratory | Wellcome Sanger Institute for the COVID-19 Genomics UK (COG-UK) Consortium | Berkshire and Surrey Pathology Services Lighthouse Laboratory and Alex Alderton, Roberto Amato, Jeffrey Barrett, Sonia Goncalves, Ewan Harrison, David K. Jackson, Ian Johnston, Dominic Kwiatkowski, Cordelia Langford, John Sillitoe on behalf of the Wellcome Sanger Institute COVID-19 Surveillance Team |
| EPI_ISL_2867214 | Lighthouse Lab in Alderley Park | Wellcome Sanger Institute for the COVID-19 Genomics UK (COG-UK) | Jacquelyn Wynn, Mairead Hyland, The Lighthouse Lab in Alderley Park and Alex Alderton, Roberto Amato, Jeffrey Barrett, Sonia Goncalves, Ewan Harrison, David K. Jackson, Ian Johnston, Dominic Kwiatkowski, Cordelia Langford, John Sillitoe on |

| | | | |
|---|---|---|---|
| | | Consortium | behalf of the Wellcome Sanger Institute COVID-19 Surveillance Team |
| EPI_ISL_2867215, EPI_ISL_2867216, EPI_ISL_2867217, EPI_ISL_2867218 | Berkshire and Surrey Pathology Services Lighthouse Laboratory | Wellcome Sanger Institute for the COVID-19 Genomics UK (COG-UK) Consortium | Berkshire and Surrey Pathology Services Lighthouse Laboratory and Alex Alderton, Roberto Amato, Jeffrey Barrett, Sonia Goncalves, Ewan Harrison, David K. Jackson, Ian Johnston, Dominic Kwiatkowski, Cordelia Langford, John Sillitoe on behalf of the Wellcome Sanger Institute COVID-19 Surveillance Team |
| EPI_ISL_2867219 | Lighthouse Lab in Alderley Park | Wellcome Sanger Institute for the COVID-19 Genomics UK (COG-UK) Consortium | Jacquelyn Wynn, Mairead Hyland, The Lighthouse Lab in Alderley Park and Alex Alderton, Roberto Amato, Jeffrey Barrett, Sonia Goncalves, Ewan Harrison, David K. Jackson, Ian Johnston, Dominic Kwiatkowski, Cordelia Langford, John Sillitoe on behalf of the Wellcome Sanger Institute COVID-19 Surveillance Team |
| EPI_ISL_2867220 | Berkshire and Surrey Pathology Services Lighthouse Laboratory | Wellcome Sanger Institute for the COVID-19 Genomics UK (COG-UK) Consortium | Berkshire and Surrey Pathology Services Lighthouse Laboratory and Alex Alderton, Roberto Amato, Jeffrey Barrett, Sonia Goncalves, Ewan Harrison, David K. Jackson, Ian Johnston, Dominic Kwiatkowski, Cordelia Langford, John Sillitoe on behalf of the Wellcome Sanger Institute COVID-19 Surveillance Team |
| EPI_ISL_2867221 | Lighthouse Lab in Alderley Park | Wellcome Sanger Institute for the COVID-19 Genomics UK (COG-UK) Consortium | Jacquelyn Wynn, Mairead Hyland, The Lighthouse Lab in Alderley Park and Alex Alderton, Roberto Amato, Jeffrey Barrett, Sonia Goncalves, Ewan Harrison, David K. Jackson, Ian Johnston, Dominic Kwiatkowski, Cordelia Langford, John Sillitoe on behalf of the Wellcome Sanger Institute COVID-19 Surveillance Team |
| EPI_ISL_2867222 | Berkshire and Surrey Pathology Services Lighthouse Laboratory | Wellcome Sanger Institute for the COVID-19 Genomics UK (COG-UK) Consortium | Berkshire and Surrey Pathology Services Lighthouse Laboratory and Alex Alderton, Roberto Amato, Jeffrey Barrett, Sonia Goncalves, Ewan Harrison, David K. Jackson, Ian Johnston, Dominic Kwiatkowski, Cordelia Langford, John Sillitoe on behalf of the Wellcome Sanger Institute COVID-19 Surveillance Team |
| EPI_ISL_2867223 | Lighthouse Lab in Alderley Park | Wellcome Sanger Institute for the COVID-19 Genomics UK (COG-UK) Consortium | Jacquelyn Wynn, Mairead Hyland, The Lighthouse Lab in Alderley Park and Alex Alderton, Roberto Amato, Jeffrey Barrett, Sonia Goncalves, Ewan Harrison, David K. Jackson, Ian Johnston, Dominic Kwiatkowski, Cordelia Langford, John Sillitoe on behalf of the Wellcome Sanger Institute COVID-19 Surveillance Team |
| EPI_ISL_2867224, EPI_ISL_2867225, EPI_ISL_2867226, EPI_ISL_2867227, EPI_ISL_2867228, EPI_ISL_2867229, EPI_ISL_2867230, EPI_ISL_2867231 | Berkshire and Surrey Pathology Services Lighthouse Laboratory | Wellcome Sanger Institute for the COVID-19 Genomics UK (COG-UK) Consortium | Berkshire and Surrey Pathology Services Lighthouse Laboratory and Alex Alderton, Roberto Amato, Jeffrey Barrett, Sonia Goncalves, Ewan Harrison, David K. Jackson, Ian Johnston, Dominic Kwiatkowski, Cordelia Langford, John Sillitoe on behalf of the Wellcome Sanger Institute COVID-19 Surveillance Team |
| EPI_ISL_2867232 | Lighthouse Laboratory Plymouth | Wellcome Sanger Institute for the COVID-19 Genomics UK (COG-UK) Consortium | Lighthouse Laboratory Plymouth and Alex Alderton, Roberto Amato, Jeffrey Barrett, Sonia Goncalves, Ewan Harrison, David K. Jackson, Ian Johnston, Dominic Kwiatkowski, Cordelia Langford, John Sillitoe on behalf of the Wellcome Sanger Institute COVID-19 Surveillance Team |
| EPI_ISL_2867233 | Lighthouse Lab in Alderley Park | Wellcome Sanger Institute for the COVID-19 Genomics UK (COG-UK) Consortium | Jacquelyn Wynn, Mairead Hyland, The Lighthouse Lab in Alderley Park and Alex Alderton, Roberto Amato, Jeffrey Barrett, Sonia Goncalves, Ewan Harrison, David K. Jackson, Ian Johnston, Dominic Kwiatkowski, Cordelia Langford, John Sillitoe on behalf of the Wellcome Sanger Institute COVID-19 Surveillance Team |
| EPI_ISL_2867234, EPI_ISL_2867235, EPI_ISL_2867236 | Berkshire and Surrey Pathology Services Lighthouse Laboratory | Wellcome Sanger Institute for the COVID-19 Genomics UK (COG-UK) Consortium | Berkshire and Surrey Pathology Services Lighthouse Laboratory and Alex Alderton, Roberto Amato, Jeffrey Barrett, Sonia Goncalves, Ewan Harrison, David K. Jackson, Ian Johnston, Dominic Kwiatkowski, Cordelia Langford, John Sillitoe on behalf of the Wellcome Sanger Institute COVID-19 Surveillance Team |
| EPI_ISL_2867237 | Lighthouse Laboratory Plymouth | Wellcome Sanger Institute for the COVID-19 Genomics UK (COG-UK) Consortium | Lighthouse Laboratory Plymouth and Alex Alderton, Roberto Amato, Jeffrey Barrett, Sonia Goncalves, Ewan Harrison, David K. Jackson, Ian Johnston, Dominic Kwiatkowski, Cordelia Langford, John Sillitoe on behalf of the Wellcome Sanger Institute COVID-19 Surveillance Team |
| EPI_ISL_2867238, EPI_ISL_2867239, EPI_ISL_2867240 | Berkshire and Surrey Pathology Services Lighthouse Laboratory | Wellcome Sanger Institute for the COVID-19 Genomics UK (COG-UK) Consortium | Berkshire and Surrey Pathology Services Lighthouse Laboratory and Alex Alderton, Roberto Amato, Jeffrey Barrett, Sonia Goncalves, Ewan Harrison, David K. Jackson, Ian Johnston, Dominic Kwiatkowski, Cordelia Langford, John Sillitoe on behalf of the Wellcome Sanger Institute COVID-19 Surveillance Team |
| EPI_ISL_2867241 | Lighthouse Lab in Alderley Park | Wellcome Sanger Institute for the COVID-19 Genomics UK (COG-UK) Consortium | Jacquelyn Wynn, Mairead Hyland, The Lighthouse Lab in Alderley Park and Alex Alderton, Roberto Amato, Jeffrey Barrett, Sonia Goncalves, Ewan Harrison, David K. Jackson, Ian Johnston, Dominic Kwiatkowski, Cordelia Langford, John Sillitoe on behalf of the Wellcome Sanger Institute COVID-19 Surveillance Team |
| EPI_ISL_2867242, EPI_ISL_2867243 | Berkshire and Surrey Pathology Services Lighthouse Laboratory | Wellcome Sanger Institute for the COVID-19 Genomics UK (COG-UK) Consortium | Berkshire and Surrey Pathology Services Lighthouse Laboratory and Alex Alderton, Roberto Amato, Jeffrey Barrett, Sonia Goncalves, Ewan Harrison, David K. Jackson, Ian Johnston, Dominic Kwiatkowski, Cordelia Langford, John Sillitoe on behalf of the Wellcome Sanger Institute COVID-19 Surveillance Team |
| EPI_ISL_2867244 | Lighthouse Lab in Alderley Park | Wellcome Sanger Institute for the COVID-19 Genomics UK (COG-UK) Consortium | Jacquelyn Wynn, Mairead Hyland, The Lighthouse Lab in Alderley Park and Alex Alderton, Roberto Amato, Jeffrey Barrett, Sonia Goncalves, Ewan Harrison, David K. Jackson, Ian Johnston, Dominic Kwiatkowski, Cordelia Langford, John Sillitoe on behalf of the Wellcome Sanger Institute COVID-19 Surveillance Team |
| EPI_ISL_2867245 | Berkshire and Surrey Pathology Services Lighthouse Laboratory | Wellcome Sanger Institute for the COVID-19 Genomics UK (COG-UK) Consortium | Berkshire and Surrey Pathology Services Lighthouse Laboratory and Alex Alderton, Roberto Amato, Jeffrey Barrett, Sonia Goncalves, Ewan Harrison, David K. Jackson, Ian Johnston, Dominic Kwiatkowski, Cordelia Langford, John Sillitoe on behalf of the Wellcome Sanger Institute COVID-19 Surveillance Team |
| EPI_ISL_2867246 | Lighthouse Lab in Alderley Park | Wellcome Sanger Institute for the COVID-19 Genomics UK (COG-UK) Consortium | Jacquelyn Wynn, Mairead Hyland, The Lighthouse Lab in Alderley Park and Alex Alderton, Roberto Amato, Jeffrey Barrett, Sonia Goncalves, Ewan Harrison, David K. Jackson, Ian Johnston, Dominic Kwiatkowski, Cordelia Langford, John Sillitoe on behalf of the Wellcome Sanger Institute COVID-19 Surveillance Team |
| EPI_ISL_2867247 | Lighthouse Laboratory Plymouth | Wellcome Sanger Institute for the COVID-19 Genomics UK (COG-UK) Consortium | Lighthouse Laboratory Plymouth and Alex Alderton, Roberto Amato, Jeffrey Barrett, Sonia Goncalves, Ewan Harrison, David K. Jackson, Ian Johnston, Dominic Kwiatkowski, Cordelia Langford, John Sillitoe on behalf of the Wellcome Sanger Institute COVID-19 Surveillance Team |
| EPI_ISL_2867248, EPI_ISL_2867249, EPI_ISL_2867250, EPI_ISL_2867251, EPI_ISL_2867252 | Berkshire and Surrey Pathology Services Lighthouse Laboratory | Wellcome Sanger Institute for the COVID-19 Genomics UK (COG-UK) Consortium | Berkshire and Surrey Pathology Services Lighthouse Laboratory and Alex Alderton, Roberto Amato, Jeffrey Barrett, Sonia Goncalves, Ewan Harrison, David K. Jackson, Ian Johnston, Dominic Kwiatkowski, Cordelia Langford, John Sillitoe on behalf of the Wellcome Sanger Institute COVID-19 Surveillance Team |
| EPI_ISL_2867253 | Lighthouse Lab in Alderley Park | Wellcome Sanger Institute for the COVID-19 Genomics UK (COG-UK) Consortium | Jacquelyn Wynn, Mairead Hyland, The Lighthouse Lab in Alderley Park and Alex Alderton, Roberto Amato, Jeffrey Barrett, Sonia Goncalves, Ewan Harrison, David K. Jackson, Ian Johnston, Dominic Kwiatkowski, Cordelia Langford, John Sillitoe on behalf of the Wellcome Sanger Institute COVID-19 Surveillance Team |
| EPI_ISL_2867254 | Lighthouse Laboratory Plymouth | Wellcome Sanger Institute for the COVID-19 Genomics UK (COG-UK) Consortium | Lighthouse Laboratory Plymouth and Alex Alderton, Roberto Amato, Jeffrey Barrett, Sonia Goncalves, Ewan Harrison, David K. Jackson, Ian Johnston, Dominic Kwiatkowski, Cordelia Langford, John Sillitoe on behalf of the Wellcome Sanger Institute COVID-19 Surveillance Team |
| EPI_ISL_2867255 | Lighthouse Lab in Alderley Park | Wellcome Sanger Institute for the COVID-19 Genomics UK (COG-UK) Consortium | Jacquelyn Wynn, Mairead Hyland, The Lighthouse Lab in Alderley Park and Alex Alderton, Roberto Amato, Jeffrey Barrett, Sonia Goncalves, Ewan Harrison, David K. Jackson, Ian Johnston, Dominic Kwiatkowski, Cordelia Langford, John Sillitoe on behalf of the Wellcome Sanger Institute COVID-19 Surveillance Team |
| EPI_ISL_2867256 | Berkshire and Surrey Pathology Services Lighthouse Laboratory | Wellcome Sanger Institute for the COVID-19 Genomics UK (COG-UK) Consortium | Berkshire and Surrey Pathology Services Lighthouse Laboratory and Alex Alderton, Roberto Amato, Jeffrey Barrett, Sonia Goncalves, Ewan Harrison, David K. Jackson, Ian Johnston, Dominic Kwiatkowski, Cordelia Langford, John Sillitoe on behalf of the Wellcome Sanger Institute COVID-19 Surveillance Team |
| EPI_ISL_2867257 | Lighthouse Laboratory Plymouth | Wellcome Sanger Institute for the COVID-19 Genomics UK (COG-UK) Consortium | Lighthouse Laboratory Plymouth and Alex Alderton, Roberto Amato, Jeffrey Barrett, Sonia Goncalves, Ewan Harrison, David K. Jackson, Ian Johnston, Dominic Kwiatkowski, Cordelia Langford, John Sillitoe on behalf of the Wellcome Sanger Institute COVID-19 Surveillance Team |
| EPI_ISL_2867258, EPI_ISL_2867259, EPI_ISL_2867260 | Berkshire and Surrey Pathology Services Lighthouse Laboratory | Wellcome Sanger Institute for the COVID-19 Genomics UK (COG-UK) Consortium | Berkshire and Surrey Pathology Services Lighthouse Laboratory and Alex Alderton, Roberto Amato, Jeffrey Barrett, Sonia Goncalves, Ewan Harrison, David K. Jackson, Ian Johnston, Dominic Kwiatkowski, Cordelia Langford, John Sillitoe on behalf of the Wellcome Sanger Institute COVID-19 Surveillance Team |
| EPI_ISL_2867261, EPI_ISL_2867262 | Lighthouse Lab in Alderley Park | Wellcome Sanger Institute for the COVID-19 Genomics UK (COG-UK) Consortium | Jacquelyn Wynn, Mairead Hyland, The Lighthouse Lab in Alderley Park and Alex Alderton, Roberto Amato, Jeffrey Barrett, Sonia Goncalves, Ewan Harrison, David K. Jackson, Ian Johnston, Dominic Kwiatkowski, Cordelia Langford, John Sillitoe on behalf of the Wellcome Sanger Institute COVID-19 Surveillance Team |
| EPI_ISL_2867263, EPI_ISL_2867264 | Berkshire and Surrey Pathology Services Lighthouse Laboratory | Wellcome Sanger Institute for the COVID-19 Genomics UK (COG-UK) Consortium | Berkshire and Surrey Pathology Services Lighthouse Laboratory and Alex Alderton, Roberto Amato, Jeffrey Barrett, Sonia Goncalves, Ewan Harrison, David K. Jackson, Ian Johnston, Dominic Kwiatkowski, Cordelia Langford, John Sillitoe on behalf of the Wellcome Sanger Institute COVID-19 Surveillance Team |
| EPI_ISL_2867265 | Lighthouse Laboratory Plymouth | Wellcome Sanger Institute for the COVID-19 Genomics UK (COG-UK) Consortium | Lighthouse Laboratory Plymouth and Alex Alderton, Roberto Amato, Jeffrey Barrett, Sonia Goncalves, Ewan Harrison, David K. Jackson, Ian Johnston, Dominic Kwiatkowski, Cordelia Langford, John Sillitoe on behalf of the Wellcome Sanger Institute COVID-19 Surveillance Team |
| EPI_ISL_2867266 | Lighthouse Lab in Alderley Park | Wellcome Sanger Institute for the COVID-19 Genomics UK (COG-UK) Consortium | Jacquelyn Wynn, Mairead Hyland, The Lighthouse Lab in Alderley Park and Alex Alderton, Roberto Amato, Jeffrey Barrett, Sonia Goncalves, Ewan Harrison, David K. Jackson, Ian Johnston, Dominic Kwiatkowski, Cordelia Langford, John Sillitoe on behalf of the Wellcome Sanger Institute COVID-19 Surveillance Team |
| EPI_ISL_2867267, EPI_ISL_2867268 | Lighthouse Laboratory Plymouth | Wellcome Sanger Institute for the COVID-19 Genomics UK (COG-UK) Consortium | Lighthouse Laboratory Plymouth and Alex Alderton, Roberto Amato, Jeffrey Barrett, Sonia Goncalves, Ewan Harrison, David K. Jackson, Ian Johnston, Dominic Kwiatkowski, Cordelia Langford, John Sillitoe on behalf of the Wellcome Sanger Institute COVID-19 Surveillance Team |
| EPI_ISL_2867269, EPI_ISL_2867270, EPI_ISL_2867271, EPI_ISL_2867272 | Lighthouse Lab in Alderley Park | Wellcome Sanger Institute for the COVID-19 Genomics UK (COG-UK) Consortium | Jacquelyn Wynn, Mairead Hyland, The Lighthouse Lab in Alderley Park and Alex Alderton, Roberto Amato, Jeffrey Barrett, Sonia Goncalves, Ewan Harrison, David K. Jackson, Ian Johnston, Dominic Kwiatkowski, Cordelia Langford, John Sillitoe on behalf of the Wellcome Sanger Institute COVID-19 Surveillance Team |
| EPI_ISL_2867273, EPI_ISL_2867274 | Berkshire and Surrey Pathology Services Lighthouse Laboratory | Wellcome Sanger Institute for the COVID-19 Genomics UK (COG-UK) Consortium | Berkshire and Surrey Pathology Services Lighthouse Laboratory and Alex Alderton, Roberto Amato, Jeffrey Barrett, Sonia Goncalves, Ewan Harrison, David K. Jackson, Ian Johnston, Dominic Kwiatkowski, Cordelia Langford, John Sillitoe on behalf of the Wellcome Sanger Institute COVID-19 Surveillance Team |
| EPI_ISL_2867275 | Lighthouse Lab in Alderley Park | Wellcome Sanger Institute for the COVID-19 Genomics UK (COG-UK) Consortium | Jacquelyn Wynn, Mairead Hyland, The Lighthouse Lab in Alderley Park and Alex Alderton, Roberto Amato, Jeffrey Barrett, Sonia Goncalves, Ewan Harrison, David K. Jackson, Ian Johnston, Dominic Kwiatkowski, Cordelia Langford, John Sillitoe on behalf of the Wellcome Sanger Institute COVID-19 Surveillance Team |
| EPI_ISL_2867276 | Berkshire and Surrey Pathology Services Lighthouse Laboratory | Wellcome Sanger Institute for the COVID-19 Genomics UK (COG-UK) Consortium | Berkshire and Surrey Pathology Services Lighthouse Laboratory and Alex Alderton, Roberto Amato, Jeffrey Barrett, Sonia Goncalves, Ewan Harrison, David K. Jackson, Ian Johnston, Dominic Kwiatkowski, Cordelia Langford, John Sillitoe on behalf of the Wellcome Sanger Institute COVID-19 Surveillance Team |
| EPI_ISL_2867277 | Lighthouse Laboratory Plymouth | Wellcome Sanger Institute for the COVID-19 Genomics UK (COG-UK) Consortium | Lighthouse Laboratory Plymouth and Alex Alderton, Roberto Amato, Jeffrey Barrett, Sonia Goncalves, Ewan Harrison, David K. Jackson, Ian Johnston, Dominic Kwiatkowski, Cordelia Langford, John Sillitoe on behalf of the Wellcome Sanger Institute COVID-19 Surveillance Team |
| EPI_ISL_2867278 | Lighthouse Lab in Alderley Park | Wellcome Sanger Institute for the COVID-19 Genomics UK (COG-UK) Consortium | Jacquelyn Wynn, Mairead Hyland, The Lighthouse Lab in Alderley Park and Alex Alderton, Roberto Amato, Jeffrey Barrett, Sonia Goncalves, Ewan Harrison, David K. Jackson, Ian Johnston, Dominic Kwiatkowski, Cordelia Langford, John Sillitoe on behalf of the Wellcome Sanger Institute COVID-19 Surveillance Team |
| EPI_ISL_2867279, EPI_ISL_2867280 | Berkshire and Surrey Pathology Services Lighthouse Laboratory | Wellcome Sanger Institute for the COVID-19 Genomics UK (COG-UK) Consortium | Berkshire and Surrey Pathology Services Lighthouse Laboratory and Alex Alderton, Roberto Amato, Jeffrey Barrett, Sonia Goncalves, Ewan Harrison, David K. Jackson, Ian Johnston, Dominic Kwiatkowski, Cordelia Langford, John Sillitoe on behalf of the Wellcome Sanger Institute COVID-19 Surveillance Team |
| EPI_ISL_2867281 | Lighthouse Laboratory Plymouth | Wellcome Sanger Institute for the COVID-19 Genomics UK (COG-UK) Consortium | Lighthouse Laboratory Plymouth and Alex Alderton, Roberto Amato, Jeffrey Barrett, Sonia Goncalves, Ewan Harrison, David K. Jackson, Ian Johnston, Dominic Kwiatkowski, Cordelia Langford, John Sillitoe on behalf of the Wellcome Sanger Institute COVID-19 Surveillance Team |

| | | | |
|---|---|---|---|
| EPI_ISL_2867282 | Berkshire and Surrey Pathology Services Lighthouse Laboratory | Wellcome Sanger Institute for the COVID-19 Genomics UK (COG-UK) Consortium | Berkshire and Surrey Pathology Services Lighthouse Laboratory and Alex Alderton, Roberto Amato, Jeffrey Barrett, Sonia Goncalves, Ewan Harrison, David K. Jackson, Ian Johnston, Dominic Kwiatkowski, Cordelia Langford, John Sillitoe on behalf of the Wellcome Sanger Institute COVID-19 Surveillance Team |
| EPI_ISL_2867283, EPI_ISL_2867284, EPI_ISL_2867285 | Lighthouse Lab in Alderley Park | Wellcome Sanger Institute for the COVID-19 Genomics UK (COG-UK) Consortium | Jacquelyn Wynn, Mairead Hyland, The Lighthouse Lab in Alderley Park and Alex Alderton, Roberto Amato, Jeffrey Barrett, Sonia Goncalves, Ewan Harrison, David K. Jackson, Ian Johnston, Dominic Kwiatkowski, Cordelia Langford, John Sillitoe on behalf of the Wellcome Sanger Institute COVID-19 Surveillance Team |
| EPI_ISL_2867286 | Berkshire and Surrey Pathology Services Lighthouse Laboratory | Wellcome Sanger Institute for the COVID-19 Genomics UK (COG-UK) Consortium | Berkshire and Surrey Pathology Services Lighthouse Laboratory and Alex Alderton, Roberto Amato, Jeffrey Barrett, Sonia Goncalves, Ewan Harrison, David K. Jackson, Ian Johnston, Dominic Kwiatkowski, Cordelia Langford, John Sillitoe on behalf of the Wellcome Sanger Institute COVID-19 Surveillance Team |
| EPI_ISL_2867287 | Lighthouse Lab in Alderley Park | Wellcome Sanger Institute for the COVID-19 Genomics UK (COG-UK) Consortium | Jacquelyn Wynn, Mairead Hyland, The Lighthouse Lab in Alderley Park and Alex Alderton, Roberto Amato, Jeffrey Barrett, Sonia Goncalves, Ewan Harrison, David K. Jackson, Ian Johnston, Dominic Kwiatkowski, Cordelia Langford, John Sillitoe on behalf of the Wellcome Sanger Institute COVID-19 Surveillance Team |
| EPI_ISL_2867288, EPI_ISL_2867289 | Berkshire and Surrey Pathology Services Lighthouse Laboratory | Wellcome Sanger Institute for the COVID-19 Genomics UK (COG-UK) Consortium | Berkshire and Surrey Pathology Services Lighthouse Laboratory and Alex Alderton, Roberto Amato, Jeffrey Barrett, Sonia Goncalves, Ewan Harrison, David K. Jackson, Ian Johnston, Dominic Kwiatkowski, Cordelia Langford, John Sillitoe on behalf of the Wellcome Sanger Institute COVID-19 Surveillance Team |
| EPI_ISL_2867290 | Lighthouse Laboratory Plymouth | Wellcome Sanger Institute for the COVID-19 Genomics UK (COG-UK) Consortium | Lighthouse Laboratory Plymouth and Alex Alderton, Roberto Amato, Jeffrey Barrett, Sonia Goncalves, Ewan Harrison, David K. Jackson, Ian Johnston, Dominic Kwiatkowski, Cordelia Langford, John Sillitoe on behalf of the Wellcome Sanger Institute COVID-19 Surveillance Team |
| EPI_ISL_2867291, EPI_ISL_2867292, EPI_ISL_2867293 | Berkshire and Surrey Pathology Services Lighthouse Laboratory | Wellcome Sanger Institute for the COVID-19 Genomics UK (COG-UK) Consortium | Berkshire and Surrey Pathology Services Lighthouse Laboratory and Alex Alderton, Roberto Amato, Jeffrey Barrett, Sonia Goncalves, Ewan Harrison, David K. Jackson, Ian Johnston, Dominic Kwiatkowski, Cordelia Langford, John Sillitoe on behalf of the Wellcome Sanger Institute COVID-19 Surveillance Team |
| EPI_ISL_2867294 | Lighthouse Lab in Alderley Park | Wellcome Sanger Institute for the COVID-19 Genomics UK (COG-UK) Consortium | Jacquelyn Wynn, Mairead Hyland, The Lighthouse Lab in Alderley Park and Alex Alderton, Roberto Amato, Jeffrey Barrett, Sonia Goncalves, Ewan Harrison, David K. Jackson, Ian Johnston, Dominic Kwiatkowski, Cordelia Langford, John Sillitoe on behalf of the Wellcome Sanger Institute COVID-19 Surveillance Team |
| EPI_ISL_2867295 | Berkshire and Surrey Pathology Services Lighthouse Laboratory | Wellcome Sanger Institute for the COVID-19 Genomics UK (COG-UK) Consortium | Berkshire and Surrey Pathology Services Lighthouse Laboratory and Alex Alderton, Roberto Amato, Jeffrey Barrett, Sonia Goncalves, Ewan Harrison, David K. Jackson, Ian Johnston, Dominic Kwiatkowski, Cordelia Langford, John Sillitoe on behalf of the Wellcome Sanger Institute COVID-19 Surveillance Team |
| EPI_ISL_2867296 | Lighthouse Lab in Alderley Park | Wellcome Sanger Institute for the COVID-19 Genomics UK (COG-UK) Consortium | Jacquelyn Wynn, Mairead Hyland, The Lighthouse Lab in Alderley Park and Alex Alderton, Roberto Amato, Jeffrey Barrett, Sonia Goncalves, Ewan Harrison, David K. Jackson, Ian Johnston, Dominic Kwiatkowski, Cordelia Langford, John Sillitoe on behalf of the Wellcome Sanger Institute COVID-19 Surveillance Team |
| EPI_ISL_2867297 | Lighthouse Laboratory Plymouth | Wellcome Sanger Institute for the COVID-19 Genomics UK (COG-UK) Consortium | Lighthouse Laboratory Plymouth and Alex Alderton, Roberto Amato, Jeffrey Barrett, Sonia Goncalves, Ewan Harrison, David K. Jackson, Ian Johnston, Dominic Kwiatkowski, Cordelia Langford, John Sillitoe on behalf of the Wellcome Sanger Institute COVID-19 Surveillance Team |
| EPI_ISL_2867298, EPI_ISL_2867299, EPI_ISL_2867300 | Berkshire and Surrey Pathology Services Lighthouse Laboratory | Wellcome Sanger Institute for the COVID-19 Genomics UK (COG-UK) Consortium | Berkshire and Surrey Pathology Services Lighthouse Laboratory and Alex Alderton, Roberto Amato, Jeffrey Barrett, Sonia Goncalves, Ewan Harrison, David K. Jackson, Ian Johnston, Dominic Kwiatkowski, Cordelia Langford, John Sillitoe on behalf of the Wellcome Sanger Institute COVID-19 Surveillance Team |
| EPI_ISL_2867301 | Lighthouse Lab in Alderley Park | Wellcome Sanger Institute for the COVID-19 Genomics UK (COG-UK) Consortium | Jacquelyn Wynn, Mairead Hyland, The Lighthouse Lab in Alderley Park and Alex Alderton, Roberto Amato, Jeffrey Barrett, Sonia Goncalves, Ewan Harrison, David K. Jackson, Ian Johnston, Dominic Kwiatkowski, Cordelia Langford, John Sillitoe on behalf of the Wellcome Sanger Institute COVID-19 Surveillance Team |
| EPI_ISL_2867302 | Lighthouse Laboratory Plymouth | Wellcome Sanger Institute for the COVID-19 Genomics UK (COG-UK) Consortium | Lighthouse Laboratory Plymouth and Alex Alderton, Roberto Amato, Jeffrey Barrett, Sonia Goncalves, Ewan Harrison, David K. Jackson, Ian Johnston, Dominic Kwiatkowski, Cordelia Langford, John Sillitoe on behalf of the Wellcome Sanger Institute COVID-19 Surveillance Team |
| EPI_ISL_2867303, EPI_ISL_2867304 | Berkshire and Surrey Pathology Services Lighthouse Laboratory | Wellcome Sanger Institute for the COVID-19 Genomics UK (COG-UK) Consortium | Berkshire and Surrey Pathology Services Lighthouse Laboratory and Alex Alderton, Roberto Amato, Jeffrey Barrett, Sonia Goncalves, Ewan Harrison, David K. Jackson, Ian Johnston, Dominic Kwiatkowski, Cordelia Langford, John Sillitoe on behalf of the Wellcome Sanger Institute COVID-19 Surveillance Team |
| EPI_ISL_2867305 | Lighthouse Laboratory Plymouth | Wellcome Sanger Institute for the COVID-19 Genomics UK (COG-UK) Consortium | Lighthouse Laboratory Plymouth and Alex Alderton, Roberto Amato, Jeffrey Barrett, Sonia Goncalves, Ewan Harrison, David K. Jackson, Ian Johnston, Dominic Kwiatkowski, Cordelia Langford, John Sillitoe on behalf of the Wellcome Sanger Institute COVID-19 Surveillance Team |
| EPI_ISL_2867306 | Lighthouse Lab in Alderley Park | Wellcome Sanger Institute for the COVID-19 Genomics UK (COG-UK) Consortium | Jacquelyn Wynn, Mairead Hyland, The Lighthouse Lab in Alderley Park and Alex Alderton, Roberto Amato, Jeffrey Barrett, Sonia Goncalves, Ewan Harrison, David K. Jackson, Ian Johnston, Dominic Kwiatkowski, Cordelia Langford, John Sillitoe on behalf of the Wellcome Sanger Institute COVID-19 Surveillance Team |
| EPI_ISL_2867307, EPI_ISL_2867308 | Berkshire and Surrey Pathology Services Lighthouse Laboratory | Wellcome Sanger Institute for the COVID-19 Genomics UK (COG-UK) Consortium | Berkshire and Surrey Pathology Services Lighthouse Laboratory and Alex Alderton, Roberto Amato, Jeffrey Barrett, Sonia Goncalves, Ewan Harrison, David K. Jackson, Ian Johnston, Dominic Kwiatkowski, Cordelia Langford, John Sillitoe on behalf of the Wellcome Sanger Institute COVID-19 Surveillance Team |
| EPI_ISL_2867309 | Lighthouse Laboratory Plymouth | Wellcome Sanger Institute for the COVID-19 Genomics UK (COG-UK) Consortium | Lighthouse Laboratory Plymouth and Alex Alderton, Roberto Amato, Jeffrey Barrett, Sonia Goncalves, Ewan Harrison, David K. Jackson, Ian Johnston, Dominic Kwiatkowski, Cordelia Langford, John Sillitoe on behalf of the Wellcome Sanger Institute COVID-19 Surveillance Team |
| EPI_ISL_2867310, EPI_ISL_2867311, EPI_ISL_2867312 | Berkshire and Surrey Pathology Services Lighthouse Laboratory | Wellcome Sanger Institute for the COVID-19 Genomics UK (COG-UK) Consortium | Berkshire and Surrey Pathology Services Lighthouse Laboratory and Alex Alderton, Roberto Amato, Jeffrey Barrett, Sonia Goncalves, Ewan Harrison, David K. Jackson, Ian Johnston, Dominic Kwiatkowski, Cordelia Langford, John Sillitoe on behalf of the Wellcome Sanger Institute COVID-19 Surveillance Team |
| EPI_ISL_2867313 | Lighthouse Lab in Alderley Park | Wellcome Sanger Institute for the COVID-19 Genomics UK (COG-UK) Consortium | Jacquelyn Wynn, Mairead Hyland, The Lighthouse Lab in Alderley Park and Alex Alderton, Roberto Amato, Jeffrey Barrett, Sonia Goncalves, Ewan Harrison, David K. Jackson, Ian Johnston, Dominic Kwiatkowski, Cordelia Langford, John Sillitoe on behalf of the Wellcome Sanger Institute COVID-19 Surveillance Team |
| EPI_ISL_2867314, EPI_ISL_2867315 | Berkshire and Surrey Pathology Services Lighthouse Laboratory | Wellcome Sanger Institute for the COVID-19 Genomics UK (COG-UK) Consortium | Berkshire and Surrey Pathology Services Lighthouse Laboratory and Alex Alderton, Roberto Amato, Jeffrey Barrett, Sonia Goncalves, Ewan Harrison, David K. Jackson, Ian Johnston, Dominic Kwiatkowski, Cordelia Langford, John Sillitoe on behalf of the Wellcome Sanger Institute COVID-19 Surveillance Team |
| EPI_ISL_2867316 | Lighthouse Lab in Alderley Park | Wellcome Sanger Institute for the COVID-19 Genomics UK (COG-UK) Consortium | Jacquelyn Wynn, Mairead Hyland, The Lighthouse Lab in Alderley Park and Alex Alderton, Roberto Amato, Jeffrey Barrett, Sonia Goncalves, Ewan Harrison, David K. Jackson, Ian Johnston, Dominic Kwiatkowski, Cordelia Langford, John Sillitoe on behalf of the Wellcome Sanger Institute COVID-19 Surveillance Team |
| EPI_ISL_2867317 | Berkshire and Surrey Pathology Services Lighthouse Laboratory | Wellcome Sanger Institute for the COVID-19 Genomics UK (COG-UK) Consortium | Berkshire and Surrey Pathology Services Lighthouse Laboratory and Alex Alderton, Roberto Amato, Jeffrey Barrett, Sonia Goncalves, Ewan Harrison, David K. Jackson, Ian Johnston, Dominic Kwiatkowski, Cordelia Langford, John Sillitoe on behalf of the Wellcome Sanger Institute COVID-19 Surveillance Team |
| EPI_ISL_2867318 | Lighthouse Lab in Alderley Park | Wellcome Sanger Institute for the COVID-19 Genomics UK (COG-UK) Consortium | Jacquelyn Wynn, Mairead Hyland, The Lighthouse Lab in Alderley Park and Alex Alderton, Roberto Amato, Jeffrey Barrett, Sonia Goncalves, Ewan Harrison, David K. Jackson, Ian Johnston, Dominic Kwiatkowski, Cordelia Langford, John Sillitoe on behalf of the Wellcome Sanger Institute COVID-19 Surveillance Team |
| EPI_ISL_2867319, EPI_ISL_2867320 | Berkshire and Surrey Pathology Services Lighthouse Laboratory | Wellcome Sanger Institute for the COVID-19 Genomics UK (COG-UK) Consortium | Berkshire and Surrey Pathology Services Lighthouse Laboratory and Alex Alderton, Roberto Amato, Jeffrey Barrett, Sonia Goncalves, Ewan Harrison, David K. Jackson, Ian Johnston, Dominic Kwiatkowski, Cordelia Langford, John Sillitoe on behalf of the Wellcome Sanger Institute COVID-19 Surveillance Team |
| EPI_ISL_2867321 | Lighthouse Laboratory Plymouth | Wellcome Sanger Institute for the COVID-19 Genomics UK (COG-UK) Consortium | Lighthouse Laboratory Plymouth and Alex Alderton, Roberto Amato, Jeffrey Barrett, Sonia Goncalves, Ewan Harrison, David K. Jackson, Ian Johnston, Dominic Kwiatkowski, Cordelia Langford, John Sillitoe on behalf of the Wellcome Sanger Institute COVID-19 Surveillance Team |
| EPI_ISL_2867322, EPI_ISL_2867323, EPI_ISL_2867324 | Lighthouse Lab in Alderley Park | Wellcome Sanger Institute for the COVID-19 Genomics UK (COG-UK) Consortium | Jacquelyn Wynn, Mairead Hyland, The Lighthouse Lab in Alderley Park and Alex Alderton, Roberto Amato, Jeffrey Barrett, Sonia Goncalves, Ewan Harrison, David K. Jackson, Ian Johnston, Dominic Kwiatkowski, Cordelia Langford, John Sillitoe on behalf of the Wellcome Sanger Institute COVID-19 Surveillance Team |
| EPI_ISL_2867325, EPI_ISL_2867326 | Berkshire and Surrey Pathology Services Lighthouse Laboratory | Wellcome Sanger Institute for the COVID-19 Genomics UK (COG-UK) Consortium | Berkshire and Surrey Pathology Services Lighthouse Laboratory and Alex Alderton, Roberto Amato, Jeffrey Barrett, Sonia Goncalves, Ewan Harrison, David K. Jackson, Ian Johnston, Dominic Kwiatkowski, Cordelia Langford, John Sillitoe on behalf of the Wellcome Sanger Institute COVID-19 Surveillance Team |
| EPI_ISL_2867327 | Lighthouse Laboratory Plymouth | Wellcome Sanger Institute for the COVID-19 Genomics UK (COG-UK) Consortium | Lighthouse Laboratory Plymouth and Alex Alderton, Roberto Amato, Jeffrey Barrett, Sonia Goncalves, Ewan Harrison, David K. Jackson, Ian Johnston, Dominic Kwiatkowski, Cordelia Langford, John Sillitoe on behalf of the Wellcome Sanger Institute COVID-19 Surveillance Team |
| EPI_ISL_2867328, EPI_ISL_2867329 | Berkshire and Surrey Pathology Services Lighthouse Laboratory | Wellcome Sanger Institute for the COVID-19 Genomics UK (COG-UK) Consortium | Berkshire and Surrey Pathology Services Lighthouse Laboratory and Alex Alderton, Roberto Amato, Jeffrey Barrett, Sonia Goncalves, Ewan Harrison, David K. Jackson, Ian Johnston, Dominic Kwiatkowski, Cordelia Langford, John Sillitoe on behalf of the Wellcome Sanger Institute COVID-19 Surveillance Team |
| EPI_ISL_2867330 | Lighthouse Lab in Alderley Park | Wellcome Sanger Institute for the COVID-19 Genomics UK (COG-UK) Consortium | Jacquelyn Wynn, Mairead Hyland, The Lighthouse Lab in Alderley Park and Alex Alderton, Roberto Amato, Jeffrey Barrett, Sonia Goncalves, Ewan Harrison, David K. Jackson, Ian Johnston, Dominic Kwiatkowski, Cordelia Langford, John Sillitoe on behalf of the Wellcome Sanger Institute COVID-19 Surveillance Team |
| EPI_ISL_2867331, EPI_ISL_2867332 | Lighthouse Laboratory Plymouth | Wellcome Sanger Institute for the COVID-19 Genomics UK (COG-UK) Consortium | Lighthouse Laboratory Plymouth and Alex Alderton, Roberto Amato, Jeffrey Barrett, Sonia Goncalves, Ewan Harrison, David K. Jackson, Ian Johnston, Dominic Kwiatkowski, Cordelia Langford, John Sillitoe on behalf of the Wellcome Sanger Institute COVID-19 Surveillance Team |
| EPI_ISL_2867333 | Lighthouse Lab in Alderley Park | Wellcome Sanger Institute for the COVID-19 Genomics UK (COG-UK) Consortium | Jacquelyn Wynn, Mairead Hyland, The Lighthouse Lab in Alderley Park and Alex Alderton, Roberto Amato, Jeffrey Barrett, Sonia Goncalves, Ewan Harrison, David K. Jackson, Ian Johnston, Dominic Kwiatkowski, Cordelia Langford, John Sillitoe on behalf of the Wellcome Sanger Institute COVID-19 Surveillance Team |
| EPI_ISL_2867334 | Lighthouse Laboratory Plymouth | Wellcome Sanger Institute for the COVID-19 Genomics UK (COG-UK) Consortium | Lighthouse Laboratory Plymouth and Alex Alderton, Roberto Amato, Jeffrey Barrett, Sonia Goncalves, Ewan Harrison, David K. Jackson, Ian Johnston, Dominic Kwiatkowski, Cordelia Langford, John Sillitoe on behalf of the Wellcome Sanger Institute COVID-19 Surveillance Team |
| EPI_ISL_2867335, EPI_ISL_2867336 | Berkshire and Surrey Pathology Services Lighthouse Laboratory | Wellcome Sanger Institute for the COVID-19 Genomics UK (COG-UK) Consortium | Berkshire and Surrey Pathology Services Lighthouse Laboratory and Alex Alderton, Roberto Amato, Jeffrey Barrett, Sonia Goncalves, Ewan Harrison, David K. Jackson, Ian Johnston, Dominic Kwiatkowski, Cordelia Langford, John Sillitoe on behalf of the Wellcome Sanger Institute COVID-19 Surveillance Team |
| EPI_ISL_2867337 | Lighthouse Laboratory Plymouth | Wellcome Sanger Institute for the COVID-19 Genomics UK (COG-UK) Consortium | Lighthouse Laboratory Plymouth and Alex Alderton, Roberto Amato, Jeffrey Barrett, Sonia Goncalves, Ewan Harrison, David K. Jackson, Ian Johnston, Dominic Kwiatkowski, Cordelia Langford, John Sillitoe on behalf of the Wellcome Sanger Institute COVID-19 Surveillance Team |
| EPI_ISL_2867338, EPI_ISL_2867339, EPI_ISL_2867340 | Lighthouse Lab in Alderley Park | Wellcome Sanger Institute for the COVID-19 Genomics UK (COG-UK) Consortium | Jacquelyn Wynn, Mairead Hyland, The Lighthouse Lab in Alderley Park and Alex Alderton, Roberto Amato, Jeffrey Barrett, Sonia Goncalves, Ewan Harrison, David K. Jackson, Ian Johnston, Dominic Kwiatkowski, Cordelia Langford, John Sillitoe on behalf of the Wellcome Sanger Institute COVID-19 Surveillance Team |
| EPI_ISL_2867341 | Lighthouse Laboratory Plymouth | Wellcome Sanger Institute for the COVID-19 Genomics UK (COG-UK) Consortium | Lighthouse Laboratory Plymouth and Alex Alderton, Roberto Amato, Jeffrey Barrett, Sonia Goncalves, Ewan Harrison, David K. Jackson, Ian Johnston, Dominic Kwiatkowski, Cordelia Langford, John Sillitoe on behalf of the Wellcome Sanger Institute COVID-19 Surveillance Team |
| EPI_ISL_2867342 | Berkshire and Surrey Pathology Services Lighthouse Laboratory | Wellcome Sanger Institute for the COVID-19 Genomics UK (COG-UK) Consortium | Berkshire and Surrey Pathology Services Lighthouse Laboratory and Alex Alderton, Roberto Amato, Jeffrey Barrett, Sonia Goncalves, Ewan Harrison, David K. Jackson, Ian Johnston, Dominic Kwiatkowski, Cordelia Langford, John Sillitoe on behalf of the Wellcome Sanger Institute COVID-19 Surveillance Team |
| EPI_ISL_2867343 | Lighthouse Laboratory Plymouth | Wellcome Sanger Institute for the COVID-19 Genomics UK (COG-UK) Consortium | Lighthouse Laboratory Plymouth and Alex Alderton, Roberto Amato, Jeffrey Barrett, Sonia Goncalves, Ewan Harrison, David K. Jackson, Ian Johnston, Dominic Kwiatkowski, Cordelia Langford, John Sillitoe on behalf of the Wellcome Sanger Institute COVID-19 Surveillance Team |
| EPI_ISL_2867344 | Lighthouse Lab in Alderley Park | Wellcome Sanger Institute for the COVID-19 Genomics UK (COG-UK) Consortium | Jacquelyn Wynn, Mairead Hyland, The Lighthouse Lab in Alderley Park and Alex Alderton, Roberto Amato, Jeffrey Barrett, Sonia Goncalves, Ewan Harrison, David K. Jackson, Ian Johnston, Dominic Kwiatkowski, Cordelia Langford, John Sillitoe on behalf of the Wellcome Sanger Institute COVID-19 Surveillance Team |

| | | | |
|---|---|---|---|
| EPI_ISL_2867345, EPI_ISL_2867346, EPI_ISL_2867347 | Berkshire and Surrey Pathology Services Lighthouse Laboratory | Wellcome Sanger Institute for the COVID-19 Genomics UK (COG-UK) Consortium | Berkshire and Surrey Pathology Services Lighthouse Laboratory and Alex Alderton, Roberto Amato, Jeffrey Barrett, Sonia Goncalves, Ewan Harrison, David K. Jackson, Ian Johnston, Dominic Kwiatkowski, Cordelia Langford, John Sillitoe on behalf of the Wellcome Sanger Institute COVID-19 Surveillance Team |
| EPI_ISL_2867348 | Lighthouse Laboratory Plymouth | Wellcome Sanger Institute for the COVID-19 Genomics UK (COG-UK) Consortium | Lighthouse Laboratory Plymouth and Alex Alderton, Roberto Amato, Jeffrey Barrett, Sonia Goncalves, Ewan Harrison, David K. Jackson, Ian Johnston, Dominic Kwiatkowski, Cordelia Langford, John Sillitoe on behalf of the Wellcome Sanger Institute COVID-19 Surveillance Team |
| EPI_ISL_2867349, EPI_ISL_2867350 | Berkshire and Surrey Pathology Services Lighthouse Laboratory | Wellcome Sanger Institute for the COVID-19 Genomics UK (COG-UK) Consortium | Berkshire and Surrey Pathology Services Lighthouse Laboratory and Alex Alderton, Roberto Amato, Jeffrey Barrett, Sonia Goncalves, Ewan Harrison, David K. Jackson, Ian Johnston, Dominic Kwiatkowski, Cordelia Langford, John Sillitoe on behalf of the Wellcome Sanger Institute COVID-19 Surveillance Team |
| EPI_ISL_2867351 | Lighthouse Laboratory Plymouth | Wellcome Sanger Institute for the COVID-19 Genomics UK (COG-UK) Consortium | Lighthouse Laboratory Plymouth and Alex Alderton, Roberto Amato, Jeffrey Barrett, Sonia Goncalves, Ewan Harrison, David K. Jackson, Ian Johnston, Dominic Kwiatkowski, Cordelia Langford, John Sillitoe on behalf of the Wellcome Sanger Institute COVID-19 Surveillance Team |
| EPI_ISL_2867352 | Lighthouse Lab in Alderley Park | Wellcome Sanger Institute for the COVID-19 Genomics UK (COG-UK) Consortium | Jacquelyn Wynn, Mairead Hyland, The Lighthouse Lab in Alderley Park and Alex Alderton, Roberto Amato, Jeffrey Barrett, Sonia Goncalves, Ewan Harrison, David K. Jackson, Ian Johnston, Dominic Kwiatkowski, Cordelia Langford, John Sillitoe on behalf of the Wellcome Sanger Institute COVID-19 Surveillance Team |
| EPI_ISL_2867353, EPI_ISL_2867354 | Lighthouse Laboratory Plymouth | Wellcome Sanger Institute for the COVID-19 Genomics UK (COG-UK) Consortium | Lighthouse Laboratory Plymouth and Alex Alderton, Roberto Amato, Jeffrey Barrett, Sonia Goncalves, Ewan Harrison, David K. Jackson, Ian Johnston, Dominic Kwiatkowski, Cordelia Langford, John Sillitoe on behalf of the Wellcome Sanger Institute COVID-19 Surveillance Team |
| EPI_ISL_2867355, EPI_ISL_2867356 | Lighthouse Lab in Alderley Park | Wellcome Sanger Institute for the COVID-19 Genomics UK (COG-UK) Consortium | Jacquelyn Wynn, Mairead Hyland, The Lighthouse Lab in Alderley Park and Alex Alderton, Roberto Amato, Jeffrey Barrett, Sonia Goncalves, Ewan Harrison, David K. Jackson, Ian Johnston, Dominic Kwiatkowski, Cordelia Langford, John Sillitoe on behalf of the Wellcome Sanger Institute COVID-19 Surveillance Team |
| EPI_ISL_2867357 | Lighthouse Laboratory Plymouth | Wellcome Sanger Institute for the COVID-19 Genomics UK (COG-UK) Consortium | Lighthouse Laboratory Plymouth and Alex Alderton, Roberto Amato, Jeffrey Barrett, Sonia Goncalves, Ewan Harrison, David K. Jackson, Ian Johnston, Dominic Kwiatkowski, Cordelia Langford, John Sillitoe on behalf of the Wellcome Sanger Institute COVID-19 Surveillance Team |
| EPI_ISL_2867358 | Lighthouse Lab in Alderley Park | Wellcome Sanger Institute for the COVID-19 Genomics UK (COG-UK) Consortium | Jacquelyn Wynn, Mairead Hyland, The Lighthouse Lab in Alderley Park and Alex Alderton, Roberto Amato, Jeffrey Barrett, Sonia Goncalves, Ewan Harrison, David K. Jackson, Ian Johnston, Dominic Kwiatkowski, Cordelia Langford, John Sillitoe on behalf of the Wellcome Sanger Institute COVID-19 Surveillance Team |
| EPI_ISL_2867359 | Berkshire and Surrey Pathology Services Lighthouse Laboratory | Wellcome Sanger Institute for the COVID-19 Genomics UK (COG-UK) Consortium | Berkshire and Surrey Pathology Services Lighthouse Laboratory and Alex Alderton, Roberto Amato, Jeffrey Barrett, Sonia Goncalves, Ewan Harrison, David K. Jackson, Ian Johnston, Dominic Kwiatkowski, Cordelia Langford, John Sillitoe on behalf of the Wellcome Sanger Institute COVID-19 Surveillance Team |
| EPI_ISL_2867360 | Lighthouse Laboratory Plymouth | Wellcome Sanger Institute for the COVID-19 Genomics UK (COG-UK) Consortium | Lighthouse Laboratory Plymouth and Alex Alderton, Roberto Amato, Jeffrey Barrett, Sonia Goncalves, Ewan Harrison, David K. Jackson, Ian Johnston, Dominic Kwiatkowski, Cordelia Langford, John Sillitoe on behalf of the Wellcome Sanger Institute COVID-19 Surveillance Team |
| EPI_ISL_2867361, EPI_ISL_2867362, EPI_ISL_2867363, EPI_ISL_2867364 | Lighthouse Lab in Milton Keynes | Wellcome Sanger Institute for the COVID-19 Genomics UK (COG-UK) Consortium | The Lighthouse Lab in Milton Keynes and Alex Alderton, Roberto Amato, Jeffrey Barrett, Sonia Goncalves, Ewan Harrison, David K. Jackson, Ian Johnston, Dominic Kwiatkowski, Cordelia Langford, John Sillitoe on behalf of the Wellcome Sanger Institute COVID-19 Surveillance Team |
| EPI_ISL_2867365, EPI_ISL_2867366, EPI_ISL_2867367 | Berkshire and Surrey Pathology Services Lighthouse Laboratory | Wellcome Sanger Institute for the COVID-19 Genomics UK (COG-UK) Consortium | Berkshire and Surrey Pathology Services Lighthouse Laboratory and Alex Alderton, Roberto Amato, Jeffrey Barrett, Sonia Goncalves, Ewan Harrison, David K. Jackson, Ian Johnston, Dominic Kwiatkowski, Cordelia Langford, John Sillitoe on behalf of the Wellcome Sanger Institute COVID-19 Surveillance Team |
| EPI_ISL_2867368, EPI_ISL_2867369, EPI_ISL_2867370 | Lighthouse Lab in Milton Keynes | Wellcome Sanger Institute for the COVID-19 Genomics UK (COG-UK) Consortium | The Lighthouse Lab in Milton Keynes and Alex Alderton, Roberto Amato, Jeffrey Barrett, Sonia Goncalves, Ewan Harrison, David K. Jackson, Ian Johnston, Dominic Kwiatkowski, Cordelia Langford, John Sillitoe on behalf of the Wellcome Sanger Institute COVID-19 Surveillance Team |
| EPI_ISL_2867371 | Berkshire and Surrey Pathology Services Lighthouse Laboratory | Wellcome Sanger Institute for the COVID-19 Genomics UK (COG-UK) Consortium | Berkshire and Surrey Pathology Services Lighthouse Laboratory and Alex Alderton, Roberto Amato, Jeffrey Barrett, Sonia Goncalves, Ewan Harrison, David K. Jackson, Ian Johnston, Dominic Kwiatkowski, Cordelia Langford, John Sillitoe on behalf of the Wellcome Sanger Institute COVID-19 Surveillance Team |
| EPI_ISL_2867372 | Lighthouse Lab in Milton Keynes | Wellcome Sanger Institute for the COVID-19 Genomics UK (COG-UK) Consortium | The Lighthouse Lab in Milton Keynes and Alex Alderton, Roberto Amato, Jeffrey Barrett, Sonia Goncalves, Ewan Harrison, David K. Jackson, Ian Johnston, Dominic Kwiatkowski, Cordelia Langford, John Sillitoe on behalf of the Wellcome Sanger Institute COVID-19 Surveillance Team |
| EPI_ISL_2867373, EPI_ISL_2867374, EPI_ISL_2867375 | Berkshire and Surrey Pathology Services Lighthouse Laboratory | Wellcome Sanger Institute for the COVID-19 Genomics UK (COG-UK) Consortium | Berkshire and Surrey Pathology Services Lighthouse Laboratory and Alex Alderton, Roberto Amato, Jeffrey Barrett, Sonia Goncalves, Ewan Harrison, David K. Jackson, Ian Johnston, Dominic Kwiatkowski, Cordelia Langford, John Sillitoe on behalf of the Wellcome Sanger Institute COVID-19 Surveillance Team |
| EPI_ISL_2867376 | Lighthouse Lab in Milton Keynes | Wellcome Sanger Institute for the COVID-19 Genomics UK (COG-UK) Consortium | The Lighthouse Lab in Milton Keynes and Alex Alderton, Roberto Amato, Jeffrey Barrett, Sonia Goncalves, Ewan Harrison, David K. Jackson, Ian Johnston, Dominic Kwiatkowski, Cordelia Langford, John Sillitoe on behalf of the Wellcome Sanger Institute COVID-19 Surveillance Team |
| EPI_ISL_2867377 | Berkshire and Surrey Pathology Services Lighthouse Laboratory | Wellcome Sanger Institute for the COVID-19 Genomics UK (COG-UK) Consortium | Berkshire and Surrey Pathology Services Lighthouse Laboratory and Alex Alderton, Roberto Amato, Jeffrey Barrett, Sonia Goncalves, Ewan Harrison, David K. Jackson, Ian Johnston, Dominic Kwiatkowski, Cordelia Langford, John Sillitoe on behalf of the Wellcome Sanger Institute COVID-19 Surveillance Team |
| EPI_ISL_2867378, EPI_ISL_2867379, EPI_ISL_2867380, EPI_ISL_2867381, EPI_ISL_2867382 | Lighthouse Lab in Milton Keynes | Wellcome Sanger Institute for the COVID-19 Genomics UK (COG-UK) Consortium | The Lighthouse Lab in Milton Keynes and Alex Alderton, Roberto Amato, Jeffrey Barrett, Sonia Goncalves, Ewan Harrison, David K. Jackson, Ian Johnston, Dominic Kwiatkowski, Cordelia Langford, John Sillitoe on behalf of the Wellcome Sanger Institute COVID-19 Surveillance Team |
| EPI_ISL_2867383 | Berkshire and Surrey Pathology Services Lighthouse Laboratory | Wellcome Sanger Institute for the COVID-19 Genomics UK (COG-UK) Consortium | Berkshire and Surrey Pathology Services Lighthouse Laboratory and Alex Alderton, Roberto Amato, Jeffrey Barrett, Sonia Goncalves, Ewan Harrison, David K. Jackson, Ian Johnston, Dominic Kwiatkowski, Cordelia Langford, John Sillitoe on behalf of the Wellcome Sanger Institute COVID-19 Surveillance Team |
| EPI_ISL_2867384, EPI_ISL_2867385, EPI_ISL_2867386, EPI_ISL_2867387 | Lighthouse Lab in Milton Keynes | Wellcome Sanger Institute for the COVID-19 Genomics UK (COG-UK) Consortium | The Lighthouse Lab in Milton Keynes and Alex Alderton, Roberto Amato, Jeffrey Barrett, Sonia Goncalves, Ewan Harrison, David K. Jackson, Ian Johnston, Dominic Kwiatkowski, Cordelia Langford, John Sillitoe on behalf of the Wellcome Sanger Institute COVID-19 Surveillance Team |
| EPI_ISL_2867388, EPI_ISL_2867389, EPI_ISL_2867390 | Berkshire and Surrey Pathology Services Lighthouse Laboratory | Wellcome Sanger Institute for the COVID-19 Genomics UK (COG-UK) Consortium | Berkshire and Surrey Pathology Services Lighthouse Laboratory and Alex Alderton, Roberto Amato, Jeffrey Barrett, Sonia Goncalves, Ewan Harrison, David K. Jackson, Ian Johnston, Dominic Kwiatkowski, Cordelia Langford, John Sillitoe on behalf of the Wellcome Sanger Institute COVID-19 Surveillance Team |
| EPI_ISL_2867391, EPI_ISL_2867392, EPI_ISL_2867393 | Lighthouse Lab in Milton Keynes | Wellcome Sanger Institute for the COVID-19 Genomics UK (COG-UK) Consortium | The Lighthouse Lab in Milton Keynes and Alex Alderton, Roberto Amato, Jeffrey Barrett, Sonia Goncalves, Ewan Harrison, David K. Jackson, Ian Johnston, Dominic Kwiatkowski, Cordelia Langford, John Sillitoe on behalf of the Wellcome Sanger Institute COVID-19 Surveillance Team |
| EPI_ISL_2867394 | Berkshire and Surrey Pathology Services Lighthouse Laboratory | Wellcome Sanger Institute for the COVID-19 Genomics UK (COG-UK) Consortium | Berkshire and Surrey Pathology Services Lighthouse Laboratory and Alex Alderton, Roberto Amato, Jeffrey Barrett, Sonia Goncalves, Ewan Harrison, David K. Jackson, Ian Johnston, Dominic Kwiatkowski, Cordelia Langford, John Sillitoe on behalf of the Wellcome Sanger Institute COVID-19 Surveillance Team |
| EPI_ISL_2867395, EPI_ISL_2867396 | Lighthouse Lab in Milton Keynes | Wellcome Sanger Institute for the COVID-19 Genomics UK (COG-UK) Consortium | The Lighthouse Lab in Milton Keynes and Alex Alderton, Roberto Amato, Jeffrey Barrett, Sonia Goncalves, Ewan Harrison, David K. Jackson, Ian Johnston, Dominic Kwiatkowski, Cordelia Langford, John Sillitoe on behalf of the Wellcome Sanger Institute COVID-19 Surveillance Team |
| EPI_ISL_2867397, EPI_ISL_2867398 | Berkshire and Surrey Pathology Services Lighthouse Laboratory | Wellcome Sanger Institute for the COVID-19 Genomics UK (COG-UK) Consortium | Berkshire and Surrey Pathology Services Lighthouse Laboratory and Alex Alderton, Roberto Amato, Jeffrey Barrett, Sonia Goncalves, Ewan Harrison, David K. Jackson, Ian Johnston, Dominic Kwiatkowski, Cordelia Langford, John Sillitoe on behalf of the Wellcome Sanger Institute COVID-19 Surveillance Team |
| EPI_ISL_2867399 | Lighthouse Lab in Milton Keynes | Wellcome Sanger Institute for the COVID-19 Genomics UK (COG-UK) Consortium | The Lighthouse Lab in Milton Keynes and Alex Alderton, Roberto Amato, Jeffrey Barrett, Sonia Goncalves, Ewan Harrison, David K. Jackson, Ian Johnston, Dominic Kwiatkowski, Cordelia Langford, John Sillitoe on behalf of the Wellcome Sanger Institute COVID-19 Surveillance Team |
| EPI_ISL_2867400 | Berkshire and Surrey Pathology Services Lighthouse Laboratory | Wellcome Sanger Institute for the COVID-19 Genomics UK (COG-UK) Consortium | Berkshire and Surrey Pathology Services Lighthouse Laboratory and Alex Alderton, Roberto Amato, Jeffrey Barrett, Sonia Goncalves, Ewan Harrison, David K. Jackson, Ian Johnston, Dominic Kwiatkowski, Cordelia Langford, John Sillitoe on behalf of the Wellcome Sanger Institute COVID-19 Surveillance Team |
| EPI_ISL_2867401 | Lighthouse Lab in Milton Keynes | Wellcome Sanger Institute for the COVID-19 Genomics UK (COG-UK) Consortium | The Lighthouse Lab in Milton Keynes and Alex Alderton, Roberto Amato, Jeffrey Barrett, Sonia Goncalves, Ewan Harrison, David K. Jackson, Ian Johnston, Dominic Kwiatkowski, Cordelia Langford, John Sillitoe on behalf of the Wellcome Sanger Institute COVID-19 Surveillance Team |
| EPI_ISL_2867402, EPI_ISL_2867403, EPI_ISL_2867404, EPI_ISL_2867405 | Berkshire and Surrey Pathology Services Lighthouse Laboratory | Wellcome Sanger Institute for the COVID-19 Genomics UK (COG-UK) Consortium | Berkshire and Surrey Pathology Services Lighthouse Laboratory and Alex Alderton, Roberto Amato, Jeffrey Barrett, Sonia Goncalves, Ewan Harrison, David K. Jackson, Ian Johnston, Dominic Kwiatkowski, Cordelia Langford, John Sillitoe on behalf of the Wellcome Sanger Institute COVID-19 Surveillance Team |
| EPI_ISL_2867406 | Lighthouse Lab in Milton Keynes | Wellcome Sanger Institute for the COVID-19 Genomics UK (COG-UK) Consortium | The Lighthouse Lab in Milton Keynes and Alex Alderton, Roberto Amato, Jeffrey Barrett, Sonia Goncalves, Ewan Harrison, David K. Jackson, Ian Johnston, Dominic Kwiatkowski, Cordelia Langford, John Sillitoe on behalf of the Wellcome Sanger Institute COVID-19 Surveillance Team |
| EPI_ISL_2867407 | Berkshire and Surrey Pathology Services Lighthouse Laboratory | Wellcome Sanger Institute for the COVID-19 Genomics UK (COG-UK) Consortium | Berkshire and Surrey Pathology Services Lighthouse Laboratory and Alex Alderton, Roberto Amato, Jeffrey Barrett, Sonia Goncalves, Ewan Harrison, David K. Jackson, Ian Johnston, Dominic Kwiatkowski, Cordelia Langford, John Sillitoe on behalf of the Wellcome Sanger Institute COVID-19 Surveillance Team |
| EPI_ISL_2867408 | Lighthouse Lab in Milton Keynes | Wellcome Sanger Institute for the COVID-19 Genomics UK (COG-UK) Consortium | The Lighthouse Lab in Milton Keynes and Alex Alderton, Roberto Amato, Jeffrey Barrett, Sonia Goncalves, Ewan Harrison, David K. Jackson, Ian Johnston, Dominic Kwiatkowski, Cordelia Langford, John Sillitoe on behalf of the Wellcome Sanger Institute COVID-19 Surveillance Team |
| EPI_ISL_2867409 | Berkshire and Surrey Pathology Services Lighthouse Laboratory | Wellcome Sanger Institute for the COVID-19 Genomics UK (COG-UK) Consortium | Berkshire and Surrey Pathology Services Lighthouse Laboratory and Alex Alderton, Roberto Amato, Jeffrey Barrett, Sonia Goncalves, Ewan Harrison, David K. Jackson, Ian Johnston, Dominic Kwiatkowski, Cordelia Langford, John Sillitoe on behalf of the Wellcome Sanger Institute COVID-19 Surveillance Team |
| EPI_ISL_2867410 | Lighthouse Lab in Milton Keynes | Wellcome Sanger Institute for the COVID-19 Genomics UK (COG-UK) Consortium | The Lighthouse Lab in Milton Keynes and Alex Alderton, Roberto Amato, Jeffrey Barrett, Sonia Goncalves, Ewan Harrison, David K. Jackson, Ian Johnston, Dominic Kwiatkowski, Cordelia Langford, John Sillitoe on behalf of the Wellcome Sanger Institute COVID-19 Surveillance Team |
| EPI_ISL_2867411, EPI_ISL_2867412 | Berkshire and Surrey Pathology Services Lighthouse Laboratory | Wellcome Sanger Institute for the COVID-19 Genomics UK (COG-UK) Consortium | Berkshire and Surrey Pathology Services Lighthouse Laboratory and Alex Alderton, Roberto Amato, Jeffrey Barrett, Sonia Goncalves, Ewan Harrison, David K. Jackson, Ian Johnston, Dominic Kwiatkowski, Cordelia Langford, John Sillitoe on behalf of the Wellcome Sanger Institute COVID-19 Surveillance Team |
| EPI_ISL_2867413, EPI_ISL_2867414, EPI_ISL_2867415 | Lighthouse Lab in Milton Keynes | Wellcome Sanger Institute for the COVID-19 Genomics UK (COG-UK) Consortium | The Lighthouse Lab in Milton Keynes and Alex Alderton, Roberto Amato, Jeffrey Barrett, Sonia Goncalves, Ewan Harrison, David K. Jackson, Ian Johnston, Dominic Kwiatkowski, Cordelia Langford, John Sillitoe on behalf of the Wellcome Sanger Institute COVID-19 Surveillance Team |
| EPI_ISL_2867416, | Berkshire and Surrey Pathology Services Lighthouse Laboratory | Wellcome Sanger Institute for the COVID-19 Genomics UK (COG-UK) | Berkshire and Surrey Pathology Services Lighthouse Laboratory and Alex Alderton, Roberto Amato, Jeffrey Barrett, Sonia Goncalves, Ewan Harrison, David K. Jackson, Ian Johnston, Dominic Kwiatkowski, Cordelia Langford, John Sillitoe on behalf |

| | | | |
|---|---|---|---|
| EPI_ISL_2867417, EPI_ISL_2867418, EPI_ISL_2867419 | | Consortium | of the Wellcome Sanger Institute COVID-19 Surveillance Team |
| EPI_ISL_2867420, EPI_ISL_2867421, EPI_ISL_2867422 | Lighthouse Lab in Milton Keynes | Wellcome Sanger Institute for the COVID-19 Genomics UK (COG-UK) Consortium | The Lighthouse Lab in Milton Keynes and Alex Alderton, Roberto Amato, Jeffrey Barrett, Sonia Goncalves, Ewan Harrison, David K. Jackson, Ian Johnston, Dominic Kwiatkowski, Cordelia Langford, John Sillitoe on behalf of the Wellcome Sanger Institute COVID-19 Surveillance Team |
| EPI_ISL_2867423 | Berkshire and Surrey Pathology Services Lighthouse Laboratory | Wellcome Sanger Institute for the COVID-19 Genomics UK (COG-UK) Consortium | Berkshire and Surrey Pathology Services Lighthouse Laboratory and Alex Alderton, Roberto Amato, Jeffrey Barrett, Sonia Goncalves, Ewan Harrison, David K. Jackson, Ian Johnston, Dominic Kwiatkowski, Cordelia Langford, John Sillitoe on behalf of the Wellcome Sanger Institute COVID-19 Surveillance Team |
| EPI_ISL_2867424, EPI_ISL_2867425, EPI_ISL_2867426 | Lighthouse Lab in Milton Keynes | Wellcome Sanger Institute for the COVID-19 Genomics UK (COG-UK) Consortium | The Lighthouse Lab in Milton Keynes and Alex Alderton, Roberto Amato, Jeffrey Barrett, Sonia Goncalves, Ewan Harrison, David K. Jackson, Ian Johnston, Dominic Kwiatkowski, Cordelia Langford, John Sillitoe on behalf of the Wellcome Sanger Institute COVID-19 Surveillance Team |
| EPI_ISL_2867427, EPI_ISL_2867428, EPI_ISL_2867429 | Berkshire and Surrey Pathology Services Lighthouse Laboratory | Wellcome Sanger Institute for the COVID-19 Genomics UK (COG-UK) Consortium | Berkshire and Surrey Pathology Services Lighthouse Laboratory and Alex Alderton, Roberto Amato, Jeffrey Barrett, Sonia Goncalves, Ewan Harrison, David K. Jackson, Ian Johnston, Dominic Kwiatkowski, Cordelia Langford, John Sillitoe on behalf of the Wellcome Sanger Institute COVID-19 Surveillance Team |
| EPI_ISL_2867430, EPI_ISL_2867431 | Lighthouse Lab in Milton Keynes | Wellcome Sanger Institute for the COVID-19 Genomics UK (COG-UK) Consortium | The Lighthouse Lab in Milton Keynes and Alex Alderton, Roberto Amato, Jeffrey Barrett, Sonia Goncalves, Ewan Harrison, David K. Jackson, Ian Johnston, Dominic Kwiatkowski, Cordelia Langford, John Sillitoe on behalf of the Wellcome Sanger Institute COVID-19 Surveillance Team |
| EPI_ISL_2867432 | Berkshire and Surrey Pathology Services Lighthouse Laboratory | Wellcome Sanger Institute for the COVID-19 Genomics UK (COG-UK) Consortium | Berkshire and Surrey Pathology Services Lighthouse Laboratory and Alex Alderton, Roberto Amato, Jeffrey Barrett, Sonia Goncalves, Ewan Harrison, David K. Jackson, Ian Johnston, Dominic Kwiatkowski, Cordelia Langford, John Sillitoe on behalf of the Wellcome Sanger Institute COVID-19 Surveillance Team |
| EPI_ISL_2867433 | Lighthouse Lab in Milton Keynes | Wellcome Sanger Institute for the COVID-19 Genomics UK (COG-UK) Consortium | The Lighthouse Lab in Milton Keynes and Alex Alderton, Roberto Amato, Jeffrey Barrett, Sonia Goncalves, Ewan Harrison, David K. Jackson, Ian Johnston, Dominic Kwiatkowski, Cordelia Langford, John Sillitoe on behalf of the Wellcome Sanger Institute COVID-19 Surveillance Team |
| EPI_ISL_2867434, EPI_ISL_2867435, EPI_ISL_2867436, EPI_ISL_2867437 | Berkshire and Surrey Pathology Services Lighthouse Laboratory | Wellcome Sanger Institute for the COVID-19 Genomics UK (COG-UK) Consortium | Berkshire and Surrey Pathology Services Lighthouse Laboratory and Alex Alderton, Roberto Amato, Jeffrey Barrett, Sonia Goncalves, Ewan Harrison, David K. Jackson, Ian Johnston, Dominic Kwiatkowski, Cordelia Langford, John Sillitoe on behalf of the Wellcome Sanger Institute COVID-19 Surveillance Team |
| EPI_ISL_2867438 | Lighthouse Lab in Milton Keynes | Wellcome Sanger Institute for the COVID-19 Genomics UK (COG-UK) Consortium | The Lighthouse Lab in Milton Keynes and Alex Alderton, Roberto Amato, Jeffrey Barrett, Sonia Goncalves, Ewan Harrison, David K. Jackson, Ian Johnston, Dominic Kwiatkowski, Cordelia Langford, John Sillitoe on behalf of the Wellcome Sanger Institute COVID-19 Surveillance Team |
| EPI_ISL_2867439, EPI_ISL_2867440 | Berkshire and Surrey Pathology Services Lighthouse Laboratory | Wellcome Sanger Institute for the COVID-19 Genomics UK (COG-UK) Consortium | Berkshire and Surrey Pathology Services Lighthouse Laboratory and Alex Alderton, Roberto Amato, Jeffrey Barrett, Sonia Goncalves, Ewan Harrison, David K. Jackson, Ian Johnston, Dominic Kwiatkowski, Cordelia Langford, John Sillitoe on behalf of the Wellcome Sanger Institute COVID-19 Surveillance Team |
| EPI_ISL_2867441 | Lighthouse Lab in Milton Keynes | Wellcome Sanger Institute for the COVID-19 Genomics UK (COG-UK) Consortium | The Lighthouse Lab in Milton Keynes and Alex Alderton, Roberto Amato, Jeffrey Barrett, Sonia Goncalves, Ewan Harrison, David K. Jackson, Ian Johnston, Dominic Kwiatkowski, Cordelia Langford, John Sillitoe on behalf of the Wellcome Sanger Institute COVID-19 Surveillance Team |
| EPI_ISL_2867442, EPI_ISL_2867443, EPI_ISL_2867444, EPI_ISL_2867445, EPI_ISL_2867446, EPI_ISL_2867447 | Berkshire and Surrey Pathology Services Lighthouse Laboratory | Wellcome Sanger Institute for the COVID-19 Genomics UK (COG-UK) Consortium | Berkshire and Surrey Pathology Services Lighthouse Laboratory and Alex Alderton, Roberto Amato, Jeffrey Barrett, Sonia Goncalves, Ewan Harrison, David K. Jackson, Ian Johnston, Dominic Kwiatkowski, Cordelia Langford, John Sillitoe on behalf of the Wellcome Sanger Institute COVID-19 Surveillance Team |
| EPI_ISL_2867448 | Lighthouse Lab in Milton Keynes | Wellcome Sanger Institute for the COVID-19 Genomics UK (COG-UK) Consortium | The Lighthouse Lab in Milton Keynes and Alex Alderton, Roberto Amato, Jeffrey Barrett, Sonia Goncalves, Ewan Harrison, David K. Jackson, Ian Johnston, Dominic Kwiatkowski, Cordelia Langford, John Sillitoe on behalf of the Wellcome Sanger Institute COVID-19 Surveillance Team |
| EPI_ISL_2867449, EPI_ISL_2867450 | Berkshire and Surrey Pathology Services Lighthouse Laboratory | Wellcome Sanger Institute for the COVID-19 Genomics UK (COG-UK) Consortium | Berkshire and Surrey Pathology Services Lighthouse Laboratory and Alex Alderton, Roberto Amato, Jeffrey Barrett, Sonia Goncalves, Ewan Harrison, David K. Jackson, Ian Johnston, Dominic Kwiatkowski, Cordelia Langford, John Sillitoe on behalf of the Wellcome Sanger Institute COVID-19 Surveillance Team |
| EPI_ISL_2867451, EPI_ISL_2867452 | Lighthouse Lab in Milton Keynes | Wellcome Sanger Institute for the COVID-19 Genomics UK (COG-UK) Consortium | The Lighthouse Lab in Milton Keynes and Alex Alderton, Roberto Amato, Jeffrey Barrett, Sonia Goncalves, Ewan Harrison, David K. Jackson, Ian Johnston, Dominic Kwiatkowski, Cordelia Langford, John Sillitoe on behalf of the Wellcome Sanger Institute COVID-19 Surveillance Team |
| EPI_ISL_2867453 | Berkshire and Surrey Pathology Services Lighthouse Laboratory | Wellcome Sanger Institute for the COVID-19 Genomics UK (COG-UK) Consortium | Berkshire and Surrey Pathology Services Lighthouse Laboratory and Alex Alderton, Roberto Amato, Jeffrey Barrett, Sonia Goncalves, Ewan Harrison, David K. Jackson, Ian Johnston, Dominic Kwiatkowski, Cordelia Langford, John Sillitoe on behalf of the Wellcome Sanger Institute COVID-19 Surveillance Team |
| EPI_ISL_2867454 | Lighthouse Lab in Milton Keynes | Wellcome Sanger Institute for the COVID-19 Genomics UK (COG-UK) Consortium | The Lighthouse Lab in Milton Keynes and Alex Alderton, Roberto Amato, Jeffrey Barrett, Sonia Goncalves, Ewan Harrison, David K. Jackson, Ian Johnston, Dominic Kwiatkowski, Cordelia Langford, John Sillitoe on behalf of the Wellcome Sanger Institute COVID-19 Surveillance Team |
| EPI_ISL_2867455, EPI_ISL_2867456 | Berkshire and Surrey Pathology Services Lighthouse Laboratory | Wellcome Sanger Institute for the COVID-19 Genomics UK (COG-UK) Consortium | Berkshire and Surrey Pathology Services Lighthouse Laboratory and Alex Alderton, Roberto Amato, Jeffrey Barrett, Sonia Goncalves, Ewan Harrison, David K. Jackson, Ian Johnston, Dominic Kwiatkowski, Cordelia Langford, John Sillitoe on behalf of the Wellcome Sanger Institute COVID-19 Surveillance Team |
| EPI_ISL_2867457 | Lighthouse Lab in Milton Keynes | Wellcome Sanger Institute for the COVID-19 Genomics UK (COG-UK) Consortium | The Lighthouse Lab in Milton Keynes and Alex Alderton, Roberto Amato, Jeffrey Barrett, Sonia Goncalves, Ewan Harrison, David K. Jackson, Ian Johnston, Dominic Kwiatkowski, Cordelia Langford, John Sillitoe on behalf of the Wellcome Sanger Institute COVID-19 Surveillance Team |
| EPI_ISL_2867458, EPI_ISL_2867459 | Berkshire and Surrey Pathology Services Lighthouse Laboratory | Wellcome Sanger Institute for the COVID-19 Genomics UK (COG-UK) Consortium | Berkshire and Surrey Pathology Services Lighthouse Laboratory and Alex Alderton, Roberto Amato, Jeffrey Barrett, Sonia Goncalves, Ewan Harrison, David K. Jackson, Ian Johnston, Dominic Kwiatkowski, Cordelia Langford, John Sillitoe on behalf of the Wellcome Sanger Institute COVID-19 Surveillance Team |
| EPI_ISL_2867460 | Lighthouse Lab in Milton Keynes | Wellcome Sanger Institute for the COVID-19 Genomics UK (COG-UK) Consortium | The Lighthouse Lab in Milton Keynes and Alex Alderton, Roberto Amato, Jeffrey Barrett, Sonia Goncalves, Ewan Harrison, David K. Jackson, Ian Johnston, Dominic Kwiatkowski, Cordelia Langford, John Sillitoe on behalf of the Wellcome Sanger Institute COVID-19 Surveillance Team |
| EPI_ISL_2867461 | Berkshire and Surrey Pathology Services Lighthouse Laboratory | Wellcome Sanger Institute for the COVID-19 Genomics UK (COG-UK) Consortium | Berkshire and Surrey Pathology Services Lighthouse Laboratory and Alex Alderton, Roberto Amato, Jeffrey Barrett, Sonia Goncalves, Ewan Harrison, David K. Jackson, Ian Johnston, Dominic Kwiatkowski, Cordelia Langford, John Sillitoe on behalf of the Wellcome Sanger Institute COVID-19 Surveillance Team |
| EPI_ISL_2867462 | Lighthouse Lab in Milton Keynes | Wellcome Sanger Institute for the COVID-19 Genomics UK (COG-UK) Consortium | The Lighthouse Lab in Milton Keynes and Alex Alderton, Roberto Amato, Jeffrey Barrett, Sonia Goncalves, Ewan Harrison, David K. Jackson, Ian Johnston, Dominic Kwiatkowski, Cordelia Langford, John Sillitoe on behalf of the Wellcome Sanger Institute COVID-19 Surveillance Team |
| EPI_ISL_2867463 | Berkshire and Surrey Pathology Services Lighthouse Laboratory | Wellcome Sanger Institute for the COVID-19 Genomics UK (COG-UK) Consortium | Berkshire and Surrey Pathology Services Lighthouse Laboratory and Alex Alderton, Roberto Amato, Jeffrey Barrett, Sonia Goncalves, Ewan Harrison, David K. Jackson, Ian Johnston, Dominic Kwiatkowski, Cordelia Langford, John Sillitoe on behalf of the Wellcome Sanger Institute COVID-19 Surveillance Team |
| EPI_ISL_2867464 | Lighthouse Lab in Milton Keynes | Wellcome Sanger Institute for the COVID-19 Genomics UK (COG-UK) Consortium | The Lighthouse Lab in Milton Keynes and Alex Alderton, Roberto Amato, Jeffrey Barrett, Sonia Goncalves, Ewan Harrison, David K. Jackson, Ian Johnston, Dominic Kwiatkowski, Cordelia Langford, John Sillitoe on behalf of the Wellcome Sanger Institute COVID-19 Surveillance Team |
| EPI_ISL_2867465, EPI_ISL_2867466 | Berkshire and Surrey Pathology Services Lighthouse Laboratory | Wellcome Sanger Institute for the COVID-19 Genomics UK (COG-UK) Consortium | Berkshire and Surrey Pathology Services Lighthouse Laboratory and Alex Alderton, Roberto Amato, Jeffrey Barrett, Sonia Goncalves, Ewan Harrison, David K. Jackson, Ian Johnston, Dominic Kwiatkowski, Cordelia Langford, John Sillitoe on behalf of the Wellcome Sanger Institute COVID-19 Surveillance Team |
| EPI_ISL_2867467, EPI_ISL_2867468, EPI_ISL_2867469 | Lighthouse Lab in Milton Keynes | Wellcome Sanger Institute for the COVID-19 Genomics UK (COG-UK) Consortium | The Lighthouse Lab in Milton Keynes and Alex Alderton, Roberto Amato, Jeffrey Barrett, Sonia Goncalves, Ewan Harrison, David K. Jackson, Ian Johnston, Dominic Kwiatkowski, Cordelia Langford, John Sillitoe on behalf of the Wellcome Sanger Institute COVID-19 Surveillance Team |
| EPI_ISL_2867470 | Berkshire and Surrey Pathology Services Lighthouse Laboratory | Wellcome Sanger Institute for the COVID-19 Genomics UK (COG-UK) Consortium | Berkshire and Surrey Pathology Services Lighthouse Laboratory and Alex Alderton, Roberto Amato, Jeffrey Barrett, Sonia Goncalves, Ewan Harrison, David K. Jackson, Ian Johnston, Dominic Kwiatkowski, Cordelia Langford, John Sillitoe on behalf of the Wellcome Sanger Institute COVID-19 Surveillance Team |
| EPI_ISL_2867471, EPI_ISL_2867472, EPI_ISL_2867473, EPI_ISL_2867474, EPI_ISL_2867475 | Lighthouse Lab in Milton Keynes | Wellcome Sanger Institute for the COVID-19 Genomics UK (COG-UK) Consortium | The Lighthouse Lab in Milton Keynes and Alex Alderton, Roberto Amato, Jeffrey Barrett, Sonia Goncalves, Ewan Harrison, David K. Jackson, Ian Johnston, Dominic Kwiatkowski, Cordelia Langford, John Sillitoe on behalf of the Wellcome Sanger Institute COVID-19 Surveillance Team |
| EPI_ISL_2867476, EPI_ISL_2867477, EPI_ISL_2867478 | Berkshire and Surrey Pathology Services Lighthouse Laboratory | Wellcome Sanger Institute for the COVID-19 Genomics UK (COG-UK) Consortium | Berkshire and Surrey Pathology Services Lighthouse Laboratory and Alex Alderton, Roberto Amato, Jeffrey Barrett, Sonia Goncalves, Ewan Harrison, David K. Jackson, Ian Johnston, Dominic Kwiatkowski, Cordelia Langford, John Sillitoe on behalf of the Wellcome Sanger Institute COVID-19 Surveillance Team |
| EPI_ISL_2867479, EPI_ISL_2867480 | Lighthouse Lab in Milton Keynes | Wellcome Sanger Institute for the COVID-19 Genomics UK (COG-UK) Consortium | The Lighthouse Lab in Milton Keynes and Alex Alderton, Roberto Amato, Jeffrey Barrett, Sonia Goncalves, Ewan Harrison, David K. Jackson, Ian Johnston, Dominic Kwiatkowski, Cordelia Langford, John Sillitoe on behalf of the Wellcome Sanger Institute COVID-19 Surveillance Team |
| EPI_ISL_2867481, EPI_ISL_2867482 | Berkshire and Surrey Pathology Services Lighthouse Laboratory | Wellcome Sanger Institute for the COVID-19 Genomics UK (COG-UK) Consortium | Berkshire and Surrey Pathology Services Lighthouse Laboratory and Alex Alderton, Roberto Amato, Jeffrey Barrett, Sonia Goncalves, Ewan Harrison, David K. Jackson, Ian Johnston, Dominic Kwiatkowski, Cordelia Langford, John Sillitoe on behalf of the Wellcome Sanger Institute COVID-19 Surveillance Team |
| EPI_ISL_2867483, EPI_ISL_2867484 | Lighthouse Lab in Milton Keynes | Wellcome Sanger Institute for the COVID-19 Genomics UK (COG-UK) Consortium | The Lighthouse Lab in Milton Keynes and Alex Alderton, Roberto Amato, Jeffrey Barrett, Sonia Goncalves, Ewan Harrison, David K. Jackson, Ian Johnston, Dominic Kwiatkowski, Cordelia Langford, John Sillitoe on behalf of the Wellcome Sanger Institute COVID-19 Surveillance Team |
| EPI_ISL_2867485 | Berkshire and Surrey Pathology Services Lighthouse Laboratory | Wellcome Sanger Institute for the COVID-19 Genomics UK (COG-UK) Consortium | Berkshire and Surrey Pathology Services Lighthouse Laboratory and Alex Alderton, Roberto Amato, Jeffrey Barrett, Sonia Goncalves, Ewan Harrison, David K. Jackson, Ian Johnston, Dominic Kwiatkowski, Cordelia Langford, John Sillitoe on behalf of the Wellcome Sanger Institute COVID-19 Surveillance Team |
| EPI_ISL_2867486 | Lighthouse Lab in Milton Keynes | Wellcome Sanger Institute for the COVID-19 Genomics UK (COG-UK) Consortium | The Lighthouse Lab in Milton Keynes and Alex Alderton, Roberto Amato, Jeffrey Barrett, Sonia Goncalves, Ewan Harrison, David K. Jackson, Ian Johnston, Dominic Kwiatkowski, Cordelia Langford, John Sillitoe on behalf of the Wellcome Sanger Institute COVID-19 Surveillance Team |
| EPI_ISL_2867487, EPI_ISL_2867488 | Berkshire and Surrey Pathology Services Lighthouse Laboratory | Wellcome Sanger Institute for the COVID-19 Genomics UK (COG-UK) Consortium | Berkshire and Surrey Pathology Services Lighthouse Laboratory and Alex Alderton, Roberto Amato, Jeffrey Barrett, Sonia Goncalves, Ewan Harrison, David K. Jackson, Ian Johnston, Dominic Kwiatkowski, Cordelia Langford, John Sillitoe on behalf of the Wellcome Sanger Institute COVID-19 Surveillance Team |
| EPI_ISL_2867489, EPI_ISL_2867490 | Lighthouse Lab in Milton Keynes | Wellcome Sanger Institute for the COVID-19 Genomics UK (COG-UK) Consortium | The Lighthouse Lab in Milton Keynes and Alex Alderton, Roberto Amato, Jeffrey Barrett, Sonia Goncalves, Ewan Harrison, David K. Jackson, Ian Johnston, Dominic Kwiatkowski, Cordelia Langford, John Sillitoe on behalf of the Wellcome Sanger Institute COVID-19 Surveillance Team |
| EPI_ISL_2867491, EPI_ISL_2867492 | Berkshire and Surrey Pathology Services Lighthouse Laboratory | Wellcome Sanger Institute for the COVID-19 Genomics UK (COG-UK) Consortium | Berkshire and Surrey Pathology Services Lighthouse Laboratory and Alex Alderton, Roberto Amato, Jeffrey Barrett, Sonia Goncalves, Ewan Harrison, David K. Jackson, Ian Johnston, Dominic Kwiatkowski, Cordelia Langford, John Sillitoe on behalf of the Wellcome Sanger Institute COVID-19 Surveillance Team |

| | | | |
|---|---|---|---|
| EPI_ISL_2867493, EPI_ISL_2867494, EPI_ISL_2867495, EPI_ISL_2867496, EPI_ISL_2867497, EPI_ISL_2867498, EPI_ISL_2867499, EPI_ISL_2867501, EPI_ISL_2867502 | Lighthouse Lab in Milton Keynes | Wellcome Sanger Institute for the COVID-19 Genomics UK (COG-UK) Consortium | The Lighthouse Lab in Milton Keynes and Alex Alderton, Roberto Amato, Jeffrey Barrett, Sonia Goncalves, Ewan Harrison, David K. Jackson, Ian Johnston, Dominic Kwiatkowski, Cordelia Langford, John Sillitoe on behalf of the Wellcome Sanger Institute COVID-19 Surveillance Team |
| EPI_ISL_2867503, EPI_ISL_2867504, EPI_ISL_2867505 | Berkshire and Surrey Pathology Services Lighthouse Laboratory | Wellcome Sanger Institute for the COVID-19 Genomics UK (COG-UK) Consortium | Berkshire and Surrey Pathology Services Lighthouse Laboratory and Alex Alderton, Roberto Amato, Jeffrey Barrett, Sonia Goncalves, Ewan Harrison, David K. Jackson, Ian Johnston, Dominic Kwiatkowski, Cordelia Langford, John Sillitoe on behalf of the Wellcome Sanger Institute COVID-19 Surveillance Team |
| EPI_ISL_2867507 | Lighthouse Lab in Milton Keynes | Wellcome Sanger Institute for the COVID-19 Genomics UK (COG-UK) Consortium | The Lighthouse Lab in Milton Keynes and Alex Alderton, Roberto Amato, Jeffrey Barrett, Sonia Goncalves, Ewan Harrison, David K. Jackson, Ian Johnston, Dominic Kwiatkowski, Cordelia Langford, John Sillitoe on behalf of the Wellcome Sanger Institute COVID-19 Surveillance Team |
| EPI_ISL_2867508, EPI_ISL_2867509 | Berkshire and Surrey Pathology Services Lighthouse Laboratory | Wellcome Sanger Institute for the COVID-19 Genomics UK (COG-UK) Consortium | Berkshire and Surrey Pathology Services Lighthouse Laboratory and Alex Alderton, Roberto Amato, Jeffrey Barrett, Sonia Goncalves, Ewan Harrison, David K. Jackson, Ian Johnston, Dominic Kwiatkowski, Cordelia Langford, John Sillitoe on behalf of the Wellcome Sanger Institute COVID-19 Surveillance Team |
| EPI_ISL_2867510 | Lighthouse Lab in Milton Keynes | Wellcome Sanger Institute for the COVID-19 Genomics UK (COG-UK) Consortium | The Lighthouse Lab in Milton Keynes and Alex Alderton, Roberto Amato, Jeffrey Barrett, Sonia Goncalves, Ewan Harrison, David K. Jackson, Ian Johnston, Dominic Kwiatkowski, Cordelia Langford, John Sillitoe on behalf of the Wellcome Sanger Institute COVID-19 Surveillance Team |
| EPI_ISL_2867511 | Berkshire and Surrey Pathology Services Lighthouse Laboratory | Wellcome Sanger Institute for the COVID-19 Genomics UK (COG-UK) Consortium | Berkshire and Surrey Pathology Services Lighthouse Laboratory and Alex Alderton, Roberto Amato, Jeffrey Barrett, Sonia Goncalves, Ewan Harrison, David K. Jackson, Ian Johnston, Dominic Kwiatkowski, Cordelia Langford, John Sillitoe on behalf of the Wellcome Sanger Institute COVID-19 Surveillance Team |
| EPI_ISL_2867512, EPI_ISL_2867513 | Lighthouse Lab in Milton Keynes | Wellcome Sanger Institute for the COVID-19 Genomics UK (COG-UK) Consortium | The Lighthouse Lab in Milton Keynes and Alex Alderton, Roberto Amato, Jeffrey Barrett, Sonia Goncalves, Ewan Harrison, David K. Jackson, Ian Johnston, Dominic Kwiatkowski, Cordelia Langford, John Sillitoe on behalf of the Wellcome Sanger Institute COVID-19 Surveillance Team |
| EPI_ISL_2867514, EPI_ISL_2867515 | Berkshire and Surrey Pathology Services Lighthouse Laboratory | Wellcome Sanger Institute for the COVID-19 Genomics UK (COG-UK) Consortium | Berkshire and Surrey Pathology Services Lighthouse Laboratory and Alex Alderton, Roberto Amato, Jeffrey Barrett, Sonia Goncalves, Ewan Harrison, David K. Jackson, Ian Johnston, Dominic Kwiatkowski, Cordelia Langford, John Sillitoe on behalf of the Wellcome Sanger Institute COVID-19 Surveillance Team |
| EPI_ISL_2867516 | Lighthouse Lab in Milton Keynes | Wellcome Sanger Institute for the COVID-19 Genomics UK (COG-UK) Consortium | The Lighthouse Lab in Milton Keynes and Alex Alderton, Roberto Amato, Jeffrey Barrett, Sonia Goncalves, Ewan Harrison, David K. Jackson, Ian Johnston, Dominic Kwiatkowski, Cordelia Langford, John Sillitoe on behalf of the Wellcome Sanger Institute COVID-19 Surveillance Team |
| EPI_ISL_2867517 | Berkshire and Surrey Pathology Services Lighthouse Laboratory | Wellcome Sanger Institute for the COVID-19 Genomics UK (COG-UK) Consortium | Berkshire and Surrey Pathology Services Lighthouse Laboratory and Alex Alderton, Roberto Amato, Jeffrey Barrett, Sonia Goncalves, Ewan Harrison, David K. Jackson, Ian Johnston, Dominic Kwiatkowski, Cordelia Langford, John Sillitoe on behalf of the Wellcome Sanger Institute COVID-19 Surveillance Team |
| EPI_ISL_2867518, EPI_ISL_2867519, EPI_ISL_2867520 | Lighthouse Lab in Milton Keynes | Wellcome Sanger Institute for the COVID-19 Genomics UK (COG-UK) Consortium | The Lighthouse Lab in Milton Keynes and Alex Alderton, Roberto Amato, Jeffrey Barrett, Sonia Goncalves, Ewan Harrison, David K. Jackson, Ian Johnston, Dominic Kwiatkowski, Cordelia Langford, John Sillitoe on behalf of the Wellcome Sanger Institute COVID-19 Surveillance Team |
| EPI_ISL_2867521 | Berkshire and Surrey Pathology Services Lighthouse Laboratory | Wellcome Sanger Institute for the COVID-19 Genomics UK (COG-UK) Consortium | Berkshire and Surrey Pathology Services Lighthouse Laboratory and Alex Alderton, Roberto Amato, Jeffrey Barrett, Sonia Goncalves, Ewan Harrison, David K. Jackson, Ian Johnston, Dominic Kwiatkowski, Cordelia Langford, John Sillitoe on behalf of the Wellcome Sanger Institute COVID-19 Surveillance Team |
| EPI_ISL_2867522 | Lighthouse Lab in Milton Keynes | Wellcome Sanger Institute for the COVID-19 Genomics UK (COG-UK) Consortium | The Lighthouse Lab in Milton Keynes and Alex Alderton, Roberto Amato, Jeffrey Barrett, Sonia Goncalves, Ewan Harrison, David K. Jackson, Ian Johnston, Dominic Kwiatkowski, Cordelia Langford, John Sillitoe on behalf of the Wellcome Sanger Institute COVID-19 Surveillance Team |
| EPI_ISL_2867523, EPI_ISL_2867524 | Berkshire and Surrey Pathology Services Lighthouse Laboratory | Wellcome Sanger Institute for the COVID-19 Genomics UK (COG-UK) Consortium | Berkshire and Surrey Pathology Services Lighthouse Laboratory and Alex Alderton, Roberto Amato, Jeffrey Barrett, Sonia Goncalves, Ewan Harrison, David K. Jackson, Ian Johnston, Dominic Kwiatkowski, Cordelia Langford, John Sillitoe on behalf of the Wellcome Sanger Institute COVID-19 Surveillance Team |
| EPI_ISL_2867525, EPI_ISL_2867526 | Lighthouse Lab in Milton Keynes | Wellcome Sanger Institute for the COVID-19 Genomics UK (COG-UK) Consortium | The Lighthouse Lab in Milton Keynes and Alex Alderton, Roberto Amato, Jeffrey Barrett, Sonia Goncalves, Ewan Harrison, David K. Jackson, Ian Johnston, Dominic Kwiatkowski, Cordelia Langford, John Sillitoe on behalf of the Wellcome Sanger Institute COVID-19 Surveillance Team |
| EPI_ISL_2867527, EPI_ISL_2867528 | Berkshire and Surrey Pathology Services Lighthouse Laboratory | Wellcome Sanger Institute for the COVID-19 Genomics UK (COG-UK) Consortium | Berkshire and Surrey Pathology Services Lighthouse Laboratory and Alex Alderton, Roberto Amato, Jeffrey Barrett, Sonia Goncalves, Ewan Harrison, David K. Jackson, Ian Johnston, Dominic Kwiatkowski, Cordelia Langford, John Sillitoe on behalf of the Wellcome Sanger Institute COVID-19 Surveillance Team |
| EPI_ISL_2867529, EPI_ISL_2867530, EPI_ISL_2867531, EPI_ISL_2867532 | Lighthouse Lab in Milton Keynes | Wellcome Sanger Institute for the COVID-19 Genomics UK (COG-UK) Consortium | The Lighthouse Lab in Milton Keynes and Alex Alderton, Roberto Amato, Jeffrey Barrett, Sonia Goncalves, Ewan Harrison, David K. Jackson, Ian Johnston, Dominic Kwiatkowski, Cordelia Langford, John Sillitoe on behalf of the Wellcome Sanger Institute COVID-19 Surveillance Team |
| EPI_ISL_2867533, EPI_ISL_2867534, EPI_ISL_2867535, EPI_ISL_2867536, EPI_ISL_2867537 | Berkshire and Surrey Pathology Services Lighthouse Laboratory | Wellcome Sanger Institute for the COVID-19 Genomics UK (COG-UK) Consortium | Berkshire and Surrey Pathology Services Lighthouse Laboratory and Alex Alderton, Roberto Amato, Jeffrey Barrett, Sonia Goncalves, Ewan Harrison, David K. Jackson, Ian Johnston, Dominic Kwiatkowski, Cordelia Langford, John Sillitoe on behalf of the Wellcome Sanger Institute COVID-19 Surveillance Team |
| EPI_ISL_2867538, EPI_ISL_2867539 | Lighthouse Lab in Milton Keynes | Wellcome Sanger Institute for the COVID-19 Genomics UK (COG-UK) Consortium | The Lighthouse Lab in Milton Keynes and Alex Alderton, Roberto Amato, Jeffrey Barrett, Sonia Goncalves, Ewan Harrison, David K. Jackson, Ian Johnston, Dominic Kwiatkowski, Cordelia Langford, John Sillitoe on behalf of the Wellcome Sanger Institute COVID-19 Surveillance Team |
| EPI_ISL_2867540, EPI_ISL_2867541 | Berkshire and Surrey Pathology Services Lighthouse Laboratory | Wellcome Sanger Institute for the COVID-19 Genomics UK (COG-UK) Consortium | Berkshire and Surrey Pathology Services Lighthouse Laboratory and Alex Alderton, Roberto Amato, Jeffrey Barrett, Sonia Goncalves, Ewan Harrison, David K. Jackson, Ian Johnston, Dominic Kwiatkowski, Cordelia Langford, John Sillitoe on behalf of the Wellcome Sanger Institute COVID-19 Surveillance Team |
| EPI_ISL_2867542, EPI_ISL_2867543, EPI_ISL_2867544, EPI_ISL_2867545, EPI_ISL_2867546, EPI_ISL_2867547 | Lighthouse Lab in Milton Keynes | Wellcome Sanger Institute for the COVID-19 Genomics UK (COG-UK) Consortium | The Lighthouse Lab in Milton Keynes and Alex Alderton, Roberto Amato, Jeffrey Barrett, Sonia Goncalves, Ewan Harrison, David K. Jackson, Ian Johnston, Dominic Kwiatkowski, Cordelia Langford, John Sillitoe on behalf of the Wellcome Sanger Institute COVID-19 Surveillance Team |
| EPI_ISL_2867548 | Berkshire and Surrey Pathology Services Lighthouse Laboratory | Wellcome Sanger Institute for the COVID-19 Genomics UK (COG-UK) Consortium | Berkshire and Surrey Pathology Services Lighthouse Laboratory and Alex Alderton, Roberto Amato, Jeffrey Barrett, Sonia Goncalves, Ewan Harrison, David K. Jackson, Ian Johnston, Dominic Kwiatkowski, Cordelia Langford, John Sillitoe on behalf of the Wellcome Sanger Institute COVID-19 Surveillance Team |
| EPI_ISL_2867549, EPI_ISL_2867550, EPI_ISL_2867551, EPI_ISL_2867552, EPI_ISL_2867553 | Lighthouse Lab in Milton Keynes | Wellcome Sanger Institute for the COVID-19 Genomics UK (COG-UK) Consortium | The Lighthouse Lab in Milton Keynes and Alex Alderton, Roberto Amato, Jeffrey Barrett, Sonia Goncalves, Ewan Harrison, David K. Jackson, Ian Johnston, Dominic Kwiatkowski, Cordelia Langford, John Sillitoe on behalf of the Wellcome Sanger Institute COVID-19 Surveillance Team |
| EPI_ISL_2867554, EPI_ISL_2867555 | Berkshire and Surrey Pathology Services Lighthouse Laboratory | Wellcome Sanger Institute for the COVID-19 Genomics UK (COG-UK) Consortium | Berkshire and Surrey Pathology Services Lighthouse Laboratory and Alex Alderton, Roberto Amato, Jeffrey Barrett, Sonia Goncalves, Ewan Harrison, David K. Jackson, Ian Johnston, Dominic Kwiatkowski, Cordelia Langford, John Sillitoe on behalf of the Wellcome Sanger Institute COVID-19 Surveillance Team |
| EPI_ISL_2867556, EPI_ISL_2867557, EPI_ISL_2867558, EPI_ISL_2867559 | Lighthouse Lab in Milton Keynes | Wellcome Sanger Institute for the COVID-19 Genomics UK (COG-UK) Consortium | The Lighthouse Lab in Milton Keynes and Alex Alderton, Roberto Amato, Jeffrey Barrett, Sonia Goncalves, Ewan Harrison, David K. Jackson, Ian Johnston, Dominic Kwiatkowski, Cordelia Langford, John Sillitoe on behalf of the Wellcome Sanger Institute COVID-19 Surveillance Team |
| EPI_ISL_2867560, EPI_ISL_2867561 | Berkshire and Surrey Pathology Services Lighthouse Laboratory | Wellcome Sanger Institute for the COVID-19 Genomics UK (COG-UK) Consortium | Berkshire and Surrey Pathology Services Lighthouse Laboratory and Alex Alderton, Roberto Amato, Jeffrey Barrett, Sonia Goncalves, Ewan Harrison, David K. Jackson, Ian Johnston, Dominic Kwiatkowski, Cordelia Langford, John Sillitoe on behalf of the Wellcome Sanger Institute COVID-19 Surveillance Team |
| EPI_ISL_2867562, EPI_ISL_2867563 | Lighthouse Lab in Milton Keynes | Wellcome Sanger Institute for the COVID-19 Genomics UK (COG-UK) Consortium | The Lighthouse Lab in Milton Keynes and Alex Alderton, Roberto Amato, Jeffrey Barrett, Sonia Goncalves, Ewan Harrison, David K. Jackson, Ian Johnston, Dominic Kwiatkowski, Cordelia Langford, John Sillitoe on behalf of the Wellcome Sanger Institute COVID-19 Surveillance Team |
| EPI_ISL_2867564, EPI_ISL_2867565, EPI_ISL_2867566 | Berkshire and Surrey Pathology Services Lighthouse Laboratory | Wellcome Sanger Institute for the COVID-19 Genomics UK (COG-UK) Consortium | Berkshire and Surrey Pathology Services Lighthouse Laboratory and Alex Alderton, Roberto Amato, Jeffrey Barrett, Sonia Goncalves, Ewan Harrison, David K. Jackson, Ian Johnston, Dominic Kwiatkowski, Cordelia Langford, John Sillitoe on behalf of the Wellcome Sanger Institute COVID-19 Surveillance Team |
| EPI_ISL_2867567 | Lighthouse Lab in Milton Keynes | Wellcome Sanger Institute for the COVID-19 Genomics UK (COG-UK) Consortium | The Lighthouse Lab in Milton Keynes and Alex Alderton, Roberto Amato, Jeffrey Barrett, Sonia Goncalves, Ewan Harrison, David K. Jackson, Ian Johnston, Dominic Kwiatkowski, Cordelia Langford, John Sillitoe on behalf of the Wellcome Sanger Institute COVID-19 Surveillance Team |
| EPI_ISL_2867568, EPI_ISL_2867569, EPI_ISL_2867570, EPI_ISL_2867571, EPI_ISL_2867572, EPI_ISL_2867573 | Berkshire and Surrey Pathology Services Lighthouse Laboratory | Wellcome Sanger Institute for the COVID-19 Genomics UK (COG-UK) Consortium | Berkshire and Surrey Pathology Services Lighthouse Laboratory and Alex Alderton, Roberto Amato, Jeffrey Barrett, Sonia Goncalves, Ewan Harrison, David K. Jackson, Ian Johnston, Dominic Kwiatkowski, Cordelia Langford, John Sillitoe on behalf of the Wellcome Sanger Institute COVID-19 Surveillance Team |
| EPI_ISL_2867574 | Lighthouse Lab in Milton Keynes | Wellcome Sanger Institute for the COVID-19 Genomics UK (COG-UK) Consortium | The Lighthouse Lab in Milton Keynes and Alex Alderton, Roberto Amato, Jeffrey Barrett, Sonia Goncalves, Ewan Harrison, David K. Jackson, Ian Johnston, Dominic Kwiatkowski, Cordelia Langford, John Sillitoe on behalf of the Wellcome Sanger Institute COVID-19 Surveillance Team |
| EPI_ISL_2867575, EPI_ISL_2867576, EPI_ISL_2867577 | Berkshire and Surrey Pathology Services Lighthouse Laboratory | Wellcome Sanger Institute for the COVID-19 Genomics UK (COG-UK) Consortium | Berkshire and Surrey Pathology Services Lighthouse Laboratory and Alex Alderton, Roberto Amato, Jeffrey Barrett, Sonia Goncalves, Ewan Harrison, David K. Jackson, Ian Johnston, Dominic Kwiatkowski, Cordelia Langford, John Sillitoe on behalf of the Wellcome Sanger Institute COVID-19 Surveillance Team |
| EPI_ISL_2867578, EPI_ISL_2867579, | Lighthouse Lab in Milton Keynes | Wellcome Sanger Institute for the COVID-19 Genomics UK (COG-UK) Consortium | The Lighthouse Lab in Milton Keynes and Alex Alderton, Roberto Amato, Jeffrey Barrett, Sonia Goncalves, Ewan Harrison, David K. Jackson, Ian Johnston, Dominic Kwiatkowski, Cordelia Langford, John Sillitoe on behalf of the Wellcome Sanger Institute COVID-19 Surveillance Team |

| | | | |
|---|---|---|---|
| EPI_ISL_2867580 | | | |
| EPI_ISL_2867581 | Berkshire and Surrey Pathology Services Lighthouse Laboratory | Wellcome Sanger Institute for the COVID-19 Genomics UK (COG-UK) Consortium | Berkshire and Surrey Pathology Services Lighthouse Laboratory and Alex Alderton, Roberto Amato, Jeffrey Barrett, Sonia Goncalves, Ewan Harrison, David K. Jackson, Ian Johnston, Dominic Kwiatkowski, Cordelia Langford, John Sillitoe on behalf of the Wellcome Sanger Institute COVID-19 Surveillance Team |
| EPI_ISL_2867582 | Lighthouse Lab in Milton Keynes | Wellcome Sanger Institute for the COVID-19 Genomics UK (COG-UK) Consortium | The Lighthouse Lab in Milton Keynes and Alex Alderton, Roberto Amato, Jeffrey Barrett, Sonia Goncalves, Ewan Harrison, David K. Jackson, Ian Johnston, Dominic Kwiatkowski, Cordelia Langford, John Sillitoe on behalf of the Wellcome Sanger Institute COVID-19 Surveillance Team |
| EPI_ISL_2867583, EPI_ISL_2867584, EPI_ISL_2867585 | Berkshire and Surrey Pathology Services Lighthouse Laboratory | Wellcome Sanger Institute for the COVID-19 Genomics UK (COG-UK) Consortium | Berkshire and Surrey Pathology Services Lighthouse Laboratory and Alex Alderton, Roberto Amato, Jeffrey Barrett, Sonia Goncalves, Ewan Harrison, David K. Jackson, Ian Johnston, Dominic Kwiatkowski, Cordelia Langford, John Sillitoe on behalf of the Wellcome Sanger Institute COVID-19 Surveillance Team |
| EPI_ISL_2867586, EPI_ISL_2867587, EPI_ISL_2867588 | Lighthouse Lab in Milton Keynes | Wellcome Sanger Institute for the COVID-19 Genomics UK (COG-UK) Consortium | The Lighthouse Lab in Milton Keynes and Alex Alderton, Roberto Amato, Jeffrey Barrett, Sonia Goncalves, Ewan Harrison, David K. Jackson, Ian Johnston, Dominic Kwiatkowski, Cordelia Langford, John Sillitoe on behalf of the Wellcome Sanger Institute COVID-19 Surveillance Team |
| EPI_ISL_2867589, EPI_ISL_2867590, EPI_ISL_2867591, EPI_ISL_2867592, EPI_ISL_2867593, EPI_ISL_2867594 | Berkshire and Surrey Pathology Services Lighthouse Laboratory | Wellcome Sanger Institute for the COVID-19 Genomics UK (COG-UK) Consortium | Berkshire and Surrey Pathology Services Lighthouse Laboratory and Alex Alderton, Roberto Amato, Jeffrey Barrett, Sonia Goncalves, Ewan Harrison, David K. Jackson, Ian Johnston, Dominic Kwiatkowski, Cordelia Langford, John Sillitoe on behalf of the Wellcome Sanger Institute COVID-19 Surveillance Team |
| EPI_ISL_2867595 | Lighthouse Lab in Milton Keynes | Wellcome Sanger Institute for the COVID-19 Genomics UK (COG-UK) Consortium | The Lighthouse Lab in Milton Keynes and Alex Alderton, Roberto Amato, Jeffrey Barrett, Sonia Goncalves, Ewan Harrison, David K. Jackson, Ian Johnston, Dominic Kwiatkowski, Cordelia Langford, John Sillitoe on behalf of the Wellcome Sanger Institute COVID-19 Surveillance Team |
| EPI_ISL_2867596 | Berkshire and Surrey Pathology Services Lighthouse Laboratory | Wellcome Sanger Institute for the COVID-19 Genomics UK (COG-UK) Consortium | Berkshire and Surrey Pathology Services Lighthouse Laboratory and Alex Alderton, Roberto Amato, Jeffrey Barrett, Sonia Goncalves, Ewan Harrison, David K. Jackson, Ian Johnston, Dominic Kwiatkowski, Cordelia Langford, John Sillitoe on behalf of the Wellcome Sanger Institute COVID-19 Surveillance Team |
| EPI_ISL_2867597 | Lighthouse Lab in Milton Keynes | Wellcome Sanger Institute for the COVID-19 Genomics UK (COG-UK) Consortium | The Lighthouse Lab in Milton Keynes and Alex Alderton, Roberto Amato, Jeffrey Barrett, Sonia Goncalves, Ewan Harrison, David K. Jackson, Ian Johnston, Dominic Kwiatkowski, Cordelia Langford, John Sillitoe on behalf of the Wellcome Sanger Institute COVID-19 Surveillance Team |
| EPI_ISL_2867598 | Berkshire and Surrey Pathology Services Lighthouse Laboratory | Wellcome Sanger Institute for the COVID-19 Genomics UK (COG-UK) Consortium | Berkshire and Surrey Pathology Services Lighthouse Laboratory and Alex Alderton, Roberto Amato, Jeffrey Barrett, Sonia Goncalves, Ewan Harrison, David K. Jackson, Ian Johnston, Dominic Kwiatkowski, Cordelia Langford, John Sillitoe on behalf of the Wellcome Sanger Institute COVID-19 Surveillance Team |
| EPI_ISL_2867599, EPI_ISL_2867600 | Lighthouse Lab in Milton Keynes | Wellcome Sanger Institute for the COVID-19 Genomics UK (COG-UK) Consortium | The Lighthouse Lab in Milton Keynes and Alex Alderton, Roberto Amato, Jeffrey Barrett, Sonia Goncalves, Ewan Harrison, David K. Jackson, Ian Johnston, Dominic Kwiatkowski, Cordelia Langford, John Sillitoe on behalf of the Wellcome Sanger Institute COVID-19 Surveillance Team |
| EPI_ISL_2867601 | Berkshire and Surrey Pathology Services Lighthouse Laboratory | Wellcome Sanger Institute for the COVID-19 Genomics UK (COG-UK) Consortium | Berkshire and Surrey Pathology Services Lighthouse Laboratory and Alex Alderton, Roberto Amato, Jeffrey Barrett, Sonia Goncalves, Ewan Harrison, David K. Jackson, Ian Johnston, Dominic Kwiatkowski, Cordelia Langford, John Sillitoe on behalf of the Wellcome Sanger Institute COVID-19 Surveillance Team |
| EPI_ISL_2867602 | Lighthouse Lab in Milton Keynes | Wellcome Sanger Institute for the COVID-19 Genomics UK (COG-UK) Consortium | The Lighthouse Lab in Milton Keynes and Alex Alderton, Roberto Amato, Jeffrey Barrett, Sonia Goncalves, Ewan Harrison, David K. Jackson, Ian Johnston, Dominic Kwiatkowski, Cordelia Langford, John Sillitoe on behalf of the Wellcome Sanger Institute COVID-19 Surveillance Team |
| EPI_ISL_2867603, EPI_ISL_2867604 | Berkshire and Surrey Pathology Services Lighthouse Laboratory | Wellcome Sanger Institute for the COVID-19 Genomics UK (COG-UK) Consortium | Berkshire and Surrey Pathology Services Lighthouse Laboratory and Alex Alderton, Roberto Amato, Jeffrey Barrett, Sonia Goncalves, Ewan Harrison, David K. Jackson, Ian Johnston, Dominic Kwiatkowski, Cordelia Langford, John Sillitoe on behalf of the Wellcome Sanger Institute COVID-19 Surveillance Team |
| EPI_ISL_2867605, EPI_ISL_2867606 | Lighthouse Lab in Milton Keynes | Wellcome Sanger Institute for the COVID-19 Genomics UK (COG-UK) Consortium | The Lighthouse Lab in Milton Keynes and Alex Alderton, Roberto Amato, Jeffrey Barrett, Sonia Goncalves, Ewan Harrison, David K. Jackson, Ian Johnston, Dominic Kwiatkowski, Cordelia Langford, John Sillitoe on behalf of the Wellcome Sanger Institute COVID-19 Surveillance Team |
| EPI_ISL_2867607 | Berkshire and Surrey Pathology Services Lighthouse Laboratory | Wellcome Sanger Institute for the COVID-19 Genomics UK (COG-UK) Consortium | Berkshire and Surrey Pathology Services Lighthouse Laboratory and Alex Alderton, Roberto Amato, Jeffrey Barrett, Sonia Goncalves, Ewan Harrison, David K. Jackson, Ian Johnston, Dominic Kwiatkowski, Cordelia Langford, John Sillitoe on behalf of the Wellcome Sanger Institute COVID-19 Surveillance Team |
| EPI_ISL_2867608, EPI_ISL_2867609 | Lighthouse Lab in Milton Keynes | Wellcome Sanger Institute for the COVID-19 Genomics UK (COG-UK) Consortium | The Lighthouse Lab in Milton Keynes and Alex Alderton, Roberto Amato, Jeffrey Barrett, Sonia Goncalves, Ewan Harrison, David K. Jackson, Ian Johnston, Dominic Kwiatkowski, Cordelia Langford, John Sillitoe on behalf of the Wellcome Sanger Institute COVID-19 Surveillance Team |
| EPI_ISL_2867610 | Berkshire and Surrey Pathology Services Lighthouse Laboratory | Wellcome Sanger Institute for the COVID-19 Genomics UK (COG-UK) Consortium | Berkshire and Surrey Pathology Services Lighthouse Laboratory and Alex Alderton, Roberto Amato, Jeffrey Barrett, Sonia Goncalves, Ewan Harrison, David K. Jackson, Ian Johnston, Dominic Kwiatkowski, Cordelia Langford, John Sillitoe on behalf of the Wellcome Sanger Institute COVID-19 Surveillance Team |
| EPI_ISL_2867611 | Lighthouse Lab in Milton Keynes | Wellcome Sanger Institute for the COVID-19 Genomics UK (COG-UK) Consortium | The Lighthouse Lab in Milton Keynes and Alex Alderton, Roberto Amato, Jeffrey Barrett, Sonia Goncalves, Ewan Harrison, David K. Jackson, Ian Johnston, Dominic Kwiatkowski, Cordelia Langford, John Sillitoe on behalf of the Wellcome Sanger Institute COVID-19 Surveillance Team |
| EPI_ISL_2867612 | Berkshire and Surrey Pathology Services Lighthouse Laboratory | Wellcome Sanger Institute for the COVID-19 Genomics UK (COG-UK) Consortium | Berkshire and Surrey Pathology Services Lighthouse Laboratory and Alex Alderton, Roberto Amato, Jeffrey Barrett, Sonia Goncalves, Ewan Harrison, David K. Jackson, Ian Johnston, Dominic Kwiatkowski, Cordelia Langford, John Sillitoe on behalf of the Wellcome Sanger Institute COVID-19 Surveillance Team |
| EPI_ISL_2867613, EPI_ISL_2867614, EPI_ISL_2867615 | Lighthouse Lab in Milton Keynes | Wellcome Sanger Institute for the COVID-19 Genomics UK (COG-UK) Consortium | The Lighthouse Lab in Milton Keynes and Alex Alderton, Roberto Amato, Jeffrey Barrett, Sonia Goncalves, Ewan Harrison, David K. Jackson, Ian Johnston, Dominic Kwiatkowski, Cordelia Langford, John Sillitoe on behalf of the Wellcome Sanger Institute COVID-19 Surveillance Team |
| EPI_ISL_2867616, EPI_ISL_2867617, EPI_ISL_2867618, EPI_ISL_2867619, EPI_ISL_2867620 | Berkshire and Surrey Pathology Services Lighthouse Laboratory | Wellcome Sanger Institute for the COVID-19 Genomics UK (COG-UK) Consortium | Berkshire and Surrey Pathology Services Lighthouse Laboratory and Alex Alderton, Roberto Amato, Jeffrey Barrett, Sonia Goncalves, Ewan Harrison, David K. Jackson, Ian Johnston, Dominic Kwiatkowski, Cordelia Langford, John Sillitoe on behalf of the Wellcome Sanger Institute COVID-19 Surveillance Team |
| EPI_ISL_2867621, EPI_ISL_2867622 | Lighthouse Lab in Milton Keynes | Wellcome Sanger Institute for the COVID-19 Genomics UK (COG-UK) Consortium | The Lighthouse Lab in Milton Keynes and Alex Alderton, Roberto Amato, Jeffrey Barrett, Sonia Goncalves, Ewan Harrison, David K. Jackson, Ian Johnston, Dominic Kwiatkowski, Cordelia Langford, John Sillitoe on behalf of the Wellcome Sanger Institute COVID-19 Surveillance Team |
| EPI_ISL_2867623, EPI_ISL_2867624, EPI_ISL_2867625, EPI_ISL_2867626, EPI_ISL_2867627 | Berkshire and Surrey Pathology Services Lighthouse Laboratory | Wellcome Sanger Institute for the COVID-19 Genomics UK (COG-UK) Consortium | Berkshire and Surrey Pathology Services Lighthouse Laboratory and Alex Alderton, Roberto Amato, Jeffrey Barrett, Sonia Goncalves, Ewan Harrison, David K. Jackson, Ian Johnston, Dominic Kwiatkowski, Cordelia Langford, John Sillitoe on behalf of the Wellcome Sanger Institute COVID-19 Surveillance Team |
| EPI_ISL_2867628 | Lighthouse Lab in Milton Keynes | Wellcome Sanger Institute for the COVID-19 Genomics UK (COG-UK) Consortium | The Lighthouse Lab in Milton Keynes and Alex Alderton, Roberto Amato, Jeffrey Barrett, Sonia Goncalves, Ewan Harrison, David K. Jackson, Ian Johnston, Dominic Kwiatkowski, Cordelia Langford, John Sillitoe on behalf of the Wellcome Sanger Institute COVID-19 Surveillance Team |
| EPI_ISL_2867629, EPI_ISL_2867630 | Berkshire and Surrey Pathology Services Lighthouse Laboratory | Wellcome Sanger Institute for the COVID-19 Genomics UK (COG-UK) Consortium | Berkshire and Surrey Pathology Services Lighthouse Laboratory and Alex Alderton, Roberto Amato, Jeffrey Barrett, Sonia Goncalves, Ewan Harrison, David K. Jackson, Ian Johnston, Dominic Kwiatkowski, Cordelia Langford, John Sillitoe on behalf of the Wellcome Sanger Institute COVID-19 Surveillance Team |
| EPI_ISL_2867631, EPI_ISL_2867632 | Lighthouse Lab in Milton Keynes | Wellcome Sanger Institute for the COVID-19 Genomics UK (COG-UK) Consortium | The Lighthouse Lab in Milton Keynes and Alex Alderton, Roberto Amato, Jeffrey Barrett, Sonia Goncalves, Ewan Harrison, David K. Jackson, Ian Johnston, Dominic Kwiatkowski, Cordelia Langford, John Sillitoe on behalf of the Wellcome Sanger Institute COVID-19 Surveillance Team |
| EPI_ISL_2867633 | Berkshire and Surrey Pathology Services Lighthouse Laboratory | Wellcome Sanger Institute for the COVID-19 Genomics UK (COG-UK) Consortium | Berkshire and Surrey Pathology Services Lighthouse Laboratory and Alex Alderton, Roberto Amato, Jeffrey Barrett, Sonia Goncalves, Ewan Harrison, David K. Jackson, Ian Johnston, Dominic Kwiatkowski, Cordelia Langford, John Sillitoe on behalf of the Wellcome Sanger Institute COVID-19 Surveillance Team |
| EPI_ISL_2867634, EPI_ISL_2867635, EPI_ISL_2867636 | Lighthouse Lab in Milton Keynes | Wellcome Sanger Institute for the COVID-19 Genomics UK (COG-UK) Consortium | The Lighthouse Lab in Milton Keynes and Alex Alderton, Roberto Amato, Jeffrey Barrett, Sonia Goncalves, Ewan Harrison, David K. Jackson, Ian Johnston, Dominic Kwiatkowski, Cordelia Langford, John Sillitoe on behalf of the Wellcome Sanger Institute COVID-19 Surveillance Team |
| EPI_ISL_2867637 | Berkshire and Surrey Pathology Services Lighthouse Laboratory | Wellcome Sanger Institute for the COVID-19 Genomics UK (COG-UK) Consortium | Berkshire and Surrey Pathology Services Lighthouse Laboratory and Alex Alderton, Roberto Amato, Jeffrey Barrett, Sonia Goncalves, Ewan Harrison, David K. Jackson, Ian Johnston, Dominic Kwiatkowski, Cordelia Langford, John Sillitoe on behalf of the Wellcome Sanger Institute COVID-19 Surveillance Team |
| EPI_ISL_2867638 | Lighthouse Lab in Milton Keynes | Wellcome Sanger Institute for the COVID-19 Genomics UK (COG-UK) Consortium | The Lighthouse Lab in Milton Keynes and Alex Alderton, Roberto Amato, Jeffrey Barrett, Sonia Goncalves, Ewan Harrison, David K. Jackson, Ian Johnston, Dominic Kwiatkowski, Cordelia Langford, John Sillitoe on behalf of the Wellcome Sanger Institute COVID-19 Surveillance Team |
| EPI_ISL_2867639, EPI_ISL_2867640 | Berkshire and Surrey Pathology Services Lighthouse Laboratory | Wellcome Sanger Institute for the COVID-19 Genomics UK (COG-UK) Consortium | Berkshire and Surrey Pathology Services Lighthouse Laboratory and Alex Alderton, Roberto Amato, Jeffrey Barrett, Sonia Goncalves, Ewan Harrison, David K. Jackson, Ian Johnston, Dominic Kwiatkowski, Cordelia Langford, John Sillitoe on behalf of the Wellcome Sanger Institute COVID-19 Surveillance Team |
| EPI_ISL_2867641, EPI_ISL_2867642 | Lighthouse Lab in Milton Keynes | Wellcome Sanger Institute for the COVID-19 Genomics UK (COG-UK) Consortium | The Lighthouse Lab in Milton Keynes and Alex Alderton, Roberto Amato, Jeffrey Barrett, Sonia Goncalves, Ewan Harrison, David K. Jackson, Ian Johnston, Dominic Kwiatkowski, Cordelia Langford, John Sillitoe on behalf of the Wellcome Sanger Institute COVID-19 Surveillance Team |
| EPI_ISL_2867643, EPI_ISL_2867644, EPI_ISL_2867645 | Berkshire and Surrey Pathology Services Lighthouse Laboratory | Wellcome Sanger Institute for the COVID-19 Genomics UK (COG-UK) Consortium | Berkshire and Surrey Pathology Services Lighthouse Laboratory and Alex Alderton, Roberto Amato, Jeffrey Barrett, Sonia Goncalves, Ewan Harrison, David K. Jackson, Ian Johnston, Dominic Kwiatkowski, Cordelia Langford, John Sillitoe on behalf of the Wellcome Sanger Institute COVID-19 Surveillance Team |
| EPI_ISL_2867646, EPI_ISL_2867647 | Lighthouse Lab in Milton Keynes | Wellcome Sanger Institute for the COVID-19 Genomics UK (COG-UK) Consortium | The Lighthouse Lab in Milton Keynes and Alex Alderton, Roberto Amato, Jeffrey Barrett, Sonia Goncalves, Ewan Harrison, David K. Jackson, Ian Johnston, Dominic Kwiatkowski, Cordelia Langford, John Sillitoe on behalf of the Wellcome Sanger Institute COVID-19 Surveillance Team |
| EPI_ISL_2867648 | Berkshire and Surrey Pathology Services Lighthouse Laboratory | Wellcome Sanger Institute for the COVID-19 Genomics UK (COG-UK) Consortium | Berkshire and Surrey Pathology Services Lighthouse Laboratory and Alex Alderton, Roberto Amato, Jeffrey Barrett, Sonia Goncalves, Ewan Harrison, David K. Jackson, Ian Johnston, Dominic Kwiatkowski, Cordelia Langford, John Sillitoe on behalf of the Wellcome Sanger Institute COVID-19 Surveillance Team |
| EPI_ISL_2867649, EPI_ISL_2867650 | Lighthouse Lab in Milton Keynes | Wellcome Sanger Institute for the COVID-19 Genomics UK (COG-UK) Consortium | The Lighthouse Lab in Milton Keynes and Alex Alderton, Roberto Amato, Jeffrey Barrett, Sonia Goncalves, Ewan Harrison, David K. Jackson, Ian Johnston, Dominic Kwiatkowski, Cordelia Langford, John Sillitoe on behalf of the Wellcome Sanger Institute COVID-19 Surveillance Team |
| EPI_ISL_2867651 | Berkshire and Surrey Pathology Services Lighthouse Laboratory | Wellcome Sanger Institute for the COVID-19 Genomics UK (COG-UK) Consortium | Berkshire and Surrey Pathology Services Lighthouse Laboratory and Alex Alderton, Roberto Amato, Jeffrey Barrett, Sonia Goncalves, Ewan Harrison, David K. Jackson, Ian Johnston, Dominic Kwiatkowski, Cordelia Langford, John Sillitoe on behalf of the Wellcome Sanger Institute COVID-19 Surveillance Team |
| EPI_ISL_2867652, EPI_ISL_2867653, EPI_ISL_2867654 | Lighthouse Lab in Milton Keynes | Wellcome Sanger Institute for the COVID-19 Genomics UK (COG-UK) Consortium | The Lighthouse Lab in Milton Keynes and Alex Alderton, Roberto Amato, Jeffrey Barrett, Sonia Goncalves, Ewan Harrison, David K. Jackson, Ian Johnston, Dominic Kwiatkowski, Cordelia Langford, John Sillitoe on behalf of the Wellcome Sanger Institute COVID-19 Surveillance Team |

| | | | |
|---|---|---|---|
| EPI_ISL_2867655, EPI_ISL_2867656 | Berkshire and Surrey Pathology Services Lighthouse Laboratory | Wellcome Sanger Institute for the COVID-19 Genomics UK (COG-UK) Consortium | Berkshire and Surrey Pathology Services Lighthouse Laboratory and Alex Alderton, Roberto Amato, Jeffrey Barrett, Sonia Goncalves, Ewan Harrison, David K. Jackson, Ian Johnston, Dominic Kwiatkowski, Cordelia Langford, John Sillitoe on behalf of the Wellcome Sanger Institute COVID-19 Surveillance Team |
| EPI_ISL_2867657, EPI_ISL_2867658, EPI_ISL_2867659, EPI_ISL_2867660 | Lighthouse Lab in Milton Keynes | Wellcome Sanger Institute for the COVID-19 Genomics UK (COG-UK) Consortium | The Lighthouse Lab in Milton Keynes and Alex Alderton, Roberto Amato, Jeffrey Barrett, Sonia Goncalves, Ewan Harrison, David K. Jackson, Ian Johnston, Dominic Kwiatkowski, Cordelia Langford, John Sillitoe on behalf of the Wellcome Sanger Institute COVID-19 Surveillance Team |
| EPI_ISL_2867661 | Berkshire and Surrey Pathology Services Lighthouse Laboratory | Wellcome Sanger Institute for the COVID-19 Genomics UK (COG-UK) Consortium | Berkshire and Surrey Pathology Services Lighthouse Laboratory and Alex Alderton, Roberto Amato, Jeffrey Barrett, Sonia Goncalves, Ewan Harrison, David K. Jackson, Ian Johnston, Dominic Kwiatkowski, Cordelia Langford, John Sillitoe on behalf of the Wellcome Sanger Institute COVID-19 Surveillance Team |
| EPI_ISL_2867662 | Lighthouse Lab in Milton Keynes | Wellcome Sanger Institute for the COVID-19 Genomics UK (COG-UK) Consortium | The Lighthouse Lab in Milton Keynes and Alex Alderton, Roberto Amato, Jeffrey Barrett, Sonia Goncalves, Ewan Harrison, David K. Jackson, Ian Johnston, Dominic Kwiatkowski, Cordelia Langford, John Sillitoe on behalf of the Wellcome Sanger Institute COVID-19 Surveillance Team |
| EPI_ISL_2867663, EPI_ISL_2867664 | Berkshire and Surrey Pathology Services Lighthouse Laboratory | Wellcome Sanger Institute for the COVID-19 Genomics UK (COG-UK) Consortium | Berkshire and Surrey Pathology Services Lighthouse Laboratory and Alex Alderton, Roberto Amato, Jeffrey Barrett, Sonia Goncalves, Ewan Harrison, David K. Jackson, Ian Johnston, Dominic Kwiatkowski, Cordelia Langford, John Sillitoe on behalf of the Wellcome Sanger Institute COVID-19 Surveillance Team |
| EPI_ISL_2867665 | Lighthouse Lab in Milton Keynes | Wellcome Sanger Institute for the COVID-19 Genomics UK (COG-UK) Consortium | The Lighthouse Lab in Milton Keynes and Alex Alderton, Roberto Amato, Jeffrey Barrett, Sonia Goncalves, Ewan Harrison, David K. Jackson, Ian Johnston, Dominic Kwiatkowski, Cordelia Langford, John Sillitoe on behalf of the Wellcome Sanger Institute COVID-19 Surveillance Team |
| EPI_ISL_2867666 | Berkshire and Surrey Pathology Services Lighthouse Laboratory | Wellcome Sanger Institute for the COVID-19 Genomics UK (COG-UK) Consortium | Berkshire and Surrey Pathology Services Lighthouse Laboratory and Alex Alderton, Roberto Amato, Jeffrey Barrett, Sonia Goncalves, Ewan Harrison, David K. Jackson, Ian Johnston, Dominic Kwiatkowski, Cordelia Langford, John Sillitoe on behalf of the Wellcome Sanger Institute COVID-19 Surveillance Team |
| EPI_ISL_2867667, EPI_ISL_2867668, EPI_ISL_2867669 | Lighthouse Lab in Milton Keynes | Wellcome Sanger Institute for the COVID-19 Genomics UK (COG-UK) Consortium | The Lighthouse Lab in Milton Keynes and Alex Alderton, Roberto Amato, Jeffrey Barrett, Sonia Goncalves, Ewan Harrison, David K. Jackson, Ian Johnston, Dominic Kwiatkowski, Cordelia Langford, John Sillitoe on behalf of the Wellcome Sanger Institute COVID-19 Surveillance Team |
| EPI_ISL_2867670 | Berkshire and Surrey Pathology Services Lighthouse Laboratory | Wellcome Sanger Institute for the COVID-19 Genomics UK (COG-UK) Consortium | Berkshire and Surrey Pathology Services Lighthouse Laboratory and Alex Alderton, Roberto Amato, Jeffrey Barrett, Sonia Goncalves, Ewan Harrison, David K. Jackson, Ian Johnston, Dominic Kwiatkowski, Cordelia Langford, John Sillitoe on behalf of the Wellcome Sanger Institute COVID-19 Surveillance Team |
| EPI_ISL_2867671 | Lighthouse Lab in Milton Keynes | Wellcome Sanger Institute for the COVID-19 Genomics UK (COG-UK) Consortium | The Lighthouse Lab in Milton Keynes and Alex Alderton, Roberto Amato, Jeffrey Barrett, Sonia Goncalves, Ewan Harrison, David K. Jackson, Ian Johnston, Dominic Kwiatkowski, Cordelia Langford, John Sillitoe on behalf of the Wellcome Sanger Institute COVID-19 Surveillance Team |
| EPI_ISL_2867672, EPI_ISL_2867673 | Berkshire and Surrey Pathology Services Lighthouse Laboratory | Wellcome Sanger Institute for the COVID-19 Genomics UK (COG-UK) Consortium | Berkshire and Surrey Pathology Services Lighthouse Laboratory and Alex Alderton, Roberto Amato, Jeffrey Barrett, Sonia Goncalves, Ewan Harrison, David K. Jackson, Ian Johnston, Dominic Kwiatkowski, Cordelia Langford, John Sillitoe on behalf of the Wellcome Sanger Institute COVID-19 Surveillance Team |
| EPI_ISL_2867674, EPI_ISL_2867675, EPI_ISL_2867676, EPI_ISL_2867677, EPI_ISL_2867678, EPI_ISL_2867679, EPI_ISL_2867680, EPI_ISL_2867681, EPI_ISL_2867682 | Lighthouse Lab in Milton Keynes | Wellcome Sanger Institute for the COVID-19 Genomics UK (COG-UK) Consortium | The Lighthouse Lab in Milton Keynes and Alex Alderton, Roberto Amato, Jeffrey Barrett, Sonia Goncalves, Ewan Harrison, David K. Jackson, Ian Johnston, Dominic Kwiatkowski, Cordelia Langford, John Sillitoe on behalf of the Wellcome Sanger Institute COVID-19 Surveillance Team |
| EPI_ISL_2867683 | Berkshire and Surrey Pathology Services Lighthouse Laboratory | Wellcome Sanger Institute for the COVID-19 Genomics UK (COG-UK) Consortium | Berkshire and Surrey Pathology Services Lighthouse Laboratory and Alex Alderton, Roberto Amato, Jeffrey Barrett, Sonia Goncalves, Ewan Harrison, David K. Jackson, Ian Johnston, Dominic Kwiatkowski, Cordelia Langford, John Sillitoe on behalf of the Wellcome Sanger Institute COVID-19 Surveillance Team |
| EPI_ISL_2867684 | Lighthouse Lab in Milton Keynes | Wellcome Sanger Institute for the COVID-19 Genomics UK (COG-UK) Consortium | The Lighthouse Lab in Milton Keynes and Alex Alderton, Roberto Amato, Jeffrey Barrett, Sonia Goncalves, Ewan Harrison, David K. Jackson, Ian Johnston, Dominic Kwiatkowski, Cordelia Langford, John Sillitoe on behalf of the Wellcome Sanger Institute COVID-19 Surveillance Team |
| EPI_ISL_2867685 | Berkshire and Surrey Pathology Services Lighthouse Laboratory | Wellcome Sanger Institute for the COVID-19 Genomics UK (COG-UK) Consortium | Berkshire and Surrey Pathology Services Lighthouse Laboratory and Alex Alderton, Roberto Amato, Jeffrey Barrett, Sonia Goncalves, Ewan Harrison, David K. Jackson, Ian Johnston, Dominic Kwiatkowski, Cordelia Langford, John Sillitoe on behalf of the Wellcome Sanger Institute COVID-19 Surveillance Team |
| EPI_ISL_2867686, EPI_ISL_2867687 | Lighthouse Lab in Milton Keynes | Wellcome Sanger Institute for the COVID-19 Genomics UK (COG-UK) Consortium | The Lighthouse Lab in Milton Keynes and Alex Alderton, Roberto Amato, Jeffrey Barrett, Sonia Goncalves, Ewan Harrison, David K. Jackson, Ian Johnston, Dominic Kwiatkowski, Cordelia Langford, John Sillitoe on behalf of the Wellcome Sanger Institute COVID-19 Surveillance Team |
| EPI_ISL_2867688 | Berkshire and Surrey Pathology Services Lighthouse Laboratory | Wellcome Sanger Institute for the COVID-19 Genomics UK (COG-UK) Consortium | Berkshire and Surrey Pathology Services Lighthouse Laboratory and Alex Alderton, Roberto Amato, Jeffrey Barrett, Sonia Goncalves, Ewan Harrison, David K. Jackson, Ian Johnston, Dominic Kwiatkowski, Cordelia Langford, John Sillitoe on behalf of the Wellcome Sanger Institute COVID-19 Surveillance Team |
| EPI_ISL_2867689, EPI_ISL_2867690 | Lighthouse Lab in Milton Keynes | Wellcome Sanger Institute for the COVID-19 Genomics UK (COG-UK) Consortium | The Lighthouse Lab in Milton Keynes and Alex Alderton, Roberto Amato, Jeffrey Barrett, Sonia Goncalves, Ewan Harrison, David K. Jackson, Ian Johnston, Dominic Kwiatkowski, Cordelia Langford, John Sillitoe on behalf of the Wellcome Sanger Institute COVID-19 Surveillance Team |
| EPI_ISL_2867691, EPI_ISL_2867692 | Berkshire and Surrey Pathology Services Lighthouse Laboratory | Wellcome Sanger Institute for the COVID-19 Genomics UK (COG-UK) Consortium | Berkshire and Surrey Pathology Services Lighthouse Laboratory and Alex Alderton, Roberto Amato, Jeffrey Barrett, Sonia Goncalves, Ewan Harrison, David K. Jackson, Ian Johnston, Dominic Kwiatkowski, Cordelia Langford, John Sillitoe on behalf of the Wellcome Sanger Institute COVID-19 Surveillance Team |
| EPI_ISL_2867693 | Lighthouse Lab in Milton Keynes | Wellcome Sanger Institute for the COVID-19 Genomics UK (COG-UK) Consortium | The Lighthouse Lab in Milton Keynes and Alex Alderton, Roberto Amato, Jeffrey Barrett, Sonia Goncalves, Ewan Harrison, David K. Jackson, Ian Johnston, Dominic Kwiatkowski, Cordelia Langford, John Sillitoe on behalf of the Wellcome Sanger Institute COVID-19 Surveillance Team |
| EPI_ISL_2867694, EPI_ISL_2867695 | Lighthouse Lab in Alderley Park | Wellcome Sanger Institute for the COVID-19 Genomics UK (COG-UK) Consortium | Jacquelyn Wynn, Mairead Hyland, The Lighthouse Lab in Alderley Park and Alex Alderton, Roberto Amato, Jeffrey Barrett, Sonia Goncalves, Ewan Harrison, David K. Jackson, Ian Johnston, Dominic Kwiatkowski, Cordelia Langford, John Sillitoe on behalf of the Wellcome Sanger Institute COVID-19 Surveillance Team |
| EPI_ISL_2867696 | Lighthouse Laboratory Plymouth | Wellcome Sanger Institute for the COVID-19 Genomics UK (COG-UK) Consortium | Lighthouse Laboratory Plymouth and Alex Alderton, Roberto Amato, Jeffrey Barrett, Sonia Goncalves, Ewan Harrison, David K. Jackson, Ian Johnston, Dominic Kwiatkowski, Cordelia Langford, John Sillitoe on behalf of the Wellcome Sanger Institute COVID-19 Surveillance Team |
| EPI_ISL_2867697 | Lighthouse Lab in Alderley Park | Wellcome Sanger Institute for the COVID-19 Genomics UK (COG-UK) Consortium | Jacquelyn Wynn, Mairead Hyland, The Lighthouse Lab in Alderley Park and Alex Alderton, Roberto Amato, Jeffrey Barrett, Sonia Goncalves, Ewan Harrison, David K. Jackson, Ian Johnston, Dominic Kwiatkowski, Cordelia Langford, John Sillitoe on behalf of the Wellcome Sanger Institute COVID-19 Surveillance Team |
| EPI_ISL_2867698 | Lighthouse Laboratory Plymouth | Wellcome Sanger Institute for the COVID-19 Genomics UK (COG-UK) Consortium | Lighthouse Laboratory Plymouth and Alex Alderton, Roberto Amato, Jeffrey Barrett, Sonia Goncalves, Ewan Harrison, David K. Jackson, Ian Johnston, Dominic Kwiatkowski, Cordelia Langford, John Sillitoe on behalf of the Wellcome Sanger Institute COVID-19 Surveillance Team |
| EPI_ISL_2867699, EPI_ISL_2867700 | Lighthouse Lab in Alderley Park | Wellcome Sanger Institute for the COVID-19 Genomics UK (COG-UK) Consortium | Jacquelyn Wynn, Mairead Hyland, The Lighthouse Lab in Alderley Park and Alex Alderton, Roberto Amato, Jeffrey Barrett, Sonia Goncalves, Ewan Harrison, David K. Jackson, Ian Johnston, Dominic Kwiatkowski, Cordelia Langford, John Sillitoe on behalf of the Wellcome Sanger Institute COVID-19 Surveillance Team |
| EPI_ISL_2867701 | Lighthouse Laboratory Plymouth | Wellcome Sanger Institute for the COVID-19 Genomics UK (COG-UK) Consortium | Lighthouse Laboratory Plymouth and Alex Alderton, Roberto Amato, Jeffrey Barrett, Sonia Goncalves, Ewan Harrison, David K. Jackson, Ian Johnston, Dominic Kwiatkowski, Cordelia Langford, John Sillitoe on behalf of the Wellcome Sanger Institute COVID-19 Surveillance Team |
| EPI_ISL_2867702 | Lighthouse Lab in Alderley Park | Wellcome Sanger Institute for the COVID-19 Genomics UK (COG-UK) Consortium | Jacquelyn Wynn, Mairead Hyland, The Lighthouse Lab in Alderley Park and Alex Alderton, Roberto Amato, Jeffrey Barrett, Sonia Goncalves, Ewan Harrison, David K. Jackson, Ian Johnston, Dominic Kwiatkowski, Cordelia Langford, John Sillitoe on behalf of the Wellcome Sanger Institute COVID-19 Surveillance Team |
| EPI_ISL_2867703 | Lighthouse Laboratory Plymouth | Wellcome Sanger Institute for the COVID-19 Genomics UK (COG-UK) Consortium | Lighthouse Laboratory Plymouth and Alex Alderton, Roberto Amato, Jeffrey Barrett, Sonia Goncalves, Ewan Harrison, David K. Jackson, Ian Johnston, Dominic Kwiatkowski, Cordelia Langford, John Sillitoe on behalf of the Wellcome Sanger Institute COVID-19 Surveillance Team |
| EPI_ISL_2867704 | Lighthouse Lab in Alderley Park | Wellcome Sanger Institute for the COVID-19 Genomics UK (COG-UK) Consortium | Jacquelyn Wynn, Mairead Hyland, The Lighthouse Lab in Alderley Park and Alex Alderton, Roberto Amato, Jeffrey Barrett, Sonia Goncalves, Ewan Harrison, David K. Jackson, Ian Johnston, Dominic Kwiatkowski, Cordelia Langford, John Sillitoe on behalf of the Wellcome Sanger Institute COVID-19 Surveillance Team |
| EPI_ISL_2867705 | Lighthouse Laboratory Plymouth | Wellcome Sanger Institute for the COVID-19 Genomics UK (COG-UK) Consortium | Lighthouse Laboratory Plymouth and Alex Alderton, Roberto Amato, Jeffrey Barrett, Sonia Goncalves, Ewan Harrison, David K. Jackson, Ian Johnston, Dominic Kwiatkowski, Cordelia Langford, John Sillitoe on behalf of the Wellcome Sanger Institute COVID-19 Surveillance Team |
| EPI_ISL_2867706 | Lighthouse Lab in Alderley Park | Wellcome Sanger Institute for the COVID-19 Genomics UK (COG-UK) Consortium | Jacquelyn Wynn, Mairead Hyland, The Lighthouse Lab in Alderley Park and Alex Alderton, Roberto Amato, Jeffrey Barrett, Sonia Goncalves, Ewan Harrison, David K. Jackson, Ian Johnston, Dominic Kwiatkowski, Cordelia Langford, John Sillitoe on behalf of the Wellcome Sanger Institute COVID-19 Surveillance Team |
| EPI_ISL_2867707, EPI_ISL_2867708 | Lighthouse Laboratory Plymouth | Wellcome Sanger Institute for the COVID-19 Genomics UK (COG-UK) Consortium | Lighthouse Laboratory Plymouth and Alex Alderton, Roberto Amato, Jeffrey Barrett, Sonia Goncalves, Ewan Harrison, David K. Jackson, Ian Johnston, Dominic Kwiatkowski, Cordelia Langford, John Sillitoe on behalf of the Wellcome Sanger Institute COVID-19 Surveillance Team |
| EPI_ISL_2867709, EPI_ISL_2867710, EPI_ISL_2867711 | Lighthouse Lab in Alderley Park | Wellcome Sanger Institute for the COVID-19 Genomics UK (COG-UK) Consortium | Jacquelyn Wynn, Mairead Hyland, The Lighthouse Lab in Alderley Park and Alex Alderton, Roberto Amato, Jeffrey Barrett, Sonia Goncalves, Ewan Harrison, David K. Jackson, Ian Johnston, Dominic Kwiatkowski, Cordelia Langford, John Sillitoe on behalf of the Wellcome Sanger Institute COVID-19 Surveillance Team |
| EPI_ISL_2867712, EPI_ISL_2867713, EPI_ISL_2867714, EPI_ISL_2867715, EPI_ISL_2867716 | Lighthouse Laboratory Plymouth | Wellcome Sanger Institute for the COVID-19 Genomics UK (COG-UK) Consortium | Lighthouse Laboratory Plymouth and Alex Alderton, Roberto Amato, Jeffrey Barrett, Sonia Goncalves, Ewan Harrison, David K. Jackson, Ian Johnston, Dominic Kwiatkowski, Cordelia Langford, John Sillitoe on behalf of the Wellcome Sanger Institute COVID-19 Surveillance Team |
| EPI_ISL_2867717 | Lighthouse Lab in Alderley Park | Wellcome Sanger Institute for the COVID-19 Genomics UK (COG-UK) Consortium | Jacquelyn Wynn, Mairead Hyland, The Lighthouse Lab in Alderley Park and Alex Alderton, Roberto Amato, Jeffrey Barrett, Sonia Goncalves, Ewan Harrison, David K. Jackson, Ian Johnston, Dominic Kwiatkowski, Cordelia Langford, John Sillitoe on behalf of the Wellcome Sanger Institute COVID-19 Surveillance Team |
| EPI_ISL_2867718, EPI_ISL_2867719 | Lighthouse Laboratory Plymouth | Wellcome Sanger Institute for the COVID-19 Genomics UK (COG-UK) Consortium | Lighthouse Laboratory Plymouth and Alex Alderton, Roberto Amato, Jeffrey Barrett, Sonia Goncalves, Ewan Harrison, David K. Jackson, Ian Johnston, Dominic Kwiatkowski, Cordelia Langford, John Sillitoe on behalf of the Wellcome Sanger Institute COVID-19 Surveillance Team |
| EPI_ISL_2867720 | Lighthouse Lab in Alderley Park | Wellcome Sanger Institute for the COVID-19 Genomics UK (COG-UK) Consortium | Jacquelyn Wynn, Mairead Hyland, The Lighthouse Lab in Alderley Park and Alex Alderton, Roberto Amato, Jeffrey Barrett, Sonia Goncalves, Ewan Harrison, David K. Jackson, Ian Johnston, Dominic Kwiatkowski, Cordelia Langford, John Sillitoe on behalf of the Wellcome Sanger Institute COVID-19 Surveillance Team |
| EPI_ISL_2867721, EPI_ISL_2867722, EPI_ISL_2867723 | Lighthouse Laboratory Plymouth | Wellcome Sanger Institute for the COVID-19 Genomics UK (COG-UK) Consortium | Lighthouse Laboratory Plymouth and Alex Alderton, Roberto Amato, Jeffrey Barrett, Sonia Goncalves, Ewan Harrison, David K. Jackson, Ian Johnston, Dominic Kwiatkowski, Cordelia Langford, John Sillitoe on behalf of the Wellcome Sanger Institute COVID-19 Surveillance Team |
| EPI_ISL_2867724, EPI_ISL_2867725, | Lighthouse Lab in Alderley Park | Wellcome Sanger Institute for the COVID-19 Genomics UK (COG-UK) Consortium | Jacquelyn Wynn, Mairead Hyland, The Lighthouse Lab in Alderley Park and Alex Alderton, Roberto Amato, Jeffrey Barrett, Sonia Goncalves, Ewan Harrison, David K. Jackson, Ian Johnston, Dominic Kwiatkowski, Cordelia Langford, John Sillitoe on behalf of the Wellcome Sanger Institute COVID-19 Surveillance Team |

| | | | |
|---|---|---|---|
| EPI_ISL_2867726 | | | |
| EPI_ISL_2867727, EPI_ISL_2867728 | Lighthouse Laboratory Plymouth | Wellcome Sanger Institute for the COVID-19 Genomics UK (COG-UK) Consortium | Lighthouse Laboratory Plymouth and Alex Alderton, Roberto Amato, Jeffrey Barrett, Sonia Goncalves, Ewan Harrison, David K. Jackson, Ian Johnston, Dominic Kwiatkowski, Cordelia Langford, John Sillitoe on behalf of the Wellcome Sanger Institute COVID-19 Surveillance Team |
| EPI_ISL_2867729, EPI_ISL_2867730, EPI_ISL_2867731, EPI_ISL_2867732, EPI_ISL_2867733, EPI_ISL_2867734 | Lighthouse Lab in Alderley Park | Wellcome Sanger Institute for the COVID-19 Genomics UK (COG-UK) Consortium | Jacquelyn Wynn, Mairead Hyland, The Lighthouse Lab in Alderley Park and Alex Alderton, Roberto Amato, Jeffrey Barrett, Sonia Goncalves, Ewan Harrison, David K. Jackson, Ian Johnston, Dominic Kwiatkowski, Cordelia Langford, John Sillitoe on behalf of the Wellcome Sanger Institute COVID-19 Surveillance Team |
| EPI_ISL_2867735, EPI_ISL_2867736 | Lighthouse Laboratory Plymouth | Wellcome Sanger Institute for the COVID-19 Genomics UK (COG-UK) Consortium | Lighthouse Laboratory Plymouth and Alex Alderton, Roberto Amato, Jeffrey Barrett, Sonia Goncalves, Ewan Harrison, David K. Jackson, Ian Johnston, Dominic Kwiatkowski, Cordelia Langford, John Sillitoe on behalf of the Wellcome Sanger Institute COVID-19 Surveillance Team |
| EPI_ISL_2867737, EPI_ISL_2867738 | Lighthouse Lab in Alderley Park | Wellcome Sanger Institute for the COVID-19 Genomics UK (COG-UK) Consortium | Jacquelyn Wynn, Mairead Hyland, The Lighthouse Lab in Alderley Park and Alex Alderton, Roberto Amato, Jeffrey Barrett, Sonia Goncalves, Ewan Harrison, David K. Jackson, Ian Johnston, Dominic Kwiatkowski, Cordelia Langford, John Sillitoe on behalf of the Wellcome Sanger Institute COVID-19 Surveillance Team |
| EPI_ISL_2867739 | Lighthouse Laboratory Plymouth | Wellcome Sanger Institute for the COVID-19 Genomics UK (COG-UK) Consortium | Lighthouse Laboratory Plymouth and Alex Alderton, Roberto Amato, Jeffrey Barrett, Sonia Goncalves, Ewan Harrison, David K. Jackson, Ian Johnston, Dominic Kwiatkowski, Cordelia Langford, John Sillitoe on behalf of the Wellcome Sanger Institute COVID-19 Surveillance Team |
| EPI_ISL_2867740 | Lighthouse Lab in Alderley Park | Wellcome Sanger Institute for the COVID-19 Genomics UK (COG-UK) Consortium | Jacquelyn Wynn, Mairead Hyland, The Lighthouse Lab in Alderley Park and Alex Alderton, Roberto Amato, Jeffrey Barrett, Sonia Goncalves, Ewan Harrison, David K. Jackson, Ian Johnston, Dominic Kwiatkowski, Cordelia Langford, John Sillitoe on behalf of the Wellcome Sanger Institute COVID-19 Surveillance Team |
| EPI_ISL_2867741, EPI_ISL_2867742 | Lighthouse Laboratory Plymouth | Wellcome Sanger Institute for the COVID-19 Genomics UK (COG-UK) Consortium | Lighthouse Laboratory Plymouth and Alex Alderton, Roberto Amato, Jeffrey Barrett, Sonia Goncalves, Ewan Harrison, David K. Jackson, Ian Johnston, Dominic Kwiatkowski, Cordelia Langford, John Sillitoe on behalf of the Wellcome Sanger Institute COVID-19 Surveillance Team |
| EPI_ISL_2867743, EPI_ISL_2867744 | Lighthouse Lab in Alderley Park | Wellcome Sanger Institute for the COVID-19 Genomics UK (COG-UK) Consortium | Jacquelyn Wynn, Mairead Hyland, The Lighthouse Lab in Alderley Park and Alex Alderton, Roberto Amato, Jeffrey Barrett, Sonia Goncalves, Ewan Harrison, David K. Jackson, Ian Johnston, Dominic Kwiatkowski, Cordelia Langford, John Sillitoe on behalf of the Wellcome Sanger Institute COVID-19 Surveillance Team |
| EPI_ISL_2867745 | Lighthouse Laboratory Plymouth | Wellcome Sanger Institute for the COVID-19 Genomics UK (COG-UK) Consortium | Lighthouse Laboratory Plymouth and Alex Alderton, Roberto Amato, Jeffrey Barrett, Sonia Goncalves, Ewan Harrison, David K. Jackson, Ian Johnston, Dominic Kwiatkowski, Cordelia Langford, John Sillitoe on behalf of the Wellcome Sanger Institute COVID-19 Surveillance Team |
| EPI_ISL_2867746 | Lighthouse Lab in Alderley Park | Wellcome Sanger Institute for the COVID-19 Genomics UK (COG-UK) Consortium | Jacquelyn Wynn, Mairead Hyland, The Lighthouse Lab in Alderley Park and Alex Alderton, Roberto Amato, Jeffrey Barrett, Sonia Goncalves, Ewan Harrison, David K. Jackson, Ian Johnston, Dominic Kwiatkowski, Cordelia Langford, John Sillitoe on behalf of the Wellcome Sanger Institute COVID-19 Surveillance Team |
| EPI_ISL_2867747 | Lighthouse Laboratory Plymouth | Wellcome Sanger Institute for the COVID-19 Genomics UK (COG-UK) Consortium | Lighthouse Laboratory Plymouth and Alex Alderton, Roberto Amato, Jeffrey Barrett, Sonia Goncalves, Ewan Harrison, David K. Jackson, Ian Johnston, Dominic Kwiatkowski, Cordelia Langford, John Sillitoe on behalf of the Wellcome Sanger Institute COVID-19 Surveillance Team |
| EPI_ISL_2867748 | Lighthouse Lab in Alderley Park | Wellcome Sanger Institute for the COVID-19 Genomics UK (COG-UK) Consortium | Jacquelyn Wynn, Mairead Hyland, The Lighthouse Lab in Alderley Park and Alex Alderton, Roberto Amato, Jeffrey Barrett, Sonia Goncalves, Ewan Harrison, David K. Jackson, Ian Johnston, Dominic Kwiatkowski, Cordelia Langford, John Sillitoe on behalf of the Wellcome Sanger Institute COVID-19 Surveillance Team |
| EPI_ISL_2867749, EPI_ISL_2867750 | Lighthouse Laboratory Plymouth | Wellcome Sanger Institute for the COVID-19 Genomics UK (COG-UK) Consortium | Lighthouse Laboratory Plymouth and Alex Alderton, Roberto Amato, Jeffrey Barrett, Sonia Goncalves, Ewan Harrison, David K. Jackson, Ian Johnston, Dominic Kwiatkowski, Cordelia Langford, John Sillitoe on behalf of the Wellcome Sanger Institute COVID-19 Surveillance Team |
| EPI_ISL_2867751, EPI_ISL_2867752, EPI_ISL_2867753, EPI_ISL_2867754 | Lighthouse Lab in Alderley Park | Wellcome Sanger Institute for the COVID-19 Genomics UK (COG-UK) Consortium | Jacquelyn Wynn, Mairead Hyland, The Lighthouse Lab in Alderley Park and Alex Alderton, Roberto Amato, Jeffrey Barrett, Sonia Goncalves, Ewan Harrison, David K. Jackson, Ian Johnston, Dominic Kwiatkowski, Cordelia Langford, John Sillitoe on behalf of the Wellcome Sanger Institute COVID-19 Surveillance Team |
| EPI_ISL_2867755 | Lighthouse Laboratory Plymouth | Wellcome Sanger Institute for the COVID-19 Genomics UK (COG-UK) Consortium | Lighthouse Laboratory Plymouth and Alex Alderton, Roberto Amato, Jeffrey Barrett, Sonia Goncalves, Ewan Harrison, David K. Jackson, Ian Johnston, Dominic Kwiatkowski, Cordelia Langford, John Sillitoe on behalf of the Wellcome Sanger Institute COVID-19 Surveillance Team |
| EPI_ISL_2867756, EPI_ISL_2867757 | Lighthouse Lab in Alderley Park | Wellcome Sanger Institute for the COVID-19 Genomics UK (COG-UK) Consortium | Jacquelyn Wynn, Mairead Hyland, The Lighthouse Lab in Alderley Park and Alex Alderton, Roberto Amato, Jeffrey Barrett, Sonia Goncalves, Ewan Harrison, David K. Jackson, Ian Johnston, Dominic Kwiatkowski, Cordelia Langford, John Sillitoe on behalf of the Wellcome Sanger Institute COVID-19 Surveillance Team |
| EPI_ISL_2867758 | Lighthouse Laboratory Plymouth | Wellcome Sanger Institute for the COVID-19 Genomics UK (COG-UK) Consortium | Lighthouse Laboratory Plymouth and Alex Alderton, Roberto Amato, Jeffrey Barrett, Sonia Goncalves, Ewan Harrison, David K. Jackson, Ian Johnston, Dominic Kwiatkowski, Cordelia Langford, John Sillitoe on behalf of the Wellcome Sanger Institute COVID-19 Surveillance Team |
| EPI_ISL_2867759 | Lighthouse Lab in Alderley Park | Wellcome Sanger Institute for the COVID-19 Genomics UK (COG-UK) Consortium | Jacquelyn Wynn, Mairead Hyland, The Lighthouse Lab in Alderley Park and Alex Alderton, Roberto Amato, Jeffrey Barrett, Sonia Goncalves, Ewan Harrison, David K. Jackson, Ian Johnston, Dominic Kwiatkowski, Cordelia Langford, John Sillitoe on behalf of the Wellcome Sanger Institute COVID-19 Surveillance Team |
| EPI_ISL_2867760, EPI_ISL_2867761 | Lighthouse Laboratory Plymouth | Wellcome Sanger Institute for the COVID-19 Genomics UK (COG-UK) Consortium | Lighthouse Laboratory Plymouth and Alex Alderton, Roberto Amato, Jeffrey Barrett, Sonia Goncalves, Ewan Harrison, David K. Jackson, Ian Johnston, Dominic Kwiatkowski, Cordelia Langford, John Sillitoe on behalf of the Wellcome Sanger Institute COVID-19 Surveillance Team |
| EPI_ISL_2867762 | Lighthouse Lab in Alderley Park | Wellcome Sanger Institute for the COVID-19 Genomics UK (COG-UK) Consortium | Jacquelyn Wynn, Mairead Hyland, The Lighthouse Lab in Alderley Park and Alex Alderton, Roberto Amato, Jeffrey Barrett, Sonia Goncalves, Ewan Harrison, David K. Jackson, Ian Johnston, Dominic Kwiatkowski, Cordelia Langford, John Sillitoe on behalf of the Wellcome Sanger Institute COVID-19 Surveillance Team |
| EPI_ISL_2867763, EPI_ISL_2867764 | Lighthouse Laboratory Plymouth | Wellcome Sanger Institute for the COVID-19 Genomics UK (COG-UK) Consortium | Lighthouse Laboratory Plymouth and Alex Alderton, Roberto Amato, Jeffrey Barrett, Sonia Goncalves, Ewan Harrison, David K. Jackson, Ian Johnston, Dominic Kwiatkowski, Cordelia Langford, John Sillitoe on behalf of the Wellcome Sanger Institute COVID-19 Surveillance Team |
| EPI_ISL_2867765, EPI_ISL_2867766 | Lighthouse Lab in Alderley Park | Wellcome Sanger Institute for the COVID-19 Genomics UK (COG-UK) Consortium | Jacquelyn Wynn, Mairead Hyland, The Lighthouse Lab in Alderley Park and Alex Alderton, Roberto Amato, Jeffrey Barrett, Sonia Goncalves, Ewan Harrison, David K. Jackson, Ian Johnston, Dominic Kwiatkowski, Cordelia Langford, John Sillitoe on behalf of the Wellcome Sanger Institute COVID-19 Surveillance Team |
| EPI_ISL_2867767 | Lighthouse Laboratory Plymouth | Wellcome Sanger Institute for the COVID-19 Genomics UK (COG-UK) Consortium | Lighthouse Laboratory Plymouth and Alex Alderton, Roberto Amato, Jeffrey Barrett, Sonia Goncalves, Ewan Harrison, David K. Jackson, Ian Johnston, Dominic Kwiatkowski, Cordelia Langford, John Sillitoe on behalf of the Wellcome Sanger Institute COVID-19 Surveillance Team |
| EPI_ISL_2867768, EPI_ISL_2867769, EPI_ISL_2867770, EPI_ISL_2867771 | Lighthouse Lab in Alderley Park | Wellcome Sanger Institute for the COVID-19 Genomics UK (COG-UK) Consortium | Jacquelyn Wynn, Mairead Hyland, The Lighthouse Lab in Alderley Park and Alex Alderton, Roberto Amato, Jeffrey Barrett, Sonia Goncalves, Ewan Harrison, David K. Jackson, Ian Johnston, Dominic Kwiatkowski, Cordelia Langford, John Sillitoe on behalf of the Wellcome Sanger Institute COVID-19 Surveillance Team |
| EPI_ISL_2867772 | Lighthouse Laboratory Plymouth | Wellcome Sanger Institute for the COVID-19 Genomics UK (COG-UK) Consortium | Lighthouse Laboratory Plymouth and Alex Alderton, Roberto Amato, Jeffrey Barrett, Sonia Goncalves, Ewan Harrison, David K. Jackson, Ian Johnston, Dominic Kwiatkowski, Cordelia Langford, John Sillitoe on behalf of the Wellcome Sanger Institute COVID-19 Surveillance Team |
| EPI_ISL_2867773 | Lighthouse Lab in Alderley Park | Wellcome Sanger Institute for the COVID-19 Genomics UK (COG-UK) Consortium | Jacquelyn Wynn, Mairead Hyland, The Lighthouse Lab in Alderley Park and Alex Alderton, Roberto Amato, Jeffrey Barrett, Sonia Goncalves, Ewan Harrison, David K. Jackson, Ian Johnston, Dominic Kwiatkowski, Cordelia Langford, John Sillitoe on behalf of the Wellcome Sanger Institute COVID-19 Surveillance Team |
| EPI_ISL_2867774, EPI_ISL_2867775, EPI_ISL_2867776, EPI_ISL_2867777, EPI_ISL_2867778 | Lighthouse Laboratory Plymouth | Wellcome Sanger Institute for the COVID-19 Genomics UK (COG-UK) Consortium | Lighthouse Laboratory Plymouth and Alex Alderton, Roberto Amato, Jeffrey Barrett, Sonia Goncalves, Ewan Harrison, David K. Jackson, Ian Johnston, Dominic Kwiatkowski, Cordelia Langford, John Sillitoe on behalf of the Wellcome Sanger Institute COVID-19 Surveillance Team |
| EPI_ISL_2867779, EPI_ISL_2867780, EPI_ISL_2867781 | Lighthouse Lab in Alderley Park | Wellcome Sanger Institute for the COVID-19 Genomics UK (COG-UK) Consortium | Jacquelyn Wynn, Mairead Hyland, The Lighthouse Lab in Alderley Park and Alex Alderton, Roberto Amato, Jeffrey Barrett, Sonia Goncalves, Ewan Harrison, David K. Jackson, Ian Johnston, Dominic Kwiatkowski, Cordelia Langford, John Sillitoe on behalf of the Wellcome Sanger Institute COVID-19 Surveillance Team |
| EPI_ISL_2867782 | Lighthouse Laboratory Plymouth | Wellcome Sanger Institute for the COVID-19 Genomics UK (COG-UK) Consortium | Lighthouse Laboratory Plymouth and Alex Alderton, Roberto Amato, Jeffrey Barrett, Sonia Goncalves, Ewan Harrison, David K. Jackson, Ian Johnston, Dominic Kwiatkowski, Cordelia Langford, John Sillitoe on behalf of the Wellcome Sanger Institute COVID-19 Surveillance Team |
| EPI_ISL_2867783 | Lighthouse Lab in Alderley Park | Wellcome Sanger Institute for the COVID-19 Genomics UK (COG-UK) Consortium | Jacquelyn Wynn, Mairead Hyland, The Lighthouse Lab in Alderley Park and Alex Alderton, Roberto Amato, Jeffrey Barrett, Sonia Goncalves, Ewan Harrison, David K. Jackson, Ian Johnston, Dominic Kwiatkowski, Cordelia Langford, John Sillitoe on behalf of the Wellcome Sanger Institute COVID-19 Surveillance Team |
| EPI_ISL_2867784, EPI_ISL_2867785, EPI_ISL_2867786, EPI_ISL_2867787, EPI_ISL_2867788 | Lighthouse Laboratory Plymouth | Wellcome Sanger Institute for the COVID-19 Genomics UK (COG-UK) Consortium | Lighthouse Laboratory Plymouth and Alex Alderton, Roberto Amato, Jeffrey Barrett, Sonia Goncalves, Ewan Harrison, David K. Jackson, Ian Johnston, Dominic Kwiatkowski, Cordelia Langford, John Sillitoe on behalf of the Wellcome Sanger Institute COVID-19 Surveillance Team |
| EPI_ISL_2867789, EPI_ISL_2867790, EPI_ISL_2867791, EPI_ISL_2867792 | Lighthouse Lab in Alderley Park | Wellcome Sanger Institute for the COVID-19 Genomics UK (COG-UK) Consortium | Jacquelyn Wynn, Mairead Hyland, The Lighthouse Lab in Alderley Park and Alex Alderton, Roberto Amato, Jeffrey Barrett, Sonia Goncalves, Ewan Harrison, David K. Jackson, Ian Johnston, Dominic Kwiatkowski, Cordelia Langford, John Sillitoe on behalf of the Wellcome Sanger Institute COVID-19 Surveillance Team |
| EPI_ISL_2867793 | Lighthouse Laboratory Plymouth | Wellcome Sanger Institute for the COVID-19 Genomics UK (COG-UK) Consortium | Lighthouse Laboratory Plymouth and Alex Alderton, Roberto Amato, Jeffrey Barrett, Sonia Goncalves, Ewan Harrison, David K. Jackson, Ian Johnston, Dominic Kwiatkowski, Cordelia Langford, John Sillitoe on behalf of the Wellcome Sanger Institute COVID-19 Surveillance Team |
| EPI_ISL_2867794 | Lighthouse Lab in Alderley Park | Wellcome Sanger Institute for the COVID-19 Genomics UK (COG-UK) Consortium | Jacquelyn Wynn, Mairead Hyland, The Lighthouse Lab in Alderley Park and Alex Alderton, Roberto Amato, Jeffrey Barrett, Sonia Goncalves, Ewan Harrison, David K. Jackson, Ian Johnston, Dominic Kwiatkowski, Cordelia Langford, John Sillitoe on behalf of the Wellcome Sanger Institute COVID-19 Surveillance Team |
| EPI_ISL_2867795, EPI_ISL_2867796, EPI_ISL_2867797, EPI_ISL_2867798, EPI_ISL_2867799, EPI_ISL_2867800 | Lighthouse Laboratory Plymouth | Wellcome Sanger Institute for the COVID-19 Genomics UK (COG-UK) Consortium | Lighthouse Laboratory Plymouth and Alex Alderton, Roberto Amato, Jeffrey Barrett, Sonia Goncalves, Ewan Harrison, David K. Jackson, Ian Johnston, Dominic Kwiatkowski, Cordelia Langford, John Sillitoe on behalf of the Wellcome Sanger Institute COVID-19 Surveillance Team |
| EPI_ISL_2867801 | Lighthouse Lab in Alderley Park | Wellcome Sanger Institute for the COVID-19 Genomics UK (COG-UK) Consortium | Jacquelyn Wynn, Mairead Hyland, The Lighthouse Lab in Alderley Park and Alex Alderton, Roberto Amato, Jeffrey Barrett, Sonia Goncalves, Ewan Harrison, David K. Jackson, Ian Johnston, Dominic Kwiatkowski, Cordelia Langford, John Sillitoe on behalf of the Wellcome Sanger Institute COVID-19 Surveillance Team |

| | | | |
|---|---|---|---|
| EPI_ISL_2867802, EPI_ISL_2867803, EPI_ISL_2867804 | Lighthouse Laboratory Plymouth | Wellcome Sanger Institute for the COVID-19 Genomics UK (COG-UK) Consortium | Lighthouse Laboratory Plymouth and Alex Alderton, Roberto Amato, Jeffrey Barrett, Sonia Goncalves, Ewan Harrison, David K. Jackson, Ian Johnston, Dominic Kwiatkowski, Cordelia Langford, John Sillitoe on behalf of the Wellcome Sanger Institute COVID-19 Surveillance Team |
| EPI_ISL_2867805 | Lighthouse Lab in Alderley Park | Wellcome Sanger Institute for the COVID-19 Genomics UK (COG-UK) Consortium | Jacquelyn Wynn, Mairead Hyland, The Lighthouse Lab in Alderley Park and Alex Alderton, Roberto Amato, Jeffrey Barrett, Sonia Goncalves, Ewan Harrison, David K. Jackson, Ian Johnston, Dominic Kwiatkowski, Cordelia Langford, John Sillitoe on behalf of the Wellcome Sanger Institute COVID-19 Surveillance Team |
| EPI_ISL_2867806 | Lighthouse Laboratory Plymouth | Wellcome Sanger Institute for the COVID-19 Genomics UK (COG-UK) Consortium | Lighthouse Laboratory Plymouth and Alex Alderton, Roberto Amato, Jeffrey Barrett, Sonia Goncalves, Ewan Harrison, David K. Jackson, Ian Johnston, Dominic Kwiatkowski, Cordelia Langford, John Sillitoe on behalf of the Wellcome Sanger Institute COVID-19 Surveillance Team |
| EPI_ISL_2867807 | Lighthouse Lab in Alderley Park | Wellcome Sanger Institute for the COVID-19 Genomics UK (COG-UK) Consortium | Jacquelyn Wynn, Mairead Hyland, The Lighthouse Lab in Alderley Park and Alex Alderton, Roberto Amato, Jeffrey Barrett, Sonia Goncalves, Ewan Harrison, David K. Jackson, Ian Johnston, Dominic Kwiatkowski, Cordelia Langford, John Sillitoe on behalf of the Wellcome Sanger Institute COVID-19 Surveillance Team |
| EPI_ISL_2867808 | Lighthouse Laboratory Plymouth | Wellcome Sanger Institute for the COVID-19 Genomics UK (COG-UK) Consortium | Lighthouse Laboratory Plymouth and Alex Alderton, Roberto Amato, Jeffrey Barrett, Sonia Goncalves, Ewan Harrison, David K. Jackson, Ian Johnston, Dominic Kwiatkowski, Cordelia Langford, John Sillitoe on behalf of the Wellcome Sanger Institute COVID-19 Surveillance Team |
| EPI_ISL_2867809, EPI_ISL_2867810 | Lighthouse Lab in Alderley Park | Wellcome Sanger Institute for the COVID-19 Genomics UK (COG-UK) Consortium | Jacquelyn Wynn, Mairead Hyland, The Lighthouse Lab in Alderley Park and Alex Alderton, Roberto Amato, Jeffrey Barrett, Sonia Goncalves, Ewan Harrison, David K. Jackson, Ian Johnston, Dominic Kwiatkowski, Cordelia Langford, John Sillitoe on behalf of the Wellcome Sanger Institute COVID-19 Surveillance Team |
| EPI_ISL_2867811 | Lighthouse Laboratory Plymouth | Wellcome Sanger Institute for the COVID-19 Genomics UK (COG-UK) Consortium | Lighthouse Laboratory Plymouth and Alex Alderton, Roberto Amato, Jeffrey Barrett, Sonia Goncalves, Ewan Harrison, David K. Jackson, Ian Johnston, Dominic Kwiatkowski, Cordelia Langford, John Sillitoe on behalf of the Wellcome Sanger Institute COVID-19 Surveillance Team |
| EPI_ISL_2867812, EPI_ISL_2867813 | Lighthouse Lab in Alderley Park | Wellcome Sanger Institute for the COVID-19 Genomics UK (COG-UK) Consortium | Jacquelyn Wynn, Mairead Hyland, The Lighthouse Lab in Alderley Park and Alex Alderton, Roberto Amato, Jeffrey Barrett, Sonia Goncalves, Ewan Harrison, David K. Jackson, Ian Johnston, Dominic Kwiatkowski, Cordelia Langford, John Sillitoe on behalf of the Wellcome Sanger Institute COVID-19 Surveillance Team |
| EPI_ISL_2867814 | Lighthouse Laboratory Plymouth | Wellcome Sanger Institute for the COVID-19 Genomics UK (COG-UK) Consortium | Lighthouse Laboratory Plymouth and Alex Alderton, Roberto Amato, Jeffrey Barrett, Sonia Goncalves, Ewan Harrison, David K. Jackson, Ian Johnston, Dominic Kwiatkowski, Cordelia Langford, John Sillitoe on behalf of the Wellcome Sanger Institute COVID-19 Surveillance Team |
| EPI_ISL_2867815 | Lighthouse Lab in Alderley Park | Wellcome Sanger Institute for the COVID-19 Genomics UK (COG-UK) Consortium | Jacquelyn Wynn, Mairead Hyland, The Lighthouse Lab in Alderley Park and Alex Alderton, Roberto Amato, Jeffrey Barrett, Sonia Goncalves, Ewan Harrison, David K. Jackson, Ian Johnston, Dominic Kwiatkowski, Cordelia Langford, John Sillitoe on behalf of the Wellcome Sanger Institute COVID-19 Surveillance Team |
| EPI_ISL_2867816, EPI_ISL_2867817 | Lighthouse Laboratory Plymouth | Wellcome Sanger Institute for the COVID-19 Genomics UK (COG-UK) Consortium | Lighthouse Laboratory Plymouth and Alex Alderton, Roberto Amato, Jeffrey Barrett, Sonia Goncalves, Ewan Harrison, David K. Jackson, Ian Johnston, Dominic Kwiatkowski, Cordelia Langford, John Sillitoe on behalf of the Wellcome Sanger Institute COVID-19 Surveillance Team |
| EPI_ISL_2867818 | Lighthouse Lab in Alderley Park | Wellcome Sanger Institute for the COVID-19 Genomics UK (COG-UK) Consortium | Jacquelyn Wynn, Mairead Hyland, The Lighthouse Lab in Alderley Park and Alex Alderton, Roberto Amato, Jeffrey Barrett, Sonia Goncalves, Ewan Harrison, David K. Jackson, Ian Johnston, Dominic Kwiatkowski, Cordelia Langford, John Sillitoe on behalf of the Wellcome Sanger Institute COVID-19 Surveillance Team |
| EPI_ISL_2867819 | Lighthouse Laboratory Plymouth | Wellcome Sanger Institute for the COVID-19 Genomics UK (COG-UK) Consortium | Lighthouse Laboratory Plymouth and Alex Alderton, Roberto Amato, Jeffrey Barrett, Sonia Goncalves, Ewan Harrison, David K. Jackson, Ian Johnston, Dominic Kwiatkowski, Cordelia Langford, John Sillitoe on behalf of the Wellcome Sanger Institute COVID-19 Surveillance Team |
| EPI_ISL_2867820 | Lighthouse Lab in Alderley Park | Wellcome Sanger Institute for the COVID-19 Genomics UK (COG-UK) Consortium | Jacquelyn Wynn, Mairead Hyland, The Lighthouse Lab in Alderley Park and Alex Alderton, Roberto Amato, Jeffrey Barrett, Sonia Goncalves, Ewan Harrison, David K. Jackson, Ian Johnston, Dominic Kwiatkowski, Cordelia Langford, John Sillitoe on behalf of the Wellcome Sanger Institute COVID-19 Surveillance Team |
| EPI_ISL_2867821 | Lighthouse Laboratory Plymouth | Wellcome Sanger Institute for the COVID-19 Genomics UK (COG-UK) Consortium | Lighthouse Laboratory Plymouth and Alex Alderton, Roberto Amato, Jeffrey Barrett, Sonia Goncalves, Ewan Harrison, David K. Jackson, Ian Johnston, Dominic Kwiatkowski, Cordelia Langford, John Sillitoe on behalf of the Wellcome Sanger Institute COVID-19 Surveillance Team |
| EPI_ISL_2867822, EPI_ISL_2867823 | Lighthouse Lab in Alderley Park | Wellcome Sanger Institute for the COVID-19 Genomics UK (COG-UK) Consortium | Jacquelyn Wynn, Mairead Hyland, The Lighthouse Lab in Alderley Park and Alex Alderton, Roberto Amato, Jeffrey Barrett, Sonia Goncalves, Ewan Harrison, David K. Jackson, Ian Johnston, Dominic Kwiatkowski, Cordelia Langford, John Sillitoe on behalf of the Wellcome Sanger Institute COVID-19 Surveillance Team |
| EPI_ISL_2867824, EPI_ISL_2867825 | Lighthouse Laboratory Plymouth | Wellcome Sanger Institute for the COVID-19 Genomics UK (COG-UK) Consortium | Lighthouse Laboratory Plymouth and Alex Alderton, Roberto Amato, Jeffrey Barrett, Sonia Goncalves, Ewan Harrison, David K. Jackson, Ian Johnston, Dominic Kwiatkowski, Cordelia Langford, John Sillitoe on behalf of the Wellcome Sanger Institute COVID-19 Surveillance Team |
| EPI_ISL_2867826, EPI_ISL_2867827 | Lighthouse Lab in Alderley Park | Wellcome Sanger Institute for the COVID-19 Genomics UK (COG-UK) Consortium | Jacquelyn Wynn, Mairead Hyland, The Lighthouse Lab in Alderley Park and Alex Alderton, Roberto Amato, Jeffrey Barrett, Sonia Goncalves, Ewan Harrison, David K. Jackson, Ian Johnston, Dominic Kwiatkowski, Cordelia Langford, John Sillitoe on behalf of the Wellcome Sanger Institute COVID-19 Surveillance Team |
| EPI_ISL_2867828, EPI_ISL_2867829 | Lighthouse Laboratory Plymouth | Wellcome Sanger Institute for the COVID-19 Genomics UK (COG-UK) Consortium | Lighthouse Laboratory Plymouth and Alex Alderton, Roberto Amato, Jeffrey Barrett, Sonia Goncalves, Ewan Harrison, David K. Jackson, Ian Johnston, Dominic Kwiatkowski, Cordelia Langford, John Sillitoe on behalf of the Wellcome Sanger Institute COVID-19 Surveillance Team |
| EPI_ISL_2867830 | Lighthouse Lab in Alderley Park | Wellcome Sanger Institute for the COVID-19 Genomics UK (COG-UK) Consortium | Jacquelyn Wynn, Mairead Hyland, The Lighthouse Lab in Alderley Park and Alex Alderton, Roberto Amato, Jeffrey Barrett, Sonia Goncalves, Ewan Harrison, David K. Jackson, Ian Johnston, Dominic Kwiatkowski, Cordelia Langford, John Sillitoe on behalf of the Wellcome Sanger Institute COVID-19 Surveillance Team |
| EPI_ISL_2867831 | Lighthouse Laboratory Plymouth | Wellcome Sanger Institute for the COVID-19 Genomics UK (COG-UK) Consortium | Lighthouse Laboratory Plymouth and Alex Alderton, Roberto Amato, Jeffrey Barrett, Sonia Goncalves, Ewan Harrison, David K. Jackson, Ian Johnston, Dominic Kwiatkowski, Cordelia Langford, John Sillitoe on behalf of the Wellcome Sanger Institute COVID-19 Surveillance Team |
| EPI_ISL_2867832 | Lighthouse Lab in Alderley Park | Wellcome Sanger Institute for the COVID-19 Genomics UK (COG-UK) Consortium | Jacquelyn Wynn, Mairead Hyland, The Lighthouse Lab in Alderley Park and Alex Alderton, Roberto Amato, Jeffrey Barrett, Sonia Goncalves, Ewan Harrison, David K. Jackson, Ian Johnston, Dominic Kwiatkowski, Cordelia Langford, John Sillitoe on behalf of the Wellcome Sanger Institute COVID-19 Surveillance Team |
| EPI_ISL_2867833, EPI_ISL_2867834, EPI_ISL_2867835 | Lighthouse Laboratory Plymouth | Wellcome Sanger Institute for the COVID-19 Genomics UK (COG-UK) Consortium | Lighthouse Laboratory Plymouth and Alex Alderton, Roberto Amato, Jeffrey Barrett, Sonia Goncalves, Ewan Harrison, David K. Jackson, Ian Johnston, Dominic Kwiatkowski, Cordelia Langford, John Sillitoe on behalf of the Wellcome Sanger Institute COVID-19 Surveillance Team |
| EPI_ISL_2867836 | Lighthouse Lab in Alderley Park | Wellcome Sanger Institute for the COVID-19 Genomics UK (COG-UK) Consortium | Jacquelyn Wynn, Mairead Hyland, The Lighthouse Lab in Alderley Park and Alex Alderton, Roberto Amato, Jeffrey Barrett, Sonia Goncalves, Ewan Harrison, David K. Jackson, Ian Johnston, Dominic Kwiatkowski, Cordelia Langford, John Sillitoe on behalf of the Wellcome Sanger Institute COVID-19 Surveillance Team |
| EPI_ISL_2867837, EPI_ISL_2867838, EPI_ISL_2867839, EPI_ISL_2867840 | Lighthouse Laboratory Plymouth | Wellcome Sanger Institute for the COVID-19 Genomics UK (COG-UK) Consortium | Lighthouse Laboratory Plymouth and Alex Alderton, Roberto Amato, Jeffrey Barrett, Sonia Goncalves, Ewan Harrison, David K. Jackson, Ian Johnston, Dominic Kwiatkowski, Cordelia Langford, John Sillitoe on behalf of the Wellcome Sanger Institute COVID-19 Surveillance Team |
| EPI_ISL_2867841, EPI_ISL_2867842, EPI_ISL_2867843, EPI_ISL_2867844, EPI_ISL_2867845 | Lighthouse Lab in Alderley Park | Wellcome Sanger Institute for the COVID-19 Genomics UK (COG-UK) Consortium | Jacquelyn Wynn, Mairead Hyland, The Lighthouse Lab in Alderley Park and Alex Alderton, Roberto Amato, Jeffrey Barrett, Sonia Goncalves, Ewan Harrison, David K. Jackson, Ian Johnston, Dominic Kwiatkowski, Cordelia Langford, John Sillitoe on behalf of the Wellcome Sanger Institute COVID-19 Surveillance Team |
| EPI_ISL_2867846, EPI_ISL_2867847 | Lighthouse Laboratory Plymouth | Wellcome Sanger Institute for the COVID-19 Genomics UK (COG-UK) Consortium | Lighthouse Laboratory Plymouth and Alex Alderton, Roberto Amato, Jeffrey Barrett, Sonia Goncalves, Ewan Harrison, David K. Jackson, Ian Johnston, Dominic Kwiatkowski, Cordelia Langford, John Sillitoe on behalf of the Wellcome Sanger Institute COVID-19 Surveillance Team |
| EPI_ISL_2867848 | Lighthouse Lab in Alderley Park | Wellcome Sanger Institute for the COVID-19 Genomics UK (COG-UK) Consortium | Jacquelyn Wynn, Mairead Hyland, The Lighthouse Lab in Alderley Park and Alex Alderton, Roberto Amato, Jeffrey Barrett, Sonia Goncalves, Ewan Harrison, David K. Jackson, Ian Johnston, Dominic Kwiatkowski, Cordelia Langford, John Sillitoe on behalf of the Wellcome Sanger Institute COVID-19 Surveillance Team |
| EPI_ISL_2867849, EPI_ISL_2867850, EPI_ISL_2867851 | Lighthouse Laboratory Plymouth | Wellcome Sanger Institute for the COVID-19 Genomics UK (COG-UK) Consortium | Lighthouse Laboratory Plymouth and Alex Alderton, Roberto Amato, Jeffrey Barrett, Sonia Goncalves, Ewan Harrison, David K. Jackson, Ian Johnston, Dominic Kwiatkowski, Cordelia Langford, John Sillitoe on behalf of the Wellcome Sanger Institute COVID-19 Surveillance Team |
| EPI_ISL_2867852, EPI_ISL_2867853 | Lighthouse Lab in Alderley Park | Wellcome Sanger Institute for the COVID-19 Genomics UK (COG-UK) Consortium | Jacquelyn Wynn, Mairead Hyland, The Lighthouse Lab in Alderley Park and Alex Alderton, Roberto Amato, Jeffrey Barrett, Sonia Goncalves, Ewan Harrison, David K. Jackson, Ian Johnston, Dominic Kwiatkowski, Cordelia Langford, John Sillitoe on behalf of the Wellcome Sanger Institute COVID-19 Surveillance Team |
| EPI_ISL_2867854, EPI_ISL_2867855 | Lighthouse Laboratory Plymouth | Wellcome Sanger Institute for the COVID-19 Genomics UK (COG-UK) Consortium | Lighthouse Laboratory Plymouth and Alex Alderton, Roberto Amato, Jeffrey Barrett, Sonia Goncalves, Ewan Harrison, David K. Jackson, Ian Johnston, Dominic Kwiatkowski, Cordelia Langford, John Sillitoe on behalf of the Wellcome Sanger Institute COVID-19 Surveillance Team |
| EPI_ISL_2867856, EPI_ISL_2867857 | Lighthouse Lab in Alderley Park | Wellcome Sanger Institute for the COVID-19 Genomics UK (COG-UK) Consortium | Jacquelyn Wynn, Mairead Hyland, The Lighthouse Lab in Alderley Park and Alex Alderton, Roberto Amato, Jeffrey Barrett, Sonia Goncalves, Ewan Harrison, David K. Jackson, Ian Johnston, Dominic Kwiatkowski, Cordelia Langford, John Sillitoe on behalf of the Wellcome Sanger Institute COVID-19 Surveillance Team |
| EPI_ISL_2867858 | Lighthouse Laboratory Plymouth | Wellcome Sanger Institute for the COVID-19 Genomics UK (COG-UK) Consortium | Lighthouse Laboratory Plymouth and Alex Alderton, Roberto Amato, Jeffrey Barrett, Sonia Goncalves, Ewan Harrison, David K. Jackson, Ian Johnston, Dominic Kwiatkowski, Cordelia Langford, John Sillitoe on behalf of the Wellcome Sanger Institute COVID-19 Surveillance Team |
| EPI_ISL_2867859 | Lighthouse Lab in Alderley Park | Wellcome Sanger Institute for the COVID-19 Genomics UK (COG-UK) Consortium | Jacquelyn Wynn, Mairead Hyland, The Lighthouse Lab in Alderley Park and Alex Alderton, Roberto Amato, Jeffrey Barrett, Sonia Goncalves, Ewan Harrison, David K. Jackson, Ian Johnston, Dominic Kwiatkowski, Cordelia Langford, John Sillitoe on behalf of the Wellcome Sanger Institute COVID-19 Surveillance Team |
| EPI_ISL_2867860 | Lighthouse Laboratory Plymouth | Wellcome Sanger Institute for the COVID-19 Genomics UK (COG-UK) Consortium | Lighthouse Laboratory Plymouth and Alex Alderton, Roberto Amato, Jeffrey Barrett, Sonia Goncalves, Ewan Harrison, David K. Jackson, Ian Johnston, Dominic Kwiatkowski, Cordelia Langford, John Sillitoe on behalf of the Wellcome Sanger Institute COVID-19 Surveillance Team |
| EPI_ISL_2867861, EPI_ISL_2867862 | Lighthouse Lab in Alderley Park | Wellcome Sanger Institute for the COVID-19 Genomics UK (COG-UK) Consortium | Jacquelyn Wynn, Mairead Hyland, The Lighthouse Lab in Alderley Park and Alex Alderton, Roberto Amato, Jeffrey Barrett, Sonia Goncalves, Ewan Harrison, David K. Jackson, Ian Johnston, Dominic Kwiatkowski, Cordelia Langford, John Sillitoe on behalf of the Wellcome Sanger Institute COVID-19 Surveillance Team |
| EPI_ISL_2867863, EPI_ISL_2867864 | Lighthouse Laboratory Plymouth | Wellcome Sanger Institute for the COVID-19 Genomics UK (COG-UK) Consortium | Lighthouse Laboratory Plymouth and Alex Alderton, Roberto Amato, Jeffrey Barrett, Sonia Goncalves, Ewan Harrison, David K. Jackson, Ian Johnston, Dominic Kwiatkowski, Cordelia Langford, John Sillitoe on behalf of the Wellcome Sanger Institute COVID-19 Surveillance Team |
| EPI_ISL_2867865 | Lighthouse Lab in Alderley Park | Wellcome Sanger Institute for the COVID-19 Genomics UK (COG-UK) Consortium | Jacquelyn Wynn, Mairead Hyland, The Lighthouse Lab in Alderley Park and Alex Alderton, Roberto Amato, Jeffrey Barrett, Sonia Goncalves, Ewan Harrison, David K. Jackson, Ian Johnston, Dominic Kwiatkowski, Cordelia Langford, John Sillitoe on behalf of the Wellcome Sanger Institute COVID-19 Surveillance Team |
| EPI_ISL_2867866 | Lighthouse Laboratory Plymouth | Wellcome Sanger Institute for the COVID-19 Genomics UK (COG-UK) Consortium | Lighthouse Laboratory Plymouth and Alex Alderton, Roberto Amato, Jeffrey Barrett, Sonia Goncalves, Ewan Harrison, David K. Jackson, Ian Johnston, Dominic Kwiatkowski, Cordelia Langford, John Sillitoe on behalf of the Wellcome Sanger Institute COVID-19 Surveillance Team |
| EPI_ISL_2867867 | Lighthouse Lab in Alderley Park | Wellcome Sanger Institute for the COVID-19 Genomics UK (COG-UK) Consortium | Jacquelyn Wynn, Mairead Hyland, The Lighthouse Lab in Alderley Park and Alex Alderton, Roberto Amato, Jeffrey Barrett, Sonia Goncalves, Ewan Harrison, David K. Jackson, Ian Johnston, Dominic Kwiatkowski, Cordelia Langford, John Sillitoe on behalf of the Wellcome Sanger Institute COVID-19 Surveillance Team |
| EPI_ISL_2867868 | Lighthouse Laboratory Plymouth | Wellcome Sanger Institute for the COVID-19 Genomics UK (COG-UK) Consortium | Lighthouse Laboratory Plymouth and Alex Alderton, Roberto Amato, Jeffrey Barrett, Sonia Goncalves, Ewan Harrison, David K. Jackson, Ian Johnston, Dominic Kwiatkowski, Cordelia Langford, John Sillitoe on behalf of the Wellcome Sanger Institute COVID-19 Surveillance Team |
| EPI_ISL_2867869 | Lighthouse Lab in Alderley Park | Wellcome Sanger Institute for the COVID-19 Genomics UK (COG-UK) Consortium | Jacquelyn Wynn, Mairead Hyland, The Lighthouse Lab in Alderley Park and Alex Alderton, Roberto Amato, Jeffrey Barrett, Sonia Goncalves, Ewan Harrison, David K. Jackson, Ian Johnston, Dominic Kwiatkowski, Cordelia Langford, John Sillitoe on behalf of the Wellcome Sanger Institute COVID-19 Surveillance Team |

| | | | |
|---|---|---|---|
| EPI_ISL_2867870, EPI_ISL_2867871, EPI_ISL_2867872 | Lighthouse Laboratory Plymouth | Wellcome Sanger Institute for the COVID-19 Genomics UK (COG-UK) Consortium | Lighthouse Laboratory Plymouth and Alex Alderton, Roberto Amato, Jeffrey Barrett, Sonia Goncalves, Ewan Harrison, David K. Jackson, Ian Johnston, Dominic Kwiatkowski, Cordelia Langford, John Sillitoe on behalf of the Wellcome Sanger Institute COVID-19 Surveillance Team |
| EPI_ISL_2867873, EPI_ISL_2867874, EPI_ISL_2867875, EPI_ISL_2867876, EPI_ISL_2867877 | Lighthouse Lab in Alderley Park | Wellcome Sanger Institute for the COVID-19 Genomics UK (COG-UK) Consortium | Jacquelyn Wynn, Mairead Hyland, The Lighthouse Lab in Alderley Park and Alex Alderton, Roberto Amato, Jeffrey Barrett, Sonia Goncalves, Ewan Harrison, David K. Jackson, Ian Johnston, Dominic Kwiatkowski, Cordelia Langford, John Sillitoe on behalf of the Wellcome Sanger Institute COVID-19 Surveillance Team |
| EPI_ISL_2867878 | Lighthouse Laboratory Plymouth | Wellcome Sanger Institute for the COVID-19 Genomics UK (COG-UK) Consortium | Lighthouse Laboratory Plymouth and Alex Alderton, Roberto Amato, Jeffrey Barrett, Sonia Goncalves, Ewan Harrison, David K. Jackson, Ian Johnston, Dominic Kwiatkowski, Cordelia Langford, John Sillitoe on behalf of the Wellcome Sanger Institute COVID-19 Surveillance Team |
| EPI_ISL_2867879, EPI_ISL_2867880, EPI_ISL_2867881 | Lighthouse Lab in Alderley Park | Wellcome Sanger Institute for the COVID-19 Genomics UK (COG-UK) Consortium | Jacquelyn Wynn, Mairead Hyland, The Lighthouse Lab in Alderley Park and Alex Alderton, Roberto Amato, Jeffrey Barrett, Sonia Goncalves, Ewan Harrison, David K. Jackson, Ian Johnston, Dominic Kwiatkowski, Cordelia Langford, John Sillitoe on behalf of the Wellcome Sanger Institute COVID-19 Surveillance Team |
| EPI_ISL_2867882 | Lighthouse Laboratory Plymouth | Wellcome Sanger Institute for the COVID-19 Genomics UK (COG-UK) Consortium | Lighthouse Laboratory Plymouth and Alex Alderton, Roberto Amato, Jeffrey Barrett, Sonia Goncalves, Ewan Harrison, David K. Jackson, Ian Johnston, Dominic Kwiatkowski, Cordelia Langford, John Sillitoe on behalf of the Wellcome Sanger Institute COVID-19 Surveillance Team |
| EPI_ISL_2867883 | Lighthouse Lab in Alderley Park | Wellcome Sanger Institute for the COVID-19 Genomics UK (COG-UK) Consortium | Jacquelyn Wynn, Mairead Hyland, The Lighthouse Lab in Alderley Park and Alex Alderton, Roberto Amato, Jeffrey Barrett, Sonia Goncalves, Ewan Harrison, David K. Jackson, Ian Johnston, Dominic Kwiatkowski, Cordelia Langford, John Sillitoe on behalf of the Wellcome Sanger Institute COVID-19 Surveillance Team |
| EPI_ISL_2867884, EPI_ISL_2867885, EPI_ISL_2867886, EPI_ISL_2867887 | Lighthouse Laboratory Plymouth | Wellcome Sanger Institute for the COVID-19 Genomics UK (COG-UK) Consortium | Lighthouse Laboratory Plymouth and Alex Alderton, Roberto Amato, Jeffrey Barrett, Sonia Goncalves, Ewan Harrison, David K. Jackson, Ian Johnston, Dominic Kwiatkowski, Cordelia Langford, John Sillitoe on behalf of the Wellcome Sanger Institute COVID-19 Surveillance Team |
| EPI_ISL_2867888, EPI_ISL_2867889, EPI_ISL_2867890 | Lighthouse Lab in Alderley Park | Wellcome Sanger Institute for the COVID-19 Genomics UK (COG-UK) Consortium | Jacquelyn Wynn, Mairead Hyland, The Lighthouse Lab in Alderley Park and Alex Alderton, Roberto Amato, Jeffrey Barrett, Sonia Goncalves, Ewan Harrison, David K. Jackson, Ian Johnston, Dominic Kwiatkowski, Cordelia Langford, John Sillitoe on behalf of the Wellcome Sanger Institute COVID-19 Surveillance Team |
| EPI_ISL_2867891 | Lighthouse Laboratory Plymouth | Wellcome Sanger Institute for the COVID-19 Genomics UK (COG-UK) Consortium | Lighthouse Laboratory Plymouth and Alex Alderton, Roberto Amato, Jeffrey Barrett, Sonia Goncalves, Ewan Harrison, David K. Jackson, Ian Johnston, Dominic Kwiatkowski, Cordelia Langford, John Sillitoe on behalf of the Wellcome Sanger Institute COVID-19 Surveillance Team |
| EPI_ISL_2867892 | Lighthouse Lab in Alderley Park | Wellcome Sanger Institute for the COVID-19 Genomics UK (COG-UK) Consortium | Jacquelyn Wynn, Mairead Hyland, The Lighthouse Lab in Alderley Park and Alex Alderton, Roberto Amato, Jeffrey Barrett, Sonia Goncalves, Ewan Harrison, David K. Jackson, Ian Johnston, Dominic Kwiatkowski, Cordelia Langford, John Sillitoe on behalf of the Wellcome Sanger Institute COVID-19 Surveillance Team |
| EPI_ISL_2867893, EPI_ISL_2867894 | Lighthouse Laboratory Plymouth | Wellcome Sanger Institute for the COVID-19 Genomics UK (COG-UK) Consortium | Lighthouse Laboratory Plymouth and Alex Alderton, Roberto Amato, Jeffrey Barrett, Sonia Goncalves, Ewan Harrison, David K. Jackson, Ian Johnston, Dominic Kwiatkowski, Cordelia Langford, John Sillitoe on behalf of the Wellcome Sanger Institute COVID-19 Surveillance Team |
| EPI_ISL_2867895 | Lighthouse Lab in Alderley Park | Wellcome Sanger Institute for the COVID-19 Genomics UK (COG-UK) Consortium | Jacquelyn Wynn, Mairead Hyland, The Lighthouse Lab in Alderley Park and Alex Alderton, Roberto Amato, Jeffrey Barrett, Sonia Goncalves, Ewan Harrison, David K. Jackson, Ian Johnston, Dominic Kwiatkowski, Cordelia Langford, John Sillitoe on behalf of the Wellcome Sanger Institute COVID-19 Surveillance Team |
| EPI_ISL_2867896 | Lighthouse Laboratory Plymouth | Wellcome Sanger Institute for the COVID-19 Genomics UK (COG-UK) Consortium | Lighthouse Laboratory Plymouth and Alex Alderton, Roberto Amato, Jeffrey Barrett, Sonia Goncalves, Ewan Harrison, David K. Jackson, Ian Johnston, Dominic Kwiatkowski, Cordelia Langford, John Sillitoe on behalf of the Wellcome Sanger Institute COVID-19 Surveillance Team |
| EPI_ISL_2867897, EPI_ISL_2867898 | Lighthouse Lab in Alderley Park | Wellcome Sanger Institute for the COVID-19 Genomics UK (COG-UK) Consortium | Jacquelyn Wynn, Mairead Hyland, The Lighthouse Lab in Alderley Park and Alex Alderton, Roberto Amato, Jeffrey Barrett, Sonia Goncalves, Ewan Harrison, David K. Jackson, Ian Johnston, Dominic Kwiatkowski, Cordelia Langford, John Sillitoe on behalf of the Wellcome Sanger Institute COVID-19 Surveillance Team |
| EPI_ISL_2867899, EPI_ISL_2867900 | Lighthouse Laboratory Plymouth | Wellcome Sanger Institute for the COVID-19 Genomics UK (COG-UK) Consortium | Lighthouse Laboratory Plymouth and Alex Alderton, Roberto Amato, Jeffrey Barrett, Sonia Goncalves, Ewan Harrison, David K. Jackson, Ian Johnston, Dominic Kwiatkowski, Cordelia Langford, John Sillitoe on behalf of the Wellcome Sanger Institute COVID-19 Surveillance Team |
| EPI_ISL_2867901 | Lighthouse Lab in Alderley Park | Wellcome Sanger Institute for the COVID-19 Genomics UK (COG-UK) Consortium | Jacquelyn Wynn, Mairead Hyland, The Lighthouse Lab in Alderley Park and Alex Alderton, Roberto Amato, Jeffrey Barrett, Sonia Goncalves, Ewan Harrison, David K. Jackson, Ian Johnston, Dominic Kwiatkowski, Cordelia Langford, John Sillitoe on behalf of the Wellcome Sanger Institute COVID-19 Surveillance Team |
| EPI_ISL_2867902 | Lighthouse Laboratory Plymouth | Wellcome Sanger Institute for the COVID-19 Genomics UK (COG-UK) Consortium | Lighthouse Laboratory Plymouth and Alex Alderton, Roberto Amato, Jeffrey Barrett, Sonia Goncalves, Ewan Harrison, David K. Jackson, Ian Johnston, Dominic Kwiatkowski, Cordelia Langford, John Sillitoe on behalf of the Wellcome Sanger Institute COVID-19 Surveillance Team |
| EPI_ISL_2867903 | Lighthouse Lab in Alderley Park | Wellcome Sanger Institute for the COVID-19 Genomics UK (COG-UK) Consortium | Jacquelyn Wynn, Mairead Hyland, The Lighthouse Lab in Alderley Park and Alex Alderton, Roberto Amato, Jeffrey Barrett, Sonia Goncalves, Ewan Harrison, David K. Jackson, Ian Johnston, Dominic Kwiatkowski, Cordelia Langford, John Sillitoe on behalf of the Wellcome Sanger Institute COVID-19 Surveillance Team |
| EPI_ISL_2867904 | Lighthouse Laboratory Plymouth | Wellcome Sanger Institute for the COVID-19 Genomics UK (COG-UK) Consortium | Lighthouse Laboratory Plymouth and Alex Alderton, Roberto Amato, Jeffrey Barrett, Sonia Goncalves, Ewan Harrison, David K. Jackson, Ian Johnston, Dominic Kwiatkowski, Cordelia Langford, John Sillitoe on behalf of the Wellcome Sanger Institute COVID-19 Surveillance Team |
| EPI_ISL_2867905, EPI_ISL_2867906 | Lighthouse Lab in Alderley Park | Wellcome Sanger Institute for the COVID-19 Genomics UK (COG-UK) Consortium | Jacquelyn Wynn, Mairead Hyland, The Lighthouse Lab in Alderley Park and Alex Alderton, Roberto Amato, Jeffrey Barrett, Sonia Goncalves, Ewan Harrison, David K. Jackson, Ian Johnston, Dominic Kwiatkowski, Cordelia Langford, John Sillitoe on behalf of the Wellcome Sanger Institute COVID-19 Surveillance Team |
| EPI_ISL_2867907 | Lighthouse Laboratory Plymouth | Wellcome Sanger Institute for the COVID-19 Genomics UK (COG-UK) Consortium | Lighthouse Laboratory Plymouth and Alex Alderton, Roberto Amato, Jeffrey Barrett, Sonia Goncalves, Ewan Harrison, David K. Jackson, Ian Johnston, Dominic Kwiatkowski, Cordelia Langford, John Sillitoe on behalf of the Wellcome Sanger Institute COVID-19 Surveillance Team |
| EPI_ISL_2867908 | Lighthouse Lab in Alderley Park | Wellcome Sanger Institute for the COVID-19 Genomics UK (COG-UK) Consortium | Jacquelyn Wynn, Mairead Hyland, The Lighthouse Lab in Alderley Park and Alex Alderton, Roberto Amato, Jeffrey Barrett, Sonia Goncalves, Ewan Harrison, David K. Jackson, Ian Johnston, Dominic Kwiatkowski, Cordelia Langford, John Sillitoe on behalf of the Wellcome Sanger Institute COVID-19 Surveillance Team |
| EPI_ISL_2867909 | Lighthouse Laboratory Plymouth | Wellcome Sanger Institute for the COVID-19 Genomics UK (COG-UK) Consortium | Lighthouse Laboratory Plymouth and Alex Alderton, Roberto Amato, Jeffrey Barrett, Sonia Goncalves, Ewan Harrison, David K. Jackson, Ian Johnston, Dominic Kwiatkowski, Cordelia Langford, John Sillitoe on behalf of the Wellcome Sanger Institute COVID-19 Surveillance Team |
| EPI_ISL_2867910, EPI_ISL_2867911, EPI_ISL_2867912, EPI_ISL_2867913 | Lighthouse Lab in Alderley Park | Wellcome Sanger Institute for the COVID-19 Genomics UK (COG-UK) Consortium | Jacquelyn Wynn, Mairead Hyland, The Lighthouse Lab in Alderley Park and Alex Alderton, Roberto Amato, Jeffrey Barrett, Sonia Goncalves, Ewan Harrison, David K. Jackson, Ian Johnston, Dominic Kwiatkowski, Cordelia Langford, John Sillitoe on behalf of the Wellcome Sanger Institute COVID-19 Surveillance Team |
| EPI_ISL_2867914 | Lighthouse Laboratory Plymouth | Wellcome Sanger Institute for the COVID-19 Genomics UK (COG-UK) Consortium | Lighthouse Laboratory Plymouth and Alex Alderton, Roberto Amato, Jeffrey Barrett, Sonia Goncalves, Ewan Harrison, David K. Jackson, Ian Johnston, Dominic Kwiatkowski, Cordelia Langford, John Sillitoe on behalf of the Wellcome Sanger Institute COVID-19 Surveillance Team |
| EPI_ISL_2867915 | Lighthouse Lab in Alderley Park | Wellcome Sanger Institute for the COVID-19 Genomics UK (COG-UK) Consortium | Jacquelyn Wynn, Mairead Hyland, The Lighthouse Lab in Alderley Park and Alex Alderton, Roberto Amato, Jeffrey Barrett, Sonia Goncalves, Ewan Harrison, David K. Jackson, Ian Johnston, Dominic Kwiatkowski, Cordelia Langford, John Sillitoe on behalf of the Wellcome Sanger Institute COVID-19 Surveillance Team |
| EPI_ISL_2867916, EPI_ISL_2867917, EPI_ISL_2867918 | Lighthouse Laboratory Plymouth | Wellcome Sanger Institute for the COVID-19 Genomics UK (COG-UK) Consortium | Lighthouse Laboratory Plymouth and Alex Alderton, Roberto Amato, Jeffrey Barrett, Sonia Goncalves, Ewan Harrison, David K. Jackson, Ian Johnston, Dominic Kwiatkowski, Cordelia Langford, John Sillitoe on behalf of the Wellcome Sanger Institute COVID-19 Surveillance Team |
| EPI_ISL_2867919, EPI_ISL_2867920, EPI_ISL_2867921, EPI_ISL_2867922 | Lighthouse Lab in Alderley Park | Wellcome Sanger Institute for the COVID-19 Genomics UK (COG-UK) Consortium | Jacquelyn Wynn, Mairead Hyland, The Lighthouse Lab in Alderley Park and Alex Alderton, Roberto Amato, Jeffrey Barrett, Sonia Goncalves, Ewan Harrison, David K. Jackson, Ian Johnston, Dominic Kwiatkowski, Cordelia Langford, John Sillitoe on behalf of the Wellcome Sanger Institute COVID-19 Surveillance Team |
| EPI_ISL_2867923 | Lighthouse Laboratory Plymouth | Wellcome Sanger Institute for the COVID-19 Genomics UK (COG-UK) Consortium | Lighthouse Laboratory Plymouth and Alex Alderton, Roberto Amato, Jeffrey Barrett, Sonia Goncalves, Ewan Harrison, David K. Jackson, Ian Johnston, Dominic Kwiatkowski, Cordelia Langford, John Sillitoe on behalf of the Wellcome Sanger Institute COVID-19 Surveillance Team |
| EPI_ISL_2867924, EPI_ISL_2867925, EPI_ISL_2867926, EPI_ISL_2867927 | Lighthouse Lab in Alderley Park | Wellcome Sanger Institute for the COVID-19 Genomics UK (COG-UK) Consortium | Jacquelyn Wynn, Mairead Hyland, The Lighthouse Lab in Alderley Park and Alex Alderton, Roberto Amato, Jeffrey Barrett, Sonia Goncalves, Ewan Harrison, David K. Jackson, Ian Johnston, Dominic Kwiatkowski, Cordelia Langford, John Sillitoe on behalf of the Wellcome Sanger Institute COVID-19 Surveillance Team |
| EPI_ISL_2867928 | Lighthouse Laboratory Plymouth | Wellcome Sanger Institute for the COVID-19 Genomics UK (COG-UK) Consortium | Lighthouse Laboratory Plymouth and Alex Alderton, Roberto Amato, Jeffrey Barrett, Sonia Goncalves, Ewan Harrison, David K. Jackson, Ian Johnston, Dominic Kwiatkowski, Cordelia Langford, John Sillitoe on behalf of the Wellcome Sanger Institute COVID-19 Surveillance Team |
| EPI_ISL_2867929 | Lighthouse Lab in Alderley Park | Wellcome Sanger Institute for the COVID-19 Genomics UK (COG-UK) Consortium | Jacquelyn Wynn, Mairead Hyland, The Lighthouse Lab in Alderley Park and Alex Alderton, Roberto Amato, Jeffrey Barrett, Sonia Goncalves, Ewan Harrison, David K. Jackson, Ian Johnston, Dominic Kwiatkowski, Cordelia Langford, John Sillitoe on behalf of the Wellcome Sanger Institute COVID-19 Surveillance Team |
| EPI_ISL_2867930, EPI_ISL_2867931, EPI_ISL_2867932, EPI_ISL_2867933, EPI_ISL_2867934 | Lighthouse Laboratory Plymouth | Wellcome Sanger Institute for the COVID-19 Genomics UK (COG-UK) Consortium | Lighthouse Laboratory Plymouth and Alex Alderton, Roberto Amato, Jeffrey Barrett, Sonia Goncalves, Ewan Harrison, David K. Jackson, Ian Johnston, Dominic Kwiatkowski, Cordelia Langford, John Sillitoe on behalf of the Wellcome Sanger Institute COVID-19 Surveillance Team |
| EPI_ISL_2867935, EPI_ISL_2867936 | Lighthouse Lab in Alderley Park | Wellcome Sanger Institute for the COVID-19 Genomics UK (COG-UK) Consortium | Jacquelyn Wynn, Mairead Hyland, The Lighthouse Lab in Alderley Park and Alex Alderton, Roberto Amato, Jeffrey Barrett, Sonia Goncalves, Ewan Harrison, David K. Jackson, Ian Johnston, Dominic Kwiatkowski, Cordelia Langford, John Sillitoe on behalf of the Wellcome Sanger Institute COVID-19 Surveillance Team |
| EPI_ISL_2867937, EPI_ISL_2867938 | Lighthouse Laboratory Plymouth | Wellcome Sanger Institute for the COVID-19 Genomics UK (COG-UK) Consortium | Lighthouse Laboratory Plymouth and Alex Alderton, Roberto Amato, Jeffrey Barrett, Sonia Goncalves, Ewan Harrison, David K. Jackson, Ian Johnston, Dominic Kwiatkowski, Cordelia Langford, John Sillitoe on behalf of the Wellcome Sanger Institute COVID-19 Surveillance Team |
| EPI_ISL_2867939, EPI_ISL_2867940, EPI_ISL_2867941 | Lighthouse Lab in Alderley Park | Wellcome Sanger Institute for the COVID-19 Genomics UK (COG-UK) Consortium | Jacquelyn Wynn, Mairead Hyland, The Lighthouse Lab in Alderley Park and Alex Alderton, Roberto Amato, Jeffrey Barrett, Sonia Goncalves, Ewan Harrison, David K. Jackson, Ian Johnston, Dominic Kwiatkowski, Cordelia Langford, John Sillitoe on behalf of the Wellcome Sanger Institute COVID-19 Surveillance Team |
| EPI_ISL_2867942, EPI_ISL_2867943, | Lighthouse Laboratory Plymouth | Wellcome Sanger Institute for the COVID-19 Genomics UK (COG-UK) Consortium | Lighthouse Laboratory Plymouth and Alex Alderton, Roberto Amato, Jeffrey Barrett, Sonia Goncalves, Ewan Harrison, David K. Jackson, Ian Johnston, Dominic Kwiatkowski, Cordelia Langford, John Sillitoe on behalf of the Wellcome Sanger Institute COVID-19 Surveillance Team |

| | | | |
|---|---|---|---|
| EPI_ISL_2867944 | | | |
| EPI_ISL_2867945 | Lighthouse Lab in Alderley Park | Wellcome Sanger Institute for the COVID-19 Genomics UK (COG-UK) Consortium | Jacquelyn Wynn, Mairead Hyland, The Lighthouse Lab in Alderley Park and Alex Alderton, Roberto Amato, Jeffrey Barrett, Sonia Goncalves, Ewan Harrison, David K. Jackson, Ian Johnston, Dominic Kwiatkowski, Cordelia Langford, John Sillitoe on behalf of the Wellcome Sanger Institute COVID-19 Surveillance Team |
| EPI_ISL_2867946 | Lighthouse Laboratory Plymouth | Wellcome Sanger Institute for the COVID-19 Genomics UK (COG-UK) Consortium | Lighthouse Laboratory Plymouth and Alex Alderton, Roberto Amato, Jeffrey Barrett, Sonia Goncalves, Ewan Harrison, David K. Jackson, Ian Johnston, Dominic Kwiatkowski, Cordelia Langford, John Sillitoe on behalf of the Wellcome Sanger Institute COVID-19 Surveillance Team |
| EPI_ISL_2867947 | Lighthouse Lab in Alderley Park | Wellcome Sanger Institute for the COVID-19 Genomics UK (COG-UK) Consortium | Jacquelyn Wynn, Mairead Hyland, The Lighthouse Lab in Alderley Park and Alex Alderton, Roberto Amato, Jeffrey Barrett, Sonia Goncalves, Ewan Harrison, David K. Jackson, Ian Johnston, Dominic Kwiatkowski, Cordelia Langford, John Sillitoe on behalf of the Wellcome Sanger Institute COVID-19 Surveillance Team |
| EPI_ISL_2867948, EPI_ISL_2867949, EPI_ISL_2867950, EPI_ISL_2867951, EPI_ISL_2867952, EPI_ISL_2867953 | Lighthouse Laboratory Plymouth | Wellcome Sanger Institute for the COVID-19 Genomics UK (COG-UK) Consortium | Lighthouse Laboratory Plymouth and Alex Alderton, Roberto Amato, Jeffrey Barrett, Sonia Goncalves, Ewan Harrison, David K. Jackson, Ian Johnston, Dominic Kwiatkowski, Cordelia Langford, John Sillitoe on behalf of the Wellcome Sanger Institute COVID-19 Surveillance Team |
| EPI_ISL_2867954, EPI_ISL_2867955, EPI_ISL_2867956, EPI_ISL_2867957, EPI_ISL_2867958 | Lighthouse Lab in Alderley Park | Wellcome Sanger Institute for the COVID-19 Genomics UK (COG-UK) Consortium | Jacquelyn Wynn, Mairead Hyland, The Lighthouse Lab in Alderley Park and Alex Alderton, Roberto Amato, Jeffrey Barrett, Sonia Goncalves, Ewan Harrison, David K. Jackson, Ian Johnston, Dominic Kwiatkowski, Cordelia Langford, John Sillitoe on behalf of the Wellcome Sanger Institute COVID-19 Surveillance Team |
| EPI_ISL_2867959 | Lighthouse Laboratory Plymouth | Wellcome Sanger Institute for the COVID-19 Genomics UK (COG-UK) Consortium | Lighthouse Laboratory Plymouth and Alex Alderton, Roberto Amato, Jeffrey Barrett, Sonia Goncalves, Ewan Harrison, David K. Jackson, Ian Johnston, Dominic Kwiatkowski, Cordelia Langford, John Sillitoe on behalf of the Wellcome Sanger Institute COVID-19 Surveillance Team |
| EPI_ISL_2867960 | Lighthouse Lab in Alderley Park | Wellcome Sanger Institute for the COVID-19 Genomics UK (COG-UK) Consortium | Jacquelyn Wynn, Mairead Hyland, The Lighthouse Lab in Alderley Park and Alex Alderton, Roberto Amato, Jeffrey Barrett, Sonia Goncalves, Ewan Harrison, David K. Jackson, Ian Johnston, Dominic Kwiatkowski, Cordelia Langford, John Sillitoe on behalf of the Wellcome Sanger Institute COVID-19 Surveillance Team |
| EPI_ISL_2867961 | Lighthouse Laboratory Plymouth | Wellcome Sanger Institute for the COVID-19 Genomics UK (COG-UK) Consortium | Lighthouse Laboratory Plymouth and Alex Alderton, Roberto Amato, Jeffrey Barrett, Sonia Goncalves, Ewan Harrison, David K. Jackson, Ian Johnston, Dominic Kwiatkowski, Cordelia Langford, John Sillitoe on behalf of the Wellcome Sanger Institute COVID-19 Surveillance Team |
| EPI_ISL_2867962, EPI_ISL_2867963, EPI_ISL_2867964, EPI_ISL_2867965, EPI_ISL_2867966, EPI_ISL_2867967 | Lighthouse Lab in Alderley Park | Wellcome Sanger Institute for the COVID-19 Genomics UK (COG-UK) Consortium | Jacquelyn Wynn, Mairead Hyland, The Lighthouse Lab in Alderley Park and Alex Alderton, Roberto Amato, Jeffrey Barrett, Sonia Goncalves, Ewan Harrison, David K. Jackson, Ian Johnston, Dominic Kwiatkowski, Cordelia Langford, John Sillitoe on behalf of the Wellcome Sanger Institute COVID-19 Surveillance Team |
| EPI_ISL_2867968 | Lighthouse Laboratory Plymouth | Wellcome Sanger Institute for the COVID-19 Genomics UK (COG-UK) Consortium | Lighthouse Laboratory Plymouth and Alex Alderton, Roberto Amato, Jeffrey Barrett, Sonia Goncalves, Ewan Harrison, David K. Jackson, Ian Johnston, Dominic Kwiatkowski, Cordelia Langford, John Sillitoe on behalf of the Wellcome Sanger Institute COVID-19 Surveillance Team |
| EPI_ISL_2867969, EPI_ISL_2867970, EPI_ISL_2867971 | Lighthouse Lab in Alderley Park | Wellcome Sanger Institute for the COVID-19 Genomics UK (COG-UK) Consortium | Jacquelyn Wynn, Mairead Hyland, The Lighthouse Lab in Alderley Park and Alex Alderton, Roberto Amato, Jeffrey Barrett, Sonia Goncalves, Ewan Harrison, David K. Jackson, Ian Johnston, Dominic Kwiatkowski, Cordelia Langford, John Sillitoe on behalf of the Wellcome Sanger Institute COVID-19 Surveillance Team |
| EPI_ISL_2867972, EPI_ISL_2867973 | Lighthouse Laboratory Plymouth | Wellcome Sanger Institute for the COVID-19 Genomics UK (COG-UK) Consortium | Lighthouse Laboratory Plymouth and Alex Alderton, Roberto Amato, Jeffrey Barrett, Sonia Goncalves, Ewan Harrison, David K. Jackson, Ian Johnston, Dominic Kwiatkowski, Cordelia Langford, John Sillitoe on behalf of the Wellcome Sanger Institute COVID-19 Surveillance Team |
| EPI_ISL_2867974 | Lighthouse Lab in Alderley Park | Wellcome Sanger Institute for the COVID-19 Genomics UK (COG-UK) Consortium | Jacquelyn Wynn, Mairead Hyland, The Lighthouse Lab in Alderley Park and Alex Alderton, Roberto Amato, Jeffrey Barrett, Sonia Goncalves, Ewan Harrison, David K. Jackson, Ian Johnston, Dominic Kwiatkowski, Cordelia Langford, John Sillitoe on behalf of the Wellcome Sanger Institute COVID-19 Surveillance Team |
| EPI_ISL_2867975, EPI_ISL_2867976, EPI_ISL_2867977, EPI_ISL_2867978 | Lighthouse Laboratory Plymouth | Wellcome Sanger Institute for the COVID-19 Genomics UK (COG-UK) Consortium | Lighthouse Laboratory Plymouth and Alex Alderton, Roberto Amato, Jeffrey Barrett, Sonia Goncalves, Ewan Harrison, David K. Jackson, Ian Johnston, Dominic Kwiatkowski, Cordelia Langford, John Sillitoe on behalf of the Wellcome Sanger Institute COVID-19 Surveillance Team |
| EPI_ISL_2867979, EPI_ISL_2867980, EPI_ISL_2867981, EPI_ISL_2867982 | Lighthouse Lab in Alderley Park | Wellcome Sanger Institute for the COVID-19 Genomics UK (COG-UK) Consortium | Jacquelyn Wynn, Mairead Hyland, The Lighthouse Lab in Alderley Park and Alex Alderton, Roberto Amato, Jeffrey Barrett, Sonia Goncalves, Ewan Harrison, David K. Jackson, Ian Johnston, Dominic Kwiatkowski, Cordelia Langford, John Sillitoe on behalf of the Wellcome Sanger Institute COVID-19 Surveillance Team |
| EPI_ISL_2867983 | Lighthouse Laboratory Plymouth | Wellcome Sanger Institute for the COVID-19 Genomics UK (COG-UK) Consortium | Lighthouse Laboratory Plymouth and Alex Alderton, Roberto Amato, Jeffrey Barrett, Sonia Goncalves, Ewan Harrison, David K. Jackson, Ian Johnston, Dominic Kwiatkowski, Cordelia Langford, John Sillitoe on behalf of the Wellcome Sanger Institute COVID-19 Surveillance Team |
| EPI_ISL_2867984 | Lighthouse Lab in Alderley Park | Wellcome Sanger Institute for the COVID-19 Genomics UK (COG-UK) Consortium | Jacquelyn Wynn, Mairead Hyland, The Lighthouse Lab in Alderley Park and Alex Alderton, Roberto Amato, Jeffrey Barrett, Sonia Goncalves, Ewan Harrison, David K. Jackson, Ian Johnston, Dominic Kwiatkowski, Cordelia Langford, John Sillitoe on behalf of the Wellcome Sanger Institute COVID-19 Surveillance Team |
| EPI_ISL_2867985, EPI_ISL_2867986 | Lighthouse Laboratory Plymouth | Wellcome Sanger Institute for the COVID-19 Genomics UK (COG-UK) Consortium | Lighthouse Laboratory Plymouth and Alex Alderton, Roberto Amato, Jeffrey Barrett, Sonia Goncalves, Ewan Harrison, David K. Jackson, Ian Johnston, Dominic Kwiatkowski, Cordelia Langford, John Sillitoe on behalf of the Wellcome Sanger Institute COVID-19 Surveillance Team |
| EPI_ISL_2867987 | Lighthouse Lab in Alderley Park | Wellcome Sanger Institute for the COVID-19 Genomics UK (COG-UK) Consortium | Jacquelyn Wynn, Mairead Hyland, The Lighthouse Lab in Alderley Park and Alex Alderton, Roberto Amato, Jeffrey Barrett, Sonia Goncalves, Ewan Harrison, David K. Jackson, Ian Johnston, Dominic Kwiatkowski, Cordelia Langford, John Sillitoe on behalf of the Wellcome Sanger Institute COVID-19 Surveillance Team |
| EPI_ISL_2867988 | Lighthouse Laboratory Plymouth | Wellcome Sanger Institute for the COVID-19 Genomics UK (COG-UK) Consortium | Lighthouse Laboratory Plymouth and Alex Alderton, Roberto Amato, Jeffrey Barrett, Sonia Goncalves, Ewan Harrison, David K. Jackson, Ian Johnston, Dominic Kwiatkowski, Cordelia Langford, John Sillitoe on behalf of the Wellcome Sanger Institute COVID-19 Surveillance Team |
| EPI_ISL_2867989, EPI_ISL_2867990, EPI_ISL_2867991, EPI_ISL_2867992, EPI_ISL_2867993, EPI_ISL_2867994, EPI_ISL_2867995 | Lighthouse Lab in Alderley Park | Wellcome Sanger Institute for the COVID-19 Genomics UK (COG-UK) Consortium | Jacquelyn Wynn, Mairead Hyland, The Lighthouse Lab in Alderley Park and Alex Alderton, Roberto Amato, Jeffrey Barrett, Sonia Goncalves, Ewan Harrison, David K. Jackson, Ian Johnston, Dominic Kwiatkowski, Cordelia Langford, John Sillitoe on behalf of the Wellcome Sanger Institute COVID-19 Surveillance Team |
| EPI_ISL_2867996 | Lighthouse Laboratory Plymouth | Wellcome Sanger Institute for the COVID-19 Genomics UK (COG-UK) Consortium | Lighthouse Laboratory Plymouth and Alex Alderton, Roberto Amato, Jeffrey Barrett, Sonia Goncalves, Ewan Harrison, David K. Jackson, Ian Johnston, Dominic Kwiatkowski, Cordelia Langford, John Sillitoe on behalf of the Wellcome Sanger Institute COVID-19 Surveillance Team |
| EPI_ISL_2867997 | Lighthouse Lab in Alderley Park | Wellcome Sanger Institute for the COVID-19 Genomics UK (COG-UK) Consortium | Jacquelyn Wynn, Mairead Hyland, The Lighthouse Lab in Alderley Park and Alex Alderton, Roberto Amato, Jeffrey Barrett, Sonia Goncalves, Ewan Harrison, David K. Jackson, Ian Johnston, Dominic Kwiatkowski, Cordelia Langford, John Sillitoe on behalf of the Wellcome Sanger Institute COVID-19 Surveillance Team |
| EPI_ISL_2867998, EPI_ISL_2867999 | Lighthouse Laboratory Plymouth | Wellcome Sanger Institute for the COVID-19 Genomics UK (COG-UK) Consortium | Lighthouse Laboratory Plymouth and Alex Alderton, Roberto Amato, Jeffrey Barrett, Sonia Goncalves, Ewan Harrison, David K. Jackson, Ian Johnston, Dominic Kwiatkowski, Cordelia Langford, John Sillitoe on behalf of the Wellcome Sanger Institute COVID-19 Surveillance Team |
| EPI_ISL_2868000, EPI_ISL_2868001 | Lighthouse Lab in Alderley Park | Wellcome Sanger Institute for the COVID-19 Genomics UK (COG-UK) Consortium | Jacquelyn Wynn, Mairead Hyland, The Lighthouse Lab in Alderley Park and Alex Alderton, Roberto Amato, Jeffrey Barrett, Sonia Goncalves, Ewan Harrison, David K. Jackson, Ian Johnston, Dominic Kwiatkowski, Cordelia Langford, John Sillitoe on behalf of the Wellcome Sanger Institute COVID-19 Surveillance Team |
| EPI_ISL_2868002, EPI_ISL_2868003, EPI_ISL_2868004, EPI_ISL_2868005, EPI_ISL_2868006 | Lighthouse Laboratory Plymouth | Wellcome Sanger Institute for the COVID-19 Genomics UK (COG-UK) Consortium | Lighthouse Laboratory Plymouth and Alex Alderton, Roberto Amato, Jeffrey Barrett, Sonia Goncalves, Ewan Harrison, David K. Jackson, Ian Johnston, Dominic Kwiatkowski, Cordelia Langford, John Sillitoe on behalf of the Wellcome Sanger Institute COVID-19 Surveillance Team |
| EPI_ISL_2868007, EPI_ISL_2868008 | Lighthouse Lab in Alderley Park | Wellcome Sanger Institute for the COVID-19 Genomics UK (COG-UK) Consortium | Jacquelyn Wynn, Mairead Hyland, The Lighthouse Lab in Alderley Park and Alex Alderton, Roberto Amato, Jeffrey Barrett, Sonia Goncalves, Ewan Harrison, David K. Jackson, Ian Johnston, Dominic Kwiatkowski, Cordelia Langford, John Sillitoe on behalf of the Wellcome Sanger Institute COVID-19 Surveillance Team |
| EPI_ISL_2868009, EPI_ISL_2868010, EPI_ISL_2868011 | Lighthouse Laboratory Plymouth | Wellcome Sanger Institute for the COVID-19 Genomics UK (COG-UK) Consortium | Lighthouse Laboratory Plymouth and Alex Alderton, Roberto Amato, Jeffrey Barrett, Sonia Goncalves, Ewan Harrison, David K. Jackson, Ian Johnston, Dominic Kwiatkowski, Cordelia Langford, John Sillitoe on behalf of the Wellcome Sanger Institute COVID-19 Surveillance Team |
| EPI_ISL_2868012 | Lighthouse Lab in Alderley Park | Wellcome Sanger Institute for the COVID-19 Genomics UK (COG-UK) Consortium | Jacquelyn Wynn, Mairead Hyland, The Lighthouse Lab in Alderley Park and Alex Alderton, Roberto Amato, Jeffrey Barrett, Sonia Goncalves, Ewan Harrison, David K. Jackson, Ian Johnston, Dominic Kwiatkowski, Cordelia Langford, John Sillitoe on behalf of the Wellcome Sanger Institute COVID-19 Surveillance Team |
| EPI_ISL_2868013 | Lighthouse Laboratory Plymouth | Wellcome Sanger Institute for the COVID-19 Genomics UK (COG-UK) Consortium | Lighthouse Laboratory Plymouth and Alex Alderton, Roberto Amato, Jeffrey Barrett, Sonia Goncalves, Ewan Harrison, David K. Jackson, Ian Johnston, Dominic Kwiatkowski, Cordelia Langford, John Sillitoe on behalf of the Wellcome Sanger Institute COVID-19 Surveillance Team |
| EPI_ISL_2868014 | Lighthouse Lab in Alderley Park | Wellcome Sanger Institute for the COVID-19 Genomics UK (COG-UK) Consortium | Jacquelyn Wynn, Mairead Hyland, The Lighthouse Lab in Alderley Park and Alex Alderton, Roberto Amato, Jeffrey Barrett, Sonia Goncalves, Ewan Harrison, David K. Jackson, Ian Johnston, Dominic Kwiatkowski, Cordelia Langford, John Sillitoe on behalf of the Wellcome Sanger Institute COVID-19 Surveillance Team |
| EPI_ISL_2868015 | Lighthouse Laboratory Plymouth | Wellcome Sanger Institute for the COVID-19 Genomics UK (COG-UK) Consortium | Lighthouse Laboratory Plymouth and Alex Alderton, Roberto Amato, Jeffrey Barrett, Sonia Goncalves, Ewan Harrison, David K. Jackson, Ian Johnston, Dominic Kwiatkowski, Cordelia Langford, John Sillitoe on behalf of the Wellcome Sanger Institute COVID-19 Surveillance Team |
| EPI_ISL_2868016, EPI_ISL_2868017 | Lighthouse Lab in Alderley Park | Wellcome Sanger Institute for the COVID-19 Genomics UK (COG-UK) Consortium | Jacquelyn Wynn, Mairead Hyland, The Lighthouse Lab in Alderley Park and Alex Alderton, Roberto Amato, Jeffrey Barrett, Sonia Goncalves, Ewan Harrison, David K. Jackson, Ian Johnston, Dominic Kwiatkowski, Cordelia Langford, John Sillitoe on behalf of the Wellcome Sanger Institute COVID-19 Surveillance Team |
| EPI_ISL_2868018 | Lighthouse Laboratory Plymouth | Wellcome Sanger Institute for the COVID-19 Genomics UK (COG-UK) Consortium | Lighthouse Laboratory Plymouth and Alex Alderton, Roberto Amato, Jeffrey Barrett, Sonia Goncalves, Ewan Harrison, David K. Jackson, Ian Johnston, Dominic Kwiatkowski, Cordelia Langford, John Sillitoe on behalf of the Wellcome Sanger Institute COVID-19 Surveillance Team |
| EPI_ISL_2868019 | Lighthouse Lab in Alderley Park | Wellcome Sanger Institute for the COVID-19 Genomics UK (COG-UK) | Jacquelyn Wynn, Mairead Hyland, The Lighthouse Lab in Alderley Park and Alex Alderton, Roberto Amato, Jeffrey Barrett, Sonia Goncalves, Ewan Harrison, David K. Jackson, Ian Johnston, Dominic Kwiatkowski, Cordelia Langford, John Sillitoe on |

| | | | |
|---|---|---|---|
| | | Consortium | behalf of the Wellcome Sanger Institute COVID-19 Surveillance Team |
| EPI_ISL_2868020, EPI_ISL_2868021 | Lighthouse Laboratory Plymouth | Wellcome Sanger Institute for the COVID-19 Genomics UK (COG-UK) Consortium | Lighthouse Laboratory Plymouth and Alex Alderton, Roberto Amato, Jeffrey Barrett, Sonia Goncalves, Ewan Harrison, David K. Jackson, Ian Johnston, Dominic Kwiatkowski, Cordelia Langford, John Sillitoe on behalf of the Wellcome Sanger Institute COVID-19 Surveillance Team |
| EPI_ISL_2868022, EPI_ISL_2868023, EPI_ISL_2868024, EPI_ISL_2868025 | Lighthouse Lab in Alderley Park | Wellcome Sanger Institute for the COVID-19 Genomics UK (COG-UK) Consortium | Jacquelyn Wynn, Mairead Hyland, The Lighthouse Lab in Alderley Park and Alex Alderton, Roberto Amato, Jeffrey Barrett, Sonia Goncalves, Ewan Harrison, David K. Jackson, Ian Johnston, Dominic Kwiatkowski, Cordelia Langford, John Sillitoe on behalf of the Wellcome Sanger Institute COVID-19 Surveillance Team |
| EPI_ISL_2868026, EPI_ISL_2868027 | Lighthouse Laboratory Plymouth | Wellcome Sanger Institute for the COVID-19 Genomics UK (COG-UK) Consortium | Lighthouse Laboratory Plymouth and Alex Alderton, Roberto Amato, Jeffrey Barrett, Sonia Goncalves, Ewan Harrison, David K. Jackson, Ian Johnston, Dominic Kwiatkowski, Cordelia Langford, John Sillitoe on behalf of the Wellcome Sanger Institute COVID-19 Surveillance Team |
| EPI_ISL_2868028, EPI_ISL_2868029 | Lighthouse Lab in Alderley Park | Wellcome Sanger Institute for the COVID-19 Genomics UK (COG-UK) Consortium | Jacquelyn Wynn, Mairead Hyland, The Lighthouse Lab in Alderley Park and Alex Alderton, Roberto Amato, Jeffrey Barrett, Sonia Goncalves, Ewan Harrison, David K. Jackson, Ian Johnston, Dominic Kwiatkowski, Cordelia Langford, John Sillitoe on behalf of the Wellcome Sanger Institute COVID-19 Surveillance Team |
| EPI_ISL_2868030, EPI_ISL_2868031, EPI_ISL_2868032 | Lighthouse Laboratory Plymouth | Wellcome Sanger Institute for the COVID-19 Genomics UK (COG-UK) Consortium | Lighthouse Laboratory Plymouth and Alex Alderton, Roberto Amato, Jeffrey Barrett, Sonia Goncalves, Ewan Harrison, David K. Jackson, Ian Johnston, Dominic Kwiatkowski, Cordelia Langford, John Sillitoe on behalf of the Wellcome Sanger Institute COVID-19 Surveillance Team |
| EPI_ISL_2868033 | Lighthouse Lab in Alderley Park | Wellcome Sanger Institute for the COVID-19 Genomics UK (COG-UK) Consortium | Jacquelyn Wynn, Mairead Hyland, The Lighthouse Lab in Alderley Park and Alex Alderton, Roberto Amato, Jeffrey Barrett, Sonia Goncalves, Ewan Harrison, David K. Jackson, Ian Johnston, Dominic Kwiatkowski, Cordelia Langford, John Sillitoe on behalf of the Wellcome Sanger Institute COVID-19 Surveillance Team |
| EPI_ISL_2868034, EPI_ISL_2868035 | Lighthouse Laboratory Plymouth | Wellcome Sanger Institute for the COVID-19 Genomics UK (COG-UK) Consortium | Lighthouse Laboratory Plymouth and Alex Alderton, Roberto Amato, Jeffrey Barrett, Sonia Goncalves, Ewan Harrison, David K. Jackson, Ian Johnston, Dominic Kwiatkowski, Cordelia Langford, John Sillitoe on behalf of the Wellcome Sanger Institute COVID-19 Surveillance Team |
| EPI_ISL_2868036, EPI_ISL_2868037, EPI_ISL_2868038 | Lighthouse Lab in Alderley Park | Wellcome Sanger Institute for the COVID-19 Genomics UK (COG-UK) Consortium | Jacquelyn Wynn, Mairead Hyland, The Lighthouse Lab in Alderley Park and Alex Alderton, Roberto Amato, Jeffrey Barrett, Sonia Goncalves, Ewan Harrison, David K. Jackson, Ian Johnston, Dominic Kwiatkowski, Cordelia Langford, John Sillitoe on behalf of the Wellcome Sanger Institute COVID-19 Surveillance Team |
| EPI_ISL_2868039, EPI_ISL_2868040, EPI_ISL_2868041, EPI_ISL_2868042 | Lighthouse Laboratory Plymouth | Wellcome Sanger Institute for the COVID-19 Genomics UK (COG-UK) Consortium | Lighthouse Laboratory Plymouth and Alex Alderton, Roberto Amato, Jeffrey Barrett, Sonia Goncalves, Ewan Harrison, David K. Jackson, Ian Johnston, Dominic Kwiatkowski, Cordelia Langford, John Sillitoe on behalf of the Wellcome Sanger Institute COVID-19 Surveillance Team |
| EPI_ISL_2868043 | Lighthouse Lab in Alderley Park | Wellcome Sanger Institute for the COVID-19 Genomics UK (COG-UK) Consortium | Jacquelyn Wynn, Mairead Hyland, The Lighthouse Lab in Alderley Park and Alex Alderton, Roberto Amato, Jeffrey Barrett, Sonia Goncalves, Ewan Harrison, David K. Jackson, Ian Johnston, Dominic Kwiatkowski, Cordelia Langford, John Sillitoe on behalf of the Wellcome Sanger Institute COVID-19 Surveillance Team |
| EPI_ISL_2868044 | Lighthouse Laboratory Plymouth | Wellcome Sanger Institute for the COVID-19 Genomics UK (COG-UK) Consortium | Lighthouse Laboratory Plymouth and Alex Alderton, Roberto Amato, Jeffrey Barrett, Sonia Goncalves, Ewan Harrison, David K. Jackson, Ian Johnston, Dominic Kwiatkowski, Cordelia Langford, John Sillitoe on behalf of the Wellcome Sanger Institute COVID-19 Surveillance Team |
| EPI_ISL_2868045 | Lighthouse Lab in Alderley Park | Wellcome Sanger Institute for the COVID-19 Genomics UK (COG-UK) Consortium | Jacquelyn Wynn, Mairead Hyland, The Lighthouse Lab in Alderley Park and Alex Alderton, Roberto Amato, Jeffrey Barrett, Sonia Goncalves, Ewan Harrison, David K. Jackson, Ian Johnston, Dominic Kwiatkowski, Cordelia Langford, John Sillitoe on behalf of the Wellcome Sanger Institute COVID-19 Surveillance Team |
| EPI_ISL_2868046 | Lighthouse Laboratory Plymouth | Wellcome Sanger Institute for the COVID-19 Genomics UK (COG-UK) Consortium | Lighthouse Laboratory Plymouth and Alex Alderton, Roberto Amato, Jeffrey Barrett, Sonia Goncalves, Ewan Harrison, David K. Jackson, Ian Johnston, Dominic Kwiatkowski, Cordelia Langford, John Sillitoe on behalf of the Wellcome Sanger Institute COVID-19 Surveillance Team |
| EPI_ISL_2868047, EPI_ISL_2868048, EPI_ISL_2868049, EPI_ISL_2868050 | Lighthouse Lab in Alderley Park | Wellcome Sanger Institute for the COVID-19 Genomics UK (COG-UK) Consortium | Jacquelyn Wynn, Mairead Hyland, The Lighthouse Lab in Alderley Park and Alex Alderton, Roberto Amato, Jeffrey Barrett, Sonia Goncalves, Ewan Harrison, David K. Jackson, Ian Johnston, Dominic Kwiatkowski, Cordelia Langford, John Sillitoe on behalf of the Wellcome Sanger Institute COVID-19 Surveillance Team |
| EPI_ISL_2868051 | Lighthouse Laboratory Plymouth | Wellcome Sanger Institute for the COVID-19 Genomics UK (COG-UK) Consortium | Lighthouse Laboratory Plymouth and Alex Alderton, Roberto Amato, Jeffrey Barrett, Sonia Goncalves, Ewan Harrison, David K. Jackson, Ian Johnston, Dominic Kwiatkowski, Cordelia Langford, John Sillitoe on behalf of the Wellcome Sanger Institute COVID-19 Surveillance Team |
| EPI_ISL_2868052 | Lighthouse Lab in Alderley Park | Wellcome Sanger Institute for the COVID-19 Genomics UK (COG-UK) Consortium | Jacquelyn Wynn, Mairead Hyland, The Lighthouse Lab in Alderley Park and Alex Alderton, Roberto Amato, Jeffrey Barrett, Sonia Goncalves, Ewan Harrison, David K. Jackson, Ian Johnston, Dominic Kwiatkowski, Cordelia Langford, John Sillitoe on behalf of the Wellcome Sanger Institute COVID-19 Surveillance Team |
| EPI_ISL_2868053 | Health Services Laboratories | Wellcome Sanger Institute for the COVID-19 Genomics UK (COG-UK) Consortium | Health Services Laboratories and Alex Alderton, Roberto Amato, Jeffrey Barrett, Sonia Goncalves, Ewan Harrison, David K. Jackson, Ian Johnston, Dominic Kwiatkowski, Cordelia Langford, John Sillitoe on behalf of the Wellcome Sanger Institute COVID-19 Surveillance Team |
| EPI_ISL_2868054, EPI_ISL_2868055 | Lighthouse Laboratory Plymouth | Wellcome Sanger Institute for the COVID-19 Genomics UK (COG-UK) Consortium | Lighthouse Laboratory Plymouth and Alex Alderton, Roberto Amato, Jeffrey Barrett, Sonia Goncalves, Ewan Harrison, David K. Jackson, Ian Johnston, Dominic Kwiatkowski, Cordelia Langford, John Sillitoe on behalf of the Wellcome Sanger Institute COVID-19 Surveillance Team |
| EPI_ISL_2868056, EPI_ISL_2868057 | Lighthouse Lab in Alderley Park | Wellcome Sanger Institute for the COVID-19 Genomics UK (COG-UK) Consortium | Jacquelyn Wynn, Mairead Hyland, The Lighthouse Lab in Alderley Park and Alex Alderton, Roberto Amato, Jeffrey Barrett, Sonia Goncalves, Ewan Harrison, David K. Jackson, Ian Johnston, Dominic Kwiatkowski, Cordelia Langford, John Sillitoe on behalf of the Wellcome Sanger Institute COVID-19 Surveillance Team |
| EPI_ISL_2868058 | Health Services Laboratories | Wellcome Sanger Institute for the COVID-19 Genomics UK (COG-UK) Consortium | Health Services Laboratories and Alex Alderton, Roberto Amato, Jeffrey Barrett, Sonia Goncalves, Ewan Harrison, David K. Jackson, Ian Johnston, Dominic Kwiatkowski, Cordelia Langford, John Sillitoe on behalf of the Wellcome Sanger Institute COVID-19 Surveillance Team |
| EPI_ISL_2868059 | Lighthouse Lab in Alderley Park | Wellcome Sanger Institute for the COVID-19 Genomics UK (COG-UK) Consortium | Jacquelyn Wynn, Mairead Hyland, The Lighthouse Lab in Alderley Park and Alex Alderton, Roberto Amato, Jeffrey Barrett, Sonia Goncalves, Ewan Harrison, David K. Jackson, Ian Johnston, Dominic Kwiatkowski, Cordelia Langford, John Sillitoe on behalf of the Wellcome Sanger Institute COVID-19 Surveillance Team |
| EPI_ISL_2868060 | Lighthouse Laboratory Plymouth | Wellcome Sanger Institute for the COVID-19 Genomics UK (COG-UK) Consortium | Lighthouse Laboratory Plymouth and Alex Alderton, Roberto Amato, Jeffrey Barrett, Sonia Goncalves, Ewan Harrison, David K. Jackson, Ian Johnston, Dominic Kwiatkowski, Cordelia Langford, John Sillitoe on behalf of the Wellcome Sanger Institute COVID-19 Surveillance Team |
| EPI_ISL_2868061 | Lighthouse Lab in Alderley Park | Wellcome Sanger Institute for the COVID-19 Genomics UK (COG-UK) Consortium | Jacquelyn Wynn, Mairead Hyland, The Lighthouse Lab in Alderley Park and Alex Alderton, Roberto Amato, Jeffrey Barrett, Sonia Goncalves, Ewan Harrison, David K. Jackson, Ian Johnston, Dominic Kwiatkowski, Cordelia Langford, John Sillitoe on behalf of the Wellcome Sanger Institute COVID-19 Surveillance Team |
| EPI_ISL_2868062 | Lighthouse Lab in Glasgow | Wellcome Sanger Institute for the COVID-19 Genomics UK (COG-UK) Consortium | Harper VanSteenhouse, Yumi Kasai, David Gray, Carol Clugston, Anna Dominiczak and Alex Alderton, Roberto Amato, Jeffrey Barrett, Sonia Goncalves, Ewan Harrison, David K. Jackson, Ian Johnston, Dominic Kwiatkowski, Cordelia Langford, John Sillitoe on behalf of the Wellcome Sanger Institute COVID-19 Surveillance Team |
| EPI_ISL_2868063, EPI_ISL_2868064 | Lighthouse Laboratory Plymouth | Wellcome Sanger Institute for the COVID-19 Genomics UK (COG-UK) Consortium | Lighthouse Laboratory Plymouth and Alex Alderton, Roberto Amato, Jeffrey Barrett, Sonia Goncalves, Ewan Harrison, David K. Jackson, Ian Johnston, Dominic Kwiatkowski, Cordelia Langford, John Sillitoe on behalf of the Wellcome Sanger Institute COVID-19 Surveillance Team |
| EPI_ISL_2868065, EPI_ISL_2868066, EPI_ISL_2868067, EPI_ISL_2868068, EPI_ISL_2868069 | Lighthouse Lab in Milton Keynes | Wellcome Sanger Institute for the COVID-19 Genomics UK (COG-UK) Consortium | The Lighthouse Lab in Milton Keynes and Alex Alderton, Roberto Amato, Jeffrey Barrett, Sonia Goncalves, Ewan Harrison, David K. Jackson, Ian Johnston, Dominic Kwiatkowski, Cordelia Langford, John Sillitoe on behalf of the Wellcome Sanger Institute COVID-19 Surveillance Team |
| EPI_ISL_2868070, EPI_ISL_2868071, EPI_ISL_2868072, EPI_ISL_2868073 | Lighthouse Laboratory Plymouth | Wellcome Sanger Institute for the COVID-19 Genomics UK (COG-UK) Consortium | Lighthouse Laboratory Plymouth and Alex Alderton, Roberto Amato, Jeffrey Barrett, Sonia Goncalves, Ewan Harrison, David K. Jackson, Ian Johnston, Dominic Kwiatkowski, Cordelia Langford, John Sillitoe on behalf of the Wellcome Sanger Institute COVID-19 Surveillance Team |
| EPI_ISL_2868074, EPI_ISL_2868075 | Lighthouse Lab in Alderley Park | Wellcome Sanger Institute for the COVID-19 Genomics UK (COG-UK) Consortium | Jacquelyn Wynn, Mairead Hyland, The Lighthouse Lab in Alderley Park and Alex Alderton, Roberto Amato, Jeffrey Barrett, Sonia Goncalves, Ewan Harrison, David K. Jackson, Ian Johnston, Dominic Kwiatkowski, Cordelia Langford, John Sillitoe on behalf of the Wellcome Sanger Institute COVID-19 Surveillance Team |
| EPI_ISL_2868076, EPI_ISL_2868077 | Lighthouse Laboratory Plymouth | Wellcome Sanger Institute for the COVID-19 Genomics UK (COG-UK) Consortium | Lighthouse Laboratory Plymouth and Alex Alderton, Roberto Amato, Jeffrey Barrett, Sonia Goncalves, Ewan Harrison, David K. Jackson, Ian Johnston, Dominic Kwiatkowski, Cordelia Langford, John Sillitoe on behalf of the Wellcome Sanger Institute COVID-19 Surveillance Team |
| EPI_ISL_2868078 | Lighthouse Lab in Milton Keynes | Wellcome Sanger Institute for the COVID-19 Genomics UK (COG-UK) Consortium | The Lighthouse Lab in Milton Keynes and Alex Alderton, Roberto Amato, Jeffrey Barrett, Sonia Goncalves, Ewan Harrison, David K. Jackson, Ian Johnston, Dominic Kwiatkowski, Cordelia Langford, John Sillitoe on behalf of the Wellcome Sanger Institute COVID-19 Surveillance Team |
| EPI_ISL_2868079 | Lighthouse Laboratory Plymouth | Wellcome Sanger Institute for the COVID-19 Genomics UK (COG-UK) Consortium | Lighthouse Laboratory Plymouth and Alex Alderton, Roberto Amato, Jeffrey Barrett, Sonia Goncalves, Ewan Harrison, David K. Jackson, Ian Johnston, Dominic Kwiatkowski, Cordelia Langford, John Sillitoe on behalf of the Wellcome Sanger Institute COVID-19 Surveillance Team |
| EPI_ISL_2868080 | Lighthouse Lab in Alderley Park | Wellcome Sanger Institute for the COVID-19 Genomics UK (COG-UK) Consortium | Jacquelyn Wynn, Mairead Hyland, The Lighthouse Lab in Alderley Park and Alex Alderton, Roberto Amato, Jeffrey Barrett, Sonia Goncalves, Ewan Harrison, David K. Jackson, Ian Johnston, Dominic Kwiatkowski, Cordelia Langford, John Sillitoe on behalf of the Wellcome Sanger Institute COVID-19 Surveillance Team |
| EPI_ISL_2868081, EPI_ISL_2868082 | Lighthouse Laboratory Plymouth | Wellcome Sanger Institute for the COVID-19 Genomics UK (COG-UK) Consortium | Lighthouse Laboratory Plymouth and Alex Alderton, Roberto Amato, Jeffrey Barrett, Sonia Goncalves, Ewan Harrison, David K. Jackson, Ian Johnston, Dominic Kwiatkowski, Cordelia Langford, John Sillitoe on behalf of the Wellcome Sanger Institute COVID-19 Surveillance Team |
| EPI_ISL_2868083 | Lighthouse Lab in Alderley Park | Wellcome Sanger Institute for the COVID-19 Genomics UK (COG-UK) Consortium | Jacquelyn Wynn, Mairead Hyland, The Lighthouse Lab in Alderley Park and Alex Alderton, Roberto Amato, Jeffrey Barrett, Sonia Goncalves, Ewan Harrison, David K. Jackson, Ian Johnston, Dominic Kwiatkowski, Cordelia Langford, John Sillitoe on behalf of the Wellcome Sanger Institute COVID-19 Surveillance Team |
| EPI_ISL_2868084 | Lighthouse Lab in Milton Keynes | Wellcome Sanger Institute for the COVID-19 Genomics UK (COG-UK) Consortium | The Lighthouse Lab in Milton Keynes and Alex Alderton, Roberto Amato, Jeffrey Barrett, Sonia Goncalves, Ewan Harrison, David K. Jackson, Ian Johnston, Dominic Kwiatkowski, Cordelia Langford, John Sillitoe on behalf of the Wellcome Sanger Institute COVID-19 Surveillance Team |
| EPI_ISL_2868085, EPI_ISL_2868086, EPI_ISL_2868087, EPI_ISL_2868088, EPI_ISL_2868089, EPI_ISL_2868090, EPI_ISL_2868091, EPI_ISL_2868092, EPI_ISL_2868093, EPI_ISL_2868094, EPI_ISL_2868095, EPI_ISL_2868096, EPI_ISL_2868097, EPI_ISL_2868098, EPI_ISL_2868099, EPI_ISL_2868100, EPI_ISL_2868101, EPI_ISL_2868102, EPI_ISL_2868103, EPI_ISL_2868104, EPI_ISL_2868105, EPI_ISL_2868106, EPI_ISL_2868107, EPI_ISL_2868108, EPI_ISL_2868109, EPI_ISL_2868110, EPI_ISL_2868111, EPI_ISL_2868112, EPI_ISL_2868113, EPI_ISL_2868114, EPI_ISL_2868115, EPI_ISL_2868116, EPI_ISL_2868117, EPI_ISL_2868118, EPI_ISL_2868119, EPI_ISL_2868120, EPI_ISL_2868121, EPI_ISL_2868122, EPI_ISL_2868123, EPI_ISL_2868124, EPI_ISL_2868125, EPI_ISL_2868126, EPI_ISL_2868127, EPI_ISL_2868128, EPI_ISL_2868129, EPI_ISL_2868130, EPI_ISL_2868131, EPI_ISL_2868132, EPI_ISL_2868133, EPI_ISL_2868134, EPI_ISL_2868135, EPI_ISL_2868136, EPI_ISL_2868137, EPI_ISL_2868138, EPI_ISL_2868139, EPI_ISL_2868140, EPI_ISL_2868141, EPI_ISL_2868142, EPI_ISL_2868143, EPI_ISL_2868144, EPI_ISL_2868145, EPI_ISL_2868146, EPI_ISL_2868147, EPI_ISL_2868148, EPI_ISL_2868149, EPI_ISL_2868150, EPI_ISL_2868151, EPI_ISL_2868152, EPI_ISL_2868153, EPI_ISL_2868154, EPI_ISL_2868155, EPI_ISL_2868156, EPI_ISL_2868157, EPI_ISL_2868158, EPI_ISL_2868159, EPI_ISL_2868160, EPI_ISL_2868161, EPI_ISL_2868162, EPI_ISL_2868163, EPI_ISL_2868164, EPI_ISL_2868165, EPI_ISL_2868166, EPI_ISL_2868167 | | | |
| see above | Lighthouse Lab in Glasgow | Wellcome Sanger Institute for the COVID-19 Genomics UK (COG-UK) Consortium | Harper VanSteenhouse, Yumi Kasai, David Gray, Carol Clugston, Anna Dominiczak and Alex Alderton, Roberto Amato, Jeffrey Barrett, Sonia Goncalves, Ewan Harrison, David K. Jackson, Ian Johnston, Dominic Kwiatkowski, Cordelia Langford, John Sillitoe on behalf of the Wellcome Sanger Institute COVID-19 Surveillance Team |
| EPI_ISL_2868168 | Lighthouse Laboratory Plymouth | Wellcome Sanger Institute for the COVID-19 Genomics UK (COG-UK) Consortium | Lighthouse Laboratory Plymouth and Alex Alderton, Roberto Amato, Jeffrey Barrett, Sonia Goncalves, Ewan Harrison, David K. Jackson, Ian Johnston, Dominic Kwiatkowski, Cordelia Langford, John Sillitoe on behalf of the Wellcome Sanger Institute COVID-19 Surveillance Team |

| | | | |
|---|---|---|---|
| EPI_ISL_2868169, EPI_ISL_2868170, EPI_ISL_2868171, EPI_ISL_2868172, EPI_ISL_2868173, EPI_ISL_2868174, EPI_ISL_2868175, EPI_ISL_2868176, EPI_ISL_2868177, EPI_ISL_2868178, EPI_ISL_2868179, EPI_ISL_2868180, EPI_ISL_2868181, EPI_ISL_2868182, EPI_ISL_2868183, EPI_ISL_2868184, EPI_ISL_2868185, EPI_ISL_2868186, EPI_ISL_2868187, EPI_ISL_2868188, EPI_ISL_2868189, EPI_ISL_2868190, EPI_ISL_2868191, EPI_ISL_2868192, EPI_ISL_2868193, EPI_ISL_2868194, EPI_ISL_2868195, EPI_ISL_2868196, EPI_ISL_2868197, EPI_ISL_2868198, EPI_ISL_2868199, EPI_ISL_2868200, EPI_ISL_2868201, EPI_ISL_2868202, EPI_ISL_2868203, EPI_ISL_2868204, EPI_ISL_2868205, EPI_ISL_2868206, EPI_ISL_2868207, EPI_ISL_2868208, EPI_ISL_2868209 | | | |
| see above | Lighthouse Lab in Glasgow | Wellcome Sanger Institute for the COVID-19 Genomics UK (COG-UK) Consortium | Harper VanSteenhouse, Yumi Kasai, David Gray, Carol Clugston, Anna Dominiczak and Alex Alderton, Roberto Amato, Jeffrey Barrett, Sonia Goncalves, Ewan Harrison, David K. Jackson, Ian Johnston, Dominic Kwiatkowski, Cordelia Langford, John Sillitoe on behalf of the Wellcome Sanger Institute COVID-19 Surveillance Team |
| EPI_ISL_2868210 | Berkshire and Surrey Pathology Services Lighthouse Laboratory | Wellcome Sanger Institute for the COVID-19 Genomics UK (COG-UK) Consortium | Berkshire and Surrey Pathology Services Lighthouse Laboratory and Alex Alderton, Roberto Amato, Jeffrey Barrett, Sonia Goncalves, Ewan Harrison, David K. Jackson, Ian Johnston, Dominic Kwiatkowski, Cordelia Langford, John Sillitoe on behalf of the Wellcome Sanger Institute COVID-19 Surveillance Team |
| EPI_ISL_2868211, EPI_ISL_2868212, EPI_ISL_2868213 | Lighthouse Lab in Glasgow | Wellcome Sanger Institute for the COVID-19 Genomics UK (COG-UK) Consortium | Harper VanSteenhouse, Yumi Kasai, David Gray, Carol Clugston, Anna Dominiczak and Alex Alderton, Roberto Amato, Jeffrey Barrett, Sonia Goncalves, Ewan Harrison, David K. Jackson, Ian Johnston, Dominic Kwiatkowski, Cordelia Langford, John Sillitoe on behalf of the Wellcome Sanger Institute COVID-19 Surveillance Team |
| EPI_ISL_2868214, EPI_ISL_2868215, EPI_ISL_2868216, EPI_ISL_2868217, EPI_ISL_2868218 | Lighthouse Lab in Alderley Park | Wellcome Sanger Institute for the COVID-19 Genomics UK (COG-UK) Consortium | Jacquelyn Wynn, Mairead Hyland, The Lighthouse Lab in Alderley Park and Alex Alderton, Roberto Amato, Jeffrey Barrett, Sonia Goncalves, Ewan Harrison, David K. Jackson, Ian Johnston, Dominic Kwiatkowski, Cordelia Langford, John Sillitoe on behalf of the Wellcome Sanger Institute COVID-19 Surveillance Team |
| EPI_ISL_2868219 | Lighthouse Lab in Glasgow | Wellcome Sanger Institute for the COVID-19 Genomics UK (COG-UK) Consortium | Harper VanSteenhouse, Yumi Kasai, David Gray, Carol Clugston, Anna Dominiczak and Alex Alderton, Roberto Amato, Jeffrey Barrett, Sonia Goncalves, Ewan Harrison, David K. Jackson, Ian Johnston, Dominic Kwiatkowski, Cordelia Langford, John Sillitoe on behalf of the Wellcome Sanger Institute COVID-19 Surveillance Team |
| EPI_ISL_2868220, EPI_ISL_2868221 | Health Services Laboratories | Wellcome Sanger Institute for the COVID-19 Genomics UK (COG-UK) Consortium | Health Services Laboratories and Alex Alderton, Roberto Amato, Jeffrey Barrett, Sonia Goncalves, Ewan Harrison, David K. Jackson, Ian Johnston, Dominic Kwiatkowski, Cordelia Langford, John Sillitoe on behalf of the Wellcome Sanger Institute COVID-19 Surveillance Team |
| EPI_ISL_2868222 | Lighthouse Lab in Alderley Park | Wellcome Sanger Institute for the COVID-19 Genomics UK (COG-UK) Consortium | Jacquelyn Wynn, Mairead Hyland, The Lighthouse Lab in Alderley Park and Alex Alderton, Roberto Amato, Jeffrey Barrett, Sonia Goncalves, Ewan Harrison, David K. Jackson, Ian Johnston, Dominic Kwiatkowski, Cordelia Langford, John Sillitoe on behalf of the Wellcome Sanger Institute COVID-19 Surveillance Team |
| EPI_ISL_2868223 | Lighthouse Lab in Milton Keynes | Wellcome Sanger Institute for the COVID-19 Genomics UK (COG-UK) Consortium | The Lighthouse Lab in Milton Keynes and Alex Alderton, Roberto Amato, Jeffrey Barrett, Sonia Goncalves, Ewan Harrison, David K. Jackson, Ian Johnston, Dominic Kwiatkowski, Cordelia Langford, John Sillitoe on behalf of the Wellcome Sanger Institute COVID-19 Surveillance Team |
| EPI_ISL_2868224, EPI_ISL_2868225, EPI_ISL_2868226 | Lighthouse Lab in Alderley Park | Wellcome Sanger Institute for the COVID-19 Genomics UK (COG-UK) Consortium | Jacquelyn Wynn, Mairead Hyland, The Lighthouse Lab in Alderley Park and Alex Alderton, Roberto Amato, Jeffrey Barrett, Sonia Goncalves, Ewan Harrison, David K. Jackson, Ian Johnston, Dominic Kwiatkowski, Cordelia Langford, John Sillitoe on behalf of the Wellcome Sanger Institute COVID-19 Surveillance Team |
| EPI_ISL_2868227, EPI_ISL_2868228 | Lighthouse Laboratory Plymouth | Wellcome Sanger Institute for the COVID-19 Genomics UK (COG-UK) Consortium | Lighthouse Laboratory Plymouth and Alex Alderton, Roberto Amato, Jeffrey Barrett, Sonia Goncalves, Ewan Harrison, David K. Jackson, Ian Johnston, Dominic Kwiatkowski, Cordelia Langford, John Sillitoe on behalf of the Wellcome Sanger Institute COVID-19 Surveillance Team |
| EPI_ISL_2868229, EPI_ISL_2868230, EPI_ISL_2868231, EPI_ISL_2868233 | Lighthouse Lab in Alderley Park | Wellcome Sanger Institute for the COVID-19 Genomics UK (COG-UK) Consortium | Jacquelyn Wynn, Mairead Hyland, The Lighthouse Lab in Alderley Park and Alex Alderton, Roberto Amato, Jeffrey Barrett, Sonia Goncalves, Ewan Harrison, David K. Jackson, Ian Johnston, Dominic Kwiatkowski, Cordelia Langford, John Sillitoe on behalf of the Wellcome Sanger Institute COVID-19 Surveillance Team |
| EPI_ISL_2868234 | Lighthouse Laboratory Plymouth | Wellcome Sanger Institute for the COVID-19 Genomics UK (COG-UK) Consortium | Lighthouse Laboratory Plymouth and Alex Alderton, Roberto Amato, Jeffrey Barrett, Sonia Goncalves, Ewan Harrison, David K. Jackson, Ian Johnston, Dominic Kwiatkowski, Cordelia Langford, John Sillitoe on behalf of the Wellcome Sanger Institute COVID-19 Surveillance Team |
| EPI_ISL_2868235 | Lighthouse Lab in Alderley Park | Wellcome Sanger Institute for the COVID-19 Genomics UK (COG-UK) Consortium | Jacquelyn Wynn, Mairead Hyland, The Lighthouse Lab in Alderley Park and Alex Alderton, Roberto Amato, Jeffrey Barrett, Sonia Goncalves, Ewan Harrison, David K. Jackson, Ian Johnston, Dominic Kwiatkowski, Cordelia Langford, John Sillitoe on behalf of the Wellcome Sanger Institute COVID-19 Surveillance Team |
| EPI_ISL_2868267, EPI_ISL_2868286, EPI_ISL_2868287, EPI_ISL_2868300, EPI_ISL_2868305, EPI_ISL_2868311, EPI_ISL_2868323 | URMC LABS | Wadsworth Center, New York State Department of Health | Kirsten St. George, Daryl M. Lamson, Alexis Russell, Matthew Shudt, Melissa A Leisner, Jonathan Plitnick, Catharine Prussing, Navjot Singh, John Kelly, Erasmus Schneider, Erica Lasek-Nesselquist |
| EPI_ISL_2868327, EPI_ISL_2868328, EPI_ISL_2868329, EPI_ISL_2868330, EPI_ISL_2868331 | Botswana Harvard HIV Reference Laboratory | Botswana Harvard HIV Reference Laboratory | Sikhulile Moyo, Wonderful T. Choga, Dorcas Maruapula, Thongbotho Mphoyakgosi, Boitumelo Zuze, Botshelo Radibe, Legodile Kooepile, Ontlametse T. Bareng, Keoratile Ntshambiwa, Modisa Motswaledi, Madisa Mine, Joseph Makhema, Roger Shapiro, Shahin Lockman, Mosepele Mosepele, Simani Gaseitsiwe |
| EPI_ISL_2868335 | Botswana Harvard HIV Reference Laboratory | Botswana Harvard HIV Reference Laboratory | Sikhulile Moyo, Wonderful T. Choga, Dorcas Maruapula, Thongbotho Mphoyakgosi, Boitumelo Zuze, Botshelo Radibe, Legodile Kooepile, Ontlametse T. Bareng, Motlalepule L. Pone, Godfrey Simoonga, Kereng Mphoyakgosi, Keoratile Ntshambiwa, Modisa Motswaledi, Madisa Mine, Joseph Makhema, Roger Shapiro, Shahin Lockman, Mosepele Mosepele, Simani Gaseitsiwe |
| EPI_ISL_2868336, EPI_ISL_2868338, EPI_ISL_2868339, EPI_ISL_2868341, EPI_ISL_2868342, EPI_ISL_2868343, EPI_ISL_2868344, EPI_ISL_2868345, EPI_ISL_2868346, EPI_ISL_2868347, EPI_ISL_2868348, EPI_ISL_2868349, EPI_ISL_2868350, EPI_ISL_2868352, EPI_ISL_2868353, EPI_ISL_2868355, EPI_ISL_2868356, EPI_ISL_2868357, EPI_ISL_2868358, EPI_ISL_2868359, EPI_ISL_2868360 | | | |
| see above | Botswana Harvard HIV Reference Laboratory | Botswana Harvard HIV Reference Laboratory | Sikhulile Moyo, Wonderful T. Choga, Dorcas Maruapula, Thongbotho Mphoyakgosi, Boitumelo Zuze, Botshelo Radibe, Legodile Kooepile, Ontlametse T. Bareng, Keoratile Ntshambiwa, Modisa Motswaledi, Madisa Mine, Joseph Makhema, Roger Shapiro, Shahin Lockman, Mosepele Mosepele, Simani Gaseitsiwe |
| EPI_ISL_2868361 | Botswana Harvard HIV Reference Laboratory | Botswana Harvard HIV Reference Laboratory | Sikhulile Moyo, Wonderful T. Choga, Dorcas Maruapula, Thongbotho Mphoyakgosi, Boitumelo Zuze, Botshelo Radibe, Legodile Kooepile, Ontlametse T. Bareng, Motlalepule L. Pone, Godfrey Simoonga, Kereng Mphoyakgosi, Keoratile Ntshambiwa, Modisa Motswaledi, Madisa Mine, Joseph Makhema, Roger Shapiro, Shahin Lockman, Mosepele Mosepele, Simani Gaseitsiwe |
| EPI_ISL_2868362, EPI_ISL_2868363, EPI_ISL_2868364, EPI_ISL_2868365, EPI_ISL_2868367, EPI_ISL_2868368, EPI_ISL_2868369, EPI_ISL_2868370, EPI_ISL_2868371 | Botswana Harvard HIV Reference Laboratory | Botswana Harvard HIV Reference Laboratory | Sikhulile Moyo, Wonderful T. Choga, Dorcas Maruapula, Thongbotho Mphoyakgosi, Boitumelo Zuze, Botshelo Radibe, Legodile Kooepile, Ontlametse T. Bareng, Keoratile Ntshambiwa, Modisa Motswaledi, Madisa Mine, Joseph Makhema, Roger Shapiro, Shahin Lockman, Mosepele Mosepele, Simani Gaseitsiwe |
| EPI_ISL_2868373 | Botswana Harvard HIV Reference Laboratory | Botswana Harvard HIV Reference Laboratory | Sikhulile Moyo, Wonderful T. Choga, Dorcas Maruapula, Thongbotho Mphoyakgosi, Boitumelo Zuze, Botshelo Radibe, Legodile Kooepile, Ontlametse T. Bareng, Motlalepule L. Pone, Godfrey Simoonga, Kereng Mphoyakgosi, Keoratile Ntshambiwa, Modisa Motswaledi, Madisa Mine, Joseph Makhema, Roger Shapiro, Shahin Lockman, Mosepele Mosepele, Simani Gaseitsiwe |
| EPI_ISL_2868374, EPI_ISL_2868375, EPI_ISL_2868376, EPI_ISL_2868378, EPI_ISL_2868379, EPI_ISL_2868381, EPI_ISL_2868383, EPI_ISL_2868384, EPI_ISL_2868386, EPI_ISL_2868387, EPI_ISL_2868388, EPI_ISL_2868389, EPI_ISL_2868391, EPI_ISL_2868393, EPI_ISL_2868394, EPI_ISL_2868395, EPI_ISL_2868399, EPI_ISL_2868400, EPI_ISL_2868401, EPI_ISL_2868403, EPI_ISL_2868405, EPI_ISL_2868406, EPI_ISL_2868407, EPI_ISL_2868408, EPI_ISL_2868409, EPI_ISL_2868413 | | | |
| see above | Botswana Harvard HIV Reference Laboratory | Botswana Harvard HIV Reference Laboratory | Sikhulile Moyo, Wonderful T. Choga, Dorcas Maruapula, Thongbotho Mphoyakgosi, Boitumelo Zuze, Botshelo Radibe, Legodile Kooepile, Ontlametse T. Bareng, Keoratile Ntshambiwa, Modisa Motswaledi, Madisa Mine, Joseph Makhema, Roger Shapiro, Shahin Lockman, Mosepele Mosepele, Simani Gaseitsiwe |
| EPI_ISL_2868422 | Biolab Diagnostic Laboratories | Biolab Diagnostic Laboratories | Issa Abu-Dayyeh, Ahmad Tibi, Lama Hussein, Shaima Ali, Badia Saddedin, Eiad Atwa, Amid Abdelnour |
| EPI_ISL_2868424, EPI_ISL_2868426, EPI_ISL_2868428, EPI_ISL_2868430, EPI_ISL_2868431, EPI_ISL_2868433, EPI_ISL_2868434, EPI_ISL_2868435, EPI_ISL_2868437, EPI_ISL_2868439, EPI_ISL_2868440, EPI_ISL_2868443, EPI_ISL_2868444, EPI_ISL_2868445, EPI_ISL_2868446, EPI_ISL_2868447, EPI_ISL_2868448, EPI_ISL_2868449, EPI_ISL_2868450, EPI_ISL_2868451, EPI_ISL_2868453, EPI_ISL_2868454, EPI_ISL_2868455, EPI_ISL_2868456, EPI_ISL_2868457, EPI_ISL_2868458, EPI_ISL_2868459 | | | |
| see above | Dept. of Microbiology and Infection Control, Akershus University Hospital HF | Dept. of Microbiology and Infection Control, Akershus University Hospital HF | Hege Vangstein Aamot, Alexander Hesselberg Løvestad |
| EPI_ISL_2868471, EPI_ISL_2868536 | URMC LABS | Wadsworth Center, New York State Department of Health | Kirsten St. George, Daryl M. Lamson, Alexis Russell, Matthew Shudt, Melissa A Leisner, Jonathan Plitnick, Catharine Prussing, Navjot Singh, John Kelly, Erasmus Schneider, Erica Lasek-Nesselquist |
| EPI_ISL_2868537 | STONY BROOK UNIVERSITY HOSPITAL | Wadsworth Center, New York State Department of Health | Kirsten St. George, Daryl M. Lamson, Alexis Russell, Matthew Shudt, Melissa A Leisner, Jonathan Plitnick, Catharine Prussing, Navjot Singh, John Kelly, Erasmus Schneider, Erica Lasek-Nesselquist |
| EPI_ISL_2868549 | URMC LABS | Wadsworth Center, New York State Department of Health | Kirsten St. George, Daryl M. Lamson, Alexis Russell, Matthew Shudt, Melissa A Leisner, Jonathan Plitnick, Catharine Prussing, Navjot Singh, John Kelly, Erasmus Schneider, Erica Lasek-Nesselquist |
| EPI_ISL_2868554 | BOSTON HEART DIAGNOSTICS CORP | Wadsworth Center, New York State Department of Health | Kirsten St. George, Daryl M. Lamson, Alexis Russell, Matthew Shudt, Melissa A Leisner, Jonathan Plitnick, Catharine Prussing, Navjot Singh, John Kelly, Erasmus Schneider, Erica Lasek-Nesselquist |
| EPI_ISL_2868558 | URMC LABS | Wadsworth Center, New York State Department of Health | Kirsten St. George, Daryl M. Lamson, Alexis Russell, Matthew Shudt, Melissa A Leisner, Jonathan Plitnick, Catharine Prussing, Navjot Singh, John Kelly, Erasmus Schneider, Erica Lasek-Nesselquist |
| EPI_ISL_2868568, EPI_ISL_2868585, EPI_ISL_2868587 | CHUV | Laboratory of genomics and metagenomics | Trestan Pillonel, Damien Jacot, Sébastien Aeby, Gilbert Greub, Claire Bertelli |
| EPI_ISL_2868591 | VIDYMED EPALINGES | Laboratory of genomics and metagenomics | Trestan Pillonel, Damien Jacot, Sébastien Aeby, Gilbert Greub, Claire Bertelli |
| EPI_ISL_2868593, EPI_ISL_2868596 | EHNV | Laboratory of genomics and metagenomics | Trestan Pillonel, Damien Jacot, Sébastien Aeby, Gilbert Greub, Claire Bertelli |
| EPI_ISL_2868600, EPI_ISL_2868601, EPI_ISL_2868603, EPI_ISL_2868606, EPI_ISL_2868608, EPI_ISL_2868612, EPI_ISL_2868614 | CHUV | Laboratory of genomics and metagenomics | Trestan Pillonel, Damien Jacot, Sébastien Aeby, Gilbert Greub, Claire Bertelli |
| EPI_ISL_2868615, | TEMPUS LABS INC | Wadsworth Center, New York State Department of Health | Kirsten St. George, Daryl M. Lamson, Alexis Russell, Matthew Shudt, Melissa A Leisner, Jonathan Plitnick, Catharine Prussing, Navjot Singh, John Kelly, Erasmus Schneider, Erica Lasek-Nesselquist |

| | | | |
|---|---|---|---|
| EPI_ISL_2868617 | | | |
| EPI_ISL_2868618 | UNILABS - LAUSANNE | Laboratory of genomics and metagenomics | Trestan Pillonel, Damien Jacot, Sébastien Aeby, Gilbert Greub, Claire Bertelli |
| EPI_ISL_2868621, EPI_ISL_2868622 | NORTHWELL HEALTH LABORATORIES | Wadsworth Center, New York State Department of Health | Kirsten St. George, Daryl M. Lamson, Alexis Russell, Matthew Shudt, Melissa A Leisner, Jonathan Plitnick, Catharine Prussing, Navjot Singh, John Kelly, Erasmus Schneider, Erica Lasek-Nesselquist |
| EPI_ISL_2868693, EPI_ISL_2868695, EPI_ISL_2868696, EPI_ISL_2868697, EPI_ISL_2868699, EPI_ISL_2868700, EPI_ISL_2868701, EPI_ISL_2868702, EPI_ISL_2868705, EPI_ISL_2868706, EPI_ISL_2868707, EPI_ISL_2868708, EPI_ISL_2868709, EPI_ISL_2868711, EPI_ISL_2868712, EPI_ISL_2868715, EPI_ISL_2868719, EPI_ISL_2868720, EPI_ISL_2868721, EPI_ISL_2868722, EPI_ISL_2868723, EPI_ISL_2868725, EPI_ISL_2868726, EPI_ISL_2868731, EPI_ISL_2868732, EPI_ISL_2868733, EPI_ISL_2868734, EPI_ISL_2868735, EPI_ISL_2868737, EPI_ISL_2868738, EPI_ISL_2868739, EPI_ISL_2868740, EPI_ISL_2868741, EPI_ISL_2868743, EPI_ISL_2868744, EPI_ISL_2868745, EPI_ISL_2868746, EPI_ISL_2868748, EPI_ISL_2868749, EPI_ISL_2868750, EPI_ISL_2868751, EPI_ISL_2868752, EPI_ISL_2868754, EPI_ISL_2868755, EPI_ISL_2868757, EPI_ISL_2868759, EPI_ISL_2868762, EPI_ISL_2868764, EPI_ISL_2868765, EPI_ISL_2868767, EPI_ISL_2868768, EPI_ISL_2868769, EPI_ISL_2868773, EPI_ISL_2868774, EPI_ISL_2868775, EPI_ISL_2868776, EPI_ISL_2868777, EPI_ISL_2868782, EPI_ISL_2868788, EPI_ISL_2868790, EPI_ISL_2868791, EPI_ISL_2868793, EPI_ISL_2868794, EPI_ISL_2868796, EPI_ISL_2868797, EPI_ISL_2868798, EPI_ISL_2868799, EPI_ISL_2868800, EPI_ISL_2868801, EPI_ISL_2868802, EPI_ISL_2868805, EPI_ISL_2868806, EPI_ISL_2868807, EPI_ISL_2868809, EPI_ISL_2868810, EPI_ISL_2868811, EPI_ISL_2868812, EPI_ISL_2868813, EPI_ISL_2868814, EPI_ISL_2868815, EPI_ISL_2868816, EPI_ISL_2868821, EPI_ISL_2868822, EPI_ISL_2868823, EPI_ISL_2868824, EPI_ISL_2868826, EPI_ISL_2868827, EPI_ISL_2868828, EPI_ISL_2868829, EPI_ISL_2868830, EPI_ISL_2868833, EPI_ISL_2868835, EPI_ISL_2868838, EPI_ISL_2868839, EPI_ISL_2868841, EPI_ISL_2868842, EPI_ISL_2868843, EPI_ISL_2868844, EPI_ISL_2868845, EPI_ISL_2868847, EPI_ISL_2868848, EPI_ISL_2868849, EPI_ISL_2868851, EPI_ISL_2868853, EPI_ISL_2868854, EPI_ISL_2868855, EPI_ISL_2868856, EPI_ISL_2868858, EPI_ISL_2868859, EPI_ISL_2868860, EPI_ISL_2868863, EPI_ISL_2868865, EPI_ISL_2868868, EPI_ISL_2868870 | | | |
| see above | Quest Diagnostics Incorporated | Centers for Disease Control and Prevention Division of Viral Diseases, Pathogen Discovery | Dakota Howard, Dhwani Batra, Peter W. Cook, Kara Moser, Adrian Paskey, Jason Caravas, Benjamin Rambo-Martin, Shatavia Morrison, Christopher Gulvick, Scott Sammons, Yvette Unoarumhi, Darlene Wagner, Matthew Schmerer, S. H. Rosenthal, A. Gerasimova, R. M. Kagan, B. Anderson, M. Hua, Y. Liu, L.E. Bernstein, K.E. Livingston, A. Perez, I. A. Shlyakhter, R. V. Rolando, R. Owen, P. Tanpaiboon, F. Lacbawan, Clinton R. Paden, Duncan MacCannell |
| EPI_ISL_2868873 | RSPI Sulianti Saroso | National Institute of Health Research and Development | Vivi Setiawaty, Hana Apsari Pawestri, Subangkit, Kartika Dewi Puspa, Arie Ardiansyah Nugraha, Hartanti Dian Ikawati, Krisna Pangesti, Triyani Soekarso, Nelly Puspandari |
| EPI_ISL_2868874 | Quest Diagnostics Incorporated | Centers for Disease Control and Prevention Division of Viral Diseases, Pathogen Discovery | Dakota Howard, Dhwani Batra, Peter W. Cook, Kara Moser, Adrian Paskey, Jason Caravas, Benjamin Rambo-Martin, Shatavia Morrison, Christopher Gulvick, Scott Sammons, Yvette Unoarumhi, Darlene Wagner, Matthew Schmerer, S. H. Rosenthal, A. Gerasimova, R. M. Kagan, B. Anderson, M. Hua, Y. Liu, L.E. Bernstein, K.E. Livingston, A. Perez, I. A. Shlyakhter, R. V. Rolando, R. Owen, P. Tanpaiboon, F. Lacbawan, Clinton R. Paden, Duncan MacCannell |
| EPI_ISL_2868875 | BTKLPP Kelas I Makassar | National Institute of Health Research and Development | Vivi Setiawaty, Hana Apsari Pawestri, Subangkit, Kartika Dewi Puspa, Arie Ardiansyah Nugraha, Hartanti Dian Ikawati, Krisna Pangesti, Triyani Soekarso, Nelly Puspandari |
| EPI_ISL_2868876 | RSPI Sulianti Saroso | National Institute of Health Research and Development | Vivi Setiawaty, Hana Apsari Pawestri, Subangkit, Kartika Dewi Puspa, Arie Ardiansyah Nugraha, Hartanti Dian Ikawati, Krisna Pangesti, Triyani Soekarso, Nelly Puspandari |
| EPI_ISL_2868877 | Quest Diagnostics Incorporated | Centers for Disease Control and Prevention Division of Viral Diseases, Pathogen Discovery | Dakota Howard, Dhwani Batra, Peter W. Cook, Kara Moser, Adrian Paskey, Jason Caravas, Benjamin Rambo-Martin, Shatavia Morrison, Christopher Gulvick, Scott Sammons, Yvette Unoarumhi, Darlene Wagner, Matthew Schmerer, S. H. Rosenthal, A. Gerasimova, R. M. Kagan, B. Anderson, M. Hua, Y. Liu, L.E. Bernstein, K.E. Livingston, A. Perez, I. A. Shlyakhter, R. V. Rolando, R. Owen, P. Tanpaiboon, F. Lacbawan, Clinton R. Paden, Duncan MacCannell |
| EPI_ISL_2868879 | RSPI Sulianti Saroso | National Institute of Health Research and Development | Vivi Setiawaty, Hana Apsari Pawestri, Subangkit, Kartika Dewi Puspa, Arie Ardiansyah Nugraha, Hartanti Dian Ikawati, Krisna Pangesti, Triyani Soekarso, Nelly Puspandari |
| EPI_ISL_2868880 | Quest Diagnostics Incorporated | Centers for Disease Control and Prevention Division of Viral Diseases, Pathogen Discovery | Dakota Howard, Dhwani Batra, Peter W. Cook, Kara Moser, Adrian Paskey, Jason Caravas, Benjamin Rambo-Martin, Shatavia Morrison, Christopher Gulvick, Scott Sammons, Yvette Unoarumhi, Darlene Wagner, Matthew Schmerer, S. H. Rosenthal, A. Gerasimova, R. M. Kagan, B. Anderson, M. Hua, Y. Liu, L.E. Bernstein, K.E. Livingston, A. Perez, I. A. Shlyakhter, R. V. Rolando, R. Owen, P. Tanpaiboon, F. Lacbawan, Clinton R. Paden, Duncan MacCannell |
| EPI_ISL_2868881, EPI_ISL_2868883, EPI_ISL_2868884, EPI_ISL_2868886 | RSPI Sulianti Saroso | National Institute of Health Research and Development | Vivi Setiawaty, Hana Apsari Pawestri, Subangkit, Kartika Dewi Puspa, Arie Ardiansyah Nugraha, Hartanti Dian Ikawati, Krisna Pangesti, Triyani Soekarso, Nelly Puspandari |
| EPI_ISL_2868887, EPI_ISL_2868889 | BTKLPP Kelas I Makassar | National Institute of Health Research and Development | Vivi Setiawaty, Hana Apsari Pawestri, Subangkit, Kartika Dewi Puspa, Arie Ardiansyah Nugraha, Hartanti Dian Ikawati, Krisna Pangesti, Triyani Soekarso, Nelly Puspandari |
| EPI_ISL_2868890 | Quest Diagnostics Incorporated | Centers for Disease Control and Prevention Division of Viral Diseases, Pathogen Discovery | Dakota Howard, Dhwani Batra, Peter W. Cook, Kara Moser, Adrian Paskey, Jason Caravas, Benjamin Rambo-Martin, Shatavia Morrison, Christopher Gulvick, Scott Sammons, Yvette Unoarumhi, Darlene Wagner, Matthew Schmerer, S. H. Rosenthal, A. Gerasimova, R. M. Kagan, B. Anderson, M. Hua, Y. Liu, L.E. Bernstein, K.E. Livingston, A. Perez, I. A. Shlyakhter, R. V. Rolando, R. Owen, P. Tanpaiboon, F. Lacbawan, Clinton R. Paden, Duncan MacCannell |
| EPI_ISL_2868891, EPI_ISL_2868892, EPI_ISL_2868894 | BTKLPP Kelas I Makassar | National Institute of Health Research and Development | Vivi Setiawaty, Hana Apsari Pawestri, Subangkit, Kartika Dewi Puspa, Arie Ardiansyah Nugraha, Hartanti Dian Ikawati, Krisna Pangesti, Triyani Soekarso, Nelly Puspandari |
| EPI_ISL_2868895 | Quest Diagnostics Incorporated | Centers for Disease Control and Prevention Division of Viral Diseases, Pathogen Discovery | Dakota Howard, Dhwani Batra, Peter W. Cook, Kara Moser, Adrian Paskey, Jason Caravas, Benjamin Rambo-Martin, Shatavia Morrison, Christopher Gulvick, Scott Sammons, Yvette Unoarumhi, Darlene Wagner, Matthew Schmerer, S. H. Rosenthal, A. Gerasimova, R. M. Kagan, B. Anderson, M. Hua, Y. Liu, L.E. Bernstein, K.E. Livingston, A. Perez, I. A. Shlyakhter, R. V. Rolando, R. Owen, P. Tanpaiboon, F. Lacbawan, Clinton R. Paden, Duncan MacCannell |
| EPI_ISL_2868896, EPI_ISL_2868897 | BTKLPP Kelas I Makassar | National Institute of Health Research and Development | Vivi Setiawaty, Hana Apsari Pawestri, Subangkit, Kartika Dewi Puspa, Arie Ardiansyah Nugraha, Hartanti Dian Ikawati, Krisna Pangesti, Triyani Soekarso, Nelly Puspandari |
| EPI_ISL_2868898 | Quest Diagnostics Incorporated | Centers for Disease Control and Prevention Division of Viral Diseases, Pathogen Discovery | Dakota Howard, Dhwani Batra, Peter W. Cook, Kara Moser, Adrian Paskey, Jason Caravas, Benjamin Rambo-Martin, Shatavia Morrison, Christopher Gulvick, Scott Sammons, Yvette Unoarumhi, Darlene Wagner, Matthew Schmerer, S. H. Rosenthal, A. Gerasimova, R. M. Kagan, B. Anderson, M. Hua, Y. Liu, L.E. Bernstein, K.E. Livingston, A. Perez, I. A. Shlyakhter, R. V. Rolando, R. Owen, P. Tanpaiboon, F. Lacbawan, Clinton R. Paden, Duncan MacCannell |
| EPI_ISL_2868899, EPI_ISL_2868901 | BTKLPP Kelas I Makassar | National Institute of Health Research and Development | Vivi Setiawaty, Hana Apsari Pawestri, Subangkit, Kartika Dewi Puspa, Arie Ardiansyah Nugraha, Hartanti Dian Ikawati, Krisna Pangesti, Triyani Soekarso, Nelly Puspandari |
| EPI_ISL_2868902 | RS Cinta Kasih Tzu Chi | National Institute of Health Research and Development | Vivi Setiawaty, Hana Apsari Pawestri, Subangkit, Kartika Dewi Puspa, Arie Ardiansyah Nugraha, Hartanti Dian Ikawati, Krisna Pangesti, Triyani Soekarso, Nelly Puspandari |
| EPI_ISL_2868903 | Quest Diagnostics Incorporated | Centers for Disease Control and Prevention Division of Viral Diseases, Pathogen Discovery | Dakota Howard, Dhwani Batra, Peter W. Cook, Kara Moser, Adrian Paskey, Jason Caravas, Benjamin Rambo-Martin, Shatavia Morrison, Christopher Gulvick, Scott Sammons, Yvette Unoarumhi, Darlene Wagner, Matthew Schmerer, S. H. Rosenthal, A. Gerasimova, R. M. Kagan, B. Anderson, M. Hua, Y. Liu, L.E. Bernstein, K.E. Livingston, A. Perez, I. A. Shlyakhter, R. V. Rolando, R. Owen, P. Tanpaiboon, F. Lacbawan, Clinton R. Paden, Duncan MacCannell |
| EPI_ISL_2868904 | RSUD Pademangan | National Institute of Health Research and Development | Vivi Setiawaty, Hana Apsari Pawestri, Subangkit, Kartika Dewi Puspa, Arie Ardiansyah Nugraha, Hartanti Dian Ikawati, Krisna Pangesti, Triyani Soekarso, Nelly Puspandari |
| EPI_ISL_2868905 | RS Kartika Pulomas | National Institute of Health Research and Development | Vivi Setiawaty, Hana Apsari Pawestri, Subangkit, Kartika Dewi Puspa, Arie Ardiansyah Nugraha, Hartanti Dian Ikawati, Krisna Pangesti, Triyani Soekarso, Nelly Puspandari |
| EPI_ISL_2868907, EPI_ISL_2868909, EPI_ISL_2868910 | RS Cinta Kasih Tzu Chi | National Institute of Health Research and Development | Vivi Setiawaty, Hana Apsari Pawestri, Subangkit, Kartika Dewi Puspa, Arie Ardiansyah Nugraha, Hartanti Dian Ikawati, Krisna Pangesti, Triyani Soekarso, Nelly Puspandari |
| EPI_ISL_2868911 | Quest Diagnostics Incorporated | Centers for Disease Control and Prevention Division of Viral Diseases, Pathogen Discovery | Dakota Howard, Dhwani Batra, Peter W. Cook, Kara Moser, Adrian Paskey, Jason Caravas, Benjamin Rambo-Martin, Shatavia Morrison, Christopher Gulvick, Scott Sammons, Yvette Unoarumhi, Darlene Wagner, Matthew Schmerer, S. H. Rosenthal, A. Gerasimova, R. M. Kagan, B. Anderson, M. Hua, Y. Liu, L.E. Bernstein, K.E. Livingston, A. Perez, I. A. Shlyakhter, R. V. Rolando, R. Owen, P. Tanpaiboon, F. Lacbawan, Clinton R. Paden, Duncan MacCannell |
| EPI_ISL_2868912, EPI_ISL_2868913 | RS Royal Progress | National Institute of Health Research and Development | Vivi Setiawaty, Hana Apsari Pawestri, Subangkit, Kartika Dewi Puspa, Arie Ardiansyah Nugraha, Hartanti Dian Ikawati, Krisna Pangesti, Triyani Soekarso, Nelly Puspandari |
| EPI_ISL_2868915 | RS Gading Pluit | National Institute of Health Research and Development | Vivi Setiawaty, Hana Apsari Pawestri, Subangkit, Kartika Dewi Puspa, Arie Ardiansyah Nugraha, Hartanti Dian Ikawati, Krisna Pangesti, Triyani Soekarso, Nelly Puspandari |
| EPI_ISL_2868916 | DINKES Kab Bogor | National Institute of Health Research and Development | Vivi Setiawaty, Hana Apsari Pawestri, Subangkit, Kartika Dewi Puspa, Arie Ardiansyah Nugraha, Hartanti Dian Ikawati, Krisna Pangesti, Triyani Soekarso, Nelly Puspandari |
| EPI_ISL_2868917 | Quest Diagnostics Incorporated | Centers for Disease Control and Prevention Division of Viral Diseases, Pathogen Discovery | Dakota Howard, Dhwani Batra, Peter W. Cook, Kara Moser, Adrian Paskey, Jason Caravas, Benjamin Rambo-Martin, Shatavia Morrison, Christopher Gulvick, Scott Sammons, Yvette Unoarumhi, Darlene Wagner, Matthew Schmerer, S. H. Rosenthal, A. Gerasimova, R. M. Kagan, B. Anderson, M. Hua, Y. Liu, L.E. Bernstein, K.E. Livingston, A. Perez, I. A. Shlyakhter, R. V. Rolando, R. Owen, P. Tanpaiboon, F. Lacbawan, Clinton R. Paden, Duncan MacCannell |
| EPI_ISL_2868918 | DINKES Kab Bogor | National Institute of Health Research and Development | Vivi Setiawaty, Hana Apsari Pawestri, Subangkit, Kartika Dewi Puspa, Arie Ardiansyah Nugraha, Hartanti Dian Ikawati, Krisna Pangesti, Triyani Soekarso, Nelly Puspandari |
| EPI_ISL_2868919, EPI_ISL_2868922, EPI_ISL_2868927, EPI_ISL_2868930, EPI_ISL_2868935, EPI_ISL_2868938, EPI_ISL_2868940, EPI_ISL_2868943, EPI_ISL_2868949, EPI_ISL_2868950, EPI_ISL_2868951, EPI_ISL_2868952, EPI_ISL_2868955, EPI_ISL_2868956, EPI_ISL_2868957, EPI_ISL_2868958, EPI_ISL_2868960, EPI_ISL_2868961, EPI_ISL_2868965, EPI_ISL_2868966, EPI_ISL_2868969, EPI_ISL_2868970, EPI_ISL_2868971, EPI_ISL_2868972, EPI_ISL_2868975, EPI_ISL_2868976, EPI_ISL_2868977, EPI_ISL_2868978, EPI_ISL_2868979, EPI_ISL_2868980, EPI_ISL_2868982, EPI_ISL_2868984, EPI_ISL_2868985, EPI_ISL_2868986, EPI_ISL_2868989, EPI_ISL_2868990, EPI_ISL_2868991, EPI_ISL_2868993, EPI_ISL_2868994, EPI_ISL_2868995, EPI_ISL_2868996 | | | |
| see above | Quest Diagnostics Incorporated | Centers for Disease Control and Prevention Division of Viral Diseases, Pathogen Discovery | Dakota Howard, Dhwani Batra, Peter W. Cook, Kara Moser, Adrian Paskey, Jason Caravas, Benjamin Rambo-Martin, Shatavia Morrison, Christopher Gulvick, Scott Sammons, Yvette Unoarumhi, Darlene Wagner, Matthew Schmerer, S. H. Rosenthal, A. Gerasimova, R. M. Kagan, B. Anderson, M. Hua, Y. Liu, L.E. Bernstein, K.E. Livingston, A. Perez, I. A. Shlyakhter, R. V. Rolando, R. Owen, P. Tanpaiboon, F. Lacbawan, Clinton R. Paden, Duncan MacCannell |
| EPI_ISL_2868997 | Heilig hart Lier | Imelda | Johan Frans, Dagmar Obbels, Hanne Valgaeren |
| EPI_ISL_2868998, EPI_ISL_2869000 | AZ Jan Portaels | Imelda | Johan Frans, Dagmar Obbels, Hanne Valgaeren |
| EPI_ISL_2869001 | Imelda | Imelda | Johan Frans, Dagmar Obbels, Hanne Valgaeren |
| EPI_ISL_2869002, EPI_ISL_2869003, EPI_ISL_2869004, EPI_ISL_2869006, EPI_ISL_2869007, EPI_ISL_2869008, EPI_ISL_2869010, EPI_ISL_2869011, EPI_ISL_2869012, EPI_ISL_2869013, EPI_ISL_2869017, EPI_ISL_2869018, EPI_ISL_2869019, EPI_ISL_2869021, EPI_ISL_2869022, EPI_ISL_2869024, EPI_ISL_2869025, EPI_ISL_2869028, EPI_ISL_2869029, EPI_ISL_2869030, EPI_ISL_2869031, EPI_ISL_2869032, EPI_ISL_2869035, EPI_ISL_2869036, EPI_ISL_2869041, EPI_ISL_2869042, EPI_ISL_2869043, EPI_ISL_2869045, EPI_ISL_2869046, EPI_ISL_2869048, EPI_ISL_2869049, EPI_ISL_2869050, EPI_ISL_2869054, EPI_ISL_2869056, EPI_ISL_2869058, EPI_ISL_2869060, EPI_ISL_2869063, EPI_ISL_2869064, EPI_ISL_2869065, EPI_ISL_2869066, EPI_ISL_2869068, EPI_ISL_2869069, EPI_ISL_2869070, EPI_ISL_2869072, EPI_ISL_2869073, EPI_ISL_2869074, EPI_ISL_2869076, EPI_ISL_2869077, EPI_ISL_2869080, EPI_ISL_2869082, EPI_ISL_2869083, EPI_ISL_2869085, EPI_ISL_2869086, EPI_ISL_2869087, EPI_ISL_2869089, EPI_ISL_2869090, EPI_ISL_2869092, EPI_ISL_2869093, EPI_ISL_2869095, EPI_ISL_2869096, EPI_ISL_2869097, EPI_ISL_2869101, EPI_ISL_2869103, EPI_ISL_2869104, EPI_ISL_2869106, EPI_ISL_2869108, EPI_ISL_2869109, EPI_ISL_2869110, EPI_ISL_2869111, EPI_ISL_2869112, EPI_ISL_2869113, EPI_ISL_2869114, EPI_ISL_2869115, EPI_ISL_2869116, EPI_ISL_2869117, EPI_ISL_2869118, EPI_ISL_2869120, EPI_ISL_2869121, EPI_ISL_2869122, EPI_ISL_2869123, EPI_ISL_2869124, EPI_ISL_2869126, EPI_ISL_2869127, EPI_ISL_2869128, EPI_ISL_2869129, EPI_ISL_2869130, EPI_ISL_2869131, EPI_ISL_2869132, EPI_ISL_2869133, EPI_ISL_2869135, EPI_ISL_2869137, EPI_ISL_2869142, EPI_ISL_2869144, EPI_ISL_2869145, EPI_ISL_2869146, EPI_ISL_2869149, EPI_ISL_2869150, EPI_ISL_2869151, EPI_ISL_2869152, EPI_ISL_2869153, EPI_ISL_2869154, EPI_ISL_2869156, EPI_ISL_2869157, EPI_ISL_2869159, EPI_ISL_2869161, EPI_ISL_2869164, EPI_ISL_2869165, EPI_ISL_2869166, EPI_ISL_2869167, EPI_ISL_2869168, EPI_ISL_2869169, EPI_ISL_2869170, EPI_ISL_2869171, EPI_ISL_2869172, EPI_ISL_2869173, EPI_ISL_2869174, EPI_ISL_2869175, EPI_ISL_2869178, EPI_ISL_2869179, EPI_ISL_2869180, EPI_ISL_2869181, EPI_ISL_2869186, EPI_ISL_2869187, EPI_ISL_2869188, EPI_ISL_2869189, EPI_ISL_2869190, EPI_ISL_2869192, EPI_ISL_2869193, EPI_ISL_2869195, EPI_ISL_2869196, EPI_ISL_2869197, EPI_ISL_2869198, EPI_ISL_2869199, EPI_ISL_2869201, EPI_ISL_2869202, EPI_ISL_2869204, EPI_ISL_2869206, EPI_ISL_2869207, EPI_ISL_2869208, EPI_ISL_2869209, EPI_ISL_2869211, EPI_ISL_2869213, EPI_ISL_2869214, EPI_ISL_2869215, EPI_ISL_2869216, EPI_ISL_2869220, EPI_ISL_2869221, EPI_ISL_2869222, EPI_ISL_2869223, EPI_ISL_2869226, EPI_ISL_2869227, EPI_ISL_2869229, EPI_ISL_2869230, EPI_ISL_2869231, EPI_ISL_2869232, EPI_ISL_2869233, EPI_ISL_2869234, EPI_ISL_2869235, EPI_ISL_2869238, EPI_ISL_2869240, EPI_ISL_2869241, EPI_ISL_2869242, EPI_ISL_2869243, EPI_ISL_2869244, EPI_ISL_2869246, EPI_ISL_2869247, EPI_ISL_2869249, EPI_ISL_2869250, EPI_ISL_2869251, EPI_ISL_2869252, EPI_ISL_2869256, EPI_ISL_2869258, EPI_ISL_2869259, EPI_ISL_2869260, EPI_ISL_2869262, EPI_ISL_2869263, EPI_ISL_2869264, EPI_ISL_2869265, EPI_ISL_2869266, EPI_ISL_2869267, EPI_ISL_2869268, EPI_ISL_2869271, EPI_ISL_2869275, EPI_ISL_2869276, EPI_ISL_2869277, EPI_ISL_2869278, EPI_ISL_2869279, EPI_ISL_2869284, EPI_ISL_2869285, EPI_ISL_2869289, EPI_ISL_2869291, EPI_ISL_2869292, EPI_ISL_2869296, EPI_ISL_2869297, EPI_ISL_2869298, EPI_ISL_2869299, EPI_ISL_2869300, EPI_ISL_2869301, EPI_ISL_2869302, EPI_ISL_2869304, EPI_ISL_2869305, EPI_ISL_2869306, EPI_ISL_2869307, EPI_ISL_2869308, EPI_ISL_2869309, EPI_ISL_2869310, EPI_ISL_2869312, EPI_ISL_2869313, EPI_ISL_2869314, EPI_ISL_2869315, EPI_ISL_2869316, EPI_ISL_2869317, EPI_ISL_2869318, EPI_ISL_2869319, EPI_ISL_2869320, EPI_ISL_2869321, EPI_ISL_2869323, EPI_ISL_2869324, EPI_ISL_2869325, EPI_ISL_2869326, EPI_ISL_2869328, EPI_ISL_2869330, EPI_ISL_2869332, EPI_ISL_2869333, EPI_ISL_2869337, EPI_ISL_2869338, EPI_ISL_2869340, EPI_ISL_2869341, EPI_ISL_2869342, EPI_ISL_2869343, EPI_ISL_2869344, EPI_ISL_2869345, EPI_ISL_2869346, EPI_ISL_2869347, EPI_ISL_2869348, EPI_ISL_2869350, EPI_ISL_2869351, EPI_ISL_2869352, EPI_ISL_2869353, EPI_ISL_2869354, EPI_ISL_2869356, EPI_ISL_2869359, EPI_ISL_2869360, EPI_ISL_2869361, EPI_ISL_2869362, EPI_ISL_2869363, EPI_ISL_2869364, EPI_ISL_2869369, EPI_ISL_2869370, EPI_ISL_2869372, EPI_ISL_2869373, EPI_ISL_2869374, EPI_ISL_2869376, EPI_ISL_2869377, EPI_ISL_2869378, EPI_ISL_2869379, EPI_ISL_2869380, EPI_ISL_2869381, EPI_ISL_2869382, EPI_ISL_2869383, EPI_ISL_2869386, EPI_ISL_2869388, EPI_ISL_2869389, EPI_ISL_2869391, EPI_ISL_2869393, EPI_ISL_2869394, EPI_ISL_2869395, EPI_ISL_2869396, EPI_ISL_2869400, EPI_ISL_2869402, EPI_ISL_2869405, EPI_ISL_2869407, EPI_ISL_2869408, EPI_ISL_2869409, EPI_ISL_2869410, EPI_ISL_2869412, EPI_ISL_2869413, EPI_ISL_2869414, EPI_ISL_2869415, EPI_ISL_2869416, EPI_ISL_2869418, EPI_ISL_2869419, EPI_ISL_2869420, EPI_ISL_2869421, EPI_ISL_2869422, EPI_ISL_2869424, EPI_ISL_2869426, EPI_ISL_2869427, EPI_ISL_2869431, EPI_ISL_2869433, EPI_ISL_2869437, EPI_ISL_2869440, EPI_ISL_2869441, EPI_ISL_2869442, EPI_ISL_2869443, EPI_ISL_2869444, EPI_ISL_2869445, EPI_ISL_2869446, EPI_ISL_2869447, EPI_ISL_2869448, EPI_ISL_2869449, EPI_ISL_2869451, EPI_ISL_2869454, EPI_ISL_2869458, EPI_ISL_2869460, EPI_ISL_2869462, EPI_ISL_2869463, EPI_ISL_2869464, EPI_ISL_2869465, EPI_ISL_2869466, EPI_ISL_2869467, EPI_ISL_2869468, EPI_ISL_2869469, EPI_ISL_2869471, EPI_ISL_2869472, EPI_ISL_2869473, EPI_ISL_2869476, EPI_ISL_2869478, EPI_ISL_2869479, EPI_ISL_2869480, EPI_ISL_2869481, EPI_ISL_2869483, EPI_ISL_2869484, EPI_ISL_2869485, EPI_ISL_2869487, EPI_ISL_2869488, EPI_ISL_2869490, EPI_ISL_2869492, EPI_ISL_2869493, EPI_ISL_2869496, EPI_ISL_2869498, EPI_ISL_2869500, EPI_ISL_2869501, EPI_ISL_2869502, EPI_ISL_2869504, EPI_ISL_2869506, EPI_ISL_2869509, EPI_ISL_2869510, EPI_ISL_2869514, EPI_ISL_2869515, EPI_ISL_2869516, EPI_ISL_2869517, EPI_ISL_2869518, EPI_ISL_2869519, EPI_ISL_2869520, EPI_ISL_2869521, EPI_ISL_2869523, EPI_ISL_2869524, EPI_ISL_2869528, EPI_ISL_2869530, EPI_ISL_2869531, EPI_ISL_2869532, EPI_ISL_2869533, EPI_ISL_2869534, EPI_ISL_2869535, EPI_ISL_2869538, EPI_ISL_2869540, EPI_ISL_2869541, EPI_ISL_2869542, EPI_ISL_2869543, EPI_ISL_2869547, EPI_ISL_2869548, EPI_ISL_2869549, EPI_ISL_2869550, EPI_ISL_2869551, EPI_ISL_2869552, EPI_ISL_2869553, EPI_ISL_2869555, EPI_ISL_2869557, EPI_ISL_2869558, EPI_ISL_2869561, EPI_ISL_2869563, EPI_ISL_2869564, EPI_ISL_2869565, EPI_ISL_2869566, EPI_ISL_2869567, EPI_ISL_2869569, EPI_ISL_2869570, EPI_ISL_2869571, EPI_ISL_2869574, EPI_ISL_2869575, EPI_ISL_2869577, EPI_ISL_2869579, EPI_ISL_2869580, EPI_ISL_2869581, EPI_ISL_2869582, EPI_ISL_2869583, EPI_ISL_2869584, EPI_ISL_2869585, EPI_ISL_2869586, EPI_ISL_2869587, EPI_ISL_2869588, EPI_ISL_2869589, EPI_ISL_2869590, EPI_ISL_2869591, EPI_ISL_2869597, EPI_ISL_2869599, EPI_ISL_2869600, EPI_ISL_2869601, EPI_ISL_2869602, EPI_ISL_2869605, EPI_ISL_2869607, EPI_ISL_2869610, EPI_ISL_2869611, EPI_ISL_2869613, EPI_ISL_2869615, EPI_ISL_2869616, EPI_ISL_2869619, EPI_ISL_2869620, EPI_ISL_2869621, EPI_ISL_2869623, EPI_ISL_2869625, EPI_ISL_2869627, EPI_ISL_2869630, EPI_ISL_2869632, EPI_ISL_2869633, EPI_ISL_2869635, EPI_ISL_2869636, EPI_ISL_2869638, EPI_ISL_2869643, EPI_ISL_2869646, EPI_ISL_2869647, EPI_ISL_2869652, EPI_ISL_2869655, EPI_ISL_2869658, EPI_ISL_2869660, EPI_ISL_2869662, EPI_ISL_2869664, EPI_ISL_2869670, EPI_ISL_2869676, EPI_ISL_2869679, EPI_ISL_2869690, EPI_ISL_2869692, EPI_ISL_2869695, EPI_ISL_2869697, EPI_ISL_2869698, EPI_ISL_2869702, EPI_ISL_2869705, EPI_ISL_2869707, EPI_ISL_2869712, EPI_ISL_2869714, EPI_ISL_2869718, EPI_ISL_2869721, EPI_ISL_2869731, EPI_ISL_2869741, EPI_ISL_2869763, EPI_ISL_2869766, EPI_ISL_2869773, EPI_ISL_2869782, EPI_ISL_2869792, EPI_ISL_2869830, EPI_ISL_2869833, EPI_ISL_2869848, EPI_ISL_2869863, EPI_ISL_2869881, EPI_ISL_2869884, EPI_ISL_2869898, EPI_ISL_2869901, EPI_ISL_2869905, EPI_ISL_2869919, EPI_ISL_2869934, EPI_ISL_2869940, EPI_ISL_2869967, EPI_ISL_2869970, EPI_ISL_2869982, EPI_ISL_2869984, EPI_ISL_2869987, EPI_ISL_2869990, EPI_ISL_2869996, EPI_ISL_2870016, EPI_ISL_2870024, EPI_ISL_2870027, EPI_ISL_2870044, EPI_ISL_2870052, EPI_ISL_2870055, EPI_ISL_2870078, EPI_ISL_2870086, EPI_ISL_2870098, EPI_ISL_2870106, EPI_ISL_2870109, EPI_ISL_2870115, EPI_ISL_2870127, EPI_ISL_2870138, EPI_ISL_2870164, EPI_ISL_2870173, EPI_ISL_2870194, EPI_ISL_2870199, EPI_ISL_2870214, EPI_ISL_2870217, EPI_ISL_2870234, EPI_ISL_2870246, EPI_ISL_2870263, EPI_ISL_2870307, EPI_ISL_2870330, EPI_ISL_2870336, EPI_ISL_2870342, EPI_ISL_2870345, EPI_ISL_2870357, EPI_ISL_2870363, EPI_ISL_2870369, EPI_ISL_2870372, EPI_ISL_2870378, EPI_ISL_2870384, EPI_ISL_2870412, EPI_ISL_2870418, EPI_ISL_2870427, EPI_ISL_2870452, EPI_ISL_2870484, EPI_ISL_2870504, EPI_ISL_2870527, EPI_ISL_2870533, EPI_ISL_2870545, EPI_ISL_2870548, EPI_ISL_2870574, EPI_ISL_2870577, EPI_ISL_2870586, EPI_ISL_2870598, EPI_ISL_2870637, EPI_ISL_2870660, EPI_ISL_2870663, EPI_ISL_2870670, EPI_ISL_2870678, EPI_ISL_2870684, EPI_ISL_2870687, EPI_ISL_2870690, EPI_ISL_2870696, EPI_ISL_2870730, EPI_ISL_2870733, EPI_ISL_2870769, EPI_ISL_2870810, EPI_ISL_2870822, EPI_ISL_2870828, EPI_ISL_2870842, EPI_ISL_2870845, EPI_ISL_2870874, EPI_ISL_2870897, EPI_ISL_2870909, EPI_ISL_2870915, EPI_ISL_2870927, EPI_ISL_2870945, EPI_ISL_2870954, EPI_ISL_2870975, EPI_ISL_2870979, EPI_ISL_2870982, EPI_ISL_2870984, EPI_ISL_2870991, EPI_ISL_2871000, EPI_ISL_2871003, EPI_ISL_2871009, EPI_ISL_2871015, EPI_ISL_2871024, EPI_ISL_2871026, EPI_ISL_2871035, EPI_ISL_2871044, EPI_ISL_2871048, EPI_ISL_2871051, EPI_ISL_2871056, EPI_ISL_2871059, EPI_ISL_2871063, EPI_ISL_2871075, EPI_ISL_2871078, EPI_ISL_2871090, EPI_ISL_2871093, EPI_ISL_2871096, EPI_ISL_2871098, EPI_ISL_2871104, EPI_ISL_2871116, EPI_ISL_2871119, EPI_ISL_2871122, EPI_ISL_2871151, EPI_ISL_2871169, EPI_ISL_2871172, EPI_ISL_2871190, EPI_ISL_2871199, EPI_ISL_2871202, EPI_ISL_2871208, EPI_ISL_2871216, EPI_ISL_2871228, EPI_ISL_2871234, EPI_ISL_2871237, EPI_ISL_2871240, EPI_ISL_2871243, EPI_ISL_2871249, EPI_ISL_2871254, EPI_ISL_2871261, EPI_ISL_2871270, EPI_ISL_2871278, EPI_ISL_2871284, EPI_ISL_2871287, EPI_ISL_2871290, EPI_ISL_2871296, EPI_ISL_2871302, EPI_ISL_2871308, EPI_ISL_2871323, EPI_ISL_2871335, EPI_ISL_2871337, EPI_ISL_2871341, EPI_ISL_2871344, EPI_ISL_2871346, EPI_ISL_2871355, EPI_ISL_2871358, EPI_ISL_2871367, EPI_ISL_2871370, EPI_ISL_2871373, EPI_ISL_2871379, EPI_ISL_2871383, EPI_ISL_2871386, EPI_ISL_2871392, EPI_ISL_2871404, EPI_ISL_2871413, EPI_ISL_2871419, EPI_ISL_2871422, EPI_ISL_2871424, EPI_ISL_2871427, EPI_ISL_2871430, EPI_ISL_2871433, | | | |

| | | | |
|---|---|---|---|
| EPI_ISL_2871439, EPI_ISL_2871442, EPI_ISL_2871454, EPI_ISL_2871462, EPI_ISL_2871465, EPI_ISL_2871468, EPI_ISL_2871474, EPI_ISL_2871482, EPI_ISL_2871485, EPI_ISL_2871509, EPI_ISL_2871515, EPI_ISL_2871532, EPI_ISL_2871535, EPI_ISL_2871541, EPI_ISL_2871551, EPI_ISL_2871553, EPI_ISL_2871560, EPI_ISL_2871563, EPI_ISL_2871566, EPI_ISL_2871574, EPI_ISL_2871580, EPI_ISL_2871583, EPI_ISL_2871601, EPI_ISL_2871610, EPI_ISL_2871616, EPI_ISL_2871620, EPI_ISL_2871627, EPI_ISL_2871630, EPI_ISL_2871638, EPI_ISL_2871658, EPI_ISL_2871661, EPI_ISL_2871678, EPI_ISL_2871681, EPI_ISL_2871690, EPI_ISL_2871693, EPI_ISL_2871696, EPI_ISL_2871713, EPI_ISL_2871719, EPI_ISL_2871725, EPI_ISL_2871737, EPI_ISL_2871748, EPI_ISL_2871751, EPI_ISL_2871760, EPI_ISL_2871764, EPI_ISL_2871766, EPI_ISL_2871773, EPI_ISL_2871779, EPI_ISL_2871782, EPI_ISL_2871791, EPI_ISL_2871793, EPI_ISL_2871796, EPI_ISL_2871805, EPI_ISL_2871808, EPI_ISL_2871831, EPI_ISL_2871861, EPI_ISL_2871864, EPI_ISL_2871881, EPI_ISL_2871904, EPI_ISL_2871914, EPI_ISL_2871931, EPI_ISL_2871937, EPI_ISL_2871946, EPI_ISL_2871951, EPI_ISL_2871963, EPI_ISL_2871972, EPI_ISL_2871980, EPI_ISL_2871983, EPI_ISL_2871989, EPI_ISL_2871992, EPI_ISL_2872009, EPI_ISL_2872024, EPI_ISL_2872027, EPI_ISL_2872030, EPI_ISL_2872039, EPI_ISL_2872042, EPI_ISL_2872062, EPI_ISL_2872065, EPI_ISL_2872085, EPI_ISL_2872088, EPI_ISL_2872091, EPI_ISL_2872098, EPI_ISL_2872100, EPI_ISL_2872103, EPI_ISL_2872109, EPI_ISL_2872117, EPI_ISL_2872120 | | | |
| see above | Quest Diagnostics Incorporated | Centers for Disease Control and Prevention Division of Viral Diseases, Pathogen Discovery | Dakota Howard, Dhwani Batra, Peter W. Cook, Kara Moser, Adrian Paskey, Jason Caravas, Benjamin Rambo-Martin, Shatavia Morrison, Christopher Gulvick, Scott Sammons, Yvette Unoarumhi, Darlene Wagner, Matthew Schmerer, S. H. Rosenthal, A. Gerasimova, R. M. Kagan, B. Anderson, M. Hua, Y. Liu, L.E. Bernstein, K.E. Livingston, A. Perez, I. A. Shlyakhter, R. V. Rolando, R. Owen, P. Tanpaiboon, F. Lacbawan, Clinton R. Paden, Duncan MacCannell |
| EPI_ISL_2872162, EPI_ISL_2872167, EPI_ISL_2872199, EPI_ISL_2872211, EPI_ISL_2872214, EPI_ISL_2872226, EPI_ISL_2872229, EPI_ISL_2872249, EPI_ISL_2872252, EPI_ISL_2872261, EPI_ISL_2872271, EPI_ISL_2872277, EPI_ISL_2872280, EPI_ISL_2872282, EPI_ISL_2872291, EPI_ISL_2872294, EPI_ISL_2872297, EPI_ISL_2872314, EPI_ISL_2872317, EPI_ISL_2872319, EPI_ISL_2872345, EPI_ISL_2872348, EPI_ISL_2872364, EPI_ISL_2872370, EPI_ISL_2872373, EPI_ISL_2872394, EPI_ISL_2872401, EPI_ISL_2872403, EPI_ISL_2872421, EPI_ISL_2872423, EPI_ISL_2872426, EPI_ISL_2872427, EPI_ISL_2872434, EPI_ISL_2872437, EPI_ISL_2872440, EPI_ISL_2872441, EPI_ISL_2872454, EPI_ISL_2872457, EPI_ISL_2872458, EPI_ISL_2872464, EPI_ISL_2872466, EPI_ISL_2872467, EPI_ISL_2872468, EPI_ISL_2872472, EPI_ISL_2872475, EPI_ISL_2872477, EPI_ISL_2872478, EPI_ISL_2872479, EPI_ISL_2872487, EPI_ISL_2872492, EPI_ISL_2872494, EPI_ISL_2872495, EPI_ISL_2872496, EPI_ISL_2872497, EPI_ISL_2872498, EPI_ISL_2872499, EPI_ISL_2872501, EPI_ISL_2872503, EPI_ISL_2872504, EPI_ISL_2872505, EPI_ISL_2872507, EPI_ISL_2872508, EPI_ISL_2872510, EPI_ISL_2872511, EPI_ISL_2872512, EPI_ISL_2872513, EPI_ISL_2872514, EPI_ISL_2872515, EPI_ISL_2872516, EPI_ISL_2872519, EPI_ISL_2872526, EPI_ISL_2872528, EPI_ISL_2872529, EPI_ISL_2872533, EPI_ISL_2872539, EPI_ISL_2872541, EPI_ISL_2872550, EPI_ISL_2872551, EPI_ISL_2872553, EPI_ISL_2872554, EPI_ISL_2872562, EPI_ISL_2872563, EPI_ISL_2872564, EPI_ISL_2872567, EPI_ISL_2872568, EPI_ISL_2872569, EPI_ISL_2872570, EPI_ISL_2872571, EPI_ISL_2872572, EPI_ISL_2872573, EPI_ISL_2872583 | | | |
| see above | Laboratory Corporation of America | Centers for Disease Control and Prevention Division of Viral Diseases, Pathogen Discovery | Dakota Howard, Dhwani Batra, Peter W. Cook, Kara Moser, Adrian Paskey, Jason Caravas, Benjamin Rambo-Martin, Shatavia Morrison, Christopher Gulvick, Scott Sammons, Yvette Unoarumhi, Darlene Wagner, Matthew Schmerer, Minoo Agarwal, Eyad Almasri, Debbie Boles, Ayla Burns, Nuthawin Charoensri, Oren Cohen, Susan Countryman, Mary Ann Cristobal, Bobbi Croy, Suzanne Dale, Hrushikesh Deshmukh, Amanda Douglas, Vincent Drouillon, Marcia Eisenberg, Howard Engler, Rama Ghatti, Prashant Gupta, Susan Hicks, Jake Humphrey, Lax Iyer, Lisa Pfefferle, Manoj Jain, Matthew Robinson, Mohan Kolli, Brian Krueger, Tim Kuphal, Stanley Letovsky, Michael Levandoski, Craig Lukasik, Jonathan Meltzer, Brian Norvell, Mindy Nye, Scott Parker, Christos Petropoulos, John Pruitt, Steven Ragan, Scott Ryan, Mike Sapeta, Jana Schroth, Suresh Babu Selvaraju, Goran Stevovic, Amanda Suchanek, Andrea Throop, Lyndon Tilson, Thomas Urban, Joe Voshell, Kimberly Wagner, Jonathan Williams, Mary Williamson, Qian Zeng, Tricia Zwiefelhofer, Clinton R. Paden, Duncan MacCannell |
| EPI_ISL_2872584, EPI_ISL_2872585, EPI_ISL_2872587 | Azienda Ospedaliero Universitaria di Sassari | AMES Centro Polidiagnostico Strumentale S.r.l. | Salvatore Rubino; Flavia Angioj; Laura Firino; Rosalba Govoni; Gabriele Ibba; Vincenzo Lai; Erica Mura; Bianca Paglietti; Claudia Piu; Anna Puggioni; Elena Rimini; Giulia Rocca; Caterina Serra; Sergio Uzzau |
| EPI_ISL_2872591, EPI_ISL_2872596 | Laboratory Corporation of America | Centers for Disease Control and Prevention Division of Viral Diseases, Pathogen Discovery | Dakota Howard, Dhwani Batra, Peter W. Cook, Kara Moser, Adrian Paskey, Jason Caravas, Benjamin Rambo-Martin, Shatavia Morrison, Christopher Gulvick, Scott Sammons, Yvette Unoarumhi, Darlene Wagner, Matthew Schmerer, Minoo Agarwal, Eyad Almasri, Debbie Boles, Ayla Burns, Nuthawin Charoensri, Oren Cohen, Susan Countryman, Mary Ann Cristobal, Bobbi Croy, Suzanne Dale, Hrushikesh Deshmukh, Amanda Douglas, Vincent Drouillon, Marcia Eisenberg, Howard Engler, Rama Ghatti, Prashant Gupta, Susan Hicks, Jake Humphrey, Lax Iyer, Lisa Pfefferle, Manoj Jain, Matthew Robinson, Mohan Kolli, Brian Krueger, Tim Kuphal, Stanley Letovsky, Michael Levandoski, Craig Lukasik, Jonathan Meltzer, Brian Norvell, Mindy Nye, Scott Parker, Christos Petropoulos, John Pruitt, Steven Ragan, Scott Ryan, Mike Sapeta, Jana Schroth, Suresh Babu Selvaraju, Goran Stevovic, Amanda Suchanek, Andrea Throop, Lyndon Tilson, Thomas Urban, Joe Voshell, Kimberly Wagner, Jonathan Williams, Mary Williamson, Qian Zeng, Tricia Zwiefelhofer, Clinton R. Paden, Duncan MacCannell |
| EPI_ISL_2872597, EPI_ISL_2872600, EPI_ISL_2872603, EPI_ISL_2872610 | Azienda Ospedaliero Universitaria di Sassari | AMES Centro Polidiagnostico Strumentale S.r.l. | Salvatore Rubino; Flavia Angioj; Laura Firino; Rosalba Govoni; Gabriele Ibba; Vincenzo Lai; Erica Mura; Bianca Paglietti; Claudia Piu; Anna Puggioni; Elena Rimini; Giulia Rocca; Caterina Serra; Sergio Uzzau |
| EPI_ISL_2872611 | Laboratory Corporation of America | Centers for Disease Control and Prevention Division of Viral Diseases, Pathogen Discovery | Dakota Howard, Dhwani Batra, Peter W. Cook, Kara Moser, Adrian Paskey, Jason Caravas, Benjamin Rambo-Martin, Shatavia Morrison, Christopher Gulvick, Scott Sammons, Yvette Unoarumhi, Darlene Wagner, Matthew Schmerer, Minoo Agarwal, Eyad Almasri, Debbie Boles, Ayla Burns, Nuthawin Charoensri, Oren Cohen, Susan Countryman, Mary Ann Cristobal, Bobbi Croy, Suzanne Dale, Hrushikesh Deshmukh, Amanda Douglas, Vincent Drouillon, Marcia Eisenberg, Howard Engler, Rama Ghatti, Prashant Gupta, Susan Hicks, Jake Humphrey, Lax Iyer, Lisa Pfefferle, Manoj Jain, Matthew Robinson, Mohan Kolli, Brian Krueger, Tim Kuphal, Stanley Letovsky, Michael Levandoski, Craig Lukasik, Jonathan Meltzer, Brian Norvell, Mindy Nye, Scott Parker, Christos Petropoulos, John Pruitt, Steven Ragan, Scott Ryan, Mike Sapeta, Jana Schroth, Suresh Babu Selvaraju, Goran Stevovic, Amanda Suchanek, Andrea Throop, Lyndon Tilson, Thomas Urban, Joe Voshell, Kimberly Wagner, Jonathan Williams, Mary Williamson, Qian Zeng, Tricia Zwiefelhofer, Clinton R. Paden, Duncan MacCannell |
| EPI_ISL_2872612 | Azienda Ospedaliero Universitaria di Sassari | AMES Centro Polidiagnostico Strumentale S.r.l. | Salvatore Rubino; Flavia Angioj; Laura Firino; Rosalba Govoni; Gabriele Ibba; Vincenzo Lai; Erica Mura; Bianca Paglietti; Claudia Piu; Anna Puggioni; Elena Rimini; Giulia Rocca; Caterina Serra; Sergio Uzzau |
| EPI_ISL_2872613 | Laboratory Corporation of America | Centers for Disease Control and Prevention Division of Viral Diseases, Pathogen Discovery | Dakota Howard, Dhwani Batra, Peter W. Cook, Kara Moser, Adrian Paskey, Jason Caravas, Benjamin Rambo-Martin, Shatavia Morrison, Christopher Gulvick, Scott Sammons, Yvette Unoarumhi, Darlene Wagner, Matthew Schmerer, Minoo Agarwal, Eyad Almasri, Debbie Boles, Ayla Burns, Nuthawin Charoensri, Oren Cohen, Susan Countryman, Mary Ann Cristobal, Bobbi Croy, Suzanne Dale, Hrushikesh Deshmukh, Amanda Douglas, Vincent Drouillon, Marcia Eisenberg, Howard Engler, Rama Ghatti, Prashant Gupta, Susan Hicks, Jake Humphrey, Lax Iyer, Lisa Pfefferle, Manoj Jain, Matthew Robinson, Mohan Kolli, Brian Krueger, Tim Kuphal, Stanley Letovsky, Michael Levandoski, Craig Lukasik, Jonathan Meltzer, Brian Norvell, Mindy Nye, Scott Parker, Christos Petropoulos, John Pruitt, Steven Ragan, Scott Ryan, Mike Sapeta, Jana Schroth, Suresh Babu Selvaraju, Goran Stevovic, Amanda Suchanek, Andrea Throop, Lyndon Tilson, Thomas Urban, Joe Voshell, Kimberly Wagner, Jonathan Williams, Mary Williamson, Qian Zeng, Tricia Zwiefelhofer, Clinton R. Paden, Duncan MacCannell |
| EPI_ISL_2872617, EPI_ISL_2872619, EPI_ISL_2872621, EPI_ISL_2872622, EPI_ISL_2872624, EPI_ISL_2872626, EPI_ISL_2872627 | Azienda Ospedaliero Universitaria di Sassari | AMES Centro Polidiagnostico Strumentale S.r.l. | Salvatore Rubino; Flavia Angioj; Laura Firino; Rosalba Govoni; Gabriele Ibba; Vincenzo Lai; Erica Mura; Bianca Paglietti; Claudia Piu; Anna Puggioni; Elena Rimini; Giulia Rocca; Caterina Serra; Sergio Uzzau |
| EPI_ISL_2872628 | Laboratory Corporation of America | Centers for Disease Control and Prevention Division of Viral Diseases, Pathogen Discovery | Dakota Howard, Dhwani Batra, Peter W. Cook, Kara Moser, Adrian Paskey, Jason Caravas, Benjamin Rambo-Martin, Shatavia Morrison, Christopher Gulvick, Scott Sammons, Yvette Unoarumhi, Darlene Wagner, Matthew Schmerer, Minoo Agarwal, Eyad Almasri, Debbie Boles, Ayla Burns, Nuthawin Charoensri, Oren Cohen, Susan Countryman, Mary Ann Cristobal, Bobbi Croy, Suzanne Dale, Hrushikesh Deshmukh, Amanda Douglas, Vincent Drouillon, Marcia Eisenberg, Howard Engler, Rama Ghatti, Prashant Gupta, Susan Hicks, Jake Humphrey, Lax Iyer, Lisa Pfefferle, Manoj Jain, Matthew Robinson, Mohan Kolli, Brian Krueger, Tim Kuphal, Stanley Letovsky, Michael Levandoski, Craig Lukasik, Jonathan Meltzer, Brian Norvell, Mindy Nye, Scott Parker, Christos Petropoulos, John Pruitt, Steven Ragan, Scott Ryan, Mike Sapeta, Jana Schroth, Suresh Babu Selvaraju, Goran Stevovic, Amanda Suchanek, Andrea Throop, Lyndon Tilson, Thomas Urban, Joe Voshell, Kimberly Wagner, Jonathan Williams, Mary Williamson, Qian Zeng, Tricia Zwiefelhofer, Clinton R. Paden, Duncan MacCannell |
| EPI_ISL_2872629 | Azienda Ospedaliero Universitaria di Sassari | AMES Centro Polidiagnostico Strumentale S.r.l. | Salvatore Rubino; Flavia Angioj; Laura Firino; Rosalba Govoni; Gabriele Ibba; Vincenzo Lai; Erica Mura; Bianca Paglietti; Claudia Piu; Anna Puggioni; Elena Rimini; Giulia Rocca; Caterina Serra; Sergio Uzzau |
| EPI_ISL_2872630, EPI_ISL_2872639, EPI_ISL_2872644, EPI_ISL_2872646 | Laboratory Corporation of America | Centers for Disease Control and Prevention Division of Viral Diseases, Pathogen Discovery | Dakota Howard, Dhwani Batra, Peter W. Cook, Kara Moser, Adrian Paskey, Jason Caravas, Benjamin Rambo-Martin, Shatavia Morrison, Christopher Gulvick, Scott Sammons, Yvette Unoarumhi, Darlene Wagner, Matthew Schmerer, Minoo Agarwal, Eyad Almasri, Debbie Boles, Ayla Burns, Nuthawin Charoensri, Oren Cohen, Susan Countryman, Mary Ann Cristobal, Bobbi Croy, Suzanne Dale, Hrushikesh Deshmukh, Amanda Douglas, Vincent Drouillon, Marcia Eisenberg, Howard Engler, Rama Ghatti, Prashant Gupta, Susan Hicks, Jake Humphrey, Lax Iyer, Lisa Pfefferle, Manoj Jain, Matthew Robinson, Mohan Kolli, Brian Krueger, Tim Kuphal, Stanley Letovsky, Michael Levandoski, Craig Lukasik, Jonathan Meltzer, Brian Norvell, Mindy Nye, Scott Parker, Christos Petropoulos, John Pruitt, Steven Ragan, Scott Ryan, Mike Sapeta, Jana Schroth, Suresh Babu Selvaraju, Goran Stevovic, Amanda Suchanek, Andrea Throop, Lyndon Tilson, Thomas Urban, Joe Voshell, Kimberly Wagner, Jonathan Williams, Mary Williamson, Qian Zeng, Tricia Zwiefelhofer, Clinton R. Paden, Duncan MacCannell |
| EPI_ISL_2872647 | Azienda Ospedaliero Universitaria di Sassari | AMES Centro Polidiagnostico Strumentale S.r.l. | Salvatore Rubino; Flavia Angioj; Laura Firino; Rosalba Govoni; Gabriele Ibba; Vincenzo Lai; Erica Mura; Bianca Paglietti; Claudia Piu; Anna Puggioni; Elena Rimini; Giulia Rocca; Caterina Serra; Sergio Uzzau |
| EPI_ISL_2872654 | Laboratory Corporation of America | Centers for Disease Control and Prevention Division of Viral Diseases, Pathogen Discovery | Dakota Howard, Dhwani Batra, Peter W. Cook, Kara Moser, Adrian Paskey, Jason Caravas, Benjamin Rambo-Martin, Shatavia Morrison, Christopher Gulvick, Scott Sammons, Yvette Unoarumhi, Darlene Wagner, Matthew Schmerer, Minoo Agarwal, Eyad Almasri, Debbie Boles, Ayla Burns, Nuthawin Charoensri, Oren Cohen, Susan Countryman, Mary Ann Cristobal, Bobbi Croy, Suzanne Dale, Hrushikesh Deshmukh, Amanda Douglas, Vincent Drouillon, Marcia Eisenberg, Howard Engler, Rama Ghatti, Prashant Gupta, Susan Hicks, Jake Humphrey, Lax Iyer, Lisa Pfefferle, Manoj Jain, Matthew Robinson, Mohan Kolli, Brian Krueger, Tim Kuphal, Stanley Letovsky, Michael Levandoski, Craig Lukasik, Jonathan Meltzer, Brian Norvell, Mindy Nye, Scott Parker, Christos Petropoulos, John Pruitt, Steven Ragan, Scott Ryan, Mike Sapeta, Jana Schroth, Suresh Babu Selvaraju, Goran Stevovic, Amanda Suchanek, Andrea Throop, Lyndon Tilson, Thomas Urban, Joe Voshell, Kimberly Wagner, Jonathan Williams, Mary Williamson, Qian Zeng, Tricia Zwiefelhofer, Clinton R. Paden, Duncan MacCannell |
| EPI_ISL_2872656 | Azienda Ospedaliero Universitaria di Sassari | AMES Centro Polidiagnostico Strumentale S.r.l. | Salvatore Rubino; Flavia Angioj; Laura Firino; Rosalba Govoni; Gabriele Ibba; Vincenzo Lai; Erica Mura; Bianca Paglietti; Claudia Piu; Anna Puggioni; Elena Rimini; Giulia Rocca; Caterina Serra; Sergio Uzzau |
| EPI_ISL_2872657 | Laboratory Corporation of America | Centers for Disease Control and Prevention Division of Viral Diseases, Pathogen Discovery | Dakota Howard, Dhwani Batra, Peter W. Cook, Kara Moser, Adrian Paskey, Jason Caravas, Benjamin Rambo-Martin, Shatavia Morrison, Christopher Gulvick, Scott Sammons, Yvette Unoarumhi, Darlene Wagner, Matthew Schmerer, Minoo Agarwal, Eyad Almasri, Debbie Boles, Ayla Burns, Nuthawin Charoensri, Oren Cohen, Susan Countryman, Mary Ann Cristobal, Bobbi Croy, Suzanne Dale, Hrushikesh Deshmukh, Amanda Douglas, Vincent Drouillon, Marcia Eisenberg, Howard Engler, Rama Ghatti, Prashant Gupta, Susan Hicks, Jake Humphrey, Lax Iyer, Lisa Pfefferle, Manoj Jain, Matthew Robinson, Mohan Kolli, Brian Krueger, Tim Kuphal, Stanley Letovsky, Michael Levandoski, Craig Lukasik, Jonathan Meltzer, Brian Norvell, Mindy Nye, Scott Parker, Christos Petropoulos, John Pruitt, Steven Ragan, Scott Ryan, Mike Sapeta, Jana Schroth, Suresh Babu Selvaraju, Goran Stevovic, Amanda Suchanek, Andrea Throop, Lyndon Tilson, Thomas Urban, Joe Voshell, Kimberly Wagner, Jonathan Williams, Mary Williamson, Qian Zeng, Tricia Zwiefelhofer, Clinton R. Paden, Duncan MacCannell |
| EPI_ISL_2872658, EPI_ISL_2872660, EPI_ISL_2872664, EPI_ISL_2872666, EPI_ISL_2872668, EPI_ISL_2872669 | Azienda Ospedaliero Universitaria di Sassari | AMES Centro Polidiagnostico Strumentale S.r.l. | Salvatore Rubino; Flavia Angioj; Laura Firino; Rosalba Govoni; Gabriele Ibba; Vincenzo Lai; Erica Mura; Bianca Paglietti; Claudia Piu; Anna Puggioni; Elena Rimini; Giulia Rocca; Caterina Serra; Sergio Uzzau |
| EPI_ISL_2872672 | Laboratory Corporation of America | Centers for Disease Control and Prevention Division of Viral Diseases, Pathogen Discovery | Dakota Howard, Dhwani Batra, Peter W. Cook, Kara Moser, Adrian Paskey, Jason Caravas, Benjamin Rambo-Martin, Shatavia Morrison, Christopher Gulvick, Scott Sammons, Yvette Unoarumhi, Darlene Wagner, Matthew Schmerer, Minoo Agarwal, Eyad Almasri, Debbie Boles, Ayla Burns, Nuthawin Charoensri, Oren Cohen, Susan Countryman, Mary Ann Cristobal, Bobbi Croy, Suzanne Dale, Hrushikesh Deshmukh, Amanda Douglas, Vincent Drouillon, Marcia Eisenberg, Howard Engler, Rama Ghatti, Prashant Gupta, Susan Hicks, Jake Humphrey, Lax Iyer, Lisa Pfefferle, Manoj Jain, Matthew Robinson, Mohan Kolli, Brian Krueger, Tim Kuphal, Stanley Letovsky, Michael Levandoski, Craig Lukasik, Jonathan Meltzer, Brian Norvell, Mindy Nye, Scott Parker, Christos Petropoulos, John Pruitt, Steven Ragan, Scott Ryan, Mike Sapeta, Jana Schroth, Suresh Babu Selvaraju, Goran Stevovic, Amanda Suchanek, Andrea Throop, Lyndon Tilson, Thomas Urban, Joe Voshell, Kimberly Wagner, Jonathan Williams, Mary Williamson, Qian Zeng, Tricia Zwiefelhofer, Clinton R. Paden, Duncan MacCannell |
| EPI_ISL_2872675, EPI_ISL_2872683, EPI_ISL_2872685 | Azienda Ospedaliero Universitaria di Sassari | AMES Centro Polidiagnostico Strumentale S.r.l. | Salvatore Rubino; Flavia Angioj; Laura Firino; Rosalba Govoni; Gabriele Ibba; Vincenzo Lai; Erica Mura; Bianca Paglietti; Claudia Piu; Anna Puggioni; Elena Rimini; Giulia Rocca; Caterina Serra; Sergio Uzzau |
| EPI_ISL_2872692, EPI_ISL_2872697, EPI_ISL_2872699, EPI_ISL_2872702, EPI_ISL_2872704, EPI_ISL_2872715, EPI_ISL_2872716, EPI_ISL_2872717, EPI_ISL_2872718, EPI_ISL_2872719, EPI_ISL_2872720, EPI_ISL_2872721, EPI_ISL_2872724, EPI_ISL_2872725, EPI_ISL_2872727, EPI_ISL_2872728, EPI_ISL_2872729, EPI_ISL_2872730, EPI_ISL_2872731, EPI_ISL_2872733, EPI_ISL_2872734, EPI_ISL_2872737, EPI_ISL_2872738, EPI_ISL_2872739, EPI_ISL_2872740, EPI_ISL_2872741, EPI_ISL_2872742, EPI_ISL_2872743, EPI_ISL_2872744, EPI_ISL_2872745, EPI_ISL_2872746, EPI_ISL_2872747, EPI_ISL_2872748, EPI_ISL_2872749, EPI_ISL_2872750, EPI_ISL_2872751, EPI_ISL_2872752, EPI_ISL_2872754, EPI_ISL_2872755, EPI_ISL_2872756, EPI_ISL_2872757, EPI_ISL_2872758, EPI_ISL_2872759, EPI_ISL_2872760, EPI_ISL_2872762, EPI_ISL_2872763, EPI_ISL_2872764, EPI_ISL_2872765, EPI_ISL_2872766, EPI_ISL_2872767, EPI_ISL_2872768, EPI_ISL_2872769, EPI_ISL_2872775, EPI_ISL_2872779, EPI_ISL_2872780, EPI_ISL_2872783, EPI_ISL_2872784, EPI_ISL_2872788, EPI_ISL_2872790, EPI_ISL_2872792, EPI_ISL_2872793, EPI_ISL_2872794, EPI_ISL_2872796, EPI_ISL_2872797, EPI_ISL_2872798, EPI_ISL_2872799, EPI_ISL_2872801, EPI_ISL_2872803, EPI_ISL_2872805, EPI_ISL_2872812, EPI_ISL_2872813, EPI_ISL_2872814, EPI_ISL_2872816, EPI_ISL_2872829, EPI_ISL_2872830, EPI_ISL_2872832, EPI_ISL_2872833, EPI_ISL_2872840, EPI_ISL_2872845, EPI_ISL_2872847, EPI_ISL_2872852, EPI_ISL_2872853, EPI_ISL_2872855, EPI_ISL_2872858, EPI_ISL_2872861, EPI_ISL_2872863, EPI_ISL_2872864, EPI_ISL_2872876, EPI_ISL_2872880, EPI_ISL_2872882, EPI_ISL_2872884, EPI_ISL_2872890, EPI_ISL_2872891, EPI_ISL_2872897, EPI_ISL_2872898, EPI_ISL_2872901, EPI_ISL_2872902, EPI_ISL_2872903, EPI_ISL_2872906, EPI_ISL_2872911, EPI_ISL_2872915, EPI_ISL_2872920, EPI_ISL_2872922, EPI_ISL_2872925, EPI_ISL_2872926, EPI_ISL_2872928, EPI_ISL_2872933, EPI_ISL_2872949, EPI_ISL_2872952, EPI_ISL_2872953, EPI_ISL_2872955, EPI_ISL_2872956, EPI_ISL_2872959, EPI_ISL_2872961, EPI_ISL_2872964, EPI_ISL_2872965, EPI_ISL_2872967, EPI_ISL_2872968, EPI_ISL_2872970, EPI_ISL_2872972, EPI_ISL_2872973, EPI_ISL_2872977, EPI_ISL_2872979, EPI_ISL_2872982, EPI_ISL_2872984, EPI_ISL_2872989, EPI_ISL_2872990, EPI_ISL_2872996, EPI_ISL_2872999, EPI_ISL_2873000, EPI_ISL_2873003, EPI_ISL_2873005, EPI_ISL_2873007, EPI_ISL_2873008, EPI_ISL_2873010, EPI_ISL_2873011, EPI_ISL_2873012, EPI_ISL_2873013, EPI_ISL_2873014, EPI_ISL_2873016, EPI_ISL_2873017, EPI_ISL_2873019, EPI_ISL_2873021, EPI_ISL_2873024, EPI_ISL_2873025, EPI_ISL_2873026, EPI_ISL_2873028, EPI_ISL_2873032, EPI_ISL_2873033, EPI_ISL_2873039, EPI_ISL_2873043, EPI_ISL_2873044, EPI_ISL_2873046, EPI_ISL_2873048, EPI_ISL_2873052, EPI_ISL_2873054, EPI_ISL_2873057, EPI_ISL_2873061 | | | |
| see above | Laboratory Corporation of America | Centers for Disease Control and Prevention Division of Viral Diseases, Pathogen Discovery | Dakota Howard, Dhwani Batra, Peter W. Cook, Kara Moser, Adrian Paskey, Jason Caravas, Benjamin Rambo-Martin, Shatavia Morrison, Christopher Gulvick, Scott Sammons, Yvette Unoarumhi, Darlene Wagner, Matthew Schmerer, Minoo Agarwal, Eyad Almasri, Debbie Boles, Ayla Burns, Nuthawin Charoensri, Oren Cohen, Susan Countryman, Mary Ann Cristobal, Bobbi Croy, Suzanne Dale, Hrushikesh Deshmukh, Amanda Douglas, Vincent Drouillon, Marcia Eisenberg, Howard Engler, Rama Ghatti, Prashant Gupta, Susan Hicks, Jake Humphrey, Lax Iyer, Lisa Pfefferle, Manoj Jain, Matthew Robinson, Mohan Kolli, Brian Krueger, Tim Kuphal, Stanley Letovsky, Michael Levandoski, Craig Lukasik, Jonathan Meltzer, Brian Norvell, Mindy Nye, Scott Parker, Christos Petropoulos, John Pruitt, Steven Ragan, Scott Ryan, Mike Sapeta, Jana Schroth, Suresh Babu Selvaraju, Goran Stevovic, Amanda Suchanek, Andrea Throop, Lyndon Tilson, Thomas Urban, Joe Voshell, Kimberly Wagner, Jonathan Williams, Mary Williamson, Qian Zeng, Tricia Zwiefelhofer, Clinton R. Paden, Duncan MacCannell |
| EPI_ISL_2873062 | ASST MONZA | ASST MONZA | Sergio Maria Ivano Malandrin |
| EPI_ISL_2873063 | Laboratory Corporation of America | Centers for Disease Control and Prevention Division of Viral Diseases, Pathogen Discovery | Dakota Howard, Dhwani Batra, Peter W. Cook, Kara Moser, Adrian Paskey, Jason Caravas, Benjamin Rambo-Martin, Shatavia Morrison, Christopher Gulvick, Scott Sammons, Yvette Unoarumhi, Darlene Wagner, Matthew Schmerer, Minoo Agarwal, Eyad Almasri, Debbie Boles, Ayla Burns, Nuthawin Charoensri, Oren Cohen, Susan Countryman, Mary Ann Cristobal, Bobbi Croy, Suzanne Dale, Hrushikesh Deshmukh, Amanda Douglas, Vincent Drouillon, Marcia Eisenberg, Howard |

| | | | |
|---|---|---|---|
| | | | Engler, Rama Ghatti, Prashant Gupta, Susan Hicks, Jake Humphrey, Lax Iyer, Lisa Pfefferle, Manoj Jain, Matthew Robinson, Mohan Kolli, Brian Krueger, Tim Kuphal, Stanley Letovsky, Michael Levandoski, Craig Lukasik, Jonathan Meltzer, Brian Norvell, Mindy Nye, Scott Parker, Christos Petropoulos, John Pruitt, Steven Ragan, Scott Ryan, Mike Sapeta, Jana Schroth, Suresh Babu Selvaraju, Goran Stevovic, Amanda Suchanek, Andrea Throop, Lyndon Tilson, Thomas Urban, Joe Voshell, Kimberly Wagner, Jonathan Williams, Mary Williamson, Qian Zeng, Tricia Zwiefelhofer, Clinton R. Paden, Duncan MacCannell |
| EPI_ISL_2873064, EPI_ISL_2873065 | ASST MONZA | ASST MONZA | Sergio Maria Ivano Malandrin |
| EPI_ISL_2873066 | Laboratory Corporation of America | Centers for Disease Control and Prevention Division of Viral Diseases, Pathogen Discovery | Dakota Howard, Dhwani Batra, Peter W. Cook, Kara Moser, Adrian Paskey, Jason Caravas, Benjamin Rambo-Martin, Shatavia Morrison, Christopher Gulvick, Scott Sammons, Yvette Unoarumhi, Darlene Wagner, Matthew Schmerer, Minoo Agarwal, Eyad Almasri, Debbie Boles, Ayla Burns, Nuthawin Charoensri, Oren Cohen, Susan Countryman, Mary Ann Cristobal, Bobbi Croy, Suzanne Dale, Hrushikesh Deshmukh, Amanda Douglas, Vincent Drouillon, Marcia Eisenberg, Howard Engler, Rama Ghatti, Prashant Gupta, Susan Hicks, Jake Humphrey, Lax Iyer, Lisa Pfefferle, Manoj Jain, Matthew Robinson, Mohan Kolli, Brian Krueger, Tim Kuphal, Stanley Letovsky, Michael Levandoski, Craig Lukasik, Jonathan Meltzer, Brian Norvell, Mindy Nye, Scott Parker, Christos Petropoulos, John Pruitt, Steven Ragan, Scott Ryan, Mike Sapeta, Jana Schroth, Suresh Babu Selvaraju, Goran Stevovic, Amanda Suchanek, Andrea Throop, Lyndon Tilson, Thomas Urban, Joe Voshell, Kimberly Wagner, Jonathan Williams, Mary Williamson, Qian Zeng, Tricia Zwiefelhofer, Clinton R. Paden, Duncan MacCannell |
| EPI_ISL_2873067 | ASST MONZA | ASST MONZA | Sergio Maria Ivano Malandrin |
| EPI_ISL_2873068 | Laboratory Corporation of America | Centers for Disease Control and Prevention Division of Viral Diseases, Pathogen Discovery | Dakota Howard, Dhwani Batra, Peter W. Cook, Kara Moser, Adrian Paskey, Jason Caravas, Benjamin Rambo-Martin, Shatavia Morrison, Christopher Gulvick, Scott Sammons, Yvette Unoarumhi, Darlene Wagner, Matthew Schmerer, Minoo Agarwal, Eyad Almasri, Debbie Boles, Ayla Burns, Nuthawin Charoensri, Oren Cohen, Susan Countryman, Mary Ann Cristobal, Bobbi Croy, Suzanne Dale, Hrushikesh Deshmukh, Amanda Douglas, Vincent Drouillon, Marcia Eisenberg, Howard Engler, Rama Ghatti, Prashant Gupta, Susan Hicks, Jake Humphrey, Lax Iyer, Lisa Pfefferle, Manoj Jain, Matthew Robinson, Mohan Kolli, Brian Krueger, Tim Kuphal, Stanley Letovsky, Michael Levandoski, Craig Lukasik, Jonathan Meltzer, Brian Norvell, Mindy Nye, Scott Parker, Christos Petropoulos, John Pruitt, Steven Ragan, Scott Ryan, Mike Sapeta, Jana Schroth, Suresh Babu Selvaraju, Goran Stevovic, Amanda Suchanek, Andrea Throop, Lyndon Tilson, Thomas Urban, Joe Voshell, Kimberly Wagner, Jonathan Williams, Mary Williamson, Qian Zeng, Tricia Zwiefelhofer, Clinton R. Paden, Duncan MacCannell |
| EPI_ISL_2873070, EPI_ISL_2873071 | ASST MONZA | ASST MONZA | Sergio Maria Ivano Malandrin |
| EPI_ISL_2873072, EPI_ISL_2873074 | Laboratory Corporation of America | Centers for Disease Control and Prevention Division of Viral Diseases, Pathogen Discovery | Dakota Howard, Dhwani Batra, Peter W. Cook, Kara Moser, Adrian Paskey, Jason Caravas, Benjamin Rambo-Martin, Shatavia Morrison, Christopher Gulvick, Scott Sammons, Yvette Unoarumhi, Darlene Wagner, Matthew Schmerer, Minoo Agarwal, Eyad Almasri, Debbie Boles, Ayla Burns, Nuthawin Charoensri, Oren Cohen, Susan Countryman, Mary Ann Cristobal, Bobbi Croy, Suzanne Dale, Hrushikesh Deshmukh, Amanda Douglas, Vincent Drouillon, Marcia Eisenberg, Howard Engler, Rama Ghatti, Prashant Gupta, Susan Hicks, Jake Humphrey, Lax Iyer, Lisa Pfefferle, Manoj Jain, Matthew Robinson, Mohan Kolli, Brian Krueger, Tim Kuphal, Stanley Letovsky, Michael Levandoski, Craig Lukasik, Jonathan Meltzer, Brian Norvell, Mindy Nye, Scott Parker, Christos Petropoulos, John Pruitt, Steven Ragan, Scott Ryan, Mike Sapeta, Jana Schroth, Suresh Babu Selvaraju, Goran Stevovic, Amanda Suchanek, Andrea Throop, Lyndon Tilson, Thomas Urban, Joe Voshell, Kimberly Wagner, Jonathan Williams, Mary Williamson, Qian Zeng, Tricia Zwiefelhofer, Clinton R. Paden, Duncan MacCannell |
| EPI_ISL_2873075 | ASST MONZA | ASST MONZA | Sergio Maria Ivano Malandrin |
| EPI_ISL_2873076 | Laboratory Corporation of America | Centers for Disease Control and Prevention Division of Viral Diseases, Pathogen Discovery | Dakota Howard, Dhwani Batra, Peter W. Cook, Kara Moser, Adrian Paskey, Jason Caravas, Benjamin Rambo-Martin, Shatavia Morrison, Christopher Gulvick, Scott Sammons, Yvette Unoarumhi, Darlene Wagner, Matthew Schmerer, Minoo Agarwal, Eyad Almasri, Debbie Boles, Ayla Burns, Nuthawin Charoensri, Oren Cohen, Susan Countryman, Mary Ann Cristobal, Bobbi Croy, Suzanne Dale, Hrushikesh Deshmukh, Amanda Douglas, Vincent Drouillon, Marcia Eisenberg, Howard Engler, Rama Ghatti, Prashant Gupta, Susan Hicks, Jake Humphrey, Lax Iyer, Lisa Pfefferle, Manoj Jain, Matthew Robinson, Mohan Kolli, Brian Krueger, Tim Kuphal, Stanley Letovsky, Michael Levandoski, Craig Lukasik, Jonathan Meltzer, Brian Norvell, Mindy Nye, Scott Parker, Christos Petropoulos, John Pruitt, Steven Ragan, Scott Ryan, Mike Sapeta, Jana Schroth, Suresh Babu Selvaraju, Goran Stevovic, Amanda Suchanek, Andrea Throop, Lyndon Tilson, Thomas Urban, Joe Voshell, Kimberly Wagner, Jonathan Williams, Mary Williamson, Qian Zeng, Tricia Zwiefelhofer, Clinton R. Paden, Duncan MacCannell |
| EPI_ISL_2873077, EPI_ISL_2873079, EPI_ISL_2873081, EPI_ISL_2873083, EPI_ISL_2873085 | ASST MONZA | ASST MONZA | Sergio Maria Ivano Malandrin |
| EPI_ISL_2873086 | Laboratory Corporation of America | Centers for Disease Control and Prevention Division of Viral Diseases, Pathogen Discovery | Dakota Howard, Dhwani Batra, Peter W. Cook, Kara Moser, Adrian Paskey, Jason Caravas, Benjamin Rambo-Martin, Shatavia Morrison, Christopher Gulvick, Scott Sammons, Yvette Unoarumhi, Darlene Wagner, Matthew Schmerer, Minoo Agarwal, Eyad Almasri, Debbie Boles, Ayla Burns, Nuthawin Charoensri, Oren Cohen, Susan Countryman, Mary Ann Cristobal, Bobbi Croy, Suzanne Dale, Hrushikesh Deshmukh, Amanda Douglas, Vincent Drouillon, Marcia Eisenberg, Howard Engler, Rama Ghatti, Prashant Gupta, Susan Hicks, Jake Humphrey, Lax Iyer, Lisa Pfefferle, Manoj Jain, Matthew Robinson, Mohan Kolli, Brian Krueger, Tim Kuphal, Stanley Letovsky, Michael Levandoski, Craig Lukasik, Jonathan Meltzer, Brian Norvell, Mindy Nye, Scott Parker, Christos Petropoulos, John Pruitt, Steven Ragan, Scott Ryan, Mike Sapeta, Jana Schroth, Suresh Babu Selvaraju, Goran Stevovic, Amanda Suchanek, Andrea Throop, Lyndon Tilson, Thomas Urban, Joe Voshell, Kimberly Wagner, Jonathan Williams, Mary Williamson, Qian Zeng, Tricia Zwiefelhofer, Clinton R. Paden, Duncan MacCannell |
| EPI_ISL_2873087, EPI_ISL_2873089 | ASST MONZA | ASST MONZA | Sergio Maria Ivano Malandrin |
| EPI_ISL_2873090 | Laboratory Corporation of America | Centers for Disease Control and Prevention Division of Viral Diseases, Pathogen Discovery | Dakota Howard, Dhwani Batra, Peter W. Cook, Kara Moser, Adrian Paskey, Jason Caravas, Benjamin Rambo-Martin, Shatavia Morrison, Christopher Gulvick, Scott Sammons, Yvette Unoarumhi, Darlene Wagner, Matthew Schmerer, Minoo Agarwal, Eyad Almasri, Debbie Boles, Ayla Burns, Nuthawin Charoensri, Oren Cohen, Susan Countryman, Mary Ann Cristobal, Bobbi Croy, Suzanne Dale, Hrushikesh Deshmukh, Amanda Douglas, Vincent Drouillon, Marcia Eisenberg, Howard Engler, Rama Ghatti, Prashant Gupta, Susan Hicks, Jake Humphrey, Lax Iyer, Lisa Pfefferle, Manoj Jain, Matthew Robinson, Mohan Kolli, Brian Krueger, Tim Kuphal, Stanley Letovsky, Michael Levandoski, Craig Lukasik, Jonathan Meltzer, Brian Norvell, Mindy Nye, Scott Parker, Christos Petropoulos, John Pruitt, Steven Ragan, Scott Ryan, Mike Sapeta, Jana Schroth, Suresh Babu Selvaraju, Goran Stevovic, Amanda Suchanek, Andrea Throop, Lyndon Tilson, Thomas Urban, Joe Voshell, Kimberly Wagner, Jonathan Williams, Mary Williamson, Qian Zeng, Tricia Zwiefelhofer, Clinton R. Paden, Duncan MacCannell |
| EPI_ISL_2873091, EPI_ISL_2873093, EPI_ISL_2873095, EPI_ISL_2873097, EPI_ISL_2873099, EPI_ISL_2873101, EPI_ISL_2873103, EPI_ISL_2873105, EPI_ISL_2873107 | ASST MONZA | ASST MONZA | Sergio Maria Ivano Malandrin |
| EPI_ISL_2873110, EPI_ISL_2873116, EPI_ISL_2873117, EPI_ISL_2873118, EPI_ISL_2873129, EPI_ISL_2873130, EPI_ISL_2873131, EPI_ISL_2873132, EPI_ISL_2873133, EPI_ISL_2873135, EPI_ISL_2873136, EPI_ISL_2873137, EPI_ISL_2873138, EPI_ISL_2873141, EPI_ISL_2873144, EPI_ISL_2873145, EPI_ISL_2873146, EPI_ISL_2873150, EPI_ISL_2873157, EPI_ISL_2873158, EPI_ISL_2873166, EPI_ISL_2873170, EPI_ISL_2873179, EPI_ISL_2873188, EPI_ISL_2873197, EPI_ISL_2873202, EPI_ISL_2873203, EPI_ISL_2873204, EPI_ISL_2873207, EPI_ISL_2873210, EPI_ISL_2873213, EPI_ISL_2873214, EPI_ISL_2873216, EPI_ISL_2873218, EPI_ISL_2873224, EPI_ISL_2873225, EPI_ISL_2873226, EPI_ISL_2873228, EPI_ISL_2873231, EPI_ISL_2873233, EPI_ISL_2873234, EPI_ISL_2873238, EPI_ISL_2873240, EPI_ISL_2873243, EPI_ISL_2873246, EPI_ISL_2873248, EPI_ISL_2873249, EPI_ISL_2873251, EPI_ISL_2873252, EPI_ISL_2873258, EPI_ISL_2873259, EPI_ISL_2873260, EPI_ISL_2873262, EPI_ISL_2873263, EPI_ISL_2873264, EPI_ISL_2873265, EPI_ISL_2873267, EPI_ISL_2873272, EPI_ISL_2873280, EPI_ISL_2873281, EPI_ISL_2873283, EPI_ISL_2873285, EPI_ISL_2873286, EPI_ISL_2873287, EPI_ISL_2873289, EPI_ISL_2873291, EPI_ISL_2873294, EPI_ISL_2873298, EPI_ISL_2873299, EPI_ISL_2873300, EPI_ISL_2873301, EPI_ISL_2873302, EPI_ISL_2873303, EPI_ISL_2873308, EPI_ISL_2873309, EPI_ISL_2873310, EPI_ISL_2873311, EPI_ISL_2873312, EPI_ISL_2873314, EPI_ISL_2873315, EPI_ISL_2873316, EPI_ISL_2873321, EPI_ISL_2873323, EPI_ISL_2873325 | | | |
| see above | Laboratory Corporation of America | Centers for Disease Control and Prevention Division of Viral Diseases, Pathogen Discovery | Dakota Howard, Dhwani Batra, Peter W. Cook, Kara Moser, Adrian Paskey, Jason Caravas, Benjamin Rambo-Martin, Shatavia Morrison, Christopher Gulvick, Scott Sammons, Yvette Unoarumhi, Darlene Wagner, Matthew Schmerer, Minoo Agarwal, Eyad Almasri, Debbie Boles, Ayla Burns, Nuthawin Charoensri, Oren Cohen, Susan Countryman, Mary Ann Cristobal, Bobbi Croy, Suzanne Dale, Hrushikesh Deshmukh, Amanda Douglas, Vincent Drouillon, Marcia Eisenberg, Howard Engler, Rama Ghatti, Prashant Gupta, Susan Hicks, Jake Humphrey, Lax Iyer, Lisa Pfefferle, Manoj Jain, Matthew Robinson, Mohan Kolli, Brian Krueger, Tim Kuphal, Stanley Letovsky, Michael Levandoski, Craig Lukasik, Jonathan Meltzer, Brian Norvell, Mindy Nye, Scott Parker, Christos Petropoulos, John Pruitt, Steven Ragan, Scott Ryan, Mike Sapeta, Jana Schroth, Suresh Babu Selvaraju, Goran Stevovic, Amanda Suchanek, Andrea Throop, Lyndon Tilson, Thomas Urban, Joe Voshell, Kimberly Wagner, Jonathan Williams, Mary Williamson, Qian Zeng, Tricia Zwiefelhofer, Clinton R. Paden, Duncan MacCannell |
| EPI_ISL_2873326, EPI_ISL_2873327, EPI_ISL_2873328, EPI_ISL_2873329, EPI_ISL_2873330, EPI_ISL_2873331, EPI_ISL_2873332, EPI_ISL_2873334, EPI_ISL_2873335, EPI_ISL_2873336, EPI_ISL_2873337, EPI_ISL_2873338, EPI_ISL_2873340, EPI_ISL_2873341, EPI_ISL_2873344, EPI_ISL_2873345, EPI_ISL_2873346, EPI_ISL_2873347, EPI_ISL_2873348, EPI_ISL_2873349, EPI_ISL_2873350, EPI_ISL_2873351, EPI_ISL_2873352, EPI_ISL_2873353, EPI_ISL_2873354, EPI_ISL_2873355, EPI_ISL_2873357, EPI_ISL_2873358, EPI_ISL_2873359, EPI_ISL_2873360, EPI_ISL_2873361, EPI_ISL_2873362, EPI_ISL_2873364, EPI_ISL_2873365, EPI_ISL_2873366, EPI_ISL_2873367, EPI_ISL_2873368, EPI_ISL_2873372, EPI_ISL_2873374, EPI_ISL_2873375, EPI_ISL_2873376, EPI_ISL_2873378, EPI_ISL_2873379, EPI_ISL_2873380, EPI_ISL_2873381, EPI_ISL_2873383, EPI_ISL_2873384, EPI_ISL_2873385, EPI_ISL_2873386, EPI_ISL_2873387, EPI_ISL_2873388, EPI_ISL_2873389, EPI_ISL_2873391, EPI_ISL_2873392, EPI_ISL_2873393, EPI_ISL_2873394, EPI_ISL_2873395, EPI_ISL_2873396, EPI_ISL_2873397, EPI_ISL_2873398, EPI_ISL_2873399, EPI_ISL_2873400, EPI_ISL_2873402, EPI_ISL_2873403, EPI_ISL_2873406, EPI_ISL_2873407, EPI_ISL_2873408, EPI_ISL_2873411, EPI_ISL_2873414, EPI_ISL_2873416, EPI_ISL_2873417, EPI_ISL_2873418, EPI_ISL_2873419, EPI_ISL_2873425, EPI_ISL_2873430, EPI_ISL_2873431, EPI_ISL_2873432, EPI_ISL_2873435, EPI_ISL_2873436, EPI_ISL_2873437, EPI_ISL_2873438, EPI_ISL_2873439, EPI_ISL_2873440, EPI_ISL_2873441, EPI_ISL_2873445, EPI_ISL_2873446, EPI_ISL_2873447, EPI_ISL_2873449, EPI_ISL_2873450, EPI_ISL_2873451, EPI_ISL_2873453, EPI_ISL_2873454, EPI_ISL_2873455, EPI_ISL_2873456, EPI_ISL_2873459, EPI_ISL_2873460, EPI_ISL_2873462, EPI_ISL_2873465, EPI_ISL_2873467, EPI_ISL_2873469, EPI_ISL_2873471, EPI_ISL_2873472, EPI_ISL_2873473, EPI_ISL_2873475, EPI_ISL_2873476, EPI_ISL_2873477, EPI_ISL_2873478, EPI_ISL_2873480, EPI_ISL_2873481, EPI_ISL_2873482, EPI_ISL_2873483, EPI_ISL_2873485, EPI_ISL_2873486, EPI_ISL_2873487, EPI_ISL_2873488, EPI_ISL_2873494, EPI_ISL_2873500, EPI_ISL_2873501, EPI_ISL_2873502, EPI_ISL_2873504, EPI_ISL_2873505, EPI_ISL_2873506, EPI_ISL_2873509, EPI_ISL_2873510, EPI_ISL_2873511, EPI_ISL_2873512, EPI_ISL_2873515, EPI_ISL_2873516, EPI_ISL_2873518, EPI_ISL_2873519, EPI_ISL_2873520, EPI_ISL_2873521, EPI_ISL_2873522, EPI_ISL_2873523, EPI_ISL_2873524, EPI_ISL_2873526, EPI_ISL_2873528, EPI_ISL_2873529, EPI_ISL_2873530, EPI_ISL_2873532, EPI_ISL_2873533, EPI_ISL_2873534, EPI_ISL_2873537, EPI_ISL_2873539, EPI_ISL_2873540, EPI_ISL_2873541, EPI_ISL_2873542, EPI_ISL_2873543, EPI_ISL_2873544, EPI_ISL_2873545, EPI_ISL_2873546, EPI_ISL_2873547, EPI_ISL_2873548, EPI_ISL_2873549, EPI_ISL_2873550, EPI_ISL_2873552, EPI_ISL_2873554, EPI_ISL_2873556, EPI_ISL_2873558, EPI_ISL_2873560, EPI_ISL_2873561, EPI_ISL_2873562, EPI_ISL_2873563, EPI_ISL_2873565, EPI_ISL_2873566, EPI_ISL_2873567, EPI_ISL_2873570, EPI_ISL_2873571, EPI_ISL_2873572, EPI_ISL_2873574, EPI_ISL_2873576, EPI_ISL_2873577, EPI_ISL_2873578, EPI_ISL_2873579, EPI_ISL_2873580, EPI_ISL_2873581, EPI_ISL_2873582, EPI_ISL_2873583, EPI_ISL_2873584, EPI_ISL_2873585, EPI_ISL_2873589, EPI_ISL_2873590, EPI_ISL_2873591, EPI_ISL_2873592, EPI_ISL_2873594, EPI_ISL_2873595, EPI_ISL_2873596, EPI_ISL_2873597, EPI_ISL_2873598, EPI_ISL_2873599, EPI_ISL_2873600, EPI_ISL_2873602, EPI_ISL_2873603, EPI_ISL_2873604, EPI_ISL_2873607, EPI_ISL_2873608, EPI_ISL_2873609, EPI_ISL_2873610, EPI_ISL_2873611, EPI_ISL_2873612, EPI_ISL_2873615, EPI_ISL_2873616, EPI_ISL_2873618, EPI_ISL_2873620, EPI_ISL_2873621, EPI_ISL_2873622, EPI_ISL_2873623, EPI_ISL_2873624, EPI_ISL_2873626, EPI_ISL_2873627, EPI_ISL_2873628, EPI_ISL_2873630, EPI_ISL_2873632, EPI_ISL_2873633, EPI_ISL_2873634, EPI_ISL_2873635, EPI_ISL_2873637, EPI_ISL_2873638, EPI_ISL_2873641, EPI_ISL_2873643, EPI_ISL_2873645, EPI_ISL_2873649, EPI_ISL_2873650, EPI_ISL_2873651, EPI_ISL_2873652, EPI_ISL_2873654, EPI_ISL_2873655, EPI_ISL_2873656, EPI_ISL_2873657, EPI_ISL_2873659, EPI_ISL_2873660, EPI_ISL_2873661, EPI_ISL_2873662, EPI_ISL_2873663, EPI_ISL_2873664, EPI_ISL_2873665, EPI_ISL_2873666, EPI_ISL_2873667, EPI_ISL_2873668, EPI_ISL_2873670, EPI_ISL_2873671, EPI_ISL_2873672, EPI_ISL_2873673, EPI_ISL_2873674, EPI_ISL_2873676, EPI_ISL_2873677, EPI_ISL_2873679, EPI_ISL_2873682, EPI_ISL_2873683, EPI_ISL_2873684, EPI_ISL_2873685, EPI_ISL_2873687, EPI_ISL_2873689, EPI_ISL_2873691, EPI_ISL_2873693, EPI_ISL_2873696, EPI_ISL_2873697, EPI_ISL_2873698, EPI_ISL_2873699, EPI_ISL_2873700, EPI_ISL_2873701, EPI_ISL_2873702, EPI_ISL_2873703, EPI_ISL_2873704, EPI_ISL_2873705, EPI_ISL_2873706, EPI_ISL_2873708, EPI_ISL_2873710, EPI_ISL_2873713, EPI_ISL_2873714, EPI_ISL_2873715, EPI_ISL_2873716, EPI_ISL_2873717, EPI_ISL_2873718, EPI_ISL_2873719, EPI_ISL_2873720, EPI_ISL_2873723, EPI_ISL_2873724, EPI_ISL_2873725, EPI_ISL_2873727, EPI_ISL_2873728, EPI_ISL_2873729, EPI_ISL_2873731, EPI_ISL_2873732, EPI_ISL_2873733, EPI_ISL_2873734, EPI_ISL_2873735, EPI_ISL_2873737, EPI_ISL_2873740, EPI_ISL_2873743, EPI_ISL_2873744, EPI_ISL_2873745, EPI_ISL_2873746, EPI_ISL_2873747, EPI_ISL_2873748, EPI_ISL_2873749, EPI_ISL_2873751, EPI_ISL_2873752, EPI_ISL_2873753, EPI_ISL_2873754, EPI_ISL_2873755, EPI_ISL_2873756, EPI_ISL_2873757, EPI_ISL_2873759, EPI_ISL_2873760, EPI_ISL_2873761, EPI_ISL_2873763, EPI_ISL_2873764, EPI_ISL_2873766, EPI_ISL_2873768, EPI_ISL_2873769, EPI_ISL_2873772, EPI_ISL_2873774, EPI_ISL_2873775, EPI_ISL_2873776, EPI_ISL_2873777, EPI_ISL_2873778, EPI_ISL_2873779, EPI_ISL_2873780, EPI_ISL_2873781, EPI_ISL_2873783, EPI_ISL_2873784, EPI_ISL_2873785, EPI_ISL_2873786, EPI_ISL_2873787, EPI_ISL_2873788, EPI_ISL_2873789, EPI_ISL_2873791, EPI_ISL_2873792, EPI_ISL_2873795, EPI_ISL_2873797, EPI_ISL_2873799, EPI_ISL_2873806, EPI_ISL_2873807, EPI_ISL_2873809, EPI_ISL_2873812, EPI_ISL_2873814 | | | |
| see above | Fulgent Genetics | Centers for Disease Control and Prevention Division of Viral Diseases, Pathogen Discovery | Dakota Howard, Dhwani Batra, Peter W. Cook, Kara Moser, Adrian Paskey, Jason Caravas, Benjamin Rambo-Martin, Shatavia Morrison, Christopher Gulvick, Scott Sammons, Yvette Unoarumhi, Darlene Wagner, Matthew Schmerer, Harry Gao, Mickey Li, John Gao, Joseph Fierro, Benafsh Sapra, Becky Tsai, Yan Meng, Doreen Ng, James Xie, Clinton R. Paden, Duncan MacCannell |
| EPI_ISL_2873828, EPI_ISL_2873829, EPI_ISL_2873830, EPI_ISL_2873831, EPI_ISL_2873833, EPI_ISL_2873834, EPI_ISL_2873835 | Iressef Genomics lab | IRESSEF | Souleymane MBOUP, Abdou PADANE , Khadim GUEYE, Diabou Diagne , Samba Ndiour, Papa Alassane DIAW, Birahim Piere NDIAYE, Barada CISSE, Aminata MBOUP, Moustapha MBOW, Ndeye Coumba Toure KANE, Nafisatou LEYE, Gora LO, Ambroise AHOUIDI , Astou Gaye GAYE, Aminata DIA, Yacine DIA,Seni Ndiaye |
| EPI_ISL_2873839 | Lafene Health Center | Kansas State Veterinary Diagnostic Laboratory | Tyler Doerksen, Andreea Lu, Lance Noll, Elizabeth Porter, Emily Cox, Jianfa Bai, Jamie Henningson, Rachel Palinski |
| EPI_ISL_2873874, EPI_ISL_2873875, EPI_ISL_2873876, EPI_ISL_2873877, EPI_ISL_2873878, EPI_ISL_2873879, EPI_ISL_2873881, EPI_ISL_2873883, EPI_ISL_2873884, EPI_ISL_2873887, EPI_ISL_2873888, EPI_ISL_2873889, EPI_ISL_2873890, EPI_ISL_2873892, EPI_ISL_2873893, EPI_ISL_2873896, EPI_ISL_2873899, EPI_ISL_2873900, EPI_ISL_2873901, EPI_ISL_2873902, EPI_ISL_2873903, EPI_ISL_2873904, EPI_ISL_2873906, EPI_ISL_2873907, EPI_ISL_2873908, EPI_ISL_2873909, EPI_ISL_2873910, EPI_ISL_2873911, EPI_ISL_2873912, EPI_ISL_2873913, EPI_ISL_2873914, EPI_ISL_2873915, EPI_ISL_2873916, EPI_ISL_2873917, EPI_ISL_2873919, EPI_ISL_2873920, EPI_ISL_2873921, EPI_ISL_2873922, EPI_ISL_2873924, EPI_ISL_2873925, EPI_ISL_2873926, EPI_ISL_2873927, EPI_ISL_2873929, EPI_ISL_2873930, EPI_ISL_2873931, EPI_ISL_2873932, EPI_ISL_2873933, EPI_ISL_2873935, EPI_ISL_2873937, EPI_ISL_2873939, EPI_ISL_2873942, EPI_ISL_2873943, EPI_ISL_2873945, EPI_ISL_2873946, EPI_ISL_2873948, EPI_ISL_2873950, EPI_ISL_2873951, EPI_ISL_2873952, EPI_ISL_2873953, EPI_ISL_2873954, EPI_ISL_2873955, EPI_ISL_2873960, EPI_ISL_2873961, EPI_ISL_2873962, EPI_ISL_2873963, EPI_ISL_2873964, EPI_ISL_2873965, EPI_ISL_2873966, EPI_ISL_2873967, EPI_ISL_2873968, EPI_ISL_2873969, EPI_ISL_2873970, EPI_ISL_2873971, EPI_ISL_2873972, EPI_ISL_2873974, EPI_ISL_2873975, EPI_ISL_2873976, EPI_ISL_2873978, EPI_ISL_2873980, EPI_ISL_2873981, EPI_ISL_2873983, EPI_ISL_2873984, EPI_ISL_2873986, EPI_ISL_2873987, EPI_ISL_2873989, EPI_ISL_2873990, EPI_ISL_2873993, EPI_ISL_2873994, EPI_ISL_2873995, EPI_ISL_2873996, EPI_ISL_2873998, EPI_ISL_2873999, EPI_ISL_2874000, EPI_ISL_2874003, EPI_ISL_2874004, EPI_ISL_2874005, EPI_ISL_2874008, EPI_ISL_2874009, EPI_ISL_2874010, EPI_ISL_2874012, EPI_ISL_2874013, EPI_ISL_2874014, EPI_ISL_2874015, EPI_ISL_2874016, EPI_ISL_2874017, EPI_ISL_2874018, EPI_ISL_2874019, EPI_ISL_2874021, EPI_ISL_2874023, EPI_ISL_2874024, EPI_ISL_2874025, EPI_ISL_2874027, EPI_ISL_2874029, EPI_ISL_2874030, EPI_ISL_2874031, EPI_ISL_2874033, EPI_ISL_2874038, EPI_ISL_2874039, EPI_ISL_2874042, EPI_ISL_2874045, EPI_ISL_2874047, EPI_ISL_2874049, EPI_ISL_2874050, EPI_ISL_2874051, EPI_ISL_2874052, EPI_ISL_2874054, EPI_ISL_2874055, EPI_ISL_2874057, EPI_ISL_2874058, EPI_ISL_2874059, EPI_ISL_2874061, EPI_ISL_2874063, EPI_ISL_2874064, EPI_ISL_2874065, EPI_ISL_2874067, EPI_ISL_2874069, EPI_ISL_2874070, EPI_ISL_2874071, EPI_ISL_2874072, EPI_ISL_2874073, EPI_ISL_2874074, EPI_ISL_2874075, EPI_ISL_2874076, EPI_ISL_2874078, EPI_ISL_2874079, EPI_ISL_2874083, EPI_ISL_2874084, EPI_ISL_2874085, EPI_ISL_2874086, EPI_ISL_2874088, EPI_ISL_2874089, EPI_ISL_2874090, EPI_ISL_2874093, EPI_ISL_2874094, EPI_ISL_2874098, EPI_ISL_2874100, EPI_ISL_2874103, EPI_ISL_2874104, EPI_ISL_2874107, EPI_ISL_2874108, EPI_ISL_2874109, EPI_ISL_2874110, EPI_ISL_2874111, EPI_ISL_2874112, EPI_ISL_2874114, EPI_ISL_2874115, EPI_ISL_2874117, EPI_ISL_2874118, EPI_ISL_2874119, EPI_ISL_2874121, EPI_ISL_2874123, EPI_ISL_2874125, EPI_ISL_2874126, EPI_ISL_2874128, EPI_ISL_2874129, EPI_ISL_2874131, EPI_ISL_2874132, EPI_ISL_2874133, EPI_ISL_2874135, EPI_ISL_2874136, EPI_ISL_2874137, EPI_ISL_2874138, EPI_ISL_2874139, EPI_ISL_2874140, EPI_ISL_2874143, EPI_ISL_2874144, EPI_ISL_2874145, EPI_ISL_2874146, EPI_ISL_2874148, EPI_ISL_2874150, EPI_ISL_2874151, EPI_ISL_2874152, EPI_ISL_2874153, EPI_ISL_2874154, EPI_ISL_2874158, EPI_ISL_2874159, EPI_ISL_2874160, EPI_ISL_2874161, EPI_ISL_2874165, EPI_ISL_2874166, EPI_ISL_2874168, EPI_ISL_2874169, EPI_ISL_2874171, EPI_ISL_2874172, EPI_ISL_2874174, EPI_ISL_2874177, EPI_ISL_2874178, EPI_ISL_2874180, EPI_ISL_2874181, EPI_ISL_2874186, EPI_ISL_2874187, EPI_ISL_2874190, EPI_ISL_2874191, EPI_ISL_2874193, EPI_ISL_2874194, EPI_ISL_2874195, EPI_ISL_2874196, EPI_ISL_2874197, EPI_ISL_2874199, EPI_ISL_2874201, EPI_ISL_2874203, EPI_ISL_2874204, EPI_ISL_2874206, EPI_ISL_2874208, EPI_ISL_2874209, EPI_ISL_2874210, EPI_ISL_2874211, EPI_ISL_2874212, EPI_ISL_2874214, EPI_ISL_2874215, EPI_ISL_2874216, EPI_ISL_2874218, EPI_ISL_2874219, EPI_ISL_2874220, EPI_ISL_2874221, EPI_ISL_2874222, EPI_ISL_2874223, EPI_ISL_2874225, EPI_ISL_2874226, EPI_ISL_2874228, EPI_ISL_2874230, EPI_ISL_2874231, | | | |

| | | | |
|---|---|---|---|
| EPI_ISL_2874233, EPI_ISL_2874234, EPI_ISL_2874235, EPI_ISL_2874237, EPI_ISL_2874240, EPI_ISL_2874242, EPI_ISL_2874244, EPI_ISL_2874245, EPI_ISL_2874246, EPI_ISL_2874247, EPI_ISL_2874248, EPI_ISL_2874249, EPI_ISL_2874251, EPI_ISL_2874252, EPI_ISL_2874253, EPI_ISL_2874254, EPI_ISL_2874255, EPI_ISL_2874256, EPI_ISL_2874257, EPI_ISL_2874259, EPI_ISL_2874261, EPI_ISL_2874263, EPI_ISL_2874264, EPI_ISL_2874265, EPI_ISL_2874266, EPI_ISL_2874267, EPI_ISL_2874268, EPI_ISL_2874269, EPI_ISL_2874270, EPI_ISL_2874272, EPI_ISL_2874273, EPI_ISL_2874274, EPI_ISL_2874276, EPI_ISL_2874277, EPI_ISL_2874278, EPI_ISL_2874279, EPI_ISL_2874281, EPI_ISL_2874284, EPI_ISL_2874285, EPI_ISL_2874286, EPI_ISL_2874287, EPI_ISL_2874288, EPI_ISL_2874289, EPI_ISL_2874290, EPI_ISL_2874291, EPI_ISL_2874292, EPI_ISL_2874294, EPI_ISL_2874295, EPI_ISL_2874296, EPI_ISL_2874297, EPI_ISL_2874298, EPI_ISL_2874299, EPI_ISL_2874300, EPI_ISL_2874302, EPI_ISL_2874303, EPI_ISL_2874304, EPI_ISL_2874305, EPI_ISL_2874306, EPI_ISL_2874307, EPI_ISL_2874308, EPI_ISL_2874309, EPI_ISL_2874310, EPI_ISL_2874311, EPI_ISL_2874312, EPI_ISL_2874313, EPI_ISL_2874314, EPI_ISL_2874316, EPI_ISL_2874317, EPI_ISL_2874318, EPI_ISL_2874320, EPI_ISL_2874321, EPI_ISL_2874322, EPI_ISL_2874324, EPI_ISL_2874325, EPI_ISL_2874326, EPI_ISL_2874328, EPI_ISL_2874329, EPI_ISL_2874331, EPI_ISL_2874332, EPI_ISL_2874336, EPI_ISL_2874338, EPI_ISL_2874339, EPI_ISL_2874341, EPI_ISL_2874342, EPI_ISL_2874344, EPI_ISL_2874345, EPI_ISL_2874346, EPI_ISL_2874347, EPI_ISL_2874348, EPI_ISL_2874349, EPI_ISL_2874350, EPI_ISL_2874351, EPI_ISL_2874353, EPI_ISL_2874354, EPI_ISL_2874355, EPI_ISL_2874356, EPI_ISL_2874357, EPI_ISL_2874358, EPI_ISL_2874359, EPI_ISL_2874360, EPI_ISL_2874361, EPI_ISL_2874362, EPI_ISL_2874363, EPI_ISL_2874364, EPI_ISL_2874365, EPI_ISL_2874366, EPI_ISL_2874367, EPI_ISL_2874368, EPI_ISL_2874371, EPI_ISL_2874373, EPI_ISL_2874374, EPI_ISL_2874375, EPI_ISL_2874376, EPI_ISL_2874377, EPI_ISL_2874378, EPI_ISL_2874379, EPI_ISL_2874380, EPI_ISL_2874381, EPI_ISL_2874383, EPI_ISL_2874384, EPI_ISL_2874385, EPI_ISL_2874386, EPI_ISL_2874387, EPI_ISL_2874390, EPI_ISL_2874391, EPI_ISL_2874392, EPI_ISL_2874393, EPI_ISL_2874394, EPI_ISL_2874396, EPI_ISL_2874397, EPI_ISL_2874399, EPI_ISL_2874400, EPI_ISL_2874401, EPI_ISL_2874403, EPI_ISL_2874405, EPI_ISL_2874406, EPI_ISL_2874407, EPI_ISL_2874409, EPI_ISL_2874411, EPI_ISL_2874412, EPI_ISL_2874413, EPI_ISL_2874414, EPI_ISL_2874415, EPI_ISL_2874416, EPI_ISL_2874418, EPI_ISL_2874419, EPI_ISL_2874420, EPI_ISL_2874422, EPI_ISL_2874423, EPI_ISL_2874424, EPI_ISL_2874425, EPI_ISL_2874426, EPI_ISL_2874427, EPI_ISL_2874428, EPI_ISL_2874430, EPI_ISL_2874431, EPI_ISL_2874432, EPI_ISL_2874433, EPI_ISL_2874434, EPI_ISL_2874436, EPI_ISL_2874437, EPI_ISL_2874438, EPI_ISL_2874439, EPI_ISL_2874440, EPI_ISL_2874441, EPI_ISL_2874442, EPI_ISL_2874444, EPI_ISL_2874445, EPI_ISL_2874448, EPI_ISL_2874449, EPI_ISL_2874451, EPI_ISL_2874452, EPI_ISL_2874453, EPI_ISL_2874454, EPI_ISL_2874455, EPI_ISL_2874456, EPI_ISL_2874459, EPI_ISL_2874460, EPI_ISL_2874461, EPI_ISL_2874464, EPI_ISL_2874465, EPI_ISL_2874467, EPI_ISL_2874470, EPI_ISL_2874471, EPI_ISL_2874472, EPI_ISL_2874473, EPI_ISL_2874474, EPI_ISL_2874475, EPI_ISL_2874476, EPI_ISL_2874477, EPI_ISL_2874479, EPI_ISL_2874480, EPI_ISL_2874481, EPI_ISL_2874483, EPI_ISL_2874484, EPI_ISL_2874485, EPI_ISL_2874486, EPI_ISL_2874487, EPI_ISL_2874488, EPI_ISL_2874490, EPI_ISL_2874491, EPI_ISL_2874492, EPI_ISL_2874495, EPI_ISL_2874496, EPI_ISL_2874497, EPI_ISL_2874498, EPI_ISL_2874499, EPI_ISL_2874501, EPI_ISL_2874502, EPI_ISL_2874503, EPI_ISL_2874504, EPI_ISL_2874505, EPI_ISL_2874506, EPI_ISL_2874507, EPI_ISL_2874509, EPI_ISL_2874511, EPI_ISL_2874513, EPI_ISL_2874515, EPI_ISL_2874516, EPI_ISL_2874517, EPI_ISL_2874520, EPI_ISL_2874521, EPI_ISL_2874522, EPI_ISL_2874525, EPI_ISL_2874526, EPI_ISL_2874529, EPI_ISL_2874530, EPI_ISL_2874531, EPI_ISL_2874532, EPI_ISL_2874533, EPI_ISL_2874534, EPI_ISL_2874536, EPI_ISL_2874540, EPI_ISL_2874541, EPI_ISL_2874542, EPI_ISL_2874543, EPI_ISL_2874544, EPI_ISL_2874545, EPI_ISL_2874546, EPI_ISL_2874547, EPI_ISL_2874548, EPI_ISL_2874549, EPI_ISL_2874551, EPI_ISL_2874552, EPI_ISL_2874555, EPI_ISL_2874557, EPI_ISL_2874558, EPI_ISL_2874559, EPI_ISL_2874560, EPI_ISL_2874561, EPI_ISL_2874563, EPI_ISL_2874564, EPI_ISL_2874565, EPI_ISL_2874566, EPI_ISL_2874567, EPI_ISL_2874568, EPI_ISL_2874571, EPI_ISL_2874572, EPI_ISL_2874574, EPI_ISL_2874575, EPI_ISL_2874576, EPI_ISL_2874578, EPI_ISL_2874579, EPI_ISL_2874581, EPI_ISL_2874582, EPI_ISL_2874583, EPI_ISL_2874584, EPI_ISL_2874585, EPI_ISL_2874587, EPI_ISL_2874588, EPI_ISL_2874589, EPI_ISL_2874590, EPI_ISL_2874591, EPI_ISL_2874592, EPI_ISL_2874593, EPI_ISL_2874594, EPI_ISL_2874595, EPI_ISL_2874597, EPI_ISL_2874598, EPI_ISL_2874599, EPI_ISL_2874600, EPI_ISL_2874601, EPI_ISL_2874602, EPI_ISL_2874603, EPI_ISL_2874605, EPI_ISL_2874606, EPI_ISL_2874608, EPI_ISL_2874609, EPI_ISL_2874610, EPI_ISL_2874611, EPI_ISL_2874612, EPI_ISL_2874613, EPI_ISL_2874615, EPI_ISL_2874618, EPI_ISL_2874619, EPI_ISL_2874620, EPI_ISL_2874624, EPI_ISL_2874625, EPI_ISL_2874626, EPI_ISL_2874628, EPI_ISL_2874629, EPI_ISL_2874630, EPI_ISL_2874631, EPI_ISL_2874632, EPI_ISL_2874633, EPI_ISL_2874635, EPI_ISL_2874636, EPI_ISL_2874637, EPI_ISL_2874638, EPI_ISL_2874639, EPI_ISL_2874640, EPI_ISL_2874641, EPI_ISL_2874642, EPI_ISL_2874643, EPI_ISL_2874644, EPI_ISL_2874645, EPI_ISL_2874646, EPI_ISL_2874647, EPI_ISL_2874648, EPI_ISL_2874650, EPI_ISL_2874651, EPI_ISL_2874652, EPI_ISL_2874653, EPI_ISL_2874654, EPI_ISL_2874656, EPI_ISL_2874657, EPI_ISL_2874658, EPI_ISL_2874659, EPI_ISL_2874661, EPI_ISL_2874662, EPI_ISL_2874663, EPI_ISL_2874665, EPI_ISL_2874666, EPI_ISL_2874667, EPI_ISL_2874668, EPI_ISL_2874669, EPI_ISL_2874670, EPI_ISL_2874671, EPI_ISL_2874672, EPI_ISL_2874673, EPI_ISL_2874674, EPI_ISL_2874675, EPI_ISL_2874679, EPI_ISL_2874680, EPI_ISL_2874682, EPI_ISL_2874684, EPI_ISL_2874685, EPI_ISL_2874686, EPI_ISL_2874688, EPI_ISL_2874689, EPI_ISL_2874690, EPI_ISL_2874691, EPI_ISL_2874692, EPI_ISL_2874693, EPI_ISL_2874695, EPI_ISL_2874697, EPI_ISL_2874698, EPI_ISL_2874699, EPI_ISL_2874700, EPI_ISL_2874702, EPI_ISL_2874704, EPI_ISL_2874705, EPI_ISL_2874706, EPI_ISL_2874708, EPI_ISL_2874709, EPI_ISL_2874710, EPI_ISL_2874711, EPI_ISL_2874712, EPI_ISL_2874715, EPI_ISL_2874716, EPI_ISL_2874717, EPI_ISL_2874718, EPI_ISL_2874719, EPI_ISL_2874721, EPI_ISL_2874722, EPI_ISL_2874723, EPI_ISL_2874724, EPI_ISL_2874725, EPI_ISL_2874726, EPI_ISL_2874727, EPI_ISL_2874728, EPI_ISL_2874729, EPI_ISL_2874730, EPI_ISL_2874731, EPI_ISL_2874732, EPI_ISL_2874733, EPI_ISL_2874734, EPI_ISL_2874738, EPI_ISL_2874739, EPI_ISL_2874740, EPI_ISL_2874743, EPI_ISL_2874749, EPI_ISL_2874750, EPI_ISL_2874751, EPI_ISL_2874752, EPI_ISL_2874754, EPI_ISL_2874755, EPI_ISL_2874756, EPI_ISL_2874757, EPI_ISL_2874758, EPI_ISL_2874759, EPI_ISL_2874761, EPI_ISL_2874762, EPI_ISL_2874766, EPI_ISL_2874769, EPI_ISL_2874770, EPI_ISL_2874771, EPI_ISL_2874772, EPI_ISL_2874774, EPI_ISL_2874775, EPI_ISL_2874776, EPI_ISL_2874777, EPI_ISL_2874778, EPI_ISL_2874779, EPI_ISL_2874780, EPI_ISL_2874781, EPI_ISL_2874782, EPI_ISL_2874783, EPI_ISL_2874784, EPI_ISL_2874785, EPI_ISL_2874786, EPI_ISL_2874787, EPI_ISL_2874788, EPI_ISL_2874790, EPI_ISL_2874791, EPI_ISL_2874793, EPI_ISL_2874794, EPI_ISL_2874795, EPI_ISL_2874796, EPI_ISL_2874797, EPI_ISL_2874798, EPI_ISL_2874799, EPI_ISL_2874800, EPI_ISL_2874801, EPI_ISL_2874802, EPI_ISL_2874803, EPI_ISL_2874804, EPI_ISL_2874806, EPI_ISL_2874807, EPI_ISL_2874808, EPI_ISL_2874809, EPI_ISL_2874810, EPI_ISL_2874812, EPI_ISL_2874814, EPI_ISL_2874815, EPI_ISL_2874816, EPI_ISL_2874817, EPI_ISL_2874818, EPI_ISL_2874819, EPI_ISL_2874820, EPI_ISL_2874821, EPI_ISL_2874822, EPI_ISL_2874823, EPI_ISL_2874824, EPI_ISL_2874825, EPI_ISL_2874827, EPI_ISL_2874829, EPI_ISL_2874831, EPI_ISL_2874832, EPI_ISL_2874833, EPI_ISL_2874834, EPI_ISL_2874835, EPI_ISL_2874836, EPI_ISL_2874839, EPI_ISL_2874840, EPI_ISL_2874841, EPI_ISL_2874842, EPI_ISL_2874843, EPI_ISL_2874845, EPI_ISL_2874846, EPI_ISL_2874847, EPI_ISL_2874848, EPI_ISL_2874849, EPI_ISL_2874850, EPI_ISL_2874851, EPI_ISL_2874854, EPI_ISL_2874855, EPI_ISL_2874857, EPI_ISL_2874858, EPI_ISL_2874860, EPI_ISL_2874861, EPI_ISL_2874862, EPI_ISL_2874864, EPI_ISL_2874865, EPI_ISL_2874867, EPI_ISL_2874868, EPI_ISL_2874869, EPI_ISL_2874870, EPI_ISL_2874871, EPI_ISL_2874873, EPI_ISL_2874875, EPI_ISL_2874876, EPI_ISL_2874877, EPI_ISL_2874879, EPI_ISL_2874881, EPI_ISL_2874882, EPI_ISL_2874884, EPI_ISL_2874885, EPI_ISL_2874890, EPI_ISL_2874891, EPI_ISL_2874892, EPI_ISL_2874894, EPI_ISL_2874895, EPI_ISL_2874896, EPI_ISL_2874897, EPI_ISL_2874899, EPI_ISL_2874900, EPI_ISL_2874901, EPI_ISL_2874903, EPI_ISL_2874904, EPI_ISL_2874906, EPI_ISL_2874909, EPI_ISL_2874911, EPI_ISL_2874912, EPI_ISL_2874913, EPI_ISL_2874914, EPI_ISL_2874915, EPI_ISL_2874916, EPI_ISL_2874917, EPI_ISL_2874919, EPI_ISL_2874920, EPI_ISL_2874924, EPI_ISL_2874926, EPI_ISL_2874927, EPI_ISL_2874928, EPI_ISL_2874929, EPI_ISL_2874931, EPI_ISL_2874932, EPI_ISL_2874933, EPI_ISL_2874935, EPI_ISL_2874936, EPI_ISL_2874939, EPI_ISL_2874940, EPI_ISL_2874941, EPI_ISL_2874942, EPI_ISL_2874944, EPI_ISL_2874946, EPI_ISL_2874949, EPI_ISL_2874950, EPI_ISL_2874951, EPI_ISL_2874953, EPI_ISL_2874954, EPI_ISL_2874957, EPI_ISL_2874958, EPI_ISL_2874959, EPI_ISL_2874960, EPI_ISL_2874961, EPI_ISL_2874962, EPI_ISL_2874963, EPI_ISL_2874964, EPI_ISL_2874965, EPI_ISL_2874966, EPI_ISL_2874967, EPI_ISL_2874968, EPI_ISL_2874970, EPI_ISL_2874971, EPI_ISL_2874972, EPI_ISL_2874973, EPI_ISL_2874978, EPI_ISL_2874979, EPI_ISL_2874980, EPI_ISL_2874981, EPI_ISL_2874983, EPI_ISL_2874984, EPI_ISL_2874986, EPI_ISL_2874987, EPI_ISL_2874988, EPI_ISL_2874989, EPI_ISL_2874990, EPI_ISL_2874993, EPI_ISL_2874994, EPI_ISL_2874995, EPI_ISL_2874999, EPI_ISL_2875000, EPI_ISL_2875001, EPI_ISL_2875004, EPI_ISL_2875006, EPI_ISL_2875007, EPI_ISL_2875011, EPI_ISL_2875014, EPI_ISL_2875015, EPI_ISL_2875016, EPI_ISL_2875017, EPI_ISL_2875020, EPI_ISL_2875022, EPI_ISL_2875024, EPI_ISL_2875026, EPI_ISL_2875027, EPI_ISL_2875028, EPI_ISL_2875030, EPI_ISL_2875031, EPI_ISL_2875033, EPI_ISL_2875036, EPI_ISL_2875037, EPI_ISL_2875038, EPI_ISL_2875039, EPI_ISL_2875040, EPI_ISL_2875042, EPI_ISL_2875043, EPI_ISL_2875044, EPI_ISL_2875046, EPI_ISL_2875047, EPI_ISL_2875048, EPI_ISL_2875049, EPI_ISL_2875050, EPI_ISL_2875051, EPI_ISL_2875052, EPI_ISL_2875053, EPI_ISL_2875054, EPI_ISL_2875058, EPI_ISL_2875060, EPI_ISL_2875061, EPI_ISL_2875062, EPI_ISL_2875064, EPI_ISL_2875066, EPI_ISL_2875067, EPI_ISL_2875069, EPI_ISL_2875070, EPI_ISL_2875072, EPI_ISL_2875073, EPI_ISL_2875074, EPI_ISL_2875075, EPI_ISL_2875076, EPI_ISL_2875077, EPI_ISL_2875079, EPI_ISL_2875081, EPI_ISL_2875082, EPI_ISL_2875086, EPI_ISL_2875089, EPI_ISL_2875090, EPI_ISL_2875091, EPI_ISL_2875094, EPI_ISL_2875096, EPI_ISL_2875099, EPI_ISL_2875100, EPI_ISL_2875102, EPI_ISL_2875104, EPI_ISL_2875106, EPI_ISL_2875107, EPI_ISL_2875109, EPI_ISL_2875110, EPI_ISL_2875112, EPI_ISL_2875113, EPI_ISL_2875116, EPI_ISL_2875117, EPI_ISL_2875118, EPI_ISL_2875119, EPI_ISL_2875120, EPI_ISL_2875121, EPI_ISL_2875126, EPI_ISL_2875127, EPI_ISL_2875128, EPI_ISL_2875129, EPI_ISL_2875132, EPI_ISL_2875133, EPI_ISL_2875135, EPI_ISL_2875136, EPI_ISL_2875137, EPI_ISL_2875138, EPI_ISL_2875139, EPI_ISL_2875140, EPI_ISL_2875141, EPI_ISL_2875143, EPI_ISL_2875144, EPI_ISL_2875145, EPI_ISL_2875148, EPI_ISL_2875150, EPI_ISL_2875151, EPI_ISL_2875156, EPI_ISL_2875157, EPI_ISL_2875159, EPI_ISL_2875160, EPI_ISL_2875161, EPI_ISL_2875162, EPI_ISL_2875163, EPI_ISL_2875164, EPI_ISL_2875165, EPI_ISL_2875166, EPI_ISL_2875167 | | | |
| see above | Quest Diagnostics Incorporated | Centers for Disease Control and Prevention Division of Viral Diseases, Pathogen Discovery | Dakota Howard, Dhwani Batra, Peter W. Cook, Kara Moser, Adrian Paskey, Jason Caravas, Benjamin Rambo-Martin, Shatavia Morrison, Christopher Gulvick, Scott Sammons, Yvette Unoarumhi, Darlene Wagner, Matthew Schmerer, S. H. Rosenthal, A. Gerasimova, R. M. Kagan, B. Anderson, M. Hua, Y. Liu, L.E. Bernstein, K.E. Livingston, A. Perez, I. A. Shlyakhter, R. V. Rolando, R. Owen, P. Tanpaiboon, F. Lacbawan, Clinton R. Paden, Duncan MacCannell |
| EPI_ISL_2875171, EPI_ISL_2875173, EPI_ISL_2875177, EPI_ISL_2875178, EPI_ISL_2875180, EPI_ISL_2875185, EPI_ISL_2875187, EPI_ISL_2875188, EPI_ISL_2875189, EPI_ISL_2875193, EPI_ISL_2875194, EPI_ISL_2875201, EPI_ISL_2875206, EPI_ISL_2875207, EPI_ISL_2875208, EPI_ISL_2875214, EPI_ISL_2875233, EPI_ISL_2875234, EPI_ISL_2875246, EPI_ISL_2875249, EPI_ISL_2875251, EPI_ISL_2875256, EPI_ISL_2875257, EPI_ISL_2875269, EPI_ISL_2875270, EPI_ISL_2875272, EPI_ISL_2875277, EPI_ISL_2875278, EPI_ISL_2875280, EPI_ISL_2875288, EPI_ISL_2875291, EPI_ISL_2875293, EPI_ISL_2875299, EPI_ISL_2875305, EPI_ISL_2875307, EPI_ISL_2875312, EPI_ISL_2875317, EPI_ISL_2875318, EPI_ISL_2875324, EPI_ISL_2875326, EPI_ISL_2875330, EPI_ISL_2875332, EPI_ISL_2875335, EPI_ISL_2875337, EPI_ISL_2875339, EPI_ISL_2875340, EPI_ISL_2875345, EPI_ISL_2875346, EPI_ISL_2875348, EPI_ISL_2875353, EPI_ISL_2875360, EPI_ISL_2875363, EPI_ISL_2875365, EPI_ISL_2875367, EPI_ISL_2875369, EPI_ISL_2875375, EPI_ISL_2875377, EPI_ISL_2875378, EPI_ISL_2875379, EPI_ISL_2875380, EPI_ISL_2875381, EPI_ISL_2875382, EPI_ISL_2875384, EPI_ISL_2875385, EPI_ISL_2875387, EPI_ISL_2875388, EPI_ISL_2875389, EPI_ISL_2875392, EPI_ISL_2875393, EPI_ISL_2875396, EPI_ISL_2875398, EPI_ISL_2875399, EPI_ISL_2875400, EPI_ISL_2875401, EPI_ISL_2875403, EPI_ISL_2875404, EPI_ISL_2875405, EPI_ISL_2875408, EPI_ISL_2875409, EPI_ISL_2875411, EPI_ISL_2875413, EPI_ISL_2875416, EPI_ISL_2875417, EPI_ISL_2875419, EPI_ISL_2875420, EPI_ISL_2875421, EPI_ISL_2875422, EPI_ISL_2875423, EPI_ISL_2875424, EPI_ISL_2875425, EPI_ISL_2875428, EPI_ISL_2875429, EPI_ISL_2875431, EPI_ISL_2875432, EPI_ISL_2875433, EPI_ISL_2875436, EPI_ISL_2875437, EPI_ISL_2875439, EPI_ISL_2875440, EPI_ISL_2875442, EPI_ISL_2875444, EPI_ISL_2875446, EPI_ISL_2875447, EPI_ISL_2875449, EPI_ISL_2875450, EPI_ISL_2875451, EPI_ISL_2875452, EPI_ISL_2875453, EPI_ISL_2875454, EPI_ISL_2875455, EPI_ISL_2875456, EPI_ISL_2875458, EPI_ISL_2875460, EPI_ISL_2875461, EPI_ISL_2875467, EPI_ISL_2875468, EPI_ISL_2875469, EPI_ISL_2875470, EPI_ISL_2875471, EPI_ISL_2875472, EPI_ISL_2875475, EPI_ISL_2875478, EPI_ISL_2875480, EPI_ISL_2875482, EPI_ISL_2875484, EPI_ISL_2875485, EPI_ISL_2875488, EPI_ISL_2875492, EPI_ISL_2875493, EPI_ISL_2875495, EPI_ISL_2875496, EPI_ISL_2875497, EPI_ISL_2875498, EPI_ISL_2875501, EPI_ISL_2875502, EPI_ISL_2875504, EPI_ISL_2875505, EPI_ISL_2875506, EPI_ISL_2875509, EPI_ISL_2875510, EPI_ISL_2875511, EPI_ISL_2875517, EPI_ISL_2875519 | | | |
| see above | Mako Medical | Centers for Disease Control and Prevention Division of Viral Diseases, Pathogen Discovery | Dakota Howard, Dhwani Batra, Peter W. Cook, Kara Moser, Adrian Paskey, Jason Caravas, Benjamin Rambo-Martin, Shatavia Morrison, Christopher Gulvick, Scott Sammons, Yvette Unoarumhi, Darlene Wagner, Matthew Schmerer,Matthew Tugwell, Lauren Moon, Clinton R. Paden, Duncan MacCannell |
| EPI_ISL_2875520, EPI_ISL_2875521, EPI_ISL_2875522, EPI_ISL_2875523, EPI_ISL_2875524, EPI_ISL_2875525, EPI_ISL_2875526, EPI_ISL_2875533, EPI_ISL_2875535, EPI_ISL_2875543, EPI_ISL_2875545, EPI_ISL_2875546, EPI_ISL_2875547, EPI_ISL_2875548, EPI_ISL_2875552, EPI_ISL_2875553, EPI_ISL_2875554, EPI_ISL_2875555, EPI_ISL_2875556, EPI_ISL_2875557, EPI_ISL_2875559, EPI_ISL_2875560, EPI_ISL_2875562, EPI_ISL_2875563, EPI_ISL_2875565, EPI_ISL_2875568, EPI_ISL_2875574, EPI_ISL_2875577, EPI_ISL_2875578, EPI_ISL_2875580, EPI_ISL_2875581, EPI_ISL_2875582, EPI_ISL_2875583, EPI_ISL_2875584, EPI_ISL_2875586, EPI_ISL_2875589, EPI_ISL_2875591, EPI_ISL_2875593, EPI_ISL_2875594, EPI_ISL_2875605, EPI_ISL_2875608, EPI_ISL_2875610, EPI_ISL_2875611, EPI_ISL_2875613, EPI_ISL_2875618, EPI_ISL_2875619, EPI_ISL_2875621, EPI_ISL_2875622, EPI_ISL_2875623, EPI_ISL_2875625, EPI_ISL_2875629, EPI_ISL_2875630, EPI_ISL_2875633 | | | |
| see above | Aegis Sciences Corporation | Centers for Disease Control and Prevention Division of Viral Diseases, Pathogen Discovery | Dakota Howard, Dhwani Batra, Peter W. Cook, Kara Moser, Adrian Paskey, Jason Caravas, Benjamin Rambo-Martin, Shatavia Morrison, Christopher Gulvick, Scott Sammons, Yvette Unoarumhi, Darlene Wagner, Matthew Schmerer, Cyndi Clark, Patrick Campbell, Rob Case, Vikramsinha Ghorpade, Holly Houdeshell, Ola Kvalvaag, Dillon Nall, Ethan Sanders, Alec Vest, Shaun Westlund, Matthew Hardison, Clinton R. Paden, Duncan MacCannell |
| EPI_ISL_2875634, EPI_ISL_2875635, EPI_ISL_2875637 | Center for Laboratory Medicine | Center for Laboratory Medicine | Yannick Gerth |
| EPI_ISL_2875638 | Aegis Sciences Corporation | Centers for Disease Control and Prevention Division of Viral Diseases, Pathogen Discovery | Dakota Howard, Dhwani Batra, Peter W. Cook, Kara Moser, Adrian Paskey, Jason Caravas, Benjamin Rambo-Martin, Shatavia Morrison, Christopher Gulvick, Scott Sammons, Yvette Unoarumhi, Darlene Wagner, Matthew Schmerer, Cyndi Clark, Patrick Campbell, Rob Case, Vikramsinha Ghorpade, Holly Houdeshell, Ola Kvalvaag, Dillon Nall, Ethan Sanders, Alec Vest, Shaun Westlund, Matthew Hardison, Clinton R. Paden, Duncan MacCannell |
| EPI_ISL_2875639, EPI_ISL_2875640 | Center for Laboratory Medicine | Center for Laboratory Medicine | Yannick Gerth |
| EPI_ISL_2875641, EPI_ISL_2875643 | Aegis Sciences Corporation | Centers for Disease Control and Prevention Division of Viral Diseases, Pathogen Discovery | Dakota Howard, Dhwani Batra, Peter W. Cook, Kara Moser, Adrian Paskey, Jason Caravas, Benjamin Rambo-Martin, Shatavia Morrison, Christopher Gulvick, Scott Sammons, Yvette Unoarumhi, Darlene Wagner, Matthew Schmerer, Cyndi Clark, Patrick Campbell, Rob Case, Vikramsinha Ghorpade, Holly Houdeshell, Ola Kvalvaag, Dillon Nall, Ethan Sanders, Alec Vest, Shaun Westlund, Matthew Hardison, Clinton R. Paden, Duncan MacCannell |
| EPI_ISL_2875644 | Center for Laboratory Medicine | Center for Laboratory Medicine | Yannick Gerth |
| EPI_ISL_2875645, EPI_ISL_2875648, EPI_ISL_2875651, EPI_ISL_2875652, EPI_ISL_2875654, EPI_ISL_2875655, EPI_ISL_2875656, EPI_ISL_2875657, EPI_ISL_2875660, EPI_ISL_2875661, EPI_ISL_2875662, EPI_ISL_2875663, EPI_ISL_2875665, EPI_ISL_2875666, EPI_ISL_2875668, EPI_ISL_2875669, EPI_ISL_2875670, EPI_ISL_2875671, EPI_ISL_2875674, EPI_ISL_2875676, EPI_ISL_2875680, EPI_ISL_2875684, EPI_ISL_2875689, EPI_ISL_2875690, EPI_ISL_2875692, EPI_ISL_2875694, EPI_ISL_2875695, EPI_ISL_2875702, EPI_ISL_2875703, EPI_ISL_2875706, EPI_ISL_2875707, EPI_ISL_2875708, EPI_ISL_2875710, EPI_ISL_2875711, EPI_ISL_2875712, EPI_ISL_2875713, EPI_ISL_2875714, EPI_ISL_2875716, EPI_ISL_2875718, EPI_ISL_2875719, EPI_ISL_2875720, EPI_ISL_2875721, EPI_ISL_2875725, EPI_ISL_2875726, EPI_ISL_2875728, EPI_ISL_2875729, EPI_ISL_2875732, EPI_ISL_2875733, EPI_ISL_2875734, EPI_ISL_2875735, EPI_ISL_2875736, EPI_ISL_2875739, EPI_ISL_2875740, EPI_ISL_2875741, EPI_ISL_2875742, EPI_ISL_2875743, EPI_ISL_2875744, EPI_ISL_2875745, EPI_ISL_2875746, EPI_ISL_2875751, EPI_ISL_2875753, EPI_ISL_2875754, EPI_ISL_2875757, EPI_ISL_2875758, EPI_ISL_2875760, EPI_ISL_2875761, EPI_ISL_2875762, EPI_ISL_2875766 | | | |
| see above | Aegis Sciences Corporation | Centers for Disease Control and Prevention Division of Viral Diseases, Pathogen Discovery | Dakota Howard, Dhwani Batra, Peter W. Cook, Kara Moser, Adrian Paskey, Jason Caravas, Benjamin Rambo-Martin, Shatavia Morrison, Christopher Gulvick, Scott Sammons, Yvette Unoarumhi, Darlene Wagner, Matthew Schmerer, Cyndi Clark, Patrick Campbell, Rob Case, Vikramsinha Ghorpade, Holly Houdeshell, Ola Kvalvaag, Dillon Nall, Ethan Sanders, Alec Vest, Shaun Westlund, Matthew Hardison, Clinton R. Paden, Duncan MacCannell |
| EPI_ISL_2875769, EPI_ISL_2875770 | Rumah Sakit Umum Daerah Kota Mataram | Genomik Solidaritas Indonesia Laboratorium | Anna Christina Brazia, Himawan Masyhuri, Normastuti Adhini Tantyo, Alfin Muhammad Abdillah, Dhahlia Agustina Cahyono, Ahmad Zoebad Foeady, Annisa Muthiah Sukirman, Gracia Felias Enos Korompis, Laely Indrayanti, Louisa Markus, Anuraj Shankar, Meutia Ayuputeri Kumaheri, Anindya Pradipta Susanto, Ariel Pradipta |
| EPI_ISL_2875771 | Aegis Sciences Corporation | Centers for Disease Control and Prevention Division of Viral Diseases, Pathogen Discovery | Dakota Howard, Dhwani Batra, Peter W. Cook, Kara Moser, Adrian Paskey, Jason Caravas, Benjamin Rambo-Martin, Shatavia Morrison, Christopher Gulvick, Scott Sammons, Yvette Unoarumhi, Darlene Wagner, Matthew Schmerer, Cyndi Clark, Patrick Campbell, Rob Case, Vikramsinha Ghorpade, Holly Houdeshell, Ola Kvalvaag, Dillon Nall, Ethan Sanders, Alec Vest, Shaun Westlund, Matthew Hardison, Clinton R. Paden, Duncan MacCannell |
| EPI_ISL_2875772, EPI_ISL_2875773 | Rumah Sakit Umum Daerah Kota Mataram | Genomik Solidaritas Indonesia Laboratorium | Himawan Masyhuri, Normastuti Adhini Tantyo, Alfin Muhammad Abdillah, Dhahlia Agustina Cahyono, Anna Christina Brazia, Ahmad Zoebad Foeady, Annisa Muthiah Sukirman, Gracia Felias Enos Korompis, Laely Indrayanti, Louisa Markus, Anuraj Shankar, Meutia Ayuputeri Kumaheri, Anindya Pradipta Susanto, Ariel Pradipta |
| EPI_ISL_2875774 | Aegis Sciences Corporation | Centers for Disease Control and Prevention Division of Viral Diseases, Pathogen Discovery | Dakota Howard, Dhwani Batra, Peter W. Cook, Kara Moser, Adrian Paskey, Jason Caravas, Benjamin Rambo-Martin, Shatavia Morrison, Christopher Gulvick, Scott Sammons, Yvette Unoarumhi, Darlene Wagner, Matthew Schmerer, Cyndi Clark, Patrick Campbell, Rob Case, Vikramsinha Ghorpade, Holly Houdeshell, Ola Kvalvaag, Dillon Nall, Ethan Sanders, Alec Vest, Shaun Westlund, Matthew Hardison, Clinton R. Paden, Duncan MacCannell |
| EPI_ISL_2875775, EPI_ISL_2875776 | Rumah Sakit Umum Daerah Kota Mataram | Genomik Solidaritas Indonesia Laboratorium | Normastuti Adhini Tantyo, Alfin Muhammad Abdillah, Dhahlia Agustina Cahyono, Anna Christina Brazia, Himawan Masyhuri, Ahmad Zoebad Foeady, Annisa Muthiah Sukirman, Gracia Felias Enos Korompis, Laely Indrayanti, Louisa Markus, Anuraj Shankar, Meutia Ayuputeri Kumaheri, Anindya Pradipta Susanto, Ariel Pradipta |
| EPI_ISL_2875777 | Aegis Sciences Corporation | Centers for Disease Control and Prevention Division of Viral Diseases, Pathogen Discovery | Dakota Howard, Dhwani Batra, Peter W. Cook, Kara Moser, Adrian Paskey, Jason Caravas, Benjamin Rambo-Martin, Shatavia Morrison, Christopher Gulvick, Scott Sammons, Yvette Unoarumhi, Darlene Wagner, Matthew Schmerer, Cyndi Clark, Patrick Campbell, Rob Case, Vikramsinha Ghorpade, Holly Houdeshell, Ola Kvalvaag, Dillon Nall, Ethan Sanders, Alec Vest, Shaun Westlund, Matthew Hardison, Clinton R. Paden, Duncan MacCannell |
| EPI_ISL_2875778 | Rumah Sakit Umum Daerah Kota Mataram | Genomik Solidaritas Indonesia Laboratorium | Alfin Muhammad Abdillah, Dhahlia Agustina Cahyono, Anna Christina Brazia, Himawan Masyhuri, Normastuti Adhini Tantyo, Ahmad Zoebad Foeady, Annisa Muthiah Sukirman, Gracia Felias Enos Korompis, Laely Indrayanti, Louisa Markus, Anuraj Shankar, Meutia Ayuputeri Kumaheri, Anindya Pradipta Susanto, Ariel Pradipta |
| EPI_ISL_2875779 | Aegis Sciences Corporation | Centers for Disease Control and Prevention Division of Viral Diseases, Pathogen Discovery | Dakota Howard, Dhwani Batra, Peter W. Cook, Kara Moser, Adrian Paskey, Jason Caravas, Benjamin Rambo-Martin, Shatavia Morrison, Christopher Gulvick, Scott Sammons, Yvette Unoarumhi, Darlene Wagner, Matthew Schmerer, Cyndi Clark, Patrick Campbell, Rob Case, Vikramsinha Ghorpade, Holly Houdeshell, Ola Kvalvaag, Dillon Nall, Ethan Sanders, Alec Vest, Shaun Westlund, Matthew Hardison, Clinton R. Paden, Duncan MacCannell |
| EPI_ISL_2875780 | Rumah Sakit Umum Daerah Kota Mataram | Genomik Solidaritas Indonesia Laboratorium | Alfin Muhammad Abdillah, Dhahlia Agustina Cahyono, Anna Christina Brazia, Himawan Masyhuri, Normastuti Adhini Tantyo, Ahmad Zoebad Foeady, Annisa Muthiah Sukirman, Gracia Felias Enos Korompis, Laely Indrayanti, Louisa Markus, Anuraj Shankar, Meutia Ayuputeri Kumaheri, Anindya Pradipta Susanto, Ariel Pradipta |
| EPI_ISL_2875781, EPI_ISL_2875783 | Rumah Sakit Umum Daerah Kota Mataram | Genomik Solidaritas Indonesia Laboratorium | Dhahlia Agustina Cahyono, Anna Christina Brazia, Himawan Masyhuri, Normastuti Adhini Tantyo, Alfin Muhammad Abdillah, Ahmad Zoebad Foeady, Annisa Muthiah Sukirman, Gracia Felias Enos Korompis, Laely Indrayanti, Louisa Markus, Anuraj Shankar, Meutia Ayuputeri Kumaheri, Anindya Pradipta Susanto, Ariel Pradipta |
| EPI_ISL_2875784 | Rumah Sakit Umum Daerah Kota Mataram | Genomik Solidaritas Indonesia Laboratorium | Anna Christina Brazia, Himawan Masyhuri, Normastuti Adhini Tantyo, Alfin Muhammad Abdillah, Dhahlia Agustina Cahyono, Ahmad Zoebad Foeady, Annisa Muthiah Sukirman, Gracia Felias Enos Korompis, Laely Indrayanti, Louisa Markus, Anuraj Shankar, Meutia Ayuputeri Kumaheri, Anindya Pradipta Susanto, Ariel Pradipta |
| EPI_ISL_2875786 | Rumah Sakit Umum Daerah Kota Mataram | Genomik Solidaritas Indonesia Laboratorium | Normastuti Adhini Tantyo, Alfin Muhammad Abdillah, Dhahlia Agustina Cahyono, Anna Christina Brazia, Himawan Masyhuri, Ahmad Zoebad Foeady, Annisa Muthiah Sukirman, Gracia Felias Enos Korompis, Laely Indrayanti, Louisa Markus, Anuraj Shankar, Meutia Ayuputeri Kumaheri, Anindya Pradipta Susanto, Ariel Pradipta |
| EPI_ISL_2875787 | Rumah Sakit Umum Daerah Kota Mataram | Genomik Solidaritas Indonesia Laboratorium | Dhahlia Agustina Cahyono, Anna Christina Brazia, Himawan Masyhuri, Normastuti Adhini Tantyo, Alfin Muhammad Abdillah, Ahmad Zoebad Foeady, Annisa Muthiah Sukirman, Gracia Felias Enos Korompis, Laely Indrayanti, Louisa Markus, Anuraj Shankar, Meutia Ayuputeri Kumaheri, Anindya Pradipta Susanto, Ariel Pradipta |
| EPI_ISL_2875788, EPI_ISL_2875791, EPI_ISL_2875792, EPI_ISL_2875793, EPI_ISL_2875794, EPI_ISL_2875796, EPI_ISL_2875800, EPI_ISL_2875802, EPI_ISL_2875803, EPI_ISL_2875805, EPI_ISL_2875806, EPI_ISL_2875807, EPI_ISL_2875808, EPI_ISL_2875809, EPI_ISL_2875810, EPI_ISL_2875811, EPI_ISL_2875813, EPI_ISL_2875814, EPI_ISL_2875815, EPI_ISL_2875816, EPI_ISL_2875818, EPI_ISL_2875819, EPI_ISL_2875820, EPI_ISL_2875821, EPI_ISL_2875822, EPI_ISL_2875823, EPI_ISL_2875824, EPI_ISL_2875825, EPI_ISL_2875826, EPI_ISL_2875827, EPI_ISL_2875828, EPI_ISL_2875829, EPI_ISL_2875830, EPI_ISL_2875831, EPI_ISL_2875833 | | | |
| see above | Aegis Sciences Corporation | Centers for Disease Control and Prevention Division of Viral Diseases, Pathogen Discovery | Dakota Howard, Dhwani Batra, Peter W. Cook, Kara Moser, Adrian Paskey, Jason Caravas, Benjamin Rambo-Martin, Shatavia Morrison, Christopher Gulvick, Scott Sammons, Yvette Unoarumhi, Darlene Wagner, Matthew Schmerer, Cyndi Clark, Patrick Campbell, Rob Case, Vikramsinha Ghorpade, Holly Houdeshell, Ola Kvalvaag, Dillon Nall, Ethan Sanders, Alec Vest, Shaun Westlund, Matthew Hardison, Clinton R. Paden, Duncan MacCannell |
| EPI_ISL_2875836 | UniversitätsSpital Zürich | Institute of Medical Virology | Daniel Ehrsam, Isabel Stürmer, Catharine Aquino, Joel Wirz, Weihong Qi, Hubert Rehrauer, Verena Kufner, Gabriela Ziltener, Maryam Zaheri, Stefan Schmutz, Annette Audigé, Maria Grünberg, Kevin Steiner, Jon Huder, Cyril Shah, Riccarda Capaul, Guido Bloemberg, Jürg Böni, Michael Huber, Alexandra Trkola |
| EPI_ISL_2875844, EPI_ISL_2875846, EPI_ISL_2875848, EPI_ISL_2875849, EPI_ISL_2875850, EPI_ISL_2875854, EPI_ISL_2875856, EPI_ISL_2875857, EPI_ISL_2875858, EPI_ISL_2875860, EPI_ISL_2875864, EPI_ISL_2875867, EPI_ISL_2875868, EPI_ISL_2875869, EPI_ISL_2875870, EPI_ISL_2875873, EPI_ISL_2875874, EPI_ISL_2875875, EPI_ISL_2875876, EPI_ISL_2875880, EPI_ISL_2875881, EPI_ISL_2875882, EPI_ISL_2875883, EPI_ISL_2875884, EPI_ISL_2875887, EPI_ISL_2875888, EPI_ISL_2875889, EPI_ISL_2875890, EPI_ISL_2875891, EPI_ISL_2875895, EPI_ISL_2875896, EPI_ISL_2875897, EPI_ISL_2875898, EPI_ISL_2875899, EPI_ISL_2875900, EPI_ISL_2875901, EPI_ISL_2875902, EPI_ISL_2875903, EPI_ISL_2875905, EPI_ISL_2875906, EPI_ISL_2875908, EPI_ISL_2875909, EPI_ISL_2875910, EPI_ISL_2875911, | | | |

| | | | |
|---|---|---|---|
| EPI_ISL_2875912, EPI_ISL_2875915, EPI_ISL_2875917, EPI_ISL_2875919, EPI_ISL_2875920, EPI_ISL_2875921, EPI_ISL_2875927, EPI_ISL_2875930, EPI_ISL_2875932, EPI_ISL_2875933, EPI_ISL_2875937 | | | |
| see above | Aegis Sciences Corporation | Centers for Disease Control and Prevention Division of Viral Diseases, Pathogen Discovery | Dakota Howard, Dhwani Batra, Peter W. Cook, Kara Moser, Adrian Paskey, Jason Caravas, Benjamin Rambo-Martin, Shatavia Morrison, Christopher Gulvick, Scott Sammons, Yvette Unoarumhi, Darlene Wagner, Matthew Schmerer, Cyndi Clark, Patrick Campbell, Rob Case, Vikramsinha Ghorpade, Holly Houdeshell, Ola Kvalvaag, Dillon Nall, Ethan Sanders, Alec Vest, Shaun Westlund, Matthew Hardison, Clinton R. Paden, Duncan MacCannell |
| EPI_ISL_2875938 | UniversitätsSpital Zürich | Institute of Medical Virology | Daniel Ehrsam, Isabel Stürmer, Catharine Aquino, Joel Wirz, Weihong Qi, Hubert Rehrauer, Verena Kufner, Gabriela Ziltener, Maryam Zaheri, Stefan Schmutz, Annette Audigé, Maria Grünberg, Kevin Steiner, Jon Huder, Cyril Shah, Riccarda Capaul, Guido Bloemberg, Jürg Böni, Michael Huber, Alexandra Trkola |
| EPI_ISL_2875940 | Aegis Sciences Corporation | Centers for Disease Control and Prevention Division of Viral Diseases, Pathogen Discovery | Dakota Howard, Dhwani Batra, Peter W. Cook, Kara Moser, Adrian Paskey, Jason Caravas, Benjamin Rambo-Martin, Shatavia Morrison, Christopher Gulvick, Scott Sammons, Yvette Unoarumhi, Darlene Wagner, Matthew Schmerer, Cyndi Clark, Patrick Campbell, Rob Case, Vikramsinha Ghorpade, Holly Houdeshell, Ola Kvalvaag, Dillon Nall, Ethan Sanders, Alec Vest, Shaun Westlund, Matthew Hardison, Clinton R. Paden, Duncan MacCannell |
| EPI_ISL_2875941, EPI_ISL_2875942 | UniversitätsSpital Zürich | Institute of Medical Virology | Daniel Ehrsam, Isabel Stürmer, Catharine Aquino, Joel Wirz, Weihong Qi, Hubert Rehrauer, Verena Kufner, Gabriela Ziltener, Maryam Zaheri, Stefan Schmutz, Annette Audigé, Maria Grünberg, Kevin Steiner, Jon Huder, Cyril Shah, Riccarda Capaul, Guido Bloemberg, Jürg Böni, Michael Huber, Alexandra Trkola |
| EPI_ISL_2875945, EPI_ISL_2875946, EPI_ISL_2875947, EPI_ISL_2875948, EPI_ISL_2875949, EPI_ISL_2875950, EPI_ISL_2875951, EPI_ISL_2875952, EPI_ISL_2875953, EPI_ISL_2875955, EPI_ISL_2875958, EPI_ISL_2875960, EPI_ISL_2875961, EPI_ISL_2875962, EPI_ISL_2875964, EPI_ISL_2875965, EPI_ISL_2875968, EPI_ISL_2875969, EPI_ISL_2875971, EPI_ISL_2875973, EPI_ISL_2875974, EPI_ISL_2875975, EPI_ISL_2875976, EPI_ISL_2875977, EPI_ISL_2875979, EPI_ISL_2875980, EPI_ISL_2875981, EPI_ISL_2875982, EPI_ISL_2875983, EPI_ISL_2875986, EPI_ISL_2875991, EPI_ISL_2875993, EPI_ISL_2875995, EPI_ISL_2875996, EPI_ISL_2875997, EPI_ISL_2875999, EPI_ISL_2876000, EPI_ISL_2876003, EPI_ISL_2876004, EPI_ISL_2876005, EPI_ISL_2876006, EPI_ISL_2876008, EPI_ISL_2876009, EPI_ISL_2876010, EPI_ISL_2876011, EPI_ISL_2876012, EPI_ISL_2876013 | | | |
| see above | Aegis Sciences Corporation | Centers for Disease Control and Prevention Division of Viral Diseases, Pathogen Discovery | Dakota Howard, Dhwani Batra, Peter W. Cook, Kara Moser, Adrian Paskey, Jason Caravas, Benjamin Rambo-Martin, Shatavia Morrison, Christopher Gulvick, Scott Sammons, Yvette Unoarumhi, Darlene Wagner, Matthew Schmerer, Cyndi Clark, Patrick Campbell, Rob Case, Vikramsinha Ghorpade, Holly Houdeshell, Ola Kvalvaag, Dillon Nall, Ethan Sanders, Alec Vest, Shaun Westlund, Matthew Hardison, Clinton R. Paden, Duncan MacCannell |
| EPI_ISL_2876015, EPI_ISL_2876016, EPI_ISL_2876017, EPI_ISL_2876018, EPI_ISL_2876019, EPI_ISL_2876020, EPI_ISL_2876021, EPI_ISL_2876022, EPI_ISL_2876023, EPI_ISL_2876024, EPI_ISL_2876025 | | | |
| see above | St Vincent's Pathology (SydPath) | NSW Health Pathology - Institute of Clinical Pathology and Medical Research; Westmead Hospital; University of Sydney | CIDM-PH et al. |
| EPI_ISL_2876026 | Medlab Pathology | NSW Health Pathology - Institute of Clinical Pathology and Medical Research; Westmead Hospital; University of Sydney | CIDM-PH et al. |
| EPI_ISL_2876027 | 4Cyte Pathology | NSW Health Pathology - Institute of Clinical Pathology and Medical Research; Westmead Hospital; University of Sydney | CIDM-PH et al. |
| EPI_ISL_2876028 | Liverpool Hospital | NSW Health Pathology - Institute of Clinical Pathology and Medical Research; Westmead Hospital; University of Sydney | CIDM-PH et al. |
| EPI_ISL_2876029, EPI_ISL_2876030 | Histopath | NSW Health Pathology - Institute of Clinical Pathology and Medical Research; Westmead Hospital; University of Sydney | CIDM-PH et al. |
| EPI_ISL_2876031 | Austech Medical Laboratories | NSW Health Pathology - Institute of Clinical Pathology and Medical Research; Westmead Hospital; University of Sydney | CIDM-PH et al. |
| EPI_ISL_2876032, EPI_ISL_2876033 | 4Cyte Pathology | NSW Health Pathology - Institute of Clinical Pathology and Medical Research; Westmead Hospital; University of Sydney | CIDM-PH et al. |
| EPI_ISL_2876034, EPI_ISL_2876035, EPI_ISL_2876036, EPI_ISL_2876037 | Laverty Pathology | NSW Health Pathology - Institute of Clinical Pathology and Medical Research; Westmead Hospital; University of Sydney | CIDM-PH et al. |
| EPI_ISL_2876038, EPI_ISL_2876039, EPI_ISL_2876040, EPI_ISL_2876041, EPI_ISL_2876042 | St Vincent's Pathology (SydPath) | NSW Health Pathology - Institute of Clinical Pathology and Medical Research; Westmead Hospital; University of Sydney | CIDM-PH et al. |
| EPI_ISL_2876043, EPI_ISL_2876044, EPI_ISL_2876045 | Sydney South West Pathology Service (SSWPS) - Liverpool Hospital - NSW Health Pathology | NSW Health Pathology - Institute of Clinical Pathology and Medical Research; Westmead Hospital; University of Sydney | CIDM-PH et al. |
| EPI_ISL_2876048, EPI_ISL_2876049, EPI_ISL_2876053, EPI_ISL_2876055, EPI_ISL_2876056, EPI_ISL_2876061, EPI_ISL_2876062, EPI_ISL_2876063, EPI_ISL_2876068, EPI_ISL_2876070, EPI_ISL_2876071, EPI_ISL_2876073, EPI_ISL_2876074, EPI_ISL_2876077, EPI_ISL_2876079, EPI_ISL_2876082, EPI_ISL_2876084, EPI_ISL_2876085 | | | |
| see above | Infinity Biologix | Centers for Disease Control and Prevention Division of Viral Diseases, Pathogen Discovery | Dakota Howard, Dhwani Batra, Peter W. Cook, Kara Moser, Adrian Paskey, Jason Caravas, Benjamin Rambo-Martin, Shatavia Morrison, Christopher Gulvick, Scott Sammons, Yvette Unoarumhi, Darlene Wagner, Matthew Schmerer, Christian Bixby, Yihe Wang, Jonathan Schultz, Chirayu Goswami, Russ Hager, Robin Grimwood, Clinton R. Paden, Duncan MacCannell |
| EPI_ISL_2876086 | A.S.L. NOVARA-BORGOMANERO | Fondazione del Piemonte per l'Oncologia IRCCS | Antonino Sottile, Silvia Brossa, Paola Marino, Giorgio Giardina |
| EPI_ISL_2876089, EPI_ISL_2876094 | A.S.L. VCO | Fondazione del Piemonte per l'Oncologia IRCCS | Antonino Sottile, Silvia Brossa, Paola Marino, Giorgio Giardina |
| EPI_ISL_2876104 | Wexner Medical Center | The Ohio State University College of Medicine | Koenig,S., Seminetta,J. |
| EPI_ISL_2876177, EPI_ISL_2876178, EPI_ISL_2876179, EPI_ISL_2876180, EPI_ISL_2876182, EPI_ISL_2876183, EPI_ISL_2876184, EPI_ISL_2876185, EPI_ISL_2876186, EPI_ISL_2876190, EPI_ISL_2876191, EPI_ISL_2876192, EPI_ISL_2876193, EPI_ISL_2876194, EPI_ISL_2876195, EPI_ISL_2876196, EPI_ISL_2876197, EPI_ISL_2876198, EPI_ISL_2876199, EPI_ISL_2876200, EPI_ISL_2876202, EPI_ISL_2876203, EPI_ISL_2876205, EPI_ISL_2876206, EPI_ISL_2876207, EPI_ISL_2876208, EPI_ISL_2876209, EPI_ISL_2876211, EPI_ISL_2876212, EPI_ISL_2876213, EPI_ISL_2876215, EPI_ISL_2876216, EPI_ISL_2876219, EPI_ISL_2876221, EPI_ISL_2876225, EPI_ISL_2876228, EPI_ISL_2876230, EPI_ISL_2876231, EPI_ISL_2876232, EPI_ISL_2876233, EPI_ISL_2876234, EPI_ISL_2876235, EPI_ISL_2876236, EPI_ISL_2876240, EPI_ISL_2876241, EPI_ISL_2876242, EPI_ISL_2876243, EPI_ISL_2876244, EPI_ISL_2876245, EPI_ISL_2876247, EPI_ISL_2876248, EPI_ISL_2876249, EPI_ISL_2876250, EPI_ISL_2876251, EPI_ISL_2876252, EPI_ISL_2876255, EPI_ISL_2876256, EPI_ISL_2876257, EPI_ISL_2876258, EPI_ISL_2876260, EPI_ISL_2876261, EPI_ISL_2876262, EPI_ISL_2876263, EPI_ISL_2876264, EPI_ISL_2876265, EPI_ISL_2876266, EPI_ISL_2876267, EPI_ISL_2876268, EPI_ISL_2876269, EPI_ISL_2876270, EPI_ISL_2876271, EPI_ISL_2876273, EPI_ISL_2876274, EPI_ISL_2876275 | | | |
| see above | Public Health Laboratory, Minnesota Department of Health | University of Minnesota Genomics Center | Daryl M. Gohl, John Garbe, Jaquelyn Kuriger-Laber, Corbin Dirkx, and Sean Wang |
| EPI_ISL_2876284, EPI_ISL_2876287, EPI_ISL_2876289 | City of Milwaukee Health Department Laboratory | City of Milwaukee Health Department Laboratory | Manjeet Khubbar, Amy Bauer, Jennifer Lentz, Samantha Scott, Sanjib Bhattacharyya |
| EPI_ISL_2876306, EPI_ISL_2876307, EPI_ISL_2876308, EPI_ISL_2876309, EPI_ISL_2876310, EPI_ISL_2876311, EPI_ISL_2876312, EPI_ISL_2876313, EPI_ISL_2876314 | Division of Medical Virology, National Health Laboratory Service (NHLS), Tygerberg Hospital / Stellenbosch University | Division of Medical Virology, National Health Laboratory Service (NHLS), Tygerberg Hospital / Stellenbosch University | Susan Engelbrecht, Tongai Maponga, Bronwyn Kleinhans, Tania Stander, Gert van Zyl, Wolfgang Preiser |
| EPI_ISL_2876316 | PathCare, Cape Town | Division of Medical Virology, National Health Laboratory Service (NHLS), Tygerberg Hospital / Stellenbosch University | Jean Maritz, Petra Raimond, Susan Engelbrecht, Tongai Maponga, Bronwyn Kleinhans, Tania Stander, Gert van Zyl, Wolfgang Preiser |
| EPI_ISL_2876317 | Division of Medical Virology, National Health Laboratory Service (NHLS), Tygerberg Hospital / Stellenbosch University | Division of Medical Virology, National Health Laboratory Service (NHLS), Tygerberg Hospital / Stellenbosch University | Jean Maritz, Petra Raimond, Susan Engelbrecht, Tongai Maponga, Bronwyn Kleinhans, Tania Stander, Gert van Zyl, Wolfgang Preiser |
| EPI_ISL_2876318, EPI_ISL_2876319, EPI_ISL_2876321, EPI_ISL_2876322, EPI_ISL_2876323, EPI_ISL_2876324, EPI_ISL_2876325, EPI_ISL_2876326 | Division of Medical Virology, National Health Laboratory Service (NHLS), Tygerberg Hospital / Stellenbosch University | Division of Medical Virology, National Health Laboratory Service (NHLS), Tygerberg Hospital / Stellenbosch University | Susan Engelbrecht, Tongai Maponga, Bronwyn Kleinhans, Tania Stander, Gert van Zyl, Wolfgang Preiser |
| EPI_ISL_2876328 | PathCare, Cape Town | Division of Medical Virology, National Health Laboratory Service (NHLS), Tygerberg Hospital / Stellenbosch University | Jean Maritz, Petra Raimond, Susan Engelbrecht, Tongai Maponga, Bronwyn Kleinhans, Tania Stander, Gert van Zyl, Wolfgang Preiser |
| EPI_ISL_2876329, EPI_ISL_2876330, EPI_ISL_2876331, EPI_ISL_2876332, EPI_ISL_2876335, EPI_ISL_2876336, EPI_ISL_2876337 | Division of Medical Virology, National Health Laboratory Service (NHLS), Tygerberg Hospital / Stellenbosch University | Division of Medical Virology, National Health Laboratory Service (NHLS), Tygerberg Hospital / Stellenbosch University | Susan Engelbrecht, Tongai Maponga, Bronwyn Kleinhans, Tania Stander, Gert van Zyl, Wolfgang Preiser |
| EPI_ISL_2876339 | PathCare, Cape Town | Division of Medical Virology, National Health Laboratory Service (NHLS), Tygerberg Hospital / Stellenbosch University | Jean Maritz, Petra Raimond, Susan Engelbrecht, Tongai Maponga, Bronwyn Kleinhans, Tania Stander, Gert van Zyl, Wolfgang Preiser |
| EPI_ISL_2876340, EPI_ISL_2876341, EPI_ISL_2876342, EPI_ISL_2876343, EPI_ISL_2876344, EPI_ISL_2876345, EPI_ISL_2876346, | Division of Medical Virology, National Health Laboratory Service (NHLS), Tygerberg Hospital / Stellenbosch University | Division of Medical Virology, National Health Laboratory Service (NHLS), Tygerberg Hospital / Stellenbosch University | Susan Engelbrecht, Tongai Maponga, Bronwyn Kleinhans, Tania Stander, Gert van Zyl, Wolfgang Preiser |

| | | | |
|---|---|---|---|
| EPI_ISL_2876347 | | | |
| EPI_ISL_2876352, EPI_ISL_2876353 | PathCare, Cape Town | Division of Medical Virology, National Health Laboratory Service (NHLS), Tygerberg Hospital / Stellenbosch University | Jean Maritz, Petra Raimond, Susan Engelbrecht, Tongai Maponga, Bronwyn Kleinhans, Tania Stander, Gert van Zyl, Wolfgang Preiser |
| EPI_ISL_2876354, EPI_ISL_2876355, EPI_ISL_2876356, EPI_ISL_2876357, EPI_ISL_2876358, EPI_ISL_2876359, EPI_ISL_2876360, EPI_ISL_2876361, EPI_ISL_2876362 | Division of Medical Virology, National Health Laboratory Service (NHLS), Tygerberg Hospital / Stellenbosch University | Division of Medical Virology, National Health Laboratory Service (NHLS), Tygerberg Hospital / Stellenbosch University | Susan Engelbrecht, Tongai Maponga, Bronwyn Kleinhans, Tania Stander, Gert van Zyl, Wolfgang Preiser |
| EPI_ISL_2876363, EPI_ISL_2876364 | PathCare, Cape Town | Division of Medical Virology, National Health Laboratory Service (NHLS), Tygerberg Hospital / Stellenbosch University | Jean Maritz, Petra Raimond, Susan Engelbrecht, Tongai Maponga, Bronwyn Kleinhans, Tania Stander, Gert van Zyl, Wolfgang Preiser |
| EPI_ISL_2876365, EPI_ISL_2876367, EPI_ISL_2876368, EPI_ISL_2876369, EPI_ISL_2876370, EPI_ISL_2876372, EPI_ISL_2876373 | Division of Medical Virology, National Health Laboratory Service (NHLS), Tygerberg Hospital / Stellenbosch University | Division of Medical Virology, National Health Laboratory Service (NHLS), Tygerberg Hospital / Stellenbosch University | Susan Engelbrecht, Tongai Maponga, Bronwyn Kleinhans, Tania Stander, Gert van Zyl, Wolfgang Preiser |
| EPI_ISL_2876374, EPI_ISL_2876375 | PathCare, Cape Town | Division of Medical Virology, National Health Laboratory Service (NHLS), Tygerberg Hospital / Stellenbosch University | Jean Maritz, Petra Raimond, Susan Engelbrecht, Tongai Maponga, Bronwyn Kleinhans, Tania Stander, Gert van Zyl, Wolfgang Preiser |
| EPI_ISL_2876376, EPI_ISL_2876378, EPI_ISL_2876379, EPI_ISL_2876380, EPI_ISL_2876381, EPI_ISL_2876383, EPI_ISL_2876384 | Division of Medical Virology, National Health Laboratory Service (NHLS), Tygerberg Hospital / Stellenbosch University | Division of Medical Virology, National Health Laboratory Service (NHLS), Tygerberg Hospital / Stellenbosch University | Susan Engelbrecht, Tongai Maponga, Bronwyn Kleinhans, Tania Stander, Gert van Zyl, Wolfgang Preiser |
| EPI_ISL_2876387 | PathCare, Cape Town | Division of Medical Virology, National Health Laboratory Service (NHLS), Tygerberg Hospital / Stellenbosch University | Jean Maritz, Petra Raimond, Susan Engelbrecht, Tongai Maponga, Bronwyn Kleinhans, Tania Stander, Gert van Zyl, Wolfgang Preiser |
| EPI_ISL_2876388, EPI_ISL_2876389, EPI_ISL_2876390, EPI_ISL_2876391, EPI_ISL_2876392, EPI_ISL_2876393, EPI_ISL_2876394, EPI_ISL_2876395 | Division of Medical Virology, National Health Laboratory Service (NHLS), Tygerberg Hospital / Stellenbosch University | Division of Medical Virology, National Health Laboratory Service (NHLS), Tygerberg Hospital / Stellenbosch University | Susan Engelbrecht, Tongai Maponga, Bronwyn Kleinhans, Tania Stander, Gert van Zyl, Wolfgang Preiser |
| EPI_ISL_2876396, EPI_ISL_2876397, EPI_ISL_2876398 | PathCare, Cape Town | Division of Medical Virology, National Health Laboratory Service (NHLS), Tygerberg Hospital / Stellenbosch University | Jean Maritz, Petra Raimond, Susan Engelbrecht, Tongai Maponga, Bronwyn Kleinhans, Tania Stander, Gert van Zyl, Wolfgang Preiser |
| EPI_ISL_2876399, EPI_ISL_2876400, EPI_ISL_2876401, EPI_ISL_2876402, EPI_ISL_2876403, EPI_ISL_2876404, EPI_ISL_2876405, EPI_ISL_2876406, EPI_ISL_2876407, EPI_ISL_2876408, EPI_ISL_2876409, EPI_ISL_2876410, EPI_ISL_2876411, EPI_ISL_2876412, EPI_ISL_2876413, EPI_ISL_2876414, EPI_ISL_2876415, EPI_ISL_2876416, EPI_ISL_2876417, EPI_ISL_2876418, EPI_ISL_2876419, EPI_ISL_2876420, EPI_ISL_2876421, EPI_ISL_2876422 | | | |
| see above | Kansas Health and Environmental Lab | Kansas Health and Environmental Lab | Mike Grose, Katherine Wiggins, Jonathan Barnell, Ben Olsen, and Phil Adam |
| EPI_ISL_2876423 | Osaka Prefecture | Clinical Laboratory Medicine, Kyoto University Graduate School of Medicine | Yasufumi Matsumura, Masaki Yamamoto, Miki Nagao |
| EPI_ISL_2876524 | DC Public Health Lab/ Dept. of Forensic Sciences | DC Public Health Lab/ Dept. of Forensic Sciences | Janis Doss, Scott Nguyen, Elizabeth Zelaya, Sarah Scott, Connie Maza, Monica Mann, Brittany Hamilton, David Payne, Jocelyn Hauser |
| EPI_ISL_2876556 | North Dakota Department of Health, Public Health Laboratory | North Dakota Department of Health, Public Health Laboratory | Lisa Wingerter |
| EPI_ISL_2876622, EPI_ISL_2876700, EPI_ISL_2876726 | Quest Diagnostics Incorporated | Centers for Disease Control and Prevention Division of Viral Diseases, Pathogen Discovery | Dakota Howard, Dhwani Batra, Peter W. Cook, Kara Moser, Adrian Paskey, Jason Caravas, Benjamin Rambo-Martin, Shatavia Morrison, Christopher Gulvick, Scott Sammons, Yvette Unoarumhi, Darlene Wagner, Matthew Schmerer, S. H. Rosenthal, A. Gerasimova, R. M. Kagan, B. Anderson, M. Hua, Y. Liu, L.E. Bernstein, K.E. Livingston, A. Perez, I. A. Shlyakhter, R. V. Rolando, R. Owen, P. Tanpaiboon, F. Lacbawan, Clinton R. Paden, Duncan MacCannell |
| EPI_ISL_2876733 | Laboratory Corporation of America | Centers for Disease Control and Prevention Division of Viral Diseases, Pathogen Discovery | Dakota Howard, Dhwani Batra, Peter W. Cook, Kara Moser, Adrian Paskey, Jason Caravas, Benjamin Rambo-Martin, Shatavia Morrison, Christopher Gulvick, Scott Sammons, Yvette Unoarumhi, Darlene Wagner, Matthew Schmerer, Minoo Agarwal, Eyad Almasri, Debbie Boles, Ayla Burns, Nuthawin Charoensri, Oren Cohen, Susan Countryman, Mary Ann Cristobal, Bobbi Croy, Suzanne Dale, Hrushikesh Deshmukh, Amanda Douglas, Vincent Drouillon, Marcia Eisenberg, Howard Engler, Rama Ghatti, Prashant Gupta, Susan Hicks, Jake Humphrey, Lax Iyer, Lisa Pfefferle, Manoj Jain, Matthew Robinson, Mohan Kolli, Brian Krueger, Tim Kuphal, Stanley Letovsky, Michael Levandoski, Craig Lukasik, Jonathan Meltzer, Brian Norvell, Mindy Nye, Scott Parker, Christos Petropoulos, John Pruitt, Steven Ragan, Scott Ryan, Mike Sapeta, Jana Schroth, Suresh Babu Selvaraju, Goran Stevovic, Amanda Suchanek, Andrea Throop, Lyndon Tilson, Thomas Urban, Joe Voshell, Kimberly Wagner, Jonathan Williams, Mary Williamson, Qian Zeng, Tricia Zwiefelhofer, Clinton R. Paden, Duncan MacCannell |
| EPI_ISL_2876857 | Quest Diagnostics Incorporated | Centers for Disease Control and Prevention Division of Viral Diseases, Pathogen Discovery | Dakota Howard, Dhwani Batra, Peter W. Cook, Kara Moser, Adrian Paskey, Jason Caravas, Benjamin Rambo-Martin, Shatavia Morrison, Christopher Gulvick, Scott Sammons, Yvette Unoarumhi, Darlene Wagner, Matthew Schmerer, S. H. Rosenthal, A. Gerasimova, R. M. Kagan, B. Anderson, M. Hua, Y. Liu, L.E. Bernstein, K.E. Livingston, A. Perez, I. A. Shlyakhter, R. V. Rolando, R. Owen, P. Tanpaiboon, F. Lacbawan, Clinton R. Paden, Duncan MacCannell |
| EPI_ISL_2876937 | Mitchells Plain Hospital wc MPH | NHLS/UCT | Arash Iranzadeh, Deelan Doolabh, Lynn Tyers, Bruna Galvao, Innocent Mudau, Marvin Hsiao, Gert Marais, Diana Hardie, Stephen Korsman, Carolyn Williamson |
| EPI_ISL_2876940 | Heidelberg Clinic wc HBC | NHLS/UCT | Arash Iranzadeh, Deelan Doolabh, Lynn Tyers, Bruna Galvao, Innocent Mudau, Marvin Hsiao, Gert Marais, Diana Hardie, Stephen Korsman, Carolyn Williamson |
| EPI_ISL_2876941 | Great Brak River Clinic wc GBC | NHLS/UCT | Arash Iranzadeh, Deelan Doolabh, Lynn Tyers, Bruna Galvao, Innocent Mudau, Marvin Hsiao, Gert Marais, Diana Hardie, Stephen Korsman, Carolyn Williamson |
| EPI_ISL_2876942 | Clinic-in-Asla | NHLS/UCT | Arash Iranzadeh, Deelan Doolabh, Lynn Tyers, Bruna Galvao, Innocent Mudau, Marvin Hsiao, Gert Marais, Diana Hardie, Stephen Korsman, Carolyn Williamson |
| EPI_ISL_2876944, EPI_ISL_2876945 | 2 Military Hospital wc MAA | NHLS/UCT | Arash Iranzadeh, Deelan Doolabh, Lynn Tyers, Bruna Galvao, Innocent Mudau, Marvin Hsiao, Gert Marais, Diana Hardie, Stephen Korsman, Carolyn Williamson |
| EPI_ISL_2876946 | Victoria Hospital wc VHW | NHLS/UCT | Arash Iranzadeh, Deelan Doolabh, Lynn Tyers, Bruna Galvao, Innocent Mudau, Marvin Hsiao, Gert Marais, Diana Hardie, Stephen Korsman, Carolyn Williamson |
| EPI_ISL_2876948 | Great Brak River Clinic wc GBC | NHLS/UCT | Arash Iranzadeh, Deelan Doolabh, Lynn Tyers, Bruna Galvao, Innocent Mudau, Marvin Hsiao, Gert Marais, Diana Hardie, Stephen Korsman, Carolyn Williamson |
| EPI_ISL_2876949 | Clinic-in-Asla | NHLS/UCT | Arash Iranzadeh, Deelan Doolabh, Lynn Tyers, Bruna Galvao, Innocent Mudau, Marvin Hsiao, Gert Marais, Diana Hardie, Stephen Korsman, Carolyn Williamson |
| EPI_ISL_2876950 | Great Brak River Clinic wc GBC | NHLS/UCT | Arash Iranzadeh, Deelan Doolabh, Lynn Tyers, Bruna Galvao, Innocent Mudau, Marvin Hsiao, Gert Marais, Diana Hardie, Stephen Korsman, Carolyn Williamson |
| EPI_ISL_2876952, EPI_ISL_2876953 | D'Almeida Clinic wc DAL | NHLS/UCT | Arash Iranzadeh, Deelan Doolabh, Lynn Tyers, Bruna Galvao, Innocent Mudau, Marvin Hsiao, Gert Marais, Diana Hardie, Stephen Korsman, Carolyn Williamson |
| EPI_ISL_2876954 | Conville CDC wc CVC | NHLS/UCT | Arash Iranzadeh, Deelan Doolabh, Lynn Tyers, Bruna Galvao, Innocent Mudau, Marvin Hsiao, Gert Marais, Diana Hardie, Stephen Korsman, Carolyn Williamson |
| EPI_ISL_2876955 | Groote Schuur Hospital wc GSH | NHLS/UCT | Arash Iranzadeh, Deelan Doolabh, Lynn Tyers, Bruna Galvao, Innocent Mudau, Marvin Hsiao, Gert Marais, Diana Hardie, Stephen Korsman, Carolyn Williamson |
| EPI_ISL_2876957 | 2 Military Hospital wc MAA | NHLS/UCT | Arash Iranzadeh, Deelan Doolabh, Lynn Tyers, Bruna Galvao, Innocent Mudau, Marvin Hsiao, Gert Marais, Diana Hardie, Stephen Korsman, Carolyn Williamson |
| EPI_ISL_2876959 | George Hospital wc GRH | NHLS/UCT | Arash Iranzadeh, Deelan Doolabh, Lynn Tyers, Bruna Galvao, Innocent Mudau, Marvin Hsiao, Gert Marais, Diana Hardie, Stephen Korsman, Carolyn Williamson |
| EPI_ISL_2876960 | Pacaltsdorp Clinic wc PAC | NHLS/UCT | Arash Iranzadeh, Deelan Doolabh, Lynn Tyers, Bruna Galvao, Innocent Mudau, Marvin Hsiao, Gert Marais, Diana Hardie, Stephen Korsman, Carolyn Williamson |
| EPI_ISL_2876961, EPI_ISL_2876962, EPI_ISL_2876963, EPI_ISL_2876964 | Great Brak River Clinic wc GBC | NHLS/UCT | Arash Iranzadeh, Deelan Doolabh, Lynn Tyers, Bruna Galvao, Innocent Mudau, Marvin Hsiao, Gert Marais, Diana Hardie, Stephen Korsman, Carolyn Williamson |
| EPI_ISL_2876965 | Alma CDC wc AHC | NHLS/UCT | Arash Iranzadeh, Deelan Doolabh, Lynn Tyers, Bruna Galvao, Innocent Mudau, Marvin Hsiao, Gert Marais, Diana Hardie, Stephen Korsman, Carolyn Williamson |
| EPI_ISL_2876966 | Pacaltsdorp Clinic wc PAC | NHLS/UCT | Arash Iranzadeh, Deelan Doolabh, Lynn Tyers, Bruna Galvao, Innocent Mudau, Marvin Hsiao, Gert Marais, Diana Hardie, Stephen Korsman, Carolyn Williamson |
| EPI_ISL_2876967, EPI_ISL_2876968, EPI_ISL_2876969 | Great Brak River Clinic wc GBC | NHLS/UCT | Arash Iranzadeh, Deelan Doolabh, Lynn Tyers, Bruna Galvao, Innocent Mudau, Marvin Hsiao, Gert Marais, Diana Hardie, Stephen Korsman, Carolyn Williamson |
| EPI_ISL_2876970 | Victoria Hospital wc VHW | NHLS/UCT | Arash Iranzadeh, Deelan Doolabh, Lynn Tyers, Bruna Galvao, Innocent Mudau, Marvin Hsiao, Gert Marais, Diana Hardie, Stephen Korsman, Carolyn Williamson |
| EPI_ISL_2876971, EPI_ISL_2876973, EPI_ISL_2876975 | Groote Schuur Hospital wc GSH | NHLS/UCT | Arash Iranzadeh, Deelan Doolabh, Lynn Tyers, Bruna Galvao, Innocent Mudau, Marvin Hsiao, Gert Marais, Diana Hardie, Stephen Korsman, Carolyn Williamson |
| EPI_ISL_2876976 | Victoria Hospital wc VHW | NHLS/UCT | Arash Iranzadeh, Deelan Doolabh, Lynn Tyers, Bruna Galvao, Innocent Mudau, Marvin Hsiao, Gert Marais, Diana Hardie, Stephen Korsman, Carolyn Williamson |
| EPI_ISL_2876977, | Groote Schuur Hospital wc GSH | NHLS/UCT | Arash Iranzadeh, Deelan Doolabh, Lynn Tyers, Bruna Galvao, Innocent Mudau, Marvin Hsiao, Gert Marais, Diana Hardie, Stephen Korsman, Carolyn Williamson |

| | | | |
|---|---|---|---|
| EPI_ISL_2876979 | | | |
| EPI_ISL_2876980 | Claremont Clinic wc CLA | NHLS/UCT | Arash Iranzadeh, Deelan Doolabh, Lynn Tyers, Bruna Galvao, Innocent Mudau, Marvin Hsiao, Gert Marais, Diana Hardie, Stephen Korsman, Carolyn Williamson |
| EPI_ISL_2876981, EPI_ISL_2876983 | Groote Schuur Hospital wc GSH | NHLS/UCT | Arash Iranzadeh, Deelan Doolabh, Lynn Tyers, Bruna Galvao, Innocent Mudau, Marvin Hsiao, Gert Marais, Diana Hardie, Stephen Korsman, Carolyn Williamson |
| EPI_ISL_2876984 | George Hospital wc GRH | NHLS/UCT | Arash Iranzadeh, Deelan Doolabh, Lynn Tyers, Bruna Galvao, Innocent Mudau, Marvin Hsiao, Gert Marais, Diana Hardie, Stephen Korsman, Carolyn Williamson |
| EPI_ISL_2876985 | Thembalethu CDC wc THC | NHLS/UCT | Arash Iranzadeh, Deelan Doolabh, Lynn Tyers, Bruna Galvao, Innocent Mudau, Marvin Hsiao, Gert Marais, Diana Hardie, Stephen Korsman, Carolyn Williamson |
| EPI_ISL_2876986 | Pacaltsdorp Clinic wc PAC | NHLS/UCT | Arash Iranzadeh, Deelan Doolabh, Lynn Tyers, Bruna Galvao, Innocent Mudau, Marvin Hsiao, Gert Marais, Diana Hardie, Stephen Korsman, Carolyn Williamson |
| EPI_ISL_2876987 | Conville CDC wc CVC | NHLS/UCT | Arash Iranzadeh, Deelan Doolabh, Lynn Tyers, Bruna Galvao, Innocent Mudau, Marvin Hsiao, Gert Marais, Diana Hardie, Stephen Korsman, Carolyn Williamson |
| EPI_ISL_2876988 | Clinic-in-Asla | NHLS/UCT | Arash Iranzadeh, Deelan Doolabh, Lynn Tyers, Bruna Galvao, Innocent Mudau, Marvin Hsiao, Gert Marais, Diana Hardie, Stephen Korsman, Carolyn Williamson |
| EPI_ISL_2876990 | D'Almeida Clinic wc DAL | NHLS/UCT | Arash Iranzadeh, Deelan Doolabh, Lynn Tyers, Bruna Galvao, Innocent Mudau, Marvin Hsiao, Gert Marais, Diana Hardie, Stephen Korsman, Carolyn Williamson |
| EPI_ISL_2876991 | Groote Schuur Hospital wc GSH | NHLS/UCT | Arash Iranzadeh, Deelan Doolabh, Lynn Tyers, Bruna Galvao, Innocent Mudau, Marvin Hsiao, Gert Marais, Diana Hardie, Stephen Korsman, Carolyn Williamson |
| EPI_ISL_2876992 | D'Almeida Clinic wc DAL | NHLS/UCT | Arash Iranzadeh, Deelan Doolabh, Lynn Tyers, Bruna Galvao, Innocent Mudau, Marvin Hsiao, Gert Marais, Diana Hardie, Stephen Korsman, Carolyn Williamson |
| EPI_ISL_2876994 | Pelican Park CDC wc PAX | NHLS/UCT | Arash Iranzadeh, Deelan Doolabh, Lynn Tyers, Bruna Galvao, Innocent Mudau, Marvin Hsiao, Gert Marais, Diana Hardie, Stephen Korsman, Carolyn Williamson |
| EPI_ISL_2876995, EPI_ISL_2876996 | Parkwood Clinic wc PRK | NHLS/UCT | Arash Iranzadeh, Deelan Doolabh, Lynn Tyers, Bruna Galvao, Innocent Mudau, Marvin Hsiao, Gert Marais, Diana Hardie, Stephen Korsman, Carolyn Williamson |
| EPI_ISL_2876997 | 2 Military Hospital wc MAA | NHLS/UCT | Arash Iranzadeh, Deelan Doolabh, Lynn Tyers, Bruna Galvao, Innocent Mudau, Marvin Hsiao, Gert Marais, Diana Hardie, Stephen Korsman, Carolyn Williamson |
| EPI_ISL_2876998 | Groote Schuur Hospital wc GSH | NHLS/UCT | Arash Iranzadeh, Deelan Doolabh, Lynn Tyers, Bruna Galvao, Innocent Mudau, Marvin Hsiao, Gert Marais, Diana Hardie, Stephen Korsman, Carolyn Williamson |
| EPI_ISL_2876999 | Red Cross Children's Hospital wc RXH | NHLS/UCT | Arash Iranzadeh, Deelan Doolabh, Lynn Tyers, Bruna Galvao, Innocent Mudau, Marvin Hsiao, Gert Marais, Diana Hardie, Stephen Korsman, Carolyn Williamson |
| EPI_ISL_2877000 | Mitchells Plain Hospital wc MPH | NHLS/UCT | Arash Iranzadeh, Deelan Doolabh, Lynn Tyers, Bruna Galvao, Innocent Mudau, Marvin Hsiao, Gert Marais, Diana Hardie, Stephen Korsman, Carolyn Williamson |
| EPI_ISL_2877001 | Heideveld Emergency Centre | NHLS/UCT | Arash Iranzadeh, Deelan Doolabh, Lynn Tyers, Bruna Galvao, Innocent Mudau, Marvin Hsiao, Gert Marais, Diana Hardie, Stephen Korsman, Carolyn Williamson |
| EPI_ISL_2877002 | Knysna Hospital wc KNY | NHLS/UCT | Arash Iranzadeh, Deelan Doolabh, Lynn Tyers, Bruna Galvao, Innocent Mudau, Marvin Hsiao, Gert Marais, Diana Hardie, Stephen Korsman, Carolyn Williamson |
| EPI_ISL_2877003 | Knysna CDC wc WLC | NHLS/UCT | Arash Iranzadeh, Deelan Doolabh, Lynn Tyers, Bruna Galvao, Innocent Mudau, Marvin Hsiao, Gert Marais, Diana Hardie, Stephen Korsman, Carolyn Williamson |
| EPI_ISL_2877004, EPI_ISL_2877005 | Kwanokuthula CDC wc KWA | NHLS/UCT | Arash Iranzadeh, Deelan Doolabh, Lynn Tyers, Bruna Galvao, Innocent Mudau, Marvin Hsiao, Gert Marais, Diana Hardie, Stephen Korsman, Carolyn Williamson |
| EPI_ISL_2877006 | Hornlee Clinic wc HLC | NHLS/UCT | Arash Iranzadeh, Deelan Doolabh, Lynn Tyers, Bruna Galvao, Innocent Mudau, Marvin Hsiao, Gert Marais, Diana Hardie, Stephen Korsman, Carolyn Williamson |
| EPI_ISL_2877030, EPI_ISL_2877031 | Chulabhorn Hospital | National Institute of Health, Department of Medical Sciences, Ministry of Public Health, Thailand | Pilailuk Okada; Siripaporn Phuygun; Nuttida Thongpramul; Sittiporn Parnmen; Ratana Tacharoenmuang; Pakorn Piromtong; Natchaya Khiadsang; Thanutsapa Thanadachakul; Warawan Wongboot; sirikanda wimol; Sunthareeya Waicharoen; |
| EPI_ISL_2877032, EPI_ISL_2877033 | Regional Medical Sciences Center 1/1 Chiangrai | National Institute of Health, Department of Medical Sciences, Ministry of Public Health, Thailand | Pilailuk Okada; Siripaporn Phuygun; Nuttida Thongpramul; Sittiporn Parnmen; Ratana Tacharoenmuang; Pakorn Piromtong; Natchaya Khiadsang; Thanutsapa Thanadachakul; Warawan Wongboot; sirikanda wimol; Sunthareeya Waicharoen; |
| EPI_ISL_2877034, EPI_ISL_2877035, EPI_ISL_2877036, EPI_ISL_2877037 | Regional Medical Sciences Center 8 Udonthani | National Institute of Health, Department of Medical Sciences, Ministry of Public Health, Thailand | Pilailuk Okada; Siripaporn Phuygun; Nuttida Thongpramul; Sittiporn Parnmen; Ratana Tacharoenmuang; Pakorn Piromtong; Natchaya Khiadsang; Thanutsapa Thanadachakul; Warawan Wongboot; sirikanda wimol; Sunthareeya Waicharoen; |
| EPI_ISL_2877038 | St Vincent's Pathology (SydPath) | NSW Health Pathology - Institute of Clinical Pathology and Medical Research; Westmead Hospital; University of Sydney | CIDM-PH et al. |
| EPI_ISL_2877039, EPI_ISL_2877040, EPI_ISL_2877041, EPI_ISL_2877042 | Laverty Pathology | NSW Health Pathology - Institute of Clinical Pathology and Medical Research; Westmead Hospital; University of Sydney | CIDM-PH et al. |
| EPI_ISL_2877043, EPI_ISL_2877044 | Medlab Pathology | NSW Health Pathology - Institute of Clinical Pathology and Medical Research; Westmead Hospital; University of Sydney | CIDM-PH et al. |
| EPI_ISL_2877050, EPI_ISL_2877052, EPI_ISL_2877053, EPI_ISL_2877055, EPI_ISL_2877056, EPI_ISL_2877362, EPI_ISL_2877426, EPI_ISL_2877428, EPI_ISL_2877430, EPI_ISL_2877432, EPI_ISL_2877434, EPI_ISL_2877436, EPI_ISL_2877515, EPI_ISL_2877516, EPI_ISL_2877533, EPI_ISL_2877534, EPI_ISL_2877543, EPI_ISL_2877553, EPI_ISL_2877559, EPI_ISL_2877560, EPI_ISL_2877567, EPI_ISL_2877580, EPI_ISL_2877582, EPI_ISL_2877583, EPI_ISL_2877624 | | | |
| see above | KU Leuven, Rega Institute, Clinical and Epidemiological Virology | KU Leuven, Rega Institute, Clinical and Epidemiological Virology | Tony Wawina-Bokalanga, Bert Vanmechelen, Joan Marti-Carerras, Piet Maes |
| EPI_ISL_2877649, EPI_ISL_2877650, EPI_ISL_2877651, EPI_ISL_2877652, EPI_ISL_2877653, EPI_ISL_2877654 | Genome Information Research Center, Research Institute for Microbial Diseases, Osaka University | Genome Information Research Center, Research Institute for Microbial Diseases, Osaka University | Daisuke Motooka, Kentaro Tanaka, Hiroya Oki, and Shota Nakamura |
| EPI_ISL_2877656, EPI_ISL_2877658, EPI_ISL_2877660, EPI_ISL_2877661, EPI_ISL_2877666, EPI_ISL_2877670, EPI_ISL_2877672, EPI_ISL_2877674, EPI_ISL_2877675, EPI_ISL_2877677, EPI_ISL_2877679, EPI_ISL_2877680, EPI_ISL_2877681, EPI_ISL_2877682, EPI_ISL_2877683, EPI_ISL_2877684, EPI_ISL_2877685, EPI_ISL_2877687, EPI_ISL_2877688, EPI_ISL_2877689, EPI_ISL_2877690, EPI_ISL_2877693, EPI_ISL_2877695 | | | |
| see above | Aegis Sciences Corporation | Centers for Disease Control and Prevention Division of Viral Diseases, Pathogen Discovery | Dakota Howard, Dhwani Batra, Peter W. Cook, Kara Moser, Adrian Paskey, Jason Caravas, Benjamin Rambo-Martin, Shatavia Morrison, Christopher Gulvick, Scott Sammons, Yvette Unoarumhi, Darlene Wagner, Matthew Schmerer, Cyndi Clark, Patrick Campbell, Rob Case, Vikramsinha Ghorpade, Holly Houdeshell, Ola Kvalvaag, Dillon Nall, Ethan Sanders, Alec Vest, Shaun Westlund, Matthew Hardison, Clinton R. Paden, Duncan MacCannell |
| EPI_ISL_2877701 | Infinity Biologix | Centers for Disease Control and Prevention Division of Viral Diseases, Pathogen Discovery | Dakota Howard, Dhwani Batra, Peter W. Cook, Kara Moser, Adrian Paskey, Jason Caravas, Benjamin Rambo-Martin, Shatavia Morrison, Christopher Gulvick, Scott Sammons, Yvette Unoarumhi, Darlene Wagner, Matthew Schmerer, Christian Bixby, Yihe Wang, Jonathan Schultz, Chirayu Goswami, Russ Hager, Robin Grimwood, Clinton R. Paden, Duncan MacCannell |
| EPI_ISL_2877706, EPI_ISL_2877708, EPI_ISL_2877709, EPI_ISL_2877711, EPI_ISL_2877712, EPI_ISL_2877713, EPI_ISL_2877714, EPI_ISL_2877715, EPI_ISL_2877717, EPI_ISL_2877721, EPI_ISL_2877723, EPI_ISL_2877725, EPI_ISL_2877726, EPI_ISL_2877727, EPI_ISL_2877728 | | | |
| see above | County Hospital of Liberec, Dept. of genetic and molecular diagnostic | Institute of Medical Microbiology and Virology, University Hospital Carl Gustav Carus, TU Dresden | Marlena Stadtmüller, Alexa Laubner, Johanna Beil, Fabian Rost, Sylvia Klemroth, Grit Mehnert, Montserrat Palau de Miquel, Susanne Reinhardt, Leo Büttner, Anett Zabzinski, Sylke Winkler, Alexander Dalpke, Iva Dolinová |
| EPI_ISL_2877729, EPI_ISL_2877730, EPI_ISL_2877734 | Oddělení klinické mikrobiologie | Institute of Medical Microbiology and Virology, University Hospital Carl Gustav Carus, TU Dresden | Marlena Stadtmüller, Alexa Laubner, Johanna Beil, Fabian Rost, Sylvia Klemroth, Grit Mehnert, Montserrat Palau de Miquel, Susanne Reinhardt, Leo Büttner, Anett Zabzinski, Sylke Winkler, Alexander Dalpke, Lenka Šimůnková, Lenka Dvořáková |
| EPI_ISL_2877746, EPI_ISL_2877747, EPI_ISL_2877748, EPI_ISL_2877749, EPI_ISL_2877750, EPI_ISL_2877751, EPI_ISL_2877752, EPI_ISL_2877753, EPI_ISL_2877754, EPI_ISL_2877755, EPI_ISL_2877756, EPI_ISL_2877757, EPI_ISL_2877758, EPI_ISL_2877759, EPI_ISL_2877760, EPI_ISL_2877761, EPI_ISL_2877762, EPI_ISL_2877763, EPI_ISL_2877764, EPI_ISL_2877765, EPI_ISL_2877766, EPI_ISL_2877767, EPI_ISL_2877768, EPI_ISL_2877769, EPI_ISL_2877770, EPI_ISL_2877771, EPI_ISL_2877772, EPI_ISL_2877773, EPI_ISL_2877774, EPI_ISL_2877775, EPI_ISL_2877776, EPI_ISL_2877777, EPI_ISL_2877778, EPI_ISL_2877779, EPI_ISL_2877780, EPI_ISL_2877781, EPI_ISL_2877782, EPI_ISL_2877783, EPI_ISL_2877784, EPI_ISL_2877785, EPI_ISL_2877786, EPI_ISL_2877787, EPI_ISL_2877788, EPI_ISL_2877789, EPI_ISL_2877790, EPI_ISL_2877791, EPI_ISL_2877792, EPI_ISL_2877793, EPI_ISL_2877794, EPI_ISL_2877795, EPI_ISL_2877797, EPI_ISL_2877798, EPI_ISL_2877799, EPI_ISL_2877800, EPI_ISL_2877801, EPI_ISL_2877802, EPI_ISL_2877803, EPI_ISL_2877804, EPI_ISL_2877805, EPI_ISL_2877806, EPI_ISL_2877807, EPI_ISL_2877808, EPI_ISL_2877809, EPI_ISL_2877810, EPI_ISL_2877811, EPI_ISL_2877812, EPI_ISL_2877813, EPI_ISL_2877814, EPI_ISL_2877815, EPI_ISL_2877816, EPI_ISL_2877817, EPI_ISL_2877818, EPI_ISL_2877819, EPI_ISL_2877820, EPI_ISL_2877821, EPI_ISL_2877822, EPI_ISL_2877823, EPI_ISL_2877824, EPI_ISL_2877825, EPI_ISL_2877826, EPI_ISL_2877827, EPI_ISL_2877828, EPI_ISL_2877829, EPI_ISL_2877830, EPI_ISL_2877831, EPI_ISL_2877832, EPI_ISL_2877833, EPI_ISL_2877834, EPI_ISL_2877835, EPI_ISL_2877836, EPI_ISL_2877837, EPI_ISL_2877838, EPI_ISL_2877839, EPI_ISL_2877840, EPI_ISL_2877841, EPI_ISL_2877842, EPI_ISL_2877843, EPI_ISL_2877844, EPI_ISL_2877845, EPI_ISL_2877846, EPI_ISL_2877847, EPI_ISL_2877848, EPI_ISL_2877849, EPI_ISL_2877850, EPI_ISL_2877851, EPI_ISL_2877852, EPI_ISL_2877853, EPI_ISL_2877854, EPI_ISL_2877855, EPI_ISL_2877856, EPI_ISL_2877857, EPI_ISL_2877858, EPI_ISL_2877859, EPI_ISL_2877860, EPI_ISL_2877861, EPI_ISL_2877862, EPI_ISL_2877863, EPI_ISL_2877864, EPI_ISL_2877865, EPI_ISL_2877866, EPI_ISL_2877867, EPI_ISL_2877868, EPI_ISL_2877869, EPI_ISL_2877870, EPI_ISL_2877871, EPI_ISL_2877872, EPI_ISL_2877873, EPI_ISL_2877874, EPI_ISL_2877875, EPI_ISL_2877876, EPI_ISL_2877877, EPI_ISL_2877878, EPI_ISL_2877879, EPI_ISL_2877880, EPI_ISL_2877881, EPI_ISL_2877882, EPI_ISL_2877883, EPI_ISL_2877884, EPI_ISL_2877885, EPI_ISL_2877886, EPI_ISL_2877887, EPI_ISL_2877888, EPI_ISL_2877889 | | | |
| see above | INSACOG-Assam | National Institute of Biomedical Genomics – INSACOG | Arindam Maitra, Ajanta Sharma, Nidhan Kumar Biswas, Lahari Saikia, Sreedhar Chinnaswamy, Kailash Chamuah, Saumitra Das |
| EPI_ISL_2877890, EPI_ISL_2877891, EPI_ISL_2877892, EPI_ISL_2877893, EPI_ISL_2877894, EPI_ISL_2877895, EPI_ISL_2877896, EPI_ISL_2877897, EPI_ISL_2877900, EPI_ISL_2877901, EPI_ISL_2877902, EPI_ISL_2877903, EPI_ISL_2877906, EPI_ISL_2877907, EPI_ISL_2877908, EPI_ISL_2877909, EPI_ISL_2877910, EPI_ISL_2877911, EPI_ISL_2877913, EPI_ISL_2877914, EPI_ISL_2877915, EPI_ISL_2877917, EPI_ISL_2877918, EPI_ISL_2877919, EPI_ISL_2877920, EPI_ISL_2877922, EPI_ISL_2877923, EPI_ISL_2877924, EPI_ISL_2877926, EPI_ISL_2877927, EPI_ISL_2877928, EPI_ISL_2877929, EPI_ISL_2877930, EPI_ISL_2877931, EPI_ISL_2877932, EPI_ISL_2877933, EPI_ISL_2877934, EPI_ISL_2877935, EPI_ISL_2877936, EPI_ISL_2877937, EPI_ISL_2877938, EPI_ISL_2877939, EPI_ISL_2877941, EPI_ISL_2877942, EPI_ISL_2877943, EPI_ISL_2877944, EPI_ISL_2877945, EPI_ISL_2877946, EPI_ISL_2877947, EPI_ISL_2877948, EPI_ISL_2877949, EPI_ISL_2877950, EPI_ISL_2877954, EPI_ISL_2877955, EPI_ISL_2877956, EPI_ISL_2877957, EPI_ISL_2877958, EPI_ISL_2877959, EPI_ISL_2877960, EPI_ISL_2877961, EPI_ISL_2877962, EPI_ISL_2877963, EPI_ISL_2877964, EPI_ISL_2877965, EPI_ISL_2877966, EPI_ISL_2877967, EPI_ISL_2877968, EPI_ISL_2877969, EPI_ISL_2877970, EPI_ISL_2877971, EPI_ISL_2877972, EPI_ISL_2877973, EPI_ISL_2877974, EPI_ISL_2877975, EPI_ISL_2877976, EPI_ISL_2877977, EPI_ISL_2877978, EPI_ISL_2877979, EPI_ISL_2877980, EPI_ISL_2877981, EPI_ISL_2877982, EPI_ISL_2877983, EPI_ISL_2877984, EPI_ISL_2877985, EPI_ISL_2877986, EPI_ISL_2877987, EPI_ISL_2877988, EPI_ISL_2877989, EPI_ISL_2877990, EPI_ISL_2877991, EPI_ISL_2877992, EPI_ISL_2877993, EPI_ISL_2877994, EPI_ISL_2877995, EPI_ISL_2877996, EPI_ISL_2877997, EPI_ISL_2877998, EPI_ISL_2877999, EPI_ISL_2878001, EPI_ISL_2878002, EPI_ISL_2878003, EPI_ISL_2878004, EPI_ISL_2878005, EPI_ISL_2878006, EPI_ISL_2878007, EPI_ISL_2878008, EPI_ISL_2878009, EPI_ISL_2878010, EPI_ISL_2878011, EPI_ISL_2878012, EPI_ISL_2878013, EPI_ISL_2878014, EPI_ISL_2878015, EPI_ISL_2878016, EPI_ISL_2878017, EPI_ISL_2878018, EPI_ISL_2878019, EPI_ISL_2878020, EPI_ISL_2878021, EPI_ISL_2878022, EPI_ISL_2878023, EPI_ISL_2878024, EPI_ISL_2878025, EPI_ISL_2878026, EPI_ISL_2878027, EPI_ISL_2878028, EPI_ISL_2878029, EPI_ISL_2878030, EPI_ISL_2878031, EPI_ISL_2878032, EPI_ISL_2878033, EPI_ISL_2878034, EPI_ISL_2878035, EPI_ISL_2878036, EPI_ISL_2878037, EPI_ISL_2878038, EPI_ISL_2878039 | | | |
| see above | INSACOG-Tripura | National Institute of Biomedical Genomics – INSACOG | Arindam Maitra, Tapan Majumdar, Nidhan Kumar Biswas, Saikat Majumder, Sreedhar Chinnaswamy, Apurba Sarkar, Saumitra Das |
| EPI_ISL_2878040, EPI_ISL_2878045, EPI_ISL_2878046, EPI_ISL_2878052, EPI_ISL_2878056 | County Hospital of Liberec, Dept. of genetic and molecular diagnostic | Institute of Medical Microbiology and Virology, University Hospital Carl Gustav Carus, TU Dresden | Marlena Stadtmüller, Alexa Laubner, Johanna Beil, Fabian Rost, Sylvia Klemroth, Grit Mehnert, Montserrat Palau de Miquel, Susanne Reinhardt, Leo Büttner, Anett Zabzinski, Sylke Winkler, Alexander Dalpke, Iva Dolinová |
| EPI_ISL_2878070 | Viešoji įstaiga Klaipėdos universitetinė ligoninė | Institute of Biotechnology, Life Sciences Center, Vilnius University | Emilija Vasiliunaite; Milda Norkiene; Danguole Ziogiene; Albertas Timinskas; Alma Gedvilaite |
| EPI_ISL_2878075 | UAB Medicina practica laboratorija | Institute of Biotechnology, Life Sciences Center, Vilnius University | Emilija Vasiliunaite; Milda Norkiene; Danguole Ziogiene; Albertas Timinskas; Alma Gedvilaite |
| EPI_ISL_2878097 | UAB Diagnostikos laboratorija | National Public Health Surveillance Laboratory | Lukas Zemaitis, Migle Gabrielaite, Jelena Razmuk, Svajune Muralyte, Ana Steponkiene, Lukas Vasionis, Danas Baksa |
| EPI_ISL_2878099 | Viešoji įstaiga Klaipėdos universitetinė ligoninė | National Public Health Surveillance Laboratory | Lukas Zemaitis, Migle Gabrielaite, Jelena Razmuk, Svajune Muralyte, Ana Steponkiene, Lukas Vasionis, Danas Baksa |
| EPI_ISL_2878104, EPI_ISL_2878129 | UAB Diagnostikos laboratorija | National Public Health Surveillance Laboratory | Lukas Zemaitis, Migle Gabrielaite, Jelena Razmuk, Svajune Muralyte, Ana Steponkiene, Lukas Vasionis, Danas Baksa |

EPI_ISL_2878165, EPI_ISL_2878167, EPI_ISL_2878168, EPI_ISL_2878169, EPI_ISL_2878170, EPI_ISL_2878171, EPI_ISL_2878172, EPI_ISL_2878173, EPI_ISL_2878179, EPI_ISL_2878182, EPI_ISL_2878183, EPI_ISL_2878184, EPI_ISL_2878185, EPI_ISL_2878187, EPI_ISL_2878188, EPI_ISL_2878189, EPI_ISL_2878190, EPI_ISL_2878192, EPI_ISL_2878194, EPI_ISL_2878195, EPI_ISL_2878196, EPI_ISL_2878197, EPI_ISL_2878200, EPI_ISL_2878201, EPI_ISL_2878202

| | | | |
|---|---|---|---|
| see above | KU Leuven, Rega Institute, Clinical and Epidemiological Virology | KU Leuven, Rega Institute, Clinical and Epidemiological Virology | Tony Wawina-Bokalanga, Bert Vanmechelen, Joan Marti-Carerras, Piet Maes |
| EPI_ISL_2878247 | Biogroup Bio Lam-LCD Saint-Denis | Department of Virology, Henri Mondor University Hospital, Assistance Publique Hôpitaux de Paris, Université Paris-Est Créteil, INSERM U955 | Christophe Rodriguez, Slim Fourati, Vanessa Demontant, Guillaume Gricourt, Melissa N'Debi, Alexandre Soulier, Elisabeth Trawinski, Jean-Michel Pawlotsky |
| EPI_ISL_2878249, EPI_ISL_2878250 | SELAS MEDILYS | Department of Virology, Henri Mondor University Hospital, Assistance Publique Hôpitaux de Paris, Université Paris-Est Créteil, INSERM U955 | Christophe Rodriguez, Slim Fourati, Vanessa Demontant, Guillaume Gricourt, Melissa N'Debi, Alexandre Soulier, Elisabeth Trawinski, Jean-Michel Pawlotsky |
| EPI_ISL_2878251 | BROUSSAIS | Department of Virology, Henri Mondor University Hospital, Assistance Publique Hôpitaux de Paris, Université Paris-Est Créteil, INSERM U955 | Christophe Rodriguez, Slim Fourati, Vanessa Demontant, Guillaume Gricourt, Melissa N'Debi, Alexandre Soulier, Elisabeth Trawinski, Jean-Michel Pawlotsky |
| EPI_ISL_2878252 | Biogroup Bio Lam-LCD Saint-Denis | Department of Virology, Henri Mondor University Hospital, Assistance Publique Hôpitaux de Paris, Université Paris-Est Créteil, INSERM U955 | Christophe Rodriguez, Slim Fourati, Vanessa Demontant, Guillaume Gricourt, Melissa N'Debi, Alexandre Soulier, Elisabeth Trawinski, Jean-Michel Pawlotsky |
| EPI_ISL_2878253 | Hôpital Avicenne | Department of Virology, Henri Mondor University Hospital, Assistance Publique Hôpitaux de Paris, Université Paris-Est Créteil, INSERM U955 | Christophe Rodriguez, Slim Fourati, Vanessa Demontant, Guillaume Gricourt, Melissa N'Debi, Alexandre Soulier, Elisabeth Trawinski, Jean-Michel Pawlotsky |
| EPI_ISL_2878254 | BROUSSAIS | Department of Virology, Henri Mondor University Hospital, Assistance Publique Hôpitaux de Paris, Université Paris-Est Créteil, INSERM U955 | Christophe Rodriguez, Slim Fourati, Vanessa Demontant, Guillaume Gricourt, Melissa N'Debi, Alexandre Soulier, Elisabeth Trawinski, Jean-Michel Pawlotsky |
| EPI_ISL_2878257 | SELAS MEDILYS | Department of Virology, Henri Mondor University Hospital, Assistance Publique Hôpitaux de Paris, Université Paris-Est Créteil, INSERM U955 | Christophe Rodriguez, Slim Fourati, Vanessa Demontant, Guillaume Gricourt, Melissa N'Debi, Alexandre Soulier, Elisabeth Trawinski, Jean-Michel Pawlotsky |
| EPI_ISL_2878258 | Biogroup Bio Lam-LCD Saint-Denis | Department of Virology, Henri Mondor University Hospital, Assistance Publique Hôpitaux de Paris, Université Paris-Est Créteil, INSERM U955 | Christophe Rodriguez, Slim Fourati, Vanessa Demontant, Guillaume Gricourt, Melissa N'Debi, Alexandre Soulier, Elisabeth Trawinski, Jean-Michel Pawlotsky |
| EPI_ISL_2878259 | SELAS MEDILYS | Department of Virology, Henri Mondor University Hospital, Assistance Publique Hôpitaux de Paris, Université Paris-Est Créteil, INSERM U955 | Christophe Rodriguez, Slim Fourati, Vanessa Demontant, Guillaume Gricourt, Melissa N'Debi, Alexandre Soulier, Elisabeth Trawinski, Jean-Michel Pawlotsky |
| EPI_ISL_2878260 | Biogroup Bio Lam-LCD Saint-Denis | Department of Virology, Henri Mondor University Hospital, Assistance Publique Hôpitaux de Paris, Université Paris-Est Créteil, INSERM U955 | Christophe Rodriguez, Slim Fourati, Vanessa Demontant, Guillaume Gricourt, Melissa N'Debi, Alexandre Soulier, Elisabeth Trawinski, Jean-Michel Pawlotsky |
| EPI_ISL_2878261 | SELAS MEDILYS | Department of Virology, Henri Mondor University Hospital, Assistance Publique Hôpitaux de Paris, Université Paris-Est Créteil, INSERM U955 | Christophe Rodriguez, Slim Fourati, Vanessa Demontant, Guillaume Gricourt, Melissa N'Debi, Alexandre Soulier, Elisabeth Trawinski, Jean-Michel Pawlotsky |
| EPI_ISL_2878262, EPI_ISL_2878263, EPI_ISL_2878264, EPI_ISL_2878265 | Biogroup Bio Lam-LCD Saint-Denis | Department of Virology, Henri Mondor University Hospital, Assistance Publique Hôpitaux de Paris, Université Paris-Est Créteil, INSERM U955 | Christophe Rodriguez, Slim Fourati, Vanessa Demontant, Guillaume Gricourt, Melissa N'Debi, Alexandre Soulier, Elisabeth Trawinski, Jean-Michel Pawlotsky |
| EPI_ISL_2878266 | SELAS MEDILYS | Department of Virology, Henri Mondor University Hospital, Assistance Publique Hôpitaux de Paris, Université Paris-Est Créteil, INSERM U955 | Christophe Rodriguez, Slim Fourati, Vanessa Demontant, Guillaume Gricourt, Melissa N'Debi, Alexandre Soulier, Elisabeth Trawinski, Jean-Michel Pawlotsky |
| EPI_ISL_2878268 | Biogroup Bio Lam-LCD Saint-Denis | Department of Virology, Henri Mondor University Hospital, Assistance Publique Hôpitaux de Paris, Université Paris-Est Créteil, INSERM U955 | Christophe Rodriguez, Slim Fourati, Vanessa Demontant, Guillaume Gricourt, Melissa N'Debi, Alexandre Soulier, Elisabeth Trawinski, Jean-Michel Pawlotsky |
| EPI_ISL_2878269 | BROUSSAIS | Department of Virology, Henri Mondor University Hospital, Assistance Publique Hôpitaux de Paris, Université Paris-Est Créteil, INSERM U955 | Christophe Rodriguez, Slim Fourati, Vanessa Demontant, Guillaume Gricourt, Melissa N'Debi, Alexandre Soulier, Elisabeth Trawinski, Jean-Michel Pawlotsky |
| EPI_ISL_2878271 | Biogroup Bio Lam-LCD Saint-Denis | Department of Virology, Henri Mondor University Hospital, Assistance Publique Hôpitaux de Paris, Université Paris-Est Créteil, INSERM U955 | Christophe Rodriguez, Slim Fourati, Vanessa Demontant, Guillaume Gricourt, Melissa N'Debi, Alexandre Soulier, Elisabeth Trawinski, Jean-Michel Pawlotsky |
| EPI_ISL_2878272 | BROUSSAIS | Department of Virology, Henri Mondor University Hospital, Assistance Publique Hôpitaux de Paris, Université Paris-Est Créteil, INSERM U955 | Christophe Rodriguez, Slim Fourati, Vanessa Demontant, Guillaume Gricourt, Melissa N'Debi, Alexandre Soulier, Elisabeth Trawinski, Jean-Michel Pawlotsky |
| EPI_ISL_2878273, EPI_ISL_2878274, EPI_ISL_2878275, EPI_ISL_2878277, EPI_ISL_2878278, EPI_ISL_2878279, EPI_ISL_2878280 | Biogroup Bio Lam-LCD Saint-Denis | Department of Virology, Henri Mondor University Hospital, Assistance Publique Hôpitaux de Paris, Université Paris-Est Créteil, INSERM U955 | Christophe Rodriguez, Slim Fourati, Vanessa Demontant, Guillaume Gricourt, Melissa N'Debi, Alexandre Soulier, Elisabeth Trawinski, Jean-Michel Pawlotsky |
| EPI_ISL_2878281 | SELAS MEDILYS | Department of Virology, Henri Mondor University Hospital, Assistance Publique Hôpitaux de Paris, Université Paris-Est Créteil, INSERM U955 | Christophe Rodriguez, Slim Fourati, Vanessa Demontant, Guillaume Gricourt, Melissa N'Debi, Alexandre Soulier, Elisabeth Trawinski, Jean-Michel Pawlotsky |
| EPI_ISL_2878283 | BROUSSAIS | Department of Virology, Henri Mondor University Hospital, Assistance Publique Hôpitaux de Paris, Université Paris-Est Créteil, INSERM U955 | Christophe Rodriguez, Slim Fourati, Vanessa Demontant, Guillaume Gricourt, Melissa N'Debi, Alexandre Soulier, Elisabeth Trawinski, Jean-Michel Pawlotsky |
| EPI_ISL_2878286 | Biogroup Bio Lam-LCD Saint-Denis | Department of Virology, Henri Mondor University Hospital, Assistance Publique Hôpitaux de Paris, Université Paris-Est Créteil, INSERM U955 | Christophe Rodriguez, Slim Fourati, Vanessa Demontant, Guillaume Gricourt, Melissa N'Debi, Alexandre Soulier, Elisabeth Trawinski, Jean-Michel Pawlotsky |
| EPI_ISL_2878287 | SELAS MEDILYS | Department of Virology, Henri Mondor University Hospital, Assistance Publique Hôpitaux de Paris, Université Paris-Est Créteil, INSERM U955 | Christophe Rodriguez, Slim Fourati, Vanessa Demontant, Guillaume Gricourt, Melissa N'Debi, Alexandre Soulier, Elisabeth Trawinski, Jean-Michel Pawlotsky |
| EPI_ISL_2878288, EPI_ISL_2878289, EPI_ISL_2878290 | Biogroup Bio Lam-LCD Saint-Denis | Department of Virology, Henri Mondor University Hospital, Assistance Publique Hôpitaux de Paris, Université Paris-Est Créteil, INSERM U955 | Christophe Rodriguez, Slim Fourati, Vanessa Demontant, Guillaume Gricourt, Melissa N'Debi, Alexandre Soulier, Elisabeth Trawinski, Jean-Michel Pawlotsky |
| EPI_ISL_2878291 | BIO67-BIOSPHERE | Department of Virology, Henri Mondor University Hospital, Assistance Publique Hôpitaux de Paris, Université Paris-Est Créteil, INSERM U955 | Christophe Rodriguez, Slim Fourati, Vanessa Demontant, Guillaume Gricourt, Melissa N'Debi, Alexandre Soulier, Elisabeth Trawinski, Jean-Michel Pawlotsky |
| EPI_ISL_2878292, EPI_ISL_2878294 | Biogroup Bio Lam-LCD Saint-Denis | Department of Virology, Henri Mondor University Hospital, Assistance Publique Hôpitaux de Paris, Université Paris-Est Créteil, INSERM U955 | Christophe Rodriguez, Slim Fourati, Vanessa Demontant, Guillaume Gricourt, Melissa N'Debi, Alexandre Soulier, Elisabeth Trawinski, Jean-Michel Pawlotsky |
| EPI_ISL_2878295 | SELAS MEDILYS | Department of Virology, Henri Mondor University Hospital, Assistance Publique Hôpitaux de Paris, Université Paris-Est Créteil, INSERM U955 | Christophe Rodriguez, Slim Fourati, Vanessa Demontant, Guillaume Gricourt, Melissa N'Debi, Alexandre Soulier, Elisabeth Trawinski, Jean-Michel Pawlotsky |
| EPI_ISL_2878296 | Biogroup Bio Lam-LCD Saint-Denis | Department of Virology, Henri Mondor University Hospital, Assistance Publique Hôpitaux de Paris, Université Paris-Est Créteil, INSERM U955 | Christophe Rodriguez, Slim Fourati, Vanessa Demontant, Guillaume Gricourt, Melissa N'Debi, Alexandre Soulier, Elisabeth Trawinski, Jean-Michel Pawlotsky |
| EPI_ISL_2878297 | SELAS MEDILYS | Department of Virology, Henri Mondor University Hospital, Assistance Publique Hôpitaux de Paris, Université Paris-Est Créteil, INSERM U955 | Christophe Rodriguez, Slim Fourati, Vanessa Demontant, Guillaume Gricourt, Melissa N'Debi, Alexandre Soulier, Elisabeth Trawinski, Jean-Michel Pawlotsky |
| EPI_ISL_2878298 | Biogroup Bio Lam-LCD Saint-Denis | Department of Virology, Henri Mondor University Hospital, Assistance Publique Hôpitaux de Paris, Université Paris-Est Créteil, INSERM U955 | Christophe Rodriguez, Slim Fourati, Vanessa Demontant, Guillaume Gricourt, Melissa N'Debi, Alexandre Soulier, Elisabeth Trawinski, Jean-Michel Pawlotsky |
| EPI_ISL_2878299 | CH. CHU DE REIMS | Department of Virology, Henri Mondor University Hospital, | Christophe Rodriguez, Slim Fourati, Vanessa Demontant, Guillaume Gricourt, Melissa N'Debi, Alexandre Soulier, Elisabeth Trawinski, Jean-Michel Pawlotsky |

| | | | |
|---|---|---|---|
| | | Assistance Publique Hôpitaux de Paris, Université Paris-Est Créteil, INSERM U955 | |
| EPI_ISL_2878300 | GH de l'Est Francilien | Department of Virology, Henri Mondor University Hospital, Assistance Publique Hôpitaux de Paris, Université Paris-Est Créteil, INSERM U955 | Christophe Rodriguez, Slim Fourati, Vanessa Demontant, Guillaume Gricourt, Melissa N'Debi, Alexandre Soulier, Elisabeth Trawinski, Jean-Michel Pawlotsky |
| EPI_ISL_2878302, EPI_ISL_2878303 | SELAS MEDILYS | Department of Virology, Henri Mondor University Hospital, Assistance Publique Hôpitaux de Paris, Université Paris-Est Créteil, INSERM U955 | Christophe Rodriguez, Slim Fourati, Vanessa Demontant, Guillaume Gricourt, Melissa N'Debi, Alexandre Soulier, Elisabeth Trawinski, Jean-Michel Pawlotsky |
| EPI_ISL_2878304, EPI_ISL_2878305, EPI_ISL_2878306 | Biogroup Bio Lam-LCD Saint-Denis | Department of Virology, Henri Mondor University Hospital, Assistance Publique Hôpitaux de Paris, Université Paris-Est Créteil, INSERM U955 | Christophe Rodriguez, Slim Fourati, Vanessa Demontant, Guillaume Gricourt, Melissa N'Debi, Alexandre Soulier, Elisabeth Trawinski, Jean-Michel Pawlotsky |
| EPI_ISL_2878307 | BROUSSAIS | Department of Virology, Henri Mondor University Hospital, Assistance Publique Hôpitaux de Paris, Université Paris-Est Créteil, INSERM U955 | Christophe Rodriguez, Slim Fourati, Vanessa Demontant, Guillaume Gricourt, Melissa N'Debi, Alexandre Soulier, Elisabeth Trawinski, Jean-Michel Pawlotsky |
| EPI_ISL_2878308 | SELAS MEDILYS | Department of Virology, Henri Mondor University Hospital, Assistance Publique Hôpitaux de Paris, Université Paris-Est Créteil, INSERM U955 | Christophe Rodriguez, Slim Fourati, Vanessa Demontant, Guillaume Gricourt, Melissa N'Debi, Alexandre Soulier, Elisabeth Trawinski, Jean-Michel Pawlotsky |
| EPI_ISL_2878309, EPI_ISL_2878310, EPI_ISL_2878311 | Biogroup Bio Lam-LCD Saint-Denis | Department of Virology, Henri Mondor University Hospital, Assistance Publique Hôpitaux de Paris, Université Paris-Est Créteil, INSERM U955 | Christophe Rodriguez, Slim Fourati, Vanessa Demontant, Guillaume Gricourt, Melissa N'Debi, Alexandre Soulier, Elisabeth Trawinski, Jean-Michel Pawlotsky |
| EPI_ISL_2878312 | BIO67-BIOSPHERE | Department of Virology, Henri Mondor University Hospital, Assistance Publique Hôpitaux de Paris, Université Paris-Est Créteil, INSERM U955 | Christophe Rodriguez, Slim Fourati, Vanessa Demontant, Guillaume Gricourt, Melissa N'Debi, Alexandre Soulier, Elisabeth Trawinski, Jean-Michel Pawlotsky |
| EPI_ISL_2878313 | SELAS MEDILYS | Department of Virology, Henri Mondor University Hospital, Assistance Publique Hôpitaux de Paris, Université Paris-Est Créteil, INSERM U955 | Christophe Rodriguez, Slim Fourati, Vanessa Demontant, Guillaume Gricourt, Melissa N'Debi, Alexandre Soulier, Elisabeth Trawinski, Jean-Michel Pawlotsky |
| EPI_ISL_2878314 | BROUSSAIS | Department of Virology, Henri Mondor University Hospital, Assistance Publique Hôpitaux de Paris, Université Paris-Est Créteil, INSERM U955 | Christophe Rodriguez, Slim Fourati, Vanessa Demontant, Guillaume Gricourt, Melissa N'Debi, Alexandre Soulier, Elisabeth Trawinski, Jean-Michel Pawlotsky |
| EPI_ISL_2878315 | Biogroup Bio Lam-LCD Saint-Denis | Department of Virology, Henri Mondor University Hospital, Assistance Publique Hôpitaux de Paris, Université Paris-Est Créteil, INSERM U955 | Christophe Rodriguez, Slim Fourati, Vanessa Demontant, Guillaume Gricourt, Melissa N'Debi, Alexandre Soulier, Elisabeth Trawinski, Jean-Michel Pawlotsky |
| EPI_ISL_2878316, EPI_ISL_2878317 | SELAS MEDILYS | Department of Virology, Henri Mondor University Hospital, Assistance Publique Hôpitaux de Paris, Université Paris-Est Créteil, INSERM U955 | Christophe Rodriguez, Slim Fourati, Vanessa Demontant, Guillaume Gricourt, Melissa N'Debi, Alexandre Soulier, Elisabeth Trawinski, Jean-Michel Pawlotsky |
| EPI_ISL_2878318, EPI_ISL_2878319 | Biogroup Bio Lam-LCD Saint-Denis | Department of Virology, Henri Mondor University Hospital, Assistance Publique Hôpitaux de Paris, Université Paris-Est Créteil, INSERM U955 | Christophe Rodriguez, Slim Fourati, Vanessa Demontant, Guillaume Gricourt, Melissa N'Debi, Alexandre Soulier, Elisabeth Trawinski, Jean-Michel Pawlotsky |
| EPI_ISL_2878321, EPI_ISL_2878322 | SELAS MEDILYS | Department of Virology, Henri Mondor University Hospital, Assistance Publique Hôpitaux de Paris, Université Paris-Est Créteil, INSERM U955 | Christophe Rodriguez, Slim Fourati, Vanessa Demontant, Guillaume Gricourt, Melissa N'Debi, Alexandre Soulier, Elisabeth Trawinski, Jean-Michel Pawlotsky |
| EPI_ISL_2878323 | Biogroup Bio Lam-LCD Saint-Denis | Department of Virology, Henri Mondor University Hospital, Assistance Publique Hôpitaux de Paris, Université Paris-Est Créteil, INSERM U955 | Christophe Rodriguez, Slim Fourati, Vanessa Demontant, Guillaume Gricourt, Melissa N'Debi, Alexandre Soulier, Elisabeth Trawinski, Jean-Michel Pawlotsky |
| EPI_ISL_2878324 | Hôpital Cochin | Department of Virology, Henri Mondor University Hospital, Assistance Publique Hôpitaux de Paris, Université Paris-Est Créteil, INSERM U955 | Christophe Rodriguez, Slim Fourati, Vanessa Demontant, Guillaume Gricourt, Melissa N'Debi, Alexandre Soulier, Elisabeth Trawinski, Jean-Michel Pawlotsky |
| EPI_ISL_2878325 | BROUSSAIS | Department of Virology, Henri Mondor University Hospital, Assistance Publique Hôpitaux de Paris, Université Paris-Est Créteil, INSERM U955 | Christophe Rodriguez, Slim Fourati, Vanessa Demontant, Guillaume Gricourt, Melissa N'Debi, Alexandre Soulier, Elisabeth Trawinski, Jean-Michel Pawlotsky |
| EPI_ISL_2878326, EPI_ISL_2878327, EPI_ISL_2878329 | SELAS MEDILYS | Department of Virology, Henri Mondor University Hospital, Assistance Publique Hôpitaux de Paris, Université Paris-Est Créteil, INSERM U955 | Christophe Rodriguez, Slim Fourati, Vanessa Demontant, Guillaume Gricourt, Melissa N'Debi, Alexandre Soulier, Elisabeth Trawinski, Jean-Michel Pawlotsky |
| EPI_ISL_2878330 | CHRU STRASBOURG | Department of Virology, Henri Mondor University Hospital, Assistance Publique Hôpitaux de Paris, Université Paris-Est Créteil, INSERM U955 | Christophe Rodriguez, Slim Fourati, Vanessa Demontant, Guillaume Gricourt, Melissa N'Debi, Alexandre Soulier, Elisabeth Trawinski, Jean-Michel Pawlotsky |
| EPI_ISL_2878331 | SELAS MEDILYS | Department of Virology, Henri Mondor University Hospital, Assistance Publique Hôpitaux de Paris, Université Paris-Est Créteil, INSERM U955 | Christophe Rodriguez, Slim Fourati, Vanessa Demontant, Guillaume Gricourt, Melissa N'Debi, Alexandre Soulier, Elisabeth Trawinski, Jean-Michel Pawlotsky |
| EPI_ISL_2878334, EPI_ISL_2878335, EPI_ISL_2878336, EPI_ISL_2878337 | Biogroup Bio Lam-LCD Saint-Denis | Department of Virology, Henri Mondor University Hospital, Assistance Publique Hôpitaux de Paris, Université Paris-Est Créteil, INSERM U955 | Christophe Rodriguez, Slim Fourati, Vanessa Demontant, Guillaume Gricourt, Melissa N'Debi, Alexandre Soulier, Elisabeth Trawinski, Jean-Michel Pawlotsky |
| EPI_ISL_2878338 | CH. ANDRE GREGOIRE | Department of Virology, Henri Mondor University Hospital, Assistance Publique Hôpitaux de Paris, Université Paris-Est Créteil, INSERM U955 | Christophe Rodriguez, Slim Fourati, Vanessa Demontant, Guillaume Gricourt, Melissa N'Debi, Alexandre Soulier, Elisabeth Trawinski, Jean-Michel Pawlotsky |
| EPI_ISL_2878339, EPI_ISL_2878341 | Biogroup Bio Lam-LCD Saint-Denis | Department of Virology, Henri Mondor University Hospital, Assistance Publique Hôpitaux de Paris, Université Paris-Est Créteil, INSERM U955 | Christophe Rodriguez, Slim Fourati, Vanessa Demontant, Guillaume Gricourt, Melissa N'Debi, Alexandre Soulier, Elisabeth Trawinski, Jean-Michel Pawlotsky |
| EPI_ISL_2878342, EPI_ISL_2878343, EPI_ISL_2878344 | BIO67-BIOSPHERE | Department of Virology, Henri Mondor University Hospital, Assistance Publique Hôpitaux de Paris, Université Paris-Est Créteil, INSERM U955 | Christophe Rodriguez, Slim Fourati, Vanessa Demontant, Guillaume Gricourt, Melissa N'Debi, Alexandre Soulier, Elisabeth Trawinski, Jean-Michel Pawlotsky |
| EPI_ISL_2878345, EPI_ISL_2878346, EPI_ISL_2878347, EPI_ISL_2878348, EPI_ISL_2878349, EPI_ISL_2878350, EPI_ISL_2878351, EPI_ISL_2878352 | Biogroup Bio Lam-LCD Saint-Denis | Department of Virology, Henri Mondor University Hospital, Assistance Publique Hôpitaux de Paris, Université Paris-Est Créteil, INSERM U955 | Christophe Rodriguez, Slim Fourati, Vanessa Demontant, Guillaume Gricourt, Melissa N'Debi, Alexandre Soulier, Elisabeth Trawinski, Jean-Michel Pawlotsky |
| EPI_ISL_2878353 | SELAS MEDILYS | Department of Virology, Henri Mondor University Hospital, Assistance Publique Hôpitaux de Paris, Université Paris-Est Créteil, INSERM U955 | Christophe Rodriguez, Slim Fourati, Vanessa Demontant, Guillaume Gricourt, Melissa N'Debi, Alexandre Soulier, Elisabeth Trawinski, Jean-Michel Pawlotsky |
| EPI_ISL_2878354, EPI_ISL_2878355 | Biogroup Bio Lam-LCD Saint-Denis | Department of Virology, Henri Mondor University Hospital, Assistance Publique Hôpitaux de Paris, Université Paris-Est Créteil, INSERM U955 | Christophe Rodriguez, Slim Fourati, Vanessa Demontant, Guillaume Gricourt, Melissa N'Debi, Alexandre Soulier, Elisabeth Trawinski, Jean-Michel Pawlotsky |
| EPI_ISL_2878356 | SELAS MEDILYS | Department of Virology, Henri Mondor University Hospital, Assistance Publique Hôpitaux de Paris, Université Paris-Est Créteil, INSERM U955 | Christophe Rodriguez, Slim Fourati, Vanessa Demontant, Guillaume Gricourt, Melissa N'Debi, Alexandre Soulier, Elisabeth Trawinski, Jean-Michel Pawlotsky |
| EPI_ISL_2878357, EPI_ISL_2878358, EPI_ISL_2878359, EPI_ISL_2878361, EPI_ISL_2878362, EPI_ISL_2878363, EPI_ISL_2878364, EPI_ISL_2878365, EPI_ISL_2878366 | Biogroup Bio Lam-LCD Saint-Denis | Department of Virology, Henri Mondor University Hospital, Assistance Publique Hôpitaux de Paris, Université Paris-Est Créteil, INSERM U955 | Christophe Rodriguez, Slim Fourati, Vanessa Demontant, Guillaume Gricourt, Melissa N'Debi, Alexandre Soulier, Elisabeth Trawinski, Jean-Michel Pawlotsky |

| | | | |
|---|---|---|---|
| EPI_ISL_2878367 | BROUSSAIS | Department of Virology, Henri Mondor University Hospital, Assistance Publique Hôpitaux de Paris, Université Paris-Est Créteil, INSERM U955 | Christophe Rodriguez, Slim Fourati, Vanessa Demontant, Guillaume Gricourt, Melissa N'Debi, Alexandre Soulier, Elisabeth Trawinski, Jean-Michel Pawlotsky |
| EPI_ISL_2878368, EPI_ISL_2878369, EPI_ISL_2878370, EPI_ISL_2878371, EPI_ISL_2878372, EPI_ISL_2878373, EPI_ISL_2878376 | Biogroup Bio Lam-LCD Saint-Denis | Department of Virology, Henri Mondor University Hospital, Assistance Publique Hôpitaux de Paris, Université Paris-Est Créteil, INSERM U955 | Christophe Rodriguez, Slim Fourati, Vanessa Demontant, Guillaume Gricourt, Melissa N'Debi, Alexandre Soulier, Elisabeth Trawinski, Jean-Michel Pawlotsky |
| EPI_ISL_2878377 | CH.INTERCOMMUNAL DE CRETEIL | Department of Virology, Henri Mondor University Hospital, Assistance Publique Hôpitaux de Paris, Université Paris-Est Créteil, INSERM U955 | Christophe Rodriguez, Slim Fourati, Vanessa Demontant, Guillaume Gricourt, Melissa N'Debi, Alexandre Soulier, Elisabeth Trawinski, Jean-Michel Pawlotsky |
| EPI_ISL_2878380 | Biogroup Bio Lam-LCD Saint-Denis | Department of Virology, Henri Mondor University Hospital, Assistance Publique Hôpitaux de Paris, Université Paris-Est Créteil, INSERM U955 | Christophe Rodriguez, Slim Fourati, Vanessa Demontant, Guillaume Gricourt, Melissa N'Debi, Alexandre Soulier, Elisabeth Trawinski, Jean-Michel Pawlotsky |
| EPI_ISL_2878381 | BIO67-BIOSPHERE | Department of Virology, Henri Mondor University Hospital, Assistance Publique Hôpitaux de Paris, Université Paris-Est Créteil, INSERM U955 | Christophe Rodriguez, Slim Fourati, Vanessa Demontant, Guillaume Gricourt, Melissa N'Debi, Alexandre Soulier, Elisabeth Trawinski, Jean-Michel Pawlotsky |
| EPI_ISL_2878382 | GH de l'Est Francilien | Department of Virology, Henri Mondor University Hospital, Assistance Publique Hôpitaux de Paris, Université Paris-Est Créteil, INSERM U955 | Christophe Rodriguez, Slim Fourati, Vanessa Demontant, Guillaume Gricourt, Melissa N'Debi, Alexandre Soulier, Elisabeth Trawinski, Jean-Michel Pawlotsky |
| EPI_ISL_2878383, EPI_ISL_2878384, EPI_ISL_2878385 | Biogroup Bio Lam-LCD Saint-Denis | Department of Virology, Henri Mondor University Hospital, Assistance Publique Hôpitaux de Paris, Université Paris-Est Créteil, INSERM U955 | Christophe Rodriguez, Slim Fourati, Vanessa Demontant, Guillaume Gricourt, Melissa N'Debi, Alexandre Soulier, Elisabeth Trawinski, Jean-Michel Pawlotsky |
| EPI_ISL_2878386 | BIO67-BIOSPHERE | Department of Virology, Henri Mondor University Hospital, Assistance Publique Hôpitaux de Paris, Université Paris-Est Créteil, INSERM U955 | Christophe Rodriguez, Slim Fourati, Vanessa Demontant, Guillaume Gricourt, Melissa N'Debi, Alexandre Soulier, Elisabeth Trawinski, Jean-Michel Pawlotsky |
| EPI_ISL_2878387 | Biogroup Bio Lam-LCD Saint-Denis | Department of Virology, Henri Mondor University Hospital, Assistance Publique Hôpitaux de Paris, Université Paris-Est Créteil, INSERM U955 | Christophe Rodriguez, Slim Fourati, Vanessa Demontant, Guillaume Gricourt, Melissa N'Debi, Alexandre Soulier, Elisabeth Trawinski, Jean-Michel Pawlotsky |
| EPI_ISL_2878388 | SELAS MEDILYS | Department of Virology, Henri Mondor University Hospital, Assistance Publique Hôpitaux de Paris, Université Paris-Est Créteil, INSERM U955 | Christophe Rodriguez, Slim Fourati, Vanessa Demontant, Guillaume Gricourt, Melissa N'Debi, Alexandre Soulier, Elisabeth Trawinski, Jean-Michel Pawlotsky |
| EPI_ISL_2878389 | Biogroup Bio Lam-LCD Saint-Denis | Department of Virology, Henri Mondor University Hospital, Assistance Publique Hôpitaux de Paris, Université Paris-Est Créteil, INSERM U955 | Christophe Rodriguez, Slim Fourati, Vanessa Demontant, Guillaume Gricourt, Melissa N'Debi, Alexandre Soulier, Elisabeth Trawinski, Jean-Michel Pawlotsky |
| EPI_ISL_2878390 | GH A.CHENEVIER-H.MONDOR | Department of Virology, Henri Mondor University Hospital, Assistance Publique Hôpitaux de Paris, Université Paris-Est Créteil, INSERM U955 | Christophe Rodriguez, Slim Fourati, Vanessa Demontant, Guillaume Gricourt, Melissa N'Debi, Alexandre Soulier, Elisabeth Trawinski, Jean-Michel Pawlotsky |
| EPI_ISL_2878391 | SELAS MEDILYS | Department of Virology, Henri Mondor University Hospital, Assistance Publique Hôpitaux de Paris, Université Paris-Est Créteil, INSERM U955 | Christophe Rodriguez, Slim Fourati, Vanessa Demontant, Guillaume Gricourt, Melissa N'Debi, Alexandre Soulier, Elisabeth Trawinski, Jean-Michel Pawlotsky |
| EPI_ISL_2878392, EPI_ISL_2878393, EPI_ISL_2878394 | Biogroup Bio Lam-LCD Saint-Denis | Department of Virology, Henri Mondor University Hospital, Assistance Publique Hôpitaux de Paris, Université Paris-Est Créteil, INSERM U955 | Christophe Rodriguez, Slim Fourati, Vanessa Demontant, Guillaume Gricourt, Melissa N'Debi, Alexandre Soulier, Elisabeth Trawinski, Jean-Michel Pawlotsky |
| EPI_ISL_2878395 | GH A.CHENEVIER-H.MONDOR | Department of Virology, Henri Mondor University Hospital, Assistance Publique Hôpitaux de Paris, Université Paris-Est Créteil, INSERM U955 | Christophe Rodriguez, Slim Fourati, Vanessa Demontant, Guillaume Gricourt, Melissa N'Debi, Alexandre Soulier, Elisabeth Trawinski, Jean-Michel Pawlotsky |
| EPI_ISL_2878397 | Cerballiance Paris IDF est Saint-Denis | Department of Virology, Henri Mondor University Hospital, Assistance Publique Hôpitaux de Paris, Université Paris-Est Créteil, INSERM U955 | Christophe Rodriguez, Slim Fourati, Vanessa Demontant, Guillaume Gricourt, Melissa N'Debi, Alexandre Soulier, Elisabeth Trawinski, Jean-Michel Pawlotsky |
| EPI_ISL_2878399, EPI_ISL_2878400 | BIO67-BIOSPHERE | Department of Virology, Henri Mondor University Hospital, Assistance Publique Hôpitaux de Paris, Université Paris-Est Créteil, INSERM U955 | Christophe Rodriguez, Slim Fourati, Vanessa Demontant, Guillaume Gricourt, Melissa N'Debi, Alexandre Soulier, Elisabeth Trawinski, Jean-Michel Pawlotsky |
| EPI_ISL_2878402, EPI_ISL_2878404 | SELAS MEDILYS | Department of Virology, Henri Mondor University Hospital, Assistance Publique Hôpitaux de Paris, Université Paris-Est Créteil, INSERM U955 | Christophe Rodriguez, Slim Fourati, Vanessa Demontant, Guillaume Gricourt, Melissa N'Debi, Alexandre Soulier, Elisabeth Trawinski, Jean-Michel Pawlotsky |
| EPI_ISL_2878405 | BIO67-BIOSPHERE | Department of Virology, Henri Mondor University Hospital, Assistance Publique Hôpitaux de Paris, Université Paris-Est Créteil, INSERM U955 | Christophe Rodriguez, Slim Fourati, Vanessa Demontant, Guillaume Gricourt, Melissa N'Debi, Alexandre Soulier, Elisabeth Trawinski, Jean-Michel Pawlotsky |
| EPI_ISL_2878407 | Biogroup Bio Lam-LCD Saint-Denis | Department of Virology, Henri Mondor University Hospital, Assistance Publique Hôpitaux de Paris, Université Paris-Est Créteil, INSERM U955 | Christophe Rodriguez, Slim Fourati, Vanessa Demontant, Guillaume Gricourt, Melissa N'Debi, Alexandre Soulier, Elisabeth Trawinski, Jean-Michel Pawlotsky |
| EPI_ISL_2878408 | BIOMNIS EUROFINS IVRY | Department of Virology, Henri Mondor University Hospital, Assistance Publique Hôpitaux de Paris, Université Paris-Est Créteil, INSERM U955 | Christophe Rodriguez, Slim Fourati, Vanessa Demontant, Guillaume Gricourt, Melissa N'Debi, Alexandre Soulier, Elisabeth Trawinski, Jean-Michel Pawlotsky |
| EPI_ISL_2878410 | BIO67-BIOSPHERE | Department of Virology, Henri Mondor University Hospital, Assistance Publique Hôpitaux de Paris, Université Paris-Est Créteil, INSERM U955 | Christophe Rodriguez, Slim Fourati, Vanessa Demontant, Guillaume Gricourt, Melissa N'Debi, Alexandre Soulier, Elisabeth Trawinski, Jean-Michel Pawlotsky |
| EPI_ISL_2878411, EPI_ISL_2878412 | Cerballiance Paris IDF est Saint-Denis | Department of Virology, Henri Mondor University Hospital, Assistance Publique Hôpitaux de Paris, Université Paris-Est Créteil, INSERM U955 | Christophe Rodriguez, Slim Fourati, Vanessa Demontant, Guillaume Gricourt, Melissa N'Debi, Alexandre Soulier, Elisabeth Trawinski, Jean-Michel Pawlotsky |
| EPI_ISL_2878414 | LISIEUX Cerballiance | Department of Virology, Henri Mondor University Hospital, Assistance Publique Hôpitaux de Paris, Université Paris-Est Créteil, INSERM U955 | Christophe Rodriguez, Slim Fourati, Vanessa Demontant, Guillaume Gricourt, Melissa N'Debi, Alexandre Soulier, Elisabeth Trawinski, Jean-Michel Pawlotsky |
| EPI_ISL_2878415 | Biogroup Bio Lam-LCD Saint-Denis | Department of Virology, Henri Mondor University Hospital, Assistance Publique Hôpitaux de Paris, Université Paris-Est Créteil, INSERM U955 | Christophe Rodriguez, Slim Fourati, Vanessa Demontant, Guillaume Gricourt, Melissa N'Debi, Alexandre Soulier, Elisabeth Trawinski, Jean-Michel Pawlotsky |
| EPI_ISL_2878417 | SELAS MEDILYS | Department of Virology, Henri Mondor University Hospital, Assistance Publique Hôpitaux de Paris, Université Paris-Est Créteil, INSERM U955 | Christophe Rodriguez, Slim Fourati, Vanessa Demontant, Guillaume Gricourt, Melissa N'Debi, Alexandre Soulier, Elisabeth Trawinski, Jean-Michel Pawlotsky |
| EPI_ISL_2878419 | Laboratoire LENYS | Department of Virology, Henri Mondor University Hospital, Assistance Publique Hôpitaux de Paris, Université Paris-Est Créteil, INSERM U955 | Christophe Rodriguez, Slim Fourati, Vanessa Demontant, Guillaume Gricourt, Melissa N'Debi, Alexandre Soulier, Elisabeth Trawinski, Jean-Michel Pawlotsky |
| EPI_ISL_2878420 | LISIEUX Cerballiance | Department of Virology, Henri Mondor University Hospital, Assistance Publique Hôpitaux de Paris, Université Paris-Est Créteil, INSERM U955 | Christophe Rodriguez, Slim Fourati, Vanessa Demontant, Guillaume Gricourt, Melissa N'Debi, Alexandre Soulier, Elisabeth Trawinski, Jean-Michel Pawlotsky |
| EPI_ISL_2878421 | Laboratoire Ana-L | Department of Virology, Henri Mondor University Hospital, Assistance Publique Hôpitaux de Paris, Université Paris-Est Créteil, INSERM U955 | Christophe Rodriguez, Slim Fourati, Vanessa Demontant, Guillaume Gricourt, Melissa N'Debi, Alexandre Soulier, Elisabeth Trawinski, Jean-Michel Pawlotsky |
| EPI_ISL_2878422 | BIOLBS Rouen | Department of Virology, Henri Mondor University Hospital, Assistance Publique Hôpitaux de Paris, Université Paris-Est Créteil, INSERM U955 | Christophe Rodriguez, Slim Fourati, Vanessa Demontant, Guillaume Gricourt, Melissa N'Debi, Alexandre Soulier, Elisabeth Trawinski, Jean-Michel Pawlotsky |
| EPI_ISL_2878424 | SELAS MEDILYS | Department of Virology, Henri Mondor University Hospital, Assistance Publique Hôpitaux de Paris, Université Paris-Est Créteil, INSERM U955 | Christophe Rodriguez, Slim Fourati, Vanessa Demontant, Guillaume Gricourt, Melissa N'Debi, Alexandre Soulier, Elisabeth Trawinski, Jean-Michel Pawlotsky |

| | | | |
|---|---|---|---|
| EPI_ISL_2878426 | Cerballiance Paris IDF est Saint-Denis | Department of Virology, Henri Mondor University Hospital, Assistance Publique Hôpitaux de Paris, Université Paris-Est Créteil, INSERM U955 | Christophe Rodriguez, Slim Fourati, Vanessa Demontant, Guillaume Gricourt, Melissa N'Debi, Alexandre Soulier, Elisabeth Trawinski, Jean-Michel Pawlotsky |
| EPI_ISL_2878427 | laboratoire Belle Epine | Department of Virology, Henri Mondor University Hospital, Assistance Publique Hôpitaux de Paris, Université Paris-Est Créteil, INSERM U955 | Christophe Rodriguez, Slim Fourati, Vanessa Demontant, Guillaume Gricourt, Melissa N'Debi, Alexandre Soulier, Elisabeth Trawinski, Jean-Michel Pawlotsky |
| EPI_ISL_2878429 | Laboratoire LENYS | Department of Virology, Henri Mondor University Hospital, Assistance Publique Hôpitaux de Paris, Université Paris-Est Créteil, INSERM U955 | Christophe Rodriguez, Slim Fourati, Vanessa Demontant, Guillaume Gricourt, Melissa N'Debi, Alexandre Soulier, Elisabeth Trawinski, Jean-Michel Pawlotsky |
| EPI_ISL_2878430 | Plateau technique Biomer Metz Verriers | Department of Virology, Henri Mondor University Hospital, Assistance Publique Hôpitaux de Paris, Université Paris-Est Créteil, INSERM U955 | Christophe Rodriguez, Slim Fourati, Vanessa Demontant, Guillaume Gricourt, Melissa N'Debi, Alexandre Soulier, Elisabeth Trawinski, Jean-Michel Pawlotsky |
| EPI_ISL_2878431 | BIO67-BIOSPHERE | Department of Virology, Henri Mondor University Hospital, Assistance Publique Hôpitaux de Paris, Université Paris-Est Créteil, INSERM U955 | Christophe Rodriguez, Slim Fourati, Vanessa Demontant, Guillaume Gricourt, Melissa N'Debi, Alexandre Soulier, Elisabeth Trawinski, Jean-Michel Pawlotsky |
| EPI_ISL_2878432 | BIOMNIS EUROFINS IVRY | Department of Virology, Henri Mondor University Hospital, Assistance Publique Hôpitaux de Paris, Université Paris-Est Créteil, INSERM U955 | Christophe Rodriguez, Slim Fourati, Vanessa Demontant, Guillaume Gricourt, Melissa N'Debi, Alexandre Soulier, Elisabeth Trawinski, Jean-Michel Pawlotsky |
| EPI_ISL_2878434 | Laboratoire Coste | Department of Virology, Henri Mondor University Hospital, Assistance Publique Hôpitaux de Paris, Université Paris-Est Créteil, INSERM U955 | Christophe Rodriguez, Slim Fourati, Vanessa Demontant, Guillaume Gricourt, Melissa N'Debi, Alexandre Soulier, Elisabeth Trawinski, Jean-Michel Pawlotsky |
| EPI_ISL_2878436 | BIO67-BIOSPHERE | Department of Virology, Henri Mondor University Hospital, Assistance Publique Hôpitaux de Paris, Université Paris-Est Créteil, INSERM U955 | Christophe Rodriguez, Slim Fourati, Vanessa Demontant, Guillaume Gricourt, Melissa N'Debi, Alexandre Soulier, Elisabeth Trawinski, Jean-Michel Pawlotsky |
| EPI_ISL_2878439 | laboratoire Belle Epine | Department of Virology, Henri Mondor University Hospital, Assistance Publique Hôpitaux de Paris, Université Paris-Est Créteil, INSERM U955 | Christophe Rodriguez, Slim Fourati, Vanessa Demontant, Guillaume Gricourt, Melissa N'Debi, Alexandre Soulier, Elisabeth Trawinski, Jean-Michel Pawlotsky |
| EPI_ISL_2878440, EPI_ISL_2878441 | Biogroup Bio Lam-LCD Saint-Denis | Department of Virology, Henri Mondor University Hospital, Assistance Publique Hôpitaux de Paris, Université Paris-Est Créteil, INSERM U955 | Christophe Rodriguez, Slim Fourati, Vanessa Demontant, Guillaume Gricourt, Melissa N'Debi, Alexandre Soulier, Elisabeth Trawinski, Jean-Michel Pawlotsky |
| EPI_ISL_2878442 | laboratoire Belle Epine | Department of Virology, Henri Mondor University Hospital, Assistance Publique Hôpitaux de Paris, Université Paris-Est Créteil, INSERM U955 | Christophe Rodriguez, Slim Fourati, Vanessa Demontant, Guillaume Gricourt, Melissa N'Debi, Alexandre Soulier, Elisabeth Trawinski, Jean-Michel Pawlotsky |
| EPI_ISL_2878443, EPI_ISL_2878445 | Cerballiance Paris IDF est Saint-Denis | Department of Virology, Henri Mondor University Hospital, Assistance Publique Hôpitaux de Paris, Université Paris-Est Créteil, INSERM U955 | Christophe Rodriguez, Slim Fourati, Vanessa Demontant, Guillaume Gricourt, Melissa N'Debi, Alexandre Soulier, Elisabeth Trawinski, Jean-Michel Pawlotsky |
| EPI_ISL_2878446 | Laboratoire ESPACEBIO | Department of Virology, Henri Mondor University Hospital, Assistance Publique Hôpitaux de Paris, Université Paris-Est Créteil, INSERM U955 | Christophe Rodriguez, Slim Fourati, Vanessa Demontant, Guillaume Gricourt, Melissa N'Debi, Alexandre Soulier, Elisabeth Trawinski, Jean-Michel Pawlotsky |
| EPI_ISL_2878450 | laboratoire Belle Epine | Department of Virology, Henri Mondor University Hospital, Assistance Publique Hôpitaux de Paris, Université Paris-Est Créteil, INSERM U955 | Christophe Rodriguez, Slim Fourati, Vanessa Demontant, Guillaume Gricourt, Melissa N'Debi, Alexandre Soulier, Elisabeth Trawinski, Jean-Michel Pawlotsky |
| EPI_ISL_2878451 | Laboratoire Ana-L | Department of Virology, Henri Mondor University Hospital, Assistance Publique Hôpitaux de Paris, Université Paris-Est Créteil, INSERM U955 | Christophe Rodriguez, Slim Fourati, Vanessa Demontant, Guillaume Gricourt, Melissa N'Debi, Alexandre Soulier, Elisabeth Trawinski, Jean-Michel Pawlotsky |
| EPI_ISL_2878452, EPI_ISL_2878454 | BIO67-BIOSPHERE | Department of Virology, Henri Mondor University Hospital, Assistance Publique Hôpitaux de Paris, Université Paris-Est Créteil, INSERM U955 | Christophe Rodriguez, Slim Fourati, Vanessa Demontant, Guillaume Gricourt, Melissa N'Debi, Alexandre Soulier, Elisabeth Trawinski, Jean-Michel Pawlotsky |
| EPI_ISL_2878456, EPI_ISL_2878458 | BIOMNIS EUROFINS IVRY | Department of Virology, Henri Mondor University Hospital, Assistance Publique Hôpitaux de Paris, Université Paris-Est Créteil, INSERM U955 | Christophe Rodriguez, Slim Fourati, Vanessa Demontant, Guillaume Gricourt, Melissa N'Debi, Alexandre Soulier, Elisabeth Trawinski, Jean-Michel Pawlotsky |
| EPI_ISL_2878459 | SELAS MEDILYS | Department of Virology, Henri Mondor University Hospital, Assistance Publique Hôpitaux de Paris, Université Paris-Est Créteil, INSERM U955 | Christophe Rodriguez, Slim Fourati, Vanessa Demontant, Guillaume Gricourt, Melissa N'Debi, Alexandre Soulier, Elisabeth Trawinski, Jean-Michel Pawlotsky |
| EPI_ISL_2878460 | Biogroup Bio Lam-LCD Saint-Denis | Department of Virology, Henri Mondor University Hospital, Assistance Publique Hôpitaux de Paris, Université Paris-Est Créteil, INSERM U955 | Christophe Rodriguez, Slim Fourati, Vanessa Demontant, Guillaume Gricourt, Melissa N'Debi, Alexandre Soulier, Elisabeth Trawinski, Jean-Michel Pawlotsky |
| EPI_ISL_2878461 | SELAS MEDILYS | Department of Virology, Henri Mondor University Hospital, Assistance Publique Hôpitaux de Paris, Université Paris-Est Créteil, INSERM U955 | Christophe Rodriguez, Slim Fourati, Vanessa Demontant, Guillaume Gricourt, Melissa N'Debi, Alexandre Soulier, Elisabeth Trawinski, Jean-Michel Pawlotsky |
| EPI_ISL_2878463 | BIOMNIS EUROFINS IVRY | Department of Virology, Henri Mondor University Hospital, Assistance Publique Hôpitaux de Paris, Université Paris-Est Créteil, INSERM U955 | Christophe Rodriguez, Slim Fourati, Vanessa Demontant, Guillaume Gricourt, Melissa N'Debi, Alexandre Soulier, Elisabeth Trawinski, Jean-Michel Pawlotsky |
| EPI_ISL_2878464 | Laboratoire CBM 25 TERRE ROUGE | Department of Virology, Henri Mondor University Hospital, Assistance Publique Hôpitaux de Paris, Université Paris-Est Créteil, INSERM U955 | Christophe Rodriguez, Slim Fourati, Vanessa Demontant, Guillaume Gricourt, Melissa N'Debi, Alexandre Soulier, Elisabeth Trawinski, Jean-Michel Pawlotsky |
| EPI_ISL_2878465 | SELAS MEDILYS | Department of Virology, Henri Mondor University Hospital, Assistance Publique Hôpitaux de Paris, Université Paris-Est Créteil, INSERM U955 | Christophe Rodriguez, Slim Fourati, Vanessa Demontant, Guillaume Gricourt, Melissa N'Debi, Alexandre Soulier, Elisabeth Trawinski, Jean-Michel Pawlotsky |
| EPI_ISL_2878466 | BIO67-BIOSPHERE | Department of Virology, Henri Mondor University Hospital, Assistance Publique Hôpitaux de Paris, Université Paris-Est Créteil, INSERM U955 | Christophe Rodriguez, Slim Fourati, Vanessa Demontant, Guillaume Gricourt, Melissa N'Debi, Alexandre Soulier, Elisabeth Trawinski, Jean-Michel Pawlotsky |
| EPI_ISL_2878467 | Laboratoire Ana-L | Department of Virology, Henri Mondor University Hospital, Assistance Publique Hôpitaux de Paris, Université Paris-Est Créteil, INSERM U955 | Christophe Rodriguez, Slim Fourati, Vanessa Demontant, Guillaume Gricourt, Melissa N'Debi, Alexandre Soulier, Elisabeth Trawinski, Jean-Michel Pawlotsky |
| EPI_ISL_2878470 | BIO67-BIOSPHERE | Department of Virology, Henri Mondor University Hospital, Assistance Publique Hôpitaux de Paris, Université Paris-Est Créteil, INSERM U955 | Christophe Rodriguez, Slim Fourati, Vanessa Demontant, Guillaume Gricourt, Melissa N'Debi, Alexandre Soulier, Elisabeth Trawinski, Jean-Michel Pawlotsky |
| EPI_ISL_2878471 | laboratoire Belle Epine | Department of Virology, Henri Mondor University Hospital, Assistance Publique Hôpitaux de Paris, Université Paris-Est Créteil, INSERM U955 | Christophe Rodriguez, Slim Fourati, Vanessa Demontant, Guillaume Gricourt, Melissa N'Debi, Alexandre Soulier, Elisabeth Trawinski, Jean-Michel Pawlotsky |
| EPI_ISL_2878472 | BIO67-BIOSPHERE | Department of Virology, Henri Mondor University Hospital, Assistance Publique Hôpitaux de Paris, Université Paris-Est Créteil, INSERM U955 | Christophe Rodriguez, Slim Fourati, Vanessa Demontant, Guillaume Gricourt, Melissa N'Debi, Alexandre Soulier, Elisabeth Trawinski, Jean-Michel Pawlotsky |
| EPI_ISL_2878473 | BIOMNIS EUROFINS IVRY | Department of Virology, Henri Mondor University Hospital, Assistance Publique Hôpitaux de Paris, Université Paris-Est Créteil, INSERM U955 | Christophe Rodriguez, Slim Fourati, Vanessa Demontant, Guillaume Gricourt, Melissa N'Debi, Alexandre Soulier, Elisabeth Trawinski, Jean-Michel Pawlotsky |
| EPI_ISL_2878474 | Cerballiance Paris IDF est Saint-Denis | Department of Virology, Henri Mondor University Hospital, Assistance Publique Hôpitaux de Paris, Université Paris-Est Créteil, INSERM U955 | Christophe Rodriguez, Slim Fourati, Vanessa Demontant, Guillaume Gricourt, Melissa N'Debi, Alexandre Soulier, Elisabeth Trawinski, Jean-Michel Pawlotsky |
| EPI_ISL_2878476, EPI_ISL_2878479 | BIOMNIS EUROFINS IVRY | Department of Virology, Henri Mondor University Hospital, Assistance Publique Hôpitaux de Paris, Université Paris-Est Créteil, INSERM U955 | Christophe Rodriguez, Slim Fourati, Vanessa Demontant, Guillaume Gricourt, Melissa N'Debi, Alexandre Soulier, Elisabeth Trawinski, Jean-Michel Pawlotsky |
| EPI_ISL_2878481 | SELAS MEDILYS | Department of Virology, Henri Mondor University Hospital, Assistance Publique Hôpitaux de Paris, Université Paris-Est Créteil, INSERM U955 | Christophe Rodriguez, Slim Fourati, Vanessa Demontant, Guillaume Gricourt, Melissa N'Debi, Alexandre Soulier, Elisabeth Trawinski, Jean-Michel Pawlotsky |

| | | | |
|---|---|---|---|
| EPI_ISL_2878482 | Laboratoire ESPACEBIO | Department of Virology, Henri Mondor University Hospital, Assistance Publique Hôpitaux de Paris, Université Paris-Est Créteil, INSERM U955 | Christophe Rodriguez, Slim Fourati, Vanessa Demontant, Guillaume Gricourt, Melissa N'Debi, Alexandre Soulier, Elisabeth Trawinski, Jean-Michel Pawlotsky |
| EPI_ISL_2878484 | Biogroup Bio Lam-LCD Saint-Denis | Department of Virology, Henri Mondor University Hospital, Assistance Publique Hôpitaux de Paris, Université Paris-Est Créteil, INSERM U955 | Christophe Rodriguez, Slim Fourati, Vanessa Demontant, Guillaume Gricourt, Melissa N'Debi, Alexandre Soulier, Elisabeth Trawinski, Jean-Michel Pawlotsky |
| EPI_ISL_2878485 | Laboratoire Ana-L | Department of Virology, Henri Mondor University Hospital, Assistance Publique Hôpitaux de Paris, Université Paris-Est Créteil, INSERM U955 | Christophe Rodriguez, Slim Fourati, Vanessa Demontant, Guillaume Gricourt, Melissa N'Debi, Alexandre Soulier, Elisabeth Trawinski, Jean-Michel Pawlotsky |
| EPI_ISL_2878488 | LISIEUX Cerballiance | Department of Virology, Henri Mondor University Hospital, Assistance Publique Hôpitaux de Paris, Université Paris-Est Créteil, INSERM U955 | Christophe Rodriguez, Slim Fourati, Vanessa Demontant, Guillaume Gricourt, Melissa N'Debi, Alexandre Soulier, Elisabeth Trawinski, Jean-Michel Pawlotsky |
| EPI_ISL_2878489 | BIO67-BIOSPHERE | Department of Virology, Henri Mondor University Hospital, Assistance Publique Hôpitaux de Paris, Université Paris-Est Créteil, INSERM U955 | Christophe Rodriguez, Slim Fourati, Vanessa Demontant, Guillaume Gricourt, Melissa N'Debi, Alexandre Soulier, Elisabeth Trawinski, Jean-Michel Pawlotsky |
| EPI_ISL_2878490 | laboratoire Belle Epine | Department of Virology, Henri Mondor University Hospital, Assistance Publique Hôpitaux de Paris, Université Paris-Est Créteil, INSERM U955 | Christophe Rodriguez, Slim Fourati, Vanessa Demontant, Guillaume Gricourt, Melissa N'Debi, Alexandre Soulier, Elisabeth Trawinski, Jean-Michel Pawlotsky |
| EPI_ISL_2878492 | SELAS BC-LAB | Department of Virology, Henri Mondor University Hospital, Assistance Publique Hôpitaux de Paris, Université Paris-Est Créteil, INSERM U955 | Christophe Rodriguez, Slim Fourati, Vanessa Demontant, Guillaume Gricourt, Melissa N'Debi, Alexandre Soulier, Elisabeth Trawinski, Jean-Michel Pawlotsky |
| EPI_ISL_2878495 | BIOMNIS EUROFINS IVRY | Department of Virology, Henri Mondor University Hospital, Assistance Publique Hôpitaux de Paris, Université Paris-Est Créteil, INSERM U955 | Christophe Rodriguez, Slim Fourati, Vanessa Demontant, Guillaume Gricourt, Melissa N'Debi, Alexandre Soulier, Elisabeth Trawinski, Jean-Michel Pawlotsky |
| EPI_ISL_2878499 | Cerballiance Paris IDF est Saint-Denis | Department of Virology, Henri Mondor University Hospital, Assistance Publique Hôpitaux de Paris, Université Paris-Est Créteil, INSERM U955 | Christophe Rodriguez, Slim Fourati, Vanessa Demontant, Guillaume Gricourt, Melissa N'Debi, Alexandre Soulier, Elisabeth Trawinski, Jean-Michel Pawlotsky |
| EPI_ISL_2878501 | Laboratoire Ana-L | Department of Virology, Henri Mondor University Hospital, Assistance Publique Hôpitaux de Paris, Université Paris-Est Créteil, INSERM U955 | Christophe Rodriguez, Slim Fourati, Vanessa Demontant, Guillaume Gricourt, Melissa N'Debi, Alexandre Soulier, Elisabeth Trawinski, Jean-Michel Pawlotsky |
| EPI_ISL_2878503 | SELAS MEDILYS | Department of Virology, Henri Mondor University Hospital, Assistance Publique Hôpitaux de Paris, Université Paris-Est Créteil, INSERM U955 | Christophe Rodriguez, Slim Fourati, Vanessa Demontant, Guillaume Gricourt, Melissa N'Debi, Alexandre Soulier, Elisabeth Trawinski, Jean-Michel Pawlotsky |
| EPI_ISL_2878506 | SELAS BC-LAB | Department of Virology, Henri Mondor University Hospital, Assistance Publique Hôpitaux de Paris, Université Paris-Est Créteil, INSERM U955 | Christophe Rodriguez, Slim Fourati, Vanessa Demontant, Guillaume Gricourt, Melissa N'Debi, Alexandre Soulier, Elisabeth Trawinski, Jean-Michel Pawlotsky |
| EPI_ISL_2878507 | LISIEUX Cerballiance | Department of Virology, Henri Mondor University Hospital, Assistance Publique Hôpitaux de Paris, Université Paris-Est Créteil, INSERM U955 | Christophe Rodriguez, Slim Fourati, Vanessa Demontant, Guillaume Gricourt, Melissa N'Debi, Alexandre Soulier, Elisabeth Trawinski, Jean-Michel Pawlotsky |
| EPI_ISL_2878509 | Laboratoire Coste | Department of Virology, Henri Mondor University Hospital, Assistance Publique Hôpitaux de Paris, Université Paris-Est Créteil, INSERM U955 | Christophe Rodriguez, Slim Fourati, Vanessa Demontant, Guillaume Gricourt, Melissa N'Debi, Alexandre Soulier, Elisabeth Trawinski, Jean-Michel Pawlotsky |
| EPI_ISL_2878510 | Cerballiance Paris IDF est Saint-Denis | Department of Virology, Henri Mondor University Hospital, Assistance Publique Hôpitaux de Paris, Université Paris-Est Créteil, INSERM U955 | Christophe Rodriguez, Slim Fourati, Vanessa Demontant, Guillaume Gricourt, Melissa N'Debi, Alexandre Soulier, Elisabeth Trawinski, Jean-Michel Pawlotsky |
| EPI_ISL_2878512 | Biogroup Bio Lam-LCD Saint-Denis | Department of Virology, Henri Mondor University Hospital, Assistance Publique Hôpitaux de Paris, Université Paris-Est Créteil, INSERM U955 | Christophe Rodriguez, Slim Fourati, Vanessa Demontant, Guillaume Gricourt, Melissa N'Debi, Alexandre Soulier, Elisabeth Trawinski, Jean-Michel Pawlotsky |
| EPI_ISL_2878514 | BIO67-BIOSPHERE | Department of Virology, Henri Mondor University Hospital, Assistance Publique Hôpitaux de Paris, Université Paris-Est Créteil, INSERM U955 | Christophe Rodriguez, Slim Fourati, Vanessa Demontant, Guillaume Gricourt, Melissa N'Debi, Alexandre Soulier, Elisabeth Trawinski, Jean-Michel Pawlotsky |
| EPI_ISL_2878517 | Cerballiance Paris IDF est Saint-Denis | Department of Virology, Henri Mondor University Hospital, Assistance Publique Hôpitaux de Paris, Université Paris-Est Créteil, INSERM U955 | Christophe Rodriguez, Slim Fourati, Vanessa Demontant, Guillaume Gricourt, Melissa N'Debi, Alexandre Soulier, Elisabeth Trawinski, Jean-Michel Pawlotsky |
| EPI_ISL_2878520 | laboratoire Belle Epine | Department of Virology, Henri Mondor University Hospital, Assistance Publique Hôpitaux de Paris, Université Paris-Est Créteil, INSERM U955 | Christophe Rodriguez, Slim Fourati, Vanessa Demontant, Guillaume Gricourt, Melissa N'Debi, Alexandre Soulier, Elisabeth Trawinski, Jean-Michel Pawlotsky |
| EPI_ISL_2878521 | BIOMNIS EUROFINS IVRY | Department of Virology, Henri Mondor University Hospital, Assistance Publique Hôpitaux de Paris, Université Paris-Est Créteil, INSERM U955 | Christophe Rodriguez, Slim Fourati, Vanessa Demontant, Guillaume Gricourt, Melissa N'Debi, Alexandre Soulier, Elisabeth Trawinski, Jean-Michel Pawlotsky |
| EPI_ISL_2878522 | Biogroup Bio Lam-LCD Saint-Denis | Department of Virology, Henri Mondor University Hospital, Assistance Publique Hôpitaux de Paris, Université Paris-Est Créteil, INSERM U955 | Christophe Rodriguez, Slim Fourati, Vanessa Demontant, Guillaume Gricourt, Melissa N'Debi, Alexandre Soulier, Elisabeth Trawinski, Jean-Michel Pawlotsky |
| EPI_ISL_2878523 | SELAS MEDILYS | Department of Virology, Henri Mondor University Hospital, Assistance Publique Hôpitaux de Paris, Université Paris-Est Créteil, INSERM U955 | Christophe Rodriguez, Slim Fourati, Vanessa Demontant, Guillaume Gricourt, Melissa N'Debi, Alexandre Soulier, Elisabeth Trawinski, Jean-Michel Pawlotsky |
| EPI_ISL_2878525 | Cerballiance Paris IDF est Saint-Denis | Department of Virology, Henri Mondor University Hospital, Assistance Publique Hôpitaux de Paris, Université Paris-Est Créteil, INSERM U955 | Christophe Rodriguez, Slim Fourati, Vanessa Demontant, Guillaume Gricourt, Melissa N'Debi, Alexandre Soulier, Elisabeth Trawinski, Jean-Michel Pawlotsky |
| EPI_ISL_2878526 | Laboratoire CBM 25 TERRE ROUGE | Department of Virology, Henri Mondor University Hospital, Assistance Publique Hôpitaux de Paris, Université Paris-Est Créteil, INSERM U955 | Christophe Rodriguez, Slim Fourati, Vanessa Demontant, Guillaume Gricourt, Melissa N'Debi, Alexandre Soulier, Elisabeth Trawinski, Jean-Michel Pawlotsky |
| EPI_ISL_2878528 | Cerballiance Paris IDF est Saint-Denis | Department of Virology, Henri Mondor University Hospital, Assistance Publique Hôpitaux de Paris, Université Paris-Est Créteil, INSERM U955 | Christophe Rodriguez, Slim Fourati, Vanessa Demontant, Guillaume Gricourt, Melissa N'Debi, Alexandre Soulier, Elisabeth Trawinski, Jean-Michel Pawlotsky |
| EPI_ISL_2878529 | Biogroup Bio Lam-LCD Saint-Denis | Department of Virology, Henri Mondor University Hospital, Assistance Publique Hôpitaux de Paris, Université Paris-Est Créteil, INSERM U955 | Christophe Rodriguez, Slim Fourati, Vanessa Demontant, Guillaume Gricourt, Melissa N'Debi, Alexandre Soulier, Elisabeth Trawinski, Jean-Michel Pawlotsky |
| EPI_ISL_2878532 | laboratoire Belle Epine | Department of Virology, Henri Mondor University Hospital, Assistance Publique Hôpitaux de Paris, Université Paris-Est Créteil, INSERM U955 | Christophe Rodriguez, Slim Fourati, Vanessa Demontant, Guillaume Gricourt, Melissa N'Debi, Alexandre Soulier, Elisabeth Trawinski, Jean-Michel Pawlotsky |
| EPI_ISL_2878534 | SELAS MEDILYS | Department of Virology, Henri Mondor University Hospital, Assistance Publique Hôpitaux de Paris, Université Paris-Est Créteil, INSERM U955 | Christophe Rodriguez, Slim Fourati, Vanessa Demontant, Guillaume Gricourt, Melissa N'Debi, Alexandre Soulier, Elisabeth Trawinski, Jean-Michel Pawlotsky |
| EPI_ISL_2878537 | BIOMNIS EUROFINS IVRY | Department of Virology, Henri Mondor University Hospital, Assistance Publique Hôpitaux de Paris, Université Paris-Est Créteil, INSERM U955 | Christophe Rodriguez, Slim Fourati, Vanessa Demontant, Guillaume Gricourt, Melissa N'Debi, Alexandre Soulier, Elisabeth Trawinski, Jean-Michel Pawlotsky |
| EPI_ISL_2878538, EPI_ISL_2878540 | SELAS MEDILYS | Department of Virology, Henri Mondor University Hospital, Assistance Publique Hôpitaux de Paris, Université Paris-Est Créteil, INSERM U955 | Christophe Rodriguez, Slim Fourati, Vanessa Demontant, Guillaume Gricourt, Melissa N'Debi, Alexandre Soulier, Elisabeth Trawinski, Jean-Michel Pawlotsky |
| EPI_ISL_2878542 | BIO67-BIOSPHERE | Department of Virology, Henri Mondor University Hospital, Assistance Publique Hôpitaux de Paris, Université Paris-Est Créteil, INSERM U955 | Christophe Rodriguez, Slim Fourati, Vanessa Demontant, Guillaume Gricourt, Melissa N'Debi, Alexandre Soulier, Elisabeth Trawinski, Jean-Michel Pawlotsky |
| EPI_ISL_2878545 | SELAS MEDILYS | Department of Virology, Henri Mondor University Hospital, Assistance Publique Hôpitaux de Paris, Université Paris-Est Créteil, INSERM U955 | Christophe Rodriguez, Slim Fourati, Vanessa Demontant, Guillaume Gricourt, Melissa N'Debi, Alexandre Soulier, Elisabeth Trawinski, Jean-Michel Pawlotsky |

| | | | |
|---|---|---|---|
| EPI_ISL_2878546 | BIOMNIS EUROFINS IVRY | Department of Virology, Henri Mondor University Hospital, Assistance Publique Hôpitaux de Paris, Université Paris-Est Créteil, INSERM U955 | Christophe Rodriguez, Slim Fourati, Vanessa Demontant, Guillaume Gricourt, Melissa N'Debi, Alexandre Soulier, Elisabeth Trawinski, Jean-Michel Pawlotsky |
| EPI_ISL_2878547 | laboratoire Belle Epine | Department of Virology, Henri Mondor University Hospital, Assistance Publique Hôpitaux de Paris, Université Paris-Est Créteil, INSERM U955 | Christophe Rodriguez, Slim Fourati, Vanessa Demontant, Guillaume Gricourt, Melissa N'Debi, Alexandre Soulier, Elisabeth Trawinski, Jean-Michel Pawlotsky |
| EPI_ISL_2878548 | BIOMNIS EUROFINS IVRY | Department of Virology, Henri Mondor University Hospital, Assistance Publique Hôpitaux de Paris, Université Paris-Est Créteil, INSERM U955 | Christophe Rodriguez, Slim Fourati, Vanessa Demontant, Guillaume Gricourt, Melissa N'Debi, Alexandre Soulier, Elisabeth Trawinski, Jean-Michel Pawlotsky |
| EPI_ISL_2878549 | Plateau technique Biomer Metz Verriers | Department of Virology, Henri Mondor University Hospital, Assistance Publique Hôpitaux de Paris, Université Paris-Est Créteil, INSERM U955 | Christophe Rodriguez, Slim Fourati, Vanessa Demontant, Guillaume Gricourt, Melissa N'Debi, Alexandre Soulier, Elisabeth Trawinski, Jean-Michel Pawlotsky |
| EPI_ISL_2878551 | Biogroup Bio Lam-LCD Saint-Denis | Department of Virology, Henri Mondor University Hospital, Assistance Publique Hôpitaux de Paris, Université Paris-Est Créteil, INSERM U955 | Christophe Rodriguez, Slim Fourati, Vanessa Demontant, Guillaume Gricourt, Melissa N'Debi, Alexandre Soulier, Elisabeth Trawinski, Jean-Michel Pawlotsky |
| EPI_ISL_2878552 | Laboratoire ESPACEBIO | Department of Virology, Henri Mondor University Hospital, Assistance Publique Hôpitaux de Paris, Université Paris-Est Créteil, INSERM U955 | Christophe Rodriguez, Slim Fourati, Vanessa Demontant, Guillaume Gricourt, Melissa N'Debi, Alexandre Soulier, Elisabeth Trawinski, Jean-Michel Pawlotsky |
| EPI_ISL_2878554 | Laboratoire LENYS | Department of Virology, Henri Mondor University Hospital, Assistance Publique Hôpitaux de Paris, Université Paris-Est Créteil, INSERM U955 | Christophe Rodriguez, Slim Fourati, Vanessa Demontant, Guillaume Gricourt, Melissa N'Debi, Alexandre Soulier, Elisabeth Trawinski, Jean-Michel Pawlotsky |
| EPI_ISL_2878555, EPI_ISL_2878558 | BIO67-BIOSPHERE | Department of Virology, Henri Mondor University Hospital, Assistance Publique Hôpitaux de Paris, Université Paris-Est Créteil, INSERM U955 | Christophe Rodriguez, Slim Fourati, Vanessa Demontant, Guillaume Gricourt, Melissa N'Debi, Alexandre Soulier, Elisabeth Trawinski, Jean-Michel Pawlotsky |

EPI_ISL_2878570, EPI_ISL_2878571, EPI_ISL_2878573, EPI_ISL_2878574, EPI_ISL_2878575, EPI_ISL_2878576, EPI_ISL_2878577, EPI_ISL_2878578, EPI_ISL_2878579, EPI_ISL_2878580, EPI_ISL_2878706, EPI_ISL_2878730, EPI_ISL_2878743, EPI_ISL_2878796, EPI_ISL_2878804, EPI_ISL_2878824, EPI_ISL_2878830, EPI_ISL_2878837, EPI_ISL_2878867, EPI_ISL_2878876, EPI_ISL_2878885, EPI_ISL_2878887, EPI_ISL_2878889, EPI_ISL_2878902, EPI_ISL_2878904, EPI_ISL_2878906, EPI_ISL_2878907, EPI_ISL_2878924, EPI_ISL_2878925, EPI_ISL_2878931, EPI_ISL_2878932, EPI_ISL_2878943, EPI_ISL_2878959, EPI_ISL_2878960, EPI_ISL_2878961, EPI_ISL_2878962, EPI_ISL_2878964, EPI_ISL_2878968, EPI_ISL_2878977, EPI_ISL_2878978, EPI_ISL_2878980, EPI_ISL_2878983, EPI_ISL_2878985, EPI_ISL_2878988, EPI_ISL_2878991, EPI_ISL_2878992, EPI_ISL_2878996, EPI_ISL_2878999, EPI_ISL_2879009, EPI_ISL_2879017, EPI_ISL_2879018, EPI_ISL_2879059, EPI_ISL_2879065, EPI_ISL_2879066, EPI_ISL_2879072, EPI_ISL_2879073, EPI_ISL_2879074, EPI_ISL_2879078, EPI_ISL_2879079, EPI_ISL_2879081, EPI_ISL_2879083, EPI_ISL_2879089, EPI_ISL_2879091, EPI_ISL_2879092, EPI_ISL_2879094, EPI_ISL_2879095, EPI_ISL_2879100, EPI_ISL_2879102, EPI_ISL_2879103, EPI_ISL_2879105, EPI_ISL_2879106, EPI_ISL_2879112, EPI_ISL_2879113, EPI_ISL_2879114, EPI_ISL_2879115, EPI_ISL_2879116, EPI_ISL_2879117, EPI_ISL_2879124, EPI_ISL_2879127, EPI_ISL_2879129, EPI_ISL_2879131, EPI_ISL_2879133, EPI_ISL_2879134, EPI_ISL_2879137, EPI_ISL_2879138, EPI_ISL_2879140, EPI_ISL_2879141, EPI_ISL_2879146, EPI_ISL_2879147, EPI_ISL_2879152, EPI_ISL_2879153, EPI_ISL_2879154, EPI_ISL_2879160, EPI_ISL_2879167, EPI_ISL_2879176, EPI_ISL_2879177, EPI_ISL_2879178, EPI_ISL_2879183, EPI_ISL_2879190, EPI_ISL_2879197, EPI_ISL_2879199, EPI_ISL_2879201, EPI_ISL_2879204, EPI_ISL_2879205, EPI_ISL_2879206, EPI_ISL_2879208, EPI_ISL_2879209, EPI_ISL_2879210, EPI_ISL_2879211, EPI_ISL_2879213, EPI_ISL_2879217, EPI_ISL_2879218, EPI_ISL_2879219, EPI_ISL_2879220, EPI_ISL_2879221, EPI_ISL_2879223, EPI_ISL_2879224, EPI_ISL_2879226, EPI_ISL_2879230, EPI_ISL_2879231, EPI_ISL_2879232, EPI_ISL_2879233, EPI_ISL_2879234, EPI_ISL_2879235, EPI_ISL_2879236, EPI_ISL_2879238, EPI_ISL_2879246, EPI_ISL_2879248, EPI_ISL_2879249, EPI_ISL_2879259, EPI_ISL_2879260, EPI_ISL_2879261, EPI_ISL_2879266, EPI_ISL_2879267, EPI_ISL_2879268, EPI_ISL_2879270, EPI_ISL_2879271, EPI_ISL_2879272, EPI_ISL_2879273, EPI_ISL_2879274, EPI_ISL_2879278, EPI_ISL_2879284, EPI_ISL_2879290, EPI_ISL_2879292, EPI_ISL_2879313, EPI_ISL_2879314, EPI_ISL_2879317, EPI_ISL_2879318, EPI_ISL_2879327, EPI_ISL_2879338, EPI_ISL_2879345, EPI_ISL_2879349, EPI_ISL_2879351, EPI_ISL_2879353, EPI_ISL_2879354, EPI_ISL_2879355, EPI_ISL_2879360, EPI_ISL_2879361, EPI_ISL_2879362, EPI_ISL_2879363, EPI_ISL_2879366, EPI_ISL_2879368, EPI_ISL_2879369, EPI_ISL_2879371, EPI_ISL_2879372, EPI_ISL_2879374, EPI_ISL_2879379, EPI_ISL_2879382, EPI_ISL_2879383, EPI_ISL_2879388, EPI_ISL_2879390, EPI_ISL_2879393, EPI_ISL_2879396, EPI_ISL_2879397, EPI_ISL_2879398, EPI_ISL_2879399, EPI_ISL_2879400, EPI_ISL_2879401, EPI_ISL_2879402, EPI_ISL_2879403, EPI_ISL_2879404, EPI_ISL_2879405, EPI_ISL_2879406, EPI_ISL_2879407, EPI_ISL_2879408, EPI_ISL_2879409, EPI_ISL_2879410, EPI_ISL_2879411, EPI_ISL_2879412, EPI_ISL_2879413, EPI_ISL_2879415, EPI_ISL_2879416, EPI_ISL_2879417, EPI_ISL_2879421, EPI_ISL_2879432, EPI_ISL_2879434, EPI_ISL_2879436, EPI_ISL_2879438, EPI_ISL_2879439, EPI_ISL_2879441, EPI_ISL_2879442, EPI_ISL_2879444, EPI_ISL_2879445, EPI_ISL_2879453, EPI_ISL_2879454, EPI_ISL_2879455, EPI_ISL_2879456, EPI_ISL_2879458, EPI_ISL_2879459, EPI_ISL_2879460, EPI_ISL_2879461, EPI_ISL_2879463, EPI_ISL_2879466, EPI_ISL_2879467, EPI_ISL_2879468, EPI_ISL_2879469, EPI_ISL_2879470, EPI_ISL_2879471, EPI_ISL_2879472, EPI_ISL_2879473, EPI_ISL_2879475, EPI_ISL_2879478, EPI_ISL_2879479, EPI_ISL_2879481, EPI_ISL_2879482, EPI_ISL_2879484, EPI_ISL_2879491, EPI_ISL_2879496, EPI_ISL_2879500, EPI_ISL_2879501, EPI_ISL_2879502, EPI_ISL_2879509, EPI_ISL_2879510, EPI_ISL_2879512, EPI_ISL_2879513, EPI_ISL_2879515, EPI_ISL_2879516, EPI_ISL_2879517, EPI_ISL_2879518, EPI_ISL_2879520, EPI_ISL_2879521, EPI_ISL_2879524, EPI_ISL_2879525, EPI_ISL_2879526, EPI_ISL_2879528, EPI_ISL_2879529, EPI_ISL_2879530, EPI_ISL_2879531, EPI_ISL_2879532, EPI_ISL_2879533, EPI_ISL_2879535, EPI_ISL_2879537, EPI_ISL_2879538, EPI_ISL_2879539, EPI_ISL_2879540, EPI_ISL_2879541, EPI_ISL_2879542, EPI_ISL_2879544, EPI_ISL_2879546, EPI_ISL_2879547, EPI_ISL_2879548, EPI_ISL_2879557, EPI_ISL_2879558, EPI_ISL_2879567, EPI_ISL_2879570, EPI_ISL_2879571, EPI_ISL_2879578, EPI_ISL_2879587, EPI_ISL_2879588, EPI_ISL_2879589, EPI_ISL_2879591, EPI_ISL_2879594, EPI_ISL_2879595, EPI_ISL_2879597, EPI_ISL_2879598, EPI_ISL_2879599, EPI_ISL_2879602, EPI_ISL_2879603, EPI_ISL_2879604, EPI_ISL_2879606, EPI_ISL_2879607, EPI_ISL_2879608, EPI_ISL_2879609, EPI_ISL_2879610, EPI_ISL_2879611, EPI_ISL_2879612, EPI_ISL_2879615, EPI_ISL_2879616, EPI_ISL_2879619, EPI_ISL_2879620, EPI_ISL_2879622, EPI_ISL_2879624, EPI_ISL_2879626, EPI_ISL_2879628, EPI_ISL_2879629, EPI_ISL_2879630, EPI_ISL_2879631, EPI_ISL_2879632, EPI_ISL_2879633, EPI_ISL_2879634, EPI_ISL_2879635, EPI_ISL_2879636, EPI_ISL_2879637, EPI_ISL_2879638, EPI_ISL_2879639, EPI_ISL_2879640, EPI_ISL_2879641, EPI_ISL_2879643, EPI_ISL_2879645, EPI_ISL_2879649, EPI_ISL_2879651, EPI_ISL_2879657, EPI_ISL_2879659, EPI_ISL_2879664, EPI_ISL_2879667, EPI_ISL_2879668, EPI_ISL_2879670, EPI_ISL_2879671, EPI_ISL_2879673, EPI_ISL_2879674, EPI_ISL_2879675, EPI_ISL_2879676, EPI_ISL_2879677, EPI_ISL_2879678, EPI_ISL_2879681, EPI_ISL_2879683, EPI_ISL_2879684, EPI_ISL_2879685, EPI_ISL_2879686, EPI_ISL_2879687, EPI_ISL_2879688, EPI_ISL_2879689, EPI_ISL_2879690, EPI_ISL_2879692, EPI_ISL_2879695, EPI_ISL_2879702, EPI_ISL_2879703, EPI_ISL_2879709, EPI_ISL_2879710, EPI_ISL_2879711, EPI_ISL_2879717, EPI_ISL_2879718, EPI_ISL_2879719, EPI_ISL_2879720, EPI_ISL_2879721, EPI_ISL_2879722, EPI_ISL_2879728, EPI_ISL_2879730, EPI_ISL_2879731, EPI_ISL_2879734, EPI_ISL_2879736, EPI_ISL_2879737, EPI_ISL_2879739, EPI_ISL_2879742, EPI_ISL_2879745, EPI_ISL_2879746, EPI_ISL_2879747, EPI_ISL_2879748, EPI_ISL_2879749, EPI_ISL_2879751, EPI_ISL_2879752, EPI_ISL_2879753, EPI_ISL_2879754, EPI_ISL_2879755, EPI_ISL_2879756, EPI_ISL_2879757, EPI_ISL_2879758, EPI_ISL_2879759, EPI_ISL_2879760, EPI_ISL_2879761, EPI_ISL_2879762, EPI_ISL_2879766, EPI_ISL_2879767, EPI_ISL_2879768, EPI_ISL_2879769, EPI_ISL_2879770, EPI_ISL_2879771, EPI_ISL_2879772, EPI_ISL_2879773, EPI_ISL_2879774, EPI_ISL_2879775, EPI_ISL_2879776, EPI_ISL_2879777, EPI_ISL_2879778, EPI_ISL_2879779, EPI_ISL_2879780, EPI_ISL_2879781, EPI_ISL_2879782, EPI_ISL_2879783, EPI_ISL_2879784, EPI_ISL_2879785, EPI_ISL_2879786, EPI_ISL_2879818, EPI_ISL_2879837, EPI_ISL_2879838, EPI_ISL_2879839, EPI_ISL_2879840, EPI_ISL_2879842, EPI_ISL_2879844, EPI_ISL_2879845, EPI_ISL_2879846, EPI_ISL_2879848, EPI_ISL_2879849, EPI_ISL_2879850, EPI_ISL_2879851, EPI_ISL_2879852, EPI_ISL_2879854, EPI_ISL_2879855, EPI_ISL_2879856, EPI_ISL_2879857, EPI_ISL_2879858, EPI_ISL_2879859, EPI_ISL_2879860, EPI_ISL_2879862, EPI_ISL_2879863, EPI_ISL_2879866, EPI_ISL_2879870, EPI_ISL_2879871, EPI_ISL_2879872, EPI_ISL_2879873, EPI_ISL_2879876, EPI_ISL_2879877, EPI_ISL_2879878, EPI_ISL_2879879, EPI_ISL_2879881, EPI_ISL_2879882, EPI_ISL_2879883, EPI_ISL_2879884, EPI_ISL_2879885, EPI_ISL_2879886, EPI_ISL_2879887, EPI_ISL_2879888, EPI_ISL_2879889, EPI_ISL_2879890, EPI_ISL_2879891, EPI_ISL_2879893, EPI_ISL_2879894, EPI_ISL_2879895, EPI_ISL_2879896, EPI_ISL_2879897, EPI_ISL_2879898, EPI_ISL_2879899, EPI_ISL_2879900, EPI_ISL_2879901, EPI_ISL_2879902, EPI_ISL_2879904, EPI_ISL_2879905, EPI_ISL_2879906, EPI_ISL_2879907, EPI_ISL_2879908, EPI_ISL_2879909, EPI_ISL_2879910, EPI_ISL_2879911, EPI_ISL_2879912, EPI_ISL_2879914, EPI_ISL_2879915, EPI_ISL_2879917, EPI_ISL_2879918, EPI_ISL_2879919, EPI_ISL_2879920, EPI_ISL_2879921, EPI_ISL_2879922, EPI_ISL_2879923, EPI_ISL_2879924, EPI_ISL_2879928, EPI_ISL_2879929, EPI_ISL_2879930, EPI_ISL_2879933, EPI_ISL_2879934, EPI_ISL_2879935, EPI_ISL_2879936, EPI_ISL_2879939, EPI_ISL_2879942, EPI_ISL_2879945, EPI_ISL_2879946, EPI_ISL_2879947, EPI_ISL_2879948, EPI_ISL_2879949, EPI_ISL_2879951, EPI_ISL_2879952, EPI_ISL_2879953, EPI_ISL_2879954, EPI_ISL_2879956, EPI_ISL_2879959, EPI_ISL_2879960, EPI_ISL_2879964, EPI_ISL_2879965, EPI_ISL_2879966, EPI_ISL_2879967, EPI_ISL_2879968, EPI_ISL_2879969, EPI_ISL_2879970, EPI_ISL_2879971, EPI_ISL_2879972, EPI_ISL_2879974, EPI_ISL_2879976, EPI_ISL_2879977, EPI_ISL_2879978, EPI_ISL_2879979, EPI_ISL_2879980, EPI_ISL_2879981, EPI_ISL_2879982, EPI_ISL_2879983, EPI_ISL_2879984, EPI_ISL_2879985, EPI_ISL_2879988, EPI_ISL_2879989, EPI_ISL_2879991, EPI_ISL_2879992, EPI_ISL_2879993, EPI_ISL_2879994, EPI_ISL_2879995, EPI_ISL_2879996, EPI_ISL_2879997, EPI_ISL_2880000, EPI_ISL_2880001, EPI_ISL_2880002, EPI_ISL_2880003, EPI_ISL_2880004, EPI_ISL_2880007, EPI_ISL_2880008, EPI_ISL_2880009, EPI_ISL_2880011, EPI_ISL_2880012, EPI_ISL_2880013, EPI_ISL_2880014, EPI_ISL_2880015, EPI_ISL_2880016, EPI_ISL_2880018, EPI_ISL_2880019, EPI_ISL_2880020, EPI_ISL_2880021, EPI_ISL_2880022, EPI_ISL_2880023, EPI_ISL_2880024, EPI_ISL_2880025, EPI_ISL_2880026, EPI_ISL_2880027, EPI_ISL_2880030, EPI_ISL_2880031, EPI_ISL_2880032, EPI_ISL_2880033, EPI_ISL_2880036, EPI_ISL_2880037, EPI_ISL_2880038, EPI_ISL_2880039, EPI_ISL_2880040, EPI_ISL_2880041, EPI_ISL_2880042, EPI_ISL_2880043, EPI_ISL_2880044, EPI_ISL_2880045, EPI_ISL_2880046, EPI_ISL_2880047, EPI_ISL_2880050, EPI_ISL_2880051, EPI_ISL_2880052, EPI_ISL_2880053, EPI_ISL_2880054, EPI_ISL_2880057, EPI_ISL_2880058, EPI_ISL_2880059, EPI_ISL_2880060, EPI_ISL_2880066, EPI_ISL_2880067, EPI_ISL_2880068, EPI_ISL_2880070, EPI_ISL_2880071, EPI_ISL_2880072, EPI_ISL_2880073, EPI_ISL_2880074, EPI_ISL_2880075, EPI_ISL_2880076, EPI_ISL_2880077, EPI_ISL_2880078, EPI_ISL_2880079, EPI_ISL_2880080, EPI_ISL_2880082, EPI_ISL_2880083, EPI_ISL_2880084, EPI_ISL_2880085, EPI_ISL_2880086, EPI_ISL_2880087, EPI_ISL_2880088, EPI_ISL_2880089, EPI_ISL_2880091, EPI_ISL_2880092, EPI_ISL_2880094, EPI_ISL_2880096, EPI_ISL_2880097, EPI_ISL_2880098, EPI_ISL_2880100, EPI_ISL_2880102, EPI_ISL_2880103, EPI_ISL_2880104, EPI_ISL_2880105, EPI_ISL_2880106, EPI_ISL_2880108, EPI_ISL_2880109, EPI_ISL_2880110, EPI_ISL_2880111, EPI_ISL_2880112, EPI_ISL_2880113, EPI_ISL_2880114, EPI_ISL_2880115, EPI_ISL_2880116, EPI_ISL_2880117, EPI_ISL_2880118, EPI_ISL_2880119, EPI_ISL_2880120, EPI_ISL_2880121, EPI_ISL_2880122, EPI_ISL_2880123, EPI_ISL_2880124, EPI_ISL_2880126, EPI_ISL_2880127, EPI_ISL_2880128, EPI_ISL_2880129, EPI_ISL_2880131, EPI_ISL_2880132, EPI_ISL_2880133, EPI_ISL_2880134, EPI_ISL_2880135, EPI_ISL_2880136, EPI_ISL_2880137, EPI_ISL_2880138, EPI_ISL_2880139, EPI_ISL_2880140, EPI_ISL_2880141, EPI_ISL_2880142, EPI_ISL_2880143, EPI_ISL_2880144, EPI_ISL_2880145, EPI_ISL_2880146, EPI_ISL_2880147, EPI_ISL_2880148, EPI_ISL_2880150, EPI_ISL_2880151, EPI_ISL_2880152, EPI_ISL_2880153, EPI_ISL_2880156, EPI_ISL_2880158, EPI_ISL_2880162, EPI_ISL_2880163, EPI_ISL_2880165, EPI_ISL_2880167, EPI_ISL_2880168, EPI_ISL_2880169, EPI_ISL_2880170, EPI_ISL_2880171, EPI_ISL_2880172, EPI_ISL_2880173, EPI_ISL_2880176, EPI_ISL_2880177, EPI_ISL_2880178, EPI_ISL_2880179, EPI_ISL_2880180, EPI_ISL_2880181, EPI_ISL_2880182, EPI_ISL_2880183, EPI_ISL_2880184, EPI_ISL_2880185, EPI_ISL_2880186, EPI_ISL_2880187, EPI_ISL_2880188, EPI_ISL_2880189, EPI_ISL_2880190, EPI_ISL_2880191, EPI_ISL_2880192, EPI_ISL_2880193, EPI_ISL_2880194, EPI_ISL_2880195, EPI_ISL_2880196, EPI_ISL_2880197, EPI_ISL_2880198, EPI_ISL_2880199, EPI_ISL_2880201, EPI_ISL_2880202, EPI_ISL_2880204, EPI_ISL_2880205, EPI_ISL_2880206, EPI_ISL_2880207, EPI_ISL_2880208, EPI_ISL_2880209, EPI_ISL_2880210, EPI_ISL_2880211, EPI_ISL_2880212, EPI_ISL_2880213, EPI_ISL_2880214, EPI_ISL_2880218, EPI_ISL_2880219, EPI_ISL_2880220, EPI_ISL_2880221, EPI_ISL_2880222, EPI_ISL_2880223, EPI_ISL_2880224, EPI_ISL_2880225, EPI_ISL_2880226, EPI_ISL_2880227, EPI_ISL_2880228, EPI_ISL_2880229, EPI_ISL_2880230, EPI_ISL_2880233, EPI_ISL_2880234, EPI_ISL_2880235, EPI_ISL_2880236, EPI_ISL_2880237, EPI_ISL_2880238, EPI_ISL_2880239, EPI_ISL_2880240, EPI_ISL_2880241, EPI_ISL_2880242, EPI_ISL_2880243, EPI_ISL_2880244, EPI_ISL_2880245, EPI_ISL_2880246, EPI_ISL_2880247, EPI_ISL_2880248, EPI_ISL_2880249, EPI_ISL_2880250, EPI_ISL_2880251, EPI_ISL_2880252, EPI_ISL_2880253, EPI_ISL_2880254, EPI_ISL_2880255, EPI_ISL_2880256, EPI_ISL_2880257, EPI_ISL_2880258, EPI_ISL_2880259, EPI_ISL_2880260, EPI_ISL_2880261, EPI_ISL_2880271, EPI_ISL_2880272, EPI_ISL_2880273, EPI_ISL_2880274, EPI_ISL_2880276, EPI_ISL_2880277, EPI_ISL_2880279, EPI_ISL_2880280, EPI_ISL_2880282, EPI_ISL_2880284, EPI_ISL_2880285, EPI_ISL_2880286, EPI_ISL_2880288, EPI_ISL_2880289, EPI_ISL_2880292, EPI_ISL_2880293, EPI_ISL_2880294, EPI_ISL_2880295, EPI_ISL_2880298, EPI_ISL_2880299, EPI_ISL_2880301, EPI_ISL_2880302, EPI_ISL_2880303, EPI_ISL_2880304, EPI_ISL_2880305, EPI_ISL_2880306, EPI_ISL_2880307, EPI_ISL_2880308, EPI_ISL_2880309, EPI_ISL_2880310, EPI_ISL_2880311, EPI_ISL_2880313, EPI_ISL_2880314, EPI_ISL_2880315, EPI_ISL_2880316, EPI_ISL_2880317, EPI_ISL_2880319, EPI_ISL_2880321, EPI_ISL_2880322, EPI_ISL_2880323, EPI_ISL_2880324, EPI_ISL_2880326, EPI_ISL_2880328, EPI_ISL_2880329, EPI_ISL_2880332, EPI_ISL_2880333, EPI_ISL_2880334, EPI_ISL_2880335, EPI_ISL_2880336, EPI_ISL_2880337, EPI_ISL_2880338, EPI_ISL_2880340, EPI_ISL_2880341, EPI_ISL_2880342, EPI_ISL_2880343, EPI_ISL_2880344, EPI_ISL_2880345, EPI_ISL_2880347, EPI_ISL_2880348, EPI_ISL_2880349, EPI_ISL_2880350, EPI_ISL_2880351, EPI_ISL_2880352, EPI_ISL_2880353, EPI_ISL_2880354, EPI_ISL_2880355, EPI_ISL_2880356, EPI_ISL_2880357, EPI_ISL_2880358, EPI_ISL_2880359, EPI_ISL_2880360, EPI_ISL_2880361, EPI_ISL_2880362, EPI_ISL_2880363, EPI_ISL_2880364, EPI_ISL_2880365, EPI_ISL_2880366, EPI_ISL_2880367, EPI_ISL_2880368, EPI_ISL_2880369, EPI_ISL_2880370, EPI_ISL_2880371, EPI_ISL_2880372, EPI_ISL_2880373, EPI_ISL_2880374, EPI_ISL_2880375, EPI_ISL_2880377, EPI_ISL_2880378, EPI_ISL_2880379, EPI_ISL_2880380, EPI_ISL_2880381, EPI_ISL_2880382, EPI_ISL_2880383, EPI_ISL_2880384, EPI_ISL_2880386, EPI_ISL_2880387, EPI_ISL_2880389, EPI_ISL_2880390, EPI_ISL_2880391, EPI_ISL_2880392, EPI_ISL_2880393, EPI_ISL_2880394, EPI_ISL_2880395, EPI_ISL_2880396, EPI_ISL_2880397, EPI_ISL_2880398, EPI_ISL_2880399, EPI_ISL_2880400, EPI_ISL_2880401, EPI_ISL_2880402, EPI_ISL_2880404, EPI_ISL_2880405, EPI_ISL_2880407, EPI_ISL_2880410, EPI_ISL_2880411, EPI_ISL_2880419, EPI_ISL_2880429, EPI_ISL_2880433, EPI_ISL_2880434, EPI_ISL_2880435, EPI_ISL_2880440, EPI_ISL_2880443, EPI_ISL_2880444, EPI_ISL_2880447, EPI_ISL_2880448, EPI_ISL_2880449, EPI_ISL_2880450, EPI_ISL_2880451, EPI_ISL_2880452, EPI_ISL_2880453, EPI_ISL_2880454, EPI_ISL_2880456, EPI_ISL_2880457, EPI_ISL_2880458, EPI_ISL_2880459, EPI_ISL_2880460, EPI_ISL_2880461, EPI_ISL_2880462, EPI_ISL_2880463, EPI_ISL_2880464, EPI_ISL_2880465, EPI_ISL_2880467, EPI_ISL_2880468, EPI_ISL_2880469, EPI_ISL_2880470, EPI_ISL_2880471, EPI_ISL_2880472, EPI_ISL_2880473, EPI_ISL_2880474, EPI_ISL_2880475, EPI_ISL_2880476, EPI_ISL_2880477, EPI_ISL_2880478, EPI_ISL_2880479, EPI_ISL_2880480, EPI_ISL_2880481, EPI_ISL_2880482, EPI_ISL_2880483, EPI_ISL_2880484, EPI_ISL_2880485, EPI_ISL_2880486, EPI_ISL_2880487, EPI_ISL_2880488, EPI_ISL_2880489, EPI_ISL_2880490, EPI_ISL_2880491, EPI_ISL_2880492, EPI_ISL_2880493, EPI_ISL_2880494, EPI_ISL_2880496, EPI_ISL_2880497, EPI_ISL_2880498, EPI_ISL_2880499, EPI_ISL_2880500, EPI_ISL_2880501, EPI_ISL_2880502, EPI_ISL_2880503, EPI_ISL_2880504, EPI_ISL_2880505, EPI_ISL_2880506, EPI_ISL_2880507, EPI_ISL_2880508, EPI_ISL_2880509, EPI_ISL_2880510, EPI_ISL_2880511, EPI_ISL_2880512, EPI_ISL_2880513, EPI_ISL_2880514, EPI_ISL_2880515, EPI_ISL_2880516, EPI_ISL_2880517, EPI_ISL_2880518, EPI_ISL_2880519, EPI_ISL_2880520, EPI_ISL_2880521, EPI_ISL_2880522, EPI_ISL_2880523, EPI_ISL_2880524, EPI_ISL_2880525, EPI_ISL_2880526, EPI_ISL_2880527, EPI_ISL_2880528, EPI_ISL_2880529, EPI_ISL_2880530, EPI_ISL_2880531, EPI_ISL_2880532, EPI_ISL_2880533, EPI_ISL_2880534, EPI_ISL_2880535, EPI_ISL_2880536, EPI_ISL_2880537, EPI_ISL_2880538, EPI_ISL_2880539, EPI_ISL_2880540, EPI_ISL_2880541, EPI_ISL_2880542, EPI_ISL_2880543, EPI_ISL_2880544, EPI_ISL_2880546, EPI_ISL_2880548, EPI_ISL_2880549, EPI_ISL_2880550, EPI_ISL_2880553, EPI_ISL_2880554, EPI_ISL_2880555, EPI_ISL_2880556, EPI_ISL_2880557, EPI_ISL_2880558, EPI_ISL_2880560, EPI_ISL_2880561, EPI_ISL_2880562, EPI_ISL_2880563, EPI_ISL_2880564, EPI_ISL_2880565, EPI_ISL_2880566, EPI_ISL_2880567, EPI_ISL_2880568, EPI_ISL_2880569, EPI_ISL_2880570, EPI_ISL_2880571, EPI_ISL_2880573, EPI_ISL_2880574, EPI_ISL_2880575, EPI_ISL_2880576, EPI_ISL_2880577, EPI_ISL_2880578, EPI_ISL_2880579, EPI_ISL_2880580, EPI_ISL_2880581, EPI_ISL_2880582, EPI_ISL_2880583, EPI_ISL_2880584, EPI_ISL_2880585, EPI_ISL_2880586, EPI_ISL_2880587, EPI_ISL_2880590, EPI_ISL_2880592, EPI_ISL_2880593, EPI_ISL_2880594, EPI_ISL_2880595, EPI_ISL_2880596, EPI_ISL_2880598, EPI_ISL_2880599, EPI_ISL_2880600, EPI_ISL_2880601, EPI_ISL_2880602, EPI_ISL_2880605, EPI_ISL_2880606, EPI_ISL_2880607, EPI_ISL_2880608, EPI_ISL_2880609, EPI_ISL_2880610, EPI_ISL_2880611, EPI_ISL_2880612, EPI_ISL_2880613, EPI_ISL_2880614, EPI_ISL_2880615, EPI_ISL_2880616, EPI_ISL_2880617, EPI_ISL_2880619, EPI_ISL_2880620, EPI_ISL_2880621, EPI_ISL_2880622, EPI_ISL_2880623, EPI_ISL_2880627, EPI_ISL_2880629, EPI_ISL_2880630, EPI_ISL_2880631, EPI_ISL_2880632, EPI_ISL_2880633, EPI_ISL_2880635, EPI_ISL_2880636, EPI_ISL_2880637, EPI_ISL_2880638, EPI_ISL_2880639, EPI_ISL_2880641, EPI_ISL_2880642, EPI_ISL_2880643, EPI_ISL_2880644, EPI_ISL_2880645, EPI_ISL_2880646, EPI_ISL_2880647, EPI_ISL_2880648, EPI_ISL_2880649, EPI_ISL_2880650, EPI_ISL_2880651, EPI_ISL_2880652, EPI_ISL_2880653, EPI_ISL_2880655, EPI_ISL_2880656, EPI_ISL_2880659, EPI_ISL_2880660, EPI_ISL_2880661, EPI_ISL_2880662, EPI_ISL_2880663, EPI_ISL_2880664, EPI_ISL_2880665, EPI_ISL_2880666, EPI_ISL_2880668, EPI_ISL_2880669, EPI_ISL_2880670, EPI_ISL_2880671, EPI_ISL_2880672, EPI_ISL_2880673, EPI_ISL_2880674, EPI_ISL_2880675, EPI_ISL_2880676, EPI_ISL_2880677, EPI_ISL_2880678, EPI_ISL_2880679, EPI_ISL_2880680, EPI_ISL_2880681, EPI_ISL_2880682, EPI_ISL_2880683, EPI_ISL_2880684, EPI_ISL_2880685, EPI_ISL_2880686, EPI_ISL_2880687, EPI_ISL_2880688, EPI_ISL_2880689, EPI_ISL_2880690, EPI_ISL_2880691, EPI_ISL_2880692, EPI_ISL_2880693, EPI_ISL_2880694, EPI_ISL_2880695, EPI_ISL_2880696, EPI_ISL_2880697, EPI_ISL_2880698, EPI_ISL_2880699, EPI_ISL_2880701, EPI_ISL_2880717, EPI_ISL_2880721, EPI_ISL_2880726, EPI_ISL_2880729, EPI_ISL_2880734, EPI_ISL_2880737, EPI_ISL_2880739, EPI_ISL_2880755, EPI_ISL_2880757, EPI_ISL_2880758, EPI_ISL_2880759, EPI_ISL_2880760, EPI_ISL_2880761, EPI_ISL_2880765, EPI_ISL_2880772, EPI_ISL_2880775, EPI_ISL_2880781, EPI_ISL_2880799, EPI_ISL_2880801, EPI_ISL_2880809, EPI_ISL_2880821, EPI_ISL_2880824, EPI_ISL_2880832, EPI_ISL_2880835, EPI_ISL_2880847, EPI_ISL_2880853, EPI_ISL_2880861, EPI_ISL_2880876, EPI_ISL_2880877, EPI_ISL_2880879, EPI_ISL_2880880, EPI_ISL_2880887, EPI_ISL_2880891, EPI_ISL_2880912, EPI_ISL_2880915, EPI_ISL_2880917, EPI_ISL_2880920, EPI_ISL_2880922, EPI_ISL_2880928, EPI_ISL_2880930, EPI_ISL_2880935, EPI_ISL_2880936, EPI_ISL_2880937, EPI_ISL_2880938, EPI_ISL_2880939, EPI_ISL_2880940, EPI_ISL_2880941, EPI_ISL_2880942, EPI_ISL_2880943, EPI_ISL_2880944, EPI_ISL_2880945, EPI_ISL_2880946, EPI_ISL_2880947, EPI_ISL_2880948, EPI_ISL_2880949, EPI_ISL_2880950, EPI_ISL_2880951, EPI_ISL_2880952, EPI_ISL_2880953, EPI_ISL_2880954, EPI_ISL_2880955, EPI_ISL_2880956, EPI_ISL_2880957, EPI_ISL_2880958, EPI_ISL_2880959, EPI_ISL_2880960, EPI_ISL_2880961, EPI_ISL_2880962, EPI_ISL_2880963, EPI_ISL_2880964, EPI_ISL_2880965, EPI_ISL_2880966, EPI_ISL_2880967, EPI_ISL_2880968, EPI_ISL_2880969, EPI_ISL_2880970, EPI_ISL_2880971, EPI_ISL_2880972, EPI_ISL_2880973, EPI_ISL_2880974, EPI_ISL_2880975, EPI_ISL_2880976, EPI_ISL_2880977, EPI_ISL_2880978, EPI_ISL_2880980, EPI_ISL_2880981, EPI_ISL_2880983, EPI_ISL_2880984, EPI_ISL_2880985, EPI_ISL_2880986, EPI_ISL_2880988, EPI_ISL_2880989, EPI_ISL_2880990, EPI_ISL_2880991, EPI_ISL_2880992, EPI_ISL_2880993, EPI_ISL_2880994, EPI_ISL_2880995, EPI_ISL_2880996, EPI_ISL_2880997, EPI_ISL_2880998, EPI_ISL_2880999, EPI_ISL_2881000, EPI_ISL_2881001, EPI_ISL_2881002, EPI_ISL_2881003, EPI_ISL_2881004, EPI_ISL_2881005, EPI_ISL_2881006, EPI_ISL_2881007, EPI_ISL_2881008, EPI_ISL_2881009, EPI_ISL_2881010, EPI_ISL_2881011, EPI_ISL_2881012, EPI_ISL_2881013, EPI_ISL_2881014, EPI_ISL_2881015, EPI_ISL_2881016, EPI_ISL_2881017, EPI_ISL_2881019, EPI_ISL_2881020, EPI_ISL_2881021, EPI_ISL_2881022, EPI_ISL_2881023, EPI_ISL_2881024, EPI_ISL_2881026, EPI_ISL_2881027, EPI_ISL_2881028, EPI_ISL_2881029, EPI_ISL_2881030, EPI_ISL_2881031, EPI_ISL_2881032, EPI_ISL_2881033, EPI_ISL_2881034, EPI_ISL_2881035, EPI_ISL_2881038, EPI_ISL_2881039, EPI_ISL_2881040, EPI_ISL_2881041, EPI_ISL_2881042, EPI_ISL_2881044, EPI_ISL_2881045, EPI_ISL_2881046, EPI_ISL_2881047, EPI_ISL_2881048, EPI_ISL_2881049, EPI_ISL_2881050, EPI_ISL_2881051, EPI_ISL_2881052, EPI_ISL_2881053, EPI_ISL_2881054, EPI_ISL_2881055, EPI_ISL_2881056, EPI_ISL_2881058, EPI_ISL_2881059, EPI_ISL_2881061, EPI_ISL_2881062, EPI_ISL_2881063, EPI_ISL_2881064, EPI_ISL_2881066, EPI_ISL_2881067, EPI_ISL_2881068, EPI_ISL_2881071, EPI_ISL_2881072, EPI_ISL_2881073, EPI_ISL_2881074, EPI_ISL_2881075, EPI_ISL_2881076, EPI_ISL_2881077, EPI_ISL_2881078, EPI_ISL_2881080, EPI_ISL_2881081, EPI_ISL_2881082, EPI_ISL_2881083, EPI_ISL_2881085, EPI_ISL_2881086, EPI_ISL_2881087, EPI_ISL_2881088, EPI_ISL_2881089, EPI_ISL_2881090, EPI_ISL_2881091, EPI_ISL_2881092, EPI_ISL_2881093, EPI_ISL_2881094, EPI_ISL_2881095, EPI_ISL_2881096, EPI_ISL_2881097, EPI_ISL_2881098, EPI_ISL_2881099, EPI_ISL_2881101, EPI_ISL_2881102, EPI_ISL_2881103, EPI_ISL_2881104, EPI_ISL_2881105, EPI_ISL_2881106, EPI_ISL_2881107, EPI_ISL_2881108, EPI_ISL_2881109, EPI_ISL_2881110, EPI_ISL_2881111, EPI_ISL_2881112, EPI_ISL_2881113, EPI_ISL_2881114, EPI_ISL_2881115, EPI_ISL_2881116, EPI_ISL_2881117, EPI_ISL_2881118, EPI_ISL_2881119, EPI_ISL_2881121, EPI_ISL_2881122, EPI_ISL_2881123, EPI_ISL_2881124, EPI_ISL_2881125, EPI_ISL_2881126, EPI_ISL_2881127, EPI_ISL_2881128, EPI_ISL_2881129, EPI_ISL_2881130, EPI_ISL_2881131, EPI_ISL_2881132, EPI_ISL_2881133, EPI_ISL_2881134, EPI_ISL_2881145, EPI_ISL_2881146, EPI_ISL_2881148, EPI_ISL_2881149, EPI_ISL_2881150, EPI_ISL_2881153, EPI_ISL_2881157, EPI_ISL_2881158, EPI_ISL_2881159, EPI_ISL_2881160, EPI_ISL_2881161, EPI_ISL_2881163, EPI_ISL_2881164, EPI_ISL_2881165, EPI_ISL_2881166, EPI_ISL_2881167, EPI_ISL_2881168, EPI_ISL_2881169, EPI_ISL_2881170, EPI_ISL_2881171, EPI_ISL_2881172, EPI_ISL_2881173, EPI_ISL_2881174, EPI_ISL_2881175, EPI_ISL_2881176, EPI_ISL_2881177, EPI_ISL_2881178, EPI_ISL_2881179, EPI_ISL_2881180, EPI_ISL_2881181, EPI_ISL_2881183, EPI_ISL_2881185, EPI_ISL_2881186, EPI_ISL_2881187, EPI_ISL_2881188, EPI_ISL_2881189, EPI_ISL_2881190, EPI_ISL_2881191, EPI_ISL_2881192, EPI_ISL_2881193, EPI_ISL_2881194, EPI_ISL_2881195, EPI_ISL_2881196, EPI_ISL_2881197, EPI_ISL_2881198, EPI_ISL_2881199, EPI_ISL_2881200, EPI_ISL_2881201, EPI_ISL_2881202, EPI_ISL_2881203, EPI_ISL_2881204, EPI_ISL_2881205, EPI_ISL_2881206, EPI_ISL_2881213, EPI_ISL_2881214, EPI_ISL_2881215, EPI_ISL_2881216, EPI_ISL_2881217, EPI_ISL_2881219, EPI_ISL_2881220, EPI_ISL_2881221, EPI_ISL_2881222, EPI_ISL_2881223, EPI_ISL_2881224, EPI_ISL_2881225, EPI_ISL_2881226, EPI_ISL_2881227, EPI_ISL_2881228, EPI_ISL_2881229, EPI_ISL_2881233, EPI_ISL_2881236, EPI_ISL_2881238, EPI_ISL_2881239, EPI_ISL_2881240, EPI_ISL_2881241, EPI_ISL_2881247, EPI_ISL_2881248, EPI_ISL_2881251, EPI_ISL_2881256, EPI_ISL_2881258, EPI_ISL_2881259, EPI_ISL_2881260, EPI_ISL_2881261, EPI_ISL_2881262, EPI_ISL_2881263, EPI_ISL_2881264, EPI_ISL_2881265, EPI_ISL_2881266, EPI_ISL_2881267, EPI_ISL_2881269, EPI_ISL_2881270, EPI_ISL_2881272, EPI_ISL_2881274, EPI_ISL_2881275, EPI_ISL_2881277, EPI_ISL_2881278, EPI_ISL_2881279, EPI_ISL_2881280, EPI_ISL_2881281, EPI_ISL_2881282, EPI_ISL_2881283, EPI_ISL_2881284, EPI_ISL_2881286, EPI_ISL_2881287, EPI_ISL_2881288, EPI_ISL_2881289, EPI_ISL_2881290, EPI_ISL_2881291, EPI_ISL_2881292, EPI_ISL_2881293, EPI_ISL_2881294, EPI_ISL_2881295, EPI_ISL_2881296, EPI_ISL_2881297, EPI_ISL_2881298, EPI_ISL_2881299, EPI_ISL_2881300, EPI_ISL_2881302, EPI_ISL_2881303, EPI_ISL_2881304, EPI_ISL_2881305, EPI_ISL_2881306, EPI_ISL_2881307, EPI_ISL_2881308, EPI_ISL_2881309, EPI_ISL_2881310, EPI_ISL_2881311, EPI_ISL_2881312, EPI_ISL_2881313, EPI_ISL_2881314, EPI_ISL_2881315, EPI_ISL_2881316, EPI_ISL_2881317, EPI_ISL_2881318, EPI_ISL_2881319, EPI_ISL_2881320, EPI_ISL_2881321, EPI_ISL_2881322, EPI_ISL_2881323, EPI_ISL_2881324, EPI_ISL_2881325, EPI_ISL_2881329, EPI_ISL_2881331, EPI_ISL_2881333, EPI_ISL_2881337, EPI_ISL_2881339, EPI_ISL_2881341, EPI_ISL_2881342, EPI_ISL_2881350, EPI_ISL_2881351, EPI_ISL_2881354, EPI_ISL_2881355, EPI_ISL_2881356, EPI_ISL_2881357, EPI_ISL_2881358, EPI_ISL_2881359, EPI_ISL_2881360, EPI_ISL_2881362, EPI_ISL_2881363, EPI_ISL_2881364, EPI_ISL_2881366, EPI_ISL_2881370, EPI_ISL_2881376, EPI_ISL_2881377, EPI_ISL_2881378, EPI_ISL_2881379, EPI_ISL_2881380, EPI_ISL_2881381, EPI_ISL_2881382, EPI_ISL_2881383, EPI_ISL_2881384, EPI_ISL_2881385, EPI_ISL_2881386, EPI_ISL_2881387, EPI_ISL_2881388, EPI_ISL_2881389, EPI_ISL_2881390, EPI_ISL_2881392, EPI_ISL_2881393, EPI_ISL_2881394, EPI_ISL_2881395, EPI_ISL_2881396, EPI_ISL_2881397, EPI_ISL_2881398, EPI_ISL_2881399, EPI_ISL_2881400, EPI_ISL_2881401, EPI_ISL_2881403, EPI_ISL_2881404, EPI_ISL_2881405, EPI_ISL_2881406, EPI_ISL_2881407, EPI_ISL_2881408, EPI_ISL_2881409, EPI_ISL_2881410, EPI_ISL_2881411, EPI_ISL_2881412, EPI_ISL_2881413, EPI_ISL_2881414, EPI_ISL_2881416, EPI_ISL_2881417, EPI_ISL_2881418, EPI_ISL_2881419, EPI_ISL_2881420, EPI_ISL_2881421, EPI_ISL_2881422, EPI_ISL_2881423, EPI_ISL_2881424, EPI_ISL_2881425, EPI_ISL_2881426, EPI_ISL_2881427, EPI_ISL_2881428, EPI_ISL_2881429, EPI_ISL_2881430, EPI_ISL_2881431, EPI_ISL_2881432, EPI_ISL_2881435, EPI_ISL_2881437, EPI_ISL_2881439, EPI_ISL_2881440, EPI_ISL_2881441, EPI_ISL_2881443, EPI_ISL_2881444, EPI_ISL_2881446, EPI_ISL_2881449, EPI_ISL_2881450, EPI_ISL_2881451, EPI_ISL_2881452, EPI_ISL_2881453, EPI_ISL_2881454, EPI_ISL_2881455, EPI_ISL_2881456, EPI_ISL_2881457, EPI_ISL_2881458, EPI_ISL_2881459, EPI_ISL_2881460, EPI_ISL_2881461, EPI_ISL_2881462, EPI_ISL_2881463, EPI_ISL_2881464, EPI_ISL_2881465, EPI_ISL_2881466, EPI_ISL_2881468, EPI_ISL_2881469, EPI_ISL_2881470, EPI_ISL_2881471, EPI_ISL_2881472, EPI_ISL_2881473, EPI_ISL_2881474, EPI_ISL_2881475, EPI_ISL_2881476, EPI_ISL_2881477, EPI_ISL_2881478, EPI_ISL_2881479, EPI_ISL_2881480, EPI_ISL_2881481, EPI_ISL_2881482, EPI_ISL_2881483, EPI_ISL_2881485, EPI_ISL_2881486, EPI_ISL_2881488, EPI_ISL_2881489, EPI_ISL_2881491, EPI_ISL_2881492, EPI_ISL_2881494, EPI_ISL_2881495, EPI_ISL_2881496, EPI_ISL_2881497, EPI_ISL_2881498, EPI_ISL_2881499, EPI_ISL_2881500, EPI_ISL_2881501, EPI_ISL_2881502, EPI_ISL_2881503, EPI_ISL_2881504, EPI_ISL_2881505, EPI_ISL_2881506, EPI_ISL_2881507, EPI_ISL_2881508, EPI_ISL_2881509, EPI_ISL_2881510, EPI_ISL_2881511, EPI_ISL_2881512, EPI_ISL_2881513, EPI_ISL_2881514, EPI_ISL_2881515, EPI_ISL_2881516, EPI_ISL_2881517, EPI_ISL_2881518, EPI_ISL_2881519, EPI_ISL_2881520, EPI_ISL_2881521, EPI_ISL_2881522, EPI_ISL_2881523, EPI_ISL_2881524, EPI_ISL_2881525, EPI_ISL_2881526, EPI_ISL_2881527, EPI_ISL_2881528, EPI_ISL_2881531, EPI_ISL_2881532, EPI_ISL_2881533, EPI_ISL_2881534, EPI_ISL_2881537, EPI_ISL_2881538, EPI_ISL_2881539, EPI_ISL_2881540, EPI_ISL_2881541, EPI_ISL_2881542, EPI_ISL_2881544, EPI_ISL_2881545, EPI_ISL_2881546, EPI_ISL_2881547, EPI_ISL_2881548, EPI_ISL_2881549, EPI_ISL_2881550, EPI_ISL_2881551, EPI_ISL_2881552, EPI_ISL_2881553, EPI_ISL_2881554, EPI_ISL_2881555, EPI_ISL_2881556, EPI_ISL_2881558, EPI_ISL_2881559, EPI_ISL_2881561, EPI_ISL_2881562, EPI_ISL_2881563, EPI_ISL_2881564, EPI_ISL_2881565, EPI_ISL_2881566, EPI_ISL_2881567, EPI_ISL_2881568, EPI_ISL_2881569, EPI_ISL_2881570, EPI_ISL_2881571, EPI_ISL_2881572, EPI_ISL_2881573, EPI_ISL_2881574, EPI_ISL_2881575, EPI_ISL_2881576, EPI_ISL_2881578, EPI_ISL_2881579, EPI_ISL_2881580, EPI_ISL_2881581, EPI_ISL_2881582, EPI_ISL_2881583, EPI_ISL_2881584, EPI_ISL_2881585, EPI_ISL_2881586, EPI_ISL_2881587, EPI_ISL_2881588, EPI_ISL_2881589, EPI_ISL_2881590, EPI_ISL_2881591, EPI_ISL_2881592, EPI_ISL_2881593, EPI_ISL_2881594, EPI_ISL_2881595, EPI_ISL_2881596, EPI_ISL_2881597, EPI_ISL_2881598, EPI_ISL_2881599, EPI_ISL_2881600, EPI_ISL_2881601, EPI_ISL_2881602, EPI_ISL_2881603, EPI_ISL_2881604, EPI_ISL_2881605, EPI_ISL_2881606, EPI_ISL_2881607, EPI_ISL_2881608, EPI_ISL_2881609, EPI_ISL_2881610, EPI_ISL_2881611, EPI_ISL_2881612, EPI_ISL_2881614, EPI_ISL_2881615, EPI_ISL_2881616, EPI_ISL_2881617, EPI_ISL_2881618, EPI_ISL_2881619, EPI_ISL_2881620, EPI_ISL_2881623, EPI_ISL_2881624, EPI_ISL_2881625, EPI_ISL_2881626, EPI_ISL_2881627, EPI_ISL_2881628, EPI_ISL_2881629, EPI_ISL_2881630, EPI_ISL_2881631, EPI_ISL_2881632, EPI_ISL_2881633, EPI_ISL_2881634, EPI_ISL_2881635, EPI_ISL_2881636, EPI_ISL_2881637, EPI_ISL_2881638, EPI_ISL_2881639, EPI_ISL_2881640, EPI_ISL_2881641, EPI_ISL_2881642, EPI_ISL_2881643, EPI_ISL_2881644, EPI_ISL_2881645, EPI_ISL_2881646, EPI_ISL_2881647, EPI_ISL_2881648, EPI_ISL_2881649, EPI_ISL_2881650, EPI_ISL_2881651, EPI_ISL_2881652, EPI_ISL_2881653, EPI_ISL_2881654, EPI_ISL_2881655, EPI_ISL_2881656, EPI_ISL_2881657, EPI_ISL_2881658, EPI_ISL_2881659, EPI_ISL_2881662, EPI_ISL_2881665, EPI_ISL_2881668, EPI_ISL_2881669, EPI_ISL_2881670, EPI_ISL_2881672, EPI_ISL_2881673, EPI_ISL_2881674, EPI_ISL_2881675, EPI_ISL_2881676, EPI_ISL_2881677, EPI_ISL_2881678, EPI_ISL_2881679, EPI_ISL_2881680, EPI_ISL_2881681, EPI_ISL_2881682, EPI_ISL_2881683, EPI_ISL_2881684, EPI_ISL_2881685, EPI_ISL_2881686, EPI_ISL_2881687, EPI_ISL_2881688, EPI_ISL_2881689, EPI_ISL_2881690, EPI_ISL_2881691, EPI_ISL_2881692, EPI_ISL_2881693, EPI_ISL_2881694, EPI_ISL_2881695, EPI_ISL_2881696, EPI_ISL_2881697, EPI_ISL_2881698, EPI_ISL_2881699, EPI_ISL_2881700, EPI_ISL_2881701, EPI_ISL_2881702, EPI_ISL_2881703, EPI_ISL_2881704, EPI_ISL_2881705, EPI_ISL_2881706, EPI_ISL_2881707, EPI_ISL_2881708, EPI_ISL_2881709, EPI_ISL_2881710, EPI_ISL_2881711, EPI_ISL_2881712, EPI_ISL_2881713, EPI_ISL_2881714, EPI_ISL_2881715, EPI_ISL_2881716, EPI_ISL_2881717, EPI_ISL_2881718, EPI_ISL_2881719, EPI_ISL_2881720

| | | | |
|---|---|---|---|
| see above | INSACOG Surveillance | INSACOG at CSIR Institute of Genomics and Integrative Biology | INSACOG |
| EPI_ISL_2881809, EPI_ISL_2881856 | Salud Digna | Instituto Nacional de Medicina Genomica | Cedro-Tanda A, Hidalgo-Miranda A, Mendoza-Vargas A, Reyes-Grajeda JP, Abraham Campos-Romero, Moreno-Camacho José Luis, Rodriguez-Gallegos Jorge, Luna-Ruiz Marco, Gonzalez-Barrera D, Rangel-DeLeon D, Munguia-Garza P, Ramirez-Vega O, Escobar-Arrazola, M, Herrera-Montalvo LA. |
| EPI_ISL_2882329, EPI_ISL_2882330, EPI_ISL_2882331, EPI_ISL_2882363, EPI_ISL_2882365, EPI_ISL_2882627 | CDPH VBL | California Department of Public Health | CDPH-COVIDNet |
| EPI_ISL_2884065, EPI_ISL_2884157, EPI_ISL_2884324, EPI_ISL_2884491, EPI_ISL_2884492, EPI_ISL_2884493, EPI_ISL_2884494, EPI_ISL_2884495, EPI_ISL_2884496, EPI_ISL_2884497, EPI_ISL_2884498, EPI_ISL_2884499, EPI_ISL_2884500, EPI_ISL_2884501, EPI_ISL_2884570, EPI_ISL_2884599, EPI_ISL_2884615, EPI_ISL_2884616, EPI_ISL_2884617, EPI_ISL_2884618, EPI_ISL_2884619, EPI_ISL_2884620, EPI_ISL_2884621, EPI_ISL_2884622, EPI_ISL_2884623, EPI_ISL_2884624, EPI_ISL_2884625, EPI_ISL_2884626, EPI_ISL_2884627, EPI_ISL_2884629, EPI_ISL_2884630 | | | |
| see above | Virology Laboratory, Scientific Department, Army Medical Center | Virology Laboratory, Scientific Department, Army Medical Center | Silvia Fillo, Riccardo De Sanctis, Antonella Fortunato, Anella Monte, Rossella Brandi, Giulia Campoli, Marzia Cavalli, Lucia Nicosia, Anna Anselmo, Vanessa Vera Fain, Francesco Giordani, Giandomenico Cerreto, Filippo Molinari, Giancarlo Petralito, Florigio Lista |
| EPI_ISL_2884880, EPI_ISL_2884881, EPI_ISL_2884884, EPI_ISL_2884907, EPI_ISL_2884908, EPI_ISL_2884971, EPI_ISL_2885150, EPI_ISL_2885151 | Johns Hopkins Hospital Department of Pathology | Johns Hopkins Hospital Department of Pathology | C. Paul Morris, Chun Huai Luo, Adannaya Amadi, Matthew Schwartz, Nicholas Gallagher, Heba H. Mostafa |
| EPI_ISL_2885266, EPI_ISL_2885268, EPI_ISL_2885270, EPI_ISL_2885271, EPI_ISL_2885272, EPI_ISL_2885273, EPI_ISL_2885274, EPI_ISL_2885275, EPI_ISL_2885276, EPI_ISL_2885278, EPI_ISL_2885280, EPI_ISL_2885281, EPI_ISL_2885282, EPI_ISL_2885283, EPI_ISL_2885284, EPI_ISL_2885285, EPI_ISL_2885286, EPI_ISL_2885287, EPI_ISL_2885288, EPI_ISL_2885289, EPI_ISL_2885290, EPI_ISL_2885291, EPI_ISL_2885292, EPI_ISL_2885293, EPI_ISL_2885294, EPI_ISL_2885295, EPI_ISL_2885297, EPI_ISL_2885298, EPI_ISL_2885299, EPI_ISL_2885300, EPI_ISL_2885301, EPI_ISL_2885303, EPI_ISL_2885304, EPI_ISL_2885305, EPI_ISL_2885307, EPI_ISL_2885308, EPI_ISL_2885309, EPI_ISL_2885310, EPI_ISL_2885311, EPI_ISL_2885312, EPI_ISL_2885313, EPI_ISL_2885314, EPI_ISL_2885315, EPI_ISL_2885316, EPI_ISL_2885317, EPI_ISL_2885318, EPI_ISL_2885319, EPI_ISL_2885320, EPI_ISL_2885321, EPI_ISL_2885323, EPI_ISL_2885324, EPI_ISL_2885325, EPI_ISL_2885326, EPI_ISL_2885327, EPI_ISL_2885328, EPI_ISL_2885329, EPI_ISL_2885330, EPI_ISL_2885331, EPI_ISL_2885332, EPI_ISL_2885334, EPI_ISL_2885335, EPI_ISL_2885336, EPI_ISL_2885338, EPI_ISL_2885341, EPI_ISL_2885343, EPI_ISL_2885344, EPI_ISL_2885346, EPI_ISL_2885347, EPI_ISL_2885348, EPI_ISL_2885349 | | | |
| see above | Hospital Universitari Vall d'Hebron - Vall d'Hebron Institut de Recerca | Hospital Universitari Vall d'Hebron - Vall d'Hebron Institut de Recerca | Cristina Andrés, Maria Piñana, Alejandra González-Sánchez, Damir Garcia-Cehic, Ariadna Rando, Juliana Esperalba, Maria Gema Codina, Carla Castillo, Karen García, Rodrigo Vásquez, Maria Carmen Martin, Tomàs Pumarola, Josep Quer, Andrés Antón |
| EPI_ISL_2885390, EPI_ISL_2885391, EPI_ISL_2885393, EPI_ISL_2885396, EPI_ISL_2885400, EPI_ISL_2885401, EPI_ISL_2885402, EPI_ISL_2885403, EPI_ISL_2885404, EPI_ISL_2885405, EPI_ISL_2885408, EPI_ISL_2885411, EPI_ISL_2885412, EPI_ISL_2885413, EPI_ISL_2885414, EPI_ISL_2885415, EPI_ISL_2885416, EPI_ISL_2885417, EPI_ISL_2885418, EPI_ISL_2885419, EPI_ISL_2885420, EPI_ISL_2885421, EPI_ISL_2885422, EPI_ISL_2885426, EPI_ISL_2885428, EPI_ISL_2885430, EPI_ISL_2885431, EPI_ISL_2885432, EPI_ISL_2885433, EPI_ISL_2885434, EPI_ISL_2885436, EPI_ISL_2885437, EPI_ISL_2885438, EPI_ISL_2885439, EPI_ISL_2885440, EPI_ISL_2885441, EPI_ISL_2885442, EPI_ISL_2885444, EPI_ISL_2885445, EPI_ISL_2885447, EPI_ISL_2885448, EPI_ISL_2885450, EPI_ISL_2885451, EPI_ISL_2885452, EPI_ISL_2885453, EPI_ISL_2885454, EPI_ISL_2885455, EPI_ISL_2885456, EPI_ISL_2885457, EPI_ISL_2885458, EPI_ISL_2885462, EPI_ISL_2885463, EPI_ISL_2885464, EPI_ISL_2885465, EPI_ISL_2885466, EPI_ISL_2885467, EPI_ISL_2885468, EPI_ISL_2885469, EPI_ISL_2885470, EPI_ISL_2885471, EPI_ISL_2885472 | | | |
| see above | Illinois Department of Public Health - Springfield Lab | Illinois Department of Public Health - Springfield Lab | Bryan Sim, Gordon McCall |
| EPI_ISL_2885473, EPI_ISL_2885476, EPI_ISL_2885480, EPI_ISL_2885487, EPI_ISL_2885488, EPI_ISL_2885489, EPI_ISL_2885497, EPI_ISL_2885498, EPI_ISL_2885499, EPI_ISL_2885502, EPI_ISL_2885503, EPI_ISL_2885504, EPI_ISL_2885506, EPI_ISL_2885507, EPI_ISL_2885509 | | | |
| see above | University Hospitals of Geneva, Laboratory of Virology | HUG, Laboratory of Virology and the Health2030 Genome Center | Samuel Cordey, Ana Rita Goncalves, Laurent Kaiser, Lorenzo Cerutti, Henri Pegeot, Melyssa Elies, Deborah Penet, Keith Harshman, Ioannis Xenarios, Emmanouil Dermitzakis |
| EPI_ISL_2885611, EPI_ISL_2885612, EPI_ISL_2885613, EPI_ISL_2885614, EPI_ISL_2885615, EPI_ISL_2885616, EPI_ISL_2885617, EPI_ISL_2885618, EPI_ISL_2885619, EPI_ISL_2885620, EPI_ISL_2885621, EPI_ISL_2885623, EPI_ISL_2885624, EPI_ISL_2885625, EPI_ISL_2885626, EPI_ISL_2885627, EPI_ISL_2885628, EPI_ISL_2885629, EPI_ISL_2885633, EPI_ISL_2885634, EPI_ISL_2885636, EPI_ISL_2885637, EPI_ISL_2885638, EPI_ISL_2885640, EPI_ISL_2885641, EPI_ISL_2885642, EPI_ISL_2885643, EPI_ISL_2885644, EPI_ISL_2885645, EPI_ISL_2885647, EPI_ISL_2885648, EPI_ISL_2885649, EPI_ISL_2885651, EPI_ISL_2885652, EPI_ISL_2885653, EPI_ISL_2885654, EPI_ISL_2885655, EPI_ISL_2885656, EPI_ISL_2885657, EPI_ISL_2885658, EPI_ISL_2885660, EPI_ISL_2885661, EPI_ISL_2885663, EPI_ISL_2885664, EPI_ISL_2885665, EPI_ISL_2885666, EPI_ISL_2885668, EPI_ISL_2885669, EPI_ISL_2885671, EPI_ISL_2885675, EPI_ISL_2885676, EPI_ISL_2885677, EPI_ISL_2885678, EPI_ISL_2885679 | | | |
| see above | Cliniques universitaires Saint-Luc | UCLouvain/IREC/MBLG | Jean Ruelle, Bertrand Bearzatto, Benoit Kabamba Mukadi |
| EPI_ISL_2885680, EPI_ISL_2885681, EPI_ISL_2885682, EPI_ISL_2885683, EPI_ISL_2885684, EPI_ISL_2885685, EPI_ISL_2885686, EPI_ISL_2885687, EPI_ISL_2885690, EPI_ISL_2885691, EPI_ISL_2885692, EPI_ISL_2885694, EPI_ISL_2885695 | | | |
| see above | Clinique Saint-Pierre Ottignies | UCLouvain/IREC/MBLG | Jean Ruelle, Bertrand Bearzatto, Benoit Kabamba Mukadi |
| EPI_ISL_2885698 | Cliniques universitaires Saint-Luc | UCLouvain/IREC/MBLG | Jean Ruelle, Bertrand Bearzatto, Benoit Kabamba Mukadi |
| EPI_ISL_2885737 | URMC LABS | URMC LABS | Ghinwa Dumyati, Dwight Hardy, Andrew Cameron, Stepan Voljehn, Kelly DeLary |
| EPI_ISL_2885743, EPI_ISL_2885746, EPI_ISL_2885747 | Laboratorio HUB -Azienda Ospedaliero Universitaria - AOU – Cagliari | Laboratorio SPOKE Biologia Molecolare -Azienda Ospedaliero Universitaria - AOU – Cagliari | Germano Orrù, Sara Fais, Valentina Medda, Alessandra Scano, Miriam Loddo, Riccardo Cappai, Ferdinando Coghe |
| EPI_ISL_2885770 | Centro de Investigación Biomédica de La Rioja - Hospital San Pedro Logroño | SeqCOVID-SPAIN consortium/IBV(CSIC) | María de Toro, José Manuel Azcona Gutiérrez, María Pilar Bea Escudero, Miriam Blasco Alberdi and SeqCOVID-SPAIN consortium |
| EPI_ISL_2885868 | Virginia Division of Consolidated Laboratory Services | Virginia Division of Consolidated Laboratory Services | Virginia Division of Consolidated Laboratory Services |
| EPI_ISL_2885869, EPI_ISL_2885870, EPI_ISL_2885871, EPI_ISL_2885872 | Kansas Health and Environmental Lab | Kansas Health and Environmental Lab | Mike Grose, Katherine Wiggins, Jonathan Barnell, Ben Olsen, and Phil Adam |
| EPI_ISL_2885873, EPI_ISL_2885874, EPI_ISL_2885875, EPI_ISL_2885876, EPI_ISL_2885877, EPI_ISL_2885878, EPI_ISL_2885880, EPI_ISL_2885881, EPI_ISL_2885882, EPI_ISL_2885883, EPI_ISL_2885884, EPI_ISL_2885885, EPI_ISL_2885887, EPI_ISL_2885888, EPI_ISL_2885889, EPI_ISL_2885890, EPI_ISL_2885891, EPI_ISL_2885892, EPI_ISL_2885893, EPI_ISL_2885894, EPI_ISL_2885895, EPI_ISL_2885896, EPI_ISL_2885898, EPI_ISL_2885899, EPI_ISL_2885900 | | | |
| see above | Wichita State University- Molecular Diagnostic Lab | Kansas Health and Environmental Lab | Mike Grose, Katherine Wiggins, Jonathan Barnell, Ben Olsen, and Phil Adam |
| EPI_ISL_2885922 | Viešoji įstaiga Klaipėdos universitetinė ligoninė | Institute of Biotechnology, Life Sciences Center, Vilnius University | Emilija Vasiliunaite; Milda Norkiene; Danguole Ziogiene; Albertas Timinskas; Alma Gedvilaite |
| EPI_ISL_2885938, EPI_ISL_2885944, EPI_ISL_2885945, EPI_ISL_2885946 | Viešoji įstaiga Vilniaus universiteto ligoninė Santaros klinikos | Institute of Biotechnology, Life Sciences Center, Vilnius University | Emilija Vasiliunaite; Milda Norkiene; Danguole Ziogiene; Albertas Timinskas; Alma Gedvilaite |
| EPI_ISL_2885952, EPI_ISL_2885957, EPI_ISL_2885960, EPI_ISL_2885961 | ADPH | ADPH | Adrianna Walker |
| EPI_ISL_2885978, EPI_ISL_2885995, EPI_ISL_2886031, EPI_ISL_2886039, EPI_ISL_2886044 | Hospital Universitario de Guadalajara | SeqCOVID-SPAIN consortium/IBV(CSIC) | Alejandro González Praetorius y Mariela Martínez Ramírez and SeqCOVID-SPAIN consortium |
| EPI_ISL_2886054, EPI_ISL_2886055, EPI_ISL_2886056, EPI_ISL_2886057, EPI_ISL_2886058, EPI_ISL_2886059, EPI_ISL_2886060, EPI_ISL_2886061, EPI_ISL_2886062, EPI_ISL_2886063, EPI_ISL_2886064, EPI_ISL_2886065, EPI_ISL_2886066, EPI_ISL_2886067, EPI_ISL_2886068, EPI_ISL_2886069, EPI_ISL_2886070, EPI_ISL_2886071, EPI_ISL_2886073, EPI_ISL_2886075, EPI_ISL_2886076, EPI_ISL_2886077, EPI_ISL_2886078, EPI_ISL_2886080, EPI_ISL_2886081, EPI_ISL_2886082, EPI_ISL_2886083, EPI_ISL_2886085, EPI_ISL_2886086, EPI_ISL_2886087, EPI_ISL_2886088, EPI_ISL_2886089 | | | |
| see above | Dept. of Microbiology and Infection Control, Akershus University Hospital HF | Dept. of Microbiology and Infection Control, Akershus University Hospital HF | Hege Vangstein Aamot, Alexander Hesselberg Løvestad |
| EPI_ISL_2886094, EPI_ISL_2886095, EPI_ISL_2886096, EPI_ISL_2886099, EPI_ISL_2886100, EPI_ISL_2886101, EPI_ISL_2886102, EPI_ISL_2886104, EPI_ISL_2886106, EPI_ISL_2886107, EPI_ISL_2886109, EPI_ISL_2886110, EPI_ISL_2886111, EPI_ISL_2886112, EPI_ISL_2886113, EPI_ISL_2886114, EPI_ISL_2886115, EPI_ISL_2886116, EPI_ISL_2886117, EPI_ISL_2886119, EPI_ISL_2886120, EPI_ISL_2886121, EPI_ISL_2886124, EPI_ISL_2886125, EPI_ISL_2886126, EPI_ISL_2886127, EPI_ISL_2886128, EPI_ISL_2886129, EPI_ISL_2886130, EPI_ISL_2886132 | | | |
| see above | UMC Utrecht | UMC Utrecht | Rob Schuurman, Anniek Tanja, Annemarie Wensing |
| EPI_ISL_2886157, EPI_ISL_2886159 | IVRCRD RSPAW | Institute for Vector and Reservoir Control, Research and Development (IVRCRD)- National Institute of Health Research and Development (NIHRD), Indonesia | Arief Mulyono, Triwibowo Ambar Garjito, Restu Choirul Saban, Ayu Pradipta Pratiwi, Dwi Susilo, Yusnita Mirna Anggraeni, Mujiyanto, Mega Tyas Prihatin, Nur Hidayati, Arif Suryo Prasetyo, Tika Fiona Sari, Sri Wahyuni Handayani, Aryo Ardanto, Wimbi Kartika Ratnasari, Kusumaningtyas Sekar Negari, Aprilia Safitri, Dimas Bagus Wicaksono Putro, Esti Rahardianingtyas, Sidiq Setyo Nugroho, Marjianto, Muhidin, Subangkit, Hana Apsari Pawestri, Siti Alfiah, Ristiyanto, Evi sulistyorini, Lulus Susanti, Vivi Setiawaty, Joko Waluyo |
| EPI_ISL_2886160, EPI_ISL_2886161 | IVRCRD Puskesmas Muntilan 1 | Institute for Vector and Reservoir Control, Research and Development (IVRCRD)- National Institute of Health Research and Development (NIHRD), Indonesia | Arief Mulyono, Triwibowo Ambar Garjito, Restu Choirul Saban, Ayu Pradipta Pratiwi, Dwi Susilo, Yusnita Mirna Anggraeni, Mujiyanto, Mega Tyas Prihatin, Nur Hidayati, Arif Suryo Prasetyo, Tika Fiona Sari, Sri Wahyuni Handayani, Aryo Ardanto, Wimbi Kartika Ratnasari, Kusumaningtyas Sekar Negari, Aprilia Safitri, Dimas Bagus Wicaksono Putro, Esti Rahardianingtyas, Sidiq Setyo Nugroho, Marjianto, Muhidin, Subangkit, Hana Apsari Pawestri, Siti Alfiah, Ristiyanto, Evi sulistyorini, Lulus Susanti, Vivi Setiawaty, Joko Waluyo |
| EPI_ISL_2886162, EPI_ISL_2886163 | IVRCRD RSUD Budi Rahayu | Institute for Vector and Reservoir Control, Research and Development (IVRCRD)- National Institute of Health Research and Development (NIHRD), Indonesia | Arief Mulyono, Triwibowo Ambar Garjito, Restu Choirul Saban, Ayu Pradipta Pratiwi, Dwi Susilo, Yusnita Mirna Anggraeni, Mujiyanto, Mega Tyas Prihatin, Nur Hidayati, Arif Suryo Prasetyo, Tika Fiona Sari, Sri Wahyuni Handayani, Aryo Ardanto, Wimbi Kartika Ratnasari, Kusumaningtyas Sekar Negari, Aprilia Safitri, Dimas Bagus Wicaksono Putro, Esti Rahardianingtyas, Sidiq Setyo Nugroho, Marjianto, Muhidin, Subangkit, Hana Apsari Pawestri, Siti Alfiah, Ristiyanto, Evi sulistyorini, Lulus Susanti, Vivi Setiawaty, Joko Waluyo |
| EPI_ISL_2886164 | IVRCRD Puskesmas Welahan 1 | Institute for Vector and Reservoir Control, Research and Development (IVRCRD)- National Institute of Health Research and Development (NIHRD), Indonesia | Arief Mulyono, Triwibowo Ambar Garjito, Restu Choirul Saban, Ayu Pradipta Pratiwi, Dwi Susilo, Yusnita Mirna Anggraeni, Mujiyanto, Mega Tyas Prihatin, Nur Hidayati, Arif Suryo Prasetyo, Tika Fiona Sari, Sri Wahyuni Handayani, Aryo Ardanto, Wimbi Kartika Ratnasari, Kusumaningtyas Sekar Negari, Aprilia Safitri, Dimas Bagus Wicaksono Putro, Esti Rahardianingtyas, Sidiq Setyo Nugroho, Marjianto, Muhidin, Subangkit, Hana Apsari Pawestri, Siti Alfiah, Ristiyanto, Evi sulistyorini, Lulus Susanti, Vivi Setiawaty, Joko Waluyo |
| EPI_ISL_2886165 | IVRCRD RSUD Budi Rahayu | Institute for Vector and Reservoir Control, Research and Development (IVRCRD)- National Institute of Health Research and Development (NIHRD), Indonesia | Arief Mulyono, Triwibowo Ambar Garjito, Restu Choirul Saban, Ayu Pradipta Pratiwi, Dwi Susilo, Yusnita Mirna Anggraeni, Mujiyanto, Mega Tyas Prihatin, Nur Hidayati, Arif Suryo Prasetyo, Tika Fiona Sari, Sri Wahyuni Handayani, Aryo Ardanto, Wimbi Kartika Ratnasari, Kusumaningtyas Sekar Negari, Aprilia Safitri, Dimas Bagus Wicaksono Putro, Esti Rahardianingtyas, Sidiq Setyo Nugroho, Marjianto, Muhidin, Subangkit, Hana Apsari Pawestri, Siti Alfiah, Ristiyanto, Evi sulistyorini, Lulus Susanti, Vivi Setiawaty, Joko Waluyo |
| EPI_ISL_2886166 | IVRCRD Puskesmas Cebongan | Institute for Vector and Reservoir Control, Research and Development (IVRCRD)- National Institute of Health Research and Development (NIHRD), Indonesia | Arief Mulyono, Triwibowo Ambar Garjito, Restu Choirul Saban, Ayu Pradipta Pratiwi, Dwi Susilo, Yusnita Mirna Anggraeni, Mujiyanto, Mega Tyas Prihatin, Nur Hidayati, Arif Suryo Prasetyo, Tika Fiona Sari, Sri Wahyuni Handayani, Aryo Ardanto, Wimbi Kartika Ratnasari, Kusumaningtyas Sekar Negari, Aprilia Safitri, Dimas Bagus Wicaksono Putro, Esti Rahardianingtyas, Sidiq Setyo Nugroho, Marjianto, Muhidin, Subangkit, Hana Apsari Pawestri, Siti Alfiah, Ristiyanto, Evi sulistyorini, Lulus Susanti, Vivi Setiawaty, Joko Waluyo |

| | | | |
|---|---|---|---|
| EPI_ISL_2886167 | IVRCRD Puskesmas Sidorejo Lor | Institute for Vector and Reservoir Control, Research and Development (IVRCRD)- National Institute of Health Research and Development (NIHRD), Indonesia | Arief Mulyono, Triwibowo Ambar Garjito, Restu Choirul Saban, Ayu Pradipta Pratiwi, Dwi Susilo, Yusnita Mirna Anggraeni, Mujiyanto, Mega Tyas Prihatin, Nur Hidayati, Arif Suryo Prasetyo, Tika Fiona Sari, Sri Wahyuni Handayani, Aryo Ardanto, Wimbi Kartika Ratnasari, Kusumaningtyas Sekar Negari, Aprilia Safitri, Dimas Bagus Wicaksono Putro, Esti Rahardianingtyas, Sidiq Setyo Nugroho, Marjianto, Muhidin, Subangkit, Hana Apsari Pawestri, Siti Alfiah, Ristiyanto, Evi sulistyorini, Lulus Susanti, Vivi Setiawaty, Joko Waluyo |
| EPI_ISL_2886168 | Institute for Vector and Reservoir Control, Research and Development (IVRCRD)- National Institute of Health Research and Development (NIHRD), Indonesia | Institute for Vector and Reservoir Control, Research and Development (IVRCRD)- National Institute of Health Research and Development (NIHRD), Indonesia | Arief Mulyono, Triwibowo Ambar Garjito, Restu Choirul Saban, Ayu Pradipta Pratiwi, Dwi Susilo, Yusnita Mirna Anggraeni, Mujiyanto, Mega Tyas Prihatin, Nur Hidayati, Arif Suryo Prasetyo, Tika Fiona Sari, Sri Wahyuni Handayani, Aryo Ardanto, Wimbi Kartika Ratnasari, Kusumaningtyas Sekar Negari, Aprilia Safitri, Dimas Bagus Wicaksono Putro, Esti Rahardianingtyas, Sidiq Setyo Nugroho, Marjianto, Muhidin, Subangkit, Hana Apsari Pawestri, Siti Alfiah, Ristiyanto, Evi sulistyorini, Lulus Susanti, Vivi Setiawaty, Joko Waluyo |
| EPI_ISL_2886172, EPI_ISL_2886173, EPI_ISL_2886175, EPI_ISL_2886176 | Dr. Gernot Walder GmbH | Dr. Gernot Walder GmbH | Sissy T. Sonnleitner |
| EPI_ISL_2886182, EPI_ISL_2886183, EPI_ISL_2886186, EPI_ISL_2886189, EPI_ISL_2886199 | Laboratorio di Riferimento Regionale della Sicilia Occidentale per l'Emergenza COVID-19 | Laboratorio di Riferimento Regionale della Sicilia Occidentale per l'Emergenza COVID-19 | Fabio Tramuto, Carmelo Massimo Maida, Daniela Di Naro, Giulia Randazzo, Walter Mazzucco, Giorgio Graziano, Vincenzo Restivo, Claudio Costantino, Francesco Vitale |
| EPI_ISL_2886215, EPI_ISL_2886216, EPI_ISL_2886217, EPI_ISL_2886218, EPI_ISL_2886219, EPI_ISL_2886220, EPI_ISL_2886221, EPI_ISL_2886232, EPI_ISL_2886237, EPI_ISL_2886241, EPI_ISL_2886250, EPI_ISL_2886256, EPI_ISL_2886322, EPI_ISL_2886323, EPI_ISL_2886325, EPI_ISL_2886334, EPI_ISL_2886339, EPI_ISL_2886356, EPI_ISL_2886357, EPI_ISL_2886359, EPI_ISL_2886360, EPI_ISL_2886361, EPI_ISL_2886374, EPI_ISL_2886382 | | | |
| see above | KU Leuven, Rega Institute, Clinical and Epidemiological Virology | KU Leuven, Rega Institute, Clinical and Epidemiological Virology | Tony Wawina-Bokalanga, Bert Vanmechelen, Joan Marti-Carerras, Piet Maes |
| EPI_ISL_2886575, EPI_ISL_2886903 | Institute of Microbiology and Immunology, Faculty of Medicine, University of Ljubljana | Institute of Microbiology and Immunology, Faculty of Medicine, University of Ljubljana | Alen Suljič, Samo Zakotnik, Tomaž Mark Zorec, Matic Brvar, Doroteja Vlaj, Andraž Celar, Dominika Šturm, Patricija Pozvek, Špela Pleh, Miša Korva, Mario Poljak, Tatjana Avšič – Županc |
| EPI_ISL_2887353, EPI_ISL_2887396, EPI_ISL_2887401, EPI_ISL_2887405, EPI_ISL_2887412 | Division of Emerging Infectious Diseases, Bureau of Infectious Diseases Diagnosis Control, Korea Disease Control and Prevention Agency | Division of Emerging Infectious Diseases, Bureau of Infectious Diseases Diagnosis Control, Korea Disease Control and Prevention Agency | Jeong-Ah Kim, Ae Kyung Park, Il-Hwan Kim, Heui Man Kim, Jeong-Min Kim, Chae Young Lee, Eun-Jin Kim |
| EPI_ISL_2887417, EPI_ISL_2887419, EPI_ISL_2887420, EPI_ISL_2887426, EPI_ISL_2887427, EPI_ISL_2887429, EPI_ISL_2887431, EPI_ISL_2887436, EPI_ISL_2887437, EPI_ISL_2887438, EPI_ISL_2887439, EPI_ISL_2887444, EPI_ISL_2887445, EPI_ISL_2887446, EPI_ISL_2887447, EPI_ISL_2887448, EPI_ISL_2887454, EPI_ISL_2887455, EPI_ISL_2887456, EPI_ISL_2887457, EPI_ISL_2887458, EPI_ISL_2887459, EPI_ISL_2887463, EPI_ISL_2887464, EPI_ISL_2887465, EPI_ISL_2887466, EPI_ISL_2887467, EPI_ISL_2887468, EPI_ISL_2887470, EPI_ISL_2887473, EPI_ISL_2887474, EPI_ISL_2887475, EPI_ISL_2887476, EPI_ISL_2887477, EPI_ISL_2887479, EPI_ISL_2887484, EPI_ISL_2887485, EPI_ISL_2887486, EPI_ISL_2887487, EPI_ISL_2887489, EPI_ISL_2887492, EPI_ISL_2887494, EPI_ISL_2887495, EPI_ISL_2887503, EPI_ISL_2887504, EPI_ISL_2887514, EPI_ISL_2887515, EPI_ISL_2887516, EPI_ISL_2887517, EPI_ISL_2887518, EPI_ISL_2887527, EPI_ISL_2887528, EPI_ISL_2887533, EPI_ISL_2887537, EPI_ISL_2887538, EPI_ISL_2887539, EPI_ISL_2887549, EPI_ISL_2887550, EPI_ISL_2887551, EPI_ISL_2887552, EPI_ISL_2887561, EPI_ISL_2887562, EPI_ISL_2887565, EPI_ISL_2887572, EPI_ISL_2887573, EPI_ISL_2887575, EPI_ISL_2887584, EPI_ISL_2887585, EPI_ISL_2887591, EPI_ISL_2887592, EPI_ISL_2887593, EPI_ISL_2887594, EPI_ISL_2887595, EPI_ISL_2887603, EPI_ISL_2887604, EPI_ISL_2887605, EPI_ISL_2887606, EPI_ISL_2887607, EPI_ISL_2887608, EPI_ISL_2887610, EPI_ISL_2887615, EPI_ISL_2887616, EPI_ISL_2887617, EPI_ISL_2887618, EPI_ISL_2887619, EPI_ISL_2887621, EPI_ISL_2887625, EPI_ISL_2887626, EPI_ISL_2887627, EPI_ISL_2887628, EPI_ISL_2887629, EPI_ISL_2887630, EPI_ISL_2887636, EPI_ISL_2887637, EPI_ISL_2887638, EPI_ISL_2887639, EPI_ISL_2887640, EPI_ISL_2887641, EPI_ISL_2887642, EPI_ISL_2887646, EPI_ISL_2887647, EPI_ISL_2887649, EPI_ISL_2887650, EPI_ISL_2887651, EPI_ISL_2887652, EPI_ISL_2887653, EPI_ISL_2887654, EPI_ISL_2887661, EPI_ISL_2887662, EPI_ISL_2887663, EPI_ISL_2887664, EPI_ISL_2887665, EPI_ISL_2887666, EPI_ISL_2887673, EPI_ISL_2887674, EPI_ISL_2887675, EPI_ISL_2887676 | | | |
| see above | Austrian Agency for Health and Food Safety (AGES) | Bergthaler laboratory, CeMM Research Center for Molecular Medicine of the Austrian Academy of Sciences | Lukas Endler, Anna Schedl, Fabian Amman, Petr Triska, Matthew Thornton, Thomas Penz, Benedikt Agerer, Maelle Le Moing, Michael Schuster, Bekir Erguner, Jan Laine, Martin Senekowitsch, Christoph Bock, Andreas Bergthaler |
| EPI_ISL_2887685 | Universität Innsbruck, Institut für Mikrobiologie | Bergthaler laboratory, CeMM Research Center for Molecular Medicine of the Austrian Academy of Sciences | Lukas Endler, Anna Schedl, Fabian Amman, Petr Triska, Matthew Thornton, Thomas Penz, Benedikt Agerer, Maelle Le Moing, Michael Schuster, Bekir Erguner, Jan Laine, Martin Senekowitsch, Christoph Bock, Andreas Bergthaler |
| EPI_ISL_2887701 | Center for Virology, Medical University of Vienna | Bergthaler laboratory, CeMM Research Center for Molecular Medicine of the Austrian Academy of Sciences | Lukas Endler, Anna Schedl, Fabian Amman, Petr Triska, Matthew Thornton, Thomas Penz, Benedikt Agerer, Maelle Le Moing, Michael Schuster, Bekir Erguner, Jan Laine, Martin Senekowitsch, Christoph Bock, Andreas Bergthaler |
| EPI_ISL_2887734 | CHU Lille | CHU Lille - Laboratoire de Virologie | AIT YAHYA Emilie, ALIDJINOU Enagnon Kazali, BOCKET Laurence, CREPIN Michel, DEMAY Christophe, ENGELMANN Ilka, GEFFROY Sandrine, GUIGON Aurélie, LAMBERT Valérie, LAZREK Mouna, NOBILLIAUX Florian, PREVOST Brigitte, THUILLIER Caroline, TINEZ Claire ,TCHANTCHOU NJOSSE YANICK |
| EPI_ISL_2887746 | CH Tourcoing | CHU Lille - Laboratoire de Virologie | AIT YAHYA Emilie, ALIDJINOU Enagnon Kazali, BOCKET Laurence, CREPIN Michel, DEMAY Christophe, ENGELMANN Ilka, GEFFROY Sandrine, GUIGON Aurélie, LAMBERT Valérie, LAZREK Mouna, NOBILLIAUX Florian, PREVOST Brigitte, THUILLIER Caroline, TINEZ Claire ,TCHANTCHOU NJOSSE YANICK |
| EPI_ISL_2887750, EPI_ISL_2887755, EPI_ISL_2887758, EPI_ISL_2887760, EPI_ISL_2887761, EPI_ISL_2887764, EPI_ISL_2887765, EPI_ISL_2887766, EPI_ISL_2887773, EPI_ISL_2887774, EPI_ISL_2887776, EPI_ISL_2887777 | | | |
| see above | Clinical Microbiology, Infection Prevention and Control | Section for Molecular Diagnostics | Björn Hallström, Jonas Björkman |
| EPI_ISL_2887781, EPI_ISL_2887782, EPI_ISL_2887783, EPI_ISL_2887784, EPI_ISL_2887785, EPI_ISL_2887786, EPI_ISL_2887787, EPI_ISL_2887788, EPI_ISL_2887789, EPI_ISL_2887790, EPI_ISL_2887791, EPI_ISL_2887792, EPI_ISL_2887793, EPI_ISL_2887794, EPI_ISL_2887795 | | | |
| see above | National Public Health Laboratory, National Centre for Infectious Diseases | National Public Health Laboratory, National Centre for Infectious Diseases | Tze Minn Mak, Katherine Ching, Zhenyang Zhou, Royce Ang, Lin Cui, Raymond Tzer Pin Lin |
| EPI_ISL_2887847 | Dutch COVID-19 response team | Erasmus Medical Center | Bas Oude Munnink, Reina Sikkema, David Nieuwenhuijse, Irina Chestakova, Anne van der Linden, Marjan Boter, Emmanuelle Munger, Corine GeurtsvanKessel, Annemiek van der Eijk, Richard Molenkamp, Marion Koopmans, on behalf of the Dutch national COVID-19 response team. |
| EPI_ISL_2887853, EPI_ISL_2887855, EPI_ISL_2887856, EPI_ISL_2887857, EPI_ISL_2887858 | Iressef Genomics lab | IRESSEF | Souleymane MBOUP, Abdou PADANE , Khadim GUEYE, Diabou Diagne , Samba Ndiour, Papa Alassane DIAW, Birahim Piere NDIAYE, Barada CISSE, Aminata MBOUP, Moustapha MBOW, Ndeye Coumba Toure KANE, Nafisatou LEYE, Gora LO, Ambroise AHOUIDI , Astou Gaye GAYE, Aminata DIA, Yacine DIA,Seni Ndiaye |
| EPI_ISL_2887964, EPI_ISL_2887965, EPI_ISL_2887966, EPI_ISL_2887967, EPI_ISL_2887969, EPI_ISL_2887972, EPI_ISL_2887976, EPI_ISL_2887979, EPI_ISL_2887980, EPI_ISL_2887981, EPI_ISL_2887982, EPI_ISL_2887983, EPI_ISL_2887985, EPI_ISL_2887986, EPI_ISL_2887988, EPI_ISL_2887990, EPI_ISL_2887991, EPI_ISL_2887993, EPI_ISL_2887996, EPI_ISL_2888003, EPI_ISL_2888004, EPI_ISL_2888006, EPI_ISL_2888008, EPI_ISL_2888011 | | | |
| see above | UMC Utrecht | UMC Utrecht | Rob Schuurman, Anniek Tanja, Annemarie Wensing, Pascal Mantiri |
| EPI_ISL_2888016, EPI_ISL_2888017, EPI_ISL_2888018, EPI_ISL_2888020 | Ospedale Sandro Pertini ASL Roma2 | Department of General Diagnostics; Department of Virology; Istituto Zooprofilattico Sperimentale del Lazio e della Toscana (IZSLT) | Patricia Alba, Giuseppe Manna, Elena L. Diaconu, Fabiola Feltrin, Raffaela Conti, Teresa Scicluna, Virginia Carfora, Antonella Cersini, Alessia Franco, Antonio Battisti. |
| EPI_ISL_2888023, EPI_ISL_2888027, EPI_ISL_2888028, EPI_ISL_2888033, EPI_ISL_2888034, EPI_ISL_2888036, EPI_ISL_2888037, EPI_ISL_2888038, EPI_ISL_2888039, EPI_ISL_2888040, EPI_ISL_2888042 | | | |
| see above | Ospedale San Filippo Neri | Department of General Diagnostics; Department of Virology; Istituto Zooprofilattico Sperimentale del Lazio e della Toscana (IZSLT) | Patricia Alba, Giuseppe Manna, Elena L. Diaconu, Fabiola Feltrin, Raffaela Conti, Teresa Scicluna, Virginia Carfora, Antonella Cersini, Alessia Franco, Antonio Battisti. |
| EPI_ISL_2888043, EPI_ISL_2888044, EPI_ISL_2888045, EPI_ISL_2888046, EPI_ISL_2888049, EPI_ISL_2888051, EPI_ISL_2888052, EPI_ISL_2888053, EPI_ISL_2888055, EPI_ISL_2888057, EPI_ISL_2888062, EPI_ISL_2888065, EPI_ISL_2888068, EPI_ISL_2888069, EPI_ISL_2888070, EPI_ISL_2888073, EPI_ISL_2888076, EPI_ISL_2888080, EPI_ISL_2888081, EPI_ISL_2888082, EPI_ISL_2888085, EPI_ISL_2888086, EPI_ISL_2888087, EPI_ISL_2888089, EPI_ISL_2888090, EPI_ISL_2888092, EPI_ISL_2888095, EPI_ISL_2888099, EPI_ISL_2888100, EPI_ISL_2888101, EPI_ISL_2888102, EPI_ISL_2888106, EPI_ISL_2888108, EPI_ISL_2888109, EPI_ISL_2888110, EPI_ISL_2888113, EPI_ISL_2888117, EPI_ISL_2888121, EPI_ISL_2888122, EPI_ISL_2888125, EPI_ISL_2888126, EPI_ISL_2888129, EPI_ISL_2888130, EPI_ISL_2888132, EPI_ISL_2888133, EPI_ISL_2888134, EPI_ISL_2888135, EPI_ISL_2888136, EPI_ISL_2888137, EPI_ISL_2888139, EPI_ISL_2888140, EPI_ISL_2888141, EPI_ISL_2888142, EPI_ISL_2888143, EPI_ISL_2888145, EPI_ISL_2888146, EPI_ISL_2888147, EPI_ISL_2888151, EPI_ISL_2888152, EPI_ISL_2888153, EPI_ISL_2888154, EPI_ISL_2888155, EPI_ISL_2888163, EPI_ISL_2888164, EPI_ISL_2888165, EPI_ISL_2888166, EPI_ISL_2888168, EPI_ISL_2888170, EPI_ISL_2888171, EPI_ISL_2888175, EPI_ISL_2888176, EPI_ISL_2888177, EPI_ISL_2888178, EPI_ISL_2888179, EPI_ISL_2888180, EPI_ISL_2888181, EPI_ISL_2888182, EPI_ISL_2888183, EPI_ISL_2888184, EPI_ISL_2888185, EPI_ISL_2888186, EPI_ISL_2888187, EPI_ISL_2888197, EPI_ISL_2888199, EPI_ISL_2888200, EPI_ISL_2888201, EPI_ISL_2888202, EPI_ISL_2888203, EPI_ISL_2888204, EPI_ISL_2888215, EPI_ISL_2888216, EPI_ISL_2888217, EPI_ISL_2888218, EPI_ISL_2888219, EPI_ISL_2888220, EPI_ISL_2888221, EPI_ISL_2888222, EPI_ISL_2888229, EPI_ISL_2888231, EPI_ISL_2888269, EPI_ISL_2888278, EPI_ISL_2888283, EPI_ISL_2888297, EPI_ISL_2888299, EPI_ISL_2888324, EPI_ISL_2888325, EPI_ISL_2888336, EPI_ISL_2888337, EPI_ISL_2888352, EPI_ISL_2888357, EPI_ISL_2888375, EPI_ISL_2888385, EPI_ISL_2888387, EPI_ISL_2888388, EPI_ISL_2888389, EPI_ISL_2888396, EPI_ISL_2888397, EPI_ISL_2888398, EPI_ISL_2888478, EPI_ISL_2888479, EPI_ISL_2888480, EPI_ISL_2888481, EPI_ISL_2888482, EPI_ISL_2888483, EPI_ISL_2888484, EPI_ISL_2888485, EPI_ISL_2888486, EPI_ISL_2888487, EPI_ISL_2888488, EPI_ISL_2888489, EPI_ISL_2888490, EPI_ISL_2888491, EPI_ISL_2888492, EPI_ISL_2888493, EPI_ISL_2888494, EPI_ISL_2888495, EPI_ISL_2888496, EPI_ISL_2888497, EPI_ISL_2888498, EPI_ISL_2888499, EPI_ISL_2888500, EPI_ISL_2888501, EPI_ISL_2888502, EPI_ISL_2888503, EPI_ISL_2888504, EPI_ISL_2888505, EPI_ISL_2888506, EPI_ISL_2888507, EPI_ISL_2888508, EPI_ISL_2888509, EPI_ISL_2888510, EPI_ISL_2888511, EPI_ISL_2888512, EPI_ISL_2888513, EPI_ISL_2888514, EPI_ISL_2888515, EPI_ISL_2888516, EPI_ISL_2888517, EPI_ISL_2888518, EPI_ISL_2888519, EPI_ISL_2888520, EPI_ISL_2888521, EPI_ISL_2888522, EPI_ISL_2888523, EPI_ISL_2888524, EPI_ISL_2888525, EPI_ISL_2888526, EPI_ISL_2888527, EPI_ISL_2888528, EPI_ISL_2888529, EPI_ISL_2888530, EPI_ISL_2888531, EPI_ISL_2888532, EPI_ISL_2888533, EPI_ISL_2888534, EPI_ISL_2888535, EPI_ISL_2888536, EPI_ISL_2888537, EPI_ISL_2888538, EPI_ISL_2888539, EPI_ISL_2888540, EPI_ISL_2888541, EPI_ISL_2888542, EPI_ISL_2888543, EPI_ISL_2888544, EPI_ISL_2888545, EPI_ISL_2888546, EPI_ISL_2888547, EPI_ISL_2888548, EPI_ISL_2888549, EPI_ISL_2888550, EPI_ISL_2888551, EPI_ISL_2888552, EPI_ISL_2888553, EPI_ISL_2888554, EPI_ISL_2888555, EPI_ISL_2888556, EPI_ISL_2888557, EPI_ISL_2888558, EPI_ISL_2888559, EPI_ISL_2888560, EPI_ISL_2888561, EPI_ISL_2888562, EPI_ISL_2888563, EPI_ISL_2888564, EPI_ISL_2888565, EPI_ISL_2888566, EPI_ISL_2888567, EPI_ISL_2888568, EPI_ISL_2888569, EPI_ISL_2888570, EPI_ISL_2888571, EPI_ISL_2888572, EPI_ISL_2888573, EPI_ISL_2888574, EPI_ISL_2888575, EPI_ISL_2888576, EPI_ISL_2888577, EPI_ISL_2888578, EPI_ISL_2888579, EPI_ISL_2888580, EPI_ISL_2888581, EPI_ISL_2888582, EPI_ISL_2888583, EPI_ISL_2888584, EPI_ISL_2888585, EPI_ISL_2888586, EPI_ISL_2888587, EPI_ISL_2888588, EPI_ISL_2888589, EPI_ISL_2888590, EPI_ISL_2888591, EPI_ISL_2888592, EPI_ISL_2888593, EPI_ISL_2888594, EPI_ISL_2888595, EPI_ISL_2888596, EPI_ISL_2888597, EPI_ISL_2888598, EPI_ISL_2888599, EPI_ISL_2888600, EPI_ISL_2888601, EPI_ISL_2888602, EPI_ISL_2888603, EPI_ISL_2888604, EPI_ISL_2888605, EPI_ISL_2888606, EPI_ISL_2888607, EPI_ISL_2888608, EPI_ISL_2888609, EPI_ISL_2888610, EPI_ISL_2888611, EPI_ISL_2888612, EPI_ISL_2888613, EPI_ISL_2888614, EPI_ISL_2888615, EPI_ISL_2888616, EPI_ISL_2888617, EPI_ISL_2888619, EPI_ISL_2888620, EPI_ISL_2888621, EPI_ISL_2888622, EPI_ISL_2888623, EPI_ISL_2888624, EPI_ISL_2888625, EPI_ISL_2888626, EPI_ISL_2888633, EPI_ISL_2888634, EPI_ISL_2888635, EPI_ISL_2888636, EPI_ISL_2888637, EPI_ISL_2888638, EPI_ISL_2888639, EPI_ISL_2888640, EPI_ISL_2888641, EPI_ISL_2888642, EPI_ISL_2888643, EPI_ISL_2888644, EPI_ISL_2888645, EPI_ISL_2888646, EPI_ISL_2888647, EPI_ISL_2888648, EPI_ISL_2888649, EPI_ISL_2888650, EPI_ISL_2888651, EPI_ISL_2888652, EPI_ISL_2888653, EPI_ISL_2888654, EPI_ISL_2888655, EPI_ISL_2888656, EPI_ISL_2888657, EPI_ISL_2888658, EPI_ISL_2888659, EPI_ISL_2888660, EPI_ISL_2888661, EPI_ISL_2888662, EPI_ISL_2888663, EPI_ISL_2888666, EPI_ISL_2888667, EPI_ISL_2888668, EPI_ISL_2888669, EPI_ISL_2888670, EPI_ISL_2888671, EPI_ISL_2888672, EPI_ISL_2888673, EPI_ISL_2888674, EPI_ISL_2888675, EPI_ISL_2888677, EPI_ISL_2888678, EPI_ISL_2888679, EPI_ISL_2888680, EPI_ISL_2888681, EPI_ISL_2888682, EPI_ISL_2888683, EPI_ISL_2888684, EPI_ISL_2888685, EPI_ISL_2888686, EPI_ISL_2888687, EPI_ISL_2888688, EPI_ISL_2888689, EPI_ISL_2888690, EPI_ISL_2888691, EPI_ISL_2888692, EPI_ISL_2888693, EPI_ISL_2888694, EPI_ISL_2888695, EPI_ISL_2888696, EPI_ISL_2888697, EPI_ISL_2888698, EPI_ISL_2888703, EPI_ISL_2888704, EPI_ISL_2888705, EPI_ISL_2888706, EPI_ISL_2888707, EPI_ISL_2888708, EPI_ISL_2888709, EPI_ISL_2888710, EPI_ISL_2888711, EPI_ISL_2888712, EPI_ISL_2888713, EPI_ISL_2888714, EPI_ISL_2888715, EPI_ISL_2888716, EPI_ISL_2888717, EPI_ISL_2888718, EPI_ISL_2888719, EPI_ISL_2888720, EPI_ISL_2888721, EPI_ISL_2888722, EPI_ISL_2888723, EPI_ISL_2888724, EPI_ISL_2888725, EPI_ISL_2888730, EPI_ISL_2888743, EPI_ISL_2888744, EPI_ISL_2888745, EPI_ISL_2888746, EPI_ISL_2888747, EPI_ISL_2888748, EPI_ISL_2888749, EPI_ISL_2888750, EPI_ISL_2888751, EPI_ISL_2888768, EPI_ISL_2888771, EPI_ISL_2888773, EPI_ISL_2888775, EPI_ISL_2888776, EPI_ISL_2888777, EPI_ISL_2888778, EPI_ISL_2888779, EPI_ISL_2888780, EPI_ISL_2888781, EPI_ISL_2888782, EPI_ISL_2888783, EPI_ISL_2888784, EPI_ISL_2888980, EPI_ISL_2888982, EPI_ISL_2889181, EPI_ISL_2889182, EPI_ISL_2889183, EPI_ISL_2889226, EPI_ISL_2889227, EPI_ISL_2889228, EPI_ISL_2889229 | | | |
| see above | Swedish national genomic surveillance program of SARS-CoV-2 | The Public Health Agency of Sweden | Maximilian Riess, Maria Lind Karlberg, Alma Brolund, Swedish national genomic surveillance program of SARS-CoV-2 |
| EPI_ISL_2889292, EPI_ISL_2889295, EPI_ISL_2889298, EPI_ISL_2889301, EPI_ISL_2889307, EPI_ISL_2889308, EPI_ISL_2889342, EPI_ISL_2889382, EPI_ISL_2889416, EPI_ISL_2889422, EPI_ISL_2889433, EPI_ISL_2889448, EPI_ISL_2889456, EPI_ISL_2889457, EPI_ISL_2889481, EPI_ISL_2889486, EPI_ISL_2889489, EPI_ISL_2889492, EPI_ISL_2889542, EPI_ISL_2889544, EPI_ISL_2889545, EPI_ISL_2889567, EPI_ISL_2889580, EPI_ISL_2889596, EPI_ISL_2889605, EPI_ISL_2889608, EPI_ISL_2889609, EPI_ISL_2889610, EPI_ISL_2889613, EPI_ISL_2889616, EPI_ISL_2889625, EPI_ISL_2889626, EPI_ISL_2889668, EPI_ISL_2889672, EPI_ISL_2889674, EPI_ISL_2889677, EPI_ISL_2889682, EPI_ISL_2889687, EPI_ISL_2889688, EPI_ISL_2889691, EPI_ISL_2889692, EPI_ISL_2889693, EPI_ISL_2889697, EPI_ISL_2889711, EPI_ISL_2889743, EPI_ISL_2889747, EPI_ISL_2889755, EPI_ISL_2889756, EPI_ISL_2889757, EPI_ISL_2889770, EPI_ISL_2889813, EPI_ISL_2889814 | | | |
| see above | SK-Roy Romanow Provincial Laboratory | National Microbiology Laboratory (NML) | Anna Majer, Shari Tyson, Grace Seo, Philip Mabon, Elsie Grudeski, Rhiannon Huzarewich, Russell Mandes, Anneliese Landgraff, Jennifer Tanner, Natalie Knox, Morag Graham, Gary Van Domselaar, Ryan McDonald, Keith MacKenzie, Meredith Faires, Kara Loos, Rachel DePaulo, Alanna Senecal, Amanda Lang, Jessica Minion, Nathalie Bastien, Yan Li, Timothy Booth, Darian Hole, Madison Chapel, Kirsten Biggar, CanCOGeN's metadata curation team, Public Health Agency of Canada CanCOGeN team |
| EPI_ISL_2889854 | Fondation Congolaise pour la recherche medicale (FCRM) | Fondation Congolaise pour la Recherche Médicale | Mfoutou Mapanguy Claujens Chastel; Batchi-Bouyou Armel Landry; Jean Claude Djontu; Abel Lissom; Thirumalaisamy P. Velavan; Francine Ntoumi |
| EPI_ISL_2890202, EPI_ISL_2891708, EPI_ISL_2891801, EPI_ISL_2891813, EPI_ISL_2891818, EPI_ISL_2891849, EPI_ISL_2891907, EPI_ISL_2891963, EPI_ISL_2891964, EPI_ISL_2891967, EPI_ISL_2892022, EPI_ISL_2892062, EPI_ISL_2892083, EPI_ISL_2892085, EPI_ISL_2892087, EPI_ISL_2892088, EPI_ISL_2892089, EPI_ISL_2892109, EPI_ISL_2892130, EPI_ISL_2892133, EPI_ISL_2892134, EPI_ISL_2892152, EPI_ISL_2892153, EPI_ISL_2892154, EPI_ISL_2892155, EPI_ISL_2892156, EPI_ISL_2892204, EPI_ISL_2892263, EPI_ISL_2892294, EPI_ISL_2892298, EPI_ISL_2892299 | | | |
| see above | CDPH VBL | California Department of Public Health | CDPH-COVIDNet |
| EPI_ISL_2892409, EPI_ISL_2892413, EPI_ISL_2892414, EPI_ISL_2892415, EPI_ISL_2892416, EPI_ISL_2892417, EPI_ISL_2892418, EPI_ISL_2892419, EPI_ISL_2892436, EPI_ISL_2892468, EPI_ISL_2892472, EPI_ISL_2892483, EPI_ISL_2892485, EPI_ISL_2892486, EPI_ISL_2892494, EPI_ISL_2892519, EPI_ISL_2892522, EPI_ISL_2892523, EPI_ISL_2892524, EPI_ISL_2892529, EPI_ISL_2892531, EPI_ISL_2892540, EPI_ISL_2892550, EPI_ISL_2892553, EPI_ISL_2892558, EPI_ISL_2892563, EPI_ISL_2892564 | | | |
| see above | California Department of Public Health | California Department of Public Health | CDPH IDLB COVIDNet et al |

| | | | |
|---|---|---|---|
| EPI_ISL_2893002 | Genetica Molecular and Subdepartamento de Virologia ISP Chile | Instituto de Salud Publica de Chile | Karen Orostica, Constanza Campano, Barbara Parra, Loredana Arata, Gisselle Barra, Patricia Bustos, Rodrigo Fasce, Javier Tognarelli, Andres Castillo, Soledad Ulloa, Jorge Fernandez |
| EPI_ISL_2893034, EPI_ISL_2893035, EPI_ISL_2893036, EPI_ISL_2893037, EPI_ISL_2893038, EPI_ISL_2893039, EPI_ISL_2893040, EPI_ISL_2893041, EPI_ISL_2893042, EPI_ISL_2893043, EPI_ISL_2893044, EPI_ISL_2893045, EPI_ISL_2893046, EPI_ISL_2893047, EPI_ISL_2893048, EPI_ISL_2893049, EPI_ISL_2893050, EPI_ISL_2893051, EPI_ISL_2893052, EPI_ISL_2893053, EPI_ISL_2893054, EPI_ISL_2893055, EPI_ISL_2893057, EPI_ISL_2893058, EPI_ISL_2893059, EPI_ISL_2893060, EPI_ISL_2893061, EPI_ISL_2893062, EPI_ISL_2893063, EPI_ISL_2893064, EPI_ISL_2893065, EPI_ISL_2893066, EPI_ISL_2893067, EPI_ISL_2893068, EPI_ISL_2893069, EPI_ISL_2893070, EPI_ISL_2893071, EPI_ISL_2893072, EPI_ISL_2893073, EPI_ISL_2893074, EPI_ISL_2893075, EPI_ISL_2893076, EPI_ISL_2893077, EPI_ISL_2893078, EPI_ISL_2893079, EPI_ISL_2893080, EPI_ISL_2893081, EPI_ISL_2893082, EPI_ISL_2893083, EPI_ISL_2893084, EPI_ISL_2893085, EPI_ISL_2893086, EPI_ISL_2893087, EPI_ISL_2893088, EPI_ISL_2893089, EPI_ISL_2893090, EPI_ISL_2893091, EPI_ISL_2893092, EPI_ISL_2893093, EPI_ISL_2893094, EPI_ISL_2893095, EPI_ISL_2893096, EPI_ISL_2893097, EPI_ISL_2893098, EPI_ISL_2893099, EPI_ISL_2893100, EPI_ISL_2893101, EPI_ISL_2893102, EPI_ISL_2893103, EPI_ISL_2893104, EPI_ISL_2893105, EPI_ISL_2893106, EPI_ISL_2893107, EPI_ISL_2893108, EPI_ISL_2893109, EPI_ISL_2893110, EPI_ISL_2893111, EPI_ISL_2893112, EPI_ISL_2893113, EPI_ISL_2893114, EPI_ISL_2893115, EPI_ISL_2893116, EPI_ISL_2893117, EPI_ISL_2893118, EPI_ISL_2893119, EPI_ISL_2893120, EPI_ISL_2893121, EPI_ISL_2893122, EPI_ISL_2893124, EPI_ISL_2893125, EPI_ISL_2893126, EPI_ISL_2893127, EPI_ISL_2893128, EPI_ISL_2893129, EPI_ISL_2893130, EPI_ISL_2893131, EPI_ISL_2893132, EPI_ISL_2893133, EPI_ISL_2893134, EPI_ISL_2893135, EPI_ISL_2893136, EPI_ISL_2893137, EPI_ISL_2893138, EPI_ISL_2893139, EPI_ISL_2893140, EPI_ISL_2893141, EPI_ISL_2893142, EPI_ISL_2893143, EPI_ISL_2893146, EPI_ISL_2893147, EPI_ISL_2893148, EPI_ISL_2893149, EPI_ISL_2893150, EPI_ISL_2893151, EPI_ISL_2893152, EPI_ISL_2893153, EPI_ISL_2893154, EPI_ISL_2893155, EPI_ISL_2893156, EPI_ISL_2893157, EPI_ISL_2893158, EPI_ISL_2893159, EPI_ISL_2893160, EPI_ISL_2893161, EPI_ISL_2893162, EPI_ISL_2893163, EPI_ISL_2893164, EPI_ISL_2893165, EPI_ISL_2893166, EPI_ISL_2893167, EPI_ISL_2893168, EPI_ISL_2893169, EPI_ISL_2893170, EPI_ISL_2893171, EPI_ISL_2893173, EPI_ISL_2893174, EPI_ISL_2893175, EPI_ISL_2893176, EPI_ISL_2893177, EPI_ISL_2893178, EPI_ISL_2893179, EPI_ISL_2893180, EPI_ISL_2893181, EPI_ISL_2893182, EPI_ISL_2893184, EPI_ISL_2893185, EPI_ISL_2893186, EPI_ISL_2893187, EPI_ISL_2893189, EPI_ISL_2893190, EPI_ISL_2893191, EPI_ISL_2893192, EPI_ISL_2893193, EPI_ISL_2893194, EPI_ISL_2893195, EPI_ISL_2893196, EPI_ISL_2893197, EPI_ISL_2893198, EPI_ISL_2893199, EPI_ISL_2893200, EPI_ISL_2893201, EPI_ISL_2893202, EPI_ISL_2893203, EPI_ISL_2893204, EPI_ISL_2893205, EPI_ISL_2893206 | | | |
| see above | National Virus Reference Laboratory | National Virus Reference Laboratory | Zoe Yandle, Charlene Bennett, Gabriel Gonzalez, Michael Carr, Jonathan Dean, Cillian F De Gascun |
| EPI_ISL_2893207, EPI_ISL_2893208, EPI_ISL_2893209, EPI_ISL_2893210, EPI_ISL_2893211, EPI_ISL_2893212, EPI_ISL_2893213, EPI_ISL_2893214, EPI_ISL_2893215, EPI_ISL_2893216, EPI_ISL_2893217, EPI_ISL_2893218, EPI_ISL_2893219, EPI_ISL_2893220, EPI_ISL_2893221, EPI_ISL_2893222, EPI_ISL_2893223 | | | |
| see above | Laboratory of Molecular Biology and Cancer Immunology, Faculty of Sciences, Lebanese University | Microbial Pathogenomics Lab - LAU | Sima tokajian, Georgi Merhi, Jad Koweyes, Bassam Badran, Fadi Abdel Sater, Rawan Makki, Hamad Hassan |
| EPI_ISL_2893224, EPI_ISL_2893225 | Laboratory of Molecular Biology and Cancer Immunology, Faculty of Sciences, Lebanese University | Microbial Pathogenomics Lab - LAU | Sima tokajian, Georgi Merhi, Jad Koweyes, Bassam Badran, Fadi Abdel Sater, Rawan Makki, Tatiana Tohmeh, Nada Ghosn, Hamad Hassan |
| EPI_ISL_2893243, EPI_ISL_2893246, EPI_ISL_2893247, EPI_ISL_2893248, EPI_ISL_2893251, EPI_ISL_2893252, EPI_ISL_2893253, EPI_ISL_2893255, EPI_ISL_2893256, EPI_ISL_2893257, EPI_ISL_2893258, EPI_ISL_2893259, EPI_ISL_2893260, EPI_ISL_2893261, EPI_ISL_2893262, EPI_ISL_2893264, EPI_ISL_2893265, EPI_ISL_2893266, EPI_ISL_2893267, EPI_ISL_2893268, EPI_ISL_2893269, EPI_ISL_2893270, EPI_ISL_2893276, EPI_ISL_2893279, EPI_ISL_2893281, EPI_ISL_2893282, EPI_ISL_2893284, EPI_ISL_2893286, EPI_ISL_2893287, EPI_ISL_2893288, EPI_ISL_2893292, EPI_ISL_2893293, EPI_ISL_2893294, EPI_ISL_2893295, EPI_ISL_2893297, EPI_ISL_2893298, EPI_ISL_2893299 | | | |
| see above | NHLS_VIRO Eastern Cape | KRISP, KZN Research Innovation and Sequencing Platform | Laguda-Akingba O, Giandhari J, Pillay S, Lessells R, Mdlalose K, York D, Khan S, Emmanuel SJ, Tegally H, Wilkinson E, de Oliveira T |
| EPI_ISL_2893300, EPI_ISL_2893301, EPI_ISL_2893302, EPI_ISL_2893304, EPI_ISL_2893305, EPI_ISL_2893306, EPI_ISL_2893308, EPI_ISL_2893309, EPI_ISL_2893311, EPI_ISL_2893313, EPI_ISL_2893316, EPI_ISL_2893318 | | | |
| see above | BARC - Sisonke | KRISP, KZN Research Innovation and Sequencing Platform | Sisonke, Giandhari Jennifer, Pillay Sureshnee, Yajna Ramphal, Naidoo Yeshnee, Tshabuila Derek,Tegally Houriiyah, San James, Wilkinson Eduan, de Oliveira Tulio |
| EPI_ISL_2893319, EPI_ISL_2893320, EPI_ISL_2893321, EPI_ISL_2893322, EPI_ISL_2893323, EPI_ISL_2893340, EPI_ISL_2893341, EPI_ISL_2893342, EPI_ISL_2893355, EPI_ISL_2893356, EPI_ISL_2893357, EPI_ISL_2893358, EPI_ISL_2893359, EPI_ISL_2893360, EPI_ISL_2893361, EPI_ISL_2893362, EPI_ISL_2893374, EPI_ISL_2893377, EPI_ISL_2893392, EPI_ISL_2893393 | | | |
| see above | Alaska State Virology Laboratory | Alaska State Virology Laboratory | Stephanie DeRonde, Elva House, Jacob Zidek, Lisa Smith, Ph.D., Jack Chen, Ph.D. |
| EPI_ISL_2893436, EPI_ISL_2893437, EPI_ISL_2893439, EPI_ISL_2893440, EPI_ISL_2893441, EPI_ISL_2893442, EPI_ISL_2893443, EPI_ISL_2893444 | Oregon State Public Health Laboratory | Oregon State Public Health Laboratory | Rafia Razzaque, Eugene Yeboah, Vanda Makris, Laura Tsaknaridis, John Fontana and Shane Sevey |
| EPI_ISL_2893453 | Jessa | Jessa | Severine Berden et al. on behalf of the Jessa_cmdLab |
| EPI_ISL_2893454, EPI_ISL_2893455, EPI_ISL_2893456, EPI_ISL_2893457, EPI_ISL_2893458, EPI_ISL_2893459, EPI_ISL_2893460, EPI_ISL_2893461 | ZOL | Jessa | Severine Berden et al. on behalf of the Jessa_cmdLab |
| EPI_ISL_2893462 | Jessa | Jessa | Severine Berden et al. on behalf of the Jessa_cmdLab |
| EPI_ISL_2893463, EPI_ISL_2893464 | ZOL | Jessa | Severine Berden et al. on behalf of the Jessa_cmdLab |
| EPI_ISL_2893465, EPI_ISL_2893466, EPI_ISL_2893467, EPI_ISL_2893468, EPI_ISL_2893469, EPI_ISL_2893470, EPI_ISL_2893471, EPI_ISL_2893472, EPI_ISL_2893473, EPI_ISL_2893474, EPI_ISL_2893475 | | | |
| see above | Jessa | Jessa | Severine Berden et al. on behalf of the Jessa_cmdLab |
| EPI_ISL_2893477, EPI_ISL_2893498, EPI_ISL_2893504, EPI_ISL_2893511, EPI_ISL_2893513, EPI_ISL_2893515, EPI_ISL_2893516, EPI_ISL_2893517, EPI_ISL_2893519 | Texas Children's Hospital | Texas Children's Microbiome Center | Ruth Ann Luna, Jennifer K. Spinler, James Dunn, James Versalovic, Ila Singh |
| EPI_ISL_2893520, EPI_ISL_2893521, EPI_ISL_2893522, EPI_ISL_2893523, EPI_ISL_2893527, EPI_ISL_2893530, EPI_ISL_2893531, EPI_ISL_2893532 | Laboratoire Hôpital Sainte Musse | Laboratoire Hôpital Sainte Musse | Lionel Chollet |
| EPI_ISL_2893668, EPI_ISL_2893669, EPI_ISL_2893670, EPI_ISL_2893671, EPI_ISL_2893672, EPI_ISL_2893673, EPI_ISL_2893674, EPI_ISL_2893675, EPI_ISL_2893676, EPI_ISL_2893677, EPI_ISL_2893681, EPI_ISL_2893682, EPI_ISL_2893683, EPI_ISL_2893684, EPI_ISL_2893685, EPI_ISL_2893686, EPI_ISL_2893687, EPI_ISL_2893688, EPI_ISL_2893689, EPI_ISL_2893690, EPI_ISL_2893691, EPI_ISL_2893692, EPI_ISL_2893693 | | | |
| see above | Kansas Health and Environmental Lab | Kansas Health and Environmental Lab | Katherine Wiggins, Mike Grose, Jonathan Barnell, Ben Olsen, and Phil Adam |
| EPI_ISL_2893698, EPI_ISL_2893699, EPI_ISL_2893700 | BIOPOLE 66 (IASCB) | GenBio | Gwladys SEVRIN |
| EPI_ISL_2893701 | MEDILAB 66 | GenBio | Gwladys SEVRIN |
| EPI_ISL_2893702 | LABOSUD | GenBio | Gwladys SEVRIN |
| EPI_ISL_2893703 | MEDILAB 66 | GenBio | Gwladys SEVRIN |
| EPI_ISL_2893704 | LABOSUD | GenBio | Gwladys SEVRIN |
| EPI_ISL_2893705 | MEDILAB 66 | GenBio | Gwladys SEVRIN |
| EPI_ISL_2893706 | BIOPOLE 66 (IASCB) | GenBio | Gwladys SEVRIN |
| EPI_ISL_2893707 | LABOSUD | GenBio | Gwladys SEVRIN |
| EPI_ISL_2893708 | MEDILAB 66 | GenBio | Gwladys SEVRIN |
| EPI_ISL_2893709, EPI_ISL_2893710 | BIOPOLE 66 (IASCB) | GenBio | Gwladys SEVRIN |
| EPI_ISL_2893712 | LABOSUD | GenBio | Gwladys SEVRIN |
| EPI_ISL_2893713 | BIOFUSION | GenBio | Gwladys SEVRIN |
| EPI_ISL_2893714, EPI_ISL_2893715 | BIOPOLE 66 (IASCB) | GenBio | Gwladys SEVRIN |
| EPI_ISL_2893716 | BIOPOLE 66 (IASCC) | GenBio | Gwladys SEVRIN |
| EPI_ISL_2893717 | BIOFUSION | GenBio | Gwladys SEVRIN |
| EPI_ISL_2893718, EPI_ISL_2893719 | BIOPOLE 66 (IASCB) | GenBio | Gwladys SEVRIN |
| EPI_ISL_2893720 | LABOSUD | GenBio | Gwladys SEVRIN |
| EPI_ISL_2893721, EPI_ISL_2893722 | BIOPOLE 66 (IASCB) | GenBio | Gwladys SEVRIN |
| EPI_ISL_2893723 | LABOSUD | GenBio | Gwladys SEVRIN |
| EPI_ISL_2893724 | MEDILAB 66 | GenBio | Gwladys SEVRIN |
| EPI_ISL_2893725 | BIOPOLE 66 (IASCB) | GenBio | Gwladys SEVRIN |

| | | | |
|---|---|---|---|
| EPI_ISL_2893726 | BIOFUSION | GenBio | Gwladys SEVRIN |
| EPI_ISL_2893727 | MEDILAB 66 | GenBio | Gwladys SEVRIN |
| EPI_ISL_2893728 | BIOMEDIVAL V | GenBio | Gwladys SEVRIN |
| EPI_ISL_2893729 | MEDILAB 66 | GenBio | Gwladys SEVRIN |
| EPI_ISL_2893730 | LABOSUD | GenBio | Gwladys SEVRIN |
| EPI_ISL_2893731, EPI_ISL_2893732, EPI_ISL_2893733, EPI_ISL_2893734 | BIOPOLE 66 (IASCB) | GenBio | Gwladys SEVRIN |
| EPI_ISL_2893735 | LABOSUD | GenBio | Gwladys SEVRIN |
| EPI_ISL_2893736 | BIOPOLE 66 (IASCB) | GenBio | Gwladys SEVRIN |
| EPI_ISL_2893737 | LABOSUD | GenBio | Gwladys SEVRIN |
| EPI_ISL_2893738 | BIOFUSION | GenBio | Gwladys SEVRIN |
| EPI_ISL_2893739 | LABOSUD | GenBio | Gwladys SEVRIN |
| EPI_ISL_2893740 | BIOFUSION | GenBio | Gwladys SEVRIN |
| EPI_ISL_2893741 | LABOSUD | GenBio | Gwladys SEVRIN |
| EPI_ISL_2893742 | BIOPOLE 66 (IASCB) | GenBio | Gwladys SEVRIN |
| EPI_ISL_2893743 | LABOSUD | GenBio | Gwladys SEVRIN |
| EPI_ISL_2893744 | BIOFUSION | GenBio | Gwladys SEVRIN |
| EPI_ISL_2893745 | BIOPOLE 66 (IASCB) | GenBio | Gwladys SEVRIN |
| EPI_ISL_2893746 | MEDILAB 66 | GenBio | Gwladys SEVRIN |
| EPI_ISL_2893747 | BIOPOLE 66 (IASCB) | GenBio | Gwladys SEVRIN |
| EPI_ISL_2893748 | BIOPOLE 66 (IASCC) | GenBio | Gwladys SEVRIN |
| EPI_ISL_2893749 | LABOSUD | GenBio | Gwladys SEVRIN |
| EPI_ISL_2893750 | BIOPOLE 66 (IASCB) | GenBio | Gwladys SEVRIN |
| EPI_ISL_2893751, EPI_ISL_2893752 | MEDILAB 66 | GenBio | Gwladys SEVRIN |
| EPI_ISL_2893753 | BIOPOLE 66 (IASCB) | GenBio | Gwladys SEVRIN |
| EPI_ISL_2893754, EPI_ISL_2893755 | LABOSUD | GenBio | Gwladys SEVRIN |
| EPI_ISL_2893756, EPI_ISL_2893757 | MEDILAB 66 | GenBio | Gwladys SEVRIN |
| EPI_ISL_2893758 | BIOPOLE 66 (IASCB) | GenBio | Gwladys SEVRIN |
| EPI_ISL_2893759 | MEDILAB 66 | GenBio | Gwladys SEVRIN |
| EPI_ISL_2893760, EPI_ISL_2893761 | LABOSUD | GenBio | Gwladys SEVRIN |
| EPI_ISL_2893762 | BIOFUSION | GenBio | Gwladys SEVRIN |
| EPI_ISL_2893763 | BIOPOLE 66 (IASCB) | GenBio | Gwladys SEVRIN |
| EPI_ISL_2893764 | BIOFUSION | GenBio | Gwladys SEVRIN |
| EPI_ISL_2893765, EPI_ISL_2893766 | MEDILAB 66 | GenBio | Gwladys SEVRIN |
| EPI_ISL_2893767 | BIOMED 34 | GenBio | Gwladys SEVRIN |
| EPI_ISL_2893768, EPI_ISL_2893769 | LABOSUD | GenBio | Gwladys SEVRIN |
| EPI_ISL_2893770 | MEDILAB 66 | GenBio | Gwladys SEVRIN |
| EPI_ISL_2893771 | BIOPOLE 66 (IASCB) | GenBio | Gwladys SEVRIN |
| EPI_ISL_2893772, EPI_ISL_2893773, EPI_ISL_2893774, EPI_ISL_2893775, EPI_ISL_2893776, EPI_ISL_2893777, EPI_ISL_2893778 | Department of Acute Infectious Diseases Control and Prevention, Yunnan Provincial Center for Disease Control and Prevention | Department of Acute Infectious Diseases Control and Prevention, Yunnan Provincial Center for Disease Control and Prevention | Meiling Zhang, Jienan Zhou, Zhaosheng Liu, Xiaoqing Fu |
| EPI_ISL_2893779, EPI_ISL_2893780 | CENTRE HOSPITALIER ST JOSEPH ST LUC | CNR Virus des Infections Respiratoires - France SUD | Antonin Bal, Gregory Destras, Gwendolyne Burfin, Hadrien Regue, Quentin Semanas, Martine Valette, Bruno Lina, Laurence Josset |
| EPI_ISL_2893781 | CHU ST ETIENNE HOPITAL NORD | CNR Virus des Infections Respiratoires - France SUD | Antonin Bal, Gregory Destras, Gwendolyne Burfin, Hadrien Regue, Quentin Semanas, Martine Valette, Bruno Lina, Laurence Josset |
| EPI_ISL_2893782 | CENTRE HOSPITALIER PIERRE OUDOT | CNR Virus des Infections Respiratoires - France SUD | Antonin Bal, Gregory Destras, Gwendolyne Burfin, Hadrien Regue, Quentin Semanas, Martine Valette, Bruno Lina, Laurence Josset |
| EPI_ISL_2893783 | CHU ST ETIENNE HOPITAL NORD | CNR Virus des Infections Respiratoires - France SUD | Antonin Bal, Gregory Destras, Gwendolyne Burfin, Hadrien Regue, Quentin Semanas, Martine Valette, Bruno Lina, Laurence Josset |
| EPI_ISL_2893784, EPI_ISL_2893785, EPI_ISL_2893786, EPI_ISL_2893787 | LABORATOIRE UNILIANS DECINES | CNR Virus des Infections Respiratoires - France SUD | Antonin Bal, Gregory Destras, Gwendolyne Burfin, Hadrien Regue, Quentin Semanas, Martine Valette, Bruno Lina, Laurence Josset |
| EPI_ISL_2893788 | MIRIALIS CLUSES BECHET | CNR Virus des Infections Respiratoires - France SUD | Antonin Bal, Gregory Destras, Gwendolyne Burfin, Hadrien Regue, Quentin Semanas, Martine Valette, Bruno Lina, Laurence Josset |
| EPI_ISL_2893789 | LBM NEOLAB | CNR Virus des Infections Respiratoires - France SUD | Antonin Bal, Gregory Destras, Gwendolyne Burfin, Hadrien Regue, Quentin Semanas, Martine Valette, Bruno Lina, Laurence Josset |
| EPI_ISL_2893790 | DYOMEDEA-LABORATOIRE DE LA SAUVEGARDE | CNR Virus des Infections Respiratoires - France SUD | Antonin Bal, Gregory Destras, Gwendolyne Burfin, Hadrien Regue, Quentin Semanas, Martine Valette, Bruno Lina, Laurence Josset |
| EPI_ISL_2893791 | LAM GEN-BIO GRAVANCHES | CNR Virus des Infections Respiratoires - France SUD | Antonin Bal, Gregory Destras, Gwendolyne Burfin, Hadrien Regue, Quentin Semanas, Martine Valette, Bruno Lina, Laurence Josset |
| EPI_ISL_2893792, EPI_ISL_2893793, EPI_ISL_2893794 | DYOMEDEA-LABORATOIRE DE LA SAUVEGARDE | CNR Virus des Infections Respiratoires - France SUD | Antonin Bal, Gregory Destras, Gwendolyne Burfin, Hadrien Regue, Quentin Semanas, Martine Valette, Bruno Lina, Laurence Josset |
| EPI_ISL_2893795, EPI_ISL_2893796, EPI_ISL_2893797 | SCM B12 | CNR Virus des Infections Respiratoires - France SUD | Antonin Bal, Gregory Destras, Gwendolyne Burfin, Hadrien Regue, Quentin Semanas, Martine Valette, Bruno Lina, Laurence Josset |
| EPI_ISL_2893798 | ORIAPOLE | CNR Virus des Infections Respiratoires - France SUD | Antonin Bal, Gregory Destras, Gwendolyne Burfin, Hadrien Regue, Quentin Semanas, Martine Valette, Bruno Lina, Laurence Josset |
| EPI_ISL_2893799 | SCM B12 | CNR Virus des Infections Respiratoires - France SUD | Antonin Bal, Gregory Destras, Gwendolyne Burfin, Hadrien Regue, Quentin Semanas, Martine Valette, Bruno Lina, Laurence Josset |
| EPI_ISL_2893800 | ORIAPOLE | CNR Virus des Infections Respiratoires - France SUD | Antonin Bal, Gregory Destras, Gwendolyne Burfin, Hadrien Regue, Quentin Semanas, Martine Valette, Bruno Lina, Laurence Josset |
| EPI_ISL_2893801, EPI_ISL_2893802 | SCM B12 | CNR Virus des Infections Respiratoires - France SUD | Antonin Bal, Gregory Destras, Gwendolyne Burfin, Hadrien Regue, Quentin Semanas, Martine Valette, Bruno Lina, Laurence Josset |
| EPI_ISL_2893803, EPI_ISL_2893804 | ORIAPOLE | CNR Virus des Infections Respiratoires - France SUD | Antonin Bal, Gregory Destras, Gwendolyne Burfin, Hadrien Regue, Quentin Semanas, Martine Valette, Bruno Lina, Laurence Josset |
| EPI_ISL_2893805 | SCM B12 | CNR Virus des Infections Respiratoires - France SUD | Antonin Bal, Gregory Destras, Gwendolyne Burfin, Hadrien Regue, Quentin Semanas, Martine Valette, Bruno Lina, Laurence Josset |
| EPI_ISL_2893806 | ORIAPOLE | CNR Virus des Infections Respiratoires - France SUD | Antonin Bal, Gregory Destras, Gwendolyne Burfin, Hadrien Regue, Quentin Semanas, Martine Valette, Bruno Lina, Laurence Josset |
| EPI_ISL_2893807 | SCM B12 | CNR Virus des Infections Respiratoires - France SUD | Antonin Bal, Gregory Destras, Gwendolyne Burfin, Hadrien Regue, Quentin Semanas, Martine Valette, Bruno Lina, Laurence Josset |
| EPI_ISL_2893808, EPI_ISL_2893809, | ORIAPOLE | CNR Virus des Infections Respiratoires - France SUD | Antonin Bal, Gregory Destras, Gwendolyne Burfin, Hadrien Regue, Quentin Semanas, Martine Valette, Bruno Lina, Laurence Josset |

| | | | |
|---|---|---|---|
| EPI_ISL_2893810, EPI_ISL_2893811 | | | |
| EPI_ISL_2893812, EPI_ISL_2893813, EPI_ISL_2893814, EPI_ISL_2893815 | LABORATOIRE CERBALLIANCE PLT VILLON | CNR Virus des Infections Respiratoires - France SUD | Antonin Bal, Gregory Destras, Gwendolyne Burfin, Hadrien Regue, Quentin Semanas, Martine Valette, Bruno Lina, Laurence Josset |
| EPI_ISL_2893816 | LABORATOIRE NOVELAB | CNR Virus des Infections Respiratoires - France SUD | Antonin Bal, Gregory Destras, Gwendolyne Burfin, Hadrien Regue, Quentin Semanas, Martine Valette, Bruno Lina, Laurence Josset |
| EPI_ISL_2893817 | CENTRE HOSPITALIER PIERRE OUDOT | CNR Virus des Infections Respiratoires - France SUD | Antonin Bal, Gregory Destras, Gwendolyne Burfin, Hadrien Regue, Quentin Semanas, Martine Valette, Bruno Lina, Laurence Josset |
| EPI_ISL_2893818 | LABAZUR BRETAGNE CHATEAULIN | CNR Virus des Infections Respiratoires - France SUD | Antonin Bal, Gregory Destras, Gwendolyne Burfin, Hadrien Regue, Quentin Semanas, Martine Valette, Bruno Lina, Laurence Josset |
| EPI_ISL_2893819, EPI_ISL_2893820, EPI_ISL_2893821 | LBM DU PARC DES POTERIES | CNR Virus des Infections Respiratoires - France SUD | Antonin Bal, Gregory Destras, Gwendolyne Burfin, Hadrien Regue, Quentin Semanas, Martine Valette, Bruno Lina, Laurence Josset |
| EPI_ISL_2893822, EPI_ISL_2893823, EPI_ISL_2893824 | LBM SEALAB DARRASSE ET ASSOCIES | CNR Virus des Infections Respiratoires - France SUD | Antonin Bal, Gregory Destras, Gwendolyne Burfin, Hadrien Regue, Quentin Semanas, Martine Valette, Bruno Lina, Laurence Josset |
| EPI_ISL_2893825, EPI_ISL_2893826, EPI_ISL_2893827, EPI_ISL_2893828, EPI_ISL_2893829, EPI_ISL_2893830, EPI_ISL_2893831 | HOPITAL MARIN HENDAYE | CNR Virus des Infections Respiratoires - France SUD | Antonin Bal, Gregory Destras, Gwendolyne Burfin, Hadrien Regue, Quentin Semanas, Martine Valette, Bruno Lina, Laurence Josset |
| EPI_ISL_2893832 | LABORATOIRE d'ANALYSES de BIOLOGIE MEDICALES | CNR Virus des Infections Respiratoires - France SUD | Antonin Bal, Gregory Destras, Gwendolyne Burfin, Hadrien Regue, Quentin Semanas, Martine Valette, Bruno Lina, Laurence Josset |
| EPI_ISL_2893833 | ASTRALAB TASSIGNY | CNR Virus des Infections Respiratoires - France SUD | Antonin Bal, Gregory Destras, Gwendolyne Burfin, Hadrien Regue, Quentin Semanas, Martine Valette, Bruno Lina, Laurence Josset |
| EPI_ISL_2893834 | LAM BIOPOLE ORTHEZ | CNR Virus des Infections Respiratoires - France SUD | Antonin Bal, Gregory Destras, Gwendolyne Burfin, Hadrien Regue, Quentin Semanas, Martine Valette, Bruno Lina, Laurence Josset |
| EPI_ISL_2893835 | LABORATOIRE FORTE BIO ST PAUL LES DAX | CNR Virus des Infections Respiratoires - France SUD | Antonin Bal, Gregory Destras, Gwendolyne Burfin, Hadrien Regue, Quentin Semanas, Martine Valette, Bruno Lina, Laurence Josset |
| EPI_ISL_2893836 | EXALAB FOCH | CNR Virus des Infections Respiratoires - France SUD | Antonin Bal, Gregory Destras, Gwendolyne Burfin, Hadrien Regue, Quentin Semanas, Martine Valette, Bruno Lina, Laurence Josset |
| EPI_ISL_2893837, EPI_ISL_2893838, EPI_ISL_2893839, EPI_ISL_2893840, EPI_ISL_2893841, EPI_ISL_2893842, EPI_ISL_2893843, EPI_ISL_2893844, EPI_ISL_2893845, EPI_ISL_2893846, EPI_ISL_2893847, EPI_ISL_2893848, EPI_ISL_2893849, EPI_ISL_2893850, EPI_ISL_2893851, EPI_ISL_2893852, EPI_ISL_2893853, EPI_ISL_2893854, EPI_ISL_2893855, EPI_ISL_2893856, EPI_ISL_2893857, EPI_ISL_2893858, EPI_ISL_2893859, EPI_ISL_2893860 | | | |
| see above | AX BIO OCEAN | CNR Virus des Infections Respiratoires - France SUD | Antonin Bal, Gregory Destras, Gwendolyne Burfin, Hadrien Regue, Quentin Semanas, Martine Valette, Bruno Lina, Laurence Josset |
| EPI_ISL_2893861 | CERBALLIANCE MIDI-PYRENEES | CNR Virus des Infections Respiratoires - France SUD | Antonin Bal, Gregory Destras, Gwendolyne Burfin, Hadrien Regue, Quentin Semanas, Martine Valette, Bruno Lina, Laurence Josset |
| EPI_ISL_2893862, EPI_ISL_2893863, EPI_ISL_2893864, EPI_ISL_2893865 | LABORATOIRE BIOESTEREL | CNR Virus des Infections Respiratoires - France SUD | Antonin Bal, Gregory Destras, Gwendolyne Burfin, Hadrien Regue, Quentin Semanas, Martine Valette, Bruno Lina, Laurence Josset |
| EPI_ISL_2893866, EPI_ISL_2893867 | LABORATOIRE- BIOESTERL | CNR Virus des Infections Respiratoires- France SUD | Antonin Bal, Gregory Destras, Gwendolyne Burfin, Hadrien Regue, Quentin Semanas, Martine Valette, Bruno Lina, Laurence Josset |
| EPI_ISL_2893868 | CERBALLIANCE - LAMAT | CNR Virus des Infections Respiratoires - France SUD | Antonin Bal, Gregory Destras, Gwendolyne Burfin, Hadrien Regue, Quentin Semanas, Martine Valette, Bruno Lina, Laurence Josset |
| EPI_ISL_2893869 | LABORATOIRE CERBALLIANCE PLT VILLON | CNR Virus des Infections Respiratoires - France SUD | Antonin Bal, Gregory Destras, Gwendolyne Burfin, Hadrien Regue, Quentin Semanas, Martine Valette, Bruno Lina, Laurence Josset |
| EPI_ISL_2893883 | LAM GEN-BIO GRAVANCHES | CNR Virus des Infections Respiratoires - France SUD | Antonin Bal, Gregory Destras, Gwendolyne Burfin, Hadrien Regue, Quentin Semanas, Martine Valette, Bruno Lina, Laurence Josset |
| EPI_ISL_2893887 | AX BIO OCEAN | CNR Virus des Infections Respiratoires - France SUD | Antonin Bal, Gregory Destras, Gwendolyne Burfin, Hadrien Regue, Quentin Semanas, Martine Valette, Bruno Lina, Laurence Josset |
| EPI_ISL_2893920 | LABORATOIRE BIOESTEREL | CNR Virus des Infections Respiratoires - France SUD | Antonin Bal, Gregory Destras, Gwendolyne Burfin, Hadrien Regue, Quentin Semanas, Martine Valette, Bruno Lina, Laurence Josset |
| EPI_ISL_2893922 | LBM DU PARC DES POTERIES | CNR Virus des Infections Respiratoires - France SUD | Antonin Bal, Gregory Destras, Gwendolyne Burfin, Hadrien Regue, Quentin Semanas, Martine Valette, Bruno Lina, Laurence Josset |
| EPI_ISL_2893935, EPI_ISL_2893936 | LABORATOIRE INGELS-VIGNON | CNR Virus des Infections Respiratoires - France SUD | Antonin Bal, Gregory Destras, Gwendolyne Burfin, Hadrien Regue, Quentin Semanas, Martine Valette, Bruno Lina, Laurence Josset |
| EPI_ISL_2893941 | LABORATOIRE XLABS | CNR Virus des Infections Respiratoires - France SUD | Antonin Bal, Gregory Destras, Gwendolyne Burfin, Hadrien Regue, Quentin Semanas, Martine Valette, Bruno Lina, Laurence Josset |
| EPI_ISL_2893945 | BIOLITTORAL BIOGROUP PLATEAU TECH | CNR Virus des Infections Respiratoires - France SUD | Antonin Bal, Gregory Destras, Gwendolyne Burfin, Hadrien Regue, Quentin Semanas, Martine Valette, Bruno Lina, Laurence Josset |
| EPI_ISL_2894078 | AX BIO OCEAN | CNR Virus des Infections Respiratoires - France SUD | Antonin Bal, Gregory Destras, Gwendolyne Burfin, Hadrien Regue, Quentin Semanas, Martine Valette, Bruno Lina, Laurence Josset |
| EPI_ISL_2894080, EPI_ISL_2894083 | ORIAPOLE | CNR Virus des Infections Respiratoires - France SUD | Antonin Bal, Gregory Destras, Gwendolyne Burfin, Hadrien Regue, Quentin Semanas, Martine Valette, Bruno Lina, Laurence Josset |
| EPI_ISL_2894085 | DYOMEDEA-LABORATOIRE DE LA SAUVEGARDE | CNR Virus des Infections Respiratoires - France SUD | Antonin Bal, Gregory Destras, Gwendolyne Burfin, Hadrien Regue, Quentin Semanas, Martine Valette, Bruno Lina, Laurence Josset |
| EPI_ISL_2894086 | LAM GEN-BIO GRAVANCHES | CNR Virus des Infections Respiratoires - France SUD | Antonin Bal, Gregory Destras, Gwendolyne Burfin, Hadrien Regue, Quentin Semanas, Martine Valette, Bruno Lina, Laurence Josset |
| EPI_ISL_2894089 | CNR Virus des Infections Respiratoires – France Sud | CNR Virus des Infections Respiratoires - France SUD | Antonin Bal, Gregory Destras, Gwendolyne Burfin, Hadrien Regue, Quentin Semanas, Martine Valette, Bruno Lina, Laurence Josset |
| EPI_ISL_2894090 | ORIAPOLE | CNR Virus des Infections Respiratoires - France SUD | Antonin Bal, Gregory Destras, Gwendolyne Burfin, Hadrien Regue, Quentin Semanas, Martine Valette, Bruno Lina, Laurence Josset |
| EPI_ISL_2894092 | LABORATOIRE LABAZUR FELIX FAURE | CNR Virus des Infections Respiratoires - France SUD | Antonin Bal, Gregory Destras, Gwendolyne Burfin, Hadrien Regue, Quentin Semanas, Martine Valette, Bruno Lina, Laurence Josset |
| EPI_ISL_2894093, EPI_ISL_2894095 | ORIAPOLE | CNR Virus des Infections Respiratoires - France SUD | Antonin Bal, Gregory Destras, Gwendolyne Burfin, Hadrien Regue, Quentin Semanas, Martine Valette, Bruno Lina, Laurence Josset |
| EPI_ISL_2894096 | AX BIO OCEAN | CNR Virus des Infections Respiratoires - France SUD | Antonin Bal, Gregory Destras, Gwendolyne Burfin, Hadrien Regue, Quentin Semanas, Martine Valette, Bruno Lina, Laurence Josset |
| EPI_ISL_2894098, EPI_ISL_2894099, EPI_ISL_2894100, EPI_ISL_2894102 | DYOMEDEA-LABORATOIRE DE LA SAUVEGARDE | CNR Virus des Infections Respiratoires - France SUD | Antonin Bal, Gregory Destras, Gwendolyne Burfin, Hadrien Regue, Quentin Semanas, Martine Valette, Bruno Lina, Laurence Josset |
| EPI_ISL_2894106 | LBM DU PARC DES POTERIES | CNR Virus des Infections Respiratoires - France SUD | Antonin Bal, Gregory Destras, Gwendolyne Burfin, Hadrien Regue, Quentin Semanas, Martine Valette, Bruno Lina, Laurence Josset |
| EPI_ISL_2894107 | LAM BIOPOLE ORTHEZ | CNR Virus des Infections Respiratoires - France SUD | Antonin Bal, Gregory Destras, Gwendolyne Burfin, Hadrien Regue, Quentin Semanas, Martine Valette, Bruno Lina, Laurence Josset |
| EPI_ISL_2894109 | DYOMEDEA-LABORATOIRE DE LA SAUVEGARDE | CNR Virus des Infections Respiratoires - France SUD | Antonin Bal, Gregory Destras, Gwendolyne Burfin, Hadrien Regue, Quentin Semanas, Martine Valette, Bruno Lina, Laurence Josset |
| EPI_ISL_2894110 | ORIAPOLE | CNR Virus des Infections Respiratoires - France SUD | Antonin Bal, Gregory Destras, Gwendolyne Burfin, Hadrien Regue, Quentin Semanas, Martine Valette, Bruno Lina, Laurence Josset |
| EPI_ISL_2894111, EPI_ISL_2894112 | AX BIO OCEAN | CNR Virus des Infections Respiratoires - France SUD | Antonin Bal, Gregory Destras, Gwendolyne Burfin, Hadrien Regue, Quentin Semanas, Martine Valette, Bruno Lina, Laurence Josset |
| EPI_ISL_2894113 | CNR Virus des Infections Respiratoires – France Sud | CNR Virus des Infections Respiratoires - France SUD | Antonin Bal, Gregory Destras, Gwendolyne Burfin, Hadrien Regue, Quentin Semanas, Martine Valette, Bruno Lina, Laurence Josset |
| EPI_ISL_2894114 | CHU ST ETIENNE HOPITAL NORD | CNR Virus des Infections Respiratoires - France SUD | Antonin Bal, Gregory Destras, Gwendolyne Burfin, Hadrien Regue, Quentin Semanas, Martine Valette, Bruno Lina, Laurence Josset |
| EPI_ISL_2894130 | MEDILAB | CHU Poitiers | Caroline MICHAUD, Maxime PICHON, Manon PRAT, Valentin BON-BARET, Birama N'DIAYE, Magali GARCIA, Agnes BEBY-DEFAUX, Nicolas LEVEQUE |
| EPI_ISL_2894131, EPI_ISL_2894132, EPI_ISL_2894133 | BIO86 GENCAY | CHU Poitiers | Caroline MICHAUD, Maxime PICHON, Manon PRAT, Valentin BON-BARET, Birama N'DIAYE, Magali GARCIA, Agnes BEBY-DEFAUX, Nicolas LEVEQUE |
| EPI_ISL_2894134 | Laverty Pathology | NSW Health Pathology - Institute of Clinical Pathology and Medical Research; Westmead Hospital; University of Sydney | CIDM-PH et al. |
| EPI_ISL_2894135 | 4Cyte Pathology | NSW Health Pathology - Institute of Clinical Pathology and Medical Research; Westmead Hospital; University of Sydney | CIDM-PH et al. |
| EPI_ISL_2894136 | Medlab Pathology | NSW Health Pathology - Institute of Clinical Pathology and Medical Research; Westmead Hospital; University of Sydney | CIDM-PH et al. |
| EPI_ISL_2894137 | Southern. IML Pathology | NSW Health Pathology - Institute of Clinical Pathology and Medical Research; Westmead Hospital; University of Sydney | CIDM-PH et al. |
| EPI_ISL_2894138 | Histopath | NSW Health Pathology - Institute of Clinical Pathology and Medical Research; Westmead Hospital; University of Sydney | CIDM-PH et al. |
| EPI_ISL_2894139, EPI_ISL_2894140 | Laverty Pathology | NSW Health Pathology - Institute of Clinical Pathology and Medical Research; Westmead Hospital; University of Sydney | CIDM-PH et al. |
| EPI_ISL_2894141, | Sydney South West Pathology Service (SSWPS) - Liverpool | NSW Health Pathology - Institute of Clinical Pathology and Medical | CIDM-PH et al. |

| | | | |
|---|---|---|---|
| EPI_ISL_2894142 | Hospital - NSW Health Pathology | Research; Westmead Hospital; University of Sydney | |
| EPI_ISL_2894143, EPI_ISL_2894144 | Histopath | NSW Health Pathology - Institute of Clinical Pathology and Medical Research; Westmead Hospital; University of Sydney | CIDM-PH et al. |
| EPI_ISL_2894145, EPI_ISL_2894146 | Laverty Pathology | NSW Health Pathology - Institute of Clinical Pathology and Medical Research; Westmead Hospital; University of Sydney | CIDM-PH et al. |
| EPI_ISL_2894147 | Medlab Pathology | NSW Health Pathology - Institute of Clinical Pathology and Medical Research; Westmead Hospital; University of Sydney | CIDM-PH et al. |
| EPI_ISL_2894148 | Pathology West - NSW Health Pathology | NSW Health Pathology - Institute of Clinical Pathology and Medical Research; Westmead Hospital; University of Sydney | CIDM-PH et al. |
| EPI_ISL_2894149, EPI_ISL_2894150 | Australian Clinical Labs (formerly Healthscope Pathology) | NSW Health Pathology - Institute of Clinical Pathology and Medical Research; Westmead Hospital; University of Sydney | CIDM-PH et al. |
| EPI_ISL_2894151 | Pathology West - NSW Health Pathology | NSW Health Pathology - Institute of Clinical Pathology and Medical Research; Westmead Hospital; University of Sydney | CIDM-PH et al. |
| EPI_ISL_2894152, EPI_ISL_2894153, EPI_ISL_2894159, EPI_ISL_2894160, EPI_ISL_2894166, EPI_ISL_2894167, EPI_ISL_2894171, EPI_ISL_2894173, EPI_ISL_2894179, EPI_ISL_2894184, EPI_ISL_2894186, EPI_ISL_2894192, EPI_ISL_2894194, EPI_ISL_2894201, EPI_ISL_2894209, EPI_ISL_2894211, EPI_ISL_2894216, EPI_ISL_2894217, EPI_ISL_2894218, EPI_ISL_2894224 | | | |
| see above | Salud Digna | Instituto Nacional de Medicina Genomica | Cedro-Tanda A, Hidalgo-Miranda A, Mendoza-Vargas A, Reyes-Grajeda JP, Abraham Campos-Romero, Moreno-Camacho José Luis, Rodriguez-Gallegos Jorge, Luna-Ruiz Marco, Gonzalez-Barrera D, Rangel-DeLeon D, Munguia-Garza P, Ramirez-Vega O, Escobar-Arrazola, M, Herrera-Montalvo LA. |
| EPI_ISL_2894238, EPI_ISL_2894239, EPI_ISL_2894240, EPI_ISL_2894241, EPI_ISL_2894242, EPI_ISL_2894243, EPI_ISL_2894244, EPI_ISL_2894245, EPI_ISL_2894246, EPI_ISL_2894247, EPI_ISL_2894248, EPI_ISL_2894249, EPI_ISL_2894250, EPI_ISL_2894251, EPI_ISL_2894252, EPI_ISL_2894253, EPI_ISL_2894254, EPI_ISL_2894256, EPI_ISL_2894257, EPI_ISL_2894258, EPI_ISL_2894259, EPI_ISL_2894260, EPI_ISL_2894262, EPI_ISL_2894263 | | | |
| see above | Instituto Nacional de Medicina Genomica | Instituto Nacional de Medicina Genomica | Hidalgo-Miranda A, Cedro-Tanda A, Mendoza-Vargas A, Reyes-Grajeda JP, Cisneros- Villanueva M, Gonzalez-Barrera D, Rangel-DeLeon D, Munguia-Garza P, Ramirez-Vega O, Escobar-Arrazola, M, Herrera-Montalvo LA. |
| EPI_ISL_2894264 | Salud Digna | Instituto Nacional de Medicina Genomica | Cedro-Tanda A, Hidalgo-Miranda A, Mendoza-Vargas A, Reyes-Grajeda JP, Abraham Campos-Romero, Moreno-Camacho José Luis, Rodriguez-Gallegos Jorge, Luna-Ruiz Marco, Gonzalez-Barrera D, Rangel-DeLeon D, Munguia-Garza P, Ramirez-Vega O, Escobar-Arrazola, M, Herrera-Montalvo LA. |
| EPI_ISL_2894266, EPI_ISL_2894267, EPI_ISL_2894270, EPI_ISL_2894271, EPI_ISL_2894272, EPI_ISL_2894275, EPI_ISL_2894276, EPI_ISL_2894277, EPI_ISL_2894278, EPI_ISL_2894279, EPI_ISL_2894280, EPI_ISL_2894281, EPI_ISL_2894282, EPI_ISL_2894283, EPI_ISL_2894285, EPI_ISL_2894286, EPI_ISL_2894287, EPI_ISL_2894288, EPI_ISL_2894289, EPI_ISL_2894290, EPI_ISL_2894291, EPI_ISL_2894292, EPI_ISL_2894293, EPI_ISL_2894294, EPI_ISL_2894295, EPI_ISL_2894296, EPI_ISL_2894297, EPI_ISL_2894298, EPI_ISL_2894300, EPI_ISL_2894301, EPI_ISL_2894302, EPI_ISL_2894303, EPI_ISL_2894305, EPI_ISL_2894307, EPI_ISL_2894308, EPI_ISL_2894309, EPI_ISL_2894310, EPI_ISL_2894313, EPI_ISL_2894314, EPI_ISL_2894315, EPI_ISL_2894316, EPI_ISL_2894317, EPI_ISL_2894318, EPI_ISL_2894319, EPI_ISL_2894320, EPI_ISL_2894321, EPI_ISL_2894322, EPI_ISL_2894323, EPI_ISL_2894325, EPI_ISL_2894327, EPI_ISL_2894328, EPI_ISL_2894330 | | | |
| see above | Instituto Nacional de Medicina Genomica | Instituto Nacional de Medicina Genomica | Hidalgo-Miranda A, Cedro-Tanda A, Mendoza-Vargas A, Reyes-Grajeda JP, Cisneros- Villanueva M, Gonzalez-Barrera D, Rangel-DeLeon D, Munguia-Garza P, Ramirez-Vega O, Escobar-Arrazola, M, Herrera-Montalvo LA. |
| EPI_ISL_2894371 | Salud Digna | Instituto Nacional de Medicina Genomica | Cedro-Tanda A, Hidalgo-Miranda A, Mendoza-Vargas A, Reyes-Grajeda JP, Abraham Campos-Romero, Moreno-Camacho José Luis, Rodriguez-Gallegos Jorge, Luna-Ruiz Marco, Gonzalez-Barrera D, Rangel-DeLeon D, Munguia-Garza P, Ramirez-Vega O, Escobar-Arrazola, M, Herrera-Montalvo LA. |
| EPI_ISL_2894395, EPI_ISL_2894396, EPI_ISL_2894397, EPI_ISL_2894398, EPI_ISL_2894399, EPI_ISL_2894400, EPI_ISL_2894401, EPI_ISL_2894402, EPI_ISL_2894403, EPI_ISL_2894404, EPI_ISL_2894405 | | | |
| see above | Instituto Nacional de Medicina Genomica | Instituto Nacional de Medicina Genomica | Hidalgo-Miranda A, Cedro-Tanda A, Mendoza-Vargas A, Reyes-Grajeda JP, Cisneros- Villanueva M, Gonzalez-Barrera D, Rangel-DeLeon D, Munguia-Garza P, Ramirez-Vega O, Escobar-Arrazola, M, Herrera-Montalvo LA. |
| EPI_ISL_2894406, EPI_ISL_2894407, EPI_ISL_2894408, EPI_ISL_2894409, EPI_ISL_2894411, EPI_ISL_2894412, EPI_ISL_2894413, EPI_ISL_2894415 | Salud Digna | Instituto Nacional de Medicina Genomica | Cedro-Tanda A, Hidalgo-Miranda A, Mendoza-Vargas A, Reyes-Grajeda JP, Abraham Campos-Romero, Moreno-Camacho José Luis, Rodriguez-Gallegos Jorge, Luna-Ruiz Marco, Gonzalez-Barrera D, Rangel-DeLeon D, Munguia-Garza P, Ramirez-Vega O, Escobar-Arrazola, M, Herrera-Montalvo LA. |
| EPI_ISL_2894423, EPI_ISL_2894425, EPI_ISL_2894428, EPI_ISL_2894436, EPI_ISL_2894439, EPI_ISL_2894448 | Institute of Virology, Biomedical Research Center of the Slovak Academy of Sciences, Bratislava | Faculty of Natural Sciences, Comenius University, Bratislava | Brona Brejova, Viktoria Cabanova, Kristina Borsova, Viktoria Hodorova, Sabina Fumacova Havlikova, Juraj Kopacek, Martina Lickova, Lubomira Lukacikova, Martina Nebohacova, Monika Slavikova, Tomas Vinar, Jozef Nosek, Boris Klempa |
| EPI_ISL_2894453 | Institute of Virology, Biomedical Research Center of the Slovak Academy of Sciences, Bratislava | Faculty of Natural Sciences, Comenius University, Bratislava | Kristina Borsova, Viktoria Cabanova, Brona Brejova, Viktoria Hodorova, Sabina Fumacova Havlikova, Juraj Kopacek, Martina Lickova, Lubomira Lukacikova, Martina Nebohacova, Monika Slavikova, Tomas Vinar, Boris Klempa, Jozef Nosek |
| EPI_ISL_2894456, EPI_ISL_2894457 | Institute of Virology, Biomedical Research Center of the Slovak Academy of Sciences, Bratislava | Faculty of Natural Sciences, Comenius University, Bratislava | Viktoria Cabanova, Kristina Borsova, Brona Brejova, Viktoria Hodorova, Sabina Fumacova Havlikova, Juraj Kopacek, Martina Lickova, Lubomira Lukacikova, Martina Nebohacova, Monika Slavikova, Tomas Vinar, Jozef Nosek, Boris Klempa |
| EPI_ISL_2894459 | Institute of Virology, Biomedical Research Center of the Slovak Academy of Sciences, Bratislava | Faculty of Natural Sciences, Comenius University, Bratislava | Brona Brejova, Viktoria Cabanova, Kristina Borsova, Viktoria Hodorova, Sabina Fumacova Havlikova, Juraj Kopacek, Martina Lickova, Lubomira Lukacikova, Martina Nebohacova, Monika Slavikova, Tomas Vinar, Jozef Nosek, Boris Klempa |
| EPI_ISL_2894463 | Institute of Virology, Biomedical Research Center of the Slovak Academy of Sciences, Bratislava | Faculty of Natural Sciences, Comenius University, Bratislava | Kristina Borsova, Viktoria Cabanova, Brona Brejova, Viktoria Hodorova, Sabina Fumacova Havlikova, Juraj Kopacek, Martina Lickova, Lubomira Lukacikova, Martina Nebohacova, Monika Slavikova, Tomas Vinar, Boris Klempa, Jozef Nosek |
| EPI_ISL_2894465 | Institute of Virology, Biomedical Research Center of the Slovak Academy of Sciences, Bratislava | Faculty of Natural Sciences, Comenius University, Bratislava | Brona Brejova, Viktoria Cabanova, Kristina Borsova, Viktoria Hodorova, Sabina Fumacova Havlikova, Juraj Kopacek, Martina Lickova, Lubomira Lukacikova, Martina Nebohacova, Monika Slavikova, Tomas Vinar, Jozef Nosek, Boris Klempa |
| EPI_ISL_2894466 | Institute of Virology, Biomedical Research Center of the Slovak Academy of Sciences, Bratislava | Faculty of Natural Sciences, Comenius University, Bratislava | Kristina Borsova, Viktoria Cabanova, Brona Brejova, Viktoria Hodorova, Sabina Fumacova Havlikova, Juraj Kopacek, Martina Lickova, Lubomira Lukacikova, Martina Nebohacova, Monika Slavikova, Tomas Vinar, Boris Klempa, Jozef Nosek |
| EPI_ISL_2894475 | Institute of Virology, Biomedical Research Center of the Slovak Academy of Sciences, Bratislava | Faculty of Natural Sciences, Comenius University, Bratislava | Brona Brejova, Viktoria Cabanova, Kristina Borsova, Viktoria Hodorova, Sabina Fumacova Havlikova, Juraj Kopacek, Martina Lickova, Lubomira Lukacikova, Martina Nebohacova, Monika Slavikova, Tomas Vinar, Jozef Nosek, Boris Klempa |
| EPI_ISL_2894476 | Institute of Virology, Biomedical Research Center of the Slovak Academy of Sciences, Bratislava | Faculty of Natural Sciences, Comenius University, Bratislava | Kristina Borsova, Viktoria Cabanova, Brona Brejova, Viktoria Hodorova, Sabina Fumacova Havlikova, Juraj Kopacek, Martina Lickova, Lubomira Lukacikova, Martina Nebohacova, Monika Slavikova, Tomas Vinar, Boris Klempa, Jozef Nosek |
| EPI_ISL_2894477 | Institute of Virology, Biomedical Research Center of the Slovak Academy of Sciences, Bratislava | Faculty of Natural Sciences, Comenius University, Bratislava | Brona Brejova, Viktoria Cabanova, Kristina Borsova, Viktoria Hodorova, Sabina Fumacova Havlikova, Juraj Kopacek, Martina Lickova, Lubomira Lukacikova, Martina Nebohacova, Monika Slavikova, Tomas Vinar, Jozef Nosek, Boris Klempa |
| EPI_ISL_2894484 | Institute of Virology, Biomedical Research Center of the Slovak Academy of Sciences, Bratislava | Faculty of Natural Sciences, Comenius University, Bratislava | Viktoria Cabanova, Kristina Borsova, Brona Brejova, Viktoria Hodorova, Sabina Fumacova Havlikova, Juraj Kopacek, Martina Lickova, Lubomira Lukacikova, Martina Nebohacova, Monika Slavikova, Tomas Vinar, Jozef Nosek, Boris Klempa |
| EPI_ISL_2894486 | Institute of Virology, Biomedical Research Center of the Slovak Academy of Sciences, Bratislava | Faculty of Natural Sciences, Comenius University, Bratislava | Brona Brejova, Viktoria Cabanova, Kristina Borsova, Viktoria Hodorova, Sabina Fumacova Havlikova, Juraj Kopacek, Martina Lickova, Lubomira Lukacikova, Martina Nebohacova, Monika Slavikova, Tomas Vinar, Jozef Nosek, Boris Klempa |
| EPI_ISL_2894493, EPI_ISL_2894494, EPI_ISL_2894495, EPI_ISL_2894496, EPI_ISL_2894497, EPI_ISL_2894498, EPI_ISL_2894499, EPI_ISL_2894500, EPI_ISL_2894501 | Wyoming Public Health Laboratory | Wyoming Public Health Laboratory | Jim Mildenberger, Taylor Fearing, Channing Weber, Ashley Norberg, Chayse Rowley, Marley Goetz, Brian Dominguez, Elliot Thomasson, Sam Britz, Cari Sloma, Robert Petit, and Rob Christensen |
| EPI_ISL_2894516 | Nucleic Acid Testing, National Reference Laboratory | GIGA Medical Genomics | Yvan Butera, Keith Durkin, Maria Artesi, Bouchra Boujemla, Robert Rutayisire, Reuben Sindayiheba, Patrick Tuyisenge, Esperence Umumararungu, Sébastien Bontems, Marie-Pierre Hayette, Nathalie Renotte, Swaibu Gatare, Jacob Souopgui, Sabin Nsanzimana, Vincent Bours, Léon Mutesa |
| EPI_ISL_2894517, EPI_ISL_2894518, EPI_ISL_2894519, EPI_ISL_2894520, EPI_ISL_2894521, EPI_ISL_2894522, EPI_ISL_2894523, EPI_ISL_2894524, EPI_ISL_2894525, EPI_ISL_2894526, EPI_ISL_2894527, EPI_ISL_2894528, EPI_ISL_2894532, EPI_ISL_2894533, EPI_ISL_2894534, EPI_ISL_2894535, EPI_ISL_2894536, EPI_ISL_2894537, EPI_ISL_2894538, EPI_ISL_2894539, EPI_ISL_2894540, EPI_ISL_2894541, EPI_ISL_2894542, EPI_ISL_2894543, EPI_ISL_2894544, EPI_ISL_2894545, EPI_ISL_2894546 | | | |
| see above | Kansas Health and Environmental Lab | Kansas Health and Environmental Lab | Katherine Wiggins, Mike Grose, Jonathan Barnell, Ben Olsen, and Phil Adam |
| EPI_ISL_2894549, EPI_ISL_2894550 | Laboratorios Delia Barraza | Laboratorio nacional-LANIIA-CIAD | Consorcio Mexicano de Vigilancia Genómica (CoViGen-Mex). Authors (in alphabetical order): Julio Elias Alvarado-Yaah, Carlos F. Arias, Santiago Ávila-Ríos, Eduardo Becerril-Vargas, Víctor Hugo Borja-Aburto, Celia Boukadida, Cristóbal Cháidez-Quiróz, Juan Bautista Chale-Dzul, Ricardo Ciria Merce, Andreu Comas-García, Célida Duque Molina, Julissa Enciso-Ibarra, José Antonio Enciso-Moreno, Gloria Elena Espinosa-Ayala, Fernando Fontove-Herrera, Daniel Fregoso-Rueda, Victor Eduardo García-Arias, Alejandra García-Gasca, Bruno Gómez-Gil, Jean Pierre González, Irvin González-López, Concepción Grajales-Muñiz, Ricardo Grande, Rosa María Gutiérrez Rios, Jesús Hernández, Alejandra Hernández-Terán, Alfredo Herrera-Estrella, Carla Ivón Herrera-Najera, Pavel Isa, Verónica Mata-Haro, Daniel Lira Morales, Susana Lopez, Antonio Loza Román, Brenda Irasema Maldonado-Meza, Bernardo Martínez-Miguel, José Arturo Martínez-Orozco, Célida Martínez-Rodríguez, Margarita Matías-Florentino, Fidencio Mejía-Nepomuceno, María Guadalupe de Jesús Mireles-Rivera, Gloria María Molina-Salinas, Hector Montoya-Fuentes, , Mario Mújica-Sánchez, José Esteban Muñoz-Medina, José de Jesús Nuñez-Contreras, Alicia Ocaña-Mondragón, Luis Alberto Ochoa-Carrera, Hector Esteban Paz-Juárez, Marissa Perez-Garcia, Francisco Pulido, Helen Haydee Fernanda Ramírez-Plascencia, Jorge Salas-Hernández, Angel Gustavo Salas-Lais, Alejandro Sánchez-Flores, Clara Esperanza Santacruz-Tinoco, María Guadalupe Santiago-Mauricio, Selene Zárate, Nelly Sélem-Mojica, Blanca Taboada, Gloria Vazquez, Joel Armando Vázquez-Pérez. Laboratorios Delia Barraza: María Delia Barraza Sámano, Claudia Soto Félix, Melissa García Angulo. |
| EPI_ISL_2894553 | Hospital Margarita Maza de Juárez | Microbial Genomics Laboratory | Consorcio Mexicano de Vigilancia Genómica (CoViGen-Mex). Authors (in alphabetical order): Julio Elias Alvarado-Yaah, Carlos F. Arias, Santiago Ávila-Ríos, Eduardo Becerril-Vargas, Víctor Hugo Borja-Aburto, Celia Boukadida, Cristóbal Cháidez-Quiróz, Juan Bautista Chale-Dzul, Ricardo Ciria Merce, Andreu Comas-García, Célida Duque Molina, Julissa Enciso-Ibarra, José Antonio Enciso-Moreno, Gloria Elena Espinosa-Ayala, Fernando Fontove-Herrera, Daniel Fregoso-Rueda, Victor Eduardo García-Arias, Alejandra García-Gasca, Bruno Gómez-Gil, Jean Pierre González, Irvin González-López, Concepción Grajales-Muñiz, Ricardo Grande, Rosa María Gutiérrez Rios, Jesús Hernández, Alejandra Hernández-Terán, Alfredo Herrera-Estrella, Carla Ivón Herrera-Najera, Pavel Isa, Verónica Mata-Haro, Daniel Lira Morales, Susana Lopez, Antonio Loza Román, Brenda Irasema Maldonado-Meza, Bernardo Martínez-Miguel, José Arturo Martínez-Orozco, Célida Martínez-Rodríguez, Margarita Matías-Florentino, Fidencio Mejía-Nepomuceno, María Guadalupe de Jesús Mireles-Rivera, Gloria María Molina-Salinas, Hector Montoya-Fuentes, , Mario Mújica-Sánchez, José Esteban Muñoz-Medina, José de Jesús Nuñez-Contreras, Alicia Ocaña-Mondragón, Luis Alberto Ochoa-Carrera, Hector Esteban Paz-Juárez, Marissa Perez-Garcia, Francisco Pulido, Helen Haydee Fernanda Ramírez-Plascencia, Jorge Salas-Hernández, Angel Gustavo Salas-Lais, Alejandro Sánchez-Flores, Clara Esperanza Santacruz-Tinoco, María Guadalupe Santiago-Mauricio, Selene Zárate, Nelly Sélem-Mojica, Blanca Taboada, Gloria Vazquez, Joel Armando Vázquez-Pérez. |

| | | | |
|---|---|---|---|
| EPI_ISL_2894563 | Hospital Sharp | Microbial Genomics Laboratory | Consorcio Mexicano de Vigilancia Genómica (CoViGen-Mex). Authors (in alphabetical order): Julio Elias Alvarado-Yaah, Carlos F. Arias, Santiago Ávila-Ríos, Eduardo Becerril-Vargas, Víctor Hugo Borja-Aburto, Celia Boukadida, Cristóbal Cháidez-Quiróz, Juan Bautista Chale-Dzul, Ricardo Ciria Merce, Andreu Comas-García, Célida Duque Molina, Julissa Enciso-Ibarra, José Antonio Enciso-Moreno, Gloria Elena Espinosa-Ayala, Fernando Fontove-Herrera, Daniel Fregoso-Rueda, Victor Eduardo García-Arias, Alejandra García-Gasca, Bruno Gómez-Gil, Jean Pierre González, Irvin González-López, Concepción Grajales-Muñiz, Ricardo Grande, Rosa María Gutiérrez Rios, Jesús Hernández, Alejandra Hernández-Terán, Alfredo Herrera-Estrella, Carla Ivón Herrera-Najera, Pavel Isa, Verónica Mata-Haro, Daniel Lira Morales, Susana Lopez, Antonio Loza Román, Brenda Irasema Maldonado-Meza, Bernardo Martínez-Miguel, José Arturo Martínez-Orozco, Célida Martínez-Rodríguez, Margarita Matías-Florentino, Fidencio Mejía-Nepomuceno, María Guadalupe de Jesús Mireles-Rivera, Gloria María Molina-Salinas, Hector Montoya-Fuentes, , Mario Mújica-Sánchez, José Esteban Muñoz-Medina, José de Jesús Nuñez-Contreras, Alicia Ocaña-Mondragón, Luis Alberto Ochoa-Carrera, Hector Esteban Paz-Juárez, Marissa Perez-Garcia, Francisco Pulido, Helen Haydee Fernanda Ramírez-Plascencia, Jorge Salas-Hernández, Angel Gustavo Salas-Lais, Alejandro Sánchez-Flores, Clara Esperanza Santacruz-Tinoco, María Guadalupe Santiago-Mauricio, Selene Zárate, Nelly Sélem-Mojica, Blanca Taboada, Gloria Vazquez, Joel Armando Vázquez-Pérez. |
| EPI_ISL_2894572 | Pomona Valley Hospital Medical Center | Los Angeles County Public Health Laboratories | P. Hemarajata et al. |
| EPI_ISL_2894579, EPI_ISL_2894580, EPI_ISL_2894581 | Olive View Medical Center | Los Angeles County Public Health Laboratories | P. Hemarajata et al. |
| EPI_ISL_2894591, EPI_ISL_2894592 | Palmdale Regional Medical Center | Los Angeles County Public Health Laboratories | P. Hemarajata et al. |
| EPI_ISL_2894609, EPI_ISL_2894610, EPI_ISL_2894611, EPI_ISL_2894612 | Wichita State University - Molecular Diagnostics Lab | Kansas Health and Environmental Lab | Katherine Wiggins, Mike Grose, Jonathan Barnell, Ben Olsen, and Phil Adam |
| EPI_ISL_2894613, EPI_ISL_2894614, EPI_ISL_2894615, EPI_ISL_2894616, EPI_ISL_2894617, EPI_ISL_2894618, EPI_ISL_2894619, EPI_ISL_2894620, EPI_ISL_2894621, EPI_ISL_2894622, EPI_ISL_2894623, EPI_ISL_2894624, EPI_ISL_2894625, EPI_ISL_2894626, EPI_ISL_2894627, EPI_ISL_2894628, EPI_ISL_2894629, EPI_ISL_2894630 | | | |
| see above | Kansas Health and Environmental Lab | Kansas Health and Environmental Lab | Katherine Wiggins, Mike Grose, Jonathan Barnell, Ben Olsen, and Phil Adam |
| EPI_ISL_2894958, EPI_ISL_2894965 | Genetica Molecular and Subdepartamento de Virologia ISP Chile | Instituto de Salud Publica de Chile | Karen Orostica, Constanza Campano, Barbara Parra, Loredana Arata, Gisselle Barra, Patricia Bustos, Rodrigo Fasce, Javier Tognarelli, Andres Castillo, Soledad Ulloa, Jorge Fernandez |
| EPI_ISL_2894974, EPI_ISL_2894977, EPI_ISL_2894982, EPI_ISL_2894983 | Department of Virology | Department of Virology | Massab Umair, Aamer Ikram, Muhammad Salman, Nazish Badar, Zaira Rehman, Muhammad Ammar, Syed Adnan Haider |
| EPI_ISL_2894997, EPI_ISL_2894998, EPI_ISL_2894999, EPI_ISL_2895000 | A. Krumbholz, Labor Dr. Krause und Kollegen MVZ GmbH, Kiel | Charité Universitätsmedizin Berlin, Institute of Virology | Victor M Corman, Barbara Mühlemann, Julia Schneider, Jörn Beheim-Schwarzbach, Tobias Bleicker, Julia Tesch, Talitha Veith, Terry Jones, Christian Drosten |
| EPI_ISL_2895001, EPI_ISL_2895002 | University College Hospital, Ibadan | African Centre of Excellence for Genomics of Infectious Diseases (ACEGID), Redeemer's University, Nigeria | Olawoye, I.B., Oluniyi, P.E., Eromon, P.E., Oguzie, J.U., Kayode, A.T., Uwanibe, J.N., Ugwu, C.A., Akano, K.O., Ajogbasile, F.V., Abechi, P.S., Olumade, T.J., Nosamiefan, I., Fowotade, A., Folarin, O., Happi, C.T. |
| EPI_ISL_2895004, EPI_ISL_2895005, EPI_ISL_2895006, EPI_ISL_2895007, EPI_ISL_2895008, EPI_ISL_2895009, EPI_ISL_2895010, EPI_ISL_2895011, EPI_ISL_2895013, EPI_ISL_2895014, EPI_ISL_2895015, EPI_ISL_2895016, EPI_ISL_2895017, EPI_ISL_2895018, EPI_ISL_2895019, EPI_ISL_2895020, EPI_ISL_2895021, EPI_ISL_2895023, EPI_ISL_2895025, EPI_ISL_2895026, EPI_ISL_2895027, EPI_ISL_2895028, EPI_ISL_2895029, EPI_ISL_2895031, EPI_ISL_2895033, EPI_ISL_2895034, EPI_ISL_2895035, EPI_ISL_2895036, EPI_ISL_2895038, EPI_ISL_2895039, EPI_ISL_2895040, EPI_ISL_2895041, EPI_ISL_2895042, EPI_ISL_2895043, EPI_ISL_2895044, EPI_ISL_2895045, EPI_ISL_2895046, EPI_ISL_2895047, EPI_ISL_2895048, EPI_ISL_2895049, EPI_ISL_2895050, EPI_ISL_2895051, EPI_ISL_2895052, EPI_ISL_2895053, EPI_ISL_2895054, EPI_ISL_2895055, EPI_ISL_2895058, EPI_ISL_2895059, EPI_ISL_2895060, EPI_ISL_2895061, EPI_ISL_2895062, EPI_ISL_2895063, EPI_ISL_2895064, EPI_ISL_2895065, EPI_ISL_2895066, EPI_ISL_2895067 | | | |
| see above | UW Virology Lab | UW Virology Lab | Pavitra Roychoudhury, Hong Xie, Lasata Shrestha, Tien V. Nguyen, Shah Mohamed Bakhash, Michelle Lin, Noah R. Baker, Ricardo Perez, Sean Ellis, Nathan Breit, Robert J. Livingston, Meei-Li Huang, Keith R Jerome, Patrick Mathias, Alexander Greninger |
| EPI_ISL_2895068, EPI_ISL_2895069, EPI_ISL_2895070, EPI_ISL_2895071, EPI_ISL_2895072, EPI_ISL_2895073, EPI_ISL_2895074, EPI_ISL_2895075, EPI_ISL_2895076, EPI_ISL_2895077, EPI_ISL_2895078, EPI_ISL_2895079, EPI_ISL_2895080, EPI_ISL_2895081, EPI_ISL_2895082, EPI_ISL_2895083 | | | |
| see above | H Fernando Fonseca | Instituto Nacional de Saude (INSA) | Borges et al |
| EPI_ISL_2895084, EPI_ISL_2895085, EPI_ISL_2895086, EPI_ISL_2895087, EPI_ISL_2895088, EPI_ISL_2895089, EPI_ISL_2895090, EPI_ISL_2895091, EPI_ISL_2895092, EPI_ISL_2895093, EPI_ISL_2895094, EPI_ISL_2895095, EPI_ISL_2895096, EPI_ISL_2895097, EPI_ISL_2895098, EPI_ISL_2895099, EPI_ISL_2895100, EPI_ISL_2895101 | | | |
| see above | ULS Norte Alentejano | Instituto Nacional de Saude (INSA) | Borges et al |
| EPI_ISL_2895103, EPI_ISL_2895104, EPI_ISL_2895105, EPI_ISL_2895106, EPI_ISL_2895107, EPI_ISL_2895108, EPI_ISL_2895109, EPI_ISL_2895110, EPI_ISL_2895111, EPI_ISL_2895112, EPI_ISL_2895113, EPI_ISL_2895115, EPI_ISL_2895116, EPI_ISL_2895117, EPI_ISL_2895118, EPI_ISL_2895119, EPI_ISL_2895120, EPI_ISL_2895121, EPI_ISL_2895122, EPI_ISL_2895123, EPI_ISL_2895124, EPI_ISL_2895125, EPI_ISL_2895126, EPI_ISL_2895128, EPI_ISL_2895129, EPI_ISL_2895130, EPI_ISL_2895131, EPI_ISL_2895132, EPI_ISL_2895133, EPI_ISL_2895134 | | | |
| see above | ABC Algarve | Instituto Nacional de Saude (INSA) | Borges et al |
| EPI_ISL_2895135, EPI_ISL_2895136, EPI_ISL_2895137, EPI_ISL_2895138, EPI_ISL_2895139, EPI_ISL_2895140, EPI_ISL_2895141, EPI_ISL_2895142, EPI_ISL_2895143, EPI_ISL_2895144, EPI_ISL_2895145, EPI_ISL_2895147, EPI_ISL_2895148, EPI_ISL_2895149, EPI_ISL_2895150, EPI_ISL_2895151, EPI_ISL_2895152, EPI_ISL_2895153, EPI_ISL_2895154, EPI_ISL_2895155, EPI_ISL_2895156, EPI_ISL_2895157, EPI_ISL_2895158 | | | |
| see above | ARS Algarve - Laboratorio Laura Ayres | Instituto Nacional de Saude (INSA) | Borges et al |
| EPI_ISL_2895159, EPI_ISL_2895160, EPI_ISL_2895161, EPI_ISL_2895162, EPI_ISL_2895164, EPI_ISL_2895165, EPI_ISL_2895166, EPI_ISL_2895167, EPI_ISL_2895170, EPI_ISL_2895171, EPI_ISL_2895172 | | | |
| see above | ULS Castelo Branco | Instituto Nacional de Saude (INSA) | Borges et al |
| EPI_ISL_2895173, EPI_ISL_2895174, EPI_ISL_2895175, EPI_ISL_2895176, EPI_ISL_2895178, EPI_ISL_2895181, EPI_ISL_2895182, EPI_ISL_2895183, EPI_ISL_2895185, EPI_ISL_2895186 | LAC UCoimbra | Instituto Nacional de Saude (INSA) | Borges et al |
| EPI_ISL_2895189, EPI_ISL_2895190, EPI_ISL_2895191, EPI_ISL_2895192, EPI_ISL_2895193, EPI_ISL_2895194, EPI_ISL_2895195, EPI_ISL_2895196, EPI_ISL_2895197, EPI_ISL_2895198, EPI_ISL_2895199, EPI_ISL_2895200, EPI_ISL_2895201, EPI_ISL_2895202, EPI_ISL_2895203, EPI_ISL_2895204, EPI_ISL_2895205, EPI_ISL_2895206 | | | |
| see above | H Garcia de Orta | Instituto Nacional de Saude (INSA) | Borges et al |
| EPI_ISL_2895207, EPI_ISL_2895208, EPI_ISL_2895209, EPI_ISL_2895210, EPI_ISL_2895211, EPI_ISL_2895212, EPI_ISL_2895213, EPI_ISL_2895214, EPI_ISL_2895215, EPI_ISL_2895216, EPI_ISL_2895217, EPI_ISL_2895218, EPI_ISL_2895219, EPI_ISL_2895220, EPI_ISL_2895221 | | | |
| see above | CHULN | Instituto Nacional de Saude (INSA) | Borges et al |
| EPI_ISL_2895222, EPI_ISL_2895223, EPI_ISL_2895224, EPI_ISL_2895225, EPI_ISL_2895226, EPI_ISL_2895227, EPI_ISL_2895228, EPI_ISL_2895229, EPI_ISL_2895230, EPI_ISL_2895231, EPI_ISL_2895232, EPI_ISL_2895233, EPI_ISL_2895234, EPI_ISL_2895235, EPI_ISL_2895236 | | | |
| see above | CHLC | Instituto Nacional de Saude (INSA) | Borges et al |
| EPI_ISL_2895238, EPI_ISL_2895239, EPI_ISL_2895240, EPI_ISL_2895242, EPI_ISL_2895243, EPI_ISL_2895244, EPI_ISL_2895245, EPI_ISL_2895246, EPI_ISL_2895247, EPI_ISL_2895248, EPI_ISL_2895249 | | | |
| see above | H Evora | Instituto Nacional de Saude (INSA) | Borges et al |
| EPI_ISL_2895250, EPI_ISL_2895251, EPI_ISL_2895252, EPI_ISL_2895253, EPI_ISL_2895254, EPI_ISL_2895255, EPI_ISL_2895256, EPI_ISL_2895257, EPI_ISL_2895258, EPI_ISL_2895259, EPI_ISL_2895260, EPI_ISL_2895261, EPI_ISL_2895262, EPI_ISL_2895263, EPI_ISL_2895264, EPI_ISL_2895265 | | | |
| see above | Labeto - CAB - Leiria | Instituto Nacional de Saude (INSA) | Borges et al |
| EPI_ISL_2895271, EPI_ISL_2895272 | H Braga | Instituto Nacional de Saude (INSA) | Borges et al |
| EPI_ISL_2895274, EPI_ISL_2895276, EPI_ISL_2895277, EPI_ISL_2895278, EPI_ISL_2895279, EPI_ISL_2895280 | CHEDV | Instituto Nacional de Saude (INSA) | Borges et al |
| EPI_ISL_2895281, EPI_ISL_2895282 | CH Tondela Viseu | Instituto Nacional de Saude (INSA) | Borges et al |
| EPI_ISL_2895283, EPI_ISL_2895284, EPI_ISL_2895285, EPI_ISL_2895286, EPI_ISL_2895287, EPI_ISL_2895288 | CHTMAD | Instituto Nacional de Saude (INSA) | Borges et al |
| EPI_ISL_2895290, EPI_ISL_2895291, EPI_ISL_2895292, EPI_ISL_2895293, EPI_ISL_2895295, EPI_ISL_2895297, EPI_ISL_2895298, EPI_ISL_2895299, EPI_ISL_2895300, EPI_ISL_2895301, EPI_ISL_2895302 | | | |
| see above | CH Porto - H Sto Antonio | Instituto Nacional de Saude (INSA) | Borges et al |
| EPI_ISL_2895303, EPI_ISL_2895304, EPI_ISL_2895305, EPI_ISL_2895306, EPI_ISL_2895307, EPI_ISL_2895308, EPI_ISL_2895309, EPI_ISL_2895310, EPI_ISL_2895311, EPI_ISL_2895312, EPI_ISL_2895313, EPI_ISL_2895314, EPI_ISL_2895315 | | | |
| see above | H Vila Franca Xira | Instituto Nacional de Saude (INSA) | Borges et al |
| EPI_ISL_2895316, EPI_ISL_2895317, EPI_ISL_2895318, | CH Tondela Viseu | Instituto Nacional de Saude (INSA) | Borges et al |

| | | | |
|---|---|---|---|
| EPI_ISL_2895319, EPI_ISL_2895320, EPI_ISL_2895321, EPI_ISL_2895322 | | | |
| EPI_ISL_2895324, EPI_ISL_2895325 | CH Setubal | Instituto Nacional de Saude (INSA) | Borges et al |
| EPI_ISL_2895326, EPI_ISL_2895327, EPI_ISL_2895328, EPI_ISL_2895329, EPI_ISL_2895330, EPI_ISL_2895331, EPI_ISL_2895332, EPI_ISL_2895333, EPI_ISL_2895334, EPI_ISL_2895335, EPI_ISL_2895336, EPI_ISL_2895337, EPI_ISL_2895339, EPI_ISL_2895340, EPI_ISL_2895341, EPI_ISL_2895342, EPI_ISL_2895343, EPI_ISL_2895344, EPI_ISL_2895345, EPI_ISL_2895346, EPI_ISL_2895347, EPI_ISL_2895348, EPI_ISL_2895349 | | | |
| see above | Avelab | Instituto Nacional de Saude (INSA) | Borges et al |
| EPI_ISL_2895351, EPI_ISL_2895355, EPI_ISL_2895356, EPI_ISL_2895357, EPI_ISL_2895358, EPI_ISL_2895359, EPI_ISL_2895360, EPI_ISL_2895361, EPI_ISL_2895362, EPI_ISL_2895363, EPI_ISL_2895365, EPI_ISL_2895366, EPI_ISL_2895367, EPI_ISL_2895369, EPI_ISL_2895370, EPI_ISL_2895371, EPI_ISL_2895374, EPI_ISL_2895375, EPI_ISL_2895379, EPI_ISL_2895380 | | | |
| see above | SEEBMO | Instituto Nacional de Saude (INSA) | Borges et al |
| EPI_ISL_2895381, EPI_ISL_2895382, EPI_ISL_2895383, EPI_ISL_2895385, EPI_ISL_2895386, EPI_ISL_2895387, EPI_ISL_2895388, EPI_ISL_2895390, EPI_ISL_2895391, EPI_ISL_2895393, EPI_ISL_2895394, EPI_ISL_2895395, EPI_ISL_2895396, EPI_ISL_2895397, EPI_ISL_2895398, EPI_ISL_2895399, EPI_ISL_2895400, EPI_ISL_2895401, EPI_ISL_2895402, EPI_ISL_2895403, EPI_ISL_2895404, EPI_ISL_2895405, EPI_ISL_2895410, EPI_ISL_2895411, EPI_ISL_2895412, EPI_ISL_2895413, EPI_ISL_2895414, EPI_ISL_2895415, EPI_ISL_2895416 | | | |
| see above | SESARAM | Instituto Nacional de Saude (INSA) | Borges et al |
| EPI_ISL_2895417, EPI_ISL_2895418, EPI_ISL_2895419 | ULS Guarda | Instituto Nacional de Saude (INSA) | Borges et al |
| EPI_ISL_2895421, EPI_ISL_2895426, EPI_ISL_2895427, EPI_ISL_2895428, EPI_ISL_2895429, EPI_ISL_2895430, EPI_ISL_2895431, EPI_ISL_2895434, EPI_ISL_2895435, EPI_ISL_2895436, EPI_ISL_2895437, EPI_ISL_2895439, EPI_ISL_2895440, EPI_ISL_2895441, EPI_ISL_2895443, EPI_ISL_2895444, EPI_ISL_2895445, EPI_ISL_2895446, EPI_ISL_2895447 | | | |
| see above | H Divino Espirito Santo - Ponta Delgada | Instituto Nacional de Saude (INSA) | Borges et al |
| EPI_ISL_2895449, EPI_ISL_2895453, EPI_ISL_2895454, EPI_ISL_2895455, EPI_ISL_2895456, EPI_ISL_2895457, EPI_ISL_2895458, EPI_ISL_2895459, EPI_ISL_2895460, EPI_ISL_2895461, EPI_ISL_2895465, EPI_ISL_2895468, EPI_ISL_2895469, EPI_ISL_2895470, EPI_ISL_2895471, EPI_ISL_2895472, EPI_ISL_2895473, EPI_ISL_2895474, EPI_ISL_2895475, EPI_ISL_2895477, EPI_ISL_2895479, EPI_ISL_2895480, EPI_ISL_2895481, EPI_ISL_2895482, EPI_ISL_2895484, EPI_ISL_2895485, EPI_ISL_2895486, EPI_ISL_2895487, EPI_ISL_2895488, EPI_ISL_2895490, EPI_ISL_2895492, EPI_ISL_2895494, EPI_ISL_2895495, EPI_ISL_2895496, EPI_ISL_2895497, EPI_ISL_2895498, EPI_ISL_2895499, EPI_ISL_2895500, EPI_ISL_2895501, EPI_ISL_2895502, EPI_ISL_2895503, EPI_ISL_2895504, EPI_ISL_2895505, EPI_ISL_2895506, EPI_ISL_2895507, EPI_ISL_2895508, EPI_ISL_2895509, EPI_ISL_2895510, EPI_ISL_2895511, EPI_ISL_2895512, EPI_ISL_2895513, EPI_ISL_2895514, EPI_ISL_2895515, EPI_ISL_2895516, EPI_ISL_2895517, EPI_ISL_2895518, EPI_ISL_2895519, EPI_ISL_2895520, EPI_ISL_2895521, EPI_ISL_2895522, EPI_ISL_2895523, EPI_ISL_2895524, EPI_ISL_2895525, EPI_ISL_2895526, EPI_ISL_2895527, EPI_ISL_2895528, EPI_ISL_2895529, EPI_ISL_2895530, EPI_ISL_2895531, EPI_ISL_2895532, EPI_ISL_2895533, EPI_ISL_2895535, EPI_ISL_2895536, EPI_ISL_2895537, EPI_ISL_2895538, EPI_ISL_2895539, EPI_ISL_2895540, EPI_ISL_2895541, EPI_ISL_2895542, EPI_ISL_2895543, EPI_ISL_2895544, EPI_ISL_2895545, EPI_ISL_2895546, EPI_ISL_2895547, EPI_ISL_2895548, EPI_ISL_2895549, EPI_ISL_2895550, EPI_ISL_2895552, EPI_ISL_2895553, EPI_ISL_2895554, EPI_ISL_2895556, EPI_ISL_2895557, EPI_ISL_2895558, EPI_ISL_2895559, EPI_ISL_2895560, EPI_ISL_2895561, EPI_ISL_2895562, EPI_ISL_2895563, EPI_ISL_2895565, EPI_ISL_2895566, EPI_ISL_2895567, EPI_ISL_2895568, EPI_ISL_2895569, EPI_ISL_2895570, EPI_ISL_2895571, EPI_ISL_2895572, EPI_ISL_2895573, EPI_ISL_2895574, EPI_ISL_2895575, EPI_ISL_2895576, EPI_ISL_2895577, EPI_ISL_2895578, EPI_ISL_2895579, EPI_ISL_2895580, EPI_ISL_2895581, EPI_ISL_2895582, EPI_ISL_2895583, EPI_ISL_2895584, EPI_ISL_2895585, EPI_ISL_2895586, EPI_ISL_2895587, EPI_ISL_2895588, EPI_ISL_2895589, EPI_ISL_2895590, EPI_ISL_2895592, EPI_ISL_2895593, EPI_ISL_2895594, EPI_ISL_2895595, EPI_ISL_2895596, EPI_ISL_2895597, EPI_ISL_2895598, EPI_ISL_2895599, EPI_ISL_2895600, EPI_ISL_2895601, EPI_ISL_2895602, EPI_ISL_2895603, EPI_ISL_2895604, EPI_ISL_2895605, EPI_ISL_2895606, EPI_ISL_2895607, EPI_ISL_2895608, EPI_ISL_2895609, EPI_ISL_2895610, EPI_ISL_2895611, EPI_ISL_2895612, EPI_ISL_2895613, EPI_ISL_2895614, EPI_ISL_2895615, EPI_ISL_2895616, EPI_ISL_2895617, EPI_ISL_2895618, EPI_ISL_2895619, EPI_ISL_2895620, EPI_ISL_2895621, EPI_ISL_2895622, EPI_ISL_2895623, EPI_ISL_2895624, EPI_ISL_2895626, EPI_ISL_2895627, EPI_ISL_2895628, EPI_ISL_2895629 | | | |
| see above | UNILABS | Instituto Nacional de Saude (INSA) | Borges et al |
| EPI_ISL_2895630, EPI_ISL_2895632, EPI_ISL_2895633, EPI_ISL_2895634, EPI_ISL_2895636, EPI_ISL_2895641, EPI_ISL_2895642, EPI_ISL_2895646, EPI_ISL_2895647, EPI_ISL_2895651, EPI_ISL_2895652, EPI_ISL_2895659 | | | |
| see above | National Platform bis UMONS/Jolimont | National Platform bis UMONS/Jolimont | François Dufrasne, Guillaume Bayon-Vicente, Florian Juszczak, Eric Tarantino, Gautier Detry, Ruddy Wattiez |
| EPI_ISL_2895665, EPI_ISL_2895666, EPI_ISL_2895667, EPI_ISL_2895668, EPI_ISL_2895669, EPI_ISL_2895670, EPI_ISL_2895671, EPI_ISL_2895672, EPI_ISL_2895673, EPI_ISL_2895674 | INSPI-CRN DE INFLUENZA Y OTROS VIRUS RESPIRATORIOS | NIC-INSPI | Alfredo Bruno , Maritza Olmedo, Michelle Páez, Jimmy Garcés, Johanna Laines, Lizbeth Patiño, Manuel Gonzalez, Domenica de Mora. |
| EPI_ISL_2895679, EPI_ISL_2895680, EPI_ISL_2895681, EPI_ISL_2895682, EPI_ISL_2895683, EPI_ISL_2895684, EPI_ISL_2895686, EPI_ISL_2895689, EPI_ISL_2895690, EPI_ISL_2895691, EPI_ISL_2895692, EPI_ISL_2895693, EPI_ISL_2895694, EPI_ISL_2895695, EPI_ISL_2895696, EPI_ISL_2895697, EPI_ISL_2895698, EPI_ISL_2895700, EPI_ISL_2895701, EPI_ISL_2895702, EPI_ISL_2895704, EPI_ISL_2895705, EPI_ISL_2895707, EPI_ISL_2895708, EPI_ISL_2895709, EPI_ISL_2895710, EPI_ISL_2895711, EPI_ISL_2895712, EPI_ISL_2895713, EPI_ISL_2895714, EPI_ISL_2895715, EPI_ISL_2895716 | | | |
| see above | RUSTENBURG LABORATORY | National Institute for Communicable Diseases of the National Health Laboratory Service | Amoako DG, Everatt J, Scheepers C, Mohale T, Ntuli N, Mahlangu B, Mnguni A, Ismail A, Bhiman JN |
| EPI_ISL_2895717, EPI_ISL_2895718, EPI_ISL_2895719, EPI_ISL_2895720, EPI_ISL_2895721, EPI_ISL_2895722, EPI_ISL_2895723, EPI_ISL_2895724, EPI_ISL_2895725, EPI_ISL_2895726, EPI_ISL_2895727, EPI_ISL_2895728, EPI_ISL_2895729, EPI_ISL_2895730, EPI_ISL_2895731, EPI_ISL_2895732, EPI_ISL_2895734, EPI_ISL_2895735, EPI_ISL_2895736, EPI_ISL_2895737, EPI_ISL_2895738, EPI_ISL_2895739, EPI_ISL_2895740, EPI_ISL_2895741, EPI_ISL_2895742, EPI_ISL_2895744, EPI_ISL_2895745, EPI_ISL_2895746, EPI_ISL_2895747 | | | |
| see above | PORT ELIZABETH LABORATORY | National Institute for Communicable Diseases of the National Health Laboratory Service | Amoako DG, Everatt J, Scheepers C, Mohale T, Ntuli N, Mahlangu B, Mnguni A, Ismail A, Bhiman JN |
| EPI_ISL_2895753, EPI_ISL_2895757, EPI_ISL_2895765 | MIDDELBURG LABORATORY | National Institute for Communicable Diseases of the National Health Laboratory Service | Amoako DG, Everatt J, Scheepers C, Mohale T, Ntuli N, Mahlangu B, Mnguni A, Ismail A, Bhiman JN |
| EPI_ISL_2895766, EPI_ISL_2895768, EPI_ISL_2895770, EPI_ISL_2895771 | KOPANONG LABORATORY | National Institute for Communicable Diseases of the National Health Laboratory Service | Amoako DG, Everatt J, Scheepers C, Mohale T, Ntuli N, Mahlangu B, Mnguni A, Ismail A, Bhiman JN |
| EPI_ISL_2895772, EPI_ISL_2895773, EPI_ISL_2895774, EPI_ISL_2895775, EPI_ISL_2895776, EPI_ISL_2895777, EPI_ISL_2895778, EPI_ISL_2895779, EPI_ISL_2895780, EPI_ISL_2895781, EPI_ISL_2895783, EPI_ISL_2895784, EPI_ISL_2895785, EPI_ISL_2895786, EPI_ISL_2895788, EPI_ISL_2895789, EPI_ISL_2895790, EPI_ISL_2895791, EPI_ISL_2895793, EPI_ISL_2895794, EPI_ISL_2895795, EPI_ISL_2895797 | | | |
| see above | HELEN JOSEPH LABORATORY | National Institute for Communicable Diseases of the National Health Laboratory Service | Amoako DG, Everatt J, Scheepers C, Mohale T, Ntuli N, Mahlangu B, Mnguni A, Ismail A, Bhiman JN |
| EPI_ISL_2895798 | PORT ELIZABETH LABORATORY | National Institute for Communicable Diseases of the National Health Laboratory Service | Amoako DG, Everatt J, Scheepers C, Mohale T, Ntuli N, Mahlangu B, Mnguni A, Ismail A, Bhiman JN |
| EPI_ISL_2896141 | OHSU-MM Lab | OHSU-MM Lab | Rebecca O'Gara |
| EPI_ISL_2896198 | RS Pusat Otak Nasional Prof dr. Mahar Mardjono | National Institute of Health Research and Development | Hana Apsari Pawestri, Subangkit, Kartika Dewi Puspa, Arie Ardiansyah Nugraha, Hartanti Dian Ikawati, Krisna Pangesti, Triyani Soekarso, Nelly Puspandari, Vivi Setiawaty |
| EPI_ISL_2896199, EPI_ISL_2896201 | Kantor Kesehatan Pelabuhan Kelas I Medan | National Institute of Health Research and Development | Hana Apsari Pawestri, Subangkit, Kartika Dewi Puspa, Arie Ardiansyah Nugraha, Hartanti Dian Ikawati, Krisna Pangesti, Triyani Soekarso, Nelly Puspandari, Vivi Setiawaty |
| EPI_ISL_2896214, EPI_ISL_2896215, EPI_ISL_2896216, EPI_ISL_2896217, EPI_ISL_2896218, EPI_ISL_2896219, EPI_ISL_2896220, EPI_ISL_2896221, EPI_ISL_2896222, EPI_ISL_2896223, EPI_ISL_2896224, EPI_ISL_2896225, EPI_ISL_2896226, EPI_ISL_2896227, EPI_ISL_2896228, EPI_ISL_2896229, EPI_ISL_2896230, EPI_ISL_2896231 | | | |
| see above | Department of Acute Infectious Diseases Control and Prevention, Yunnan Provincial Center for Disease Control and Prevention | Department of Acute Infectious Diseases Control and Prevention, Yunnan Provincial Center for Disease Control and Prevention | Meiling Zhang, Jienan Zhou, Zhaosheng Liu, Xiaoqing Fu |
| EPI_ISL_2896244, EPI_ISL_2896245, EPI_ISL_2896246, EPI_ISL_2896247, EPI_ISL_2896248, EPI_ISL_2896249 | Queensland Health Forensic and Scientific Services | Queensland Health Forensic and Scientific Services | Son Nguyen |
| EPI_ISL_2896288, EPI_ISL_2896290, EPI_ISL_2896291, EPI_ISL_2896293, EPI_ISL_2896294, EPI_ISL_2896295, EPI_ISL_2896296, EPI_ISL_2896298, EPI_ISL_2896299, EPI_ISL_2896301 | Department of Bacteria, Parasites and Fungi, Statens Serum Institut, Copenhagen, Denmark | Statens Serum Institut Bioinformatics and Microbial Genomics | Danish Covid-19 Genome Consortium |
| EPI_ISL_2896303 | Molekylær Medicinsk Afdeling, Aarhus University Hospital, Aarhus, Denmark | Statens Serum Institut Bioinformatics and Microbial Genomics | Danish Covid-19 Genome Consortium |
| EPI_ISL_2896305, EPI_ISL_2896306, EPI_ISL_2896307, EPI_ISL_2896308, EPI_ISL_2896309, EPI_ISL_2896310, EPI_ISL_2896311, EPI_ISL_2896312, EPI_ISL_2896313, EPI_ISL_2896314, EPI_ISL_2896315, EPI_ISL_2896316, EPI_ISL_2896317, EPI_ISL_2896318, EPI_ISL_2896319, EPI_ISL_2896320, EPI_ISL_2896321, EPI_ISL_2896322, EPI_ISL_2896323, EPI_ISL_2896324, EPI_ISL_2896325, EPI_ISL_2896326, EPI_ISL_2896327, EPI_ISL_2896328, EPI_ISL_2896329, EPI_ISL_2896330, EPI_ISL_2896332, EPI_ISL_2896333, EPI_ISL_2896334, EPI_ISL_2896336, EPI_ISL_2896337, EPI_ISL_2896338, EPI_ISL_2896339, EPI_ISL_2896340, EPI_ISL_2896341, EPI_ISL_2896343, EPI_ISL_2896344, EPI_ISL_2896345, EPI_ISL_2896346, EPI_ISL_2896347, EPI_ISL_2896348, EPI_ISL_2896349, EPI_ISL_2896350, EPI_ISL_2896351, EPI_ISL_2896352, EPI_ISL_2896354, EPI_ISL_2896356, EPI_ISL_2896357, EPI_ISL_2896358, EPI_ISL_2896359, EPI_ISL_2896361, EPI_ISL_2896362, EPI_ISL_2896363, EPI_ISL_2896364, EPI_ISL_2896365, EPI_ISL_2896366, EPI_ISL_2896367, EPI_ISL_2896368, EPI_ISL_2896369, EPI_ISL_2896370, EPI_ISL_2896371, EPI_ISL_2896372, EPI_ISL_2896373, EPI_ISL_2896374, EPI_ISL_2896375, EPI_ISL_2896376, EPI_ISL_2896378, EPI_ISL_2896380, EPI_ISL_2896381, EPI_ISL_2896382, EPI_ISL_2896383, EPI_ISL_2896386, EPI_ISL_2896388, EPI_ISL_2896389, EPI_ISL_2896390, EPI_ISL_2896391, EPI_ISL_2896392, EPI_ISL_2896394, EPI_ISL_2896395, EPI_ISL_2896396, EPI_ISL_2896397, EPI_ISL_2896398, EPI_ISL_2896399, EPI_ISL_2896400, EPI_ISL_2896401, EPI_ISL_2896402, EPI_ISL_2896403, EPI_ISL_2896404, EPI_ISL_2896405, EPI_ISL_2896406, EPI_ISL_2896407, EPI_ISL_2896408, EPI_ISL_2896409, EPI_ISL_2896410, EPI_ISL_2896411, EPI_ISL_2896412, EPI_ISL_2896413, EPI_ISL_2896415 | | | |
| see above | Department of Bacteria, Parasites and Fungi, Statens Serum Institut, Copenhagen, Denmark | Statens Serum Institut Bioinformatics and Microbial Genomics | Danish Covid-19 Genome Consortium |
| EPI_ISL_2896416 | Molekylær Medicinsk Afdeling, Aarhus University Hospital, Aarhus, Denmark | Statens Serum Institut Bioinformatics and Microbial Genomics | Danish Covid-19 Genome Consortium |

EPI_ISL_2896418, EPI_ISL_2896419, EPI_ISL_2896420, EPI_ISL_2896421, EPI_ISL_2896422, EPI_ISL_2896423, EPI_ISL_2896424, EPI_ISL_2896425, EPI_ISL_2896426, EPI_ISL_2896427, EPI_ISL_2896429, EPI_ISL_2896430, EPI_ISL_2896431, EPI_ISL_2896432, EPI_ISL_2896433, EPI_ISL_2896434, EPI_ISL_2896435, EPI_ISL_2896436, EPI_ISL_2896437, EPI_ISL_2896438, EPI_ISL_2896439, EPI_ISL_2896440, EPI_ISL_2896441

| | | | |
|---|---|---|---|
| see above | Department of Bacteria, Parasites and Fungi, Statens Serum Institut, Copenhagen, Denmark | Statens Serum Institut Bioinformatics and Microbial Genomics | Danish Covid-19 Genome Consortium |
| EPI_ISL_2896442 | Molekylær Medicinsk Afdeling, Aarhus University Hospital, Aarhus, Denmark | Statens Serum Institut Bioinformatics and Microbial Genomics | Danish Covid-19 Genome Consortium |

EPI_ISL_2896443, EPI_ISL_2896444, EPI_ISL_2896445, EPI_ISL_2896446, EPI_ISL_2896447, EPI_ISL_2896448, EPI_ISL_2896449, EPI_ISL_2896451, EPI_ISL_2896452, EPI_ISL_2896453, EPI_ISL_2896454, EPI_ISL_2896455, EPI_ISL_2896456, EPI_ISL_2896457, EPI_ISL_2896458, EPI_ISL_2896459, EPI_ISL_2896461, EPI_ISL_2896462, EPI_ISL_2896463, EPI_ISL_2896464, EPI_ISL_2896465, EPI_ISL_2896466, EPI_ISL_2896467, EPI_ISL_2896468, EPI_ISL_2896469, EPI_ISL_2896470, EPI_ISL_2896471, EPI_ISL_2896472, EPI_ISL_2896473, EPI_ISL_2896474, EPI_ISL_2896475, EPI_ISL_2896476, EPI_ISL_2896477, EPI_ISL_2896478, EPI_ISL_2896479, EPI_ISL_2896480, EPI_ISL_2896481, EPI_ISL_2896482, EPI_ISL_2896484, EPI_ISL_2896486, EPI_ISL_2896487, EPI_ISL_2896488

| | | | |
|---|---|---|---|
| see above | Department of Bacteria, Parasites and Fungi, Statens Serum Institut, Copenhagen, Denmark | Statens Serum Institut Bioinformatics and Microbial Genomics | Danish Covid-19 Genome Consortium |
| EPI_ISL_2896489 | Molekylær Medicinsk Afdeling, Aarhus University Hospital, Aarhus, Denmark | Statens Serum Institut Bioinformatics and Microbial Genomics | Danish Covid-19 Genome Consortium |

EPI_ISL_2896493, EPI_ISL_2896494, EPI_ISL_2896496, EPI_ISL_2896497, EPI_ISL_2896498, EPI_ISL_2896499, EPI_ISL_2896500, EPI_ISL_2896501, EPI_ISL_2896502, EPI_ISL_2896503, EPI_ISL_2896504, EPI_ISL_2896506, EPI_ISL_2896507, EPI_ISL_2896508, EPI_ISL_2896509, EPI_ISL_2896510, EPI_ISL_2896511

| | | | |
|---|---|---|---|
| see above | Department of Bacteria, Parasites and Fungi, Statens Serum Institut, Copenhagen, Denmark | Statens Serum Institut Bioinformatics and Microbial Genomics | Danish Covid-19 Genome Consortium |
| EPI_ISL_2896512 | Department of Clinical Microbiology and Center for Genomic Medicine, Rigshospitalet, Copenhagen, Denmark | Statens Serum Institut Bioinformatics and Microbial Genomics | Danish Covid-19 Genome Consortium |

EPI_ISL_2896513, EPI_ISL_2896515, EPI_ISL_2896517, EPI_ISL_2896518, EPI_ISL_2896519, EPI_ISL_2896520, EPI_ISL_2896521, EPI_ISL_2896522, EPI_ISL_2896523, EPI_ISL_2896524, EPI_ISL_2896526, EPI_ISL_2896528, EPI_ISL_2896529, EPI_ISL_2896530, EPI_ISL_2896531, EPI_ISL_2896532, EPI_ISL_2896533, EPI_ISL_2896534, EPI_ISL_2896535, EPI_ISL_2896536, EPI_ISL_2896537, EPI_ISL_2896539, EPI_ISL_2896541, EPI_ISL_2896542, EPI_ISL_2896543, EPI_ISL_2896544, EPI_ISL_2896545, EPI_ISL_2896546, EPI_ISL_2896547, EPI_ISL_2896548, EPI_ISL_2896549, EPI_ISL_2896550

| | | | |
|---|---|---|---|
| see above | Department of Bacteria, Parasites and Fungi, Statens Serum Institut, Copenhagen, Denmark | Statens Serum Institut Bioinformatics and Microbial Genomics | Danish Covid-19 Genome Consortium |
| EPI_ISL_2896551 | Molekylær Medicinsk Afdeling, Aarhus University Hospital, Aarhus, Denmark | Statens Serum Institut Bioinformatics and Microbial Genomics | Danish Covid-19 Genome Consortium |

EPI_ISL_2896552, EPI_ISL_2896553, EPI_ISL_2896554, EPI_ISL_2896555, EPI_ISL_2896556, EPI_ISL_2896557, EPI_ISL_2896558, EPI_ISL_2896559, EPI_ISL_2896560, EPI_ISL_2896561, EPI_ISL_2896563, EPI_ISL_2896564, EPI_ISL_2896565, EPI_ISL_2896566, EPI_ISL_2896568, EPI_ISL_2896569, EPI_ISL_2896570

| | | | |
|---|---|---|---|
| see above | Department of Bacteria, Parasites and Fungi, Statens Serum Institut, Copenhagen, Denmark | Statens Serum Institut Bioinformatics and Microbial Genomics | Danish Covid-19 Genome Consortium |
| EPI_ISL_2896571 | Department of Clinical Microbiology and Center for Genomic Medicine, Rigshospitalet, Copenhagen, Denmark | Statens Serum Institut Bioinformatics and Microbial Genomics | Danish Covid-19 Genome Consortium |

EPI_ISL_2896572, EPI_ISL_2896573, EPI_ISL_2896574, EPI_ISL_2896575, EPI_ISL_2896576, EPI_ISL_2896577, EPI_ISL_2896578, EPI_ISL_2896579, EPI_ISL_2896580, EPI_ISL_2896581, EPI_ISL_2896582, EPI_ISL_2896583, EPI_ISL_2896584

| | | | |
|---|---|---|---|
| see above | Department of Bacteria, Parasites and Fungi, Statens Serum Institut, Copenhagen, Denmark | Statens Serum Institut Bioinformatics and Microbial Genomics | Danish Covid-19 Genome Consortium |
| EPI_ISL_2896585 | Molekylær Medicinsk Afdeling, Aarhus University Hospital, Aarhus, Denmark | Statens Serum Institut Bioinformatics and Microbial Genomics | Danish Covid-19 Genome Consortium |

EPI_ISL_2896586, EPI_ISL_2896587, EPI_ISL_2896588, EPI_ISL_2896589, EPI_ISL_2896590, EPI_ISL_2896591, EPI_ISL_2896592, EPI_ISL_2896593, EPI_ISL_2896594, EPI_ISL_2896595, EPI_ISL_2896596, EPI_ISL_2896597

| | | | |
|---|---|---|---|
| see above | Department of Bacteria, Parasites and Fungi, Statens Serum Institut, Copenhagen, Denmark | Statens Serum Institut Bioinformatics and Microbial Genomics | Danish Covid-19 Genome Consortium |
| EPI_ISL_2896598 | Molekylær Medicinsk Afdeling, Aarhus University Hospital, Aarhus, Denmark | Statens Serum Institut Bioinformatics and Microbial Genomics | Danish Covid-19 Genome Consortium |

EPI_ISL_2896599, EPI_ISL_2896600, EPI_ISL_2896601, EPI_ISL_2896602, EPI_ISL_2896603, EPI_ISL_2896604, EPI_ISL_2896605, EPI_ISL_2896606, EPI_ISL_2896607, EPI_ISL_2896608, EPI_ISL_2896610, EPI_ISL_2896611, EPI_ISL_2896612, EPI_ISL_2896613, EPI_ISL_2896614, EPI_ISL_2896615, EPI_ISL_2896616, EPI_ISL_2896617, EPI_ISL_2896618, EPI_ISL_2896619, EPI_ISL_2896621, EPI_ISL_2896622, EPI_ISL_2896623, EPI_ISL_2896624, EPI_ISL_2896625, EPI_ISL_2896626, EPI_ISL_2896627, EPI_ISL_2896628, EPI_ISL_2896629, EPI_ISL_2896630, EPI_ISL_2896631, EPI_ISL_2896632, EPI_ISL_2896633, EPI_ISL_2896634, EPI_ISL_2896635, EPI_ISL_2896636, EPI_ISL_2896637, EPI_ISL_2896638, EPI_ISL_2896640, EPI_ISL_2896641, EPI_ISL_2896642, EPI_ISL_2896643, EPI_ISL_2896644, EPI_ISL_2896645, EPI_ISL_2896646, EPI_ISL_2896647, EPI_ISL_2896648, EPI_ISL_2896649, EPI_ISL_2896650, EPI_ISL_2896651, EPI_ISL_2896653, EPI_ISL_2896654, EPI_ISL_2896655, EPI_ISL_2896657, EPI_ISL_2896658, EPI_ISL_2896659, EPI_ISL_2896660, EPI_ISL_2896661, EPI_ISL_2896662, EPI_ISL_2896663, EPI_ISL_2896664, EPI_ISL_2896665, EPI_ISL_2896669, EPI_ISL_2896670, EPI_ISL_2896671, EPI_ISL_2896672

| | | | |
|---|---|---|---|
| see above | Department of Bacteria, Parasites and Fungi, Statens Serum Institut, Copenhagen, Denmark | Statens Serum Institut Bioinformatics and Microbial Genomics | Danish Covid-19 Genome Consortium |
| EPI_ISL_2896674 | Molekylær Medicinsk Afdeling, Aarhus University Hospital, Aarhus, Denmark | Statens Serum Institut Bioinformatics and Microbial Genomics | Danish Covid-19 Genome Consortium |
| EPI_ISL_2896675 | Department of Clinical Microbiology, Odense University Hospital, Odense, Denmark | Statens Serum Institut Bioinformatics and Microbial Genomics | Danish Covid-19 Genome Consortium |

EPI_ISL_2896677, EPI_ISL_2896678, EPI_ISL_2896679, EPI_ISL_2896680, EPI_ISL_2896681, EPI_ISL_2896682, EPI_ISL_2896683, EPI_ISL_2896684, EPI_ISL_2896685, EPI_ISL_2896686, EPI_ISL_2896687, EPI_ISL_2896688, EPI_ISL_2896689, EPI_ISL_2896691, EPI_ISL_2896692, EPI_ISL_2896693, EPI_ISL_2896694, EPI_ISL_2896695, EPI_ISL_2896696, EPI_ISL_2896697, EPI_ISL_2896698, EPI_ISL_2896699, EPI_ISL_2896700, EPI_ISL_2896701, EPI_ISL_2896702, EPI_ISL_2896703, EPI_ISL_2896704, EPI_ISL_2896706, EPI_ISL_2896707, EPI_ISL_2896708, EPI_ISL_2896709, EPI_ISL_2896710, EPI_ISL_2896711, EPI_ISL_2896712, EPI_ISL_2896713, EPI_ISL_2896714, EPI_ISL_2896715, EPI_ISL_2896716, EPI_ISL_2896717, EPI_ISL_2896718, EPI_ISL_2896721, EPI_ISL_2896722, EPI_ISL_2896723, EPI_ISL_2896724, EPI_ISL_2896725, EPI_ISL_2896726, EPI_ISL_2896727, EPI_ISL_2896728, EPI_ISL_2896729, EPI_ISL_2896730, EPI_ISL_2896731, EPI_ISL_2896732, EPI_ISL_2896733, EPI_ISL_2896734, EPI_ISL_2896735, EPI_ISL_2896736, EPI_ISL_2896737, EPI_ISL_2896738, EPI_ISL_2896739, EPI_ISL_2896740, EPI_ISL_2896741, EPI_ISL_2896742, EPI_ISL_2896743, EPI_ISL_2896744, EPI_ISL_2896745, EPI_ISL_2896746, EPI_ISL_2896747, EPI_ISL_2896749, EPI_ISL_2896750, EPI_ISL_2896751, EPI_ISL_2896752, EPI_ISL_2896753, EPI_ISL_2896754, EPI_ISL_2896755, EPI_ISL_2896756

| | | | |
|---|---|---|---|
| see above | Department of Bacteria, Parasites and Fungi, Statens Serum Institut, Copenhagen, Denmark | Statens Serum Institut Bioinformatics and Microbial Genomics | Danish Covid-19 Genome Consortium |
| EPI_ISL_2896757 | Department of Clinical Microbiology, Odense University Hospital, Odense, Denmark | Statens Serum Institut Bioinformatics and Microbial Genomics | Danish Covid-19 Genome Consortium |

EPI_ISL_2896758, EPI_ISL_2896759, EPI_ISL_2896760, EPI_ISL_2896761, EPI_ISL_2896764, EPI_ISL_2896765, EPI_ISL_2896766, EPI_ISL_2896768, EPI_ISL_2896769, EPI_ISL_2896770, EPI_ISL_2896771, EPI_ISL_2896772, EPI_ISL_2896773, EPI_ISL_2896774, EPI_ISL_2896775, EPI_ISL_2896776, EPI_ISL_2896777, EPI_ISL_2896779, EPI_ISL_2896780, EPI_ISL_2896782, EPI_ISL_2896783, EPI_ISL_2896784, EPI_ISL_2896785, EPI_ISL_2896786, EPI_ISL_2896787, EPI_ISL_2896788, EPI_ISL_2896789, EPI_ISL_2896790, EPI_ISL_2896791, EPI_ISL_2896792, EPI_ISL_2896793, EPI_ISL_2896794, EPI_ISL_2896795, EPI_ISL_2896797, EPI_ISL_2896798, EPI_ISL_2896799, EPI_ISL_2896800, EPI_ISL_2896802, EPI_ISL_2896803, EPI_ISL_2896805, EPI_ISL_2896807, EPI_ISL_2896809, EPI_ISL_2896811, EPI_ISL_2896813, EPI_ISL_2896816, EPI_ISL_2896818, EPI_ISL_2896820, EPI_ISL_2896825, EPI_ISL_2896827, EPI_ISL_2896832, EPI_ISL_2896836, EPI_ISL_2896837, EPI_ISL_2896838, EPI_ISL_2896839, EPI_ISL_2896840, EPI_ISL_2896842, EPI_ISL_2896844, EPI_ISL_2896845, EPI_ISL_2896846, EPI_ISL_2896847, EPI_ISL_2896848, EPI_ISL_2896849, EPI_ISL_2896850, EPI_ISL_2896851, EPI_ISL_2896852, EPI_ISL_2896853

| | | | |
|---|---|---|---|
| see above | Department of Bacteria, Parasites and Fungi, Statens Serum Institut, Copenhagen, Denmark | Statens Serum Institut Bioinformatics and Microbial Genomics | Danish Covid-19 Genome Consortium |
| EPI_ISL_2896854 | Molekylær Medicinsk Afdeling, Aarhus University Hospital, Aarhus, Denmark | Statens Serum Institut Bioinformatics and Microbial Genomics | Danish Covid-19 Genome Consortium |

EPI_ISL_2896855, EPI_ISL_2896856, EPI_ISL_2896857, EPI_ISL_2896858, EPI_ISL_2896859, EPI_ISL_2896860, EPI_ISL_2896861, EPI_ISL_2896862, EPI_ISL_2896863, EPI_ISL_2896864, EPI_ISL_2896865, EPI_ISL_2896866, EPI_ISL_2896867, EPI_ISL_2896868, EPI_ISL_2896869, EPI_ISL_2896871, EPI_ISL_2896873, EPI_ISL_2896875, EPI_ISL_2896876, EPI_ISL_2896877, EPI_ISL_2896878, EPI_ISL_2896879, EPI_ISL_2896880, EPI_ISL_2896882, EPI_ISL_2896883, EPI_ISL_2896884, EPI_ISL_2896885, EPI_ISL_2896886, EPI_ISL_2896887, EPI_ISL_2896888, EPI_ISL_2896889, EPI_ISL_2896890, EPI_ISL_2896891, EPI_ISL_2896892, EPI_ISL_2896893, EPI_ISL_2896894, EPI_ISL_2896895, EPI_ISL_2896896, EPI_ISL_2896897, EPI_ISL_2896898, EPI_ISL_2896899, EPI_ISL_2896900, EPI_ISL_2896901, EPI_ISL_2896902, EPI_ISL_2896903, EPI_ISL_2896904, EPI_ISL_2896906, EPI_ISL_2896907, EPI_ISL_2896908, EPI_ISL_2896909, EPI_ISL_2896910, EPI_ISL_2896911, EPI_ISL_2896912, EPI_ISL_2896914, EPI_ISL_2896915, EPI_ISL_2896916, EPI_ISL_2896917, EPI_ISL_2896918, EPI_ISL_2896919, EPI_ISL_2896920, EPI_ISL_2896921, EPI_ISL_2896922, EPI_ISL_2896924, EPI_ISL_2896925, EPI_ISL_2896926, EPI_ISL_2896928, EPI_ISL_2896929, EPI_ISL_2896930, EPI_ISL_2896932, EPI_ISL_2896933, EPI_ISL_2896934, EPI_ISL_2896935, EPI_ISL_2896936, EPI_ISL_2896937, EPI_ISL_2896938, EPI_ISL_2896939, EPI_ISL_2896940, EPI_ISL_2896941, EPI_ISL_2896942

| | | | |
|---|---|---|---|
| see above | Department of Bacteria, Parasites and Fungi, Statens Serum Institut, Copenhagen, Denmark | Statens Serum Institut Bioinformatics and Microbial Genomics | Danish Covid-19 Genome Consortium |
| EPI_ISL_2896983 | Microbiology Department, Laboratori Clínic Metropolitana Nord. Hospital Universitari Germans Trias i Pujol | Can Ruti SARS-CoV-2 Sequencing Hub (HUGTiP/IrsiCaixa/IGTP) | Marc Noguera-Julian, Pilar Armengol, Ignacio Blanco, Antoni E Bordoy, Francesc Catala-Moll, Pere-Joan Cardona, Maria Casadellà, Cristina Casañ, Gemma Clara, Bonaventura Clotet, Montserrat Giménez, Mercedes Guerrero, Anna Not, David Panisello, Roger Paredes, Mariona Parera, Alexia París, Verónica Saludes, Alba Sánchez, Laia Soler, Lauro Sumoy, and Elisa Martró on behalf of the Can Ruti SARS-CoV-2 Sequencing Hub |
| EPI_ISL_2896984, EPI_ISL_2896985 | Parc Taulí Hospital Universitari | Can Ruti SARS-CoV-2 Sequencing Hub (HUGTiP/IrsiCaixa/IGTP) | Marc Noguera-Julian, Pilar Armengol, Ignacio Blanco, Antoni E Bordoy, Francesc Catala-Moll, Pere-Joan Cardona, Maria Casadellà, Cristina Casañ, Gemma Clara, Bonaventura Clotet, Montserrat Giménez, Mercedes Guerrero, Anna Not, David Panisello, Roger Paredes, Mariona Parera, Alexia París, Verónica Saludes, Alba Sánchez, Laia Soler, Lauro Sumoy, and Elisa Martró on behalf of the Can Ruti SARS-CoV-2 Sequencing Hub |
| EPI_ISL_2896986, EPI_ISL_2896987, EPI_ISL_2896989, EPI_ISL_2896990, EPI_ISL_2896991, EPI_ISL_2896994 | Microbiology Department, Laboratori Clínic Metropolitana Nord. Hospital Universitari Germans Trias i Pujol | Can Ruti SARS-CoV-2 Sequencing Hub (HUGTiP/IrsiCaixa/IGTP) | Marc Noguera-Julian, Pilar Armengol, Ignacio Blanco, Antoni E Bordoy, Francesc Catala-Moll, Pere-Joan Cardona, Maria Casadellà, Cristina Casañ, Gemma Clara, Bonaventura Clotet, Montserrat Giménez, Mercedes Guerrero, Anna Not, David Panisello, Roger Paredes, Mariona Parera, Alexia París, Verónica Saludes, Alba Sánchez, Laia Soler, Lauro Sumoy, and Elisa Martró on behalf of the Can Ruti SARS-CoV-2 Sequencing Hub |
| EPI_ISL_2896998, EPI_ISL_2896999 | Parc Taulí Hospital Universitari | Can Ruti SARS-CoV-2 Sequencing Hub (HUGTiP/IrsiCaixa/IGTP) | Marc Noguera-Julian, Pilar Armengol, Ignacio Blanco, Antoni E Bordoy, Francesc Catala-Moll, Pere-Joan Cardona, Maria Casadellà, Cristina Casañ, Gemma Clara, Bonaventura Clotet, Montserrat Giménez, Mercedes Guerrero, Anna Not, David Panisello, Roger Paredes, Mariona Parera, Alexia París, Verónica Saludes, Alba Sánchez, Laia Soler, Lauro Sumoy, and Elisa Martró on behalf of the Can Ruti SARS-CoV-2 Sequencing Hub |
| EPI_ISL_2897000, EPI_ISL_2897001, EPI_ISL_2897002, EPI_ISL_2897004, EPI_ISL_2897005, EPI_ISL_2897007, EPI_ISL_2897008, EPI_ISL_2897009 | Microbiology Department, Laboratori Clínic Metropolitana Nord. Hospital Universitari Germans Trias i Pujol | Can Ruti SARS-CoV-2 Sequencing Hub (HUGTiP/IrsiCaixa/IGTP) | Marc Noguera-Julian, Pilar Armengol, Ignacio Blanco, Antoni E Bordoy, Francesc Catala-Moll, Pere-Joan Cardona, Maria Casadellà, Cristina Casañ, Gemma Clara, Bonaventura Clotet, Montserrat Giménez, Mercedes Guerrero, Anna Not, David Panisello, Roger Paredes, Mariona Parera, Alexia París, Verónica Saludes, Alba Sánchez, Laia Soler, Lauro Sumoy, and Elisa Martró on behalf of the Can Ruti SARS-CoV-2 Sequencing Hub |
| EPI_ISL_2897010, EPI_ISL_2897011 | Parc Taulí Hospital Universitari | Can Ruti SARS-CoV-2 Sequencing Hub (HUGTiP/IrsiCaixa/IGTP) | Marc Noguera-Julian, Pilar Armengol, Ignacio Blanco, Antoni E Bordoy, Francesc Catala-Moll, Pere-Joan Cardona, Maria Casadellà, Cristina Casañ, Gemma Clara, Bonaventura Clotet, Montserrat Giménez, Mercedes Guerrero, Anna Not, David Panisello, Roger Paredes, Mariona Parera, Alexia París, Verónica Saludes, Alba Sánchez, Laia Soler, Lauro Sumoy, and Elisa Martró on behalf of the Can Ruti SARS-CoV-2 Sequencing Hub |
| EPI_ISL_2897012, EPI_ISL_2897013, | Microbiology Department, Laboratori Clínic Metropolitana Nord. Hospital Universitari Germans Trias i Pujol | Can Ruti SARS-CoV-2 Sequencing Hub (HUGTiP/IrsiCaixa/IGTP) | Marc Noguera-Julian, Pilar Armengol, Ignacio Blanco, Antoni E Bordoy, Francesc Catala-Moll, Pere-Joan Cardona, Maria Casadellà, Cristina Casañ, Gemma Clara, Bonaventura Clotet, Montserrat Giménez, Mercedes Guerrero, Anna Not, David Panisello, Roger Paredes, Mariona Parera, Alexia París, Verónica Saludes, Alba Sánchez, Laia Soler, Lauro Sumoy, and Elisa Martró on behalf of the Can Ruti SARS-CoV-2 Sequencing Hub |

| | | | |
|---|---|---|---|
| EPI_ISL_2897014, EPI_ISL_2897015, EPI_ISL_2897016, EPI_ISL_2897017, EPI_ISL_2897018, EPI_ISL_2897019, EPI_ISL_2897020, EPI_ISL_2897021 | | | |
| EPI_ISL_2897022 | Parc Taulí Hospital Universitari | Can Ruti SARS-CoV-2 Sequencing Hub (HUGTiP/IrsiCaixa/IGTP) | Marc Noguera-Julian, Pilar Armengol, Ignacio Blanco, Antoni E Bordoy, Francesc Catala-Moll, Pere-Joan Cardona, Maria Casadellà, Cristina Casañ, Gemma Clara, Bonaventura Clotet, Montserrat Giménez, Mercedes Guerrero, Anna Not, David Panisello, Roger Paredes, Mariona Parera, Alexia París, Verónica Saludes, Alba Sánchez, Laia Soler, Lauro Sumoy, and Elisa Martró on behalf of the Can Ruti SARS-CoV-2 Sequencing Hub |
| EPI_ISL_2897023, EPI_ISL_2897024, EPI_ISL_2897025, EPI_ISL_2897026, EPI_ISL_2897027, EPI_ISL_2897028, EPI_ISL_2897029, EPI_ISL_2897031 | Microbiology Department, Laboratori Clínic Metropolitana Nord. Hospital Universitari Germans Trias i Pujol | Can Ruti SARS-CoV-2 Sequencing Hub (HUGTiP/IrsiCaixa/IGTP) | Marc Noguera-Julian, Pilar Armengol, Ignacio Blanco, Antoni E Bordoy, Francesc Catala-Moll, Pere-Joan Cardona, Maria Casadellà, Cristina Casañ, Gemma Clara, Bonaventura Clotet, Montserrat Giménez, Mercedes Guerrero, Anna Not, David Panisello, Roger Paredes, Mariona Parera, Alexia París, Verónica Saludes, Alba Sánchez, Laia Soler, Lauro Sumoy, and Elisa Martró on behalf of the Can Ruti SARS-CoV-2 Sequencing Hub |
| EPI_ISL_2897032 | Microbiologia CATLAB | Can Ruti SARS-CoV-2 Sequencing Hub (HUGTiP/IrsiCaixa/IGTP) | Marc Noguera-Julian, Pilar Armengol, Ignacio Blanco, Antoni E Bordoy, Francesc Catala-Moll, Pere-Joan Cardona, Maria Casadellà, Cristina Casañ, Gemma Clara, Bonaventura Clotet, Montserrat Giménez, Mercedes Guerrero, Anna Not, David Panisello, Roger Paredes, Mariona Parera, Alexia París, Verónica Saludes, Alba Sánchez, Laia Soler, Lauro Sumoy, and Elisa Martró on behalf of the Can Ruti SARS-CoV-2 Sequencing Hub |
| EPI_ISL_2897034 | Parc Taulí Hospital Universitari | Can Ruti SARS-CoV-2 Sequencing Hub (HUGTiP/IrsiCaixa/IGTP) | Marc Noguera-Julian, Pilar Armengol, Ignacio Blanco, Antoni E Bordoy, Francesc Catala-Moll, Pere-Joan Cardona, Maria Casadellà, Cristina Casañ, Gemma Clara, Bonaventura Clotet, Montserrat Giménez, Mercedes Guerrero, Anna Not, David Panisello, Roger Paredes, Mariona Parera, Alexia París, Verónica Saludes, Alba Sánchez, Laia Soler, Lauro Sumoy, and Elisa Martró on behalf of the Can Ruti SARS-CoV-2 Sequencing Hub |
| EPI_ISL_2897035, EPI_ISL_2897036, EPI_ISL_2897037, EPI_ISL_2897038, EPI_ISL_2897040, EPI_ISL_2897041, EPI_ISL_2897042, EPI_ISL_2897043 | Microbiology Department, Laboratori Clínic Metropolitana Nord. Hospital Universitari Germans Trias i Pujol | Can Ruti SARS-CoV-2 Sequencing Hub (HUGTiP/IrsiCaixa/IGTP) | Marc Noguera-Julian, Pilar Armengol, Ignacio Blanco, Antoni E Bordoy, Francesc Catala-Moll, Pere-Joan Cardona, Maria Casadellà, Cristina Casañ, Gemma Clara, Bonaventura Clotet, Montserrat Giménez, Mercedes Guerrero, Anna Not, David Panisello, Roger Paredes, Mariona Parera, Alexia París, Verónica Saludes, Alba Sánchez, Laia Soler, Lauro Sumoy, and Elisa Martró on behalf of the Can Ruti SARS-CoV-2 Sequencing Hub |
| EPI_ISL_2897045, EPI_ISL_2897046 | Parc Taulí Hospital Universitari | Can Ruti SARS-CoV-2 Sequencing Hub (HUGTiP/IrsiCaixa/IGTP) | Marc Noguera-Julian, Pilar Armengol, Ignacio Blanco, Antoni E Bordoy, Francesc Catala-Moll, Pere-Joan Cardona, Maria Casadellà, Cristina Casañ, Gemma Clara, Bonaventura Clotet, Montserrat Giménez, Mercedes Guerrero, Anna Not, David Panisello, Roger Paredes, Mariona Parera, Alexia París, Verónica Saludes, Alba Sánchez, Laia Soler, Lauro Sumoy, and Elisa Martró on behalf of the Can Ruti SARS-CoV-2 Sequencing Hub |
| EPI_ISL_2897047, EPI_ISL_2897049, EPI_ISL_2897050, EPI_ISL_2897052, EPI_ISL_2897053, EPI_ISL_2897054 | Microbiology Department, Laboratori Clínic Metropolitana Nord. Hospital Universitari Germans Trias i Pujol | Can Ruti SARS-CoV-2 Sequencing Hub (HUGTiP/IrsiCaixa/IGTP) | Marc Noguera-Julian, Pilar Armengol, Ignacio Blanco, Antoni E Bordoy, Francesc Catala-Moll, Pere-Joan Cardona, Maria Casadellà, Cristina Casañ, Gemma Clara, Bonaventura Clotet, Montserrat Giménez, Mercedes Guerrero, Anna Not, David Panisello, Roger Paredes, Mariona Parera, Alexia París, Verónica Saludes, Alba Sánchez, Laia Soler, Lauro Sumoy, and Elisa Martró on behalf of the Can Ruti SARS-CoV-2 Sequencing Hub |
| EPI_ISL_2897057 | Parc Taulí Hospital Universitari | Can Ruti SARS-CoV-2 Sequencing Hub (HUGTiP/IrsiCaixa/IGTP) | Marc Noguera-Julian, Pilar Armengol, Ignacio Blanco, Antoni E Bordoy, Francesc Catala-Moll, Pere-Joan Cardona, Maria Casadellà, Cristina Casañ, Gemma Clara, Bonaventura Clotet, Montserrat Giménez, Mercedes Guerrero, Anna Not, David Panisello, Roger Paredes, Mariona Parera, Alexia París, Verónica Saludes, Alba Sánchez, Laia Soler, Lauro Sumoy, and Elisa Martró on behalf of the Can Ruti SARS-CoV-2 Sequencing Hub |
| EPI_ISL_2897058, EPI_ISL_2897060, EPI_ISL_2897061, EPI_ISL_2897062, EPI_ISL_2897063, EPI_ISL_2897064, EPI_ISL_2897065, EPI_ISL_2897069, EPI_ISL_2897070, EPI_ISL_2897071, EPI_ISL_2897072, EPI_ISL_2897073, EPI_ISL_2897074, EPI_ISL_2897075, EPI_ISL_2897076, EPI_ISL_2897077, EPI_ISL_2897078 | | | |
| see above | Microbiology Department, Laboratori Clínic Metropolitana Nord. Hospital Universitari Germans Trias i Pujol | Can Ruti SARS-CoV-2 Sequencing Hub (HUGTiP/IrsiCaixa/IGTP) | Marc Noguera-Julian, Pilar Armengol, Ignacio Blanco, Antoni E Bordoy, Francesc Catala-Moll, Pere-Joan Cardona, Maria Casadellà, Cristina Casañ, Gemma Clara, Bonaventura Clotet, Montserrat Giménez, Mercedes Guerrero, Anna Not, David Panisello, Roger Paredes, Mariona Parera, Alexia París, Verónica Saludes, Alba Sánchez, Laia Soler, Lauro Sumoy, and Elisa Martró on behalf of the Can Ruti SARS-CoV-2 Sequencing Hub |
| EPI_ISL_2897079, EPI_ISL_2897080, EPI_ISL_2897081, EPI_ISL_2897083, EPI_ISL_2897084, EPI_ISL_2897085, EPI_ISL_2897086, EPI_ISL_2897087, EPI_ISL_2897088, EPI_ISL_2897090, EPI_ISL_2897091, EPI_ISL_2897093 | | | |
| see above | Althaia. Xarxa Assistencial Universitària de Manresa | IrsiCaixa | Marc Noguera-Julian, Mariona Parera, Maria Casadellà, Pilar Armengol, Francesc Catala-Moll, Roger Paredes, Bonaventura Clotet Gloria Trujillo, Rafel Perez Vidal, Jaume Trapé, Carolina Gonzalez-Fernandez, Miquel Micó, Antonia Flor, Roger Paredes, Eulalia Grau, Bonaventura Clotet |
| EPI_ISL_2897096 | DL OBM WSSE w Lublinie | 1. Tricity SARS-CoV-2 sequencing consortium: University of Gdansk, Medical University of Gdansk, Vaxican Ltd., Invicta Ltd. 2. National Institute of Public Health - National Institute of Hygiene, Warsaw, Poland | Maciej Kosinski, Celina Cybulska, Krystyna Bienkowska Szewczyk, Maciej Grzybek, Karolina Gackowska, Marcin Lubocki, Katarzyna Groth, Lukasz Rabalski, Katarzyna Zacharczuk, Magdalena Nowakowska, Małgorzata Sadkowska-Todys, Tomasz Wołkowicz |
| EPI_ISL_2897097, EPI_ISL_2897098, EPI_ISL_2897099 | Laboratorium Diagnostyczne Centrum Medycznego Luxmed | 1. Tricity SARS-CoV-2 sequencing consortium: University of Gdansk, Medical University of Gdansk, Vaxican Ltd., Invicta Ltd. 2. National Institute of Public Health - National Institute of Hygiene, Warsaw, Poland | Maciej Kosinski, Celina Cybulska, Krystyna Bienkowska Szewczyk, Maciej Grzybek, Karolina Gackowska, Marcin Lubocki, Katarzyna Groth, Lukasz Rabalski, Katarzyna Zacharczuk, Magdalena Nowakowska, Małgorzata Sadkowska-Todys, Tomasz Wołkowicz |
| EPI_ISL_2897101, EPI_ISL_2897102, EPI_ISL_2897105, EPI_ISL_2897106, EPI_ISL_2897107, EPI_ISL_2897108, EPI_ISL_2897109, EPI_ISL_2897110 | Zakład Diagnostyki Laboratoryjnej SPWSZ w Szczecinie | 1. Tricity SARS-CoV-2 sequencing consortium: University of Gdansk, Medical University of Gdansk, Vaxican Ltd., Invicta Ltd. 2. National Institute of Public Health - National Institute of Hygiene, Warsaw, Poland | Maciej Kosinski, Celina Cybulska, Krystyna Bienkowska Szewczyk, Maciej Grzybek, Karolina Gackowska, Marcin Lubocki, Katarzyna Groth, Lukasz Rabalski, Katarzyna Zacharczuk, Magdalena Nowakowska, Małgorzata Sadkowska-Todys, Tomasz Wołkowicz |
| EPI_ISL_2897111 | Diagnostyka Sp. z o. o. Szczecin | 1. Tricity SARS-CoV-2 sequencing consortium: University of Gdansk, Medical University of Gdansk, Vaxican Ltd., Invicta Ltd. 2. National Institute of Public Health - National Institute of Hygiene, Warsaw, Poland | Maciej Kosinski, Celina Cybulska, Krystyna Bienkowska Szewczyk, Maciej Grzybek, Karolina Gackowska, Marcin Lubocki, Katarzyna Groth, Lukasz Rabalski, Katarzyna Zacharczuk, Magdalena Nowakowska, Małgorzata Sadkowska-Todys, Tomasz Wołkowicz |
| EPI_ISL_2897113 | NZOZ Meditest Diagnostyka Medyczna | 1. Tricity SARS-CoV-2 sequencing consortium: University of Gdansk, Medical University of Gdansk, Vaxican Ltd., Invicta Ltd. 2. National Institute of Public Health - National Institute of Hygiene, Warsaw, Poland | Maciej Kosinski, Celina Cybulska, Krystyna Bienkowska Szewczyk, Maciej Grzybek, Karolina Gackowska, Marcin Lubocki, Katarzyna Groth, Lukasz Rabalski, Katarzyna Zacharczuk, Magdalena Nowakowska, Małgorzata Sadkowska-Todys, Tomasz Wołkowicz |
| EPI_ISL_2897116, EPI_ISL_2897121 | Laboratorium Epidemiologii WSSE w Szczecinie | 1. Tricity SARS-CoV-2 sequencing consortium: University of Gdansk, Medical University of Gdansk, Vaxican Ltd., Invicta Ltd. 2. National Institute of Public Health - National Institute of Hygiene, Warsaw, Poland | Maciej Kosinski, Celina Cybulska, Krystyna Bienkowska Szewczyk, Maciej Grzybek, Karolina Gackowska, Marcin Lubocki, Katarzyna Groth, Lukasz Rabalski, Katarzyna Zacharczuk, Magdalena Nowakowska, Małgorzata Sadkowska-Todys, Tomasz Wołkowicz |
| EPI_ISL_2897123 | Vitomed Sp. z o.o. | 1. Tricity SARS-CoV-2 sequencing consortium: University of Gdansk, Medical University of Gdansk, Vaxican Ltd., Invicta Ltd. 2. National Institute of Public Health - National Institute of Hygiene, Warsaw, Poland | Maciej Kosinski, Celina Cybulska, Krystyna Bienkowska Szewczyk, Maciej Grzybek, Karolina Gackowska, Marcin Lubocki, Katarzyna Groth, Lukasz Rabalski, Katarzyna Zacharczuk, Magdalena Nowakowska, Małgorzata Sadkowska-Todys, Tomasz Wołkowicz |
| EPI_ISL_2897129 | WSSE w Krakowie | 1. Tricity SARS-CoV-2 sequencing consortium: University of Gdansk, Medical University of Gdansk, Vaxican Ltd., Invicta Ltd. 2. National Institute of Public Health - National Institute of Hygiene, Warsaw, Poland | Maciej Kosinski, Celina Cybulska, Krystyna Bienkowska Szewczyk, Maciej Grzybek, Karolina Gackowska, Marcin Lubocki, Katarzyna Groth, Lukasz Rabalski, Katarzyna Zacharczuk, Magdalena Nowakowska, Małgorzata Sadkowska-Todys, Tomasz Wołkowicz |
| EPI_ISL_2897132 | Szpital Uniwersytecki w Krakowie | 1. Tricity SARS-CoV-2 sequencing consortium: University of Gdansk, Medical University of Gdansk, Vaxican Ltd., Invicta Ltd. 2. National Institute of Public Health - National Institute of Hygiene, Warsaw, Poland | Maciej Kosinski, Celina Cybulska, Krystyna Bienkowska Szewczyk, Maciej Grzybek, Karolina Gackowska, Marcin Lubocki, Katarzyna Groth, Lukasz Rabalski, Katarzyna Zacharczuk, Magdalena Nowakowska, Małgorzata Sadkowska-Todys, Tomasz Wołkowicz |
| EPI_ISL_2897134 | Laboratorium Diagnostyczno-Analityczne Ośrodek Profilaktyki i Epidemiologii Nowotworów im. A. Pieńkowskiej | 1. Tricity SARS-CoV-2 sequencing consortium: University of Gdansk, Medical University of Gdansk, Vaxican Ltd., Invicta Ltd. 2. National Institute of Public Health - National Institute of Hygiene, Warsaw, Poland | Maciej Kosinski, Celina Cybulska, Krystyna Bienkowska Szewczyk, Maciej Grzybek, Karolina Gackowska, Marcin Lubocki, Katarzyna Groth, Lukasz Rabalski, Katarzyna Zacharczuk, Magdalena Nowakowska, Małgorzata Sadkowska-Todys, Tomasz Wołkowicz |
| EPI_ISL_2897137, EPI_ISL_2897139, EPI_ISL_2897141, EPI_ISL_2897142 | UM im. Karola Marcinkowskiego w Poznaniu Laboratorium UCA_Covid-19 Centrum Biologii Medycznej | 1. Tricity SARS-CoV-2 sequencing consortium: University of Gdansk, Medical University of Gdansk, Vaxican Ltd., Invicta Ltd. 2. National Institute of Public Health - National Institute of Hygiene, Warsaw, Poland | Maciej Kosinski, Celina Cybulska, Krystyna Bienkowska Szewczyk, Maciej Grzybek, Karolina Gackowska, Marcin Lubocki, Katarzyna Groth, Lukasz Rabalski, Katarzyna Zacharczuk, Magdalena Nowakowska, Małgorzata Sadkowska-Todys, Tomasz Wołkowicz |
| EPI_ISL_2897155 | NZOZ Medyczne Laboratorium Diagnostyczne | 1. Tricity SARS-CoV-2 sequencing consortium: University of Gdansk, Medical University of Gdansk, Vaxican Ltd., Invicta Ltd. 2. National Institute of Public Health - National Institute of Hygiene, Warsaw, Poland | Maciej Kosinski, Celina Cybulska, Krystyna Bienkowska Szewczyk, Maciej Grzybek, Karolina Gackowska, Marcin Lubocki, Katarzyna Groth, Lukasz Rabalski, Katarzyna Zacharczuk, Magdalena Nowakowska, Małgorzata Sadkowska-Todys, Tomasz Wołkowicz |
| EPI_ISL_2897156 | Laboratorium Mikrobiologiczne R.Sz.S | 1. Tricity SARS-CoV-2 sequencing consortium: University of Gdansk, | Maciej Kosinski, Celina Cybulska, Krystyna Bienkowska Szewczyk, Maciej Grzybek, Karolina Gackowska, Marcin Lubocki, Katarzyna Groth, Lukasz Rabalski, Katarzyna Zacharczuk, Magdalena Nowakowska, Małgorzata Sadkowska-Todys, Tomasz |

| | | | |
|---|---|---|---|
| | | Medical University of Gdansk, Vaxican Ltd., Invicta Ltd. 2. National Institute of Public Health - National Institute of Hygiene, Warsaw, Poland | Wołkowicz |
| EPI_ISL_2897158, EPI_ISL_2897159 | National Institute of Public Health - National Institute of Hygiene | 1. Tricity SARS-CoV-2 sequencing consortium: University of Gdansk, Medical University of Gdansk, Vaxican Ltd., Invicta Ltd. 2. National Institute of Public Health - National Institute of Hygiene, Warsaw, Poland | Maciej Kosinski, Celina Cybulska, Krystyna Bienkowska Szewczyk, Maciej Grzybek, Karolina Gackowska, Marcin Lubocki, Katarzyna Groth, Lukasz Rabalski, Katarzyna Zacharczuk, Magdalena Nowakowska, Małgorzata Sadkowska-Todys, Tomasz Wołkowicz |
| EPI_ISL_2897363 | Unilabs | Karolinska University Hospital | Jan Albert ,Tobias Allander ,Annelie Bjerkner ,Sandra Broddesson ,Robert Dyrdak ,Martin Ekman ,Lynda Eneh ,Lina Guerra Blomqvist ,Karolina Ininbergs ,Tanja Normark ,Isak Sylvin ,Zhibing Yun ,Martina Wahlund ,Valtteri Wirta |
| EPI_ISL_2897364 | Karolinska University Hospital Solna | Karolinska University Hospital | Jan Albert ,Tobias Allander ,Annelie Bjerkner ,Sandra Broddesson ,Robert Dyrdak ,Martin Ekman ,Lynda Eneh ,Lina Guerra Blomqvist ,Karolina Ininbergs ,Tanja Normark ,Isak Sylvin ,Zhibing Yun ,Martina Wahlund ,Valtteri Wirta |
| EPI_ISL_2897366 | Unilabs | Karolinska University Hospital | Jan Albert ,Tobias Allander ,Annelie Bjerkner ,Sandra Broddesson ,Robert Dyrdak ,Martin Ekman ,Lynda Eneh ,Lina Guerra Blomqvist ,Karolina Ininbergs ,Tanja Normark ,Isak Sylvin ,Zhibing Yun ,Martina Wahlund ,Valtteri Wirta |
| EPI_ISL_2897368 | Karolinska University Hospital Solna | Karolinska University Hospital | Jan Albert ,Tobias Allander ,Annelie Bjerkner ,Sandra Broddesson ,Robert Dyrdak ,Martin Ekman ,Lynda Eneh ,Lina Guerra Blomqvist ,Karolina Ininbergs ,Tanja Normark ,Isak Sylvin ,Zhibing Yun ,Martina Wahlund ,Valtteri Wirta |
| EPI_ISL_2897369 | Unilabs | Karolinska University Hospital | Jan Albert ,Tobias Allander ,Annelie Bjerkner ,Sandra Broddesson ,Robert Dyrdak ,Martin Ekman ,Lynda Eneh ,Lina Guerra Blomqvist ,Karolina Ininbergs ,Tanja Normark ,Isak Sylvin ,Zhibing Yun ,Martina Wahlund ,Valtteri Wirta |
| EPI_ISL_2897370 | Karolinska University Hospital Solna | Karolinska University Hospital | Jan Albert ,Tobias Allander ,Annelie Bjerkner ,Sandra Broddesson ,Robert Dyrdak ,Martin Ekman ,Lynda Eneh ,Lina Guerra Blomqvist ,Karolina Ininbergs ,Tanja Normark ,Isak Sylvin ,Zhibing Yun ,Martina Wahlund ,Valtteri Wirta |
| EPI_ISL_2897372 | Unilabs | Karolinska University Hospital | Jan Albert ,Tobias Allander ,Annelie Bjerkner ,Sandra Broddesson ,Robert Dyrdak ,Martin Ekman ,Lynda Eneh ,Lina Guerra Blomqvist ,Karolina Ininbergs ,Tanja Normark ,Isak Sylvin ,Zhibing Yun ,Martina Wahlund ,Valtteri Wirta |
| EPI_ISL_2897374 | Karolinska University Hospital Solna | Karolinska University Hospital | Jan Albert ,Tobias Allander ,Annelie Bjerkner ,Sandra Broddesson ,Robert Dyrdak ,Martin Ekman ,Lynda Eneh ,Lina Guerra Blomqvist ,Karolina Ininbergs ,Tanja Normark ,Isak Sylvin ,Zhibing Yun ,Martina Wahlund ,Valtteri Wirta |
| EPI_ISL_2897375 | Synlab Medilab | Karolinska University Hospital | Jan Albert ,Tobias Allander ,Annelie Bjerkner ,Sandra Broddesson ,Robert Dyrdak ,Martin Ekman ,Lynda Eneh ,Lina Guerra Blomqvist ,Karolina Ininbergs ,Tanja Normark ,Isak Sylvin ,Zhibing Yun ,Martina Wahlund ,Valtteri Wirta |
| EPI_ISL_2897378, EPI_ISL_2897382, EPI_ISL_2897383 | Karolinska University Hospital Solna | Karolinska University Hospital | Jan Albert ,Tobias Allander ,Annelie Bjerkner ,Sandra Broddesson ,Robert Dyrdak ,Martin Ekman ,Lynda Eneh ,Lina Guerra Blomqvist ,Karolina Ininbergs ,Tanja Normark ,Isak Sylvin ,Zhibing Yun ,Martina Wahlund ,Valtteri Wirta |
| EPI_ISL_2897385 | Unilabs | Karolinska University Hospital | Jan Albert ,Tobias Allander ,Annelie Bjerkner ,Sandra Broddesson ,Robert Dyrdak ,Martin Ekman ,Lynda Eneh ,Lina Guerra Blomqvist ,Karolina Ininbergs ,Tanja Normark ,Isak Sylvin ,Zhibing Yun ,Martina Wahlund ,Valtteri Wirta |
| EPI_ISL_2897387 | Karolinska University Hospital Solna | Karolinska University Hospital | Jan Albert ,Tobias Allander ,Annelie Bjerkner ,Sandra Broddesson ,Robert Dyrdak ,Martin Ekman ,Lynda Eneh ,Lina Guerra Blomqvist ,Karolina Ininbergs ,Tanja Normark ,Isak Sylvin ,Zhibing Yun ,Martina Wahlund ,Valtteri Wirta |
| EPI_ISL_2897388, EPI_ISL_2897389, EPI_ISL_2897391 | Unilabs | Karolinska University Hospital | Jan Albert ,Tobias Allander ,Annelie Bjerkner ,Sandra Broddesson ,Robert Dyrdak ,Martin Ekman ,Lynda Eneh ,Lina Guerra Blomqvist ,Karolina Ininbergs ,Tanja Normark ,Isak Sylvin ,Zhibing Yun ,Martina Wahlund ,Valtteri Wirta |
| EPI_ISL_2897394, EPI_ISL_2897396 | Karolinska University Hospital Solna | Karolinska University Hospital | Jan Albert ,Tobias Allander ,Annelie Bjerkner ,Sandra Broddesson ,Robert Dyrdak ,Martin Ekman ,Lynda Eneh ,Lina Guerra Blomqvist ,Karolina Ininbergs ,Tanja Normark ,Isak Sylvin ,Zhibing Yun ,Martina Wahlund ,Valtteri Wirta |
| EPI_ISL_2897397 | Unilabs | Karolinska University Hospital | Jan Albert ,Tobias Allander ,Annelie Bjerkner ,Sandra Broddesson ,Robert Dyrdak ,Martin Ekman ,Lynda Eneh ,Lina Guerra Blomqvist ,Karolina Ininbergs ,Tanja Normark ,Isak Sylvin ,Zhibing Yun ,Martina Wahlund ,Valtteri Wirta |
| EPI_ISL_2897398 | Synlab Medilab | Karolinska University Hospital | Jan Albert ,Tobias Allander ,Annelie Bjerkner ,Sandra Broddesson ,Robert Dyrdak ,Martin Ekman ,Lynda Eneh ,Lina Guerra Blomqvist ,Karolina Ininbergs ,Tanja Normark ,Isak Sylvin ,Zhibing Yun ,Martina Wahlund ,Valtteri Wirta |
| EPI_ISL_2897400, EPI_ISL_2897401 | Unilabs | Karolinska University Hospital | Jan Albert ,Tobias Allander ,Annelie Bjerkner ,Sandra Broddesson ,Robert Dyrdak ,Martin Ekman ,Lynda Eneh ,Lina Guerra Blomqvist ,Karolina Ininbergs ,Tanja Normark ,Isak Sylvin ,Zhibing Yun ,Martina Wahlund ,Valtteri Wirta |
| EPI_ISL_2897404, EPI_ISL_2897405 | Karolinska University Hospital Solna | Karolinska University Hospital | Jan Albert ,Tobias Allander ,Annelie Bjerkner ,Sandra Broddesson ,Robert Dyrdak ,Martin Ekman ,Lynda Eneh ,Lina Guerra Blomqvist ,Karolina Ininbergs ,Tanja Normark ,Isak Sylvin ,Zhibing Yun ,Martina Wahlund ,Valtteri Wirta |
| EPI_ISL_2897406, EPI_ISL_2897407, EPI_ISL_2897408, EPI_ISL_2897414, EPI_ISL_2897415, EPI_ISL_2897416, EPI_ISL_2897417, EPI_ISL_2897418, EPI_ISL_2897420, EPI_ISL_2897421, EPI_ISL_2897422, EPI_ISL_2897427 | | | |
| see above | Unilabs | Karolinska University Hospital | Jan Albert ,Tobias Allander ,Annelie Bjerkner ,Sandra Broddesson ,Robert Dyrdak ,Martin Ekman ,Lynda Eneh ,Lina Guerra Blomqvist ,Karolina Ininbergs ,Tanja Normark ,Isak Sylvin ,Zhibing Yun ,Martina Wahlund ,Valtteri Wirta |
| EPI_ISL_2897430 | Karolinska University Hospital Solna | Karolinska University Hospital | Jan Albert ,Tobias Allander ,Annelie Bjerkner ,Sandra Broddesson ,Robert Dyrdak ,Martin Ekman ,Lynda Eneh ,Lina Guerra Blomqvist ,Karolina Ininbergs ,Tanja Normark ,Isak Sylvin ,Zhibing Yun ,Martina Wahlund ,Valtteri Wirta |
| EPI_ISL_2897431, EPI_ISL_2897433, EPI_ISL_2897438, EPI_ISL_2897439 | Unilabs | Karolinska University Hospital | Jan Albert ,Tobias Allander ,Annelie Bjerkner ,Sandra Broddesson ,Robert Dyrdak ,Martin Ekman ,Lynda Eneh ,Lina Guerra Blomqvist ,Karolina Ininbergs ,Tanja Normark ,Isak Sylvin ,Zhibing Yun ,Martina Wahlund ,Valtteri Wirta |
| EPI_ISL_2897440, EPI_ISL_2897443, EPI_ISL_2897445, EPI_ISL_2897447, EPI_ISL_2897449, EPI_ISL_2897451, EPI_ISL_2897454, EPI_ISL_2897455, EPI_ISL_2897456, EPI_ISL_2897457, EPI_ISL_2897458, EPI_ISL_2897459, EPI_ISL_2897460, EPI_ISL_2897461, EPI_ISL_2897462, EPI_ISL_2897463, EPI_ISL_2897464 | | | |
| see above | Karolinska University Hospital Huddinge | Karolinska University Hospital | Jan Albert ,Tobias Allander ,Annelie Bjerkner ,Sandra Broddesson ,Robert Dyrdak ,Martin Ekman ,Lynda Eneh ,Lina Guerra Blomqvist ,Karolina Ininbergs ,Tanja Normark ,Isak Sylvin ,Zhibing Yun ,Martina Wahlund ,Valtteri Wirta |
| EPI_ISL_2897472, EPI_ISL_2897475, EPI_ISL_2897482 | Unilabs | Karolinska University Hospital | Jan Albert ,Tobias Allander ,Annelie Bjerkner ,Sandra Broddesson ,Robert Dyrdak ,Martin Ekman ,Lynda Eneh ,Lina Guerra Blomqvist ,Karolina Ininbergs ,Tanja Normark ,Isak Sylvin ,Zhibing Yun ,Martina Wahlund ,Valtteri Wirta |
| EPI_ISL_2897483, EPI_ISL_2897489 | Karolinska University Hospital Solna | Karolinska University Hospital | Jan Albert ,Tobias Allander ,Annelie Bjerkner ,Sandra Broddesson ,Robert Dyrdak ,Martin Ekman ,Lynda Eneh ,Lina Guerra Blomqvist ,Karolina Ininbergs ,Tanja Normark ,Isak Sylvin ,Zhibing Yun ,Martina Wahlund ,Valtteri Wirta |
| EPI_ISL_2897494, EPI_ISL_2897499, EPI_ISL_2897502, EPI_ISL_2897503, EPI_ISL_2897504, EPI_ISL_2897506, EPI_ISL_2897510 | Unilabs | Karolinska University Hospital | Jan Albert ,Tobias Allander ,Annelie Bjerkner ,Sandra Broddesson ,Robert Dyrdak ,Martin Ekman ,Lynda Eneh ,Lina Guerra Blomqvist ,Karolina Ininbergs ,Tanja Normark ,Isak Sylvin ,Zhibing Yun ,Martina Wahlund ,Valtteri Wirta |
| EPI_ISL_2897511 | Karolinska University Hospital Solna | Karolinska University Hospital | Jan Albert ,Tobias Allander ,Annelie Bjerkner ,Sandra Broddesson ,Robert Dyrdak ,Martin Ekman ,Lynda Eneh ,Lina Guerra Blomqvist ,Karolina Ininbergs ,Tanja Normark ,Isak Sylvin ,Zhibing Yun ,Martina Wahlund ,Valtteri Wirta |
| EPI_ISL_2897517 | Unilabs | Karolinska University Hospital | Jan Albert ,Tobias Allander ,Annelie Bjerkner ,Sandra Broddesson ,Robert Dyrdak ,Martin Ekman ,Lynda Eneh ,Lina Guerra Blomqvist ,Karolina Ininbergs ,Tanja Normark ,Isak Sylvin ,Zhibing Yun ,Martina Wahlund ,Valtteri Wirta |
| EPI_ISL_2897525 | Synlab Medilab | Karolinska University Hospital | Jan Albert ,Tobias Allander ,Annelie Bjerkner ,Sandra Broddesson ,Robert Dyrdak ,Martin Ekman ,Lynda Eneh ,Lina Guerra Blomqvist ,Karolina Ininbergs ,Tanja Normark ,Isak Sylvin ,Zhibing Yun ,Martina Wahlund ,Valtteri Wirta |
| EPI_ISL_2897536 | Unilabs | Karolinska University Hospital | Jan Albert ,Tobias Allander ,Annelie Bjerkner ,Sandra Broddesson ,Robert Dyrdak ,Martin Ekman ,Lynda Eneh ,Lina Guerra Blomqvist ,Karolina Ininbergs ,Tanja Normark ,Isak Sylvin ,Zhibing Yun ,Martina Wahlund ,Valtteri Wirta |
| EPI_ISL_2897563, EPI_ISL_2897564, EPI_ISL_2897565, EPI_ISL_2897570, EPI_ISL_2897571, EPI_ISL_2897575, EPI_ISL_2897578, EPI_ISL_2897579, EPI_ISL_2897580, EPI_ISL_2897581, EPI_ISL_2897582, EPI_ISL_2897583, EPI_ISL_2897585, EPI_ISL_2897586, EPI_ISL_2897587, EPI_ISL_2897588, EPI_ISL_2897589, EPI_ISL_2897591, EPI_ISL_2897592, EPI_ISL_2897594, EPI_ISL_2897595, EPI_ISL_2897597, EPI_ISL_2897598, EPI_ISL_2897599, EPI_ISL_2897600, EPI_ISL_2897601, EPI_ISL_2897603, EPI_ISL_2897604, EPI_ISL_2897605, EPI_ISL_2897606, EPI_ISL_2897607, EPI_ISL_2897609, EPI_ISL_2897610, EPI_ISL_2897611, EPI_ISL_2897612, EPI_ISL_2897613, EPI_ISL_2897614, EPI_ISL_2897615, EPI_ISL_2897616, EPI_ISL_2897617, EPI_ISL_2897618, EPI_ISL_2897619, EPI_ISL_2897620, EPI_ISL_2897621, EPI_ISL_2897622, EPI_ISL_2897623, EPI_ISL_2897626, EPI_ISL_2897627, EPI_ISL_2897628, EPI_ISL_2897629, EPI_ISL_2897630, EPI_ISL_2897633, EPI_ISL_2897636, EPI_ISL_2897637, EPI_ISL_2897639, EPI_ISL_2897640, EPI_ISL_2897641, EPI_ISL_2897642, EPI_ISL_2897643, EPI_ISL_2897644, EPI_ISL_2897645, EPI_ISL_2897646, EPI_ISL_2897647, EPI_ISL_2897648, EPI_ISL_2897650, EPI_ISL_2897651, EPI_ISL_2897652, EPI_ISL_2897653, EPI_ISL_2897654, EPI_ISL_2897655, EPI_ISL_2897656, EPI_ISL_2897657, EPI_ISL_2897659, EPI_ISL_2897660, EPI_ISL_2897661, EPI_ISL_2897662, EPI_ISL_2897663, EPI_ISL_2897664, EPI_ISL_2897665, EPI_ISL_2897666, EPI_ISL_2897667, EPI_ISL_2897673, EPI_ISL_2897675, EPI_ISL_2897676, EPI_ISL_2897680, EPI_ISL_2897681, EPI_ISL_2897682, EPI_ISL_2897683, EPI_ISL_2897684, EPI_ISL_2897685, EPI_ISL_2897686, EPI_ISL_2897687, EPI_ISL_2897688, EPI_ISL_2897689, EPI_ISL_2897690, EPI_ISL_2897692, EPI_ISL_2897693, EPI_ISL_2897694, EPI_ISL_2897695, EPI_ISL_2897697, EPI_ISL_2897698, EPI_ISL_2897700, EPI_ISL_2897701, EPI_ISL_2897702, EPI_ISL_2897703, EPI_ISL_2897704, EPI_ISL_2897705, EPI_ISL_2897708, EPI_ISL_2897709, EPI_ISL_2897710, EPI_ISL_2897711, EPI_ISL_2897713, EPI_ISL_2897714, EPI_ISL_2897717, EPI_ISL_2897718, EPI_ISL_2897719, EPI_ISL_2897721, EPI_ISL_2897723, EPI_ISL_2897724, EPI_ISL_2897725, EPI_ISL_2897726, EPI_ISL_2897728, EPI_ISL_2897730, EPI_ISL_2897732, EPI_ISL_2897733, EPI_ISL_2897734, EPI_ISL_2897735, EPI_ISL_2897736, EPI_ISL_2897737, EPI_ISL_2897738, EPI_ISL_2897739, EPI_ISL_2897740, EPI_ISL_2897741, EPI_ISL_2897742, EPI_ISL_2897743, EPI_ISL_2897744, EPI_ISL_2897745, EPI_ISL_2897746, EPI_ISL_2897747, EPI_ISL_2897748, EPI_ISL_2897749, EPI_ISL_2897750, EPI_ISL_2897751, EPI_ISL_2897752, EPI_ISL_2897753, EPI_ISL_2897754, EPI_ISL_2897755, EPI_ISL_2897756, EPI_ISL_2897757, EPI_ISL_2897758, EPI_ISL_2897759, EPI_ISL_2897760, EPI_ISL_2897761, EPI_ISL_2897762, EPI_ISL_2897763, EPI_ISL_2897764, EPI_ISL_2897765, EPI_ISL_2897767, EPI_ISL_2897768, EPI_ISL_2897769, EPI_ISL_2897770, EPI_ISL_2897771, EPI_ISL_2897772, EPI_ISL_2897773, EPI_ISL_2897774, EPI_ISL_2897775, EPI_ISL_2897776, EPI_ISL_2897777, EPI_ISL_2897778, EPI_ISL_2897779, EPI_ISL_2897780, EPI_ISL_2897781, EPI_ISL_2897782, EPI_ISL_2897783, EPI_ISL_2897784, EPI_ISL_2897785, EPI_ISL_2897786, EPI_ISL_2897787, EPI_ISL_2897788, EPI_ISL_2897789, EPI_ISL_2897790, EPI_ISL_2897791, EPI_ISL_2897792, EPI_ISL_2897793, EPI_ISL_2897794, EPI_ISL_2897796, EPI_ISL_2897797, EPI_ISL_2897798, EPI_ISL_2897799, EPI_ISL_2897800, EPI_ISL_2897801, EPI_ISL_2897802, EPI_ISL_2897804, EPI_ISL_2897805, EPI_ISL_2897806, EPI_ISL_2897807, EPI_ISL_2897808, EPI_ISL_2897809, EPI_ISL_2897811, EPI_ISL_2897812, EPI_ISL_2897813, EPI_ISL_2897814, EPI_ISL_2897815, EPI_ISL_2897816, EPI_ISL_2897817, EPI_ISL_2897818, EPI_ISL_2897819, EPI_ISL_2897820, EPI_ISL_2897821, EPI_ISL_2897822, EPI_ISL_2897823, EPI_ISL_2897824, EPI_ISL_2897826, EPI_ISL_2897832, EPI_ISL_2897833, EPI_ISL_2897834, EPI_ISL_2897837 | | | |
| see above | National Centre for Cell Science | CDFD | Murali Bashyam, Asmita Gupta, Reelina Basu, Vinay Donipadi, Divya Vashisht, Ashwin Dalal |
| EPI_ISL_2897840, EPI_ISL_2897841, EPI_ISL_2897842, EPI_ISL_2897843, EPI_ISL_2897844, EPI_ISL_2897845, EPI_ISL_2897846, EPI_ISL_2897847, EPI_ISL_2897848, EPI_ISL_2897849, EPI_ISL_2897850 | | | |
| see above | CDFD | CDFD | Murali Bashyam, Asmita Gupta, Reelina Basu, Vinay Donipadi, Nagamani Kammili, Divya Vashisht, Ashwin Dalal |
| EPI_ISL_2897851, EPI_ISL_2897852, EPI_ISL_2897853, EPI_ISL_2897854, EPI_ISL_2897855, EPI_ISL_2897856, EPI_ISL_2897857, EPI_ISL_2897858, EPI_ISL_2897859, EPI_ISL_2897860, EPI_ISL_2897861, EPI_ISL_2897862, EPI_ISL_2897863, EPI_ISL_2897864, EPI_ISL_2897865, EPI_ISL_2897866, EPI_ISL_2897867, EPI_ISL_2897868, EPI_ISL_2897869, EPI_ISL_2897870, EPI_ISL_2897871, EPI_ISL_2897872, EPI_ISL_2897873, EPI_ISL_2897875 | | | |
| see above | Government Fever Hospital | CDFD | Murali Bashyam, Asmita Gupta, Reelina Basu, Vinay Donipadi, Nagamani Kammili, Divya Vashisht, Ashwin Dalal |
| EPI_ISL_2897876, EPI_ISL_2897877, EPI_ISL_2897878, EPI_ISL_2897879, EPI_ISL_2897880, EPI_ISL_2897881, EPI_ISL_2897882, EPI_ISL_2897883, EPI_ISL_2897884, EPI_ISL_2897885, EPI_ISL_2897886, EPI_ISL_2897888, EPI_ISL_2897889, EPI_ISL_2897890, EPI_ISL_2897891, EPI_ISL_2897892 | | | |
| see above | CDFD | CDFD | Murali Bashyam, Asmita Gupta, Reelina Basu, Vinay Donipadi, Nagamani Kammili, Divya Vashisht, Ashwin Dalal |
| EPI_ISL_2897893, EPI_ISL_2897894, EPI_ISL_2897895, EPI_ISL_2897896, EPI_ISL_2897897, EPI_ISL_2897898, EPI_ISL_2897899, EPI_ISL_2897900, EPI_ISL_2897901, EPI_ISL_2897902, EPI_ISL_2897903, EPI_ISL_2897904, EPI_ISL_2897905 | | | |
| see above | Institute of Preventive Medicine | CDFD | Murali Bashyam, Asmita Gupta, Reelina Basu, Vinay Donipadi, Nagamani Kammili, Divya Vashisht, Ashwin Dalal |
| EPI_ISL_2897906 | Gandhi Hospital | CDFD | Murali Bashyam, Asmita Gupta, Reelina Basu, Vinay Donipadi, Nagamani Kammili, Divya Vashisht, Ashwin Dalal |
| EPI_ISL_2897907, EPI_ISL_2897908 | CDFD | CDFD | Murali Bashyam, Asmita Gupta, Reelina Basu, Vinay Donipadi, Nagamani Kammili, Divya Vashisht, Ashwin Dalal |
| EPI_ISL_2897910, EPI_ISL_2897911, EPI_ISL_2897912, EPI_ISL_2897913, EPI_ISL_2897914, EPI_ISL_2897915, EPI_ISL_2897916, EPI_ISL_2897917, EPI_ISL_2897918, EPI_ISL_2897919, EPI_ISL_2897920, EPI_ISL_2897921, EPI_ISL_2897922, EPI_ISL_2897923, EPI_ISL_2897924, EPI_ISL_2897925, EPI_ISL_2897926, EPI_ISL_2897927, EPI_ISL_2897928, EPI_ISL_2897929, EPI_ISL_2897930, EPI_ISL_2897931, EPI_ISL_2897932, EPI_ISL_2897933, EPI_ISL_2897934, EPI_ISL_2897935 | | | |
| see above | Gandhi Hospital | CDFD | Murali Bashyam, Asmita Gupta, Reelina Basu, Vinay Donipadi, Nagamani Kammili, Divya Vashisht, Ashwin Dalal |
| EPI_ISL_2897936, EPI_ISL_2897938, EPI_ISL_2897939, EPI_ISL_2897940, EPI_ISL_2897941, EPI_ISL_2897942 | Government Medical College | CDFD | Murali Bashyam, Asmita Gupta, Reelina Basu, Vinay Donipadi, Nagamani Kammili, Divya Vashisht, Ashwin Dalal |

| | | | |
|---|---|---|---|
| EPI_ISL_2897943, EPI_ISL_2897944, EPI_ISL_2897945, EPI_ISL_2897946, EPI_ISL_2897947, EPI_ISL_2897948, EPI_ISL_2897949 | Government General Hospital | CDFD | Murali Bashyam, Asmita Gupta, Reelina Basu, Vinay Donipadi, Nagamani Kammili, Divya Vashisht, Ashwin Dalal |
| EPI_ISL_2897950, EPI_ISL_2897951, EPI_ISL_2897952, EPI_ISL_2897953, EPI_ISL_2897954, EPI_ISL_2897955, EPI_ISL_2897956, EPI_ISL_2897957, EPI_ISL_2897958, EPI_ISL_2897959 | Government Maternity Hospital | CDFD | Murali Bashyam, Asmita Gupta, Reelina Basu, Vinay Donipadi, Nagamani Kammili, Divya Vashisht, Ashwin Dalal |
| EPI_ISL_2897960, EPI_ISL_2897961, EPI_ISL_2897963, EPI_ISL_2897964, EPI_ISL_2897965, EPI_ISL_2897966, EPI_ISL_2897967, EPI_ISL_2897968, EPI_ISL_2897969, EPI_ISL_2897972, EPI_ISL_2897973, EPI_ISL_2897974, EPI_ISL_2897975, EPI_ISL_2897976, EPI_ISL_2897977, EPI_ISL_2897978, EPI_ISL_2897980, EPI_ISL_2897981, EPI_ISL_2897983, EPI_ISL_2897984, EPI_ISL_2897986, EPI_ISL_2897987, EPI_ISL_2897988, EPI_ISL_2897989, EPI_ISL_2897990, EPI_ISL_2897991 | | | |
| see above | District Hospital | CDFD | Murali Bashyam, Asmita Gupta, Reelina Basu, Vinay Donipadi, Nagamani Kammili, Divya Vashisht, Ashwin Dalal |
| EPI_ISL_2898084 | Karolinska University Hospital Solna | Karolinska University Hospital | Jan Albert ,Tobias Allander ,Annelie Bjerkner ,Sandra Broddesson ,Robert Dyrdak ,Martin Ekman ,Lynda Eneh ,Lina Guerra Blomqvist ,Karolina Ininbergs ,Tanja Normark ,Isak Sylvin ,Zhibing Yun ,Martina Wahlund ,Valtteri Wirta |
| EPI_ISL_2898159 | Unilabs | Karolinska University Hospital | Jan Albert ,Tobias Allander ,Annelie Bjerkner ,Sandra Broddesson ,Robert Dyrdak ,Martin Ekman ,Lynda Eneh ,Lina Guerra Blomqvist ,Karolina Ininbergs ,Tanja Normark ,Isak Sylvin ,Zhibing Yun ,Martina Wahlund ,Valtteri Wirta |
| EPI_ISL_2898189 | Karolinska University Hospital Solna | Karolinska University Hospital | Jan Albert ,Tobias Allander ,Annelie Bjerkner ,Sandra Broddesson ,Robert Dyrdak ,Martin Ekman ,Lynda Eneh ,Lina Guerra Blomqvist ,Karolina Ininbergs ,Tanja Normark ,Isak Sylvin ,Zhibing Yun ,Martina Wahlund ,Valtteri Wirta |
| EPI_ISL_2898227, EPI_ISL_2898245, EPI_ISL_2898271 | Karolinska University Hospital Huddinge | Karolinska University Hospital | Jan Albert ,Tobias Allander ,Annelie Bjerkner ,Sandra Broddesson ,Robert Dyrdak ,Martin Ekman ,Lynda Eneh ,Lina Guerra Blomqvist ,Karolina Ininbergs ,Tanja Normark ,Isak Sylvin ,Zhibing Yun ,Martina Wahlund ,Valtteri Wirta |
| EPI_ISL_2898384, EPI_ISL_2898731 | Karolinska University Hospital Solna | Karolinska University Hospital | Jan Albert ,Tobias Allander ,Annelie Bjerkner ,Sandra Broddesson ,Robert Dyrdak ,Martin Ekman ,Lynda Eneh ,Lina Guerra Blomqvist ,Karolina Ininbergs ,Tanja Normark ,Isak Sylvin ,Zhibing Yun ,Martina Wahlund ,Valtteri Wirta |
| EPI_ISL_2898777, EPI_ISL_2898784, EPI_ISL_2898785 | Karolinska University Hospital Huddinge | Karolinska University Hospital | Jan Albert ,Tobias Allander ,Annelie Bjerkner ,Sandra Broddesson ,Robert Dyrdak ,Martin Ekman ,Lynda Eneh ,Lina Guerra Blomqvist ,Karolina Ininbergs ,Tanja Normark ,Isak Sylvin ,Zhibing Yun ,Martina Wahlund ,Valtteri Wirta |
| EPI_ISL_2898822 | Unilabs | Karolinska University Hospital | Jan Albert ,Tobias Allander ,Annelie Bjerkner ,Sandra Broddesson ,Robert Dyrdak ,Martin Ekman ,Lynda Eneh ,Lina Guerra Blomqvist ,Karolina Ininbergs ,Tanja Normark ,Isak Sylvin ,Zhibing Yun ,Martina Wahlund ,Valtteri Wirta |
| EPI_ISL_2898878 | Karolinska University Hospital Solna | Karolinska University Hospital | Jan Albert ,Tobias Allander ,Annelie Bjerkner ,Sandra Broddesson ,Robert Dyrdak ,Martin Ekman ,Lynda Eneh ,Lina Guerra Blomqvist ,Karolina Ininbergs ,Tanja Normark ,Isak Sylvin ,Zhibing Yun ,Martina Wahlund ,Valtteri Wirta |
| EPI_ISL_2898898 | Unilabs | Karolinska University Hospital | Jan Albert ,Tobias Allander ,Annelie Bjerkner ,Sandra Broddesson ,Robert Dyrdak ,Martin Ekman ,Lynda Eneh ,Lina Guerra Blomqvist ,Karolina Ininbergs ,Tanja Normark ,Isak Sylvin ,Zhibing Yun ,Martina Wahlund ,Valtteri Wirta |
| EPI_ISL_2898906 | Karolinska University Hospital Solna | Karolinska University Hospital | Jan Albert ,Tobias Allander ,Annelie Bjerkner ,Sandra Broddesson ,Robert Dyrdak ,Martin Ekman ,Lynda Eneh ,Lina Guerra Blomqvist ,Karolina Ininbergs ,Tanja Normark ,Isak Sylvin ,Zhibing Yun ,Martina Wahlund ,Valtteri Wirta |
| EPI_ISL_2898911 | Unilabs | Karolinska University Hospital | Jan Albert ,Tobias Allander ,Annelie Bjerkner ,Sandra Broddesson ,Robert Dyrdak ,Martin Ekman ,Lynda Eneh ,Lina Guerra Blomqvist ,Karolina Ininbergs ,Tanja Normark ,Isak Sylvin ,Zhibing Yun ,Martina Wahlund ,Valtteri Wirta |
| EPI_ISL_2898925 | Karolinska University Hospital Solna | Karolinska University Hospital | Jan Albert ,Tobias Allander ,Annelie Bjerkner ,Sandra Broddesson ,Robert Dyrdak ,Martin Ekman ,Lynda Eneh ,Lina Guerra Blomqvist ,Karolina Ininbergs ,Tanja Normark ,Isak Sylvin ,Zhibing Yun ,Martina Wahlund ,Valtteri Wirta |
| EPI_ISL_2898933 | Unilabs | Karolinska University Hospital | Jan Albert ,Tobias Allander ,Annelie Bjerkner ,Sandra Broddesson ,Robert Dyrdak ,Martin Ekman ,Lynda Eneh ,Lina Guerra Blomqvist ,Karolina Ininbergs ,Tanja Normark ,Isak Sylvin ,Zhibing Yun ,Martina Wahlund ,Valtteri Wirta |
| EPI_ISL_2898977, EPI_ISL_2898981, EPI_ISL_2898986, EPI_ISL_2898988, EPI_ISL_2898990, EPI_ISL_2898991 | Karolinska University Hospital Huddinge | Karolinska University Hospital | Jan Albert ,Tobias Allander ,Annelie Bjerkner ,Sandra Broddesson ,Robert Dyrdak ,Martin Ekman ,Lynda Eneh ,Lina Guerra Blomqvist ,Karolina Ininbergs ,Tanja Normark ,Isak Sylvin ,Zhibing Yun ,Martina Wahlund ,Valtteri Wirta |
| EPI_ISL_2899007 | Karolinska University Hospital Solna | Karolinska University Hospital | Jan Albert ,Tobias Allander ,Annelie Bjerkner ,Sandra Broddesson ,Robert Dyrdak ,Martin Ekman ,Lynda Eneh ,Lina Guerra Blomqvist ,Karolina Ininbergs ,Tanja Normark ,Isak Sylvin ,Zhibing Yun ,Martina Wahlund ,Valtteri Wirta |
| EPI_ISL_2899014, EPI_ISL_2899033 | Unilabs | Karolinska University Hospital | Jan Albert ,Tobias Allander ,Annelie Bjerkner ,Sandra Broddesson ,Robert Dyrdak ,Martin Ekman ,Lynda Eneh ,Lina Guerra Blomqvist ,Karolina Ininbergs ,Tanja Normark ,Isak Sylvin ,Zhibing Yun ,Martina Wahlund ,Valtteri Wirta |
| EPI_ISL_2899047 | Synlab Medilab | Karolinska University Hospital | Jan Albert ,Tobias Allander ,Annelie Bjerkner ,Sandra Broddesson ,Robert Dyrdak ,Martin Ekman ,Lynda Eneh ,Lina Guerra Blomqvist ,Karolina Ininbergs ,Tanja Normark ,Isak Sylvin ,Zhibing Yun ,Martina Wahlund ,Valtteri Wirta |
| EPI_ISL_2899050, EPI_ISL_2899053 | Unilabs | Karolinska University Hospital | Jan Albert ,Tobias Allander ,Annelie Bjerkner ,Sandra Broddesson ,Robert Dyrdak ,Martin Ekman ,Lynda Eneh ,Lina Guerra Blomqvist ,Karolina Ininbergs ,Tanja Normark ,Isak Sylvin ,Zhibing Yun ,Martina Wahlund ,Valtteri Wirta |
| EPI_ISL_2899066, EPI_ISL_2899077 | Karolinska University Hospital Solna | Karolinska University Hospital | Jan Albert ,Tobias Allander ,Annelie Bjerkner ,Sandra Broddesson ,Robert Dyrdak ,Martin Ekman ,Lynda Eneh ,Lina Guerra Blomqvist ,Karolina Ininbergs ,Tanja Normark ,Isak Sylvin ,Zhibing Yun ,Martina Wahlund ,Valtteri Wirta |
| EPI_ISL_2899112, EPI_ISL_2899118, EPI_ISL_2899126, EPI_ISL_2899146, EPI_ISL_2899151, EPI_ISL_2899160, EPI_ISL_2899162, EPI_ISL_2899191, EPI_ISL_2899197, EPI_ISL_2899198, EPI_ISL_2899210, EPI_ISL_2899217, EPI_ISL_2899250 | | | |
| see above | Unilabs | Karolinska University Hospital | Jan Albert ,Tobias Allander ,Annelie Bjerkner ,Sandra Broddesson ,Robert Dyrdak ,Martin Ekman ,Lynda Eneh ,Lina Guerra Blomqvist ,Karolina Ininbergs ,Tanja Normark ,Isak Sylvin ,Zhibing Yun ,Martina Wahlund ,Valtteri Wirta |
| EPI_ISL_2899253 | Karolinska University Hospital Solna | Karolinska University Hospital | Jan Albert ,Tobias Allander ,Annelie Bjerkner ,Sandra Broddesson ,Robert Dyrdak ,Martin Ekman ,Lynda Eneh ,Lina Guerra Blomqvist ,Karolina Ininbergs ,Tanja Normark ,Isak Sylvin ,Zhibing Yun ,Martina Wahlund ,Valtteri Wirta |
| EPI_ISL_2899277 | Unilabs | Karolinska University Hospital | Jan Albert ,Tobias Allander ,Annelie Bjerkner ,Sandra Broddesson ,Robert Dyrdak ,Martin Ekman ,Lynda Eneh ,Lina Guerra Blomqvist ,Karolina Ininbergs ,Tanja Normark ,Isak Sylvin ,Zhibing Yun ,Martina Wahlund ,Valtteri Wirta |
| EPI_ISL_2899282 | Karolinska University Hospital Solna | Karolinska University Hospital | Jan Albert ,Tobias Allander ,Annelie Bjerkner ,Sandra Broddesson ,Robert Dyrdak ,Martin Ekman ,Lynda Eneh ,Lina Guerra Blomqvist ,Karolina Ininbergs ,Tanja Normark ,Isak Sylvin ,Zhibing Yun ,Martina Wahlund ,Valtteri Wirta |
| EPI_ISL_2899300, EPI_ISL_2899304, EPI_ISL_2899306, EPI_ISL_2899308, EPI_ISL_2899315, EPI_ISL_2899319, EPI_ISL_2899320, EPI_ISL_2899321, EPI_ISL_2899328, EPI_ISL_2899330, EPI_ISL_2899334 | | | |
| see above | Karolinska University Hospital Huddinge | Karolinska University Hospital | Jan Albert ,Tobias Allander ,Annelie Bjerkner ,Sandra Broddesson ,Robert Dyrdak ,Martin Ekman ,Lynda Eneh ,Lina Guerra Blomqvist ,Karolina Ininbergs ,Tanja Normark ,Isak Sylvin ,Zhibing Yun ,Martina Wahlund ,Valtteri Wirta |
| EPI_ISL_2899344, EPI_ISL_2899362, EPI_ISL_2899364, EPI_ISL_2899366, EPI_ISL_2899368, EPI_ISL_2899369 | Unilabs | Karolinska University Hospital | Jan Albert ,Tobias Allander ,Annelie Bjerkner ,Sandra Broddesson ,Robert Dyrdak ,Martin Ekman ,Lynda Eneh ,Lina Guerra Blomqvist ,Karolina Ininbergs ,Tanja Normark ,Isak Sylvin ,Zhibing Yun ,Martina Wahlund ,Valtteri Wirta |
| EPI_ISL_2899384 | Synlab Medilab | Karolinska University Hospital | Jan Albert ,Tobias Allander ,Annelie Bjerkner ,Sandra Broddesson ,Robert Dyrdak ,Martin Ekman ,Lynda Eneh ,Lina Guerra Blomqvist ,Karolina Ininbergs ,Tanja Normark ,Isak Sylvin ,Zhibing Yun ,Martina Wahlund ,Valtteri Wirta |
| EPI_ISL_2899405, EPI_ISL_2899413 | Unilabs | Karolinska University Hospital | Jan Albert ,Tobias Allander ,Annelie Bjerkner ,Sandra Broddesson ,Robert Dyrdak ,Martin Ekman ,Lynda Eneh ,Lina Guerra Blomqvist ,Karolina Ininbergs ,Tanja Normark ,Isak Sylvin ,Zhibing Yun ,Martina Wahlund ,Valtteri Wirta |
| EPI_ISL_2899415, EPI_ISL_2899417 | Synlab Medilab | Karolinska University Hospital | Jan Albert ,Tobias Allander ,Annelie Bjerkner ,Sandra Broddesson ,Robert Dyrdak ,Martin Ekman ,Lynda Eneh ,Lina Guerra Blomqvist ,Karolina Ininbergs ,Tanja Normark ,Isak Sylvin ,Zhibing Yun ,Martina Wahlund ,Valtteri Wirta |
| EPI_ISL_2899420 | Karolinska University Hospital Solna | Karolinska University Hospital | Jan Albert ,Tobias Allander ,Annelie Bjerkner ,Sandra Broddesson ,Robert Dyrdak ,Martin Ekman ,Lynda Eneh ,Lina Guerra Blomqvist ,Karolina Ininbergs ,Tanja Normark ,Isak Sylvin ,Zhibing Yun ,Martina Wahlund ,Valtteri Wirta |
| EPI_ISL_2899425, EPI_ISL_2899426 | Synlab Medilab | Karolinska University Hospital | Jan Albert ,Tobias Allander ,Annelie Bjerkner ,Sandra Broddesson ,Robert Dyrdak ,Martin Ekman ,Lynda Eneh ,Lina Guerra Blomqvist ,Karolina Ininbergs ,Tanja Normark ,Isak Sylvin ,Zhibing Yun ,Martina Wahlund ,Valtteri Wirta |
| EPI_ISL_2899427 | Karolinska University Hospital Solna | Karolinska University Hospital | Jan Albert ,Tobias Allander ,Annelie Bjerkner ,Sandra Broddesson ,Robert Dyrdak ,Martin Ekman ,Lynda Eneh ,Lina Guerra Blomqvist ,Karolina Ininbergs ,Tanja Normark ,Isak Sylvin ,Zhibing Yun ,Martina Wahlund ,Valtteri Wirta |
| EPI_ISL_2899430, EPI_ISL_2899437, EPI_ISL_2899440, EPI_ISL_2899447, EPI_ISL_2899451, EPI_ISL_2899457 | Unilabs | Karolinska University Hospital | Jan Albert ,Tobias Allander ,Annelie Bjerkner ,Sandra Broddesson ,Robert Dyrdak ,Martin Ekman ,Lynda Eneh ,Lina Guerra Blomqvist ,Karolina Ininbergs ,Tanja Normark ,Isak Sylvin ,Zhibing Yun ,Martina Wahlund ,Valtteri Wirta |
| EPI_ISL_2899462 | Karolinska University Hospital Solna | Karolinska University Hospital | Jan Albert ,Tobias Allander ,Annelie Bjerkner ,Sandra Broddesson ,Robert Dyrdak ,Martin Ekman ,Lynda Eneh ,Lina Guerra Blomqvist ,Karolina Ininbergs ,Tanja Normark ,Isak Sylvin ,Zhibing Yun ,Martina Wahlund ,Valtteri Wirta |
| EPI_ISL_2899467 | Unilabs | Karolinska University Hospital | Jan Albert ,Tobias Allander ,Annelie Bjerkner ,Sandra Broddesson ,Robert Dyrdak ,Martin Ekman ,Lynda Eneh ,Lina Guerra Blomqvist ,Karolina Ininbergs ,Tanja Normark ,Isak Sylvin ,Zhibing Yun ,Martina Wahlund ,Valtteri Wirta |
| EPI_ISL_2899469 | Karolinska University Hospital Solna | Karolinska University Hospital | Jan Albert ,Tobias Allander ,Annelie Bjerkner ,Sandra Broddesson ,Robert Dyrdak ,Martin Ekman ,Lynda Eneh ,Lina Guerra Blomqvist ,Karolina Ininbergs ,Tanja Normark ,Isak Sylvin ,Zhibing Yun ,Martina Wahlund ,Valtteri Wirta |
| EPI_ISL_2899491 | Unilabs | Karolinska University Hospital | Jan Albert ,Tobias Allander ,Annelie Bjerkner ,Sandra Broddesson ,Robert Dyrdak ,Martin Ekman ,Lynda Eneh ,Lina Guerra Blomqvist ,Karolina Ininbergs ,Tanja Normark ,Isak Sylvin ,Zhibing Yun ,Martina Wahlund ,Valtteri Wirta |
| EPI_ISL_2899511, EPI_ISL_2899515, EPI_ISL_2899516, EPI_ISL_2899517, EPI_ISL_2899520, EPI_ISL_2899526, EPI_ISL_2899527 | Karolinska University Hospital Huddinge | Karolinska University Hospital | Jan Albert ,Tobias Allander ,Annelie Bjerkner ,Sandra Broddesson ,Robert Dyrdak ,Martin Ekman ,Lynda Eneh ,Lina Guerra Blomqvist ,Karolina Ininbergs ,Tanja Normark ,Isak Sylvin ,Zhibing Yun ,Martina Wahlund ,Valtteri Wirta |

| | | | |
|---|---|---|---|
| EPI_ISL_2899528, EPI_ISL_2899529, EPI_ISL_2899531, EPI_ISL_2899532, EPI_ISL_2899534, EPI_ISL_2899537, EPI_ISL_2899539, EPI_ISL_2899540, EPI_ISL_2899542, EPI_ISL_2899553, EPI_ISL_2899557, EPI_ISL_2899561, EPI_ISL_2899569, EPI_ISL_2899574, EPI_ISL_2899575, EPI_ISL_2899576, EPI_ISL_2899582, EPI_ISL_2899586, EPI_ISL_2899591 | | | |
| see above | Unilabs | Karolinska University Hospital | Jan Albert ,Tobias Allander ,Annelie Bjerkner ,Sandra Broddesson ,Robert Dyrdak ,Martin Ekman ,Lynda Eneh ,Lina Guerra Blomqvist ,Karolina Ininbergs ,Tanja Normark ,Isak Sylvin ,Zhibing Yun ,Martina Wahlund ,Valtteri Wirta |
| EPI_ISL_2899666, EPI_ISL_2899674 | Karolinska University Hospital Huddinge | Karolinska University Hospital | Jan Albert ,Tobias Allander ,Annelie Bjerkner ,Sandra Broddesson ,Robert Dyrdak ,Martin Ekman ,Lynda Eneh ,Lina Guerra Blomqvist ,Karolina Ininbergs ,Tanja Normark ,Isak Sylvin ,Zhibing Yun ,Martina Wahlund ,Valtteri Wirta |
| EPI_ISL_2899687, EPI_ISL_2899688, EPI_ISL_2899689, EPI_ISL_2899690, EPI_ISL_2899691, EPI_ISL_2899692, EPI_ISL_2899693, EPI_ISL_2899695, EPI_ISL_2899696, EPI_ISL_2899697, EPI_ISL_2899698, EPI_ISL_2899699, EPI_ISL_2899700, EPI_ISL_2899701, EPI_ISL_2899703, EPI_ISL_2899704, EPI_ISL_2899705, EPI_ISL_2899706, EPI_ISL_2899707, EPI_ISL_2899708, EPI_ISL_2899709, EPI_ISL_2899710, EPI_ISL_2899711, EPI_ISL_2899712, EPI_ISL_2899713, EPI_ISL_2899714, EPI_ISL_2899715, EPI_ISL_2899716, EPI_ISL_2899717, EPI_ISL_2899718, EPI_ISL_2899719, EPI_ISL_2899720, EPI_ISL_2899721, EPI_ISL_2899722, EPI_ISL_2899723, EPI_ISL_2899724, EPI_ISL_2899725, EPI_ISL_2899726, EPI_ISL_2899727, EPI_ISL_2899728, EPI_ISL_2899729, EPI_ISL_2899730, EPI_ISL_2899731, EPI_ISL_2899732, EPI_ISL_2899733, EPI_ISL_2899734, EPI_ISL_2899735, EPI_ISL_2899736, EPI_ISL_2899738, EPI_ISL_2899739, EPI_ISL_2899740 | | | |
| see above | National Health Laboratory Service, South Africa | KRISP, KZn Research Innovation and Sequencing Platform | Giandhari Jennifer, Pillay Sureshnee, Yajna Ramphal, Naidoo Yeshnee, Tshabuila Derek,Tegally Houriiyah, San James, Wilkinson Eduan, de Oliveira Tulio |
| EPI_ISL_2899741, EPI_ISL_2899742, EPI_ISL_2899743, EPI_ISL_2899744, EPI_ISL_2899745, EPI_ISL_2899746, EPI_ISL_2899747, EPI_ISL_2899748, EPI_ISL_2899749, EPI_ISL_2899750, EPI_ISL_2899751, EPI_ISL_2899752, EPI_ISL_2899753, EPI_ISL_2899754, EPI_ISL_2899755, EPI_ISL_2899756, EPI_ISL_2899757 | | | |
| see above | ZARV/NHLS, Department Medical Virology, University of Pretoria | KRISP, KZn Research Innovation and Sequencing Platform | Amy Strydom, Adriano Mendes, Micheala Davids, Sim Mayaphi and Marietjie Venter, Giandhari J, Pillay S, Naidoo Y, Ramphal U, Lessells R, Emmanuel SJ, Tegally H, Wilkinson E, de Oliveira T |
| EPI_ISL_2899759, EPI_ISL_2899760, EPI_ISL_2899761, EPI_ISL_2899762, EPI_ISL_2899763, EPI_ISL_2899764, EPI_ISL_2899765, EPI_ISL_2899767, EPI_ISL_2899768, EPI_ISL_2899769, EPI_ISL_2899771, EPI_ISL_2899772, EPI_ISL_2899773, EPI_ISL_2899775, EPI_ISL_2899777, EPI_ISL_2899778, EPI_ISL_2899779, EPI_ISL_2899780, EPI_ISL_2899781, EPI_ISL_2899782, EPI_ISL_2899783, EPI_ISL_2899784, EPI_ISL_2899785, EPI_ISL_2899786, EPI_ISL_2899787, EPI_ISL_2899788, EPI_ISL_2899789, EPI_ISL_2899790 | | | |
| see above | HELIX LLC | WHO National Influenza Centre Russian Federation | Andrey Komissarov, Artem Fadeev, Kseniya Komissarova, Oula Mansour, Kirill Varchenko, Mikhail Bakaev, Tamila Musaeva, Veronika Eder, Maria Pisareva, Nikita Yolshin, Daria Danilenko, Ksenia Safina, Elena Nabieva, Georgii Bazykin, Dmitry Lioznov |
| EPI_ISL_2899791, EPI_ISL_2899792 | City Hospital No 40 | WHO National Influenza Centre Russian Federation | Andrey Komissarov, Artem Fadeev, Kseniya Komissarova, Oula Mansour, Kirill Varchenko, Mikhail Bakaev, Tamila Musaeva, Veronika Eder, Maria Pisareva, Nikita Yolshin, Daria Danilenko, Olga Shneider, Sergey Scherbak, Ksenia Safina, Elena Nabieva, Georgii Bazykin, Dmitry Lioznov |
| EPI_ISL_2899794, EPI_ISL_2899796, EPI_ISL_2899797 | D. I. Ivanovskiy Institute of Virology | WHO National Influenza Centre Russian Federation | Andrey Komissarov, Artem Fadeev, Kseniya Komissarova, Oula Mansour, Kirill Varchenko, Mikhail Bakaev, Tamila Musaeva, Veronika Eder, Maria Pisareva, Nikita Yolshin, Daria Danilenko, Elena Burtseva, Kirill Krasnoslobodtsev, Ksenia Safina, Elena Nabieva, Georgii Bazykin, Dmitry Lioznov |
| EPI_ISL_2899801, EPI_ISL_2899802, EPI_ISL_2899804, EPI_ISL_2899805, EPI_ISL_2899806, EPI_ISL_2899807, EPI_ISL_2899808, EPI_ISL_2899809, EPI_ISL_2899811, EPI_ISL_2899812, EPI_ISL_2899813, EPI_ISL_2899815, EPI_ISL_2899816, EPI_ISL_2899817, EPI_ISL_2899818, EPI_ISL_2899819, EPI_ISL_2899820, EPI_ISL_2899821, EPI_ISL_2899822, EPI_ISL_2899823, EPI_ISL_2899824, EPI_ISL_2899825, EPI_ISL_2899826, EPI_ISL_2899827, EPI_ISL_2899828, EPI_ISL_2899829, EPI_ISL_2899830, EPI_ISL_2899831, EPI_ISL_2899832, EPI_ISL_2899833, EPI_ISL_2899834, EPI_ISL_2899835, EPI_ISL_2899836, EPI_ISL_2899837, EPI_ISL_2899838, EPI_ISL_2899839, EPI_ISL_2899840, EPI_ISL_2899841, EPI_ISL_2899842, EPI_ISL_2899843, EPI_ISL_2899844, EPI_ISL_2899845, EPI_ISL_2899846, EPI_ISL_2899847, EPI_ISL_2899848, EPI_ISL_2899849, EPI_ISL_2899850, EPI_ISL_2899851, EPI_ISL_2899852, EPI_ISL_2899853, EPI_ISL_2899854, EPI_ISL_2899855, EPI_ISL_2899856, EPI_ISL_2899857, EPI_ISL_2899858, EPI_ISL_2899859, EPI_ISL_2899860, EPI_ISL_2899861, EPI_ISL_2899862, EPI_ISL_2899863, EPI_ISL_2899864, EPI_ISL_2899865, EPI_ISL_2899866, EPI_ISL_2899867, EPI_ISL_2899868, EPI_ISL_2899869, EPI_ISL_2899870, EPI_ISL_2899871, EPI_ISL_2899872, EPI_ISL_2899873, EPI_ISL_2899874, EPI_ISL_2899875, EPI_ISL_2899876, EPI_ISL_2899877, EPI_ISL_2899878, EPI_ISL_2899879, EPI_ISL_2899880, EPI_ISL_2899881, EPI_ISL_2899882, EPI_ISL_2899883, EPI_ISL_2899884, EPI_ISL_2899885, EPI_ISL_2899886, EPI_ISL_2899887, EPI_ISL_2899888, EPI_ISL_2899889, EPI_ISL_2899890, EPI_ISL_2899891, EPI_ISL_2899892, EPI_ISL_2899893, EPI_ISL_2899894, EPI_ISL_2899895, EPI_ISL_2899896, EPI_ISL_2899897, EPI_ISL_2899898, EPI_ISL_2899901, EPI_ISL_2899902, EPI_ISL_2899903, EPI_ISL_2899904, EPI_ISL_2899905, EPI_ISL_2899906, EPI_ISL_2899907, EPI_ISL_2899908, EPI_ISL_2899909, EPI_ISL_2899910, EPI_ISL_2899911, EPI_ISL_2899912, EPI_ISL_2899913, EPI_ISL_2899914, EPI_ISL_2899915, EPI_ISL_2899916, EPI_ISL_2899917, EPI_ISL_2899918, EPI_ISL_2899919, EPI_ISL_2899920, EPI_ISL_2899921, EPI_ISL_2899922, EPI_ISL_2899923, EPI_ISL_2899924, EPI_ISL_2899925 | | | |
| see above | N.F. Gamaleya Research Center for Epidemiology and Microbiology | WHO National Influenza Centre Russian Federation | Andrey Komissarov, Artem Fadeev, Kseniya Komissarova, Oula Mansour, Kirill Varchenko, Mikhail Bakaev, Tamila Musaeva, Veronika Eder, Maria Pisareva, Nikita Yolshin, Daria Danilenko, Elizaveta Divisenko, Alina Odintsova, Elena Shidlovskaya, Ludmila Vasilchenko, Nadezhda Kuznetsova, Maria Nikiforova, Alexey Shchetinin, Artem Tkachuk, Andrei Pochtovyi, Andrei Siniavin, Denis Kleymenov, Evgeniia Bykonia, Liubov Popova, Elena Mazunina, Inna Dolzhikova, Anna Kovyrshina, Andrei Botikov, Dmitry Shcheblyakov, Vladimir Gushchin, Denis Logunov, Alexander Gintsburg, Ksenia Safina, Elena Nabieva, Georgii Bazykin, Dmitry Lioznov |
| EPI_ISL_2900247, EPI_ISL_2900253 | CHUGA-IBP-laboratoire de Virologie | IBP-laboratoire de virologie | Sylvie Larrat; Patrice Morand; Pascal Poignard; Raphaële Germi; Julien Lupo; Anne Signori -Schmuck; Anne-Karen Faure; Benjamin Nemoz; Aurélie Truffot; Léa Ponderand; Julien Andréani; Alban Caporossi; Hugo Jardin |
| EPI_ISL_2900365, EPI_ISL_2900366, EPI_ISL_2900367, EPI_ISL_2900368, EPI_ISL_2900370, EPI_ISL_2900371, EPI_ISL_2900372, EPI_ISL_2900373, EPI_ISL_2900374, EPI_ISL_2900375, EPI_ISL_2900376, EPI_ISL_2900377, EPI_ISL_2900378, EPI_ISL_2900379, EPI_ISL_2900380, EPI_ISL_2900381, EPI_ISL_2900382, EPI_ISL_2900383, EPI_ISL_2900384, EPI_ISL_2900385, EPI_ISL_2900386, EPI_ISL_2900387, EPI_ISL_2900388, EPI_ISL_2900389, EPI_ISL_2900390, EPI_ISL_2900391, EPI_ISL_2900392, EPI_ISL_2900393, EPI_ISL_2900394, EPI_ISL_2900395, EPI_ISL_2900396, EPI_ISL_2900397, EPI_ISL_2900398, EPI_ISL_2900399, EPI_ISL_2900400, EPI_ISL_2900401, EPI_ISL_2900402, EPI_ISL_2900403, EPI_ISL_2900404, EPI_ISL_2900405, EPI_ISL_2900406, EPI_ISL_2900407, EPI_ISL_2900408, EPI_ISL_2900409, EPI_ISL_2900410, EPI_ISL_2900411, EPI_ISL_2900412, EPI_ISL_2900413, EPI_ISL_2900414, EPI_ISL_2900415, EPI_ISL_2900416, EPI_ISL_2900417, EPI_ISL_2900418, EPI_ISL_2900419, EPI_ISL_2900420, EPI_ISL_2900421, EPI_ISL_2900422, EPI_ISL_2900423, EPI_ISL_2900424, EPI_ISL_2900425, EPI_ISL_2900426, EPI_ISL_2900427, EPI_ISL_2900428, EPI_ISL_2900429, EPI_ISL_2900430, EPI_ISL_2900431, EPI_ISL_2900432, EPI_ISL_2900433, EPI_ISL_2900434, EPI_ISL_2900435, EPI_ISL_2900436, EPI_ISL_2900437, EPI_ISL_2900438, EPI_ISL_2900439, EPI_ISL_2900440, EPI_ISL_2900441, EPI_ISL_2900442, EPI_ISL_2900443, EPI_ISL_2900444, EPI_ISL_2900445, EPI_ISL_2900446, EPI_ISL_2900447, EPI_ISL_2900448, EPI_ISL_2900449, EPI_ISL_2900450, EPI_ISL_2900451, EPI_ISL_2900452, EPI_ISL_2900453, EPI_ISL_2900454, EPI_ISL_2900455, EPI_ISL_2900456, EPI_ISL_2900457, EPI_ISL_2900458, EPI_ISL_2900459, EPI_ISL_2900460, EPI_ISL_2900462, EPI_ISL_2900463, EPI_ISL_2900464, EPI_ISL_2900465, EPI_ISL_2900466, EPI_ISL_2900467, EPI_ISL_2900468, EPI_ISL_2900469, EPI_ISL_2900470, EPI_ISL_2900471, EPI_ISL_2900472, EPI_ISL_2900473, EPI_ISL_2900474, EPI_ISL_2900475, EPI_ISL_2900476, EPI_ISL_2900477, EPI_ISL_2900478, EPI_ISL_2900479, EPI_ISL_2900480, EPI_ISL_2900481, EPI_ISL_2900482, EPI_ISL_2900483, EPI_ISL_2900484, EPI_ISL_2900485, EPI_ISL_2900486, EPI_ISL_2900487, EPI_ISL_2900488, EPI_ISL_2900489, EPI_ISL_2900490, EPI_ISL_2900491, EPI_ISL_2900492, EPI_ISL_2900493, EPI_ISL_2900494, EPI_ISL_2900495, EPI_ISL_2900496, EPI_ISL_2900497, EPI_ISL_2900498, EPI_ISL_2900499, EPI_ISL_2900500, EPI_ISL_2900501, EPI_ISL_2900502, EPI_ISL_2900503, EPI_ISL_2900504, EPI_ISL_2900505, EPI_ISL_2900506, EPI_ISL_2900507, EPI_ISL_2900508, EPI_ISL_2900509, EPI_ISL_2900510, EPI_ISL_2900511, EPI_ISL_2900512, EPI_ISL_2900513, EPI_ISL_2900514, EPI_ISL_2900516, EPI_ISL_2900517, EPI_ISL_2900518, EPI_ISL_2900519, EPI_ISL_2900520, EPI_ISL_2900521, EPI_ISL_2900522, EPI_ISL_2900523, EPI_ISL_2900524, EPI_ISL_2900525, EPI_ISL_2900526, EPI_ISL_2900527, EPI_ISL_2900528, EPI_ISL_2900529, EPI_ISL_2900530, EPI_ISL_2900531, EPI_ISL_2900532, EPI_ISL_2900533, EPI_ISL_2900535, EPI_ISL_2900536, EPI_ISL_2900537, EPI_ISL_2900538, EPI_ISL_2900539, EPI_ISL_2900540, EPI_ISL_2900541, EPI_ISL_2900542, EPI_ISL_2900543, EPI_ISL_2900544, EPI_ISL_2900545, EPI_ISL_2900546, EPI_ISL_2900547, EPI_ISL_2900548, EPI_ISL_2900549, EPI_ISL_2900550, EPI_ISL_2900551, EPI_ISL_2900552, EPI_ISL_2900553, EPI_ISL_2900554, EPI_ISL_2900555, EPI_ISL_2900556, EPI_ISL_2900557, EPI_ISL_2900558, EPI_ISL_2900559, EPI_ISL_2900560, EPI_ISL_2900561, EPI_ISL_2900562, EPI_ISL_2900563, EPI_ISL_2900564, EPI_ISL_2900565, EPI_ISL_2900566, EPI_ISL_2900567, EPI_ISL_2900568, EPI_ISL_2900569, EPI_ISL_2900570, EPI_ISL_2900571, EPI_ISL_2900572, EPI_ISL_2900573, EPI_ISL_2900574, EPI_ISL_2900575, EPI_ISL_2900576, EPI_ISL_2900577, EPI_ISL_2900578, EPI_ISL_2900579, EPI_ISL_2900580, EPI_ISL_2900581, EPI_ISL_2900582, EPI_ISL_2900583, EPI_ISL_2900584, EPI_ISL_2900585, EPI_ISL_2900586, EPI_ISL_2900587, EPI_ISL_2900588, EPI_ISL_2900589, EPI_ISL_2900590, EPI_ISL_2900591, EPI_ISL_2900592, EPI_ISL_2900593, EPI_ISL_2900594, EPI_ISL_2900595, EPI_ISL_2900596, EPI_ISL_2900597, EPI_ISL_2900598, EPI_ISL_2900599, EPI_ISL_2900600, EPI_ISL_2900601, EPI_ISL_2900602, EPI_ISL_2900603, EPI_ISL_2900604, EPI_ISL_2900605, EPI_ISL_2900606, EPI_ISL_2900607, EPI_ISL_2900608, EPI_ISL_2900609, EPI_ISL_2900610, EPI_ISL_2900611, EPI_ISL_2900612, EPI_ISL_2900613, EPI_ISL_2900614, EPI_ISL_2900615, EPI_ISL_2900616, EPI_ISL_2900617, EPI_ISL_2900618, EPI_ISL_2900619, EPI_ISL_2900620, EPI_ISL_2900621, EPI_ISL_2900622, EPI_ISL_2900623, EPI_ISL_2900624, EPI_ISL_2900625, EPI_ISL_2900626, EPI_ISL_2900627, EPI_ISL_2900628, EPI_ISL_2900629, EPI_ISL_2900630, EPI_ISL_2900631, EPI_ISL_2900632, EPI_ISL_2900633, EPI_ISL_2900634, EPI_ISL_2900635, EPI_ISL_2900636, EPI_ISL_2900637, EPI_ISL_2900638, EPI_ISL_2900639, EPI_ISL_2900640, EPI_ISL_2900641, EPI_ISL_2900642, EPI_ISL_2900643, EPI_ISL_2900644, EPI_ISL_2900645, EPI_ISL_2900646, EPI_ISL_2900647, EPI_ISL_2900648, EPI_ISL_2900649, EPI_ISL_2900650, EPI_ISL_2900651, EPI_ISL_2900652, EPI_ISL_2900653, EPI_ISL_2900654, EPI_ISL_2900655, EPI_ISL_2900656, EPI_ISL_2900657, EPI_ISL_2900658, EPI_ISL_2900659, EPI_ISL_2900660, EPI_ISL_2900661, EPI_ISL_2900662, EPI_ISL_2900663, EPI_ISL_2900664, EPI_ISL_2900665, EPI_ISL_2900666, EPI_ISL_2900667, EPI_ISL_2900668, EPI_ISL_2900669, EPI_ISL_2900670, EPI_ISL_2900671, EPI_ISL_2900672, EPI_ISL_2900673, EPI_ISL_2900674, EPI_ISL_2900675, EPI_ISL_2900676, EPI_ISL_2900677, EPI_ISL_2900678, EPI_ISL_2900679, EPI_ISL_2900680, EPI_ISL_2900681, EPI_ISL_2900682, EPI_ISL_2900683, EPI_ISL_2900684, EPI_ISL_2900685, EPI_ISL_2900686, EPI_ISL_2900687, EPI_ISL_2900688, EPI_ISL_2900689, EPI_ISL_2900690 | | | |
| see above | Department of Pathology, University of Cambridge | COVID-19 Genomics UK (COG-UK) Consortium | Aminu S. Jahun, Yasmin Chaudhry, Iliana Georgana, Myra Hosmillo, Rhys Izuagbe, William L. Hamilton, Martin D. Curran, Surendra Parmar, Ian Goodfellow |
| EPI_ISL_2900691, EPI_ISL_2900692, EPI_ISL_2900693, EPI_ISL_2900694, EPI_ISL_2900695, EPI_ISL_2900696, EPI_ISL_2900697, EPI_ISL_2900698, EPI_ISL_2900699, EPI_ISL_2900700, EPI_ISL_2900701, EPI_ISL_2900702, EPI_ISL_2900703, EPI_ISL_2900704, EPI_ISL_2900705, EPI_ISL_2900706, EPI_ISL_2900707, EPI_ISL_2900708, EPI_ISL_2900709, EPI_ISL_2900710, EPI_ISL_2900711, EPI_ISL_2900712, EPI_ISL_2900713, EPI_ISL_2900714, EPI_ISL_2900715, EPI_ISL_2900716, EPI_ISL_2900717, EPI_ISL_2900719, EPI_ISL_2900720, EPI_ISL_2900721, EPI_ISL_2900722, EPI_ISL_2900724, EPI_ISL_2900725, EPI_ISL_2900726, EPI_ISL_2900727, EPI_ISL_2900728, EPI_ISL_2900729, EPI_ISL_2900730, EPI_ISL_2900731, EPI_ISL_2900732, EPI_ISL_2900735, EPI_ISL_2900736, EPI_ISL_2900737, EPI_ISL_2900738, EPI_ISL_2900739, EPI_ISL_2900740, EPI_ISL_2900742, EPI_ISL_2900743, EPI_ISL_2900744, EPI_ISL_2900745, EPI_ISL_2900746 | | | |
| see above | West of Scotland Specialist Virology Centre, NHSGGC / MRC-University of Glasgow Centre for Virus Research | COVID-19 Genomics UK (COG-UK) Consortium | Ana da Silva Filipe, Natasha Johnson, Kathy Smollett, Daniel Mair, Stephen Carmichael, Alice Broos, Lily Tong, Jenna Nichols, Kyriaki Nomikou; Sarah McDonald; Ioulia Tsatsani, Richard Orton, Joseph Hughes, Sreenu Vattipally, David L Robertson; Alasdair MacLean, Rory Gunson; Sharif Shaaban, Matthew Holden; Rachel Blacow, Guy Mollett, Kathy Li, James Shepherd, Antonia Ho, Emma Thomson |
| EPI_ISL_2900747, EPI_ISL_2900748, EPI_ISL_2900749, EPI_ISL_2900750, EPI_ISL_2900751, EPI_ISL_2900752, EPI_ISL_2900753, EPI_ISL_2900754, EPI_ISL_2900755, EPI_ISL_2900756, EPI_ISL_2900757, EPI_ISL_2900758, EPI_ISL_2900759, EPI_ISL_2900760, EPI_ISL_2900761, EPI_ISL_2900762, EPI_ISL_2900763, EPI_ISL_2900764, EPI_ISL_2900765, EPI_ISL_2900766, EPI_ISL_2900767, EPI_ISL_2900769, EPI_ISL_2900770, EPI_ISL_2900771, EPI_ISL_2900772, EPI_ISL_2900774, EPI_ISL_2900775, EPI_ISL_2900776, EPI_ISL_2900777, EPI_ISL_2900778, EPI_ISL_2900779, EPI_ISL_2900780, EPI_ISL_2900781, EPI_ISL_2900782, EPI_ISL_2900783, EPI_ISL_2900784, EPI_ISL_2900785, EPI_ISL_2900786, EPI_ISL_2900787, EPI_ISL_2900788, EPI_ISL_2900789, EPI_ISL_2900790, EPI_ISL_2900791, EPI_ISL_2900792, EPI_ISL_2900793, EPI_ISL_2900794, EPI_ISL_2900795, EPI_ISL_2900796, EPI_ISL_2900797, EPI_ISL_2900798, EPI_ISL_2900799, EPI_ISL_2900800, EPI_ISL_2900801, EPI_ISL_2900802, EPI_ISL_2900803, EPI_ISL_2900804, EPI_ISL_2900805, EPI_ISL_2900806, EPI_ISL_2900807, EPI_ISL_2900808, EPI_ISL_2900809, EPI_ISL_2900810, EPI_ISL_2900811, EPI_ISL_2900812, EPI_ISL_2900813, EPI_ISL_2900814, EPI_ISL_2900815, EPI_ISL_2900816, EPI_ISL_2900817, EPI_ISL_2900818, EPI_ISL_2900819, EPI_ISL_2900820, EPI_ISL_2900821, EPI_ISL_2900822, EPI_ISL_2900823, EPI_ISL_2900824, EPI_ISL_2900825, EPI_ISL_2900826, EPI_ISL_2900827, EPI_ISL_2900828, EPI_ISL_2900829, EPI_ISL_2900830, EPI_ISL_2900831, EPI_ISL_2900832, EPI_ISL_2900833, EPI_ISL_2900834, EPI_ISL_2900835, EPI_ISL_2900836, EPI_ISL_2900837, EPI_ISL_2900838, EPI_ISL_2900839, EPI_ISL_2900840, EPI_ISL_2900841, EPI_ISL_2900842, EPI_ISL_2900843, EPI_ISL_2900844 | | | |
| see above | Virology Department, Royal Infirmary of Edinburgh, NHS Lothian / School of Biological Sciences, University of Edinburgh | COVID-19 Genomics UK (COG-UK) Consortium | McHugh M, Dewar R, Cotton S, Gallagher A, Maloney D, Rooke S, O'Toole Á, Scher E, Hill V, McCrone JT, Colquhoun R, Yu X, Jackson B, Rambaut A, Templeton K |
| EPI_ISL_2900845, EPI_ISL_2900846, EPI_ISL_2900847, EPI_ISL_2900848, EPI_ISL_2900849, EPI_ISL_2900850, EPI_ISL_2900851, EPI_ISL_2900852, EPI_ISL_2900853, EPI_ISL_2900854, EPI_ISL_2900855, EPI_ISL_2900856, EPI_ISL_2900857, EPI_ISL_2900858, EPI_ISL_2900859, EPI_ISL_2900860, EPI_ISL_2900861, EPI_ISL_2900862, EPI_ISL_2900863, EPI_ISL_2900864, EPI_ISL_2900865, EPI_ISL_2900866, EPI_ISL_2900867, EPI_ISL_2900868, EPI_ISL_2900869, EPI_ISL_2900870, EPI_ISL_2900871, EPI_ISL_2900872, EPI_ISL_2900873, EPI_ISL_2900874, EPI_ISL_2900875, EPI_ISL_2900876, EPI_ISL_2900877, EPI_ISL_2900878, EPI_ISL_2900879, EPI_ISL_2900880, EPI_ISL_2900881, EPI_ISL_2900882, EPI_ISL_2900883, EPI_ISL_2900884, EPI_ISL_2900885, EPI_ISL_2900886, EPI_ISL_2900887, EPI_ISL_2900888, EPI_ISL_2900889, EPI_ISL_2900890, EPI_ISL_2900891, EPI_ISL_2900892, EPI_ISL_2900893, EPI_ISL_2900894, EPI_ISL_2900895, EPI_ISL_2900896, EPI_ISL_2900897, EPI_ISL_2900898, EPI_ISL_2900899, EPI_ISL_2900900, EPI_ISL_2900901, EPI_ISL_2900902, EPI_ISL_2900903, EPI_ISL_2900904, EPI_ISL_2900905, EPI_ISL_2900906, EPI_ISL_2900907, EPI_ISL_2900908, EPI_ISL_2900909, EPI_ISL_2900910, EPI_ISL_2900911, EPI_ISL_2900912, EPI_ISL_2900913, EPI_ISL_2900914, EPI_ISL_2900915, EPI_ISL_2900916, EPI_ISL_2900917, EPI_ISL_2900918, EPI_ISL_2900919, EPI_ISL_2900920, EPI_ISL_2900921, EPI_ISL_2900922, EPI_ISL_2900923, EPI_ISL_2900924, EPI_ISL_2900925, EPI_ISL_2900926, EPI_ISL_2900927, EPI_ISL_2900928, EPI_ISL_2900929, EPI_ISL_2900930, EPI_ISL_2900931, EPI_ISL_2900932, EPI_ISL_2900933, EPI_ISL_2900934, EPI_ISL_2900935, EPI_ISL_2900936, EPI_ISL_2900937, EPI_ISL_2900938, EPI_ISL_2900939, EPI_ISL_2900940, EPI_ISL_2900941, EPI_ISL_2900942, EPI_ISL_2900943, EPI_ISL_2900944, EPI_ISL_2900945, EPI_ISL_2900946, EPI_ISL_2900947, EPI_ISL_2900948, EPI_ISL_2900949, EPI_ISL_2900950, EPI_ISL_2900951, EPI_ISL_2900952, EPI_ISL_2900953, EPI_ISL_2900954, EPI_ISL_2900955, EPI_ISL_2900956, EPI_ISL_2900957, EPI_ISL_2900958, EPI_ISL_2900959, EPI_ISL_2900960, EPI_ISL_2900961, EPI_ISL_2900962, EPI_ISL_2900963, EPI_ISL_2900964, EPI_ISL_2900965, EPI_ISL_2900966, EPI_ISL_2900967, EPI_ISL_2900968, EPI_ISL_2900969, EPI_ISL_2900970, EPI_ISL_2900971, EPI_ISL_2900972, EPI_ISL_2900973, EPI_ISL_2900974, EPI_ISL_2900975, EPI_ISL_2900976, EPI_ISL_2900977, EPI_ISL_2900978, EPI_ISL_2900979, EPI_ISL_2900980, EPI_ISL_2900981, EPI_ISL_2900982, EPI_ISL_2900983, EPI_ISL_2900984, EPI_ISL_2900985, EPI_ISL_2900986, EPI_ISL_2900987, EPI_ISL_2900988, EPI_ISL_2900989, EPI_ISL_2900990, EPI_ISL_2900991, EPI_ISL_2900992, EPI_ISL_2900993, EPI_ISL_2900994, EPI_ISL_2900995, EPI_ISL_2900996, EPI_ISL_2900997, EPI_ISL_2900998, EPI_ISL_2900999, EPI_ISL_2901000, EPI_ISL_2901001, EPI_ISL_2901002, EPI_ISL_2901003, EPI_ISL_2901004, EPI_ISL_2901005, EPI_ISL_2901006, EPI_ISL_2901007, EPI_ISL_2901008, EPI_ISL_2901009, EPI_ISL_2901010, EPI_ISL_2901011, EPI_ISL_2901012, EPI_ISL_2901013, EPI_ISL_2901014, EPI_ISL_2901015, EPI_ISL_2901016, EPI_ISL_2901017, EPI_ISL_2901018, EPI_ISL_2901019, EPI_ISL_2901020, EPI_ISL_2901021, EPI_ISL_2901022, EPI_ISL_2901023, EPI_ISL_2901024, EPI_ISL_2901025, EPI_ISL_2901026, EPI_ISL_2901027, EPI_ISL_2901028, EPI_ISL_2901029, EPI_ISL_2901030, EPI_ISL_2901031, EPI_ISL_2901032, EPI_ISL_2901033, EPI_ISL_2901034, EPI_ISL_2901035, EPI_ISL_2901036, EPI_ISL_2901037, EPI_ISL_2901038, EPI_ISL_2901039, EPI_ISL_2901040, EPI_ISL_2901041, EPI_ISL_2901043, EPI_ISL_2901044, EPI_ISL_2901045, EPI_ISL_2901046, EPI_ISL_2901047, EPI_ISL_2901048, EPI_ISL_2901049, EPI_ISL_2901050, EPI_ISL_2901051, EPI_ISL_2901052, EPI_ISL_2901053, EPI_ISL_2901055, EPI_ISL_2901056, EPI_ISL_2901057, EPI_ISL_2901058, EPI_ISL_2901059, EPI_ISL_2901060, EPI_ISL_2901061, EPI_ISL_2901062, EPI_ISL_2901063, EPI_ISL_2901064, EPI_ISL_2901065, EPI_ISL_2901066, EPI_ISL_2901067, EPI_ISL_2901068, EPI_ISL_2901069, EPI_ISL_2901070, EPI_ISL_2901071, EPI_ISL_2901072, EPI_ISL_2901073, EPI_ISL_2901075, EPI_ISL_2901076, EPI_ISL_2901077, EPI_ISL_2901078, EPI_ISL_2901079, EPI_ISL_2901080, EPI_ISL_2901081, EPI_ISL_2901082, EPI_ISL_2901083, EPI_ISL_2901084, EPI_ISL_2901085, EPI_ISL_2901086, EPI_ISL_2901087, EPI_ISL_2901088, EPI_ISL_2901089, EPI_ISL_2901090, EPI_ISL_2901091, EPI_ISL_2901092, EPI_ISL_2901093, EPI_ISL_2901094, EPI_ISL_2901095, EPI_ISL_2901096, EPI_ISL_2901097, EPI_ISL_2901098, EPI_ISL_2901099, EPI_ISL_2901100, EPI_ISL_2901101, EPI_ISL_2901102, EPI_ISL_2901103, EPI_ISL_2901104, EPI_ISL_2901105, EPI_ISL_2901106, EPI_ISL_2901107, EPI_ISL_2901108, EPI_ISL_2901109, EPI_ISL_2901110, EPI_ISL_2901111, EPI_ISL_2901112, EPI_ISL_2901113, EPI_ISL_2901114, EPI_ISL_2901115, EPI_ISL_2901116, EPI_ISL_2901117, EPI_ISL_2901118, EPI_ISL_2901119, EPI_ISL_2901120, EPI_ISL_2901121, EPI_ISL_2901122, EPI_ISL_2901123, EPI_ISL_2901124, EPI_ISL_2901125, EPI_ISL_2901126, EPI_ISL_2901127, EPI_ISL_2901128, EPI_ISL_2901129, EPI_ISL_2901130, EPI_ISL_2901131, EPI_ISL_2901132, EPI_ISL_2901133, EPI_ISL_2901134, EPI_ISL_2901135, EPI_ISL_2901136, EPI_ISL_2901137, EPI_ISL_2901138, EPI_ISL_2901139, EPI_ISL_2901140, EPI_ISL_2901141, EPI_ISL_2901142, EPI_ISL_2901143, EPI_ISL_2901144, EPI_ISL_2901145, EPI_ISL_2901146, EPI_ISL_2901147, EPI_ISL_2901148, EPI_ISL_2901149, EPI_ISL_2901150, EPI_ISL_2901151, EPI_ISL_2901152, EPI_ISL_2901153, EPI_ISL_2901154, EPI_ISL_2901155, EPI_ISL_2901156, EPI_ISL_2901157, EPI_ISL_2901158, EPI_ISL_2901159, EPI_ISL_2901160, EPI_ISL_2901161, EPI_ISL_2901162, EPI_ISL_2901163, EPI_ISL_2901164, EPI_ISL_2901166, EPI_ISL_2901167, EPI_ISL_2901168, EPI_ISL_2901169, EPI_ISL_2901170, EPI_ISL_2901171, EPI_ISL_2901172, EPI_ISL_2901173, EPI_ISL_2901174, EPI_ISL_2901175, EPI_ISL_2901176, EPI_ISL_2901177, EPI_ISL_2901178, EPI_ISL_2901179, EPI_ISL_2901180, EPI_ISL_2901181, EPI_ISL_2901182, EPI_ISL_2901183, EPI_ISL_2901184, EPI_ISL_2901185 | | | |
| see above | Liverpool Clinical Laboratories | COVID-19 Genomics UK (COG-UK) Consortium | Sam Haldenby, Alistair Darby, Steve Paterson, Anita Lucaci, Julian Hiscox, M Almsaud, A Alrezaihi, Muhannad Alruwaili, Stuart D Armstrong, Jones Benjamin, Eleanor G Bentley, Anu Chawla, Jordan J Clark, Angela Cowell, Richard Eccles, Isabel García-Dorival, Matthew Gemmell, Alessandro Gerada, PKF Gilmore, Richard Gregory, Ximeng Han, Catherine Hartley, Margaret Hughes, Miren Iturriza-Gomara, James Johnson, L Luu, Jenifer Manson, Charlotte Nelson, Elaine O'Toole, Cassie Olateju, Rebekah Penrice-Randal , Lucille Rainbow, N.P Randle, Trevor Ian Robinson, Parul Sharma, Ghada T Shawli, James P Stewart, Neil Swainston, Ecaterina Vamos, Joanne Watts, Mark Whitehead, Hermione Webster |
| EPI_ISL_2901186, EPI_ISL_2901187, EPI_ISL_2901188, EPI_ISL_2901189, EPI_ISL_2901190, EPI_ISL_2901191, EPI_ISL_2901192, EPI_ISL_2901193, EPI_ISL_2901194, EPI_ISL_2901195, EPI_ISL_2901196, EPI_ISL_2901197, EPI_ISL_2901198, EPI_ISL_2901199, EPI_ISL_2901200, EPI_ISL_2901201, EPI_ISL_2901202, EPI_ISL_2901203, EPI_ISL_2901204, EPI_ISL_2901205, EPI_ISL_2901206, EPI_ISL_2901207, EPI_ISL_2901208, EPI_ISL_2901209, EPI_ISL_2901210, EPI_ISL_2901211, EPI_ISL_2901212, EPI_ISL_2901213, EPI_ISL_2901214, EPI_ISL_2901215, EPI_ISL_2901216, EPI_ISL_2901217, EPI_ISL_2901218, EPI_ISL_2901219, EPI_ISL_2901220, EPI_ISL_2901221, EPI_ISL_2901222, EPI_ISL_2901223, EPI_ISL_2901224, EPI_ISL_2901225, EPI_ISL_2901226, EPI_ISL_2901227, EPI_ISL_2901228, EPI_ISL_2901229, EPI_ISL_2901230, EPI_ISL_2901231, EPI_ISL_2901232, EPI_ISL_2901233, EPI_ISL_2901235, EPI_ISL_2901236, EPI_ISL_2901237, EPI_ISL_2901238, EPI_ISL_2901239, EPI_ISL_2901240, EPI_ISL_2901241, EPI_ISL_2901242, EPI_ISL_2901243, EPI_ISL_2901244, EPI_ISL_2901245, EPI_ISL_2901246, EPI_ISL_2901247, EPI_ISL_2901248, EPI_ISL_2901249, EPI_ISL_2901250, EPI_ISL_2901251, EPI_ISL_2901252, EPI_ISL_2901253, EPI_ISL_2901254, EPI_ISL_2901255, EPI_ISL_2901256, EPI_ISL_2901257, EPI_ISL_2901258, EPI_ISL_2901259, EPI_ISL_2901260, EPI_ISL_2901261, EPI_ISL_2901262, EPI_ISL_2901263, EPI_ISL_2901264, EPI_ISL_2901265, EPI_ISL_2901266, EPI_ISL_2901267, EPI_ISL_2901268, EPI_ISL_2901269, EPI_ISL_2901270, EPI_ISL_2901271, EPI_ISL_2901272, EPI_ISL_2901273, EPI_ISL_2901274, EPI_ISL_2901275, EPI_ISL_2901276, EPI_ISL_2901277, EPI_ISL_2901278, EPI_ISL_2901279, EPI_ISL_2901280, EPI_ISL_2901281, EPI_ISL_2901282, EPI_ISL_2901283, EPI_ISL_2901284, EPI_ISL_2901285, EPI_ISL_2901286, EPI_ISL_2901287, EPI_ISL_2901288, EPI_ISL_2901289, EPI_ISL_2901290, EPI_ISL_2901291, EPI_ISL_2901292, EPI_ISL_2901293, EPI_ISL_2901294, EPI_ISL_2901295, EPI_ISL_2901296, EPI_ISL_2901297, EPI_ISL_2901298, EPI_ISL_2901299, EPI_ISL_2901300, EPI_ISL_2901301, EPI_ISL_2901302, EPI_ISL_2901303, EPI_ISL_2901304, EPI_ISL_2901305, EPI_ISL_2901306, EPI_ISL_2901307, EPI_ISL_2901308, EPI_ISL_2901309, EPI_ISL_2901310, EPI_ISL_2901311, EPI_ISL_2901312, EPI_ISL_2901313, EPI_ISL_2901314, EPI_ISL_2901315, EPI_ISL_2901316, EPI_ISL_2901317, EPI_ISL_2901318, EPI_ISL_2901319, EPI_ISL_2901320, EPI_ISL_2901321, EPI_ISL_2901322, EPI_ISL_2901323, EPI_ISL_2901324, EPI_ISL_2901325, EPI_ISL_2901326, EPI_ISL_2901327, EPI_ISL_2901328, EPI_ISL_2901329, EPI_ISL_2901330, EPI_ISL_2901331, EPI_ISL_2901332, EPI_ISL_2901333, EPI_ISL_2901334, EPI_ISL_2901335, EPI_ISL_2901336, EPI_ISL_2901337, EPI_ISL_2901338, EPI_ISL_2901339, EPI_ISL_2901340, EPI_ISL_2901341, EPI_ISL_2901342, EPI_ISL_2901343, EPI_ISL_2901344, EPI_ISL_2901345, EPI_ISL_2901346, EPI_ISL_2901347, EPI_ISL_2901348, EPI_ISL_2901349, EPI_ISL_2901350, EPI_ISL_2901351, EPI_ISL_2901352, EPI_ISL_2901353, EPI_ISL_2901354, EPI_ISL_2901355, EPI_ISL_2901356, EPI_ISL_2901357, EPI_ISL_2901358, EPI_ISL_2901359, EPI_ISL_2901360, EPI_ISL_2901361, EPI_ISL_2901362, EPI_ISL_2901363, EPI_ISL_2901364, EPI_ISL_2901365, EPI_ISL_2901366, EPI_ISL_2901367, EPI_ISL_2901368, EPI_ISL_2901369, EPI_ISL_2901370, EPI_ISL_2901371, EPI_ISL_2901372, EPI_ISL_2901373, EPI_ISL_2901374, EPI_ISL_2901375, EPI_ISL_2901376, EPI_ISL_2901377, EPI_ISL_2901378, EPI_ISL_2901379, EPI_ISL_2901380, EPI_ISL_2901381, EPI_ISL_2901382, EPI_ISL_2901383, EPI_ISL_2901384, EPI_ISL_2901385, EPI_ISL_2901386, EPI_ISL_2901387, EPI_ISL_2901388, EPI_ISL_2901389, EPI_ISL_2901390, EPI_ISL_2901391, EPI_ISL_2901392, EPI_ISL_2901393, EPI_ISL_2901394, EPI_ISL_2901395, EPI_ISL_2901396, EPI_ISL_2901397, EPI_ISL_2901398, EPI_ISL_2901399, EPI_ISL_2901400, EPI_ISL_2901401, EPI_ISL_2901402, EPI_ISL_2901403, EPI_ISL_2901404, EPI_ISL_2901405, EPI_ISL_2901406, EPI_ISL_2901407, EPI_ISL_2901408, EPI_ISL_2901409, EPI_ISL_2901410, EPI_ISL_2901411, EPI_ISL_2901412, EPI_ISL_2901413, EPI_ISL_2901414, EPI_ISL_2901415, EPI_ISL_2901416, EPI_ISL_2901417, EPI_ISL_2901418, EPI_ISL_2901419, EPI_ISL_2901420, EPI_ISL_2901421, EPI_ISL_2901422, EPI_ISL_2901423, EPI_ISL_2901424, EPI_ISL_2901425, EPI_ISL_2901426, EPI_ISL_2901427, EPI_ISL_2901428, EPI_ISL_2901429, EPI_ISL_2901430, EPI_ISL_2901431, EPI_ISL_2901432, EPI_ISL_2901433, EPI_ISL_2901434, EPI_ISL_2901435, EPI_ISL_2901436, EPI_ISL_2901437, EPI_ISL_2901438, EPI_ISL_2901439, EPI_ISL_2901440, EPI_ISL_2901441, EPI_ISL_2901442, EPI_ISL_2901443, EPI_ISL_2901444, EPI_ISL_2901445, EPI_ISL_2901446, EPI_ISL_2901447, EPI_ISL_2901448, EPI_ISL_2901449, EPI_ISL_2901450, EPI_ISL_2901451, EPI_ISL_2901452, EPI_ISL_2901453, EPI_ISL_2901454, EPI_ISL_2901455, EPI_ISL_2901456, EPI_ISL_2901457, EPI_ISL_2901458, EPI_ISL_2901459, EPI_ISL_2901460, EPI_ISL_2901461, EPI_ISL_2901462, EPI_ISL_2901463, EPI_ISL_2901464, EPI_ISL_2901465, EPI_ISL_2901466, EPI_ISL_2901467, EPI_ISL_2901468, EPI_ISL_2901469, EPI_ISL_2901470, EPI_ISL_2901471, EPI_ISL_2901472, EPI_ISL_2901473, EPI_ISL_2901474, EPI_ISL_2901475, EPI_ISL_2901476, EPI_ISL_2901477, EPI_ISL_2901478, EPI_ISL_2901479, EPI_ISL_2901480, EPI_ISL_2901481, EPI_ISL_2901482, EPI_ISL_2901483, EPI_ISL_2901484, EPI_ISL_2901485, EPI_ISL_2901486, EPI_ISL_2901487, EPI_ISL_2901488, EPI_ISL_2901489, EPI_ISL_2901490, EPI_ISL_2901491, EPI_ISL_2901492, EPI_ISL_2901493, EPI_ISL_2901494, EPI_ISL_2901495, EPI_ISL_2901496, EPI_ISL_2901497, EPI_ISL_2901498, EPI_ISL_2901499, EPI_ISL_2901500, EPI_ISL_2901501, EPI_ISL_2901502, EPI_ISL_2901503, EPI_ISL_2901504, EPI_ISL_2901505, EPI_ISL_2901506, EPI_ISL_2901507, EPI_ISL_2901508, EPI_ISL_2901509, EPI_ISL_2901510, EPI_ISL_2901511, EPI_ISL_2901512, EPI_ISL_2901513, EPI_ISL_2901514, EPI_ISL_2901515, EPI_ISL_2901516, EPI_ISL_2901517, EPI_ISL_2901518, EPI_ISL_2901519, EPI_ISL_2901520, EPI_ISL_2901521, EPI_ISL_2901522, EPI_ISL_2901523, EPI_ISL_2901524, EPI_ISL_2901525, EPI_ISL_2901526, EPI_ISL_2901527, EPI_ISL_2901528, EPI_ISL_2901529, EPI_ISL_2901530, EPI_ISL_2901531, EPI_ISL_2901532, EPI_ISL_2901533, EPI_ISL_2901534, EPI_ISL_2901535, EPI_ISL_2901536, EPI_ISL_2901537, EPI_ISL_2901538, EPI_ISL_2901539, EPI_ISL_2901540, EPI_ISL_2901541, EPI_ISL_2901542, EPI_ISL_2901543, EPI_ISL_2901544, EPI_ISL_2901545, EPI_ISL_2901546, EPI_ISL_2901547, EPI_ISL_2901548, EPI_ISL_2901549, EPI_ISL_2901550, EPI_ISL_2901551, EPI_ISL_2901552, EPI_ISL_2901553, EPI_ISL_2901554, EPI_ISL_2901555, EPI_ISL_2901556, EPI_ISL_2901557, EPI_ISL_2901558, EPI_ISL_2901559, EPI_ISL_2901560, EPI_ISL_2901561, EPI_ISL_2901562 | | | |
| see above | University College London, Great Ormond Street Hospital for Children NHS Foundation Trust, Imperial College Healthcare NHS Trust | COVID-19 Genomics UK (COG-UK) Consortium | Sergi Castellano, Rachel Williams, Mark Kristiansen, Sunando Roy, Charlotte Williams, Nadua Bayzid, Marius Cotic, Helena Tutill, Patricia Dyal, Judith Breuer |
| EPI_ISL_2901563, EPI_ISL_2901564, EPI_ISL_2901565, EPI_ISL_2901566, EPI_ISL_2901567, EPI_ISL_2901568, EPI_ISL_2901569, EPI_ISL_2901570, EPI_ISL_2901571, EPI_ISL_2901572, EPI_ISL_2901573, EPI_ISL_2901574, EPI_ISL_2901575, EPI_ISL_2901576, EPI_ISL_2901577, EPI_ISL_2901578, EPI_ISL_2901579, EPI_ISL_2901580, EPI_ISL_2901581, EPI_ISL_2901582, EPI_ISL_2901583, EPI_ISL_2901584, | | | |

EPI_ISL_2901585, EPI_ISL_2901586, EPI_ISL_2901587, EPI_ISL_2901588, EPI_ISL_2901589, EPI_ISL_2901590, EPI_ISL_2901591, EPI_ISL_2901592, EPI_ISL_2901593, EPI_ISL_2901594, EPI_ISL_2901595, EPI_ISL_2901596, EPI_ISL_2901597, EPI_ISL_2901598, EPI_ISL_2901599, EPI_ISL_2901600, EPI_ISL_2901601, EPI_ISL_2901602, EPI_ISL_2901603, EPI_ISL_2901604, EPI_ISL_2901605, EPI_ISL_2901606,
EPI_ISL_2901607, EPI_ISL_2901608, EPI_ISL_2901609, EPI_ISL_2901610, EPI_ISL_2901611, EPI_ISL_2901612, EPI_ISL_2901613, EPI_ISL_2901614, EPI_ISL_2901615, EPI_ISL_2901616, EPI_ISL_2901617, EPI_ISL_2901618, EPI_ISL_2901619, EPI_ISL_2901620, EPI_ISL_2901621, EPI_ISL_2901622, EPI_ISL_2901623, EPI_ISL_2901624, EPI_ISL_2901625, EPI_ISL_2901626, EPI_ISL_2901627, EPI_ISL_2901628,
EPI_ISL_2901629, EPI_ISL_2901630, EPI_ISL_2901631, EPI_ISL_2901632, EPI_ISL_2901633, EPI_ISL_2901634, EPI_ISL_2901635, EPI_ISL_2901636, EPI_ISL_2901637, EPI_ISL_2901638, EPI_ISL_2901639, EPI_ISL_2901640, EPI_ISL_2901641, EPI_ISL_2901642, EPI_ISL_2901643, EPI_ISL_2901644, EPI_ISL_2901645, EPI_ISL_2901646, EPI_ISL_2901647, EPI_ISL_2901648, EPI_ISL_2901649, EPI_ISL_2901650,
EPI_ISL_2901651, EPI_ISL_2901652, EPI_ISL_2901653, EPI_ISL_2901654, EPI_ISL_2901655, EPI_ISL_2901656, EPI_ISL_2901657, EPI_ISL_2901658, EPI_ISL_2901659, EPI_ISL_2901660, EPI_ISL_2901661, EPI_ISL_2901662, EPI_ISL_2901663, EPI_ISL_2901664, EPI_ISL_2901665, EPI_ISL_2901666, EPI_ISL_2901667, EPI_ISL_2901668, EPI_ISL_2901669, EPI_ISL_2901671, EPI_ISL_2901672, EPI_ISL_2901673,
EPI_ISL_2901674, EPI_ISL_2901675, EPI_ISL_2901676, EPI_ISL_2901677, EPI_ISL_2901678, EPI_ISL_2901679, EPI_ISL_2901680, EPI_ISL_2901681, EPI_ISL_2901682, EPI_ISL_2901683, EPI_ISL_2901684, EPI_ISL_2901685, EPI_ISL_2901686, EPI_ISL_2901687, EPI_ISL_2901688, EPI_ISL_2901689, EPI_ISL_2901690, EPI_ISL_2901691, EPI_ISL_2901692, EPI_ISL_2901693, EPI_ISL_2901694, EPI_ISL_2901695,
EPI_ISL_2901696, EPI_ISL_2901697, EPI_ISL_2901698, EPI_ISL_2901699, EPI_ISL_2901700, EPI_ISL_2901701, EPI_ISL_2901702, EPI_ISL_2901703, EPI_ISL_2901704, EPI_ISL_2901705, EPI_ISL_2901706, EPI_ISL_2901707, EPI_ISL_2901708, EPI_ISL_2901709, EPI_ISL_2901710, EPI_ISL_2901711, EPI_ISL_2901712, EPI_ISL_2901713, EPI_ISL_2901714, EPI_ISL_2901715, EPI_ISL_2901716, EPI_ISL_2901717,
EPI_ISL_2901718, EPI_ISL_2901719, EPI_ISL_2901720, EPI_ISL_2901721, EPI_ISL_2901722, EPI_ISL_2901723, EPI_ISL_2901724, EPI_ISL_2901725, EPI_ISL_2901726, EPI_ISL_2901727, EPI_ISL_2901728, EPI_ISL_2901729, EPI_ISL_2901730, EPI_ISL_2901731, EPI_ISL_2901732, EPI_ISL_2901733, EPI_ISL_2901734, EPI_ISL_2901735, EPI_ISL_2901736, EPI_ISL_2901737, EPI_ISL_2901738, EPI_ISL_2901739,
EPI_ISL_2901740, EPI_ISL_2901741, EPI_ISL_2901742, EPI_ISL_2901743, EPI_ISL_2901744, EPI_ISL_2901745, EPI_ISL_2901746, EPI_ISL_2901747, EPI_ISL_2901748, EPI_ISL_2901749, EPI_ISL_2901750, EPI_ISL_2901751, EPI_ISL_2901752, EPI_ISL_2901753, EPI_ISL_2901754, EPI_ISL_2901755, EPI_ISL_2901756, EPI_ISL_2901757, EPI_ISL_2901758, EPI_ISL_2901759, EPI_ISL_2901760, EPI_ISL_2901761,
EPI_ISL_2901762, EPI_ISL_2901763, EPI_ISL_2901764, EPI_ISL_2901765, EPI_ISL_2901766, EPI_ISL_2901767, EPI_ISL_2901768, EPI_ISL_2901769, EPI_ISL_2901770, EPI_ISL_2901771, EPI_ISL_2901772, EPI_ISL_2901773, EPI_ISL_2901774, EPI_ISL_2901775, EPI_ISL_2901776, EPI_ISL_2901777, EPI_ISL_2901778, EPI_ISL_2901779, EPI_ISL_2901780, EPI_ISL_2901781, EPI_ISL_2901782, EPI_ISL_2901783,
EPI_ISL_2901784, EPI_ISL_2901785, EPI_ISL_2901786, EPI_ISL_2901787, EPI_ISL_2901788, EPI_ISL_2901789, EPI_ISL_2901790, EPI_ISL_2901791, EPI_ISL_2901792, EPI_ISL_2901793, EPI_ISL_2901794, EPI_ISL_2901795, EPI_ISL_2901796, EPI_ISL_2901797, EPI_ISL_2901798, EPI_ISL_2901799, EPI_ISL_2901800, EPI_ISL_2901801, EPI_ISL_2901802, EPI_ISL_2901803, EPI_ISL_2901804, EPI_ISL_2901805,
EPI_ISL_2901806, EPI_ISL_2901807, EPI_ISL_2901808, EPI_ISL_2901809, EPI_ISL_2901810, EPI_ISL_2901811, EPI_ISL_2901812, EPI_ISL_2901813, EPI_ISL_2901814, EPI_ISL_2901815, EPI_ISL_2901816, EPI_ISL_2901817, EPI_ISL_2901818, EPI_ISL_2901819, EPI_ISL_2901820, EPI_ISL_2901821, EPI_ISL_2901822, EPI_ISL_2901823, EPI_ISL_2901824, EPI_ISL_2901825, EPI_ISL_2901826, EPI_ISL_2901827,
EPI_ISL_2901828, EPI_ISL_2901829, EPI_ISL_2901830, EPI_ISL_2901831, EPI_ISL_2901832, EPI_ISL_2901833, EPI_ISL_2901834, EPI_ISL_2901835, EPI_ISL_2901836, EPI_ISL_2901837, EPI_ISL_2901838, EPI_ISL_2901839, EPI_ISL_2901841, EPI_ISL_2901842, EPI_ISL_2901843, EPI_ISL_2901844, EPI_ISL_2901845, EPI_ISL_2901846, EPI_ISL_2901847, EPI_ISL_2901848, EPI_ISL_2901849, EPI_ISL_2901850,
EPI_ISL_2901851, EPI_ISL_2901852, EPI_ISL_2901853, EPI_ISL_2901854, EPI_ISL_2901855, EPI_ISL_2901856, EPI_ISL_2901857, EPI_ISL_2901858, EPI_ISL_2901859, EPI_ISL_2901860, EPI_ISL_2901861, EPI_ISL_2901862, EPI_ISL_2901863, EPI_ISL_2901864, EPI_ISL_2901865, EPI_ISL_2901866, EPI_ISL_2901867, EPI_ISL_2901868, EPI_ISL_2901869, EPI_ISL_2901870, EPI_ISL_2901871, EPI_ISL_2901872,
EPI_ISL_2901873, EPI_ISL_2901874, EPI_ISL_2901875, EPI_ISL_2901876, EPI_ISL_2901877, EPI_ISL_2901878, EPI_ISL_2901879, EPI_ISL_2901880, EPI_ISL_2901881, EPI_ISL_2901882, EPI_ISL_2901883, EPI_ISL_2901884, EPI_ISL_2901885, EPI_ISL_2901886, EPI_ISL_2901887, EPI_ISL_2901888, EPI_ISL_2901889, EPI_ISL_2901890, EPI_ISL_2901891, EPI_ISL_2901892, EPI_ISL_2901893, EPI_ISL_2901894,
EPI_ISL_2901895, EPI_ISL_2901896, EPI_ISL_2901897, EPI_ISL_2901898, EPI_ISL_2901899, EPI_ISL_2901900, EPI_ISL_2901901, EPI_ISL_2901902, EPI_ISL_2901903, EPI_ISL_2901904, EPI_ISL_2901905, EPI_ISL_2901906, EPI_ISL_2901907, EPI_ISL_2901908, EPI_ISL_2901909, EPI_ISL_2901910, EPI_ISL_2901911, EPI_ISL_2901912, EPI_ISL_2901913, EPI_ISL_2901914, EPI_ISL_2901915, EPI_ISL_2901916,
EPI_ISL_2901917, EPI_ISL_2901918, EPI_ISL_2901919, EPI_ISL_2901920, EPI_ISL_2901921, EPI_ISL_2901922, EPI_ISL_2901923, EPI_ISL_2901924, EPI_ISL_2901925, EPI_ISL_2901926, EPI_ISL_2901927, EPI_ISL_2901928, EPI_ISL_2901929, EPI_ISL_2901930, EPI_ISL_2901931, EPI_ISL_2901932, EPI_ISL_2901933, EPI_ISL_2901934, EPI_ISL_2901935, EPI_ISL_2901936, EPI_ISL_2901937, EPI_ISL_2901938,
EPI_ISL_2901939, EPI_ISL_2901940, EPI_ISL_2901941, EPI_ISL_2901942, EPI_ISL_2901943, EPI_ISL_2901944, EPI_ISL_2901945, EPI_ISL_2901946, EPI_ISL_2901947, EPI_ISL_2901948, EPI_ISL_2901949, EPI_ISL_2901950, EPI_ISL_2901951, EPI_ISL_2901952, EPI_ISL_2901953, EPI_ISL_2901954, EPI_ISL_2901955, EPI_ISL_2901956, EPI_ISL_2901957, EPI_ISL_2901958, EPI_ISL_2901959, EPI_ISL_2901960,
EPI_ISL_2901961, EPI_ISL_2901962, EPI_ISL_2901963, EPI_ISL_2901964, EPI_ISL_2901965, EPI_ISL_2901966, EPI_ISL_2901967, EPI_ISL_2901968, EPI_ISL_2901969, EPI_ISL_2901970, EPI_ISL_2901971, EPI_ISL_2901972, EPI_ISL_2901973, EPI_ISL_2901974, EPI_ISL_2901975, EPI_ISL_2901976, EPI_ISL_2901977, EPI_ISL_2901978, EPI_ISL_2901979, EPI_ISL_2901980, EPI_ISL_2901981, EPI_ISL_2901982,
EPI_ISL_2901983, EPI_ISL_2901984, EPI_ISL_2901985, EPI_ISL_2901986, EPI_ISL_2901987, EPI_ISL_2901988, EPI_ISL_2901989, EPI_ISL_2901990, EPI_ISL_2901991, EPI_ISL_2901992, EPI_ISL_2901993, EPI_ISL_2901994, EPI_ISL_2901995, EPI_ISL_2901996, EPI_ISL_2901997, EPI_ISL_2901998, EPI_ISL_2901999, EPI_ISL_2902000, EPI_ISL_2902001, EPI_ISL_2902002, EPI_ISL_2902003, EPI_ISL_2902004,
EPI_ISL_2902005, EPI_ISL_2902006, EPI_ISL_2902007, EPI_ISL_2902008, EPI_ISL_2902009, EPI_ISL_2902010, EPI_ISL_2902011, EPI_ISL_2902012, EPI_ISL_2902013, EPI_ISL_2902014, EPI_ISL_2902015, EPI_ISL_2902016, EPI_ISL_2902017, EPI_ISL_2902018, EPI_ISL_2902019, EPI_ISL_2902020, EPI_ISL_2902021, EPI_ISL_2902022, EPI_ISL_2902023, EPI_ISL_2902024, EPI_ISL_2902025, EPI_ISL_2902026,
EPI_ISL_2902027, EPI_ISL_2902028, EPI_ISL_2902029, EPI_ISL_2902030, EPI_ISL_2902031, EPI_ISL_2902032, EPI_ISL_2902033, EPI_ISL_2902034, EPI_ISL_2902035, EPI_ISL_2902036, EPI_ISL_2902037, EPI_ISL_2902038, EPI_ISL_2902039, EPI_ISL_2902040, EPI_ISL_2902041, EPI_ISL_2902042, EPI_ISL_2902043, EPI_ISL_2902044, EPI_ISL_2902045, EPI_ISL_2902046, EPI_ISL_2902047, EPI_ISL_2902048,
EPI_ISL_2902049, EPI_ISL_2902050, EPI_ISL_2902051, EPI_ISL_2902052, EPI_ISL_2902053, EPI_ISL_2902054, EPI_ISL_2902055, EPI_ISL_2902056, EPI_ISL_2902057, EPI_ISL_2902058, EPI_ISL_2902059, EPI_ISL_2902060, EPI_ISL_2902061, EPI_ISL_2902062, EPI_ISL_2902063, EPI_ISL_2902064, EPI_ISL_2902065, EPI_ISL_2902066, EPI_ISL_2902067, EPI_ISL_2902068, EPI_ISL_2902069, EPI_ISL_2902070,
EPI_ISL_2902071, EPI_ISL_2902072, EPI_ISL_2902073, EPI_ISL_2902074, EPI_ISL_2902075, EPI_ISL_2902076, EPI_ISL_2902077, EPI_ISL_2902078, EPI_ISL_2902079, EPI_ISL_2902080, EPI_ISL_2902081, EPI_ISL_2902082, EPI_ISL_2902083, EPI_ISL_2902084, EPI_ISL_2902085, EPI_ISL_2902086, EPI_ISL_2902087, EPI_ISL_2902088, EPI_ISL_2902089, EPI_ISL_2902090, EPI_ISL_2902091, EPI_ISL_2902092,
EPI_ISL_2902093, EPI_ISL_2902094, EPI_ISL_2902095, EPI_ISL_2902096, EPI_ISL_2902097, EPI_ISL_2902098, EPI_ISL_2902099, EPI_ISL_2902100, EPI_ISL_2902101, EPI_ISL_2902102, EPI_ISL_2902103, EPI_ISL_2902104, EPI_ISL_2902105, EPI_ISL_2902106, EPI_ISL_2902107, EPI_ISL_2902108, EPI_ISL_2902109, EPI_ISL_2902110, EPI_ISL_2902111, EPI_ISL_2902112, EPI_ISL_2902113, EPI_ISL_2902114,
EPI_ISL_2902115, EPI_ISL_2902116, EPI_ISL_2902117, EPI_ISL_2902118, EPI_ISL_2902119, EPI_ISL_2902120, EPI_ISL_2902121, EPI_ISL_2902122, EPI_ISL_2902123, EPI_ISL_2902124, EPI_ISL_2902125, EPI_ISL_2902126, EPI_ISL_2902127, EPI_ISL_2902128, EPI_ISL_2902129, EPI_ISL_2902130, EPI_ISL_2902131, EPI_ISL_2902132, EPI_ISL_2902133, EPI_ISL_2902134, EPI_ISL_2902135, EPI_ISL_2902136,
EPI_ISL_2902137, EPI_ISL_2902138, EPI_ISL_2902139, EPI_ISL_2902140, EPI_ISL_2902141, EPI_ISL_2902142, EPI_ISL_2902143, EPI_ISL_2902144, EPI_ISL_2902145, EPI_ISL_2902146, EPI_ISL_2902147, EPI_ISL_2902148, EPI_ISL_2902149, EPI_ISL_2902150, EPI_ISL_2902151, EPI_ISL_2902152, EPI_ISL_2902153, EPI_ISL_2902154, EPI_ISL_2902155, EPI_ISL_2902156, EPI_ISL_2902157, EPI_ISL_2902158,
EPI_ISL_2902159, EPI_ISL_2902160, EPI_ISL_2902161, EPI_ISL_2902162, EPI_ISL_2902163, EPI_ISL_2902164, EPI_ISL_2902165, EPI_ISL_2902166, EPI_ISL_2902167, EPI_ISL_2902168, EPI_ISL_2902169, EPI_ISL_2902170, EPI_ISL_2902171, EPI_ISL_2902172, EPI_ISL_2902173, EPI_ISL_2902174, EPI_ISL_2902175, EPI_ISL_2902176, EPI_ISL_2902177, EPI_ISL_2902178, EPI_ISL_2902179, EPI_ISL_2902180,
EPI_ISL_2902181, EPI_ISL_2902182, EPI_ISL_2902183, EPI_ISL_2902184, EPI_ISL_2902185, EPI_ISL_2902186, EPI_ISL_2902187, EPI_ISL_2902188, EPI_ISL_2902189, EPI_ISL_2902190, EPI_ISL_2902191, EPI_ISL_2902192, EPI_ISL_2902193, EPI_ISL_2902194, EPI_ISL_2902195, EPI_ISL_2902196, EPI_ISL_2902197, EPI_ISL_2902198, EPI_ISL_2902199, EPI_ISL_2902200, EPI_ISL_2902201, EPI_ISL_2902202,
EPI_ISL_2902203, EPI_ISL_2902204, EPI_ISL_2902205, EPI_ISL_2902206, EPI_ISL_2902207, EPI_ISL_2902208, EPI_ISL_2902209, EPI_ISL_2902210, EPI_ISL_2902211, EPI_ISL_2902212, EPI_ISL_2902213, EPI_ISL_2902214, EPI_ISL_2902215, EPI_ISL_2902216, EPI_ISL_2902217, EPI_ISL_2902218, EPI_ISL_2902219, EPI_ISL_2902220, EPI_ISL_2902221, EPI_ISL_2902222, EPI_ISL_2902223, EPI_ISL_2902224,
EPI_ISL_2902225, EPI_ISL_2902226, EPI_ISL_2902227, EPI_ISL_2902228, EPI_ISL_2902229, EPI_ISL_2902230, EPI_ISL_2902231, EPI_ISL_2902232, EPI_ISL_2902233, EPI_ISL_2902234, EPI_ISL_2902235, EPI_ISL_2902236, EPI_ISL_2902237, EPI_ISL_2902238, EPI_ISL_2902239, EPI_ISL_2902240, EPI_ISL_2902241, EPI_ISL_2902242, EPI_ISL_2902243, EPI_ISL_2902244, EPI_ISL_2902245, EPI_ISL_2902246,
EPI_ISL_2902247, EPI_ISL_2902248, EPI_ISL_2902249, EPI_ISL_2902250, EPI_ISL_2902251, EPI_ISL_2902252, EPI_ISL_2902253, EPI_ISL_2902254, EPI_ISL_2902255, EPI_ISL_2902256, EPI_ISL_2902257, EPI_ISL_2902258, EPI_ISL_2902259, EPI_ISL_2902260, EPI_ISL_2902261, EPI_ISL_2902262, EPI_ISL_2902263, EPI_ISL_2902264, EPI_ISL_2902265, EPI_ISL_2902266, EPI_ISL_2902267, EPI_ISL_2902268,
EPI_ISL_2902269, EPI_ISL_2902270, EPI_ISL_2902271, EPI_ISL_2902272, EPI_ISL_2902273, EPI_ISL_2902274, EPI_ISL_2902275, EPI_ISL_2902276, EPI_ISL_2902277, EPI_ISL_2902278, EPI_ISL_2902279, EPI_ISL_2902280, EPI_ISL_2902281, EPI_ISL_2902282, EPI_ISL_2902283, EPI_ISL_2902284, EPI_ISL_2902285, EPI_ISL_2902286, EPI_ISL_2902287, EPI_ISL_2902288, EPI_ISL_2902289, EPI_ISL_2902290,
EPI_ISL_2902291, EPI_ISL_2902292, EPI_ISL_2902293, EPI_ISL_2902294, EPI_ISL_2902295, EPI_ISL_2902296, EPI_ISL_2902297, EPI_ISL_2902298, EPI_ISL_2902299, EPI_ISL_2902300, EPI_ISL_2902301, EPI_ISL_2902302, EPI_ISL_2902303, EPI_ISL_2902304, EPI_ISL_2902305, EPI_ISL_2902306, EPI_ISL_2902307, EPI_ISL_2902308, EPI_ISL_2902309, EPI_ISL_2902310, EPI_ISL_2902311, EPI_ISL_2902312,
EPI_ISL_2902313, EPI_ISL_2902314, EPI_ISL_2902315, EPI_ISL_2902316, EPI_ISL_2902317, EPI_ISL_2902318, EPI_ISL_2902319, EPI_ISL_2902320, EPI_ISL_2902321, EPI_ISL_2902322, EPI_ISL_2902323, EPI_ISL_2902324, EPI_ISL_2902325, EPI_ISL_2902326, EPI_ISL_2902327, EPI_ISL_2902328, EPI_ISL_2902329, EPI_ISL_2902330, EPI_ISL_2902331, EPI_ISL_2902332, EPI_ISL_2902333, EPI_ISL_2902334,
EPI_ISL_2902335, EPI_ISL_2902336, EPI_ISL_2902337, EPI_ISL_2902338, EPI_ISL_2902339, EPI_ISL_2902340, EPI_ISL_2902341, EPI_ISL_2902342, EPI_ISL_2902343, EPI_ISL_2902344, EPI_ISL_2902345, EPI_ISL_2902346, EPI_ISL_2902347, EPI_ISL_2902348, EPI_ISL_2902349, EPI_ISL_2902350, EPI_ISL_2902351, EPI_ISL_2902352, EPI_ISL_2902353, EPI_ISL_2902354, EPI_ISL_2902355, EPI_ISL_2902356,
EPI_ISL_2902357, EPI_ISL_2902358, EPI_ISL_2902359, EPI_ISL_2902360, EPI_ISL_2902361, EPI_ISL_2902362, EPI_ISL_2902363, EPI_ISL_2902364, EPI_ISL_2902365, EPI_ISL_2902366, EPI_ISL_2902367, EPI_ISL_2902368, EPI_ISL_2902369, EPI_ISL_2902370, EPI_ISL_2902371, EPI_ISL_2902372, EPI_ISL_2902373, EPI_ISL_2902374, EPI_ISL_2902375, EPI_ISL_2902376, EPI_ISL_2902377, EPI_ISL_2902378,
EPI_ISL_2902379, EPI_ISL_2902380, EPI_ISL_2902381, EPI_ISL_2902382, EPI_ISL_2902383, EPI_ISL_2902384, EPI_ISL_2902385, EPI_ISL_2902386, EPI_ISL_2902387, EPI_ISL_2902388, EPI_ISL_2902389, EPI_ISL_2902390, EPI_ISL_2902391, EPI_ISL_2902392, EPI_ISL_2902393, EPI_ISL_2902394, EPI_ISL_2902395, EPI_ISL_2902396, EPI_ISL_2902397, EPI_ISL_2902398, EPI_ISL_2902399, EPI_ISL_2902400,
EPI_ISL_2902401, EPI_ISL_2902402, EPI_ISL_2902403, EPI_ISL_2902404, EPI_ISL_2902405, EPI_ISL_2902406, EPI_ISL_2902407, EPI_ISL_2902408, EPI_ISL_2902409

| | | | |
|---|---|---|---|
| see above | Northumbria University / South Tees Hospitals NHS Foundation Trust / North Cumbria Integrated Care NHS Foundation Trust / North Tees and Hartlepool NHS Foundation Trust / Newcastle Hospitals NHS Foundation Trust | COVID-19 Genomics UK (COG-UK) Consortium | Darren L Smith,Andrew Nelson,Matthew Bashton,Greg R Young,Joshua Loh,John Allan,Mohammad A Tariq,Giles S Holt,Gary Black,Wen C Yew,Lynn Dover,Paul Baker,Steve Liggett,Sarah Essex,Jane Greenaway,Debra Padgett,Clive Graham,Garren Scott,Edward Barton,Emma Swindells,Brendan Payne,Jennifer Collins,Yusri Taha,Gary Eltringham |

EPI_ISL_2902410, EPI_ISL_2902411, EPI_ISL_2902412, EPI_ISL_2902413, EPI_ISL_2902414, EPI_ISL_2902415, EPI_ISL_2902416, EPI_ISL_2902417, EPI_ISL_2902418, EPI_ISL_2902419, EPI_ISL_2902420, EPI_ISL_2902421, EPI_ISL_2902422, EPI_ISL_2902424, EPI_ISL_2902425, EPI_ISL_2902426, EPI_ISL_2902427, EPI_ISL_2902428, EPI_ISL_2902429, EPI_ISL_2902430, EPI_ISL_2902431, EPI_ISL_2902432,
EPI_ISL_2902433, EPI_ISL_2902434, EPI_ISL_2902435, EPI_ISL_2902436, EPI_ISL_2902437, EPI_ISL_2902438, EPI_ISL_2902439, EPI_ISL_2902440, EPI_ISL_2902441, EPI_ISL_2902442, EPI_ISL_2902443, EPI_ISL_2902444, EPI_ISL_2902445, EPI_ISL_2902446, EPI_ISL_2902447, EPI_ISL_2902448, EPI_ISL_2902449, EPI_ISL_2902450, EPI_ISL_2902451, EPI_ISL_2902452, EPI_ISL_2902453, EPI_ISL_2902454,
EPI_ISL_2902455, EPI_ISL_2902456, EPI_ISL_2902457, EPI_ISL_2902458, EPI_ISL_2902459, EPI_ISL_2902460, EPI_ISL_2902461, EPI_ISL_2902462, EPI_ISL_2902463, EPI_ISL_2902464, EPI_ISL_2902465, EPI_ISL_2902466, EPI_ISL_2902467, EPI_ISL_2902468, EPI_ISL_2902469, EPI_ISL_2902470, EPI_ISL_2902471, EPI_ISL_2902473, EPI_ISL_2902474, EPI_ISL_2902475, EPI_ISL_2902476, EPI_ISL_2902477,
EPI_ISL_2902478, EPI_ISL_2902479, EPI_ISL_2902480, EPI_ISL_2902481, EPI_ISL_2902482, EPI_ISL_2902483, EPI_ISL_2902484, EPI_ISL_2902485, EPI_ISL_2902486, EPI_ISL_2902487, EPI_ISL_2902488, EPI_ISL_2902489, EPI_ISL_2902490, EPI_ISL_2902491, EPI_ISL_2902492, EPI_ISL_2902493, EPI_ISL_2902494, EPI_ISL_2902495, EPI_ISL_2902496, EPI_ISL_2902497, EPI_ISL_2902498, EPI_ISL_2902499,
EPI_ISL_2902500, EPI_ISL_2902501, EPI_ISL_2902502, EPI_ISL_2902503, EPI_ISL_2902504, EPI_ISL_2902505, EPI_ISL_2902506, EPI_ISL_2902507, EPI_ISL_2902508, EPI_ISL_2902509, EPI_ISL_2902510, EPI_ISL_2902511, EPI_ISL_2902512, EPI_ISL_2902513, EPI_ISL_2902514, EPI_ISL_2902515, EPI_ISL_2902516, EPI_ISL_2902517, EPI_ISL_2902518, EPI_ISL_2902519, EPI_ISL_2902520, EPI_ISL_2902521,
EPI_ISL_2902522, EPI_ISL_2902523, EPI_ISL_2902524, EPI_ISL_2902525, EPI_ISL_2902526, EPI_ISL_2902527, EPI_ISL_2902528, EPI_ISL_2902529, EPI_ISL_2902530, EPI_ISL_2902531, EPI_ISL_2902532, EPI_ISL_2902533, EPI_ISL_2902534, EPI_ISL_2902535, EPI_ISL_2902536, EPI_ISL_2902537, EPI_ISL_2902538, EPI_ISL_2902539, EPI_ISL_2902540, EPI_ISL_2902541, EPI_ISL_2902542, EPI_ISL_2902543,
EPI_ISL_2902544, EPI_ISL_2902545, EPI_ISL_2902546, EPI_ISL_2902547, EPI_ISL_2902548, EPI_ISL_2902549, EPI_ISL_2902550, EPI_ISL_2902551, EPI_ISL_2902552, EPI_ISL_2902553, EPI_ISL_2902554, EPI_ISL_2902555, EPI_ISL_2902556, EPI_ISL_2902557, EPI_ISL_2902558, EPI_ISL_2902559, EPI_ISL_2902560, EPI_ISL_2902561, EPI_ISL_2902562, EPI_ISL_2902563, EPI_ISL_2902564, EPI_ISL_2902565,
EPI_ISL_2902566, EPI_ISL_2902567, EPI_ISL_2902568, EPI_ISL_2902569, EPI_ISL_2902570, EPI_ISL_2902571, EPI_ISL_2902572, EPI_ISL_2902573, EPI_ISL_2902574, EPI_ISL_2902575, EPI_ISL_2902576, EPI_ISL_2902577, EPI_ISL_2902578, EPI_ISL_2902579, EPI_ISL_2902580, EPI_ISL_2902581, EPI_ISL_2902582, EPI_ISL_2902583, EPI_ISL_2902584, EPI_ISL_2902585, EPI_ISL_2902586, EPI_ISL_2902587,
EPI_ISL_2902588, EPI_ISL_2902589, EPI_ISL_2902590, EPI_ISL_2902591, EPI_ISL_2902592, EPI_ISL_2902593, EPI_ISL_2902594, EPI_ISL_2902595, EPI_ISL_2902596, EPI_ISL_2902597, EPI_ISL_2902598, EPI_ISL_2902599, EPI_ISL_2902600, EPI_ISL_2902601, EPI_ISL_2902602, EPI_ISL_2902603, EPI_ISL_2902604, EPI_ISL_2902605, EPI_ISL_2902606, EPI_ISL_2902607, EPI_ISL_2902608, EPI_ISL_2902609,
EPI_ISL_2902610, EPI_ISL_2902611, EPI_ISL_2902612, EPI_ISL_2902613, EPI_ISL_2902614, EPI_ISL_2902615, EPI_ISL_2902616, EPI_ISL_2902617, EPI_ISL_2902618, EPI_ISL_2902619, EPI_ISL_2902620, EPI_ISL_2902621, EPI_ISL_2902622, EPI_ISL_2902623, EPI_ISL_2902624, EPI_ISL_2902625, EPI_ISL_2902626, EPI_ISL_2902627, EPI_ISL_2902628, EPI_ISL_2902629, EPI_ISL_2902630, EPI_ISL_2902631,
EPI_ISL_2902632, EPI_ISL_2902633, EPI_ISL_2902634, EPI_ISL_2902635, EPI_ISL_2902636, EPI_ISL_2902637, EPI_ISL_2902638, EPI_ISL_2902639, EPI_ISL_2902640, EPI_ISL_2902641, EPI_ISL_2902642, EPI_ISL_2902643, EPI_ISL_2902644, EPI_ISL_2902645, EPI_ISL_2902646, EPI_ISL_2902647, EPI_ISL_2902648, EPI_ISL_2902649, EPI_ISL_2902650, EPI_ISL_2902651, EPI_ISL_2902652, EPI_ISL_2902653,
EPI_ISL_2902654, EPI_ISL_2902655, EPI_ISL_2902656, EPI_ISL_2902657, EPI_ISL_2902658, EPI_ISL_2902659, EPI_ISL_2902660, EPI_ISL_2902661, EPI_ISL_2902662, EPI_ISL_2902663, EPI_ISL_2902664, EPI_ISL_2902665, EPI_ISL_2902666, EPI_ISL_2902667, EPI_ISL_2902668, EPI_ISL_2902669, EPI_ISL_2902670, EPI_ISL_2902671, EPI_ISL_2902672, EPI_ISL_2902673, EPI_ISL_2902674, EPI_ISL_2902675,
EPI_ISL_2902676, EPI_ISL_2902677, EPI_ISL_2902678, EPI_ISL_2902679, EPI_ISL_2902680, EPI_ISL_2902681, EPI_ISL_2902682, EPI_ISL_2902683, EPI_ISL_2902684, EPI_ISL_2902685, EPI_ISL_2902686, EPI_ISL_2902687, EPI_ISL_2902688, EPI_ISL_2902689, EPI_ISL_2902690, EPI_ISL_2902691, EPI_ISL_2902692, EPI_ISL_2902693, EPI_ISL_2902694, EPI_ISL_2902695, EPI_ISL_2902696, EPI_ISL_2902697,
EPI_ISL_2902698, EPI_ISL_2902699, EPI_ISL_2902700, EPI_ISL_2902701, EPI_ISL_2902702, EPI_ISL_2902703, EPI_ISL_2902704, EPI_ISL_2902705, EPI_ISL_2902706, EPI_ISL_2902707, EPI_ISL_2902708, EPI_ISL_2902709, EPI_ISL_2902710, EPI_ISL_2902711, EPI_ISL_2902712, EPI_ISL_2902713, EPI_ISL_2902714, EPI_ISL_2902715, EPI_ISL_2902716, EPI_ISL_2902717, EPI_ISL_2902718, EPI_ISL_2902719,
EPI_ISL_2902720, EPI_ISL_2902721, EPI_ISL_2902722, EPI_ISL_2902723, EPI_ISL_2902724, EPI_ISL_2902725, EPI_ISL_2902726, EPI_ISL_2902727, EPI_ISL_2902728, EPI_ISL_2902729, EPI_ISL_2902730, EPI_ISL_2902731, EPI_ISL_2902732, EPI_ISL_2902733, EPI_ISL_2902734, EPI_ISL_2902735, EPI_ISL_2902736, EPI_ISL_2902737, EPI_ISL_2902738, EPI_ISL_2902739, EPI_ISL_2902740, EPI_ISL_2902741,
EPI_ISL_2902742, EPI_ISL_2902743, EPI_ISL_2902744, EPI_ISL_2902745, EPI_ISL_2902746, EPI_ISL_2902747, EPI_ISL_2902748, EPI_ISL_2902749, EPI_ISL_2902750, EPI_ISL_2902751, EPI_ISL_2902752, EPI_ISL_2902753, EPI_ISL_2902754, EPI_ISL_2902755, EPI_ISL_2902756, EPI_ISL_2902757, EPI_ISL_2902758, EPI_ISL_2902759, EPI_ISL_2902760, EPI_ISL_2902761, EPI_ISL_2902762, EPI_ISL_2902763,
EPI_ISL_2902764, EPI_ISL_2902765, EPI_ISL_2902766, EPI_ISL_2902767, EPI_ISL_2902768, EPI_ISL_2902769, EPI_ISL_2902770, EPI_ISL_2902771, EPI_ISL_2902772, EPI_ISL_2902773, EPI_ISL_2902774, EPI_ISL_2902775, EPI_ISL_2902776, EPI_ISL_2902777, EPI_ISL_2902778, EPI_ISL_2902779, EPI_ISL_2902780, EPI_ISL_2902781, EPI_ISL_2902782, EPI_ISL_2902783, EPI_ISL_2902784, EPI_ISL_2902785,
EPI_ISL_2902786, EPI_ISL_2902787, EPI_ISL_2902788, EPI_ISL_2902789, EPI_ISL_2902790, EPI_ISL_2902791, EPI_ISL_2902792, EPI_ISL_2902793, EPI_ISL_2902794, EPI_ISL_2902795, EPI_ISL_2902796, EPI_ISL_2902797, EPI_ISL_2902798, EPI_ISL_2902799, EPI_ISL_2902800, EPI_ISL_2902801, EPI_ISL_2902802, EPI_ISL_2902803, EPI_ISL_2902804, EPI_ISL_2902805, EPI_ISL_2902806, EPI_ISL_2902807,
EPI_ISL_2902808, EPI_ISL_2902809, EPI_ISL_2902810, EPI_ISL_2902811, EPI_ISL_2902812, EPI_ISL_2902813, EPI_ISL_2902814, EPI_ISL_2902815, EPI_ISL_2902816, EPI_ISL_2902817, EPI_ISL_2902818, EPI_ISL_2902819, EPI_ISL_2902820, EPI_ISL_2902821, EPI_ISL_2902822, EPI_ISL_2902823, EPI_ISL_2902824, EPI_ISL_2902825, EPI_ISL_2902826, EPI_ISL_2902827, EPI_ISL_2902828, EPI_ISL_2902829,
EPI_ISL_2902830, EPI_ISL_2902831, EPI_ISL_2902832, EPI_ISL_2902833, EPI_ISL_2902834, EPI_ISL_2902835, EPI_ISL_2902836, EPI_ISL_2902837, EPI_ISL_2902838, EPI_ISL_2902839, EPI_ISL_2902840, EPI_ISL_2902841, EPI_ISL_2902842, EPI_ISL_2902843, EPI_ISL_2902844, EPI_ISL_2902845, EPI_ISL_2902846, EPI_ISL_2902847, EPI_ISL_2902848, EPI_ISL_2902849, EPI_ISL_2902850, EPI_ISL_2902851,
EPI_ISL_2902852, EPI_ISL_2902853, EPI_ISL_2902854, EPI_ISL_2902855, EPI_ISL_2902856, EPI_ISL_2902857, EPI_ISL_2902858, EPI_ISL_2902859, EPI_ISL_2902860, EPI_ISL_2902861, EPI_ISL_2902862, EPI_ISL_2902863, EPI_ISL_2902864, EPI_ISL_2902865, EPI_ISL_2902866, EPI_ISL_2902867, EPI_ISL_2902868, EPI_ISL_2902869, EPI_ISL_2902870, EPI_ISL_2902871, EPI_ISL_2902872, EPI_ISL_2902873,
EPI_ISL_2902874, EPI_ISL_2902875, EPI_ISL_2902876, EPI_ISL_2902877, EPI_ISL_2902878, EPI_ISL_2902879, EPI_ISL_2902880, EPI_ISL_2902881, EPI_ISL_2902882, EPI_ISL_2902883, EPI_ISL_2902884, EPI_ISL_2902885, EPI_ISL_2902886, EPI_ISL_2902887, EPI_ISL_2902888, EPI_ISL_2902889, EPI_ISL_2902890, EPI_ISL_2902891, EPI_ISL_2902892, EPI_ISL_2902893, EPI_ISL_2902894, EPI_ISL_2902895,
EPI_ISL_2902896, EPI_ISL_2902897, EPI_ISL_2902898, EPI_ISL_2902899, EPI_ISL_2902900, EPI_ISL_2902901, EPI_ISL_2902902, EPI_ISL_2902903, EPI_ISL_2902904, EPI_ISL_2902905, EPI_ISL_2902906, EPI_ISL_2902907, EPI_ISL_2902908, EPI_ISL_2902909, EPI_ISL_2902910, EPI_ISL_2902911, EPI_ISL_2902912, EPI_ISL_2902913, EPI_ISL_2902914, EPI_ISL_2902915, EPI_ISL_2902916, EPI_ISL_2902917,
EPI_ISL_2902918, EPI_ISL_2902919, EPI_ISL_2902920, EPI_ISL_2902921, EPI_ISL_2902922, EPI_ISL_2902923, EPI_ISL_2902924, EPI_ISL_2902925, EPI_ISL_2902926, EPI_ISL_2902927, EPI_ISL_2902928, EPI_ISL_2902929, EPI_ISL_2902930, EPI_ISL_2902931, EPI_ISL_2902932, EPI_ISL_2902933, EPI_ISL_2902934, EPI_ISL_2902935, EPI_ISL_2902936, EPI_ISL_2902937, EPI_ISL_2902938, EPI_ISL_2902939,
EPI_ISL_2902940, EPI_ISL_2902941, EPI_ISL_2902942, EPI_ISL_2902943, EPI_ISL_2902944, EPI_ISL_2902945, EPI_ISL_2902946, EPI_ISL_2902947, EPI_ISL_2902948, EPI_ISL_2902949, EPI_ISL_2902950, EPI_ISL_2902951, EPI_ISL_2902952, EPI_ISL_2902953, EPI_ISL_2902954, EPI_ISL_2902955, EPI_ISL_2902956, EPI_ISL_2902957, EPI_ISL_2902958, EPI_ISL_2902959, EPI_ISL_2902960, EPI_ISL_2902961,
EPI_ISL_2902962, EPI_ISL_2902963, EPI_ISL_2902964, EPI_ISL_2902965, EPI_ISL_2902966, EPI_ISL_2902967, EPI_ISL_2902968, EPI_ISL_2902969, EPI_ISL_2902970, EPI_ISL_2902971, EPI_ISL_2902972, EPI_ISL_2902973, EPI_ISL_2902974, EPI_ISL_2902975, EPI_ISL_2902976, EPI_ISL_2902977, EPI_ISL_2902978, EPI_ISL_2902979, EPI_ISL_2902980, EPI_ISL_2902981, EPI_ISL_2902982, EPI_ISL_2902983,
EPI_ISL_2902984, EPI_ISL_2902985, EPI_ISL_2902986, EPI_ISL_2902987, EPI_ISL_2902988, EPI_ISL_2902989, EPI_ISL_2902990, EPI_ISL_2902991, EPI_ISL_2902992, EPI_ISL_2902993, EPI_ISL_2902994, EPI_ISL_2902995, EPI_ISL_2902996, EPI_ISL_2902997, EPI_ISL_2902998, EPI_ISL_2902999, EPI_ISL_2903000

| | | | |
|---|---|---|---|
| see above | Quadram Institute Bioscience | COVID-19 Genomics UK (COG-UK) Consortium | Dave J. Baker, Gemma L. Kay, Alp Aydin, Thanh Le-Viet, Steven Rudder, Ana P. Tedim, Anastasia Kolyva, Maria Diaz, Leonardo de Oliveira Martins, Nabil-Fareed Alikhan, Lizzie Meadows, Rachael Stanley, Rose K Davidson, Ngozi Elumogo, Muhammed Yasir, Sophie J. Prosolek, Ebenezer Foster-Nyarko, Martin Lott, Christopher Jeanes, Lewis G. Spurgin, Nicholas M. Thomson, Alexander J Trotter, Rachel Gilroy, Luke Bedford, Samuel Bloomfield, Claire Stuart, Andrew Bell, Reenesh Prakash, Lindsay Coupland, Samir Dervisevic, Alison E. Mather, John Wain, Mark Webber, Andrew J. Page, Justin O'Grady |

EPI_ISL_2903040, EPI_ISL_2903042, EPI_ISL_2903043, EPI_ISL_2903045, EPI_ISL_2903046, EPI_ISL_2903047, EPI_ISL_2903048, EPI_ISL_2903049, EPI_ISL_2903050, EPI_ISL_2903053, EPI_ISL_2903056, EPI_ISL_2903059, EPI_ISL_2903060, EPI_ISL_2903061, EPI_ISL_2903062, EPI_ISL_2903063, EPI_ISL_2903064, EPI_ISL_2903066, EPI_ISL_2903068, EPI_ISL_2903069, EPI_ISL_2903070, EPI_ISL_2903071,
EPI_ISL_2903072, EPI_ISL_2903073, EPI_ISL_2903074, EPI_ISL_2903075, EPI_ISL_2903077, EPI_ISL_2903079, EPI_ISL_2903080, EPI_ISL_2903081, EPI_ISL_2903082, EPI_ISL_2903084, EPI_ISL_2903085, EPI_ISL_2903088, EPI_ISL_2903089, EPI_ISL_2903090, EPI_ISL_2903091, EPI_ISL_2903093, EPI_ISL_2903094, EPI_ISL_2903095, EPI_ISL_2903096, EPI_ISL_2903097, EPI_ISL_2903098, EPI_ISL_2903099,
EPI_ISL_2903100, EPI_ISL_2903101, EPI_ISL_2903102, EPI_ISL_2903103, EPI_ISL_2903105, EPI_ISL_2903106, EPI_ISL_2903107, EPI_ISL_2903108, EPI_ISL_2903109, EPI_ISL_2903110, EPI_ISL_2903111, EPI_ISL_2903113, EPI_ISL_2903116, EPI_ISL_2903117, EPI_ISL_2903118, EPI_ISL_2903119, EPI_ISL_2903120, EPI_ISL_2903121, EPI_ISL_2903122, EPI_ISL_2903124, EPI_ISL_2903125, EPI_ISL_2903126,
EPI_ISL_2903127, EPI_ISL_2903128, EPI_ISL_2903130, EPI_ISL_2903131, EPI_ISL_2903133, EPI_ISL_2903134, EPI_ISL_2903136, EPI_ISL_2903137, EPI_ISL_2903138, EPI_ISL_2903139, EPI_ISL_2903141, EPI_ISL_2903142, EPI_ISL_2903143, EPI_ISL_2903144, EPI_ISL_2903145, EPI_ISL_2903147, EPI_ISL_2903148, EPI_ISL_2903150, EPI_ISL_2903151, EPI_ISL_2903152, EPI_ISL_2903153, EPI_ISL_2903154,
EPI_ISL_2903155, EPI_ISL_2903156, EPI_ISL_2903157, EPI_ISL_2903158, EPI_ISL_2903159, EPI_ISL_2903160, EPI_ISL_2903162, EPI_ISL_2903163, EPI_ISL_2903164, EPI_ISL_2903165, EPI_ISL_2903167, EPI_ISL_2903168, EPI_ISL_2903169

| | | | |
|---|---|---|---|
| see above | Oxford Viromics, NDM, University of Oxford; Oxford University Hospitals; Basingstoke and North Hampshire Hospital | COVID-19 Genomics UK (COG-UK) Consortium | Tanya Golubchik, David Bonsall, George Macintyre, Amy Trebes, Mariateresa de Cesare, Catrin Moore, Alex Mobbs, Anita Justice, Robert Shaw, Monique Andersson, Timothy Peto, Emma Wise, Nathan Moore, Jessica Lynch, Nick Cortes, Matilde Mori, Stephen Kidd, David Buck, John Todd, Christophe Fraser |

EPI_ISL_2903412, EPI_ISL_2903413, EPI_ISL_2903414, EPI_ISL_2903415, EPI_ISL_2903416, EPI_ISL_2903417, EPI_ISL_2903418, EPI_ISL_2903419, EPI_ISL_2903420, EPI_ISL_2903421, EPI_ISL_2903422, EPI_ISL_2903423, EPI_ISL_2903424, EPI_ISL_2903425, EPI_ISL_2903426, EPI_ISL_2903427, EPI_ISL_2903428, EPI_ISL_2903429, EPI_ISL_2903430, EPI_ISL_2903431, EPI_ISL_2903432, EPI_ISL_2903433,
EPI_ISL_2903434, EPI_ISL_2903435, EPI_ISL_2903436, EPI_ISL_2903437, EPI_ISL_2903439, EPI_ISL_2903440, EPI_ISL_2903441, EPI_ISL_2903442, EPI_ISL_2903443, EPI_ISL_2903444, EPI_ISL_2903445, EPI_ISL_2903446, EPI_ISL_2903447, EPI_ISL_2903448, EPI_ISL_2903449, EPI_ISL_2903450, EPI_ISL_2903451, EPI_ISL_2903453, EPI_ISL_2903454, EPI_ISL_2903455, EPI_ISL_2903456, EPI_ISL_2903457,
EPI_ISL_2903458, EPI_ISL_2903460, EPI_ISL_2903461, EPI_ISL_2903462, EPI_ISL_2903463, EPI_ISL_2903464, EPI_ISL_2903465, EPI_ISL_2903466, EPI_ISL_2903467, EPI_ISL_2903468, EPI_ISL_2903469, EPI_ISL_2903470, EPI_ISL_2903471, EPI_ISL_2903472, EPI_ISL_2903473, EPI_ISL_2903474, EPI_ISL_2903475, EPI_ISL_2903476, EPI_ISL_2903477, EPI_ISL_2903479, EPI_ISL_2903480, EPI_ISL_2903481,
EPI_ISL_2903482, EPI_ISL_2903483, EPI_ISL_2903484, EPI_ISL_2903485, EPI_ISL_2903486, EPI_ISL_2903487, EPI_ISL_2903488, EPI_ISL_2903489, EPI_ISL_2903490, EPI_ISL_2903491, EPI_ISL_2903492, EPI_ISL_2903493, EPI_ISL_2903494, EPI_ISL_2903496, EPI_ISL_2903497, EPI_ISL_2903498, EPI_ISL_2903499, EPI_ISL_2903500, EPI_ISL_2903501, EPI_ISL_2903502, EPI_ISL_2903503, EPI_ISL_2903504,
EPI_ISL_2903505, EPI_ISL_2903506, EPI_ISL_2903507, EPI_ISL_2903508, EPI_ISL_2903509, EPI_ISL_2903510, EPI_ISL_2903511, EPI_ISL_2903512, EPI_ISL_2903513, EPI_ISL_2903514, EPI_ISL_2903515, EPI_ISL_2903516, EPI_ISL_2903517, EPI_ISL_2903518, EPI_ISL_2903519, EPI_ISL_2903520, EPI_ISL_2903521, EPI_ISL_2903522, EPI_ISL_2903523, EPI_ISL_2903524, EPI_ISL_2903525, EPI_ISL_2903526,
EPI_ISL_2903527, EPI_ISL_2903528, EPI_ISL_2903529, EPI_ISL_2903530, EPI_ISL_2903531, EPI_ISL_2903532, EPI_ISL_2903533, EPI_ISL_2903534, EPI_ISL_2903535, EPI_ISL_2903536, EPI_ISL_2903537, EPI_ISL_2903538, EPI_ISL_2903539, EPI_ISL_2903540, EPI_ISL_2903541, EPI_ISL_2903542, EPI_ISL_2903543, EPI_ISL_2903544, EPI_ISL_2903545, EPI_ISL_2903546, EPI_ISL_2903547, EPI_ISL_2903548,
EPI_ISL_2903549, EPI_ISL_2903550, EPI_ISL_2903551, EPI_ISL_2903552, EPI_ISL_2903553, EPI_ISL_2903554, EPI_ISL_2903555, EPI_ISL_2903556, EPI_ISL_2903557, EPI_ISL_2903558, EPI_ISL_2903559, EPI_ISL_2903560, EPI_ISL_2903561, EPI_ISL_2903562, EPI_ISL_2903563, EPI_ISL_2903564, EPI_ISL_2903565, EPI_ISL_2903566, EPI_ISL_2903567, EPI_ISL_2903568, EPI_ISL_2903569, EPI_ISL_2903570,
EPI_ISL_2903571, EPI_ISL_2903572, EPI_ISL_2903573, EPI_ISL_2903574, EPI_ISL_2903575, EPI_ISL_2903576, EPI_ISL_2903577, EPI_ISL_2903578, EPI_ISL_2903579, EPI_ISL_2903580, EPI_ISL_2903581, EPI_ISL_2903582, EPI_ISL_2903583, EPI_ISL_2903584, EPI_ISL_2903585, EPI_ISL_2903586, EPI_ISL_2903587, EPI_ISL_2903588, EPI_ISL_2903589, EPI_ISL_2903590, EPI_ISL_2903591, EPI_ISL_2903592,
EPI_ISL_2903593, EPI_ISL_2903594, EPI_ISL_2903595, EPI_ISL_2903596, EPI_ISL_2903597, EPI_ISL_2903598, EPI_ISL_2903599, EPI_ISL_2903600, EPI_ISL_2903601, EPI_ISL_2903603, EPI_ISL_2903604, EPI_ISL_2903605, EPI_ISL_2903606, EPI_ISL_2903607, EPI_ISL_2903608, EPI_ISL_2903609, EPI_ISL_2903610, EPI_ISL_2903611, EPI_ISL_2903612, EPI_ISL_2903613, EPI_ISL_2903615, EPI_ISL_2903616,
EPI_ISL_2903617, EPI_ISL_2903618, EPI_ISL_2903619, EPI_ISL_2903620, EPI_ISL_2903621, EPI_ISL_2903622, EPI_ISL_2903623, EPI_ISL_2903624, EPI_ISL_2903625, EPI_ISL_2903626, EPI_ISL_2903627, EPI_ISL_2903628, EPI_ISL_2903629, EPI_ISL_2903630, EPI_ISL_2903631, EPI_ISL_2903632, EPI_ISL_2903633, EPI_ISL_2903634, EPI_ISL_2903635, EPI_ISL_2903636, EPI_ISL_2903637, EPI_ISL_2903638,
EPI_ISL_2903639, EPI_ISL_2903640, EPI_ISL_2903641, EPI_ISL_2903642, EPI_ISL_2903643, EPI_ISL_2903644, EPI_ISL_2903645, EPI_ISL_2903646, EPI_ISL_2903647, EPI_ISL_2903648, EPI_ISL_2903653, EPI_ISL_2903654, EPI_ISL_2903655, EPI_ISL_2903656, EPI_ISL_2903657, EPI_ISL_2903658, EPI_ISL_2903659, EPI_ISL_2903660, EPI_ISL_2903661, EPI_ISL_2903662, EPI_ISL_2903663, EPI_ISL_2903664,
EPI_ISL_2903665, EPI_ISL_2903666, EPI_ISL_2903667, EPI_ISL_2903668, EPI_ISL_2903669, EPI_ISL_2903670, EPI_ISL_2903671, EPI_ISL_2903672, EPI_ISL_2903673, EPI_ISL_2903674, EPI_ISL_2903675, EPI_ISL_2903676, EPI_ISL_2903677, EPI_ISL_2903678, EPI_ISL_2903679, EPI_ISL_2903680, EPI_ISL_2903681, EPI_ISL_2903682, EPI_ISL_2903683, EPI_ISL_2903684, EPI_ISL_2903685, EPI_ISL_2903686,
EPI_ISL_2903687, EPI_ISL_2903688, EPI_ISL_2903689, EPI_ISL_2903690, EPI_ISL_2903691, EPI_ISL_2903692, EPI_ISL_2903693, EPI_ISL_2903694, EPI_ISL_2903695, EPI_ISL_2903696, EPI_ISL_2903697, EPI_ISL_2903698, EPI_ISL_2903699, EPI_ISL_2903700, EPI_ISL_2903702, EPI_ISL_2903703, EPI_ISL_2903704, EPI_ISL_2903705, EPI_ISL_2903706, EPI_ISL_2903707, EPI_ISL_2903708, EPI_ISL_2903709,
EPI_ISL_2903710, EPI_ISL_2903711, EPI_ISL_2903712, EPI_ISL_2903713, EPI_ISL_2903714, EPI_ISL_2903715, EPI_ISL_2903716, EPI_ISL_2903717, EPI_ISL_2903718, EPI_ISL_2903719, EPI_ISL_2903720, EPI_ISL_2903721, EPI_ISL_2903722, EPI_ISL_2903723, EPI_ISL_2903724, EPI_ISL_2903725, EPI_ISL_2903726, EPI_ISL_2903727, EPI_ISL_2903728, EPI_ISL_2903729, EPI_ISL_2903730, EPI_ISL_2903731,
EPI_ISL_2903732, EPI_ISL_2903733, EPI_ISL_2903734, EPI_ISL_2903735, EPI_ISL_2903736, EPI_ISL_2903737, EPI_ISL_2903738, EPI_ISL_2903739, EPI_ISL_2903740, EPI_ISL_2903741, EPI_ISL_2903742, EPI_ISL_2903743, EPI_ISL_2903744, EPI_ISL_2903745, EPI_ISL_2903746, EPI_ISL_2903747, EPI_ISL_2903748, EPI_ISL_2903749, EPI_ISL_2903750, EPI_ISL_2903751, EPI_ISL_2903752, EPI_ISL_2903753,
EPI_ISL_2903754, EPI_ISL_2903755, EPI_ISL_2903756, EPI_ISL_2903757, EPI_ISL_2903758, EPI_ISL_2903759, EPI_ISL_2903760, EPI_ISL_2903761, EPI_ISL_2903762, EPI_ISL_2903763, EPI_ISL_2903764, EPI_ISL_2903765, EPI_ISL_2903766, EPI_ISL_2903767, EPI_ISL_2903768, EPI_ISL_2903769, EPI_ISL_2903770, EPI_ISL_2903771, EPI_ISL_2903772, EPI_ISL_2903773, EPI_ISL_2903774, EPI_ISL_2903775,
EPI_ISL_2903776, EPI_ISL_2903777, EPI_ISL_2903778, EPI_ISL_2903779, EPI_ISL_2903780, EPI_ISL_2903781, EPI_ISL_2903782, EPI_ISL_2903783, EPI_ISL_2903784, EPI_ISL_2903785, EPI_ISL_2903786, EPI_ISL_2903787, EPI_ISL_2903788, EPI_ISL_2903789, EPI_ISL_2903792, EPI_ISL_2903793, EPI_ISL_2903794, EPI_ISL_2903795, EPI_ISL_2903796, EPI_ISL_2903797, EPI_ISL_2903798, EPI_ISL_2903799,
EPI_ISL_2903800, EPI_ISL_2903801, EPI_ISL_2903802, EPI_ISL_2903803, EPI_ISL_2903805, EPI_ISL_2903806, EPI_ISL_2903807, EPI_ISL_2903809, EPI_ISL_2903810, EPI_ISL_2903811, EPI_ISL_2903812, EPI_ISL_2903813, EPI_ISL_2903814, EPI_ISL_2903815, EPI_ISL_2903816, EPI_ISL_2903817, EPI_ISL_2903818, EPI_ISL_2903819, EPI_ISL_2903820, EPI_ISL_2903821, EPI_ISL_2903822, EPI_ISL_2903823,
EPI_ISL_2903824, EPI_ISL_2903825, EPI_ISL_2903828, EPI_ISL_2903829, EPI_ISL_2903830, EPI_ISL_2903831, EPI_ISL_2903832, EPI_ISL_2903833, EPI_ISL_2903834, EPI_ISL_2903835, EPI_ISL_2903836, EPI_ISL_2903837, EPI_ISL_2903838, EPI_ISL_2903839, EPI_ISL_2903840, EPI_ISL_2903841, EPI_ISL_2903842, EPI_ISL_2903843, EPI_ISL_2903844, EPI_ISL_2903845, EPI_ISL_2903846, EPI_ISL_2903847,
EPI_ISL_2903848, EPI_ISL_2903850, EPI_ISL_2903851, EPI_ISL_2903852, EPI_ISL_2903853, EPI_ISL_2903854, EPI_ISL_2903855, EPI_ISL_2903856, EPI_ISL_2903857, EPI_ISL_2903858, EPI_ISL_2903859, EPI_ISL_2903860, EPI_ISL_2903861, EPI_ISL_2903862, EPI_ISL_2903863, EPI_ISL_2903864, EPI_ISL_2903865, EPI_ISL_2903866, EPI_ISL_2903867, EPI_ISL_2903868, EPI_ISL_2903869, EPI_ISL_2903870,
EPI_ISL_2903871, EPI_ISL_2903872, EPI_ISL_2903873, EPI_ISL_2903874, EPI_ISL_2903875, EPI_ISL_2903876, EPI_ISL_2903877, EPI_ISL_2903878, EPI_ISL_2903879, EPI_ISL_2903880, EPI_ISL_2903881, EPI_ISL_2903882, EPI_ISL_2903883, EPI_ISL_2903884, EPI_ISL_2903885, EPI_ISL_2903886, EPI_ISL_2903887, EPI_ISL_2903888, EPI_ISL_2903889, EPI_ISL_2903890, EPI_ISL_2903891, EPI_ISL_2903892,
EPI_ISL_2903894, EPI_ISL_2903895, EPI_ISL_2903896, EPI_ISL_2903897, EPI_ISL_2903898, EPI_ISL_2903899, EPI_ISL_2903900, EPI_ISL_2903902, EPI_ISL_2903903, EPI_ISL_2903904, EPI_ISL_2903905, EPI_ISL_2903906, EPI_ISL_2903907, EPI_ISL_2903908, EPI_ISL_2903909, EPI_ISL_2903910, EPI_ISL_2903911, EPI_ISL_2903913, EPI_ISL_2903916, EPI_ISL_2903918, EPI_ISL_2903919, EPI_ISL_2903921,
EPI_ISL_2903924, EPI_ISL_2903926, EPI_ISL_2903927, EPI_ISL_2903928, EPI_ISL_2903929, EPI_ISL_2903930, EPI_ISL_2903931, EPI_ISL_2903932, EPI_ISL_2903933, EPI_ISL_2903934, EPI_ISL_2903935, EPI_ISL_2903936, EPI_ISL_2903937, EPI_ISL_2903938, EPI_ISL_2903939, EPI_ISL_2903942, EPI_ISL_2903943, EPI_ISL_2903945, EPI_ISL_2903946, EPI_ISL_2903947, EPI_ISL_2903949, EPI_ISL_2903950,
EPI_ISL_2903951, EPI_ISL_2903952, EPI_ISL_2903953, EPI_ISL_2903954, EPI_ISL_2903955, EPI_ISL_2903956, EPI_ISL_2903957, EPI_ISL_2903958, EPI_ISL_2903959, EPI_ISL_2903960, EPI_ISL_2903962, EPI_ISL_2903963, EPI_ISL_2903964, EPI_ISL_2903965, EPI_ISL_2903966, EPI_ISL_2903968, EPI_ISL_2903969, EPI_ISL_2903970, EPI_ISL_2903971, EPI_ISL_2903972, EPI_ISL_2903973, EPI_ISL_2903974,
EPI_ISL_2903975, EPI_ISL_2903976, EPI_ISL_2903977, EPI_ISL_2903978, EPI_ISL_2903979, EPI_ISL_2903980, EPI_ISL_2903981, EPI_ISL_2903982, EPI_ISL_2903983, EPI_ISL_2903984, EPI_ISL_2903985, EPI_ISL_2903986, EPI_ISL_2903987, EPI_ISL_2903989, EPI_ISL_2903990, EPI_ISL_2903991, EPI_ISL_2903992, EPI_ISL_2903993, EPI_ISL_2903994, EPI_ISL_2903995, EPI_ISL_2903996, EPI_ISL_2903997,
EPI_ISL_2903998, EPI_ISL_2904000, EPI_ISL_2904001, EPI_ISL_2904002, EPI_ISL_2904003, EPI_ISL_2904004, EPI_ISL_2904005, EPI_ISL_2904006, EPI_ISL_2904007, EPI_ISL_2904008, EPI_ISL_2904009, EPI_ISL_2904010, EPI_ISL_2904011, EPI_ISL_2904012, EPI_ISL_2904013, EPI_ISL_2904014, EPI_ISL_2904015, EPI_ISL_2904016, EPI_ISL_2904017, EPI_ISL_2904018, EPI_ISL_2904019, EPI_ISL_2904020,
EPI_ISL_2904021, EPI_ISL_2904022, EPI_ISL_2904023, EPI_ISL_2904024, EPI_ISL_2904025, EPI_ISL_2904026, EPI_ISL_2904027, EPI_ISL_2904029, EPI_ISL_2904030, EPI_ISL_2904031, EPI_ISL_2904032, EPI_ISL_2904033, EPI_ISL_2904034, EPI_ISL_2904035, EPI_ISL_2904036, EPI_ISL_2904037, EPI_ISL_2904038, EPI_ISL_2904039, EPI_ISL_2904040, EPI_ISL_2904041, EPI_ISL_2904042, EPI_ISL_2904043,
EPI_ISL_2904044, EPI_ISL_2904045, EPI_ISL_2904046, EPI_ISL_2904047, EPI_ISL_2904048, EPI_ISL_2904049, EPI_ISL_2904050, EPI_ISL_2904051, EPI_ISL_2904053, EPI_ISL_2904054, EPI_ISL_2904055, EPI_ISL_2904056, EPI_ISL_2904057, EPI_ISL_2904058, EPI_ISL_2904059, EPI_ISL_2904060, EPI_ISL_2904061, EPI_ISL_2904062, EPI_ISL_2904063, EPI_ISL_2904064, EPI_ISL_2904065, EPI_ISL_2904066,
EPI_ISL_2904067, EPI_ISL_2904068, EPI_ISL_2904069, EPI_ISL_2904070, EPI_ISL_2904071, EPI_ISL_2904072, EPI_ISL_2904073, EPI_ISL_2904074, EPI_ISL_2904075, EPI_ISL_2904076, EPI_ISL_2904077, EPI_ISL_2904078, EPI_ISL_2904079, EPI_ISL_2904080, EPI_ISL_2904081, EPI_ISL_2904082, EPI_ISL_2904083, EPI_ISL_2904084, EPI_ISL_2904085, EPI_ISL_2904086, EPI_ISL_2904087, EPI_ISL_2904088,

EPI_ISL_2904089, EPI_ISL_2904090, EPI_ISL_2904091, EPI_ISL_2904092, EPI_ISL_2904093, EPI_ISL_2904094, EPI_ISL_2904095, EPI_ISL_2904096, EPI_ISL_2904097, EPI_ISL_2904098, EPI_ISL_2904099, EPI_ISL_2904100, EPI_ISL_2904101, EPI_ISL_2904102, EPI_ISL_2904103, EPI_ISL_2904104, EPI_ISL_2904105, EPI_ISL_2904106, EPI_ISL_2904107, EPI_ISL_2904108, EPI_ISL_2904109, EPI_ISL_2904110, EPI_ISL_2904111, EPI_ISL_2904112, EPI_ISL_2904113, EPI_ISL_2904114, EPI_ISL_2904115, EPI_ISL_2904116, EPI_ISL_2904117, EPI_ISL_2904118, EPI_ISL_2904119, EPI_ISL_2904120, EPI_ISL_2904121, EPI_ISL_2904122, EPI_ISL_2904123, EPI_ISL_2904124, EPI_ISL_2904125, EPI_ISL_2904126, EPI_ISL_2904127, EPI_ISL_2904128, EPI_ISL_2904129, EPI_ISL_2904130, EPI_ISL_2904131, EPI_ISL_2904132, EPI_ISL_2904133, EPI_ISL_2904134, EPI_ISL_2904135, EPI_ISL_2904136, EPI_ISL_2904137, EPI_ISL_2904138, EPI_ISL_2904139, EPI_ISL_2904140, EPI_ISL_2904141, EPI_ISL_2904142, EPI_ISL_2904143, EPI_ISL_2904144, EPI_ISL_2904145, EPI_ISL_2904146, EPI_ISL_2904147, EPI_ISL_2904148, EPI_ISL_2904149, EPI_ISL_2904150, EPI_ISL_2904151, EPI_ISL_2904152, EPI_ISL_2904154, EPI_ISL_2904155, EPI_ISL_2904156, EPI_ISL_2904157, EPI_ISL_2904158, EPI_ISL_2904159, EPI_ISL_2904160, EPI_ISL_2904161, EPI_ISL_2904162, EPI_ISL_2904163, EPI_ISL_2904164, EPI_ISL_2904165, EPI_ISL_2904166, EPI_ISL_2904167, EPI_ISL_2904168, EPI_ISL_2904169, EPI_ISL_2904170, EPI_ISL_2904171, EPI_ISL_2904172, EPI_ISL_2904173, EPI_ISL_2904174, EPI_ISL_2904175, EPI_ISL_2904176, EPI_ISL_2904177, EPI_ISL_2904178, EPI_ISL_2904179, EPI_ISL_2904180, EPI_ISL_2904181, EPI_ISL_2904182, EPI_ISL_2904183, EPI_ISL_2904184, EPI_ISL_2904185, EPI_ISL_2904186, EPI_ISL_2904187, EPI_ISL_2904188, EPI_ISL_2904189, EPI_ISL_2904190, EPI_ISL_2904191, EPI_ISL_2904192, EPI_ISL_2904193, EPI_ISL_2904194, EPI_ISL_2904195, EPI_ISL_2904196, EPI_ISL_2904197, EPI_ISL_2904198, EPI_ISL_2904199, EPI_ISL_2904200, EPI_ISL_2904201, EPI_ISL_2904202, EPI_ISL_2904203, EPI_ISL_2904204, EPI_ISL_2904205, EPI_ISL_2904206, EPI_ISL_2904207, EPI_ISL_2904208, EPI_ISL_2904209, EPI_ISL_2904210, EPI_ISL_2904211, EPI_ISL_2904212, EPI_ISL_2904213, EPI_ISL_2904214, EPI_ISL_2904215, EPI_ISL_2904216, EPI_ISL_2904217, EPI_ISL_2904218, EPI_ISL_2904219, EPI_ISL_2904220, EPI_ISL_2904221, EPI_ISL_2904222, EPI_ISL_2904223, EPI_ISL_2904224, EPI_ISL_2904225, EPI_ISL_2904226, EPI_ISL_2904227, EPI_ISL_2904228, EPI_ISL_2904229, EPI_ISL_2904230, EPI_ISL_2904231, EPI_ISL_2904232, EPI_ISL_2904233, EPI_ISL_2904234, EPI_ISL_2904235, EPI_ISL_2904236, EPI_ISL_2904237, EPI_ISL_2904238, EPI_ISL_2904239, EPI_ISL_2904240, EPI_ISL_2904241, EPI_ISL_2904242, EPI_ISL_2904243, EPI_ISL_2904244, EPI_ISL_2904245, EPI_ISL_2904246, EPI_ISL_2904247, EPI_ISL_2904248, EPI_ISL_2904249, EPI_ISL_2904250, EPI_ISL_2904251, EPI_ISL_2904252, EPI_ISL_2904253, EPI_ISL_2904254, EPI_ISL_2904255, EPI_ISL_2904256, EPI_ISL_2904257, EPI_ISL_2904258, EPI_ISL_2904259, EPI_ISL_2904260, EPI_ISL_2904261, EPI_ISL_2904262, EPI_ISL_2904263, EPI_ISL_2904264, EPI_ISL_2904266, EPI_ISL_2904270, EPI_ISL_2904274, EPI_ISL_2904275, EPI_ISL_2904276, EPI_ISL_2904281, EPI_ISL_2904282, EPI_ISL_2904283, EPI_ISL_2904285, EPI_ISL_2904306, EPI_ISL_2904308, EPI_ISL_2904311, EPI_ISL_2904312, EPI_ISL_2904315, EPI_ISL_2904318, EPI_ISL_2904320, EPI_ISL_2904327, EPI_ISL_2904328, EPI_ISL_2904329, EPI_ISL_2904330, EPI_ISL_2904332, EPI_ISL_2904333, EPI_ISL_2904334, EPI_ISL_2904335, EPI_ISL_2904336, EPI_ISL_2904338, EPI_ISL_2904339, EPI_ISL_2904340, EPI_ISL_2904341, EPI_ISL_2904342, EPI_ISL_2904343, EPI_ISL_2904344, EPI_ISL_2904345, EPI_ISL_2904346, EPI_ISL_2904347, EPI_ISL_2904348, EPI_ISL_2904349, EPI_ISL_2904350, EPI_ISL_2904352, EPI_ISL_2904353, EPI_ISL_2904354, EPI_ISL_2904355, EPI_ISL_2904356, EPI_ISL_2904357, EPI_ISL_2904358, EPI_ISL_2904359, EPI_ISL_2904360, EPI_ISL_2904361, EPI_ISL_2904362, EPI_ISL_2904363, EPI_ISL_2904364, EPI_ISL_2904365, EPI_ISL_2904367, EPI_ISL_2904368, EPI_ISL_2904369, EPI_ISL_2904370, EPI_ISL_2904371, EPI_ISL_2904372, EPI_ISL_2904373, EPI_ISL_2904375, EPI_ISL_2904377, EPI_ISL_2904378, EPI_ISL_2904379, EPI_ISL_2904380, EPI_ISL_2904381, EPI_ISL_2904383, EPI_ISL_2904384, EPI_ISL_2904385, EPI_ISL_2904386, EPI_ISL_2904387, EPI_ISL_2904388, EPI_ISL_2904389, EPI_ISL_2904390, EPI_ISL_2904391, EPI_ISL_2904393, EPI_ISL_2904394, EPI_ISL_2904395, EPI_ISL_2904396, EPI_ISL_2904397, EPI_ISL_2904398, EPI_ISL_2904399, EPI_ISL_2904400, EPI_ISL_2904401, EPI_ISL_2904402, EPI_ISL_2904404, EPI_ISL_2904405, EPI_ISL_2904407, EPI_ISL_2904408, EPI_ISL_2904409, EPI_ISL_2904410, EPI_ISL_2904411, EPI_ISL_2904412, EPI_ISL_2904414, EPI_ISL_2904415, EPI_ISL_2904416, EPI_ISL_2904418, EPI_ISL_2904419, EPI_ISL_2904420, EPI_ISL_2904422, EPI_ISL_2904424, EPI_ISL_2904425, EPI_ISL_2904426, EPI_ISL_2904428, EPI_ISL_2904429, EPI_ISL_2904430, EPI_ISL_2904432, EPI_ISL_2904433, EPI_ISL_2904434, EPI_ISL_2904435, EPI_ISL_2904436, EPI_ISL_2904437, EPI_ISL_2904439, EPI_ISL_2904441, EPI_ISL_2904442, EPI_ISL_2904443, EPI_ISL_2904444, EPI_ISL_2904445, EPI_ISL_2904446, EPI_ISL_2904448, EPI_ISL_2904449, EPI_ISL_2904450, EPI_ISL_2904451, EPI_ISL_2904453, EPI_ISL_2904455, EPI_ISL_2904456, EPI_ISL_2904457, EPI_ISL_2904458, EPI_ISL_2904459, EPI_ISL_2904460, EPI_ISL_2904461, EPI_ISL_2904462, EPI_ISL_2904463, EPI_ISL_2904464, EPI_ISL_2904465, EPI_ISL_2904466, EPI_ISL_2904467, EPI_ISL_2904469, EPI_ISL_2904470, EPI_ISL_2904473

| | | | |
|---|---|---|---|
| see above | Respiratory Virus Unit, Microbiology Services Colindale, Public Health England | COVID-19 Genomics UK (COG-UK) Consortium | PHE Covid Sequencing Team |

EPI_ISL_2904475, EPI_ISL_2904476, EPI_ISL_2904477, EPI_ISL_2904478, EPI_ISL_2904479, EPI_ISL_2904480, EPI_ISL_2904481, EPI_ISL_2904482, EPI_ISL_2904483, EPI_ISL_2904484, EPI_ISL_2904485, EPI_ISL_2904486, EPI_ISL_2904487, EPI_ISL_2904488, EPI_ISL_2904489, EPI_ISL_2904490, EPI_ISL_2904491, EPI_ISL_2904492, EPI_ISL_2904493, EPI_ISL_2904494, EPI_ISL_2904495, EPI_ISL_2904496, EPI_ISL_2904497, EPI_ISL_2904498, EPI_ISL_2904499, EPI_ISL_2904500, EPI_ISL_2904501, EPI_ISL_2904502, EPI_ISL_2904503, EPI_ISL_2904504, EPI_ISL_2904505, EPI_ISL_2904506, EPI_ISL_2904507, EPI_ISL_2904508, EPI_ISL_2904509, EPI_ISL_2904510, EPI_ISL_2904511, EPI_ISL_2904512, EPI_ISL_2904513, EPI_ISL_2904514, EPI_ISL_2904515, EPI_ISL_2904516, EPI_ISL_2904517, EPI_ISL_2904518, EPI_ISL_2904519, EPI_ISL_2904520, EPI_ISL_2904521, EPI_ISL_2904522, EPI_ISL_2904523, EPI_ISL_2904524, EPI_ISL_2904525, EPI_ISL_2904526, EPI_ISL_2904527, EPI_ISL_2904528, EPI_ISL_2904529, EPI_ISL_2904530, EPI_ISL_2904531, EPI_ISL_2904532, EPI_ISL_2904533, EPI_ISL_2904534, EPI_ISL_2904535, EPI_ISL_2904536, EPI_ISL_2904537, EPI_ISL_2904538, EPI_ISL_2904539, EPI_ISL_2904540, EPI_ISL_2904541, EPI_ISL_2904542, EPI_ISL_2904543, EPI_ISL_2904544, EPI_ISL_2904545, EPI_ISL_2904546, EPI_ISL_2904547, EPI_ISL_2904548, EPI_ISL_2904549, EPI_ISL_2904550, EPI_ISL_2904551, EPI_ISL_2904552, EPI_ISL_2904553, EPI_ISL_2904554, EPI_ISL_2904555, EPI_ISL_2904556, EPI_ISL_2904557, EPI_ISL_2904558, EPI_ISL_2904559, EPI_ISL_2904560, EPI_ISL_2904561, EPI_ISL_2904562, EPI_ISL_2904563, EPI_ISL_2904564, EPI_ISL_2904565, EPI_ISL_2904566, EPI_ISL_2904567, EPI_ISL_2904568, EPI_ISL_2904569, EPI_ISL_2904570, EPI_ISL_2904571, EPI_ISL_2904572, EPI_ISL_2904573, EPI_ISL_2904574, EPI_ISL_2904575, EPI_ISL_2904576, EPI_ISL_2904577, EPI_ISL_2904578, EPI_ISL_2904579, EPI_ISL_2904580, EPI_ISL_2904581, EPI_ISL_2904582, EPI_ISL_2904583, EPI_ISL_2904584, EPI_ISL_2904585, EPI_ISL_2904586, EPI_ISL_2904587, EPI_ISL_2904588, EPI_ISL_2904589, EPI_ISL_2904590, EPI_ISL_2904591, EPI_ISL_2904592, EPI_ISL_2904593, EPI_ISL_2904594, EPI_ISL_2904595, EPI_ISL_2904596, EPI_ISL_2904597, EPI_ISL_2904598, EPI_ISL_2904599, EPI_ISL_2904600, EPI_ISL_2904601, EPI_ISL_2904602, EPI_ISL_2904603, EPI_ISL_2904604, EPI_ISL_2904605, EPI_ISL_2904606, EPI_ISL_2904607, EPI_ISL_2904608, EPI_ISL_2904609, EPI_ISL_2904610, EPI_ISL_2904611, EPI_ISL_2904612, EPI_ISL_2904613, EPI_ISL_2904614, EPI_ISL_2904615, EPI_ISL_2904616, EPI_ISL_2904617, EPI_ISL_2904618, EPI_ISL_2904619, EPI_ISL_2904620, EPI_ISL_2904621, EPI_ISL_2904622, EPI_ISL_2904623, EPI_ISL_2904624, EPI_ISL_2904625, EPI_ISL_2904626, EPI_ISL_2904627, EPI_ISL_2904628, EPI_ISL_2904629, EPI_ISL_2904630, EPI_ISL_2904631, EPI_ISL_2904632, EPI_ISL_2904633, EPI_ISL_2904634, EPI_ISL_2904635, EPI_ISL_2904636, EPI_ISL_2904637, EPI_ISL_2904638, EPI_ISL_2904639, EPI_ISL_2904640, EPI_ISL_2904641, EPI_ISL_2904642, EPI_ISL_2904643, EPI_ISL_2904644, EPI_ISL_2904645, EPI_ISL_2904646, EPI_ISL_2904647, EPI_ISL_2904648, EPI_ISL_2904649, EPI_ISL_2904650, EPI_ISL_2904651, EPI_ISL_2904652, EPI_ISL_2904653, EPI_ISL_2904654, EPI_ISL_2904655, EPI_ISL_2904656, EPI_ISL_2904657, EPI_ISL_2904658, EPI_ISL_2904659, EPI_ISL_2904660, EPI_ISL_2904661, EPI_ISL_2904662, EPI_ISL_2904663, EPI_ISL_2904664, EPI_ISL_2904665, EPI_ISL_2904666, EPI_ISL_2904667, EPI_ISL_2904668, EPI_ISL_2904669, EPI_ISL_2904670, EPI_ISL_2904671, EPI_ISL_2904672, EPI_ISL_2904673, EPI_ISL_2904674, EPI_ISL_2904675, EPI_ISL_2904676, EPI_ISL_2904677, EPI_ISL_2904678, EPI_ISL_2904679, EPI_ISL_2904680, EPI_ISL_2904681, EPI_ISL_2904682, EPI_ISL_2904683, EPI_ISL_2904684, EPI_ISL_2904685, EPI_ISL_2904686, EPI_ISL_2904687, EPI_ISL_2904688, EPI_ISL_2904689, EPI_ISL_2904690, EPI_ISL_2904691, EPI_ISL_2904692, EPI_ISL_2904693, EPI_ISL_2904694, EPI_ISL_2904695, EPI_ISL_2904696, EPI_ISL_2904697, EPI_ISL_2904698, EPI_ISL_2904699, EPI_ISL_2904700, EPI_ISL_2904701, EPI_ISL_2904702, EPI_ISL_2904703, EPI_ISL_2904704, EPI_ISL_2904705, EPI_ISL_2904706, EPI_ISL_2904707, EPI_ISL_2904708, EPI_ISL_2904709, EPI_ISL_2904710, EPI_ISL_2904711, EPI_ISL_2904712, EPI_ISL_2904713, EPI_ISL_2904714, EPI_ISL_2904715, EPI_ISL_2904716, EPI_ISL_2904717, EPI_ISL_2904718, EPI_ISL_2904719, EPI_ISL_2904720, EPI_ISL_2904721, EPI_ISL_2904722, EPI_ISL_2904723, EPI_ISL_2904724, EPI_ISL_2904725, EPI_ISL_2904726, EPI_ISL_2904727, EPI_ISL_2904728, EPI_ISL_2904729, EPI_ISL_2904730, EPI_ISL_2904731, EPI_ISL_2904732, EPI_ISL_2904733, EPI_ISL_2904734, EPI_ISL_2904735, EPI_ISL_2904736, EPI_ISL_2904737, EPI_ISL_2904738, EPI_ISL_2904739, EPI_ISL_2904740, EPI_ISL_2904741, EPI_ISL_2904742, EPI_ISL_2904743, EPI_ISL_2904744, EPI_ISL_2904745, EPI_ISL_2904746, EPI_ISL_2904747, EPI_ISL_2904748, EPI_ISL_2904749, EPI_ISL_2904750, EPI_ISL_2904751, EPI_ISL_2904752, EPI_ISL_2904753, EPI_ISL_2904754, EPI_ISL_2904755, EPI_ISL_2904756, EPI_ISL_2904757, EPI_ISL_2904758, EPI_ISL_2904759, EPI_ISL_2904760, EPI_ISL_2904761, EPI_ISL_2904762, EPI_ISL_2904764, EPI_ISL_2904765, EPI_ISL_2904766, EPI_ISL_2904767, EPI_ISL_2904768, EPI_ISL_2904769, EPI_ISL_2904770, EPI_ISL_2904771, EPI_ISL_2904772, EPI_ISL_2904773, EPI_ISL_2904774, EPI_ISL_2904775, EPI_ISL_2904776, EPI_ISL_2904777, EPI_ISL_2904778, EPI_ISL_2904779, EPI_ISL_2904780, EPI_ISL_2904781, EPI_ISL_2904782, EPI_ISL_2904783, EPI_ISL_2904784, EPI_ISL_2904785, EPI_ISL_2904786, EPI_ISL_2904787, EPI_ISL_2904788, EPI_ISL_2904789, EPI_ISL_2904790, EPI_ISL_2904791, EPI_ISL_2904792, EPI_ISL_2904793, EPI_ISL_2904794, EPI_ISL_2904795, EPI_ISL_2904796, EPI_ISL_2904797, EPI_ISL_2904798, EPI_ISL_2904799, EPI_ISL_2904800, EPI_ISL_2904801, EPI_ISL_2904802, EPI_ISL_2904803, EPI_ISL_2904804, EPI_ISL_2904805, EPI_ISL_2904806, EPI_ISL_2904807, EPI_ISL_2904808, EPI_ISL_2904809, EPI_ISL_2904810, EPI_ISL_2904811, EPI_ISL_2904812, EPI_ISL_2904813, EPI_ISL_2904814, EPI_ISL_2904815, EPI_ISL_2904816, EPI_ISL_2904817, EPI_ISL_2904818, EPI_ISL_2904819, EPI_ISL_2904820, EPI_ISL_2904821, EPI_ISL_2904822, EPI_ISL_2904823, EPI_ISL_2904824, EPI_ISL_2904825, EPI_ISL_2904826, EPI_ISL_2904827, EPI_ISL_2904828, EPI_ISL_2904829, EPI_ISL_2904830, EPI_ISL_2904831, EPI_ISL_2904832, EPI_ISL_2904833, EPI_ISL_2904834, EPI_ISL_2904835, EPI_ISL_2904836, EPI_ISL_2904837, EPI_ISL_2904838, EPI_ISL_2904839, EPI_ISL_2904840, EPI_ISL_2904841, EPI_ISL_2904842, EPI_ISL_2904843, EPI_ISL_2904844, EPI_ISL_2904845, EPI_ISL_2904846, EPI_ISL_2904847, EPI_ISL_2904848, EPI_ISL_2904849, EPI_ISL_2904850, EPI_ISL_2904851, EPI_ISL_2904852, EPI_ISL_2904853, EPI_ISL_2904854, EPI_ISL_2904855, EPI_ISL_2904856, EPI_ISL_2904857, EPI_ISL_2904858, EPI_ISL_2904859, EPI_ISL_2904860, EPI_ISL_2904861, EPI_ISL_2904862, EPI_ISL_2904863, EPI_ISL_2904864, EPI_ISL_2904866, EPI_ISL_2904867, EPI_ISL_2904868, EPI_ISL_2904869, EPI_ISL_2904870, EPI_ISL_2904871, EPI_ISL_2904872, EPI_ISL_2904873, EPI_ISL_2904874, EPI_ISL_2904875, EPI_ISL_2904876, EPI_ISL_2904878, EPI_ISL_2904879, EPI_ISL_2904880, EPI_ISL_2904881, EPI_ISL_2904882, EPI_ISL_2904883, EPI_ISL_2904884, EPI_ISL_2904885, EPI_ISL_2904886, EPI_ISL_2904887, EPI_ISL_2904888, EPI_ISL_2904889, EPI_ISL_2904890, EPI_ISL_2904891, EPI_ISL_2904892, EPI_ISL_2904893, EPI_ISL_2904894, EPI_ISL_2904895, EPI_ISL_2904896, EPI_ISL_2904897, EPI_ISL_2904898, EPI_ISL_2904899, EPI_ISL_2904900, EPI_ISL_2904901, EPI_ISL_2904902, EPI_ISL_2904903, EPI_ISL_2904904, EPI_ISL_2904905, EPI_ISL_2904906, EPI_ISL_2904907, EPI_ISL_2904908, EPI_ISL_2904909, EPI_ISL_2904910, EPI_ISL_2904911, EPI_ISL_2904912, EPI_ISL_2904913, EPI_ISL_2904914, EPI_ISL_2904915, EPI_ISL_2904917, EPI_ISL_2904918, EPI_ISL_2904919, EPI_ISL_2904921, EPI_ISL_2904922, EPI_ISL_2904923, EPI_ISL_2904924, EPI_ISL_2904925, EPI_ISL_2904926, EPI_ISL_2904927, EPI_ISL_2904928, EPI_ISL_2904929, EPI_ISL_2904930, EPI_ISL_2904931, EPI_ISL_2904932, EPI_ISL_2904933, EPI_ISL_2904934, EPI_ISL_2904935, EPI_ISL_2904936, EPI_ISL_2904937, EPI_ISL_2904938, EPI_ISL_2904939, EPI_ISL_2904940, EPI_ISL_2904941, EPI_ISL_2904942, EPI_ISL_2904943, EPI_ISL_2904944, EPI_ISL_2904945, EPI_ISL_2904946, EPI_ISL_2904947, EPI_ISL_2904948, EPI_ISL_2904949, EPI_ISL_2904950, EPI_ISL_2904951, EPI_ISL_2904952, EPI_ISL_2904953, EPI_ISL_2904954, EPI_ISL_2904955, EPI_ISL_2904956, EPI_ISL_2904957, EPI_ISL_2904958, EPI_ISL_2904959, EPI_ISL_2904960, EPI_ISL_2904961, EPI_ISL_2904962, EPI_ISL_2904963, EPI_ISL_2904964, EPI_ISL_2904965, EPI_ISL_2904966, EPI_ISL_2904967, EPI_ISL_2904968, EPI_ISL_2904969, EPI_ISL_2904970, EPI_ISL_2904971, EPI_ISL_2904972, EPI_ISL_2904973, EPI_ISL_2904974, EPI_ISL_2904975, EPI_ISL_2904976, EPI_ISL_2904977, EPI_ISL_2904978, EPI_ISL_2904979, EPI_ISL_2904980, EPI_ISL_2904981, EPI_ISL_2904982, EPI_ISL_2904983, EPI_ISL_2904984, EPI_ISL_2904985, EPI_ISL_2904986, EPI_ISL_2904987, EPI_ISL_2904988, EPI_ISL_2904989, EPI_ISL_2904990, EPI_ISL_2904991, EPI_ISL_2904992, EPI_ISL_2904993, EPI_ISL_2904994, EPI_ISL_2904995, EPI_ISL_2904996, EPI_ISL_2904997, EPI_ISL_2904998, EPI_ISL_2904999, EPI_ISL_2905000, EPI_ISL_2905001, EPI_ISL_2905002, EPI_ISL_2905003, EPI_ISL_2905004, EPI_ISL_2905005, EPI_ISL_2905006, EPI_ISL_2905007, EPI_ISL_2905008, EPI_ISL_2905009, EPI_ISL_2905010, EPI_ISL_2905011, EPI_ISL_2905012, EPI_ISL_2905013, EPI_ISL_2905014, EPI_ISL_2905015, EPI_ISL_2905016, EPI_ISL_2905017, EPI_ISL_2905018, EPI_ISL_2905019, EPI_ISL_2905020, EPI_ISL_2905021, EPI_ISL_2905022, EPI_ISL_2905023, EPI_ISL_2905024, EPI_ISL_2905025, EPI_ISL_2905026, EPI_ISL_2905027, EPI_ISL_2905028, EPI_ISL_2905029, EPI_ISL_2905030, EPI_ISL_2905031, EPI_ISL_2905032, EPI_ISL_2905033, EPI_ISL_2905034, EPI_ISL_2905035, EPI_ISL_2905036, EPI_ISL_2905037, EPI_ISL_2905038, EPI_ISL_2905039, EPI_ISL_2905040, EPI_ISL_2905041, EPI_ISL_2905042, EPI_ISL_2905043, EPI_ISL_2905044, EPI_ISL_2905045, EPI_ISL_2905046, EPI_ISL_2905047, EPI_ISL_2905048, EPI_ISL_2905049, EPI_ISL_2905050, EPI_ISL_2905051, EPI_ISL_2905052, EPI_ISL_2905053, EPI_ISL_2905054, EPI_ISL_2905055, EPI_ISL_2905056, EPI_ISL_2905057, EPI_ISL_2905058, EPI_ISL_2905059, EPI_ISL_2905060, EPI_ISL_2905061, EPI_ISL_2905062, EPI_ISL_2905063, EPI_ISL_2905064, EPI_ISL_2905065, EPI_ISL_2905066, EPI_ISL_2905067, EPI_ISL_2905068, EPI_ISL_2905069, EPI_ISL_2905070, EPI_ISL_2905071, EPI_ISL_2905072, EPI_ISL_2905073, EPI_ISL_2905074, EPI_ISL_2905075, EPI_ISL_2905076, EPI_ISL_2905077, EPI_ISL_2905078, EPI_ISL_2905079, EPI_ISL_2905080, EPI_ISL_2905081, EPI_ISL_2905082, EPI_ISL_2905083, EPI_ISL_2905084, EPI_ISL_2905085, EPI_ISL_2905086, EPI_ISL_2905087, EPI_ISL_2905088, EPI_ISL_2905089, EPI_ISL_2905090, EPI_ISL_2905091, EPI_ISL_2905092, EPI_ISL_2905093, EPI_ISL_2905094, EPI_ISL_2905095, EPI_ISL_2905096, EPI_ISL_2905097, EPI_ISL_2905098, EPI_ISL_2905099, EPI_ISL_2905100, EPI_ISL_2905101, EPI_ISL_2905102, EPI_ISL_2905103, EPI_ISL_2905104, EPI_ISL_2905105, EPI_ISL_2905106, EPI_ISL_2905107, EPI_ISL_2905108, EPI_ISL_2905109, EPI_ISL_2905110, EPI_ISL_2905111, EPI_ISL_2905112, EPI_ISL_2905113, EPI_ISL_2905114, EPI_ISL_2905115, EPI_ISL_2905116, EPI_ISL_2905117, EPI_ISL_2905118, EPI_ISL_2905119, EPI_ISL_2905120, EPI_ISL_2905121, EPI_ISL_2905122, EPI_ISL_2905123, EPI_ISL_2905124, EPI_ISL_2905125, EPI_ISL_2905126, EPI_ISL_2905127, EPI_ISL_2905128, EPI_ISL_2905129, EPI_ISL_2905130, EPI_ISL_2905131, EPI_ISL_2905132, EPI_ISL_2905133, EPI_ISL_2905134, EPI_ISL_2905135, EPI_ISL_2905136, EPI_ISL_2905137, EPI_ISL_2905138, EPI_ISL_2905139, EPI_ISL_2905140, EPI_ISL_2905141, EPI_ISL_2905142, EPI_ISL_2905143, EPI_ISL_2905144, EPI_ISL_2905145, EPI_ISL_2905146, EPI_ISL_2905147, EPI_ISL_2905148, EPI_ISL_2905149, EPI_ISL_2905150, EPI_ISL_2905151, EPI_ISL_2905152, EPI_ISL_2905153, EPI_ISL_2905154, EPI_ISL_2905155, EPI_ISL_2905156, EPI_ISL_2905158, EPI_ISL_2905159, EPI_ISL_2905160, EPI_ISL_2905161, EPI_ISL_2905162, EPI_ISL_2905163, EPI_ISL_2905164, EPI_ISL_2905165, EPI_ISL_2905166, EPI_ISL_2905167, EPI_ISL_2905168, EPI_ISL_2905169, EPI_ISL_2905170, EPI_ISL_2905171, EPI_ISL_2905172, EPI_ISL_2905173, EPI_ISL_2905174, EPI_ISL_2905176, EPI_ISL_2905177, EPI_ISL_2905178, EPI_ISL_2905179, EPI_ISL_2905181, EPI_ISL_2905182, EPI_ISL_2905183, EPI_ISL_2905184, EPI_ISL_2905185, EPI_ISL_2905186, EPI_ISL_2905187, EPI_ISL_2905188, EPI_ISL_2905189, EPI_ISL_2905190, EPI_ISL_2905191, EPI_ISL_2905192, EPI_ISL_2905193, EPI_ISL_2905194, EPI_ISL_2905195, EPI_ISL_2905196, EPI_ISL_2905198, EPI_ISL_2905199, EPI_ISL_2905200, EPI_ISL_2905201, EPI_ISL_2905202, EPI_ISL_2905203, EPI_ISL_2905204, EPI_ISL_2905205, EPI_ISL_2905206, EPI_ISL_2905207, EPI_ISL_2905208, EPI_ISL_2905209, EPI_ISL_2905210, EPI_ISL_2905211, EPI_ISL_2905212, EPI_ISL_2905213, EPI_ISL_2905214, EPI_ISL_2905215, EPI_ISL_2905216, EPI_ISL_2905217, EPI_ISL_2905218, EPI_ISL_2905219, EPI_ISL_2905220, EPI_ISL_2905221, EPI_ISL_2905222, EPI_ISL_2905223, EPI_ISL_2905224, EPI_ISL_2905225, EPI_ISL_2905226, EPI_ISL_2905227, EPI_ISL_2905228, EPI_ISL_2905229, EPI_ISL_2905230, EPI_ISL_2905231, EPI_ISL_2905232, EPI_ISL_2905233, EPI_ISL_2905234, EPI_ISL_2905235, EPI_ISL_2905236, EPI_ISL_2905237, EPI_ISL_2905238, EPI_ISL_2905239, EPI_ISL_2905240, EPI_ISL_2905241, EPI_ISL_2905242, EPI_ISL_2905243, EPI_ISL_2905244, EPI_ISL_2905245, EPI_ISL_2905246, EPI_ISL_2905247, EPI_ISL_2905248, EPI_ISL_2905249, EPI_ISL_2905250, EPI_ISL_2905251, EPI_ISL_2905252, EPI_ISL_2905253, EPI_ISL_2905254, EPI_ISL_2905256, EPI_ISL_2905257, EPI_ISL_2905258, EPI_ISL_2905259, EPI_ISL_2905260, EPI_ISL_2905261, EPI_ISL_2905262, EPI_ISL_2905263, EPI_ISL_2905264, EPI_ISL_2905265, EPI_ISL_2905266, EPI_ISL_2905267, EPI_ISL_2905268, EPI_ISL_2905269, EPI_ISL_2905270, EPI_ISL_2905271, EPI_ISL_2905272, EPI_ISL_2905273, EPI_ISL_2905274, EPI_ISL_2905275, EPI_ISL_2905276, EPI_ISL_2905277, EPI_ISL_2905278, EPI_ISL_2905279, EPI_ISL_2905280, EPI_ISL_2905281, EPI_ISL_2905282, EPI_ISL_2905283, EPI_ISL_2905284, EPI_ISL_2905285, EPI_ISL_2905286, EPI_ISL_2905287, EPI_ISL_2905288, EPI_ISL_2905289, EPI_ISL_2905290, EPI_ISL_2905291, EPI_ISL_2905292, EPI_ISL_2905293, EPI_ISL_2905294, EPI_ISL_2905295, EPI_ISL_2905296, EPI_ISL_2905297, EPI_ISL_2905298, EPI_ISL_2905299, EPI_ISL_2905300, EPI_ISL_2905301, EPI_ISL_2905302, EPI_ISL_2905303, EPI_ISL_2905304, EPI_ISL_2905305, EPI_ISL_2905306, EPI_ISL_2905307, EPI_ISL_2905308, EPI_ISL_2905309, EPI_ISL_2905310, EPI_ISL_2905311, EPI_ISL_2905312, EPI_ISL_2905313, EPI_ISL_2905314, EPI_ISL_2905315, EPI_ISL_2905316, EPI_ISL_2905317, EPI_ISL_2905318, EPI_ISL_2905319, EPI_ISL_2905320, EPI_ISL_2905321, EPI_ISL_2905322, EPI_ISL_2905323, EPI_ISL_2905324, EPI_ISL_2905325, EPI_ISL_2905326, EPI_ISL_2905327, EPI_ISL_2905328, EPI_ISL_2905329, EPI_ISL_2905330, EPI_ISL_2905331, EPI_ISL_2905332, EPI_ISL_2905333, EPI_ISL_2905334, EPI_ISL_2905335, EPI_ISL_2905336, EPI_ISL_2905337, EPI_ISL_2905338, EPI_ISL_2905339, EPI_ISL_2905340, EPI_ISL_2905341, EPI_ISL_2905342, EPI_ISL_2905343, EPI_ISL_2905344, EPI_ISL_2905345, EPI_ISL_2905346, EPI_ISL_2905347, EPI_ISL_2905348, EPI_ISL_2905349, EPI_ISL_2905350, EPI_ISL_2905351, EPI_ISL_2905352, EPI_ISL_2905353, EPI_ISL_2905354, EPI_ISL_2905355, EPI_ISL_2905356, EPI_ISL_2905357, EPI_ISL_2905358, EPI_ISL_2905359, EPI_ISL_2905360, EPI_ISL_2905361, EPI_ISL_2905362, EPI_ISL_2905363, EPI_ISL_2905364, EPI_ISL_2905365, EPI_ISL_2905366, EPI_ISL_2905367, EPI_ISL_2905368, EPI_ISL_2905369, EPI_ISL_2905370, EPI_ISL_2905371, EPI_ISL_2905372, EPI_ISL_2905373, EPI_ISL_2905374, EPI_ISL_2905375, EPI_ISL_2905376, EPI_ISL_2905377, EPI_ISL_2905378, EPI_ISL_2905379, EPI_ISL_2905380, EPI_ISL_2905381, EPI_ISL_2905382, EPI_ISL_2905383, EPI_ISL_2905384, EPI_ISL_2905385, EPI_ISL_2905386, EPI_ISL_2905387, EPI_ISL_2905388, EPI_ISL_2905389, EPI_ISL_2905390, EPI_ISL_2905391, EPI_ISL_2905392, EPI_ISL_2905393, EPI_ISL_2905394, EPI_ISL_2905395, EPI_ISL_2905396, EPI_ISL_2905397, EPI_ISL_2905398, EPI_ISL_2905399, EPI_ISL_2905400, EPI_ISL_2905401, EPI_ISL_2905402, EPI_ISL_2905403, EPI_ISL_2905404, EPI_ISL_2905405, EPI_ISL_2905406, EPI_ISL_2905407, EPI_ISL_2905408, EPI_ISL_2905409, EPI_ISL_2905410, EPI_ISL_2905411, EPI_ISL_2905412, EPI_ISL_2905413, EPI_ISL_2905414, EPI_ISL_2905415, EPI_ISL_2905416, EPI_ISL_2905417, EPI_ISL_2905418, EPI_ISL_2905419, EPI_ISL_2905420, EPI_ISL_2905421, EPI_ISL_2905422, EPI_ISL_2905423, EPI_ISL_2905424, EPI_ISL_2905425, EPI_ISL_2905426, EPI_ISL_2905427, EPI_ISL_2905428, EPI_ISL_2905429, EPI_ISL_2905430, EPI_ISL_2905431, EPI_ISL_2905432, EPI_ISL_2905433, EPI_ISL_2905434, EPI_ISL_2905435, EPI_ISL_2905436, EPI_ISL_2905437, EPI_ISL_2905438, EPI_ISL_2905439, EPI_ISL_2905440, EPI_ISL_2905441, EPI_ISL_2905442, EPI_ISL_2905443, EPI_ISL_2905444, EPI_ISL_2905445, EPI_ISL_2905446, EPI_ISL_2905447, EPI_ISL_2905448, EPI_ISL_2905449, EPI_ISL_2905450, EPI_ISL_2905451, EPI_ISL_2905452, EPI_ISL_2905453, EPI_ISL_2905454, EPI_ISL_2905455, EPI_ISL_2905456, EPI_ISL_2905457, EPI_ISL_2905458, EPI_ISL_2905459, EPI_ISL_2905460, EPI_ISL_2905461, EPI_ISL_2905462, EPI_ISL_2905463, EPI_ISL_2905464, EPI_ISL_2905465, EPI_ISL_2905466, EPI_ISL_2905467, EPI_ISL_2905468, EPI_ISL_2905469, EPI_ISL_2905470, EPI_ISL_2905471, EPI_ISL_2905472, EPI_ISL_2905473, EPI_ISL_2905474, EPI_ISL_2905475, EPI_ISL_2905476, EPI_ISL_2905478, EPI_ISL_2905479, EPI_ISL_2905480, EPI_ISL_2905481, EPI_ISL_2905482, EPI_ISL_2905483, EPI_ISL_2905484, EPI_ISL_2905485, EPI_ISL_2905486, EPI_ISL_2905487, EPI_ISL_2905488, EPI_ISL_2905489, EPI_ISL_2905490, EPI_ISL_2905491, EPI_ISL_2905492, EPI_ISL_2905493, EPI_ISL_2905494, EPI_ISL_2905495, EPI_ISL_2905496, EPI_ISL_2905497, EPI_ISL_2905498, EPI_ISL_2905499, EPI_ISL_2905500, EPI_ISL_2905501, EPI_ISL_2905502, EPI_ISL_2905503, EPI_ISL_2905504, EPI_ISL_2905505, EPI_ISL_2905506, EPI_ISL_2905507, EPI_ISL_2905508, EPI_ISL_2905509, EPI_ISL_2905510, EPI_ISL_2905511, EPI_ISL_2905512, EPI_ISL_2905513, EPI_ISL_2905514, EPI_ISL_2905515, EPI_ISL_2905516, EPI_ISL_2905517, EPI_ISL_2905518, EPI_ISL_2905519, EPI_ISL_2905520, EPI_ISL_2905521, EPI_ISL_2905522, EPI_ISL_2905523, EPI_ISL_2905524, EPI_ISL_2905525, EPI_ISL_2905526, EPI_ISL_2905527, EPI_ISL_2905528, EPI_ISL_2905529, EPI_ISL_2905530, EPI_ISL_2905531, EPI_ISL_2905532, EPI_ISL_2905533, EPI_ISL_2905534, EPI_ISL_2905535, EPI_ISL_2905536, EPI_ISL_2905537, EPI_ISL_2905538, EPI_ISL_2905539, EPI_ISL_2905540, EPI_ISL_2905541, EPI_ISL_2905542, EPI_ISL_2905543, EPI_ISL_2905544, EPI_ISL_2905545, EPI_ISL_2905546, EPI_ISL_2905547, EPI_ISL_2905548, EPI_ISL_2905549, EPI_ISL_2905550, EPI_ISL_2905551, EPI_ISL_2905552, EPI_ISL_2905553, EPI_ISL_2905554, EPI_ISL_2905555, EPI_ISL_2905556, EPI_ISL_2905557, EPI_ISL_2905558, EPI_ISL_2905559, EPI_ISL_2905560, EPI_ISL_2905561, EPI_ISL_2905562, EPI_ISL_2905563, EPI_ISL_2905564, EPI_ISL_2905565, EPI_ISL_2905566, EPI_ISL_2905568, EPI_ISL_2905569, EPI_ISL_2905571, EPI_ISL_2905572, EPI_ISL_2905573, EPI_ISL_2905574, EPI_ISL_2905575, EPI_ISL_2905576, EPI_ISL_2905577, EPI_ISL_2905578, EPI_ISL_2905579, EPI_ISL_2905580, EPI_ISL_2905581, EPI_ISL_2905582, EPI_ISL_2905583, EPI_ISL_2905584, EPI_ISL_2905585, EPI_ISL_2905586, EPI_ISL_2905587, EPI_ISL_2905588, EPI_ISL_2905589, EPI_ISL_2905590, EPI_ISL_2905591, EPI_ISL_2905592, EPI_ISL_2905593, EPI_ISL_2905594, EPI_ISL_2905595, EPI_ISL_2905596, EPI_ISL_2905597, EPI_ISL_2905598, EPI_ISL_2905599, EPI_ISL_2905600, EPI_ISL_2905601, EPI_ISL_2905602, EPI_ISL_2905603, EPI_ISL_2905604, EPI_ISL_2905605, EPI_ISL_2905606, EPI_ISL_2905607, EPI_ISL_2905608, EPI_ISL_2905609, EPI_ISL_2905610, EPI_ISL_2905611, EPI_ISL_2905612, EPI_ISL_2905613, EPI_ISL_2905614, EPI_ISL_2905616, EPI_ISL_2905617, EPI_ISL_2905618, EPI_ISL_2905619, EPI_ISL_2905620, EPI_ISL_2905621, EPI_ISL_2905622, EPI_ISL_2905623, EPI_ISL_2905624, EPI_ISL_2905625, EPI_ISL_2905626, EPI_ISL_2905627, EPI_ISL_2905628, EPI_ISL_2905629, EPI_ISL_2905630, EPI_ISL_2905631, EPI_ISL_2905632, EPI_ISL_2905633, EPI_ISL_2905634, EPI_ISL_2905635, EPI_ISL_2905636, EPI_ISL_2905637, EPI_ISL_2905638, EPI_ISL_2905639, EPI_ISL_2905640, EPI_ISL_2905641, EPI_ISL_2905642, EPI_ISL_2905643, EPI_ISL_2905644, EPI_ISL_2905645, EPI_ISL_2905646, EPI_ISL_2905647, EPI_ISL_2905648, EPI_ISL_2905649, EPI_ISL_2905650, EPI_ISL_2905651, EPI_ISL_2905652, EPI_ISL_2905653, EPI_ISL_2905654, EPI_ISL_2905655, EPI_ISL_2905656, EPI_ISL_2905657, EPI_ISL_2905658, EPI_ISL_2905659, EPI_ISL_2905660, EPI_ISL_2905661, EPI_ISL_2905662, EPI_ISL_2905663, EPI_ISL_2905664, EPI_ISL_2905665, EPI_ISL_2905666, EPI_ISL_2905667, EPI_ISL_2905668, EPI_ISL_2905669, EPI_ISL_2905670, EPI_ISL_2905671, EPI_ISL_2905672, EPI_ISL_2905673, EPI_ISL_2905674, EPI_ISL_2905675, EPI_ISL_2905676, EPI_ISL_2905677, EPI_ISL_2905678, EPI_ISL_2905679, EPI_ISL_2905680, EPI_ISL_2905681, EPI_ISL_2905682, EPI_ISL_2905683, EPI_ISL_2905684, EPI_ISL_2905685, EPI_ISL_2905686, EPI_ISL_2905687, EPI_ISL_2905688, EPI_ISL_2905689, EPI_ISL_2905690, EPI_ISL_2905691, EPI_ISL_2905692, EPI_ISL_2905693, EPI_ISL_2905694, EPI_ISL_2905696, EPI_ISL_2905697, EPI_ISL_2905698, EPI_ISL_2905699, EPI_ISL_2905700, EPI_ISL_2905701, EPI_ISL_2905702, EPI_ISL_2905703, EPI_ISL_2905704, EPI_ISL_2905705, EPI_ISL_2905706, EPI_ISL_2905707, EPI_ISL_2905708, EPI_ISL_2905709, EPI_ISL_2905710, EPI_ISL_2905711, EPI_ISL_2905712, EPI_ISL_2905713, EPI_ISL_2905714, EPI_ISL_2905715, EPI_ISL_2905716, EPI_ISL_2905717, EPI_ISL_2905718, EPI_ISL_2905719, EPI_ISL_2905720, EPI_ISL_2905721, EPI_ISL_2905722, EPI_ISL_2905723, EPI_ISL_2905724, EPI_ISL_2905725, EPI_ISL_2905726, EPI_ISL_2905727, EPI_ISL_2905728, EPI_ISL_2905729, EPI_ISL_2905730, EPI_ISL_2905731, EPI_ISL_2905732, EPI_ISL_2905733, EPI_ISL_2905734, EPI_ISL_2905735, EPI_ISL_2905736, EPI_ISL_2905737, EPI_ISL_2905739, EPI_ISL_2905740, EPI_ISL_2905741, EPI_ISL_2905742, EPI_ISL_2905743, EPI_ISL_2905744, EPI_ISL_2905745, EPI_ISL_2905746, EPI_ISL_2905747, EPI_ISL_2905748, EPI_ISL_2905749, EPI_ISL_2905750, EPI_ISL_2905751, EPI_ISL_2905752, EPI_ISL_2905753, EPI_ISL_2905754, EPI_ISL_2905755, EPI_ISL_2905756, EPI_ISL_2905757, EPI_ISL_2905758, EPI_ISL_2905759, EPI_ISL_2905760, EPI_ISL_2905761, EPI_ISL_2905762, EPI_ISL_2905763, EPI_ISL_2905764, EPI_ISL_2905765, EPI_ISL_2905766, EPI_ISL_2905767, EPI_ISL_2905768, EPI_ISL_2905769, EPI_ISL_2905770, EPI_ISL_2905771, EPI_ISL_2905772, EPI_ISL_2905773, EPI_ISL_2905774, EPI_ISL_2905775, EPI_ISL_2905776, EPI_ISL_2905777, EPI_ISL_2905778, EPI_ISL_2905779, EPI_ISL_2905780, EPI_ISL_2905781, EPI_ISL_2905782, EPI_ISL_2905783, EPI_ISL_2905784, EPI_ISL_2905785, EPI_ISL_2905786, EPI_ISL_2905787, EPI_ISL_2905788, EPI_ISL_2905789, EPI_ISL_2905790, EPI_ISL_2905791, EPI_ISL_2905792, EPI_ISL_2905793, EPI_ISL_2905794, EPI_ISL_2905795, EPI_ISL_2905796, EPI_ISL_2905797, EPI_ISL_2905798, EPI_ISL_2905799, EPI_ISL_2905800, EPI_ISL_2905801, EPI_ISL_2905802, EPI_ISL_2905803, EPI_ISL_2905804, EPI_ISL_2905806, EPI_ISL_2905807, EPI_ISL_2905808, EPI_ISL_2905809, EPI_ISL_2905810, EPI_ISL_2905811, EPI_ISL_2905812, EPI_ISL_2905813, EPI_ISL_2905814, EPI_ISL_2905815, EPI_ISL_2905816, EPI_ISL_2905817, EPI_ISL_2905820, EPI_ISL_2905821, EPI_ISL_2905822, EPI_ISL_2905823, EPI_ISL_2905824, EPI_ISL_2905825, EPI_ISL_2905826, EPI_ISL_2905827, EPI_ISL_2905828, EPI_ISL_2905829, EPI_ISL_2905830, EPI_ISL_2905831, EPI_ISL_2905832, EPI_ISL_2905833, EPI_ISL_2905834, EPI_ISL_2905835, EPI_ISL_2905836, EPI_ISL_2905837, EPI_ISL_2905838, EPI_ISL_2905839, EPI_ISL_2905840, EPI_ISL_2905841, EPI_ISL_2905842, EPI_ISL_2905843, EPI_ISL_2905844, EPI_ISL_2905845, EPI_ISL_2905846, EPI_ISL_2905847, EPI_ISL_2905848, EPI_ISL_2905849, EPI_ISL_2905850, EPI_ISL_2905851, EPI_ISL_2905852, EPI_ISL_2905853, EPI_ISL_2905854, EPI_ISL_2905855, EPI_ISL_2905856, EPI_ISL_2905857, EPI_ISL_2905858, EPI_ISL_2905859, EPI_ISL_2905860, EPI_ISL_2905861, EPI_ISL_2905862, EPI_ISL_2905863, EPI_ISL_2905864, EPI_ISL_2905865, EPI_ISL_2905866, EPI_ISL_2905867, EPI_ISL_2905868, EPI_ISL_2905869, EPI_ISL_2905870, EPI_ISL_2905871, EPI_ISL_2905872, EPI_ISL_2905873, EPI_ISL_2905874, EPI_ISL_2905875, EPI_ISL_2905876, EPI_ISL_2905877, EPI_ISL_2905878, EPI_ISL_2905879, EPI_ISL_2905880, EPI_ISL_2905881, EPI_ISL_2905882, EPI_ISL_2905883, EPI_ISL_2905884, EPI_ISL_2905885, EPI_ISL_2905886, EPI_ISL_2905887, EPI_ISL_2905888, EPI_ISL_2905889, EPI_ISL_2905890, EPI_ISL_2905891, EPI_ISL_2905892, EPI_ISL_2905893, EPI_ISL_2905894, EPI_ISL_2905895, EPI_ISL_2905896, EPI_ISL_2905897, EPI_ISL_2905898, EPI_ISL_2905899, EPI_ISL_2905900, EPI_ISL_2905901, EPI_ISL_2905902, EPI_ISL_2905903, EPI_ISL_2905904, EPI_ISL_2905905, EPI_ISL_2905906, EPI_ISL_2905907, EPI_ISL_2905908, EPI_ISL_2905909, EPI_ISL_2905910, EPI_ISL_2905911, EPI_ISL_2905912, EPI_ISL_2905913, EPI_ISL_2905914, EPI_ISL_2905915, EPI_ISL_2905916, EPI_ISL_2905917, EPI_ISL_2905918, EPI_ISL_2905919, EPI_ISL_2905920, EPI_ISL_2905921, EPI_ISL_2905922, EPI_ISL_2905923, EPI_ISL_2905924, EPI_ISL_2905925, EPI_ISL_2905926, EPI_ISL_2905927, EPI_ISL_2905928, EPI_ISL_2905929, EPI_ISL_2905930, EPI_ISL_2905931, EPI_ISL_2905932, EPI_ISL_2905933, EPI_ISL_2905934, EPI_ISL_2905935, EPI_ISL_2905936, EPI_ISL_2905937, EPI_ISL_2905938, EPI_ISL_2905939, EPI_ISL_2905940, EPI_ISL_2905941, EPI_ISL_2905942, EPI_ISL_2905943, EPI_ISL_2905944, EPI_ISL_2905945, EPI_ISL_2905946, EPI_ISL_2905947, EPI_ISL_2905948, EPI_ISL_2905949, EPI_ISL_2905950, EPI_ISL_2905951, EPI_ISL_2905952, EPI_ISL_2905953, EPI_ISL_2905954, EPI_ISL_2905955, EPI_ISL_2905956, EPI_ISL_2905957, EPI_ISL_2905958, EPI_ISL_2905959, EPI_ISL_2905960, EPI_ISL_2905961, EPI_ISL_2905962, EPI_ISL_2905963, EPI_ISL_2905964, EPI_ISL_2905965, EPI_ISL_2905967, EPI_ISL_2905968, EPI_ISL_2905969, EPI_ISL_2905970, EPI_ISL_2905971, EPI_ISL_2905972, EPI_ISL_2905973, EPI_ISL_2905974, EPI_ISL_2905975, EPI_ISL_2905976, EPI_ISL_2905977, EPI_ISL_2905978, EPI_ISL_2905979, EPI_ISL_2905980, EPI_ISL_2905981, EPI_ISL_2905982, EPI_ISL_2905983, EPI_ISL_2905984, EPI_ISL_2905985, EPI_ISL_2905986, EPI_ISL_2905987, EPI_ISL_2905988, EPI_ISL_2905989, EPI_ISL_2905990, EPI_ISL_2905991, EPI_ISL_2905992, EPI_ISL_2905993, EPI_ISL_2905994, EPI_ISL_2905995, EPI_ISL_2905996, EPI_ISL_2905997, EPI_ISL_2905998, EPI_ISL_2905999, EPI_ISL_2906000, EPI_ISL_2906001, EPI_ISL_2906002, EPI_ISL_2906003, EPI_ISL_2906004, EPI_ISL_2906005, EPI_ISL_2906006, EPI_ISL_2906007, EPI_ISL_2906008, EPI_ISL_2906009, EPI_ISL_2906010, EPI_ISL_2906011, EPI_ISL_2906012, EPI_ISL_2906013, EPI_ISL_2906014, EPI_ISL_2906015, EPI_ISL_2906016, EPI_ISL_2906017, EPI_ISL_2906018, EPI_ISL_2906019, EPI_ISL_2906020, EPI_ISL_2906021, EPI_ISL_2906022, EPI_ISL_2906023, EPI_ISL_2906024, EPI_ISL_2906025, EPI_ISL_2906026, EPI_ISL_2906027, EPI_ISL_2906028, EPI_ISL_2906029, EPI_ISL_2906030, EPI_ISL_2906031, EPI_ISL_2906032, EPI_ISL_2906033, EPI_ISL_2906034, EPI_ISL_2906035, EPI_ISL_2906036, EPI_ISL_2906037, EPI_ISL_2906038, EPI_ISL_2906039, EPI_ISL_2906040, EPI_ISL_2906041, EPI_ISL_2906042, EPI_ISL_2906043, EPI_ISL_2906044, EPI_ISL_2906045, EPI_ISL_2906046, EPI_ISL_2906047, EPI_ISL_2906048, EPI_ISL_2906049, EPI_ISL_2906050, EPI_ISL_2906051, EPI_ISL_2906052, EPI_ISL_2906053, EPI_ISL_2906054, EPI_ISL_2906055, EPI_ISL_2906056, EPI_ISL_2906057, EPI_ISL_2906058, EPI_ISL_2906059, EPI_ISL_2906060, EPI_ISL_2906061, EPI_ISL_2906062, EPI_ISL_2906063, EPI_ISL_2906064, EPI_ISL_2906065, EPI_ISL_2906066, EPI_ISL_2906067, EPI_ISL_2906068, EPI_ISL_2906069, EPI_ISL_2906070, EPI_ISL_2906071, EPI_ISL_2906072, EPI_ISL_2906073, EPI_ISL_2906074, EPI_ISL_2906075, EPI_ISL_2906076, EPI_ISL_2906077, EPI_ISL_2906078, EPI_ISL_2906079, EPI_ISL_2906080, EPI_ISL_2906081, EPI_ISL_2906082, EPI_ISL_2906083, EPI_ISL_2906084, EPI_ISL_2906085, EPI_ISL_2906086, EPI_ISL_2906087, EPI_ISL_2906088, EPI_ISL_2906089, EPI_ISL_2906090, EPI_ISL_2906091, EPI_ISL_2906092, EPI_ISL_2906093, EPI_ISL_2906094, EPI_ISL_2906095, EPI_ISL_2906096, EPI_ISL_2906097, EPI_ISL_2906098, EPI_ISL_2906099, EPI_ISL_2906100, EPI_ISL_2906101, EPI_ISL_2906102, EPI_ISL_2906103, EPI_ISL_2906104, EPI_ISL_2906105, EPI_ISL_2906106, EPI_ISL_2906107, EPI_ISL_2906108, EPI_ISL_2906109, EPI_ISL_2906110, EPI_ISL_2906111, EPI_ISL_2906112, EPI_ISL_2906113, EPI_ISL_2906114, EPI_ISL_2906115, EPI_ISL_2906116, EPI_ISL_2906117, EPI_ISL_2906118, EPI_ISL_2906119, EPI_ISL_2906120, EPI_ISL_2906121, EPI_ISL_2906122, EPI_ISL_2906123, EPI_ISL_2906124, EPI_ISL_2906125, EPI_ISL_2906126, EPI_ISL_2906127, EPI_ISL_2906128, EPI_ISL_2906129, EPI_ISL_2906130, EPI_ISL_2906131, EPI_ISL_2906132, EPI_ISL_2906133, EPI_ISL_2906134, EPI_ISL_2906135, EPI_ISL_2906136, EPI_ISL_2906137, EPI_ISL_2906138, EPI_ISL_2906139, EPI_ISL_2906140, EPI_ISL_2906141, EPI_ISL_2906142, EPI_ISL_2906143, EPI_ISL_2906144, EPI_ISL_2906145, EPI_ISL_2906146, EPI_ISL_2906147, EPI_ISL_2906148, EPI_ISL_2906149, EPI_ISL_2906150, EPI_ISL_2906151, EPI_ISL_2906152, EPI_ISL_2906153, EPI_ISL_2906154, EPI_ISL_2906155, EPI_ISL_2906156, EPI_ISL_2906157, EPI_ISL_2906158, EPI_ISL_2906159, EPI_ISL_2906160, EPI_ISL_2906161, EPI_ISL_2906162, EPI_ISL_2906163, EPI_ISL_2906164, EPI_ISL_2906165, EPI_ISL_2906166, EPI_ISL_2906167, EPI_ISL_2906168, EPI_ISL_2906169, EPI_ISL_2906170, EPI_ISL_2906171, EPI_ISL_2906172, EPI_ISL_2906173, EPI_ISL_2906174, EPI_ISL_2906175, EPI_ISL_2906176, EPI_ISL_2906177, EPI_ISL_2906178, EPI_ISL_2906179, EPI_ISL_2906180, EPI_ISL_2906182, EPI_ISL_2906183, EPI_ISL_2906184, EPI_ISL_2906185, EPI_ISL_2906186, EPI_ISL_2906187, EPI_ISL_2906188, EPI_ISL_2906189, EPI_ISL_2906190, EPI_ISL_2906191, EPI_ISL_2906192, EPI_ISL_2906193, EPI_ISL_2906194, EPI_ISL_2906195, EPI_ISL_2906196, EPI_ISL_2906197, EPI_ISL_2906198, EPI_ISL_2906199, EPI_ISL_2906200, EPI_ISL_2906201, EPI_ISL_2906202, EPI_ISL_2906203, EPI_ISL_2906204, EPI_ISL_2906205, EPI_ISL_2906206, EPI_ISL_2906207, EPI_ISL_2906208, EPI_ISL_2906209, EPI_ISL_2906210, EPI_ISL_2906211, EPI_ISL_2906212, EPI_ISL_2906213, EPI_ISL_2906214, EPI_ISL_2906215, EPI_ISL_2906216, EPI_ISL_2906217, EPI_ISL_2906218, EPI_ISL_2906219, EPI_ISL_2906220, EPI_ISL_2906221, EPI_ISL_2906222, EPI_ISL_2906223, EPI_ISL_2906224, EPI_ISL_2906225, EPI_ISL_2906226, EPI_ISL_2906227, EPI_ISL_2906228, EPI_ISL_2906229, EPI_ISL_2906230, EPI_ISL_2906231, EPI_ISL_2906232, EPI_ISL_2906233, EPI_ISL_2906234, EPI_ISL_2906235, EPI_ISL_2906236, EPI_ISL_2906237, EPI_ISL_2906238, EPI_ISL_2906239, EPI_ISL_2906240, EPI_ISL_2906241, EPI_ISL_2906242, EPI_ISL_2906243, EPI_ISL_2906244, EPI_ISL_2906245, EPI_ISL_2906246, EPI_ISL_2906247, EPI_ISL_2906248, EPI_ISL_2906249, EPI_ISL_2906250, EPI_ISL_2906251, EPI_ISL_2906252, EPI_ISL_2906253, EPI_ISL_2906254, EPI_ISL_2906255, EPI_ISL_2906256, EPI_ISL_2906257, EPI_ISL_2906258, EPI_ISL_2906259, EPI_ISL_2906260, EPI_ISL_2906261, EPI_ISL_2906262, EPI_ISL_2906263, EPI_ISL_2906264, EPI_ISL_2906265, EPI_ISL_2906266, EPI_ISL_2906267, EPI_ISL_2906268, EPI_ISL_2906269, EPI_ISL_2906270, EPI_ISL_2906271, EPI_ISL_2906272, EPI_ISL_2906273, EPI_ISL_2906274, EPI_ISL_2906275, EPI_ISL_2906276, EPI_ISL_2906277, EPI_ISL_2906278, EPI_ISL_2906279, EPI_ISL_2906280, EPI_ISL_2906281, EPI_ISL_2906282, EPI_ISL_2906283, EPI_ISL_2906284, EPI_ISL_2906285, EPI_ISL_2906286, EPI_ISL_2906287, EPI_ISL_2906288, EPI_ISL_2906289, EPI_ISL_2906290, EPI_ISL_2906291, EPI_ISL_2906292, EPI_ISL_2906293, EPI_ISL_2906294, EPI_ISL_2906295, EPI_ISL_2906296, EPI_ISL_2906297, EPI_ISL_2906298, EPI_ISL_2906299, EPI_ISL_2906300, EPI_ISL_2906301, EPI_ISL_2906302, EPI_ISL_2906303, EPI_ISL_2906304, EPI_ISL_2906305, EPI_ISL_2906306, EPI_ISL_2906307, EPI_ISL_2906308, EPI_ISL_2906309, EPI_ISL_2906310, EPI_ISL_2906311, EPI_ISL_2906312, EPI_ISL_2906313, EPI_ISL_2906314, EPI_ISL_2906315, EPI_ISL_2906316, EPI_ISL_2906317, EPI_ISL_2906318, EPI_ISL_2906319, EPI_ISL_2906320, EPI_ISL_2906321, EPI_ISL_2906322, EPI_ISL_2906323, EPI_ISL_2906324, EPI_ISL_2906325, EPI_ISL_2906326, EPI_ISL_2906327, EPI_ISL_2906328, EPI_ISL_2906329, EPI_ISL_2906330, EPI_ISL_2906331, EPI_ISL_2906332, EPI_ISL_2906333, EPI_ISL_2906334, EPI_ISL_2906335, EPI_ISL_2906336, EPI_ISL_2906337, EPI_ISL_2906338, EPI_ISL_2906339, EPI_ISL_2906340, EPI_ISL_2906341, EPI_ISL_2906342, EPI_ISL_2906343, EPI_ISL_2906344, EPI_ISL_2906345, EPI_ISL_2906346, EPI_ISL_2906347, EPI_ISL_2906348, EPI_ISL_2906349, EPI_ISL_2906350, EPI_ISL_2906351, EPI_ISL_2906352, EPI_ISL_2906353, EPI_ISL_2906354, EPI_ISL_2906355, EPI_ISL_2906356, EPI_ISL_2906357, EPI_ISL_2906358, EPI_ISL_2906359, EPI_ISL_2906360, EPI_ISL_2906361, EPI_ISL_2906362, EPI_ISL_2906363, EPI_ISL_2906364, EPI_ISL_2906365, EPI_ISL_2906366, EPI_ISL_2906367, EPI_ISL_2906368, EPI_ISL_2906369, EPI_ISL_2906370, EPI_ISL_2906371, EPI_ISL_2906372, EPI_ISL_2906373, EPI_ISL_2906374, EPI_ISL_2906375, EPI_ISL_2906376, EPI_ISL_2906377, EPI_ISL_2906378, EPI_ISL_2906379, EPI_ISL_2906380, EPI_ISL_2906381, EPI_ISL_2906382, EPI_ISL_2906383, EPI_ISL_2906384, EPI_ISL_2906385, EPI_ISL_2906386, EPI_ISL_2906387, EPI_ISL_2906388, EPI_ISL_2906389, EPI_ISL_2906390, EPI_ISL_2906391, EPI_ISL_2906392, EPI_ISL_2906393, EPI_ISL_2906395, EPI_ISL_2906397, EPI_ISL_2906398, EPI_ISL_2906399, EPI_ISL_2906400, EPI_ISL_2906401, EPI_ISL_2906402, EPI_ISL_2906403, EPI_ISL_2906404, EPI_ISL_2906405, EPI_ISL_2906406, EPI_ISL_2906407, EPI_ISL_2906408, EPI_ISL_2906409, EPI_ISL_2906410, EPI_ISL_2906411, EPI_ISL_2906412, EPI_ISL_2906413, EPI_ISL_2906414, EPI_ISL_2906415, EPI_ISL_2906416, EPI_ISL_2906417, EPI_ISL_2906418, EPI_ISL_2906419, EPI_ISL_2906420, EPI_ISL_2906421, EPI_ISL_2906422, EPI_ISL_2906423, EPI_ISL_2906424, EPI_ISL_2906425, EPI_ISL_2906426, EPI_ISL_2906427, EPI_ISL_2906428, EPI_ISL_2906429, EPI_ISL_2906430, EPI_ISL_2906431, EPI_ISL_2906432, EPI_ISL_2906433, EPI_ISL_2906434, EPI_ISL_2906435, EPI_ISL_2906436, EPI_ISL_2906437, EPI_ISL_2906438, EPI_ISL_2906439, EPI_ISL_2906440, EPI_ISL_2906441, EPI_ISL_2906442, EPI_ISL_2906443, EPI_ISL_2906444, EPI_ISL_2906445, EPI_ISL_2906446, EPI_ISL_2906447, EPI_ISL_2906448, EPI_ISL_2906449, EPI_ISL_2906450, EPI_ISL_2906451, EPI_ISL_2906452, EPI_ISL_2906453, EPI_ISL_2906454, EPI_ISL_2906455, EPI_ISL_2906456, EPI_ISL_2906457, EPI_ISL_2906458, EPI_ISL_2906459, EPI_ISL_2906460, EPI_ISL_2906461, EPI_ISL_2906462, EPI_ISL_2906463, EPI_ISL_2906464, EPI_ISL_2906465, EPI_ISL_2906466, EPI_ISL_2906467, EPI_ISL_2906468, EPI_ISL_2906469, EPI_ISL_2906470, EPI_ISL_2906471, EPI_ISL_2906472, EPI_ISL_2906473, EPI_ISL_2906474, EPI_ISL_2906475, EPI_ISL_2906476, EPI_ISL_2906477, EPI_ISL_2906479, EPI_ISL_2906480, EPI_ISL_2906481, EPI_ISL_2906482, EPI_ISL_2906483, EPI_ISL_2906484, EPI_ISL_2906485, EPI_ISL_2906486, EPI_ISL_2906487, EPI_ISL_2906488, EPI_ISL_2906489, EPI_ISL_2906490, EPI_ISL_2906491, EPI_ISL_2906492, EPI_ISL_2906493, EPI_ISL_2906494, EPI_ISL_2906495, EPI_ISL_2906496, EPI_ISL_2906497, EPI_ISL_2906498, EPI_ISL_2906500, EPI_ISL_2906501, EPI_ISL_2906502, EPI_ISL_2906503, EPI_ISL_2906504, EPI_ISL_2906505, EPI_ISL_2906506, EPI_ISL_2906507, EPI_ISL_2906508, EPI_ISL_2906509, EPI_ISL_2906510, EPI_ISL_2906511, EPI_ISL_2906512, EPI_ISL_2906513, EPI_ISL_2906514, EPI_ISL_2906515, EPI_ISL_2906516, EPI_ISL_2906517, EPI_ISL_2906518, EPI_ISL_2906519, EPI_ISL_2906520, EPI_ISL_2906521, EPI_ISL_2906522, EPI_ISL_2906523,

| | | | |
|---|---|---|---|
| EPI_ISL_2906524, EPI_ISL_2906525, EPI_ISL_2906526, EPI_ISL_2906527, EPI_ISL_2906528, EPI_ISL_2906529, EPI_ISL_2906530, EPI_ISL_2906531, EPI_ISL_2906532, EPI_ISL_2906533, EPI_ISL_2906534, EPI_ISL_2906535, EPI_ISL_2906536, EPI_ISL_2906537, EPI_ISL_2906538, EPI_ISL_2906539, EPI_ISL_2906540, EPI_ISL_2906541, EPI_ISL_2906542, EPI_ISL_2906543, EPI_ISL_2906544, EPI_ISL_2906545, EPI_ISL_2906546, EPI_ISL_2906547, EPI_ISL_2906548, EPI_ISL_2906549, EPI_ISL_2906550, EPI_ISL_2906551, EPI_ISL_2906552, EPI_ISL_2906553, EPI_ISL_2906554, EPI_ISL_2906555, EPI_ISL_2906556, EPI_ISL_2906557, EPI_ISL_2906558, EPI_ISL_2906559, EPI_ISL_2906560, EPI_ISL_2906561, EPI_ISL_2906562, EPI_ISL_2906563, EPI_ISL_2906564, EPI_ISL_2906565, EPI_ISL_2906566, EPI_ISL_2906567, EPI_ISL_2906568, EPI_ISL_2906569, EPI_ISL_2906570, EPI_ISL_2906571, EPI_ISL_2906572, EPI_ISL_2906573, EPI_ISL_2906574, EPI_ISL_2906575, EPI_ISL_2906576, EPI_ISL_2906577, EPI_ISL_2906578, EPI_ISL_2906579, EPI_ISL_2906580, EPI_ISL_2906581, EPI_ISL_2906582, EPI_ISL_2906583, EPI_ISL_2906584, EPI_ISL_2906585, EPI_ISL_2906586, EPI_ISL_2906587, EPI_ISL_2906588, EPI_ISL_2906589, EPI_ISL_2906590, EPI_ISL_2906591, EPI_ISL_2906592, EPI_ISL_2906593, EPI_ISL_2906594, EPI_ISL_2906595, EPI_ISL_2906596, EPI_ISL_2906597, EPI_ISL_2906598, EPI_ISL_2906599, EPI_ISL_2906600, EPI_ISL_2906601, EPI_ISL_2906602, EPI_ISL_2906603, EPI_ISL_2906604, EPI_ISL_2906605, EPI_ISL_2906606, EPI_ISL_2906607, EPI_ISL_2906608, EPI_ISL_2906609, EPI_ISL_2906610, EPI_ISL_2906611, EPI_ISL_2906612, EPI_ISL_2906613, EPI_ISL_2906614, EPI_ISL_2906615, EPI_ISL_2906616, EPI_ISL_2906617, EPI_ISL_2906618, EPI_ISL_2906619, EPI_ISL_2906620, EPI_ISL_2906622, EPI_ISL_2906623, EPI_ISL_2906624, EPI_ISL_2906625, EPI_ISL_2906626, EPI_ISL_2906627, EPI_ISL_2906628, EPI_ISL_2906629, EPI_ISL_2906630, EPI_ISL_2906631, EPI_ISL_2906632, EPI_ISL_2906633, EPI_ISL_2906634, EPI_ISL_2906635, EPI_ISL_2906636, EPI_ISL_2906637, EPI_ISL_2906638, EPI_ISL_2906639, EPI_ISL_2906640, EPI_ISL_2906641, EPI_ISL_2906642, EPI_ISL_2906643, EPI_ISL_2906644, EPI_ISL_2906645, EPI_ISL_2906646, EPI_ISL_2906647, EPI_ISL_2906648, EPI_ISL_2906649, EPI_ISL_2906650, EPI_ISL_2906651, EPI_ISL_2906652, EPI_ISL_2906653, EPI_ISL_2906654, EPI_ISL_2906655, EPI_ISL_2906656, EPI_ISL_2906657, EPI_ISL_2906658, EPI_ISL_2906659, EPI_ISL_2906660, EPI_ISL_2906661, EPI_ISL_2906662, EPI_ISL_2906663, EPI_ISL_2906664, EPI_ISL_2906665, EPI_ISL_2906666, EPI_ISL_2906667, EPI_ISL_2906668, EPI_ISL_2906669, EPI_ISL_2906670, EPI_ISL_2906671, EPI_ISL_2906672, EPI_ISL_2906673, EPI_ISL_2906674, EPI_ISL_2906675, EPI_ISL_2906676, EPI_ISL_2906677, EPI_ISL_2906678, EPI_ISL_2906679, EPI_ISL_2906680, EPI_ISL_2906681, EPI_ISL_2906682, EPI_ISL_2906683, EPI_ISL_2906684, EPI_ISL_2906685, EPI_ISL_2906686, EPI_ISL_2906687, EPI_ISL_2906688, EPI_ISL_2906689, EPI_ISL_2906690, EPI_ISL_2906691, EPI_ISL_2906692, EPI_ISL_2906693, EPI_ISL_2906694, EPI_ISL_2906695, EPI_ISL_2906696, EPI_ISL_2906697, EPI_ISL_2906698, EPI_ISL_2906699, EPI_ISL_2906700, EPI_ISL_2906701, EPI_ISL_2906702, EPI_ISL_2906703, EPI_ISL_2906704, EPI_ISL_2906705, EPI_ISL_2906706, EPI_ISL_2906707, EPI_ISL_2906708, EPI_ISL_2906709, EPI_ISL_2906710, EPI_ISL_2906711, EPI_ISL_2906712, EPI_ISL_2906713, EPI_ISL_2906714, EPI_ISL_2906715, EPI_ISL_2906716, EPI_ISL_2906717, EPI_ISL_2906718, EPI_ISL_2906719, EPI_ISL_2906720, EPI_ISL_2906721, EPI_ISL_2906722, EPI_ISL_2906723, EPI_ISL_2906724, EPI_ISL_2906725, EPI_ISL_2906726, EPI_ISL_2906727, EPI_ISL_2906728, EPI_ISL_2906729, EPI_ISL_2906730 | | | |
| see above | Originating lab: Wales Specialist Virology Centre Sequencing lab: Pathogen Genomics Unit | Public Health Wales Microbiology Cardiff Wales Specialist Virology Centre | Catherine Moore, Johnathan Evans, Laura Gifford, Malorie Perry, Simon Cottrell, Angela Marchbank, Alec Birchley, Alexander Adams, Amy Gaskin, Bree Gatica-Wilcox, Jason Coombes, Joel Southgate, Lauren Gilbert, Lee Graham, Nicole Pacchiarini, Sara Kumziene-Summerhayes, Sarah Taylor, Sophie Jones, Sara Rey, Matthew Bull, Joanne Watkins, Sally Corden, Tom Connor |
| EPI_ISL_2906731, EPI_ISL_2906732, EPI_ISL_2906733, EPI_ISL_2906734, EPI_ISL_2906735, EPI_ISL_2906736, EPI_ISL_2906737, EPI_ISL_2906738, EPI_ISL_2906739, EPI_ISL_2906740, EPI_ISL_2906741, EPI_ISL_2906742, EPI_ISL_2906743, EPI_ISL_2906744, EPI_ISL_2906745, EPI_ISL_2906746, EPI_ISL_2906747, EPI_ISL_2906748, EPI_ISL_2906749, EPI_ISL_2906750, EPI_ISL_2906751, EPI_ISL_2906752, EPI_ISL_2906753, EPI_ISL_2906754, EPI_ISL_2906755, EPI_ISL_2906756, EPI_ISL_2906757, EPI_ISL_2906758, EPI_ISL_2906759, EPI_ISL_2906760, EPI_ISL_2906761, EPI_ISL_2906762, EPI_ISL_2906763, EPI_ISL_2906764, EPI_ISL_2906765, EPI_ISL_2906766, EPI_ISL_2906767, EPI_ISL_2906768, EPI_ISL_2906769, EPI_ISL_2906770, EPI_ISL_2906771, EPI_ISL_2906772, EPI_ISL_2906773, EPI_ISL_2906774, EPI_ISL_2906775, EPI_ISL_2906776, EPI_ISL_2906777, EPI_ISL_2906778, EPI_ISL_2906779, EPI_ISL_2906780, EPI_ISL_2906781, EPI_ISL_2906782, EPI_ISL_2906783, EPI_ISL_2906784, EPI_ISL_2906785, EPI_ISL_2906786, EPI_ISL_2906787, EPI_ISL_2906788, EPI_ISL_2906789, EPI_ISL_2906790, EPI_ISL_2906791, EPI_ISL_2906792, EPI_ISL_2906793, EPI_ISL_2906794, EPI_ISL_2906795, EPI_ISL_2906796, EPI_ISL_2906797, EPI_ISL_2906798, EPI_ISL_2906799, EPI_ISL_2906800, EPI_ISL_2906801, EPI_ISL_2906802, EPI_ISL_2906803, EPI_ISL_2906804, EPI_ISL_2906805, EPI_ISL_2906806, EPI_ISL_2906807, EPI_ISL_2906808, EPI_ISL_2906809, EPI_ISL_2906810, EPI_ISL_2906811, EPI_ISL_2906812, EPI_ISL_2906813, EPI_ISL_2906814, EPI_ISL_2906815, EPI_ISL_2906816, EPI_ISL_2906817, EPI_ISL_2906818, EPI_ISL_2906819, EPI_ISL_2906820, EPI_ISL_2906821, EPI_ISL_2906822, EPI_ISL_2906823, EPI_ISL_2906824, EPI_ISL_2906825, EPI_ISL_2906826, EPI_ISL_2906827, EPI_ISL_2906828, EPI_ISL_2906829, EPI_ISL_2906830, EPI_ISL_2906831, EPI_ISL_2906832, EPI_ISL_2906833, EPI_ISL_2906834, EPI_ISL_2906835, EPI_ISL_2906836, EPI_ISL_2906837, EPI_ISL_2906838, EPI_ISL_2906839, EPI_ISL_2906840, EPI_ISL_2906841, EPI_ISL_2906842, EPI_ISL_2906843, EPI_ISL_2906844, EPI_ISL_2906845, EPI_ISL_2906846, EPI_ISL_2906847, EPI_ISL_2906848, EPI_ISL_2906849, EPI_ISL_2906850, EPI_ISL_2906851, EPI_ISL_2906852, EPI_ISL_2906853, EPI_ISL_2906854, EPI_ISL_2906855, EPI_ISL_2906856, EPI_ISL_2906857, EPI_ISL_2906858, EPI_ISL_2906859, EPI_ISL_2906860, EPI_ISL_2906861, EPI_ISL_2906862, EPI_ISL_2906863, EPI_ISL_2906864, EPI_ISL_2906865, EPI_ISL_2906866, EPI_ISL_2906867, EPI_ISL_2906868, EPI_ISL_2906869, EPI_ISL_2906870, EPI_ISL_2906871, EPI_ISL_2906872, EPI_ISL_2906873, EPI_ISL_2906874, EPI_ISL_2906875, EPI_ISL_2906876, EPI_ISL_2906877, EPI_ISL_2906878, EPI_ISL_2906879, EPI_ISL_2906880, EPI_ISL_2906881, EPI_ISL_2906882, EPI_ISL_2906883, EPI_ISL_2906884, EPI_ISL_2906885, EPI_ISL_2906886, EPI_ISL_2906887, EPI_ISL_2906888, EPI_ISL_2906889, EPI_ISL_2906890, EPI_ISL_2906891, EPI_ISL_2906892, EPI_ISL_2906893, EPI_ISL_2906894, EPI_ISL_2906895, EPI_ISL_2906896, EPI_ISL_2906897, EPI_ISL_2906898, EPI_ISL_2906899, EPI_ISL_2906900, EPI_ISL_2906901, EPI_ISL_2906902, EPI_ISL_2906903, EPI_ISL_2906904, EPI_ISL_2906905, EPI_ISL_2906906, EPI_ISL_2906907, EPI_ISL_2906908, EPI_ISL_2906909, EPI_ISL_2906910, EPI_ISL_2906911, EPI_ISL_2906912, EPI_ISL_2906913, EPI_ISL_2906914, EPI_ISL_2906915, EPI_ISL_2906916, EPI_ISL_2906917, EPI_ISL_2906918, EPI_ISL_2906919, EPI_ISL_2906920, EPI_ISL_2906921, EPI_ISL_2906922, EPI_ISL_2906923, EPI_ISL_2906924, EPI_ISL_2906925, EPI_ISL_2906926, EPI_ISL_2906927, EPI_ISL_2906928, EPI_ISL_2906929, EPI_ISL_2906930, EPI_ISL_2906931, EPI_ISL_2906932, EPI_ISL_2906933, EPI_ISL_2906934, EPI_ISL_2906935, EPI_ISL_2906936, EPI_ISL_2906937, EPI_ISL_2906938, EPI_ISL_2906939, EPI_ISL_2906940, EPI_ISL_2906941, EPI_ISL_2906942, EPI_ISL_2906943, EPI_ISL_2906944, EPI_ISL_2906945, EPI_ISL_2906946, EPI_ISL_2906947, EPI_ISL_2906948, EPI_ISL_2906949, EPI_ISL_2906950, EPI_ISL_2906951, EPI_ISL_2906952, EPI_ISL_2906953, EPI_ISL_2906954, EPI_ISL_2906955, EPI_ISL_2906956, EPI_ISL_2906957, EPI_ISL_2906958, EPI_ISL_2906959, EPI_ISL_2906960, EPI_ISL_2906961, EPI_ISL_2906962, EPI_ISL_2906963, EPI_ISL_2906964, EPI_ISL_2906965, EPI_ISL_2906966, EPI_ISL_2906967, EPI_ISL_2906968, EPI_ISL_2906969, EPI_ISL_2906970, EPI_ISL_2906971, EPI_ISL_2906972, EPI_ISL_2906973, EPI_ISL_2906974, EPI_ISL_2906975, EPI_ISL_2906976, EPI_ISL_2906977, EPI_ISL_2906978, EPI_ISL_2906979, EPI_ISL_2906980, EPI_ISL_2906981, EPI_ISL_2906982, EPI_ISL_2906983, EPI_ISL_2906984, EPI_ISL_2906985, EPI_ISL_2906986, EPI_ISL_2906987, EPI_ISL_2906988, EPI_ISL_2906989, EPI_ISL_2906990, EPI_ISL_2906991, EPI_ISL_2906992, EPI_ISL_2906993, EPI_ISL_2906994, EPI_ISL_2906995, EPI_ISL_2906996, EPI_ISL_2906997, EPI_ISL_2906998, EPI_ISL_2906999, EPI_ISL_2907000, EPI_ISL_2907001, EPI_ISL_2907002, EPI_ISL_2907003, EPI_ISL_2907004, EPI_ISL_2907005, EPI_ISL_2907006, EPI_ISL_2907007, EPI_ISL_2907008, EPI_ISL_2907009, EPI_ISL_2907010, EPI_ISL_2907011, EPI_ISL_2907012, EPI_ISL_2907013, EPI_ISL_2907014, EPI_ISL_2907015, EPI_ISL_2907016, EPI_ISL_2907017, EPI_ISL_2907018, EPI_ISL_2907019, EPI_ISL_2907020, EPI_ISL_2907021, EPI_ISL_2907022, EPI_ISL_2907023, EPI_ISL_2907024, EPI_ISL_2907025, EPI_ISL_2907026, EPI_ISL_2907027, EPI_ISL_2907028, EPI_ISL_2907029, EPI_ISL_2907030, EPI_ISL_2907031, EPI_ISL_2907032, EPI_ISL_2907033, EPI_ISL_2907034, EPI_ISL_2907035, EPI_ISL_2907036, EPI_ISL_2907037, EPI_ISL_2907038, EPI_ISL_2907039, EPI_ISL_2907040, EPI_ISL_2907041, EPI_ISL_2907042, EPI_ISL_2907043, EPI_ISL_2907044, EPI_ISL_2907045, EPI_ISL_2907046, EPI_ISL_2907047 | | | |
| see above | Centre for Enzyme Innovation, University of Portsmouth / Translational Research Laboratory, Portsmouth Hospitals NHS Trust / Hampshire Hospitals NHS Foundation Trust / Bournemouth University / University Hospitals Dorset NHS Foundation Trust / Brighton and Sussex University Hospitals NHS Trust / Dartford and Gravesham NHS Trust / Isle of Wight NHS Trust / Maidstone and Tunbridge Wells NHS Trust / University Hospital Southampton NHS Foundation Trust | COVID-19 Genomics UK (COG-UK) Consortium | Samuel Robson, Angela Beckett, Salman Goudarzi, Christopher Fearn, Kate Cook, Katie Loveson, Sharon Glaysher, Scott Elliott, Kelly Bicknell, Sarah Wyllie, Allyson Lloyd, Robert Impey, Anoop Chauhan, Stephen Kidd, Nathan Moore, Nick Cortes, Claire Thomas, Anna Mantzaouratou, Sarah Buchan, Magdalena Barrow, Andrew Butt, Liz Sheridan, Jonnie Seymour, Dorian Crudgington, Ben Macklin, Mohammed Hassan-Ibrahim, Cassandra Malone, Benjamin Cogger, Kevin Tucker, Samirakhon Raupova, Rachael Jeremiah, Anibolina Castigador, Emily Macnaughton, Karen Withell, Kordo Saeed, Jacqui Prieto, Adhyana Mahanama, Buddhini Samaraweera, Siona Silvieri, Emanuela Pelosi, Eleri Wilson-Davies, Sarah Jeremiah, Helen Wheeler, Matthew Harvey, Thea Sass, Helen Umpleby, Stephen Aplin |
| EPI_ISL_2907048, EPI_ISL_2907049, EPI_ISL_2907050, EPI_ISL_2907051, EPI_ISL_2907052, EPI_ISL_2907053, EPI_ISL_2907054, EPI_ISL_2907055, EPI_ISL_2907056, EPI_ISL_2907057, EPI_ISL_2907058, EPI_ISL_2907059, EPI_ISL_2907060, EPI_ISL_2907061, EPI_ISL_2907062, EPI_ISL_2907063, EPI_ISL_2907064, EPI_ISL_2907065, EPI_ISL_2907066, EPI_ISL_2907067, EPI_ISL_2907068, EPI_ISL_2907069, EPI_ISL_2907070, EPI_ISL_2907071, EPI_ISL_2907072, EPI_ISL_2907073, EPI_ISL_2907074, EPI_ISL_2907075, EPI_ISL_2907076, EPI_ISL_2907077, EPI_ISL_2907078, EPI_ISL_2907079, EPI_ISL_2907080, EPI_ISL_2907081, EPI_ISL_2907082, EPI_ISL_2907083, EPI_ISL_2907084, EPI_ISL_2907085, EPI_ISL_2907086, EPI_ISL_2907087, EPI_ISL_2907088, EPI_ISL_2907089, EPI_ISL_2907090, EPI_ISL_2907091, EPI_ISL_2907092, EPI_ISL_2907093, EPI_ISL_2907094, EPI_ISL_2907095, EPI_ISL_2907096, EPI_ISL_2907097, EPI_ISL_2907098, EPI_ISL_2907099, EPI_ISL_2907100, EPI_ISL_2907101, EPI_ISL_2907102, EPI_ISL_2907103, EPI_ISL_2907104, EPI_ISL_2907105, EPI_ISL_2907106, EPI_ISL_2907107, EPI_ISL_2907108, EPI_ISL_2907109, EPI_ISL_2907110, EPI_ISL_2907111, EPI_ISL_2907112, EPI_ISL_2907113, EPI_ISL_2907114, EPI_ISL_2907115, EPI_ISL_2907116, EPI_ISL_2907117, EPI_ISL_2907118, EPI_ISL_2907119, EPI_ISL_2907120, EPI_ISL_2907121, EPI_ISL_2907122, EPI_ISL_2907123, EPI_ISL_2907124, EPI_ISL_2907125, EPI_ISL_2907126, EPI_ISL_2907127, EPI_ISL_2907128, EPI_ISL_2907129, EPI_ISL_2907130, EPI_ISL_2907131, EPI_ISL_2907132, EPI_ISL_2907133, EPI_ISL_2907135, EPI_ISL_2907136, EPI_ISL_2907137, EPI_ISL_2907138, EPI_ISL_2907139, EPI_ISL_2907140, EPI_ISL_2907141, EPI_ISL_2907142, EPI_ISL_2907143, EPI_ISL_2907144, EPI_ISL_2907145, EPI_ISL_2907146, EPI_ISL_2907147, EPI_ISL_2907148, EPI_ISL_2907149, EPI_ISL_2907150, EPI_ISL_2907151, EPI_ISL_2907152, EPI_ISL_2907153, EPI_ISL_2907154, EPI_ISL_2907155, EPI_ISL_2907156, EPI_ISL_2907157, EPI_ISL_2907158, EPI_ISL_2907159, EPI_ISL_2907160, EPI_ISL_2907161, EPI_ISL_2907162, EPI_ISL_2907163, EPI_ISL_2907164, EPI_ISL_2907165, EPI_ISL_2907166, EPI_ISL_2907167, EPI_ISL_2907168, EPI_ISL_2907169, EPI_ISL_2907170, EPI_ISL_2907171, EPI_ISL_2907173, EPI_ISL_2907174, EPI_ISL_2907176, EPI_ISL_2907177, EPI_ISL_2907178, EPI_ISL_2907179, EPI_ISL_2907180, EPI_ISL_2907181, EPI_ISL_2907182, EPI_ISL_2907184, EPI_ISL_2907185, EPI_ISL_2907186, EPI_ISL_2907187, EPI_ISL_2907188, EPI_ISL_2907189, EPI_ISL_2907190, EPI_ISL_2907191, EPI_ISL_2907192, EPI_ISL_2907193, EPI_ISL_2907194, EPI_ISL_2907195, EPI_ISL_2907196, EPI_ISL_2907197, EPI_ISL_2907198, EPI_ISL_2907199, EPI_ISL_2907200, EPI_ISL_2907201, EPI_ISL_2907202, EPI_ISL_2907203, EPI_ISL_2907204, EPI_ISL_2907205, EPI_ISL_2907206, EPI_ISL_2907207, EPI_ISL_2907208, EPI_ISL_2907209, EPI_ISL_2907210, EPI_ISL_2907211, EPI_ISL_2907212, EPI_ISL_2907213, EPI_ISL_2907214, EPI_ISL_2907215, EPI_ISL_2907216, EPI_ISL_2907217, EPI_ISL_2907218, EPI_ISL_2907219, EPI_ISL_2907220, EPI_ISL_2907221, EPI_ISL_2907222, EPI_ISL_2907223, EPI_ISL_2907224, EPI_ISL_2907225, EPI_ISL_2907226, EPI_ISL_2907227, EPI_ISL_2907228, EPI_ISL_2907229, EPI_ISL_2907230, EPI_ISL_2907231, EPI_ISL_2907232, EPI_ISL_2907233, EPI_ISL_2907234, EPI_ISL_2907235, EPI_ISL_2907236, EPI_ISL_2907237, EPI_ISL_2907238, EPI_ISL_2907240, EPI_ISL_2907241, EPI_ISL_2907242, EPI_ISL_2907243, EPI_ISL_2907244, EPI_ISL_2907245, EPI_ISL_2907246, EPI_ISL_2907247, EPI_ISL_2907248, EPI_ISL_2907249, EPI_ISL_2907250, EPI_ISL_2907251, EPI_ISL_2907252, EPI_ISL_2907253, EPI_ISL_2907254, EPI_ISL_2907255, EPI_ISL_2907256, EPI_ISL_2907257, EPI_ISL_2907258, EPI_ISL_2907259, EPI_ISL_2907260, EPI_ISL_2907261, EPI_ISL_2907262, EPI_ISL_2907263, EPI_ISL_2907264, EPI_ISL_2907265, EPI_ISL_2907266, EPI_ISL_2907267, EPI_ISL_2907268, EPI_ISL_2907269, EPI_ISL_2907270, EPI_ISL_2907271, EPI_ISL_2907272, EPI_ISL_2907273, EPI_ISL_2907275, EPI_ISL_2907276, EPI_ISL_2907277, EPI_ISL_2907278, EPI_ISL_2907279, EPI_ISL_2907280, EPI_ISL_2907281, EPI_ISL_2907282, EPI_ISL_2907283, EPI_ISL_2907284, EPI_ISL_2907285, EPI_ISL_2907286, EPI_ISL_2907287, EPI_ISL_2907288, EPI_ISL_2907289, EPI_ISL_2907290, EPI_ISL_2907291, EPI_ISL_2907292, EPI_ISL_2907293, EPI_ISL_2907294, EPI_ISL_2907295, EPI_ISL_2907296, EPI_ISL_2907297, EPI_ISL_2907298, EPI_ISL_2907299, EPI_ISL_2907300, EPI_ISL_2907301, EPI_ISL_2907302, EPI_ISL_2907303, EPI_ISL_2907304, EPI_ISL_2907305, EPI_ISL_2907306, EPI_ISL_2907307, EPI_ISL_2907308, EPI_ISL_2907309, EPI_ISL_2907310, EPI_ISL_2907311, EPI_ISL_2907312, EPI_ISL_2907313, EPI_ISL_2907314, EPI_ISL_2907315, EPI_ISL_2907316, EPI_ISL_2907317, EPI_ISL_2907318, EPI_ISL_2907319, EPI_ISL_2907320, EPI_ISL_2907321, EPI_ISL_2907322, EPI_ISL_2907323, EPI_ISL_2907324, EPI_ISL_2907325, EPI_ISL_2907326, EPI_ISL_2907327, EPI_ISL_2907328, EPI_ISL_2907329, EPI_ISL_2907331, EPI_ISL_2907332, EPI_ISL_2907333, EPI_ISL_2907334, EPI_ISL_2907335, EPI_ISL_2907336, EPI_ISL_2907337, EPI_ISL_2907338, EPI_ISL_2907340, EPI_ISL_2907341, EPI_ISL_2907342, EPI_ISL_2907343, EPI_ISL_2907344, EPI_ISL_2907345, EPI_ISL_2907346, EPI_ISL_2907347, EPI_ISL_2907348, EPI_ISL_2907349, EPI_ISL_2907350, EPI_ISL_2907351, EPI_ISL_2907352, EPI_ISL_2907353, EPI_ISL_2907354, EPI_ISL_2907355, EPI_ISL_2907356, EPI_ISL_2907357, EPI_ISL_2907358, EPI_ISL_2907359, EPI_ISL_2907360, EPI_ISL_2907361, EPI_ISL_2907362, EPI_ISL_2907363, EPI_ISL_2907364, EPI_ISL_2907365, EPI_ISL_2907366, EPI_ISL_2907367, EPI_ISL_2907368, EPI_ISL_2907369, EPI_ISL_2907370, EPI_ISL_2907371, EPI_ISL_2907372, EPI_ISL_2907373, EPI_ISL_2907374, EPI_ISL_2907375, EPI_ISL_2907376, EPI_ISL_2907377, EPI_ISL_2907378, EPI_ISL_2907379, EPI_ISL_2907380, EPI_ISL_2907381, EPI_ISL_2907382, EPI_ISL_2907383, EPI_ISL_2907384, EPI_ISL_2907385, EPI_ISL_2907386, EPI_ISL_2907387, EPI_ISL_2907388, EPI_ISL_2907389, EPI_ISL_2907390, EPI_ISL_2907391, EPI_ISL_2907392, EPI_ISL_2907393, EPI_ISL_2907394, EPI_ISL_2907395, EPI_ISL_2907396, EPI_ISL_2907397, EPI_ISL_2907398, EPI_ISL_2907399, EPI_ISL_2907400, EPI_ISL_2907401, EPI_ISL_2907402, EPI_ISL_2907403, EPI_ISL_2907404, EPI_ISL_2907405, EPI_ISL_2907406, EPI_ISL_2907407, EPI_ISL_2907408, EPI_ISL_2907409, EPI_ISL_2907410, EPI_ISL_2907411, EPI_ISL_2907412, EPI_ISL_2907413, EPI_ISL_2907414, EPI_ISL_2907415, EPI_ISL_2907416, EPI_ISL_2907417, EPI_ISL_2907418, EPI_ISL_2907419, EPI_ISL_2907420, EPI_ISL_2907421, EPI_ISL_2907422, EPI_ISL_2907423, EPI_ISL_2907424, EPI_ISL_2907425, EPI_ISL_2907427, EPI_ISL_2907428, EPI_ISL_2907429, EPI_ISL_2907430, EPI_ISL_2907431, EPI_ISL_2907432, EPI_ISL_2907433, EPI_ISL_2907434, EPI_ISL_2907435, EPI_ISL_2907436, EPI_ISL_2907437, EPI_ISL_2907438, EPI_ISL_2907439, EPI_ISL_2907440, EPI_ISL_2907441, EPI_ISL_2907442, EPI_ISL_2907443, EPI_ISL_2907444, EPI_ISL_2907445, EPI_ISL_2907446, EPI_ISL_2907447, EPI_ISL_2907448, EPI_ISL_2907449, EPI_ISL_2907450, EPI_ISL_2907451, EPI_ISL_2907452, EPI_ISL_2907453, EPI_ISL_2907454, EPI_ISL_2907455, EPI_ISL_2907456, EPI_ISL_2907457, EPI_ISL_2907458, EPI_ISL_2907459, EPI_ISL_2907460, EPI_ISL_2907461, EPI_ISL_2907462, EPI_ISL_2907463, EPI_ISL_2907464, EPI_ISL_2907465, EPI_ISL_2907466, EPI_ISL_2907467, EPI_ISL_2907468, EPI_ISL_2907469, EPI_ISL_2907470, EPI_ISL_2907472, EPI_ISL_2907473, EPI_ISL_2907474, EPI_ISL_2907475, EPI_ISL_2907476, EPI_ISL_2907477, EPI_ISL_2907478, EPI_ISL_2907479, EPI_ISL_2907480, EPI_ISL_2907481 | | | |
| see above | Virology Department, Sheffield Teaching Hospitals NHS Foundation Trust/Department of Infection, Immunity and Cardiovascular Disease, The Medical School, University of Sheffield | COVID-19 Genomics UK (COG-UK) Consortium | Thushan de Silva, Matthew Parker, Nikki Smith, Adri Angyal, Rebecca Brown, Luke Green, Rachel Tucker, Paul Parsons, Danielle Groves, Katie Johnson, Laura Carrilero, Alex Keeley, Dave Partridge, Matthew Wyles, Benjamin Lindsey, Mehmet Yavuz, Mohammad Raza, Cariad Evans |
| EPI_ISL_2907488 | Department of Microbiology, Yamanashi Institute for Public Health and Environment | Genome Analysis Center, Yamanashi Central Hospital | Yosuke Hirotsu |
| EPI_ISL_2907518, EPI_ISL_2907520, EPI_ISL_2907522, EPI_ISL_2907523, EPI_ISL_2907524, EPI_ISL_2907526, EPI_ISL_2907527, EPI_ISL_2907528, EPI_ISL_2907529, EPI_ISL_2907530, EPI_ISL_2907531, EPI_ISL_2907532, EPI_ISL_2907533, EPI_ISL_2907534, EPI_ISL_2907535, EPI_ISL_2907536, EPI_ISL_2907537, EPI_ISL_2907538, EPI_ISL_2907540, EPI_ISL_2907541, EPI_ISL_2907542, EPI_ISL_2907543 | | | |
| see above | Azienda Sanitaria dell'Alto Adige - Laboratorio Aziendale di Microbiologia e Virologia | Azienda Sanitaria dell'Alto Adige | Irene Bianconi |
| EPI_ISL_2907544, EPI_ISL_2907545, EPI_ISL_2907546 | Royal Darwin Hospital Pathology | MDU-PHL | Meumann, E., Caly L., Seemann T., Sait, M.L., Druce J., Sherry, N.L. |
| EPI_ISL_2907549, EPI_ISL_2907550 | Microbiological Diagnostic Unit - Public Health Laboratory (MDU-PHL) | MDU-PHL | Seemann T., Sait, M.L., Sherry, N.L. |
| EPI_ISL_2907555, EPI_ISL_2907556, EPI_ISL_2907557, EPI_ISL_2907559, EPI_ISL_2907560 | Centinela Hospital Medical Center | Los Angeles County Public Health Laboratories | P. Hemarajata et al. |
| EPI_ISL_2907562 | Martin Luther King, Jr. Outpatient Center | Los Angeles County Public Health Laboratories | P. Hemarajata et al. |
| EPI_ISL_2907565, EPI_ISL_2907566 | Saint Francis Medical Center | Los Angeles County Public Health Laboratories | P. Hemarajata et al. |
| EPI_ISL_2907567, EPI_ISL_2907568, EPI_ISL_2907569, EPI_ISL_2907570, EPI_ISL_2907571, EPI_ISL_2907572 | Providence Saint John's Health Center | Los Angeles County Public Health Laboratories | P. Hemarajata et al. |
| EPI_ISL_2907573, EPI_ISL_2907574, EPI_ISL_2907575 | USC Clinical Lab | Los Angeles County Public Health Laboratories | P. Hemarajata et al. |
| EPI_ISL_2907576 | Centinela Hospital Medical Center | Los Angeles County Public Health Laboratories | P. Hemarajata et al. |
| EPI_ISL_2907577 | Martin Luther King Jr. Community Hospital | Los Angeles County Public Health Laboratories | P. Hemarajata et al. |
| EPI_ISL_2907578 | Laboratory of Microbiology, National Reference Lab, Charles Nicolle Hospital; 2-University of Tunis ElManar, Faculty of Medicine of Tunis, LR99ES09, Tunis, Tunisia | 1-Clinical and Experimental Pharmacology Lab, LR16SP02, National Center of Pharmacovigilance, University of Tunis El Manar, Tunis, Tunisia. 2-Neurodegenerative diseases and psychiatric troubles, LR18SP03, Razi Hospital, University of Tunis El Manar, Tunis, Tunisia. 3- Ministry of Health, National Observatory of New and Emerging Diseases, 1006, Tunis, Tunisia | Sameh Trabelsi, Nissaf Ben Alaya, Mouna Ben Sassi, Sana Ferjani, Salma Abid, Mouna Safer, Roua Ben Othman, Sarra Chammam, Ines Mdini, Imen Mkada, Imen Kacem, Maher Kharrat, Alia Benkahla, Jalila Ben Khelil, Emna Gaies, Riadh Daghfous, Riadh Gouider, Ilhem Boutiba-Ben Boubaker. |
| EPI_ISL_2907579, EPI_ISL_2907580 | Lighthouse Lab in Alderley Park | Wellcome Sanger Institute for the COVID-19 Genomics UK (COG-UK) Consortium | Jacquelyn Wynn, Mairead Hyland, The Lighthouse Lab in Alderley Park and Alex Alderton, Roberto Amato, Jeffrey Barrett, Sonia Goncalves, Ewan Harrison, David K. Jackson, Ian Johnston, Dominic Kwiatkowski, Cordelia Langford, John Sillitoe on behalf of the Wellcome Sanger Institute COVID-19 Surveillance Team |
| EPI_ISL_2907581, EPI_ISL_2907582, EPI_ISL_2907583 | Lighthouse Lab in Milton Keynes | Wellcome Sanger Institute for the COVID-19 Genomics UK (COG-UK) Consortium | The Lighthouse Lab in Milton Keynes and Alex Alderton, Roberto Amato, Jeffrey Barrett, Sonia Goncalves, Ewan Harrison, David K. Jackson, Ian Johnston, Dominic Kwiatkowski, Cordelia Langford, John Sillitoe on behalf of the Wellcome Sanger Institute COVID-19 Surveillance Team |
| EPI_ISL_2907584 | Lighthouse Lab in Alderley Park | Wellcome Sanger Institute for the COVID-19 Genomics UK (COG-UK) | Jacquelyn Wynn, Mairead Hyland, The Lighthouse Lab in Alderley Park and Alex Alderton, Roberto Amato, Jeffrey Barrett, Sonia Goncalves, Ewan Harrison, David K. Jackson, Ian Johnston, Dominic Kwiatkowski, Cordelia Langford, John Sillitoe on |

| | | | |
|---|---|---|---|
| | | Consortium | behalf of the Wellcome Sanger Institute COVID-19 Surveillance Team |
| EPI_ISL_2907585 | Lighthouse Lab in Milton Keynes | Wellcome Sanger Institute for the COVID-19 Genomics UK (COG-UK) Consortium | The Lighthouse Lab in Milton Keynes and Alex Alderton, Roberto Amato, Jeffrey Barrett, Sonia Goncalves, Ewan Harrison, David K. Jackson, Ian Johnston, Dominic Kwiatkowski, Cordelia Langford, John Sillitoe on behalf of the Wellcome Sanger Institute COVID-19 Surveillance Team |
| EPI_ISL_2907587 | Lighthouse Lab in Alderley Park | Wellcome Sanger Institute for the COVID-19 Genomics UK (COG-UK) Consortium | Jacquelyn Wynn, Mairead Hyland, The Lighthouse Lab in Alderley Park and Alex Alderton, Roberto Amato, Jeffrey Barrett, Sonia Goncalves, Ewan Harrison, David K. Jackson, Ian Johnston, Dominic Kwiatkowski, Cordelia Langford, John Sillitoe on behalf of the Wellcome Sanger Institute COVID-19 Surveillance Team |
| EPI_ISL_2907588, EPI_ISL_2907589 | Lighthouse Lab in Milton Keynes | Wellcome Sanger Institute for the COVID-19 Genomics UK (COG-UK) Consortium | The Lighthouse Lab in Milton Keynes and Alex Alderton, Roberto Amato, Jeffrey Barrett, Sonia Goncalves, Ewan Harrison, David K. Jackson, Ian Johnston, Dominic Kwiatkowski, Cordelia Langford, John Sillitoe on behalf of the Wellcome Sanger Institute COVID-19 Surveillance Team |
| EPI_ISL_2907590 | Lighthouse Lab in Alderley Park | Wellcome Sanger Institute for the COVID-19 Genomics UK (COG-UK) Consortium | Jacquelyn Wynn, Mairead Hyland, The Lighthouse Lab in Alderley Park and Alex Alderton, Roberto Amato, Jeffrey Barrett, Sonia Goncalves, Ewan Harrison, David K. Jackson, Ian Johnston, Dominic Kwiatkowski, Cordelia Langford, John Sillitoe on behalf of the Wellcome Sanger Institute COVID-19 Surveillance Team |
| EPI_ISL_2907591, EPI_ISL_2907592, EPI_ISL_2907593, EPI_ISL_2907594 | Lighthouse Lab in Milton Keynes | Wellcome Sanger Institute for the COVID-19 Genomics UK (COG-UK) Consortium | The Lighthouse Lab in Milton Keynes and Alex Alderton, Roberto Amato, Jeffrey Barrett, Sonia Goncalves, Ewan Harrison, David K. Jackson, Ian Johnston, Dominic Kwiatkowski, Cordelia Langford, John Sillitoe on behalf of the Wellcome Sanger Institute COVID-19 Surveillance Team |
| EPI_ISL_2907595, EPI_ISL_2907596, EPI_ISL_2907597, EPI_ISL_2907598, EPI_ISL_2907599 | Lighthouse Lab in Alderley Park | Wellcome Sanger Institute for the COVID-19 Genomics UK (COG-UK) Consortium | Jacquelyn Wynn, Mairead Hyland, The Lighthouse Lab in Alderley Park and Alex Alderton, Roberto Amato, Jeffrey Barrett, Sonia Goncalves, Ewan Harrison, David K. Jackson, Ian Johnston, Dominic Kwiatkowski, Cordelia Langford, John Sillitoe on behalf of the Wellcome Sanger Institute COVID-19 Surveillance Team |
| EPI_ISL_2907600 | Lighthouse Lab in Milton Keynes | Wellcome Sanger Institute for the COVID-19 Genomics UK (COG-UK) Consortium | The Lighthouse Lab in Milton Keynes and Alex Alderton, Roberto Amato, Jeffrey Barrett, Sonia Goncalves, Ewan Harrison, David K. Jackson, Ian Johnston, Dominic Kwiatkowski, Cordelia Langford, John Sillitoe on behalf of the Wellcome Sanger Institute COVID-19 Surveillance Team |
| EPI_ISL_2907601 | Lighthouse Lab in Alderley Park | Wellcome Sanger Institute for the COVID-19 Genomics UK (COG-UK) Consortium | Jacquelyn Wynn, Mairead Hyland, The Lighthouse Lab in Alderley Park and Alex Alderton, Roberto Amato, Jeffrey Barrett, Sonia Goncalves, Ewan Harrison, David K. Jackson, Ian Johnston, Dominic Kwiatkowski, Cordelia Langford, John Sillitoe on behalf of the Wellcome Sanger Institute COVID-19 Surveillance Team |
| EPI_ISL_2907602 | Lighthouse Lab in Milton Keynes | Wellcome Sanger Institute for the COVID-19 Genomics UK (COG-UK) Consortium | The Lighthouse Lab in Milton Keynes and Alex Alderton, Roberto Amato, Jeffrey Barrett, Sonia Goncalves, Ewan Harrison, David K. Jackson, Ian Johnston, Dominic Kwiatkowski, Cordelia Langford, John Sillitoe on behalf of the Wellcome Sanger Institute COVID-19 Surveillance Team |
| EPI_ISL_2907603 | Lighthouse Lab in Alderley Park | Wellcome Sanger Institute for the COVID-19 Genomics UK (COG-UK) Consortium | Jacquelyn Wynn, Mairead Hyland, The Lighthouse Lab in Alderley Park and Alex Alderton, Roberto Amato, Jeffrey Barrett, Sonia Goncalves, Ewan Harrison, David K. Jackson, Ian Johnston, Dominic Kwiatkowski, Cordelia Langford, John Sillitoe on behalf of the Wellcome Sanger Institute COVID-19 Surveillance Team |
| EPI_ISL_2907604 | Lighthouse Lab in Milton Keynes | Wellcome Sanger Institute for the COVID-19 Genomics UK (COG-UK) Consortium | The Lighthouse Lab in Milton Keynes and Alex Alderton, Roberto Amato, Jeffrey Barrett, Sonia Goncalves, Ewan Harrison, David K. Jackson, Ian Johnston, Dominic Kwiatkowski, Cordelia Langford, John Sillitoe on behalf of the Wellcome Sanger Institute COVID-19 Surveillance Team |
| EPI_ISL_2907605 | Lighthouse Lab in Alderley Park | Wellcome Sanger Institute for the COVID-19 Genomics UK (COG-UK) Consortium | Jacquelyn Wynn, Mairead Hyland, The Lighthouse Lab in Alderley Park and Alex Alderton, Roberto Amato, Jeffrey Barrett, Sonia Goncalves, Ewan Harrison, David K. Jackson, Ian Johnston, Dominic Kwiatkowski, Cordelia Langford, John Sillitoe on behalf of the Wellcome Sanger Institute COVID-19 Surveillance Team |
| EPI_ISL_2907606, EPI_ISL_2907607, EPI_ISL_2907608 | Lighthouse Lab in Milton Keynes | Wellcome Sanger Institute for the COVID-19 Genomics UK (COG-UK) Consortium | The Lighthouse Lab in Milton Keynes and Alex Alderton, Roberto Amato, Jeffrey Barrett, Sonia Goncalves, Ewan Harrison, David K. Jackson, Ian Johnston, Dominic Kwiatkowski, Cordelia Langford, John Sillitoe on behalf of the Wellcome Sanger Institute COVID-19 Surveillance Team |
| EPI_ISL_2907609 | Lighthouse Lab in Alderley Park | Wellcome Sanger Institute for the COVID-19 Genomics UK (COG-UK) Consortium | Jacquelyn Wynn, Mairead Hyland, The Lighthouse Lab in Alderley Park and Alex Alderton, Roberto Amato, Jeffrey Barrett, Sonia Goncalves, Ewan Harrison, David K. Jackson, Ian Johnston, Dominic Kwiatkowski, Cordelia Langford, John Sillitoe on behalf of the Wellcome Sanger Institute COVID-19 Surveillance Team |
| EPI_ISL_2907610, EPI_ISL_2907611 | Lighthouse Lab in Milton Keynes | Wellcome Sanger Institute for the COVID-19 Genomics UK (COG-UK) Consortium | The Lighthouse Lab in Milton Keynes and Alex Alderton, Roberto Amato, Jeffrey Barrett, Sonia Goncalves, Ewan Harrison, David K. Jackson, Ian Johnston, Dominic Kwiatkowski, Cordelia Langford, John Sillitoe on behalf of the Wellcome Sanger Institute COVID-19 Surveillance Team |
| EPI_ISL_2907612 | Lighthouse Lab in Alderley Park | Wellcome Sanger Institute for the COVID-19 Genomics UK (COG-UK) Consortium | Jacquelyn Wynn, Mairead Hyland, The Lighthouse Lab in Alderley Park and Alex Alderton, Roberto Amato, Jeffrey Barrett, Sonia Goncalves, Ewan Harrison, David K. Jackson, Ian Johnston, Dominic Kwiatkowski, Cordelia Langford, John Sillitoe on behalf of the Wellcome Sanger Institute COVID-19 Surveillance Team |
| EPI_ISL_2907613, EPI_ISL_2907614 | Lighthouse Lab in Milton Keynes | Wellcome Sanger Institute for the COVID-19 Genomics UK (COG-UK) Consortium | The Lighthouse Lab in Milton Keynes and Alex Alderton, Roberto Amato, Jeffrey Barrett, Sonia Goncalves, Ewan Harrison, David K. Jackson, Ian Johnston, Dominic Kwiatkowski, Cordelia Langford, John Sillitoe on behalf of the Wellcome Sanger Institute COVID-19 Surveillance Team |
| EPI_ISL_2907615 | Lighthouse Lab in Alderley Park | Wellcome Sanger Institute for the COVID-19 Genomics UK (COG-UK) Consortium | Jacquelyn Wynn, Mairead Hyland, The Lighthouse Lab in Alderley Park and Alex Alderton, Roberto Amato, Jeffrey Barrett, Sonia Goncalves, Ewan Harrison, David K. Jackson, Ian Johnston, Dominic Kwiatkowski, Cordelia Langford, John Sillitoe on behalf of the Wellcome Sanger Institute COVID-19 Surveillance Team |
| EPI_ISL_2907616, EPI_ISL_2907617, EPI_ISL_2907618 | Lighthouse Lab in Milton Keynes | Wellcome Sanger Institute for the COVID-19 Genomics UK (COG-UK) Consortium | The Lighthouse Lab in Milton Keynes and Alex Alderton, Roberto Amato, Jeffrey Barrett, Sonia Goncalves, Ewan Harrison, David K. Jackson, Ian Johnston, Dominic Kwiatkowski, Cordelia Langford, John Sillitoe on behalf of the Wellcome Sanger Institute COVID-19 Surveillance Team |
| EPI_ISL_2907619 | Lighthouse Lab in Alderley Park | Wellcome Sanger Institute for the COVID-19 Genomics UK (COG-UK) Consortium | Jacquelyn Wynn, Mairead Hyland, The Lighthouse Lab in Alderley Park and Alex Alderton, Roberto Amato, Jeffrey Barrett, Sonia Goncalves, Ewan Harrison, David K. Jackson, Ian Johnston, Dominic Kwiatkowski, Cordelia Langford, John Sillitoe on behalf of the Wellcome Sanger Institute COVID-19 Surveillance Team |
| EPI_ISL_2907620 | Lighthouse Lab in Milton Keynes | Wellcome Sanger Institute for the COVID-19 Genomics UK (COG-UK) Consortium | The Lighthouse Lab in Milton Keynes and Alex Alderton, Roberto Amato, Jeffrey Barrett, Sonia Goncalves, Ewan Harrison, David K. Jackson, Ian Johnston, Dominic Kwiatkowski, Cordelia Langford, John Sillitoe on behalf of the Wellcome Sanger Institute COVID-19 Surveillance Team |
| EPI_ISL_2907621 | Lighthouse Lab in Alderley Park | Wellcome Sanger Institute for the COVID-19 Genomics UK (COG-UK) Consortium | Jacquelyn Wynn, Mairead Hyland, The Lighthouse Lab in Alderley Park and Alex Alderton, Roberto Amato, Jeffrey Barrett, Sonia Goncalves, Ewan Harrison, David K. Jackson, Ian Johnston, Dominic Kwiatkowski, Cordelia Langford, John Sillitoe on behalf of the Wellcome Sanger Institute COVID-19 Surveillance Team |
| EPI_ISL_2907622, EPI_ISL_2907623 | Lighthouse Lab in Milton Keynes | Wellcome Sanger Institute for the COVID-19 Genomics UK (COG-UK) Consortium | The Lighthouse Lab in Milton Keynes and Alex Alderton, Roberto Amato, Jeffrey Barrett, Sonia Goncalves, Ewan Harrison, David K. Jackson, Ian Johnston, Dominic Kwiatkowski, Cordelia Langford, John Sillitoe on behalf of the Wellcome Sanger Institute COVID-19 Surveillance Team |
| EPI_ISL_2907624, EPI_ISL_2907625 | Lighthouse Lab in Alderley Park | Wellcome Sanger Institute for the COVID-19 Genomics UK (COG-UK) Consortium | Jacquelyn Wynn, Mairead Hyland, The Lighthouse Lab in Alderley Park and Alex Alderton, Roberto Amato, Jeffrey Barrett, Sonia Goncalves, Ewan Harrison, David K. Jackson, Ian Johnston, Dominic Kwiatkowski, Cordelia Langford, John Sillitoe on behalf of the Wellcome Sanger Institute COVID-19 Surveillance Team |
| EPI_ISL_2907626 | Lighthouse Lab in Milton Keynes | Wellcome Sanger Institute for the COVID-19 Genomics UK (COG-UK) Consortium | The Lighthouse Lab in Milton Keynes and Alex Alderton, Roberto Amato, Jeffrey Barrett, Sonia Goncalves, Ewan Harrison, David K. Jackson, Ian Johnston, Dominic Kwiatkowski, Cordelia Langford, John Sillitoe on behalf of the Wellcome Sanger Institute COVID-19 Surveillance Team |
| EPI_ISL_2907627, EPI_ISL_2907628, EPI_ISL_2907629, EPI_ISL_2907630, EPI_ISL_2907631 | Lighthouse Lab in Alderley Park | Wellcome Sanger Institute for the COVID-19 Genomics UK (COG-UK) Consortium | Jacquelyn Wynn, Mairead Hyland, The Lighthouse Lab in Alderley Park and Alex Alderton, Roberto Amato, Jeffrey Barrett, Sonia Goncalves, Ewan Harrison, David K. Jackson, Ian Johnston, Dominic Kwiatkowski, Cordelia Langford, John Sillitoe on behalf of the Wellcome Sanger Institute COVID-19 Surveillance Team |
| EPI_ISL_2907632, EPI_ISL_2907633 | Lighthouse Lab in Milton Keynes | Wellcome Sanger Institute for the COVID-19 Genomics UK (COG-UK) Consortium | The Lighthouse Lab in Milton Keynes and Alex Alderton, Roberto Amato, Jeffrey Barrett, Sonia Goncalves, Ewan Harrison, David K. Jackson, Ian Johnston, Dominic Kwiatkowski, Cordelia Langford, John Sillitoe on behalf of the Wellcome Sanger Institute COVID-19 Surveillance Team |
| EPI_ISL_2907634 | Lighthouse Lab in Alderley Park | Wellcome Sanger Institute for the COVID-19 Genomics UK (COG-UK) Consortium | Jacquelyn Wynn, Mairead Hyland, The Lighthouse Lab in Alderley Park and Alex Alderton, Roberto Amato, Jeffrey Barrett, Sonia Goncalves, Ewan Harrison, David K. Jackson, Ian Johnston, Dominic Kwiatkowski, Cordelia Langford, John Sillitoe on behalf of the Wellcome Sanger Institute COVID-19 Surveillance Team |
| EPI_ISL_2907635 | Lighthouse Lab in Milton Keynes | Wellcome Sanger Institute for the COVID-19 Genomics UK (COG-UK) Consortium | The Lighthouse Lab in Milton Keynes and Alex Alderton, Roberto Amato, Jeffrey Barrett, Sonia Goncalves, Ewan Harrison, David K. Jackson, Ian Johnston, Dominic Kwiatkowski, Cordelia Langford, John Sillitoe on behalf of the Wellcome Sanger Institute COVID-19 Surveillance Team |
| EPI_ISL_2907636 | Lighthouse Lab in Alderley Park | Wellcome Sanger Institute for the COVID-19 Genomics UK (COG-UK) Consortium | Jacquelyn Wynn, Mairead Hyland, The Lighthouse Lab in Alderley Park and Alex Alderton, Roberto Amato, Jeffrey Barrett, Sonia Goncalves, Ewan Harrison, David K. Jackson, Ian Johnston, Dominic Kwiatkowski, Cordelia Langford, John Sillitoe on behalf of the Wellcome Sanger Institute COVID-19 Surveillance Team |
| EPI_ISL_2907637, EPI_ISL_2907638 | Lighthouse Lab in Milton Keynes | Wellcome Sanger Institute for the COVID-19 Genomics UK (COG-UK) Consortium | The Lighthouse Lab in Milton Keynes and Alex Alderton, Roberto Amato, Jeffrey Barrett, Sonia Goncalves, Ewan Harrison, David K. Jackson, Ian Johnston, Dominic Kwiatkowski, Cordelia Langford, John Sillitoe on behalf of the Wellcome Sanger Institute COVID-19 Surveillance Team |
| EPI_ISL_2907639 | Lighthouse Lab in Alderley Park | Wellcome Sanger Institute for the COVID-19 Genomics UK (COG-UK) Consortium | Jacquelyn Wynn, Mairead Hyland, The Lighthouse Lab in Alderley Park and Alex Alderton, Roberto Amato, Jeffrey Barrett, Sonia Goncalves, Ewan Harrison, David K. Jackson, Ian Johnston, Dominic Kwiatkowski, Cordelia Langford, John Sillitoe on behalf of the Wellcome Sanger Institute COVID-19 Surveillance Team |
| EPI_ISL_2907640, EPI_ISL_2907641, EPI_ISL_2907642, EPI_ISL_2907643, EPI_ISL_2907644 | Lighthouse Lab in Milton Keynes | Wellcome Sanger Institute for the COVID-19 Genomics UK (COG-UK) Consortium | The Lighthouse Lab in Milton Keynes and Alex Alderton, Roberto Amato, Jeffrey Barrett, Sonia Goncalves, Ewan Harrison, David K. Jackson, Ian Johnston, Dominic Kwiatkowski, Cordelia Langford, John Sillitoe on behalf of the Wellcome Sanger Institute COVID-19 Surveillance Team |
| EPI_ISL_2907645, EPI_ISL_2907646 | Lighthouse Lab in Alderley Park | Wellcome Sanger Institute for the COVID-19 Genomics UK (COG-UK) Consortium | Jacquelyn Wynn, Mairead Hyland, The Lighthouse Lab in Alderley Park and Alex Alderton, Roberto Amato, Jeffrey Barrett, Sonia Goncalves, Ewan Harrison, David K. Jackson, Ian Johnston, Dominic Kwiatkowski, Cordelia Langford, John Sillitoe on behalf of the Wellcome Sanger Institute COVID-19 Surveillance Team |
| EPI_ISL_2907647, EPI_ISL_2907648, EPI_ISL_2907649 | Lighthouse Lab in Milton Keynes | Wellcome Sanger Institute for the COVID-19 Genomics UK (COG-UK) Consortium | The Lighthouse Lab in Milton Keynes and Alex Alderton, Roberto Amato, Jeffrey Barrett, Sonia Goncalves, Ewan Harrison, David K. Jackson, Ian Johnston, Dominic Kwiatkowski, Cordelia Langford, John Sillitoe on behalf of the Wellcome Sanger Institute COVID-19 Surveillance Team |
| EPI_ISL_2907650, EPI_ISL_2907651, EPI_ISL_2907652, EPI_ISL_2907653 | Lighthouse Lab in Alderley Park | Wellcome Sanger Institute for the COVID-19 Genomics UK (COG-UK) Consortium | Jacquelyn Wynn, Mairead Hyland, The Lighthouse Lab in Alderley Park and Alex Alderton, Roberto Amato, Jeffrey Barrett, Sonia Goncalves, Ewan Harrison, David K. Jackson, Ian Johnston, Dominic Kwiatkowski, Cordelia Langford, John Sillitoe on behalf of the Wellcome Sanger Institute COVID-19 Surveillance Team |
| EPI_ISL_2907654 | Lighthouse Lab in Milton Keynes | Wellcome Sanger Institute for the COVID-19 Genomics UK (COG-UK) Consortium | The Lighthouse Lab in Milton Keynes and Alex Alderton, Roberto Amato, Jeffrey Barrett, Sonia Goncalves, Ewan Harrison, David K. Jackson, Ian Johnston, Dominic Kwiatkowski, Cordelia Langford, John Sillitoe on behalf of the Wellcome Sanger Institute COVID-19 Surveillance Team |
| EPI_ISL_2907655 | Lighthouse Lab in Alderley Park | Wellcome Sanger Institute for the COVID-19 Genomics UK (COG-UK) Consortium | Jacquelyn Wynn, Mairead Hyland, The Lighthouse Lab in Alderley Park and Alex Alderton, Roberto Amato, Jeffrey Barrett, Sonia Goncalves, Ewan Harrison, David K. Jackson, Ian Johnston, Dominic Kwiatkowski, Cordelia Langford, John Sillitoe on behalf of the Wellcome Sanger Institute COVID-19 Surveillance Team |

| | | | |
|---|---|---|---|
| EPI_ISL_2907656, EPI_ISL_2907657 | Lighthouse Lab in Milton Keynes | Wellcome Sanger Institute for the COVID-19 Genomics UK (COG-UK) Consortium | The Lighthouse Lab in Milton Keynes and Alex Alderton, Roberto Amato, Jeffrey Barrett, Sonia Goncalves, Ewan Harrison, David K. Jackson, Ian Johnston, Dominic Kwiatkowski, Cordelia Langford, John Sillitoe on behalf of the Wellcome Sanger Institute COVID-19 Surveillance Team |
| EPI_ISL_2907658 | Lighthouse Lab in Alderley Park | Wellcome Sanger Institute for the COVID-19 Genomics UK (COG-UK) Consortium | Jacquelyn Wynn, Mairead Hyland, The Lighthouse Lab in Alderley Park and Alex Alderton, Roberto Amato, Jeffrey Barrett, Sonia Goncalves, Ewan Harrison, David K. Jackson, Ian Johnston, Dominic Kwiatkowski, Cordelia Langford, John Sillitoe on behalf of the Wellcome Sanger Institute COVID-19 Surveillance Team |
| EPI_ISL_2907659, EPI_ISL_2907660 | Lighthouse Lab in Milton Keynes | Wellcome Sanger Institute for the COVID-19 Genomics UK (COG-UK) Consortium | The Lighthouse Lab in Milton Keynes and Alex Alderton, Roberto Amato, Jeffrey Barrett, Sonia Goncalves, Ewan Harrison, David K. Jackson, Ian Johnston, Dominic Kwiatkowski, Cordelia Langford, John Sillitoe on behalf of the Wellcome Sanger Institute COVID-19 Surveillance Team |
| EPI_ISL_2907661 | Lighthouse Lab in Alderley Park | Wellcome Sanger Institute for the COVID-19 Genomics UK (COG-UK) Consortium | Jacquelyn Wynn, Mairead Hyland, The Lighthouse Lab in Alderley Park and Alex Alderton, Roberto Amato, Jeffrey Barrett, Sonia Goncalves, Ewan Harrison, David K. Jackson, Ian Johnston, Dominic Kwiatkowski, Cordelia Langford, John Sillitoe on behalf of the Wellcome Sanger Institute COVID-19 Surveillance Team |
| EPI_ISL_2907662 | Lighthouse Lab in Milton Keynes | Wellcome Sanger Institute for the COVID-19 Genomics UK (COG-UK) Consortium | The Lighthouse Lab in Milton Keynes and Alex Alderton, Roberto Amato, Jeffrey Barrett, Sonia Goncalves, Ewan Harrison, David K. Jackson, Ian Johnston, Dominic Kwiatkowski, Cordelia Langford, John Sillitoe on behalf of the Wellcome Sanger Institute COVID-19 Surveillance Team |
| EPI_ISL_2907663 | Lighthouse Lab in Alderley Park | Wellcome Sanger Institute for the COVID-19 Genomics UK (COG-UK) Consortium | Jacquelyn Wynn, Mairead Hyland, The Lighthouse Lab in Alderley Park and Alex Alderton, Roberto Amato, Jeffrey Barrett, Sonia Goncalves, Ewan Harrison, David K. Jackson, Ian Johnston, Dominic Kwiatkowski, Cordelia Langford, John Sillitoe on behalf of the Wellcome Sanger Institute COVID-19 Surveillance Team |
| EPI_ISL_2907664, EPI_ISL_2907665 | Lighthouse Lab in Milton Keynes | Wellcome Sanger Institute for the COVID-19 Genomics UK (COG-UK) Consortium | The Lighthouse Lab in Milton Keynes and Alex Alderton, Roberto Amato, Jeffrey Barrett, Sonia Goncalves, Ewan Harrison, David K. Jackson, Ian Johnston, Dominic Kwiatkowski, Cordelia Langford, John Sillitoe on behalf of the Wellcome Sanger Institute COVID-19 Surveillance Team |
| EPI_ISL_2907666 | Lighthouse Lab in Alderley Park | Wellcome Sanger Institute for the COVID-19 Genomics UK (COG-UK) Consortium | Jacquelyn Wynn, Mairead Hyland, The Lighthouse Lab in Alderley Park and Alex Alderton, Roberto Amato, Jeffrey Barrett, Sonia Goncalves, Ewan Harrison, David K. Jackson, Ian Johnston, Dominic Kwiatkowski, Cordelia Langford, John Sillitoe on behalf of the Wellcome Sanger Institute COVID-19 Surveillance Team |
| EPI_ISL_2907667 | Lighthouse Lab in Milton Keynes | Wellcome Sanger Institute for the COVID-19 Genomics UK (COG-UK) Consortium | The Lighthouse Lab in Milton Keynes and Alex Alderton, Roberto Amato, Jeffrey Barrett, Sonia Goncalves, Ewan Harrison, David K. Jackson, Ian Johnston, Dominic Kwiatkowski, Cordelia Langford, John Sillitoe on behalf of the Wellcome Sanger Institute COVID-19 Surveillance Team |
| EPI_ISL_2907668 | Lighthouse Lab in Alderley Park | Wellcome Sanger Institute for the COVID-19 Genomics UK (COG-UK) Consortium | Jacquelyn Wynn, Mairead Hyland, The Lighthouse Lab in Alderley Park and Alex Alderton, Roberto Amato, Jeffrey Barrett, Sonia Goncalves, Ewan Harrison, David K. Jackson, Ian Johnston, Dominic Kwiatkowski, Cordelia Langford, John Sillitoe on behalf of the Wellcome Sanger Institute COVID-19 Surveillance Team |
| EPI_ISL_2907669, EPI_ISL_2907670 | Lighthouse Lab in Milton Keynes | Wellcome Sanger Institute for the COVID-19 Genomics UK (COG-UK) Consortium | The Lighthouse Lab in Milton Keynes and Alex Alderton, Roberto Amato, Jeffrey Barrett, Sonia Goncalves, Ewan Harrison, David K. Jackson, Ian Johnston, Dominic Kwiatkowski, Cordelia Langford, John Sillitoe on behalf of the Wellcome Sanger Institute COVID-19 Surveillance Team |
| EPI_ISL_2907671 | Lighthouse Lab in Alderley Park | Wellcome Sanger Institute for the COVID-19 Genomics UK (COG-UK) Consortium | Jacquelyn Wynn, Mairead Hyland, The Lighthouse Lab in Alderley Park and Alex Alderton, Roberto Amato, Jeffrey Barrett, Sonia Goncalves, Ewan Harrison, David K. Jackson, Ian Johnston, Dominic Kwiatkowski, Cordelia Langford, John Sillitoe on behalf of the Wellcome Sanger Institute COVID-19 Surveillance Team |
| EPI_ISL_2907672 | Lighthouse Lab in Milton Keynes | Wellcome Sanger Institute for the COVID-19 Genomics UK (COG-UK) Consortium | The Lighthouse Lab in Milton Keynes and Alex Alderton, Roberto Amato, Jeffrey Barrett, Sonia Goncalves, Ewan Harrison, David K. Jackson, Ian Johnston, Dominic Kwiatkowski, Cordelia Langford, John Sillitoe on behalf of the Wellcome Sanger Institute COVID-19 Surveillance Team |
| EPI_ISL_2907673 | Lighthouse Lab in Alderley Park | Wellcome Sanger Institute for the COVID-19 Genomics UK (COG-UK) Consortium | Jacquelyn Wynn, Mairead Hyland, The Lighthouse Lab in Alderley Park and Alex Alderton, Roberto Amato, Jeffrey Barrett, Sonia Goncalves, Ewan Harrison, David K. Jackson, Ian Johnston, Dominic Kwiatkowski, Cordelia Langford, John Sillitoe on behalf of the Wellcome Sanger Institute COVID-19 Surveillance Team |
| EPI_ISL_2907674, EPI_ISL_2907675 | Lighthouse Lab in Milton Keynes | Wellcome Sanger Institute for the COVID-19 Genomics UK (COG-UK) Consortium | The Lighthouse Lab in Milton Keynes and Alex Alderton, Roberto Amato, Jeffrey Barrett, Sonia Goncalves, Ewan Harrison, David K. Jackson, Ian Johnston, Dominic Kwiatkowski, Cordelia Langford, John Sillitoe on behalf of the Wellcome Sanger Institute COVID-19 Surveillance Team |
| EPI_ISL_2907676 | Lighthouse Lab in Alderley Park | Wellcome Sanger Institute for the COVID-19 Genomics UK (COG-UK) Consortium | Jacquelyn Wynn, Mairead Hyland, The Lighthouse Lab in Alderley Park and Alex Alderton, Roberto Amato, Jeffrey Barrett, Sonia Goncalves, Ewan Harrison, David K. Jackson, Ian Johnston, Dominic Kwiatkowski, Cordelia Langford, John Sillitoe on behalf of the Wellcome Sanger Institute COVID-19 Surveillance Team |
| EPI_ISL_2907677 | Lighthouse Lab in Milton Keynes | Wellcome Sanger Institute for the COVID-19 Genomics UK (COG-UK) Consortium | The Lighthouse Lab in Milton Keynes and Alex Alderton, Roberto Amato, Jeffrey Barrett, Sonia Goncalves, Ewan Harrison, David K. Jackson, Ian Johnston, Dominic Kwiatkowski, Cordelia Langford, John Sillitoe on behalf of the Wellcome Sanger Institute COVID-19 Surveillance Team |
| EPI_ISL_2907678 | Lighthouse Lab in Alderley Park | Wellcome Sanger Institute for the COVID-19 Genomics UK (COG-UK) Consortium | Jacquelyn Wynn, Mairead Hyland, The Lighthouse Lab in Alderley Park and Alex Alderton, Roberto Amato, Jeffrey Barrett, Sonia Goncalves, Ewan Harrison, David K. Jackson, Ian Johnston, Dominic Kwiatkowski, Cordelia Langford, John Sillitoe on behalf of the Wellcome Sanger Institute COVID-19 Surveillance Team |
| EPI_ISL_2907679, EPI_ISL_2907680, EPI_ISL_2907681, EPI_ISL_2907682, EPI_ISL_2907683 | Lighthouse Lab in Milton Keynes | Wellcome Sanger Institute for the COVID-19 Genomics UK (COG-UK) Consortium | The Lighthouse Lab in Milton Keynes and Alex Alderton, Roberto Amato, Jeffrey Barrett, Sonia Goncalves, Ewan Harrison, David K. Jackson, Ian Johnston, Dominic Kwiatkowski, Cordelia Langford, John Sillitoe on behalf of the Wellcome Sanger Institute COVID-19 Surveillance Team |
| EPI_ISL_2907684 | Lighthouse Lab in Alderley Park | Wellcome Sanger Institute for the COVID-19 Genomics UK (COG-UK) Consortium | Jacquelyn Wynn, Mairead Hyland, The Lighthouse Lab in Alderley Park and Alex Alderton, Roberto Amato, Jeffrey Barrett, Sonia Goncalves, Ewan Harrison, David K. Jackson, Ian Johnston, Dominic Kwiatkowski, Cordelia Langford, John Sillitoe on behalf of the Wellcome Sanger Institute COVID-19 Surveillance Team |
| EPI_ISL_2907685 | Lighthouse Lab in Milton Keynes | Wellcome Sanger Institute for the COVID-19 Genomics UK (COG-UK) Consortium | The Lighthouse Lab in Milton Keynes and Alex Alderton, Roberto Amato, Jeffrey Barrett, Sonia Goncalves, Ewan Harrison, David K. Jackson, Ian Johnston, Dominic Kwiatkowski, Cordelia Langford, John Sillitoe on behalf of the Wellcome Sanger Institute COVID-19 Surveillance Team |
| EPI_ISL_2907686 | Lighthouse Lab in Alderley Park | Wellcome Sanger Institute for the COVID-19 Genomics UK (COG-UK) Consortium | Jacquelyn Wynn, Mairead Hyland, The Lighthouse Lab in Alderley Park and Alex Alderton, Roberto Amato, Jeffrey Barrett, Sonia Goncalves, Ewan Harrison, David K. Jackson, Ian Johnston, Dominic Kwiatkowski, Cordelia Langford, John Sillitoe on behalf of the Wellcome Sanger Institute COVID-19 Surveillance Team |
| EPI_ISL_2907687, EPI_ISL_2907688, EPI_ISL_2907689 | Lighthouse Lab in Milton Keynes | Wellcome Sanger Institute for the COVID-19 Genomics UK (COG-UK) Consortium | The Lighthouse Lab in Milton Keynes and Alex Alderton, Roberto Amato, Jeffrey Barrett, Sonia Goncalves, Ewan Harrison, David K. Jackson, Ian Johnston, Dominic Kwiatkowski, Cordelia Langford, John Sillitoe on behalf of the Wellcome Sanger Institute COVID-19 Surveillance Team |
| EPI_ISL_2907690, EPI_ISL_2907691, EPI_ISL_2907692, EPI_ISL_2907693, EPI_ISL_2907694, EPI_ISL_2907695 | Lighthouse Lab in Alderley Park | Wellcome Sanger Institute for the COVID-19 Genomics UK (COG-UK) Consortium | Jacquelyn Wynn, Mairead Hyland, The Lighthouse Lab in Alderley Park and Alex Alderton, Roberto Amato, Jeffrey Barrett, Sonia Goncalves, Ewan Harrison, David K. Jackson, Ian Johnston, Dominic Kwiatkowski, Cordelia Langford, John Sillitoe on behalf of the Wellcome Sanger Institute COVID-19 Surveillance Team |
| EPI_ISL_2907696, EPI_ISL_2907697, EPI_ISL_2907698, EPI_ISL_2907699 | Lighthouse Lab in Milton Keynes | Wellcome Sanger Institute for the COVID-19 Genomics UK (COG-UK) Consortium | The Lighthouse Lab in Milton Keynes and Alex Alderton, Roberto Amato, Jeffrey Barrett, Sonia Goncalves, Ewan Harrison, David K. Jackson, Ian Johnston, Dominic Kwiatkowski, Cordelia Langford, John Sillitoe on behalf of the Wellcome Sanger Institute COVID-19 Surveillance Team |
| EPI_ISL_2907700, EPI_ISL_2907701 | Lighthouse Lab in Alderley Park | Wellcome Sanger Institute for the COVID-19 Genomics UK (COG-UK) Consortium | Jacquelyn Wynn, Mairead Hyland, The Lighthouse Lab in Alderley Park and Alex Alderton, Roberto Amato, Jeffrey Barrett, Sonia Goncalves, Ewan Harrison, David K. Jackson, Ian Johnston, Dominic Kwiatkowski, Cordelia Langford, John Sillitoe on behalf of the Wellcome Sanger Institute COVID-19 Surveillance Team |
| EPI_ISL_2907702 | Lighthouse Lab in Milton Keynes | Wellcome Sanger Institute for the COVID-19 Genomics UK (COG-UK) Consortium | The Lighthouse Lab in Milton Keynes and Alex Alderton, Roberto Amato, Jeffrey Barrett, Sonia Goncalves, Ewan Harrison, David K. Jackson, Ian Johnston, Dominic Kwiatkowski, Cordelia Langford, John Sillitoe on behalf of the Wellcome Sanger Institute COVID-19 Surveillance Team |
| EPI_ISL_2907703 | Lighthouse Lab in Alderley Park | Wellcome Sanger Institute for the COVID-19 Genomics UK (COG-UK) Consortium | Jacquelyn Wynn, Mairead Hyland, The Lighthouse Lab in Alderley Park and Alex Alderton, Roberto Amato, Jeffrey Barrett, Sonia Goncalves, Ewan Harrison, David K. Jackson, Ian Johnston, Dominic Kwiatkowski, Cordelia Langford, John Sillitoe on behalf of the Wellcome Sanger Institute COVID-19 Surveillance Team |
| EPI_ISL_2907704, EPI_ISL_2907705 | Lighthouse Lab in Milton Keynes | Wellcome Sanger Institute for the COVID-19 Genomics UK (COG-UK) Consortium | The Lighthouse Lab in Milton Keynes and Alex Alderton, Roberto Amato, Jeffrey Barrett, Sonia Goncalves, Ewan Harrison, David K. Jackson, Ian Johnston, Dominic Kwiatkowski, Cordelia Langford, John Sillitoe on behalf of the Wellcome Sanger Institute COVID-19 Surveillance Team |
| EPI_ISL_2907706, EPI_ISL_2907707, EPI_ISL_2907708, EPI_ISL_2907709 | Lighthouse Lab in Alderley Park | Wellcome Sanger Institute for the COVID-19 Genomics UK (COG-UK) Consortium | Jacquelyn Wynn, Mairead Hyland, The Lighthouse Lab in Alderley Park and Alex Alderton, Roberto Amato, Jeffrey Barrett, Sonia Goncalves, Ewan Harrison, David K. Jackson, Ian Johnston, Dominic Kwiatkowski, Cordelia Langford, John Sillitoe on behalf of the Wellcome Sanger Institute COVID-19 Surveillance Team |
| EPI_ISL_2907710, EPI_ISL_2907711 | Lighthouse Lab in Milton Keynes | Wellcome Sanger Institute for the COVID-19 Genomics UK (COG-UK) Consortium | The Lighthouse Lab in Milton Keynes and Alex Alderton, Roberto Amato, Jeffrey Barrett, Sonia Goncalves, Ewan Harrison, David K. Jackson, Ian Johnston, Dominic Kwiatkowski, Cordelia Langford, John Sillitoe on behalf of the Wellcome Sanger Institute COVID-19 Surveillance Team |
| EPI_ISL_2907712, EPI_ISL_2907713 | Lighthouse Lab in Alderley Park | Wellcome Sanger Institute for the COVID-19 Genomics UK (COG-UK) Consortium | Jacquelyn Wynn, Mairead Hyland, The Lighthouse Lab in Alderley Park and Alex Alderton, Roberto Amato, Jeffrey Barrett, Sonia Goncalves, Ewan Harrison, David K. Jackson, Ian Johnston, Dominic Kwiatkowski, Cordelia Langford, John Sillitoe on behalf of the Wellcome Sanger Institute COVID-19 Surveillance Team |
| EPI_ISL_2907714 | Lighthouse Lab in Milton Keynes | Wellcome Sanger Institute for the COVID-19 Genomics UK (COG-UK) Consortium | The Lighthouse Lab in Milton Keynes and Alex Alderton, Roberto Amato, Jeffrey Barrett, Sonia Goncalves, Ewan Harrison, David K. Jackson, Ian Johnston, Dominic Kwiatkowski, Cordelia Langford, John Sillitoe on behalf of the Wellcome Sanger Institute COVID-19 Surveillance Team |
| EPI_ISL_2907715 | Lighthouse Lab in Alderley Park | Wellcome Sanger Institute for the COVID-19 Genomics UK (COG-UK) Consortium | Jacquelyn Wynn, Mairead Hyland, The Lighthouse Lab in Alderley Park and Alex Alderton, Roberto Amato, Jeffrey Barrett, Sonia Goncalves, Ewan Harrison, David K. Jackson, Ian Johnston, Dominic Kwiatkowski, Cordelia Langford, John Sillitoe on behalf of the Wellcome Sanger Institute COVID-19 Surveillance Team |
| EPI_ISL_2907716, EPI_ISL_2907717, EPI_ISL_2907718, EPI_ISL_2907719 | Lighthouse Lab in Milton Keynes | Wellcome Sanger Institute for the COVID-19 Genomics UK (COG-UK) Consortium | The Lighthouse Lab in Milton Keynes and Alex Alderton, Roberto Amato, Jeffrey Barrett, Sonia Goncalves, Ewan Harrison, David K. Jackson, Ian Johnston, Dominic Kwiatkowski, Cordelia Langford, John Sillitoe on behalf of the Wellcome Sanger Institute COVID-19 Surveillance Team |
| EPI_ISL_2907720, EPI_ISL_2907721 | Lighthouse Lab in Alderley Park | Wellcome Sanger Institute for the COVID-19 Genomics UK (COG-UK) Consortium | Jacquelyn Wynn, Mairead Hyland, The Lighthouse Lab in Alderley Park and Alex Alderton, Roberto Amato, Jeffrey Barrett, Sonia Goncalves, Ewan Harrison, David K. Jackson, Ian Johnston, Dominic Kwiatkowski, Cordelia Langford, John Sillitoe on behalf of the Wellcome Sanger Institute COVID-19 Surveillance Team |
| EPI_ISL_2907722 | Lighthouse Lab in Milton Keynes | Wellcome Sanger Institute for the COVID-19 Genomics UK (COG-UK) Consortium | The Lighthouse Lab in Milton Keynes and Alex Alderton, Roberto Amato, Jeffrey Barrett, Sonia Goncalves, Ewan Harrison, David K. Jackson, Ian Johnston, Dominic Kwiatkowski, Cordelia Langford, John Sillitoe on behalf of the Wellcome Sanger Institute COVID-19 Surveillance Team |
| EPI_ISL_2907723, EPI_ISL_2907724, EPI_ISL_2907725 | Lighthouse Lab in Alderley Park | Wellcome Sanger Institute for the COVID-19 Genomics UK (COG-UK) Consortium | Jacquelyn Wynn, Mairead Hyland, The Lighthouse Lab in Alderley Park and Alex Alderton, Roberto Amato, Jeffrey Barrett, Sonia Goncalves, Ewan Harrison, David K. Jackson, Ian Johnston, Dominic Kwiatkowski, Cordelia Langford, John Sillitoe on behalf of the Wellcome Sanger Institute COVID-19 Surveillance Team |
| EPI_ISL_2907726, EPI_ISL_2907727 | Lighthouse Lab in Milton Keynes | Wellcome Sanger Institute for the COVID-19 Genomics UK (COG-UK) Consortium | The Lighthouse Lab in Milton Keynes and Alex Alderton, Roberto Amato, Jeffrey Barrett, Sonia Goncalves, Ewan Harrison, David K. Jackson, Ian Johnston, Dominic Kwiatkowski, Cordelia Langford, John Sillitoe on behalf of the Wellcome Sanger Institute COVID-19 Surveillance Team |

| | | | |
|---|---|---|---|
| EPI_ISL_2907728 | Lighthouse Lab in Alderley Park | Wellcome Sanger Institute for the COVID-19 Genomics UK (COG-UK) Consortium | Jacquelyn Wynn, Mairead Hyland, The Lighthouse Lab in Alderley Park and Alex Alderton, Roberto Amato, Jeffrey Barrett, Sonia Goncalves, Ewan Harrison, David K. Jackson, Ian Johnston, Dominic Kwiatkowski, Cordelia Langford, John Sillitoe on behalf of the Wellcome Sanger Institute COVID-19 Surveillance Team |
| EPI_ISL_2907729 | Lighthouse Lab in Milton Keynes | Wellcome Sanger Institute for the COVID-19 Genomics UK (COG-UK) Consortium | The Lighthouse Lab in Milton Keynes and Alex Alderton, Roberto Amato, Jeffrey Barrett, Sonia Goncalves, Ewan Harrison, David K. Jackson, Ian Johnston, Dominic Kwiatkowski, Cordelia Langford, John Sillitoe on behalf of the Wellcome Sanger Institute COVID-19 Surveillance Team |
| EPI_ISL_2907730 | Lighthouse Lab in Alderley Park | Wellcome Sanger Institute for the COVID-19 Genomics UK (COG-UK) Consortium | Jacquelyn Wynn, Mairead Hyland, The Lighthouse Lab in Alderley Park and Alex Alderton, Roberto Amato, Jeffrey Barrett, Sonia Goncalves, Ewan Harrison, David K. Jackson, Ian Johnston, Dominic Kwiatkowski, Cordelia Langford, John Sillitoe on behalf of the Wellcome Sanger Institute COVID-19 Surveillance Team |
| EPI_ISL_2907731 | Lighthouse Lab in Milton Keynes | Wellcome Sanger Institute for the COVID-19 Genomics UK (COG-UK) Consortium | The Lighthouse Lab in Milton Keynes and Alex Alderton, Roberto Amato, Jeffrey Barrett, Sonia Goncalves, Ewan Harrison, David K. Jackson, Ian Johnston, Dominic Kwiatkowski, Cordelia Langford, John Sillitoe on behalf of the Wellcome Sanger Institute COVID-19 Surveillance Team |
| EPI_ISL_2907732, EPI_ISL_2907733 | Lighthouse Lab in Alderley Park | Wellcome Sanger Institute for the COVID-19 Genomics UK (COG-UK) Consortium | Jacquelyn Wynn, Mairead Hyland, The Lighthouse Lab in Alderley Park and Alex Alderton, Roberto Amato, Jeffrey Barrett, Sonia Goncalves, Ewan Harrison, David K. Jackson, Ian Johnston, Dominic Kwiatkowski, Cordelia Langford, John Sillitoe on behalf of the Wellcome Sanger Institute COVID-19 Surveillance Team |
| EPI_ISL_2907734 | Lighthouse Lab in Milton Keynes | Wellcome Sanger Institute for the COVID-19 Genomics UK (COG-UK) Consortium | The Lighthouse Lab in Milton Keynes and Alex Alderton, Roberto Amato, Jeffrey Barrett, Sonia Goncalves, Ewan Harrison, David K. Jackson, Ian Johnston, Dominic Kwiatkowski, Cordelia Langford, John Sillitoe on behalf of the Wellcome Sanger Institute COVID-19 Surveillance Team |
| EPI_ISL_2907735, EPI_ISL_2907736 | Lighthouse Lab in Alderley Park | Wellcome Sanger Institute for the COVID-19 Genomics UK (COG-UK) Consortium | Jacquelyn Wynn, Mairead Hyland, The Lighthouse Lab in Alderley Park and Alex Alderton, Roberto Amato, Jeffrey Barrett, Sonia Goncalves, Ewan Harrison, David K. Jackson, Ian Johnston, Dominic Kwiatkowski, Cordelia Langford, John Sillitoe on behalf of the Wellcome Sanger Institute COVID-19 Surveillance Team |
| EPI_ISL_2907737 | Lighthouse Lab in Milton Keynes | Wellcome Sanger Institute for the COVID-19 Genomics UK (COG-UK) Consortium | The Lighthouse Lab in Milton Keynes and Alex Alderton, Roberto Amato, Jeffrey Barrett, Sonia Goncalves, Ewan Harrison, David K. Jackson, Ian Johnston, Dominic Kwiatkowski, Cordelia Langford, John Sillitoe on behalf of the Wellcome Sanger Institute COVID-19 Surveillance Team |
| EPI_ISL_2907738, EPI_ISL_2907739 | Lighthouse Lab in Alderley Park | Wellcome Sanger Institute for the COVID-19 Genomics UK (COG-UK) Consortium | Jacquelyn Wynn, Mairead Hyland, The Lighthouse Lab in Alderley Park and Alex Alderton, Roberto Amato, Jeffrey Barrett, Sonia Goncalves, Ewan Harrison, David K. Jackson, Ian Johnston, Dominic Kwiatkowski, Cordelia Langford, John Sillitoe on behalf of the Wellcome Sanger Institute COVID-19 Surveillance Team |
| EPI_ISL_2907740, EPI_ISL_2907741 | Lighthouse Lab in Milton Keynes | Wellcome Sanger Institute for the COVID-19 Genomics UK (COG-UK) Consortium | The Lighthouse Lab in Milton Keynes and Alex Alderton, Roberto Amato, Jeffrey Barrett, Sonia Goncalves, Ewan Harrison, David K. Jackson, Ian Johnston, Dominic Kwiatkowski, Cordelia Langford, John Sillitoe on behalf of the Wellcome Sanger Institute COVID-19 Surveillance Team |
| EPI_ISL_2907742, EPI_ISL_2907743 | Lighthouse Lab in Alderley Park | Wellcome Sanger Institute for the COVID-19 Genomics UK (COG-UK) Consortium | Jacquelyn Wynn, Mairead Hyland, The Lighthouse Lab in Alderley Park and Alex Alderton, Roberto Amato, Jeffrey Barrett, Sonia Goncalves, Ewan Harrison, David K. Jackson, Ian Johnston, Dominic Kwiatkowski, Cordelia Langford, John Sillitoe on behalf of the Wellcome Sanger Institute COVID-19 Surveillance Team |
| EPI_ISL_2907744 | Lighthouse Lab in Milton Keynes | Wellcome Sanger Institute for the COVID-19 Genomics UK (COG-UK) Consortium | The Lighthouse Lab in Milton Keynes and Alex Alderton, Roberto Amato, Jeffrey Barrett, Sonia Goncalves, Ewan Harrison, David K. Jackson, Ian Johnston, Dominic Kwiatkowski, Cordelia Langford, John Sillitoe on behalf of the Wellcome Sanger Institute COVID-19 Surveillance Team |
| EPI_ISL_2907745, EPI_ISL_2907746, EPI_ISL_2907747, EPI_ISL_2907748, EPI_ISL_2907749, EPI_ISL_2907750, EPI_ISL_2907751, EPI_ISL_2907752 | Lighthouse Lab in Alderley Park | Wellcome Sanger Institute for the COVID-19 Genomics UK (COG-UK) Consortium | Jacquelyn Wynn, Mairead Hyland, The Lighthouse Lab in Alderley Park and Alex Alderton, Roberto Amato, Jeffrey Barrett, Sonia Goncalves, Ewan Harrison, David K. Jackson, Ian Johnston, Dominic Kwiatkowski, Cordelia Langford, John Sillitoe on behalf of the Wellcome Sanger Institute COVID-19 Surveillance Team |
| EPI_ISL_2907753 | Lighthouse Lab in Milton Keynes | Wellcome Sanger Institute for the COVID-19 Genomics UK (COG-UK) Consortium | The Lighthouse Lab in Milton Keynes and Alex Alderton, Roberto Amato, Jeffrey Barrett, Sonia Goncalves, Ewan Harrison, David K. Jackson, Ian Johnston, Dominic Kwiatkowski, Cordelia Langford, John Sillitoe on behalf of the Wellcome Sanger Institute COVID-19 Surveillance Team |
| EPI_ISL_2907754, EPI_ISL_2907755, EPI_ISL_2907756, EPI_ISL_2907757, EPI_ISL_2907758 | Lighthouse Lab in Alderley Park | Wellcome Sanger Institute for the COVID-19 Genomics UK (COG-UK) Consortium | Jacquelyn Wynn, Mairead Hyland, The Lighthouse Lab in Alderley Park and Alex Alderton, Roberto Amato, Jeffrey Barrett, Sonia Goncalves, Ewan Harrison, David K. Jackson, Ian Johnston, Dominic Kwiatkowski, Cordelia Langford, John Sillitoe on behalf of the Wellcome Sanger Institute COVID-19 Surveillance Team |
| EPI_ISL_2907759 | Lighthouse Lab in Milton Keynes | Wellcome Sanger Institute for the COVID-19 Genomics UK (COG-UK) Consortium | The Lighthouse Lab in Milton Keynes and Alex Alderton, Roberto Amato, Jeffrey Barrett, Sonia Goncalves, Ewan Harrison, David K. Jackson, Ian Johnston, Dominic Kwiatkowski, Cordelia Langford, John Sillitoe on behalf of the Wellcome Sanger Institute COVID-19 Surveillance Team |
| EPI_ISL_2907760 | Lighthouse Lab in Alderley Park | Wellcome Sanger Institute for the COVID-19 Genomics UK (COG-UK) Consortium | Jacquelyn Wynn, Mairead Hyland, The Lighthouse Lab in Alderley Park and Alex Alderton, Roberto Amato, Jeffrey Barrett, Sonia Goncalves, Ewan Harrison, David K. Jackson, Ian Johnston, Dominic Kwiatkowski, Cordelia Langford, John Sillitoe on behalf of the Wellcome Sanger Institute COVID-19 Surveillance Team |
| EPI_ISL_2907761, EPI_ISL_2907762, EPI_ISL_2907763, EPI_ISL_2907764, EPI_ISL_2907765, EPI_ISL_2907766 | Lighthouse Lab in Milton Keynes | Wellcome Sanger Institute for the COVID-19 Genomics UK (COG-UK) Consortium | The Lighthouse Lab in Milton Keynes and Alex Alderton, Roberto Amato, Jeffrey Barrett, Sonia Goncalves, Ewan Harrison, David K. Jackson, Ian Johnston, Dominic Kwiatkowski, Cordelia Langford, John Sillitoe on behalf of the Wellcome Sanger Institute COVID-19 Surveillance Team |
| EPI_ISL_2907767, EPI_ISL_2907768 | Lighthouse Lab in Alderley Park | Wellcome Sanger Institute for the COVID-19 Genomics UK (COG-UK) Consortium | Jacquelyn Wynn, Mairead Hyland, The Lighthouse Lab in Alderley Park and Alex Alderton, Roberto Amato, Jeffrey Barrett, Sonia Goncalves, Ewan Harrison, David K. Jackson, Ian Johnston, Dominic Kwiatkowski, Cordelia Langford, John Sillitoe on behalf of the Wellcome Sanger Institute COVID-19 Surveillance Team |
| EPI_ISL_2907769, EPI_ISL_2907770 | Lighthouse Lab in Milton Keynes | Wellcome Sanger Institute for the COVID-19 Genomics UK (COG-UK) Consortium | The Lighthouse Lab in Milton Keynes and Alex Alderton, Roberto Amato, Jeffrey Barrett, Sonia Goncalves, Ewan Harrison, David K. Jackson, Ian Johnston, Dominic Kwiatkowski, Cordelia Langford, John Sillitoe on behalf of the Wellcome Sanger Institute COVID-19 Surveillance Team |
| EPI_ISL_2907771 | Lighthouse Lab in Alderley Park | Wellcome Sanger Institute for the COVID-19 Genomics UK (COG-UK) Consortium | Jacquelyn Wynn, Mairead Hyland, The Lighthouse Lab in Alderley Park and Alex Alderton, Roberto Amato, Jeffrey Barrett, Sonia Goncalves, Ewan Harrison, David K. Jackson, Ian Johnston, Dominic Kwiatkowski, Cordelia Langford, John Sillitoe on behalf of the Wellcome Sanger Institute COVID-19 Surveillance Team |
| EPI_ISL_2907772, EPI_ISL_2907773, EPI_ISL_2907774 | Lighthouse Lab in Milton Keynes | Wellcome Sanger Institute for the COVID-19 Genomics UK (COG-UK) Consortium | The Lighthouse Lab in Milton Keynes and Alex Alderton, Roberto Amato, Jeffrey Barrett, Sonia Goncalves, Ewan Harrison, David K. Jackson, Ian Johnston, Dominic Kwiatkowski, Cordelia Langford, John Sillitoe on behalf of the Wellcome Sanger Institute COVID-19 Surveillance Team |
| EPI_ISL_2907775 | Lighthouse Lab in Alderley Park | Wellcome Sanger Institute for the COVID-19 Genomics UK (COG-UK) Consortium | Jacquelyn Wynn, Mairead Hyland, The Lighthouse Lab in Alderley Park and Alex Alderton, Roberto Amato, Jeffrey Barrett, Sonia Goncalves, Ewan Harrison, David K. Jackson, Ian Johnston, Dominic Kwiatkowski, Cordelia Langford, John Sillitoe on behalf of the Wellcome Sanger Institute COVID-19 Surveillance Team |
| EPI_ISL_2907776 | Lighthouse Lab in Milton Keynes | Wellcome Sanger Institute for the COVID-19 Genomics UK (COG-UK) Consortium | The Lighthouse Lab in Milton Keynes and Alex Alderton, Roberto Amato, Jeffrey Barrett, Sonia Goncalves, Ewan Harrison, David K. Jackson, Ian Johnston, Dominic Kwiatkowski, Cordelia Langford, John Sillitoe on behalf of the Wellcome Sanger Institute COVID-19 Surveillance Team |
| EPI_ISL_2907777, EPI_ISL_2907778, EPI_ISL_2907779, EPI_ISL_2907780, EPI_ISL_2907781, EPI_ISL_2907782, EPI_ISL_2907783, EPI_ISL_2907784 | Lighthouse Lab in Alderley Park | Wellcome Sanger Institute for the COVID-19 Genomics UK (COG-UK) Consortium | Jacquelyn Wynn, Mairead Hyland, The Lighthouse Lab in Alderley Park and Alex Alderton, Roberto Amato, Jeffrey Barrett, Sonia Goncalves, Ewan Harrison, David K. Jackson, Ian Johnston, Dominic Kwiatkowski, Cordelia Langford, John Sillitoe on behalf of the Wellcome Sanger Institute COVID-19 Surveillance Team |
| EPI_ISL_2907785, EPI_ISL_2907786 | Lighthouse Lab in Milton Keynes | Wellcome Sanger Institute for the COVID-19 Genomics UK (COG-UK) Consortium | The Lighthouse Lab in Milton Keynes and Alex Alderton, Roberto Amato, Jeffrey Barrett, Sonia Goncalves, Ewan Harrison, David K. Jackson, Ian Johnston, Dominic Kwiatkowski, Cordelia Langford, John Sillitoe on behalf of the Wellcome Sanger Institute COVID-19 Surveillance Team |
| EPI_ISL_2907787 | Lighthouse Lab in Alderley Park | Wellcome Sanger Institute for the COVID-19 Genomics UK (COG-UK) Consortium | Jacquelyn Wynn, Mairead Hyland, The Lighthouse Lab in Alderley Park and Alex Alderton, Roberto Amato, Jeffrey Barrett, Sonia Goncalves, Ewan Harrison, David K. Jackson, Ian Johnston, Dominic Kwiatkowski, Cordelia Langford, John Sillitoe on behalf of the Wellcome Sanger Institute COVID-19 Surveillance Team |
| EPI_ISL_2907788 | Lighthouse Lab in Milton Keynes | Wellcome Sanger Institute for the COVID-19 Genomics UK (COG-UK) Consortium | The Lighthouse Lab in Milton Keynes and Alex Alderton, Roberto Amato, Jeffrey Barrett, Sonia Goncalves, Ewan Harrison, David K. Jackson, Ian Johnston, Dominic Kwiatkowski, Cordelia Langford, John Sillitoe on behalf of the Wellcome Sanger Institute COVID-19 Surveillance Team |
| EPI_ISL_2907789 | Lighthouse Lab in Alderley Park | Wellcome Sanger Institute for the COVID-19 Genomics UK (COG-UK) Consortium | Jacquelyn Wynn, Mairead Hyland, The Lighthouse Lab in Alderley Park and Alex Alderton, Roberto Amato, Jeffrey Barrett, Sonia Goncalves, Ewan Harrison, David K. Jackson, Ian Johnston, Dominic Kwiatkowski, Cordelia Langford, John Sillitoe on behalf of the Wellcome Sanger Institute COVID-19 Surveillance Team |
| EPI_ISL_2907790, EPI_ISL_2907791, EPI_ISL_2907792, EPI_ISL_2907793 | Lighthouse Lab in Milton Keynes | Wellcome Sanger Institute for the COVID-19 Genomics UK (COG-UK) Consortium | The Lighthouse Lab in Milton Keynes and Alex Alderton, Roberto Amato, Jeffrey Barrett, Sonia Goncalves, Ewan Harrison, David K. Jackson, Ian Johnston, Dominic Kwiatkowski, Cordelia Langford, John Sillitoe on behalf of the Wellcome Sanger Institute COVID-19 Surveillance Team |
| EPI_ISL_2907794 | Lighthouse Lab in Alderley Park | Wellcome Sanger Institute for the COVID-19 Genomics UK (COG-UK) Consortium | Jacquelyn Wynn, Mairead Hyland, The Lighthouse Lab in Alderley Park and Alex Alderton, Roberto Amato, Jeffrey Barrett, Sonia Goncalves, Ewan Harrison, David K. Jackson, Ian Johnston, Dominic Kwiatkowski, Cordelia Langford, John Sillitoe on behalf of the Wellcome Sanger Institute COVID-19 Surveillance Team |
| EPI_ISL_2907795 | Lighthouse Lab in Milton Keynes | Wellcome Sanger Institute for the COVID-19 Genomics UK (COG-UK) Consortium | The Lighthouse Lab in Milton Keynes and Alex Alderton, Roberto Amato, Jeffrey Barrett, Sonia Goncalves, Ewan Harrison, David K. Jackson, Ian Johnston, Dominic Kwiatkowski, Cordelia Langford, John Sillitoe on behalf of the Wellcome Sanger Institute COVID-19 Surveillance Team |
| EPI_ISL_2907796 | Lighthouse Lab in Alderley Park | Wellcome Sanger Institute for the COVID-19 Genomics UK (COG-UK) Consortium | Jacquelyn Wynn, Mairead Hyland, The Lighthouse Lab in Alderley Park and Alex Alderton, Roberto Amato, Jeffrey Barrett, Sonia Goncalves, Ewan Harrison, David K. Jackson, Ian Johnston, Dominic Kwiatkowski, Cordelia Langford, John Sillitoe on behalf of the Wellcome Sanger Institute COVID-19 Surveillance Team |
| EPI_ISL_2907797 | Lighthouse Lab in Milton Keynes | Wellcome Sanger Institute for the COVID-19 Genomics UK (COG-UK) Consortium | The Lighthouse Lab in Milton Keynes and Alex Alderton, Roberto Amato, Jeffrey Barrett, Sonia Goncalves, Ewan Harrison, David K. Jackson, Ian Johnston, Dominic Kwiatkowski, Cordelia Langford, John Sillitoe on behalf of the Wellcome Sanger Institute COVID-19 Surveillance Team |
| EPI_ISL_2907798, EPI_ISL_2907799, EPI_ISL_2907800 | Lighthouse Lab in Alderley Park | Wellcome Sanger Institute for the COVID-19 Genomics UK (COG-UK) Consortium | Jacquelyn Wynn, Mairead Hyland, The Lighthouse Lab in Alderley Park and Alex Alderton, Roberto Amato, Jeffrey Barrett, Sonia Goncalves, Ewan Harrison, David K. Jackson, Ian Johnston, Dominic Kwiatkowski, Cordelia Langford, John Sillitoe on behalf of the Wellcome Sanger Institute COVID-19 Surveillance Team |
| EPI_ISL_2907801, | Lighthouse Lab in Milton Keynes | Wellcome Sanger Institute for the COVID-19 Genomics UK (COG-UK) | The Lighthouse Lab in Milton Keynes and Alex Alderton, Roberto Amato, Jeffrey Barrett, Sonia Goncalves, Ewan Harrison, David K. Jackson, Ian Johnston, Dominic Kwiatkowski, Cordelia Langford, John Sillitoe on behalf of the Wellcome Sanger |

| | | | |
|---|---|---|---|
| EPI_ISL_2907802 | | Consortium | Institute COVID-19 Surveillance Team |
| EPI_ISL_2907803, EPI_ISL_2907804, EPI_ISL_2907805, EPI_ISL_2907806, EPI_ISL_2907807 | Lighthouse Lab in Alderley Park | Wellcome Sanger Institute for the COVID-19 Genomics UK (COG-UK) Consortium | Jacquelyn Wynn, Mairead Hyland, The Lighthouse Lab in Alderley Park and Alex Alderton, Roberto Amato, Jeffrey Barrett, Sonia Goncalves, Ewan Harrison, David K. Jackson, Ian Johnston, Dominic Kwiatkowski, Cordelia Langford, John Sillitoe on behalf of the Wellcome Sanger Institute COVID-19 Surveillance Team |
| EPI_ISL_2907808, EPI_ISL_2907809, EPI_ISL_2907810 | Lighthouse Lab in Milton Keynes | Wellcome Sanger Institute for the COVID-19 Genomics UK (COG-UK) Consortium | The Lighthouse Lab in Milton Keynes and Alex Alderton, Roberto Amato, Jeffrey Barrett, Sonia Goncalves, Ewan Harrison, David K. Jackson, Ian Johnston, Dominic Kwiatkowski, Cordelia Langford, John Sillitoe on behalf of the Wellcome Sanger Institute COVID-19 Surveillance Team |
| EPI_ISL_2907811, EPI_ISL_2907812, EPI_ISL_2907813 | Lighthouse Lab in Alderley Park | Wellcome Sanger Institute for the COVID-19 Genomics UK (COG-UK) Consortium | Jacquelyn Wynn, Mairead Hyland, The Lighthouse Lab in Alderley Park and Alex Alderton, Roberto Amato, Jeffrey Barrett, Sonia Goncalves, Ewan Harrison, David K. Jackson, Ian Johnston, Dominic Kwiatkowski, Cordelia Langford, John Sillitoe on behalf of the Wellcome Sanger Institute COVID-19 Surveillance Team |
| EPI_ISL_2907814, EPI_ISL_2907815, EPI_ISL_2907816, EPI_ISL_2907817, EPI_ISL_2907818, EPI_ISL_2907819 | Lighthouse Lab in Milton Keynes | Wellcome Sanger Institute for the COVID-19 Genomics UK (COG-UK) Consortium | The Lighthouse Lab in Milton Keynes and Alex Alderton, Roberto Amato, Jeffrey Barrett, Sonia Goncalves, Ewan Harrison, David K. Jackson, Ian Johnston, Dominic Kwiatkowski, Cordelia Langford, John Sillitoe on behalf of the Wellcome Sanger Institute COVID-19 Surveillance Team |
| EPI_ISL_2907820, EPI_ISL_2907821 | Lighthouse Lab in Alderley Park | Wellcome Sanger Institute for the COVID-19 Genomics UK (COG-UK) Consortium | Jacquelyn Wynn, Mairead Hyland, The Lighthouse Lab in Alderley Park and Alex Alderton, Roberto Amato, Jeffrey Barrett, Sonia Goncalves, Ewan Harrison, David K. Jackson, Ian Johnston, Dominic Kwiatkowski, Cordelia Langford, John Sillitoe on behalf of the Wellcome Sanger Institute COVID-19 Surveillance Team |
| EPI_ISL_2907822 | Lighthouse Lab in Milton Keynes | Wellcome Sanger Institute for the COVID-19 Genomics UK (COG-UK) Consortium | The Lighthouse Lab in Milton Keynes and Alex Alderton, Roberto Amato, Jeffrey Barrett, Sonia Goncalves, Ewan Harrison, David K. Jackson, Ian Johnston, Dominic Kwiatkowski, Cordelia Langford, John Sillitoe on behalf of the Wellcome Sanger Institute COVID-19 Surveillance Team |
| EPI_ISL_2907823 | Lighthouse Lab in Alderley Park | Wellcome Sanger Institute for the COVID-19 Genomics UK (COG-UK) Consortium | Jacquelyn Wynn, Mairead Hyland, The Lighthouse Lab in Alderley Park and Alex Alderton, Roberto Amato, Jeffrey Barrett, Sonia Goncalves, Ewan Harrison, David K. Jackson, Ian Johnston, Dominic Kwiatkowski, Cordelia Langford, John Sillitoe on behalf of the Wellcome Sanger Institute COVID-19 Surveillance Team |
| EPI_ISL_2907824 | Lighthouse Lab in Milton Keynes | Wellcome Sanger Institute for the COVID-19 Genomics UK (COG-UK) Consortium | The Lighthouse Lab in Milton Keynes and Alex Alderton, Roberto Amato, Jeffrey Barrett, Sonia Goncalves, Ewan Harrison, David K. Jackson, Ian Johnston, Dominic Kwiatkowski, Cordelia Langford, John Sillitoe on behalf of the Wellcome Sanger Institute COVID-19 Surveillance Team |
| EPI_ISL_2907825 | Lighthouse Lab in Alderley Park | Wellcome Sanger Institute for the COVID-19 Genomics UK (COG-UK) Consortium | Jacquelyn Wynn, Mairead Hyland, The Lighthouse Lab in Alderley Park and Alex Alderton, Roberto Amato, Jeffrey Barrett, Sonia Goncalves, Ewan Harrison, David K. Jackson, Ian Johnston, Dominic Kwiatkowski, Cordelia Langford, John Sillitoe on behalf of the Wellcome Sanger Institute COVID-19 Surveillance Team |
| EPI_ISL_2907826, EPI_ISL_2907827, EPI_ISL_2907828 | Lighthouse Lab in Milton Keynes | Wellcome Sanger Institute for the COVID-19 Genomics UK (COG-UK) Consortium | The Lighthouse Lab in Milton Keynes and Alex Alderton, Roberto Amato, Jeffrey Barrett, Sonia Goncalves, Ewan Harrison, David K. Jackson, Ian Johnston, Dominic Kwiatkowski, Cordelia Langford, John Sillitoe on behalf of the Wellcome Sanger Institute COVID-19 Surveillance Team |
| EPI_ISL_2907829 | Lighthouse Lab in Alderley Park | Wellcome Sanger Institute for the COVID-19 Genomics UK (COG-UK) Consortium | Jacquelyn Wynn, Mairead Hyland, The Lighthouse Lab in Alderley Park and Alex Alderton, Roberto Amato, Jeffrey Barrett, Sonia Goncalves, Ewan Harrison, David K. Jackson, Ian Johnston, Dominic Kwiatkowski, Cordelia Langford, John Sillitoe on behalf of the Wellcome Sanger Institute COVID-19 Surveillance Team |
| EPI_ISL_2907830, EPI_ISL_2907831, EPI_ISL_2907832 | Lighthouse Lab in Milton Keynes | Wellcome Sanger Institute for the COVID-19 Genomics UK (COG-UK) Consortium | The Lighthouse Lab in Milton Keynes and Alex Alderton, Roberto Amato, Jeffrey Barrett, Sonia Goncalves, Ewan Harrison, David K. Jackson, Ian Johnston, Dominic Kwiatkowski, Cordelia Langford, John Sillitoe on behalf of the Wellcome Sanger Institute COVID-19 Surveillance Team |
| EPI_ISL_2907833, EPI_ISL_2907834, EPI_ISL_2907835, EPI_ISL_2907836, EPI_ISL_2907837, EPI_ISL_2907838, EPI_ISL_2907839 | Lighthouse Lab in Alderley Park | Wellcome Sanger Institute for the COVID-19 Genomics UK (COG-UK) Consortium | Jacquelyn Wynn, Mairead Hyland, The Lighthouse Lab in Alderley Park and Alex Alderton, Roberto Amato, Jeffrey Barrett, Sonia Goncalves, Ewan Harrison, David K. Jackson, Ian Johnston, Dominic Kwiatkowski, Cordelia Langford, John Sillitoe on behalf of the Wellcome Sanger Institute COVID-19 Surveillance Team |
| EPI_ISL_2907840, EPI_ISL_2907841, EPI_ISL_2907842 | Lighthouse Lab in Milton Keynes | Wellcome Sanger Institute for the COVID-19 Genomics UK (COG-UK) Consortium | The Lighthouse Lab in Milton Keynes and Alex Alderton, Roberto Amato, Jeffrey Barrett, Sonia Goncalves, Ewan Harrison, David K. Jackson, Ian Johnston, Dominic Kwiatkowski, Cordelia Langford, John Sillitoe on behalf of the Wellcome Sanger Institute COVID-19 Surveillance Team |
| EPI_ISL_2907843, EPI_ISL_2907844 | Lighthouse Lab in Alderley Park | Wellcome Sanger Institute for the COVID-19 Genomics UK (COG-UK) Consortium | Jacquelyn Wynn, Mairead Hyland, The Lighthouse Lab in Alderley Park and Alex Alderton, Roberto Amato, Jeffrey Barrett, Sonia Goncalves, Ewan Harrison, David K. Jackson, Ian Johnston, Dominic Kwiatkowski, Cordelia Langford, John Sillitoe on behalf of the Wellcome Sanger Institute COVID-19 Surveillance Team |
| EPI_ISL_2907845, EPI_ISL_2907846, EPI_ISL_2907847 | Lighthouse Lab in Milton Keynes | Wellcome Sanger Institute for the COVID-19 Genomics UK (COG-UK) Consortium | The Lighthouse Lab in Milton Keynes and Alex Alderton, Roberto Amato, Jeffrey Barrett, Sonia Goncalves, Ewan Harrison, David K. Jackson, Ian Johnston, Dominic Kwiatkowski, Cordelia Langford, John Sillitoe on behalf of the Wellcome Sanger Institute COVID-19 Surveillance Team |
| EPI_ISL_2907848, EPI_ISL_2907849, EPI_ISL_2907850, EPI_ISL_2907851, EPI_ISL_2907852, EPI_ISL_2907853, EPI_ISL_2907854, EPI_ISL_2907855, EPI_ISL_2907856 | Lighthouse Lab in Alderley Park | Wellcome Sanger Institute for the COVID-19 Genomics UK (COG-UK) Consortium | Jacquelyn Wynn, Mairead Hyland, The Lighthouse Lab in Alderley Park and Alex Alderton, Roberto Amato, Jeffrey Barrett, Sonia Goncalves, Ewan Harrison, David K. Jackson, Ian Johnston, Dominic Kwiatkowski, Cordelia Langford, John Sillitoe on behalf of the Wellcome Sanger Institute COVID-19 Surveillance Team |
| EPI_ISL_2907857 | Lighthouse Lab in Milton Keynes | Wellcome Sanger Institute for the COVID-19 Genomics UK (COG-UK) Consortium | The Lighthouse Lab in Milton Keynes and Alex Alderton, Roberto Amato, Jeffrey Barrett, Sonia Goncalves, Ewan Harrison, David K. Jackson, Ian Johnston, Dominic Kwiatkowski, Cordelia Langford, John Sillitoe on behalf of the Wellcome Sanger Institute COVID-19 Surveillance Team |
| EPI_ISL_2907858 | Lighthouse Lab in Alderley Park | Wellcome Sanger Institute for the COVID-19 Genomics UK (COG-UK) Consortium | Jacquelyn Wynn, Mairead Hyland, The Lighthouse Lab in Alderley Park and Alex Alderton, Roberto Amato, Jeffrey Barrett, Sonia Goncalves, Ewan Harrison, David K. Jackson, Ian Johnston, Dominic Kwiatkowski, Cordelia Langford, John Sillitoe on behalf of the Wellcome Sanger Institute COVID-19 Surveillance Team |
| EPI_ISL_2907859 | Lighthouse Lab in Milton Keynes | Wellcome Sanger Institute for the COVID-19 Genomics UK (COG-UK) Consortium | The Lighthouse Lab in Milton Keynes and Alex Alderton, Roberto Amato, Jeffrey Barrett, Sonia Goncalves, Ewan Harrison, David K. Jackson, Ian Johnston, Dominic Kwiatkowski, Cordelia Langford, John Sillitoe on behalf of the Wellcome Sanger Institute COVID-19 Surveillance Team |
| EPI_ISL_2907860, EPI_ISL_2907861, EPI_ISL_2907862 | Lighthouse Lab in Alderley Park | Wellcome Sanger Institute for the COVID-19 Genomics UK (COG-UK) Consortium | Jacquelyn Wynn, Mairead Hyland, The Lighthouse Lab in Alderley Park and Alex Alderton, Roberto Amato, Jeffrey Barrett, Sonia Goncalves, Ewan Harrison, David K. Jackson, Ian Johnston, Dominic Kwiatkowski, Cordelia Langford, John Sillitoe on behalf of the Wellcome Sanger Institute COVID-19 Surveillance Team |
| EPI_ISL_2907863, EPI_ISL_2907864, EPI_ISL_2907865 | Lighthouse Lab in Milton Keynes | Wellcome Sanger Institute for the COVID-19 Genomics UK (COG-UK) Consortium | The Lighthouse Lab in Milton Keynes and Alex Alderton, Roberto Amato, Jeffrey Barrett, Sonia Goncalves, Ewan Harrison, David K. Jackson, Ian Johnston, Dominic Kwiatkowski, Cordelia Langford, John Sillitoe on behalf of the Wellcome Sanger Institute COVID-19 Surveillance Team |
| EPI_ISL_2907866, EPI_ISL_2907867 | Lighthouse Lab in Alderley Park | Wellcome Sanger Institute for the COVID-19 Genomics UK (COG-UK) Consortium | Jacquelyn Wynn, Mairead Hyland, The Lighthouse Lab in Alderley Park and Alex Alderton, Roberto Amato, Jeffrey Barrett, Sonia Goncalves, Ewan Harrison, David K. Jackson, Ian Johnston, Dominic Kwiatkowski, Cordelia Langford, John Sillitoe on behalf of the Wellcome Sanger Institute COVID-19 Surveillance Team |
| EPI_ISL_2907868, EPI_ISL_2907869, EPI_ISL_2907870, EPI_ISL_2907871 | Lighthouse Lab in Milton Keynes | Wellcome Sanger Institute for the COVID-19 Genomics UK (COG-UK) Consortium | The Lighthouse Lab in Milton Keynes and Alex Alderton, Roberto Amato, Jeffrey Barrett, Sonia Goncalves, Ewan Harrison, David K. Jackson, Ian Johnston, Dominic Kwiatkowski, Cordelia Langford, John Sillitoe on behalf of the Wellcome Sanger Institute COVID-19 Surveillance Team |
| EPI_ISL_2907872, EPI_ISL_2907873 | Lighthouse Lab in Alderley Park | Wellcome Sanger Institute for the COVID-19 Genomics UK (COG-UK) Consortium | Jacquelyn Wynn, Mairead Hyland, The Lighthouse Lab in Alderley Park and Alex Alderton, Roberto Amato, Jeffrey Barrett, Sonia Goncalves, Ewan Harrison, David K. Jackson, Ian Johnston, Dominic Kwiatkowski, Cordelia Langford, John Sillitoe on behalf of the Wellcome Sanger Institute COVID-19 Surveillance Team |
| EPI_ISL_2907874 | Lighthouse Lab in Milton Keynes | Wellcome Sanger Institute for the COVID-19 Genomics UK (COG-UK) Consortium | The Lighthouse Lab in Milton Keynes and Alex Alderton, Roberto Amato, Jeffrey Barrett, Sonia Goncalves, Ewan Harrison, David K. Jackson, Ian Johnston, Dominic Kwiatkowski, Cordelia Langford, John Sillitoe on behalf of the Wellcome Sanger Institute COVID-19 Surveillance Team |
| EPI_ISL_2907875 | Lighthouse Lab in Alderley Park | Wellcome Sanger Institute for the COVID-19 Genomics UK (COG-UK) Consortium | Jacquelyn Wynn, Mairead Hyland, The Lighthouse Lab in Alderley Park and Alex Alderton, Roberto Amato, Jeffrey Barrett, Sonia Goncalves, Ewan Harrison, David K. Jackson, Ian Johnston, Dominic Kwiatkowski, Cordelia Langford, John Sillitoe on behalf of the Wellcome Sanger Institute COVID-19 Surveillance Team |
| EPI_ISL_2907876 | Lighthouse Lab in Milton Keynes | Wellcome Sanger Institute for the COVID-19 Genomics UK (COG-UK) Consortium | The Lighthouse Lab in Milton Keynes and Alex Alderton, Roberto Amato, Jeffrey Barrett, Sonia Goncalves, Ewan Harrison, David K. Jackson, Ian Johnston, Dominic Kwiatkowski, Cordelia Langford, John Sillitoe on behalf of the Wellcome Sanger Institute COVID-19 Surveillance Team |
| EPI_ISL_2907877 | Lighthouse Lab in Alderley Park | Wellcome Sanger Institute for the COVID-19 Genomics UK (COG-UK) Consortium | Jacquelyn Wynn, Mairead Hyland, The Lighthouse Lab in Alderley Park and Alex Alderton, Roberto Amato, Jeffrey Barrett, Sonia Goncalves, Ewan Harrison, David K. Jackson, Ian Johnston, Dominic Kwiatkowski, Cordelia Langford, John Sillitoe on behalf of the Wellcome Sanger Institute COVID-19 Surveillance Team |
| EPI_ISL_2907878, EPI_ISL_2907879 | Lighthouse Lab in Milton Keynes | Wellcome Sanger Institute for the COVID-19 Genomics UK (COG-UK) Consortium | The Lighthouse Lab in Milton Keynes and Alex Alderton, Roberto Amato, Jeffrey Barrett, Sonia Goncalves, Ewan Harrison, David K. Jackson, Ian Johnston, Dominic Kwiatkowski, Cordelia Langford, John Sillitoe on behalf of the Wellcome Sanger Institute COVID-19 Surveillance Team |
| EPI_ISL_2907880 | Lighthouse Lab in Alderley Park | Wellcome Sanger Institute for the COVID-19 Genomics UK (COG-UK) Consortium | Jacquelyn Wynn, Mairead Hyland, The Lighthouse Lab in Alderley Park and Alex Alderton, Roberto Amato, Jeffrey Barrett, Sonia Goncalves, Ewan Harrison, David K. Jackson, Ian Johnston, Dominic Kwiatkowski, Cordelia Langford, John Sillitoe on behalf of the Wellcome Sanger Institute COVID-19 Surveillance Team |
| EPI_ISL_2907881 | Lighthouse Lab in Milton Keynes | Wellcome Sanger Institute for the COVID-19 Genomics UK (COG-UK) Consortium | The Lighthouse Lab in Milton Keynes and Alex Alderton, Roberto Amato, Jeffrey Barrett, Sonia Goncalves, Ewan Harrison, David K. Jackson, Ian Johnston, Dominic Kwiatkowski, Cordelia Langford, John Sillitoe on behalf of the Wellcome Sanger Institute COVID-19 Surveillance Team |
| EPI_ISL_2907882, | Lighthouse Lab in Alderley Park | Wellcome Sanger Institute for the COVID-19 Genomics UK (COG-UK) | Jacquelyn Wynn, Mairead Hyland, The Lighthouse Lab in Alderley Park and Alex Alderton, Roberto Amato, Jeffrey Barrett, Sonia Goncalves, Ewan Harrison, David K. Jackson, Ian Johnston, Dominic Kwiatkowski, Cordelia Langford, John Sillitoe on |

| | | | |
|---|---|---|---|
| EPI_ISL_2907883, EPI_ISL_2907884 | | Consortium | behalf of the Wellcome Sanger Institute COVID-19 Surveillance Team |
| EPI_ISL_2907885, EPI_ISL_2907886, EPI_ISL_2907887 | Lighthouse Lab in Milton Keynes | Wellcome Sanger Institute for the COVID-19 Genomics UK (COG-UK) Consortium | The Lighthouse Lab in Milton Keynes and Alex Alderton, Roberto Amato, Jeffrey Barrett, Sonia Goncalves, Ewan Harrison, David K. Jackson, Ian Johnston, Dominic Kwiatkowski, Cordelia Langford, John Sillitoe on behalf of the Wellcome Sanger Institute COVID-19 Surveillance Team |
| EPI_ISL_2907888 | Lighthouse Lab in Alderley Park | Wellcome Sanger Institute for the COVID-19 Genomics UK (COG-UK) Consortium | Jacquelyn Wynn, Mairead Hyland, The Lighthouse Lab in Alderley Park and Alex Alderton, Roberto Amato, Jeffrey Barrett, Sonia Goncalves, Ewan Harrison, David K. Jackson, Ian Johnston, Dominic Kwiatkowski, Cordelia Langford, John Sillitoe on behalf of the Wellcome Sanger Institute COVID-19 Surveillance Team |
| EPI_ISL_2907889 | Lighthouse Lab in Milton Keynes | Wellcome Sanger Institute for the COVID-19 Genomics UK (COG-UK) Consortium | The Lighthouse Lab in Milton Keynes and Alex Alderton, Roberto Amato, Jeffrey Barrett, Sonia Goncalves, Ewan Harrison, David K. Jackson, Ian Johnston, Dominic Kwiatkowski, Cordelia Langford, John Sillitoe on behalf of the Wellcome Sanger Institute COVID-19 Surveillance Team |
| EPI_ISL_2907890 | Lighthouse Lab in Alderley Park | Wellcome Sanger Institute for the COVID-19 Genomics UK (COG-UK) Consortium | Jacquelyn Wynn, Mairead Hyland, The Lighthouse Lab in Alderley Park and Alex Alderton, Roberto Amato, Jeffrey Barrett, Sonia Goncalves, Ewan Harrison, David K. Jackson, Ian Johnston, Dominic Kwiatkowski, Cordelia Langford, John Sillitoe on behalf of the Wellcome Sanger Institute COVID-19 Surveillance Team |
| EPI_ISL_2907891 | Lighthouse Lab in Milton Keynes | Wellcome Sanger Institute for the COVID-19 Genomics UK (COG-UK) Consortium | The Lighthouse Lab in Milton Keynes and Alex Alderton, Roberto Amato, Jeffrey Barrett, Sonia Goncalves, Ewan Harrison, David K. Jackson, Ian Johnston, Dominic Kwiatkowski, Cordelia Langford, John Sillitoe on behalf of the Wellcome Sanger Institute COVID-19 Surveillance Team |
| EPI_ISL_2907892, EPI_ISL_2907893 | Lighthouse Lab in Alderley Park | Wellcome Sanger Institute for the COVID-19 Genomics UK (COG-UK) Consortium | Jacquelyn Wynn, Mairead Hyland, The Lighthouse Lab in Alderley Park and Alex Alderton, Roberto Amato, Jeffrey Barrett, Sonia Goncalves, Ewan Harrison, David K. Jackson, Ian Johnston, Dominic Kwiatkowski, Cordelia Langford, John Sillitoe on behalf of the Wellcome Sanger Institute COVID-19 Surveillance Team |
| EPI_ISL_2907894, EPI_ISL_2907895, EPI_ISL_2907896 | Lighthouse Lab in Milton Keynes | Wellcome Sanger Institute for the COVID-19 Genomics UK (COG-UK) Consortium | The Lighthouse Lab in Milton Keynes and Alex Alderton, Roberto Amato, Jeffrey Barrett, Sonia Goncalves, Ewan Harrison, David K. Jackson, Ian Johnston, Dominic Kwiatkowski, Cordelia Langford, John Sillitoe on behalf of the Wellcome Sanger Institute COVID-19 Surveillance Team |
| EPI_ISL_2907897 | Lighthouse Lab in Alderley Park | Wellcome Sanger Institute for the COVID-19 Genomics UK (COG-UK) Consortium | Jacquelyn Wynn, Mairead Hyland, The Lighthouse Lab in Alderley Park and Alex Alderton, Roberto Amato, Jeffrey Barrett, Sonia Goncalves, Ewan Harrison, David K. Jackson, Ian Johnston, Dominic Kwiatkowski, Cordelia Langford, John Sillitoe on behalf of the Wellcome Sanger Institute COVID-19 Surveillance Team |
| EPI_ISL_2907898, EPI_ISL_2907899, EPI_ISL_2907900, EPI_ISL_2907901, EPI_ISL_2907902, EPI_ISL_2907903, EPI_ISL_2907904 | Lighthouse Lab in Milton Keynes | Wellcome Sanger Institute for the COVID-19 Genomics UK (COG-UK) Consortium | The Lighthouse Lab in Milton Keynes and Alex Alderton, Roberto Amato, Jeffrey Barrett, Sonia Goncalves, Ewan Harrison, David K. Jackson, Ian Johnston, Dominic Kwiatkowski, Cordelia Langford, John Sillitoe on behalf of the Wellcome Sanger Institute COVID-19 Surveillance Team |
| EPI_ISL_2907905 | Lighthouse Lab in Alderley Park | Wellcome Sanger Institute for the COVID-19 Genomics UK (COG-UK) Consortium | Jacquelyn Wynn, Mairead Hyland, The Lighthouse Lab in Alderley Park and Alex Alderton, Roberto Amato, Jeffrey Barrett, Sonia Goncalves, Ewan Harrison, David K. Jackson, Ian Johnston, Dominic Kwiatkowski, Cordelia Langford, John Sillitoe on behalf of the Wellcome Sanger Institute COVID-19 Surveillance Team |
| EPI_ISL_2907906, EPI_ISL_2907907 | Lighthouse Lab in Milton Keynes | Wellcome Sanger Institute for the COVID-19 Genomics UK (COG-UK) Consortium | The Lighthouse Lab in Milton Keynes and Alex Alderton, Roberto Amato, Jeffrey Barrett, Sonia Goncalves, Ewan Harrison, David K. Jackson, Ian Johnston, Dominic Kwiatkowski, Cordelia Langford, John Sillitoe on behalf of the Wellcome Sanger Institute COVID-19 Surveillance Team |
| EPI_ISL_2907908, EPI_ISL_2907909, EPI_ISL_2907910, EPI_ISL_2907911 | Lighthouse Lab in Alderley Park | Wellcome Sanger Institute for the COVID-19 Genomics UK (COG-UK) Consortium | Jacquelyn Wynn, Mairead Hyland, The Lighthouse Lab in Alderley Park and Alex Alderton, Roberto Amato, Jeffrey Barrett, Sonia Goncalves, Ewan Harrison, David K. Jackson, Ian Johnston, Dominic Kwiatkowski, Cordelia Langford, John Sillitoe on behalf of the Wellcome Sanger Institute COVID-19 Surveillance Team |
| EPI_ISL_2907912, EPI_ISL_2907913 | Lighthouse Lab in Milton Keynes | Wellcome Sanger Institute for the COVID-19 Genomics UK (COG-UK) Consortium | The Lighthouse Lab in Milton Keynes and Alex Alderton, Roberto Amato, Jeffrey Barrett, Sonia Goncalves, Ewan Harrison, David K. Jackson, Ian Johnston, Dominic Kwiatkowski, Cordelia Langford, John Sillitoe on behalf of the Wellcome Sanger Institute COVID-19 Surveillance Team |
| EPI_ISL_2907914, EPI_ISL_2907915 | Lighthouse Lab in Alderley Park | Wellcome Sanger Institute for the COVID-19 Genomics UK (COG-UK) Consortium | Jacquelyn Wynn, Mairead Hyland, The Lighthouse Lab in Alderley Park and Alex Alderton, Roberto Amato, Jeffrey Barrett, Sonia Goncalves, Ewan Harrison, David K. Jackson, Ian Johnston, Dominic Kwiatkowski, Cordelia Langford, John Sillitoe on behalf of the Wellcome Sanger Institute COVID-19 Surveillance Team |
| EPI_ISL_2907916 | Lighthouse Lab in Milton Keynes | Wellcome Sanger Institute for the COVID-19 Genomics UK (COG-UK) Consortium | The Lighthouse Lab in Milton Keynes and Alex Alderton, Roberto Amato, Jeffrey Barrett, Sonia Goncalves, Ewan Harrison, David K. Jackson, Ian Johnston, Dominic Kwiatkowski, Cordelia Langford, John Sillitoe on behalf of the Wellcome Sanger Institute COVID-19 Surveillance Team |
| EPI_ISL_2907917, EPI_ISL_2907918 | Lighthouse Lab in Alderley Park | Wellcome Sanger Institute for the COVID-19 Genomics UK (COG-UK) Consortium | Jacquelyn Wynn, Mairead Hyland, The Lighthouse Lab in Alderley Park and Alex Alderton, Roberto Amato, Jeffrey Barrett, Sonia Goncalves, Ewan Harrison, David K. Jackson, Ian Johnston, Dominic Kwiatkowski, Cordelia Langford, John Sillitoe on behalf of the Wellcome Sanger Institute COVID-19 Surveillance Team |
| EPI_ISL_2907919, EPI_ISL_2907920, EPI_ISL_2907921 | Lighthouse Lab in Milton Keynes | Wellcome Sanger Institute for the COVID-19 Genomics UK (COG-UK) Consortium | The Lighthouse Lab in Milton Keynes and Alex Alderton, Roberto Amato, Jeffrey Barrett, Sonia Goncalves, Ewan Harrison, David K. Jackson, Ian Johnston, Dominic Kwiatkowski, Cordelia Langford, John Sillitoe on behalf of the Wellcome Sanger Institute COVID-19 Surveillance Team |
| EPI_ISL_2907922 | Lighthouse Lab in Alderley Park | Wellcome Sanger Institute for the COVID-19 Genomics UK (COG-UK) Consortium | Jacquelyn Wynn, Mairead Hyland, The Lighthouse Lab in Alderley Park and Alex Alderton, Roberto Amato, Jeffrey Barrett, Sonia Goncalves, Ewan Harrison, David K. Jackson, Ian Johnston, Dominic Kwiatkowski, Cordelia Langford, John Sillitoe on behalf of the Wellcome Sanger Institute COVID-19 Surveillance Team |
| EPI_ISL_2907923 | Lighthouse Lab in Milton Keynes | Wellcome Sanger Institute for the COVID-19 Genomics UK (COG-UK) Consortium | The Lighthouse Lab in Milton Keynes and Alex Alderton, Roberto Amato, Jeffrey Barrett, Sonia Goncalves, Ewan Harrison, David K. Jackson, Ian Johnston, Dominic Kwiatkowski, Cordelia Langford, John Sillitoe on behalf of the Wellcome Sanger Institute COVID-19 Surveillance Team |
| EPI_ISL_2907924, EPI_ISL_2907925 | Lighthouse Lab in Alderley Park | Wellcome Sanger Institute for the COVID-19 Genomics UK (COG-UK) Consortium | Jacquelyn Wynn, Mairead Hyland, The Lighthouse Lab in Alderley Park and Alex Alderton, Roberto Amato, Jeffrey Barrett, Sonia Goncalves, Ewan Harrison, David K. Jackson, Ian Johnston, Dominic Kwiatkowski, Cordelia Langford, John Sillitoe on behalf of the Wellcome Sanger Institute COVID-19 Surveillance Team |
| EPI_ISL_2907926 | Lighthouse Lab in Milton Keynes | Wellcome Sanger Institute for the COVID-19 Genomics UK (COG-UK) Consortium | The Lighthouse Lab in Milton Keynes and Alex Alderton, Roberto Amato, Jeffrey Barrett, Sonia Goncalves, Ewan Harrison, David K. Jackson, Ian Johnston, Dominic Kwiatkowski, Cordelia Langford, John Sillitoe on behalf of the Wellcome Sanger Institute COVID-19 Surveillance Team |
| EPI_ISL_2907927, EPI_ISL_2907928 | Lighthouse Lab in Alderley Park | Wellcome Sanger Institute for the COVID-19 Genomics UK (COG-UK) Consortium | Jacquelyn Wynn, Mairead Hyland, The Lighthouse Lab in Alderley Park and Alex Alderton, Roberto Amato, Jeffrey Barrett, Sonia Goncalves, Ewan Harrison, David K. Jackson, Ian Johnston, Dominic Kwiatkowski, Cordelia Langford, John Sillitoe on behalf of the Wellcome Sanger Institute COVID-19 Surveillance Team |
| EPI_ISL_2907929 | Lighthouse Lab in Milton Keynes | Wellcome Sanger Institute for the COVID-19 Genomics UK (COG-UK) Consortium | The Lighthouse Lab in Milton Keynes and Alex Alderton, Roberto Amato, Jeffrey Barrett, Sonia Goncalves, Ewan Harrison, David K. Jackson, Ian Johnston, Dominic Kwiatkowski, Cordelia Langford, John Sillitoe on behalf of the Wellcome Sanger Institute COVID-19 Surveillance Team |
| EPI_ISL_2907930, EPI_ISL_2907931 | Lighthouse Lab in Alderley Park | Wellcome Sanger Institute for the COVID-19 Genomics UK (COG-UK) Consortium | Jacquelyn Wynn, Mairead Hyland, The Lighthouse Lab in Alderley Park and Alex Alderton, Roberto Amato, Jeffrey Barrett, Sonia Goncalves, Ewan Harrison, David K. Jackson, Ian Johnston, Dominic Kwiatkowski, Cordelia Langford, John Sillitoe on behalf of the Wellcome Sanger Institute COVID-19 Surveillance Team |
| EPI_ISL_2907932 | Lighthouse Lab in Milton Keynes | Wellcome Sanger Institute for the COVID-19 Genomics UK (COG-UK) Consortium | The Lighthouse Lab in Milton Keynes and Alex Alderton, Roberto Amato, Jeffrey Barrett, Sonia Goncalves, Ewan Harrison, David K. Jackson, Ian Johnston, Dominic Kwiatkowski, Cordelia Langford, John Sillitoe on behalf of the Wellcome Sanger Institute COVID-19 Surveillance Team |
| EPI_ISL_2907933, EPI_ISL_2907934 | Lighthouse Lab in Alderley Park | Wellcome Sanger Institute for the COVID-19 Genomics UK (COG-UK) Consortium | Jacquelyn Wynn, Mairead Hyland, The Lighthouse Lab in Alderley Park and Alex Alderton, Roberto Amato, Jeffrey Barrett, Sonia Goncalves, Ewan Harrison, David K. Jackson, Ian Johnston, Dominic Kwiatkowski, Cordelia Langford, John Sillitoe on behalf of the Wellcome Sanger Institute COVID-19 Surveillance Team |
| EPI_ISL_2907935 | Lighthouse Lab in Milton Keynes | Wellcome Sanger Institute for the COVID-19 Genomics UK (COG-UK) Consortium | The Lighthouse Lab in Milton Keynes and Alex Alderton, Roberto Amato, Jeffrey Barrett, Sonia Goncalves, Ewan Harrison, David K. Jackson, Ian Johnston, Dominic Kwiatkowski, Cordelia Langford, John Sillitoe on behalf of the Wellcome Sanger Institute COVID-19 Surveillance Team |
| EPI_ISL_2907936, EPI_ISL_2907937 | Lighthouse Lab in Alderley Park | Wellcome Sanger Institute for the COVID-19 Genomics UK (COG-UK) Consortium | Jacquelyn Wynn, Mairead Hyland, The Lighthouse Lab in Alderley Park and Alex Alderton, Roberto Amato, Jeffrey Barrett, Sonia Goncalves, Ewan Harrison, David K. Jackson, Ian Johnston, Dominic Kwiatkowski, Cordelia Langford, John Sillitoe on behalf of the Wellcome Sanger Institute COVID-19 Surveillance Team |
| EPI_ISL_2907938, EPI_ISL_2907939 | Lighthouse Lab in Milton Keynes | Wellcome Sanger Institute for the COVID-19 Genomics UK (COG-UK) Consortium | The Lighthouse Lab in Milton Keynes and Alex Alderton, Roberto Amato, Jeffrey Barrett, Sonia Goncalves, Ewan Harrison, David K. Jackson, Ian Johnston, Dominic Kwiatkowski, Cordelia Langford, John Sillitoe on behalf of the Wellcome Sanger Institute COVID-19 Surveillance Team |
| EPI_ISL_2907940 | Lighthouse Lab in Alderley Park | Wellcome Sanger Institute for the COVID-19 Genomics UK (COG-UK) Consortium | Jacquelyn Wynn, Mairead Hyland, The Lighthouse Lab in Alderley Park and Alex Alderton, Roberto Amato, Jeffrey Barrett, Sonia Goncalves, Ewan Harrison, David K. Jackson, Ian Johnston, Dominic Kwiatkowski, Cordelia Langford, John Sillitoe on behalf of the Wellcome Sanger Institute COVID-19 Surveillance Team |
| EPI_ISL_2907941, EPI_ISL_2907942 | Lighthouse Lab in Milton Keynes | Wellcome Sanger Institute for the COVID-19 Genomics UK (COG-UK) Consortium | The Lighthouse Lab in Milton Keynes and Alex Alderton, Roberto Amato, Jeffrey Barrett, Sonia Goncalves, Ewan Harrison, David K. Jackson, Ian Johnston, Dominic Kwiatkowski, Cordelia Langford, John Sillitoe on behalf of the Wellcome Sanger Institute COVID-19 Surveillance Team |
| EPI_ISL_2907943 | Lighthouse Lab in Glasgow | Wellcome Sanger Institute for the COVID-19 Genomics UK (COG-UK) Consortium | Harper VanSteenhouse, Yumi Kasai, David Gray, Carol Clugston, Anna Dominiczak and Alex Alderton, Roberto Amato, Jeffrey Barrett, Sonia Goncalves, Ewan Harrison, David K. Jackson, Ian Johnston, Dominic Kwiatkowski, Cordelia Langford, John Sillitoe on behalf of the Wellcome Sanger Institute COVID-19 Surveillance Team |
| EPI_ISL_2907944, EPI_ISL_2907945, EPI_ISL_2907946, EPI_ISL_2907947, EPI_ISL_2907948, EPI_ISL_2907949, EPI_ISL_2907950, EPI_ISL_2907951, EPI_ISL_2907952, EPI_ISL_2907953, EPI_ISL_2907954, EPI_ISL_2907955 | | | |
| see above | Lighthouse Lab in Milton Keynes | Wellcome Sanger Institute for the COVID-19 Genomics UK (COG-UK) Consortium | The Lighthouse Lab in Milton Keynes and Alex Alderton, Roberto Amato, Jeffrey Barrett, Sonia Goncalves, Ewan Harrison, David K. Jackson, Ian Johnston, Dominic Kwiatkowski, Cordelia Langford, John Sillitoe on behalf of the Wellcome Sanger Institute COVID-19 Surveillance Team |
| EPI_ISL_2907956 | Lighthouse Lab in Glasgow | Wellcome Sanger Institute for the COVID-19 Genomics UK (COG-UK) Consortium | Harper VanSteenhouse, Yumi Kasai, David Gray, Carol Clugston, Anna Dominiczak and Alex Alderton, Roberto Amato, Jeffrey Barrett, Sonia Goncalves, Ewan Harrison, David K. Jackson, Ian Johnston, Dominic Kwiatkowski, Cordelia Langford, John Sillitoe on behalf of the Wellcome Sanger Institute COVID-19 Surveillance Team |
| EPI_ISL_2907957 | Lighthouse Lab in Milton Keynes | Wellcome Sanger Institute for the COVID-19 Genomics UK (COG-UK) Consortium | The Lighthouse Lab in Milton Keynes and Alex Alderton, Roberto Amato, Jeffrey Barrett, Sonia Goncalves, Ewan Harrison, David K. Jackson, Ian Johnston, Dominic Kwiatkowski, Cordelia Langford, John Sillitoe on behalf of the Wellcome Sanger Institute COVID-19 Surveillance Team |
| EPI_ISL_2907958, EPI_ISL_2907959 | Lighthouse Lab in Glasgow | Wellcome Sanger Institute for the COVID-19 Genomics UK (COG-UK) Consortium | Harper VanSteenhouse, Yumi Kasai, David Gray, Carol Clugston, Anna Dominiczak and Alex Alderton, Roberto Amato, Jeffrey Barrett, Sonia Goncalves, Ewan Harrison, David K. Jackson, Ian Johnston, Dominic Kwiatkowski, Cordelia Langford, John Sillitoe on behalf of the Wellcome Sanger Institute COVID-19 Surveillance Team |
| EPI_ISL_2907960, EPI_ISL_2907961 | Lighthouse Lab in Milton Keynes | Wellcome Sanger Institute for the COVID-19 Genomics UK (COG-UK) Consortium | The Lighthouse Lab in Milton Keynes and Alex Alderton, Roberto Amato, Jeffrey Barrett, Sonia Goncalves, Ewan Harrison, David K. Jackson, Ian Johnston, Dominic Kwiatkowski, Cordelia Langford, John Sillitoe on behalf of the Wellcome Sanger Institute COVID-19 Surveillance Team |
| EPI_ISL_2907962 | Lighthouse Lab in Glasgow | Wellcome Sanger Institute for the COVID-19 Genomics UK (COG-UK) Consortium | Harper VanSteenhouse, Yumi Kasai, David Gray, Carol Clugston, Anna Dominiczak and Alex Alderton, Roberto Amato, Jeffrey Barrett, Sonia Goncalves, Ewan Harrison, David K. Jackson, Ian Johnston, Dominic Kwiatkowski, Cordelia Langford, John Sillitoe on behalf of the Wellcome Sanger Institute COVID-19 Surveillance Team |
| EPI_ISL_2907963, EPI_ISL_2907964 | Lighthouse Lab in Milton Keynes | Wellcome Sanger Institute for the COVID-19 Genomics UK (COG-UK) Consortium | The Lighthouse Lab in Milton Keynes and Alex Alderton, Roberto Amato, Jeffrey Barrett, Sonia Goncalves, Ewan Harrison, David K. Jackson, Ian Johnston, Dominic Kwiatkowski, Cordelia Langford, John Sillitoe on behalf of the Wellcome Sanger Institute COVID-19 Surveillance Team |
| EPI_ISL_2907965 | Lighthouse Lab in Glasgow | Wellcome Sanger Institute for the COVID-19 Genomics UK (COG-UK) | Harper VanSteenhouse, Yumi Kasai, David Gray, Carol Clugston, Anna Dominiczak and Alex Alderton, Roberto Amato, Jeffrey Barrett, Sonia Goncalves, Ewan Harrison, David K. Jackson, Ian Johnston, Dominic Kwiatkowski, Cordelia Langford, |

| | | | |
|---|---|---|---|
| | | Consortium | John Sillitoe on behalf of the Wellcome Sanger Institute COVID-19 Surveillance Team |
| EPI_ISL_2907966, EPI_ISL_2907967, EPI_ISL_2907968, EPI_ISL_2907969 | Lighthouse Lab in Milton Keynes | Wellcome Sanger Institute for the COVID-19 Genomics UK (COG-UK) Consortium | The Lighthouse Lab in Milton Keynes and Alex Alderton, Roberto Amato, Jeffrey Barrett, Sonia Goncalves, Ewan Harrison, David K. Jackson, Ian Johnston, Dominic Kwiatkowski, Cordelia Langford, John Sillitoe on behalf of the Wellcome Sanger Institute COVID-19 Surveillance Team |
| EPI_ISL_2907970 | Lighthouse Lab in Glasgow | Wellcome Sanger Institute for the COVID-19 Genomics UK (COG-UK) Consortium | Harper VanSteenhouse, Yumi Kasai, David Gray, Carol Clugston, Anna Dominiczak and Alex Alderton, Roberto Amato, Jeffrey Barrett, Sonia Goncalves, Ewan Harrison, David K. Jackson, Ian Johnston, Dominic Kwiatkowski, Cordelia Langford, John Sillitoe on behalf of the Wellcome Sanger Institute COVID-19 Surveillance Team |
| EPI_ISL_2907971, EPI_ISL_2907972, EPI_ISL_2907973 | Lighthouse Lab in Milton Keynes | Wellcome Sanger Institute for the COVID-19 Genomics UK (COG-UK) Consortium | The Lighthouse Lab in Milton Keynes and Alex Alderton, Roberto Amato, Jeffrey Barrett, Sonia Goncalves, Ewan Harrison, David K. Jackson, Ian Johnston, Dominic Kwiatkowski, Cordelia Langford, John Sillitoe on behalf of the Wellcome Sanger Institute COVID-19 Surveillance Team |
| EPI_ISL_2907974, EPI_ISL_2907975, EPI_ISL_2907976 | Lighthouse Lab in Glasgow | Wellcome Sanger Institute for the COVID-19 Genomics UK (COG-UK) Consortium | Harper VanSteenhouse, Yumi Kasai, David Gray, Carol Clugston, Anna Dominiczak and Alex Alderton, Roberto Amato, Jeffrey Barrett, Sonia Goncalves, Ewan Harrison, David K. Jackson, Ian Johnston, Dominic Kwiatkowski, Cordelia Langford, John Sillitoe on behalf of the Wellcome Sanger Institute COVID-19 Surveillance Team |
| EPI_ISL_2907977, EPI_ISL_2907978, EPI_ISL_2907979, EPI_ISL_2907980, EPI_ISL_2907981, EPI_ISL_2907982, EPI_ISL_2907983 | Lighthouse Lab in Milton Keynes | Wellcome Sanger Institute for the COVID-19 Genomics UK (COG-UK) Consortium | The Lighthouse Lab in Milton Keynes and Alex Alderton, Roberto Amato, Jeffrey Barrett, Sonia Goncalves, Ewan Harrison, David K. Jackson, Ian Johnston, Dominic Kwiatkowski, Cordelia Langford, John Sillitoe on behalf of the Wellcome Sanger Institute COVID-19 Surveillance Team |
| EPI_ISL_2907984 | Lighthouse Lab in Glasgow | Wellcome Sanger Institute for the COVID-19 Genomics UK (COG-UK) Consortium | Harper VanSteenhouse, Yumi Kasai, David Gray, Carol Clugston, Anna Dominiczak and Alex Alderton, Roberto Amato, Jeffrey Barrett, Sonia Goncalves, Ewan Harrison, David K. Jackson, Ian Johnston, Dominic Kwiatkowski, Cordelia Langford, John Sillitoe on behalf of the Wellcome Sanger Institute COVID-19 Surveillance Team |
| EPI_ISL_2907985, EPI_ISL_2907986, EPI_ISL_2907987, EPI_ISL_2907988 | Lighthouse Lab in Milton Keynes | Wellcome Sanger Institute for the COVID-19 Genomics UK (COG-UK) Consortium | The Lighthouse Lab in Milton Keynes and Alex Alderton, Roberto Amato, Jeffrey Barrett, Sonia Goncalves, Ewan Harrison, David K. Jackson, Ian Johnston, Dominic Kwiatkowski, Cordelia Langford, John Sillitoe on behalf of the Wellcome Sanger Institute COVID-19 Surveillance Team |
| EPI_ISL_2907989 | Lighthouse Lab in Glasgow | Wellcome Sanger Institute for the COVID-19 Genomics UK (COG-UK) Consortium | Harper VanSteenhouse, Yumi Kasai, David Gray, Carol Clugston, Anna Dominiczak and Alex Alderton, Roberto Amato, Jeffrey Barrett, Sonia Goncalves, Ewan Harrison, David K. Jackson, Ian Johnston, Dominic Kwiatkowski, Cordelia Langford, John Sillitoe on behalf of the Wellcome Sanger Institute COVID-19 Surveillance Team |
| EPI_ISL_2907990, EPI_ISL_2907991, EPI_ISL_2907992, EPI_ISL_2907993, EPI_ISL_2907994, EPI_ISL_2907995, EPI_ISL_2907996, EPI_ISL_2907997, EPI_ISL_2907998, EPI_ISL_2907999, EPI_ISL_2908000, EPI_ISL_2908001, EPI_ISL_2908002, EPI_ISL_2908003, EPI_ISL_2908004, EPI_ISL_2908005, EPI_ISL_2908006, EPI_ISL_2908007, EPI_ISL_2908008, EPI_ISL_2908009, EPI_ISL_2908010, EPI_ISL_2908011, EPI_ISL_2908012 | | | |
| see above | Lighthouse Lab in Milton Keynes | Wellcome Sanger Institute for the COVID-19 Genomics UK (COG-UK) Consortium | The Lighthouse Lab in Milton Keynes and Alex Alderton, Roberto Amato, Jeffrey Barrett, Sonia Goncalves, Ewan Harrison, David K. Jackson, Ian Johnston, Dominic Kwiatkowski, Cordelia Langford, John Sillitoe on behalf of the Wellcome Sanger Institute COVID-19 Surveillance Team |
| EPI_ISL_2908013 | Lighthouse Lab in Glasgow | Wellcome Sanger Institute for the COVID-19 Genomics UK (COG-UK) Consortium | Harper VanSteenhouse, Yumi Kasai, David Gray, Carol Clugston, Anna Dominiczak and Alex Alderton, Roberto Amato, Jeffrey Barrett, Sonia Goncalves, Ewan Harrison, David K. Jackson, Ian Johnston, Dominic Kwiatkowski, Cordelia Langford, John Sillitoe on behalf of the Wellcome Sanger Institute COVID-19 Surveillance Team |
| EPI_ISL_2908014 | Lighthouse Lab in Milton Keynes | Wellcome Sanger Institute for the COVID-19 Genomics UK (COG-UK) Consortium | The Lighthouse Lab in Milton Keynes and Alex Alderton, Roberto Amato, Jeffrey Barrett, Sonia Goncalves, Ewan Harrison, David K. Jackson, Ian Johnston, Dominic Kwiatkowski, Cordelia Langford, John Sillitoe on behalf of the Wellcome Sanger Institute COVID-19 Surveillance Team |
| EPI_ISL_2908015, EPI_ISL_2908016, EPI_ISL_2908017 | Lighthouse Lab in Glasgow | Wellcome Sanger Institute for the COVID-19 Genomics UK (COG-UK) Consortium | Harper VanSteenhouse, Yumi Kasai, David Gray, Carol Clugston, Anna Dominiczak and Alex Alderton, Roberto Amato, Jeffrey Barrett, Sonia Goncalves, Ewan Harrison, David K. Jackson, Ian Johnston, Dominic Kwiatkowski, Cordelia Langford, John Sillitoe on behalf of the Wellcome Sanger Institute COVID-19 Surveillance Team |
| EPI_ISL_2908018 | Lighthouse Lab in Milton Keynes | Wellcome Sanger Institute for the COVID-19 Genomics UK (COG-UK) Consortium | The Lighthouse Lab in Milton Keynes and Alex Alderton, Roberto Amato, Jeffrey Barrett, Sonia Goncalves, Ewan Harrison, David K. Jackson, Ian Johnston, Dominic Kwiatkowski, Cordelia Langford, John Sillitoe on behalf of the Wellcome Sanger Institute COVID-19 Surveillance Team |
| EPI_ISL_2908019 | Lighthouse Lab in Glasgow | Wellcome Sanger Institute for the COVID-19 Genomics UK (COG-UK) Consortium | Harper VanSteenhouse, Yumi Kasai, David Gray, Carol Clugston, Anna Dominiczak and Alex Alderton, Roberto Amato, Jeffrey Barrett, Sonia Goncalves, Ewan Harrison, David K. Jackson, Ian Johnston, Dominic Kwiatkowski, Cordelia Langford, John Sillitoe on behalf of the Wellcome Sanger Institute COVID-19 Surveillance Team |
| EPI_ISL_2908020, EPI_ISL_2908021, EPI_ISL_2908022, EPI_ISL_2908023 | Lighthouse Lab in Milton Keynes | Wellcome Sanger Institute for the COVID-19 Genomics UK (COG-UK) Consortium | The Lighthouse Lab in Milton Keynes and Alex Alderton, Roberto Amato, Jeffrey Barrett, Sonia Goncalves, Ewan Harrison, David K. Jackson, Ian Johnston, Dominic Kwiatkowski, Cordelia Langford, John Sillitoe on behalf of the Wellcome Sanger Institute COVID-19 Surveillance Team |
| EPI_ISL_2908024 | Lighthouse Lab in Glasgow | Wellcome Sanger Institute for the COVID-19 Genomics UK (COG-UK) Consortium | Harper VanSteenhouse, Yumi Kasai, David Gray, Carol Clugston, Anna Dominiczak and Alex Alderton, Roberto Amato, Jeffrey Barrett, Sonia Goncalves, Ewan Harrison, David K. Jackson, Ian Johnston, Dominic Kwiatkowski, Cordelia Langford, John Sillitoe on behalf of the Wellcome Sanger Institute COVID-19 Surveillance Team |
| EPI_ISL_2908026, EPI_ISL_2908027 | Lighthouse Lab in Milton Keynes | Wellcome Sanger Institute for the COVID-19 Genomics UK (COG-UK) Consortium | The Lighthouse Lab in Milton Keynes and Alex Alderton, Roberto Amato, Jeffrey Barrett, Sonia Goncalves, Ewan Harrison, David K. Jackson, Ian Johnston, Dominic Kwiatkowski, Cordelia Langford, John Sillitoe on behalf of the Wellcome Sanger Institute COVID-19 Surveillance Team |
| EPI_ISL_2908028 | Lighthouse Lab in Glasgow | Wellcome Sanger Institute for the COVID-19 Genomics UK (COG-UK) Consortium | Harper VanSteenhouse, Yumi Kasai, David Gray, Carol Clugston, Anna Dominiczak and Alex Alderton, Roberto Amato, Jeffrey Barrett, Sonia Goncalves, Ewan Harrison, David K. Jackson, Ian Johnston, Dominic Kwiatkowski, Cordelia Langford, John Sillitoe on behalf of the Wellcome Sanger Institute COVID-19 Surveillance Team |
| EPI_ISL_2908029 | Lighthouse Lab in Milton Keynes | Wellcome Sanger Institute for the COVID-19 Genomics UK (COG-UK) Consortium | The Lighthouse Lab in Milton Keynes and Alex Alderton, Roberto Amato, Jeffrey Barrett, Sonia Goncalves, Ewan Harrison, David K. Jackson, Ian Johnston, Dominic Kwiatkowski, Cordelia Langford, John Sillitoe on behalf of the Wellcome Sanger Institute COVID-19 Surveillance Team |
| EPI_ISL_2908030 | Lighthouse Lab in Glasgow | Wellcome Sanger Institute for the COVID-19 Genomics UK (COG-UK) Consortium | Harper VanSteenhouse, Yumi Kasai, David Gray, Carol Clugston, Anna Dominiczak and Alex Alderton, Roberto Amato, Jeffrey Barrett, Sonia Goncalves, Ewan Harrison, David K. Jackson, Ian Johnston, Dominic Kwiatkowski, Cordelia Langford, John Sillitoe on behalf of the Wellcome Sanger Institute COVID-19 Surveillance Team |
| EPI_ISL_2908031, EPI_ISL_2908032 | Lighthouse Lab in Milton Keynes | Wellcome Sanger Institute for the COVID-19 Genomics UK (COG-UK) Consortium | The Lighthouse Lab in Milton Keynes and Alex Alderton, Roberto Amato, Jeffrey Barrett, Sonia Goncalves, Ewan Harrison, David K. Jackson, Ian Johnston, Dominic Kwiatkowski, Cordelia Langford, John Sillitoe on behalf of the Wellcome Sanger Institute COVID-19 Surveillance Team |
| EPI_ISL_2908033, EPI_ISL_2908034 | Lighthouse Lab in Glasgow | Wellcome Sanger Institute for the COVID-19 Genomics UK (COG-UK) Consortium | Harper VanSteenhouse, Yumi Kasai, David Gray, Carol Clugston, Anna Dominiczak and Alex Alderton, Roberto Amato, Jeffrey Barrett, Sonia Goncalves, Ewan Harrison, David K. Jackson, Ian Johnston, Dominic Kwiatkowski, Cordelia Langford, John Sillitoe on behalf of the Wellcome Sanger Institute COVID-19 Surveillance Team |
| EPI_ISL_2908035, EPI_ISL_2908036, EPI_ISL_2908037, EPI_ISL_2908038, EPI_ISL_2908039, EPI_ISL_2908040 | Lighthouse Lab in Milton Keynes | Wellcome Sanger Institute for the COVID-19 Genomics UK (COG-UK) Consortium | The Lighthouse Lab in Milton Keynes and Alex Alderton, Roberto Amato, Jeffrey Barrett, Sonia Goncalves, Ewan Harrison, David K. Jackson, Ian Johnston, Dominic Kwiatkowski, Cordelia Langford, John Sillitoe on behalf of the Wellcome Sanger Institute COVID-19 Surveillance Team |
| EPI_ISL_2908041 | Lighthouse Lab in Glasgow | Wellcome Sanger Institute for the COVID-19 Genomics UK (COG-UK) Consortium | Harper VanSteenhouse, Yumi Kasai, David Gray, Carol Clugston, Anna Dominiczak and Alex Alderton, Roberto Amato, Jeffrey Barrett, Sonia Goncalves, Ewan Harrison, David K. Jackson, Ian Johnston, Dominic Kwiatkowski, Cordelia Langford, John Sillitoe on behalf of the Wellcome Sanger Institute COVID-19 Surveillance Team |
| EPI_ISL_2908042, EPI_ISL_2908043, EPI_ISL_2908044 | Lighthouse Lab in Milton Keynes | Wellcome Sanger Institute for the COVID-19 Genomics UK (COG-UK) Consortium | The Lighthouse Lab in Milton Keynes and Alex Alderton, Roberto Amato, Jeffrey Barrett, Sonia Goncalves, Ewan Harrison, David K. Jackson, Ian Johnston, Dominic Kwiatkowski, Cordelia Langford, John Sillitoe on behalf of the Wellcome Sanger Institute COVID-19 Surveillance Team |
| EPI_ISL_2908045 | Lighthouse Lab in Glasgow | Wellcome Sanger Institute for the COVID-19 Genomics UK (COG-UK) Consortium | Harper VanSteenhouse, Yumi Kasai, David Gray, Carol Clugston, Anna Dominiczak and Alex Alderton, Roberto Amato, Jeffrey Barrett, Sonia Goncalves, Ewan Harrison, David K. Jackson, Ian Johnston, Dominic Kwiatkowski, Cordelia Langford, John Sillitoe on behalf of the Wellcome Sanger Institute COVID-19 Surveillance Team |
| EPI_ISL_2908046, EPI_ISL_2908047 | Lighthouse Lab in Milton Keynes | Wellcome Sanger Institute for the COVID-19 Genomics UK (COG-UK) Consortium | The Lighthouse Lab in Milton Keynes and Alex Alderton, Roberto Amato, Jeffrey Barrett, Sonia Goncalves, Ewan Harrison, David K. Jackson, Ian Johnston, Dominic Kwiatkowski, Cordelia Langford, John Sillitoe on behalf of the Wellcome Sanger Institute COVID-19 Surveillance Team |
| EPI_ISL_2908048 | Lighthouse Lab in Glasgow | Wellcome Sanger Institute for the COVID-19 Genomics UK (COG-UK) Consortium | Harper VanSteenhouse, Yumi Kasai, David Gray, Carol Clugston, Anna Dominiczak and Alex Alderton, Roberto Amato, Jeffrey Barrett, Sonia Goncalves, Ewan Harrison, David K. Jackson, Ian Johnston, Dominic Kwiatkowski, Cordelia Langford, John Sillitoe on behalf of the Wellcome Sanger Institute COVID-19 Surveillance Team |
| EPI_ISL_2908049, EPI_ISL_2908050, EPI_ISL_2908051, EPI_ISL_2908052, EPI_ISL_2908053, EPI_ISL_2908054, EPI_ISL_2908055, EPI_ISL_2908056 | Lighthouse Lab in Milton Keynes | Wellcome Sanger Institute for the COVID-19 Genomics UK (COG-UK) Consortium | The Lighthouse Lab in Milton Keynes and Alex Alderton, Roberto Amato, Jeffrey Barrett, Sonia Goncalves, Ewan Harrison, David K. Jackson, Ian Johnston, Dominic Kwiatkowski, Cordelia Langford, John Sillitoe on behalf of the Wellcome Sanger Institute COVID-19 Surveillance Team |
| EPI_ISL_2908057 | Lighthouse Lab in Glasgow | Wellcome Sanger Institute for the COVID-19 Genomics UK (COG-UK) Consortium | Harper VanSteenhouse, Yumi Kasai, David Gray, Carol Clugston, Anna Dominiczak and Alex Alderton, Roberto Amato, Jeffrey Barrett, Sonia Goncalves, Ewan Harrison, David K. Jackson, Ian Johnston, Dominic Kwiatkowski, Cordelia Langford, John Sillitoe on behalf of the Wellcome Sanger Institute COVID-19 Surveillance Team |
| EPI_ISL_2908058, EPI_ISL_2908059, EPI_ISL_2908060, EPI_ISL_2908061, EPI_ISL_2908062, EPI_ISL_2908063, EPI_ISL_2908064, | Lighthouse Lab in Milton Keynes | Wellcome Sanger Institute for the COVID-19 Genomics UK (COG-UK) Consortium | The Lighthouse Lab in Milton Keynes and Alex Alderton, Roberto Amato, Jeffrey Barrett, Sonia Goncalves, Ewan Harrison, David K. Jackson, Ian Johnston, Dominic Kwiatkowski, Cordelia Langford, John Sillitoe on behalf of the Wellcome Sanger Institute COVID-19 Surveillance Team |

| | | | |
|---|---|---|---|
| EPI_ISL_2908065 | | | |
| EPI_ISL_2908066 | Lighthouse Lab in Glasgow | Wellcome Sanger Institute for the COVID-19 Genomics UK (COG-UK) Consortium | Harper VanSteenhouse, Yumi Kasai, David Gray, Carol Clugston, Anna Dominiczak and Alex Alderton, Roberto Amato, Jeffrey Barrett, Sonia Goncalves, Ewan Harrison, David K. Jackson, Ian Johnston, Dominic Kwiatkowski, Cordelia Langford, John Sillitoe on behalf of the Wellcome Sanger Institute COVID-19 Surveillance Team |
| EPI_ISL_2908067, EPI_ISL_2908068 | Lighthouse Lab in Milton Keynes | Wellcome Sanger Institute for the COVID-19 Genomics UK (COG-UK) Consortium | The Lighthouse Lab in Milton Keynes and Alex Alderton, Roberto Amato, Jeffrey Barrett, Sonia Goncalves, Ewan Harrison, David K. Jackson, Ian Johnston, Dominic Kwiatkowski, Cordelia Langford, John Sillitoe on behalf of the Wellcome Sanger Institute COVID-19 Surveillance Team |
| EPI_ISL_2908069, EPI_ISL_2908070 | Lighthouse Lab in Glasgow | Wellcome Sanger Institute for the COVID-19 Genomics UK (COG-UK) Consortium | Harper VanSteenhouse, Yumi Kasai, David Gray, Carol Clugston, Anna Dominiczak and Alex Alderton, Roberto Amato, Jeffrey Barrett, Sonia Goncalves, Ewan Harrison, David K. Jackson, Ian Johnston, Dominic Kwiatkowski, Cordelia Langford, John Sillitoe on behalf of the Wellcome Sanger Institute COVID-19 Surveillance Team |
| EPI_ISL_2908071, EPI_ISL_2908072, EPI_ISL_2908073 | Lighthouse Lab in Milton Keynes | Wellcome Sanger Institute for the COVID-19 Genomics UK (COG-UK) Consortium | The Lighthouse Lab in Milton Keynes and Alex Alderton, Roberto Amato, Jeffrey Barrett, Sonia Goncalves, Ewan Harrison, David K. Jackson, Ian Johnston, Dominic Kwiatkowski, Cordelia Langford, John Sillitoe on behalf of the Wellcome Sanger Institute COVID-19 Surveillance Team |
| EPI_ISL_2908074 | Lighthouse Lab in Glasgow | Wellcome Sanger Institute for the COVID-19 Genomics UK (COG-UK) Consortium | Harper VanSteenhouse, Yumi Kasai, David Gray, Carol Clugston, Anna Dominiczak and Alex Alderton, Roberto Amato, Jeffrey Barrett, Sonia Goncalves, Ewan Harrison, David K. Jackson, Ian Johnston, Dominic Kwiatkowski, Cordelia Langford, John Sillitoe on behalf of the Wellcome Sanger Institute COVID-19 Surveillance Team |
| EPI_ISL_2908075 | Lighthouse Lab in Milton Keynes | Wellcome Sanger Institute for the COVID-19 Genomics UK (COG-UK) Consortium | The Lighthouse Lab in Milton Keynes and Alex Alderton, Roberto Amato, Jeffrey Barrett, Sonia Goncalves, Ewan Harrison, David K. Jackson, Ian Johnston, Dominic Kwiatkowski, Cordelia Langford, John Sillitoe on behalf of the Wellcome Sanger Institute COVID-19 Surveillance Team |
| EPI_ISL_2908076, EPI_ISL_2908077 | Lighthouse Lab in Glasgow | Wellcome Sanger Institute for the COVID-19 Genomics UK (COG-UK) Consortium | Harper VanSteenhouse, Yumi Kasai, David Gray, Carol Clugston, Anna Dominiczak and Alex Alderton, Roberto Amato, Jeffrey Barrett, Sonia Goncalves, Ewan Harrison, David K. Jackson, Ian Johnston, Dominic Kwiatkowski, Cordelia Langford, John Sillitoe on behalf of the Wellcome Sanger Institute COVID-19 Surveillance Team |
| EPI_ISL_2908078 | Lighthouse Lab in Milton Keynes | Wellcome Sanger Institute for the COVID-19 Genomics UK (COG-UK) Consortium | The Lighthouse Lab in Milton Keynes and Alex Alderton, Roberto Amato, Jeffrey Barrett, Sonia Goncalves, Ewan Harrison, David K. Jackson, Ian Johnston, Dominic Kwiatkowski, Cordelia Langford, John Sillitoe on behalf of the Wellcome Sanger Institute COVID-19 Surveillance Team |
| EPI_ISL_2908079 | Lighthouse Lab in Glasgow | Wellcome Sanger Institute for the COVID-19 Genomics UK (COG-UK) Consortium | Harper VanSteenhouse, Yumi Kasai, David Gray, Carol Clugston, Anna Dominiczak and Alex Alderton, Roberto Amato, Jeffrey Barrett, Sonia Goncalves, Ewan Harrison, David K. Jackson, Ian Johnston, Dominic Kwiatkowski, Cordelia Langford, John Sillitoe on behalf of the Wellcome Sanger Institute COVID-19 Surveillance Team |
| EPI_ISL_2908080, EPI_ISL_2908081 | Lighthouse Lab in Milton Keynes | Wellcome Sanger Institute for the COVID-19 Genomics UK (COG-UK) Consortium | The Lighthouse Lab in Milton Keynes and Alex Alderton, Roberto Amato, Jeffrey Barrett, Sonia Goncalves, Ewan Harrison, David K. Jackson, Ian Johnston, Dominic Kwiatkowski, Cordelia Langford, John Sillitoe on behalf of the Wellcome Sanger Institute COVID-19 Surveillance Team |
| EPI_ISL_2908082 | Lighthouse Lab in Glasgow | Wellcome Sanger Institute for the COVID-19 Genomics UK (COG-UK) Consortium | Harper VanSteenhouse, Yumi Kasai, David Gray, Carol Clugston, Anna Dominiczak and Alex Alderton, Roberto Amato, Jeffrey Barrett, Sonia Goncalves, Ewan Harrison, David K. Jackson, Ian Johnston, Dominic Kwiatkowski, Cordelia Langford, John Sillitoe on behalf of the Wellcome Sanger Institute COVID-19 Surveillance Team |
| EPI_ISL_2908083, EPI_ISL_2908084 | Lighthouse Lab in Milton Keynes | Wellcome Sanger Institute for the COVID-19 Genomics UK (COG-UK) Consortium | The Lighthouse Lab in Milton Keynes and Alex Alderton, Roberto Amato, Jeffrey Barrett, Sonia Goncalves, Ewan Harrison, David K. Jackson, Ian Johnston, Dominic Kwiatkowski, Cordelia Langford, John Sillitoe on behalf of the Wellcome Sanger Institute COVID-19 Surveillance Team |
| EPI_ISL_2908085, EPI_ISL_2908086 | Lighthouse Lab in Glasgow | Wellcome Sanger Institute for the COVID-19 Genomics UK (COG-UK) Consortium | Harper VanSteenhouse, Yumi Kasai, David Gray, Carol Clugston, Anna Dominiczak and Alex Alderton, Roberto Amato, Jeffrey Barrett, Sonia Goncalves, Ewan Harrison, David K. Jackson, Ian Johnston, Dominic Kwiatkowski, Cordelia Langford, John Sillitoe on behalf of the Wellcome Sanger Institute COVID-19 Surveillance Team |
| EPI_ISL_2908087, EPI_ISL_2908088, EPI_ISL_2908089, EPI_ISL_2908090, EPI_ISL_2908091, EPI_ISL_2908092 | Lighthouse Lab in Milton Keynes | Wellcome Sanger Institute for the COVID-19 Genomics UK (COG-UK) Consortium | The Lighthouse Lab in Milton Keynes and Alex Alderton, Roberto Amato, Jeffrey Barrett, Sonia Goncalves, Ewan Harrison, David K. Jackson, Ian Johnston, Dominic Kwiatkowski, Cordelia Langford, John Sillitoe on behalf of the Wellcome Sanger Institute COVID-19 Surveillance Team |
| EPI_ISL_2908093 | Lighthouse Lab in Glasgow | Wellcome Sanger Institute for the COVID-19 Genomics UK (COG-UK) Consortium | Harper VanSteenhouse, Yumi Kasai, David Gray, Carol Clugston, Anna Dominiczak and Alex Alderton, Roberto Amato, Jeffrey Barrett, Sonia Goncalves, Ewan Harrison, David K. Jackson, Ian Johnston, Dominic Kwiatkowski, Cordelia Langford, John Sillitoe on behalf of the Wellcome Sanger Institute COVID-19 Surveillance Team |
| EPI_ISL_2908094 | Lighthouse Lab in Milton Keynes | Wellcome Sanger Institute for the COVID-19 Genomics UK (COG-UK) Consortium | The Lighthouse Lab in Milton Keynes and Alex Alderton, Roberto Amato, Jeffrey Barrett, Sonia Goncalves, Ewan Harrison, David K. Jackson, Ian Johnston, Dominic Kwiatkowski, Cordelia Langford, John Sillitoe on behalf of the Wellcome Sanger Institute COVID-19 Surveillance Team |
| EPI_ISL_2908095 | Lighthouse Lab in Glasgow | Wellcome Sanger Institute for the COVID-19 Genomics UK (COG-UK) Consortium | Harper VanSteenhouse, Yumi Kasai, David Gray, Carol Clugston, Anna Dominiczak and Alex Alderton, Roberto Amato, Jeffrey Barrett, Sonia Goncalves, Ewan Harrison, David K. Jackson, Ian Johnston, Dominic Kwiatkowski, Cordelia Langford, John Sillitoe on behalf of the Wellcome Sanger Institute COVID-19 Surveillance Team |
| EPI_ISL_2908096 | Lighthouse Lab in Milton Keynes | Wellcome Sanger Institute for the COVID-19 Genomics UK (COG-UK) Consortium | The Lighthouse Lab in Milton Keynes and Alex Alderton, Roberto Amato, Jeffrey Barrett, Sonia Goncalves, Ewan Harrison, David K. Jackson, Ian Johnston, Dominic Kwiatkowski, Cordelia Langford, John Sillitoe on behalf of the Wellcome Sanger Institute COVID-19 Surveillance Team |
| EPI_ISL_2908097 | Lighthouse Lab in Glasgow | Wellcome Sanger Institute for the COVID-19 Genomics UK (COG-UK) Consortium | Harper VanSteenhouse, Yumi Kasai, David Gray, Carol Clugston, Anna Dominiczak and Alex Alderton, Roberto Amato, Jeffrey Barrett, Sonia Goncalves, Ewan Harrison, David K. Jackson, Ian Johnston, Dominic Kwiatkowski, Cordelia Langford, John Sillitoe on behalf of the Wellcome Sanger Institute COVID-19 Surveillance Team |
| EPI_ISL_2908098 | Lighthouse Lab in Milton Keynes | Wellcome Sanger Institute for the COVID-19 Genomics UK (COG-UK) Consortium | The Lighthouse Lab in Milton Keynes and Alex Alderton, Roberto Amato, Jeffrey Barrett, Sonia Goncalves, Ewan Harrison, David K. Jackson, Ian Johnston, Dominic Kwiatkowski, Cordelia Langford, John Sillitoe on behalf of the Wellcome Sanger Institute COVID-19 Surveillance Team |
| EPI_ISL_2908099 | Lighthouse Lab in Glasgow | Wellcome Sanger Institute for the COVID-19 Genomics UK (COG-UK) Consortium | Harper VanSteenhouse, Yumi Kasai, David Gray, Carol Clugston, Anna Dominiczak and Alex Alderton, Roberto Amato, Jeffrey Barrett, Sonia Goncalves, Ewan Harrison, David K. Jackson, Ian Johnston, Dominic Kwiatkowski, Cordelia Langford, John Sillitoe on behalf of the Wellcome Sanger Institute COVID-19 Surveillance Team |
| EPI_ISL_2908100 | Lighthouse Lab in Milton Keynes | Wellcome Sanger Institute for the COVID-19 Genomics UK (COG-UK) Consortium | The Lighthouse Lab in Milton Keynes and Alex Alderton, Roberto Amato, Jeffrey Barrett, Sonia Goncalves, Ewan Harrison, David K. Jackson, Ian Johnston, Dominic Kwiatkowski, Cordelia Langford, John Sillitoe on behalf of the Wellcome Sanger Institute COVID-19 Surveillance Team |
| EPI_ISL_2908101 | Lighthouse Lab in Glasgow | Wellcome Sanger Institute for the COVID-19 Genomics UK (COG-UK) Consortium | Harper VanSteenhouse, Yumi Kasai, David Gray, Carol Clugston, Anna Dominiczak and Alex Alderton, Roberto Amato, Jeffrey Barrett, Sonia Goncalves, Ewan Harrison, David K. Jackson, Ian Johnston, Dominic Kwiatkowski, Cordelia Langford, John Sillitoe on behalf of the Wellcome Sanger Institute COVID-19 Surveillance Team |
| EPI_ISL_2908102, EPI_ISL_2908103, EPI_ISL_2908104, EPI_ISL_2908105, EPI_ISL_2908106, EPI_ISL_2908107, EPI_ISL_2908108 | Lighthouse Lab in Milton Keynes | Wellcome Sanger Institute for the COVID-19 Genomics UK (COG-UK) Consortium | The Lighthouse Lab in Milton Keynes and Alex Alderton, Roberto Amato, Jeffrey Barrett, Sonia Goncalves, Ewan Harrison, David K. Jackson, Ian Johnston, Dominic Kwiatkowski, Cordelia Langford, John Sillitoe on behalf of the Wellcome Sanger Institute COVID-19 Surveillance Team |
| EPI_ISL_2908109 | Lighthouse Lab in Glasgow | Wellcome Sanger Institute for the COVID-19 Genomics UK (COG-UK) Consortium | Harper VanSteenhouse, Yumi Kasai, David Gray, Carol Clugston, Anna Dominiczak and Alex Alderton, Roberto Amato, Jeffrey Barrett, Sonia Goncalves, Ewan Harrison, David K. Jackson, Ian Johnston, Dominic Kwiatkowski, Cordelia Langford, John Sillitoe on behalf of the Wellcome Sanger Institute COVID-19 Surveillance Team |
| EPI_ISL_2908110, EPI_ISL_2908111, EPI_ISL_2908112, EPI_ISL_2908113, EPI_ISL_2908114, EPI_ISL_2908115, EPI_ISL_2908116, EPI_ISL_2908117, EPI_ISL_2908119, EPI_ISL_2908120, EPI_ISL_2908121, EPI_ISL_2908122, EPI_ISL_2908123, EPI_ISL_2908124, EPI_ISL_2908125, EPI_ISL_2908126, EPI_ISL_2908127 | | | |
| see above | Lighthouse Lab in Milton Keynes | Wellcome Sanger Institute for the COVID-19 Genomics UK (COG-UK) Consortium | The Lighthouse Lab in Milton Keynes and Alex Alderton, Roberto Amato, Jeffrey Barrett, Sonia Goncalves, Ewan Harrison, David K. Jackson, Ian Johnston, Dominic Kwiatkowski, Cordelia Langford, John Sillitoe on behalf of the Wellcome Sanger Institute COVID-19 Surveillance Team |
| EPI_ISL_2908128, EPI_ISL_2908129, EPI_ISL_2908130 | Lighthouse Lab in Glasgow | Wellcome Sanger Institute for the COVID-19 Genomics UK (COG-UK) Consortium | Harper VanSteenhouse, Yumi Kasai, David Gray, Carol Clugston, Anna Dominiczak and Alex Alderton, Roberto Amato, Jeffrey Barrett, Sonia Goncalves, Ewan Harrison, David K. Jackson, Ian Johnston, Dominic Kwiatkowski, Cordelia Langford, John Sillitoe on behalf of the Wellcome Sanger Institute COVID-19 Surveillance Team |
| EPI_ISL_2908131, EPI_ISL_2908132, EPI_ISL_2908133, EPI_ISL_2908134, EPI_ISL_2908135 | Lighthouse Lab in Milton Keynes | Wellcome Sanger Institute for the COVID-19 Genomics UK (COG-UK) Consortium | The Lighthouse Lab in Milton Keynes and Alex Alderton, Roberto Amato, Jeffrey Barrett, Sonia Goncalves, Ewan Harrison, David K. Jackson, Ian Johnston, Dominic Kwiatkowski, Cordelia Langford, John Sillitoe on behalf of the Wellcome Sanger Institute COVID-19 Surveillance Team |
| EPI_ISL_2908136 | Lighthouse Lab in Glasgow | Wellcome Sanger Institute for the COVID-19 Genomics UK (COG-UK) Consortium | Harper VanSteenhouse, Yumi Kasai, David Gray, Carol Clugston, Anna Dominiczak and Alex Alderton, Roberto Amato, Jeffrey Barrett, Sonia Goncalves, Ewan Harrison, David K. Jackson, Ian Johnston, Dominic Kwiatkowski, Cordelia Langford, John Sillitoe on behalf of the Wellcome Sanger Institute COVID-19 Surveillance Team |
| EPI_ISL_2908137, EPI_ISL_2908138, EPI_ISL_2908139, EPI_ISL_2908140, EPI_ISL_2908141, EPI_ISL_2908142 | Lighthouse Lab in Milton Keynes | Wellcome Sanger Institute for the COVID-19 Genomics UK (COG-UK) Consortium | The Lighthouse Lab in Milton Keynes and Alex Alderton, Roberto Amato, Jeffrey Barrett, Sonia Goncalves, Ewan Harrison, David K. Jackson, Ian Johnston, Dominic Kwiatkowski, Cordelia Langford, John Sillitoe on behalf of the Wellcome Sanger Institute COVID-19 Surveillance Team |
| EPI_ISL_2908143, EPI_ISL_2908144 | Lighthouse Lab in Glasgow | Wellcome Sanger Institute for the COVID-19 Genomics UK (COG-UK) Consortium | Harper VanSteenhouse, Yumi Kasai, David Gray, Carol Clugston, Anna Dominiczak and Alex Alderton, Roberto Amato, Jeffrey Barrett, Sonia Goncalves, Ewan Harrison, David K. Jackson, Ian Johnston, Dominic Kwiatkowski, Cordelia Langford, John Sillitoe on behalf of the Wellcome Sanger Institute COVID-19 Surveillance Team |
| EPI_ISL_2908145, EPI_ISL_2908146, EPI_ISL_2908147, EPI_ISL_2908148, EPI_ISL_2908149, EPI_ISL_2908150 | Lighthouse Lab in Milton Keynes | Wellcome Sanger Institute for the COVID-19 Genomics UK (COG-UK) Consortium | The Lighthouse Lab in Milton Keynes and Alex Alderton, Roberto Amato, Jeffrey Barrett, Sonia Goncalves, Ewan Harrison, David K. Jackson, Ian Johnston, Dominic Kwiatkowski, Cordelia Langford, John Sillitoe on behalf of the Wellcome Sanger Institute COVID-19 Surveillance Team |
| EPI_ISL_2908151, EPI_ISL_2908152 | Lighthouse Lab in Glasgow | Wellcome Sanger Institute for the COVID-19 Genomics UK (COG-UK) Consortium | Harper VanSteenhouse, Yumi Kasai, David Gray, Carol Clugston, Anna Dominiczak and Alex Alderton, Roberto Amato, Jeffrey Barrett, Sonia Goncalves, Ewan Harrison, David K. Jackson, Ian Johnston, Dominic Kwiatkowski, Cordelia Langford, John Sillitoe on behalf of the Wellcome Sanger Institute COVID-19 Surveillance Team |
| EPI_ISL_2908153, EPI_ISL_2908154, EPI_ISL_2908155, EPI_ISL_2908156 | Lighthouse Lab in Milton Keynes | Wellcome Sanger Institute for the COVID-19 Genomics UK (COG-UK) Consortium | The Lighthouse Lab in Milton Keynes and Alex Alderton, Roberto Amato, Jeffrey Barrett, Sonia Goncalves, Ewan Harrison, David K. Jackson, Ian Johnston, Dominic Kwiatkowski, Cordelia Langford, John Sillitoe on behalf of the Wellcome Sanger Institute COVID-19 Surveillance Team |

| | | | |
|---|---|---|---|
| EPI_ISL_2908157 | Lighthouse Lab in Glasgow | Wellcome Sanger Institute for the COVID-19 Genomics UK (COG-UK) Consortium | Harper VanSteenhouse, Yumi Kasai, David Gray, Carol Clugston, Anna Dominiczak and Alex Alderton, Roberto Amato, Jeffrey Barrett, Sonia Goncalves, Ewan Harrison, David K. Jackson, Ian Johnston, Dominic Kwiatkowski, Cordelia Langford, John Sillitoe on behalf of the Wellcome Sanger Institute COVID-19 Surveillance Team |
| EPI_ISL_2908158 | Lighthouse Lab in Milton Keynes | Wellcome Sanger Institute for the COVID-19 Genomics UK (COG-UK) Consortium | The Lighthouse Lab in Milton Keynes and Alex Alderton, Roberto Amato, Jeffrey Barrett, Sonia Goncalves, Ewan Harrison, David K. Jackson, Ian Johnston, Dominic Kwiatkowski, Cordelia Langford, John Sillitoe on behalf of the Wellcome Sanger Institute COVID-19 Surveillance Team |
| EPI_ISL_2908159 | Lighthouse Lab in Glasgow | Wellcome Sanger Institute for the COVID-19 Genomics UK (COG-UK) Consortium | Harper VanSteenhouse, Yumi Kasai, David Gray, Carol Clugston, Anna Dominiczak and Alex Alderton, Roberto Amato, Jeffrey Barrett, Sonia Goncalves, Ewan Harrison, David K. Jackson, Ian Johnston, Dominic Kwiatkowski, Cordelia Langford, John Sillitoe on behalf of the Wellcome Sanger Institute COVID-19 Surveillance Team |
| EPI_ISL_2908160, EPI_ISL_2908161, EPI_ISL_2908162, EPI_ISL_2908163, EPI_ISL_2908164, EPI_ISL_2908165, EPI_ISL_2908166, EPI_ISL_2908167, EPI_ISL_2908168, EPI_ISL_2908169, EPI_ISL_2908170, EPI_ISL_2908171, EPI_ISL_2908172, EPI_ISL_2908173, EPI_ISL_2908174 | | | |
| see above | Lighthouse Lab in Milton Keynes | Wellcome Sanger Institute for the COVID-19 Genomics UK (COG-UK) Consortium | The Lighthouse Lab in Milton Keynes and Alex Alderton, Roberto Amato, Jeffrey Barrett, Sonia Goncalves, Ewan Harrison, David K. Jackson, Ian Johnston, Dominic Kwiatkowski, Cordelia Langford, John Sillitoe on behalf of the Wellcome Sanger Institute COVID-19 Surveillance Team |
| EPI_ISL_2908175 | Lighthouse Lab in Glasgow | Wellcome Sanger Institute for the COVID-19 Genomics UK (COG-UK) Consortium | Harper VanSteenhouse, Yumi Kasai, David Gray, Carol Clugston, Anna Dominiczak and Alex Alderton, Roberto Amato, Jeffrey Barrett, Sonia Goncalves, Ewan Harrison, David K. Jackson, Ian Johnston, Dominic Kwiatkowski, Cordelia Langford, John Sillitoe on behalf of the Wellcome Sanger Institute COVID-19 Surveillance Team |
| EPI_ISL_2908176 | Lighthouse Lab in Milton Keynes | Wellcome Sanger Institute for the COVID-19 Genomics UK (COG-UK) Consortium | The Lighthouse Lab in Milton Keynes and Alex Alderton, Roberto Amato, Jeffrey Barrett, Sonia Goncalves, Ewan Harrison, David K. Jackson, Ian Johnston, Dominic Kwiatkowski, Cordelia Langford, John Sillitoe on behalf of the Wellcome Sanger Institute COVID-19 Surveillance Team |
| EPI_ISL_2908177, EPI_ISL_2908178 | Lighthouse Lab in Glasgow | Wellcome Sanger Institute for the COVID-19 Genomics UK (COG-UK) Consortium | Harper VanSteenhouse, Yumi Kasai, David Gray, Carol Clugston, Anna Dominiczak and Alex Alderton, Roberto Amato, Jeffrey Barrett, Sonia Goncalves, Ewan Harrison, David K. Jackson, Ian Johnston, Dominic Kwiatkowski, Cordelia Langford, John Sillitoe on behalf of the Wellcome Sanger Institute COVID-19 Surveillance Team |
| EPI_ISL_2908179 | Lighthouse Lab in Milton Keynes | Wellcome Sanger Institute for the COVID-19 Genomics UK (COG-UK) Consortium | The Lighthouse Lab in Milton Keynes and Alex Alderton, Roberto Amato, Jeffrey Barrett, Sonia Goncalves, Ewan Harrison, David K. Jackson, Ian Johnston, Dominic Kwiatkowski, Cordelia Langford, John Sillitoe on behalf of the Wellcome Sanger Institute COVID-19 Surveillance Team |
| EPI_ISL_2908180, EPI_ISL_2908181, EPI_ISL_2908182 | Lighthouse Lab in Glasgow | Wellcome Sanger Institute for the COVID-19 Genomics UK (COG-UK) Consortium | Harper VanSteenhouse, Yumi Kasai, David Gray, Carol Clugston, Anna Dominiczak and Alex Alderton, Roberto Amato, Jeffrey Barrett, Sonia Goncalves, Ewan Harrison, David K. Jackson, Ian Johnston, Dominic Kwiatkowski, Cordelia Langford, John Sillitoe on behalf of the Wellcome Sanger Institute COVID-19 Surveillance Team |
| EPI_ISL_2908183 | Lighthouse Lab in Milton Keynes | Wellcome Sanger Institute for the COVID-19 Genomics UK (COG-UK) Consortium | The Lighthouse Lab in Milton Keynes and Alex Alderton, Roberto Amato, Jeffrey Barrett, Sonia Goncalves, Ewan Harrison, David K. Jackson, Ian Johnston, Dominic Kwiatkowski, Cordelia Langford, John Sillitoe on behalf of the Wellcome Sanger Institute COVID-19 Surveillance Team |
| EPI_ISL_2908184, EPI_ISL_2908185, EPI_ISL_2908186 | Lighthouse Lab in Glasgow | Wellcome Sanger Institute for the COVID-19 Genomics UK (COG-UK) Consortium | Harper VanSteenhouse, Yumi Kasai, David Gray, Carol Clugston, Anna Dominiczak and Alex Alderton, Roberto Amato, Jeffrey Barrett, Sonia Goncalves, Ewan Harrison, David K. Jackson, Ian Johnston, Dominic Kwiatkowski, Cordelia Langford, John Sillitoe on behalf of the Wellcome Sanger Institute COVID-19 Surveillance Team |
| EPI_ISL_2908187 | Lighthouse Lab in Milton Keynes | Wellcome Sanger Institute for the COVID-19 Genomics UK (COG-UK) Consortium | The Lighthouse Lab in Milton Keynes and Alex Alderton, Roberto Amato, Jeffrey Barrett, Sonia Goncalves, Ewan Harrison, David K. Jackson, Ian Johnston, Dominic Kwiatkowski, Cordelia Langford, John Sillitoe on behalf of the Wellcome Sanger Institute COVID-19 Surveillance Team |
| EPI_ISL_2908188 | Lighthouse Lab in Glasgow | Wellcome Sanger Institute for the COVID-19 Genomics UK (COG-UK) Consortium | Harper VanSteenhouse, Yumi Kasai, David Gray, Carol Clugston, Anna Dominiczak and Alex Alderton, Roberto Amato, Jeffrey Barrett, Sonia Goncalves, Ewan Harrison, David K. Jackson, Ian Johnston, Dominic Kwiatkowski, Cordelia Langford, John Sillitoe on behalf of the Wellcome Sanger Institute COVID-19 Surveillance Team |
| EPI_ISL_2908189, EPI_ISL_2908190 | Lighthouse Lab in Milton Keynes | Wellcome Sanger Institute for the COVID-19 Genomics UK (COG-UK) Consortium | The Lighthouse Lab in Milton Keynes and Alex Alderton, Roberto Amato, Jeffrey Barrett, Sonia Goncalves, Ewan Harrison, David K. Jackson, Ian Johnston, Dominic Kwiatkowski, Cordelia Langford, John Sillitoe on behalf of the Wellcome Sanger Institute COVID-19 Surveillance Team |
| EPI_ISL_2908191, EPI_ISL_2908192, EPI_ISL_2908193, EPI_ISL_2908194 | Lighthouse Lab in Glasgow | Wellcome Sanger Institute for the COVID-19 Genomics UK (COG-UK) Consortium | Harper VanSteenhouse, Yumi Kasai, David Gray, Carol Clugston, Anna Dominiczak and Alex Alderton, Roberto Amato, Jeffrey Barrett, Sonia Goncalves, Ewan Harrison, David K. Jackson, Ian Johnston, Dominic Kwiatkowski, Cordelia Langford, John Sillitoe on behalf of the Wellcome Sanger Institute COVID-19 Surveillance Team |
| EPI_ISL_2908195, EPI_ISL_2908196 | Lighthouse Lab in Milton Keynes | Wellcome Sanger Institute for the COVID-19 Genomics UK (COG-UK) Consortium | The Lighthouse Lab in Milton Keynes and Alex Alderton, Roberto Amato, Jeffrey Barrett, Sonia Goncalves, Ewan Harrison, David K. Jackson, Ian Johnston, Dominic Kwiatkowski, Cordelia Langford, John Sillitoe on behalf of the Wellcome Sanger Institute COVID-19 Surveillance Team |
| EPI_ISL_2908197 | Lighthouse Lab in Glasgow | Wellcome Sanger Institute for the COVID-19 Genomics UK (COG-UK) Consortium | Harper VanSteenhouse, Yumi Kasai, David Gray, Carol Clugston, Anna Dominiczak and Alex Alderton, Roberto Amato, Jeffrey Barrett, Sonia Goncalves, Ewan Harrison, David K. Jackson, Ian Johnston, Dominic Kwiatkowski, Cordelia Langford, John Sillitoe on behalf of the Wellcome Sanger Institute COVID-19 Surveillance Team |
| EPI_ISL_2908198, EPI_ISL_2908199 | Lighthouse Lab in Milton Keynes | Wellcome Sanger Institute for the COVID-19 Genomics UK (COG-UK) Consortium | The Lighthouse Lab in Milton Keynes and Alex Alderton, Roberto Amato, Jeffrey Barrett, Sonia Goncalves, Ewan Harrison, David K. Jackson, Ian Johnston, Dominic Kwiatkowski, Cordelia Langford, John Sillitoe on behalf of the Wellcome Sanger Institute COVID-19 Surveillance Team |
| EPI_ISL_2908200 | Lighthouse Lab in Glasgow | Wellcome Sanger Institute for the COVID-19 Genomics UK (COG-UK) Consortium | Harper VanSteenhouse, Yumi Kasai, David Gray, Carol Clugston, Anna Dominiczak and Alex Alderton, Roberto Amato, Jeffrey Barrett, Sonia Goncalves, Ewan Harrison, David K. Jackson, Ian Johnston, Dominic Kwiatkowski, Cordelia Langford, John Sillitoe on behalf of the Wellcome Sanger Institute COVID-19 Surveillance Team |
| EPI_ISL_2908201 | Lighthouse Lab in Milton Keynes | Wellcome Sanger Institute for the COVID-19 Genomics UK (COG-UK) Consortium | The Lighthouse Lab in Milton Keynes and Alex Alderton, Roberto Amato, Jeffrey Barrett, Sonia Goncalves, Ewan Harrison, David K. Jackson, Ian Johnston, Dominic Kwiatkowski, Cordelia Langford, John Sillitoe on behalf of the Wellcome Sanger Institute COVID-19 Surveillance Team |
| EPI_ISL_2908202, EPI_ISL_2908203, EPI_ISL_2908204, EPI_ISL_2908205, EPI_ISL_2908206 | Lighthouse Lab in Glasgow | Wellcome Sanger Institute for the COVID-19 Genomics UK (COG-UK) Consortium | Harper VanSteenhouse, Yumi Kasai, David Gray, Carol Clugston, Anna Dominiczak and Alex Alderton, Roberto Amato, Jeffrey Barrett, Sonia Goncalves, Ewan Harrison, David K. Jackson, Ian Johnston, Dominic Kwiatkowski, Cordelia Langford, John Sillitoe on behalf of the Wellcome Sanger Institute COVID-19 Surveillance Team |
| EPI_ISL_2908207, EPI_ISL_2908208, EPI_ISL_2908209 | Lighthouse Lab in Milton Keynes | Wellcome Sanger Institute for the COVID-19 Genomics UK (COG-UK) Consortium | The Lighthouse Lab in Milton Keynes and Alex Alderton, Roberto Amato, Jeffrey Barrett, Sonia Goncalves, Ewan Harrison, David K. Jackson, Ian Johnston, Dominic Kwiatkowski, Cordelia Langford, John Sillitoe on behalf of the Wellcome Sanger Institute COVID-19 Surveillance Team |
| EPI_ISL_2908210, EPI_ISL_2908211, EPI_ISL_2908212, EPI_ISL_2908213 | Lighthouse Lab in Glasgow | Wellcome Sanger Institute for the COVID-19 Genomics UK (COG-UK) Consortium | Harper VanSteenhouse, Yumi Kasai, David Gray, Carol Clugston, Anna Dominiczak and Alex Alderton, Roberto Amato, Jeffrey Barrett, Sonia Goncalves, Ewan Harrison, David K. Jackson, Ian Johnston, Dominic Kwiatkowski, Cordelia Langford, John Sillitoe on behalf of the Wellcome Sanger Institute COVID-19 Surveillance Team |
| EPI_ISL_2908214 | Lighthouse Lab in Milton Keynes | Wellcome Sanger Institute for the COVID-19 Genomics UK (COG-UK) Consortium | The Lighthouse Lab in Milton Keynes and Alex Alderton, Roberto Amato, Jeffrey Barrett, Sonia Goncalves, Ewan Harrison, David K. Jackson, Ian Johnston, Dominic Kwiatkowski, Cordelia Langford, John Sillitoe on behalf of the Wellcome Sanger Institute COVID-19 Surveillance Team |
| EPI_ISL_2908215 | Lighthouse Lab in Glasgow | Wellcome Sanger Institute for the COVID-19 Genomics UK (COG-UK) Consortium | Harper VanSteenhouse, Yumi Kasai, David Gray, Carol Clugston, Anna Dominiczak and Alex Alderton, Roberto Amato, Jeffrey Barrett, Sonia Goncalves, Ewan Harrison, David K. Jackson, Ian Johnston, Dominic Kwiatkowski, Cordelia Langford, John Sillitoe on behalf of the Wellcome Sanger Institute COVID-19 Surveillance Team |
| EPI_ISL_2908216 | Lighthouse Lab in Milton Keynes | Wellcome Sanger Institute for the COVID-19 Genomics UK (COG-UK) Consortium | The Lighthouse Lab in Milton Keynes and Alex Alderton, Roberto Amato, Jeffrey Barrett, Sonia Goncalves, Ewan Harrison, David K. Jackson, Ian Johnston, Dominic Kwiatkowski, Cordelia Langford, John Sillitoe on behalf of the Wellcome Sanger Institute COVID-19 Surveillance Team |
| EPI_ISL_2908217 | Lighthouse Lab in Glasgow | Wellcome Sanger Institute for the COVID-19 Genomics UK (COG-UK) Consortium | Harper VanSteenhouse, Yumi Kasai, David Gray, Carol Clugston, Anna Dominiczak and Alex Alderton, Roberto Amato, Jeffrey Barrett, Sonia Goncalves, Ewan Harrison, David K. Jackson, Ian Johnston, Dominic Kwiatkowski, Cordelia Langford, John Sillitoe on behalf of the Wellcome Sanger Institute COVID-19 Surveillance Team |
| EPI_ISL_2908218 | Lighthouse Lab in Milton Keynes | Wellcome Sanger Institute for the COVID-19 Genomics UK (COG-UK) Consortium | The Lighthouse Lab in Milton Keynes and Alex Alderton, Roberto Amato, Jeffrey Barrett, Sonia Goncalves, Ewan Harrison, David K. Jackson, Ian Johnston, Dominic Kwiatkowski, Cordelia Langford, John Sillitoe on behalf of the Wellcome Sanger Institute COVID-19 Surveillance Team |
| EPI_ISL_2908219, EPI_ISL_2908220, EPI_ISL_2908221, EPI_ISL_2908222 | Lighthouse Lab in Glasgow | Wellcome Sanger Institute for the COVID-19 Genomics UK (COG-UK) Consortium | Harper VanSteenhouse, Yumi Kasai, David Gray, Carol Clugston, Anna Dominiczak and Alex Alderton, Roberto Amato, Jeffrey Barrett, Sonia Goncalves, Ewan Harrison, David K. Jackson, Ian Johnston, Dominic Kwiatkowski, Cordelia Langford, John Sillitoe on behalf of the Wellcome Sanger Institute COVID-19 Surveillance Team |
| EPI_ISL_2908223 | Lighthouse Lab in Milton Keynes | Wellcome Sanger Institute for the COVID-19 Genomics UK (COG-UK) Consortium | The Lighthouse Lab in Milton Keynes and Alex Alderton, Roberto Amato, Jeffrey Barrett, Sonia Goncalves, Ewan Harrison, David K. Jackson, Ian Johnston, Dominic Kwiatkowski, Cordelia Langford, John Sillitoe on behalf of the Wellcome Sanger Institute COVID-19 Surveillance Team |
| EPI_ISL_2908224, EPI_ISL_2908225 | Lighthouse Lab in Glasgow | Wellcome Sanger Institute for the COVID-19 Genomics UK (COG-UK) Consortium | Harper VanSteenhouse, Yumi Kasai, David Gray, Carol Clugston, Anna Dominiczak and Alex Alderton, Roberto Amato, Jeffrey Barrett, Sonia Goncalves, Ewan Harrison, David K. Jackson, Ian Johnston, Dominic Kwiatkowski, Cordelia Langford, John Sillitoe on behalf of the Wellcome Sanger Institute COVID-19 Surveillance Team |
| EPI_ISL_2908226 | Lighthouse Lab in Milton Keynes | Wellcome Sanger Institute for the COVID-19 Genomics UK (COG-UK) Consortium | The Lighthouse Lab in Milton Keynes and Alex Alderton, Roberto Amato, Jeffrey Barrett, Sonia Goncalves, Ewan Harrison, David K. Jackson, Ian Johnston, Dominic Kwiatkowski, Cordelia Langford, John Sillitoe on behalf of the Wellcome Sanger Institute COVID-19 Surveillance Team |
| EPI_ISL_2908227, EPI_ISL_2908228 | Lighthouse Lab in Glasgow | Wellcome Sanger Institute for the COVID-19 Genomics UK (COG-UK) Consortium | Harper VanSteenhouse, Yumi Kasai, David Gray, Carol Clugston, Anna Dominiczak and Alex Alderton, Roberto Amato, Jeffrey Barrett, Sonia Goncalves, Ewan Harrison, David K. Jackson, Ian Johnston, Dominic Kwiatkowski, Cordelia Langford, John Sillitoe on behalf of the Wellcome Sanger Institute COVID-19 Surveillance Team |
| EPI_ISL_2908229 | Lighthouse Lab in Milton Keynes | Wellcome Sanger Institute for the COVID-19 Genomics UK (COG-UK) Consortium | The Lighthouse Lab in Milton Keynes and Alex Alderton, Roberto Amato, Jeffrey Barrett, Sonia Goncalves, Ewan Harrison, David K. Jackson, Ian Johnston, Dominic Kwiatkowski, Cordelia Langford, John Sillitoe on behalf of the Wellcome Sanger Institute COVID-19 Surveillance Team |
| EPI_ISL_2908230 | Lighthouse Lab in Glasgow | Wellcome Sanger Institute for the COVID-19 Genomics UK (COG-UK) Consortium | Harper VanSteenhouse, Yumi Kasai, David Gray, Carol Clugston, Anna Dominiczak and Alex Alderton, Roberto Amato, Jeffrey Barrett, Sonia Goncalves, Ewan Harrison, David K. Jackson, Ian Johnston, Dominic Kwiatkowski, Cordelia Langford, John Sillitoe on behalf of the Wellcome Sanger Institute COVID-19 Surveillance Team |
| EPI_ISL_2908231, EPI_ISL_2908232 | Lighthouse Lab in Milton Keynes | Wellcome Sanger Institute for the COVID-19 Genomics UK (COG-UK) Consortium | The Lighthouse Lab in Milton Keynes and Alex Alderton, Roberto Amato, Jeffrey Barrett, Sonia Goncalves, Ewan Harrison, David K. Jackson, Ian Johnston, Dominic Kwiatkowski, Cordelia Langford, John Sillitoe on behalf of the Wellcome Sanger Institute COVID-19 Surveillance Team |
| EPI_ISL_2908233, EPI_ISL_2908234, EPI_ISL_2908235 | Lighthouse Lab in Glasgow | Wellcome Sanger Institute for the COVID-19 Genomics UK (COG-UK) Consortium | Harper VanSteenhouse, Yumi Kasai, David Gray, Carol Clugston, Anna Dominiczak and Alex Alderton, Roberto Amato, Jeffrey Barrett, Sonia Goncalves, Ewan Harrison, David K. Jackson, Ian Johnston, Dominic Kwiatkowski, Cordelia Langford, John Sillitoe on behalf of the Wellcome Sanger Institute COVID-19 Surveillance Team |
| EPI_ISL_2908236 | Lighthouse Lab in Milton Keynes | Wellcome Sanger Institute for the COVID-19 Genomics UK (COG-UK) Consortium | The Lighthouse Lab in Milton Keynes and Alex Alderton, Roberto Amato, Jeffrey Barrett, Sonia Goncalves, Ewan Harrison, David K. Jackson, Ian Johnston, Dominic Kwiatkowski, Cordelia Langford, John Sillitoe on behalf of the Wellcome Sanger Institute COVID-19 Surveillance Team |
| EPI_ISL_2908237, EPI_ISL_2908238 | Lighthouse Lab in Glasgow | Wellcome Sanger Institute for the COVID-19 Genomics UK (COG-UK) Consortium | Harper VanSteenhouse, Yumi Kasai, David Gray, Carol Clugston, Anna Dominiczak and Alex Alderton, Roberto Amato, Jeffrey Barrett, Sonia Goncalves, Ewan Harrison, David K. Jackson, Ian Johnston, Dominic Kwiatkowski, Cordelia Langford, John Sillitoe on behalf of the Wellcome Sanger Institute COVID-19 Surveillance Team |
| EPI_ISL_2908239 | Lighthouse Lab in Milton Keynes | Wellcome Sanger Institute for the COVID-19 Genomics UK (COG-UK) | The Lighthouse Lab in Milton Keynes and Alex Alderton, Roberto Amato, Jeffrey Barrett, Sonia Goncalves, Ewan Harrison, David K. Jackson, Ian Johnston, Dominic Kwiatkowski, Cordelia Langford, John Sillitoe on behalf of the Wellcome Sanger |

| | | | |
|---|---|---|---|
| | | Consortium | Institute COVID-19 Surveillance Team |
| EPI_ISL_2908240 | Lighthouse Lab in Glasgow | Wellcome Sanger Institute for the COVID-19 Genomics UK (COG-UK) Consortium | Harper VanSteenhouse, Yumi Kasai, David Gray, Carol Clugston, Anna Dominiczak and Alex Alderton, Roberto Amato, Jeffrey Barrett, Sonia Goncalves, Ewan Harrison, David K. Jackson, Ian Johnston, Dominic Kwiatkowski, Cordelia Langford, John Sillitoe on behalf of the Wellcome Sanger Institute COVID-19 Surveillance Team |
| EPI_ISL_2908241, EPI_ISL_2908242, EPI_ISL_2908243, EPI_ISL_2908244, EPI_ISL_2908245, EPI_ISL_2908246, EPI_ISL_2908247, EPI_ISL_2908248 | Lighthouse Lab in Milton Keynes | Wellcome Sanger Institute for the COVID-19 Genomics UK (COG-UK) Consortium | The Lighthouse Lab in Milton Keynes and Alex Alderton, Roberto Amato, Jeffrey Barrett, Sonia Goncalves, Ewan Harrison, David K. Jackson, Ian Johnston, Dominic Kwiatkowski, Cordelia Langford, John Sillitoe on behalf of the Wellcome Sanger Institute COVID-19 Surveillance Team |
| EPI_ISL_2908249 | Lighthouse Lab in Glasgow | Wellcome Sanger Institute for the COVID-19 Genomics UK (COG-UK) Consortium | Harper VanSteenhouse, Yumi Kasai, David Gray, Carol Clugston, Anna Dominiczak and Alex Alderton, Roberto Amato, Jeffrey Barrett, Sonia Goncalves, Ewan Harrison, David K. Jackson, Ian Johnston, Dominic Kwiatkowski, Cordelia Langford, John Sillitoe on behalf of the Wellcome Sanger Institute COVID-19 Surveillance Team |
| EPI_ISL_2908250, EPI_ISL_2908251, EPI_ISL_2908252, EPI_ISL_2908253, EPI_ISL_2908254 | Lighthouse Lab in Milton Keynes | Wellcome Sanger Institute for the COVID-19 Genomics UK (COG-UK) Consortium | The Lighthouse Lab in Milton Keynes and Alex Alderton, Roberto Amato, Jeffrey Barrett, Sonia Goncalves, Ewan Harrison, David K. Jackson, Ian Johnston, Dominic Kwiatkowski, Cordelia Langford, John Sillitoe on behalf of the Wellcome Sanger Institute COVID-19 Surveillance Team |
| EPI_ISL_2908255, EPI_ISL_2908256, EPI_ISL_2908257, EPI_ISL_2908258 | Lighthouse Lab in Glasgow | Wellcome Sanger Institute for the COVID-19 Genomics UK (COG-UK) Consortium | Harper VanSteenhouse, Yumi Kasai, David Gray, Carol Clugston, Anna Dominiczak and Alex Alderton, Roberto Amato, Jeffrey Barrett, Sonia Goncalves, Ewan Harrison, David K. Jackson, Ian Johnston, Dominic Kwiatkowski, Cordelia Langford, John Sillitoe on behalf of the Wellcome Sanger Institute COVID-19 Surveillance Team |
| EPI_ISL_2908259, EPI_ISL_2908260, EPI_ISL_2908261, EPI_ISL_2908262 | Lighthouse Lab in Milton Keynes | Wellcome Sanger Institute for the COVID-19 Genomics UK (COG-UK) Consortium | The Lighthouse Lab in Milton Keynes and Alex Alderton, Roberto Amato, Jeffrey Barrett, Sonia Goncalves, Ewan Harrison, David K. Jackson, Ian Johnston, Dominic Kwiatkowski, Cordelia Langford, John Sillitoe on behalf of the Wellcome Sanger Institute COVID-19 Surveillance Team |
| EPI_ISL_2908263, EPI_ISL_2908264 | Lighthouse Lab in Glasgow | Wellcome Sanger Institute for the COVID-19 Genomics UK (COG-UK) Consortium | Harper VanSteenhouse, Yumi Kasai, David Gray, Carol Clugston, Anna Dominiczak and Alex Alderton, Roberto Amato, Jeffrey Barrett, Sonia Goncalves, Ewan Harrison, David K. Jackson, Ian Johnston, Dominic Kwiatkowski, Cordelia Langford, John Sillitoe on behalf of the Wellcome Sanger Institute COVID-19 Surveillance Team |
| EPI_ISL_2908265 | Lighthouse Lab in Milton Keynes | Wellcome Sanger Institute for the COVID-19 Genomics UK (COG-UK) Consortium | The Lighthouse Lab in Milton Keynes and Alex Alderton, Roberto Amato, Jeffrey Barrett, Sonia Goncalves, Ewan Harrison, David K. Jackson, Ian Johnston, Dominic Kwiatkowski, Cordelia Langford, John Sillitoe on behalf of the Wellcome Sanger Institute COVID-19 Surveillance Team |
| EPI_ISL_2908266, EPI_ISL_2908267 | Lighthouse Lab in Glasgow | Wellcome Sanger Institute for the COVID-19 Genomics UK (COG-UK) Consortium | Harper VanSteenhouse, Yumi Kasai, David Gray, Carol Clugston, Anna Dominiczak and Alex Alderton, Roberto Amato, Jeffrey Barrett, Sonia Goncalves, Ewan Harrison, David K. Jackson, Ian Johnston, Dominic Kwiatkowski, Cordelia Langford, John Sillitoe on behalf of the Wellcome Sanger Institute COVID-19 Surveillance Team |
| EPI_ISL_2908268 | Lighthouse Lab in Milton Keynes | Wellcome Sanger Institute for the COVID-19 Genomics UK (COG-UK) Consortium | The Lighthouse Lab in Milton Keynes and Alex Alderton, Roberto Amato, Jeffrey Barrett, Sonia Goncalves, Ewan Harrison, David K. Jackson, Ian Johnston, Dominic Kwiatkowski, Cordelia Langford, John Sillitoe on behalf of the Wellcome Sanger Institute COVID-19 Surveillance Team |
| EPI_ISL_2908269 | Lighthouse Lab in Glasgow | Wellcome Sanger Institute for the COVID-19 Genomics UK (COG-UK) Consortium | Harper VanSteenhouse, Yumi Kasai, David Gray, Carol Clugston, Anna Dominiczak and Alex Alderton, Roberto Amato, Jeffrey Barrett, Sonia Goncalves, Ewan Harrison, David K. Jackson, Ian Johnston, Dominic Kwiatkowski, Cordelia Langford, John Sillitoe on behalf of the Wellcome Sanger Institute COVID-19 Surveillance Team |
| EPI_ISL_2908270, EPI_ISL_2908271, EPI_ISL_2908272 | Lighthouse Lab in Milton Keynes | Wellcome Sanger Institute for the COVID-19 Genomics UK (COG-UK) Consortium | The Lighthouse Lab in Milton Keynes and Alex Alderton, Roberto Amato, Jeffrey Barrett, Sonia Goncalves, Ewan Harrison, David K. Jackson, Ian Johnston, Dominic Kwiatkowski, Cordelia Langford, John Sillitoe on behalf of the Wellcome Sanger Institute COVID-19 Surveillance Team |
| EPI_ISL_2908273 | Lighthouse Lab in Glasgow | Wellcome Sanger Institute for the COVID-19 Genomics UK (COG-UK) Consortium | Harper VanSteenhouse, Yumi Kasai, David Gray, Carol Clugston, Anna Dominiczak and Alex Alderton, Roberto Amato, Jeffrey Barrett, Sonia Goncalves, Ewan Harrison, David K. Jackson, Ian Johnston, Dominic Kwiatkowski, Cordelia Langford, John Sillitoe on behalf of the Wellcome Sanger Institute COVID-19 Surveillance Team |
| EPI_ISL_2908274, EPI_ISL_2908275, EPI_ISL_2908276, EPI_ISL_2908277, EPI_ISL_2908278, EPI_ISL_2908279, EPI_ISL_2908280, EPI_ISL_2908281 | Lighthouse Lab in Milton Keynes | Wellcome Sanger Institute for the COVID-19 Genomics UK (COG-UK) Consortium | The Lighthouse Lab in Milton Keynes and Alex Alderton, Roberto Amato, Jeffrey Barrett, Sonia Goncalves, Ewan Harrison, David K. Jackson, Ian Johnston, Dominic Kwiatkowski, Cordelia Langford, John Sillitoe on behalf of the Wellcome Sanger Institute COVID-19 Surveillance Team |
| EPI_ISL_2908282 | Lighthouse Lab in Glasgow | Wellcome Sanger Institute for the COVID-19 Genomics UK (COG-UK) Consortium | Harper VanSteenhouse, Yumi Kasai, David Gray, Carol Clugston, Anna Dominiczak and Alex Alderton, Roberto Amato, Jeffrey Barrett, Sonia Goncalves, Ewan Harrison, David K. Jackson, Ian Johnston, Dominic Kwiatkowski, Cordelia Langford, John Sillitoe on behalf of the Wellcome Sanger Institute COVID-19 Surveillance Team |
| EPI_ISL_2908283, EPI_ISL_2908284, EPI_ISL_2908285, EPI_ISL_2908286 | Lighthouse Lab in Milton Keynes | Wellcome Sanger Institute for the COVID-19 Genomics UK (COG-UK) Consortium | The Lighthouse Lab in Milton Keynes and Alex Alderton, Roberto Amato, Jeffrey Barrett, Sonia Goncalves, Ewan Harrison, David K. Jackson, Ian Johnston, Dominic Kwiatkowski, Cordelia Langford, John Sillitoe on behalf of the Wellcome Sanger Institute COVID-19 Surveillance Team |
| EPI_ISL_2908287 | Lighthouse Lab in Glasgow | Wellcome Sanger Institute for the COVID-19 Genomics UK (COG-UK) Consortium | Harper VanSteenhouse, Yumi Kasai, David Gray, Carol Clugston, Anna Dominiczak and Alex Alderton, Roberto Amato, Jeffrey Barrett, Sonia Goncalves, Ewan Harrison, David K. Jackson, Ian Johnston, Dominic Kwiatkowski, Cordelia Langford, John Sillitoe on behalf of the Wellcome Sanger Institute COVID-19 Surveillance Team |
| EPI_ISL_2908288 | Lighthouse Lab in Milton Keynes | Wellcome Sanger Institute for the COVID-19 Genomics UK (COG-UK) Consortium | The Lighthouse Lab in Milton Keynes and Alex Alderton, Roberto Amato, Jeffrey Barrett, Sonia Goncalves, Ewan Harrison, David K. Jackson, Ian Johnston, Dominic Kwiatkowski, Cordelia Langford, John Sillitoe on behalf of the Wellcome Sanger Institute COVID-19 Surveillance Team |
| EPI_ISL_2908289, EPI_ISL_2908290 | Lighthouse Lab in Glasgow | Wellcome Sanger Institute for the COVID-19 Genomics UK (COG-UK) Consortium | Harper VanSteenhouse, Yumi Kasai, David Gray, Carol Clugston, Anna Dominiczak and Alex Alderton, Roberto Amato, Jeffrey Barrett, Sonia Goncalves, Ewan Harrison, David K. Jackson, Ian Johnston, Dominic Kwiatkowski, Cordelia Langford, John Sillitoe on behalf of the Wellcome Sanger Institute COVID-19 Surveillance Team |
| EPI_ISL_2908291, EPI_ISL_2908292 | Lighthouse Lab in Milton Keynes | Wellcome Sanger Institute for the COVID-19 Genomics UK (COG-UK) Consortium | The Lighthouse Lab in Milton Keynes and Alex Alderton, Roberto Amato, Jeffrey Barrett, Sonia Goncalves, Ewan Harrison, David K. Jackson, Ian Johnston, Dominic Kwiatkowski, Cordelia Langford, John Sillitoe on behalf of the Wellcome Sanger Institute COVID-19 Surveillance Team |
| EPI_ISL_2908293 | Lighthouse Lab in Glasgow | Wellcome Sanger Institute for the COVID-19 Genomics UK (COG-UK) Consortium | Harper VanSteenhouse, Yumi Kasai, David Gray, Carol Clugston, Anna Dominiczak and Alex Alderton, Roberto Amato, Jeffrey Barrett, Sonia Goncalves, Ewan Harrison, David K. Jackson, Ian Johnston, Dominic Kwiatkowski, Cordelia Langford, John Sillitoe on behalf of the Wellcome Sanger Institute COVID-19 Surveillance Team |
| EPI_ISL_2908294, EPI_ISL_2908295 | Lighthouse Lab in Milton Keynes | Wellcome Sanger Institute for the COVID-19 Genomics UK (COG-UK) Consortium | The Lighthouse Lab in Milton Keynes and Alex Alderton, Roberto Amato, Jeffrey Barrett, Sonia Goncalves, Ewan Harrison, David K. Jackson, Ian Johnston, Dominic Kwiatkowski, Cordelia Langford, John Sillitoe on behalf of the Wellcome Sanger Institute COVID-19 Surveillance Team |
| EPI_ISL_2908296, EPI_ISL_2908297 | Lighthouse Lab in Glasgow | Wellcome Sanger Institute for the COVID-19 Genomics UK (COG-UK) Consortium | Harper VanSteenhouse, Yumi Kasai, David Gray, Carol Clugston, Anna Dominiczak and Alex Alderton, Roberto Amato, Jeffrey Barrett, Sonia Goncalves, Ewan Harrison, David K. Jackson, Ian Johnston, Dominic Kwiatkowski, Cordelia Langford, John Sillitoe on behalf of the Wellcome Sanger Institute COVID-19 Surveillance Team |
| EPI_ISL_2908298 | Lighthouse Lab in Milton Keynes | Wellcome Sanger Institute for the COVID-19 Genomics UK (COG-UK) Consortium | The Lighthouse Lab in Milton Keynes and Alex Alderton, Roberto Amato, Jeffrey Barrett, Sonia Goncalves, Ewan Harrison, David K. Jackson, Ian Johnston, Dominic Kwiatkowski, Cordelia Langford, John Sillitoe on behalf of the Wellcome Sanger Institute COVID-19 Surveillance Team |
| EPI_ISL_2908299, EPI_ISL_2908300, EPI_ISL_2908301 | Lighthouse Lab in Glasgow | Wellcome Sanger Institute for the COVID-19 Genomics UK (COG-UK) Consortium | Harper VanSteenhouse, Yumi Kasai, David Gray, Carol Clugston, Anna Dominiczak and Alex Alderton, Roberto Amato, Jeffrey Barrett, Sonia Goncalves, Ewan Harrison, David K. Jackson, Ian Johnston, Dominic Kwiatkowski, Cordelia Langford, John Sillitoe on behalf of the Wellcome Sanger Institute COVID-19 Surveillance Team |
| EPI_ISL_2908302, EPI_ISL_2908303, EPI_ISL_2908304, EPI_ISL_2908305 | Lighthouse Lab in Milton Keynes | Wellcome Sanger Institute for the COVID-19 Genomics UK (COG-UK) Consortium | The Lighthouse Lab in Milton Keynes and Alex Alderton, Roberto Amato, Jeffrey Barrett, Sonia Goncalves, Ewan Harrison, David K. Jackson, Ian Johnston, Dominic Kwiatkowski, Cordelia Langford, John Sillitoe on behalf of the Wellcome Sanger Institute COVID-19 Surveillance Team |
| EPI_ISL_2908306, EPI_ISL_2908307 | Lighthouse Lab in Glasgow | Wellcome Sanger Institute for the COVID-19 Genomics UK (COG-UK) Consortium | Harper VanSteenhouse, Yumi Kasai, David Gray, Carol Clugston, Anna Dominiczak and Alex Alderton, Roberto Amato, Jeffrey Barrett, Sonia Goncalves, Ewan Harrison, David K. Jackson, Ian Johnston, Dominic Kwiatkowski, Cordelia Langford, John Sillitoe on behalf of the Wellcome Sanger Institute COVID-19 Surveillance Team |
| EPI_ISL_2908308, EPI_ISL_2908309 | Lighthouse Lab in Milton Keynes | Wellcome Sanger Institute for the COVID-19 Genomics UK (COG-UK) Consortium | The Lighthouse Lab in Milton Keynes and Alex Alderton, Roberto Amato, Jeffrey Barrett, Sonia Goncalves, Ewan Harrison, David K. Jackson, Ian Johnston, Dominic Kwiatkowski, Cordelia Langford, John Sillitoe on behalf of the Wellcome Sanger Institute COVID-19 Surveillance Team |
| EPI_ISL_2908310, EPI_ISL_2908311 | Lighthouse Lab in Glasgow | Wellcome Sanger Institute for the COVID-19 Genomics UK (COG-UK) Consortium | Harper VanSteenhouse, Yumi Kasai, David Gray, Carol Clugston, Anna Dominiczak and Alex Alderton, Roberto Amato, Jeffrey Barrett, Sonia Goncalves, Ewan Harrison, David K. Jackson, Ian Johnston, Dominic Kwiatkowski, Cordelia Langford, John Sillitoe on behalf of the Wellcome Sanger Institute COVID-19 Surveillance Team |
| EPI_ISL_2908312, EPI_ISL_2908313, EPI_ISL_2908314 | Lighthouse Lab in Milton Keynes | Wellcome Sanger Institute for the COVID-19 Genomics UK (COG-UK) Consortium | The Lighthouse Lab in Milton Keynes and Alex Alderton, Roberto Amato, Jeffrey Barrett, Sonia Goncalves, Ewan Harrison, David K. Jackson, Ian Johnston, Dominic Kwiatkowski, Cordelia Langford, John Sillitoe on behalf of the Wellcome Sanger Institute COVID-19 Surveillance Team |
| EPI_ISL_2908315, EPI_ISL_2908316 | Lighthouse Lab in Glasgow | Wellcome Sanger Institute for the COVID-19 Genomics UK (COG-UK) Consortium | Harper VanSteenhouse, Yumi Kasai, David Gray, Carol Clugston, Anna Dominiczak and Alex Alderton, Roberto Amato, Jeffrey Barrett, Sonia Goncalves, Ewan Harrison, David K. Jackson, Ian Johnston, Dominic Kwiatkowski, Cordelia Langford, John Sillitoe on behalf of the Wellcome Sanger Institute COVID-19 Surveillance Team |
| EPI_ISL_2908317, EPI_ISL_2908318 | Lighthouse Lab in Milton Keynes | Wellcome Sanger Institute for the COVID-19 Genomics UK (COG-UK) Consortium | The Lighthouse Lab in Milton Keynes and Alex Alderton, Roberto Amato, Jeffrey Barrett, Sonia Goncalves, Ewan Harrison, David K. Jackson, Ian Johnston, Dominic Kwiatkowski, Cordelia Langford, John Sillitoe on behalf of the Wellcome Sanger Institute COVID-19 Surveillance Team |
| EPI_ISL_2908319, EPI_ISL_2908320, EPI_ISL_2908321, EPI_ISL_2908322 | Lighthouse Lab in Glasgow | Wellcome Sanger Institute for the COVID-19 Genomics UK (COG-UK) Consortium | Harper VanSteenhouse, Yumi Kasai, David Gray, Carol Clugston, Anna Dominiczak and Alex Alderton, Roberto Amato, Jeffrey Barrett, Sonia Goncalves, Ewan Harrison, David K. Jackson, Ian Johnston, Dominic Kwiatkowski, Cordelia Langford, John Sillitoe on behalf of the Wellcome Sanger Institute COVID-19 Surveillance Team |

| | | | |
|---|---|---|---|
| EPI_ISL_2908323 | Lighthouse Lab in Milton Keynes | Wellcome Sanger Institute for the COVID-19 Genomics UK (COG-UK) Consortium | The Lighthouse Lab in Milton Keynes and Alex Alderton, Roberto Amato, Jeffrey Barrett, Sonia Goncalves, Ewan Harrison, David K. Jackson, Ian Johnston, Dominic Kwiatkowski, Cordelia Langford, John Sillitoe on behalf of the Wellcome Sanger Institute COVID-19 Surveillance Team |
| EPI_ISL_2908324 | Lighthouse Lab in Glasgow | Wellcome Sanger Institute for the COVID-19 Genomics UK (COG-UK) Consortium | Harper VanSteenhouse, Yumi Kasai, David Gray, Carol Clugston, Anna Dominiczak and Alex Alderton, Roberto Amato, Jeffrey Barrett, Sonia Goncalves, Ewan Harrison, David K. Jackson, Ian Johnston, Dominic Kwiatkowski, Cordelia Langford, John Sillitoe on behalf of the Wellcome Sanger Institute COVID-19 Surveillance Team |
| EPI_ISL_2908325 | Lighthouse Lab in Milton Keynes | Wellcome Sanger Institute for the COVID-19 Genomics UK (COG-UK) Consortium | The Lighthouse Lab in Milton Keynes and Alex Alderton, Roberto Amato, Jeffrey Barrett, Sonia Goncalves, Ewan Harrison, David K. Jackson, Ian Johnston, Dominic Kwiatkowski, Cordelia Langford, John Sillitoe on behalf of the Wellcome Sanger Institute COVID-19 Surveillance Team |
| EPI_ISL_2908326 | Lighthouse Lab in Glasgow | Wellcome Sanger Institute for the COVID-19 Genomics UK (COG-UK) Consortium | Harper VanSteenhouse, Yumi Kasai, David Gray, Carol Clugston, Anna Dominiczak and Alex Alderton, Roberto Amato, Jeffrey Barrett, Sonia Goncalves, Ewan Harrison, David K. Jackson, Ian Johnston, Dominic Kwiatkowski, Cordelia Langford, John Sillitoe on behalf of the Wellcome Sanger Institute COVID-19 Surveillance Team |
| EPI_ISL_2908327, EPI_ISL_2908328, EPI_ISL_2908329, EPI_ISL_2908330, EPI_ISL_2908331, EPI_ISL_2908332, EPI_ISL_2908333, EPI_ISL_2908334 | Lighthouse Lab in Milton Keynes | Wellcome Sanger Institute for the COVID-19 Genomics UK (COG-UK) Consortium | The Lighthouse Lab in Milton Keynes and Alex Alderton, Roberto Amato, Jeffrey Barrett, Sonia Goncalves, Ewan Harrison, David K. Jackson, Ian Johnston, Dominic Kwiatkowski, Cordelia Langford, John Sillitoe on behalf of the Wellcome Sanger Institute COVID-19 Surveillance Team |
| EPI_ISL_2908335 | Lighthouse Lab in Glasgow | Wellcome Sanger Institute for the COVID-19 Genomics UK (COG-UK) Consortium | Harper VanSteenhouse, Yumi Kasai, David Gray, Carol Clugston, Anna Dominiczak and Alex Alderton, Roberto Amato, Jeffrey Barrett, Sonia Goncalves, Ewan Harrison, David K. Jackson, Ian Johnston, Dominic Kwiatkowski, Cordelia Langford, John Sillitoe on behalf of the Wellcome Sanger Institute COVID-19 Surveillance Team |
| EPI_ISL_2908336 | Lighthouse Lab in Milton Keynes | Wellcome Sanger Institute for the COVID-19 Genomics UK (COG-UK) Consortium | The Lighthouse Lab in Milton Keynes and Alex Alderton, Roberto Amato, Jeffrey Barrett, Sonia Goncalves, Ewan Harrison, David K. Jackson, Ian Johnston, Dominic Kwiatkowski, Cordelia Langford, John Sillitoe on behalf of the Wellcome Sanger Institute COVID-19 Surveillance Team |
| EPI_ISL_2908337 | Lighthouse Lab in Glasgow | Wellcome Sanger Institute for the COVID-19 Genomics UK (COG-UK) Consortium | Harper VanSteenhouse, Yumi Kasai, David Gray, Carol Clugston, Anna Dominiczak and Alex Alderton, Roberto Amato, Jeffrey Barrett, Sonia Goncalves, Ewan Harrison, David K. Jackson, Ian Johnston, Dominic Kwiatkowski, Cordelia Langford, John Sillitoe on behalf of the Wellcome Sanger Institute COVID-19 Surveillance Team |
| EPI_ISL_2908338 | Lighthouse Lab in Milton Keynes | Wellcome Sanger Institute for the COVID-19 Genomics UK (COG-UK) Consortium | The Lighthouse Lab in Milton Keynes and Alex Alderton, Roberto Amato, Jeffrey Barrett, Sonia Goncalves, Ewan Harrison, David K. Jackson, Ian Johnston, Dominic Kwiatkowski, Cordelia Langford, John Sillitoe on behalf of the Wellcome Sanger Institute COVID-19 Surveillance Team |
| EPI_ISL_2908339, EPI_ISL_2908340 | Lighthouse Lab in Glasgow | Wellcome Sanger Institute for the COVID-19 Genomics UK (COG-UK) Consortium | Harper VanSteenhouse, Yumi Kasai, David Gray, Carol Clugston, Anna Dominiczak and Alex Alderton, Roberto Amato, Jeffrey Barrett, Sonia Goncalves, Ewan Harrison, David K. Jackson, Ian Johnston, Dominic Kwiatkowski, Cordelia Langford, John Sillitoe on behalf of the Wellcome Sanger Institute COVID-19 Surveillance Team |
| EPI_ISL_2908341, EPI_ISL_2908342, EPI_ISL_2908343 | Lighthouse Lab in Milton Keynes | Wellcome Sanger Institute for the COVID-19 Genomics UK (COG-UK) Consortium | The Lighthouse Lab in Milton Keynes and Alex Alderton, Roberto Amato, Jeffrey Barrett, Sonia Goncalves, Ewan Harrison, David K. Jackson, Ian Johnston, Dominic Kwiatkowski, Cordelia Langford, John Sillitoe on behalf of the Wellcome Sanger Institute COVID-19 Surveillance Team |
| EPI_ISL_2908344 | Lighthouse Lab in Glasgow | Wellcome Sanger Institute for the COVID-19 Genomics UK (COG-UK) Consortium | Harper VanSteenhouse, Yumi Kasai, David Gray, Carol Clugston, Anna Dominiczak and Alex Alderton, Roberto Amato, Jeffrey Barrett, Sonia Goncalves, Ewan Harrison, David K. Jackson, Ian Johnston, Dominic Kwiatkowski, Cordelia Langford, John Sillitoe on behalf of the Wellcome Sanger Institute COVID-19 Surveillance Team |
| EPI_ISL_2908345 | Lighthouse Lab in Milton Keynes | Wellcome Sanger Institute for the COVID-19 Genomics UK (COG-UK) Consortium | The Lighthouse Lab in Milton Keynes and Alex Alderton, Roberto Amato, Jeffrey Barrett, Sonia Goncalves, Ewan Harrison, David K. Jackson, Ian Johnston, Dominic Kwiatkowski, Cordelia Langford, John Sillitoe on behalf of the Wellcome Sanger Institute COVID-19 Surveillance Team |
| EPI_ISL_2908346 | Lighthouse Lab in Glasgow | Wellcome Sanger Institute for the COVID-19 Genomics UK (COG-UK) Consortium | Harper VanSteenhouse, Yumi Kasai, David Gray, Carol Clugston, Anna Dominiczak and Alex Alderton, Roberto Amato, Jeffrey Barrett, Sonia Goncalves, Ewan Harrison, David K. Jackson, Ian Johnston, Dominic Kwiatkowski, Cordelia Langford, John Sillitoe on behalf of the Wellcome Sanger Institute COVID-19 Surveillance Team |
| EPI_ISL_2908347, EPI_ISL_2908348, EPI_ISL_2908349, EPI_ISL_2908350, EPI_ISL_2908351 | Lighthouse Lab in Milton Keynes | Wellcome Sanger Institute for the COVID-19 Genomics UK (COG-UK) Consortium | The Lighthouse Lab in Milton Keynes and Alex Alderton, Roberto Amato, Jeffrey Barrett, Sonia Goncalves, Ewan Harrison, David K. Jackson, Ian Johnston, Dominic Kwiatkowski, Cordelia Langford, John Sillitoe on behalf of the Wellcome Sanger Institute COVID-19 Surveillance Team |
| EPI_ISL_2908352 | Lighthouse Lab in Glasgow | Wellcome Sanger Institute for the COVID-19 Genomics UK (COG-UK) Consortium | Harper VanSteenhouse, Yumi Kasai, David Gray, Carol Clugston, Anna Dominiczak and Alex Alderton, Roberto Amato, Jeffrey Barrett, Sonia Goncalves, Ewan Harrison, David K. Jackson, Ian Johnston, Dominic Kwiatkowski, Cordelia Langford, John Sillitoe on behalf of the Wellcome Sanger Institute COVID-19 Surveillance Team |
| EPI_ISL_2908353, EPI_ISL_2908354, EPI_ISL_2908355 | Lighthouse Lab in Milton Keynes | Wellcome Sanger Institute for the COVID-19 Genomics UK (COG-UK) Consortium | The Lighthouse Lab in Milton Keynes and Alex Alderton, Roberto Amato, Jeffrey Barrett, Sonia Goncalves, Ewan Harrison, David K. Jackson, Ian Johnston, Dominic Kwiatkowski, Cordelia Langford, John Sillitoe on behalf of the Wellcome Sanger Institute COVID-19 Surveillance Team |
| EPI_ISL_2908356, EPI_ISL_2908357 | Lighthouse Lab in Glasgow | Wellcome Sanger Institute for the COVID-19 Genomics UK (COG-UK) Consortium | Harper VanSteenhouse, Yumi Kasai, David Gray, Carol Clugston, Anna Dominiczak and Alex Alderton, Roberto Amato, Jeffrey Barrett, Sonia Goncalves, Ewan Harrison, David K. Jackson, Ian Johnston, Dominic Kwiatkowski, Cordelia Langford, John Sillitoe on behalf of the Wellcome Sanger Institute COVID-19 Surveillance Team |
| EPI_ISL_2908358 | Lighthouse Lab in Milton Keynes | Wellcome Sanger Institute for the COVID-19 Genomics UK (COG-UK) Consortium | The Lighthouse Lab in Milton Keynes and Alex Alderton, Roberto Amato, Jeffrey Barrett, Sonia Goncalves, Ewan Harrison, David K. Jackson, Ian Johnston, Dominic Kwiatkowski, Cordelia Langford, John Sillitoe on behalf of the Wellcome Sanger Institute COVID-19 Surveillance Team |
| EPI_ISL_2908359, EPI_ISL_2908360 | Lighthouse Lab in Glasgow | Wellcome Sanger Institute for the COVID-19 Genomics UK (COG-UK) Consortium | Harper VanSteenhouse, Yumi Kasai, David Gray, Carol Clugston, Anna Dominiczak and Alex Alderton, Roberto Amato, Jeffrey Barrett, Sonia Goncalves, Ewan Harrison, David K. Jackson, Ian Johnston, Dominic Kwiatkowski, Cordelia Langford, John Sillitoe on behalf of the Wellcome Sanger Institute COVID-19 Surveillance Team |
| EPI_ISL_2908361, EPI_ISL_2908362 | Lighthouse Lab in Milton Keynes | Wellcome Sanger Institute for the COVID-19 Genomics UK (COG-UK) Consortium | The Lighthouse Lab in Milton Keynes and Alex Alderton, Roberto Amato, Jeffrey Barrett, Sonia Goncalves, Ewan Harrison, David K. Jackson, Ian Johnston, Dominic Kwiatkowski, Cordelia Langford, John Sillitoe on behalf of the Wellcome Sanger Institute COVID-19 Surveillance Team |
| EPI_ISL_2908363 | Lighthouse Lab in Glasgow | Wellcome Sanger Institute for the COVID-19 Genomics UK (COG-UK) Consortium | Harper VanSteenhouse, Yumi Kasai, David Gray, Carol Clugston, Anna Dominiczak and Alex Alderton, Roberto Amato, Jeffrey Barrett, Sonia Goncalves, Ewan Harrison, David K. Jackson, Ian Johnston, Dominic Kwiatkowski, Cordelia Langford, John Sillitoe on behalf of the Wellcome Sanger Institute COVID-19 Surveillance Team |
| EPI_ISL_2908364 | Lighthouse Lab in Milton Keynes | Wellcome Sanger Institute for the COVID-19 Genomics UK (COG-UK) Consortium | The Lighthouse Lab in Milton Keynes and Alex Alderton, Roberto Amato, Jeffrey Barrett, Sonia Goncalves, Ewan Harrison, David K. Jackson, Ian Johnston, Dominic Kwiatkowski, Cordelia Langford, John Sillitoe on behalf of the Wellcome Sanger Institute COVID-19 Surveillance Team |
| EPI_ISL_2908365 | Lighthouse Lab in Glasgow | Wellcome Sanger Institute for the COVID-19 Genomics UK (COG-UK) Consortium | Harper VanSteenhouse, Yumi Kasai, David Gray, Carol Clugston, Anna Dominiczak and Alex Alderton, Roberto Amato, Jeffrey Barrett, Sonia Goncalves, Ewan Harrison, David K. Jackson, Ian Johnston, Dominic Kwiatkowski, Cordelia Langford, John Sillitoe on behalf of the Wellcome Sanger Institute COVID-19 Surveillance Team |
| EPI_ISL_2908366, EPI_ISL_2908367 | Lighthouse Lab in Milton Keynes | Wellcome Sanger Institute for the COVID-19 Genomics UK (COG-UK) Consortium | The Lighthouse Lab in Milton Keynes and Alex Alderton, Roberto Amato, Jeffrey Barrett, Sonia Goncalves, Ewan Harrison, David K. Jackson, Ian Johnston, Dominic Kwiatkowski, Cordelia Langford, John Sillitoe on behalf of the Wellcome Sanger Institute COVID-19 Surveillance Team |
| EPI_ISL_2908368, EPI_ISL_2908369, EPI_ISL_2908370 | Lighthouse Lab in Glasgow | Wellcome Sanger Institute for the COVID-19 Genomics UK (COG-UK) Consortium | Harper VanSteenhouse, Yumi Kasai, David Gray, Carol Clugston, Anna Dominiczak and Alex Alderton, Roberto Amato, Jeffrey Barrett, Sonia Goncalves, Ewan Harrison, David K. Jackson, Ian Johnston, Dominic Kwiatkowski, Cordelia Langford, John Sillitoe on behalf of the Wellcome Sanger Institute COVID-19 Surveillance Team |
| EPI_ISL_2908371 | Lighthouse Lab in Milton Keynes | Wellcome Sanger Institute for the COVID-19 Genomics UK (COG-UK) Consortium | The Lighthouse Lab in Milton Keynes and Alex Alderton, Roberto Amato, Jeffrey Barrett, Sonia Goncalves, Ewan Harrison, David K. Jackson, Ian Johnston, Dominic Kwiatkowski, Cordelia Langford, John Sillitoe on behalf of the Wellcome Sanger Institute COVID-19 Surveillance Team |
| EPI_ISL_2908372, EPI_ISL_2908373, EPI_ISL_2908374 | Lighthouse Lab in Glasgow | Wellcome Sanger Institute for the COVID-19 Genomics UK (COG-UK) Consortium | Harper VanSteenhouse, Yumi Kasai, David Gray, Carol Clugston, Anna Dominiczak and Alex Alderton, Roberto Amato, Jeffrey Barrett, Sonia Goncalves, Ewan Harrison, David K. Jackson, Ian Johnston, Dominic Kwiatkowski, Cordelia Langford, John Sillitoe on behalf of the Wellcome Sanger Institute COVID-19 Surveillance Team |
| EPI_ISL_2908375, EPI_ISL_2908376, EPI_ISL_2908377, EPI_ISL_2908378, EPI_ISL_2908379 | Lighthouse Lab in Milton Keynes | Wellcome Sanger Institute for the COVID-19 Genomics UK (COG-UK) Consortium | The Lighthouse Lab in Milton Keynes and Alex Alderton, Roberto Amato, Jeffrey Barrett, Sonia Goncalves, Ewan Harrison, David K. Jackson, Ian Johnston, Dominic Kwiatkowski, Cordelia Langford, John Sillitoe on behalf of the Wellcome Sanger Institute COVID-19 Surveillance Team |
| EPI_ISL_2908380, EPI_ISL_2908381, EPI_ISL_2908382 | Lighthouse Lab in Glasgow | Wellcome Sanger Institute for the COVID-19 Genomics UK (COG-UK) Consortium | Harper VanSteenhouse, Yumi Kasai, David Gray, Carol Clugston, Anna Dominiczak and Alex Alderton, Roberto Amato, Jeffrey Barrett, Sonia Goncalves, Ewan Harrison, David K. Jackson, Ian Johnston, Dominic Kwiatkowski, Cordelia Langford, John Sillitoe on behalf of the Wellcome Sanger Institute COVID-19 Surveillance Team |
| EPI_ISL_2908383 | Lighthouse Lab in Milton Keynes | Wellcome Sanger Institute for the COVID-19 Genomics UK (COG-UK) Consortium | The Lighthouse Lab in Milton Keynes and Alex Alderton, Roberto Amato, Jeffrey Barrett, Sonia Goncalves, Ewan Harrison, David K. Jackson, Ian Johnston, Dominic Kwiatkowski, Cordelia Langford, John Sillitoe on behalf of the Wellcome Sanger Institute COVID-19 Surveillance Team |
| EPI_ISL_2908384, EPI_ISL_2908385, EPI_ISL_2908386 | Lighthouse Lab in Glasgow | Wellcome Sanger Institute for the COVID-19 Genomics UK (COG-UK) Consortium | Harper VanSteenhouse, Yumi Kasai, David Gray, Carol Clugston, Anna Dominiczak and Alex Alderton, Roberto Amato, Jeffrey Barrett, Sonia Goncalves, Ewan Harrison, David K. Jackson, Ian Johnston, Dominic Kwiatkowski, Cordelia Langford, John Sillitoe on behalf of the Wellcome Sanger Institute COVID-19 Surveillance Team |
| EPI_ISL_2908387 | Lighthouse Lab in Milton Keynes | Wellcome Sanger Institute for the COVID-19 Genomics UK (COG-UK) Consortium | The Lighthouse Lab in Milton Keynes and Alex Alderton, Roberto Amato, Jeffrey Barrett, Sonia Goncalves, Ewan Harrison, David K. Jackson, Ian Johnston, Dominic Kwiatkowski, Cordelia Langford, John Sillitoe on behalf of the Wellcome Sanger Institute COVID-19 Surveillance Team |
| EPI_ISL_2908388, EPI_ISL_2908389 | Lighthouse Lab in Glasgow | Wellcome Sanger Institute for the COVID-19 Genomics UK (COG-UK) Consortium | Harper VanSteenhouse, Yumi Kasai, David Gray, Carol Clugston, Anna Dominiczak and Alex Alderton, Roberto Amato, Jeffrey Barrett, Sonia Goncalves, Ewan Harrison, David K. Jackson, Ian Johnston, Dominic Kwiatkowski, Cordelia Langford, John Sillitoe on behalf of the Wellcome Sanger Institute COVID-19 Surveillance Team |
| EPI_ISL_2908390 | Lighthouse Lab in Milton Keynes | Wellcome Sanger Institute for the COVID-19 Genomics UK (COG-UK) Consortium | The Lighthouse Lab in Milton Keynes and Alex Alderton, Roberto Amato, Jeffrey Barrett, Sonia Goncalves, Ewan Harrison, David K. Jackson, Ian Johnston, Dominic Kwiatkowski, Cordelia Langford, John Sillitoe on behalf of the Wellcome Sanger Institute COVID-19 Surveillance Team |
| EPI_ISL_2908391, EPI_ISL_2908392 | Lighthouse Lab in Glasgow | Wellcome Sanger Institute for the COVID-19 Genomics UK (COG-UK) Consortium | Harper VanSteenhouse, Yumi Kasai, David Gray, Carol Clugston, Anna Dominiczak and Alex Alderton, Roberto Amato, Jeffrey Barrett, Sonia Goncalves, Ewan Harrison, David K. Jackson, Ian Johnston, Dominic Kwiatkowski, Cordelia Langford, John Sillitoe on behalf of the Wellcome Sanger Institute COVID-19 Surveillance Team |
| EPI_ISL_2908393, EPI_ISL_2908394 | Lighthouse Lab in Milton Keynes | Wellcome Sanger Institute for the COVID-19 Genomics UK (COG-UK) Consortium | The Lighthouse Lab in Milton Keynes and Alex Alderton, Roberto Amato, Jeffrey Barrett, Sonia Goncalves, Ewan Harrison, David K. Jackson, Ian Johnston, Dominic Kwiatkowski, Cordelia Langford, John Sillitoe on behalf of the Wellcome Sanger Institute COVID-19 Surveillance Team |
| EPI_ISL_2908395 | Lighthouse Lab in Glasgow | Wellcome Sanger Institute for the COVID-19 Genomics UK (COG-UK) Consortium | Harper VanSteenhouse, Yumi Kasai, David Gray, Carol Clugston, Anna Dominiczak and Alex Alderton, Roberto Amato, Jeffrey Barrett, Sonia Goncalves, Ewan Harrison, David K. Jackson, Ian Johnston, Dominic Kwiatkowski, Cordelia Langford, John Sillitoe on behalf of the Wellcome Sanger Institute COVID-19 Surveillance Team |

| | | | |
|---|---|---|---|
| EPI_ISL_2908396 | Lighthouse Lab in Milton Keynes | Wellcome Sanger Institute for the COVID-19 Genomics UK (COG-UK) Consortium | The Lighthouse Lab in Milton Keynes and Alex Alderton, Roberto Amato, Jeffrey Barrett, Sonia Goncalves, Ewan Harrison, David K. Jackson, Ian Johnston, Dominic Kwiatkowski, Cordelia Langford, John Sillitoe on behalf of the Wellcome Sanger Institute COVID-19 Surveillance Team |
| EPI_ISL_2908397, EPI_ISL_2908398 | Lighthouse Lab in Glasgow | Wellcome Sanger Institute for the COVID-19 Genomics UK (COG-UK) Consortium | Harper VanSteenhouse, Yumi Kasai, David Gray, Carol Clugston, Anna Dominiczak and Alex Alderton, Roberto Amato, Jeffrey Barrett, Sonia Goncalves, Ewan Harrison, David K. Jackson, Ian Johnston, Dominic Kwiatkowski, Cordelia Langford, John Sillitoe on behalf of the Wellcome Sanger Institute COVID-19 Surveillance Team |
| EPI_ISL_2908399, EPI_ISL_2908400, EPI_ISL_2908401 | Lighthouse Lab in Milton Keynes | Wellcome Sanger Institute for the COVID-19 Genomics UK (COG-UK) Consortium | The Lighthouse Lab in Milton Keynes and Alex Alderton, Roberto Amato, Jeffrey Barrett, Sonia Goncalves, Ewan Harrison, David K. Jackson, Ian Johnston, Dominic Kwiatkowski, Cordelia Langford, John Sillitoe on behalf of the Wellcome Sanger Institute COVID-19 Surveillance Team |
| EPI_ISL_2908402, EPI_ISL_2908403, EPI_ISL_2908404 | Lighthouse Lab in Glasgow | Wellcome Sanger Institute for the COVID-19 Genomics UK (COG-UK) Consortium | Harper VanSteenhouse, Yumi Kasai, David Gray, Carol Clugston, Anna Dominiczak and Alex Alderton, Roberto Amato, Jeffrey Barrett, Sonia Goncalves, Ewan Harrison, David K. Jackson, Ian Johnston, Dominic Kwiatkowski, Cordelia Langford, John Sillitoe on behalf of the Wellcome Sanger Institute COVID-19 Surveillance Team |
| EPI_ISL_2908405, EPI_ISL_2908406, EPI_ISL_2908407, EPI_ISL_2908408, EPI_ISL_2908409, EPI_ISL_2908410 | Lighthouse Lab in Milton Keynes | Wellcome Sanger Institute for the COVID-19 Genomics UK (COG-UK) Consortium | The Lighthouse Lab in Milton Keynes and Alex Alderton, Roberto Amato, Jeffrey Barrett, Sonia Goncalves, Ewan Harrison, David K. Jackson, Ian Johnston, Dominic Kwiatkowski, Cordelia Langford, John Sillitoe on behalf of the Wellcome Sanger Institute COVID-19 Surveillance Team |
| EPI_ISL_2908411, EPI_ISL_2908412 | Lighthouse Lab in Glasgow | Wellcome Sanger Institute for the COVID-19 Genomics UK (COG-UK) Consortium | Harper VanSteenhouse, Yumi Kasai, David Gray, Carol Clugston, Anna Dominiczak and Alex Alderton, Roberto Amato, Jeffrey Barrett, Sonia Goncalves, Ewan Harrison, David K. Jackson, Ian Johnston, Dominic Kwiatkowski, Cordelia Langford, John Sillitoe on behalf of the Wellcome Sanger Institute COVID-19 Surveillance Team |
| EPI_ISL_2908413, EPI_ISL_2908414, EPI_ISL_2908415, EPI_ISL_2908416, EPI_ISL_2908417 | Lighthouse Lab in Milton Keynes | Wellcome Sanger Institute for the COVID-19 Genomics UK (COG-UK) Consortium | The Lighthouse Lab in Milton Keynes and Alex Alderton, Roberto Amato, Jeffrey Barrett, Sonia Goncalves, Ewan Harrison, David K. Jackson, Ian Johnston, Dominic Kwiatkowski, Cordelia Langford, John Sillitoe on behalf of the Wellcome Sanger Institute COVID-19 Surveillance Team |
| EPI_ISL_2908418 | Lighthouse Lab in Glasgow | Wellcome Sanger Institute for the COVID-19 Genomics UK (COG-UK) Consortium | Harper VanSteenhouse, Yumi Kasai, David Gray, Carol Clugston, Anna Dominiczak and Alex Alderton, Roberto Amato, Jeffrey Barrett, Sonia Goncalves, Ewan Harrison, David K. Jackson, Ian Johnston, Dominic Kwiatkowski, Cordelia Langford, John Sillitoe on behalf of the Wellcome Sanger Institute COVID-19 Surveillance Team |
| EPI_ISL_2908419 | Lighthouse Lab in Milton Keynes | Wellcome Sanger Institute for the COVID-19 Genomics UK (COG-UK) Consortium | The Lighthouse Lab in Milton Keynes and Alex Alderton, Roberto Amato, Jeffrey Barrett, Sonia Goncalves, Ewan Harrison, David K. Jackson, Ian Johnston, Dominic Kwiatkowski, Cordelia Langford, John Sillitoe on behalf of the Wellcome Sanger Institute COVID-19 Surveillance Team |
| EPI_ISL_2908420, EPI_ISL_2908421 | Lighthouse Lab in Glasgow | Wellcome Sanger Institute for the COVID-19 Genomics UK (COG-UK) Consortium | Harper VanSteenhouse, Yumi Kasai, David Gray, Carol Clugston, Anna Dominiczak and Alex Alderton, Roberto Amato, Jeffrey Barrett, Sonia Goncalves, Ewan Harrison, David K. Jackson, Ian Johnston, Dominic Kwiatkowski, Cordelia Langford, John Sillitoe on behalf of the Wellcome Sanger Institute COVID-19 Surveillance Team |
| EPI_ISL_2908422, EPI_ISL_2908423 | Lighthouse Lab in Milton Keynes | Wellcome Sanger Institute for the COVID-19 Genomics UK (COG-UK) Consortium | The Lighthouse Lab in Milton Keynes and Alex Alderton, Roberto Amato, Jeffrey Barrett, Sonia Goncalves, Ewan Harrison, David K. Jackson, Ian Johnston, Dominic Kwiatkowski, Cordelia Langford, John Sillitoe on behalf of the Wellcome Sanger Institute COVID-19 Surveillance Team |
| EPI_ISL_2908424, EPI_ISL_2908425 | Lighthouse Lab in Glasgow | Wellcome Sanger Institute for the COVID-19 Genomics UK (COG-UK) Consortium | Harper VanSteenhouse, Yumi Kasai, David Gray, Carol Clugston, Anna Dominiczak and Alex Alderton, Roberto Amato, Jeffrey Barrett, Sonia Goncalves, Ewan Harrison, David K. Jackson, Ian Johnston, Dominic Kwiatkowski, Cordelia Langford, John Sillitoe on behalf of the Wellcome Sanger Institute COVID-19 Surveillance Team |
| EPI_ISL_2908426, EPI_ISL_2908427 | Lighthouse Lab in Milton Keynes | Wellcome Sanger Institute for the COVID-19 Genomics UK (COG-UK) Consortium | The Lighthouse Lab in Milton Keynes and Alex Alderton, Roberto Amato, Jeffrey Barrett, Sonia Goncalves, Ewan Harrison, David K. Jackson, Ian Johnston, Dominic Kwiatkowski, Cordelia Langford, John Sillitoe on behalf of the Wellcome Sanger Institute COVID-19 Surveillance Team |
| EPI_ISL_2908428 | Lighthouse Lab in Glasgow | Wellcome Sanger Institute for the COVID-19 Genomics UK (COG-UK) Consortium | Harper VanSteenhouse, Yumi Kasai, David Gray, Carol Clugston, Anna Dominiczak and Alex Alderton, Roberto Amato, Jeffrey Barrett, Sonia Goncalves, Ewan Harrison, David K. Jackson, Ian Johnston, Dominic Kwiatkowski, Cordelia Langford, John Sillitoe on behalf of the Wellcome Sanger Institute COVID-19 Surveillance Team |
| EPI_ISL_2908429, EPI_ISL_2908430 | Lighthouse Lab in Milton Keynes | Wellcome Sanger Institute for the COVID-19 Genomics UK (COG-UK) Consortium | The Lighthouse Lab in Milton Keynes and Alex Alderton, Roberto Amato, Jeffrey Barrett, Sonia Goncalves, Ewan Harrison, David K. Jackson, Ian Johnston, Dominic Kwiatkowski, Cordelia Langford, John Sillitoe on behalf of the Wellcome Sanger Institute COVID-19 Surveillance Team |
| EPI_ISL_2908431 | Lighthouse Lab in Glasgow | Wellcome Sanger Institute for the COVID-19 Genomics UK (COG-UK) Consortium | Harper VanSteenhouse, Yumi Kasai, David Gray, Carol Clugston, Anna Dominiczak and Alex Alderton, Roberto Amato, Jeffrey Barrett, Sonia Goncalves, Ewan Harrison, David K. Jackson, Ian Johnston, Dominic Kwiatkowski, Cordelia Langford, John Sillitoe on behalf of the Wellcome Sanger Institute COVID-19 Surveillance Team |
| EPI_ISL_2908432 | Lighthouse Lab in Milton Keynes | Wellcome Sanger Institute for the COVID-19 Genomics UK (COG-UK) Consortium | The Lighthouse Lab in Milton Keynes and Alex Alderton, Roberto Amato, Jeffrey Barrett, Sonia Goncalves, Ewan Harrison, David K. Jackson, Ian Johnston, Dominic Kwiatkowski, Cordelia Langford, John Sillitoe on behalf of the Wellcome Sanger Institute COVID-19 Surveillance Team |
| EPI_ISL_2908433 | Lighthouse Lab in Glasgow | Wellcome Sanger Institute for the COVID-19 Genomics UK (COG-UK) Consortium | Harper VanSteenhouse, Yumi Kasai, David Gray, Carol Clugston, Anna Dominiczak and Alex Alderton, Roberto Amato, Jeffrey Barrett, Sonia Goncalves, Ewan Harrison, David K. Jackson, Ian Johnston, Dominic Kwiatkowski, Cordelia Langford, John Sillitoe on behalf of the Wellcome Sanger Institute COVID-19 Surveillance Team |
| EPI_ISL_2908434, EPI_ISL_2908435 | Lighthouse Lab in Milton Keynes | Wellcome Sanger Institute for the COVID-19 Genomics UK (COG-UK) Consortium | The Lighthouse Lab in Milton Keynes and Alex Alderton, Roberto Amato, Jeffrey Barrett, Sonia Goncalves, Ewan Harrison, David K. Jackson, Ian Johnston, Dominic Kwiatkowski, Cordelia Langford, John Sillitoe on behalf of the Wellcome Sanger Institute COVID-19 Surveillance Team |
| EPI_ISL_2908436 | Lighthouse Lab in Glasgow | Wellcome Sanger Institute for the COVID-19 Genomics UK (COG-UK) Consortium | Harper VanSteenhouse, Yumi Kasai, David Gray, Carol Clugston, Anna Dominiczak and Alex Alderton, Roberto Amato, Jeffrey Barrett, Sonia Goncalves, Ewan Harrison, David K. Jackson, Ian Johnston, Dominic Kwiatkowski, Cordelia Langford, John Sillitoe on behalf of the Wellcome Sanger Institute COVID-19 Surveillance Team |
| EPI_ISL_2908437 | Lighthouse Lab in Milton Keynes | Wellcome Sanger Institute for the COVID-19 Genomics UK (COG-UK) Consortium | The Lighthouse Lab in Milton Keynes and Alex Alderton, Roberto Amato, Jeffrey Barrett, Sonia Goncalves, Ewan Harrison, David K. Jackson, Ian Johnston, Dominic Kwiatkowski, Cordelia Langford, John Sillitoe on behalf of the Wellcome Sanger Institute COVID-19 Surveillance Team |
| EPI_ISL_2908438, EPI_ISL_2908439, EPI_ISL_2908440, EPI_ISL_2908441 | Lighthouse Lab in Glasgow | Wellcome Sanger Institute for the COVID-19 Genomics UK (COG-UK) Consortium | Harper VanSteenhouse, Yumi Kasai, David Gray, Carol Clugston, Anna Dominiczak and Alex Alderton, Roberto Amato, Jeffrey Barrett, Sonia Goncalves, Ewan Harrison, David K. Jackson, Ian Johnston, Dominic Kwiatkowski, Cordelia Langford, John Sillitoe on behalf of the Wellcome Sanger Institute COVID-19 Surveillance Team |
| EPI_ISL_2908442, EPI_ISL_2908443 | Lighthouse Lab in Milton Keynes | Wellcome Sanger Institute for the COVID-19 Genomics UK (COG-UK) Consortium | The Lighthouse Lab in Milton Keynes and Alex Alderton, Roberto Amato, Jeffrey Barrett, Sonia Goncalves, Ewan Harrison, David K. Jackson, Ian Johnston, Dominic Kwiatkowski, Cordelia Langford, John Sillitoe on behalf of the Wellcome Sanger Institute COVID-19 Surveillance Team |
| EPI_ISL_2908444, EPI_ISL_2908445, EPI_ISL_2908446, EPI_ISL_2908447 | Lighthouse Lab in Glasgow | Wellcome Sanger Institute for the COVID-19 Genomics UK (COG-UK) Consortium | Harper VanSteenhouse, Yumi Kasai, David Gray, Carol Clugston, Anna Dominiczak and Alex Alderton, Roberto Amato, Jeffrey Barrett, Sonia Goncalves, Ewan Harrison, David K. Jackson, Ian Johnston, Dominic Kwiatkowski, Cordelia Langford, John Sillitoe on behalf of the Wellcome Sanger Institute COVID-19 Surveillance Team |
| EPI_ISL_2908448, EPI_ISL_2908449, EPI_ISL_2908450 | Lighthouse Lab in Milton Keynes | Wellcome Sanger Institute for the COVID-19 Genomics UK (COG-UK) Consortium | The Lighthouse Lab in Milton Keynes and Alex Alderton, Roberto Amato, Jeffrey Barrett, Sonia Goncalves, Ewan Harrison, David K. Jackson, Ian Johnston, Dominic Kwiatkowski, Cordelia Langford, John Sillitoe on behalf of the Wellcome Sanger Institute COVID-19 Surveillance Team |
| EPI_ISL_2908451 | Lighthouse Lab in Glasgow | Wellcome Sanger Institute for the COVID-19 Genomics UK (COG-UK) Consortium | Harper VanSteenhouse, Yumi Kasai, David Gray, Carol Clugston, Anna Dominiczak and Alex Alderton, Roberto Amato, Jeffrey Barrett, Sonia Goncalves, Ewan Harrison, David K. Jackson, Ian Johnston, Dominic Kwiatkowski, Cordelia Langford, John Sillitoe on behalf of the Wellcome Sanger Institute COVID-19 Surveillance Team |
| EPI_ISL_2908452, EPI_ISL_2908453 | Lighthouse Lab in Milton Keynes | Wellcome Sanger Institute for the COVID-19 Genomics UK (COG-UK) Consortium | The Lighthouse Lab in Milton Keynes and Alex Alderton, Roberto Amato, Jeffrey Barrett, Sonia Goncalves, Ewan Harrison, David K. Jackson, Ian Johnston, Dominic Kwiatkowski, Cordelia Langford, John Sillitoe on behalf of the Wellcome Sanger Institute COVID-19 Surveillance Team |
| EPI_ISL_2908454 | Lighthouse Lab in Glasgow | Wellcome Sanger Institute for the COVID-19 Genomics UK (COG-UK) Consortium | Harper VanSteenhouse, Yumi Kasai, David Gray, Carol Clugston, Anna Dominiczak and Alex Alderton, Roberto Amato, Jeffrey Barrett, Sonia Goncalves, Ewan Harrison, David K. Jackson, Ian Johnston, Dominic Kwiatkowski, Cordelia Langford, John Sillitoe on behalf of the Wellcome Sanger Institute COVID-19 Surveillance Team |
| EPI_ISL_2908455, EPI_ISL_2908456 | Lighthouse Lab in Milton Keynes | Wellcome Sanger Institute for the COVID-19 Genomics UK (COG-UK) Consortium | The Lighthouse Lab in Milton Keynes and Alex Alderton, Roberto Amato, Jeffrey Barrett, Sonia Goncalves, Ewan Harrison, David K. Jackson, Ian Johnston, Dominic Kwiatkowski, Cordelia Langford, John Sillitoe on behalf of the Wellcome Sanger Institute COVID-19 Surveillance Team |
| EPI_ISL_2908457 | Lighthouse Lab in Glasgow | Wellcome Sanger Institute for the COVID-19 Genomics UK (COG-UK) Consortium | Harper VanSteenhouse, Yumi Kasai, David Gray, Carol Clugston, Anna Dominiczak and Alex Alderton, Roberto Amato, Jeffrey Barrett, Sonia Goncalves, Ewan Harrison, David K. Jackson, Ian Johnston, Dominic Kwiatkowski, Cordelia Langford, John Sillitoe on behalf of the Wellcome Sanger Institute COVID-19 Surveillance Team |
| EPI_ISL_2908458, EPI_ISL_2908459, EPI_ISL_2908460, EPI_ISL_2908461 | Lighthouse Lab in Milton Keynes | Wellcome Sanger Institute for the COVID-19 Genomics UK (COG-UK) Consortium | The Lighthouse Lab in Milton Keynes and Alex Alderton, Roberto Amato, Jeffrey Barrett, Sonia Goncalves, Ewan Harrison, David K. Jackson, Ian Johnston, Dominic Kwiatkowski, Cordelia Langford, John Sillitoe on behalf of the Wellcome Sanger Institute COVID-19 Surveillance Team |
| EPI_ISL_2908462 | Lighthouse Lab in Glasgow | Wellcome Sanger Institute for the COVID-19 Genomics UK (COG-UK) Consortium | Harper VanSteenhouse, Yumi Kasai, David Gray, Carol Clugston, Anna Dominiczak and Alex Alderton, Roberto Amato, Jeffrey Barrett, Sonia Goncalves, Ewan Harrison, David K. Jackson, Ian Johnston, Dominic Kwiatkowski, Cordelia Langford, John Sillitoe on behalf of the Wellcome Sanger Institute COVID-19 Surveillance Team |
| EPI_ISL_2908463, EPI_ISL_2908464, EPI_ISL_2908465 | Lighthouse Lab in Milton Keynes | Wellcome Sanger Institute for the COVID-19 Genomics UK (COG-UK) Consortium | The Lighthouse Lab in Milton Keynes and Alex Alderton, Roberto Amato, Jeffrey Barrett, Sonia Goncalves, Ewan Harrison, David K. Jackson, Ian Johnston, Dominic Kwiatkowski, Cordelia Langford, John Sillitoe on behalf of the Wellcome Sanger Institute COVID-19 Surveillance Team |
| EPI_ISL_2908466 | Lighthouse Lab in Glasgow | Wellcome Sanger Institute for the COVID-19 Genomics UK (COG-UK) Consortium | Harper VanSteenhouse, Yumi Kasai, David Gray, Carol Clugston, Anna Dominiczak and Alex Alderton, Roberto Amato, Jeffrey Barrett, Sonia Goncalves, Ewan Harrison, David K. Jackson, Ian Johnston, Dominic Kwiatkowski, Cordelia Langford, John Sillitoe on behalf of the Wellcome Sanger Institute COVID-19 Surveillance Team |
| EPI_ISL_2908467, EPI_ISL_2908468, EPI_ISL_2908469 | Lighthouse Lab in Milton Keynes | Wellcome Sanger Institute for the COVID-19 Genomics UK (COG-UK) Consortium | The Lighthouse Lab in Milton Keynes and Alex Alderton, Roberto Amato, Jeffrey Barrett, Sonia Goncalves, Ewan Harrison, David K. Jackson, Ian Johnston, Dominic Kwiatkowski, Cordelia Langford, John Sillitoe on behalf of the Wellcome Sanger Institute COVID-19 Surveillance Team |
| EPI_ISL_2908470, EPI_ISL_2908471 | Lighthouse Lab in Glasgow | Wellcome Sanger Institute for the COVID-19 Genomics UK (COG-UK) Consortium | Harper VanSteenhouse, Yumi Kasai, David Gray, Carol Clugston, Anna Dominiczak and Alex Alderton, Roberto Amato, Jeffrey Barrett, Sonia Goncalves, Ewan Harrison, David K. Jackson, Ian Johnston, Dominic Kwiatkowski, Cordelia Langford, John Sillitoe on behalf of the Wellcome Sanger Institute COVID-19 Surveillance Team |
| EPI_ISL_2908472 | Lighthouse Lab in Milton Keynes | Wellcome Sanger Institute for the COVID-19 Genomics UK (COG-UK) Consortium | The Lighthouse Lab in Milton Keynes and Alex Alderton, Roberto Amato, Jeffrey Barrett, Sonia Goncalves, Ewan Harrison, David K. Jackson, Ian Johnston, Dominic Kwiatkowski, Cordelia Langford, John Sillitoe on behalf of the Wellcome Sanger Institute COVID-19 Surveillance Team |
| EPI_ISL_2908473 | Lighthouse Lab in Glasgow | Wellcome Sanger Institute for the COVID-19 Genomics UK (COG-UK) Consortium | Harper VanSteenhouse, Yumi Kasai, David Gray, Carol Clugston, Anna Dominiczak and Alex Alderton, Roberto Amato, Jeffrey Barrett, Sonia Goncalves, Ewan Harrison, David K. Jackson, Ian Johnston, Dominic Kwiatkowski, Cordelia Langford, John Sillitoe on behalf of the Wellcome Sanger Institute COVID-19 Surveillance Team |

| | | | |
|---|---|---|---|
| EPI_ISL_2908474, EPI_ISL_2908475, EPI_ISL_2908476 | Lighthouse Lab in Milton Keynes | Wellcome Sanger Institute for the COVID-19 Genomics UK (COG-UK) Consortium | The Lighthouse Lab in Milton Keynes and Alex Alderton, Roberto Amato, Jeffrey Barrett, Sonia Goncalves, Ewan Harrison, David K. Jackson, Ian Johnston, Dominic Kwiatkowski, Cordelia Langford, John Sillitoe on behalf of the Wellcome Sanger Institute COVID-19 Surveillance Team |
| EPI_ISL_2908477 | Lighthouse Lab in Glasgow | Wellcome Sanger Institute for the COVID-19 Genomics UK (COG-UK) Consortium | Harper VanSteenhouse, Yumi Kasai, David Gray, Carol Clugston, Anna Dominiczak and Alex Alderton, Roberto Amato, Jeffrey Barrett, Sonia Goncalves, Ewan Harrison, David K. Jackson, Ian Johnston, Dominic Kwiatkowski, Cordelia Langford, John Sillitoe on behalf of the Wellcome Sanger Institute COVID-19 Surveillance Team |
| EPI_ISL_2908478, EPI_ISL_2908479 | Lighthouse Lab in Milton Keynes | Wellcome Sanger Institute for the COVID-19 Genomics UK (COG-UK) Consortium | The Lighthouse Lab in Milton Keynes and Alex Alderton, Roberto Amato, Jeffrey Barrett, Sonia Goncalves, Ewan Harrison, David K. Jackson, Ian Johnston, Dominic Kwiatkowski, Cordelia Langford, John Sillitoe on behalf of the Wellcome Sanger Institute COVID-19 Surveillance Team |
| EPI_ISL_2908480, EPI_ISL_2908481 | Lighthouse Lab in Glasgow | Wellcome Sanger Institute for the COVID-19 Genomics UK (COG-UK) Consortium | Harper VanSteenhouse, Yumi Kasai, David Gray, Carol Clugston, Anna Dominiczak and Alex Alderton, Roberto Amato, Jeffrey Barrett, Sonia Goncalves, Ewan Harrison, David K. Jackson, Ian Johnston, Dominic Kwiatkowski, Cordelia Langford, John Sillitoe on behalf of the Wellcome Sanger Institute COVID-19 Surveillance Team |
| EPI_ISL_2908482 | Lighthouse Lab in Milton Keynes | Wellcome Sanger Institute for the COVID-19 Genomics UK (COG-UK) Consortium | The Lighthouse Lab in Milton Keynes and Alex Alderton, Roberto Amato, Jeffrey Barrett, Sonia Goncalves, Ewan Harrison, David K. Jackson, Ian Johnston, Dominic Kwiatkowski, Cordelia Langford, John Sillitoe on behalf of the Wellcome Sanger Institute COVID-19 Surveillance Team |
| EPI_ISL_2908483, EPI_ISL_2908484 | Lighthouse Lab in Glasgow | Wellcome Sanger Institute for the COVID-19 Genomics UK (COG-UK) Consortium | Harper VanSteenhouse, Yumi Kasai, David Gray, Carol Clugston, Anna Dominiczak and Alex Alderton, Roberto Amato, Jeffrey Barrett, Sonia Goncalves, Ewan Harrison, David K. Jackson, Ian Johnston, Dominic Kwiatkowski, Cordelia Langford, John Sillitoe on behalf of the Wellcome Sanger Institute COVID-19 Surveillance Team |
| EPI_ISL_2908485 | Lighthouse Lab in Milton Keynes | Wellcome Sanger Institute for the COVID-19 Genomics UK (COG-UK) Consortium | The Lighthouse Lab in Milton Keynes and Alex Alderton, Roberto Amato, Jeffrey Barrett, Sonia Goncalves, Ewan Harrison, David K. Jackson, Ian Johnston, Dominic Kwiatkowski, Cordelia Langford, John Sillitoe on behalf of the Wellcome Sanger Institute COVID-19 Surveillance Team |
| EPI_ISL_2908486 | Lighthouse Lab in Glasgow | Wellcome Sanger Institute for the COVID-19 Genomics UK (COG-UK) Consortium | Harper VanSteenhouse, Yumi Kasai, David Gray, Carol Clugston, Anna Dominiczak and Alex Alderton, Roberto Amato, Jeffrey Barrett, Sonia Goncalves, Ewan Harrison, David K. Jackson, Ian Johnston, Dominic Kwiatkowski, Cordelia Langford, John Sillitoe on behalf of the Wellcome Sanger Institute COVID-19 Surveillance Team |
| EPI_ISL_2908487, EPI_ISL_2908488 | Lighthouse Lab in Milton Keynes | Wellcome Sanger Institute for the COVID-19 Genomics UK (COG-UK) Consortium | The Lighthouse Lab in Milton Keynes and Alex Alderton, Roberto Amato, Jeffrey Barrett, Sonia Goncalves, Ewan Harrison, David K. Jackson, Ian Johnston, Dominic Kwiatkowski, Cordelia Langford, John Sillitoe on behalf of the Wellcome Sanger Institute COVID-19 Surveillance Team |
| EPI_ISL_2908489 | Lighthouse Lab in Glasgow | Wellcome Sanger Institute for the COVID-19 Genomics UK (COG-UK) Consortium | Harper VanSteenhouse, Yumi Kasai, David Gray, Carol Clugston, Anna Dominiczak and Alex Alderton, Roberto Amato, Jeffrey Barrett, Sonia Goncalves, Ewan Harrison, David K. Jackson, Ian Johnston, Dominic Kwiatkowski, Cordelia Langford, John Sillitoe on behalf of the Wellcome Sanger Institute COVID-19 Surveillance Team |
| EPI_ISL_2908490, EPI_ISL_2908491 | Lighthouse Lab in Milton Keynes | Wellcome Sanger Institute for the COVID-19 Genomics UK (COG-UK) Consortium | The Lighthouse Lab in Milton Keynes and Alex Alderton, Roberto Amato, Jeffrey Barrett, Sonia Goncalves, Ewan Harrison, David K. Jackson, Ian Johnston, Dominic Kwiatkowski, Cordelia Langford, John Sillitoe on behalf of the Wellcome Sanger Institute COVID-19 Surveillance Team |
| EPI_ISL_2908492 | Lighthouse Lab in Glasgow | Wellcome Sanger Institute for the COVID-19 Genomics UK (COG-UK) Consortium | Harper VanSteenhouse, Yumi Kasai, David Gray, Carol Clugston, Anna Dominiczak and Alex Alderton, Roberto Amato, Jeffrey Barrett, Sonia Goncalves, Ewan Harrison, David K. Jackson, Ian Johnston, Dominic Kwiatkowski, Cordelia Langford, John Sillitoe on behalf of the Wellcome Sanger Institute COVID-19 Surveillance Team |
| EPI_ISL_2908493, EPI_ISL_2908494, EPI_ISL_2908495, EPI_ISL_2908496, EPI_ISL_2908497 | Lighthouse Lab in Milton Keynes | Wellcome Sanger Institute for the COVID-19 Genomics UK (COG-UK) Consortium | The Lighthouse Lab in Milton Keynes and Alex Alderton, Roberto Amato, Jeffrey Barrett, Sonia Goncalves, Ewan Harrison, David K. Jackson, Ian Johnston, Dominic Kwiatkowski, Cordelia Langford, John Sillitoe on behalf of the Wellcome Sanger Institute COVID-19 Surveillance Team |
| EPI_ISL_2908498, EPI_ISL_2908499 | Lighthouse Laboratory Plymouth | Wellcome Sanger Institute for the COVID-19 Genomics UK (COG-UK) Consortium | Lighthouse Laboratory Plymouth and Alex Alderton, Roberto Amato, Jeffrey Barrett, Sonia Goncalves, Ewan Harrison, David K. Jackson, Ian Johnston, Dominic Kwiatkowski, Cordelia Langford, John Sillitoe on behalf of the Wellcome Sanger Institute COVID-19 Surveillance Team |
| EPI_ISL_2908500, EPI_ISL_2908501 | Lighthouse Lab in Milton Keynes | Wellcome Sanger Institute for the COVID-19 Genomics UK (COG-UK) Consortium | The Lighthouse Lab in Milton Keynes and Alex Alderton, Roberto Amato, Jeffrey Barrett, Sonia Goncalves, Ewan Harrison, David K. Jackson, Ian Johnston, Dominic Kwiatkowski, Cordelia Langford, John Sillitoe on behalf of the Wellcome Sanger Institute COVID-19 Surveillance Team |
| EPI_ISL_2908502 | Lighthouse Laboratory Plymouth | Wellcome Sanger Institute for the COVID-19 Genomics UK (COG-UK) Consortium | Lighthouse Laboratory Plymouth and Alex Alderton, Roberto Amato, Jeffrey Barrett, Sonia Goncalves, Ewan Harrison, David K. Jackson, Ian Johnston, Dominic Kwiatkowski, Cordelia Langford, John Sillitoe on behalf of the Wellcome Sanger Institute COVID-19 Surveillance Team |
| EPI_ISL_2908503, EPI_ISL_2908504, EPI_ISL_2908505 | Lighthouse Lab in Milton Keynes | Wellcome Sanger Institute for the COVID-19 Genomics UK (COG-UK) Consortium | The Lighthouse Lab in Milton Keynes and Alex Alderton, Roberto Amato, Jeffrey Barrett, Sonia Goncalves, Ewan Harrison, David K. Jackson, Ian Johnston, Dominic Kwiatkowski, Cordelia Langford, John Sillitoe on behalf of the Wellcome Sanger Institute COVID-19 Surveillance Team |
| EPI_ISL_2908506 | Lighthouse Laboratory Plymouth | Wellcome Sanger Institute for the COVID-19 Genomics UK (COG-UK) Consortium | Lighthouse Laboratory Plymouth and Alex Alderton, Roberto Amato, Jeffrey Barrett, Sonia Goncalves, Ewan Harrison, David K. Jackson, Ian Johnston, Dominic Kwiatkowski, Cordelia Langford, John Sillitoe on behalf of the Wellcome Sanger Institute COVID-19 Surveillance Team |
| EPI_ISL_2908507, EPI_ISL_2908508, EPI_ISL_2908509, EPI_ISL_2908510, EPI_ISL_2908511 | Lighthouse Lab in Milton Keynes | Wellcome Sanger Institute for the COVID-19 Genomics UK (COG-UK) Consortium | The Lighthouse Lab in Milton Keynes and Alex Alderton, Roberto Amato, Jeffrey Barrett, Sonia Goncalves, Ewan Harrison, David K. Jackson, Ian Johnston, Dominic Kwiatkowski, Cordelia Langford, John Sillitoe on behalf of the Wellcome Sanger Institute COVID-19 Surveillance Team |
| EPI_ISL_2908512, EPI_ISL_2908513 | Lighthouse Laboratory Plymouth | Wellcome Sanger Institute for the COVID-19 Genomics UK (COG-UK) Consortium | Lighthouse Laboratory Plymouth and Alex Alderton, Roberto Amato, Jeffrey Barrett, Sonia Goncalves, Ewan Harrison, David K. Jackson, Ian Johnston, Dominic Kwiatkowski, Cordelia Langford, John Sillitoe on behalf of the Wellcome Sanger Institute COVID-19 Surveillance Team |
| EPI_ISL_2908514 | Lighthouse Lab in Milton Keynes | Wellcome Sanger Institute for the COVID-19 Genomics UK (COG-UK) Consortium | The Lighthouse Lab in Milton Keynes and Alex Alderton, Roberto Amato, Jeffrey Barrett, Sonia Goncalves, Ewan Harrison, David K. Jackson, Ian Johnston, Dominic Kwiatkowski, Cordelia Langford, John Sillitoe on behalf of the Wellcome Sanger Institute COVID-19 Surveillance Team |
| EPI_ISL_2908515 | Lighthouse Laboratory Plymouth | Wellcome Sanger Institute for the COVID-19 Genomics UK (COG-UK) Consortium | Lighthouse Laboratory Plymouth and Alex Alderton, Roberto Amato, Jeffrey Barrett, Sonia Goncalves, Ewan Harrison, David K. Jackson, Ian Johnston, Dominic Kwiatkowski, Cordelia Langford, John Sillitoe on behalf of the Wellcome Sanger Institute COVID-19 Surveillance Team |
| EPI_ISL_2908516, EPI_ISL_2908517, EPI_ISL_2908518 | Lighthouse Lab in Milton Keynes | Wellcome Sanger Institute for the COVID-19 Genomics UK (COG-UK) Consortium | The Lighthouse Lab in Milton Keynes and Alex Alderton, Roberto Amato, Jeffrey Barrett, Sonia Goncalves, Ewan Harrison, David K. Jackson, Ian Johnston, Dominic Kwiatkowski, Cordelia Langford, John Sillitoe on behalf of the Wellcome Sanger Institute COVID-19 Surveillance Team |
| EPI_ISL_2908519 | Lighthouse Laboratory Plymouth | Wellcome Sanger Institute for the COVID-19 Genomics UK (COG-UK) Consortium | Lighthouse Laboratory Plymouth and Alex Alderton, Roberto Amato, Jeffrey Barrett, Sonia Goncalves, Ewan Harrison, David K. Jackson, Ian Johnston, Dominic Kwiatkowski, Cordelia Langford, John Sillitoe on behalf of the Wellcome Sanger Institute COVID-19 Surveillance Team |
| EPI_ISL_2908520, EPI_ISL_2908521, EPI_ISL_2908522 | Lighthouse Lab in Milton Keynes | Wellcome Sanger Institute for the COVID-19 Genomics UK (COG-UK) Consortium | The Lighthouse Lab in Milton Keynes and Alex Alderton, Roberto Amato, Jeffrey Barrett, Sonia Goncalves, Ewan Harrison, David K. Jackson, Ian Johnston, Dominic Kwiatkowski, Cordelia Langford, John Sillitoe on behalf of the Wellcome Sanger Institute COVID-19 Surveillance Team |
| EPI_ISL_2908523, EPI_ISL_2908524, EPI_ISL_2908525, EPI_ISL_2908526 | Lighthouse Laboratory Plymouth | Wellcome Sanger Institute for the COVID-19 Genomics UK (COG-UK) Consortium | Lighthouse Laboratory Plymouth and Alex Alderton, Roberto Amato, Jeffrey Barrett, Sonia Goncalves, Ewan Harrison, David K. Jackson, Ian Johnston, Dominic Kwiatkowski, Cordelia Langford, John Sillitoe on behalf of the Wellcome Sanger Institute COVID-19 Surveillance Team |
| EPI_ISL_2908527, EPI_ISL_2908528 | Lighthouse Lab in Milton Keynes | Wellcome Sanger Institute for the COVID-19 Genomics UK (COG-UK) Consortium | The Lighthouse Lab in Milton Keynes and Alex Alderton, Roberto Amato, Jeffrey Barrett, Sonia Goncalves, Ewan Harrison, David K. Jackson, Ian Johnston, Dominic Kwiatkowski, Cordelia Langford, John Sillitoe on behalf of the Wellcome Sanger Institute COVID-19 Surveillance Team |
| EPI_ISL_2908529 | Lighthouse Laboratory Plymouth | Wellcome Sanger Institute for the COVID-19 Genomics UK (COG-UK) Consortium | Lighthouse Laboratory Plymouth and Alex Alderton, Roberto Amato, Jeffrey Barrett, Sonia Goncalves, Ewan Harrison, David K. Jackson, Ian Johnston, Dominic Kwiatkowski, Cordelia Langford, John Sillitoe on behalf of the Wellcome Sanger Institute COVID-19 Surveillance Team |
| EPI_ISL_2908530 | Lighthouse Lab in Milton Keynes | Wellcome Sanger Institute for the COVID-19 Genomics UK (COG-UK) Consortium | The Lighthouse Lab in Milton Keynes and Alex Alderton, Roberto Amato, Jeffrey Barrett, Sonia Goncalves, Ewan Harrison, David K. Jackson, Ian Johnston, Dominic Kwiatkowski, Cordelia Langford, John Sillitoe on behalf of the Wellcome Sanger Institute COVID-19 Surveillance Team |
| EPI_ISL_2908531, EPI_ISL_2908532 | Lighthouse Laboratory Plymouth | Wellcome Sanger Institute for the COVID-19 Genomics UK (COG-UK) Consortium | Lighthouse Laboratory Plymouth and Alex Alderton, Roberto Amato, Jeffrey Barrett, Sonia Goncalves, Ewan Harrison, David K. Jackson, Ian Johnston, Dominic Kwiatkowski, Cordelia Langford, John Sillitoe on behalf of the Wellcome Sanger Institute COVID-19 Surveillance Team |
| EPI_ISL_2908533 | Lighthouse Lab in Milton Keynes | Wellcome Sanger Institute for the COVID-19 Genomics UK (COG-UK) Consortium | The Lighthouse Lab in Milton Keynes and Alex Alderton, Roberto Amato, Jeffrey Barrett, Sonia Goncalves, Ewan Harrison, David K. Jackson, Ian Johnston, Dominic Kwiatkowski, Cordelia Langford, John Sillitoe on behalf of the Wellcome Sanger Institute COVID-19 Surveillance Team |
| EPI_ISL_2908534 | Lighthouse Laboratory Plymouth | Wellcome Sanger Institute for the COVID-19 Genomics UK (COG-UK) Consortium | Lighthouse Laboratory Plymouth and Alex Alderton, Roberto Amato, Jeffrey Barrett, Sonia Goncalves, Ewan Harrison, David K. Jackson, Ian Johnston, Dominic Kwiatkowski, Cordelia Langford, John Sillitoe on behalf of the Wellcome Sanger Institute COVID-19 Surveillance Team |
| EPI_ISL_2908535, EPI_ISL_2908536, EPI_ISL_2908537, EPI_ISL_2908538, EPI_ISL_2908539, EPI_ISL_2908540, EPI_ISL_2908541, EPI_ISL_2908542, EPI_ISL_2908543 | Lighthouse Lab in Milton Keynes | Wellcome Sanger Institute for the COVID-19 Genomics UK (COG-UK) Consortium | The Lighthouse Lab in Milton Keynes and Alex Alderton, Roberto Amato, Jeffrey Barrett, Sonia Goncalves, Ewan Harrison, David K. Jackson, Ian Johnston, Dominic Kwiatkowski, Cordelia Langford, John Sillitoe on behalf of the Wellcome Sanger Institute COVID-19 Surveillance Team |
| EPI_ISL_2908544 | Lighthouse Laboratory Plymouth | Wellcome Sanger Institute for the COVID-19 Genomics UK (COG-UK) Consortium | Lighthouse Laboratory Plymouth and Alex Alderton, Roberto Amato, Jeffrey Barrett, Sonia Goncalves, Ewan Harrison, David K. Jackson, Ian Johnston, Dominic Kwiatkowski, Cordelia Langford, John Sillitoe on behalf of the Wellcome Sanger Institute COVID-19 Surveillance Team |
| EPI_ISL_2908545, EPI_ISL_2908546 | Lighthouse Lab in Milton Keynes | Wellcome Sanger Institute for the COVID-19 Genomics UK (COG-UK) Consortium | The Lighthouse Lab in Milton Keynes and Alex Alderton, Roberto Amato, Jeffrey Barrett, Sonia Goncalves, Ewan Harrison, David K. Jackson, Ian Johnston, Dominic Kwiatkowski, Cordelia Langford, John Sillitoe on behalf of the Wellcome Sanger Institute COVID-19 Surveillance Team |
| EPI_ISL_2908547 | Lighthouse Laboratory Plymouth | Wellcome Sanger Institute for the COVID-19 Genomics UK (COG-UK) Consortium | Lighthouse Laboratory Plymouth and Alex Alderton, Roberto Amato, Jeffrey Barrett, Sonia Goncalves, Ewan Harrison, David K. Jackson, Ian Johnston, Dominic Kwiatkowski, Cordelia Langford, John Sillitoe on behalf of the Wellcome Sanger Institute COVID-19 Surveillance Team |
| EPI_ISL_2908548, EPI_ISL_2908549 | Lighthouse Lab in Milton Keynes | Wellcome Sanger Institute for the COVID-19 Genomics UK (COG-UK) Consortium | The Lighthouse Lab in Milton Keynes and Alex Alderton, Roberto Amato, Jeffrey Barrett, Sonia Goncalves, Ewan Harrison, David K. Jackson, Ian Johnston, Dominic Kwiatkowski, Cordelia Langford, John Sillitoe on behalf of the Wellcome Sanger Institute COVID-19 Surveillance Team |

| | | | |
|---|---|---|---|
| EPI_ISL_2908550, EPI_ISL_2908551, EPI_ISL_2908552 | Lighthouse Laboratory Plymouth | Wellcome Sanger Institute for the COVID-19 Genomics UK (COG-UK) Consortium | Lighthouse Laboratory Plymouth and Alex Alderton, Roberto Amato, Jeffrey Barrett, Sonia Goncalves, Ewan Harrison, David K. Jackson, Ian Johnston, Dominic Kwiatkowski, Cordelia Langford, John Sillitoe on behalf of the Wellcome Sanger Institute COVID-19 Surveillance Team |
| EPI_ISL_2908553 | Lighthouse Lab in Milton Keynes | Wellcome Sanger Institute for the COVID-19 Genomics UK (COG-UK) Consortium | The Lighthouse Lab in Milton Keynes and Alex Alderton, Roberto Amato, Jeffrey Barrett, Sonia Goncalves, Ewan Harrison, David K. Jackson, Ian Johnston, Dominic Kwiatkowski, Cordelia Langford, John Sillitoe on behalf of the Wellcome Sanger Institute COVID-19 Surveillance Team |
| EPI_ISL_2908554, EPI_ISL_2908555, EPI_ISL_2908556 | Lighthouse Laboratory Plymouth | Wellcome Sanger Institute for the COVID-19 Genomics UK (COG-UK) Consortium | Lighthouse Laboratory Plymouth and Alex Alderton, Roberto Amato, Jeffrey Barrett, Sonia Goncalves, Ewan Harrison, David K. Jackson, Ian Johnston, Dominic Kwiatkowski, Cordelia Langford, John Sillitoe on behalf of the Wellcome Sanger Institute COVID-19 Surveillance Team |
| EPI_ISL_2908557 | Lighthouse Lab in Milton Keynes | Wellcome Sanger Institute for the COVID-19 Genomics UK (COG-UK) Consortium | The Lighthouse Lab in Milton Keynes and Alex Alderton, Roberto Amato, Jeffrey Barrett, Sonia Goncalves, Ewan Harrison, David K. Jackson, Ian Johnston, Dominic Kwiatkowski, Cordelia Langford, John Sillitoe on behalf of the Wellcome Sanger Institute COVID-19 Surveillance Team |
| EPI_ISL_2908558, EPI_ISL_2908559 | Lighthouse Laboratory Plymouth | Wellcome Sanger Institute for the COVID-19 Genomics UK (COG-UK) Consortium | Lighthouse Laboratory Plymouth and Alex Alderton, Roberto Amato, Jeffrey Barrett, Sonia Goncalves, Ewan Harrison, David K. Jackson, Ian Johnston, Dominic Kwiatkowski, Cordelia Langford, John Sillitoe on behalf of the Wellcome Sanger Institute COVID-19 Surveillance Team |
| EPI_ISL_2908560 | Lighthouse Lab in Milton Keynes | Wellcome Sanger Institute for the COVID-19 Genomics UK (COG-UK) Consortium | The Lighthouse Lab in Milton Keynes and Alex Alderton, Roberto Amato, Jeffrey Barrett, Sonia Goncalves, Ewan Harrison, David K. Jackson, Ian Johnston, Dominic Kwiatkowski, Cordelia Langford, John Sillitoe on behalf of the Wellcome Sanger Institute COVID-19 Surveillance Team |
| EPI_ISL_2908561 | Lighthouse Laboratory Plymouth | Wellcome Sanger Institute for the COVID-19 Genomics UK (COG-UK) Consortium | Lighthouse Laboratory Plymouth and Alex Alderton, Roberto Amato, Jeffrey Barrett, Sonia Goncalves, Ewan Harrison, David K. Jackson, Ian Johnston, Dominic Kwiatkowski, Cordelia Langford, John Sillitoe on behalf of the Wellcome Sanger Institute COVID-19 Surveillance Team |
| EPI_ISL_2908562, EPI_ISL_2908563 | Lighthouse Lab in Milton Keynes | Wellcome Sanger Institute for the COVID-19 Genomics UK (COG-UK) Consortium | The Lighthouse Lab in Milton Keynes and Alex Alderton, Roberto Amato, Jeffrey Barrett, Sonia Goncalves, Ewan Harrison, David K. Jackson, Ian Johnston, Dominic Kwiatkowski, Cordelia Langford, John Sillitoe on behalf of the Wellcome Sanger Institute COVID-19 Surveillance Team |
| EPI_ISL_2908564, EPI_ISL_2908565, EPI_ISL_2908566, EPI_ISL_2908567 | Lighthouse Laboratory Plymouth | Wellcome Sanger Institute for the COVID-19 Genomics UK (COG-UK) Consortium | Lighthouse Laboratory Plymouth and Alex Alderton, Roberto Amato, Jeffrey Barrett, Sonia Goncalves, Ewan Harrison, David K. Jackson, Ian Johnston, Dominic Kwiatkowski, Cordelia Langford, John Sillitoe on behalf of the Wellcome Sanger Institute COVID-19 Surveillance Team |
| EPI_ISL_2908568, EPI_ISL_2908569, EPI_ISL_2908570 | Lighthouse Lab in Milton Keynes | Wellcome Sanger Institute for the COVID-19 Genomics UK (COG-UK) Consortium | The Lighthouse Lab in Milton Keynes and Alex Alderton, Roberto Amato, Jeffrey Barrett, Sonia Goncalves, Ewan Harrison, David K. Jackson, Ian Johnston, Dominic Kwiatkowski, Cordelia Langford, John Sillitoe on behalf of the Wellcome Sanger Institute COVID-19 Surveillance Team |
| EPI_ISL_2908571, EPI_ISL_2908572, EPI_ISL_2908573 | Lighthouse Laboratory Plymouth | Wellcome Sanger Institute for the COVID-19 Genomics UK (COG-UK) Consortium | Lighthouse Laboratory Plymouth and Alex Alderton, Roberto Amato, Jeffrey Barrett, Sonia Goncalves, Ewan Harrison, David K. Jackson, Ian Johnston, Dominic Kwiatkowski, Cordelia Langford, John Sillitoe on behalf of the Wellcome Sanger Institute COVID-19 Surveillance Team |
| EPI_ISL_2908574, EPI_ISL_2908575 | Lighthouse Lab in Milton Keynes | Wellcome Sanger Institute for the COVID-19 Genomics UK (COG-UK) Consortium | The Lighthouse Lab in Milton Keynes and Alex Alderton, Roberto Amato, Jeffrey Barrett, Sonia Goncalves, Ewan Harrison, David K. Jackson, Ian Johnston, Dominic Kwiatkowski, Cordelia Langford, John Sillitoe on behalf of the Wellcome Sanger Institute COVID-19 Surveillance Team |
| EPI_ISL_2908576, EPI_ISL_2908577, EPI_ISL_2908578, EPI_ISL_2908579, EPI_ISL_2908580, EPI_ISL_2908581, EPI_ISL_2908582 | Lighthouse Laboratory Plymouth | Wellcome Sanger Institute for the COVID-19 Genomics UK (COG-UK) Consortium | Lighthouse Laboratory Plymouth and Alex Alderton, Roberto Amato, Jeffrey Barrett, Sonia Goncalves, Ewan Harrison, David K. Jackson, Ian Johnston, Dominic Kwiatkowski, Cordelia Langford, John Sillitoe on behalf of the Wellcome Sanger Institute COVID-19 Surveillance Team |
| EPI_ISL_2908583, EPI_ISL_2908584, EPI_ISL_2908585, EPI_ISL_2908586, EPI_ISL_2908587 | Lighthouse Lab in Milton Keynes | Wellcome Sanger Institute for the COVID-19 Genomics UK (COG-UK) Consortium | The Lighthouse Lab in Milton Keynes and Alex Alderton, Roberto Amato, Jeffrey Barrett, Sonia Goncalves, Ewan Harrison, David K. Jackson, Ian Johnston, Dominic Kwiatkowski, Cordelia Langford, John Sillitoe on behalf of the Wellcome Sanger Institute COVID-19 Surveillance Team |
| EPI_ISL_2908588, EPI_ISL_2908589 | Lighthouse Laboratory Plymouth | Wellcome Sanger Institute for the COVID-19 Genomics UK (COG-UK) Consortium | Lighthouse Laboratory Plymouth and Alex Alderton, Roberto Amato, Jeffrey Barrett, Sonia Goncalves, Ewan Harrison, David K. Jackson, Ian Johnston, Dominic Kwiatkowski, Cordelia Langford, John Sillitoe on behalf of the Wellcome Sanger Institute COVID-19 Surveillance Team |
| EPI_ISL_2908590, EPI_ISL_2908591, EPI_ISL_2908592 | Lighthouse Lab in Milton Keynes | Wellcome Sanger Institute for the COVID-19 Genomics UK (COG-UK) Consortium | The Lighthouse Lab in Milton Keynes and Alex Alderton, Roberto Amato, Jeffrey Barrett, Sonia Goncalves, Ewan Harrison, David K. Jackson, Ian Johnston, Dominic Kwiatkowski, Cordelia Langford, John Sillitoe on behalf of the Wellcome Sanger Institute COVID-19 Surveillance Team |
| EPI_ISL_2908593, EPI_ISL_2908594, EPI_ISL_2908595, EPI_ISL_2908596 | Lighthouse Laboratory Plymouth | Wellcome Sanger Institute for the COVID-19 Genomics UK (COG-UK) Consortium | Lighthouse Laboratory Plymouth and Alex Alderton, Roberto Amato, Jeffrey Barrett, Sonia Goncalves, Ewan Harrison, David K. Jackson, Ian Johnston, Dominic Kwiatkowski, Cordelia Langford, John Sillitoe on behalf of the Wellcome Sanger Institute COVID-19 Surveillance Team |
| EPI_ISL_2908597, EPI_ISL_2908598 | Lighthouse Lab in Milton Keynes | Wellcome Sanger Institute for the COVID-19 Genomics UK (COG-UK) Consortium | The Lighthouse Lab in Milton Keynes and Alex Alderton, Roberto Amato, Jeffrey Barrett, Sonia Goncalves, Ewan Harrison, David K. Jackson, Ian Johnston, Dominic Kwiatkowski, Cordelia Langford, John Sillitoe on behalf of the Wellcome Sanger Institute COVID-19 Surveillance Team |
| EPI_ISL_2908599 | Lighthouse Laboratory Plymouth | Wellcome Sanger Institute for the COVID-19 Genomics UK (COG-UK) Consortium | Lighthouse Laboratory Plymouth and Alex Alderton, Roberto Amato, Jeffrey Barrett, Sonia Goncalves, Ewan Harrison, David K. Jackson, Ian Johnston, Dominic Kwiatkowski, Cordelia Langford, John Sillitoe on behalf of the Wellcome Sanger Institute COVID-19 Surveillance Team |
| EPI_ISL_2908600 | Lighthouse Lab in Milton Keynes | Wellcome Sanger Institute for the COVID-19 Genomics UK (COG-UK) Consortium | The Lighthouse Lab in Milton Keynes and Alex Alderton, Roberto Amato, Jeffrey Barrett, Sonia Goncalves, Ewan Harrison, David K. Jackson, Ian Johnston, Dominic Kwiatkowski, Cordelia Langford, John Sillitoe on behalf of the Wellcome Sanger Institute COVID-19 Surveillance Team |
| EPI_ISL_2908601 | Lighthouse Laboratory Plymouth | Wellcome Sanger Institute for the COVID-19 Genomics UK (COG-UK) Consortium | Lighthouse Laboratory Plymouth and Alex Alderton, Roberto Amato, Jeffrey Barrett, Sonia Goncalves, Ewan Harrison, David K. Jackson, Ian Johnston, Dominic Kwiatkowski, Cordelia Langford, John Sillitoe on behalf of the Wellcome Sanger Institute COVID-19 Surveillance Team |
| EPI_ISL_2908602 | Lighthouse Lab in Milton Keynes | Wellcome Sanger Institute for the COVID-19 Genomics UK (COG-UK) Consortium | The Lighthouse Lab in Milton Keynes and Alex Alderton, Roberto Amato, Jeffrey Barrett, Sonia Goncalves, Ewan Harrison, David K. Jackson, Ian Johnston, Dominic Kwiatkowski, Cordelia Langford, John Sillitoe on behalf of the Wellcome Sanger Institute COVID-19 Surveillance Team |
| EPI_ISL_2908603, EPI_ISL_2908604 | Lighthouse Laboratory Plymouth | Wellcome Sanger Institute for the COVID-19 Genomics UK (COG-UK) Consortium | Lighthouse Laboratory Plymouth and Alex Alderton, Roberto Amato, Jeffrey Barrett, Sonia Goncalves, Ewan Harrison, David K. Jackson, Ian Johnston, Dominic Kwiatkowski, Cordelia Langford, John Sillitoe on behalf of the Wellcome Sanger Institute COVID-19 Surveillance Team |
| EPI_ISL_2908605, EPI_ISL_2908606 | Lighthouse Lab in Milton Keynes | Wellcome Sanger Institute for the COVID-19 Genomics UK (COG-UK) Consortium | The Lighthouse Lab in Milton Keynes and Alex Alderton, Roberto Amato, Jeffrey Barrett, Sonia Goncalves, Ewan Harrison, David K. Jackson, Ian Johnston, Dominic Kwiatkowski, Cordelia Langford, John Sillitoe on behalf of the Wellcome Sanger Institute COVID-19 Surveillance Team |
| EPI_ISL_2908607, EPI_ISL_2908608 | Lighthouse Laboratory Plymouth | Wellcome Sanger Institute for the COVID-19 Genomics UK (COG-UK) Consortium | Lighthouse Laboratory Plymouth and Alex Alderton, Roberto Amato, Jeffrey Barrett, Sonia Goncalves, Ewan Harrison, David K. Jackson, Ian Johnston, Dominic Kwiatkowski, Cordelia Langford, John Sillitoe on behalf of the Wellcome Sanger Institute COVID-19 Surveillance Team |
| EPI_ISL_2908609 | Lighthouse Lab in Milton Keynes | Wellcome Sanger Institute for the COVID-19 Genomics UK (COG-UK) Consortium | The Lighthouse Lab in Milton Keynes and Alex Alderton, Roberto Amato, Jeffrey Barrett, Sonia Goncalves, Ewan Harrison, David K. Jackson, Ian Johnston, Dominic Kwiatkowski, Cordelia Langford, John Sillitoe on behalf of the Wellcome Sanger Institute COVID-19 Surveillance Team |
| EPI_ISL_2908610, EPI_ISL_2908611 | Lighthouse Laboratory Plymouth | Wellcome Sanger Institute for the COVID-19 Genomics UK (COG-UK) Consortium | Lighthouse Laboratory Plymouth and Alex Alderton, Roberto Amato, Jeffrey Barrett, Sonia Goncalves, Ewan Harrison, David K. Jackson, Ian Johnston, Dominic Kwiatkowski, Cordelia Langford, John Sillitoe on behalf of the Wellcome Sanger Institute COVID-19 Surveillance Team |
| EPI_ISL_2908612 | Lighthouse Lab in Milton Keynes | Wellcome Sanger Institute for the COVID-19 Genomics UK (COG-UK) Consortium | The Lighthouse Lab in Milton Keynes and Alex Alderton, Roberto Amato, Jeffrey Barrett, Sonia Goncalves, Ewan Harrison, David K. Jackson, Ian Johnston, Dominic Kwiatkowski, Cordelia Langford, John Sillitoe on behalf of the Wellcome Sanger Institute COVID-19 Surveillance Team |
| EPI_ISL_2908613 | Lighthouse Laboratory Plymouth | Wellcome Sanger Institute for the COVID-19 Genomics UK (COG-UK) Consortium | Lighthouse Laboratory Plymouth and Alex Alderton, Roberto Amato, Jeffrey Barrett, Sonia Goncalves, Ewan Harrison, David K. Jackson, Ian Johnston, Dominic Kwiatkowski, Cordelia Langford, John Sillitoe on behalf of the Wellcome Sanger Institute COVID-19 Surveillance Team |
| EPI_ISL_2908614 | Lighthouse Lab in Milton Keynes | Wellcome Sanger Institute for the COVID-19 Genomics UK (COG-UK) Consortium | The Lighthouse Lab in Milton Keynes and Alex Alderton, Roberto Amato, Jeffrey Barrett, Sonia Goncalves, Ewan Harrison, David K. Jackson, Ian Johnston, Dominic Kwiatkowski, Cordelia Langford, John Sillitoe on behalf of the Wellcome Sanger Institute COVID-19 Surveillance Team |
| EPI_ISL_2908615 | Lighthouse Laboratory Plymouth | Wellcome Sanger Institute for the COVID-19 Genomics UK (COG-UK) Consortium | Lighthouse Laboratory Plymouth and Alex Alderton, Roberto Amato, Jeffrey Barrett, Sonia Goncalves, Ewan Harrison, David K. Jackson, Ian Johnston, Dominic Kwiatkowski, Cordelia Langford, John Sillitoe on behalf of the Wellcome Sanger Institute COVID-19 Surveillance Team |
| EPI_ISL_2908616 | Lighthouse Lab in Milton Keynes | Wellcome Sanger Institute for the COVID-19 Genomics UK (COG-UK) Consortium | The Lighthouse Lab in Milton Keynes and Alex Alderton, Roberto Amato, Jeffrey Barrett, Sonia Goncalves, Ewan Harrison, David K. Jackson, Ian Johnston, Dominic Kwiatkowski, Cordelia Langford, John Sillitoe on behalf of the Wellcome Sanger Institute COVID-19 Surveillance Team |
| EPI_ISL_2908617, EPI_ISL_2908618, EPI_ISL_2908619, EPI_ISL_2908620 | Lighthouse Laboratory Plymouth | Wellcome Sanger Institute for the COVID-19 Genomics UK (COG-UK) Consortium | Lighthouse Laboratory Plymouth and Alex Alderton, Roberto Amato, Jeffrey Barrett, Sonia Goncalves, Ewan Harrison, David K. Jackson, Ian Johnston, Dominic Kwiatkowski, Cordelia Langford, John Sillitoe on behalf of the Wellcome Sanger Institute COVID-19 Surveillance Team |
| EPI_ISL_2908621 | Lighthouse Lab in Milton Keynes | Wellcome Sanger Institute for the COVID-19 Genomics UK (COG-UK) Consortium | The Lighthouse Lab in Milton Keynes and Alex Alderton, Roberto Amato, Jeffrey Barrett, Sonia Goncalves, Ewan Harrison, David K. Jackson, Ian Johnston, Dominic Kwiatkowski, Cordelia Langford, John Sillitoe on behalf of the Wellcome Sanger Institute COVID-19 Surveillance Team |
| EPI_ISL_2908622 | Lighthouse Laboratory Plymouth | Wellcome Sanger Institute for the COVID-19 Genomics UK (COG-UK) Consortium | Lighthouse Laboratory Plymouth and Alex Alderton, Roberto Amato, Jeffrey Barrett, Sonia Goncalves, Ewan Harrison, David K. Jackson, Ian Johnston, Dominic Kwiatkowski, Cordelia Langford, John Sillitoe on behalf of the Wellcome Sanger Institute COVID-19 Surveillance Team |
| EPI_ISL_2908623 | Lighthouse Lab in Milton Keynes | Wellcome Sanger Institute for the COVID-19 Genomics UK (COG-UK) Consortium | The Lighthouse Lab in Milton Keynes and Alex Alderton, Roberto Amato, Jeffrey Barrett, Sonia Goncalves, Ewan Harrison, David K. Jackson, Ian Johnston, Dominic Kwiatkowski, Cordelia Langford, John Sillitoe on behalf of the Wellcome Sanger Institute COVID-19 Surveillance Team |
| EPI_ISL_2908624, EPI_ISL_2908625 | Lighthouse Laboratory Plymouth | Wellcome Sanger Institute for the COVID-19 Genomics UK (COG-UK) Consortium | Lighthouse Laboratory Plymouth and Alex Alderton, Roberto Amato, Jeffrey Barrett, Sonia Goncalves, Ewan Harrison, David K. Jackson, Ian Johnston, Dominic Kwiatkowski, Cordelia Langford, John Sillitoe on behalf of the Wellcome Sanger Institute COVID-19 Surveillance Team |
| EPI_ISL_2908626, | Lighthouse Lab in Milton Keynes | Wellcome Sanger Institute for the COVID-19 Genomics UK (COG-UK) | The Lighthouse Lab in Milton Keynes and Alex Alderton, Roberto Amato, Jeffrey Barrett, Sonia Goncalves, Ewan Harrison, David K. Jackson, Ian Johnston, Dominic Kwiatkowski, Cordelia Langford, John Sillitoe on behalf of the Wellcome Sanger |

| | | | |
|---|---|---|---|
| EPI_ISL_2908627, EPI_ISL_2908628 | | Consortium | Institute COVID-19 Surveillance Team |
| EPI_ISL_2908629 | Lighthouse Laboratory Plymouth | Wellcome Sanger Institute for the COVID-19 Genomics UK (COG-UK) Consortium | Lighthouse Laboratory Plymouth and Alex Alderton, Roberto Amato, Jeffrey Barrett, Sonia Goncalves, Ewan Harrison, David K. Jackson, Ian Johnston, Dominic Kwiatkowski, Cordelia Langford, John Sillitoe on behalf of the Wellcome Sanger Institute COVID-19 Surveillance Team |
| EPI_ISL_2908630 | Lighthouse Lab in Milton Keynes | Wellcome Sanger Institute for the COVID-19 Genomics UK (COG-UK) Consortium | The Lighthouse Lab in Milton Keynes and Alex Alderton, Roberto Amato, Jeffrey Barrett, Sonia Goncalves, Ewan Harrison, David K. Jackson, Ian Johnston, Dominic Kwiatkowski, Cordelia Langford, John Sillitoe on behalf of the Wellcome Sanger Institute COVID-19 Surveillance Team |
| EPI_ISL_2908631 | Lighthouse Laboratory Plymouth | Wellcome Sanger Institute for the COVID-19 Genomics UK (COG-UK) Consortium | Lighthouse Laboratory Plymouth and Alex Alderton, Roberto Amato, Jeffrey Barrett, Sonia Goncalves, Ewan Harrison, David K. Jackson, Ian Johnston, Dominic Kwiatkowski, Cordelia Langford, John Sillitoe on behalf of the Wellcome Sanger Institute COVID-19 Surveillance Team |
| EPI_ISL_2908632 | Lighthouse Lab in Milton Keynes | Wellcome Sanger Institute for the COVID-19 Genomics UK (COG-UK) Consortium | The Lighthouse Lab in Milton Keynes and Alex Alderton, Roberto Amato, Jeffrey Barrett, Sonia Goncalves, Ewan Harrison, David K. Jackson, Ian Johnston, Dominic Kwiatkowski, Cordelia Langford, John Sillitoe on behalf of the Wellcome Sanger Institute COVID-19 Surveillance Team |
| EPI_ISL_2908633, EPI_ISL_2908634, EPI_ISL_2908635 | Lighthouse Laboratory Plymouth | Wellcome Sanger Institute for the COVID-19 Genomics UK (COG-UK) Consortium | Lighthouse Laboratory Plymouth and Alex Alderton, Roberto Amato, Jeffrey Barrett, Sonia Goncalves, Ewan Harrison, David K. Jackson, Ian Johnston, Dominic Kwiatkowski, Cordelia Langford, John Sillitoe on behalf of the Wellcome Sanger Institute COVID-19 Surveillance Team |
| EPI_ISL_2908636 | Lighthouse Lab in Milton Keynes | Wellcome Sanger Institute for the COVID-19 Genomics UK (COG-UK) Consortium | The Lighthouse Lab in Milton Keynes and Alex Alderton, Roberto Amato, Jeffrey Barrett, Sonia Goncalves, Ewan Harrison, David K. Jackson, Ian Johnston, Dominic Kwiatkowski, Cordelia Langford, John Sillitoe on behalf of the Wellcome Sanger Institute COVID-19 Surveillance Team |
| EPI_ISL_2908637 | Lighthouse Laboratory Plymouth | Wellcome Sanger Institute for the COVID-19 Genomics UK (COG-UK) Consortium | Lighthouse Laboratory Plymouth and Alex Alderton, Roberto Amato, Jeffrey Barrett, Sonia Goncalves, Ewan Harrison, David K. Jackson, Ian Johnston, Dominic Kwiatkowski, Cordelia Langford, John Sillitoe on behalf of the Wellcome Sanger Institute COVID-19 Surveillance Team |
| EPI_ISL_2908638, EPI_ISL_2908639 | Lighthouse Lab in Milton Keynes | Wellcome Sanger Institute for the COVID-19 Genomics UK (COG-UK) Consortium | The Lighthouse Lab in Milton Keynes and Alex Alderton, Roberto Amato, Jeffrey Barrett, Sonia Goncalves, Ewan Harrison, David K. Jackson, Ian Johnston, Dominic Kwiatkowski, Cordelia Langford, John Sillitoe on behalf of the Wellcome Sanger Institute COVID-19 Surveillance Team |
| EPI_ISL_2908640 | Lighthouse Laboratory Plymouth | Wellcome Sanger Institute for the COVID-19 Genomics UK (COG-UK) Consortium | Lighthouse Laboratory Plymouth and Alex Alderton, Roberto Amato, Jeffrey Barrett, Sonia Goncalves, Ewan Harrison, David K. Jackson, Ian Johnston, Dominic Kwiatkowski, Cordelia Langford, John Sillitoe on behalf of the Wellcome Sanger Institute COVID-19 Surveillance Team |
| EPI_ISL_2908641 | Lighthouse Lab in Milton Keynes | Wellcome Sanger Institute for the COVID-19 Genomics UK (COG-UK) Consortium | The Lighthouse Lab in Milton Keynes and Alex Alderton, Roberto Amato, Jeffrey Barrett, Sonia Goncalves, Ewan Harrison, David K. Jackson, Ian Johnston, Dominic Kwiatkowski, Cordelia Langford, John Sillitoe on behalf of the Wellcome Sanger Institute COVID-19 Surveillance Team |
| EPI_ISL_2908642, EPI_ISL_2908643, EPI_ISL_2908644 | Lighthouse Laboratory Plymouth | Wellcome Sanger Institute for the COVID-19 Genomics UK (COG-UK) Consortium | Lighthouse Laboratory Plymouth and Alex Alderton, Roberto Amato, Jeffrey Barrett, Sonia Goncalves, Ewan Harrison, David K. Jackson, Ian Johnston, Dominic Kwiatkowski, Cordelia Langford, John Sillitoe on behalf of the Wellcome Sanger Institute COVID-19 Surveillance Team |
| EPI_ISL_2908645, EPI_ISL_2908646 | Lighthouse Lab in Milton Keynes | Wellcome Sanger Institute for the COVID-19 Genomics UK (COG-UK) Consortium | The Lighthouse Lab in Milton Keynes and Alex Alderton, Roberto Amato, Jeffrey Barrett, Sonia Goncalves, Ewan Harrison, David K. Jackson, Ian Johnston, Dominic Kwiatkowski, Cordelia Langford, John Sillitoe on behalf of the Wellcome Sanger Institute COVID-19 Surveillance Team |
| EPI_ISL_2908647 | Lighthouse Laboratory Plymouth | Wellcome Sanger Institute for the COVID-19 Genomics UK (COG-UK) Consortium | Lighthouse Laboratory Plymouth and Alex Alderton, Roberto Amato, Jeffrey Barrett, Sonia Goncalves, Ewan Harrison, David K. Jackson, Ian Johnston, Dominic Kwiatkowski, Cordelia Langford, John Sillitoe on behalf of the Wellcome Sanger Institute COVID-19 Surveillance Team |
| EPI_ISL_2908648 | Lighthouse Lab in Milton Keynes | Wellcome Sanger Institute for the COVID-19 Genomics UK (COG-UK) Consortium | The Lighthouse Lab in Milton Keynes and Alex Alderton, Roberto Amato, Jeffrey Barrett, Sonia Goncalves, Ewan Harrison, David K. Jackson, Ian Johnston, Dominic Kwiatkowski, Cordelia Langford, John Sillitoe on behalf of the Wellcome Sanger Institute COVID-19 Surveillance Team |
| EPI_ISL_2908649, EPI_ISL_2908650 | Lighthouse Laboratory Plymouth | Wellcome Sanger Institute for the COVID-19 Genomics UK (COG-UK) Consortium | Lighthouse Laboratory Plymouth and Alex Alderton, Roberto Amato, Jeffrey Barrett, Sonia Goncalves, Ewan Harrison, David K. Jackson, Ian Johnston, Dominic Kwiatkowski, Cordelia Langford, John Sillitoe on behalf of the Wellcome Sanger Institute COVID-19 Surveillance Team |
| EPI_ISL_2908651, EPI_ISL_2908652 | Lighthouse Lab in Milton Keynes | Wellcome Sanger Institute for the COVID-19 Genomics UK (COG-UK) Consortium | The Lighthouse Lab in Milton Keynes and Alex Alderton, Roberto Amato, Jeffrey Barrett, Sonia Goncalves, Ewan Harrison, David K. Jackson, Ian Johnston, Dominic Kwiatkowski, Cordelia Langford, John Sillitoe on behalf of the Wellcome Sanger Institute COVID-19 Surveillance Team |
| EPI_ISL_2908653 | Lighthouse Laboratory Plymouth | Wellcome Sanger Institute for the COVID-19 Genomics UK (COG-UK) Consortium | Lighthouse Laboratory Plymouth and Alex Alderton, Roberto Amato, Jeffrey Barrett, Sonia Goncalves, Ewan Harrison, David K. Jackson, Ian Johnston, Dominic Kwiatkowski, Cordelia Langford, John Sillitoe on behalf of the Wellcome Sanger Institute COVID-19 Surveillance Team |
| EPI_ISL_2908654, EPI_ISL_2908655, EPI_ISL_2908656, EPI_ISL_2908657, EPI_ISL_2908658 | Lighthouse Lab in Milton Keynes | Wellcome Sanger Institute for the COVID-19 Genomics UK (COG-UK) Consortium | The Lighthouse Lab in Milton Keynes and Alex Alderton, Roberto Amato, Jeffrey Barrett, Sonia Goncalves, Ewan Harrison, David K. Jackson, Ian Johnston, Dominic Kwiatkowski, Cordelia Langford, John Sillitoe on behalf of the Wellcome Sanger Institute COVID-19 Surveillance Team |
| EPI_ISL_2908659 | Lighthouse Laboratory Plymouth | Wellcome Sanger Institute for the COVID-19 Genomics UK (COG-UK) Consortium | Lighthouse Laboratory Plymouth and Alex Alderton, Roberto Amato, Jeffrey Barrett, Sonia Goncalves, Ewan Harrison, David K. Jackson, Ian Johnston, Dominic Kwiatkowski, Cordelia Langford, John Sillitoe on behalf of the Wellcome Sanger Institute COVID-19 Surveillance Team |
| EPI_ISL_2908660 | Lighthouse Lab in Milton Keynes | Wellcome Sanger Institute for the COVID-19 Genomics UK (COG-UK) Consortium | The Lighthouse Lab in Milton Keynes and Alex Alderton, Roberto Amato, Jeffrey Barrett, Sonia Goncalves, Ewan Harrison, David K. Jackson, Ian Johnston, Dominic Kwiatkowski, Cordelia Langford, John Sillitoe on behalf of the Wellcome Sanger Institute COVID-19 Surveillance Team |
| EPI_ISL_2908661 | Lighthouse Laboratory Plymouth | Wellcome Sanger Institute for the COVID-19 Genomics UK (COG-UK) Consortium | Lighthouse Laboratory Plymouth and Alex Alderton, Roberto Amato, Jeffrey Barrett, Sonia Goncalves, Ewan Harrison, David K. Jackson, Ian Johnston, Dominic Kwiatkowski, Cordelia Langford, John Sillitoe on behalf of the Wellcome Sanger Institute COVID-19 Surveillance Team |
| EPI_ISL_2908662, EPI_ISL_2908663 | Lighthouse Lab in Milton Keynes | Wellcome Sanger Institute for the COVID-19 Genomics UK (COG-UK) Consortium | The Lighthouse Lab in Milton Keynes and Alex Alderton, Roberto Amato, Jeffrey Barrett, Sonia Goncalves, Ewan Harrison, David K. Jackson, Ian Johnston, Dominic Kwiatkowski, Cordelia Langford, John Sillitoe on behalf of the Wellcome Sanger Institute COVID-19 Surveillance Team |
| EPI_ISL_2908664 | Lighthouse Laboratory Plymouth | Wellcome Sanger Institute for the COVID-19 Genomics UK (COG-UK) Consortium | Lighthouse Laboratory Plymouth and Alex Alderton, Roberto Amato, Jeffrey Barrett, Sonia Goncalves, Ewan Harrison, David K. Jackson, Ian Johnston, Dominic Kwiatkowski, Cordelia Langford, John Sillitoe on behalf of the Wellcome Sanger Institute COVID-19 Surveillance Team |
| EPI_ISL_2908665 | Lighthouse Lab in Milton Keynes | Wellcome Sanger Institute for the COVID-19 Genomics UK (COG-UK) Consortium | The Lighthouse Lab in Milton Keynes and Alex Alderton, Roberto Amato, Jeffrey Barrett, Sonia Goncalves, Ewan Harrison, David K. Jackson, Ian Johnston, Dominic Kwiatkowski, Cordelia Langford, John Sillitoe on behalf of the Wellcome Sanger Institute COVID-19 Surveillance Team |
| EPI_ISL_2908666, EPI_ISL_2908667 | Lighthouse Laboratory Plymouth | Wellcome Sanger Institute for the COVID-19 Genomics UK (COG-UK) Consortium | Lighthouse Laboratory Plymouth and Alex Alderton, Roberto Amato, Jeffrey Barrett, Sonia Goncalves, Ewan Harrison, David K. Jackson, Ian Johnston, Dominic Kwiatkowski, Cordelia Langford, John Sillitoe on behalf of the Wellcome Sanger Institute COVID-19 Surveillance Team |
| EPI_ISL_2908668, EPI_ISL_2908669, EPI_ISL_2908670 | Lighthouse Lab in Milton Keynes | Wellcome Sanger Institute for the COVID-19 Genomics UK (COG-UK) Consortium | The Lighthouse Lab in Milton Keynes and Alex Alderton, Roberto Amato, Jeffrey Barrett, Sonia Goncalves, Ewan Harrison, David K. Jackson, Ian Johnston, Dominic Kwiatkowski, Cordelia Langford, John Sillitoe on behalf of the Wellcome Sanger Institute COVID-19 Surveillance Team |
| EPI_ISL_2908671, EPI_ISL_2908672, EPI_ISL_2908673, EPI_ISL_2908674 | Lighthouse Laboratory Plymouth | Wellcome Sanger Institute for the COVID-19 Genomics UK (COG-UK) Consortium | Lighthouse Laboratory Plymouth and Alex Alderton, Roberto Amato, Jeffrey Barrett, Sonia Goncalves, Ewan Harrison, David K. Jackson, Ian Johnston, Dominic Kwiatkowski, Cordelia Langford, John Sillitoe on behalf of the Wellcome Sanger Institute COVID-19 Surveillance Team |
| EPI_ISL_2908675, EPI_ISL_2908676 | Lighthouse Lab in Milton Keynes | Wellcome Sanger Institute for the COVID-19 Genomics UK (COG-UK) Consortium | The Lighthouse Lab in Milton Keynes and Alex Alderton, Roberto Amato, Jeffrey Barrett, Sonia Goncalves, Ewan Harrison, David K. Jackson, Ian Johnston, Dominic Kwiatkowski, Cordelia Langford, John Sillitoe on behalf of the Wellcome Sanger Institute COVID-19 Surveillance Team |
| EPI_ISL_2908677 | Lighthouse Laboratory Plymouth | Wellcome Sanger Institute for the COVID-19 Genomics UK (COG-UK) Consortium | Lighthouse Laboratory Plymouth and Alex Alderton, Roberto Amato, Jeffrey Barrett, Sonia Goncalves, Ewan Harrison, David K. Jackson, Ian Johnston, Dominic Kwiatkowski, Cordelia Langford, John Sillitoe on behalf of the Wellcome Sanger Institute COVID-19 Surveillance Team |
| EPI_ISL_2908678 | Lighthouse Lab in Milton Keynes | Wellcome Sanger Institute for the COVID-19 Genomics UK (COG-UK) Consortium | The Lighthouse Lab in Milton Keynes and Alex Alderton, Roberto Amato, Jeffrey Barrett, Sonia Goncalves, Ewan Harrison, David K. Jackson, Ian Johnston, Dominic Kwiatkowski, Cordelia Langford, John Sillitoe on behalf of the Wellcome Sanger Institute COVID-19 Surveillance Team |
| EPI_ISL_2908679 | Lighthouse Laboratory Plymouth | Wellcome Sanger Institute for the COVID-19 Genomics UK (COG-UK) Consortium | Lighthouse Laboratory Plymouth and Alex Alderton, Roberto Amato, Jeffrey Barrett, Sonia Goncalves, Ewan Harrison, David K. Jackson, Ian Johnston, Dominic Kwiatkowski, Cordelia Langford, John Sillitoe on behalf of the Wellcome Sanger Institute COVID-19 Surveillance Team |
| EPI_ISL_2908680, EPI_ISL_2908681, EPI_ISL_2908682 | Lighthouse Lab in Milton Keynes | Wellcome Sanger Institute for the COVID-19 Genomics UK (COG-UK) Consortium | The Lighthouse Lab in Milton Keynes and Alex Alderton, Roberto Amato, Jeffrey Barrett, Sonia Goncalves, Ewan Harrison, David K. Jackson, Ian Johnston, Dominic Kwiatkowski, Cordelia Langford, John Sillitoe on behalf of the Wellcome Sanger Institute COVID-19 Surveillance Team |
| EPI_ISL_2908683, EPI_ISL_2908684 | Lighthouse Laboratory Plymouth | Wellcome Sanger Institute for the COVID-19 Genomics UK (COG-UK) Consortium | Lighthouse Laboratory Plymouth and Alex Alderton, Roberto Amato, Jeffrey Barrett, Sonia Goncalves, Ewan Harrison, David K. Jackson, Ian Johnston, Dominic Kwiatkowski, Cordelia Langford, John Sillitoe on behalf of the Wellcome Sanger Institute COVID-19 Surveillance Team |
| EPI_ISL_2908685 | Lighthouse Lab in Milton Keynes | Wellcome Sanger Institute for the COVID-19 Genomics UK (COG-UK) Consortium | The Lighthouse Lab in Milton Keynes and Alex Alderton, Roberto Amato, Jeffrey Barrett, Sonia Goncalves, Ewan Harrison, David K. Jackson, Ian Johnston, Dominic Kwiatkowski, Cordelia Langford, John Sillitoe on behalf of the Wellcome Sanger Institute COVID-19 Surveillance Team |
| EPI_ISL_2908686, EPI_ISL_2908687 | Lighthouse Laboratory Plymouth | Wellcome Sanger Institute for the COVID-19 Genomics UK (COG-UK) Consortium | Lighthouse Laboratory Plymouth and Alex Alderton, Roberto Amato, Jeffrey Barrett, Sonia Goncalves, Ewan Harrison, David K. Jackson, Ian Johnston, Dominic Kwiatkowski, Cordelia Langford, John Sillitoe on behalf of the Wellcome Sanger Institute COVID-19 Surveillance Team |
| EPI_ISL_2908688 | Lighthouse Lab in Milton Keynes | Wellcome Sanger Institute for the COVID-19 Genomics UK (COG-UK) Consortium | The Lighthouse Lab in Milton Keynes and Alex Alderton, Roberto Amato, Jeffrey Barrett, Sonia Goncalves, Ewan Harrison, David K. Jackson, Ian Johnston, Dominic Kwiatkowski, Cordelia Langford, John Sillitoe on behalf of the Wellcome Sanger Institute COVID-19 Surveillance Team |
| EPI_ISL_2908689 | Lighthouse Laboratory Plymouth | Wellcome Sanger Institute for the COVID-19 Genomics UK (COG-UK) Consortium | Lighthouse Laboratory Plymouth and Alex Alderton, Roberto Amato, Jeffrey Barrett, Sonia Goncalves, Ewan Harrison, David K. Jackson, Ian Johnston, Dominic Kwiatkowski, Cordelia Langford, John Sillitoe on behalf of the Wellcome Sanger Institute COVID-19 Surveillance Team |
| EPI_ISL_2908690 | Lighthouse Lab in Milton Keynes | Wellcome Sanger Institute for the COVID-19 Genomics UK (COG-UK) Consortium | The Lighthouse Lab in Milton Keynes and Alex Alderton, Roberto Amato, Jeffrey Barrett, Sonia Goncalves, Ewan Harrison, David K. Jackson, Ian Johnston, Dominic Kwiatkowski, Cordelia Langford, John Sillitoe on behalf of the Wellcome Sanger Institute COVID-19 Surveillance Team |
| EPI_ISL_2908691 | Lighthouse Laboratory Plymouth | Wellcome Sanger Institute for the COVID-19 Genomics UK (COG-UK) Consortium | Lighthouse Laboratory Plymouth and Alex Alderton, Roberto Amato, Jeffrey Barrett, Sonia Goncalves, Ewan Harrison, David K. Jackson, Ian Johnston, Dominic Kwiatkowski, Cordelia Langford, John Sillitoe on behalf of the Wellcome Sanger Institute COVID-19 Surveillance Team |
| EPI_ISL_2908692 | Lighthouse Lab in Milton Keynes | Wellcome Sanger Institute for the COVID-19 Genomics UK (COG-UK) Consortium | The Lighthouse Lab in Milton Keynes and Alex Alderton, Roberto Amato, Jeffrey Barrett, Sonia Goncalves, Ewan Harrison, David K. Jackson, Ian Johnston, Dominic Kwiatkowski, Cordelia Langford, John Sillitoe on behalf of the Wellcome Sanger Institute COVID-19 Surveillance Team |
| EPI_ISL_2908693 | Lighthouse Laboratory Plymouth | Wellcome Sanger Institute for the COVID-19 Genomics UK (COG-UK) | Lighthouse Laboratory Plymouth and Alex Alderton, Roberto Amato, Jeffrey Barrett, Sonia Goncalves, Ewan Harrison, David K. Jackson, Ian Johnston, Dominic Kwiatkowski, Cordelia Langford, John Sillitoe on behalf of the Wellcome Sanger |

| | | | |
|---|---|---|---|
| | | Consortium | Institute COVID-19 Surveillance Team |
| EPI_ISL_2908694, EPI_ISL_2908695, EPI_ISL_2908696, EPI_ISL_2908697 | Lighthouse Lab in Milton Keynes | Wellcome Sanger Institute for the COVID-19 Genomics UK (COG-UK) Consortium | The Lighthouse Lab in Milton Keynes and Alex Alderton, Roberto Amato, Jeffrey Barrett, Sonia Goncalves, Ewan Harrison, David K. Jackson, Ian Johnston, Dominic Kwiatkowski, Cordelia Langford, John Sillitoe on behalf of the Wellcome Sanger Institute COVID-19 Surveillance Team |
| EPI_ISL_2908698, EPI_ISL_2908699, EPI_ISL_2908700 | Lighthouse Laboratory Plymouth | Wellcome Sanger Institute for the COVID-19 Genomics UK (COG-UK) Consortium | Lighthouse Laboratory Plymouth and Alex Alderton, Roberto Amato, Jeffrey Barrett, Sonia Goncalves, Ewan Harrison, David K. Jackson, Ian Johnston, Dominic Kwiatkowski, Cordelia Langford, John Sillitoe on behalf of the Wellcome Sanger Institute COVID-19 Surveillance Team |
| EPI_ISL_2908701 | Lighthouse Lab in Milton Keynes | Wellcome Sanger Institute for the COVID-19 Genomics UK (COG-UK) Consortium | The Lighthouse Lab in Milton Keynes and Alex Alderton, Roberto Amato, Jeffrey Barrett, Sonia Goncalves, Ewan Harrison, David K. Jackson, Ian Johnston, Dominic Kwiatkowski, Cordelia Langford, John Sillitoe on behalf of the Wellcome Sanger Institute COVID-19 Surveillance Team |
| EPI_ISL_2908702 | Lighthouse Laboratory Plymouth | Wellcome Sanger Institute for the COVID-19 Genomics UK (COG-UK) Consortium | Lighthouse Laboratory Plymouth and Alex Alderton, Roberto Amato, Jeffrey Barrett, Sonia Goncalves, Ewan Harrison, David K. Jackson, Ian Johnston, Dominic Kwiatkowski, Cordelia Langford, John Sillitoe on behalf of the Wellcome Sanger Institute COVID-19 Surveillance Team |
| EPI_ISL_2908703, EPI_ISL_2908704 | Lighthouse Lab in Milton Keynes | Wellcome Sanger Institute for the COVID-19 Genomics UK (COG-UK) Consortium | The Lighthouse Lab in Milton Keynes and Alex Alderton, Roberto Amato, Jeffrey Barrett, Sonia Goncalves, Ewan Harrison, David K. Jackson, Ian Johnston, Dominic Kwiatkowski, Cordelia Langford, John Sillitoe on behalf of the Wellcome Sanger Institute COVID-19 Surveillance Team |
| EPI_ISL_2908705 | Lighthouse Laboratory Plymouth | Wellcome Sanger Institute for the COVID-19 Genomics UK (COG-UK) Consortium | Lighthouse Laboratory Plymouth and Alex Alderton, Roberto Amato, Jeffrey Barrett, Sonia Goncalves, Ewan Harrison, David K. Jackson, Ian Johnston, Dominic Kwiatkowski, Cordelia Langford, John Sillitoe on behalf of the Wellcome Sanger Institute COVID-19 Surveillance Team |
| EPI_ISL_2908706, EPI_ISL_2908707, EPI_ISL_2908708, EPI_ISL_2908709 | Lighthouse Lab in Milton Keynes | Wellcome Sanger Institute for the COVID-19 Genomics UK (COG-UK) Consortium | The Lighthouse Lab in Milton Keynes and Alex Alderton, Roberto Amato, Jeffrey Barrett, Sonia Goncalves, Ewan Harrison, David K. Jackson, Ian Johnston, Dominic Kwiatkowski, Cordelia Langford, John Sillitoe on behalf of the Wellcome Sanger Institute COVID-19 Surveillance Team |
| EPI_ISL_2908710 | Lighthouse Laboratory Plymouth | Wellcome Sanger Institute for the COVID-19 Genomics UK (COG-UK) Consortium | Lighthouse Laboratory Plymouth and Alex Alderton, Roberto Amato, Jeffrey Barrett, Sonia Goncalves, Ewan Harrison, David K. Jackson, Ian Johnston, Dominic Kwiatkowski, Cordelia Langford, John Sillitoe on behalf of the Wellcome Sanger Institute COVID-19 Surveillance Team |
| EPI_ISL_2908711, EPI_ISL_2908712 | Lighthouse Lab in Milton Keynes | Wellcome Sanger Institute for the COVID-19 Genomics UK (COG-UK) Consortium | The Lighthouse Lab in Milton Keynes and Alex Alderton, Roberto Amato, Jeffrey Barrett, Sonia Goncalves, Ewan Harrison, David K. Jackson, Ian Johnston, Dominic Kwiatkowski, Cordelia Langford, John Sillitoe on behalf of the Wellcome Sanger Institute COVID-19 Surveillance Team |
| EPI_ISL_2908713, EPI_ISL_2908714 | Lighthouse Laboratory Plymouth | Wellcome Sanger Institute for the COVID-19 Genomics UK (COG-UK) Consortium | Lighthouse Laboratory Plymouth and Alex Alderton, Roberto Amato, Jeffrey Barrett, Sonia Goncalves, Ewan Harrison, David K. Jackson, Ian Johnston, Dominic Kwiatkowski, Cordelia Langford, John Sillitoe on behalf of the Wellcome Sanger Institute COVID-19 Surveillance Team |
| EPI_ISL_2908715, EPI_ISL_2908716 | Lighthouse Lab in Milton Keynes | Wellcome Sanger Institute for the COVID-19 Genomics UK (COG-UK) Consortium | The Lighthouse Lab in Milton Keynes and Alex Alderton, Roberto Amato, Jeffrey Barrett, Sonia Goncalves, Ewan Harrison, David K. Jackson, Ian Johnston, Dominic Kwiatkowski, Cordelia Langford, John Sillitoe on behalf of the Wellcome Sanger Institute COVID-19 Surveillance Team |
| EPI_ISL_2908717, EPI_ISL_2908718, EPI_ISL_2908719, EPI_ISL_2908720, EPI_ISL_2908721 | Lighthouse Laboratory Plymouth | Wellcome Sanger Institute for the COVID-19 Genomics UK (COG-UK) Consortium | Lighthouse Laboratory Plymouth and Alex Alderton, Roberto Amato, Jeffrey Barrett, Sonia Goncalves, Ewan Harrison, David K. Jackson, Ian Johnston, Dominic Kwiatkowski, Cordelia Langford, John Sillitoe on behalf of the Wellcome Sanger Institute COVID-19 Surveillance Team |
| EPI_ISL_2908722 | Lighthouse Lab in Milton Keynes | Wellcome Sanger Institute for the COVID-19 Genomics UK (COG-UK) Consortium | The Lighthouse Lab in Milton Keynes and Alex Alderton, Roberto Amato, Jeffrey Barrett, Sonia Goncalves, Ewan Harrison, David K. Jackson, Ian Johnston, Dominic Kwiatkowski, Cordelia Langford, John Sillitoe on behalf of the Wellcome Sanger Institute COVID-19 Surveillance Team |
| EPI_ISL_2908723 | Lighthouse Laboratory Plymouth | Wellcome Sanger Institute for the COVID-19 Genomics UK (COG-UK) Consortium | Lighthouse Laboratory Plymouth and Alex Alderton, Roberto Amato, Jeffrey Barrett, Sonia Goncalves, Ewan Harrison, David K. Jackson, Ian Johnston, Dominic Kwiatkowski, Cordelia Langford, John Sillitoe on behalf of the Wellcome Sanger Institute COVID-19 Surveillance Team |
| EPI_ISL_2908724 | Lighthouse Lab in Milton Keynes | Wellcome Sanger Institute for the COVID-19 Genomics UK (COG-UK) Consortium | The Lighthouse Lab in Milton Keynes and Alex Alderton, Roberto Amato, Jeffrey Barrett, Sonia Goncalves, Ewan Harrison, David K. Jackson, Ian Johnston, Dominic Kwiatkowski, Cordelia Langford, John Sillitoe on behalf of the Wellcome Sanger Institute COVID-19 Surveillance Team |
| EPI_ISL_2908725, EPI_ISL_2908726 | Lighthouse Laboratory Plymouth | Wellcome Sanger Institute for the COVID-19 Genomics UK (COG-UK) Consortium | Lighthouse Laboratory Plymouth and Alex Alderton, Roberto Amato, Jeffrey Barrett, Sonia Goncalves, Ewan Harrison, David K. Jackson, Ian Johnston, Dominic Kwiatkowski, Cordelia Langford, John Sillitoe on behalf of the Wellcome Sanger Institute COVID-19 Surveillance Team |
| EPI_ISL_2908727, EPI_ISL_2908728 | Lighthouse Lab in Milton Keynes | Wellcome Sanger Institute for the COVID-19 Genomics UK (COG-UK) Consortium | The Lighthouse Lab in Milton Keynes and Alex Alderton, Roberto Amato, Jeffrey Barrett, Sonia Goncalves, Ewan Harrison, David K. Jackson, Ian Johnston, Dominic Kwiatkowski, Cordelia Langford, John Sillitoe on behalf of the Wellcome Sanger Institute COVID-19 Surveillance Team |
| EPI_ISL_2908729, EPI_ISL_2908730 | Lighthouse Laboratory Plymouth | Wellcome Sanger Institute for the COVID-19 Genomics UK (COG-UK) Consortium | Lighthouse Laboratory Plymouth and Alex Alderton, Roberto Amato, Jeffrey Barrett, Sonia Goncalves, Ewan Harrison, David K. Jackson, Ian Johnston, Dominic Kwiatkowski, Cordelia Langford, John Sillitoe on behalf of the Wellcome Sanger Institute COVID-19 Surveillance Team |
| EPI_ISL_2908731, EPI_ISL_2908732 | Lighthouse Lab in Milton Keynes | Wellcome Sanger Institute for the COVID-19 Genomics UK (COG-UK) Consortium | The Lighthouse Lab in Milton Keynes and Alex Alderton, Roberto Amato, Jeffrey Barrett, Sonia Goncalves, Ewan Harrison, David K. Jackson, Ian Johnston, Dominic Kwiatkowski, Cordelia Langford, John Sillitoe on behalf of the Wellcome Sanger Institute COVID-19 Surveillance Team |
| EPI_ISL_2908733, EPI_ISL_2908734 | Lighthouse Laboratory Plymouth | Wellcome Sanger Institute for the COVID-19 Genomics UK (COG-UK) Consortium | Lighthouse Laboratory Plymouth and Alex Alderton, Roberto Amato, Jeffrey Barrett, Sonia Goncalves, Ewan Harrison, David K. Jackson, Ian Johnston, Dominic Kwiatkowski, Cordelia Langford, John Sillitoe on behalf of the Wellcome Sanger Institute COVID-19 Surveillance Team |
| EPI_ISL_2908735, EPI_ISL_2908736, EPI_ISL_2908737, EPI_ISL_2908738, EPI_ISL_2908739, EPI_ISL_2908740, EPI_ISL_2908741 | Lighthouse Lab in Milton Keynes | Wellcome Sanger Institute for the COVID-19 Genomics UK (COG-UK) Consortium | The Lighthouse Lab in Milton Keynes and Alex Alderton, Roberto Amato, Jeffrey Barrett, Sonia Goncalves, Ewan Harrison, David K. Jackson, Ian Johnston, Dominic Kwiatkowski, Cordelia Langford, John Sillitoe on behalf of the Wellcome Sanger Institute COVID-19 Surveillance Team |
| EPI_ISL_2908742 | Lighthouse Laboratory Plymouth | Wellcome Sanger Institute for the COVID-19 Genomics UK (COG-UK) Consortium | Lighthouse Laboratory Plymouth and Alex Alderton, Roberto Amato, Jeffrey Barrett, Sonia Goncalves, Ewan Harrison, David K. Jackson, Ian Johnston, Dominic Kwiatkowski, Cordelia Langford, John Sillitoe on behalf of the Wellcome Sanger Institute COVID-19 Surveillance Team |
| EPI_ISL_2908743, EPI_ISL_2908744 | Lighthouse Lab in Milton Keynes | Wellcome Sanger Institute for the COVID-19 Genomics UK (COG-UK) Consortium | The Lighthouse Lab in Milton Keynes and Alex Alderton, Roberto Amato, Jeffrey Barrett, Sonia Goncalves, Ewan Harrison, David K. Jackson, Ian Johnston, Dominic Kwiatkowski, Cordelia Langford, John Sillitoe on behalf of the Wellcome Sanger Institute COVID-19 Surveillance Team |
| EPI_ISL_2908745 | Lighthouse Laboratory Plymouth | Wellcome Sanger Institute for the COVID-19 Genomics UK (COG-UK) Consortium | Lighthouse Laboratory Plymouth and Alex Alderton, Roberto Amato, Jeffrey Barrett, Sonia Goncalves, Ewan Harrison, David K. Jackson, Ian Johnston, Dominic Kwiatkowski, Cordelia Langford, John Sillitoe on behalf of the Wellcome Sanger Institute COVID-19 Surveillance Team |
| EPI_ISL_2908746 | Lighthouse Lab in Milton Keynes | Wellcome Sanger Institute for the COVID-19 Genomics UK (COG-UK) Consortium | The Lighthouse Lab in Milton Keynes and Alex Alderton, Roberto Amato, Jeffrey Barrett, Sonia Goncalves, Ewan Harrison, David K. Jackson, Ian Johnston, Dominic Kwiatkowski, Cordelia Langford, John Sillitoe on behalf of the Wellcome Sanger Institute COVID-19 Surveillance Team |
| EPI_ISL_2908747, EPI_ISL_2908748, EPI_ISL_2908749, EPI_ISL_2908750 | Lighthouse Laboratory Plymouth | Wellcome Sanger Institute for the COVID-19 Genomics UK (COG-UK) Consortium | Lighthouse Laboratory Plymouth and Alex Alderton, Roberto Amato, Jeffrey Barrett, Sonia Goncalves, Ewan Harrison, David K. Jackson, Ian Johnston, Dominic Kwiatkowski, Cordelia Langford, John Sillitoe on behalf of the Wellcome Sanger Institute COVID-19 Surveillance Team |
| EPI_ISL_2908751, EPI_ISL_2908752 | Lighthouse Lab in Milton Keynes | Wellcome Sanger Institute for the COVID-19 Genomics UK (COG-UK) Consortium | The Lighthouse Lab in Milton Keynes and Alex Alderton, Roberto Amato, Jeffrey Barrett, Sonia Goncalves, Ewan Harrison, David K. Jackson, Ian Johnston, Dominic Kwiatkowski, Cordelia Langford, John Sillitoe on behalf of the Wellcome Sanger Institute COVID-19 Surveillance Team |
| EPI_ISL_2908753, EPI_ISL_2908754 | Lighthouse Laboratory Plymouth | Wellcome Sanger Institute for the COVID-19 Genomics UK (COG-UK) Consortium | Lighthouse Laboratory Plymouth and Alex Alderton, Roberto Amato, Jeffrey Barrett, Sonia Goncalves, Ewan Harrison, David K. Jackson, Ian Johnston, Dominic Kwiatkowski, Cordelia Langford, John Sillitoe on behalf of the Wellcome Sanger Institute COVID-19 Surveillance Team |
| EPI_ISL_2908755 | Lighthouse Lab in Milton Keynes | Wellcome Sanger Institute for the COVID-19 Genomics UK (COG-UK) Consortium | The Lighthouse Lab in Milton Keynes and Alex Alderton, Roberto Amato, Jeffrey Barrett, Sonia Goncalves, Ewan Harrison, David K. Jackson, Ian Johnston, Dominic Kwiatkowski, Cordelia Langford, John Sillitoe on behalf of the Wellcome Sanger Institute COVID-19 Surveillance Team |
| EPI_ISL_2908756, EPI_ISL_2908757 | Lighthouse Laboratory Plymouth | Wellcome Sanger Institute for the COVID-19 Genomics UK (COG-UK) Consortium | Lighthouse Laboratory Plymouth and Alex Alderton, Roberto Amato, Jeffrey Barrett, Sonia Goncalves, Ewan Harrison, David K. Jackson, Ian Johnston, Dominic Kwiatkowski, Cordelia Langford, John Sillitoe on behalf of the Wellcome Sanger Institute COVID-19 Surveillance Team |
| EPI_ISL_2908758 | Lighthouse Lab in Milton Keynes | Wellcome Sanger Institute for the COVID-19 Genomics UK (COG-UK) Consortium | The Lighthouse Lab in Milton Keynes and Alex Alderton, Roberto Amato, Jeffrey Barrett, Sonia Goncalves, Ewan Harrison, David K. Jackson, Ian Johnston, Dominic Kwiatkowski, Cordelia Langford, John Sillitoe on behalf of the Wellcome Sanger Institute COVID-19 Surveillance Team |
| EPI_ISL_2908759, EPI_ISL_2908760, EPI_ISL_2908761, EPI_ISL_2908762, EPI_ISL_2908763, EPI_ISL_2908764, EPI_ISL_2908765 | Lighthouse Laboratory Plymouth | Wellcome Sanger Institute for the COVID-19 Genomics UK (COG-UK) Consortium | Lighthouse Laboratory Plymouth and Alex Alderton, Roberto Amato, Jeffrey Barrett, Sonia Goncalves, Ewan Harrison, David K. Jackson, Ian Johnston, Dominic Kwiatkowski, Cordelia Langford, John Sillitoe on behalf of the Wellcome Sanger Institute COVID-19 Surveillance Team |
| EPI_ISL_2908766 | Lighthouse Lab in Milton Keynes | Wellcome Sanger Institute for the COVID-19 Genomics UK (COG-UK) Consortium | The Lighthouse Lab in Milton Keynes and Alex Alderton, Roberto Amato, Jeffrey Barrett, Sonia Goncalves, Ewan Harrison, David K. Jackson, Ian Johnston, Dominic Kwiatkowski, Cordelia Langford, John Sillitoe on behalf of the Wellcome Sanger Institute COVID-19 Surveillance Team |
| EPI_ISL_2908767, EPI_ISL_2908768 | Lighthouse Laboratory Plymouth | Wellcome Sanger Institute for the COVID-19 Genomics UK (COG-UK) Consortium | Lighthouse Laboratory Plymouth and Alex Alderton, Roberto Amato, Jeffrey Barrett, Sonia Goncalves, Ewan Harrison, David K. Jackson, Ian Johnston, Dominic Kwiatkowski, Cordelia Langford, John Sillitoe on behalf of the Wellcome Sanger Institute COVID-19 Surveillance Team |
| EPI_ISL_2908769 | Lighthouse Lab in Milton Keynes | Wellcome Sanger Institute for the COVID-19 Genomics UK (COG-UK) Consortium | The Lighthouse Lab in Milton Keynes and Alex Alderton, Roberto Amato, Jeffrey Barrett, Sonia Goncalves, Ewan Harrison, David K. Jackson, Ian Johnston, Dominic Kwiatkowski, Cordelia Langford, John Sillitoe on behalf of the Wellcome Sanger Institute COVID-19 Surveillance Team |
| EPI_ISL_2908770 | Lighthouse Laboratory Plymouth | Wellcome Sanger Institute for the COVID-19 Genomics UK (COG-UK) Consortium | Lighthouse Laboratory Plymouth and Alex Alderton, Roberto Amato, Jeffrey Barrett, Sonia Goncalves, Ewan Harrison, David K. Jackson, Ian Johnston, Dominic Kwiatkowski, Cordelia Langford, John Sillitoe on behalf of the Wellcome Sanger Institute COVID-19 Surveillance Team |
| EPI_ISL_2908771 | Lighthouse Lab in Milton Keynes | Wellcome Sanger Institute for the COVID-19 Genomics UK (COG-UK) | The Lighthouse Lab in Milton Keynes and Alex Alderton, Roberto Amato, Jeffrey Barrett, Sonia Goncalves, Ewan Harrison, David K. Jackson, Ian Johnston, Dominic Kwiatkowski, Cordelia Langford, John Sillitoe on behalf of the Wellcome Sanger |

| | | | |
|---|---|---|---|
| | | Consortium | Institute COVID-19 Surveillance Team |
| EPI_ISL_2908772 | Lighthouse Laboratory Plymouth | Wellcome Sanger Institute for the COVID-19 Genomics UK (COG-UK) Consortium | Lighthouse Laboratory Plymouth and Alex Alderton, Roberto Amato, Jeffrey Barrett, Sonia Goncalves, Ewan Harrison, David K. Jackson, Ian Johnston, Dominic Kwiatkowski, Cordelia Langford, John Sillitoe on behalf of the Wellcome Sanger Institute COVID-19 Surveillance Team |
| EPI_ISL_2908773 | Lighthouse Lab in Milton Keynes | Wellcome Sanger Institute for the COVID-19 Genomics UK (COG-UK) Consortium | The Lighthouse Lab in Milton Keynes and Alex Alderton, Roberto Amato, Jeffrey Barrett, Sonia Goncalves, Ewan Harrison, David K. Jackson, Ian Johnston, Dominic Kwiatkowski, Cordelia Langford, John Sillitoe on behalf of the Wellcome Sanger Institute COVID-19 Surveillance Team |
| EPI_ISL_2908774, EPI_ISL_2908775, EPI_ISL_2908776 | Lighthouse Laboratory Plymouth | Wellcome Sanger Institute for the COVID-19 Genomics UK (COG-UK) Consortium | Lighthouse Laboratory Plymouth and Alex Alderton, Roberto Amato, Jeffrey Barrett, Sonia Goncalves, Ewan Harrison, David K. Jackson, Ian Johnston, Dominic Kwiatkowski, Cordelia Langford, John Sillitoe on behalf of the Wellcome Sanger Institute COVID-19 Surveillance Team |
| EPI_ISL_2908777, EPI_ISL_2908778, EPI_ISL_2908779 | Lighthouse Lab in Milton Keynes | Wellcome Sanger Institute for the COVID-19 Genomics UK (COG-UK) Consortium | The Lighthouse Lab in Milton Keynes and Alex Alderton, Roberto Amato, Jeffrey Barrett, Sonia Goncalves, Ewan Harrison, David K. Jackson, Ian Johnston, Dominic Kwiatkowski, Cordelia Langford, John Sillitoe on behalf of the Wellcome Sanger Institute COVID-19 Surveillance Team |
| EPI_ISL_2908780 | Lighthouse Laboratory Plymouth | Wellcome Sanger Institute for the COVID-19 Genomics UK (COG-UK) Consortium | Lighthouse Laboratory Plymouth and Alex Alderton, Roberto Amato, Jeffrey Barrett, Sonia Goncalves, Ewan Harrison, David K. Jackson, Ian Johnston, Dominic Kwiatkowski, Cordelia Langford, John Sillitoe on behalf of the Wellcome Sanger Institute COVID-19 Surveillance Team |
| EPI_ISL_2908781, EPI_ISL_2908782, EPI_ISL_2908783 | Lighthouse Lab in Milton Keynes | Wellcome Sanger Institute for the COVID-19 Genomics UK (COG-UK) Consortium | The Lighthouse Lab in Milton Keynes and Alex Alderton, Roberto Amato, Jeffrey Barrett, Sonia Goncalves, Ewan Harrison, David K. Jackson, Ian Johnston, Dominic Kwiatkowski, Cordelia Langford, John Sillitoe on behalf of the Wellcome Sanger Institute COVID-19 Surveillance Team |
| EPI_ISL_2908784, EPI_ISL_2908785 | Lighthouse Laboratory Plymouth | Wellcome Sanger Institute for the COVID-19 Genomics UK (COG-UK) Consortium | Lighthouse Laboratory Plymouth and Alex Alderton, Roberto Amato, Jeffrey Barrett, Sonia Goncalves, Ewan Harrison, David K. Jackson, Ian Johnston, Dominic Kwiatkowski, Cordelia Langford, John Sillitoe on behalf of the Wellcome Sanger Institute COVID-19 Surveillance Team |
| EPI_ISL_2908786, EPI_ISL_2908787 | Lighthouse Lab in Milton Keynes | Wellcome Sanger Institute for the COVID-19 Genomics UK (COG-UK) Consortium | The Lighthouse Lab in Milton Keynes and Alex Alderton, Roberto Amato, Jeffrey Barrett, Sonia Goncalves, Ewan Harrison, David K. Jackson, Ian Johnston, Dominic Kwiatkowski, Cordelia Langford, John Sillitoe on behalf of the Wellcome Sanger Institute COVID-19 Surveillance Team |
| EPI_ISL_2908788 | Lighthouse Laboratory Plymouth | Wellcome Sanger Institute for the COVID-19 Genomics UK (COG-UK) Consortium | Lighthouse Laboratory Plymouth and Alex Alderton, Roberto Amato, Jeffrey Barrett, Sonia Goncalves, Ewan Harrison, David K. Jackson, Ian Johnston, Dominic Kwiatkowski, Cordelia Langford, John Sillitoe on behalf of the Wellcome Sanger Institute COVID-19 Surveillance Team |
| EPI_ISL_2908789, EPI_ISL_2908790, EPI_ISL_2908791, EPI_ISL_2908792 | Lighthouse Lab in Milton Keynes | Wellcome Sanger Institute for the COVID-19 Genomics UK (COG-UK) Consortium | The Lighthouse Lab in Milton Keynes and Alex Alderton, Roberto Amato, Jeffrey Barrett, Sonia Goncalves, Ewan Harrison, David K. Jackson, Ian Johnston, Dominic Kwiatkowski, Cordelia Langford, John Sillitoe on behalf of the Wellcome Sanger Institute COVID-19 Surveillance Team |
| EPI_ISL_2908793 | Lighthouse Laboratory Plymouth | Wellcome Sanger Institute for the COVID-19 Genomics UK (COG-UK) Consortium | Lighthouse Laboratory Plymouth and Alex Alderton, Roberto Amato, Jeffrey Barrett, Sonia Goncalves, Ewan Harrison, David K. Jackson, Ian Johnston, Dominic Kwiatkowski, Cordelia Langford, John Sillitoe on behalf of the Wellcome Sanger Institute COVID-19 Surveillance Team |
| EPI_ISL_2908794, EPI_ISL_2908795, EPI_ISL_2908796 | Lighthouse Lab in Milton Keynes | Wellcome Sanger Institute for the COVID-19 Genomics UK (COG-UK) Consortium | The Lighthouse Lab in Milton Keynes and Alex Alderton, Roberto Amato, Jeffrey Barrett, Sonia Goncalves, Ewan Harrison, David K. Jackson, Ian Johnston, Dominic Kwiatkowski, Cordelia Langford, John Sillitoe on behalf of the Wellcome Sanger Institute COVID-19 Surveillance Team |
| EPI_ISL_2908797, EPI_ISL_2908798, EPI_ISL_2908799 | Lighthouse Laboratory Plymouth | Wellcome Sanger Institute for the COVID-19 Genomics UK (COG-UK) Consortium | Lighthouse Laboratory Plymouth and Alex Alderton, Roberto Amato, Jeffrey Barrett, Sonia Goncalves, Ewan Harrison, David K. Jackson, Ian Johnston, Dominic Kwiatkowski, Cordelia Langford, John Sillitoe on behalf of the Wellcome Sanger Institute COVID-19 Surveillance Team |
| EPI_ISL_2908800 | Lighthouse Lab in Milton Keynes | Wellcome Sanger Institute for the COVID-19 Genomics UK (COG-UK) Consortium | The Lighthouse Lab in Milton Keynes and Alex Alderton, Roberto Amato, Jeffrey Barrett, Sonia Goncalves, Ewan Harrison, David K. Jackson, Ian Johnston, Dominic Kwiatkowski, Cordelia Langford, John Sillitoe on behalf of the Wellcome Sanger Institute COVID-19 Surveillance Team |
| EPI_ISL_2908801, EPI_ISL_2908802 | Lighthouse Laboratory Plymouth | Wellcome Sanger Institute for the COVID-19 Genomics UK (COG-UK) Consortium | Lighthouse Laboratory Plymouth and Alex Alderton, Roberto Amato, Jeffrey Barrett, Sonia Goncalves, Ewan Harrison, David K. Jackson, Ian Johnston, Dominic Kwiatkowski, Cordelia Langford, John Sillitoe on behalf of the Wellcome Sanger Institute COVID-19 Surveillance Team |
| EPI_ISL_2908803 | Lighthouse Lab in Milton Keynes | Wellcome Sanger Institute for the COVID-19 Genomics UK (COG-UK) Consortium | The Lighthouse Lab in Milton Keynes and Alex Alderton, Roberto Amato, Jeffrey Barrett, Sonia Goncalves, Ewan Harrison, David K. Jackson, Ian Johnston, Dominic Kwiatkowski, Cordelia Langford, John Sillitoe on behalf of the Wellcome Sanger Institute COVID-19 Surveillance Team |
| EPI_ISL_2908804, EPI_ISL_2908805, EPI_ISL_2908806, EPI_ISL_2908807 | Lighthouse Laboratory Plymouth | Wellcome Sanger Institute for the COVID-19 Genomics UK (COG-UK) Consortium | Lighthouse Laboratory Plymouth and Alex Alderton, Roberto Amato, Jeffrey Barrett, Sonia Goncalves, Ewan Harrison, David K. Jackson, Ian Johnston, Dominic Kwiatkowski, Cordelia Langford, John Sillitoe on behalf of the Wellcome Sanger Institute COVID-19 Surveillance Team |
| EPI_ISL_2908808 | Lighthouse Lab in Milton Keynes | Wellcome Sanger Institute for the COVID-19 Genomics UK (COG-UK) Consortium | The Lighthouse Lab in Milton Keynes and Alex Alderton, Roberto Amato, Jeffrey Barrett, Sonia Goncalves, Ewan Harrison, David K. Jackson, Ian Johnston, Dominic Kwiatkowski, Cordelia Langford, John Sillitoe on behalf of the Wellcome Sanger Institute COVID-19 Surveillance Team |
| EPI_ISL_2908809 | Lighthouse Laboratory Plymouth | Wellcome Sanger Institute for the COVID-19 Genomics UK (COG-UK) Consortium | Lighthouse Laboratory Plymouth and Alex Alderton, Roberto Amato, Jeffrey Barrett, Sonia Goncalves, Ewan Harrison, David K. Jackson, Ian Johnston, Dominic Kwiatkowski, Cordelia Langford, John Sillitoe on behalf of the Wellcome Sanger Institute COVID-19 Surveillance Team |
| EPI_ISL_2908810 | Lighthouse Lab in Milton Keynes | Wellcome Sanger Institute for the COVID-19 Genomics UK (COG-UK) Consortium | The Lighthouse Lab in Milton Keynes and Alex Alderton, Roberto Amato, Jeffrey Barrett, Sonia Goncalves, Ewan Harrison, David K. Jackson, Ian Johnston, Dominic Kwiatkowski, Cordelia Langford, John Sillitoe on behalf of the Wellcome Sanger Institute COVID-19 Surveillance Team |
| EPI_ISL_2908811 | Lighthouse Laboratory Plymouth | Wellcome Sanger Institute for the COVID-19 Genomics UK (COG-UK) Consortium | Lighthouse Laboratory Plymouth and Alex Alderton, Roberto Amato, Jeffrey Barrett, Sonia Goncalves, Ewan Harrison, David K. Jackson, Ian Johnston, Dominic Kwiatkowski, Cordelia Langford, John Sillitoe on behalf of the Wellcome Sanger Institute COVID-19 Surveillance Team |
| EPI_ISL_2908812 | Lighthouse Lab in Milton Keynes | Wellcome Sanger Institute for the COVID-19 Genomics UK (COG-UK) Consortium | The Lighthouse Lab in Milton Keynes and Alex Alderton, Roberto Amato, Jeffrey Barrett, Sonia Goncalves, Ewan Harrison, David K. Jackson, Ian Johnston, Dominic Kwiatkowski, Cordelia Langford, John Sillitoe on behalf of the Wellcome Sanger Institute COVID-19 Surveillance Team |
| EPI_ISL_2908813 | Lighthouse Laboratory Plymouth | Wellcome Sanger Institute for the COVID-19 Genomics UK (COG-UK) Consortium | Lighthouse Laboratory Plymouth and Alex Alderton, Roberto Amato, Jeffrey Barrett, Sonia Goncalves, Ewan Harrison, David K. Jackson, Ian Johnston, Dominic Kwiatkowski, Cordelia Langford, John Sillitoe on behalf of the Wellcome Sanger Institute COVID-19 Surveillance Team |
| EPI_ISL_2908814, EPI_ISL_2908815 | Lighthouse Lab in Milton Keynes | Wellcome Sanger Institute for the COVID-19 Genomics UK (COG-UK) Consortium | The Lighthouse Lab in Milton Keynes and Alex Alderton, Roberto Amato, Jeffrey Barrett, Sonia Goncalves, Ewan Harrison, David K. Jackson, Ian Johnston, Dominic Kwiatkowski, Cordelia Langford, John Sillitoe on behalf of the Wellcome Sanger Institute COVID-19 Surveillance Team |
| EPI_ISL_2908816, EPI_ISL_2908817, EPI_ISL_2908818 | Lighthouse Laboratory Plymouth | Wellcome Sanger Institute for the COVID-19 Genomics UK (COG-UK) Consortium | Lighthouse Laboratory Plymouth and Alex Alderton, Roberto Amato, Jeffrey Barrett, Sonia Goncalves, Ewan Harrison, David K. Jackson, Ian Johnston, Dominic Kwiatkowski, Cordelia Langford, John Sillitoe on behalf of the Wellcome Sanger Institute COVID-19 Surveillance Team |
| EPI_ISL_2908819, EPI_ISL_2908820 | Lighthouse Lab in Milton Keynes | Wellcome Sanger Institute for the COVID-19 Genomics UK (COG-UK) Consortium | The Lighthouse Lab in Milton Keynes and Alex Alderton, Roberto Amato, Jeffrey Barrett, Sonia Goncalves, Ewan Harrison, David K. Jackson, Ian Johnston, Dominic Kwiatkowski, Cordelia Langford, John Sillitoe on behalf of the Wellcome Sanger Institute COVID-19 Surveillance Team |
| EPI_ISL_2908821 | Lighthouse Laboratory Plymouth | Wellcome Sanger Institute for the COVID-19 Genomics UK (COG-UK) Consortium | Lighthouse Laboratory Plymouth and Alex Alderton, Roberto Amato, Jeffrey Barrett, Sonia Goncalves, Ewan Harrison, David K. Jackson, Ian Johnston, Dominic Kwiatkowski, Cordelia Langford, John Sillitoe on behalf of the Wellcome Sanger Institute COVID-19 Surveillance Team |
| EPI_ISL_2908822 | Lighthouse Lab in Milton Keynes | Wellcome Sanger Institute for the COVID-19 Genomics UK (COG-UK) Consortium | The Lighthouse Lab in Milton Keynes and Alex Alderton, Roberto Amato, Jeffrey Barrett, Sonia Goncalves, Ewan Harrison, David K. Jackson, Ian Johnston, Dominic Kwiatkowski, Cordelia Langford, John Sillitoe on behalf of the Wellcome Sanger Institute COVID-19 Surveillance Team |
| EPI_ISL_2908823 | Lighthouse Laboratory Plymouth | Wellcome Sanger Institute for the COVID-19 Genomics UK (COG-UK) Consortium | Lighthouse Laboratory Plymouth and Alex Alderton, Roberto Amato, Jeffrey Barrett, Sonia Goncalves, Ewan Harrison, David K. Jackson, Ian Johnston, Dominic Kwiatkowski, Cordelia Langford, John Sillitoe on behalf of the Wellcome Sanger Institute COVID-19 Surveillance Team |
| EPI_ISL_2908824, EPI_ISL_2908825 | Lighthouse Lab in Milton Keynes | Wellcome Sanger Institute for the COVID-19 Genomics UK (COG-UK) Consortium | The Lighthouse Lab in Milton Keynes and Alex Alderton, Roberto Amato, Jeffrey Barrett, Sonia Goncalves, Ewan Harrison, David K. Jackson, Ian Johnston, Dominic Kwiatkowski, Cordelia Langford, John Sillitoe on behalf of the Wellcome Sanger Institute COVID-19 Surveillance Team |
| EPI_ISL_2908826 | Lighthouse Laboratory Plymouth | Wellcome Sanger Institute for the COVID-19 Genomics UK (COG-UK) Consortium | Lighthouse Laboratory Plymouth and Alex Alderton, Roberto Amato, Jeffrey Barrett, Sonia Goncalves, Ewan Harrison, David K. Jackson, Ian Johnston, Dominic Kwiatkowski, Cordelia Langford, John Sillitoe on behalf of the Wellcome Sanger Institute COVID-19 Surveillance Team |
| EPI_ISL_2908827 | Lighthouse Lab in Milton Keynes | Wellcome Sanger Institute for the COVID-19 Genomics UK (COG-UK) Consortium | The Lighthouse Lab in Milton Keynes and Alex Alderton, Roberto Amato, Jeffrey Barrett, Sonia Goncalves, Ewan Harrison, David K. Jackson, Ian Johnston, Dominic Kwiatkowski, Cordelia Langford, John Sillitoe on behalf of the Wellcome Sanger Institute COVID-19 Surveillance Team |
| EPI_ISL_2908828 | Lighthouse Laboratory Plymouth | Wellcome Sanger Institute for the COVID-19 Genomics UK (COG-UK) Consortium | Lighthouse Laboratory Plymouth and Alex Alderton, Roberto Amato, Jeffrey Barrett, Sonia Goncalves, Ewan Harrison, David K. Jackson, Ian Johnston, Dominic Kwiatkowski, Cordelia Langford, John Sillitoe on behalf of the Wellcome Sanger Institute COVID-19 Surveillance Team |
| EPI_ISL_2908829, EPI_ISL_2908830, EPI_ISL_2908831, EPI_ISL_2908832 | Lighthouse Lab in Milton Keynes | Wellcome Sanger Institute for the COVID-19 Genomics UK (COG-UK) Consortium | The Lighthouse Lab in Milton Keynes and Alex Alderton, Roberto Amato, Jeffrey Barrett, Sonia Goncalves, Ewan Harrison, David K. Jackson, Ian Johnston, Dominic Kwiatkowski, Cordelia Langford, John Sillitoe on behalf of the Wellcome Sanger Institute COVID-19 Surveillance Team |
| EPI_ISL_2908833, EPI_ISL_2908834 | Lighthouse Laboratory Plymouth | Wellcome Sanger Institute for the COVID-19 Genomics UK (COG-UK) Consortium | Lighthouse Laboratory Plymouth and Alex Alderton, Roberto Amato, Jeffrey Barrett, Sonia Goncalves, Ewan Harrison, David K. Jackson, Ian Johnston, Dominic Kwiatkowski, Cordelia Langford, John Sillitoe on behalf of the Wellcome Sanger Institute COVID-19 Surveillance Team |
| EPI_ISL_2908835 | Lighthouse Lab in Milton Keynes | Wellcome Sanger Institute for the COVID-19 Genomics UK (COG-UK) Consortium | The Lighthouse Lab in Milton Keynes and Alex Alderton, Roberto Amato, Jeffrey Barrett, Sonia Goncalves, Ewan Harrison, David K. Jackson, Ian Johnston, Dominic Kwiatkowski, Cordelia Langford, John Sillitoe on behalf of the Wellcome Sanger Institute COVID-19 Surveillance Team |
| EPI_ISL_2908836 | Lighthouse Laboratory Plymouth | Wellcome Sanger Institute for the COVID-19 Genomics UK (COG-UK) Consortium | Lighthouse Laboratory Plymouth and Alex Alderton, Roberto Amato, Jeffrey Barrett, Sonia Goncalves, Ewan Harrison, David K. Jackson, Ian Johnston, Dominic Kwiatkowski, Cordelia Langford, John Sillitoe on behalf of the Wellcome Sanger Institute COVID-19 Surveillance Team |
| EPI_ISL_2908837 | Lighthouse Lab in Milton Keynes | Wellcome Sanger Institute for the COVID-19 Genomics UK (COG-UK) Consortium | The Lighthouse Lab in Milton Keynes and Alex Alderton, Roberto Amato, Jeffrey Barrett, Sonia Goncalves, Ewan Harrison, David K. Jackson, Ian Johnston, Dominic Kwiatkowski, Cordelia Langford, John Sillitoe on behalf of the Wellcome Sanger Institute COVID-19 Surveillance Team |
| EPI_ISL_2908838, EPI_ISL_2908839, EPI_ISL_2908840 | Lighthouse Laboratory Plymouth | Wellcome Sanger Institute for the COVID-19 Genomics UK (COG-UK) Consortium | Lighthouse Laboratory Plymouth and Alex Alderton, Roberto Amato, Jeffrey Barrett, Sonia Goncalves, Ewan Harrison, David K. Jackson, Ian Johnston, Dominic Kwiatkowski, Cordelia Langford, John Sillitoe on behalf of the Wellcome Sanger Institute COVID-19 Surveillance Team |

| | | | |
|---|---|---|---|
| EPI_ISL_2908841, EPI_ISL_2908842 | Lighthouse Lab in Milton Keynes | Wellcome Sanger Institute for the COVID-19 Genomics UK (COG-UK) Consortium | The Lighthouse Lab in Milton Keynes and Alex Alderton, Roberto Amato, Jeffrey Barrett, Sonia Goncalves, Ewan Harrison, David K. Jackson, Ian Johnston, Dominic Kwiatkowski, Cordelia Langford, John Sillitoe on behalf of the Wellcome Sanger Institute COVID-19 Surveillance Team |
| EPI_ISL_2908843, EPI_ISL_2908844, EPI_ISL_2908845, EPI_ISL_2908846, EPI_ISL_2908847, EPI_ISL_2908848, EPI_ISL_2908849, EPI_ISL_2908850, EPI_ISL_2908851, EPI_ISL_2908852, EPI_ISL_2908853, EPI_ISL_2908854, EPI_ISL_2908855, EPI_ISL_2908856 | | | |
| see above | Lighthouse Laboratory Plymouth | Wellcome Sanger Institute for the COVID-19 Genomics UK (COG-UK) Consortium | Lighthouse Laboratory Plymouth and Alex Alderton, Roberto Amato, Jeffrey Barrett, Sonia Goncalves, Ewan Harrison, David K. Jackson, Ian Johnston, Dominic Kwiatkowski, Cordelia Langford, John Sillitoe on behalf of the Wellcome Sanger Institute COVID-19 Surveillance Team |
| EPI_ISL_2908857, EPI_ISL_2908858 | Lighthouse Lab in Milton Keynes | Wellcome Sanger Institute for the COVID-19 Genomics UK (COG-UK) Consortium | The Lighthouse Lab in Milton Keynes and Alex Alderton, Roberto Amato, Jeffrey Barrett, Sonia Goncalves, Ewan Harrison, David K. Jackson, Ian Johnston, Dominic Kwiatkowski, Cordelia Langford, John Sillitoe on behalf of the Wellcome Sanger Institute COVID-19 Surveillance Team |
| EPI_ISL_2908859, EPI_ISL_2908860, EPI_ISL_2908861, EPI_ISL_2908862 | Lighthouse Laboratory Plymouth | Wellcome Sanger Institute for the COVID-19 Genomics UK (COG-UK) Consortium | Lighthouse Laboratory Plymouth and Alex Alderton, Roberto Amato, Jeffrey Barrett, Sonia Goncalves, Ewan Harrison, David K. Jackson, Ian Johnston, Dominic Kwiatkowski, Cordelia Langford, John Sillitoe on behalf of the Wellcome Sanger Institute COVID-19 Surveillance Team |
| EPI_ISL_2908863 | Lighthouse Lab in Milton Keynes | Wellcome Sanger Institute for the COVID-19 Genomics UK (COG-UK) Consortium | The Lighthouse Lab in Milton Keynes and Alex Alderton, Roberto Amato, Jeffrey Barrett, Sonia Goncalves, Ewan Harrison, David K. Jackson, Ian Johnston, Dominic Kwiatkowski, Cordelia Langford, John Sillitoe on behalf of the Wellcome Sanger Institute COVID-19 Surveillance Team |
| EPI_ISL_2908864, EPI_ISL_2908865, EPI_ISL_2908866, EPI_ISL_2908867 | Lighthouse Laboratory Plymouth | Wellcome Sanger Institute for the COVID-19 Genomics UK (COG-UK) Consortium | Lighthouse Laboratory Plymouth and Alex Alderton, Roberto Amato, Jeffrey Barrett, Sonia Goncalves, Ewan Harrison, David K. Jackson, Ian Johnston, Dominic Kwiatkowski, Cordelia Langford, John Sillitoe on behalf of the Wellcome Sanger Institute COVID-19 Surveillance Team |
| EPI_ISL_2908868 | Lighthouse Lab in Milton Keynes | Wellcome Sanger Institute for the COVID-19 Genomics UK (COG-UK) Consortium | The Lighthouse Lab in Milton Keynes and Alex Alderton, Roberto Amato, Jeffrey Barrett, Sonia Goncalves, Ewan Harrison, David K. Jackson, Ian Johnston, Dominic Kwiatkowski, Cordelia Langford, John Sillitoe on behalf of the Wellcome Sanger Institute COVID-19 Surveillance Team |
| EPI_ISL_2908869, EPI_ISL_2908870 | Lighthouse Laboratory Plymouth | Wellcome Sanger Institute for the COVID-19 Genomics UK (COG-UK) Consortium | Lighthouse Laboratory Plymouth and Alex Alderton, Roberto Amato, Jeffrey Barrett, Sonia Goncalves, Ewan Harrison, David K. Jackson, Ian Johnston, Dominic Kwiatkowski, Cordelia Langford, John Sillitoe on behalf of the Wellcome Sanger Institute COVID-19 Surveillance Team |
| EPI_ISL_2908871 | Lighthouse Lab in Milton Keynes | Wellcome Sanger Institute for the COVID-19 Genomics UK (COG-UK) Consortium | The Lighthouse Lab in Milton Keynes and Alex Alderton, Roberto Amato, Jeffrey Barrett, Sonia Goncalves, Ewan Harrison, David K. Jackson, Ian Johnston, Dominic Kwiatkowski, Cordelia Langford, John Sillitoe on behalf of the Wellcome Sanger Institute COVID-19 Surveillance Team |
| EPI_ISL_2908872, EPI_ISL_2908873, EPI_ISL_2908874, EPI_ISL_2908875, EPI_ISL_2908876, EPI_ISL_2908877, EPI_ISL_2908878, EPI_ISL_2908879 | Lighthouse Laboratory Plymouth | Wellcome Sanger Institute for the COVID-19 Genomics UK (COG-UK) Consortium | Lighthouse Laboratory Plymouth and Alex Alderton, Roberto Amato, Jeffrey Barrett, Sonia Goncalves, Ewan Harrison, David K. Jackson, Ian Johnston, Dominic Kwiatkowski, Cordelia Langford, John Sillitoe on behalf of the Wellcome Sanger Institute COVID-19 Surveillance Team |
| EPI_ISL_2908880 | Lighthouse Lab in Milton Keynes | Wellcome Sanger Institute for the COVID-19 Genomics UK (COG-UK) Consortium | The Lighthouse Lab in Milton Keynes and Alex Alderton, Roberto Amato, Jeffrey Barrett, Sonia Goncalves, Ewan Harrison, David K. Jackson, Ian Johnston, Dominic Kwiatkowski, Cordelia Langford, John Sillitoe on behalf of the Wellcome Sanger Institute COVID-19 Surveillance Team |
| EPI_ISL_2908881, EPI_ISL_2908882, EPI_ISL_2908883 | Lighthouse Laboratory Plymouth | Wellcome Sanger Institute for the COVID-19 Genomics UK (COG-UK) Consortium | Lighthouse Laboratory Plymouth and Alex Alderton, Roberto Amato, Jeffrey Barrett, Sonia Goncalves, Ewan Harrison, David K. Jackson, Ian Johnston, Dominic Kwiatkowski, Cordelia Langford, John Sillitoe on behalf of the Wellcome Sanger Institute COVID-19 Surveillance Team |
| EPI_ISL_2908884 | Lighthouse Lab in Milton Keynes | Wellcome Sanger Institute for the COVID-19 Genomics UK (COG-UK) Consortium | The Lighthouse Lab in Milton Keynes and Alex Alderton, Roberto Amato, Jeffrey Barrett, Sonia Goncalves, Ewan Harrison, David K. Jackson, Ian Johnston, Dominic Kwiatkowski, Cordelia Langford, John Sillitoe on behalf of the Wellcome Sanger Institute COVID-19 Surveillance Team |
| EPI_ISL_2908885 | Lighthouse Laboratory Plymouth | Wellcome Sanger Institute for the COVID-19 Genomics UK (COG-UK) Consortium | Lighthouse Laboratory Plymouth and Alex Alderton, Roberto Amato, Jeffrey Barrett, Sonia Goncalves, Ewan Harrison, David K. Jackson, Ian Johnston, Dominic Kwiatkowski, Cordelia Langford, John Sillitoe on behalf of the Wellcome Sanger Institute COVID-19 Surveillance Team |
| EPI_ISL_2908886 | Lighthouse Lab in Milton Keynes | Wellcome Sanger Institute for the COVID-19 Genomics UK (COG-UK) Consortium | The Lighthouse Lab in Milton Keynes and Alex Alderton, Roberto Amato, Jeffrey Barrett, Sonia Goncalves, Ewan Harrison, David K. Jackson, Ian Johnston, Dominic Kwiatkowski, Cordelia Langford, John Sillitoe on behalf of the Wellcome Sanger Institute COVID-19 Surveillance Team |
| EPI_ISL_2908887 | Lighthouse Laboratory Plymouth | Wellcome Sanger Institute for the COVID-19 Genomics UK (COG-UK) Consortium | Lighthouse Laboratory Plymouth and Alex Alderton, Roberto Amato, Jeffrey Barrett, Sonia Goncalves, Ewan Harrison, David K. Jackson, Ian Johnston, Dominic Kwiatkowski, Cordelia Langford, John Sillitoe on behalf of the Wellcome Sanger Institute COVID-19 Surveillance Team |
| EPI_ISL_2908888 | Lighthouse Lab in Milton Keynes | Wellcome Sanger Institute for the COVID-19 Genomics UK (COG-UK) Consortium | The Lighthouse Lab in Milton Keynes and Alex Alderton, Roberto Amato, Jeffrey Barrett, Sonia Goncalves, Ewan Harrison, David K. Jackson, Ian Johnston, Dominic Kwiatkowski, Cordelia Langford, John Sillitoe on behalf of the Wellcome Sanger Institute COVID-19 Surveillance Team |
| EPI_ISL_2908889, EPI_ISL_2908890 | Lighthouse Laboratory Plymouth | Wellcome Sanger Institute for the COVID-19 Genomics UK (COG-UK) Consortium | Lighthouse Laboratory Plymouth and Alex Alderton, Roberto Amato, Jeffrey Barrett, Sonia Goncalves, Ewan Harrison, David K. Jackson, Ian Johnston, Dominic Kwiatkowski, Cordelia Langford, John Sillitoe on behalf of the Wellcome Sanger Institute COVID-19 Surveillance Team |
| EPI_ISL_2908891 | Lighthouse Lab in Milton Keynes | Wellcome Sanger Institute for the COVID-19 Genomics UK (COG-UK) Consortium | The Lighthouse Lab in Milton Keynes and Alex Alderton, Roberto Amato, Jeffrey Barrett, Sonia Goncalves, Ewan Harrison, David K. Jackson, Ian Johnston, Dominic Kwiatkowski, Cordelia Langford, John Sillitoe on behalf of the Wellcome Sanger Institute COVID-19 Surveillance Team |
| EPI_ISL_2908892, EPI_ISL_2908893 | Lighthouse Laboratory Plymouth | Wellcome Sanger Institute for the COVID-19 Genomics UK (COG-UK) Consortium | Lighthouse Laboratory Plymouth and Alex Alderton, Roberto Amato, Jeffrey Barrett, Sonia Goncalves, Ewan Harrison, David K. Jackson, Ian Johnston, Dominic Kwiatkowski, Cordelia Langford, John Sillitoe on behalf of the Wellcome Sanger Institute COVID-19 Surveillance Team |
| EPI_ISL_2908894 | Lighthouse Lab in Milton Keynes | Wellcome Sanger Institute for the COVID-19 Genomics UK (COG-UK) Consortium | The Lighthouse Lab in Milton Keynes and Alex Alderton, Roberto Amato, Jeffrey Barrett, Sonia Goncalves, Ewan Harrison, David K. Jackson, Ian Johnston, Dominic Kwiatkowski, Cordelia Langford, John Sillitoe on behalf of the Wellcome Sanger Institute COVID-19 Surveillance Team |
| EPI_ISL_2908895, EPI_ISL_2908896 | Lighthouse Laboratory Plymouth | Wellcome Sanger Institute for the COVID-19 Genomics UK (COG-UK) Consortium | Lighthouse Laboratory Plymouth and Alex Alderton, Roberto Amato, Jeffrey Barrett, Sonia Goncalves, Ewan Harrison, David K. Jackson, Ian Johnston, Dominic Kwiatkowski, Cordelia Langford, John Sillitoe on behalf of the Wellcome Sanger Institute COVID-19 Surveillance Team |
| EPI_ISL_2908897 | Lighthouse Lab in Milton Keynes | Wellcome Sanger Institute for the COVID-19 Genomics UK (COG-UK) Consortium | The Lighthouse Lab in Milton Keynes and Alex Alderton, Roberto Amato, Jeffrey Barrett, Sonia Goncalves, Ewan Harrison, David K. Jackson, Ian Johnston, Dominic Kwiatkowski, Cordelia Langford, John Sillitoe on behalf of the Wellcome Sanger Institute COVID-19 Surveillance Team |
| EPI_ISL_2908898, EPI_ISL_2908899, EPI_ISL_2908900, EPI_ISL_2908901, EPI_ISL_2908902, EPI_ISL_2908903, EPI_ISL_2908904, EPI_ISL_2908905, EPI_ISL_2908906 | Lighthouse Laboratory Plymouth | Wellcome Sanger Institute for the COVID-19 Genomics UK (COG-UK) Consortium | Lighthouse Laboratory Plymouth and Alex Alderton, Roberto Amato, Jeffrey Barrett, Sonia Goncalves, Ewan Harrison, David K. Jackson, Ian Johnston, Dominic Kwiatkowski, Cordelia Langford, John Sillitoe on behalf of the Wellcome Sanger Institute COVID-19 Surveillance Team |
| EPI_ISL_2908907 | Lighthouse Lab in Milton Keynes | Wellcome Sanger Institute for the COVID-19 Genomics UK (COG-UK) Consortium | The Lighthouse Lab in Milton Keynes and Alex Alderton, Roberto Amato, Jeffrey Barrett, Sonia Goncalves, Ewan Harrison, David K. Jackson, Ian Johnston, Dominic Kwiatkowski, Cordelia Langford, John Sillitoe on behalf of the Wellcome Sanger Institute COVID-19 Surveillance Team |
| EPI_ISL_2908908, EPI_ISL_2908909, EPI_ISL_2908910, EPI_ISL_2908911, EPI_ISL_2908912, EPI_ISL_2908913, EPI_ISL_2908914, EPI_ISL_2908915, EPI_ISL_2908916 | Lighthouse Laboratory Plymouth | Wellcome Sanger Institute for the COVID-19 Genomics UK (COG-UK) Consortium | Lighthouse Laboratory Plymouth and Alex Alderton, Roberto Amato, Jeffrey Barrett, Sonia Goncalves, Ewan Harrison, David K. Jackson, Ian Johnston, Dominic Kwiatkowski, Cordelia Langford, John Sillitoe on behalf of the Wellcome Sanger Institute COVID-19 Surveillance Team |
| EPI_ISL_2908917 | Lighthouse Lab in Milton Keynes | Wellcome Sanger Institute for the COVID-19 Genomics UK (COG-UK) Consortium | The Lighthouse Lab in Milton Keynes and Alex Alderton, Roberto Amato, Jeffrey Barrett, Sonia Goncalves, Ewan Harrison, David K. Jackson, Ian Johnston, Dominic Kwiatkowski, Cordelia Langford, John Sillitoe on behalf of the Wellcome Sanger Institute COVID-19 Surveillance Team |
| EPI_ISL_2908918, EPI_ISL_2908919, EPI_ISL_2908920 | Lighthouse Laboratory Plymouth | Wellcome Sanger Institute for the COVID-19 Genomics UK (COG-UK) Consortium | Lighthouse Laboratory Plymouth and Alex Alderton, Roberto Amato, Jeffrey Barrett, Sonia Goncalves, Ewan Harrison, David K. Jackson, Ian Johnston, Dominic Kwiatkowski, Cordelia Langford, John Sillitoe on behalf of the Wellcome Sanger Institute COVID-19 Surveillance Team |
| EPI_ISL_2908921 | Lighthouse Lab in Milton Keynes | Wellcome Sanger Institute for the COVID-19 Genomics UK (COG-UK) Consortium | The Lighthouse Lab in Milton Keynes and Alex Alderton, Roberto Amato, Jeffrey Barrett, Sonia Goncalves, Ewan Harrison, David K. Jackson, Ian Johnston, Dominic Kwiatkowski, Cordelia Langford, John Sillitoe on behalf of the Wellcome Sanger Institute COVID-19 Surveillance Team |
| EPI_ISL_2908922, EPI_ISL_2908923, EPI_ISL_2908924, EPI_ISL_2908925 | Lighthouse Laboratory Plymouth | Wellcome Sanger Institute for the COVID-19 Genomics UK (COG-UK) Consortium | Lighthouse Laboratory Plymouth and Alex Alderton, Roberto Amato, Jeffrey Barrett, Sonia Goncalves, Ewan Harrison, David K. Jackson, Ian Johnston, Dominic Kwiatkowski, Cordelia Langford, John Sillitoe on behalf of the Wellcome Sanger Institute COVID-19 Surveillance Team |
| EPI_ISL_2908926 | Lighthouse Lab in Milton Keynes | Wellcome Sanger Institute for the COVID-19 Genomics UK (COG-UK) Consortium | The Lighthouse Lab in Milton Keynes and Alex Alderton, Roberto Amato, Jeffrey Barrett, Sonia Goncalves, Ewan Harrison, David K. Jackson, Ian Johnston, Dominic Kwiatkowski, Cordelia Langford, John Sillitoe on behalf of the Wellcome Sanger Institute COVID-19 Surveillance Team |
| EPI_ISL_2908927, EPI_ISL_2908928, EPI_ISL_2908929 | Lighthouse Laboratory Plymouth | Wellcome Sanger Institute for the COVID-19 Genomics UK (COG-UK) Consortium | Lighthouse Laboratory Plymouth and Alex Alderton, Roberto Amato, Jeffrey Barrett, Sonia Goncalves, Ewan Harrison, David K. Jackson, Ian Johnston, Dominic Kwiatkowski, Cordelia Langford, John Sillitoe on behalf of the Wellcome Sanger Institute COVID-19 Surveillance Team |
| EPI_ISL_2908930 | Lighthouse Lab in Milton Keynes | Wellcome Sanger Institute for the COVID-19 Genomics UK (COG-UK) | The Lighthouse Lab in Milton Keynes and Alex Alderton, Roberto Amato, Jeffrey Barrett, Sonia Goncalves, Ewan Harrison, David K. Jackson, Ian Johnston, Dominic Kwiatkowski, Cordelia Langford, John Sillitoe on behalf of the Wellcome Sanger |

| | | | |
|---|---|---|---|
| | | Consortium | Institute COVID-19 Surveillance Team |
| EPI_ISL_2908931, EPI_ISL_2908932, EPI_ISL_2908933 | Lighthouse Laboratory Plymouth | Wellcome Sanger Institute for the COVID-19 Genomics UK (COG-UK) Consortium | Lighthouse Laboratory Plymouth and Alex Alderton, Roberto Amato, Jeffrey Barrett, Sonia Goncalves, Ewan Harrison, David K. Jackson, Ian Johnston, Dominic Kwiatkowski, Cordelia Langford, John Sillitoe on behalf of the Wellcome Sanger Institute COVID-19 Surveillance Team |
| EPI_ISL_2908934 | Lighthouse Lab in Milton Keynes | Wellcome Sanger Institute for the COVID-19 Genomics UK (COG-UK) Consortium | The Lighthouse Lab in Milton Keynes and Alex Alderton, Roberto Amato, Jeffrey Barrett, Sonia Goncalves, Ewan Harrison, David K. Jackson, Ian Johnston, Dominic Kwiatkowski, Cordelia Langford, John Sillitoe on behalf of the Wellcome Sanger Institute COVID-19 Surveillance Team |
| EPI_ISL_2908935, EPI_ISL_2908936 | Lighthouse Laboratory Plymouth | Wellcome Sanger Institute for the COVID-19 Genomics UK (COG-UK) Consortium | Lighthouse Laboratory Plymouth and Alex Alderton, Roberto Amato, Jeffrey Barrett, Sonia Goncalves, Ewan Harrison, David K. Jackson, Ian Johnston, Dominic Kwiatkowski, Cordelia Langford, John Sillitoe on behalf of the Wellcome Sanger Institute COVID-19 Surveillance Team |
| EPI_ISL_2908937 | Lighthouse Lab in Milton Keynes | Wellcome Sanger Institute for the COVID-19 Genomics UK (COG-UK) Consortium | The Lighthouse Lab in Milton Keynes and Alex Alderton, Roberto Amato, Jeffrey Barrett, Sonia Goncalves, Ewan Harrison, David K. Jackson, Ian Johnston, Dominic Kwiatkowski, Cordelia Langford, John Sillitoe on behalf of the Wellcome Sanger Institute COVID-19 Surveillance Team |
| EPI_ISL_2908938, EPI_ISL_2908939, EPI_ISL_2908940, EPI_ISL_2908941, EPI_ISL_2908942, EPI_ISL_2908943, EPI_ISL_2908944 | Lighthouse Laboratory Plymouth | Wellcome Sanger Institute for the COVID-19 Genomics UK (COG-UK) Consortium | Lighthouse Laboratory Plymouth and Alex Alderton, Roberto Amato, Jeffrey Barrett, Sonia Goncalves, Ewan Harrison, David K. Jackson, Ian Johnston, Dominic Kwiatkowski, Cordelia Langford, John Sillitoe on behalf of the Wellcome Sanger Institute COVID-19 Surveillance Team |
| EPI_ISL_2908945 | Lighthouse Lab in Milton Keynes | Wellcome Sanger Institute for the COVID-19 Genomics UK (COG-UK) Consortium | The Lighthouse Lab in Milton Keynes and Alex Alderton, Roberto Amato, Jeffrey Barrett, Sonia Goncalves, Ewan Harrison, David K. Jackson, Ian Johnston, Dominic Kwiatkowski, Cordelia Langford, John Sillitoe on behalf of the Wellcome Sanger Institute COVID-19 Surveillance Team |
| EPI_ISL_2908946, EPI_ISL_2908947, EPI_ISL_2908948, EPI_ISL_2908949, EPI_ISL_2908950, EPI_ISL_2908951, EPI_ISL_2908952, EPI_ISL_2908953 | Lighthouse Laboratory Plymouth | Wellcome Sanger Institute for the COVID-19 Genomics UK (COG-UK) Consortium | Lighthouse Laboratory Plymouth and Alex Alderton, Roberto Amato, Jeffrey Barrett, Sonia Goncalves, Ewan Harrison, David K. Jackson, Ian Johnston, Dominic Kwiatkowski, Cordelia Langford, John Sillitoe on behalf of the Wellcome Sanger Institute COVID-19 Surveillance Team |
| EPI_ISL_2908954 | Lighthouse Lab in Milton Keynes | Wellcome Sanger Institute for the COVID-19 Genomics UK (COG-UK) Consortium | The Lighthouse Lab in Milton Keynes and Alex Alderton, Roberto Amato, Jeffrey Barrett, Sonia Goncalves, Ewan Harrison, David K. Jackson, Ian Johnston, Dominic Kwiatkowski, Cordelia Langford, John Sillitoe on behalf of the Wellcome Sanger Institute COVID-19 Surveillance Team |
| EPI_ISL_2908955, EPI_ISL_2908956 | Lighthouse Laboratory Plymouth | Wellcome Sanger Institute for the COVID-19 Genomics UK (COG-UK) Consortium | Lighthouse Laboratory Plymouth and Alex Alderton, Roberto Amato, Jeffrey Barrett, Sonia Goncalves, Ewan Harrison, David K. Jackson, Ian Johnston, Dominic Kwiatkowski, Cordelia Langford, John Sillitoe on behalf of the Wellcome Sanger Institute COVID-19 Surveillance Team |
| EPI_ISL_2908957 | Lighthouse Lab in Milton Keynes | Wellcome Sanger Institute for the COVID-19 Genomics UK (COG-UK) Consortium | The Lighthouse Lab in Milton Keynes and Alex Alderton, Roberto Amato, Jeffrey Barrett, Sonia Goncalves, Ewan Harrison, David K. Jackson, Ian Johnston, Dominic Kwiatkowski, Cordelia Langford, John Sillitoe on behalf of the Wellcome Sanger Institute COVID-19 Surveillance Team |
| EPI_ISL_2908958 | Lighthouse Laboratory Plymouth | Wellcome Sanger Institute for the COVID-19 Genomics UK (COG-UK) Consortium | Lighthouse Laboratory Plymouth and Alex Alderton, Roberto Amato, Jeffrey Barrett, Sonia Goncalves, Ewan Harrison, David K. Jackson, Ian Johnston, Dominic Kwiatkowski, Cordelia Langford, John Sillitoe on behalf of the Wellcome Sanger Institute COVID-19 Surveillance Team |
| EPI_ISL_2908959, EPI_ISL_2908960 | Lighthouse Lab in Milton Keynes | Wellcome Sanger Institute for the COVID-19 Genomics UK (COG-UK) Consortium | The Lighthouse Lab in Milton Keynes and Alex Alderton, Roberto Amato, Jeffrey Barrett, Sonia Goncalves, Ewan Harrison, David K. Jackson, Ian Johnston, Dominic Kwiatkowski, Cordelia Langford, John Sillitoe on behalf of the Wellcome Sanger Institute COVID-19 Surveillance Team |
| EPI_ISL_2908961 | Lighthouse Laboratory Plymouth | Wellcome Sanger Institute for the COVID-19 Genomics UK (COG-UK) Consortium | Lighthouse Laboratory Plymouth and Alex Alderton, Roberto Amato, Jeffrey Barrett, Sonia Goncalves, Ewan Harrison, David K. Jackson, Ian Johnston, Dominic Kwiatkowski, Cordelia Langford, John Sillitoe on behalf of the Wellcome Sanger Institute COVID-19 Surveillance Team |
| EPI_ISL_2908962 | Lighthouse Lab in Milton Keynes | Wellcome Sanger Institute for the COVID-19 Genomics UK (COG-UK) Consortium | The Lighthouse Lab in Milton Keynes and Alex Alderton, Roberto Amato, Jeffrey Barrett, Sonia Goncalves, Ewan Harrison, David K. Jackson, Ian Johnston, Dominic Kwiatkowski, Cordelia Langford, John Sillitoe on behalf of the Wellcome Sanger Institute COVID-19 Surveillance Team |
| EPI_ISL_2908963, EPI_ISL_2908964 | Lighthouse Laboratory Plymouth | Wellcome Sanger Institute for the COVID-19 Genomics UK (COG-UK) Consortium | Lighthouse Laboratory Plymouth and Alex Alderton, Roberto Amato, Jeffrey Barrett, Sonia Goncalves, Ewan Harrison, David K. Jackson, Ian Johnston, Dominic Kwiatkowski, Cordelia Langford, John Sillitoe on behalf of the Wellcome Sanger Institute COVID-19 Surveillance Team |
| EPI_ISL_2908965 | Lighthouse Lab in Milton Keynes | Wellcome Sanger Institute for the COVID-19 Genomics UK (COG-UK) Consortium | The Lighthouse Lab in Milton Keynes and Alex Alderton, Roberto Amato, Jeffrey Barrett, Sonia Goncalves, Ewan Harrison, David K. Jackson, Ian Johnston, Dominic Kwiatkowski, Cordelia Langford, John Sillitoe on behalf of the Wellcome Sanger Institute COVID-19 Surveillance Team |
| EPI_ISL_2908966, EPI_ISL_2908967 | Lighthouse Laboratory Plymouth | Wellcome Sanger Institute for the COVID-19 Genomics UK (COG-UK) Consortium | Lighthouse Laboratory Plymouth and Alex Alderton, Roberto Amato, Jeffrey Barrett, Sonia Goncalves, Ewan Harrison, David K. Jackson, Ian Johnston, Dominic Kwiatkowski, Cordelia Langford, John Sillitoe on behalf of the Wellcome Sanger Institute COVID-19 Surveillance Team |
| EPI_ISL_2908968 | Lighthouse Lab in Milton Keynes | Wellcome Sanger Institute for the COVID-19 Genomics UK (COG-UK) Consortium | The Lighthouse Lab in Milton Keynes and Alex Alderton, Roberto Amato, Jeffrey Barrett, Sonia Goncalves, Ewan Harrison, David K. Jackson, Ian Johnston, Dominic Kwiatkowski, Cordelia Langford, John Sillitoe on behalf of the Wellcome Sanger Institute COVID-19 Surveillance Team |
| EPI_ISL_2908969, EPI_ISL_2908970, EPI_ISL_2908971 | Lighthouse Laboratory Plymouth | Wellcome Sanger Institute for the COVID-19 Genomics UK (COG-UK) Consortium | Lighthouse Laboratory Plymouth and Alex Alderton, Roberto Amato, Jeffrey Barrett, Sonia Goncalves, Ewan Harrison, David K. Jackson, Ian Johnston, Dominic Kwiatkowski, Cordelia Langford, John Sillitoe on behalf of the Wellcome Sanger Institute COVID-19 Surveillance Team |
| EPI_ISL_2908972 | Lighthouse Lab in Milton Keynes | Wellcome Sanger Institute for the COVID-19 Genomics UK (COG-UK) Consortium | The Lighthouse Lab in Milton Keynes and Alex Alderton, Roberto Amato, Jeffrey Barrett, Sonia Goncalves, Ewan Harrison, David K. Jackson, Ian Johnston, Dominic Kwiatkowski, Cordelia Langford, John Sillitoe on behalf of the Wellcome Sanger Institute COVID-19 Surveillance Team |
| EPI_ISL_2908973 | Lighthouse Laboratory Plymouth | Wellcome Sanger Institute for the COVID-19 Genomics UK (COG-UK) Consortium | Lighthouse Laboratory Plymouth and Alex Alderton, Roberto Amato, Jeffrey Barrett, Sonia Goncalves, Ewan Harrison, David K. Jackson, Ian Johnston, Dominic Kwiatkowski, Cordelia Langford, John Sillitoe on behalf of the Wellcome Sanger Institute COVID-19 Surveillance Team |
| EPI_ISL_2908974 | Lighthouse Lab in Milton Keynes | Wellcome Sanger Institute for the COVID-19 Genomics UK (COG-UK) Consortium | The Lighthouse Lab in Milton Keynes and Alex Alderton, Roberto Amato, Jeffrey Barrett, Sonia Goncalves, Ewan Harrison, David K. Jackson, Ian Johnston, Dominic Kwiatkowski, Cordelia Langford, John Sillitoe on behalf of the Wellcome Sanger Institute COVID-19 Surveillance Team |
| EPI_ISL_2908975, EPI_ISL_2908976 | Lighthouse Laboratory Plymouth | Wellcome Sanger Institute for the COVID-19 Genomics UK (COG-UK) Consortium | Lighthouse Laboratory Plymouth and Alex Alderton, Roberto Amato, Jeffrey Barrett, Sonia Goncalves, Ewan Harrison, David K. Jackson, Ian Johnston, Dominic Kwiatkowski, Cordelia Langford, John Sillitoe on behalf of the Wellcome Sanger Institute COVID-19 Surveillance Team |
| EPI_ISL_2908977, EPI_ISL_2908978 | Lighthouse Lab in Milton Keynes | Wellcome Sanger Institute for the COVID-19 Genomics UK (COG-UK) Consortium | The Lighthouse Lab in Milton Keynes and Alex Alderton, Roberto Amato, Jeffrey Barrett, Sonia Goncalves, Ewan Harrison, David K. Jackson, Ian Johnston, Dominic Kwiatkowski, Cordelia Langford, John Sillitoe on behalf of the Wellcome Sanger Institute COVID-19 Surveillance Team |
| EPI_ISL_2908979, EPI_ISL_2908980 | Lighthouse Laboratory Plymouth | Wellcome Sanger Institute for the COVID-19 Genomics UK (COG-UK) Consortium | Lighthouse Laboratory Plymouth and Alex Alderton, Roberto Amato, Jeffrey Barrett, Sonia Goncalves, Ewan Harrison, David K. Jackson, Ian Johnston, Dominic Kwiatkowski, Cordelia Langford, John Sillitoe on behalf of the Wellcome Sanger Institute COVID-19 Surveillance Team |
| EPI_ISL_2908981 | Lighthouse Lab in Milton Keynes | Wellcome Sanger Institute for the COVID-19 Genomics UK (COG-UK) Consortium | The Lighthouse Lab in Milton Keynes and Alex Alderton, Roberto Amato, Jeffrey Barrett, Sonia Goncalves, Ewan Harrison, David K. Jackson, Ian Johnston, Dominic Kwiatkowski, Cordelia Langford, John Sillitoe on behalf of the Wellcome Sanger Institute COVID-19 Surveillance Team |
| EPI_ISL_2908982, EPI_ISL_2908983, EPI_ISL_2908984, EPI_ISL_2908985, EPI_ISL_2908986 | Lighthouse Laboratory Plymouth | Wellcome Sanger Institute for the COVID-19 Genomics UK (COG-UK) Consortium | Lighthouse Laboratory Plymouth and Alex Alderton, Roberto Amato, Jeffrey Barrett, Sonia Goncalves, Ewan Harrison, David K. Jackson, Ian Johnston, Dominic Kwiatkowski, Cordelia Langford, John Sillitoe on behalf of the Wellcome Sanger Institute COVID-19 Surveillance Team |
| EPI_ISL_2908987 | Lighthouse Lab in Milton Keynes | Wellcome Sanger Institute for the COVID-19 Genomics UK (COG-UK) Consortium | The Lighthouse Lab in Milton Keynes and Alex Alderton, Roberto Amato, Jeffrey Barrett, Sonia Goncalves, Ewan Harrison, David K. Jackson, Ian Johnston, Dominic Kwiatkowski, Cordelia Langford, John Sillitoe on behalf of the Wellcome Sanger Institute COVID-19 Surveillance Team |
| EPI_ISL_2908988, EPI_ISL_2908989, EPI_ISL_2908990, EPI_ISL_2908991, EPI_ISL_2908992, EPI_ISL_2908993, EPI_ISL_2908994, EPI_ISL_2908995, EPI_ISL_2908996, EPI_ISL_2908997, EPI_ISL_2908998 | | | |
| see above | Lighthouse Laboratory Plymouth | Wellcome Sanger Institute for the COVID-19 Genomics UK (COG-UK) Consortium | Lighthouse Laboratory Plymouth and Alex Alderton, Roberto Amato, Jeffrey Barrett, Sonia Goncalves, Ewan Harrison, David K. Jackson, Ian Johnston, Dominic Kwiatkowski, Cordelia Langford, John Sillitoe on behalf of the Wellcome Sanger Institute COVID-19 Surveillance Team |
| EPI_ISL_2908999 | Lighthouse Lab in Milton Keynes | Wellcome Sanger Institute for the COVID-19 Genomics UK (COG-UK) Consortium | The Lighthouse Lab in Milton Keynes and Alex Alderton, Roberto Amato, Jeffrey Barrett, Sonia Goncalves, Ewan Harrison, David K. Jackson, Ian Johnston, Dominic Kwiatkowski, Cordelia Langford, John Sillitoe on behalf of the Wellcome Sanger Institute COVID-19 Surveillance Team |
| EPI_ISL_2909000, EPI_ISL_2909001, EPI_ISL_2909002 | Lighthouse Laboratory Plymouth | Wellcome Sanger Institute for the COVID-19 Genomics UK (COG-UK) Consortium | Lighthouse Laboratory Plymouth and Alex Alderton, Roberto Amato, Jeffrey Barrett, Sonia Goncalves, Ewan Harrison, David K. Jackson, Ian Johnston, Dominic Kwiatkowski, Cordelia Langford, John Sillitoe on behalf of the Wellcome Sanger Institute COVID-19 Surveillance Team |
| EPI_ISL_2909003 | Lighthouse Lab in Milton Keynes | Wellcome Sanger Institute for the COVID-19 Genomics UK (COG-UK) Consortium | The Lighthouse Lab in Milton Keynes and Alex Alderton, Roberto Amato, Jeffrey Barrett, Sonia Goncalves, Ewan Harrison, David K. Jackson, Ian Johnston, Dominic Kwiatkowski, Cordelia Langford, John Sillitoe on behalf of the Wellcome Sanger Institute COVID-19 Surveillance Team |
| EPI_ISL_2909004 | Lighthouse Laboratory Plymouth | Wellcome Sanger Institute for the COVID-19 Genomics UK (COG-UK) Consortium | Lighthouse Laboratory Plymouth and Alex Alderton, Roberto Amato, Jeffrey Barrett, Sonia Goncalves, Ewan Harrison, David K. Jackson, Ian Johnston, Dominic Kwiatkowski, Cordelia Langford, John Sillitoe on behalf of the Wellcome Sanger Institute COVID-19 Surveillance Team |
| EPI_ISL_2909005, EPI_ISL_2909006 | Lighthouse Lab in Milton Keynes | Wellcome Sanger Institute for the COVID-19 Genomics UK (COG-UK) Consortium | The Lighthouse Lab in Milton Keynes and Alex Alderton, Roberto Amato, Jeffrey Barrett, Sonia Goncalves, Ewan Harrison, David K. Jackson, Ian Johnston, Dominic Kwiatkowski, Cordelia Langford, John Sillitoe on behalf of the Wellcome Sanger Institute COVID-19 Surveillance Team |
| EPI_ISL_2909007 | Lighthouse Laboratory Plymouth | Wellcome Sanger Institute for the COVID-19 Genomics UK (COG-UK) Consortium | Lighthouse Laboratory Plymouth and Alex Alderton, Roberto Amato, Jeffrey Barrett, Sonia Goncalves, Ewan Harrison, David K. Jackson, Ian Johnston, Dominic Kwiatkowski, Cordelia Langford, John Sillitoe on behalf of the Wellcome Sanger Institute COVID-19 Surveillance Team |
| EPI_ISL_2909008 | Lighthouse Lab in Milton Keynes | Wellcome Sanger Institute for the COVID-19 Genomics UK (COG-UK) Consortium | The Lighthouse Lab in Milton Keynes and Alex Alderton, Roberto Amato, Jeffrey Barrett, Sonia Goncalves, Ewan Harrison, David K. Jackson, Ian Johnston, Dominic Kwiatkowski, Cordelia Langford, John Sillitoe on behalf of the Wellcome Sanger Institute COVID-19 Surveillance Team |
| EPI_ISL_2909009, EPI_ISL_2909010, | Lighthouse Laboratory Plymouth | Wellcome Sanger Institute for the COVID-19 Genomics UK (COG-UK) Consortium | Lighthouse Laboratory Plymouth and Alex Alderton, Roberto Amato, Jeffrey Barrett, Sonia Goncalves, Ewan Harrison, David K. Jackson, Ian Johnston, Dominic Kwiatkowski, Cordelia Langford, John Sillitoe on behalf of the Wellcome Sanger Institute COVID-19 Surveillance Team |

| | | | |
|---|---|---|---|
| EPI_ISL_2909011 | | | |
| EPI_ISL_2909012 | Lighthouse Lab in Milton Keynes | Wellcome Sanger Institute for the COVID-19 Genomics UK (COG-UK) Consortium | The Lighthouse Lab in Milton Keynes and Alex Alderton, Roberto Amato, Jeffrey Barrett, Sonia Goncalves, Ewan Harrison, David K. Jackson, Ian Johnston, Dominic Kwiatkowski, Cordelia Langford, John Sillitoe on behalf of the Wellcome Sanger Institute COVID-19 Surveillance Team |
| EPI_ISL_2909013, EPI_ISL_2909014, EPI_ISL_2909015 | Lighthouse Laboratory Plymouth | Wellcome Sanger Institute for the COVID-19 Genomics UK (COG-UK) Consortium | Lighthouse Laboratory Plymouth and Alex Alderton, Roberto Amato, Jeffrey Barrett, Sonia Goncalves, Ewan Harrison, David K. Jackson, Ian Johnston, Dominic Kwiatkowski, Cordelia Langford, John Sillitoe on behalf of the Wellcome Sanger Institute COVID-19 Surveillance Team |
| EPI_ISL_2909016 | Lighthouse Lab in Milton Keynes | Wellcome Sanger Institute for the COVID-19 Genomics UK (COG-UK) Consortium | The Lighthouse Lab in Milton Keynes and Alex Alderton, Roberto Amato, Jeffrey Barrett, Sonia Goncalves, Ewan Harrison, David K. Jackson, Ian Johnston, Dominic Kwiatkowski, Cordelia Langford, John Sillitoe on behalf of the Wellcome Sanger Institute COVID-19 Surveillance Team |
| EPI_ISL_2909017 | Lighthouse Laboratory Plymouth | Wellcome Sanger Institute for the COVID-19 Genomics UK (COG-UK) Consortium | Lighthouse Laboratory Plymouth and Alex Alderton, Roberto Amato, Jeffrey Barrett, Sonia Goncalves, Ewan Harrison, David K. Jackson, Ian Johnston, Dominic Kwiatkowski, Cordelia Langford, John Sillitoe on behalf of the Wellcome Sanger Institute COVID-19 Surveillance Team |
| EPI_ISL_2909018 | Lighthouse Lab in Milton Keynes | Wellcome Sanger Institute for the COVID-19 Genomics UK (COG-UK) Consortium | The Lighthouse Lab in Milton Keynes and Alex Alderton, Roberto Amato, Jeffrey Barrett, Sonia Goncalves, Ewan Harrison, David K. Jackson, Ian Johnston, Dominic Kwiatkowski, Cordelia Langford, John Sillitoe on behalf of the Wellcome Sanger Institute COVID-19 Surveillance Team |
| EPI_ISL_2909019 | Lighthouse Laboratory Plymouth | Wellcome Sanger Institute for the COVID-19 Genomics UK (COG-UK) Consortium | Lighthouse Laboratory Plymouth and Alex Alderton, Roberto Amato, Jeffrey Barrett, Sonia Goncalves, Ewan Harrison, David K. Jackson, Ian Johnston, Dominic Kwiatkowski, Cordelia Langford, John Sillitoe on behalf of the Wellcome Sanger Institute COVID-19 Surveillance Team |
| EPI_ISL_2909020 | Lighthouse Lab in Milton Keynes | Wellcome Sanger Institute for the COVID-19 Genomics UK (COG-UK) Consortium | The Lighthouse Lab in Milton Keynes and Alex Alderton, Roberto Amato, Jeffrey Barrett, Sonia Goncalves, Ewan Harrison, David K. Jackson, Ian Johnston, Dominic Kwiatkowski, Cordelia Langford, John Sillitoe on behalf of the Wellcome Sanger Institute COVID-19 Surveillance Team |
| EPI_ISL_2909021, EPI_ISL_2909022 | Lighthouse Laboratory Plymouth | Wellcome Sanger Institute for the COVID-19 Genomics UK (COG-UK) Consortium | Lighthouse Laboratory Plymouth and Alex Alderton, Roberto Amato, Jeffrey Barrett, Sonia Goncalves, Ewan Harrison, David K. Jackson, Ian Johnston, Dominic Kwiatkowski, Cordelia Langford, John Sillitoe on behalf of the Wellcome Sanger Institute COVID-19 Surveillance Team |
| EPI_ISL_2909023 | Lighthouse Lab in Milton Keynes | Wellcome Sanger Institute for the COVID-19 Genomics UK (COG-UK) Consortium | The Lighthouse Lab in Milton Keynes and Alex Alderton, Roberto Amato, Jeffrey Barrett, Sonia Goncalves, Ewan Harrison, David K. Jackson, Ian Johnston, Dominic Kwiatkowski, Cordelia Langford, John Sillitoe on behalf of the Wellcome Sanger Institute COVID-19 Surveillance Team |
| EPI_ISL_2909024, EPI_ISL_2909025, EPI_ISL_2909026 | Lighthouse Laboratory Plymouth | Wellcome Sanger Institute for the COVID-19 Genomics UK (COG-UK) Consortium | Lighthouse Laboratory Plymouth and Alex Alderton, Roberto Amato, Jeffrey Barrett, Sonia Goncalves, Ewan Harrison, David K. Jackson, Ian Johnston, Dominic Kwiatkowski, Cordelia Langford, John Sillitoe on behalf of the Wellcome Sanger Institute COVID-19 Surveillance Team |
| EPI_ISL_2909027 | Lighthouse Lab in Milton Keynes | Wellcome Sanger Institute for the COVID-19 Genomics UK (COG-UK) Consortium | The Lighthouse Lab in Milton Keynes and Alex Alderton, Roberto Amato, Jeffrey Barrett, Sonia Goncalves, Ewan Harrison, David K. Jackson, Ian Johnston, Dominic Kwiatkowski, Cordelia Langford, John Sillitoe on behalf of the Wellcome Sanger Institute COVID-19 Surveillance Team |
| EPI_ISL_2909028, EPI_ISL_2909029, EPI_ISL_2909030 | Lighthouse Laboratory Plymouth | Wellcome Sanger Institute for the COVID-19 Genomics UK (COG-UK) Consortium | Lighthouse Laboratory Plymouth and Alex Alderton, Roberto Amato, Jeffrey Barrett, Sonia Goncalves, Ewan Harrison, David K. Jackson, Ian Johnston, Dominic Kwiatkowski, Cordelia Langford, John Sillitoe on behalf of the Wellcome Sanger Institute COVID-19 Surveillance Team |
| EPI_ISL_2909031 | Lighthouse Lab in Milton Keynes | Wellcome Sanger Institute for the COVID-19 Genomics UK (COG-UK) Consortium | The Lighthouse Lab in Milton Keynes and Alex Alderton, Roberto Amato, Jeffrey Barrett, Sonia Goncalves, Ewan Harrison, David K. Jackson, Ian Johnston, Dominic Kwiatkowski, Cordelia Langford, John Sillitoe on behalf of the Wellcome Sanger Institute COVID-19 Surveillance Team |
| EPI_ISL_2909032, EPI_ISL_2909033, EPI_ISL_2909034, EPI_ISL_2909035, EPI_ISL_2909036, EPI_ISL_2909037, EPI_ISL_2909038, EPI_ISL_2909039, EPI_ISL_2909041 | Lighthouse Laboratory Plymouth | Wellcome Sanger Institute for the COVID-19 Genomics UK (COG-UK) Consortium | Lighthouse Laboratory Plymouth and Alex Alderton, Roberto Amato, Jeffrey Barrett, Sonia Goncalves, Ewan Harrison, David K. Jackson, Ian Johnston, Dominic Kwiatkowski, Cordelia Langford, John Sillitoe on behalf of the Wellcome Sanger Institute COVID-19 Surveillance Team |
| EPI_ISL_2909042 | Lighthouse Lab in Milton Keynes | Wellcome Sanger Institute for the COVID-19 Genomics UK (COG-UK) Consortium | The Lighthouse Lab in Milton Keynes and Alex Alderton, Roberto Amato, Jeffrey Barrett, Sonia Goncalves, Ewan Harrison, David K. Jackson, Ian Johnston, Dominic Kwiatkowski, Cordelia Langford, John Sillitoe on behalf of the Wellcome Sanger Institute COVID-19 Surveillance Team |
| EPI_ISL_2909043, EPI_ISL_2909044, EPI_ISL_2909045, EPI_ISL_2909046 | Lighthouse Laboratory Plymouth | Wellcome Sanger Institute for the COVID-19 Genomics UK (COG-UK) Consortium | Lighthouse Laboratory Plymouth and Alex Alderton, Roberto Amato, Jeffrey Barrett, Sonia Goncalves, Ewan Harrison, David K. Jackson, Ian Johnston, Dominic Kwiatkowski, Cordelia Langford, John Sillitoe on behalf of the Wellcome Sanger Institute COVID-19 Surveillance Team |
| EPI_ISL_2909047 | Lighthouse Lab in Milton Keynes | Wellcome Sanger Institute for the COVID-19 Genomics UK (COG-UK) Consortium | The Lighthouse Lab in Milton Keynes and Alex Alderton, Roberto Amato, Jeffrey Barrett, Sonia Goncalves, Ewan Harrison, David K. Jackson, Ian Johnston, Dominic Kwiatkowski, Cordelia Langford, John Sillitoe on behalf of the Wellcome Sanger Institute COVID-19 Surveillance Team |
| EPI_ISL_2909048, EPI_ISL_2909049, EPI_ISL_2909050, EPI_ISL_2909051, EPI_ISL_2909052, EPI_ISL_2909053, EPI_ISL_2909054 | Lighthouse Laboratory Plymouth | Wellcome Sanger Institute for the COVID-19 Genomics UK (COG-UK) Consortium | Lighthouse Laboratory Plymouth and Alex Alderton, Roberto Amato, Jeffrey Barrett, Sonia Goncalves, Ewan Harrison, David K. Jackson, Ian Johnston, Dominic Kwiatkowski, Cordelia Langford, John Sillitoe on behalf of the Wellcome Sanger Institute COVID-19 Surveillance Team |
| EPI_ISL_2909055 | Lighthouse Lab in Milton Keynes | Wellcome Sanger Institute for the COVID-19 Genomics UK (COG-UK) Consortium | The Lighthouse Lab in Milton Keynes and Alex Alderton, Roberto Amato, Jeffrey Barrett, Sonia Goncalves, Ewan Harrison, David K. Jackson, Ian Johnston, Dominic Kwiatkowski, Cordelia Langford, John Sillitoe on behalf of the Wellcome Sanger Institute COVID-19 Surveillance Team |
| EPI_ISL_2909056, EPI_ISL_2909057, EPI_ISL_2909058, EPI_ISL_2909059, EPI_ISL_2909060, EPI_ISL_2909061 | Lighthouse Laboratory Plymouth | Wellcome Sanger Institute for the COVID-19 Genomics UK (COG-UK) Consortium | Lighthouse Laboratory Plymouth and Alex Alderton, Roberto Amato, Jeffrey Barrett, Sonia Goncalves, Ewan Harrison, David K. Jackson, Ian Johnston, Dominic Kwiatkowski, Cordelia Langford, John Sillitoe on behalf of the Wellcome Sanger Institute COVID-19 Surveillance Team |
| EPI_ISL_2909062, EPI_ISL_2909063 | Lighthouse Lab in Milton Keynes | Wellcome Sanger Institute for the COVID-19 Genomics UK (COG-UK) Consortium | The Lighthouse Lab in Milton Keynes and Alex Alderton, Roberto Amato, Jeffrey Barrett, Sonia Goncalves, Ewan Harrison, David K. Jackson, Ian Johnston, Dominic Kwiatkowski, Cordelia Langford, John Sillitoe on behalf of the Wellcome Sanger Institute COVID-19 Surveillance Team |
| EPI_ISL_2909064, EPI_ISL_2909065, EPI_ISL_2909066 | Lighthouse Laboratory Plymouth | Wellcome Sanger Institute for the COVID-19 Genomics UK (COG-UK) Consortium | Lighthouse Laboratory Plymouth and Alex Alderton, Roberto Amato, Jeffrey Barrett, Sonia Goncalves, Ewan Harrison, David K. Jackson, Ian Johnston, Dominic Kwiatkowski, Cordelia Langford, John Sillitoe on behalf of the Wellcome Sanger Institute COVID-19 Surveillance Team |
| EPI_ISL_2909067 | Lighthouse Lab in Milton Keynes | Wellcome Sanger Institute for the COVID-19 Genomics UK (COG-UK) Consortium | The Lighthouse Lab in Milton Keynes and Alex Alderton, Roberto Amato, Jeffrey Barrett, Sonia Goncalves, Ewan Harrison, David K. Jackson, Ian Johnston, Dominic Kwiatkowski, Cordelia Langford, John Sillitoe on behalf of the Wellcome Sanger Institute COVID-19 Surveillance Team |
| EPI_ISL_2909068, EPI_ISL_2909069 | Lighthouse Laboratory Plymouth | Wellcome Sanger Institute for the COVID-19 Genomics UK (COG-UK) Consortium | Lighthouse Laboratory Plymouth and Alex Alderton, Roberto Amato, Jeffrey Barrett, Sonia Goncalves, Ewan Harrison, David K. Jackson, Ian Johnston, Dominic Kwiatkowski, Cordelia Langford, John Sillitoe on behalf of the Wellcome Sanger Institute COVID-19 Surveillance Team |
| EPI_ISL_2909070, EPI_ISL_2909071 | Lighthouse Lab in Milton Keynes | Wellcome Sanger Institute for the COVID-19 Genomics UK (COG-UK) Consortium | The Lighthouse Lab in Milton Keynes and Alex Alderton, Roberto Amato, Jeffrey Barrett, Sonia Goncalves, Ewan Harrison, David K. Jackson, Ian Johnston, Dominic Kwiatkowski, Cordelia Langford, John Sillitoe on behalf of the Wellcome Sanger Institute COVID-19 Surveillance Team |
| EPI_ISL_2909072, EPI_ISL_2909073, EPI_ISL_2909074, EPI_ISL_2909075, EPI_ISL_2909076, EPI_ISL_2909077 | Lighthouse Laboratory Plymouth | Wellcome Sanger Institute for the COVID-19 Genomics UK (COG-UK) Consortium | Lighthouse Laboratory Plymouth and Alex Alderton, Roberto Amato, Jeffrey Barrett, Sonia Goncalves, Ewan Harrison, David K. Jackson, Ian Johnston, Dominic Kwiatkowski, Cordelia Langford, John Sillitoe on behalf of the Wellcome Sanger Institute COVID-19 Surveillance Team |
| EPI_ISL_2909078 | Lighthouse Lab in Milton Keynes | Wellcome Sanger Institute for the COVID-19 Genomics UK (COG-UK) Consortium | The Lighthouse Lab in Milton Keynes and Alex Alderton, Roberto Amato, Jeffrey Barrett, Sonia Goncalves, Ewan Harrison, David K. Jackson, Ian Johnston, Dominic Kwiatkowski, Cordelia Langford, John Sillitoe on behalf of the Wellcome Sanger Institute COVID-19 Surveillance Team |
| EPI_ISL_2909079, EPI_ISL_2909080 | Lighthouse Laboratory Plymouth | Wellcome Sanger Institute for the COVID-19 Genomics UK (COG-UK) Consortium | Lighthouse Laboratory Plymouth and Alex Alderton, Roberto Amato, Jeffrey Barrett, Sonia Goncalves, Ewan Harrison, David K. Jackson, Ian Johnston, Dominic Kwiatkowski, Cordelia Langford, John Sillitoe on behalf of the Wellcome Sanger Institute COVID-19 Surveillance Team |
| EPI_ISL_2909081 | Lighthouse Lab in Milton Keynes | Wellcome Sanger Institute for the COVID-19 Genomics UK (COG-UK) Consortium | The Lighthouse Lab in Milton Keynes and Alex Alderton, Roberto Amato, Jeffrey Barrett, Sonia Goncalves, Ewan Harrison, David K. Jackson, Ian Johnston, Dominic Kwiatkowski, Cordelia Langford, John Sillitoe on behalf of the Wellcome Sanger Institute COVID-19 Surveillance Team |
| EPI_ISL_2909082 | Lighthouse Laboratory Plymouth | Wellcome Sanger Institute for the COVID-19 Genomics UK (COG-UK) Consortium | Lighthouse Laboratory Plymouth and Alex Alderton, Roberto Amato, Jeffrey Barrett, Sonia Goncalves, Ewan Harrison, David K. Jackson, Ian Johnston, Dominic Kwiatkowski, Cordelia Langford, John Sillitoe on behalf of the Wellcome Sanger Institute COVID-19 Surveillance Team |
| EPI_ISL_2909083 | Lighthouse Lab in Milton Keynes | Wellcome Sanger Institute for the COVID-19 Genomics UK (COG-UK) Consortium | The Lighthouse Lab in Milton Keynes and Alex Alderton, Roberto Amato, Jeffrey Barrett, Sonia Goncalves, Ewan Harrison, David K. Jackson, Ian Johnston, Dominic Kwiatkowski, Cordelia Langford, John Sillitoe on behalf of the Wellcome Sanger Institute COVID-19 Surveillance Team |
| EPI_ISL_2909084 | Lighthouse Laboratory Plymouth | Wellcome Sanger Institute for the COVID-19 Genomics UK (COG-UK) Consortium | Lighthouse Laboratory Plymouth and Alex Alderton, Roberto Amato, Jeffrey Barrett, Sonia Goncalves, Ewan Harrison, David K. Jackson, Ian Johnston, Dominic Kwiatkowski, Cordelia Langford, John Sillitoe on behalf of the Wellcome Sanger Institute COVID-19 Surveillance Team |
| EPI_ISL_2909085 | Lighthouse Lab in Milton Keynes | Wellcome Sanger Institute for the COVID-19 Genomics UK (COG-UK) Consortium | The Lighthouse Lab in Milton Keynes and Alex Alderton, Roberto Amato, Jeffrey Barrett, Sonia Goncalves, Ewan Harrison, David K. Jackson, Ian Johnston, Dominic Kwiatkowski, Cordelia Langford, John Sillitoe on behalf of the Wellcome Sanger Institute COVID-19 Surveillance Team |
| EPI_ISL_2909087, | Lighthouse Laboratory Plymouth | Wellcome Sanger Institute for the COVID-19 Genomics UK (COG-UK) | Lighthouse Laboratory Plymouth and Alex Alderton, Roberto Amato, Jeffrey Barrett, Sonia Goncalves, Ewan Harrison, David K. Jackson, Ian Johnston, Dominic Kwiatkowski, Cordelia Langford, John Sillitoe on behalf of the Wellcome Sanger |

| | | | |
|---|---|---|---|
| EPI_ISL_2909088, EPI_ISL_2909089, EPI_ISL_2909090 | | Consortium | Institute COVID-19 Surveillance Team |
| EPI_ISL_2909091 | Lighthouse Lab in Milton Keynes | Wellcome Sanger Institute for the COVID-19 Genomics UK (COG-UK) Consortium | The Lighthouse Lab in Milton Keynes and Alex Alderton, Roberto Amato, Jeffrey Barrett, Sonia Goncalves, Ewan Harrison, David K. Jackson, Ian Johnston, Dominic Kwiatkowski, Cordelia Langford, John Sillitoe on behalf of the Wellcome Sanger Institute COVID-19 Surveillance Team |
| EPI_ISL_2909092 | Lighthouse Laboratory Plymouth | Wellcome Sanger Institute for the COVID-19 Genomics UK (COG-UK) Consortium | Lighthouse Laboratory Plymouth and Alex Alderton, Roberto Amato, Jeffrey Barrett, Sonia Goncalves, Ewan Harrison, David K. Jackson, Ian Johnston, Dominic Kwiatkowski, Cordelia Langford, John Sillitoe on behalf of the Wellcome Sanger Institute COVID-19 Surveillance Team |
| EPI_ISL_2909093 | Lighthouse Lab in Milton Keynes | Wellcome Sanger Institute for the COVID-19 Genomics UK (COG-UK) Consortium | The Lighthouse Lab in Milton Keynes and Alex Alderton, Roberto Amato, Jeffrey Barrett, Sonia Goncalves, Ewan Harrison, David K. Jackson, Ian Johnston, Dominic Kwiatkowski, Cordelia Langford, John Sillitoe on behalf of the Wellcome Sanger Institute COVID-19 Surveillance Team |
| EPI_ISL_2909094, EPI_ISL_2909095 | Lighthouse Laboratory Plymouth | Wellcome Sanger Institute for the COVID-19 Genomics UK (COG-UK) Consortium | Lighthouse Laboratory Plymouth and Alex Alderton, Roberto Amato, Jeffrey Barrett, Sonia Goncalves, Ewan Harrison, David K. Jackson, Ian Johnston, Dominic Kwiatkowski, Cordelia Langford, John Sillitoe on behalf of the Wellcome Sanger Institute COVID-19 Surveillance Team |
| EPI_ISL_2909096 | Lighthouse Lab in Milton Keynes | Wellcome Sanger Institute for the COVID-19 Genomics UK (COG-UK) Consortium | The Lighthouse Lab in Milton Keynes and Alex Alderton, Roberto Amato, Jeffrey Barrett, Sonia Goncalves, Ewan Harrison, David K. Jackson, Ian Johnston, Dominic Kwiatkowski, Cordelia Langford, John Sillitoe on behalf of the Wellcome Sanger Institute COVID-19 Surveillance Team |
| EPI_ISL_2909097 | Lighthouse Laboratory Plymouth | Wellcome Sanger Institute for the COVID-19 Genomics UK (COG-UK) Consortium | Lighthouse Laboratory Plymouth and Alex Alderton, Roberto Amato, Jeffrey Barrett, Sonia Goncalves, Ewan Harrison, David K. Jackson, Ian Johnston, Dominic Kwiatkowski, Cordelia Langford, John Sillitoe on behalf of the Wellcome Sanger Institute COVID-19 Surveillance Team |
| EPI_ISL_2909098 | Lighthouse Lab in Milton Keynes | Wellcome Sanger Institute for the COVID-19 Genomics UK (COG-UK) Consortium | The Lighthouse Lab in Milton Keynes and Alex Alderton, Roberto Amato, Jeffrey Barrett, Sonia Goncalves, Ewan Harrison, David K. Jackson, Ian Johnston, Dominic Kwiatkowski, Cordelia Langford, John Sillitoe on behalf of the Wellcome Sanger Institute COVID-19 Surveillance Team |
| EPI_ISL_2909099 | Lighthouse Laboratory Plymouth | Wellcome Sanger Institute for the COVID-19 Genomics UK (COG-UK) Consortium | Lighthouse Laboratory Plymouth and Alex Alderton, Roberto Amato, Jeffrey Barrett, Sonia Goncalves, Ewan Harrison, David K. Jackson, Ian Johnston, Dominic Kwiatkowski, Cordelia Langford, John Sillitoe on behalf of the Wellcome Sanger Institute COVID-19 Surveillance Team |
| EPI_ISL_2909100 | Lighthouse Lab in Milton Keynes | Wellcome Sanger Institute for the COVID-19 Genomics UK (COG-UK) Consortium | The Lighthouse Lab in Milton Keynes and Alex Alderton, Roberto Amato, Jeffrey Barrett, Sonia Goncalves, Ewan Harrison, David K. Jackson, Ian Johnston, Dominic Kwiatkowski, Cordelia Langford, John Sillitoe on behalf of the Wellcome Sanger Institute COVID-19 Surveillance Team |
| EPI_ISL_2909101, EPI_ISL_2909102, EPI_ISL_2909103, EPI_ISL_2909104, EPI_ISL_2909105 | Lighthouse Laboratory Plymouth | Wellcome Sanger Institute for the COVID-19 Genomics UK (COG-UK) Consortium | Lighthouse Laboratory Plymouth and Alex Alderton, Roberto Amato, Jeffrey Barrett, Sonia Goncalves, Ewan Harrison, David K. Jackson, Ian Johnston, Dominic Kwiatkowski, Cordelia Langford, John Sillitoe on behalf of the Wellcome Sanger Institute COVID-19 Surveillance Team |
| EPI_ISL_2909106, EPI_ISL_2909107, EPI_ISL_2909108, EPI_ISL_2909109 | Lighthouse Lab in Milton Keynes | Wellcome Sanger Institute for the COVID-19 Genomics UK (COG-UK) Consortium | The Lighthouse Lab in Milton Keynes and Alex Alderton, Roberto Amato, Jeffrey Barrett, Sonia Goncalves, Ewan Harrison, David K. Jackson, Ian Johnston, Dominic Kwiatkowski, Cordelia Langford, John Sillitoe on behalf of the Wellcome Sanger Institute COVID-19 Surveillance Team |
| EPI_ISL_2909110 | Lighthouse Laboratory Plymouth | Wellcome Sanger Institute for the COVID-19 Genomics UK (COG-UK) Consortium | Lighthouse Laboratory Plymouth and Alex Alderton, Roberto Amato, Jeffrey Barrett, Sonia Goncalves, Ewan Harrison, David K. Jackson, Ian Johnston, Dominic Kwiatkowski, Cordelia Langford, John Sillitoe on behalf of the Wellcome Sanger Institute COVID-19 Surveillance Team |
| EPI_ISL_2909111 | Lighthouse Lab in Milton Keynes | Wellcome Sanger Institute for the COVID-19 Genomics UK (COG-UK) Consortium | The Lighthouse Lab in Milton Keynes and Alex Alderton, Roberto Amato, Jeffrey Barrett, Sonia Goncalves, Ewan Harrison, David K. Jackson, Ian Johnston, Dominic Kwiatkowski, Cordelia Langford, John Sillitoe on behalf of the Wellcome Sanger Institute COVID-19 Surveillance Team |
| EPI_ISL_2909112 | Lighthouse Laboratory Plymouth | Wellcome Sanger Institute for the COVID-19 Genomics UK (COG-UK) Consortium | Lighthouse Laboratory Plymouth and Alex Alderton, Roberto Amato, Jeffrey Barrett, Sonia Goncalves, Ewan Harrison, David K. Jackson, Ian Johnston, Dominic Kwiatkowski, Cordelia Langford, John Sillitoe on behalf of the Wellcome Sanger Institute COVID-19 Surveillance Team |
| EPI_ISL_2909113 | Lighthouse Lab in Milton Keynes | Wellcome Sanger Institute for the COVID-19 Genomics UK (COG-UK) Consortium | The Lighthouse Lab in Milton Keynes and Alex Alderton, Roberto Amato, Jeffrey Barrett, Sonia Goncalves, Ewan Harrison, David K. Jackson, Ian Johnston, Dominic Kwiatkowski, Cordelia Langford, John Sillitoe on behalf of the Wellcome Sanger Institute COVID-19 Surveillance Team |
| EPI_ISL_2909114, EPI_ISL_2909115, EPI_ISL_2909116 | Lighthouse Laboratory Plymouth | Wellcome Sanger Institute for the COVID-19 Genomics UK (COG-UK) Consortium | Lighthouse Laboratory Plymouth and Alex Alderton, Roberto Amato, Jeffrey Barrett, Sonia Goncalves, Ewan Harrison, David K. Jackson, Ian Johnston, Dominic Kwiatkowski, Cordelia Langford, John Sillitoe on behalf of the Wellcome Sanger Institute COVID-19 Surveillance Team |
| EPI_ISL_2909117, EPI_ISL_2909118 | Lighthouse Lab in Milton Keynes | Wellcome Sanger Institute for the COVID-19 Genomics UK (COG-UK) Consortium | The Lighthouse Lab in Milton Keynes and Alex Alderton, Roberto Amato, Jeffrey Barrett, Sonia Goncalves, Ewan Harrison, David K. Jackson, Ian Johnston, Dominic Kwiatkowski, Cordelia Langford, John Sillitoe on behalf of the Wellcome Sanger Institute COVID-19 Surveillance Team |
| EPI_ISL_2909119, EPI_ISL_2909120, EPI_ISL_2909121, EPI_ISL_2909122, EPI_ISL_2909123 | Lighthouse Laboratory Plymouth | Wellcome Sanger Institute for the COVID-19 Genomics UK (COG-UK) Consortium | Lighthouse Laboratory Plymouth and Alex Alderton, Roberto Amato, Jeffrey Barrett, Sonia Goncalves, Ewan Harrison, David K. Jackson, Ian Johnston, Dominic Kwiatkowski, Cordelia Langford, John Sillitoe on behalf of the Wellcome Sanger Institute COVID-19 Surveillance Team |
| EPI_ISL_2909124 | Lighthouse Lab in Milton Keynes | Wellcome Sanger Institute for the COVID-19 Genomics UK (COG-UK) Consortium | The Lighthouse Lab in Milton Keynes and Alex Alderton, Roberto Amato, Jeffrey Barrett, Sonia Goncalves, Ewan Harrison, David K. Jackson, Ian Johnston, Dominic Kwiatkowski, Cordelia Langford, John Sillitoe on behalf of the Wellcome Sanger Institute COVID-19 Surveillance Team |
| EPI_ISL_2909125, EPI_ISL_2909126 | Lighthouse Laboratory Plymouth | Wellcome Sanger Institute for the COVID-19 Genomics UK (COG-UK) Consortium | Lighthouse Laboratory Plymouth and Alex Alderton, Roberto Amato, Jeffrey Barrett, Sonia Goncalves, Ewan Harrison, David K. Jackson, Ian Johnston, Dominic Kwiatkowski, Cordelia Langford, John Sillitoe on behalf of the Wellcome Sanger Institute COVID-19 Surveillance Team |
| EPI_ISL_2909127 | Lighthouse Lab in Milton Keynes | Wellcome Sanger Institute for the COVID-19 Genomics UK (COG-UK) Consortium | The Lighthouse Lab in Milton Keynes and Alex Alderton, Roberto Amato, Jeffrey Barrett, Sonia Goncalves, Ewan Harrison, David K. Jackson, Ian Johnston, Dominic Kwiatkowski, Cordelia Langford, John Sillitoe on behalf of the Wellcome Sanger Institute COVID-19 Surveillance Team |
| EPI_ISL_2909128, EPI_ISL_2909129, EPI_ISL_2909130 | Lighthouse Laboratory Plymouth | Wellcome Sanger Institute for the COVID-19 Genomics UK (COG-UK) Consortium | Lighthouse Laboratory Plymouth and Alex Alderton, Roberto Amato, Jeffrey Barrett, Sonia Goncalves, Ewan Harrison, David K. Jackson, Ian Johnston, Dominic Kwiatkowski, Cordelia Langford, John Sillitoe on behalf of the Wellcome Sanger Institute COVID-19 Surveillance Team |
| EPI_ISL_2909131 | Lighthouse Lab in Milton Keynes | Wellcome Sanger Institute for the COVID-19 Genomics UK (COG-UK) Consortium | The Lighthouse Lab in Milton Keynes and Alex Alderton, Roberto Amato, Jeffrey Barrett, Sonia Goncalves, Ewan Harrison, David K. Jackson, Ian Johnston, Dominic Kwiatkowski, Cordelia Langford, John Sillitoe on behalf of the Wellcome Sanger Institute COVID-19 Surveillance Team |
| EPI_ISL_2909132, EPI_ISL_2909133 | Lighthouse Laboratory Plymouth | Wellcome Sanger Institute for the COVID-19 Genomics UK (COG-UK) Consortium | Lighthouse Laboratory Plymouth and Alex Alderton, Roberto Amato, Jeffrey Barrett, Sonia Goncalves, Ewan Harrison, David K. Jackson, Ian Johnston, Dominic Kwiatkowski, Cordelia Langford, John Sillitoe on behalf of the Wellcome Sanger Institute COVID-19 Surveillance Team |
| EPI_ISL_2909134, EPI_ISL_2909135 | Lighthouse Lab in Milton Keynes | Wellcome Sanger Institute for the COVID-19 Genomics UK (COG-UK) Consortium | The Lighthouse Lab in Milton Keynes and Alex Alderton, Roberto Amato, Jeffrey Barrett, Sonia Goncalves, Ewan Harrison, David K. Jackson, Ian Johnston, Dominic Kwiatkowski, Cordelia Langford, John Sillitoe on behalf of the Wellcome Sanger Institute COVID-19 Surveillance Team |
| EPI_ISL_2909136, EPI_ISL_2909137, EPI_ISL_2909138, EPI_ISL_2909139 | Lighthouse Laboratory Plymouth | Wellcome Sanger Institute for the COVID-19 Genomics UK (COG-UK) Consortium | Lighthouse Laboratory Plymouth and Alex Alderton, Roberto Amato, Jeffrey Barrett, Sonia Goncalves, Ewan Harrison, David K. Jackson, Ian Johnston, Dominic Kwiatkowski, Cordelia Langford, John Sillitoe on behalf of the Wellcome Sanger Institute COVID-19 Surveillance Team |
| EPI_ISL_2909140 | Lighthouse Lab in Milton Keynes | Wellcome Sanger Institute for the COVID-19 Genomics UK (COG-UK) Consortium | The Lighthouse Lab in Milton Keynes and Alex Alderton, Roberto Amato, Jeffrey Barrett, Sonia Goncalves, Ewan Harrison, David K. Jackson, Ian Johnston, Dominic Kwiatkowski, Cordelia Langford, John Sillitoe on behalf of the Wellcome Sanger Institute COVID-19 Surveillance Team |
| EPI_ISL_2909141 | Lighthouse Laboratory Plymouth | Wellcome Sanger Institute for the COVID-19 Genomics UK (COG-UK) Consortium | Lighthouse Laboratory Plymouth and Alex Alderton, Roberto Amato, Jeffrey Barrett, Sonia Goncalves, Ewan Harrison, David K. Jackson, Ian Johnston, Dominic Kwiatkowski, Cordelia Langford, John Sillitoe on behalf of the Wellcome Sanger Institute COVID-19 Surveillance Team |
| EPI_ISL_2909142 | Lighthouse Lab in Milton Keynes | Wellcome Sanger Institute for the COVID-19 Genomics UK (COG-UK) Consortium | The Lighthouse Lab in Milton Keynes and Alex Alderton, Roberto Amato, Jeffrey Barrett, Sonia Goncalves, Ewan Harrison, David K. Jackson, Ian Johnston, Dominic Kwiatkowski, Cordelia Langford, John Sillitoe on behalf of the Wellcome Sanger Institute COVID-19 Surveillance Team |
| EPI_ISL_2909143, EPI_ISL_2909144, EPI_ISL_2909145 | Lighthouse Laboratory Plymouth | Wellcome Sanger Institute for the COVID-19 Genomics UK (COG-UK) Consortium | Lighthouse Laboratory Plymouth and Alex Alderton, Roberto Amato, Jeffrey Barrett, Sonia Goncalves, Ewan Harrison, David K. Jackson, Ian Johnston, Dominic Kwiatkowski, Cordelia Langford, John Sillitoe on behalf of the Wellcome Sanger Institute COVID-19 Surveillance Team |
| EPI_ISL_2909146 | Lighthouse Lab in Milton Keynes | Wellcome Sanger Institute for the COVID-19 Genomics UK (COG-UK) Consortium | The Lighthouse Lab in Milton Keynes and Alex Alderton, Roberto Amato, Jeffrey Barrett, Sonia Goncalves, Ewan Harrison, David K. Jackson, Ian Johnston, Dominic Kwiatkowski, Cordelia Langford, John Sillitoe on behalf of the Wellcome Sanger Institute COVID-19 Surveillance Team |
| EPI_ISL_2909147, EPI_ISL_2909148, EPI_ISL_2909149, EPI_ISL_2909150, EPI_ISL_2909151, EPI_ISL_2909152 | Lighthouse Laboratory Plymouth | Wellcome Sanger Institute for the COVID-19 Genomics UK (COG-UK) Consortium | Lighthouse Laboratory Plymouth and Alex Alderton, Roberto Amato, Jeffrey Barrett, Sonia Goncalves, Ewan Harrison, David K. Jackson, Ian Johnston, Dominic Kwiatkowski, Cordelia Langford, John Sillitoe on behalf of the Wellcome Sanger Institute COVID-19 Surveillance Team |
| EPI_ISL_2909153, EPI_ISL_2909154 | Lighthouse Lab in Milton Keynes | Wellcome Sanger Institute for the COVID-19 Genomics UK (COG-UK) Consortium | The Lighthouse Lab in Milton Keynes and Alex Alderton, Roberto Amato, Jeffrey Barrett, Sonia Goncalves, Ewan Harrison, David K. Jackson, Ian Johnston, Dominic Kwiatkowski, Cordelia Langford, John Sillitoe on behalf of the Wellcome Sanger Institute COVID-19 Surveillance Team |
| EPI_ISL_2909156, EPI_ISL_2909157 | Lighthouse Laboratory Plymouth | Wellcome Sanger Institute for the COVID-19 Genomics UK (COG-UK) Consortium | Lighthouse Laboratory Plymouth and Alex Alderton, Roberto Amato, Jeffrey Barrett, Sonia Goncalves, Ewan Harrison, David K. Jackson, Ian Johnston, Dominic Kwiatkowski, Cordelia Langford, John Sillitoe on behalf of the Wellcome Sanger Institute COVID-19 Surveillance Team |
| EPI_ISL_2909158 | Lighthouse Lab in Milton Keynes | Wellcome Sanger Institute for the COVID-19 Genomics UK (COG-UK) Consortium | The Lighthouse Lab in Milton Keynes and Alex Alderton, Roberto Amato, Jeffrey Barrett, Sonia Goncalves, Ewan Harrison, David K. Jackson, Ian Johnston, Dominic Kwiatkowski, Cordelia Langford, John Sillitoe on behalf of the Wellcome Sanger Institute COVID-19 Surveillance Team |
| EPI_ISL_2909159 | Lighthouse Laboratory Plymouth | Wellcome Sanger Institute for the COVID-19 Genomics UK (COG-UK) Consortium | Lighthouse Laboratory Plymouth and Alex Alderton, Roberto Amato, Jeffrey Barrett, Sonia Goncalves, Ewan Harrison, David K. Jackson, Ian Johnston, Dominic Kwiatkowski, Cordelia Langford, John Sillitoe on behalf of the Wellcome Sanger Institute COVID-19 Surveillance Team |
| EPI_ISL_2909160 | Lighthouse Lab in Milton Keynes | Wellcome Sanger Institute for the COVID-19 Genomics UK (COG-UK) Consortium | The Lighthouse Lab in Milton Keynes and Alex Alderton, Roberto Amato, Jeffrey Barrett, Sonia Goncalves, Ewan Harrison, David K. Jackson, Ian Johnston, Dominic Kwiatkowski, Cordelia Langford, John Sillitoe on behalf of the Wellcome Sanger Institute COVID-19 Surveillance Team |

| | | | |
|---|---|---|---|
| EPI_ISL_2909161 | Lighthouse Laboratory Plymouth | Wellcome Sanger Institute for the COVID-19 Genomics UK (COG-UK) Consortium | Lighthouse Laboratory Plymouth and Alex Alderton, Roberto Amato, Jeffrey Barrett, Sonia Goncalves, Ewan Harrison, David K. Jackson, Ian Johnston, Dominic Kwiatkowski, Cordelia Langford, John Sillitoe on behalf of the Wellcome Sanger Institute COVID-19 Surveillance Team |
| EPI_ISL_2909162, EPI_ISL_2909163 | Lighthouse Lab in Milton Keynes | Wellcome Sanger Institute for the COVID-19 Genomics UK (COG-UK) Consortium | The Lighthouse Lab in Milton Keynes and Alex Alderton, Roberto Amato, Jeffrey Barrett, Sonia Goncalves, Ewan Harrison, David K. Jackson, Ian Johnston, Dominic Kwiatkowski, Cordelia Langford, John Sillitoe on behalf of the Wellcome Sanger Institute COVID-19 Surveillance Team |
| EPI_ISL_2909164, EPI_ISL_2909165, EPI_ISL_2909166, EPI_ISL_2909167, EPI_ISL_2909168, EPI_ISL_2909169, EPI_ISL_2909170 | Lighthouse Laboratory Plymouth | Wellcome Sanger Institute for the COVID-19 Genomics UK (COG-UK) Consortium | Lighthouse Laboratory Plymouth and Alex Alderton, Roberto Amato, Jeffrey Barrett, Sonia Goncalves, Ewan Harrison, David K. Jackson, Ian Johnston, Dominic Kwiatkowski, Cordelia Langford, John Sillitoe on behalf of the Wellcome Sanger Institute COVID-19 Surveillance Team |
| EPI_ISL_2909171 | Lighthouse Lab in Milton Keynes | Wellcome Sanger Institute for the COVID-19 Genomics UK (COG-UK) Consortium | The Lighthouse Lab in Milton Keynes and Alex Alderton, Roberto Amato, Jeffrey Barrett, Sonia Goncalves, Ewan Harrison, David K. Jackson, Ian Johnston, Dominic Kwiatkowski, Cordelia Langford, John Sillitoe on behalf of the Wellcome Sanger Institute COVID-19 Surveillance Team |
| EPI_ISL_2909172 | Lighthouse Laboratory Plymouth | Wellcome Sanger Institute for the COVID-19 Genomics UK (COG-UK) Consortium | Lighthouse Laboratory Plymouth and Alex Alderton, Roberto Amato, Jeffrey Barrett, Sonia Goncalves, Ewan Harrison, David K. Jackson, Ian Johnston, Dominic Kwiatkowski, Cordelia Langford, John Sillitoe on behalf of the Wellcome Sanger Institute COVID-19 Surveillance Team |
| EPI_ISL_2909173 | Lighthouse Lab in Milton Keynes | Wellcome Sanger Institute for the COVID-19 Genomics UK (COG-UK) Consortium | The Lighthouse Lab in Milton Keynes and Alex Alderton, Roberto Amato, Jeffrey Barrett, Sonia Goncalves, Ewan Harrison, David K. Jackson, Ian Johnston, Dominic Kwiatkowski, Cordelia Langford, John Sillitoe on behalf of the Wellcome Sanger Institute COVID-19 Surveillance Team |
| EPI_ISL_2909174, EPI_ISL_2909175, EPI_ISL_2909176, EPI_ISL_2909177, EPI_ISL_2909178, EPI_ISL_2909179, EPI_ISL_2909180 | Lighthouse Laboratory Plymouth | Wellcome Sanger Institute for the COVID-19 Genomics UK (COG-UK) Consortium | Lighthouse Laboratory Plymouth and Alex Alderton, Roberto Amato, Jeffrey Barrett, Sonia Goncalves, Ewan Harrison, David K. Jackson, Ian Johnston, Dominic Kwiatkowski, Cordelia Langford, John Sillitoe on behalf of the Wellcome Sanger Institute COVID-19 Surveillance Team |
| EPI_ISL_2909181 | Lighthouse Lab in Milton Keynes | Wellcome Sanger Institute for the COVID-19 Genomics UK (COG-UK) Consortium | The Lighthouse Lab in Milton Keynes and Alex Alderton, Roberto Amato, Jeffrey Barrett, Sonia Goncalves, Ewan Harrison, David K. Jackson, Ian Johnston, Dominic Kwiatkowski, Cordelia Langford, John Sillitoe on behalf of the Wellcome Sanger Institute COVID-19 Surveillance Team |
| EPI_ISL_2909182, EPI_ISL_2909183, EPI_ISL_2909184, EPI_ISL_2909185 | Lighthouse Laboratory Plymouth | Wellcome Sanger Institute for the COVID-19 Genomics UK (COG-UK) Consortium | Lighthouse Laboratory Plymouth and Alex Alderton, Roberto Amato, Jeffrey Barrett, Sonia Goncalves, Ewan Harrison, David K. Jackson, Ian Johnston, Dominic Kwiatkowski, Cordelia Langford, John Sillitoe on behalf of the Wellcome Sanger Institute COVID-19 Surveillance Team |
| EPI_ISL_2909186, EPI_ISL_2909187, EPI_ISL_2909188, EPI_ISL_2909189 | Lighthouse Lab in Milton Keynes | Wellcome Sanger Institute for the COVID-19 Genomics UK (COG-UK) Consortium | The Lighthouse Lab in Milton Keynes and Alex Alderton, Roberto Amato, Jeffrey Barrett, Sonia Goncalves, Ewan Harrison, David K. Jackson, Ian Johnston, Dominic Kwiatkowski, Cordelia Langford, John Sillitoe on behalf of the Wellcome Sanger Institute COVID-19 Surveillance Team |
| EPI_ISL_2909190, EPI_ISL_2909191, EPI_ISL_2909192, EPI_ISL_2909193, EPI_ISL_2909194, EPI_ISL_2909195 | Lighthouse Laboratory Plymouth | Wellcome Sanger Institute for the COVID-19 Genomics UK (COG-UK) Consortium | Lighthouse Laboratory Plymouth and Alex Alderton, Roberto Amato, Jeffrey Barrett, Sonia Goncalves, Ewan Harrison, David K. Jackson, Ian Johnston, Dominic Kwiatkowski, Cordelia Langford, John Sillitoe on behalf of the Wellcome Sanger Institute COVID-19 Surveillance Team |
| EPI_ISL_2909196, EPI_ISL_2909197, EPI_ISL_2909198, EPI_ISL_2909199, EPI_ISL_2909200, EPI_ISL_2909201, EPI_ISL_2909202, EPI_ISL_2909203 | Lighthouse Lab in Glasgow | Wellcome Sanger Institute for the COVID-19 Genomics UK (COG-UK) Consortium | Harper VanSteenhouse, Yumi Kasai, David Gray, Carol Clugston, Anna Dominiczak and Alex Alderton, Roberto Amato, Jeffrey Barrett, Sonia Goncalves, Ewan Harrison, David K. Jackson, Ian Johnston, Dominic Kwiatkowski, Cordelia Langford, John Sillitoe on behalf of the Wellcome Sanger Institute COVID-19 Surveillance Team |
| EPI_ISL_2909204 | Lighthouse Lab in Milton Keynes | Wellcome Sanger Institute for the COVID-19 Genomics UK (COG-UK) Consortium | The Lighthouse Lab in Milton Keynes and Alex Alderton, Roberto Amato, Jeffrey Barrett, Sonia Goncalves, Ewan Harrison, David K. Jackson, Ian Johnston, Dominic Kwiatkowski, Cordelia Langford, John Sillitoe on behalf of the Wellcome Sanger Institute COVID-19 Surveillance Team |
| EPI_ISL_2909205, EPI_ISL_2909206, EPI_ISL_2909207 | Lighthouse Lab in Glasgow | Wellcome Sanger Institute for the COVID-19 Genomics UK (COG-UK) Consortium | Harper VanSteenhouse, Yumi Kasai, David Gray, Carol Clugston, Anna Dominiczak and Alex Alderton, Roberto Amato, Jeffrey Barrett, Sonia Goncalves, Ewan Harrison, David K. Jackson, Ian Johnston, Dominic Kwiatkowski, Cordelia Langford, John Sillitoe on behalf of the Wellcome Sanger Institute COVID-19 Surveillance Team |
| EPI_ISL_2909208 | Lighthouse Lab in Milton Keynes | Wellcome Sanger Institute for the COVID-19 Genomics UK (COG-UK) Consortium | The Lighthouse Lab in Milton Keynes and Alex Alderton, Roberto Amato, Jeffrey Barrett, Sonia Goncalves, Ewan Harrison, David K. Jackson, Ian Johnston, Dominic Kwiatkowski, Cordelia Langford, John Sillitoe on behalf of the Wellcome Sanger Institute COVID-19 Surveillance Team |
| EPI_ISL_2909209, EPI_ISL_2909210 | Lighthouse Lab in Glasgow | Wellcome Sanger Institute for the COVID-19 Genomics UK (COG-UK) Consortium | Harper VanSteenhouse, Yumi Kasai, David Gray, Carol Clugston, Anna Dominiczak and Alex Alderton, Roberto Amato, Jeffrey Barrett, Sonia Goncalves, Ewan Harrison, David K. Jackson, Ian Johnston, Dominic Kwiatkowski, Cordelia Langford, John Sillitoe on behalf of the Wellcome Sanger Institute COVID-19 Surveillance Team |
| EPI_ISL_2909211 | Lighthouse Lab in Milton Keynes | Wellcome Sanger Institute for the COVID-19 Genomics UK (COG-UK) Consortium | The Lighthouse Lab in Milton Keynes and Alex Alderton, Roberto Amato, Jeffrey Barrett, Sonia Goncalves, Ewan Harrison, David K. Jackson, Ian Johnston, Dominic Kwiatkowski, Cordelia Langford, John Sillitoe on behalf of the Wellcome Sanger Institute COVID-19 Surveillance Team |
| EPI_ISL_2909212, EPI_ISL_2909213, EPI_ISL_2909214, EPI_ISL_2909215, EPI_ISL_2909216, EPI_ISL_2909217, EPI_ISL_2909218 | Lighthouse Lab in Glasgow | Wellcome Sanger Institute for the COVID-19 Genomics UK (COG-UK) Consortium | Harper VanSteenhouse, Yumi Kasai, David Gray, Carol Clugston, Anna Dominiczak and Alex Alderton, Roberto Amato, Jeffrey Barrett, Sonia Goncalves, Ewan Harrison, David K. Jackson, Ian Johnston, Dominic Kwiatkowski, Cordelia Langford, John Sillitoe on behalf of the Wellcome Sanger Institute COVID-19 Surveillance Team |
| EPI_ISL_2909219, EPI_ISL_2909220 | Lighthouse Lab in Milton Keynes | Wellcome Sanger Institute for the COVID-19 Genomics UK (COG-UK) Consortium | The Lighthouse Lab in Milton Keynes and Alex Alderton, Roberto Amato, Jeffrey Barrett, Sonia Goncalves, Ewan Harrison, David K. Jackson, Ian Johnston, Dominic Kwiatkowski, Cordelia Langford, John Sillitoe on behalf of the Wellcome Sanger Institute COVID-19 Surveillance Team |
| EPI_ISL_2909221, EPI_ISL_2909222, EPI_ISL_2909223 | Lighthouse Lab in Glasgow | Wellcome Sanger Institute for the COVID-19 Genomics UK (COG-UK) Consortium | Harper VanSteenhouse, Yumi Kasai, David Gray, Carol Clugston, Anna Dominiczak and Alex Alderton, Roberto Amato, Jeffrey Barrett, Sonia Goncalves, Ewan Harrison, David K. Jackson, Ian Johnston, Dominic Kwiatkowski, Cordelia Langford, John Sillitoe on behalf of the Wellcome Sanger Institute COVID-19 Surveillance Team |
| EPI_ISL_2909224 | Lighthouse Lab in Milton Keynes | Wellcome Sanger Institute for the COVID-19 Genomics UK (COG-UK) Consortium | The Lighthouse Lab in Milton Keynes and Alex Alderton, Roberto Amato, Jeffrey Barrett, Sonia Goncalves, Ewan Harrison, David K. Jackson, Ian Johnston, Dominic Kwiatkowski, Cordelia Langford, John Sillitoe on behalf of the Wellcome Sanger Institute COVID-19 Surveillance Team |
| EPI_ISL_2909225, EPI_ISL_2909226, EPI_ISL_2909227 | Lighthouse Lab in Glasgow | Wellcome Sanger Institute for the COVID-19 Genomics UK (COG-UK) Consortium | Harper VanSteenhouse, Yumi Kasai, David Gray, Carol Clugston, Anna Dominiczak and Alex Alderton, Roberto Amato, Jeffrey Barrett, Sonia Goncalves, Ewan Harrison, David K. Jackson, Ian Johnston, Dominic Kwiatkowski, Cordelia Langford, John Sillitoe on behalf of the Wellcome Sanger Institute COVID-19 Surveillance Team |
| EPI_ISL_2909228, EPI_ISL_2909229 | Lighthouse Lab in Milton Keynes | Wellcome Sanger Institute for the COVID-19 Genomics UK (COG-UK) Consortium | The Lighthouse Lab in Milton Keynes and Alex Alderton, Roberto Amato, Jeffrey Barrett, Sonia Goncalves, Ewan Harrison, David K. Jackson, Ian Johnston, Dominic Kwiatkowski, Cordelia Langford, John Sillitoe on behalf of the Wellcome Sanger Institute COVID-19 Surveillance Team |
| EPI_ISL_2909230, EPI_ISL_2909231, EPI_ISL_2909232 | Lighthouse Lab in Glasgow | Wellcome Sanger Institute for the COVID-19 Genomics UK (COG-UK) Consortium | Harper VanSteenhouse, Yumi Kasai, David Gray, Carol Clugston, Anna Dominiczak and Alex Alderton, Roberto Amato, Jeffrey Barrett, Sonia Goncalves, Ewan Harrison, David K. Jackson, Ian Johnston, Dominic Kwiatkowski, Cordelia Langford, John Sillitoe on behalf of the Wellcome Sanger Institute COVID-19 Surveillance Team |
| EPI_ISL_2909233 | Lighthouse Lab in Milton Keynes | Wellcome Sanger Institute for the COVID-19 Genomics UK (COG-UK) Consortium | The Lighthouse Lab in Milton Keynes and Alex Alderton, Roberto Amato, Jeffrey Barrett, Sonia Goncalves, Ewan Harrison, David K. Jackson, Ian Johnston, Dominic Kwiatkowski, Cordelia Langford, John Sillitoe on behalf of the Wellcome Sanger Institute COVID-19 Surveillance Team |
| EPI_ISL_2909234 | Lighthouse Lab in Glasgow | Wellcome Sanger Institute for the COVID-19 Genomics UK (COG-UK) Consortium | Harper VanSteenhouse, Yumi Kasai, David Gray, Carol Clugston, Anna Dominiczak and Alex Alderton, Roberto Amato, Jeffrey Barrett, Sonia Goncalves, Ewan Harrison, David K. Jackson, Ian Johnston, Dominic Kwiatkowski, Cordelia Langford, John Sillitoe on behalf of the Wellcome Sanger Institute COVID-19 Surveillance Team |
| EPI_ISL_2909235 | Lighthouse Lab in Milton Keynes | Wellcome Sanger Institute for the COVID-19 Genomics UK (COG-UK) Consortium | The Lighthouse Lab in Milton Keynes and Alex Alderton, Roberto Amato, Jeffrey Barrett, Sonia Goncalves, Ewan Harrison, David K. Jackson, Ian Johnston, Dominic Kwiatkowski, Cordelia Langford, John Sillitoe on behalf of the Wellcome Sanger Institute COVID-19 Surveillance Team |
| EPI_ISL_2909236, EPI_ISL_2909237, EPI_ISL_2909238, EPI_ISL_2909239 | Lighthouse Lab in Glasgow | Wellcome Sanger Institute for the COVID-19 Genomics UK (COG-UK) Consortium | Harper VanSteenhouse, Yumi Kasai, David Gray, Carol Clugston, Anna Dominiczak and Alex Alderton, Roberto Amato, Jeffrey Barrett, Sonia Goncalves, Ewan Harrison, David K. Jackson, Ian Johnston, Dominic Kwiatkowski, Cordelia Langford, John Sillitoe on behalf of the Wellcome Sanger Institute COVID-19 Surveillance Team |
| EPI_ISL_2909240 | Lighthouse Lab in Milton Keynes | Wellcome Sanger Institute for the COVID-19 Genomics UK (COG-UK) Consortium | The Lighthouse Lab in Milton Keynes and Alex Alderton, Roberto Amato, Jeffrey Barrett, Sonia Goncalves, Ewan Harrison, David K. Jackson, Ian Johnston, Dominic Kwiatkowski, Cordelia Langford, John Sillitoe on behalf of the Wellcome Sanger Institute COVID-19 Surveillance Team |
| EPI_ISL_2909241, EPI_ISL_2909242 | Lighthouse Lab in Glasgow | Wellcome Sanger Institute for the COVID-19 Genomics UK (COG-UK) Consortium | Harper VanSteenhouse, Yumi Kasai, David Gray, Carol Clugston, Anna Dominiczak and Alex Alderton, Roberto Amato, Jeffrey Barrett, Sonia Goncalves, Ewan Harrison, David K. Jackson, Ian Johnston, Dominic Kwiatkowski, Cordelia Langford, John Sillitoe on behalf of the Wellcome Sanger Institute COVID-19 Surveillance Team |
| EPI_ISL_2909243 | Lighthouse Lab in Milton Keynes | Wellcome Sanger Institute for the COVID-19 Genomics UK (COG-UK) | The Lighthouse Lab in Milton Keynes and Alex Alderton, Roberto Amato, Jeffrey Barrett, Sonia Goncalves, Ewan Harrison, David K. Jackson, Ian Johnston, Dominic Kwiatkowski, Cordelia Langford, John Sillitoe on behalf of the Wellcome Sanger |

| | | | |
|---|---|---|---|
| | | Consortium | Institute COVID-19 Surveillance Team |
| EPI_ISL_2909244, EPI_ISL_2909245, EPI_ISL_2909246 | Lighthouse Lab in Glasgow | Wellcome Sanger Institute for the COVID-19 Genomics UK (COG-UK) Consortium | Harper VanSteenhouse, Yumi Kasai, David Gray, Carol Clugston, Anna Dominiczak and Alex Alderton, Roberto Amato, Jeffrey Barrett, Sonia Goncalves, Ewan Harrison, David K. Jackson, Ian Johnston, Dominic Kwiatkowski, Cordelia Langford, John Sillitoe on behalf of the Wellcome Sanger Institute COVID-19 Surveillance Team |
| EPI_ISL_2909247 | Lighthouse Lab in Milton Keynes | Wellcome Sanger Institute for the COVID-19 Genomics UK (COG-UK) Consortium | The Lighthouse Lab in Milton Keynes and Alex Alderton, Roberto Amato, Jeffrey Barrett, Sonia Goncalves, Ewan Harrison, David K. Jackson, Ian Johnston, Dominic Kwiatkowski, Cordelia Langford, John Sillitoe on behalf of the Wellcome Sanger Institute COVID-19 Surveillance Team |
| EPI_ISL_2909248 | Lighthouse Lab in Glasgow | Wellcome Sanger Institute for the COVID-19 Genomics UK (COG-UK) Consortium | Harper VanSteenhouse, Yumi Kasai, David Gray, Carol Clugston, Anna Dominiczak and Alex Alderton, Roberto Amato, Jeffrey Barrett, Sonia Goncalves, Ewan Harrison, David K. Jackson, Ian Johnston, Dominic Kwiatkowski, Cordelia Langford, John Sillitoe on behalf of the Wellcome Sanger Institute COVID-19 Surveillance Team |
| EPI_ISL_2909249 | Lighthouse Lab in Milton Keynes | Wellcome Sanger Institute for the COVID-19 Genomics UK (COG-UK) Consortium | The Lighthouse Lab in Milton Keynes and Alex Alderton, Roberto Amato, Jeffrey Barrett, Sonia Goncalves, Ewan Harrison, David K. Jackson, Ian Johnston, Dominic Kwiatkowski, Cordelia Langford, John Sillitoe on behalf of the Wellcome Sanger Institute COVID-19 Surveillance Team |
| EPI_ISL_2909250, EPI_ISL_2909251, EPI_ISL_2909252 | Lighthouse Lab in Glasgow | Wellcome Sanger Institute for the COVID-19 Genomics UK (COG-UK) Consortium | Harper VanSteenhouse, Yumi Kasai, David Gray, Carol Clugston, Anna Dominiczak and Alex Alderton, Roberto Amato, Jeffrey Barrett, Sonia Goncalves, Ewan Harrison, David K. Jackson, Ian Johnston, Dominic Kwiatkowski, Cordelia Langford, John Sillitoe on behalf of the Wellcome Sanger Institute COVID-19 Surveillance Team |
| EPI_ISL_2909253 | Lighthouse Lab in Milton Keynes | Wellcome Sanger Institute for the COVID-19 Genomics UK (COG-UK) Consortium | The Lighthouse Lab in Milton Keynes and Alex Alderton, Roberto Amato, Jeffrey Barrett, Sonia Goncalves, Ewan Harrison, David K. Jackson, Ian Johnston, Dominic Kwiatkowski, Cordelia Langford, John Sillitoe on behalf of the Wellcome Sanger Institute COVID-19 Surveillance Team |
| EPI_ISL_2909254, EPI_ISL_2909255, EPI_ISL_2909256, EPI_ISL_2909257, EPI_ISL_2909258 | Lighthouse Lab in Glasgow | Wellcome Sanger Institute for the COVID-19 Genomics UK (COG-UK) Consortium | Harper VanSteenhouse, Yumi Kasai, David Gray, Carol Clugston, Anna Dominiczak and Alex Alderton, Roberto Amato, Jeffrey Barrett, Sonia Goncalves, Ewan Harrison, David K. Jackson, Ian Johnston, Dominic Kwiatkowski, Cordelia Langford, John Sillitoe on behalf of the Wellcome Sanger Institute COVID-19 Surveillance Team |
| EPI_ISL_2909259 | Lighthouse Lab in Milton Keynes | Wellcome Sanger Institute for the COVID-19 Genomics UK (COG-UK) Consortium | The Lighthouse Lab in Milton Keynes and Alex Alderton, Roberto Amato, Jeffrey Barrett, Sonia Goncalves, Ewan Harrison, David K. Jackson, Ian Johnston, Dominic Kwiatkowski, Cordelia Langford, John Sillitoe on behalf of the Wellcome Sanger Institute COVID-19 Surveillance Team |
| EPI_ISL_2909260, EPI_ISL_2909261, EPI_ISL_2909262 | Lighthouse Lab in Glasgow | Wellcome Sanger Institute for the COVID-19 Genomics UK (COG-UK) Consortium | Harper VanSteenhouse, Yumi Kasai, David Gray, Carol Clugston, Anna Dominiczak and Alex Alderton, Roberto Amato, Jeffrey Barrett, Sonia Goncalves, Ewan Harrison, David K. Jackson, Ian Johnston, Dominic Kwiatkowski, Cordelia Langford, John Sillitoe on behalf of the Wellcome Sanger Institute COVID-19 Surveillance Team |
| EPI_ISL_2909263 | Lighthouse Lab in Milton Keynes | Wellcome Sanger Institute for the COVID-19 Genomics UK (COG-UK) Consortium | The Lighthouse Lab in Milton Keynes and Alex Alderton, Roberto Amato, Jeffrey Barrett, Sonia Goncalves, Ewan Harrison, David K. Jackson, Ian Johnston, Dominic Kwiatkowski, Cordelia Langford, John Sillitoe on behalf of the Wellcome Sanger Institute COVID-19 Surveillance Team |
| EPI_ISL_2909264 | Lighthouse Lab in Glasgow | Wellcome Sanger Institute for the COVID-19 Genomics UK (COG-UK) Consortium | Harper VanSteenhouse, Yumi Kasai, David Gray, Carol Clugston, Anna Dominiczak and Alex Alderton, Roberto Amato, Jeffrey Barrett, Sonia Goncalves, Ewan Harrison, David K. Jackson, Ian Johnston, Dominic Kwiatkowski, Cordelia Langford, John Sillitoe on behalf of the Wellcome Sanger Institute COVID-19 Surveillance Team |
| EPI_ISL_2909265 | Lighthouse Lab in Milton Keynes | Wellcome Sanger Institute for the COVID-19 Genomics UK (COG-UK) Consortium | The Lighthouse Lab in Milton Keynes and Alex Alderton, Roberto Amato, Jeffrey Barrett, Sonia Goncalves, Ewan Harrison, David K. Jackson, Ian Johnston, Dominic Kwiatkowski, Cordelia Langford, John Sillitoe on behalf of the Wellcome Sanger Institute COVID-19 Surveillance Team |
| EPI_ISL_2909266 | Lighthouse Lab in Glasgow | Wellcome Sanger Institute for the COVID-19 Genomics UK (COG-UK) Consortium | Harper VanSteenhouse, Yumi Kasai, David Gray, Carol Clugston, Anna Dominiczak and Alex Alderton, Roberto Amato, Jeffrey Barrett, Sonia Goncalves, Ewan Harrison, David K. Jackson, Ian Johnston, Dominic Kwiatkowski, Cordelia Langford, John Sillitoe on behalf of the Wellcome Sanger Institute COVID-19 Surveillance Team |
| EPI_ISL_2909267 | Lighthouse Lab in Milton Keynes | Wellcome Sanger Institute for the COVID-19 Genomics UK (COG-UK) Consortium | The Lighthouse Lab in Milton Keynes and Alex Alderton, Roberto Amato, Jeffrey Barrett, Sonia Goncalves, Ewan Harrison, David K. Jackson, Ian Johnston, Dominic Kwiatkowski, Cordelia Langford, John Sillitoe on behalf of the Wellcome Sanger Institute COVID-19 Surveillance Team |
| EPI_ISL_2909268, EPI_ISL_2909269, EPI_ISL_2909270, EPI_ISL_2909271 | Lighthouse Lab in Glasgow | Wellcome Sanger Institute for the COVID-19 Genomics UK (COG-UK) Consortium | Harper VanSteenhouse, Yumi Kasai, David Gray, Carol Clugston, Anna Dominiczak and Alex Alderton, Roberto Amato, Jeffrey Barrett, Sonia Goncalves, Ewan Harrison, David K. Jackson, Ian Johnston, Dominic Kwiatkowski, Cordelia Langford, John Sillitoe on behalf of the Wellcome Sanger Institute COVID-19 Surveillance Team |
| EPI_ISL_2909272 | Lighthouse Lab in Milton Keynes | Wellcome Sanger Institute for the COVID-19 Genomics UK (COG-UK) Consortium | The Lighthouse Lab in Milton Keynes and Alex Alderton, Roberto Amato, Jeffrey Barrett, Sonia Goncalves, Ewan Harrison, David K. Jackson, Ian Johnston, Dominic Kwiatkowski, Cordelia Langford, John Sillitoe on behalf of the Wellcome Sanger Institute COVID-19 Surveillance Team |
| EPI_ISL_2909273, EPI_ISL_2909274, EPI_ISL_2909275 | Lighthouse Lab in Glasgow | Wellcome Sanger Institute for the COVID-19 Genomics UK (COG-UK) Consortium | Harper VanSteenhouse, Yumi Kasai, David Gray, Carol Clugston, Anna Dominiczak and Alex Alderton, Roberto Amato, Jeffrey Barrett, Sonia Goncalves, Ewan Harrison, David K. Jackson, Ian Johnston, Dominic Kwiatkowski, Cordelia Langford, John Sillitoe on behalf of the Wellcome Sanger Institute COVID-19 Surveillance Team |
| EPI_ISL_2909276, EPI_ISL_2909277 | Lighthouse Lab in Milton Keynes | Wellcome Sanger Institute for the COVID-19 Genomics UK (COG-UK) Consortium | The Lighthouse Lab in Milton Keynes and Alex Alderton, Roberto Amato, Jeffrey Barrett, Sonia Goncalves, Ewan Harrison, David K. Jackson, Ian Johnston, Dominic Kwiatkowski, Cordelia Langford, John Sillitoe on behalf of the Wellcome Sanger Institute COVID-19 Surveillance Team |
| EPI_ISL_2909278, EPI_ISL_2909279, EPI_ISL_2909280, EPI_ISL_2909281 | Lighthouse Lab in Glasgow | Wellcome Sanger Institute for the COVID-19 Genomics UK (COG-UK) Consortium | Harper VanSteenhouse, Yumi Kasai, David Gray, Carol Clugston, Anna Dominiczak and Alex Alderton, Roberto Amato, Jeffrey Barrett, Sonia Goncalves, Ewan Harrison, David K. Jackson, Ian Johnston, Dominic Kwiatkowski, Cordelia Langford, John Sillitoe on behalf of the Wellcome Sanger Institute COVID-19 Surveillance Team |
| EPI_ISL_2909282 | Lighthouse Lab in Milton Keynes | Wellcome Sanger Institute for the COVID-19 Genomics UK (COG-UK) Consortium | The Lighthouse Lab in Milton Keynes and Alex Alderton, Roberto Amato, Jeffrey Barrett, Sonia Goncalves, Ewan Harrison, David K. Jackson, Ian Johnston, Dominic Kwiatkowski, Cordelia Langford, John Sillitoe on behalf of the Wellcome Sanger Institute COVID-19 Surveillance Team |
| EPI_ISL_2909283 | Lighthouse Lab in Glasgow | Wellcome Sanger Institute for the COVID-19 Genomics UK (COG-UK) Consortium | Harper VanSteenhouse, Yumi Kasai, David Gray, Carol Clugston, Anna Dominiczak and Alex Alderton, Roberto Amato, Jeffrey Barrett, Sonia Goncalves, Ewan Harrison, David K. Jackson, Ian Johnston, Dominic Kwiatkowski, Cordelia Langford, John Sillitoe on behalf of the Wellcome Sanger Institute COVID-19 Surveillance Team |
| EPI_ISL_2909284 | Lighthouse Lab in Milton Keynes | Wellcome Sanger Institute for the COVID-19 Genomics UK (COG-UK) Consortium | The Lighthouse Lab in Milton Keynes and Alex Alderton, Roberto Amato, Jeffrey Barrett, Sonia Goncalves, Ewan Harrison, David K. Jackson, Ian Johnston, Dominic Kwiatkowski, Cordelia Langford, John Sillitoe on behalf of the Wellcome Sanger Institute COVID-19 Surveillance Team |
| EPI_ISL_2909285, EPI_ISL_2909286, EPI_ISL_2909287, EPI_ISL_2909288, EPI_ISL_2909289, EPI_ISL_2909290, EPI_ISL_2909291, EPI_ISL_2909292, EPI_ISL_2909293, EPI_ISL_2909294, EPI_ISL_2909295, EPI_ISL_2909296, EPI_ISL_2909297, EPI_ISL_2909298, EPI_ISL_2909299, EPI_ISL_2909300, EPI_ISL_2909301, EPI_ISL_2909302, EPI_ISL_2909303 | | | |
| see above | Lighthouse Lab in Glasgow | Wellcome Sanger Institute for the COVID-19 Genomics UK (COG-UK) Consortium | Harper VanSteenhouse, Yumi Kasai, David Gray, Carol Clugston, Anna Dominiczak and Alex Alderton, Roberto Amato, Jeffrey Barrett, Sonia Goncalves, Ewan Harrison, David K. Jackson, Ian Johnston, Dominic Kwiatkowski, Cordelia Langford, John Sillitoe on behalf of the Wellcome Sanger Institute COVID-19 Surveillance Team |
| EPI_ISL_2909304 | Lighthouse Lab in Milton Keynes | Wellcome Sanger Institute for the COVID-19 Genomics UK (COG-UK) Consortium | The Lighthouse Lab in Milton Keynes and Alex Alderton, Roberto Amato, Jeffrey Barrett, Sonia Goncalves, Ewan Harrison, David K. Jackson, Ian Johnston, Dominic Kwiatkowski, Cordelia Langford, John Sillitoe on behalf of the Wellcome Sanger Institute COVID-19 Surveillance Team |
| EPI_ISL_2909305 | Lighthouse Lab in Glasgow | Wellcome Sanger Institute for the COVID-19 Genomics UK (COG-UK) Consortium | Harper VanSteenhouse, Yumi Kasai, David Gray, Carol Clugston, Anna Dominiczak and Alex Alderton, Roberto Amato, Jeffrey Barrett, Sonia Goncalves, Ewan Harrison, David K. Jackson, Ian Johnston, Dominic Kwiatkowski, Cordelia Langford, John Sillitoe on behalf of the Wellcome Sanger Institute COVID-19 Surveillance Team |
| EPI_ISL_2909306 | Lighthouse Lab in Milton Keynes | Wellcome Sanger Institute for the COVID-19 Genomics UK (COG-UK) Consortium | The Lighthouse Lab in Milton Keynes and Alex Alderton, Roberto Amato, Jeffrey Barrett, Sonia Goncalves, Ewan Harrison, David K. Jackson, Ian Johnston, Dominic Kwiatkowski, Cordelia Langford, John Sillitoe on behalf of the Wellcome Sanger Institute COVID-19 Surveillance Team |
| EPI_ISL_2909307, EPI_ISL_2909308 | Lighthouse Lab in Glasgow | Wellcome Sanger Institute for the COVID-19 Genomics UK (COG-UK) Consortium | Harper VanSteenhouse, Yumi Kasai, David Gray, Carol Clugston, Anna Dominiczak and Alex Alderton, Roberto Amato, Jeffrey Barrett, Sonia Goncalves, Ewan Harrison, David K. Jackson, Ian Johnston, Dominic Kwiatkowski, Cordelia Langford, John Sillitoe on behalf of the Wellcome Sanger Institute COVID-19 Surveillance Team |
| EPI_ISL_2909309 | Lighthouse Lab in Milton Keynes | Wellcome Sanger Institute for the COVID-19 Genomics UK (COG-UK) Consortium | The Lighthouse Lab in Milton Keynes and Alex Alderton, Roberto Amato, Jeffrey Barrett, Sonia Goncalves, Ewan Harrison, David K. Jackson, Ian Johnston, Dominic Kwiatkowski, Cordelia Langford, John Sillitoe on behalf of the Wellcome Sanger Institute COVID-19 Surveillance Team |
| EPI_ISL_2909310, EPI_ISL_2909311, EPI_ISL_2909312, EPI_ISL_2909313, EPI_ISL_2909314, EPI_ISL_2909315 | Lighthouse Lab in Glasgow | Wellcome Sanger Institute for the COVID-19 Genomics UK (COG-UK) Consortium | Harper VanSteenhouse, Yumi Kasai, David Gray, Carol Clugston, Anna Dominiczak and Alex Alderton, Roberto Amato, Jeffrey Barrett, Sonia Goncalves, Ewan Harrison, David K. Jackson, Ian Johnston, Dominic Kwiatkowski, Cordelia Langford, John Sillitoe on behalf of the Wellcome Sanger Institute COVID-19 Surveillance Team |
| EPI_ISL_2909316, EPI_ISL_2909317 | Lighthouse Lab in Milton Keynes | Wellcome Sanger Institute for the COVID-19 Genomics UK (COG-UK) Consortium | The Lighthouse Lab in Milton Keynes and Alex Alderton, Roberto Amato, Jeffrey Barrett, Sonia Goncalves, Ewan Harrison, David K. Jackson, Ian Johnston, Dominic Kwiatkowski, Cordelia Langford, John Sillitoe on behalf of the Wellcome Sanger Institute COVID-19 Surveillance Team |
| EPI_ISL_2909318, EPI_ISL_2909319, EPI_ISL_2909320 | Lighthouse Lab in Glasgow | Wellcome Sanger Institute for the COVID-19 Genomics UK (COG-UK) Consortium | Harper VanSteenhouse, Yumi Kasai, David Gray, Carol Clugston, Anna Dominiczak and Alex Alderton, Roberto Amato, Jeffrey Barrett, Sonia Goncalves, Ewan Harrison, David K. Jackson, Ian Johnston, Dominic Kwiatkowski, Cordelia Langford, John Sillitoe on behalf of the Wellcome Sanger Institute COVID-19 Surveillance Team |
| EPI_ISL_2909321 | Lighthouse Lab in Milton Keynes | Wellcome Sanger Institute for the COVID-19 Genomics UK (COG-UK) Consortium | The Lighthouse Lab in Milton Keynes and Alex Alderton, Roberto Amato, Jeffrey Barrett, Sonia Goncalves, Ewan Harrison, David K. Jackson, Ian Johnston, Dominic Kwiatkowski, Cordelia Langford, John Sillitoe on behalf of the Wellcome Sanger Institute COVID-19 Surveillance Team |
| EPI_ISL_2909322, EPI_ISL_2909323, EPI_ISL_2909324, EPI_ISL_2909325 | Lighthouse Lab in Glasgow | Wellcome Sanger Institute for the COVID-19 Genomics UK (COG-UK) Consortium | Harper VanSteenhouse, Yumi Kasai, David Gray, Carol Clugston, Anna Dominiczak and Alex Alderton, Roberto Amato, Jeffrey Barrett, Sonia Goncalves, Ewan Harrison, David K. Jackson, Ian Johnston, Dominic Kwiatkowski, Cordelia Langford, John Sillitoe on behalf of the Wellcome Sanger Institute COVID-19 Surveillance Team |
| EPI_ISL_2909326 | Lighthouse Lab in Milton Keynes | Wellcome Sanger Institute for the COVID-19 Genomics UK (COG-UK) Consortium | The Lighthouse Lab in Milton Keynes and Alex Alderton, Roberto Amato, Jeffrey Barrett, Sonia Goncalves, Ewan Harrison, David K. Jackson, Ian Johnston, Dominic Kwiatkowski, Cordelia Langford, John Sillitoe on behalf of the Wellcome Sanger Institute COVID-19 Surveillance Team |
| EPI_ISL_2909327, EPI_ISL_2909328 | Lighthouse Lab in Glasgow | Wellcome Sanger Institute for the COVID-19 Genomics UK (COG-UK) Consortium | Harper VanSteenhouse, Yumi Kasai, David Gray, Carol Clugston, Anna Dominiczak and Alex Alderton, Roberto Amato, Jeffrey Barrett, Sonia Goncalves, Ewan Harrison, David K. Jackson, Ian Johnston, Dominic Kwiatkowski, Cordelia Langford, John Sillitoe on behalf of the Wellcome Sanger Institute COVID-19 Surveillance Team |
| EPI_ISL_2909329 | Lighthouse Lab in Milton Keynes | Wellcome Sanger Institute for the COVID-19 Genomics UK (COG-UK) Consortium | The Lighthouse Lab in Milton Keynes and Alex Alderton, Roberto Amato, Jeffrey Barrett, Sonia Goncalves, Ewan Harrison, David K. Jackson, Ian Johnston, Dominic Kwiatkowski, Cordelia Langford, John Sillitoe on behalf of the Wellcome Sanger Institute COVID-19 Surveillance Team |
| EPI_ISL_2909330, EPI_ISL_2909331, EPI_ISL_2909332, EPI_ISL_2909333, EPI_ISL_2909334, EPI_ISL_2909335, EPI_ISL_2909336, EPI_ISL_2909337, EPI_ISL_2909338, EPI_ISL_2909339, EPI_ISL_2909340, EPI_ISL_2909341, EPI_ISL_2909342 | | | |

| | | | |
|---|---|---|---|
| see above | Lighthouse Lab in Glasgow | Wellcome Sanger Institute for the COVID-19 Genomics UK (COG-UK) Consortium | Harper VanSteenhouse, Yumi Kasai, David Gray, Carol Clugston, Anna Dominiczak and Alex Alderton, Roberto Amato, Jeffrey Barrett, Sonia Goncalves, Ewan Harrison, David K. Jackson, Ian Johnston, Dominic Kwiatkowski, Cordelia Langford, John Sillitoe on behalf of the Wellcome Sanger Institute COVID-19 Surveillance Team |
| EPI_ISL_2909343 | Lighthouse Lab in Milton Keynes | Wellcome Sanger Institute for the COVID-19 Genomics UK (COG-UK) Consortium | The Lighthouse Lab in Milton Keynes and Alex Alderton, Roberto Amato, Jeffrey Barrett, Sonia Goncalves, Ewan Harrison, David K. Jackson, Ian Johnston, Dominic Kwiatkowski, Cordelia Langford, John Sillitoe on behalf of the Wellcome Sanger Institute COVID-19 Surveillance Team |
| EPI_ISL_2909344, EPI_ISL_2909345 | Lighthouse Lab in Glasgow | Wellcome Sanger Institute for the COVID-19 Genomics UK (COG-UK) Consortium | Harper VanSteenhouse, Yumi Kasai, David Gray, Carol Clugston, Anna Dominiczak and Alex Alderton, Roberto Amato, Jeffrey Barrett, Sonia Goncalves, Ewan Harrison, David K. Jackson, Ian Johnston, Dominic Kwiatkowski, Cordelia Langford, John Sillitoe on behalf of the Wellcome Sanger Institute COVID-19 Surveillance Team |
| EPI_ISL_2909346 | Lighthouse Lab in Milton Keynes | Wellcome Sanger Institute for the COVID-19 Genomics UK (COG-UK) Consortium | The Lighthouse Lab in Milton Keynes and Alex Alderton, Roberto Amato, Jeffrey Barrett, Sonia Goncalves, Ewan Harrison, David K. Jackson, Ian Johnston, Dominic Kwiatkowski, Cordelia Langford, John Sillitoe on behalf of the Wellcome Sanger Institute COVID-19 Surveillance Team |
| EPI_ISL_2909347, EPI_ISL_2909348, EPI_ISL_2909349 | Lighthouse Lab in Glasgow | Wellcome Sanger Institute for the COVID-19 Genomics UK (COG-UK) Consortium | Harper VanSteenhouse, Yumi Kasai, David Gray, Carol Clugston, Anna Dominiczak and Alex Alderton, Roberto Amato, Jeffrey Barrett, Sonia Goncalves, Ewan Harrison, David K. Jackson, Ian Johnston, Dominic Kwiatkowski, Cordelia Langford, John Sillitoe on behalf of the Wellcome Sanger Institute COVID-19 Surveillance Team |
| EPI_ISL_2909350, EPI_ISL_2909351 | Lighthouse Lab in Milton Keynes | Wellcome Sanger Institute for the COVID-19 Genomics UK (COG-UK) Consortium | The Lighthouse Lab in Milton Keynes and Alex Alderton, Roberto Amato, Jeffrey Barrett, Sonia Goncalves, Ewan Harrison, David K. Jackson, Ian Johnston, Dominic Kwiatkowski, Cordelia Langford, John Sillitoe on behalf of the Wellcome Sanger Institute COVID-19 Surveillance Team |
| EPI_ISL_2909352, EPI_ISL_2909353, EPI_ISL_2909354, EPI_ISL_2909355, EPI_ISL_2909356 | Lighthouse Lab in Glasgow | Wellcome Sanger Institute for the COVID-19 Genomics UK (COG-UK) Consortium | Harper VanSteenhouse, Yumi Kasai, David Gray, Carol Clugston, Anna Dominiczak and Alex Alderton, Roberto Amato, Jeffrey Barrett, Sonia Goncalves, Ewan Harrison, David K. Jackson, Ian Johnston, Dominic Kwiatkowski, Cordelia Langford, John Sillitoe on behalf of the Wellcome Sanger Institute COVID-19 Surveillance Team |
| EPI_ISL_2909357 | Lighthouse Lab in Milton Keynes | Wellcome Sanger Institute for the COVID-19 Genomics UK (COG-UK) Consortium | The Lighthouse Lab in Milton Keynes and Alex Alderton, Roberto Amato, Jeffrey Barrett, Sonia Goncalves, Ewan Harrison, David K. Jackson, Ian Johnston, Dominic Kwiatkowski, Cordelia Langford, John Sillitoe on behalf of the Wellcome Sanger Institute COVID-19 Surveillance Team |
| EPI_ISL_2909358, EPI_ISL_2909359, EPI_ISL_2909360, EPI_ISL_2909361, EPI_ISL_2909362, EPI_ISL_2909363, EPI_ISL_2909364, EPI_ISL_2909365, EPI_ISL_2909366 | Lighthouse Lab in Glasgow | Wellcome Sanger Institute for the COVID-19 Genomics UK (COG-UK) Consortium | Harper VanSteenhouse, Yumi Kasai, David Gray, Carol Clugston, Anna Dominiczak and Alex Alderton, Roberto Amato, Jeffrey Barrett, Sonia Goncalves, Ewan Harrison, David K. Jackson, Ian Johnston, Dominic Kwiatkowski, Cordelia Langford, John Sillitoe on behalf of the Wellcome Sanger Institute COVID-19 Surveillance Team |
| EPI_ISL_2909367, EPI_ISL_2909368 | Lighthouse Lab in Milton Keynes | Wellcome Sanger Institute for the COVID-19 Genomics UK (COG-UK) Consortium | The Lighthouse Lab in Milton Keynes and Alex Alderton, Roberto Amato, Jeffrey Barrett, Sonia Goncalves, Ewan Harrison, David K. Jackson, Ian Johnston, Dominic Kwiatkowski, Cordelia Langford, John Sillitoe on behalf of the Wellcome Sanger Institute COVID-19 Surveillance Team |
| EPI_ISL_2909369, EPI_ISL_2909370, EPI_ISL_2909371, EPI_ISL_2909372, EPI_ISL_2909373, EPI_ISL_2909374, EPI_ISL_2909375, EPI_ISL_2909376, EPI_ISL_2909377, EPI_ISL_2909378 | Lighthouse Lab in Glasgow | Wellcome Sanger Institute for the COVID-19 Genomics UK (COG-UK) Consortium | Harper VanSteenhouse, Yumi Kasai, David Gray, Carol Clugston, Anna Dominiczak and Alex Alderton, Roberto Amato, Jeffrey Barrett, Sonia Goncalves, Ewan Harrison, David K. Jackson, Ian Johnston, Dominic Kwiatkowski, Cordelia Langford, John Sillitoe on behalf of the Wellcome Sanger Institute COVID-19 Surveillance Team |
| EPI_ISL_2909379 | Lighthouse Lab in Milton Keynes | Wellcome Sanger Institute for the COVID-19 Genomics UK (COG-UK) Consortium | The Lighthouse Lab in Milton Keynes and Alex Alderton, Roberto Amato, Jeffrey Barrett, Sonia Goncalves, Ewan Harrison, David K. Jackson, Ian Johnston, Dominic Kwiatkowski, Cordelia Langford, John Sillitoe on behalf of the Wellcome Sanger Institute COVID-19 Surveillance Team |
| EPI_ISL_2909380, EPI_ISL_2909381, EPI_ISL_2909382 | Lighthouse Lab in Glasgow | Wellcome Sanger Institute for the COVID-19 Genomics UK (COG-UK) Consortium | Harper VanSteenhouse, Yumi Kasai, David Gray, Carol Clugston, Anna Dominiczak and Alex Alderton, Roberto Amato, Jeffrey Barrett, Sonia Goncalves, Ewan Harrison, David K. Jackson, Ian Johnston, Dominic Kwiatkowski, Cordelia Langford, John Sillitoe on behalf of the Wellcome Sanger Institute COVID-19 Surveillance Team |
| EPI_ISL_2909383, EPI_ISL_2909384, EPI_ISL_2909385 | Lighthouse Lab in Milton Keynes | Wellcome Sanger Institute for the COVID-19 Genomics UK (COG-UK) Consortium | The Lighthouse Lab in Milton Keynes and Alex Alderton, Roberto Amato, Jeffrey Barrett, Sonia Goncalves, Ewan Harrison, David K. Jackson, Ian Johnston, Dominic Kwiatkowski, Cordelia Langford, John Sillitoe on behalf of the Wellcome Sanger Institute COVID-19 Surveillance Team |
| EPI_ISL_2909386 | Lighthouse Lab in Glasgow | Wellcome Sanger Institute for the COVID-19 Genomics UK (COG-UK) Consortium | Harper VanSteenhouse, Yumi Kasai, David Gray, Carol Clugston, Anna Dominiczak and Alex Alderton, Roberto Amato, Jeffrey Barrett, Sonia Goncalves, Ewan Harrison, David K. Jackson, Ian Johnston, Dominic Kwiatkowski, Cordelia Langford, John Sillitoe on behalf of the Wellcome Sanger Institute COVID-19 Surveillance Team |
| EPI_ISL_2909387 | Lighthouse Lab in Milton Keynes | Wellcome Sanger Institute for the COVID-19 Genomics UK (COG-UK) Consortium | The Lighthouse Lab in Milton Keynes and Alex Alderton, Roberto Amato, Jeffrey Barrett, Sonia Goncalves, Ewan Harrison, David K. Jackson, Ian Johnston, Dominic Kwiatkowski, Cordelia Langford, John Sillitoe on behalf of the Wellcome Sanger Institute COVID-19 Surveillance Team |
| EPI_ISL_2909388, EPI_ISL_2909389, EPI_ISL_2909390 | Lighthouse Lab in Glasgow | Wellcome Sanger Institute for the COVID-19 Genomics UK (COG-UK) Consortium | Harper VanSteenhouse, Yumi Kasai, David Gray, Carol Clugston, Anna Dominiczak and Alex Alderton, Roberto Amato, Jeffrey Barrett, Sonia Goncalves, Ewan Harrison, David K. Jackson, Ian Johnston, Dominic Kwiatkowski, Cordelia Langford, John Sillitoe on behalf of the Wellcome Sanger Institute COVID-19 Surveillance Team |
| EPI_ISL_2909391, EPI_ISL_2909392 | Lighthouse Lab in Milton Keynes | Wellcome Sanger Institute for the COVID-19 Genomics UK (COG-UK) Consortium | The Lighthouse Lab in Milton Keynes and Alex Alderton, Roberto Amato, Jeffrey Barrett, Sonia Goncalves, Ewan Harrison, David K. Jackson, Ian Johnston, Dominic Kwiatkowski, Cordelia Langford, John Sillitoe on behalf of the Wellcome Sanger Institute COVID-19 Surveillance Team |
| EPI_ISL_2909393 | Lighthouse Lab in Glasgow | Wellcome Sanger Institute for the COVID-19 Genomics UK (COG-UK) Consortium | Harper VanSteenhouse, Yumi Kasai, David Gray, Carol Clugston, Anna Dominiczak and Alex Alderton, Roberto Amato, Jeffrey Barrett, Sonia Goncalves, Ewan Harrison, David K. Jackson, Ian Johnston, Dominic Kwiatkowski, Cordelia Langford, John Sillitoe on behalf of the Wellcome Sanger Institute COVID-19 Surveillance Team |
| EPI_ISL_2909394 | Lighthouse Lab in Milton Keynes | Wellcome Sanger Institute for the COVID-19 Genomics UK (COG-UK) Consortium | The Lighthouse Lab in Milton Keynes and Alex Alderton, Roberto Amato, Jeffrey Barrett, Sonia Goncalves, Ewan Harrison, David K. Jackson, Ian Johnston, Dominic Kwiatkowski, Cordelia Langford, John Sillitoe on behalf of the Wellcome Sanger Institute COVID-19 Surveillance Team |
| EPI_ISL_2909395, EPI_ISL_2909396, EPI_ISL_2909397, EPI_ISL_2909398, EPI_ISL_2909399, EPI_ISL_2909400, EPI_ISL_2909401, EPI_ISL_2909402 | Lighthouse Lab in Glasgow | Wellcome Sanger Institute for the COVID-19 Genomics UK (COG-UK) Consortium | Harper VanSteenhouse, Yumi Kasai, David Gray, Carol Clugston, Anna Dominiczak and Alex Alderton, Roberto Amato, Jeffrey Barrett, Sonia Goncalves, Ewan Harrison, David K. Jackson, Ian Johnston, Dominic Kwiatkowski, Cordelia Langford, John Sillitoe on behalf of the Wellcome Sanger Institute COVID-19 Surveillance Team |
| EPI_ISL_2909403 | Lighthouse Lab in Milton Keynes | Wellcome Sanger Institute for the COVID-19 Genomics UK (COG-UK) Consortium | The Lighthouse Lab in Milton Keynes and Alex Alderton, Roberto Amato, Jeffrey Barrett, Sonia Goncalves, Ewan Harrison, David K. Jackson, Ian Johnston, Dominic Kwiatkowski, Cordelia Langford, John Sillitoe on behalf of the Wellcome Sanger Institute COVID-19 Surveillance Team |
| EPI_ISL_2909404, EPI_ISL_2909405, EPI_ISL_2909406, EPI_ISL_2909407 | Lighthouse Lab in Glasgow | Wellcome Sanger Institute for the COVID-19 Genomics UK (COG-UK) Consortium | Harper VanSteenhouse, Yumi Kasai, David Gray, Carol Clugston, Anna Dominiczak and Alex Alderton, Roberto Amato, Jeffrey Barrett, Sonia Goncalves, Ewan Harrison, David K. Jackson, Ian Johnston, Dominic Kwiatkowski, Cordelia Langford, John Sillitoe on behalf of the Wellcome Sanger Institute COVID-19 Surveillance Team |
| EPI_ISL_2909408 | Lighthouse Lab in Milton Keynes | Wellcome Sanger Institute for the COVID-19 Genomics UK (COG-UK) Consortium | The Lighthouse Lab in Milton Keynes and Alex Alderton, Roberto Amato, Jeffrey Barrett, Sonia Goncalves, Ewan Harrison, David K. Jackson, Ian Johnston, Dominic Kwiatkowski, Cordelia Langford, John Sillitoe on behalf of the Wellcome Sanger Institute COVID-19 Surveillance Team |
| EPI_ISL_2909409, EPI_ISL_2909410, EPI_ISL_2909411, EPI_ISL_2909412 | Lighthouse Lab in Glasgow | Wellcome Sanger Institute for the COVID-19 Genomics UK (COG-UK) Consortium | Harper VanSteenhouse, Yumi Kasai, David Gray, Carol Clugston, Anna Dominiczak and Alex Alderton, Roberto Amato, Jeffrey Barrett, Sonia Goncalves, Ewan Harrison, David K. Jackson, Ian Johnston, Dominic Kwiatkowski, Cordelia Langford, John Sillitoe on behalf of the Wellcome Sanger Institute COVID-19 Surveillance Team |
| EPI_ISL_2909413 | Lighthouse Lab in Milton Keynes | Wellcome Sanger Institute for the COVID-19 Genomics UK (COG-UK) Consortium | The Lighthouse Lab in Milton Keynes and Alex Alderton, Roberto Amato, Jeffrey Barrett, Sonia Goncalves, Ewan Harrison, David K. Jackson, Ian Johnston, Dominic Kwiatkowski, Cordelia Langford, John Sillitoe on behalf of the Wellcome Sanger Institute COVID-19 Surveillance Team |
| EPI_ISL_2909414, EPI_ISL_2909415, EPI_ISL_2909416 | Lighthouse Lab in Glasgow | Wellcome Sanger Institute for the COVID-19 Genomics UK (COG-UK) Consortium | Harper VanSteenhouse, Yumi Kasai, David Gray, Carol Clugston, Anna Dominiczak and Alex Alderton, Roberto Amato, Jeffrey Barrett, Sonia Goncalves, Ewan Harrison, David K. Jackson, Ian Johnston, Dominic Kwiatkowski, Cordelia Langford, John Sillitoe on behalf of the Wellcome Sanger Institute COVID-19 Surveillance Team |
| EPI_ISL_2909417 | Lighthouse Lab in Milton Keynes | Wellcome Sanger Institute for the COVID-19 Genomics UK (COG-UK) Consortium | The Lighthouse Lab in Milton Keynes and Alex Alderton, Roberto Amato, Jeffrey Barrett, Sonia Goncalves, Ewan Harrison, David K. Jackson, Ian Johnston, Dominic Kwiatkowski, Cordelia Langford, John Sillitoe on behalf of the Wellcome Sanger Institute COVID-19 Surveillance Team |
| EPI_ISL_2909418, EPI_ISL_2909419, EPI_ISL_2909420, EPI_ISL_2909421, EPI_ISL_2909422, EPI_ISL_2909423, EPI_ISL_2909424 | Lighthouse Lab in Glasgow | Wellcome Sanger Institute for the COVID-19 Genomics UK (COG-UK) Consortium | Harper VanSteenhouse, Yumi Kasai, David Gray, Carol Clugston, Anna Dominiczak and Alex Alderton, Roberto Amato, Jeffrey Barrett, Sonia Goncalves, Ewan Harrison, David K. Jackson, Ian Johnston, Dominic Kwiatkowski, Cordelia Langford, John Sillitoe on behalf of the Wellcome Sanger Institute COVID-19 Surveillance Team |
| EPI_ISL_2909425 | Lighthouse Lab in Milton Keynes | Wellcome Sanger Institute for the COVID-19 Genomics UK (COG-UK) | The Lighthouse Lab in Milton Keynes and Alex Alderton, Roberto Amato, Jeffrey Barrett, Sonia Goncalves, Ewan Harrison, David K. Jackson, Ian Johnston, Dominic Kwiatkowski, Cordelia Langford, John Sillitoe on behalf of the Wellcome Sanger |

| | | | |
|---|---|---|---|
| | | Consortium | Institute COVID-19 Surveillance Team |
| EPI_ISL_2909426, EPI_ISL_2909427, EPI_ISL_2909428, EPI_ISL_2909429, EPI_ISL_2909430, EPI_ISL_2909431 | Lighthouse Lab in Glasgow | Wellcome Sanger Institute for the COVID-19 Genomics UK (COG-UK) Consortium | Harper VanSteenhouse, Yumi Kasai, David Gray, Carol Clugston, Anna Dominiczak and Alex Alderton, Roberto Amato, Jeffrey Barrett, Sonia Goncalves, Ewan Harrison, David K. Jackson, Ian Johnston, Dominic Kwiatkowski, Cordelia Langford, John Sillitoe on behalf of the Wellcome Sanger Institute COVID-19 Surveillance Team |
| EPI_ISL_2909432 | Lighthouse Lab in Milton Keynes | Wellcome Sanger Institute for the COVID-19 Genomics UK (COG-UK) Consortium | The Lighthouse Lab in Milton Keynes and Alex Alderton, Roberto Amato, Jeffrey Barrett, Sonia Goncalves, Ewan Harrison, David K. Jackson, Ian Johnston, Dominic Kwiatkowski, Cordelia Langford, John Sillitoe on behalf of the Wellcome Sanger Institute COVID-19 Surveillance Team |
| EPI_ISL_2909433 | Lighthouse Lab in Glasgow | Wellcome Sanger Institute for the COVID-19 Genomics UK (COG-UK) Consortium | Harper VanSteenhouse, Yumi Kasai, David Gray, Carol Clugston, Anna Dominiczak and Alex Alderton, Roberto Amato, Jeffrey Barrett, Sonia Goncalves, Ewan Harrison, David K. Jackson, Ian Johnston, Dominic Kwiatkowski, Cordelia Langford, John Sillitoe on behalf of the Wellcome Sanger Institute COVID-19 Surveillance Team |
| EPI_ISL_2909434 | Lighthouse Lab in Milton Keynes | Wellcome Sanger Institute for the COVID-19 Genomics UK (COG-UK) Consortium | The Lighthouse Lab in Milton Keynes and Alex Alderton, Roberto Amato, Jeffrey Barrett, Sonia Goncalves, Ewan Harrison, David K. Jackson, Ian Johnston, Dominic Kwiatkowski, Cordelia Langford, John Sillitoe on behalf of the Wellcome Sanger Institute COVID-19 Surveillance Team |
| EPI_ISL_2909435, EPI_ISL_2909436, EPI_ISL_2909437 | Lighthouse Lab in Glasgow | Wellcome Sanger Institute for the COVID-19 Genomics UK (COG-UK) Consortium | Harper VanSteenhouse, Yumi Kasai, David Gray, Carol Clugston, Anna Dominiczak and Alex Alderton, Roberto Amato, Jeffrey Barrett, Sonia Goncalves, Ewan Harrison, David K. Jackson, Ian Johnston, Dominic Kwiatkowski, Cordelia Langford, John Sillitoe on behalf of the Wellcome Sanger Institute COVID-19 Surveillance Team |
| EPI_ISL_2909438 | Lighthouse Lab in Milton Keynes | Wellcome Sanger Institute for the COVID-19 Genomics UK (COG-UK) Consortium | The Lighthouse Lab in Milton Keynes and Alex Alderton, Roberto Amato, Jeffrey Barrett, Sonia Goncalves, Ewan Harrison, David K. Jackson, Ian Johnston, Dominic Kwiatkowski, Cordelia Langford, John Sillitoe on behalf of the Wellcome Sanger Institute COVID-19 Surveillance Team |
| EPI_ISL_2909439, EPI_ISL_2909440, EPI_ISL_2909441, EPI_ISL_2909442 | Lighthouse Lab in Glasgow | Wellcome Sanger Institute for the COVID-19 Genomics UK (COG-UK) Consortium | Harper VanSteenhouse, Yumi Kasai, David Gray, Carol Clugston, Anna Dominiczak and Alex Alderton, Roberto Amato, Jeffrey Barrett, Sonia Goncalves, Ewan Harrison, David K. Jackson, Ian Johnston, Dominic Kwiatkowski, Cordelia Langford, John Sillitoe on behalf of the Wellcome Sanger Institute COVID-19 Surveillance Team |
| EPI_ISL_2909443 | Lighthouse Lab in Milton Keynes | Wellcome Sanger Institute for the COVID-19 Genomics UK (COG-UK) Consortium | The Lighthouse Lab in Milton Keynes and Alex Alderton, Roberto Amato, Jeffrey Barrett, Sonia Goncalves, Ewan Harrison, David K. Jackson, Ian Johnston, Dominic Kwiatkowski, Cordelia Langford, John Sillitoe on behalf of the Wellcome Sanger Institute COVID-19 Surveillance Team |
| EPI_ISL_2909444, EPI_ISL_2909445, EPI_ISL_2909446 | Lighthouse Lab in Glasgow | Wellcome Sanger Institute for the COVID-19 Genomics UK (COG-UK) Consortium | Harper VanSteenhouse, Yumi Kasai, David Gray, Carol Clugston, Anna Dominiczak and Alex Alderton, Roberto Amato, Jeffrey Barrett, Sonia Goncalves, Ewan Harrison, David K. Jackson, Ian Johnston, Dominic Kwiatkowski, Cordelia Langford, John Sillitoe on behalf of the Wellcome Sanger Institute COVID-19 Surveillance Team |
| EPI_ISL_2909447 | Lighthouse Lab in Milton Keynes | Wellcome Sanger Institute for the COVID-19 Genomics UK (COG-UK) Consortium | The Lighthouse Lab in Milton Keynes and Alex Alderton, Roberto Amato, Jeffrey Barrett, Sonia Goncalves, Ewan Harrison, David K. Jackson, Ian Johnston, Dominic Kwiatkowski, Cordelia Langford, John Sillitoe on behalf of the Wellcome Sanger Institute COVID-19 Surveillance Team |
| EPI_ISL_2909448, EPI_ISL_2909449, EPI_ISL_2909450, EPI_ISL_2909451, EPI_ISL_2909452, EPI_ISL_2909453, EPI_ISL_2909454, EPI_ISL_2909455 | Lighthouse Lab in Glasgow | Wellcome Sanger Institute for the COVID-19 Genomics UK (COG-UK) Consortium | Harper VanSteenhouse, Yumi Kasai, David Gray, Carol Clugston, Anna Dominiczak and Alex Alderton, Roberto Amato, Jeffrey Barrett, Sonia Goncalves, Ewan Harrison, David K. Jackson, Ian Johnston, Dominic Kwiatkowski, Cordelia Langford, John Sillitoe on behalf of the Wellcome Sanger Institute COVID-19 Surveillance Team |
| EPI_ISL_2909456, EPI_ISL_2909457, EPI_ISL_2909458 | Lighthouse Lab in Milton Keynes | Wellcome Sanger Institute for the COVID-19 Genomics UK (COG-UK) Consortium | The Lighthouse Lab in Milton Keynes and Alex Alderton, Roberto Amato, Jeffrey Barrett, Sonia Goncalves, Ewan Harrison, David K. Jackson, Ian Johnston, Dominic Kwiatkowski, Cordelia Langford, John Sillitoe on behalf of the Wellcome Sanger Institute COVID-19 Surveillance Team |
| EPI_ISL_2909459, EPI_ISL_2909460, EPI_ISL_2909461, EPI_ISL_2909462 | Lighthouse Lab in Glasgow | Wellcome Sanger Institute for the COVID-19 Genomics UK (COG-UK) Consortium | Harper VanSteenhouse, Yumi Kasai, David Gray, Carol Clugston, Anna Dominiczak and Alex Alderton, Roberto Amato, Jeffrey Barrett, Sonia Goncalves, Ewan Harrison, David K. Jackson, Ian Johnston, Dominic Kwiatkowski, Cordelia Langford, John Sillitoe on behalf of the Wellcome Sanger Institute COVID-19 Surveillance Team |
| EPI_ISL_2909463, EPI_ISL_2909464 | Lighthouse Lab in Milton Keynes | Wellcome Sanger Institute for the COVID-19 Genomics UK (COG-UK) Consortium | The Lighthouse Lab in Milton Keynes and Alex Alderton, Roberto Amato, Jeffrey Barrett, Sonia Goncalves, Ewan Harrison, David K. Jackson, Ian Johnston, Dominic Kwiatkowski, Cordelia Langford, John Sillitoe on behalf of the Wellcome Sanger Institute COVID-19 Surveillance Team |
| EPI_ISL_2909465, EPI_ISL_2909466, EPI_ISL_2909467, EPI_ISL_2909468 | Lighthouse Lab in Glasgow | Wellcome Sanger Institute for the COVID-19 Genomics UK (COG-UK) Consortium | Harper VanSteenhouse, Yumi Kasai, David Gray, Carol Clugston, Anna Dominiczak and Alex Alderton, Roberto Amato, Jeffrey Barrett, Sonia Goncalves, Ewan Harrison, David K. Jackson, Ian Johnston, Dominic Kwiatkowski, Cordelia Langford, John Sillitoe on behalf of the Wellcome Sanger Institute COVID-19 Surveillance Team |
| EPI_ISL_2909469 | Lighthouse Lab in Milton Keynes | Wellcome Sanger Institute for the COVID-19 Genomics UK (COG-UK) Consortium | The Lighthouse Lab in Milton Keynes and Alex Alderton, Roberto Amato, Jeffrey Barrett, Sonia Goncalves, Ewan Harrison, David K. Jackson, Ian Johnston, Dominic Kwiatkowski, Cordelia Langford, John Sillitoe on behalf of the Wellcome Sanger Institute COVID-19 Surveillance Team |
| EPI_ISL_2909470, EPI_ISL_2909471 | Lighthouse Lab in Glasgow | Wellcome Sanger Institute for the COVID-19 Genomics UK (COG-UK) Consortium | Harper VanSteenhouse, Yumi Kasai, David Gray, Carol Clugston, Anna Dominiczak and Alex Alderton, Roberto Amato, Jeffrey Barrett, Sonia Goncalves, Ewan Harrison, David K. Jackson, Ian Johnston, Dominic Kwiatkowski, Cordelia Langford, John Sillitoe on behalf of the Wellcome Sanger Institute COVID-19 Surveillance Team |
| EPI_ISL_2909472 | Lighthouse Lab in Milton Keynes | Wellcome Sanger Institute for the COVID-19 Genomics UK (COG-UK) Consortium | The Lighthouse Lab in Milton Keynes and Alex Alderton, Roberto Amato, Jeffrey Barrett, Sonia Goncalves, Ewan Harrison, David K. Jackson, Ian Johnston, Dominic Kwiatkowski, Cordelia Langford, John Sillitoe on behalf of the Wellcome Sanger Institute COVID-19 Surveillance Team |
| EPI_ISL_2909473 | Lighthouse Lab in Glasgow | Wellcome Sanger Institute for the COVID-19 Genomics UK (COG-UK) Consortium | Harper VanSteenhouse, Yumi Kasai, David Gray, Carol Clugston, Anna Dominiczak and Alex Alderton, Roberto Amato, Jeffrey Barrett, Sonia Goncalves, Ewan Harrison, David K. Jackson, Ian Johnston, Dominic Kwiatkowski, Cordelia Langford, John Sillitoe on behalf of the Wellcome Sanger Institute COVID-19 Surveillance Team |
| EPI_ISL_2909474 | Lighthouse Lab in Milton Keynes | Wellcome Sanger Institute for the COVID-19 Genomics UK (COG-UK) Consortium | The Lighthouse Lab in Milton Keynes and Alex Alderton, Roberto Amato, Jeffrey Barrett, Sonia Goncalves, Ewan Harrison, David K. Jackson, Ian Johnston, Dominic Kwiatkowski, Cordelia Langford, John Sillitoe on behalf of the Wellcome Sanger Institute COVID-19 Surveillance Team |
| EPI_ISL_2909475 | Lighthouse Lab in Glasgow | Wellcome Sanger Institute for the COVID-19 Genomics UK (COG-UK) Consortium | Harper VanSteenhouse, Yumi Kasai, David Gray, Carol Clugston, Anna Dominiczak and Alex Alderton, Roberto Amato, Jeffrey Barrett, Sonia Goncalves, Ewan Harrison, David K. Jackson, Ian Johnston, Dominic Kwiatkowski, Cordelia Langford, John Sillitoe on behalf of the Wellcome Sanger Institute COVID-19 Surveillance Team |
| EPI_ISL_2909476 | Lighthouse Lab in Milton Keynes | Wellcome Sanger Institute for the COVID-19 Genomics UK (COG-UK) Consortium | The Lighthouse Lab in Milton Keynes and Alex Alderton, Roberto Amato, Jeffrey Barrett, Sonia Goncalves, Ewan Harrison, David K. Jackson, Ian Johnston, Dominic Kwiatkowski, Cordelia Langford, John Sillitoe on behalf of the Wellcome Sanger Institute COVID-19 Surveillance Team |
| EPI_ISL_2909477, EPI_ISL_2909478, EPI_ISL_2909479, EPI_ISL_2909480 | Lighthouse Lab in Glasgow | Wellcome Sanger Institute for the COVID-19 Genomics UK (COG-UK) Consortium | Harper VanSteenhouse, Yumi Kasai, David Gray, Carol Clugston, Anna Dominiczak and Alex Alderton, Roberto Amato, Jeffrey Barrett, Sonia Goncalves, Ewan Harrison, David K. Jackson, Ian Johnston, Dominic Kwiatkowski, Cordelia Langford, John Sillitoe on behalf of the Wellcome Sanger Institute COVID-19 Surveillance Team |
| EPI_ISL_2909481 | Lighthouse Lab in Milton Keynes | Wellcome Sanger Institute for the COVID-19 Genomics UK (COG-UK) Consortium | The Lighthouse Lab in Milton Keynes and Alex Alderton, Roberto Amato, Jeffrey Barrett, Sonia Goncalves, Ewan Harrison, David K. Jackson, Ian Johnston, Dominic Kwiatkowski, Cordelia Langford, John Sillitoe on behalf of the Wellcome Sanger Institute COVID-19 Surveillance Team |
| EPI_ISL_2909482, EPI_ISL_2909483, EPI_ISL_2909484, EPI_ISL_2909485, EPI_ISL_2909486, EPI_ISL_2909487, EPI_ISL_2909488, EPI_ISL_2909489, EPI_ISL_2909490, EPI_ISL_2909491, EPI_ISL_2909492, EPI_ISL_2909493, EPI_ISL_2909494, EPI_ISL_2909495 | | | |
| see above | Lighthouse Lab in Glasgow | Wellcome Sanger Institute for the COVID-19 Genomics UK (COG-UK) Consortium | Harper VanSteenhouse, Yumi Kasai, David Gray, Carol Clugston, Anna Dominiczak and Alex Alderton, Roberto Amato, Jeffrey Barrett, Sonia Goncalves, Ewan Harrison, David K. Jackson, Ian Johnston, Dominic Kwiatkowski, Cordelia Langford, John Sillitoe on behalf of the Wellcome Sanger Institute COVID-19 Surveillance Team |
| EPI_ISL_2909496 | Lighthouse Lab in Milton Keynes | Wellcome Sanger Institute for the COVID-19 Genomics UK (COG-UK) Consortium | The Lighthouse Lab in Milton Keynes and Alex Alderton, Roberto Amato, Jeffrey Barrett, Sonia Goncalves, Ewan Harrison, David K. Jackson, Ian Johnston, Dominic Kwiatkowski, Cordelia Langford, John Sillitoe on behalf of the Wellcome Sanger Institute COVID-19 Surveillance Team |
| EPI_ISL_2909497, EPI_ISL_2909498, EPI_ISL_2909499, EPI_ISL_2909500, EPI_ISL_2909501 | Lighthouse Lab in Glasgow | Wellcome Sanger Institute for the COVID-19 Genomics UK (COG-UK) Consortium | Harper VanSteenhouse, Yumi Kasai, David Gray, Carol Clugston, Anna Dominiczak and Alex Alderton, Roberto Amato, Jeffrey Barrett, Sonia Goncalves, Ewan Harrison, David K. Jackson, Ian Johnston, Dominic Kwiatkowski, Cordelia Langford, John Sillitoe on behalf of the Wellcome Sanger Institute COVID-19 Surveillance Team |
| EPI_ISL_2909502 | Lighthouse Lab in Milton Keynes | Wellcome Sanger Institute for the COVID-19 Genomics UK (COG-UK) Consortium | The Lighthouse Lab in Milton Keynes and Alex Alderton, Roberto Amato, Jeffrey Barrett, Sonia Goncalves, Ewan Harrison, David K. Jackson, Ian Johnston, Dominic Kwiatkowski, Cordelia Langford, John Sillitoe on behalf of the Wellcome Sanger Institute COVID-19 Surveillance Team |
| EPI_ISL_2909503, EPI_ISL_2909504, EPI_ISL_2909505, EPI_ISL_2909506, EPI_ISL_2909507 | Lighthouse Lab in Glasgow | Wellcome Sanger Institute for the COVID-19 Genomics UK (COG-UK) Consortium | Harper VanSteenhouse, Yumi Kasai, David Gray, Carol Clugston, Anna Dominiczak and Alex Alderton, Roberto Amato, Jeffrey Barrett, Sonia Goncalves, Ewan Harrison, David K. Jackson, Ian Johnston, Dominic Kwiatkowski, Cordelia Langford, John Sillitoe on behalf of the Wellcome Sanger Institute COVID-19 Surveillance Team |
| EPI_ISL_2909508, EPI_ISL_2909509 | Lighthouse Lab in Milton Keynes | Wellcome Sanger Institute for the COVID-19 Genomics UK (COG-UK) Consortium | The Lighthouse Lab in Milton Keynes and Alex Alderton, Roberto Amato, Jeffrey Barrett, Sonia Goncalves, Ewan Harrison, David K. Jackson, Ian Johnston, Dominic Kwiatkowski, Cordelia Langford, John Sillitoe on behalf of the Wellcome Sanger Institute COVID-19 Surveillance Team |
| EPI_ISL_2909510, EPI_ISL_2909511, EPI_ISL_2909512 | Lighthouse Lab in Glasgow | Wellcome Sanger Institute for the COVID-19 Genomics UK (COG-UK) Consortium | Harper VanSteenhouse, Yumi Kasai, David Gray, Carol Clugston, Anna Dominiczak and Alex Alderton, Roberto Amato, Jeffrey Barrett, Sonia Goncalves, Ewan Harrison, David K. Jackson, Ian Johnston, Dominic Kwiatkowski, Cordelia Langford, John Sillitoe on behalf of the Wellcome Sanger Institute COVID-19 Surveillance Team |
| EPI_ISL_2909513 | Lighthouse Lab in Milton Keynes | Wellcome Sanger Institute for the COVID-19 Genomics UK (COG-UK) Consortium | The Lighthouse Lab in Milton Keynes and Alex Alderton, Roberto Amato, Jeffrey Barrett, Sonia Goncalves, Ewan Harrison, David K. Jackson, Ian Johnston, Dominic Kwiatkowski, Cordelia Langford, John Sillitoe on behalf of the Wellcome Sanger Institute COVID-19 Surveillance Team |
| EPI_ISL_2909514, | Lighthouse Lab in Glasgow | Wellcome Sanger Institute for the COVID-19 Genomics UK (COG-UK) | Harper VanSteenhouse, Yumi Kasai, David Gray, Carol Clugston, Anna Dominiczak and Alex Alderton, Roberto Amato, Jeffrey Barrett, Sonia Goncalves, Ewan Harrison, David K. Jackson, Ian Johnston, Dominic Kwiatkowski, Cordelia Langford, |

| | | | |
|---|---|---|---|
| EPI_ISL_2909515 | | Consortium | John Sillitoe on behalf of the Wellcome Sanger Institute COVID-19 Surveillance Team |
| EPI_ISL_2909516 | Lighthouse Lab in Milton Keynes | Wellcome Sanger Institute for the COVID-19 Genomics UK (COG-UK) Consortium | The Lighthouse Lab in Milton Keynes and Alex Alderton, Roberto Amato, Jeffrey Barrett, Sonia Goncalves, Ewan Harrison, David K. Jackson, Ian Johnston, Dominic Kwiatkowski, Cordelia Langford, John Sillitoe on behalf of the Wellcome Sanger Institute COVID-19 Surveillance Team |
| EPI_ISL_2909517, EPI_ISL_2909518, EPI_ISL_2909519 | Lighthouse Lab in Glasgow | Wellcome Sanger Institute for the COVID-19 Genomics UK (COG-UK) Consortium | Harper VanSteenhouse, Yumi Kasai, David Gray, Carol Clugston, Anna Dominiczak and Alex Alderton, Roberto Amato, Jeffrey Barrett, Sonia Goncalves, Ewan Harrison, David K. Jackson, Ian Johnston, Dominic Kwiatkowski, Cordelia Langford, John Sillitoe on behalf of the Wellcome Sanger Institute COVID-19 Surveillance Team |
| EPI_ISL_2909520 | Lighthouse Lab in Milton Keynes | Wellcome Sanger Institute for the COVID-19 Genomics UK (COG-UK) Consortium | The Lighthouse Lab in Milton Keynes and Alex Alderton, Roberto Amato, Jeffrey Barrett, Sonia Goncalves, Ewan Harrison, David K. Jackson, Ian Johnston, Dominic Kwiatkowski, Cordelia Langford, John Sillitoe on behalf of the Wellcome Sanger Institute COVID-19 Surveillance Team |
| EPI_ISL_2909521, EPI_ISL_2909522, EPI_ISL_2909523, EPI_ISL_2909524, EPI_ISL_2909525, EPI_ISL_2909526, EPI_ISL_2909527, EPI_ISL_2909528, EPI_ISL_2909529 | Lighthouse Lab in Glasgow | Wellcome Sanger Institute for the COVID-19 Genomics UK (COG-UK) Consortium | Harper VanSteenhouse, Yumi Kasai, David Gray, Carol Clugston, Anna Dominiczak and Alex Alderton, Roberto Amato, Jeffrey Barrett, Sonia Goncalves, Ewan Harrison, David K. Jackson, Ian Johnston, Dominic Kwiatkowski, Cordelia Langford, John Sillitoe on behalf of the Wellcome Sanger Institute COVID-19 Surveillance Team |
| EPI_ISL_2909530 | Lighthouse Lab in Milton Keynes | Wellcome Sanger Institute for the COVID-19 Genomics UK (COG-UK) Consortium | The Lighthouse Lab in Milton Keynes and Alex Alderton, Roberto Amato, Jeffrey Barrett, Sonia Goncalves, Ewan Harrison, David K. Jackson, Ian Johnston, Dominic Kwiatkowski, Cordelia Langford, John Sillitoe on behalf of the Wellcome Sanger Institute COVID-19 Surveillance Team |
| EPI_ISL_2909531 | Lighthouse Lab in Glasgow | Wellcome Sanger Institute for the COVID-19 Genomics UK (COG-UK) Consortium | Harper VanSteenhouse, Yumi Kasai, David Gray, Carol Clugston, Anna Dominiczak and Alex Alderton, Roberto Amato, Jeffrey Barrett, Sonia Goncalves, Ewan Harrison, David K. Jackson, Ian Johnston, Dominic Kwiatkowski, Cordelia Langford, John Sillitoe on behalf of the Wellcome Sanger Institute COVID-19 Surveillance Team |
| EPI_ISL_2909532 | Lighthouse Lab in Milton Keynes | Wellcome Sanger Institute for the COVID-19 Genomics UK (COG-UK) Consortium | The Lighthouse Lab in Milton Keynes and Alex Alderton, Roberto Amato, Jeffrey Barrett, Sonia Goncalves, Ewan Harrison, David K. Jackson, Ian Johnston, Dominic Kwiatkowski, Cordelia Langford, John Sillitoe on behalf of the Wellcome Sanger Institute COVID-19 Surveillance Team |
| EPI_ISL_2909533, EPI_ISL_2909534, EPI_ISL_2909535, EPI_ISL_2909536, EPI_ISL_2909537, EPI_ISL_2909538 | Lighthouse Lab in Glasgow | Wellcome Sanger Institute for the COVID-19 Genomics UK (COG-UK) Consortium | Harper VanSteenhouse, Yumi Kasai, David Gray, Carol Clugston, Anna Dominiczak and Alex Alderton, Roberto Amato, Jeffrey Barrett, Sonia Goncalves, Ewan Harrison, David K. Jackson, Ian Johnston, Dominic Kwiatkowski, Cordelia Langford, John Sillitoe on behalf of the Wellcome Sanger Institute COVID-19 Surveillance Team |
| EPI_ISL_2909539 | Lighthouse Lab in Milton Keynes | Wellcome Sanger Institute for the COVID-19 Genomics UK (COG-UK) Consortium | The Lighthouse Lab in Milton Keynes and Alex Alderton, Roberto Amato, Jeffrey Barrett, Sonia Goncalves, Ewan Harrison, David K. Jackson, Ian Johnston, Dominic Kwiatkowski, Cordelia Langford, John Sillitoe on behalf of the Wellcome Sanger Institute COVID-19 Surveillance Team |
| EPI_ISL_2909540, EPI_ISL_2909541, EPI_ISL_2909542, EPI_ISL_2909543 | Lighthouse Lab in Glasgow | Wellcome Sanger Institute for the COVID-19 Genomics UK (COG-UK) Consortium | Harper VanSteenhouse, Yumi Kasai, David Gray, Carol Clugston, Anna Dominiczak and Alex Alderton, Roberto Amato, Jeffrey Barrett, Sonia Goncalves, Ewan Harrison, David K. Jackson, Ian Johnston, Dominic Kwiatkowski, Cordelia Langford, John Sillitoe on behalf of the Wellcome Sanger Institute COVID-19 Surveillance Team |
| EPI_ISL_2909544 | Lighthouse Lab in Milton Keynes | Wellcome Sanger Institute for the COVID-19 Genomics UK (COG-UK) Consortium | The Lighthouse Lab in Milton Keynes and Alex Alderton, Roberto Amato, Jeffrey Barrett, Sonia Goncalves, Ewan Harrison, David K. Jackson, Ian Johnston, Dominic Kwiatkowski, Cordelia Langford, John Sillitoe on behalf of the Wellcome Sanger Institute COVID-19 Surveillance Team |
| EPI_ISL_2909545 | Lighthouse Lab in Glasgow | Wellcome Sanger Institute for the COVID-19 Genomics UK (COG-UK) Consortium | Harper VanSteenhouse, Yumi Kasai, David Gray, Carol Clugston, Anna Dominiczak and Alex Alderton, Roberto Amato, Jeffrey Barrett, Sonia Goncalves, Ewan Harrison, David K. Jackson, Ian Johnston, Dominic Kwiatkowski, Cordelia Langford, John Sillitoe on behalf of the Wellcome Sanger Institute COVID-19 Surveillance Team |
| EPI_ISL_2909546 | Lighthouse Lab in Alderley Park | Wellcome Sanger Institute for the COVID-19 Genomics UK (COG-UK) Consortium | Jacquelyn Wynn, Mairead Hyland, The Lighthouse Lab in Alderley Park and Alex Alderton, Roberto Amato, Jeffrey Barrett, Sonia Goncalves, Ewan Harrison, David K. Jackson, Ian Johnston, Dominic Kwiatkowski, Cordelia Langford, John Sillitoe on behalf of the Wellcome Sanger Institute COVID-19 Surveillance Team |
| EPI_ISL_2909547 | Berkshire and Surrey Pathology Services Lighthouse Laboratory | Wellcome Sanger Institute for the COVID-19 Genomics UK (COG-UK) Consortium | Berkshire and Surrey Pathology Services Lighthouse Laboratory and Alex Alderton, Roberto Amato, Jeffrey Barrett, Sonia Goncalves, Ewan Harrison, David K. Jackson, Ian Johnston, Dominic Kwiatkowski, Cordelia Langford, John Sillitoe on behalf of the Wellcome Sanger Institute COVID-19 Surveillance Team |
| EPI_ISL_2909548 | Lighthouse Laboratory Plymouth | Wellcome Sanger Institute for the COVID-19 Genomics UK (COG-UK) Consortium | Lighthouse Laboratory Plymouth and Alex Alderton, Roberto Amato, Jeffrey Barrett, Sonia Goncalves, Ewan Harrison, David K. Jackson, Ian Johnston, Dominic Kwiatkowski, Cordelia Langford, John Sillitoe on behalf of the Wellcome Sanger Institute COVID-19 Surveillance Team |
| EPI_ISL_2909549, EPI_ISL_2909550 | Lighthouse Lab in Alderley Park | Wellcome Sanger Institute for the COVID-19 Genomics UK (COG-UK) Consortium | Jacquelyn Wynn, Mairead Hyland, The Lighthouse Lab in Alderley Park and Alex Alderton, Roberto Amato, Jeffrey Barrett, Sonia Goncalves, Ewan Harrison, David K. Jackson, Ian Johnston, Dominic Kwiatkowski, Cordelia Langford, John Sillitoe on behalf of the Wellcome Sanger Institute COVID-19 Surveillance Team |
| EPI_ISL_2909551 | Berkshire and Surrey Pathology Services Lighthouse Laboratory | Wellcome Sanger Institute for the COVID-19 Genomics UK (COG-UK) Consortium | Berkshire and Surrey Pathology Services Lighthouse Laboratory and Alex Alderton, Roberto Amato, Jeffrey Barrett, Sonia Goncalves, Ewan Harrison, David K. Jackson, Ian Johnston, Dominic Kwiatkowski, Cordelia Langford, John Sillitoe on behalf of the Wellcome Sanger Institute COVID-19 Surveillance Team |
| EPI_ISL_2909552, EPI_ISL_2909553 | Lighthouse Lab in Alderley Park | Wellcome Sanger Institute for the COVID-19 Genomics UK (COG-UK) Consortium | Jacquelyn Wynn, Mairead Hyland, The Lighthouse Lab in Alderley Park and Alex Alderton, Roberto Amato, Jeffrey Barrett, Sonia Goncalves, Ewan Harrison, David K. Jackson, Ian Johnston, Dominic Kwiatkowski, Cordelia Langford, John Sillitoe on behalf of the Wellcome Sanger Institute COVID-19 Surveillance Team |
| EPI_ISL_2909554, EPI_ISL_2909555, EPI_ISL_2909556 | Lighthouse Laboratory Plymouth | Wellcome Sanger Institute for the COVID-19 Genomics UK (COG-UK) Consortium | Lighthouse Laboratory Plymouth and Alex Alderton, Roberto Amato, Jeffrey Barrett, Sonia Goncalves, Ewan Harrison, David K. Jackson, Ian Johnston, Dominic Kwiatkowski, Cordelia Langford, John Sillitoe on behalf of the Wellcome Sanger Institute COVID-19 Surveillance Team |
| EPI_ISL_2909557, EPI_ISL_2909558 | Lighthouse Lab in Alderley Park | Wellcome Sanger Institute for the COVID-19 Genomics UK (COG-UK) Consortium | Jacquelyn Wynn, Mairead Hyland, The Lighthouse Lab in Alderley Park and Alex Alderton, Roberto Amato, Jeffrey Barrett, Sonia Goncalves, Ewan Harrison, David K. Jackson, Ian Johnston, Dominic Kwiatkowski, Cordelia Langford, John Sillitoe on behalf of the Wellcome Sanger Institute COVID-19 Surveillance Team |
| EPI_ISL_2909559 | Berkshire and Surrey Pathology Services Lighthouse Laboratory | Wellcome Sanger Institute for the COVID-19 Genomics UK (COG-UK) Consortium | Berkshire and Surrey Pathology Services Lighthouse Laboratory and Alex Alderton, Roberto Amato, Jeffrey Barrett, Sonia Goncalves, Ewan Harrison, David K. Jackson, Ian Johnston, Dominic Kwiatkowski, Cordelia Langford, John Sillitoe on behalf of the Wellcome Sanger Institute COVID-19 Surveillance Team |
| EPI_ISL_2909560, EPI_ISL_2909561, EPI_ISL_2909562, EPI_ISL_2909563, EPI_ISL_2909564, EPI_ISL_2909565 | Lighthouse Lab in Alderley Park | Wellcome Sanger Institute for the COVID-19 Genomics UK (COG-UK) Consortium | Jacquelyn Wynn, Mairead Hyland, The Lighthouse Lab in Alderley Park and Alex Alderton, Roberto Amato, Jeffrey Barrett, Sonia Goncalves, Ewan Harrison, David K. Jackson, Ian Johnston, Dominic Kwiatkowski, Cordelia Langford, John Sillitoe on behalf of the Wellcome Sanger Institute COVID-19 Surveillance Team |
| EPI_ISL_2909566 | Lighthouse Laboratory Plymouth | Wellcome Sanger Institute for the COVID-19 Genomics UK (COG-UK) Consortium | Lighthouse Laboratory Plymouth and Alex Alderton, Roberto Amato, Jeffrey Barrett, Sonia Goncalves, Ewan Harrison, David K. Jackson, Ian Johnston, Dominic Kwiatkowski, Cordelia Langford, John Sillitoe on behalf of the Wellcome Sanger Institute COVID-19 Surveillance Team |
| EPI_ISL_2909567 | Lighthouse Lab in Alderley Park | Wellcome Sanger Institute for the COVID-19 Genomics UK (COG-UK) Consortium | Jacquelyn Wynn, Mairead Hyland, The Lighthouse Lab in Alderley Park and Alex Alderton, Roberto Amato, Jeffrey Barrett, Sonia Goncalves, Ewan Harrison, David K. Jackson, Ian Johnston, Dominic Kwiatkowski, Cordelia Langford, John Sillitoe on behalf of the Wellcome Sanger Institute COVID-19 Surveillance Team |
| EPI_ISL_2909568 | Lighthouse Laboratory Plymouth | Wellcome Sanger Institute for the COVID-19 Genomics UK (COG-UK) Consortium | Lighthouse Laboratory Plymouth and Alex Alderton, Roberto Amato, Jeffrey Barrett, Sonia Goncalves, Ewan Harrison, David K. Jackson, Ian Johnston, Dominic Kwiatkowski, Cordelia Langford, John Sillitoe on behalf of the Wellcome Sanger Institute COVID-19 Surveillance Team |
| EPI_ISL_2909569, EPI_ISL_2909570 | Lighthouse Lab in Alderley Park | Wellcome Sanger Institute for the COVID-19 Genomics UK (COG-UK) Consortium | Jacquelyn Wynn, Mairead Hyland, The Lighthouse Lab in Alderley Park and Alex Alderton, Roberto Amato, Jeffrey Barrett, Sonia Goncalves, Ewan Harrison, David K. Jackson, Ian Johnston, Dominic Kwiatkowski, Cordelia Langford, John Sillitoe on behalf of the Wellcome Sanger Institute COVID-19 Surveillance Team |
| EPI_ISL_2909571, EPI_ISL_2909572 | Lighthouse Laboratory Plymouth | Wellcome Sanger Institute for the COVID-19 Genomics UK (COG-UK) Consortium | Lighthouse Laboratory Plymouth and Alex Alderton, Roberto Amato, Jeffrey Barrett, Sonia Goncalves, Ewan Harrison, David K. Jackson, Ian Johnston, Dominic Kwiatkowski, Cordelia Langford, John Sillitoe on behalf of the Wellcome Sanger Institute COVID-19 Surveillance Team |
| EPI_ISL_2909573, EPI_ISL_2909574, EPI_ISL_2909575 | Lighthouse Lab in Alderley Park | Wellcome Sanger Institute for the COVID-19 Genomics UK (COG-UK) Consortium | Jacquelyn Wynn, Mairead Hyland, The Lighthouse Lab in Alderley Park and Alex Alderton, Roberto Amato, Jeffrey Barrett, Sonia Goncalves, Ewan Harrison, David K. Jackson, Ian Johnston, Dominic Kwiatkowski, Cordelia Langford, John Sillitoe on behalf of the Wellcome Sanger Institute COVID-19 Surveillance Team |
| EPI_ISL_2909576, EPI_ISL_2909577 | Lighthouse Laboratory Plymouth | Wellcome Sanger Institute for the COVID-19 Genomics UK (COG-UK) Consortium | Lighthouse Laboratory Plymouth and Alex Alderton, Roberto Amato, Jeffrey Barrett, Sonia Goncalves, Ewan Harrison, David K. Jackson, Ian Johnston, Dominic Kwiatkowski, Cordelia Langford, John Sillitoe on behalf of the Wellcome Sanger Institute COVID-19 Surveillance Team |
| EPI_ISL_2909578, EPI_ISL_2909579, EPI_ISL_2909580 | Lighthouse Lab in Alderley Park | Wellcome Sanger Institute for the COVID-19 Genomics UK (COG-UK) Consortium | Jacquelyn Wynn, Mairead Hyland, The Lighthouse Lab in Alderley Park and Alex Alderton, Roberto Amato, Jeffrey Barrett, Sonia Goncalves, Ewan Harrison, David K. Jackson, Ian Johnston, Dominic Kwiatkowski, Cordelia Langford, John Sillitoe on behalf of the Wellcome Sanger Institute COVID-19 Surveillance Team |
| EPI_ISL_2909581 | Lighthouse Laboratory Plymouth | Wellcome Sanger Institute for the COVID-19 Genomics UK (COG-UK) Consortium | Lighthouse Laboratory Plymouth and Alex Alderton, Roberto Amato, Jeffrey Barrett, Sonia Goncalves, Ewan Harrison, David K. Jackson, Ian Johnston, Dominic Kwiatkowski, Cordelia Langford, John Sillitoe on behalf of the Wellcome Sanger Institute COVID-19 Surveillance Team |
| EPI_ISL_2909582, EPI_ISL_2909583 | Lighthouse Lab in Alderley Park | Wellcome Sanger Institute for the COVID-19 Genomics UK (COG-UK) Consortium | Jacquelyn Wynn, Mairead Hyland, The Lighthouse Lab in Alderley Park and Alex Alderton, Roberto Amato, Jeffrey Barrett, Sonia Goncalves, Ewan Harrison, David K. Jackson, Ian Johnston, Dominic Kwiatkowski, Cordelia Langford, John Sillitoe on behalf of the Wellcome Sanger Institute COVID-19 Surveillance Team |
| EPI_ISL_2909584 | Berkshire and Surrey Pathology Services Lighthouse Laboratory | Wellcome Sanger Institute for the COVID-19 Genomics UK (COG-UK) Consortium | Berkshire and Surrey Pathology Services Lighthouse Laboratory and Alex Alderton, Roberto Amato, Jeffrey Barrett, Sonia Goncalves, Ewan Harrison, David K. Jackson, Ian Johnston, Dominic Kwiatkowski, Cordelia Langford, John Sillitoe on behalf of the Wellcome Sanger Institute COVID-19 Surveillance Team |
| EPI_ISL_2909585, EPI_ISL_2909586, EPI_ISL_2909587, EPI_ISL_2909588 | Lighthouse Lab in Alderley Park | Wellcome Sanger Institute for the COVID-19 Genomics UK (COG-UK) Consortium | Jacquelyn Wynn, Mairead Hyland, The Lighthouse Lab in Alderley Park and Alex Alderton, Roberto Amato, Jeffrey Barrett, Sonia Goncalves, Ewan Harrison, David K. Jackson, Ian Johnston, Dominic Kwiatkowski, Cordelia Langford, John Sillitoe on behalf of the Wellcome Sanger Institute COVID-19 Surveillance Team |
| EPI_ISL_2909589, | Lighthouse Laboratory Plymouth | Wellcome Sanger Institute for the COVID-19 Genomics UK (COG-UK) | Lighthouse Laboratory Plymouth and Alex Alderton, Roberto Amato, Jeffrey Barrett, Sonia Goncalves, Ewan Harrison, David K. Jackson, Ian Johnston, Dominic Kwiatkowski, Cordelia Langford, John Sillitoe on behalf of the Wellcome Sanger |

| | | | |
|---|---|---|---|
| EPI_ISL_2909590 | | Consortium | Institute COVID-19 Surveillance Team |
| EPI_ISL_2909591 | Lighthouse Lab in Alderley Park | Wellcome Sanger Institute for the COVID-19 Genomics UK (COG-UK) Consortium | Jacquelyn Wynn, Mairead Hyland, The Lighthouse Lab in Alderley Park and Alex Alderton, Roberto Amato, Jeffrey Barrett, Sonia Goncalves, Ewan Harrison, David K. Jackson, Ian Johnston, Dominic Kwiatkowski, Cordelia Langford, John Sillitoe on behalf of the Wellcome Sanger Institute COVID-19 Surveillance Team |
| EPI_ISL_2909592, EPI_ISL_2909593 | Lighthouse Laboratory Plymouth | Wellcome Sanger Institute for the COVID-19 Genomics UK (COG-UK) Consortium | Lighthouse Laboratory Plymouth and Alex Alderton, Roberto Amato, Jeffrey Barrett, Sonia Goncalves, Ewan Harrison, David K. Jackson, Ian Johnston, Dominic Kwiatkowski, Cordelia Langford, John Sillitoe on behalf of the Wellcome Sanger Institute COVID-19 Surveillance Team |
| EPI_ISL_2909594, EPI_ISL_2909595 | Lighthouse Lab in Alderley Park | Wellcome Sanger Institute for the COVID-19 Genomics UK (COG-UK) Consortium | Jacquelyn Wynn, Mairead Hyland, The Lighthouse Lab in Alderley Park and Alex Alderton, Roberto Amato, Jeffrey Barrett, Sonia Goncalves, Ewan Harrison, David K. Jackson, Ian Johnston, Dominic Kwiatkowski, Cordelia Langford, John Sillitoe on behalf of the Wellcome Sanger Institute COVID-19 Surveillance Team |
| EPI_ISL_2909596 | Lighthouse Laboratory Plymouth | Wellcome Sanger Institute for the COVID-19 Genomics UK (COG-UK) Consortium | Lighthouse Laboratory Plymouth and Alex Alderton, Roberto Amato, Jeffrey Barrett, Sonia Goncalves, Ewan Harrison, David K. Jackson, Ian Johnston, Dominic Kwiatkowski, Cordelia Langford, John Sillitoe on behalf of the Wellcome Sanger Institute COVID-19 Surveillance Team |
| EPI_ISL_2909597, EPI_ISL_2909598 | Lighthouse Lab in Alderley Park | Wellcome Sanger Institute for the COVID-19 Genomics UK (COG-UK) Consortium | Jacquelyn Wynn, Mairead Hyland, The Lighthouse Lab in Alderley Park and Alex Alderton, Roberto Amato, Jeffrey Barrett, Sonia Goncalves, Ewan Harrison, David K. Jackson, Ian Johnston, Dominic Kwiatkowski, Cordelia Langford, John Sillitoe on behalf of the Wellcome Sanger Institute COVID-19 Surveillance Team |
| EPI_ISL_2909599 | Lighthouse Laboratory Plymouth | Wellcome Sanger Institute for the COVID-19 Genomics UK (COG-UK) Consortium | Lighthouse Laboratory Plymouth and Alex Alderton, Roberto Amato, Jeffrey Barrett, Sonia Goncalves, Ewan Harrison, David K. Jackson, Ian Johnston, Dominic Kwiatkowski, Cordelia Langford, John Sillitoe on behalf of the Wellcome Sanger Institute COVID-19 Surveillance Team |
| EPI_ISL_2909600, EPI_ISL_2909601, EPI_ISL_2909602 | Lighthouse Lab in Alderley Park | Wellcome Sanger Institute for the COVID-19 Genomics UK (COG-UK) Consortium | Jacquelyn Wynn, Mairead Hyland, The Lighthouse Lab in Alderley Park and Alex Alderton, Roberto Amato, Jeffrey Barrett, Sonia Goncalves, Ewan Harrison, David K. Jackson, Ian Johnston, Dominic Kwiatkowski, Cordelia Langford, John Sillitoe on behalf of the Wellcome Sanger Institute COVID-19 Surveillance Team |
| EPI_ISL_2909603 | Lighthouse Laboratory Plymouth | Wellcome Sanger Institute for the COVID-19 Genomics UK (COG-UK) Consortium | Lighthouse Laboratory Plymouth and Alex Alderton, Roberto Amato, Jeffrey Barrett, Sonia Goncalves, Ewan Harrison, David K. Jackson, Ian Johnston, Dominic Kwiatkowski, Cordelia Langford, John Sillitoe on behalf of the Wellcome Sanger Institute COVID-19 Surveillance Team |
| EPI_ISL_2909604 | Berkshire and Surrey Pathology Services Lighthouse Laboratory | Wellcome Sanger Institute for the COVID-19 Genomics UK (COG-UK) Consortium | Berkshire and Surrey Pathology Services Lighthouse Laboratory and Alex Alderton, Roberto Amato, Jeffrey Barrett, Sonia Goncalves, Ewan Harrison, David K. Jackson, Ian Johnston, Dominic Kwiatkowski, Cordelia Langford, John Sillitoe on behalf of the Wellcome Sanger Institute COVID-19 Surveillance Team |
| EPI_ISL_2909605, EPI_ISL_2909606 | Lighthouse Laboratory Plymouth | Wellcome Sanger Institute for the COVID-19 Genomics UK (COG-UK) Consortium | Lighthouse Laboratory Plymouth and Alex Alderton, Roberto Amato, Jeffrey Barrett, Sonia Goncalves, Ewan Harrison, David K. Jackson, Ian Johnston, Dominic Kwiatkowski, Cordelia Langford, John Sillitoe on behalf of the Wellcome Sanger Institute COVID-19 Surveillance Team |
| EPI_ISL_2909607 | Berkshire and Surrey Pathology Services Lighthouse Laboratory | Wellcome Sanger Institute for the COVID-19 Genomics UK (COG-UK) Consortium | Berkshire and Surrey Pathology Services Lighthouse Laboratory and Alex Alderton, Roberto Amato, Jeffrey Barrett, Sonia Goncalves, Ewan Harrison, David K. Jackson, Ian Johnston, Dominic Kwiatkowski, Cordelia Langford, John Sillitoe on behalf of the Wellcome Sanger Institute COVID-19 Surveillance Team |
| EPI_ISL_2909608 | Lighthouse Lab in Alderley Park | Wellcome Sanger Institute for the COVID-19 Genomics UK (COG-UK) Consortium | Jacquelyn Wynn, Mairead Hyland, The Lighthouse Lab in Alderley Park and Alex Alderton, Roberto Amato, Jeffrey Barrett, Sonia Goncalves, Ewan Harrison, David K. Jackson, Ian Johnston, Dominic Kwiatkowski, Cordelia Langford, John Sillitoe on behalf of the Wellcome Sanger Institute COVID-19 Surveillance Team |
| EPI_ISL_2909609 | Lighthouse Laboratory Plymouth | Wellcome Sanger Institute for the COVID-19 Genomics UK (COG-UK) Consortium | Lighthouse Laboratory Plymouth and Alex Alderton, Roberto Amato, Jeffrey Barrett, Sonia Goncalves, Ewan Harrison, David K. Jackson, Ian Johnston, Dominic Kwiatkowski, Cordelia Langford, John Sillitoe on behalf of the Wellcome Sanger Institute COVID-19 Surveillance Team |
| EPI_ISL_2909610 | Lighthouse Lab in Alderley Park | Wellcome Sanger Institute for the COVID-19 Genomics UK (COG-UK) Consortium | Jacquelyn Wynn, Mairead Hyland, The Lighthouse Lab in Alderley Park and Alex Alderton, Roberto Amato, Jeffrey Barrett, Sonia Goncalves, Ewan Harrison, David K. Jackson, Ian Johnston, Dominic Kwiatkowski, Cordelia Langford, John Sillitoe on behalf of the Wellcome Sanger Institute COVID-19 Surveillance Team |
| EPI_ISL_2909611 | Lighthouse Laboratory Plymouth | Wellcome Sanger Institute for the COVID-19 Genomics UK (COG-UK) Consortium | Lighthouse Laboratory Plymouth and Alex Alderton, Roberto Amato, Jeffrey Barrett, Sonia Goncalves, Ewan Harrison, David K. Jackson, Ian Johnston, Dominic Kwiatkowski, Cordelia Langford, John Sillitoe on behalf of the Wellcome Sanger Institute COVID-19 Surveillance Team |
| EPI_ISL_2909612, EPI_ISL_2909613 | Lighthouse Lab in Alderley Park | Wellcome Sanger Institute for the COVID-19 Genomics UK (COG-UK) Consortium | Jacquelyn Wynn, Mairead Hyland, The Lighthouse Lab in Alderley Park and Alex Alderton, Roberto Amato, Jeffrey Barrett, Sonia Goncalves, Ewan Harrison, David K. Jackson, Ian Johnston, Dominic Kwiatkowski, Cordelia Langford, John Sillitoe on behalf of the Wellcome Sanger Institute COVID-19 Surveillance Team |
| EPI_ISL_2909614 | Berkshire and Surrey Pathology Services Lighthouse Laboratory | Wellcome Sanger Institute for the COVID-19 Genomics UK (COG-UK) Consortium | Berkshire and Surrey Pathology Services Lighthouse Laboratory and Alex Alderton, Roberto Amato, Jeffrey Barrett, Sonia Goncalves, Ewan Harrison, David K. Jackson, Ian Johnston, Dominic Kwiatkowski, Cordelia Langford, John Sillitoe on behalf of the Wellcome Sanger Institute COVID-19 Surveillance Team |
| EPI_ISL_2909615, EPI_ISL_2909616 | Lighthouse Lab in Alderley Park | Wellcome Sanger Institute for the COVID-19 Genomics UK (COG-UK) Consortium | Jacquelyn Wynn, Mairead Hyland, The Lighthouse Lab in Alderley Park and Alex Alderton, Roberto Amato, Jeffrey Barrett, Sonia Goncalves, Ewan Harrison, David K. Jackson, Ian Johnston, Dominic Kwiatkowski, Cordelia Langford, John Sillitoe on behalf of the Wellcome Sanger Institute COVID-19 Surveillance Team |
| EPI_ISL_2909617 | Lighthouse Laboratory Plymouth | Wellcome Sanger Institute for the COVID-19 Genomics UK (COG-UK) Consortium | Lighthouse Laboratory Plymouth and Alex Alderton, Roberto Amato, Jeffrey Barrett, Sonia Goncalves, Ewan Harrison, David K. Jackson, Ian Johnston, Dominic Kwiatkowski, Cordelia Langford, John Sillitoe on behalf of the Wellcome Sanger Institute COVID-19 Surveillance Team |
| EPI_ISL_2909618, EPI_ISL_2909619, EPI_ISL_2909620, EPI_ISL_2909621, EPI_ISL_2909622, EPI_ISL_2909623 | Lighthouse Lab in Alderley Park | Wellcome Sanger Institute for the COVID-19 Genomics UK (COG-UK) Consortium | Jacquelyn Wynn, Mairead Hyland, The Lighthouse Lab in Alderley Park and Alex Alderton, Roberto Amato, Jeffrey Barrett, Sonia Goncalves, Ewan Harrison, David K. Jackson, Ian Johnston, Dominic Kwiatkowski, Cordelia Langford, John Sillitoe on behalf of the Wellcome Sanger Institute COVID-19 Surveillance Team |
| EPI_ISL_2909624 | Lighthouse Laboratory Plymouth | Wellcome Sanger Institute for the COVID-19 Genomics UK (COG-UK) Consortium | Lighthouse Laboratory Plymouth and Alex Alderton, Roberto Amato, Jeffrey Barrett, Sonia Goncalves, Ewan Harrison, David K. Jackson, Ian Johnston, Dominic Kwiatkowski, Cordelia Langford, John Sillitoe on behalf of the Wellcome Sanger Institute COVID-19 Surveillance Team |
| EPI_ISL_2909625, EPI_ISL_2909626, EPI_ISL_2909627, EPI_ISL_2909628 | Lighthouse Lab in Alderley Park | Wellcome Sanger Institute for the COVID-19 Genomics UK (COG-UK) Consortium | Jacquelyn Wynn, Mairead Hyland, The Lighthouse Lab in Alderley Park and Alex Alderton, Roberto Amato, Jeffrey Barrett, Sonia Goncalves, Ewan Harrison, David K. Jackson, Ian Johnston, Dominic Kwiatkowski, Cordelia Langford, John Sillitoe on behalf of the Wellcome Sanger Institute COVID-19 Surveillance Team |
| EPI_ISL_2909629, EPI_ISL_2909631 | Lighthouse Laboratory Plymouth | Wellcome Sanger Institute for the COVID-19 Genomics UK (COG-UK) Consortium | Lighthouse Laboratory Plymouth and Alex Alderton, Roberto Amato, Jeffrey Barrett, Sonia Goncalves, Ewan Harrison, David K. Jackson, Ian Johnston, Dominic Kwiatkowski, Cordelia Langford, John Sillitoe on behalf of the Wellcome Sanger Institute COVID-19 Surveillance Team |
| EPI_ISL_2909632, EPI_ISL_2909633 | Lighthouse Lab in Alderley Park | Wellcome Sanger Institute for the COVID-19 Genomics UK (COG-UK) Consortium | Jacquelyn Wynn, Mairead Hyland, The Lighthouse Lab in Alderley Park and Alex Alderton, Roberto Amato, Jeffrey Barrett, Sonia Goncalves, Ewan Harrison, David K. Jackson, Ian Johnston, Dominic Kwiatkowski, Cordelia Langford, John Sillitoe on behalf of the Wellcome Sanger Institute COVID-19 Surveillance Team |
| EPI_ISL_2909634 | Lighthouse Laboratory Plymouth | Wellcome Sanger Institute for the COVID-19 Genomics UK (COG-UK) Consortium | Lighthouse Laboratory Plymouth and Alex Alderton, Roberto Amato, Jeffrey Barrett, Sonia Goncalves, Ewan Harrison, David K. Jackson, Ian Johnston, Dominic Kwiatkowski, Cordelia Langford, John Sillitoe on behalf of the Wellcome Sanger Institute COVID-19 Surveillance Team |
| EPI_ISL_2909635, EPI_ISL_2909636, EPI_ISL_2909637, EPI_ISL_2909638, EPI_ISL_2909639, EPI_ISL_2909640, EPI_ISL_2909641 | Lighthouse Lab in Alderley Park | Wellcome Sanger Institute for the COVID-19 Genomics UK (COG-UK) Consortium | Jacquelyn Wynn, Mairead Hyland, The Lighthouse Lab in Alderley Park and Alex Alderton, Roberto Amato, Jeffrey Barrett, Sonia Goncalves, Ewan Harrison, David K. Jackson, Ian Johnston, Dominic Kwiatkowski, Cordelia Langford, John Sillitoe on behalf of the Wellcome Sanger Institute COVID-19 Surveillance Team |
| EPI_ISL_2909642 | Lighthouse Laboratory Plymouth | Wellcome Sanger Institute for the COVID-19 Genomics UK (COG-UK) Consortium | Lighthouse Laboratory Plymouth and Alex Alderton, Roberto Amato, Jeffrey Barrett, Sonia Goncalves, Ewan Harrison, David K. Jackson, Ian Johnston, Dominic Kwiatkowski, Cordelia Langford, John Sillitoe on behalf of the Wellcome Sanger Institute COVID-19 Surveillance Team |
| EPI_ISL_2909643, EPI_ISL_2909644, EPI_ISL_2909645 | Lighthouse Lab in Alderley Park | Wellcome Sanger Institute for the COVID-19 Genomics UK (COG-UK) Consortium | Jacquelyn Wynn, Mairead Hyland, The Lighthouse Lab in Alderley Park and Alex Alderton, Roberto Amato, Jeffrey Barrett, Sonia Goncalves, Ewan Harrison, David K. Jackson, Ian Johnston, Dominic Kwiatkowski, Cordelia Langford, John Sillitoe on behalf of the Wellcome Sanger Institute COVID-19 Surveillance Team |
| EPI_ISL_2909646, EPI_ISL_2909647 | Lighthouse Laboratory Plymouth | Wellcome Sanger Institute for the COVID-19 Genomics UK (COG-UK) Consortium | Lighthouse Laboratory Plymouth and Alex Alderton, Roberto Amato, Jeffrey Barrett, Sonia Goncalves, Ewan Harrison, David K. Jackson, Ian Johnston, Dominic Kwiatkowski, Cordelia Langford, John Sillitoe on behalf of the Wellcome Sanger Institute COVID-19 Surveillance Team |
| EPI_ISL_2909648, EPI_ISL_2909649 | Lighthouse Lab in Alderley Park | Wellcome Sanger Institute for the COVID-19 Genomics UK (COG-UK) Consortium | Jacquelyn Wynn, Mairead Hyland, The Lighthouse Lab in Alderley Park and Alex Alderton, Roberto Amato, Jeffrey Barrett, Sonia Goncalves, Ewan Harrison, David K. Jackson, Ian Johnston, Dominic Kwiatkowski, Cordelia Langford, John Sillitoe on behalf of the Wellcome Sanger Institute COVID-19 Surveillance Team |
| EPI_ISL_2909650 | Berkshire and Surrey Pathology Services Lighthouse Laboratory | Wellcome Sanger Institute for the COVID-19 Genomics UK (COG-UK) Consortium | Berkshire and Surrey Pathology Services Lighthouse Laboratory and Alex Alderton, Roberto Amato, Jeffrey Barrett, Sonia Goncalves, Ewan Harrison, David K. Jackson, Ian Johnston, Dominic Kwiatkowski, Cordelia Langford, John Sillitoe on behalf of the Wellcome Sanger Institute COVID-19 Surveillance Team |
| EPI_ISL_2909651 | Lighthouse Lab in Alderley Park | Wellcome Sanger Institute for the COVID-19 Genomics UK (COG-UK) Consortium | Jacquelyn Wynn, Mairead Hyland, The Lighthouse Lab in Alderley Park and Alex Alderton, Roberto Amato, Jeffrey Barrett, Sonia Goncalves, Ewan Harrison, David K. Jackson, Ian Johnston, Dominic Kwiatkowski, Cordelia Langford, John Sillitoe on behalf of the Wellcome Sanger Institute COVID-19 Surveillance Team |
| EPI_ISL_2909652 | Lighthouse Laboratory Plymouth | Wellcome Sanger Institute for the COVID-19 Genomics UK (COG-UK) Consortium | Lighthouse Laboratory Plymouth and Alex Alderton, Roberto Amato, Jeffrey Barrett, Sonia Goncalves, Ewan Harrison, David K. Jackson, Ian Johnston, Dominic Kwiatkowski, Cordelia Langford, John Sillitoe on behalf of the Wellcome Sanger Institute COVID-19 Surveillance Team |
| EPI_ISL_2909653, EPI_ISL_2909654, EPI_ISL_2909655, EPI_ISL_2909656, EPI_ISL_2909657 | Lighthouse Lab in Alderley Park | Wellcome Sanger Institute for the COVID-19 Genomics UK (COG-UK) Consortium | Jacquelyn Wynn, Mairead Hyland, The Lighthouse Lab in Alderley Park and Alex Alderton, Roberto Amato, Jeffrey Barrett, Sonia Goncalves, Ewan Harrison, David K. Jackson, Ian Johnston, Dominic Kwiatkowski, Cordelia Langford, John Sillitoe on behalf of the Wellcome Sanger Institute COVID-19 Surveillance Team |
| EPI_ISL_2909658, EPI_ISL_2909659 | Lighthouse Laboratory Plymouth | Wellcome Sanger Institute for the COVID-19 Genomics UK (COG-UK) Consortium | Lighthouse Laboratory Plymouth and Alex Alderton, Roberto Amato, Jeffrey Barrett, Sonia Goncalves, Ewan Harrison, David K. Jackson, Ian Johnston, Dominic Kwiatkowski, Cordelia Langford, John Sillitoe on behalf of the Wellcome Sanger Institute COVID-19 Surveillance Team |
| EPI_ISL_2909660, EPI_ISL_2909661 | Lighthouse Lab in Alderley Park | Wellcome Sanger Institute for the COVID-19 Genomics UK (COG-UK) Consortium | Jacquelyn Wynn, Mairead Hyland, The Lighthouse Lab in Alderley Park and Alex Alderton, Roberto Amato, Jeffrey Barrett, Sonia Goncalves, Ewan Harrison, David K. Jackson, Ian Johnston, Dominic Kwiatkowski, Cordelia Langford, John Sillitoe on behalf of the Wellcome Sanger Institute COVID-19 Surveillance Team |
| EPI_ISL_2909662 | Lighthouse Laboratory Plymouth | Wellcome Sanger Institute for the COVID-19 Genomics UK (COG-UK) Consortium | Lighthouse Laboratory Plymouth and Alex Alderton, Roberto Amato, Jeffrey Barrett, Sonia Goncalves, Ewan Harrison, David K. Jackson, Ian Johnston, Dominic Kwiatkowski, Cordelia Langford, John Sillitoe on behalf of the Wellcome Sanger Institute COVID-19 Surveillance Team |
| EPI_ISL_2909663, EPI_ISL_2909664 | Lighthouse Lab in Alderley Park | Wellcome Sanger Institute for the COVID-19 Genomics UK (COG-UK) Consortium | Jacquelyn Wynn, Mairead Hyland, The Lighthouse Lab in Alderley Park and Alex Alderton, Roberto Amato, Jeffrey Barrett, Sonia Goncalves, Ewan Harrison, David K. Jackson, Ian Johnston, Dominic Kwiatkowski, Cordelia Langford, John Sillitoe on behalf of the Wellcome Sanger Institute COVID-19 Surveillance Team |

| | | | |
|---|---|---|---|
| EPI_ISL_2909665, EPI_ISL_2909666 | Lighthouse Laboratory Plymouth | Wellcome Sanger Institute for the COVID-19 Genomics UK (COG-UK) Consortium | Lighthouse Laboratory Plymouth and Alex Alderton, Roberto Amato, Jeffrey Barrett, Sonia Goncalves, Ewan Harrison, David K. Jackson, Ian Johnston, Dominic Kwiatkowski, Cordelia Langford, John Sillitoe on behalf of the Wellcome Sanger Institute COVID-19 Surveillance Team |
| EPI_ISL_2909667, EPI_ISL_2909668, EPI_ISL_2909669, EPI_ISL_2909670, EPI_ISL_2909671, EPI_ISL_2909672 | Lighthouse Lab in Alderley Park | Wellcome Sanger Institute for the COVID-19 Genomics UK (COG-UK) Consortium | Jacquelyn Wynn, Mairead Hyland, The Lighthouse Lab in Alderley Park and Alex Alderton, Roberto Amato, Jeffrey Barrett, Sonia Goncalves, Ewan Harrison, David K. Jackson, Ian Johnston, Dominic Kwiatkowski, Cordelia Langford, John Sillitoe on behalf of the Wellcome Sanger Institute COVID-19 Surveillance Team |
| EPI_ISL_2909673 | Berkshire and Surrey Pathology Services Lighthouse Laboratory | Wellcome Sanger Institute for the COVID-19 Genomics UK (COG-UK) Consortium | Berkshire and Surrey Pathology Services Lighthouse Laboratory and Alex Alderton, Roberto Amato, Jeffrey Barrett, Sonia Goncalves, Ewan Harrison, David K. Jackson, Ian Johnston, Dominic Kwiatkowski, Cordelia Langford, John Sillitoe on behalf of the Wellcome Sanger Institute COVID-19 Surveillance Team |
| EPI_ISL_2909674, EPI_ISL_2909675, EPI_ISL_2909676, EPI_ISL_2909677 | Lighthouse Lab in Alderley Park | Wellcome Sanger Institute for the COVID-19 Genomics UK (COG-UK) Consortium | Jacquelyn Wynn, Mairead Hyland, The Lighthouse Lab in Alderley Park and Alex Alderton, Roberto Amato, Jeffrey Barrett, Sonia Goncalves, Ewan Harrison, David K. Jackson, Ian Johnston, Dominic Kwiatkowski, Cordelia Langford, John Sillitoe on behalf of the Wellcome Sanger Institute COVID-19 Surveillance Team |
| EPI_ISL_2909678 | Lighthouse Laboratory Plymouth | Wellcome Sanger Institute for the COVID-19 Genomics UK (COG-UK) Consortium | Lighthouse Laboratory Plymouth and Alex Alderton, Roberto Amato, Jeffrey Barrett, Sonia Goncalves, Ewan Harrison, David K. Jackson, Ian Johnston, Dominic Kwiatkowski, Cordelia Langford, John Sillitoe on behalf of the Wellcome Sanger Institute COVID-19 Surveillance Team |
| EPI_ISL_2909679, EPI_ISL_2909680, EPI_ISL_2909681, EPI_ISL_2909682, EPI_ISL_2909683, EPI_ISL_2909684, EPI_ISL_2909685, EPI_ISL_2909686, EPI_ISL_2909687, EPI_ISL_2909688, EPI_ISL_2909689, EPI_ISL_2909690, EPI_ISL_2909691, EPI_ISL_2909692, EPI_ISL_2909693, EPI_ISL_2909694 | | | |
| see above | Lighthouse Lab in Alderley Park | Wellcome Sanger Institute for the COVID-19 Genomics UK (COG-UK) Consortium | Jacquelyn Wynn, Mairead Hyland, The Lighthouse Lab in Alderley Park and Alex Alderton, Roberto Amato, Jeffrey Barrett, Sonia Goncalves, Ewan Harrison, David K. Jackson, Ian Johnston, Dominic Kwiatkowski, Cordelia Langford, John Sillitoe on behalf of the Wellcome Sanger Institute COVID-19 Surveillance Team |
| EPI_ISL_2909695, EPI_ISL_2909696, EPI_ISL_2909697 | Lighthouse Laboratory Plymouth | Wellcome Sanger Institute for the COVID-19 Genomics UK (COG-UK) Consortium | Lighthouse Laboratory Plymouth and Alex Alderton, Roberto Amato, Jeffrey Barrett, Sonia Goncalves, Ewan Harrison, David K. Jackson, Ian Johnston, Dominic Kwiatkowski, Cordelia Langford, John Sillitoe on behalf of the Wellcome Sanger Institute COVID-19 Surveillance Team |
| EPI_ISL_2909698 | Lighthouse Lab in Alderley Park | Wellcome Sanger Institute for the COVID-19 Genomics UK (COG-UK) Consortium | Jacquelyn Wynn, Mairead Hyland, The Lighthouse Lab in Alderley Park and Alex Alderton, Roberto Amato, Jeffrey Barrett, Sonia Goncalves, Ewan Harrison, David K. Jackson, Ian Johnston, Dominic Kwiatkowski, Cordelia Langford, John Sillitoe on behalf of the Wellcome Sanger Institute COVID-19 Surveillance Team |
| EPI_ISL_2909699, EPI_ISL_2909700 | Lighthouse Laboratory Plymouth | Wellcome Sanger Institute for the COVID-19 Genomics UK (COG-UK) Consortium | Lighthouse Laboratory Plymouth and Alex Alderton, Roberto Amato, Jeffrey Barrett, Sonia Goncalves, Ewan Harrison, David K. Jackson, Ian Johnston, Dominic Kwiatkowski, Cordelia Langford, John Sillitoe on behalf of the Wellcome Sanger Institute COVID-19 Surveillance Team |
| EPI_ISL_2909701, EPI_ISL_2909702 | Lighthouse Lab in Alderley Park | Wellcome Sanger Institute for the COVID-19 Genomics UK (COG-UK) Consortium | Jacquelyn Wynn, Mairead Hyland, The Lighthouse Lab in Alderley Park and Alex Alderton, Roberto Amato, Jeffrey Barrett, Sonia Goncalves, Ewan Harrison, David K. Jackson, Ian Johnston, Dominic Kwiatkowski, Cordelia Langford, John Sillitoe on behalf of the Wellcome Sanger Institute COVID-19 Surveillance Team |
| EPI_ISL_2909703 | Lighthouse Laboratory Plymouth | Wellcome Sanger Institute for the COVID-19 Genomics UK (COG-UK) Consortium | Lighthouse Laboratory Plymouth and Alex Alderton, Roberto Amato, Jeffrey Barrett, Sonia Goncalves, Ewan Harrison, David K. Jackson, Ian Johnston, Dominic Kwiatkowski, Cordelia Langford, John Sillitoe on behalf of the Wellcome Sanger Institute COVID-19 Surveillance Team |
| EPI_ISL_2909704, EPI_ISL_2909705, EPI_ISL_2909706, EPI_ISL_2909707, EPI_ISL_2909708, EPI_ISL_2909709, EPI_ISL_2909710, EPI_ISL_2909711 | Lighthouse Lab in Alderley Park | Wellcome Sanger Institute for the COVID-19 Genomics UK (COG-UK) Consortium | Jacquelyn Wynn, Mairead Hyland, The Lighthouse Lab in Alderley Park and Alex Alderton, Roberto Amato, Jeffrey Barrett, Sonia Goncalves, Ewan Harrison, David K. Jackson, Ian Johnston, Dominic Kwiatkowski, Cordelia Langford, John Sillitoe on behalf of the Wellcome Sanger Institute COVID-19 Surveillance Team |
| EPI_ISL_2909712 | Lighthouse Laboratory Plymouth | Wellcome Sanger Institute for the COVID-19 Genomics UK (COG-UK) Consortium | Lighthouse Laboratory Plymouth and Alex Alderton, Roberto Amato, Jeffrey Barrett, Sonia Goncalves, Ewan Harrison, David K. Jackson, Ian Johnston, Dominic Kwiatkowski, Cordelia Langford, John Sillitoe on behalf of the Wellcome Sanger Institute COVID-19 Surveillance Team |
| EPI_ISL_2909713, EPI_ISL_2909714, EPI_ISL_2909715, EPI_ISL_2909716, EPI_ISL_2909717 | Lighthouse Lab in Alderley Park | Wellcome Sanger Institute for the COVID-19 Genomics UK (COG-UK) Consortium | Jacquelyn Wynn, Mairead Hyland, The Lighthouse Lab in Alderley Park and Alex Alderton, Roberto Amato, Jeffrey Barrett, Sonia Goncalves, Ewan Harrison, David K. Jackson, Ian Johnston, Dominic Kwiatkowski, Cordelia Langford, John Sillitoe on behalf of the Wellcome Sanger Institute COVID-19 Surveillance Team |
| EPI_ISL_2909718 | Lighthouse Laboratory Plymouth | Wellcome Sanger Institute for the COVID-19 Genomics UK (COG-UK) Consortium | Lighthouse Laboratory Plymouth and Alex Alderton, Roberto Amato, Jeffrey Barrett, Sonia Goncalves, Ewan Harrison, David K. Jackson, Ian Johnston, Dominic Kwiatkowski, Cordelia Langford, John Sillitoe on behalf of the Wellcome Sanger Institute COVID-19 Surveillance Team |
| EPI_ISL_2909719, EPI_ISL_2909720, EPI_ISL_2909721, EPI_ISL_2909722, EPI_ISL_2909723 | Lighthouse Lab in Alderley Park | Wellcome Sanger Institute for the COVID-19 Genomics UK (COG-UK) Consortium | Jacquelyn Wynn, Mairead Hyland, The Lighthouse Lab in Alderley Park and Alex Alderton, Roberto Amato, Jeffrey Barrett, Sonia Goncalves, Ewan Harrison, David K. Jackson, Ian Johnston, Dominic Kwiatkowski, Cordelia Langford, John Sillitoe on behalf of the Wellcome Sanger Institute COVID-19 Surveillance Team |
| EPI_ISL_2909724 | Lighthouse Laboratory Plymouth | Wellcome Sanger Institute for the COVID-19 Genomics UK (COG-UK) Consortium | Lighthouse Laboratory Plymouth and Alex Alderton, Roberto Amato, Jeffrey Barrett, Sonia Goncalves, Ewan Harrison, David K. Jackson, Ian Johnston, Dominic Kwiatkowski, Cordelia Langford, John Sillitoe on behalf of the Wellcome Sanger Institute COVID-19 Surveillance Team |
| EPI_ISL_2909725 | Lighthouse Lab in Alderley Park | Wellcome Sanger Institute for the COVID-19 Genomics UK (COG-UK) Consortium | Jacquelyn Wynn, Mairead Hyland, The Lighthouse Lab in Alderley Park and Alex Alderton, Roberto Amato, Jeffrey Barrett, Sonia Goncalves, Ewan Harrison, David K. Jackson, Ian Johnston, Dominic Kwiatkowski, Cordelia Langford, John Sillitoe on behalf of the Wellcome Sanger Institute COVID-19 Surveillance Team |
| EPI_ISL_2909726 | Lighthouse Laboratory Plymouth | Wellcome Sanger Institute for the COVID-19 Genomics UK (COG-UK) Consortium | Lighthouse Laboratory Plymouth and Alex Alderton, Roberto Amato, Jeffrey Barrett, Sonia Goncalves, Ewan Harrison, David K. Jackson, Ian Johnston, Dominic Kwiatkowski, Cordelia Langford, John Sillitoe on behalf of the Wellcome Sanger Institute COVID-19 Surveillance Team |
| EPI_ISL_2909727, EPI_ISL_2909728, EPI_ISL_2909729, EPI_ISL_2909730, EPI_ISL_2909731, EPI_ISL_2909732 | Lighthouse Lab in Alderley Park | Wellcome Sanger Institute for the COVID-19 Genomics UK (COG-UK) Consortium | Jacquelyn Wynn, Mairead Hyland, The Lighthouse Lab in Alderley Park and Alex Alderton, Roberto Amato, Jeffrey Barrett, Sonia Goncalves, Ewan Harrison, David K. Jackson, Ian Johnston, Dominic Kwiatkowski, Cordelia Langford, John Sillitoe on behalf of the Wellcome Sanger Institute COVID-19 Surveillance Team |
| EPI_ISL_2909733 | Lighthouse Laboratory Plymouth | Wellcome Sanger Institute for the COVID-19 Genomics UK (COG-UK) Consortium | Lighthouse Laboratory Plymouth and Alex Alderton, Roberto Amato, Jeffrey Barrett, Sonia Goncalves, Ewan Harrison, David K. Jackson, Ian Johnston, Dominic Kwiatkowski, Cordelia Langford, John Sillitoe on behalf of the Wellcome Sanger Institute COVID-19 Surveillance Team |
| EPI_ISL_2909734, EPI_ISL_2909735, EPI_ISL_2909736 | Lighthouse Lab in Alderley Park | Wellcome Sanger Institute for the COVID-19 Genomics UK (COG-UK) Consortium | Jacquelyn Wynn, Mairead Hyland, The Lighthouse Lab in Alderley Park and Alex Alderton, Roberto Amato, Jeffrey Barrett, Sonia Goncalves, Ewan Harrison, David K. Jackson, Ian Johnston, Dominic Kwiatkowski, Cordelia Langford, John Sillitoe on behalf of the Wellcome Sanger Institute COVID-19 Surveillance Team |
| EPI_ISL_2909737 | Berkshire and Surrey Pathology Services Lighthouse Laboratory | Wellcome Sanger Institute for the COVID-19 Genomics UK (COG-UK) Consortium | Berkshire and Surrey Pathology Services Lighthouse Laboratory and Alex Alderton, Roberto Amato, Jeffrey Barrett, Sonia Goncalves, Ewan Harrison, David K. Jackson, Ian Johnston, Dominic Kwiatkowski, Cordelia Langford, John Sillitoe on behalf of the Wellcome Sanger Institute COVID-19 Surveillance Team |
| EPI_ISL_2909738 | Lighthouse Laboratory Plymouth | Wellcome Sanger Institute for the COVID-19 Genomics UK (COG-UK) Consortium | Lighthouse Laboratory Plymouth and Alex Alderton, Roberto Amato, Jeffrey Barrett, Sonia Goncalves, Ewan Harrison, David K. Jackson, Ian Johnston, Dominic Kwiatkowski, Cordelia Langford, John Sillitoe on behalf of the Wellcome Sanger Institute COVID-19 Surveillance Team |
| EPI_ISL_2909739, EPI_ISL_2909740, EPI_ISL_2909741, EPI_ISL_2909742, EPI_ISL_2909743 | Lighthouse Lab in Alderley Park | Wellcome Sanger Institute for the COVID-19 Genomics UK (COG-UK) Consortium | Jacquelyn Wynn, Mairead Hyland, The Lighthouse Lab in Alderley Park and Alex Alderton, Roberto Amato, Jeffrey Barrett, Sonia Goncalves, Ewan Harrison, David K. Jackson, Ian Johnston, Dominic Kwiatkowski, Cordelia Langford, John Sillitoe on behalf of the Wellcome Sanger Institute COVID-19 Surveillance Team |
| EPI_ISL_2909744 | Lighthouse Laboratory Plymouth | Wellcome Sanger Institute for the COVID-19 Genomics UK (COG-UK) Consortium | Lighthouse Laboratory Plymouth and Alex Alderton, Roberto Amato, Jeffrey Barrett, Sonia Goncalves, Ewan Harrison, David K. Jackson, Ian Johnston, Dominic Kwiatkowski, Cordelia Langford, John Sillitoe on behalf of the Wellcome Sanger Institute COVID-19 Surveillance Team |
| EPI_ISL_2909745, EPI_ISL_2909746, EPI_ISL_2909747 | Lighthouse Lab in Alderley Park | Wellcome Sanger Institute for the COVID-19 Genomics UK (COG-UK) Consortium | Jacquelyn Wynn, Mairead Hyland, The Lighthouse Lab in Alderley Park and Alex Alderton, Roberto Amato, Jeffrey Barrett, Sonia Goncalves, Ewan Harrison, David K. Jackson, Ian Johnston, Dominic Kwiatkowski, Cordelia Langford, John Sillitoe on behalf of the Wellcome Sanger Institute COVID-19 Surveillance Team |
| EPI_ISL_2909748 | Berkshire and Surrey Pathology Services Lighthouse Laboratory | Wellcome Sanger Institute for the COVID-19 Genomics UK (COG-UK) Consortium | Berkshire and Surrey Pathology Services Lighthouse Laboratory and Alex Alderton, Roberto Amato, Jeffrey Barrett, Sonia Goncalves, Ewan Harrison, David K. Jackson, Ian Johnston, Dominic Kwiatkowski, Cordelia Langford, John Sillitoe on behalf of the Wellcome Sanger Institute COVID-19 Surveillance Team |
| EPI_ISL_2909749 | Lighthouse Lab in Alderley Park | Wellcome Sanger Institute for the COVID-19 Genomics UK (COG-UK) Consortium | Jacquelyn Wynn, Mairead Hyland, The Lighthouse Lab in Alderley Park and Alex Alderton, Roberto Amato, Jeffrey Barrett, Sonia Goncalves, Ewan Harrison, David K. Jackson, Ian Johnston, Dominic Kwiatkowski, Cordelia Langford, John Sillitoe on behalf of the Wellcome Sanger Institute COVID-19 Surveillance Team |
| EPI_ISL_2909750 | Berkshire and Surrey Pathology Services Lighthouse Laboratory | Wellcome Sanger Institute for the COVID-19 Genomics UK (COG-UK) Consortium | Berkshire and Surrey Pathology Services Lighthouse Laboratory and Alex Alderton, Roberto Amato, Jeffrey Barrett, Sonia Goncalves, Ewan Harrison, David K. Jackson, Ian Johnston, Dominic Kwiatkowski, Cordelia Langford, John Sillitoe on behalf of the Wellcome Sanger Institute COVID-19 Surveillance Team |
| EPI_ISL_2909751 | Lighthouse Lab in Alderley Park | Wellcome Sanger Institute for the COVID-19 Genomics UK (COG-UK) Consortium | Jacquelyn Wynn, Mairead Hyland, The Lighthouse Lab in Alderley Park and Alex Alderton, Roberto Amato, Jeffrey Barrett, Sonia Goncalves, Ewan Harrison, David K. Jackson, Ian Johnston, Dominic Kwiatkowski, Cordelia Langford, John Sillitoe on behalf of the Wellcome Sanger Institute COVID-19 Surveillance Team |
| EPI_ISL_2909752 | Berkshire and Surrey Pathology Services Lighthouse Laboratory | Wellcome Sanger Institute for the COVID-19 Genomics UK (COG-UK) Consortium | Berkshire and Surrey Pathology Services Lighthouse Laboratory and Alex Alderton, Roberto Amato, Jeffrey Barrett, Sonia Goncalves, Ewan Harrison, David K. Jackson, Ian Johnston, Dominic Kwiatkowski, Cordelia Langford, John Sillitoe on behalf of the Wellcome Sanger Institute COVID-19 Surveillance Team |
| EPI_ISL_2909754, EPI_ISL_2909755, EPI_ISL_2909756 | Lighthouse Lab in Alderley Park | Wellcome Sanger Institute for the COVID-19 Genomics UK (COG-UK) Consortium | Jacquelyn Wynn, Mairead Hyland, The Lighthouse Lab in Alderley Park and Alex Alderton, Roberto Amato, Jeffrey Barrett, Sonia Goncalves, Ewan Harrison, David K. Jackson, Ian Johnston, Dominic Kwiatkowski, Cordelia Langford, John Sillitoe on behalf of the Wellcome Sanger Institute COVID-19 Surveillance Team |

| | | | |
|---|---|---|---|
| EPI_ISL_2909757 | Berkshire and Surrey Pathology Services Lighthouse Laboratory | Wellcome Sanger Institute for the COVID-19 Genomics UK (COG-UK) Consortium | Berkshire and Surrey Pathology Services Lighthouse Laboratory and Alex Alderton, Roberto Amato, Jeffrey Barrett, Sonia Goncalves, Ewan Harrison, David K. Jackson, Ian Johnston, Dominic Kwiatkowski, Cordelia Langford, John Sillitoe on behalf of the Wellcome Sanger Institute COVID-19 Surveillance Team |
| EPI_ISL_2909758, EPI_ISL_2909759 | Lighthouse Laboratory Plymouth | Wellcome Sanger Institute for the COVID-19 Genomics UK (COG-UK) Consortium | Lighthouse Laboratory Plymouth and Alex Alderton, Roberto Amato, Jeffrey Barrett, Sonia Goncalves, Ewan Harrison, David K. Jackson, Ian Johnston, Dominic Kwiatkowski, Cordelia Langford, John Sillitoe on behalf of the Wellcome Sanger Institute COVID-19 Surveillance Team |
| EPI_ISL_2909760, EPI_ISL_2909761, EPI_ISL_2909762 | Lighthouse Lab in Alderley Park | Wellcome Sanger Institute for the COVID-19 Genomics UK (COG-UK) Consortium | Jacquelyn Wynn, Mairead Hyland, The Lighthouse Lab in Alderley Park and Alex Alderton, Roberto Amato, Jeffrey Barrett, Sonia Goncalves, Ewan Harrison, David K. Jackson, Ian Johnston, Dominic Kwiatkowski, Cordelia Langford, John Sillitoe on behalf of the Wellcome Sanger Institute COVID-19 Surveillance Team |
| EPI_ISL_2909763 | Lighthouse Laboratory Plymouth | Wellcome Sanger Institute for the COVID-19 Genomics UK (COG-UK) Consortium | Lighthouse Laboratory Plymouth and Alex Alderton, Roberto Amato, Jeffrey Barrett, Sonia Goncalves, Ewan Harrison, David K. Jackson, Ian Johnston, Dominic Kwiatkowski, Cordelia Langford, John Sillitoe on behalf of the Wellcome Sanger Institute COVID-19 Surveillance Team |
| EPI_ISL_2909764, EPI_ISL_2909765, EPI_ISL_2909766, EPI_ISL_2909767, EPI_ISL_2909768 | Lighthouse Lab in Alderley Park | Wellcome Sanger Institute for the COVID-19 Genomics UK (COG-UK) Consortium | Jacquelyn Wynn, Mairead Hyland, The Lighthouse Lab in Alderley Park and Alex Alderton, Roberto Amato, Jeffrey Barrett, Sonia Goncalves, Ewan Harrison, David K. Jackson, Ian Johnston, Dominic Kwiatkowski, Cordelia Langford, John Sillitoe on behalf of the Wellcome Sanger Institute COVID-19 Surveillance Team |
| EPI_ISL_2909769 | Berkshire and Surrey Pathology Services Lighthouse Laboratory | Wellcome Sanger Institute for the COVID-19 Genomics UK (COG-UK) Consortium | Berkshire and Surrey Pathology Services Lighthouse Laboratory and Alex Alderton, Roberto Amato, Jeffrey Barrett, Sonia Goncalves, Ewan Harrison, David K. Jackson, Ian Johnston, Dominic Kwiatkowski, Cordelia Langford, John Sillitoe on behalf of the Wellcome Sanger Institute COVID-19 Surveillance Team |
| EPI_ISL_2909770 | Lighthouse Lab in Alderley Park | Wellcome Sanger Institute for the COVID-19 Genomics UK (COG-UK) Consortium | Jacquelyn Wynn, Mairead Hyland, The Lighthouse Lab in Alderley Park and Alex Alderton, Roberto Amato, Jeffrey Barrett, Sonia Goncalves, Ewan Harrison, David K. Jackson, Ian Johnston, Dominic Kwiatkowski, Cordelia Langford, John Sillitoe on behalf of the Wellcome Sanger Institute COVID-19 Surveillance Team |
| EPI_ISL_2909771, EPI_ISL_2909772 | Lighthouse Laboratory Plymouth | Wellcome Sanger Institute for the COVID-19 Genomics UK (COG-UK) Consortium | Lighthouse Laboratory Plymouth and Alex Alderton, Roberto Amato, Jeffrey Barrett, Sonia Goncalves, Ewan Harrison, David K. Jackson, Ian Johnston, Dominic Kwiatkowski, Cordelia Langford, John Sillitoe on behalf of the Wellcome Sanger Institute COVID-19 Surveillance Team |
| EPI_ISL_2909773, EPI_ISL_2909774, EPI_ISL_2909775 | Lighthouse Lab in Alderley Park | Wellcome Sanger Institute for the COVID-19 Genomics UK (COG-UK) Consortium | Jacquelyn Wynn, Mairead Hyland, The Lighthouse Lab in Alderley Park and Alex Alderton, Roberto Amato, Jeffrey Barrett, Sonia Goncalves, Ewan Harrison, David K. Jackson, Ian Johnston, Dominic Kwiatkowski, Cordelia Langford, John Sillitoe on behalf of the Wellcome Sanger Institute COVID-19 Surveillance Team |
| EPI_ISL_2909776 | Lighthouse Laboratory Plymouth | Wellcome Sanger Institute for the COVID-19 Genomics UK (COG-UK) Consortium | Lighthouse Laboratory Plymouth and Alex Alderton, Roberto Amato, Jeffrey Barrett, Sonia Goncalves, Ewan Harrison, David K. Jackson, Ian Johnston, Dominic Kwiatkowski, Cordelia Langford, John Sillitoe on behalf of the Wellcome Sanger Institute COVID-19 Surveillance Team |
| EPI_ISL_2909777, EPI_ISL_2909778 | Lighthouse Lab in Alderley Park | Wellcome Sanger Institute for the COVID-19 Genomics UK (COG-UK) Consortium | Jacquelyn Wynn, Mairead Hyland, The Lighthouse Lab in Alderley Park and Alex Alderton, Roberto Amato, Jeffrey Barrett, Sonia Goncalves, Ewan Harrison, David K. Jackson, Ian Johnston, Dominic Kwiatkowski, Cordelia Langford, John Sillitoe on behalf of the Wellcome Sanger Institute COVID-19 Surveillance Team |
| EPI_ISL_2909779 | Lighthouse Laboratory Plymouth | Wellcome Sanger Institute for the COVID-19 Genomics UK (COG-UK) Consortium | Lighthouse Laboratory Plymouth and Alex Alderton, Roberto Amato, Jeffrey Barrett, Sonia Goncalves, Ewan Harrison, David K. Jackson, Ian Johnston, Dominic Kwiatkowski, Cordelia Langford, John Sillitoe on behalf of the Wellcome Sanger Institute COVID-19 Surveillance Team |
| EPI_ISL_2909780, EPI_ISL_2909781, EPI_ISL_2909782, EPI_ISL_2909783 | Lighthouse Lab in Alderley Park | Wellcome Sanger Institute for the COVID-19 Genomics UK (COG-UK) Consortium | Jacquelyn Wynn, Mairead Hyland, The Lighthouse Lab in Alderley Park and Alex Alderton, Roberto Amato, Jeffrey Barrett, Sonia Goncalves, Ewan Harrison, David K. Jackson, Ian Johnston, Dominic Kwiatkowski, Cordelia Langford, John Sillitoe on behalf of the Wellcome Sanger Institute COVID-19 Surveillance Team |
| EPI_ISL_2909784 | Lighthouse Laboratory Plymouth | Wellcome Sanger Institute for the COVID-19 Genomics UK (COG-UK) Consortium | Lighthouse Laboratory Plymouth and Alex Alderton, Roberto Amato, Jeffrey Barrett, Sonia Goncalves, Ewan Harrison, David K. Jackson, Ian Johnston, Dominic Kwiatkowski, Cordelia Langford, John Sillitoe on behalf of the Wellcome Sanger Institute COVID-19 Surveillance Team |
| EPI_ISL_2909785, EPI_ISL_2909786, EPI_ISL_2909787, EPI_ISL_2909788, EPI_ISL_2909789 | Lighthouse Lab in Alderley Park | Wellcome Sanger Institute for the COVID-19 Genomics UK (COG-UK) Consortium | Jacquelyn Wynn, Mairead Hyland, The Lighthouse Lab in Alderley Park and Alex Alderton, Roberto Amato, Jeffrey Barrett, Sonia Goncalves, Ewan Harrison, David K. Jackson, Ian Johnston, Dominic Kwiatkowski, Cordelia Langford, John Sillitoe on behalf of the Wellcome Sanger Institute COVID-19 Surveillance Team |
| EPI_ISL_2909790 | Lighthouse Laboratory Plymouth | Wellcome Sanger Institute for the COVID-19 Genomics UK (COG-UK) Consortium | Lighthouse Laboratory Plymouth and Alex Alderton, Roberto Amato, Jeffrey Barrett, Sonia Goncalves, Ewan Harrison, David K. Jackson, Ian Johnston, Dominic Kwiatkowski, Cordelia Langford, John Sillitoe on behalf of the Wellcome Sanger Institute COVID-19 Surveillance Team |
| EPI_ISL_2909791 | Lighthouse Lab in Alderley Park | Wellcome Sanger Institute for the COVID-19 Genomics UK (COG-UK) Consortium | Jacquelyn Wynn, Mairead Hyland, The Lighthouse Lab in Alderley Park and Alex Alderton, Roberto Amato, Jeffrey Barrett, Sonia Goncalves, Ewan Harrison, David K. Jackson, Ian Johnston, Dominic Kwiatkowski, Cordelia Langford, John Sillitoe on behalf of the Wellcome Sanger Institute COVID-19 Surveillance Team |
| EPI_ISL_2909792 | Berkshire and Surrey Pathology Services Lighthouse Laboratory | Wellcome Sanger Institute for the COVID-19 Genomics UK (COG-UK) Consortium | Berkshire and Surrey Pathology Services Lighthouse Laboratory and Alex Alderton, Roberto Amato, Jeffrey Barrett, Sonia Goncalves, Ewan Harrison, David K. Jackson, Ian Johnston, Dominic Kwiatkowski, Cordelia Langford, John Sillitoe on behalf of the Wellcome Sanger Institute COVID-19 Surveillance Team |
| EPI_ISL_2909793, EPI_ISL_2909794, EPI_ISL_2909795, EPI_ISL_2909796, EPI_ISL_2909797, EPI_ISL_2909798, EPI_ISL_2909799 | Lighthouse Lab in Alderley Park | Wellcome Sanger Institute for the COVID-19 Genomics UK (COG-UK) Consortium | Jacquelyn Wynn, Mairead Hyland, The Lighthouse Lab in Alderley Park and Alex Alderton, Roberto Amato, Jeffrey Barrett, Sonia Goncalves, Ewan Harrison, David K. Jackson, Ian Johnston, Dominic Kwiatkowski, Cordelia Langford, John Sillitoe on behalf of the Wellcome Sanger Institute COVID-19 Surveillance Team |
| EPI_ISL_2909800 | Lighthouse Laboratory Plymouth | Wellcome Sanger Institute for the COVID-19 Genomics UK (COG-UK) Consortium | Lighthouse Laboratory Plymouth and Alex Alderton, Roberto Amato, Jeffrey Barrett, Sonia Goncalves, Ewan Harrison, David K. Jackson, Ian Johnston, Dominic Kwiatkowski, Cordelia Langford, John Sillitoe on behalf of the Wellcome Sanger Institute COVID-19 Surveillance Team |
| EPI_ISL_2909801 | Lighthouse Lab in Alderley Park | Wellcome Sanger Institute for the COVID-19 Genomics UK (COG-UK) Consortium | Jacquelyn Wynn, Mairead Hyland, The Lighthouse Lab in Alderley Park and Alex Alderton, Roberto Amato, Jeffrey Barrett, Sonia Goncalves, Ewan Harrison, David K. Jackson, Ian Johnston, Dominic Kwiatkowski, Cordelia Langford, John Sillitoe on behalf of the Wellcome Sanger Institute COVID-19 Surveillance Team |
| EPI_ISL_2909802 | Lighthouse Laboratory Plymouth | Wellcome Sanger Institute for the COVID-19 Genomics UK (COG-UK) Consortium | Lighthouse Laboratory Plymouth and Alex Alderton, Roberto Amato, Jeffrey Barrett, Sonia Goncalves, Ewan Harrison, David K. Jackson, Ian Johnston, Dominic Kwiatkowski, Cordelia Langford, John Sillitoe on behalf of the Wellcome Sanger Institute COVID-19 Surveillance Team |
| EPI_ISL_2909803, EPI_ISL_2909804, EPI_ISL_2909805, EPI_ISL_2909806, EPI_ISL_2909807, EPI_ISL_2909808, EPI_ISL_2909809, EPI_ISL_2909810 | Lighthouse Lab in Alderley Park | Wellcome Sanger Institute for the COVID-19 Genomics UK (COG-UK) Consortium | Jacquelyn Wynn, Mairead Hyland, The Lighthouse Lab in Alderley Park and Alex Alderton, Roberto Amato, Jeffrey Barrett, Sonia Goncalves, Ewan Harrison, David K. Jackson, Ian Johnston, Dominic Kwiatkowski, Cordelia Langford, John Sillitoe on behalf of the Wellcome Sanger Institute COVID-19 Surveillance Team |
| EPI_ISL_2909811 | Lighthouse Laboratory Plymouth | Wellcome Sanger Institute for the COVID-19 Genomics UK (COG-UK) Consortium | Lighthouse Laboratory Plymouth and Alex Alderton, Roberto Amato, Jeffrey Barrett, Sonia Goncalves, Ewan Harrison, David K. Jackson, Ian Johnston, Dominic Kwiatkowski, Cordelia Langford, John Sillitoe on behalf of the Wellcome Sanger Institute COVID-19 Surveillance Team |
| EPI_ISL_2909812 | Lighthouse Lab in Alderley Park | Wellcome Sanger Institute for the COVID-19 Genomics UK (COG-UK) Consortium | Jacquelyn Wynn, Mairead Hyland, The Lighthouse Lab in Alderley Park and Alex Alderton, Roberto Amato, Jeffrey Barrett, Sonia Goncalves, Ewan Harrison, David K. Jackson, Ian Johnston, Dominic Kwiatkowski, Cordelia Langford, John Sillitoe on behalf of the Wellcome Sanger Institute COVID-19 Surveillance Team |
| EPI_ISL_2909813 | Lighthouse Laboratory Plymouth | Wellcome Sanger Institute for the COVID-19 Genomics UK (COG-UK) Consortium | Lighthouse Laboratory Plymouth and Alex Alderton, Roberto Amato, Jeffrey Barrett, Sonia Goncalves, Ewan Harrison, David K. Jackson, Ian Johnston, Dominic Kwiatkowski, Cordelia Langford, John Sillitoe on behalf of the Wellcome Sanger Institute COVID-19 Surveillance Team |
| EPI_ISL_2909814 | Berkshire and Surrey Pathology Services Lighthouse Laboratory | Wellcome Sanger Institute for the COVID-19 Genomics UK (COG-UK) Consortium | Berkshire and Surrey Pathology Services Lighthouse Laboratory and Alex Alderton, Roberto Amato, Jeffrey Barrett, Sonia Goncalves, Ewan Harrison, David K. Jackson, Ian Johnston, Dominic Kwiatkowski, Cordelia Langford, John Sillitoe on behalf of the Wellcome Sanger Institute COVID-19 Surveillance Team |
| EPI_ISL_2909815, EPI_ISL_2909816, EPI_ISL_2909817, EPI_ISL_2909818 | Lighthouse Laboratory Plymouth | Wellcome Sanger Institute for the COVID-19 Genomics UK (COG-UK) Consortium | Lighthouse Laboratory Plymouth and Alex Alderton, Roberto Amato, Jeffrey Barrett, Sonia Goncalves, Ewan Harrison, David K. Jackson, Ian Johnston, Dominic Kwiatkowski, Cordelia Langford, John Sillitoe on behalf of the Wellcome Sanger Institute COVID-19 Surveillance Team |
| EPI_ISL_2909819, EPI_ISL_2909820 | Lighthouse Lab in Alderley Park | Wellcome Sanger Institute for the COVID-19 Genomics UK (COG-UK) Consortium | Jacquelyn Wynn, Mairead Hyland, The Lighthouse Lab in Alderley Park and Alex Alderton, Roberto Amato, Jeffrey Barrett, Sonia Goncalves, Ewan Harrison, David K. Jackson, Ian Johnston, Dominic Kwiatkowski, Cordelia Langford, John Sillitoe on behalf of the Wellcome Sanger Institute COVID-19 Surveillance Team |
| EPI_ISL_2909821 | Lighthouse Laboratory Plymouth | Wellcome Sanger Institute for the COVID-19 Genomics UK (COG-UK) Consortium | Lighthouse Laboratory Plymouth and Alex Alderton, Roberto Amato, Jeffrey Barrett, Sonia Goncalves, Ewan Harrison, David K. Jackson, Ian Johnston, Dominic Kwiatkowski, Cordelia Langford, John Sillitoe on behalf of the Wellcome Sanger Institute COVID-19 Surveillance Team |
| EPI_ISL_2909822, EPI_ISL_2909823 | Lighthouse Lab in Alderley Park | Wellcome Sanger Institute for the COVID-19 Genomics UK (COG-UK) Consortium | Jacquelyn Wynn, Mairead Hyland, The Lighthouse Lab in Alderley Park and Alex Alderton, Roberto Amato, Jeffrey Barrett, Sonia Goncalves, Ewan Harrison, David K. Jackson, Ian Johnston, Dominic Kwiatkowski, Cordelia Langford, John Sillitoe on behalf of the Wellcome Sanger Institute COVID-19 Surveillance Team |
| EPI_ISL_2909824 | Lighthouse Laboratory Plymouth | Wellcome Sanger Institute for the COVID-19 Genomics UK (COG-UK) Consortium | Lighthouse Laboratory Plymouth and Alex Alderton, Roberto Amato, Jeffrey Barrett, Sonia Goncalves, Ewan Harrison, David K. Jackson, Ian Johnston, Dominic Kwiatkowski, Cordelia Langford, John Sillitoe on behalf of the Wellcome Sanger Institute COVID-19 Surveillance Team |
| EPI_ISL_2909825, EPI_ISL_2909826, EPI_ISL_2909827 | Lighthouse Lab in Alderley Park | Wellcome Sanger Institute for the COVID-19 Genomics UK (COG-UK) Consortium | Jacquelyn Wynn, Mairead Hyland, The Lighthouse Lab in Alderley Park and Alex Alderton, Roberto Amato, Jeffrey Barrett, Sonia Goncalves, Ewan Harrison, David K. Jackson, Ian Johnston, Dominic Kwiatkowski, Cordelia Langford, John Sillitoe on behalf of the Wellcome Sanger Institute COVID-19 Surveillance Team |
| EPI_ISL_2909828 | Berkshire and Surrey Pathology Services Lighthouse Laboratory | Wellcome Sanger Institute for the COVID-19 Genomics UK (COG-UK) Consortium | Berkshire and Surrey Pathology Services Lighthouse Laboratory and Alex Alderton, Roberto Amato, Jeffrey Barrett, Sonia Goncalves, Ewan Harrison, David K. Jackson, Ian Johnston, Dominic Kwiatkowski, Cordelia Langford, John Sillitoe on behalf of the Wellcome Sanger Institute COVID-19 Surveillance Team |
| EPI_ISL_2909829 | Lighthouse Laboratory Plymouth | Wellcome Sanger Institute for the COVID-19 Genomics UK (COG-UK) Consortium | Lighthouse Laboratory Plymouth and Alex Alderton, Roberto Amato, Jeffrey Barrett, Sonia Goncalves, Ewan Harrison, David K. Jackson, Ian Johnston, Dominic Kwiatkowski, Cordelia Langford, John Sillitoe on behalf of the Wellcome Sanger Institute COVID-19 Surveillance Team |

| | | | |
|---|---|---|---|
| EPI_ISL_2909830, EPI_ISL_2909831 | Lighthouse Lab in Alderley Park | Wellcome Sanger Institute for the COVID-19 Genomics UK (COG-UK) Consortium | Jacquelyn Wynn, Mairead Hyland, The Lighthouse Lab in Alderley Park and Alex Alderton, Roberto Amato, Jeffrey Barrett, Sonia Goncalves, Ewan Harrison, David K. Jackson, Ian Johnston, Dominic Kwiatkowski, Cordelia Langford, John Sillitoe on behalf of the Wellcome Sanger Institute COVID-19 Surveillance Team |
| EPI_ISL_2909832, EPI_ISL_2909833 | Lighthouse Laboratory Plymouth | Wellcome Sanger Institute for the COVID-19 Genomics UK (COG-UK) Consortium | Lighthouse Laboratory Plymouth and Alex Alderton, Roberto Amato, Jeffrey Barrett, Sonia Goncalves, Ewan Harrison, David K. Jackson, Ian Johnston, Dominic Kwiatkowski, Cordelia Langford, John Sillitoe on behalf of the Wellcome Sanger Institute COVID-19 Surveillance Team |
| EPI_ISL_2909834, EPI_ISL_2909835, EPI_ISL_2909836, EPI_ISL_2909837, EPI_ISL_2909838, EPI_ISL_2909839 | Lighthouse Lab in Alderley Park | Wellcome Sanger Institute for the COVID-19 Genomics UK (COG-UK) Consortium | Jacquelyn Wynn, Mairead Hyland, The Lighthouse Lab in Alderley Park and Alex Alderton, Roberto Amato, Jeffrey Barrett, Sonia Goncalves, Ewan Harrison, David K. Jackson, Ian Johnston, Dominic Kwiatkowski, Cordelia Langford, John Sillitoe on behalf of the Wellcome Sanger Institute COVID-19 Surveillance Team |
| EPI_ISL_2909840 | Lighthouse Laboratory Plymouth | Wellcome Sanger Institute for the COVID-19 Genomics UK (COG-UK) Consortium | Lighthouse Laboratory Plymouth and Alex Alderton, Roberto Amato, Jeffrey Barrett, Sonia Goncalves, Ewan Harrison, David K. Jackson, Ian Johnston, Dominic Kwiatkowski, Cordelia Langford, John Sillitoe on behalf of the Wellcome Sanger Institute COVID-19 Surveillance Team |
| EPI_ISL_2909841, EPI_ISL_2909842, EPI_ISL_2909843, EPI_ISL_2909844, EPI_ISL_2909845 | Lighthouse Lab in Alderley Park | Wellcome Sanger Institute for the COVID-19 Genomics UK (COG-UK) Consortium | Jacquelyn Wynn, Mairead Hyland, The Lighthouse Lab in Alderley Park and Alex Alderton, Roberto Amato, Jeffrey Barrett, Sonia Goncalves, Ewan Harrison, David K. Jackson, Ian Johnston, Dominic Kwiatkowski, Cordelia Langford, John Sillitoe on behalf of the Wellcome Sanger Institute COVID-19 Surveillance Team |
| EPI_ISL_2909846, EPI_ISL_2909847, EPI_ISL_2909848 | Lighthouse Laboratory Plymouth | Wellcome Sanger Institute for the COVID-19 Genomics UK (COG-UK) Consortium | Lighthouse Laboratory Plymouth and Alex Alderton, Roberto Amato, Jeffrey Barrett, Sonia Goncalves, Ewan Harrison, David K. Jackson, Ian Johnston, Dominic Kwiatkowski, Cordelia Langford, John Sillitoe on behalf of the Wellcome Sanger Institute COVID-19 Surveillance Team |
| EPI_ISL_2909849 | Lighthouse Lab in Alderley Park | Wellcome Sanger Institute for the COVID-19 Genomics UK (COG-UK) Consortium | Jacquelyn Wynn, Mairead Hyland, The Lighthouse Lab in Alderley Park and Alex Alderton, Roberto Amato, Jeffrey Barrett, Sonia Goncalves, Ewan Harrison, David K. Jackson, Ian Johnston, Dominic Kwiatkowski, Cordelia Langford, John Sillitoe on behalf of the Wellcome Sanger Institute COVID-19 Surveillance Team |
| EPI_ISL_2909850, EPI_ISL_2909851 | Lighthouse Laboratory Plymouth | Wellcome Sanger Institute for the COVID-19 Genomics UK (COG-UK) Consortium | Lighthouse Laboratory Plymouth and Alex Alderton, Roberto Amato, Jeffrey Barrett, Sonia Goncalves, Ewan Harrison, David K. Jackson, Ian Johnston, Dominic Kwiatkowski, Cordelia Langford, John Sillitoe on behalf of the Wellcome Sanger Institute COVID-19 Surveillance Team |
| EPI_ISL_2909852, EPI_ISL_2909853 | Lighthouse Lab in Alderley Park | Wellcome Sanger Institute for the COVID-19 Genomics UK (COG-UK) Consortium | Jacquelyn Wynn, Mairead Hyland, The Lighthouse Lab in Alderley Park and Alex Alderton, Roberto Amato, Jeffrey Barrett, Sonia Goncalves, Ewan Harrison, David K. Jackson, Ian Johnston, Dominic Kwiatkowski, Cordelia Langford, John Sillitoe on behalf of the Wellcome Sanger Institute COVID-19 Surveillance Team |
| EPI_ISL_2909854 | Lighthouse Laboratory Plymouth | Wellcome Sanger Institute for the COVID-19 Genomics UK (COG-UK) Consortium | Lighthouse Laboratory Plymouth and Alex Alderton, Roberto Amato, Jeffrey Barrett, Sonia Goncalves, Ewan Harrison, David K. Jackson, Ian Johnston, Dominic Kwiatkowski, Cordelia Langford, John Sillitoe on behalf of the Wellcome Sanger Institute COVID-19 Surveillance Team |
| EPI_ISL_2909855 | Lighthouse Lab in Alderley Park | Wellcome Sanger Institute for the COVID-19 Genomics UK (COG-UK) Consortium | Jacquelyn Wynn, Mairead Hyland, The Lighthouse Lab in Alderley Park and Alex Alderton, Roberto Amato, Jeffrey Barrett, Sonia Goncalves, Ewan Harrison, David K. Jackson, Ian Johnston, Dominic Kwiatkowski, Cordelia Langford, John Sillitoe on behalf of the Wellcome Sanger Institute COVID-19 Surveillance Team |
| EPI_ISL_2909856 | Lighthouse Laboratory Plymouth | Wellcome Sanger Institute for the COVID-19 Genomics UK (COG-UK) Consortium | Lighthouse Laboratory Plymouth and Alex Alderton, Roberto Amato, Jeffrey Barrett, Sonia Goncalves, Ewan Harrison, David K. Jackson, Ian Johnston, Dominic Kwiatkowski, Cordelia Langford, John Sillitoe on behalf of the Wellcome Sanger Institute COVID-19 Surveillance Team |
| EPI_ISL_2909857, EPI_ISL_2909858, EPI_ISL_2909859, EPI_ISL_2909860, EPI_ISL_2909861 | Lighthouse Lab in Alderley Park | Wellcome Sanger Institute for the COVID-19 Genomics UK (COG-UK) Consortium | Jacquelyn Wynn, Mairead Hyland, The Lighthouse Lab in Alderley Park and Alex Alderton, Roberto Amato, Jeffrey Barrett, Sonia Goncalves, Ewan Harrison, David K. Jackson, Ian Johnston, Dominic Kwiatkowski, Cordelia Langford, John Sillitoe on behalf of the Wellcome Sanger Institute COVID-19 Surveillance Team |
| EPI_ISL_2909862 | Lighthouse Laboratory Plymouth | Wellcome Sanger Institute for the COVID-19 Genomics UK (COG-UK) Consortium | Lighthouse Laboratory Plymouth and Alex Alderton, Roberto Amato, Jeffrey Barrett, Sonia Goncalves, Ewan Harrison, David K. Jackson, Ian Johnston, Dominic Kwiatkowski, Cordelia Langford, John Sillitoe on behalf of the Wellcome Sanger Institute COVID-19 Surveillance Team |
| EPI_ISL_2909863, EPI_ISL_2909864 | Lighthouse Lab in Alderley Park | Wellcome Sanger Institute for the COVID-19 Genomics UK (COG-UK) Consortium | Jacquelyn Wynn, Mairead Hyland, The Lighthouse Lab in Alderley Park and Alex Alderton, Roberto Amato, Jeffrey Barrett, Sonia Goncalves, Ewan Harrison, David K. Jackson, Ian Johnston, Dominic Kwiatkowski, Cordelia Langford, John Sillitoe on behalf of the Wellcome Sanger Institute COVID-19 Surveillance Team |
| EPI_ISL_2909865, EPI_ISL_2909866 | Berkshire and Surrey Pathology Services Lighthouse Laboratory | Wellcome Sanger Institute for the COVID-19 Genomics UK (COG-UK) Consortium | Berkshire and Surrey Pathology Services Lighthouse Laboratory and Alex Alderton, Roberto Amato, Jeffrey Barrett, Sonia Goncalves, Ewan Harrison, David K. Jackson, Ian Johnston, Dominic Kwiatkowski, Cordelia Langford, John Sillitoe on behalf of the Wellcome Sanger Institute COVID-19 Surveillance Team |
| EPI_ISL_2909867 | Lighthouse Lab in Alderley Park | Wellcome Sanger Institute for the COVID-19 Genomics UK (COG-UK) Consortium | Jacquelyn Wynn, Mairead Hyland, The Lighthouse Lab in Alderley Park and Alex Alderton, Roberto Amato, Jeffrey Barrett, Sonia Goncalves, Ewan Harrison, David K. Jackson, Ian Johnston, Dominic Kwiatkowski, Cordelia Langford, John Sillitoe on behalf of the Wellcome Sanger Institute COVID-19 Surveillance Team |
| EPI_ISL_2909868, EPI_ISL_2909869 | Berkshire and Surrey Pathology Services Lighthouse Laboratory | Wellcome Sanger Institute for the COVID-19 Genomics UK (COG-UK) Consortium | Berkshire and Surrey Pathology Services Lighthouse Laboratory and Alex Alderton, Roberto Amato, Jeffrey Barrett, Sonia Goncalves, Ewan Harrison, David K. Jackson, Ian Johnston, Dominic Kwiatkowski, Cordelia Langford, John Sillitoe on behalf of the Wellcome Sanger Institute COVID-19 Surveillance Team |
| EPI_ISL_2909870, EPI_ISL_2909871, EPI_ISL_2909872 | Lighthouse Lab in Alderley Park | Wellcome Sanger Institute for the COVID-19 Genomics UK (COG-UK) Consortium | Jacquelyn Wynn, Mairead Hyland, The Lighthouse Lab in Alderley Park and Alex Alderton, Roberto Amato, Jeffrey Barrett, Sonia Goncalves, Ewan Harrison, David K. Jackson, Ian Johnston, Dominic Kwiatkowski, Cordelia Langford, John Sillitoe on behalf of the Wellcome Sanger Institute COVID-19 Surveillance Team |
| EPI_ISL_2909873, EPI_ISL_2909874, EPI_ISL_2909875 | Lighthouse Laboratory Plymouth | Wellcome Sanger Institute for the COVID-19 Genomics UK (COG-UK) Consortium | Lighthouse Laboratory Plymouth and Alex Alderton, Roberto Amato, Jeffrey Barrett, Sonia Goncalves, Ewan Harrison, David K. Jackson, Ian Johnston, Dominic Kwiatkowski, Cordelia Langford, John Sillitoe on behalf of the Wellcome Sanger Institute COVID-19 Surveillance Team |
| EPI_ISL_2909876, EPI_ISL_2909877 | Lighthouse Lab in Alderley Park | Wellcome Sanger Institute for the COVID-19 Genomics UK (COG-UK) Consortium | Jacquelyn Wynn, Mairead Hyland, The Lighthouse Lab in Alderley Park and Alex Alderton, Roberto Amato, Jeffrey Barrett, Sonia Goncalves, Ewan Harrison, David K. Jackson, Ian Johnston, Dominic Kwiatkowski, Cordelia Langford, John Sillitoe on behalf of the Wellcome Sanger Institute COVID-19 Surveillance Team |
| EPI_ISL_2909878 | Lighthouse Laboratory Plymouth | Wellcome Sanger Institute for the COVID-19 Genomics UK (COG-UK) Consortium | Lighthouse Laboratory Plymouth and Alex Alderton, Roberto Amato, Jeffrey Barrett, Sonia Goncalves, Ewan Harrison, David K. Jackson, Ian Johnston, Dominic Kwiatkowski, Cordelia Langford, John Sillitoe on behalf of the Wellcome Sanger Institute COVID-19 Surveillance Team |
| EPI_ISL_2909879 | Lighthouse Lab in Alderley Park | Wellcome Sanger Institute for the COVID-19 Genomics UK (COG-UK) Consortium | Jacquelyn Wynn, Mairead Hyland, The Lighthouse Lab in Alderley Park and Alex Alderton, Roberto Amato, Jeffrey Barrett, Sonia Goncalves, Ewan Harrison, David K. Jackson, Ian Johnston, Dominic Kwiatkowski, Cordelia Langford, John Sillitoe on behalf of the Wellcome Sanger Institute COVID-19 Surveillance Team |
| EPI_ISL_2909880 | Lighthouse Laboratory Plymouth | Wellcome Sanger Institute for the COVID-19 Genomics UK (COG-UK) Consortium | Lighthouse Laboratory Plymouth and Alex Alderton, Roberto Amato, Jeffrey Barrett, Sonia Goncalves, Ewan Harrison, David K. Jackson, Ian Johnston, Dominic Kwiatkowski, Cordelia Langford, John Sillitoe on behalf of the Wellcome Sanger Institute COVID-19 Surveillance Team |
| EPI_ISL_2909881 | Lighthouse Lab in Alderley Park | Wellcome Sanger Institute for the COVID-19 Genomics UK (COG-UK) Consortium | Jacquelyn Wynn, Mairead Hyland, The Lighthouse Lab in Alderley Park and Alex Alderton, Roberto Amato, Jeffrey Barrett, Sonia Goncalves, Ewan Harrison, David K. Jackson, Ian Johnston, Dominic Kwiatkowski, Cordelia Langford, John Sillitoe on behalf of the Wellcome Sanger Institute COVID-19 Surveillance Team |
| EPI_ISL_2909882 | Lighthouse Laboratory Plymouth | Wellcome Sanger Institute for the COVID-19 Genomics UK (COG-UK) Consortium | Lighthouse Laboratory Plymouth and Alex Alderton, Roberto Amato, Jeffrey Barrett, Sonia Goncalves, Ewan Harrison, David K. Jackson, Ian Johnston, Dominic Kwiatkowski, Cordelia Langford, John Sillitoe on behalf of the Wellcome Sanger Institute COVID-19 Surveillance Team |
| EPI_ISL_2909883, EPI_ISL_2909884, EPI_ISL_2909885, EPI_ISL_2909886 | Lighthouse Lab in Alderley Park | Wellcome Sanger Institute for the COVID-19 Genomics UK (COG-UK) Consortium | Jacquelyn Wynn, Mairead Hyland, The Lighthouse Lab in Alderley Park and Alex Alderton, Roberto Amato, Jeffrey Barrett, Sonia Goncalves, Ewan Harrison, David K. Jackson, Ian Johnston, Dominic Kwiatkowski, Cordelia Langford, John Sillitoe on behalf of the Wellcome Sanger Institute COVID-19 Surveillance Team |
| EPI_ISL_2909887 | Berkshire and Surrey Pathology Services Lighthouse Laboratory | Wellcome Sanger Institute for the COVID-19 Genomics UK (COG-UK) Consortium | Berkshire and Surrey Pathology Services Lighthouse Laboratory and Alex Alderton, Roberto Amato, Jeffrey Barrett, Sonia Goncalves, Ewan Harrison, David K. Jackson, Ian Johnston, Dominic Kwiatkowski, Cordelia Langford, John Sillitoe on behalf of the Wellcome Sanger Institute COVID-19 Surveillance Team |
| EPI_ISL_2909888, EPI_ISL_2909889, EPI_ISL_2909890, EPI_ISL_2909891 | Lighthouse Lab in Alderley Park | Wellcome Sanger Institute for the COVID-19 Genomics UK (COG-UK) Consortium | Jacquelyn Wynn, Mairead Hyland, The Lighthouse Lab in Alderley Park and Alex Alderton, Roberto Amato, Jeffrey Barrett, Sonia Goncalves, Ewan Harrison, David K. Jackson, Ian Johnston, Dominic Kwiatkowski, Cordelia Langford, John Sillitoe on behalf of the Wellcome Sanger Institute COVID-19 Surveillance Team |
| EPI_ISL_2909892 | Lighthouse Laboratory Plymouth | Wellcome Sanger Institute for the COVID-19 Genomics UK (COG-UK) Consortium | Lighthouse Laboratory Plymouth and Alex Alderton, Roberto Amato, Jeffrey Barrett, Sonia Goncalves, Ewan Harrison, David K. Jackson, Ian Johnston, Dominic Kwiatkowski, Cordelia Langford, John Sillitoe on behalf of the Wellcome Sanger Institute COVID-19 Surveillance Team |
| EPI_ISL_2909893, EPI_ISL_2909894, EPI_ISL_2909895, EPI_ISL_2909896, EPI_ISL_2909897, EPI_ISL_2909898, EPI_ISL_2909899 | Lighthouse Lab in Alderley Park | Wellcome Sanger Institute for the COVID-19 Genomics UK (COG-UK) Consortium | Jacquelyn Wynn, Mairead Hyland, The Lighthouse Lab in Alderley Park and Alex Alderton, Roberto Amato, Jeffrey Barrett, Sonia Goncalves, Ewan Harrison, David K. Jackson, Ian Johnston, Dominic Kwiatkowski, Cordelia Langford, John Sillitoe on behalf of the Wellcome Sanger Institute COVID-19 Surveillance Team |
| EPI_ISL_2909900 | Lighthouse Lab in Milton Keynes | Wellcome Sanger Institute for the COVID-19 Genomics UK (COG-UK) Consortium | The Lighthouse Lab in Milton Keynes and Alex Alderton, Roberto Amato, Jeffrey Barrett, Sonia Goncalves, Ewan Harrison, David K. Jackson, Ian Johnston, Dominic Kwiatkowski, Cordelia Langford, John Sillitoe on behalf of the Wellcome Sanger Institute COVID-19 Surveillance Team |
| EPI_ISL_2909901 | Lighthouse Laboratory Plymouth | Wellcome Sanger Institute for the COVID-19 Genomics UK (COG-UK) Consortium | Lighthouse Laboratory Plymouth and Alex Alderton, Roberto Amato, Jeffrey Barrett, Sonia Goncalves, Ewan Harrison, David K. Jackson, Ian Johnston, Dominic Kwiatkowski, Cordelia Langford, John Sillitoe on behalf of the Wellcome Sanger Institute COVID-19 Surveillance Team |
| EPI_ISL_2909902 | Lighthouse Lab in Milton Keynes | Wellcome Sanger Institute for the COVID-19 Genomics UK (COG-UK) Consortium | The Lighthouse Lab in Milton Keynes and Alex Alderton, Roberto Amato, Jeffrey Barrett, Sonia Goncalves, Ewan Harrison, David K. Jackson, Ian Johnston, Dominic Kwiatkowski, Cordelia Langford, John Sillitoe on behalf of the Wellcome Sanger Institute COVID-19 Surveillance Team |
| EPI_ISL_2909903 | Lighthouse Laboratory Plymouth | Wellcome Sanger Institute for the COVID-19 Genomics UK (COG-UK) Consortium | Lighthouse Laboratory Plymouth and Alex Alderton, Roberto Amato, Jeffrey Barrett, Sonia Goncalves, Ewan Harrison, David K. Jackson, Ian Johnston, Dominic Kwiatkowski, Cordelia Langford, John Sillitoe on behalf of the Wellcome Sanger Institute COVID-19 Surveillance Team |
| EPI_ISL_2909904, EPI_ISL_2909905, EPI_ISL_2909906, | Lighthouse Lab in Alderley Park | Wellcome Sanger Institute for the COVID-19 Genomics UK (COG-UK) Consortium | Jacquelyn Wynn, Mairead Hyland, The Lighthouse Lab in Alderley Park and Alex Alderton, Roberto Amato, Jeffrey Barrett, Sonia Goncalves, Ewan Harrison, David K. Jackson, Ian Johnston, Dominic Kwiatkowski, Cordelia Langford, John Sillitoe on behalf of the Wellcome Sanger Institute COVID-19 Surveillance Team |

| | | | |
|---|---|---|---|
| EPI_ISL_2909907, EPI_ISL_2909908, EPI_ISL_2909909, EPI_ISL_2909910, EPI_ISL_2909911 | | | |
| EPI_ISL_2909912 | Lighthouse Laboratory Plymouth | Wellcome Sanger Institute for the COVID-19 Genomics UK (COG-UK) Consortium | Lighthouse Laboratory Plymouth and Alex Alderton, Roberto Amato, Jeffrey Barrett, Sonia Goncalves, Ewan Harrison, David K. Jackson, Ian Johnston, Dominic Kwiatkowski, Cordelia Langford, John Sillitoe on behalf of the Wellcome Sanger Institute COVID-19 Surveillance Team |
| EPI_ISL_2909913 | Lighthouse Lab in Alderley Park | Wellcome Sanger Institute for the COVID-19 Genomics UK (COG-UK) Consortium | Jacquelyn Wynn, Mairead Hyland, The Lighthouse Lab in Alderley Park and Alex Alderton, Roberto Amato, Jeffrey Barrett, Sonia Goncalves, Ewan Harrison, David K. Jackson, Ian Johnston, Dominic Kwiatkowski, Cordelia Langford, John Sillitoe on behalf of the Wellcome Sanger Institute COVID-19 Surveillance Team |
| EPI_ISL_2909914 | Lighthouse Laboratory Plymouth | Wellcome Sanger Institute for the COVID-19 Genomics UK (COG-UK) Consortium | Lighthouse Laboratory Plymouth and Alex Alderton, Roberto Amato, Jeffrey Barrett, Sonia Goncalves, Ewan Harrison, David K. Jackson, Ian Johnston, Dominic Kwiatkowski, Cordelia Langford, John Sillitoe on behalf of the Wellcome Sanger Institute COVID-19 Surveillance Team |
| EPI_ISL_2909915 | Lighthouse Lab in Alderley Park | Wellcome Sanger Institute for the COVID-19 Genomics UK (COG-UK) Consortium | Jacquelyn Wynn, Mairead Hyland, The Lighthouse Lab in Alderley Park and Alex Alderton, Roberto Amato, Jeffrey Barrett, Sonia Goncalves, Ewan Harrison, David K. Jackson, Ian Johnston, Dominic Kwiatkowski, Cordelia Langford, John Sillitoe on behalf of the Wellcome Sanger Institute COVID-19 Surveillance Team |
| EPI_ISL_2909916, EPI_ISL_2909917 | Lighthouse Lab in Milton Keynes | Wellcome Sanger Institute for the COVID-19 Genomics UK (COG-UK) Consortium | The Lighthouse Lab in Milton Keynes and Alex Alderton, Roberto Amato, Jeffrey Barrett, Sonia Goncalves, Ewan Harrison, David K. Jackson, Ian Johnston, Dominic Kwiatkowski, Cordelia Langford, John Sillitoe on behalf of the Wellcome Sanger Institute COVID-19 Surveillance Team |
| EPI_ISL_2909918, EPI_ISL_2909919, EPI_ISL_2909920, EPI_ISL_2909921 | Lighthouse Lab in Alderley Park | Wellcome Sanger Institute for the COVID-19 Genomics UK (COG-UK) Consortium | Jacquelyn Wynn, Mairead Hyland, The Lighthouse Lab in Alderley Park and Alex Alderton, Roberto Amato, Jeffrey Barrett, Sonia Goncalves, Ewan Harrison, David K. Jackson, Ian Johnston, Dominic Kwiatkowski, Cordelia Langford, John Sillitoe on behalf of the Wellcome Sanger Institute COVID-19 Surveillance Team |
| EPI_ISL_2909922 | Lighthouse Lab in Milton Keynes | Wellcome Sanger Institute for the COVID-19 Genomics UK (COG-UK) Consortium | The Lighthouse Lab in Milton Keynes and Alex Alderton, Roberto Amato, Jeffrey Barrett, Sonia Goncalves, Ewan Harrison, David K. Jackson, Ian Johnston, Dominic Kwiatkowski, Cordelia Langford, John Sillitoe on behalf of the Wellcome Sanger Institute COVID-19 Surveillance Team |
| EPI_ISL_2909923 | Lighthouse Lab in Alderley Park | Wellcome Sanger Institute for the COVID-19 Genomics UK (COG-UK) Consortium | Jacquelyn Wynn, Mairead Hyland, The Lighthouse Lab in Alderley Park and Alex Alderton, Roberto Amato, Jeffrey Barrett, Sonia Goncalves, Ewan Harrison, David K. Jackson, Ian Johnston, Dominic Kwiatkowski, Cordelia Langford, John Sillitoe on behalf of the Wellcome Sanger Institute COVID-19 Surveillance Team |
| EPI_ISL_2909924 | Lighthouse Laboratory Plymouth | Wellcome Sanger Institute for the COVID-19 Genomics UK (COG-UK) Consortium | Lighthouse Laboratory Plymouth and Alex Alderton, Roberto Amato, Jeffrey Barrett, Sonia Goncalves, Ewan Harrison, David K. Jackson, Ian Johnston, Dominic Kwiatkowski, Cordelia Langford, John Sillitoe on behalf of the Wellcome Sanger Institute COVID-19 Surveillance Team |
| EPI_ISL_2909925, EPI_ISL_2909926 | Lighthouse Lab in Alderley Park | Wellcome Sanger Institute for the COVID-19 Genomics UK (COG-UK) Consortium | Jacquelyn Wynn, Mairead Hyland, The Lighthouse Lab in Alderley Park and Alex Alderton, Roberto Amato, Jeffrey Barrett, Sonia Goncalves, Ewan Harrison, David K. Jackson, Ian Johnston, Dominic Kwiatkowski, Cordelia Langford, John Sillitoe on behalf of the Wellcome Sanger Institute COVID-19 Surveillance Team |
| EPI_ISL_2909927 | Lighthouse Lab in Milton Keynes | Wellcome Sanger Institute for the COVID-19 Genomics UK (COG-UK) Consortium | The Lighthouse Lab in Milton Keynes and Alex Alderton, Roberto Amato, Jeffrey Barrett, Sonia Goncalves, Ewan Harrison, David K. Jackson, Ian Johnston, Dominic Kwiatkowski, Cordelia Langford, John Sillitoe on behalf of the Wellcome Sanger Institute COVID-19 Surveillance Team |
| EPI_ISL_2909928 | Lighthouse Lab in Alderley Park | Wellcome Sanger Institute for the COVID-19 Genomics UK (COG-UK) Consortium | Jacquelyn Wynn, Mairead Hyland, The Lighthouse Lab in Alderley Park and Alex Alderton, Roberto Amato, Jeffrey Barrett, Sonia Goncalves, Ewan Harrison, David K. Jackson, Ian Johnston, Dominic Kwiatkowski, Cordelia Langford, John Sillitoe on behalf of the Wellcome Sanger Institute COVID-19 Surveillance Team |
| EPI_ISL_2909929 | Lighthouse Lab in Milton Keynes | Wellcome Sanger Institute for the COVID-19 Genomics UK (COG-UK) Consortium | The Lighthouse Lab in Milton Keynes and Alex Alderton, Roberto Amato, Jeffrey Barrett, Sonia Goncalves, Ewan Harrison, David K. Jackson, Ian Johnston, Dominic Kwiatkowski, Cordelia Langford, John Sillitoe on behalf of the Wellcome Sanger Institute COVID-19 Surveillance Team |
| EPI_ISL_2909930 | Lighthouse Lab in Alderley Park | Wellcome Sanger Institute for the COVID-19 Genomics UK (COG-UK) Consortium | Jacquelyn Wynn, Mairead Hyland, The Lighthouse Lab in Alderley Park and Alex Alderton, Roberto Amato, Jeffrey Barrett, Sonia Goncalves, Ewan Harrison, David K. Jackson, Ian Johnston, Dominic Kwiatkowski, Cordelia Langford, John Sillitoe on behalf of the Wellcome Sanger Institute COVID-19 Surveillance Team |
| EPI_ISL_2909931, EPI_ISL_2909932 | Lighthouse Lab in Milton Keynes | Wellcome Sanger Institute for the COVID-19 Genomics UK (COG-UK) Consortium | The Lighthouse Lab in Milton Keynes and Alex Alderton, Roberto Amato, Jeffrey Barrett, Sonia Goncalves, Ewan Harrison, David K. Jackson, Ian Johnston, Dominic Kwiatkowski, Cordelia Langford, John Sillitoe on behalf of the Wellcome Sanger Institute COVID-19 Surveillance Team |
| EPI_ISL_2909933 | Lighthouse Laboratory Plymouth | Wellcome Sanger Institute for the COVID-19 Genomics UK (COG-UK) Consortium | Lighthouse Laboratory Plymouth and Alex Alderton, Roberto Amato, Jeffrey Barrett, Sonia Goncalves, Ewan Harrison, David K. Jackson, Ian Johnston, Dominic Kwiatkowski, Cordelia Langford, John Sillitoe on behalf of the Wellcome Sanger Institute COVID-19 Surveillance Team |
| EPI_ISL_2909934, EPI_ISL_2909935, EPI_ISL_2909936 | Lighthouse Lab in Alderley Park | Wellcome Sanger Institute for the COVID-19 Genomics UK (COG-UK) Consortium | Jacquelyn Wynn, Mairead Hyland, The Lighthouse Lab in Alderley Park and Alex Alderton, Roberto Amato, Jeffrey Barrett, Sonia Goncalves, Ewan Harrison, David K. Jackson, Ian Johnston, Dominic Kwiatkowski, Cordelia Langford, John Sillitoe on behalf of the Wellcome Sanger Institute COVID-19 Surveillance Team |
| EPI_ISL_2909937, EPI_ISL_2909938, EPI_ISL_2909939 | Lighthouse Laboratory Plymouth | Wellcome Sanger Institute for the COVID-19 Genomics UK (COG-UK) Consortium | Lighthouse Laboratory Plymouth and Alex Alderton, Roberto Amato, Jeffrey Barrett, Sonia Goncalves, Ewan Harrison, David K. Jackson, Ian Johnston, Dominic Kwiatkowski, Cordelia Langford, John Sillitoe on behalf of the Wellcome Sanger Institute COVID-19 Surveillance Team |
| EPI_ISL_2909940 | Lighthouse Lab in Alderley Park | Wellcome Sanger Institute for the COVID-19 Genomics UK (COG-UK) Consortium | Jacquelyn Wynn, Mairead Hyland, The Lighthouse Lab in Alderley Park and Alex Alderton, Roberto Amato, Jeffrey Barrett, Sonia Goncalves, Ewan Harrison, David K. Jackson, Ian Johnston, Dominic Kwiatkowski, Cordelia Langford, John Sillitoe on behalf of the Wellcome Sanger Institute COVID-19 Surveillance Team |
| EPI_ISL_2909941, EPI_ISL_2909942, EPI_ISL_2909943 | Lighthouse Laboratory Plymouth | Wellcome Sanger Institute for the COVID-19 Genomics UK (COG-UK) Consortium | Lighthouse Laboratory Plymouth and Alex Alderton, Roberto Amato, Jeffrey Barrett, Sonia Goncalves, Ewan Harrison, David K. Jackson, Ian Johnston, Dominic Kwiatkowski, Cordelia Langford, John Sillitoe on behalf of the Wellcome Sanger Institute COVID-19 Surveillance Team |
| EPI_ISL_2909944 | Lighthouse Lab in Alderley Park | Wellcome Sanger Institute for the COVID-19 Genomics UK (COG-UK) Consortium | Jacquelyn Wynn, Mairead Hyland, The Lighthouse Lab in Alderley Park and Alex Alderton, Roberto Amato, Jeffrey Barrett, Sonia Goncalves, Ewan Harrison, David K. Jackson, Ian Johnston, Dominic Kwiatkowski, Cordelia Langford, John Sillitoe on behalf of the Wellcome Sanger Institute COVID-19 Surveillance Team |
| EPI_ISL_2909945 | Lighthouse Lab in Milton Keynes | Wellcome Sanger Institute for the COVID-19 Genomics UK (COG-UK) Consortium | The Lighthouse Lab in Milton Keynes and Alex Alderton, Roberto Amato, Jeffrey Barrett, Sonia Goncalves, Ewan Harrison, David K. Jackson, Ian Johnston, Dominic Kwiatkowski, Cordelia Langford, John Sillitoe on behalf of the Wellcome Sanger Institute COVID-19 Surveillance Team |
| EPI_ISL_2909946 | Lighthouse Lab in Alderley Park | Wellcome Sanger Institute for the COVID-19 Genomics UK (COG-UK) Consortium | Jacquelyn Wynn, Mairead Hyland, The Lighthouse Lab in Alderley Park and Alex Alderton, Roberto Amato, Jeffrey Barrett, Sonia Goncalves, Ewan Harrison, David K. Jackson, Ian Johnston, Dominic Kwiatkowski, Cordelia Langford, John Sillitoe on behalf of the Wellcome Sanger Institute COVID-19 Surveillance Team |
| EPI_ISL_2909947, EPI_ISL_2909948 | Lighthouse Laboratory Plymouth | Wellcome Sanger Institute for the COVID-19 Genomics UK (COG-UK) Consortium | Lighthouse Laboratory Plymouth and Alex Alderton, Roberto Amato, Jeffrey Barrett, Sonia Goncalves, Ewan Harrison, David K. Jackson, Ian Johnston, Dominic Kwiatkowski, Cordelia Langford, John Sillitoe on behalf of the Wellcome Sanger Institute COVID-19 Surveillance Team |
| EPI_ISL_2909949 | Lighthouse Lab in Alderley Park | Wellcome Sanger Institute for the COVID-19 Genomics UK (COG-UK) Consortium | Jacquelyn Wynn, Mairead Hyland, The Lighthouse Lab in Alderley Park and Alex Alderton, Roberto Amato, Jeffrey Barrett, Sonia Goncalves, Ewan Harrison, David K. Jackson, Ian Johnston, Dominic Kwiatkowski, Cordelia Langford, John Sillitoe on behalf of the Wellcome Sanger Institute COVID-19 Surveillance Team |
| EPI_ISL_2909950 | Lighthouse Laboratory Plymouth | Wellcome Sanger Institute for the COVID-19 Genomics UK (COG-UK) Consortium | Lighthouse Laboratory Plymouth and Alex Alderton, Roberto Amato, Jeffrey Barrett, Sonia Goncalves, Ewan Harrison, David K. Jackson, Ian Johnston, Dominic Kwiatkowski, Cordelia Langford, John Sillitoe on behalf of the Wellcome Sanger Institute COVID-19 Surveillance Team |
| EPI_ISL_2909951, EPI_ISL_2909952 | Lighthouse Lab in Milton Keynes | Wellcome Sanger Institute for the COVID-19 Genomics UK (COG-UK) Consortium | The Lighthouse Lab in Milton Keynes and Alex Alderton, Roberto Amato, Jeffrey Barrett, Sonia Goncalves, Ewan Harrison, David K. Jackson, Ian Johnston, Dominic Kwiatkowski, Cordelia Langford, John Sillitoe on behalf of the Wellcome Sanger Institute COVID-19 Surveillance Team |
| EPI_ISL_2909953 | Lighthouse Lab in Alderley Park | Wellcome Sanger Institute for the COVID-19 Genomics UK (COG-UK) Consortium | Jacquelyn Wynn, Mairead Hyland, The Lighthouse Lab in Alderley Park and Alex Alderton, Roberto Amato, Jeffrey Barrett, Sonia Goncalves, Ewan Harrison, David K. Jackson, Ian Johnston, Dominic Kwiatkowski, Cordelia Langford, John Sillitoe on behalf of the Wellcome Sanger Institute COVID-19 Surveillance Team |
| EPI_ISL_2909954 | Lighthouse Laboratory Plymouth | Wellcome Sanger Institute for the COVID-19 Genomics UK (COG-UK) Consortium | Lighthouse Laboratory Plymouth and Alex Alderton, Roberto Amato, Jeffrey Barrett, Sonia Goncalves, Ewan Harrison, David K. Jackson, Ian Johnston, Dominic Kwiatkowski, Cordelia Langford, John Sillitoe on behalf of the Wellcome Sanger Institute COVID-19 Surveillance Team |
| EPI_ISL_2909955 | Lighthouse Lab in Alderley Park | Wellcome Sanger Institute for the COVID-19 Genomics UK (COG-UK) Consortium | Jacquelyn Wynn, Mairead Hyland, The Lighthouse Lab in Alderley Park and Alex Alderton, Roberto Amato, Jeffrey Barrett, Sonia Goncalves, Ewan Harrison, David K. Jackson, Ian Johnston, Dominic Kwiatkowski, Cordelia Langford, John Sillitoe on behalf of the Wellcome Sanger Institute COVID-19 Surveillance Team |
| EPI_ISL_2909956 | Lighthouse Laboratory Plymouth | Wellcome Sanger Institute for the COVID-19 Genomics UK (COG-UK) Consortium | Lighthouse Laboratory Plymouth and Alex Alderton, Roberto Amato, Jeffrey Barrett, Sonia Goncalves, Ewan Harrison, David K. Jackson, Ian Johnston, Dominic Kwiatkowski, Cordelia Langford, John Sillitoe on behalf of the Wellcome Sanger Institute COVID-19 Surveillance Team |
| EPI_ISL_2909957, EPI_ISL_2909958 | Lighthouse Lab in Milton Keynes | Wellcome Sanger Institute for the COVID-19 Genomics UK (COG-UK) Consortium | The Lighthouse Lab in Milton Keynes and Alex Alderton, Roberto Amato, Jeffrey Barrett, Sonia Goncalves, Ewan Harrison, David K. Jackson, Ian Johnston, Dominic Kwiatkowski, Cordelia Langford, John Sillitoe on behalf of the Wellcome Sanger Institute COVID-19 Surveillance Team |
| EPI_ISL_2909959 | Lighthouse Lab in Alderley Park | Wellcome Sanger Institute for the COVID-19 Genomics UK (COG-UK) Consortium | Jacquelyn Wynn, Mairead Hyland, The Lighthouse Lab in Alderley Park and Alex Alderton, Roberto Amato, Jeffrey Barrett, Sonia Goncalves, Ewan Harrison, David K. Jackson, Ian Johnston, Dominic Kwiatkowski, Cordelia Langford, John Sillitoe on behalf of the Wellcome Sanger Institute COVID-19 Surveillance Team |
| EPI_ISL_2909960 | Lighthouse Lab in Milton Keynes | Wellcome Sanger Institute for the COVID-19 Genomics UK (COG-UK) Consortium | The Lighthouse Lab in Milton Keynes and Alex Alderton, Roberto Amato, Jeffrey Barrett, Sonia Goncalves, Ewan Harrison, David K. Jackson, Ian Johnston, Dominic Kwiatkowski, Cordelia Langford, John Sillitoe on behalf of the Wellcome Sanger Institute COVID-19 Surveillance Team |
| EPI_ISL_2909961 | Lighthouse Lab in Alderley Park | Wellcome Sanger Institute for the COVID-19 Genomics UK (COG-UK) Consortium | Jacquelyn Wynn, Mairead Hyland, The Lighthouse Lab in Alderley Park and Alex Alderton, Roberto Amato, Jeffrey Barrett, Sonia Goncalves, Ewan Harrison, David K. Jackson, Ian Johnston, Dominic Kwiatkowski, Cordelia Langford, John Sillitoe on behalf of the Wellcome Sanger Institute COVID-19 Surveillance Team |
| EPI_ISL_2909962 | Lighthouse Lab in Milton Keynes | Wellcome Sanger Institute for the COVID-19 Genomics UK (COG-UK) Consortium | The Lighthouse Lab in Milton Keynes and Alex Alderton, Roberto Amato, Jeffrey Barrett, Sonia Goncalves, Ewan Harrison, David K. Jackson, Ian Johnston, Dominic Kwiatkowski, Cordelia Langford, John Sillitoe on behalf of the Wellcome Sanger Institute COVID-19 Surveillance Team |
| EPI_ISL_2909963 | Lighthouse Lab in Alderley Park | Wellcome Sanger Institute for the COVID-19 Genomics UK (COG-UK) Consortium | Jacquelyn Wynn, Mairead Hyland, The Lighthouse Lab in Alderley Park and Alex Alderton, Roberto Amato, Jeffrey Barrett, Sonia Goncalves, Ewan Harrison, David K. Jackson, Ian Johnston, Dominic Kwiatkowski, Cordelia Langford, John Sillitoe on behalf of the Wellcome Sanger Institute COVID-19 Surveillance Team |
| EPI_ISL_2909964 | Lighthouse Lab in Milton Keynes | Wellcome Sanger Institute for the COVID-19 Genomics UK (COG-UK) Consortium | The Lighthouse Lab in Milton Keynes and Alex Alderton, Roberto Amato, Jeffrey Barrett, Sonia Goncalves, Ewan Harrison, David K. Jackson, Ian Johnston, Dominic Kwiatkowski, Cordelia Langford, John Sillitoe on behalf of the Wellcome Sanger Institute COVID-19 Surveillance Team |
| EPI_ISL_2909965 | Lighthouse Laboratory Plymouth | Wellcome Sanger Institute for the COVID-19 Genomics UK (COG-UK) Consortium | Lighthouse Laboratory Plymouth and Alex Alderton, Roberto Amato, Jeffrey Barrett, Sonia Goncalves, Ewan Harrison, David K. Jackson, Ian Johnston, Dominic Kwiatkowski, Cordelia Langford, John Sillitoe on behalf of the Wellcome Sanger Institute COVID-19 Surveillance Team |
| EPI_ISL_2909966, EPI_ISL_2909967 | Lighthouse Lab in Alderley Park | Wellcome Sanger Institute for the COVID-19 Genomics UK (COG-UK) Consortium | Jacquelyn Wynn, Mairead Hyland, The Lighthouse Lab in Alderley Park and Alex Alderton, Roberto Amato, Jeffrey Barrett, Sonia Goncalves, Ewan Harrison, David K. Jackson, Ian Johnston, Dominic Kwiatkowski, Cordelia Langford, John Sillitoe on behalf of the Wellcome Sanger Institute COVID-19 Surveillance Team |
| EPI_ISL_2909968 | Lighthouse Lab in Milton Keynes | Wellcome Sanger Institute for the COVID-19 Genomics UK (COG-UK) | The Lighthouse Lab in Milton Keynes and Alex Alderton, Roberto Amato, Jeffrey Barrett, Sonia Goncalves, Ewan Harrison, David K. Jackson, Ian Johnston, Dominic Kwiatkowski, Cordelia Langford, John Sillitoe on behalf of the Wellcome Sanger |

| | | | |
|---|---|---|---|
| | | Consortium | Institute COVID-19 Surveillance Team |
| EPI_ISL_2909969, EPI_ISL_2909970 | Lighthouse Laboratory Plymouth | Wellcome Sanger Institute for the COVID-19 Genomics UK (COG-UK) Consortium | Lighthouse Laboratory Plymouth and Alex Alderton, Roberto Amato, Jeffrey Barrett, Sonia Goncalves, Ewan Harrison, David K. Jackson, Ian Johnston, Dominic Kwiatkowski, Cordelia Langford, John Sillitoe on behalf of the Wellcome Sanger Institute COVID-19 Surveillance Team |
| EPI_ISL_2909971, EPI_ISL_2909972, EPI_ISL_2909973 | Lighthouse Lab in Alderley Park | Wellcome Sanger Institute for the COVID-19 Genomics UK (COG-UK) Consortium | Jacquelyn Wynn, Mairead Hyland, The Lighthouse Lab in Alderley Park and Alex Alderton, Roberto Amato, Jeffrey Barrett, Sonia Goncalves, Ewan Harrison, David K. Jackson, Ian Johnston, Dominic Kwiatkowski, Cordelia Langford, John Sillitoe on behalf of the Wellcome Sanger Institute COVID-19 Surveillance Team |
| EPI_ISL_2909974 | Lighthouse Laboratory Plymouth | Wellcome Sanger Institute for the COVID-19 Genomics UK (COG-UK) Consortium | Lighthouse Laboratory Plymouth and Alex Alderton, Roberto Amato, Jeffrey Barrett, Sonia Goncalves, Ewan Harrison, David K. Jackson, Ian Johnston, Dominic Kwiatkowski, Cordelia Langford, John Sillitoe on behalf of the Wellcome Sanger Institute COVID-19 Surveillance Team |
| EPI_ISL_2909975, EPI_ISL_2909976 | Lighthouse Lab in Milton Keynes | Wellcome Sanger Institute for the COVID-19 Genomics UK (COG-UK) Consortium | The Lighthouse Lab in Milton Keynes and Alex Alderton, Roberto Amato, Jeffrey Barrett, Sonia Goncalves, Ewan Harrison, David K. Jackson, Ian Johnston, Dominic Kwiatkowski, Cordelia Langford, John Sillitoe on behalf of the Wellcome Sanger Institute COVID-19 Surveillance Team |
| EPI_ISL_2909977, EPI_ISL_2909978, EPI_ISL_2909979 | Lighthouse Laboratory Plymouth | Wellcome Sanger Institute for the COVID-19 Genomics UK (COG-UK) Consortium | Lighthouse Laboratory Plymouth and Alex Alderton, Roberto Amato, Jeffrey Barrett, Sonia Goncalves, Ewan Harrison, David K. Jackson, Ian Johnston, Dominic Kwiatkowski, Cordelia Langford, John Sillitoe on behalf of the Wellcome Sanger Institute COVID-19 Surveillance Team |
| EPI_ISL_2909980 | Lighthouse Lab in Alderley Park | Wellcome Sanger Institute for the COVID-19 Genomics UK (COG-UK) Consortium | Jacquelyn Wynn, Mairead Hyland, The Lighthouse Lab in Alderley Park and Alex Alderton, Roberto Amato, Jeffrey Barrett, Sonia Goncalves, Ewan Harrison, David K. Jackson, Ian Johnston, Dominic Kwiatkowski, Cordelia Langford, John Sillitoe on behalf of the Wellcome Sanger Institute COVID-19 Surveillance Team |
| EPI_ISL_2909981 | Lighthouse Lab in Milton Keynes | Wellcome Sanger Institute for the COVID-19 Genomics UK (COG-UK) Consortium | The Lighthouse Lab in Milton Keynes and Alex Alderton, Roberto Amato, Jeffrey Barrett, Sonia Goncalves, Ewan Harrison, David K. Jackson, Ian Johnston, Dominic Kwiatkowski, Cordelia Langford, John Sillitoe on behalf of the Wellcome Sanger Institute COVID-19 Surveillance Team |
| EPI_ISL_2909982 | Lighthouse Lab in Alderley Park | Wellcome Sanger Institute for the COVID-19 Genomics UK (COG-UK) Consortium | Jacquelyn Wynn, Mairead Hyland, The Lighthouse Lab in Alderley Park and Alex Alderton, Roberto Amato, Jeffrey Barrett, Sonia Goncalves, Ewan Harrison, David K. Jackson, Ian Johnston, Dominic Kwiatkowski, Cordelia Langford, John Sillitoe on behalf of the Wellcome Sanger Institute COVID-19 Surveillance Team |
| EPI_ISL_2909983 | Lighthouse Lab in Milton Keynes | Wellcome Sanger Institute for the COVID-19 Genomics UK (COG-UK) Consortium | The Lighthouse Lab in Milton Keynes and Alex Alderton, Roberto Amato, Jeffrey Barrett, Sonia Goncalves, Ewan Harrison, David K. Jackson, Ian Johnston, Dominic Kwiatkowski, Cordelia Langford, John Sillitoe on behalf of the Wellcome Sanger Institute COVID-19 Surveillance Team |
| EPI_ISL_2909984 | Lighthouse Lab in Alderley Park | Wellcome Sanger Institute for the COVID-19 Genomics UK (COG-UK) Consortium | Jacquelyn Wynn, Mairead Hyland, The Lighthouse Lab in Alderley Park and Alex Alderton, Roberto Amato, Jeffrey Barrett, Sonia Goncalves, Ewan Harrison, David K. Jackson, Ian Johnston, Dominic Kwiatkowski, Cordelia Langford, John Sillitoe on behalf of the Wellcome Sanger Institute COVID-19 Surveillance Team |
| EPI_ISL_2909985 | Lighthouse Laboratory Plymouth | Wellcome Sanger Institute for the COVID-19 Genomics UK (COG-UK) Consortium | Lighthouse Laboratory Plymouth and Alex Alderton, Roberto Amato, Jeffrey Barrett, Sonia Goncalves, Ewan Harrison, David K. Jackson, Ian Johnston, Dominic Kwiatkowski, Cordelia Langford, John Sillitoe on behalf of the Wellcome Sanger Institute COVID-19 Surveillance Team |
| EPI_ISL_2909986, EPI_ISL_2909987, EPI_ISL_2909988, EPI_ISL_2909989, EPI_ISL_2909990, EPI_ISL_2909991, EPI_ISL_2909992, EPI_ISL_2909993 | Lighthouse Lab in Alderley Park | Wellcome Sanger Institute for the COVID-19 Genomics UK (COG-UK) Consortium | Jacquelyn Wynn, Mairead Hyland, The Lighthouse Lab in Alderley Park and Alex Alderton, Roberto Amato, Jeffrey Barrett, Sonia Goncalves, Ewan Harrison, David K. Jackson, Ian Johnston, Dominic Kwiatkowski, Cordelia Langford, John Sillitoe on behalf of the Wellcome Sanger Institute COVID-19 Surveillance Team |
| EPI_ISL_2909994 | Lighthouse Laboratory Plymouth | Wellcome Sanger Institute for the COVID-19 Genomics UK (COG-UK) Consortium | Lighthouse Laboratory Plymouth and Alex Alderton, Roberto Amato, Jeffrey Barrett, Sonia Goncalves, Ewan Harrison, David K. Jackson, Ian Johnston, Dominic Kwiatkowski, Cordelia Langford, John Sillitoe on behalf of the Wellcome Sanger Institute COVID-19 Surveillance Team |
| EPI_ISL_2909995, EPI_ISL_2909996, EPI_ISL_2909997 | Lighthouse Lab in Alderley Park | Wellcome Sanger Institute for the COVID-19 Genomics UK (COG-UK) Consortium | Jacquelyn Wynn, Mairead Hyland, The Lighthouse Lab in Alderley Park and Alex Alderton, Roberto Amato, Jeffrey Barrett, Sonia Goncalves, Ewan Harrison, David K. Jackson, Ian Johnston, Dominic Kwiatkowski, Cordelia Langford, John Sillitoe on behalf of the Wellcome Sanger Institute COVID-19 Surveillance Team |
| EPI_ISL_2909998 | Lighthouse Lab in Milton Keynes | Wellcome Sanger Institute for the COVID-19 Genomics UK (COG-UK) Consortium | The Lighthouse Lab in Milton Keynes and Alex Alderton, Roberto Amato, Jeffrey Barrett, Sonia Goncalves, Ewan Harrison, David K. Jackson, Ian Johnston, Dominic Kwiatkowski, Cordelia Langford, John Sillitoe on behalf of the Wellcome Sanger Institute COVID-19 Surveillance Team |
| EPI_ISL_2909999 | Lighthouse Laboratory Plymouth | Wellcome Sanger Institute for the COVID-19 Genomics UK (COG-UK) Consortium | Lighthouse Laboratory Plymouth and Alex Alderton, Roberto Amato, Jeffrey Barrett, Sonia Goncalves, Ewan Harrison, David K. Jackson, Ian Johnston, Dominic Kwiatkowski, Cordelia Langford, John Sillitoe on behalf of the Wellcome Sanger Institute COVID-19 Surveillance Team |
| EPI_ISL_2910000, EPI_ISL_2910001, EPI_ISL_2910002 | Lighthouse Lab in Alderley Park | Wellcome Sanger Institute for the COVID-19 Genomics UK (COG-UK) Consortium | Jacquelyn Wynn, Mairead Hyland, The Lighthouse Lab in Alderley Park and Alex Alderton, Roberto Amato, Jeffrey Barrett, Sonia Goncalves, Ewan Harrison, David K. Jackson, Ian Johnston, Dominic Kwiatkowski, Cordelia Langford, John Sillitoe on behalf of the Wellcome Sanger Institute COVID-19 Surveillance Team |
| EPI_ISL_2910003 | Lighthouse Lab in Milton Keynes | Wellcome Sanger Institute for the COVID-19 Genomics UK (COG-UK) Consortium | The Lighthouse Lab in Milton Keynes and Alex Alderton, Roberto Amato, Jeffrey Barrett, Sonia Goncalves, Ewan Harrison, David K. Jackson, Ian Johnston, Dominic Kwiatkowski, Cordelia Langford, John Sillitoe on behalf of the Wellcome Sanger Institute COVID-19 Surveillance Team |
| EPI_ISL_2910004, EPI_ISL_2910005 | Lighthouse Lab in Alderley Park | Wellcome Sanger Institute for the COVID-19 Genomics UK (COG-UK) Consortium | Jacquelyn Wynn, Mairead Hyland, The Lighthouse Lab in Alderley Park and Alex Alderton, Roberto Amato, Jeffrey Barrett, Sonia Goncalves, Ewan Harrison, David K. Jackson, Ian Johnston, Dominic Kwiatkowski, Cordelia Langford, John Sillitoe on behalf of the Wellcome Sanger Institute COVID-19 Surveillance Team |
| EPI_ISL_2910006, EPI_ISL_2910007 | Lighthouse Lab in Milton Keynes | Wellcome Sanger Institute for the COVID-19 Genomics UK (COG-UK) Consortium | The Lighthouse Lab in Milton Keynes and Alex Alderton, Roberto Amato, Jeffrey Barrett, Sonia Goncalves, Ewan Harrison, David K. Jackson, Ian Johnston, Dominic Kwiatkowski, Cordelia Langford, John Sillitoe on behalf of the Wellcome Sanger Institute COVID-19 Surveillance Team |
| EPI_ISL_2910008 | Lighthouse Lab in Alderley Park | Wellcome Sanger Institute for the COVID-19 Genomics UK (COG-UK) Consortium | Jacquelyn Wynn, Mairead Hyland, The Lighthouse Lab in Alderley Park and Alex Alderton, Roberto Amato, Jeffrey Barrett, Sonia Goncalves, Ewan Harrison, David K. Jackson, Ian Johnston, Dominic Kwiatkowski, Cordelia Langford, John Sillitoe on behalf of the Wellcome Sanger Institute COVID-19 Surveillance Team |
| EPI_ISL_2910009, EPI_ISL_2910010 | Lighthouse Lab in Milton Keynes | Wellcome Sanger Institute for the COVID-19 Genomics UK (COG-UK) Consortium | The Lighthouse Lab in Milton Keynes and Alex Alderton, Roberto Amato, Jeffrey Barrett, Sonia Goncalves, Ewan Harrison, David K. Jackson, Ian Johnston, Dominic Kwiatkowski, Cordelia Langford, John Sillitoe on behalf of the Wellcome Sanger Institute COVID-19 Surveillance Team |
| EPI_ISL_2910011, EPI_ISL_2910012, EPI_ISL_2910013, EPI_ISL_2910014 | Lighthouse Lab in Alderley Park | Wellcome Sanger Institute for the COVID-19 Genomics UK (COG-UK) Consortium | Jacquelyn Wynn, Mairead Hyland, The Lighthouse Lab in Alderley Park and Alex Alderton, Roberto Amato, Jeffrey Barrett, Sonia Goncalves, Ewan Harrison, David K. Jackson, Ian Johnston, Dominic Kwiatkowski, Cordelia Langford, John Sillitoe on behalf of the Wellcome Sanger Institute COVID-19 Surveillance Team |
| EPI_ISL_2910015 | Lighthouse Laboratory Plymouth | Wellcome Sanger Institute for the COVID-19 Genomics UK (COG-UK) Consortium | Lighthouse Laboratory Plymouth and Alex Alderton, Roberto Amato, Jeffrey Barrett, Sonia Goncalves, Ewan Harrison, David K. Jackson, Ian Johnston, Dominic Kwiatkowski, Cordelia Langford, John Sillitoe on behalf of the Wellcome Sanger Institute COVID-19 Surveillance Team |
| EPI_ISL_2910016 | Lighthouse Lab in Alderley Park | Wellcome Sanger Institute for the COVID-19 Genomics UK (COG-UK) Consortium | Jacquelyn Wynn, Mairead Hyland, The Lighthouse Lab in Alderley Park and Alex Alderton, Roberto Amato, Jeffrey Barrett, Sonia Goncalves, Ewan Harrison, David K. Jackson, Ian Johnston, Dominic Kwiatkowski, Cordelia Langford, John Sillitoe on behalf of the Wellcome Sanger Institute COVID-19 Surveillance Team |
| EPI_ISL_2910017, EPI_ISL_2910019 | Lighthouse Laboratory Plymouth | Wellcome Sanger Institute for the COVID-19 Genomics UK (COG-UK) Consortium | Lighthouse Laboratory Plymouth and Alex Alderton, Roberto Amato, Jeffrey Barrett, Sonia Goncalves, Ewan Harrison, David K. Jackson, Ian Johnston, Dominic Kwiatkowski, Cordelia Langford, John Sillitoe on behalf of the Wellcome Sanger Institute COVID-19 Surveillance Team |
| EPI_ISL_2910020, EPI_ISL_2910021, EPI_ISL_2910022 | Lighthouse Lab in Alderley Park | Wellcome Sanger Institute for the COVID-19 Genomics UK (COG-UK) Consortium | Jacquelyn Wynn, Mairead Hyland, The Lighthouse Lab in Alderley Park and Alex Alderton, Roberto Amato, Jeffrey Barrett, Sonia Goncalves, Ewan Harrison, David K. Jackson, Ian Johnston, Dominic Kwiatkowski, Cordelia Langford, John Sillitoe on behalf of the Wellcome Sanger Institute COVID-19 Surveillance Team |
| EPI_ISL_2910023, EPI_ISL_2910024 | Lighthouse Lab in Milton Keynes | Wellcome Sanger Institute for the COVID-19 Genomics UK (COG-UK) Consortium | The Lighthouse Lab in Milton Keynes and Alex Alderton, Roberto Amato, Jeffrey Barrett, Sonia Goncalves, Ewan Harrison, David K. Jackson, Ian Johnston, Dominic Kwiatkowski, Cordelia Langford, John Sillitoe on behalf of the Wellcome Sanger Institute COVID-19 Surveillance Team |
| EPI_ISL_2910025 | Lighthouse Lab in Alderley Park | Wellcome Sanger Institute for the COVID-19 Genomics UK (COG-UK) Consortium | Jacquelyn Wynn, Mairead Hyland, The Lighthouse Lab in Alderley Park and Alex Alderton, Roberto Amato, Jeffrey Barrett, Sonia Goncalves, Ewan Harrison, David K. Jackson, Ian Johnston, Dominic Kwiatkowski, Cordelia Langford, John Sillitoe on behalf of the Wellcome Sanger Institute COVID-19 Surveillance Team |
| EPI_ISL_2910026, EPI_ISL_2910027 | Lighthouse Laboratory Plymouth | Wellcome Sanger Institute for the COVID-19 Genomics UK (COG-UK) Consortium | Lighthouse Laboratory Plymouth and Alex Alderton, Roberto Amato, Jeffrey Barrett, Sonia Goncalves, Ewan Harrison, David K. Jackson, Ian Johnston, Dominic Kwiatkowski, Cordelia Langford, John Sillitoe on behalf of the Wellcome Sanger Institute COVID-19 Surveillance Team |
| EPI_ISL_2910028, EPI_ISL_2910029 | Lighthouse Lab in Milton Keynes | Wellcome Sanger Institute for the COVID-19 Genomics UK (COG-UK) Consortium | The Lighthouse Lab in Milton Keynes and Alex Alderton, Roberto Amato, Jeffrey Barrett, Sonia Goncalves, Ewan Harrison, David K. Jackson, Ian Johnston, Dominic Kwiatkowski, Cordelia Langford, John Sillitoe on behalf of the Wellcome Sanger Institute COVID-19 Surveillance Team |
| EPI_ISL_2910030 | Lighthouse Laboratory Plymouth | Wellcome Sanger Institute for the COVID-19 Genomics UK (COG-UK) Consortium | Lighthouse Laboratory Plymouth and Alex Alderton, Roberto Amato, Jeffrey Barrett, Sonia Goncalves, Ewan Harrison, David K. Jackson, Ian Johnston, Dominic Kwiatkowski, Cordelia Langford, John Sillitoe on behalf of the Wellcome Sanger Institute COVID-19 Surveillance Team |
| EPI_ISL_2910031, EPI_ISL_2910032, EPI_ISL_2910033 | Lighthouse Lab in Alderley Park | Wellcome Sanger Institute for the COVID-19 Genomics UK (COG-UK) Consortium | Jacquelyn Wynn, Mairead Hyland, The Lighthouse Lab in Alderley Park and Alex Alderton, Roberto Amato, Jeffrey Barrett, Sonia Goncalves, Ewan Harrison, David K. Jackson, Ian Johnston, Dominic Kwiatkowski, Cordelia Langford, John Sillitoe on behalf of the Wellcome Sanger Institute COVID-19 Surveillance Team |
| EPI_ISL_2910034, EPI_ISL_2910035 | Lighthouse Lab in Milton Keynes | Wellcome Sanger Institute for the COVID-19 Genomics UK (COG-UK) Consortium | The Lighthouse Lab in Milton Keynes and Alex Alderton, Roberto Amato, Jeffrey Barrett, Sonia Goncalves, Ewan Harrison, David K. Jackson, Ian Johnston, Dominic Kwiatkowski, Cordelia Langford, John Sillitoe on behalf of the Wellcome Sanger Institute COVID-19 Surveillance Team |
| EPI_ISL_2910036 | Lighthouse Laboratory Plymouth | Wellcome Sanger Institute for the COVID-19 Genomics UK (COG-UK) Consortium | Lighthouse Laboratory Plymouth and Alex Alderton, Roberto Amato, Jeffrey Barrett, Sonia Goncalves, Ewan Harrison, David K. Jackson, Ian Johnston, Dominic Kwiatkowski, Cordelia Langford, John Sillitoe on behalf of the Wellcome Sanger Institute COVID-19 Surveillance Team |
| EPI_ISL_2910037, EPI_ISL_2910038, EPI_ISL_2910039 | Lighthouse Lab in Alderley Park | Wellcome Sanger Institute for the COVID-19 Genomics UK (COG-UK) Consortium | Jacquelyn Wynn, Mairead Hyland, The Lighthouse Lab in Alderley Park and Alex Alderton, Roberto Amato, Jeffrey Barrett, Sonia Goncalves, Ewan Harrison, David K. Jackson, Ian Johnston, Dominic Kwiatkowski, Cordelia Langford, John Sillitoe on behalf of the Wellcome Sanger Institute COVID-19 Surveillance Team |
| EPI_ISL_2910040 | Lighthouse Laboratory Plymouth | Wellcome Sanger Institute for the COVID-19 Genomics UK (COG-UK) Consortium | Lighthouse Laboratory Plymouth and Alex Alderton, Roberto Amato, Jeffrey Barrett, Sonia Goncalves, Ewan Harrison, David K. Jackson, Ian Johnston, Dominic Kwiatkowski, Cordelia Langford, John Sillitoe on behalf of the Wellcome Sanger Institute COVID-19 Surveillance Team |
| EPI_ISL_2910041 | Lighthouse Lab in Alderley Park | Wellcome Sanger Institute for the COVID-19 Genomics UK (COG-UK) Consortium | Jacquelyn Wynn, Mairead Hyland, The Lighthouse Lab in Alderley Park and Alex Alderton, Roberto Amato, Jeffrey Barrett, Sonia Goncalves, Ewan Harrison, David K. Jackson, Ian Johnston, Dominic Kwiatkowski, Cordelia Langford, John Sillitoe on behalf of the Wellcome Sanger Institute COVID-19 Surveillance Team |

| | | | |
|---|---|---|---|
| EPI_ISL_2910042 | Lighthouse Lab in Milton Keynes | Wellcome Sanger Institute for the COVID-19 Genomics UK (COG-UK) Consortium | The Lighthouse Lab in Milton Keynes and Alex Alderton, Roberto Amato, Jeffrey Barrett, Sonia Goncalves, Ewan Harrison, David K. Jackson, Ian Johnston, Dominic Kwiatkowski, Cordelia Langford, John Sillitoe on behalf of the Wellcome Sanger Institute COVID-19 Surveillance Team |
| EPI_ISL_2910043 | Lighthouse Lab in Alderley Park | Wellcome Sanger Institute for the COVID-19 Genomics UK (COG-UK) Consortium | Jacquelyn Wynn, Mairead Hyland, The Lighthouse Lab in Alderley Park and Alex Alderton, Roberto Amato, Jeffrey Barrett, Sonia Goncalves, Ewan Harrison, David K. Jackson, Ian Johnston, Dominic Kwiatkowski, Cordelia Langford, John Sillitoe on behalf of the Wellcome Sanger Institute COVID-19 Surveillance Team |
| EPI_ISL_2910044 | Lighthouse Lab in Milton Keynes | Wellcome Sanger Institute for the COVID-19 Genomics UK (COG-UK) Consortium | The Lighthouse Lab in Milton Keynes and Alex Alderton, Roberto Amato, Jeffrey Barrett, Sonia Goncalves, Ewan Harrison, David K. Jackson, Ian Johnston, Dominic Kwiatkowski, Cordelia Langford, John Sillitoe on behalf of the Wellcome Sanger Institute COVID-19 Surveillance Team |
| EPI_ISL_2910045 | Lighthouse Lab in Alderley Park | Wellcome Sanger Institute for the COVID-19 Genomics UK (COG-UK) Consortium | Jacquelyn Wynn, Mairead Hyland, The Lighthouse Lab in Alderley Park and Alex Alderton, Roberto Amato, Jeffrey Barrett, Sonia Goncalves, Ewan Harrison, David K. Jackson, Ian Johnston, Dominic Kwiatkowski, Cordelia Langford, John Sillitoe on behalf of the Wellcome Sanger Institute COVID-19 Surveillance Team |
| EPI_ISL_2910046 | Lighthouse Lab in Milton Keynes | Wellcome Sanger Institute for the COVID-19 Genomics UK (COG-UK) Consortium | The Lighthouse Lab in Milton Keynes and Alex Alderton, Roberto Amato, Jeffrey Barrett, Sonia Goncalves, Ewan Harrison, David K. Jackson, Ian Johnston, Dominic Kwiatkowski, Cordelia Langford, John Sillitoe on behalf of the Wellcome Sanger Institute COVID-19 Surveillance Team |
| EPI_ISL_2910047 | Lighthouse Lab in Alderley Park | Wellcome Sanger Institute for the COVID-19 Genomics UK (COG-UK) Consortium | Jacquelyn Wynn, Mairead Hyland, The Lighthouse Lab in Alderley Park and Alex Alderton, Roberto Amato, Jeffrey Barrett, Sonia Goncalves, Ewan Harrison, David K. Jackson, Ian Johnston, Dominic Kwiatkowski, Cordelia Langford, John Sillitoe on behalf of the Wellcome Sanger Institute COVID-19 Surveillance Team |
| EPI_ISL_2910048 | Lighthouse Laboratory Plymouth | Wellcome Sanger Institute for the COVID-19 Genomics UK (COG-UK) Consortium | Lighthouse Laboratory Plymouth and Alex Alderton, Roberto Amato, Jeffrey Barrett, Sonia Goncalves, Ewan Harrison, David K. Jackson, Ian Johnston, Dominic Kwiatkowski, Cordelia Langford, John Sillitoe on behalf of the Wellcome Sanger Institute COVID-19 Surveillance Team |
| EPI_ISL_2910049 | Lighthouse Lab in Milton Keynes | Wellcome Sanger Institute for the COVID-19 Genomics UK (COG-UK) Consortium | The Lighthouse Lab in Milton Keynes and Alex Alderton, Roberto Amato, Jeffrey Barrett, Sonia Goncalves, Ewan Harrison, David K. Jackson, Ian Johnston, Dominic Kwiatkowski, Cordelia Langford, John Sillitoe on behalf of the Wellcome Sanger Institute COVID-19 Surveillance Team |
| EPI_ISL_2910050 | Lighthouse Lab in Alderley Park | Wellcome Sanger Institute for the COVID-19 Genomics UK (COG-UK) Consortium | Jacquelyn Wynn, Mairead Hyland, The Lighthouse Lab in Alderley Park and Alex Alderton, Roberto Amato, Jeffrey Barrett, Sonia Goncalves, Ewan Harrison, David K. Jackson, Ian Johnston, Dominic Kwiatkowski, Cordelia Langford, John Sillitoe on behalf of the Wellcome Sanger Institute COVID-19 Surveillance Team |
| EPI_ISL_2910051 | Lighthouse Lab in Milton Keynes | Wellcome Sanger Institute for the COVID-19 Genomics UK (COG-UK) Consortium | The Lighthouse Lab in Milton Keynes and Alex Alderton, Roberto Amato, Jeffrey Barrett, Sonia Goncalves, Ewan Harrison, David K. Jackson, Ian Johnston, Dominic Kwiatkowski, Cordelia Langford, John Sillitoe on behalf of the Wellcome Sanger Institute COVID-19 Surveillance Team |
| EPI_ISL_2910052, EPI_ISL_2910053, EPI_ISL_2910054 | Lighthouse Laboratory Plymouth | Wellcome Sanger Institute for the COVID-19 Genomics UK (COG-UK) Consortium | Lighthouse Laboratory Plymouth and Alex Alderton, Roberto Amato, Jeffrey Barrett, Sonia Goncalves, Ewan Harrison, David K. Jackson, Ian Johnston, Dominic Kwiatkowski, Cordelia Langford, John Sillitoe on behalf of the Wellcome Sanger Institute COVID-19 Surveillance Team |
| EPI_ISL_2910055, EPI_ISL_2910056 | Lighthouse Lab in Alderley Park | Wellcome Sanger Institute for the COVID-19 Genomics UK (COG-UK) Consortium | Jacquelyn Wynn, Mairead Hyland, The Lighthouse Lab in Alderley Park and Alex Alderton, Roberto Amato, Jeffrey Barrett, Sonia Goncalves, Ewan Harrison, David K. Jackson, Ian Johnston, Dominic Kwiatkowski, Cordelia Langford, John Sillitoe on behalf of the Wellcome Sanger Institute COVID-19 Surveillance Team |
| EPI_ISL_2910057 | Lighthouse Laboratory Plymouth | Wellcome Sanger Institute for the COVID-19 Genomics UK (COG-UK) Consortium | Lighthouse Laboratory Plymouth and Alex Alderton, Roberto Amato, Jeffrey Barrett, Sonia Goncalves, Ewan Harrison, David K. Jackson, Ian Johnston, Dominic Kwiatkowski, Cordelia Langford, John Sillitoe on behalf of the Wellcome Sanger Institute COVID-19 Surveillance Team |
| EPI_ISL_2910058 | Lighthouse Lab in Milton Keynes | Wellcome Sanger Institute for the COVID-19 Genomics UK (COG-UK) Consortium | The Lighthouse Lab in Milton Keynes and Alex Alderton, Roberto Amato, Jeffrey Barrett, Sonia Goncalves, Ewan Harrison, David K. Jackson, Ian Johnston, Dominic Kwiatkowski, Cordelia Langford, John Sillitoe on behalf of the Wellcome Sanger Institute COVID-19 Surveillance Team |
| EPI_ISL_2910059, EPI_ISL_2910060 | Lighthouse Lab in Alderley Park | Wellcome Sanger Institute for the COVID-19 Genomics UK (COG-UK) Consortium | Jacquelyn Wynn, Mairead Hyland, The Lighthouse Lab in Alderley Park and Alex Alderton, Roberto Amato, Jeffrey Barrett, Sonia Goncalves, Ewan Harrison, David K. Jackson, Ian Johnston, Dominic Kwiatkowski, Cordelia Langford, John Sillitoe on behalf of the Wellcome Sanger Institute COVID-19 Surveillance Team |
| EPI_ISL_2910061, EPI_ISL_2910062 | Lighthouse Lab in Milton Keynes | Wellcome Sanger Institute for the COVID-19 Genomics UK (COG-UK) Consortium | The Lighthouse Lab in Milton Keynes and Alex Alderton, Roberto Amato, Jeffrey Barrett, Sonia Goncalves, Ewan Harrison, David K. Jackson, Ian Johnston, Dominic Kwiatkowski, Cordelia Langford, John Sillitoe on behalf of the Wellcome Sanger Institute COVID-19 Surveillance Team |
| EPI_ISL_2910063, EPI_ISL_2910064, EPI_ISL_2910065, EPI_ISL_2910066, EPI_ISL_2910067 | Lighthouse Lab in Alderley Park | Wellcome Sanger Institute for the COVID-19 Genomics UK (COG-UK) Consortium | Jacquelyn Wynn, Mairead Hyland, The Lighthouse Lab in Alderley Park and Alex Alderton, Roberto Amato, Jeffrey Barrett, Sonia Goncalves, Ewan Harrison, David K. Jackson, Ian Johnston, Dominic Kwiatkowski, Cordelia Langford, John Sillitoe on behalf of the Wellcome Sanger Institute COVID-19 Surveillance Team |
| EPI_ISL_2910068 | Lighthouse Lab in Milton Keynes | Wellcome Sanger Institute for the COVID-19 Genomics UK (COG-UK) Consortium | The Lighthouse Lab in Milton Keynes and Alex Alderton, Roberto Amato, Jeffrey Barrett, Sonia Goncalves, Ewan Harrison, David K. Jackson, Ian Johnston, Dominic Kwiatkowski, Cordelia Langford, John Sillitoe on behalf of the Wellcome Sanger Institute COVID-19 Surveillance Team |
| EPI_ISL_2910069, EPI_ISL_2910070, EPI_ISL_2910071, EPI_ISL_2910072, EPI_ISL_2910073, EPI_ISL_2910074, EPI_ISL_2910075, EPI_ISL_2910076, EPI_ISL_2910077 | Lighthouse Lab in Alderley Park | Wellcome Sanger Institute for the COVID-19 Genomics UK (COG-UK) Consortium | Jacquelyn Wynn, Mairead Hyland, The Lighthouse Lab in Alderley Park and Alex Alderton, Roberto Amato, Jeffrey Barrett, Sonia Goncalves, Ewan Harrison, David K. Jackson, Ian Johnston, Dominic Kwiatkowski, Cordelia Langford, John Sillitoe on behalf of the Wellcome Sanger Institute COVID-19 Surveillance Team |
| EPI_ISL_2910078 | Lighthouse Lab in Milton Keynes | Wellcome Sanger Institute for the COVID-19 Genomics UK (COG-UK) Consortium | The Lighthouse Lab in Milton Keynes and Alex Alderton, Roberto Amato, Jeffrey Barrett, Sonia Goncalves, Ewan Harrison, David K. Jackson, Ian Johnston, Dominic Kwiatkowski, Cordelia Langford, John Sillitoe on behalf of the Wellcome Sanger Institute COVID-19 Surveillance Team |
| EPI_ISL_2910079 | Lighthouse Lab in Alderley Park | Wellcome Sanger Institute for the COVID-19 Genomics UK (COG-UK) Consortium | Jacquelyn Wynn, Mairead Hyland, The Lighthouse Lab in Alderley Park and Alex Alderton, Roberto Amato, Jeffrey Barrett, Sonia Goncalves, Ewan Harrison, David K. Jackson, Ian Johnston, Dominic Kwiatkowski, Cordelia Langford, John Sillitoe on behalf of the Wellcome Sanger Institute COVID-19 Surveillance Team |
| EPI_ISL_2910080 | Lighthouse Lab in Milton Keynes | Wellcome Sanger Institute for the COVID-19 Genomics UK (COG-UK) Consortium | The Lighthouse Lab in Milton Keynes and Alex Alderton, Roberto Amato, Jeffrey Barrett, Sonia Goncalves, Ewan Harrison, David K. Jackson, Ian Johnston, Dominic Kwiatkowski, Cordelia Langford, John Sillitoe on behalf of the Wellcome Sanger Institute COVID-19 Surveillance Team |
| EPI_ISL_2910081 | Lighthouse Laboratory Plymouth | Wellcome Sanger Institute for the COVID-19 Genomics UK (COG-UK) Consortium | Lighthouse Laboratory Plymouth and Alex Alderton, Roberto Amato, Jeffrey Barrett, Sonia Goncalves, Ewan Harrison, David K. Jackson, Ian Johnston, Dominic Kwiatkowski, Cordelia Langford, John Sillitoe on behalf of the Wellcome Sanger Institute COVID-19 Surveillance Team |
| EPI_ISL_2910082 | Lighthouse Lab in Milton Keynes | Wellcome Sanger Institute for the COVID-19 Genomics UK (COG-UK) Consortium | The Lighthouse Lab in Milton Keynes and Alex Alderton, Roberto Amato, Jeffrey Barrett, Sonia Goncalves, Ewan Harrison, David K. Jackson, Ian Johnston, Dominic Kwiatkowski, Cordelia Langford, John Sillitoe on behalf of the Wellcome Sanger Institute COVID-19 Surveillance Team |
| EPI_ISL_2910083 | Lighthouse Laboratory Plymouth | Wellcome Sanger Institute for the COVID-19 Genomics UK (COG-UK) Consortium | Lighthouse Laboratory Plymouth and Alex Alderton, Roberto Amato, Jeffrey Barrett, Sonia Goncalves, Ewan Harrison, David K. Jackson, Ian Johnston, Dominic Kwiatkowski, Cordelia Langford, John Sillitoe on behalf of the Wellcome Sanger Institute COVID-19 Surveillance Team |
| EPI_ISL_2910084, EPI_ISL_2910085, EPI_ISL_2910086 | Lighthouse Lab in Alderley Park | Wellcome Sanger Institute for the COVID-19 Genomics UK (COG-UK) Consortium | Jacquelyn Wynn, Mairead Hyland, The Lighthouse Lab in Alderley Park and Alex Alderton, Roberto Amato, Jeffrey Barrett, Sonia Goncalves, Ewan Harrison, David K. Jackson, Ian Johnston, Dominic Kwiatkowski, Cordelia Langford, John Sillitoe on behalf of the Wellcome Sanger Institute COVID-19 Surveillance Team |
| EPI_ISL_2910087 | Lighthouse Lab in Milton Keynes | Wellcome Sanger Institute for the COVID-19 Genomics UK (COG-UK) Consortium | The Lighthouse Lab in Milton Keynes and Alex Alderton, Roberto Amato, Jeffrey Barrett, Sonia Goncalves, Ewan Harrison, David K. Jackson, Ian Johnston, Dominic Kwiatkowski, Cordelia Langford, John Sillitoe on behalf of the Wellcome Sanger Institute COVID-19 Surveillance Team |
| EPI_ISL_2910088 | Lighthouse Lab in Alderley Park | Wellcome Sanger Institute for the COVID-19 Genomics UK (COG-UK) Consortium | Jacquelyn Wynn, Mairead Hyland, The Lighthouse Lab in Alderley Park and Alex Alderton, Roberto Amato, Jeffrey Barrett, Sonia Goncalves, Ewan Harrison, David K. Jackson, Ian Johnston, Dominic Kwiatkowski, Cordelia Langford, John Sillitoe on behalf of the Wellcome Sanger Institute COVID-19 Surveillance Team |
| EPI_ISL_2910089 | Lighthouse Laboratory Plymouth | Wellcome Sanger Institute for the COVID-19 Genomics UK (COG-UK) Consortium | Lighthouse Laboratory Plymouth and Alex Alderton, Roberto Amato, Jeffrey Barrett, Sonia Goncalves, Ewan Harrison, David K. Jackson, Ian Johnston, Dominic Kwiatkowski, Cordelia Langford, John Sillitoe on behalf of the Wellcome Sanger Institute COVID-19 Surveillance Team |
| EPI_ISL_2910090, EPI_ISL_2910091 | Lighthouse Lab in Alderley Park | Wellcome Sanger Institute for the COVID-19 Genomics UK (COG-UK) Consortium | Jacquelyn Wynn, Mairead Hyland, The Lighthouse Lab in Alderley Park and Alex Alderton, Roberto Amato, Jeffrey Barrett, Sonia Goncalves, Ewan Harrison, David K. Jackson, Ian Johnston, Dominic Kwiatkowski, Cordelia Langford, John Sillitoe on behalf of the Wellcome Sanger Institute COVID-19 Surveillance Team |
| EPI_ISL_2910092 | Lighthouse Laboratory Plymouth | Wellcome Sanger Institute for the COVID-19 Genomics UK (COG-UK) Consortium | Lighthouse Laboratory Plymouth and Alex Alderton, Roberto Amato, Jeffrey Barrett, Sonia Goncalves, Ewan Harrison, David K. Jackson, Ian Johnston, Dominic Kwiatkowski, Cordelia Langford, John Sillitoe on behalf of the Wellcome Sanger Institute COVID-19 Surveillance Team |
| EPI_ISL_2910093 | Lighthouse Lab in Milton Keynes | Wellcome Sanger Institute for the COVID-19 Genomics UK (COG-UK) Consortium | The Lighthouse Lab in Milton Keynes and Alex Alderton, Roberto Amato, Jeffrey Barrett, Sonia Goncalves, Ewan Harrison, David K. Jackson, Ian Johnston, Dominic Kwiatkowski, Cordelia Langford, John Sillitoe on behalf of the Wellcome Sanger Institute COVID-19 Surveillance Team |
| EPI_ISL_2910094, EPI_ISL_2910095 | Lighthouse Lab in Alderley Park | Wellcome Sanger Institute for the COVID-19 Genomics UK (COG-UK) Consortium | Jacquelyn Wynn, Mairead Hyland, The Lighthouse Lab in Alderley Park and Alex Alderton, Roberto Amato, Jeffrey Barrett, Sonia Goncalves, Ewan Harrison, David K. Jackson, Ian Johnston, Dominic Kwiatkowski, Cordelia Langford, John Sillitoe on behalf of the Wellcome Sanger Institute COVID-19 Surveillance Team |
| EPI_ISL_2910096 | Lighthouse Lab in Milton Keynes | Wellcome Sanger Institute for the COVID-19 Genomics UK (COG-UK) Consortium | The Lighthouse Lab in Milton Keynes and Alex Alderton, Roberto Amato, Jeffrey Barrett, Sonia Goncalves, Ewan Harrison, David K. Jackson, Ian Johnston, Dominic Kwiatkowski, Cordelia Langford, John Sillitoe on behalf of the Wellcome Sanger Institute COVID-19 Surveillance Team |
| EPI_ISL_2910097, EPI_ISL_2910098 | Lighthouse Lab in Alderley Park | Wellcome Sanger Institute for the COVID-19 Genomics UK (COG-UK) Consortium | Jacquelyn Wynn, Mairead Hyland, The Lighthouse Lab in Alderley Park and Alex Alderton, Roberto Amato, Jeffrey Barrett, Sonia Goncalves, Ewan Harrison, David K. Jackson, Ian Johnston, Dominic Kwiatkowski, Cordelia Langford, John Sillitoe on behalf of the Wellcome Sanger Institute COVID-19 Surveillance Team |
| EPI_ISL_2910099, EPI_ISL_2910100, EPI_ISL_2910101 | Lighthouse Laboratory Plymouth | Wellcome Sanger Institute for the COVID-19 Genomics UK (COG-UK) Consortium | Lighthouse Laboratory Plymouth and Alex Alderton, Roberto Amato, Jeffrey Barrett, Sonia Goncalves, Ewan Harrison, David K. Jackson, Ian Johnston, Dominic Kwiatkowski, Cordelia Langford, John Sillitoe on behalf of the Wellcome Sanger Institute COVID-19 Surveillance Team |
| EPI_ISL_2910102 | Lighthouse Lab in Alderley Park | Wellcome Sanger Institute for the COVID-19 Genomics UK (COG-UK) Consortium | Jacquelyn Wynn, Mairead Hyland, The Lighthouse Lab in Alderley Park and Alex Alderton, Roberto Amato, Jeffrey Barrett, Sonia Goncalves, Ewan Harrison, David K. Jackson, Ian Johnston, Dominic Kwiatkowski, Cordelia Langford, John Sillitoe on behalf of the Wellcome Sanger Institute COVID-19 Surveillance Team |
| EPI_ISL_2910103 | Lighthouse Laboratory Plymouth | Wellcome Sanger Institute for the COVID-19 Genomics UK (COG-UK) Consortium | Lighthouse Laboratory Plymouth and Alex Alderton, Roberto Amato, Jeffrey Barrett, Sonia Goncalves, Ewan Harrison, David K. Jackson, Ian Johnston, Dominic Kwiatkowski, Cordelia Langford, John Sillitoe on behalf of the Wellcome Sanger Institute COVID-19 Surveillance Team |
| EPI_ISL_2910104 | Lighthouse Lab in Milton Keynes | Wellcome Sanger Institute for the COVID-19 Genomics UK (COG-UK) Consortium | The Lighthouse Lab in Milton Keynes and Alex Alderton, Roberto Amato, Jeffrey Barrett, Sonia Goncalves, Ewan Harrison, David K. Jackson, Ian Johnston, Dominic Kwiatkowski, Cordelia Langford, John Sillitoe on behalf of the Wellcome Sanger Institute COVID-19 Surveillance Team |
| EPI_ISL_2910105 | Lighthouse Laboratory Plymouth | Wellcome Sanger Institute for the COVID-19 Genomics UK (COG-UK) Consortium | Lighthouse Laboratory Plymouth and Alex Alderton, Roberto Amato, Jeffrey Barrett, Sonia Goncalves, Ewan Harrison, David K. Jackson, Ian Johnston, Dominic Kwiatkowski, Cordelia Langford, John Sillitoe on behalf of the Wellcome Sanger Institute COVID-19 Surveillance Team |

| | | | |
|---|---|---|---|
| EPI_ISL_2910106, EPI_ISL_2910107 | Lighthouse Lab in Alderley Park | Wellcome Sanger Institute for the COVID-19 Genomics UK (COG-UK) Consortium | Jacquelyn Wynn, Mairead Hyland, The Lighthouse Lab in Alderley Park and Alex Alderton, Roberto Amato, Jeffrey Barrett, Sonia Goncalves, Ewan Harrison, David K. Jackson, Ian Johnston, Dominic Kwiatkowski, Cordelia Langford, John Sillitoe on behalf of the Wellcome Sanger Institute COVID-19 Surveillance Team |
| EPI_ISL_2910108 | Lighthouse Lab in Milton Keynes | Wellcome Sanger Institute for the COVID-19 Genomics UK (COG-UK) Consortium | The Lighthouse Lab in Milton Keynes and Alex Alderton, Roberto Amato, Jeffrey Barrett, Sonia Goncalves, Ewan Harrison, David K. Jackson, Ian Johnston, Dominic Kwiatkowski, Cordelia Langford, John Sillitoe on behalf of the Wellcome Sanger Institute COVID-19 Surveillance Team |
| EPI_ISL_2910109 | Lighthouse Lab in Alderley Park | Wellcome Sanger Institute for the COVID-19 Genomics UK (COG-UK) Consortium | Jacquelyn Wynn, Mairead Hyland, The Lighthouse Lab in Alderley Park and Alex Alderton, Roberto Amato, Jeffrey Barrett, Sonia Goncalves, Ewan Harrison, David K. Jackson, Ian Johnston, Dominic Kwiatkowski, Cordelia Langford, John Sillitoe on behalf of the Wellcome Sanger Institute COVID-19 Surveillance Team |
| EPI_ISL_2910110 | Lighthouse Laboratory Plymouth | Wellcome Sanger Institute for the COVID-19 Genomics UK (COG-UK) Consortium | Lighthouse Laboratory Plymouth and Alex Alderton, Roberto Amato, Jeffrey Barrett, Sonia Goncalves, Ewan Harrison, David K. Jackson, Ian Johnston, Dominic Kwiatkowski, Cordelia Langford, John Sillitoe on behalf of the Wellcome Sanger Institute COVID-19 Surveillance Team |
| EPI_ISL_2910111, EPI_ISL_2910112 | Lighthouse Lab in Milton Keynes | Wellcome Sanger Institute for the COVID-19 Genomics UK (COG-UK) Consortium | The Lighthouse Lab in Milton Keynes and Alex Alderton, Roberto Amato, Jeffrey Barrett, Sonia Goncalves, Ewan Harrison, David K. Jackson, Ian Johnston, Dominic Kwiatkowski, Cordelia Langford, John Sillitoe on behalf of the Wellcome Sanger Institute COVID-19 Surveillance Team |
| EPI_ISL_2910113, EPI_ISL_2910114, EPI_ISL_2910115 | Lighthouse Lab in Alderley Park | Wellcome Sanger Institute for the COVID-19 Genomics UK (COG-UK) Consortium | Jacquelyn Wynn, Mairead Hyland, The Lighthouse Lab in Alderley Park and Alex Alderton, Roberto Amato, Jeffrey Barrett, Sonia Goncalves, Ewan Harrison, David K. Jackson, Ian Johnston, Dominic Kwiatkowski, Cordelia Langford, John Sillitoe on behalf of the Wellcome Sanger Institute COVID-19 Surveillance Team |
| EPI_ISL_2910116, EPI_ISL_2910117, EPI_ISL_2910118 | Lighthouse Lab in Milton Keynes | Wellcome Sanger Institute for the COVID-19 Genomics UK (COG-UK) Consortium | The Lighthouse Lab in Milton Keynes and Alex Alderton, Roberto Amato, Jeffrey Barrett, Sonia Goncalves, Ewan Harrison, David K. Jackson, Ian Johnston, Dominic Kwiatkowski, Cordelia Langford, John Sillitoe on behalf of the Wellcome Sanger Institute COVID-19 Surveillance Team |
| EPI_ISL_2910119, EPI_ISL_2910120, EPI_ISL_2910121, EPI_ISL_2910122 | Lighthouse Lab in Alderley Park | Wellcome Sanger Institute for the COVID-19 Genomics UK (COG-UK) Consortium | Jacquelyn Wynn, Mairead Hyland, The Lighthouse Lab in Alderley Park and Alex Alderton, Roberto Amato, Jeffrey Barrett, Sonia Goncalves, Ewan Harrison, David K. Jackson, Ian Johnston, Dominic Kwiatkowski, Cordelia Langford, John Sillitoe on behalf of the Wellcome Sanger Institute COVID-19 Surveillance Team |
| EPI_ISL_2910123 | Lighthouse Laboratory Plymouth | Wellcome Sanger Institute for the COVID-19 Genomics UK (COG-UK) Consortium | Lighthouse Laboratory Plymouth and Alex Alderton, Roberto Amato, Jeffrey Barrett, Sonia Goncalves, Ewan Harrison, David K. Jackson, Ian Johnston, Dominic Kwiatkowski, Cordelia Langford, John Sillitoe on behalf of the Wellcome Sanger Institute COVID-19 Surveillance Team |
| EPI_ISL_2910124 | Lighthouse Lab in Alderley Park | Wellcome Sanger Institute for the COVID-19 Genomics UK (COG-UK) Consortium | Jacquelyn Wynn, Mairead Hyland, The Lighthouse Lab in Alderley Park and Alex Alderton, Roberto Amato, Jeffrey Barrett, Sonia Goncalves, Ewan Harrison, David K. Jackson, Ian Johnston, Dominic Kwiatkowski, Cordelia Langford, John Sillitoe on behalf of the Wellcome Sanger Institute COVID-19 Surveillance Team |
| EPI_ISL_2910125 | Lighthouse Laboratory Plymouth | Wellcome Sanger Institute for the COVID-19 Genomics UK (COG-UK) Consortium | Lighthouse Laboratory Plymouth and Alex Alderton, Roberto Amato, Jeffrey Barrett, Sonia Goncalves, Ewan Harrison, David K. Jackson, Ian Johnston, Dominic Kwiatkowski, Cordelia Langford, John Sillitoe on behalf of the Wellcome Sanger Institute COVID-19 Surveillance Team |
| EPI_ISL_2910126 | Lighthouse Lab in Alderley Park | Wellcome Sanger Institute for the COVID-19 Genomics UK (COG-UK) Consortium | Jacquelyn Wynn, Mairead Hyland, The Lighthouse Lab in Alderley Park and Alex Alderton, Roberto Amato, Jeffrey Barrett, Sonia Goncalves, Ewan Harrison, David K. Jackson, Ian Johnston, Dominic Kwiatkowski, Cordelia Langford, John Sillitoe on behalf of the Wellcome Sanger Institute COVID-19 Surveillance Team |
| EPI_ISL_2910127 | Lighthouse Laboratory Plymouth | Wellcome Sanger Institute for the COVID-19 Genomics UK (COG-UK) Consortium | Lighthouse Laboratory Plymouth and Alex Alderton, Roberto Amato, Jeffrey Barrett, Sonia Goncalves, Ewan Harrison, David K. Jackson, Ian Johnston, Dominic Kwiatkowski, Cordelia Langford, John Sillitoe on behalf of the Wellcome Sanger Institute COVID-19 Surveillance Team |
| EPI_ISL_2910128 | Lighthouse Lab in Milton Keynes | Wellcome Sanger Institute for the COVID-19 Genomics UK (COG-UK) Consortium | The Lighthouse Lab in Milton Keynes and Alex Alderton, Roberto Amato, Jeffrey Barrett, Sonia Goncalves, Ewan Harrison, David K. Jackson, Ian Johnston, Dominic Kwiatkowski, Cordelia Langford, John Sillitoe on behalf of the Wellcome Sanger Institute COVID-19 Surveillance Team |
| EPI_ISL_2910129 | Lighthouse Lab in Alderley Park | Wellcome Sanger Institute for the COVID-19 Genomics UK (COG-UK) Consortium | Jacquelyn Wynn, Mairead Hyland, The Lighthouse Lab in Alderley Park and Alex Alderton, Roberto Amato, Jeffrey Barrett, Sonia Goncalves, Ewan Harrison, David K. Jackson, Ian Johnston, Dominic Kwiatkowski, Cordelia Langford, John Sillitoe on behalf of the Wellcome Sanger Institute COVID-19 Surveillance Team |
| EPI_ISL_2910130 | Lighthouse Laboratory Plymouth | Wellcome Sanger Institute for the COVID-19 Genomics UK (COG-UK) Consortium | Lighthouse Laboratory Plymouth and Alex Alderton, Roberto Amato, Jeffrey Barrett, Sonia Goncalves, Ewan Harrison, David K. Jackson, Ian Johnston, Dominic Kwiatkowski, Cordelia Langford, John Sillitoe on behalf of the Wellcome Sanger Institute COVID-19 Surveillance Team |
| EPI_ISL_2910131 | Lighthouse Lab in Alderley Park | Wellcome Sanger Institute for the COVID-19 Genomics UK (COG-UK) Consortium | Jacquelyn Wynn, Mairead Hyland, The Lighthouse Lab in Alderley Park and Alex Alderton, Roberto Amato, Jeffrey Barrett, Sonia Goncalves, Ewan Harrison, David K. Jackson, Ian Johnston, Dominic Kwiatkowski, Cordelia Langford, John Sillitoe on behalf of the Wellcome Sanger Institute COVID-19 Surveillance Team |
| EPI_ISL_2910132 | Lighthouse Lab in Milton Keynes | Wellcome Sanger Institute for the COVID-19 Genomics UK (COG-UK) Consortium | The Lighthouse Lab in Milton Keynes and Alex Alderton, Roberto Amato, Jeffrey Barrett, Sonia Goncalves, Ewan Harrison, David K. Jackson, Ian Johnston, Dominic Kwiatkowski, Cordelia Langford, John Sillitoe on behalf of the Wellcome Sanger Institute COVID-19 Surveillance Team |
| EPI_ISL_2910133, EPI_ISL_2910134, EPI_ISL_2910135 | Lighthouse Lab in Alderley Park | Wellcome Sanger Institute for the COVID-19 Genomics UK (COG-UK) Consortium | Jacquelyn Wynn, Mairead Hyland, The Lighthouse Lab in Alderley Park and Alex Alderton, Roberto Amato, Jeffrey Barrett, Sonia Goncalves, Ewan Harrison, David K. Jackson, Ian Johnston, Dominic Kwiatkowski, Cordelia Langford, John Sillitoe on behalf of the Wellcome Sanger Institute COVID-19 Surveillance Team |
| EPI_ISL_2910136 | Lighthouse Laboratory Plymouth | Wellcome Sanger Institute for the COVID-19 Genomics UK (COG-UK) Consortium | Lighthouse Laboratory Plymouth and Alex Alderton, Roberto Amato, Jeffrey Barrett, Sonia Goncalves, Ewan Harrison, David K. Jackson, Ian Johnston, Dominic Kwiatkowski, Cordelia Langford, John Sillitoe on behalf of the Wellcome Sanger Institute COVID-19 Surveillance Team |
| EPI_ISL_2910137, EPI_ISL_2910138, EPI_ISL_2910139 | Lighthouse Lab in Alderley Park | Wellcome Sanger Institute for the COVID-19 Genomics UK (COG-UK) Consortium | Jacquelyn Wynn, Mairead Hyland, The Lighthouse Lab in Alderley Park and Alex Alderton, Roberto Amato, Jeffrey Barrett, Sonia Goncalves, Ewan Harrison, David K. Jackson, Ian Johnston, Dominic Kwiatkowski, Cordelia Langford, John Sillitoe on behalf of the Wellcome Sanger Institute COVID-19 Surveillance Team |
| EPI_ISL_2910140, EPI_ISL_2910141 | Lighthouse Lab in Milton Keynes | Wellcome Sanger Institute for the COVID-19 Genomics UK (COG-UK) Consortium | The Lighthouse Lab in Milton Keynes and Alex Alderton, Roberto Amato, Jeffrey Barrett, Sonia Goncalves, Ewan Harrison, David K. Jackson, Ian Johnston, Dominic Kwiatkowski, Cordelia Langford, John Sillitoe on behalf of the Wellcome Sanger Institute COVID-19 Surveillance Team |
| EPI_ISL_2910142 | Lighthouse Lab in Alderley Park | Wellcome Sanger Institute for the COVID-19 Genomics UK (COG-UK) Consortium | Jacquelyn Wynn, Mairead Hyland, The Lighthouse Lab in Alderley Park and Alex Alderton, Roberto Amato, Jeffrey Barrett, Sonia Goncalves, Ewan Harrison, David K. Jackson, Ian Johnston, Dominic Kwiatkowski, Cordelia Langford, John Sillitoe on behalf of the Wellcome Sanger Institute COVID-19 Surveillance Team |
| EPI_ISL_2910143 | Lighthouse Laboratory Plymouth | Wellcome Sanger Institute for the COVID-19 Genomics UK (COG-UK) Consortium | Lighthouse Laboratory Plymouth and Alex Alderton, Roberto Amato, Jeffrey Barrett, Sonia Goncalves, Ewan Harrison, David K. Jackson, Ian Johnston, Dominic Kwiatkowski, Cordelia Langford, John Sillitoe on behalf of the Wellcome Sanger Institute COVID-19 Surveillance Team |
| EPI_ISL_2910144, EPI_ISL_2910145, EPI_ISL_2910146, EPI_ISL_2910147, EPI_ISL_2910148, EPI_ISL_2910149 | Lighthouse Lab in Alderley Park | Wellcome Sanger Institute for the COVID-19 Genomics UK (COG-UK) Consortium | Jacquelyn Wynn, Mairead Hyland, The Lighthouse Lab in Alderley Park and Alex Alderton, Roberto Amato, Jeffrey Barrett, Sonia Goncalves, Ewan Harrison, David K. Jackson, Ian Johnston, Dominic Kwiatkowski, Cordelia Langford, John Sillitoe on behalf of the Wellcome Sanger Institute COVID-19 Surveillance Team |
| EPI_ISL_2910150 | Lighthouse Lab in Milton Keynes | Wellcome Sanger Institute for the COVID-19 Genomics UK (COG-UK) Consortium | The Lighthouse Lab in Milton Keynes and Alex Alderton, Roberto Amato, Jeffrey Barrett, Sonia Goncalves, Ewan Harrison, David K. Jackson, Ian Johnston, Dominic Kwiatkowski, Cordelia Langford, John Sillitoe on behalf of the Wellcome Sanger Institute COVID-19 Surveillance Team |
| EPI_ISL_2910151 | Lighthouse Lab in Alderley Park | Wellcome Sanger Institute for the COVID-19 Genomics UK (COG-UK) Consortium | Jacquelyn Wynn, Mairead Hyland, The Lighthouse Lab in Alderley Park and Alex Alderton, Roberto Amato, Jeffrey Barrett, Sonia Goncalves, Ewan Harrison, David K. Jackson, Ian Johnston, Dominic Kwiatkowski, Cordelia Langford, John Sillitoe on behalf of the Wellcome Sanger Institute COVID-19 Surveillance Team |
| EPI_ISL_2910152 | Lighthouse Laboratory Plymouth | Wellcome Sanger Institute for the COVID-19 Genomics UK (COG-UK) Consortium | Lighthouse Laboratory Plymouth and Alex Alderton, Roberto Amato, Jeffrey Barrett, Sonia Goncalves, Ewan Harrison, David K. Jackson, Ian Johnston, Dominic Kwiatkowski, Cordelia Langford, John Sillitoe on behalf of the Wellcome Sanger Institute COVID-19 Surveillance Team |
| EPI_ISL_2910153 | Lighthouse Lab in Alderley Park | Wellcome Sanger Institute for the COVID-19 Genomics UK (COG-UK) Consortium | Jacquelyn Wynn, Mairead Hyland, The Lighthouse Lab in Alderley Park and Alex Alderton, Roberto Amato, Jeffrey Barrett, Sonia Goncalves, Ewan Harrison, David K. Jackson, Ian Johnston, Dominic Kwiatkowski, Cordelia Langford, John Sillitoe on behalf of the Wellcome Sanger Institute COVID-19 Surveillance Team |
| EPI_ISL_2910154, EPI_ISL_2910155 | Lighthouse Lab in Milton Keynes | Wellcome Sanger Institute for the COVID-19 Genomics UK (COG-UK) Consortium | The Lighthouse Lab in Milton Keynes and Alex Alderton, Roberto Amato, Jeffrey Barrett, Sonia Goncalves, Ewan Harrison, David K. Jackson, Ian Johnston, Dominic Kwiatkowski, Cordelia Langford, John Sillitoe on behalf of the Wellcome Sanger Institute COVID-19 Surveillance Team |
| EPI_ISL_2910156 | Lighthouse Laboratory Plymouth | Wellcome Sanger Institute for the COVID-19 Genomics UK (COG-UK) Consortium | Lighthouse Laboratory Plymouth and Alex Alderton, Roberto Amato, Jeffrey Barrett, Sonia Goncalves, Ewan Harrison, David K. Jackson, Ian Johnston, Dominic Kwiatkowski, Cordelia Langford, John Sillitoe on behalf of the Wellcome Sanger Institute COVID-19 Surveillance Team |
| EPI_ISL_2910157 | Lighthouse Lab in Alderley Park | Wellcome Sanger Institute for the COVID-19 Genomics UK (COG-UK) Consortium | Jacquelyn Wynn, Mairead Hyland, The Lighthouse Lab in Alderley Park and Alex Alderton, Roberto Amato, Jeffrey Barrett, Sonia Goncalves, Ewan Harrison, David K. Jackson, Ian Johnston, Dominic Kwiatkowski, Cordelia Langford, John Sillitoe on behalf of the Wellcome Sanger Institute COVID-19 Surveillance Team |
| EPI_ISL_2910158, EPI_ISL_2910159, EPI_ISL_2910160, EPI_ISL_2910161 | Lighthouse Lab in Milton Keynes | Wellcome Sanger Institute for the COVID-19 Genomics UK (COG-UK) Consortium | The Lighthouse Lab in Milton Keynes and Alex Alderton, Roberto Amato, Jeffrey Barrett, Sonia Goncalves, Ewan Harrison, David K. Jackson, Ian Johnston, Dominic Kwiatkowski, Cordelia Langford, John Sillitoe on behalf of the Wellcome Sanger Institute COVID-19 Surveillance Team |
| EPI_ISL_2910162, EPI_ISL_2910163, EPI_ISL_2910164 | Lighthouse Laboratory Plymouth | Wellcome Sanger Institute for the COVID-19 Genomics UK (COG-UK) Consortium | Lighthouse Laboratory Plymouth and Alex Alderton, Roberto Amato, Jeffrey Barrett, Sonia Goncalves, Ewan Harrison, David K. Jackson, Ian Johnston, Dominic Kwiatkowski, Cordelia Langford, John Sillitoe on behalf of the Wellcome Sanger Institute COVID-19 Surveillance Team |
| EPI_ISL_2910165, EPI_ISL_2910166, EPI_ISL_2910167 | Lighthouse Lab in Alderley Park | Wellcome Sanger Institute for the COVID-19 Genomics UK (COG-UK) Consortium | Jacquelyn Wynn, Mairead Hyland, The Lighthouse Lab in Alderley Park and Alex Alderton, Roberto Amato, Jeffrey Barrett, Sonia Goncalves, Ewan Harrison, David K. Jackson, Ian Johnston, Dominic Kwiatkowski, Cordelia Langford, John Sillitoe on behalf of the Wellcome Sanger Institute COVID-19 Surveillance Team |
| EPI_ISL_2910168 | Lighthouse Lab in Milton Keynes | Wellcome Sanger Institute for the COVID-19 Genomics UK (COG-UK) Consortium | The Lighthouse Lab in Milton Keynes and Alex Alderton, Roberto Amato, Jeffrey Barrett, Sonia Goncalves, Ewan Harrison, David K. Jackson, Ian Johnston, Dominic Kwiatkowski, Cordelia Langford, John Sillitoe on behalf of the Wellcome Sanger Institute COVID-19 Surveillance Team |
| EPI_ISL_2910169, EPI_ISL_2910170, EPI_ISL_2910171, EPI_ISL_2910172 | Lighthouse Laboratory Plymouth | Wellcome Sanger Institute for the COVID-19 Genomics UK (COG-UK) Consortium | Lighthouse Laboratory Plymouth and Alex Alderton, Roberto Amato, Jeffrey Barrett, Sonia Goncalves, Ewan Harrison, David K. Jackson, Ian Johnston, Dominic Kwiatkowski, Cordelia Langford, John Sillitoe on behalf of the Wellcome Sanger Institute COVID-19 Surveillance Team |
| EPI_ISL_2910173 | Lighthouse Lab in Milton Keynes | Wellcome Sanger Institute for the COVID-19 Genomics UK (COG-UK) Consortium | The Lighthouse Lab in Milton Keynes and Alex Alderton, Roberto Amato, Jeffrey Barrett, Sonia Goncalves, Ewan Harrison, David K. Jackson, Ian Johnston, Dominic Kwiatkowski, Cordelia Langford, John Sillitoe on behalf of the Wellcome Sanger Institute COVID-19 Surveillance Team |

| | | | |
|---|---|---|---|
| EPI_ISL_2910174, EPI_ISL_2910175 | Lighthouse Laboratory Plymouth | Wellcome Sanger Institute for the COVID-19 Genomics UK (COG-UK) Consortium | Lighthouse Laboratory Plymouth and Alex Alderton, Roberto Amato, Jeffrey Barrett, Sonia Goncalves, Ewan Harrison, David K. Jackson, Ian Johnston, Dominic Kwiatkowski, Cordelia Langford, John Sillitoe on behalf of the Wellcome Sanger Institute COVID-19 Surveillance Team |
| EPI_ISL_2910176, EPI_ISL_2910177, EPI_ISL_2910178, EPI_ISL_2910179 | Lighthouse Lab in Alderley Park | Wellcome Sanger Institute for the COVID-19 Genomics UK (COG-UK) Consortium | Jacquelyn Wynn, Mairead Hyland, The Lighthouse Lab in Alderley Park and Alex Alderton, Roberto Amato, Jeffrey Barrett, Sonia Goncalves, Ewan Harrison, David K. Jackson, Ian Johnston, Dominic Kwiatkowski, Cordelia Langford, John Sillitoe on behalf of the Wellcome Sanger Institute COVID-19 Surveillance Team |
| EPI_ISL_2910180, EPI_ISL_2910181 | Lighthouse Lab in Milton Keynes | Wellcome Sanger Institute for the COVID-19 Genomics UK (COG-UK) Consortium | The Lighthouse Lab in Milton Keynes and Alex Alderton, Roberto Amato, Jeffrey Barrett, Sonia Goncalves, Ewan Harrison, David K. Jackson, Ian Johnston, Dominic Kwiatkowski, Cordelia Langford, John Sillitoe on behalf of the Wellcome Sanger Institute COVID-19 Surveillance Team |
| EPI_ISL_2910182 | Lighthouse Lab in Alderley Park | Wellcome Sanger Institute for the COVID-19 Genomics UK (COG-UK) Consortium | Jacquelyn Wynn, Mairead Hyland, The Lighthouse Lab in Alderley Park and Alex Alderton, Roberto Amato, Jeffrey Barrett, Sonia Goncalves, Ewan Harrison, David K. Jackson, Ian Johnston, Dominic Kwiatkowski, Cordelia Langford, John Sillitoe on behalf of the Wellcome Sanger Institute COVID-19 Surveillance Team |
| EPI_ISL_2910183 | Lighthouse Lab in Milton Keynes | Wellcome Sanger Institute for the COVID-19 Genomics UK (COG-UK) Consortium | The Lighthouse Lab in Milton Keynes and Alex Alderton, Roberto Amato, Jeffrey Barrett, Sonia Goncalves, Ewan Harrison, David K. Jackson, Ian Johnston, Dominic Kwiatkowski, Cordelia Langford, John Sillitoe on behalf of the Wellcome Sanger Institute COVID-19 Surveillance Team |
| EPI_ISL_2910184 | Lighthouse Lab in Alderley Park | Wellcome Sanger Institute for the COVID-19 Genomics UK (COG-UK) Consortium | Jacquelyn Wynn, Mairead Hyland, The Lighthouse Lab in Alderley Park and Alex Alderton, Roberto Amato, Jeffrey Barrett, Sonia Goncalves, Ewan Harrison, David K. Jackson, Ian Johnston, Dominic Kwiatkowski, Cordelia Langford, John Sillitoe on behalf of the Wellcome Sanger Institute COVID-19 Surveillance Team |
| EPI_ISL_2910185 | Lighthouse Lab in Milton Keynes | Wellcome Sanger Institute for the COVID-19 Genomics UK (COG-UK) Consortium | The Lighthouse Lab in Milton Keynes and Alex Alderton, Roberto Amato, Jeffrey Barrett, Sonia Goncalves, Ewan Harrison, David K. Jackson, Ian Johnston, Dominic Kwiatkowski, Cordelia Langford, John Sillitoe on behalf of the Wellcome Sanger Institute COVID-19 Surveillance Team |
| EPI_ISL_2910186 | Lighthouse Lab in Alderley Park | Wellcome Sanger Institute for the COVID-19 Genomics UK (COG-UK) Consortium | Jacquelyn Wynn, Mairead Hyland, The Lighthouse Lab in Alderley Park and Alex Alderton, Roberto Amato, Jeffrey Barrett, Sonia Goncalves, Ewan Harrison, David K. Jackson, Ian Johnston, Dominic Kwiatkowski, Cordelia Langford, John Sillitoe on behalf of the Wellcome Sanger Institute COVID-19 Surveillance Team |
| EPI_ISL_2910187 | Lighthouse Laboratory Plymouth | Wellcome Sanger Institute for the COVID-19 Genomics UK (COG-UK) Consortium | Lighthouse Laboratory Plymouth and Alex Alderton, Roberto Amato, Jeffrey Barrett, Sonia Goncalves, Ewan Harrison, David K. Jackson, Ian Johnston, Dominic Kwiatkowski, Cordelia Langford, John Sillitoe on behalf of the Wellcome Sanger Institute COVID-19 Surveillance Team |
| EPI_ISL_2910188 | Lighthouse Lab in Alderley Park | Wellcome Sanger Institute for the COVID-19 Genomics UK (COG-UK) Consortium | Jacquelyn Wynn, Mairead Hyland, The Lighthouse Lab in Alderley Park and Alex Alderton, Roberto Amato, Jeffrey Barrett, Sonia Goncalves, Ewan Harrison, David K. Jackson, Ian Johnston, Dominic Kwiatkowski, Cordelia Langford, John Sillitoe on behalf of the Wellcome Sanger Institute COVID-19 Surveillance Team |
| EPI_ISL_2910189 | Lighthouse Laboratory Plymouth | Wellcome Sanger Institute for the COVID-19 Genomics UK (COG-UK) Consortium | Lighthouse Laboratory Plymouth and Alex Alderton, Roberto Amato, Jeffrey Barrett, Sonia Goncalves, Ewan Harrison, David K. Jackson, Ian Johnston, Dominic Kwiatkowski, Cordelia Langford, John Sillitoe on behalf of the Wellcome Sanger Institute COVID-19 Surveillance Team |
| EPI_ISL_2910190, EPI_ISL_2910191 | Lighthouse Lab in Alderley Park | Wellcome Sanger Institute for the COVID-19 Genomics UK (COG-UK) Consortium | Jacquelyn Wynn, Mairead Hyland, The Lighthouse Lab in Alderley Park and Alex Alderton, Roberto Amato, Jeffrey Barrett, Sonia Goncalves, Ewan Harrison, David K. Jackson, Ian Johnston, Dominic Kwiatkowski, Cordelia Langford, John Sillitoe on behalf of the Wellcome Sanger Institute COVID-19 Surveillance Team |
| EPI_ISL_2910192 | Lighthouse Laboratory Plymouth | Wellcome Sanger Institute for the COVID-19 Genomics UK (COG-UK) Consortium | Lighthouse Laboratory Plymouth and Alex Alderton, Roberto Amato, Jeffrey Barrett, Sonia Goncalves, Ewan Harrison, David K. Jackson, Ian Johnston, Dominic Kwiatkowski, Cordelia Langford, John Sillitoe on behalf of the Wellcome Sanger Institute COVID-19 Surveillance Team |
| EPI_ISL_2910193 | Lighthouse Lab in Milton Keynes | Wellcome Sanger Institute for the COVID-19 Genomics UK (COG-UK) Consortium | The Lighthouse Lab in Milton Keynes and Alex Alderton, Roberto Amato, Jeffrey Barrett, Sonia Goncalves, Ewan Harrison, David K. Jackson, Ian Johnston, Dominic Kwiatkowski, Cordelia Langford, John Sillitoe on behalf of the Wellcome Sanger Institute COVID-19 Surveillance Team |
| EPI_ISL_2910194, EPI_ISL_2910195 | Lighthouse Lab in Alderley Park | Wellcome Sanger Institute for the COVID-19 Genomics UK (COG-UK) Consortium | Jacquelyn Wynn, Mairead Hyland, The Lighthouse Lab in Alderley Park and Alex Alderton, Roberto Amato, Jeffrey Barrett, Sonia Goncalves, Ewan Harrison, David K. Jackson, Ian Johnston, Dominic Kwiatkowski, Cordelia Langford, John Sillitoe on behalf of the Wellcome Sanger Institute COVID-19 Surveillance Team |
| EPI_ISL_2910196 | Lighthouse Laboratory Plymouth | Wellcome Sanger Institute for the COVID-19 Genomics UK (COG-UK) Consortium | Lighthouse Laboratory Plymouth and Alex Alderton, Roberto Amato, Jeffrey Barrett, Sonia Goncalves, Ewan Harrison, David K. Jackson, Ian Johnston, Dominic Kwiatkowski, Cordelia Langford, John Sillitoe on behalf of the Wellcome Sanger Institute COVID-19 Surveillance Team |
| EPI_ISL_2910197, EPI_ISL_2910198 | Lighthouse Lab in Milton Keynes | Wellcome Sanger Institute for the COVID-19 Genomics UK (COG-UK) Consortium | The Lighthouse Lab in Milton Keynes and Alex Alderton, Roberto Amato, Jeffrey Barrett, Sonia Goncalves, Ewan Harrison, David K. Jackson, Ian Johnston, Dominic Kwiatkowski, Cordelia Langford, John Sillitoe on behalf of the Wellcome Sanger Institute COVID-19 Surveillance Team |
| EPI_ISL_2910199 | Lighthouse Lab in Alderley Park | Wellcome Sanger Institute for the COVID-19 Genomics UK (COG-UK) Consortium | Jacquelyn Wynn, Mairead Hyland, The Lighthouse Lab in Alderley Park and Alex Alderton, Roberto Amato, Jeffrey Barrett, Sonia Goncalves, Ewan Harrison, David K. Jackson, Ian Johnston, Dominic Kwiatkowski, Cordelia Langford, John Sillitoe on behalf of the Wellcome Sanger Institute COVID-19 Surveillance Team |
| EPI_ISL_2910200 | Lighthouse Laboratory Plymouth | Wellcome Sanger Institute for the COVID-19 Genomics UK (COG-UK) Consortium | Lighthouse Laboratory Plymouth and Alex Alderton, Roberto Amato, Jeffrey Barrett, Sonia Goncalves, Ewan Harrison, David K. Jackson, Ian Johnston, Dominic Kwiatkowski, Cordelia Langford, John Sillitoe on behalf of the Wellcome Sanger Institute COVID-19 Surveillance Team |
| EPI_ISL_2910201 | Lighthouse Lab in Alderley Park | Wellcome Sanger Institute for the COVID-19 Genomics UK (COG-UK) Consortium | Jacquelyn Wynn, Mairead Hyland, The Lighthouse Lab in Alderley Park and Alex Alderton, Roberto Amato, Jeffrey Barrett, Sonia Goncalves, Ewan Harrison, David K. Jackson, Ian Johnston, Dominic Kwiatkowski, Cordelia Langford, John Sillitoe on behalf of the Wellcome Sanger Institute COVID-19 Surveillance Team |
| EPI_ISL_2910202, EPI_ISL_2910203 | Lighthouse Laboratory Plymouth | Wellcome Sanger Institute for the COVID-19 Genomics UK (COG-UK) Consortium | Lighthouse Laboratory Plymouth and Alex Alderton, Roberto Amato, Jeffrey Barrett, Sonia Goncalves, Ewan Harrison, David K. Jackson, Ian Johnston, Dominic Kwiatkowski, Cordelia Langford, John Sillitoe on behalf of the Wellcome Sanger Institute COVID-19 Surveillance Team |
| EPI_ISL_2910204, EPI_ISL_2910205 | Lighthouse Lab in Milton Keynes | Wellcome Sanger Institute for the COVID-19 Genomics UK (COG-UK) Consortium | The Lighthouse Lab in Milton Keynes and Alex Alderton, Roberto Amato, Jeffrey Barrett, Sonia Goncalves, Ewan Harrison, David K. Jackson, Ian Johnston, Dominic Kwiatkowski, Cordelia Langford, John Sillitoe on behalf of the Wellcome Sanger Institute COVID-19 Surveillance Team |
| EPI_ISL_2910206, EPI_ISL_2910207, EPI_ISL_2910208, EPI_ISL_2910209 | Lighthouse Lab in Alderley Park | Wellcome Sanger Institute for the COVID-19 Genomics UK (COG-UK) Consortium | Jacquelyn Wynn, Mairead Hyland, The Lighthouse Lab in Alderley Park and Alex Alderton, Roberto Amato, Jeffrey Barrett, Sonia Goncalves, Ewan Harrison, David K. Jackson, Ian Johnston, Dominic Kwiatkowski, Cordelia Langford, John Sillitoe on behalf of the Wellcome Sanger Institute COVID-19 Surveillance Team |
| EPI_ISL_2910210 | Lighthouse Laboratory Plymouth | Wellcome Sanger Institute for the COVID-19 Genomics UK (COG-UK) Consortium | Lighthouse Laboratory Plymouth and Alex Alderton, Roberto Amato, Jeffrey Barrett, Sonia Goncalves, Ewan Harrison, David K. Jackson, Ian Johnston, Dominic Kwiatkowski, Cordelia Langford, John Sillitoe on behalf of the Wellcome Sanger Institute COVID-19 Surveillance Team |
| EPI_ISL_2910211, EPI_ISL_2910212 | Lighthouse Lab in Alderley Park | Wellcome Sanger Institute for the COVID-19 Genomics UK (COG-UK) Consortium | Jacquelyn Wynn, Mairead Hyland, The Lighthouse Lab in Alderley Park and Alex Alderton, Roberto Amato, Jeffrey Barrett, Sonia Goncalves, Ewan Harrison, David K. Jackson, Ian Johnston, Dominic Kwiatkowski, Cordelia Langford, John Sillitoe on behalf of the Wellcome Sanger Institute COVID-19 Surveillance Team |
| EPI_ISL_2910213 | Lighthouse Lab in Milton Keynes | Wellcome Sanger Institute for the COVID-19 Genomics UK (COG-UK) Consortium | The Lighthouse Lab in Milton Keynes and Alex Alderton, Roberto Amato, Jeffrey Barrett, Sonia Goncalves, Ewan Harrison, David K. Jackson, Ian Johnston, Dominic Kwiatkowski, Cordelia Langford, John Sillitoe on behalf of the Wellcome Sanger Institute COVID-19 Surveillance Team |
| EPI_ISL_2910214 | Lighthouse Lab in Alderley Park | Wellcome Sanger Institute for the COVID-19 Genomics UK (COG-UK) Consortium | Jacquelyn Wynn, Mairead Hyland, The Lighthouse Lab in Alderley Park and Alex Alderton, Roberto Amato, Jeffrey Barrett, Sonia Goncalves, Ewan Harrison, David K. Jackson, Ian Johnston, Dominic Kwiatkowski, Cordelia Langford, John Sillitoe on behalf of the Wellcome Sanger Institute COVID-19 Surveillance Team |
| EPI_ISL_2910215 | Lighthouse Lab in Milton Keynes | Wellcome Sanger Institute for the COVID-19 Genomics UK (COG-UK) Consortium | The Lighthouse Lab in Milton Keynes and Alex Alderton, Roberto Amato, Jeffrey Barrett, Sonia Goncalves, Ewan Harrison, David K. Jackson, Ian Johnston, Dominic Kwiatkowski, Cordelia Langford, John Sillitoe on behalf of the Wellcome Sanger Institute COVID-19 Surveillance Team |
| EPI_ISL_2910216 | Lighthouse Lab in Alderley Park | Wellcome Sanger Institute for the COVID-19 Genomics UK (COG-UK) Consortium | Jacquelyn Wynn, Mairead Hyland, The Lighthouse Lab in Alderley Park and Alex Alderton, Roberto Amato, Jeffrey Barrett, Sonia Goncalves, Ewan Harrison, David K. Jackson, Ian Johnston, Dominic Kwiatkowski, Cordelia Langford, John Sillitoe on behalf of the Wellcome Sanger Institute COVID-19 Surveillance Team |
| EPI_ISL_2910217 | Lighthouse Laboratory Plymouth | Wellcome Sanger Institute for the COVID-19 Genomics UK (COG-UK) Consortium | Lighthouse Laboratory Plymouth and Alex Alderton, Roberto Amato, Jeffrey Barrett, Sonia Goncalves, Ewan Harrison, David K. Jackson, Ian Johnston, Dominic Kwiatkowski, Cordelia Langford, John Sillitoe on behalf of the Wellcome Sanger Institute COVID-19 Surveillance Team |
| EPI_ISL_2910218, EPI_ISL_2910219 | Lighthouse Lab in Alderley Park | Wellcome Sanger Institute for the COVID-19 Genomics UK (COG-UK) Consortium | Jacquelyn Wynn, Mairead Hyland, The Lighthouse Lab in Alderley Park and Alex Alderton, Roberto Amato, Jeffrey Barrett, Sonia Goncalves, Ewan Harrison, David K. Jackson, Ian Johnston, Dominic Kwiatkowski, Cordelia Langford, John Sillitoe on behalf of the Wellcome Sanger Institute COVID-19 Surveillance Team |
| EPI_ISL_2910220, EPI_ISL_2910221, EPI_ISL_2910222 | Lighthouse Lab in Milton Keynes | Wellcome Sanger Institute for the COVID-19 Genomics UK (COG-UK) Consortium | The Lighthouse Lab in Milton Keynes and Alex Alderton, Roberto Amato, Jeffrey Barrett, Sonia Goncalves, Ewan Harrison, David K. Jackson, Ian Johnston, Dominic Kwiatkowski, Cordelia Langford, John Sillitoe on behalf of the Wellcome Sanger Institute COVID-19 Surveillance Team |
| EPI_ISL_2910223 | Lighthouse Lab in Alderley Park | Wellcome Sanger Institute for the COVID-19 Genomics UK (COG-UK) Consortium | Jacquelyn Wynn, Mairead Hyland, The Lighthouse Lab in Alderley Park and Alex Alderton, Roberto Amato, Jeffrey Barrett, Sonia Goncalves, Ewan Harrison, David K. Jackson, Ian Johnston, Dominic Kwiatkowski, Cordelia Langford, John Sillitoe on behalf of the Wellcome Sanger Institute COVID-19 Surveillance Team |
| EPI_ISL_2910224 | Lighthouse Lab in Milton Keynes | Wellcome Sanger Institute for the COVID-19 Genomics UK (COG-UK) Consortium | The Lighthouse Lab in Milton Keynes and Alex Alderton, Roberto Amato, Jeffrey Barrett, Sonia Goncalves, Ewan Harrison, David K. Jackson, Ian Johnston, Dominic Kwiatkowski, Cordelia Langford, John Sillitoe on behalf of the Wellcome Sanger Institute COVID-19 Surveillance Team |
| EPI_ISL_2910225 | Lighthouse Lab in Alderley Park | Wellcome Sanger Institute for the COVID-19 Genomics UK (COG-UK) Consortium | Jacquelyn Wynn, Mairead Hyland, The Lighthouse Lab in Alderley Park and Alex Alderton, Roberto Amato, Jeffrey Barrett, Sonia Goncalves, Ewan Harrison, David K. Jackson, Ian Johnston, Dominic Kwiatkowski, Cordelia Langford, John Sillitoe on behalf of the Wellcome Sanger Institute COVID-19 Surveillance Team |
| EPI_ISL_2910226 | Lighthouse Lab in Milton Keynes | Wellcome Sanger Institute for the COVID-19 Genomics UK (COG-UK) Consortium | The Lighthouse Lab in Milton Keynes and Alex Alderton, Roberto Amato, Jeffrey Barrett, Sonia Goncalves, Ewan Harrison, David K. Jackson, Ian Johnston, Dominic Kwiatkowski, Cordelia Langford, John Sillitoe on behalf of the Wellcome Sanger Institute COVID-19 Surveillance Team |
| EPI_ISL_2910227, EPI_ISL_2910228, EPI_ISL_2910229 | Lighthouse Laboratory Plymouth | Wellcome Sanger Institute for the COVID-19 Genomics UK (COG-UK) Consortium | Lighthouse Laboratory Plymouth and Alex Alderton, Roberto Amato, Jeffrey Barrett, Sonia Goncalves, Ewan Harrison, David K. Jackson, Ian Johnston, Dominic Kwiatkowski, Cordelia Langford, John Sillitoe on behalf of the Wellcome Sanger Institute COVID-19 Surveillance Team |
| EPI_ISL_2910230 | Lighthouse Lab in Alderley Park | Wellcome Sanger Institute for the COVID-19 Genomics UK (COG-UK) Consortium | Jacquelyn Wynn, Mairead Hyland, The Lighthouse Lab in Alderley Park and Alex Alderton, Roberto Amato, Jeffrey Barrett, Sonia Goncalves, Ewan Harrison, David K. Jackson, Ian Johnston, Dominic Kwiatkowski, Cordelia Langford, John Sillitoe on behalf of the Wellcome Sanger Institute COVID-19 Surveillance Team |
| EPI_ISL_2910231 | Lighthouse Laboratory Plymouth | Wellcome Sanger Institute for the COVID-19 Genomics UK (COG-UK) Consortium | Lighthouse Laboratory Plymouth and Alex Alderton, Roberto Amato, Jeffrey Barrett, Sonia Goncalves, Ewan Harrison, David K. Jackson, Ian Johnston, Dominic Kwiatkowski, Cordelia Langford, John Sillitoe on behalf of the Wellcome Sanger Institute COVID-19 Surveillance Team |
| EPI_ISL_2910232 | Lighthouse Lab in Alderley Park | Wellcome Sanger Institute for the COVID-19 Genomics UK (COG-UK) Consortium | Jacquelyn Wynn, Mairead Hyland, The Lighthouse Lab in Alderley Park and Alex Alderton, Roberto Amato, Jeffrey Barrett, Sonia Goncalves, Ewan Harrison, David K. Jackson, Ian Johnston, Dominic Kwiatkowski, Cordelia Langford, John Sillitoe on behalf of the Wellcome Sanger Institute COVID-19 Surveillance Team |
| EPI_ISL_2910233 | Lighthouse Laboratory Plymouth | Wellcome Sanger Institute for the COVID-19 Genomics UK (COG-UK) Consortium | Lighthouse Laboratory Plymouth and Alex Alderton, Roberto Amato, Jeffrey Barrett, Sonia Goncalves, Ewan Harrison, David K. Jackson, Ian Johnston, Dominic Kwiatkowski, Cordelia Langford, John Sillitoe on behalf of the Wellcome Sanger Institute COVID-19 Surveillance Team |
| EPI_ISL_2910234 | Lighthouse Lab in Alderley Park | Wellcome Sanger Institute for the COVID-19 Genomics UK (COG-UK) Consortium | Jacquelyn Wynn, Mairead Hyland, The Lighthouse Lab in Alderley Park and Alex Alderton, Roberto Amato, Jeffrey Barrett, Sonia Goncalves, Ewan Harrison, David K. Jackson, Ian Johnston, Dominic Kwiatkowski, Cordelia Langford, John Sillitoe on behalf of the Wellcome Sanger Institute COVID-19 Surveillance Team |
| EPI_ISL_2910235 | Lighthouse Lab in Milton Keynes | Wellcome Sanger Institute for the COVID-19 Genomics UK (COG-UK) Consortium | The Lighthouse Lab in Milton Keynes and Alex Alderton, Roberto Amato, Jeffrey Barrett, Sonia Goncalves, Ewan Harrison, David K. Jackson, Ian Johnston, Dominic Kwiatkowski, Cordelia Langford, John Sillitoe on behalf of the Wellcome Sanger Institute COVID-19 Surveillance Team |

| | | | |
|---|---|---|---|
| EPI_ISL_2910236, EPI_ISL_2910238 | Lighthouse Lab in Alderley Park | Wellcome Sanger Institute for the COVID-19 Genomics UK (COG-UK) Consortium | Jacquelyn Wynn, Mairead Hyland, The Lighthouse Lab in Alderley Park and Alex Alderton, Roberto Amato, Jeffrey Barrett, Sonia Goncalves, Ewan Harrison, David K. Jackson, Ian Johnston, Dominic Kwiatkowski, Cordelia Langford, John Sillitoe on behalf of the Wellcome Sanger Institute COVID-19 Surveillance Team |
| EPI_ISL_2910239 | Lighthouse Laboratory Plymouth | Wellcome Sanger Institute for the COVID-19 Genomics UK (COG-UK) Consortium | Lighthouse Laboratory Plymouth and Alex Alderton, Roberto Amato, Jeffrey Barrett, Sonia Goncalves, Ewan Harrison, David K. Jackson, Ian Johnston, Dominic Kwiatkowski, Cordelia Langford, John Sillitoe on behalf of the Wellcome Sanger Institute COVID-19 Surveillance Team |
| EPI_ISL_2910240, EPI_ISL_2910241, EPI_ISL_2910242, EPI_ISL_2910243 | Lighthouse Lab in Alderley Park | Wellcome Sanger Institute for the COVID-19 Genomics UK (COG-UK) Consortium | Jacquelyn Wynn, Mairead Hyland, The Lighthouse Lab in Alderley Park and Alex Alderton, Roberto Amato, Jeffrey Barrett, Sonia Goncalves, Ewan Harrison, David K. Jackson, Ian Johnston, Dominic Kwiatkowski, Cordelia Langford, John Sillitoe on behalf of the Wellcome Sanger Institute COVID-19 Surveillance Team |
| EPI_ISL_2910244 | Lighthouse Lab in Milton Keynes | Wellcome Sanger Institute for the COVID-19 Genomics UK (COG-UK) Consortium | The Lighthouse Lab in Milton Keynes and Alex Alderton, Roberto Amato, Jeffrey Barrett, Sonia Goncalves, Ewan Harrison, David K. Jackson, Ian Johnston, Dominic Kwiatkowski, Cordelia Langford, John Sillitoe on behalf of the Wellcome Sanger Institute COVID-19 Surveillance Team |
| EPI_ISL_2910245 | Lighthouse Laboratory Plymouth | Wellcome Sanger Institute for the COVID-19 Genomics UK (COG-UK) Consortium | Lighthouse Laboratory Plymouth and Alex Alderton, Roberto Amato, Jeffrey Barrett, Sonia Goncalves, Ewan Harrison, David K. Jackson, Ian Johnston, Dominic Kwiatkowski, Cordelia Langford, John Sillitoe on behalf of the Wellcome Sanger Institute COVID-19 Surveillance Team |
| EPI_ISL_2910246, EPI_ISL_2910247 | Lighthouse Lab in Milton Keynes | Wellcome Sanger Institute for the COVID-19 Genomics UK (COG-UK) Consortium | The Lighthouse Lab in Milton Keynes and Alex Alderton, Roberto Amato, Jeffrey Barrett, Sonia Goncalves, Ewan Harrison, David K. Jackson, Ian Johnston, Dominic Kwiatkowski, Cordelia Langford, John Sillitoe on behalf of the Wellcome Sanger Institute COVID-19 Surveillance Team |
| EPI_ISL_2910248, EPI_ISL_2910249, EPI_ISL_2910250, EPI_ISL_2910251 | Lighthouse Laboratory Plymouth | Wellcome Sanger Institute for the COVID-19 Genomics UK (COG-UK) Consortium | Lighthouse Laboratory Plymouth and Alex Alderton, Roberto Amato, Jeffrey Barrett, Sonia Goncalves, Ewan Harrison, David K. Jackson, Ian Johnston, Dominic Kwiatkowski, Cordelia Langford, John Sillitoe on behalf of the Wellcome Sanger Institute COVID-19 Surveillance Team |
| EPI_ISL_2910252, EPI_ISL_2910253 | Lighthouse Lab in Alderley Park | Wellcome Sanger Institute for the COVID-19 Genomics UK (COG-UK) Consortium | Jacquelyn Wynn, Mairead Hyland, The Lighthouse Lab in Alderley Park and Alex Alderton, Roberto Amato, Jeffrey Barrett, Sonia Goncalves, Ewan Harrison, David K. Jackson, Ian Johnston, Dominic Kwiatkowski, Cordelia Langford, John Sillitoe on behalf of the Wellcome Sanger Institute COVID-19 Surveillance Team |
| EPI_ISL_2910254 | Lighthouse Laboratory Plymouth | Wellcome Sanger Institute for the COVID-19 Genomics UK (COG-UK) Consortium | Lighthouse Laboratory Plymouth and Alex Alderton, Roberto Amato, Jeffrey Barrett, Sonia Goncalves, Ewan Harrison, David K. Jackson, Ian Johnston, Dominic Kwiatkowski, Cordelia Langford, John Sillitoe on behalf of the Wellcome Sanger Institute COVID-19 Surveillance Team |
| EPI_ISL_2910255 | Lighthouse Lab in Alderley Park | Wellcome Sanger Institute for the COVID-19 Genomics UK (COG-UK) Consortium | Jacquelyn Wynn, Mairead Hyland, The Lighthouse Lab in Alderley Park and Alex Alderton, Roberto Amato, Jeffrey Barrett, Sonia Goncalves, Ewan Harrison, David K. Jackson, Ian Johnston, Dominic Kwiatkowski, Cordelia Langford, John Sillitoe on behalf of the Wellcome Sanger Institute COVID-19 Surveillance Team |
| EPI_ISL_2910256, EPI_ISL_2910257 | Lighthouse Laboratory Plymouth | Wellcome Sanger Institute for the COVID-19 Genomics UK (COG-UK) Consortium | Lighthouse Laboratory Plymouth and Alex Alderton, Roberto Amato, Jeffrey Barrett, Sonia Goncalves, Ewan Harrison, David K. Jackson, Ian Johnston, Dominic Kwiatkowski, Cordelia Langford, John Sillitoe on behalf of the Wellcome Sanger Institute COVID-19 Surveillance Team |
| EPI_ISL_2910258, EPI_ISL_2910259 | Lighthouse Lab in Alderley Park | Wellcome Sanger Institute for the COVID-19 Genomics UK (COG-UK) Consortium | Jacquelyn Wynn, Mairead Hyland, The Lighthouse Lab in Alderley Park and Alex Alderton, Roberto Amato, Jeffrey Barrett, Sonia Goncalves, Ewan Harrison, David K. Jackson, Ian Johnston, Dominic Kwiatkowski, Cordelia Langford, John Sillitoe on behalf of the Wellcome Sanger Institute COVID-19 Surveillance Team |
| EPI_ISL_2910260 | Lighthouse Lab in Glasgow | Wellcome Sanger Institute for the COVID-19 Genomics UK (COG-UK) Consortium | Harper VanSteenhouse, Yumi Kasai, David Gray, Carol Clugston, Anna Dominiczak and Alex Alderton, Roberto Amato, Jeffrey Barrett, Sonia Goncalves, Ewan Harrison, David K. Jackson, Ian Johnston, Dominic Kwiatkowski, Cordelia Langford, John Sillitoe on behalf of the Wellcome Sanger Institute COVID-19 Surveillance Team |
| EPI_ISL_2910261 | Lighthouse Lab in Alderley Park | Wellcome Sanger Institute for the COVID-19 Genomics UK (COG-UK) Consortium | Jacquelyn Wynn, Mairead Hyland, The Lighthouse Lab in Alderley Park and Alex Alderton, Roberto Amato, Jeffrey Barrett, Sonia Goncalves, Ewan Harrison, David K. Jackson, Ian Johnston, Dominic Kwiatkowski, Cordelia Langford, John Sillitoe on behalf of the Wellcome Sanger Institute COVID-19 Surveillance Team |
| EPI_ISL_2910262, EPI_ISL_2910263 | Lighthouse Lab in Milton Keynes | Wellcome Sanger Institute for the COVID-19 Genomics UK (COG-UK) Consortium | The Lighthouse Lab in Milton Keynes and Alex Alderton, Roberto Amato, Jeffrey Barrett, Sonia Goncalves, Ewan Harrison, David K. Jackson, Ian Johnston, Dominic Kwiatkowski, Cordelia Langford, John Sillitoe on behalf of the Wellcome Sanger Institute COVID-19 Surveillance Team |
| EPI_ISL_2910264, EPI_ISL_2910265 | Lighthouse Lab in Alderley Park | Wellcome Sanger Institute for the COVID-19 Genomics UK (COG-UK) Consortium | Jacquelyn Wynn, Mairead Hyland, The Lighthouse Lab in Alderley Park and Alex Alderton, Roberto Amato, Jeffrey Barrett, Sonia Goncalves, Ewan Harrison, David K. Jackson, Ian Johnston, Dominic Kwiatkowski, Cordelia Langford, John Sillitoe on behalf of the Wellcome Sanger Institute COVID-19 Surveillance Team |
| EPI_ISL_2910266, EPI_ISL_2910267 | Lighthouse Lab in Milton Keynes | Wellcome Sanger Institute for the COVID-19 Genomics UK (COG-UK) Consortium | The Lighthouse Lab in Milton Keynes and Alex Alderton, Roberto Amato, Jeffrey Barrett, Sonia Goncalves, Ewan Harrison, David K. Jackson, Ian Johnston, Dominic Kwiatkowski, Cordelia Langford, John Sillitoe on behalf of the Wellcome Sanger Institute COVID-19 Surveillance Team |
| EPI_ISL_2910268 | Lighthouse Lab in Glasgow | Wellcome Sanger Institute for the COVID-19 Genomics UK (COG-UK) Consortium | Harper VanSteenhouse, Yumi Kasai, David Gray, Carol Clugston, Anna Dominiczak and Alex Alderton, Roberto Amato, Jeffrey Barrett, Sonia Goncalves, Ewan Harrison, David K. Jackson, Ian Johnston, Dominic Kwiatkowski, Cordelia Langford, John Sillitoe on behalf of the Wellcome Sanger Institute COVID-19 Surveillance Team |
| EPI_ISL_2910269 | Lighthouse Lab in Alderley Park | Wellcome Sanger Institute for the COVID-19 Genomics UK (COG-UK) Consortium | Jacquelyn Wynn, Mairead Hyland, The Lighthouse Lab in Alderley Park and Alex Alderton, Roberto Amato, Jeffrey Barrett, Sonia Goncalves, Ewan Harrison, David K. Jackson, Ian Johnston, Dominic Kwiatkowski, Cordelia Langford, John Sillitoe on behalf of the Wellcome Sanger Institute COVID-19 Surveillance Team |
| EPI_ISL_2910270 | Lighthouse Lab in Glasgow | Wellcome Sanger Institute for the COVID-19 Genomics UK (COG-UK) Consortium | Harper VanSteenhouse, Yumi Kasai, David Gray, Carol Clugston, Anna Dominiczak and Alex Alderton, Roberto Amato, Jeffrey Barrett, Sonia Goncalves, Ewan Harrison, David K. Jackson, Ian Johnston, Dominic Kwiatkowski, Cordelia Langford, John Sillitoe on behalf of the Wellcome Sanger Institute COVID-19 Surveillance Team |
| EPI_ISL_2910271, EPI_ISL_2910272 | Lighthouse Lab in Alderley Park | Wellcome Sanger Institute for the COVID-19 Genomics UK (COG-UK) Consortium | Jacquelyn Wynn, Mairead Hyland, The Lighthouse Lab in Alderley Park and Alex Alderton, Roberto Amato, Jeffrey Barrett, Sonia Goncalves, Ewan Harrison, David K. Jackson, Ian Johnston, Dominic Kwiatkowski, Cordelia Langford, John Sillitoe on behalf of the Wellcome Sanger Institute COVID-19 Surveillance Team |
| EPI_ISL_2910273 | Lighthouse Lab in Glasgow | Wellcome Sanger Institute for the COVID-19 Genomics UK (COG-UK) Consortium | Harper VanSteenhouse, Yumi Kasai, David Gray, Carol Clugston, Anna Dominiczak and Alex Alderton, Roberto Amato, Jeffrey Barrett, Sonia Goncalves, Ewan Harrison, David K. Jackson, Ian Johnston, Dominic Kwiatkowski, Cordelia Langford, John Sillitoe on behalf of the Wellcome Sanger Institute COVID-19 Surveillance Team |
| EPI_ISL_2910274 | Lighthouse Lab in Alderley Park | Wellcome Sanger Institute for the COVID-19 Genomics UK (COG-UK) Consortium | Jacquelyn Wynn, Mairead Hyland, The Lighthouse Lab in Alderley Park and Alex Alderton, Roberto Amato, Jeffrey Barrett, Sonia Goncalves, Ewan Harrison, David K. Jackson, Ian Johnston, Dominic Kwiatkowski, Cordelia Langford, John Sillitoe on behalf of the Wellcome Sanger Institute COVID-19 Surveillance Team |
| EPI_ISL_2910275 | Lighthouse Lab in Milton Keynes | Wellcome Sanger Institute for the COVID-19 Genomics UK (COG-UK) Consortium | The Lighthouse Lab in Milton Keynes and Alex Alderton, Roberto Amato, Jeffrey Barrett, Sonia Goncalves, Ewan Harrison, David K. Jackson, Ian Johnston, Dominic Kwiatkowski, Cordelia Langford, John Sillitoe on behalf of the Wellcome Sanger Institute COVID-19 Surveillance Team |
| EPI_ISL_2910276, EPI_ISL_2910277 | Lighthouse Lab in Glasgow | Wellcome Sanger Institute for the COVID-19 Genomics UK (COG-UK) Consortium | Harper VanSteenhouse, Yumi Kasai, David Gray, Carol Clugston, Anna Dominiczak and Alex Alderton, Roberto Amato, Jeffrey Barrett, Sonia Goncalves, Ewan Harrison, David K. Jackson, Ian Johnston, Dominic Kwiatkowski, Cordelia Langford, John Sillitoe on behalf of the Wellcome Sanger Institute COVID-19 Surveillance Team |
| EPI_ISL_2910278 | Lighthouse Lab in Milton Keynes | Wellcome Sanger Institute for the COVID-19 Genomics UK (COG-UK) Consortium | The Lighthouse Lab in Milton Keynes and Alex Alderton, Roberto Amato, Jeffrey Barrett, Sonia Goncalves, Ewan Harrison, David K. Jackson, Ian Johnston, Dominic Kwiatkowski, Cordelia Langford, John Sillitoe on behalf of the Wellcome Sanger Institute COVID-19 Surveillance Team |
| EPI_ISL_2910279 | Lighthouse Lab in Alderley Park | Wellcome Sanger Institute for the COVID-19 Genomics UK (COG-UK) Consortium | Jacquelyn Wynn, Mairead Hyland, The Lighthouse Lab in Alderley Park and Alex Alderton, Roberto Amato, Jeffrey Barrett, Sonia Goncalves, Ewan Harrison, David K. Jackson, Ian Johnston, Dominic Kwiatkowski, Cordelia Langford, John Sillitoe on behalf of the Wellcome Sanger Institute COVID-19 Surveillance Team |
| EPI_ISL_2910280 | Lighthouse Lab in Milton Keynes | Wellcome Sanger Institute for the COVID-19 Genomics UK (COG-UK) Consortium | The Lighthouse Lab in Milton Keynes and Alex Alderton, Roberto Amato, Jeffrey Barrett, Sonia Goncalves, Ewan Harrison, David K. Jackson, Ian Johnston, Dominic Kwiatkowski, Cordelia Langford, John Sillitoe on behalf of the Wellcome Sanger Institute COVID-19 Surveillance Team |
| EPI_ISL_2910281 | Lighthouse Lab in Alderley Park | Wellcome Sanger Institute for the COVID-19 Genomics UK (COG-UK) Consortium | Jacquelyn Wynn, Mairead Hyland, The Lighthouse Lab in Alderley Park and Alex Alderton, Roberto Amato, Jeffrey Barrett, Sonia Goncalves, Ewan Harrison, David K. Jackson, Ian Johnston, Dominic Kwiatkowski, Cordelia Langford, John Sillitoe on behalf of the Wellcome Sanger Institute COVID-19 Surveillance Team |
| EPI_ISL_2910282 | Lighthouse Lab in Glasgow | Wellcome Sanger Institute for the COVID-19 Genomics UK (COG-UK) Consortium | Harper VanSteenhouse, Yumi Kasai, David Gray, Carol Clugston, Anna Dominiczak and Alex Alderton, Roberto Amato, Jeffrey Barrett, Sonia Goncalves, Ewan Harrison, David K. Jackson, Ian Johnston, Dominic Kwiatkowski, Cordelia Langford, John Sillitoe on behalf of the Wellcome Sanger Institute COVID-19 Surveillance Team |
| EPI_ISL_2910283, EPI_ISL_2910284, EPI_ISL_2910285 | Lighthouse Lab in Alderley Park | Wellcome Sanger Institute for the COVID-19 Genomics UK (COG-UK) Consortium | Jacquelyn Wynn, Mairead Hyland, The Lighthouse Lab in Alderley Park and Alex Alderton, Roberto Amato, Jeffrey Barrett, Sonia Goncalves, Ewan Harrison, David K. Jackson, Ian Johnston, Dominic Kwiatkowski, Cordelia Langford, John Sillitoe on behalf of the Wellcome Sanger Institute COVID-19 Surveillance Team |
| EPI_ISL_2910286, EPI_ISL_2910287 | Lighthouse Lab in Milton Keynes | Wellcome Sanger Institute for the COVID-19 Genomics UK (COG-UK) Consortium | The Lighthouse Lab in Milton Keynes and Alex Alderton, Roberto Amato, Jeffrey Barrett, Sonia Goncalves, Ewan Harrison, David K. Jackson, Ian Johnston, Dominic Kwiatkowski, Cordelia Langford, John Sillitoe on behalf of the Wellcome Sanger Institute COVID-19 Surveillance Team |
| EPI_ISL_2910288 | Lighthouse Lab in Alderley Park | Wellcome Sanger Institute for the COVID-19 Genomics UK (COG-UK) Consortium | Jacquelyn Wynn, Mairead Hyland, The Lighthouse Lab in Alderley Park and Alex Alderton, Roberto Amato, Jeffrey Barrett, Sonia Goncalves, Ewan Harrison, David K. Jackson, Ian Johnston, Dominic Kwiatkowski, Cordelia Langford, John Sillitoe on behalf of the Wellcome Sanger Institute COVID-19 Surveillance Team |
| EPI_ISL_2910289 | Lighthouse Lab in Glasgow | Wellcome Sanger Institute for the COVID-19 Genomics UK (COG-UK) Consortium | Harper VanSteenhouse, Yumi Kasai, David Gray, Carol Clugston, Anna Dominiczak and Alex Alderton, Roberto Amato, Jeffrey Barrett, Sonia Goncalves, Ewan Harrison, David K. Jackson, Ian Johnston, Dominic Kwiatkowski, Cordelia Langford, John Sillitoe on behalf of the Wellcome Sanger Institute COVID-19 Surveillance Team |
| EPI_ISL_2910290, EPI_ISL_2910291, EPI_ISL_2910292, EPI_ISL_2910293, EPI_ISL_2910294, EPI_ISL_2910295, EPI_ISL_2910296 | Lighthouse Lab in Alderley Park | Wellcome Sanger Institute for the COVID-19 Genomics UK (COG-UK) Consortium | Jacquelyn Wynn, Mairead Hyland, The Lighthouse Lab in Alderley Park and Alex Alderton, Roberto Amato, Jeffrey Barrett, Sonia Goncalves, Ewan Harrison, David K. Jackson, Ian Johnston, Dominic Kwiatkowski, Cordelia Langford, John Sillitoe on behalf of the Wellcome Sanger Institute COVID-19 Surveillance Team |
| EPI_ISL_2910297 | Lighthouse Lab in Glasgow | Wellcome Sanger Institute for the COVID-19 Genomics UK (COG-UK) Consortium | Harper VanSteenhouse, Yumi Kasai, David Gray, Carol Clugston, Anna Dominiczak and Alex Alderton, Roberto Amato, Jeffrey Barrett, Sonia Goncalves, Ewan Harrison, David K. Jackson, Ian Johnston, Dominic Kwiatkowski, Cordelia Langford, John Sillitoe on behalf of the Wellcome Sanger Institute COVID-19 Surveillance Team |
| EPI_ISL_2910298 | Lighthouse Lab in Alderley Park | Wellcome Sanger Institute for the COVID-19 Genomics UK (COG-UK) Consortium | Jacquelyn Wynn, Mairead Hyland, The Lighthouse Lab in Alderley Park and Alex Alderton, Roberto Amato, Jeffrey Barrett, Sonia Goncalves, Ewan Harrison, David K. Jackson, Ian Johnston, Dominic Kwiatkowski, Cordelia Langford, John Sillitoe on behalf of the Wellcome Sanger Institute COVID-19 Surveillance Team |
| EPI_ISL_2910299 | Lighthouse Lab in Milton Keynes | Wellcome Sanger Institute for the COVID-19 Genomics UK (COG-UK) Consortium | The Lighthouse Lab in Milton Keynes and Alex Alderton, Roberto Amato, Jeffrey Barrett, Sonia Goncalves, Ewan Harrison, David K. Jackson, Ian Johnston, Dominic Kwiatkowski, Cordelia Langford, John Sillitoe on behalf of the Wellcome Sanger Institute COVID-19 Surveillance Team |
| EPI_ISL_2910300, EPI_ISL_2910301, EPI_ISL_2910302 | Lighthouse Lab in Alderley Park | Wellcome Sanger Institute for the COVID-19 Genomics UK (COG-UK) Consortium | Jacquelyn Wynn, Mairead Hyland, The Lighthouse Lab in Alderley Park and Alex Alderton, Roberto Amato, Jeffrey Barrett, Sonia Goncalves, Ewan Harrison, David K. Jackson, Ian Johnston, Dominic Kwiatkowski, Cordelia Langford, John Sillitoe on behalf of the Wellcome Sanger Institute COVID-19 Surveillance Team |
| EPI_ISL_2910303 | Lighthouse Lab in Glasgow | Wellcome Sanger Institute for the COVID-19 Genomics UK (COG-UK) Consortium | Harper VanSteenhouse, Yumi Kasai, David Gray, Carol Clugston, Anna Dominiczak and Alex Alderton, Roberto Amato, Jeffrey Barrett, Sonia Goncalves, Ewan Harrison, David K. Jackson, Ian Johnston, Dominic Kwiatkowski, Cordelia Langford, John Sillitoe on behalf of the Wellcome Sanger Institute COVID-19 Surveillance Team |

| | | | |
|---|---|---|---|
| EPI_ISL_2910304, EPI_ISL_2910305 | Lighthouse Lab in Alderley Park | Wellcome Sanger Institute for the COVID-19 Genomics UK (COG-UK) Consortium | Jacquelyn Wynn, Mairead Hyland, The Lighthouse Lab in Alderley Park and Alex Alderton, Roberto Amato, Jeffrey Barrett, Sonia Goncalves, Ewan Harrison, David K. Jackson, Ian Johnston, Dominic Kwiatkowski, Cordelia Langford, John Sillitoe on behalf of the Wellcome Sanger Institute COVID-19 Surveillance Team |
| EPI_ISL_2910306 | Lighthouse Lab in Glasgow | Wellcome Sanger Institute for the COVID-19 Genomics UK (COG-UK) Consortium | Harper VanSteenhouse, Yumi Kasai, David Gray, Carol Clugston, Anna Dominiczak and Alex Alderton, Roberto Amato, Jeffrey Barrett, Sonia Goncalves, Ewan Harrison, David K. Jackson, Ian Johnston, Dominic Kwiatkowski, Cordelia Langford, John Sillitoe on behalf of the Wellcome Sanger Institute COVID-19 Surveillance Team |
| EPI_ISL_2910307, EPI_ISL_2910308 | Lighthouse Lab in Alderley Park | Wellcome Sanger Institute for the COVID-19 Genomics UK (COG-UK) Consortium | Jacquelyn Wynn, Mairead Hyland, The Lighthouse Lab in Alderley Park and Alex Alderton, Roberto Amato, Jeffrey Barrett, Sonia Goncalves, Ewan Harrison, David K. Jackson, Ian Johnston, Dominic Kwiatkowski, Cordelia Langford, John Sillitoe on behalf of the Wellcome Sanger Institute COVID-19 Surveillance Team |
| EPI_ISL_2910309, EPI_ISL_2910310, EPI_ISL_2910311 | Lighthouse Lab in Milton Keynes | Wellcome Sanger Institute for the COVID-19 Genomics UK (COG-UK) Consortium | The Lighthouse Lab in Milton Keynes and Alex Alderton, Roberto Amato, Jeffrey Barrett, Sonia Goncalves, Ewan Harrison, David K. Jackson, Ian Johnston, Dominic Kwiatkowski, Cordelia Langford, John Sillitoe on behalf of the Wellcome Sanger Institute COVID-19 Surveillance Team |
| EPI_ISL_2910312 | Lighthouse Lab in Alderley Park | Wellcome Sanger Institute for the COVID-19 Genomics UK (COG-UK) Consortium | Jacquelyn Wynn, Mairead Hyland, The Lighthouse Lab in Alderley Park and Alex Alderton, Roberto Amato, Jeffrey Barrett, Sonia Goncalves, Ewan Harrison, David K. Jackson, Ian Johnston, Dominic Kwiatkowski, Cordelia Langford, John Sillitoe on behalf of the Wellcome Sanger Institute COVID-19 Surveillance Team |
| EPI_ISL_2910313 | Lighthouse Lab in Milton Keynes | Wellcome Sanger Institute for the COVID-19 Genomics UK (COG-UK) Consortium | The Lighthouse Lab in Milton Keynes and Alex Alderton, Roberto Amato, Jeffrey Barrett, Sonia Goncalves, Ewan Harrison, David K. Jackson, Ian Johnston, Dominic Kwiatkowski, Cordelia Langford, John Sillitoe on behalf of the Wellcome Sanger Institute COVID-19 Surveillance Team |
| EPI_ISL_2910314 | Lighthouse Lab in Glasgow | Wellcome Sanger Institute for the COVID-19 Genomics UK (COG-UK) Consortium | Harper VanSteenhouse, Yumi Kasai, David Gray, Carol Clugston, Anna Dominiczak and Alex Alderton, Roberto Amato, Jeffrey Barrett, Sonia Goncalves, Ewan Harrison, David K. Jackson, Ian Johnston, Dominic Kwiatkowski, Cordelia Langford, John Sillitoe on behalf of the Wellcome Sanger Institute COVID-19 Surveillance Team |
| EPI_ISL_2910315 | Lighthouse Lab in Alderley Park | Wellcome Sanger Institute for the COVID-19 Genomics UK (COG-UK) Consortium | Jacquelyn Wynn, Mairead Hyland, The Lighthouse Lab in Alderley Park and Alex Alderton, Roberto Amato, Jeffrey Barrett, Sonia Goncalves, Ewan Harrison, David K. Jackson, Ian Johnston, Dominic Kwiatkowski, Cordelia Langford, John Sillitoe on behalf of the Wellcome Sanger Institute COVID-19 Surveillance Team |
| EPI_ISL_2910316 | Lighthouse Lab in Glasgow | Wellcome Sanger Institute for the COVID-19 Genomics UK (COG-UK) Consortium | Harper VanSteenhouse, Yumi Kasai, David Gray, Carol Clugston, Anna Dominiczak and Alex Alderton, Roberto Amato, Jeffrey Barrett, Sonia Goncalves, Ewan Harrison, David K. Jackson, Ian Johnston, Dominic Kwiatkowski, Cordelia Langford, John Sillitoe on behalf of the Wellcome Sanger Institute COVID-19 Surveillance Team |
| EPI_ISL_2910317, EPI_ISL_2910318, EPI_ISL_2910319, EPI_ISL_2910320, EPI_ISL_2910321, EPI_ISL_2910322, EPI_ISL_2910323 | Lighthouse Lab in Alderley Park | Wellcome Sanger Institute for the COVID-19 Genomics UK (COG-UK) Consortium | Jacquelyn Wynn, Mairead Hyland, The Lighthouse Lab in Alderley Park and Alex Alderton, Roberto Amato, Jeffrey Barrett, Sonia Goncalves, Ewan Harrison, David K. Jackson, Ian Johnston, Dominic Kwiatkowski, Cordelia Langford, John Sillitoe on behalf of the Wellcome Sanger Institute COVID-19 Surveillance Team |
| EPI_ISL_2910324 | Lighthouse Lab in Milton Keynes | Wellcome Sanger Institute for the COVID-19 Genomics UK (COG-UK) Consortium | The Lighthouse Lab in Milton Keynes and Alex Alderton, Roberto Amato, Jeffrey Barrett, Sonia Goncalves, Ewan Harrison, David K. Jackson, Ian Johnston, Dominic Kwiatkowski, Cordelia Langford, John Sillitoe on behalf of the Wellcome Sanger Institute COVID-19 Surveillance Team |
| EPI_ISL_2910325, EPI_ISL_2910326 | Lighthouse Lab in Alderley Park | Wellcome Sanger Institute for the COVID-19 Genomics UK (COG-UK) Consortium | Jacquelyn Wynn, Mairead Hyland, The Lighthouse Lab in Alderley Park and Alex Alderton, Roberto Amato, Jeffrey Barrett, Sonia Goncalves, Ewan Harrison, David K. Jackson, Ian Johnston, Dominic Kwiatkowski, Cordelia Langford, John Sillitoe on behalf of the Wellcome Sanger Institute COVID-19 Surveillance Team |
| EPI_ISL_2910327, EPI_ISL_2910328 | Lighthouse Lab in Milton Keynes | Wellcome Sanger Institute for the COVID-19 Genomics UK (COG-UK) Consortium | The Lighthouse Lab in Milton Keynes and Alex Alderton, Roberto Amato, Jeffrey Barrett, Sonia Goncalves, Ewan Harrison, David K. Jackson, Ian Johnston, Dominic Kwiatkowski, Cordelia Langford, John Sillitoe on behalf of the Wellcome Sanger Institute COVID-19 Surveillance Team |
| EPI_ISL_2910329 | Lighthouse Lab in Alderley Park | Wellcome Sanger Institute for the COVID-19 Genomics UK (COG-UK) Consortium | Jacquelyn Wynn, Mairead Hyland, The Lighthouse Lab in Alderley Park and Alex Alderton, Roberto Amato, Jeffrey Barrett, Sonia Goncalves, Ewan Harrison, David K. Jackson, Ian Johnston, Dominic Kwiatkowski, Cordelia Langford, John Sillitoe on behalf of the Wellcome Sanger Institute COVID-19 Surveillance Team |
| EPI_ISL_2910330 | Lighthouse Lab in Milton Keynes | Wellcome Sanger Institute for the COVID-19 Genomics UK (COG-UK) Consortium | The Lighthouse Lab in Milton Keynes and Alex Alderton, Roberto Amato, Jeffrey Barrett, Sonia Goncalves, Ewan Harrison, David K. Jackson, Ian Johnston, Dominic Kwiatkowski, Cordelia Langford, John Sillitoe on behalf of the Wellcome Sanger Institute COVID-19 Surveillance Team |
| EPI_ISL_2910331, EPI_ISL_2910332 | Lighthouse Lab in Alderley Park | Wellcome Sanger Institute for the COVID-19 Genomics UK (COG-UK) Consortium | Jacquelyn Wynn, Mairead Hyland, The Lighthouse Lab in Alderley Park and Alex Alderton, Roberto Amato, Jeffrey Barrett, Sonia Goncalves, Ewan Harrison, David K. Jackson, Ian Johnston, Dominic Kwiatkowski, Cordelia Langford, John Sillitoe on behalf of the Wellcome Sanger Institute COVID-19 Surveillance Team |
| EPI_ISL_2910333, EPI_ISL_2910334 | Lighthouse Lab in Milton Keynes | Wellcome Sanger Institute for the COVID-19 Genomics UK (COG-UK) Consortium | The Lighthouse Lab in Milton Keynes and Alex Alderton, Roberto Amato, Jeffrey Barrett, Sonia Goncalves, Ewan Harrison, David K. Jackson, Ian Johnston, Dominic Kwiatkowski, Cordelia Langford, John Sillitoe on behalf of the Wellcome Sanger Institute COVID-19 Surveillance Team |
| EPI_ISL_2910335 | Lighthouse Lab in Alderley Park | Wellcome Sanger Institute for the COVID-19 Genomics UK (COG-UK) Consortium | Jacquelyn Wynn, Mairead Hyland, The Lighthouse Lab in Alderley Park and Alex Alderton, Roberto Amato, Jeffrey Barrett, Sonia Goncalves, Ewan Harrison, David K. Jackson, Ian Johnston, Dominic Kwiatkowski, Cordelia Langford, John Sillitoe on behalf of the Wellcome Sanger Institute COVID-19 Surveillance Team |
| EPI_ISL_2910336 | Lighthouse Lab in Glasgow | Wellcome Sanger Institute for the COVID-19 Genomics UK (COG-UK) Consortium | Harper VanSteenhouse, Yumi Kasai, David Gray, Carol Clugston, Anna Dominiczak and Alex Alderton, Roberto Amato, Jeffrey Barrett, Sonia Goncalves, Ewan Harrison, David K. Jackson, Ian Johnston, Dominic Kwiatkowski, Cordelia Langford, John Sillitoe on behalf of the Wellcome Sanger Institute COVID-19 Surveillance Team |
| EPI_ISL_2910337 | Lighthouse Lab in Alderley Park | Wellcome Sanger Institute for the COVID-19 Genomics UK (COG-UK) Consortium | Jacquelyn Wynn, Mairead Hyland, The Lighthouse Lab in Alderley Park and Alex Alderton, Roberto Amato, Jeffrey Barrett, Sonia Goncalves, Ewan Harrison, David K. Jackson, Ian Johnston, Dominic Kwiatkowski, Cordelia Langford, John Sillitoe on behalf of the Wellcome Sanger Institute COVID-19 Surveillance Team |
| EPI_ISL_2910338 | Lighthouse Lab in Milton Keynes | Wellcome Sanger Institute for the COVID-19 Genomics UK (COG-UK) Consortium | The Lighthouse Lab in Milton Keynes and Alex Alderton, Roberto Amato, Jeffrey Barrett, Sonia Goncalves, Ewan Harrison, David K. Jackson, Ian Johnston, Dominic Kwiatkowski, Cordelia Langford, John Sillitoe on behalf of the Wellcome Sanger Institute COVID-19 Surveillance Team |
| EPI_ISL_2910339, EPI_ISL_2910340 | Lighthouse Lab in Alderley Park | Wellcome Sanger Institute for the COVID-19 Genomics UK (COG-UK) Consortium | Jacquelyn Wynn, Mairead Hyland, The Lighthouse Lab in Alderley Park and Alex Alderton, Roberto Amato, Jeffrey Barrett, Sonia Goncalves, Ewan Harrison, David K. Jackson, Ian Johnston, Dominic Kwiatkowski, Cordelia Langford, John Sillitoe on behalf of the Wellcome Sanger Institute COVID-19 Surveillance Team |
| EPI_ISL_2910341 | Lighthouse Lab in Milton Keynes | Wellcome Sanger Institute for the COVID-19 Genomics UK (COG-UK) Consortium | The Lighthouse Lab in Milton Keynes and Alex Alderton, Roberto Amato, Jeffrey Barrett, Sonia Goncalves, Ewan Harrison, David K. Jackson, Ian Johnston, Dominic Kwiatkowski, Cordelia Langford, John Sillitoe on behalf of the Wellcome Sanger Institute COVID-19 Surveillance Team |
| EPI_ISL_2910342 | Lighthouse Lab in Alderley Park | Wellcome Sanger Institute for the COVID-19 Genomics UK (COG-UK) Consortium | Jacquelyn Wynn, Mairead Hyland, The Lighthouse Lab in Alderley Park and Alex Alderton, Roberto Amato, Jeffrey Barrett, Sonia Goncalves, Ewan Harrison, David K. Jackson, Ian Johnston, Dominic Kwiatkowski, Cordelia Langford, John Sillitoe on behalf of the Wellcome Sanger Institute COVID-19 Surveillance Team |
| EPI_ISL_2910343 | Lighthouse Lab in Glasgow | Wellcome Sanger Institute for the COVID-19 Genomics UK (COG-UK) Consortium | Harper VanSteenhouse, Yumi Kasai, David Gray, Carol Clugston, Anna Dominiczak and Alex Alderton, Roberto Amato, Jeffrey Barrett, Sonia Goncalves, Ewan Harrison, David K. Jackson, Ian Johnston, Dominic Kwiatkowski, Cordelia Langford, John Sillitoe on behalf of the Wellcome Sanger Institute COVID-19 Surveillance Team |
| EPI_ISL_2910344, EPI_ISL_2910345, EPI_ISL_2910346 | Lighthouse Lab in Alderley Park | Wellcome Sanger Institute for the COVID-19 Genomics UK (COG-UK) Consortium | Jacquelyn Wynn, Mairead Hyland, The Lighthouse Lab in Alderley Park and Alex Alderton, Roberto Amato, Jeffrey Barrett, Sonia Goncalves, Ewan Harrison, David K. Jackson, Ian Johnston, Dominic Kwiatkowski, Cordelia Langford, John Sillitoe on behalf of the Wellcome Sanger Institute COVID-19 Surveillance Team |
| EPI_ISL_2910347 | Lighthouse Lab in Milton Keynes | Wellcome Sanger Institute for the COVID-19 Genomics UK (COG-UK) Consortium | The Lighthouse Lab in Milton Keynes and Alex Alderton, Roberto Amato, Jeffrey Barrett, Sonia Goncalves, Ewan Harrison, David K. Jackson, Ian Johnston, Dominic Kwiatkowski, Cordelia Langford, John Sillitoe on behalf of the Wellcome Sanger Institute COVID-19 Surveillance Team |
| EPI_ISL_2910348, EPI_ISL_2910349, EPI_ISL_2910350 | Lighthouse Lab in Alderley Park | Wellcome Sanger Institute for the COVID-19 Genomics UK (COG-UK) Consortium | Jacquelyn Wynn, Mairead Hyland, The Lighthouse Lab in Alderley Park and Alex Alderton, Roberto Amato, Jeffrey Barrett, Sonia Goncalves, Ewan Harrison, David K. Jackson, Ian Johnston, Dominic Kwiatkowski, Cordelia Langford, John Sillitoe on behalf of the Wellcome Sanger Institute COVID-19 Surveillance Team |
| EPI_ISL_2910351 | Lighthouse Lab in Glasgow | Wellcome Sanger Institute for the COVID-19 Genomics UK (COG-UK) Consortium | Harper VanSteenhouse, Yumi Kasai, David Gray, Carol Clugston, Anna Dominiczak and Alex Alderton, Roberto Amato, Jeffrey Barrett, Sonia Goncalves, Ewan Harrison, David K. Jackson, Ian Johnston, Dominic Kwiatkowski, Cordelia Langford, John Sillitoe on behalf of the Wellcome Sanger Institute COVID-19 Surveillance Team |
| EPI_ISL_2910352, EPI_ISL_2910353 | Lighthouse Lab in Alderley Park | Wellcome Sanger Institute for the COVID-19 Genomics UK (COG-UK) Consortium | Jacquelyn Wynn, Mairead Hyland, The Lighthouse Lab in Alderley Park and Alex Alderton, Roberto Amato, Jeffrey Barrett, Sonia Goncalves, Ewan Harrison, David K. Jackson, Ian Johnston, Dominic Kwiatkowski, Cordelia Langford, John Sillitoe on behalf of the Wellcome Sanger Institute COVID-19 Surveillance Team |
| EPI_ISL_2910354 | Lighthouse Lab in Glasgow | Wellcome Sanger Institute for the COVID-19 Genomics UK (COG-UK) Consortium | Harper VanSteenhouse, Yumi Kasai, David Gray, Carol Clugston, Anna Dominiczak and Alex Alderton, Roberto Amato, Jeffrey Barrett, Sonia Goncalves, Ewan Harrison, David K. Jackson, Ian Johnston, Dominic Kwiatkowski, Cordelia Langford, John Sillitoe on behalf of the Wellcome Sanger Institute COVID-19 Surveillance Team |
| EPI_ISL_2910355, EPI_ISL_2910356 | Lighthouse Lab in Milton Keynes | Wellcome Sanger Institute for the COVID-19 Genomics UK (COG-UK) Consortium | The Lighthouse Lab in Milton Keynes and Alex Alderton, Roberto Amato, Jeffrey Barrett, Sonia Goncalves, Ewan Harrison, David K. Jackson, Ian Johnston, Dominic Kwiatkowski, Cordelia Langford, John Sillitoe on behalf of the Wellcome Sanger Institute COVID-19 Surveillance Team |
| EPI_ISL_2910357 | Lighthouse Lab in Glasgow | Wellcome Sanger Institute for the COVID-19 Genomics UK (COG-UK) Consortium | Harper VanSteenhouse, Yumi Kasai, David Gray, Carol Clugston, Anna Dominiczak and Alex Alderton, Roberto Amato, Jeffrey Barrett, Sonia Goncalves, Ewan Harrison, David K. Jackson, Ian Johnston, Dominic Kwiatkowski, Cordelia Langford, John Sillitoe on behalf of the Wellcome Sanger Institute COVID-19 Surveillance Team |
| EPI_ISL_2910358, EPI_ISL_2910360 | Lighthouse Lab in Milton Keynes | Wellcome Sanger Institute for the COVID-19 Genomics UK (COG-UK) Consortium | The Lighthouse Lab in Milton Keynes and Alex Alderton, Roberto Amato, Jeffrey Barrett, Sonia Goncalves, Ewan Harrison, David K. Jackson, Ian Johnston, Dominic Kwiatkowski, Cordelia Langford, John Sillitoe on behalf of the Wellcome Sanger Institute COVID-19 Surveillance Team |
| EPI_ISL_2910361, EPI_ISL_2910362, EPI_ISL_2910363 | Lighthouse Lab in Alderley Park | Wellcome Sanger Institute for the COVID-19 Genomics UK (COG-UK) Consortium | Jacquelyn Wynn, Mairead Hyland, The Lighthouse Lab in Alderley Park and Alex Alderton, Roberto Amato, Jeffrey Barrett, Sonia Goncalves, Ewan Harrison, David K. Jackson, Ian Johnston, Dominic Kwiatkowski, Cordelia Langford, John Sillitoe on behalf of the Wellcome Sanger Institute COVID-19 Surveillance Team |
| EPI_ISL_2910364 | Lighthouse Lab in Milton Keynes | Wellcome Sanger Institute for the COVID-19 Genomics UK (COG-UK) Consortium | The Lighthouse Lab in Milton Keynes and Alex Alderton, Roberto Amato, Jeffrey Barrett, Sonia Goncalves, Ewan Harrison, David K. Jackson, Ian Johnston, Dominic Kwiatkowski, Cordelia Langford, John Sillitoe on behalf of the Wellcome Sanger Institute COVID-19 Surveillance Team |
| EPI_ISL_2910365, EPI_ISL_2910366, EPI_ISL_2910367, EPI_ISL_2910368 | Lighthouse Lab in Alderley Park | Wellcome Sanger Institute for the COVID-19 Genomics UK (COG-UK) Consortium | Jacquelyn Wynn, Mairead Hyland, The Lighthouse Lab in Alderley Park and Alex Alderton, Roberto Amato, Jeffrey Barrett, Sonia Goncalves, Ewan Harrison, David K. Jackson, Ian Johnston, Dominic Kwiatkowski, Cordelia Langford, John Sillitoe on behalf of the Wellcome Sanger Institute COVID-19 Surveillance Team |
| EPI_ISL_2910369 | Lighthouse Lab in Milton Keynes | Wellcome Sanger Institute for the COVID-19 Genomics UK (COG-UK) Consortium | The Lighthouse Lab in Milton Keynes and Alex Alderton, Roberto Amato, Jeffrey Barrett, Sonia Goncalves, Ewan Harrison, David K. Jackson, Ian Johnston, Dominic Kwiatkowski, Cordelia Langford, John Sillitoe on behalf of the Wellcome Sanger Institute COVID-19 Surveillance Team |
| EPI_ISL_2910370 | Lighthouse Lab in Alderley Park | Wellcome Sanger Institute for the COVID-19 Genomics UK (COG-UK) Consortium | Jacquelyn Wynn, Mairead Hyland, The Lighthouse Lab in Alderley Park and Alex Alderton, Roberto Amato, Jeffrey Barrett, Sonia Goncalves, Ewan Harrison, David K. Jackson, Ian Johnston, Dominic Kwiatkowski, Cordelia Langford, John Sillitoe on behalf of the Wellcome Sanger Institute COVID-19 Surveillance Team |
| EPI_ISL_2910371 | Lighthouse Lab in Glasgow | Wellcome Sanger Institute for the COVID-19 Genomics UK (COG-UK) Consortium | Harper VanSteenhouse, Yumi Kasai, David Gray, Carol Clugston, Anna Dominiczak and Alex Alderton, Roberto Amato, Jeffrey Barrett, Sonia Goncalves, Ewan Harrison, David K. Jackson, Ian Johnston, Dominic Kwiatkowski, Cordelia Langford, John Sillitoe on behalf of the Wellcome Sanger Institute COVID-19 Surveillance Team |
| EPI_ISL_2910372, EPI_ISL_2910373 | Lighthouse Lab in Alderley Park | Wellcome Sanger Institute for the COVID-19 Genomics UK (COG-UK) Consortium | Jacquelyn Wynn, Mairead Hyland, The Lighthouse Lab in Alderley Park and Alex Alderton, Roberto Amato, Jeffrey Barrett, Sonia Goncalves, Ewan Harrison, David K. Jackson, Ian Johnston, Dominic Kwiatkowski, Cordelia Langford, John Sillitoe on behalf of the Wellcome Sanger Institute COVID-19 Surveillance Team |

| | | | |
|---|---|---|---|
| EPI_ISL_2910374 | Lighthouse Lab in Milton Keynes | Wellcome Sanger Institute for the COVID-19 Genomics UK (COG-UK) Consortium | The Lighthouse Lab in Milton Keynes and Alex Alderton, Roberto Amato, Jeffrey Barrett, Sonia Goncalves, Ewan Harrison, David K. Jackson, Ian Johnston, Dominic Kwiatkowski, Cordelia Langford, John Sillitoe on behalf of the Wellcome Sanger Institute COVID-19 Surveillance Team |
| EPI_ISL_2910375 | Lighthouse Lab in Alderley Park | Wellcome Sanger Institute for the COVID-19 Genomics UK (COG-UK) Consortium | Jacquelyn Wynn, Mairead Hyland, The Lighthouse Lab in Alderley Park and Alex Alderton, Roberto Amato, Jeffrey Barrett, Sonia Goncalves, Ewan Harrison, David K. Jackson, Ian Johnston, Dominic Kwiatkowski, Cordelia Langford, John Sillitoe on behalf of the Wellcome Sanger Institute COVID-19 Surveillance Team |
| EPI_ISL_2910376, EPI_ISL_2910377 | Lighthouse Lab in Glasgow | Wellcome Sanger Institute for the COVID-19 Genomics UK (COG-UK) Consortium | Harper VanSteenhouse, Yumi Kasai, David Gray, Carol Clugston, Anna Dominiczak and Alex Alderton, Roberto Amato, Jeffrey Barrett, Sonia Goncalves, Ewan Harrison, David K. Jackson, Ian Johnston, Dominic Kwiatkowski, Cordelia Langford, John Sillitoe on behalf of the Wellcome Sanger Institute COVID-19 Surveillance Team |
| EPI_ISL_2910378, EPI_ISL_2910379, EPI_ISL_2910380 | Lighthouse Lab in Alderley Park | Wellcome Sanger Institute for the COVID-19 Genomics UK (COG-UK) Consortium | Jacquelyn Wynn, Mairead Hyland, The Lighthouse Lab in Alderley Park and Alex Alderton, Roberto Amato, Jeffrey Barrett, Sonia Goncalves, Ewan Harrison, David K. Jackson, Ian Johnston, Dominic Kwiatkowski, Cordelia Langford, John Sillitoe on behalf of the Wellcome Sanger Institute COVID-19 Surveillance Team |
| EPI_ISL_2910381 | Lighthouse Lab in Glasgow | Wellcome Sanger Institute for the COVID-19 Genomics UK (COG-UK) Consortium | Harper VanSteenhouse, Yumi Kasai, David Gray, Carol Clugston, Anna Dominiczak and Alex Alderton, Roberto Amato, Jeffrey Barrett, Sonia Goncalves, Ewan Harrison, David K. Jackson, Ian Johnston, Dominic Kwiatkowski, Cordelia Langford, John Sillitoe on behalf of the Wellcome Sanger Institute COVID-19 Surveillance Team |
| EPI_ISL_2910382 | Lighthouse Lab in Alderley Park | Wellcome Sanger Institute for the COVID-19 Genomics UK (COG-UK) Consortium | Jacquelyn Wynn, Mairead Hyland, The Lighthouse Lab in Alderley Park and Alex Alderton, Roberto Amato, Jeffrey Barrett, Sonia Goncalves, Ewan Harrison, David K. Jackson, Ian Johnston, Dominic Kwiatkowski, Cordelia Langford, John Sillitoe on behalf of the Wellcome Sanger Institute COVID-19 Surveillance Team |
| EPI_ISL_2910383 | Lighthouse Lab in Glasgow | Wellcome Sanger Institute for the COVID-19 Genomics UK (COG-UK) Consortium | Harper VanSteenhouse, Yumi Kasai, David Gray, Carol Clugston, Anna Dominiczak and Alex Alderton, Roberto Amato, Jeffrey Barrett, Sonia Goncalves, Ewan Harrison, David K. Jackson, Ian Johnston, Dominic Kwiatkowski, Cordelia Langford, John Sillitoe on behalf of the Wellcome Sanger Institute COVID-19 Surveillance Team |
| EPI_ISL_2910384 | Lighthouse Lab in Milton Keynes | Wellcome Sanger Institute for the COVID-19 Genomics UK (COG-UK) Consortium | The Lighthouse Lab in Milton Keynes and Alex Alderton, Roberto Amato, Jeffrey Barrett, Sonia Goncalves, Ewan Harrison, David K. Jackson, Ian Johnston, Dominic Kwiatkowski, Cordelia Langford, John Sillitoe on behalf of the Wellcome Sanger Institute COVID-19 Surveillance Team |
| EPI_ISL_2910385, EPI_ISL_2910386, EPI_ISL_2910387 | Lighthouse Lab in Alderley Park | Wellcome Sanger Institute for the COVID-19 Genomics UK (COG-UK) Consortium | Jacquelyn Wynn, Mairead Hyland, The Lighthouse Lab in Alderley Park and Alex Alderton, Roberto Amato, Jeffrey Barrett, Sonia Goncalves, Ewan Harrison, David K. Jackson, Ian Johnston, Dominic Kwiatkowski, Cordelia Langford, John Sillitoe on behalf of the Wellcome Sanger Institute COVID-19 Surveillance Team |
| EPI_ISL_2910388 | Lighthouse Lab in Glasgow | Wellcome Sanger Institute for the COVID-19 Genomics UK (COG-UK) Consortium | Harper VanSteenhouse, Yumi Kasai, David Gray, Carol Clugston, Anna Dominiczak and Alex Alderton, Roberto Amato, Jeffrey Barrett, Sonia Goncalves, Ewan Harrison, David K. Jackson, Ian Johnston, Dominic Kwiatkowski, Cordelia Langford, John Sillitoe on behalf of the Wellcome Sanger Institute COVID-19 Surveillance Team |
| EPI_ISL_2910389 | Lighthouse Lab in Alderley Park | Wellcome Sanger Institute for the COVID-19 Genomics UK (COG-UK) Consortium | Jacquelyn Wynn, Mairead Hyland, The Lighthouse Lab in Alderley Park and Alex Alderton, Roberto Amato, Jeffrey Barrett, Sonia Goncalves, Ewan Harrison, David K. Jackson, Ian Johnston, Dominic Kwiatkowski, Cordelia Langford, John Sillitoe on behalf of the Wellcome Sanger Institute COVID-19 Surveillance Team |
| EPI_ISL_2910390, EPI_ISL_2910391 | Lighthouse Lab in Milton Keynes | Wellcome Sanger Institute for the COVID-19 Genomics UK (COG-UK) Consortium | The Lighthouse Lab in Milton Keynes and Alex Alderton, Roberto Amato, Jeffrey Barrett, Sonia Goncalves, Ewan Harrison, David K. Jackson, Ian Johnston, Dominic Kwiatkowski, Cordelia Langford, John Sillitoe on behalf of the Wellcome Sanger Institute COVID-19 Surveillance Team |
| EPI_ISL_2910392 | Lighthouse Lab in Glasgow | Wellcome Sanger Institute for the COVID-19 Genomics UK (COG-UK) Consortium | Harper VanSteenhouse, Yumi Kasai, David Gray, Carol Clugston, Anna Dominiczak and Alex Alderton, Roberto Amato, Jeffrey Barrett, Sonia Goncalves, Ewan Harrison, David K. Jackson, Ian Johnston, Dominic Kwiatkowski, Cordelia Langford, John Sillitoe on behalf of the Wellcome Sanger Institute COVID-19 Surveillance Team |
| EPI_ISL_2910393 | Lighthouse Lab in Alderley Park | Wellcome Sanger Institute for the COVID-19 Genomics UK (COG-UK) Consortium | Jacquelyn Wynn, Mairead Hyland, The Lighthouse Lab in Alderley Park and Alex Alderton, Roberto Amato, Jeffrey Barrett, Sonia Goncalves, Ewan Harrison, David K. Jackson, Ian Johnston, Dominic Kwiatkowski, Cordelia Langford, John Sillitoe on behalf of the Wellcome Sanger Institute COVID-19 Surveillance Team |
| EPI_ISL_2910394 | Lighthouse Lab in Milton Keynes | Wellcome Sanger Institute for the COVID-19 Genomics UK (COG-UK) Consortium | The Lighthouse Lab in Milton Keynes and Alex Alderton, Roberto Amato, Jeffrey Barrett, Sonia Goncalves, Ewan Harrison, David K. Jackson, Ian Johnston, Dominic Kwiatkowski, Cordelia Langford, John Sillitoe on behalf of the Wellcome Sanger Institute COVID-19 Surveillance Team |
| EPI_ISL_2910395, EPI_ISL_2910396 | Lighthouse Lab in Alderley Park | Wellcome Sanger Institute for the COVID-19 Genomics UK (COG-UK) Consortium | Jacquelyn Wynn, Mairead Hyland, The Lighthouse Lab in Alderley Park and Alex Alderton, Roberto Amato, Jeffrey Barrett, Sonia Goncalves, Ewan Harrison, David K. Jackson, Ian Johnston, Dominic Kwiatkowski, Cordelia Langford, John Sillitoe on behalf of the Wellcome Sanger Institute COVID-19 Surveillance Team |
| EPI_ISL_2910397 | Lighthouse Lab in Glasgow | Wellcome Sanger Institute for the COVID-19 Genomics UK (COG-UK) Consortium | Harper VanSteenhouse, Yumi Kasai, David Gray, Carol Clugston, Anna Dominiczak and Alex Alderton, Roberto Amato, Jeffrey Barrett, Sonia Goncalves, Ewan Harrison, David K. Jackson, Ian Johnston, Dominic Kwiatkowski, Cordelia Langford, John Sillitoe on behalf of the Wellcome Sanger Institute COVID-19 Surveillance Team |
| EPI_ISL_2910398, EPI_ISL_2910399, EPI_ISL_2910400 | Lighthouse Lab in Alderley Park | Wellcome Sanger Institute for the COVID-19 Genomics UK (COG-UK) Consortium | Jacquelyn Wynn, Mairead Hyland, The Lighthouse Lab in Alderley Park and Alex Alderton, Roberto Amato, Jeffrey Barrett, Sonia Goncalves, Ewan Harrison, David K. Jackson, Ian Johnston, Dominic Kwiatkowski, Cordelia Langford, John Sillitoe on behalf of the Wellcome Sanger Institute COVID-19 Surveillance Team |
| EPI_ISL_2910401 | Lighthouse Lab in Milton Keynes | Wellcome Sanger Institute for the COVID-19 Genomics UK (COG-UK) Consortium | The Lighthouse Lab in Milton Keynes and Alex Alderton, Roberto Amato, Jeffrey Barrett, Sonia Goncalves, Ewan Harrison, David K. Jackson, Ian Johnston, Dominic Kwiatkowski, Cordelia Langford, John Sillitoe on behalf of the Wellcome Sanger Institute COVID-19 Surveillance Team |
| EPI_ISL_2910402, EPI_ISL_2910403 | Lighthouse Lab in Glasgow | Wellcome Sanger Institute for the COVID-19 Genomics UK (COG-UK) Consortium | Harper VanSteenhouse, Yumi Kasai, David Gray, Carol Clugston, Anna Dominiczak and Alex Alderton, Roberto Amato, Jeffrey Barrett, Sonia Goncalves, Ewan Harrison, David K. Jackson, Ian Johnston, Dominic Kwiatkowski, Cordelia Langford, John Sillitoe on behalf of the Wellcome Sanger Institute COVID-19 Surveillance Team |
| EPI_ISL_2910404 | Lighthouse Lab in Milton Keynes | Wellcome Sanger Institute for the COVID-19 Genomics UK (COG-UK) Consortium | The Lighthouse Lab in Milton Keynes and Alex Alderton, Roberto Amato, Jeffrey Barrett, Sonia Goncalves, Ewan Harrison, David K. Jackson, Ian Johnston, Dominic Kwiatkowski, Cordelia Langford, John Sillitoe on behalf of the Wellcome Sanger Institute COVID-19 Surveillance Team |
| EPI_ISL_2910405 | Lighthouse Lab in Glasgow | Wellcome Sanger Institute for the COVID-19 Genomics UK (COG-UK) Consortium | Harper VanSteenhouse, Yumi Kasai, David Gray, Carol Clugston, Anna Dominiczak and Alex Alderton, Roberto Amato, Jeffrey Barrett, Sonia Goncalves, Ewan Harrison, David K. Jackson, Ian Johnston, Dominic Kwiatkowski, Cordelia Langford, John Sillitoe on behalf of the Wellcome Sanger Institute COVID-19 Surveillance Team |
| EPI_ISL_2910406, EPI_ISL_2910407 | Lighthouse Lab in Milton Keynes | Wellcome Sanger Institute for the COVID-19 Genomics UK (COG-UK) Consortium | The Lighthouse Lab in Milton Keynes and Alex Alderton, Roberto Amato, Jeffrey Barrett, Sonia Goncalves, Ewan Harrison, David K. Jackson, Ian Johnston, Dominic Kwiatkowski, Cordelia Langford, John Sillitoe on behalf of the Wellcome Sanger Institute COVID-19 Surveillance Team |
| EPI_ISL_2910408 | Lighthouse Lab in Alderley Park | Wellcome Sanger Institute for the COVID-19 Genomics UK (COG-UK) Consortium | Jacquelyn Wynn, Mairead Hyland, The Lighthouse Lab in Alderley Park and Alex Alderton, Roberto Amato, Jeffrey Barrett, Sonia Goncalves, Ewan Harrison, David K. Jackson, Ian Johnston, Dominic Kwiatkowski, Cordelia Langford, John Sillitoe on behalf of the Wellcome Sanger Institute COVID-19 Surveillance Team |
| EPI_ISL_2910409 | Lighthouse Lab in Glasgow | Wellcome Sanger Institute for the COVID-19 Genomics UK (COG-UK) Consortium | Harper VanSteenhouse, Yumi Kasai, David Gray, Carol Clugston, Anna Dominiczak and Alex Alderton, Roberto Amato, Jeffrey Barrett, Sonia Goncalves, Ewan Harrison, David K. Jackson, Ian Johnston, Dominic Kwiatkowski, Cordelia Langford, John Sillitoe on behalf of the Wellcome Sanger Institute COVID-19 Surveillance Team |
| EPI_ISL_2910410 | Lighthouse Lab in Milton Keynes | Wellcome Sanger Institute for the COVID-19 Genomics UK (COG-UK) Consortium | The Lighthouse Lab in Milton Keynes and Alex Alderton, Roberto Amato, Jeffrey Barrett, Sonia Goncalves, Ewan Harrison, David K. Jackson, Ian Johnston, Dominic Kwiatkowski, Cordelia Langford, John Sillitoe on behalf of the Wellcome Sanger Institute COVID-19 Surveillance Team |
| EPI_ISL_2910411, EPI_ISL_2910412 | Lighthouse Lab in Alderley Park | Wellcome Sanger Institute for the COVID-19 Genomics UK (COG-UK) Consortium | Jacquelyn Wynn, Mairead Hyland, The Lighthouse Lab in Alderley Park and Alex Alderton, Roberto Amato, Jeffrey Barrett, Sonia Goncalves, Ewan Harrison, David K. Jackson, Ian Johnston, Dominic Kwiatkowski, Cordelia Langford, John Sillitoe on behalf of the Wellcome Sanger Institute COVID-19 Surveillance Team |
| EPI_ISL_2910413 | Lighthouse Lab in Milton Keynes | Wellcome Sanger Institute for the COVID-19 Genomics UK (COG-UK) Consortium | The Lighthouse Lab in Milton Keynes and Alex Alderton, Roberto Amato, Jeffrey Barrett, Sonia Goncalves, Ewan Harrison, David K. Jackson, Ian Johnston, Dominic Kwiatkowski, Cordelia Langford, John Sillitoe on behalf of the Wellcome Sanger Institute COVID-19 Surveillance Team |
| EPI_ISL_2910414, EPI_ISL_2910415 | Lighthouse Lab in Alderley Park | Wellcome Sanger Institute for the COVID-19 Genomics UK (COG-UK) Consortium | Jacquelyn Wynn, Mairead Hyland, The Lighthouse Lab in Alderley Park and Alex Alderton, Roberto Amato, Jeffrey Barrett, Sonia Goncalves, Ewan Harrison, David K. Jackson, Ian Johnston, Dominic Kwiatkowski, Cordelia Langford, John Sillitoe on behalf of the Wellcome Sanger Institute COVID-19 Surveillance Team |
| EPI_ISL_2910416 | Lighthouse Lab in Glasgow | Wellcome Sanger Institute for the COVID-19 Genomics UK (COG-UK) Consortium | Harper VanSteenhouse, Yumi Kasai, David Gray, Carol Clugston, Anna Dominiczak and Alex Alderton, Roberto Amato, Jeffrey Barrett, Sonia Goncalves, Ewan Harrison, David K. Jackson, Ian Johnston, Dominic Kwiatkowski, Cordelia Langford, John Sillitoe on behalf of the Wellcome Sanger Institute COVID-19 Surveillance Team |
| EPI_ISL_2910417, EPI_ISL_2910418 | Lighthouse Lab in Alderley Park | Wellcome Sanger Institute for the COVID-19 Genomics UK (COG-UK) Consortium | Jacquelyn Wynn, Mairead Hyland, The Lighthouse Lab in Alderley Park and Alex Alderton, Roberto Amato, Jeffrey Barrett, Sonia Goncalves, Ewan Harrison, David K. Jackson, Ian Johnston, Dominic Kwiatkowski, Cordelia Langford, John Sillitoe on behalf of the Wellcome Sanger Institute COVID-19 Surveillance Team |
| EPI_ISL_2910419, EPI_ISL_2910420 | Lighthouse Lab in Glasgow | Wellcome Sanger Institute for the COVID-19 Genomics UK (COG-UK) Consortium | Harper VanSteenhouse, Yumi Kasai, David Gray, Carol Clugston, Anna Dominiczak and Alex Alderton, Roberto Amato, Jeffrey Barrett, Sonia Goncalves, Ewan Harrison, David K. Jackson, Ian Johnston, Dominic Kwiatkowski, Cordelia Langford, John Sillitoe on behalf of the Wellcome Sanger Institute COVID-19 Surveillance Team |
| EPI_ISL_2910421, EPI_ISL_2910422 | Lighthouse Lab in Alderley Park | Wellcome Sanger Institute for the COVID-19 Genomics UK (COG-UK) Consortium | Jacquelyn Wynn, Mairead Hyland, The Lighthouse Lab in Alderley Park and Alex Alderton, Roberto Amato, Jeffrey Barrett, Sonia Goncalves, Ewan Harrison, David K. Jackson, Ian Johnston, Dominic Kwiatkowski, Cordelia Langford, John Sillitoe on behalf of the Wellcome Sanger Institute COVID-19 Surveillance Team |
| EPI_ISL_2910423 | Lighthouse Lab in Milton Keynes | Wellcome Sanger Institute for the COVID-19 Genomics UK (COG-UK) Consortium | The Lighthouse Lab in Milton Keynes and Alex Alderton, Roberto Amato, Jeffrey Barrett, Sonia Goncalves, Ewan Harrison, David K. Jackson, Ian Johnston, Dominic Kwiatkowski, Cordelia Langford, John Sillitoe on behalf of the Wellcome Sanger Institute COVID-19 Surveillance Team |
| EPI_ISL_2910424, EPI_ISL_2910425 | Lighthouse Lab in Glasgow | Wellcome Sanger Institute for the COVID-19 Genomics UK (COG-UK) Consortium | Harper VanSteenhouse, Yumi Kasai, David Gray, Carol Clugston, Anna Dominiczak and Alex Alderton, Roberto Amato, Jeffrey Barrett, Sonia Goncalves, Ewan Harrison, David K. Jackson, Ian Johnston, Dominic Kwiatkowski, Cordelia Langford, John Sillitoe on behalf of the Wellcome Sanger Institute COVID-19 Surveillance Team |
| EPI_ISL_2910426 | Lighthouse Lab in Alderley Park | Wellcome Sanger Institute for the COVID-19 Genomics UK (COG-UK) Consortium | Jacquelyn Wynn, Mairead Hyland, The Lighthouse Lab in Alderley Park and Alex Alderton, Roberto Amato, Jeffrey Barrett, Sonia Goncalves, Ewan Harrison, David K. Jackson, Ian Johnston, Dominic Kwiatkowski, Cordelia Langford, John Sillitoe on behalf of the Wellcome Sanger Institute COVID-19 Surveillance Team |
| EPI_ISL_2910427 | Lighthouse Lab in Glasgow | Wellcome Sanger Institute for the COVID-19 Genomics UK (COG-UK) Consortium | Harper VanSteenhouse, Yumi Kasai, David Gray, Carol Clugston, Anna Dominiczak and Alex Alderton, Roberto Amato, Jeffrey Barrett, Sonia Goncalves, Ewan Harrison, David K. Jackson, Ian Johnston, Dominic Kwiatkowski, Cordelia Langford, John Sillitoe on behalf of the Wellcome Sanger Institute COVID-19 Surveillance Team |
| EPI_ISL_2910428 | Lighthouse Lab in Alderley Park | Wellcome Sanger Institute for the COVID-19 Genomics UK (COG-UK) Consortium | Jacquelyn Wynn, Mairead Hyland, The Lighthouse Lab in Alderley Park and Alex Alderton, Roberto Amato, Jeffrey Barrett, Sonia Goncalves, Ewan Harrison, David K. Jackson, Ian Johnston, Dominic Kwiatkowski, Cordelia Langford, John Sillitoe on behalf of the Wellcome Sanger Institute COVID-19 Surveillance Team |
| EPI_ISL_2910429 | Lighthouse Lab in Glasgow | Wellcome Sanger Institute for the COVID-19 Genomics UK (COG-UK) Consortium | Harper VanSteenhouse, Yumi Kasai, David Gray, Carol Clugston, Anna Dominiczak and Alex Alderton, Roberto Amato, Jeffrey Barrett, Sonia Goncalves, Ewan Harrison, David K. Jackson, Ian Johnston, Dominic Kwiatkowski, Cordelia Langford, John Sillitoe on behalf of the Wellcome Sanger Institute COVID-19 Surveillance Team |
| EPI_ISL_2910430, EPI_ISL_2910431, EPI_ISL_2910432 | Lighthouse Lab in Milton Keynes | Wellcome Sanger Institute for the COVID-19 Genomics UK (COG-UK) Consortium | The Lighthouse Lab in Milton Keynes and Alex Alderton, Roberto Amato, Jeffrey Barrett, Sonia Goncalves, Ewan Harrison, David K. Jackson, Ian Johnston, Dominic Kwiatkowski, Cordelia Langford, John Sillitoe on behalf of the Wellcome Sanger Institute COVID-19 Surveillance Team |
| EPI_ISL_2910433 | Lighthouse Lab in Glasgow | Wellcome Sanger Institute for the COVID-19 Genomics UK (COG-UK) Consortium | Harper VanSteenhouse, Yumi Kasai, David Gray, Carol Clugston, Anna Dominiczak and Alex Alderton, Roberto Amato, Jeffrey Barrett, Sonia Goncalves, Ewan Harrison, David K. Jackson, Ian Johnston, Dominic Kwiatkowski, Cordelia Langford, John Sillitoe on behalf of the Wellcome Sanger Institute COVID-19 Surveillance Team |
| EPI_ISL_2910434 | Lighthouse Lab in Milton Keynes | Wellcome Sanger Institute for the COVID-19 Genomics UK (COG-UK) Consortium | The Lighthouse Lab in Milton Keynes and Alex Alderton, Roberto Amato, Jeffrey Barrett, Sonia Goncalves, Ewan Harrison, David K. Jackson, Ian Johnston, Dominic Kwiatkowski, Cordelia Langford, John Sillitoe on behalf of the Wellcome Sanger Institute COVID-19 Surveillance Team |
| EPI_ISL_2910435 | Lighthouse Lab in Glasgow | Wellcome Sanger Institute for the COVID-19 Genomics UK (COG-UK) Consortium | Harper VanSteenhouse, Yumi Kasai, David Gray, Carol Clugston, Anna Dominiczak and Alex Alderton, Roberto Amato, Jeffrey Barrett, Sonia Goncalves, Ewan Harrison, David K. Jackson, Ian Johnston, Dominic Kwiatkowski, Cordelia Langford, John Sillitoe on behalf of the Wellcome Sanger Institute COVID-19 Surveillance Team |
| EPI_ISL_2910436, | Lighthouse Lab in Milton Keynes | Wellcome Sanger Institute for the COVID-19 Genomics UK (COG-UK) | The Lighthouse Lab in Milton Keynes and Alex Alderton, Roberto Amato, Jeffrey Barrett, Sonia Goncalves, Ewan Harrison, David K. Jackson, Ian Johnston, Dominic Kwiatkowski, Cordelia Langford, John Sillitoe on behalf of the Wellcome Sanger |

| | | | |
|---|---|---|---|
| EPI_ISL_2910437, EPI_ISL_2910438 | | Consortium | Institute COVID-19 Surveillance Team |
| EPI_ISL_2910439 | Lighthouse Lab in Glasgow | Wellcome Sanger Institute for the COVID-19 Genomics UK (COG-UK) Consortium | Harper VanSteenhouse, Yumi Kasai, David Gray, Carol Clugston, Anna Dominiczak and Alex Alderton, Roberto Amato, Jeffrey Barrett, Sonia Goncalves, Ewan Harrison, David K. Jackson, Ian Johnston, Dominic Kwiatkowski, Cordelia Langford, John Sillitoe on behalf of the Wellcome Sanger Institute COVID-19 Surveillance Team |
| EPI_ISL_2910440 | Lighthouse Lab in Milton Keynes | Wellcome Sanger Institute for the COVID-19 Genomics UK (COG-UK) Consortium | The Lighthouse Lab in Milton Keynes and Alex Alderton, Roberto Amato, Jeffrey Barrett, Sonia Goncalves, Ewan Harrison, David K. Jackson, Ian Johnston, Dominic Kwiatkowski, Cordelia Langford, John Sillitoe on behalf of the Wellcome Sanger Institute COVID-19 Surveillance Team |
| EPI_ISL_2910441 | Lighthouse Lab in Glasgow | Wellcome Sanger Institute for the COVID-19 Genomics UK (COG-UK) Consortium | Harper VanSteenhouse, Yumi Kasai, David Gray, Carol Clugston, Anna Dominiczak and Alex Alderton, Roberto Amato, Jeffrey Barrett, Sonia Goncalves, Ewan Harrison, David K. Jackson, Ian Johnston, Dominic Kwiatkowski, Cordelia Langford, John Sillitoe on behalf of the Wellcome Sanger Institute COVID-19 Surveillance Team |
| EPI_ISL_2910442 | Lighthouse Lab in Alderley Park | Wellcome Sanger Institute for the COVID-19 Genomics UK (COG-UK) Consortium | Jacquelyn Wynn, Mairead Hyland, The Lighthouse Lab in Alderley Park and Alex Alderton, Roberto Amato, Jeffrey Barrett, Sonia Goncalves, Ewan Harrison, David K. Jackson, Ian Johnston, Dominic Kwiatkowski, Cordelia Langford, John Sillitoe on behalf of the Wellcome Sanger Institute COVID-19 Surveillance Team |
| EPI_ISL_2910443 | Lighthouse Lab in Milton Keynes | Wellcome Sanger Institute for the COVID-19 Genomics UK (COG-UK) Consortium | The Lighthouse Lab in Milton Keynes and Alex Alderton, Roberto Amato, Jeffrey Barrett, Sonia Goncalves, Ewan Harrison, David K. Jackson, Ian Johnston, Dominic Kwiatkowski, Cordelia Langford, John Sillitoe on behalf of the Wellcome Sanger Institute COVID-19 Surveillance Team |
| EPI_ISL_2910444 | Lighthouse Lab in Glasgow | Wellcome Sanger Institute for the COVID-19 Genomics UK (COG-UK) Consortium | Harper VanSteenhouse, Yumi Kasai, David Gray, Carol Clugston, Anna Dominiczak and Alex Alderton, Roberto Amato, Jeffrey Barrett, Sonia Goncalves, Ewan Harrison, David K. Jackson, Ian Johnston, Dominic Kwiatkowski, Cordelia Langford, John Sillitoe on behalf of the Wellcome Sanger Institute COVID-19 Surveillance Team |
| EPI_ISL_2910445 | Lighthouse Lab in Milton Keynes | Wellcome Sanger Institute for the COVID-19 Genomics UK (COG-UK) Consortium | The Lighthouse Lab in Milton Keynes and Alex Alderton, Roberto Amato, Jeffrey Barrett, Sonia Goncalves, Ewan Harrison, David K. Jackson, Ian Johnston, Dominic Kwiatkowski, Cordelia Langford, John Sillitoe on behalf of the Wellcome Sanger Institute COVID-19 Surveillance Team |
| EPI_ISL_2910446, EPI_ISL_2910447 | Lighthouse Lab in Glasgow | Wellcome Sanger Institute for the COVID-19 Genomics UK (COG-UK) Consortium | Harper VanSteenhouse, Yumi Kasai, David Gray, Carol Clugston, Anna Dominiczak and Alex Alderton, Roberto Amato, Jeffrey Barrett, Sonia Goncalves, Ewan Harrison, David K. Jackson, Ian Johnston, Dominic Kwiatkowski, Cordelia Langford, John Sillitoe on behalf of the Wellcome Sanger Institute COVID-19 Surveillance Team |
| EPI_ISL_2910448 | Lighthouse Lab in Alderley Park | Wellcome Sanger Institute for the COVID-19 Genomics UK (COG-UK) Consortium | Jacquelyn Wynn, Mairead Hyland, The Lighthouse Lab in Alderley Park and Alex Alderton, Roberto Amato, Jeffrey Barrett, Sonia Goncalves, Ewan Harrison, David K. Jackson, Ian Johnston, Dominic Kwiatkowski, Cordelia Langford, John Sillitoe on behalf of the Wellcome Sanger Institute COVID-19 Surveillance Team |
| EPI_ISL_2910449 | Lighthouse Lab in Milton Keynes | Wellcome Sanger Institute for the COVID-19 Genomics UK (COG-UK) Consortium | The Lighthouse Lab in Milton Keynes and Alex Alderton, Roberto Amato, Jeffrey Barrett, Sonia Goncalves, Ewan Harrison, David K. Jackson, Ian Johnston, Dominic Kwiatkowski, Cordelia Langford, John Sillitoe on behalf of the Wellcome Sanger Institute COVID-19 Surveillance Team |
| EPI_ISL_2910450 | Lighthouse Lab in Alderley Park | Wellcome Sanger Institute for the COVID-19 Genomics UK (COG-UK) Consortium | Jacquelyn Wynn, Mairead Hyland, The Lighthouse Lab in Alderley Park and Alex Alderton, Roberto Amato, Jeffrey Barrett, Sonia Goncalves, Ewan Harrison, David K. Jackson, Ian Johnston, Dominic Kwiatkowski, Cordelia Langford, John Sillitoe on behalf of the Wellcome Sanger Institute COVID-19 Surveillance Team |
| EPI_ISL_2910451, EPI_ISL_2910452 | Lighthouse Lab in Glasgow | Wellcome Sanger Institute for the COVID-19 Genomics UK (COG-UK) Consortium | Harper VanSteenhouse, Yumi Kasai, David Gray, Carol Clugston, Anna Dominiczak and Alex Alderton, Roberto Amato, Jeffrey Barrett, Sonia Goncalves, Ewan Harrison, David K. Jackson, Ian Johnston, Dominic Kwiatkowski, Cordelia Langford, John Sillitoe on behalf of the Wellcome Sanger Institute COVID-19 Surveillance Team |
| EPI_ISL_2910453 | Lighthouse Lab in Milton Keynes | Wellcome Sanger Institute for the COVID-19 Genomics UK (COG-UK) Consortium | The Lighthouse Lab in Milton Keynes and Alex Alderton, Roberto Amato, Jeffrey Barrett, Sonia Goncalves, Ewan Harrison, David K. Jackson, Ian Johnston, Dominic Kwiatkowski, Cordelia Langford, John Sillitoe on behalf of the Wellcome Sanger Institute COVID-19 Surveillance Team |
| EPI_ISL_2910454 | Lighthouse Lab in Alderley Park | Wellcome Sanger Institute for the COVID-19 Genomics UK (COG-UK) Consortium | Jacquelyn Wynn, Mairead Hyland, The Lighthouse Lab in Alderley Park and Alex Alderton, Roberto Amato, Jeffrey Barrett, Sonia Goncalves, Ewan Harrison, David K. Jackson, Ian Johnston, Dominic Kwiatkowski, Cordelia Langford, John Sillitoe on behalf of the Wellcome Sanger Institute COVID-19 Surveillance Team |
| EPI_ISL_2910455, EPI_ISL_2910456 | Lighthouse Lab in Milton Keynes | Wellcome Sanger Institute for the COVID-19 Genomics UK (COG-UK) Consortium | The Lighthouse Lab in Milton Keynes and Alex Alderton, Roberto Amato, Jeffrey Barrett, Sonia Goncalves, Ewan Harrison, David K. Jackson, Ian Johnston, Dominic Kwiatkowski, Cordelia Langford, John Sillitoe on behalf of the Wellcome Sanger Institute COVID-19 Surveillance Team |
| EPI_ISL_2910457 | Lighthouse Lab in Alderley Park | Wellcome Sanger Institute for the COVID-19 Genomics UK (COG-UK) Consortium | Jacquelyn Wynn, Mairead Hyland, The Lighthouse Lab in Alderley Park and Alex Alderton, Roberto Amato, Jeffrey Barrett, Sonia Goncalves, Ewan Harrison, David K. Jackson, Ian Johnston, Dominic Kwiatkowski, Cordelia Langford, John Sillitoe on behalf of the Wellcome Sanger Institute COVID-19 Surveillance Team |
| EPI_ISL_2910458 | Lighthouse Lab in Milton Keynes | Wellcome Sanger Institute for the COVID-19 Genomics UK (COG-UK) Consortium | The Lighthouse Lab in Milton Keynes and Alex Alderton, Roberto Amato, Jeffrey Barrett, Sonia Goncalves, Ewan Harrison, David K. Jackson, Ian Johnston, Dominic Kwiatkowski, Cordelia Langford, John Sillitoe on behalf of the Wellcome Sanger Institute COVID-19 Surveillance Team |
| EPI_ISL_2910459 | Lighthouse Lab in Alderley Park | Wellcome Sanger Institute for the COVID-19 Genomics UK (COG-UK) Consortium | Jacquelyn Wynn, Mairead Hyland, The Lighthouse Lab in Alderley Park and Alex Alderton, Roberto Amato, Jeffrey Barrett, Sonia Goncalves, Ewan Harrison, David K. Jackson, Ian Johnston, Dominic Kwiatkowski, Cordelia Langford, John Sillitoe on behalf of the Wellcome Sanger Institute COVID-19 Surveillance Team |
| EPI_ISL_2910460 | Lighthouse Lab in Glasgow | Wellcome Sanger Institute for the COVID-19 Genomics UK (COG-UK) Consortium | Harper VanSteenhouse, Yumi Kasai, David Gray, Carol Clugston, Anna Dominiczak and Alex Alderton, Roberto Amato, Jeffrey Barrett, Sonia Goncalves, Ewan Harrison, David K. Jackson, Ian Johnston, Dominic Kwiatkowski, Cordelia Langford, John Sillitoe on behalf of the Wellcome Sanger Institute COVID-19 Surveillance Team |
| EPI_ISL_2910461, EPI_ISL_2910462 | Lighthouse Lab in Alderley Park | Wellcome Sanger Institute for the COVID-19 Genomics UK (COG-UK) Consortium | Jacquelyn Wynn, Mairead Hyland, The Lighthouse Lab in Alderley Park and Alex Alderton, Roberto Amato, Jeffrey Barrett, Sonia Goncalves, Ewan Harrison, David K. Jackson, Ian Johnston, Dominic Kwiatkowski, Cordelia Langford, John Sillitoe on behalf of the Wellcome Sanger Institute COVID-19 Surveillance Team |
| EPI_ISL_2910463 | Lighthouse Lab in Glasgow | Wellcome Sanger Institute for the COVID-19 Genomics UK (COG-UK) Consortium | Harper VanSteenhouse, Yumi Kasai, David Gray, Carol Clugston, Anna Dominiczak and Alex Alderton, Roberto Amato, Jeffrey Barrett, Sonia Goncalves, Ewan Harrison, David K. Jackson, Ian Johnston, Dominic Kwiatkowski, Cordelia Langford, John Sillitoe on behalf of the Wellcome Sanger Institute COVID-19 Surveillance Team |
| EPI_ISL_2910464 | Lighthouse Lab in Alderley Park | Wellcome Sanger Institute for the COVID-19 Genomics UK (COG-UK) Consortium | Jacquelyn Wynn, Mairead Hyland, The Lighthouse Lab in Alderley Park and Alex Alderton, Roberto Amato, Jeffrey Barrett, Sonia Goncalves, Ewan Harrison, David K. Jackson, Ian Johnston, Dominic Kwiatkowski, Cordelia Langford, John Sillitoe on behalf of the Wellcome Sanger Institute COVID-19 Surveillance Team |
| EPI_ISL_2910465 | Lighthouse Lab in Milton Keynes | Wellcome Sanger Institute for the COVID-19 Genomics UK (COG-UK) Consortium | The Lighthouse Lab in Milton Keynes and Alex Alderton, Roberto Amato, Jeffrey Barrett, Sonia Goncalves, Ewan Harrison, David K. Jackson, Ian Johnston, Dominic Kwiatkowski, Cordelia Langford, John Sillitoe on behalf of the Wellcome Sanger Institute COVID-19 Surveillance Team |
| EPI_ISL_2910466, EPI_ISL_2910467 | Lighthouse Lab in Alderley Park | Wellcome Sanger Institute for the COVID-19 Genomics UK (COG-UK) Consortium | Jacquelyn Wynn, Mairead Hyland, The Lighthouse Lab in Alderley Park and Alex Alderton, Roberto Amato, Jeffrey Barrett, Sonia Goncalves, Ewan Harrison, David K. Jackson, Ian Johnston, Dominic Kwiatkowski, Cordelia Langford, John Sillitoe on behalf of the Wellcome Sanger Institute COVID-19 Surveillance Team |
| EPI_ISL_2910468, EPI_ISL_2910469 | Lighthouse Lab in Glasgow | Wellcome Sanger Institute for the COVID-19 Genomics UK (COG-UK) Consortium | Harper VanSteenhouse, Yumi Kasai, David Gray, Carol Clugston, Anna Dominiczak and Alex Alderton, Roberto Amato, Jeffrey Barrett, Sonia Goncalves, Ewan Harrison, David K. Jackson, Ian Johnston, Dominic Kwiatkowski, Cordelia Langford, John Sillitoe on behalf of the Wellcome Sanger Institute COVID-19 Surveillance Team |
| EPI_ISL_2910470, EPI_ISL_2910471 | Lighthouse Lab in Alderley Park | Wellcome Sanger Institute for the COVID-19 Genomics UK (COG-UK) Consortium | Jacquelyn Wynn, Mairead Hyland, The Lighthouse Lab in Alderley Park and Alex Alderton, Roberto Amato, Jeffrey Barrett, Sonia Goncalves, Ewan Harrison, David K. Jackson, Ian Johnston, Dominic Kwiatkowski, Cordelia Langford, John Sillitoe on behalf of the Wellcome Sanger Institute COVID-19 Surveillance Team |
| EPI_ISL_2910472 | Lighthouse Lab in Milton Keynes | Wellcome Sanger Institute for the COVID-19 Genomics UK (COG-UK) Consortium | The Lighthouse Lab in Milton Keynes and Alex Alderton, Roberto Amato, Jeffrey Barrett, Sonia Goncalves, Ewan Harrison, David K. Jackson, Ian Johnston, Dominic Kwiatkowski, Cordelia Langford, John Sillitoe on behalf of the Wellcome Sanger Institute COVID-19 Surveillance Team |
| EPI_ISL_2910473, EPI_ISL_2910474 | Lighthouse Lab in Alderley Park | Wellcome Sanger Institute for the COVID-19 Genomics UK (COG-UK) Consortium | Jacquelyn Wynn, Mairead Hyland, The Lighthouse Lab in Alderley Park and Alex Alderton, Roberto Amato, Jeffrey Barrett, Sonia Goncalves, Ewan Harrison, David K. Jackson, Ian Johnston, Dominic Kwiatkowski, Cordelia Langford, John Sillitoe on behalf of the Wellcome Sanger Institute COVID-19 Surveillance Team |
| EPI_ISL_2910475 | Lighthouse Lab in Glasgow | Wellcome Sanger Institute for the COVID-19 Genomics UK (COG-UK) Consortium | Harper VanSteenhouse, Yumi Kasai, David Gray, Carol Clugston, Anna Dominiczak and Alex Alderton, Roberto Amato, Jeffrey Barrett, Sonia Goncalves, Ewan Harrison, David K. Jackson, Ian Johnston, Dominic Kwiatkowski, Cordelia Langford, John Sillitoe on behalf of the Wellcome Sanger Institute COVID-19 Surveillance Team |
| EPI_ISL_2910476 | Lighthouse Lab in Alderley Park | Wellcome Sanger Institute for the COVID-19 Genomics UK (COG-UK) Consortium | Jacquelyn Wynn, Mairead Hyland, The Lighthouse Lab in Alderley Park and Alex Alderton, Roberto Amato, Jeffrey Barrett, Sonia Goncalves, Ewan Harrison, David K. Jackson, Ian Johnston, Dominic Kwiatkowski, Cordelia Langford, John Sillitoe on behalf of the Wellcome Sanger Institute COVID-19 Surveillance Team |
| EPI_ISL_2910477 | Lighthouse Lab in Glasgow | Wellcome Sanger Institute for the COVID-19 Genomics UK (COG-UK) Consortium | Harper VanSteenhouse, Yumi Kasai, David Gray, Carol Clugston, Anna Dominiczak and Alex Alderton, Roberto Amato, Jeffrey Barrett, Sonia Goncalves, Ewan Harrison, David K. Jackson, Ian Johnston, Dominic Kwiatkowski, Cordelia Langford, John Sillitoe on behalf of the Wellcome Sanger Institute COVID-19 Surveillance Team |
| EPI_ISL_2910478, EPI_ISL_2910479 | Lighthouse Lab in Milton Keynes | Wellcome Sanger Institute for the COVID-19 Genomics UK (COG-UK) Consortium | The Lighthouse Lab in Milton Keynes and Alex Alderton, Roberto Amato, Jeffrey Barrett, Sonia Goncalves, Ewan Harrison, David K. Jackson, Ian Johnston, Dominic Kwiatkowski, Cordelia Langford, John Sillitoe on behalf of the Wellcome Sanger Institute COVID-19 Surveillance Team |
| EPI_ISL_2910480, EPI_ISL_2910481, EPI_ISL_2910482, EPI_ISL_2910483 | Lighthouse Lab in Alderley Park | Wellcome Sanger Institute for the COVID-19 Genomics UK (COG-UK) Consortium | Jacquelyn Wynn, Mairead Hyland, The Lighthouse Lab in Alderley Park and Alex Alderton, Roberto Amato, Jeffrey Barrett, Sonia Goncalves, Ewan Harrison, David K. Jackson, Ian Johnston, Dominic Kwiatkowski, Cordelia Langford, John Sillitoe on behalf of the Wellcome Sanger Institute COVID-19 Surveillance Team |
| EPI_ISL_2910484 | Lighthouse Lab in Milton Keynes | Wellcome Sanger Institute for the COVID-19 Genomics UK (COG-UK) Consortium | The Lighthouse Lab in Milton Keynes and Alex Alderton, Roberto Amato, Jeffrey Barrett, Sonia Goncalves, Ewan Harrison, David K. Jackson, Ian Johnston, Dominic Kwiatkowski, Cordelia Langford, John Sillitoe on behalf of the Wellcome Sanger Institute COVID-19 Surveillance Team |
| EPI_ISL_2910485 | Lighthouse Lab in Alderley Park | Wellcome Sanger Institute for the COVID-19 Genomics UK (COG-UK) Consortium | Jacquelyn Wynn, Mairead Hyland, The Lighthouse Lab in Alderley Park and Alex Alderton, Roberto Amato, Jeffrey Barrett, Sonia Goncalves, Ewan Harrison, David K. Jackson, Ian Johnston, Dominic Kwiatkowski, Cordelia Langford, John Sillitoe on behalf of the Wellcome Sanger Institute COVID-19 Surveillance Team |
| EPI_ISL_2910486 | Lighthouse Lab in Glasgow | Wellcome Sanger Institute for the COVID-19 Genomics UK (COG-UK) Consortium | Harper VanSteenhouse, Yumi Kasai, David Gray, Carol Clugston, Anna Dominiczak and Alex Alderton, Roberto Amato, Jeffrey Barrett, Sonia Goncalves, Ewan Harrison, David K. Jackson, Ian Johnston, Dominic Kwiatkowski, Cordelia Langford, John Sillitoe on behalf of the Wellcome Sanger Institute COVID-19 Surveillance Team |
| EPI_ISL_2910487 | Lighthouse Lab in Alderley Park | Wellcome Sanger Institute for the COVID-19 Genomics UK (COG-UK) Consortium | Jacquelyn Wynn, Mairead Hyland, The Lighthouse Lab in Alderley Park and Alex Alderton, Roberto Amato, Jeffrey Barrett, Sonia Goncalves, Ewan Harrison, David K. Jackson, Ian Johnston, Dominic Kwiatkowski, Cordelia Langford, John Sillitoe on behalf of the Wellcome Sanger Institute COVID-19 Surveillance Team |
| EPI_ISL_2910488, EPI_ISL_2910489, EPI_ISL_2910490 | Lighthouse Lab in Milton Keynes | Wellcome Sanger Institute for the COVID-19 Genomics UK (COG-UK) Consortium | The Lighthouse Lab in Milton Keynes and Alex Alderton, Roberto Amato, Jeffrey Barrett, Sonia Goncalves, Ewan Harrison, David K. Jackson, Ian Johnston, Dominic Kwiatkowski, Cordelia Langford, John Sillitoe on behalf of the Wellcome Sanger Institute COVID-19 Surveillance Team |
| EPI_ISL_2910491, EPI_ISL_2910492, EPI_ISL_2910493 | Lighthouse Lab in Alderley Park | Wellcome Sanger Institute for the COVID-19 Genomics UK (COG-UK) Consortium | Jacquelyn Wynn, Mairead Hyland, The Lighthouse Lab in Alderley Park and Alex Alderton, Roberto Amato, Jeffrey Barrett, Sonia Goncalves, Ewan Harrison, David K. Jackson, Ian Johnston, Dominic Kwiatkowski, Cordelia Langford, John Sillitoe on behalf of the Wellcome Sanger Institute COVID-19 Surveillance Team |
| EPI_ISL_2910494 | Lighthouse Lab in Milton Keynes | Wellcome Sanger Institute for the COVID-19 Genomics UK (COG-UK) Consortium | The Lighthouse Lab in Milton Keynes and Alex Alderton, Roberto Amato, Jeffrey Barrett, Sonia Goncalves, Ewan Harrison, David K. Jackson, Ian Johnston, Dominic Kwiatkowski, Cordelia Langford, John Sillitoe on behalf of the Wellcome Sanger Institute COVID-19 Surveillance Team |
| EPI_ISL_2910495, EPI_ISL_2910496, EPI_ISL_2910497 | Lighthouse Lab in Alderley Park | Wellcome Sanger Institute for the COVID-19 Genomics UK (COG-UK) Consortium | Jacquelyn Wynn, Mairead Hyland, The Lighthouse Lab in Alderley Park and Alex Alderton, Roberto Amato, Jeffrey Barrett, Sonia Goncalves, Ewan Harrison, David K. Jackson, Ian Johnston, Dominic Kwiatkowski, Cordelia Langford, John Sillitoe on behalf of the Wellcome Sanger Institute COVID-19 Surveillance Team |
| EPI_ISL_2910498 | Lighthouse Lab in Glasgow | Wellcome Sanger Institute for the COVID-19 Genomics UK (COG-UK) Consortium | Harper VanSteenhouse, Yumi Kasai, David Gray, Carol Clugston, Anna Dominiczak and Alex Alderton, Roberto Amato, Jeffrey Barrett, Sonia Goncalves, Ewan Harrison, David K. Jackson, Ian Johnston, Dominic Kwiatkowski, Cordelia Langford, John Sillitoe on behalf of the Wellcome Sanger Institute COVID-19 Surveillance Team |

| | | | |
|---|---|---|---|
| EPI_ISL_2910499 | Lighthouse Lab in Alderley Park | Wellcome Sanger Institute for the COVID-19 Genomics UK (COG-UK) Consortium | Jacquelyn Wynn, Mairead Hyland, The Lighthouse Lab in Alderley Park and Alex Alderton, Roberto Amato, Jeffrey Barrett, Sonia Goncalves, Ewan Harrison, David K. Jackson, Ian Johnston, Dominic Kwiatkowski, Cordelia Langford, John Sillitoe on behalf of the Wellcome Sanger Institute COVID-19 Surveillance Team |
| EPI_ISL_2910500, EPI_ISL_2910501 | Lighthouse Lab in Glasgow | Wellcome Sanger Institute for the COVID-19 Genomics UK (COG-UK) Consortium | Harper VanSteenhouse, Yumi Kasai, David Gray, Carol Clugston, Anna Dominiczak and Alex Alderton, Roberto Amato, Jeffrey Barrett, Sonia Goncalves, Ewan Harrison, David K. Jackson, Ian Johnston, Dominic Kwiatkowski, Cordelia Langford, John Sillitoe on behalf of the Wellcome Sanger Institute COVID-19 Surveillance Team |
| EPI_ISL_2910502 | Lighthouse Lab in Alderley Park | Wellcome Sanger Institute for the COVID-19 Genomics UK (COG-UK) Consortium | Jacquelyn Wynn, Mairead Hyland, The Lighthouse Lab in Alderley Park and Alex Alderton, Roberto Amato, Jeffrey Barrett, Sonia Goncalves, Ewan Harrison, David K. Jackson, Ian Johnston, Dominic Kwiatkowski, Cordelia Langford, John Sillitoe on behalf of the Wellcome Sanger Institute COVID-19 Surveillance Team |
| EPI_ISL_2910503, EPI_ISL_2910504 | Lighthouse Lab in Milton Keynes | Wellcome Sanger Institute for the COVID-19 Genomics UK (COG-UK) Consortium | The Lighthouse Lab in Milton Keynes and Alex Alderton, Roberto Amato, Jeffrey Barrett, Sonia Goncalves, Ewan Harrison, David K. Jackson, Ian Johnston, Dominic Kwiatkowski, Cordelia Langford, John Sillitoe on behalf of the Wellcome Sanger Institute COVID-19 Surveillance Team |
| EPI_ISL_2910505, EPI_ISL_2910506, EPI_ISL_2910507, EPI_ISL_2910508 | Lighthouse Lab in Alderley Park | Wellcome Sanger Institute for the COVID-19 Genomics UK (COG-UK) Consortium | Jacquelyn Wynn, Mairead Hyland, The Lighthouse Lab in Alderley Park and Alex Alderton, Roberto Amato, Jeffrey Barrett, Sonia Goncalves, Ewan Harrison, David K. Jackson, Ian Johnston, Dominic Kwiatkowski, Cordelia Langford, John Sillitoe on behalf of the Wellcome Sanger Institute COVID-19 Surveillance Team |
| EPI_ISL_2910509, EPI_ISL_2910510, EPI_ISL_2910511 | Lighthouse Lab in Glasgow | Wellcome Sanger Institute for the COVID-19 Genomics UK (COG-UK) Consortium | Harper VanSteenhouse, Yumi Kasai, David Gray, Carol Clugston, Anna Dominiczak and Alex Alderton, Roberto Amato, Jeffrey Barrett, Sonia Goncalves, Ewan Harrison, David K. Jackson, Ian Johnston, Dominic Kwiatkowski, Cordelia Langford, John Sillitoe on behalf of the Wellcome Sanger Institute COVID-19 Surveillance Team |
| EPI_ISL_2910512, EPI_ISL_2910513 | Lighthouse Lab in Alderley Park | Wellcome Sanger Institute for the COVID-19 Genomics UK (COG-UK) Consortium | Jacquelyn Wynn, Mairead Hyland, The Lighthouse Lab in Alderley Park and Alex Alderton, Roberto Amato, Jeffrey Barrett, Sonia Goncalves, Ewan Harrison, David K. Jackson, Ian Johnston, Dominic Kwiatkowski, Cordelia Langford, John Sillitoe on behalf of the Wellcome Sanger Institute COVID-19 Surveillance Team |
| EPI_ISL_2910514 | Lighthouse Lab in Milton Keynes | Wellcome Sanger Institute for the COVID-19 Genomics UK (COG-UK) Consortium | The Lighthouse Lab in Milton Keynes and Alex Alderton, Roberto Amato, Jeffrey Barrett, Sonia Goncalves, Ewan Harrison, David K. Jackson, Ian Johnston, Dominic Kwiatkowski, Cordelia Langford, John Sillitoe on behalf of the Wellcome Sanger Institute COVID-19 Surveillance Team |
| EPI_ISL_2910515 | Lighthouse Lab in Alderley Park | Wellcome Sanger Institute for the COVID-19 Genomics UK (COG-UK) Consortium | Jacquelyn Wynn, Mairead Hyland, The Lighthouse Lab in Alderley Park and Alex Alderton, Roberto Amato, Jeffrey Barrett, Sonia Goncalves, Ewan Harrison, David K. Jackson, Ian Johnston, Dominic Kwiatkowski, Cordelia Langford, John Sillitoe on behalf of the Wellcome Sanger Institute COVID-19 Surveillance Team |
| EPI_ISL_2910516 | Lighthouse Lab in Milton Keynes | Wellcome Sanger Institute for the COVID-19 Genomics UK (COG-UK) Consortium | The Lighthouse Lab in Milton Keynes and Alex Alderton, Roberto Amato, Jeffrey Barrett, Sonia Goncalves, Ewan Harrison, David K. Jackson, Ian Johnston, Dominic Kwiatkowski, Cordelia Langford, John Sillitoe on behalf of the Wellcome Sanger Institute COVID-19 Surveillance Team |
| EPI_ISL_2910517, EPI_ISL_2910518, EPI_ISL_2910519, EPI_ISL_2910520, EPI_ISL_2910521 | Lighthouse Lab in Alderley Park | Wellcome Sanger Institute for the COVID-19 Genomics UK (COG-UK) Consortium | Jacquelyn Wynn, Mairead Hyland, The Lighthouse Lab in Alderley Park and Alex Alderton, Roberto Amato, Jeffrey Barrett, Sonia Goncalves, Ewan Harrison, David K. Jackson, Ian Johnston, Dominic Kwiatkowski, Cordelia Langford, John Sillitoe on behalf of the Wellcome Sanger Institute COVID-19 Surveillance Team |
| EPI_ISL_2910522, EPI_ISL_2910523, EPI_ISL_2910525 | Lighthouse Lab in Glasgow | Wellcome Sanger Institute for the COVID-19 Genomics UK (COG-UK) Consortium | Harper VanSteenhouse, Yumi Kasai, David Gray, Carol Clugston, Anna Dominiczak and Alex Alderton, Roberto Amato, Jeffrey Barrett, Sonia Goncalves, Ewan Harrison, David K. Jackson, Ian Johnston, Dominic Kwiatkowski, Cordelia Langford, John Sillitoe on behalf of the Wellcome Sanger Institute COVID-19 Surveillance Team |
| EPI_ISL_2910526, EPI_ISL_2910527 | Lighthouse Lab in Alderley Park | Wellcome Sanger Institute for the COVID-19 Genomics UK (COG-UK) Consortium | Jacquelyn Wynn, Mairead Hyland, The Lighthouse Lab in Alderley Park and Alex Alderton, Roberto Amato, Jeffrey Barrett, Sonia Goncalves, Ewan Harrison, David K. Jackson, Ian Johnston, Dominic Kwiatkowski, Cordelia Langford, John Sillitoe on behalf of the Wellcome Sanger Institute COVID-19 Surveillance Team |
| EPI_ISL_2910528 | Lighthouse Lab in Glasgow | Wellcome Sanger Institute for the COVID-19 Genomics UK (COG-UK) Consortium | Harper VanSteenhouse, Yumi Kasai, David Gray, Carol Clugston, Anna Dominiczak and Alex Alderton, Roberto Amato, Jeffrey Barrett, Sonia Goncalves, Ewan Harrison, David K. Jackson, Ian Johnston, Dominic Kwiatkowski, Cordelia Langford, John Sillitoe on behalf of the Wellcome Sanger Institute COVID-19 Surveillance Team |
| EPI_ISL_2910529 | Lighthouse Lab in Alderley Park | Wellcome Sanger Institute for the COVID-19 Genomics UK (COG-UK) Consortium | Jacquelyn Wynn, Mairead Hyland, The Lighthouse Lab in Alderley Park and Alex Alderton, Roberto Amato, Jeffrey Barrett, Sonia Goncalves, Ewan Harrison, David K. Jackson, Ian Johnston, Dominic Kwiatkowski, Cordelia Langford, John Sillitoe on behalf of the Wellcome Sanger Institute COVID-19 Surveillance Team |
| EPI_ISL_2910530 | Lighthouse Lab in Milton Keynes | Wellcome Sanger Institute for the COVID-19 Genomics UK (COG-UK) Consortium | The Lighthouse Lab in Milton Keynes and Alex Alderton, Roberto Amato, Jeffrey Barrett, Sonia Goncalves, Ewan Harrison, David K. Jackson, Ian Johnston, Dominic Kwiatkowski, Cordelia Langford, John Sillitoe on behalf of the Wellcome Sanger Institute COVID-19 Surveillance Team |
| EPI_ISL_2910531, EPI_ISL_2910532 | Lighthouse Lab in Alderley Park | Wellcome Sanger Institute for the COVID-19 Genomics UK (COG-UK) Consortium | Jacquelyn Wynn, Mairead Hyland, The Lighthouse Lab in Alderley Park and Alex Alderton, Roberto Amato, Jeffrey Barrett, Sonia Goncalves, Ewan Harrison, David K. Jackson, Ian Johnston, Dominic Kwiatkowski, Cordelia Langford, John Sillitoe on behalf of the Wellcome Sanger Institute COVID-19 Surveillance Team |
| EPI_ISL_2910533 | Lighthouse Lab in Milton Keynes | Wellcome Sanger Institute for the COVID-19 Genomics UK (COG-UK) Consortium | The Lighthouse Lab in Milton Keynes and Alex Alderton, Roberto Amato, Jeffrey Barrett, Sonia Goncalves, Ewan Harrison, David K. Jackson, Ian Johnston, Dominic Kwiatkowski, Cordelia Langford, John Sillitoe on behalf of the Wellcome Sanger Institute COVID-19 Surveillance Team |
| EPI_ISL_2910534 | Lighthouse Lab in Glasgow | Wellcome Sanger Institute for the COVID-19 Genomics UK (COG-UK) Consortium | Harper VanSteenhouse, Yumi Kasai, David Gray, Carol Clugston, Anna Dominiczak and Alex Alderton, Roberto Amato, Jeffrey Barrett, Sonia Goncalves, Ewan Harrison, David K. Jackson, Ian Johnston, Dominic Kwiatkowski, Cordelia Langford, John Sillitoe on behalf of the Wellcome Sanger Institute COVID-19 Surveillance Team |
| EPI_ISL_2910535 | Lighthouse Lab in Alderley Park | Wellcome Sanger Institute for the COVID-19 Genomics UK (COG-UK) Consortium | Jacquelyn Wynn, Mairead Hyland, The Lighthouse Lab in Alderley Park and Alex Alderton, Roberto Amato, Jeffrey Barrett, Sonia Goncalves, Ewan Harrison, David K. Jackson, Ian Johnston, Dominic Kwiatkowski, Cordelia Langford, John Sillitoe on behalf of the Wellcome Sanger Institute COVID-19 Surveillance Team |
| EPI_ISL_2910536, EPI_ISL_2910537 | Lighthouse Lab in Glasgow | Wellcome Sanger Institute for the COVID-19 Genomics UK (COG-UK) Consortium | Harper VanSteenhouse, Yumi Kasai, David Gray, Carol Clugston, Anna Dominiczak and Alex Alderton, Roberto Amato, Jeffrey Barrett, Sonia Goncalves, Ewan Harrison, David K. Jackson, Ian Johnston, Dominic Kwiatkowski, Cordelia Langford, John Sillitoe on behalf of the Wellcome Sanger Institute COVID-19 Surveillance Team |
| EPI_ISL_2910538 | Lighthouse Lab in Alderley Park | Wellcome Sanger Institute for the COVID-19 Genomics UK (COG-UK) Consortium | Jacquelyn Wynn, Mairead Hyland, The Lighthouse Lab in Alderley Park and Alex Alderton, Roberto Amato, Jeffrey Barrett, Sonia Goncalves, Ewan Harrison, David K. Jackson, Ian Johnston, Dominic Kwiatkowski, Cordelia Langford, John Sillitoe on behalf of the Wellcome Sanger Institute COVID-19 Surveillance Team |
| EPI_ISL_2910539 | Lighthouse Lab in Glasgow | Wellcome Sanger Institute for the COVID-19 Genomics UK (COG-UK) Consortium | Harper VanSteenhouse, Yumi Kasai, David Gray, Carol Clugston, Anna Dominiczak and Alex Alderton, Roberto Amato, Jeffrey Barrett, Sonia Goncalves, Ewan Harrison, David K. Jackson, Ian Johnston, Dominic Kwiatkowski, Cordelia Langford, John Sillitoe on behalf of the Wellcome Sanger Institute COVID-19 Surveillance Team |
| EPI_ISL_2910540 | Lighthouse Lab in Alderley Park | Wellcome Sanger Institute for the COVID-19 Genomics UK (COG-UK) Consortium | Jacquelyn Wynn, Mairead Hyland, The Lighthouse Lab in Alderley Park and Alex Alderton, Roberto Amato, Jeffrey Barrett, Sonia Goncalves, Ewan Harrison, David K. Jackson, Ian Johnston, Dominic Kwiatkowski, Cordelia Langford, John Sillitoe on behalf of the Wellcome Sanger Institute COVID-19 Surveillance Team |
| EPI_ISL_2910541, EPI_ISL_2910542 | Lighthouse Lab in Milton Keynes | Wellcome Sanger Institute for the COVID-19 Genomics UK (COG-UK) Consortium | The Lighthouse Lab in Milton Keynes and Alex Alderton, Roberto Amato, Jeffrey Barrett, Sonia Goncalves, Ewan Harrison, David K. Jackson, Ian Johnston, Dominic Kwiatkowski, Cordelia Langford, John Sillitoe on behalf of the Wellcome Sanger Institute COVID-19 Surveillance Team |
| EPI_ISL_2910543 | Lighthouse Lab in Alderley Park | Wellcome Sanger Institute for the COVID-19 Genomics UK (COG-UK) Consortium | Jacquelyn Wynn, Mairead Hyland, The Lighthouse Lab in Alderley Park and Alex Alderton, Roberto Amato, Jeffrey Barrett, Sonia Goncalves, Ewan Harrison, David K. Jackson, Ian Johnston, Dominic Kwiatkowski, Cordelia Langford, John Sillitoe on behalf of the Wellcome Sanger Institute COVID-19 Surveillance Team |
| EPI_ISL_2910544 | Lighthouse Lab in Glasgow | Wellcome Sanger Institute for the COVID-19 Genomics UK (COG-UK) Consortium | Harper VanSteenhouse, Yumi Kasai, David Gray, Carol Clugston, Anna Dominiczak and Alex Alderton, Roberto Amato, Jeffrey Barrett, Sonia Goncalves, Ewan Harrison, David K. Jackson, Ian Johnston, Dominic Kwiatkowski, Cordelia Langford, John Sillitoe on behalf of the Wellcome Sanger Institute COVID-19 Surveillance Team |
| EPI_ISL_2910545 | Lighthouse Lab in Alderley Park | Wellcome Sanger Institute for the COVID-19 Genomics UK (COG-UK) Consortium | Jacquelyn Wynn, Mairead Hyland, The Lighthouse Lab in Alderley Park and Alex Alderton, Roberto Amato, Jeffrey Barrett, Sonia Goncalves, Ewan Harrison, David K. Jackson, Ian Johnston, Dominic Kwiatkowski, Cordelia Langford, John Sillitoe on behalf of the Wellcome Sanger Institute COVID-19 Surveillance Team |
| EPI_ISL_2910546 | Lighthouse Lab in Milton Keynes | Wellcome Sanger Institute for the COVID-19 Genomics UK (COG-UK) Consortium | The Lighthouse Lab in Milton Keynes and Alex Alderton, Roberto Amato, Jeffrey Barrett, Sonia Goncalves, Ewan Harrison, David K. Jackson, Ian Johnston, Dominic Kwiatkowski, Cordelia Langford, John Sillitoe on behalf of the Wellcome Sanger Institute COVID-19 Surveillance Team |
| EPI_ISL_2910547, EPI_ISL_2910548 | Lighthouse Lab in Alderley Park | Wellcome Sanger Institute for the COVID-19 Genomics UK (COG-UK) Consortium | Jacquelyn Wynn, Mairead Hyland, The Lighthouse Lab in Alderley Park and Alex Alderton, Roberto Amato, Jeffrey Barrett, Sonia Goncalves, Ewan Harrison, David K. Jackson, Ian Johnston, Dominic Kwiatkowski, Cordelia Langford, John Sillitoe on behalf of the Wellcome Sanger Institute COVID-19 Surveillance Team |
| EPI_ISL_2910549, EPI_ISL_2910550 | Lighthouse Lab in Milton Keynes | Wellcome Sanger Institute for the COVID-19 Genomics UK (COG-UK) Consortium | The Lighthouse Lab in Milton Keynes and Alex Alderton, Roberto Amato, Jeffrey Barrett, Sonia Goncalves, Ewan Harrison, David K. Jackson, Ian Johnston, Dominic Kwiatkowski, Cordelia Langford, John Sillitoe on behalf of the Wellcome Sanger Institute COVID-19 Surveillance Team |
| EPI_ISL_2910551, EPI_ISL_2910552 | Lighthouse Lab in Alderley Park | Wellcome Sanger Institute for the COVID-19 Genomics UK (COG-UK) Consortium | Jacquelyn Wynn, Mairead Hyland, The Lighthouse Lab in Alderley Park and Alex Alderton, Roberto Amato, Jeffrey Barrett, Sonia Goncalves, Ewan Harrison, David K. Jackson, Ian Johnston, Dominic Kwiatkowski, Cordelia Langford, John Sillitoe on behalf of the Wellcome Sanger Institute COVID-19 Surveillance Team |
| EPI_ISL_2910553 | Lighthouse Lab in Glasgow | Wellcome Sanger Institute for the COVID-19 Genomics UK (COG-UK) Consortium | Harper VanSteenhouse, Yumi Kasai, David Gray, Carol Clugston, Anna Dominiczak and Alex Alderton, Roberto Amato, Jeffrey Barrett, Sonia Goncalves, Ewan Harrison, David K. Jackson, Ian Johnston, Dominic Kwiatkowski, Cordelia Langford, John Sillitoe on behalf of the Wellcome Sanger Institute COVID-19 Surveillance Team |
| EPI_ISL_2910554 | Lighthouse Lab in Alderley Park | Wellcome Sanger Institute for the COVID-19 Genomics UK (COG-UK) Consortium | Jacquelyn Wynn, Mairead Hyland, The Lighthouse Lab in Alderley Park and Alex Alderton, Roberto Amato, Jeffrey Barrett, Sonia Goncalves, Ewan Harrison, David K. Jackson, Ian Johnston, Dominic Kwiatkowski, Cordelia Langford, John Sillitoe on behalf of the Wellcome Sanger Institute COVID-19 Surveillance Team |
| EPI_ISL_2910555, EPI_ISL_2910556 | Lighthouse Lab in Milton Keynes | Wellcome Sanger Institute for the COVID-19 Genomics UK (COG-UK) Consortium | The Lighthouse Lab in Milton Keynes and Alex Alderton, Roberto Amato, Jeffrey Barrett, Sonia Goncalves, Ewan Harrison, David K. Jackson, Ian Johnston, Dominic Kwiatkowski, Cordelia Langford, John Sillitoe on behalf of the Wellcome Sanger Institute COVID-19 Surveillance Team |
| EPI_ISL_2910557 | Lighthouse Lab in Glasgow | Wellcome Sanger Institute for the COVID-19 Genomics UK (COG-UK) Consortium | Harper VanSteenhouse, Yumi Kasai, David Gray, Carol Clugston, Anna Dominiczak and Alex Alderton, Roberto Amato, Jeffrey Barrett, Sonia Goncalves, Ewan Harrison, David K. Jackson, Ian Johnston, Dominic Kwiatkowski, Cordelia Langford, John Sillitoe on behalf of the Wellcome Sanger Institute COVID-19 Surveillance Team |
| EPI_ISL_2910558 | Lighthouse Lab in Milton Keynes | Wellcome Sanger Institute for the COVID-19 Genomics UK (COG-UK) Consortium | The Lighthouse Lab in Milton Keynes and Alex Alderton, Roberto Amato, Jeffrey Barrett, Sonia Goncalves, Ewan Harrison, David K. Jackson, Ian Johnston, Dominic Kwiatkowski, Cordelia Langford, John Sillitoe on behalf of the Wellcome Sanger Institute COVID-19 Surveillance Team |
| EPI_ISL_2910559 | Lighthouse Lab in Glasgow | Wellcome Sanger Institute for the COVID-19 Genomics UK (COG-UK) Consortium | Harper VanSteenhouse, Yumi Kasai, David Gray, Carol Clugston, Anna Dominiczak and Alex Alderton, Roberto Amato, Jeffrey Barrett, Sonia Goncalves, Ewan Harrison, David K. Jackson, Ian Johnston, Dominic Kwiatkowski, Cordelia Langford, John Sillitoe on behalf of the Wellcome Sanger Institute COVID-19 Surveillance Team |
| EPI_ISL_2910560, EPI_ISL_2910561, EPI_ISL_2910562 | Lighthouse Lab in Milton Keynes | Wellcome Sanger Institute for the COVID-19 Genomics UK (COG-UK) Consortium | The Lighthouse Lab in Milton Keynes and Alex Alderton, Roberto Amato, Jeffrey Barrett, Sonia Goncalves, Ewan Harrison, David K. Jackson, Ian Johnston, Dominic Kwiatkowski, Cordelia Langford, John Sillitoe on behalf of the Wellcome Sanger Institute COVID-19 Surveillance Team |
| EPI_ISL_2910563 | Lighthouse Lab in Glasgow | Wellcome Sanger Institute for the COVID-19 Genomics UK (COG-UK) Consortium | Harper VanSteenhouse, Yumi Kasai, David Gray, Carol Clugston, Anna Dominiczak and Alex Alderton, Roberto Amato, Jeffrey Barrett, Sonia Goncalves, Ewan Harrison, David K. Jackson, Ian Johnston, Dominic Kwiatkowski, Cordelia Langford, John Sillitoe on behalf of the Wellcome Sanger Institute COVID-19 Surveillance Team |
| EPI_ISL_2910564 | Lighthouse Lab in Milton Keynes | Wellcome Sanger Institute for the COVID-19 Genomics UK (COG-UK) Consortium | The Lighthouse Lab in Milton Keynes and Alex Alderton, Roberto Amato, Jeffrey Barrett, Sonia Goncalves, Ewan Harrison, David K. Jackson, Ian Johnston, Dominic Kwiatkowski, Cordelia Langford, John Sillitoe on behalf of the Wellcome Sanger Institute COVID-19 Surveillance Team |
| EPI_ISL_2910565 | Lighthouse Lab in Glasgow | Wellcome Sanger Institute for the COVID-19 Genomics UK (COG-UK) Consortium | Harper VanSteenhouse, Yumi Kasai, David Gray, Carol Clugston, Anna Dominiczak and Alex Alderton, Roberto Amato, Jeffrey Barrett, Sonia Goncalves, Ewan Harrison, David K. Jackson, Ian Johnston, Dominic Kwiatkowski, Cordelia Langford, John Sillitoe on behalf of the Wellcome Sanger Institute COVID-19 Surveillance Team |

| | | | |
|---|---|---|---|
| EPI_ISL_2910566 | Lighthouse Lab in Milton Keynes | Wellcome Sanger Institute for the COVID-19 Genomics UK (COG-UK) Consortium | The Lighthouse Lab in Milton Keynes and Alex Alderton, Roberto Amato, Jeffrey Barrett, Sonia Goncalves, Ewan Harrison, David K. Jackson, Ian Johnston, Dominic Kwiatkowski, Cordelia Langford, John Sillitoe on behalf of the Wellcome Sanger Institute COVID-19 Surveillance Team |
| EPI_ISL_2910567 | Lighthouse Lab in Glasgow | Wellcome Sanger Institute for the COVID-19 Genomics UK (COG-UK) Consortium | Harper VanSteenhouse, Yumi Kasai, David Gray, Carol Clugston, Anna Dominiczak and Alex Alderton, Roberto Amato, Jeffrey Barrett, Sonia Goncalves, Ewan Harrison, David K. Jackson, Ian Johnston, Dominic Kwiatkowski, Cordelia Langford, John Sillitoe on behalf of the Wellcome Sanger Institute COVID-19 Surveillance Team |
| EPI_ISL_2910568, EPI_ISL_2910569, EPI_ISL_2910570, EPI_ISL_2910571, EPI_ISL_2910572, EPI_ISL_2910573 | Lighthouse Lab in Alderley Park | Wellcome Sanger Institute for the COVID-19 Genomics UK (COG-UK) Consortium | Jacquelyn Wynn, Mairead Hyland, The Lighthouse Lab in Alderley Park and Alex Alderton, Roberto Amato, Jeffrey Barrett, Sonia Goncalves, Ewan Harrison, David K. Jackson, Ian Johnston, Dominic Kwiatkowski, Cordelia Langford, John Sillitoe on behalf of the Wellcome Sanger Institute COVID-19 Surveillance Team |
| EPI_ISL_2910574 | Lighthouse Lab in Milton Keynes | Wellcome Sanger Institute for the COVID-19 Genomics UK (COG-UK) Consortium | The Lighthouse Lab in Milton Keynes and Alex Alderton, Roberto Amato, Jeffrey Barrett, Sonia Goncalves, Ewan Harrison, David K. Jackson, Ian Johnston, Dominic Kwiatkowski, Cordelia Langford, John Sillitoe on behalf of the Wellcome Sanger Institute COVID-19 Surveillance Team |
| EPI_ISL_2910575 | Lighthouse Lab in Glasgow | Wellcome Sanger Institute for the COVID-19 Genomics UK (COG-UK) Consortium | Harper VanSteenhouse, Yumi Kasai, David Gray, Carol Clugston, Anna Dominiczak and Alex Alderton, Roberto Amato, Jeffrey Barrett, Sonia Goncalves, Ewan Harrison, David K. Jackson, Ian Johnston, Dominic Kwiatkowski, Cordelia Langford, John Sillitoe on behalf of the Wellcome Sanger Institute COVID-19 Surveillance Team |
| EPI_ISL_2910576 | Lighthouse Lab in Milton Keynes | Wellcome Sanger Institute for the COVID-19 Genomics UK (COG-UK) Consortium | The Lighthouse Lab in Milton Keynes and Alex Alderton, Roberto Amato, Jeffrey Barrett, Sonia Goncalves, Ewan Harrison, David K. Jackson, Ian Johnston, Dominic Kwiatkowski, Cordelia Langford, John Sillitoe on behalf of the Wellcome Sanger Institute COVID-19 Surveillance Team |
| EPI_ISL_2910577, EPI_ISL_2910578, EPI_ISL_2910579, EPI_ISL_2910580, EPI_ISL_2910581 | Lighthouse Lab in Alderley Park | Wellcome Sanger Institute for the COVID-19 Genomics UK (COG-UK) Consortium | Jacquelyn Wynn, Mairead Hyland, The Lighthouse Lab in Alderley Park and Alex Alderton, Roberto Amato, Jeffrey Barrett, Sonia Goncalves, Ewan Harrison, David K. Jackson, Ian Johnston, Dominic Kwiatkowski, Cordelia Langford, John Sillitoe on behalf of the Wellcome Sanger Institute COVID-19 Surveillance Team |
| EPI_ISL_2910582 | Lighthouse Lab in Milton Keynes | Wellcome Sanger Institute for the COVID-19 Genomics UK (COG-UK) Consortium | The Lighthouse Lab in Milton Keynes and Alex Alderton, Roberto Amato, Jeffrey Barrett, Sonia Goncalves, Ewan Harrison, David K. Jackson, Ian Johnston, Dominic Kwiatkowski, Cordelia Langford, John Sillitoe on behalf of the Wellcome Sanger Institute COVID-19 Surveillance Team |
| EPI_ISL_2910583 | Lighthouse Lab in Glasgow | Wellcome Sanger Institute for the COVID-19 Genomics UK (COG-UK) Consortium | Harper VanSteenhouse, Yumi Kasai, David Gray, Carol Clugston, Anna Dominiczak and Alex Alderton, Roberto Amato, Jeffrey Barrett, Sonia Goncalves, Ewan Harrison, David K. Jackson, Ian Johnston, Dominic Kwiatkowski, Cordelia Langford, John Sillitoe on behalf of the Wellcome Sanger Institute COVID-19 Surveillance Team |
| EPI_ISL_2910584 | Lighthouse Lab in Milton Keynes | Wellcome Sanger Institute for the COVID-19 Genomics UK (COG-UK) Consortium | The Lighthouse Lab in Milton Keynes and Alex Alderton, Roberto Amato, Jeffrey Barrett, Sonia Goncalves, Ewan Harrison, David K. Jackson, Ian Johnston, Dominic Kwiatkowski, Cordelia Langford, John Sillitoe on behalf of the Wellcome Sanger Institute COVID-19 Surveillance Team |
| EPI_ISL_2910585 | Lighthouse Lab in Alderley Park | Wellcome Sanger Institute for the COVID-19 Genomics UK (COG-UK) Consortium | Jacquelyn Wynn, Mairead Hyland, The Lighthouse Lab in Alderley Park and Alex Alderton, Roberto Amato, Jeffrey Barrett, Sonia Goncalves, Ewan Harrison, David K. Jackson, Ian Johnston, Dominic Kwiatkowski, Cordelia Langford, John Sillitoe on behalf of the Wellcome Sanger Institute COVID-19 Surveillance Team |
| EPI_ISL_2910586 | Lighthouse Lab in Glasgow | Wellcome Sanger Institute for the COVID-19 Genomics UK (COG-UK) Consortium | Harper VanSteenhouse, Yumi Kasai, David Gray, Carol Clugston, Anna Dominiczak and Alex Alderton, Roberto Amato, Jeffrey Barrett, Sonia Goncalves, Ewan Harrison, David K. Jackson, Ian Johnston, Dominic Kwiatkowski, Cordelia Langford, John Sillitoe on behalf of the Wellcome Sanger Institute COVID-19 Surveillance Team |
| EPI_ISL_2910587, EPI_ISL_2910588, EPI_ISL_2910589, EPI_ISL_2910590, EPI_ISL_2910591, EPI_ISL_2910592, EPI_ISL_2910593, EPI_ISL_2910594 | Lighthouse Lab in Alderley Park | Wellcome Sanger Institute for the COVID-19 Genomics UK (COG-UK) Consortium | Jacquelyn Wynn, Mairead Hyland, The Lighthouse Lab in Alderley Park and Alex Alderton, Roberto Amato, Jeffrey Barrett, Sonia Goncalves, Ewan Harrison, David K. Jackson, Ian Johnston, Dominic Kwiatkowski, Cordelia Langford, John Sillitoe on behalf of the Wellcome Sanger Institute COVID-19 Surveillance Team |
| EPI_ISL_2910595 | Lighthouse Lab in Glasgow | Wellcome Sanger Institute for the COVID-19 Genomics UK (COG-UK) Consortium | Harper VanSteenhouse, Yumi Kasai, David Gray, Carol Clugston, Anna Dominiczak and Alex Alderton, Roberto Amato, Jeffrey Barrett, Sonia Goncalves, Ewan Harrison, David K. Jackson, Ian Johnston, Dominic Kwiatkowski, Cordelia Langford, John Sillitoe on behalf of the Wellcome Sanger Institute COVID-19 Surveillance Team |
| EPI_ISL_2910596, EPI_ISL_2910597, EPI_ISL_2910598, EPI_ISL_2910599, EPI_ISL_2910600 | Lighthouse Lab in Alderley Park | Wellcome Sanger Institute for the COVID-19 Genomics UK (COG-UK) Consortium | Jacquelyn Wynn, Mairead Hyland, The Lighthouse Lab in Alderley Park and Alex Alderton, Roberto Amato, Jeffrey Barrett, Sonia Goncalves, Ewan Harrison, David K. Jackson, Ian Johnston, Dominic Kwiatkowski, Cordelia Langford, John Sillitoe on behalf of the Wellcome Sanger Institute COVID-19 Surveillance Team |
| EPI_ISL_2910601 | Lighthouse Lab in Milton Keynes | Wellcome Sanger Institute for the COVID-19 Genomics UK (COG-UK) Consortium | The Lighthouse Lab in Milton Keynes and Alex Alderton, Roberto Amato, Jeffrey Barrett, Sonia Goncalves, Ewan Harrison, David K. Jackson, Ian Johnston, Dominic Kwiatkowski, Cordelia Langford, John Sillitoe on behalf of the Wellcome Sanger Institute COVID-19 Surveillance Team |
| EPI_ISL_2910602, EPI_ISL_2910603 | Lighthouse Lab in Alderley Park | Wellcome Sanger Institute for the COVID-19 Genomics UK (COG-UK) Consortium | Jacquelyn Wynn, Mairead Hyland, The Lighthouse Lab in Alderley Park and Alex Alderton, Roberto Amato, Jeffrey Barrett, Sonia Goncalves, Ewan Harrison, David K. Jackson, Ian Johnston, Dominic Kwiatkowski, Cordelia Langford, John Sillitoe on behalf of the Wellcome Sanger Institute COVID-19 Surveillance Team |
| EPI_ISL_2910604, EPI_ISL_2910605, EPI_ISL_2910606 | Lighthouse Lab in Milton Keynes | Wellcome Sanger Institute for the COVID-19 Genomics UK (COG-UK) Consortium | The Lighthouse Lab in Milton Keynes and Alex Alderton, Roberto Amato, Jeffrey Barrett, Sonia Goncalves, Ewan Harrison, David K. Jackson, Ian Johnston, Dominic Kwiatkowski, Cordelia Langford, John Sillitoe on behalf of the Wellcome Sanger Institute COVID-19 Surveillance Team |
| EPI_ISL_2910607 | Lighthouse Lab in Glasgow | Wellcome Sanger Institute for the COVID-19 Genomics UK (COG-UK) Consortium | Harper VanSteenhouse, Yumi Kasai, David Gray, Carol Clugston, Anna Dominiczak and Alex Alderton, Roberto Amato, Jeffrey Barrett, Sonia Goncalves, Ewan Harrison, David K. Jackson, Ian Johnston, Dominic Kwiatkowski, Cordelia Langford, John Sillitoe on behalf of the Wellcome Sanger Institute COVID-19 Surveillance Team |
| EPI_ISL_2910608 | Lighthouse Lab in Alderley Park | Wellcome Sanger Institute for the COVID-19 Genomics UK (COG-UK) Consortium | Jacquelyn Wynn, Mairead Hyland, The Lighthouse Lab in Alderley Park and Alex Alderton, Roberto Amato, Jeffrey Barrett, Sonia Goncalves, Ewan Harrison, David K. Jackson, Ian Johnston, Dominic Kwiatkowski, Cordelia Langford, John Sillitoe on behalf of the Wellcome Sanger Institute COVID-19 Surveillance Team |
| EPI_ISL_2910609 | Lighthouse Lab in Glasgow | Wellcome Sanger Institute for the COVID-19 Genomics UK (COG-UK) Consortium | Harper VanSteenhouse, Yumi Kasai, David Gray, Carol Clugston, Anna Dominiczak and Alex Alderton, Roberto Amato, Jeffrey Barrett, Sonia Goncalves, Ewan Harrison, David K. Jackson, Ian Johnston, Dominic Kwiatkowski, Cordelia Langford, John Sillitoe on behalf of the Wellcome Sanger Institute COVID-19 Surveillance Team |
| EPI_ISL_2910610 | Lighthouse Lab in Milton Keynes | Wellcome Sanger Institute for the COVID-19 Genomics UK (COG-UK) Consortium | The Lighthouse Lab in Milton Keynes and Alex Alderton, Roberto Amato, Jeffrey Barrett, Sonia Goncalves, Ewan Harrison, David K. Jackson, Ian Johnston, Dominic Kwiatkowski, Cordelia Langford, John Sillitoe on behalf of the Wellcome Sanger Institute COVID-19 Surveillance Team |
| EPI_ISL_2910611, EPI_ISL_2910612 | Lighthouse Lab in Glasgow | Wellcome Sanger Institute for the COVID-19 Genomics UK (COG-UK) Consortium | Harper VanSteenhouse, Yumi Kasai, David Gray, Carol Clugston, Anna Dominiczak and Alex Alderton, Roberto Amato, Jeffrey Barrett, Sonia Goncalves, Ewan Harrison, David K. Jackson, Ian Johnston, Dominic Kwiatkowski, Cordelia Langford, John Sillitoe on behalf of the Wellcome Sanger Institute COVID-19 Surveillance Team |
| EPI_ISL_2910613, EPI_ISL_2910614, EPI_ISL_2910615, EPI_ISL_2910616, EPI_ISL_2910617 | Lighthouse Lab in Milton Keynes | Wellcome Sanger Institute for the COVID-19 Genomics UK (COG-UK) Consortium | The Lighthouse Lab in Milton Keynes and Alex Alderton, Roberto Amato, Jeffrey Barrett, Sonia Goncalves, Ewan Harrison, David K. Jackson, Ian Johnston, Dominic Kwiatkowski, Cordelia Langford, John Sillitoe on behalf of the Wellcome Sanger Institute COVID-19 Surveillance Team |
| EPI_ISL_2910618 | Lighthouse Lab in Alderley Park | Wellcome Sanger Institute for the COVID-19 Genomics UK (COG-UK) Consortium | Jacquelyn Wynn, Mairead Hyland, The Lighthouse Lab in Alderley Park and Alex Alderton, Roberto Amato, Jeffrey Barrett, Sonia Goncalves, Ewan Harrison, David K. Jackson, Ian Johnston, Dominic Kwiatkowski, Cordelia Langford, John Sillitoe on behalf of the Wellcome Sanger Institute COVID-19 Surveillance Team |
| EPI_ISL_2910619, EPI_ISL_2910620, EPI_ISL_2910621 | Lighthouse Lab in Milton Keynes | Wellcome Sanger Institute for the COVID-19 Genomics UK (COG-UK) Consortium | The Lighthouse Lab in Milton Keynes and Alex Alderton, Roberto Amato, Jeffrey Barrett, Sonia Goncalves, Ewan Harrison, David K. Jackson, Ian Johnston, Dominic Kwiatkowski, Cordelia Langford, John Sillitoe on behalf of the Wellcome Sanger Institute COVID-19 Surveillance Team |
| EPI_ISL_2910622 | Lighthouse Lab in Glasgow | Wellcome Sanger Institute for the COVID-19 Genomics UK (COG-UK) Consortium | Harper VanSteenhouse, Yumi Kasai, David Gray, Carol Clugston, Anna Dominiczak and Alex Alderton, Roberto Amato, Jeffrey Barrett, Sonia Goncalves, Ewan Harrison, David K. Jackson, Ian Johnston, Dominic Kwiatkowski, Cordelia Langford, John Sillitoe on behalf of the Wellcome Sanger Institute COVID-19 Surveillance Team |
| EPI_ISL_2910623 | Lighthouse Lab in Milton Keynes | Wellcome Sanger Institute for the COVID-19 Genomics UK (COG-UK) Consortium | The Lighthouse Lab in Milton Keynes and Alex Alderton, Roberto Amato, Jeffrey Barrett, Sonia Goncalves, Ewan Harrison, David K. Jackson, Ian Johnston, Dominic Kwiatkowski, Cordelia Langford, John Sillitoe on behalf of the Wellcome Sanger Institute COVID-19 Surveillance Team |
| EPI_ISL_2910624 | Lighthouse Lab in Alderley Park | Wellcome Sanger Institute for the COVID-19 Genomics UK (COG-UK) Consortium | Jacquelyn Wynn, Mairead Hyland, The Lighthouse Lab in Alderley Park and Alex Alderton, Roberto Amato, Jeffrey Barrett, Sonia Goncalves, Ewan Harrison, David K. Jackson, Ian Johnston, Dominic Kwiatkowski, Cordelia Langford, John Sillitoe on behalf of the Wellcome Sanger Institute COVID-19 Surveillance Team |
| EPI_ISL_2910625 | Lighthouse Lab in Glasgow | Wellcome Sanger Institute for the COVID-19 Genomics UK (COG-UK) Consortium | Harper VanSteenhouse, Yumi Kasai, David Gray, Carol Clugston, Anna Dominiczak and Alex Alderton, Roberto Amato, Jeffrey Barrett, Sonia Goncalves, Ewan Harrison, David K. Jackson, Ian Johnston, Dominic Kwiatkowski, Cordelia Langford, John Sillitoe on behalf of the Wellcome Sanger Institute COVID-19 Surveillance Team |
| EPI_ISL_2910626, EPI_ISL_2910627, EPI_ISL_2910628, EPI_ISL_2910629, EPI_ISL_2910630, EPI_ISL_2910631, EPI_ISL_2910632, EPI_ISL_2910633 | Lighthouse Lab in Milton Keynes | Wellcome Sanger Institute for the COVID-19 Genomics UK (COG-UK) Consortium | The Lighthouse Lab in Milton Keynes and Alex Alderton, Roberto Amato, Jeffrey Barrett, Sonia Goncalves, Ewan Harrison, David K. Jackson, Ian Johnston, Dominic Kwiatkowski, Cordelia Langford, John Sillitoe on behalf of the Wellcome Sanger Institute COVID-19 Surveillance Team |
| EPI_ISL_2910634 | Lighthouse Lab in Alderley Park | Wellcome Sanger Institute for the COVID-19 Genomics UK (COG-UK) Consortium | Jacquelyn Wynn, Mairead Hyland, The Lighthouse Lab in Alderley Park and Alex Alderton, Roberto Amato, Jeffrey Barrett, Sonia Goncalves, Ewan Harrison, David K. Jackson, Ian Johnston, Dominic Kwiatkowski, Cordelia Langford, John Sillitoe on behalf of the Wellcome Sanger Institute COVID-19 Surveillance Team |
| EPI_ISL_2910635, EPI_ISL_2910636, EPI_ISL_2910637 | Lighthouse Lab in Milton Keynes | Wellcome Sanger Institute for the COVID-19 Genomics UK (COG-UK) Consortium | The Lighthouse Lab in Milton Keynes and Alex Alderton, Roberto Amato, Jeffrey Barrett, Sonia Goncalves, Ewan Harrison, David K. Jackson, Ian Johnston, Dominic Kwiatkowski, Cordelia Langford, John Sillitoe on behalf of the Wellcome Sanger Institute COVID-19 Surveillance Team |

| | | | |
|---|---|---|---|
| EPI_ISL_2910638 | Lighthouse Lab in Alderley Park | Wellcome Sanger Institute for the COVID-19 Genomics UK (COG-UK) Consortium | Jacquelyn Wynn, Mairead Hyland, The Lighthouse Lab in Alderley Park and Alex Alderton, Roberto Amato, Jeffrey Barrett, Sonia Goncalves, Ewan Harrison, David K. Jackson, Ian Johnston, Dominic Kwiatkowski, Cordelia Langford, John Sillitoe on behalf of the Wellcome Sanger Institute COVID-19 Surveillance Team |
| EPI_ISL_2910639 | Lighthouse Lab in Glasgow | Wellcome Sanger Institute for the COVID-19 Genomics UK (COG-UK) Consortium | Harper VanSteenhouse, Yumi Kasai, David Gray, Carol Clugston, Anna Dominiczak and Alex Alderton, Roberto Amato, Jeffrey Barrett, Sonia Goncalves, Ewan Harrison, David K. Jackson, Ian Johnston, Dominic Kwiatkowski, Cordelia Langford, John Sillitoe on behalf of the Wellcome Sanger Institute COVID-19 Surveillance Team |
| EPI_ISL_2910640 | Lighthouse Lab in Milton Keynes | Wellcome Sanger Institute for the COVID-19 Genomics UK (COG-UK) Consortium | The Lighthouse Lab in Milton Keynes and Alex Alderton, Roberto Amato, Jeffrey Barrett, Sonia Goncalves, Ewan Harrison, David K. Jackson, Ian Johnston, Dominic Kwiatkowski, Cordelia Langford, John Sillitoe on behalf of the Wellcome Sanger Institute COVID-19 Surveillance Team |
| EPI_ISL_2910641, EPI_ISL_2910642 | Lighthouse Lab in Glasgow | Wellcome Sanger Institute for the COVID-19 Genomics UK (COG-UK) Consortium | Harper VanSteenhouse, Yumi Kasai, David Gray, Carol Clugston, Anna Dominiczak and Alex Alderton, Roberto Amato, Jeffrey Barrett, Sonia Goncalves, Ewan Harrison, David K. Jackson, Ian Johnston, Dominic Kwiatkowski, Cordelia Langford, John Sillitoe on behalf of the Wellcome Sanger Institute COVID-19 Surveillance Team |
| EPI_ISL_2910643 | Lighthouse Lab in Milton Keynes | Wellcome Sanger Institute for the COVID-19 Genomics UK (COG-UK) Consortium | The Lighthouse Lab in Milton Keynes and Alex Alderton, Roberto Amato, Jeffrey Barrett, Sonia Goncalves, Ewan Harrison, David K. Jackson, Ian Johnston, Dominic Kwiatkowski, Cordelia Langford, John Sillitoe on behalf of the Wellcome Sanger Institute COVID-19 Surveillance Team |
| EPI_ISL_2910644 | Lighthouse Lab in Glasgow | Wellcome Sanger Institute for the COVID-19 Genomics UK (COG-UK) Consortium | Harper VanSteenhouse, Yumi Kasai, David Gray, Carol Clugston, Anna Dominiczak and Alex Alderton, Roberto Amato, Jeffrey Barrett, Sonia Goncalves, Ewan Harrison, David K. Jackson, Ian Johnston, Dominic Kwiatkowski, Cordelia Langford, John Sillitoe on behalf of the Wellcome Sanger Institute COVID-19 Surveillance Team |
| EPI_ISL_2910645, EPI_ISL_2910646, EPI_ISL_2910647, EPI_ISL_2910648 | Lighthouse Lab in Milton Keynes | Wellcome Sanger Institute for the COVID-19 Genomics UK (COG-UK) Consortium | The Lighthouse Lab in Milton Keynes and Alex Alderton, Roberto Amato, Jeffrey Barrett, Sonia Goncalves, Ewan Harrison, David K. Jackson, Ian Johnston, Dominic Kwiatkowski, Cordelia Langford, John Sillitoe on behalf of the Wellcome Sanger Institute COVID-19 Surveillance Team |
| EPI_ISL_2910649 | Lighthouse Lab in Glasgow | Wellcome Sanger Institute for the COVID-19 Genomics UK (COG-UK) Consortium | Harper VanSteenhouse, Yumi Kasai, David Gray, Carol Clugston, Anna Dominiczak and Alex Alderton, Roberto Amato, Jeffrey Barrett, Sonia Goncalves, Ewan Harrison, David K. Jackson, Ian Johnston, Dominic Kwiatkowski, Cordelia Langford, John Sillitoe on behalf of the Wellcome Sanger Institute COVID-19 Surveillance Team |
| EPI_ISL_2910650 | Lighthouse Lab in Alderley Park | Wellcome Sanger Institute for the COVID-19 Genomics UK (COG-UK) Consortium | Jacquelyn Wynn, Mairead Hyland, The Lighthouse Lab in Alderley Park and Alex Alderton, Roberto Amato, Jeffrey Barrett, Sonia Goncalves, Ewan Harrison, David K. Jackson, Ian Johnston, Dominic Kwiatkowski, Cordelia Langford, John Sillitoe on behalf of the Wellcome Sanger Institute COVID-19 Surveillance Team |
| EPI_ISL_2910651 | Lighthouse Lab in Glasgow | Wellcome Sanger Institute for the COVID-19 Genomics UK (COG-UK) Consortium | Harper VanSteenhouse, Yumi Kasai, David Gray, Carol Clugston, Anna Dominiczak and Alex Alderton, Roberto Amato, Jeffrey Barrett, Sonia Goncalves, Ewan Harrison, David K. Jackson, Ian Johnston, Dominic Kwiatkowski, Cordelia Langford, John Sillitoe on behalf of the Wellcome Sanger Institute COVID-19 Surveillance Team |
| EPI_ISL_2910652, EPI_ISL_2910653, EPI_ISL_2910654, EPI_ISL_2910655, EPI_ISL_2910656, EPI_ISL_2910657, EPI_ISL_2910658, EPI_ISL_2910659, EPI_ISL_2910660, EPI_ISL_2910661 | Lighthouse Lab in Milton Keynes | Wellcome Sanger Institute for the COVID-19 Genomics UK (COG-UK) Consortium | The Lighthouse Lab in Milton Keynes and Alex Alderton, Roberto Amato, Jeffrey Barrett, Sonia Goncalves, Ewan Harrison, David K. Jackson, Ian Johnston, Dominic Kwiatkowski, Cordelia Langford, John Sillitoe on behalf of the Wellcome Sanger Institute COVID-19 Surveillance Team |
| EPI_ISL_2910662 | Lighthouse Lab in Alderley Park | Wellcome Sanger Institute for the COVID-19 Genomics UK (COG-UK) Consortium | Jacquelyn Wynn, Mairead Hyland, The Lighthouse Lab in Alderley Park and Alex Alderton, Roberto Amato, Jeffrey Barrett, Sonia Goncalves, Ewan Harrison, David K. Jackson, Ian Johnston, Dominic Kwiatkowski, Cordelia Langford, John Sillitoe on behalf of the Wellcome Sanger Institute COVID-19 Surveillance Team |
| EPI_ISL_2910663 | Lighthouse Lab in Milton Keynes | Wellcome Sanger Institute for the COVID-19 Genomics UK (COG-UK) Consortium | The Lighthouse Lab in Milton Keynes and Alex Alderton, Roberto Amato, Jeffrey Barrett, Sonia Goncalves, Ewan Harrison, David K. Jackson, Ian Johnston, Dominic Kwiatkowski, Cordelia Langford, John Sillitoe on behalf of the Wellcome Sanger Institute COVID-19 Surveillance Team |
| EPI_ISL_2910664 | Lighthouse Lab in Glasgow | Wellcome Sanger Institute for the COVID-19 Genomics UK (COG-UK) Consortium | Harper VanSteenhouse, Yumi Kasai, David Gray, Carol Clugston, Anna Dominiczak and Alex Alderton, Roberto Amato, Jeffrey Barrett, Sonia Goncalves, Ewan Harrison, David K. Jackson, Ian Johnston, Dominic Kwiatkowski, Cordelia Langford, John Sillitoe on behalf of the Wellcome Sanger Institute COVID-19 Surveillance Team |
| EPI_ISL_2910665 | Lighthouse Lab in Alderley Park | Wellcome Sanger Institute for the COVID-19 Genomics UK (COG-UK) Consortium | Jacquelyn Wynn, Mairead Hyland, The Lighthouse Lab in Alderley Park and Alex Alderton, Roberto Amato, Jeffrey Barrett, Sonia Goncalves, Ewan Harrison, David K. Jackson, Ian Johnston, Dominic Kwiatkowski, Cordelia Langford, John Sillitoe on behalf of the Wellcome Sanger Institute COVID-19 Surveillance Team |
| EPI_ISL_2910666 | Lighthouse Lab in Glasgow | Wellcome Sanger Institute for the COVID-19 Genomics UK (COG-UK) Consortium | Harper VanSteenhouse, Yumi Kasai, David Gray, Carol Clugston, Anna Dominiczak and Alex Alderton, Roberto Amato, Jeffrey Barrett, Sonia Goncalves, Ewan Harrison, David K. Jackson, Ian Johnston, Dominic Kwiatkowski, Cordelia Langford, John Sillitoe on behalf of the Wellcome Sanger Institute COVID-19 Surveillance Team |
| EPI_ISL_2910667 | Lighthouse Lab in Milton Keynes | Wellcome Sanger Institute for the COVID-19 Genomics UK (COG-UK) Consortium | The Lighthouse Lab in Milton Keynes and Alex Alderton, Roberto Amato, Jeffrey Barrett, Sonia Goncalves, Ewan Harrison, David K. Jackson, Ian Johnston, Dominic Kwiatkowski, Cordelia Langford, John Sillitoe on behalf of the Wellcome Sanger Institute COVID-19 Surveillance Team |
| EPI_ISL_2910668 | Lighthouse Lab in Alderley Park | Wellcome Sanger Institute for the COVID-19 Genomics UK (COG-UK) Consortium | Jacquelyn Wynn, Mairead Hyland, The Lighthouse Lab in Alderley Park and Alex Alderton, Roberto Amato, Jeffrey Barrett, Sonia Goncalves, Ewan Harrison, David K. Jackson, Ian Johnston, Dominic Kwiatkowski, Cordelia Langford, John Sillitoe on behalf of the Wellcome Sanger Institute COVID-19 Surveillance Team |
| EPI_ISL_2910669 | Lighthouse Lab in Glasgow | Wellcome Sanger Institute for the COVID-19 Genomics UK (COG-UK) Consortium | Harper VanSteenhouse, Yumi Kasai, David Gray, Carol Clugston, Anna Dominiczak and Alex Alderton, Roberto Amato, Jeffrey Barrett, Sonia Goncalves, Ewan Harrison, David K. Jackson, Ian Johnston, Dominic Kwiatkowski, Cordelia Langford, John Sillitoe on behalf of the Wellcome Sanger Institute COVID-19 Surveillance Team |
| EPI_ISL_2910670 | Lighthouse Lab in Milton Keynes | Wellcome Sanger Institute for the COVID-19 Genomics UK (COG-UK) Consortium | The Lighthouse Lab in Milton Keynes and Alex Alderton, Roberto Amato, Jeffrey Barrett, Sonia Goncalves, Ewan Harrison, David K. Jackson, Ian Johnston, Dominic Kwiatkowski, Cordelia Langford, John Sillitoe on behalf of the Wellcome Sanger Institute COVID-19 Surveillance Team |
| EPI_ISL_2910672 | Lighthouse Lab in Glasgow | Wellcome Sanger Institute for the COVID-19 Genomics UK (COG-UK) Consortium | Harper VanSteenhouse, Yumi Kasai, David Gray, Carol Clugston, Anna Dominiczak and Alex Alderton, Roberto Amato, Jeffrey Barrett, Sonia Goncalves, Ewan Harrison, David K. Jackson, Ian Johnston, Dominic Kwiatkowski, Cordelia Langford, John Sillitoe on behalf of the Wellcome Sanger Institute COVID-19 Surveillance Team |
| EPI_ISL_2910673, EPI_ISL_2910674, EPI_ISL_2910675, EPI_ISL_2910676, EPI_ISL_2910677 | Lighthouse Lab in Milton Keynes | Wellcome Sanger Institute for the COVID-19 Genomics UK (COG-UK) Consortium | The Lighthouse Lab in Milton Keynes and Alex Alderton, Roberto Amato, Jeffrey Barrett, Sonia Goncalves, Ewan Harrison, David K. Jackson, Ian Johnston, Dominic Kwiatkowski, Cordelia Langford, John Sillitoe on behalf of the Wellcome Sanger Institute COVID-19 Surveillance Team |
| EPI_ISL_2910678 | Lighthouse Lab in Glasgow | Wellcome Sanger Institute for the COVID-19 Genomics UK (COG-UK) Consortium | Harper VanSteenhouse, Yumi Kasai, David Gray, Carol Clugston, Anna Dominiczak and Alex Alderton, Roberto Amato, Jeffrey Barrett, Sonia Goncalves, Ewan Harrison, David K. Jackson, Ian Johnston, Dominic Kwiatkowski, Cordelia Langford, John Sillitoe on behalf of the Wellcome Sanger Institute COVID-19 Surveillance Team |
| EPI_ISL_2910679 | Lighthouse Lab in Alderley Park | Wellcome Sanger Institute for the COVID-19 Genomics UK (COG-UK) Consortium | Jacquelyn Wynn, Mairead Hyland, The Lighthouse Lab in Alderley Park and Alex Alderton, Roberto Amato, Jeffrey Barrett, Sonia Goncalves, Ewan Harrison, David K. Jackson, Ian Johnston, Dominic Kwiatkowski, Cordelia Langford, John Sillitoe on behalf of the Wellcome Sanger Institute COVID-19 Surveillance Team |
| EPI_ISL_2910680, EPI_ISL_2910681, EPI_ISL_2910682, EPI_ISL_2910683 | Lighthouse Lab in Milton Keynes | Wellcome Sanger Institute for the COVID-19 Genomics UK (COG-UK) Consortium | The Lighthouse Lab in Milton Keynes and Alex Alderton, Roberto Amato, Jeffrey Barrett, Sonia Goncalves, Ewan Harrison, David K. Jackson, Ian Johnston, Dominic Kwiatkowski, Cordelia Langford, John Sillitoe on behalf of the Wellcome Sanger Institute COVID-19 Surveillance Team |
| EPI_ISL_2910684, EPI_ISL_2910685, EPI_ISL_2910686 | Lighthouse Lab in Glasgow | Wellcome Sanger Institute for the COVID-19 Genomics UK (COG-UK) Consortium | Harper VanSteenhouse, Yumi Kasai, David Gray, Carol Clugston, Anna Dominiczak and Alex Alderton, Roberto Amato, Jeffrey Barrett, Sonia Goncalves, Ewan Harrison, David K. Jackson, Ian Johnston, Dominic Kwiatkowski, Cordelia Langford, John Sillitoe on behalf of the Wellcome Sanger Institute COVID-19 Surveillance Team |
| EPI_ISL_2910687, EPI_ISL_2910688, EPI_ISL_2910689 | Lighthouse Lab in Alderley Park | Wellcome Sanger Institute for the COVID-19 Genomics UK (COG-UK) Consortium | Jacquelyn Wynn, Mairead Hyland, The Lighthouse Lab in Alderley Park and Alex Alderton, Roberto Amato, Jeffrey Barrett, Sonia Goncalves, Ewan Harrison, David K. Jackson, Ian Johnston, Dominic Kwiatkowski, Cordelia Langford, John Sillitoe on behalf of the Wellcome Sanger Institute COVID-19 Surveillance Team |
| EPI_ISL_2910690 | Lighthouse Lab in Glasgow | Wellcome Sanger Institute for the COVID-19 Genomics UK (COG-UK) Consortium | Harper VanSteenhouse, Yumi Kasai, David Gray, Carol Clugston, Anna Dominiczak and Alex Alderton, Roberto Amato, Jeffrey Barrett, Sonia Goncalves, Ewan Harrison, David K. Jackson, Ian Johnston, Dominic Kwiatkowski, Cordelia Langford, John Sillitoe on behalf of the Wellcome Sanger Institute COVID-19 Surveillance Team |
| EPI_ISL_2910691 | Lighthouse Lab in Alderley Park | Wellcome Sanger Institute for the COVID-19 Genomics UK (COG-UK) Consortium | Jacquelyn Wynn, Mairead Hyland, The Lighthouse Lab in Alderley Park and Alex Alderton, Roberto Amato, Jeffrey Barrett, Sonia Goncalves, Ewan Harrison, David K. Jackson, Ian Johnston, Dominic Kwiatkowski, Cordelia Langford, John Sillitoe on behalf of the Wellcome Sanger Institute COVID-19 Surveillance Team |
| EPI_ISL_2910692, EPI_ISL_2910693, EPI_ISL_2910694, EPI_ISL_2910695 | Lighthouse Lab in Milton Keynes | Wellcome Sanger Institute for the COVID-19 Genomics UK (COG-UK) Consortium | The Lighthouse Lab in Milton Keynes and Alex Alderton, Roberto Amato, Jeffrey Barrett, Sonia Goncalves, Ewan Harrison, David K. Jackson, Ian Johnston, Dominic Kwiatkowski, Cordelia Langford, John Sillitoe on behalf of the Wellcome Sanger Institute COVID-19 Surveillance Team |
| EPI_ISL_2910696, EPI_ISL_2910697 | Lighthouse Lab in Alderley Park | Wellcome Sanger Institute for the COVID-19 Genomics UK (COG-UK) Consortium | Jacquelyn Wynn, Mairead Hyland, The Lighthouse Lab in Alderley Park and Alex Alderton, Roberto Amato, Jeffrey Barrett, Sonia Goncalves, Ewan Harrison, David K. Jackson, Ian Johnston, Dominic Kwiatkowski, Cordelia Langford, John Sillitoe on behalf of the Wellcome Sanger Institute COVID-19 Surveillance Team |
| EPI_ISL_2910698, EPI_ISL_2910699 | Lighthouse Lab in Milton Keynes | Wellcome Sanger Institute for the COVID-19 Genomics UK (COG-UK) Consortium | The Lighthouse Lab in Milton Keynes and Alex Alderton, Roberto Amato, Jeffrey Barrett, Sonia Goncalves, Ewan Harrison, David K. Jackson, Ian Johnston, Dominic Kwiatkowski, Cordelia Langford, John Sillitoe on behalf of the Wellcome Sanger Institute COVID-19 Surveillance Team |
| EPI_ISL_2910700 | Lighthouse Lab in Alderley Park | Wellcome Sanger Institute for the COVID-19 Genomics UK (COG-UK) Consortium | Jacquelyn Wynn, Mairead Hyland, The Lighthouse Lab in Alderley Park and Alex Alderton, Roberto Amato, Jeffrey Barrett, Sonia Goncalves, Ewan Harrison, David K. Jackson, Ian Johnston, Dominic Kwiatkowski, Cordelia Langford, John Sillitoe on behalf of the Wellcome Sanger Institute COVID-19 Surveillance Team |
| EPI_ISL_2910701 | Lighthouse Lab in Milton Keynes | Wellcome Sanger Institute for the COVID-19 Genomics UK (COG-UK) Consortium | The Lighthouse Lab in Milton Keynes and Alex Alderton, Roberto Amato, Jeffrey Barrett, Sonia Goncalves, Ewan Harrison, David K. Jackson, Ian Johnston, Dominic Kwiatkowski, Cordelia Langford, John Sillitoe on behalf of the Wellcome Sanger Institute COVID-19 Surveillance Team |
| EPI_ISL_2910702, EPI_ISL_2910703 | Lighthouse Lab in Glasgow | Wellcome Sanger Institute for the COVID-19 Genomics UK (COG-UK) Consortium | Harper VanSteenhouse, Yumi Kasai, David Gray, Carol Clugston, Anna Dominiczak and Alex Alderton, Roberto Amato, Jeffrey Barrett, Sonia Goncalves, Ewan Harrison, David K. Jackson, Ian Johnston, Dominic Kwiatkowski, Cordelia Langford, John Sillitoe on behalf of the Wellcome Sanger Institute COVID-19 Surveillance Team |
| EPI_ISL_2910704, EPI_ISL_2910705, EPI_ISL_2910706, EPI_ISL_2910707, EPI_ISL_2910708, EPI_ISL_2910709 | Lighthouse Lab in Milton Keynes | Wellcome Sanger Institute for the COVID-19 Genomics UK (COG-UK) Consortium | The Lighthouse Lab in Milton Keynes and Alex Alderton, Roberto Amato, Jeffrey Barrett, Sonia Goncalves, Ewan Harrison, David K. Jackson, Ian Johnston, Dominic Kwiatkowski, Cordelia Langford, John Sillitoe on behalf of the Wellcome Sanger Institute COVID-19 Surveillance Team |

| | | | |
|---|---|---|---|
| EPI_ISL_2910710 | Lighthouse Lab in Glasgow | Wellcome Sanger Institute for the COVID-19 Genomics UK (COG-UK) Consortium | Harper VanSteenhouse, Yumi Kasai, David Gray, Carol Clugston, Anna Dominiczak and Alex Alderton, Roberto Amato, Jeffrey Barrett, Sonia Goncalves, Ewan Harrison, David K. Jackson, Ian Johnston, Dominic Kwiatkowski, Cordelia Langford, John Sillitoe on behalf of the Wellcome Sanger Institute COVID-19 Surveillance Team |
| EPI_ISL_2910711 | Lighthouse Lab in Milton Keynes | Wellcome Sanger Institute for the COVID-19 Genomics UK (COG-UK) Consortium | The Lighthouse Lab in Milton Keynes and Alex Alderton, Roberto Amato, Jeffrey Barrett, Sonia Goncalves, Ewan Harrison, David K. Jackson, Ian Johnston, Dominic Kwiatkowski, Cordelia Langford, John Sillitoe on behalf of the Wellcome Sanger Institute COVID-19 Surveillance Team |
| EPI_ISL_2910712 | Lighthouse Lab in Alderley Park | Wellcome Sanger Institute for the COVID-19 Genomics UK (COG-UK) Consortium | Jacquelyn Wynn, Mairead Hyland, The Lighthouse Lab in Alderley Park and Alex Alderton, Roberto Amato, Jeffrey Barrett, Sonia Goncalves, Ewan Harrison, David K. Jackson, Ian Johnston, Dominic Kwiatkowski, Cordelia Langford, John Sillitoe on behalf of the Wellcome Sanger Institute COVID-19 Surveillance Team |
| EPI_ISL_2910713 | Lighthouse Lab in Milton Keynes | Wellcome Sanger Institute for the COVID-19 Genomics UK (COG-UK) Consortium | The Lighthouse Lab in Milton Keynes and Alex Alderton, Roberto Amato, Jeffrey Barrett, Sonia Goncalves, Ewan Harrison, David K. Jackson, Ian Johnston, Dominic Kwiatkowski, Cordelia Langford, John Sillitoe on behalf of the Wellcome Sanger Institute COVID-19 Surveillance Team |
| EPI_ISL_2910714 | Lighthouse Lab in Glasgow | Wellcome Sanger Institute for the COVID-19 Genomics UK (COG-UK) Consortium | Harper VanSteenhouse, Yumi Kasai, David Gray, Carol Clugston, Anna Dominiczak and Alex Alderton, Roberto Amato, Jeffrey Barrett, Sonia Goncalves, Ewan Harrison, David K. Jackson, Ian Johnston, Dominic Kwiatkowski, Cordelia Langford, John Sillitoe on behalf of the Wellcome Sanger Institute COVID-19 Surveillance Team |
| EPI_ISL_2910715, EPI_ISL_2910716, EPI_ISL_2910717, EPI_ISL_2910718, EPI_ISL_2910719 | Lighthouse Lab in Milton Keynes | Wellcome Sanger Institute for the COVID-19 Genomics UK (COG-UK) Consortium | The Lighthouse Lab in Milton Keynes and Alex Alderton, Roberto Amato, Jeffrey Barrett, Sonia Goncalves, Ewan Harrison, David K. Jackson, Ian Johnston, Dominic Kwiatkowski, Cordelia Langford, John Sillitoe on behalf of the Wellcome Sanger Institute COVID-19 Surveillance Team |
| EPI_ISL_2910720 | Lighthouse Lab in Glasgow | Wellcome Sanger Institute for the COVID-19 Genomics UK (COG-UK) Consortium | Harper VanSteenhouse, Yumi Kasai, David Gray, Carol Clugston, Anna Dominiczak and Alex Alderton, Roberto Amato, Jeffrey Barrett, Sonia Goncalves, Ewan Harrison, David K. Jackson, Ian Johnston, Dominic Kwiatkowski, Cordelia Langford, John Sillitoe on behalf of the Wellcome Sanger Institute COVID-19 Surveillance Team |
| EPI_ISL_2910721, EPI_ISL_2910722 | Lighthouse Lab in Alderley Park | Wellcome Sanger Institute for the COVID-19 Genomics UK (COG-UK) Consortium | Jacquelyn Wynn, Mairead Hyland, The Lighthouse Lab in Alderley Park and Alex Alderton, Roberto Amato, Jeffrey Barrett, Sonia Goncalves, Ewan Harrison, David K. Jackson, Ian Johnston, Dominic Kwiatkowski, Cordelia Langford, John Sillitoe on behalf of the Wellcome Sanger Institute COVID-19 Surveillance Team |
| EPI_ISL_2910723, EPI_ISL_2910724 | Lighthouse Lab in Milton Keynes | Wellcome Sanger Institute for the COVID-19 Genomics UK (COG-UK) Consortium | The Lighthouse Lab in Milton Keynes and Alex Alderton, Roberto Amato, Jeffrey Barrett, Sonia Goncalves, Ewan Harrison, David K. Jackson, Ian Johnston, Dominic Kwiatkowski, Cordelia Langford, John Sillitoe on behalf of the Wellcome Sanger Institute COVID-19 Surveillance Team |
| EPI_ISL_2910725 | Lighthouse Lab in Alderley Park | Wellcome Sanger Institute for the COVID-19 Genomics UK (COG-UK) Consortium | Jacquelyn Wynn, Mairead Hyland, The Lighthouse Lab in Alderley Park and Alex Alderton, Roberto Amato, Jeffrey Barrett, Sonia Goncalves, Ewan Harrison, David K. Jackson, Ian Johnston, Dominic Kwiatkowski, Cordelia Langford, John Sillitoe on behalf of the Wellcome Sanger Institute COVID-19 Surveillance Team |
| EPI_ISL_2910726, EPI_ISL_2910727, EPI_ISL_2910728, EPI_ISL_2910729, EPI_ISL_2910730 | Lighthouse Lab in Milton Keynes | Wellcome Sanger Institute for the COVID-19 Genomics UK (COG-UK) Consortium | The Lighthouse Lab in Milton Keynes and Alex Alderton, Roberto Amato, Jeffrey Barrett, Sonia Goncalves, Ewan Harrison, David K. Jackson, Ian Johnston, Dominic Kwiatkowski, Cordelia Langford, John Sillitoe on behalf of the Wellcome Sanger Institute COVID-19 Surveillance Team |
| EPI_ISL_2910731 | Lighthouse Lab in Glasgow | Wellcome Sanger Institute for the COVID-19 Genomics UK (COG-UK) Consortium | Harper VanSteenhouse, Yumi Kasai, David Gray, Carol Clugston, Anna Dominiczak and Alex Alderton, Roberto Amato, Jeffrey Barrett, Sonia Goncalves, Ewan Harrison, David K. Jackson, Ian Johnston, Dominic Kwiatkowski, Cordelia Langford, John Sillitoe on behalf of the Wellcome Sanger Institute COVID-19 Surveillance Team |
| EPI_ISL_2910732 | Lighthouse Lab in Milton Keynes | Wellcome Sanger Institute for the COVID-19 Genomics UK (COG-UK) Consortium | The Lighthouse Lab in Milton Keynes and Alex Alderton, Roberto Amato, Jeffrey Barrett, Sonia Goncalves, Ewan Harrison, David K. Jackson, Ian Johnston, Dominic Kwiatkowski, Cordelia Langford, John Sillitoe on behalf of the Wellcome Sanger Institute COVID-19 Surveillance Team |
| EPI_ISL_2910733 | Lighthouse Lab in Alderley Park | Wellcome Sanger Institute for the COVID-19 Genomics UK (COG-UK) Consortium | Jacquelyn Wynn, Mairead Hyland, The Lighthouse Lab in Alderley Park and Alex Alderton, Roberto Amato, Jeffrey Barrett, Sonia Goncalves, Ewan Harrison, David K. Jackson, Ian Johnston, Dominic Kwiatkowski, Cordelia Langford, John Sillitoe on behalf of the Wellcome Sanger Institute COVID-19 Surveillance Team |
| EPI_ISL_2910734 | Lighthouse Lab in Milton Keynes | Wellcome Sanger Institute for the COVID-19 Genomics UK (COG-UK) Consortium | The Lighthouse Lab in Milton Keynes and Alex Alderton, Roberto Amato, Jeffrey Barrett, Sonia Goncalves, Ewan Harrison, David K. Jackson, Ian Johnston, Dominic Kwiatkowski, Cordelia Langford, John Sillitoe on behalf of the Wellcome Sanger Institute COVID-19 Surveillance Team |
| EPI_ISL_2910735 | Lighthouse Lab in Alderley Park | Wellcome Sanger Institute for the COVID-19 Genomics UK (COG-UK) Consortium | Jacquelyn Wynn, Mairead Hyland, The Lighthouse Lab in Alderley Park and Alex Alderton, Roberto Amato, Jeffrey Barrett, Sonia Goncalves, Ewan Harrison, David K. Jackson, Ian Johnston, Dominic Kwiatkowski, Cordelia Langford, John Sillitoe on behalf of the Wellcome Sanger Institute COVID-19 Surveillance Team |
| EPI_ISL_2910736, EPI_ISL_2910737 | Lighthouse Lab in Milton Keynes | Wellcome Sanger Institute for the COVID-19 Genomics UK (COG-UK) Consortium | The Lighthouse Lab in Milton Keynes and Alex Alderton, Roberto Amato, Jeffrey Barrett, Sonia Goncalves, Ewan Harrison, David K. Jackson, Ian Johnston, Dominic Kwiatkowski, Cordelia Langford, John Sillitoe on behalf of the Wellcome Sanger Institute COVID-19 Surveillance Team |
| EPI_ISL_2910738 | Lighthouse Lab in Alderley Park | Wellcome Sanger Institute for the COVID-19 Genomics UK (COG-UK) Consortium | Jacquelyn Wynn, Mairead Hyland, The Lighthouse Lab in Alderley Park and Alex Alderton, Roberto Amato, Jeffrey Barrett, Sonia Goncalves, Ewan Harrison, David K. Jackson, Ian Johnston, Dominic Kwiatkowski, Cordelia Langford, John Sillitoe on behalf of the Wellcome Sanger Institute COVID-19 Surveillance Team |
| EPI_ISL_2910739, EPI_ISL_2910740 | Lighthouse Lab in Milton Keynes | Wellcome Sanger Institute for the COVID-19 Genomics UK (COG-UK) Consortium | The Lighthouse Lab in Milton Keynes and Alex Alderton, Roberto Amato, Jeffrey Barrett, Sonia Goncalves, Ewan Harrison, David K. Jackson, Ian Johnston, Dominic Kwiatkowski, Cordelia Langford, John Sillitoe on behalf of the Wellcome Sanger Institute COVID-19 Surveillance Team |
| EPI_ISL_2910741 | Lighthouse Lab in Alderley Park | Wellcome Sanger Institute for the COVID-19 Genomics UK (COG-UK) Consortium | Jacquelyn Wynn, Mairead Hyland, The Lighthouse Lab in Alderley Park and Alex Alderton, Roberto Amato, Jeffrey Barrett, Sonia Goncalves, Ewan Harrison, David K. Jackson, Ian Johnston, Dominic Kwiatkowski, Cordelia Langford, John Sillitoe on behalf of the Wellcome Sanger Institute COVID-19 Surveillance Team |
| EPI_ISL_2910742, EPI_ISL_2910743, EPI_ISL_2910744, EPI_ISL_2910745, EPI_ISL_2910746 | Lighthouse Lab in Milton Keynes | Wellcome Sanger Institute for the COVID-19 Genomics UK (COG-UK) Consortium | The Lighthouse Lab in Milton Keynes and Alex Alderton, Roberto Amato, Jeffrey Barrett, Sonia Goncalves, Ewan Harrison, David K. Jackson, Ian Johnston, Dominic Kwiatkowski, Cordelia Langford, John Sillitoe on behalf of the Wellcome Sanger Institute COVID-19 Surveillance Team |
| EPI_ISL_2910747 | Lighthouse Lab in Glasgow | Wellcome Sanger Institute for the COVID-19 Genomics UK (COG-UK) Consortium | Harper VanSteenhouse, Yumi Kasai, David Gray, Carol Clugston, Anna Dominiczak and Alex Alderton, Roberto Amato, Jeffrey Barrett, Sonia Goncalves, Ewan Harrison, David K. Jackson, Ian Johnston, Dominic Kwiatkowski, Cordelia Langford, John Sillitoe on behalf of the Wellcome Sanger Institute COVID-19 Surveillance Team |
| EPI_ISL_2910748, EPI_ISL_2910749 | Lighthouse Lab in Milton Keynes | Wellcome Sanger Institute for the COVID-19 Genomics UK (COG-UK) Consortium | The Lighthouse Lab in Milton Keynes and Alex Alderton, Roberto Amato, Jeffrey Barrett, Sonia Goncalves, Ewan Harrison, David K. Jackson, Ian Johnston, Dominic Kwiatkowski, Cordelia Langford, John Sillitoe on behalf of the Wellcome Sanger Institute COVID-19 Surveillance Team |
| EPI_ISL_2910750 | Lighthouse Lab in Alderley Park | Wellcome Sanger Institute for the COVID-19 Genomics UK (COG-UK) Consortium | Jacquelyn Wynn, Mairead Hyland, The Lighthouse Lab in Alderley Park and Alex Alderton, Roberto Amato, Jeffrey Barrett, Sonia Goncalves, Ewan Harrison, David K. Jackson, Ian Johnston, Dominic Kwiatkowski, Cordelia Langford, John Sillitoe on behalf of the Wellcome Sanger Institute COVID-19 Surveillance Team |
| EPI_ISL_2910751 | Lighthouse Lab in Milton Keynes | Wellcome Sanger Institute for the COVID-19 Genomics UK (COG-UK) Consortium | The Lighthouse Lab in Milton Keynes and Alex Alderton, Roberto Amato, Jeffrey Barrett, Sonia Goncalves, Ewan Harrison, David K. Jackson, Ian Johnston, Dominic Kwiatkowski, Cordelia Langford, John Sillitoe on behalf of the Wellcome Sanger Institute COVID-19 Surveillance Team |
| EPI_ISL_2910752 | Lighthouse Lab in Alderley Park | Wellcome Sanger Institute for the COVID-19 Genomics UK (COG-UK) Consortium | Jacquelyn Wynn, Mairead Hyland, The Lighthouse Lab in Alderley Park and Alex Alderton, Roberto Amato, Jeffrey Barrett, Sonia Goncalves, Ewan Harrison, David K. Jackson, Ian Johnston, Dominic Kwiatkowski, Cordelia Langford, John Sillitoe on behalf of the Wellcome Sanger Institute COVID-19 Surveillance Team |
| EPI_ISL_2910753, EPI_ISL_2910754 | Lighthouse Lab in Milton Keynes | Wellcome Sanger Institute for the COVID-19 Genomics UK (COG-UK) Consortium | The Lighthouse Lab in Milton Keynes and Alex Alderton, Roberto Amato, Jeffrey Barrett, Sonia Goncalves, Ewan Harrison, David K. Jackson, Ian Johnston, Dominic Kwiatkowski, Cordelia Langford, John Sillitoe on behalf of the Wellcome Sanger Institute COVID-19 Surveillance Team |
| EPI_ISL_2910755 | Lighthouse Lab in Glasgow | Wellcome Sanger Institute for the COVID-19 Genomics UK (COG-UK) Consortium | Harper VanSteenhouse, Yumi Kasai, David Gray, Carol Clugston, Anna Dominiczak and Alex Alderton, Roberto Amato, Jeffrey Barrett, Sonia Goncalves, Ewan Harrison, David K. Jackson, Ian Johnston, Dominic Kwiatkowski, Cordelia Langford, John Sillitoe on behalf of the Wellcome Sanger Institute COVID-19 Surveillance Team |
| EPI_ISL_2910756 | Lighthouse Lab in Milton Keynes | Wellcome Sanger Institute for the COVID-19 Genomics UK (COG-UK) Consortium | The Lighthouse Lab in Milton Keynes and Alex Alderton, Roberto Amato, Jeffrey Barrett, Sonia Goncalves, Ewan Harrison, David K. Jackson, Ian Johnston, Dominic Kwiatkowski, Cordelia Langford, John Sillitoe on behalf of the Wellcome Sanger Institute COVID-19 Surveillance Team |
| EPI_ISL_2910757 | Lighthouse Lab in Alderley Park | Wellcome Sanger Institute for the COVID-19 Genomics UK (COG-UK) Consortium | Jacquelyn Wynn, Mairead Hyland, The Lighthouse Lab in Alderley Park and Alex Alderton, Roberto Amato, Jeffrey Barrett, Sonia Goncalves, Ewan Harrison, David K. Jackson, Ian Johnston, Dominic Kwiatkowski, Cordelia Langford, John Sillitoe on behalf of the Wellcome Sanger Institute COVID-19 Surveillance Team |
| EPI_ISL_2910758 | Lighthouse Lab in Glasgow | Wellcome Sanger Institute for the COVID-19 Genomics UK (COG-UK) Consortium | Harper VanSteenhouse, Yumi Kasai, David Gray, Carol Clugston, Anna Dominiczak and Alex Alderton, Roberto Amato, Jeffrey Barrett, Sonia Goncalves, Ewan Harrison, David K. Jackson, Ian Johnston, Dominic Kwiatkowski, Cordelia Langford, John Sillitoe on behalf of the Wellcome Sanger Institute COVID-19 Surveillance Team |
| EPI_ISL_2910759, EPI_ISL_2910760, EPI_ISL_2910761, EPI_ISL_2910762 | Lighthouse Lab in Milton Keynes | Wellcome Sanger Institute for the COVID-19 Genomics UK (COG-UK) Consortium | The Lighthouse Lab in Milton Keynes and Alex Alderton, Roberto Amato, Jeffrey Barrett, Sonia Goncalves, Ewan Harrison, David K. Jackson, Ian Johnston, Dominic Kwiatkowski, Cordelia Langford, John Sillitoe on behalf of the Wellcome Sanger Institute COVID-19 Surveillance Team |
| EPI_ISL_2910763 | Lighthouse Lab in Alderley Park | Wellcome Sanger Institute for the COVID-19 Genomics UK (COG-UK) Consortium | Jacquelyn Wynn, Mairead Hyland, The Lighthouse Lab in Alderley Park and Alex Alderton, Roberto Amato, Jeffrey Barrett, Sonia Goncalves, Ewan Harrison, David K. Jackson, Ian Johnston, Dominic Kwiatkowski, Cordelia Langford, John Sillitoe on behalf of the Wellcome Sanger Institute COVID-19 Surveillance Team |
| EPI_ISL_2910764 | Lighthouse Lab in Milton Keynes | Wellcome Sanger Institute for the COVID-19 Genomics UK (COG-UK) Consortium | The Lighthouse Lab in Milton Keynes and Alex Alderton, Roberto Amato, Jeffrey Barrett, Sonia Goncalves, Ewan Harrison, David K. Jackson, Ian Johnston, Dominic Kwiatkowski, Cordelia Langford, John Sillitoe on behalf of the Wellcome Sanger Institute COVID-19 Surveillance Team |
| EPI_ISL_2910765 | Lighthouse Lab in Alderley Park | Wellcome Sanger Institute for the COVID-19 Genomics UK (COG-UK) Consortium | Jacquelyn Wynn, Mairead Hyland, The Lighthouse Lab in Alderley Park and Alex Alderton, Roberto Amato, Jeffrey Barrett, Sonia Goncalves, Ewan Harrison, David K. Jackson, Ian Johnston, Dominic Kwiatkowski, Cordelia Langford, John Sillitoe on behalf of the Wellcome Sanger Institute COVID-19 Surveillance Team |
| EPI_ISL_2910766, EPI_ISL_2910767, EPI_ISL_2910768, EPI_ISL_2910769, EPI_ISL_2910770, EPI_ISL_2910771 | Lighthouse Lab in Milton Keynes | Wellcome Sanger Institute for the COVID-19 Genomics UK (COG-UK) Consortium | The Lighthouse Lab in Milton Keynes and Alex Alderton, Roberto Amato, Jeffrey Barrett, Sonia Goncalves, Ewan Harrison, David K. Jackson, Ian Johnston, Dominic Kwiatkowski, Cordelia Langford, John Sillitoe on behalf of the Wellcome Sanger Institute COVID-19 Surveillance Team |
| EPI_ISL_2910772 | Lighthouse Lab in Alderley Park | Wellcome Sanger Institute for the COVID-19 Genomics UK (COG-UK) Consortium | Jacquelyn Wynn, Mairead Hyland, The Lighthouse Lab in Alderley Park and Alex Alderton, Roberto Amato, Jeffrey Barrett, Sonia Goncalves, Ewan Harrison, David K. Jackson, Ian Johnston, Dominic Kwiatkowski, Cordelia Langford, John Sillitoe on behalf of the Wellcome Sanger Institute COVID-19 Surveillance Team |
| EPI_ISL_2910773 | Lighthouse Lab in Milton Keynes | Wellcome Sanger Institute for the COVID-19 Genomics UK (COG-UK) Consortium | The Lighthouse Lab in Milton Keynes and Alex Alderton, Roberto Amato, Jeffrey Barrett, Sonia Goncalves, Ewan Harrison, David K. Jackson, Ian Johnston, Dominic Kwiatkowski, Cordelia Langford, John Sillitoe on behalf of the Wellcome Sanger Institute COVID-19 Surveillance Team |
| EPI_ISL_2910774, EPI_ISL_2910775 | Lighthouse Lab in Alderley Park | Wellcome Sanger Institute for the COVID-19 Genomics UK (COG-UK) Consortium | Jacquelyn Wynn, Mairead Hyland, The Lighthouse Lab in Alderley Park and Alex Alderton, Roberto Amato, Jeffrey Barrett, Sonia Goncalves, Ewan Harrison, David K. Jackson, Ian Johnston, Dominic Kwiatkowski, Cordelia Langford, John Sillitoe on behalf of the Wellcome Sanger Institute COVID-19 Surveillance Team |

| | | | |
|---|---|---|---|
| EPI_ISL_2910776, EPI_ISL_2910777 | Lighthouse Lab in Milton Keynes | Wellcome Sanger Institute for the COVID-19 Genomics UK (COG-UK) Consortium | The Lighthouse Lab in Milton Keynes and Alex Alderton, Roberto Amato, Jeffrey Barrett, Sonia Goncalves, Ewan Harrison, David K. Jackson, Ian Johnston, Dominic Kwiatkowski, Cordelia Langford, John Sillitoe on behalf of the Wellcome Sanger Institute COVID-19 Surveillance Team |
| EPI_ISL_2910778 | Lighthouse Lab in Glasgow | Wellcome Sanger Institute for the COVID-19 Genomics UK (COG-UK) Consortium | Harper VanSteenhouse, Yumi Kasai, David Gray, Carol Clugston, Anna Dominiczak and Alex Alderton, Roberto Amato, Jeffrey Barrett, Sonia Goncalves, Ewan Harrison, David K. Jackson, Ian Johnston, Dominic Kwiatkowski, Cordelia Langford, John Sillitoe on behalf of the Wellcome Sanger Institute COVID-19 Surveillance Team |
| EPI_ISL_2910779, EPI_ISL_2910780 | Lighthouse Lab in Alderley Park | Wellcome Sanger Institute for the COVID-19 Genomics UK (COG-UK) Consortium | Jacquelyn Wynn, Mairead Hyland, The Lighthouse Lab in Alderley Park and Alex Alderton, Roberto Amato, Jeffrey Barrett, Sonia Goncalves, Ewan Harrison, David K. Jackson, Ian Johnston, Dominic Kwiatkowski, Cordelia Langford, John Sillitoe on behalf of the Wellcome Sanger Institute COVID-19 Surveillance Team |
| EPI_ISL_2910781, EPI_ISL_2910782 | Lighthouse Lab in Milton Keynes | Wellcome Sanger Institute for the COVID-19 Genomics UK (COG-UK) Consortium | The Lighthouse Lab in Milton Keynes and Alex Alderton, Roberto Amato, Jeffrey Barrett, Sonia Goncalves, Ewan Harrison, David K. Jackson, Ian Johnston, Dominic Kwiatkowski, Cordelia Langford, John Sillitoe on behalf of the Wellcome Sanger Institute COVID-19 Surveillance Team |
| EPI_ISL_2910783 | Lighthouse Lab in Alderley Park | Wellcome Sanger Institute for the COVID-19 Genomics UK (COG-UK) Consortium | Jacquelyn Wynn, Mairead Hyland, The Lighthouse Lab in Alderley Park and Alex Alderton, Roberto Amato, Jeffrey Barrett, Sonia Goncalves, Ewan Harrison, David K. Jackson, Ian Johnston, Dominic Kwiatkowski, Cordelia Langford, John Sillitoe on behalf of the Wellcome Sanger Institute COVID-19 Surveillance Team |
| EPI_ISL_2910784, EPI_ISL_2910785, EPI_ISL_2910786, EPI_ISL_2910787, EPI_ISL_2910788, EPI_ISL_2910789, EPI_ISL_2910790, EPI_ISL_2910791 | Lighthouse Lab in Milton Keynes | Wellcome Sanger Institute for the COVID-19 Genomics UK (COG-UK) Consortium | The Lighthouse Lab in Milton Keynes and Alex Alderton, Roberto Amato, Jeffrey Barrett, Sonia Goncalves, Ewan Harrison, David K. Jackson, Ian Johnston, Dominic Kwiatkowski, Cordelia Langford, John Sillitoe on behalf of the Wellcome Sanger Institute COVID-19 Surveillance Team |
| EPI_ISL_2910792 | Lighthouse Lab in Glasgow | Wellcome Sanger Institute for the COVID-19 Genomics UK (COG-UK) Consortium | Harper VanSteenhouse, Yumi Kasai, David Gray, Carol Clugston, Anna Dominiczak and Alex Alderton, Roberto Amato, Jeffrey Barrett, Sonia Goncalves, Ewan Harrison, David K. Jackson, Ian Johnston, Dominic Kwiatkowski, Cordelia Langford, John Sillitoe on behalf of the Wellcome Sanger Institute COVID-19 Surveillance Team |
| EPI_ISL_2910793, EPI_ISL_2910794, EPI_ISL_2910795 | Lighthouse Lab in Milton Keynes | Wellcome Sanger Institute for the COVID-19 Genomics UK (COG-UK) Consortium | The Lighthouse Lab in Milton Keynes and Alex Alderton, Roberto Amato, Jeffrey Barrett, Sonia Goncalves, Ewan Harrison, David K. Jackson, Ian Johnston, Dominic Kwiatkowski, Cordelia Langford, John Sillitoe on behalf of the Wellcome Sanger Institute COVID-19 Surveillance Team |
| EPI_ISL_2910796 | Lighthouse Lab in Alderley Park | Wellcome Sanger Institute for the COVID-19 Genomics UK (COG-UK) Consortium | Jacquelyn Wynn, Mairead Hyland, The Lighthouse Lab in Alderley Park and Alex Alderton, Roberto Amato, Jeffrey Barrett, Sonia Goncalves, Ewan Harrison, David K. Jackson, Ian Johnston, Dominic Kwiatkowski, Cordelia Langford, John Sillitoe on behalf of the Wellcome Sanger Institute COVID-19 Surveillance Team |
| EPI_ISL_2910797, EPI_ISL_2910798, EPI_ISL_2910799 | Lighthouse Lab in Milton Keynes | Wellcome Sanger Institute for the COVID-19 Genomics UK (COG-UK) Consortium | The Lighthouse Lab in Milton Keynes and Alex Alderton, Roberto Amato, Jeffrey Barrett, Sonia Goncalves, Ewan Harrison, David K. Jackson, Ian Johnston, Dominic Kwiatkowski, Cordelia Langford, John Sillitoe on behalf of the Wellcome Sanger Institute COVID-19 Surveillance Team |
| EPI_ISL_2910800 | Lighthouse Lab in Glasgow | Wellcome Sanger Institute for the COVID-19 Genomics UK (COG-UK) Consortium | Harper VanSteenhouse, Yumi Kasai, David Gray, Carol Clugston, Anna Dominiczak and Alex Alderton, Roberto Amato, Jeffrey Barrett, Sonia Goncalves, Ewan Harrison, David K. Jackson, Ian Johnston, Dominic Kwiatkowski, Cordelia Langford, John Sillitoe on behalf of the Wellcome Sanger Institute COVID-19 Surveillance Team |
| EPI_ISL_2910801 | Lighthouse Lab in Milton Keynes | Wellcome Sanger Institute for the COVID-19 Genomics UK (COG-UK) Consortium | The Lighthouse Lab in Milton Keynes and Alex Alderton, Roberto Amato, Jeffrey Barrett, Sonia Goncalves, Ewan Harrison, David K. Jackson, Ian Johnston, Dominic Kwiatkowski, Cordelia Langford, John Sillitoe on behalf of the Wellcome Sanger Institute COVID-19 Surveillance Team |
| EPI_ISL_2910802, EPI_ISL_2910803 | Lighthouse Lab in Alderley Park | Wellcome Sanger Institute for the COVID-19 Genomics UK (COG-UK) Consortium | Jacquelyn Wynn, Mairead Hyland, The Lighthouse Lab in Alderley Park and Alex Alderton, Roberto Amato, Jeffrey Barrett, Sonia Goncalves, Ewan Harrison, David K. Jackson, Ian Johnston, Dominic Kwiatkowski, Cordelia Langford, John Sillitoe on behalf of the Wellcome Sanger Institute COVID-19 Surveillance Team |
| EPI_ISL_2910804, EPI_ISL_2910805 | Lighthouse Lab in Milton Keynes | Wellcome Sanger Institute for the COVID-19 Genomics UK (COG-UK) Consortium | The Lighthouse Lab in Milton Keynes and Alex Alderton, Roberto Amato, Jeffrey Barrett, Sonia Goncalves, Ewan Harrison, David K. Jackson, Ian Johnston, Dominic Kwiatkowski, Cordelia Langford, John Sillitoe on behalf of the Wellcome Sanger Institute COVID-19 Surveillance Team |
| EPI_ISL_2910806 | Lighthouse Lab in Alderley Park | Wellcome Sanger Institute for the COVID-19 Genomics UK (COG-UK) Consortium | Jacquelyn Wynn, Mairead Hyland, The Lighthouse Lab in Alderley Park and Alex Alderton, Roberto Amato, Jeffrey Barrett, Sonia Goncalves, Ewan Harrison, David K. Jackson, Ian Johnston, Dominic Kwiatkowski, Cordelia Langford, John Sillitoe on behalf of the Wellcome Sanger Institute COVID-19 Surveillance Team |
| EPI_ISL_2910808 | Lighthouse Lab in Milton Keynes | Wellcome Sanger Institute for the COVID-19 Genomics UK (COG-UK) Consortium | The Lighthouse Lab in Milton Keynes and Alex Alderton, Roberto Amato, Jeffrey Barrett, Sonia Goncalves, Ewan Harrison, David K. Jackson, Ian Johnston, Dominic Kwiatkowski, Cordelia Langford, John Sillitoe on behalf of the Wellcome Sanger Institute COVID-19 Surveillance Team |
| EPI_ISL_2910809 | Lighthouse Lab in Alderley Park | Wellcome Sanger Institute for the COVID-19 Genomics UK (COG-UK) Consortium | Jacquelyn Wynn, Mairead Hyland, The Lighthouse Lab in Alderley Park and Alex Alderton, Roberto Amato, Jeffrey Barrett, Sonia Goncalves, Ewan Harrison, David K. Jackson, Ian Johnston, Dominic Kwiatkowski, Cordelia Langford, John Sillitoe on behalf of the Wellcome Sanger Institute COVID-19 Surveillance Team |
| EPI_ISL_2910810, EPI_ISL_2910811, EPI_ISL_2910812 | Lighthouse Lab in Milton Keynes | Wellcome Sanger Institute for the COVID-19 Genomics UK (COG-UK) Consortium | The Lighthouse Lab in Milton Keynes and Alex Alderton, Roberto Amato, Jeffrey Barrett, Sonia Goncalves, Ewan Harrison, David K. Jackson, Ian Johnston, Dominic Kwiatkowski, Cordelia Langford, John Sillitoe on behalf of the Wellcome Sanger Institute COVID-19 Surveillance Team |
| EPI_ISL_2910813 | Lighthouse Lab in Alderley Park | Wellcome Sanger Institute for the COVID-19 Genomics UK (COG-UK) Consortium | Jacquelyn Wynn, Mairead Hyland, The Lighthouse Lab in Alderley Park and Alex Alderton, Roberto Amato, Jeffrey Barrett, Sonia Goncalves, Ewan Harrison, David K. Jackson, Ian Johnston, Dominic Kwiatkowski, Cordelia Langford, John Sillitoe on behalf of the Wellcome Sanger Institute COVID-19 Surveillance Team |
| EPI_ISL_2910814, EPI_ISL_2910815 | Lighthouse Lab in Milton Keynes | Wellcome Sanger Institute for the COVID-19 Genomics UK (COG-UK) Consortium | The Lighthouse Lab in Milton Keynes and Alex Alderton, Roberto Amato, Jeffrey Barrett, Sonia Goncalves, Ewan Harrison, David K. Jackson, Ian Johnston, Dominic Kwiatkowski, Cordelia Langford, John Sillitoe on behalf of the Wellcome Sanger Institute COVID-19 Surveillance Team |
| EPI_ISL_2910816 | Lighthouse Lab in Glasgow | Wellcome Sanger Institute for the COVID-19 Genomics UK (COG-UK) Consortium | Harper VanSteenhouse, Yumi Kasai, David Gray, Carol Clugston, Anna Dominiczak and Alex Alderton, Roberto Amato, Jeffrey Barrett, Sonia Goncalves, Ewan Harrison, David K. Jackson, Ian Johnston, Dominic Kwiatkowski, Cordelia Langford, John Sillitoe on behalf of the Wellcome Sanger Institute COVID-19 Surveillance Team |
| EPI_ISL_2910817 | Lighthouse Lab in Alderley Park | Wellcome Sanger Institute for the COVID-19 Genomics UK (COG-UK) Consortium | Jacquelyn Wynn, Mairead Hyland, The Lighthouse Lab in Alderley Park and Alex Alderton, Roberto Amato, Jeffrey Barrett, Sonia Goncalves, Ewan Harrison, David K. Jackson, Ian Johnston, Dominic Kwiatkowski, Cordelia Langford, John Sillitoe on behalf of the Wellcome Sanger Institute COVID-19 Surveillance Team |
| EPI_ISL_2910818 | Lighthouse Lab in Milton Keynes | Wellcome Sanger Institute for the COVID-19 Genomics UK (COG-UK) Consortium | The Lighthouse Lab in Milton Keynes and Alex Alderton, Roberto Amato, Jeffrey Barrett, Sonia Goncalves, Ewan Harrison, David K. Jackson, Ian Johnston, Dominic Kwiatkowski, Cordelia Langford, John Sillitoe on behalf of the Wellcome Sanger Institute COVID-19 Surveillance Team |
| EPI_ISL_2910819, EPI_ISL_2910820 | Lighthouse Lab in Glasgow | Wellcome Sanger Institute for the COVID-19 Genomics UK (COG-UK) Consortium | Harper VanSteenhouse, Yumi Kasai, David Gray, Carol Clugston, Anna Dominiczak and Alex Alderton, Roberto Amato, Jeffrey Barrett, Sonia Goncalves, Ewan Harrison, David K. Jackson, Ian Johnston, Dominic Kwiatkowski, Cordelia Langford, John Sillitoe on behalf of the Wellcome Sanger Institute COVID-19 Surveillance Team |
| EPI_ISL_2910821, EPI_ISL_2910822 | Lighthouse Lab in Milton Keynes | Wellcome Sanger Institute for the COVID-19 Genomics UK (COG-UK) Consortium | The Lighthouse Lab in Milton Keynes and Alex Alderton, Roberto Amato, Jeffrey Barrett, Sonia Goncalves, Ewan Harrison, David K. Jackson, Ian Johnston, Dominic Kwiatkowski, Cordelia Langford, John Sillitoe on behalf of the Wellcome Sanger Institute COVID-19 Surveillance Team |
| EPI_ISL_2910823 | Lighthouse Lab in Alderley Park | Wellcome Sanger Institute for the COVID-19 Genomics UK (COG-UK) Consortium | Jacquelyn Wynn, Mairead Hyland, The Lighthouse Lab in Alderley Park and Alex Alderton, Roberto Amato, Jeffrey Barrett, Sonia Goncalves, Ewan Harrison, David K. Jackson, Ian Johnston, Dominic Kwiatkowski, Cordelia Langford, John Sillitoe on behalf of the Wellcome Sanger Institute COVID-19 Surveillance Team |
| EPI_ISL_2910824 | Lighthouse Lab in Milton Keynes | Wellcome Sanger Institute for the COVID-19 Genomics UK (COG-UK) Consortium | The Lighthouse Lab in Milton Keynes and Alex Alderton, Roberto Amato, Jeffrey Barrett, Sonia Goncalves, Ewan Harrison, David K. Jackson, Ian Johnston, Dominic Kwiatkowski, Cordelia Langford, John Sillitoe on behalf of the Wellcome Sanger Institute COVID-19 Surveillance Team |
| EPI_ISL_2910825 | Lighthouse Lab in Glasgow | Wellcome Sanger Institute for the COVID-19 Genomics UK (COG-UK) Consortium | Harper VanSteenhouse, Yumi Kasai, David Gray, Carol Clugston, Anna Dominiczak and Alex Alderton, Roberto Amato, Jeffrey Barrett, Sonia Goncalves, Ewan Harrison, David K. Jackson, Ian Johnston, Dominic Kwiatkowski, Cordelia Langford, John Sillitoe on behalf of the Wellcome Sanger Institute COVID-19 Surveillance Team |
| EPI_ISL_2910826 | Lighthouse Lab in Milton Keynes | Wellcome Sanger Institute for the COVID-19 Genomics UK (COG-UK) Consortium | The Lighthouse Lab in Milton Keynes and Alex Alderton, Roberto Amato, Jeffrey Barrett, Sonia Goncalves, Ewan Harrison, David K. Jackson, Ian Johnston, Dominic Kwiatkowski, Cordelia Langford, John Sillitoe on behalf of the Wellcome Sanger Institute COVID-19 Surveillance Team |
| EPI_ISL_2910827, EPI_ISL_2910828 | Lighthouse Lab in Glasgow | Wellcome Sanger Institute for the COVID-19 Genomics UK (COG-UK) Consortium | Harper VanSteenhouse, Yumi Kasai, David Gray, Carol Clugston, Anna Dominiczak and Alex Alderton, Roberto Amato, Jeffrey Barrett, Sonia Goncalves, Ewan Harrison, David K. Jackson, Ian Johnston, Dominic Kwiatkowski, Cordelia Langford, John Sillitoe on behalf of the Wellcome Sanger Institute COVID-19 Surveillance Team |
| EPI_ISL_2910829, EPI_ISL_2910830, EPI_ISL_2910831, EPI_ISL_2910832, EPI_ISL_2910833 | Lighthouse Lab in Milton Keynes | Wellcome Sanger Institute for the COVID-19 Genomics UK (COG-UK) Consortium | The Lighthouse Lab in Milton Keynes and Alex Alderton, Roberto Amato, Jeffrey Barrett, Sonia Goncalves, Ewan Harrison, David K. Jackson, Ian Johnston, Dominic Kwiatkowski, Cordelia Langford, John Sillitoe on behalf of the Wellcome Sanger Institute COVID-19 Surveillance Team |
| EPI_ISL_2910834, EPI_ISL_2910835 | Lighthouse Lab in Glasgow | Wellcome Sanger Institute for the COVID-19 Genomics UK (COG-UK) Consortium | Harper VanSteenhouse, Yumi Kasai, David Gray, Carol Clugston, Anna Dominiczak and Alex Alderton, Roberto Amato, Jeffrey Barrett, Sonia Goncalves, Ewan Harrison, David K. Jackson, Ian Johnston, Dominic Kwiatkowski, Cordelia Langford, John Sillitoe on behalf of the Wellcome Sanger Institute COVID-19 Surveillance Team |
| EPI_ISL_2910836 | Lighthouse Lab in Alderley Park | Wellcome Sanger Institute for the COVID-19 Genomics UK (COG-UK) Consortium | Jacquelyn Wynn, Mairead Hyland, The Lighthouse Lab in Alderley Park and Alex Alderton, Roberto Amato, Jeffrey Barrett, Sonia Goncalves, Ewan Harrison, David K. Jackson, Ian Johnston, Dominic Kwiatkowski, Cordelia Langford, John Sillitoe on behalf of the Wellcome Sanger Institute COVID-19 Surveillance Team |
| EPI_ISL_2910837 | Lighthouse Lab in Milton Keynes | Wellcome Sanger Institute for the COVID-19 Genomics UK (COG-UK) Consortium | The Lighthouse Lab in Milton Keynes and Alex Alderton, Roberto Amato, Jeffrey Barrett, Sonia Goncalves, Ewan Harrison, David K. Jackson, Ian Johnston, Dominic Kwiatkowski, Cordelia Langford, John Sillitoe on behalf of the Wellcome Sanger Institute COVID-19 Surveillance Team |
| EPI_ISL_2910838 | Lighthouse Lab in Alderley Park | Wellcome Sanger Institute for the COVID-19 Genomics UK (COG-UK) Consortium | Jacquelyn Wynn, Mairead Hyland, The Lighthouse Lab in Alderley Park and Alex Alderton, Roberto Amato, Jeffrey Barrett, Sonia Goncalves, Ewan Harrison, David K. Jackson, Ian Johnston, Dominic Kwiatkowski, Cordelia Langford, John Sillitoe on behalf of the Wellcome Sanger Institute COVID-19 Surveillance Team |
| EPI_ISL_2910839, EPI_ISL_2910840 | Lighthouse Lab in Milton Keynes | Wellcome Sanger Institute for the COVID-19 Genomics UK (COG-UK) Consortium | The Lighthouse Lab in Milton Keynes and Alex Alderton, Roberto Amato, Jeffrey Barrett, Sonia Goncalves, Ewan Harrison, David K. Jackson, Ian Johnston, Dominic Kwiatkowski, Cordelia Langford, John Sillitoe on behalf of the Wellcome Sanger Institute COVID-19 Surveillance Team |
| EPI_ISL_2910841, EPI_ISL_2910842, EPI_ISL_2910843 | Lighthouse Lab in Glasgow | Wellcome Sanger Institute for the COVID-19 Genomics UK (COG-UK) Consortium | Harper VanSteenhouse, Yumi Kasai, David Gray, Carol Clugston, Anna Dominiczak and Alex Alderton, Roberto Amato, Jeffrey Barrett, Sonia Goncalves, Ewan Harrison, David K. Jackson, Ian Johnston, Dominic Kwiatkowski, Cordelia Langford, John Sillitoe on behalf of the Wellcome Sanger Institute COVID-19 Surveillance Team |
| EPI_ISL_2910844 | Lighthouse Lab in Alderley Park | Wellcome Sanger Institute for the COVID-19 Genomics UK (COG-UK) Consortium | Jacquelyn Wynn, Mairead Hyland, The Lighthouse Lab in Alderley Park and Alex Alderton, Roberto Amato, Jeffrey Barrett, Sonia Goncalves, Ewan Harrison, David K. Jackson, Ian Johnston, Dominic Kwiatkowski, Cordelia Langford, John Sillitoe on behalf of the Wellcome Sanger Institute COVID-19 Surveillance Team |
| EPI_ISL_2910845, EPI_ISL_2910846, EPI_ISL_2910847, EPI_ISL_2910848, EPI_ISL_2910849, EPI_ISL_2910850, EPI_ISL_2910851, EPI_ISL_2910852, EPI_ISL_2910853, EPI_ISL_2910854, EPI_ISL_2910855, EPI_ISL_2910856, EPI_ISL_2910857, EPI_ISL_2910858, EPI_ISL_2910859, EPI_ISL_2910860, EPI_ISL_2910861, EPI_ISL_2910862, EPI_ISL_2910863 | | | |
| see above | Lighthouse Lab in Milton Keynes | Wellcome Sanger Institute for the COVID-19 Genomics UK (COG-UK) Consortium | The Lighthouse Lab in Milton Keynes and Alex Alderton, Roberto Amato, Jeffrey Barrett, Sonia Goncalves, Ewan Harrison, David K. Jackson, Ian Johnston, Dominic Kwiatkowski, Cordelia Langford, John Sillitoe on behalf of the Wellcome Sanger Institute COVID-19 Surveillance Team |

| | | | |
|---|---|---|---|
| EPI_ISL_2910864, EPI_ISL_2910865, EPI_ISL_2910866 | Lighthouse Laboratory Plymouth | Wellcome Sanger Institute for the COVID-19 Genomics UK (COG-UK) Consortium | Lighthouse Laboratory Plymouth and Alex Alderton, Roberto Amato, Jeffrey Barrett, Sonia Goncalves, Ewan Harrison, David K. Jackson, Ian Johnston, Dominic Kwiatkowski, Cordelia Langford, John Sillitoe on behalf of the Wellcome Sanger Institute COVID-19 Surveillance Team |
| EPI_ISL_2910867 | Lighthouse Lab in Milton Keynes | Wellcome Sanger Institute for the COVID-19 Genomics UK (COG-UK) Consortium | The Lighthouse Lab in Milton Keynes and Alex Alderton, Roberto Amato, Jeffrey Barrett, Sonia Goncalves, Ewan Harrison, David K. Jackson, Ian Johnston, Dominic Kwiatkowski, Cordelia Langford, John Sillitoe on behalf of the Wellcome Sanger Institute COVID-19 Surveillance Team |
| EPI_ISL_2910869 | Lighthouse Laboratory Plymouth | Wellcome Sanger Institute for the COVID-19 Genomics UK (COG-UK) Consortium | Lighthouse Laboratory Plymouth and Alex Alderton, Roberto Amato, Jeffrey Barrett, Sonia Goncalves, Ewan Harrison, David K. Jackson, Ian Johnston, Dominic Kwiatkowski, Cordelia Langford, John Sillitoe on behalf of the Wellcome Sanger Institute COVID-19 Surveillance Team |
| EPI_ISL_2910870, EPI_ISL_2910871 | Lighthouse Lab in Milton Keynes | Wellcome Sanger Institute for the COVID-19 Genomics UK (COG-UK) Consortium | The Lighthouse Lab in Milton Keynes and Alex Alderton, Roberto Amato, Jeffrey Barrett, Sonia Goncalves, Ewan Harrison, David K. Jackson, Ian Johnston, Dominic Kwiatkowski, Cordelia Langford, John Sillitoe on behalf of the Wellcome Sanger Institute COVID-19 Surveillance Team |
| EPI_ISL_2910872 | Lighthouse Laboratory Plymouth | Wellcome Sanger Institute for the COVID-19 Genomics UK (COG-UK) Consortium | Lighthouse Laboratory Plymouth and Alex Alderton, Roberto Amato, Jeffrey Barrett, Sonia Goncalves, Ewan Harrison, David K. Jackson, Ian Johnston, Dominic Kwiatkowski, Cordelia Langford, John Sillitoe on behalf of the Wellcome Sanger Institute COVID-19 Surveillance Team |
| EPI_ISL_2910873, EPI_ISL_2910874, EPI_ISL_2910875, EPI_ISL_2910876, EPI_ISL_2910877, EPI_ISL_2910878, EPI_ISL_2910879, EPI_ISL_2910880, EPI_ISL_2910881, EPI_ISL_2910882 | Lighthouse Lab in Milton Keynes | Wellcome Sanger Institute for the COVID-19 Genomics UK (COG-UK) Consortium | The Lighthouse Lab in Milton Keynes and Alex Alderton, Roberto Amato, Jeffrey Barrett, Sonia Goncalves, Ewan Harrison, David K. Jackson, Ian Johnston, Dominic Kwiatkowski, Cordelia Langford, John Sillitoe on behalf of the Wellcome Sanger Institute COVID-19 Surveillance Team |
| EPI_ISL_2910883, EPI_ISL_2910884, EPI_ISL_2910885 | Lighthouse Laboratory Plymouth | Wellcome Sanger Institute for the COVID-19 Genomics UK (COG-UK) Consortium | Lighthouse Laboratory Plymouth and Alex Alderton, Roberto Amato, Jeffrey Barrett, Sonia Goncalves, Ewan Harrison, David K. Jackson, Ian Johnston, Dominic Kwiatkowski, Cordelia Langford, John Sillitoe on behalf of the Wellcome Sanger Institute COVID-19 Surveillance Team |
| EPI_ISL_2910886, EPI_ISL_2910887, EPI_ISL_2910888 | Lighthouse Lab in Milton Keynes | Wellcome Sanger Institute for the COVID-19 Genomics UK (COG-UK) Consortium | The Lighthouse Lab in Milton Keynes and Alex Alderton, Roberto Amato, Jeffrey Barrett, Sonia Goncalves, Ewan Harrison, David K. Jackson, Ian Johnston, Dominic Kwiatkowski, Cordelia Langford, John Sillitoe on behalf of the Wellcome Sanger Institute COVID-19 Surveillance Team |
| EPI_ISL_2910889 | Lighthouse Laboratory Plymouth | Wellcome Sanger Institute for the COVID-19 Genomics UK (COG-UK) Consortium | Lighthouse Laboratory Plymouth and Alex Alderton, Roberto Amato, Jeffrey Barrett, Sonia Goncalves, Ewan Harrison, David K. Jackson, Ian Johnston, Dominic Kwiatkowski, Cordelia Langford, John Sillitoe on behalf of the Wellcome Sanger Institute COVID-19 Surveillance Team |
| EPI_ISL_2910890, EPI_ISL_2910891, EPI_ISL_2910892, EPI_ISL_2910893, EPI_ISL_2910894, EPI_ISL_2910895, EPI_ISL_2910896, EPI_ISL_2910897 | Lighthouse Lab in Milton Keynes | Wellcome Sanger Institute for the COVID-19 Genomics UK (COG-UK) Consortium | The Lighthouse Lab in Milton Keynes and Alex Alderton, Roberto Amato, Jeffrey Barrett, Sonia Goncalves, Ewan Harrison, David K. Jackson, Ian Johnston, Dominic Kwiatkowski, Cordelia Langford, John Sillitoe on behalf of the Wellcome Sanger Institute COVID-19 Surveillance Team |
| EPI_ISL_2910898 | Lighthouse Laboratory Plymouth | Wellcome Sanger Institute for the COVID-19 Genomics UK (COG-UK) Consortium | Lighthouse Laboratory Plymouth and Alex Alderton, Roberto Amato, Jeffrey Barrett, Sonia Goncalves, Ewan Harrison, David K. Jackson, Ian Johnston, Dominic Kwiatkowski, Cordelia Langford, John Sillitoe on behalf of the Wellcome Sanger Institute COVID-19 Surveillance Team |
| EPI_ISL_2910899, EPI_ISL_2910900 | Lighthouse Lab in Milton Keynes | Wellcome Sanger Institute for the COVID-19 Genomics UK (COG-UK) Consortium | The Lighthouse Lab in Milton Keynes and Alex Alderton, Roberto Amato, Jeffrey Barrett, Sonia Goncalves, Ewan Harrison, David K. Jackson, Ian Johnston, Dominic Kwiatkowski, Cordelia Langford, John Sillitoe on behalf of the Wellcome Sanger Institute COVID-19 Surveillance Team |
| EPI_ISL_2910901 | Lighthouse Laboratory Plymouth | Wellcome Sanger Institute for the COVID-19 Genomics UK (COG-UK) Consortium | Lighthouse Laboratory Plymouth and Alex Alderton, Roberto Amato, Jeffrey Barrett, Sonia Goncalves, Ewan Harrison, David K. Jackson, Ian Johnston, Dominic Kwiatkowski, Cordelia Langford, John Sillitoe on behalf of the Wellcome Sanger Institute COVID-19 Surveillance Team |
| EPI_ISL_2910902, EPI_ISL_2910903, EPI_ISL_2910904, EPI_ISL_2910905, EPI_ISL_2910906, EPI_ISL_2910907 | Lighthouse Lab in Milton Keynes | Wellcome Sanger Institute for the COVID-19 Genomics UK (COG-UK) Consortium | The Lighthouse Lab in Milton Keynes and Alex Alderton, Roberto Amato, Jeffrey Barrett, Sonia Goncalves, Ewan Harrison, David K. Jackson, Ian Johnston, Dominic Kwiatkowski, Cordelia Langford, John Sillitoe on behalf of the Wellcome Sanger Institute COVID-19 Surveillance Team |
| EPI_ISL_2910908 | Lighthouse Laboratory Plymouth | Wellcome Sanger Institute for the COVID-19 Genomics UK (COG-UK) Consortium | Lighthouse Laboratory Plymouth and Alex Alderton, Roberto Amato, Jeffrey Barrett, Sonia Goncalves, Ewan Harrison, David K. Jackson, Ian Johnston, Dominic Kwiatkowski, Cordelia Langford, John Sillitoe on behalf of the Wellcome Sanger Institute COVID-19 Surveillance Team |
| EPI_ISL_2910909, EPI_ISL_2910910, EPI_ISL_2910911, EPI_ISL_2910912, EPI_ISL_2910913, EPI_ISL_2910914, EPI_ISL_2910915, EPI_ISL_2910916, EPI_ISL_2910917, EPI_ISL_2910918, EPI_ISL_2910919 | | | |
| see above | Lighthouse Lab in Milton Keynes | Wellcome Sanger Institute for the COVID-19 Genomics UK (COG-UK) Consortium | The Lighthouse Lab in Milton Keynes and Alex Alderton, Roberto Amato, Jeffrey Barrett, Sonia Goncalves, Ewan Harrison, David K. Jackson, Ian Johnston, Dominic Kwiatkowski, Cordelia Langford, John Sillitoe on behalf of the Wellcome Sanger Institute COVID-19 Surveillance Team |
| EPI_ISL_2910920 | Lighthouse Laboratory Plymouth | Wellcome Sanger Institute for the COVID-19 Genomics UK (COG-UK) Consortium | Lighthouse Laboratory Plymouth and Alex Alderton, Roberto Amato, Jeffrey Barrett, Sonia Goncalves, Ewan Harrison, David K. Jackson, Ian Johnston, Dominic Kwiatkowski, Cordelia Langford, John Sillitoe on behalf of the Wellcome Sanger Institute COVID-19 Surveillance Team |
| EPI_ISL_2910921, EPI_ISL_2910922 | Lighthouse Lab in Milton Keynes | Wellcome Sanger Institute for the COVID-19 Genomics UK (COG-UK) Consortium | The Lighthouse Lab in Milton Keynes and Alex Alderton, Roberto Amato, Jeffrey Barrett, Sonia Goncalves, Ewan Harrison, David K. Jackson, Ian Johnston, Dominic Kwiatkowski, Cordelia Langford, John Sillitoe on behalf of the Wellcome Sanger Institute COVID-19 Surveillance Team |
| EPI_ISL_2910923 | Lighthouse Laboratory Plymouth | Wellcome Sanger Institute for the COVID-19 Genomics UK (COG-UK) Consortium | Lighthouse Laboratory Plymouth and Alex Alderton, Roberto Amato, Jeffrey Barrett, Sonia Goncalves, Ewan Harrison, David K. Jackson, Ian Johnston, Dominic Kwiatkowski, Cordelia Langford, John Sillitoe on behalf of the Wellcome Sanger Institute COVID-19 Surveillance Team |
| EPI_ISL_2910924 | Lighthouse Lab in Milton Keynes | Wellcome Sanger Institute for the COVID-19 Genomics UK (COG-UK) Consortium | The Lighthouse Lab in Milton Keynes and Alex Alderton, Roberto Amato, Jeffrey Barrett, Sonia Goncalves, Ewan Harrison, David K. Jackson, Ian Johnston, Dominic Kwiatkowski, Cordelia Langford, John Sillitoe on behalf of the Wellcome Sanger Institute COVID-19 Surveillance Team |
| EPI_ISL_2910925, EPI_ISL_2910926 | Lighthouse Laboratory Plymouth | Wellcome Sanger Institute for the COVID-19 Genomics UK (COG-UK) Consortium | Lighthouse Laboratory Plymouth and Alex Alderton, Roberto Amato, Jeffrey Barrett, Sonia Goncalves, Ewan Harrison, David K. Jackson, Ian Johnston, Dominic Kwiatkowski, Cordelia Langford, John Sillitoe on behalf of the Wellcome Sanger Institute COVID-19 Surveillance Team |
| EPI_ISL_2910927, EPI_ISL_2910928, EPI_ISL_2910929, EPI_ISL_2910930, EPI_ISL_2910931, EPI_ISL_2910932, EPI_ISL_2910933 | Lighthouse Lab in Milton Keynes | Wellcome Sanger Institute for the COVID-19 Genomics UK (COG-UK) Consortium | The Lighthouse Lab in Milton Keynes and Alex Alderton, Roberto Amato, Jeffrey Barrett, Sonia Goncalves, Ewan Harrison, David K. Jackson, Ian Johnston, Dominic Kwiatkowski, Cordelia Langford, John Sillitoe on behalf of the Wellcome Sanger Institute COVID-19 Surveillance Team |
| EPI_ISL_2910934, EPI_ISL_2910935 | Lighthouse Laboratory Plymouth | Wellcome Sanger Institute for the COVID-19 Genomics UK (COG-UK) Consortium | Lighthouse Laboratory Plymouth and Alex Alderton, Roberto Amato, Jeffrey Barrett, Sonia Goncalves, Ewan Harrison, David K. Jackson, Ian Johnston, Dominic Kwiatkowski, Cordelia Langford, John Sillitoe on behalf of the Wellcome Sanger Institute COVID-19 Surveillance Team |
| EPI_ISL_2910936 | Lighthouse Lab in Milton Keynes | Wellcome Sanger Institute for the COVID-19 Genomics UK (COG-UK) Consortium | The Lighthouse Lab in Milton Keynes and Alex Alderton, Roberto Amato, Jeffrey Barrett, Sonia Goncalves, Ewan Harrison, David K. Jackson, Ian Johnston, Dominic Kwiatkowski, Cordelia Langford, John Sillitoe on behalf of the Wellcome Sanger Institute COVID-19 Surveillance Team |
| EPI_ISL_2910937, EPI_ISL_2910938, EPI_ISL_2910939 | Lighthouse Laboratory Plymouth | Wellcome Sanger Institute for the COVID-19 Genomics UK (COG-UK) Consortium | Lighthouse Laboratory Plymouth and Alex Alderton, Roberto Amato, Jeffrey Barrett, Sonia Goncalves, Ewan Harrison, David K. Jackson, Ian Johnston, Dominic Kwiatkowski, Cordelia Langford, John Sillitoe on behalf of the Wellcome Sanger Institute COVID-19 Surveillance Team |
| EPI_ISL_2910940, EPI_ISL_2910941, EPI_ISL_2910942, EPI_ISL_2910943, EPI_ISL_2910944, EPI_ISL_2910945, EPI_ISL_2910946 | Lighthouse Lab in Milton Keynes | Wellcome Sanger Institute for the COVID-19 Genomics UK (COG-UK) Consortium | The Lighthouse Lab in Milton Keynes and Alex Alderton, Roberto Amato, Jeffrey Barrett, Sonia Goncalves, Ewan Harrison, David K. Jackson, Ian Johnston, Dominic Kwiatkowski, Cordelia Langford, John Sillitoe on behalf of the Wellcome Sanger Institute COVID-19 Surveillance Team |
| EPI_ISL_2910947 | Lighthouse Laboratory Plymouth | Wellcome Sanger Institute for the COVID-19 Genomics UK (COG-UK) Consortium | Lighthouse Laboratory Plymouth and Alex Alderton, Roberto Amato, Jeffrey Barrett, Sonia Goncalves, Ewan Harrison, David K. Jackson, Ian Johnston, Dominic Kwiatkowski, Cordelia Langford, John Sillitoe on behalf of the Wellcome Sanger Institute COVID-19 Surveillance Team |
| EPI_ISL_2910948, EPI_ISL_2910949, EPI_ISL_2910950, EPI_ISL_2910951, EPI_ISL_2910952 | Lighthouse Lab in Milton Keynes | Wellcome Sanger Institute for the COVID-19 Genomics UK (COG-UK) Consortium | The Lighthouse Lab in Milton Keynes and Alex Alderton, Roberto Amato, Jeffrey Barrett, Sonia Goncalves, Ewan Harrison, David K. Jackson, Ian Johnston, Dominic Kwiatkowski, Cordelia Langford, John Sillitoe on behalf of the Wellcome Sanger Institute COVID-19 Surveillance Team |
| EPI_ISL_2910953 | Lighthouse Laboratory Plymouth | Wellcome Sanger Institute for the COVID-19 Genomics UK (COG-UK) Consortium | Lighthouse Laboratory Plymouth and Alex Alderton, Roberto Amato, Jeffrey Barrett, Sonia Goncalves, Ewan Harrison, David K. Jackson, Ian Johnston, Dominic Kwiatkowski, Cordelia Langford, John Sillitoe on behalf of the Wellcome Sanger Institute COVID-19 Surveillance Team |
| EPI_ISL_2910954 | Lighthouse Lab in Milton Keynes | Wellcome Sanger Institute for the COVID-19 Genomics UK (COG-UK) Consortium | The Lighthouse Lab in Milton Keynes and Alex Alderton, Roberto Amato, Jeffrey Barrett, Sonia Goncalves, Ewan Harrison, David K. Jackson, Ian Johnston, Dominic Kwiatkowski, Cordelia Langford, John Sillitoe on behalf of the Wellcome Sanger Institute COVID-19 Surveillance Team |

| | | | |
|---|---|---|---|
| EPI_ISL_2910955 | Lighthouse Laboratory Plymouth | Wellcome Sanger Institute for the COVID-19 Genomics UK (COG-UK) Consortium | Lighthouse Laboratory Plymouth and Alex Alderton, Roberto Amato, Jeffrey Barrett, Sonia Goncalves, Ewan Harrison, David K. Jackson, Ian Johnston, Dominic Kwiatkowski, Cordelia Langford, John Sillitoe on behalf of the Wellcome Sanger Institute COVID-19 Surveillance Team |
| EPI_ISL_2910956 | Lighthouse Lab in Milton Keynes | Wellcome Sanger Institute for the COVID-19 Genomics UK (COG-UK) Consortium | The Lighthouse Lab in Milton Keynes and Alex Alderton, Roberto Amato, Jeffrey Barrett, Sonia Goncalves, Ewan Harrison, David K. Jackson, Ian Johnston, Dominic Kwiatkowski, Cordelia Langford, John Sillitoe on behalf of the Wellcome Sanger Institute COVID-19 Surveillance Team |
| EPI_ISL_2910957, EPI_ISL_2910958 | Lighthouse Laboratory Plymouth | Wellcome Sanger Institute for the COVID-19 Genomics UK (COG-UK) Consortium | Lighthouse Laboratory Plymouth and Alex Alderton, Roberto Amato, Jeffrey Barrett, Sonia Goncalves, Ewan Harrison, David K. Jackson, Ian Johnston, Dominic Kwiatkowski, Cordelia Langford, John Sillitoe on behalf of the Wellcome Sanger Institute COVID-19 Surveillance Team |
| EPI_ISL_2910959, EPI_ISL_2910960, EPI_ISL_2910961 | Lighthouse Lab in Milton Keynes | Wellcome Sanger Institute for the COVID-19 Genomics UK (COG-UK) Consortium | The Lighthouse Lab in Milton Keynes and Alex Alderton, Roberto Amato, Jeffrey Barrett, Sonia Goncalves, Ewan Harrison, David K. Jackson, Ian Johnston, Dominic Kwiatkowski, Cordelia Langford, John Sillitoe on behalf of the Wellcome Sanger Institute COVID-19 Surveillance Team |
| EPI_ISL_2910962 | Lighthouse Laboratory Plymouth | Wellcome Sanger Institute for the COVID-19 Genomics UK (COG-UK) Consortium | Lighthouse Laboratory Plymouth and Alex Alderton, Roberto Amato, Jeffrey Barrett, Sonia Goncalves, Ewan Harrison, David K. Jackson, Ian Johnston, Dominic Kwiatkowski, Cordelia Langford, John Sillitoe on behalf of the Wellcome Sanger Institute COVID-19 Surveillance Team |
| EPI_ISL_2910963, EPI_ISL_2910964 | Lighthouse Lab in Milton Keynes | Wellcome Sanger Institute for the COVID-19 Genomics UK (COG-UK) Consortium | The Lighthouse Lab in Milton Keynes and Alex Alderton, Roberto Amato, Jeffrey Barrett, Sonia Goncalves, Ewan Harrison, David K. Jackson, Ian Johnston, Dominic Kwiatkowski, Cordelia Langford, John Sillitoe on behalf of the Wellcome Sanger Institute COVID-19 Surveillance Team |
| EPI_ISL_2910965, EPI_ISL_2910966 | Lighthouse Laboratory Plymouth | Wellcome Sanger Institute for the COVID-19 Genomics UK (COG-UK) Consortium | Lighthouse Laboratory Plymouth and Alex Alderton, Roberto Amato, Jeffrey Barrett, Sonia Goncalves, Ewan Harrison, David K. Jackson, Ian Johnston, Dominic Kwiatkowski, Cordelia Langford, John Sillitoe on behalf of the Wellcome Sanger Institute COVID-19 Surveillance Team |
| EPI_ISL_2910967, EPI_ISL_2910968 | Lighthouse Lab in Milton Keynes | Wellcome Sanger Institute for the COVID-19 Genomics UK (COG-UK) Consortium | The Lighthouse Lab in Milton Keynes and Alex Alderton, Roberto Amato, Jeffrey Barrett, Sonia Goncalves, Ewan Harrison, David K. Jackson, Ian Johnston, Dominic Kwiatkowski, Cordelia Langford, John Sillitoe on behalf of the Wellcome Sanger Institute COVID-19 Surveillance Team |
| EPI_ISL_2910969, EPI_ISL_2910970 | Lighthouse Laboratory Plymouth | Wellcome Sanger Institute for the COVID-19 Genomics UK (COG-UK) Consortium | Lighthouse Laboratory Plymouth and Alex Alderton, Roberto Amato, Jeffrey Barrett, Sonia Goncalves, Ewan Harrison, David K. Jackson, Ian Johnston, Dominic Kwiatkowski, Cordelia Langford, John Sillitoe on behalf of the Wellcome Sanger Institute COVID-19 Surveillance Team |
| EPI_ISL_2910971, EPI_ISL_2910972, EPI_ISL_2910973, EPI_ISL_2910974, EPI_ISL_2910975, EPI_ISL_2910976 | Lighthouse Lab in Milton Keynes | Wellcome Sanger Institute for the COVID-19 Genomics UK (COG-UK) Consortium | The Lighthouse Lab in Milton Keynes and Alex Alderton, Roberto Amato, Jeffrey Barrett, Sonia Goncalves, Ewan Harrison, David K. Jackson, Ian Johnston, Dominic Kwiatkowski, Cordelia Langford, John Sillitoe on behalf of the Wellcome Sanger Institute COVID-19 Surveillance Team |
| EPI_ISL_2910977, EPI_ISL_2910978 | Lighthouse Laboratory Plymouth | Wellcome Sanger Institute for the COVID-19 Genomics UK (COG-UK) Consortium | Lighthouse Laboratory Plymouth and Alex Alderton, Roberto Amato, Jeffrey Barrett, Sonia Goncalves, Ewan Harrison, David K. Jackson, Ian Johnston, Dominic Kwiatkowski, Cordelia Langford, John Sillitoe on behalf of the Wellcome Sanger Institute COVID-19 Surveillance Team |
| EPI_ISL_2910979, EPI_ISL_2910980, EPI_ISL_2910981, EPI_ISL_2910982 | Lighthouse Lab in Milton Keynes | Wellcome Sanger Institute for the COVID-19 Genomics UK (COG-UK) Consortium | The Lighthouse Lab in Milton Keynes and Alex Alderton, Roberto Amato, Jeffrey Barrett, Sonia Goncalves, Ewan Harrison, David K. Jackson, Ian Johnston, Dominic Kwiatkowski, Cordelia Langford, John Sillitoe on behalf of the Wellcome Sanger Institute COVID-19 Surveillance Team |
| EPI_ISL_2910983 | Lighthouse Laboratory Plymouth | Wellcome Sanger Institute for the COVID-19 Genomics UK (COG-UK) Consortium | Lighthouse Laboratory Plymouth and Alex Alderton, Roberto Amato, Jeffrey Barrett, Sonia Goncalves, Ewan Harrison, David K. Jackson, Ian Johnston, Dominic Kwiatkowski, Cordelia Langford, John Sillitoe on behalf of the Wellcome Sanger Institute COVID-19 Surveillance Team |
| EPI_ISL_2910984 | Lighthouse Lab in Milton Keynes | Wellcome Sanger Institute for the COVID-19 Genomics UK (COG-UK) Consortium | The Lighthouse Lab in Milton Keynes and Alex Alderton, Roberto Amato, Jeffrey Barrett, Sonia Goncalves, Ewan Harrison, David K. Jackson, Ian Johnston, Dominic Kwiatkowski, Cordelia Langford, John Sillitoe on behalf of the Wellcome Sanger Institute COVID-19 Surveillance Team |
| EPI_ISL_2910985 | Lighthouse Laboratory Plymouth | Wellcome Sanger Institute for the COVID-19 Genomics UK (COG-UK) Consortium | Lighthouse Laboratory Plymouth and Alex Alderton, Roberto Amato, Jeffrey Barrett, Sonia Goncalves, Ewan Harrison, David K. Jackson, Ian Johnston, Dominic Kwiatkowski, Cordelia Langford, John Sillitoe on behalf of the Wellcome Sanger Institute COVID-19 Surveillance Team |
| EPI_ISL_2910986 | Lighthouse Lab in Milton Keynes | Wellcome Sanger Institute for the COVID-19 Genomics UK (COG-UK) Consortium | The Lighthouse Lab in Milton Keynes and Alex Alderton, Roberto Amato, Jeffrey Barrett, Sonia Goncalves, Ewan Harrison, David K. Jackson, Ian Johnston, Dominic Kwiatkowski, Cordelia Langford, John Sillitoe on behalf of the Wellcome Sanger Institute COVID-19 Surveillance Team |
| EPI_ISL_2910987, EPI_ISL_2910988 | Lighthouse Laboratory Plymouth | Wellcome Sanger Institute for the COVID-19 Genomics UK (COG-UK) Consortium | Lighthouse Laboratory Plymouth and Alex Alderton, Roberto Amato, Jeffrey Barrett, Sonia Goncalves, Ewan Harrison, David K. Jackson, Ian Johnston, Dominic Kwiatkowski, Cordelia Langford, John Sillitoe on behalf of the Wellcome Sanger Institute COVID-19 Surveillance Team |
| EPI_ISL_2910989, EPI_ISL_2910990 | Lighthouse Lab in Milton Keynes | Wellcome Sanger Institute for the COVID-19 Genomics UK (COG-UK) Consortium | The Lighthouse Lab in Milton Keynes and Alex Alderton, Roberto Amato, Jeffrey Barrett, Sonia Goncalves, Ewan Harrison, David K. Jackson, Ian Johnston, Dominic Kwiatkowski, Cordelia Langford, John Sillitoe on behalf of the Wellcome Sanger Institute COVID-19 Surveillance Team |
| EPI_ISL_2910991, EPI_ISL_2910992 | Lighthouse Laboratory Plymouth | Wellcome Sanger Institute for the COVID-19 Genomics UK (COG-UK) Consortium | Lighthouse Laboratory Plymouth and Alex Alderton, Roberto Amato, Jeffrey Barrett, Sonia Goncalves, Ewan Harrison, David K. Jackson, Ian Johnston, Dominic Kwiatkowski, Cordelia Langford, John Sillitoe on behalf of the Wellcome Sanger Institute COVID-19 Surveillance Team |
| EPI_ISL_2910993, EPI_ISL_2910994, EPI_ISL_2910995, EPI_ISL_2910996, EPI_ISL_2910997, EPI_ISL_2910998, EPI_ISL_2910999 | Lighthouse Lab in Milton Keynes | Wellcome Sanger Institute for the COVID-19 Genomics UK (COG-UK) Consortium | The Lighthouse Lab in Milton Keynes and Alex Alderton, Roberto Amato, Jeffrey Barrett, Sonia Goncalves, Ewan Harrison, David K. Jackson, Ian Johnston, Dominic Kwiatkowski, Cordelia Langford, John Sillitoe on behalf of the Wellcome Sanger Institute COVID-19 Surveillance Team |
| EPI_ISL_2911000 | Lighthouse Laboratory Plymouth | Wellcome Sanger Institute for the COVID-19 Genomics UK (COG-UK) Consortium | Lighthouse Laboratory Plymouth and Alex Alderton, Roberto Amato, Jeffrey Barrett, Sonia Goncalves, Ewan Harrison, David K. Jackson, Ian Johnston, Dominic Kwiatkowski, Cordelia Langford, John Sillitoe on behalf of the Wellcome Sanger Institute COVID-19 Surveillance Team |
| EPI_ISL_2911001, EPI_ISL_2911002, EPI_ISL_2911003, EPI_ISL_2911004 | Lighthouse Lab in Milton Keynes | Wellcome Sanger Institute for the COVID-19 Genomics UK (COG-UK) Consortium | The Lighthouse Lab in Milton Keynes and Alex Alderton, Roberto Amato, Jeffrey Barrett, Sonia Goncalves, Ewan Harrison, David K. Jackson, Ian Johnston, Dominic Kwiatkowski, Cordelia Langford, John Sillitoe on behalf of the Wellcome Sanger Institute COVID-19 Surveillance Team |
| EPI_ISL_2911005 | Lighthouse Laboratory Plymouth | Wellcome Sanger Institute for the COVID-19 Genomics UK (COG-UK) Consortium | Lighthouse Laboratory Plymouth and Alex Alderton, Roberto Amato, Jeffrey Barrett, Sonia Goncalves, Ewan Harrison, David K. Jackson, Ian Johnston, Dominic Kwiatkowski, Cordelia Langford, John Sillitoe on behalf of the Wellcome Sanger Institute COVID-19 Surveillance Team |
| EPI_ISL_2911006, EPI_ISL_2911007 | Lighthouse Lab in Milton Keynes | Wellcome Sanger Institute for the COVID-19 Genomics UK (COG-UK) Consortium | The Lighthouse Lab in Milton Keynes and Alex Alderton, Roberto Amato, Jeffrey Barrett, Sonia Goncalves, Ewan Harrison, David K. Jackson, Ian Johnston, Dominic Kwiatkowski, Cordelia Langford, John Sillitoe on behalf of the Wellcome Sanger Institute COVID-19 Surveillance Team |
| EPI_ISL_2911008 | Lighthouse Laboratory Plymouth | Wellcome Sanger Institute for the COVID-19 Genomics UK (COG-UK) Consortium | Lighthouse Laboratory Plymouth and Alex Alderton, Roberto Amato, Jeffrey Barrett, Sonia Goncalves, Ewan Harrison, David K. Jackson, Ian Johnston, Dominic Kwiatkowski, Cordelia Langford, John Sillitoe on behalf of the Wellcome Sanger Institute COVID-19 Surveillance Team |
| EPI_ISL_2911009, EPI_ISL_2911010, EPI_ISL_2911011, EPI_ISL_2911012, EPI_ISL_2911013, EPI_ISL_2911014, EPI_ISL_2911015, EPI_ISL_2911016, EPI_ISL_2911017, EPI_ISL_2911019 | Lighthouse Lab in Milton Keynes | Wellcome Sanger Institute for the COVID-19 Genomics UK (COG-UK) Consortium | The Lighthouse Lab in Milton Keynes and Alex Alderton, Roberto Amato, Jeffrey Barrett, Sonia Goncalves, Ewan Harrison, David K. Jackson, Ian Johnston, Dominic Kwiatkowski, Cordelia Langford, John Sillitoe on behalf of the Wellcome Sanger Institute COVID-19 Surveillance Team |
| EPI_ISL_2911020, EPI_ISL_2911021 | Lighthouse Laboratory Plymouth | Wellcome Sanger Institute for the COVID-19 Genomics UK (COG-UK) Consortium | Lighthouse Laboratory Plymouth and Alex Alderton, Roberto Amato, Jeffrey Barrett, Sonia Goncalves, Ewan Harrison, David K. Jackson, Ian Johnston, Dominic Kwiatkowski, Cordelia Langford, John Sillitoe on behalf of the Wellcome Sanger Institute COVID-19 Surveillance Team |
| EPI_ISL_2911022 | Lighthouse Lab in Milton Keynes | Wellcome Sanger Institute for the COVID-19 Genomics UK (COG-UK) Consortium | The Lighthouse Lab in Milton Keynes and Alex Alderton, Roberto Amato, Jeffrey Barrett, Sonia Goncalves, Ewan Harrison, David K. Jackson, Ian Johnston, Dominic Kwiatkowski, Cordelia Langford, John Sillitoe on behalf of the Wellcome Sanger Institute COVID-19 Surveillance Team |
| EPI_ISL_2911023, EPI_ISL_2911024 | Lighthouse Laboratory Plymouth | Wellcome Sanger Institute for the COVID-19 Genomics UK (COG-UK) Consortium | Lighthouse Laboratory Plymouth and Alex Alderton, Roberto Amato, Jeffrey Barrett, Sonia Goncalves, Ewan Harrison, David K. Jackson, Ian Johnston, Dominic Kwiatkowski, Cordelia Langford, John Sillitoe on behalf of the Wellcome Sanger Institute COVID-19 Surveillance Team |
| EPI_ISL_2911025, EPI_ISL_2911026 | Lighthouse Lab in Milton Keynes | Wellcome Sanger Institute for the COVID-19 Genomics UK (COG-UK) Consortium | The Lighthouse Lab in Milton Keynes and Alex Alderton, Roberto Amato, Jeffrey Barrett, Sonia Goncalves, Ewan Harrison, David K. Jackson, Ian Johnston, Dominic Kwiatkowski, Cordelia Langford, John Sillitoe on behalf of the Wellcome Sanger Institute COVID-19 Surveillance Team |
| EPI_ISL_2911027, EPI_ISL_2911028 | Lighthouse Laboratory Plymouth | Wellcome Sanger Institute for the COVID-19 Genomics UK (COG-UK) Consortium | Lighthouse Laboratory Plymouth and Alex Alderton, Roberto Amato, Jeffrey Barrett, Sonia Goncalves, Ewan Harrison, David K. Jackson, Ian Johnston, Dominic Kwiatkowski, Cordelia Langford, John Sillitoe on behalf of the Wellcome Sanger Institute COVID-19 Surveillance Team |
| EPI_ISL_2911029, EPI_ISL_2911030 | Lighthouse Lab in Milton Keynes | Wellcome Sanger Institute for the COVID-19 Genomics UK (COG-UK) Consortium | The Lighthouse Lab in Milton Keynes and Alex Alderton, Roberto Amato, Jeffrey Barrett, Sonia Goncalves, Ewan Harrison, David K. Jackson, Ian Johnston, Dominic Kwiatkowski, Cordelia Langford, John Sillitoe on behalf of the Wellcome Sanger Institute COVID-19 Surveillance Team |
| EPI_ISL_2911031 | Lighthouse Laboratory Plymouth | Wellcome Sanger Institute for the COVID-19 Genomics UK (COG-UK) Consortium | Lighthouse Laboratory Plymouth and Alex Alderton, Roberto Amato, Jeffrey Barrett, Sonia Goncalves, Ewan Harrison, David K. Jackson, Ian Johnston, Dominic Kwiatkowski, Cordelia Langford, John Sillitoe on behalf of the Wellcome Sanger Institute COVID-19 Surveillance Team |
| EPI_ISL_2911032 | Lighthouse Lab in Milton Keynes | Wellcome Sanger Institute for the COVID-19 Genomics UK (COG-UK) Consortium | The Lighthouse Lab in Milton Keynes and Alex Alderton, Roberto Amato, Jeffrey Barrett, Sonia Goncalves, Ewan Harrison, David K. Jackson, Ian Johnston, Dominic Kwiatkowski, Cordelia Langford, John Sillitoe on behalf of the Wellcome Sanger Institute COVID-19 Surveillance Team |
| EPI_ISL_2911033 | Lighthouse Laboratory Plymouth | Wellcome Sanger Institute for the COVID-19 Genomics UK (COG-UK) Consortium | Lighthouse Laboratory Plymouth and Alex Alderton, Roberto Amato, Jeffrey Barrett, Sonia Goncalves, Ewan Harrison, David K. Jackson, Ian Johnston, Dominic Kwiatkowski, Cordelia Langford, John Sillitoe on behalf of the Wellcome Sanger Institute COVID-19 Surveillance Team |
| EPI_ISL_2911034 | Lighthouse Lab in Milton Keynes | Wellcome Sanger Institute for the COVID-19 Genomics UK (COG-UK) Consortium | The Lighthouse Lab in Milton Keynes and Alex Alderton, Roberto Amato, Jeffrey Barrett, Sonia Goncalves, Ewan Harrison, David K. Jackson, Ian Johnston, Dominic Kwiatkowski, Cordelia Langford, John Sillitoe on behalf of the Wellcome Sanger Institute COVID-19 Surveillance Team |

| | | | |
|---|---|---|---|
| EPI_ISL_2911035 | Lighthouse Laboratory Plymouth | Wellcome Sanger Institute for the COVID-19 Genomics UK (COG-UK) Consortium | Lighthouse Laboratory Plymouth and Alex Alderton, Roberto Amato, Jeffrey Barrett, Sonia Goncalves, Ewan Harrison, David K. Jackson, Ian Johnston, Dominic Kwiatkowski, Cordelia Langford, John Sillitoe on behalf of the Wellcome Sanger Institute COVID-19 Surveillance Team |
| EPI_ISL_2911036, EPI_ISL_2911037, EPI_ISL_2911038 | Lighthouse Lab in Milton Keynes | Wellcome Sanger Institute for the COVID-19 Genomics UK (COG-UK) Consortium | The Lighthouse Lab in Milton Keynes and Alex Alderton, Roberto Amato, Jeffrey Barrett, Sonia Goncalves, Ewan Harrison, David K. Jackson, Ian Johnston, Dominic Kwiatkowski, Cordelia Langford, John Sillitoe on behalf of the Wellcome Sanger Institute COVID-19 Surveillance Team |
| EPI_ISL_2911039 | Lighthouse Laboratory Plymouth | Wellcome Sanger Institute for the COVID-19 Genomics UK (COG-UK) Consortium | Lighthouse Laboratory Plymouth and Alex Alderton, Roberto Amato, Jeffrey Barrett, Sonia Goncalves, Ewan Harrison, David K. Jackson, Ian Johnston, Dominic Kwiatkowski, Cordelia Langford, John Sillitoe on behalf of the Wellcome Sanger Institute COVID-19 Surveillance Team |
| EPI_ISL_2911041, EPI_ISL_2911042 | Lighthouse Lab in Milton Keynes | Wellcome Sanger Institute for the COVID-19 Genomics UK (COG-UK) Consortium | The Lighthouse Lab in Milton Keynes and Alex Alderton, Roberto Amato, Jeffrey Barrett, Sonia Goncalves, Ewan Harrison, David K. Jackson, Ian Johnston, Dominic Kwiatkowski, Cordelia Langford, John Sillitoe on behalf of the Wellcome Sanger Institute COVID-19 Surveillance Team |
| EPI_ISL_2911043, EPI_ISL_2911044 | Lighthouse Laboratory Plymouth | Wellcome Sanger Institute for the COVID-19 Genomics UK (COG-UK) Consortium | Lighthouse Laboratory Plymouth and Alex Alderton, Roberto Amato, Jeffrey Barrett, Sonia Goncalves, Ewan Harrison, David K. Jackson, Ian Johnston, Dominic Kwiatkowski, Cordelia Langford, John Sillitoe on behalf of the Wellcome Sanger Institute COVID-19 Surveillance Team |
| EPI_ISL_2911045, EPI_ISL_2911046 | Lighthouse Lab in Milton Keynes | Wellcome Sanger Institute for the COVID-19 Genomics UK (COG-UK) Consortium | The Lighthouse Lab in Milton Keynes and Alex Alderton, Roberto Amato, Jeffrey Barrett, Sonia Goncalves, Ewan Harrison, David K. Jackson, Ian Johnston, Dominic Kwiatkowski, Cordelia Langford, John Sillitoe on behalf of the Wellcome Sanger Institute COVID-19 Surveillance Team |
| EPI_ISL_2911047, EPI_ISL_2911048 | Lighthouse Laboratory Plymouth | Wellcome Sanger Institute for the COVID-19 Genomics UK (COG-UK) Consortium | Lighthouse Laboratory Plymouth and Alex Alderton, Roberto Amato, Jeffrey Barrett, Sonia Goncalves, Ewan Harrison, David K. Jackson, Ian Johnston, Dominic Kwiatkowski, Cordelia Langford, John Sillitoe on behalf of the Wellcome Sanger Institute COVID-19 Surveillance Team |
| EPI_ISL_2911049, EPI_ISL_2911050 | Lighthouse Lab in Milton Keynes | Wellcome Sanger Institute for the COVID-19 Genomics UK (COG-UK) Consortium | The Lighthouse Lab in Milton Keynes and Alex Alderton, Roberto Amato, Jeffrey Barrett, Sonia Goncalves, Ewan Harrison, David K. Jackson, Ian Johnston, Dominic Kwiatkowski, Cordelia Langford, John Sillitoe on behalf of the Wellcome Sanger Institute COVID-19 Surveillance Team |
| EPI_ISL_2911051 | Lighthouse Laboratory Plymouth | Wellcome Sanger Institute for the COVID-19 Genomics UK (COG-UK) Consortium | Lighthouse Laboratory Plymouth and Alex Alderton, Roberto Amato, Jeffrey Barrett, Sonia Goncalves, Ewan Harrison, David K. Jackson, Ian Johnston, Dominic Kwiatkowski, Cordelia Langford, John Sillitoe on behalf of the Wellcome Sanger Institute COVID-19 Surveillance Team |
| EPI_ISL_2911052, EPI_ISL_2911053, EPI_ISL_2911054, EPI_ISL_2911055, EPI_ISL_2911056, EPI_ISL_2911057, EPI_ISL_2911058, EPI_ISL_2911059, EPI_ISL_2911060 | Lighthouse Lab in Milton Keynes | Wellcome Sanger Institute for the COVID-19 Genomics UK (COG-UK) Consortium | The Lighthouse Lab in Milton Keynes and Alex Alderton, Roberto Amato, Jeffrey Barrett, Sonia Goncalves, Ewan Harrison, David K. Jackson, Ian Johnston, Dominic Kwiatkowski, Cordelia Langford, John Sillitoe on behalf of the Wellcome Sanger Institute COVID-19 Surveillance Team |
| EPI_ISL_2911061 | Lighthouse Laboratory Plymouth | Wellcome Sanger Institute for the COVID-19 Genomics UK (COG-UK) Consortium | Lighthouse Laboratory Plymouth and Alex Alderton, Roberto Amato, Jeffrey Barrett, Sonia Goncalves, Ewan Harrison, David K. Jackson, Ian Johnston, Dominic Kwiatkowski, Cordelia Langford, John Sillitoe on behalf of the Wellcome Sanger Institute COVID-19 Surveillance Team |
| EPI_ISL_2911062, EPI_ISL_2911063, EPI_ISL_2911064, EPI_ISL_2911065, EPI_ISL_2911066, EPI_ISL_2911067 | Lighthouse Lab in Milton Keynes | Wellcome Sanger Institute for the COVID-19 Genomics UK (COG-UK) Consortium | The Lighthouse Lab in Milton Keynes and Alex Alderton, Roberto Amato, Jeffrey Barrett, Sonia Goncalves, Ewan Harrison, David K. Jackson, Ian Johnston, Dominic Kwiatkowski, Cordelia Langford, John Sillitoe on behalf of the Wellcome Sanger Institute COVID-19 Surveillance Team |
| EPI_ISL_2911068 | Lighthouse Laboratory Plymouth | Wellcome Sanger Institute for the COVID-19 Genomics UK (COG-UK) Consortium | Lighthouse Laboratory Plymouth and Alex Alderton, Roberto Amato, Jeffrey Barrett, Sonia Goncalves, Ewan Harrison, David K. Jackson, Ian Johnston, Dominic Kwiatkowski, Cordelia Langford, John Sillitoe on behalf of the Wellcome Sanger Institute COVID-19 Surveillance Team |
| EPI_ISL_2911069, EPI_ISL_2911070, EPI_ISL_2911071, EPI_ISL_2911072, EPI_ISL_2911073, EPI_ISL_2911074, EPI_ISL_2911075, EPI_ISL_2911076 | Lighthouse Lab in Milton Keynes | Wellcome Sanger Institute for the COVID-19 Genomics UK (COG-UK) Consortium | The Lighthouse Lab in Milton Keynes and Alex Alderton, Roberto Amato, Jeffrey Barrett, Sonia Goncalves, Ewan Harrison, David K. Jackson, Ian Johnston, Dominic Kwiatkowski, Cordelia Langford, John Sillitoe on behalf of the Wellcome Sanger Institute COVID-19 Surveillance Team |
| EPI_ISL_2911077, EPI_ISL_2911078 | Lighthouse Laboratory Plymouth | Wellcome Sanger Institute for the COVID-19 Genomics UK (COG-UK) Consortium | Lighthouse Laboratory Plymouth and Alex Alderton, Roberto Amato, Jeffrey Barrett, Sonia Goncalves, Ewan Harrison, David K. Jackson, Ian Johnston, Dominic Kwiatkowski, Cordelia Langford, John Sillitoe on behalf of the Wellcome Sanger Institute COVID-19 Surveillance Team |
| EPI_ISL_2911079 | Lighthouse Lab in Milton Keynes | Wellcome Sanger Institute for the COVID-19 Genomics UK (COG-UK) Consortium | The Lighthouse Lab in Milton Keynes and Alex Alderton, Roberto Amato, Jeffrey Barrett, Sonia Goncalves, Ewan Harrison, David K. Jackson, Ian Johnston, Dominic Kwiatkowski, Cordelia Langford, John Sillitoe on behalf of the Wellcome Sanger Institute COVID-19 Surveillance Team |
| EPI_ISL_2911080 | Lighthouse Laboratory Plymouth | Wellcome Sanger Institute for the COVID-19 Genomics UK (COG-UK) Consortium | Lighthouse Laboratory Plymouth and Alex Alderton, Roberto Amato, Jeffrey Barrett, Sonia Goncalves, Ewan Harrison, David K. Jackson, Ian Johnston, Dominic Kwiatkowski, Cordelia Langford, John Sillitoe on behalf of the Wellcome Sanger Institute COVID-19 Surveillance Team |
| EPI_ISL_2911081 | Lighthouse Lab in Milton Keynes | Wellcome Sanger Institute for the COVID-19 Genomics UK (COG-UK) Consortium | The Lighthouse Lab in Milton Keynes and Alex Alderton, Roberto Amato, Jeffrey Barrett, Sonia Goncalves, Ewan Harrison, David K. Jackson, Ian Johnston, Dominic Kwiatkowski, Cordelia Langford, John Sillitoe on behalf of the Wellcome Sanger Institute COVID-19 Surveillance Team |
| EPI_ISL_2911082 | Lighthouse Laboratory Plymouth | Wellcome Sanger Institute for the COVID-19 Genomics UK (COG-UK) Consortium | Lighthouse Laboratory Plymouth and Alex Alderton, Roberto Amato, Jeffrey Barrett, Sonia Goncalves, Ewan Harrison, David K. Jackson, Ian Johnston, Dominic Kwiatkowski, Cordelia Langford, John Sillitoe on behalf of the Wellcome Sanger Institute COVID-19 Surveillance Team |
| EPI_ISL_2911083 | Lighthouse Lab in Milton Keynes | Wellcome Sanger Institute for the COVID-19 Genomics UK (COG-UK) Consortium | The Lighthouse Lab in Milton Keynes and Alex Alderton, Roberto Amato, Jeffrey Barrett, Sonia Goncalves, Ewan Harrison, David K. Jackson, Ian Johnston, Dominic Kwiatkowski, Cordelia Langford, John Sillitoe on behalf of the Wellcome Sanger Institute COVID-19 Surveillance Team |
| EPI_ISL_2911084 | Lighthouse Laboratory Plymouth | Wellcome Sanger Institute for the COVID-19 Genomics UK (COG-UK) Consortium | Lighthouse Laboratory Plymouth and Alex Alderton, Roberto Amato, Jeffrey Barrett, Sonia Goncalves, Ewan Harrison, David K. Jackson, Ian Johnston, Dominic Kwiatkowski, Cordelia Langford, John Sillitoe on behalf of the Wellcome Sanger Institute COVID-19 Surveillance Team |
| EPI_ISL_2911085, EPI_ISL_2911086, EPI_ISL_2911087, EPI_ISL_2911088, EPI_ISL_2911089 | Lighthouse Lab in Milton Keynes | Wellcome Sanger Institute for the COVID-19 Genomics UK (COG-UK) Consortium | The Lighthouse Lab in Milton Keynes and Alex Alderton, Roberto Amato, Jeffrey Barrett, Sonia Goncalves, Ewan Harrison, David K. Jackson, Ian Johnston, Dominic Kwiatkowski, Cordelia Langford, John Sillitoe on behalf of the Wellcome Sanger Institute COVID-19 Surveillance Team |
| EPI_ISL_2911090 | Lighthouse Laboratory Plymouth | Wellcome Sanger Institute for the COVID-19 Genomics UK (COG-UK) Consortium | Lighthouse Laboratory Plymouth and Alex Alderton, Roberto Amato, Jeffrey Barrett, Sonia Goncalves, Ewan Harrison, David K. Jackson, Ian Johnston, Dominic Kwiatkowski, Cordelia Langford, John Sillitoe on behalf of the Wellcome Sanger Institute COVID-19 Surveillance Team |
| EPI_ISL_2911091, EPI_ISL_2911092, EPI_ISL_2911093 | Lighthouse Lab in Milton Keynes | Wellcome Sanger Institute for the COVID-19 Genomics UK (COG-UK) Consortium | The Lighthouse Lab in Milton Keynes and Alex Alderton, Roberto Amato, Jeffrey Barrett, Sonia Goncalves, Ewan Harrison, David K. Jackson, Ian Johnston, Dominic Kwiatkowski, Cordelia Langford, John Sillitoe on behalf of the Wellcome Sanger Institute COVID-19 Surveillance Team |
| EPI_ISL_2911094 | Lighthouse Laboratory Plymouth | Wellcome Sanger Institute for the COVID-19 Genomics UK (COG-UK) Consortium | Lighthouse Laboratory Plymouth and Alex Alderton, Roberto Amato, Jeffrey Barrett, Sonia Goncalves, Ewan Harrison, David K. Jackson, Ian Johnston, Dominic Kwiatkowski, Cordelia Langford, John Sillitoe on behalf of the Wellcome Sanger Institute COVID-19 Surveillance Team |
| EPI_ISL_2911095, EPI_ISL_2911096, EPI_ISL_2911097, EPI_ISL_2911098, EPI_ISL_2911099 | Lighthouse Lab in Milton Keynes | Wellcome Sanger Institute for the COVID-19 Genomics UK (COG-UK) Consortium | The Lighthouse Lab in Milton Keynes and Alex Alderton, Roberto Amato, Jeffrey Barrett, Sonia Goncalves, Ewan Harrison, David K. Jackson, Ian Johnston, Dominic Kwiatkowski, Cordelia Langford, John Sillitoe on behalf of the Wellcome Sanger Institute COVID-19 Surveillance Team |
| EPI_ISL_2911100, EPI_ISL_2911101 | Lighthouse Laboratory Plymouth | Wellcome Sanger Institute for the COVID-19 Genomics UK (COG-UK) Consortium | Lighthouse Laboratory Plymouth and Alex Alderton, Roberto Amato, Jeffrey Barrett, Sonia Goncalves, Ewan Harrison, David K. Jackson, Ian Johnston, Dominic Kwiatkowski, Cordelia Langford, John Sillitoe on behalf of the Wellcome Sanger Institute COVID-19 Surveillance Team |
| EPI_ISL_2911102, EPI_ISL_2911103 | Lighthouse Lab in Milton Keynes | Wellcome Sanger Institute for the COVID-19 Genomics UK (COG-UK) Consortium | The Lighthouse Lab in Milton Keynes and Alex Alderton, Roberto Amato, Jeffrey Barrett, Sonia Goncalves, Ewan Harrison, David K. Jackson, Ian Johnston, Dominic Kwiatkowski, Cordelia Langford, John Sillitoe on behalf of the Wellcome Sanger Institute COVID-19 Surveillance Team |
| EPI_ISL_2911104 | Lighthouse Laboratory Plymouth | Wellcome Sanger Institute for the COVID-19 Genomics UK (COG-UK) Consortium | Lighthouse Laboratory Plymouth and Alex Alderton, Roberto Amato, Jeffrey Barrett, Sonia Goncalves, Ewan Harrison, David K. Jackson, Ian Johnston, Dominic Kwiatkowski, Cordelia Langford, John Sillitoe on behalf of the Wellcome Sanger Institute COVID-19 Surveillance Team |
| EPI_ISL_2911105, EPI_ISL_2911106, EPI_ISL_2911107 | Lighthouse Lab in Milton Keynes | Wellcome Sanger Institute for the COVID-19 Genomics UK (COG-UK) Consortium | The Lighthouse Lab in Milton Keynes and Alex Alderton, Roberto Amato, Jeffrey Barrett, Sonia Goncalves, Ewan Harrison, David K. Jackson, Ian Johnston, Dominic Kwiatkowski, Cordelia Langford, John Sillitoe on behalf of the Wellcome Sanger Institute COVID-19 Surveillance Team |
| EPI_ISL_2911108 | Lighthouse Laboratory Plymouth | Wellcome Sanger Institute for the COVID-19 Genomics UK (COG-UK) Consortium | Lighthouse Laboratory Plymouth and Alex Alderton, Roberto Amato, Jeffrey Barrett, Sonia Goncalves, Ewan Harrison, David K. Jackson, Ian Johnston, Dominic Kwiatkowski, Cordelia Langford, John Sillitoe on behalf of the Wellcome Sanger Institute COVID-19 Surveillance Team |
| EPI_ISL_2911109, EPI_ISL_2911110 | Lighthouse Lab in Milton Keynes | Wellcome Sanger Institute for the COVID-19 Genomics UK (COG-UK) Consortium | The Lighthouse Lab in Milton Keynes and Alex Alderton, Roberto Amato, Jeffrey Barrett, Sonia Goncalves, Ewan Harrison, David K. Jackson, Ian Johnston, Dominic Kwiatkowski, Cordelia Langford, John Sillitoe on behalf of the Wellcome Sanger Institute COVID-19 Surveillance Team |
| EPI_ISL_2911111 | Lighthouse Laboratory Plymouth | Wellcome Sanger Institute for the COVID-19 Genomics UK (COG-UK) Consortium | Lighthouse Laboratory Plymouth and Alex Alderton, Roberto Amato, Jeffrey Barrett, Sonia Goncalves, Ewan Harrison, David K. Jackson, Ian Johnston, Dominic Kwiatkowski, Cordelia Langford, John Sillitoe on behalf of the Wellcome Sanger Institute COVID-19 Surveillance Team |
| EPI_ISL_2911112, EPI_ISL_2911113 | Lighthouse Lab in Milton Keynes | Wellcome Sanger Institute for the COVID-19 Genomics UK (COG-UK) Consortium | The Lighthouse Lab in Milton Keynes and Alex Alderton, Roberto Amato, Jeffrey Barrett, Sonia Goncalves, Ewan Harrison, David K. Jackson, Ian Johnston, Dominic Kwiatkowski, Cordelia Langford, John Sillitoe on behalf of the Wellcome Sanger Institute COVID-19 Surveillance Team |

| | | | |
|---|---|---|---|
| EPI_ISL_2911114, EPI_ISL_2911115 | Lighthouse Laboratory Plymouth | Wellcome Sanger Institute for the COVID-19 Genomics UK (COG-UK) Consortium | Lighthouse Laboratory Plymouth and Alex Alderton, Roberto Amato, Jeffrey Barrett, Sonia Goncalves, Ewan Harrison, David K. Jackson, Ian Johnston, Dominic Kwiatkowski, Cordelia Langford, John Sillitoe on behalf of the Wellcome Sanger Institute COVID-19 Surveillance Team |
| EPI_ISL_2911116, EPI_ISL_2911117, EPI_ISL_2911118, EPI_ISL_2911119, EPI_ISL_2911120, EPI_ISL_2911121, EPI_ISL_2911122, EPI_ISL_2911123, EPI_ISL_2911124, EPI_ISL_2911125, EPI_ISL_2911126 | | | |
| see above | Lighthouse Lab in Milton Keynes | Wellcome Sanger Institute for the COVID-19 Genomics UK (COG-UK) Consortium | The Lighthouse Lab in Milton Keynes and Alex Alderton, Roberto Amato, Jeffrey Barrett, Sonia Goncalves, Ewan Harrison, David K. Jackson, Ian Johnston, Dominic Kwiatkowski, Cordelia Langford, John Sillitoe on behalf of the Wellcome Sanger Institute COVID-19 Surveillance Team |
| EPI_ISL_2911127 | Lighthouse Laboratory Plymouth | Wellcome Sanger Institute for the COVID-19 Genomics UK (COG-UK) Consortium | Lighthouse Laboratory Plymouth and Alex Alderton, Roberto Amato, Jeffrey Barrett, Sonia Goncalves, Ewan Harrison, David K. Jackson, Ian Johnston, Dominic Kwiatkowski, Cordelia Langford, John Sillitoe on behalf of the Wellcome Sanger Institute COVID-19 Surveillance Team |
| EPI_ISL_2911128, EPI_ISL_2911129, EPI_ISL_2911130, EPI_ISL_2911131, EPI_ISL_2911132, EPI_ISL_2911133, EPI_ISL_2911134, EPI_ISL_2911135 | Lighthouse Lab in Milton Keynes | Wellcome Sanger Institute for the COVID-19 Genomics UK (COG-UK) Consortium | The Lighthouse Lab in Milton Keynes and Alex Alderton, Roberto Amato, Jeffrey Barrett, Sonia Goncalves, Ewan Harrison, David K. Jackson, Ian Johnston, Dominic Kwiatkowski, Cordelia Langford, John Sillitoe on behalf of the Wellcome Sanger Institute COVID-19 Surveillance Team |
| EPI_ISL_2911136 | Lighthouse Laboratory Plymouth | Wellcome Sanger Institute for the COVID-19 Genomics UK (COG-UK) Consortium | Lighthouse Laboratory Plymouth and Alex Alderton, Roberto Amato, Jeffrey Barrett, Sonia Goncalves, Ewan Harrison, David K. Jackson, Ian Johnston, Dominic Kwiatkowski, Cordelia Langford, John Sillitoe on behalf of the Wellcome Sanger Institute COVID-19 Surveillance Team |
| EPI_ISL_2911137, EPI_ISL_2911138 | Lighthouse Lab in Milton Keynes | Wellcome Sanger Institute for the COVID-19 Genomics UK (COG-UK) Consortium | The Lighthouse Lab in Milton Keynes and Alex Alderton, Roberto Amato, Jeffrey Barrett, Sonia Goncalves, Ewan Harrison, David K. Jackson, Ian Johnston, Dominic Kwiatkowski, Cordelia Langford, John Sillitoe on behalf of the Wellcome Sanger Institute COVID-19 Surveillance Team |
| EPI_ISL_2911139 | Lighthouse Laboratory Plymouth | Wellcome Sanger Institute for the COVID-19 Genomics UK (COG-UK) Consortium | Lighthouse Laboratory Plymouth and Alex Alderton, Roberto Amato, Jeffrey Barrett, Sonia Goncalves, Ewan Harrison, David K. Jackson, Ian Johnston, Dominic Kwiatkowski, Cordelia Langford, John Sillitoe on behalf of the Wellcome Sanger Institute COVID-19 Surveillance Team |
| EPI_ISL_2911140, EPI_ISL_2911141, EPI_ISL_2911142, EPI_ISL_2911143, EPI_ISL_2911144, EPI_ISL_2911145, EPI_ISL_2911146, EPI_ISL_2911147 | Lighthouse Lab in Milton Keynes | Wellcome Sanger Institute for the COVID-19 Genomics UK (COG-UK) Consortium | The Lighthouse Lab in Milton Keynes and Alex Alderton, Roberto Amato, Jeffrey Barrett, Sonia Goncalves, Ewan Harrison, David K. Jackson, Ian Johnston, Dominic Kwiatkowski, Cordelia Langford, John Sillitoe on behalf of the Wellcome Sanger Institute COVID-19 Surveillance Team |
| EPI_ISL_2911148 | Lighthouse Laboratory Plymouth | Wellcome Sanger Institute for the COVID-19 Genomics UK (COG-UK) Consortium | Lighthouse Laboratory Plymouth and Alex Alderton, Roberto Amato, Jeffrey Barrett, Sonia Goncalves, Ewan Harrison, David K. Jackson, Ian Johnston, Dominic Kwiatkowski, Cordelia Langford, John Sillitoe on behalf of the Wellcome Sanger Institute COVID-19 Surveillance Team |
| EPI_ISL_2911149 | Lighthouse Lab in Milton Keynes | Wellcome Sanger Institute for the COVID-19 Genomics UK (COG-UK) Consortium | The Lighthouse Lab in Milton Keynes and Alex Alderton, Roberto Amato, Jeffrey Barrett, Sonia Goncalves, Ewan Harrison, David K. Jackson, Ian Johnston, Dominic Kwiatkowski, Cordelia Langford, John Sillitoe on behalf of the Wellcome Sanger Institute COVID-19 Surveillance Team |
| EPI_ISL_2911150 | Lighthouse Laboratory Plymouth | Wellcome Sanger Institute for the COVID-19 Genomics UK (COG-UK) Consortium | Lighthouse Laboratory Plymouth and Alex Alderton, Roberto Amato, Jeffrey Barrett, Sonia Goncalves, Ewan Harrison, David K. Jackson, Ian Johnston, Dominic Kwiatkowski, Cordelia Langford, John Sillitoe on behalf of the Wellcome Sanger Institute COVID-19 Surveillance Team |
| EPI_ISL_2911151, EPI_ISL_2911152, EPI_ISL_2911153, EPI_ISL_2911154, EPI_ISL_2911155 | Lighthouse Lab in Milton Keynes | Wellcome Sanger Institute for the COVID-19 Genomics UK (COG-UK) Consortium | The Lighthouse Lab in Milton Keynes and Alex Alderton, Roberto Amato, Jeffrey Barrett, Sonia Goncalves, Ewan Harrison, David K. Jackson, Ian Johnston, Dominic Kwiatkowski, Cordelia Langford, John Sillitoe on behalf of the Wellcome Sanger Institute COVID-19 Surveillance Team |
| EPI_ISL_2911156 | Lighthouse Laboratory Plymouth | Wellcome Sanger Institute for the COVID-19 Genomics UK (COG-UK) Consortium | Lighthouse Laboratory Plymouth and Alex Alderton, Roberto Amato, Jeffrey Barrett, Sonia Goncalves, Ewan Harrison, David K. Jackson, Ian Johnston, Dominic Kwiatkowski, Cordelia Langford, John Sillitoe on behalf of the Wellcome Sanger Institute COVID-19 Surveillance Team |
| EPI_ISL_2911157, EPI_ISL_2911158, EPI_ISL_2911159 | Lighthouse Lab in Milton Keynes | Wellcome Sanger Institute for the COVID-19 Genomics UK (COG-UK) Consortium | The Lighthouse Lab in Milton Keynes and Alex Alderton, Roberto Amato, Jeffrey Barrett, Sonia Goncalves, Ewan Harrison, David K. Jackson, Ian Johnston, Dominic Kwiatkowski, Cordelia Langford, John Sillitoe on behalf of the Wellcome Sanger Institute COVID-19 Surveillance Team |
| EPI_ISL_2911160 | Lighthouse Laboratory Plymouth | Wellcome Sanger Institute for the COVID-19 Genomics UK (COG-UK) Consortium | Lighthouse Laboratory Plymouth and Alex Alderton, Roberto Amato, Jeffrey Barrett, Sonia Goncalves, Ewan Harrison, David K. Jackson, Ian Johnston, Dominic Kwiatkowski, Cordelia Langford, John Sillitoe on behalf of the Wellcome Sanger Institute COVID-19 Surveillance Team |
| EPI_ISL_2911161 | Lighthouse Lab in Milton Keynes | Wellcome Sanger Institute for the COVID-19 Genomics UK (COG-UK) Consortium | The Lighthouse Lab in Milton Keynes and Alex Alderton, Roberto Amato, Jeffrey Barrett, Sonia Goncalves, Ewan Harrison, David K. Jackson, Ian Johnston, Dominic Kwiatkowski, Cordelia Langford, John Sillitoe on behalf of the Wellcome Sanger Institute COVID-19 Surveillance Team |
| EPI_ISL_2911162, EPI_ISL_2911163, EPI_ISL_2911164 | Lighthouse Laboratory Plymouth | Wellcome Sanger Institute for the COVID-19 Genomics UK (COG-UK) Consortium | Lighthouse Laboratory Plymouth and Alex Alderton, Roberto Amato, Jeffrey Barrett, Sonia Goncalves, Ewan Harrison, David K. Jackson, Ian Johnston, Dominic Kwiatkowski, Cordelia Langford, John Sillitoe on behalf of the Wellcome Sanger Institute COVID-19 Surveillance Team |
| EPI_ISL_2911165 | Lighthouse Lab in Milton Keynes | Wellcome Sanger Institute for the COVID-19 Genomics UK (COG-UK) Consortium | The Lighthouse Lab in Milton Keynes and Alex Alderton, Roberto Amato, Jeffrey Barrett, Sonia Goncalves, Ewan Harrison, David K. Jackson, Ian Johnston, Dominic Kwiatkowski, Cordelia Langford, John Sillitoe on behalf of the Wellcome Sanger Institute COVID-19 Surveillance Team |
| EPI_ISL_2911166 | Lighthouse Laboratory Plymouth | Wellcome Sanger Institute for the COVID-19 Genomics UK (COG-UK) Consortium | Lighthouse Laboratory Plymouth and Alex Alderton, Roberto Amato, Jeffrey Barrett, Sonia Goncalves, Ewan Harrison, David K. Jackson, Ian Johnston, Dominic Kwiatkowski, Cordelia Langford, John Sillitoe on behalf of the Wellcome Sanger Institute COVID-19 Surveillance Team |
| EPI_ISL_2911167, EPI_ISL_2911168, EPI_ISL_2911169, EPI_ISL_2911170, EPI_ISL_2911171, EPI_ISL_2911172 | Lighthouse Lab in Milton Keynes | Wellcome Sanger Institute for the COVID-19 Genomics UK (COG-UK) Consortium | The Lighthouse Lab in Milton Keynes and Alex Alderton, Roberto Amato, Jeffrey Barrett, Sonia Goncalves, Ewan Harrison, David K. Jackson, Ian Johnston, Dominic Kwiatkowski, Cordelia Langford, John Sillitoe on behalf of the Wellcome Sanger Institute COVID-19 Surveillance Team |
| EPI_ISL_2911173, EPI_ISL_2911174, EPI_ISL_2911175, EPI_ISL_2911176 | Lighthouse Laboratory Plymouth | Wellcome Sanger Institute for the COVID-19 Genomics UK (COG-UK) Consortium | Lighthouse Laboratory Plymouth and Alex Alderton, Roberto Amato, Jeffrey Barrett, Sonia Goncalves, Ewan Harrison, David K. Jackson, Ian Johnston, Dominic Kwiatkowski, Cordelia Langford, John Sillitoe on behalf of the Wellcome Sanger Institute COVID-19 Surveillance Team |
| EPI_ISL_2911177, EPI_ISL_2911178, EPI_ISL_2911179, EPI_ISL_2911180, EPI_ISL_2911181, EPI_ISL_2911182, EPI_ISL_2911183 | Lighthouse Lab in Milton Keynes | Wellcome Sanger Institute for the COVID-19 Genomics UK (COG-UK) Consortium | The Lighthouse Lab in Milton Keynes and Alex Alderton, Roberto Amato, Jeffrey Barrett, Sonia Goncalves, Ewan Harrison, David K. Jackson, Ian Johnston, Dominic Kwiatkowski, Cordelia Langford, John Sillitoe on behalf of the Wellcome Sanger Institute COVID-19 Surveillance Team |
| EPI_ISL_2911184 | Lighthouse Laboratory Plymouth | Wellcome Sanger Institute for the COVID-19 Genomics UK (COG-UK) Consortium | Lighthouse Laboratory Plymouth and Alex Alderton, Roberto Amato, Jeffrey Barrett, Sonia Goncalves, Ewan Harrison, David K. Jackson, Ian Johnston, Dominic Kwiatkowski, Cordelia Langford, John Sillitoe on behalf of the Wellcome Sanger Institute COVID-19 Surveillance Team |
| EPI_ISL_2911185, EPI_ISL_2911186, EPI_ISL_2911187, EPI_ISL_2911188 | Lighthouse Lab in Milton Keynes | Wellcome Sanger Institute for the COVID-19 Genomics UK (COG-UK) Consortium | The Lighthouse Lab in Milton Keynes and Alex Alderton, Roberto Amato, Jeffrey Barrett, Sonia Goncalves, Ewan Harrison, David K. Jackson, Ian Johnston, Dominic Kwiatkowski, Cordelia Langford, John Sillitoe on behalf of the Wellcome Sanger Institute COVID-19 Surveillance Team |
| EPI_ISL_2911189 | Lighthouse Laboratory Plymouth | Wellcome Sanger Institute for the COVID-19 Genomics UK (COG-UK) Consortium | Lighthouse Laboratory Plymouth and Alex Alderton, Roberto Amato, Jeffrey Barrett, Sonia Goncalves, Ewan Harrison, David K. Jackson, Ian Johnston, Dominic Kwiatkowski, Cordelia Langford, John Sillitoe on behalf of the Wellcome Sanger Institute COVID-19 Surveillance Team |
| EPI_ISL_2911190 | Lighthouse Lab in Milton Keynes | Wellcome Sanger Institute for the COVID-19 Genomics UK (COG-UK) Consortium | The Lighthouse Lab in Milton Keynes and Alex Alderton, Roberto Amato, Jeffrey Barrett, Sonia Goncalves, Ewan Harrison, David K. Jackson, Ian Johnston, Dominic Kwiatkowski, Cordelia Langford, John Sillitoe on behalf of the Wellcome Sanger Institute COVID-19 Surveillance Team |
| EPI_ISL_2911191, EPI_ISL_2911192 | Lighthouse Laboratory Plymouth | Wellcome Sanger Institute for the COVID-19 Genomics UK (COG-UK) Consortium | Lighthouse Laboratory Plymouth and Alex Alderton, Roberto Amato, Jeffrey Barrett, Sonia Goncalves, Ewan Harrison, David K. Jackson, Ian Johnston, Dominic Kwiatkowski, Cordelia Langford, John Sillitoe on behalf of the Wellcome Sanger Institute COVID-19 Surveillance Team |
| EPI_ISL_2911193, EPI_ISL_2911194, EPI_ISL_2911195, EPI_ISL_2911196 | Lighthouse Lab in Milton Keynes | Wellcome Sanger Institute for the COVID-19 Genomics UK (COG-UK) Consortium | The Lighthouse Lab in Milton Keynes and Alex Alderton, Roberto Amato, Jeffrey Barrett, Sonia Goncalves, Ewan Harrison, David K. Jackson, Ian Johnston, Dominic Kwiatkowski, Cordelia Langford, John Sillitoe on behalf of the Wellcome Sanger Institute COVID-19 Surveillance Team |
| EPI_ISL_2911197 | Lighthouse Lab in Glasgow | Wellcome Sanger Institute for the COVID-19 Genomics UK (COG-UK) Consortium | Harper VanSteenhouse, Yumi Kasai, David Gray, Carol Clugston, Anna Dominiczak and Alex Alderton, Roberto Amato, Jeffrey Barrett, Sonia Goncalves, Ewan Harrison, David K. Jackson, Ian Johnston, Dominic Kwiatkowski, Cordelia Langford, John Sillitoe on behalf of the Wellcome Sanger Institute COVID-19 Surveillance Team |
| EPI_ISL_2911198, EPI_ISL_2911199 | Lighthouse Lab in Milton Keynes | Wellcome Sanger Institute for the COVID-19 Genomics UK (COG-UK) Consortium | The Lighthouse Lab in Milton Keynes and Alex Alderton, Roberto Amato, Jeffrey Barrett, Sonia Goncalves, Ewan Harrison, David K. Jackson, Ian Johnston, Dominic Kwiatkowski, Cordelia Langford, John Sillitoe on behalf of the Wellcome Sanger Institute COVID-19 Surveillance Team |
| EPI_ISL_2911200 | Lighthouse Lab in Glasgow | Wellcome Sanger Institute for the COVID-19 Genomics UK (COG-UK) Consortium | Harper VanSteenhouse, Yumi Kasai, David Gray, Carol Clugston, Anna Dominiczak and Alex Alderton, Roberto Amato, Jeffrey Barrett, Sonia Goncalves, Ewan Harrison, David K. Jackson, Ian Johnston, Dominic Kwiatkowski, Cordelia Langford, John Sillitoe on behalf of the Wellcome Sanger Institute COVID-19 Surveillance Team |
| EPI_ISL_2911201, | Lighthouse Lab in Milton Keynes | Wellcome Sanger Institute for the COVID-19 Genomics UK (COG-UK) | The Lighthouse Lab in Milton Keynes and Alex Alderton, Roberto Amato, Jeffrey Barrett, Sonia Goncalves, Ewan Harrison, David K. Jackson, Ian Johnston, Dominic Kwiatkowski, Cordelia Langford, John Sillitoe on behalf of the Wellcome Sanger |

| | | | |
|---|---|---|---|
| EPI_ISL_2911202, EPI_ISL_2911203, EPI_ISL_2911204, EPI_ISL_2911205, EPI_ISL_2911206, EPI_ISL_2911207, EPI_ISL_2911208 | | Consortium | Institute COVID-19 Surveillance Team |
| EPI_ISL_2911209 | Lighthouse Lab in Glasgow | Wellcome Sanger Institute for the COVID-19 Genomics UK (COG-UK) Consortium | Harper VanSteenhouse, Yumi Kasai, David Gray, Carol Clugston, Anna Dominiczak and Alex Alderton, Roberto Amato, Jeffrey Barrett, Sonia Goncalves, Ewan Harrison, David K. Jackson, Ian Johnston, Dominic Kwiatkowski, Cordelia Langford, John Sillitoe on behalf of the Wellcome Sanger Institute COVID-19 Surveillance Team |
| EPI_ISL_2911210, EPI_ISL_2911211 | Lighthouse Lab in Milton Keynes | Wellcome Sanger Institute for the COVID-19 Genomics UK (COG-UK) Consortium | The Lighthouse Lab in Milton Keynes and Alex Alderton, Roberto Amato, Jeffrey Barrett, Sonia Goncalves, Ewan Harrison, David K. Jackson, Ian Johnston, Dominic Kwiatkowski, Cordelia Langford, John Sillitoe on behalf of the Wellcome Sanger Institute COVID-19 Surveillance Team |
| EPI_ISL_2911212 | Lighthouse Lab in Glasgow | Wellcome Sanger Institute for the COVID-19 Genomics UK (COG-UK) Consortium | Harper VanSteenhouse, Yumi Kasai, David Gray, Carol Clugston, Anna Dominiczak and Alex Alderton, Roberto Amato, Jeffrey Barrett, Sonia Goncalves, Ewan Harrison, David K. Jackson, Ian Johnston, Dominic Kwiatkowski, Cordelia Langford, John Sillitoe on behalf of the Wellcome Sanger Institute COVID-19 Surveillance Team |
| EPI_ISL_2911213, EPI_ISL_2911214, EPI_ISL_2911215, EPI_ISL_2911216 | Lighthouse Lab in Milton Keynes | Wellcome Sanger Institute for the COVID-19 Genomics UK (COG-UK) Consortium | The Lighthouse Lab in Milton Keynes and Alex Alderton, Roberto Amato, Jeffrey Barrett, Sonia Goncalves, Ewan Harrison, David K. Jackson, Ian Johnston, Dominic Kwiatkowski, Cordelia Langford, John Sillitoe on behalf of the Wellcome Sanger Institute COVID-19 Surveillance Team |
| EPI_ISL_2911217 | Lighthouse Lab in Glasgow | Wellcome Sanger Institute for the COVID-19 Genomics UK (COG-UK) Consortium | Harper VanSteenhouse, Yumi Kasai, David Gray, Carol Clugston, Anna Dominiczak and Alex Alderton, Roberto Amato, Jeffrey Barrett, Sonia Goncalves, Ewan Harrison, David K. Jackson, Ian Johnston, Dominic Kwiatkowski, Cordelia Langford, John Sillitoe on behalf of the Wellcome Sanger Institute COVID-19 Surveillance Team |
| EPI_ISL_2911218, EPI_ISL_2911219, EPI_ISL_2911220, EPI_ISL_2911221, EPI_ISL_2911222, EPI_ISL_2911223, EPI_ISL_2911224 | Lighthouse Lab in Milton Keynes | Wellcome Sanger Institute for the COVID-19 Genomics UK (COG-UK) Consortium | The Lighthouse Lab in Milton Keynes and Alex Alderton, Roberto Amato, Jeffrey Barrett, Sonia Goncalves, Ewan Harrison, David K. Jackson, Ian Johnston, Dominic Kwiatkowski, Cordelia Langford, John Sillitoe on behalf of the Wellcome Sanger Institute COVID-19 Surveillance Team |
| EPI_ISL_2911225 | Lighthouse Lab in Glasgow | Wellcome Sanger Institute for the COVID-19 Genomics UK (COG-UK) Consortium | Harper VanSteenhouse, Yumi Kasai, David Gray, Carol Clugston, Anna Dominiczak and Alex Alderton, Roberto Amato, Jeffrey Barrett, Sonia Goncalves, Ewan Harrison, David K. Jackson, Ian Johnston, Dominic Kwiatkowski, Cordelia Langford, John Sillitoe on behalf of the Wellcome Sanger Institute COVID-19 Surveillance Team |
| EPI_ISL_2911226, EPI_ISL_2911227, EPI_ISL_2911228, EPI_ISL_2911229 | Lighthouse Lab in Milton Keynes | Wellcome Sanger Institute for the COVID-19 Genomics UK (COG-UK) Consortium | The Lighthouse Lab in Milton Keynes and Alex Alderton, Roberto Amato, Jeffrey Barrett, Sonia Goncalves, Ewan Harrison, David K. Jackson, Ian Johnston, Dominic Kwiatkowski, Cordelia Langford, John Sillitoe on behalf of the Wellcome Sanger Institute COVID-19 Surveillance Team |
| EPI_ISL_2911230 | Lighthouse Lab in Glasgow | Wellcome Sanger Institute for the COVID-19 Genomics UK (COG-UK) Consortium | Harper VanSteenhouse, Yumi Kasai, David Gray, Carol Clugston, Anna Dominiczak and Alex Alderton, Roberto Amato, Jeffrey Barrett, Sonia Goncalves, Ewan Harrison, David K. Jackson, Ian Johnston, Dominic Kwiatkowski, Cordelia Langford, John Sillitoe on behalf of the Wellcome Sanger Institute COVID-19 Surveillance Team |
| EPI_ISL_2911231, EPI_ISL_2911232, EPI_ISL_2911233 | Lighthouse Lab in Milton Keynes | Wellcome Sanger Institute for the COVID-19 Genomics UK (COG-UK) Consortium | The Lighthouse Lab in Milton Keynes and Alex Alderton, Roberto Amato, Jeffrey Barrett, Sonia Goncalves, Ewan Harrison, David K. Jackson, Ian Johnston, Dominic Kwiatkowski, Cordelia Langford, John Sillitoe on behalf of the Wellcome Sanger Institute COVID-19 Surveillance Team |
| EPI_ISL_2911234, EPI_ISL_2911235 | Lighthouse Lab in Glasgow | Wellcome Sanger Institute for the COVID-19 Genomics UK (COG-UK) Consortium | Harper VanSteenhouse, Yumi Kasai, David Gray, Carol Clugston, Anna Dominiczak and Alex Alderton, Roberto Amato, Jeffrey Barrett, Sonia Goncalves, Ewan Harrison, David K. Jackson, Ian Johnston, Dominic Kwiatkowski, Cordelia Langford, John Sillitoe on behalf of the Wellcome Sanger Institute COVID-19 Surveillance Team |
| EPI_ISL_2911236 | Lighthouse Lab in Milton Keynes | Wellcome Sanger Institute for the COVID-19 Genomics UK (COG-UK) Consortium | The Lighthouse Lab in Milton Keynes and Alex Alderton, Roberto Amato, Jeffrey Barrett, Sonia Goncalves, Ewan Harrison, David K. Jackson, Ian Johnston, Dominic Kwiatkowski, Cordelia Langford, John Sillitoe on behalf of the Wellcome Sanger Institute COVID-19 Surveillance Team |
| EPI_ISL_2911237, EPI_ISL_2911238 | Lighthouse Lab in Glasgow | Wellcome Sanger Institute for the COVID-19 Genomics UK (COG-UK) Consortium | Harper VanSteenhouse, Yumi Kasai, David Gray, Carol Clugston, Anna Dominiczak and Alex Alderton, Roberto Amato, Jeffrey Barrett, Sonia Goncalves, Ewan Harrison, David K. Jackson, Ian Johnston, Dominic Kwiatkowski, Cordelia Langford, John Sillitoe on behalf of the Wellcome Sanger Institute COVID-19 Surveillance Team |
| EPI_ISL_2911239, EPI_ISL_2911240, EPI_ISL_2911241, EPI_ISL_2911242, EPI_ISL_2911243, EPI_ISL_2911244, EPI_ISL_2911245, EPI_ISL_2911246, EPI_ISL_2911247, EPI_ISL_2911248, EPI_ISL_2911249 | | | |
| see above | Lighthouse Lab in Milton Keynes | Wellcome Sanger Institute for the COVID-19 Genomics UK (COG-UK) Consortium | The Lighthouse Lab in Milton Keynes and Alex Alderton, Roberto Amato, Jeffrey Barrett, Sonia Goncalves, Ewan Harrison, David K. Jackson, Ian Johnston, Dominic Kwiatkowski, Cordelia Langford, John Sillitoe on behalf of the Wellcome Sanger Institute COVID-19 Surveillance Team |
| EPI_ISL_2911250 | Lighthouse Lab in Glasgow | Wellcome Sanger Institute for the COVID-19 Genomics UK (COG-UK) Consortium | Harper VanSteenhouse, Yumi Kasai, David Gray, Carol Clugston, Anna Dominiczak and Alex Alderton, Roberto Amato, Jeffrey Barrett, Sonia Goncalves, Ewan Harrison, David K. Jackson, Ian Johnston, Dominic Kwiatkowski, Cordelia Langford, John Sillitoe on behalf of the Wellcome Sanger Institute COVID-19 Surveillance Team |
| EPI_ISL_2911251, EPI_ISL_2911252 | Lighthouse Lab in Milton Keynes | Wellcome Sanger Institute for the COVID-19 Genomics UK (COG-UK) Consortium | The Lighthouse Lab in Milton Keynes and Alex Alderton, Roberto Amato, Jeffrey Barrett, Sonia Goncalves, Ewan Harrison, David K. Jackson, Ian Johnston, Dominic Kwiatkowski, Cordelia Langford, John Sillitoe on behalf of the Wellcome Sanger Institute COVID-19 Surveillance Team |
| EPI_ISL_2911253 | Lighthouse Lab in Glasgow | Wellcome Sanger Institute for the COVID-19 Genomics UK (COG-UK) Consortium | Harper VanSteenhouse, Yumi Kasai, David Gray, Carol Clugston, Anna Dominiczak and Alex Alderton, Roberto Amato, Jeffrey Barrett, Sonia Goncalves, Ewan Harrison, David K. Jackson, Ian Johnston, Dominic Kwiatkowski, Cordelia Langford, John Sillitoe on behalf of the Wellcome Sanger Institute COVID-19 Surveillance Team |
| EPI_ISL_2911254, EPI_ISL_2911255, EPI_ISL_2911256 | Lighthouse Lab in Milton Keynes | Wellcome Sanger Institute for the COVID-19 Genomics UK (COG-UK) Consortium | The Lighthouse Lab in Milton Keynes and Alex Alderton, Roberto Amato, Jeffrey Barrett, Sonia Goncalves, Ewan Harrison, David K. Jackson, Ian Johnston, Dominic Kwiatkowski, Cordelia Langford, John Sillitoe on behalf of the Wellcome Sanger Institute COVID-19 Surveillance Team |
| EPI_ISL_2911257, EPI_ISL_2911258 | Lighthouse Lab in Glasgow | Wellcome Sanger Institute for the COVID-19 Genomics UK (COG-UK) Consortium | Harper VanSteenhouse, Yumi Kasai, David Gray, Carol Clugston, Anna Dominiczak and Alex Alderton, Roberto Amato, Jeffrey Barrett, Sonia Goncalves, Ewan Harrison, David K. Jackson, Ian Johnston, Dominic Kwiatkowski, Cordelia Langford, John Sillitoe on behalf of the Wellcome Sanger Institute COVID-19 Surveillance Team |
| EPI_ISL_2911259, EPI_ISL_2911260 | Lighthouse Lab in Milton Keynes | Wellcome Sanger Institute for the COVID-19 Genomics UK (COG-UK) Consortium | The Lighthouse Lab in Milton Keynes and Alex Alderton, Roberto Amato, Jeffrey Barrett, Sonia Goncalves, Ewan Harrison, David K. Jackson, Ian Johnston, Dominic Kwiatkowski, Cordelia Langford, John Sillitoe on behalf of the Wellcome Sanger Institute COVID-19 Surveillance Team |
| EPI_ISL_2911261 | Lighthouse Lab in Glasgow | Wellcome Sanger Institute for the COVID-19 Genomics UK (COG-UK) Consortium | Harper VanSteenhouse, Yumi Kasai, David Gray, Carol Clugston, Anna Dominiczak and Alex Alderton, Roberto Amato, Jeffrey Barrett, Sonia Goncalves, Ewan Harrison, David K. Jackson, Ian Johnston, Dominic Kwiatkowski, Cordelia Langford, John Sillitoe on behalf of the Wellcome Sanger Institute COVID-19 Surveillance Team |
| EPI_ISL_2911262, EPI_ISL_2911263, EPI_ISL_2911264, EPI_ISL_2911265, EPI_ISL_2911266, EPI_ISL_2911267, EPI_ISL_2911268, EPI_ISL_2911269 | Lighthouse Lab in Milton Keynes | Wellcome Sanger Institute for the COVID-19 Genomics UK (COG-UK) Consortium | The Lighthouse Lab in Milton Keynes and Alex Alderton, Roberto Amato, Jeffrey Barrett, Sonia Goncalves, Ewan Harrison, David K. Jackson, Ian Johnston, Dominic Kwiatkowski, Cordelia Langford, John Sillitoe on behalf of the Wellcome Sanger Institute COVID-19 Surveillance Team |
| EPI_ISL_2911270, EPI_ISL_2911271 | Lighthouse Lab in Glasgow | Wellcome Sanger Institute for the COVID-19 Genomics UK (COG-UK) Consortium | Harper VanSteenhouse, Yumi Kasai, David Gray, Carol Clugston, Anna Dominiczak and Alex Alderton, Roberto Amato, Jeffrey Barrett, Sonia Goncalves, Ewan Harrison, David K. Jackson, Ian Johnston, Dominic Kwiatkowski, Cordelia Langford, John Sillitoe on behalf of the Wellcome Sanger Institute COVID-19 Surveillance Team |
| EPI_ISL_2911272, EPI_ISL_2911273, EPI_ISL_2911274, EPI_ISL_2911275, EPI_ISL_2911276, EPI_ISL_2911277 | Lighthouse Lab in Milton Keynes | Wellcome Sanger Institute for the COVID-19 Genomics UK (COG-UK) Consortium | The Lighthouse Lab in Milton Keynes and Alex Alderton, Roberto Amato, Jeffrey Barrett, Sonia Goncalves, Ewan Harrison, David K. Jackson, Ian Johnston, Dominic Kwiatkowski, Cordelia Langford, John Sillitoe on behalf of the Wellcome Sanger Institute COVID-19 Surveillance Team |
| EPI_ISL_2911278 | Lighthouse Lab in Glasgow | Wellcome Sanger Institute for the COVID-19 Genomics UK (COG-UK) Consortium | Harper VanSteenhouse, Yumi Kasai, David Gray, Carol Clugston, Anna Dominiczak and Alex Alderton, Roberto Amato, Jeffrey Barrett, Sonia Goncalves, Ewan Harrison, David K. Jackson, Ian Johnston, Dominic Kwiatkowski, Cordelia Langford, John Sillitoe on behalf of the Wellcome Sanger Institute COVID-19 Surveillance Team |
| EPI_ISL_2911279, EPI_ISL_2911280, EPI_ISL_2911281, EPI_ISL_2911282, EPI_ISL_2911283, EPI_ISL_2911284, EPI_ISL_2911285, EPI_ISL_2911286, EPI_ISL_2911287, EPI_ISL_2911288, EPI_ISL_2911289, EPI_ISL_2911290 | | | |
| see above | Lighthouse Lab in Milton Keynes | Wellcome Sanger Institute for the COVID-19 Genomics UK (COG-UK) Consortium | The Lighthouse Lab in Milton Keynes and Alex Alderton, Roberto Amato, Jeffrey Barrett, Sonia Goncalves, Ewan Harrison, David K. Jackson, Ian Johnston, Dominic Kwiatkowski, Cordelia Langford, John Sillitoe on behalf of the Wellcome Sanger Institute COVID-19 Surveillance Team |
| EPI_ISL_2911291 | Lighthouse Lab in Glasgow | Wellcome Sanger Institute for the COVID-19 Genomics UK (COG-UK) Consortium | Harper VanSteenhouse, Yumi Kasai, David Gray, Carol Clugston, Anna Dominiczak and Alex Alderton, Roberto Amato, Jeffrey Barrett, Sonia Goncalves, Ewan Harrison, David K. Jackson, Ian Johnston, Dominic Kwiatkowski, Cordelia Langford, John Sillitoe on behalf of the Wellcome Sanger Institute COVID-19 Surveillance Team |
| EPI_ISL_2911292 | Lighthouse Lab in Milton Keynes | Wellcome Sanger Institute for the COVID-19 Genomics UK (COG-UK) Consortium | The Lighthouse Lab in Milton Keynes and Alex Alderton, Roberto Amato, Jeffrey Barrett, Sonia Goncalves, Ewan Harrison, David K. Jackson, Ian Johnston, Dominic Kwiatkowski, Cordelia Langford, John Sillitoe on behalf of the Wellcome Sanger Institute COVID-19 Surveillance Team |
| EPI_ISL_2911293 | Lighthouse Lab in Glasgow | Wellcome Sanger Institute for the COVID-19 Genomics UK (COG-UK) Consortium | Harper VanSteenhouse, Yumi Kasai, David Gray, Carol Clugston, Anna Dominiczak and Alex Alderton, Roberto Amato, Jeffrey Barrett, Sonia Goncalves, Ewan Harrison, David K. Jackson, Ian Johnston, Dominic Kwiatkowski, Cordelia Langford, John Sillitoe on behalf of the Wellcome Sanger Institute COVID-19 Surveillance Team |
| EPI_ISL_2911294, EPI_ISL_2911295, EPI_ISL_2911296, EPI_ISL_2911297, EPI_ISL_2911298 | Lighthouse Lab in Milton Keynes | Wellcome Sanger Institute for the COVID-19 Genomics UK (COG-UK) Consortium | The Lighthouse Lab in Milton Keynes and Alex Alderton, Roberto Amato, Jeffrey Barrett, Sonia Goncalves, Ewan Harrison, David K. Jackson, Ian Johnston, Dominic Kwiatkowski, Cordelia Langford, John Sillitoe on behalf of the Wellcome Sanger Institute COVID-19 Surveillance Team |
| EPI_ISL_2911299 | Lighthouse Lab in Glasgow | Wellcome Sanger Institute for the COVID-19 Genomics UK (COG-UK) Consortium | Harper VanSteenhouse, Yumi Kasai, David Gray, Carol Clugston, Anna Dominiczak and Alex Alderton, Roberto Amato, Jeffrey Barrett, Sonia Goncalves, Ewan Harrison, David K. Jackson, Ian Johnston, Dominic Kwiatkowski, Cordelia Langford, John Sillitoe on behalf of the Wellcome Sanger Institute COVID-19 Surveillance Team |

| | | | |
|---|---|---|---|
| EPI_ISL_2911300, EPI_ISL_2911301 | Lighthouse Lab in Milton Keynes | Wellcome Sanger Institute for the COVID-19 Genomics UK (COG-UK) Consortium | The Lighthouse Lab in Milton Keynes and Alex Alderton, Roberto Amato, Jeffrey Barrett, Sonia Goncalves, Ewan Harrison, David K. Jackson, Ian Johnston, Dominic Kwiatkowski, Cordelia Langford, John Sillitoe on behalf of the Wellcome Sanger Institute COVID-19 Surveillance Team |
| EPI_ISL_2911302, EPI_ISL_2911303, EPI_ISL_2911304 | Lighthouse Lab in Glasgow | Wellcome Sanger Institute for the COVID-19 Genomics UK (COG-UK) Consortium | Harper VanSteenhouse, Yumi Kasai, David Gray, Carol Clugston, Anna Dominiczak and Alex Alderton, Roberto Amato, Jeffrey Barrett, Sonia Goncalves, Ewan Harrison, David K. Jackson, Ian Johnston, Dominic Kwiatkowski, Cordelia Langford, John Sillitoe on behalf of the Wellcome Sanger Institute COVID-19 Surveillance Team |
| EPI_ISL_2911305, EPI_ISL_2911306, EPI_ISL_2911307, EPI_ISL_2911308, EPI_ISL_2911309, EPI_ISL_2911310, EPI_ISL_2911311 | Lighthouse Lab in Milton Keynes | Wellcome Sanger Institute for the COVID-19 Genomics UK (COG-UK) Consortium | The Lighthouse Lab in Milton Keynes and Alex Alderton, Roberto Amato, Jeffrey Barrett, Sonia Goncalves, Ewan Harrison, David K. Jackson, Ian Johnston, Dominic Kwiatkowski, Cordelia Langford, John Sillitoe on behalf of the Wellcome Sanger Institute COVID-19 Surveillance Team |
| EPI_ISL_2911312 | Lighthouse Lab in Glasgow | Wellcome Sanger Institute for the COVID-19 Genomics UK (COG-UK) Consortium | Harper VanSteenhouse, Yumi Kasai, David Gray, Carol Clugston, Anna Dominiczak and Alex Alderton, Roberto Amato, Jeffrey Barrett, Sonia Goncalves, Ewan Harrison, David K. Jackson, Ian Johnston, Dominic Kwiatkowski, Cordelia Langford, John Sillitoe on behalf of the Wellcome Sanger Institute COVID-19 Surveillance Team |
| EPI_ISL_2911313, EPI_ISL_2911314, EPI_ISL_2911315 | Lighthouse Lab in Milton Keynes | Wellcome Sanger Institute for the COVID-19 Genomics UK (COG-UK) Consortium | The Lighthouse Lab in Milton Keynes and Alex Alderton, Roberto Amato, Jeffrey Barrett, Sonia Goncalves, Ewan Harrison, David K. Jackson, Ian Johnston, Dominic Kwiatkowski, Cordelia Langford, John Sillitoe on behalf of the Wellcome Sanger Institute COVID-19 Surveillance Team |
| EPI_ISL_2911316 | Lighthouse Lab in Glasgow | Wellcome Sanger Institute for the COVID-19 Genomics UK (COG-UK) Consortium | Harper VanSteenhouse, Yumi Kasai, David Gray, Carol Clugston, Anna Dominiczak and Alex Alderton, Roberto Amato, Jeffrey Barrett, Sonia Goncalves, Ewan Harrison, David K. Jackson, Ian Johnston, Dominic Kwiatkowski, Cordelia Langford, John Sillitoe on behalf of the Wellcome Sanger Institute COVID-19 Surveillance Team |
| EPI_ISL_2911317, EPI_ISL_2911318 | Lighthouse Lab in Milton Keynes | Wellcome Sanger Institute for the COVID-19 Genomics UK (COG-UK) Consortium | The Lighthouse Lab in Milton Keynes and Alex Alderton, Roberto Amato, Jeffrey Barrett, Sonia Goncalves, Ewan Harrison, David K. Jackson, Ian Johnston, Dominic Kwiatkowski, Cordelia Langford, John Sillitoe on behalf of the Wellcome Sanger Institute COVID-19 Surveillance Team |
| EPI_ISL_2911319 | Lighthouse Lab in Glasgow | Wellcome Sanger Institute for the COVID-19 Genomics UK (COG-UK) Consortium | Harper VanSteenhouse, Yumi Kasai, David Gray, Carol Clugston, Anna Dominiczak and Alex Alderton, Roberto Amato, Jeffrey Barrett, Sonia Goncalves, Ewan Harrison, David K. Jackson, Ian Johnston, Dominic Kwiatkowski, Cordelia Langford, John Sillitoe on behalf of the Wellcome Sanger Institute COVID-19 Surveillance Team |
| EPI_ISL_2911320, EPI_ISL_2911321, EPI_ISL_2911322, EPI_ISL_2911323, EPI_ISL_2911324, EPI_ISL_2911325, EPI_ISL_2911326, EPI_ISL_2911327, EPI_ISL_2911328, EPI_ISL_2911329, EPI_ISL_2911330, EPI_ISL_2911331, EPI_ISL_2911332 | | | |
| see above | Lighthouse Lab in Milton Keynes | Wellcome Sanger Institute for the COVID-19 Genomics UK (COG-UK) Consortium | The Lighthouse Lab in Milton Keynes and Alex Alderton, Roberto Amato, Jeffrey Barrett, Sonia Goncalves, Ewan Harrison, David K. Jackson, Ian Johnston, Dominic Kwiatkowski, Cordelia Langford, John Sillitoe on behalf of the Wellcome Sanger Institute COVID-19 Surveillance Team |
| EPI_ISL_2911333 | Lighthouse Lab in Glasgow | Wellcome Sanger Institute for the COVID-19 Genomics UK (COG-UK) Consortium | Harper VanSteenhouse, Yumi Kasai, David Gray, Carol Clugston, Anna Dominiczak and Alex Alderton, Roberto Amato, Jeffrey Barrett, Sonia Goncalves, Ewan Harrison, David K. Jackson, Ian Johnston, Dominic Kwiatkowski, Cordelia Langford, John Sillitoe on behalf of the Wellcome Sanger Institute COVID-19 Surveillance Team |
| EPI_ISL_2911334, EPI_ISL_2911335, EPI_ISL_2911336, EPI_ISL_2911337, EPI_ISL_2911338, EPI_ISL_2911339, EPI_ISL_2911340, EPI_ISL_2911341, EPI_ISL_2911342 | Lighthouse Lab in Milton Keynes | Wellcome Sanger Institute for the COVID-19 Genomics UK (COG-UK) Consortium | The Lighthouse Lab in Milton Keynes and Alex Alderton, Roberto Amato, Jeffrey Barrett, Sonia Goncalves, Ewan Harrison, David K. Jackson, Ian Johnston, Dominic Kwiatkowski, Cordelia Langford, John Sillitoe on behalf of the Wellcome Sanger Institute COVID-19 Surveillance Team |
| EPI_ISL_2911343 | Lighthouse Lab in Glasgow | Wellcome Sanger Institute for the COVID-19 Genomics UK (COG-UK) Consortium | Harper VanSteenhouse, Yumi Kasai, David Gray, Carol Clugston, Anna Dominiczak and Alex Alderton, Roberto Amato, Jeffrey Barrett, Sonia Goncalves, Ewan Harrison, David K. Jackson, Ian Johnston, Dominic Kwiatkowski, Cordelia Langford, John Sillitoe on behalf of the Wellcome Sanger Institute COVID-19 Surveillance Team |
| EPI_ISL_2911344, EPI_ISL_2911345, EPI_ISL_2911346 | Lighthouse Lab in Milton Keynes | Wellcome Sanger Institute for the COVID-19 Genomics UK (COG-UK) Consortium | The Lighthouse Lab in Milton Keynes and Alex Alderton, Roberto Amato, Jeffrey Barrett, Sonia Goncalves, Ewan Harrison, David K. Jackson, Ian Johnston, Dominic Kwiatkowski, Cordelia Langford, John Sillitoe on behalf of the Wellcome Sanger Institute COVID-19 Surveillance Team |
| EPI_ISL_2911347, EPI_ISL_2911348 | Lighthouse Lab in Glasgow | Wellcome Sanger Institute for the COVID-19 Genomics UK (COG-UK) Consortium | Harper VanSteenhouse, Yumi Kasai, David Gray, Carol Clugston, Anna Dominiczak and Alex Alderton, Roberto Amato, Jeffrey Barrett, Sonia Goncalves, Ewan Harrison, David K. Jackson, Ian Johnston, Dominic Kwiatkowski, Cordelia Langford, John Sillitoe on behalf of the Wellcome Sanger Institute COVID-19 Surveillance Team |
| EPI_ISL_2911349, EPI_ISL_2911350, EPI_ISL_2911351 | Lighthouse Lab in Milton Keynes | Wellcome Sanger Institute for the COVID-19 Genomics UK (COG-UK) Consortium | The Lighthouse Lab in Milton Keynes and Alex Alderton, Roberto Amato, Jeffrey Barrett, Sonia Goncalves, Ewan Harrison, David K. Jackson, Ian Johnston, Dominic Kwiatkowski, Cordelia Langford, John Sillitoe on behalf of the Wellcome Sanger Institute COVID-19 Surveillance Team |
| EPI_ISL_2911352 | Lighthouse Lab in Glasgow | Wellcome Sanger Institute for the COVID-19 Genomics UK (COG-UK) Consortium | Harper VanSteenhouse, Yumi Kasai, David Gray, Carol Clugston, Anna Dominiczak and Alex Alderton, Roberto Amato, Jeffrey Barrett, Sonia Goncalves, Ewan Harrison, David K. Jackson, Ian Johnston, Dominic Kwiatkowski, Cordelia Langford, John Sillitoe on behalf of the Wellcome Sanger Institute COVID-19 Surveillance Team |
| EPI_ISL_2911353, EPI_ISL_2911354 | Lighthouse Lab in Milton Keynes | Wellcome Sanger Institute for the COVID-19 Genomics UK (COG-UK) Consortium | The Lighthouse Lab in Milton Keynes and Alex Alderton, Roberto Amato, Jeffrey Barrett, Sonia Goncalves, Ewan Harrison, David K. Jackson, Ian Johnston, Dominic Kwiatkowski, Cordelia Langford, John Sillitoe on behalf of the Wellcome Sanger Institute COVID-19 Surveillance Team |
| EPI_ISL_2911355 | Lighthouse Lab in Glasgow | Wellcome Sanger Institute for the COVID-19 Genomics UK (COG-UK) Consortium | Harper VanSteenhouse, Yumi Kasai, David Gray, Carol Clugston, Anna Dominiczak and Alex Alderton, Roberto Amato, Jeffrey Barrett, Sonia Goncalves, Ewan Harrison, David K. Jackson, Ian Johnston, Dominic Kwiatkowski, Cordelia Langford, John Sillitoe on behalf of the Wellcome Sanger Institute COVID-19 Surveillance Team |
| EPI_ISL_2911356, EPI_ISL_2911357, EPI_ISL_2911358, EPI_ISL_2911359, EPI_ISL_2911360, EPI_ISL_2911361, EPI_ISL_2911362, EPI_ISL_2911363, EPI_ISL_2911364, EPI_ISL_2911365, EPI_ISL_2911366, EPI_ISL_2911367, EPI_ISL_2911368, EPI_ISL_2911369, EPI_ISL_2911370, EPI_ISL_2911371 | | | |
| see above | Lighthouse Lab in Milton Keynes | Wellcome Sanger Institute for the COVID-19 Genomics UK (COG-UK) Consortium | The Lighthouse Lab in Milton Keynes and Alex Alderton, Roberto Amato, Jeffrey Barrett, Sonia Goncalves, Ewan Harrison, David K. Jackson, Ian Johnston, Dominic Kwiatkowski, Cordelia Langford, John Sillitoe on behalf of the Wellcome Sanger Institute COVID-19 Surveillance Team |
| EPI_ISL_2911372 | Lighthouse Lab in Glasgow | Wellcome Sanger Institute for the COVID-19 Genomics UK (COG-UK) Consortium | Harper VanSteenhouse, Yumi Kasai, David Gray, Carol Clugston, Anna Dominiczak and Alex Alderton, Roberto Amato, Jeffrey Barrett, Sonia Goncalves, Ewan Harrison, David K. Jackson, Ian Johnston, Dominic Kwiatkowski, Cordelia Langford, John Sillitoe on behalf of the Wellcome Sanger Institute COVID-19 Surveillance Team |
| EPI_ISL_2911373 | Lighthouse Lab in Milton Keynes | Wellcome Sanger Institute for the COVID-19 Genomics UK (COG-UK) Consortium | The Lighthouse Lab in Milton Keynes and Alex Alderton, Roberto Amato, Jeffrey Barrett, Sonia Goncalves, Ewan Harrison, David K. Jackson, Ian Johnston, Dominic Kwiatkowski, Cordelia Langford, John Sillitoe on behalf of the Wellcome Sanger Institute COVID-19 Surveillance Team |
| EPI_ISL_2911374 | Lighthouse Lab in Glasgow | Wellcome Sanger Institute for the COVID-19 Genomics UK (COG-UK) Consortium | Harper VanSteenhouse, Yumi Kasai, David Gray, Carol Clugston, Anna Dominiczak and Alex Alderton, Roberto Amato, Jeffrey Barrett, Sonia Goncalves, Ewan Harrison, David K. Jackson, Ian Johnston, Dominic Kwiatkowski, Cordelia Langford, John Sillitoe on behalf of the Wellcome Sanger Institute COVID-19 Surveillance Team |
| EPI_ISL_2911375, EPI_ISL_2911376, EPI_ISL_2911377, EPI_ISL_2911378 | Lighthouse Lab in Milton Keynes | Wellcome Sanger Institute for the COVID-19 Genomics UK (COG-UK) Consortium | The Lighthouse Lab in Milton Keynes and Alex Alderton, Roberto Amato, Jeffrey Barrett, Sonia Goncalves, Ewan Harrison, David K. Jackson, Ian Johnston, Dominic Kwiatkowski, Cordelia Langford, John Sillitoe on behalf of the Wellcome Sanger Institute COVID-19 Surveillance Team |
| EPI_ISL_2911379 | Lighthouse Lab in Glasgow | Wellcome Sanger Institute for the COVID-19 Genomics UK (COG-UK) Consortium | Harper VanSteenhouse, Yumi Kasai, David Gray, Carol Clugston, Anna Dominiczak and Alex Alderton, Roberto Amato, Jeffrey Barrett, Sonia Goncalves, Ewan Harrison, David K. Jackson, Ian Johnston, Dominic Kwiatkowski, Cordelia Langford, John Sillitoe on behalf of the Wellcome Sanger Institute COVID-19 Surveillance Team |
| EPI_ISL_2911380, EPI_ISL_2911381, EPI_ISL_2911382, EPI_ISL_2911383 | Lighthouse Lab in Milton Keynes | Wellcome Sanger Institute for the COVID-19 Genomics UK (COG-UK) Consortium | The Lighthouse Lab in Milton Keynes and Alex Alderton, Roberto Amato, Jeffrey Barrett, Sonia Goncalves, Ewan Harrison, David K. Jackson, Ian Johnston, Dominic Kwiatkowski, Cordelia Langford, John Sillitoe on behalf of the Wellcome Sanger Institute COVID-19 Surveillance Team |
| EPI_ISL_2911384 | Lighthouse Lab in Glasgow | Wellcome Sanger Institute for the COVID-19 Genomics UK (COG-UK) Consortium | Harper VanSteenhouse, Yumi Kasai, David Gray, Carol Clugston, Anna Dominiczak and Alex Alderton, Roberto Amato, Jeffrey Barrett, Sonia Goncalves, Ewan Harrison, David K. Jackson, Ian Johnston, Dominic Kwiatkowski, Cordelia Langford, John Sillitoe on behalf of the Wellcome Sanger Institute COVID-19 Surveillance Team |
| EPI_ISL_2911385 | Lighthouse Lab in Milton Keynes | Wellcome Sanger Institute for the COVID-19 Genomics UK (COG-UK) Consortium | The Lighthouse Lab in Milton Keynes and Alex Alderton, Roberto Amato, Jeffrey Barrett, Sonia Goncalves, Ewan Harrison, David K. Jackson, Ian Johnston, Dominic Kwiatkowski, Cordelia Langford, John Sillitoe on behalf of the Wellcome Sanger Institute COVID-19 Surveillance Team |
| EPI_ISL_2911386 | Lighthouse Lab in Glasgow | Wellcome Sanger Institute for the COVID-19 Genomics UK (COG-UK) Consortium | Harper VanSteenhouse, Yumi Kasai, David Gray, Carol Clugston, Anna Dominiczak and Alex Alderton, Roberto Amato, Jeffrey Barrett, Sonia Goncalves, Ewan Harrison, David K. Jackson, Ian Johnston, Dominic Kwiatkowski, Cordelia Langford, John Sillitoe on behalf of the Wellcome Sanger Institute COVID-19 Surveillance Team |
| EPI_ISL_2911387 | Lighthouse Lab in Milton Keynes | Wellcome Sanger Institute for the COVID-19 Genomics UK (COG-UK) Consortium | The Lighthouse Lab in Milton Keynes and Alex Alderton, Roberto Amato, Jeffrey Barrett, Sonia Goncalves, Ewan Harrison, David K. Jackson, Ian Johnston, Dominic Kwiatkowski, Cordelia Langford, John Sillitoe on behalf of the Wellcome Sanger Institute COVID-19 Surveillance Team |
| EPI_ISL_2911388 | Lighthouse Lab in Glasgow | Wellcome Sanger Institute for the COVID-19 Genomics UK (COG-UK) Consortium | Harper VanSteenhouse, Yumi Kasai, David Gray, Carol Clugston, Anna Dominiczak and Alex Alderton, Roberto Amato, Jeffrey Barrett, Sonia Goncalves, Ewan Harrison, David K. Jackson, Ian Johnston, Dominic Kwiatkowski, Cordelia Langford, John Sillitoe on behalf of the Wellcome Sanger Institute COVID-19 Surveillance Team |
| EPI_ISL_2911389, EPI_ISL_2911390, EPI_ISL_2911391 | Lighthouse Lab in Milton Keynes | Wellcome Sanger Institute for the COVID-19 Genomics UK (COG-UK) Consortium | The Lighthouse Lab in Milton Keynes and Alex Alderton, Roberto Amato, Jeffrey Barrett, Sonia Goncalves, Ewan Harrison, David K. Jackson, Ian Johnston, Dominic Kwiatkowski, Cordelia Langford, John Sillitoe on behalf of the Wellcome Sanger Institute COVID-19 Surveillance Team |
| EPI_ISL_2911392 | Lighthouse Lab in Glasgow | Wellcome Sanger Institute for the COVID-19 Genomics UK (COG-UK) Consortium | Harper VanSteenhouse, Yumi Kasai, David Gray, Carol Clugston, Anna Dominiczak and Alex Alderton, Roberto Amato, Jeffrey Barrett, Sonia Goncalves, Ewan Harrison, David K. Jackson, Ian Johnston, Dominic Kwiatkowski, Cordelia Langford, John Sillitoe on behalf of the Wellcome Sanger Institute COVID-19 Surveillance Team |
| EPI_ISL_2911393, EPI_ISL_2911394, EPI_ISL_2911395, EPI_ISL_2911396, EPI_ISL_2911397, EPI_ISL_2911398, EPI_ISL_2911399 | Lighthouse Lab in Milton Keynes | Wellcome Sanger Institute for the COVID-19 Genomics UK (COG-UK) Consortium | The Lighthouse Lab in Milton Keynes and Alex Alderton, Roberto Amato, Jeffrey Barrett, Sonia Goncalves, Ewan Harrison, David K. Jackson, Ian Johnston, Dominic Kwiatkowski, Cordelia Langford, John Sillitoe on behalf of the Wellcome Sanger Institute COVID-19 Surveillance Team |

| | | | |
|---|---|---|---|
| EPI_ISL_2911400 | Lighthouse Lab in Glasgow | Wellcome Sanger Institute for the COVID-19 Genomics UK (COG-UK) Consortium | Harper VanSteenhouse, Yumi Kasai, David Gray, Carol Clugston, Anna Dominiczak and Alex Alderton, Roberto Amato, Jeffrey Barrett, Sonia Goncalves, Ewan Harrison, David K. Jackson, Ian Johnston, Dominic Kwiatkowski, Cordelia Langford, John Sillitoe on behalf of the Wellcome Sanger Institute COVID-19 Surveillance Team |
| EPI_ISL_2911401, EPI_ISL_2911402, EPI_ISL_2911403, EPI_ISL_2911404, EPI_ISL_2911405, EPI_ISL_2911406, EPI_ISL_2911407 | Lighthouse Lab in Milton Keynes | Wellcome Sanger Institute for the COVID-19 Genomics UK (COG-UK) Consortium | The Lighthouse Lab in Milton Keynes and Alex Alderton, Roberto Amato, Jeffrey Barrett, Sonia Goncalves, Ewan Harrison, David K. Jackson, Ian Johnston, Dominic Kwiatkowski, Cordelia Langford, John Sillitoe on behalf of the Wellcome Sanger Institute COVID-19 Surveillance Team |
| EPI_ISL_2911408 | Lighthouse Lab in Glasgow | Wellcome Sanger Institute for the COVID-19 Genomics UK (COG-UK) Consortium | Harper VanSteenhouse, Yumi Kasai, David Gray, Carol Clugston, Anna Dominiczak and Alex Alderton, Roberto Amato, Jeffrey Barrett, Sonia Goncalves, Ewan Harrison, David K. Jackson, Ian Johnston, Dominic Kwiatkowski, Cordelia Langford, John Sillitoe on behalf of the Wellcome Sanger Institute COVID-19 Surveillance Team |
| EPI_ISL_2911409, EPI_ISL_2911410, EPI_ISL_2911411 | Lighthouse Lab in Milton Keynes | Wellcome Sanger Institute for the COVID-19 Genomics UK (COG-UK) Consortium | The Lighthouse Lab in Milton Keynes and Alex Alderton, Roberto Amato, Jeffrey Barrett, Sonia Goncalves, Ewan Harrison, David K. Jackson, Ian Johnston, Dominic Kwiatkowski, Cordelia Langford, John Sillitoe on behalf of the Wellcome Sanger Institute COVID-19 Surveillance Team |
| EPI_ISL_2911412 | Lighthouse Lab in Glasgow | Wellcome Sanger Institute for the COVID-19 Genomics UK (COG-UK) Consortium | Harper VanSteenhouse, Yumi Kasai, David Gray, Carol Clugston, Anna Dominiczak and Alex Alderton, Roberto Amato, Jeffrey Barrett, Sonia Goncalves, Ewan Harrison, David K. Jackson, Ian Johnston, Dominic Kwiatkowski, Cordelia Langford, John Sillitoe on behalf of the Wellcome Sanger Institute COVID-19 Surveillance Team |
| EPI_ISL_2911413, EPI_ISL_2911414, EPI_ISL_2911415, EPI_ISL_2911416, EPI_ISL_2911417, EPI_ISL_2911418 | Lighthouse Lab in Milton Keynes | Wellcome Sanger Institute for the COVID-19 Genomics UK (COG-UK) Consortium | The Lighthouse Lab in Milton Keynes and Alex Alderton, Roberto Amato, Jeffrey Barrett, Sonia Goncalves, Ewan Harrison, David K. Jackson, Ian Johnston, Dominic Kwiatkowski, Cordelia Langford, John Sillitoe on behalf of the Wellcome Sanger Institute COVID-19 Surveillance Team |
| EPI_ISL_2911419 | Lighthouse Lab in Glasgow | Wellcome Sanger Institute for the COVID-19 Genomics UK (COG-UK) Consortium | Harper VanSteenhouse, Yumi Kasai, David Gray, Carol Clugston, Anna Dominiczak and Alex Alderton, Roberto Amato, Jeffrey Barrett, Sonia Goncalves, Ewan Harrison, David K. Jackson, Ian Johnston, Dominic Kwiatkowski, Cordelia Langford, John Sillitoe on behalf of the Wellcome Sanger Institute COVID-19 Surveillance Team |
| EPI_ISL_2911420, EPI_ISL_2911421, EPI_ISL_2911422 | Lighthouse Lab in Milton Keynes | Wellcome Sanger Institute for the COVID-19 Genomics UK (COG-UK) Consortium | The Lighthouse Lab in Milton Keynes and Alex Alderton, Roberto Amato, Jeffrey Barrett, Sonia Goncalves, Ewan Harrison, David K. Jackson, Ian Johnston, Dominic Kwiatkowski, Cordelia Langford, John Sillitoe on behalf of the Wellcome Sanger Institute COVID-19 Surveillance Team |
| EPI_ISL_2911423 | Lighthouse Lab in Glasgow | Wellcome Sanger Institute for the COVID-19 Genomics UK (COG-UK) Consortium | Harper VanSteenhouse, Yumi Kasai, David Gray, Carol Clugston, Anna Dominiczak and Alex Alderton, Roberto Amato, Jeffrey Barrett, Sonia Goncalves, Ewan Harrison, David K. Jackson, Ian Johnston, Dominic Kwiatkowski, Cordelia Langford, John Sillitoe on behalf of the Wellcome Sanger Institute COVID-19 Surveillance Team |
| EPI_ISL_2911424, EPI_ISL_2911425, EPI_ISL_2911426 | Lighthouse Lab in Milton Keynes | Wellcome Sanger Institute for the COVID-19 Genomics UK (COG-UK) Consortium | The Lighthouse Lab in Milton Keynes and Alex Alderton, Roberto Amato, Jeffrey Barrett, Sonia Goncalves, Ewan Harrison, David K. Jackson, Ian Johnston, Dominic Kwiatkowski, Cordelia Langford, John Sillitoe on behalf of the Wellcome Sanger Institute COVID-19 Surveillance Team |
| EPI_ISL_2911427, EPI_ISL_2911428 | Lighthouse Lab in Glasgow | Wellcome Sanger Institute for the COVID-19 Genomics UK (COG-UK) Consortium | Harper VanSteenhouse, Yumi Kasai, David Gray, Carol Clugston, Anna Dominiczak and Alex Alderton, Roberto Amato, Jeffrey Barrett, Sonia Goncalves, Ewan Harrison, David K. Jackson, Ian Johnston, Dominic Kwiatkowski, Cordelia Langford, John Sillitoe on behalf of the Wellcome Sanger Institute COVID-19 Surveillance Team |
| EPI_ISL_2911429 | Lighthouse Lab in Milton Keynes | Wellcome Sanger Institute for the COVID-19 Genomics UK (COG-UK) Consortium | The Lighthouse Lab in Milton Keynes and Alex Alderton, Roberto Amato, Jeffrey Barrett, Sonia Goncalves, Ewan Harrison, David K. Jackson, Ian Johnston, Dominic Kwiatkowski, Cordelia Langford, John Sillitoe on behalf of the Wellcome Sanger Institute COVID-19 Surveillance Team |
| EPI_ISL_2911430 | Lighthouse Lab in Glasgow | Wellcome Sanger Institute for the COVID-19 Genomics UK (COG-UK) Consortium | Harper VanSteenhouse, Yumi Kasai, David Gray, Carol Clugston, Anna Dominiczak and Alex Alderton, Roberto Amato, Jeffrey Barrett, Sonia Goncalves, Ewan Harrison, David K. Jackson, Ian Johnston, Dominic Kwiatkowski, Cordelia Langford, John Sillitoe on behalf of the Wellcome Sanger Institute COVID-19 Surveillance Team |
| EPI_ISL_2911431, EPI_ISL_2911432 | Lighthouse Lab in Milton Keynes | Wellcome Sanger Institute for the COVID-19 Genomics UK (COG-UK) Consortium | The Lighthouse Lab in Milton Keynes and Alex Alderton, Roberto Amato, Jeffrey Barrett, Sonia Goncalves, Ewan Harrison, David K. Jackson, Ian Johnston, Dominic Kwiatkowski, Cordelia Langford, John Sillitoe on behalf of the Wellcome Sanger Institute COVID-19 Surveillance Team |
| EPI_ISL_2911433 | Lighthouse Lab in Glasgow | Wellcome Sanger Institute for the COVID-19 Genomics UK (COG-UK) Consortium | Harper VanSteenhouse, Yumi Kasai, David Gray, Carol Clugston, Anna Dominiczak and Alex Alderton, Roberto Amato, Jeffrey Barrett, Sonia Goncalves, Ewan Harrison, David K. Jackson, Ian Johnston, Dominic Kwiatkowski, Cordelia Langford, John Sillitoe on behalf of the Wellcome Sanger Institute COVID-19 Surveillance Team |
| EPI_ISL_2911434 | Lighthouse Lab in Milton Keynes | Wellcome Sanger Institute for the COVID-19 Genomics UK (COG-UK) Consortium | The Lighthouse Lab in Milton Keynes and Alex Alderton, Roberto Amato, Jeffrey Barrett, Sonia Goncalves, Ewan Harrison, David K. Jackson, Ian Johnston, Dominic Kwiatkowski, Cordelia Langford, John Sillitoe on behalf of the Wellcome Sanger Institute COVID-19 Surveillance Team |
| EPI_ISL_2911435 | Lighthouse Lab in Glasgow | Wellcome Sanger Institute for the COVID-19 Genomics UK (COG-UK) Consortium | Harper VanSteenhouse, Yumi Kasai, David Gray, Carol Clugston, Anna Dominiczak and Alex Alderton, Roberto Amato, Jeffrey Barrett, Sonia Goncalves, Ewan Harrison, David K. Jackson, Ian Johnston, Dominic Kwiatkowski, Cordelia Langford, John Sillitoe on behalf of the Wellcome Sanger Institute COVID-19 Surveillance Team |
| EPI_ISL_2911436, EPI_ISL_2911437, EPI_ISL_2911438, EPI_ISL_2911439, EPI_ISL_2911440, EPI_ISL_2911441, EPI_ISL_2911442, EPI_ISL_2911443 | Lighthouse Lab in Milton Keynes | Wellcome Sanger Institute for the COVID-19 Genomics UK (COG-UK) Consortium | The Lighthouse Lab in Milton Keynes and Alex Alderton, Roberto Amato, Jeffrey Barrett, Sonia Goncalves, Ewan Harrison, David K. Jackson, Ian Johnston, Dominic Kwiatkowski, Cordelia Langford, John Sillitoe on behalf of the Wellcome Sanger Institute COVID-19 Surveillance Team |
| EPI_ISL_2911444 | Lighthouse Lab in Glasgow | Wellcome Sanger Institute for the COVID-19 Genomics UK (COG-UK) Consortium | Harper VanSteenhouse, Yumi Kasai, David Gray, Carol Clugston, Anna Dominiczak and Alex Alderton, Roberto Amato, Jeffrey Barrett, Sonia Goncalves, Ewan Harrison, David K. Jackson, Ian Johnston, Dominic Kwiatkowski, Cordelia Langford, John Sillitoe on behalf of the Wellcome Sanger Institute COVID-19 Surveillance Team |
| EPI_ISL_2911445, EPI_ISL_2911446, EPI_ISL_2911447 | Lighthouse Lab in Milton Keynes | Wellcome Sanger Institute for the COVID-19 Genomics UK (COG-UK) Consortium | The Lighthouse Lab in Milton Keynes and Alex Alderton, Roberto Amato, Jeffrey Barrett, Sonia Goncalves, Ewan Harrison, David K. Jackson, Ian Johnston, Dominic Kwiatkowski, Cordelia Langford, John Sillitoe on behalf of the Wellcome Sanger Institute COVID-19 Surveillance Team |
| EPI_ISL_2911448 | Lighthouse Lab in Glasgow | Wellcome Sanger Institute for the COVID-19 Genomics UK (COG-UK) Consortium | Harper VanSteenhouse, Yumi Kasai, David Gray, Carol Clugston, Anna Dominiczak and Alex Alderton, Roberto Amato, Jeffrey Barrett, Sonia Goncalves, Ewan Harrison, David K. Jackson, Ian Johnston, Dominic Kwiatkowski, Cordelia Langford, John Sillitoe on behalf of the Wellcome Sanger Institute COVID-19 Surveillance Team |
| EPI_ISL_2911449, EPI_ISL_2911450, EPI_ISL_2911451, EPI_ISL_2911452, EPI_ISL_2911453, EPI_ISL_2911454, EPI_ISL_2911455, EPI_ISL_2911456, EPI_ISL_2911457, EPI_ISL_2911458, EPI_ISL_2911459, EPI_ISL_2911460, EPI_ISL_2911461 | | | |
| see above | Lighthouse Lab in Milton Keynes | Wellcome Sanger Institute for the COVID-19 Genomics UK (COG-UK) Consortium | The Lighthouse Lab in Milton Keynes and Alex Alderton, Roberto Amato, Jeffrey Barrett, Sonia Goncalves, Ewan Harrison, David K. Jackson, Ian Johnston, Dominic Kwiatkowski, Cordelia Langford, John Sillitoe on behalf of the Wellcome Sanger Institute COVID-19 Surveillance Team |
| EPI_ISL_2911462, EPI_ISL_2911463 | Lighthouse Lab in Glasgow | Wellcome Sanger Institute for the COVID-19 Genomics UK (COG-UK) Consortium | Harper VanSteenhouse, Yumi Kasai, David Gray, Carol Clugston, Anna Dominiczak and Alex Alderton, Roberto Amato, Jeffrey Barrett, Sonia Goncalves, Ewan Harrison, David K. Jackson, Ian Johnston, Dominic Kwiatkowski, Cordelia Langford, John Sillitoe on behalf of the Wellcome Sanger Institute COVID-19 Surveillance Team |
| EPI_ISL_2911464, EPI_ISL_2911465, EPI_ISL_2911466, EPI_ISL_2911467, EPI_ISL_2911468, EPI_ISL_2911469, EPI_ISL_2911470, EPI_ISL_2911471, EPI_ISL_2911472, EPI_ISL_2911473, EPI_ISL_2911474 | | | |
| see above | Lighthouse Lab in Milton Keynes | Wellcome Sanger Institute for the COVID-19 Genomics UK (COG-UK) Consortium | The Lighthouse Lab in Milton Keynes and Alex Alderton, Roberto Amato, Jeffrey Barrett, Sonia Goncalves, Ewan Harrison, David K. Jackson, Ian Johnston, Dominic Kwiatkowski, Cordelia Langford, John Sillitoe on behalf of the Wellcome Sanger Institute COVID-19 Surveillance Team |
| EPI_ISL_2911475 | Lighthouse Lab in Glasgow | Wellcome Sanger Institute for the COVID-19 Genomics UK (COG-UK) Consortium | Harper VanSteenhouse, Yumi Kasai, David Gray, Carol Clugston, Anna Dominiczak and Alex Alderton, Roberto Amato, Jeffrey Barrett, Sonia Goncalves, Ewan Harrison, David K. Jackson, Ian Johnston, Dominic Kwiatkowski, Cordelia Langford, John Sillitoe on behalf of the Wellcome Sanger Institute COVID-19 Surveillance Team |
| EPI_ISL_2911476 | Lighthouse Lab in Milton Keynes | Wellcome Sanger Institute for the COVID-19 Genomics UK (COG-UK) Consortium | The Lighthouse Lab in Milton Keynes and Alex Alderton, Roberto Amato, Jeffrey Barrett, Sonia Goncalves, Ewan Harrison, David K. Jackson, Ian Johnston, Dominic Kwiatkowski, Cordelia Langford, John Sillitoe on behalf of the Wellcome Sanger Institute COVID-19 Surveillance Team |
| EPI_ISL_2911477 | Lighthouse Lab in Glasgow | Wellcome Sanger Institute for the COVID-19 Genomics UK (COG-UK) Consortium | Harper VanSteenhouse, Yumi Kasai, David Gray, Carol Clugston, Anna Dominiczak and Alex Alderton, Roberto Amato, Jeffrey Barrett, Sonia Goncalves, Ewan Harrison, David K. Jackson, Ian Johnston, Dominic Kwiatkowski, Cordelia Langford, John Sillitoe on behalf of the Wellcome Sanger Institute COVID-19 Surveillance Team |
| EPI_ISL_2911478, EPI_ISL_2911479, EPI_ISL_2911480, EPI_ISL_2911481 | Lighthouse Lab in Milton Keynes | Wellcome Sanger Institute for the COVID-19 Genomics UK (COG-UK) Consortium | The Lighthouse Lab in Milton Keynes and Alex Alderton, Roberto Amato, Jeffrey Barrett, Sonia Goncalves, Ewan Harrison, David K. Jackson, Ian Johnston, Dominic Kwiatkowski, Cordelia Langford, John Sillitoe on behalf of the Wellcome Sanger Institute COVID-19 Surveillance Team |
| EPI_ISL_2911482 | Lighthouse Lab in Glasgow | Wellcome Sanger Institute for the COVID-19 Genomics UK (COG-UK) Consortium | Harper VanSteenhouse, Yumi Kasai, David Gray, Carol Clugston, Anna Dominiczak and Alex Alderton, Roberto Amato, Jeffrey Barrett, Sonia Goncalves, Ewan Harrison, David K. Jackson, Ian Johnston, Dominic Kwiatkowski, Cordelia Langford, John Sillitoe on behalf of the Wellcome Sanger Institute COVID-19 Surveillance Team |
| EPI_ISL_2911483, EPI_ISL_2911484 | Lighthouse Lab in Milton Keynes | Wellcome Sanger Institute for the COVID-19 Genomics UK (COG-UK) Consortium | The Lighthouse Lab in Milton Keynes and Alex Alderton, Roberto Amato, Jeffrey Barrett, Sonia Goncalves, Ewan Harrison, David K. Jackson, Ian Johnston, Dominic Kwiatkowski, Cordelia Langford, John Sillitoe on behalf of the Wellcome Sanger Institute COVID-19 Surveillance Team |
| EPI_ISL_2911485 | Lighthouse Lab in Glasgow | Wellcome Sanger Institute for the COVID-19 Genomics UK (COG-UK) Consortium | Harper VanSteenhouse, Yumi Kasai, David Gray, Carol Clugston, Anna Dominiczak and Alex Alderton, Roberto Amato, Jeffrey Barrett, Sonia Goncalves, Ewan Harrison, David K. Jackson, Ian Johnston, Dominic Kwiatkowski, Cordelia Langford, John Sillitoe on behalf of the Wellcome Sanger Institute COVID-19 Surveillance Team |
| EPI_ISL_2911486 | Lighthouse Lab in Milton Keynes | Wellcome Sanger Institute for the COVID-19 Genomics UK (COG-UK) Consortium | The Lighthouse Lab in Milton Keynes and Alex Alderton, Roberto Amato, Jeffrey Barrett, Sonia Goncalves, Ewan Harrison, David K. Jackson, Ian Johnston, Dominic Kwiatkowski, Cordelia Langford, John Sillitoe on behalf of the Wellcome Sanger Institute COVID-19 Surveillance Team |
| EPI_ISL_2911487 | Lighthouse Lab in Glasgow | Wellcome Sanger Institute for the COVID-19 Genomics UK (COG-UK) Consortium | Harper VanSteenhouse, Yumi Kasai, David Gray, Carol Clugston, Anna Dominiczak and Alex Alderton, Roberto Amato, Jeffrey Barrett, Sonia Goncalves, Ewan Harrison, David K. Jackson, Ian Johnston, Dominic Kwiatkowski, Cordelia Langford, John Sillitoe on behalf of the Wellcome Sanger Institute COVID-19 Surveillance Team |
| EPI_ISL_2911488, EPI_ISL_2911489 | Lighthouse Lab in Milton Keynes | Wellcome Sanger Institute for the COVID-19 Genomics UK (COG-UK) Consortium | The Lighthouse Lab in Milton Keynes and Alex Alderton, Roberto Amato, Jeffrey Barrett, Sonia Goncalves, Ewan Harrison, David K. Jackson, Ian Johnston, Dominic Kwiatkowski, Cordelia Langford, John Sillitoe on behalf of the Wellcome Sanger Institute COVID-19 Surveillance Team |
| EPI_ISL_2911490, EPI_ISL_2911491 | Lighthouse Lab in Glasgow | Wellcome Sanger Institute for the COVID-19 Genomics UK (COG-UK) Consortium | Harper VanSteenhouse, Yumi Kasai, David Gray, Carol Clugston, Anna Dominiczak and Alex Alderton, Roberto Amato, Jeffrey Barrett, Sonia Goncalves, Ewan Harrison, David K. Jackson, Ian Johnston, Dominic Kwiatkowski, Cordelia Langford, John Sillitoe on behalf of the Wellcome Sanger Institute COVID-19 Surveillance Team |
| EPI_ISL_2911492, | Lighthouse Lab in Milton Keynes | Wellcome Sanger Institute for the COVID-19 Genomics UK (COG-UK) | The Lighthouse Lab in Milton Keynes and Alex Alderton, Roberto Amato, Jeffrey Barrett, Sonia Goncalves, Ewan Harrison, David K. Jackson, Ian Johnston, Dominic Kwiatkowski, Cordelia Langford, John Sillitoe on behalf of the Wellcome Sanger |

| | | | |
|---|---|---|---|
| EPI_ISL_2911493, EPI_ISL_2911494, EPI_ISL_2911495, EPI_ISL_2911496 | | Consortium | Institute COVID-19 Surveillance Team |
| EPI_ISL_2911497 | Lighthouse Lab in Glasgow | Wellcome Sanger Institute for the COVID-19 Genomics UK (COG-UK) Consortium | Harper VanSteenhouse, Yumi Kasai, David Gray, Carol Clugston, Anna Dominiczak and Alex Alderton, Roberto Amato, Jeffrey Barrett, Sonia Goncalves, Ewan Harrison, David K. Jackson, Ian Johnston, Dominic Kwiatkowski, Cordelia Langford, John Sillitoe on behalf of the Wellcome Sanger Institute COVID-19 Surveillance Team |
| EPI_ISL_2911498, EPI_ISL_2911499, EPI_ISL_2911500, EPI_ISL_2911501, EPI_ISL_2911502, EPI_ISL_2911503, EPI_ISL_2911504, EPI_ISL_2911505, EPI_ISL_2911506, EPI_ISL_2911507 | Lighthouse Lab in Milton Keynes | Wellcome Sanger Institute for the COVID-19 Genomics UK (COG-UK) Consortium | The Lighthouse Lab in Milton Keynes and Alex Alderton, Roberto Amato, Jeffrey Barrett, Sonia Goncalves, Ewan Harrison, David K. Jackson, Ian Johnston, Dominic Kwiatkowski, Cordelia Langford, John Sillitoe on behalf of the Wellcome Sanger Institute COVID-19 Surveillance Team |
| EPI_ISL_2911508 | Lighthouse Lab in Glasgow | Wellcome Sanger Institute for the COVID-19 Genomics UK (COG-UK) Consortium | Harper VanSteenhouse, Yumi Kasai, David Gray, Carol Clugston, Anna Dominiczak and Alex Alderton, Roberto Amato, Jeffrey Barrett, Sonia Goncalves, Ewan Harrison, David K. Jackson, Ian Johnston, Dominic Kwiatkowski, Cordelia Langford, John Sillitoe on behalf of the Wellcome Sanger Institute COVID-19 Surveillance Team |
| EPI_ISL_2911509, EPI_ISL_2911510, EPI_ISL_2911511, EPI_ISL_2911512, EPI_ISL_2911513, EPI_ISL_2911514, EPI_ISL_2911515, EPI_ISL_2911516, EPI_ISL_2911517, EPI_ISL_2911518, EPI_ISL_2911519, EPI_ISL_2911520, EPI_ISL_2911521, EPI_ISL_2911522, EPI_ISL_2911523, EPI_ISL_2911524, EPI_ISL_2911525, EPI_ISL_2911526 | | | |
| see above | Lighthouse Lab in Milton Keynes | Wellcome Sanger Institute for the COVID-19 Genomics UK (COG-UK) Consortium | The Lighthouse Lab in Milton Keynes and Alex Alderton, Roberto Amato, Jeffrey Barrett, Sonia Goncalves, Ewan Harrison, David K. Jackson, Ian Johnston, Dominic Kwiatkowski, Cordelia Langford, John Sillitoe on behalf of the Wellcome Sanger Institute COVID-19 Surveillance Team |
| EPI_ISL_2911527, EPI_ISL_2911528, EPI_ISL_2911529, EPI_ISL_2911530, EPI_ISL_2911531, EPI_ISL_2911532, EPI_ISL_2911533, EPI_ISL_2911534, EPI_ISL_2911535, EPI_ISL_2911536, EPI_ISL_2911537, EPI_ISL_2911538, EPI_ISL_2911539, EPI_ISL_2911540, EPI_ISL_2911541, EPI_ISL_2911542, EPI_ISL_2911543, EPI_ISL_2911544, EPI_ISL_2911545, EPI_ISL_2911546, EPI_ISL_2911547, EPI_ISL_2911548, EPI_ISL_2911549, EPI_ISL_2911550, EPI_ISL_2911551, EPI_ISL_2911552, EPI_ISL_2911553, EPI_ISL_2911554, EPI_ISL_2911555, EPI_ISL_2911556, EPI_ISL_2911557, EPI_ISL_2911558, EPI_ISL_2911559, EPI_ISL_2911560, EPI_ISL_2911561, EPI_ISL_2911562, EPI_ISL_2911563, EPI_ISL_2911564, EPI_ISL_2911565, EPI_ISL_2911566, EPI_ISL_2911567, EPI_ISL_2911568, EPI_ISL_2911569, EPI_ISL_2911570, EPI_ISL_2911571, EPI_ISL_2911572, EPI_ISL_2911573, EPI_ISL_2911574, EPI_ISL_2911575, EPI_ISL_2911576, EPI_ISL_2911577, EPI_ISL_2911578, EPI_ISL_2911579, EPI_ISL_2911580, EPI_ISL_2911582, EPI_ISL_2911583, EPI_ISL_2911584, EPI_ISL_2911585, EPI_ISL_2911586, EPI_ISL_2911587, EPI_ISL_2911588, EPI_ISL_2911589, EPI_ISL_2911590, EPI_ISL_2911591, EPI_ISL_2911592, EPI_ISL_2911593, EPI_ISL_2911594, EPI_ISL_2911595, EPI_ISL_2911596, EPI_ISL_2911597, EPI_ISL_2911598, EPI_ISL_2911599, EPI_ISL_2911600, EPI_ISL_2911601, EPI_ISL_2911602, EPI_ISL_2911603, EPI_ISL_2911604, EPI_ISL_2911605, EPI_ISL_2911606, EPI_ISL_2911607, EPI_ISL_2911608, EPI_ISL_2911609, EPI_ISL_2911610, EPI_ISL_2911611, EPI_ISL_2911612, EPI_ISL_2911613, EPI_ISL_2911614, EPI_ISL_2911615 | | | |
| see above | Lighthouse Lab in Glasgow | Wellcome Sanger Institute for the COVID-19 Genomics UK (COG-UK) Consortium | Harper VanSteenhouse, Yumi Kasai, David Gray, Carol Clugston, Anna Dominiczak and Alex Alderton, Roberto Amato, Jeffrey Barrett, Sonia Goncalves, Ewan Harrison, David K. Jackson, Ian Johnston, Dominic Kwiatkowski, Cordelia Langford, John Sillitoe on behalf of the Wellcome Sanger Institute COVID-19 Surveillance Team |
| EPI_ISL_2911616, EPI_ISL_2911617 | Lighthouse Lab in Milton Keynes | Wellcome Sanger Institute for the COVID-19 Genomics UK (COG-UK) Consortium | The Lighthouse Lab in Milton Keynes and Alex Alderton, Roberto Amato, Jeffrey Barrett, Sonia Goncalves, Ewan Harrison, David K. Jackson, Ian Johnston, Dominic Kwiatkowski, Cordelia Langford, John Sillitoe on behalf of the Wellcome Sanger Institute COVID-19 Surveillance Team |
| EPI_ISL_2911618, EPI_ISL_2911619, EPI_ISL_2911620, EPI_ISL_2911621 | Lighthouse Lab in Alderley Park | Wellcome Sanger Institute for the COVID-19 Genomics UK (COG-UK) Consortium | Jacquelyn Wynn, Mairead Hyland, The Lighthouse Lab in Alderley Park and Alex Alderton, Roberto Amato, Jeffrey Barrett, Sonia Goncalves, Ewan Harrison, David K. Jackson, Ian Johnston, Dominic Kwiatkowski, Cordelia Langford, John Sillitoe on behalf of the Wellcome Sanger Institute COVID-19 Surveillance Team |
| EPI_ISL_2911622 | Lighthouse Lab in Milton Keynes | Wellcome Sanger Institute for the COVID-19 Genomics UK (COG-UK) Consortium | The Lighthouse Lab in Milton Keynes and Alex Alderton, Roberto Amato, Jeffrey Barrett, Sonia Goncalves, Ewan Harrison, David K. Jackson, Ian Johnston, Dominic Kwiatkowski, Cordelia Langford, John Sillitoe on behalf of the Wellcome Sanger Institute COVID-19 Surveillance Team |
| EPI_ISL_2911623 | Lighthouse Lab in Alderley Park | Wellcome Sanger Institute for the COVID-19 Genomics UK (COG-UK) Consortium | Jacquelyn Wynn, Mairead Hyland, The Lighthouse Lab in Alderley Park and Alex Alderton, Roberto Amato, Jeffrey Barrett, Sonia Goncalves, Ewan Harrison, David K. Jackson, Ian Johnston, Dominic Kwiatkowski, Cordelia Langford, John Sillitoe on behalf of the Wellcome Sanger Institute COVID-19 Surveillance Team |
| EPI_ISL_2911624 | Lighthouse Lab in Glasgow | Wellcome Sanger Institute for the COVID-19 Genomics UK (COG-UK) Consortium | Harper VanSteenhouse, Yumi Kasai, David Gray, Carol Clugston, Anna Dominiczak and Alex Alderton, Roberto Amato, Jeffrey Barrett, Sonia Goncalves, Ewan Harrison, David K. Jackson, Ian Johnston, Dominic Kwiatkowski, Cordelia Langford, John Sillitoe on behalf of the Wellcome Sanger Institute COVID-19 Surveillance Team |
| EPI_ISL_2911625, EPI_ISL_2911626 | Lighthouse Lab in Alderley Park | Wellcome Sanger Institute for the COVID-19 Genomics UK (COG-UK) Consortium | Jacquelyn Wynn, Mairead Hyland, The Lighthouse Lab in Alderley Park and Alex Alderton, Roberto Amato, Jeffrey Barrett, Sonia Goncalves, Ewan Harrison, David K. Jackson, Ian Johnston, Dominic Kwiatkowski, Cordelia Langford, John Sillitoe on behalf of the Wellcome Sanger Institute COVID-19 Surveillance Team |
| EPI_ISL_2911627 | Lighthouse Lab in Milton Keynes | Wellcome Sanger Institute for the COVID-19 Genomics UK (COG-UK) Consortium | The Lighthouse Lab in Milton Keynes and Alex Alderton, Roberto Amato, Jeffrey Barrett, Sonia Goncalves, Ewan Harrison, David K. Jackson, Ian Johnston, Dominic Kwiatkowski, Cordelia Langford, John Sillitoe on behalf of the Wellcome Sanger Institute COVID-19 Surveillance Team |
| EPI_ISL_2911628 | Lighthouse Lab in Alderley Park | Wellcome Sanger Institute for the COVID-19 Genomics UK (COG-UK) Consortium | Jacquelyn Wynn, Mairead Hyland, The Lighthouse Lab in Alderley Park and Alex Alderton, Roberto Amato, Jeffrey Barrett, Sonia Goncalves, Ewan Harrison, David K. Jackson, Ian Johnston, Dominic Kwiatkowski, Cordelia Langford, John Sillitoe on behalf of the Wellcome Sanger Institute COVID-19 Surveillance Team |
| EPI_ISL_2911629, EPI_ISL_2911630, EPI_ISL_2911631 | Lighthouse Lab in Milton Keynes | Wellcome Sanger Institute for the COVID-19 Genomics UK (COG-UK) Consortium | The Lighthouse Lab in Milton Keynes and Alex Alderton, Roberto Amato, Jeffrey Barrett, Sonia Goncalves, Ewan Harrison, David K. Jackson, Ian Johnston, Dominic Kwiatkowski, Cordelia Langford, John Sillitoe on behalf of the Wellcome Sanger Institute COVID-19 Surveillance Team |
| EPI_ISL_2911632, EPI_ISL_2911633, EPI_ISL_2911634 | Lighthouse Lab in Alderley Park | Wellcome Sanger Institute for the COVID-19 Genomics UK (COG-UK) Consortium | Jacquelyn Wynn, Mairead Hyland, The Lighthouse Lab in Alderley Park and Alex Alderton, Roberto Amato, Jeffrey Barrett, Sonia Goncalves, Ewan Harrison, David K. Jackson, Ian Johnston, Dominic Kwiatkowski, Cordelia Langford, John Sillitoe on behalf of the Wellcome Sanger Institute COVID-19 Surveillance Team |
| EPI_ISL_2911635 | Lighthouse Lab in Milton Keynes | Wellcome Sanger Institute for the COVID-19 Genomics UK (COG-UK) Consortium | The Lighthouse Lab in Milton Keynes and Alex Alderton, Roberto Amato, Jeffrey Barrett, Sonia Goncalves, Ewan Harrison, David K. Jackson, Ian Johnston, Dominic Kwiatkowski, Cordelia Langford, John Sillitoe on behalf of the Wellcome Sanger Institute COVID-19 Surveillance Team |
| EPI_ISL_2911636, EPI_ISL_2911637, EPI_ISL_2911638 | Lighthouse Lab in Alderley Park | Wellcome Sanger Institute for the COVID-19 Genomics UK (COG-UK) Consortium | Jacquelyn Wynn, Mairead Hyland, The Lighthouse Lab in Alderley Park and Alex Alderton, Roberto Amato, Jeffrey Barrett, Sonia Goncalves, Ewan Harrison, David K. Jackson, Ian Johnston, Dominic Kwiatkowski, Cordelia Langford, John Sillitoe on behalf of the Wellcome Sanger Institute COVID-19 Surveillance Team |
| EPI_ISL_2911639 | Lighthouse Lab in Milton Keynes | Wellcome Sanger Institute for the COVID-19 Genomics UK (COG-UK) Consortium | The Lighthouse Lab in Milton Keynes and Alex Alderton, Roberto Amato, Jeffrey Barrett, Sonia Goncalves, Ewan Harrison, David K. Jackson, Ian Johnston, Dominic Kwiatkowski, Cordelia Langford, John Sillitoe on behalf of the Wellcome Sanger Institute COVID-19 Surveillance Team |
| EPI_ISL_2911640, EPI_ISL_2911641, EPI_ISL_2911642, EPI_ISL_2911643, EPI_ISL_2911644, EPI_ISL_2911645, EPI_ISL_2911646 | Lighthouse Lab in Alderley Park | Wellcome Sanger Institute for the COVID-19 Genomics UK (COG-UK) Consortium | Jacquelyn Wynn, Mairead Hyland, The Lighthouse Lab in Alderley Park and Alex Alderton, Roberto Amato, Jeffrey Barrett, Sonia Goncalves, Ewan Harrison, David K. Jackson, Ian Johnston, Dominic Kwiatkowski, Cordelia Langford, John Sillitoe on behalf of the Wellcome Sanger Institute COVID-19 Surveillance Team |
| EPI_ISL_2911647 | Lighthouse Lab in Milton Keynes | Wellcome Sanger Institute for the COVID-19 Genomics UK (COG-UK) Consortium | The Lighthouse Lab in Milton Keynes and Alex Alderton, Roberto Amato, Jeffrey Barrett, Sonia Goncalves, Ewan Harrison, David K. Jackson, Ian Johnston, Dominic Kwiatkowski, Cordelia Langford, John Sillitoe on behalf of the Wellcome Sanger Institute COVID-19 Surveillance Team |
| EPI_ISL_2911649, EPI_ISL_2911650, EPI_ISL_2911651, EPI_ISL_2911652, EPI_ISL_2911654 | Lighthouse Lab in Alderley Park | Wellcome Sanger Institute for the COVID-19 Genomics UK (COG-UK) Consortium | Jacquelyn Wynn, Mairead Hyland, The Lighthouse Lab in Alderley Park and Alex Alderton, Roberto Amato, Jeffrey Barrett, Sonia Goncalves, Ewan Harrison, David K. Jackson, Ian Johnston, Dominic Kwiatkowski, Cordelia Langford, John Sillitoe on behalf of the Wellcome Sanger Institute COVID-19 Surveillance Team |
| EPI_ISL_2911655 | Lighthouse Lab in Milton Keynes | Wellcome Sanger Institute for the COVID-19 Genomics UK (COG-UK) Consortium | The Lighthouse Lab in Milton Keynes and Alex Alderton, Roberto Amato, Jeffrey Barrett, Sonia Goncalves, Ewan Harrison, David K. Jackson, Ian Johnston, Dominic Kwiatkowski, Cordelia Langford, John Sillitoe on behalf of the Wellcome Sanger Institute COVID-19 Surveillance Team |
| EPI_ISL_2911656, EPI_ISL_2911657, EPI_ISL_2911658, EPI_ISL_2911659, EPI_ISL_2911660 | Lighthouse Lab in Alderley Park | Wellcome Sanger Institute for the COVID-19 Genomics UK (COG-UK) Consortium | Jacquelyn Wynn, Mairead Hyland, The Lighthouse Lab in Alderley Park and Alex Alderton, Roberto Amato, Jeffrey Barrett, Sonia Goncalves, Ewan Harrison, David K. Jackson, Ian Johnston, Dominic Kwiatkowski, Cordelia Langford, John Sillitoe on behalf of the Wellcome Sanger Institute COVID-19 Surveillance Team |
| EPI_ISL_2911661, EPI_ISL_2911662 | Lighthouse Lab in Milton Keynes | Wellcome Sanger Institute for the COVID-19 Genomics UK (COG-UK) Consortium | The Lighthouse Lab in Milton Keynes and Alex Alderton, Roberto Amato, Jeffrey Barrett, Sonia Goncalves, Ewan Harrison, David K. Jackson, Ian Johnston, Dominic Kwiatkowski, Cordelia Langford, John Sillitoe on behalf of the Wellcome Sanger Institute COVID-19 Surveillance Team |
| EPI_ISL_2911663, EPI_ISL_2911664, EPI_ISL_2911665, EPI_ISL_2911666, EPI_ISL_2911667 | Lighthouse Lab in Alderley Park | Wellcome Sanger Institute for the COVID-19 Genomics UK (COG-UK) Consortium | Jacquelyn Wynn, Mairead Hyland, The Lighthouse Lab in Alderley Park and Alex Alderton, Roberto Amato, Jeffrey Barrett, Sonia Goncalves, Ewan Harrison, David K. Jackson, Ian Johnston, Dominic Kwiatkowski, Cordelia Langford, John Sillitoe on behalf of the Wellcome Sanger Institute COVID-19 Surveillance Team |
| EPI_ISL_2911668 | Lighthouse Lab in Milton Keynes | Wellcome Sanger Institute for the COVID-19 Genomics UK (COG-UK) Consortium | The Lighthouse Lab in Milton Keynes and Alex Alderton, Roberto Amato, Jeffrey Barrett, Sonia Goncalves, Ewan Harrison, David K. Jackson, Ian Johnston, Dominic Kwiatkowski, Cordelia Langford, John Sillitoe on behalf of the Wellcome Sanger Institute COVID-19 Surveillance Team |
| EPI_ISL_2911669, EPI_ISL_2911670, EPI_ISL_2911671, EPI_ISL_2911672, EPI_ISL_2911673, EPI_ISL_2911674, EPI_ISL_2911675, EPI_ISL_2911676, EPI_ISL_2911677, | Lighthouse Lab in Alderley Park | Wellcome Sanger Institute for the COVID-19 Genomics UK (COG-UK) Consortium | Jacquelyn Wynn, Mairead Hyland, The Lighthouse Lab in Alderley Park and Alex Alderton, Roberto Amato, Jeffrey Barrett, Sonia Goncalves, Ewan Harrison, David K. Jackson, Ian Johnston, Dominic Kwiatkowski, Cordelia Langford, John Sillitoe on behalf of the Wellcome Sanger Institute COVID-19 Surveillance Team |

| | | | |
|---|---|---|---|
| EPI_ISL_2911678 | | | |
| EPI_ISL_2911679 | Lighthouse Lab in Milton Keynes | Wellcome Sanger Institute for the COVID-19 Genomics UK (COG-UK) Consortium | The Lighthouse Lab in Milton Keynes and Alex Alderton, Roberto Amato, Jeffrey Barrett, Sonia Goncalves, Ewan Harrison, David K. Jackson, Ian Johnston, Dominic Kwiatkowski, Cordelia Langford, John Sillitoe on behalf of the Wellcome Sanger Institute COVID-19 Surveillance Team |
| EPI_ISL_2911680 | Lighthouse Lab in Alderley Park | Wellcome Sanger Institute for the COVID-19 Genomics UK (COG-UK) Consortium | Jacquelyn Wynn, Mairead Hyland, The Lighthouse Lab in Alderley Park and Alex Alderton, Roberto Amato, Jeffrey Barrett, Sonia Goncalves, Ewan Harrison, David K. Jackson, Ian Johnston, Dominic Kwiatkowski, Cordelia Langford, John Sillitoe on behalf of the Wellcome Sanger Institute COVID-19 Surveillance Team |
| EPI_ISL_2911681 | Lighthouse Lab in Milton Keynes | Wellcome Sanger Institute for the COVID-19 Genomics UK (COG-UK) Consortium | The Lighthouse Lab in Milton Keynes and Alex Alderton, Roberto Amato, Jeffrey Barrett, Sonia Goncalves, Ewan Harrison, David K. Jackson, Ian Johnston, Dominic Kwiatkowski, Cordelia Langford, John Sillitoe on behalf of the Wellcome Sanger Institute COVID-19 Surveillance Team |
| EPI_ISL_2911682 | Lighthouse Lab in Glasgow | Wellcome Sanger Institute for the COVID-19 Genomics UK (COG-UK) Consortium | Harper VanSteenhouse, Yumi Kasai, David Gray, Carol Clugston, Anna Dominiczak and Alex Alderton, Roberto Amato, Jeffrey Barrett, Sonia Goncalves, Ewan Harrison, David K. Jackson, Ian Johnston, Dominic Kwiatkowski, Cordelia Langford, John Sillitoe on behalf of the Wellcome Sanger Institute COVID-19 Surveillance Team |
| EPI_ISL_2911683 | Lighthouse Lab in Alderley Park | Wellcome Sanger Institute for the COVID-19 Genomics UK (COG-UK) Consortium | Jacquelyn Wynn, Mairead Hyland, The Lighthouse Lab in Alderley Park and Alex Alderton, Roberto Amato, Jeffrey Barrett, Sonia Goncalves, Ewan Harrison, David K. Jackson, Ian Johnston, Dominic Kwiatkowski, Cordelia Langford, John Sillitoe on behalf of the Wellcome Sanger Institute COVID-19 Surveillance Team |
| EPI_ISL_2911684 | Lighthouse Lab in Milton Keynes | Wellcome Sanger Institute for the COVID-19 Genomics UK (COG-UK) Consortium | The Lighthouse Lab in Milton Keynes and Alex Alderton, Roberto Amato, Jeffrey Barrett, Sonia Goncalves, Ewan Harrison, David K. Jackson, Ian Johnston, Dominic Kwiatkowski, Cordelia Langford, John Sillitoe on behalf of the Wellcome Sanger Institute COVID-19 Surveillance Team |
| EPI_ISL_2911685, EPI_ISL_2911686, EPI_ISL_2911687, EPI_ISL_2911688, EPI_ISL_2911689, EPI_ISL_2911690, EPI_ISL_2911691, EPI_ISL_2911692, EPI_ISL_2911693, EPI_ISL_2911694, EPI_ISL_2911695 | | | |
| see above | Lighthouse Lab in Alderley Park | Wellcome Sanger Institute for the COVID-19 Genomics UK (COG-UK) Consortium | Jacquelyn Wynn, Mairead Hyland, The Lighthouse Lab in Alderley Park and Alex Alderton, Roberto Amato, Jeffrey Barrett, Sonia Goncalves, Ewan Harrison, David K. Jackson, Ian Johnston, Dominic Kwiatkowski, Cordelia Langford, John Sillitoe on behalf of the Wellcome Sanger Institute COVID-19 Surveillance Team |
| EPI_ISL_2911696, EPI_ISL_2911697 | Lighthouse Lab in Milton Keynes | Wellcome Sanger Institute for the COVID-19 Genomics UK (COG-UK) Consortium | The Lighthouse Lab in Milton Keynes and Alex Alderton, Roberto Amato, Jeffrey Barrett, Sonia Goncalves, Ewan Harrison, David K. Jackson, Ian Johnston, Dominic Kwiatkowski, Cordelia Langford, John Sillitoe on behalf of the Wellcome Sanger Institute COVID-19 Surveillance Team |
| EPI_ISL_2911698, EPI_ISL_2911699, EPI_ISL_2911700, EPI_ISL_2911701, EPI_ISL_2911702, EPI_ISL_2911703 | Lighthouse Lab in Alderley Park | Wellcome Sanger Institute for the COVID-19 Genomics UK (COG-UK) Consortium | Jacquelyn Wynn, Mairead Hyland, The Lighthouse Lab in Alderley Park and Alex Alderton, Roberto Amato, Jeffrey Barrett, Sonia Goncalves, Ewan Harrison, David K. Jackson, Ian Johnston, Dominic Kwiatkowski, Cordelia Langford, John Sillitoe on behalf of the Wellcome Sanger Institute COVID-19 Surveillance Team |
| EPI_ISL_2911704 | Lighthouse Lab in Milton Keynes | Wellcome Sanger Institute for the COVID-19 Genomics UK (COG-UK) Consortium | The Lighthouse Lab in Milton Keynes and Alex Alderton, Roberto Amato, Jeffrey Barrett, Sonia Goncalves, Ewan Harrison, David K. Jackson, Ian Johnston, Dominic Kwiatkowski, Cordelia Langford, John Sillitoe on behalf of the Wellcome Sanger Institute COVID-19 Surveillance Team |
| EPI_ISL_2911705 | Lighthouse Lab in Alderley Park | Wellcome Sanger Institute for the COVID-19 Genomics UK (COG-UK) Consortium | Jacquelyn Wynn, Mairead Hyland, The Lighthouse Lab in Alderley Park and Alex Alderton, Roberto Amato, Jeffrey Barrett, Sonia Goncalves, Ewan Harrison, David K. Jackson, Ian Johnston, Dominic Kwiatkowski, Cordelia Langford, John Sillitoe on behalf of the Wellcome Sanger Institute COVID-19 Surveillance Team |
| EPI_ISL_2911706 | Lighthouse Lab in Glasgow | Wellcome Sanger Institute for the COVID-19 Genomics UK (COG-UK) Consortium | Harper VanSteenhouse, Yumi Kasai, David Gray, Carol Clugston, Anna Dominiczak and Alex Alderton, Roberto Amato, Jeffrey Barrett, Sonia Goncalves, Ewan Harrison, David K. Jackson, Ian Johnston, Dominic Kwiatkowski, Cordelia Langford, John Sillitoe on behalf of the Wellcome Sanger Institute COVID-19 Surveillance Team |
| EPI_ISL_2911707 | Lighthouse Lab in Alderley Park | Wellcome Sanger Institute for the COVID-19 Genomics UK (COG-UK) Consortium | Jacquelyn Wynn, Mairead Hyland, The Lighthouse Lab in Alderley Park and Alex Alderton, Roberto Amato, Jeffrey Barrett, Sonia Goncalves, Ewan Harrison, David K. Jackson, Ian Johnston, Dominic Kwiatkowski, Cordelia Langford, John Sillitoe on behalf of the Wellcome Sanger Institute COVID-19 Surveillance Team |
| EPI_ISL_2911708 | Lighthouse Lab in Glasgow | Wellcome Sanger Institute for the COVID-19 Genomics UK (COG-UK) Consortium | Harper VanSteenhouse, Yumi Kasai, David Gray, Carol Clugston, Anna Dominiczak and Alex Alderton, Roberto Amato, Jeffrey Barrett, Sonia Goncalves, Ewan Harrison, David K. Jackson, Ian Johnston, Dominic Kwiatkowski, Cordelia Langford, John Sillitoe on behalf of the Wellcome Sanger Institute COVID-19 Surveillance Team |
| EPI_ISL_2911709, EPI_ISL_2911710 | Lighthouse Lab in Alderley Park | Wellcome Sanger Institute for the COVID-19 Genomics UK (COG-UK) Consortium | Jacquelyn Wynn, Mairead Hyland, The Lighthouse Lab in Alderley Park and Alex Alderton, Roberto Amato, Jeffrey Barrett, Sonia Goncalves, Ewan Harrison, David K. Jackson, Ian Johnston, Dominic Kwiatkowski, Cordelia Langford, John Sillitoe on behalf of the Wellcome Sanger Institute COVID-19 Surveillance Team |
| EPI_ISL_2911711 | Lighthouse Lab in Milton Keynes | Wellcome Sanger Institute for the COVID-19 Genomics UK (COG-UK) Consortium | The Lighthouse Lab in Milton Keynes and Alex Alderton, Roberto Amato, Jeffrey Barrett, Sonia Goncalves, Ewan Harrison, David K. Jackson, Ian Johnston, Dominic Kwiatkowski, Cordelia Langford, John Sillitoe on behalf of the Wellcome Sanger Institute COVID-19 Surveillance Team |
| EPI_ISL_2911712 | Lighthouse Lab in Alderley Park | Wellcome Sanger Institute for the COVID-19 Genomics UK (COG-UK) Consortium | Jacquelyn Wynn, Mairead Hyland, The Lighthouse Lab in Alderley Park and Alex Alderton, Roberto Amato, Jeffrey Barrett, Sonia Goncalves, Ewan Harrison, David K. Jackson, Ian Johnston, Dominic Kwiatkowski, Cordelia Langford, John Sillitoe on behalf of the Wellcome Sanger Institute COVID-19 Surveillance Team |
| EPI_ISL_2911713 | Lighthouse Lab in Milton Keynes | Wellcome Sanger Institute for the COVID-19 Genomics UK (COG-UK) Consortium | The Lighthouse Lab in Milton Keynes and Alex Alderton, Roberto Amato, Jeffrey Barrett, Sonia Goncalves, Ewan Harrison, David K. Jackson, Ian Johnston, Dominic Kwiatkowski, Cordelia Langford, John Sillitoe on behalf of the Wellcome Sanger Institute COVID-19 Surveillance Team |
| EPI_ISL_2911714, EPI_ISL_2911715 | Lighthouse Lab in Alderley Park | Wellcome Sanger Institute for the COVID-19 Genomics UK (COG-UK) Consortium | Jacquelyn Wynn, Mairead Hyland, The Lighthouse Lab in Alderley Park and Alex Alderton, Roberto Amato, Jeffrey Barrett, Sonia Goncalves, Ewan Harrison, David K. Jackson, Ian Johnston, Dominic Kwiatkowski, Cordelia Langford, John Sillitoe on behalf of the Wellcome Sanger Institute COVID-19 Surveillance Team |
| EPI_ISL_2911716, EPI_ISL_2911717 | Lighthouse Lab in Milton Keynes | Wellcome Sanger Institute for the COVID-19 Genomics UK (COG-UK) Consortium | The Lighthouse Lab in Milton Keynes and Alex Alderton, Roberto Amato, Jeffrey Barrett, Sonia Goncalves, Ewan Harrison, David K. Jackson, Ian Johnston, Dominic Kwiatkowski, Cordelia Langford, John Sillitoe on behalf of the Wellcome Sanger Institute COVID-19 Surveillance Team |
| EPI_ISL_2911718 | Lighthouse Lab in Alderley Park | Wellcome Sanger Institute for the COVID-19 Genomics UK (COG-UK) Consortium | Jacquelyn Wynn, Mairead Hyland, The Lighthouse Lab in Alderley Park and Alex Alderton, Roberto Amato, Jeffrey Barrett, Sonia Goncalves, Ewan Harrison, David K. Jackson, Ian Johnston, Dominic Kwiatkowski, Cordelia Langford, John Sillitoe on behalf of the Wellcome Sanger Institute COVID-19 Surveillance Team |
| EPI_ISL_2911719 | Lighthouse Lab in Milton Keynes | Wellcome Sanger Institute for the COVID-19 Genomics UK (COG-UK) Consortium | The Lighthouse Lab in Milton Keynes and Alex Alderton, Roberto Amato, Jeffrey Barrett, Sonia Goncalves, Ewan Harrison, David K. Jackson, Ian Johnston, Dominic Kwiatkowski, Cordelia Langford, John Sillitoe on behalf of the Wellcome Sanger Institute COVID-19 Surveillance Team |
| EPI_ISL_2911720 | Lighthouse Lab in Alderley Park | Wellcome Sanger Institute for the COVID-19 Genomics UK (COG-UK) Consortium | Jacquelyn Wynn, Mairead Hyland, The Lighthouse Lab in Alderley Park and Alex Alderton, Roberto Amato, Jeffrey Barrett, Sonia Goncalves, Ewan Harrison, David K. Jackson, Ian Johnston, Dominic Kwiatkowski, Cordelia Langford, John Sillitoe on behalf of the Wellcome Sanger Institute COVID-19 Surveillance Team |
| EPI_ISL_2911721 | Lighthouse Lab in Milton Keynes | Wellcome Sanger Institute for the COVID-19 Genomics UK (COG-UK) Consortium | The Lighthouse Lab in Milton Keynes and Alex Alderton, Roberto Amato, Jeffrey Barrett, Sonia Goncalves, Ewan Harrison, David K. Jackson, Ian Johnston, Dominic Kwiatkowski, Cordelia Langford, John Sillitoe on behalf of the Wellcome Sanger Institute COVID-19 Surveillance Team |
| EPI_ISL_2911722 | Lighthouse Lab in Alderley Park | Wellcome Sanger Institute for the COVID-19 Genomics UK (COG-UK) Consortium | Jacquelyn Wynn, Mairead Hyland, The Lighthouse Lab in Alderley Park and Alex Alderton, Roberto Amato, Jeffrey Barrett, Sonia Goncalves, Ewan Harrison, David K. Jackson, Ian Johnston, Dominic Kwiatkowski, Cordelia Langford, John Sillitoe on behalf of the Wellcome Sanger Institute COVID-19 Surveillance Team |
| EPI_ISL_2911723 | Lighthouse Lab in Milton Keynes | Wellcome Sanger Institute for the COVID-19 Genomics UK (COG-UK) Consortium | The Lighthouse Lab in Milton Keynes and Alex Alderton, Roberto Amato, Jeffrey Barrett, Sonia Goncalves, Ewan Harrison, David K. Jackson, Ian Johnston, Dominic Kwiatkowski, Cordelia Langford, John Sillitoe on behalf of the Wellcome Sanger Institute COVID-19 Surveillance Team |
| EPI_ISL_2911724, EPI_ISL_2911725 | Lighthouse Lab in Alderley Park | Wellcome Sanger Institute for the COVID-19 Genomics UK (COG-UK) Consortium | Jacquelyn Wynn, Mairead Hyland, The Lighthouse Lab in Alderley Park and Alex Alderton, Roberto Amato, Jeffrey Barrett, Sonia Goncalves, Ewan Harrison, David K. Jackson, Ian Johnston, Dominic Kwiatkowski, Cordelia Langford, John Sillitoe on behalf of the Wellcome Sanger Institute COVID-19 Surveillance Team |
| EPI_ISL_2911726 | Lighthouse Lab in Glasgow | Wellcome Sanger Institute for the COVID-19 Genomics UK (COG-UK) Consortium | Harper VanSteenhouse, Yumi Kasai, David Gray, Carol Clugston, Anna Dominiczak and Alex Alderton, Roberto Amato, Jeffrey Barrett, Sonia Goncalves, Ewan Harrison, David K. Jackson, Ian Johnston, Dominic Kwiatkowski, Cordelia Langford, John Sillitoe on behalf of the Wellcome Sanger Institute COVID-19 Surveillance Team |
| EPI_ISL_2911727, EPI_ISL_2911728 | Lighthouse Lab in Alderley Park | Wellcome Sanger Institute for the COVID-19 Genomics UK (COG-UK) Consortium | Jacquelyn Wynn, Mairead Hyland, The Lighthouse Lab in Alderley Park and Alex Alderton, Roberto Amato, Jeffrey Barrett, Sonia Goncalves, Ewan Harrison, David K. Jackson, Ian Johnston, Dominic Kwiatkowski, Cordelia Langford, John Sillitoe on behalf of the Wellcome Sanger Institute COVID-19 Surveillance Team |
| EPI_ISL_2911729, EPI_ISL_2911730 | Lighthouse Lab in Milton Keynes | Wellcome Sanger Institute for the COVID-19 Genomics UK (COG-UK) Consortium | The Lighthouse Lab in Milton Keynes and Alex Alderton, Roberto Amato, Jeffrey Barrett, Sonia Goncalves, Ewan Harrison, David K. Jackson, Ian Johnston, Dominic Kwiatkowski, Cordelia Langford, John Sillitoe on behalf of the Wellcome Sanger Institute COVID-19 Surveillance Team |
| EPI_ISL_2911731 | Lighthouse Lab in Alderley Park | Wellcome Sanger Institute for the COVID-19 Genomics UK (COG-UK) Consortium | Jacquelyn Wynn, Mairead Hyland, The Lighthouse Lab in Alderley Park and Alex Alderton, Roberto Amato, Jeffrey Barrett, Sonia Goncalves, Ewan Harrison, David K. Jackson, Ian Johnston, Dominic Kwiatkowski, Cordelia Langford, John Sillitoe on behalf of the Wellcome Sanger Institute COVID-19 Surveillance Team |
| EPI_ISL_2911732 | Lighthouse Lab in Glasgow | Wellcome Sanger Institute for the COVID-19 Genomics UK (COG-UK) Consortium | Harper VanSteenhouse, Yumi Kasai, David Gray, Carol Clugston, Anna Dominiczak and Alex Alderton, Roberto Amato, Jeffrey Barrett, Sonia Goncalves, Ewan Harrison, David K. Jackson, Ian Johnston, Dominic Kwiatkowski, Cordelia Langford, John Sillitoe on behalf of the Wellcome Sanger Institute COVID-19 Surveillance Team |
| EPI_ISL_2911733 | Lighthouse Lab in Alderley Park | Wellcome Sanger Institute for the COVID-19 Genomics UK (COG-UK) Consortium | Jacquelyn Wynn, Mairead Hyland, The Lighthouse Lab in Alderley Park and Alex Alderton, Roberto Amato, Jeffrey Barrett, Sonia Goncalves, Ewan Harrison, David K. Jackson, Ian Johnston, Dominic Kwiatkowski, Cordelia Langford, John Sillitoe on behalf of the Wellcome Sanger Institute COVID-19 Surveillance Team |
| EPI_ISL_2911734, EPI_ISL_2911735, EPI_ISL_2911736 | Lighthouse Lab in Milton Keynes | Wellcome Sanger Institute for the COVID-19 Genomics UK (COG-UK) Consortium | The Lighthouse Lab in Milton Keynes and Alex Alderton, Roberto Amato, Jeffrey Barrett, Sonia Goncalves, Ewan Harrison, David K. Jackson, Ian Johnston, Dominic Kwiatkowski, Cordelia Langford, John Sillitoe on behalf of the Wellcome Sanger Institute COVID-19 Surveillance Team |
| EPI_ISL_2911737 | Lighthouse Lab in Alderley Park | Wellcome Sanger Institute for the COVID-19 Genomics UK (COG-UK) Consortium | Jacquelyn Wynn, Mairead Hyland, The Lighthouse Lab in Alderley Park and Alex Alderton, Roberto Amato, Jeffrey Barrett, Sonia Goncalves, Ewan Harrison, David K. Jackson, Ian Johnston, Dominic Kwiatkowski, Cordelia Langford, John Sillitoe on behalf of the Wellcome Sanger Institute COVID-19 Surveillance Team |
| EPI_ISL_2911738, EPI_ISL_2911739 | Lighthouse Lab in Milton Keynes | Wellcome Sanger Institute for the COVID-19 Genomics UK (COG-UK) Consortium | The Lighthouse Lab in Milton Keynes and Alex Alderton, Roberto Amato, Jeffrey Barrett, Sonia Goncalves, Ewan Harrison, David K. Jackson, Ian Johnston, Dominic Kwiatkowski, Cordelia Langford, John Sillitoe on behalf of the Wellcome Sanger Institute COVID-19 Surveillance Team |
| EPI_ISL_2911740 | Lighthouse Lab in Alderley Park | Wellcome Sanger Institute for the COVID-19 Genomics UK (COG-UK) Consortium | Jacquelyn Wynn, Mairead Hyland, The Lighthouse Lab in Alderley Park and Alex Alderton, Roberto Amato, Jeffrey Barrett, Sonia Goncalves, Ewan Harrison, David K. Jackson, Ian Johnston, Dominic Kwiatkowski, Cordelia Langford, John Sillitoe on behalf of the Wellcome Sanger Institute COVID-19 Surveillance Team |
| EPI_ISL_2911741 | Lighthouse Lab in Glasgow | Wellcome Sanger Institute for the COVID-19 Genomics UK (COG-UK) Consortium | Harper VanSteenhouse, Yumi Kasai, David Gray, Carol Clugston, Anna Dominiczak and Alex Alderton, Roberto Amato, Jeffrey Barrett, Sonia Goncalves, Ewan Harrison, David K. Jackson, Ian Johnston, Dominic Kwiatkowski, Cordelia Langford, John Sillitoe on behalf of the Wellcome Sanger Institute COVID-19 Surveillance Team |
| EPI_ISL_2911742, EPI_ISL_2911743, EPI_ISL_2911744 | Lighthouse Lab in Milton Keynes | Wellcome Sanger Institute for the COVID-19 Genomics UK (COG-UK) Consortium | The Lighthouse Lab in Milton Keynes and Alex Alderton, Roberto Amato, Jeffrey Barrett, Sonia Goncalves, Ewan Harrison, David K. Jackson, Ian Johnston, Dominic Kwiatkowski, Cordelia Langford, John Sillitoe on behalf of the Wellcome Sanger Institute COVID-19 Surveillance Team |
| EPI_ISL_2911745, EPI_ISL_2911746, EPI_ISL_2911747, EPI_ISL_2911748 | Lighthouse Lab in Alderley Park | Wellcome Sanger Institute for the COVID-19 Genomics UK (COG-UK) Consortium | Jacquelyn Wynn, Mairead Hyland, The Lighthouse Lab in Alderley Park and Alex Alderton, Roberto Amato, Jeffrey Barrett, Sonia Goncalves, Ewan Harrison, David K. Jackson, Ian Johnston, Dominic Kwiatkowski, Cordelia Langford, John Sillitoe on behalf of the Wellcome Sanger Institute COVID-19 Surveillance Team |
| EPI_ISL_2911749, | Lighthouse Lab in Milton Keynes | Wellcome Sanger Institute for the COVID-19 Genomics UK (COG-UK) | The Lighthouse Lab in Milton Keynes and Alex Alderton, Roberto Amato, Jeffrey Barrett, Sonia Goncalves, Ewan Harrison, David K. Jackson, Ian Johnston, Dominic Kwiatkowski, Cordelia Langford, John Sillitoe on behalf of the Wellcome Sanger |

| | | | |
|---|---|---|---|
| EPI_ISL_2911750 | | Consortium | Institute COVID-19 Surveillance Team |
| EPI_ISL_2911751, EPI_ISL_2911752 | Lighthouse Lab in Alderley Park | Wellcome Sanger Institute for the COVID-19 Genomics UK (COG-UK) Consortium | Jacquelyn Wynn, Mairead Hyland, The Lighthouse Lab in Alderley Park and Alex Alderton, Roberto Amato, Jeffrey Barrett, Sonia Goncalves, Ewan Harrison, David K. Jackson, Ian Johnston, Dominic Kwiatkowski, Cordelia Langford, John Sillitoe on behalf of the Wellcome Sanger Institute COVID-19 Surveillance Team |
| EPI_ISL_2911753 | Lighthouse Lab in Milton Keynes | Wellcome Sanger Institute for the COVID-19 Genomics UK (COG-UK) Consortium | The Lighthouse Lab in Milton Keynes and Alex Alderton, Roberto Amato, Jeffrey Barrett, Sonia Goncalves, Ewan Harrison, David K. Jackson, Ian Johnston, Dominic Kwiatkowski, Cordelia Langford, John Sillitoe on behalf of the Wellcome Sanger Institute COVID-19 Surveillance Team |
| EPI_ISL_2911754, EPI_ISL_2911755 | Lighthouse Lab in Alderley Park | Wellcome Sanger Institute for the COVID-19 Genomics UK (COG-UK) Consortium | Jacquelyn Wynn, Mairead Hyland, The Lighthouse Lab in Alderley Park and Alex Alderton, Roberto Amato, Jeffrey Barrett, Sonia Goncalves, Ewan Harrison, David K. Jackson, Ian Johnston, Dominic Kwiatkowski, Cordelia Langford, John Sillitoe on behalf of the Wellcome Sanger Institute COVID-19 Surveillance Team |
| EPI_ISL_2911756 | Lighthouse Lab in Glasgow | Wellcome Sanger Institute for the COVID-19 Genomics UK (COG-UK) Consortium | Harper VanSteenhouse, Yumi Kasai, David Gray, Carol Clugston, Anna Dominiczak and Alex Alderton, Roberto Amato, Jeffrey Barrett, Sonia Goncalves, Ewan Harrison, David K. Jackson, Ian Johnston, Dominic Kwiatkowski, Cordelia Langford, John Sillitoe on behalf of the Wellcome Sanger Institute COVID-19 Surveillance Team |
| EPI_ISL_2911757, EPI_ISL_2911758, EPI_ISL_2911759, EPI_ISL_2911760, EPI_ISL_2911761, EPI_ISL_2911762, EPI_ISL_2911763, EPI_ISL_2911764 | Lighthouse Lab in Alderley Park | Wellcome Sanger Institute for the COVID-19 Genomics UK (COG-UK) Consortium | Jacquelyn Wynn, Mairead Hyland, The Lighthouse Lab in Alderley Park and Alex Alderton, Roberto Amato, Jeffrey Barrett, Sonia Goncalves, Ewan Harrison, David K. Jackson, Ian Johnston, Dominic Kwiatkowski, Cordelia Langford, John Sillitoe on behalf of the Wellcome Sanger Institute COVID-19 Surveillance Team |
| EPI_ISL_2911765 | Lighthouse Lab in Milton Keynes | Wellcome Sanger Institute for the COVID-19 Genomics UK (COG-UK) Consortium | The Lighthouse Lab in Milton Keynes and Alex Alderton, Roberto Amato, Jeffrey Barrett, Sonia Goncalves, Ewan Harrison, David K. Jackson, Ian Johnston, Dominic Kwiatkowski, Cordelia Langford, John Sillitoe on behalf of the Wellcome Sanger Institute COVID-19 Surveillance Team |
| EPI_ISL_2911766, EPI_ISL_2911767, EPI_ISL_2911768 | Lighthouse Lab in Alderley Park | Wellcome Sanger Institute for the COVID-19 Genomics UK (COG-UK) Consortium | Jacquelyn Wynn, Mairead Hyland, The Lighthouse Lab in Alderley Park and Alex Alderton, Roberto Amato, Jeffrey Barrett, Sonia Goncalves, Ewan Harrison, David K. Jackson, Ian Johnston, Dominic Kwiatkowski, Cordelia Langford, John Sillitoe on behalf of the Wellcome Sanger Institute COVID-19 Surveillance Team |
| EPI_ISL_2911769 | Lighthouse Lab in Milton Keynes | Wellcome Sanger Institute for the COVID-19 Genomics UK (COG-UK) Consortium | The Lighthouse Lab in Milton Keynes and Alex Alderton, Roberto Amato, Jeffrey Barrett, Sonia Goncalves, Ewan Harrison, David K. Jackson, Ian Johnston, Dominic Kwiatkowski, Cordelia Langford, John Sillitoe on behalf of the Wellcome Sanger Institute COVID-19 Surveillance Team |
| EPI_ISL_2911770 | Lighthouse Lab in Glasgow | Wellcome Sanger Institute for the COVID-19 Genomics UK (COG-UK) Consortium | Harper VanSteenhouse, Yumi Kasai, David Gray, Carol Clugston, Anna Dominiczak and Alex Alderton, Roberto Amato, Jeffrey Barrett, Sonia Goncalves, Ewan Harrison, David K. Jackson, Ian Johnston, Dominic Kwiatkowski, Cordelia Langford, John Sillitoe on behalf of the Wellcome Sanger Institute COVID-19 Surveillance Team |
| EPI_ISL_2911771 | Lighthouse Lab in Alderley Park | Wellcome Sanger Institute for the COVID-19 Genomics UK (COG-UK) Consortium | Jacquelyn Wynn, Mairead Hyland, The Lighthouse Lab in Alderley Park and Alex Alderton, Roberto Amato, Jeffrey Barrett, Sonia Goncalves, Ewan Harrison, David K. Jackson, Ian Johnston, Dominic Kwiatkowski, Cordelia Langford, John Sillitoe on behalf of the Wellcome Sanger Institute COVID-19 Surveillance Team |
| EPI_ISL_2911772 | Lighthouse Lab in Glasgow | Wellcome Sanger Institute for the COVID-19 Genomics UK (COG-UK) Consortium | Harper VanSteenhouse, Yumi Kasai, David Gray, Carol Clugston, Anna Dominiczak and Alex Alderton, Roberto Amato, Jeffrey Barrett, Sonia Goncalves, Ewan Harrison, David K. Jackson, Ian Johnston, Dominic Kwiatkowski, Cordelia Langford, John Sillitoe on behalf of the Wellcome Sanger Institute COVID-19 Surveillance Team |
| EPI_ISL_2911773, EPI_ISL_2911774, EPI_ISL_2911775 | Lighthouse Lab in Alderley Park | Wellcome Sanger Institute for the COVID-19 Genomics UK (COG-UK) Consortium | Jacquelyn Wynn, Mairead Hyland, The Lighthouse Lab in Alderley Park and Alex Alderton, Roberto Amato, Jeffrey Barrett, Sonia Goncalves, Ewan Harrison, David K. Jackson, Ian Johnston, Dominic Kwiatkowski, Cordelia Langford, John Sillitoe on behalf of the Wellcome Sanger Institute COVID-19 Surveillance Team |
| EPI_ISL_2911776 | Lighthouse Lab in Milton Keynes | Wellcome Sanger Institute for the COVID-19 Genomics UK (COG-UK) Consortium | The Lighthouse Lab in Milton Keynes and Alex Alderton, Roberto Amato, Jeffrey Barrett, Sonia Goncalves, Ewan Harrison, David K. Jackson, Ian Johnston, Dominic Kwiatkowski, Cordelia Langford, John Sillitoe on behalf of the Wellcome Sanger Institute COVID-19 Surveillance Team |
| EPI_ISL_2911777, EPI_ISL_2911778, EPI_ISL_2911779, EPI_ISL_2911780, EPI_ISL_2911781, EPI_ISL_2911782 | Lighthouse Lab in Alderley Park | Wellcome Sanger Institute for the COVID-19 Genomics UK (COG-UK) Consortium | Jacquelyn Wynn, Mairead Hyland, The Lighthouse Lab in Alderley Park and Alex Alderton, Roberto Amato, Jeffrey Barrett, Sonia Goncalves, Ewan Harrison, David K. Jackson, Ian Johnston, Dominic Kwiatkowski, Cordelia Langford, John Sillitoe on behalf of the Wellcome Sanger Institute COVID-19 Surveillance Team |
| EPI_ISL_2911783, EPI_ISL_2911784 | Lighthouse Lab in Milton Keynes | Wellcome Sanger Institute for the COVID-19 Genomics UK (COG-UK) Consortium | The Lighthouse Lab in Milton Keynes and Alex Alderton, Roberto Amato, Jeffrey Barrett, Sonia Goncalves, Ewan Harrison, David K. Jackson, Ian Johnston, Dominic Kwiatkowski, Cordelia Langford, John Sillitoe on behalf of the Wellcome Sanger Institute COVID-19 Surveillance Team |
| EPI_ISL_2911785, EPI_ISL_2911786, EPI_ISL_2911787, EPI_ISL_2911788 | Lighthouse Lab in Alderley Park | Wellcome Sanger Institute for the COVID-19 Genomics UK (COG-UK) Consortium | Jacquelyn Wynn, Mairead Hyland, The Lighthouse Lab in Alderley Park and Alex Alderton, Roberto Amato, Jeffrey Barrett, Sonia Goncalves, Ewan Harrison, David K. Jackson, Ian Johnston, Dominic Kwiatkowski, Cordelia Langford, John Sillitoe on behalf of the Wellcome Sanger Institute COVID-19 Surveillance Team |
| EPI_ISL_2911789 | Lighthouse Lab in Milton Keynes | Wellcome Sanger Institute for the COVID-19 Genomics UK (COG-UK) Consortium | The Lighthouse Lab in Milton Keynes and Alex Alderton, Roberto Amato, Jeffrey Barrett, Sonia Goncalves, Ewan Harrison, David K. Jackson, Ian Johnston, Dominic Kwiatkowski, Cordelia Langford, John Sillitoe on behalf of the Wellcome Sanger Institute COVID-19 Surveillance Team |
| EPI_ISL_2911790, EPI_ISL_2911791, EPI_ISL_2911792, EPI_ISL_2911793, EPI_ISL_2911794 | Lighthouse Lab in Alderley Park | Wellcome Sanger Institute for the COVID-19 Genomics UK (COG-UK) Consortium | Jacquelyn Wynn, Mairead Hyland, The Lighthouse Lab in Alderley Park and Alex Alderton, Roberto Amato, Jeffrey Barrett, Sonia Goncalves, Ewan Harrison, David K. Jackson, Ian Johnston, Dominic Kwiatkowski, Cordelia Langford, John Sillitoe on behalf of the Wellcome Sanger Institute COVID-19 Surveillance Team |
| EPI_ISL_2911795 | Lighthouse Lab in Glasgow | Wellcome Sanger Institute for the COVID-19 Genomics UK (COG-UK) Consortium | Harper VanSteenhouse, Yumi Kasai, David Gray, Carol Clugston, Anna Dominiczak and Alex Alderton, Roberto Amato, Jeffrey Barrett, Sonia Goncalves, Ewan Harrison, David K. Jackson, Ian Johnston, Dominic Kwiatkowski, Cordelia Langford, John Sillitoe on behalf of the Wellcome Sanger Institute COVID-19 Surveillance Team |
| EPI_ISL_2911796 | Lighthouse Lab in Milton Keynes | Wellcome Sanger Institute for the COVID-19 Genomics UK (COG-UK) Consortium | The Lighthouse Lab in Milton Keynes and Alex Alderton, Roberto Amato, Jeffrey Barrett, Sonia Goncalves, Ewan Harrison, David K. Jackson, Ian Johnston, Dominic Kwiatkowski, Cordelia Langford, John Sillitoe on behalf of the Wellcome Sanger Institute COVID-19 Surveillance Team |
| EPI_ISL_2911797 | Lighthouse Lab in Glasgow | Wellcome Sanger Institute for the COVID-19 Genomics UK (COG-UK) Consortium | Harper VanSteenhouse, Yumi Kasai, David Gray, Carol Clugston, Anna Dominiczak and Alex Alderton, Roberto Amato, Jeffrey Barrett, Sonia Goncalves, Ewan Harrison, David K. Jackson, Ian Johnston, Dominic Kwiatkowski, Cordelia Langford, John Sillitoe on behalf of the Wellcome Sanger Institute COVID-19 Surveillance Team |
| EPI_ISL_2911798, EPI_ISL_2911799, EPI_ISL_2911800, EPI_ISL_2911801 | Lighthouse Lab in Alderley Park | Wellcome Sanger Institute for the COVID-19 Genomics UK (COG-UK) Consortium | Jacquelyn Wynn, Mairead Hyland, The Lighthouse Lab in Alderley Park and Alex Alderton, Roberto Amato, Jeffrey Barrett, Sonia Goncalves, Ewan Harrison, David K. Jackson, Ian Johnston, Dominic Kwiatkowski, Cordelia Langford, John Sillitoe on behalf of the Wellcome Sanger Institute COVID-19 Surveillance Team |
| EPI_ISL_2911802 | Lighthouse Lab in Milton Keynes | Wellcome Sanger Institute for the COVID-19 Genomics UK (COG-UK) Consortium | The Lighthouse Lab in Milton Keynes and Alex Alderton, Roberto Amato, Jeffrey Barrett, Sonia Goncalves, Ewan Harrison, David K. Jackson, Ian Johnston, Dominic Kwiatkowski, Cordelia Langford, John Sillitoe on behalf of the Wellcome Sanger Institute COVID-19 Surveillance Team |
| EPI_ISL_2911803 | Lighthouse Lab in Glasgow | Wellcome Sanger Institute for the COVID-19 Genomics UK (COG-UK) Consortium | Harper VanSteenhouse, Yumi Kasai, David Gray, Carol Clugston, Anna Dominiczak and Alex Alderton, Roberto Amato, Jeffrey Barrett, Sonia Goncalves, Ewan Harrison, David K. Jackson, Ian Johnston, Dominic Kwiatkowski, Cordelia Langford, John Sillitoe on behalf of the Wellcome Sanger Institute COVID-19 Surveillance Team |
| EPI_ISL_2911804, EPI_ISL_2911805 | Lighthouse Lab in Alderley Park | Wellcome Sanger Institute for the COVID-19 Genomics UK (COG-UK) Consortium | Jacquelyn Wynn, Mairead Hyland, The Lighthouse Lab in Alderley Park and Alex Alderton, Roberto Amato, Jeffrey Barrett, Sonia Goncalves, Ewan Harrison, David K. Jackson, Ian Johnston, Dominic Kwiatkowski, Cordelia Langford, John Sillitoe on behalf of the Wellcome Sanger Institute COVID-19 Surveillance Team |
| EPI_ISL_2911806, EPI_ISL_2911807 | Lighthouse Lab in Milton Keynes | Wellcome Sanger Institute for the COVID-19 Genomics UK (COG-UK) Consortium | The Lighthouse Lab in Milton Keynes and Alex Alderton, Roberto Amato, Jeffrey Barrett, Sonia Goncalves, Ewan Harrison, David K. Jackson, Ian Johnston, Dominic Kwiatkowski, Cordelia Langford, John Sillitoe on behalf of the Wellcome Sanger Institute COVID-19 Surveillance Team |
| EPI_ISL_2911808, EPI_ISL_2911809 | Lighthouse Lab in Alderley Park | Wellcome Sanger Institute for the COVID-19 Genomics UK (COG-UK) Consortium | Jacquelyn Wynn, Mairead Hyland, The Lighthouse Lab in Alderley Park and Alex Alderton, Roberto Amato, Jeffrey Barrett, Sonia Goncalves, Ewan Harrison, David K. Jackson, Ian Johnston, Dominic Kwiatkowski, Cordelia Langford, John Sillitoe on behalf of the Wellcome Sanger Institute COVID-19 Surveillance Team |
| EPI_ISL_2911810 | Lighthouse Lab in Milton Keynes | Wellcome Sanger Institute for the COVID-19 Genomics UK (COG-UK) Consortium | The Lighthouse Lab in Milton Keynes and Alex Alderton, Roberto Amato, Jeffrey Barrett, Sonia Goncalves, Ewan Harrison, David K. Jackson, Ian Johnston, Dominic Kwiatkowski, Cordelia Langford, John Sillitoe on behalf of the Wellcome Sanger Institute COVID-19 Surveillance Team |
| EPI_ISL_2911811 | Lighthouse Lab in Alderley Park | Wellcome Sanger Institute for the COVID-19 Genomics UK (COG-UK) Consortium | Jacquelyn Wynn, Mairead Hyland, The Lighthouse Lab in Alderley Park and Alex Alderton, Roberto Amato, Jeffrey Barrett, Sonia Goncalves, Ewan Harrison, David K. Jackson, Ian Johnston, Dominic Kwiatkowski, Cordelia Langford, John Sillitoe on behalf of the Wellcome Sanger Institute COVID-19 Surveillance Team |
| EPI_ISL_2911812 | Lighthouse Lab in Glasgow | Wellcome Sanger Institute for the COVID-19 Genomics UK (COG-UK) Consortium | Harper VanSteenhouse, Yumi Kasai, David Gray, Carol Clugston, Anna Dominiczak and Alex Alderton, Roberto Amato, Jeffrey Barrett, Sonia Goncalves, Ewan Harrison, David K. Jackson, Ian Johnston, Dominic Kwiatkowski, Cordelia Langford, John Sillitoe on behalf of the Wellcome Sanger Institute COVID-19 Surveillance Team |
| EPI_ISL_2911813, EPI_ISL_2911814, EPI_ISL_2911815, EPI_ISL_2911816 | Lighthouse Lab in Milton Keynes | Wellcome Sanger Institute for the COVID-19 Genomics UK (COG-UK) Consortium | The Lighthouse Lab in Milton Keynes and Alex Alderton, Roberto Amato, Jeffrey Barrett, Sonia Goncalves, Ewan Harrison, David K. Jackson, Ian Johnston, Dominic Kwiatkowski, Cordelia Langford, John Sillitoe on behalf of the Wellcome Sanger Institute COVID-19 Surveillance Team |
| EPI_ISL_2911817, EPI_ISL_2911818, EPI_ISL_2911819 | Lighthouse Lab in Alderley Park | Wellcome Sanger Institute for the COVID-19 Genomics UK (COG-UK) Consortium | Jacquelyn Wynn, Mairead Hyland, The Lighthouse Lab in Alderley Park and Alex Alderton, Roberto Amato, Jeffrey Barrett, Sonia Goncalves, Ewan Harrison, David K. Jackson, Ian Johnston, Dominic Kwiatkowski, Cordelia Langford, John Sillitoe on behalf of the Wellcome Sanger Institute COVID-19 Surveillance Team |
| EPI_ISL_2911820 | Lighthouse Lab in Glasgow | Wellcome Sanger Institute for the COVID-19 Genomics UK (COG-UK) Consortium | Harper VanSteenhouse, Yumi Kasai, David Gray, Carol Clugston, Anna Dominiczak and Alex Alderton, Roberto Amato, Jeffrey Barrett, Sonia Goncalves, Ewan Harrison, David K. Jackson, Ian Johnston, Dominic Kwiatkowski, Cordelia Langford, John Sillitoe on behalf of the Wellcome Sanger Institute COVID-19 Surveillance Team |
| EPI_ISL_2911821 | Lighthouse Lab in Milton Keynes | Wellcome Sanger Institute for the COVID-19 Genomics UK (COG-UK) Consortium | The Lighthouse Lab in Milton Keynes and Alex Alderton, Roberto Amato, Jeffrey Barrett, Sonia Goncalves, Ewan Harrison, David K. Jackson, Ian Johnston, Dominic Kwiatkowski, Cordelia Langford, John Sillitoe on behalf of the Wellcome Sanger Institute COVID-19 Surveillance Team |
| EPI_ISL_2911822 | Lighthouse Lab in Glasgow | Wellcome Sanger Institute for the COVID-19 Genomics UK (COG-UK) Consortium | Harper VanSteenhouse, Yumi Kasai, David Gray, Carol Clugston, Anna Dominiczak and Alex Alderton, Roberto Amato, Jeffrey Barrett, Sonia Goncalves, Ewan Harrison, David K. Jackson, Ian Johnston, Dominic Kwiatkowski, Cordelia Langford, John Sillitoe on behalf of the Wellcome Sanger Institute COVID-19 Surveillance Team |
| EPI_ISL_2911823 | Lighthouse Lab in Milton Keynes | Wellcome Sanger Institute for the COVID-19 Genomics UK (COG-UK) | The Lighthouse Lab in Milton Keynes and Alex Alderton, Roberto Amato, Jeffrey Barrett, Sonia Goncalves, Ewan Harrison, David K. Jackson, Ian Johnston, Dominic Kwiatkowski, Cordelia Langford, John Sillitoe on behalf of the Wellcome Sanger |

| | | | |
|---|---|---|---|
| | | Consortium | Institute COVID-19 Surveillance Team |
| EPI_ISL_2911824, EPI_ISL_2911825 | Lighthouse Lab in Alderley Park | Wellcome Sanger Institute for the COVID-19 Genomics UK (COG-UK) Consortium | Jacquelyn Wynn, Mairead Hyland, The Lighthouse Lab in Alderley Park and Alex Alderton, Roberto Amato, Jeffrey Barrett, Sonia Goncalves, Ewan Harrison, David K. Jackson, Ian Johnston, Dominic Kwiatkowski, Cordelia Langford, John Sillitoe on behalf of the Wellcome Sanger Institute COVID-19 Surveillance Team |
| EPI_ISL_2911826, EPI_ISL_2911827 | Lighthouse Lab in Milton Keynes | Wellcome Sanger Institute for the COVID-19 Genomics UK (COG-UK) Consortium | The Lighthouse Lab in Milton Keynes and Alex Alderton, Roberto Amato, Jeffrey Barrett, Sonia Goncalves, Ewan Harrison, David K. Jackson, Ian Johnston, Dominic Kwiatkowski, Cordelia Langford, John Sillitoe on behalf of the Wellcome Sanger Institute COVID-19 Surveillance Team |
| EPI_ISL_2911828, EPI_ISL_2911829, EPI_ISL_2911830 | Lighthouse Lab in Alderley Park | Wellcome Sanger Institute for the COVID-19 Genomics UK (COG-UK) Consortium | Jacquelyn Wynn, Mairead Hyland, The Lighthouse Lab in Alderley Park and Alex Alderton, Roberto Amato, Jeffrey Barrett, Sonia Goncalves, Ewan Harrison, David K. Jackson, Ian Johnston, Dominic Kwiatkowski, Cordelia Langford, John Sillitoe on behalf of the Wellcome Sanger Institute COVID-19 Surveillance Team |
| EPI_ISL_2911831, EPI_ISL_2911832 | Lighthouse Lab in Milton Keynes | Wellcome Sanger Institute for the COVID-19 Genomics UK (COG-UK) Consortium | The Lighthouse Lab in Milton Keynes and Alex Alderton, Roberto Amato, Jeffrey Barrett, Sonia Goncalves, Ewan Harrison, David K. Jackson, Ian Johnston, Dominic Kwiatkowski, Cordelia Langford, John Sillitoe on behalf of the Wellcome Sanger Institute COVID-19 Surveillance Team |
| EPI_ISL_2911833, EPI_ISL_2911834 | Lighthouse Lab in Alderley Park | Wellcome Sanger Institute for the COVID-19 Genomics UK (COG-UK) Consortium | Jacquelyn Wynn, Mairead Hyland, The Lighthouse Lab in Alderley Park and Alex Alderton, Roberto Amato, Jeffrey Barrett, Sonia Goncalves, Ewan Harrison, David K. Jackson, Ian Johnston, Dominic Kwiatkowski, Cordelia Langford, John Sillitoe on behalf of the Wellcome Sanger Institute COVID-19 Surveillance Team |
| EPI_ISL_2911835 | Lighthouse Lab in Milton Keynes | Wellcome Sanger Institute for the COVID-19 Genomics UK (COG-UK) Consortium | The Lighthouse Lab in Milton Keynes and Alex Alderton, Roberto Amato, Jeffrey Barrett, Sonia Goncalves, Ewan Harrison, David K. Jackson, Ian Johnston, Dominic Kwiatkowski, Cordelia Langford, John Sillitoe on behalf of the Wellcome Sanger Institute COVID-19 Surveillance Team |
| EPI_ISL_2911836, EPI_ISL_2911837, EPI_ISL_2911838 | Lighthouse Lab in Alderley Park | Wellcome Sanger Institute for the COVID-19 Genomics UK (COG-UK) Consortium | Jacquelyn Wynn, Mairead Hyland, The Lighthouse Lab in Alderley Park and Alex Alderton, Roberto Amato, Jeffrey Barrett, Sonia Goncalves, Ewan Harrison, David K. Jackson, Ian Johnston, Dominic Kwiatkowski, Cordelia Langford, John Sillitoe on behalf of the Wellcome Sanger Institute COVID-19 Surveillance Team |
| EPI_ISL_2911839, EPI_ISL_2911840 | Lighthouse Lab in Milton Keynes | Wellcome Sanger Institute for the COVID-19 Genomics UK (COG-UK) Consortium | The Lighthouse Lab in Milton Keynes and Alex Alderton, Roberto Amato, Jeffrey Barrett, Sonia Goncalves, Ewan Harrison, David K. Jackson, Ian Johnston, Dominic Kwiatkowski, Cordelia Langford, John Sillitoe on behalf of the Wellcome Sanger Institute COVID-19 Surveillance Team |
| EPI_ISL_2911841 | Lighthouse Lab in Glasgow | Wellcome Sanger Institute for the COVID-19 Genomics UK (COG-UK) Consortium | Harper VanSteenhouse, Yumi Kasai, David Gray, Carol Clugston, Anna Dominiczak and Alex Alderton, Roberto Amato, Jeffrey Barrett, Sonia Goncalves, Ewan Harrison, David K. Jackson, Ian Johnston, Dominic Kwiatkowski, Cordelia Langford, John Sillitoe on behalf of the Wellcome Sanger Institute COVID-19 Surveillance Team |
| EPI_ISL_2911842 | Lighthouse Lab in Alderley Park | Wellcome Sanger Institute for the COVID-19 Genomics UK (COG-UK) Consortium | Jacquelyn Wynn, Mairead Hyland, The Lighthouse Lab in Alderley Park and Alex Alderton, Roberto Amato, Jeffrey Barrett, Sonia Goncalves, Ewan Harrison, David K. Jackson, Ian Johnston, Dominic Kwiatkowski, Cordelia Langford, John Sillitoe on behalf of the Wellcome Sanger Institute COVID-19 Surveillance Team |
| EPI_ISL_2911843 | Lighthouse Lab in Glasgow | Wellcome Sanger Institute for the COVID-19 Genomics UK (COG-UK) Consortium | Harper VanSteenhouse, Yumi Kasai, David Gray, Carol Clugston, Anna Dominiczak and Alex Alderton, Roberto Amato, Jeffrey Barrett, Sonia Goncalves, Ewan Harrison, David K. Jackson, Ian Johnston, Dominic Kwiatkowski, Cordelia Langford, John Sillitoe on behalf of the Wellcome Sanger Institute COVID-19 Surveillance Team |
| EPI_ISL_2911844, EPI_ISL_2911845 | Lighthouse Lab in Alderley Park | Wellcome Sanger Institute for the COVID-19 Genomics UK (COG-UK) Consortium | Jacquelyn Wynn, Mairead Hyland, The Lighthouse Lab in Alderley Park and Alex Alderton, Roberto Amato, Jeffrey Barrett, Sonia Goncalves, Ewan Harrison, David K. Jackson, Ian Johnston, Dominic Kwiatkowski, Cordelia Langford, John Sillitoe on behalf of the Wellcome Sanger Institute COVID-19 Surveillance Team |
| EPI_ISL_2911846 | Lighthouse Lab in Milton Keynes | Wellcome Sanger Institute for the COVID-19 Genomics UK (COG-UK) Consortium | The Lighthouse Lab in Milton Keynes and Alex Alderton, Roberto Amato, Jeffrey Barrett, Sonia Goncalves, Ewan Harrison, David K. Jackson, Ian Johnston, Dominic Kwiatkowski, Cordelia Langford, John Sillitoe on behalf of the Wellcome Sanger Institute COVID-19 Surveillance Team |
| EPI_ISL_2911847, EPI_ISL_2911848 | Lighthouse Lab in Alderley Park | Wellcome Sanger Institute for the COVID-19 Genomics UK (COG-UK) Consortium | Jacquelyn Wynn, Mairead Hyland, The Lighthouse Lab in Alderley Park and Alex Alderton, Roberto Amato, Jeffrey Barrett, Sonia Goncalves, Ewan Harrison, David K. Jackson, Ian Johnston, Dominic Kwiatkowski, Cordelia Langford, John Sillitoe on behalf of the Wellcome Sanger Institute COVID-19 Surveillance Team |
| EPI_ISL_2911849 | Lighthouse Lab in Milton Keynes | Wellcome Sanger Institute for the COVID-19 Genomics UK (COG-UK) Consortium | The Lighthouse Lab in Milton Keynes and Alex Alderton, Roberto Amato, Jeffrey Barrett, Sonia Goncalves, Ewan Harrison, David K. Jackson, Ian Johnston, Dominic Kwiatkowski, Cordelia Langford, John Sillitoe on behalf of the Wellcome Sanger Institute COVID-19 Surveillance Team |
| EPI_ISL_2911850 | Lighthouse Lab in Alderley Park | Wellcome Sanger Institute for the COVID-19 Genomics UK (COG-UK) Consortium | Jacquelyn Wynn, Mairead Hyland, The Lighthouse Lab in Alderley Park and Alex Alderton, Roberto Amato, Jeffrey Barrett, Sonia Goncalves, Ewan Harrison, David K. Jackson, Ian Johnston, Dominic Kwiatkowski, Cordelia Langford, John Sillitoe on behalf of the Wellcome Sanger Institute COVID-19 Surveillance Team |
| EPI_ISL_2911851 | Lighthouse Lab in Milton Keynes | Wellcome Sanger Institute for the COVID-19 Genomics UK (COG-UK) Consortium | The Lighthouse Lab in Milton Keynes and Alex Alderton, Roberto Amato, Jeffrey Barrett, Sonia Goncalves, Ewan Harrison, David K. Jackson, Ian Johnston, Dominic Kwiatkowski, Cordelia Langford, John Sillitoe on behalf of the Wellcome Sanger Institute COVID-19 Surveillance Team |
| EPI_ISL_2911852, EPI_ISL_2911853, EPI_ISL_2911854 | Lighthouse Lab in Alderley Park | Wellcome Sanger Institute for the COVID-19 Genomics UK (COG-UK) Consortium | Jacquelyn Wynn, Mairead Hyland, The Lighthouse Lab in Alderley Park and Alex Alderton, Roberto Amato, Jeffrey Barrett, Sonia Goncalves, Ewan Harrison, David K. Jackson, Ian Johnston, Dominic Kwiatkowski, Cordelia Langford, John Sillitoe on behalf of the Wellcome Sanger Institute COVID-19 Surveillance Team |
| EPI_ISL_2911855 | Lighthouse Lab in Milton Keynes | Wellcome Sanger Institute for the COVID-19 Genomics UK (COG-UK) Consortium | The Lighthouse Lab in Milton Keynes and Alex Alderton, Roberto Amato, Jeffrey Barrett, Sonia Goncalves, Ewan Harrison, David K. Jackson, Ian Johnston, Dominic Kwiatkowski, Cordelia Langford, John Sillitoe on behalf of the Wellcome Sanger Institute COVID-19 Surveillance Team |
| EPI_ISL_2911856 | Lighthouse Lab in Alderley Park | Wellcome Sanger Institute for the COVID-19 Genomics UK (COG-UK) Consortium | Jacquelyn Wynn, Mairead Hyland, The Lighthouse Lab in Alderley Park and Alex Alderton, Roberto Amato, Jeffrey Barrett, Sonia Goncalves, Ewan Harrison, David K. Jackson, Ian Johnston, Dominic Kwiatkowski, Cordelia Langford, John Sillitoe on behalf of the Wellcome Sanger Institute COVID-19 Surveillance Team |
| EPI_ISL_2911857, EPI_ISL_2911858 | Lighthouse Lab in Glasgow | Wellcome Sanger Institute for the COVID-19 Genomics UK (COG-UK) Consortium | Harper VanSteenhouse, Yumi Kasai, David Gray, Carol Clugston, Anna Dominiczak and Alex Alderton, Roberto Amato, Jeffrey Barrett, Sonia Goncalves, Ewan Harrison, David K. Jackson, Ian Johnston, Dominic Kwiatkowski, Cordelia Langford, John Sillitoe on behalf of the Wellcome Sanger Institute COVID-19 Surveillance Team |
| EPI_ISL_2911859, EPI_ISL_2911860 | Lighthouse Lab in Alderley Park | Wellcome Sanger Institute for the COVID-19 Genomics UK (COG-UK) Consortium | Jacquelyn Wynn, Mairead Hyland, The Lighthouse Lab in Alderley Park and Alex Alderton, Roberto Amato, Jeffrey Barrett, Sonia Goncalves, Ewan Harrison, David K. Jackson, Ian Johnston, Dominic Kwiatkowski, Cordelia Langford, John Sillitoe on behalf of the Wellcome Sanger Institute COVID-19 Surveillance Team |
| EPI_ISL_2911861 | Lighthouse Lab in Milton Keynes | Wellcome Sanger Institute for the COVID-19 Genomics UK (COG-UK) Consortium | The Lighthouse Lab in Milton Keynes and Alex Alderton, Roberto Amato, Jeffrey Barrett, Sonia Goncalves, Ewan Harrison, David K. Jackson, Ian Johnston, Dominic Kwiatkowski, Cordelia Langford, John Sillitoe on behalf of the Wellcome Sanger Institute COVID-19 Surveillance Team |
| EPI_ISL_2911862 | Lighthouse Lab in Alderley Park | Wellcome Sanger Institute for the COVID-19 Genomics UK (COG-UK) Consortium | Jacquelyn Wynn, Mairead Hyland, The Lighthouse Lab in Alderley Park and Alex Alderton, Roberto Amato, Jeffrey Barrett, Sonia Goncalves, Ewan Harrison, David K. Jackson, Ian Johnston, Dominic Kwiatkowski, Cordelia Langford, John Sillitoe on behalf of the Wellcome Sanger Institute COVID-19 Surveillance Team |
| EPI_ISL_2911863, EPI_ISL_2911864 | Lighthouse Lab in Milton Keynes | Wellcome Sanger Institute for the COVID-19 Genomics UK (COG-UK) Consortium | The Lighthouse Lab in Milton Keynes and Alex Alderton, Roberto Amato, Jeffrey Barrett, Sonia Goncalves, Ewan Harrison, David K. Jackson, Ian Johnston, Dominic Kwiatkowski, Cordelia Langford, John Sillitoe on behalf of the Wellcome Sanger Institute COVID-19 Surveillance Team |
| EPI_ISL_2911865, EPI_ISL_2911866 | Lighthouse Lab in Alderley Park | Wellcome Sanger Institute for the COVID-19 Genomics UK (COG-UK) Consortium | Jacquelyn Wynn, Mairead Hyland, The Lighthouse Lab in Alderley Park and Alex Alderton, Roberto Amato, Jeffrey Barrett, Sonia Goncalves, Ewan Harrison, David K. Jackson, Ian Johnston, Dominic Kwiatkowski, Cordelia Langford, John Sillitoe on behalf of the Wellcome Sanger Institute COVID-19 Surveillance Team |
| EPI_ISL_2911867, EPI_ISL_2911868 | Lighthouse Lab in Milton Keynes | Wellcome Sanger Institute for the COVID-19 Genomics UK (COG-UK) Consortium | The Lighthouse Lab in Milton Keynes and Alex Alderton, Roberto Amato, Jeffrey Barrett, Sonia Goncalves, Ewan Harrison, David K. Jackson, Ian Johnston, Dominic Kwiatkowski, Cordelia Langford, John Sillitoe on behalf of the Wellcome Sanger Institute COVID-19 Surveillance Team |
| EPI_ISL_2911869, EPI_ISL_2911870, EPI_ISL_2911871, EPI_ISL_2911872, EPI_ISL_2911873 | Lighthouse Lab in Alderley Park | Wellcome Sanger Institute for the COVID-19 Genomics UK (COG-UK) Consortium | Jacquelyn Wynn, Mairead Hyland, The Lighthouse Lab in Alderley Park and Alex Alderton, Roberto Amato, Jeffrey Barrett, Sonia Goncalves, Ewan Harrison, David K. Jackson, Ian Johnston, Dominic Kwiatkowski, Cordelia Langford, John Sillitoe on behalf of the Wellcome Sanger Institute COVID-19 Surveillance Team |
| EPI_ISL_2911874 | Lighthouse Lab in Milton Keynes | Wellcome Sanger Institute for the COVID-19 Genomics UK (COG-UK) Consortium | The Lighthouse Lab in Milton Keynes and Alex Alderton, Roberto Amato, Jeffrey Barrett, Sonia Goncalves, Ewan Harrison, David K. Jackson, Ian Johnston, Dominic Kwiatkowski, Cordelia Langford, John Sillitoe on behalf of the Wellcome Sanger Institute COVID-19 Surveillance Team |
| EPI_ISL_2911875, EPI_ISL_2911876, EPI_ISL_2911877 | Lighthouse Lab in Alderley Park | Wellcome Sanger Institute for the COVID-19 Genomics UK (COG-UK) Consortium | Jacquelyn Wynn, Mairead Hyland, The Lighthouse Lab in Alderley Park and Alex Alderton, Roberto Amato, Jeffrey Barrett, Sonia Goncalves, Ewan Harrison, David K. Jackson, Ian Johnston, Dominic Kwiatkowski, Cordelia Langford, John Sillitoe on behalf of the Wellcome Sanger Institute COVID-19 Surveillance Team |
| EPI_ISL_2911878 | Lighthouse Lab in Milton Keynes | Wellcome Sanger Institute for the COVID-19 Genomics UK (COG-UK) Consortium | The Lighthouse Lab in Milton Keynes and Alex Alderton, Roberto Amato, Jeffrey Barrett, Sonia Goncalves, Ewan Harrison, David K. Jackson, Ian Johnston, Dominic Kwiatkowski, Cordelia Langford, John Sillitoe on behalf of the Wellcome Sanger Institute COVID-19 Surveillance Team |
| EPI_ISL_2911879 | Lighthouse Lab in Alderley Park | Wellcome Sanger Institute for the COVID-19 Genomics UK (COG-UK) Consortium | Jacquelyn Wynn, Mairead Hyland, The Lighthouse Lab in Alderley Park and Alex Alderton, Roberto Amato, Jeffrey Barrett, Sonia Goncalves, Ewan Harrison, David K. Jackson, Ian Johnston, Dominic Kwiatkowski, Cordelia Langford, John Sillitoe on behalf of the Wellcome Sanger Institute COVID-19 Surveillance Team |
| EPI_ISL_2911880 | Lighthouse Lab in Milton Keynes | Wellcome Sanger Institute for the COVID-19 Genomics UK (COG-UK) Consortium | The Lighthouse Lab in Milton Keynes and Alex Alderton, Roberto Amato, Jeffrey Barrett, Sonia Goncalves, Ewan Harrison, David K. Jackson, Ian Johnston, Dominic Kwiatkowski, Cordelia Langford, John Sillitoe on behalf of the Wellcome Sanger Institute COVID-19 Surveillance Team |
| EPI_ISL_2911881 | Lighthouse Lab in Alderley Park | Wellcome Sanger Institute for the COVID-19 Genomics UK (COG-UK) Consortium | Jacquelyn Wynn, Mairead Hyland, The Lighthouse Lab in Alderley Park and Alex Alderton, Roberto Amato, Jeffrey Barrett, Sonia Goncalves, Ewan Harrison, David K. Jackson, Ian Johnston, Dominic Kwiatkowski, Cordelia Langford, John Sillitoe on behalf of the Wellcome Sanger Institute COVID-19 Surveillance Team |
| EPI_ISL_2911882 | Lighthouse Lab in Milton Keynes | Wellcome Sanger Institute for the COVID-19 Genomics UK (COG-UK) Consortium | The Lighthouse Lab in Milton Keynes and Alex Alderton, Roberto Amato, Jeffrey Barrett, Sonia Goncalves, Ewan Harrison, David K. Jackson, Ian Johnston, Dominic Kwiatkowski, Cordelia Langford, John Sillitoe on behalf of the Wellcome Sanger Institute COVID-19 Surveillance Team |
| EPI_ISL_2911883, EPI_ISL_2911884, EPI_ISL_2911885 | Lighthouse Lab in Alderley Park | Wellcome Sanger Institute for the COVID-19 Genomics UK (COG-UK) Consortium | Jacquelyn Wynn, Mairead Hyland, The Lighthouse Lab in Alderley Park and Alex Alderton, Roberto Amato, Jeffrey Barrett, Sonia Goncalves, Ewan Harrison, David K. Jackson, Ian Johnston, Dominic Kwiatkowski, Cordelia Langford, John Sillitoe on behalf of the Wellcome Sanger Institute COVID-19 Surveillance Team |
| EPI_ISL_2911886 | Lighthouse Lab in Glasgow | Wellcome Sanger Institute for the COVID-19 Genomics UK (COG-UK) Consortium | Harper VanSteenhouse, Yumi Kasai, David Gray, Carol Clugston, Anna Dominiczak and Alex Alderton, Roberto Amato, Jeffrey Barrett, Sonia Goncalves, Ewan Harrison, David K. Jackson, Ian Johnston, Dominic Kwiatkowski, Cordelia Langford, John Sillitoe on behalf of the Wellcome Sanger Institute COVID-19 Surveillance Team |
| EPI_ISL_2911887 | Lighthouse Lab in Alderley Park | Wellcome Sanger Institute for the COVID-19 Genomics UK (COG-UK) Consortium | Jacquelyn Wynn, Mairead Hyland, The Lighthouse Lab in Alderley Park and Alex Alderton, Roberto Amato, Jeffrey Barrett, Sonia Goncalves, Ewan Harrison, David K. Jackson, Ian Johnston, Dominic Kwiatkowski, Cordelia Langford, John Sillitoe on behalf of the Wellcome Sanger Institute COVID-19 Surveillance Team |
| EPI_ISL_2911888 | Lighthouse Lab in Milton Keynes | Wellcome Sanger Institute for the COVID-19 Genomics UK (COG-UK) Consortium | The Lighthouse Lab in Milton Keynes and Alex Alderton, Roberto Amato, Jeffrey Barrett, Sonia Goncalves, Ewan Harrison, David K. Jackson, Ian Johnston, Dominic Kwiatkowski, Cordelia Langford, John Sillitoe on behalf of the Wellcome Sanger Institute COVID-19 Surveillance Team |
| EPI_ISL_2911889 | Lighthouse Lab in Alderley Park | Wellcome Sanger Institute for the COVID-19 Genomics UK (COG-UK) Consortium | Jacquelyn Wynn, Mairead Hyland, The Lighthouse Lab in Alderley Park and Alex Alderton, Roberto Amato, Jeffrey Barrett, Sonia Goncalves, Ewan Harrison, David K. Jackson, Ian Johnston, Dominic Kwiatkowski, Cordelia Langford, John Sillitoe on behalf of the Wellcome Sanger Institute COVID-19 Surveillance Team |
| EPI_ISL_2911890, EPI_ISL_2911891 | Lighthouse Lab in Milton Keynes | Wellcome Sanger Institute for the COVID-19 Genomics UK (COG-UK) Consortium | The Lighthouse Lab in Milton Keynes and Alex Alderton, Roberto Amato, Jeffrey Barrett, Sonia Goncalves, Ewan Harrison, David K. Jackson, Ian Johnston, Dominic Kwiatkowski, Cordelia Langford, John Sillitoe on behalf of the Wellcome Sanger Institute COVID-19 Surveillance Team |

| | | | |
|---|---|---|---|
| EPI_ISL_2911892, EPI_ISL_2911893, EPI_ISL_2911894, EPI_ISL_2911895 | Lighthouse Lab in Alderley Park | Wellcome Sanger Institute for the COVID-19 Genomics UK (COG-UK) Consortium | Jacquelyn Wynn, Mairead Hyland, The Lighthouse Lab in Alderley Park and Alex Alderton, Roberto Amato, Jeffrey Barrett, Sonia Goncalves, Ewan Harrison, David K. Jackson, Ian Johnston, Dominic Kwiatkowski, Cordelia Langford, John Sillitoe on behalf of the Wellcome Sanger Institute COVID-19 Surveillance Team |
| EPI_ISL_2911896 | Lighthouse Lab in Milton Keynes | Wellcome Sanger Institute for the COVID-19 Genomics UK (COG-UK) Consortium | The Lighthouse Lab in Milton Keynes and Alex Alderton, Roberto Amato, Jeffrey Barrett, Sonia Goncalves, Ewan Harrison, David K. Jackson, Ian Johnston, Dominic Kwiatkowski, Cordelia Langford, John Sillitoe on behalf of the Wellcome Sanger Institute COVID-19 Surveillance Team |
| EPI_ISL_2911897, EPI_ISL_2911898, EPI_ISL_2911899 | Lighthouse Lab in Alderley Park | Wellcome Sanger Institute for the COVID-19 Genomics UK (COG-UK) Consortium | Jacquelyn Wynn, Mairead Hyland, The Lighthouse Lab in Alderley Park and Alex Alderton, Roberto Amato, Jeffrey Barrett, Sonia Goncalves, Ewan Harrison, David K. Jackson, Ian Johnston, Dominic Kwiatkowski, Cordelia Langford, John Sillitoe on behalf of the Wellcome Sanger Institute COVID-19 Surveillance Team |
| EPI_ISL_2911900, EPI_ISL_2911901 | Berkshire and Surrey Pathology Services Lighthouse Laboratory | Wellcome Sanger Institute for the COVID-19 Genomics UK (COG-UK) Consortium | Berkshire and Surrey Pathology Services Lighthouse Laboratory and Alex Alderton, Roberto Amato, Jeffrey Barrett, Sonia Goncalves, Ewan Harrison, David K. Jackson, Ian Johnston, Dominic Kwiatkowski, Cordelia Langford, John Sillitoe on behalf of the Wellcome Sanger Institute COVID-19 Surveillance Team |
| EPI_ISL_2911902, EPI_ISL_2911903 | Lighthouse Laboratory Plymouth | Wellcome Sanger Institute for the COVID-19 Genomics UK (COG-UK) Consortium | Lighthouse Laboratory Plymouth and Alex Alderton, Roberto Amato, Jeffrey Barrett, Sonia Goncalves, Ewan Harrison, David K. Jackson, Ian Johnston, Dominic Kwiatkowski, Cordelia Langford, John Sillitoe on behalf of the Wellcome Sanger Institute COVID-19 Surveillance Team |
| EPI_ISL_2911904, EPI_ISL_2911905, EPI_ISL_2911906 | Berkshire and Surrey Pathology Services Lighthouse Laboratory | Wellcome Sanger Institute for the COVID-19 Genomics UK (COG-UK) Consortium | Berkshire and Surrey Pathology Services Lighthouse Laboratory and Alex Alderton, Roberto Amato, Jeffrey Barrett, Sonia Goncalves, Ewan Harrison, David K. Jackson, Ian Johnston, Dominic Kwiatkowski, Cordelia Langford, John Sillitoe on behalf of the Wellcome Sanger Institute COVID-19 Surveillance Team |
| EPI_ISL_2911907 | Lighthouse Laboratory Plymouth | Wellcome Sanger Institute for the COVID-19 Genomics UK (COG-UK) Consortium | Lighthouse Laboratory Plymouth and Alex Alderton, Roberto Amato, Jeffrey Barrett, Sonia Goncalves, Ewan Harrison, David K. Jackson, Ian Johnston, Dominic Kwiatkowski, Cordelia Langford, John Sillitoe on behalf of the Wellcome Sanger Institute COVID-19 Surveillance Team |
| EPI_ISL_2911908 | Berkshire and Surrey Pathology Services Lighthouse Laboratory | Wellcome Sanger Institute for the COVID-19 Genomics UK (COG-UK) Consortium | Berkshire and Surrey Pathology Services Lighthouse Laboratory and Alex Alderton, Roberto Amato, Jeffrey Barrett, Sonia Goncalves, Ewan Harrison, David K. Jackson, Ian Johnston, Dominic Kwiatkowski, Cordelia Langford, John Sillitoe on behalf of the Wellcome Sanger Institute COVID-19 Surveillance Team |
| EPI_ISL_2911909 | Lighthouse Lab in Milton Keynes | Wellcome Sanger Institute for the COVID-19 Genomics UK (COG-UK) Consortium | The Lighthouse Lab in Milton Keynes and Alex Alderton, Roberto Amato, Jeffrey Barrett, Sonia Goncalves, Ewan Harrison, David K. Jackson, Ian Johnston, Dominic Kwiatkowski, Cordelia Langford, John Sillitoe on behalf of the Wellcome Sanger Institute COVID-19 Surveillance Team |
| EPI_ISL_2911910, EPI_ISL_2911911 | Berkshire and Surrey Pathology Services Lighthouse Laboratory | Wellcome Sanger Institute for the COVID-19 Genomics UK (COG-UK) Consortium | Berkshire and Surrey Pathology Services Lighthouse Laboratory and Alex Alderton, Roberto Amato, Jeffrey Barrett, Sonia Goncalves, Ewan Harrison, David K. Jackson, Ian Johnston, Dominic Kwiatkowski, Cordelia Langford, John Sillitoe on behalf of the Wellcome Sanger Institute COVID-19 Surveillance Team |
| EPI_ISL_2911912 | Lighthouse Laboratory Plymouth | Wellcome Sanger Institute for the COVID-19 Genomics UK (COG-UK) Consortium | Lighthouse Laboratory Plymouth and Alex Alderton, Roberto Amato, Jeffrey Barrett, Sonia Goncalves, Ewan Harrison, David K. Jackson, Ian Johnston, Dominic Kwiatkowski, Cordelia Langford, John Sillitoe on behalf of the Wellcome Sanger Institute COVID-19 Surveillance Team |
| EPI_ISL_2911913 | Berkshire and Surrey Pathology Services Lighthouse Laboratory | Wellcome Sanger Institute for the COVID-19 Genomics UK (COG-UK) Consortium | Berkshire and Surrey Pathology Services Lighthouse Laboratory and Alex Alderton, Roberto Amato, Jeffrey Barrett, Sonia Goncalves, Ewan Harrison, David K. Jackson, Ian Johnston, Dominic Kwiatkowski, Cordelia Langford, John Sillitoe on behalf of the Wellcome Sanger Institute COVID-19 Surveillance Team |
| EPI_ISL_2911914 | Lighthouse Lab in Milton Keynes | Wellcome Sanger Institute for the COVID-19 Genomics UK (COG-UK) Consortium | The Lighthouse Lab in Milton Keynes and Alex Alderton, Roberto Amato, Jeffrey Barrett, Sonia Goncalves, Ewan Harrison, David K. Jackson, Ian Johnston, Dominic Kwiatkowski, Cordelia Langford, John Sillitoe on behalf of the Wellcome Sanger Institute COVID-19 Surveillance Team |
| EPI_ISL_2911915 | Lighthouse Laboratory Plymouth | Wellcome Sanger Institute for the COVID-19 Genomics UK (COG-UK) Consortium | Lighthouse Laboratory Plymouth and Alex Alderton, Roberto Amato, Jeffrey Barrett, Sonia Goncalves, Ewan Harrison, David K. Jackson, Ian Johnston, Dominic Kwiatkowski, Cordelia Langford, John Sillitoe on behalf of the Wellcome Sanger Institute COVID-19 Surveillance Team |
| EPI_ISL_2911916 | Lighthouse Lab in Milton Keynes | Wellcome Sanger Institute for the COVID-19 Genomics UK (COG-UK) Consortium | The Lighthouse Lab in Milton Keynes and Alex Alderton, Roberto Amato, Jeffrey Barrett, Sonia Goncalves, Ewan Harrison, David K. Jackson, Ian Johnston, Dominic Kwiatkowski, Cordelia Langford, John Sillitoe on behalf of the Wellcome Sanger Institute COVID-19 Surveillance Team |
| EPI_ISL_2911917, EPI_ISL_2911918, EPI_ISL_2911919 | Lighthouse Laboratory Plymouth | Wellcome Sanger Institute for the COVID-19 Genomics UK (COG-UK) Consortium | Lighthouse Laboratory Plymouth and Alex Alderton, Roberto Amato, Jeffrey Barrett, Sonia Goncalves, Ewan Harrison, David K. Jackson, Ian Johnston, Dominic Kwiatkowski, Cordelia Langford, John Sillitoe on behalf of the Wellcome Sanger Institute COVID-19 Surveillance Team |
| EPI_ISL_2911920, EPI_ISL_2911921, EPI_ISL_2911922 | Berkshire and Surrey Pathology Services Lighthouse Laboratory | Wellcome Sanger Institute for the COVID-19 Genomics UK (COG-UK) Consortium | Berkshire and Surrey Pathology Services Lighthouse Laboratory and Alex Alderton, Roberto Amato, Jeffrey Barrett, Sonia Goncalves, Ewan Harrison, David K. Jackson, Ian Johnston, Dominic Kwiatkowski, Cordelia Langford, John Sillitoe on behalf of the Wellcome Sanger Institute COVID-19 Surveillance Team |
| EPI_ISL_2911923 | Lighthouse Laboratory Plymouth | Wellcome Sanger Institute for the COVID-19 Genomics UK (COG-UK) Consortium | Lighthouse Laboratory Plymouth and Alex Alderton, Roberto Amato, Jeffrey Barrett, Sonia Goncalves, Ewan Harrison, David K. Jackson, Ian Johnston, Dominic Kwiatkowski, Cordelia Langford, John Sillitoe on behalf of the Wellcome Sanger Institute COVID-19 Surveillance Team |
| EPI_ISL_2911924 | Lighthouse Lab in Milton Keynes | Wellcome Sanger Institute for the COVID-19 Genomics UK (COG-UK) Consortium | The Lighthouse Lab in Milton Keynes and Alex Alderton, Roberto Amato, Jeffrey Barrett, Sonia Goncalves, Ewan Harrison, David K. Jackson, Ian Johnston, Dominic Kwiatkowski, Cordelia Langford, John Sillitoe on behalf of the Wellcome Sanger Institute COVID-19 Surveillance Team |
| EPI_ISL_2911925, EPI_ISL_2911926 | Berkshire and Surrey Pathology Services Lighthouse Laboratory | Wellcome Sanger Institute for the COVID-19 Genomics UK (COG-UK) Consortium | Berkshire and Surrey Pathology Services Lighthouse Laboratory and Alex Alderton, Roberto Amato, Jeffrey Barrett, Sonia Goncalves, Ewan Harrison, David K. Jackson, Ian Johnston, Dominic Kwiatkowski, Cordelia Langford, John Sillitoe on behalf of the Wellcome Sanger Institute COVID-19 Surveillance Team |
| EPI_ISL_2911927 | Lighthouse Lab in Milton Keynes | Wellcome Sanger Institute for the COVID-19 Genomics UK (COG-UK) Consortium | The Lighthouse Lab in Milton Keynes and Alex Alderton, Roberto Amato, Jeffrey Barrett, Sonia Goncalves, Ewan Harrison, David K. Jackson, Ian Johnston, Dominic Kwiatkowski, Cordelia Langford, John Sillitoe on behalf of the Wellcome Sanger Institute COVID-19 Surveillance Team |
| EPI_ISL_2911928, EPI_ISL_2911929 | Lighthouse Laboratory Plymouth | Wellcome Sanger Institute for the COVID-19 Genomics UK (COG-UK) Consortium | Lighthouse Laboratory Plymouth and Alex Alderton, Roberto Amato, Jeffrey Barrett, Sonia Goncalves, Ewan Harrison, David K. Jackson, Ian Johnston, Dominic Kwiatkowski, Cordelia Langford, John Sillitoe on behalf of the Wellcome Sanger Institute COVID-19 Surveillance Team |
| EPI_ISL_2911930 | Berkshire and Surrey Pathology Services Lighthouse Laboratory | Wellcome Sanger Institute for the COVID-19 Genomics UK (COG-UK) Consortium | Berkshire and Surrey Pathology Services Lighthouse Laboratory and Alex Alderton, Roberto Amato, Jeffrey Barrett, Sonia Goncalves, Ewan Harrison, David K. Jackson, Ian Johnston, Dominic Kwiatkowski, Cordelia Langford, John Sillitoe on behalf of the Wellcome Sanger Institute COVID-19 Surveillance Team |
| EPI_ISL_2911931 | Lighthouse Laboratory Plymouth | Wellcome Sanger Institute for the COVID-19 Genomics UK (COG-UK) Consortium | Lighthouse Laboratory Plymouth and Alex Alderton, Roberto Amato, Jeffrey Barrett, Sonia Goncalves, Ewan Harrison, David K. Jackson, Ian Johnston, Dominic Kwiatkowski, Cordelia Langford, John Sillitoe on behalf of the Wellcome Sanger Institute COVID-19 Surveillance Team |
| EPI_ISL_2911932, EPI_ISL_2911933, EPI_ISL_2911934 | Berkshire and Surrey Pathology Services Lighthouse Laboratory | Wellcome Sanger Institute for the COVID-19 Genomics UK (COG-UK) Consortium | Berkshire and Surrey Pathology Services Lighthouse Laboratory and Alex Alderton, Roberto Amato, Jeffrey Barrett, Sonia Goncalves, Ewan Harrison, David K. Jackson, Ian Johnston, Dominic Kwiatkowski, Cordelia Langford, John Sillitoe on behalf of the Wellcome Sanger Institute COVID-19 Surveillance Team |
| EPI_ISL_2911935, EPI_ISL_2911936, EPI_ISL_2911937, EPI_ISL_2911938 | Lighthouse Laboratory Plymouth | Wellcome Sanger Institute for the COVID-19 Genomics UK (COG-UK) Consortium | Lighthouse Laboratory Plymouth and Alex Alderton, Roberto Amato, Jeffrey Barrett, Sonia Goncalves, Ewan Harrison, David K. Jackson, Ian Johnston, Dominic Kwiatkowski, Cordelia Langford, John Sillitoe on behalf of the Wellcome Sanger Institute COVID-19 Surveillance Team |
| EPI_ISL_2911939, EPI_ISL_2911940 | Berkshire and Surrey Pathology Services Lighthouse Laboratory | Wellcome Sanger Institute for the COVID-19 Genomics UK (COG-UK) Consortium | Berkshire and Surrey Pathology Services Lighthouse Laboratory and Alex Alderton, Roberto Amato, Jeffrey Barrett, Sonia Goncalves, Ewan Harrison, David K. Jackson, Ian Johnston, Dominic Kwiatkowski, Cordelia Langford, John Sillitoe on behalf of the Wellcome Sanger Institute COVID-19 Surveillance Team |
| EPI_ISL_2911941 | Lighthouse Laboratory Plymouth | Wellcome Sanger Institute for the COVID-19 Genomics UK (COG-UK) Consortium | Lighthouse Laboratory Plymouth and Alex Alderton, Roberto Amato, Jeffrey Barrett, Sonia Goncalves, Ewan Harrison, David K. Jackson, Ian Johnston, Dominic Kwiatkowski, Cordelia Langford, John Sillitoe on behalf of the Wellcome Sanger Institute COVID-19 Surveillance Team |
| EPI_ISL_2911942, EPI_ISL_2911943 | Berkshire and Surrey Pathology Services Lighthouse Laboratory | Wellcome Sanger Institute for the COVID-19 Genomics UK (COG-UK) Consortium | Berkshire and Surrey Pathology Services Lighthouse Laboratory and Alex Alderton, Roberto Amato, Jeffrey Barrett, Sonia Goncalves, Ewan Harrison, David K. Jackson, Ian Johnston, Dominic Kwiatkowski, Cordelia Langford, John Sillitoe on behalf of the Wellcome Sanger Institute COVID-19 Surveillance Team |
| EPI_ISL_2911944 | Lighthouse Lab in Milton Keynes | Wellcome Sanger Institute for the COVID-19 Genomics UK (COG-UK) Consortium | The Lighthouse Lab in Milton Keynes and Alex Alderton, Roberto Amato, Jeffrey Barrett, Sonia Goncalves, Ewan Harrison, David K. Jackson, Ian Johnston, Dominic Kwiatkowski, Cordelia Langford, John Sillitoe on behalf of the Wellcome Sanger Institute COVID-19 Surveillance Team |
| EPI_ISL_2911945 | Berkshire and Surrey Pathology Services Lighthouse Laboratory | Wellcome Sanger Institute for the COVID-19 Genomics UK (COG-UK) Consortium | Berkshire and Surrey Pathology Services Lighthouse Laboratory and Alex Alderton, Roberto Amato, Jeffrey Barrett, Sonia Goncalves, Ewan Harrison, David K. Jackson, Ian Johnston, Dominic Kwiatkowski, Cordelia Langford, John Sillitoe on behalf of the Wellcome Sanger Institute COVID-19 Surveillance Team |
| EPI_ISL_2911946 | Lighthouse Laboratory Plymouth | Wellcome Sanger Institute for the COVID-19 Genomics UK (COG-UK) Consortium | Lighthouse Laboratory Plymouth and Alex Alderton, Roberto Amato, Jeffrey Barrett, Sonia Goncalves, Ewan Harrison, David K. Jackson, Ian Johnston, Dominic Kwiatkowski, Cordelia Langford, John Sillitoe on behalf of the Wellcome Sanger Institute COVID-19 Surveillance Team |
| EPI_ISL_2911947, EPI_ISL_2911948 | Berkshire and Surrey Pathology Services Lighthouse Laboratory | Wellcome Sanger Institute for the COVID-19 Genomics UK (COG-UK) Consortium | Berkshire and Surrey Pathology Services Lighthouse Laboratory and Alex Alderton, Roberto Amato, Jeffrey Barrett, Sonia Goncalves, Ewan Harrison, David K. Jackson, Ian Johnston, Dominic Kwiatkowski, Cordelia Langford, John Sillitoe on behalf of the Wellcome Sanger Institute COVID-19 Surveillance Team |
| EPI_ISL_2911949, EPI_ISL_2911950 | Lighthouse Lab in Milton Keynes | Wellcome Sanger Institute for the COVID-19 Genomics UK (COG-UK) Consortium | The Lighthouse Lab in Milton Keynes and Alex Alderton, Roberto Amato, Jeffrey Barrett, Sonia Goncalves, Ewan Harrison, David K. Jackson, Ian Johnston, Dominic Kwiatkowski, Cordelia Langford, John Sillitoe on behalf of the Wellcome Sanger Institute COVID-19 Surveillance Team |
| EPI_ISL_2911951 | Berkshire and Surrey Pathology Services Lighthouse Laboratory | Wellcome Sanger Institute for the COVID-19 Genomics UK (COG-UK) Consortium | Berkshire and Surrey Pathology Services Lighthouse Laboratory and Alex Alderton, Roberto Amato, Jeffrey Barrett, Sonia Goncalves, Ewan Harrison, David K. Jackson, Ian Johnston, Dominic Kwiatkowski, Cordelia Langford, John Sillitoe on behalf of the Wellcome Sanger Institute COVID-19 Surveillance Team |
| EPI_ISL_2911952, EPI_ISL_2911953, EPI_ISL_2911954 | Lighthouse Lab in Milton Keynes | Wellcome Sanger Institute for the COVID-19 Genomics UK (COG-UK) Consortium | The Lighthouse Lab in Milton Keynes and Alex Alderton, Roberto Amato, Jeffrey Barrett, Sonia Goncalves, Ewan Harrison, David K. Jackson, Ian Johnston, Dominic Kwiatkowski, Cordelia Langford, John Sillitoe on behalf of the Wellcome Sanger Institute COVID-19 Surveillance Team |
| EPI_ISL_2911955 | Lighthouse Laboratory Plymouth | Wellcome Sanger Institute for the COVID-19 Genomics UK (COG-UK) Consortium | Lighthouse Laboratory Plymouth and Alex Alderton, Roberto Amato, Jeffrey Barrett, Sonia Goncalves, Ewan Harrison, David K. Jackson, Ian Johnston, Dominic Kwiatkowski, Cordelia Langford, John Sillitoe on behalf of the Wellcome Sanger Institute COVID-19 Surveillance Team |
| EPI_ISL_2911956 | Lighthouse Lab in Milton Keynes | Wellcome Sanger Institute for the COVID-19 Genomics UK (COG-UK) Consortium | The Lighthouse Lab in Milton Keynes and Alex Alderton, Roberto Amato, Jeffrey Barrett, Sonia Goncalves, Ewan Harrison, David K. Jackson, Ian Johnston, Dominic Kwiatkowski, Cordelia Langford, John Sillitoe on behalf of the Wellcome Sanger Institute COVID-19 Surveillance Team |
| EPI_ISL_2911957, EPI_ISL_2911958 | Berkshire and Surrey Pathology Services Lighthouse Laboratory | Wellcome Sanger Institute for the COVID-19 Genomics UK (COG-UK) Consortium | Berkshire and Surrey Pathology Services Lighthouse Laboratory and Alex Alderton, Roberto Amato, Jeffrey Barrett, Sonia Goncalves, Ewan Harrison, David K. Jackson, Ian Johnston, Dominic Kwiatkowski, Cordelia Langford, John Sillitoe on behalf of the Wellcome Sanger Institute COVID-19 Surveillance Team |
| EPI_ISL_2911959, EPI_ISL_2911960 | Lighthouse Laboratory Plymouth | Wellcome Sanger Institute for the COVID-19 Genomics UK (COG-UK) Consortium | Lighthouse Laboratory Plymouth and Alex Alderton, Roberto Amato, Jeffrey Barrett, Sonia Goncalves, Ewan Harrison, David K. Jackson, Ian Johnston, Dominic Kwiatkowski, Cordelia Langford, John Sillitoe on behalf of the Wellcome Sanger Institute COVID-19 Surveillance Team |
| EPI_ISL_2911961, EPI_ISL_2911962 | Berkshire and Surrey Pathology Services Lighthouse Laboratory | Wellcome Sanger Institute for the COVID-19 Genomics UK (COG-UK) Consortium | Berkshire and Surrey Pathology Services Lighthouse Laboratory and Alex Alderton, Roberto Amato, Jeffrey Barrett, Sonia Goncalves, Ewan Harrison, David K. Jackson, Ian Johnston, Dominic Kwiatkowski, Cordelia Langford, John Sillitoe on behalf of the Wellcome Sanger Institute COVID-19 Surveillance Team |

| | | | |
|---|---|---|---|
| EPI_ISL_2911963 | Lighthouse Lab in Milton Keynes | Wellcome Sanger Institute for the COVID-19 Genomics UK (COG-UK) Consortium | The Lighthouse Lab in Milton Keynes and Alex Alderton, Roberto Amato, Jeffrey Barrett, Sonia Goncalves, Ewan Harrison, David K. Jackson, Ian Johnston, Dominic Kwiatkowski, Cordelia Langford, John Sillitoe on behalf of the Wellcome Sanger Institute COVID-19 Surveillance Team |
| EPI_ISL_2911964 | Berkshire and Surrey Pathology Services Lighthouse Laboratory | Wellcome Sanger Institute for the COVID-19 Genomics UK (COG-UK) Consortium | Berkshire and Surrey Pathology Services Lighthouse Laboratory and Alex Alderton, Roberto Amato, Jeffrey Barrett, Sonia Goncalves, Ewan Harrison, David K. Jackson, Ian Johnston, Dominic Kwiatkowski, Cordelia Langford, John Sillitoe on behalf of the Wellcome Sanger Institute COVID-19 Surveillance Team |
| EPI_ISL_2911965 | Lighthouse Lab in Milton Keynes | Wellcome Sanger Institute for the COVID-19 Genomics UK (COG-UK) Consortium | The Lighthouse Lab in Milton Keynes and Alex Alderton, Roberto Amato, Jeffrey Barrett, Sonia Goncalves, Ewan Harrison, David K. Jackson, Ian Johnston, Dominic Kwiatkowski, Cordelia Langford, John Sillitoe on behalf of the Wellcome Sanger Institute COVID-19 Surveillance Team |
| EPI_ISL_2911966 | Berkshire and Surrey Pathology Services Lighthouse Laboratory | Wellcome Sanger Institute for the COVID-19 Genomics UK (COG-UK) Consortium | Berkshire and Surrey Pathology Services Lighthouse Laboratory and Alex Alderton, Roberto Amato, Jeffrey Barrett, Sonia Goncalves, Ewan Harrison, David K. Jackson, Ian Johnston, Dominic Kwiatkowski, Cordelia Langford, John Sillitoe on behalf of the Wellcome Sanger Institute COVID-19 Surveillance Team |
| EPI_ISL_2911967 | Lighthouse Lab in Milton Keynes | Wellcome Sanger Institute for the COVID-19 Genomics UK (COG-UK) Consortium | The Lighthouse Lab in Milton Keynes and Alex Alderton, Roberto Amato, Jeffrey Barrett, Sonia Goncalves, Ewan Harrison, David K. Jackson, Ian Johnston, Dominic Kwiatkowski, Cordelia Langford, John Sillitoe on behalf of the Wellcome Sanger Institute COVID-19 Surveillance Team |
| EPI_ISL_2911968 | Berkshire and Surrey Pathology Services Lighthouse Laboratory | Wellcome Sanger Institute for the COVID-19 Genomics UK (COG-UK) Consortium | Berkshire and Surrey Pathology Services Lighthouse Laboratory and Alex Alderton, Roberto Amato, Jeffrey Barrett, Sonia Goncalves, Ewan Harrison, David K. Jackson, Ian Johnston, Dominic Kwiatkowski, Cordelia Langford, John Sillitoe on behalf of the Wellcome Sanger Institute COVID-19 Surveillance Team |
| EPI_ISL_2911969, EPI_ISL_2911970 | Lighthouse Laboratory Plymouth | Wellcome Sanger Institute for the COVID-19 Genomics UK (COG-UK) Consortium | Lighthouse Laboratory Plymouth and Alex Alderton, Roberto Amato, Jeffrey Barrett, Sonia Goncalves, Ewan Harrison, David K. Jackson, Ian Johnston, Dominic Kwiatkowski, Cordelia Langford, John Sillitoe on behalf of the Wellcome Sanger Institute COVID-19 Surveillance Team |
| EPI_ISL_2911971, EPI_ISL_2911972 | Berkshire and Surrey Pathology Services Lighthouse Laboratory | Wellcome Sanger Institute for the COVID-19 Genomics UK (COG-UK) Consortium | Berkshire and Surrey Pathology Services Lighthouse Laboratory and Alex Alderton, Roberto Amato, Jeffrey Barrett, Sonia Goncalves, Ewan Harrison, David K. Jackson, Ian Johnston, Dominic Kwiatkowski, Cordelia Langford, John Sillitoe on behalf of the Wellcome Sanger Institute COVID-19 Surveillance Team |
| EPI_ISL_2911973, EPI_ISL_2911974, EPI_ISL_2911975 | Lighthouse Lab in Milton Keynes | Wellcome Sanger Institute for the COVID-19 Genomics UK (COG-UK) Consortium | The Lighthouse Lab in Milton Keynes and Alex Alderton, Roberto Amato, Jeffrey Barrett, Sonia Goncalves, Ewan Harrison, David K. Jackson, Ian Johnston, Dominic Kwiatkowski, Cordelia Langford, John Sillitoe on behalf of the Wellcome Sanger Institute COVID-19 Surveillance Team |
| EPI_ISL_2911976 | Berkshire and Surrey Pathology Services Lighthouse Laboratory | Wellcome Sanger Institute for the COVID-19 Genomics UK (COG-UK) Consortium | Berkshire and Surrey Pathology Services Lighthouse Laboratory and Alex Alderton, Roberto Amato, Jeffrey Barrett, Sonia Goncalves, Ewan Harrison, David K. Jackson, Ian Johnston, Dominic Kwiatkowski, Cordelia Langford, John Sillitoe on behalf of the Wellcome Sanger Institute COVID-19 Surveillance Team |
| EPI_ISL_2911977, EPI_ISL_2911978 | Lighthouse Laboratory Plymouth | Wellcome Sanger Institute for the COVID-19 Genomics UK (COG-UK) Consortium | Lighthouse Laboratory Plymouth and Alex Alderton, Roberto Amato, Jeffrey Barrett, Sonia Goncalves, Ewan Harrison, David K. Jackson, Ian Johnston, Dominic Kwiatkowski, Cordelia Langford, John Sillitoe on behalf of the Wellcome Sanger Institute COVID-19 Surveillance Team |
| EPI_ISL_2911979 | Lighthouse Lab in Milton Keynes | Wellcome Sanger Institute for the COVID-19 Genomics UK (COG-UK) Consortium | The Lighthouse Lab in Milton Keynes and Alex Alderton, Roberto Amato, Jeffrey Barrett, Sonia Goncalves, Ewan Harrison, David K. Jackson, Ian Johnston, Dominic Kwiatkowski, Cordelia Langford, John Sillitoe on behalf of the Wellcome Sanger Institute COVID-19 Surveillance Team |
| EPI_ISL_2911980, EPI_ISL_2911981, EPI_ISL_2911982 | Berkshire and Surrey Pathology Services Lighthouse Laboratory | Wellcome Sanger Institute for the COVID-19 Genomics UK (COG-UK) Consortium | Berkshire and Surrey Pathology Services Lighthouse Laboratory and Alex Alderton, Roberto Amato, Jeffrey Barrett, Sonia Goncalves, Ewan Harrison, David K. Jackson, Ian Johnston, Dominic Kwiatkowski, Cordelia Langford, John Sillitoe on behalf of the Wellcome Sanger Institute COVID-19 Surveillance Team |
| EPI_ISL_2911983, EPI_ISL_2911984 | Lighthouse Lab in Milton Keynes | Wellcome Sanger Institute for the COVID-19 Genomics UK (COG-UK) Consortium | The Lighthouse Lab in Milton Keynes and Alex Alderton, Roberto Amato, Jeffrey Barrett, Sonia Goncalves, Ewan Harrison, David K. Jackson, Ian Johnston, Dominic Kwiatkowski, Cordelia Langford, John Sillitoe on behalf of the Wellcome Sanger Institute COVID-19 Surveillance Team |
| EPI_ISL_2911985 | Lighthouse Laboratory Plymouth | Wellcome Sanger Institute for the COVID-19 Genomics UK (COG-UK) Consortium | Lighthouse Laboratory Plymouth and Alex Alderton, Roberto Amato, Jeffrey Barrett, Sonia Goncalves, Ewan Harrison, David K. Jackson, Ian Johnston, Dominic Kwiatkowski, Cordelia Langford, John Sillitoe on behalf of the Wellcome Sanger Institute COVID-19 Surveillance Team |
| EPI_ISL_2911986 | Berkshire and Surrey Pathology Services Lighthouse Laboratory | Wellcome Sanger Institute for the COVID-19 Genomics UK (COG-UK) Consortium | Berkshire and Surrey Pathology Services Lighthouse Laboratory and Alex Alderton, Roberto Amato, Jeffrey Barrett, Sonia Goncalves, Ewan Harrison, David K. Jackson, Ian Johnston, Dominic Kwiatkowski, Cordelia Langford, John Sillitoe on behalf of the Wellcome Sanger Institute COVID-19 Surveillance Team |
| EPI_ISL_2911987 | Lighthouse Laboratory Plymouth | Wellcome Sanger Institute for the COVID-19 Genomics UK (COG-UK) Consortium | Lighthouse Laboratory Plymouth and Alex Alderton, Roberto Amato, Jeffrey Barrett, Sonia Goncalves, Ewan Harrison, David K. Jackson, Ian Johnston, Dominic Kwiatkowski, Cordelia Langford, John Sillitoe on behalf of the Wellcome Sanger Institute COVID-19 Surveillance Team |
| EPI_ISL_2911988 | Berkshire and Surrey Pathology Services Lighthouse Laboratory | Wellcome Sanger Institute for the COVID-19 Genomics UK (COG-UK) Consortium | Berkshire and Surrey Pathology Services Lighthouse Laboratory and Alex Alderton, Roberto Amato, Jeffrey Barrett, Sonia Goncalves, Ewan Harrison, David K. Jackson, Ian Johnston, Dominic Kwiatkowski, Cordelia Langford, John Sillitoe on behalf of the Wellcome Sanger Institute COVID-19 Surveillance Team |
| EPI_ISL_2911989 | Lighthouse Lab in Milton Keynes | Wellcome Sanger Institute for the COVID-19 Genomics UK (COG-UK) Consortium | The Lighthouse Lab in Milton Keynes and Alex Alderton, Roberto Amato, Jeffrey Barrett, Sonia Goncalves, Ewan Harrison, David K. Jackson, Ian Johnston, Dominic Kwiatkowski, Cordelia Langford, John Sillitoe on behalf of the Wellcome Sanger Institute COVID-19 Surveillance Team |
| EPI_ISL_2911990 | Lighthouse Laboratory Plymouth | Wellcome Sanger Institute for the COVID-19 Genomics UK (COG-UK) Consortium | Lighthouse Laboratory Plymouth and Alex Alderton, Roberto Amato, Jeffrey Barrett, Sonia Goncalves, Ewan Harrison, David K. Jackson, Ian Johnston, Dominic Kwiatkowski, Cordelia Langford, John Sillitoe on behalf of the Wellcome Sanger Institute COVID-19 Surveillance Team |
| EPI_ISL_2911991, EPI_ISL_2911992, EPI_ISL_2911993, EPI_ISL_2911994 | Berkshire and Surrey Pathology Services Lighthouse Laboratory | Wellcome Sanger Institute for the COVID-19 Genomics UK (COG-UK) Consortium | Berkshire and Surrey Pathology Services Lighthouse Laboratory and Alex Alderton, Roberto Amato, Jeffrey Barrett, Sonia Goncalves, Ewan Harrison, David K. Jackson, Ian Johnston, Dominic Kwiatkowski, Cordelia Langford, John Sillitoe on behalf of the Wellcome Sanger Institute COVID-19 Surveillance Team |
| EPI_ISL_2911995 | Lighthouse Laboratory Plymouth | Wellcome Sanger Institute for the COVID-19 Genomics UK (COG-UK) Consortium | Lighthouse Laboratory Plymouth and Alex Alderton, Roberto Amato, Jeffrey Barrett, Sonia Goncalves, Ewan Harrison, David K. Jackson, Ian Johnston, Dominic Kwiatkowski, Cordelia Langford, John Sillitoe on behalf of the Wellcome Sanger Institute COVID-19 Surveillance Team |
| EPI_ISL_2911996 | Berkshire and Surrey Pathology Services Lighthouse Laboratory | Wellcome Sanger Institute for the COVID-19 Genomics UK (COG-UK) Consortium | Berkshire and Surrey Pathology Services Lighthouse Laboratory and Alex Alderton, Roberto Amato, Jeffrey Barrett, Sonia Goncalves, Ewan Harrison, David K. Jackson, Ian Johnston, Dominic Kwiatkowski, Cordelia Langford, John Sillitoe on behalf of the Wellcome Sanger Institute COVID-19 Surveillance Team |
| EPI_ISL_2911997 | Lighthouse Laboratory Plymouth | Wellcome Sanger Institute for the COVID-19 Genomics UK (COG-UK) Consortium | Lighthouse Laboratory Plymouth and Alex Alderton, Roberto Amato, Jeffrey Barrett, Sonia Goncalves, Ewan Harrison, David K. Jackson, Ian Johnston, Dominic Kwiatkowski, Cordelia Langford, John Sillitoe on behalf of the Wellcome Sanger Institute COVID-19 Surveillance Team |
| EPI_ISL_2911998, EPI_ISL_2911999, EPI_ISL_2912000, EPI_ISL_2912001 | Berkshire and Surrey Pathology Services Lighthouse Laboratory | Wellcome Sanger Institute for the COVID-19 Genomics UK (COG-UK) Consortium | Berkshire and Surrey Pathology Services Lighthouse Laboratory and Alex Alderton, Roberto Amato, Jeffrey Barrett, Sonia Goncalves, Ewan Harrison, David K. Jackson, Ian Johnston, Dominic Kwiatkowski, Cordelia Langford, John Sillitoe on behalf of the Wellcome Sanger Institute COVID-19 Surveillance Team |
| EPI_ISL_2912002, EPI_ISL_2912003 | Lighthouse Laboratory Plymouth | Wellcome Sanger Institute for the COVID-19 Genomics UK (COG-UK) Consortium | Lighthouse Laboratory Plymouth and Alex Alderton, Roberto Amato, Jeffrey Barrett, Sonia Goncalves, Ewan Harrison, David K. Jackson, Ian Johnston, Dominic Kwiatkowski, Cordelia Langford, John Sillitoe on behalf of the Wellcome Sanger Institute COVID-19 Surveillance Team |
| EPI_ISL_2912004, EPI_ISL_2912005 | Berkshire and Surrey Pathology Services Lighthouse Laboratory | Wellcome Sanger Institute for the COVID-19 Genomics UK (COG-UK) Consortium | Berkshire and Surrey Pathology Services Lighthouse Laboratory and Alex Alderton, Roberto Amato, Jeffrey Barrett, Sonia Goncalves, Ewan Harrison, David K. Jackson, Ian Johnston, Dominic Kwiatkowski, Cordelia Langford, John Sillitoe on behalf of the Wellcome Sanger Institute COVID-19 Surveillance Team |
| EPI_ISL_2912006, EPI_ISL_2912007 | Lighthouse Lab in Milton Keynes | Wellcome Sanger Institute for the COVID-19 Genomics UK (COG-UK) Consortium | The Lighthouse Lab in Milton Keynes and Alex Alderton, Roberto Amato, Jeffrey Barrett, Sonia Goncalves, Ewan Harrison, David K. Jackson, Ian Johnston, Dominic Kwiatkowski, Cordelia Langford, John Sillitoe on behalf of the Wellcome Sanger Institute COVID-19 Surveillance Team |
| EPI_ISL_2912008, EPI_ISL_2912009 | Berkshire and Surrey Pathology Services Lighthouse Laboratory | Wellcome Sanger Institute for the COVID-19 Genomics UK (COG-UK) Consortium | Berkshire and Surrey Pathology Services Lighthouse Laboratory and Alex Alderton, Roberto Amato, Jeffrey Barrett, Sonia Goncalves, Ewan Harrison, David K. Jackson, Ian Johnston, Dominic Kwiatkowski, Cordelia Langford, John Sillitoe on behalf of the Wellcome Sanger Institute COVID-19 Surveillance Team |
| EPI_ISL_2912010 | Lighthouse Laboratory Plymouth | Wellcome Sanger Institute for the COVID-19 Genomics UK (COG-UK) Consortium | Lighthouse Laboratory Plymouth and Alex Alderton, Roberto Amato, Jeffrey Barrett, Sonia Goncalves, Ewan Harrison, David K. Jackson, Ian Johnston, Dominic Kwiatkowski, Cordelia Langford, John Sillitoe on behalf of the Wellcome Sanger Institute COVID-19 Surveillance Team |
| EPI_ISL_2912011, EPI_ISL_2912012, EPI_ISL_2912013, EPI_ISL_2912014 | Lighthouse Lab in Milton Keynes | Wellcome Sanger Institute for the COVID-19 Genomics UK (COG-UK) Consortium | The Lighthouse Lab in Milton Keynes and Alex Alderton, Roberto Amato, Jeffrey Barrett, Sonia Goncalves, Ewan Harrison, David K. Jackson, Ian Johnston, Dominic Kwiatkowski, Cordelia Langford, John Sillitoe on behalf of the Wellcome Sanger Institute COVID-19 Surveillance Team |
| EPI_ISL_2912015, EPI_ISL_2912016 | Lighthouse Laboratory Plymouth | Wellcome Sanger Institute for the COVID-19 Genomics UK (COG-UK) Consortium | Lighthouse Laboratory Plymouth and Alex Alderton, Roberto Amato, Jeffrey Barrett, Sonia Goncalves, Ewan Harrison, David K. Jackson, Ian Johnston, Dominic Kwiatkowski, Cordelia Langford, John Sillitoe on behalf of the Wellcome Sanger Institute COVID-19 Surveillance Team |
| EPI_ISL_2912017 | Berkshire and Surrey Pathology Services Lighthouse Laboratory | Wellcome Sanger Institute for the COVID-19 Genomics UK (COG-UK) Consortium | Berkshire and Surrey Pathology Services Lighthouse Laboratory and Alex Alderton, Roberto Amato, Jeffrey Barrett, Sonia Goncalves, Ewan Harrison, David K. Jackson, Ian Johnston, Dominic Kwiatkowski, Cordelia Langford, John Sillitoe on behalf of the Wellcome Sanger Institute COVID-19 Surveillance Team |
| EPI_ISL_2912018 | Lighthouse Lab in Milton Keynes | Wellcome Sanger Institute for the COVID-19 Genomics UK (COG-UK) Consortium | The Lighthouse Lab in Milton Keynes and Alex Alderton, Roberto Amato, Jeffrey Barrett, Sonia Goncalves, Ewan Harrison, David K. Jackson, Ian Johnston, Dominic Kwiatkowski, Cordelia Langford, John Sillitoe on behalf of the Wellcome Sanger Institute COVID-19 Surveillance Team |
| EPI_ISL_2912019, EPI_ISL_2912020 | Berkshire and Surrey Pathology Services Lighthouse Laboratory | Wellcome Sanger Institute for the COVID-19 Genomics UK (COG-UK) Consortium | Berkshire and Surrey Pathology Services Lighthouse Laboratory and Alex Alderton, Roberto Amato, Jeffrey Barrett, Sonia Goncalves, Ewan Harrison, David K. Jackson, Ian Johnston, Dominic Kwiatkowski, Cordelia Langford, John Sillitoe on behalf of the Wellcome Sanger Institute COVID-19 Surveillance Team |
| EPI_ISL_2912021, EPI_ISL_2912022, EPI_ISL_2912023, EPI_ISL_2912024 | Lighthouse Lab in Milton Keynes | Wellcome Sanger Institute for the COVID-19 Genomics UK (COG-UK) Consortium | The Lighthouse Lab in Milton Keynes and Alex Alderton, Roberto Amato, Jeffrey Barrett, Sonia Goncalves, Ewan Harrison, David K. Jackson, Ian Johnston, Dominic Kwiatkowski, Cordelia Langford, John Sillitoe on behalf of the Wellcome Sanger Institute COVID-19 Surveillance Team |
| EPI_ISL_2912025, EPI_ISL_2912026 | Berkshire and Surrey Pathology Services Lighthouse Laboratory | Wellcome Sanger Institute for the COVID-19 Genomics UK (COG-UK) Consortium | Berkshire and Surrey Pathology Services Lighthouse Laboratory and Alex Alderton, Roberto Amato, Jeffrey Barrett, Sonia Goncalves, Ewan Harrison, David K. Jackson, Ian Johnston, Dominic Kwiatkowski, Cordelia Langford, John Sillitoe on behalf of the Wellcome Sanger Institute COVID-19 Surveillance Team |
| EPI_ISL_2912027, EPI_ISL_2912028, EPI_ISL_2912029 | Lighthouse Lab in Milton Keynes | Wellcome Sanger Institute for the COVID-19 Genomics UK (COG-UK) Consortium | The Lighthouse Lab in Milton Keynes and Alex Alderton, Roberto Amato, Jeffrey Barrett, Sonia Goncalves, Ewan Harrison, David K. Jackson, Ian Johnston, Dominic Kwiatkowski, Cordelia Langford, John Sillitoe on behalf of the Wellcome Sanger Institute COVID-19 Surveillance Team |
| EPI_ISL_2912030, EPI_ISL_2912031 | Berkshire and Surrey Pathology Services Lighthouse Laboratory | Wellcome Sanger Institute for the COVID-19 Genomics UK (COG-UK) Consortium | Berkshire and Surrey Pathology Services Lighthouse Laboratory and Alex Alderton, Roberto Amato, Jeffrey Barrett, Sonia Goncalves, Ewan Harrison, David K. Jackson, Ian Johnston, Dominic Kwiatkowski, Cordelia Langford, John Sillitoe on behalf of the Wellcome Sanger Institute COVID-19 Surveillance Team |
| EPI_ISL_2912032 | Lighthouse Laboratory Plymouth | Wellcome Sanger Institute for the COVID-19 Genomics UK (COG-UK) Consortium | Lighthouse Laboratory Plymouth and Alex Alderton, Roberto Amato, Jeffrey Barrett, Sonia Goncalves, Ewan Harrison, David K. Jackson, Ian Johnston, Dominic Kwiatkowski, Cordelia Langford, John Sillitoe on behalf of the Wellcome Sanger Institute COVID-19 Surveillance Team |
| EPI_ISL_2912033, | Berkshire and Surrey Pathology Services Lighthouse Laboratory | Wellcome Sanger Institute for the COVID-19 Genomics UK (COG-UK) | Berkshire and Surrey Pathology Services Lighthouse Laboratory and Alex Alderton, Roberto Amato, Jeffrey Barrett, Sonia Goncalves, Ewan Harrison, David K. Jackson, Ian Johnston, Dominic Kwiatkowski, Cordelia Langford, John Sillitoe on behalf |

| | | | |
|---|---|---|---|
| EPI_ISL_2912034 | | Consortium | of the Wellcome Sanger Institute COVID-19 Surveillance Team |
| EPI_ISL_2912035 | Lighthouse Laboratory Plymouth | Wellcome Sanger Institute for the COVID-19 Genomics UK (COG-UK) Consortium | Lighthouse Laboratory Plymouth and Alex Alderton, Roberto Amato, Jeffrey Barrett, Sonia Goncalves, Ewan Harrison, David K. Jackson, Ian Johnston, Dominic Kwiatkowski, Cordelia Langford, John Sillitoe on behalf of the Wellcome Sanger Institute COVID-19 Surveillance Team |
| EPI_ISL_2912036 | Lighthouse Lab in Milton Keynes | Wellcome Sanger Institute for the COVID-19 Genomics UK (COG-UK) Consortium | The Lighthouse Lab in Milton Keynes and Alex Alderton, Roberto Amato, Jeffrey Barrett, Sonia Goncalves, Ewan Harrison, David K. Jackson, Ian Johnston, Dominic Kwiatkowski, Cordelia Langford, John Sillitoe on behalf of the Wellcome Sanger Institute COVID-19 Surveillance Team |
| EPI_ISL_2912037, EPI_ISL_2912038, EPI_ISL_2912039, EPI_ISL_2912040, EPI_ISL_2912041, EPI_ISL_2912042 | Berkshire and Surrey Pathology Services Lighthouse Laboratory | Wellcome Sanger Institute for the COVID-19 Genomics UK (COG-UK) Consortium | Berkshire and Surrey Pathology Services Lighthouse Laboratory and Alex Alderton, Roberto Amato, Jeffrey Barrett, Sonia Goncalves, Ewan Harrison, David K. Jackson, Ian Johnston, Dominic Kwiatkowski, Cordelia Langford, John Sillitoe on behalf of the Wellcome Sanger Institute COVID-19 Surveillance Team |
| EPI_ISL_2912043 | Lighthouse Laboratory Plymouth | Wellcome Sanger Institute for the COVID-19 Genomics UK (COG-UK) Consortium | Lighthouse Laboratory Plymouth and Alex Alderton, Roberto Amato, Jeffrey Barrett, Sonia Goncalves, Ewan Harrison, David K. Jackson, Ian Johnston, Dominic Kwiatkowski, Cordelia Langford, John Sillitoe on behalf of the Wellcome Sanger Institute COVID-19 Surveillance Team |
| EPI_ISL_2912044 | Berkshire and Surrey Pathology Services Lighthouse Laboratory | Wellcome Sanger Institute for the COVID-19 Genomics UK (COG-UK) Consortium | Berkshire and Surrey Pathology Services Lighthouse Laboratory and Alex Alderton, Roberto Amato, Jeffrey Barrett, Sonia Goncalves, Ewan Harrison, David K. Jackson, Ian Johnston, Dominic Kwiatkowski, Cordelia Langford, John Sillitoe on behalf of the Wellcome Sanger Institute COVID-19 Surveillance Team |
| EPI_ISL_2912045 | Lighthouse Laboratory Plymouth | Wellcome Sanger Institute for the COVID-19 Genomics UK (COG-UK) Consortium | Lighthouse Laboratory Plymouth and Alex Alderton, Roberto Amato, Jeffrey Barrett, Sonia Goncalves, Ewan Harrison, David K. Jackson, Ian Johnston, Dominic Kwiatkowski, Cordelia Langford, John Sillitoe on behalf of the Wellcome Sanger Institute COVID-19 Surveillance Team |
| EPI_ISL_2912046, EPI_ISL_2912047 | Berkshire and Surrey Pathology Services Lighthouse Laboratory | Wellcome Sanger Institute for the COVID-19 Genomics UK (COG-UK) Consortium | Berkshire and Surrey Pathology Services Lighthouse Laboratory and Alex Alderton, Roberto Amato, Jeffrey Barrett, Sonia Goncalves, Ewan Harrison, David K. Jackson, Ian Johnston, Dominic Kwiatkowski, Cordelia Langford, John Sillitoe on behalf of the Wellcome Sanger Institute COVID-19 Surveillance Team |
| EPI_ISL_2912048 | Lighthouse Laboratory Plymouth | Wellcome Sanger Institute for the COVID-19 Genomics UK (COG-UK) Consortium | Lighthouse Laboratory Plymouth and Alex Alderton, Roberto Amato, Jeffrey Barrett, Sonia Goncalves, Ewan Harrison, David K. Jackson, Ian Johnston, Dominic Kwiatkowski, Cordelia Langford, John Sillitoe on behalf of the Wellcome Sanger Institute COVID-19 Surveillance Team |
| EPI_ISL_2912049 | Berkshire and Surrey Pathology Services Lighthouse Laboratory | Wellcome Sanger Institute for the COVID-19 Genomics UK (COG-UK) Consortium | Berkshire and Surrey Pathology Services Lighthouse Laboratory and Alex Alderton, Roberto Amato, Jeffrey Barrett, Sonia Goncalves, Ewan Harrison, David K. Jackson, Ian Johnston, Dominic Kwiatkowski, Cordelia Langford, John Sillitoe on behalf of the Wellcome Sanger Institute COVID-19 Surveillance Team |
| EPI_ISL_2912050 | Lighthouse Laboratory Plymouth | Wellcome Sanger Institute for the COVID-19 Genomics UK (COG-UK) Consortium | Lighthouse Laboratory Plymouth and Alex Alderton, Roberto Amato, Jeffrey Barrett, Sonia Goncalves, Ewan Harrison, David K. Jackson, Ian Johnston, Dominic Kwiatkowski, Cordelia Langford, John Sillitoe on behalf of the Wellcome Sanger Institute COVID-19 Surveillance Team |
| EPI_ISL_2912051 | Berkshire and Surrey Pathology Services Lighthouse Laboratory | Wellcome Sanger Institute for the COVID-19 Genomics UK (COG-UK) Consortium | Berkshire and Surrey Pathology Services Lighthouse Laboratory and Alex Alderton, Roberto Amato, Jeffrey Barrett, Sonia Goncalves, Ewan Harrison, David K. Jackson, Ian Johnston, Dominic Kwiatkowski, Cordelia Langford, John Sillitoe on behalf of the Wellcome Sanger Institute COVID-19 Surveillance Team |
| EPI_ISL_2912052 | Lighthouse Laboratory Plymouth | Wellcome Sanger Institute for the COVID-19 Genomics UK (COG-UK) Consortium | Lighthouse Laboratory Plymouth and Alex Alderton, Roberto Amato, Jeffrey Barrett, Sonia Goncalves, Ewan Harrison, David K. Jackson, Ian Johnston, Dominic Kwiatkowski, Cordelia Langford, John Sillitoe on behalf of the Wellcome Sanger Institute COVID-19 Surveillance Team |
| EPI_ISL_2912053, EPI_ISL_2912054 | Berkshire and Surrey Pathology Services Lighthouse Laboratory | Wellcome Sanger Institute for the COVID-19 Genomics UK (COG-UK) Consortium | Berkshire and Surrey Pathology Services Lighthouse Laboratory and Alex Alderton, Roberto Amato, Jeffrey Barrett, Sonia Goncalves, Ewan Harrison, David K. Jackson, Ian Johnston, Dominic Kwiatkowski, Cordelia Langford, John Sillitoe on behalf of the Wellcome Sanger Institute COVID-19 Surveillance Team |
| EPI_ISL_2912055 | Lighthouse Lab in Milton Keynes | Wellcome Sanger Institute for the COVID-19 Genomics UK (COG-UK) Consortium | The Lighthouse Lab in Milton Keynes and Alex Alderton, Roberto Amato, Jeffrey Barrett, Sonia Goncalves, Ewan Harrison, David K. Jackson, Ian Johnston, Dominic Kwiatkowski, Cordelia Langford, John Sillitoe on behalf of the Wellcome Sanger Institute COVID-19 Surveillance Team |
| EPI_ISL_2912056 | Berkshire and Surrey Pathology Services Lighthouse Laboratory | Wellcome Sanger Institute for the COVID-19 Genomics UK (COG-UK) Consortium | Berkshire and Surrey Pathology Services Lighthouse Laboratory and Alex Alderton, Roberto Amato, Jeffrey Barrett, Sonia Goncalves, Ewan Harrison, David K. Jackson, Ian Johnston, Dominic Kwiatkowski, Cordelia Langford, John Sillitoe on behalf of the Wellcome Sanger Institute COVID-19 Surveillance Team |
| EPI_ISL_2912057 | Lighthouse Lab in Milton Keynes | Wellcome Sanger Institute for the COVID-19 Genomics UK (COG-UK) Consortium | The Lighthouse Lab in Milton Keynes and Alex Alderton, Roberto Amato, Jeffrey Barrett, Sonia Goncalves, Ewan Harrison, David K. Jackson, Ian Johnston, Dominic Kwiatkowski, Cordelia Langford, John Sillitoe on behalf of the Wellcome Sanger Institute COVID-19 Surveillance Team |
| EPI_ISL_2912058, EPI_ISL_2912059, EPI_ISL_2912060 | Berkshire and Surrey Pathology Services Lighthouse Laboratory | Wellcome Sanger Institute for the COVID-19 Genomics UK (COG-UK) Consortium | Berkshire and Surrey Pathology Services Lighthouse Laboratory and Alex Alderton, Roberto Amato, Jeffrey Barrett, Sonia Goncalves, Ewan Harrison, David K. Jackson, Ian Johnston, Dominic Kwiatkowski, Cordelia Langford, John Sillitoe on behalf of the Wellcome Sanger Institute COVID-19 Surveillance Team |
| EPI_ISL_2912061 | Lighthouse Lab in Milton Keynes | Wellcome Sanger Institute for the COVID-19 Genomics UK (COG-UK) Consortium | The Lighthouse Lab in Milton Keynes and Alex Alderton, Roberto Amato, Jeffrey Barrett, Sonia Goncalves, Ewan Harrison, David K. Jackson, Ian Johnston, Dominic Kwiatkowski, Cordelia Langford, John Sillitoe on behalf of the Wellcome Sanger Institute COVID-19 Surveillance Team |
| EPI_ISL_2912062 | Berkshire and Surrey Pathology Services Lighthouse Laboratory | Wellcome Sanger Institute for the COVID-19 Genomics UK (COG-UK) Consortium | Berkshire and Surrey Pathology Services Lighthouse Laboratory and Alex Alderton, Roberto Amato, Jeffrey Barrett, Sonia Goncalves, Ewan Harrison, David K. Jackson, Ian Johnston, Dominic Kwiatkowski, Cordelia Langford, John Sillitoe on behalf of the Wellcome Sanger Institute COVID-19 Surveillance Team |
| EPI_ISL_2912063 | Lighthouse Laboratory Plymouth | Wellcome Sanger Institute for the COVID-19 Genomics UK (COG-UK) Consortium | Lighthouse Laboratory Plymouth and Alex Alderton, Roberto Amato, Jeffrey Barrett, Sonia Goncalves, Ewan Harrison, David K. Jackson, Ian Johnston, Dominic Kwiatkowski, Cordelia Langford, John Sillitoe on behalf of the Wellcome Sanger Institute COVID-19 Surveillance Team |
| EPI_ISL_2912064 | Berkshire and Surrey Pathology Services Lighthouse Laboratory | Wellcome Sanger Institute for the COVID-19 Genomics UK (COG-UK) Consortium | Berkshire and Surrey Pathology Services Lighthouse Laboratory and Alex Alderton, Roberto Amato, Jeffrey Barrett, Sonia Goncalves, Ewan Harrison, David K. Jackson, Ian Johnston, Dominic Kwiatkowski, Cordelia Langford, John Sillitoe on behalf of the Wellcome Sanger Institute COVID-19 Surveillance Team |
| EPI_ISL_2912065 | Lighthouse Laboratory Plymouth | Wellcome Sanger Institute for the COVID-19 Genomics UK (COG-UK) Consortium | Lighthouse Laboratory Plymouth and Alex Alderton, Roberto Amato, Jeffrey Barrett, Sonia Goncalves, Ewan Harrison, David K. Jackson, Ian Johnston, Dominic Kwiatkowski, Cordelia Langford, John Sillitoe on behalf of the Wellcome Sanger Institute COVID-19 Surveillance Team |
| EPI_ISL_2912066 | Berkshire and Surrey Pathology Services Lighthouse Laboratory | Wellcome Sanger Institute for the COVID-19 Genomics UK (COG-UK) Consortium | Berkshire and Surrey Pathology Services Lighthouse Laboratory and Alex Alderton, Roberto Amato, Jeffrey Barrett, Sonia Goncalves, Ewan Harrison, David K. Jackson, Ian Johnston, Dominic Kwiatkowski, Cordelia Langford, John Sillitoe on behalf of the Wellcome Sanger Institute COVID-19 Surveillance Team |
| EPI_ISL_2912067 | Lighthouse Lab in Milton Keynes | Wellcome Sanger Institute for the COVID-19 Genomics UK (COG-UK) Consortium | The Lighthouse Lab in Milton Keynes and Alex Alderton, Roberto Amato, Jeffrey Barrett, Sonia Goncalves, Ewan Harrison, David K. Jackson, Ian Johnston, Dominic Kwiatkowski, Cordelia Langford, John Sillitoe on behalf of the Wellcome Sanger Institute COVID-19 Surveillance Team |
| EPI_ISL_2912068, EPI_ISL_2912069 | Berkshire and Surrey Pathology Services Lighthouse Laboratory | Wellcome Sanger Institute for the COVID-19 Genomics UK (COG-UK) Consortium | Berkshire and Surrey Pathology Services Lighthouse Laboratory and Alex Alderton, Roberto Amato, Jeffrey Barrett, Sonia Goncalves, Ewan Harrison, David K. Jackson, Ian Johnston, Dominic Kwiatkowski, Cordelia Langford, John Sillitoe on behalf of the Wellcome Sanger Institute COVID-19 Surveillance Team |
| EPI_ISL_2912070 | Lighthouse Laboratory Plymouth | Wellcome Sanger Institute for the COVID-19 Genomics UK (COG-UK) Consortium | Lighthouse Laboratory Plymouth and Alex Alderton, Roberto Amato, Jeffrey Barrett, Sonia Goncalves, Ewan Harrison, David K. Jackson, Ian Johnston, Dominic Kwiatkowski, Cordelia Langford, John Sillitoe on behalf of the Wellcome Sanger Institute COVID-19 Surveillance Team |
| EPI_ISL_2912071 | Lighthouse Lab in Milton Keynes | Wellcome Sanger Institute for the COVID-19 Genomics UK (COG-UK) Consortium | The Lighthouse Lab in Milton Keynes and Alex Alderton, Roberto Amato, Jeffrey Barrett, Sonia Goncalves, Ewan Harrison, David K. Jackson, Ian Johnston, Dominic Kwiatkowski, Cordelia Langford, John Sillitoe on behalf of the Wellcome Sanger Institute COVID-19 Surveillance Team |
| EPI_ISL_2912072 | Lighthouse Laboratory Plymouth | Wellcome Sanger Institute for the COVID-19 Genomics UK (COG-UK) Consortium | Lighthouse Laboratory Plymouth and Alex Alderton, Roberto Amato, Jeffrey Barrett, Sonia Goncalves, Ewan Harrison, David K. Jackson, Ian Johnston, Dominic Kwiatkowski, Cordelia Langford, John Sillitoe on behalf of the Wellcome Sanger Institute COVID-19 Surveillance Team |
| EPI_ISL_2912073 | Lighthouse Lab in Milton Keynes | Wellcome Sanger Institute for the COVID-19 Genomics UK (COG-UK) Consortium | The Lighthouse Lab in Milton Keynes and Alex Alderton, Roberto Amato, Jeffrey Barrett, Sonia Goncalves, Ewan Harrison, David K. Jackson, Ian Johnston, Dominic Kwiatkowski, Cordelia Langford, John Sillitoe on behalf of the Wellcome Sanger Institute COVID-19 Surveillance Team |
| EPI_ISL_2912074, EPI_ISL_2912075 | Lighthouse Laboratory Plymouth | Wellcome Sanger Institute for the COVID-19 Genomics UK (COG-UK) Consortium | Lighthouse Laboratory Plymouth and Alex Alderton, Roberto Amato, Jeffrey Barrett, Sonia Goncalves, Ewan Harrison, David K. Jackson, Ian Johnston, Dominic Kwiatkowski, Cordelia Langford, John Sillitoe on behalf of the Wellcome Sanger Institute COVID-19 Surveillance Team |
| EPI_ISL_2912076, EPI_ISL_2912077, EPI_ISL_2912078, EPI_ISL_2912079, EPI_ISL_2912080 | Berkshire and Surrey Pathology Services Lighthouse Laboratory | Wellcome Sanger Institute for the COVID-19 Genomics UK (COG-UK) Consortium | Berkshire and Surrey Pathology Services Lighthouse Laboratory and Alex Alderton, Roberto Amato, Jeffrey Barrett, Sonia Goncalves, Ewan Harrison, David K. Jackson, Ian Johnston, Dominic Kwiatkowski, Cordelia Langford, John Sillitoe on behalf of the Wellcome Sanger Institute COVID-19 Surveillance Team |
| EPI_ISL_2912081, EPI_ISL_2912082 | Lighthouse Lab in Milton Keynes | Wellcome Sanger Institute for the COVID-19 Genomics UK (COG-UK) Consortium | The Lighthouse Lab in Milton Keynes and Alex Alderton, Roberto Amato, Jeffrey Barrett, Sonia Goncalves, Ewan Harrison, David K. Jackson, Ian Johnston, Dominic Kwiatkowski, Cordelia Langford, John Sillitoe on behalf of the Wellcome Sanger Institute COVID-19 Surveillance Team |
| EPI_ISL_2912083 | Berkshire and Surrey Pathology Services Lighthouse Laboratory | Wellcome Sanger Institute for the COVID-19 Genomics UK (COG-UK) Consortium | Berkshire and Surrey Pathology Services Lighthouse Laboratory and Alex Alderton, Roberto Amato, Jeffrey Barrett, Sonia Goncalves, Ewan Harrison, David K. Jackson, Ian Johnston, Dominic Kwiatkowski, Cordelia Langford, John Sillitoe on behalf of the Wellcome Sanger Institute COVID-19 Surveillance Team |
| EPI_ISL_2912084, EPI_ISL_2912085, EPI_ISL_2912086 | Lighthouse Lab in Milton Keynes | Wellcome Sanger Institute for the COVID-19 Genomics UK (COG-UK) Consortium | The Lighthouse Lab in Milton Keynes and Alex Alderton, Roberto Amato, Jeffrey Barrett, Sonia Goncalves, Ewan Harrison, David K. Jackson, Ian Johnston, Dominic Kwiatkowski, Cordelia Langford, John Sillitoe on behalf of the Wellcome Sanger Institute COVID-19 Surveillance Team |
| EPI_ISL_2912087 | Lighthouse Laboratory Plymouth | Wellcome Sanger Institute for the COVID-19 Genomics UK (COG-UK) Consortium | Lighthouse Laboratory Plymouth and Alex Alderton, Roberto Amato, Jeffrey Barrett, Sonia Goncalves, Ewan Harrison, David K. Jackson, Ian Johnston, Dominic Kwiatkowski, Cordelia Langford, John Sillitoe on behalf of the Wellcome Sanger Institute COVID-19 Surveillance Team |
| EPI_ISL_2912088, EPI_ISL_2912089 | Lighthouse Lab in Milton Keynes | Wellcome Sanger Institute for the COVID-19 Genomics UK (COG-UK) Consortium | The Lighthouse Lab in Milton Keynes and Alex Alderton, Roberto Amato, Jeffrey Barrett, Sonia Goncalves, Ewan Harrison, David K. Jackson, Ian Johnston, Dominic Kwiatkowski, Cordelia Langford, John Sillitoe on behalf of the Wellcome Sanger Institute COVID-19 Surveillance Team |
| EPI_ISL_2912090, EPI_ISL_2912091, EPI_ISL_2912092, EPI_ISL_2912093, EPI_ISL_2912094, EPI_ISL_2912095 | Berkshire and Surrey Pathology Services Lighthouse Laboratory | Wellcome Sanger Institute for the COVID-19 Genomics UK (COG-UK) Consortium | Berkshire and Surrey Pathology Services Lighthouse Laboratory and Alex Alderton, Roberto Amato, Jeffrey Barrett, Sonia Goncalves, Ewan Harrison, David K. Jackson, Ian Johnston, Dominic Kwiatkowski, Cordelia Langford, John Sillitoe on behalf of the Wellcome Sanger Institute COVID-19 Surveillance Team |
| EPI_ISL_2912096 | Lighthouse Lab in Milton Keynes | Wellcome Sanger Institute for the COVID-19 Genomics UK (COG-UK) Consortium | The Lighthouse Lab in Milton Keynes and Alex Alderton, Roberto Amato, Jeffrey Barrett, Sonia Goncalves, Ewan Harrison, David K. Jackson, Ian Johnston, Dominic Kwiatkowski, Cordelia Langford, John Sillitoe on behalf of the Wellcome Sanger Institute COVID-19 Surveillance Team |
| EPI_ISL_2912097 | Lighthouse Laboratory Plymouth | Wellcome Sanger Institute for the COVID-19 Genomics UK (COG-UK) Consortium | Lighthouse Laboratory Plymouth and Alex Alderton, Roberto Amato, Jeffrey Barrett, Sonia Goncalves, Ewan Harrison, David K. Jackson, Ian Johnston, Dominic Kwiatkowski, Cordelia Langford, John Sillitoe on behalf of the Wellcome Sanger Institute COVID-19 Surveillance Team |

| | | | |
|---|---|---|---|
| EPI_ISL_2912098, EPI_ISL_2912099 | Berkshire and Surrey Pathology Services Lighthouse Laboratory | Wellcome Sanger Institute for the COVID-19 Genomics UK (COG-UK) Consortium | Berkshire and Surrey Pathology Services Lighthouse Laboratory and Alex Alderton, Roberto Amato, Jeffrey Barrett, Sonia Goncalves, Ewan Harrison, David K. Jackson, Ian Johnston, Dominic Kwiatkowski, Cordelia Langford, John Sillitoe on behalf of the Wellcome Sanger Institute COVID-19 Surveillance Team |
| EPI_ISL_2912100, EPI_ISL_2912101 | Lighthouse Laboratory Plymouth | Wellcome Sanger Institute for the COVID-19 Genomics UK (COG-UK) Consortium | Lighthouse Laboratory Plymouth and Alex Alderton, Roberto Amato, Jeffrey Barrett, Sonia Goncalves, Ewan Harrison, David K. Jackson, Ian Johnston, Dominic Kwiatkowski, Cordelia Langford, John Sillitoe on behalf of the Wellcome Sanger Institute COVID-19 Surveillance Team |
| EPI_ISL_2912102 | Lighthouse Lab in Milton Keynes | Wellcome Sanger Institute for the COVID-19 Genomics UK (COG-UK) Consortium | The Lighthouse Lab in Milton Keynes and Alex Alderton, Roberto Amato, Jeffrey Barrett, Sonia Goncalves, Ewan Harrison, David K. Jackson, Ian Johnston, Dominic Kwiatkowski, Cordelia Langford, John Sillitoe on behalf of the Wellcome Sanger Institute COVID-19 Surveillance Team |
| EPI_ISL_2912103 | Lighthouse Laboratory Plymouth | Wellcome Sanger Institute for the COVID-19 Genomics UK (COG-UK) Consortium | Lighthouse Laboratory Plymouth and Alex Alderton, Roberto Amato, Jeffrey Barrett, Sonia Goncalves, Ewan Harrison, David K. Jackson, Ian Johnston, Dominic Kwiatkowski, Cordelia Langford, John Sillitoe on behalf of the Wellcome Sanger Institute COVID-19 Surveillance Team |
| EPI_ISL_2912104 | Lighthouse Lab in Milton Keynes | Wellcome Sanger Institute for the COVID-19 Genomics UK (COG-UK) Consortium | The Lighthouse Lab in Milton Keynes and Alex Alderton, Roberto Amato, Jeffrey Barrett, Sonia Goncalves, Ewan Harrison, David K. Jackson, Ian Johnston, Dominic Kwiatkowski, Cordelia Langford, John Sillitoe on behalf of the Wellcome Sanger Institute COVID-19 Surveillance Team |
| EPI_ISL_2912105 | Berkshire and Surrey Pathology Services Lighthouse Laboratory | Wellcome Sanger Institute for the COVID-19 Genomics UK (COG-UK) Consortium | Berkshire and Surrey Pathology Services Lighthouse Laboratory and Alex Alderton, Roberto Amato, Jeffrey Barrett, Sonia Goncalves, Ewan Harrison, David K. Jackson, Ian Johnston, Dominic Kwiatkowski, Cordelia Langford, John Sillitoe on behalf of the Wellcome Sanger Institute COVID-19 Surveillance Team |
| EPI_ISL_2912106 | Lighthouse Lab in Milton Keynes | Wellcome Sanger Institute for the COVID-19 Genomics UK (COG-UK) Consortium | The Lighthouse Lab in Milton Keynes and Alex Alderton, Roberto Amato, Jeffrey Barrett, Sonia Goncalves, Ewan Harrison, David K. Jackson, Ian Johnston, Dominic Kwiatkowski, Cordelia Langford, John Sillitoe on behalf of the Wellcome Sanger Institute COVID-19 Surveillance Team |
| EPI_ISL_2912107 | Lighthouse Laboratory Plymouth | Wellcome Sanger Institute for the COVID-19 Genomics UK (COG-UK) Consortium | Lighthouse Laboratory Plymouth and Alex Alderton, Roberto Amato, Jeffrey Barrett, Sonia Goncalves, Ewan Harrison, David K. Jackson, Ian Johnston, Dominic Kwiatkowski, Cordelia Langford, John Sillitoe on behalf of the Wellcome Sanger Institute COVID-19 Surveillance Team |
| EPI_ISL_2912108 | Lighthouse Lab in Milton Keynes | Wellcome Sanger Institute for the COVID-19 Genomics UK (COG-UK) Consortium | The Lighthouse Lab in Milton Keynes and Alex Alderton, Roberto Amato, Jeffrey Barrett, Sonia Goncalves, Ewan Harrison, David K. Jackson, Ian Johnston, Dominic Kwiatkowski, Cordelia Langford, John Sillitoe on behalf of the Wellcome Sanger Institute COVID-19 Surveillance Team |
| EPI_ISL_2912109 | Berkshire and Surrey Pathology Services Lighthouse Laboratory | Wellcome Sanger Institute for the COVID-19 Genomics UK (COG-UK) Consortium | Berkshire and Surrey Pathology Services Lighthouse Laboratory and Alex Alderton, Roberto Amato, Jeffrey Barrett, Sonia Goncalves, Ewan Harrison, David K. Jackson, Ian Johnston, Dominic Kwiatkowski, Cordelia Langford, John Sillitoe on behalf of the Wellcome Sanger Institute COVID-19 Surveillance Team |
| EPI_ISL_2912110 | Lighthouse Lab in Milton Keynes | Wellcome Sanger Institute for the COVID-19 Genomics UK (COG-UK) Consortium | The Lighthouse Lab in Milton Keynes and Alex Alderton, Roberto Amato, Jeffrey Barrett, Sonia Goncalves, Ewan Harrison, David K. Jackson, Ian Johnston, Dominic Kwiatkowski, Cordelia Langford, John Sillitoe on behalf of the Wellcome Sanger Institute COVID-19 Surveillance Team |
| EPI_ISL_2912111, EPI_ISL_2912112, EPI_ISL_2912113 | Berkshire and Surrey Pathology Services Lighthouse Laboratory | Wellcome Sanger Institute for the COVID-19 Genomics UK (COG-UK) Consortium | Berkshire and Surrey Pathology Services Lighthouse Laboratory and Alex Alderton, Roberto Amato, Jeffrey Barrett, Sonia Goncalves, Ewan Harrison, David K. Jackson, Ian Johnston, Dominic Kwiatkowski, Cordelia Langford, John Sillitoe on behalf of the Wellcome Sanger Institute COVID-19 Surveillance Team |
| EPI_ISL_2912114, EPI_ISL_2912115 | Lighthouse Lab in Milton Keynes | Wellcome Sanger Institute for the COVID-19 Genomics UK (COG-UK) Consortium | The Lighthouse Lab in Milton Keynes and Alex Alderton, Roberto Amato, Jeffrey Barrett, Sonia Goncalves, Ewan Harrison, David K. Jackson, Ian Johnston, Dominic Kwiatkowski, Cordelia Langford, John Sillitoe on behalf of the Wellcome Sanger Institute COVID-19 Surveillance Team |
| EPI_ISL_2912116, EPI_ISL_2912117 | Berkshire and Surrey Pathology Services Lighthouse Laboratory | Wellcome Sanger Institute for the COVID-19 Genomics UK (COG-UK) Consortium | Berkshire and Surrey Pathology Services Lighthouse Laboratory and Alex Alderton, Roberto Amato, Jeffrey Barrett, Sonia Goncalves, Ewan Harrison, David K. Jackson, Ian Johnston, Dominic Kwiatkowski, Cordelia Langford, John Sillitoe on behalf of the Wellcome Sanger Institute COVID-19 Surveillance Team |
| EPI_ISL_2912118 | Lighthouse Laboratory Plymouth | Wellcome Sanger Institute for the COVID-19 Genomics UK (COG-UK) Consortium | Lighthouse Laboratory Plymouth and Alex Alderton, Roberto Amato, Jeffrey Barrett, Sonia Goncalves, Ewan Harrison, David K. Jackson, Ian Johnston, Dominic Kwiatkowski, Cordelia Langford, John Sillitoe on behalf of the Wellcome Sanger Institute COVID-19 Surveillance Team |
| EPI_ISL_2912119 | Lighthouse Lab in Milton Keynes | Wellcome Sanger Institute for the COVID-19 Genomics UK (COG-UK) Consortium | The Lighthouse Lab in Milton Keynes and Alex Alderton, Roberto Amato, Jeffrey Barrett, Sonia Goncalves, Ewan Harrison, David K. Jackson, Ian Johnston, Dominic Kwiatkowski, Cordelia Langford, John Sillitoe on behalf of the Wellcome Sanger Institute COVID-19 Surveillance Team |
| EPI_ISL_2912120, EPI_ISL_2912121 | Berkshire and Surrey Pathology Services Lighthouse Laboratory | Wellcome Sanger Institute for the COVID-19 Genomics UK (COG-UK) Consortium | Berkshire and Surrey Pathology Services Lighthouse Laboratory and Alex Alderton, Roberto Amato, Jeffrey Barrett, Sonia Goncalves, Ewan Harrison, David K. Jackson, Ian Johnston, Dominic Kwiatkowski, Cordelia Langford, John Sillitoe on behalf of the Wellcome Sanger Institute COVID-19 Surveillance Team |
| EPI_ISL_2912122, EPI_ISL_2912123 | Lighthouse Lab in Milton Keynes | Wellcome Sanger Institute for the COVID-19 Genomics UK (COG-UK) Consortium | The Lighthouse Lab in Milton Keynes and Alex Alderton, Roberto Amato, Jeffrey Barrett, Sonia Goncalves, Ewan Harrison, David K. Jackson, Ian Johnston, Dominic Kwiatkowski, Cordelia Langford, John Sillitoe on behalf of the Wellcome Sanger Institute COVID-19 Surveillance Team |
| EPI_ISL_2912124 | Lighthouse Laboratory Plymouth | Wellcome Sanger Institute for the COVID-19 Genomics UK (COG-UK) Consortium | Lighthouse Laboratory Plymouth and Alex Alderton, Roberto Amato, Jeffrey Barrett, Sonia Goncalves, Ewan Harrison, David K. Jackson, Ian Johnston, Dominic Kwiatkowski, Cordelia Langford, John Sillitoe on behalf of the Wellcome Sanger Institute COVID-19 Surveillance Team |
| EPI_ISL_2912125, EPI_ISL_2912126 | Berkshire and Surrey Pathology Services Lighthouse Laboratory | Wellcome Sanger Institute for the COVID-19 Genomics UK (COG-UK) Consortium | Berkshire and Surrey Pathology Services Lighthouse Laboratory and Alex Alderton, Roberto Amato, Jeffrey Barrett, Sonia Goncalves, Ewan Harrison, David K. Jackson, Ian Johnston, Dominic Kwiatkowski, Cordelia Langford, John Sillitoe on behalf of the Wellcome Sanger Institute COVID-19 Surveillance Team |
| EPI_ISL_2912127 | Lighthouse Laboratory Plymouth | Wellcome Sanger Institute for the COVID-19 Genomics UK (COG-UK) Consortium | Lighthouse Laboratory Plymouth and Alex Alderton, Roberto Amato, Jeffrey Barrett, Sonia Goncalves, Ewan Harrison, David K. Jackson, Ian Johnston, Dominic Kwiatkowski, Cordelia Langford, John Sillitoe on behalf of the Wellcome Sanger Institute COVID-19 Surveillance Team |
| EPI_ISL_2912128 | Berkshire and Surrey Pathology Services Lighthouse Laboratory | Wellcome Sanger Institute for the COVID-19 Genomics UK (COG-UK) Consortium | Berkshire and Surrey Pathology Services Lighthouse Laboratory and Alex Alderton, Roberto Amato, Jeffrey Barrett, Sonia Goncalves, Ewan Harrison, David K. Jackson, Ian Johnston, Dominic Kwiatkowski, Cordelia Langford, John Sillitoe on behalf of the Wellcome Sanger Institute COVID-19 Surveillance Team |
| EPI_ISL_2912129 | Lighthouse Laboratory Plymouth | Wellcome Sanger Institute for the COVID-19 Genomics UK (COG-UK) Consortium | Lighthouse Laboratory Plymouth and Alex Alderton, Roberto Amato, Jeffrey Barrett, Sonia Goncalves, Ewan Harrison, David K. Jackson, Ian Johnston, Dominic Kwiatkowski, Cordelia Langford, John Sillitoe on behalf of the Wellcome Sanger Institute COVID-19 Surveillance Team |
| EPI_ISL_2912130 | Lighthouse Lab in Milton Keynes | Wellcome Sanger Institute for the COVID-19 Genomics UK (COG-UK) Consortium | The Lighthouse Lab in Milton Keynes and Alex Alderton, Roberto Amato, Jeffrey Barrett, Sonia Goncalves, Ewan Harrison, David K. Jackson, Ian Johnston, Dominic Kwiatkowski, Cordelia Langford, John Sillitoe on behalf of the Wellcome Sanger Institute COVID-19 Surveillance Team |
| EPI_ISL_2912131 | Berkshire and Surrey Pathology Services Lighthouse Laboratory | Wellcome Sanger Institute for the COVID-19 Genomics UK (COG-UK) Consortium | Berkshire and Surrey Pathology Services Lighthouse Laboratory and Alex Alderton, Roberto Amato, Jeffrey Barrett, Sonia Goncalves, Ewan Harrison, David K. Jackson, Ian Johnston, Dominic Kwiatkowski, Cordelia Langford, John Sillitoe on behalf of the Wellcome Sanger Institute COVID-19 Surveillance Team |
| EPI_ISL_2912132, EPI_ISL_2912133 | Lighthouse Laboratory Plymouth | Wellcome Sanger Institute for the COVID-19 Genomics UK (COG-UK) Consortium | Lighthouse Laboratory Plymouth and Alex Alderton, Roberto Amato, Jeffrey Barrett, Sonia Goncalves, Ewan Harrison, David K. Jackson, Ian Johnston, Dominic Kwiatkowski, Cordelia Langford, John Sillitoe on behalf of the Wellcome Sanger Institute COVID-19 Surveillance Team |
| EPI_ISL_2912134 | Lighthouse Lab in Milton Keynes | Wellcome Sanger Institute for the COVID-19 Genomics UK (COG-UK) Consortium | The Lighthouse Lab in Milton Keynes and Alex Alderton, Roberto Amato, Jeffrey Barrett, Sonia Goncalves, Ewan Harrison, David K. Jackson, Ian Johnston, Dominic Kwiatkowski, Cordelia Langford, John Sillitoe on behalf of the Wellcome Sanger Institute COVID-19 Surveillance Team |
| EPI_ISL_2912135, EPI_ISL_2912136 | Berkshire and Surrey Pathology Services Lighthouse Laboratory | Wellcome Sanger Institute for the COVID-19 Genomics UK (COG-UK) Consortium | Berkshire and Surrey Pathology Services Lighthouse Laboratory and Alex Alderton, Roberto Amato, Jeffrey Barrett, Sonia Goncalves, Ewan Harrison, David K. Jackson, Ian Johnston, Dominic Kwiatkowski, Cordelia Langford, John Sillitoe on behalf of the Wellcome Sanger Institute COVID-19 Surveillance Team |
| EPI_ISL_2912137 | Lighthouse Lab in Milton Keynes | Wellcome Sanger Institute for the COVID-19 Genomics UK (COG-UK) Consortium | The Lighthouse Lab in Milton Keynes and Alex Alderton, Roberto Amato, Jeffrey Barrett, Sonia Goncalves, Ewan Harrison, David K. Jackson, Ian Johnston, Dominic Kwiatkowski, Cordelia Langford, John Sillitoe on behalf of the Wellcome Sanger Institute COVID-19 Surveillance Team |
| EPI_ISL_2912138 | Lighthouse Laboratory Plymouth | Wellcome Sanger Institute for the COVID-19 Genomics UK (COG-UK) Consortium | Lighthouse Laboratory Plymouth and Alex Alderton, Roberto Amato, Jeffrey Barrett, Sonia Goncalves, Ewan Harrison, David K. Jackson, Ian Johnston, Dominic Kwiatkowski, Cordelia Langford, John Sillitoe on behalf of the Wellcome Sanger Institute COVID-19 Surveillance Team |
| EPI_ISL_2912139 | Berkshire and Surrey Pathology Services Lighthouse Laboratory | Wellcome Sanger Institute for the COVID-19 Genomics UK (COG-UK) Consortium | Berkshire and Surrey Pathology Services Lighthouse Laboratory and Alex Alderton, Roberto Amato, Jeffrey Barrett, Sonia Goncalves, Ewan Harrison, David K. Jackson, Ian Johnston, Dominic Kwiatkowski, Cordelia Langford, John Sillitoe on behalf of the Wellcome Sanger Institute COVID-19 Surveillance Team |
| EPI_ISL_2912140 | Lighthouse Lab in Milton Keynes | Wellcome Sanger Institute for the COVID-19 Genomics UK (COG-UK) Consortium | The Lighthouse Lab in Milton Keynes and Alex Alderton, Roberto Amato, Jeffrey Barrett, Sonia Goncalves, Ewan Harrison, David K. Jackson, Ian Johnston, Dominic Kwiatkowski, Cordelia Langford, John Sillitoe on behalf of the Wellcome Sanger Institute COVID-19 Surveillance Team |
| EPI_ISL_2912141 | Berkshire and Surrey Pathology Services Lighthouse Laboratory | Wellcome Sanger Institute for the COVID-19 Genomics UK (COG-UK) Consortium | Berkshire and Surrey Pathology Services Lighthouse Laboratory and Alex Alderton, Roberto Amato, Jeffrey Barrett, Sonia Goncalves, Ewan Harrison, David K. Jackson, Ian Johnston, Dominic Kwiatkowski, Cordelia Langford, John Sillitoe on behalf of the Wellcome Sanger Institute COVID-19 Surveillance Team |
| EPI_ISL_2912142 | Lighthouse Lab in Milton Keynes | Wellcome Sanger Institute for the COVID-19 Genomics UK (COG-UK) Consortium | The Lighthouse Lab in Milton Keynes and Alex Alderton, Roberto Amato, Jeffrey Barrett, Sonia Goncalves, Ewan Harrison, David K. Jackson, Ian Johnston, Dominic Kwiatkowski, Cordelia Langford, John Sillitoe on behalf of the Wellcome Sanger Institute COVID-19 Surveillance Team |
| EPI_ISL_2912143 | Lighthouse Laboratory Plymouth | Wellcome Sanger Institute for the COVID-19 Genomics UK (COG-UK) Consortium | Lighthouse Laboratory Plymouth and Alex Alderton, Roberto Amato, Jeffrey Barrett, Sonia Goncalves, Ewan Harrison, David K. Jackson, Ian Johnston, Dominic Kwiatkowski, Cordelia Langford, John Sillitoe on behalf of the Wellcome Sanger Institute COVID-19 Surveillance Team |
| EPI_ISL_2912144 | Lighthouse Lab in Milton Keynes | Wellcome Sanger Institute for the COVID-19 Genomics UK (COG-UK) Consortium | The Lighthouse Lab in Milton Keynes and Alex Alderton, Roberto Amato, Jeffrey Barrett, Sonia Goncalves, Ewan Harrison, David K. Jackson, Ian Johnston, Dominic Kwiatkowski, Cordelia Langford, John Sillitoe on behalf of the Wellcome Sanger Institute COVID-19 Surveillance Team |
| EPI_ISL_2912145, EPI_ISL_2912146, EPI_ISL_2912147 | Berkshire and Surrey Pathology Services Lighthouse Laboratory | Wellcome Sanger Institute for the COVID-19 Genomics UK (COG-UK) Consortium | Berkshire and Surrey Pathology Services Lighthouse Laboratory and Alex Alderton, Roberto Amato, Jeffrey Barrett, Sonia Goncalves, Ewan Harrison, David K. Jackson, Ian Johnston, Dominic Kwiatkowski, Cordelia Langford, John Sillitoe on behalf of the Wellcome Sanger Institute COVID-19 Surveillance Team |
| EPI_ISL_2912148 | Lighthouse Laboratory Plymouth | Wellcome Sanger Institute for the COVID-19 Genomics UK (COG-UK) Consortium | Lighthouse Laboratory Plymouth and Alex Alderton, Roberto Amato, Jeffrey Barrett, Sonia Goncalves, Ewan Harrison, David K. Jackson, Ian Johnston, Dominic Kwiatkowski, Cordelia Langford, John Sillitoe on behalf of the Wellcome Sanger Institute COVID-19 Surveillance Team |
| EPI_ISL_2912149, EPI_ISL_2912150, EPI_ISL_2912151 | Berkshire and Surrey Pathology Services Lighthouse Laboratory | Wellcome Sanger Institute for the COVID-19 Genomics UK (COG-UK) Consortium | Berkshire and Surrey Pathology Services Lighthouse Laboratory and Alex Alderton, Roberto Amato, Jeffrey Barrett, Sonia Goncalves, Ewan Harrison, David K. Jackson, Ian Johnston, Dominic Kwiatkowski, Cordelia Langford, John Sillitoe on behalf of the Wellcome Sanger Institute COVID-19 Surveillance Team |
| EPI_ISL_2912152, EPI_ISL_2912153 | Lighthouse Laboratory Plymouth | Wellcome Sanger Institute for the COVID-19 Genomics UK (COG-UK) Consortium | Lighthouse Laboratory Plymouth and Alex Alderton, Roberto Amato, Jeffrey Barrett, Sonia Goncalves, Ewan Harrison, David K. Jackson, Ian Johnston, Dominic Kwiatkowski, Cordelia Langford, John Sillitoe on behalf of the Wellcome Sanger Institute COVID-19 Surveillance Team |
| EPI_ISL_2912154, EPI_ISL_2912155 | Lighthouse Lab in Milton Keynes | Wellcome Sanger Institute for the COVID-19 Genomics UK (COG-UK) Consortium | The Lighthouse Lab in Milton Keynes and Alex Alderton, Roberto Amato, Jeffrey Barrett, Sonia Goncalves, Ewan Harrison, David K. Jackson, Ian Johnston, Dominic Kwiatkowski, Cordelia Langford, John Sillitoe on behalf of the Wellcome Sanger Institute COVID-19 Surveillance Team |
| EPI_ISL_2912156 | Lighthouse Laboratory Plymouth | Wellcome Sanger Institute for the COVID-19 Genomics UK (COG-UK) Consortium | Lighthouse Laboratory Plymouth and Alex Alderton, Roberto Amato, Jeffrey Barrett, Sonia Goncalves, Ewan Harrison, David K. Jackson, Ian Johnston, Dominic Kwiatkowski, Cordelia Langford, John Sillitoe on behalf of the Wellcome Sanger Institute COVID-19 Surveillance Team |
| EPI_ISL_2912157 | Berkshire and Surrey Pathology Services Lighthouse Laboratory | Wellcome Sanger Institute for the COVID-19 Genomics UK (COG-UK) Consortium | Berkshire and Surrey Pathology Services Lighthouse Laboratory and Alex Alderton, Roberto Amato, Jeffrey Barrett, Sonia Goncalves, Ewan Harrison, David K. Jackson, Ian Johnston, Dominic Kwiatkowski, Cordelia Langford, John Sillitoe on behalf of the Wellcome Sanger Institute COVID-19 Surveillance Team |
| EPI_ISL_2912158 | Lighthouse Laboratory Plymouth | Wellcome Sanger Institute for the COVID-19 Genomics UK (COG-UK) Consortium | Lighthouse Laboratory Plymouth and Alex Alderton, Roberto Amato, Jeffrey Barrett, Sonia Goncalves, Ewan Harrison, David K. Jackson, Ian Johnston, Dominic Kwiatkowski, Cordelia Langford, John Sillitoe on behalf of the Wellcome Sanger Institute COVID-19 Surveillance Team |

| | | | |
|---|---|---|---|
| EPI_ISL_2912159 | Lighthouse Lab in Milton Keynes | Wellcome Sanger Institute for the COVID-19 Genomics UK (COG-UK) Consortium | The Lighthouse Lab in Milton Keynes and Alex Alderton, Roberto Amato, Jeffrey Barrett, Sonia Goncalves, Ewan Harrison, David K. Jackson, Ian Johnston, Dominic Kwiatkowski, Cordelia Langford, John Sillitoe on behalf of the Wellcome Sanger Institute COVID-19 Surveillance Team |
| EPI_ISL_2912160 | Berkshire and Surrey Pathology Services Lighthouse Laboratory | Wellcome Sanger Institute for the COVID-19 Genomics UK (COG-UK) Consortium | Berkshire and Surrey Pathology Services Lighthouse Laboratory and Alex Alderton, Roberto Amato, Jeffrey Barrett, Sonia Goncalves, Ewan Harrison, David K. Jackson, Ian Johnston, Dominic Kwiatkowski, Cordelia Langford, John Sillitoe on behalf of the Wellcome Sanger Institute COVID-19 Surveillance Team |
| EPI_ISL_2912161, EPI_ISL_2912162 | Lighthouse Laboratory Plymouth | Wellcome Sanger Institute for the COVID-19 Genomics UK (COG-UK) Consortium | Lighthouse Laboratory Plymouth and Alex Alderton, Roberto Amato, Jeffrey Barrett, Sonia Goncalves, Ewan Harrison, David K. Jackson, Ian Johnston, Dominic Kwiatkowski, Cordelia Langford, John Sillitoe on behalf of the Wellcome Sanger Institute COVID-19 Surveillance Team |
| EPI_ISL_2912163 | Berkshire and Surrey Pathology Services Lighthouse Laboratory | Wellcome Sanger Institute for the COVID-19 Genomics UK (COG-UK) Consortium | Berkshire and Surrey Pathology Services Lighthouse Laboratory and Alex Alderton, Roberto Amato, Jeffrey Barrett, Sonia Goncalves, Ewan Harrison, David K. Jackson, Ian Johnston, Dominic Kwiatkowski, Cordelia Langford, John Sillitoe on behalf of the Wellcome Sanger Institute COVID-19 Surveillance Team |
| EPI_ISL_2912164 | Lighthouse Laboratory Plymouth | Wellcome Sanger Institute for the COVID-19 Genomics UK (COG-UK) Consortium | Lighthouse Laboratory Plymouth and Alex Alderton, Roberto Amato, Jeffrey Barrett, Sonia Goncalves, Ewan Harrison, David K. Jackson, Ian Johnston, Dominic Kwiatkowski, Cordelia Langford, John Sillitoe on behalf of the Wellcome Sanger Institute COVID-19 Surveillance Team |
| EPI_ISL_2912165 | Berkshire and Surrey Pathology Services Lighthouse Laboratory | Wellcome Sanger Institute for the COVID-19 Genomics UK (COG-UK) Consortium | Berkshire and Surrey Pathology Services Lighthouse Laboratory and Alex Alderton, Roberto Amato, Jeffrey Barrett, Sonia Goncalves, Ewan Harrison, David K. Jackson, Ian Johnston, Dominic Kwiatkowski, Cordelia Langford, John Sillitoe on behalf of the Wellcome Sanger Institute COVID-19 Surveillance Team |
| EPI_ISL_2912166 | Lighthouse Laboratory Plymouth | Wellcome Sanger Institute for the COVID-19 Genomics UK (COG-UK) Consortium | Lighthouse Laboratory Plymouth and Alex Alderton, Roberto Amato, Jeffrey Barrett, Sonia Goncalves, Ewan Harrison, David K. Jackson, Ian Johnston, Dominic Kwiatkowski, Cordelia Langford, John Sillitoe on behalf of the Wellcome Sanger Institute COVID-19 Surveillance Team |
| EPI_ISL_2912167 | Berkshire and Surrey Pathology Services Lighthouse Laboratory | Wellcome Sanger Institute for the COVID-19 Genomics UK (COG-UK) Consortium | Berkshire and Surrey Pathology Services Lighthouse Laboratory and Alex Alderton, Roberto Amato, Jeffrey Barrett, Sonia Goncalves, Ewan Harrison, David K. Jackson, Ian Johnston, Dominic Kwiatkowski, Cordelia Langford, John Sillitoe on behalf of the Wellcome Sanger Institute COVID-19 Surveillance Team |
| EPI_ISL_2912168 | Lighthouse Lab in Milton Keynes | Wellcome Sanger Institute for the COVID-19 Genomics UK (COG-UK) Consortium | The Lighthouse Lab in Milton Keynes and Alex Alderton, Roberto Amato, Jeffrey Barrett, Sonia Goncalves, Ewan Harrison, David K. Jackson, Ian Johnston, Dominic Kwiatkowski, Cordelia Langford, John Sillitoe on behalf of the Wellcome Sanger Institute COVID-19 Surveillance Team |
| EPI_ISL_2912169, EPI_ISL_2912170, EPI_ISL_2912171 | Berkshire and Surrey Pathology Services Lighthouse Laboratory | Wellcome Sanger Institute for the COVID-19 Genomics UK (COG-UK) Consortium | Berkshire and Surrey Pathology Services Lighthouse Laboratory and Alex Alderton, Roberto Amato, Jeffrey Barrett, Sonia Goncalves, Ewan Harrison, David K. Jackson, Ian Johnston, Dominic Kwiatkowski, Cordelia Langford, John Sillitoe on behalf of the Wellcome Sanger Institute COVID-19 Surveillance Team |
| EPI_ISL_2912172 | Lighthouse Laboratory Plymouth | Wellcome Sanger Institute for the COVID-19 Genomics UK (COG-UK) Consortium | Lighthouse Laboratory Plymouth and Alex Alderton, Roberto Amato, Jeffrey Barrett, Sonia Goncalves, Ewan Harrison, David K. Jackson, Ian Johnston, Dominic Kwiatkowski, Cordelia Langford, John Sillitoe on behalf of the Wellcome Sanger Institute COVID-19 Surveillance Team |
| EPI_ISL_2912173, EPI_ISL_2912174, EPI_ISL_2912175, EPI_ISL_2912176 | Berkshire and Surrey Pathology Services Lighthouse Laboratory | Wellcome Sanger Institute for the COVID-19 Genomics UK (COG-UK) Consortium | Berkshire and Surrey Pathology Services Lighthouse Laboratory and Alex Alderton, Roberto Amato, Jeffrey Barrett, Sonia Goncalves, Ewan Harrison, David K. Jackson, Ian Johnston, Dominic Kwiatkowski, Cordelia Langford, John Sillitoe on behalf of the Wellcome Sanger Institute COVID-19 Surveillance Team |
| EPI_ISL_2912177 | Lighthouse Lab in Milton Keynes | Wellcome Sanger Institute for the COVID-19 Genomics UK (COG-UK) Consortium | The Lighthouse Lab in Milton Keynes and Alex Alderton, Roberto Amato, Jeffrey Barrett, Sonia Goncalves, Ewan Harrison, David K. Jackson, Ian Johnston, Dominic Kwiatkowski, Cordelia Langford, John Sillitoe on behalf of the Wellcome Sanger Institute COVID-19 Surveillance Team |
| EPI_ISL_2912178, EPI_ISL_2912179 | Berkshire and Surrey Pathology Services Lighthouse Laboratory | Wellcome Sanger Institute for the COVID-19 Genomics UK (COG-UK) Consortium | Berkshire and Surrey Pathology Services Lighthouse Laboratory and Alex Alderton, Roberto Amato, Jeffrey Barrett, Sonia Goncalves, Ewan Harrison, David K. Jackson, Ian Johnston, Dominic Kwiatkowski, Cordelia Langford, John Sillitoe on behalf of the Wellcome Sanger Institute COVID-19 Surveillance Team |
| EPI_ISL_2912180 | Lighthouse Laboratory Plymouth | Wellcome Sanger Institute for the COVID-19 Genomics UK (COG-UK) Consortium | Lighthouse Laboratory Plymouth and Alex Alderton, Roberto Amato, Jeffrey Barrett, Sonia Goncalves, Ewan Harrison, David K. Jackson, Ian Johnston, Dominic Kwiatkowski, Cordelia Langford, John Sillitoe on behalf of the Wellcome Sanger Institute COVID-19 Surveillance Team |
| EPI_ISL_2912181 | Lighthouse Lab in Milton Keynes | Wellcome Sanger Institute for the COVID-19 Genomics UK (COG-UK) Consortium | The Lighthouse Lab in Milton Keynes and Alex Alderton, Roberto Amato, Jeffrey Barrett, Sonia Goncalves, Ewan Harrison, David K. Jackson, Ian Johnston, Dominic Kwiatkowski, Cordelia Langford, John Sillitoe on behalf of the Wellcome Sanger Institute COVID-19 Surveillance Team |
| EPI_ISL_2912182 | Berkshire and Surrey Pathology Services Lighthouse Laboratory | Wellcome Sanger Institute for the COVID-19 Genomics UK (COG-UK) Consortium | Berkshire and Surrey Pathology Services Lighthouse Laboratory and Alex Alderton, Roberto Amato, Jeffrey Barrett, Sonia Goncalves, Ewan Harrison, David K. Jackson, Ian Johnston, Dominic Kwiatkowski, Cordelia Langford, John Sillitoe on behalf of the Wellcome Sanger Institute COVID-19 Surveillance Team |
| EPI_ISL_2912183 | Lighthouse Laboratory Plymouth | Wellcome Sanger Institute for the COVID-19 Genomics UK (COG-UK) Consortium | Lighthouse Laboratory Plymouth and Alex Alderton, Roberto Amato, Jeffrey Barrett, Sonia Goncalves, Ewan Harrison, David K. Jackson, Ian Johnston, Dominic Kwiatkowski, Cordelia Langford, John Sillitoe on behalf of the Wellcome Sanger Institute COVID-19 Surveillance Team |
| EPI_ISL_2912184 | Lighthouse Lab in Milton Keynes | Wellcome Sanger Institute for the COVID-19 Genomics UK (COG-UK) Consortium | The Lighthouse Lab in Milton Keynes and Alex Alderton, Roberto Amato, Jeffrey Barrett, Sonia Goncalves, Ewan Harrison, David K. Jackson, Ian Johnston, Dominic Kwiatkowski, Cordelia Langford, John Sillitoe on behalf of the Wellcome Sanger Institute COVID-19 Surveillance Team |
| EPI_ISL_2912185 | Berkshire and Surrey Pathology Services Lighthouse Laboratory | Wellcome Sanger Institute for the COVID-19 Genomics UK (COG-UK) Consortium | Berkshire and Surrey Pathology Services Lighthouse Laboratory and Alex Alderton, Roberto Amato, Jeffrey Barrett, Sonia Goncalves, Ewan Harrison, David K. Jackson, Ian Johnston, Dominic Kwiatkowski, Cordelia Langford, John Sillitoe on behalf of the Wellcome Sanger Institute COVID-19 Surveillance Team |
| EPI_ISL_2912186 | Lighthouse Laboratory Plymouth | Wellcome Sanger Institute for the COVID-19 Genomics UK (COG-UK) Consortium | Lighthouse Laboratory Plymouth and Alex Alderton, Roberto Amato, Jeffrey Barrett, Sonia Goncalves, Ewan Harrison, David K. Jackson, Ian Johnston, Dominic Kwiatkowski, Cordelia Langford, John Sillitoe on behalf of the Wellcome Sanger Institute COVID-19 Surveillance Team |
| EPI_ISL_2912187 | Berkshire and Surrey Pathology Services Lighthouse Laboratory | Wellcome Sanger Institute for the COVID-19 Genomics UK (COG-UK) Consortium | Berkshire and Surrey Pathology Services Lighthouse Laboratory and Alex Alderton, Roberto Amato, Jeffrey Barrett, Sonia Goncalves, Ewan Harrison, David K. Jackson, Ian Johnston, Dominic Kwiatkowski, Cordelia Langford, John Sillitoe on behalf of the Wellcome Sanger Institute COVID-19 Surveillance Team |
| EPI_ISL_2912188 | Lighthouse Lab in Milton Keynes | Wellcome Sanger Institute for the COVID-19 Genomics UK (COG-UK) Consortium | The Lighthouse Lab in Milton Keynes and Alex Alderton, Roberto Amato, Jeffrey Barrett, Sonia Goncalves, Ewan Harrison, David K. Jackson, Ian Johnston, Dominic Kwiatkowski, Cordelia Langford, John Sillitoe on behalf of the Wellcome Sanger Institute COVID-19 Surveillance Team |
| EPI_ISL_2912189 | Berkshire and Surrey Pathology Services Lighthouse Laboratory | Wellcome Sanger Institute for the COVID-19 Genomics UK (COG-UK) Consortium | Berkshire and Surrey Pathology Services Lighthouse Laboratory and Alex Alderton, Roberto Amato, Jeffrey Barrett, Sonia Goncalves, Ewan Harrison, David K. Jackson, Ian Johnston, Dominic Kwiatkowski, Cordelia Langford, John Sillitoe on behalf of the Wellcome Sanger Institute COVID-19 Surveillance Team |
| EPI_ISL_2912190 | Lighthouse Lab in Milton Keynes | Wellcome Sanger Institute for the COVID-19 Genomics UK (COG-UK) Consortium | The Lighthouse Lab in Milton Keynes and Alex Alderton, Roberto Amato, Jeffrey Barrett, Sonia Goncalves, Ewan Harrison, David K. Jackson, Ian Johnston, Dominic Kwiatkowski, Cordelia Langford, John Sillitoe on behalf of the Wellcome Sanger Institute COVID-19 Surveillance Team |
| EPI_ISL_2912191, EPI_ISL_2912192 | Berkshire and Surrey Pathology Services Lighthouse Laboratory | Wellcome Sanger Institute for the COVID-19 Genomics UK (COG-UK) Consortium | Berkshire and Surrey Pathology Services Lighthouse Laboratory and Alex Alderton, Roberto Amato, Jeffrey Barrett, Sonia Goncalves, Ewan Harrison, David K. Jackson, Ian Johnston, Dominic Kwiatkowski, Cordelia Langford, John Sillitoe on behalf of the Wellcome Sanger Institute COVID-19 Surveillance Team |
| EPI_ISL_2912193 | Lighthouse Lab in Milton Keynes | Wellcome Sanger Institute for the COVID-19 Genomics UK (COG-UK) Consortium | The Lighthouse Lab in Milton Keynes and Alex Alderton, Roberto Amato, Jeffrey Barrett, Sonia Goncalves, Ewan Harrison, David K. Jackson, Ian Johnston, Dominic Kwiatkowski, Cordelia Langford, John Sillitoe on behalf of the Wellcome Sanger Institute COVID-19 Surveillance Team |
| EPI_ISL_2912194 | Berkshire and Surrey Pathology Services Lighthouse Laboratory | Wellcome Sanger Institute for the COVID-19 Genomics UK (COG-UK) Consortium | Berkshire and Surrey Pathology Services Lighthouse Laboratory and Alex Alderton, Roberto Amato, Jeffrey Barrett, Sonia Goncalves, Ewan Harrison, David K. Jackson, Ian Johnston, Dominic Kwiatkowski, Cordelia Langford, John Sillitoe on behalf of the Wellcome Sanger Institute COVID-19 Surveillance Team |
| EPI_ISL_2912195, EPI_ISL_2912196 | Lighthouse Laboratory Plymouth | Wellcome Sanger Institute for the COVID-19 Genomics UK (COG-UK) Consortium | Lighthouse Laboratory Plymouth and Alex Alderton, Roberto Amato, Jeffrey Barrett, Sonia Goncalves, Ewan Harrison, David K. Jackson, Ian Johnston, Dominic Kwiatkowski, Cordelia Langford, John Sillitoe on behalf of the Wellcome Sanger Institute COVID-19 Surveillance Team |
| EPI_ISL_2912197, EPI_ISL_2912198 | Berkshire and Surrey Pathology Services Lighthouse Laboratory | Wellcome Sanger Institute for the COVID-19 Genomics UK (COG-UK) Consortium | Berkshire and Surrey Pathology Services Lighthouse Laboratory and Alex Alderton, Roberto Amato, Jeffrey Barrett, Sonia Goncalves, Ewan Harrison, David K. Jackson, Ian Johnston, Dominic Kwiatkowski, Cordelia Langford, John Sillitoe on behalf of the Wellcome Sanger Institute COVID-19 Surveillance Team |
| EPI_ISL_2912199 | Lighthouse Laboratory Plymouth | Wellcome Sanger Institute for the COVID-19 Genomics UK (COG-UK) Consortium | Lighthouse Laboratory Plymouth and Alex Alderton, Roberto Amato, Jeffrey Barrett, Sonia Goncalves, Ewan Harrison, David K. Jackson, Ian Johnston, Dominic Kwiatkowski, Cordelia Langford, John Sillitoe on behalf of the Wellcome Sanger Institute COVID-19 Surveillance Team |
| EPI_ISL_2912200, EPI_ISL_2912201 | Berkshire and Surrey Pathology Services Lighthouse Laboratory | Wellcome Sanger Institute for the COVID-19 Genomics UK (COG-UK) Consortium | Berkshire and Surrey Pathology Services Lighthouse Laboratory and Alex Alderton, Roberto Amato, Jeffrey Barrett, Sonia Goncalves, Ewan Harrison, David K. Jackson, Ian Johnston, Dominic Kwiatkowski, Cordelia Langford, John Sillitoe on behalf of the Wellcome Sanger Institute COVID-19 Surveillance Team |
| EPI_ISL_2912202 | Lighthouse Laboratory Plymouth | Wellcome Sanger Institute for the COVID-19 Genomics UK (COG-UK) Consortium | Lighthouse Laboratory Plymouth and Alex Alderton, Roberto Amato, Jeffrey Barrett, Sonia Goncalves, Ewan Harrison, David K. Jackson, Ian Johnston, Dominic Kwiatkowski, Cordelia Langford, John Sillitoe on behalf of the Wellcome Sanger Institute COVID-19 Surveillance Team |
| EPI_ISL_2912203 | Berkshire and Surrey Pathology Services Lighthouse Laboratory | Wellcome Sanger Institute for the COVID-19 Genomics UK (COG-UK) Consortium | Berkshire and Surrey Pathology Services Lighthouse Laboratory and Alex Alderton, Roberto Amato, Jeffrey Barrett, Sonia Goncalves, Ewan Harrison, David K. Jackson, Ian Johnston, Dominic Kwiatkowski, Cordelia Langford, John Sillitoe on behalf of the Wellcome Sanger Institute COVID-19 Surveillance Team |
| EPI_ISL_2912204, EPI_ISL_2912205, EPI_ISL_2912206 | Lighthouse Lab in Milton Keynes | Wellcome Sanger Institute for the COVID-19 Genomics UK (COG-UK) Consortium | The Lighthouse Lab in Milton Keynes and Alex Alderton, Roberto Amato, Jeffrey Barrett, Sonia Goncalves, Ewan Harrison, David K. Jackson, Ian Johnston, Dominic Kwiatkowski, Cordelia Langford, John Sillitoe on behalf of the Wellcome Sanger Institute COVID-19 Surveillance Team |
| EPI_ISL_2912207 | Berkshire and Surrey Pathology Services Lighthouse Laboratory | Wellcome Sanger Institute for the COVID-19 Genomics UK (COG-UK) Consortium | Berkshire and Surrey Pathology Services Lighthouse Laboratory and Alex Alderton, Roberto Amato, Jeffrey Barrett, Sonia Goncalves, Ewan Harrison, David K. Jackson, Ian Johnston, Dominic Kwiatkowski, Cordelia Langford, John Sillitoe on behalf of the Wellcome Sanger Institute COVID-19 Surveillance Team |
| EPI_ISL_2912208, EPI_ISL_2912209 | Lighthouse Laboratory Plymouth | Wellcome Sanger Institute for the COVID-19 Genomics UK (COG-UK) Consortium | Lighthouse Laboratory Plymouth and Alex Alderton, Roberto Amato, Jeffrey Barrett, Sonia Goncalves, Ewan Harrison, David K. Jackson, Ian Johnston, Dominic Kwiatkowski, Cordelia Langford, John Sillitoe on behalf of the Wellcome Sanger Institute COVID-19 Surveillance Team |
| EPI_ISL_2912210, EPI_ISL_2912211, EPI_ISL_2912212 | Berkshire and Surrey Pathology Services Lighthouse Laboratory | Wellcome Sanger Institute for the COVID-19 Genomics UK (COG-UK) Consortium | Berkshire and Surrey Pathology Services Lighthouse Laboratory and Alex Alderton, Roberto Amato, Jeffrey Barrett, Sonia Goncalves, Ewan Harrison, David K. Jackson, Ian Johnston, Dominic Kwiatkowski, Cordelia Langford, John Sillitoe on behalf of the Wellcome Sanger Institute COVID-19 Surveillance Team |
| EPI_ISL_2912213 | Lighthouse Lab in Milton Keynes | Wellcome Sanger Institute for the COVID-19 Genomics UK (COG-UK) Consortium | The Lighthouse Lab in Milton Keynes and Alex Alderton, Roberto Amato, Jeffrey Barrett, Sonia Goncalves, Ewan Harrison, David K. Jackson, Ian Johnston, Dominic Kwiatkowski, Cordelia Langford, John Sillitoe on behalf of the Wellcome Sanger Institute COVID-19 Surveillance Team |
| EPI_ISL_2912214 | Berkshire and Surrey Pathology Services Lighthouse Laboratory | Wellcome Sanger Institute for the COVID-19 Genomics UK (COG-UK) Consortium | Berkshire and Surrey Pathology Services Lighthouse Laboratory and Alex Alderton, Roberto Amato, Jeffrey Barrett, Sonia Goncalves, Ewan Harrison, David K. Jackson, Ian Johnston, Dominic Kwiatkowski, Cordelia Langford, John Sillitoe on behalf of the Wellcome Sanger Institute COVID-19 Surveillance Team |
| EPI_ISL_2912215 | Lighthouse Laboratory Plymouth | Wellcome Sanger Institute for the COVID-19 Genomics UK (COG-UK) Consortium | Lighthouse Laboratory Plymouth and Alex Alderton, Roberto Amato, Jeffrey Barrett, Sonia Goncalves, Ewan Harrison, David K. Jackson, Ian Johnston, Dominic Kwiatkowski, Cordelia Langford, John Sillitoe on behalf of the Wellcome Sanger Institute COVID-19 Surveillance Team |
| EPI_ISL_2912216 | Lighthouse Lab in Milton Keynes | Wellcome Sanger Institute for the COVID-19 Genomics UK (COG-UK) Consortium | The Lighthouse Lab in Milton Keynes and Alex Alderton, Roberto Amato, Jeffrey Barrett, Sonia Goncalves, Ewan Harrison, David K. Jackson, Ian Johnston, Dominic Kwiatkowski, Cordelia Langford, John Sillitoe on behalf of the Wellcome Sanger Institute COVID-19 Surveillance Team |
| EPI_ISL_2912217 | Berkshire and Surrey Pathology Services Lighthouse Laboratory | Wellcome Sanger Institute for the COVID-19 Genomics UK (COG-UK) Consortium | Berkshire and Surrey Pathology Services Lighthouse Laboratory and Alex Alderton, Roberto Amato, Jeffrey Barrett, Sonia Goncalves, Ewan Harrison, David K. Jackson, Ian Johnston, Dominic Kwiatkowski, Cordelia Langford, John Sillitoe on behalf of the Wellcome Sanger Institute COVID-19 Surveillance Team |

| | | | |
|---|---|---|---|
| EPI_ISL_2912218 | Lighthouse Laboratory Plymouth | Wellcome Sanger Institute for the COVID-19 Genomics UK (COG-UK) Consortium | Lighthouse Laboratory Plymouth and Alex Alderton, Roberto Amato, Jeffrey Barrett, Sonia Goncalves, Ewan Harrison, David K. Jackson, Ian Johnston, Dominic Kwiatkowski, Cordelia Langford, John Sillitoe on behalf of the Wellcome Sanger Institute COVID-19 Surveillance Team |
| EPI_ISL_2912219 | Berkshire and Surrey Pathology Services Lighthouse Laboratory | Wellcome Sanger Institute for the COVID-19 Genomics UK (COG-UK) Consortium | Berkshire and Surrey Pathology Services Lighthouse Laboratory and Alex Alderton, Roberto Amato, Jeffrey Barrett, Sonia Goncalves, Ewan Harrison, David K. Jackson, Ian Johnston, Dominic Kwiatkowski, Cordelia Langford, John Sillitoe on behalf of the Wellcome Sanger Institute COVID-19 Surveillance Team |
| EPI_ISL_2912220 | Lighthouse Lab in Milton Keynes | Wellcome Sanger Institute for the COVID-19 Genomics UK (COG-UK) Consortium | The Lighthouse Lab in Milton Keynes and Alex Alderton, Roberto Amato, Jeffrey Barrett, Sonia Goncalves, Ewan Harrison, David K. Jackson, Ian Johnston, Dominic Kwiatkowski, Cordelia Langford, John Sillitoe on behalf of the Wellcome Sanger Institute COVID-19 Surveillance Team |
| EPI_ISL_2912221 | Berkshire and Surrey Pathology Services Lighthouse Laboratory | Wellcome Sanger Institute for the COVID-19 Genomics UK (COG-UK) Consortium | Berkshire and Surrey Pathology Services Lighthouse Laboratory and Alex Alderton, Roberto Amato, Jeffrey Barrett, Sonia Goncalves, Ewan Harrison, David K. Jackson, Ian Johnston, Dominic Kwiatkowski, Cordelia Langford, John Sillitoe on behalf of the Wellcome Sanger Institute COVID-19 Surveillance Team |
| EPI_ISL_2912222 | Lighthouse Lab in Milton Keynes | Wellcome Sanger Institute for the COVID-19 Genomics UK (COG-UK) Consortium | The Lighthouse Lab in Milton Keynes and Alex Alderton, Roberto Amato, Jeffrey Barrett, Sonia Goncalves, Ewan Harrison, David K. Jackson, Ian Johnston, Dominic Kwiatkowski, Cordelia Langford, John Sillitoe on behalf of the Wellcome Sanger Institute COVID-19 Surveillance Team |
| EPI_ISL_2912223 | Berkshire and Surrey Pathology Services Lighthouse Laboratory | Wellcome Sanger Institute for the COVID-19 Genomics UK (COG-UK) Consortium | Berkshire and Surrey Pathology Services Lighthouse Laboratory and Alex Alderton, Roberto Amato, Jeffrey Barrett, Sonia Goncalves, Ewan Harrison, David K. Jackson, Ian Johnston, Dominic Kwiatkowski, Cordelia Langford, John Sillitoe on behalf of the Wellcome Sanger Institute COVID-19 Surveillance Team |
| EPI_ISL_2912224, EPI_ISL_2912225 | Lighthouse Laboratory Plymouth | Wellcome Sanger Institute for the COVID-19 Genomics UK (COG-UK) Consortium | Lighthouse Laboratory Plymouth and Alex Alderton, Roberto Amato, Jeffrey Barrett, Sonia Goncalves, Ewan Harrison, David K. Jackson, Ian Johnston, Dominic Kwiatkowski, Cordelia Langford, John Sillitoe on behalf of the Wellcome Sanger Institute COVID-19 Surveillance Team |
| EPI_ISL_2912226 | Berkshire and Surrey Pathology Services Lighthouse Laboratory | Wellcome Sanger Institute for the COVID-19 Genomics UK (COG-UK) Consortium | Berkshire and Surrey Pathology Services Lighthouse Laboratory and Alex Alderton, Roberto Amato, Jeffrey Barrett, Sonia Goncalves, Ewan Harrison, David K. Jackson, Ian Johnston, Dominic Kwiatkowski, Cordelia Langford, John Sillitoe on behalf of the Wellcome Sanger Institute COVID-19 Surveillance Team |
| EPI_ISL_2912227 | Lighthouse Laboratory Plymouth | Wellcome Sanger Institute for the COVID-19 Genomics UK (COG-UK) Consortium | Lighthouse Laboratory Plymouth and Alex Alderton, Roberto Amato, Jeffrey Barrett, Sonia Goncalves, Ewan Harrison, David K. Jackson, Ian Johnston, Dominic Kwiatkowski, Cordelia Langford, John Sillitoe on behalf of the Wellcome Sanger Institute COVID-19 Surveillance Team |
| EPI_ISL_2912228, EPI_ISL_2912229 | Berkshire and Surrey Pathology Services Lighthouse Laboratory | Wellcome Sanger Institute for the COVID-19 Genomics UK (COG-UK) Consortium | Berkshire and Surrey Pathology Services Lighthouse Laboratory and Alex Alderton, Roberto Amato, Jeffrey Barrett, Sonia Goncalves, Ewan Harrison, David K. Jackson, Ian Johnston, Dominic Kwiatkowski, Cordelia Langford, John Sillitoe on behalf of the Wellcome Sanger Institute COVID-19 Surveillance Team |
| EPI_ISL_2912230 | Lighthouse Lab in Milton Keynes | Wellcome Sanger Institute for the COVID-19 Genomics UK (COG-UK) Consortium | The Lighthouse Lab in Milton Keynes and Alex Alderton, Roberto Amato, Jeffrey Barrett, Sonia Goncalves, Ewan Harrison, David K. Jackson, Ian Johnston, Dominic Kwiatkowski, Cordelia Langford, John Sillitoe on behalf of the Wellcome Sanger Institute COVID-19 Surveillance Team |
| EPI_ISL_2912231 | Lighthouse Laboratory Plymouth | Wellcome Sanger Institute for the COVID-19 Genomics UK (COG-UK) Consortium | Lighthouse Laboratory Plymouth and Alex Alderton, Roberto Amato, Jeffrey Barrett, Sonia Goncalves, Ewan Harrison, David K. Jackson, Ian Johnston, Dominic Kwiatkowski, Cordelia Langford, John Sillitoe on behalf of the Wellcome Sanger Institute COVID-19 Surveillance Team |
| EPI_ISL_2912232 | Berkshire and Surrey Pathology Services Lighthouse Laboratory | Wellcome Sanger Institute for the COVID-19 Genomics UK (COG-UK) Consortium | Berkshire and Surrey Pathology Services Lighthouse Laboratory and Alex Alderton, Roberto Amato, Jeffrey Barrett, Sonia Goncalves, Ewan Harrison, David K. Jackson, Ian Johnston, Dominic Kwiatkowski, Cordelia Langford, John Sillitoe on behalf of the Wellcome Sanger Institute COVID-19 Surveillance Team |
| EPI_ISL_2912233 | Lighthouse Laboratory Plymouth | Wellcome Sanger Institute for the COVID-19 Genomics UK (COG-UK) Consortium | Lighthouse Laboratory Plymouth and Alex Alderton, Roberto Amato, Jeffrey Barrett, Sonia Goncalves, Ewan Harrison, David K. Jackson, Ian Johnston, Dominic Kwiatkowski, Cordelia Langford, John Sillitoe on behalf of the Wellcome Sanger Institute COVID-19 Surveillance Team |
| EPI_ISL_2912234 | Lighthouse Lab in Milton Keynes | Wellcome Sanger Institute for the COVID-19 Genomics UK (COG-UK) Consortium | The Lighthouse Lab in Milton Keynes and Alex Alderton, Roberto Amato, Jeffrey Barrett, Sonia Goncalves, Ewan Harrison, David K. Jackson, Ian Johnston, Dominic Kwiatkowski, Cordelia Langford, John Sillitoe on behalf of the Wellcome Sanger Institute COVID-19 Surveillance Team |
| EPI_ISL_2912235, EPI_ISL_2912236 | Berkshire and Surrey Pathology Services Lighthouse Laboratory | Wellcome Sanger Institute for the COVID-19 Genomics UK (COG-UK) Consortium | Berkshire and Surrey Pathology Services Lighthouse Laboratory and Alex Alderton, Roberto Amato, Jeffrey Barrett, Sonia Goncalves, Ewan Harrison, David K. Jackson, Ian Johnston, Dominic Kwiatkowski, Cordelia Langford, John Sillitoe on behalf of the Wellcome Sanger Institute COVID-19 Surveillance Team |
| EPI_ISL_2912237, EPI_ISL_2912238, EPI_ISL_2912239 | Lighthouse Laboratory Plymouth | Wellcome Sanger Institute for the COVID-19 Genomics UK (COG-UK) Consortium | Lighthouse Laboratory Plymouth and Alex Alderton, Roberto Amato, Jeffrey Barrett, Sonia Goncalves, Ewan Harrison, David K. Jackson, Ian Johnston, Dominic Kwiatkowski, Cordelia Langford, John Sillitoe on behalf of the Wellcome Sanger Institute COVID-19 Surveillance Team |
| EPI_ISL_2912240 | Berkshire and Surrey Pathology Services Lighthouse Laboratory | Wellcome Sanger Institute for the COVID-19 Genomics UK (COG-UK) Consortium | Berkshire and Surrey Pathology Services Lighthouse Laboratory and Alex Alderton, Roberto Amato, Jeffrey Barrett, Sonia Goncalves, Ewan Harrison, David K. Jackson, Ian Johnston, Dominic Kwiatkowski, Cordelia Langford, John Sillitoe on behalf of the Wellcome Sanger Institute COVID-19 Surveillance Team |
| EPI_ISL_2912241, EPI_ISL_2912242 | Lighthouse Lab in Milton Keynes | Wellcome Sanger Institute for the COVID-19 Genomics UK (COG-UK) Consortium | The Lighthouse Lab in Milton Keynes and Alex Alderton, Roberto Amato, Jeffrey Barrett, Sonia Goncalves, Ewan Harrison, David K. Jackson, Ian Johnston, Dominic Kwiatkowski, Cordelia Langford, John Sillitoe on behalf of the Wellcome Sanger Institute COVID-19 Surveillance Team |
| EPI_ISL_2912243 | Lighthouse Laboratory Plymouth | Wellcome Sanger Institute for the COVID-19 Genomics UK (COG-UK) Consortium | Lighthouse Laboratory Plymouth and Alex Alderton, Roberto Amato, Jeffrey Barrett, Sonia Goncalves, Ewan Harrison, David K. Jackson, Ian Johnston, Dominic Kwiatkowski, Cordelia Langford, John Sillitoe on behalf of the Wellcome Sanger Institute COVID-19 Surveillance Team |
| EPI_ISL_2912244, EPI_ISL_2912245 | Berkshire and Surrey Pathology Services Lighthouse Laboratory | Wellcome Sanger Institute for the COVID-19 Genomics UK (COG-UK) Consortium | Berkshire and Surrey Pathology Services Lighthouse Laboratory and Alex Alderton, Roberto Amato, Jeffrey Barrett, Sonia Goncalves, Ewan Harrison, David K. Jackson, Ian Johnston, Dominic Kwiatkowski, Cordelia Langford, John Sillitoe on behalf of the Wellcome Sanger Institute COVID-19 Surveillance Team |
| EPI_ISL_2912246 | Lighthouse Lab in Glasgow | Wellcome Sanger Institute for the COVID-19 Genomics UK (COG-UK) Consortium | Harper VanSteenhouse, Yumi Kasai, David Gray, Carol Clugston, Anna Dominiczak and Alex Alderton, Roberto Amato, Jeffrey Barrett, Sonia Goncalves, Ewan Harrison, David K. Jackson, Ian Johnston, Dominic Kwiatkowski, Cordelia Langford, John Sillitoe on behalf of the Wellcome Sanger Institute COVID-19 Surveillance Team |
| EPI_ISL_2912247, EPI_ISL_2912248, EPI_ISL_2912249 | Berkshire and Surrey Pathology Services Lighthouse Laboratory | Wellcome Sanger Institute for the COVID-19 Genomics UK (COG-UK) Consortium | Berkshire and Surrey Pathology Services Lighthouse Laboratory and Alex Alderton, Roberto Amato, Jeffrey Barrett, Sonia Goncalves, Ewan Harrison, David K. Jackson, Ian Johnston, Dominic Kwiatkowski, Cordelia Langford, John Sillitoe on behalf of the Wellcome Sanger Institute COVID-19 Surveillance Team |
| EPI_ISL_2912250 | Lighthouse Lab in Milton Keynes | Wellcome Sanger Institute for the COVID-19 Genomics UK (COG-UK) Consortium | The Lighthouse Lab in Milton Keynes and Alex Alderton, Roberto Amato, Jeffrey Barrett, Sonia Goncalves, Ewan Harrison, David K. Jackson, Ian Johnston, Dominic Kwiatkowski, Cordelia Langford, John Sillitoe on behalf of the Wellcome Sanger Institute COVID-19 Surveillance Team |
| EPI_ISL_2912251 | Lighthouse Lab in Glasgow | Wellcome Sanger Institute for the COVID-19 Genomics UK (COG-UK) Consortium | Harper VanSteenhouse, Yumi Kasai, David Gray, Carol Clugston, Anna Dominiczak and Alex Alderton, Roberto Amato, Jeffrey Barrett, Sonia Goncalves, Ewan Harrison, David K. Jackson, Ian Johnston, Dominic Kwiatkowski, Cordelia Langford, John Sillitoe on behalf of the Wellcome Sanger Institute COVID-19 Surveillance Team |
| EPI_ISL_2912252 | Berkshire and Surrey Pathology Services Lighthouse Laboratory | Wellcome Sanger Institute for the COVID-19 Genomics UK (COG-UK) Consortium | Berkshire and Surrey Pathology Services Lighthouse Laboratory and Alex Alderton, Roberto Amato, Jeffrey Barrett, Sonia Goncalves, Ewan Harrison, David K. Jackson, Ian Johnston, Dominic Kwiatkowski, Cordelia Langford, John Sillitoe on behalf of the Wellcome Sanger Institute COVID-19 Surveillance Team |
| EPI_ISL_2912253, EPI_ISL_2912254 | Lighthouse Lab in Glasgow | Wellcome Sanger Institute for the COVID-19 Genomics UK (COG-UK) Consortium | Harper VanSteenhouse, Yumi Kasai, David Gray, Carol Clugston, Anna Dominiczak and Alex Alderton, Roberto Amato, Jeffrey Barrett, Sonia Goncalves, Ewan Harrison, David K. Jackson, Ian Johnston, Dominic Kwiatkowski, Cordelia Langford, John Sillitoe on behalf of the Wellcome Sanger Institute COVID-19 Surveillance Team |
| EPI_ISL_2912255, EPI_ISL_2912256 | Berkshire and Surrey Pathology Services Lighthouse Laboratory | Wellcome Sanger Institute for the COVID-19 Genomics UK (COG-UK) Consortium | Berkshire and Surrey Pathology Services Lighthouse Laboratory and Alex Alderton, Roberto Amato, Jeffrey Barrett, Sonia Goncalves, Ewan Harrison, David K. Jackson, Ian Johnston, Dominic Kwiatkowski, Cordelia Langford, John Sillitoe on behalf of the Wellcome Sanger Institute COVID-19 Surveillance Team |
| EPI_ISL_2912257, EPI_ISL_2912258, EPI_ISL_2912259 | Lighthouse Lab in Milton Keynes | Wellcome Sanger Institute for the COVID-19 Genomics UK (COG-UK) Consortium | The Lighthouse Lab in Milton Keynes and Alex Alderton, Roberto Amato, Jeffrey Barrett, Sonia Goncalves, Ewan Harrison, David K. Jackson, Ian Johnston, Dominic Kwiatkowski, Cordelia Langford, John Sillitoe on behalf of the Wellcome Sanger Institute COVID-19 Surveillance Team |
| EPI_ISL_2912260 | Berkshire and Surrey Pathology Services Lighthouse Laboratory | Wellcome Sanger Institute for the COVID-19 Genomics UK (COG-UK) Consortium | Berkshire and Surrey Pathology Services Lighthouse Laboratory and Alex Alderton, Roberto Amato, Jeffrey Barrett, Sonia Goncalves, Ewan Harrison, David K. Jackson, Ian Johnston, Dominic Kwiatkowski, Cordelia Langford, John Sillitoe on behalf of the Wellcome Sanger Institute COVID-19 Surveillance Team |
| EPI_ISL_2912261 | Lighthouse Lab in Milton Keynes | Wellcome Sanger Institute for the COVID-19 Genomics UK (COG-UK) Consortium | The Lighthouse Lab in Milton Keynes and Alex Alderton, Roberto Amato, Jeffrey Barrett, Sonia Goncalves, Ewan Harrison, David K. Jackson, Ian Johnston, Dominic Kwiatkowski, Cordelia Langford, John Sillitoe on behalf of the Wellcome Sanger Institute COVID-19 Surveillance Team |
| EPI_ISL_2912262 | Lighthouse Lab in Glasgow | Wellcome Sanger Institute for the COVID-19 Genomics UK (COG-UK) Consortium | Harper VanSteenhouse, Yumi Kasai, David Gray, Carol Clugston, Anna Dominiczak and Alex Alderton, Roberto Amato, Jeffrey Barrett, Sonia Goncalves, Ewan Harrison, David K. Jackson, Ian Johnston, Dominic Kwiatkowski, Cordelia Langford, John Sillitoe on behalf of the Wellcome Sanger Institute COVID-19 Surveillance Team |
| EPI_ISL_2912263 | Lighthouse Lab in Milton Keynes | Wellcome Sanger Institute for the COVID-19 Genomics UK (COG-UK) Consortium | The Lighthouse Lab in Milton Keynes and Alex Alderton, Roberto Amato, Jeffrey Barrett, Sonia Goncalves, Ewan Harrison, David K. Jackson, Ian Johnston, Dominic Kwiatkowski, Cordelia Langford, John Sillitoe on behalf of the Wellcome Sanger Institute COVID-19 Surveillance Team |
| EPI_ISL_2912264, EPI_ISL_2912265 | Berkshire and Surrey Pathology Services Lighthouse Laboratory | Wellcome Sanger Institute for the COVID-19 Genomics UK (COG-UK) Consortium | Berkshire and Surrey Pathology Services Lighthouse Laboratory and Alex Alderton, Roberto Amato, Jeffrey Barrett, Sonia Goncalves, Ewan Harrison, David K. Jackson, Ian Johnston, Dominic Kwiatkowski, Cordelia Langford, John Sillitoe on behalf of the Wellcome Sanger Institute COVID-19 Surveillance Team |
| EPI_ISL_2912266 | Lighthouse Lab in Milton Keynes | Wellcome Sanger Institute for the COVID-19 Genomics UK (COG-UK) Consortium | The Lighthouse Lab in Milton Keynes and Alex Alderton, Roberto Amato, Jeffrey Barrett, Sonia Goncalves, Ewan Harrison, David K. Jackson, Ian Johnston, Dominic Kwiatkowski, Cordelia Langford, John Sillitoe on behalf of the Wellcome Sanger Institute COVID-19 Surveillance Team |
| EPI_ISL_2912267 | Lighthouse Lab in Glasgow | Wellcome Sanger Institute for the COVID-19 Genomics UK (COG-UK) Consortium | Harper VanSteenhouse, Yumi Kasai, David Gray, Carol Clugston, Anna Dominiczak and Alex Alderton, Roberto Amato, Jeffrey Barrett, Sonia Goncalves, Ewan Harrison, David K. Jackson, Ian Johnston, Dominic Kwiatkowski, Cordelia Langford, John Sillitoe on behalf of the Wellcome Sanger Institute COVID-19 Surveillance Team |
| EPI_ISL_2912268, EPI_ISL_2912269 | Lighthouse Lab in Milton Keynes | Wellcome Sanger Institute for the COVID-19 Genomics UK (COG-UK) Consortium | The Lighthouse Lab in Milton Keynes and Alex Alderton, Roberto Amato, Jeffrey Barrett, Sonia Goncalves, Ewan Harrison, David K. Jackson, Ian Johnston, Dominic Kwiatkowski, Cordelia Langford, John Sillitoe on behalf of the Wellcome Sanger Institute COVID-19 Surveillance Team |
| EPI_ISL_2912270 | Berkshire and Surrey Pathology Services Lighthouse Laboratory | Wellcome Sanger Institute for the COVID-19 Genomics UK (COG-UK) Consortium | Berkshire and Surrey Pathology Services Lighthouse Laboratory and Alex Alderton, Roberto Amato, Jeffrey Barrett, Sonia Goncalves, Ewan Harrison, David K. Jackson, Ian Johnston, Dominic Kwiatkowski, Cordelia Langford, John Sillitoe on behalf of the Wellcome Sanger Institute COVID-19 Surveillance Team |
| EPI_ISL_2912271 | Lighthouse Lab in Glasgow | Wellcome Sanger Institute for the COVID-19 Genomics UK (COG-UK) Consortium | Harper VanSteenhouse, Yumi Kasai, David Gray, Carol Clugston, Anna Dominiczak and Alex Alderton, Roberto Amato, Jeffrey Barrett, Sonia Goncalves, Ewan Harrison, David K. Jackson, Ian Johnston, Dominic Kwiatkowski, Cordelia Langford, John Sillitoe on behalf of the Wellcome Sanger Institute COVID-19 Surveillance Team |
| EPI_ISL_2912272 | Lighthouse Lab in Milton Keynes | Wellcome Sanger Institute for the COVID-19 Genomics UK (COG-UK) Consortium | The Lighthouse Lab in Milton Keynes and Alex Alderton, Roberto Amato, Jeffrey Barrett, Sonia Goncalves, Ewan Harrison, David K. Jackson, Ian Johnston, Dominic Kwiatkowski, Cordelia Langford, John Sillitoe on behalf of the Wellcome Sanger Institute COVID-19 Surveillance Team |
| EPI_ISL_2912273, EPI_ISL_2912274 | Berkshire and Surrey Pathology Services Lighthouse Laboratory | Wellcome Sanger Institute for the COVID-19 Genomics UK (COG-UK) Consortium | Berkshire and Surrey Pathology Services Lighthouse Laboratory and Alex Alderton, Roberto Amato, Jeffrey Barrett, Sonia Goncalves, Ewan Harrison, David K. Jackson, Ian Johnston, Dominic Kwiatkowski, Cordelia Langford, John Sillitoe on behalf of the Wellcome Sanger Institute COVID-19 Surveillance Team |
| EPI_ISL_2912275 | Lighthouse Lab in Glasgow | Wellcome Sanger Institute for the COVID-19 Genomics UK (COG-UK) Consortium | Harper VanSteenhouse, Yumi Kasai, David Gray, Carol Clugston, Anna Dominiczak and Alex Alderton, Roberto Amato, Jeffrey Barrett, Sonia Goncalves, Ewan Harrison, David K. Jackson, Ian Johnston, Dominic Kwiatkowski, Cordelia Langford, John Sillitoe on behalf of the Wellcome Sanger Institute COVID-19 Surveillance Team |
| EPI_ISL_2912276, EPI_ISL_2912277, EPI_ISL_2912278, EPI_ISL_2912279 | Berkshire and Surrey Pathology Services Lighthouse Laboratory | Wellcome Sanger Institute for the COVID-19 Genomics UK (COG-UK) Consortium | Berkshire and Surrey Pathology Services Lighthouse Laboratory and Alex Alderton, Roberto Amato, Jeffrey Barrett, Sonia Goncalves, Ewan Harrison, David K. Jackson, Ian Johnston, Dominic Kwiatkowski, Cordelia Langford, John Sillitoe on behalf of the Wellcome Sanger Institute COVID-19 Surveillance Team |

| | | | |
|---|---|---|---|
| EPI_ISL_2912280, EPI_ISL_2912281, EPI_ISL_2912282 | Lighthouse Lab in Glasgow | Wellcome Sanger Institute for the COVID-19 Genomics UK (COG-UK) Consortium | Harper VanSteenhouse, Yumi Kasai, David Gray, Carol Clugston, Anna Dominiczak and Alex Alderton, Roberto Amato, Jeffrey Barrett, Sonia Goncalves, Ewan Harrison, David K. Jackson, Ian Johnston, Dominic Kwiatkowski, Cordelia Langford, John Sillitoe on behalf of the Wellcome Sanger Institute COVID-19 Surveillance Team |
| EPI_ISL_2912283, EPI_ISL_2912284 | Berkshire and Surrey Pathology Services Lighthouse Laboratory | Wellcome Sanger Institute for the COVID-19 Genomics UK (COG-UK) Consortium | Berkshire and Surrey Pathology Services Lighthouse Laboratory and Alex Alderton, Roberto Amato, Jeffrey Barrett, Sonia Goncalves, Ewan Harrison, David K. Jackson, Ian Johnston, Dominic Kwiatkowski, Cordelia Langford, John Sillitoe on behalf of the Wellcome Sanger Institute COVID-19 Surveillance Team |
| EPI_ISL_2912285 | Lighthouse Lab in Milton Keynes | Wellcome Sanger Institute for the COVID-19 Genomics UK (COG-UK) Consortium | The Lighthouse Lab in Milton Keynes and Alex Alderton, Roberto Amato, Jeffrey Barrett, Sonia Goncalves, Ewan Harrison, David K. Jackson, Ian Johnston, Dominic Kwiatkowski, Cordelia Langford, John Sillitoe on behalf of the Wellcome Sanger Institute COVID-19 Surveillance Team |
| EPI_ISL_2912286 | Berkshire and Surrey Pathology Services Lighthouse Laboratory | Wellcome Sanger Institute for the COVID-19 Genomics UK (COG-UK) Consortium | Berkshire and Surrey Pathology Services Lighthouse Laboratory and Alex Alderton, Roberto Amato, Jeffrey Barrett, Sonia Goncalves, Ewan Harrison, David K. Jackson, Ian Johnston, Dominic Kwiatkowski, Cordelia Langford, John Sillitoe on behalf of the Wellcome Sanger Institute COVID-19 Surveillance Team |
| EPI_ISL_2912287 | Lighthouse Lab in Glasgow | Wellcome Sanger Institute for the COVID-19 Genomics UK (COG-UK) Consortium | Harper VanSteenhouse, Yumi Kasai, David Gray, Carol Clugston, Anna Dominiczak and Alex Alderton, Roberto Amato, Jeffrey Barrett, Sonia Goncalves, Ewan Harrison, David K. Jackson, Ian Johnston, Dominic Kwiatkowski, Cordelia Langford, John Sillitoe on behalf of the Wellcome Sanger Institute COVID-19 Surveillance Team |
| EPI_ISL_2912288 | Berkshire and Surrey Pathology Services Lighthouse Laboratory | Wellcome Sanger Institute for the COVID-19 Genomics UK (COG-UK) Consortium | Berkshire and Surrey Pathology Services Lighthouse Laboratory and Alex Alderton, Roberto Amato, Jeffrey Barrett, Sonia Goncalves, Ewan Harrison, David K. Jackson, Ian Johnston, Dominic Kwiatkowski, Cordelia Langford, John Sillitoe on behalf of the Wellcome Sanger Institute COVID-19 Surveillance Team |
| EPI_ISL_2912289 | Lighthouse Lab in Milton Keynes | Wellcome Sanger Institute for the COVID-19 Genomics UK (COG-UK) Consortium | The Lighthouse Lab in Milton Keynes and Alex Alderton, Roberto Amato, Jeffrey Barrett, Sonia Goncalves, Ewan Harrison, David K. Jackson, Ian Johnston, Dominic Kwiatkowski, Cordelia Langford, John Sillitoe on behalf of the Wellcome Sanger Institute COVID-19 Surveillance Team |
| EPI_ISL_2912290, EPI_ISL_2912291 | Berkshire and Surrey Pathology Services Lighthouse Laboratory | Wellcome Sanger Institute for the COVID-19 Genomics UK (COG-UK) Consortium | Berkshire and Surrey Pathology Services Lighthouse Laboratory and Alex Alderton, Roberto Amato, Jeffrey Barrett, Sonia Goncalves, Ewan Harrison, David K. Jackson, Ian Johnston, Dominic Kwiatkowski, Cordelia Langford, John Sillitoe on behalf of the Wellcome Sanger Institute COVID-19 Surveillance Team |
| EPI_ISL_2912292, EPI_ISL_2912293, EPI_ISL_2912294, EPI_ISL_2912295 | Lighthouse Lab in Glasgow | Wellcome Sanger Institute for the COVID-19 Genomics UK (COG-UK) Consortium | Harper VanSteenhouse, Yumi Kasai, David Gray, Carol Clugston, Anna Dominiczak and Alex Alderton, Roberto Amato, Jeffrey Barrett, Sonia Goncalves, Ewan Harrison, David K. Jackson, Ian Johnston, Dominic Kwiatkowski, Cordelia Langford, John Sillitoe on behalf of the Wellcome Sanger Institute COVID-19 Surveillance Team |
| EPI_ISL_2912296, EPI_ISL_2912297 | Berkshire and Surrey Pathology Services Lighthouse Laboratory | Wellcome Sanger Institute for the COVID-19 Genomics UK (COG-UK) Consortium | Berkshire and Surrey Pathology Services Lighthouse Laboratory and Alex Alderton, Roberto Amato, Jeffrey Barrett, Sonia Goncalves, Ewan Harrison, David K. Jackson, Ian Johnston, Dominic Kwiatkowski, Cordelia Langford, John Sillitoe on behalf of the Wellcome Sanger Institute COVID-19 Surveillance Team |
| EPI_ISL_2912298, EPI_ISL_2912299, EPI_ISL_2912300 | Lighthouse Lab in Glasgow | Wellcome Sanger Institute for the COVID-19 Genomics UK (COG-UK) Consortium | Harper VanSteenhouse, Yumi Kasai, David Gray, Carol Clugston, Anna Dominiczak and Alex Alderton, Roberto Amato, Jeffrey Barrett, Sonia Goncalves, Ewan Harrison, David K. Jackson, Ian Johnston, Dominic Kwiatkowski, Cordelia Langford, John Sillitoe on behalf of the Wellcome Sanger Institute COVID-19 Surveillance Team |
| EPI_ISL_2912301, EPI_ISL_2912302 | Berkshire and Surrey Pathology Services Lighthouse Laboratory | Wellcome Sanger Institute for the COVID-19 Genomics UK (COG-UK) Consortium | Berkshire and Surrey Pathology Services Lighthouse Laboratory and Alex Alderton, Roberto Amato, Jeffrey Barrett, Sonia Goncalves, Ewan Harrison, David K. Jackson, Ian Johnston, Dominic Kwiatkowski, Cordelia Langford, John Sillitoe on behalf of the Wellcome Sanger Institute COVID-19 Surveillance Team |
| EPI_ISL_2912303, EPI_ISL_2912304 | Lighthouse Lab in Glasgow | Wellcome Sanger Institute for the COVID-19 Genomics UK (COG-UK) Consortium | Harper VanSteenhouse, Yumi Kasai, David Gray, Carol Clugston, Anna Dominiczak and Alex Alderton, Roberto Amato, Jeffrey Barrett, Sonia Goncalves, Ewan Harrison, David K. Jackson, Ian Johnston, Dominic Kwiatkowski, Cordelia Langford, John Sillitoe on behalf of the Wellcome Sanger Institute COVID-19 Surveillance Team |
| EPI_ISL_2912305 | Lighthouse Lab in Milton Keynes | Wellcome Sanger Institute for the COVID-19 Genomics UK (COG-UK) Consortium | The Lighthouse Lab in Milton Keynes and Alex Alderton, Roberto Amato, Jeffrey Barrett, Sonia Goncalves, Ewan Harrison, David K. Jackson, Ian Johnston, Dominic Kwiatkowski, Cordelia Langford, John Sillitoe on behalf of the Wellcome Sanger Institute COVID-19 Surveillance Team |
| EPI_ISL_2912306, EPI_ISL_2912307, EPI_ISL_2912308 | Lighthouse Lab in Glasgow | Wellcome Sanger Institute for the COVID-19 Genomics UK (COG-UK) Consortium | Harper VanSteenhouse, Yumi Kasai, David Gray, Carol Clugston, Anna Dominiczak and Alex Alderton, Roberto Amato, Jeffrey Barrett, Sonia Goncalves, Ewan Harrison, David K. Jackson, Ian Johnston, Dominic Kwiatkowski, Cordelia Langford, John Sillitoe on behalf of the Wellcome Sanger Institute COVID-19 Surveillance Team |
| EPI_ISL_2912309, EPI_ISL_2912310, EPI_ISL_2912311, EPI_ISL_2912312, EPI_ISL_2912313, EPI_ISL_2912314, EPI_ISL_2912315, EPI_ISL_2912316 | Berkshire and Surrey Pathology Services Lighthouse Laboratory | Wellcome Sanger Institute for the COVID-19 Genomics UK (COG-UK) Consortium | Berkshire and Surrey Pathology Services Lighthouse Laboratory and Alex Alderton, Roberto Amato, Jeffrey Barrett, Sonia Goncalves, Ewan Harrison, David K. Jackson, Ian Johnston, Dominic Kwiatkowski, Cordelia Langford, John Sillitoe on behalf of the Wellcome Sanger Institute COVID-19 Surveillance Team |
| EPI_ISL_2912317 | Lighthouse Lab in Milton Keynes | Wellcome Sanger Institute for the COVID-19 Genomics UK (COG-UK) Consortium | The Lighthouse Lab in Milton Keynes and Alex Alderton, Roberto Amato, Jeffrey Barrett, Sonia Goncalves, Ewan Harrison, David K. Jackson, Ian Johnston, Dominic Kwiatkowski, Cordelia Langford, John Sillitoe on behalf of the Wellcome Sanger Institute COVID-19 Surveillance Team |
| EPI_ISL_2912318 | Berkshire and Surrey Pathology Services Lighthouse Laboratory | Wellcome Sanger Institute for the COVID-19 Genomics UK (COG-UK) Consortium | Berkshire and Surrey Pathology Services Lighthouse Laboratory and Alex Alderton, Roberto Amato, Jeffrey Barrett, Sonia Goncalves, Ewan Harrison, David K. Jackson, Ian Johnston, Dominic Kwiatkowski, Cordelia Langford, John Sillitoe on behalf of the Wellcome Sanger Institute COVID-19 Surveillance Team |
| EPI_ISL_2912319 | Lighthouse Lab in Milton Keynes | Wellcome Sanger Institute for the COVID-19 Genomics UK (COG-UK) Consortium | The Lighthouse Lab in Milton Keynes and Alex Alderton, Roberto Amato, Jeffrey Barrett, Sonia Goncalves, Ewan Harrison, David K. Jackson, Ian Johnston, Dominic Kwiatkowski, Cordelia Langford, John Sillitoe on behalf of the Wellcome Sanger Institute COVID-19 Surveillance Team |
| EPI_ISL_2912320, EPI_ISL_2912321, EPI_ISL_2912322, EPI_ISL_2912323 | Berkshire and Surrey Pathology Services Lighthouse Laboratory | Wellcome Sanger Institute for the COVID-19 Genomics UK (COG-UK) Consortium | Berkshire and Surrey Pathology Services Lighthouse Laboratory and Alex Alderton, Roberto Amato, Jeffrey Barrett, Sonia Goncalves, Ewan Harrison, David K. Jackson, Ian Johnston, Dominic Kwiatkowski, Cordelia Langford, John Sillitoe on behalf of the Wellcome Sanger Institute COVID-19 Surveillance Team |
| EPI_ISL_2912324 | Lighthouse Lab in Milton Keynes | Wellcome Sanger Institute for the COVID-19 Genomics UK (COG-UK) Consortium | The Lighthouse Lab in Milton Keynes and Alex Alderton, Roberto Amato, Jeffrey Barrett, Sonia Goncalves, Ewan Harrison, David K. Jackson, Ian Johnston, Dominic Kwiatkowski, Cordelia Langford, John Sillitoe on behalf of the Wellcome Sanger Institute COVID-19 Surveillance Team |
| EPI_ISL_2912325 | Lighthouse Lab in Glasgow | Wellcome Sanger Institute for the COVID-19 Genomics UK (COG-UK) Consortium | Harper VanSteenhouse, Yumi Kasai, David Gray, Carol Clugston, Anna Dominiczak and Alex Alderton, Roberto Amato, Jeffrey Barrett, Sonia Goncalves, Ewan Harrison, David K. Jackson, Ian Johnston, Dominic Kwiatkowski, Cordelia Langford, John Sillitoe on behalf of the Wellcome Sanger Institute COVID-19 Surveillance Team |
| EPI_ISL_2912326 | Berkshire and Surrey Pathology Services Lighthouse Laboratory | Wellcome Sanger Institute for the COVID-19 Genomics UK (COG-UK) Consortium | Berkshire and Surrey Pathology Services Lighthouse Laboratory and Alex Alderton, Roberto Amato, Jeffrey Barrett, Sonia Goncalves, Ewan Harrison, David K. Jackson, Ian Johnston, Dominic Kwiatkowski, Cordelia Langford, John Sillitoe on behalf of the Wellcome Sanger Institute COVID-19 Surveillance Team |
| EPI_ISL_2912327 | Lighthouse Lab in Milton Keynes | Wellcome Sanger Institute for the COVID-19 Genomics UK (COG-UK) Consortium | The Lighthouse Lab in Milton Keynes and Alex Alderton, Roberto Amato, Jeffrey Barrett, Sonia Goncalves, Ewan Harrison, David K. Jackson, Ian Johnston, Dominic Kwiatkowski, Cordelia Langford, John Sillitoe on behalf of the Wellcome Sanger Institute COVID-19 Surveillance Team |
| EPI_ISL_2912328, EPI_ISL_2912329 | Berkshire and Surrey Pathology Services Lighthouse Laboratory | Wellcome Sanger Institute for the COVID-19 Genomics UK (COG-UK) Consortium | Berkshire and Surrey Pathology Services Lighthouse Laboratory and Alex Alderton, Roberto Amato, Jeffrey Barrett, Sonia Goncalves, Ewan Harrison, David K. Jackson, Ian Johnston, Dominic Kwiatkowski, Cordelia Langford, John Sillitoe on behalf of the Wellcome Sanger Institute COVID-19 Surveillance Team |
| EPI_ISL_2912330 | Lighthouse Lab in Milton Keynes | Wellcome Sanger Institute for the COVID-19 Genomics UK (COG-UK) Consortium | The Lighthouse Lab in Milton Keynes and Alex Alderton, Roberto Amato, Jeffrey Barrett, Sonia Goncalves, Ewan Harrison, David K. Jackson, Ian Johnston, Dominic Kwiatkowski, Cordelia Langford, John Sillitoe on behalf of the Wellcome Sanger Institute COVID-19 Surveillance Team |
| EPI_ISL_2912331 | Berkshire and Surrey Pathology Services Lighthouse Laboratory | Wellcome Sanger Institute for the COVID-19 Genomics UK (COG-UK) Consortium | Berkshire and Surrey Pathology Services Lighthouse Laboratory and Alex Alderton, Roberto Amato, Jeffrey Barrett, Sonia Goncalves, Ewan Harrison, David K. Jackson, Ian Johnston, Dominic Kwiatkowski, Cordelia Langford, John Sillitoe on behalf of the Wellcome Sanger Institute COVID-19 Surveillance Team |
| EPI_ISL_2912332 | Lighthouse Lab in Milton Keynes | Wellcome Sanger Institute for the COVID-19 Genomics UK (COG-UK) Consortium | The Lighthouse Lab in Milton Keynes and Alex Alderton, Roberto Amato, Jeffrey Barrett, Sonia Goncalves, Ewan Harrison, David K. Jackson, Ian Johnston, Dominic Kwiatkowski, Cordelia Langford, John Sillitoe on behalf of the Wellcome Sanger Institute COVID-19 Surveillance Team |
| EPI_ISL_2912333, EPI_ISL_2912334, EPI_ISL_2912335 | Berkshire and Surrey Pathology Services Lighthouse Laboratory | Wellcome Sanger Institute for the COVID-19 Genomics UK (COG-UK) Consortium | Berkshire and Surrey Pathology Services Lighthouse Laboratory and Alex Alderton, Roberto Amato, Jeffrey Barrett, Sonia Goncalves, Ewan Harrison, David K. Jackson, Ian Johnston, Dominic Kwiatkowski, Cordelia Langford, John Sillitoe on behalf of the Wellcome Sanger Institute COVID-19 Surveillance Team |
| EPI_ISL_2912336 | Lighthouse Lab in Milton Keynes | Wellcome Sanger Institute for the COVID-19 Genomics UK (COG-UK) Consortium | The Lighthouse Lab in Milton Keynes and Alex Alderton, Roberto Amato, Jeffrey Barrett, Sonia Goncalves, Ewan Harrison, David K. Jackson, Ian Johnston, Dominic Kwiatkowski, Cordelia Langford, John Sillitoe on behalf of the Wellcome Sanger Institute COVID-19 Surveillance Team |
| EPI_ISL_2912337 | Berkshire and Surrey Pathology Services Lighthouse Laboratory | Wellcome Sanger Institute for the COVID-19 Genomics UK (COG-UK) Consortium | Berkshire and Surrey Pathology Services Lighthouse Laboratory and Alex Alderton, Roberto Amato, Jeffrey Barrett, Sonia Goncalves, Ewan Harrison, David K. Jackson, Ian Johnston, Dominic Kwiatkowski, Cordelia Langford, John Sillitoe on behalf of the Wellcome Sanger Institute COVID-19 Surveillance Team |
| EPI_ISL_2912338 | Lighthouse Lab in Milton Keynes | Wellcome Sanger Institute for the COVID-19 Genomics UK (COG-UK) Consortium | The Lighthouse Lab in Milton Keynes and Alex Alderton, Roberto Amato, Jeffrey Barrett, Sonia Goncalves, Ewan Harrison, David K. Jackson, Ian Johnston, Dominic Kwiatkowski, Cordelia Langford, John Sillitoe on behalf of the Wellcome Sanger Institute COVID-19 Surveillance Team |
| EPI_ISL_2912339, EPI_ISL_2912340 | Lighthouse Lab in Glasgow | Wellcome Sanger Institute for the COVID-19 Genomics UK (COG-UK) Consortium | Harper VanSteenhouse, Yumi Kasai, David Gray, Carol Clugston, Anna Dominiczak and Alex Alderton, Roberto Amato, Jeffrey Barrett, Sonia Goncalves, Ewan Harrison, David K. Jackson, Ian Johnston, Dominic Kwiatkowski, Cordelia Langford, John Sillitoe on behalf of the Wellcome Sanger Institute COVID-19 Surveillance Team |
| EPI_ISL_2912341, EPI_ISL_2912342, EPI_ISL_2912343, EPI_ISL_2912344 | Berkshire and Surrey Pathology Services Lighthouse Laboratory | Wellcome Sanger Institute for the COVID-19 Genomics UK (COG-UK) Consortium | Berkshire and Surrey Pathology Services Lighthouse Laboratory and Alex Alderton, Roberto Amato, Jeffrey Barrett, Sonia Goncalves, Ewan Harrison, David K. Jackson, Ian Johnston, Dominic Kwiatkowski, Cordelia Langford, John Sillitoe on behalf of the Wellcome Sanger Institute COVID-19 Surveillance Team |
| EPI_ISL_2912345, EPI_ISL_2912346 | Lighthouse Lab in Milton Keynes | Wellcome Sanger Institute for the COVID-19 Genomics UK (COG-UK) Consortium | The Lighthouse Lab in Milton Keynes and Alex Alderton, Roberto Amato, Jeffrey Barrett, Sonia Goncalves, Ewan Harrison, David K. Jackson, Ian Johnston, Dominic Kwiatkowski, Cordelia Langford, John Sillitoe on behalf of the Wellcome Sanger Institute COVID-19 Surveillance Team |
| EPI_ISL_2912347, EPI_ISL_2912348 | Berkshire and Surrey Pathology Services Lighthouse Laboratory | Wellcome Sanger Institute for the COVID-19 Genomics UK (COG-UK) Consortium | Berkshire and Surrey Pathology Services Lighthouse Laboratory and Alex Alderton, Roberto Amato, Jeffrey Barrett, Sonia Goncalves, Ewan Harrison, David K. Jackson, Ian Johnston, Dominic Kwiatkowski, Cordelia Langford, John Sillitoe on behalf of the Wellcome Sanger Institute COVID-19 Surveillance Team |
| EPI_ISL_2912349, EPI_ISL_2912350 | Lighthouse Lab in Milton Keynes | Wellcome Sanger Institute for the COVID-19 Genomics UK (COG-UK) Consortium | The Lighthouse Lab in Milton Keynes and Alex Alderton, Roberto Amato, Jeffrey Barrett, Sonia Goncalves, Ewan Harrison, David K. Jackson, Ian Johnston, Dominic Kwiatkowski, Cordelia Langford, John Sillitoe on behalf of the Wellcome Sanger Institute COVID-19 Surveillance Team |
| EPI_ISL_2912351 | Berkshire and Surrey Pathology Services Lighthouse Laboratory | Wellcome Sanger Institute for the COVID-19 Genomics UK (COG-UK) Consortium | Berkshire and Surrey Pathology Services Lighthouse Laboratory and Alex Alderton, Roberto Amato, Jeffrey Barrett, Sonia Goncalves, Ewan Harrison, David K. Jackson, Ian Johnston, Dominic Kwiatkowski, Cordelia Langford, John Sillitoe on behalf of the Wellcome Sanger Institute COVID-19 Surveillance Team |
| EPI_ISL_2912352 | Lighthouse Lab in Milton Keynes | Wellcome Sanger Institute for the COVID-19 Genomics UK (COG-UK) | The Lighthouse Lab in Milton Keynes and Alex Alderton, Roberto Amato, Jeffrey Barrett, Sonia Goncalves, Ewan Harrison, David K. Jackson, Ian Johnston, Dominic Kwiatkowski, Cordelia Langford, John Sillitoe on behalf of the Wellcome Sanger |

| | | | |
|---|---|---|---|
| | | Consortium | Institute COVID-19 Surveillance Team |
| EPI_ISL_2912353 | Berkshire and Surrey Pathology Services Lighthouse Laboratory | Wellcome Sanger Institute for the COVID-19 Genomics UK (COG-UK) Consortium | Berkshire and Surrey Pathology Services Lighthouse Laboratory and Alex Alderton, Roberto Amato, Jeffrey Barrett, Sonia Goncalves, Ewan Harrison, David K. Jackson, Ian Johnston, Dominic Kwiatkowski, Cordelia Langford, John Sillitoe on behalf of the Wellcome Sanger Institute COVID-19 Surveillance Team |
| EPI_ISL_2912354 | Lighthouse Lab in Milton Keynes | Wellcome Sanger Institute for the COVID-19 Genomics UK (COG-UK) Consortium | The Lighthouse Lab in Milton Keynes and Alex Alderton, Roberto Amato, Jeffrey Barrett, Sonia Goncalves, Ewan Harrison, David K. Jackson, Ian Johnston, Dominic Kwiatkowski, Cordelia Langford, John Sillitoe on behalf of the Wellcome Sanger Institute COVID-19 Surveillance Team |
| EPI_ISL_2912355, EPI_ISL_2912356 | Lighthouse Lab in Glasgow | Wellcome Sanger Institute for the COVID-19 Genomics UK (COG-UK) Consortium | Harper VanSteenhouse, Yumi Kasai, David Gray, Carol Clugston, Anna Dominiczak and Alex Alderton, Roberto Amato, Jeffrey Barrett, Sonia Goncalves, Ewan Harrison, David K. Jackson, Ian Johnston, Dominic Kwiatkowski, Cordelia Langford, John Sillitoe on behalf of the Wellcome Sanger Institute COVID-19 Surveillance Team |
| EPI_ISL_2912357, EPI_ISL_2912358, EPI_ISL_2912359 | Berkshire and Surrey Pathology Services Lighthouse Laboratory | Wellcome Sanger Institute for the COVID-19 Genomics UK (COG-UK) Consortium | Berkshire and Surrey Pathology Services Lighthouse Laboratory and Alex Alderton, Roberto Amato, Jeffrey Barrett, Sonia Goncalves, Ewan Harrison, David K. Jackson, Ian Johnston, Dominic Kwiatkowski, Cordelia Langford, John Sillitoe on behalf of the Wellcome Sanger Institute COVID-19 Surveillance Team |
| EPI_ISL_2912360, EPI_ISL_2912361, EPI_ISL_2912362 | Lighthouse Lab in Milton Keynes | Wellcome Sanger Institute for the COVID-19 Genomics UK (COG-UK) Consortium | The Lighthouse Lab in Milton Keynes and Alex Alderton, Roberto Amato, Jeffrey Barrett, Sonia Goncalves, Ewan Harrison, David K. Jackson, Ian Johnston, Dominic Kwiatkowski, Cordelia Langford, John Sillitoe on behalf of the Wellcome Sanger Institute COVID-19 Surveillance Team |
| EPI_ISL_2912363, EPI_ISL_2912364 | Berkshire and Surrey Pathology Services Lighthouse Laboratory | Wellcome Sanger Institute for the COVID-19 Genomics UK (COG-UK) Consortium | Berkshire and Surrey Pathology Services Lighthouse Laboratory and Alex Alderton, Roberto Amato, Jeffrey Barrett, Sonia Goncalves, Ewan Harrison, David K. Jackson, Ian Johnston, Dominic Kwiatkowski, Cordelia Langford, John Sillitoe on behalf of the Wellcome Sanger Institute COVID-19 Surveillance Team |
| EPI_ISL_2912365, EPI_ISL_2912366 | Lighthouse Lab in Milton Keynes | Wellcome Sanger Institute for the COVID-19 Genomics UK (COG-UK) Consortium | The Lighthouse Lab in Milton Keynes and Alex Alderton, Roberto Amato, Jeffrey Barrett, Sonia Goncalves, Ewan Harrison, David K. Jackson, Ian Johnston, Dominic Kwiatkowski, Cordelia Langford, John Sillitoe on behalf of the Wellcome Sanger Institute COVID-19 Surveillance Team |
| EPI_ISL_2912367, EPI_ISL_2912368 | Berkshire and Surrey Pathology Services Lighthouse Laboratory | Wellcome Sanger Institute for the COVID-19 Genomics UK (COG-UK) Consortium | Berkshire and Surrey Pathology Services Lighthouse Laboratory and Alex Alderton, Roberto Amato, Jeffrey Barrett, Sonia Goncalves, Ewan Harrison, David K. Jackson, Ian Johnston, Dominic Kwiatkowski, Cordelia Langford, John Sillitoe on behalf of the Wellcome Sanger Institute COVID-19 Surveillance Team |
| EPI_ISL_2912369 | Lighthouse Lab in Milton Keynes | Wellcome Sanger Institute for the COVID-19 Genomics UK (COG-UK) Consortium | The Lighthouse Lab in Milton Keynes and Alex Alderton, Roberto Amato, Jeffrey Barrett, Sonia Goncalves, Ewan Harrison, David K. Jackson, Ian Johnston, Dominic Kwiatkowski, Cordelia Langford, John Sillitoe on behalf of the Wellcome Sanger Institute COVID-19 Surveillance Team |
| EPI_ISL_2912370 | Berkshire and Surrey Pathology Services Lighthouse Laboratory | Wellcome Sanger Institute for the COVID-19 Genomics UK (COG-UK) Consortium | Berkshire and Surrey Pathology Services Lighthouse Laboratory and Alex Alderton, Roberto Amato, Jeffrey Barrett, Sonia Goncalves, Ewan Harrison, David K. Jackson, Ian Johnston, Dominic Kwiatkowski, Cordelia Langford, John Sillitoe on behalf of the Wellcome Sanger Institute COVID-19 Surveillance Team |
| EPI_ISL_2912371 | Lighthouse Lab in Milton Keynes | Wellcome Sanger Institute for the COVID-19 Genomics UK (COG-UK) Consortium | The Lighthouse Lab in Milton Keynes and Alex Alderton, Roberto Amato, Jeffrey Barrett, Sonia Goncalves, Ewan Harrison, David K. Jackson, Ian Johnston, Dominic Kwiatkowski, Cordelia Langford, John Sillitoe on behalf of the Wellcome Sanger Institute COVID-19 Surveillance Team |
| EPI_ISL_2912372 | Berkshire and Surrey Pathology Services Lighthouse Laboratory | Wellcome Sanger Institute for the COVID-19 Genomics UK (COG-UK) Consortium | Berkshire and Surrey Pathology Services Lighthouse Laboratory and Alex Alderton, Roberto Amato, Jeffrey Barrett, Sonia Goncalves, Ewan Harrison, David K. Jackson, Ian Johnston, Dominic Kwiatkowski, Cordelia Langford, John Sillitoe on behalf of the Wellcome Sanger Institute COVID-19 Surveillance Team |
| EPI_ISL_2912373 | Lighthouse Lab in Milton Keynes | Wellcome Sanger Institute for the COVID-19 Genomics UK (COG-UK) Consortium | The Lighthouse Lab in Milton Keynes and Alex Alderton, Roberto Amato, Jeffrey Barrett, Sonia Goncalves, Ewan Harrison, David K. Jackson, Ian Johnston, Dominic Kwiatkowski, Cordelia Langford, John Sillitoe on behalf of the Wellcome Sanger Institute COVID-19 Surveillance Team |
| EPI_ISL_2912374, EPI_ISL_2912375 | Berkshire and Surrey Pathology Services Lighthouse Laboratory | Wellcome Sanger Institute for the COVID-19 Genomics UK (COG-UK) Consortium | Berkshire and Surrey Pathology Services Lighthouse Laboratory and Alex Alderton, Roberto Amato, Jeffrey Barrett, Sonia Goncalves, Ewan Harrison, David K. Jackson, Ian Johnston, Dominic Kwiatkowski, Cordelia Langford, John Sillitoe on behalf of the Wellcome Sanger Institute COVID-19 Surveillance Team |
| EPI_ISL_2912376, EPI_ISL_2912377 | Lighthouse Lab in Glasgow | Wellcome Sanger Institute for the COVID-19 Genomics UK (COG-UK) Consortium | Harper VanSteenhouse, Yumi Kasai, David Gray, Carol Clugston, Anna Dominiczak and Alex Alderton, Roberto Amato, Jeffrey Barrett, Sonia Goncalves, Ewan Harrison, David K. Jackson, Ian Johnston, Dominic Kwiatkowski, Cordelia Langford, John Sillitoe on behalf of the Wellcome Sanger Institute COVID-19 Surveillance Team |
| EPI_ISL_2912378 | Berkshire and Surrey Pathology Services Lighthouse Laboratory | Wellcome Sanger Institute for the COVID-19 Genomics UK (COG-UK) Consortium | Berkshire and Surrey Pathology Services Lighthouse Laboratory and Alex Alderton, Roberto Amato, Jeffrey Barrett, Sonia Goncalves, Ewan Harrison, David K. Jackson, Ian Johnston, Dominic Kwiatkowski, Cordelia Langford, John Sillitoe on behalf of the Wellcome Sanger Institute COVID-19 Surveillance Team |
| EPI_ISL_2912379 | Lighthouse Lab in Milton Keynes | Wellcome Sanger Institute for the COVID-19 Genomics UK (COG-UK) Consortium | The Lighthouse Lab in Milton Keynes and Alex Alderton, Roberto Amato, Jeffrey Barrett, Sonia Goncalves, Ewan Harrison, David K. Jackson, Ian Johnston, Dominic Kwiatkowski, Cordelia Langford, John Sillitoe on behalf of the Wellcome Sanger Institute COVID-19 Surveillance Team |
| EPI_ISL_2912380, EPI_ISL_2912381 | Lighthouse Lab in Glasgow | Wellcome Sanger Institute for the COVID-19 Genomics UK (COG-UK) Consortium | Harper VanSteenhouse, Yumi Kasai, David Gray, Carol Clugston, Anna Dominiczak and Alex Alderton, Roberto Amato, Jeffrey Barrett, Sonia Goncalves, Ewan Harrison, David K. Jackson, Ian Johnston, Dominic Kwiatkowski, Cordelia Langford, John Sillitoe on behalf of the Wellcome Sanger Institute COVID-19 Surveillance Team |
| EPI_ISL_2912382 | Lighthouse Lab in Milton Keynes | Wellcome Sanger Institute for the COVID-19 Genomics UK (COG-UK) Consortium | The Lighthouse Lab in Milton Keynes and Alex Alderton, Roberto Amato, Jeffrey Barrett, Sonia Goncalves, Ewan Harrison, David K. Jackson, Ian Johnston, Dominic Kwiatkowski, Cordelia Langford, John Sillitoe on behalf of the Wellcome Sanger Institute COVID-19 Surveillance Team |
| EPI_ISL_2912383 | Berkshire and Surrey Pathology Services Lighthouse Laboratory | Wellcome Sanger Institute for the COVID-19 Genomics UK (COG-UK) Consortium | Berkshire and Surrey Pathology Services Lighthouse Laboratory and Alex Alderton, Roberto Amato, Jeffrey Barrett, Sonia Goncalves, Ewan Harrison, David K. Jackson, Ian Johnston, Dominic Kwiatkowski, Cordelia Langford, John Sillitoe on behalf of the Wellcome Sanger Institute COVID-19 Surveillance Team |
| EPI_ISL_2912384 | Lighthouse Lab in Milton Keynes | Wellcome Sanger Institute for the COVID-19 Genomics UK (COG-UK) Consortium | The Lighthouse Lab in Milton Keynes and Alex Alderton, Roberto Amato, Jeffrey Barrett, Sonia Goncalves, Ewan Harrison, David K. Jackson, Ian Johnston, Dominic Kwiatkowski, Cordelia Langford, John Sillitoe on behalf of the Wellcome Sanger Institute COVID-19 Surveillance Team |
| EPI_ISL_2912385 | Berkshire and Surrey Pathology Services Lighthouse Laboratory | Wellcome Sanger Institute for the COVID-19 Genomics UK (COG-UK) Consortium | Berkshire and Surrey Pathology Services Lighthouse Laboratory and Alex Alderton, Roberto Amato, Jeffrey Barrett, Sonia Goncalves, Ewan Harrison, David K. Jackson, Ian Johnston, Dominic Kwiatkowski, Cordelia Langford, John Sillitoe on behalf of the Wellcome Sanger Institute COVID-19 Surveillance Team |
| EPI_ISL_2912386 | Lighthouse Lab in Glasgow | Wellcome Sanger Institute for the COVID-19 Genomics UK (COG-UK) Consortium | Harper VanSteenhouse, Yumi Kasai, David Gray, Carol Clugston, Anna Dominiczak and Alex Alderton, Roberto Amato, Jeffrey Barrett, Sonia Goncalves, Ewan Harrison, David K. Jackson, Ian Johnston, Dominic Kwiatkowski, Cordelia Langford, John Sillitoe on behalf of the Wellcome Sanger Institute COVID-19 Surveillance Team |
| EPI_ISL_2912387 | Berkshire and Surrey Pathology Services Lighthouse Laboratory | Wellcome Sanger Institute for the COVID-19 Genomics UK (COG-UK) Consortium | Berkshire and Surrey Pathology Services Lighthouse Laboratory and Alex Alderton, Roberto Amato, Jeffrey Barrett, Sonia Goncalves, Ewan Harrison, David K. Jackson, Ian Johnston, Dominic Kwiatkowski, Cordelia Langford, John Sillitoe on behalf of the Wellcome Sanger Institute COVID-19 Surveillance Team |
| EPI_ISL_2912388 | Lighthouse Lab in Glasgow | Wellcome Sanger Institute for the COVID-19 Genomics UK (COG-UK) Consortium | Harper VanSteenhouse, Yumi Kasai, David Gray, Carol Clugston, Anna Dominiczak and Alex Alderton, Roberto Amato, Jeffrey Barrett, Sonia Goncalves, Ewan Harrison, David K. Jackson, Ian Johnston, Dominic Kwiatkowski, Cordelia Langford, John Sillitoe on behalf of the Wellcome Sanger Institute COVID-19 Surveillance Team |
| EPI_ISL_2912389, EPI_ISL_2912390, EPI_ISL_2912391 | Berkshire and Surrey Pathology Services Lighthouse Laboratory | Wellcome Sanger Institute for the COVID-19 Genomics UK (COG-UK) Consortium | Berkshire and Surrey Pathology Services Lighthouse Laboratory and Alex Alderton, Roberto Amato, Jeffrey Barrett, Sonia Goncalves, Ewan Harrison, David K. Jackson, Ian Johnston, Dominic Kwiatkowski, Cordelia Langford, John Sillitoe on behalf of the Wellcome Sanger Institute COVID-19 Surveillance Team |
| EPI_ISL_2912392 | Lighthouse Lab in Milton Keynes | Wellcome Sanger Institute for the COVID-19 Genomics UK (COG-UK) Consortium | The Lighthouse Lab in Milton Keynes and Alex Alderton, Roberto Amato, Jeffrey Barrett, Sonia Goncalves, Ewan Harrison, David K. Jackson, Ian Johnston, Dominic Kwiatkowski, Cordelia Langford, John Sillitoe on behalf of the Wellcome Sanger Institute COVID-19 Surveillance Team |
| EPI_ISL_2912393 | Berkshire and Surrey Pathology Services Lighthouse Laboratory | Wellcome Sanger Institute for the COVID-19 Genomics UK (COG-UK) Consortium | Berkshire and Surrey Pathology Services Lighthouse Laboratory and Alex Alderton, Roberto Amato, Jeffrey Barrett, Sonia Goncalves, Ewan Harrison, David K. Jackson, Ian Johnston, Dominic Kwiatkowski, Cordelia Langford, John Sillitoe on behalf of the Wellcome Sanger Institute COVID-19 Surveillance Team |
| EPI_ISL_2912394 | Lighthouse Lab in Milton Keynes | Wellcome Sanger Institute for the COVID-19 Genomics UK (COG-UK) Consortium | The Lighthouse Lab in Milton Keynes and Alex Alderton, Roberto Amato, Jeffrey Barrett, Sonia Goncalves, Ewan Harrison, David K. Jackson, Ian Johnston, Dominic Kwiatkowski, Cordelia Langford, John Sillitoe on behalf of the Wellcome Sanger Institute COVID-19 Surveillance Team |
| EPI_ISL_2912395, EPI_ISL_2912396 | Berkshire and Surrey Pathology Services Lighthouse Laboratory | Wellcome Sanger Institute for the COVID-19 Genomics UK (COG-UK) Consortium | Berkshire and Surrey Pathology Services Lighthouse Laboratory and Alex Alderton, Roberto Amato, Jeffrey Barrett, Sonia Goncalves, Ewan Harrison, David K. Jackson, Ian Johnston, Dominic Kwiatkowski, Cordelia Langford, John Sillitoe on behalf of the Wellcome Sanger Institute COVID-19 Surveillance Team |
| EPI_ISL_2912397 | Lighthouse Lab in Milton Keynes | Wellcome Sanger Institute for the COVID-19 Genomics UK (COG-UK) Consortium | The Lighthouse Lab in Milton Keynes and Alex Alderton, Roberto Amato, Jeffrey Barrett, Sonia Goncalves, Ewan Harrison, David K. Jackson, Ian Johnston, Dominic Kwiatkowski, Cordelia Langford, John Sillitoe on behalf of the Wellcome Sanger Institute COVID-19 Surveillance Team |
| EPI_ISL_2912398, EPI_ISL_2912399, EPI_ISL_2912400 | Berkshire and Surrey Pathology Services Lighthouse Laboratory | Wellcome Sanger Institute for the COVID-19 Genomics UK (COG-UK) Consortium | Berkshire and Surrey Pathology Services Lighthouse Laboratory and Alex Alderton, Roberto Amato, Jeffrey Barrett, Sonia Goncalves, Ewan Harrison, David K. Jackson, Ian Johnston, Dominic Kwiatkowski, Cordelia Langford, John Sillitoe on behalf of the Wellcome Sanger Institute COVID-19 Surveillance Team |
| EPI_ISL_2912401, EPI_ISL_2912402, EPI_ISL_2912403 | Lighthouse Lab in Glasgow | Wellcome Sanger Institute for the COVID-19 Genomics UK (COG-UK) Consortium | Harper VanSteenhouse, Yumi Kasai, David Gray, Carol Clugston, Anna Dominiczak and Alex Alderton, Roberto Amato, Jeffrey Barrett, Sonia Goncalves, Ewan Harrison, David K. Jackson, Ian Johnston, Dominic Kwiatkowski, Cordelia Langford, John Sillitoe on behalf of the Wellcome Sanger Institute COVID-19 Surveillance Team |
| EPI_ISL_2912404 | Lighthouse Lab in Milton Keynes | Wellcome Sanger Institute for the COVID-19 Genomics UK (COG-UK) Consortium | The Lighthouse Lab in Milton Keynes and Alex Alderton, Roberto Amato, Jeffrey Barrett, Sonia Goncalves, Ewan Harrison, David K. Jackson, Ian Johnston, Dominic Kwiatkowski, Cordelia Langford, John Sillitoe on behalf of the Wellcome Sanger Institute COVID-19 Surveillance Team |
| EPI_ISL_2912405 | Lighthouse Lab in Glasgow | Wellcome Sanger Institute for the COVID-19 Genomics UK (COG-UK) Consortium | Harper VanSteenhouse, Yumi Kasai, David Gray, Carol Clugston, Anna Dominiczak and Alex Alderton, Roberto Amato, Jeffrey Barrett, Sonia Goncalves, Ewan Harrison, David K. Jackson, Ian Johnston, Dominic Kwiatkowski, Cordelia Langford, John Sillitoe on behalf of the Wellcome Sanger Institute COVID-19 Surveillance Team |
| EPI_ISL_2912406, EPI_ISL_2912407, EPI_ISL_2912408, EPI_ISL_2912409 | Lighthouse Lab in Milton Keynes | Wellcome Sanger Institute for the COVID-19 Genomics UK (COG-UK) Consortium | The Lighthouse Lab in Milton Keynes and Alex Alderton, Roberto Amato, Jeffrey Barrett, Sonia Goncalves, Ewan Harrison, David K. Jackson, Ian Johnston, Dominic Kwiatkowski, Cordelia Langford, John Sillitoe on behalf of the Wellcome Sanger Institute COVID-19 Surveillance Team |
| EPI_ISL_2912410, EPI_ISL_2912411 | Berkshire and Surrey Pathology Services Lighthouse Laboratory | Wellcome Sanger Institute for the COVID-19 Genomics UK (COG-UK) Consortium | Berkshire and Surrey Pathology Services Lighthouse Laboratory and Alex Alderton, Roberto Amato, Jeffrey Barrett, Sonia Goncalves, Ewan Harrison, David K. Jackson, Ian Johnston, Dominic Kwiatkowski, Cordelia Langford, John Sillitoe on behalf of the Wellcome Sanger Institute COVID-19 Surveillance Team |
| EPI_ISL_2912412 | Lighthouse Lab in Glasgow | Wellcome Sanger Institute for the COVID-19 Genomics UK (COG-UK) Consortium | Harper VanSteenhouse, Yumi Kasai, David Gray, Carol Clugston, Anna Dominiczak and Alex Alderton, Roberto Amato, Jeffrey Barrett, Sonia Goncalves, Ewan Harrison, David K. Jackson, Ian Johnston, Dominic Kwiatkowski, Cordelia Langford, John Sillitoe on behalf of the Wellcome Sanger Institute COVID-19 Surveillance Team |
| EPI_ISL_2912413, EPI_ISL_2912414, EPI_ISL_2912415 | Berkshire and Surrey Pathology Services Lighthouse Laboratory | Wellcome Sanger Institute for the COVID-19 Genomics UK (COG-UK) Consortium | Berkshire and Surrey Pathology Services Lighthouse Laboratory and Alex Alderton, Roberto Amato, Jeffrey Barrett, Sonia Goncalves, Ewan Harrison, David K. Jackson, Ian Johnston, Dominic Kwiatkowski, Cordelia Langford, John Sillitoe on behalf of the Wellcome Sanger Institute COVID-19 Surveillance Team |
| EPI_ISL_2912416 | Lighthouse Lab in Milton Keynes | Wellcome Sanger Institute for the COVID-19 Genomics UK (COG-UK) Consortium | The Lighthouse Lab in Milton Keynes and Alex Alderton, Roberto Amato, Jeffrey Barrett, Sonia Goncalves, Ewan Harrison, David K. Jackson, Ian Johnston, Dominic Kwiatkowski, Cordelia Langford, John Sillitoe on behalf of the Wellcome Sanger Institute COVID-19 Surveillance Team |
| EPI_ISL_2912417 | Berkshire and Surrey Pathology Services Lighthouse Laboratory | Wellcome Sanger Institute for the COVID-19 Genomics UK (COG-UK) Consortium | Berkshire and Surrey Pathology Services Lighthouse Laboratory and Alex Alderton, Roberto Amato, Jeffrey Barrett, Sonia Goncalves, Ewan Harrison, David K. Jackson, Ian Johnston, Dominic Kwiatkowski, Cordelia Langford, John Sillitoe on behalf of the Wellcome Sanger Institute COVID-19 Surveillance Team |
| EPI_ISL_2912418 | Lighthouse Lab in Glasgow | Wellcome Sanger Institute for the COVID-19 Genomics UK (COG-UK) | Harper VanSteenhouse, Yumi Kasai, David Gray, Carol Clugston, Anna Dominiczak and Alex Alderton, Roberto Amato, Jeffrey Barrett, Sonia Goncalves, Ewan Harrison, David K. Jackson, Ian Johnston, Dominic Kwiatkowski, Cordelia Langford, |

| | | | |
|---|---|---|---|
| | | Consortium | John Sillitoe on behalf of the Wellcome Sanger Institute COVID-19 Surveillance Team |
| EPI_ISL_2912419, EPI_ISL_2912420 | Berkshire and Surrey Pathology Services Lighthouse Laboratory | Wellcome Sanger Institute for the COVID-19 Genomics UK (COG-UK) Consortium | Berkshire and Surrey Pathology Services Lighthouse Laboratory and Alex Alderton, Roberto Amato, Jeffrey Barrett, Sonia Goncalves, Ewan Harrison, David K. Jackson, Ian Johnston, Dominic Kwiatkowski, Cordelia Langford, John Sillitoe on behalf of the Wellcome Sanger Institute COVID-19 Surveillance Team |
| EPI_ISL_2912421 | Lighthouse Lab in Milton Keynes | Wellcome Sanger Institute for the COVID-19 Genomics UK (COG-UK) Consortium | The Lighthouse Lab in Milton Keynes and Alex Alderton, Roberto Amato, Jeffrey Barrett, Sonia Goncalves, Ewan Harrison, David K. Jackson, Ian Johnston, Dominic Kwiatkowski, Cordelia Langford, John Sillitoe on behalf of the Wellcome Sanger Institute COVID-19 Surveillance Team |
| EPI_ISL_2912422 | Berkshire and Surrey Pathology Services Lighthouse Laboratory | Wellcome Sanger Institute for the COVID-19 Genomics UK (COG-UK) Consortium | Berkshire and Surrey Pathology Services Lighthouse Laboratory and Alex Alderton, Roberto Amato, Jeffrey Barrett, Sonia Goncalves, Ewan Harrison, David K. Jackson, Ian Johnston, Dominic Kwiatkowski, Cordelia Langford, John Sillitoe on behalf of the Wellcome Sanger Institute COVID-19 Surveillance Team |
| EPI_ISL_2912423, EPI_ISL_2912424, EPI_ISL_2912425 | Lighthouse Lab in Milton Keynes | Wellcome Sanger Institute for the COVID-19 Genomics UK (COG-UK) Consortium | The Lighthouse Lab in Milton Keynes and Alex Alderton, Roberto Amato, Jeffrey Barrett, Sonia Goncalves, Ewan Harrison, David K. Jackson, Ian Johnston, Dominic Kwiatkowski, Cordelia Langford, John Sillitoe on behalf of the Wellcome Sanger Institute COVID-19 Surveillance Team |
| EPI_ISL_2912426 | Berkshire and Surrey Pathology Services Lighthouse Laboratory | Wellcome Sanger Institute for the COVID-19 Genomics UK (COG-UK) Consortium | Berkshire and Surrey Pathology Services Lighthouse Laboratory and Alex Alderton, Roberto Amato, Jeffrey Barrett, Sonia Goncalves, Ewan Harrison, David K. Jackson, Ian Johnston, Dominic Kwiatkowski, Cordelia Langford, John Sillitoe on behalf of the Wellcome Sanger Institute COVID-19 Surveillance Team |
| EPI_ISL_2912427 | Lighthouse Lab in Milton Keynes | Wellcome Sanger Institute for the COVID-19 Genomics UK (COG-UK) Consortium | The Lighthouse Lab in Milton Keynes and Alex Alderton, Roberto Amato, Jeffrey Barrett, Sonia Goncalves, Ewan Harrison, David K. Jackson, Ian Johnston, Dominic Kwiatkowski, Cordelia Langford, John Sillitoe on behalf of the Wellcome Sanger Institute COVID-19 Surveillance Team |
| EPI_ISL_2912428 | Lighthouse Lab in Glasgow | Wellcome Sanger Institute for the COVID-19 Genomics UK (COG-UK) Consortium | Harper VanSteenhouse, Yumi Kasai, David Gray, Carol Clugston, Anna Dominiczak and Alex Alderton, Roberto Amato, Jeffrey Barrett, Sonia Goncalves, Ewan Harrison, David K. Jackson, Ian Johnston, Dominic Kwiatkowski, Cordelia Langford, John Sillitoe on behalf of the Wellcome Sanger Institute COVID-19 Surveillance Team |
| EPI_ISL_2912429 | Lighthouse Lab in Milton Keynes | Wellcome Sanger Institute for the COVID-19 Genomics UK (COG-UK) Consortium | The Lighthouse Lab in Milton Keynes and Alex Alderton, Roberto Amato, Jeffrey Barrett, Sonia Goncalves, Ewan Harrison, David K. Jackson, Ian Johnston, Dominic Kwiatkowski, Cordelia Langford, John Sillitoe on behalf of the Wellcome Sanger Institute COVID-19 Surveillance Team |
| EPI_ISL_2912430, EPI_ISL_2912431, EPI_ISL_2912432, EPI_ISL_2912433 | Berkshire and Surrey Pathology Services Lighthouse Laboratory | Wellcome Sanger Institute for the COVID-19 Genomics UK (COG-UK) Consortium | Berkshire and Surrey Pathology Services Lighthouse Laboratory and Alex Alderton, Roberto Amato, Jeffrey Barrett, Sonia Goncalves, Ewan Harrison, David K. Jackson, Ian Johnston, Dominic Kwiatkowski, Cordelia Langford, John Sillitoe on behalf of the Wellcome Sanger Institute COVID-19 Surveillance Team |
| EPI_ISL_2912434, EPI_ISL_2912435 | Lighthouse Lab in Milton Keynes | Wellcome Sanger Institute for the COVID-19 Genomics UK (COG-UK) Consortium | The Lighthouse Lab in Milton Keynes and Alex Alderton, Roberto Amato, Jeffrey Barrett, Sonia Goncalves, Ewan Harrison, David K. Jackson, Ian Johnston, Dominic Kwiatkowski, Cordelia Langford, John Sillitoe on behalf of the Wellcome Sanger Institute COVID-19 Surveillance Team |
| EPI_ISL_2912436, EPI_ISL_2912437 | Lighthouse Lab in Glasgow | Wellcome Sanger Institute for the COVID-19 Genomics UK (COG-UK) Consortium | Harper VanSteenhouse, Yumi Kasai, David Gray, Carol Clugston, Anna Dominiczak and Alex Alderton, Roberto Amato, Jeffrey Barrett, Sonia Goncalves, Ewan Harrison, David K. Jackson, Ian Johnston, Dominic Kwiatkowski, Cordelia Langford, John Sillitoe on behalf of the Wellcome Sanger Institute COVID-19 Surveillance Team |
| EPI_ISL_2912438 | Lighthouse Lab in Milton Keynes | Wellcome Sanger Institute for the COVID-19 Genomics UK (COG-UK) Consortium | The Lighthouse Lab in Milton Keynes and Alex Alderton, Roberto Amato, Jeffrey Barrett, Sonia Goncalves, Ewan Harrison, David K. Jackson, Ian Johnston, Dominic Kwiatkowski, Cordelia Langford, John Sillitoe on behalf of the Wellcome Sanger Institute COVID-19 Surveillance Team |
| EPI_ISL_2912439, EPI_ISL_2912440, EPI_ISL_2912441 | Berkshire and Surrey Pathology Services Lighthouse Laboratory | Wellcome Sanger Institute for the COVID-19 Genomics UK (COG-UK) Consortium | Berkshire and Surrey Pathology Services Lighthouse Laboratory and Alex Alderton, Roberto Amato, Jeffrey Barrett, Sonia Goncalves, Ewan Harrison, David K. Jackson, Ian Johnston, Dominic Kwiatkowski, Cordelia Langford, John Sillitoe on behalf of the Wellcome Sanger Institute COVID-19 Surveillance Team |
| EPI_ISL_2912442, EPI_ISL_2912443, EPI_ISL_2912444 | Lighthouse Lab in Glasgow | Wellcome Sanger Institute for the COVID-19 Genomics UK (COG-UK) Consortium | Harper VanSteenhouse, Yumi Kasai, David Gray, Carol Clugston, Anna Dominiczak and Alex Alderton, Roberto Amato, Jeffrey Barrett, Sonia Goncalves, Ewan Harrison, David K. Jackson, Ian Johnston, Dominic Kwiatkowski, Cordelia Langford, John Sillitoe on behalf of the Wellcome Sanger Institute COVID-19 Surveillance Team |
| EPI_ISL_2912445 | Lighthouse Lab in Milton Keynes | Wellcome Sanger Institute for the COVID-19 Genomics UK (COG-UK) Consortium | The Lighthouse Lab in Milton Keynes and Alex Alderton, Roberto Amato, Jeffrey Barrett, Sonia Goncalves, Ewan Harrison, David K. Jackson, Ian Johnston, Dominic Kwiatkowski, Cordelia Langford, John Sillitoe on behalf of the Wellcome Sanger Institute COVID-19 Surveillance Team |
| EPI_ISL_2912446, EPI_ISL_2912447, EPI_ISL_2912448, EPI_ISL_2912449 | Berkshire and Surrey Pathology Services Lighthouse Laboratory | Wellcome Sanger Institute for the COVID-19 Genomics UK (COG-UK) Consortium | Berkshire and Surrey Pathology Services Lighthouse Laboratory and Alex Alderton, Roberto Amato, Jeffrey Barrett, Sonia Goncalves, Ewan Harrison, David K. Jackson, Ian Johnston, Dominic Kwiatkowski, Cordelia Langford, John Sillitoe on behalf of the Wellcome Sanger Institute COVID-19 Surveillance Team |
| EPI_ISL_2912450, EPI_ISL_2912451 | Lighthouse Lab in Glasgow | Wellcome Sanger Institute for the COVID-19 Genomics UK (COG-UK) Consortium | Harper VanSteenhouse, Yumi Kasai, David Gray, Carol Clugston, Anna Dominiczak and Alex Alderton, Roberto Amato, Jeffrey Barrett, Sonia Goncalves, Ewan Harrison, David K. Jackson, Ian Johnston, Dominic Kwiatkowski, Cordelia Langford, John Sillitoe on behalf of the Wellcome Sanger Institute COVID-19 Surveillance Team |
| EPI_ISL_2912452, EPI_ISL_2912453, EPI_ISL_2912454, EPI_ISL_2912455 | Berkshire and Surrey Pathology Services Lighthouse Laboratory | Wellcome Sanger Institute for the COVID-19 Genomics UK (COG-UK) Consortium | Berkshire and Surrey Pathology Services Lighthouse Laboratory and Alex Alderton, Roberto Amato, Jeffrey Barrett, Sonia Goncalves, Ewan Harrison, David K. Jackson, Ian Johnston, Dominic Kwiatkowski, Cordelia Langford, John Sillitoe on behalf of the Wellcome Sanger Institute COVID-19 Surveillance Team |
| EPI_ISL_2912456 | Lighthouse Lab in Milton Keynes | Wellcome Sanger Institute for the COVID-19 Genomics UK (COG-UK) Consortium | The Lighthouse Lab in Milton Keynes and Alex Alderton, Roberto Amato, Jeffrey Barrett, Sonia Goncalves, Ewan Harrison, David K. Jackson, Ian Johnston, Dominic Kwiatkowski, Cordelia Langford, John Sillitoe on behalf of the Wellcome Sanger Institute COVID-19 Surveillance Team |
| EPI_ISL_2912457, EPI_ISL_2912458 | Lighthouse Lab in Glasgow | Wellcome Sanger Institute for the COVID-19 Genomics UK (COG-UK) Consortium | Harper VanSteenhouse, Yumi Kasai, David Gray, Carol Clugston, Anna Dominiczak and Alex Alderton, Roberto Amato, Jeffrey Barrett, Sonia Goncalves, Ewan Harrison, David K. Jackson, Ian Johnston, Dominic Kwiatkowski, Cordelia Langford, John Sillitoe on behalf of the Wellcome Sanger Institute COVID-19 Surveillance Team |
| EPI_ISL_2912459 | Lighthouse Lab in Milton Keynes | Wellcome Sanger Institute for the COVID-19 Genomics UK (COG-UK) Consortium | The Lighthouse Lab in Milton Keynes and Alex Alderton, Roberto Amato, Jeffrey Barrett, Sonia Goncalves, Ewan Harrison, David K. Jackson, Ian Johnston, Dominic Kwiatkowski, Cordelia Langford, John Sillitoe on behalf of the Wellcome Sanger Institute COVID-19 Surveillance Team |
| EPI_ISL_2912460, EPI_ISL_2912461 | Berkshire and Surrey Pathology Services Lighthouse Laboratory | Wellcome Sanger Institute for the COVID-19 Genomics UK (COG-UK) Consortium | Berkshire and Surrey Pathology Services Lighthouse Laboratory and Alex Alderton, Roberto Amato, Jeffrey Barrett, Sonia Goncalves, Ewan Harrison, David K. Jackson, Ian Johnston, Dominic Kwiatkowski, Cordelia Langford, John Sillitoe on behalf of the Wellcome Sanger Institute COVID-19 Surveillance Team |
| EPI_ISL_2912462 | Lighthouse Lab in Glasgow | Wellcome Sanger Institute for the COVID-19 Genomics UK (COG-UK) Consortium | Harper VanSteenhouse, Yumi Kasai, David Gray, Carol Clugston, Anna Dominiczak and Alex Alderton, Roberto Amato, Jeffrey Barrett, Sonia Goncalves, Ewan Harrison, David K. Jackson, Ian Johnston, Dominic Kwiatkowski, Cordelia Langford, John Sillitoe on behalf of the Wellcome Sanger Institute COVID-19 Surveillance Team |
| EPI_ISL_2912463, EPI_ISL_2912464, EPI_ISL_2912465 | Berkshire and Surrey Pathology Services Lighthouse Laboratory | Wellcome Sanger Institute for the COVID-19 Genomics UK (COG-UK) Consortium | Berkshire and Surrey Pathology Services Lighthouse Laboratory and Alex Alderton, Roberto Amato, Jeffrey Barrett, Sonia Goncalves, Ewan Harrison, David K. Jackson, Ian Johnston, Dominic Kwiatkowski, Cordelia Langford, John Sillitoe on behalf of the Wellcome Sanger Institute COVID-19 Surveillance Team |
| EPI_ISL_2912466 | Lighthouse Lab in Glasgow | Wellcome Sanger Institute for the COVID-19 Genomics UK (COG-UK) Consortium | Harper VanSteenhouse, Yumi Kasai, David Gray, Carol Clugston, Anna Dominiczak and Alex Alderton, Roberto Amato, Jeffrey Barrett, Sonia Goncalves, Ewan Harrison, David K. Jackson, Ian Johnston, Dominic Kwiatkowski, Cordelia Langford, John Sillitoe on behalf of the Wellcome Sanger Institute COVID-19 Surveillance Team |
| EPI_ISL_2912467 | Berkshire and Surrey Pathology Services Lighthouse Laboratory | Wellcome Sanger Institute for the COVID-19 Genomics UK (COG-UK) Consortium | Berkshire and Surrey Pathology Services Lighthouse Laboratory and Alex Alderton, Roberto Amato, Jeffrey Barrett, Sonia Goncalves, Ewan Harrison, David K. Jackson, Ian Johnston, Dominic Kwiatkowski, Cordelia Langford, John Sillitoe on behalf of the Wellcome Sanger Institute COVID-19 Surveillance Team |
| EPI_ISL_2912468, EPI_ISL_2912469 | Lighthouse Lab in Milton Keynes | Wellcome Sanger Institute for the COVID-19 Genomics UK (COG-UK) Consortium | The Lighthouse Lab in Milton Keynes and Alex Alderton, Roberto Amato, Jeffrey Barrett, Sonia Goncalves, Ewan Harrison, David K. Jackson, Ian Johnston, Dominic Kwiatkowski, Cordelia Langford, John Sillitoe on behalf of the Wellcome Sanger Institute COVID-19 Surveillance Team |
| EPI_ISL_2912470 | Berkshire and Surrey Pathology Services Lighthouse Laboratory | Wellcome Sanger Institute for the COVID-19 Genomics UK (COG-UK) Consortium | Berkshire and Surrey Pathology Services Lighthouse Laboratory and Alex Alderton, Roberto Amato, Jeffrey Barrett, Sonia Goncalves, Ewan Harrison, David K. Jackson, Ian Johnston, Dominic Kwiatkowski, Cordelia Langford, John Sillitoe on behalf of the Wellcome Sanger Institute COVID-19 Surveillance Team |
| EPI_ISL_2912471 | Lighthouse Lab in Milton Keynes | Wellcome Sanger Institute for the COVID-19 Genomics UK (COG-UK) Consortium | The Lighthouse Lab in Milton Keynes and Alex Alderton, Roberto Amato, Jeffrey Barrett, Sonia Goncalves, Ewan Harrison, David K. Jackson, Ian Johnston, Dominic Kwiatkowski, Cordelia Langford, John Sillitoe on behalf of the Wellcome Sanger Institute COVID-19 Surveillance Team |
| EPI_ISL_2912472 | Berkshire and Surrey Pathology Services Lighthouse Laboratory | Wellcome Sanger Institute for the COVID-19 Genomics UK (COG-UK) Consortium | Berkshire and Surrey Pathology Services Lighthouse Laboratory and Alex Alderton, Roberto Amato, Jeffrey Barrett, Sonia Goncalves, Ewan Harrison, David K. Jackson, Ian Johnston, Dominic Kwiatkowski, Cordelia Langford, John Sillitoe on behalf of the Wellcome Sanger Institute COVID-19 Surveillance Team |
| EPI_ISL_2912473 | Lighthouse Lab in Glasgow | Wellcome Sanger Institute for the COVID-19 Genomics UK (COG-UK) Consortium | Harper VanSteenhouse, Yumi Kasai, David Gray, Carol Clugston, Anna Dominiczak and Alex Alderton, Roberto Amato, Jeffrey Barrett, Sonia Goncalves, Ewan Harrison, David K. Jackson, Ian Johnston, Dominic Kwiatkowski, Cordelia Langford, John Sillitoe on behalf of the Wellcome Sanger Institute COVID-19 Surveillance Team |
| EPI_ISL_2912474 | Lighthouse Lab in Milton Keynes | Wellcome Sanger Institute for the COVID-19 Genomics UK (COG-UK) Consortium | The Lighthouse Lab in Milton Keynes and Alex Alderton, Roberto Amato, Jeffrey Barrett, Sonia Goncalves, Ewan Harrison, David K. Jackson, Ian Johnston, Dominic Kwiatkowski, Cordelia Langford, John Sillitoe on behalf of the Wellcome Sanger Institute COVID-19 Surveillance Team |
| EPI_ISL_2912475 | Lighthouse Lab in Glasgow | Wellcome Sanger Institute for the COVID-19 Genomics UK (COG-UK) Consortium | Harper VanSteenhouse, Yumi Kasai, David Gray, Carol Clugston, Anna Dominiczak and Alex Alderton, Roberto Amato, Jeffrey Barrett, Sonia Goncalves, Ewan Harrison, David K. Jackson, Ian Johnston, Dominic Kwiatkowski, Cordelia Langford, John Sillitoe on behalf of the Wellcome Sanger Institute COVID-19 Surveillance Team |
| EPI_ISL_2912476 | Berkshire and Surrey Pathology Services Lighthouse Laboratory | Wellcome Sanger Institute for the COVID-19 Genomics UK (COG-UK) Consortium | Berkshire and Surrey Pathology Services Lighthouse Laboratory and Alex Alderton, Roberto Amato, Jeffrey Barrett, Sonia Goncalves, Ewan Harrison, David K. Jackson, Ian Johnston, Dominic Kwiatkowski, Cordelia Langford, John Sillitoe on behalf of the Wellcome Sanger Institute COVID-19 Surveillance Team |
| EPI_ISL_2912477 | Lighthouse Lab in Glasgow | Wellcome Sanger Institute for the COVID-19 Genomics UK (COG-UK) Consortium | Harper VanSteenhouse, Yumi Kasai, David Gray, Carol Clugston, Anna Dominiczak and Alex Alderton, Roberto Amato, Jeffrey Barrett, Sonia Goncalves, Ewan Harrison, David K. Jackson, Ian Johnston, Dominic Kwiatkowski, Cordelia Langford, John Sillitoe on behalf of the Wellcome Sanger Institute COVID-19 Surveillance Team |
| EPI_ISL_2912478 | Berkshire and Surrey Pathology Services Lighthouse Laboratory | Wellcome Sanger Institute for the COVID-19 Genomics UK (COG-UK) Consortium | Berkshire and Surrey Pathology Services Lighthouse Laboratory and Alex Alderton, Roberto Amato, Jeffrey Barrett, Sonia Goncalves, Ewan Harrison, David K. Jackson, Ian Johnston, Dominic Kwiatkowski, Cordelia Langford, John Sillitoe on behalf of the Wellcome Sanger Institute COVID-19 Surveillance Team |
| EPI_ISL_2912479, EPI_ISL_2912480 | Lighthouse Lab in Glasgow | Wellcome Sanger Institute for the COVID-19 Genomics UK (COG-UK) Consortium | Harper VanSteenhouse, Yumi Kasai, David Gray, Carol Clugston, Anna Dominiczak and Alex Alderton, Roberto Amato, Jeffrey Barrett, Sonia Goncalves, Ewan Harrison, David K. Jackson, Ian Johnston, Dominic Kwiatkowski, Cordelia Langford, John Sillitoe on behalf of the Wellcome Sanger Institute COVID-19 Surveillance Team |
| EPI_ISL_2912481, EPI_ISL_2912482 | Berkshire and Surrey Pathology Services Lighthouse Laboratory | Wellcome Sanger Institute for the COVID-19 Genomics UK (COG-UK) Consortium | Berkshire and Surrey Pathology Services Lighthouse Laboratory and Alex Alderton, Roberto Amato, Jeffrey Barrett, Sonia Goncalves, Ewan Harrison, David K. Jackson, Ian Johnston, Dominic Kwiatkowski, Cordelia Langford, John Sillitoe on behalf of the Wellcome Sanger Institute COVID-19 Surveillance Team |
| EPI_ISL_2912483, EPI_ISL_2912484, EPI_ISL_2912485, EPI_ISL_2912486 | Lighthouse Lab in Milton Keynes | Wellcome Sanger Institute for the COVID-19 Genomics UK (COG-UK) Consortium | The Lighthouse Lab in Milton Keynes and Alex Alderton, Roberto Amato, Jeffrey Barrett, Sonia Goncalves, Ewan Harrison, David K. Jackson, Ian Johnston, Dominic Kwiatkowski, Cordelia Langford, John Sillitoe on behalf of the Wellcome Sanger Institute COVID-19 Surveillance Team |

| | | | |
|---|---|---|---|
| EPI_ISL_2912487 | Berkshire and Surrey Pathology Services Lighthouse Laboratory | Wellcome Sanger Institute for the COVID-19 Genomics UK (COG-UK) Consortium | Berkshire and Surrey Pathology Services Lighthouse Laboratory and Alex Alderton, Roberto Amato, Jeffrey Barrett, Sonia Goncalves, Ewan Harrison, David K. Jackson, Ian Johnston, Dominic Kwiatkowski, Cordelia Langford, John Sillitoe on behalf of the Wellcome Sanger Institute COVID-19 Surveillance Team |
| EPI_ISL_2912488 | Lighthouse Lab in Glasgow | Wellcome Sanger Institute for the COVID-19 Genomics UK (COG-UK) Consortium | Harper VanSteenhouse, Yumi Kasai, David Gray, Carol Clugston, Anna Dominiczak and Alex Alderton, Roberto Amato, Jeffrey Barrett, Sonia Goncalves, Ewan Harrison, David K. Jackson, Ian Johnston, Dominic Kwiatkowski, Cordelia Langford, John Sillitoe on behalf of the Wellcome Sanger Institute COVID-19 Surveillance Team |
| EPI_ISL_2912489, EPI_ISL_2912490, EPI_ISL_2912491, EPI_ISL_2912492 | Berkshire and Surrey Pathology Services Lighthouse Laboratory | Wellcome Sanger Institute for the COVID-19 Genomics UK (COG-UK) Consortium | Berkshire and Surrey Pathology Services Lighthouse Laboratory and Alex Alderton, Roberto Amato, Jeffrey Barrett, Sonia Goncalves, Ewan Harrison, David K. Jackson, Ian Johnston, Dominic Kwiatkowski, Cordelia Langford, John Sillitoe on behalf of the Wellcome Sanger Institute COVID-19 Surveillance Team |
| EPI_ISL_2912493 | Lighthouse Lab in Glasgow | Wellcome Sanger Institute for the COVID-19 Genomics UK (COG-UK) Consortium | Harper VanSteenhouse, Yumi Kasai, David Gray, Carol Clugston, Anna Dominiczak and Alex Alderton, Roberto Amato, Jeffrey Barrett, Sonia Goncalves, Ewan Harrison, David K. Jackson, Ian Johnston, Dominic Kwiatkowski, Cordelia Langford, John Sillitoe on behalf of the Wellcome Sanger Institute COVID-19 Surveillance Team |
| EPI_ISL_2912494, EPI_ISL_2912495 | Berkshire and Surrey Pathology Services Lighthouse Laboratory | Wellcome Sanger Institute for the COVID-19 Genomics UK (COG-UK) Consortium | Berkshire and Surrey Pathology Services Lighthouse Laboratory and Alex Alderton, Roberto Amato, Jeffrey Barrett, Sonia Goncalves, Ewan Harrison, David K. Jackson, Ian Johnston, Dominic Kwiatkowski, Cordelia Langford, John Sillitoe on behalf of the Wellcome Sanger Institute COVID-19 Surveillance Team |
| EPI_ISL_2912496, EPI_ISL_2912497, EPI_ISL_2912498, EPI_ISL_2912499 | Lighthouse Lab in Glasgow | Wellcome Sanger Institute for the COVID-19 Genomics UK (COG-UK) Consortium | Harper VanSteenhouse, Yumi Kasai, David Gray, Carol Clugston, Anna Dominiczak and Alex Alderton, Roberto Amato, Jeffrey Barrett, Sonia Goncalves, Ewan Harrison, David K. Jackson, Ian Johnston, Dominic Kwiatkowski, Cordelia Langford, John Sillitoe on behalf of the Wellcome Sanger Institute COVID-19 Surveillance Team |
| EPI_ISL_2912500 | Lighthouse Lab in Milton Keynes | Wellcome Sanger Institute for the COVID-19 Genomics UK (COG-UK) Consortium | The Lighthouse Lab in Milton Keynes and Alex Alderton, Roberto Amato, Jeffrey Barrett, Sonia Goncalves, Ewan Harrison, David K. Jackson, Ian Johnston, Dominic Kwiatkowski, Cordelia Langford, John Sillitoe on behalf of the Wellcome Sanger Institute COVID-19 Surveillance Team |
| EPI_ISL_2912501 | Berkshire and Surrey Pathology Services Lighthouse Laboratory | Wellcome Sanger Institute for the COVID-19 Genomics UK (COG-UK) Consortium | Berkshire and Surrey Pathology Services Lighthouse Laboratory and Alex Alderton, Roberto Amato, Jeffrey Barrett, Sonia Goncalves, Ewan Harrison, David K. Jackson, Ian Johnston, Dominic Kwiatkowski, Cordelia Langford, John Sillitoe on behalf of the Wellcome Sanger Institute COVID-19 Surveillance Team |
| EPI_ISL_2912502, EPI_ISL_2912503 | Lighthouse Lab in Glasgow | Wellcome Sanger Institute for the COVID-19 Genomics UK (COG-UK) Consortium | Harper VanSteenhouse, Yumi Kasai, David Gray, Carol Clugston, Anna Dominiczak and Alex Alderton, Roberto Amato, Jeffrey Barrett, Sonia Goncalves, Ewan Harrison, David K. Jackson, Ian Johnston, Dominic Kwiatkowski, Cordelia Langford, John Sillitoe on behalf of the Wellcome Sanger Institute COVID-19 Surveillance Team |
| EPI_ISL_2912504 | Berkshire and Surrey Pathology Services Lighthouse Laboratory | Wellcome Sanger Institute for the COVID-19 Genomics UK (COG-UK) Consortium | Berkshire and Surrey Pathology Services Lighthouse Laboratory and Alex Alderton, Roberto Amato, Jeffrey Barrett, Sonia Goncalves, Ewan Harrison, David K. Jackson, Ian Johnston, Dominic Kwiatkowski, Cordelia Langford, John Sillitoe on behalf of the Wellcome Sanger Institute COVID-19 Surveillance Team |
| EPI_ISL_2912505 | Lighthouse Lab in Glasgow | Wellcome Sanger Institute for the COVID-19 Genomics UK (COG-UK) Consortium | Harper VanSteenhouse, Yumi Kasai, David Gray, Carol Clugston, Anna Dominiczak and Alex Alderton, Roberto Amato, Jeffrey Barrett, Sonia Goncalves, Ewan Harrison, David K. Jackson, Ian Johnston, Dominic Kwiatkowski, Cordelia Langford, John Sillitoe on behalf of the Wellcome Sanger Institute COVID-19 Surveillance Team |
| EPI_ISL_2912506 | Berkshire and Surrey Pathology Services Lighthouse Laboratory | Wellcome Sanger Institute for the COVID-19 Genomics UK (COG-UK) Consortium | Berkshire and Surrey Pathology Services Lighthouse Laboratory and Alex Alderton, Roberto Amato, Jeffrey Barrett, Sonia Goncalves, Ewan Harrison, David K. Jackson, Ian Johnston, Dominic Kwiatkowski, Cordelia Langford, John Sillitoe on behalf of the Wellcome Sanger Institute COVID-19 Surveillance Team |
| EPI_ISL_2912507, EPI_ISL_2912508 | Lighthouse Lab in Milton Keynes | Wellcome Sanger Institute for the COVID-19 Genomics UK (COG-UK) Consortium | The Lighthouse Lab in Milton Keynes and Alex Alderton, Roberto Amato, Jeffrey Barrett, Sonia Goncalves, Ewan Harrison, David K. Jackson, Ian Johnston, Dominic Kwiatkowski, Cordelia Langford, John Sillitoe on behalf of the Wellcome Sanger Institute COVID-19 Surveillance Team |
| EPI_ISL_2912509 | Berkshire and Surrey Pathology Services Lighthouse Laboratory | Wellcome Sanger Institute for the COVID-19 Genomics UK (COG-UK) Consortium | Berkshire and Surrey Pathology Services Lighthouse Laboratory and Alex Alderton, Roberto Amato, Jeffrey Barrett, Sonia Goncalves, Ewan Harrison, David K. Jackson, Ian Johnston, Dominic Kwiatkowski, Cordelia Langford, John Sillitoe on behalf of the Wellcome Sanger Institute COVID-19 Surveillance Team |
| EPI_ISL_2912510 | Lighthouse Lab in Milton Keynes | Wellcome Sanger Institute for the COVID-19 Genomics UK (COG-UK) Consortium | The Lighthouse Lab in Milton Keynes and Alex Alderton, Roberto Amato, Jeffrey Barrett, Sonia Goncalves, Ewan Harrison, David K. Jackson, Ian Johnston, Dominic Kwiatkowski, Cordelia Langford, John Sillitoe on behalf of the Wellcome Sanger Institute COVID-19 Surveillance Team |
| EPI_ISL_2912511 | Lighthouse Lab in Glasgow | Wellcome Sanger Institute for the COVID-19 Genomics UK (COG-UK) Consortium | Harper VanSteenhouse, Yumi Kasai, David Gray, Carol Clugston, Anna Dominiczak and Alex Alderton, Roberto Amato, Jeffrey Barrett, Sonia Goncalves, Ewan Harrison, David K. Jackson, Ian Johnston, Dominic Kwiatkowski, Cordelia Langford, John Sillitoe on behalf of the Wellcome Sanger Institute COVID-19 Surveillance Team |
| EPI_ISL_2912513, EPI_ISL_2912514, EPI_ISL_2912515, EPI_ISL_2912516 | Berkshire and Surrey Pathology Services Lighthouse Laboratory | Wellcome Sanger Institute for the COVID-19 Genomics UK (COG-UK) Consortium | Berkshire and Surrey Pathology Services Lighthouse Laboratory and Alex Alderton, Roberto Amato, Jeffrey Barrett, Sonia Goncalves, Ewan Harrison, David K. Jackson, Ian Johnston, Dominic Kwiatkowski, Cordelia Langford, John Sillitoe on behalf of the Wellcome Sanger Institute COVID-19 Surveillance Team |
| EPI_ISL_2912517 | Lighthouse Lab in Glasgow | Wellcome Sanger Institute for the COVID-19 Genomics UK (COG-UK) Consortium | Harper VanSteenhouse, Yumi Kasai, David Gray, Carol Clugston, Anna Dominiczak and Alex Alderton, Roberto Amato, Jeffrey Barrett, Sonia Goncalves, Ewan Harrison, David K. Jackson, Ian Johnston, Dominic Kwiatkowski, Cordelia Langford, John Sillitoe on behalf of the Wellcome Sanger Institute COVID-19 Surveillance Team |
| EPI_ISL_2912518, EPI_ISL_2912519 | Lighthouse Lab in Milton Keynes | Wellcome Sanger Institute for the COVID-19 Genomics UK (COG-UK) Consortium | The Lighthouse Lab in Milton Keynes and Alex Alderton, Roberto Amato, Jeffrey Barrett, Sonia Goncalves, Ewan Harrison, David K. Jackson, Ian Johnston, Dominic Kwiatkowski, Cordelia Langford, John Sillitoe on behalf of the Wellcome Sanger Institute COVID-19 Surveillance Team |
| EPI_ISL_2912520 | Berkshire and Surrey Pathology Services Lighthouse Laboratory | Wellcome Sanger Institute for the COVID-19 Genomics UK (COG-UK) Consortium | Berkshire and Surrey Pathology Services Lighthouse Laboratory and Alex Alderton, Roberto Amato, Jeffrey Barrett, Sonia Goncalves, Ewan Harrison, David K. Jackson, Ian Johnston, Dominic Kwiatkowski, Cordelia Langford, John Sillitoe on behalf of the Wellcome Sanger Institute COVID-19 Surveillance Team |
| EPI_ISL_2912521 | Lighthouse Lab in Milton Keynes | Wellcome Sanger Institute for the COVID-19 Genomics UK (COG-UK) Consortium | The Lighthouse Lab in Milton Keynes and Alex Alderton, Roberto Amato, Jeffrey Barrett, Sonia Goncalves, Ewan Harrison, David K. Jackson, Ian Johnston, Dominic Kwiatkowski, Cordelia Langford, John Sillitoe on behalf of the Wellcome Sanger Institute COVID-19 Surveillance Team |
| EPI_ISL_2912522 | Lighthouse Lab in Glasgow | Wellcome Sanger Institute for the COVID-19 Genomics UK (COG-UK) Consortium | Harper VanSteenhouse, Yumi Kasai, David Gray, Carol Clugston, Anna Dominiczak and Alex Alderton, Roberto Amato, Jeffrey Barrett, Sonia Goncalves, Ewan Harrison, David K. Jackson, Ian Johnston, Dominic Kwiatkowski, Cordelia Langford, John Sillitoe on behalf of the Wellcome Sanger Institute COVID-19 Surveillance Team |
| EPI_ISL_2912523, EPI_ISL_2912524 | Berkshire and Surrey Pathology Services Lighthouse Laboratory | Wellcome Sanger Institute for the COVID-19 Genomics UK (COG-UK) Consortium | Berkshire and Surrey Pathology Services Lighthouse Laboratory and Alex Alderton, Roberto Amato, Jeffrey Barrett, Sonia Goncalves, Ewan Harrison, David K. Jackson, Ian Johnston, Dominic Kwiatkowski, Cordelia Langford, John Sillitoe on behalf of the Wellcome Sanger Institute COVID-19 Surveillance Team |
| EPI_ISL_2912525, EPI_ISL_2912526 | Lighthouse Lab in Glasgow | Wellcome Sanger Institute for the COVID-19 Genomics UK (COG-UK) Consortium | Harper VanSteenhouse, Yumi Kasai, David Gray, Carol Clugston, Anna Dominiczak and Alex Alderton, Roberto Amato, Jeffrey Barrett, Sonia Goncalves, Ewan Harrison, David K. Jackson, Ian Johnston, Dominic Kwiatkowski, Cordelia Langford, John Sillitoe on behalf of the Wellcome Sanger Institute COVID-19 Surveillance Team |
| EPI_ISL_2912527, EPI_ISL_2912528 | Berkshire and Surrey Pathology Services Lighthouse Laboratory | Wellcome Sanger Institute for the COVID-19 Genomics UK (COG-UK) Consortium | Berkshire and Surrey Pathology Services Lighthouse Laboratory and Alex Alderton, Roberto Amato, Jeffrey Barrett, Sonia Goncalves, Ewan Harrison, David K. Jackson, Ian Johnston, Dominic Kwiatkowski, Cordelia Langford, John Sillitoe on behalf of the Wellcome Sanger Institute COVID-19 Surveillance Team |
| EPI_ISL_2912529 | Lighthouse Lab in Glasgow | Wellcome Sanger Institute for the COVID-19 Genomics UK (COG-UK) Consortium | Harper VanSteenhouse, Yumi Kasai, David Gray, Carol Clugston, Anna Dominiczak and Alex Alderton, Roberto Amato, Jeffrey Barrett, Sonia Goncalves, Ewan Harrison, David K. Jackson, Ian Johnston, Dominic Kwiatkowski, Cordelia Langford, John Sillitoe on behalf of the Wellcome Sanger Institute COVID-19 Surveillance Team |
| EPI_ISL_2912530, EPI_ISL_2912531, EPI_ISL_2912532 | Lighthouse Lab in Milton Keynes | Wellcome Sanger Institute for the COVID-19 Genomics UK (COG-UK) Consortium | The Lighthouse Lab in Milton Keynes and Alex Alderton, Roberto Amato, Jeffrey Barrett, Sonia Goncalves, Ewan Harrison, David K. Jackson, Ian Johnston, Dominic Kwiatkowski, Cordelia Langford, John Sillitoe on behalf of the Wellcome Sanger Institute COVID-19 Surveillance Team |
| EPI_ISL_2912533 | Lighthouse Lab in Glasgow | Wellcome Sanger Institute for the COVID-19 Genomics UK (COG-UK) Consortium | Harper VanSteenhouse, Yumi Kasai, David Gray, Carol Clugston, Anna Dominiczak and Alex Alderton, Roberto Amato, Jeffrey Barrett, Sonia Goncalves, Ewan Harrison, David K. Jackson, Ian Johnston, Dominic Kwiatkowski, Cordelia Langford, John Sillitoe on behalf of the Wellcome Sanger Institute COVID-19 Surveillance Team |
| EPI_ISL_2912534 | Lighthouse Lab in Milton Keynes | Wellcome Sanger Institute for the COVID-19 Genomics UK (COG-UK) Consortium | The Lighthouse Lab in Milton Keynes and Alex Alderton, Roberto Amato, Jeffrey Barrett, Sonia Goncalves, Ewan Harrison, David K. Jackson, Ian Johnston, Dominic Kwiatkowski, Cordelia Langford, John Sillitoe on behalf of the Wellcome Sanger Institute COVID-19 Surveillance Team |
| EPI_ISL_2912535, EPI_ISL_2912536, EPI_ISL_2912537 | Lighthouse Lab in Glasgow | Wellcome Sanger Institute for the COVID-19 Genomics UK (COG-UK) Consortium | Harper VanSteenhouse, Yumi Kasai, David Gray, Carol Clugston, Anna Dominiczak and Alex Alderton, Roberto Amato, Jeffrey Barrett, Sonia Goncalves, Ewan Harrison, David K. Jackson, Ian Johnston, Dominic Kwiatkowski, Cordelia Langford, John Sillitoe on behalf of the Wellcome Sanger Institute COVID-19 Surveillance Team |
| EPI_ISL_2912538 | Lighthouse Lab in Milton Keynes | Wellcome Sanger Institute for the COVID-19 Genomics UK (COG-UK) Consortium | The Lighthouse Lab in Milton Keynes and Alex Alderton, Roberto Amato, Jeffrey Barrett, Sonia Goncalves, Ewan Harrison, David K. Jackson, Ian Johnston, Dominic Kwiatkowski, Cordelia Langford, John Sillitoe on behalf of the Wellcome Sanger Institute COVID-19 Surveillance Team |
| EPI_ISL_2912539 | Lighthouse Lab in Glasgow | Wellcome Sanger Institute for the COVID-19 Genomics UK (COG-UK) Consortium | Harper VanSteenhouse, Yumi Kasai, David Gray, Carol Clugston, Anna Dominiczak and Alex Alderton, Roberto Amato, Jeffrey Barrett, Sonia Goncalves, Ewan Harrison, David K. Jackson, Ian Johnston, Dominic Kwiatkowski, Cordelia Langford, John Sillitoe on behalf of the Wellcome Sanger Institute COVID-19 Surveillance Team |
| EPI_ISL_2912540, EPI_ISL_2912541 | Lighthouse Lab in Milton Keynes | Wellcome Sanger Institute for the COVID-19 Genomics UK (COG-UK) Consortium | The Lighthouse Lab in Milton Keynes and Alex Alderton, Roberto Amato, Jeffrey Barrett, Sonia Goncalves, Ewan Harrison, David K. Jackson, Ian Johnston, Dominic Kwiatkowski, Cordelia Langford, John Sillitoe on behalf of the Wellcome Sanger Institute COVID-19 Surveillance Team |
| EPI_ISL_2912542 | Berkshire and Surrey Pathology Services Lighthouse Laboratory | Wellcome Sanger Institute for the COVID-19 Genomics UK (COG-UK) Consortium | Berkshire and Surrey Pathology Services Lighthouse Laboratory and Alex Alderton, Roberto Amato, Jeffrey Barrett, Sonia Goncalves, Ewan Harrison, David K. Jackson, Ian Johnston, Dominic Kwiatkowski, Cordelia Langford, John Sillitoe on behalf of the Wellcome Sanger Institute COVID-19 Surveillance Team |
| EPI_ISL_2912543 | Lighthouse Lab in Milton Keynes | Wellcome Sanger Institute for the COVID-19 Genomics UK (COG-UK) Consortium | The Lighthouse Lab in Milton Keynes and Alex Alderton, Roberto Amato, Jeffrey Barrett, Sonia Goncalves, Ewan Harrison, David K. Jackson, Ian Johnston, Dominic Kwiatkowski, Cordelia Langford, John Sillitoe on behalf of the Wellcome Sanger Institute COVID-19 Surveillance Team |
| EPI_ISL_2912544, EPI_ISL_2912545 | Berkshire and Surrey Pathology Services Lighthouse Laboratory | Wellcome Sanger Institute for the COVID-19 Genomics UK (COG-UK) Consortium | Berkshire and Surrey Pathology Services Lighthouse Laboratory and Alex Alderton, Roberto Amato, Jeffrey Barrett, Sonia Goncalves, Ewan Harrison, David K. Jackson, Ian Johnston, Dominic Kwiatkowski, Cordelia Langford, John Sillitoe on behalf of the Wellcome Sanger Institute COVID-19 Surveillance Team |
| EPI_ISL_2912546, EPI_ISL_2912547 | Lighthouse Lab in Glasgow | Wellcome Sanger Institute for the COVID-19 Genomics UK (COG-UK) Consortium | Harper VanSteenhouse, Yumi Kasai, David Gray, Carol Clugston, Anna Dominiczak and Alex Alderton, Roberto Amato, Jeffrey Barrett, Sonia Goncalves, Ewan Harrison, David K. Jackson, Ian Johnston, Dominic Kwiatkowski, Cordelia Langford, John Sillitoe on behalf of the Wellcome Sanger Institute COVID-19 Surveillance Team |
| EPI_ISL_2912548 | Lighthouse Lab in Milton Keynes | Wellcome Sanger Institute for the COVID-19 Genomics UK (COG-UK) Consortium | The Lighthouse Lab in Milton Keynes and Alex Alderton, Roberto Amato, Jeffrey Barrett, Sonia Goncalves, Ewan Harrison, David K. Jackson, Ian Johnston, Dominic Kwiatkowski, Cordelia Langford, John Sillitoe on behalf of the Wellcome Sanger Institute COVID-19 Surveillance Team |
| EPI_ISL_2912549, EPI_ISL_2912550 | Berkshire and Surrey Pathology Services Lighthouse Laboratory | Wellcome Sanger Institute for the COVID-19 Genomics UK (COG-UK) Consortium | Berkshire and Surrey Pathology Services Lighthouse Laboratory and Alex Alderton, Roberto Amato, Jeffrey Barrett, Sonia Goncalves, Ewan Harrison, David K. Jackson, Ian Johnston, Dominic Kwiatkowski, Cordelia Langford, John Sillitoe on behalf of the Wellcome Sanger Institute COVID-19 Surveillance Team |
| EPI_ISL_2912551, EPI_ISL_2912552 | Lighthouse Lab in Glasgow | Wellcome Sanger Institute for the COVID-19 Genomics UK (COG-UK) Consortium | Harper VanSteenhouse, Yumi Kasai, David Gray, Carol Clugston, Anna Dominiczak and Alex Alderton, Roberto Amato, Jeffrey Barrett, Sonia Goncalves, Ewan Harrison, David K. Jackson, Ian Johnston, Dominic Kwiatkowski, Cordelia Langford, John Sillitoe on behalf of the Wellcome Sanger Institute COVID-19 Surveillance Team |
| EPI_ISL_2912553 | Berkshire and Surrey Pathology Services Lighthouse Laboratory | Wellcome Sanger Institute for the COVID-19 Genomics UK (COG-UK) Consortium | Berkshire and Surrey Pathology Services Lighthouse Laboratory and Alex Alderton, Roberto Amato, Jeffrey Barrett, Sonia Goncalves, Ewan Harrison, David K. Jackson, Ian Johnston, Dominic Kwiatkowski, Cordelia Langford, John Sillitoe on behalf of the Wellcome Sanger Institute COVID-19 Surveillance Team |
| EPI_ISL_2912554 | Lighthouse Lab in Glasgow | Wellcome Sanger Institute for the COVID-19 Genomics UK (COG-UK) | Harper VanSteenhouse, Yumi Kasai, David Gray, Carol Clugston, Anna Dominiczak and Alex Alderton, Roberto Amato, Jeffrey Barrett, Sonia Goncalves, Ewan Harrison, David K. Jackson, Ian Johnston, Dominic Kwiatkowski, Cordelia Langford, |

| | | | |
|---|---|---|---|
| | | Consortium | John Sillitoe on behalf of the Wellcome Sanger Institute COVID-19 Surveillance Team |
| EPI_ISL_2912555 | Berkshire and Surrey Pathology Services Lighthouse Laboratory | Wellcome Sanger Institute for the COVID-19 Genomics UK (COG-UK) Consortium | Berkshire and Surrey Pathology Services Lighthouse Laboratory and Alex Alderton, Roberto Amato, Jeffrey Barrett, Sonia Goncalves, Ewan Harrison, David K. Jackson, Ian Johnston, Dominic Kwiatkowski, Cordelia Langford, John Sillitoe on behalf of the Wellcome Sanger Institute COVID-19 Surveillance Team |
| EPI_ISL_2912556 | Lighthouse Lab in Glasgow | Wellcome Sanger Institute for the COVID-19 Genomics UK (COG-UK) Consortium | Harper VanSteenhouse, Yumi Kasai, David Gray, Carol Clugston, Anna Dominiczak and Alex Alderton, Roberto Amato, Jeffrey Barrett, Sonia Goncalves, Ewan Harrison, David K. Jackson, Ian Johnston, Dominic Kwiatkowski, Cordelia Langford, John Sillitoe on behalf of the Wellcome Sanger Institute COVID-19 Surveillance Team |
| EPI_ISL_2912557, EPI_ISL_2912558, EPI_ISL_2912559 | Lighthouse Lab in Milton Keynes | Wellcome Sanger Institute for the COVID-19 Genomics UK (COG-UK) Consortium | The Lighthouse Lab in Milton Keynes and Alex Alderton, Roberto Amato, Jeffrey Barrett, Sonia Goncalves, Ewan Harrison, David K. Jackson, Ian Johnston, Dominic Kwiatkowski, Cordelia Langford, John Sillitoe on behalf of the Wellcome Sanger Institute COVID-19 Surveillance Team |
| EPI_ISL_2912560 | Berkshire and Surrey Pathology Services Lighthouse Laboratory | Wellcome Sanger Institute for the COVID-19 Genomics UK (COG-UK) Consortium | Berkshire and Surrey Pathology Services Lighthouse Laboratory and Alex Alderton, Roberto Amato, Jeffrey Barrett, Sonia Goncalves, Ewan Harrison, David K. Jackson, Ian Johnston, Dominic Kwiatkowski, Cordelia Langford, John Sillitoe on behalf of the Wellcome Sanger Institute COVID-19 Surveillance Team |
| EPI_ISL_2912561 | Lighthouse Lab in Glasgow | Wellcome Sanger Institute for the COVID-19 Genomics UK (COG-UK) Consortium | Harper VanSteenhouse, Yumi Kasai, David Gray, Carol Clugston, Anna Dominiczak and Alex Alderton, Roberto Amato, Jeffrey Barrett, Sonia Goncalves, Ewan Harrison, David K. Jackson, Ian Johnston, Dominic Kwiatkowski, Cordelia Langford, John Sillitoe on behalf of the Wellcome Sanger Institute COVID-19 Surveillance Team |
| EPI_ISL_2912562 | Berkshire and Surrey Pathology Services Lighthouse Laboratory | Wellcome Sanger Institute for the COVID-19 Genomics UK (COG-UK) Consortium | Berkshire and Surrey Pathology Services Lighthouse Laboratory and Alex Alderton, Roberto Amato, Jeffrey Barrett, Sonia Goncalves, Ewan Harrison, David K. Jackson, Ian Johnston, Dominic Kwiatkowski, Cordelia Langford, John Sillitoe on behalf of the Wellcome Sanger Institute COVID-19 Surveillance Team |
| EPI_ISL_2912563, EPI_ISL_2912564, EPI_ISL_2912565 | Lighthouse Lab in Glasgow | Wellcome Sanger Institute for the COVID-19 Genomics UK (COG-UK) Consortium | Harper VanSteenhouse, Yumi Kasai, David Gray, Carol Clugston, Anna Dominiczak and Alex Alderton, Roberto Amato, Jeffrey Barrett, Sonia Goncalves, Ewan Harrison, David K. Jackson, Ian Johnston, Dominic Kwiatkowski, Cordelia Langford, John Sillitoe on behalf of the Wellcome Sanger Institute COVID-19 Surveillance Team |
| EPI_ISL_2912566 | Berkshire and Surrey Pathology Services Lighthouse Laboratory | Wellcome Sanger Institute for the COVID-19 Genomics UK (COG-UK) Consortium | Berkshire and Surrey Pathology Services Lighthouse Laboratory and Alex Alderton, Roberto Amato, Jeffrey Barrett, Sonia Goncalves, Ewan Harrison, David K. Jackson, Ian Johnston, Dominic Kwiatkowski, Cordelia Langford, John Sillitoe on behalf of the Wellcome Sanger Institute COVID-19 Surveillance Team |
| EPI_ISL_2912567 | Lighthouse Lab in Milton Keynes | Wellcome Sanger Institute for the COVID-19 Genomics UK (COG-UK) Consortium | The Lighthouse Lab in Milton Keynes and Alex Alderton, Roberto Amato, Jeffrey Barrett, Sonia Goncalves, Ewan Harrison, David K. Jackson, Ian Johnston, Dominic Kwiatkowski, Cordelia Langford, John Sillitoe on behalf of the Wellcome Sanger Institute COVID-19 Surveillance Team |
| EPI_ISL_2912568 | Lighthouse Lab in Glasgow | Wellcome Sanger Institute for the COVID-19 Genomics UK (COG-UK) Consortium | Harper VanSteenhouse, Yumi Kasai, David Gray, Carol Clugston, Anna Dominiczak and Alex Alderton, Roberto Amato, Jeffrey Barrett, Sonia Goncalves, Ewan Harrison, David K. Jackson, Ian Johnston, Dominic Kwiatkowski, Cordelia Langford, John Sillitoe on behalf of the Wellcome Sanger Institute COVID-19 Surveillance Team |
| EPI_ISL_2912569, EPI_ISL_2912570 | Berkshire and Surrey Pathology Services Lighthouse Laboratory | Wellcome Sanger Institute for the COVID-19 Genomics UK (COG-UK) Consortium | Berkshire and Surrey Pathology Services Lighthouse Laboratory and Alex Alderton, Roberto Amato, Jeffrey Barrett, Sonia Goncalves, Ewan Harrison, David K. Jackson, Ian Johnston, Dominic Kwiatkowski, Cordelia Langford, John Sillitoe on behalf of the Wellcome Sanger Institute COVID-19 Surveillance Team |
| EPI_ISL_2912571 | Lighthouse Lab in Milton Keynes | Wellcome Sanger Institute for the COVID-19 Genomics UK (COG-UK) Consortium | The Lighthouse Lab in Milton Keynes and Alex Alderton, Roberto Amato, Jeffrey Barrett, Sonia Goncalves, Ewan Harrison, David K. Jackson, Ian Johnston, Dominic Kwiatkowski, Cordelia Langford, John Sillitoe on behalf of the Wellcome Sanger Institute COVID-19 Surveillance Team |
| EPI_ISL_2912572 | Berkshire and Surrey Pathology Services Lighthouse Laboratory | Wellcome Sanger Institute for the COVID-19 Genomics UK (COG-UK) Consortium | Berkshire and Surrey Pathology Services Lighthouse Laboratory and Alex Alderton, Roberto Amato, Jeffrey Barrett, Sonia Goncalves, Ewan Harrison, David K. Jackson, Ian Johnston, Dominic Kwiatkowski, Cordelia Langford, John Sillitoe on behalf of the Wellcome Sanger Institute COVID-19 Surveillance Team |
| EPI_ISL_2912573 | Lighthouse Lab in Milton Keynes | Wellcome Sanger Institute for the COVID-19 Genomics UK (COG-UK) Consortium | The Lighthouse Lab in Milton Keynes and Alex Alderton, Roberto Amato, Jeffrey Barrett, Sonia Goncalves, Ewan Harrison, David K. Jackson, Ian Johnston, Dominic Kwiatkowski, Cordelia Langford, John Sillitoe on behalf of the Wellcome Sanger Institute COVID-19 Surveillance Team |
| EPI_ISL_2912574 | Berkshire and Surrey Pathology Services Lighthouse Laboratory | Wellcome Sanger Institute for the COVID-19 Genomics UK (COG-UK) Consortium | Berkshire and Surrey Pathology Services Lighthouse Laboratory and Alex Alderton, Roberto Amato, Jeffrey Barrett, Sonia Goncalves, Ewan Harrison, David K. Jackson, Ian Johnston, Dominic Kwiatkowski, Cordelia Langford, John Sillitoe on behalf of the Wellcome Sanger Institute COVID-19 Surveillance Team |
| EPI_ISL_2912575 | Lighthouse Lab in Milton Keynes | Wellcome Sanger Institute for the COVID-19 Genomics UK (COG-UK) Consortium | The Lighthouse Lab in Milton Keynes and Alex Alderton, Roberto Amato, Jeffrey Barrett, Sonia Goncalves, Ewan Harrison, David K. Jackson, Ian Johnston, Dominic Kwiatkowski, Cordelia Langford, John Sillitoe on behalf of the Wellcome Sanger Institute COVID-19 Surveillance Team |
| EPI_ISL_2912576, EPI_ISL_2912577, EPI_ISL_2912578, EPI_ISL_2912579, EPI_ISL_2912580 | Berkshire and Surrey Pathology Services Lighthouse Laboratory | Wellcome Sanger Institute for the COVID-19 Genomics UK (COG-UK) Consortium | Berkshire and Surrey Pathology Services Lighthouse Laboratory and Alex Alderton, Roberto Amato, Jeffrey Barrett, Sonia Goncalves, Ewan Harrison, David K. Jackson, Ian Johnston, Dominic Kwiatkowski, Cordelia Langford, John Sillitoe on behalf of the Wellcome Sanger Institute COVID-19 Surveillance Team |
| EPI_ISL_2912581 | Lighthouse Lab in Glasgow | Wellcome Sanger Institute for the COVID-19 Genomics UK (COG-UK) Consortium | Harper VanSteenhouse, Yumi Kasai, David Gray, Carol Clugston, Anna Dominiczak and Alex Alderton, Roberto Amato, Jeffrey Barrett, Sonia Goncalves, Ewan Harrison, David K. Jackson, Ian Johnston, Dominic Kwiatkowski, Cordelia Langford, John Sillitoe on behalf of the Wellcome Sanger Institute COVID-19 Surveillance Team |
| EPI_ISL_2912582 | Lighthouse Lab in Milton Keynes | Wellcome Sanger Institute for the COVID-19 Genomics UK (COG-UK) Consortium | The Lighthouse Lab in Milton Keynes and Alex Alderton, Roberto Amato, Jeffrey Barrett, Sonia Goncalves, Ewan Harrison, David K. Jackson, Ian Johnston, Dominic Kwiatkowski, Cordelia Langford, John Sillitoe on behalf of the Wellcome Sanger Institute COVID-19 Surveillance Team |
| EPI_ISL_2912583, EPI_ISL_2912584, EPI_ISL_2912585 | Berkshire and Surrey Pathology Services Lighthouse Laboratory | Wellcome Sanger Institute for the COVID-19 Genomics UK (COG-UK) Consortium | Berkshire and Surrey Pathology Services Lighthouse Laboratory and Alex Alderton, Roberto Amato, Jeffrey Barrett, Sonia Goncalves, Ewan Harrison, David K. Jackson, Ian Johnston, Dominic Kwiatkowski, Cordelia Langford, John Sillitoe on behalf of the Wellcome Sanger Institute COVID-19 Surveillance Team |
| EPI_ISL_2912586 | Lighthouse Lab in Milton Keynes | Wellcome Sanger Institute for the COVID-19 Genomics UK (COG-UK) Consortium | The Lighthouse Lab in Milton Keynes and Alex Alderton, Roberto Amato, Jeffrey Barrett, Sonia Goncalves, Ewan Harrison, David K. Jackson, Ian Johnston, Dominic Kwiatkowski, Cordelia Langford, John Sillitoe on behalf of the Wellcome Sanger Institute COVID-19 Surveillance Team |
| EPI_ISL_2912587 | Berkshire and Surrey Pathology Services Lighthouse Laboratory | Wellcome Sanger Institute for the COVID-19 Genomics UK (COG-UK) Consortium | Berkshire and Surrey Pathology Services Lighthouse Laboratory and Alex Alderton, Roberto Amato, Jeffrey Barrett, Sonia Goncalves, Ewan Harrison, David K. Jackson, Ian Johnston, Dominic Kwiatkowski, Cordelia Langford, John Sillitoe on behalf of the Wellcome Sanger Institute COVID-19 Surveillance Team |
| EPI_ISL_2912588, EPI_ISL_2912589, EPI_ISL_2912590, EPI_ISL_2912591, EPI_ISL_2912592, EPI_ISL_2912593, EPI_ISL_2912594, EPI_ISL_2912595, EPI_ISL_2912596, EPI_ISL_2912597, EPI_ISL_2912598, EPI_ISL_2912599, EPI_ISL_2912600, EPI_ISL_2912601, EPI_ISL_2912602, EPI_ISL_2912603, EPI_ISL_2912604, EPI_ISL_2912605, EPI_ISL_2912606, EPI_ISL_2912607, EPI_ISL_2912608, EPI_ISL_2912609, EPI_ISL_2912610, EPI_ISL_2912611, EPI_ISL_2912612, EPI_ISL_2912613, EPI_ISL_2912614, EPI_ISL_2912615, EPI_ISL_2912616, EPI_ISL_2912617, EPI_ISL_2912618, EPI_ISL_2912619, EPI_ISL_2912620, EPI_ISL_2912621, EPI_ISL_2912622, EPI_ISL_2912623, EPI_ISL_2912624, EPI_ISL_2912625, EPI_ISL_2912626, EPI_ISL_2912627, EPI_ISL_2912628, EPI_ISL_2912629, EPI_ISL_2912630, EPI_ISL_2912631, EPI_ISL_2912632, EPI_ISL_2912634, EPI_ISL_2912635, EPI_ISL_2912636, EPI_ISL_2912637, EPI_ISL_2912638, EPI_ISL_2912639, EPI_ISL_2912640, EPI_ISL_2912641, EPI_ISL_2912642, EPI_ISL_2912643, EPI_ISL_2912644, EPI_ISL_2912645, EPI_ISL_2912646, EPI_ISL_2912647, EPI_ISL_2912648, EPI_ISL_2912649, EPI_ISL_2912650, EPI_ISL_2912651, EPI_ISL_2912652, EPI_ISL_2912653, EPI_ISL_2912654, EPI_ISL_2912655, EPI_ISL_2912656, EPI_ISL_2912657, EPI_ISL_2912658, EPI_ISL_2912659, EPI_ISL_2912660, EPI_ISL_2912661, EPI_ISL_2912662, EPI_ISL_2912663, EPI_ISL_2912664, EPI_ISL_2912665, EPI_ISL_2912667, EPI_ISL_2912668, EPI_ISL_2912669, EPI_ISL_2912670, EPI_ISL_2912671, EPI_ISL_2912672, EPI_ISL_2912673, EPI_ISL_2912674, EPI_ISL_2912675, EPI_ISL_2912676, EPI_ISL_2912677, EPI_ISL_2912678, EPI_ISL_2912679, EPI_ISL_2912680, EPI_ISL_2912681, EPI_ISL_2912682, EPI_ISL_2912683, EPI_ISL_2912684, EPI_ISL_2912685, EPI_ISL_2912686, EPI_ISL_2912687, EPI_ISL_2912688, EPI_ISL_2912689, EPI_ISL_2912690, EPI_ISL_2912691, EPI_ISL_2912692, EPI_ISL_2912693, EPI_ISL_2912694, EPI_ISL_2912695, EPI_ISL_2912696, EPI_ISL_2912697, EPI_ISL_2912698, EPI_ISL_2912699, EPI_ISL_2912700, EPI_ISL_2912701, EPI_ISL_2912702, EPI_ISL_2912703, EPI_ISL_2912704, EPI_ISL_2912705, EPI_ISL_2912706, EPI_ISL_2912707, EPI_ISL_2912708, EPI_ISL_2912709, EPI_ISL_2912710, EPI_ISL_2912711, EPI_ISL_2912712, EPI_ISL_2912713, EPI_ISL_2912714, EPI_ISL_2912715, EPI_ISL_2912716, EPI_ISL_2912717, EPI_ISL_2912718, EPI_ISL_2912719, EPI_ISL_2912720, EPI_ISL_2912721, EPI_ISL_2912722, EPI_ISL_2912723, EPI_ISL_2912724, EPI_ISL_2912725, EPI_ISL_2912726, EPI_ISL_2912727, EPI_ISL_2912728, EPI_ISL_2912730, EPI_ISL_2912731, EPI_ISL_2912732, EPI_ISL_2912733, EPI_ISL_2912734, EPI_ISL_2912735, EPI_ISL_2912736, EPI_ISL_2912737, EPI_ISL_2912738, EPI_ISL_2912739, EPI_ISL_2912740, EPI_ISL_2912741, EPI_ISL_2912742, EPI_ISL_2912743, EPI_ISL_2912744, EPI_ISL_2912745, EPI_ISL_2912746, EPI_ISL_2912747, EPI_ISL_2912748, EPI_ISL_2912749, EPI_ISL_2912750, EPI_ISL_2912751, EPI_ISL_2912752, EPI_ISL_2912753, EPI_ISL_2912754, EPI_ISL_2912755, EPI_ISL_2912756, EPI_ISL_2912757, EPI_ISL_2912758, EPI_ISL_2912759, EPI_ISL_2912760, EPI_ISL_2912761, EPI_ISL_2912762, EPI_ISL_2912763, EPI_ISL_2912764, EPI_ISL_2912765, EPI_ISL_2912766, EPI_ISL_2912767, EPI_ISL_2912768, EPI_ISL_2912769, EPI_ISL_2912770, EPI_ISL_2912771, EPI_ISL_2912772, EPI_ISL_2912773, EPI_ISL_2912774, EPI_ISL_2912775, EPI_ISL_2912776, EPI_ISL_2912777, EPI_ISL_2912778, EPI_ISL_2912779, EPI_ISL_2912780, EPI_ISL_2912781, EPI_ISL_2912782, EPI_ISL_2912783, EPI_ISL_2912784, EPI_ISL_2912785, EPI_ISL_2912786, EPI_ISL_2912787, EPI_ISL_2912788, EPI_ISL_2912789, EPI_ISL_2912790, EPI_ISL_2912791, EPI_ISL_2912792, EPI_ISL_2912793, EPI_ISL_2912795, EPI_ISL_2912796, EPI_ISL_2912797, EPI_ISL_2912798, EPI_ISL_2912799, EPI_ISL_2912800, EPI_ISL_2912801, EPI_ISL_2912802, EPI_ISL_2912803, EPI_ISL_2912804, EPI_ISL_2912805, EPI_ISL_2912806, EPI_ISL_2912807, EPI_ISL_2912808, EPI_ISL_2912809, EPI_ISL_2912810, EPI_ISL_2912811, EPI_ISL_2912812, EPI_ISL_2912813, EPI_ISL_2912814, EPI_ISL_2912815, EPI_ISL_2912816, EPI_ISL_2912817, EPI_ISL_2912818, EPI_ISL_2912819, EPI_ISL_2912820, EPI_ISL_2912821, EPI_ISL_2912823, EPI_ISL_2912824, EPI_ISL_2912825, EPI_ISL_2912826, EPI_ISL_2912827, EPI_ISL_2912828, EPI_ISL_2912829, EPI_ISL_2912830, EPI_ISL_2912831, EPI_ISL_2912832, EPI_ISL_2912833, EPI_ISL_2912834, EPI_ISL_2912835, EPI_ISL_2912836, EPI_ISL_2912837, EPI_ISL_2912838, EPI_ISL_2912839, EPI_ISL_2912840, EPI_ISL_2912841, EPI_ISL_2912842, EPI_ISL_2912843, EPI_ISL_2912844, EPI_ISL_2912845, EPI_ISL_2912846, EPI_ISL_2912847, EPI_ISL_2912848, EPI_ISL_2912849, EPI_ISL_2912850, EPI_ISL_2912851, EPI_ISL_2912852, EPI_ISL_2912853, EPI_ISL_2912854, EPI_ISL_2912855, EPI_ISL_2912856, EPI_ISL_2912857, EPI_ISL_2912858, EPI_ISL_2912859, EPI_ISL_2912860, EPI_ISL_2912861, EPI_ISL_2912862, EPI_ISL_2912863, EPI_ISL_2912864, EPI_ISL_2912865, EPI_ISL_2912866, EPI_ISL_2912867, EPI_ISL_2912868, EPI_ISL_2912869, EPI_ISL_2912870, EPI_ISL_2912871, EPI_ISL_2912872, EPI_ISL_2912873, EPI_ISL_2912874, EPI_ISL_2912875, EPI_ISL_2912876, EPI_ISL_2912877, EPI_ISL_2912878, EPI_ISL_2912879, EPI_ISL_2912880, EPI_ISL_2912881, EPI_ISL_2912882, EPI_ISL_2912883, EPI_ISL_2912884, EPI_ISL_2912885, EPI_ISL_2912886, EPI_ISL_2912887, EPI_ISL_2912888, EPI_ISL_2912889, EPI_ISL_2912890, EPI_ISL_2912891, EPI_ISL_2912892, EPI_ISL_2912893, EPI_ISL_2912894, EPI_ISL_2912895, EPI_ISL_2912896, EPI_ISL_2912897, EPI_ISL_2912899, EPI_ISL_2912900, EPI_ISL_2912901, EPI_ISL_2912902, EPI_ISL_2912903, EPI_ISL_2912904, EPI_ISL_2912905, EPI_ISL_2912906, EPI_ISL_2912907, EPI_ISL_2912908, EPI_ISL_2912909, EPI_ISL_2912910, EPI_ISL_2912911, EPI_ISL_2912912, EPI_ISL_2912913, EPI_ISL_2912914, EPI_ISL_2912915, EPI_ISL_2912916, EPI_ISL_2912918, EPI_ISL_2912919, EPI_ISL_2912920, EPI_ISL_2912921, EPI_ISL_2912922, EPI_ISL_2912923, EPI_ISL_2912924, EPI_ISL_2912926, EPI_ISL_2912927, EPI_ISL_2912928, EPI_ISL_2912929, EPI_ISL_2912930, EPI_ISL_2912931, EPI_ISL_2912932, EPI_ISL_2912933, EPI_ISL_2912934, EPI_ISL_2912935, EPI_ISL_2912936, EPI_ISL_2912937, EPI_ISL_2912938, EPI_ISL_2912939, EPI_ISL_2912940, EPI_ISL_2912941, EPI_ISL_2912942, EPI_ISL_2912943, EPI_ISL_2912944, EPI_ISL_2912945, EPI_ISL_2912946, EPI_ISL_2912947, EPI_ISL_2912948, EPI_ISL_2912949, EPI_ISL_2912950, EPI_ISL_2912951, EPI_ISL_2912952, EPI_ISL_2912953, EPI_ISL_2912954, EPI_ISL_2912955, EPI_ISL_2912956, EPI_ISL_2912957, EPI_ISL_2912958, EPI_ISL_2912959, EPI_ISL_2912960, EPI_ISL_2912961 | | | |
| see above | Lighthouse Lab in Glasgow | Wellcome Sanger Institute for the COVID-19 Genomics UK (COG-UK) Consortium | Harper VanSteenhouse, Yumi Kasai, David Gray, Carol Clugston, Anna Dominiczak and Alex Alderton, Roberto Amato, Jeffrey Barrett, Sonia Goncalves, Ewan Harrison, David K. Jackson, Ian Johnston, Dominic Kwiatkowski, Cordelia Langford, John Sillitoe on behalf of the Wellcome Sanger Institute COVID-19 Surveillance Team |
| EPI_ISL_2912962 | Lighthouse Lab in Milton Keynes | Wellcome Sanger Institute for the COVID-19 Genomics UK (COG-UK) Consortium | The Lighthouse Lab in Milton Keynes and Alex Alderton, Roberto Amato, Jeffrey Barrett, Sonia Goncalves, Ewan Harrison, David K. Jackson, Ian Johnston, Dominic Kwiatkowski, Cordelia Langford, John Sillitoe on behalf of the Wellcome Sanger Institute COVID-19 Surveillance Team |
| EPI_ISL_2912963 | Berkshire and Surrey Pathology Services Lighthouse Laboratory | Wellcome Sanger Institute for the COVID-19 Genomics UK (COG-UK) Consortium | Berkshire and Surrey Pathology Services Lighthouse Laboratory and Alex Alderton, Roberto Amato, Jeffrey Barrett, Sonia Goncalves, Ewan Harrison, David K. Jackson, Ian Johnston, Dominic Kwiatkowski, Cordelia Langford, John Sillitoe on behalf of the Wellcome Sanger Institute COVID-19 Surveillance Team |
| EPI_ISL_2912964 | Lighthouse Lab in Milton Keynes | Wellcome Sanger Institute for the COVID-19 Genomics UK (COG-UK) Consortium | The Lighthouse Lab in Milton Keynes and Alex Alderton, Roberto Amato, Jeffrey Barrett, Sonia Goncalves, Ewan Harrison, David K. Jackson, Ian Johnston, Dominic Kwiatkowski, Cordelia Langford, John Sillitoe on behalf of the Wellcome Sanger Institute COVID-19 Surveillance Team |
| EPI_ISL_2912965, EPI_ISL_2912966 | Berkshire and Surrey Pathology Services Lighthouse Laboratory | Wellcome Sanger Institute for the COVID-19 Genomics UK (COG-UK) Consortium | Berkshire and Surrey Pathology Services Lighthouse Laboratory and Alex Alderton, Roberto Amato, Jeffrey Barrett, Sonia Goncalves, Ewan Harrison, David K. Jackson, Ian Johnston, Dominic Kwiatkowski, Cordelia Langford, John Sillitoe on behalf of the Wellcome Sanger Institute COVID-19 Surveillance Team |
| EPI_ISL_2912967 | Lighthouse Lab in Glasgow | Wellcome Sanger Institute for the COVID-19 Genomics UK (COG-UK) Consortium | Harper VanSteenhouse, Yumi Kasai, David Gray, Carol Clugston, Anna Dominiczak and Alex Alderton, Roberto Amato, Jeffrey Barrett, Sonia Goncalves, Ewan Harrison, David K. Jackson, Ian Johnston, Dominic Kwiatkowski, Cordelia Langford, John Sillitoe on behalf of the Wellcome Sanger Institute COVID-19 Surveillance Team |
| EPI_ISL_2912968, EPI_ISL_2912969 | Lighthouse Lab in Milton Keynes | Wellcome Sanger Institute for the COVID-19 Genomics UK (COG-UK) Consortium | The Lighthouse Lab in Milton Keynes and Alex Alderton, Roberto Amato, Jeffrey Barrett, Sonia Goncalves, Ewan Harrison, David K. Jackson, Ian Johnston, Dominic Kwiatkowski, Cordelia Langford, John Sillitoe on behalf of the Wellcome Sanger Institute COVID-19 Surveillance Team |
| EPI_ISL_2912970 | Lighthouse Lab in Glasgow | Wellcome Sanger Institute for the COVID-19 Genomics UK (COG-UK) Consortium | Harper VanSteenhouse, Yumi Kasai, David Gray, Carol Clugston, Anna Dominiczak and Alex Alderton, Roberto Amato, Jeffrey Barrett, Sonia Goncalves, Ewan Harrison, David K. Jackson, Ian Johnston, Dominic Kwiatkowski, Cordelia Langford, John Sillitoe on behalf of the Wellcome Sanger Institute COVID-19 Surveillance Team |
| EPI_ISL_2912971 | Lighthouse Lab in Milton Keynes | Wellcome Sanger Institute for the COVID-19 Genomics UK (COG-UK) Consortium | The Lighthouse Lab in Milton Keynes and Alex Alderton, Roberto Amato, Jeffrey Barrett, Sonia Goncalves, Ewan Harrison, David K. Jackson, Ian Johnston, Dominic Kwiatkowski, Cordelia Langford, John Sillitoe on behalf of the Wellcome Sanger Institute COVID-19 Surveillance Team |
| EPI_ISL_2912972 | Berkshire and Surrey Pathology Services Lighthouse Laboratory | Wellcome Sanger Institute for the COVID-19 Genomics UK (COG-UK) Consortium | Berkshire and Surrey Pathology Services Lighthouse Laboratory and Alex Alderton, Roberto Amato, Jeffrey Barrett, Sonia Goncalves, Ewan Harrison, David K. Jackson, Ian Johnston, Dominic Kwiatkowski, Cordelia Langford, John Sillitoe on behalf of the Wellcome Sanger Institute COVID-19 Surveillance Team |
| EPI_ISL_2912973, EPI_ISL_2912974 | Lighthouse Lab in Milton Keynes | Wellcome Sanger Institute for the COVID-19 Genomics UK (COG-UK) Consortium | The Lighthouse Lab in Milton Keynes and Alex Alderton, Roberto Amato, Jeffrey Barrett, Sonia Goncalves, Ewan Harrison, David K. Jackson, Ian Johnston, Dominic Kwiatkowski, Cordelia Langford, John Sillitoe on behalf of the Wellcome Sanger Institute COVID-19 Surveillance Team |
| EPI_ISL_2912975, EPI_ISL_2912976 | Lighthouse Lab in Glasgow | Wellcome Sanger Institute for the COVID-19 Genomics UK (COG-UK) Consortium | Harper VanSteenhouse, Yumi Kasai, David Gray, Carol Clugston, Anna Dominiczak and Alex Alderton, Roberto Amato, Jeffrey Barrett, Sonia Goncalves, Ewan Harrison, David K. Jackson, Ian Johnston, Dominic Kwiatkowski, Cordelia Langford, John Sillitoe on behalf of the Wellcome Sanger Institute COVID-19 Surveillance Team |
| EPI_ISL_2912977, EPI_ISL_2912978 | Lighthouse Lab in Milton Keynes | Wellcome Sanger Institute for the COVID-19 Genomics UK (COG-UK) Consortium | The Lighthouse Lab in Milton Keynes and Alex Alderton, Roberto Amato, Jeffrey Barrett, Sonia Goncalves, Ewan Harrison, David K. Jackson, Ian Johnston, Dominic Kwiatkowski, Cordelia Langford, John Sillitoe on behalf of the Wellcome Sanger Institute COVID-19 Surveillance Team |

| | | | |
|---|---|---|---|
| EPI_ISL_2912979, EPI_ISL_2912980 | Lighthouse Lab in Glasgow | Wellcome Sanger Institute for the COVID-19 Genomics UK (COG-UK) Consortium | Harper VanSteenhouse, Yumi Kasai, David Gray, Carol Clugston, Anna Dominiczak and Alex Alderton, Roberto Amato, Jeffrey Barrett, Sonia Goncalves, Ewan Harrison, David K. Jackson, Ian Johnston, Dominic Kwiatkowski, Cordelia Langford, John Sillitoe on behalf of the Wellcome Sanger Institute COVID-19 Surveillance Team |
| EPI_ISL_2912981, EPI_ISL_2912982 | Lighthouse Lab in Milton Keynes | Wellcome Sanger Institute for the COVID-19 Genomics UK (COG-UK) Consortium | The Lighthouse Lab in Milton Keynes and Alex Alderton, Roberto Amato, Jeffrey Barrett, Sonia Goncalves, Ewan Harrison, David K. Jackson, Ian Johnston, Dominic Kwiatkowski, Cordelia Langford, John Sillitoe on behalf of the Wellcome Sanger Institute COVID-19 Surveillance Team |
| EPI_ISL_2912983 | Lighthouse Lab in Glasgow | Wellcome Sanger Institute for the COVID-19 Genomics UK (COG-UK) Consortium | Harper VanSteenhouse, Yumi Kasai, David Gray, Carol Clugston, Anna Dominiczak and Alex Alderton, Roberto Amato, Jeffrey Barrett, Sonia Goncalves, Ewan Harrison, David K. Jackson, Ian Johnston, Dominic Kwiatkowski, Cordelia Langford, John Sillitoe on behalf of the Wellcome Sanger Institute COVID-19 Surveillance Team |
| EPI_ISL_2912984 | Berkshire and Surrey Pathology Services Lighthouse Laboratory | Wellcome Sanger Institute for the COVID-19 Genomics UK (COG-UK) Consortium | Berkshire and Surrey Pathology Services Lighthouse Laboratory and Alex Alderton, Roberto Amato, Jeffrey Barrett, Sonia Goncalves, Ewan Harrison, David K. Jackson, Ian Johnston, Dominic Kwiatkowski, Cordelia Langford, John Sillitoe on behalf of the Wellcome Sanger Institute COVID-19 Surveillance Team |
| EPI_ISL_2912985, EPI_ISL_2912986 | Lighthouse Lab in Milton Keynes | Wellcome Sanger Institute for the COVID-19 Genomics UK (COG-UK) Consortium | The Lighthouse Lab in Milton Keynes and Alex Alderton, Roberto Amato, Jeffrey Barrett, Sonia Goncalves, Ewan Harrison, David K. Jackson, Ian Johnston, Dominic Kwiatkowski, Cordelia Langford, John Sillitoe on behalf of the Wellcome Sanger Institute COVID-19 Surveillance Team |
| EPI_ISL_2912987 | Lighthouse Lab in Glasgow | Wellcome Sanger Institute for the COVID-19 Genomics UK (COG-UK) Consortium | Harper VanSteenhouse, Yumi Kasai, David Gray, Carol Clugston, Anna Dominiczak and Alex Alderton, Roberto Amato, Jeffrey Barrett, Sonia Goncalves, Ewan Harrison, David K. Jackson, Ian Johnston, Dominic Kwiatkowski, Cordelia Langford, John Sillitoe on behalf of the Wellcome Sanger Institute COVID-19 Surveillance Team |
| EPI_ISL_2912988 | Berkshire and Surrey Pathology Services Lighthouse Laboratory | Wellcome Sanger Institute for the COVID-19 Genomics UK (COG-UK) Consortium | Berkshire and Surrey Pathology Services Lighthouse Laboratory and Alex Alderton, Roberto Amato, Jeffrey Barrett, Sonia Goncalves, Ewan Harrison, David K. Jackson, Ian Johnston, Dominic Kwiatkowski, Cordelia Langford, John Sillitoe on behalf of the Wellcome Sanger Institute COVID-19 Surveillance Team |
| EPI_ISL_2912989, EPI_ISL_2912990 | Lighthouse Lab in Glasgow | Wellcome Sanger Institute for the COVID-19 Genomics UK (COG-UK) Consortium | Harper VanSteenhouse, Yumi Kasai, David Gray, Carol Clugston, Anna Dominiczak and Alex Alderton, Roberto Amato, Jeffrey Barrett, Sonia Goncalves, Ewan Harrison, David K. Jackson, Ian Johnston, Dominic Kwiatkowski, Cordelia Langford, John Sillitoe on behalf of the Wellcome Sanger Institute COVID-19 Surveillance Team |
| EPI_ISL_2912991 | Berkshire and Surrey Pathology Services Lighthouse Laboratory | Wellcome Sanger Institute for the COVID-19 Genomics UK (COG-UK) Consortium | Berkshire and Surrey Pathology Services Lighthouse Laboratory and Alex Alderton, Roberto Amato, Jeffrey Barrett, Sonia Goncalves, Ewan Harrison, David K. Jackson, Ian Johnston, Dominic Kwiatkowski, Cordelia Langford, John Sillitoe on behalf of the Wellcome Sanger Institute COVID-19 Surveillance Team |
| EPI_ISL_2912992, EPI_ISL_2912993, EPI_ISL_2912994 | Lighthouse Lab in Milton Keynes | Wellcome Sanger Institute for the COVID-19 Genomics UK (COG-UK) Consortium | The Lighthouse Lab in Milton Keynes and Alex Alderton, Roberto Amato, Jeffrey Barrett, Sonia Goncalves, Ewan Harrison, David K. Jackson, Ian Johnston, Dominic Kwiatkowski, Cordelia Langford, John Sillitoe on behalf of the Wellcome Sanger Institute COVID-19 Surveillance Team |
| EPI_ISL_2912995, EPI_ISL_2912996 | Berkshire and Surrey Pathology Services Lighthouse Laboratory | Wellcome Sanger Institute for the COVID-19 Genomics UK (COG-UK) Consortium | Berkshire and Surrey Pathology Services Lighthouse Laboratory and Alex Alderton, Roberto Amato, Jeffrey Barrett, Sonia Goncalves, Ewan Harrison, David K. Jackson, Ian Johnston, Dominic Kwiatkowski, Cordelia Langford, John Sillitoe on behalf of the Wellcome Sanger Institute COVID-19 Surveillance Team |
| EPI_ISL_2912997, EPI_ISL_2912998 | Lighthouse Lab in Milton Keynes | Wellcome Sanger Institute for the COVID-19 Genomics UK (COG-UK) Consortium | The Lighthouse Lab in Milton Keynes and Alex Alderton, Roberto Amato, Jeffrey Barrett, Sonia Goncalves, Ewan Harrison, David K. Jackson, Ian Johnston, Dominic Kwiatkowski, Cordelia Langford, John Sillitoe on behalf of the Wellcome Sanger Institute COVID-19 Surveillance Team |
| EPI_ISL_2912999 | Berkshire and Surrey Pathology Services Lighthouse Laboratory | Wellcome Sanger Institute for the COVID-19 Genomics UK (COG-UK) Consortium | Berkshire and Surrey Pathology Services Lighthouse Laboratory and Alex Alderton, Roberto Amato, Jeffrey Barrett, Sonia Goncalves, Ewan Harrison, David K. Jackson, Ian Johnston, Dominic Kwiatkowski, Cordelia Langford, John Sillitoe on behalf of the Wellcome Sanger Institute COVID-19 Surveillance Team |
| EPI_ISL_2913000, EPI_ISL_2913001 | Lighthouse Lab in Glasgow | Wellcome Sanger Institute for the COVID-19 Genomics UK (COG-UK) Consortium | Harper VanSteenhouse, Yumi Kasai, David Gray, Carol Clugston, Anna Dominiczak and Alex Alderton, Roberto Amato, Jeffrey Barrett, Sonia Goncalves, Ewan Harrison, David K. Jackson, Ian Johnston, Dominic Kwiatkowski, Cordelia Langford, John Sillitoe on behalf of the Wellcome Sanger Institute COVID-19 Surveillance Team |
| EPI_ISL_2913002, EPI_ISL_2913003 | Lighthouse Lab in Milton Keynes | Wellcome Sanger Institute for the COVID-19 Genomics UK (COG-UK) Consortium | The Lighthouse Lab in Milton Keynes and Alex Alderton, Roberto Amato, Jeffrey Barrett, Sonia Goncalves, Ewan Harrison, David K. Jackson, Ian Johnston, Dominic Kwiatkowski, Cordelia Langford, John Sillitoe on behalf of the Wellcome Sanger Institute COVID-19 Surveillance Team |
| EPI_ISL_2913004, EPI_ISL_2913005 | Berkshire and Surrey Pathology Services Lighthouse Laboratory | Wellcome Sanger Institute for the COVID-19 Genomics UK (COG-UK) Consortium | Berkshire and Surrey Pathology Services Lighthouse Laboratory and Alex Alderton, Roberto Amato, Jeffrey Barrett, Sonia Goncalves, Ewan Harrison, David K. Jackson, Ian Johnston, Dominic Kwiatkowski, Cordelia Langford, John Sillitoe on behalf of the Wellcome Sanger Institute COVID-19 Surveillance Team |
| EPI_ISL_2913006, EPI_ISL_2913007 | Lighthouse Lab in Milton Keynes | Wellcome Sanger Institute for the COVID-19 Genomics UK (COG-UK) Consortium | The Lighthouse Lab in Milton Keynes and Alex Alderton, Roberto Amato, Jeffrey Barrett, Sonia Goncalves, Ewan Harrison, David K. Jackson, Ian Johnston, Dominic Kwiatkowski, Cordelia Langford, John Sillitoe on behalf of the Wellcome Sanger Institute COVID-19 Surveillance Team |
| EPI_ISL_2913008 | Lighthouse Lab in Glasgow | Wellcome Sanger Institute for the COVID-19 Genomics UK (COG-UK) Consortium | Harper VanSteenhouse, Yumi Kasai, David Gray, Carol Clugston, Anna Dominiczak and Alex Alderton, Roberto Amato, Jeffrey Barrett, Sonia Goncalves, Ewan Harrison, David K. Jackson, Ian Johnston, Dominic Kwiatkowski, Cordelia Langford, John Sillitoe on behalf of the Wellcome Sanger Institute COVID-19 Surveillance Team |
| EPI_ISL_2913009, EPI_ISL_2913010 | Lighthouse Lab in Milton Keynes | Wellcome Sanger Institute for the COVID-19 Genomics UK (COG-UK) Consortium | The Lighthouse Lab in Milton Keynes and Alex Alderton, Roberto Amato, Jeffrey Barrett, Sonia Goncalves, Ewan Harrison, David K. Jackson, Ian Johnston, Dominic Kwiatkowski, Cordelia Langford, John Sillitoe on behalf of the Wellcome Sanger Institute COVID-19 Surveillance Team |
| EPI_ISL_2913011 | Lighthouse Lab in Glasgow | Wellcome Sanger Institute for the COVID-19 Genomics UK (COG-UK) Consortium | Harper VanSteenhouse, Yumi Kasai, David Gray, Carol Clugston, Anna Dominiczak and Alex Alderton, Roberto Amato, Jeffrey Barrett, Sonia Goncalves, Ewan Harrison, David K. Jackson, Ian Johnston, Dominic Kwiatkowski, Cordelia Langford, John Sillitoe on behalf of the Wellcome Sanger Institute COVID-19 Surveillance Team |
| EPI_ISL_2913012 | Lighthouse Lab in Milton Keynes | Wellcome Sanger Institute for the COVID-19 Genomics UK (COG-UK) Consortium | The Lighthouse Lab in Milton Keynes and Alex Alderton, Roberto Amato, Jeffrey Barrett, Sonia Goncalves, Ewan Harrison, David K. Jackson, Ian Johnston, Dominic Kwiatkowski, Cordelia Langford, John Sillitoe on behalf of the Wellcome Sanger Institute COVID-19 Surveillance Team |
| EPI_ISL_2913013 | Lighthouse Lab in Glasgow | Wellcome Sanger Institute for the COVID-19 Genomics UK (COG-UK) Consortium | Harper VanSteenhouse, Yumi Kasai, David Gray, Carol Clugston, Anna Dominiczak and Alex Alderton, Roberto Amato, Jeffrey Barrett, Sonia Goncalves, Ewan Harrison, David K. Jackson, Ian Johnston, Dominic Kwiatkowski, Cordelia Langford, John Sillitoe on behalf of the Wellcome Sanger Institute COVID-19 Surveillance Team |
| EPI_ISL_2913014, EPI_ISL_2913015, EPI_ISL_2913016, EPI_ISL_2913017, EPI_ISL_2913018, EPI_ISL_2913019 | Lighthouse Lab in Milton Keynes | Wellcome Sanger Institute for the COVID-19 Genomics UK (COG-UK) Consortium | The Lighthouse Lab in Milton Keynes and Alex Alderton, Roberto Amato, Jeffrey Barrett, Sonia Goncalves, Ewan Harrison, David K. Jackson, Ian Johnston, Dominic Kwiatkowski, Cordelia Langford, John Sillitoe on behalf of the Wellcome Sanger Institute COVID-19 Surveillance Team |
| EPI_ISL_2913020 | Lighthouse Lab in Glasgow | Wellcome Sanger Institute for the COVID-19 Genomics UK (COG-UK) Consortium | Harper VanSteenhouse, Yumi Kasai, David Gray, Carol Clugston, Anna Dominiczak and Alex Alderton, Roberto Amato, Jeffrey Barrett, Sonia Goncalves, Ewan Harrison, David K. Jackson, Ian Johnston, Dominic Kwiatkowski, Cordelia Langford, John Sillitoe on behalf of the Wellcome Sanger Institute COVID-19 Surveillance Team |
| EPI_ISL_2913021 | Lighthouse Lab in Milton Keynes | Wellcome Sanger Institute for the COVID-19 Genomics UK (COG-UK) Consortium | The Lighthouse Lab in Milton Keynes and Alex Alderton, Roberto Amato, Jeffrey Barrett, Sonia Goncalves, Ewan Harrison, David K. Jackson, Ian Johnston, Dominic Kwiatkowski, Cordelia Langford, John Sillitoe on behalf of the Wellcome Sanger Institute COVID-19 Surveillance Team |
| EPI_ISL_2913022, EPI_ISL_2913023, EPI_ISL_2913024, EPI_ISL_2913025, EPI_ISL_2913026, EPI_ISL_2913027 | Berkshire and Surrey Pathology Services Lighthouse Laboratory | Wellcome Sanger Institute for the COVID-19 Genomics UK (COG-UK) Consortium | Berkshire and Surrey Pathology Services Lighthouse Laboratory and Alex Alderton, Roberto Amato, Jeffrey Barrett, Sonia Goncalves, Ewan Harrison, David K. Jackson, Ian Johnston, Dominic Kwiatkowski, Cordelia Langford, John Sillitoe on behalf of the Wellcome Sanger Institute COVID-19 Surveillance Team |
| EPI_ISL_2913028 | Lighthouse Lab in Glasgow | Wellcome Sanger Institute for the COVID-19 Genomics UK (COG-UK) Consortium | Harper VanSteenhouse, Yumi Kasai, David Gray, Carol Clugston, Anna Dominiczak and Alex Alderton, Roberto Amato, Jeffrey Barrett, Sonia Goncalves, Ewan Harrison, David K. Jackson, Ian Johnston, Dominic Kwiatkowski, Cordelia Langford, John Sillitoe on behalf of the Wellcome Sanger Institute COVID-19 Surveillance Team |
| EPI_ISL_2913029, EPI_ISL_2913030, EPI_ISL_2913031 | Lighthouse Lab in Milton Keynes | Wellcome Sanger Institute for the COVID-19 Genomics UK (COG-UK) Consortium | The Lighthouse Lab in Milton Keynes and Alex Alderton, Roberto Amato, Jeffrey Barrett, Sonia Goncalves, Ewan Harrison, David K. Jackson, Ian Johnston, Dominic Kwiatkowski, Cordelia Langford, John Sillitoe on behalf of the Wellcome Sanger Institute COVID-19 Surveillance Team |
| EPI_ISL_2913032 | Lighthouse Lab in Glasgow | Wellcome Sanger Institute for the COVID-19 Genomics UK (COG-UK) Consortium | Harper VanSteenhouse, Yumi Kasai, David Gray, Carol Clugston, Anna Dominiczak and Alex Alderton, Roberto Amato, Jeffrey Barrett, Sonia Goncalves, Ewan Harrison, David K. Jackson, Ian Johnston, Dominic Kwiatkowski, Cordelia Langford, John Sillitoe on behalf of the Wellcome Sanger Institute COVID-19 Surveillance Team |
| EPI_ISL_2913033, EPI_ISL_2913034, EPI_ISL_2913035, EPI_ISL_2913036 | Lighthouse Lab in Milton Keynes | Wellcome Sanger Institute for the COVID-19 Genomics UK (COG-UK) Consortium | The Lighthouse Lab in Milton Keynes and Alex Alderton, Roberto Amato, Jeffrey Barrett, Sonia Goncalves, Ewan Harrison, David K. Jackson, Ian Johnston, Dominic Kwiatkowski, Cordelia Langford, John Sillitoe on behalf of the Wellcome Sanger Institute COVID-19 Surveillance Team |
| EPI_ISL_2913037, EPI_ISL_2913038 | Lighthouse Lab in Glasgow | Wellcome Sanger Institute for the COVID-19 Genomics UK (COG-UK) Consortium | Harper VanSteenhouse, Yumi Kasai, David Gray, Carol Clugston, Anna Dominiczak and Alex Alderton, Roberto Amato, Jeffrey Barrett, Sonia Goncalves, Ewan Harrison, David K. Jackson, Ian Johnston, Dominic Kwiatkowski, Cordelia Langford, John Sillitoe on behalf of the Wellcome Sanger Institute COVID-19 Surveillance Team |
| EPI_ISL_2913039, EPI_ISL_2913040 | Lighthouse Lab in Milton Keynes | Wellcome Sanger Institute for the COVID-19 Genomics UK (COG-UK) Consortium | The Lighthouse Lab in Milton Keynes and Alex Alderton, Roberto Amato, Jeffrey Barrett, Sonia Goncalves, Ewan Harrison, David K. Jackson, Ian Johnston, Dominic Kwiatkowski, Cordelia Langford, John Sillitoe on behalf of the Wellcome Sanger Institute COVID-19 Surveillance Team |
| EPI_ISL_2913041 | Lighthouse Lab in Glasgow | Wellcome Sanger Institute for the COVID-19 Genomics UK (COG-UK) Consortium | Harper VanSteenhouse, Yumi Kasai, David Gray, Carol Clugston, Anna Dominiczak and Alex Alderton, Roberto Amato, Jeffrey Barrett, Sonia Goncalves, Ewan Harrison, David K. Jackson, Ian Johnston, Dominic Kwiatkowski, Cordelia Langford, John Sillitoe on behalf of the Wellcome Sanger Institute COVID-19 Surveillance Team |
| EPI_ISL_2913042 | Berkshire and Surrey Pathology Services Lighthouse Laboratory | Wellcome Sanger Institute for the COVID-19 Genomics UK (COG-UK) Consortium | Berkshire and Surrey Pathology Services Lighthouse Laboratory and Alex Alderton, Roberto Amato, Jeffrey Barrett, Sonia Goncalves, Ewan Harrison, David K. Jackson, Ian Johnston, Dominic Kwiatkowski, Cordelia Langford, John Sillitoe on behalf of the Wellcome Sanger Institute COVID-19 Surveillance Team |
| EPI_ISL_2913043 | Lighthouse Lab in Milton Keynes | Wellcome Sanger Institute for the COVID-19 Genomics UK (COG-UK) Consortium | The Lighthouse Lab in Milton Keynes and Alex Alderton, Roberto Amato, Jeffrey Barrett, Sonia Goncalves, Ewan Harrison, David K. Jackson, Ian Johnston, Dominic Kwiatkowski, Cordelia Langford, John Sillitoe on behalf of the Wellcome Sanger Institute COVID-19 Surveillance Team |
| EPI_ISL_2913044 | Lighthouse Lab in Glasgow | Wellcome Sanger Institute for the COVID-19 Genomics UK (COG-UK) Consortium | Harper VanSteenhouse, Yumi Kasai, David Gray, Carol Clugston, Anna Dominiczak and Alex Alderton, Roberto Amato, Jeffrey Barrett, Sonia Goncalves, Ewan Harrison, David K. Jackson, Ian Johnston, Dominic Kwiatkowski, Cordelia Langford, John Sillitoe on behalf of the Wellcome Sanger Institute COVID-19 Surveillance Team |
| EPI_ISL_2913045 | Lighthouse Lab in Milton Keynes | Wellcome Sanger Institute for the COVID-19 Genomics UK (COG-UK) Consortium | The Lighthouse Lab in Milton Keynes and Alex Alderton, Roberto Amato, Jeffrey Barrett, Sonia Goncalves, Ewan Harrison, David K. Jackson, Ian Johnston, Dominic Kwiatkowski, Cordelia Langford, John Sillitoe on behalf of the Wellcome Sanger Institute COVID-19 Surveillance Team |
| EPI_ISL_2913046 | Berkshire and Surrey Pathology Services Lighthouse Laboratory | Wellcome Sanger Institute for the COVID-19 Genomics UK (COG-UK) Consortium | Berkshire and Surrey Pathology Services Lighthouse Laboratory and Alex Alderton, Roberto Amato, Jeffrey Barrett, Sonia Goncalves, Ewan Harrison, David K. Jackson, Ian Johnston, Dominic Kwiatkowski, Cordelia Langford, John Sillitoe on behalf of the Wellcome Sanger Institute COVID-19 Surveillance Team |
| EPI_ISL_2913047 | Lighthouse Lab in Glasgow | Wellcome Sanger Institute for the COVID-19 Genomics UK (COG-UK) Consortium | Harper VanSteenhouse, Yumi Kasai, David Gray, Carol Clugston, Anna Dominiczak and Alex Alderton, Roberto Amato, Jeffrey Barrett, Sonia Goncalves, Ewan Harrison, David K. Jackson, Ian Johnston, Dominic Kwiatkowski, Cordelia Langford, John Sillitoe on behalf of the Wellcome Sanger Institute COVID-19 Surveillance Team |
| EPI_ISL_2913048, EPI_ISL_2913049, | Lighthouse Lab in Milton Keynes | Wellcome Sanger Institute for the COVID-19 Genomics UK (COG-UK) Consortium | The Lighthouse Lab in Milton Keynes and Alex Alderton, Roberto Amato, Jeffrey Barrett, Sonia Goncalves, Ewan Harrison, David K. Jackson, Ian Johnston, Dominic Kwiatkowski, Cordelia Langford, John Sillitoe on behalf of the Wellcome Sanger Institute COVID-19 Surveillance Team |

| | | | |
|---|---|---|---|
| EPI_ISL_2913050, EPI_ISL_2913051, EPI_ISL_2913052 | | | |
| EPI_ISL_2913053, EPI_ISL_2913054 | Berkshire and Surrey Pathology Services Lighthouse Laboratory | Wellcome Sanger Institute for the COVID-19 Genomics UK (COG-UK) Consortium | Berkshire and Surrey Pathology Services Lighthouse Laboratory and Alex Alderton, Roberto Amato, Jeffrey Barrett, Sonia Goncalves, Ewan Harrison, David K. Jackson, Ian Johnston, Dominic Kwiatkowski, Cordelia Langford, John Sillitoe on behalf of the Wellcome Sanger Institute COVID-19 Surveillance Team |
| EPI_ISL_2913055 | Lighthouse Lab in Milton Keynes | Wellcome Sanger Institute for the COVID-19 Genomics UK (COG-UK) Consortium | The Lighthouse Lab in Milton Keynes and Alex Alderton, Roberto Amato, Jeffrey Barrett, Sonia Goncalves, Ewan Harrison, David K. Jackson, Ian Johnston, Dominic Kwiatkowski, Cordelia Langford, John Sillitoe on behalf of the Wellcome Sanger Institute COVID-19 Surveillance Team |
| EPI_ISL_2913056 | Lighthouse Lab in Glasgow | Wellcome Sanger Institute for the COVID-19 Genomics UK (COG-UK) Consortium | Harper VanSteenhouse, Yumi Kasai, David Gray, Carol Clugston, Anna Dominiczak and Alex Alderton, Roberto Amato, Jeffrey Barrett, Sonia Goncalves, Ewan Harrison, David K. Jackson, Ian Johnston, Dominic Kwiatkowski, Cordelia Langford, John Sillitoe on behalf of the Wellcome Sanger Institute COVID-19 Surveillance Team |
| EPI_ISL_2913057, EPI_ISL_2913058 | Berkshire and Surrey Pathology Services Lighthouse Laboratory | Wellcome Sanger Institute for the COVID-19 Genomics UK (COG-UK) Consortium | Berkshire and Surrey Pathology Services Lighthouse Laboratory and Alex Alderton, Roberto Amato, Jeffrey Barrett, Sonia Goncalves, Ewan Harrison, David K. Jackson, Ian Johnston, Dominic Kwiatkowski, Cordelia Langford, John Sillitoe on behalf of the Wellcome Sanger Institute COVID-19 Surveillance Team |
| EPI_ISL_2913060 | Lighthouse Lab in Glasgow | Wellcome Sanger Institute for the COVID-19 Genomics UK (COG-UK) Consortium | Harper VanSteenhouse, Yumi Kasai, David Gray, Carol Clugston, Anna Dominiczak and Alex Alderton, Roberto Amato, Jeffrey Barrett, Sonia Goncalves, Ewan Harrison, David K. Jackson, Ian Johnston, Dominic Kwiatkowski, Cordelia Langford, John Sillitoe on behalf of the Wellcome Sanger Institute COVID-19 Surveillance Team |
| EPI_ISL_2913061, EPI_ISL_2913062, EPI_ISL_2913063, EPI_ISL_2913064, EPI_ISL_2913065, EPI_ISL_2913066 | Lighthouse Lab in Milton Keynes | Wellcome Sanger Institute for the COVID-19 Genomics UK (COG-UK) Consortium | The Lighthouse Lab in Milton Keynes and Alex Alderton, Roberto Amato, Jeffrey Barrett, Sonia Goncalves, Ewan Harrison, David K. Jackson, Ian Johnston, Dominic Kwiatkowski, Cordelia Langford, John Sillitoe on behalf of the Wellcome Sanger Institute COVID-19 Surveillance Team |
| EPI_ISL_2913067, EPI_ISL_2913068 | Lighthouse Lab in Glasgow | Wellcome Sanger Institute for the COVID-19 Genomics UK (COG-UK) Consortium | Harper VanSteenhouse, Yumi Kasai, David Gray, Carol Clugston, Anna Dominiczak and Alex Alderton, Roberto Amato, Jeffrey Barrett, Sonia Goncalves, Ewan Harrison, David K. Jackson, Ian Johnston, Dominic Kwiatkowski, Cordelia Langford, John Sillitoe on behalf of the Wellcome Sanger Institute COVID-19 Surveillance Team |
| EPI_ISL_2913069, EPI_ISL_2913070 | Berkshire and Surrey Pathology Services Lighthouse Laboratory | Wellcome Sanger Institute for the COVID-19 Genomics UK (COG-UK) Consortium | Berkshire and Surrey Pathology Services Lighthouse Laboratory and Alex Alderton, Roberto Amato, Jeffrey Barrett, Sonia Goncalves, Ewan Harrison, David K. Jackson, Ian Johnston, Dominic Kwiatkowski, Cordelia Langford, John Sillitoe on behalf of the Wellcome Sanger Institute COVID-19 Surveillance Team |
| EPI_ISL_2913071 | Lighthouse Lab in Glasgow | Wellcome Sanger Institute for the COVID-19 Genomics UK (COG-UK) Consortium | Harper VanSteenhouse, Yumi Kasai, David Gray, Carol Clugston, Anna Dominiczak and Alex Alderton, Roberto Amato, Jeffrey Barrett, Sonia Goncalves, Ewan Harrison, David K. Jackson, Ian Johnston, Dominic Kwiatkowski, Cordelia Langford, John Sillitoe on behalf of the Wellcome Sanger Institute COVID-19 Surveillance Team |
| EPI_ISL_2913072 | Lighthouse Lab in Milton Keynes | Wellcome Sanger Institute for the COVID-19 Genomics UK (COG-UK) Consortium | The Lighthouse Lab in Milton Keynes and Alex Alderton, Roberto Amato, Jeffrey Barrett, Sonia Goncalves, Ewan Harrison, David K. Jackson, Ian Johnston, Dominic Kwiatkowski, Cordelia Langford, John Sillitoe on behalf of the Wellcome Sanger Institute COVID-19 Surveillance Team |
| EPI_ISL_2913073, EPI_ISL_2913074 | Lighthouse Lab in Glasgow | Wellcome Sanger Institute for the COVID-19 Genomics UK (COG-UK) Consortium | Harper VanSteenhouse, Yumi Kasai, David Gray, Carol Clugston, Anna Dominiczak and Alex Alderton, Roberto Amato, Jeffrey Barrett, Sonia Goncalves, Ewan Harrison, David K. Jackson, Ian Johnston, Dominic Kwiatkowski, Cordelia Langford, John Sillitoe on behalf of the Wellcome Sanger Institute COVID-19 Surveillance Team |
| EPI_ISL_2913075, EPI_ISL_2913076 | Lighthouse Lab in Milton Keynes | Wellcome Sanger Institute for the COVID-19 Genomics UK (COG-UK) Consortium | The Lighthouse Lab in Milton Keynes and Alex Alderton, Roberto Amato, Jeffrey Barrett, Sonia Goncalves, Ewan Harrison, David K. Jackson, Ian Johnston, Dominic Kwiatkowski, Cordelia Langford, John Sillitoe on behalf of the Wellcome Sanger Institute COVID-19 Surveillance Team |
| EPI_ISL_2913077 | Berkshire and Surrey Pathology Services Lighthouse Laboratory | Wellcome Sanger Institute for the COVID-19 Genomics UK (COG-UK) Consortium | Berkshire and Surrey Pathology Services Lighthouse Laboratory and Alex Alderton, Roberto Amato, Jeffrey Barrett, Sonia Goncalves, Ewan Harrison, David K. Jackson, Ian Johnston, Dominic Kwiatkowski, Cordelia Langford, John Sillitoe on behalf of the Wellcome Sanger Institute COVID-19 Surveillance Team |
| EPI_ISL_2913078 | Lighthouse Lab in Glasgow | Wellcome Sanger Institute for the COVID-19 Genomics UK (COG-UK) Consortium | Harper VanSteenhouse, Yumi Kasai, David Gray, Carol Clugston, Anna Dominiczak and Alex Alderton, Roberto Amato, Jeffrey Barrett, Sonia Goncalves, Ewan Harrison, David K. Jackson, Ian Johnston, Dominic Kwiatkowski, Cordelia Langford, John Sillitoe on behalf of the Wellcome Sanger Institute COVID-19 Surveillance Team |
| EPI_ISL_2913079, EPI_ISL_2913080 | Lighthouse Lab in Milton Keynes | Wellcome Sanger Institute for the COVID-19 Genomics UK (COG-UK) Consortium | The Lighthouse Lab in Milton Keynes and Alex Alderton, Roberto Amato, Jeffrey Barrett, Sonia Goncalves, Ewan Harrison, David K. Jackson, Ian Johnston, Dominic Kwiatkowski, Cordelia Langford, John Sillitoe on behalf of the Wellcome Sanger Institute COVID-19 Surveillance Team |
| EPI_ISL_2913081 | Berkshire and Surrey Pathology Services Lighthouse Laboratory | Wellcome Sanger Institute for the COVID-19 Genomics UK (COG-UK) Consortium | Berkshire and Surrey Pathology Services Lighthouse Laboratory and Alex Alderton, Roberto Amato, Jeffrey Barrett, Sonia Goncalves, Ewan Harrison, David K. Jackson, Ian Johnston, Dominic Kwiatkowski, Cordelia Langford, John Sillitoe on behalf of the Wellcome Sanger Institute COVID-19 Surveillance Team |
| EPI_ISL_2913082, EPI_ISL_2913083 | Lighthouse Lab in Milton Keynes | Wellcome Sanger Institute for the COVID-19 Genomics UK (COG-UK) Consortium | The Lighthouse Lab in Milton Keynes and Alex Alderton, Roberto Amato, Jeffrey Barrett, Sonia Goncalves, Ewan Harrison, David K. Jackson, Ian Johnston, Dominic Kwiatkowski, Cordelia Langford, John Sillitoe on behalf of the Wellcome Sanger Institute COVID-19 Surveillance Team |
| EPI_ISL_2913084 | Lighthouse Lab in Glasgow | Wellcome Sanger Institute for the COVID-19 Genomics UK (COG-UK) Consortium | Harper VanSteenhouse, Yumi Kasai, David Gray, Carol Clugston, Anna Dominiczak and Alex Alderton, Roberto Amato, Jeffrey Barrett, Sonia Goncalves, Ewan Harrison, David K. Jackson, Ian Johnston, Dominic Kwiatkowski, Cordelia Langford, John Sillitoe on behalf of the Wellcome Sanger Institute COVID-19 Surveillance Team |
| EPI_ISL_2913085, EPI_ISL_2913086, EPI_ISL_2913087, EPI_ISL_2913088, EPI_ISL_2913089 | Lighthouse Lab in Milton Keynes | Wellcome Sanger Institute for the COVID-19 Genomics UK (COG-UK) Consortium | The Lighthouse Lab in Milton Keynes and Alex Alderton, Roberto Amato, Jeffrey Barrett, Sonia Goncalves, Ewan Harrison, David K. Jackson, Ian Johnston, Dominic Kwiatkowski, Cordelia Langford, John Sillitoe on behalf of the Wellcome Sanger Institute COVID-19 Surveillance Team |
| EPI_ISL_2913090 | Lighthouse Lab in Glasgow | Wellcome Sanger Institute for the COVID-19 Genomics UK (COG-UK) Consortium | Harper VanSteenhouse, Yumi Kasai, David Gray, Carol Clugston, Anna Dominiczak and Alex Alderton, Roberto Amato, Jeffrey Barrett, Sonia Goncalves, Ewan Harrison, David K. Jackson, Ian Johnston, Dominic Kwiatkowski, Cordelia Langford, John Sillitoe on behalf of the Wellcome Sanger Institute COVID-19 Surveillance Team |
| EPI_ISL_2913091, EPI_ISL_2913092, EPI_ISL_2913093, EPI_ISL_2913094 | Lighthouse Lab in Milton Keynes | Wellcome Sanger Institute for the COVID-19 Genomics UK (COG-UK) Consortium | The Lighthouse Lab in Milton Keynes and Alex Alderton, Roberto Amato, Jeffrey Barrett, Sonia Goncalves, Ewan Harrison, David K. Jackson, Ian Johnston, Dominic Kwiatkowski, Cordelia Langford, John Sillitoe on behalf of the Wellcome Sanger Institute COVID-19 Surveillance Team |
| EPI_ISL_2913095 | Berkshire and Surrey Pathology Services Lighthouse Laboratory | Wellcome Sanger Institute for the COVID-19 Genomics UK (COG-UK) Consortium | Berkshire and Surrey Pathology Services Lighthouse Laboratory and Alex Alderton, Roberto Amato, Jeffrey Barrett, Sonia Goncalves, Ewan Harrison, David K. Jackson, Ian Johnston, Dominic Kwiatkowski, Cordelia Langford, John Sillitoe on behalf of the Wellcome Sanger Institute COVID-19 Surveillance Team |
| EPI_ISL_2913096 | Lighthouse Lab in Glasgow | Wellcome Sanger Institute for the COVID-19 Genomics UK (COG-UK) Consortium | Harper VanSteenhouse, Yumi Kasai, David Gray, Carol Clugston, Anna Dominiczak and Alex Alderton, Roberto Amato, Jeffrey Barrett, Sonia Goncalves, Ewan Harrison, David K. Jackson, Ian Johnston, Dominic Kwiatkowski, Cordelia Langford, John Sillitoe on behalf of the Wellcome Sanger Institute COVID-19 Surveillance Team |
| EPI_ISL_2913097, EPI_ISL_2913098 | Lighthouse Lab in Milton Keynes | Wellcome Sanger Institute for the COVID-19 Genomics UK (COG-UK) Consortium | The Lighthouse Lab in Milton Keynes and Alex Alderton, Roberto Amato, Jeffrey Barrett, Sonia Goncalves, Ewan Harrison, David K. Jackson, Ian Johnston, Dominic Kwiatkowski, Cordelia Langford, John Sillitoe on behalf of the Wellcome Sanger Institute COVID-19 Surveillance Team |
| EPI_ISL_2913099 | Berkshire and Surrey Pathology Services Lighthouse Laboratory | Wellcome Sanger Institute for the COVID-19 Genomics UK (COG-UK) Consortium | Berkshire and Surrey Pathology Services Lighthouse Laboratory and Alex Alderton, Roberto Amato, Jeffrey Barrett, Sonia Goncalves, Ewan Harrison, David K. Jackson, Ian Johnston, Dominic Kwiatkowski, Cordelia Langford, John Sillitoe on behalf of the Wellcome Sanger Institute COVID-19 Surveillance Team |
| EPI_ISL_2913100 | Lighthouse Lab in Glasgow | Wellcome Sanger Institute for the COVID-19 Genomics UK (COG-UK) Consortium | Harper VanSteenhouse, Yumi Kasai, David Gray, Carol Clugston, Anna Dominiczak and Alex Alderton, Roberto Amato, Jeffrey Barrett, Sonia Goncalves, Ewan Harrison, David K. Jackson, Ian Johnston, Dominic Kwiatkowski, Cordelia Langford, John Sillitoe on behalf of the Wellcome Sanger Institute COVID-19 Surveillance Team |
| EPI_ISL_2913101, EPI_ISL_2913102, EPI_ISL_2913103 | Lighthouse Lab in Milton Keynes | Wellcome Sanger Institute for the COVID-19 Genomics UK (COG-UK) Consortium | The Lighthouse Lab in Milton Keynes and Alex Alderton, Roberto Amato, Jeffrey Barrett, Sonia Goncalves, Ewan Harrison, David K. Jackson, Ian Johnston, Dominic Kwiatkowski, Cordelia Langford, John Sillitoe on behalf of the Wellcome Sanger Institute COVID-19 Surveillance Team |
| EPI_ISL_2913104 | Lighthouse Lab in Glasgow | Wellcome Sanger Institute for the COVID-19 Genomics UK (COG-UK) Consortium | Harper VanSteenhouse, Yumi Kasai, David Gray, Carol Clugston, Anna Dominiczak and Alex Alderton, Roberto Amato, Jeffrey Barrett, Sonia Goncalves, Ewan Harrison, David K. Jackson, Ian Johnston, Dominic Kwiatkowski, Cordelia Langford, John Sillitoe on behalf of the Wellcome Sanger Institute COVID-19 Surveillance Team |
| EPI_ISL_2913105 | Lighthouse Lab in Milton Keynes | Wellcome Sanger Institute for the COVID-19 Genomics UK (COG-UK) Consortium | The Lighthouse Lab in Milton Keynes and Alex Alderton, Roberto Amato, Jeffrey Barrett, Sonia Goncalves, Ewan Harrison, David K. Jackson, Ian Johnston, Dominic Kwiatkowski, Cordelia Langford, John Sillitoe on behalf of the Wellcome Sanger Institute COVID-19 Surveillance Team |
| EPI_ISL_2913106, EPI_ISL_2913107 | Lighthouse Lab in Glasgow | Wellcome Sanger Institute for the COVID-19 Genomics UK (COG-UK) Consortium | Harper VanSteenhouse, Yumi Kasai, David Gray, Carol Clugston, Anna Dominiczak and Alex Alderton, Roberto Amato, Jeffrey Barrett, Sonia Goncalves, Ewan Harrison, David K. Jackson, Ian Johnston, Dominic Kwiatkowski, Cordelia Langford, John Sillitoe on behalf of the Wellcome Sanger Institute COVID-19 Surveillance Team |
| EPI_ISL_2913108, EPI_ISL_2913109, EPI_ISL_2913110 | Lighthouse Lab in Milton Keynes | Wellcome Sanger Institute for the COVID-19 Genomics UK (COG-UK) Consortium | The Lighthouse Lab in Milton Keynes and Alex Alderton, Roberto Amato, Jeffrey Barrett, Sonia Goncalves, Ewan Harrison, David K. Jackson, Ian Johnston, Dominic Kwiatkowski, Cordelia Langford, John Sillitoe on behalf of the Wellcome Sanger Institute COVID-19 Surveillance Team |
| EPI_ISL_2913111 | Lighthouse Lab in Glasgow | Wellcome Sanger Institute for the COVID-19 Genomics UK (COG-UK) Consortium | Harper VanSteenhouse, Yumi Kasai, David Gray, Carol Clugston, Anna Dominiczak and Alex Alderton, Roberto Amato, Jeffrey Barrett, Sonia Goncalves, Ewan Harrison, David K. Jackson, Ian Johnston, Dominic Kwiatkowski, Cordelia Langford, John Sillitoe on behalf of the Wellcome Sanger Institute COVID-19 Surveillance Team |
| EPI_ISL_2913112, EPI_ISL_2913113 | Lighthouse Lab in Milton Keynes | Wellcome Sanger Institute for the COVID-19 Genomics UK (COG-UK) Consortium | The Lighthouse Lab in Milton Keynes and Alex Alderton, Roberto Amato, Jeffrey Barrett, Sonia Goncalves, Ewan Harrison, David K. Jackson, Ian Johnston, Dominic Kwiatkowski, Cordelia Langford, John Sillitoe on behalf of the Wellcome Sanger Institute COVID-19 Surveillance Team |
| EPI_ISL_2913114 | Lighthouse Lab in Glasgow | Wellcome Sanger Institute for the COVID-19 Genomics UK (COG-UK) Consortium | Harper VanSteenhouse, Yumi Kasai, David Gray, Carol Clugston, Anna Dominiczak and Alex Alderton, Roberto Amato, Jeffrey Barrett, Sonia Goncalves, Ewan Harrison, David K. Jackson, Ian Johnston, Dominic Kwiatkowski, Cordelia Langford, John Sillitoe on behalf of the Wellcome Sanger Institute COVID-19 Surveillance Team |
| EPI_ISL_2913115, EPI_ISL_2913116, EPI_ISL_2913117, EPI_ISL_2913118, EPI_ISL_2913119 | Lighthouse Lab in Milton Keynes | Wellcome Sanger Institute for the COVID-19 Genomics UK (COG-UK) Consortium | The Lighthouse Lab in Milton Keynes and Alex Alderton, Roberto Amato, Jeffrey Barrett, Sonia Goncalves, Ewan Harrison, David K. Jackson, Ian Johnston, Dominic Kwiatkowski, Cordelia Langford, John Sillitoe on behalf of the Wellcome Sanger Institute COVID-19 Surveillance Team |
| EPI_ISL_2913120 | Lighthouse Lab in Glasgow | Wellcome Sanger Institute for the COVID-19 Genomics UK (COG-UK) Consortium | Harper VanSteenhouse, Yumi Kasai, David Gray, Carol Clugston, Anna Dominiczak and Alex Alderton, Roberto Amato, Jeffrey Barrett, Sonia Goncalves, Ewan Harrison, David K. Jackson, Ian Johnston, Dominic Kwiatkowski, Cordelia Langford, John Sillitoe on behalf of the Wellcome Sanger Institute COVID-19 Surveillance Team |
| EPI_ISL_2913121, EPI_ISL_2913122, EPI_ISL_2913123, EPI_ISL_2913124, | Lighthouse Lab in Milton Keynes | Wellcome Sanger Institute for the COVID-19 Genomics UK (COG-UK) Consortium | The Lighthouse Lab in Milton Keynes and Alex Alderton, Roberto Amato, Jeffrey Barrett, Sonia Goncalves, Ewan Harrison, David K. Jackson, Ian Johnston, Dominic Kwiatkowski, Cordelia Langford, John Sillitoe on behalf of the Wellcome Sanger Institute COVID-19 Surveillance Team |

| | | | |
|---|---|---|---|
| EPI_ISL_2913125 | | | |
| EPI_ISL_2913126, EPI_ISL_2913128 | Lighthouse Lab in Glasgow | Wellcome Sanger Institute for the COVID-19 Genomics UK (COG-UK) Consortium | Harper VanSteenhouse, Yumi Kasai, David Gray, Carol Clugston, Anna Dominiczak and Alex Alderton, Roberto Amato, Jeffrey Barrett, Sonia Goncalves, Ewan Harrison, David K. Jackson, Ian Johnston, Dominic Kwiatkowski, Cordelia Langford, John Sillitoe on behalf of the Wellcome Sanger Institute COVID-19 Surveillance Team |
| EPI_ISL_2913129, EPI_ISL_2913130 | Lighthouse Lab in Milton Keynes | Wellcome Sanger Institute for the COVID-19 Genomics UK (COG-UK) Consortium | The Lighthouse Lab in Milton Keynes and Alex Alderton, Roberto Amato, Jeffrey Barrett, Sonia Goncalves, Ewan Harrison, David K. Jackson, Ian Johnston, Dominic Kwiatkowski, Cordelia Langford, John Sillitoe on behalf of the Wellcome Sanger Institute COVID-19 Surveillance Team |
| EPI_ISL_2913131, EPI_ISL_2913132 | Berkshire and Surrey Pathology Services Lighthouse Laboratory | Wellcome Sanger Institute for the COVID-19 Genomics UK (COG-UK) Consortium | Berkshire and Surrey Pathology Services Lighthouse Laboratory and Alex Alderton, Roberto Amato, Jeffrey Barrett, Sonia Goncalves, Ewan Harrison, David K. Jackson, Ian Johnston, Dominic Kwiatkowski, Cordelia Langford, John Sillitoe on behalf of the Wellcome Sanger Institute COVID-19 Surveillance Team |
| EPI_ISL_2913133, EPI_ISL_2913134, EPI_ISL_2913135, EPI_ISL_2913136, EPI_ISL_2913137 | Lighthouse Lab in Milton Keynes | Wellcome Sanger Institute for the COVID-19 Genomics UK (COG-UK) Consortium | The Lighthouse Lab in Milton Keynes and Alex Alderton, Roberto Amato, Jeffrey Barrett, Sonia Goncalves, Ewan Harrison, David K. Jackson, Ian Johnston, Dominic Kwiatkowski, Cordelia Langford, John Sillitoe on behalf of the Wellcome Sanger Institute COVID-19 Surveillance Team |
| EPI_ISL_2913138 | Lighthouse Lab in Glasgow | Wellcome Sanger Institute for the COVID-19 Genomics UK (COG-UK) Consortium | Harper VanSteenhouse, Yumi Kasai, David Gray, Carol Clugston, Anna Dominiczak and Alex Alderton, Roberto Amato, Jeffrey Barrett, Sonia Goncalves, Ewan Harrison, David K. Jackson, Ian Johnston, Dominic Kwiatkowski, Cordelia Langford, John Sillitoe on behalf of the Wellcome Sanger Institute COVID-19 Surveillance Team |
| EPI_ISL_2913139, EPI_ISL_2913140, EPI_ISL_2913141 | Lighthouse Lab in Milton Keynes | Wellcome Sanger Institute for the COVID-19 Genomics UK (COG-UK) Consortium | The Lighthouse Lab in Milton Keynes and Alex Alderton, Roberto Amato, Jeffrey Barrett, Sonia Goncalves, Ewan Harrison, David K. Jackson, Ian Johnston, Dominic Kwiatkowski, Cordelia Langford, John Sillitoe on behalf of the Wellcome Sanger Institute COVID-19 Surveillance Team |
| EPI_ISL_2913142 | Lighthouse Lab in Glasgow | Wellcome Sanger Institute for the COVID-19 Genomics UK (COG-UK) Consortium | Harper VanSteenhouse, Yumi Kasai, David Gray, Carol Clugston, Anna Dominiczak and Alex Alderton, Roberto Amato, Jeffrey Barrett, Sonia Goncalves, Ewan Harrison, David K. Jackson, Ian Johnston, Dominic Kwiatkowski, Cordelia Langford, John Sillitoe on behalf of the Wellcome Sanger Institute COVID-19 Surveillance Team |
| EPI_ISL_2913143 | Berkshire and Surrey Pathology Services Lighthouse Laboratory | Wellcome Sanger Institute for the COVID-19 Genomics UK (COG-UK) Consortium | Berkshire and Surrey Pathology Services Lighthouse Laboratory and Alex Alderton, Roberto Amato, Jeffrey Barrett, Sonia Goncalves, Ewan Harrison, David K. Jackson, Ian Johnston, Dominic Kwiatkowski, Cordelia Langford, John Sillitoe on behalf of the Wellcome Sanger Institute COVID-19 Surveillance Team |
| EPI_ISL_2913144, EPI_ISL_2913145, EPI_ISL_2913146, EPI_ISL_2913147 | Lighthouse Lab in Milton Keynes | Wellcome Sanger Institute for the COVID-19 Genomics UK (COG-UK) Consortium | The Lighthouse Lab in Milton Keynes and Alex Alderton, Roberto Amato, Jeffrey Barrett, Sonia Goncalves, Ewan Harrison, David K. Jackson, Ian Johnston, Dominic Kwiatkowski, Cordelia Langford, John Sillitoe on behalf of the Wellcome Sanger Institute COVID-19 Surveillance Team |
| EPI_ISL_2913148 | Berkshire and Surrey Pathology Services Lighthouse Laboratory | Wellcome Sanger Institute for the COVID-19 Genomics UK (COG-UK) Consortium | Berkshire and Surrey Pathology Services Lighthouse Laboratory and Alex Alderton, Roberto Amato, Jeffrey Barrett, Sonia Goncalves, Ewan Harrison, David K. Jackson, Ian Johnston, Dominic Kwiatkowski, Cordelia Langford, John Sillitoe on behalf of the Wellcome Sanger Institute COVID-19 Surveillance Team |
| EPI_ISL_2913149 | Lighthouse Lab in Milton Keynes | Wellcome Sanger Institute for the COVID-19 Genomics UK (COG-UK) Consortium | The Lighthouse Lab in Milton Keynes and Alex Alderton, Roberto Amato, Jeffrey Barrett, Sonia Goncalves, Ewan Harrison, David K. Jackson, Ian Johnston, Dominic Kwiatkowski, Cordelia Langford, John Sillitoe on behalf of the Wellcome Sanger Institute COVID-19 Surveillance Team |
| EPI_ISL_2913150, EPI_ISL_2913151 | Lighthouse Lab in Glasgow | Wellcome Sanger Institute for the COVID-19 Genomics UK (COG-UK) Consortium | Harper VanSteenhouse, Yumi Kasai, David Gray, Carol Clugston, Anna Dominiczak and Alex Alderton, Roberto Amato, Jeffrey Barrett, Sonia Goncalves, Ewan Harrison, David K. Jackson, Ian Johnston, Dominic Kwiatkowski, Cordelia Langford, John Sillitoe on behalf of the Wellcome Sanger Institute COVID-19 Surveillance Team |
| EPI_ISL_2913152 | Lighthouse Lab in Milton Keynes | Wellcome Sanger Institute for the COVID-19 Genomics UK (COG-UK) Consortium | The Lighthouse Lab in Milton Keynes and Alex Alderton, Roberto Amato, Jeffrey Barrett, Sonia Goncalves, Ewan Harrison, David K. Jackson, Ian Johnston, Dominic Kwiatkowski, Cordelia Langford, John Sillitoe on behalf of the Wellcome Sanger Institute COVID-19 Surveillance Team |
| EPI_ISL_2913153, EPI_ISL_2913154 | Berkshire and Surrey Pathology Services Lighthouse Laboratory | Wellcome Sanger Institute for the COVID-19 Genomics UK (COG-UK) Consortium | Berkshire and Surrey Pathology Services Lighthouse Laboratory and Alex Alderton, Roberto Amato, Jeffrey Barrett, Sonia Goncalves, Ewan Harrison, David K. Jackson, Ian Johnston, Dominic Kwiatkowski, Cordelia Langford, John Sillitoe on behalf of the Wellcome Sanger Institute COVID-19 Surveillance Team |
| EPI_ISL_2913155 | Lighthouse Lab in Glasgow | Wellcome Sanger Institute for the COVID-19 Genomics UK (COG-UK) Consortium | Harper VanSteenhouse, Yumi Kasai, David Gray, Carol Clugston, Anna Dominiczak and Alex Alderton, Roberto Amato, Jeffrey Barrett, Sonia Goncalves, Ewan Harrison, David K. Jackson, Ian Johnston, Dominic Kwiatkowski, Cordelia Langford, John Sillitoe on behalf of the Wellcome Sanger Institute COVID-19 Surveillance Team |
| EPI_ISL_2913156 | Lighthouse Lab in Milton Keynes | Wellcome Sanger Institute for the COVID-19 Genomics UK (COG-UK) Consortium | The Lighthouse Lab in Milton Keynes and Alex Alderton, Roberto Amato, Jeffrey Barrett, Sonia Goncalves, Ewan Harrison, David K. Jackson, Ian Johnston, Dominic Kwiatkowski, Cordelia Langford, John Sillitoe on behalf of the Wellcome Sanger Institute COVID-19 Surveillance Team |
| EPI_ISL_2913157, EPI_ISL_2913158 | Berkshire and Surrey Pathology Services Lighthouse Laboratory | Wellcome Sanger Institute for the COVID-19 Genomics UK (COG-UK) Consortium | Berkshire and Surrey Pathology Services Lighthouse Laboratory and Alex Alderton, Roberto Amato, Jeffrey Barrett, Sonia Goncalves, Ewan Harrison, David K. Jackson, Ian Johnston, Dominic Kwiatkowski, Cordelia Langford, John Sillitoe on behalf of the Wellcome Sanger Institute COVID-19 Surveillance Team |
| EPI_ISL_2913159 | Lighthouse Lab in Milton Keynes | Wellcome Sanger Institute for the COVID-19 Genomics UK (COG-UK) Consortium | The Lighthouse Lab in Milton Keynes and Alex Alderton, Roberto Amato, Jeffrey Barrett, Sonia Goncalves, Ewan Harrison, David K. Jackson, Ian Johnston, Dominic Kwiatkowski, Cordelia Langford, John Sillitoe on behalf of the Wellcome Sanger Institute COVID-19 Surveillance Team |
| EPI_ISL_2913160, EPI_ISL_2913161 | Berkshire and Surrey Pathology Services Lighthouse Laboratory | Wellcome Sanger Institute for the COVID-19 Genomics UK (COG-UK) Consortium | Berkshire and Surrey Pathology Services Lighthouse Laboratory and Alex Alderton, Roberto Amato, Jeffrey Barrett, Sonia Goncalves, Ewan Harrison, David K. Jackson, Ian Johnston, Dominic Kwiatkowski, Cordelia Langford, John Sillitoe on behalf of the Wellcome Sanger Institute COVID-19 Surveillance Team |
| EPI_ISL_2913162 | Lighthouse Lab in Milton Keynes | Wellcome Sanger Institute for the COVID-19 Genomics UK (COG-UK) Consortium | The Lighthouse Lab in Milton Keynes and Alex Alderton, Roberto Amato, Jeffrey Barrett, Sonia Goncalves, Ewan Harrison, David K. Jackson, Ian Johnston, Dominic Kwiatkowski, Cordelia Langford, John Sillitoe on behalf of the Wellcome Sanger Institute COVID-19 Surveillance Team |
| EPI_ISL_2913163 | Berkshire and Surrey Pathology Services Lighthouse Laboratory | Wellcome Sanger Institute for the COVID-19 Genomics UK (COG-UK) Consortium | Berkshire and Surrey Pathology Services Lighthouse Laboratory and Alex Alderton, Roberto Amato, Jeffrey Barrett, Sonia Goncalves, Ewan Harrison, David K. Jackson, Ian Johnston, Dominic Kwiatkowski, Cordelia Langford, John Sillitoe on behalf of the Wellcome Sanger Institute COVID-19 Surveillance Team |
| EPI_ISL_2913164 | Lighthouse Lab in Milton Keynes | Wellcome Sanger Institute for the COVID-19 Genomics UK (COG-UK) Consortium | The Lighthouse Lab in Milton Keynes and Alex Alderton, Roberto Amato, Jeffrey Barrett, Sonia Goncalves, Ewan Harrison, David K. Jackson, Ian Johnston, Dominic Kwiatkowski, Cordelia Langford, John Sillitoe on behalf of the Wellcome Sanger Institute COVID-19 Surveillance Team |
| EPI_ISL_2913165 | Lighthouse Lab in Glasgow | Wellcome Sanger Institute for the COVID-19 Genomics UK (COG-UK) Consortium | Harper VanSteenhouse, Yumi Kasai, David Gray, Carol Clugston, Anna Dominiczak and Alex Alderton, Roberto Amato, Jeffrey Barrett, Sonia Goncalves, Ewan Harrison, David K. Jackson, Ian Johnston, Dominic Kwiatkowski, Cordelia Langford, John Sillitoe on behalf of the Wellcome Sanger Institute COVID-19 Surveillance Team |
| EPI_ISL_2913166 | Berkshire and Surrey Pathology Services Lighthouse Laboratory | Wellcome Sanger Institute for the COVID-19 Genomics UK (COG-UK) Consortium | Berkshire and Surrey Pathology Services Lighthouse Laboratory and Alex Alderton, Roberto Amato, Jeffrey Barrett, Sonia Goncalves, Ewan Harrison, David K. Jackson, Ian Johnston, Dominic Kwiatkowski, Cordelia Langford, John Sillitoe on behalf of the Wellcome Sanger Institute COVID-19 Surveillance Team |
| EPI_ISL_2913167 | Lighthouse Lab in Glasgow | Wellcome Sanger Institute for the COVID-19 Genomics UK (COG-UK) Consortium | Harper VanSteenhouse, Yumi Kasai, David Gray, Carol Clugston, Anna Dominiczak and Alex Alderton, Roberto Amato, Jeffrey Barrett, Sonia Goncalves, Ewan Harrison, David K. Jackson, Ian Johnston, Dominic Kwiatkowski, Cordelia Langford, John Sillitoe on behalf of the Wellcome Sanger Institute COVID-19 Surveillance Team |
| EPI_ISL_2913168 | Berkshire and Surrey Pathology Services Lighthouse Laboratory | Wellcome Sanger Institute for the COVID-19 Genomics UK (COG-UK) Consortium | Berkshire and Surrey Pathology Services Lighthouse Laboratory and Alex Alderton, Roberto Amato, Jeffrey Barrett, Sonia Goncalves, Ewan Harrison, David K. Jackson, Ian Johnston, Dominic Kwiatkowski, Cordelia Langford, John Sillitoe on behalf of the Wellcome Sanger Institute COVID-19 Surveillance Team |
| EPI_ISL_2913169, EPI_ISL_2913170 | Lighthouse Lab in Milton Keynes | Wellcome Sanger Institute for the COVID-19 Genomics UK (COG-UK) Consortium | The Lighthouse Lab in Milton Keynes and Alex Alderton, Roberto Amato, Jeffrey Barrett, Sonia Goncalves, Ewan Harrison, David K. Jackson, Ian Johnston, Dominic Kwiatkowski, Cordelia Langford, John Sillitoe on behalf of the Wellcome Sanger Institute COVID-19 Surveillance Team |
| EPI_ISL_2913171, EPI_ISL_2913172 | Berkshire and Surrey Pathology Services Lighthouse Laboratory | Wellcome Sanger Institute for the COVID-19 Genomics UK (COG-UK) Consortium | Berkshire and Surrey Pathology Services Lighthouse Laboratory and Alex Alderton, Roberto Amato, Jeffrey Barrett, Sonia Goncalves, Ewan Harrison, David K. Jackson, Ian Johnston, Dominic Kwiatkowski, Cordelia Langford, John Sillitoe on behalf of the Wellcome Sanger Institute COVID-19 Surveillance Team |
| EPI_ISL_2913173 | Lighthouse Lab in Milton Keynes | Wellcome Sanger Institute for the COVID-19 Genomics UK (COG-UK) Consortium | The Lighthouse Lab in Milton Keynes and Alex Alderton, Roberto Amato, Jeffrey Barrett, Sonia Goncalves, Ewan Harrison, David K. Jackson, Ian Johnston, Dominic Kwiatkowski, Cordelia Langford, John Sillitoe on behalf of the Wellcome Sanger Institute COVID-19 Surveillance Team |
| EPI_ISL_2913174 | Berkshire and Surrey Pathology Services Lighthouse Laboratory | Wellcome Sanger Institute for the COVID-19 Genomics UK (COG-UK) Consortium | Berkshire and Surrey Pathology Services Lighthouse Laboratory and Alex Alderton, Roberto Amato, Jeffrey Barrett, Sonia Goncalves, Ewan Harrison, David K. Jackson, Ian Johnston, Dominic Kwiatkowski, Cordelia Langford, John Sillitoe on behalf of the Wellcome Sanger Institute COVID-19 Surveillance Team |
| EPI_ISL_2913175, EPI_ISL_2913176, EPI_ISL_2913177, EPI_ISL_2913178, EPI_ISL_2913179, EPI_ISL_2913180 | Lighthouse Lab in Milton Keynes | Wellcome Sanger Institute for the COVID-19 Genomics UK (COG-UK) Consortium | The Lighthouse Lab in Milton Keynes and Alex Alderton, Roberto Amato, Jeffrey Barrett, Sonia Goncalves, Ewan Harrison, David K. Jackson, Ian Johnston, Dominic Kwiatkowski, Cordelia Langford, John Sillitoe on behalf of the Wellcome Sanger Institute COVID-19 Surveillance Team |
| EPI_ISL_2913181 | Berkshire and Surrey Pathology Services Lighthouse Laboratory | Wellcome Sanger Institute for the COVID-19 Genomics UK (COG-UK) Consortium | Berkshire and Surrey Pathology Services Lighthouse Laboratory and Alex Alderton, Roberto Amato, Jeffrey Barrett, Sonia Goncalves, Ewan Harrison, David K. Jackson, Ian Johnston, Dominic Kwiatkowski, Cordelia Langford, John Sillitoe on behalf of the Wellcome Sanger Institute COVID-19 Surveillance Team |
| EPI_ISL_2913182, EPI_ISL_2913183 | Lighthouse Lab in Glasgow | Wellcome Sanger Institute for the COVID-19 Genomics UK (COG-UK) Consortium | Harper VanSteenhouse, Yumi Kasai, David Gray, Carol Clugston, Anna Dominiczak and Alex Alderton, Roberto Amato, Jeffrey Barrett, Sonia Goncalves, Ewan Harrison, David K. Jackson, Ian Johnston, Dominic Kwiatkowski, Cordelia Langford, John Sillitoe on behalf of the Wellcome Sanger Institute COVID-19 Surveillance Team |
| EPI_ISL_2913184 | Lighthouse Lab in Milton Keynes | Wellcome Sanger Institute for the COVID-19 Genomics UK (COG-UK) Consortium | The Lighthouse Lab in Milton Keynes and Alex Alderton, Roberto Amato, Jeffrey Barrett, Sonia Goncalves, Ewan Harrison, David K. Jackson, Ian Johnston, Dominic Kwiatkowski, Cordelia Langford, John Sillitoe on behalf of the Wellcome Sanger Institute COVID-19 Surveillance Team |
| EPI_ISL_2913185, EPI_ISL_2913186, EPI_ISL_2913187 | Lighthouse Lab in Glasgow | Wellcome Sanger Institute for the COVID-19 Genomics UK (COG-UK) Consortium | Harper VanSteenhouse, Yumi Kasai, David Gray, Carol Clugston, Anna Dominiczak and Alex Alderton, Roberto Amato, Jeffrey Barrett, Sonia Goncalves, Ewan Harrison, David K. Jackson, Ian Johnston, Dominic Kwiatkowski, Cordelia Langford, John Sillitoe on behalf of the Wellcome Sanger Institute COVID-19 Surveillance Team |
| EPI_ISL_2913188 | Berkshire and Surrey Pathology Services Lighthouse Laboratory | Wellcome Sanger Institute for the COVID-19 Genomics UK (COG-UK) Consortium | Berkshire and Surrey Pathology Services Lighthouse Laboratory and Alex Alderton, Roberto Amato, Jeffrey Barrett, Sonia Goncalves, Ewan Harrison, David K. Jackson, Ian Johnston, Dominic Kwiatkowski, Cordelia Langford, John Sillitoe on behalf of the Wellcome Sanger Institute COVID-19 Surveillance Team |
| EPI_ISL_2913189 | Lighthouse Lab in Milton Keynes | Wellcome Sanger Institute for the COVID-19 Genomics UK (COG-UK) Consortium | The Lighthouse Lab in Milton Keynes and Alex Alderton, Roberto Amato, Jeffrey Barrett, Sonia Goncalves, Ewan Harrison, David K. Jackson, Ian Johnston, Dominic Kwiatkowski, Cordelia Langford, John Sillitoe on behalf of the Wellcome Sanger Institute COVID-19 Surveillance Team |
| EPI_ISL_2913190 | Berkshire and Surrey Pathology Services Lighthouse Laboratory | Wellcome Sanger Institute for the COVID-19 Genomics UK (COG-UK) Consortium | Berkshire and Surrey Pathology Services Lighthouse Laboratory and Alex Alderton, Roberto Amato, Jeffrey Barrett, Sonia Goncalves, Ewan Harrison, David K. Jackson, Ian Johnston, Dominic Kwiatkowski, Cordelia Langford, John Sillitoe on behalf of the Wellcome Sanger Institute COVID-19 Surveillance Team |
| EPI_ISL_2913191, EPI_ISL_2913192, EPI_ISL_2913193 | Lighthouse Lab in Milton Keynes | Wellcome Sanger Institute for the COVID-19 Genomics UK (COG-UK) Consortium | The Lighthouse Lab in Milton Keynes and Alex Alderton, Roberto Amato, Jeffrey Barrett, Sonia Goncalves, Ewan Harrison, David K. Jackson, Ian Johnston, Dominic Kwiatkowski, Cordelia Langford, John Sillitoe on behalf of the Wellcome Sanger Institute COVID-19 Surveillance Team |
| EPI_ISL_2913194 | Lighthouse Lab in Glasgow | Wellcome Sanger Institute for the COVID-19 Genomics UK (COG-UK) Consortium | Harper VanSteenhouse, Yumi Kasai, David Gray, Carol Clugston, Anna Dominiczak and Alex Alderton, Roberto Amato, Jeffrey Barrett, Sonia Goncalves, Ewan Harrison, David K. Jackson, Ian Johnston, Dominic Kwiatkowski, Cordelia Langford, John Sillitoe on behalf of the Wellcome Sanger Institute COVID-19 Surveillance Team |

| | | | |
|---|---|---|---|
| EPI_ISL_2913195, EPI_ISL_2913196, EPI_ISL_2913197, EPI_ISL_2913198, EPI_ISL_2913199 | Lighthouse Lab in Milton Keynes | Wellcome Sanger Institute for the COVID-19 Genomics UK (COG-UK) Consortium | The Lighthouse Lab in Milton Keynes and Alex Alderton, Roberto Amato, Jeffrey Barrett, Sonia Goncalves, Ewan Harrison, David K. Jackson, Ian Johnston, Dominic Kwiatkowski, Cordelia Langford, John Sillitoe on behalf of the Wellcome Sanger Institute COVID-19 Surveillance Team |
| EPI_ISL_2913200 | Berkshire and Surrey Pathology Services Lighthouse Laboratory | Wellcome Sanger Institute for the COVID-19 Genomics UK (COG-UK) Consortium | Berkshire and Surrey Pathology Services Lighthouse Laboratory and Alex Alderton, Roberto Amato, Jeffrey Barrett, Sonia Goncalves, Ewan Harrison, David K. Jackson, Ian Johnston, Dominic Kwiatkowski, Cordelia Langford, John Sillitoe on behalf of the Wellcome Sanger Institute COVID-19 Surveillance Team |
| EPI_ISL_2913201 | Lighthouse Lab in Glasgow | Wellcome Sanger Institute for the COVID-19 Genomics UK (COG-UK) Consortium | Harper VanSteenhouse, Yumi Kasai, David Gray, Carol Clugston, Anna Dominiczak and Alex Alderton, Roberto Amato, Jeffrey Barrett, Sonia Goncalves, Ewan Harrison, David K. Jackson, Ian Johnston, Dominic Kwiatkowski, Cordelia Langford, John Sillitoe on behalf of the Wellcome Sanger Institute COVID-19 Surveillance Team |
| EPI_ISL_2913202, EPI_ISL_2913203, EPI_ISL_2913204 | Lighthouse Lab in Milton Keynes | Wellcome Sanger Institute for the COVID-19 Genomics UK (COG-UK) Consortium | The Lighthouse Lab in Milton Keynes and Alex Alderton, Roberto Amato, Jeffrey Barrett, Sonia Goncalves, Ewan Harrison, David K. Jackson, Ian Johnston, Dominic Kwiatkowski, Cordelia Langford, John Sillitoe on behalf of the Wellcome Sanger Institute COVID-19 Surveillance Team |
| EPI_ISL_2913205, EPI_ISL_2913206 | Berkshire and Surrey Pathology Services Lighthouse Laboratory | Wellcome Sanger Institute for the COVID-19 Genomics UK (COG-UK) Consortium | Berkshire and Surrey Pathology Services Lighthouse Laboratory and Alex Alderton, Roberto Amato, Jeffrey Barrett, Sonia Goncalves, Ewan Harrison, David K. Jackson, Ian Johnston, Dominic Kwiatkowski, Cordelia Langford, John Sillitoe on behalf of the Wellcome Sanger Institute COVID-19 Surveillance Team |
| EPI_ISL_2913207, EPI_ISL_2913208, EPI_ISL_2913209 | Lighthouse Lab in Glasgow | Wellcome Sanger Institute for the COVID-19 Genomics UK (COG-UK) Consortium | Harper VanSteenhouse, Yumi Kasai, David Gray, Carol Clugston, Anna Dominiczak and Alex Alderton, Roberto Amato, Jeffrey Barrett, Sonia Goncalves, Ewan Harrison, David K. Jackson, Ian Johnston, Dominic Kwiatkowski, Cordelia Langford, John Sillitoe on behalf of the Wellcome Sanger Institute COVID-19 Surveillance Team |
| EPI_ISL_2913210 | Berkshire and Surrey Pathology Services Lighthouse Laboratory | Wellcome Sanger Institute for the COVID-19 Genomics UK (COG-UK) Consortium | Berkshire and Surrey Pathology Services Lighthouse Laboratory and Alex Alderton, Roberto Amato, Jeffrey Barrett, Sonia Goncalves, Ewan Harrison, David K. Jackson, Ian Johnston, Dominic Kwiatkowski, Cordelia Langford, John Sillitoe on behalf of the Wellcome Sanger Institute COVID-19 Surveillance Team |
| EPI_ISL_2913211 | Lighthouse Lab in Milton Keynes | Wellcome Sanger Institute for the COVID-19 Genomics UK (COG-UK) Consortium | The Lighthouse Lab in Milton Keynes and Alex Alderton, Roberto Amato, Jeffrey Barrett, Sonia Goncalves, Ewan Harrison, David K. Jackson, Ian Johnston, Dominic Kwiatkowski, Cordelia Langford, John Sillitoe on behalf of the Wellcome Sanger Institute COVID-19 Surveillance Team |
| EPI_ISL_2913212, EPI_ISL_2913213 | Lighthouse Lab in Glasgow | Wellcome Sanger Institute for the COVID-19 Genomics UK (COG-UK) Consortium | Harper VanSteenhouse, Yumi Kasai, David Gray, Carol Clugston, Anna Dominiczak and Alex Alderton, Roberto Amato, Jeffrey Barrett, Sonia Goncalves, Ewan Harrison, David K. Jackson, Ian Johnston, Dominic Kwiatkowski, Cordelia Langford, John Sillitoe on behalf of the Wellcome Sanger Institute COVID-19 Surveillance Team |
| EPI_ISL_2913214, EPI_ISL_2913215 | Lighthouse Lab in Milton Keynes | Wellcome Sanger Institute for the COVID-19 Genomics UK (COG-UK) Consortium | The Lighthouse Lab in Milton Keynes and Alex Alderton, Roberto Amato, Jeffrey Barrett, Sonia Goncalves, Ewan Harrison, David K. Jackson, Ian Johnston, Dominic Kwiatkowski, Cordelia Langford, John Sillitoe on behalf of the Wellcome Sanger Institute COVID-19 Surveillance Team |
| EPI_ISL_2913216, EPI_ISL_2913217 | Berkshire and Surrey Pathology Services Lighthouse Laboratory | Wellcome Sanger Institute for the COVID-19 Genomics UK (COG-UK) Consortium | Berkshire and Surrey Pathology Services Lighthouse Laboratory and Alex Alderton, Roberto Amato, Jeffrey Barrett, Sonia Goncalves, Ewan Harrison, David K. Jackson, Ian Johnston, Dominic Kwiatkowski, Cordelia Langford, John Sillitoe on behalf of the Wellcome Sanger Institute COVID-19 Surveillance Team |
| EPI_ISL_2913218 | Lighthouse Lab in Milton Keynes | Wellcome Sanger Institute for the COVID-19 Genomics UK (COG-UK) Consortium | The Lighthouse Lab in Milton Keynes and Alex Alderton, Roberto Amato, Jeffrey Barrett, Sonia Goncalves, Ewan Harrison, David K. Jackson, Ian Johnston, Dominic Kwiatkowski, Cordelia Langford, John Sillitoe on behalf of the Wellcome Sanger Institute COVID-19 Surveillance Team |
| EPI_ISL_2913219, EPI_ISL_2913220 | Lighthouse Lab in Glasgow | Wellcome Sanger Institute for the COVID-19 Genomics UK (COG-UK) Consortium | Harper VanSteenhouse, Yumi Kasai, David Gray, Carol Clugston, Anna Dominiczak and Alex Alderton, Roberto Amato, Jeffrey Barrett, Sonia Goncalves, Ewan Harrison, David K. Jackson, Ian Johnston, Dominic Kwiatkowski, Cordelia Langford, John Sillitoe on behalf of the Wellcome Sanger Institute COVID-19 Surveillance Team |
| EPI_ISL_2913221, EPI_ISL_2913222, EPI_ISL_2913223, EPI_ISL_2913224 | Lighthouse Lab in Milton Keynes | Wellcome Sanger Institute for the COVID-19 Genomics UK (COG-UK) Consortium | The Lighthouse Lab in Milton Keynes and Alex Alderton, Roberto Amato, Jeffrey Barrett, Sonia Goncalves, Ewan Harrison, David K. Jackson, Ian Johnston, Dominic Kwiatkowski, Cordelia Langford, John Sillitoe on behalf of the Wellcome Sanger Institute COVID-19 Surveillance Team |
| EPI_ISL_2913225 | Lighthouse Lab in Glasgow | Wellcome Sanger Institute for the COVID-19 Genomics UK (COG-UK) Consortium | Harper VanSteenhouse, Yumi Kasai, David Gray, Carol Clugston, Anna Dominiczak and Alex Alderton, Roberto Amato, Jeffrey Barrett, Sonia Goncalves, Ewan Harrison, David K. Jackson, Ian Johnston, Dominic Kwiatkowski, Cordelia Langford, John Sillitoe on behalf of the Wellcome Sanger Institute COVID-19 Surveillance Team |
| EPI_ISL_2913226 | Lighthouse Lab in Milton Keynes | Wellcome Sanger Institute for the COVID-19 Genomics UK (COG-UK) Consortium | The Lighthouse Lab in Milton Keynes and Alex Alderton, Roberto Amato, Jeffrey Barrett, Sonia Goncalves, Ewan Harrison, David K. Jackson, Ian Johnston, Dominic Kwiatkowski, Cordelia Langford, John Sillitoe on behalf of the Wellcome Sanger Institute COVID-19 Surveillance Team |
| EPI_ISL_2913227, EPI_ISL_2913228 | Lighthouse Lab in Glasgow | Wellcome Sanger Institute for the COVID-19 Genomics UK (COG-UK) Consortium | Harper VanSteenhouse, Yumi Kasai, David Gray, Carol Clugston, Anna Dominiczak and Alex Alderton, Roberto Amato, Jeffrey Barrett, Sonia Goncalves, Ewan Harrison, David K. Jackson, Ian Johnston, Dominic Kwiatkowski, Cordelia Langford, John Sillitoe on behalf of the Wellcome Sanger Institute COVID-19 Surveillance Team |
| EPI_ISL_2913229 | Lighthouse Lab in Milton Keynes | Wellcome Sanger Institute for the COVID-19 Genomics UK (COG-UK) Consortium | The Lighthouse Lab in Milton Keynes and Alex Alderton, Roberto Amato, Jeffrey Barrett, Sonia Goncalves, Ewan Harrison, David K. Jackson, Ian Johnston, Dominic Kwiatkowski, Cordelia Langford, John Sillitoe on behalf of the Wellcome Sanger Institute COVID-19 Surveillance Team |
| EPI_ISL_2913230 | Berkshire and Surrey Pathology Services Lighthouse Laboratory | Wellcome Sanger Institute for the COVID-19 Genomics UK (COG-UK) Consortium | Berkshire and Surrey Pathology Services Lighthouse Laboratory and Alex Alderton, Roberto Amato, Jeffrey Barrett, Sonia Goncalves, Ewan Harrison, David K. Jackson, Ian Johnston, Dominic Kwiatkowski, Cordelia Langford, John Sillitoe on behalf of the Wellcome Sanger Institute COVID-19 Surveillance Team |
| EPI_ISL_2913231 | Lighthouse Lab in Glasgow | Wellcome Sanger Institute for the COVID-19 Genomics UK (COG-UK) Consortium | Harper VanSteenhouse, Yumi Kasai, David Gray, Carol Clugston, Anna Dominiczak and Alex Alderton, Roberto Amato, Jeffrey Barrett, Sonia Goncalves, Ewan Harrison, David K. Jackson, Ian Johnston, Dominic Kwiatkowski, Cordelia Langford, John Sillitoe on behalf of the Wellcome Sanger Institute COVID-19 Surveillance Team |
| EPI_ISL_2913232 | Lighthouse Lab in Milton Keynes | Wellcome Sanger Institute for the COVID-19 Genomics UK (COG-UK) Consortium | The Lighthouse Lab in Milton Keynes and Alex Alderton, Roberto Amato, Jeffrey Barrett, Sonia Goncalves, Ewan Harrison, David K. Jackson, Ian Johnston, Dominic Kwiatkowski, Cordelia Langford, John Sillitoe on behalf of the Wellcome Sanger Institute COVID-19 Surveillance Team |
| EPI_ISL_2913233 | Lighthouse Lab in Glasgow | Wellcome Sanger Institute for the COVID-19 Genomics UK (COG-UK) Consortium | Harper VanSteenhouse, Yumi Kasai, David Gray, Carol Clugston, Anna Dominiczak and Alex Alderton, Roberto Amato, Jeffrey Barrett, Sonia Goncalves, Ewan Harrison, David K. Jackson, Ian Johnston, Dominic Kwiatkowski, Cordelia Langford, John Sillitoe on behalf of the Wellcome Sanger Institute COVID-19 Surveillance Team |
| EPI_ISL_2913234 | Lighthouse Lab in Milton Keynes | Wellcome Sanger Institute for the COVID-19 Genomics UK (COG-UK) Consortium | The Lighthouse Lab in Milton Keynes and Alex Alderton, Roberto Amato, Jeffrey Barrett, Sonia Goncalves, Ewan Harrison, David K. Jackson, Ian Johnston, Dominic Kwiatkowski, Cordelia Langford, John Sillitoe on behalf of the Wellcome Sanger Institute COVID-19 Surveillance Team |
| EPI_ISL_2913235 | Berkshire and Surrey Pathology Services Lighthouse Laboratory | Wellcome Sanger Institute for the COVID-19 Genomics UK (COG-UK) Consortium | Berkshire and Surrey Pathology Services Lighthouse Laboratory and Alex Alderton, Roberto Amato, Jeffrey Barrett, Sonia Goncalves, Ewan Harrison, David K. Jackson, Ian Johnston, Dominic Kwiatkowski, Cordelia Langford, John Sillitoe on behalf of the Wellcome Sanger Institute COVID-19 Surveillance Team |
| EPI_ISL_2913236 | Lighthouse Lab in Milton Keynes | Wellcome Sanger Institute for the COVID-19 Genomics UK (COG-UK) Consortium | The Lighthouse Lab in Milton Keynes and Alex Alderton, Roberto Amato, Jeffrey Barrett, Sonia Goncalves, Ewan Harrison, David K. Jackson, Ian Johnston, Dominic Kwiatkowski, Cordelia Langford, John Sillitoe on behalf of the Wellcome Sanger Institute COVID-19 Surveillance Team |
| EPI_ISL_2913237, EPI_ISL_2913238 | Berkshire and Surrey Pathology Services Lighthouse Laboratory | Wellcome Sanger Institute for the COVID-19 Genomics UK (COG-UK) Consortium | Berkshire and Surrey Pathology Services Lighthouse Laboratory and Alex Alderton, Roberto Amato, Jeffrey Barrett, Sonia Goncalves, Ewan Harrison, David K. Jackson, Ian Johnston, Dominic Kwiatkowski, Cordelia Langford, John Sillitoe on behalf of the Wellcome Sanger Institute COVID-19 Surveillance Team |
| EPI_ISL_2913239 | Lighthouse Lab in Glasgow | Wellcome Sanger Institute for the COVID-19 Genomics UK (COG-UK) Consortium | Harper VanSteenhouse, Yumi Kasai, David Gray, Carol Clugston, Anna Dominiczak and Alex Alderton, Roberto Amato, Jeffrey Barrett, Sonia Goncalves, Ewan Harrison, David K. Jackson, Ian Johnston, Dominic Kwiatkowski, Cordelia Langford, John Sillitoe on behalf of the Wellcome Sanger Institute COVID-19 Surveillance Team |
| EPI_ISL_2913240, EPI_ISL_2913241 | Berkshire and Surrey Pathology Services Lighthouse Laboratory | Wellcome Sanger Institute for the COVID-19 Genomics UK (COG-UK) Consortium | Berkshire and Surrey Pathology Services Lighthouse Laboratory and Alex Alderton, Roberto Amato, Jeffrey Barrett, Sonia Goncalves, Ewan Harrison, David K. Jackson, Ian Johnston, Dominic Kwiatkowski, Cordelia Langford, John Sillitoe on behalf of the Wellcome Sanger Institute COVID-19 Surveillance Team |
| EPI_ISL_2913242 | Lighthouse Lab in Milton Keynes | Wellcome Sanger Institute for the COVID-19 Genomics UK (COG-UK) Consortium | The Lighthouse Lab in Milton Keynes and Alex Alderton, Roberto Amato, Jeffrey Barrett, Sonia Goncalves, Ewan Harrison, David K. Jackson, Ian Johnston, Dominic Kwiatkowski, Cordelia Langford, John Sillitoe on behalf of the Wellcome Sanger Institute COVID-19 Surveillance Team |
| EPI_ISL_2913243 | Berkshire and Surrey Pathology Services Lighthouse Laboratory | Wellcome Sanger Institute for the COVID-19 Genomics UK (COG-UK) Consortium | Berkshire and Surrey Pathology Services Lighthouse Laboratory and Alex Alderton, Roberto Amato, Jeffrey Barrett, Sonia Goncalves, Ewan Harrison, David K. Jackson, Ian Johnston, Dominic Kwiatkowski, Cordelia Langford, John Sillitoe on behalf of the Wellcome Sanger Institute COVID-19 Surveillance Team |
| EPI_ISL_2913244, EPI_ISL_2913245 | Lighthouse Lab in Milton Keynes | Wellcome Sanger Institute for the COVID-19 Genomics UK (COG-UK) Consortium | The Lighthouse Lab in Milton Keynes and Alex Alderton, Roberto Amato, Jeffrey Barrett, Sonia Goncalves, Ewan Harrison, David K. Jackson, Ian Johnston, Dominic Kwiatkowski, Cordelia Langford, John Sillitoe on behalf of the Wellcome Sanger Institute COVID-19 Surveillance Team |
| EPI_ISL_2913246 | Berkshire and Surrey Pathology Services Lighthouse Laboratory | Wellcome Sanger Institute for the COVID-19 Genomics UK (COG-UK) Consortium | Berkshire and Surrey Pathology Services Lighthouse Laboratory and Alex Alderton, Roberto Amato, Jeffrey Barrett, Sonia Goncalves, Ewan Harrison, David K. Jackson, Ian Johnston, Dominic Kwiatkowski, Cordelia Langford, John Sillitoe on behalf of the Wellcome Sanger Institute COVID-19 Surveillance Team |
| EPI_ISL_2913247, EPI_ISL_2913248 | Lighthouse Lab in Glasgow | Wellcome Sanger Institute for the COVID-19 Genomics UK (COG-UK) Consortium | Harper VanSteenhouse, Yumi Kasai, David Gray, Carol Clugston, Anna Dominiczak and Alex Alderton, Roberto Amato, Jeffrey Barrett, Sonia Goncalves, Ewan Harrison, David K. Jackson, Ian Johnston, Dominic Kwiatkowski, Cordelia Langford, John Sillitoe on behalf of the Wellcome Sanger Institute COVID-19 Surveillance Team |
| EPI_ISL_2913249 | Lighthouse Lab in Milton Keynes | Wellcome Sanger Institute for the COVID-19 Genomics UK (COG-UK) Consortium | The Lighthouse Lab in Milton Keynes and Alex Alderton, Roberto Amato, Jeffrey Barrett, Sonia Goncalves, Ewan Harrison, David K. Jackson, Ian Johnston, Dominic Kwiatkowski, Cordelia Langford, John Sillitoe on behalf of the Wellcome Sanger Institute COVID-19 Surveillance Team |
| EPI_ISL_2913250, EPI_ISL_2913251 | Berkshire and Surrey Pathology Services Lighthouse Laboratory | Wellcome Sanger Institute for the COVID-19 Genomics UK (COG-UK) Consortium | Berkshire and Surrey Pathology Services Lighthouse Laboratory and Alex Alderton, Roberto Amato, Jeffrey Barrett, Sonia Goncalves, Ewan Harrison, David K. Jackson, Ian Johnston, Dominic Kwiatkowski, Cordelia Langford, John Sillitoe on behalf of the Wellcome Sanger Institute COVID-19 Surveillance Team |
| EPI_ISL_2913252 | Lighthouse Lab in Glasgow | Wellcome Sanger Institute for the COVID-19 Genomics UK (COG-UK) Consortium | Harper VanSteenhouse, Yumi Kasai, David Gray, Carol Clugston, Anna Dominiczak and Alex Alderton, Roberto Amato, Jeffrey Barrett, Sonia Goncalves, Ewan Harrison, David K. Jackson, Ian Johnston, Dominic Kwiatkowski, Cordelia Langford, John Sillitoe on behalf of the Wellcome Sanger Institute COVID-19 Surveillance Team |
| EPI_ISL_2913253 | Lighthouse Lab in Milton Keynes | Wellcome Sanger Institute for the COVID-19 Genomics UK (COG-UK) Consortium | The Lighthouse Lab in Milton Keynes and Alex Alderton, Roberto Amato, Jeffrey Barrett, Sonia Goncalves, Ewan Harrison, David K. Jackson, Ian Johnston, Dominic Kwiatkowski, Cordelia Langford, John Sillitoe on behalf of the Wellcome Sanger Institute COVID-19 Surveillance Team |
| EPI_ISL_2913254 | Lighthouse Lab in Glasgow | Wellcome Sanger Institute for the COVID-19 Genomics UK (COG-UK) Consortium | Harper VanSteenhouse, Yumi Kasai, David Gray, Carol Clugston, Anna Dominiczak and Alex Alderton, Roberto Amato, Jeffrey Barrett, Sonia Goncalves, Ewan Harrison, David K. Jackson, Ian Johnston, Dominic Kwiatkowski, Cordelia Langford, John Sillitoe on behalf of the Wellcome Sanger Institute COVID-19 Surveillance Team |
| EPI_ISL_2913255 | Lighthouse Lab in Milton Keynes | Wellcome Sanger Institute for the COVID-19 Genomics UK (COG-UK) Consortium | The Lighthouse Lab in Milton Keynes and Alex Alderton, Roberto Amato, Jeffrey Barrett, Sonia Goncalves, Ewan Harrison, David K. Jackson, Ian Johnston, Dominic Kwiatkowski, Cordelia Langford, John Sillitoe on behalf of the Wellcome Sanger Institute COVID-19 Surveillance Team |
| EPI_ISL_2913256 | Lighthouse Lab in Glasgow | Wellcome Sanger Institute for the COVID-19 Genomics UK (COG-UK) Consortium | Harper VanSteenhouse, Yumi Kasai, David Gray, Carol Clugston, Anna Dominiczak and Alex Alderton, Roberto Amato, Jeffrey Barrett, Sonia Goncalves, Ewan Harrison, David K. Jackson, Ian Johnston, Dominic Kwiatkowski, Cordelia Langford, John Sillitoe on behalf of the Wellcome Sanger Institute COVID-19 Surveillance Team |
| EPI_ISL_2913257, EPI_ISL_2913258 | Lighthouse Lab in Milton Keynes | Wellcome Sanger Institute for the COVID-19 Genomics UK (COG-UK) Consortium | The Lighthouse Lab in Milton Keynes and Alex Alderton, Roberto Amato, Jeffrey Barrett, Sonia Goncalves, Ewan Harrison, David K. Jackson, Ian Johnston, Dominic Kwiatkowski, Cordelia Langford, John Sillitoe on behalf of the Wellcome Sanger Institute COVID-19 Surveillance Team |
| EPI_ISL_2913259 | Lighthouse Lab in Glasgow | Wellcome Sanger Institute for the COVID-19 Genomics UK (COG-UK) Consortium | Harper VanSteenhouse, Yumi Kasai, David Gray, Carol Clugston, Anna Dominiczak and Alex Alderton, Roberto Amato, Jeffrey Barrett, Sonia Goncalves, Ewan Harrison, David K. Jackson, Ian Johnston, Dominic Kwiatkowski, Cordelia Langford, John Sillitoe on behalf of the Wellcome Sanger Institute COVID-19 Surveillance Team |

| | | | |
|---|---|---|---|
| EPI_ISL_2913260 | Lighthouse Lab in Milton Keynes | Wellcome Sanger Institute for the COVID-19 Genomics UK (COG-UK) Consortium | The Lighthouse Lab in Milton Keynes and Alex Alderton, Roberto Amato, Jeffrey Barrett, Sonia Goncalves, Ewan Harrison, David K. Jackson, Ian Johnston, Dominic Kwiatkowski, Cordelia Langford, John Sillitoe on behalf of the Wellcome Sanger Institute COVID-19 Surveillance Team |
| EPI_ISL_2913261 | Lighthouse Lab in Glasgow | Wellcome Sanger Institute for the COVID-19 Genomics UK (COG-UK) Consortium | Harper VanSteenhouse, Yumi Kasai, David Gray, Carol Clugston, Anna Dominiczak and Alex Alderton, Roberto Amato, Jeffrey Barrett, Sonia Goncalves, Ewan Harrison, David K. Jackson, Ian Johnston, Dominic Kwiatkowski, Cordelia Langford, John Sillitoe on behalf of the Wellcome Sanger Institute COVID-19 Surveillance Team |
| EPI_ISL_2913262 | Berkshire and Surrey Pathology Services Lighthouse Laboratory | Wellcome Sanger Institute for the COVID-19 Genomics UK (COG-UK) Consortium | Berkshire and Surrey Pathology Services Lighthouse Laboratory and Alex Alderton, Roberto Amato, Jeffrey Barrett, Sonia Goncalves, Ewan Harrison, David K. Jackson, Ian Johnston, Dominic Kwiatkowski, Cordelia Langford, John Sillitoe on behalf of the Wellcome Sanger Institute COVID-19 Surveillance Team |
| EPI_ISL_2913263 | Lighthouse Lab in Milton Keynes | Wellcome Sanger Institute for the COVID-19 Genomics UK (COG-UK) Consortium | The Lighthouse Lab in Milton Keynes and Alex Alderton, Roberto Amato, Jeffrey Barrett, Sonia Goncalves, Ewan Harrison, David K. Jackson, Ian Johnston, Dominic Kwiatkowski, Cordelia Langford, John Sillitoe on behalf of the Wellcome Sanger Institute COVID-19 Surveillance Team |
| EPI_ISL_2913264, EPI_ISL_2913265 | Berkshire and Surrey Pathology Services Lighthouse Laboratory | Wellcome Sanger Institute for the COVID-19 Genomics UK (COG-UK) Consortium | Berkshire and Surrey Pathology Services Lighthouse Laboratory and Alex Alderton, Roberto Amato, Jeffrey Barrett, Sonia Goncalves, Ewan Harrison, David K. Jackson, Ian Johnston, Dominic Kwiatkowski, Cordelia Langford, John Sillitoe on behalf of the Wellcome Sanger Institute COVID-19 Surveillance Team |
| EPI_ISL_2913266 | Lighthouse Lab in Milton Keynes | Wellcome Sanger Institute for the COVID-19 Genomics UK (COG-UK) Consortium | The Lighthouse Lab in Milton Keynes and Alex Alderton, Roberto Amato, Jeffrey Barrett, Sonia Goncalves, Ewan Harrison, David K. Jackson, Ian Johnston, Dominic Kwiatkowski, Cordelia Langford, John Sillitoe on behalf of the Wellcome Sanger Institute COVID-19 Surveillance Team |
| EPI_ISL_2913267, EPI_ISL_2913268 | Lighthouse Lab in Glasgow | Wellcome Sanger Institute for the COVID-19 Genomics UK (COG-UK) Consortium | Harper VanSteenhouse, Yumi Kasai, David Gray, Carol Clugston, Anna Dominiczak and Alex Alderton, Roberto Amato, Jeffrey Barrett, Sonia Goncalves, Ewan Harrison, David K. Jackson, Ian Johnston, Dominic Kwiatkowski, Cordelia Langford, John Sillitoe on behalf of the Wellcome Sanger Institute COVID-19 Surveillance Team |
| EPI_ISL_2913269 | Lighthouse Lab in Milton Keynes | Wellcome Sanger Institute for the COVID-19 Genomics UK (COG-UK) Consortium | The Lighthouse Lab in Milton Keynes and Alex Alderton, Roberto Amato, Jeffrey Barrett, Sonia Goncalves, Ewan Harrison, David K. Jackson, Ian Johnston, Dominic Kwiatkowski, Cordelia Langford, John Sillitoe on behalf of the Wellcome Sanger Institute COVID-19 Surveillance Team |
| EPI_ISL_2913270 | Berkshire and Surrey Pathology Services Lighthouse Laboratory | Wellcome Sanger Institute for the COVID-19 Genomics UK (COG-UK) Consortium | Berkshire and Surrey Pathology Services Lighthouse Laboratory and Alex Alderton, Roberto Amato, Jeffrey Barrett, Sonia Goncalves, Ewan Harrison, David K. Jackson, Ian Johnston, Dominic Kwiatkowski, Cordelia Langford, John Sillitoe on behalf of the Wellcome Sanger Institute COVID-19 Surveillance Team |
| EPI_ISL_2913271 | Lighthouse Lab in Glasgow | Wellcome Sanger Institute for the COVID-19 Genomics UK (COG-UK) Consortium | Harper VanSteenhouse, Yumi Kasai, David Gray, Carol Clugston, Anna Dominiczak and Alex Alderton, Roberto Amato, Jeffrey Barrett, Sonia Goncalves, Ewan Harrison, David K. Jackson, Ian Johnston, Dominic Kwiatkowski, Cordelia Langford, John Sillitoe on behalf of the Wellcome Sanger Institute COVID-19 Surveillance Team |
| EPI_ISL_2913272 | Berkshire and Surrey Pathology Services Lighthouse Laboratory | Wellcome Sanger Institute for the COVID-19 Genomics UK (COG-UK) Consortium | Berkshire and Surrey Pathology Services Lighthouse Laboratory and Alex Alderton, Roberto Amato, Jeffrey Barrett, Sonia Goncalves, Ewan Harrison, David K. Jackson, Ian Johnston, Dominic Kwiatkowski, Cordelia Langford, John Sillitoe on behalf of the Wellcome Sanger Institute COVID-19 Surveillance Team |
| EPI_ISL_2913273, EPI_ISL_2913274 | Lighthouse Lab in Milton Keynes | Wellcome Sanger Institute for the COVID-19 Genomics UK (COG-UK) Consortium | The Lighthouse Lab in Milton Keynes and Alex Alderton, Roberto Amato, Jeffrey Barrett, Sonia Goncalves, Ewan Harrison, David K. Jackson, Ian Johnston, Dominic Kwiatkowski, Cordelia Langford, John Sillitoe on behalf of the Wellcome Sanger Institute COVID-19 Surveillance Team |
| EPI_ISL_2913275 | Berkshire and Surrey Pathology Services Lighthouse Laboratory | Wellcome Sanger Institute for the COVID-19 Genomics UK (COG-UK) Consortium | Berkshire and Surrey Pathology Services Lighthouse Laboratory and Alex Alderton, Roberto Amato, Jeffrey Barrett, Sonia Goncalves, Ewan Harrison, David K. Jackson, Ian Johnston, Dominic Kwiatkowski, Cordelia Langford, John Sillitoe on behalf of the Wellcome Sanger Institute COVID-19 Surveillance Team |
| EPI_ISL_2913276 | Lighthouse Lab in Milton Keynes | Wellcome Sanger Institute for the COVID-19 Genomics UK (COG-UK) Consortium | The Lighthouse Lab in Milton Keynes and Alex Alderton, Roberto Amato, Jeffrey Barrett, Sonia Goncalves, Ewan Harrison, David K. Jackson, Ian Johnston, Dominic Kwiatkowski, Cordelia Langford, John Sillitoe on behalf of the Wellcome Sanger Institute COVID-19 Surveillance Team |
| EPI_ISL_2913277 | Berkshire and Surrey Pathology Services Lighthouse Laboratory | Wellcome Sanger Institute for the COVID-19 Genomics UK (COG-UK) Consortium | Berkshire and Surrey Pathology Services Lighthouse Laboratory and Alex Alderton, Roberto Amato, Jeffrey Barrett, Sonia Goncalves, Ewan Harrison, David K. Jackson, Ian Johnston, Dominic Kwiatkowski, Cordelia Langford, John Sillitoe on behalf of the Wellcome Sanger Institute COVID-19 Surveillance Team |
| EPI_ISL_2913279, EPI_ISL_2913280 | Lighthouse Lab in Milton Keynes | Wellcome Sanger Institute for the COVID-19 Genomics UK (COG-UK) Consortium | The Lighthouse Lab in Milton Keynes and Alex Alderton, Roberto Amato, Jeffrey Barrett, Sonia Goncalves, Ewan Harrison, David K. Jackson, Ian Johnston, Dominic Kwiatkowski, Cordelia Langford, John Sillitoe on behalf of the Wellcome Sanger Institute COVID-19 Surveillance Team |
| EPI_ISL_2913281 | Lighthouse Lab in Glasgow | Wellcome Sanger Institute for the COVID-19 Genomics UK (COG-UK) Consortium | Harper VanSteenhouse, Yumi Kasai, David Gray, Carol Clugston, Anna Dominiczak and Alex Alderton, Roberto Amato, Jeffrey Barrett, Sonia Goncalves, Ewan Harrison, David K. Jackson, Ian Johnston, Dominic Kwiatkowski, Cordelia Langford, John Sillitoe on behalf of the Wellcome Sanger Institute COVID-19 Surveillance Team |
| EPI_ISL_2913282 | Berkshire and Surrey Pathology Services Lighthouse Laboratory | Wellcome Sanger Institute for the COVID-19 Genomics UK (COG-UK) Consortium | Berkshire and Surrey Pathology Services Lighthouse Laboratory and Alex Alderton, Roberto Amato, Jeffrey Barrett, Sonia Goncalves, Ewan Harrison, David K. Jackson, Ian Johnston, Dominic Kwiatkowski, Cordelia Langford, John Sillitoe on behalf of the Wellcome Sanger Institute COVID-19 Surveillance Team |
| EPI_ISL_2913283 | Lighthouse Lab in Milton Keynes | Wellcome Sanger Institute for the COVID-19 Genomics UK (COG-UK) Consortium | The Lighthouse Lab in Milton Keynes and Alex Alderton, Roberto Amato, Jeffrey Barrett, Sonia Goncalves, Ewan Harrison, David K. Jackson, Ian Johnston, Dominic Kwiatkowski, Cordelia Langford, John Sillitoe on behalf of the Wellcome Sanger Institute COVID-19 Surveillance Team |
| EPI_ISL_2913284 | Berkshire and Surrey Pathology Services Lighthouse Laboratory | Wellcome Sanger Institute for the COVID-19 Genomics UK (COG-UK) Consortium | Berkshire and Surrey Pathology Services Lighthouse Laboratory and Alex Alderton, Roberto Amato, Jeffrey Barrett, Sonia Goncalves, Ewan Harrison, David K. Jackson, Ian Johnston, Dominic Kwiatkowski, Cordelia Langford, John Sillitoe on behalf of the Wellcome Sanger Institute COVID-19 Surveillance Team |
| EPI_ISL_2913285 | Lighthouse Lab in Milton Keynes | Wellcome Sanger Institute for the COVID-19 Genomics UK (COG-UK) Consortium | The Lighthouse Lab in Milton Keynes and Alex Alderton, Roberto Amato, Jeffrey Barrett, Sonia Goncalves, Ewan Harrison, David K. Jackson, Ian Johnston, Dominic Kwiatkowski, Cordelia Langford, John Sillitoe on behalf of the Wellcome Sanger Institute COVID-19 Surveillance Team |
| EPI_ISL_2913286, EPI_ISL_2913287, EPI_ISL_2913288 | Lighthouse Lab in Glasgow | Wellcome Sanger Institute for the COVID-19 Genomics UK (COG-UK) Consortium | Harper VanSteenhouse, Yumi Kasai, David Gray, Carol Clugston, Anna Dominiczak and Alex Alderton, Roberto Amato, Jeffrey Barrett, Sonia Goncalves, Ewan Harrison, David K. Jackson, Ian Johnston, Dominic Kwiatkowski, Cordelia Langford, John Sillitoe on behalf of the Wellcome Sanger Institute COVID-19 Surveillance Team |
| EPI_ISL_2913289, EPI_ISL_2913290 | Lighthouse Lab in Milton Keynes | Wellcome Sanger Institute for the COVID-19 Genomics UK (COG-UK) Consortium | The Lighthouse Lab in Milton Keynes and Alex Alderton, Roberto Amato, Jeffrey Barrett, Sonia Goncalves, Ewan Harrison, David K. Jackson, Ian Johnston, Dominic Kwiatkowski, Cordelia Langford, John Sillitoe on behalf of the Wellcome Sanger Institute COVID-19 Surveillance Team |
| EPI_ISL_2913291 | Berkshire and Surrey Pathology Services Lighthouse Laboratory | Wellcome Sanger Institute for the COVID-19 Genomics UK (COG-UK) Consortium | Berkshire and Surrey Pathology Services Lighthouse Laboratory and Alex Alderton, Roberto Amato, Jeffrey Barrett, Sonia Goncalves, Ewan Harrison, David K. Jackson, Ian Johnston, Dominic Kwiatkowski, Cordelia Langford, John Sillitoe on behalf of the Wellcome Sanger Institute COVID-19 Surveillance Team |
| EPI_ISL_2913292 | Lighthouse Lab in Glasgow | Wellcome Sanger Institute for the COVID-19 Genomics UK (COG-UK) Consortium | Harper VanSteenhouse, Yumi Kasai, David Gray, Carol Clugston, Anna Dominiczak and Alex Alderton, Roberto Amato, Jeffrey Barrett, Sonia Goncalves, Ewan Harrison, David K. Jackson, Ian Johnston, Dominic Kwiatkowski, Cordelia Langford, John Sillitoe on behalf of the Wellcome Sanger Institute COVID-19 Surveillance Team |
| EPI_ISL_2913293 | Berkshire and Surrey Pathology Services Lighthouse Laboratory | Wellcome Sanger Institute for the COVID-19 Genomics UK (COG-UK) Consortium | Berkshire and Surrey Pathology Services Lighthouse Laboratory and Alex Alderton, Roberto Amato, Jeffrey Barrett, Sonia Goncalves, Ewan Harrison, David K. Jackson, Ian Johnston, Dominic Kwiatkowski, Cordelia Langford, John Sillitoe on behalf of the Wellcome Sanger Institute COVID-19 Surveillance Team |
| EPI_ISL_2913294 | Lighthouse Lab in Glasgow | Wellcome Sanger Institute for the COVID-19 Genomics UK (COG-UK) Consortium | Harper VanSteenhouse, Yumi Kasai, David Gray, Carol Clugston, Anna Dominiczak and Alex Alderton, Roberto Amato, Jeffrey Barrett, Sonia Goncalves, Ewan Harrison, David K. Jackson, Ian Johnston, Dominic Kwiatkowski, Cordelia Langford, John Sillitoe on behalf of the Wellcome Sanger Institute COVID-19 Surveillance Team |
| EPI_ISL_2913295 | Berkshire and Surrey Pathology Services Lighthouse Laboratory | Wellcome Sanger Institute for the COVID-19 Genomics UK (COG-UK) Consortium | Berkshire and Surrey Pathology Services Lighthouse Laboratory and Alex Alderton, Roberto Amato, Jeffrey Barrett, Sonia Goncalves, Ewan Harrison, David K. Jackson, Ian Johnston, Dominic Kwiatkowski, Cordelia Langford, John Sillitoe on behalf of the Wellcome Sanger Institute COVID-19 Surveillance Team |
| EPI_ISL_2913296 | Lighthouse Lab in Glasgow | Wellcome Sanger Institute for the COVID-19 Genomics UK (COG-UK) Consortium | Harper VanSteenhouse, Yumi Kasai, David Gray, Carol Clugston, Anna Dominiczak and Alex Alderton, Roberto Amato, Jeffrey Barrett, Sonia Goncalves, Ewan Harrison, David K. Jackson, Ian Johnston, Dominic Kwiatkowski, Cordelia Langford, John Sillitoe on behalf of the Wellcome Sanger Institute COVID-19 Surveillance Team |
| EPI_ISL_2913297, EPI_ISL_2913298 | Lighthouse Lab in Milton Keynes | Wellcome Sanger Institute for the COVID-19 Genomics UK (COG-UK) Consortium | The Lighthouse Lab in Milton Keynes and Alex Alderton, Roberto Amato, Jeffrey Barrett, Sonia Goncalves, Ewan Harrison, David K. Jackson, Ian Johnston, Dominic Kwiatkowski, Cordelia Langford, John Sillitoe on behalf of the Wellcome Sanger Institute COVID-19 Surveillance Team |
| EPI_ISL_2913299, EPI_ISL_2913300, EPI_ISL_2913301 | Berkshire and Surrey Pathology Services Lighthouse Laboratory | Wellcome Sanger Institute for the COVID-19 Genomics UK (COG-UK) Consortium | Berkshire and Surrey Pathology Services Lighthouse Laboratory and Alex Alderton, Roberto Amato, Jeffrey Barrett, Sonia Goncalves, Ewan Harrison, David K. Jackson, Ian Johnston, Dominic Kwiatkowski, Cordelia Langford, John Sillitoe on behalf of the Wellcome Sanger Institute COVID-19 Surveillance Team |
| EPI_ISL_2913302, EPI_ISL_2913303 | Lighthouse Lab in Milton Keynes | Wellcome Sanger Institute for the COVID-19 Genomics UK (COG-UK) Consortium | The Lighthouse Lab in Milton Keynes and Alex Alderton, Roberto Amato, Jeffrey Barrett, Sonia Goncalves, Ewan Harrison, David K. Jackson, Ian Johnston, Dominic Kwiatkowski, Cordelia Langford, John Sillitoe on behalf of the Wellcome Sanger Institute COVID-19 Surveillance Team |
| EPI_ISL_2913304, EPI_ISL_2913305 | Berkshire and Surrey Pathology Services Lighthouse Laboratory | Wellcome Sanger Institute for the COVID-19 Genomics UK (COG-UK) Consortium | Berkshire and Surrey Pathology Services Lighthouse Laboratory and Alex Alderton, Roberto Amato, Jeffrey Barrett, Sonia Goncalves, Ewan Harrison, David K. Jackson, Ian Johnston, Dominic Kwiatkowski, Cordelia Langford, John Sillitoe on behalf of the Wellcome Sanger Institute COVID-19 Surveillance Team |
| EPI_ISL_2913306, EPI_ISL_2913307, EPI_ISL_2913308, EPI_ISL_2913309 | Lighthouse Lab in Glasgow | Wellcome Sanger Institute for the COVID-19 Genomics UK (COG-UK) Consortium | Harper VanSteenhouse, Yumi Kasai, David Gray, Carol Clugston, Anna Dominiczak and Alex Alderton, Roberto Amato, Jeffrey Barrett, Sonia Goncalves, Ewan Harrison, David K. Jackson, Ian Johnston, Dominic Kwiatkowski, Cordelia Langford, John Sillitoe on behalf of the Wellcome Sanger Institute COVID-19 Surveillance Team |
| EPI_ISL_2913310 | Lighthouse Lab in Milton Keynes | Wellcome Sanger Institute for the COVID-19 Genomics UK (COG-UK) Consortium | The Lighthouse Lab in Milton Keynes and Alex Alderton, Roberto Amato, Jeffrey Barrett, Sonia Goncalves, Ewan Harrison, David K. Jackson, Ian Johnston, Dominic Kwiatkowski, Cordelia Langford, John Sillitoe on behalf of the Wellcome Sanger Institute COVID-19 Surveillance Team |
| EPI_ISL_2913311 | Berkshire and Surrey Pathology Services Lighthouse Laboratory | Wellcome Sanger Institute for the COVID-19 Genomics UK (COG-UK) Consortium | Berkshire and Surrey Pathology Services Lighthouse Laboratory and Alex Alderton, Roberto Amato, Jeffrey Barrett, Sonia Goncalves, Ewan Harrison, David K. Jackson, Ian Johnston, Dominic Kwiatkowski, Cordelia Langford, John Sillitoe on behalf of the Wellcome Sanger Institute COVID-19 Surveillance Team |
| EPI_ISL_2913312, EPI_ISL_2913313 | Lighthouse Lab in Milton Keynes | Wellcome Sanger Institute for the COVID-19 Genomics UK (COG-UK) Consortium | The Lighthouse Lab in Milton Keynes and Alex Alderton, Roberto Amato, Jeffrey Barrett, Sonia Goncalves, Ewan Harrison, David K. Jackson, Ian Johnston, Dominic Kwiatkowski, Cordelia Langford, John Sillitoe on behalf of the Wellcome Sanger Institute COVID-19 Surveillance Team |
| EPI_ISL_2913314 | Berkshire and Surrey Pathology Services Lighthouse Laboratory | Wellcome Sanger Institute for the COVID-19 Genomics UK (COG-UK) Consortium | Berkshire and Surrey Pathology Services Lighthouse Laboratory and Alex Alderton, Roberto Amato, Jeffrey Barrett, Sonia Goncalves, Ewan Harrison, David K. Jackson, Ian Johnston, Dominic Kwiatkowski, Cordelia Langford, John Sillitoe on behalf of the Wellcome Sanger Institute COVID-19 Surveillance Team |
| EPI_ISL_2913315 | Lighthouse Lab in Glasgow | Wellcome Sanger Institute for the COVID-19 Genomics UK (COG-UK) Consortium | Harper VanSteenhouse, Yumi Kasai, David Gray, Carol Clugston, Anna Dominiczak and Alex Alderton, Roberto Amato, Jeffrey Barrett, Sonia Goncalves, Ewan Harrison, David K. Jackson, Ian Johnston, Dominic Kwiatkowski, Cordelia Langford, John Sillitoe on behalf of the Wellcome Sanger Institute COVID-19 Surveillance Team |
| EPI_ISL_2913316 | Berkshire and Surrey Pathology Services Lighthouse Laboratory | Wellcome Sanger Institute for the COVID-19 Genomics UK (COG-UK) Consortium | Berkshire and Surrey Pathology Services Lighthouse Laboratory and Alex Alderton, Roberto Amato, Jeffrey Barrett, Sonia Goncalves, Ewan Harrison, David K. Jackson, Ian Johnston, Dominic Kwiatkowski, Cordelia Langford, John Sillitoe on behalf of the Wellcome Sanger Institute COVID-19 Surveillance Team |
| EPI_ISL_2913317, EPI_ISL_2913318, EPI_ISL_2913319, EPI_ISL_2913320, EPI_ISL_2913321, EPI_ISL_2913322, EPI_ISL_2913323 | Lighthouse Lab in Milton Keynes | Wellcome Sanger Institute for the COVID-19 Genomics UK (COG-UK) Consortium | The Lighthouse Lab in Milton Keynes and Alex Alderton, Roberto Amato, Jeffrey Barrett, Sonia Goncalves, Ewan Harrison, David K. Jackson, Ian Johnston, Dominic Kwiatkowski, Cordelia Langford, John Sillitoe on behalf of the Wellcome Sanger Institute COVID-19 Surveillance Team |
| EPI_ISL_2913324, | Berkshire and Surrey Pathology Services Lighthouse Laboratory | Wellcome Sanger Institute for the COVID-19 Genomics UK (COG-UK) | Berkshire and Surrey Pathology Services Lighthouse Laboratory and Alex Alderton, Roberto Amato, Jeffrey Barrett, Sonia Goncalves, Ewan Harrison, David K. Jackson, Ian Johnston, Dominic Kwiatkowski, Cordelia Langford, John Sillitoe on behalf |

| | | | |
|---|---|---|---|
| EPI_ISL_2913325 | | Consortium | of the Wellcome Sanger Institute COVID-19 Surveillance Team |
| EPI_ISL_2913326 | Lighthouse Lab in Alderley Park | Wellcome Sanger Institute for the COVID-19 Genomics UK (COG-UK) Consortium | Jacquelyn Wynn, Mairead Hyland, The Lighthouse Lab in Alderley Park and Alex Alderton, Roberto Amato, Jeffrey Barrett, Sonia Goncalves, Ewan Harrison, David K. Jackson, Ian Johnston, Dominic Kwiatkowski, Cordelia Langford, John Sillitoe on behalf of the Wellcome Sanger Institute COVID-19 Surveillance Team |
| EPI_ISL_2913327 | Berkshire and Surrey Pathology Services Lighthouse Laboratory | Wellcome Sanger Institute for the COVID-19 Genomics UK (COG-UK) Consortium | Berkshire and Surrey Pathology Services Lighthouse Laboratory and Alex Alderton, Roberto Amato, Jeffrey Barrett, Sonia Goncalves, Ewan Harrison, David K. Jackson, Ian Johnston, Dominic Kwiatkowski, Cordelia Langford, John Sillitoe on behalf of the Wellcome Sanger Institute COVID-19 Surveillance Team |
| EPI_ISL_2913328 | Lighthouse Lab in Milton Keynes | Wellcome Sanger Institute for the COVID-19 Genomics UK (COG-UK) Consortium | The Lighthouse Lab in Milton Keynes and Alex Alderton, Roberto Amato, Jeffrey Barrett, Sonia Goncalves, Ewan Harrison, David K. Jackson, Ian Johnston, Dominic Kwiatkowski, Cordelia Langford, John Sillitoe on behalf of the Wellcome Sanger Institute COVID-19 Surveillance Team |
| EPI_ISL_2913329 | Lighthouse Lab in Alderley Park | Wellcome Sanger Institute for the COVID-19 Genomics UK (COG-UK) Consortium | Jacquelyn Wynn, Mairead Hyland, The Lighthouse Lab in Alderley Park and Alex Alderton, Roberto Amato, Jeffrey Barrett, Sonia Goncalves, Ewan Harrison, David K. Jackson, Ian Johnston, Dominic Kwiatkowski, Cordelia Langford, John Sillitoe on behalf of the Wellcome Sanger Institute COVID-19 Surveillance Team |
| EPI_ISL_2913330 | Berkshire and Surrey Pathology Services Lighthouse Laboratory | Wellcome Sanger Institute for the COVID-19 Genomics UK (COG-UK) Consortium | Berkshire and Surrey Pathology Services Lighthouse Laboratory and Alex Alderton, Roberto Amato, Jeffrey Barrett, Sonia Goncalves, Ewan Harrison, David K. Jackson, Ian Johnston, Dominic Kwiatkowski, Cordelia Langford, John Sillitoe on behalf of the Wellcome Sanger Institute COVID-19 Surveillance Team |
| EPI_ISL_2913331, EPI_ISL_2913332 | Lighthouse Lab in Milton Keynes | Wellcome Sanger Institute for the COVID-19 Genomics UK (COG-UK) Consortium | The Lighthouse Lab in Milton Keynes and Alex Alderton, Roberto Amato, Jeffrey Barrett, Sonia Goncalves, Ewan Harrison, David K. Jackson, Ian Johnston, Dominic Kwiatkowski, Cordelia Langford, John Sillitoe on behalf of the Wellcome Sanger Institute COVID-19 Surveillance Team |
| EPI_ISL_2913333 | Lighthouse Lab in Alderley Park | Wellcome Sanger Institute for the COVID-19 Genomics UK (COG-UK) Consortium | Jacquelyn Wynn, Mairead Hyland, The Lighthouse Lab in Alderley Park and Alex Alderton, Roberto Amato, Jeffrey Barrett, Sonia Goncalves, Ewan Harrison, David K. Jackson, Ian Johnston, Dominic Kwiatkowski, Cordelia Langford, John Sillitoe on behalf of the Wellcome Sanger Institute COVID-19 Surveillance Team |
| EPI_ISL_2913334, EPI_ISL_2913335, EPI_ISL_2913336, EPI_ISL_2913337, EPI_ISL_2913338, EPI_ISL_2913339, EPI_ISL_2913340 | Lighthouse Lab in Milton Keynes | Wellcome Sanger Institute for the COVID-19 Genomics UK (COG-UK) Consortium | The Lighthouse Lab in Milton Keynes and Alex Alderton, Roberto Amato, Jeffrey Barrett, Sonia Goncalves, Ewan Harrison, David K. Jackson, Ian Johnston, Dominic Kwiatkowski, Cordelia Langford, John Sillitoe on behalf of the Wellcome Sanger Institute COVID-19 Surveillance Team |
| EPI_ISL_2913341, EPI_ISL_2913342 | Berkshire and Surrey Pathology Services Lighthouse Laboratory | Wellcome Sanger Institute for the COVID-19 Genomics UK (COG-UK) Consortium | Berkshire and Surrey Pathology Services Lighthouse Laboratory and Alex Alderton, Roberto Amato, Jeffrey Barrett, Sonia Goncalves, Ewan Harrison, David K. Jackson, Ian Johnston, Dominic Kwiatkowski, Cordelia Langford, John Sillitoe on behalf of the Wellcome Sanger Institute COVID-19 Surveillance Team |
| EPI_ISL_2913343, EPI_ISL_2913344 | Lighthouse Lab in Milton Keynes | Wellcome Sanger Institute for the COVID-19 Genomics UK (COG-UK) Consortium | The Lighthouse Lab in Milton Keynes and Alex Alderton, Roberto Amato, Jeffrey Barrett, Sonia Goncalves, Ewan Harrison, David K. Jackson, Ian Johnston, Dominic Kwiatkowski, Cordelia Langford, John Sillitoe on behalf of the Wellcome Sanger Institute COVID-19 Surveillance Team |
| EPI_ISL_2913345 | Lighthouse Lab in Alderley Park | Wellcome Sanger Institute for the COVID-19 Genomics UK (COG-UK) Consortium | Jacquelyn Wynn, Mairead Hyland, The Lighthouse Lab in Alderley Park and Alex Alderton, Roberto Amato, Jeffrey Barrett, Sonia Goncalves, Ewan Harrison, David K. Jackson, Ian Johnston, Dominic Kwiatkowski, Cordelia Langford, John Sillitoe on behalf of the Wellcome Sanger Institute COVID-19 Surveillance Team |
| EPI_ISL_2913346, EPI_ISL_2913347, EPI_ISL_2913348 | Berkshire and Surrey Pathology Services Lighthouse Laboratory | Wellcome Sanger Institute for the COVID-19 Genomics UK (COG-UK) Consortium | Berkshire and Surrey Pathology Services Lighthouse Laboratory and Alex Alderton, Roberto Amato, Jeffrey Barrett, Sonia Goncalves, Ewan Harrison, David K. Jackson, Ian Johnston, Dominic Kwiatkowski, Cordelia Langford, John Sillitoe on behalf of the Wellcome Sanger Institute COVID-19 Surveillance Team |
| EPI_ISL_2913349 | Lighthouse Lab in Alderley Park | Wellcome Sanger Institute for the COVID-19 Genomics UK (COG-UK) Consortium | Jacquelyn Wynn, Mairead Hyland, The Lighthouse Lab in Alderley Park and Alex Alderton, Roberto Amato, Jeffrey Barrett, Sonia Goncalves, Ewan Harrison, David K. Jackson, Ian Johnston, Dominic Kwiatkowski, Cordelia Langford, John Sillitoe on behalf of the Wellcome Sanger Institute COVID-19 Surveillance Team |
| EPI_ISL_2913350 | Lighthouse Lab in Milton Keynes | Wellcome Sanger Institute for the COVID-19 Genomics UK (COG-UK) Consortium | The Lighthouse Lab in Milton Keynes and Alex Alderton, Roberto Amato, Jeffrey Barrett, Sonia Goncalves, Ewan Harrison, David K. Jackson, Ian Johnston, Dominic Kwiatkowski, Cordelia Langford, John Sillitoe on behalf of the Wellcome Sanger Institute COVID-19 Surveillance Team |
| EPI_ISL_2913351 | Berkshire and Surrey Pathology Services Lighthouse Laboratory | Wellcome Sanger Institute for the COVID-19 Genomics UK (COG-UK) Consortium | Berkshire and Surrey Pathology Services Lighthouse Laboratory and Alex Alderton, Roberto Amato, Jeffrey Barrett, Sonia Goncalves, Ewan Harrison, David K. Jackson, Ian Johnston, Dominic Kwiatkowski, Cordelia Langford, John Sillitoe on behalf of the Wellcome Sanger Institute COVID-19 Surveillance Team |
| EPI_ISL_2913352 | Lighthouse Lab in Milton Keynes | Wellcome Sanger Institute for the COVID-19 Genomics UK (COG-UK) Consortium | The Lighthouse Lab in Milton Keynes and Alex Alderton, Roberto Amato, Jeffrey Barrett, Sonia Goncalves, Ewan Harrison, David K. Jackson, Ian Johnston, Dominic Kwiatkowski, Cordelia Langford, John Sillitoe on behalf of the Wellcome Sanger Institute COVID-19 Surveillance Team |
| EPI_ISL_2913353, EPI_ISL_2913354, EPI_ISL_2913355 | Berkshire and Surrey Pathology Services Lighthouse Laboratory | Wellcome Sanger Institute for the COVID-19 Genomics UK (COG-UK) Consortium | Berkshire and Surrey Pathology Services Lighthouse Laboratory and Alex Alderton, Roberto Amato, Jeffrey Barrett, Sonia Goncalves, Ewan Harrison, David K. Jackson, Ian Johnston, Dominic Kwiatkowski, Cordelia Langford, John Sillitoe on behalf of the Wellcome Sanger Institute COVID-19 Surveillance Team |
| EPI_ISL_2913356 | Lighthouse Lab in Alderley Park | Wellcome Sanger Institute for the COVID-19 Genomics UK (COG-UK) Consortium | Jacquelyn Wynn, Mairead Hyland, The Lighthouse Lab in Alderley Park and Alex Alderton, Roberto Amato, Jeffrey Barrett, Sonia Goncalves, Ewan Harrison, David K. Jackson, Ian Johnston, Dominic Kwiatkowski, Cordelia Langford, John Sillitoe on behalf of the Wellcome Sanger Institute COVID-19 Surveillance Team |
| EPI_ISL_2913357, EPI_ISL_2913358 | Berkshire and Surrey Pathology Services Lighthouse Laboratory | Wellcome Sanger Institute for the COVID-19 Genomics UK (COG-UK) Consortium | Berkshire and Surrey Pathology Services Lighthouse Laboratory and Alex Alderton, Roberto Amato, Jeffrey Barrett, Sonia Goncalves, Ewan Harrison, David K. Jackson, Ian Johnston, Dominic Kwiatkowski, Cordelia Langford, John Sillitoe on behalf of the Wellcome Sanger Institute COVID-19 Surveillance Team |
| EPI_ISL_2913359 | Lighthouse Lab in Milton Keynes | Wellcome Sanger Institute for the COVID-19 Genomics UK (COG-UK) Consortium | The Lighthouse Lab in Milton Keynes and Alex Alderton, Roberto Amato, Jeffrey Barrett, Sonia Goncalves, Ewan Harrison, David K. Jackson, Ian Johnston, Dominic Kwiatkowski, Cordelia Langford, John Sillitoe on behalf of the Wellcome Sanger Institute COVID-19 Surveillance Team |
| EPI_ISL_2913360 | Lighthouse Lab in Alderley Park | Wellcome Sanger Institute for the COVID-19 Genomics UK (COG-UK) Consortium | Jacquelyn Wynn, Mairead Hyland, The Lighthouse Lab in Alderley Park and Alex Alderton, Roberto Amato, Jeffrey Barrett, Sonia Goncalves, Ewan Harrison, David K. Jackson, Ian Johnston, Dominic Kwiatkowski, Cordelia Langford, John Sillitoe on behalf of the Wellcome Sanger Institute COVID-19 Surveillance Team |
| EPI_ISL_2913361 | Berkshire and Surrey Pathology Services Lighthouse Laboratory | Wellcome Sanger Institute for the COVID-19 Genomics UK (COG-UK) Consortium | Berkshire and Surrey Pathology Services Lighthouse Laboratory and Alex Alderton, Roberto Amato, Jeffrey Barrett, Sonia Goncalves, Ewan Harrison, David K. Jackson, Ian Johnston, Dominic Kwiatkowski, Cordelia Langford, John Sillitoe on behalf of the Wellcome Sanger Institute COVID-19 Surveillance Team |
| EPI_ISL_2913362, EPI_ISL_2913363 | Lighthouse Lab in Alderley Park | Wellcome Sanger Institute for the COVID-19 Genomics UK (COG-UK) Consortium | Jacquelyn Wynn, Mairead Hyland, The Lighthouse Lab in Alderley Park and Alex Alderton, Roberto Amato, Jeffrey Barrett, Sonia Goncalves, Ewan Harrison, David K. Jackson, Ian Johnston, Dominic Kwiatkowski, Cordelia Langford, John Sillitoe on behalf of the Wellcome Sanger Institute COVID-19 Surveillance Team |
| EPI_ISL_2913364, EPI_ISL_2913365 | Berkshire and Surrey Pathology Services Lighthouse Laboratory | Wellcome Sanger Institute for the COVID-19 Genomics UK (COG-UK) Consortium | Berkshire and Surrey Pathology Services Lighthouse Laboratory and Alex Alderton, Roberto Amato, Jeffrey Barrett, Sonia Goncalves, Ewan Harrison, David K. Jackson, Ian Johnston, Dominic Kwiatkowski, Cordelia Langford, John Sillitoe on behalf of the Wellcome Sanger Institute COVID-19 Surveillance Team |
| EPI_ISL_2913366 | Lighthouse Lab in Alderley Park | Wellcome Sanger Institute for the COVID-19 Genomics UK (COG-UK) Consortium | Jacquelyn Wynn, Mairead Hyland, The Lighthouse Lab in Alderley Park and Alex Alderton, Roberto Amato, Jeffrey Barrett, Sonia Goncalves, Ewan Harrison, David K. Jackson, Ian Johnston, Dominic Kwiatkowski, Cordelia Langford, John Sillitoe on behalf of the Wellcome Sanger Institute COVID-19 Surveillance Team |
| EPI_ISL_2913367, EPI_ISL_2913368 | Berkshire and Surrey Pathology Services Lighthouse Laboratory | Wellcome Sanger Institute for the COVID-19 Genomics UK (COG-UK) Consortium | Berkshire and Surrey Pathology Services Lighthouse Laboratory and Alex Alderton, Roberto Amato, Jeffrey Barrett, Sonia Goncalves, Ewan Harrison, David K. Jackson, Ian Johnston, Dominic Kwiatkowski, Cordelia Langford, John Sillitoe on behalf of the Wellcome Sanger Institute COVID-19 Surveillance Team |
| EPI_ISL_2913369 | Lighthouse Lab in Milton Keynes | Wellcome Sanger Institute for the COVID-19 Genomics UK (COG-UK) Consortium | The Lighthouse Lab in Milton Keynes and Alex Alderton, Roberto Amato, Jeffrey Barrett, Sonia Goncalves, Ewan Harrison, David K. Jackson, Ian Johnston, Dominic Kwiatkowski, Cordelia Langford, John Sillitoe on behalf of the Wellcome Sanger Institute COVID-19 Surveillance Team |
| EPI_ISL_2913370, EPI_ISL_2913371 | Lighthouse Lab in Alderley Park | Wellcome Sanger Institute for the COVID-19 Genomics UK (COG-UK) Consortium | Jacquelyn Wynn, Mairead Hyland, The Lighthouse Lab in Alderley Park and Alex Alderton, Roberto Amato, Jeffrey Barrett, Sonia Goncalves, Ewan Harrison, David K. Jackson, Ian Johnston, Dominic Kwiatkowski, Cordelia Langford, John Sillitoe on behalf of the Wellcome Sanger Institute COVID-19 Surveillance Team |
| EPI_ISL_2913372 | Berkshire and Surrey Pathology Services Lighthouse Laboratory | Wellcome Sanger Institute for the COVID-19 Genomics UK (COG-UK) Consortium | Berkshire and Surrey Pathology Services Lighthouse Laboratory and Alex Alderton, Roberto Amato, Jeffrey Barrett, Sonia Goncalves, Ewan Harrison, David K. Jackson, Ian Johnston, Dominic Kwiatkowski, Cordelia Langford, John Sillitoe on behalf of the Wellcome Sanger Institute COVID-19 Surveillance Team |
| EPI_ISL_2913373 | Lighthouse Lab in Alderley Park | Wellcome Sanger Institute for the COVID-19 Genomics UK (COG-UK) Consortium | Jacquelyn Wynn, Mairead Hyland, The Lighthouse Lab in Alderley Park and Alex Alderton, Roberto Amato, Jeffrey Barrett, Sonia Goncalves, Ewan Harrison, David K. Jackson, Ian Johnston, Dominic Kwiatkowski, Cordelia Langford, John Sillitoe on behalf of the Wellcome Sanger Institute COVID-19 Surveillance Team |
| EPI_ISL_2913374, EPI_ISL_2913375, EPI_ISL_2913376 | Berkshire and Surrey Pathology Services Lighthouse Laboratory | Wellcome Sanger Institute for the COVID-19 Genomics UK (COG-UK) Consortium | Berkshire and Surrey Pathology Services Lighthouse Laboratory and Alex Alderton, Roberto Amato, Jeffrey Barrett, Sonia Goncalves, Ewan Harrison, David K. Jackson, Ian Johnston, Dominic Kwiatkowski, Cordelia Langford, John Sillitoe on behalf of the Wellcome Sanger Institute COVID-19 Surveillance Team |
| EPI_ISL_2913377 | Lighthouse Lab in Alderley Park | Wellcome Sanger Institute for the COVID-19 Genomics UK (COG-UK) Consortium | Jacquelyn Wynn, Mairead Hyland, The Lighthouse Lab in Alderley Park and Alex Alderton, Roberto Amato, Jeffrey Barrett, Sonia Goncalves, Ewan Harrison, David K. Jackson, Ian Johnston, Dominic Kwiatkowski, Cordelia Langford, John Sillitoe on behalf of the Wellcome Sanger Institute COVID-19 Surveillance Team |
| EPI_ISL_2913378 | Lighthouse Lab in Milton Keynes | Wellcome Sanger Institute for the COVID-19 Genomics UK (COG-UK) Consortium | The Lighthouse Lab in Milton Keynes and Alex Alderton, Roberto Amato, Jeffrey Barrett, Sonia Goncalves, Ewan Harrison, David K. Jackson, Ian Johnston, Dominic Kwiatkowski, Cordelia Langford, John Sillitoe on behalf of the Wellcome Sanger Institute COVID-19 Surveillance Team |
| EPI_ISL_2913379 | Lighthouse Lab in Alderley Park | Wellcome Sanger Institute for the COVID-19 Genomics UK (COG-UK) Consortium | Jacquelyn Wynn, Mairead Hyland, The Lighthouse Lab in Alderley Park and Alex Alderton, Roberto Amato, Jeffrey Barrett, Sonia Goncalves, Ewan Harrison, David K. Jackson, Ian Johnston, Dominic Kwiatkowski, Cordelia Langford, John Sillitoe on behalf of the Wellcome Sanger Institute COVID-19 Surveillance Team |
| EPI_ISL_2913380 | Lighthouse Lab in Milton Keynes | Wellcome Sanger Institute for the COVID-19 Genomics UK (COG-UK) Consortium | The Lighthouse Lab in Milton Keynes and Alex Alderton, Roberto Amato, Jeffrey Barrett, Sonia Goncalves, Ewan Harrison, David K. Jackson, Ian Johnston, Dominic Kwiatkowski, Cordelia Langford, John Sillitoe on behalf of the Wellcome Sanger Institute COVID-19 Surveillance Team |
| EPI_ISL_2913381 | Berkshire and Surrey Pathology Services Lighthouse Laboratory | Wellcome Sanger Institute for the COVID-19 Genomics UK (COG-UK) Consortium | Berkshire and Surrey Pathology Services Lighthouse Laboratory and Alex Alderton, Roberto Amato, Jeffrey Barrett, Sonia Goncalves, Ewan Harrison, David K. Jackson, Ian Johnston, Dominic Kwiatkowski, Cordelia Langford, John Sillitoe on behalf of the Wellcome Sanger Institute COVID-19 Surveillance Team |
| EPI_ISL_2913382, EPI_ISL_2913383, EPI_ISL_2913384 | Lighthouse Lab in Alderley Park | Wellcome Sanger Institute for the COVID-19 Genomics UK (COG-UK) Consortium | Jacquelyn Wynn, Mairead Hyland, The Lighthouse Lab in Alderley Park and Alex Alderton, Roberto Amato, Jeffrey Barrett, Sonia Goncalves, Ewan Harrison, David K. Jackson, Ian Johnston, Dominic Kwiatkowski, Cordelia Langford, John Sillitoe on behalf of the Wellcome Sanger Institute COVID-19 Surveillance Team |
| EPI_ISL_2913385, EPI_ISL_2913386, EPI_ISL_2913387 | Lighthouse Lab in Milton Keynes | Wellcome Sanger Institute for the COVID-19 Genomics UK (COG-UK) Consortium | The Lighthouse Lab in Milton Keynes and Alex Alderton, Roberto Amato, Jeffrey Barrett, Sonia Goncalves, Ewan Harrison, David K. Jackson, Ian Johnston, Dominic Kwiatkowski, Cordelia Langford, John Sillitoe on behalf of the Wellcome Sanger Institute COVID-19 Surveillance Team |
| EPI_ISL_2913388 | Lighthouse Lab in Alderley Park | Wellcome Sanger Institute for the COVID-19 Genomics UK (COG-UK) Consortium | Jacquelyn Wynn, Mairead Hyland, The Lighthouse Lab in Alderley Park and Alex Alderton, Roberto Amato, Jeffrey Barrett, Sonia Goncalves, Ewan Harrison, David K. Jackson, Ian Johnston, Dominic Kwiatkowski, Cordelia Langford, John Sillitoe on behalf of the Wellcome Sanger Institute COVID-19 Surveillance Team |
| EPI_ISL_2913389 | Lighthouse Lab in Milton Keynes | Wellcome Sanger Institute for the COVID-19 Genomics UK (COG-UK) Consortium | The Lighthouse Lab in Milton Keynes and Alex Alderton, Roberto Amato, Jeffrey Barrett, Sonia Goncalves, Ewan Harrison, David K. Jackson, Ian Johnston, Dominic Kwiatkowski, Cordelia Langford, John Sillitoe on behalf of the Wellcome Sanger Institute COVID-19 Surveillance Team |
| EPI_ISL_2913390 | Lighthouse Lab in Alderley Park | Wellcome Sanger Institute for the COVID-19 Genomics UK (COG-UK) | Jacquelyn Wynn, Mairead Hyland, The Lighthouse Lab in Alderley Park and Alex Alderton, Roberto Amato, Jeffrey Barrett, Sonia Goncalves, Ewan Harrison, David K. Jackson, Ian Johnston, Dominic Kwiatkowski, Cordelia Langford, John Sillitoe on |

| | | | |
|---|---|---|---|
| | | Consortium | behalf of the Wellcome Sanger Institute COVID-19 Surveillance Team |
| EPI_ISL_2913391 | Berkshire and Surrey Pathology Services Lighthouse Laboratory | Wellcome Sanger Institute for the COVID-19 Genomics UK (COG-UK) Consortium | Berkshire and Surrey Pathology Services Lighthouse Laboratory and Alex Alderton, Roberto Amato, Jeffrey Barrett, Sonia Goncalves, Ewan Harrison, David K. Jackson, Ian Johnston, Dominic Kwiatkowski, Cordelia Langford, John Sillitoe on behalf of the Wellcome Sanger Institute COVID-19 Surveillance Team |
| EPI_ISL_2913392 | Lighthouse Lab in Milton Keynes | Wellcome Sanger Institute for the COVID-19 Genomics UK (COG-UK) Consortium | The Lighthouse Lab in Milton Keynes and Alex Alderton, Roberto Amato, Jeffrey Barrett, Sonia Goncalves, Ewan Harrison, David K. Jackson, Ian Johnston, Dominic Kwiatkowski, Cordelia Langford, John Sillitoe on behalf of the Wellcome Sanger Institute COVID-19 Surveillance Team |
| EPI_ISL_2913393, EPI_ISL_2913394 | Berkshire and Surrey Pathology Services Lighthouse Laboratory | Wellcome Sanger Institute for the COVID-19 Genomics UK (COG-UK) Consortium | Berkshire and Surrey Pathology Services Lighthouse Laboratory and Alex Alderton, Roberto Amato, Jeffrey Barrett, Sonia Goncalves, Ewan Harrison, David K. Jackson, Ian Johnston, Dominic Kwiatkowski, Cordelia Langford, John Sillitoe on behalf of the Wellcome Sanger Institute COVID-19 Surveillance Team |
| EPI_ISL_2913395, EPI_ISL_2913396 | Lighthouse Lab in Milton Keynes | Wellcome Sanger Institute for the COVID-19 Genomics UK (COG-UK) Consortium | The Lighthouse Lab in Milton Keynes and Alex Alderton, Roberto Amato, Jeffrey Barrett, Sonia Goncalves, Ewan Harrison, David K. Jackson, Ian Johnston, Dominic Kwiatkowski, Cordelia Langford, John Sillitoe on behalf of the Wellcome Sanger Institute COVID-19 Surveillance Team |
| EPI_ISL_2913397 | Berkshire and Surrey Pathology Services Lighthouse Laboratory | Wellcome Sanger Institute for the COVID-19 Genomics UK (COG-UK) Consortium | Berkshire and Surrey Pathology Services Lighthouse Laboratory and Alex Alderton, Roberto Amato, Jeffrey Barrett, Sonia Goncalves, Ewan Harrison, David K. Jackson, Ian Johnston, Dominic Kwiatkowski, Cordelia Langford, John Sillitoe on behalf of the Wellcome Sanger Institute COVID-19 Surveillance Team |
| EPI_ISL_2913398 | Lighthouse Lab in Alderley Park | Wellcome Sanger Institute for the COVID-19 Genomics UK (COG-UK) Consortium | Jacquelyn Wynn, Mairead Hyland, The Lighthouse Lab in Alderley Park and Alex Alderton, Roberto Amato, Jeffrey Barrett, Sonia Goncalves, Ewan Harrison, David K. Jackson, Ian Johnston, Dominic Kwiatkowski, Cordelia Langford, John Sillitoe on behalf of the Wellcome Sanger Institute COVID-19 Surveillance Team |
| EPI_ISL_2913399, EPI_ISL_2913400 | Lighthouse Lab in Milton Keynes | Wellcome Sanger Institute for the COVID-19 Genomics UK (COG-UK) Consortium | The Lighthouse Lab in Milton Keynes and Alex Alderton, Roberto Amato, Jeffrey Barrett, Sonia Goncalves, Ewan Harrison, David K. Jackson, Ian Johnston, Dominic Kwiatkowski, Cordelia Langford, John Sillitoe on behalf of the Wellcome Sanger Institute COVID-19 Surveillance Team |
| EPI_ISL_2913401 | Berkshire and Surrey Pathology Services Lighthouse Laboratory | Wellcome Sanger Institute for the COVID-19 Genomics UK (COG-UK) Consortium | Berkshire and Surrey Pathology Services Lighthouse Laboratory and Alex Alderton, Roberto Amato, Jeffrey Barrett, Sonia Goncalves, Ewan Harrison, David K. Jackson, Ian Johnston, Dominic Kwiatkowski, Cordelia Langford, John Sillitoe on behalf of the Wellcome Sanger Institute COVID-19 Surveillance Team |
| EPI_ISL_2913402 | Lighthouse Lab in Milton Keynes | Wellcome Sanger Institute for the COVID-19 Genomics UK (COG-UK) Consortium | The Lighthouse Lab in Milton Keynes and Alex Alderton, Roberto Amato, Jeffrey Barrett, Sonia Goncalves, Ewan Harrison, David K. Jackson, Ian Johnston, Dominic Kwiatkowski, Cordelia Langford, John Sillitoe on behalf of the Wellcome Sanger Institute COVID-19 Surveillance Team |
| EPI_ISL_2913403 | Berkshire and Surrey Pathology Services Lighthouse Laboratory | Wellcome Sanger Institute for the COVID-19 Genomics UK (COG-UK) Consortium | Berkshire and Surrey Pathology Services Lighthouse Laboratory and Alex Alderton, Roberto Amato, Jeffrey Barrett, Sonia Goncalves, Ewan Harrison, David K. Jackson, Ian Johnston, Dominic Kwiatkowski, Cordelia Langford, John Sillitoe on behalf of the Wellcome Sanger Institute COVID-19 Surveillance Team |
| EPI_ISL_2913404, EPI_ISL_2913405, EPI_ISL_2913406, EPI_ISL_2913407 | Lighthouse Lab in Milton Keynes | Wellcome Sanger Institute for the COVID-19 Genomics UK (COG-UK) Consortium | The Lighthouse Lab in Milton Keynes and Alex Alderton, Roberto Amato, Jeffrey Barrett, Sonia Goncalves, Ewan Harrison, David K. Jackson, Ian Johnston, Dominic Kwiatkowski, Cordelia Langford, John Sillitoe on behalf of the Wellcome Sanger Institute COVID-19 Surveillance Team |
| EPI_ISL_2913408 | Lighthouse Lab in Alderley Park | Wellcome Sanger Institute for the COVID-19 Genomics UK (COG-UK) Consortium | Jacquelyn Wynn, Mairead Hyland, The Lighthouse Lab in Alderley Park and Alex Alderton, Roberto Amato, Jeffrey Barrett, Sonia Goncalves, Ewan Harrison, David K. Jackson, Ian Johnston, Dominic Kwiatkowski, Cordelia Langford, John Sillitoe on behalf of the Wellcome Sanger Institute COVID-19 Surveillance Team |
| EPI_ISL_2913409 | Lighthouse Lab in Milton Keynes | Wellcome Sanger Institute for the COVID-19 Genomics UK (COG-UK) Consortium | The Lighthouse Lab in Milton Keynes and Alex Alderton, Roberto Amato, Jeffrey Barrett, Sonia Goncalves, Ewan Harrison, David K. Jackson, Ian Johnston, Dominic Kwiatkowski, Cordelia Langford, John Sillitoe on behalf of the Wellcome Sanger Institute COVID-19 Surveillance Team |
| EPI_ISL_2913410 | Berkshire and Surrey Pathology Services Lighthouse Laboratory | Wellcome Sanger Institute for the COVID-19 Genomics UK (COG-UK) Consortium | Berkshire and Surrey Pathology Services Lighthouse Laboratory and Alex Alderton, Roberto Amato, Jeffrey Barrett, Sonia Goncalves, Ewan Harrison, David K. Jackson, Ian Johnston, Dominic Kwiatkowski, Cordelia Langford, John Sillitoe on behalf of the Wellcome Sanger Institute COVID-19 Surveillance Team |
| EPI_ISL_2913411, EPI_ISL_2913412 | Lighthouse Lab in Milton Keynes | Wellcome Sanger Institute for the COVID-19 Genomics UK (COG-UK) Consortium | The Lighthouse Lab in Milton Keynes and Alex Alderton, Roberto Amato, Jeffrey Barrett, Sonia Goncalves, Ewan Harrison, David K. Jackson, Ian Johnston, Dominic Kwiatkowski, Cordelia Langford, John Sillitoe on behalf of the Wellcome Sanger Institute COVID-19 Surveillance Team |
| EPI_ISL_2913413, EPI_ISL_2913414 | Lighthouse Lab in Alderley Park | Wellcome Sanger Institute for the COVID-19 Genomics UK (COG-UK) Consortium | Jacquelyn Wynn, Mairead Hyland, The Lighthouse Lab in Alderley Park and Alex Alderton, Roberto Amato, Jeffrey Barrett, Sonia Goncalves, Ewan Harrison, David K. Jackson, Ian Johnston, Dominic Kwiatkowski, Cordelia Langford, John Sillitoe on behalf of the Wellcome Sanger Institute COVID-19 Surveillance Team |
| EPI_ISL_2913415 | Berkshire and Surrey Pathology Services Lighthouse Laboratory | Wellcome Sanger Institute for the COVID-19 Genomics UK (COG-UK) Consortium | Berkshire and Surrey Pathology Services Lighthouse Laboratory and Alex Alderton, Roberto Amato, Jeffrey Barrett, Sonia Goncalves, Ewan Harrison, David K. Jackson, Ian Johnston, Dominic Kwiatkowski, Cordelia Langford, John Sillitoe on behalf of the Wellcome Sanger Institute COVID-19 Surveillance Team |
| EPI_ISL_2913416 | Lighthouse Lab in Alderley Park | Wellcome Sanger Institute for the COVID-19 Genomics UK (COG-UK) Consortium | Jacquelyn Wynn, Mairead Hyland, The Lighthouse Lab in Alderley Park and Alex Alderton, Roberto Amato, Jeffrey Barrett, Sonia Goncalves, Ewan Harrison, David K. Jackson, Ian Johnston, Dominic Kwiatkowski, Cordelia Langford, John Sillitoe on behalf of the Wellcome Sanger Institute COVID-19 Surveillance Team |
| EPI_ISL_2913417 | Lighthouse Lab in Milton Keynes | Wellcome Sanger Institute for the COVID-19 Genomics UK (COG-UK) Consortium | The Lighthouse Lab in Milton Keynes and Alex Alderton, Roberto Amato, Jeffrey Barrett, Sonia Goncalves, Ewan Harrison, David K. Jackson, Ian Johnston, Dominic Kwiatkowski, Cordelia Langford, John Sillitoe on behalf of the Wellcome Sanger Institute COVID-19 Surveillance Team |
| EPI_ISL_2913418 | Lighthouse Lab in Alderley Park | Wellcome Sanger Institute for the COVID-19 Genomics UK (COG-UK) Consortium | Jacquelyn Wynn, Mairead Hyland, The Lighthouse Lab in Alderley Park and Alex Alderton, Roberto Amato, Jeffrey Barrett, Sonia Goncalves, Ewan Harrison, David K. Jackson, Ian Johnston, Dominic Kwiatkowski, Cordelia Langford, John Sillitoe on behalf of the Wellcome Sanger Institute COVID-19 Surveillance Team |
| EPI_ISL_2913419, EPI_ISL_2913420 | Lighthouse Lab in Milton Keynes | Wellcome Sanger Institute for the COVID-19 Genomics UK (COG-UK) Consortium | The Lighthouse Lab in Milton Keynes and Alex Alderton, Roberto Amato, Jeffrey Barrett, Sonia Goncalves, Ewan Harrison, David K. Jackson, Ian Johnston, Dominic Kwiatkowski, Cordelia Langford, John Sillitoe on behalf of the Wellcome Sanger Institute COVID-19 Surveillance Team |
| EPI_ISL_2913421, EPI_ISL_2913422 | Berkshire and Surrey Pathology Services Lighthouse Laboratory | Wellcome Sanger Institute for the COVID-19 Genomics UK (COG-UK) Consortium | Berkshire and Surrey Pathology Services Lighthouse Laboratory and Alex Alderton, Roberto Amato, Jeffrey Barrett, Sonia Goncalves, Ewan Harrison, David K. Jackson, Ian Johnston, Dominic Kwiatkowski, Cordelia Langford, John Sillitoe on behalf of the Wellcome Sanger Institute COVID-19 Surveillance Team |
| EPI_ISL_2913423 | Lighthouse Lab in Alderley Park | Wellcome Sanger Institute for the COVID-19 Genomics UK (COG-UK) Consortium | Jacquelyn Wynn, Mairead Hyland, The Lighthouse Lab in Alderley Park and Alex Alderton, Roberto Amato, Jeffrey Barrett, Sonia Goncalves, Ewan Harrison, David K. Jackson, Ian Johnston, Dominic Kwiatkowski, Cordelia Langford, John Sillitoe on behalf of the Wellcome Sanger Institute COVID-19 Surveillance Team |
| EPI_ISL_2913424 | Lighthouse Lab in Milton Keynes | Wellcome Sanger Institute for the COVID-19 Genomics UK (COG-UK) Consortium | The Lighthouse Lab in Milton Keynes and Alex Alderton, Roberto Amato, Jeffrey Barrett, Sonia Goncalves, Ewan Harrison, David K. Jackson, Ian Johnston, Dominic Kwiatkowski, Cordelia Langford, John Sillitoe on behalf of the Wellcome Sanger Institute COVID-19 Surveillance Team |
| EPI_ISL_2913425 | Berkshire and Surrey Pathology Services Lighthouse Laboratory | Wellcome Sanger Institute for the COVID-19 Genomics UK (COG-UK) Consortium | Berkshire and Surrey Pathology Services Lighthouse Laboratory and Alex Alderton, Roberto Amato, Jeffrey Barrett, Sonia Goncalves, Ewan Harrison, David K. Jackson, Ian Johnston, Dominic Kwiatkowski, Cordelia Langford, John Sillitoe on behalf of the Wellcome Sanger Institute COVID-19 Surveillance Team |
| EPI_ISL_2913426 | Lighthouse Lab in Milton Keynes | Wellcome Sanger Institute for the COVID-19 Genomics UK (COG-UK) Consortium | The Lighthouse Lab in Milton Keynes and Alex Alderton, Roberto Amato, Jeffrey Barrett, Sonia Goncalves, Ewan Harrison, David K. Jackson, Ian Johnston, Dominic Kwiatkowski, Cordelia Langford, John Sillitoe on behalf of the Wellcome Sanger Institute COVID-19 Surveillance Team |
| EPI_ISL_2913427 | Lighthouse Lab in Alderley Park | Wellcome Sanger Institute for the COVID-19 Genomics UK (COG-UK) Consortium | Jacquelyn Wynn, Mairead Hyland, The Lighthouse Lab in Alderley Park and Alex Alderton, Roberto Amato, Jeffrey Barrett, Sonia Goncalves, Ewan Harrison, David K. Jackson, Ian Johnston, Dominic Kwiatkowski, Cordelia Langford, John Sillitoe on behalf of the Wellcome Sanger Institute COVID-19 Surveillance Team |
| EPI_ISL_2913428, EPI_ISL_2913429, EPI_ISL_2913430 | Berkshire and Surrey Pathology Services Lighthouse Laboratory | Wellcome Sanger Institute for the COVID-19 Genomics UK (COG-UK) Consortium | Berkshire and Surrey Pathology Services Lighthouse Laboratory and Alex Alderton, Roberto Amato, Jeffrey Barrett, Sonia Goncalves, Ewan Harrison, David K. Jackson, Ian Johnston, Dominic Kwiatkowski, Cordelia Langford, John Sillitoe on behalf of the Wellcome Sanger Institute COVID-19 Surveillance Team |
| EPI_ISL_2913431, EPI_ISL_2913432 | Lighthouse Lab in Milton Keynes | Wellcome Sanger Institute for the COVID-19 Genomics UK (COG-UK) Consortium | The Lighthouse Lab in Milton Keynes and Alex Alderton, Roberto Amato, Jeffrey Barrett, Sonia Goncalves, Ewan Harrison, David K. Jackson, Ian Johnston, Dominic Kwiatkowski, Cordelia Langford, John Sillitoe on behalf of the Wellcome Sanger Institute COVID-19 Surveillance Team |
| EPI_ISL_2913433 | Berkshire and Surrey Pathology Services Lighthouse Laboratory | Wellcome Sanger Institute for the COVID-19 Genomics UK (COG-UK) Consortium | Berkshire and Surrey Pathology Services Lighthouse Laboratory and Alex Alderton, Roberto Amato, Jeffrey Barrett, Sonia Goncalves, Ewan Harrison, David K. Jackson, Ian Johnston, Dominic Kwiatkowski, Cordelia Langford, John Sillitoe on behalf of the Wellcome Sanger Institute COVID-19 Surveillance Team |
| EPI_ISL_2913435 | Lighthouse Lab in Milton Keynes | Wellcome Sanger Institute for the COVID-19 Genomics UK (COG-UK) Consortium | The Lighthouse Lab in Milton Keynes and Alex Alderton, Roberto Amato, Jeffrey Barrett, Sonia Goncalves, Ewan Harrison, David K. Jackson, Ian Johnston, Dominic Kwiatkowski, Cordelia Langford, John Sillitoe on behalf of the Wellcome Sanger Institute COVID-19 Surveillance Team |
| EPI_ISL_2913436 | Lighthouse Lab in Alderley Park | Wellcome Sanger Institute for the COVID-19 Genomics UK (COG-UK) Consortium | Jacquelyn Wynn, Mairead Hyland, The Lighthouse Lab in Alderley Park and Alex Alderton, Roberto Amato, Jeffrey Barrett, Sonia Goncalves, Ewan Harrison, David K. Jackson, Ian Johnston, Dominic Kwiatkowski, Cordelia Langford, John Sillitoe on behalf of the Wellcome Sanger Institute COVID-19 Surveillance Team |
| EPI_ISL_2913437, EPI_ISL_2913438, EPI_ISL_2913439 | Berkshire and Surrey Pathology Services Lighthouse Laboratory | Wellcome Sanger Institute for the COVID-19 Genomics UK (COG-UK) Consortium | Berkshire and Surrey Pathology Services Lighthouse Laboratory and Alex Alderton, Roberto Amato, Jeffrey Barrett, Sonia Goncalves, Ewan Harrison, David K. Jackson, Ian Johnston, Dominic Kwiatkowski, Cordelia Langford, John Sillitoe on behalf of the Wellcome Sanger Institute COVID-19 Surveillance Team |
| EPI_ISL_2913440 | Lighthouse Lab in Milton Keynes | Wellcome Sanger Institute for the COVID-19 Genomics UK (COG-UK) Consortium | The Lighthouse Lab in Milton Keynes and Alex Alderton, Roberto Amato, Jeffrey Barrett, Sonia Goncalves, Ewan Harrison, David K. Jackson, Ian Johnston, Dominic Kwiatkowski, Cordelia Langford, John Sillitoe on behalf of the Wellcome Sanger Institute COVID-19 Surveillance Team |
| EPI_ISL_2913441 | Berkshire and Surrey Pathology Services Lighthouse Laboratory | Wellcome Sanger Institute for the COVID-19 Genomics UK (COG-UK) Consortium | Berkshire and Surrey Pathology Services Lighthouse Laboratory and Alex Alderton, Roberto Amato, Jeffrey Barrett, Sonia Goncalves, Ewan Harrison, David K. Jackson, Ian Johnston, Dominic Kwiatkowski, Cordelia Langford, John Sillitoe on behalf of the Wellcome Sanger Institute COVID-19 Surveillance Team |
| EPI_ISL_2913442 | Lighthouse Lab in Alderley Park | Wellcome Sanger Institute for the COVID-19 Genomics UK (COG-UK) Consortium | Jacquelyn Wynn, Mairead Hyland, The Lighthouse Lab in Alderley Park and Alex Alderton, Roberto Amato, Jeffrey Barrett, Sonia Goncalves, Ewan Harrison, David K. Jackson, Ian Johnston, Dominic Kwiatkowski, Cordelia Langford, John Sillitoe on behalf of the Wellcome Sanger Institute COVID-19 Surveillance Team |
| EPI_ISL_2913443 | Lighthouse Lab in Milton Keynes | Wellcome Sanger Institute for the COVID-19 Genomics UK (COG-UK) Consortium | The Lighthouse Lab in Milton Keynes and Alex Alderton, Roberto Amato, Jeffrey Barrett, Sonia Goncalves, Ewan Harrison, David K. Jackson, Ian Johnston, Dominic Kwiatkowski, Cordelia Langford, John Sillitoe on behalf of the Wellcome Sanger Institute COVID-19 Surveillance Team |
| EPI_ISL_2913444, EPI_ISL_2913445 | Berkshire and Surrey Pathology Services Lighthouse Laboratory | Wellcome Sanger Institute for the COVID-19 Genomics UK (COG-UK) Consortium | Berkshire and Surrey Pathology Services Lighthouse Laboratory and Alex Alderton, Roberto Amato, Jeffrey Barrett, Sonia Goncalves, Ewan Harrison, David K. Jackson, Ian Johnston, Dominic Kwiatkowski, Cordelia Langford, John Sillitoe on behalf of the Wellcome Sanger Institute COVID-19 Surveillance Team |
| EPI_ISL_2913446 | Lighthouse Lab in Alderley Park | Wellcome Sanger Institute for the COVID-19 Genomics UK (COG-UK) Consortium | Jacquelyn Wynn, Mairead Hyland, The Lighthouse Lab in Alderley Park and Alex Alderton, Roberto Amato, Jeffrey Barrett, Sonia Goncalves, Ewan Harrison, David K. Jackson, Ian Johnston, Dominic Kwiatkowski, Cordelia Langford, John Sillitoe on behalf of the Wellcome Sanger Institute COVID-19 Surveillance Team |
| EPI_ISL_2913447, EPI_ISL_2913448, EPI_ISL_2913449, EPI_ISL_2913450 | Lighthouse Lab in Milton Keynes | Wellcome Sanger Institute for the COVID-19 Genomics UK (COG-UK) Consortium | The Lighthouse Lab in Milton Keynes and Alex Alderton, Roberto Amato, Jeffrey Barrett, Sonia Goncalves, Ewan Harrison, David K. Jackson, Ian Johnston, Dominic Kwiatkowski, Cordelia Langford, John Sillitoe on behalf of the Wellcome Sanger Institute COVID-19 Surveillance Team |
| EPI_ISL_2913451 | Berkshire and Surrey Pathology Services Lighthouse Laboratory | Wellcome Sanger Institute for the COVID-19 Genomics UK (COG-UK) Consortium | Berkshire and Surrey Pathology Services Lighthouse Laboratory and Alex Alderton, Roberto Amato, Jeffrey Barrett, Sonia Goncalves, Ewan Harrison, David K. Jackson, Ian Johnston, Dominic Kwiatkowski, Cordelia Langford, John Sillitoe on behalf of the Wellcome Sanger Institute COVID-19 Surveillance Team |
| EPI_ISL_2913452 | Lighthouse Lab in Milton Keynes | Wellcome Sanger Institute for the COVID-19 Genomics UK (COG-UK) Consortium | The Lighthouse Lab in Milton Keynes and Alex Alderton, Roberto Amato, Jeffrey Barrett, Sonia Goncalves, Ewan Harrison, David K. Jackson, Ian Johnston, Dominic Kwiatkowski, Cordelia Langford, John Sillitoe on behalf of the Wellcome Sanger Institute COVID-19 Surveillance Team |

| | | | |
|---|---|---|---|
| EPI_ISL_2913453 | Lighthouse Lab in Alderley Park | Wellcome Sanger Institute for the COVID-19 Genomics UK (COG-UK) Consortium | Jacquelyn Wynn, Mairead Hyland, The Lighthouse Lab in Alderley Park and Alex Alderton, Roberto Amato, Jeffrey Barrett, Sonia Goncalves, Ewan Harrison, David K. Jackson, Ian Johnston, Dominic Kwiatkowski, Cordelia Langford, John Sillitoe on behalf of the Wellcome Sanger Institute COVID-19 Surveillance Team |
| EPI_ISL_2913454, EPI_ISL_2913455 | Berkshire and Surrey Pathology Services Lighthouse Laboratory | Wellcome Sanger Institute for the COVID-19 Genomics UK (COG-UK) Consortium | Berkshire and Surrey Pathology Services Lighthouse Laboratory and Alex Alderton, Roberto Amato, Jeffrey Barrett, Sonia Goncalves, Ewan Harrison, David K. Jackson, Ian Johnston, Dominic Kwiatkowski, Cordelia Langford, John Sillitoe on behalf of the Wellcome Sanger Institute COVID-19 Surveillance Team |
| EPI_ISL_2913456 | Lighthouse Lab in Alderley Park | Wellcome Sanger Institute for the COVID-19 Genomics UK (COG-UK) Consortium | Jacquelyn Wynn, Mairead Hyland, The Lighthouse Lab in Alderley Park and Alex Alderton, Roberto Amato, Jeffrey Barrett, Sonia Goncalves, Ewan Harrison, David K. Jackson, Ian Johnston, Dominic Kwiatkowski, Cordelia Langford, John Sillitoe on behalf of the Wellcome Sanger Institute COVID-19 Surveillance Team |
| EPI_ISL_2913457 | Lighthouse Lab in Milton Keynes | Wellcome Sanger Institute for the COVID-19 Genomics UK (COG-UK) Consortium | The Lighthouse Lab in Milton Keynes and Alex Alderton, Roberto Amato, Jeffrey Barrett, Sonia Goncalves, Ewan Harrison, David K. Jackson, Ian Johnston, Dominic Kwiatkowski, Cordelia Langford, John Sillitoe on behalf of the Wellcome Sanger Institute COVID-19 Surveillance Team |
| EPI_ISL_2913458 | Lighthouse Lab in Alderley Park | Wellcome Sanger Institute for the COVID-19 Genomics UK (COG-UK) Consortium | Jacquelyn Wynn, Mairead Hyland, The Lighthouse Lab in Alderley Park and Alex Alderton, Roberto Amato, Jeffrey Barrett, Sonia Goncalves, Ewan Harrison, David K. Jackson, Ian Johnston, Dominic Kwiatkowski, Cordelia Langford, John Sillitoe on behalf of the Wellcome Sanger Institute COVID-19 Surveillance Team |
| EPI_ISL_2913459 | Berkshire and Surrey Pathology Services Lighthouse Laboratory | Wellcome Sanger Institute for the COVID-19 Genomics UK (COG-UK) Consortium | Berkshire and Surrey Pathology Services Lighthouse Laboratory and Alex Alderton, Roberto Amato, Jeffrey Barrett, Sonia Goncalves, Ewan Harrison, David K. Jackson, Ian Johnston, Dominic Kwiatkowski, Cordelia Langford, John Sillitoe on behalf of the Wellcome Sanger Institute COVID-19 Surveillance Team |
| EPI_ISL_2913460, EPI_ISL_2913461 | Lighthouse Lab in Alderley Park | Wellcome Sanger Institute for the COVID-19 Genomics UK (COG-UK) Consortium | Jacquelyn Wynn, Mairead Hyland, The Lighthouse Lab in Alderley Park and Alex Alderton, Roberto Amato, Jeffrey Barrett, Sonia Goncalves, Ewan Harrison, David K. Jackson, Ian Johnston, Dominic Kwiatkowski, Cordelia Langford, John Sillitoe on behalf of the Wellcome Sanger Institute COVID-19 Surveillance Team |
| EPI_ISL_2913462 | Berkshire and Surrey Pathology Services Lighthouse Laboratory | Wellcome Sanger Institute for the COVID-19 Genomics UK (COG-UK) Consortium | Berkshire and Surrey Pathology Services Lighthouse Laboratory and Alex Alderton, Roberto Amato, Jeffrey Barrett, Sonia Goncalves, Ewan Harrison, David K. Jackson, Ian Johnston, Dominic Kwiatkowski, Cordelia Langford, John Sillitoe on behalf of the Wellcome Sanger Institute COVID-19 Surveillance Team |
| EPI_ISL_2913463 | Lighthouse Lab in Milton Keynes | Wellcome Sanger Institute for the COVID-19 Genomics UK (COG-UK) Consortium | The Lighthouse Lab in Milton Keynes and Alex Alderton, Roberto Amato, Jeffrey Barrett, Sonia Goncalves, Ewan Harrison, David K. Jackson, Ian Johnston, Dominic Kwiatkowski, Cordelia Langford, John Sillitoe on behalf of the Wellcome Sanger Institute COVID-19 Surveillance Team |
| EPI_ISL_2913464 | Lighthouse Lab in Alderley Park | Wellcome Sanger Institute for the COVID-19 Genomics UK (COG-UK) Consortium | Jacquelyn Wynn, Mairead Hyland, The Lighthouse Lab in Alderley Park and Alex Alderton, Roberto Amato, Jeffrey Barrett, Sonia Goncalves, Ewan Harrison, David K. Jackson, Ian Johnston, Dominic Kwiatkowski, Cordelia Langford, John Sillitoe on behalf of the Wellcome Sanger Institute COVID-19 Surveillance Team |
| EPI_ISL_2913465 | Lighthouse Lab in Milton Keynes | Wellcome Sanger Institute for the COVID-19 Genomics UK (COG-UK) Consortium | The Lighthouse Lab in Milton Keynes and Alex Alderton, Roberto Amato, Jeffrey Barrett, Sonia Goncalves, Ewan Harrison, David K. Jackson, Ian Johnston, Dominic Kwiatkowski, Cordelia Langford, John Sillitoe on behalf of the Wellcome Sanger Institute COVID-19 Surveillance Team |
| EPI_ISL_2913466 | Lighthouse Lab in Alderley Park | Wellcome Sanger Institute for the COVID-19 Genomics UK (COG-UK) Consortium | Jacquelyn Wynn, Mairead Hyland, The Lighthouse Lab in Alderley Park and Alex Alderton, Roberto Amato, Jeffrey Barrett, Sonia Goncalves, Ewan Harrison, David K. Jackson, Ian Johnston, Dominic Kwiatkowski, Cordelia Langford, John Sillitoe on behalf of the Wellcome Sanger Institute COVID-19 Surveillance Team |
| EPI_ISL_2913467, EPI_ISL_2913468, EPI_ISL_2913469 | Berkshire and Surrey Pathology Services Lighthouse Laboratory | Wellcome Sanger Institute for the COVID-19 Genomics UK (COG-UK) Consortium | Berkshire and Surrey Pathology Services Lighthouse Laboratory and Alex Alderton, Roberto Amato, Jeffrey Barrett, Sonia Goncalves, Ewan Harrison, David K. Jackson, Ian Johnston, Dominic Kwiatkowski, Cordelia Langford, John Sillitoe on behalf of the Wellcome Sanger Institute COVID-19 Surveillance Team |
| EPI_ISL_2913470, EPI_ISL_2913471, EPI_ISL_2913472 | Lighthouse Lab in Alderley Park | Wellcome Sanger Institute for the COVID-19 Genomics UK (COG-UK) Consortium | Jacquelyn Wynn, Mairead Hyland, The Lighthouse Lab in Alderley Park and Alex Alderton, Roberto Amato, Jeffrey Barrett, Sonia Goncalves, Ewan Harrison, David K. Jackson, Ian Johnston, Dominic Kwiatkowski, Cordelia Langford, John Sillitoe on behalf of the Wellcome Sanger Institute COVID-19 Surveillance Team |
| EPI_ISL_2913473 | Berkshire and Surrey Pathology Services Lighthouse Laboratory | Wellcome Sanger Institute for the COVID-19 Genomics UK (COG-UK) Consortium | Berkshire and Surrey Pathology Services Lighthouse Laboratory and Alex Alderton, Roberto Amato, Jeffrey Barrett, Sonia Goncalves, Ewan Harrison, David K. Jackson, Ian Johnston, Dominic Kwiatkowski, Cordelia Langford, John Sillitoe on behalf of the Wellcome Sanger Institute COVID-19 Surveillance Team |
| EPI_ISL_2913474 | Lighthouse Lab in Milton Keynes | Wellcome Sanger Institute for the COVID-19 Genomics UK (COG-UK) Consortium | The Lighthouse Lab in Milton Keynes and Alex Alderton, Roberto Amato, Jeffrey Barrett, Sonia Goncalves, Ewan Harrison, David K. Jackson, Ian Johnston, Dominic Kwiatkowski, Cordelia Langford, John Sillitoe on behalf of the Wellcome Sanger Institute COVID-19 Surveillance Team |
| EPI_ISL_2913475 | Berkshire and Surrey Pathology Services Lighthouse Laboratory | Wellcome Sanger Institute for the COVID-19 Genomics UK (COG-UK) Consortium | Berkshire and Surrey Pathology Services Lighthouse Laboratory and Alex Alderton, Roberto Amato, Jeffrey Barrett, Sonia Goncalves, Ewan Harrison, David K. Jackson, Ian Johnston, Dominic Kwiatkowski, Cordelia Langford, John Sillitoe on behalf of the Wellcome Sanger Institute COVID-19 Surveillance Team |
| EPI_ISL_2913476, EPI_ISL_2913477, EPI_ISL_2913478, EPI_ISL_2913479 | Lighthouse Lab in Milton Keynes | Wellcome Sanger Institute for the COVID-19 Genomics UK (COG-UK) Consortium | The Lighthouse Lab in Milton Keynes and Alex Alderton, Roberto Amato, Jeffrey Barrett, Sonia Goncalves, Ewan Harrison, David K. Jackson, Ian Johnston, Dominic Kwiatkowski, Cordelia Langford, John Sillitoe on behalf of the Wellcome Sanger Institute COVID-19 Surveillance Team |
| EPI_ISL_2913480 | Lighthouse Lab in Alderley Park | Wellcome Sanger Institute for the COVID-19 Genomics UK (COG-UK) Consortium | Jacquelyn Wynn, Mairead Hyland, The Lighthouse Lab in Alderley Park and Alex Alderton, Roberto Amato, Jeffrey Barrett, Sonia Goncalves, Ewan Harrison, David K. Jackson, Ian Johnston, Dominic Kwiatkowski, Cordelia Langford, John Sillitoe on behalf of the Wellcome Sanger Institute COVID-19 Surveillance Team |
| EPI_ISL_2913481 | Lighthouse Lab in Milton Keynes | Wellcome Sanger Institute for the COVID-19 Genomics UK (COG-UK) Consortium | The Lighthouse Lab in Milton Keynes and Alex Alderton, Roberto Amato, Jeffrey Barrett, Sonia Goncalves, Ewan Harrison, David K. Jackson, Ian Johnston, Dominic Kwiatkowski, Cordelia Langford, John Sillitoe on behalf of the Wellcome Sanger Institute COVID-19 Surveillance Team |
| EPI_ISL_2913482 | Berkshire and Surrey Pathology Services Lighthouse Laboratory | Wellcome Sanger Institute for the COVID-19 Genomics UK (COG-UK) Consortium | Berkshire and Surrey Pathology Services Lighthouse Laboratory and Alex Alderton, Roberto Amato, Jeffrey Barrett, Sonia Goncalves, Ewan Harrison, David K. Jackson, Ian Johnston, Dominic Kwiatkowski, Cordelia Langford, John Sillitoe on behalf of the Wellcome Sanger Institute COVID-19 Surveillance Team |
| EPI_ISL_2913483 | Lighthouse Lab in Alderley Park | Wellcome Sanger Institute for the COVID-19 Genomics UK (COG-UK) Consortium | Jacquelyn Wynn, Mairead Hyland, The Lighthouse Lab in Alderley Park and Alex Alderton, Roberto Amato, Jeffrey Barrett, Sonia Goncalves, Ewan Harrison, David K. Jackson, Ian Johnston, Dominic Kwiatkowski, Cordelia Langford, John Sillitoe on behalf of the Wellcome Sanger Institute COVID-19 Surveillance Team |
| EPI_ISL_2913484, EPI_ISL_2913485 | Lighthouse Lab in Milton Keynes | Wellcome Sanger Institute for the COVID-19 Genomics UK (COG-UK) Consortium | The Lighthouse Lab in Milton Keynes and Alex Alderton, Roberto Amato, Jeffrey Barrett, Sonia Goncalves, Ewan Harrison, David K. Jackson, Ian Johnston, Dominic Kwiatkowski, Cordelia Langford, John Sillitoe on behalf of the Wellcome Sanger Institute COVID-19 Surveillance Team |
| EPI_ISL_2913486, EPI_ISL_2913487, EPI_ISL_2913488, EPI_ISL_2913489, EPI_ISL_2913490, EPI_ISL_2913491, EPI_ISL_2913492 | Lighthouse Lab in Alderley Park | Wellcome Sanger Institute for the COVID-19 Genomics UK (COG-UK) Consortium | Jacquelyn Wynn, Mairead Hyland, The Lighthouse Lab in Alderley Park and Alex Alderton, Roberto Amato, Jeffrey Barrett, Sonia Goncalves, Ewan Harrison, David K. Jackson, Ian Johnston, Dominic Kwiatkowski, Cordelia Langford, John Sillitoe on behalf of the Wellcome Sanger Institute COVID-19 Surveillance Team |
| EPI_ISL_2913493 | Lighthouse Lab in Milton Keynes | Wellcome Sanger Institute for the COVID-19 Genomics UK (COG-UK) Consortium | The Lighthouse Lab in Milton Keynes and Alex Alderton, Roberto Amato, Jeffrey Barrett, Sonia Goncalves, Ewan Harrison, David K. Jackson, Ian Johnston, Dominic Kwiatkowski, Cordelia Langford, John Sillitoe on behalf of the Wellcome Sanger Institute COVID-19 Surveillance Team |
| EPI_ISL_2913494 | Berkshire and Surrey Pathology Services Lighthouse Laboratory | Wellcome Sanger Institute for the COVID-19 Genomics UK (COG-UK) Consortium | Berkshire and Surrey Pathology Services Lighthouse Laboratory and Alex Alderton, Roberto Amato, Jeffrey Barrett, Sonia Goncalves, Ewan Harrison, David K. Jackson, Ian Johnston, Dominic Kwiatkowski, Cordelia Langford, John Sillitoe on behalf of the Wellcome Sanger Institute COVID-19 Surveillance Team |
| EPI_ISL_2913495, EPI_ISL_2913496 | Lighthouse Lab in Milton Keynes | Wellcome Sanger Institute for the COVID-19 Genomics UK (COG-UK) Consortium | The Lighthouse Lab in Milton Keynes and Alex Alderton, Roberto Amato, Jeffrey Barrett, Sonia Goncalves, Ewan Harrison, David K. Jackson, Ian Johnston, Dominic Kwiatkowski, Cordelia Langford, John Sillitoe on behalf of the Wellcome Sanger Institute COVID-19 Surveillance Team |
| EPI_ISL_2913497, EPI_ISL_2913498, EPI_ISL_2913499 | Berkshire and Surrey Pathology Services Lighthouse Laboratory | Wellcome Sanger Institute for the COVID-19 Genomics UK (COG-UK) Consortium | Berkshire and Surrey Pathology Services Lighthouse Laboratory and Alex Alderton, Roberto Amato, Jeffrey Barrett, Sonia Goncalves, Ewan Harrison, David K. Jackson, Ian Johnston, Dominic Kwiatkowski, Cordelia Langford, John Sillitoe on behalf of the Wellcome Sanger Institute COVID-19 Surveillance Team |
| EPI_ISL_2913500 | Lighthouse Lab in Alderley Park | Wellcome Sanger Institute for the COVID-19 Genomics UK (COG-UK) Consortium | Jacquelyn Wynn, Mairead Hyland, The Lighthouse Lab in Alderley Park and Alex Alderton, Roberto Amato, Jeffrey Barrett, Sonia Goncalves, Ewan Harrison, David K. Jackson, Ian Johnston, Dominic Kwiatkowski, Cordelia Langford, John Sillitoe on behalf of the Wellcome Sanger Institute COVID-19 Surveillance Team |
| EPI_ISL_2913501, EPI_ISL_2913502, EPI_ISL_2913503 | Lighthouse Lab in Milton Keynes | Wellcome Sanger Institute for the COVID-19 Genomics UK (COG-UK) Consortium | The Lighthouse Lab in Milton Keynes and Alex Alderton, Roberto Amato, Jeffrey Barrett, Sonia Goncalves, Ewan Harrison, David K. Jackson, Ian Johnston, Dominic Kwiatkowski, Cordelia Langford, John Sillitoe on behalf of the Wellcome Sanger Institute COVID-19 Surveillance Team |
| EPI_ISL_2913504 | Berkshire and Surrey Pathology Services Lighthouse Laboratory | Wellcome Sanger Institute for the COVID-19 Genomics UK (COG-UK) Consortium | Berkshire and Surrey Pathology Services Lighthouse Laboratory and Alex Alderton, Roberto Amato, Jeffrey Barrett, Sonia Goncalves, Ewan Harrison, David K. Jackson, Ian Johnston, Dominic Kwiatkowski, Cordelia Langford, John Sillitoe on behalf of the Wellcome Sanger Institute COVID-19 Surveillance Team |
| EPI_ISL_2913505, EPI_ISL_2913506 | Lighthouse Lab in Alderley Park | Wellcome Sanger Institute for the COVID-19 Genomics UK (COG-UK) Consortium | Jacquelyn Wynn, Mairead Hyland, The Lighthouse Lab in Alderley Park and Alex Alderton, Roberto Amato, Jeffrey Barrett, Sonia Goncalves, Ewan Harrison, David K. Jackson, Ian Johnston, Dominic Kwiatkowski, Cordelia Langford, John Sillitoe on behalf of the Wellcome Sanger Institute COVID-19 Surveillance Team |
| EPI_ISL_2913507 | Berkshire and Surrey Pathology Services Lighthouse Laboratory | Wellcome Sanger Institute for the COVID-19 Genomics UK (COG-UK) Consortium | Berkshire and Surrey Pathology Services Lighthouse Laboratory and Alex Alderton, Roberto Amato, Jeffrey Barrett, Sonia Goncalves, Ewan Harrison, David K. Jackson, Ian Johnston, Dominic Kwiatkowski, Cordelia Langford, John Sillitoe on behalf of the Wellcome Sanger Institute COVID-19 Surveillance Team |
| EPI_ISL_2913508 | Lighthouse Lab in Milton Keynes | Wellcome Sanger Institute for the COVID-19 Genomics UK (COG-UK) Consortium | The Lighthouse Lab in Milton Keynes and Alex Alderton, Roberto Amato, Jeffrey Barrett, Sonia Goncalves, Ewan Harrison, David K. Jackson, Ian Johnston, Dominic Kwiatkowski, Cordelia Langford, John Sillitoe on behalf of the Wellcome Sanger Institute COVID-19 Surveillance Team |
| EPI_ISL_2913509 | Lighthouse Lab in Alderley Park | Wellcome Sanger Institute for the COVID-19 Genomics UK (COG-UK) Consortium | Jacquelyn Wynn, Mairead Hyland, The Lighthouse Lab in Alderley Park and Alex Alderton, Roberto Amato, Jeffrey Barrett, Sonia Goncalves, Ewan Harrison, David K. Jackson, Ian Johnston, Dominic Kwiatkowski, Cordelia Langford, John Sillitoe on behalf of the Wellcome Sanger Institute COVID-19 Surveillance Team |
| EPI_ISL_2913510 | Berkshire and Surrey Pathology Services Lighthouse Laboratory | Wellcome Sanger Institute for the COVID-19 Genomics UK (COG-UK) Consortium | Berkshire and Surrey Pathology Services Lighthouse Laboratory and Alex Alderton, Roberto Amato, Jeffrey Barrett, Sonia Goncalves, Ewan Harrison, David K. Jackson, Ian Johnston, Dominic Kwiatkowski, Cordelia Langford, John Sillitoe on behalf of the Wellcome Sanger Institute COVID-19 Surveillance Team |
| EPI_ISL_2913511 | Lighthouse Lab in Alderley Park | Wellcome Sanger Institute for the COVID-19 Genomics UK (COG-UK) Consortium | Jacquelyn Wynn, Mairead Hyland, The Lighthouse Lab in Alderley Park and Alex Alderton, Roberto Amato, Jeffrey Barrett, Sonia Goncalves, Ewan Harrison, David K. Jackson, Ian Johnston, Dominic Kwiatkowski, Cordelia Langford, John Sillitoe on behalf of the Wellcome Sanger Institute COVID-19 Surveillance Team |
| EPI_ISL_2913512, EPI_ISL_2913513 | Berkshire and Surrey Pathology Services Lighthouse Laboratory | Wellcome Sanger Institute for the COVID-19 Genomics UK (COG-UK) Consortium | Berkshire and Surrey Pathology Services Lighthouse Laboratory and Alex Alderton, Roberto Amato, Jeffrey Barrett, Sonia Goncalves, Ewan Harrison, David K. Jackson, Ian Johnston, Dominic Kwiatkowski, Cordelia Langford, John Sillitoe on behalf of the Wellcome Sanger Institute COVID-19 Surveillance Team |
| EPI_ISL_2913514, EPI_ISL_2913515 | Lighthouse Lab in Alderley Park | Wellcome Sanger Institute for the COVID-19 Genomics UK (COG-UK) Consortium | Jacquelyn Wynn, Mairead Hyland, The Lighthouse Lab in Alderley Park and Alex Alderton, Roberto Amato, Jeffrey Barrett, Sonia Goncalves, Ewan Harrison, David K. Jackson, Ian Johnston, Dominic Kwiatkowski, Cordelia Langford, John Sillitoe on behalf of the Wellcome Sanger Institute COVID-19 Surveillance Team |
| EPI_ISL_2913516 | Berkshire and Surrey Pathology Services Lighthouse Laboratory | Wellcome Sanger Institute for the COVID-19 Genomics UK (COG-UK) Consortium | Berkshire and Surrey Pathology Services Lighthouse Laboratory and Alex Alderton, Roberto Amato, Jeffrey Barrett, Sonia Goncalves, Ewan Harrison, David K. Jackson, Ian Johnston, Dominic Kwiatkowski, Cordelia Langford, John Sillitoe on behalf of the Wellcome Sanger Institute COVID-19 Surveillance Team |
| EPI_ISL_2913517, | Lighthouse Lab in Alderley Park | Wellcome Sanger Institute for the COVID-19 Genomics UK (COG-UK) | Jacquelyn Wynn, Mairead Hyland, The Lighthouse Lab in Alderley Park and Alex Alderton, Roberto Amato, Jeffrey Barrett, Sonia Goncalves, Ewan Harrison, David K. Jackson, Ian Johnston, Dominic Kwiatkowski, Cordelia Langford, John Sillitoe on |

| | | | |
|---|---|---|---|
| EPI_ISL_2913518, EPI_ISL_2913519 | | Consortium | behalf of the Wellcome Sanger Institute COVID-19 Surveillance Team |
| EPI_ISL_2913520 | Berkshire and Surrey Pathology Services Lighthouse Laboratory | Wellcome Sanger Institute for the COVID-19 Genomics UK (COG-UK) Consortium | Berkshire and Surrey Pathology Services Lighthouse Laboratory and Alex Alderton, Roberto Amato, Jeffrey Barrett, Sonia Goncalves, Ewan Harrison, David K. Jackson, Ian Johnston, Dominic Kwiatkowski, Cordelia Langford, John Sillitoe on behalf of the Wellcome Sanger Institute COVID-19 Surveillance Team |
| EPI_ISL_2913521 | Lighthouse Lab in Milton Keynes | Wellcome Sanger Institute for the COVID-19 Genomics UK (COG-UK) Consortium | The Lighthouse Lab in Milton Keynes and Alex Alderton, Roberto Amato, Jeffrey Barrett, Sonia Goncalves, Ewan Harrison, David K. Jackson, Ian Johnston, Dominic Kwiatkowski, Cordelia Langford, John Sillitoe on behalf of the Wellcome Sanger Institute COVID-19 Surveillance Team |
| EPI_ISL_2913522 | Berkshire and Surrey Pathology Services Lighthouse Laboratory | Wellcome Sanger Institute for the COVID-19 Genomics UK (COG-UK) Consortium | Berkshire and Surrey Pathology Services Lighthouse Laboratory and Alex Alderton, Roberto Amato, Jeffrey Barrett, Sonia Goncalves, Ewan Harrison, David K. Jackson, Ian Johnston, Dominic Kwiatkowski, Cordelia Langford, John Sillitoe on behalf of the Wellcome Sanger Institute COVID-19 Surveillance Team |
| EPI_ISL_2913523 | Lighthouse Lab in Alderley Park | Wellcome Sanger Institute for the COVID-19 Genomics UK (COG-UK) Consortium | Jacquelyn Wynn, Mairead Hyland, The Lighthouse Lab in Alderley Park and Alex Alderton, Roberto Amato, Jeffrey Barrett, Sonia Goncalves, Ewan Harrison, David K. Jackson, Ian Johnston, Dominic Kwiatkowski, Cordelia Langford, John Sillitoe on behalf of the Wellcome Sanger Institute COVID-19 Surveillance Team |
| EPI_ISL_2913524 | Berkshire and Surrey Pathology Services Lighthouse Laboratory | Wellcome Sanger Institute for the COVID-19 Genomics UK (COG-UK) Consortium | Berkshire and Surrey Pathology Services Lighthouse Laboratory and Alex Alderton, Roberto Amato, Jeffrey Barrett, Sonia Goncalves, Ewan Harrison, David K. Jackson, Ian Johnston, Dominic Kwiatkowski, Cordelia Langford, John Sillitoe on behalf of the Wellcome Sanger Institute COVID-19 Surveillance Team |
| EPI_ISL_2913525 | Lighthouse Lab in Alderley Park | Wellcome Sanger Institute for the COVID-19 Genomics UK (COG-UK) Consortium | Jacquelyn Wynn, Mairead Hyland, The Lighthouse Lab in Alderley Park and Alex Alderton, Roberto Amato, Jeffrey Barrett, Sonia Goncalves, Ewan Harrison, David K. Jackson, Ian Johnston, Dominic Kwiatkowski, Cordelia Langford, John Sillitoe on behalf of the Wellcome Sanger Institute COVID-19 Surveillance Team |
| EPI_ISL_2913526 | Berkshire and Surrey Pathology Services Lighthouse Laboratory | Wellcome Sanger Institute for the COVID-19 Genomics UK (COG-UK) Consortium | Berkshire and Surrey Pathology Services Lighthouse Laboratory and Alex Alderton, Roberto Amato, Jeffrey Barrett, Sonia Goncalves, Ewan Harrison, David K. Jackson, Ian Johnston, Dominic Kwiatkowski, Cordelia Langford, John Sillitoe on behalf of the Wellcome Sanger Institute COVID-19 Surveillance Team |
| EPI_ISL_2913527 | Lighthouse Lab in Alderley Park | Wellcome Sanger Institute for the COVID-19 Genomics UK (COG-UK) Consortium | Jacquelyn Wynn, Mairead Hyland, The Lighthouse Lab in Alderley Park and Alex Alderton, Roberto Amato, Jeffrey Barrett, Sonia Goncalves, Ewan Harrison, David K. Jackson, Ian Johnston, Dominic Kwiatkowski, Cordelia Langford, John Sillitoe on behalf of the Wellcome Sanger Institute COVID-19 Surveillance Team |
| EPI_ISL_2913528, EPI_ISL_2913529 | Lighthouse Lab in Milton Keynes | Wellcome Sanger Institute for the COVID-19 Genomics UK (COG-UK) Consortium | The Lighthouse Lab in Milton Keynes and Alex Alderton, Roberto Amato, Jeffrey Barrett, Sonia Goncalves, Ewan Harrison, David K. Jackson, Ian Johnston, Dominic Kwiatkowski, Cordelia Langford, John Sillitoe on behalf of the Wellcome Sanger Institute COVID-19 Surveillance Team |
| EPI_ISL_2913530, EPI_ISL_2913531 | Berkshire and Surrey Pathology Services Lighthouse Laboratory | Wellcome Sanger Institute for the COVID-19 Genomics UK (COG-UK) Consortium | Berkshire and Surrey Pathology Services Lighthouse Laboratory and Alex Alderton, Roberto Amato, Jeffrey Barrett, Sonia Goncalves, Ewan Harrison, David K. Jackson, Ian Johnston, Dominic Kwiatkowski, Cordelia Langford, John Sillitoe on behalf of the Wellcome Sanger Institute COVID-19 Surveillance Team |
| EPI_ISL_2913532 | Lighthouse Lab in Alderley Park | Wellcome Sanger Institute for the COVID-19 Genomics UK (COG-UK) Consortium | Jacquelyn Wynn, Mairead Hyland, The Lighthouse Lab in Alderley Park and Alex Alderton, Roberto Amato, Jeffrey Barrett, Sonia Goncalves, Ewan Harrison, David K. Jackson, Ian Johnston, Dominic Kwiatkowski, Cordelia Langford, John Sillitoe on behalf of the Wellcome Sanger Institute COVID-19 Surveillance Team |
| EPI_ISL_2913533 | Berkshire and Surrey Pathology Services Lighthouse Laboratory | Wellcome Sanger Institute for the COVID-19 Genomics UK (COG-UK) Consortium | Berkshire and Surrey Pathology Services Lighthouse Laboratory and Alex Alderton, Roberto Amato, Jeffrey Barrett, Sonia Goncalves, Ewan Harrison, David K. Jackson, Ian Johnston, Dominic Kwiatkowski, Cordelia Langford, John Sillitoe on behalf of the Wellcome Sanger Institute COVID-19 Surveillance Team |
| EPI_ISL_2913534 | Lighthouse Lab in Alderley Park | Wellcome Sanger Institute for the COVID-19 Genomics UK (COG-UK) Consortium | Jacquelyn Wynn, Mairead Hyland, The Lighthouse Lab in Alderley Park and Alex Alderton, Roberto Amato, Jeffrey Barrett, Sonia Goncalves, Ewan Harrison, David K. Jackson, Ian Johnston, Dominic Kwiatkowski, Cordelia Langford, John Sillitoe on behalf of the Wellcome Sanger Institute COVID-19 Surveillance Team |
| EPI_ISL_2913535 | Berkshire and Surrey Pathology Services Lighthouse Laboratory | Wellcome Sanger Institute for the COVID-19 Genomics UK (COG-UK) Consortium | Berkshire and Surrey Pathology Services Lighthouse Laboratory and Alex Alderton, Roberto Amato, Jeffrey Barrett, Sonia Goncalves, Ewan Harrison, David K. Jackson, Ian Johnston, Dominic Kwiatkowski, Cordelia Langford, John Sillitoe on behalf of the Wellcome Sanger Institute COVID-19 Surveillance Team |
| EPI_ISL_2913536 | Lighthouse Lab in Milton Keynes | Wellcome Sanger Institute for the COVID-19 Genomics UK (COG-UK) Consortium | The Lighthouse Lab in Milton Keynes and Alex Alderton, Roberto Amato, Jeffrey Barrett, Sonia Goncalves, Ewan Harrison, David K. Jackson, Ian Johnston, Dominic Kwiatkowski, Cordelia Langford, John Sillitoe on behalf of the Wellcome Sanger Institute COVID-19 Surveillance Team |
| EPI_ISL_2913537 | Berkshire and Surrey Pathology Services Lighthouse Laboratory | Wellcome Sanger Institute for the COVID-19 Genomics UK (COG-UK) Consortium | Berkshire and Surrey Pathology Services Lighthouse Laboratory and Alex Alderton, Roberto Amato, Jeffrey Barrett, Sonia Goncalves, Ewan Harrison, David K. Jackson, Ian Johnston, Dominic Kwiatkowski, Cordelia Langford, John Sillitoe on behalf of the Wellcome Sanger Institute COVID-19 Surveillance Team |
| EPI_ISL_2913538 | Lighthouse Lab in Milton Keynes | Wellcome Sanger Institute for the COVID-19 Genomics UK (COG-UK) Consortium | The Lighthouse Lab in Milton Keynes and Alex Alderton, Roberto Amato, Jeffrey Barrett, Sonia Goncalves, Ewan Harrison, David K. Jackson, Ian Johnston, Dominic Kwiatkowski, Cordelia Langford, John Sillitoe on behalf of the Wellcome Sanger Institute COVID-19 Surveillance Team |
| EPI_ISL_2913539 | Berkshire and Surrey Pathology Services Lighthouse Laboratory | Wellcome Sanger Institute for the COVID-19 Genomics UK (COG-UK) Consortium | Berkshire and Surrey Pathology Services Lighthouse Laboratory and Alex Alderton, Roberto Amato, Jeffrey Barrett, Sonia Goncalves, Ewan Harrison, David K. Jackson, Ian Johnston, Dominic Kwiatkowski, Cordelia Langford, John Sillitoe on behalf of the Wellcome Sanger Institute COVID-19 Surveillance Team |
| EPI_ISL_2913540 | Lighthouse Lab in Milton Keynes | Wellcome Sanger Institute for the COVID-19 Genomics UK (COG-UK) Consortium | The Lighthouse Lab in Milton Keynes and Alex Alderton, Roberto Amato, Jeffrey Barrett, Sonia Goncalves, Ewan Harrison, David K. Jackson, Ian Johnston, Dominic Kwiatkowski, Cordelia Langford, John Sillitoe on behalf of the Wellcome Sanger Institute COVID-19 Surveillance Team |
| EPI_ISL_2913541 | Lighthouse Lab in Alderley Park | Wellcome Sanger Institute for the COVID-19 Genomics UK (COG-UK) Consortium | Jacquelyn Wynn, Mairead Hyland, The Lighthouse Lab in Alderley Park and Alex Alderton, Roberto Amato, Jeffrey Barrett, Sonia Goncalves, Ewan Harrison, David K. Jackson, Ian Johnston, Dominic Kwiatkowski, Cordelia Langford, John Sillitoe on behalf of the Wellcome Sanger Institute COVID-19 Surveillance Team |
| EPI_ISL_2913542 | Berkshire and Surrey Pathology Services Lighthouse Laboratory | Wellcome Sanger Institute for the COVID-19 Genomics UK (COG-UK) Consortium | Berkshire and Surrey Pathology Services Lighthouse Laboratory and Alex Alderton, Roberto Amato, Jeffrey Barrett, Sonia Goncalves, Ewan Harrison, David K. Jackson, Ian Johnston, Dominic Kwiatkowski, Cordelia Langford, John Sillitoe on behalf of the Wellcome Sanger Institute COVID-19 Surveillance Team |
| EPI_ISL_2913543 | Lighthouse Lab in Alderley Park | Wellcome Sanger Institute for the COVID-19 Genomics UK (COG-UK) Consortium | Jacquelyn Wynn, Mairead Hyland, The Lighthouse Lab in Alderley Park and Alex Alderton, Roberto Amato, Jeffrey Barrett, Sonia Goncalves, Ewan Harrison, David K. Jackson, Ian Johnston, Dominic Kwiatkowski, Cordelia Langford, John Sillitoe on behalf of the Wellcome Sanger Institute COVID-19 Surveillance Team |
| EPI_ISL_2913544 | Lighthouse Lab in Milton Keynes | Wellcome Sanger Institute for the COVID-19 Genomics UK (COG-UK) Consortium | The Lighthouse Lab in Milton Keynes and Alex Alderton, Roberto Amato, Jeffrey Barrett, Sonia Goncalves, Ewan Harrison, David K. Jackson, Ian Johnston, Dominic Kwiatkowski, Cordelia Langford, John Sillitoe on behalf of the Wellcome Sanger Institute COVID-19 Surveillance Team |
| EPI_ISL_2913545 | Berkshire and Surrey Pathology Services Lighthouse Laboratory | Wellcome Sanger Institute for the COVID-19 Genomics UK (COG-UK) Consortium | Berkshire and Surrey Pathology Services Lighthouse Laboratory and Alex Alderton, Roberto Amato, Jeffrey Barrett, Sonia Goncalves, Ewan Harrison, David K. Jackson, Ian Johnston, Dominic Kwiatkowski, Cordelia Langford, John Sillitoe on behalf of the Wellcome Sanger Institute COVID-19 Surveillance Team |
| EPI_ISL_2913546 | Lighthouse Lab in Alderley Park | Wellcome Sanger Institute for the COVID-19 Genomics UK (COG-UK) Consortium | Jacquelyn Wynn, Mairead Hyland, The Lighthouse Lab in Alderley Park and Alex Alderton, Roberto Amato, Jeffrey Barrett, Sonia Goncalves, Ewan Harrison, David K. Jackson, Ian Johnston, Dominic Kwiatkowski, Cordelia Langford, John Sillitoe on behalf of the Wellcome Sanger Institute COVID-19 Surveillance Team |
| EPI_ISL_2913547 | Lighthouse Lab in Milton Keynes | Wellcome Sanger Institute for the COVID-19 Genomics UK (COG-UK) Consortium | The Lighthouse Lab in Milton Keynes and Alex Alderton, Roberto Amato, Jeffrey Barrett, Sonia Goncalves, Ewan Harrison, David K. Jackson, Ian Johnston, Dominic Kwiatkowski, Cordelia Langford, John Sillitoe on behalf of the Wellcome Sanger Institute COVID-19 Surveillance Team |
| EPI_ISL_2913548, EPI_ISL_2913549 | Lighthouse Lab in Alderley Park | Wellcome Sanger Institute for the COVID-19 Genomics UK (COG-UK) Consortium | Jacquelyn Wynn, Mairead Hyland, The Lighthouse Lab in Alderley Park and Alex Alderton, Roberto Amato, Jeffrey Barrett, Sonia Goncalves, Ewan Harrison, David K. Jackson, Ian Johnston, Dominic Kwiatkowski, Cordelia Langford, John Sillitoe on behalf of the Wellcome Sanger Institute COVID-19 Surveillance Team |
| EPI_ISL_2913550 | Lighthouse Lab in Milton Keynes | Wellcome Sanger Institute for the COVID-19 Genomics UK (COG-UK) Consortium | The Lighthouse Lab in Milton Keynes and Alex Alderton, Roberto Amato, Jeffrey Barrett, Sonia Goncalves, Ewan Harrison, David K. Jackson, Ian Johnston, Dominic Kwiatkowski, Cordelia Langford, John Sillitoe on behalf of the Wellcome Sanger Institute COVID-19 Surveillance Team |
| EPI_ISL_2913551, EPI_ISL_2913552, EPI_ISL_2913553 | Lighthouse Lab in Alderley Park | Wellcome Sanger Institute for the COVID-19 Genomics UK (COG-UK) Consortium | Jacquelyn Wynn, Mairead Hyland, The Lighthouse Lab in Alderley Park and Alex Alderton, Roberto Amato, Jeffrey Barrett, Sonia Goncalves, Ewan Harrison, David K. Jackson, Ian Johnston, Dominic Kwiatkowski, Cordelia Langford, John Sillitoe on behalf of the Wellcome Sanger Institute COVID-19 Surveillance Team |
| EPI_ISL_2913554 | Berkshire and Surrey Pathology Services Lighthouse Laboratory | Wellcome Sanger Institute for the COVID-19 Genomics UK (COG-UK) Consortium | Berkshire and Surrey Pathology Services Lighthouse Laboratory and Alex Alderton, Roberto Amato, Jeffrey Barrett, Sonia Goncalves, Ewan Harrison, David K. Jackson, Ian Johnston, Dominic Kwiatkowski, Cordelia Langford, John Sillitoe on behalf of the Wellcome Sanger Institute COVID-19 Surveillance Team |
| EPI_ISL_2913555 | Lighthouse Lab in Milton Keynes | Wellcome Sanger Institute for the COVID-19 Genomics UK (COG-UK) Consortium | The Lighthouse Lab in Milton Keynes and Alex Alderton, Roberto Amato, Jeffrey Barrett, Sonia Goncalves, Ewan Harrison, David K. Jackson, Ian Johnston, Dominic Kwiatkowski, Cordelia Langford, John Sillitoe on behalf of the Wellcome Sanger Institute COVID-19 Surveillance Team |
| EPI_ISL_2913556, EPI_ISL_2913557, EPI_ISL_2913558 | Lighthouse Lab in Alderley Park | Wellcome Sanger Institute for the COVID-19 Genomics UK (COG-UK) Consortium | Jacquelyn Wynn, Mairead Hyland, The Lighthouse Lab in Alderley Park and Alex Alderton, Roberto Amato, Jeffrey Barrett, Sonia Goncalves, Ewan Harrison, David K. Jackson, Ian Johnston, Dominic Kwiatkowski, Cordelia Langford, John Sillitoe on behalf of the Wellcome Sanger Institute COVID-19 Surveillance Team |
| EPI_ISL_2913559, EPI_ISL_2913560 | Berkshire and Surrey Pathology Services Lighthouse Laboratory | Wellcome Sanger Institute for the COVID-19 Genomics UK (COG-UK) Consortium | Berkshire and Surrey Pathology Services Lighthouse Laboratory and Alex Alderton, Roberto Amato, Jeffrey Barrett, Sonia Goncalves, Ewan Harrison, David K. Jackson, Ian Johnston, Dominic Kwiatkowski, Cordelia Langford, John Sillitoe on behalf of the Wellcome Sanger Institute COVID-19 Surveillance Team |
| EPI_ISL_2913561, EPI_ISL_2913562 | Lighthouse Lab in Alderley Park | Wellcome Sanger Institute for the COVID-19 Genomics UK (COG-UK) Consortium | Jacquelyn Wynn, Mairead Hyland, The Lighthouse Lab in Alderley Park and Alex Alderton, Roberto Amato, Jeffrey Barrett, Sonia Goncalves, Ewan Harrison, David K. Jackson, Ian Johnston, Dominic Kwiatkowski, Cordelia Langford, John Sillitoe on behalf of the Wellcome Sanger Institute COVID-19 Surveillance Team |
| EPI_ISL_2913563, EPI_ISL_2913564, EPI_ISL_2913565 | Lighthouse Lab in Milton Keynes | Wellcome Sanger Institute for the COVID-19 Genomics UK (COG-UK) Consortium | The Lighthouse Lab in Milton Keynes and Alex Alderton, Roberto Amato, Jeffrey Barrett, Sonia Goncalves, Ewan Harrison, David K. Jackson, Ian Johnston, Dominic Kwiatkowski, Cordelia Langford, John Sillitoe on behalf of the Wellcome Sanger Institute COVID-19 Surveillance Team |
| EPI_ISL_2913566 | Berkshire and Surrey Pathology Services Lighthouse Laboratory | Wellcome Sanger Institute for the COVID-19 Genomics UK (COG-UK) Consortium | Berkshire and Surrey Pathology Services Lighthouse Laboratory and Alex Alderton, Roberto Amato, Jeffrey Barrett, Sonia Goncalves, Ewan Harrison, David K. Jackson, Ian Johnston, Dominic Kwiatkowski, Cordelia Langford, John Sillitoe on behalf of the Wellcome Sanger Institute COVID-19 Surveillance Team |
| EPI_ISL_2913567 | Lighthouse Lab in Milton Keynes | Wellcome Sanger Institute for the COVID-19 Genomics UK (COG-UK) Consortium | The Lighthouse Lab in Milton Keynes and Alex Alderton, Roberto Amato, Jeffrey Barrett, Sonia Goncalves, Ewan Harrison, David K. Jackson, Ian Johnston, Dominic Kwiatkowski, Cordelia Langford, John Sillitoe on behalf of the Wellcome Sanger Institute COVID-19 Surveillance Team |
| EPI_ISL_2913568 | Berkshire and Surrey Pathology Services Lighthouse Laboratory | Wellcome Sanger Institute for the COVID-19 Genomics UK (COG-UK) Consortium | Berkshire and Surrey Pathology Services Lighthouse Laboratory and Alex Alderton, Roberto Amato, Jeffrey Barrett, Sonia Goncalves, Ewan Harrison, David K. Jackson, Ian Johnston, Dominic Kwiatkowski, Cordelia Langford, John Sillitoe on behalf of the Wellcome Sanger Institute COVID-19 Surveillance Team |
| EPI_ISL_2913569 | Lighthouse Lab in Alderley Park | Wellcome Sanger Institute for the COVID-19 Genomics UK (COG-UK) Consortium | Jacquelyn Wynn, Mairead Hyland, The Lighthouse Lab in Alderley Park and Alex Alderton, Roberto Amato, Jeffrey Barrett, Sonia Goncalves, Ewan Harrison, David K. Jackson, Ian Johnston, Dominic Kwiatkowski, Cordelia Langford, John Sillitoe on behalf of the Wellcome Sanger Institute COVID-19 Surveillance Team |
| EPI_ISL_2913570 | Lighthouse Lab in Milton Keynes | Wellcome Sanger Institute for the COVID-19 Genomics UK (COG-UK) Consortium | The Lighthouse Lab in Milton Keynes and Alex Alderton, Roberto Amato, Jeffrey Barrett, Sonia Goncalves, Ewan Harrison, David K. Jackson, Ian Johnston, Dominic Kwiatkowski, Cordelia Langford, John Sillitoe on behalf of the Wellcome Sanger Institute COVID-19 Surveillance Team |
| EPI_ISL_2913571 | Lighthouse Lab in Alderley Park | Wellcome Sanger Institute for the COVID-19 Genomics UK (COG-UK) Consortium | Jacquelyn Wynn, Mairead Hyland, The Lighthouse Lab in Alderley Park and Alex Alderton, Roberto Amato, Jeffrey Barrett, Sonia Goncalves, Ewan Harrison, David K. Jackson, Ian Johnston, Dominic Kwiatkowski, Cordelia Langford, John Sillitoe on behalf of the Wellcome Sanger Institute COVID-19 Surveillance Team |
| EPI_ISL_2913572 | Lighthouse Lab in Milton Keynes | Wellcome Sanger Institute for the COVID-19 Genomics UK (COG-UK) Consortium | The Lighthouse Lab in Milton Keynes and Alex Alderton, Roberto Amato, Jeffrey Barrett, Sonia Goncalves, Ewan Harrison, David K. Jackson, Ian Johnston, Dominic Kwiatkowski, Cordelia Langford, John Sillitoe on behalf of the Wellcome Sanger Institute COVID-19 Surveillance Team |
| EPI_ISL_2913573 | Lighthouse Lab in Alderley Park | Wellcome Sanger Institute for the COVID-19 Genomics UK (COG-UK) Consortium | Jacquelyn Wynn, Mairead Hyland, The Lighthouse Lab in Alderley Park and Alex Alderton, Roberto Amato, Jeffrey Barrett, Sonia Goncalves, Ewan Harrison, David K. Jackson, Ian Johnston, Dominic Kwiatkowski, Cordelia Langford, John Sillitoe on behalf of the Wellcome Sanger Institute COVID-19 Surveillance Team |

| | | | |
|---|---|---|---|
| EPI_ISL_2913574 | Berkshire and Surrey Pathology Services Lighthouse Laboratory | Wellcome Sanger Institute for the COVID-19 Genomics UK (COG-UK) Consortium | Berkshire and Surrey Pathology Services Lighthouse Laboratory and Alex Alderton, Roberto Amato, Jeffrey Barrett, Sonia Goncalves, Ewan Harrison, David K. Jackson, Ian Johnston, Dominic Kwiatkowski, Cordelia Langford, John Sillitoe on behalf of the Wellcome Sanger Institute COVID-19 Surveillance Team |
| EPI_ISL_2913575, EPI_ISL_2913576, EPI_ISL_2913577, EPI_ISL_2913578 | Lighthouse Lab in Alderley Park | Wellcome Sanger Institute for the COVID-19 Genomics UK (COG-UK) Consortium | Jacquelyn Wynn, Mairead Hyland, The Lighthouse Lab in Alderley Park and Alex Alderton, Roberto Amato, Jeffrey Barrett, Sonia Goncalves, Ewan Harrison, David K. Jackson, Ian Johnston, Dominic Kwiatkowski, Cordelia Langford, John Sillitoe on behalf of the Wellcome Sanger Institute COVID-19 Surveillance Team |
| EPI_ISL_2913579 | Berkshire and Surrey Pathology Services Lighthouse Laboratory | Wellcome Sanger Institute for the COVID-19 Genomics UK (COG-UK) Consortium | Berkshire and Surrey Pathology Services Lighthouse Laboratory and Alex Alderton, Roberto Amato, Jeffrey Barrett, Sonia Goncalves, Ewan Harrison, David K. Jackson, Ian Johnston, Dominic Kwiatkowski, Cordelia Langford, John Sillitoe on behalf of the Wellcome Sanger Institute COVID-19 Surveillance Team |
| EPI_ISL_2913580 | Lighthouse Lab in Milton Keynes | Wellcome Sanger Institute for the COVID-19 Genomics UK (COG-UK) Consortium | The Lighthouse Lab in Milton Keynes and Alex Alderton, Roberto Amato, Jeffrey Barrett, Sonia Goncalves, Ewan Harrison, David K. Jackson, Ian Johnston, Dominic Kwiatkowski, Cordelia Langford, John Sillitoe on behalf of the Wellcome Sanger Institute COVID-19 Surveillance Team |
| EPI_ISL_2913581 | Berkshire and Surrey Pathology Services Lighthouse Laboratory | Wellcome Sanger Institute for the COVID-19 Genomics UK (COG-UK) Consortium | Berkshire and Surrey Pathology Services Lighthouse Laboratory and Alex Alderton, Roberto Amato, Jeffrey Barrett, Sonia Goncalves, Ewan Harrison, David K. Jackson, Ian Johnston, Dominic Kwiatkowski, Cordelia Langford, John Sillitoe on behalf of the Wellcome Sanger Institute COVID-19 Surveillance Team |
| EPI_ISL_2913582 | Lighthouse Lab in Alderley Park | Wellcome Sanger Institute for the COVID-19 Genomics UK (COG-UK) Consortium | Jacquelyn Wynn, Mairead Hyland, The Lighthouse Lab in Alderley Park and Alex Alderton, Roberto Amato, Jeffrey Barrett, Sonia Goncalves, Ewan Harrison, David K. Jackson, Ian Johnston, Dominic Kwiatkowski, Cordelia Langford, John Sillitoe on behalf of the Wellcome Sanger Institute COVID-19 Surveillance Team |
| EPI_ISL_2913583 | Lighthouse Lab in Milton Keynes | Wellcome Sanger Institute for the COVID-19 Genomics UK (COG-UK) Consortium | The Lighthouse Lab in Milton Keynes and Alex Alderton, Roberto Amato, Jeffrey Barrett, Sonia Goncalves, Ewan Harrison, David K. Jackson, Ian Johnston, Dominic Kwiatkowski, Cordelia Langford, John Sillitoe on behalf of the Wellcome Sanger Institute COVID-19 Surveillance Team |
| EPI_ISL_2913584 | Lighthouse Lab in Alderley Park | Wellcome Sanger Institute for the COVID-19 Genomics UK (COG-UK) Consortium | Jacquelyn Wynn, Mairead Hyland, The Lighthouse Lab in Alderley Park and Alex Alderton, Roberto Amato, Jeffrey Barrett, Sonia Goncalves, Ewan Harrison, David K. Jackson, Ian Johnston, Dominic Kwiatkowski, Cordelia Langford, John Sillitoe on behalf of the Wellcome Sanger Institute COVID-19 Surveillance Team |
| EPI_ISL_2913585, EPI_ISL_2913586 | Lighthouse Lab in Milton Keynes | Wellcome Sanger Institute for the COVID-19 Genomics UK (COG-UK) Consortium | The Lighthouse Lab in Milton Keynes and Alex Alderton, Roberto Amato, Jeffrey Barrett, Sonia Goncalves, Ewan Harrison, David K. Jackson, Ian Johnston, Dominic Kwiatkowski, Cordelia Langford, John Sillitoe on behalf of the Wellcome Sanger Institute COVID-19 Surveillance Team |
| EPI_ISL_2913587, EPI_ISL_2913588 | Lighthouse Lab in Alderley Park | Wellcome Sanger Institute for the COVID-19 Genomics UK (COG-UK) Consortium | Jacquelyn Wynn, Mairead Hyland, The Lighthouse Lab in Alderley Park and Alex Alderton, Roberto Amato, Jeffrey Barrett, Sonia Goncalves, Ewan Harrison, David K. Jackson, Ian Johnston, Dominic Kwiatkowski, Cordelia Langford, John Sillitoe on behalf of the Wellcome Sanger Institute COVID-19 Surveillance Team |
| EPI_ISL_2913589 | Berkshire and Surrey Pathology Services Lighthouse Laboratory | Wellcome Sanger Institute for the COVID-19 Genomics UK (COG-UK) Consortium | Berkshire and Surrey Pathology Services Lighthouse Laboratory and Alex Alderton, Roberto Amato, Jeffrey Barrett, Sonia Goncalves, Ewan Harrison, David K. Jackson, Ian Johnston, Dominic Kwiatkowski, Cordelia Langford, John Sillitoe on behalf of the Wellcome Sanger Institute COVID-19 Surveillance Team |
| EPI_ISL_2913590, EPI_ISL_2913591 | Lighthouse Lab in Alderley Park | Wellcome Sanger Institute for the COVID-19 Genomics UK (COG-UK) Consortium | Jacquelyn Wynn, Mairead Hyland, The Lighthouse Lab in Alderley Park and Alex Alderton, Roberto Amato, Jeffrey Barrett, Sonia Goncalves, Ewan Harrison, David K. Jackson, Ian Johnston, Dominic Kwiatkowski, Cordelia Langford, John Sillitoe on behalf of the Wellcome Sanger Institute COVID-19 Surveillance Team |
| EPI_ISL_2913592 | Berkshire and Surrey Pathology Services Lighthouse Laboratory | Wellcome Sanger Institute for the COVID-19 Genomics UK (COG-UK) Consortium | Berkshire and Surrey Pathology Services Lighthouse Laboratory and Alex Alderton, Roberto Amato, Jeffrey Barrett, Sonia Goncalves, Ewan Harrison, David K. Jackson, Ian Johnston, Dominic Kwiatkowski, Cordelia Langford, John Sillitoe on behalf of the Wellcome Sanger Institute COVID-19 Surveillance Team |
| EPI_ISL_2913593, EPI_ISL_2913594 | Lighthouse Lab in Milton Keynes | Wellcome Sanger Institute for the COVID-19 Genomics UK (COG-UK) Consortium | The Lighthouse Lab in Milton Keynes and Alex Alderton, Roberto Amato, Jeffrey Barrett, Sonia Goncalves, Ewan Harrison, David K. Jackson, Ian Johnston, Dominic Kwiatkowski, Cordelia Langford, John Sillitoe on behalf of the Wellcome Sanger Institute COVID-19 Surveillance Team |
| EPI_ISL_2913595 | Lighthouse Lab in Alderley Park | Wellcome Sanger Institute for the COVID-19 Genomics UK (COG-UK) Consortium | Jacquelyn Wynn, Mairead Hyland, The Lighthouse Lab in Alderley Park and Alex Alderton, Roberto Amato, Jeffrey Barrett, Sonia Goncalves, Ewan Harrison, David K. Jackson, Ian Johnston, Dominic Kwiatkowski, Cordelia Langford, John Sillitoe on behalf of the Wellcome Sanger Institute COVID-19 Surveillance Team |
| EPI_ISL_2913596, EPI_ISL_2913597, EPI_ISL_2913598 | Lighthouse Laboratory Plymouth | Wellcome Sanger Institute for the COVID-19 Genomics UK (COG-UK) Consortium | Lighthouse Laboratory Plymouth and Alex Alderton, Roberto Amato, Jeffrey Barrett, Sonia Goncalves, Ewan Harrison, David K. Jackson, Ian Johnston, Dominic Kwiatkowski, Cordelia Langford, John Sillitoe on behalf of the Wellcome Sanger Institute COVID-19 Surveillance Team |
| EPI_ISL_2913599 | Lighthouse Lab in Glasgow | Wellcome Sanger Institute for the COVID-19 Genomics UK (COG-UK) Consortium | Harper VanSteenhouse, Yumi Kasai, David Gray, Carol Clugston, Anna Dominiczak and Alex Alderton, Roberto Amato, Jeffrey Barrett, Sonia Goncalves, Ewan Harrison, David K. Jackson, Ian Johnston, Dominic Kwiatkowski, Cordelia Langford, John Sillitoe on behalf of the Wellcome Sanger Institute COVID-19 Surveillance Team |
| EPI_ISL_2913600 | Lighthouse Laboratory Plymouth | Wellcome Sanger Institute for the COVID-19 Genomics UK (COG-UK) Consortium | Lighthouse Laboratory Plymouth and Alex Alderton, Roberto Amato, Jeffrey Barrett, Sonia Goncalves, Ewan Harrison, David K. Jackson, Ian Johnston, Dominic Kwiatkowski, Cordelia Langford, John Sillitoe on behalf of the Wellcome Sanger Institute COVID-19 Surveillance Team |
| EPI_ISL_2913601 | Lighthouse Lab in Glasgow | Wellcome Sanger Institute for the COVID-19 Genomics UK (COG-UK) Consortium | Harper VanSteenhouse, Yumi Kasai, David Gray, Carol Clugston, Anna Dominiczak and Alex Alderton, Roberto Amato, Jeffrey Barrett, Sonia Goncalves, Ewan Harrison, David K. Jackson, Ian Johnston, Dominic Kwiatkowski, Cordelia Langford, John Sillitoe on behalf of the Wellcome Sanger Institute COVID-19 Surveillance Team |
| EPI_ISL_2913602, EPI_ISL_2913603, EPI_ISL_2913604 | Lighthouse Laboratory Plymouth | Wellcome Sanger Institute for the COVID-19 Genomics UK (COG-UK) Consortium | Lighthouse Laboratory Plymouth and Alex Alderton, Roberto Amato, Jeffrey Barrett, Sonia Goncalves, Ewan Harrison, David K. Jackson, Ian Johnston, Dominic Kwiatkowski, Cordelia Langford, John Sillitoe on behalf of the Wellcome Sanger Institute COVID-19 Surveillance Team |
| EPI_ISL_2913605 | Berkshire and Surrey Pathology Services Lighthouse Laboratory | Wellcome Sanger Institute for the COVID-19 Genomics UK (COG-UK) Consortium | Berkshire and Surrey Pathology Services Lighthouse Laboratory and Alex Alderton, Roberto Amato, Jeffrey Barrett, Sonia Goncalves, Ewan Harrison, David K. Jackson, Ian Johnston, Dominic Kwiatkowski, Cordelia Langford, John Sillitoe on behalf of the Wellcome Sanger Institute COVID-19 Surveillance Team |
| EPI_ISL_2913606 | Lighthouse Laboratory Plymouth | Wellcome Sanger Institute for the COVID-19 Genomics UK (COG-UK) Consortium | Lighthouse Laboratory Plymouth and Alex Alderton, Roberto Amato, Jeffrey Barrett, Sonia Goncalves, Ewan Harrison, David K. Jackson, Ian Johnston, Dominic Kwiatkowski, Cordelia Langford, John Sillitoe on behalf of the Wellcome Sanger Institute COVID-19 Surveillance Team |
| EPI_ISL_2913607, EPI_ISL_2913608, EPI_ISL_2913609 | Lighthouse Lab in Glasgow | Wellcome Sanger Institute for the COVID-19 Genomics UK (COG-UK) Consortium | Harper VanSteenhouse, Yumi Kasai, David Gray, Carol Clugston, Anna Dominiczak and Alex Alderton, Roberto Amato, Jeffrey Barrett, Sonia Goncalves, Ewan Harrison, David K. Jackson, Ian Johnston, Dominic Kwiatkowski, Cordelia Langford, John Sillitoe on behalf of the Wellcome Sanger Institute COVID-19 Surveillance Team |
| EPI_ISL_2913610, EPI_ISL_2913611 | Lighthouse Laboratory Plymouth | Wellcome Sanger Institute for the COVID-19 Genomics UK (COG-UK) Consortium | Lighthouse Laboratory Plymouth and Alex Alderton, Roberto Amato, Jeffrey Barrett, Sonia Goncalves, Ewan Harrison, David K. Jackson, Ian Johnston, Dominic Kwiatkowski, Cordelia Langford, John Sillitoe on behalf of the Wellcome Sanger Institute COVID-19 Surveillance Team |
| EPI_ISL_2913612, EPI_ISL_2913613, EPI_ISL_2913614 | Lighthouse Lab in Glasgow | Wellcome Sanger Institute for the COVID-19 Genomics UK (COG-UK) Consortium | Harper VanSteenhouse, Yumi Kasai, David Gray, Carol Clugston, Anna Dominiczak and Alex Alderton, Roberto Amato, Jeffrey Barrett, Sonia Goncalves, Ewan Harrison, David K. Jackson, Ian Johnston, Dominic Kwiatkowski, Cordelia Langford, John Sillitoe on behalf of the Wellcome Sanger Institute COVID-19 Surveillance Team |
| EPI_ISL_2913615 | Lighthouse Laboratory Plymouth | Wellcome Sanger Institute for the COVID-19 Genomics UK (COG-UK) Consortium | Lighthouse Laboratory Plymouth and Alex Alderton, Roberto Amato, Jeffrey Barrett, Sonia Goncalves, Ewan Harrison, David K. Jackson, Ian Johnston, Dominic Kwiatkowski, Cordelia Langford, John Sillitoe on behalf of the Wellcome Sanger Institute COVID-19 Surveillance Team |
| EPI_ISL_2913616 | Berkshire and Surrey Pathology Services Lighthouse Laboratory | Wellcome Sanger Institute for the COVID-19 Genomics UK (COG-UK) Consortium | Berkshire and Surrey Pathology Services Lighthouse Laboratory and Alex Alderton, Roberto Amato, Jeffrey Barrett, Sonia Goncalves, Ewan Harrison, David K. Jackson, Ian Johnston, Dominic Kwiatkowski, Cordelia Langford, John Sillitoe on behalf of the Wellcome Sanger Institute COVID-19 Surveillance Team |
| EPI_ISL_2913617 | Lighthouse Laboratory Plymouth | Wellcome Sanger Institute for the COVID-19 Genomics UK (COG-UK) Consortium | Lighthouse Laboratory Plymouth and Alex Alderton, Roberto Amato, Jeffrey Barrett, Sonia Goncalves, Ewan Harrison, David K. Jackson, Ian Johnston, Dominic Kwiatkowski, Cordelia Langford, John Sillitoe on behalf of the Wellcome Sanger Institute COVID-19 Surveillance Team |
| EPI_ISL_2913618 | Lighthouse Lab in Glasgow | Wellcome Sanger Institute for the COVID-19 Genomics UK (COG-UK) Consortium | Harper VanSteenhouse, Yumi Kasai, David Gray, Carol Clugston, Anna Dominiczak and Alex Alderton, Roberto Amato, Jeffrey Barrett, Sonia Goncalves, Ewan Harrison, David K. Jackson, Ian Johnston, Dominic Kwiatkowski, Cordelia Langford, John Sillitoe on behalf of the Wellcome Sanger Institute COVID-19 Surveillance Team |
| EPI_ISL_2913619 | Lighthouse Laboratory Plymouth | Wellcome Sanger Institute for the COVID-19 Genomics UK (COG-UK) Consortium | Lighthouse Laboratory Plymouth and Alex Alderton, Roberto Amato, Jeffrey Barrett, Sonia Goncalves, Ewan Harrison, David K. Jackson, Ian Johnston, Dominic Kwiatkowski, Cordelia Langford, John Sillitoe on behalf of the Wellcome Sanger Institute COVID-19 Surveillance Team |
| EPI_ISL_2913620 | Lighthouse Lab in Glasgow | Wellcome Sanger Institute for the COVID-19 Genomics UK (COG-UK) Consortium | Harper VanSteenhouse, Yumi Kasai, David Gray, Carol Clugston, Anna Dominiczak and Alex Alderton, Roberto Amato, Jeffrey Barrett, Sonia Goncalves, Ewan Harrison, David K. Jackson, Ian Johnston, Dominic Kwiatkowski, Cordelia Langford, John Sillitoe on behalf of the Wellcome Sanger Institute COVID-19 Surveillance Team |
| EPI_ISL_2913621 | Lighthouse Laboratory Plymouth | Wellcome Sanger Institute for the COVID-19 Genomics UK (COG-UK) Consortium | Lighthouse Laboratory Plymouth and Alex Alderton, Roberto Amato, Jeffrey Barrett, Sonia Goncalves, Ewan Harrison, David K. Jackson, Ian Johnston, Dominic Kwiatkowski, Cordelia Langford, John Sillitoe on behalf of the Wellcome Sanger Institute COVID-19 Surveillance Team |
| EPI_ISL_2913622 | Lighthouse Lab in Glasgow | Wellcome Sanger Institute for the COVID-19 Genomics UK (COG-UK) Consortium | Harper VanSteenhouse, Yumi Kasai, David Gray, Carol Clugston, Anna Dominiczak and Alex Alderton, Roberto Amato, Jeffrey Barrett, Sonia Goncalves, Ewan Harrison, David K. Jackson, Ian Johnston, Dominic Kwiatkowski, Cordelia Langford, John Sillitoe on behalf of the Wellcome Sanger Institute COVID-19 Surveillance Team |
| EPI_ISL_2913623 | Berkshire and Surrey Pathology Services Lighthouse Laboratory | Wellcome Sanger Institute for the COVID-19 Genomics UK (COG-UK) Consortium | Berkshire and Surrey Pathology Services Lighthouse Laboratory and Alex Alderton, Roberto Amato, Jeffrey Barrett, Sonia Goncalves, Ewan Harrison, David K. Jackson, Ian Johnston, Dominic Kwiatkowski, Cordelia Langford, John Sillitoe on behalf of the Wellcome Sanger Institute COVID-19 Surveillance Team |
| EPI_ISL_2913624, EPI_ISL_2913625, EPI_ISL_2913626, EPI_ISL_2913627 | Lighthouse Lab in Glasgow | Wellcome Sanger Institute for the COVID-19 Genomics UK (COG-UK) Consortium | Harper VanSteenhouse, Yumi Kasai, David Gray, Carol Clugston, Anna Dominiczak and Alex Alderton, Roberto Amato, Jeffrey Barrett, Sonia Goncalves, Ewan Harrison, David K. Jackson, Ian Johnston, Dominic Kwiatkowski, Cordelia Langford, John Sillitoe on behalf of the Wellcome Sanger Institute COVID-19 Surveillance Team |
| EPI_ISL_2913628, EPI_ISL_2913629, EPI_ISL_2913630 | Lighthouse Laboratory Plymouth | Wellcome Sanger Institute for the COVID-19 Genomics UK (COG-UK) Consortium | Lighthouse Laboratory Plymouth and Alex Alderton, Roberto Amato, Jeffrey Barrett, Sonia Goncalves, Ewan Harrison, David K. Jackson, Ian Johnston, Dominic Kwiatkowski, Cordelia Langford, John Sillitoe on behalf of the Wellcome Sanger Institute COVID-19 Surveillance Team |
| EPI_ISL_2913631 | Lighthouse Lab in Glasgow | Wellcome Sanger Institute for the COVID-19 Genomics UK (COG-UK) Consortium | Harper VanSteenhouse, Yumi Kasai, David Gray, Carol Clugston, Anna Dominiczak and Alex Alderton, Roberto Amato, Jeffrey Barrett, Sonia Goncalves, Ewan Harrison, David K. Jackson, Ian Johnston, Dominic Kwiatkowski, Cordelia Langford, John Sillitoe on behalf of the Wellcome Sanger Institute COVID-19 Surveillance Team |
| EPI_ISL_2913632 | Berkshire and Surrey Pathology Services Lighthouse Laboratory | Wellcome Sanger Institute for the COVID-19 Genomics UK (COG-UK) Consortium | Berkshire and Surrey Pathology Services Lighthouse Laboratory and Alex Alderton, Roberto Amato, Jeffrey Barrett, Sonia Goncalves, Ewan Harrison, David K. Jackson, Ian Johnston, Dominic Kwiatkowski, Cordelia Langford, John Sillitoe on behalf of the Wellcome Sanger Institute COVID-19 Surveillance Team |
| EPI_ISL_2913633 | Lighthouse Laboratory Plymouth | Wellcome Sanger Institute for the COVID-19 Genomics UK (COG-UK) Consortium | Lighthouse Laboratory Plymouth and Alex Alderton, Roberto Amato, Jeffrey Barrett, Sonia Goncalves, Ewan Harrison, David K. Jackson, Ian Johnston, Dominic Kwiatkowski, Cordelia Langford, John Sillitoe on behalf of the Wellcome Sanger Institute COVID-19 Surveillance Team |
| EPI_ISL_2913634 | Berkshire and Surrey Pathology Services Lighthouse Laboratory | Wellcome Sanger Institute for the COVID-19 Genomics UK (COG-UK) Consortium | Berkshire and Surrey Pathology Services Lighthouse Laboratory and Alex Alderton, Roberto Amato, Jeffrey Barrett, Sonia Goncalves, Ewan Harrison, David K. Jackson, Ian Johnston, Dominic Kwiatkowski, Cordelia Langford, John Sillitoe on behalf of the Wellcome Sanger Institute COVID-19 Surveillance Team |
| EPI_ISL_2913635 | Lighthouse Laboratory Plymouth | Wellcome Sanger Institute for the COVID-19 Genomics UK (COG-UK) Consortium | Lighthouse Laboratory Plymouth and Alex Alderton, Roberto Amato, Jeffrey Barrett, Sonia Goncalves, Ewan Harrison, David K. Jackson, Ian Johnston, Dominic Kwiatkowski, Cordelia Langford, John Sillitoe on behalf of the Wellcome Sanger Institute COVID-19 Surveillance Team |

| | | | |
|---|---|---|---|
| EPI_ISL_2913636 | Berkshire and Surrey Pathology Services Lighthouse Laboratory | Wellcome Sanger Institute for the COVID-19 Genomics UK (COG-UK) Consortium | Berkshire and Surrey Pathology Services Lighthouse Laboratory and Alex Alderton, Roberto Amato, Jeffrey Barrett, Sonia Goncalves, Ewan Harrison, David K. Jackson, Ian Johnston, Dominic Kwiatkowski, Cordelia Langford, John Sillitoe on behalf of the Wellcome Sanger Institute COVID-19 Surveillance Team |
| EPI_ISL_2913637, EPI_ISL_2913638 | Lighthouse Laboratory Plymouth | Wellcome Sanger Institute for the COVID-19 Genomics UK (COG-UK) Consortium | Lighthouse Laboratory Plymouth and Alex Alderton, Roberto Amato, Jeffrey Barrett, Sonia Goncalves, Ewan Harrison, David K. Jackson, Ian Johnston, Dominic Kwiatkowski, Cordelia Langford, John Sillitoe on behalf of the Wellcome Sanger Institute COVID-19 Surveillance Team |
| EPI_ISL_2913639 | Lighthouse Lab in Glasgow | Wellcome Sanger Institute for the COVID-19 Genomics UK (COG-UK) Consortium | Harper VanSteenhouse, Yumi Kasai, David Gray, Carol Clugston, Anna Dominiczak and Alex Alderton, Roberto Amato, Jeffrey Barrett, Sonia Goncalves, Ewan Harrison, David K. Jackson, Ian Johnston, Dominic Kwiatkowski, Cordelia Langford, John Sillitoe on behalf of the Wellcome Sanger Institute COVID-19 Surveillance Team |
| EPI_ISL_2913640, EPI_ISL_2913641, EPI_ISL_2913642, EPI_ISL_2913643, EPI_ISL_2913645 | Lighthouse Laboratory Plymouth | Wellcome Sanger Institute for the COVID-19 Genomics UK (COG-UK) Consortium | Lighthouse Laboratory Plymouth and Alex Alderton, Roberto Amato, Jeffrey Barrett, Sonia Goncalves, Ewan Harrison, David K. Jackson, Ian Johnston, Dominic Kwiatkowski, Cordelia Langford, John Sillitoe on behalf of the Wellcome Sanger Institute COVID-19 Surveillance Team |
| EPI_ISL_2913646 | Lighthouse Lab in Glasgow | Wellcome Sanger Institute for the COVID-19 Genomics UK (COG-UK) Consortium | Harper VanSteenhouse, Yumi Kasai, David Gray, Carol Clugston, Anna Dominiczak and Alex Alderton, Roberto Amato, Jeffrey Barrett, Sonia Goncalves, Ewan Harrison, David K. Jackson, Ian Johnston, Dominic Kwiatkowski, Cordelia Langford, John Sillitoe on behalf of the Wellcome Sanger Institute COVID-19 Surveillance Team |
| EPI_ISL_2913647 | Lighthouse Laboratory Plymouth | Wellcome Sanger Institute for the COVID-19 Genomics UK (COG-UK) Consortium | Lighthouse Laboratory Plymouth and Alex Alderton, Roberto Amato, Jeffrey Barrett, Sonia Goncalves, Ewan Harrison, David K. Jackson, Ian Johnston, Dominic Kwiatkowski, Cordelia Langford, John Sillitoe on behalf of the Wellcome Sanger Institute COVID-19 Surveillance Team |
| EPI_ISL_2913648 | Berkshire and Surrey Pathology Services Lighthouse Laboratory | Wellcome Sanger Institute for the COVID-19 Genomics UK (COG-UK) Consortium | Berkshire and Surrey Pathology Services Lighthouse Laboratory and Alex Alderton, Roberto Amato, Jeffrey Barrett, Sonia Goncalves, Ewan Harrison, David K. Jackson, Ian Johnston, Dominic Kwiatkowski, Cordelia Langford, John Sillitoe on behalf of the Wellcome Sanger Institute COVID-19 Surveillance Team |
| EPI_ISL_2913649 | Lighthouse Laboratory Plymouth | Wellcome Sanger Institute for the COVID-19 Genomics UK (COG-UK) Consortium | Lighthouse Laboratory Plymouth and Alex Alderton, Roberto Amato, Jeffrey Barrett, Sonia Goncalves, Ewan Harrison, David K. Jackson, Ian Johnston, Dominic Kwiatkowski, Cordelia Langford, John Sillitoe on behalf of the Wellcome Sanger Institute COVID-19 Surveillance Team |
| EPI_ISL_2913650 | Lighthouse Lab in Glasgow | Wellcome Sanger Institute for the COVID-19 Genomics UK (COG-UK) Consortium | Harper VanSteenhouse, Yumi Kasai, David Gray, Carol Clugston, Anna Dominiczak and Alex Alderton, Roberto Amato, Jeffrey Barrett, Sonia Goncalves, Ewan Harrison, David K. Jackson, Ian Johnston, Dominic Kwiatkowski, Cordelia Langford, John Sillitoe on behalf of the Wellcome Sanger Institute COVID-19 Surveillance Team |
| EPI_ISL_2913651 | Berkshire and Surrey Pathology Services Lighthouse Laboratory | Wellcome Sanger Institute for the COVID-19 Genomics UK (COG-UK) Consortium | Berkshire and Surrey Pathology Services Lighthouse Laboratory and Alex Alderton, Roberto Amato, Jeffrey Barrett, Sonia Goncalves, Ewan Harrison, David K. Jackson, Ian Johnston, Dominic Kwiatkowski, Cordelia Langford, John Sillitoe on behalf of the Wellcome Sanger Institute COVID-19 Surveillance Team |
| EPI_ISL_2913652, EPI_ISL_2913653, EPI_ISL_2913654 | Lighthouse Laboratory Plymouth | Wellcome Sanger Institute for the COVID-19 Genomics UK (COG-UK) Consortium | Lighthouse Laboratory Plymouth and Alex Alderton, Roberto Amato, Jeffrey Barrett, Sonia Goncalves, Ewan Harrison, David K. Jackson, Ian Johnston, Dominic Kwiatkowski, Cordelia Langford, John Sillitoe on behalf of the Wellcome Sanger Institute COVID-19 Surveillance Team |
| EPI_ISL_2913655, EPI_ISL_2913656 | Lighthouse Lab in Glasgow | Wellcome Sanger Institute for the COVID-19 Genomics UK (COG-UK) Consortium | Harper VanSteenhouse, Yumi Kasai, David Gray, Carol Clugston, Anna Dominiczak and Alex Alderton, Roberto Amato, Jeffrey Barrett, Sonia Goncalves, Ewan Harrison, David K. Jackson, Ian Johnston, Dominic Kwiatkowski, Cordelia Langford, John Sillitoe on behalf of the Wellcome Sanger Institute COVID-19 Surveillance Team |
| EPI_ISL_2913657 | Lighthouse Laboratory Plymouth | Wellcome Sanger Institute for the COVID-19 Genomics UK (COG-UK) Consortium | Lighthouse Laboratory Plymouth and Alex Alderton, Roberto Amato, Jeffrey Barrett, Sonia Goncalves, Ewan Harrison, David K. Jackson, Ian Johnston, Dominic Kwiatkowski, Cordelia Langford, John Sillitoe on behalf of the Wellcome Sanger Institute COVID-19 Surveillance Team |
| EPI_ISL_2913658, EPI_ISL_2913659 | Berkshire and Surrey Pathology Services Lighthouse Laboratory | Wellcome Sanger Institute for the COVID-19 Genomics UK (COG-UK) Consortium | Berkshire and Surrey Pathology Services Lighthouse Laboratory and Alex Alderton, Roberto Amato, Jeffrey Barrett, Sonia Goncalves, Ewan Harrison, David K. Jackson, Ian Johnston, Dominic Kwiatkowski, Cordelia Langford, John Sillitoe on behalf of the Wellcome Sanger Institute COVID-19 Surveillance Team |
| EPI_ISL_2913660, EPI_ISL_2913661 | Lighthouse Laboratory Plymouth | Wellcome Sanger Institute for the COVID-19 Genomics UK (COG-UK) Consortium | Lighthouse Laboratory Plymouth and Alex Alderton, Roberto Amato, Jeffrey Barrett, Sonia Goncalves, Ewan Harrison, David K. Jackson, Ian Johnston, Dominic Kwiatkowski, Cordelia Langford, John Sillitoe on behalf of the Wellcome Sanger Institute COVID-19 Surveillance Team |
| EPI_ISL_2913662, EPI_ISL_2913663 | Lighthouse Lab in Glasgow | Wellcome Sanger Institute for the COVID-19 Genomics UK (COG-UK) Consortium | Harper VanSteenhouse, Yumi Kasai, David Gray, Carol Clugston, Anna Dominiczak and Alex Alderton, Roberto Amato, Jeffrey Barrett, Sonia Goncalves, Ewan Harrison, David K. Jackson, Ian Johnston, Dominic Kwiatkowski, Cordelia Langford, John Sillitoe on behalf of the Wellcome Sanger Institute COVID-19 Surveillance Team |
| EPI_ISL_2913664 | Berkshire and Surrey Pathology Services Lighthouse Laboratory | Wellcome Sanger Institute for the COVID-19 Genomics UK (COG-UK) Consortium | Berkshire and Surrey Pathology Services Lighthouse Laboratory and Alex Alderton, Roberto Amato, Jeffrey Barrett, Sonia Goncalves, Ewan Harrison, David K. Jackson, Ian Johnston, Dominic Kwiatkowski, Cordelia Langford, John Sillitoe on behalf of the Wellcome Sanger Institute COVID-19 Surveillance Team |
| EPI_ISL_2913665, EPI_ISL_2913666, EPI_ISL_2913667 | Lighthouse Laboratory Plymouth | Wellcome Sanger Institute for the COVID-19 Genomics UK (COG-UK) Consortium | Lighthouse Laboratory Plymouth and Alex Alderton, Roberto Amato, Jeffrey Barrett, Sonia Goncalves, Ewan Harrison, David K. Jackson, Ian Johnston, Dominic Kwiatkowski, Cordelia Langford, John Sillitoe on behalf of the Wellcome Sanger Institute COVID-19 Surveillance Team |
| EPI_ISL_2913668 | Lighthouse Lab in Glasgow | Wellcome Sanger Institute for the COVID-19 Genomics UK (COG-UK) Consortium | Harper VanSteenhouse, Yumi Kasai, David Gray, Carol Clugston, Anna Dominiczak and Alex Alderton, Roberto Amato, Jeffrey Barrett, Sonia Goncalves, Ewan Harrison, David K. Jackson, Ian Johnston, Dominic Kwiatkowski, Cordelia Langford, John Sillitoe on behalf of the Wellcome Sanger Institute COVID-19 Surveillance Team |
| EPI_ISL_2913669, EPI_ISL_2913670, EPI_ISL_2913671, EPI_ISL_2913672 | Lighthouse Laboratory Plymouth | Wellcome Sanger Institute for the COVID-19 Genomics UK (COG-UK) Consortium | Lighthouse Laboratory Plymouth and Alex Alderton, Roberto Amato, Jeffrey Barrett, Sonia Goncalves, Ewan Harrison, David K. Jackson, Ian Johnston, Dominic Kwiatkowski, Cordelia Langford, John Sillitoe on behalf of the Wellcome Sanger Institute COVID-19 Surveillance Team |
| EPI_ISL_2913673 | Berkshire and Surrey Pathology Services Lighthouse Laboratory | Wellcome Sanger Institute for the COVID-19 Genomics UK (COG-UK) Consortium | Berkshire and Surrey Pathology Services Lighthouse Laboratory and Alex Alderton, Roberto Amato, Jeffrey Barrett, Sonia Goncalves, Ewan Harrison, David K. Jackson, Ian Johnston, Dominic Kwiatkowski, Cordelia Langford, John Sillitoe on behalf of the Wellcome Sanger Institute COVID-19 Surveillance Team |
| EPI_ISL_2913674, EPI_ISL_2913675, EPI_ISL_2913676 | Lighthouse Laboratory Plymouth | Wellcome Sanger Institute for the COVID-19 Genomics UK (COG-UK) Consortium | Lighthouse Laboratory Plymouth and Alex Alderton, Roberto Amato, Jeffrey Barrett, Sonia Goncalves, Ewan Harrison, David K. Jackson, Ian Johnston, Dominic Kwiatkowski, Cordelia Langford, John Sillitoe on behalf of the Wellcome Sanger Institute COVID-19 Surveillance Team |
| EPI_ISL_2913677, EPI_ISL_2913678, EPI_ISL_2913679 | Lighthouse Lab in Glasgow | Wellcome Sanger Institute for the COVID-19 Genomics UK (COG-UK) Consortium | Harper VanSteenhouse, Yumi Kasai, David Gray, Carol Clugston, Anna Dominiczak and Alex Alderton, Roberto Amato, Jeffrey Barrett, Sonia Goncalves, Ewan Harrison, David K. Jackson, Ian Johnston, Dominic Kwiatkowski, Cordelia Langford, John Sillitoe on behalf of the Wellcome Sanger Institute COVID-19 Surveillance Team |
| EPI_ISL_2913680 | Berkshire and Surrey Pathology Services Lighthouse Laboratory | Wellcome Sanger Institute for the COVID-19 Genomics UK (COG-UK) Consortium | Berkshire and Surrey Pathology Services Lighthouse Laboratory and Alex Alderton, Roberto Amato, Jeffrey Barrett, Sonia Goncalves, Ewan Harrison, David K. Jackson, Ian Johnston, Dominic Kwiatkowski, Cordelia Langford, John Sillitoe on behalf of the Wellcome Sanger Institute COVID-19 Surveillance Team |
| EPI_ISL_2913681 | Lighthouse Laboratory Plymouth | Wellcome Sanger Institute for the COVID-19 Genomics UK (COG-UK) Consortium | Lighthouse Laboratory Plymouth and Alex Alderton, Roberto Amato, Jeffrey Barrett, Sonia Goncalves, Ewan Harrison, David K. Jackson, Ian Johnston, Dominic Kwiatkowski, Cordelia Langford, John Sillitoe on behalf of the Wellcome Sanger Institute COVID-19 Surveillance Team |
| EPI_ISL_2913682 | Berkshire and Surrey Pathology Services Lighthouse Laboratory | Wellcome Sanger Institute for the COVID-19 Genomics UK (COG-UK) Consortium | Berkshire and Surrey Pathology Services Lighthouse Laboratory and Alex Alderton, Roberto Amato, Jeffrey Barrett, Sonia Goncalves, Ewan Harrison, David K. Jackson, Ian Johnston, Dominic Kwiatkowski, Cordelia Langford, John Sillitoe on behalf of the Wellcome Sanger Institute COVID-19 Surveillance Team |
| EPI_ISL_2913683 | Lighthouse Lab in Glasgow | Wellcome Sanger Institute for the COVID-19 Genomics UK (COG-UK) Consortium | Harper VanSteenhouse, Yumi Kasai, David Gray, Carol Clugston, Anna Dominiczak and Alex Alderton, Roberto Amato, Jeffrey Barrett, Sonia Goncalves, Ewan Harrison, David K. Jackson, Ian Johnston, Dominic Kwiatkowski, Cordelia Langford, John Sillitoe on behalf of the Wellcome Sanger Institute COVID-19 Surveillance Team |
| EPI_ISL_2913684 | Lighthouse Laboratory Plymouth | Wellcome Sanger Institute for the COVID-19 Genomics UK (COG-UK) Consortium | Lighthouse Laboratory Plymouth and Alex Alderton, Roberto Amato, Jeffrey Barrett, Sonia Goncalves, Ewan Harrison, David K. Jackson, Ian Johnston, Dominic Kwiatkowski, Cordelia Langford, John Sillitoe on behalf of the Wellcome Sanger Institute COVID-19 Surveillance Team |
| EPI_ISL_2913685 | Berkshire and Surrey Pathology Services Lighthouse Laboratory | Wellcome Sanger Institute for the COVID-19 Genomics UK (COG-UK) Consortium | Berkshire and Surrey Pathology Services Lighthouse Laboratory and Alex Alderton, Roberto Amato, Jeffrey Barrett, Sonia Goncalves, Ewan Harrison, David K. Jackson, Ian Johnston, Dominic Kwiatkowski, Cordelia Langford, John Sillitoe on behalf of the Wellcome Sanger Institute COVID-19 Surveillance Team |
| EPI_ISL_2913686 | Lighthouse Laboratory Plymouth | Wellcome Sanger Institute for the COVID-19 Genomics UK (COG-UK) Consortium | Lighthouse Laboratory Plymouth and Alex Alderton, Roberto Amato, Jeffrey Barrett, Sonia Goncalves, Ewan Harrison, David K. Jackson, Ian Johnston, Dominic Kwiatkowski, Cordelia Langford, John Sillitoe on behalf of the Wellcome Sanger Institute COVID-19 Surveillance Team |
| EPI_ISL_2913687 | Lighthouse Lab in Glasgow | Wellcome Sanger Institute for the COVID-19 Genomics UK (COG-UK) Consortium | Harper VanSteenhouse, Yumi Kasai, David Gray, Carol Clugston, Anna Dominiczak and Alex Alderton, Roberto Amato, Jeffrey Barrett, Sonia Goncalves, Ewan Harrison, David K. Jackson, Ian Johnston, Dominic Kwiatkowski, Cordelia Langford, John Sillitoe on behalf of the Wellcome Sanger Institute COVID-19 Surveillance Team |
| EPI_ISL_2913688 | Berkshire and Surrey Pathology Services Lighthouse Laboratory | Wellcome Sanger Institute for the COVID-19 Genomics UK (COG-UK) Consortium | Berkshire and Surrey Pathology Services Lighthouse Laboratory and Alex Alderton, Roberto Amato, Jeffrey Barrett, Sonia Goncalves, Ewan Harrison, David K. Jackson, Ian Johnston, Dominic Kwiatkowski, Cordelia Langford, John Sillitoe on behalf of the Wellcome Sanger Institute COVID-19 Surveillance Team |
| EPI_ISL_2913689, EPI_ISL_2913690 | Lighthouse Laboratory Plymouth | Wellcome Sanger Institute for the COVID-19 Genomics UK (COG-UK) Consortium | Lighthouse Laboratory Plymouth and Alex Alderton, Roberto Amato, Jeffrey Barrett, Sonia Goncalves, Ewan Harrison, David K. Jackson, Ian Johnston, Dominic Kwiatkowski, Cordelia Langford, John Sillitoe on behalf of the Wellcome Sanger Institute COVID-19 Surveillance Team |
| EPI_ISL_2913691 | Berkshire and Surrey Pathology Services Lighthouse Laboratory | Wellcome Sanger Institute for the COVID-19 Genomics UK (COG-UK) Consortium | Berkshire and Surrey Pathology Services Lighthouse Laboratory and Alex Alderton, Roberto Amato, Jeffrey Barrett, Sonia Goncalves, Ewan Harrison, David K. Jackson, Ian Johnston, Dominic Kwiatkowski, Cordelia Langford, John Sillitoe on behalf of the Wellcome Sanger Institute COVID-19 Surveillance Team |
| EPI_ISL_2913692, EPI_ISL_2913693 | Lighthouse Laboratory Plymouth | Wellcome Sanger Institute for the COVID-19 Genomics UK (COG-UK) Consortium | Lighthouse Laboratory Plymouth and Alex Alderton, Roberto Amato, Jeffrey Barrett, Sonia Goncalves, Ewan Harrison, David K. Jackson, Ian Johnston, Dominic Kwiatkowski, Cordelia Langford, John Sillitoe on behalf of the Wellcome Sanger Institute COVID-19 Surveillance Team |
| EPI_ISL_2913694 | Lighthouse Lab in Glasgow | Wellcome Sanger Institute for the COVID-19 Genomics UK (COG-UK) Consortium | Harper VanSteenhouse, Yumi Kasai, David Gray, Carol Clugston, Anna Dominiczak and Alex Alderton, Roberto Amato, Jeffrey Barrett, Sonia Goncalves, Ewan Harrison, David K. Jackson, Ian Johnston, Dominic Kwiatkowski, Cordelia Langford, John Sillitoe on behalf of the Wellcome Sanger Institute COVID-19 Surveillance Team |
| EPI_ISL_2913695 | Berkshire and Surrey Pathology Services Lighthouse Laboratory | Wellcome Sanger Institute for the COVID-19 Genomics UK (COG-UK) Consortium | Berkshire and Surrey Pathology Services Lighthouse Laboratory and Alex Alderton, Roberto Amato, Jeffrey Barrett, Sonia Goncalves, Ewan Harrison, David K. Jackson, Ian Johnston, Dominic Kwiatkowski, Cordelia Langford, John Sillitoe on behalf of the Wellcome Sanger Institute COVID-19 Surveillance Team |
| EPI_ISL_2913696, EPI_ISL_2913697, EPI_ISL_2913698 | Lighthouse Laboratory Plymouth | Wellcome Sanger Institute for the COVID-19 Genomics UK (COG-UK) Consortium | Lighthouse Laboratory Plymouth and Alex Alderton, Roberto Amato, Jeffrey Barrett, Sonia Goncalves, Ewan Harrison, David K. Jackson, Ian Johnston, Dominic Kwiatkowski, Cordelia Langford, John Sillitoe on behalf of the Wellcome Sanger Institute COVID-19 Surveillance Team |
| EPI_ISL_2913699 | Lighthouse Lab in Glasgow | Wellcome Sanger Institute for the COVID-19 Genomics UK (COG-UK) Consortium | Harper VanSteenhouse, Yumi Kasai, David Gray, Carol Clugston, Anna Dominiczak and Alex Alderton, Roberto Amato, Jeffrey Barrett, Sonia Goncalves, Ewan Harrison, David K. Jackson, Ian Johnston, Dominic Kwiatkowski, Cordelia Langford, John Sillitoe on behalf of the Wellcome Sanger Institute COVID-19 Surveillance Team |
| EPI_ISL_2913700 | Lighthouse Laboratory Plymouth | Wellcome Sanger Institute for the COVID-19 Genomics UK (COG-UK) Consortium | Lighthouse Laboratory Plymouth and Alex Alderton, Roberto Amato, Jeffrey Barrett, Sonia Goncalves, Ewan Harrison, David K. Jackson, Ian Johnston, Dominic Kwiatkowski, Cordelia Langford, John Sillitoe on behalf of the Wellcome Sanger Institute COVID-19 Surveillance Team |
| EPI_ISL_2913701 | Berkshire and Surrey Pathology Services Lighthouse Laboratory | Wellcome Sanger Institute for the COVID-19 Genomics UK (COG-UK) Consortium | Berkshire and Surrey Pathology Services Lighthouse Laboratory and Alex Alderton, Roberto Amato, Jeffrey Barrett, Sonia Goncalves, Ewan Harrison, David K. Jackson, Ian Johnston, Dominic Kwiatkowski, Cordelia Langford, John Sillitoe on behalf of the Wellcome Sanger Institute COVID-19 Surveillance Team |

| | | | |
|---|---|---|---|
| EPI_ISL_2913702 | Lighthouse Lab in Glasgow | Wellcome Sanger Institute for the COVID-19 Genomics UK (COG-UK) Consortium | Harper VanSteenhouse, Yumi Kasai, David Gray, Carol Clugston, Anna Dominiczak and Alex Alderton, Roberto Amato, Jeffrey Barrett, Sonia Goncalves, Ewan Harrison, David K. Jackson, Ian Johnston, Dominic Kwiatkowski, Cordelia Langford, John Sillitoe on behalf of the Wellcome Sanger Institute COVID-19 Surveillance Team |
| EPI_ISL_2913703, EPI_ISL_2913704 | Lighthouse Laboratory Plymouth | Wellcome Sanger Institute for the COVID-19 Genomics UK (COG-UK) Consortium | Lighthouse Laboratory Plymouth and Alex Alderton, Roberto Amato, Jeffrey Barrett, Sonia Goncalves, Ewan Harrison, David K. Jackson, Ian Johnston, Dominic Kwiatkowski, Cordelia Langford, John Sillitoe on behalf of the Wellcome Sanger Institute COVID-19 Surveillance Team |
| EPI_ISL_2913705, EPI_ISL_2913706, EPI_ISL_2913707 | Berkshire and Surrey Pathology Services Lighthouse Laboratory | Wellcome Sanger Institute for the COVID-19 Genomics UK (COG-UK) Consortium | Berkshire and Surrey Pathology Services Lighthouse Laboratory and Alex Alderton, Roberto Amato, Jeffrey Barrett, Sonia Goncalves, Ewan Harrison, David K. Jackson, Ian Johnston, Dominic Kwiatkowski, Cordelia Langford, John Sillitoe on behalf of the Wellcome Sanger Institute COVID-19 Surveillance Team |
| EPI_ISL_2913708 | Lighthouse Lab in Glasgow | Wellcome Sanger Institute for the COVID-19 Genomics UK (COG-UK) Consortium | Harper VanSteenhouse, Yumi Kasai, David Gray, Carol Clugston, Anna Dominiczak and Alex Alderton, Roberto Amato, Jeffrey Barrett, Sonia Goncalves, Ewan Harrison, David K. Jackson, Ian Johnston, Dominic Kwiatkowski, Cordelia Langford, John Sillitoe on behalf of the Wellcome Sanger Institute COVID-19 Surveillance Team |
| EPI_ISL_2913709 | Lighthouse Laboratory Plymouth | Wellcome Sanger Institute for the COVID-19 Genomics UK (COG-UK) Consortium | Lighthouse Laboratory Plymouth and Alex Alderton, Roberto Amato, Jeffrey Barrett, Sonia Goncalves, Ewan Harrison, David K. Jackson, Ian Johnston, Dominic Kwiatkowski, Cordelia Langford, John Sillitoe on behalf of the Wellcome Sanger Institute COVID-19 Surveillance Team |
| EPI_ISL_2913710 | Berkshire and Surrey Pathology Services Lighthouse Laboratory | Wellcome Sanger Institute for the COVID-19 Genomics UK (COG-UK) Consortium | Berkshire and Surrey Pathology Services Lighthouse Laboratory and Alex Alderton, Roberto Amato, Jeffrey Barrett, Sonia Goncalves, Ewan Harrison, David K. Jackson, Ian Johnston, Dominic Kwiatkowski, Cordelia Langford, John Sillitoe on behalf of the Wellcome Sanger Institute COVID-19 Surveillance Team |
| EPI_ISL_2913711 | Lighthouse Lab in Glasgow | Wellcome Sanger Institute for the COVID-19 Genomics UK (COG-UK) Consortium | Harper VanSteenhouse, Yumi Kasai, David Gray, Carol Clugston, Anna Dominiczak and Alex Alderton, Roberto Amato, Jeffrey Barrett, Sonia Goncalves, Ewan Harrison, David K. Jackson, Ian Johnston, Dominic Kwiatkowski, Cordelia Langford, John Sillitoe on behalf of the Wellcome Sanger Institute COVID-19 Surveillance Team |
| EPI_ISL_2913712 | Lighthouse Laboratory Plymouth | Wellcome Sanger Institute for the COVID-19 Genomics UK (COG-UK) Consortium | Lighthouse Laboratory Plymouth and Alex Alderton, Roberto Amato, Jeffrey Barrett, Sonia Goncalves, Ewan Harrison, David K. Jackson, Ian Johnston, Dominic Kwiatkowski, Cordelia Langford, John Sillitoe on behalf of the Wellcome Sanger Institute COVID-19 Surveillance Team |
| EPI_ISL_2913713, EPI_ISL_2913714 | Berkshire and Surrey Pathology Services Lighthouse Laboratory | Wellcome Sanger Institute for the COVID-19 Genomics UK (COG-UK) Consortium | Berkshire and Surrey Pathology Services Lighthouse Laboratory and Alex Alderton, Roberto Amato, Jeffrey Barrett, Sonia Goncalves, Ewan Harrison, David K. Jackson, Ian Johnston, Dominic Kwiatkowski, Cordelia Langford, John Sillitoe on behalf of the Wellcome Sanger Institute COVID-19 Surveillance Team |
| EPI_ISL_2913715, EPI_ISL_2913716 | Lighthouse Laboratory Plymouth | Wellcome Sanger Institute for the COVID-19 Genomics UK (COG-UK) Consortium | Lighthouse Laboratory Plymouth and Alex Alderton, Roberto Amato, Jeffrey Barrett, Sonia Goncalves, Ewan Harrison, David K. Jackson, Ian Johnston, Dominic Kwiatkowski, Cordelia Langford, John Sillitoe on behalf of the Wellcome Sanger Institute COVID-19 Surveillance Team |
| EPI_ISL_2913717, EPI_ISL_2913718 | Lighthouse Lab in Glasgow | Wellcome Sanger Institute for the COVID-19 Genomics UK (COG-UK) Consortium | Harper VanSteenhouse, Yumi Kasai, David Gray, Carol Clugston, Anna Dominiczak and Alex Alderton, Roberto Amato, Jeffrey Barrett, Sonia Goncalves, Ewan Harrison, David K. Jackson, Ian Johnston, Dominic Kwiatkowski, Cordelia Langford, John Sillitoe on behalf of the Wellcome Sanger Institute COVID-19 Surveillance Team |
| EPI_ISL_2913719 | Lighthouse Laboratory Plymouth | Wellcome Sanger Institute for the COVID-19 Genomics UK (COG-UK) Consortium | Lighthouse Laboratory Plymouth and Alex Alderton, Roberto Amato, Jeffrey Barrett, Sonia Goncalves, Ewan Harrison, David K. Jackson, Ian Johnston, Dominic Kwiatkowski, Cordelia Langford, John Sillitoe on behalf of the Wellcome Sanger Institute COVID-19 Surveillance Team |
| EPI_ISL_2913720 | Berkshire and Surrey Pathology Services Lighthouse Laboratory | Wellcome Sanger Institute for the COVID-19 Genomics UK (COG-UK) Consortium | Berkshire and Surrey Pathology Services Lighthouse Laboratory and Alex Alderton, Roberto Amato, Jeffrey Barrett, Sonia Goncalves, Ewan Harrison, David K. Jackson, Ian Johnston, Dominic Kwiatkowski, Cordelia Langford, John Sillitoe on behalf of the Wellcome Sanger Institute COVID-19 Surveillance Team |
| EPI_ISL_2913721 | Lighthouse Laboratory Plymouth | Wellcome Sanger Institute for the COVID-19 Genomics UK (COG-UK) Consortium | Lighthouse Laboratory Plymouth and Alex Alderton, Roberto Amato, Jeffrey Barrett, Sonia Goncalves, Ewan Harrison, David K. Jackson, Ian Johnston, Dominic Kwiatkowski, Cordelia Langford, John Sillitoe on behalf of the Wellcome Sanger Institute COVID-19 Surveillance Team |
| EPI_ISL_2913722 | Lighthouse Lab in Glasgow | Wellcome Sanger Institute for the COVID-19 Genomics UK (COG-UK) Consortium | Harper VanSteenhouse, Yumi Kasai, David Gray, Carol Clugston, Anna Dominiczak and Alex Alderton, Roberto Amato, Jeffrey Barrett, Sonia Goncalves, Ewan Harrison, David K. Jackson, Ian Johnston, Dominic Kwiatkowski, Cordelia Langford, John Sillitoe on behalf of the Wellcome Sanger Institute COVID-19 Surveillance Team |
| EPI_ISL_2913723, EPI_ISL_2913724 | Lighthouse Laboratory Plymouth | Wellcome Sanger Institute for the COVID-19 Genomics UK (COG-UK) Consortium | Lighthouse Laboratory Plymouth and Alex Alderton, Roberto Amato, Jeffrey Barrett, Sonia Goncalves, Ewan Harrison, David K. Jackson, Ian Johnston, Dominic Kwiatkowski, Cordelia Langford, John Sillitoe on behalf of the Wellcome Sanger Institute COVID-19 Surveillance Team |
| EPI_ISL_2913725 | Berkshire and Surrey Pathology Services Lighthouse Laboratory | Wellcome Sanger Institute for the COVID-19 Genomics UK (COG-UK) Consortium | Berkshire and Surrey Pathology Services Lighthouse Laboratory and Alex Alderton, Roberto Amato, Jeffrey Barrett, Sonia Goncalves, Ewan Harrison, David K. Jackson, Ian Johnston, Dominic Kwiatkowski, Cordelia Langford, John Sillitoe on behalf of the Wellcome Sanger Institute COVID-19 Surveillance Team |
| EPI_ISL_2913726 | Lighthouse Lab in Glasgow | Wellcome Sanger Institute for the COVID-19 Genomics UK (COG-UK) Consortium | Harper VanSteenhouse, Yumi Kasai, David Gray, Carol Clugston, Anna Dominiczak and Alex Alderton, Roberto Amato, Jeffrey Barrett, Sonia Goncalves, Ewan Harrison, David K. Jackson, Ian Johnston, Dominic Kwiatkowski, Cordelia Langford, John Sillitoe on behalf of the Wellcome Sanger Institute COVID-19 Surveillance Team |
| EPI_ISL_2913727 | Berkshire and Surrey Pathology Services Lighthouse Laboratory | Wellcome Sanger Institute for the COVID-19 Genomics UK (COG-UK) Consortium | Berkshire and Surrey Pathology Services Lighthouse Laboratory and Alex Alderton, Roberto Amato, Jeffrey Barrett, Sonia Goncalves, Ewan Harrison, David K. Jackson, Ian Johnston, Dominic Kwiatkowski, Cordelia Langford, John Sillitoe on behalf of the Wellcome Sanger Institute COVID-19 Surveillance Team |
| EPI_ISL_2913728 | Lighthouse Laboratory Plymouth | Wellcome Sanger Institute for the COVID-19 Genomics UK (COG-UK) Consortium | Lighthouse Laboratory Plymouth and Alex Alderton, Roberto Amato, Jeffrey Barrett, Sonia Goncalves, Ewan Harrison, David K. Jackson, Ian Johnston, Dominic Kwiatkowski, Cordelia Langford, John Sillitoe on behalf of the Wellcome Sanger Institute COVID-19 Surveillance Team |
| EPI_ISL_2913729 | Berkshire and Surrey Pathology Services Lighthouse Laboratory | Wellcome Sanger Institute for the COVID-19 Genomics UK (COG-UK) Consortium | Berkshire and Surrey Pathology Services Lighthouse Laboratory and Alex Alderton, Roberto Amato, Jeffrey Barrett, Sonia Goncalves, Ewan Harrison, David K. Jackson, Ian Johnston, Dominic Kwiatkowski, Cordelia Langford, John Sillitoe on behalf of the Wellcome Sanger Institute COVID-19 Surveillance Team |
| EPI_ISL_2913730, EPI_ISL_2913731 | Lighthouse Laboratory Plymouth | Wellcome Sanger Institute for the COVID-19 Genomics UK (COG-UK) Consortium | Lighthouse Laboratory Plymouth and Alex Alderton, Roberto Amato, Jeffrey Barrett, Sonia Goncalves, Ewan Harrison, David K. Jackson, Ian Johnston, Dominic Kwiatkowski, Cordelia Langford, John Sillitoe on behalf of the Wellcome Sanger Institute COVID-19 Surveillance Team |
| EPI_ISL_2913732, EPI_ISL_2913733, EPI_ISL_2913734 | Berkshire and Surrey Pathology Services Lighthouse Laboratory | Wellcome Sanger Institute for the COVID-19 Genomics UK (COG-UK) Consortium | Berkshire and Surrey Pathology Services Lighthouse Laboratory and Alex Alderton, Roberto Amato, Jeffrey Barrett, Sonia Goncalves, Ewan Harrison, David K. Jackson, Ian Johnston, Dominic Kwiatkowski, Cordelia Langford, John Sillitoe on behalf of the Wellcome Sanger Institute COVID-19 Surveillance Team |
| EPI_ISL_2913735 | Lighthouse Laboratory Plymouth | Wellcome Sanger Institute for the COVID-19 Genomics UK (COG-UK) Consortium | Lighthouse Laboratory Plymouth and Alex Alderton, Roberto Amato, Jeffrey Barrett, Sonia Goncalves, Ewan Harrison, David K. Jackson, Ian Johnston, Dominic Kwiatkowski, Cordelia Langford, John Sillitoe on behalf of the Wellcome Sanger Institute COVID-19 Surveillance Team |
| EPI_ISL_2913736 | Lighthouse Lab in Glasgow | Wellcome Sanger Institute for the COVID-19 Genomics UK (COG-UK) Consortium | Harper VanSteenhouse, Yumi Kasai, David Gray, Carol Clugston, Anna Dominiczak and Alex Alderton, Roberto Amato, Jeffrey Barrett, Sonia Goncalves, Ewan Harrison, David K. Jackson, Ian Johnston, Dominic Kwiatkowski, Cordelia Langford, John Sillitoe on behalf of the Wellcome Sanger Institute COVID-19 Surveillance Team |
| EPI_ISL_2913737, EPI_ISL_2913738, EPI_ISL_2913739 | Lighthouse Laboratory Plymouth | Wellcome Sanger Institute for the COVID-19 Genomics UK (COG-UK) Consortium | Lighthouse Laboratory Plymouth and Alex Alderton, Roberto Amato, Jeffrey Barrett, Sonia Goncalves, Ewan Harrison, David K. Jackson, Ian Johnston, Dominic Kwiatkowski, Cordelia Langford, John Sillitoe on behalf of the Wellcome Sanger Institute COVID-19 Surveillance Team |
| EPI_ISL_2913740, EPI_ISL_2913741 | Lighthouse Lab in Glasgow | Wellcome Sanger Institute for the COVID-19 Genomics UK (COG-UK) Consortium | Harper VanSteenhouse, Yumi Kasai, David Gray, Carol Clugston, Anna Dominiczak and Alex Alderton, Roberto Amato, Jeffrey Barrett, Sonia Goncalves, Ewan Harrison, David K. Jackson, Ian Johnston, Dominic Kwiatkowski, Cordelia Langford, John Sillitoe on behalf of the Wellcome Sanger Institute COVID-19 Surveillance Team |
| EPI_ISL_2913742 | Lighthouse Laboratory Plymouth | Wellcome Sanger Institute for the COVID-19 Genomics UK (COG-UK) Consortium | Lighthouse Laboratory Plymouth and Alex Alderton, Roberto Amato, Jeffrey Barrett, Sonia Goncalves, Ewan Harrison, David K. Jackson, Ian Johnston, Dominic Kwiatkowski, Cordelia Langford, John Sillitoe on behalf of the Wellcome Sanger Institute COVID-19 Surveillance Team |
| EPI_ISL_2913743 | Berkshire and Surrey Pathology Services Lighthouse Laboratory | Wellcome Sanger Institute for the COVID-19 Genomics UK (COG-UK) Consortium | Berkshire and Surrey Pathology Services Lighthouse Laboratory and Alex Alderton, Roberto Amato, Jeffrey Barrett, Sonia Goncalves, Ewan Harrison, David K. Jackson, Ian Johnston, Dominic Kwiatkowski, Cordelia Langford, John Sillitoe on behalf of the Wellcome Sanger Institute COVID-19 Surveillance Team |
| EPI_ISL_2913744 | Lighthouse Lab in Glasgow | Wellcome Sanger Institute for the COVID-19 Genomics UK (COG-UK) Consortium | Harper VanSteenhouse, Yumi Kasai, David Gray, Carol Clugston, Anna Dominiczak and Alex Alderton, Roberto Amato, Jeffrey Barrett, Sonia Goncalves, Ewan Harrison, David K. Jackson, Ian Johnston, Dominic Kwiatkowski, Cordelia Langford, John Sillitoe on behalf of the Wellcome Sanger Institute COVID-19 Surveillance Team |
| EPI_ISL_2913745, EPI_ISL_2913746, EPI_ISL_2913747, EPI_ISL_2913748 | Lighthouse Laboratory Plymouth | Wellcome Sanger Institute for the COVID-19 Genomics UK (COG-UK) Consortium | Lighthouse Laboratory Plymouth and Alex Alderton, Roberto Amato, Jeffrey Barrett, Sonia Goncalves, Ewan Harrison, David K. Jackson, Ian Johnston, Dominic Kwiatkowski, Cordelia Langford, John Sillitoe on behalf of the Wellcome Sanger Institute COVID-19 Surveillance Team |
| EPI_ISL_2913749 | Lighthouse Lab in Glasgow | Wellcome Sanger Institute for the COVID-19 Genomics UK (COG-UK) Consortium | Harper VanSteenhouse, Yumi Kasai, David Gray, Carol Clugston, Anna Dominiczak and Alex Alderton, Roberto Amato, Jeffrey Barrett, Sonia Goncalves, Ewan Harrison, David K. Jackson, Ian Johnston, Dominic Kwiatkowski, Cordelia Langford, John Sillitoe on behalf of the Wellcome Sanger Institute COVID-19 Surveillance Team |
| EPI_ISL_2913750 | Berkshire and Surrey Pathology Services Lighthouse Laboratory | Wellcome Sanger Institute for the COVID-19 Genomics UK (COG-UK) Consortium | Berkshire and Surrey Pathology Services Lighthouse Laboratory and Alex Alderton, Roberto Amato, Jeffrey Barrett, Sonia Goncalves, Ewan Harrison, David K. Jackson, Ian Johnston, Dominic Kwiatkowski, Cordelia Langford, John Sillitoe on behalf of the Wellcome Sanger Institute COVID-19 Surveillance Team |
| EPI_ISL_2913751, EPI_ISL_2913752 | Lighthouse Laboratory Plymouth | Wellcome Sanger Institute for the COVID-19 Genomics UK (COG-UK) Consortium | Lighthouse Laboratory Plymouth and Alex Alderton, Roberto Amato, Jeffrey Barrett, Sonia Goncalves, Ewan Harrison, David K. Jackson, Ian Johnston, Dominic Kwiatkowski, Cordelia Langford, John Sillitoe on behalf of the Wellcome Sanger Institute COVID-19 Surveillance Team |
| EPI_ISL_2913753 | Berkshire and Surrey Pathology Services Lighthouse Laboratory | Wellcome Sanger Institute for the COVID-19 Genomics UK (COG-UK) Consortium | Berkshire and Surrey Pathology Services Lighthouse Laboratory and Alex Alderton, Roberto Amato, Jeffrey Barrett, Sonia Goncalves, Ewan Harrison, David K. Jackson, Ian Johnston, Dominic Kwiatkowski, Cordelia Langford, John Sillitoe on behalf of the Wellcome Sanger Institute COVID-19 Surveillance Team |
| EPI_ISL_2913754 | Lighthouse Laboratory Plymouth | Wellcome Sanger Institute for the COVID-19 Genomics UK (COG-UK) Consortium | Lighthouse Laboratory Plymouth and Alex Alderton, Roberto Amato, Jeffrey Barrett, Sonia Goncalves, Ewan Harrison, David K. Jackson, Ian Johnston, Dominic Kwiatkowski, Cordelia Langford, John Sillitoe on behalf of the Wellcome Sanger Institute COVID-19 Surveillance Team |
| EPI_ISL_2913755 | Lighthouse Lab in Glasgow | Wellcome Sanger Institute for the COVID-19 Genomics UK (COG-UK) Consortium | Harper VanSteenhouse, Yumi Kasai, David Gray, Carol Clugston, Anna Dominiczak and Alex Alderton, Roberto Amato, Jeffrey Barrett, Sonia Goncalves, Ewan Harrison, David K. Jackson, Ian Johnston, Dominic Kwiatkowski, Cordelia Langford, John Sillitoe on behalf of the Wellcome Sanger Institute COVID-19 Surveillance Team |
| EPI_ISL_2913756 | Lighthouse Laboratory Plymouth | Wellcome Sanger Institute for the COVID-19 Genomics UK (COG-UK) Consortium | Lighthouse Laboratory Plymouth and Alex Alderton, Roberto Amato, Jeffrey Barrett, Sonia Goncalves, Ewan Harrison, David K. Jackson, Ian Johnston, Dominic Kwiatkowski, Cordelia Langford, John Sillitoe on behalf of the Wellcome Sanger Institute COVID-19 Surveillance Team |
| EPI_ISL_2913757 | Lighthouse Lab in Glasgow | Wellcome Sanger Institute for the COVID-19 Genomics UK (COG-UK) Consortium | Harper VanSteenhouse, Yumi Kasai, David Gray, Carol Clugston, Anna Dominiczak and Alex Alderton, Roberto Amato, Jeffrey Barrett, Sonia Goncalves, Ewan Harrison, David K. Jackson, Ian Johnston, Dominic Kwiatkowski, Cordelia Langford, John Sillitoe on behalf of the Wellcome Sanger Institute COVID-19 Surveillance Team |
| EPI_ISL_2913758 | Berkshire and Surrey Pathology Services Lighthouse Laboratory | Wellcome Sanger Institute for the COVID-19 Genomics UK (COG-UK) Consortium | Berkshire and Surrey Pathology Services Lighthouse Laboratory and Alex Alderton, Roberto Amato, Jeffrey Barrett, Sonia Goncalves, Ewan Harrison, David K. Jackson, Ian Johnston, Dominic Kwiatkowski, Cordelia Langford, John Sillitoe on behalf of the Wellcome Sanger Institute COVID-19 Surveillance Team |
| EPI_ISL_2913759 | Lighthouse Laboratory Plymouth | Wellcome Sanger Institute for the COVID-19 Genomics UK (COG-UK) Consortium | Lighthouse Laboratory Plymouth and Alex Alderton, Roberto Amato, Jeffrey Barrett, Sonia Goncalves, Ewan Harrison, David K. Jackson, Ian Johnston, Dominic Kwiatkowski, Cordelia Langford, John Sillitoe on behalf of the Wellcome Sanger Institute COVID-19 Surveillance Team |
| EPI_ISL_2913760 | Lighthouse Lab in Glasgow | Wellcome Sanger Institute for the COVID-19 Genomics UK (COG-UK) Consortium | Harper VanSteenhouse, Yumi Kasai, David Gray, Carol Clugston, Anna Dominiczak and Alex Alderton, Roberto Amato, Jeffrey Barrett, Sonia Goncalves, Ewan Harrison, David K. Jackson, Ian Johnston, Dominic Kwiatkowski, Cordelia Langford, John Sillitoe on behalf of the Wellcome Sanger Institute COVID-19 Surveillance Team |
| EPI_ISL_2913761 | Berkshire and Surrey Pathology Services Lighthouse Laboratory | Wellcome Sanger Institute for the COVID-19 Genomics UK (COG-UK) Consortium | Berkshire and Surrey Pathology Services Lighthouse Laboratory and Alex Alderton, Roberto Amato, Jeffrey Barrett, Sonia Goncalves, Ewan Harrison, David K. Jackson, Ian Johnston, Dominic Kwiatkowski, Cordelia Langford, John Sillitoe on behalf of the Wellcome Sanger Institute COVID-19 Surveillance Team |

| | | | |
|---|---|---|---|
| EPI_ISL_2913762 | Lighthouse Laboratory Plymouth | Wellcome Sanger Institute for the COVID-19 Genomics UK (COG-UK) Consortium | Lighthouse Laboratory Plymouth and Alex Alderton, Roberto Amato, Jeffrey Barrett, Sonia Goncalves, Ewan Harrison, David K. Jackson, Ian Johnston, Dominic Kwiatkowski, Cordelia Langford, John Sillitoe on behalf of the Wellcome Sanger Institute COVID-19 Surveillance Team |
| EPI_ISL_2913763 | Lighthouse Lab in Glasgow | Wellcome Sanger Institute for the COVID-19 Genomics UK (COG-UK) Consortium | Harper VanSteenhouse, Yumi Kasai, David Gray, Carol Clugston, Anna Dominiczak and Alex Alderton, Roberto Amato, Jeffrey Barrett, Sonia Goncalves, Ewan Harrison, David K. Jackson, Ian Johnston, Dominic Kwiatkowski, Cordelia Langford, John Sillitoe on behalf of the Wellcome Sanger Institute COVID-19 Surveillance Team |
| EPI_ISL_2913764 | Lighthouse Laboratory Plymouth | Wellcome Sanger Institute for the COVID-19 Genomics UK (COG-UK) Consortium | Lighthouse Laboratory Plymouth and Alex Alderton, Roberto Amato, Jeffrey Barrett, Sonia Goncalves, Ewan Harrison, David K. Jackson, Ian Johnston, Dominic Kwiatkowski, Cordelia Langford, John Sillitoe on behalf of the Wellcome Sanger Institute COVID-19 Surveillance Team |
| EPI_ISL_2913765 | Lighthouse Lab in Glasgow | Wellcome Sanger Institute for the COVID-19 Genomics UK (COG-UK) Consortium | Harper VanSteenhouse, Yumi Kasai, David Gray, Carol Clugston, Anna Dominiczak and Alex Alderton, Roberto Amato, Jeffrey Barrett, Sonia Goncalves, Ewan Harrison, David K. Jackson, Ian Johnston, Dominic Kwiatkowski, Cordelia Langford, John Sillitoe on behalf of the Wellcome Sanger Institute COVID-19 Surveillance Team |
| EPI_ISL_2913766, EPI_ISL_2913767 | Berkshire and Surrey Pathology Services Lighthouse Laboratory | Wellcome Sanger Institute for the COVID-19 Genomics UK (COG-UK) Consortium | Berkshire and Surrey Pathology Services Lighthouse Laboratory and Alex Alderton, Roberto Amato, Jeffrey Barrett, Sonia Goncalves, Ewan Harrison, David K. Jackson, Ian Johnston, Dominic Kwiatkowski, Cordelia Langford, John Sillitoe on behalf of the Wellcome Sanger Institute COVID-19 Surveillance Team |
| EPI_ISL_2913768, EPI_ISL_2913769 | Lighthouse Laboratory Plymouth | Wellcome Sanger Institute for the COVID-19 Genomics UK (COG-UK) Consortium | Lighthouse Laboratory Plymouth and Alex Alderton, Roberto Amato, Jeffrey Barrett, Sonia Goncalves, Ewan Harrison, David K. Jackson, Ian Johnston, Dominic Kwiatkowski, Cordelia Langford, John Sillitoe on behalf of the Wellcome Sanger Institute COVID-19 Surveillance Team |
| EPI_ISL_2913770 | Lighthouse Lab in Glasgow | Wellcome Sanger Institute for the COVID-19 Genomics UK (COG-UK) Consortium | Harper VanSteenhouse, Yumi Kasai, David Gray, Carol Clugston, Anna Dominiczak and Alex Alderton, Roberto Amato, Jeffrey Barrett, Sonia Goncalves, Ewan Harrison, David K. Jackson, Ian Johnston, Dominic Kwiatkowski, Cordelia Langford, John Sillitoe on behalf of the Wellcome Sanger Institute COVID-19 Surveillance Team |
| EPI_ISL_2913771 | Berkshire and Surrey Pathology Services Lighthouse Laboratory | Wellcome Sanger Institute for the COVID-19 Genomics UK (COG-UK) Consortium | Berkshire and Surrey Pathology Services Lighthouse Laboratory and Alex Alderton, Roberto Amato, Jeffrey Barrett, Sonia Goncalves, Ewan Harrison, David K. Jackson, Ian Johnston, Dominic Kwiatkowski, Cordelia Langford, John Sillitoe on behalf of the Wellcome Sanger Institute COVID-19 Surveillance Team |
| EPI_ISL_2913772 | Lighthouse Laboratory Plymouth | Wellcome Sanger Institute for the COVID-19 Genomics UK (COG-UK) Consortium | Lighthouse Laboratory Plymouth and Alex Alderton, Roberto Amato, Jeffrey Barrett, Sonia Goncalves, Ewan Harrison, David K. Jackson, Ian Johnston, Dominic Kwiatkowski, Cordelia Langford, John Sillitoe on behalf of the Wellcome Sanger Institute COVID-19 Surveillance Team |
| EPI_ISL_2913773, EPI_ISL_2913774 | Lighthouse Lab in Glasgow | Wellcome Sanger Institute for the COVID-19 Genomics UK (COG-UK) Consortium | Harper VanSteenhouse, Yumi Kasai, David Gray, Carol Clugston, Anna Dominiczak and Alex Alderton, Roberto Amato, Jeffrey Barrett, Sonia Goncalves, Ewan Harrison, David K. Jackson, Ian Johnston, Dominic Kwiatkowski, Cordelia Langford, John Sillitoe on behalf of the Wellcome Sanger Institute COVID-19 Surveillance Team |
| EPI_ISL_2913775, EPI_ISL_2913776 | Lighthouse Laboratory Plymouth | Wellcome Sanger Institute for the COVID-19 Genomics UK (COG-UK) Consortium | Lighthouse Laboratory Plymouth and Alex Alderton, Roberto Amato, Jeffrey Barrett, Sonia Goncalves, Ewan Harrison, David K. Jackson, Ian Johnston, Dominic Kwiatkowski, Cordelia Langford, John Sillitoe on behalf of the Wellcome Sanger Institute COVID-19 Surveillance Team |
| EPI_ISL_2913777 | Berkshire and Surrey Pathology Services Lighthouse Laboratory | Wellcome Sanger Institute for the COVID-19 Genomics UK (COG-UK) Consortium | Berkshire and Surrey Pathology Services Lighthouse Laboratory and Alex Alderton, Roberto Amato, Jeffrey Barrett, Sonia Goncalves, Ewan Harrison, David K. Jackson, Ian Johnston, Dominic Kwiatkowski, Cordelia Langford, John Sillitoe on behalf of the Wellcome Sanger Institute COVID-19 Surveillance Team |
| EPI_ISL_2913778 | Lighthouse Lab in Glasgow | Wellcome Sanger Institute for the COVID-19 Genomics UK (COG-UK) Consortium | Harper VanSteenhouse, Yumi Kasai, David Gray, Carol Clugston, Anna Dominiczak and Alex Alderton, Roberto Amato, Jeffrey Barrett, Sonia Goncalves, Ewan Harrison, David K. Jackson, Ian Johnston, Dominic Kwiatkowski, Cordelia Langford, John Sillitoe on behalf of the Wellcome Sanger Institute COVID-19 Surveillance Team |
| EPI_ISL_2913779 | Berkshire and Surrey Pathology Services Lighthouse Laboratory | Wellcome Sanger Institute for the COVID-19 Genomics UK (COG-UK) Consortium | Berkshire and Surrey Pathology Services Lighthouse Laboratory and Alex Alderton, Roberto Amato, Jeffrey Barrett, Sonia Goncalves, Ewan Harrison, David K. Jackson, Ian Johnston, Dominic Kwiatkowski, Cordelia Langford, John Sillitoe on behalf of the Wellcome Sanger Institute COVID-19 Surveillance Team |
| EPI_ISL_2913780 | Lighthouse Laboratory Plymouth | Wellcome Sanger Institute for the COVID-19 Genomics UK (COG-UK) Consortium | Lighthouse Laboratory Plymouth and Alex Alderton, Roberto Amato, Jeffrey Barrett, Sonia Goncalves, Ewan Harrison, David K. Jackson, Ian Johnston, Dominic Kwiatkowski, Cordelia Langford, John Sillitoe on behalf of the Wellcome Sanger Institute COVID-19 Surveillance Team |
| EPI_ISL_2913781 | Lighthouse Lab in Glasgow | Wellcome Sanger Institute for the COVID-19 Genomics UK (COG-UK) Consortium | Harper VanSteenhouse, Yumi Kasai, David Gray, Carol Clugston, Anna Dominiczak and Alex Alderton, Roberto Amato, Jeffrey Barrett, Sonia Goncalves, Ewan Harrison, David K. Jackson, Ian Johnston, Dominic Kwiatkowski, Cordelia Langford, John Sillitoe on behalf of the Wellcome Sanger Institute COVID-19 Surveillance Team |
| EPI_ISL_2913782, EPI_ISL_2913783, EPI_ISL_2913784, EPI_ISL_2913785, EPI_ISL_2913786, EPI_ISL_2913787, EPI_ISL_2913788, EPI_ISL_2913789 | Lighthouse Laboratory Plymouth | Wellcome Sanger Institute for the COVID-19 Genomics UK (COG-UK) Consortium | Lighthouse Laboratory Plymouth and Alex Alderton, Roberto Amato, Jeffrey Barrett, Sonia Goncalves, Ewan Harrison, David K. Jackson, Ian Johnston, Dominic Kwiatkowski, Cordelia Langford, John Sillitoe on behalf of the Wellcome Sanger Institute COVID-19 Surveillance Team |
| EPI_ISL_2913790 | Lighthouse Lab in Glasgow | Wellcome Sanger Institute for the COVID-19 Genomics UK (COG-UK) Consortium | Harper VanSteenhouse, Yumi Kasai, David Gray, Carol Clugston, Anna Dominiczak and Alex Alderton, Roberto Amato, Jeffrey Barrett, Sonia Goncalves, Ewan Harrison, David K. Jackson, Ian Johnston, Dominic Kwiatkowski, Cordelia Langford, John Sillitoe on behalf of the Wellcome Sanger Institute COVID-19 Surveillance Team |
| EPI_ISL_2913791 | Lighthouse Laboratory Plymouth | Wellcome Sanger Institute for the COVID-19 Genomics UK (COG-UK) Consortium | Lighthouse Laboratory Plymouth and Alex Alderton, Roberto Amato, Jeffrey Barrett, Sonia Goncalves, Ewan Harrison, David K. Jackson, Ian Johnston, Dominic Kwiatkowski, Cordelia Langford, John Sillitoe on behalf of the Wellcome Sanger Institute COVID-19 Surveillance Team |
| EPI_ISL_2913792 | Berkshire and Surrey Pathology Services Lighthouse Laboratory | Wellcome Sanger Institute for the COVID-19 Genomics UK (COG-UK) Consortium | Berkshire and Surrey Pathology Services Lighthouse Laboratory and Alex Alderton, Roberto Amato, Jeffrey Barrett, Sonia Goncalves, Ewan Harrison, David K. Jackson, Ian Johnston, Dominic Kwiatkowski, Cordelia Langford, John Sillitoe on behalf of the Wellcome Sanger Institute COVID-19 Surveillance Team |
| EPI_ISL_2913793 | Lighthouse Laboratory Plymouth | Wellcome Sanger Institute for the COVID-19 Genomics UK (COG-UK) Consortium | Lighthouse Laboratory Plymouth and Alex Alderton, Roberto Amato, Jeffrey Barrett, Sonia Goncalves, Ewan Harrison, David K. Jackson, Ian Johnston, Dominic Kwiatkowski, Cordelia Langford, John Sillitoe on behalf of the Wellcome Sanger Institute COVID-19 Surveillance Team |
| EPI_ISL_2913794 | Berkshire and Surrey Pathology Services Lighthouse Laboratory | Wellcome Sanger Institute for the COVID-19 Genomics UK (COG-UK) Consortium | Berkshire and Surrey Pathology Services Lighthouse Laboratory and Alex Alderton, Roberto Amato, Jeffrey Barrett, Sonia Goncalves, Ewan Harrison, David K. Jackson, Ian Johnston, Dominic Kwiatkowski, Cordelia Langford, John Sillitoe on behalf of the Wellcome Sanger Institute COVID-19 Surveillance Team |
| EPI_ISL_2913795, EPI_ISL_2913796 | Lighthouse Laboratory Plymouth | Wellcome Sanger Institute for the COVID-19 Genomics UK (COG-UK) Consortium | Lighthouse Laboratory Plymouth and Alex Alderton, Roberto Amato, Jeffrey Barrett, Sonia Goncalves, Ewan Harrison, David K. Jackson, Ian Johnston, Dominic Kwiatkowski, Cordelia Langford, John Sillitoe on behalf of the Wellcome Sanger Institute COVID-19 Surveillance Team |
| EPI_ISL_2913797, EPI_ISL_2913798 | Lighthouse Lab in Glasgow | Wellcome Sanger Institute for the COVID-19 Genomics UK (COG-UK) Consortium | Harper VanSteenhouse, Yumi Kasai, David Gray, Carol Clugston, Anna Dominiczak and Alex Alderton, Roberto Amato, Jeffrey Barrett, Sonia Goncalves, Ewan Harrison, David K. Jackson, Ian Johnston, Dominic Kwiatkowski, Cordelia Langford, John Sillitoe on behalf of the Wellcome Sanger Institute COVID-19 Surveillance Team |
| EPI_ISL_2913799 | Berkshire and Surrey Pathology Services Lighthouse Laboratory | Wellcome Sanger Institute for the COVID-19 Genomics UK (COG-UK) Consortium | Berkshire and Surrey Pathology Services Lighthouse Laboratory and Alex Alderton, Roberto Amato, Jeffrey Barrett, Sonia Goncalves, Ewan Harrison, David K. Jackson, Ian Johnston, Dominic Kwiatkowski, Cordelia Langford, John Sillitoe on behalf of the Wellcome Sanger Institute COVID-19 Surveillance Team |
| EPI_ISL_2913800, EPI_ISL_2913801, EPI_ISL_2913802 | Lighthouse Laboratory Plymouth | Wellcome Sanger Institute for the COVID-19 Genomics UK (COG-UK) Consortium | Lighthouse Laboratory Plymouth and Alex Alderton, Roberto Amato, Jeffrey Barrett, Sonia Goncalves, Ewan Harrison, David K. Jackson, Ian Johnston, Dominic Kwiatkowski, Cordelia Langford, John Sillitoe on behalf of the Wellcome Sanger Institute COVID-19 Surveillance Team |
| EPI_ISL_2913803, EPI_ISL_2913804, EPI_ISL_2913805 | Berkshire and Surrey Pathology Services Lighthouse Laboratory | Wellcome Sanger Institute for the COVID-19 Genomics UK (COG-UK) Consortium | Berkshire and Surrey Pathology Services Lighthouse Laboratory and Alex Alderton, Roberto Amato, Jeffrey Barrett, Sonia Goncalves, Ewan Harrison, David K. Jackson, Ian Johnston, Dominic Kwiatkowski, Cordelia Langford, John Sillitoe on behalf of the Wellcome Sanger Institute COVID-19 Surveillance Team |
| EPI_ISL_2913806, EPI_ISL_2913807 | Lighthouse Laboratory Plymouth | Wellcome Sanger Institute for the COVID-19 Genomics UK (COG-UK) Consortium | Lighthouse Laboratory Plymouth and Alex Alderton, Roberto Amato, Jeffrey Barrett, Sonia Goncalves, Ewan Harrison, David K. Jackson, Ian Johnston, Dominic Kwiatkowski, Cordelia Langford, John Sillitoe on behalf of the Wellcome Sanger Institute COVID-19 Surveillance Team |
| EPI_ISL_2913808 | Lighthouse Lab in Glasgow | Wellcome Sanger Institute for the COVID-19 Genomics UK (COG-UK) Consortium | Harper VanSteenhouse, Yumi Kasai, David Gray, Carol Clugston, Anna Dominiczak and Alex Alderton, Roberto Amato, Jeffrey Barrett, Sonia Goncalves, Ewan Harrison, David K. Jackson, Ian Johnston, Dominic Kwiatkowski, Cordelia Langford, John Sillitoe on behalf of the Wellcome Sanger Institute COVID-19 Surveillance Team |
| EPI_ISL_2913809 | Lighthouse Laboratory Plymouth | Wellcome Sanger Institute for the COVID-19 Genomics UK (COG-UK) Consortium | Lighthouse Laboratory Plymouth and Alex Alderton, Roberto Amato, Jeffrey Barrett, Sonia Goncalves, Ewan Harrison, David K. Jackson, Ian Johnston, Dominic Kwiatkowski, Cordelia Langford, John Sillitoe on behalf of the Wellcome Sanger Institute COVID-19 Surveillance Team |
| EPI_ISL_2913810 | Berkshire and Surrey Pathology Services Lighthouse Laboratory | Wellcome Sanger Institute for the COVID-19 Genomics UK (COG-UK) Consortium | Berkshire and Surrey Pathology Services Lighthouse Laboratory and Alex Alderton, Roberto Amato, Jeffrey Barrett, Sonia Goncalves, Ewan Harrison, David K. Jackson, Ian Johnston, Dominic Kwiatkowski, Cordelia Langford, John Sillitoe on behalf of the Wellcome Sanger Institute COVID-19 Surveillance Team |
| EPI_ISL_2913811 | Lighthouse Lab in Glasgow | Wellcome Sanger Institute for the COVID-19 Genomics UK (COG-UK) Consortium | Harper VanSteenhouse, Yumi Kasai, David Gray, Carol Clugston, Anna Dominiczak and Alex Alderton, Roberto Amato, Jeffrey Barrett, Sonia Goncalves, Ewan Harrison, David K. Jackson, Ian Johnston, Dominic Kwiatkowski, Cordelia Langford, John Sillitoe on behalf of the Wellcome Sanger Institute COVID-19 Surveillance Team |
| EPI_ISL_2913812 | Berkshire and Surrey Pathology Services Lighthouse Laboratory | Wellcome Sanger Institute for the COVID-19 Genomics UK (COG-UK) Consortium | Berkshire and Surrey Pathology Services Lighthouse Laboratory and Alex Alderton, Roberto Amato, Jeffrey Barrett, Sonia Goncalves, Ewan Harrison, David K. Jackson, Ian Johnston, Dominic Kwiatkowski, Cordelia Langford, John Sillitoe on behalf of the Wellcome Sanger Institute COVID-19 Surveillance Team |
| EPI_ISL_2913813, EPI_ISL_2913814, EPI_ISL_2913815 | Lighthouse Laboratory Plymouth | Wellcome Sanger Institute for the COVID-19 Genomics UK (COG-UK) Consortium | Lighthouse Laboratory Plymouth and Alex Alderton, Roberto Amato, Jeffrey Barrett, Sonia Goncalves, Ewan Harrison, David K. Jackson, Ian Johnston, Dominic Kwiatkowski, Cordelia Langford, John Sillitoe on behalf of the Wellcome Sanger Institute COVID-19 Surveillance Team |
| EPI_ISL_2913816 | Lighthouse Lab in Glasgow | Wellcome Sanger Institute for the COVID-19 Genomics UK (COG-UK) Consortium | Harper VanSteenhouse, Yumi Kasai, David Gray, Carol Clugston, Anna Dominiczak and Alex Alderton, Roberto Amato, Jeffrey Barrett, Sonia Goncalves, Ewan Harrison, David K. Jackson, Ian Johnston, Dominic Kwiatkowski, Cordelia Langford, John Sillitoe on behalf of the Wellcome Sanger Institute COVID-19 Surveillance Team |
| EPI_ISL_2913817, EPI_ISL_2913818 | Lighthouse Laboratory Plymouth | Wellcome Sanger Institute for the COVID-19 Genomics UK (COG-UK) Consortium | Lighthouse Laboratory Plymouth and Alex Alderton, Roberto Amato, Jeffrey Barrett, Sonia Goncalves, Ewan Harrison, David K. Jackson, Ian Johnston, Dominic Kwiatkowski, Cordelia Langford, John Sillitoe on behalf of the Wellcome Sanger Institute COVID-19 Surveillance Team |
| EPI_ISL_2913819 | Lighthouse Lab in Glasgow | Wellcome Sanger Institute for the COVID-19 Genomics UK (COG-UK) Consortium | Harper VanSteenhouse, Yumi Kasai, David Gray, Carol Clugston, Anna Dominiczak and Alex Alderton, Roberto Amato, Jeffrey Barrett, Sonia Goncalves, Ewan Harrison, David K. Jackson, Ian Johnston, Dominic Kwiatkowski, Cordelia Langford, John Sillitoe on behalf of the Wellcome Sanger Institute COVID-19 Surveillance Team |
| EPI_ISL_2913820, EPI_ISL_2913821 | Berkshire and Surrey Pathology Services Lighthouse Laboratory | Wellcome Sanger Institute for the COVID-19 Genomics UK (COG-UK) Consortium | Berkshire and Surrey Pathology Services Lighthouse Laboratory and Alex Alderton, Roberto Amato, Jeffrey Barrett, Sonia Goncalves, Ewan Harrison, David K. Jackson, Ian Johnston, Dominic Kwiatkowski, Cordelia Langford, John Sillitoe on behalf of the Wellcome Sanger Institute COVID-19 Surveillance Team |
| EPI_ISL_2913822 | Lighthouse Laboratory Plymouth | Wellcome Sanger Institute for the COVID-19 Genomics UK (COG-UK) Consortium | Lighthouse Laboratory Plymouth and Alex Alderton, Roberto Amato, Jeffrey Barrett, Sonia Goncalves, Ewan Harrison, David K. Jackson, Ian Johnston, Dominic Kwiatkowski, Cordelia Langford, John Sillitoe on behalf of the Wellcome Sanger Institute COVID-19 Surveillance Team |
| EPI_ISL_2913823, EPI_ISL_2913824 | Berkshire and Surrey Pathology Services Lighthouse Laboratory | Wellcome Sanger Institute for the COVID-19 Genomics UK (COG-UK) Consortium | Berkshire and Surrey Pathology Services Lighthouse Laboratory and Alex Alderton, Roberto Amato, Jeffrey Barrett, Sonia Goncalves, Ewan Harrison, David K. Jackson, Ian Johnston, Dominic Kwiatkowski, Cordelia Langford, John Sillitoe on behalf of the Wellcome Sanger Institute COVID-19 Surveillance Team |
| EPI_ISL_2913825, EPI_ISL_2913826, EPI_ISL_2913827 | Lighthouse Laboratory Plymouth | Wellcome Sanger Institute for the COVID-19 Genomics UK (COG-UK) Consortium | Lighthouse Laboratory Plymouth and Alex Alderton, Roberto Amato, Jeffrey Barrett, Sonia Goncalves, Ewan Harrison, David K. Jackson, Ian Johnston, Dominic Kwiatkowski, Cordelia Langford, John Sillitoe on behalf of the Wellcome Sanger Institute COVID-19 Surveillance Team |

| | | | |
|---|---|---|---|
| EPI_ISL_2913828 | Berkshire and Surrey Pathology Services Lighthouse Laboratory | Wellcome Sanger Institute for the COVID-19 Genomics UK (COG-UK) Consortium | Berkshire and Surrey Pathology Services Lighthouse Laboratory and Alex Alderton, Roberto Amato, Jeffrey Barrett, Sonia Goncalves, Ewan Harrison, David K. Jackson, Ian Johnston, Dominic Kwiatkowski, Cordelia Langford, John Sillitoe on behalf of the Wellcome Sanger Institute COVID-19 Surveillance Team |
| EPI_ISL_2913829 | Lighthouse Laboratory Plymouth | Wellcome Sanger Institute for the COVID-19 Genomics UK (COG-UK) Consortium | Lighthouse Laboratory Plymouth and Alex Alderton, Roberto Amato, Jeffrey Barrett, Sonia Goncalves, Ewan Harrison, David K. Jackson, Ian Johnston, Dominic Kwiatkowski, Cordelia Langford, John Sillitoe on behalf of the Wellcome Sanger Institute COVID-19 Surveillance Team |
| EPI_ISL_2913830 | Berkshire and Surrey Pathology Services Lighthouse Laboratory | Wellcome Sanger Institute for the COVID-19 Genomics UK (COG-UK) Consortium | Berkshire and Surrey Pathology Services Lighthouse Laboratory and Alex Alderton, Roberto Amato, Jeffrey Barrett, Sonia Goncalves, Ewan Harrison, David K. Jackson, Ian Johnston, Dominic Kwiatkowski, Cordelia Langford, John Sillitoe on behalf of the Wellcome Sanger Institute COVID-19 Surveillance Team |
| EPI_ISL_2913831, EPI_ISL_2913832, EPI_ISL_2913833, EPI_ISL_2913834 | Lighthouse Laboratory Plymouth | Wellcome Sanger Institute for the COVID-19 Genomics UK (COG-UK) Consortium | Lighthouse Laboratory Plymouth and Alex Alderton, Roberto Amato, Jeffrey Barrett, Sonia Goncalves, Ewan Harrison, David K. Jackson, Ian Johnston, Dominic Kwiatkowski, Cordelia Langford, John Sillitoe on behalf of the Wellcome Sanger Institute COVID-19 Surveillance Team |
| EPI_ISL_2913835 | Berkshire and Surrey Pathology Services Lighthouse Laboratory | Wellcome Sanger Institute for the COVID-19 Genomics UK (COG-UK) Consortium | Berkshire and Surrey Pathology Services Lighthouse Laboratory and Alex Alderton, Roberto Amato, Jeffrey Barrett, Sonia Goncalves, Ewan Harrison, David K. Jackson, Ian Johnston, Dominic Kwiatkowski, Cordelia Langford, John Sillitoe on behalf of the Wellcome Sanger Institute COVID-19 Surveillance Team |
| EPI_ISL_2913836 | Lighthouse Laboratory Plymouth | Wellcome Sanger Institute for the COVID-19 Genomics UK (COG-UK) Consortium | Lighthouse Laboratory Plymouth and Alex Alderton, Roberto Amato, Jeffrey Barrett, Sonia Goncalves, Ewan Harrison, David K. Jackson, Ian Johnston, Dominic Kwiatkowski, Cordelia Langford, John Sillitoe on behalf of the Wellcome Sanger Institute COVID-19 Surveillance Team |
| EPI_ISL_2913837 | Berkshire and Surrey Pathology Services Lighthouse Laboratory | Wellcome Sanger Institute for the COVID-19 Genomics UK (COG-UK) Consortium | Berkshire and Surrey Pathology Services Lighthouse Laboratory and Alex Alderton, Roberto Amato, Jeffrey Barrett, Sonia Goncalves, Ewan Harrison, David K. Jackson, Ian Johnston, Dominic Kwiatkowski, Cordelia Langford, John Sillitoe on behalf of the Wellcome Sanger Institute COVID-19 Surveillance Team |
| EPI_ISL_2913838, EPI_ISL_2913839, EPI_ISL_2913840 | Lighthouse Laboratory Plymouth | Wellcome Sanger Institute for the COVID-19 Genomics UK (COG-UK) Consortium | Lighthouse Laboratory Plymouth and Alex Alderton, Roberto Amato, Jeffrey Barrett, Sonia Goncalves, Ewan Harrison, David K. Jackson, Ian Johnston, Dominic Kwiatkowski, Cordelia Langford, John Sillitoe on behalf of the Wellcome Sanger Institute COVID-19 Surveillance Team |
| EPI_ISL_2913841, EPI_ISL_2913842, EPI_ISL_2913843 | Lighthouse Lab in Glasgow | Wellcome Sanger Institute for the COVID-19 Genomics UK (COG-UK) Consortium | Harper VanSteenhouse, Yumi Kasai, David Gray, Carol Clugston, Anna Dominiczak and Alex Alderton, Roberto Amato, Jeffrey Barrett, Sonia Goncalves, Ewan Harrison, David K. Jackson, Ian Johnston, Dominic Kwiatkowski, Cordelia Langford, John Sillitoe on behalf of the Wellcome Sanger Institute COVID-19 Surveillance Team |
| EPI_ISL_2913844, EPI_ISL_2913845 | Berkshire and Surrey Pathology Services Lighthouse Laboratory | Wellcome Sanger Institute for the COVID-19 Genomics UK (COG-UK) Consortium | Berkshire and Surrey Pathology Services Lighthouse Laboratory and Alex Alderton, Roberto Amato, Jeffrey Barrett, Sonia Goncalves, Ewan Harrison, David K. Jackson, Ian Johnston, Dominic Kwiatkowski, Cordelia Langford, John Sillitoe on behalf of the Wellcome Sanger Institute COVID-19 Surveillance Team |
| EPI_ISL_2913846 | Lighthouse Laboratory Plymouth | Wellcome Sanger Institute for the COVID-19 Genomics UK (COG-UK) Consortium | Lighthouse Laboratory Plymouth and Alex Alderton, Roberto Amato, Jeffrey Barrett, Sonia Goncalves, Ewan Harrison, David K. Jackson, Ian Johnston, Dominic Kwiatkowski, Cordelia Langford, John Sillitoe on behalf of the Wellcome Sanger Institute COVID-19 Surveillance Team |
| EPI_ISL_2913847 | Berkshire and Surrey Pathology Services Lighthouse Laboratory | Wellcome Sanger Institute for the COVID-19 Genomics UK (COG-UK) Consortium | Berkshire and Surrey Pathology Services Lighthouse Laboratory and Alex Alderton, Roberto Amato, Jeffrey Barrett, Sonia Goncalves, Ewan Harrison, David K. Jackson, Ian Johnston, Dominic Kwiatkowski, Cordelia Langford, John Sillitoe on behalf of the Wellcome Sanger Institute COVID-19 Surveillance Team |
| EPI_ISL_2913848 | Lighthouse Laboratory Plymouth | Wellcome Sanger Institute for the COVID-19 Genomics UK (COG-UK) Consortium | Lighthouse Laboratory Plymouth and Alex Alderton, Roberto Amato, Jeffrey Barrett, Sonia Goncalves, Ewan Harrison, David K. Jackson, Ian Johnston, Dominic Kwiatkowski, Cordelia Langford, John Sillitoe on behalf of the Wellcome Sanger Institute COVID-19 Surveillance Team |
| EPI_ISL_2913849 | Berkshire and Surrey Pathology Services Lighthouse Laboratory | Wellcome Sanger Institute for the COVID-19 Genomics UK (COG-UK) Consortium | Berkshire and Surrey Pathology Services Lighthouse Laboratory and Alex Alderton, Roberto Amato, Jeffrey Barrett, Sonia Goncalves, Ewan Harrison, David K. Jackson, Ian Johnston, Dominic Kwiatkowski, Cordelia Langford, John Sillitoe on behalf of the Wellcome Sanger Institute COVID-19 Surveillance Team |
| EPI_ISL_2913850, EPI_ISL_2913851 | Lighthouse Laboratory Plymouth | Wellcome Sanger Institute for the COVID-19 Genomics UK (COG-UK) Consortium | Lighthouse Laboratory Plymouth and Alex Alderton, Roberto Amato, Jeffrey Barrett, Sonia Goncalves, Ewan Harrison, David K. Jackson, Ian Johnston, Dominic Kwiatkowski, Cordelia Langford, John Sillitoe on behalf of the Wellcome Sanger Institute COVID-19 Surveillance Team |
| EPI_ISL_2913852, EPI_ISL_2913853 | Berkshire and Surrey Pathology Services Lighthouse Laboratory | Wellcome Sanger Institute for the COVID-19 Genomics UK (COG-UK) Consortium | Berkshire and Surrey Pathology Services Lighthouse Laboratory and Alex Alderton, Roberto Amato, Jeffrey Barrett, Sonia Goncalves, Ewan Harrison, David K. Jackson, Ian Johnston, Dominic Kwiatkowski, Cordelia Langford, John Sillitoe on behalf of the Wellcome Sanger Institute COVID-19 Surveillance Team |
| EPI_ISL_2913854 | Lighthouse Lab in Glasgow | Wellcome Sanger Institute for the COVID-19 Genomics UK (COG-UK) Consortium | Harper VanSteenhouse, Yumi Kasai, David Gray, Carol Clugston, Anna Dominiczak and Alex Alderton, Roberto Amato, Jeffrey Barrett, Sonia Goncalves, Ewan Harrison, David K. Jackson, Ian Johnston, Dominic Kwiatkowski, Cordelia Langford, John Sillitoe on behalf of the Wellcome Sanger Institute COVID-19 Surveillance Team |
| EPI_ISL_2913855 | Berkshire and Surrey Pathology Services Lighthouse Laboratory | Wellcome Sanger Institute for the COVID-19 Genomics UK (COG-UK) Consortium | Berkshire and Surrey Pathology Services Lighthouse Laboratory and Alex Alderton, Roberto Amato, Jeffrey Barrett, Sonia Goncalves, Ewan Harrison, David K. Jackson, Ian Johnston, Dominic Kwiatkowski, Cordelia Langford, John Sillitoe on behalf of the Wellcome Sanger Institute COVID-19 Surveillance Team |
| EPI_ISL_2913856, EPI_ISL_2913857, EPI_ISL_2913858 | Lighthouse Lab in Glasgow | Wellcome Sanger Institute for the COVID-19 Genomics UK (COG-UK) Consortium | Harper VanSteenhouse, Yumi Kasai, David Gray, Carol Clugston, Anna Dominiczak and Alex Alderton, Roberto Amato, Jeffrey Barrett, Sonia Goncalves, Ewan Harrison, David K. Jackson, Ian Johnston, Dominic Kwiatkowski, Cordelia Langford, John Sillitoe on behalf of the Wellcome Sanger Institute COVID-19 Surveillance Team |
| EPI_ISL_2913859 | Lighthouse Laboratory Plymouth | Wellcome Sanger Institute for the COVID-19 Genomics UK (COG-UK) Consortium | Lighthouse Laboratory Plymouth and Alex Alderton, Roberto Amato, Jeffrey Barrett, Sonia Goncalves, Ewan Harrison, David K. Jackson, Ian Johnston, Dominic Kwiatkowski, Cordelia Langford, John Sillitoe on behalf of the Wellcome Sanger Institute COVID-19 Surveillance Team |
| EPI_ISL_2913860, EPI_ISL_2913861, EPI_ISL_2913862 | Lighthouse Lab in Glasgow | Wellcome Sanger Institute for the COVID-19 Genomics UK (COG-UK) Consortium | Harper VanSteenhouse, Yumi Kasai, David Gray, Carol Clugston, Anna Dominiczak and Alex Alderton, Roberto Amato, Jeffrey Barrett, Sonia Goncalves, Ewan Harrison, David K. Jackson, Ian Johnston, Dominic Kwiatkowski, Cordelia Langford, John Sillitoe on behalf of the Wellcome Sanger Institute COVID-19 Surveillance Team |
| EPI_ISL_2913863 | Berkshire and Surrey Pathology Services Lighthouse Laboratory | Wellcome Sanger Institute for the COVID-19 Genomics UK (COG-UK) Consortium | Berkshire and Surrey Pathology Services Lighthouse Laboratory and Alex Alderton, Roberto Amato, Jeffrey Barrett, Sonia Goncalves, Ewan Harrison, David K. Jackson, Ian Johnston, Dominic Kwiatkowski, Cordelia Langford, John Sillitoe on behalf of the Wellcome Sanger Institute COVID-19 Surveillance Team |
| EPI_ISL_2913864 | Lighthouse Lab in Glasgow | Wellcome Sanger Institute for the COVID-19 Genomics UK (COG-UK) Consortium | Harper VanSteenhouse, Yumi Kasai, David Gray, Carol Clugston, Anna Dominiczak and Alex Alderton, Roberto Amato, Jeffrey Barrett, Sonia Goncalves, Ewan Harrison, David K. Jackson, Ian Johnston, Dominic Kwiatkowski, Cordelia Langford, John Sillitoe on behalf of the Wellcome Sanger Institute COVID-19 Surveillance Team |
| EPI_ISL_2913865 | Lighthouse Laboratory Plymouth | Wellcome Sanger Institute for the COVID-19 Genomics UK (COG-UK) Consortium | Lighthouse Laboratory Plymouth and Alex Alderton, Roberto Amato, Jeffrey Barrett, Sonia Goncalves, Ewan Harrison, David K. Jackson, Ian Johnston, Dominic Kwiatkowski, Cordelia Langford, John Sillitoe on behalf of the Wellcome Sanger Institute COVID-19 Surveillance Team |
| EPI_ISL_2913866 | Lighthouse Lab in Glasgow | Wellcome Sanger Institute for the COVID-19 Genomics UK (COG-UK) Consortium | Harper VanSteenhouse, Yumi Kasai, David Gray, Carol Clugston, Anna Dominiczak and Alex Alderton, Roberto Amato, Jeffrey Barrett, Sonia Goncalves, Ewan Harrison, David K. Jackson, Ian Johnston, Dominic Kwiatkowski, Cordelia Langford, John Sillitoe on behalf of the Wellcome Sanger Institute COVID-19 Surveillance Team |
| EPI_ISL_2913867 | Lighthouse Laboratory Plymouth | Wellcome Sanger Institute for the COVID-19 Genomics UK (COG-UK) Consortium | Lighthouse Laboratory Plymouth and Alex Alderton, Roberto Amato, Jeffrey Barrett, Sonia Goncalves, Ewan Harrison, David K. Jackson, Ian Johnston, Dominic Kwiatkowski, Cordelia Langford, John Sillitoe on behalf of the Wellcome Sanger Institute COVID-19 Surveillance Team |
| EPI_ISL_2913868 | Lighthouse Lab in Glasgow | Wellcome Sanger Institute for the COVID-19 Genomics UK (COG-UK) Consortium | Harper VanSteenhouse, Yumi Kasai, David Gray, Carol Clugston, Anna Dominiczak and Alex Alderton, Roberto Amato, Jeffrey Barrett, Sonia Goncalves, Ewan Harrison, David K. Jackson, Ian Johnston, Dominic Kwiatkowski, Cordelia Langford, John Sillitoe on behalf of the Wellcome Sanger Institute COVID-19 Surveillance Team |
| EPI_ISL_2913869, EPI_ISL_2913870, EPI_ISL_2913871, EPI_ISL_2913872, EPI_ISL_2913873, EPI_ISL_2913874 | Lighthouse Laboratory Plymouth | Wellcome Sanger Institute for the COVID-19 Genomics UK (COG-UK) Consortium | Lighthouse Laboratory Plymouth and Alex Alderton, Roberto Amato, Jeffrey Barrett, Sonia Goncalves, Ewan Harrison, David K. Jackson, Ian Johnston, Dominic Kwiatkowski, Cordelia Langford, John Sillitoe on behalf of the Wellcome Sanger Institute COVID-19 Surveillance Team |
| EPI_ISL_2913875 | Lighthouse Lab in Glasgow | Wellcome Sanger Institute for the COVID-19 Genomics UK (COG-UK) Consortium | Harper VanSteenhouse, Yumi Kasai, David Gray, Carol Clugston, Anna Dominiczak and Alex Alderton, Roberto Amato, Jeffrey Barrett, Sonia Goncalves, Ewan Harrison, David K. Jackson, Ian Johnston, Dominic Kwiatkowski, Cordelia Langford, John Sillitoe on behalf of the Wellcome Sanger Institute COVID-19 Surveillance Team |
| EPI_ISL_2913876, EPI_ISL_2913877 | Lighthouse Laboratory Plymouth | Wellcome Sanger Institute for the COVID-19 Genomics UK (COG-UK) Consortium | Lighthouse Laboratory Plymouth and Alex Alderton, Roberto Amato, Jeffrey Barrett, Sonia Goncalves, Ewan Harrison, David K. Jackson, Ian Johnston, Dominic Kwiatkowski, Cordelia Langford, John Sillitoe on behalf of the Wellcome Sanger Institute COVID-19 Surveillance Team |
| EPI_ISL_2913878 | Lighthouse Lab in Glasgow | Wellcome Sanger Institute for the COVID-19 Genomics UK (COG-UK) Consortium | Harper VanSteenhouse, Yumi Kasai, David Gray, Carol Clugston, Anna Dominiczak and Alex Alderton, Roberto Amato, Jeffrey Barrett, Sonia Goncalves, Ewan Harrison, David K. Jackson, Ian Johnston, Dominic Kwiatkowski, Cordelia Langford, John Sillitoe on behalf of the Wellcome Sanger Institute COVID-19 Surveillance Team |
| EPI_ISL_2913879 | Lighthouse Laboratory Plymouth | Wellcome Sanger Institute for the COVID-19 Genomics UK (COG-UK) Consortium | Lighthouse Laboratory Plymouth and Alex Alderton, Roberto Amato, Jeffrey Barrett, Sonia Goncalves, Ewan Harrison, David K. Jackson, Ian Johnston, Dominic Kwiatkowski, Cordelia Langford, John Sillitoe on behalf of the Wellcome Sanger Institute COVID-19 Surveillance Team |
| EPI_ISL_2913880, EPI_ISL_2913881 | Lighthouse Lab in Glasgow | Wellcome Sanger Institute for the COVID-19 Genomics UK (COG-UK) Consortium | Harper VanSteenhouse, Yumi Kasai, David Gray, Carol Clugston, Anna Dominiczak and Alex Alderton, Roberto Amato, Jeffrey Barrett, Sonia Goncalves, Ewan Harrison, David K. Jackson, Ian Johnston, Dominic Kwiatkowski, Cordelia Langford, John Sillitoe on behalf of the Wellcome Sanger Institute COVID-19 Surveillance Team |
| EPI_ISL_2913882 | Berkshire and Surrey Pathology Services Lighthouse Laboratory | Wellcome Sanger Institute for the COVID-19 Genomics UK (COG-UK) Consortium | Berkshire and Surrey Pathology Services Lighthouse Laboratory and Alex Alderton, Roberto Amato, Jeffrey Barrett, Sonia Goncalves, Ewan Harrison, David K. Jackson, Ian Johnston, Dominic Kwiatkowski, Cordelia Langford, John Sillitoe on behalf of the Wellcome Sanger Institute COVID-19 Surveillance Team |
| EPI_ISL_2913883 | Lighthouse Laboratory Plymouth | Wellcome Sanger Institute for the COVID-19 Genomics UK (COG-UK) Consortium | Lighthouse Laboratory Plymouth and Alex Alderton, Roberto Amato, Jeffrey Barrett, Sonia Goncalves, Ewan Harrison, David K. Jackson, Ian Johnston, Dominic Kwiatkowski, Cordelia Langford, John Sillitoe on behalf of the Wellcome Sanger Institute COVID-19 Surveillance Team |
| EPI_ISL_2913884, EPI_ISL_2913885 | Berkshire and Surrey Pathology Services Lighthouse Laboratory | Wellcome Sanger Institute for the COVID-19 Genomics UK (COG-UK) Consortium | Berkshire and Surrey Pathology Services Lighthouse Laboratory and Alex Alderton, Roberto Amato, Jeffrey Barrett, Sonia Goncalves, Ewan Harrison, David K. Jackson, Ian Johnston, Dominic Kwiatkowski, Cordelia Langford, John Sillitoe on behalf of the Wellcome Sanger Institute COVID-19 Surveillance Team |
| EPI_ISL_2913886 | Lighthouse Lab in Glasgow | Wellcome Sanger Institute for the COVID-19 Genomics UK (COG-UK) Consortium | Harper VanSteenhouse, Yumi Kasai, David Gray, Carol Clugston, Anna Dominiczak and Alex Alderton, Roberto Amato, Jeffrey Barrett, Sonia Goncalves, Ewan Harrison, David K. Jackson, Ian Johnston, Dominic Kwiatkowski, Cordelia Langford, John Sillitoe on behalf of the Wellcome Sanger Institute COVID-19 Surveillance Team |
| EPI_ISL_2913887, EPI_ISL_2913888 | Lighthouse Laboratory Plymouth | Wellcome Sanger Institute for the COVID-19 Genomics UK (COG-UK) Consortium | Lighthouse Laboratory Plymouth and Alex Alderton, Roberto Amato, Jeffrey Barrett, Sonia Goncalves, Ewan Harrison, David K. Jackson, Ian Johnston, Dominic Kwiatkowski, Cordelia Langford, John Sillitoe on behalf of the Wellcome Sanger Institute COVID-19 Surveillance Team |
| EPI_ISL_2913889 | Berkshire and Surrey Pathology Services Lighthouse Laboratory | Wellcome Sanger Institute for the COVID-19 Genomics UK (COG-UK) Consortium | Berkshire and Surrey Pathology Services Lighthouse Laboratory and Alex Alderton, Roberto Amato, Jeffrey Barrett, Sonia Goncalves, Ewan Harrison, David K. Jackson, Ian Johnston, Dominic Kwiatkowski, Cordelia Langford, John Sillitoe on behalf of the Wellcome Sanger Institute COVID-19 Surveillance Team |
| EPI_ISL_2913890, EPI_ISL_2913891, EPI_ISL_2913892 | Lighthouse Laboratory Plymouth | Wellcome Sanger Institute for the COVID-19 Genomics UK (COG-UK) Consortium | Lighthouse Laboratory Plymouth and Alex Alderton, Roberto Amato, Jeffrey Barrett, Sonia Goncalves, Ewan Harrison, David K. Jackson, Ian Johnston, Dominic Kwiatkowski, Cordelia Langford, John Sillitoe on behalf of the Wellcome Sanger Institute COVID-19 Surveillance Team |
| EPI_ISL_2913893 | Berkshire and Surrey Pathology Services Lighthouse Laboratory | Wellcome Sanger Institute for the COVID-19 Genomics UK (COG-UK) | Berkshire and Surrey Pathology Services Lighthouse Laboratory and Alex Alderton, Roberto Amato, Jeffrey Barrett, Sonia Goncalves, Ewan Harrison, David K. Jackson, Ian Johnston, Dominic Kwiatkowski, Cordelia Langford, John Sillitoe on behalf of |

| | | | |
|---|---|---|---|
| | | Consortium | of the Wellcome Sanger Institute COVID-19 Surveillance Team |
| EPI_ISL_2913894 | Lighthouse Lab in Glasgow | Wellcome Sanger Institute for the COVID-19 Genomics UK (COG-UK) Consortium | Harper VanSteenhouse, Yumi Kasai, David Gray, Carol Clugston, Anna Dominiczak and Alex Alderton, Roberto Amato, Jeffrey Barrett, Sonia Goncalves, Ewan Harrison, David K. Jackson, Ian Johnston, Dominic Kwiatkowski, Cordelia Langford, John Sillitoe on behalf of the Wellcome Sanger Institute COVID-19 Surveillance Team |
| EPI_ISL_2913895 | Lighthouse Laboratory Plymouth | Wellcome Sanger Institute for the COVID-19 Genomics UK (COG-UK) Consortium | Lighthouse Laboratory Plymouth and Alex Alderton, Roberto Amato, Jeffrey Barrett, Sonia Goncalves, Ewan Harrison, David K. Jackson, Ian Johnston, Dominic Kwiatkowski, Cordelia Langford, John Sillitoe on behalf of the Wellcome Sanger Institute COVID-19 Surveillance Team |
| EPI_ISL_2913896, EPI_ISL_2913897 | Berkshire and Surrey Pathology Services Lighthouse Laboratory | Wellcome Sanger Institute for the COVID-19 Genomics UK (COG-UK) Consortium | Berkshire and Surrey Pathology Services Lighthouse Laboratory and Alex Alderton, Roberto Amato, Jeffrey Barrett, Sonia Goncalves, Ewan Harrison, David K. Jackson, Ian Johnston, Dominic Kwiatkowski, Cordelia Langford, John Sillitoe on behalf of the Wellcome Sanger Institute COVID-19 Surveillance Team |
| EPI_ISL_2913898 | Lighthouse Lab in Glasgow | Wellcome Sanger Institute for the COVID-19 Genomics UK (COG-UK) Consortium | Harper VanSteenhouse, Yumi Kasai, David Gray, Carol Clugston, Anna Dominiczak and Alex Alderton, Roberto Amato, Jeffrey Barrett, Sonia Goncalves, Ewan Harrison, David K. Jackson, Ian Johnston, Dominic Kwiatkowski, Cordelia Langford, John Sillitoe on behalf of the Wellcome Sanger Institute COVID-19 Surveillance Team |
| EPI_ISL_2913899 | Lighthouse Laboratory Plymouth | Wellcome Sanger Institute for the COVID-19 Genomics UK (COG-UK) Consortium | Lighthouse Laboratory Plymouth and Alex Alderton, Roberto Amato, Jeffrey Barrett, Sonia Goncalves, Ewan Harrison, David K. Jackson, Ian Johnston, Dominic Kwiatkowski, Cordelia Langford, John Sillitoe on behalf of the Wellcome Sanger Institute COVID-19 Surveillance Team |
| EPI_ISL_2913900 | Lighthouse Lab in Glasgow | Wellcome Sanger Institute for the COVID-19 Genomics UK (COG-UK) Consortium | Harper VanSteenhouse, Yumi Kasai, David Gray, Carol Clugston, Anna Dominiczak and Alex Alderton, Roberto Amato, Jeffrey Barrett, Sonia Goncalves, Ewan Harrison, David K. Jackson, Ian Johnston, Dominic Kwiatkowski, Cordelia Langford, John Sillitoe on behalf of the Wellcome Sanger Institute COVID-19 Surveillance Team |
| EPI_ISL_2913901, EPI_ISL_2913902, EPI_ISL_2913903, EPI_ISL_2913904, EPI_ISL_2913905, EPI_ISL_2913906, EPI_ISL_2913907, EPI_ISL_2913908 | Lighthouse Laboratory Plymouth | Wellcome Sanger Institute for the COVID-19 Genomics UK (COG-UK) Consortium | Lighthouse Laboratory Plymouth and Alex Alderton, Roberto Amato, Jeffrey Barrett, Sonia Goncalves, Ewan Harrison, David K. Jackson, Ian Johnston, Dominic Kwiatkowski, Cordelia Langford, John Sillitoe on behalf of the Wellcome Sanger Institute COVID-19 Surveillance Team |
| EPI_ISL_2913909 | Berkshire and Surrey Pathology Services Lighthouse Laboratory | Wellcome Sanger Institute for the COVID-19 Genomics UK (COG-UK) Consortium | Berkshire and Surrey Pathology Services Lighthouse Laboratory and Alex Alderton, Roberto Amato, Jeffrey Barrett, Sonia Goncalves, Ewan Harrison, David K. Jackson, Ian Johnston, Dominic Kwiatkowski, Cordelia Langford, John Sillitoe on behalf of the Wellcome Sanger Institute COVID-19 Surveillance Team |
| EPI_ISL_2913910 | Lighthouse Laboratory Plymouth | Wellcome Sanger Institute for the COVID-19 Genomics UK (COG-UK) Consortium | Lighthouse Laboratory Plymouth and Alex Alderton, Roberto Amato, Jeffrey Barrett, Sonia Goncalves, Ewan Harrison, David K. Jackson, Ian Johnston, Dominic Kwiatkowski, Cordelia Langford, John Sillitoe on behalf of the Wellcome Sanger Institute COVID-19 Surveillance Team |
| EPI_ISL_2913911 | Berkshire and Surrey Pathology Services Lighthouse Laboratory | Wellcome Sanger Institute for the COVID-19 Genomics UK (COG-UK) Consortium | Berkshire and Surrey Pathology Services Lighthouse Laboratory and Alex Alderton, Roberto Amato, Jeffrey Barrett, Sonia Goncalves, Ewan Harrison, David K. Jackson, Ian Johnston, Dominic Kwiatkowski, Cordelia Langford, John Sillitoe on behalf of the Wellcome Sanger Institute COVID-19 Surveillance Team |
| EPI_ISL_2913912 | Lighthouse Laboratory Plymouth | Wellcome Sanger Institute for the COVID-19 Genomics UK (COG-UK) Consortium | Lighthouse Laboratory Plymouth and Alex Alderton, Roberto Amato, Jeffrey Barrett, Sonia Goncalves, Ewan Harrison, David K. Jackson, Ian Johnston, Dominic Kwiatkowski, Cordelia Langford, John Sillitoe on behalf of the Wellcome Sanger Institute COVID-19 Surveillance Team |
| EPI_ISL_2913913, EPI_ISL_2913914 | Lighthouse Lab in Glasgow | Wellcome Sanger Institute for the COVID-19 Genomics UK (COG-UK) Consortium | Harper VanSteenhouse, Yumi Kasai, David Gray, Carol Clugston, Anna Dominiczak and Alex Alderton, Roberto Amato, Jeffrey Barrett, Sonia Goncalves, Ewan Harrison, David K. Jackson, Ian Johnston, Dominic Kwiatkowski, Cordelia Langford, John Sillitoe on behalf of the Wellcome Sanger Institute COVID-19 Surveillance Team |
| EPI_ISL_2913915 | Lighthouse Laboratory Plymouth | Wellcome Sanger Institute for the COVID-19 Genomics UK (COG-UK) Consortium | Lighthouse Laboratory Plymouth and Alex Alderton, Roberto Amato, Jeffrey Barrett, Sonia Goncalves, Ewan Harrison, David K. Jackson, Ian Johnston, Dominic Kwiatkowski, Cordelia Langford, John Sillitoe on behalf of the Wellcome Sanger Institute COVID-19 Surveillance Team |
| EPI_ISL_2913916 | Berkshire and Surrey Pathology Services Lighthouse Laboratory | Wellcome Sanger Institute for the COVID-19 Genomics UK (COG-UK) Consortium | Berkshire and Surrey Pathology Services Lighthouse Laboratory and Alex Alderton, Roberto Amato, Jeffrey Barrett, Sonia Goncalves, Ewan Harrison, David K. Jackson, Ian Johnston, Dominic Kwiatkowski, Cordelia Langford, John Sillitoe on behalf of the Wellcome Sanger Institute COVID-19 Surveillance Team |
| EPI_ISL_2913917 | Lighthouse Laboratory Plymouth | Wellcome Sanger Institute for the COVID-19 Genomics UK (COG-UK) Consortium | Lighthouse Laboratory Plymouth and Alex Alderton, Roberto Amato, Jeffrey Barrett, Sonia Goncalves, Ewan Harrison, David K. Jackson, Ian Johnston, Dominic Kwiatkowski, Cordelia Langford, John Sillitoe on behalf of the Wellcome Sanger Institute COVID-19 Surveillance Team |
| EPI_ISL_2913918 | Lighthouse Lab in Glasgow | Wellcome Sanger Institute for the COVID-19 Genomics UK (COG-UK) Consortium | Harper VanSteenhouse, Yumi Kasai, David Gray, Carol Clugston, Anna Dominiczak and Alex Alderton, Roberto Amato, Jeffrey Barrett, Sonia Goncalves, Ewan Harrison, David K. Jackson, Ian Johnston, Dominic Kwiatkowski, Cordelia Langford, John Sillitoe on behalf of the Wellcome Sanger Institute COVID-19 Surveillance Team |
| EPI_ISL_2913919, EPI_ISL_2913920, EPI_ISL_2913921, EPI_ISL_2913922, EPI_ISL_2913923, EPI_ISL_2913924, EPI_ISL_2913925, EPI_ISL_2913926, EPI_ISL_2913927 | Lighthouse Laboratory Plymouth | Wellcome Sanger Institute for the COVID-19 Genomics UK (COG-UK) Consortium | Lighthouse Laboratory Plymouth and Alex Alderton, Roberto Amato, Jeffrey Barrett, Sonia Goncalves, Ewan Harrison, David K. Jackson, Ian Johnston, Dominic Kwiatkowski, Cordelia Langford, John Sillitoe on behalf of the Wellcome Sanger Institute COVID-19 Surveillance Team |
| EPI_ISL_2913928 | Berkshire and Surrey Pathology Services Lighthouse Laboratory | Wellcome Sanger Institute for the COVID-19 Genomics UK (COG-UK) Consortium | Berkshire and Surrey Pathology Services Lighthouse Laboratory and Alex Alderton, Roberto Amato, Jeffrey Barrett, Sonia Goncalves, Ewan Harrison, David K. Jackson, Ian Johnston, Dominic Kwiatkowski, Cordelia Langford, John Sillitoe on behalf of the Wellcome Sanger Institute COVID-19 Surveillance Team |
| EPI_ISL_2913929 | Lighthouse Laboratory Plymouth | Wellcome Sanger Institute for the COVID-19 Genomics UK (COG-UK) Consortium | Lighthouse Laboratory Plymouth and Alex Alderton, Roberto Amato, Jeffrey Barrett, Sonia Goncalves, Ewan Harrison, David K. Jackson, Ian Johnston, Dominic Kwiatkowski, Cordelia Langford, John Sillitoe on behalf of the Wellcome Sanger Institute COVID-19 Surveillance Team |
| EPI_ISL_2913930 | Berkshire and Surrey Pathology Services Lighthouse Laboratory | Wellcome Sanger Institute for the COVID-19 Genomics UK (COG-UK) Consortium | Berkshire and Surrey Pathology Services Lighthouse Laboratory and Alex Alderton, Roberto Amato, Jeffrey Barrett, Sonia Goncalves, Ewan Harrison, David K. Jackson, Ian Johnston, Dominic Kwiatkowski, Cordelia Langford, John Sillitoe on behalf of the Wellcome Sanger Institute COVID-19 Surveillance Team |
| EPI_ISL_2913931 | Lighthouse Lab in Glasgow | Wellcome Sanger Institute for the COVID-19 Genomics UK (COG-UK) Consortium | Harper VanSteenhouse, Yumi Kasai, David Gray, Carol Clugston, Anna Dominiczak and Alex Alderton, Roberto Amato, Jeffrey Barrett, Sonia Goncalves, Ewan Harrison, David K. Jackson, Ian Johnston, Dominic Kwiatkowski, Cordelia Langford, John Sillitoe on behalf of the Wellcome Sanger Institute COVID-19 Surveillance Team |
| EPI_ISL_2913932, EPI_ISL_2913933, EPI_ISL_2913934, EPI_ISL_2913935 | Lighthouse Laboratory Plymouth | Wellcome Sanger Institute for the COVID-19 Genomics UK (COG-UK) Consortium | Lighthouse Laboratory Plymouth and Alex Alderton, Roberto Amato, Jeffrey Barrett, Sonia Goncalves, Ewan Harrison, David K. Jackson, Ian Johnston, Dominic Kwiatkowski, Cordelia Langford, John Sillitoe on behalf of the Wellcome Sanger Institute COVID-19 Surveillance Team |
| EPI_ISL_2913936 | Lighthouse Lab in Glasgow | Wellcome Sanger Institute for the COVID-19 Genomics UK (COG-UK) Consortium | Harper VanSteenhouse, Yumi Kasai, David Gray, Carol Clugston, Anna Dominiczak and Alex Alderton, Roberto Amato, Jeffrey Barrett, Sonia Goncalves, Ewan Harrison, David K. Jackson, Ian Johnston, Dominic Kwiatkowski, Cordelia Langford, John Sillitoe on behalf of the Wellcome Sanger Institute COVID-19 Surveillance Team |
| EPI_ISL_2913938, EPI_ISL_2913940 | Lighthouse Laboratory Plymouth | Wellcome Sanger Institute for the COVID-19 Genomics UK (COG-UK) Consortium | Lighthouse Laboratory Plymouth and Alex Alderton, Roberto Amato, Jeffrey Barrett, Sonia Goncalves, Ewan Harrison, David K. Jackson, Ian Johnston, Dominic Kwiatkowski, Cordelia Langford, John Sillitoe on behalf of the Wellcome Sanger Institute COVID-19 Surveillance Team |
| EPI_ISL_2913941 | Lighthouse Lab in Glasgow | Wellcome Sanger Institute for the COVID-19 Genomics UK (COG-UK) Consortium | Harper VanSteenhouse, Yumi Kasai, David Gray, Carol Clugston, Anna Dominiczak and Alex Alderton, Roberto Amato, Jeffrey Barrett, Sonia Goncalves, Ewan Harrison, David K. Jackson, Ian Johnston, Dominic Kwiatkowski, Cordelia Langford, John Sillitoe on behalf of the Wellcome Sanger Institute COVID-19 Surveillance Team |
| EPI_ISL_2913942 | Lighthouse Laboratory Plymouth | Wellcome Sanger Institute for the COVID-19 Genomics UK (COG-UK) Consortium | Lighthouse Laboratory Plymouth and Alex Alderton, Roberto Amato, Jeffrey Barrett, Sonia Goncalves, Ewan Harrison, David K. Jackson, Ian Johnston, Dominic Kwiatkowski, Cordelia Langford, John Sillitoe on behalf of the Wellcome Sanger Institute COVID-19 Surveillance Team |
| EPI_ISL_2913943, EPI_ISL_2913944, EPI_ISL_2913945 | Lighthouse Lab in Glasgow | Wellcome Sanger Institute for the COVID-19 Genomics UK (COG-UK) Consortium | Harper VanSteenhouse, Yumi Kasai, David Gray, Carol Clugston, Anna Dominiczak and Alex Alderton, Roberto Amato, Jeffrey Barrett, Sonia Goncalves, Ewan Harrison, David K. Jackson, Ian Johnston, Dominic Kwiatkowski, Cordelia Langford, John Sillitoe on behalf of the Wellcome Sanger Institute COVID-19 Surveillance Team |
| EPI_ISL_2913946 | Lighthouse Laboratory Plymouth | Wellcome Sanger Institute for the COVID-19 Genomics UK (COG-UK) Consortium | Lighthouse Laboratory Plymouth and Alex Alderton, Roberto Amato, Jeffrey Barrett, Sonia Goncalves, Ewan Harrison, David K. Jackson, Ian Johnston, Dominic Kwiatkowski, Cordelia Langford, John Sillitoe on behalf of the Wellcome Sanger Institute COVID-19 Surveillance Team |
| EPI_ISL_2913947, EPI_ISL_2913948 | Lighthouse Lab in Glasgow | Wellcome Sanger Institute for the COVID-19 Genomics UK (COG-UK) Consortium | Harper VanSteenhouse, Yumi Kasai, David Gray, Carol Clugston, Anna Dominiczak and Alex Alderton, Roberto Amato, Jeffrey Barrett, Sonia Goncalves, Ewan Harrison, David K. Jackson, Ian Johnston, Dominic Kwiatkowski, Cordelia Langford, John Sillitoe on behalf of the Wellcome Sanger Institute COVID-19 Surveillance Team |
| EPI_ISL_2913949 | Lighthouse Laboratory Plymouth | Wellcome Sanger Institute for the COVID-19 Genomics UK (COG-UK) Consortium | Lighthouse Laboratory Plymouth and Alex Alderton, Roberto Amato, Jeffrey Barrett, Sonia Goncalves, Ewan Harrison, David K. Jackson, Ian Johnston, Dominic Kwiatkowski, Cordelia Langford, John Sillitoe on behalf of the Wellcome Sanger Institute COVID-19 Surveillance Team |
| EPI_ISL_2913950 | Lighthouse Lab in Glasgow | Wellcome Sanger Institute for the COVID-19 Genomics UK (COG-UK) Consortium | Harper VanSteenhouse, Yumi Kasai, David Gray, Carol Clugston, Anna Dominiczak and Alex Alderton, Roberto Amato, Jeffrey Barrett, Sonia Goncalves, Ewan Harrison, David K. Jackson, Ian Johnston, Dominic Kwiatkowski, Cordelia Langford, John Sillitoe on behalf of the Wellcome Sanger Institute COVID-19 Surveillance Team |
| EPI_ISL_2913951 | Lighthouse Laboratory Plymouth | Wellcome Sanger Institute for the COVID-19 Genomics UK (COG-UK) Consortium | Lighthouse Laboratory Plymouth and Alex Alderton, Roberto Amato, Jeffrey Barrett, Sonia Goncalves, Ewan Harrison, David K. Jackson, Ian Johnston, Dominic Kwiatkowski, Cordelia Langford, John Sillitoe on behalf of the Wellcome Sanger Institute COVID-19 Surveillance Team |
| EPI_ISL_2913952, EPI_ISL_2913953 | Lighthouse Lab in Glasgow | Wellcome Sanger Institute for the COVID-19 Genomics UK (COG-UK) Consortium | Harper VanSteenhouse, Yumi Kasai, David Gray, Carol Clugston, Anna Dominiczak and Alex Alderton, Roberto Amato, Jeffrey Barrett, Sonia Goncalves, Ewan Harrison, David K. Jackson, Ian Johnston, Dominic Kwiatkowski, Cordelia Langford, John Sillitoe on behalf of the Wellcome Sanger Institute COVID-19 Surveillance Team |
| EPI_ISL_2913954 | Lighthouse Laboratory Plymouth | Wellcome Sanger Institute for the COVID-19 Genomics UK (COG-UK) Consortium | Lighthouse Laboratory Plymouth and Alex Alderton, Roberto Amato, Jeffrey Barrett, Sonia Goncalves, Ewan Harrison, David K. Jackson, Ian Johnston, Dominic Kwiatkowski, Cordelia Langford, John Sillitoe on behalf of the Wellcome Sanger Institute COVID-19 Surveillance Team |
| EPI_ISL_2913955 | Lighthouse Lab in Glasgow | Wellcome Sanger Institute for the COVID-19 Genomics UK (COG-UK) Consortium | Harper VanSteenhouse, Yumi Kasai, David Gray, Carol Clugston, Anna Dominiczak and Alex Alderton, Roberto Amato, Jeffrey Barrett, Sonia Goncalves, Ewan Harrison, David K. Jackson, Ian Johnston, Dominic Kwiatkowski, Cordelia Langford, John Sillitoe on behalf of the Wellcome Sanger Institute COVID-19 Surveillance Team |
| EPI_ISL_2913956, EPI_ISL_2913957 | Lighthouse Laboratory Plymouth | Wellcome Sanger Institute for the COVID-19 Genomics UK (COG-UK) Consortium | Lighthouse Laboratory Plymouth and Alex Alderton, Roberto Amato, Jeffrey Barrett, Sonia Goncalves, Ewan Harrison, David K. Jackson, Ian Johnston, Dominic Kwiatkowski, Cordelia Langford, John Sillitoe on behalf of the Wellcome Sanger Institute COVID-19 Surveillance Team |
| EPI_ISL_2913959 | Lighthouse Lab in Glasgow | Wellcome Sanger Institute for the COVID-19 Genomics UK (COG-UK) Consortium | Harper VanSteenhouse, Yumi Kasai, David Gray, Carol Clugston, Anna Dominiczak and Alex Alderton, Roberto Amato, Jeffrey Barrett, Sonia Goncalves, Ewan Harrison, David K. Jackson, Ian Johnston, Dominic Kwiatkowski, Cordelia Langford, John Sillitoe on behalf of the Wellcome Sanger Institute COVID-19 Surveillance Team |
| EPI_ISL_2913960, EPI_ISL_2913961, | Lighthouse Laboratory Plymouth | Wellcome Sanger Institute for the COVID-19 Genomics UK (COG-UK) Consortium | Lighthouse Laboratory Plymouth and Alex Alderton, Roberto Amato, Jeffrey Barrett, Sonia Goncalves, Ewan Harrison, David K. Jackson, Ian Johnston, Dominic Kwiatkowski, Cordelia Langford, John Sillitoe on behalf of the Wellcome Sanger Institute COVID-19 Surveillance Team |

| | | | |
|---|---|---|---|
| EPI_ISL_2913962 | | | |
| EPI_ISL_2913963 | Lighthouse Lab in Glasgow | Wellcome Sanger Institute for the COVID-19 Genomics UK (COG-UK) Consortium | Harper VanSteenhouse, Yumi Kasai, David Gray, Carol Clugston, Anna Dominiczak and Alex Alderton, Roberto Amato, Jeffrey Barrett, Sonia Goncalves, Ewan Harrison, David K. Jackson, Ian Johnston, Dominic Kwiatkowski, Cordelia Langford, John Sillitoe on behalf of the Wellcome Sanger Institute COVID-19 Surveillance Team |
| EPI_ISL_2913964, EPI_ISL_2913965, EPI_ISL_2913966 | Lighthouse Laboratory Plymouth | Wellcome Sanger Institute for the COVID-19 Genomics UK (COG-UK) Consortium | Lighthouse Laboratory Plymouth and Alex Alderton, Roberto Amato, Jeffrey Barrett, Sonia Goncalves, Ewan Harrison, David K. Jackson, Ian Johnston, Dominic Kwiatkowski, Cordelia Langford, John Sillitoe on behalf of the Wellcome Sanger Institute COVID-19 Surveillance Team |
| EPI_ISL_2913967 | Lighthouse Lab in Glasgow | Wellcome Sanger Institute for the COVID-19 Genomics UK (COG-UK) Consortium | Harper VanSteenhouse, Yumi Kasai, David Gray, Carol Clugston, Anna Dominiczak and Alex Alderton, Roberto Amato, Jeffrey Barrett, Sonia Goncalves, Ewan Harrison, David K. Jackson, Ian Johnston, Dominic Kwiatkowski, Cordelia Langford, John Sillitoe on behalf of the Wellcome Sanger Institute COVID-19 Surveillance Team |
| EPI_ISL_2913968, EPI_ISL_2913969, EPI_ISL_2913970 | Lighthouse Laboratory Plymouth | Wellcome Sanger Institute for the COVID-19 Genomics UK (COG-UK) Consortium | Lighthouse Laboratory Plymouth and Alex Alderton, Roberto Amato, Jeffrey Barrett, Sonia Goncalves, Ewan Harrison, David K. Jackson, Ian Johnston, Dominic Kwiatkowski, Cordelia Langford, John Sillitoe on behalf of the Wellcome Sanger Institute COVID-19 Surveillance Team |
| EPI_ISL_2913971, EPI_ISL_2913972 | Lighthouse Lab in Glasgow | Wellcome Sanger Institute for the COVID-19 Genomics UK (COG-UK) Consortium | Harper VanSteenhouse, Yumi Kasai, David Gray, Carol Clugston, Anna Dominiczak and Alex Alderton, Roberto Amato, Jeffrey Barrett, Sonia Goncalves, Ewan Harrison, David K. Jackson, Ian Johnston, Dominic Kwiatkowski, Cordelia Langford, John Sillitoe on behalf of the Wellcome Sanger Institute COVID-19 Surveillance Team |
| EPI_ISL_2913973 | Lighthouse Laboratory Plymouth | Wellcome Sanger Institute for the COVID-19 Genomics UK (COG-UK) Consortium | Lighthouse Laboratory Plymouth and Alex Alderton, Roberto Amato, Jeffrey Barrett, Sonia Goncalves, Ewan Harrison, David K. Jackson, Ian Johnston, Dominic Kwiatkowski, Cordelia Langford, John Sillitoe on behalf of the Wellcome Sanger Institute COVID-19 Surveillance Team |
| EPI_ISL_2913974 | Lighthouse Lab in Glasgow | Wellcome Sanger Institute for the COVID-19 Genomics UK (COG-UK) Consortium | Harper VanSteenhouse, Yumi Kasai, David Gray, Carol Clugston, Anna Dominiczak and Alex Alderton, Roberto Amato, Jeffrey Barrett, Sonia Goncalves, Ewan Harrison, David K. Jackson, Ian Johnston, Dominic Kwiatkowski, Cordelia Langford, John Sillitoe on behalf of the Wellcome Sanger Institute COVID-19 Surveillance Team |
| EPI_ISL_2913975 | Lighthouse Laboratory Plymouth | Wellcome Sanger Institute for the COVID-19 Genomics UK (COG-UK) Consortium | Lighthouse Laboratory Plymouth and Alex Alderton, Roberto Amato, Jeffrey Barrett, Sonia Goncalves, Ewan Harrison, David K. Jackson, Ian Johnston, Dominic Kwiatkowski, Cordelia Langford, John Sillitoe on behalf of the Wellcome Sanger Institute COVID-19 Surveillance Team |
| EPI_ISL_2913976 | Lighthouse Lab in Glasgow | Wellcome Sanger Institute for the COVID-19 Genomics UK (COG-UK) Consortium | Harper VanSteenhouse, Yumi Kasai, David Gray, Carol Clugston, Anna Dominiczak and Alex Alderton, Roberto Amato, Jeffrey Barrett, Sonia Goncalves, Ewan Harrison, David K. Jackson, Ian Johnston, Dominic Kwiatkowski, Cordelia Langford, John Sillitoe on behalf of the Wellcome Sanger Institute COVID-19 Surveillance Team |
| EPI_ISL_2913977 | Lighthouse Laboratory Plymouth | Wellcome Sanger Institute for the COVID-19 Genomics UK (COG-UK) Consortium | Lighthouse Laboratory Plymouth and Alex Alderton, Roberto Amato, Jeffrey Barrett, Sonia Goncalves, Ewan Harrison, David K. Jackson, Ian Johnston, Dominic Kwiatkowski, Cordelia Langford, John Sillitoe on behalf of the Wellcome Sanger Institute COVID-19 Surveillance Team |
| EPI_ISL_2913978 | Lighthouse Lab in Glasgow | Wellcome Sanger Institute for the COVID-19 Genomics UK (COG-UK) Consortium | Harper VanSteenhouse, Yumi Kasai, David Gray, Carol Clugston, Anna Dominiczak and Alex Alderton, Roberto Amato, Jeffrey Barrett, Sonia Goncalves, Ewan Harrison, David K. Jackson, Ian Johnston, Dominic Kwiatkowski, Cordelia Langford, John Sillitoe on behalf of the Wellcome Sanger Institute COVID-19 Surveillance Team |
| EPI_ISL_2913979 | Lighthouse Laboratory Plymouth | Wellcome Sanger Institute for the COVID-19 Genomics UK (COG-UK) Consortium | Lighthouse Laboratory Plymouth and Alex Alderton, Roberto Amato, Jeffrey Barrett, Sonia Goncalves, Ewan Harrison, David K. Jackson, Ian Johnston, Dominic Kwiatkowski, Cordelia Langford, John Sillitoe on behalf of the Wellcome Sanger Institute COVID-19 Surveillance Team |
| EPI_ISL_2913980 | Lighthouse Lab in Glasgow | Wellcome Sanger Institute for the COVID-19 Genomics UK (COG-UK) Consortium | Harper VanSteenhouse, Yumi Kasai, David Gray, Carol Clugston, Anna Dominiczak and Alex Alderton, Roberto Amato, Jeffrey Barrett, Sonia Goncalves, Ewan Harrison, David K. Jackson, Ian Johnston, Dominic Kwiatkowski, Cordelia Langford, John Sillitoe on behalf of the Wellcome Sanger Institute COVID-19 Surveillance Team |
| EPI_ISL_2913981 | Lighthouse Laboratory Plymouth | Wellcome Sanger Institute for the COVID-19 Genomics UK (COG-UK) Consortium | Lighthouse Laboratory Plymouth and Alex Alderton, Roberto Amato, Jeffrey Barrett, Sonia Goncalves, Ewan Harrison, David K. Jackson, Ian Johnston, Dominic Kwiatkowski, Cordelia Langford, John Sillitoe on behalf of the Wellcome Sanger Institute COVID-19 Surveillance Team |
| EPI_ISL_2913982 | Lighthouse Lab in Glasgow | Wellcome Sanger Institute for the COVID-19 Genomics UK (COG-UK) Consortium | Harper VanSteenhouse, Yumi Kasai, David Gray, Carol Clugston, Anna Dominiczak and Alex Alderton, Roberto Amato, Jeffrey Barrett, Sonia Goncalves, Ewan Harrison, David K. Jackson, Ian Johnston, Dominic Kwiatkowski, Cordelia Langford, John Sillitoe on behalf of the Wellcome Sanger Institute COVID-19 Surveillance Team |
| EPI_ISL_2913983 | Lighthouse Laboratory Plymouth | Wellcome Sanger Institute for the COVID-19 Genomics UK (COG-UK) Consortium | Lighthouse Laboratory Plymouth and Alex Alderton, Roberto Amato, Jeffrey Barrett, Sonia Goncalves, Ewan Harrison, David K. Jackson, Ian Johnston, Dominic Kwiatkowski, Cordelia Langford, John Sillitoe on behalf of the Wellcome Sanger Institute COVID-19 Surveillance Team |
| EPI_ISL_2913984 | Lighthouse Lab in Glasgow | Wellcome Sanger Institute for the COVID-19 Genomics UK (COG-UK) Consortium | Harper VanSteenhouse, Yumi Kasai, David Gray, Carol Clugston, Anna Dominiczak and Alex Alderton, Roberto Amato, Jeffrey Barrett, Sonia Goncalves, Ewan Harrison, David K. Jackson, Ian Johnston, Dominic Kwiatkowski, Cordelia Langford, John Sillitoe on behalf of the Wellcome Sanger Institute COVID-19 Surveillance Team |
| EPI_ISL_2913985, EPI_ISL_2913986, EPI_ISL_2913987, EPI_ISL_2913988, EPI_ISL_2913989 | Lighthouse Laboratory Plymouth | Wellcome Sanger Institute for the COVID-19 Genomics UK (COG-UK) Consortium | Lighthouse Laboratory Plymouth and Alex Alderton, Roberto Amato, Jeffrey Barrett, Sonia Goncalves, Ewan Harrison, David K. Jackson, Ian Johnston, Dominic Kwiatkowski, Cordelia Langford, John Sillitoe on behalf of the Wellcome Sanger Institute COVID-19 Surveillance Team |
| EPI_ISL_2913990 | Lighthouse Lab in Glasgow | Wellcome Sanger Institute for the COVID-19 Genomics UK (COG-UK) Consortium | Harper VanSteenhouse, Yumi Kasai, David Gray, Carol Clugston, Anna Dominiczak and Alex Alderton, Roberto Amato, Jeffrey Barrett, Sonia Goncalves, Ewan Harrison, David K. Jackson, Ian Johnston, Dominic Kwiatkowski, Cordelia Langford, John Sillitoe on behalf of the Wellcome Sanger Institute COVID-19 Surveillance Team |
| EPI_ISL_2913991 | Lighthouse Laboratory Plymouth | Wellcome Sanger Institute for the COVID-19 Genomics UK (COG-UK) Consortium | Lighthouse Laboratory Plymouth and Alex Alderton, Roberto Amato, Jeffrey Barrett, Sonia Goncalves, Ewan Harrison, David K. Jackson, Ian Johnston, Dominic Kwiatkowski, Cordelia Langford, John Sillitoe on behalf of the Wellcome Sanger Institute COVID-19 Surveillance Team |
| EPI_ISL_2913992 | Lighthouse Lab in Glasgow | Wellcome Sanger Institute for the COVID-19 Genomics UK (COG-UK) Consortium | Harper VanSteenhouse, Yumi Kasai, David Gray, Carol Clugston, Anna Dominiczak and Alex Alderton, Roberto Amato, Jeffrey Barrett, Sonia Goncalves, Ewan Harrison, David K. Jackson, Ian Johnston, Dominic Kwiatkowski, Cordelia Langford, John Sillitoe on behalf of the Wellcome Sanger Institute COVID-19 Surveillance Team |
| EPI_ISL_2913993, EPI_ISL_2913994 | Lighthouse Laboratory Plymouth | Wellcome Sanger Institute for the COVID-19 Genomics UK (COG-UK) Consortium | Lighthouse Laboratory Plymouth and Alex Alderton, Roberto Amato, Jeffrey Barrett, Sonia Goncalves, Ewan Harrison, David K. Jackson, Ian Johnston, Dominic Kwiatkowski, Cordelia Langford, John Sillitoe on behalf of the Wellcome Sanger Institute COVID-19 Surveillance Team |
| EPI_ISL_2913995 | Lighthouse Lab in Glasgow | Wellcome Sanger Institute for the COVID-19 Genomics UK (COG-UK) Consortium | Harper VanSteenhouse, Yumi Kasai, David Gray, Carol Clugston, Anna Dominiczak and Alex Alderton, Roberto Amato, Jeffrey Barrett, Sonia Goncalves, Ewan Harrison, David K. Jackson, Ian Johnston, Dominic Kwiatkowski, Cordelia Langford, John Sillitoe on behalf of the Wellcome Sanger Institute COVID-19 Surveillance Team |
| EPI_ISL_2913996 | Lighthouse Laboratory Plymouth | Wellcome Sanger Institute for the COVID-19 Genomics UK (COG-UK) Consortium | Lighthouse Laboratory Plymouth and Alex Alderton, Roberto Amato, Jeffrey Barrett, Sonia Goncalves, Ewan Harrison, David K. Jackson, Ian Johnston, Dominic Kwiatkowski, Cordelia Langford, John Sillitoe on behalf of the Wellcome Sanger Institute COVID-19 Surveillance Team |
| EPI_ISL_2913997 | Lighthouse Lab in Glasgow | Wellcome Sanger Institute for the COVID-19 Genomics UK (COG-UK) Consortium | Harper VanSteenhouse, Yumi Kasai, David Gray, Carol Clugston, Anna Dominiczak and Alex Alderton, Roberto Amato, Jeffrey Barrett, Sonia Goncalves, Ewan Harrison, David K. Jackson, Ian Johnston, Dominic Kwiatkowski, Cordelia Langford, John Sillitoe on behalf of the Wellcome Sanger Institute COVID-19 Surveillance Team |
| EPI_ISL_2913998 | Lighthouse Laboratory Plymouth | Wellcome Sanger Institute for the COVID-19 Genomics UK (COG-UK) Consortium | Lighthouse Laboratory Plymouth and Alex Alderton, Roberto Amato, Jeffrey Barrett, Sonia Goncalves, Ewan Harrison, David K. Jackson, Ian Johnston, Dominic Kwiatkowski, Cordelia Langford, John Sillitoe on behalf of the Wellcome Sanger Institute COVID-19 Surveillance Team |
| EPI_ISL_2913999 | Lighthouse Lab in Glasgow | Wellcome Sanger Institute for the COVID-19 Genomics UK (COG-UK) Consortium | Harper VanSteenhouse, Yumi Kasai, David Gray, Carol Clugston, Anna Dominiczak and Alex Alderton, Roberto Amato, Jeffrey Barrett, Sonia Goncalves, Ewan Harrison, David K. Jackson, Ian Johnston, Dominic Kwiatkowski, Cordelia Langford, John Sillitoe on behalf of the Wellcome Sanger Institute COVID-19 Surveillance Team |
| EPI_ISL_2914000, EPI_ISL_2914001 | Lighthouse Laboratory Plymouth | Wellcome Sanger Institute for the COVID-19 Genomics UK (COG-UK) Consortium | Lighthouse Laboratory Plymouth and Alex Alderton, Roberto Amato, Jeffrey Barrett, Sonia Goncalves, Ewan Harrison, David K. Jackson, Ian Johnston, Dominic Kwiatkowski, Cordelia Langford, John Sillitoe on behalf of the Wellcome Sanger Institute COVID-19 Surveillance Team |
| EPI_ISL_2914002 | Lighthouse Lab in Glasgow | Wellcome Sanger Institute for the COVID-19 Genomics UK (COG-UK) Consortium | Harper VanSteenhouse, Yumi Kasai, David Gray, Carol Clugston, Anna Dominiczak and Alex Alderton, Roberto Amato, Jeffrey Barrett, Sonia Goncalves, Ewan Harrison, David K. Jackson, Ian Johnston, Dominic Kwiatkowski, Cordelia Langford, John Sillitoe on behalf of the Wellcome Sanger Institute COVID-19 Surveillance Team |
| EPI_ISL_2914003, EPI_ISL_2914004 | Lighthouse Laboratory Plymouth | Wellcome Sanger Institute for the COVID-19 Genomics UK (COG-UK) Consortium | Lighthouse Laboratory Plymouth and Alex Alderton, Roberto Amato, Jeffrey Barrett, Sonia Goncalves, Ewan Harrison, David K. Jackson, Ian Johnston, Dominic Kwiatkowski, Cordelia Langford, John Sillitoe on behalf of the Wellcome Sanger Institute COVID-19 Surveillance Team |
| EPI_ISL_2914005 | Lighthouse Lab in Glasgow | Wellcome Sanger Institute for the COVID-19 Genomics UK (COG-UK) Consortium | Harper VanSteenhouse, Yumi Kasai, David Gray, Carol Clugston, Anna Dominiczak and Alex Alderton, Roberto Amato, Jeffrey Barrett, Sonia Goncalves, Ewan Harrison, David K. Jackson, Ian Johnston, Dominic Kwiatkowski, Cordelia Langford, John Sillitoe on behalf of the Wellcome Sanger Institute COVID-19 Surveillance Team |
| EPI_ISL_2914006, EPI_ISL_2914007, EPI_ISL_2914008, EPI_ISL_2914009, EPI_ISL_2914010 | Lighthouse Laboratory Plymouth | Wellcome Sanger Institute for the COVID-19 Genomics UK (COG-UK) Consortium | Lighthouse Laboratory Plymouth and Alex Alderton, Roberto Amato, Jeffrey Barrett, Sonia Goncalves, Ewan Harrison, David K. Jackson, Ian Johnston, Dominic Kwiatkowski, Cordelia Langford, John Sillitoe on behalf of the Wellcome Sanger Institute COVID-19 Surveillance Team |
| EPI_ISL_2914011, EPI_ISL_2914012, EPI_ISL_2914013, EPI_ISL_2914014, EPI_ISL_2914015, EPI_ISL_2914016 | Lighthouse Lab in Glasgow | Wellcome Sanger Institute for the COVID-19 Genomics UK (COG-UK) Consortium | Harper VanSteenhouse, Yumi Kasai, David Gray, Carol Clugston, Anna Dominiczak and Alex Alderton, Roberto Amato, Jeffrey Barrett, Sonia Goncalves, Ewan Harrison, David K. Jackson, Ian Johnston, Dominic Kwiatkowski, Cordelia Langford, John Sillitoe on behalf of the Wellcome Sanger Institute COVID-19 Surveillance Team |
| EPI_ISL_2914017, EPI_ISL_2914018, EPI_ISL_2914019, EPI_ISL_2914020, EPI_ISL_2914021, EPI_ISL_2914022 | Lighthouse Laboratory Plymouth | Wellcome Sanger Institute for the COVID-19 Genomics UK (COG-UK) Consortium | Lighthouse Laboratory Plymouth and Alex Alderton, Roberto Amato, Jeffrey Barrett, Sonia Goncalves, Ewan Harrison, David K. Jackson, Ian Johnston, Dominic Kwiatkowski, Cordelia Langford, John Sillitoe on behalf of the Wellcome Sanger Institute COVID-19 Surveillance Team |
| EPI_ISL_2914023 | Lighthouse Lab in Glasgow | Wellcome Sanger Institute for the COVID-19 Genomics UK (COG-UK) Consortium | Harper VanSteenhouse, Yumi Kasai, David Gray, Carol Clugston, Anna Dominiczak and Alex Alderton, Roberto Amato, Jeffrey Barrett, Sonia Goncalves, Ewan Harrison, David K. Jackson, Ian Johnston, Dominic Kwiatkowski, Cordelia Langford, John Sillitoe on behalf of the Wellcome Sanger Institute COVID-19 Surveillance Team |
| EPI_ISL_2914024, EPI_ISL_2914025, EPI_ISL_2914026, EPI_ISL_2914027, EPI_ISL_2914029, EPI_ISL_2914030 | Lighthouse Laboratory Plymouth | Wellcome Sanger Institute for the COVID-19 Genomics UK (COG-UK) Consortium | Lighthouse Laboratory Plymouth and Alex Alderton, Roberto Amato, Jeffrey Barrett, Sonia Goncalves, Ewan Harrison, David K. Jackson, Ian Johnston, Dominic Kwiatkowski, Cordelia Langford, John Sillitoe on behalf of the Wellcome Sanger Institute COVID-19 Surveillance Team |

| | | | |
|---|---|---|---|
| EPI_ISL_2914031, EPI_ISL_2914032 | Lighthouse Lab in Glasgow | Wellcome Sanger Institute for the COVID-19 Genomics UK (COG-UK) Consortium | Harper VanSteenhouse, Yumi Kasai, David Gray, Carol Clugston, Anna Dominiczak and Alex Alderton, Roberto Amato, Jeffrey Barrett, Sonia Goncalves, Ewan Harrison, David K. Jackson, Ian Johnston, Dominic Kwiatkowski, Cordelia Langford, John Sillitoe on behalf of the Wellcome Sanger Institute COVID-19 Surveillance Team |
| EPI_ISL_2914033, EPI_ISL_2914034, EPI_ISL_2914035, EPI_ISL_2914036, EPI_ISL_2914037, EPI_ISL_2914038, EPI_ISL_2914039, EPI_ISL_2914040 | Lighthouse Laboratory Plymouth | Wellcome Sanger Institute for the COVID-19 Genomics UK (COG-UK) Consortium | Lighthouse Laboratory Plymouth and Alex Alderton, Roberto Amato, Jeffrey Barrett, Sonia Goncalves, Ewan Harrison, David K. Jackson, Ian Johnston, Dominic Kwiatkowski, Cordelia Langford, John Sillitoe on behalf of the Wellcome Sanger Institute COVID-19 Surveillance Team |
| EPI_ISL_2914041 | Lighthouse Lab in Glasgow | Wellcome Sanger Institute for the COVID-19 Genomics UK (COG-UK) Consortium | Harper VanSteenhouse, Yumi Kasai, David Gray, Carol Clugston, Anna Dominiczak and Alex Alderton, Roberto Amato, Jeffrey Barrett, Sonia Goncalves, Ewan Harrison, David K. Jackson, Ian Johnston, Dominic Kwiatkowski, Cordelia Langford, John Sillitoe on behalf of the Wellcome Sanger Institute COVID-19 Surveillance Team |
| EPI_ISL_2914042, EPI_ISL_2914043 | Lighthouse Laboratory Plymouth | Wellcome Sanger Institute for the COVID-19 Genomics UK (COG-UK) Consortium | Lighthouse Laboratory Plymouth and Alex Alderton, Roberto Amato, Jeffrey Barrett, Sonia Goncalves, Ewan Harrison, David K. Jackson, Ian Johnston, Dominic Kwiatkowski, Cordelia Langford, John Sillitoe on behalf of the Wellcome Sanger Institute COVID-19 Surveillance Team |
| EPI_ISL_2914044, EPI_ISL_2914045 | Lighthouse Lab in Glasgow | Wellcome Sanger Institute for the COVID-19 Genomics UK (COG-UK) Consortium | Harper VanSteenhouse, Yumi Kasai, David Gray, Carol Clugston, Anna Dominiczak and Alex Alderton, Roberto Amato, Jeffrey Barrett, Sonia Goncalves, Ewan Harrison, David K. Jackson, Ian Johnston, Dominic Kwiatkowski, Cordelia Langford, John Sillitoe on behalf of the Wellcome Sanger Institute COVID-19 Surveillance Team |
| EPI_ISL_2914046 | Lighthouse Laboratory Plymouth | Wellcome Sanger Institute for the COVID-19 Genomics UK (COG-UK) Consortium | Lighthouse Laboratory Plymouth and Alex Alderton, Roberto Amato, Jeffrey Barrett, Sonia Goncalves, Ewan Harrison, David K. Jackson, Ian Johnston, Dominic Kwiatkowski, Cordelia Langford, John Sillitoe on behalf of the Wellcome Sanger Institute COVID-19 Surveillance Team |
| EPI_ISL_2914047 | Lighthouse Lab in Glasgow | Wellcome Sanger Institute for the COVID-19 Genomics UK (COG-UK) Consortium | Harper VanSteenhouse, Yumi Kasai, David Gray, Carol Clugston, Anna Dominiczak and Alex Alderton, Roberto Amato, Jeffrey Barrett, Sonia Goncalves, Ewan Harrison, David K. Jackson, Ian Johnston, Dominic Kwiatkowski, Cordelia Langford, John Sillitoe on behalf of the Wellcome Sanger Institute COVID-19 Surveillance Team |
| EPI_ISL_2914048, EPI_ISL_2914049 | Lighthouse Laboratory Plymouth | Wellcome Sanger Institute for the COVID-19 Genomics UK (COG-UK) Consortium | Lighthouse Laboratory Plymouth and Alex Alderton, Roberto Amato, Jeffrey Barrett, Sonia Goncalves, Ewan Harrison, David K. Jackson, Ian Johnston, Dominic Kwiatkowski, Cordelia Langford, John Sillitoe on behalf of the Wellcome Sanger Institute COVID-19 Surveillance Team |
| EPI_ISL_2914050 | Lighthouse Lab in Glasgow | Wellcome Sanger Institute for the COVID-19 Genomics UK (COG-UK) Consortium | Harper VanSteenhouse, Yumi Kasai, David Gray, Carol Clugston, Anna Dominiczak and Alex Alderton, Roberto Amato, Jeffrey Barrett, Sonia Goncalves, Ewan Harrison, David K. Jackson, Ian Johnston, Dominic Kwiatkowski, Cordelia Langford, John Sillitoe on behalf of the Wellcome Sanger Institute COVID-19 Surveillance Team |
| EPI_ISL_2914051, EPI_ISL_2914052, EPI_ISL_2914053, EPI_ISL_2914054, EPI_ISL_2914055, EPI_ISL_2914056 | Lighthouse Laboratory Plymouth | Wellcome Sanger Institute for the COVID-19 Genomics UK (COG-UK) Consortium | Lighthouse Laboratory Plymouth and Alex Alderton, Roberto Amato, Jeffrey Barrett, Sonia Goncalves, Ewan Harrison, David K. Jackson, Ian Johnston, Dominic Kwiatkowski, Cordelia Langford, John Sillitoe on behalf of the Wellcome Sanger Institute COVID-19 Surveillance Team |
| EPI_ISL_2914057 | Lighthouse Lab in Glasgow | Wellcome Sanger Institute for the COVID-19 Genomics UK (COG-UK) Consortium | Harper VanSteenhouse, Yumi Kasai, David Gray, Carol Clugston, Anna Dominiczak and Alex Alderton, Roberto Amato, Jeffrey Barrett, Sonia Goncalves, Ewan Harrison, David K. Jackson, Ian Johnston, Dominic Kwiatkowski, Cordelia Langford, John Sillitoe on behalf of the Wellcome Sanger Institute COVID-19 Surveillance Team |
| EPI_ISL_2914058 | Lighthouse Laboratory Plymouth | Wellcome Sanger Institute for the COVID-19 Genomics UK (COG-UK) Consortium | Lighthouse Laboratory Plymouth and Alex Alderton, Roberto Amato, Jeffrey Barrett, Sonia Goncalves, Ewan Harrison, David K. Jackson, Ian Johnston, Dominic Kwiatkowski, Cordelia Langford, John Sillitoe on behalf of the Wellcome Sanger Institute COVID-19 Surveillance Team |
| EPI_ISL_2914059 | Lighthouse Lab in Glasgow | Wellcome Sanger Institute for the COVID-19 Genomics UK (COG-UK) Consortium | Harper VanSteenhouse, Yumi Kasai, David Gray, Carol Clugston, Anna Dominiczak and Alex Alderton, Roberto Amato, Jeffrey Barrett, Sonia Goncalves, Ewan Harrison, David K. Jackson, Ian Johnston, Dominic Kwiatkowski, Cordelia Langford, John Sillitoe on behalf of the Wellcome Sanger Institute COVID-19 Surveillance Team |
| EPI_ISL_2914060, EPI_ISL_2914061, EPI_ISL_2914062, EPI_ISL_2914063, EPI_ISL_2914064, EPI_ISL_2914065, EPI_ISL_2914066, EPI_ISL_2914067 | Lighthouse Laboratory Plymouth | Wellcome Sanger Institute for the COVID-19 Genomics UK (COG-UK) Consortium | Lighthouse Laboratory Plymouth and Alex Alderton, Roberto Amato, Jeffrey Barrett, Sonia Goncalves, Ewan Harrison, David K. Jackson, Ian Johnston, Dominic Kwiatkowski, Cordelia Langford, John Sillitoe on behalf of the Wellcome Sanger Institute COVID-19 Surveillance Team |
| EPI_ISL_2914068 | Lighthouse Lab in Glasgow | Wellcome Sanger Institute for the COVID-19 Genomics UK (COG-UK) Consortium | Harper VanSteenhouse, Yumi Kasai, David Gray, Carol Clugston, Anna Dominiczak and Alex Alderton, Roberto Amato, Jeffrey Barrett, Sonia Goncalves, Ewan Harrison, David K. Jackson, Ian Johnston, Dominic Kwiatkowski, Cordelia Langford, John Sillitoe on behalf of the Wellcome Sanger Institute COVID-19 Surveillance Team |
| EPI_ISL_2914069 | Lighthouse Laboratory Plymouth | Wellcome Sanger Institute for the COVID-19 Genomics UK (COG-UK) Consortium | Lighthouse Laboratory Plymouth and Alex Alderton, Roberto Amato, Jeffrey Barrett, Sonia Goncalves, Ewan Harrison, David K. Jackson, Ian Johnston, Dominic Kwiatkowski, Cordelia Langford, John Sillitoe on behalf of the Wellcome Sanger Institute COVID-19 Surveillance Team |
| EPI_ISL_2914070 | Lighthouse Lab in Glasgow | Wellcome Sanger Institute for the COVID-19 Genomics UK (COG-UK) Consortium | Harper VanSteenhouse, Yumi Kasai, David Gray, Carol Clugston, Anna Dominiczak and Alex Alderton, Roberto Amato, Jeffrey Barrett, Sonia Goncalves, Ewan Harrison, David K. Jackson, Ian Johnston, Dominic Kwiatkowski, Cordelia Langford, John Sillitoe on behalf of the Wellcome Sanger Institute COVID-19 Surveillance Team |
| EPI_ISL_2914071, EPI_ISL_2914072 | Lighthouse Laboratory Plymouth | Wellcome Sanger Institute for the COVID-19 Genomics UK (COG-UK) Consortium | Lighthouse Laboratory Plymouth and Alex Alderton, Roberto Amato, Jeffrey Barrett, Sonia Goncalves, Ewan Harrison, David K. Jackson, Ian Johnston, Dominic Kwiatkowski, Cordelia Langford, John Sillitoe on behalf of the Wellcome Sanger Institute COVID-19 Surveillance Team |
| EPI_ISL_2914073 | Lighthouse Lab in Glasgow | Wellcome Sanger Institute for the COVID-19 Genomics UK (COG-UK) Consortium | Harper VanSteenhouse, Yumi Kasai, David Gray, Carol Clugston, Anna Dominiczak and Alex Alderton, Roberto Amato, Jeffrey Barrett, Sonia Goncalves, Ewan Harrison, David K. Jackson, Ian Johnston, Dominic Kwiatkowski, Cordelia Langford, John Sillitoe on behalf of the Wellcome Sanger Institute COVID-19 Surveillance Team |
| EPI_ISL_2914074, EPI_ISL_2914075, EPI_ISL_2914076, EPI_ISL_2914077, EPI_ISL_2914078, EPI_ISL_2914079, EPI_ISL_2914080, EPI_ISL_2914081 | Lighthouse Laboratory Plymouth | Wellcome Sanger Institute for the COVID-19 Genomics UK (COG-UK) Consortium | Lighthouse Laboratory Plymouth and Alex Alderton, Roberto Amato, Jeffrey Barrett, Sonia Goncalves, Ewan Harrison, David K. Jackson, Ian Johnston, Dominic Kwiatkowski, Cordelia Langford, John Sillitoe on behalf of the Wellcome Sanger Institute COVID-19 Surveillance Team |
| EPI_ISL_2914082 | Lighthouse Lab in Glasgow | Wellcome Sanger Institute for the COVID-19 Genomics UK (COG-UK) Consortium | Harper VanSteenhouse, Yumi Kasai, David Gray, Carol Clugston, Anna Dominiczak and Alex Alderton, Roberto Amato, Jeffrey Barrett, Sonia Goncalves, Ewan Harrison, David K. Jackson, Ian Johnston, Dominic Kwiatkowski, Cordelia Langford, John Sillitoe on behalf of the Wellcome Sanger Institute COVID-19 Surveillance Team |
| EPI_ISL_2914083 | Lighthouse Laboratory Plymouth | Wellcome Sanger Institute for the COVID-19 Genomics UK (COG-UK) Consortium | Lighthouse Laboratory Plymouth and Alex Alderton, Roberto Amato, Jeffrey Barrett, Sonia Goncalves, Ewan Harrison, David K. Jackson, Ian Johnston, Dominic Kwiatkowski, Cordelia Langford, John Sillitoe on behalf of the Wellcome Sanger Institute COVID-19 Surveillance Team |
| EPI_ISL_2914084, EPI_ISL_2914085 | Lighthouse Lab in Glasgow | Wellcome Sanger Institute for the COVID-19 Genomics UK (COG-UK) Consortium | Harper VanSteenhouse, Yumi Kasai, David Gray, Carol Clugston, Anna Dominiczak and Alex Alderton, Roberto Amato, Jeffrey Barrett, Sonia Goncalves, Ewan Harrison, David K. Jackson, Ian Johnston, Dominic Kwiatkowski, Cordelia Langford, John Sillitoe on behalf of the Wellcome Sanger Institute COVID-19 Surveillance Team |
| EPI_ISL_2914086, EPI_ISL_2914087, EPI_ISL_2914088 | Lighthouse Laboratory Plymouth | Wellcome Sanger Institute for the COVID-19 Genomics UK (COG-UK) Consortium | Lighthouse Laboratory Plymouth and Alex Alderton, Roberto Amato, Jeffrey Barrett, Sonia Goncalves, Ewan Harrison, David K. Jackson, Ian Johnston, Dominic Kwiatkowski, Cordelia Langford, John Sillitoe on behalf of the Wellcome Sanger Institute COVID-19 Surveillance Team |
| EPI_ISL_2914089 | Lighthouse Lab in Glasgow | Wellcome Sanger Institute for the COVID-19 Genomics UK (COG-UK) Consortium | Harper VanSteenhouse, Yumi Kasai, David Gray, Carol Clugston, Anna Dominiczak and Alex Alderton, Roberto Amato, Jeffrey Barrett, Sonia Goncalves, Ewan Harrison, David K. Jackson, Ian Johnston, Dominic Kwiatkowski, Cordelia Langford, John Sillitoe on behalf of the Wellcome Sanger Institute COVID-19 Surveillance Team |
| EPI_ISL_2914090, EPI_ISL_2914091, EPI_ISL_2914092 | Lighthouse Laboratory Plymouth | Wellcome Sanger Institute for the COVID-19 Genomics UK (COG-UK) Consortium | Lighthouse Laboratory Plymouth and Alex Alderton, Roberto Amato, Jeffrey Barrett, Sonia Goncalves, Ewan Harrison, David K. Jackson, Ian Johnston, Dominic Kwiatkowski, Cordelia Langford, John Sillitoe on behalf of the Wellcome Sanger Institute COVID-19 Surveillance Team |
| EPI_ISL_2914093 | Lighthouse Lab in Glasgow | Wellcome Sanger Institute for the COVID-19 Genomics UK (COG-UK) Consortium | Harper VanSteenhouse, Yumi Kasai, David Gray, Carol Clugston, Anna Dominiczak and Alex Alderton, Roberto Amato, Jeffrey Barrett, Sonia Goncalves, Ewan Harrison, David K. Jackson, Ian Johnston, Dominic Kwiatkowski, Cordelia Langford, John Sillitoe on behalf of the Wellcome Sanger Institute COVID-19 Surveillance Team |
| EPI_ISL_2914094 | Lighthouse Laboratory Plymouth | Wellcome Sanger Institute for the COVID-19 Genomics UK (COG-UK) Consortium | Lighthouse Laboratory Plymouth and Alex Alderton, Roberto Amato, Jeffrey Barrett, Sonia Goncalves, Ewan Harrison, David K. Jackson, Ian Johnston, Dominic Kwiatkowski, Cordelia Langford, John Sillitoe on behalf of the Wellcome Sanger Institute COVID-19 Surveillance Team |
| EPI_ISL_2914095 | Lighthouse Lab in Glasgow | Wellcome Sanger Institute for the COVID-19 Genomics UK (COG-UK) Consortium | Harper VanSteenhouse, Yumi Kasai, David Gray, Carol Clugston, Anna Dominiczak and Alex Alderton, Roberto Amato, Jeffrey Barrett, Sonia Goncalves, Ewan Harrison, David K. Jackson, Ian Johnston, Dominic Kwiatkowski, Cordelia Langford, John Sillitoe on behalf of the Wellcome Sanger Institute COVID-19 Surveillance Team |
| EPI_ISL_2914096 | Lighthouse Laboratory Plymouth | Wellcome Sanger Institute for the COVID-19 Genomics UK (COG-UK) Consortium | Lighthouse Laboratory Plymouth and Alex Alderton, Roberto Amato, Jeffrey Barrett, Sonia Goncalves, Ewan Harrison, David K. Jackson, Ian Johnston, Dominic Kwiatkowski, Cordelia Langford, John Sillitoe on behalf of the Wellcome Sanger Institute COVID-19 Surveillance Team |
| EPI_ISL_2914097 | Lighthouse Lab in Glasgow | Wellcome Sanger Institute for the COVID-19 Genomics UK (COG-UK) Consortium | Harper VanSteenhouse, Yumi Kasai, David Gray, Carol Clugston, Anna Dominiczak and Alex Alderton, Roberto Amato, Jeffrey Barrett, Sonia Goncalves, Ewan Harrison, David K. Jackson, Ian Johnston, Dominic Kwiatkowski, Cordelia Langford, John Sillitoe on behalf of the Wellcome Sanger Institute COVID-19 Surveillance Team |
| EPI_ISL_2914098, EPI_ISL_2914099 | Lighthouse Laboratory Plymouth | Wellcome Sanger Institute for the COVID-19 Genomics UK (COG-UK) Consortium | Lighthouse Laboratory Plymouth and Alex Alderton, Roberto Amato, Jeffrey Barrett, Sonia Goncalves, Ewan Harrison, David K. Jackson, Ian Johnston, Dominic Kwiatkowski, Cordelia Langford, John Sillitoe on behalf of the Wellcome Sanger Institute COVID-19 Surveillance Team |
| EPI_ISL_2914100 | Lighthouse Lab in Glasgow | Wellcome Sanger Institute for the COVID-19 Genomics UK (COG-UK) Consortium | Harper VanSteenhouse, Yumi Kasai, David Gray, Carol Clugston, Anna Dominiczak and Alex Alderton, Roberto Amato, Jeffrey Barrett, Sonia Goncalves, Ewan Harrison, David K. Jackson, Ian Johnston, Dominic Kwiatkowski, Cordelia Langford, John Sillitoe on behalf of the Wellcome Sanger Institute COVID-19 Surveillance Team |
| EPI_ISL_2914101, EPI_ISL_2914102, EPI_ISL_2914103 | Lighthouse Laboratory Plymouth | Wellcome Sanger Institute for the COVID-19 Genomics UK (COG-UK) Consortium | Lighthouse Laboratory Plymouth and Alex Alderton, Roberto Amato, Jeffrey Barrett, Sonia Goncalves, Ewan Harrison, David K. Jackson, Ian Johnston, Dominic Kwiatkowski, Cordelia Langford, John Sillitoe on behalf of the Wellcome Sanger Institute COVID-19 Surveillance Team |
| EPI_ISL_2914104, EPI_ISL_2914105 | Lighthouse Lab in Glasgow | Wellcome Sanger Institute for the COVID-19 Genomics UK (COG-UK) Consortium | Harper VanSteenhouse, Yumi Kasai, David Gray, Carol Clugston, Anna Dominiczak and Alex Alderton, Roberto Amato, Jeffrey Barrett, Sonia Goncalves, Ewan Harrison, David K. Jackson, Ian Johnston, Dominic Kwiatkowski, Cordelia Langford, John Sillitoe on behalf of the Wellcome Sanger Institute COVID-19 Surveillance Team |

| | | | |
|---|---|---|---|
| EPI_ISL_2914106, EPI_ISL_2914107 | Lighthouse Laboratory Plymouth | Wellcome Sanger Institute for the COVID-19 Genomics UK (COG-UK) Consortium | Lighthouse Laboratory Plymouth and Alex Alderton, Roberto Amato, Jeffrey Barrett, Sonia Goncalves, Ewan Harrison, David K. Jackson, Ian Johnston, Dominic Kwiatkowski, Cordelia Langford, John Sillitoe on behalf of the Wellcome Sanger Institute COVID-19 Surveillance Team |
| EPI_ISL_2914108, EPI_ISL_2914109, EPI_ISL_2914110 | Lighthouse Lab in Glasgow | Wellcome Sanger Institute for the COVID-19 Genomics UK (COG-UK) Consortium | Harper VanSteenhouse, Yumi Kasai, David Gray, Carol Clugston, Anna Dominiczak and Alex Alderton, Roberto Amato, Jeffrey Barrett, Sonia Goncalves, Ewan Harrison, David K. Jackson, Ian Johnston, Dominic Kwiatkowski, Cordelia Langford, John Sillitoe on behalf of the Wellcome Sanger Institute COVID-19 Surveillance Team |
| EPI_ISL_2914111 | Lighthouse Laboratory Plymouth | Wellcome Sanger Institute for the COVID-19 Genomics UK (COG-UK) Consortium | Lighthouse Laboratory Plymouth and Alex Alderton, Roberto Amato, Jeffrey Barrett, Sonia Goncalves, Ewan Harrison, David K. Jackson, Ian Johnston, Dominic Kwiatkowski, Cordelia Langford, John Sillitoe on behalf of the Wellcome Sanger Institute COVID-19 Surveillance Team |
| EPI_ISL_2914112 | Lighthouse Lab in Glasgow | Wellcome Sanger Institute for the COVID-19 Genomics UK (COG-UK) Consortium | Harper VanSteenhouse, Yumi Kasai, David Gray, Carol Clugston, Anna Dominiczak and Alex Alderton, Roberto Amato, Jeffrey Barrett, Sonia Goncalves, Ewan Harrison, David K. Jackson, Ian Johnston, Dominic Kwiatkowski, Cordelia Langford, John Sillitoe on behalf of the Wellcome Sanger Institute COVID-19 Surveillance Team |
| EPI_ISL_2914113 | Lighthouse Laboratory Plymouth | Wellcome Sanger Institute for the COVID-19 Genomics UK (COG-UK) Consortium | Lighthouse Laboratory Plymouth and Alex Alderton, Roberto Amato, Jeffrey Barrett, Sonia Goncalves, Ewan Harrison, David K. Jackson, Ian Johnston, Dominic Kwiatkowski, Cordelia Langford, John Sillitoe on behalf of the Wellcome Sanger Institute COVID-19 Surveillance Team |
| EPI_ISL_2914114 | Lighthouse Lab in Glasgow | Wellcome Sanger Institute for the COVID-19 Genomics UK (COG-UK) Consortium | Harper VanSteenhouse, Yumi Kasai, David Gray, Carol Clugston, Anna Dominiczak and Alex Alderton, Roberto Amato, Jeffrey Barrett, Sonia Goncalves, Ewan Harrison, David K. Jackson, Ian Johnston, Dominic Kwiatkowski, Cordelia Langford, John Sillitoe on behalf of the Wellcome Sanger Institute COVID-19 Surveillance Team |
| EPI_ISL_2914115, EPI_ISL_2914116, EPI_ISL_2914117 | Lighthouse Laboratory Plymouth | Wellcome Sanger Institute for the COVID-19 Genomics UK (COG-UK) Consortium | Lighthouse Laboratory Plymouth and Alex Alderton, Roberto Amato, Jeffrey Barrett, Sonia Goncalves, Ewan Harrison, David K. Jackson, Ian Johnston, Dominic Kwiatkowski, Cordelia Langford, John Sillitoe on behalf of the Wellcome Sanger Institute COVID-19 Surveillance Team |
| EPI_ISL_2914118 | Lighthouse Lab in Glasgow | Wellcome Sanger Institute for the COVID-19 Genomics UK (COG-UK) Consortium | Harper VanSteenhouse, Yumi Kasai, David Gray, Carol Clugston, Anna Dominiczak and Alex Alderton, Roberto Amato, Jeffrey Barrett, Sonia Goncalves, Ewan Harrison, David K. Jackson, Ian Johnston, Dominic Kwiatkowski, Cordelia Langford, John Sillitoe on behalf of the Wellcome Sanger Institute COVID-19 Surveillance Team |
| EPI_ISL_2914119, EPI_ISL_2914120, EPI_ISL_2914121, EPI_ISL_2914122 | Lighthouse Laboratory Plymouth | Wellcome Sanger Institute for the COVID-19 Genomics UK (COG-UK) Consortium | Lighthouse Laboratory Plymouth and Alex Alderton, Roberto Amato, Jeffrey Barrett, Sonia Goncalves, Ewan Harrison, David K. Jackson, Ian Johnston, Dominic Kwiatkowski, Cordelia Langford, John Sillitoe on behalf of the Wellcome Sanger Institute COVID-19 Surveillance Team |
| EPI_ISL_2914123 | Lighthouse Lab in Glasgow | Wellcome Sanger Institute for the COVID-19 Genomics UK (COG-UK) Consortium | Harper VanSteenhouse, Yumi Kasai, David Gray, Carol Clugston, Anna Dominiczak and Alex Alderton, Roberto Amato, Jeffrey Barrett, Sonia Goncalves, Ewan Harrison, David K. Jackson, Ian Johnston, Dominic Kwiatkowski, Cordelia Langford, John Sillitoe on behalf of the Wellcome Sanger Institute COVID-19 Surveillance Team |
| EPI_ISL_2914124 | Lighthouse Laboratory Plymouth | Wellcome Sanger Institute for the COVID-19 Genomics UK (COG-UK) Consortium | Lighthouse Laboratory Plymouth and Alex Alderton, Roberto Amato, Jeffrey Barrett, Sonia Goncalves, Ewan Harrison, David K. Jackson, Ian Johnston, Dominic Kwiatkowski, Cordelia Langford, John Sillitoe on behalf of the Wellcome Sanger Institute COVID-19 Surveillance Team |
| EPI_ISL_2914125 | Lighthouse Lab in Glasgow | Wellcome Sanger Institute for the COVID-19 Genomics UK (COG-UK) Consortium | Harper VanSteenhouse, Yumi Kasai, David Gray, Carol Clugston, Anna Dominiczak and Alex Alderton, Roberto Amato, Jeffrey Barrett, Sonia Goncalves, Ewan Harrison, David K. Jackson, Ian Johnston, Dominic Kwiatkowski, Cordelia Langford, John Sillitoe on behalf of the Wellcome Sanger Institute COVID-19 Surveillance Team |
| EPI_ISL_2914126, EPI_ISL_2914127, EPI_ISL_2914128 | Lighthouse Laboratory Plymouth | Wellcome Sanger Institute for the COVID-19 Genomics UK (COG-UK) Consortium | Lighthouse Laboratory Plymouth and Alex Alderton, Roberto Amato, Jeffrey Barrett, Sonia Goncalves, Ewan Harrison, David K. Jackson, Ian Johnston, Dominic Kwiatkowski, Cordelia Langford, John Sillitoe on behalf of the Wellcome Sanger Institute COVID-19 Surveillance Team |
| EPI_ISL_2914129 | Lighthouse Lab in Glasgow | Wellcome Sanger Institute for the COVID-19 Genomics UK (COG-UK) Consortium | Harper VanSteenhouse, Yumi Kasai, David Gray, Carol Clugston, Anna Dominiczak and Alex Alderton, Roberto Amato, Jeffrey Barrett, Sonia Goncalves, Ewan Harrison, David K. Jackson, Ian Johnston, Dominic Kwiatkowski, Cordelia Langford, John Sillitoe on behalf of the Wellcome Sanger Institute COVID-19 Surveillance Team |

EPI_ISL_2914130, EPI_ISL_2914131, EPI_ISL_2914132, EPI_ISL_2914133, EPI_ISL_2914134, EPI_ISL_2914135, EPI_ISL_2914136, EPI_ISL_2914137, EPI_ISL_2914138, EPI_ISL_2914139, EPI_ISL_2914140, EPI_ISL_2914141, EPI_ISL_2914142, EPI_ISL_2914143

| | | | |
|---|---|---|---|
| see above | Lighthouse Laboratory Plymouth | Wellcome Sanger Institute for the COVID-19 Genomics UK (COG-UK) Consortium | Lighthouse Laboratory Plymouth and Alex Alderton, Roberto Amato, Jeffrey Barrett, Sonia Goncalves, Ewan Harrison, David K. Jackson, Ian Johnston, Dominic Kwiatkowski, Cordelia Langford, John Sillitoe on behalf of the Wellcome Sanger Institute COVID-19 Surveillance Team |
| EPI_ISL_2914144 | Lighthouse Lab in Glasgow | Wellcome Sanger Institute for the COVID-19 Genomics UK (COG-UK) Consortium | Harper VanSteenhouse, Yumi Kasai, David Gray, Carol Clugston, Anna Dominiczak and Alex Alderton, Roberto Amato, Jeffrey Barrett, Sonia Goncalves, Ewan Harrison, David K. Jackson, Ian Johnston, Dominic Kwiatkowski, Cordelia Langford, John Sillitoe on behalf of the Wellcome Sanger Institute COVID-19 Surveillance Team |
| EPI_ISL_2914145, EPI_ISL_2914146, EPI_ISL_2914147, EPI_ISL_2914148, EPI_ISL_2914149, EPI_ISL_2914150, EPI_ISL_2914151 | Lighthouse Laboratory Plymouth | Wellcome Sanger Institute for the COVID-19 Genomics UK (COG-UK) Consortium | Lighthouse Laboratory Plymouth and Alex Alderton, Roberto Amato, Jeffrey Barrett, Sonia Goncalves, Ewan Harrison, David K. Jackson, Ian Johnston, Dominic Kwiatkowski, Cordelia Langford, John Sillitoe on behalf of the Wellcome Sanger Institute COVID-19 Surveillance Team |
| EPI_ISL_2914152 | Lighthouse Lab in Glasgow | Wellcome Sanger Institute for the COVID-19 Genomics UK (COG-UK) Consortium | Harper VanSteenhouse, Yumi Kasai, David Gray, Carol Clugston, Anna Dominiczak and Alex Alderton, Roberto Amato, Jeffrey Barrett, Sonia Goncalves, Ewan Harrison, David K. Jackson, Ian Johnston, Dominic Kwiatkowski, Cordelia Langford, John Sillitoe on behalf of the Wellcome Sanger Institute COVID-19 Surveillance Team |
| EPI_ISL_2914153, EPI_ISL_2914154, EPI_ISL_2914155, EPI_ISL_2914156, EPI_ISL_2914157 | Lighthouse Laboratory Plymouth | Wellcome Sanger Institute for the COVID-19 Genomics UK (COG-UK) Consortium | Lighthouse Laboratory Plymouth and Alex Alderton, Roberto Amato, Jeffrey Barrett, Sonia Goncalves, Ewan Harrison, David K. Jackson, Ian Johnston, Dominic Kwiatkowski, Cordelia Langford, John Sillitoe on behalf of the Wellcome Sanger Institute COVID-19 Surveillance Team |
| EPI_ISL_2914158 | Lighthouse Lab in Glasgow | Wellcome Sanger Institute for the COVID-19 Genomics UK (COG-UK) Consortium | Harper VanSteenhouse, Yumi Kasai, David Gray, Carol Clugston, Anna Dominiczak and Alex Alderton, Roberto Amato, Jeffrey Barrett, Sonia Goncalves, Ewan Harrison, David K. Jackson, Ian Johnston, Dominic Kwiatkowski, Cordelia Langford, John Sillitoe on behalf of the Wellcome Sanger Institute COVID-19 Surveillance Team |
| EPI_ISL_2914159 | Lighthouse Laboratory Plymouth | Wellcome Sanger Institute for the COVID-19 Genomics UK (COG-UK) Consortium | Lighthouse Laboratory Plymouth and Alex Alderton, Roberto Amato, Jeffrey Barrett, Sonia Goncalves, Ewan Harrison, David K. Jackson, Ian Johnston, Dominic Kwiatkowski, Cordelia Langford, John Sillitoe on behalf of the Wellcome Sanger Institute COVID-19 Surveillance Team |
| EPI_ISL_2914160 | Lighthouse Lab in Glasgow | Wellcome Sanger Institute for the COVID-19 Genomics UK (COG-UK) Consortium | Harper VanSteenhouse, Yumi Kasai, David Gray, Carol Clugston, Anna Dominiczak and Alex Alderton, Roberto Amato, Jeffrey Barrett, Sonia Goncalves, Ewan Harrison, David K. Jackson, Ian Johnston, Dominic Kwiatkowski, Cordelia Langford, John Sillitoe on behalf of the Wellcome Sanger Institute COVID-19 Surveillance Team |
| EPI_ISL_2914161, EPI_ISL_2914162, EPI_ISL_2914163, EPI_ISL_2914164, EPI_ISL_2914165, EPI_ISL_2914166, EPI_ISL_2914167, EPI_ISL_2914168, EPI_ISL_2914169, EPI_ISL_2914170 | Lighthouse Laboratory Plymouth | Wellcome Sanger Institute for the COVID-19 Genomics UK (COG-UK) Consortium | Lighthouse Laboratory Plymouth and Alex Alderton, Roberto Amato, Jeffrey Barrett, Sonia Goncalves, Ewan Harrison, David K. Jackson, Ian Johnston, Dominic Kwiatkowski, Cordelia Langford, John Sillitoe on behalf of the Wellcome Sanger Institute COVID-19 Surveillance Team |
| EPI_ISL_2914171 | Lighthouse Lab in Glasgow | Wellcome Sanger Institute for the COVID-19 Genomics UK (COG-UK) Consortium | Harper VanSteenhouse, Yumi Kasai, David Gray, Carol Clugston, Anna Dominiczak and Alex Alderton, Roberto Amato, Jeffrey Barrett, Sonia Goncalves, Ewan Harrison, David K. Jackson, Ian Johnston, Dominic Kwiatkowski, Cordelia Langford, John Sillitoe on behalf of the Wellcome Sanger Institute COVID-19 Surveillance Team |
| EPI_ISL_2914172, EPI_ISL_2914173, EPI_ISL_2914174, EPI_ISL_2914175 | Lighthouse Laboratory Plymouth | Wellcome Sanger Institute for the COVID-19 Genomics UK (COG-UK) Consortium | Lighthouse Laboratory Plymouth and Alex Alderton, Roberto Amato, Jeffrey Barrett, Sonia Goncalves, Ewan Harrison, David K. Jackson, Ian Johnston, Dominic Kwiatkowski, Cordelia Langford, John Sillitoe on behalf of the Wellcome Sanger Institute COVID-19 Surveillance Team |
| EPI_ISL_2914176 | Lighthouse Lab in Glasgow | Wellcome Sanger Institute for the COVID-19 Genomics UK (COG-UK) Consortium | Harper VanSteenhouse, Yumi Kasai, David Gray, Carol Clugston, Anna Dominiczak and Alex Alderton, Roberto Amato, Jeffrey Barrett, Sonia Goncalves, Ewan Harrison, David K. Jackson, Ian Johnston, Dominic Kwiatkowski, Cordelia Langford, John Sillitoe on behalf of the Wellcome Sanger Institute COVID-19 Surveillance Team |
| EPI_ISL_2914177 | Lighthouse Laboratory Plymouth | Wellcome Sanger Institute for the COVID-19 Genomics UK (COG-UK) Consortium | Lighthouse Laboratory Plymouth and Alex Alderton, Roberto Amato, Jeffrey Barrett, Sonia Goncalves, Ewan Harrison, David K. Jackson, Ian Johnston, Dominic Kwiatkowski, Cordelia Langford, John Sillitoe on behalf of the Wellcome Sanger Institute COVID-19 Surveillance Team |
| EPI_ISL_2914178 | Lighthouse Lab in Glasgow | Wellcome Sanger Institute for the COVID-19 Genomics UK (COG-UK) Consortium | Harper VanSteenhouse, Yumi Kasai, David Gray, Carol Clugston, Anna Dominiczak and Alex Alderton, Roberto Amato, Jeffrey Barrett, Sonia Goncalves, Ewan Harrison, David K. Jackson, Ian Johnston, Dominic Kwiatkowski, Cordelia Langford, John Sillitoe on behalf of the Wellcome Sanger Institute COVID-19 Surveillance Team |
| EPI_ISL_2914179 | Lighthouse Laboratory Plymouth | Wellcome Sanger Institute for the COVID-19 Genomics UK (COG-UK) Consortium | Lighthouse Laboratory Plymouth and Alex Alderton, Roberto Amato, Jeffrey Barrett, Sonia Goncalves, Ewan Harrison, David K. Jackson, Ian Johnston, Dominic Kwiatkowski, Cordelia Langford, John Sillitoe on behalf of the Wellcome Sanger Institute COVID-19 Surveillance Team |
| EPI_ISL_2914180 | Lighthouse Lab in Glasgow | Wellcome Sanger Institute for the COVID-19 Genomics UK (COG-UK) Consortium | Harper VanSteenhouse, Yumi Kasai, David Gray, Carol Clugston, Anna Dominiczak and Alex Alderton, Roberto Amato, Jeffrey Barrett, Sonia Goncalves, Ewan Harrison, David K. Jackson, Ian Johnston, Dominic Kwiatkowski, Cordelia Langford, John Sillitoe on behalf of the Wellcome Sanger Institute COVID-19 Surveillance Team |
| EPI_ISL_2914181, EPI_ISL_2914182 | Lighthouse Laboratory Plymouth | Wellcome Sanger Institute for the COVID-19 Genomics UK (COG-UK) Consortium | Lighthouse Laboratory Plymouth and Alex Alderton, Roberto Amato, Jeffrey Barrett, Sonia Goncalves, Ewan Harrison, David K. Jackson, Ian Johnston, Dominic Kwiatkowski, Cordelia Langford, John Sillitoe on behalf of the Wellcome Sanger Institute COVID-19 Surveillance Team |
| EPI_ISL_2914183 | Lighthouse Lab in Glasgow | Wellcome Sanger Institute for the COVID-19 Genomics UK (COG-UK) Consortium | Harper VanSteenhouse, Yumi Kasai, David Gray, Carol Clugston, Anna Dominiczak and Alex Alderton, Roberto Amato, Jeffrey Barrett, Sonia Goncalves, Ewan Harrison, David K. Jackson, Ian Johnston, Dominic Kwiatkowski, Cordelia Langford, John Sillitoe on behalf of the Wellcome Sanger Institute COVID-19 Surveillance Team |
| EPI_ISL_2914184, EPI_ISL_2914185, EPI_ISL_2914186, EPI_ISL_2914187, EPI_ISL_2914188 | Lighthouse Laboratory Plymouth | Wellcome Sanger Institute for the COVID-19 Genomics UK (COG-UK) Consortium | Lighthouse Laboratory Plymouth and Alex Alderton, Roberto Amato, Jeffrey Barrett, Sonia Goncalves, Ewan Harrison, David K. Jackson, Ian Johnston, Dominic Kwiatkowski, Cordelia Langford, John Sillitoe on behalf of the Wellcome Sanger Institute COVID-19 Surveillance Team |
| EPI_ISL_2914189, | Lighthouse Lab in Glasgow | Wellcome Sanger Institute for the COVID-19 Genomics UK (COG-UK) | Harper VanSteenhouse, Yumi Kasai, David Gray, Carol Clugston, Anna Dominiczak and Alex Alderton, Roberto Amato, Jeffrey Barrett, Sonia Goncalves, Ewan Harrison, David K. Jackson, Ian Johnston, Dominic Kwiatkowski, Cordelia Langford, |

| | | | |
|---|---|---|---|
| EPI_ISL_2914190 | | Consortium | John Sillitoe on behalf of the Wellcome Sanger Institute COVID-19 Surveillance Team |
| EPI_ISL_2914191, EPI_ISL_2914192 | Lighthouse Laboratory Plymouth | Wellcome Sanger Institute for the COVID-19 Genomics UK (COG-UK) Consortium | Lighthouse Laboratory Plymouth and Alex Alderton, Roberto Amato, Jeffrey Barrett, Sonia Goncalves, Ewan Harrison, David K. Jackson, Ian Johnston, Dominic Kwiatkowski, Cordelia Langford, John Sillitoe on behalf of the Wellcome Sanger Institute COVID-19 Surveillance Team |
| EPI_ISL_2914193 | Lighthouse Lab in Glasgow | Wellcome Sanger Institute for the COVID-19 Genomics UK (COG-UK) Consortium | Harper VanSteenhouse, Yumi Kasai, David Gray, Carol Clugston, Anna Dominiczak and Alex Alderton, Roberto Amato, Jeffrey Barrett, Sonia Goncalves, Ewan Harrison, David K. Jackson, Ian Johnston, Dominic Kwiatkowski, Cordelia Langford, John Sillitoe on behalf of the Wellcome Sanger Institute COVID-19 Surveillance Team |
| EPI_ISL_2914194 | Lighthouse Laboratory Plymouth | Wellcome Sanger Institute for the COVID-19 Genomics UK (COG-UK) Consortium | Lighthouse Laboratory Plymouth and Alex Alderton, Roberto Amato, Jeffrey Barrett, Sonia Goncalves, Ewan Harrison, David K. Jackson, Ian Johnston, Dominic Kwiatkowski, Cordelia Langford, John Sillitoe on behalf of the Wellcome Sanger Institute COVID-19 Surveillance Team |
| EPI_ISL_2914195, EPI_ISL_2914196, EPI_ISL_2914197 | Lighthouse Lab in Glasgow | Wellcome Sanger Institute for the COVID-19 Genomics UK (COG-UK) Consortium | Harper VanSteenhouse, Yumi Kasai, David Gray, Carol Clugston, Anna Dominiczak and Alex Alderton, Roberto Amato, Jeffrey Barrett, Sonia Goncalves, Ewan Harrison, David K. Jackson, Ian Johnston, Dominic Kwiatkowski, Cordelia Langford, John Sillitoe on behalf of the Wellcome Sanger Institute COVID-19 Surveillance Team |
| EPI_ISL_2914198, EPI_ISL_2914199, EPI_ISL_2914200, EPI_ISL_2914201, EPI_ISL_2914202, EPI_ISL_2914203, EPI_ISL_2914204, EPI_ISL_2914205, EPI_ISL_2914206 | Lighthouse Laboratory Plymouth | Wellcome Sanger Institute for the COVID-19 Genomics UK (COG-UK) Consortium | Lighthouse Laboratory Plymouth and Alex Alderton, Roberto Amato, Jeffrey Barrett, Sonia Goncalves, Ewan Harrison, David K. Jackson, Ian Johnston, Dominic Kwiatkowski, Cordelia Langford, John Sillitoe on behalf of the Wellcome Sanger Institute COVID-19 Surveillance Team |
| EPI_ISL_2914207 | Lighthouse Lab in Glasgow | Wellcome Sanger Institute for the COVID-19 Genomics UK (COG-UK) Consortium | Harper VanSteenhouse, Yumi Kasai, David Gray, Carol Clugston, Anna Dominiczak and Alex Alderton, Roberto Amato, Jeffrey Barrett, Sonia Goncalves, Ewan Harrison, David K. Jackson, Ian Johnston, Dominic Kwiatkowski, Cordelia Langford, John Sillitoe on behalf of the Wellcome Sanger Institute COVID-19 Surveillance Team |
| EPI_ISL_2914208, EPI_ISL_2914209, EPI_ISL_2914210 | Lighthouse Laboratory Plymouth | Wellcome Sanger Institute for the COVID-19 Genomics UK (COG-UK) Consortium | Lighthouse Laboratory Plymouth and Alex Alderton, Roberto Amato, Jeffrey Barrett, Sonia Goncalves, Ewan Harrison, David K. Jackson, Ian Johnston, Dominic Kwiatkowski, Cordelia Langford, John Sillitoe on behalf of the Wellcome Sanger Institute COVID-19 Surveillance Team |
| EPI_ISL_2914211, EPI_ISL_2914212 | Lighthouse Lab in Glasgow | Wellcome Sanger Institute for the COVID-19 Genomics UK (COG-UK) Consortium | Harper VanSteenhouse, Yumi Kasai, David Gray, Carol Clugston, Anna Dominiczak and Alex Alderton, Roberto Amato, Jeffrey Barrett, Sonia Goncalves, Ewan Harrison, David K. Jackson, Ian Johnston, Dominic Kwiatkowski, Cordelia Langford, John Sillitoe on behalf of the Wellcome Sanger Institute COVID-19 Surveillance Team |
| EPI_ISL_2914213, EPI_ISL_2914214, EPI_ISL_2914215, EPI_ISL_2914216, EPI_ISL_2914217, EPI_ISL_2914218, EPI_ISL_2914219, EPI_ISL_2914220 | Lighthouse Laboratory Plymouth | Wellcome Sanger Institute for the COVID-19 Genomics UK (COG-UK) Consortium | Lighthouse Laboratory Plymouth and Alex Alderton, Roberto Amato, Jeffrey Barrett, Sonia Goncalves, Ewan Harrison, David K. Jackson, Ian Johnston, Dominic Kwiatkowski, Cordelia Langford, John Sillitoe on behalf of the Wellcome Sanger Institute COVID-19 Surveillance Team |
| EPI_ISL_2914221 | Lighthouse Lab in Glasgow | Wellcome Sanger Institute for the COVID-19 Genomics UK (COG-UK) Consortium | Harper VanSteenhouse, Yumi Kasai, David Gray, Carol Clugston, Anna Dominiczak and Alex Alderton, Roberto Amato, Jeffrey Barrett, Sonia Goncalves, Ewan Harrison, David K. Jackson, Ian Johnston, Dominic Kwiatkowski, Cordelia Langford, John Sillitoe on behalf of the Wellcome Sanger Institute COVID-19 Surveillance Team |
| EPI_ISL_2914222, EPI_ISL_2914223, EPI_ISL_2914224, EPI_ISL_2914225 | Lighthouse Laboratory Plymouth | Wellcome Sanger Institute for the COVID-19 Genomics UK (COG-UK) Consortium | Lighthouse Laboratory Plymouth and Alex Alderton, Roberto Amato, Jeffrey Barrett, Sonia Goncalves, Ewan Harrison, David K. Jackson, Ian Johnston, Dominic Kwiatkowski, Cordelia Langford, John Sillitoe on behalf of the Wellcome Sanger Institute COVID-19 Surveillance Team |
| EPI_ISL_2914226, EPI_ISL_2914227 | Lighthouse Lab in Glasgow | Wellcome Sanger Institute for the COVID-19 Genomics UK (COG-UK) Consortium | Harper VanSteenhouse, Yumi Kasai, David Gray, Carol Clugston, Anna Dominiczak and Alex Alderton, Roberto Amato, Jeffrey Barrett, Sonia Goncalves, Ewan Harrison, David K. Jackson, Ian Johnston, Dominic Kwiatkowski, Cordelia Langford, John Sillitoe on behalf of the Wellcome Sanger Institute COVID-19 Surveillance Team |
| EPI_ISL_2914228, EPI_ISL_2914229, EPI_ISL_2914230, EPI_ISL_2914231, EPI_ISL_2914233 | Lighthouse Laboratory Plymouth | Wellcome Sanger Institute for the COVID-19 Genomics UK (COG-UK) Consortium | Lighthouse Laboratory Plymouth and Alex Alderton, Roberto Amato, Jeffrey Barrett, Sonia Goncalves, Ewan Harrison, David K. Jackson, Ian Johnston, Dominic Kwiatkowski, Cordelia Langford, John Sillitoe on behalf of the Wellcome Sanger Institute COVID-19 Surveillance Team |
| EPI_ISL_2914234 | Lighthouse Lab in Glasgow | Wellcome Sanger Institute for the COVID-19 Genomics UK (COG-UK) Consortium | Harper VanSteenhouse, Yumi Kasai, David Gray, Carol Clugston, Anna Dominiczak and Alex Alderton, Roberto Amato, Jeffrey Barrett, Sonia Goncalves, Ewan Harrison, David K. Jackson, Ian Johnston, Dominic Kwiatkowski, Cordelia Langford, John Sillitoe on behalf of the Wellcome Sanger Institute COVID-19 Surveillance Team |
| EPI_ISL_2914235, EPI_ISL_2914236, EPI_ISL_2914237, EPI_ISL_2914238, EPI_ISL_2914239, EPI_ISL_2914240, EPI_ISL_2914241, EPI_ISL_2914242 | Lighthouse Laboratory Plymouth | Wellcome Sanger Institute for the COVID-19 Genomics UK (COG-UK) Consortium | Lighthouse Laboratory Plymouth and Alex Alderton, Roberto Amato, Jeffrey Barrett, Sonia Goncalves, Ewan Harrison, David K. Jackson, Ian Johnston, Dominic Kwiatkowski, Cordelia Langford, John Sillitoe on behalf of the Wellcome Sanger Institute COVID-19 Surveillance Team |
| EPI_ISL_2914243 | Lighthouse Lab in Glasgow | Wellcome Sanger Institute for the COVID-19 Genomics UK (COG-UK) Consortium | Harper VanSteenhouse, Yumi Kasai, David Gray, Carol Clugston, Anna Dominiczak and Alex Alderton, Roberto Amato, Jeffrey Barrett, Sonia Goncalves, Ewan Harrison, David K. Jackson, Ian Johnston, Dominic Kwiatkowski, Cordelia Langford, John Sillitoe on behalf of the Wellcome Sanger Institute COVID-19 Surveillance Team |
| EPI_ISL_2914244, EPI_ISL_2914245, EPI_ISL_2914246, EPI_ISL_2914247, EPI_ISL_2914248, EPI_ISL_2914249, EPI_ISL_2914250, EPI_ISL_2914251, EPI_ISL_2914252, EPI_ISL_2914253, EPI_ISL_2914254, EPI_ISL_2914255, EPI_ISL_2914256, EPI_ISL_2914257, EPI_ISL_2914258, EPI_ISL_2914259, EPI_ISL_2914260, EPI_ISL_2914261 | | | |
| see above | Lighthouse Laboratory Plymouth | Wellcome Sanger Institute for the COVID-19 Genomics UK (COG-UK) Consortium | Lighthouse Laboratory Plymouth and Alex Alderton, Roberto Amato, Jeffrey Barrett, Sonia Goncalves, Ewan Harrison, David K. Jackson, Ian Johnston, Dominic Kwiatkowski, Cordelia Langford, John Sillitoe on behalf of the Wellcome Sanger Institute COVID-19 Surveillance Team |
| EPI_ISL_2914262, EPI_ISL_2914263 | Lighthouse Lab in Glasgow | Wellcome Sanger Institute for the COVID-19 Genomics UK (COG-UK) Consortium | Harper VanSteenhouse, Yumi Kasai, David Gray, Carol Clugston, Anna Dominiczak and Alex Alderton, Roberto Amato, Jeffrey Barrett, Sonia Goncalves, Ewan Harrison, David K. Jackson, Ian Johnston, Dominic Kwiatkowski, Cordelia Langford, John Sillitoe on behalf of the Wellcome Sanger Institute COVID-19 Surveillance Team |
| EPI_ISL_2914264, EPI_ISL_2914265, EPI_ISL_2914266 | Lighthouse Laboratory Plymouth | Wellcome Sanger Institute for the COVID-19 Genomics UK (COG-UK) Consortium | Lighthouse Laboratory Plymouth and Alex Alderton, Roberto Amato, Jeffrey Barrett, Sonia Goncalves, Ewan Harrison, David K. Jackson, Ian Johnston, Dominic Kwiatkowski, Cordelia Langford, John Sillitoe on behalf of the Wellcome Sanger Institute COVID-19 Surveillance Team |
| EPI_ISL_2914267 | Lighthouse Lab in Glasgow | Wellcome Sanger Institute for the COVID-19 Genomics UK (COG-UK) Consortium | Harper VanSteenhouse, Yumi Kasai, David Gray, Carol Clugston, Anna Dominiczak and Alex Alderton, Roberto Amato, Jeffrey Barrett, Sonia Goncalves, Ewan Harrison, David K. Jackson, Ian Johnston, Dominic Kwiatkowski, Cordelia Langford, John Sillitoe on behalf of the Wellcome Sanger Institute COVID-19 Surveillance Team |
| EPI_ISL_2914268, EPI_ISL_2914269, EPI_ISL_2914270, EPI_ISL_2914271, EPI_ISL_2914272, EPI_ISL_2914273, EPI_ISL_2914274, EPI_ISL_2914275, EPI_ISL_2914276, EPI_ISL_2914277, EPI_ISL_2914278 | | | |
| see above | Lighthouse Laboratory Plymouth | Wellcome Sanger Institute for the COVID-19 Genomics UK (COG-UK) Consortium | Lighthouse Laboratory Plymouth and Alex Alderton, Roberto Amato, Jeffrey Barrett, Sonia Goncalves, Ewan Harrison, David K. Jackson, Ian Johnston, Dominic Kwiatkowski, Cordelia Langford, John Sillitoe on behalf of the Wellcome Sanger Institute COVID-19 Surveillance Team |
| EPI_ISL_2914279 | Berkshire and Surrey Pathology Services Lighthouse Laboratory | Wellcome Sanger Institute for the COVID-19 Genomics UK (COG-UK) Consortium | Berkshire and Surrey Pathology Services Lighthouse Laboratory and Alex Alderton, Roberto Amato, Jeffrey Barrett, Sonia Goncalves, Ewan Harrison, David K. Jackson, Ian Johnston, Dominic Kwiatkowski, Cordelia Langford, John Sillitoe on behalf of the Wellcome Sanger Institute COVID-19 Surveillance Team |
| EPI_ISL_2914280, EPI_ISL_2914281, EPI_ISL_2914282, EPI_ISL_2914283 | Lighthouse Lab in Alderley Park | Wellcome Sanger Institute for the COVID-19 Genomics UK (COG-UK) Consortium | Jacquelyn Wynn, Mairead Hyland, The Lighthouse Lab in Alderley Park and Alex Alderton, Roberto Amato, Jeffrey Barrett, Sonia Goncalves, Ewan Harrison, David K. Jackson, Ian Johnston, Dominic Kwiatkowski, Cordelia Langford, John Sillitoe on behalf of the Wellcome Sanger Institute COVID-19 Surveillance Team |
| EPI_ISL_2914284 | Berkshire and Surrey Pathology Services Lighthouse Laboratory | Wellcome Sanger Institute for the COVID-19 Genomics UK (COG-UK) Consortium | Berkshire and Surrey Pathology Services Lighthouse Laboratory and Alex Alderton, Roberto Amato, Jeffrey Barrett, Sonia Goncalves, Ewan Harrison, David K. Jackson, Ian Johnston, Dominic Kwiatkowski, Cordelia Langford, John Sillitoe on behalf of the Wellcome Sanger Institute COVID-19 Surveillance Team |
| EPI_ISL_2914285, EPI_ISL_2914286 | Lighthouse Lab in Glasgow | Wellcome Sanger Institute for the COVID-19 Genomics UK (COG-UK) Consortium | Harper VanSteenhouse, Yumi Kasai, David Gray, Carol Clugston, Anna Dominiczak and Alex Alderton, Roberto Amato, Jeffrey Barrett, Sonia Goncalves, Ewan Harrison, David K. Jackson, Ian Johnston, Dominic Kwiatkowski, Cordelia Langford, John Sillitoe on behalf of the Wellcome Sanger Institute COVID-19 Surveillance Team |
| EPI_ISL_2914287 | Berkshire and Surrey Pathology Services Lighthouse Laboratory | Wellcome Sanger Institute for the COVID-19 Genomics UK (COG-UK) Consortium | Berkshire and Surrey Pathology Services Lighthouse Laboratory and Alex Alderton, Roberto Amato, Jeffrey Barrett, Sonia Goncalves, Ewan Harrison, David K. Jackson, Ian Johnston, Dominic Kwiatkowski, Cordelia Langford, John Sillitoe on behalf of the Wellcome Sanger Institute COVID-19 Surveillance Team |
| EPI_ISL_2914288 | Lighthouse Lab in Glasgow | Wellcome Sanger Institute for the COVID-19 Genomics UK (COG-UK) Consortium | Harper VanSteenhouse, Yumi Kasai, David Gray, Carol Clugston, Anna Dominiczak and Alex Alderton, Roberto Amato, Jeffrey Barrett, Sonia Goncalves, Ewan Harrison, David K. Jackson, Ian Johnston, Dominic Kwiatkowski, Cordelia Langford, John Sillitoe on behalf of the Wellcome Sanger Institute COVID-19 Surveillance Team |
| EPI_ISL_2914289 | Berkshire and Surrey Pathology Services Lighthouse Laboratory | Wellcome Sanger Institute for the COVID-19 Genomics UK (COG-UK) Consortium | Berkshire and Surrey Pathology Services Lighthouse Laboratory and Alex Alderton, Roberto Amato, Jeffrey Barrett, Sonia Goncalves, Ewan Harrison, David K. Jackson, Ian Johnston, Dominic Kwiatkowski, Cordelia Langford, John Sillitoe on behalf of the Wellcome Sanger Institute COVID-19 Surveillance Team |
| EPI_ISL_2914290, EPI_ISL_2914291, EPI_ISL_2914292 | Lighthouse Lab in Glasgow | Wellcome Sanger Institute for the COVID-19 Genomics UK (COG-UK) Consortium | Harper VanSteenhouse, Yumi Kasai, David Gray, Carol Clugston, Anna Dominiczak and Alex Alderton, Roberto Amato, Jeffrey Barrett, Sonia Goncalves, Ewan Harrison, David K. Jackson, Ian Johnston, Dominic Kwiatkowski, Cordelia Langford, John Sillitoe on behalf of the Wellcome Sanger Institute COVID-19 Surveillance Team |
| EPI_ISL_2914293 | Berkshire and Surrey Pathology Services Lighthouse Laboratory | Wellcome Sanger Institute for the COVID-19 Genomics UK (COG-UK) Consortium | Berkshire and Surrey Pathology Services Lighthouse Laboratory and Alex Alderton, Roberto Amato, Jeffrey Barrett, Sonia Goncalves, Ewan Harrison, David K. Jackson, Ian Johnston, Dominic Kwiatkowski, Cordelia Langford, John Sillitoe on behalf of the Wellcome Sanger Institute COVID-19 Surveillance Team |
| EPI_ISL_2914294, EPI_ISL_2914295 | Lighthouse Lab in Glasgow | Wellcome Sanger Institute for the COVID-19 Genomics UK (COG-UK) Consortium | Harper VanSteenhouse, Yumi Kasai, David Gray, Carol Clugston, Anna Dominiczak and Alex Alderton, Roberto Amato, Jeffrey Barrett, Sonia Goncalves, Ewan Harrison, David K. Jackson, Ian Johnston, Dominic Kwiatkowski, Cordelia Langford, John Sillitoe on behalf of the Wellcome Sanger Institute COVID-19 Surveillance Team |

| | | | |
|---|---|---|---|
| EPI_ISL_2914296 | Lighthouse Lab in Alderley Park | Wellcome Sanger Institute for the COVID-19 Genomics UK (COG-UK) Consortium | Jacquelyn Wynn, Mairead Hyland, The Lighthouse Lab in Alderley Park and Alex Alderton, Roberto Amato, Jeffrey Barrett, Sonia Goncalves, Ewan Harrison, David K. Jackson, Ian Johnston, Dominic Kwiatkowski, Cordelia Langford, John Sillitoe on behalf of the Wellcome Sanger Institute COVID-19 Surveillance Team |
| EPI_ISL_2914297 | Berkshire and Surrey Pathology Services Lighthouse Laboratory | Wellcome Sanger Institute for the COVID-19 Genomics UK (COG-UK) Consortium | Berkshire and Surrey Pathology Services Lighthouse Laboratory and Alex Alderton, Roberto Amato, Jeffrey Barrett, Sonia Goncalves, Ewan Harrison, David K. Jackson, Ian Johnston, Dominic Kwiatkowski, Cordelia Langford, John Sillitoe on behalf of the Wellcome Sanger Institute COVID-19 Surveillance Team |
| EPI_ISL_2914298, EPI_ISL_2914299, EPI_ISL_2914301, EPI_ISL_2914302, EPI_ISL_2914303, EPI_ISL_2914304 | Lighthouse Lab in Glasgow | Wellcome Sanger Institute for the COVID-19 Genomics UK (COG-UK) Consortium | Harper VanSteenhouse, Yumi Kasai, David Gray, Carol Clugston, Anna Dominiczak and Alex Alderton, Roberto Amato, Jeffrey Barrett, Sonia Goncalves, Ewan Harrison, David K. Jackson, Ian Johnston, Dominic Kwiatkowski, Cordelia Langford, John Sillitoe on behalf of the Wellcome Sanger Institute COVID-19 Surveillance Team |
| EPI_ISL_2914305, EPI_ISL_2914306 | Berkshire and Surrey Pathology Services Lighthouse Laboratory | Wellcome Sanger Institute for the COVID-19 Genomics UK (COG-UK) Consortium | Berkshire and Surrey Pathology Services Lighthouse Laboratory and Alex Alderton, Roberto Amato, Jeffrey Barrett, Sonia Goncalves, Ewan Harrison, David K. Jackson, Ian Johnston, Dominic Kwiatkowski, Cordelia Langford, John Sillitoe on behalf of the Wellcome Sanger Institute COVID-19 Surveillance Team |
| EPI_ISL_2914307 | Lighthouse Lab in Glasgow | Wellcome Sanger Institute for the COVID-19 Genomics UK (COG-UK) Consortium | Harper VanSteenhouse, Yumi Kasai, David Gray, Carol Clugston, Anna Dominiczak and Alex Alderton, Roberto Amato, Jeffrey Barrett, Sonia Goncalves, Ewan Harrison, David K. Jackson, Ian Johnston, Dominic Kwiatkowski, Cordelia Langford, John Sillitoe on behalf of the Wellcome Sanger Institute COVID-19 Surveillance Team |
| EPI_ISL_2914308 | Lighthouse Lab in Alderley Park | Wellcome Sanger Institute for the COVID-19 Genomics UK (COG-UK) Consortium | Jacquelyn Wynn, Mairead Hyland, The Lighthouse Lab in Alderley Park and Alex Alderton, Roberto Amato, Jeffrey Barrett, Sonia Goncalves, Ewan Harrison, David K. Jackson, Ian Johnston, Dominic Kwiatkowski, Cordelia Langford, John Sillitoe on behalf of the Wellcome Sanger Institute COVID-19 Surveillance Team |
| EPI_ISL_2914309, EPI_ISL_2914310 | Lighthouse Lab in Glasgow | Wellcome Sanger Institute for the COVID-19 Genomics UK (COG-UK) Consortium | Harper VanSteenhouse, Yumi Kasai, David Gray, Carol Clugston, Anna Dominiczak and Alex Alderton, Roberto Amato, Jeffrey Barrett, Sonia Goncalves, Ewan Harrison, David K. Jackson, Ian Johnston, Dominic Kwiatkowski, Cordelia Langford, John Sillitoe on behalf of the Wellcome Sanger Institute COVID-19 Surveillance Team |
| EPI_ISL_2914311 | Lighthouse Lab in Alderley Park | Wellcome Sanger Institute for the COVID-19 Genomics UK (COG-UK) Consortium | Jacquelyn Wynn, Mairead Hyland, The Lighthouse Lab in Alderley Park and Alex Alderton, Roberto Amato, Jeffrey Barrett, Sonia Goncalves, Ewan Harrison, David K. Jackson, Ian Johnston, Dominic Kwiatkowski, Cordelia Langford, John Sillitoe on behalf of the Wellcome Sanger Institute COVID-19 Surveillance Team |
| EPI_ISL_2914312 | Lighthouse Lab in Glasgow | Wellcome Sanger Institute for the COVID-19 Genomics UK (COG-UK) Consortium | Harper VanSteenhouse, Yumi Kasai, David Gray, Carol Clugston, Anna Dominiczak and Alex Alderton, Roberto Amato, Jeffrey Barrett, Sonia Goncalves, Ewan Harrison, David K. Jackson, Ian Johnston, Dominic Kwiatkowski, Cordelia Langford, John Sillitoe on behalf of the Wellcome Sanger Institute COVID-19 Surveillance Team |
| EPI_ISL_2914313 | Berkshire and Surrey Pathology Services Lighthouse Laboratory | Wellcome Sanger Institute for the COVID-19 Genomics UK (COG-UK) Consortium | Berkshire and Surrey Pathology Services Lighthouse Laboratory and Alex Alderton, Roberto Amato, Jeffrey Barrett, Sonia Goncalves, Ewan Harrison, David K. Jackson, Ian Johnston, Dominic Kwiatkowski, Cordelia Langford, John Sillitoe on behalf of the Wellcome Sanger Institute COVID-19 Surveillance Team |
| EPI_ISL_2914314, EPI_ISL_2914315 | Lighthouse Lab in Glasgow | Wellcome Sanger Institute for the COVID-19 Genomics UK (COG-UK) Consortium | Harper VanSteenhouse, Yumi Kasai, David Gray, Carol Clugston, Anna Dominiczak and Alex Alderton, Roberto Amato, Jeffrey Barrett, Sonia Goncalves, Ewan Harrison, David K. Jackson, Ian Johnston, Dominic Kwiatkowski, Cordelia Langford, John Sillitoe on behalf of the Wellcome Sanger Institute COVID-19 Surveillance Team |
| EPI_ISL_2914316, EPI_ISL_2914317 | Lighthouse Lab in Alderley Park | Wellcome Sanger Institute for the COVID-19 Genomics UK (COG-UK) Consortium | Jacquelyn Wynn, Mairead Hyland, The Lighthouse Lab in Alderley Park and Alex Alderton, Roberto Amato, Jeffrey Barrett, Sonia Goncalves, Ewan Harrison, David K. Jackson, Ian Johnston, Dominic Kwiatkowski, Cordelia Langford, John Sillitoe on behalf of the Wellcome Sanger Institute COVID-19 Surveillance Team |
| EPI_ISL_2914318 | Berkshire and Surrey Pathology Services Lighthouse Laboratory | Wellcome Sanger Institute for the COVID-19 Genomics UK (COG-UK) Consortium | Berkshire and Surrey Pathology Services Lighthouse Laboratory and Alex Alderton, Roberto Amato, Jeffrey Barrett, Sonia Goncalves, Ewan Harrison, David K. Jackson, Ian Johnston, Dominic Kwiatkowski, Cordelia Langford, John Sillitoe on behalf of the Wellcome Sanger Institute COVID-19 Surveillance Team |
| EPI_ISL_2914319 | Lighthouse Lab in Glasgow | Wellcome Sanger Institute for the COVID-19 Genomics UK (COG-UK) Consortium | Harper VanSteenhouse, Yumi Kasai, David Gray, Carol Clugston, Anna Dominiczak and Alex Alderton, Roberto Amato, Jeffrey Barrett, Sonia Goncalves, Ewan Harrison, David K. Jackson, Ian Johnston, Dominic Kwiatkowski, Cordelia Langford, John Sillitoe on behalf of the Wellcome Sanger Institute COVID-19 Surveillance Team |
| EPI_ISL_2914320 | Berkshire and Surrey Pathology Services Lighthouse Laboratory | Wellcome Sanger Institute for the COVID-19 Genomics UK (COG-UK) Consortium | Berkshire and Surrey Pathology Services Lighthouse Laboratory and Alex Alderton, Roberto Amato, Jeffrey Barrett, Sonia Goncalves, Ewan Harrison, David K. Jackson, Ian Johnston, Dominic Kwiatkowski, Cordelia Langford, John Sillitoe on behalf of the Wellcome Sanger Institute COVID-19 Surveillance Team |
| EPI_ISL_2914321, EPI_ISL_2914322, EPI_ISL_2914323, EPI_ISL_2914324, EPI_ISL_2914325, EPI_ISL_2914326 | Lighthouse Lab in Glasgow | Wellcome Sanger Institute for the COVID-19 Genomics UK (COG-UK) Consortium | Harper VanSteenhouse, Yumi Kasai, David Gray, Carol Clugston, Anna Dominiczak and Alex Alderton, Roberto Amato, Jeffrey Barrett, Sonia Goncalves, Ewan Harrison, David K. Jackson, Ian Johnston, Dominic Kwiatkowski, Cordelia Langford, John Sillitoe on behalf of the Wellcome Sanger Institute COVID-19 Surveillance Team |
| EPI_ISL_2914327 | Lighthouse Lab in Alderley Park | Wellcome Sanger Institute for the COVID-19 Genomics UK (COG-UK) Consortium | Jacquelyn Wynn, Mairead Hyland, The Lighthouse Lab in Alderley Park and Alex Alderton, Roberto Amato, Jeffrey Barrett, Sonia Goncalves, Ewan Harrison, David K. Jackson, Ian Johnston, Dominic Kwiatkowski, Cordelia Langford, John Sillitoe on behalf of the Wellcome Sanger Institute COVID-19 Surveillance Team |
| EPI_ISL_2914328 | Berkshire and Surrey Pathology Services Lighthouse Laboratory | Wellcome Sanger Institute for the COVID-19 Genomics UK (COG-UK) Consortium | Berkshire and Surrey Pathology Services Lighthouse Laboratory and Alex Alderton, Roberto Amato, Jeffrey Barrett, Sonia Goncalves, Ewan Harrison, David K. Jackson, Ian Johnston, Dominic Kwiatkowski, Cordelia Langford, John Sillitoe on behalf of the Wellcome Sanger Institute COVID-19 Surveillance Team |
| EPI_ISL_2914329, EPI_ISL_2914330, EPI_ISL_2914331, EPI_ISL_2914332, EPI_ISL_2914333, EPI_ISL_2914334, EPI_ISL_2914335 | Lighthouse Lab in Glasgow | Wellcome Sanger Institute for the COVID-19 Genomics UK (COG-UK) Consortium | Harper VanSteenhouse, Yumi Kasai, David Gray, Carol Clugston, Anna Dominiczak and Alex Alderton, Roberto Amato, Jeffrey Barrett, Sonia Goncalves, Ewan Harrison, David K. Jackson, Ian Johnston, Dominic Kwiatkowski, Cordelia Langford, John Sillitoe on behalf of the Wellcome Sanger Institute COVID-19 Surveillance Team |
| EPI_ISL_2914336 | Berkshire and Surrey Pathology Services Lighthouse Laboratory | Wellcome Sanger Institute for the COVID-19 Genomics UK (COG-UK) Consortium | Berkshire and Surrey Pathology Services Lighthouse Laboratory and Alex Alderton, Roberto Amato, Jeffrey Barrett, Sonia Goncalves, Ewan Harrison, David K. Jackson, Ian Johnston, Dominic Kwiatkowski, Cordelia Langford, John Sillitoe on behalf of the Wellcome Sanger Institute COVID-19 Surveillance Team |
| EPI_ISL_2914337 | Lighthouse Lab in Alderley Park | Wellcome Sanger Institute for the COVID-19 Genomics UK (COG-UK) Consortium | Jacquelyn Wynn, Mairead Hyland, The Lighthouse Lab in Alderley Park and Alex Alderton, Roberto Amato, Jeffrey Barrett, Sonia Goncalves, Ewan Harrison, David K. Jackson, Ian Johnston, Dominic Kwiatkowski, Cordelia Langford, John Sillitoe on behalf of the Wellcome Sanger Institute COVID-19 Surveillance Team |
| EPI_ISL_2914338 | Lighthouse Lab in Glasgow | Wellcome Sanger Institute for the COVID-19 Genomics UK (COG-UK) Consortium | Harper VanSteenhouse, Yumi Kasai, David Gray, Carol Clugston, Anna Dominiczak and Alex Alderton, Roberto Amato, Jeffrey Barrett, Sonia Goncalves, Ewan Harrison, David K. Jackson, Ian Johnston, Dominic Kwiatkowski, Cordelia Langford, John Sillitoe on behalf of the Wellcome Sanger Institute COVID-19 Surveillance Team |
| EPI_ISL_2914339 | Lighthouse Lab in Alderley Park | Wellcome Sanger Institute for the COVID-19 Genomics UK (COG-UK) Consortium | Jacquelyn Wynn, Mairead Hyland, The Lighthouse Lab in Alderley Park and Alex Alderton, Roberto Amato, Jeffrey Barrett, Sonia Goncalves, Ewan Harrison, David K. Jackson, Ian Johnston, Dominic Kwiatkowski, Cordelia Langford, John Sillitoe on behalf of the Wellcome Sanger Institute COVID-19 Surveillance Team |
| EPI_ISL_2914340 | Lighthouse Lab in Glasgow | Wellcome Sanger Institute for the COVID-19 Genomics UK (COG-UK) Consortium | Harper VanSteenhouse, Yumi Kasai, David Gray, Carol Clugston, Anna Dominiczak and Alex Alderton, Roberto Amato, Jeffrey Barrett, Sonia Goncalves, Ewan Harrison, David K. Jackson, Ian Johnston, Dominic Kwiatkowski, Cordelia Langford, John Sillitoe on behalf of the Wellcome Sanger Institute COVID-19 Surveillance Team |
| EPI_ISL_2914341, EPI_ISL_2914342 | Lighthouse Lab in Alderley Park | Wellcome Sanger Institute for the COVID-19 Genomics UK (COG-UK) Consortium | Jacquelyn Wynn, Mairead Hyland, The Lighthouse Lab in Alderley Park and Alex Alderton, Roberto Amato, Jeffrey Barrett, Sonia Goncalves, Ewan Harrison, David K. Jackson, Ian Johnston, Dominic Kwiatkowski, Cordelia Langford, John Sillitoe on behalf of the Wellcome Sanger Institute COVID-19 Surveillance Team |
| EPI_ISL_2914343, EPI_ISL_2914344 | Lighthouse Lab in Glasgow | Wellcome Sanger Institute for the COVID-19 Genomics UK (COG-UK) Consortium | Harper VanSteenhouse, Yumi Kasai, David Gray, Carol Clugston, Anna Dominiczak and Alex Alderton, Roberto Amato, Jeffrey Barrett, Sonia Goncalves, Ewan Harrison, David K. Jackson, Ian Johnston, Dominic Kwiatkowski, Cordelia Langford, John Sillitoe on behalf of the Wellcome Sanger Institute COVID-19 Surveillance Team |
| EPI_ISL_2914345, EPI_ISL_2914346 | Lighthouse Lab in Alderley Park | Wellcome Sanger Institute for the COVID-19 Genomics UK (COG-UK) Consortium | Jacquelyn Wynn, Mairead Hyland, The Lighthouse Lab in Alderley Park and Alex Alderton, Roberto Amato, Jeffrey Barrett, Sonia Goncalves, Ewan Harrison, David K. Jackson, Ian Johnston, Dominic Kwiatkowski, Cordelia Langford, John Sillitoe on behalf of the Wellcome Sanger Institute COVID-19 Surveillance Team |
| EPI_ISL_2914347 | Lighthouse Lab in Glasgow | Wellcome Sanger Institute for the COVID-19 Genomics UK (COG-UK) Consortium | Harper VanSteenhouse, Yumi Kasai, David Gray, Carol Clugston, Anna Dominiczak and Alex Alderton, Roberto Amato, Jeffrey Barrett, Sonia Goncalves, Ewan Harrison, David K. Jackson, Ian Johnston, Dominic Kwiatkowski, Cordelia Langford, John Sillitoe on behalf of the Wellcome Sanger Institute COVID-19 Surveillance Team |
| EPI_ISL_2914348 | Berkshire and Surrey Pathology Services Lighthouse Laboratory | Wellcome Sanger Institute for the COVID-19 Genomics UK (COG-UK) Consortium | Berkshire and Surrey Pathology Services Lighthouse Laboratory and Alex Alderton, Roberto Amato, Jeffrey Barrett, Sonia Goncalves, Ewan Harrison, David K. Jackson, Ian Johnston, Dominic Kwiatkowski, Cordelia Langford, John Sillitoe on behalf of the Wellcome Sanger Institute COVID-19 Surveillance Team |
| EPI_ISL_2914349 | Lighthouse Lab in Glasgow | Wellcome Sanger Institute for the COVID-19 Genomics UK (COG-UK) Consortium | Harper VanSteenhouse, Yumi Kasai, David Gray, Carol Clugston, Anna Dominiczak and Alex Alderton, Roberto Amato, Jeffrey Barrett, Sonia Goncalves, Ewan Harrison, David K. Jackson, Ian Johnston, Dominic Kwiatkowski, Cordelia Langford, John Sillitoe on behalf of the Wellcome Sanger Institute COVID-19 Surveillance Team |
| EPI_ISL_2914350, EPI_ISL_2914351, EPI_ISL_2914352 | Lighthouse Lab in Alderley Park | Wellcome Sanger Institute for the COVID-19 Genomics UK (COG-UK) Consortium | Jacquelyn Wynn, Mairead Hyland, The Lighthouse Lab in Alderley Park and Alex Alderton, Roberto Amato, Jeffrey Barrett, Sonia Goncalves, Ewan Harrison, David K. Jackson, Ian Johnston, Dominic Kwiatkowski, Cordelia Langford, John Sillitoe on behalf of the Wellcome Sanger Institute COVID-19 Surveillance Team |
| EPI_ISL_2914353, EPI_ISL_2914354, EPI_ISL_2914355, EPI_ISL_2914356 | Lighthouse Lab in Glasgow | Wellcome Sanger Institute for the COVID-19 Genomics UK (COG-UK) Consortium | Harper VanSteenhouse, Yumi Kasai, David Gray, Carol Clugston, Anna Dominiczak and Alex Alderton, Roberto Amato, Jeffrey Barrett, Sonia Goncalves, Ewan Harrison, David K. Jackson, Ian Johnston, Dominic Kwiatkowski, Cordelia Langford, John Sillitoe on behalf of the Wellcome Sanger Institute COVID-19 Surveillance Team |
| EPI_ISL_2914357, EPI_ISL_2914358 | Berkshire and Surrey Pathology Services Lighthouse Laboratory | Wellcome Sanger Institute for the COVID-19 Genomics UK (COG-UK) Consortium | Berkshire and Surrey Pathology Services Lighthouse Laboratory and Alex Alderton, Roberto Amato, Jeffrey Barrett, Sonia Goncalves, Ewan Harrison, David K. Jackson, Ian Johnston, Dominic Kwiatkowski, Cordelia Langford, John Sillitoe on behalf of the Wellcome Sanger Institute COVID-19 Surveillance Team |
| EPI_ISL_2914359 | Lighthouse Lab in Glasgow | Wellcome Sanger Institute for the COVID-19 Genomics UK (COG-UK) Consortium | Harper VanSteenhouse, Yumi Kasai, David Gray, Carol Clugston, Anna Dominiczak and Alex Alderton, Roberto Amato, Jeffrey Barrett, Sonia Goncalves, Ewan Harrison, David K. Jackson, Ian Johnston, Dominic Kwiatkowski, Cordelia Langford, John Sillitoe on behalf of the Wellcome Sanger Institute COVID-19 Surveillance Team |
| EPI_ISL_2914360 | Lighthouse Lab in Alderley Park | Wellcome Sanger Institute for the COVID-19 Genomics UK (COG-UK) Consortium | Jacquelyn Wynn, Mairead Hyland, The Lighthouse Lab in Alderley Park and Alex Alderton, Roberto Amato, Jeffrey Barrett, Sonia Goncalves, Ewan Harrison, David K. Jackson, Ian Johnston, Dominic Kwiatkowski, Cordelia Langford, John Sillitoe on behalf of the Wellcome Sanger Institute COVID-19 Surveillance Team |
| EPI_ISL_2914361 | Lighthouse Lab in Glasgow | Wellcome Sanger Institute for the COVID-19 Genomics UK (COG-UK) Consortium | Harper VanSteenhouse, Yumi Kasai, David Gray, Carol Clugston, Anna Dominiczak and Alex Alderton, Roberto Amato, Jeffrey Barrett, Sonia Goncalves, Ewan Harrison, David K. Jackson, Ian Johnston, Dominic Kwiatkowski, Cordelia Langford, John Sillitoe on behalf of the Wellcome Sanger Institute COVID-19 Surveillance Team |
| EPI_ISL_2914362 | Lighthouse Lab in Alderley Park | Wellcome Sanger Institute for the COVID-19 Genomics UK (COG-UK) Consortium | Jacquelyn Wynn, Mairead Hyland, The Lighthouse Lab in Alderley Park and Alex Alderton, Roberto Amato, Jeffrey Barrett, Sonia Goncalves, Ewan Harrison, David K. Jackson, Ian Johnston, Dominic Kwiatkowski, Cordelia Langford, John Sillitoe on behalf of the Wellcome Sanger Institute COVID-19 Surveillance Team |
| EPI_ISL_2914363, EPI_ISL_2914364, EPI_ISL_2914365, EPI_ISL_2914366 | Lighthouse Lab in Glasgow | Wellcome Sanger Institute for the COVID-19 Genomics UK (COG-UK) Consortium | Harper VanSteenhouse, Yumi Kasai, David Gray, Carol Clugston, Anna Dominiczak and Alex Alderton, Roberto Amato, Jeffrey Barrett, Sonia Goncalves, Ewan Harrison, David K. Jackson, Ian Johnston, Dominic Kwiatkowski, Cordelia Langford, John Sillitoe on behalf of the Wellcome Sanger Institute COVID-19 Surveillance Team |

| | | | |
|---|---|---|---|
| EPI_ISL_2914367, EPI_ISL_2914368 | Berkshire and Surrey Pathology Services Lighthouse Laboratory | Wellcome Sanger Institute for the COVID-19 Genomics UK (COG-UK) Consortium | Berkshire and Surrey Pathology Services Lighthouse Laboratory and Alex Alderton, Roberto Amato, Jeffrey Barrett, Sonia Goncalves, Ewan Harrison, David K. Jackson, Ian Johnston, Dominic Kwiatkowski, Cordelia Langford, John Sillitoe on behalf of the Wellcome Sanger Institute COVID-19 Surveillance Team |
| EPI_ISL_2914369 | Lighthouse Lab in Glasgow | Wellcome Sanger Institute for the COVID-19 Genomics UK (COG-UK) Consortium | Harper VanSteenhouse, Yumi Kasai, David Gray, Carol Clugston, Anna Dominiczak and Alex Alderton, Roberto Amato, Jeffrey Barrett, Sonia Goncalves, Ewan Harrison, David K. Jackson, Ian Johnston, Dominic Kwiatkowski, Cordelia Langford, John Sillitoe on behalf of the Wellcome Sanger Institute COVID-19 Surveillance Team |
| EPI_ISL_2914370 | Berkshire and Surrey Pathology Services Lighthouse Laboratory | Wellcome Sanger Institute for the COVID-19 Genomics UK (COG-UK) Consortium | Berkshire and Surrey Pathology Services Lighthouse Laboratory and Alex Alderton, Roberto Amato, Jeffrey Barrett, Sonia Goncalves, Ewan Harrison, David K. Jackson, Ian Johnston, Dominic Kwiatkowski, Cordelia Langford, John Sillitoe on behalf of the Wellcome Sanger Institute COVID-19 Surveillance Team |
| EPI_ISL_2914371 | Lighthouse Lab in Glasgow | Wellcome Sanger Institute for the COVID-19 Genomics UK (COG-UK) Consortium | Harper VanSteenhouse, Yumi Kasai, David Gray, Carol Clugston, Anna Dominiczak and Alex Alderton, Roberto Amato, Jeffrey Barrett, Sonia Goncalves, Ewan Harrison, David K. Jackson, Ian Johnston, Dominic Kwiatkowski, Cordelia Langford, John Sillitoe on behalf of the Wellcome Sanger Institute COVID-19 Surveillance Team |
| EPI_ISL_2914372 | Lighthouse Lab in Alderley Park | Wellcome Sanger Institute for the COVID-19 Genomics UK (COG-UK) Consortium | Jacquelyn Wynn, Mairead Hyland, The Lighthouse Lab in Alderley Park and Alex Alderton, Roberto Amato, Jeffrey Barrett, Sonia Goncalves, Ewan Harrison, David K. Jackson, Ian Johnston, Dominic Kwiatkowski, Cordelia Langford, John Sillitoe on behalf of the Wellcome Sanger Institute COVID-19 Surveillance Team |
| EPI_ISL_2914373, EPI_ISL_2914374, EPI_ISL_2914375 | Lighthouse Lab in Glasgow | Wellcome Sanger Institute for the COVID-19 Genomics UK (COG-UK) Consortium | Harper VanSteenhouse, Yumi Kasai, David Gray, Carol Clugston, Anna Dominiczak and Alex Alderton, Roberto Amato, Jeffrey Barrett, Sonia Goncalves, Ewan Harrison, David K. Jackson, Ian Johnston, Dominic Kwiatkowski, Cordelia Langford, John Sillitoe on behalf of the Wellcome Sanger Institute COVID-19 Surveillance Team |
| EPI_ISL_2914376 | Lighthouse Lab in Alderley Park | Wellcome Sanger Institute for the COVID-19 Genomics UK (COG-UK) Consortium | Jacquelyn Wynn, Mairead Hyland, The Lighthouse Lab in Alderley Park and Alex Alderton, Roberto Amato, Jeffrey Barrett, Sonia Goncalves, Ewan Harrison, David K. Jackson, Ian Johnston, Dominic Kwiatkowski, Cordelia Langford, John Sillitoe on behalf of the Wellcome Sanger Institute COVID-19 Surveillance Team |
| EPI_ISL_2914377, EPI_ISL_2914378, EPI_ISL_2914379, EPI_ISL_2914380 | Lighthouse Lab in Glasgow | Wellcome Sanger Institute for the COVID-19 Genomics UK (COG-UK) Consortium | Harper VanSteenhouse, Yumi Kasai, David Gray, Carol Clugston, Anna Dominiczak and Alex Alderton, Roberto Amato, Jeffrey Barrett, Sonia Goncalves, Ewan Harrison, David K. Jackson, Ian Johnston, Dominic Kwiatkowski, Cordelia Langford, John Sillitoe on behalf of the Wellcome Sanger Institute COVID-19 Surveillance Team |
| EPI_ISL_2914381, EPI_ISL_2914382 | Berkshire and Surrey Pathology Services Lighthouse Laboratory | Wellcome Sanger Institute for the COVID-19 Genomics UK (COG-UK) Consortium | Berkshire and Surrey Pathology Services Lighthouse Laboratory and Alex Alderton, Roberto Amato, Jeffrey Barrett, Sonia Goncalves, Ewan Harrison, David K. Jackson, Ian Johnston, Dominic Kwiatkowski, Cordelia Langford, John Sillitoe on behalf of the Wellcome Sanger Institute COVID-19 Surveillance Team |
| EPI_ISL_2914383, EPI_ISL_2914384 | Lighthouse Lab in Alderley Park | Wellcome Sanger Institute for the COVID-19 Genomics UK (COG-UK) Consortium | Jacquelyn Wynn, Mairead Hyland, The Lighthouse Lab in Alderley Park and Alex Alderton, Roberto Amato, Jeffrey Barrett, Sonia Goncalves, Ewan Harrison, David K. Jackson, Ian Johnston, Dominic Kwiatkowski, Cordelia Langford, John Sillitoe on behalf of the Wellcome Sanger Institute COVID-19 Surveillance Team |
| EPI_ISL_2914385, EPI_ISL_2914386, EPI_ISL_2914387 | Lighthouse Lab in Glasgow | Wellcome Sanger Institute for the COVID-19 Genomics UK (COG-UK) Consortium | Harper VanSteenhouse, Yumi Kasai, David Gray, Carol Clugston, Anna Dominiczak and Alex Alderton, Roberto Amato, Jeffrey Barrett, Sonia Goncalves, Ewan Harrison, David K. Jackson, Ian Johnston, Dominic Kwiatkowski, Cordelia Langford, John Sillitoe on behalf of the Wellcome Sanger Institute COVID-19 Surveillance Team |
| EPI_ISL_2914388 | Berkshire and Surrey Pathology Services Lighthouse Laboratory | Wellcome Sanger Institute for the COVID-19 Genomics UK (COG-UK) Consortium | Berkshire and Surrey Pathology Services Lighthouse Laboratory and Alex Alderton, Roberto Amato, Jeffrey Barrett, Sonia Goncalves, Ewan Harrison, David K. Jackson, Ian Johnston, Dominic Kwiatkowski, Cordelia Langford, John Sillitoe on behalf of the Wellcome Sanger Institute COVID-19 Surveillance Team |
| EPI_ISL_2914389 | Lighthouse Lab in Alderley Park | Wellcome Sanger Institute for the COVID-19 Genomics UK (COG-UK) Consortium | Jacquelyn Wynn, Mairead Hyland, The Lighthouse Lab in Alderley Park and Alex Alderton, Roberto Amato, Jeffrey Barrett, Sonia Goncalves, Ewan Harrison, David K. Jackson, Ian Johnston, Dominic Kwiatkowski, Cordelia Langford, John Sillitoe on behalf of the Wellcome Sanger Institute COVID-19 Surveillance Team |
| EPI_ISL_2914390, EPI_ISL_2914391, EPI_ISL_2914392, EPI_ISL_2914393 | Lighthouse Lab in Glasgow | Wellcome Sanger Institute for the COVID-19 Genomics UK (COG-UK) Consortium | Harper VanSteenhouse, Yumi Kasai, David Gray, Carol Clugston, Anna Dominiczak and Alex Alderton, Roberto Amato, Jeffrey Barrett, Sonia Goncalves, Ewan Harrison, David K. Jackson, Ian Johnston, Dominic Kwiatkowski, Cordelia Langford, John Sillitoe on behalf of the Wellcome Sanger Institute COVID-19 Surveillance Team |
| EPI_ISL_2914394, EPI_ISL_2914395 | Lighthouse Lab in Alderley Park | Wellcome Sanger Institute for the COVID-19 Genomics UK (COG-UK) Consortium | Jacquelyn Wynn, Mairead Hyland, The Lighthouse Lab in Alderley Park and Alex Alderton, Roberto Amato, Jeffrey Barrett, Sonia Goncalves, Ewan Harrison, David K. Jackson, Ian Johnston, Dominic Kwiatkowski, Cordelia Langford, John Sillitoe on behalf of the Wellcome Sanger Institute COVID-19 Surveillance Team |
| EPI_ISL_2914396 | Berkshire and Surrey Pathology Services Lighthouse Laboratory | Wellcome Sanger Institute for the COVID-19 Genomics UK (COG-UK) Consortium | Berkshire and Surrey Pathology Services Lighthouse Laboratory and Alex Alderton, Roberto Amato, Jeffrey Barrett, Sonia Goncalves, Ewan Harrison, David K. Jackson, Ian Johnston, Dominic Kwiatkowski, Cordelia Langford, John Sillitoe on behalf of the Wellcome Sanger Institute COVID-19 Surveillance Team |
| EPI_ISL_2914397, EPI_ISL_2914398 | Lighthouse Lab in Alderley Park | Wellcome Sanger Institute for the COVID-19 Genomics UK (COG-UK) Consortium | Jacquelyn Wynn, Mairead Hyland, The Lighthouse Lab in Alderley Park and Alex Alderton, Roberto Amato, Jeffrey Barrett, Sonia Goncalves, Ewan Harrison, David K. Jackson, Ian Johnston, Dominic Kwiatkowski, Cordelia Langford, John Sillitoe on behalf of the Wellcome Sanger Institute COVID-19 Surveillance Team |
| EPI_ISL_2914399, EPI_ISL_2914400 | Lighthouse Lab in Glasgow | Wellcome Sanger Institute for the COVID-19 Genomics UK (COG-UK) Consortium | Harper VanSteenhouse, Yumi Kasai, David Gray, Carol Clugston, Anna Dominiczak and Alex Alderton, Roberto Amato, Jeffrey Barrett, Sonia Goncalves, Ewan Harrison, David K. Jackson, Ian Johnston, Dominic Kwiatkowski, Cordelia Langford, John Sillitoe on behalf of the Wellcome Sanger Institute COVID-19 Surveillance Team |
| EPI_ISL_2914401 | Lighthouse Lab in Alderley Park | Wellcome Sanger Institute for the COVID-19 Genomics UK (COG-UK) Consortium | Jacquelyn Wynn, Mairead Hyland, The Lighthouse Lab in Alderley Park and Alex Alderton, Roberto Amato, Jeffrey Barrett, Sonia Goncalves, Ewan Harrison, David K. Jackson, Ian Johnston, Dominic Kwiatkowski, Cordelia Langford, John Sillitoe on behalf of the Wellcome Sanger Institute COVID-19 Surveillance Team |
| EPI_ISL_2914402 | Berkshire and Surrey Pathology Services Lighthouse Laboratory | Wellcome Sanger Institute for the COVID-19 Genomics UK (COG-UK) Consortium | Berkshire and Surrey Pathology Services Lighthouse Laboratory and Alex Alderton, Roberto Amato, Jeffrey Barrett, Sonia Goncalves, Ewan Harrison, David K. Jackson, Ian Johnston, Dominic Kwiatkowski, Cordelia Langford, John Sillitoe on behalf of the Wellcome Sanger Institute COVID-19 Surveillance Team |
| EPI_ISL_2914403, EPI_ISL_2914404 | Lighthouse Lab in Glasgow | Wellcome Sanger Institute for the COVID-19 Genomics UK (COG-UK) Consortium | Harper VanSteenhouse, Yumi Kasai, David Gray, Carol Clugston, Anna Dominiczak and Alex Alderton, Roberto Amato, Jeffrey Barrett, Sonia Goncalves, Ewan Harrison, David K. Jackson, Ian Johnston, Dominic Kwiatkowski, Cordelia Langford, John Sillitoe on behalf of the Wellcome Sanger Institute COVID-19 Surveillance Team |
| EPI_ISL_2914405, EPI_ISL_2914406 | Lighthouse Lab in Alderley Park | Wellcome Sanger Institute for the COVID-19 Genomics UK (COG-UK) Consortium | Jacquelyn Wynn, Mairead Hyland, The Lighthouse Lab in Alderley Park and Alex Alderton, Roberto Amato, Jeffrey Barrett, Sonia Goncalves, Ewan Harrison, David K. Jackson, Ian Johnston, Dominic Kwiatkowski, Cordelia Langford, John Sillitoe on behalf of the Wellcome Sanger Institute COVID-19 Surveillance Team |
| EPI_ISL_2914407, EPI_ISL_2914408 | Berkshire and Surrey Pathology Services Lighthouse Laboratory | Wellcome Sanger Institute for the COVID-19 Genomics UK (COG-UK) Consortium | Berkshire and Surrey Pathology Services Lighthouse Laboratory and Alex Alderton, Roberto Amato, Jeffrey Barrett, Sonia Goncalves, Ewan Harrison, David K. Jackson, Ian Johnston, Dominic Kwiatkowski, Cordelia Langford, John Sillitoe on behalf of the Wellcome Sanger Institute COVID-19 Surveillance Team |
| EPI_ISL_2914409, EPI_ISL_2914410 | Lighthouse Lab in Glasgow | Wellcome Sanger Institute for the COVID-19 Genomics UK (COG-UK) Consortium | Harper VanSteenhouse, Yumi Kasai, David Gray, Carol Clugston, Anna Dominiczak and Alex Alderton, Roberto Amato, Jeffrey Barrett, Sonia Goncalves, Ewan Harrison, David K. Jackson, Ian Johnston, Dominic Kwiatkowski, Cordelia Langford, John Sillitoe on behalf of the Wellcome Sanger Institute COVID-19 Surveillance Team |
| EPI_ISL_2914411, EPI_ISL_2914412, EPI_ISL_2914413, EPI_ISL_2914414 | Lighthouse Lab in Alderley Park | Wellcome Sanger Institute for the COVID-19 Genomics UK (COG-UK) Consortium | Jacquelyn Wynn, Mairead Hyland, The Lighthouse Lab in Alderley Park and Alex Alderton, Roberto Amato, Jeffrey Barrett, Sonia Goncalves, Ewan Harrison, David K. Jackson, Ian Johnston, Dominic Kwiatkowski, Cordelia Langford, John Sillitoe on behalf of the Wellcome Sanger Institute COVID-19 Surveillance Team |
| EPI_ISL_2914415 | Berkshire and Surrey Pathology Services Lighthouse Laboratory | Wellcome Sanger Institute for the COVID-19 Genomics UK (COG-UK) Consortium | Berkshire and Surrey Pathology Services Lighthouse Laboratory and Alex Alderton, Roberto Amato, Jeffrey Barrett, Sonia Goncalves, Ewan Harrison, David K. Jackson, Ian Johnston, Dominic Kwiatkowski, Cordelia Langford, John Sillitoe on behalf of the Wellcome Sanger Institute COVID-19 Surveillance Team |
| EPI_ISL_2914416 | Lighthouse Lab in Glasgow | Wellcome Sanger Institute for the COVID-19 Genomics UK (COG-UK) Consortium | Harper VanSteenhouse, Yumi Kasai, David Gray, Carol Clugston, Anna Dominiczak and Alex Alderton, Roberto Amato, Jeffrey Barrett, Sonia Goncalves, Ewan Harrison, David K. Jackson, Ian Johnston, Dominic Kwiatkowski, Cordelia Langford, John Sillitoe on behalf of the Wellcome Sanger Institute COVID-19 Surveillance Team |
| EPI_ISL_2914417 | Berkshire and Surrey Pathology Services Lighthouse Laboratory | Wellcome Sanger Institute for the COVID-19 Genomics UK (COG-UK) Consortium | Berkshire and Surrey Pathology Services Lighthouse Laboratory and Alex Alderton, Roberto Amato, Jeffrey Barrett, Sonia Goncalves, Ewan Harrison, David K. Jackson, Ian Johnston, Dominic Kwiatkowski, Cordelia Langford, John Sillitoe on behalf of the Wellcome Sanger Institute COVID-19 Surveillance Team |
| EPI_ISL_2914418, EPI_ISL_2914419 | Lighthouse Lab in Glasgow | Wellcome Sanger Institute for the COVID-19 Genomics UK (COG-UK) Consortium | Harper VanSteenhouse, Yumi Kasai, David Gray, Carol Clugston, Anna Dominiczak and Alex Alderton, Roberto Amato, Jeffrey Barrett, Sonia Goncalves, Ewan Harrison, David K. Jackson, Ian Johnston, Dominic Kwiatkowski, Cordelia Langford, John Sillitoe on behalf of the Wellcome Sanger Institute COVID-19 Surveillance Team |
| EPI_ISL_2914420 | Lighthouse Lab in Alderley Park | Wellcome Sanger Institute for the COVID-19 Genomics UK (COG-UK) Consortium | Jacquelyn Wynn, Mairead Hyland, The Lighthouse Lab in Alderley Park and Alex Alderton, Roberto Amato, Jeffrey Barrett, Sonia Goncalves, Ewan Harrison, David K. Jackson, Ian Johnston, Dominic Kwiatkowski, Cordelia Langford, John Sillitoe on behalf of the Wellcome Sanger Institute COVID-19 Surveillance Team |
| EPI_ISL_2914421, EPI_ISL_2914422 | Lighthouse Lab in Glasgow | Wellcome Sanger Institute for the COVID-19 Genomics UK (COG-UK) Consortium | Harper VanSteenhouse, Yumi Kasai, David Gray, Carol Clugston, Anna Dominiczak and Alex Alderton, Roberto Amato, Jeffrey Barrett, Sonia Goncalves, Ewan Harrison, David K. Jackson, Ian Johnston, Dominic Kwiatkowski, Cordelia Langford, John Sillitoe on behalf of the Wellcome Sanger Institute COVID-19 Surveillance Team |
| EPI_ISL_2914423, EPI_ISL_2914424 | Lighthouse Lab in Alderley Park | Wellcome Sanger Institute for the COVID-19 Genomics UK (COG-UK) Consortium | Jacquelyn Wynn, Mairead Hyland, The Lighthouse Lab in Alderley Park and Alex Alderton, Roberto Amato, Jeffrey Barrett, Sonia Goncalves, Ewan Harrison, David K. Jackson, Ian Johnston, Dominic Kwiatkowski, Cordelia Langford, John Sillitoe on behalf of the Wellcome Sanger Institute COVID-19 Surveillance Team |
| EPI_ISL_2914425, EPI_ISL_2914426, EPI_ISL_2914427 | Lighthouse Lab in Glasgow | Wellcome Sanger Institute for the COVID-19 Genomics UK (COG-UK) Consortium | Harper VanSteenhouse, Yumi Kasai, David Gray, Carol Clugston, Anna Dominiczak and Alex Alderton, Roberto Amato, Jeffrey Barrett, Sonia Goncalves, Ewan Harrison, David K. Jackson, Ian Johnston, Dominic Kwiatkowski, Cordelia Langford, John Sillitoe on behalf of the Wellcome Sanger Institute COVID-19 Surveillance Team |
| EPI_ISL_2914428 | Berkshire and Surrey Pathology Services Lighthouse Laboratory | Wellcome Sanger Institute for the COVID-19 Genomics UK (COG-UK) Consortium | Berkshire and Surrey Pathology Services Lighthouse Laboratory and Alex Alderton, Roberto Amato, Jeffrey Barrett, Sonia Goncalves, Ewan Harrison, David K. Jackson, Ian Johnston, Dominic Kwiatkowski, Cordelia Langford, John Sillitoe on behalf of the Wellcome Sanger Institute COVID-19 Surveillance Team |
| EPI_ISL_2914429 | Lighthouse Lab in Alderley Park | Wellcome Sanger Institute for the COVID-19 Genomics UK (COG-UK) Consortium | Jacquelyn Wynn, Mairead Hyland, The Lighthouse Lab in Alderley Park and Alex Alderton, Roberto Amato, Jeffrey Barrett, Sonia Goncalves, Ewan Harrison, David K. Jackson, Ian Johnston, Dominic Kwiatkowski, Cordelia Langford, John Sillitoe on behalf of the Wellcome Sanger Institute COVID-19 Surveillance Team |
| EPI_ISL_2914430, EPI_ISL_2914431 | Berkshire and Surrey Pathology Services Lighthouse Laboratory | Wellcome Sanger Institute for the COVID-19 Genomics UK (COG-UK) Consortium | Berkshire and Surrey Pathology Services Lighthouse Laboratory and Alex Alderton, Roberto Amato, Jeffrey Barrett, Sonia Goncalves, Ewan Harrison, David K. Jackson, Ian Johnston, Dominic Kwiatkowski, Cordelia Langford, John Sillitoe on behalf of the Wellcome Sanger Institute COVID-19 Surveillance Team |
| EPI_ISL_2914432, EPI_ISL_2914433 | Lighthouse Lab in Glasgow | Wellcome Sanger Institute for the COVID-19 Genomics UK (COG-UK) Consortium | Harper VanSteenhouse, Yumi Kasai, David Gray, Carol Clugston, Anna Dominiczak and Alex Alderton, Roberto Amato, Jeffrey Barrett, Sonia Goncalves, Ewan Harrison, David K. Jackson, Ian Johnston, Dominic Kwiatkowski, Cordelia Langford, John Sillitoe on behalf of the Wellcome Sanger Institute COVID-19 Surveillance Team |
| EPI_ISL_2914434 | Berkshire and Surrey Pathology Services Lighthouse Laboratory | Wellcome Sanger Institute for the COVID-19 Genomics UK (COG-UK) Consortium | Berkshire and Surrey Pathology Services Lighthouse Laboratory and Alex Alderton, Roberto Amato, Jeffrey Barrett, Sonia Goncalves, Ewan Harrison, David K. Jackson, Ian Johnston, Dominic Kwiatkowski, Cordelia Langford, John Sillitoe on behalf of the Wellcome Sanger Institute COVID-19 Surveillance Team |
| EPI_ISL_2914435, EPI_ISL_2914436, EPI_ISL_2914437 | Lighthouse Lab in Glasgow | Wellcome Sanger Institute for the COVID-19 Genomics UK (COG-UK) Consortium | Harper VanSteenhouse, Yumi Kasai, David Gray, Carol Clugston, Anna Dominiczak and Alex Alderton, Roberto Amato, Jeffrey Barrett, Sonia Goncalves, Ewan Harrison, David K. Jackson, Ian Johnston, Dominic Kwiatkowski, Cordelia Langford, John Sillitoe on behalf of the Wellcome Sanger Institute COVID-19 Surveillance Team |
| EPI_ISL_2914438, EPI_ISL_2914439 | Lighthouse Lab in Alderley Park | Wellcome Sanger Institute for the COVID-19 Genomics UK (COG-UK) Consortium | Jacquelyn Wynn, Mairead Hyland, The Lighthouse Lab in Alderley Park and Alex Alderton, Roberto Amato, Jeffrey Barrett, Sonia Goncalves, Ewan Harrison, David K. Jackson, Ian Johnston, Dominic Kwiatkowski, Cordelia Langford, John Sillitoe on behalf of the Wellcome Sanger Institute COVID-19 Surveillance Team |
| EPI_ISL_2914440, EPI_ISL_2914441, | Lighthouse Lab in Glasgow | Wellcome Sanger Institute for the COVID-19 Genomics UK (COG-UK) Consortium | Harper VanSteenhouse, Yumi Kasai, David Gray, Carol Clugston, Anna Dominiczak and Alex Alderton, Roberto Amato, Jeffrey Barrett, Sonia Goncalves, Ewan Harrison, David K. Jackson, Ian Johnston, Dominic Kwiatkowski, Cordelia Langford, John Sillitoe on behalf of the Wellcome Sanger Institute COVID-19 Surveillance Team |

| | | | |
|---|---|---|---|
| EPI_ISL_2914442, EPI_ISL_2914443, EPI_ISL_2914444 | | | |
| EPI_ISL_2914445 | Lighthouse Lab in Alderley Park | Wellcome Sanger Institute for the COVID-19 Genomics UK (COG-UK) Consortium | Jacquelyn Wynn, Mairead Hyland, The Lighthouse Lab in Alderley Park and Alex Alderton, Roberto Amato, Jeffrey Barrett, Sonia Goncalves, Ewan Harrison, David K. Jackson, Ian Johnston, Dominic Kwiatkowski, Cordelia Langford, John Sillitoe on behalf of the Wellcome Sanger Institute COVID-19 Surveillance Team |
| EPI_ISL_2914446 | Lighthouse Lab in Glasgow | Wellcome Sanger Institute for the COVID-19 Genomics UK (COG-UK) Consortium | Harper VanSteenhouse, Yumi Kasai, David Gray, Carol Clugston, Anna Dominiczak and Alex Alderton, Roberto Amato, Jeffrey Barrett, Sonia Goncalves, Ewan Harrison, David K. Jackson, Ian Johnston, Dominic Kwiatkowski, Cordelia Langford, John Sillitoe on behalf of the Wellcome Sanger Institute COVID-19 Surveillance Team |
| EPI_ISL_2914447 | Berkshire and Surrey Pathology Services Lighthouse Laboratory | Wellcome Sanger Institute for the COVID-19 Genomics UK (COG-UK) Consortium | Berkshire and Surrey Pathology Services Lighthouse Laboratory and Alex Alderton, Roberto Amato, Jeffrey Barrett, Sonia Goncalves, Ewan Harrison, David K. Jackson, Ian Johnston, Dominic Kwiatkowski, Cordelia Langford, John Sillitoe on behalf of the Wellcome Sanger Institute COVID-19 Surveillance Team |
| EPI_ISL_2914448 | Lighthouse Lab in Glasgow | Wellcome Sanger Institute for the COVID-19 Genomics UK (COG-UK) Consortium | Harper VanSteenhouse, Yumi Kasai, David Gray, Carol Clugston, Anna Dominiczak and Alex Alderton, Roberto Amato, Jeffrey Barrett, Sonia Goncalves, Ewan Harrison, David K. Jackson, Ian Johnston, Dominic Kwiatkowski, Cordelia Langford, John Sillitoe on behalf of the Wellcome Sanger Institute COVID-19 Surveillance Team |
| EPI_ISL_2914449 | Lighthouse Lab in Alderley Park | Wellcome Sanger Institute for the COVID-19 Genomics UK (COG-UK) Consortium | Jacquelyn Wynn, Mairead Hyland, The Lighthouse Lab in Alderley Park and Alex Alderton, Roberto Amato, Jeffrey Barrett, Sonia Goncalves, Ewan Harrison, David K. Jackson, Ian Johnston, Dominic Kwiatkowski, Cordelia Langford, John Sillitoe on behalf of the Wellcome Sanger Institute COVID-19 Surveillance Team |
| EPI_ISL_2914450, EPI_ISL_2914451 | Lighthouse Lab in Glasgow | Wellcome Sanger Institute for the COVID-19 Genomics UK (COG-UK) Consortium | Harper VanSteenhouse, Yumi Kasai, David Gray, Carol Clugston, Anna Dominiczak and Alex Alderton, Roberto Amato, Jeffrey Barrett, Sonia Goncalves, Ewan Harrison, David K. Jackson, Ian Johnston, Dominic Kwiatkowski, Cordelia Langford, John Sillitoe on behalf of the Wellcome Sanger Institute COVID-19 Surveillance Team |
| EPI_ISL_2914452 | Lighthouse Lab in Alderley Park | Wellcome Sanger Institute for the COVID-19 Genomics UK (COG-UK) Consortium | Jacquelyn Wynn, Mairead Hyland, The Lighthouse Lab in Alderley Park and Alex Alderton, Roberto Amato, Jeffrey Barrett, Sonia Goncalves, Ewan Harrison, David K. Jackson, Ian Johnston, Dominic Kwiatkowski, Cordelia Langford, John Sillitoe on behalf of the Wellcome Sanger Institute COVID-19 Surveillance Team |
| EPI_ISL_2914453 | Lighthouse Lab in Glasgow | Wellcome Sanger Institute for the COVID-19 Genomics UK (COG-UK) Consortium | Harper VanSteenhouse, Yumi Kasai, David Gray, Carol Clugston, Anna Dominiczak and Alex Alderton, Roberto Amato, Jeffrey Barrett, Sonia Goncalves, Ewan Harrison, David K. Jackson, Ian Johnston, Dominic Kwiatkowski, Cordelia Langford, John Sillitoe on behalf of the Wellcome Sanger Institute COVID-19 Surveillance Team |
| EPI_ISL_2914454 | Lighthouse Lab in Alderley Park | Wellcome Sanger Institute for the COVID-19 Genomics UK (COG-UK) Consortium | Jacquelyn Wynn, Mairead Hyland, The Lighthouse Lab in Alderley Park and Alex Alderton, Roberto Amato, Jeffrey Barrett, Sonia Goncalves, Ewan Harrison, David K. Jackson, Ian Johnston, Dominic Kwiatkowski, Cordelia Langford, John Sillitoe on behalf of the Wellcome Sanger Institute COVID-19 Surveillance Team |
| EPI_ISL_2914455 | Berkshire and Surrey Pathology Services Lighthouse Laboratory | Wellcome Sanger Institute for the COVID-19 Genomics UK (COG-UK) Consortium | Berkshire and Surrey Pathology Services Lighthouse Laboratory and Alex Alderton, Roberto Amato, Jeffrey Barrett, Sonia Goncalves, Ewan Harrison, David K. Jackson, Ian Johnston, Dominic Kwiatkowski, Cordelia Langford, John Sillitoe on behalf of the Wellcome Sanger Institute COVID-19 Surveillance Team |
| EPI_ISL_2914456, EPI_ISL_2914457, EPI_ISL_2914458 | Lighthouse Lab in Glasgow | Wellcome Sanger Institute for the COVID-19 Genomics UK (COG-UK) Consortium | Harper VanSteenhouse, Yumi Kasai, David Gray, Carol Clugston, Anna Dominiczak and Alex Alderton, Roberto Amato, Jeffrey Barrett, Sonia Goncalves, Ewan Harrison, David K. Jackson, Ian Johnston, Dominic Kwiatkowski, Cordelia Langford, John Sillitoe on behalf of the Wellcome Sanger Institute COVID-19 Surveillance Team |
| EPI_ISL_2914459, EPI_ISL_2914460 | Berkshire and Surrey Pathology Services Lighthouse Laboratory | Wellcome Sanger Institute for the COVID-19 Genomics UK (COG-UK) Consortium | Berkshire and Surrey Pathology Services Lighthouse Laboratory and Alex Alderton, Roberto Amato, Jeffrey Barrett, Sonia Goncalves, Ewan Harrison, David K. Jackson, Ian Johnston, Dominic Kwiatkowski, Cordelia Langford, John Sillitoe on behalf of the Wellcome Sanger Institute COVID-19 Surveillance Team |
| EPI_ISL_2914461, EPI_ISL_2914462 | Lighthouse Lab in Alderley Park | Wellcome Sanger Institute for the COVID-19 Genomics UK (COG-UK) Consortium | Jacquelyn Wynn, Mairead Hyland, The Lighthouse Lab in Alderley Park and Alex Alderton, Roberto Amato, Jeffrey Barrett, Sonia Goncalves, Ewan Harrison, David K. Jackson, Ian Johnston, Dominic Kwiatkowski, Cordelia Langford, John Sillitoe on behalf of the Wellcome Sanger Institute COVID-19 Surveillance Team |
| EPI_ISL_2914463, EPI_ISL_2914464, EPI_ISL_2914465 | Lighthouse Lab in Glasgow | Wellcome Sanger Institute for the COVID-19 Genomics UK (COG-UK) Consortium | Harper VanSteenhouse, Yumi Kasai, David Gray, Carol Clugston, Anna Dominiczak and Alex Alderton, Roberto Amato, Jeffrey Barrett, Sonia Goncalves, Ewan Harrison, David K. Jackson, Ian Johnston, Dominic Kwiatkowski, Cordelia Langford, John Sillitoe on behalf of the Wellcome Sanger Institute COVID-19 Surveillance Team |
| EPI_ISL_2914466 | Lighthouse Lab in Alderley Park | Wellcome Sanger Institute for the COVID-19 Genomics UK (COG-UK) Consortium | Jacquelyn Wynn, Mairead Hyland, The Lighthouse Lab in Alderley Park and Alex Alderton, Roberto Amato, Jeffrey Barrett, Sonia Goncalves, Ewan Harrison, David K. Jackson, Ian Johnston, Dominic Kwiatkowski, Cordelia Langford, John Sillitoe on behalf of the Wellcome Sanger Institute COVID-19 Surveillance Team |
| EPI_ISL_2914467, EPI_ISL_2914468 | Lighthouse Lab in Glasgow | Wellcome Sanger Institute for the COVID-19 Genomics UK (COG-UK) Consortium | Harper VanSteenhouse, Yumi Kasai, David Gray, Carol Clugston, Anna Dominiczak and Alex Alderton, Roberto Amato, Jeffrey Barrett, Sonia Goncalves, Ewan Harrison, David K. Jackson, Ian Johnston, Dominic Kwiatkowski, Cordelia Langford, John Sillitoe on behalf of the Wellcome Sanger Institute COVID-19 Surveillance Team |
| EPI_ISL_2914469, EPI_ISL_2914470 | Berkshire and Surrey Pathology Services Lighthouse Laboratory | Wellcome Sanger Institute for the COVID-19 Genomics UK (COG-UK) Consortium | Berkshire and Surrey Pathology Services Lighthouse Laboratory and Alex Alderton, Roberto Amato, Jeffrey Barrett, Sonia Goncalves, Ewan Harrison, David K. Jackson, Ian Johnston, Dominic Kwiatkowski, Cordelia Langford, John Sillitoe on behalf of the Wellcome Sanger Institute COVID-19 Surveillance Team |
| EPI_ISL_2914471 | Lighthouse Lab in Alderley Park | Wellcome Sanger Institute for the COVID-19 Genomics UK (COG-UK) Consortium | Jacquelyn Wynn, Mairead Hyland, The Lighthouse Lab in Alderley Park and Alex Alderton, Roberto Amato, Jeffrey Barrett, Sonia Goncalves, Ewan Harrison, David K. Jackson, Ian Johnston, Dominic Kwiatkowski, Cordelia Langford, John Sillitoe on behalf of the Wellcome Sanger Institute COVID-19 Surveillance Team |
| EPI_ISL_2914472, EPI_ISL_2914473, EPI_ISL_2914474 | Berkshire and Surrey Pathology Services Lighthouse Laboratory | Wellcome Sanger Institute for the COVID-19 Genomics UK (COG-UK) Consortium | Berkshire and Surrey Pathology Services Lighthouse Laboratory and Alex Alderton, Roberto Amato, Jeffrey Barrett, Sonia Goncalves, Ewan Harrison, David K. Jackson, Ian Johnston, Dominic Kwiatkowski, Cordelia Langford, John Sillitoe on behalf of the Wellcome Sanger Institute COVID-19 Surveillance Team |
| EPI_ISL_2914475 | Lighthouse Lab in Alderley Park | Wellcome Sanger Institute for the COVID-19 Genomics UK (COG-UK) Consortium | Jacquelyn Wynn, Mairead Hyland, The Lighthouse Lab in Alderley Park and Alex Alderton, Roberto Amato, Jeffrey Barrett, Sonia Goncalves, Ewan Harrison, David K. Jackson, Ian Johnston, Dominic Kwiatkowski, Cordelia Langford, John Sillitoe on behalf of the Wellcome Sanger Institute COVID-19 Surveillance Team |
| EPI_ISL_2914476, EPI_ISL_2914477 | Lighthouse Lab in Glasgow | Wellcome Sanger Institute for the COVID-19 Genomics UK (COG-UK) Consortium | Harper VanSteenhouse, Yumi Kasai, David Gray, Carol Clugston, Anna Dominiczak and Alex Alderton, Roberto Amato, Jeffrey Barrett, Sonia Goncalves, Ewan Harrison, David K. Jackson, Ian Johnston, Dominic Kwiatkowski, Cordelia Langford, John Sillitoe on behalf of the Wellcome Sanger Institute COVID-19 Surveillance Team |
| EPI_ISL_2914478 | Berkshire and Surrey Pathology Services Lighthouse Laboratory | Wellcome Sanger Institute for the COVID-19 Genomics UK (COG-UK) Consortium | Berkshire and Surrey Pathology Services Lighthouse Laboratory and Alex Alderton, Roberto Amato, Jeffrey Barrett, Sonia Goncalves, Ewan Harrison, David K. Jackson, Ian Johnston, Dominic Kwiatkowski, Cordelia Langford, John Sillitoe on behalf of the Wellcome Sanger Institute COVID-19 Surveillance Team |
| EPI_ISL_2914479 | Lighthouse Lab in Glasgow | Wellcome Sanger Institute for the COVID-19 Genomics UK (COG-UK) Consortium | Harper VanSteenhouse, Yumi Kasai, David Gray, Carol Clugston, Anna Dominiczak and Alex Alderton, Roberto Amato, Jeffrey Barrett, Sonia Goncalves, Ewan Harrison, David K. Jackson, Ian Johnston, Dominic Kwiatkowski, Cordelia Langford, John Sillitoe on behalf of the Wellcome Sanger Institute COVID-19 Surveillance Team |
| EPI_ISL_2914480 | Berkshire and Surrey Pathology Services Lighthouse Laboratory | Wellcome Sanger Institute for the COVID-19 Genomics UK (COG-UK) Consortium | Berkshire and Surrey Pathology Services Lighthouse Laboratory and Alex Alderton, Roberto Amato, Jeffrey Barrett, Sonia Goncalves, Ewan Harrison, David K. Jackson, Ian Johnston, Dominic Kwiatkowski, Cordelia Langford, John Sillitoe on behalf of the Wellcome Sanger Institute COVID-19 Surveillance Team |
| EPI_ISL_2914481 | Lighthouse Lab in Glasgow | Wellcome Sanger Institute for the COVID-19 Genomics UK (COG-UK) Consortium | Harper VanSteenhouse, Yumi Kasai, David Gray, Carol Clugston, Anna Dominiczak and Alex Alderton, Roberto Amato, Jeffrey Barrett, Sonia Goncalves, Ewan Harrison, David K. Jackson, Ian Johnston, Dominic Kwiatkowski, Cordelia Langford, John Sillitoe on behalf of the Wellcome Sanger Institute COVID-19 Surveillance Team |
| EPI_ISL_2914482 | Berkshire and Surrey Pathology Services Lighthouse Laboratory | Wellcome Sanger Institute for the COVID-19 Genomics UK (COG-UK) Consortium | Berkshire and Surrey Pathology Services Lighthouse Laboratory and Alex Alderton, Roberto Amato, Jeffrey Barrett, Sonia Goncalves, Ewan Harrison, David K. Jackson, Ian Johnston, Dominic Kwiatkowski, Cordelia Langford, John Sillitoe on behalf of the Wellcome Sanger Institute COVID-19 Surveillance Team |
| EPI_ISL_2914483 | Lighthouse Lab in Alderley Park | Wellcome Sanger Institute for the COVID-19 Genomics UK (COG-UK) Consortium | Jacquelyn Wynn, Mairead Hyland, The Lighthouse Lab in Alderley Park and Alex Alderton, Roberto Amato, Jeffrey Barrett, Sonia Goncalves, Ewan Harrison, David K. Jackson, Ian Johnston, Dominic Kwiatkowski, Cordelia Langford, John Sillitoe on behalf of the Wellcome Sanger Institute COVID-19 Surveillance Team |
| EPI_ISL_2914484 | Lighthouse Lab in Glasgow | Wellcome Sanger Institute for the COVID-19 Genomics UK (COG-UK) Consortium | Harper VanSteenhouse, Yumi Kasai, David Gray, Carol Clugston, Anna Dominiczak and Alex Alderton, Roberto Amato, Jeffrey Barrett, Sonia Goncalves, Ewan Harrison, David K. Jackson, Ian Johnston, Dominic Kwiatkowski, Cordelia Langford, John Sillitoe on behalf of the Wellcome Sanger Institute COVID-19 Surveillance Team |
| EPI_ISL_2914485 | Lighthouse Lab in Alderley Park | Wellcome Sanger Institute for the COVID-19 Genomics UK (COG-UK) Consortium | Jacquelyn Wynn, Mairead Hyland, The Lighthouse Lab in Alderley Park and Alex Alderton, Roberto Amato, Jeffrey Barrett, Sonia Goncalves, Ewan Harrison, David K. Jackson, Ian Johnston, Dominic Kwiatkowski, Cordelia Langford, John Sillitoe on behalf of the Wellcome Sanger Institute COVID-19 Surveillance Team |
| EPI_ISL_2914486 | Lighthouse Lab in Glasgow | Wellcome Sanger Institute for the COVID-19 Genomics UK (COG-UK) Consortium | Harper VanSteenhouse, Yumi Kasai, David Gray, Carol Clugston, Anna Dominiczak and Alex Alderton, Roberto Amato, Jeffrey Barrett, Sonia Goncalves, Ewan Harrison, David K. Jackson, Ian Johnston, Dominic Kwiatkowski, Cordelia Langford, John Sillitoe on behalf of the Wellcome Sanger Institute COVID-19 Surveillance Team |
| EPI_ISL_2914487 | Berkshire and Surrey Pathology Services Lighthouse Laboratory | Wellcome Sanger Institute for the COVID-19 Genomics UK (COG-UK) Consortium | Berkshire and Surrey Pathology Services Lighthouse Laboratory and Alex Alderton, Roberto Amato, Jeffrey Barrett, Sonia Goncalves, Ewan Harrison, David K. Jackson, Ian Johnston, Dominic Kwiatkowski, Cordelia Langford, John Sillitoe on behalf of the Wellcome Sanger Institute COVID-19 Surveillance Team |
| EPI_ISL_2914488, EPI_ISL_2914489 | Lighthouse Lab in Glasgow | Wellcome Sanger Institute for the COVID-19 Genomics UK (COG-UK) Consortium | Harper VanSteenhouse, Yumi Kasai, David Gray, Carol Clugston, Anna Dominiczak and Alex Alderton, Roberto Amato, Jeffrey Barrett, Sonia Goncalves, Ewan Harrison, David K. Jackson, Ian Johnston, Dominic Kwiatkowski, Cordelia Langford, John Sillitoe on behalf of the Wellcome Sanger Institute COVID-19 Surveillance Team |
| EPI_ISL_2914490 | Lighthouse Lab in Alderley Park | Wellcome Sanger Institute for the COVID-19 Genomics UK (COG-UK) Consortium | Jacquelyn Wynn, Mairead Hyland, The Lighthouse Lab in Alderley Park and Alex Alderton, Roberto Amato, Jeffrey Barrett, Sonia Goncalves, Ewan Harrison, David K. Jackson, Ian Johnston, Dominic Kwiatkowski, Cordelia Langford, John Sillitoe on behalf of the Wellcome Sanger Institute COVID-19 Surveillance Team |
| EPI_ISL_2914491, EPI_ISL_2914492 | Berkshire and Surrey Pathology Services Lighthouse Laboratory | Wellcome Sanger Institute for the COVID-19 Genomics UK (COG-UK) Consortium | Berkshire and Surrey Pathology Services Lighthouse Laboratory and Alex Alderton, Roberto Amato, Jeffrey Barrett, Sonia Goncalves, Ewan Harrison, David K. Jackson, Ian Johnston, Dominic Kwiatkowski, Cordelia Langford, John Sillitoe on behalf of the Wellcome Sanger Institute COVID-19 Surveillance Team |
| EPI_ISL_2914493, EPI_ISL_2914494 | Lighthouse Lab in Glasgow | Wellcome Sanger Institute for the COVID-19 Genomics UK (COG-UK) Consortium | Harper VanSteenhouse, Yumi Kasai, David Gray, Carol Clugston, Anna Dominiczak and Alex Alderton, Roberto Amato, Jeffrey Barrett, Sonia Goncalves, Ewan Harrison, David K. Jackson, Ian Johnston, Dominic Kwiatkowski, Cordelia Langford, John Sillitoe on behalf of the Wellcome Sanger Institute COVID-19 Surveillance Team |
| EPI_ISL_2914495, EPI_ISL_2914496 | Berkshire and Surrey Pathology Services Lighthouse Laboratory | Wellcome Sanger Institute for the COVID-19 Genomics UK (COG-UK) Consortium | Berkshire and Surrey Pathology Services Lighthouse Laboratory and Alex Alderton, Roberto Amato, Jeffrey Barrett, Sonia Goncalves, Ewan Harrison, David K. Jackson, Ian Johnston, Dominic Kwiatkowski, Cordelia Langford, John Sillitoe on behalf of the Wellcome Sanger Institute COVID-19 Surveillance Team |
| EPI_ISL_2914497, EPI_ISL_2914498 | Lighthouse Lab in Alderley Park | Wellcome Sanger Institute for the COVID-19 Genomics UK (COG-UK) Consortium | Jacquelyn Wynn, Mairead Hyland, The Lighthouse Lab in Alderley Park and Alex Alderton, Roberto Amato, Jeffrey Barrett, Sonia Goncalves, Ewan Harrison, David K. Jackson, Ian Johnston, Dominic Kwiatkowski, Cordelia Langford, John Sillitoe on behalf of the Wellcome Sanger Institute COVID-19 Surveillance Team |
| EPI_ISL_2914499 | Berkshire and Surrey Pathology Services Lighthouse Laboratory | Wellcome Sanger Institute for the COVID-19 Genomics UK (COG-UK) Consortium | Berkshire and Surrey Pathology Services Lighthouse Laboratory and Alex Alderton, Roberto Amato, Jeffrey Barrett, Sonia Goncalves, Ewan Harrison, David K. Jackson, Ian Johnston, Dominic Kwiatkowski, Cordelia Langford, John Sillitoe on behalf of the Wellcome Sanger Institute COVID-19 Surveillance Team |
| EPI_ISL_2914500, EPI_ISL_2914501, EPI_ISL_2914502, EPI_ISL_2914503, EPI_ISL_2914504, EPI_ISL_2914505, EPI_ISL_2914506 | Lighthouse Lab in Glasgow | Wellcome Sanger Institute for the COVID-19 Genomics UK (COG-UK) Consortium | Harper VanSteenhouse, Yumi Kasai, David Gray, Carol Clugston, Anna Dominiczak and Alex Alderton, Roberto Amato, Jeffrey Barrett, Sonia Goncalves, Ewan Harrison, David K. Jackson, Ian Johnston, Dominic Kwiatkowski, Cordelia Langford, John Sillitoe on behalf of the Wellcome Sanger Institute COVID-19 Surveillance Team |
| EPI_ISL_2914507, EPI_ISL_2914508 | Lighthouse Lab in Alderley Park | Wellcome Sanger Institute for the COVID-19 Genomics UK (COG-UK) Consortium | Jacquelyn Wynn, Mairead Hyland, The Lighthouse Lab in Alderley Park and Alex Alderton, Roberto Amato, Jeffrey Barrett, Sonia Goncalves, Ewan Harrison, David K. Jackson, Ian Johnston, Dominic Kwiatkowski, Cordelia Langford, John Sillitoe on behalf of the Wellcome Sanger Institute COVID-19 Surveillance Team |

| | | | |
|---|---|---|---|
| EPI_ISL_2914509, EPI_ISL_2914510, EPI_ISL_2914511 | Lighthouse Lab in Glasgow | Wellcome Sanger Institute for the COVID-19 Genomics UK (COG-UK) Consortium | Harper VanSteenhouse, Yumi Kasai, David Gray, Carol Clugston, Anna Dominiczak and Alex Alderton, Roberto Amato, Jeffrey Barrett, Sonia Goncalves, Ewan Harrison, David K. Jackson, Ian Johnston, Dominic Kwiatkowski, Cordelia Langford, John Sillitoe on behalf of the Wellcome Sanger Institute COVID-19 Surveillance Team |
| EPI_ISL_2914512 | Berkshire and Surrey Pathology Services Lighthouse Laboratory | Wellcome Sanger Institute for the COVID-19 Genomics UK (COG-UK) Consortium | Berkshire and Surrey Pathology Services Lighthouse Laboratory and Alex Alderton, Roberto Amato, Jeffrey Barrett, Sonia Goncalves, Ewan Harrison, David K. Jackson, Ian Johnston, Dominic Kwiatkowski, Cordelia Langford, John Sillitoe on behalf of the Wellcome Sanger Institute COVID-19 Surveillance Team |
| EPI_ISL_2914513 | Lighthouse Lab in Alderley Park | Wellcome Sanger Institute for the COVID-19 Genomics UK (COG-UK) Consortium | Jacquelyn Wynn, Mairead Hyland, The Lighthouse Lab in Alderley Park and Alex Alderton, Roberto Amato, Jeffrey Barrett, Sonia Goncalves, Ewan Harrison, David K. Jackson, Ian Johnston, Dominic Kwiatkowski, Cordelia Langford, John Sillitoe on behalf of the Wellcome Sanger Institute COVID-19 Surveillance Team |
| EPI_ISL_2914514 | Lighthouse Lab in Glasgow | Wellcome Sanger Institute for the COVID-19 Genomics UK (COG-UK) Consortium | Harper VanSteenhouse, Yumi Kasai, David Gray, Carol Clugston, Anna Dominiczak and Alex Alderton, Roberto Amato, Jeffrey Barrett, Sonia Goncalves, Ewan Harrison, David K. Jackson, Ian Johnston, Dominic Kwiatkowski, Cordelia Langford, John Sillitoe on behalf of the Wellcome Sanger Institute COVID-19 Surveillance Team |
| EPI_ISL_2914515 | Berkshire and Surrey Pathology Services Lighthouse Laboratory | Wellcome Sanger Institute for the COVID-19 Genomics UK (COG-UK) Consortium | Berkshire and Surrey Pathology Services Lighthouse Laboratory and Alex Alderton, Roberto Amato, Jeffrey Barrett, Sonia Goncalves, Ewan Harrison, David K. Jackson, Ian Johnston, Dominic Kwiatkowski, Cordelia Langford, John Sillitoe on behalf of the Wellcome Sanger Institute COVID-19 Surveillance Team |
| EPI_ISL_2914516 | Lighthouse Lab in Alderley Park | Wellcome Sanger Institute for the COVID-19 Genomics UK (COG-UK) Consortium | Jacquelyn Wynn, Mairead Hyland, The Lighthouse Lab in Alderley Park and Alex Alderton, Roberto Amato, Jeffrey Barrett, Sonia Goncalves, Ewan Harrison, David K. Jackson, Ian Johnston, Dominic Kwiatkowski, Cordelia Langford, John Sillitoe on behalf of the Wellcome Sanger Institute COVID-19 Surveillance Team |
| EPI_ISL_2914517 | Berkshire and Surrey Pathology Services Lighthouse Laboratory | Wellcome Sanger Institute for the COVID-19 Genomics UK (COG-UK) Consortium | Berkshire and Surrey Pathology Services Lighthouse Laboratory and Alex Alderton, Roberto Amato, Jeffrey Barrett, Sonia Goncalves, Ewan Harrison, David K. Jackson, Ian Johnston, Dominic Kwiatkowski, Cordelia Langford, John Sillitoe on behalf of the Wellcome Sanger Institute COVID-19 Surveillance Team |
| EPI_ISL_2914518, EPI_ISL_2914519 | Lighthouse Lab in Glasgow | Wellcome Sanger Institute for the COVID-19 Genomics UK (COG-UK) Consortium | Harper VanSteenhouse, Yumi Kasai, David Gray, Carol Clugston, Anna Dominiczak and Alex Alderton, Roberto Amato, Jeffrey Barrett, Sonia Goncalves, Ewan Harrison, David K. Jackson, Ian Johnston, Dominic Kwiatkowski, Cordelia Langford, John Sillitoe on behalf of the Wellcome Sanger Institute COVID-19 Surveillance Team |
| EPI_ISL_2914520 | Lighthouse Lab in Alderley Park | Wellcome Sanger Institute for the COVID-19 Genomics UK (COG-UK) Consortium | Jacquelyn Wynn, Mairead Hyland, The Lighthouse Lab in Alderley Park and Alex Alderton, Roberto Amato, Jeffrey Barrett, Sonia Goncalves, Ewan Harrison, David K. Jackson, Ian Johnston, Dominic Kwiatkowski, Cordelia Langford, John Sillitoe on behalf of the Wellcome Sanger Institute COVID-19 Surveillance Team |
| EPI_ISL_2914521, EPI_ISL_2914522, EPI_ISL_2914523, EPI_ISL_2914524, EPI_ISL_2914525 | Lighthouse Lab in Glasgow | Wellcome Sanger Institute for the COVID-19 Genomics UK (COG-UK) Consortium | Harper VanSteenhouse, Yumi Kasai, David Gray, Carol Clugston, Anna Dominiczak and Alex Alderton, Roberto Amato, Jeffrey Barrett, Sonia Goncalves, Ewan Harrison, David K. Jackson, Ian Johnston, Dominic Kwiatkowski, Cordelia Langford, John Sillitoe on behalf of the Wellcome Sanger Institute COVID-19 Surveillance Team |
| EPI_ISL_2914527 | Berkshire and Surrey Pathology Services Lighthouse Laboratory | Wellcome Sanger Institute for the COVID-19 Genomics UK (COG-UK) Consortium | Berkshire and Surrey Pathology Services Lighthouse Laboratory and Alex Alderton, Roberto Amato, Jeffrey Barrett, Sonia Goncalves, Ewan Harrison, David K. Jackson, Ian Johnston, Dominic Kwiatkowski, Cordelia Langford, John Sillitoe on behalf of the Wellcome Sanger Institute COVID-19 Surveillance Team |
| EPI_ISL_2914528, EPI_ISL_2914529, EPI_ISL_2914530 | Lighthouse Lab in Glasgow | Wellcome Sanger Institute for the COVID-19 Genomics UK (COG-UK) Consortium | Harper VanSteenhouse, Yumi Kasai, David Gray, Carol Clugston, Anna Dominiczak and Alex Alderton, Roberto Amato, Jeffrey Barrett, Sonia Goncalves, Ewan Harrison, David K. Jackson, Ian Johnston, Dominic Kwiatkowski, Cordelia Langford, John Sillitoe on behalf of the Wellcome Sanger Institute COVID-19 Surveillance Team |
| EPI_ISL_2914531 | Lighthouse Lab in Alderley Park | Wellcome Sanger Institute for the COVID-19 Genomics UK (COG-UK) Consortium | Jacquelyn Wynn, Mairead Hyland, The Lighthouse Lab in Alderley Park and Alex Alderton, Roberto Amato, Jeffrey Barrett, Sonia Goncalves, Ewan Harrison, David K. Jackson, Ian Johnston, Dominic Kwiatkowski, Cordelia Langford, John Sillitoe on behalf of the Wellcome Sanger Institute COVID-19 Surveillance Team |
| EPI_ISL_2914532 | Lighthouse Lab in Glasgow | Wellcome Sanger Institute for the COVID-19 Genomics UK (COG-UK) Consortium | Harper VanSteenhouse, Yumi Kasai, David Gray, Carol Clugston, Anna Dominiczak and Alex Alderton, Roberto Amato, Jeffrey Barrett, Sonia Goncalves, Ewan Harrison, David K. Jackson, Ian Johnston, Dominic Kwiatkowski, Cordelia Langford, John Sillitoe on behalf of the Wellcome Sanger Institute COVID-19 Surveillance Team |
| EPI_ISL_2914534 | Berkshire and Surrey Pathology Services Lighthouse Laboratory | Wellcome Sanger Institute for the COVID-19 Genomics UK (COG-UK) Consortium | Berkshire and Surrey Pathology Services Lighthouse Laboratory and Alex Alderton, Roberto Amato, Jeffrey Barrett, Sonia Goncalves, Ewan Harrison, David K. Jackson, Ian Johnston, Dominic Kwiatkowski, Cordelia Langford, John Sillitoe on behalf of the Wellcome Sanger Institute COVID-19 Surveillance Team |
| EPI_ISL_2914535 | Lighthouse Lab in Glasgow | Wellcome Sanger Institute for the COVID-19 Genomics UK (COG-UK) Consortium | Harper VanSteenhouse, Yumi Kasai, David Gray, Carol Clugston, Anna Dominiczak and Alex Alderton, Roberto Amato, Jeffrey Barrett, Sonia Goncalves, Ewan Harrison, David K. Jackson, Ian Johnston, Dominic Kwiatkowski, Cordelia Langford, John Sillitoe on behalf of the Wellcome Sanger Institute COVID-19 Surveillance Team |
| EPI_ISL_2914536 | Berkshire and Surrey Pathology Services Lighthouse Laboratory | Wellcome Sanger Institute for the COVID-19 Genomics UK (COG-UK) Consortium | Berkshire and Surrey Pathology Services Lighthouse Laboratory and Alex Alderton, Roberto Amato, Jeffrey Barrett, Sonia Goncalves, Ewan Harrison, David K. Jackson, Ian Johnston, Dominic Kwiatkowski, Cordelia Langford, John Sillitoe on behalf of the Wellcome Sanger Institute COVID-19 Surveillance Team |
| EPI_ISL_2914537 | Lighthouse Lab in Glasgow | Wellcome Sanger Institute for the COVID-19 Genomics UK (COG-UK) Consortium | Harper VanSteenhouse, Yumi Kasai, David Gray, Carol Clugston, Anna Dominiczak and Alex Alderton, Roberto Amato, Jeffrey Barrett, Sonia Goncalves, Ewan Harrison, David K. Jackson, Ian Johnston, Dominic Kwiatkowski, Cordelia Langford, John Sillitoe on behalf of the Wellcome Sanger Institute COVID-19 Surveillance Team |
| EPI_ISL_2914538 | Lighthouse Lab in Alderley Park | Wellcome Sanger Institute for the COVID-19 Genomics UK (COG-UK) Consortium | Jacquelyn Wynn, Mairead Hyland, The Lighthouse Lab in Alderley Park and Alex Alderton, Roberto Amato, Jeffrey Barrett, Sonia Goncalves, Ewan Harrison, David K. Jackson, Ian Johnston, Dominic Kwiatkowski, Cordelia Langford, John Sillitoe on behalf of the Wellcome Sanger Institute COVID-19 Surveillance Team |
| EPI_ISL_2914539 | Lighthouse Lab in Glasgow | Wellcome Sanger Institute for the COVID-19 Genomics UK (COG-UK) Consortium | Harper VanSteenhouse, Yumi Kasai, David Gray, Carol Clugston, Anna Dominiczak and Alex Alderton, Roberto Amato, Jeffrey Barrett, Sonia Goncalves, Ewan Harrison, David K. Jackson, Ian Johnston, Dominic Kwiatkowski, Cordelia Langford, John Sillitoe on behalf of the Wellcome Sanger Institute COVID-19 Surveillance Team |
| EPI_ISL_2914540 | Lighthouse Lab in Alderley Park | Wellcome Sanger Institute for the COVID-19 Genomics UK (COG-UK) Consortium | Jacquelyn Wynn, Mairead Hyland, The Lighthouse Lab in Alderley Park and Alex Alderton, Roberto Amato, Jeffrey Barrett, Sonia Goncalves, Ewan Harrison, David K. Jackson, Ian Johnston, Dominic Kwiatkowski, Cordelia Langford, John Sillitoe on behalf of the Wellcome Sanger Institute COVID-19 Surveillance Team |
| EPI_ISL_2914541, EPI_ISL_2914542 | Lighthouse Lab in Glasgow | Wellcome Sanger Institute for the COVID-19 Genomics UK (COG-UK) Consortium | Harper VanSteenhouse, Yumi Kasai, David Gray, Carol Clugston, Anna Dominiczak and Alex Alderton, Roberto Amato, Jeffrey Barrett, Sonia Goncalves, Ewan Harrison, David K. Jackson, Ian Johnston, Dominic Kwiatkowski, Cordelia Langford, John Sillitoe on behalf of the Wellcome Sanger Institute COVID-19 Surveillance Team |
| EPI_ISL_2914543 | Lighthouse Lab in Alderley Park | Wellcome Sanger Institute for the COVID-19 Genomics UK (COG-UK) Consortium | Jacquelyn Wynn, Mairead Hyland, The Lighthouse Lab in Alderley Park and Alex Alderton, Roberto Amato, Jeffrey Barrett, Sonia Goncalves, Ewan Harrison, David K. Jackson, Ian Johnston, Dominic Kwiatkowski, Cordelia Langford, John Sillitoe on behalf of the Wellcome Sanger Institute COVID-19 Surveillance Team |
| EPI_ISL_2914544 | Lighthouse Lab in Glasgow | Wellcome Sanger Institute for the COVID-19 Genomics UK (COG-UK) Consortium | Harper VanSteenhouse, Yumi Kasai, David Gray, Carol Clugston, Anna Dominiczak and Alex Alderton, Roberto Amato, Jeffrey Barrett, Sonia Goncalves, Ewan Harrison, David K. Jackson, Ian Johnston, Dominic Kwiatkowski, Cordelia Langford, John Sillitoe on behalf of the Wellcome Sanger Institute COVID-19 Surveillance Team |
| EPI_ISL_2914545 | Lighthouse Lab in Alderley Park | Wellcome Sanger Institute for the COVID-19 Genomics UK (COG-UK) Consortium | Jacquelyn Wynn, Mairead Hyland, The Lighthouse Lab in Alderley Park and Alex Alderton, Roberto Amato, Jeffrey Barrett, Sonia Goncalves, Ewan Harrison, David K. Jackson, Ian Johnston, Dominic Kwiatkowski, Cordelia Langford, John Sillitoe on behalf of the Wellcome Sanger Institute COVID-19 Surveillance Team |
| EPI_ISL_2914546, EPI_ISL_2914547 | Berkshire and Surrey Pathology Services Lighthouse Laboratory | Wellcome Sanger Institute for the COVID-19 Genomics UK (COG-UK) Consortium | Berkshire and Surrey Pathology Services Lighthouse Laboratory and Alex Alderton, Roberto Amato, Jeffrey Barrett, Sonia Goncalves, Ewan Harrison, David K. Jackson, Ian Johnston, Dominic Kwiatkowski, Cordelia Langford, John Sillitoe on behalf of the Wellcome Sanger Institute COVID-19 Surveillance Team |
| EPI_ISL_2914548, EPI_ISL_2914549 | Lighthouse Lab in Alderley Park | Wellcome Sanger Institute for the COVID-19 Genomics UK (COG-UK) Consortium | Jacquelyn Wynn, Mairead Hyland, The Lighthouse Lab in Alderley Park and Alex Alderton, Roberto Amato, Jeffrey Barrett, Sonia Goncalves, Ewan Harrison, David K. Jackson, Ian Johnston, Dominic Kwiatkowski, Cordelia Langford, John Sillitoe on behalf of the Wellcome Sanger Institute COVID-19 Surveillance Team |
| EPI_ISL_2914550, EPI_ISL_2914551 | Lighthouse Lab in Glasgow | Wellcome Sanger Institute for the COVID-19 Genomics UK (COG-UK) Consortium | Harper VanSteenhouse, Yumi Kasai, David Gray, Carol Clugston, Anna Dominiczak and Alex Alderton, Roberto Amato, Jeffrey Barrett, Sonia Goncalves, Ewan Harrison, David K. Jackson, Ian Johnston, Dominic Kwiatkowski, Cordelia Langford, John Sillitoe on behalf of the Wellcome Sanger Institute COVID-19 Surveillance Team |
| EPI_ISL_2914552 | Lighthouse Lab in Alderley Park | Wellcome Sanger Institute for the COVID-19 Genomics UK (COG-UK) Consortium | Jacquelyn Wynn, Mairead Hyland, The Lighthouse Lab in Alderley Park and Alex Alderton, Roberto Amato, Jeffrey Barrett, Sonia Goncalves, Ewan Harrison, David K. Jackson, Ian Johnston, Dominic Kwiatkowski, Cordelia Langford, John Sillitoe on behalf of the Wellcome Sanger Institute COVID-19 Surveillance Team |
| EPI_ISL_2914553, EPI_ISL_2914554 | Lighthouse Lab in Glasgow | Wellcome Sanger Institute for the COVID-19 Genomics UK (COG-UK) Consortium | Harper VanSteenhouse, Yumi Kasai, David Gray, Carol Clugston, Anna Dominiczak and Alex Alderton, Roberto Amato, Jeffrey Barrett, Sonia Goncalves, Ewan Harrison, David K. Jackson, Ian Johnston, Dominic Kwiatkowski, Cordelia Langford, John Sillitoe on behalf of the Wellcome Sanger Institute COVID-19 Surveillance Team |
| EPI_ISL_2914555, EPI_ISL_2914556, EPI_ISL_2914557 | Lighthouse Lab in Alderley Park | Wellcome Sanger Institute for the COVID-19 Genomics UK (COG-UK) Consortium | Jacquelyn Wynn, Mairead Hyland, The Lighthouse Lab in Alderley Park and Alex Alderton, Roberto Amato, Jeffrey Barrett, Sonia Goncalves, Ewan Harrison, David K. Jackson, Ian Johnston, Dominic Kwiatkowski, Cordelia Langford, John Sillitoe on behalf of the Wellcome Sanger Institute COVID-19 Surveillance Team |
| EPI_ISL_2914558 | Lighthouse Lab in Glasgow | Wellcome Sanger Institute for the COVID-19 Genomics UK (COG-UK) Consortium | Harper VanSteenhouse, Yumi Kasai, David Gray, Carol Clugston, Anna Dominiczak and Alex Alderton, Roberto Amato, Jeffrey Barrett, Sonia Goncalves, Ewan Harrison, David K. Jackson, Ian Johnston, Dominic Kwiatkowski, Cordelia Langford, John Sillitoe on behalf of the Wellcome Sanger Institute COVID-19 Surveillance Team |
| EPI_ISL_2914559 | Lighthouse Lab in Alderley Park | Wellcome Sanger Institute for the COVID-19 Genomics UK (COG-UK) Consortium | Jacquelyn Wynn, Mairead Hyland, The Lighthouse Lab in Alderley Park and Alex Alderton, Roberto Amato, Jeffrey Barrett, Sonia Goncalves, Ewan Harrison, David K. Jackson, Ian Johnston, Dominic Kwiatkowski, Cordelia Langford, John Sillitoe on behalf of the Wellcome Sanger Institute COVID-19 Surveillance Team |
| EPI_ISL_2914560 | Lighthouse Lab in Glasgow | Wellcome Sanger Institute for the COVID-19 Genomics UK (COG-UK) Consortium | Harper VanSteenhouse, Yumi Kasai, David Gray, Carol Clugston, Anna Dominiczak and Alex Alderton, Roberto Amato, Jeffrey Barrett, Sonia Goncalves, Ewan Harrison, David K. Jackson, Ian Johnston, Dominic Kwiatkowski, Cordelia Langford, John Sillitoe on behalf of the Wellcome Sanger Institute COVID-19 Surveillance Team |
| EPI_ISL_2914561 | Lighthouse Lab in Alderley Park | Wellcome Sanger Institute for the COVID-19 Genomics UK (COG-UK) Consortium | Jacquelyn Wynn, Mairead Hyland, The Lighthouse Lab in Alderley Park and Alex Alderton, Roberto Amato, Jeffrey Barrett, Sonia Goncalves, Ewan Harrison, David K. Jackson, Ian Johnston, Dominic Kwiatkowski, Cordelia Langford, John Sillitoe on behalf of the Wellcome Sanger Institute COVID-19 Surveillance Team |
| EPI_ISL_2914562, EPI_ISL_2914563, EPI_ISL_2914564 | Lighthouse Lab in Glasgow | Wellcome Sanger Institute for the COVID-19 Genomics UK (COG-UK) Consortium | Harper VanSteenhouse, Yumi Kasai, David Gray, Carol Clugston, Anna Dominiczak and Alex Alderton, Roberto Amato, Jeffrey Barrett, Sonia Goncalves, Ewan Harrison, David K. Jackson, Ian Johnston, Dominic Kwiatkowski, Cordelia Langford, John Sillitoe on behalf of the Wellcome Sanger Institute COVID-19 Surveillance Team |
| EPI_ISL_2914565 | Berkshire and Surrey Pathology Services Lighthouse Laboratory | Wellcome Sanger Institute for the COVID-19 Genomics UK (COG-UK) Consortium | Berkshire and Surrey Pathology Services Lighthouse Laboratory and Alex Alderton, Roberto Amato, Jeffrey Barrett, Sonia Goncalves, Ewan Harrison, David K. Jackson, Ian Johnston, Dominic Kwiatkowski, Cordelia Langford, John Sillitoe on behalf of the Wellcome Sanger Institute COVID-19 Surveillance Team |
| EPI_ISL_2914566, EPI_ISL_2914567, EPI_ISL_2914568 | Lighthouse Lab in Glasgow | Wellcome Sanger Institute for the COVID-19 Genomics UK (COG-UK) Consortium | Harper VanSteenhouse, Yumi Kasai, David Gray, Carol Clugston, Anna Dominiczak and Alex Alderton, Roberto Amato, Jeffrey Barrett, Sonia Goncalves, Ewan Harrison, David K. Jackson, Ian Johnston, Dominic Kwiatkowski, Cordelia Langford, John Sillitoe on behalf of the Wellcome Sanger Institute COVID-19 Surveillance Team |
| EPI_ISL_2914569, EPI_ISL_2914570 | Berkshire and Surrey Pathology Services Lighthouse Laboratory | Wellcome Sanger Institute for the COVID-19 Genomics UK (COG-UK) Consortium | Berkshire and Surrey Pathology Services Lighthouse Laboratory and Alex Alderton, Roberto Amato, Jeffrey Barrett, Sonia Goncalves, Ewan Harrison, David K. Jackson, Ian Johnston, Dominic Kwiatkowski, Cordelia Langford, John Sillitoe on behalf of the Wellcome Sanger Institute COVID-19 Surveillance Team |
| EPI_ISL_2914571 | Lighthouse Lab in Glasgow | Wellcome Sanger Institute for the COVID-19 Genomics UK (COG-UK) Consortium | Harper VanSteenhouse, Yumi Kasai, David Gray, Carol Clugston, Anna Dominiczak and Alex Alderton, Roberto Amato, Jeffrey Barrett, Sonia Goncalves, Ewan Harrison, David K. Jackson, Ian Johnston, Dominic Kwiatkowski, Cordelia Langford, John Sillitoe on behalf of the Wellcome Sanger Institute COVID-19 Surveillance Team |
| EPI_ISL_2914572 | Berkshire and Surrey Pathology Services Lighthouse Laboratory | Wellcome Sanger Institute for the COVID-19 Genomics UK (COG-UK) Consortium | Berkshire and Surrey Pathology Services Lighthouse Laboratory and Alex Alderton, Roberto Amato, Jeffrey Barrett, Sonia Goncalves, Ewan Harrison, David K. Jackson, Ian Johnston, Dominic Kwiatkowski, Cordelia Langford, John Sillitoe on behalf of the Wellcome Sanger Institute COVID-19 Surveillance Team |
| EPI_ISL_2914573, EPI_ISL_2914574 | Lighthouse Lab in Glasgow | Wellcome Sanger Institute for the COVID-19 Genomics UK (COG-UK) Consortium | Harper VanSteenhouse, Yumi Kasai, David Gray, Carol Clugston, Anna Dominiczak and Alex Alderton, Roberto Amato, Jeffrey Barrett, Sonia Goncalves, Ewan Harrison, David K. Jackson, Ian Johnston, Dominic Kwiatkowski, Cordelia Langford, John Sillitoe on behalf of the Wellcome Sanger Institute COVID-19 Surveillance Team |

| | | | |
|---|---|---|---|
| EPI_ISL_2914575 | Lighthouse Lab in Alderley Park | Wellcome Sanger Institute for the COVID-19 Genomics UK (COG-UK) Consortium | Jacquelyn Wynn, Mairead Hyland, The Lighthouse Lab in Alderley Park and Alex Alderton, Roberto Amato, Jeffrey Barrett, Sonia Goncalves, Ewan Harrison, David K. Jackson, Ian Johnston, Dominic Kwiatkowski, Cordelia Langford, John Sillitoe on behalf of the Wellcome Sanger Institute COVID-19 Surveillance Team |
| EPI_ISL_2914576, EPI_ISL_2914577 | Lighthouse Lab in Glasgow | Wellcome Sanger Institute for the COVID-19 Genomics UK (COG-UK) Consortium | Harper VanSteenhouse, Yumi Kasai, David Gray, Carol Clugston, Anna Dominiczak and Alex Alderton, Roberto Amato, Jeffrey Barrett, Sonia Goncalves, Ewan Harrison, David K. Jackson, Ian Johnston, Dominic Kwiatkowski, Cordelia Langford, John Sillitoe on behalf of the Wellcome Sanger Institute COVID-19 Surveillance Team |
| EPI_ISL_2914578, EPI_ISL_2914579 | Lighthouse Lab in Alderley Park | Wellcome Sanger Institute for the COVID-19 Genomics UK (COG-UK) Consortium | Jacquelyn Wynn, Mairead Hyland, The Lighthouse Lab in Alderley Park and Alex Alderton, Roberto Amato, Jeffrey Barrett, Sonia Goncalves, Ewan Harrison, David K. Jackson, Ian Johnston, Dominic Kwiatkowski, Cordelia Langford, John Sillitoe on behalf of the Wellcome Sanger Institute COVID-19 Surveillance Team |
| EPI_ISL_2914580 | Berkshire and Surrey Pathology Services Lighthouse Laboratory | Wellcome Sanger Institute for the COVID-19 Genomics UK (COG-UK) Consortium | Berkshire and Surrey Pathology Services Lighthouse Laboratory and Alex Alderton, Roberto Amato, Jeffrey Barrett, Sonia Goncalves, Ewan Harrison, David K. Jackson, Ian Johnston, Dominic Kwiatkowski, Cordelia Langford, John Sillitoe on behalf of the Wellcome Sanger Institute COVID-19 Surveillance Team |
| EPI_ISL_2914581 | Lighthouse Lab in Alderley Park | Wellcome Sanger Institute for the COVID-19 Genomics UK (COG-UK) Consortium | Jacquelyn Wynn, Mairead Hyland, The Lighthouse Lab in Alderley Park and Alex Alderton, Roberto Amato, Jeffrey Barrett, Sonia Goncalves, Ewan Harrison, David K. Jackson, Ian Johnston, Dominic Kwiatkowski, Cordelia Langford, John Sillitoe on behalf of the Wellcome Sanger Institute COVID-19 Surveillance Team |
| EPI_ISL_2914582 | Lighthouse Lab in Glasgow | Wellcome Sanger Institute for the COVID-19 Genomics UK (COG-UK) Consortium | Harper VanSteenhouse, Yumi Kasai, David Gray, Carol Clugston, Anna Dominiczak and Alex Alderton, Roberto Amato, Jeffrey Barrett, Sonia Goncalves, Ewan Harrison, David K. Jackson, Ian Johnston, Dominic Kwiatkowski, Cordelia Langford, John Sillitoe on behalf of the Wellcome Sanger Institute COVID-19 Surveillance Team |
| EPI_ISL_2914583, EPI_ISL_2914584 | Berkshire and Surrey Pathology Services Lighthouse Laboratory | Wellcome Sanger Institute for the COVID-19 Genomics UK (COG-UK) Consortium | Berkshire and Surrey Pathology Services Lighthouse Laboratory and Alex Alderton, Roberto Amato, Jeffrey Barrett, Sonia Goncalves, Ewan Harrison, David K. Jackson, Ian Johnston, Dominic Kwiatkowski, Cordelia Langford, John Sillitoe on behalf of the Wellcome Sanger Institute COVID-19 Surveillance Team |
| EPI_ISL_2914586 | Lighthouse Lab in Alderley Park | Wellcome Sanger Institute for the COVID-19 Genomics UK (COG-UK) Consortium | Jacquelyn Wynn, Mairead Hyland, The Lighthouse Lab in Alderley Park and Alex Alderton, Roberto Amato, Jeffrey Barrett, Sonia Goncalves, Ewan Harrison, David K. Jackson, Ian Johnston, Dominic Kwiatkowski, Cordelia Langford, John Sillitoe on behalf of the Wellcome Sanger Institute COVID-19 Surveillance Team |
| EPI_ISL_2914587 | Lighthouse Lab in Glasgow | Wellcome Sanger Institute for the COVID-19 Genomics UK (COG-UK) Consortium | Harper VanSteenhouse, Yumi Kasai, David Gray, Carol Clugston, Anna Dominiczak and Alex Alderton, Roberto Amato, Jeffrey Barrett, Sonia Goncalves, Ewan Harrison, David K. Jackson, Ian Johnston, Dominic Kwiatkowski, Cordelia Langford, John Sillitoe on behalf of the Wellcome Sanger Institute COVID-19 Surveillance Team |
| EPI_ISL_2914588 | Berkshire and Surrey Pathology Services Lighthouse Laboratory | Wellcome Sanger Institute for the COVID-19 Genomics UK (COG-UK) Consortium | Berkshire and Surrey Pathology Services Lighthouse Laboratory and Alex Alderton, Roberto Amato, Jeffrey Barrett, Sonia Goncalves, Ewan Harrison, David K. Jackson, Ian Johnston, Dominic Kwiatkowski, Cordelia Langford, John Sillitoe on behalf of the Wellcome Sanger Institute COVID-19 Surveillance Team |
| EPI_ISL_2914589, EPI_ISL_2914590, EPI_ISL_2914591, EPI_ISL_2914592, EPI_ISL_2914593, EPI_ISL_2914594, EPI_ISL_2914595 | Lighthouse Lab in Glasgow | Wellcome Sanger Institute for the COVID-19 Genomics UK (COG-UK) Consortium | Harper VanSteenhouse, Yumi Kasai, David Gray, Carol Clugston, Anna Dominiczak and Alex Alderton, Roberto Amato, Jeffrey Barrett, Sonia Goncalves, Ewan Harrison, David K. Jackson, Ian Johnston, Dominic Kwiatkowski, Cordelia Langford, John Sillitoe on behalf of the Wellcome Sanger Institute COVID-19 Surveillance Team |
| EPI_ISL_2914596 | Berkshire and Surrey Pathology Services Lighthouse Laboratory | Wellcome Sanger Institute for the COVID-19 Genomics UK (COG-UK) Consortium | Berkshire and Surrey Pathology Services Lighthouse Laboratory and Alex Alderton, Roberto Amato, Jeffrey Barrett, Sonia Goncalves, Ewan Harrison, David K. Jackson, Ian Johnston, Dominic Kwiatkowski, Cordelia Langford, John Sillitoe on behalf of the Wellcome Sanger Institute COVID-19 Surveillance Team |
| EPI_ISL_2914597 | Lighthouse Lab in Alderley Park | Wellcome Sanger Institute for the COVID-19 Genomics UK (COG-UK) Consortium | Jacquelyn Wynn, Mairead Hyland, The Lighthouse Lab in Alderley Park and Alex Alderton, Roberto Amato, Jeffrey Barrett, Sonia Goncalves, Ewan Harrison, David K. Jackson, Ian Johnston, Dominic Kwiatkowski, Cordelia Langford, John Sillitoe on behalf of the Wellcome Sanger Institute COVID-19 Surveillance Team |
| EPI_ISL_2914598 | Berkshire and Surrey Pathology Services Lighthouse Laboratory | Wellcome Sanger Institute for the COVID-19 Genomics UK (COG-UK) Consortium | Berkshire and Surrey Pathology Services Lighthouse Laboratory and Alex Alderton, Roberto Amato, Jeffrey Barrett, Sonia Goncalves, Ewan Harrison, David K. Jackson, Ian Johnston, Dominic Kwiatkowski, Cordelia Langford, John Sillitoe on behalf of the Wellcome Sanger Institute COVID-19 Surveillance Team |
| EPI_ISL_2914599 | Lighthouse Lab in Alderley Park | Wellcome Sanger Institute for the COVID-19 Genomics UK (COG-UK) Consortium | Jacquelyn Wynn, Mairead Hyland, The Lighthouse Lab in Alderley Park and Alex Alderton, Roberto Amato, Jeffrey Barrett, Sonia Goncalves, Ewan Harrison, David K. Jackson, Ian Johnston, Dominic Kwiatkowski, Cordelia Langford, John Sillitoe on behalf of the Wellcome Sanger Institute COVID-19 Surveillance Team |
| EPI_ISL_2914600, EPI_ISL_2914601, EPI_ISL_2914602, EPI_ISL_2914603 | Lighthouse Lab in Glasgow | Wellcome Sanger Institute for the COVID-19 Genomics UK (COG-UK) Consortium | Harper VanSteenhouse, Yumi Kasai, David Gray, Carol Clugston, Anna Dominiczak and Alex Alderton, Roberto Amato, Jeffrey Barrett, Sonia Goncalves, Ewan Harrison, David K. Jackson, Ian Johnston, Dominic Kwiatkowski, Cordelia Langford, John Sillitoe on behalf of the Wellcome Sanger Institute COVID-19 Surveillance Team |
| EPI_ISL_2914604, EPI_ISL_2914605, EPI_ISL_2914606 | Lighthouse Lab in Alderley Park | Wellcome Sanger Institute for the COVID-19 Genomics UK (COG-UK) Consortium | Jacquelyn Wynn, Mairead Hyland, The Lighthouse Lab in Alderley Park and Alex Alderton, Roberto Amato, Jeffrey Barrett, Sonia Goncalves, Ewan Harrison, David K. Jackson, Ian Johnston, Dominic Kwiatkowski, Cordelia Langford, John Sillitoe on behalf of the Wellcome Sanger Institute COVID-19 Surveillance Team |
| EPI_ISL_2914607, EPI_ISL_2914608 | Lighthouse Lab in Glasgow | Wellcome Sanger Institute for the COVID-19 Genomics UK (COG-UK) Consortium | Harper VanSteenhouse, Yumi Kasai, David Gray, Carol Clugston, Anna Dominiczak and Alex Alderton, Roberto Amato, Jeffrey Barrett, Sonia Goncalves, Ewan Harrison, David K. Jackson, Ian Johnston, Dominic Kwiatkowski, Cordelia Langford, John Sillitoe on behalf of the Wellcome Sanger Institute COVID-19 Surveillance Team |
| EPI_ISL_2914609 | Berkshire and Surrey Pathology Services Lighthouse Laboratory | Wellcome Sanger Institute for the COVID-19 Genomics UK (COG-UK) Consortium | Berkshire and Surrey Pathology Services Lighthouse Laboratory and Alex Alderton, Roberto Amato, Jeffrey Barrett, Sonia Goncalves, Ewan Harrison, David K. Jackson, Ian Johnston, Dominic Kwiatkowski, Cordelia Langford, John Sillitoe on behalf of the Wellcome Sanger Institute COVID-19 Surveillance Team |
| EPI_ISL_2914610, EPI_ISL_2914611 | Lighthouse Lab in Glasgow | Wellcome Sanger Institute for the COVID-19 Genomics UK (COG-UK) Consortium | Harper VanSteenhouse, Yumi Kasai, David Gray, Carol Clugston, Anna Dominiczak and Alex Alderton, Roberto Amato, Jeffrey Barrett, Sonia Goncalves, Ewan Harrison, David K. Jackson, Ian Johnston, Dominic Kwiatkowski, Cordelia Langford, John Sillitoe on behalf of the Wellcome Sanger Institute COVID-19 Surveillance Team |
| EPI_ISL_2914612 | Lighthouse Lab in Alderley Park | Wellcome Sanger Institute for the COVID-19 Genomics UK (COG-UK) Consortium | Jacquelyn Wynn, Mairead Hyland, The Lighthouse Lab in Alderley Park and Alex Alderton, Roberto Amato, Jeffrey Barrett, Sonia Goncalves, Ewan Harrison, David K. Jackson, Ian Johnston, Dominic Kwiatkowski, Cordelia Langford, John Sillitoe on behalf of the Wellcome Sanger Institute COVID-19 Surveillance Team |
| EPI_ISL_2914613 | Lighthouse Lab in Glasgow | Wellcome Sanger Institute for the COVID-19 Genomics UK (COG-UK) Consortium | Harper VanSteenhouse, Yumi Kasai, David Gray, Carol Clugston, Anna Dominiczak and Alex Alderton, Roberto Amato, Jeffrey Barrett, Sonia Goncalves, Ewan Harrison, David K. Jackson, Ian Johnston, Dominic Kwiatkowski, Cordelia Langford, John Sillitoe on behalf of the Wellcome Sanger Institute COVID-19 Surveillance Team |
| EPI_ISL_2914614 | Berkshire and Surrey Pathology Services Lighthouse Laboratory | Wellcome Sanger Institute for the COVID-19 Genomics UK (COG-UK) Consortium | Berkshire and Surrey Pathology Services Lighthouse Laboratory and Alex Alderton, Roberto Amato, Jeffrey Barrett, Sonia Goncalves, Ewan Harrison, David K. Jackson, Ian Johnston, Dominic Kwiatkowski, Cordelia Langford, John Sillitoe on behalf of the Wellcome Sanger Institute COVID-19 Surveillance Team |
| EPI_ISL_2914615 | Lighthouse Lab in Alderley Park | Wellcome Sanger Institute for the COVID-19 Genomics UK (COG-UK) Consortium | Jacquelyn Wynn, Mairead Hyland, The Lighthouse Lab in Alderley Park and Alex Alderton, Roberto Amato, Jeffrey Barrett, Sonia Goncalves, Ewan Harrison, David K. Jackson, Ian Johnston, Dominic Kwiatkowski, Cordelia Langford, John Sillitoe on behalf of the Wellcome Sanger Institute COVID-19 Surveillance Team |
| EPI_ISL_2914616, EPI_ISL_2914617 | Lighthouse Lab in Glasgow | Wellcome Sanger Institute for the COVID-19 Genomics UK (COG-UK) Consortium | Harper VanSteenhouse, Yumi Kasai, David Gray, Carol Clugston, Anna Dominiczak and Alex Alderton, Roberto Amato, Jeffrey Barrett, Sonia Goncalves, Ewan Harrison, David K. Jackson, Ian Johnston, Dominic Kwiatkowski, Cordelia Langford, John Sillitoe on behalf of the Wellcome Sanger Institute COVID-19 Surveillance Team |
| EPI_ISL_2914618, EPI_ISL_2914619 | Berkshire and Surrey Pathology Services Lighthouse Laboratory | Wellcome Sanger Institute for the COVID-19 Genomics UK (COG-UK) Consortium | Berkshire and Surrey Pathology Services Lighthouse Laboratory and Alex Alderton, Roberto Amato, Jeffrey Barrett, Sonia Goncalves, Ewan Harrison, David K. Jackson, Ian Johnston, Dominic Kwiatkowski, Cordelia Langford, John Sillitoe on behalf of the Wellcome Sanger Institute COVID-19 Surveillance Team |
| EPI_ISL_2914620 | Lighthouse Lab in Glasgow | Wellcome Sanger Institute for the COVID-19 Genomics UK (COG-UK) Consortium | Harper VanSteenhouse, Yumi Kasai, David Gray, Carol Clugston, Anna Dominiczak and Alex Alderton, Roberto Amato, Jeffrey Barrett, Sonia Goncalves, Ewan Harrison, David K. Jackson, Ian Johnston, Dominic Kwiatkowski, Cordelia Langford, John Sillitoe on behalf of the Wellcome Sanger Institute COVID-19 Surveillance Team |
| EPI_ISL_2914621, EPI_ISL_2914622 | Berkshire and Surrey Pathology Services Lighthouse Laboratory | Wellcome Sanger Institute for the COVID-19 Genomics UK (COG-UK) Consortium | Berkshire and Surrey Pathology Services Lighthouse Laboratory and Alex Alderton, Roberto Amato, Jeffrey Barrett, Sonia Goncalves, Ewan Harrison, David K. Jackson, Ian Johnston, Dominic Kwiatkowski, Cordelia Langford, John Sillitoe on behalf of the Wellcome Sanger Institute COVID-19 Surveillance Team |
| EPI_ISL_2914623 | Lighthouse Lab in Milton Keynes | Wellcome Sanger Institute for the COVID-19 Genomics UK (COG-UK) Consortium | The Lighthouse Lab in Milton Keynes and Alex Alderton, Roberto Amato, Jeffrey Barrett, Sonia Goncalves, Ewan Harrison, David K. Jackson, Ian Johnston, Dominic Kwiatkowski, Cordelia Langford, John Sillitoe on behalf of the Wellcome Sanger Institute COVID-19 Surveillance Team |
| EPI_ISL_2914624, EPI_ISL_2914625 | Berkshire and Surrey Pathology Services Lighthouse Laboratory | Wellcome Sanger Institute for the COVID-19 Genomics UK (COG-UK) Consortium | Berkshire and Surrey Pathology Services Lighthouse Laboratory and Alex Alderton, Roberto Amato, Jeffrey Barrett, Sonia Goncalves, Ewan Harrison, David K. Jackson, Ian Johnston, Dominic Kwiatkowski, Cordelia Langford, John Sillitoe on behalf of the Wellcome Sanger Institute COVID-19 Surveillance Team |
| EPI_ISL_2914626, EPI_ISL_2914627 | Lighthouse Lab in Milton Keynes | Wellcome Sanger Institute for the COVID-19 Genomics UK (COG-UK) Consortium | The Lighthouse Lab in Milton Keynes and Alex Alderton, Roberto Amato, Jeffrey Barrett, Sonia Goncalves, Ewan Harrison, David K. Jackson, Ian Johnston, Dominic Kwiatkowski, Cordelia Langford, John Sillitoe on behalf of the Wellcome Sanger Institute COVID-19 Surveillance Team |
| EPI_ISL_2914628, EPI_ISL_2914629, EPI_ISL_2914630 | Berkshire and Surrey Pathology Services Lighthouse Laboratory | Wellcome Sanger Institute for the COVID-19 Genomics UK (COG-UK) Consortium | Berkshire and Surrey Pathology Services Lighthouse Laboratory and Alex Alderton, Roberto Amato, Jeffrey Barrett, Sonia Goncalves, Ewan Harrison, David K. Jackson, Ian Johnston, Dominic Kwiatkowski, Cordelia Langford, John Sillitoe on behalf of the Wellcome Sanger Institute COVID-19 Surveillance Team |
| EPI_ISL_2914631 | Lighthouse Lab in Milton Keynes | Wellcome Sanger Institute for the COVID-19 Genomics UK (COG-UK) Consortium | The Lighthouse Lab in Milton Keynes and Alex Alderton, Roberto Amato, Jeffrey Barrett, Sonia Goncalves, Ewan Harrison, David K. Jackson, Ian Johnston, Dominic Kwiatkowski, Cordelia Langford, John Sillitoe on behalf of the Wellcome Sanger Institute COVID-19 Surveillance Team |
| EPI_ISL_2914632 | Berkshire and Surrey Pathology Services Lighthouse Laboratory | Wellcome Sanger Institute for the COVID-19 Genomics UK (COG-UK) Consortium | Berkshire and Surrey Pathology Services Lighthouse Laboratory and Alex Alderton, Roberto Amato, Jeffrey Barrett, Sonia Goncalves, Ewan Harrison, David K. Jackson, Ian Johnston, Dominic Kwiatkowski, Cordelia Langford, John Sillitoe on behalf of the Wellcome Sanger Institute COVID-19 Surveillance Team |
| EPI_ISL_2914633, EPI_ISL_2914634 | Lighthouse Lab in Glasgow | Wellcome Sanger Institute for the COVID-19 Genomics UK (COG-UK) Consortium | Harper VanSteenhouse, Yumi Kasai, David Gray, Carol Clugston, Anna Dominiczak and Alex Alderton, Roberto Amato, Jeffrey Barrett, Sonia Goncalves, Ewan Harrison, David K. Jackson, Ian Johnston, Dominic Kwiatkowski, Cordelia Langford, John Sillitoe on behalf of the Wellcome Sanger Institute COVID-19 Surveillance Team |
| EPI_ISL_2914635 | Berkshire and Surrey Pathology Services Lighthouse Laboratory | Wellcome Sanger Institute for the COVID-19 Genomics UK (COG-UK) Consortium | Berkshire and Surrey Pathology Services Lighthouse Laboratory and Alex Alderton, Roberto Amato, Jeffrey Barrett, Sonia Goncalves, Ewan Harrison, David K. Jackson, Ian Johnston, Dominic Kwiatkowski, Cordelia Langford, John Sillitoe on behalf of the Wellcome Sanger Institute COVID-19 Surveillance Team |
| EPI_ISL_2914636 | Lighthouse Lab in Milton Keynes | Wellcome Sanger Institute for the COVID-19 Genomics UK (COG-UK) Consortium | The Lighthouse Lab in Milton Keynes and Alex Alderton, Roberto Amato, Jeffrey Barrett, Sonia Goncalves, Ewan Harrison, David K. Jackson, Ian Johnston, Dominic Kwiatkowski, Cordelia Langford, John Sillitoe on behalf of the Wellcome Sanger Institute COVID-19 Surveillance Team |
| EPI_ISL_2914637, EPI_ISL_2914638, EPI_ISL_2914639, EPI_ISL_2914640 | Berkshire and Surrey Pathology Services Lighthouse Laboratory | Wellcome Sanger Institute for the COVID-19 Genomics UK (COG-UK) Consortium | Berkshire and Surrey Pathology Services Lighthouse Laboratory and Alex Alderton, Roberto Amato, Jeffrey Barrett, Sonia Goncalves, Ewan Harrison, David K. Jackson, Ian Johnston, Dominic Kwiatkowski, Cordelia Langford, John Sillitoe on behalf of the Wellcome Sanger Institute COVID-19 Surveillance Team |
| EPI_ISL_2914641, EPI_ISL_2914642 | Lighthouse Lab in Milton Keynes | Wellcome Sanger Institute for the COVID-19 Genomics UK (COG-UK) Consortium | The Lighthouse Lab in Milton Keynes and Alex Alderton, Roberto Amato, Jeffrey Barrett, Sonia Goncalves, Ewan Harrison, David K. Jackson, Ian Johnston, Dominic Kwiatkowski, Cordelia Langford, John Sillitoe on behalf of the Wellcome Sanger Institute COVID-19 Surveillance Team |
| EPI_ISL_2914643 | Berkshire and Surrey Pathology Services Lighthouse Laboratory | Wellcome Sanger Institute for the COVID-19 Genomics UK (COG-UK) Consortium | Berkshire and Surrey Pathology Services Lighthouse Laboratory and Alex Alderton, Roberto Amato, Jeffrey Barrett, Sonia Goncalves, Ewan Harrison, David K. Jackson, Ian Johnston, Dominic Kwiatkowski, Cordelia Langford, John Sillitoe on behalf of the Wellcome Sanger Institute COVID-19 Surveillance Team |
| EPI_ISL_2914644 | Lighthouse Lab in Glasgow | Wellcome Sanger Institute for the COVID-19 Genomics UK (COG-UK) | Harper VanSteenhouse, Yumi Kasai, David Gray, Carol Clugston, Anna Dominiczak and Alex Alderton, Roberto Amato, Jeffrey Barrett, Sonia Goncalves, Ewan Harrison, David K. Jackson, Ian Johnston, Dominic Kwiatkowski, Cordelia Langford, |

| | | | |
|---|---|---|---|
| | | Consortium | John Sillitoe on behalf of the Wellcome Sanger Institute COVID-19 Surveillance Team |
| EPI_ISL_2914645 | Lighthouse Lab in Milton Keynes | Wellcome Sanger Institute for the COVID-19 Genomics UK (COG-UK) Consortium | The Lighthouse Lab in Milton Keynes and Alex Alderton, Roberto Amato, Jeffrey Barrett, Sonia Goncalves, Ewan Harrison, David K. Jackson, Ian Johnston, Dominic Kwiatkowski, Cordelia Langford, John Sillitoe on behalf of the Wellcome Sanger Institute COVID-19 Surveillance Team |
| EPI_ISL_2914646 | Berkshire and Surrey Pathology Services Lighthouse Laboratory | Wellcome Sanger Institute for the COVID-19 Genomics UK (COG-UK) Consortium | Berkshire and Surrey Pathology Services Lighthouse Laboratory and Alex Alderton, Roberto Amato, Jeffrey Barrett, Sonia Goncalves, Ewan Harrison, David K. Jackson, Ian Johnston, Dominic Kwiatkowski, Cordelia Langford, John Sillitoe on behalf of the Wellcome Sanger Institute COVID-19 Surveillance Team |
| EPI_ISL_2914647 | Lighthouse Lab in Glasgow | Wellcome Sanger Institute for the COVID-19 Genomics UK (COG-UK) Consortium | Harper VanSteenhouse, Yumi Kasai, David Gray, Carol Clugston, Anna Dominiczak and Alex Alderton, Roberto Amato, Jeffrey Barrett, Sonia Goncalves, Ewan Harrison, David K. Jackson, Ian Johnston, Dominic Kwiatkowski, Cordelia Langford, John Sillitoe on behalf of the Wellcome Sanger Institute COVID-19 Surveillance Team |
| EPI_ISL_2914648 | Lighthouse Lab in Milton Keynes | Wellcome Sanger Institute for the COVID-19 Genomics UK (COG-UK) Consortium | The Lighthouse Lab in Milton Keynes and Alex Alderton, Roberto Amato, Jeffrey Barrett, Sonia Goncalves, Ewan Harrison, David K. Jackson, Ian Johnston, Dominic Kwiatkowski, Cordelia Langford, John Sillitoe on behalf of the Wellcome Sanger Institute COVID-19 Surveillance Team |
| EPI_ISL_2914649 | Lighthouse Lab in Glasgow | Wellcome Sanger Institute for the COVID-19 Genomics UK (COG-UK) Consortium | Harper VanSteenhouse, Yumi Kasai, David Gray, Carol Clugston, Anna Dominiczak and Alex Alderton, Roberto Amato, Jeffrey Barrett, Sonia Goncalves, Ewan Harrison, David K. Jackson, Ian Johnston, Dominic Kwiatkowski, Cordelia Langford, John Sillitoe on behalf of the Wellcome Sanger Institute COVID-19 Surveillance Team |
| EPI_ISL_2914650, EPI_ISL_2914651, EPI_ISL_2914652 | Berkshire and Surrey Pathology Services Lighthouse Laboratory | Wellcome Sanger Institute for the COVID-19 Genomics UK (COG-UK) Consortium | Berkshire and Surrey Pathology Services Lighthouse Laboratory and Alex Alderton, Roberto Amato, Jeffrey Barrett, Sonia Goncalves, Ewan Harrison, David K. Jackson, Ian Johnston, Dominic Kwiatkowski, Cordelia Langford, John Sillitoe on behalf of the Wellcome Sanger Institute COVID-19 Surveillance Team |
| EPI_ISL_2914653 | Lighthouse Lab in Glasgow | Wellcome Sanger Institute for the COVID-19 Genomics UK (COG-UK) Consortium | Harper VanSteenhouse, Yumi Kasai, David Gray, Carol Clugston, Anna Dominiczak and Alex Alderton, Roberto Amato, Jeffrey Barrett, Sonia Goncalves, Ewan Harrison, David K. Jackson, Ian Johnston, Dominic Kwiatkowski, Cordelia Langford, John Sillitoe on behalf of the Wellcome Sanger Institute COVID-19 Surveillance Team |
| EPI_ISL_2914654, EPI_ISL_2914655 | Berkshire and Surrey Pathology Services Lighthouse Laboratory | Wellcome Sanger Institute for the COVID-19 Genomics UK (COG-UK) Consortium | Berkshire and Surrey Pathology Services Lighthouse Laboratory and Alex Alderton, Roberto Amato, Jeffrey Barrett, Sonia Goncalves, Ewan Harrison, David K. Jackson, Ian Johnston, Dominic Kwiatkowski, Cordelia Langford, John Sillitoe on behalf of the Wellcome Sanger Institute COVID-19 Surveillance Team |
| EPI_ISL_2914656 | Lighthouse Lab in Milton Keynes | Wellcome Sanger Institute for the COVID-19 Genomics UK (COG-UK) Consortium | The Lighthouse Lab in Milton Keynes and Alex Alderton, Roberto Amato, Jeffrey Barrett, Sonia Goncalves, Ewan Harrison, David K. Jackson, Ian Johnston, Dominic Kwiatkowski, Cordelia Langford, John Sillitoe on behalf of the Wellcome Sanger Institute COVID-19 Surveillance Team |
| EPI_ISL_2914657 | Berkshire and Surrey Pathology Services Lighthouse Laboratory | Wellcome Sanger Institute for the COVID-19 Genomics UK (COG-UK) Consortium | Berkshire and Surrey Pathology Services Lighthouse Laboratory and Alex Alderton, Roberto Amato, Jeffrey Barrett, Sonia Goncalves, Ewan Harrison, David K. Jackson, Ian Johnston, Dominic Kwiatkowski, Cordelia Langford, John Sillitoe on behalf of the Wellcome Sanger Institute COVID-19 Surveillance Team |
| EPI_ISL_2914658 | Lighthouse Lab in Glasgow | Wellcome Sanger Institute for the COVID-19 Genomics UK (COG-UK) Consortium | Harper VanSteenhouse, Yumi Kasai, David Gray, Carol Clugston, Anna Dominiczak and Alex Alderton, Roberto Amato, Jeffrey Barrett, Sonia Goncalves, Ewan Harrison, David K. Jackson, Ian Johnston, Dominic Kwiatkowski, Cordelia Langford, John Sillitoe on behalf of the Wellcome Sanger Institute COVID-19 Surveillance Team |
| EPI_ISL_2914659, EPI_ISL_2914660, EPI_ISL_2914661, EPI_ISL_2914662 | Berkshire and Surrey Pathology Services Lighthouse Laboratory | Wellcome Sanger Institute for the COVID-19 Genomics UK (COG-UK) Consortium | Berkshire and Surrey Pathology Services Lighthouse Laboratory and Alex Alderton, Roberto Amato, Jeffrey Barrett, Sonia Goncalves, Ewan Harrison, David K. Jackson, Ian Johnston, Dominic Kwiatkowski, Cordelia Langford, John Sillitoe on behalf of the Wellcome Sanger Institute COVID-19 Surveillance Team |
| EPI_ISL_2914663 | Lighthouse Lab in Glasgow | Wellcome Sanger Institute for the COVID-19 Genomics UK (COG-UK) Consortium | Harper VanSteenhouse, Yumi Kasai, David Gray, Carol Clugston, Anna Dominiczak and Alex Alderton, Roberto Amato, Jeffrey Barrett, Sonia Goncalves, Ewan Harrison, David K. Jackson, Ian Johnston, Dominic Kwiatkowski, Cordelia Langford, John Sillitoe on behalf of the Wellcome Sanger Institute COVID-19 Surveillance Team |
| EPI_ISL_2914664 | Lighthouse Lab in Milton Keynes | Wellcome Sanger Institute for the COVID-19 Genomics UK (COG-UK) Consortium | The Lighthouse Lab in Milton Keynes and Alex Alderton, Roberto Amato, Jeffrey Barrett, Sonia Goncalves, Ewan Harrison, David K. Jackson, Ian Johnston, Dominic Kwiatkowski, Cordelia Langford, John Sillitoe on behalf of the Wellcome Sanger Institute COVID-19 Surveillance Team |
| EPI_ISL_2914665, EPI_ISL_2914666, EPI_ISL_2914667, EPI_ISL_2914668, EPI_ISL_2914669, EPI_ISL_2914670, EPI_ISL_2914671, EPI_ISL_2914672, EPI_ISL_2914673, EPI_ISL_2914674 | Berkshire and Surrey Pathology Services Lighthouse Laboratory | Wellcome Sanger Institute for the COVID-19 Genomics UK (COG-UK) Consortium | Berkshire and Surrey Pathology Services Lighthouse Laboratory and Alex Alderton, Roberto Amato, Jeffrey Barrett, Sonia Goncalves, Ewan Harrison, David K. Jackson, Ian Johnston, Dominic Kwiatkowski, Cordelia Langford, John Sillitoe on behalf of the Wellcome Sanger Institute COVID-19 Surveillance Team |
| EPI_ISL_2914675 | Lighthouse Lab in Milton Keynes | Wellcome Sanger Institute for the COVID-19 Genomics UK (COG-UK) Consortium | The Lighthouse Lab in Milton Keynes and Alex Alderton, Roberto Amato, Jeffrey Barrett, Sonia Goncalves, Ewan Harrison, David K. Jackson, Ian Johnston, Dominic Kwiatkowski, Cordelia Langford, John Sillitoe on behalf of the Wellcome Sanger Institute COVID-19 Surveillance Team |
| EPI_ISL_2914677 | Berkshire and Surrey Pathology Services Lighthouse Laboratory | Wellcome Sanger Institute for the COVID-19 Genomics UK (COG-UK) Consortium | Berkshire and Surrey Pathology Services Lighthouse Laboratory and Alex Alderton, Roberto Amato, Jeffrey Barrett, Sonia Goncalves, Ewan Harrison, David K. Jackson, Ian Johnston, Dominic Kwiatkowski, Cordelia Langford, John Sillitoe on behalf of the Wellcome Sanger Institute COVID-19 Surveillance Team |
| EPI_ISL_2914678 | Lighthouse Lab in Glasgow | Wellcome Sanger Institute for the COVID-19 Genomics UK (COG-UK) Consortium | Harper VanSteenhouse, Yumi Kasai, David Gray, Carol Clugston, Anna Dominiczak and Alex Alderton, Roberto Amato, Jeffrey Barrett, Sonia Goncalves, Ewan Harrison, David K. Jackson, Ian Johnston, Dominic Kwiatkowski, Cordelia Langford, John Sillitoe on behalf of the Wellcome Sanger Institute COVID-19 Surveillance Team |
| EPI_ISL_2914679 | Lighthouse Lab in Milton Keynes | Wellcome Sanger Institute for the COVID-19 Genomics UK (COG-UK) Consortium | The Lighthouse Lab in Milton Keynes and Alex Alderton, Roberto Amato, Jeffrey Barrett, Sonia Goncalves, Ewan Harrison, David K. Jackson, Ian Johnston, Dominic Kwiatkowski, Cordelia Langford, John Sillitoe on behalf of the Wellcome Sanger Institute COVID-19 Surveillance Team |
| EPI_ISL_2914680 | Berkshire and Surrey Pathology Services Lighthouse Laboratory | Wellcome Sanger Institute for the COVID-19 Genomics UK (COG-UK) Consortium | Berkshire and Surrey Pathology Services Lighthouse Laboratory and Alex Alderton, Roberto Amato, Jeffrey Barrett, Sonia Goncalves, Ewan Harrison, David K. Jackson, Ian Johnston, Dominic Kwiatkowski, Cordelia Langford, John Sillitoe on behalf of the Wellcome Sanger Institute COVID-19 Surveillance Team |
| EPI_ISL_2914681, EPI_ISL_2914682 | Lighthouse Lab in Glasgow | Wellcome Sanger Institute for the COVID-19 Genomics UK (COG-UK) Consortium | Harper VanSteenhouse, Yumi Kasai, David Gray, Carol Clugston, Anna Dominiczak and Alex Alderton, Roberto Amato, Jeffrey Barrett, Sonia Goncalves, Ewan Harrison, David K. Jackson, Ian Johnston, Dominic Kwiatkowski, Cordelia Langford, John Sillitoe on behalf of the Wellcome Sanger Institute COVID-19 Surveillance Team |
| EPI_ISL_2914683, EPI_ISL_2914684, EPI_ISL_2914685 | Berkshire and Surrey Pathology Services Lighthouse Laboratory | Wellcome Sanger Institute for the COVID-19 Genomics UK (COG-UK) Consortium | Berkshire and Surrey Pathology Services Lighthouse Laboratory and Alex Alderton, Roberto Amato, Jeffrey Barrett, Sonia Goncalves, Ewan Harrison, David K. Jackson, Ian Johnston, Dominic Kwiatkowski, Cordelia Langford, John Sillitoe on behalf of the Wellcome Sanger Institute COVID-19 Surveillance Team |
| EPI_ISL_2914686 | Lighthouse Lab in Milton Keynes | Wellcome Sanger Institute for the COVID-19 Genomics UK (COG-UK) Consortium | The Lighthouse Lab in Milton Keynes and Alex Alderton, Roberto Amato, Jeffrey Barrett, Sonia Goncalves, Ewan Harrison, David K. Jackson, Ian Johnston, Dominic Kwiatkowski, Cordelia Langford, John Sillitoe on behalf of the Wellcome Sanger Institute COVID-19 Surveillance Team |
| EPI_ISL_2914687, EPI_ISL_2914688 | Berkshire and Surrey Pathology Services Lighthouse Laboratory | Wellcome Sanger Institute for the COVID-19 Genomics UK (COG-UK) Consortium | Berkshire and Surrey Pathology Services Lighthouse Laboratory and Alex Alderton, Roberto Amato, Jeffrey Barrett, Sonia Goncalves, Ewan Harrison, David K. Jackson, Ian Johnston, Dominic Kwiatkowski, Cordelia Langford, John Sillitoe on behalf of the Wellcome Sanger Institute COVID-19 Surveillance Team |
| EPI_ISL_2914689 | Lighthouse Lab in Milton Keynes | Wellcome Sanger Institute for the COVID-19 Genomics UK (COG-UK) Consortium | The Lighthouse Lab in Milton Keynes and Alex Alderton, Roberto Amato, Jeffrey Barrett, Sonia Goncalves, Ewan Harrison, David K. Jackson, Ian Johnston, Dominic Kwiatkowski, Cordelia Langford, John Sillitoe on behalf of the Wellcome Sanger Institute COVID-19 Surveillance Team |
| EPI_ISL_2914690 | Lighthouse Lab in Glasgow | Wellcome Sanger Institute for the COVID-19 Genomics UK (COG-UK) Consortium | Harper VanSteenhouse, Yumi Kasai, David Gray, Carol Clugston, Anna Dominiczak and Alex Alderton, Roberto Amato, Jeffrey Barrett, Sonia Goncalves, Ewan Harrison, David K. Jackson, Ian Johnston, Dominic Kwiatkowski, Cordelia Langford, John Sillitoe on behalf of the Wellcome Sanger Institute COVID-19 Surveillance Team |
| EPI_ISL_2914691, EPI_ISL_2914692 | Berkshire and Surrey Pathology Services Lighthouse Laboratory | Wellcome Sanger Institute for the COVID-19 Genomics UK (COG-UK) Consortium | Berkshire and Surrey Pathology Services Lighthouse Laboratory and Alex Alderton, Roberto Amato, Jeffrey Barrett, Sonia Goncalves, Ewan Harrison, David K. Jackson, Ian Johnston, Dominic Kwiatkowski, Cordelia Langford, John Sillitoe on behalf of the Wellcome Sanger Institute COVID-19 Surveillance Team |
| EPI_ISL_2914693 | Lighthouse Lab in Glasgow | Wellcome Sanger Institute for the COVID-19 Genomics UK (COG-UK) Consortium | Harper VanSteenhouse, Yumi Kasai, David Gray, Carol Clugston, Anna Dominiczak and Alex Alderton, Roberto Amato, Jeffrey Barrett, Sonia Goncalves, Ewan Harrison, David K. Jackson, Ian Johnston, Dominic Kwiatkowski, Cordelia Langford, John Sillitoe on behalf of the Wellcome Sanger Institute COVID-19 Surveillance Team |
| EPI_ISL_2914694 | Berkshire and Surrey Pathology Services Lighthouse Laboratory | Wellcome Sanger Institute for the COVID-19 Genomics UK (COG-UK) Consortium | Berkshire and Surrey Pathology Services Lighthouse Laboratory and Alex Alderton, Roberto Amato, Jeffrey Barrett, Sonia Goncalves, Ewan Harrison, David K. Jackson, Ian Johnston, Dominic Kwiatkowski, Cordelia Langford, John Sillitoe on behalf of the Wellcome Sanger Institute COVID-19 Surveillance Team |
| EPI_ISL_2914695, EPI_ISL_2914696 | Lighthouse Lab in Milton Keynes | Wellcome Sanger Institute for the COVID-19 Genomics UK (COG-UK) Consortium | The Lighthouse Lab in Milton Keynes and Alex Alderton, Roberto Amato, Jeffrey Barrett, Sonia Goncalves, Ewan Harrison, David K. Jackson, Ian Johnston, Dominic Kwiatkowski, Cordelia Langford, John Sillitoe on behalf of the Wellcome Sanger Institute COVID-19 Surveillance Team |
| EPI_ISL_2914697, EPI_ISL_2914698, EPI_ISL_2914699 | Berkshire and Surrey Pathology Services Lighthouse Laboratory | Wellcome Sanger Institute for the COVID-19 Genomics UK (COG-UK) Consortium | Berkshire and Surrey Pathology Services Lighthouse Laboratory and Alex Alderton, Roberto Amato, Jeffrey Barrett, Sonia Goncalves, Ewan Harrison, David K. Jackson, Ian Johnston, Dominic Kwiatkowski, Cordelia Langford, John Sillitoe on behalf of the Wellcome Sanger Institute COVID-19 Surveillance Team |
| EPI_ISL_2914700 | Lighthouse Lab in Glasgow | Wellcome Sanger Institute for the COVID-19 Genomics UK (COG-UK) Consortium | Harper VanSteenhouse, Yumi Kasai, David Gray, Carol Clugston, Anna Dominiczak and Alex Alderton, Roberto Amato, Jeffrey Barrett, Sonia Goncalves, Ewan Harrison, David K. Jackson, Ian Johnston, Dominic Kwiatkowski, Cordelia Langford, John Sillitoe on behalf of the Wellcome Sanger Institute COVID-19 Surveillance Team |
| EPI_ISL_2914701, EPI_ISL_2914702, EPI_ISL_2914703, EPI_ISL_2914704, EPI_ISL_2914705, EPI_ISL_2914706, EPI_ISL_2914707 | Berkshire and Surrey Pathology Services Lighthouse Laboratory | Wellcome Sanger Institute for the COVID-19 Genomics UK (COG-UK) Consortium | Berkshire and Surrey Pathology Services Lighthouse Laboratory and Alex Alderton, Roberto Amato, Jeffrey Barrett, Sonia Goncalves, Ewan Harrison, David K. Jackson, Ian Johnston, Dominic Kwiatkowski, Cordelia Langford, John Sillitoe on behalf of the Wellcome Sanger Institute COVID-19 Surveillance Team |
| EPI_ISL_2914708 | Lighthouse Lab in Milton Keynes | Wellcome Sanger Institute for the COVID-19 Genomics UK (COG-UK) Consortium | The Lighthouse Lab in Milton Keynes and Alex Alderton, Roberto Amato, Jeffrey Barrett, Sonia Goncalves, Ewan Harrison, David K. Jackson, Ian Johnston, Dominic Kwiatkowski, Cordelia Langford, John Sillitoe on behalf of the Wellcome Sanger Institute COVID-19 Surveillance Team |
| EPI_ISL_2914709, EPI_ISL_2914710 | Lighthouse Lab in Glasgow | Wellcome Sanger Institute for the COVID-19 Genomics UK (COG-UK) Consortium | Harper VanSteenhouse, Yumi Kasai, David Gray, Carol Clugston, Anna Dominiczak and Alex Alderton, Roberto Amato, Jeffrey Barrett, Sonia Goncalves, Ewan Harrison, David K. Jackson, Ian Johnston, Dominic Kwiatkowski, Cordelia Langford, John Sillitoe on behalf of the Wellcome Sanger Institute COVID-19 Surveillance Team |
| EPI_ISL_2914711 | Berkshire and Surrey Pathology Services Lighthouse Laboratory | Wellcome Sanger Institute for the COVID-19 Genomics UK (COG-UK) Consortium | Berkshire and Surrey Pathology Services Lighthouse Laboratory and Alex Alderton, Roberto Amato, Jeffrey Barrett, Sonia Goncalves, Ewan Harrison, David K. Jackson, Ian Johnston, Dominic Kwiatkowski, Cordelia Langford, John Sillitoe on behalf of the Wellcome Sanger Institute COVID-19 Surveillance Team |
| EPI_ISL_2914712, EPI_ISL_2914713, | Lighthouse Lab in Milton Keynes | Wellcome Sanger Institute for the COVID-19 Genomics UK (COG-UK) Consortium | The Lighthouse Lab in Milton Keynes and Alex Alderton, Roberto Amato, Jeffrey Barrett, Sonia Goncalves, Ewan Harrison, David K. Jackson, Ian Johnston, Dominic Kwiatkowski, Cordelia Langford, John Sillitoe on behalf of the Wellcome Sanger Institute COVID-19 Surveillance Team |

| | | | |
|---|---|---|---|
| EPI_ISL_2914714 | | | |
| EPI_ISL_2914715, EPI_ISL_2914716, EPI_ISL_2914717, EPI_ISL_2914718 | Berkshire and Surrey Pathology Services Lighthouse Laboratory | Wellcome Sanger Institute for the COVID-19 Genomics UK (COG-UK) Consortium | Berkshire and Surrey Pathology Services Lighthouse Laboratory and Alex Alderton, Roberto Amato, Jeffrey Barrett, Sonia Goncalves, Ewan Harrison, David K. Jackson, Ian Johnston, Dominic Kwiatkowski, Cordelia Langford, John Sillitoe on behalf of the Wellcome Sanger Institute COVID-19 Surveillance Team |
| EPI_ISL_2914719 | Lighthouse Lab in Glasgow | Wellcome Sanger Institute for the COVID-19 Genomics UK (COG-UK) Consortium | Harper VanSteenhouse, Yumi Kasai, David Gray, Carol Clugston, Anna Dominiczak and Alex Alderton, Roberto Amato, Jeffrey Barrett, Sonia Goncalves, Ewan Harrison, David K. Jackson, Ian Johnston, Dominic Kwiatkowski, Cordelia Langford, John Sillitoe on behalf of the Wellcome Sanger Institute COVID-19 Surveillance Team |
| EPI_ISL_2914720 | Lighthouse Lab in Milton Keynes | Wellcome Sanger Institute for the COVID-19 Genomics UK (COG-UK) Consortium | The Lighthouse Lab in Milton Keynes and Alex Alderton, Roberto Amato, Jeffrey Barrett, Sonia Goncalves, Ewan Harrison, David K. Jackson, Ian Johnston, Dominic Kwiatkowski, Cordelia Langford, John Sillitoe on behalf of the Wellcome Sanger Institute COVID-19 Surveillance Team |
| EPI_ISL_2914721, EPI_ISL_2914722 | Berkshire and Surrey Pathology Services Lighthouse Laboratory | Wellcome Sanger Institute for the COVID-19 Genomics UK (COG-UK) Consortium | Berkshire and Surrey Pathology Services Lighthouse Laboratory and Alex Alderton, Roberto Amato, Jeffrey Barrett, Sonia Goncalves, Ewan Harrison, David K. Jackson, Ian Johnston, Dominic Kwiatkowski, Cordelia Langford, John Sillitoe on behalf of the Wellcome Sanger Institute COVID-19 Surveillance Team |
| EPI_ISL_2914723 | Lighthouse Lab in Milton Keynes | Wellcome Sanger Institute for the COVID-19 Genomics UK (COG-UK) Consortium | The Lighthouse Lab in Milton Keynes and Alex Alderton, Roberto Amato, Jeffrey Barrett, Sonia Goncalves, Ewan Harrison, David K. Jackson, Ian Johnston, Dominic Kwiatkowski, Cordelia Langford, John Sillitoe on behalf of the Wellcome Sanger Institute COVID-19 Surveillance Team |
| EPI_ISL_2914724, EPI_ISL_2914725, EPI_ISL_2914726, EPI_ISL_2914727 | Lighthouse Lab in Glasgow | Wellcome Sanger Institute for the COVID-19 Genomics UK (COG-UK) Consortium | Harper VanSteenhouse, Yumi Kasai, David Gray, Carol Clugston, Anna Dominiczak and Alex Alderton, Roberto Amato, Jeffrey Barrett, Sonia Goncalves, Ewan Harrison, David K. Jackson, Ian Johnston, Dominic Kwiatkowski, Cordelia Langford, John Sillitoe on behalf of the Wellcome Sanger Institute COVID-19 Surveillance Team |
| EPI_ISL_2914728, EPI_ISL_2914729, EPI_ISL_2914730 | Berkshire and Surrey Pathology Services Lighthouse Laboratory | Wellcome Sanger Institute for the COVID-19 Genomics UK (COG-UK) Consortium | Berkshire and Surrey Pathology Services Lighthouse Laboratory and Alex Alderton, Roberto Amato, Jeffrey Barrett, Sonia Goncalves, Ewan Harrison, David K. Jackson, Ian Johnston, Dominic Kwiatkowski, Cordelia Langford, John Sillitoe on behalf of the Wellcome Sanger Institute COVID-19 Surveillance Team |
| EPI_ISL_2914731 | Lighthouse Lab in Milton Keynes | Wellcome Sanger Institute for the COVID-19 Genomics UK (COG-UK) Consortium | The Lighthouse Lab in Milton Keynes and Alex Alderton, Roberto Amato, Jeffrey Barrett, Sonia Goncalves, Ewan Harrison, David K. Jackson, Ian Johnston, Dominic Kwiatkowski, Cordelia Langford, John Sillitoe on behalf of the Wellcome Sanger Institute COVID-19 Surveillance Team |
| EPI_ISL_2914732 | Lighthouse Lab in Glasgow | Wellcome Sanger Institute for the COVID-19 Genomics UK (COG-UK) Consortium | Harper VanSteenhouse, Yumi Kasai, David Gray, Carol Clugston, Anna Dominiczak and Alex Alderton, Roberto Amato, Jeffrey Barrett, Sonia Goncalves, Ewan Harrison, David K. Jackson, Ian Johnston, Dominic Kwiatkowski, Cordelia Langford, John Sillitoe on behalf of the Wellcome Sanger Institute COVID-19 Surveillance Team |
| EPI_ISL_2914733, EPI_ISL_2914734 | Berkshire and Surrey Pathology Services Lighthouse Laboratory | Wellcome Sanger Institute for the COVID-19 Genomics UK (COG-UK) Consortium | Berkshire and Surrey Pathology Services Lighthouse Laboratory and Alex Alderton, Roberto Amato, Jeffrey Barrett, Sonia Goncalves, Ewan Harrison, David K. Jackson, Ian Johnston, Dominic Kwiatkowski, Cordelia Langford, John Sillitoe on behalf of the Wellcome Sanger Institute COVID-19 Surveillance Team |
| EPI_ISL_2914735 | Lighthouse Lab in Glasgow | Wellcome Sanger Institute for the COVID-19 Genomics UK (COG-UK) Consortium | Harper VanSteenhouse, Yumi Kasai, David Gray, Carol Clugston, Anna Dominiczak and Alex Alderton, Roberto Amato, Jeffrey Barrett, Sonia Goncalves, Ewan Harrison, David K. Jackson, Ian Johnston, Dominic Kwiatkowski, Cordelia Langford, John Sillitoe on behalf of the Wellcome Sanger Institute COVID-19 Surveillance Team |
| EPI_ISL_2914736 | Lighthouse Lab in Milton Keynes | Wellcome Sanger Institute for the COVID-19 Genomics UK (COG-UK) Consortium | The Lighthouse Lab in Milton Keynes and Alex Alderton, Roberto Amato, Jeffrey Barrett, Sonia Goncalves, Ewan Harrison, David K. Jackson, Ian Johnston, Dominic Kwiatkowski, Cordelia Langford, John Sillitoe on behalf of the Wellcome Sanger Institute COVID-19 Surveillance Team |
| EPI_ISL_2914737, EPI_ISL_2914738, EPI_ISL_2914739 | Lighthouse Lab in Glasgow | Wellcome Sanger Institute for the COVID-19 Genomics UK (COG-UK) Consortium | Harper VanSteenhouse, Yumi Kasai, David Gray, Carol Clugston, Anna Dominiczak and Alex Alderton, Roberto Amato, Jeffrey Barrett, Sonia Goncalves, Ewan Harrison, David K. Jackson, Ian Johnston, Dominic Kwiatkowski, Cordelia Langford, John Sillitoe on behalf of the Wellcome Sanger Institute COVID-19 Surveillance Team |
| EPI_ISL_2914740, EPI_ISL_2914741 | Berkshire and Surrey Pathology Services Lighthouse Laboratory | Wellcome Sanger Institute for the COVID-19 Genomics UK (COG-UK) Consortium | Berkshire and Surrey Pathology Services Lighthouse Laboratory and Alex Alderton, Roberto Amato, Jeffrey Barrett, Sonia Goncalves, Ewan Harrison, David K. Jackson, Ian Johnston, Dominic Kwiatkowski, Cordelia Langford, John Sillitoe on behalf of the Wellcome Sanger Institute COVID-19 Surveillance Team |
| EPI_ISL_2914742 | Lighthouse Lab in Milton Keynes | Wellcome Sanger Institute for the COVID-19 Genomics UK (COG-UK) Consortium | The Lighthouse Lab in Milton Keynes and Alex Alderton, Roberto Amato, Jeffrey Barrett, Sonia Goncalves, Ewan Harrison, David K. Jackson, Ian Johnston, Dominic Kwiatkowski, Cordelia Langford, John Sillitoe on behalf of the Wellcome Sanger Institute COVID-19 Surveillance Team |
| EPI_ISL_2914743 | Lighthouse Lab in Glasgow | Wellcome Sanger Institute for the COVID-19 Genomics UK (COG-UK) Consortium | Harper VanSteenhouse, Yumi Kasai, David Gray, Carol Clugston, Anna Dominiczak and Alex Alderton, Roberto Amato, Jeffrey Barrett, Sonia Goncalves, Ewan Harrison, David K. Jackson, Ian Johnston, Dominic Kwiatkowski, Cordelia Langford, John Sillitoe on behalf of the Wellcome Sanger Institute COVID-19 Surveillance Team |
| EPI_ISL_2914744 | Berkshire and Surrey Pathology Services Lighthouse Laboratory | Wellcome Sanger Institute for the COVID-19 Genomics UK (COG-UK) Consortium | Berkshire and Surrey Pathology Services Lighthouse Laboratory and Alex Alderton, Roberto Amato, Jeffrey Barrett, Sonia Goncalves, Ewan Harrison, David K. Jackson, Ian Johnston, Dominic Kwiatkowski, Cordelia Langford, John Sillitoe on behalf of the Wellcome Sanger Institute COVID-19 Surveillance Team |
| EPI_ISL_2914745 | Lighthouse Lab in Glasgow | Wellcome Sanger Institute for the COVID-19 Genomics UK (COG-UK) Consortium | Harper VanSteenhouse, Yumi Kasai, David Gray, Carol Clugston, Anna Dominiczak and Alex Alderton, Roberto Amato, Jeffrey Barrett, Sonia Goncalves, Ewan Harrison, David K. Jackson, Ian Johnston, Dominic Kwiatkowski, Cordelia Langford, John Sillitoe on behalf of the Wellcome Sanger Institute COVID-19 Surveillance Team |
| EPI_ISL_2914746, EPI_ISL_2914747 | Lighthouse Lab in Milton Keynes | Wellcome Sanger Institute for the COVID-19 Genomics UK (COG-UK) Consortium | The Lighthouse Lab in Milton Keynes and Alex Alderton, Roberto Amato, Jeffrey Barrett, Sonia Goncalves, Ewan Harrison, David K. Jackson, Ian Johnston, Dominic Kwiatkowski, Cordelia Langford, John Sillitoe on behalf of the Wellcome Sanger Institute COVID-19 Surveillance Team |
| EPI_ISL_2914748 | Lighthouse Lab in Glasgow | Wellcome Sanger Institute for the COVID-19 Genomics UK (COG-UK) Consortium | Harper VanSteenhouse, Yumi Kasai, David Gray, Carol Clugston, Anna Dominiczak and Alex Alderton, Roberto Amato, Jeffrey Barrett, Sonia Goncalves, Ewan Harrison, David K. Jackson, Ian Johnston, Dominic Kwiatkowski, Cordelia Langford, John Sillitoe on behalf of the Wellcome Sanger Institute COVID-19 Surveillance Team |
| EPI_ISL_2914749 | Lighthouse Lab in Milton Keynes | Wellcome Sanger Institute for the COVID-19 Genomics UK (COG-UK) Consortium | The Lighthouse Lab in Milton Keynes and Alex Alderton, Roberto Amato, Jeffrey Barrett, Sonia Goncalves, Ewan Harrison, David K. Jackson, Ian Johnston, Dominic Kwiatkowski, Cordelia Langford, John Sillitoe on behalf of the Wellcome Sanger Institute COVID-19 Surveillance Team |
| EPI_ISL_2914750 | Berkshire and Surrey Pathology Services Lighthouse Laboratory | Wellcome Sanger Institute for the COVID-19 Genomics UK (COG-UK) Consortium | Berkshire and Surrey Pathology Services Lighthouse Laboratory and Alex Alderton, Roberto Amato, Jeffrey Barrett, Sonia Goncalves, Ewan Harrison, David K. Jackson, Ian Johnston, Dominic Kwiatkowski, Cordelia Langford, John Sillitoe on behalf of the Wellcome Sanger Institute COVID-19 Surveillance Team |
| EPI_ISL_2914751 | Lighthouse Lab in Milton Keynes | Wellcome Sanger Institute for the COVID-19 Genomics UK (COG-UK) Consortium | The Lighthouse Lab in Milton Keynes and Alex Alderton, Roberto Amato, Jeffrey Barrett, Sonia Goncalves, Ewan Harrison, David K. Jackson, Ian Johnston, Dominic Kwiatkowski, Cordelia Langford, John Sillitoe on behalf of the Wellcome Sanger Institute COVID-19 Surveillance Team |
| EPI_ISL_2914752 | Berkshire and Surrey Pathology Services Lighthouse Laboratory | Wellcome Sanger Institute for the COVID-19 Genomics UK (COG-UK) Consortium | Berkshire and Surrey Pathology Services Lighthouse Laboratory and Alex Alderton, Roberto Amato, Jeffrey Barrett, Sonia Goncalves, Ewan Harrison, David K. Jackson, Ian Johnston, Dominic Kwiatkowski, Cordelia Langford, John Sillitoe on behalf of the Wellcome Sanger Institute COVID-19 Surveillance Team |
| EPI_ISL_2914753 | Lighthouse Lab in Milton Keynes | Wellcome Sanger Institute for the COVID-19 Genomics UK (COG-UK) Consortium | The Lighthouse Lab in Milton Keynes and Alex Alderton, Roberto Amato, Jeffrey Barrett, Sonia Goncalves, Ewan Harrison, David K. Jackson, Ian Johnston, Dominic Kwiatkowski, Cordelia Langford, John Sillitoe on behalf of the Wellcome Sanger Institute COVID-19 Surveillance Team |
| EPI_ISL_2914754, EPI_ISL_2914755 | Berkshire and Surrey Pathology Services Lighthouse Laboratory | Wellcome Sanger Institute for the COVID-19 Genomics UK (COG-UK) Consortium | Berkshire and Surrey Pathology Services Lighthouse Laboratory and Alex Alderton, Roberto Amato, Jeffrey Barrett, Sonia Goncalves, Ewan Harrison, David K. Jackson, Ian Johnston, Dominic Kwiatkowski, Cordelia Langford, John Sillitoe on behalf of the Wellcome Sanger Institute COVID-19 Surveillance Team |
| EPI_ISL_2914756 | Lighthouse Lab in Milton Keynes | Wellcome Sanger Institute for the COVID-19 Genomics UK (COG-UK) Consortium | The Lighthouse Lab in Milton Keynes and Alex Alderton, Roberto Amato, Jeffrey Barrett, Sonia Goncalves, Ewan Harrison, David K. Jackson, Ian Johnston, Dominic Kwiatkowski, Cordelia Langford, John Sillitoe on behalf of the Wellcome Sanger Institute COVID-19 Surveillance Team |
| EPI_ISL_2914757, EPI_ISL_2914758, EPI_ISL_2914759, EPI_ISL_2914760 | Lighthouse Lab in Glasgow | Wellcome Sanger Institute for the COVID-19 Genomics UK (COG-UK) Consortium | Harper VanSteenhouse, Yumi Kasai, David Gray, Carol Clugston, Anna Dominiczak and Alex Alderton, Roberto Amato, Jeffrey Barrett, Sonia Goncalves, Ewan Harrison, David K. Jackson, Ian Johnston, Dominic Kwiatkowski, Cordelia Langford, John Sillitoe on behalf of the Wellcome Sanger Institute COVID-19 Surveillance Team |
| EPI_ISL_2914761, EPI_ISL_2914762 | Berkshire and Surrey Pathology Services Lighthouse Laboratory | Wellcome Sanger Institute for the COVID-19 Genomics UK (COG-UK) Consortium | Berkshire and Surrey Pathology Services Lighthouse Laboratory and Alex Alderton, Roberto Amato, Jeffrey Barrett, Sonia Goncalves, Ewan Harrison, David K. Jackson, Ian Johnston, Dominic Kwiatkowski, Cordelia Langford, John Sillitoe on behalf of the Wellcome Sanger Institute COVID-19 Surveillance Team |
| EPI_ISL_2914763 | Lighthouse Lab in Glasgow | Wellcome Sanger Institute for the COVID-19 Genomics UK (COG-UK) Consortium | Harper VanSteenhouse, Yumi Kasai, David Gray, Carol Clugston, Anna Dominiczak and Alex Alderton, Roberto Amato, Jeffrey Barrett, Sonia Goncalves, Ewan Harrison, David K. Jackson, Ian Johnston, Dominic Kwiatkowski, Cordelia Langford, John Sillitoe on behalf of the Wellcome Sanger Institute COVID-19 Surveillance Team |
| EPI_ISL_2914764 | Lighthouse Lab in Milton Keynes | Wellcome Sanger Institute for the COVID-19 Genomics UK (COG-UK) Consortium | The Lighthouse Lab in Milton Keynes and Alex Alderton, Roberto Amato, Jeffrey Barrett, Sonia Goncalves, Ewan Harrison, David K. Jackson, Ian Johnston, Dominic Kwiatkowski, Cordelia Langford, John Sillitoe on behalf of the Wellcome Sanger Institute COVID-19 Surveillance Team |
| EPI_ISL_2914765 | Berkshire and Surrey Pathology Services Lighthouse Laboratory | Wellcome Sanger Institute for the COVID-19 Genomics UK (COG-UK) Consortium | Berkshire and Surrey Pathology Services Lighthouse Laboratory and Alex Alderton, Roberto Amato, Jeffrey Barrett, Sonia Goncalves, Ewan Harrison, David K. Jackson, Ian Johnston, Dominic Kwiatkowski, Cordelia Langford, John Sillitoe on behalf of the Wellcome Sanger Institute COVID-19 Surveillance Team |
| EPI_ISL_2914766 | Lighthouse Lab in Glasgow | Wellcome Sanger Institute for the COVID-19 Genomics UK (COG-UK) Consortium | Harper VanSteenhouse, Yumi Kasai, David Gray, Carol Clugston, Anna Dominiczak and Alex Alderton, Roberto Amato, Jeffrey Barrett, Sonia Goncalves, Ewan Harrison, David K. Jackson, Ian Johnston, Dominic Kwiatkowski, Cordelia Langford, John Sillitoe on behalf of the Wellcome Sanger Institute COVID-19 Surveillance Team |
| EPI_ISL_2914767 | Berkshire and Surrey Pathology Services Lighthouse Laboratory | Wellcome Sanger Institute for the COVID-19 Genomics UK (COG-UK) Consortium | Berkshire and Surrey Pathology Services Lighthouse Laboratory and Alex Alderton, Roberto Amato, Jeffrey Barrett, Sonia Goncalves, Ewan Harrison, David K. Jackson, Ian Johnston, Dominic Kwiatkowski, Cordelia Langford, John Sillitoe on behalf of the Wellcome Sanger Institute COVID-19 Surveillance Team |
| EPI_ISL_2914768 | Lighthouse Lab in Glasgow | Wellcome Sanger Institute for the COVID-19 Genomics UK (COG-UK) Consortium | Harper VanSteenhouse, Yumi Kasai, David Gray, Carol Clugston, Anna Dominiczak and Alex Alderton, Roberto Amato, Jeffrey Barrett, Sonia Goncalves, Ewan Harrison, David K. Jackson, Ian Johnston, Dominic Kwiatkowski, Cordelia Langford, John Sillitoe on behalf of the Wellcome Sanger Institute COVID-19 Surveillance Team |
| EPI_ISL_2914769, EPI_ISL_2914770 | Lighthouse Lab in Milton Keynes | Wellcome Sanger Institute for the COVID-19 Genomics UK (COG-UK) Consortium | The Lighthouse Lab in Milton Keynes and Alex Alderton, Roberto Amato, Jeffrey Barrett, Sonia Goncalves, Ewan Harrison, David K. Jackson, Ian Johnston, Dominic Kwiatkowski, Cordelia Langford, John Sillitoe on behalf of the Wellcome Sanger Institute COVID-19 Surveillance Team |
| EPI_ISL_2914771 | Berkshire and Surrey Pathology Services Lighthouse Laboratory | Wellcome Sanger Institute for the COVID-19 Genomics UK (COG-UK) Consortium | Berkshire and Surrey Pathology Services Lighthouse Laboratory and Alex Alderton, Roberto Amato, Jeffrey Barrett, Sonia Goncalves, Ewan Harrison, David K. Jackson, Ian Johnston, Dominic Kwiatkowski, Cordelia Langford, John Sillitoe on behalf of the Wellcome Sanger Institute COVID-19 Surveillance Team |
| EPI_ISL_2914772 | Lighthouse Lab in Glasgow | Wellcome Sanger Institute for the COVID-19 Genomics UK (COG-UK) Consortium | Harper VanSteenhouse, Yumi Kasai, David Gray, Carol Clugston, Anna Dominiczak and Alex Alderton, Roberto Amato, Jeffrey Barrett, Sonia Goncalves, Ewan Harrison, David K. Jackson, Ian Johnston, Dominic Kwiatkowski, Cordelia Langford, John Sillitoe on behalf of the Wellcome Sanger Institute COVID-19 Surveillance Team |
| EPI_ISL_2914773 | Lighthouse Lab in Milton Keynes | Wellcome Sanger Institute for the COVID-19 Genomics UK (COG-UK) Consortium | The Lighthouse Lab in Milton Keynes and Alex Alderton, Roberto Amato, Jeffrey Barrett, Sonia Goncalves, Ewan Harrison, David K. Jackson, Ian Johnston, Dominic Kwiatkowski, Cordelia Langford, John Sillitoe on behalf of the Wellcome Sanger Institute COVID-19 Surveillance Team |
| EPI_ISL_2914774, EPI_ISL_2914775 | Lighthouse Lab in Glasgow | Wellcome Sanger Institute for the COVID-19 Genomics UK (COG-UK) Consortium | Harper VanSteenhouse, Yumi Kasai, David Gray, Carol Clugston, Anna Dominiczak and Alex Alderton, Roberto Amato, Jeffrey Barrett, Sonia Goncalves, Ewan Harrison, David K. Jackson, Ian Johnston, Dominic Kwiatkowski, Cordelia Langford, John Sillitoe on behalf of the Wellcome Sanger Institute COVID-19 Surveillance Team |
| EPI_ISL_2914776, EPI_ISL_2914777 | Berkshire and Surrey Pathology Services Lighthouse Laboratory | Wellcome Sanger Institute for the COVID-19 Genomics UK (COG-UK) Consortium | Berkshire and Surrey Pathology Services Lighthouse Laboratory and Alex Alderton, Roberto Amato, Jeffrey Barrett, Sonia Goncalves, Ewan Harrison, David K. Jackson, Ian Johnston, Dominic Kwiatkowski, Cordelia Langford, John Sillitoe on behalf of the Wellcome Sanger Institute COVID-19 Surveillance Team |

| | | | |
|---|---|---|---|
| EPI_ISL_2914778 | Lighthouse Lab in Milton Keynes | Wellcome Sanger Institute for the COVID-19 Genomics UK (COG-UK) Consortium | The Lighthouse Lab in Milton Keynes and Alex Alderton, Roberto Amato, Jeffrey Barrett, Sonia Goncalves, Ewan Harrison, David K. Jackson, Ian Johnston, Dominic Kwiatkowski, Cordelia Langford, John Sillitoe on behalf of the Wellcome Sanger Institute COVID-19 Surveillance Team |
| EPI_ISL_2914779, EPI_ISL_2914780, EPI_ISL_2914781 | Lighthouse Lab in Glasgow | Wellcome Sanger Institute for the COVID-19 Genomics UK (COG-UK) Consortium | Harper VanSteenhouse, Yumi Kasai, David Gray, Carol Clugston, Anna Dominiczak and Alex Alderton, Roberto Amato, Jeffrey Barrett, Sonia Goncalves, Ewan Harrison, David K. Jackson, Ian Johnston, Dominic Kwiatkowski, Cordelia Langford, John Sillitoe on behalf of the Wellcome Sanger Institute COVID-19 Surveillance Team |
| EPI_ISL_2914782, EPI_ISL_2914783 | Lighthouse Lab in Milton Keynes | Wellcome Sanger Institute for the COVID-19 Genomics UK (COG-UK) Consortium | The Lighthouse Lab in Milton Keynes and Alex Alderton, Roberto Amato, Jeffrey Barrett, Sonia Goncalves, Ewan Harrison, David K. Jackson, Ian Johnston, Dominic Kwiatkowski, Cordelia Langford, John Sillitoe on behalf of the Wellcome Sanger Institute COVID-19 Surveillance Team |
| EPI_ISL_2914784, EPI_ISL_2914785, EPI_ISL_2914786 | Berkshire and Surrey Pathology Services Lighthouse Laboratory | Wellcome Sanger Institute for the COVID-19 Genomics UK (COG-UK) Consortium | Berkshire and Surrey Pathology Services Lighthouse Laboratory and Alex Alderton, Roberto Amato, Jeffrey Barrett, Sonia Goncalves, Ewan Harrison, David K. Jackson, Ian Johnston, Dominic Kwiatkowski, Cordelia Langford, John Sillitoe on behalf of the Wellcome Sanger Institute COVID-19 Surveillance Team |
| EPI_ISL_2914787 | Lighthouse Lab in Milton Keynes | Wellcome Sanger Institute for the COVID-19 Genomics UK (COG-UK) Consortium | The Lighthouse Lab in Milton Keynes and Alex Alderton, Roberto Amato, Jeffrey Barrett, Sonia Goncalves, Ewan Harrison, David K. Jackson, Ian Johnston, Dominic Kwiatkowski, Cordelia Langford, John Sillitoe on behalf of the Wellcome Sanger Institute COVID-19 Surveillance Team |
| EPI_ISL_2914788, EPI_ISL_2914789 | Berkshire and Surrey Pathology Services Lighthouse Laboratory | Wellcome Sanger Institute for the COVID-19 Genomics UK (COG-UK) Consortium | Berkshire and Surrey Pathology Services Lighthouse Laboratory and Alex Alderton, Roberto Amato, Jeffrey Barrett, Sonia Goncalves, Ewan Harrison, David K. Jackson, Ian Johnston, Dominic Kwiatkowski, Cordelia Langford, John Sillitoe on behalf of the Wellcome Sanger Institute COVID-19 Surveillance Team |
| EPI_ISL_2914790 | Lighthouse Lab in Glasgow | Wellcome Sanger Institute for the COVID-19 Genomics UK (COG-UK) Consortium | Harper VanSteenhouse, Yumi Kasai, David Gray, Carol Clugston, Anna Dominiczak and Alex Alderton, Roberto Amato, Jeffrey Barrett, Sonia Goncalves, Ewan Harrison, David K. Jackson, Ian Johnston, Dominic Kwiatkowski, Cordelia Langford, John Sillitoe on behalf of the Wellcome Sanger Institute COVID-19 Surveillance Team |
| EPI_ISL_2914791, EPI_ISL_2914792 | Lighthouse Lab in Milton Keynes | Wellcome Sanger Institute for the COVID-19 Genomics UK (COG-UK) Consortium | The Lighthouse Lab in Milton Keynes and Alex Alderton, Roberto Amato, Jeffrey Barrett, Sonia Goncalves, Ewan Harrison, David K. Jackson, Ian Johnston, Dominic Kwiatkowski, Cordelia Langford, John Sillitoe on behalf of the Wellcome Sanger Institute COVID-19 Surveillance Team |
| EPI_ISL_2914793, EPI_ISL_2914794 | Lighthouse Lab in Glasgow | Wellcome Sanger Institute for the COVID-19 Genomics UK (COG-UK) Consortium | Harper VanSteenhouse, Yumi Kasai, David Gray, Carol Clugston, Anna Dominiczak and Alex Alderton, Roberto Amato, Jeffrey Barrett, Sonia Goncalves, Ewan Harrison, David K. Jackson, Ian Johnston, Dominic Kwiatkowski, Cordelia Langford, John Sillitoe on behalf of the Wellcome Sanger Institute COVID-19 Surveillance Team |
| EPI_ISL_2914795, EPI_ISL_2914796 | Berkshire and Surrey Pathology Services Lighthouse Laboratory | Wellcome Sanger Institute for the COVID-19 Genomics UK (COG-UK) Consortium | Berkshire and Surrey Pathology Services Lighthouse Laboratory and Alex Alderton, Roberto Amato, Jeffrey Barrett, Sonia Goncalves, Ewan Harrison, David K. Jackson, Ian Johnston, Dominic Kwiatkowski, Cordelia Langford, John Sillitoe on behalf of the Wellcome Sanger Institute COVID-19 Surveillance Team |
| EPI_ISL_2914797, EPI_ISL_2914798 | Lighthouse Lab in Milton Keynes | Wellcome Sanger Institute for the COVID-19 Genomics UK (COG-UK) Consortium | The Lighthouse Lab in Milton Keynes and Alex Alderton, Roberto Amato, Jeffrey Barrett, Sonia Goncalves, Ewan Harrison, David K. Jackson, Ian Johnston, Dominic Kwiatkowski, Cordelia Langford, John Sillitoe on behalf of the Wellcome Sanger Institute COVID-19 Surveillance Team |
| EPI_ISL_2914799 | Berkshire and Surrey Pathology Services Lighthouse Laboratory | Wellcome Sanger Institute for the COVID-19 Genomics UK (COG-UK) Consortium | Berkshire and Surrey Pathology Services Lighthouse Laboratory and Alex Alderton, Roberto Amato, Jeffrey Barrett, Sonia Goncalves, Ewan Harrison, David K. Jackson, Ian Johnston, Dominic Kwiatkowski, Cordelia Langford, John Sillitoe on behalf of the Wellcome Sanger Institute COVID-19 Surveillance Team |
| EPI_ISL_2914800 | Lighthouse Lab in Milton Keynes | Wellcome Sanger Institute for the COVID-19 Genomics UK (COG-UK) Consortium | The Lighthouse Lab in Milton Keynes and Alex Alderton, Roberto Amato, Jeffrey Barrett, Sonia Goncalves, Ewan Harrison, David K. Jackson, Ian Johnston, Dominic Kwiatkowski, Cordelia Langford, John Sillitoe on behalf of the Wellcome Sanger Institute COVID-19 Surveillance Team |
| EPI_ISL_2914801, EPI_ISL_2914802, EPI_ISL_2914803 | Berkshire and Surrey Pathology Services Lighthouse Laboratory | Wellcome Sanger Institute for the COVID-19 Genomics UK (COG-UK) Consortium | Berkshire and Surrey Pathology Services Lighthouse Laboratory and Alex Alderton, Roberto Amato, Jeffrey Barrett, Sonia Goncalves, Ewan Harrison, David K. Jackson, Ian Johnston, Dominic Kwiatkowski, Cordelia Langford, John Sillitoe on behalf of the Wellcome Sanger Institute COVID-19 Surveillance Team |
| EPI_ISL_2914804 | Lighthouse Lab in Milton Keynes | Wellcome Sanger Institute for the COVID-19 Genomics UK (COG-UK) Consortium | The Lighthouse Lab in Milton Keynes and Alex Alderton, Roberto Amato, Jeffrey Barrett, Sonia Goncalves, Ewan Harrison, David K. Jackson, Ian Johnston, Dominic Kwiatkowski, Cordelia Langford, John Sillitoe on behalf of the Wellcome Sanger Institute COVID-19 Surveillance Team |
| EPI_ISL_2914805 | Berkshire and Surrey Pathology Services Lighthouse Laboratory | Wellcome Sanger Institute for the COVID-19 Genomics UK (COG-UK) Consortium | Berkshire and Surrey Pathology Services Lighthouse Laboratory and Alex Alderton, Roberto Amato, Jeffrey Barrett, Sonia Goncalves, Ewan Harrison, David K. Jackson, Ian Johnston, Dominic Kwiatkowski, Cordelia Langford, John Sillitoe on behalf of the Wellcome Sanger Institute COVID-19 Surveillance Team |
| EPI_ISL_2914806 | Lighthouse Lab in Milton Keynes | Wellcome Sanger Institute for the COVID-19 Genomics UK (COG-UK) Consortium | The Lighthouse Lab in Milton Keynes and Alex Alderton, Roberto Amato, Jeffrey Barrett, Sonia Goncalves, Ewan Harrison, David K. Jackson, Ian Johnston, Dominic Kwiatkowski, Cordelia Langford, John Sillitoe on behalf of the Wellcome Sanger Institute COVID-19 Surveillance Team |
| EPI_ISL_2914807, EPI_ISL_2914808 | Berkshire and Surrey Pathology Services Lighthouse Laboratory | Wellcome Sanger Institute for the COVID-19 Genomics UK (COG-UK) Consortium | Berkshire and Surrey Pathology Services Lighthouse Laboratory and Alex Alderton, Roberto Amato, Jeffrey Barrett, Sonia Goncalves, Ewan Harrison, David K. Jackson, Ian Johnston, Dominic Kwiatkowski, Cordelia Langford, John Sillitoe on behalf of the Wellcome Sanger Institute COVID-19 Surveillance Team |
| EPI_ISL_2914809 | Lighthouse Lab in Glasgow | Wellcome Sanger Institute for the COVID-19 Genomics UK (COG-UK) Consortium | Harper VanSteenhouse, Yumi Kasai, David Gray, Carol Clugston, Anna Dominiczak and Alex Alderton, Roberto Amato, Jeffrey Barrett, Sonia Goncalves, Ewan Harrison, David K. Jackson, Ian Johnston, Dominic Kwiatkowski, Cordelia Langford, John Sillitoe on behalf of the Wellcome Sanger Institute COVID-19 Surveillance Team |
| EPI_ISL_2914810 | Berkshire and Surrey Pathology Services Lighthouse Laboratory | Wellcome Sanger Institute for the COVID-19 Genomics UK (COG-UK) Consortium | Berkshire and Surrey Pathology Services Lighthouse Laboratory and Alex Alderton, Roberto Amato, Jeffrey Barrett, Sonia Goncalves, Ewan Harrison, David K. Jackson, Ian Johnston, Dominic Kwiatkowski, Cordelia Langford, John Sillitoe on behalf of the Wellcome Sanger Institute COVID-19 Surveillance Team |
| EPI_ISL_2914811 | Lighthouse Lab in Glasgow | Wellcome Sanger Institute for the COVID-19 Genomics UK (COG-UK) Consortium | Harper VanSteenhouse, Yumi Kasai, David Gray, Carol Clugston, Anna Dominiczak and Alex Alderton, Roberto Amato, Jeffrey Barrett, Sonia Goncalves, Ewan Harrison, David K. Jackson, Ian Johnston, Dominic Kwiatkowski, Cordelia Langford, John Sillitoe on behalf of the Wellcome Sanger Institute COVID-19 Surveillance Team |
| EPI_ISL_2914812, EPI_ISL_2914813, EPI_ISL_2914814 | Berkshire and Surrey Pathology Services Lighthouse Laboratory | Wellcome Sanger Institute for the COVID-19 Genomics UK (COG-UK) Consortium | Berkshire and Surrey Pathology Services Lighthouse Laboratory and Alex Alderton, Roberto Amato, Jeffrey Barrett, Sonia Goncalves, Ewan Harrison, David K. Jackson, Ian Johnston, Dominic Kwiatkowski, Cordelia Langford, John Sillitoe on behalf of the Wellcome Sanger Institute COVID-19 Surveillance Team |
| EPI_ISL_2914815, EPI_ISL_2914816 | Lighthouse Lab in Glasgow | Wellcome Sanger Institute for the COVID-19 Genomics UK (COG-UK) Consortium | Harper VanSteenhouse, Yumi Kasai, David Gray, Carol Clugston, Anna Dominiczak and Alex Alderton, Roberto Amato, Jeffrey Barrett, Sonia Goncalves, Ewan Harrison, David K. Jackson, Ian Johnston, Dominic Kwiatkowski, Cordelia Langford, John Sillitoe on behalf of the Wellcome Sanger Institute COVID-19 Surveillance Team |
| EPI_ISL_2914817 | Lighthouse Lab in Milton Keynes | Wellcome Sanger Institute for the COVID-19 Genomics UK (COG-UK) Consortium | The Lighthouse Lab in Milton Keynes and Alex Alderton, Roberto Amato, Jeffrey Barrett, Sonia Goncalves, Ewan Harrison, David K. Jackson, Ian Johnston, Dominic Kwiatkowski, Cordelia Langford, John Sillitoe on behalf of the Wellcome Sanger Institute COVID-19 Surveillance Team |
| EPI_ISL_2914818, EPI_ISL_2914819 | Berkshire and Surrey Pathology Services Lighthouse Laboratory | Wellcome Sanger Institute for the COVID-19 Genomics UK (COG-UK) Consortium | Berkshire and Surrey Pathology Services Lighthouse Laboratory and Alex Alderton, Roberto Amato, Jeffrey Barrett, Sonia Goncalves, Ewan Harrison, David K. Jackson, Ian Johnston, Dominic Kwiatkowski, Cordelia Langford, John Sillitoe on behalf of the Wellcome Sanger Institute COVID-19 Surveillance Team |
| EPI_ISL_2914820 | Lighthouse Lab in Glasgow | Wellcome Sanger Institute for the COVID-19 Genomics UK (COG-UK) Consortium | Harper VanSteenhouse, Yumi Kasai, David Gray, Carol Clugston, Anna Dominiczak and Alex Alderton, Roberto Amato, Jeffrey Barrett, Sonia Goncalves, Ewan Harrison, David K. Jackson, Ian Johnston, Dominic Kwiatkowski, Cordelia Langford, John Sillitoe on behalf of the Wellcome Sanger Institute COVID-19 Surveillance Team |
| EPI_ISL_2914821, EPI_ISL_2914822 | Berkshire and Surrey Pathology Services Lighthouse Laboratory | Wellcome Sanger Institute for the COVID-19 Genomics UK (COG-UK) Consortium | Berkshire and Surrey Pathology Services Lighthouse Laboratory and Alex Alderton, Roberto Amato, Jeffrey Barrett, Sonia Goncalves, Ewan Harrison, David K. Jackson, Ian Johnston, Dominic Kwiatkowski, Cordelia Langford, John Sillitoe on behalf of the Wellcome Sanger Institute COVID-19 Surveillance Team |
| EPI_ISL_2914823 | Lighthouse Lab in Milton Keynes | Wellcome Sanger Institute for the COVID-19 Genomics UK (COG-UK) Consortium | The Lighthouse Lab in Milton Keynes and Alex Alderton, Roberto Amato, Jeffrey Barrett, Sonia Goncalves, Ewan Harrison, David K. Jackson, Ian Johnston, Dominic Kwiatkowski, Cordelia Langford, John Sillitoe on behalf of the Wellcome Sanger Institute COVID-19 Surveillance Team |
| EPI_ISL_2914824 | Berkshire and Surrey Pathology Services Lighthouse Laboratory | Wellcome Sanger Institute for the COVID-19 Genomics UK (COG-UK) Consortium | Berkshire and Surrey Pathology Services Lighthouse Laboratory and Alex Alderton, Roberto Amato, Jeffrey Barrett, Sonia Goncalves, Ewan Harrison, David K. Jackson, Ian Johnston, Dominic Kwiatkowski, Cordelia Langford, John Sillitoe on behalf of the Wellcome Sanger Institute COVID-19 Surveillance Team |
| EPI_ISL_2914825 | Lighthouse Lab in Glasgow | Wellcome Sanger Institute for the COVID-19 Genomics UK (COG-UK) Consortium | Harper VanSteenhouse, Yumi Kasai, David Gray, Carol Clugston, Anna Dominiczak and Alex Alderton, Roberto Amato, Jeffrey Barrett, Sonia Goncalves, Ewan Harrison, David K. Jackson, Ian Johnston, Dominic Kwiatkowski, Cordelia Langford, John Sillitoe on behalf of the Wellcome Sanger Institute COVID-19 Surveillance Team |
| EPI_ISL_2914826 | Lighthouse Lab in Milton Keynes | Wellcome Sanger Institute for the COVID-19 Genomics UK (COG-UK) Consortium | The Lighthouse Lab in Milton Keynes and Alex Alderton, Roberto Amato, Jeffrey Barrett, Sonia Goncalves, Ewan Harrison, David K. Jackson, Ian Johnston, Dominic Kwiatkowski, Cordelia Langford, John Sillitoe on behalf of the Wellcome Sanger Institute COVID-19 Surveillance Team |
| EPI_ISL_2914827 | Berkshire and Surrey Pathology Services Lighthouse Laboratory | Wellcome Sanger Institute for the COVID-19 Genomics UK (COG-UK) Consortium | Berkshire and Surrey Pathology Services Lighthouse Laboratory and Alex Alderton, Roberto Amato, Jeffrey Barrett, Sonia Goncalves, Ewan Harrison, David K. Jackson, Ian Johnston, Dominic Kwiatkowski, Cordelia Langford, John Sillitoe on behalf of the Wellcome Sanger Institute COVID-19 Surveillance Team |
| EPI_ISL_2914828, EPI_ISL_2914829, EPI_ISL_2914830, EPI_ISL_2914831 | Lighthouse Lab in Glasgow | Wellcome Sanger Institute for the COVID-19 Genomics UK (COG-UK) Consortium | Harper VanSteenhouse, Yumi Kasai, David Gray, Carol Clugston, Anna Dominiczak and Alex Alderton, Roberto Amato, Jeffrey Barrett, Sonia Goncalves, Ewan Harrison, David K. Jackson, Ian Johnston, Dominic Kwiatkowski, Cordelia Langford, John Sillitoe on behalf of the Wellcome Sanger Institute COVID-19 Surveillance Team |
| EPI_ISL_2914832 | Berkshire and Surrey Pathology Services Lighthouse Laboratory | Wellcome Sanger Institute for the COVID-19 Genomics UK (COG-UK) Consortium | Berkshire and Surrey Pathology Services Lighthouse Laboratory and Alex Alderton, Roberto Amato, Jeffrey Barrett, Sonia Goncalves, Ewan Harrison, David K. Jackson, Ian Johnston, Dominic Kwiatkowski, Cordelia Langford, John Sillitoe on behalf of the Wellcome Sanger Institute COVID-19 Surveillance Team |
| EPI_ISL_2914833 | Lighthouse Lab in Milton Keynes | Wellcome Sanger Institute for the COVID-19 Genomics UK (COG-UK) Consortium | The Lighthouse Lab in Milton Keynes and Alex Alderton, Roberto Amato, Jeffrey Barrett, Sonia Goncalves, Ewan Harrison, David K. Jackson, Ian Johnston, Dominic Kwiatkowski, Cordelia Langford, John Sillitoe on behalf of the Wellcome Sanger Institute COVID-19 Surveillance Team |
| EPI_ISL_2914834, EPI_ISL_2914835 | Berkshire and Surrey Pathology Services Lighthouse Laboratory | Wellcome Sanger Institute for the COVID-19 Genomics UK (COG-UK) Consortium | Berkshire and Surrey Pathology Services Lighthouse Laboratory and Alex Alderton, Roberto Amato, Jeffrey Barrett, Sonia Goncalves, Ewan Harrison, David K. Jackson, Ian Johnston, Dominic Kwiatkowski, Cordelia Langford, John Sillitoe on behalf of the Wellcome Sanger Institute COVID-19 Surveillance Team |
| EPI_ISL_2914836 | Lighthouse Lab in Glasgow | Wellcome Sanger Institute for the COVID-19 Genomics UK (COG-UK) Consortium | Harper VanSteenhouse, Yumi Kasai, David Gray, Carol Clugston, Anna Dominiczak and Alex Alderton, Roberto Amato, Jeffrey Barrett, Sonia Goncalves, Ewan Harrison, David K. Jackson, Ian Johnston, Dominic Kwiatkowski, Cordelia Langford, John Sillitoe on behalf of the Wellcome Sanger Institute COVID-19 Surveillance Team |
| EPI_ISL_2914837 | Berkshire and Surrey Pathology Services Lighthouse Laboratory | Wellcome Sanger Institute for the COVID-19 Genomics UK (COG-UK) Consortium | Berkshire and Surrey Pathology Services Lighthouse Laboratory and Alex Alderton, Roberto Amato, Jeffrey Barrett, Sonia Goncalves, Ewan Harrison, David K. Jackson, Ian Johnston, Dominic Kwiatkowski, Cordelia Langford, John Sillitoe on behalf of the Wellcome Sanger Institute COVID-19 Surveillance Team |
| EPI_ISL_2914838 | Lighthouse Lab in Milton Keynes | Wellcome Sanger Institute for the COVID-19 Genomics UK (COG-UK) Consortium | The Lighthouse Lab in Milton Keynes and Alex Alderton, Roberto Amato, Jeffrey Barrett, Sonia Goncalves, Ewan Harrison, David K. Jackson, Ian Johnston, Dominic Kwiatkowski, Cordelia Langford, John Sillitoe on behalf of the Wellcome Sanger Institute COVID-19 Surveillance Team |
| EPI_ISL_2914839 | Berkshire and Surrey Pathology Services Lighthouse Laboratory | Wellcome Sanger Institute for the COVID-19 Genomics UK (COG-UK) Consortium | Berkshire and Surrey Pathology Services Lighthouse Laboratory and Alex Alderton, Roberto Amato, Jeffrey Barrett, Sonia Goncalves, Ewan Harrison, David K. Jackson, Ian Johnston, Dominic Kwiatkowski, Cordelia Langford, John Sillitoe on behalf of the Wellcome Sanger Institute COVID-19 Surveillance Team |
| EPI_ISL_2914840 | Lighthouse Lab in Glasgow | Wellcome Sanger Institute for the COVID-19 Genomics UK (COG-UK) Consortium | Harper VanSteenhouse, Yumi Kasai, David Gray, Carol Clugston, Anna Dominiczak and Alex Alderton, Roberto Amato, Jeffrey Barrett, Sonia Goncalves, Ewan Harrison, David K. Jackson, Ian Johnston, Dominic Kwiatkowski, Cordelia Langford, John Sillitoe on behalf of the Wellcome Sanger Institute COVID-19 Surveillance Team |
| EPI_ISL_2914841, EPI_ISL_2914842 | Berkshire and Surrey Pathology Services Lighthouse Laboratory | Wellcome Sanger Institute for the COVID-19 Genomics UK (COG-UK) Consortium | Berkshire and Surrey Pathology Services Lighthouse Laboratory and Alex Alderton, Roberto Amato, Jeffrey Barrett, Sonia Goncalves, Ewan Harrison, David K. Jackson, Ian Johnston, Dominic Kwiatkowski, Cordelia Langford, John Sillitoe on behalf of the Wellcome Sanger Institute COVID-19 Surveillance Team |
| EPI_ISL_2914843 | Lighthouse Lab in Glasgow | Wellcome Sanger Institute for the COVID-19 Genomics UK (COG-UK) Consortium | Harper VanSteenhouse, Yumi Kasai, David Gray, Carol Clugston, Anna Dominiczak and Alex Alderton, Roberto Amato, Jeffrey Barrett, Sonia Goncalves, Ewan Harrison, David K. Jackson, Ian Johnston, Dominic Kwiatkowski, Cordelia Langford, John Sillitoe on behalf of the Wellcome Sanger Institute COVID-19 Surveillance Team |

| | | | |
|---|---|---|---|
| EPI_ISL_2914844 | Berkshire and Surrey Pathology Services Lighthouse Laboratory | Wellcome Sanger Institute for the COVID-19 Genomics UK (COG-UK) Consortium | Berkshire and Surrey Pathology Services Lighthouse Laboratory and Alex Alderton, Roberto Amato, Jeffrey Barrett, Sonia Goncalves, Ewan Harrison, David K. Jackson, Ian Johnston, Dominic Kwiatkowski, Cordelia Langford, John Sillitoe on behalf of the Wellcome Sanger Institute COVID-19 Surveillance Team |
| EPI_ISL_2914845, EPI_ISL_2914846 | Lighthouse Lab in Milton Keynes | Wellcome Sanger Institute for the COVID-19 Genomics UK (COG-UK) Consortium | The Lighthouse Lab in Milton Keynes and Alex Alderton, Roberto Amato, Jeffrey Barrett, Sonia Goncalves, Ewan Harrison, David K. Jackson, Ian Johnston, Dominic Kwiatkowski, Cordelia Langford, John Sillitoe on behalf of the Wellcome Sanger Institute COVID-19 Surveillance Team |
| EPI_ISL_2914847, EPI_ISL_2914848, EPI_ISL_2914849, EPI_ISL_2914850, EPI_ISL_2914851 | Berkshire and Surrey Pathology Services Lighthouse Laboratory | Wellcome Sanger Institute for the COVID-19 Genomics UK (COG-UK) Consortium | Berkshire and Surrey Pathology Services Lighthouse Laboratory and Alex Alderton, Roberto Amato, Jeffrey Barrett, Sonia Goncalves, Ewan Harrison, David K. Jackson, Ian Johnston, Dominic Kwiatkowski, Cordelia Langford, John Sillitoe on behalf of the Wellcome Sanger Institute COVID-19 Surveillance Team |
| EPI_ISL_2914852 | Lighthouse Lab in Glasgow | Wellcome Sanger Institute for the COVID-19 Genomics UK (COG-UK) Consortium | Harper VanSteenhouse, Yumi Kasai, David Gray, Carol Clugston, Anna Dominiczak and Alex Alderton, Roberto Amato, Jeffrey Barrett, Sonia Goncalves, Ewan Harrison, David K. Jackson, Ian Johnston, Dominic Kwiatkowski, Cordelia Langford, John Sillitoe on behalf of the Wellcome Sanger Institute COVID-19 Surveillance Team |
| EPI_ISL_2914853, EPI_ISL_2914854 | Lighthouse Lab in Milton Keynes | Wellcome Sanger Institute for the COVID-19 Genomics UK (COG-UK) Consortium | The Lighthouse Lab in Milton Keynes and Alex Alderton, Roberto Amato, Jeffrey Barrett, Sonia Goncalves, Ewan Harrison, David K. Jackson, Ian Johnston, Dominic Kwiatkowski, Cordelia Langford, John Sillitoe on behalf of the Wellcome Sanger Institute COVID-19 Surveillance Team |
| EPI_ISL_2914855 | Lighthouse Lab in Glasgow | Wellcome Sanger Institute for the COVID-19 Genomics UK (COG-UK) Consortium | Harper VanSteenhouse, Yumi Kasai, David Gray, Carol Clugston, Anna Dominiczak and Alex Alderton, Roberto Amato, Jeffrey Barrett, Sonia Goncalves, Ewan Harrison, David K. Jackson, Ian Johnston, Dominic Kwiatkowski, Cordelia Langford, John Sillitoe on behalf of the Wellcome Sanger Institute COVID-19 Surveillance Team |
| EPI_ISL_2914856 | Lighthouse Lab in Milton Keynes | Wellcome Sanger Institute for the COVID-19 Genomics UK (COG-UK) Consortium | The Lighthouse Lab in Milton Keynes and Alex Alderton, Roberto Amato, Jeffrey Barrett, Sonia Goncalves, Ewan Harrison, David K. Jackson, Ian Johnston, Dominic Kwiatkowski, Cordelia Langford, John Sillitoe on behalf of the Wellcome Sanger Institute COVID-19 Surveillance Team |
| EPI_ISL_2914857, EPI_ISL_2914858, EPI_ISL_2914859 | Berkshire and Surrey Pathology Services Lighthouse Laboratory | Wellcome Sanger Institute for the COVID-19 Genomics UK (COG-UK) Consortium | Berkshire and Surrey Pathology Services Lighthouse Laboratory and Alex Alderton, Roberto Amato, Jeffrey Barrett, Sonia Goncalves, Ewan Harrison, David K. Jackson, Ian Johnston, Dominic Kwiatkowski, Cordelia Langford, John Sillitoe on behalf of the Wellcome Sanger Institute COVID-19 Surveillance Team |
| EPI_ISL_2914860, EPI_ISL_2914861 | Lighthouse Lab in Glasgow | Wellcome Sanger Institute for the COVID-19 Genomics UK (COG-UK) Consortium | Harper VanSteenhouse, Yumi Kasai, David Gray, Carol Clugston, Anna Dominiczak and Alex Alderton, Roberto Amato, Jeffrey Barrett, Sonia Goncalves, Ewan Harrison, David K. Jackson, Ian Johnston, Dominic Kwiatkowski, Cordelia Langford, John Sillitoe on behalf of the Wellcome Sanger Institute COVID-19 Surveillance Team |
| EPI_ISL_2914862, EPI_ISL_2914863 | Lighthouse Lab in Milton Keynes | Wellcome Sanger Institute for the COVID-19 Genomics UK (COG-UK) Consortium | The Lighthouse Lab in Milton Keynes and Alex Alderton, Roberto Amato, Jeffrey Barrett, Sonia Goncalves, Ewan Harrison, David K. Jackson, Ian Johnston, Dominic Kwiatkowski, Cordelia Langford, John Sillitoe on behalf of the Wellcome Sanger Institute COVID-19 Surveillance Team |
| EPI_ISL_2914864 | Berkshire and Surrey Pathology Services Lighthouse Laboratory | Wellcome Sanger Institute for the COVID-19 Genomics UK (COG-UK) Consortium | Berkshire and Surrey Pathology Services Lighthouse Laboratory and Alex Alderton, Roberto Amato, Jeffrey Barrett, Sonia Goncalves, Ewan Harrison, David K. Jackson, Ian Johnston, Dominic Kwiatkowski, Cordelia Langford, John Sillitoe on behalf of the Wellcome Sanger Institute COVID-19 Surveillance Team |
| EPI_ISL_2914865 | Lighthouse Lab in Glasgow | Wellcome Sanger Institute for the COVID-19 Genomics UK (COG-UK) Consortium | Harper VanSteenhouse, Yumi Kasai, David Gray, Carol Clugston, Anna Dominiczak and Alex Alderton, Roberto Amato, Jeffrey Barrett, Sonia Goncalves, Ewan Harrison, David K. Jackson, Ian Johnston, Dominic Kwiatkowski, Cordelia Langford, John Sillitoe on behalf of the Wellcome Sanger Institute COVID-19 Surveillance Team |
| EPI_ISL_2914866, EPI_ISL_2914867 | Berkshire and Surrey Pathology Services Lighthouse Laboratory | Wellcome Sanger Institute for the COVID-19 Genomics UK (COG-UK) Consortium | Berkshire and Surrey Pathology Services Lighthouse Laboratory and Alex Alderton, Roberto Amato, Jeffrey Barrett, Sonia Goncalves, Ewan Harrison, David K. Jackson, Ian Johnston, Dominic Kwiatkowski, Cordelia Langford, John Sillitoe on behalf of the Wellcome Sanger Institute COVID-19 Surveillance Team |
| EPI_ISL_2914868 | Lighthouse Lab in Milton Keynes | Wellcome Sanger Institute for the COVID-19 Genomics UK (COG-UK) Consortium | The Lighthouse Lab in Milton Keynes and Alex Alderton, Roberto Amato, Jeffrey Barrett, Sonia Goncalves, Ewan Harrison, David K. Jackson, Ian Johnston, Dominic Kwiatkowski, Cordelia Langford, John Sillitoe on behalf of the Wellcome Sanger Institute COVID-19 Surveillance Team |
| EPI_ISL_2914869 | Berkshire and Surrey Pathology Services Lighthouse Laboratory | Wellcome Sanger Institute for the COVID-19 Genomics UK (COG-UK) Consortium | Berkshire and Surrey Pathology Services Lighthouse Laboratory and Alex Alderton, Roberto Amato, Jeffrey Barrett, Sonia Goncalves, Ewan Harrison, David K. Jackson, Ian Johnston, Dominic Kwiatkowski, Cordelia Langford, John Sillitoe on behalf of the Wellcome Sanger Institute COVID-19 Surveillance Team |
| EPI_ISL_2914870, EPI_ISL_2914871 | Lighthouse Lab in Milton Keynes | Wellcome Sanger Institute for the COVID-19 Genomics UK (COG-UK) Consortium | The Lighthouse Lab in Milton Keynes and Alex Alderton, Roberto Amato, Jeffrey Barrett, Sonia Goncalves, Ewan Harrison, David K. Jackson, Ian Johnston, Dominic Kwiatkowski, Cordelia Langford, John Sillitoe on behalf of the Wellcome Sanger Institute COVID-19 Surveillance Team |
| EPI_ISL_2914872, EPI_ISL_2914873, EPI_ISL_2914874 | Berkshire and Surrey Pathology Services Lighthouse Laboratory | Wellcome Sanger Institute for the COVID-19 Genomics UK (COG-UK) Consortium | Berkshire and Surrey Pathology Services Lighthouse Laboratory and Alex Alderton, Roberto Amato, Jeffrey Barrett, Sonia Goncalves, Ewan Harrison, David K. Jackson, Ian Johnston, Dominic Kwiatkowski, Cordelia Langford, John Sillitoe on behalf of the Wellcome Sanger Institute COVID-19 Surveillance Team |
| EPI_ISL_2914875, EPI_ISL_2914876, EPI_ISL_2914877 | Lighthouse Lab in Milton Keynes | Wellcome Sanger Institute for the COVID-19 Genomics UK (COG-UK) Consortium | The Lighthouse Lab in Milton Keynes and Alex Alderton, Roberto Amato, Jeffrey Barrett, Sonia Goncalves, Ewan Harrison, David K. Jackson, Ian Johnston, Dominic Kwiatkowski, Cordelia Langford, John Sillitoe on behalf of the Wellcome Sanger Institute COVID-19 Surveillance Team |
| EPI_ISL_2914878 | Berkshire and Surrey Pathology Services Lighthouse Laboratory | Wellcome Sanger Institute for the COVID-19 Genomics UK (COG-UK) Consortium | Berkshire and Surrey Pathology Services Lighthouse Laboratory and Alex Alderton, Roberto Amato, Jeffrey Barrett, Sonia Goncalves, Ewan Harrison, David K. Jackson, Ian Johnston, Dominic Kwiatkowski, Cordelia Langford, John Sillitoe on behalf of the Wellcome Sanger Institute COVID-19 Surveillance Team |
| EPI_ISL_2914879, EPI_ISL_2914880, EPI_ISL_2914881 | Lighthouse Lab in Glasgow | Wellcome Sanger Institute for the COVID-19 Genomics UK (COG-UK) Consortium | Harper VanSteenhouse, Yumi Kasai, David Gray, Carol Clugston, Anna Dominiczak and Alex Alderton, Roberto Amato, Jeffrey Barrett, Sonia Goncalves, Ewan Harrison, David K. Jackson, Ian Johnston, Dominic Kwiatkowski, Cordelia Langford, John Sillitoe on behalf of the Wellcome Sanger Institute COVID-19 Surveillance Team |
| EPI_ISL_2914882 | Berkshire and Surrey Pathology Services Lighthouse Laboratory | Wellcome Sanger Institute for the COVID-19 Genomics UK (COG-UK) Consortium | Berkshire and Surrey Pathology Services Lighthouse Laboratory and Alex Alderton, Roberto Amato, Jeffrey Barrett, Sonia Goncalves, Ewan Harrison, David K. Jackson, Ian Johnston, Dominic Kwiatkowski, Cordelia Langford, John Sillitoe on behalf of the Wellcome Sanger Institute COVID-19 Surveillance Team |
| EPI_ISL_2914883 | Lighthouse Lab in Glasgow | Wellcome Sanger Institute for the COVID-19 Genomics UK (COG-UK) Consortium | Harper VanSteenhouse, Yumi Kasai, David Gray, Carol Clugston, Anna Dominiczak and Alex Alderton, Roberto Amato, Jeffrey Barrett, Sonia Goncalves, Ewan Harrison, David K. Jackson, Ian Johnston, Dominic Kwiatkowski, Cordelia Langford, John Sillitoe on behalf of the Wellcome Sanger Institute COVID-19 Surveillance Team |
| EPI_ISL_2914884, EPI_ISL_2914885, EPI_ISL_2914886 | Berkshire and Surrey Pathology Services Lighthouse Laboratory | Wellcome Sanger Institute for the COVID-19 Genomics UK (COG-UK) Consortium | Berkshire and Surrey Pathology Services Lighthouse Laboratory and Alex Alderton, Roberto Amato, Jeffrey Barrett, Sonia Goncalves, Ewan Harrison, David K. Jackson, Ian Johnston, Dominic Kwiatkowski, Cordelia Langford, John Sillitoe on behalf of the Wellcome Sanger Institute COVID-19 Surveillance Team |
| EPI_ISL_2914887 | Lighthouse Lab in Milton Keynes | Wellcome Sanger Institute for the COVID-19 Genomics UK (COG-UK) Consortium | The Lighthouse Lab in Milton Keynes and Alex Alderton, Roberto Amato, Jeffrey Barrett, Sonia Goncalves, Ewan Harrison, David K. Jackson, Ian Johnston, Dominic Kwiatkowski, Cordelia Langford, John Sillitoe on behalf of the Wellcome Sanger Institute COVID-19 Surveillance Team |
| EPI_ISL_2914888, EPI_ISL_2914889, EPI_ISL_2914890, EPI_ISL_2914891 | Berkshire and Surrey Pathology Services Lighthouse Laboratory | Wellcome Sanger Institute for the COVID-19 Genomics UK (COG-UK) Consortium | Berkshire and Surrey Pathology Services Lighthouse Laboratory and Alex Alderton, Roberto Amato, Jeffrey Barrett, Sonia Goncalves, Ewan Harrison, David K. Jackson, Ian Johnston, Dominic Kwiatkowski, Cordelia Langford, John Sillitoe on behalf of the Wellcome Sanger Institute COVID-19 Surveillance Team |
| EPI_ISL_2914892 | Lighthouse Lab in Milton Keynes | Wellcome Sanger Institute for the COVID-19 Genomics UK (COG-UK) Consortium | The Lighthouse Lab in Milton Keynes and Alex Alderton, Roberto Amato, Jeffrey Barrett, Sonia Goncalves, Ewan Harrison, David K. Jackson, Ian Johnston, Dominic Kwiatkowski, Cordelia Langford, John Sillitoe on behalf of the Wellcome Sanger Institute COVID-19 Surveillance Team |
| EPI_ISL_2914893 | Berkshire and Surrey Pathology Services Lighthouse Laboratory | Wellcome Sanger Institute for the COVID-19 Genomics UK (COG-UK) Consortium | Berkshire and Surrey Pathology Services Lighthouse Laboratory and Alex Alderton, Roberto Amato, Jeffrey Barrett, Sonia Goncalves, Ewan Harrison, David K. Jackson, Ian Johnston, Dominic Kwiatkowski, Cordelia Langford, John Sillitoe on behalf of the Wellcome Sanger Institute COVID-19 Surveillance Team |
| EPI_ISL_2914894, EPI_ISL_2914895 | Lighthouse Lab in Glasgow | Wellcome Sanger Institute for the COVID-19 Genomics UK (COG-UK) Consortium | Harper VanSteenhouse, Yumi Kasai, David Gray, Carol Clugston, Anna Dominiczak and Alex Alderton, Roberto Amato, Jeffrey Barrett, Sonia Goncalves, Ewan Harrison, David K. Jackson, Ian Johnston, Dominic Kwiatkowski, Cordelia Langford, John Sillitoe on behalf of the Wellcome Sanger Institute COVID-19 Surveillance Team |
| EPI_ISL_2914896, EPI_ISL_2914897 | Lighthouse Lab in Milton Keynes | Wellcome Sanger Institute for the COVID-19 Genomics UK (COG-UK) Consortium | The Lighthouse Lab in Milton Keynes and Alex Alderton, Roberto Amato, Jeffrey Barrett, Sonia Goncalves, Ewan Harrison, David K. Jackson, Ian Johnston, Dominic Kwiatkowski, Cordelia Langford, John Sillitoe on behalf of the Wellcome Sanger Institute COVID-19 Surveillance Team |
| EPI_ISL_2914898 | Lighthouse Lab in Glasgow | Wellcome Sanger Institute for the COVID-19 Genomics UK (COG-UK) Consortium | Harper VanSteenhouse, Yumi Kasai, David Gray, Carol Clugston, Anna Dominiczak and Alex Alderton, Roberto Amato, Jeffrey Barrett, Sonia Goncalves, Ewan Harrison, David K. Jackson, Ian Johnston, Dominic Kwiatkowski, Cordelia Langford, John Sillitoe on behalf of the Wellcome Sanger Institute COVID-19 Surveillance Team |
| EPI_ISL_2914899 | Berkshire and Surrey Pathology Services Lighthouse Laboratory | Wellcome Sanger Institute for the COVID-19 Genomics UK (COG-UK) Consortium | Berkshire and Surrey Pathology Services Lighthouse Laboratory and Alex Alderton, Roberto Amato, Jeffrey Barrett, Sonia Goncalves, Ewan Harrison, David K. Jackson, Ian Johnston, Dominic Kwiatkowski, Cordelia Langford, John Sillitoe on behalf of the Wellcome Sanger Institute COVID-19 Surveillance Team |
| EPI_ISL_2914900, EPI_ISL_2914901 | Lighthouse Lab in Glasgow | Wellcome Sanger Institute for the COVID-19 Genomics UK (COG-UK) Consortium | Harper VanSteenhouse, Yumi Kasai, David Gray, Carol Clugston, Anna Dominiczak and Alex Alderton, Roberto Amato, Jeffrey Barrett, Sonia Goncalves, Ewan Harrison, David K. Jackson, Ian Johnston, Dominic Kwiatkowski, Cordelia Langford, John Sillitoe on behalf of the Wellcome Sanger Institute COVID-19 Surveillance Team |
| EPI_ISL_2914902 | Lighthouse Lab in Milton Keynes | Wellcome Sanger Institute for the COVID-19 Genomics UK (COG-UK) Consortium | The Lighthouse Lab in Milton Keynes and Alex Alderton, Roberto Amato, Jeffrey Barrett, Sonia Goncalves, Ewan Harrison, David K. Jackson, Ian Johnston, Dominic Kwiatkowski, Cordelia Langford, John Sillitoe on behalf of the Wellcome Sanger Institute COVID-19 Surveillance Team |
| EPI_ISL_2914903 | Lighthouse Lab in Glasgow | Wellcome Sanger Institute for the COVID-19 Genomics UK (COG-UK) Consortium | Harper VanSteenhouse, Yumi Kasai, David Gray, Carol Clugston, Anna Dominiczak and Alex Alderton, Roberto Amato, Jeffrey Barrett, Sonia Goncalves, Ewan Harrison, David K. Jackson, Ian Johnston, Dominic Kwiatkowski, Cordelia Langford, John Sillitoe on behalf of the Wellcome Sanger Institute COVID-19 Surveillance Team |
| EPI_ISL_2914904 | Lighthouse Lab in Milton Keynes | Wellcome Sanger Institute for the COVID-19 Genomics UK (COG-UK) Consortium | The Lighthouse Lab in Milton Keynes and Alex Alderton, Roberto Amato, Jeffrey Barrett, Sonia Goncalves, Ewan Harrison, David K. Jackson, Ian Johnston, Dominic Kwiatkowski, Cordelia Langford, John Sillitoe on behalf of the Wellcome Sanger Institute COVID-19 Surveillance Team |
| EPI_ISL_2914905 | Berkshire and Surrey Pathology Services Lighthouse Laboratory | Wellcome Sanger Institute for the COVID-19 Genomics UK (COG-UK) Consortium | Berkshire and Surrey Pathology Services Lighthouse Laboratory and Alex Alderton, Roberto Amato, Jeffrey Barrett, Sonia Goncalves, Ewan Harrison, David K. Jackson, Ian Johnston, Dominic Kwiatkowski, Cordelia Langford, John Sillitoe on behalf of the Wellcome Sanger Institute COVID-19 Surveillance Team |
| EPI_ISL_2914906 | Lighthouse Lab in Milton Keynes | Wellcome Sanger Institute for the COVID-19 Genomics UK (COG-UK) Consortium | The Lighthouse Lab in Milton Keynes and Alex Alderton, Roberto Amato, Jeffrey Barrett, Sonia Goncalves, Ewan Harrison, David K. Jackson, Ian Johnston, Dominic Kwiatkowski, Cordelia Langford, John Sillitoe on behalf of the Wellcome Sanger Institute COVID-19 Surveillance Team |
| EPI_ISL_2914907 | Lighthouse Lab in Glasgow | Wellcome Sanger Institute for the COVID-19 Genomics UK (COG-UK) Consortium | Harper VanSteenhouse, Yumi Kasai, David Gray, Carol Clugston, Anna Dominiczak and Alex Alderton, Roberto Amato, Jeffrey Barrett, Sonia Goncalves, Ewan Harrison, David K. Jackson, Ian Johnston, Dominic Kwiatkowski, Cordelia Langford, John Sillitoe on behalf of the Wellcome Sanger Institute COVID-19 Surveillance Team |
| EPI_ISL_2914908, EPI_ISL_2914909, EPI_ISL_2914910 | Berkshire and Surrey Pathology Services Lighthouse Laboratory | Wellcome Sanger Institute for the COVID-19 Genomics UK (COG-UK) Consortium | Berkshire and Surrey Pathology Services Lighthouse Laboratory and Alex Alderton, Roberto Amato, Jeffrey Barrett, Sonia Goncalves, Ewan Harrison, David K. Jackson, Ian Johnston, Dominic Kwiatkowski, Cordelia Langford, John Sillitoe on behalf of the Wellcome Sanger Institute COVID-19 Surveillance Team |
| EPI_ISL_2914911 | Lighthouse Lab in Milton Keynes | Wellcome Sanger Institute for the COVID-19 Genomics UK (COG-UK) Consortium | The Lighthouse Lab in Milton Keynes and Alex Alderton, Roberto Amato, Jeffrey Barrett, Sonia Goncalves, Ewan Harrison, David K. Jackson, Ian Johnston, Dominic Kwiatkowski, Cordelia Langford, John Sillitoe on behalf of the Wellcome Sanger Institute COVID-19 Surveillance Team |
| EPI_ISL_2914912, | Berkshire and Surrey Pathology Services Lighthouse Laboratory | Wellcome Sanger Institute for the COVID-19 Genomics UK (COG-UK) | Berkshire and Surrey Pathology Services Lighthouse Laboratory and Alex Alderton, Roberto Amato, Jeffrey Barrett, Sonia Goncalves, Ewan Harrison, David K. Jackson, Ian Johnston, Dominic Kwiatkowski, Cordelia Langford, John Sillitoe on behalf |

| | | | |
|---|---|---|---|
| EPI_ISL_2914913, EPI_ISL_2914914 | | Consortium | of the Wellcome Sanger Institute COVID-19 Surveillance Team |
| EPI_ISL_2914915 | Lighthouse Lab in Milton Keynes | Wellcome Sanger Institute for the COVID-19 Genomics UK (COG-UK) Consortium | The Lighthouse Lab in Milton Keynes and Alex Alderton, Roberto Amato, Jeffrey Barrett, Sonia Goncalves, Ewan Harrison, David K. Jackson, Ian Johnston, Dominic Kwiatkowski, Cordelia Langford, John Sillitoe on behalf of the Wellcome Sanger Institute COVID-19 Surveillance Team |
| EPI_ISL_2914916, EPI_ISL_2914917 | Lighthouse Lab in Glasgow | Wellcome Sanger Institute for the COVID-19 Genomics UK (COG-UK) Consortium | Harper VanSteenhouse, Yumi Kasai, David Gray, Carol Clugston, Anna Dominiczak and Alex Alderton, Roberto Amato, Jeffrey Barrett, Sonia Goncalves, Ewan Harrison, David K. Jackson, Ian Johnston, Dominic Kwiatkowski, Cordelia Langford, John Sillitoe on behalf of the Wellcome Sanger Institute COVID-19 Surveillance Team |
| EPI_ISL_2914918, EPI_ISL_2914919 | Lighthouse Lab in Milton Keynes | Wellcome Sanger Institute for the COVID-19 Genomics UK (COG-UK) Consortium | The Lighthouse Lab in Milton Keynes and Alex Alderton, Roberto Amato, Jeffrey Barrett, Sonia Goncalves, Ewan Harrison, David K. Jackson, Ian Johnston, Dominic Kwiatkowski, Cordelia Langford, John Sillitoe on behalf of the Wellcome Sanger Institute COVID-19 Surveillance Team |
| EPI_ISL_2914920 | Berkshire and Surrey Pathology Services Lighthouse Laboratory | Wellcome Sanger Institute for the COVID-19 Genomics UK (COG-UK) Consortium | Berkshire and Surrey Pathology Services Lighthouse Laboratory and Alex Alderton, Roberto Amato, Jeffrey Barrett, Sonia Goncalves, Ewan Harrison, David K. Jackson, Ian Johnston, Dominic Kwiatkowski, Cordelia Langford, John Sillitoe on behalf of the Wellcome Sanger Institute COVID-19 Surveillance Team |
| EPI_ISL_2914921 | Lighthouse Lab in Milton Keynes | Wellcome Sanger Institute for the COVID-19 Genomics UK (COG-UK) Consortium | The Lighthouse Lab in Milton Keynes and Alex Alderton, Roberto Amato, Jeffrey Barrett, Sonia Goncalves, Ewan Harrison, David K. Jackson, Ian Johnston, Dominic Kwiatkowski, Cordelia Langford, John Sillitoe on behalf of the Wellcome Sanger Institute COVID-19 Surveillance Team |
| EPI_ISL_2914922 | Lighthouse Lab in Glasgow | Wellcome Sanger Institute for the COVID-19 Genomics UK (COG-UK) Consortium | Harper VanSteenhouse, Yumi Kasai, David Gray, Carol Clugston, Anna Dominiczak and Alex Alderton, Roberto Amato, Jeffrey Barrett, Sonia Goncalves, Ewan Harrison, David K. Jackson, Ian Johnston, Dominic Kwiatkowski, Cordelia Langford, John Sillitoe on behalf of the Wellcome Sanger Institute COVID-19 Surveillance Team |
| EPI_ISL_2914923, EPI_ISL_2914924 | Berkshire and Surrey Pathology Services Lighthouse Laboratory | Wellcome Sanger Institute for the COVID-19 Genomics UK (COG-UK) Consortium | Berkshire and Surrey Pathology Services Lighthouse Laboratory and Alex Alderton, Roberto Amato, Jeffrey Barrett, Sonia Goncalves, Ewan Harrison, David K. Jackson, Ian Johnston, Dominic Kwiatkowski, Cordelia Langford, John Sillitoe on behalf of the Wellcome Sanger Institute COVID-19 Surveillance Team |
| EPI_ISL_2914925 | Lighthouse Lab in Milton Keynes | Wellcome Sanger Institute for the COVID-19 Genomics UK (COG-UK) Consortium | The Lighthouse Lab in Milton Keynes and Alex Alderton, Roberto Amato, Jeffrey Barrett, Sonia Goncalves, Ewan Harrison, David K. Jackson, Ian Johnston, Dominic Kwiatkowski, Cordelia Langford, John Sillitoe on behalf of the Wellcome Sanger Institute COVID-19 Surveillance Team |
| EPI_ISL_2914926, EPI_ISL_2914927 | Berkshire and Surrey Pathology Services Lighthouse Laboratory | Wellcome Sanger Institute for the COVID-19 Genomics UK (COG-UK) Consortium | Berkshire and Surrey Pathology Services Lighthouse Laboratory and Alex Alderton, Roberto Amato, Jeffrey Barrett, Sonia Goncalves, Ewan Harrison, David K. Jackson, Ian Johnston, Dominic Kwiatkowski, Cordelia Langford, John Sillitoe on behalf of the Wellcome Sanger Institute COVID-19 Surveillance Team |
| EPI_ISL_2914928 | Lighthouse Lab in Milton Keynes | Wellcome Sanger Institute for the COVID-19 Genomics UK (COG-UK) Consortium | The Lighthouse Lab in Milton Keynes and Alex Alderton, Roberto Amato, Jeffrey Barrett, Sonia Goncalves, Ewan Harrison, David K. Jackson, Ian Johnston, Dominic Kwiatkowski, Cordelia Langford, John Sillitoe on behalf of the Wellcome Sanger Institute COVID-19 Surveillance Team |
| EPI_ISL_2914929 | Lighthouse Lab in Glasgow | Wellcome Sanger Institute for the COVID-19 Genomics UK (COG-UK) Consortium | Harper VanSteenhouse, Yumi Kasai, David Gray, Carol Clugston, Anna Dominiczak and Alex Alderton, Roberto Amato, Jeffrey Barrett, Sonia Goncalves, Ewan Harrison, David K. Jackson, Ian Johnston, Dominic Kwiatkowski, Cordelia Langford, John Sillitoe on behalf of the Wellcome Sanger Institute COVID-19 Surveillance Team |
| EPI_ISL_2914930 | Berkshire and Surrey Pathology Services Lighthouse Laboratory | Wellcome Sanger Institute for the COVID-19 Genomics UK (COG-UK) Consortium | Berkshire and Surrey Pathology Services Lighthouse Laboratory and Alex Alderton, Roberto Amato, Jeffrey Barrett, Sonia Goncalves, Ewan Harrison, David K. Jackson, Ian Johnston, Dominic Kwiatkowski, Cordelia Langford, John Sillitoe on behalf of the Wellcome Sanger Institute COVID-19 Surveillance Team |
| EPI_ISL_2914931 | Lighthouse Lab in Milton Keynes | Wellcome Sanger Institute for the COVID-19 Genomics UK (COG-UK) Consortium | The Lighthouse Lab in Milton Keynes and Alex Alderton, Roberto Amato, Jeffrey Barrett, Sonia Goncalves, Ewan Harrison, David K. Jackson, Ian Johnston, Dominic Kwiatkowski, Cordelia Langford, John Sillitoe on behalf of the Wellcome Sanger Institute COVID-19 Surveillance Team |
| EPI_ISL_2914932 | Lighthouse Lab in Glasgow | Wellcome Sanger Institute for the COVID-19 Genomics UK (COG-UK) Consortium | Harper VanSteenhouse, Yumi Kasai, David Gray, Carol Clugston, Anna Dominiczak and Alex Alderton, Roberto Amato, Jeffrey Barrett, Sonia Goncalves, Ewan Harrison, David K. Jackson, Ian Johnston, Dominic Kwiatkowski, Cordelia Langford, John Sillitoe on behalf of the Wellcome Sanger Institute COVID-19 Surveillance Team |
| EPI_ISL_2914933 | Lighthouse Lab in Milton Keynes | Wellcome Sanger Institute for the COVID-19 Genomics UK (COG-UK) Consortium | The Lighthouse Lab in Milton Keynes and Alex Alderton, Roberto Amato, Jeffrey Barrett, Sonia Goncalves, Ewan Harrison, David K. Jackson, Ian Johnston, Dominic Kwiatkowski, Cordelia Langford, John Sillitoe on behalf of the Wellcome Sanger Institute COVID-19 Surveillance Team |
| EPI_ISL_2914934, EPI_ISL_2914935, EPI_ISL_2914936, EPI_ISL_2914937, EPI_ISL_2914938 | Berkshire and Surrey Pathology Services Lighthouse Laboratory | Wellcome Sanger Institute for the COVID-19 Genomics UK (COG-UK) Consortium | Berkshire and Surrey Pathology Services Lighthouse Laboratory and Alex Alderton, Roberto Amato, Jeffrey Barrett, Sonia Goncalves, Ewan Harrison, David K. Jackson, Ian Johnston, Dominic Kwiatkowski, Cordelia Langford, John Sillitoe on behalf of the Wellcome Sanger Institute COVID-19 Surveillance Team |
| EPI_ISL_2914939, EPI_ISL_2914940 | Lighthouse Lab in Glasgow | Wellcome Sanger Institute for the COVID-19 Genomics UK (COG-UK) Consortium | Harper VanSteenhouse, Yumi Kasai, David Gray, Carol Clugston, Anna Dominiczak and Alex Alderton, Roberto Amato, Jeffrey Barrett, Sonia Goncalves, Ewan Harrison, David K. Jackson, Ian Johnston, Dominic Kwiatkowski, Cordelia Langford, John Sillitoe on behalf of the Wellcome Sanger Institute COVID-19 Surveillance Team |
| EPI_ISL_2914941, EPI_ISL_2914942 | Berkshire and Surrey Pathology Services Lighthouse Laboratory | Wellcome Sanger Institute for the COVID-19 Genomics UK (COG-UK) Consortium | Berkshire and Surrey Pathology Services Lighthouse Laboratory and Alex Alderton, Roberto Amato, Jeffrey Barrett, Sonia Goncalves, Ewan Harrison, David K. Jackson, Ian Johnston, Dominic Kwiatkowski, Cordelia Langford, John Sillitoe on behalf of the Wellcome Sanger Institute COVID-19 Surveillance Team |
| EPI_ISL_2914943 | Lighthouse Lab in Milton Keynes | Wellcome Sanger Institute for the COVID-19 Genomics UK (COG-UK) Consortium | The Lighthouse Lab in Milton Keynes and Alex Alderton, Roberto Amato, Jeffrey Barrett, Sonia Goncalves, Ewan Harrison, David K. Jackson, Ian Johnston, Dominic Kwiatkowski, Cordelia Langford, John Sillitoe on behalf of the Wellcome Sanger Institute COVID-19 Surveillance Team |
| EPI_ISL_2914944 | Berkshire and Surrey Pathology Services Lighthouse Laboratory | Wellcome Sanger Institute for the COVID-19 Genomics UK (COG-UK) Consortium | Berkshire and Surrey Pathology Services Lighthouse Laboratory and Alex Alderton, Roberto Amato, Jeffrey Barrett, Sonia Goncalves, Ewan Harrison, David K. Jackson, Ian Johnston, Dominic Kwiatkowski, Cordelia Langford, John Sillitoe on behalf of the Wellcome Sanger Institute COVID-19 Surveillance Team |
| EPI_ISL_2914945 | Lighthouse Lab in Milton Keynes | Wellcome Sanger Institute for the COVID-19 Genomics UK (COG-UK) Consortium | The Lighthouse Lab in Milton Keynes and Alex Alderton, Roberto Amato, Jeffrey Barrett, Sonia Goncalves, Ewan Harrison, David K. Jackson, Ian Johnston, Dominic Kwiatkowski, Cordelia Langford, John Sillitoe on behalf of the Wellcome Sanger Institute COVID-19 Surveillance Team |
| EPI_ISL_2914946, EPI_ISL_2914947, EPI_ISL_2914948 | Berkshire and Surrey Pathology Services Lighthouse Laboratory | Wellcome Sanger Institute for the COVID-19 Genomics UK (COG-UK) Consortium | Berkshire and Surrey Pathology Services Lighthouse Laboratory and Alex Alderton, Roberto Amato, Jeffrey Barrett, Sonia Goncalves, Ewan Harrison, David K. Jackson, Ian Johnston, Dominic Kwiatkowski, Cordelia Langford, John Sillitoe on behalf of the Wellcome Sanger Institute COVID-19 Surveillance Team |
| EPI_ISL_2914949, EPI_ISL_2914950, EPI_ISL_2914951, EPI_ISL_2914952, EPI_ISL_2914953, EPI_ISL_2914954, EPI_ISL_2914955 | Lighthouse Lab in Glasgow | Wellcome Sanger Institute for the COVID-19 Genomics UK (COG-UK) Consortium | Harper VanSteenhouse, Yumi Kasai, David Gray, Carol Clugston, Anna Dominiczak and Alex Alderton, Roberto Amato, Jeffrey Barrett, Sonia Goncalves, Ewan Harrison, David K. Jackson, Ian Johnston, Dominic Kwiatkowski, Cordelia Langford, John Sillitoe on behalf of the Wellcome Sanger Institute COVID-19 Surveillance Team |
| EPI_ISL_2914956, EPI_ISL_2914957, EPI_ISL_2914958 | Lighthouse Lab in Milton Keynes | Wellcome Sanger Institute for the COVID-19 Genomics UK (COG-UK) Consortium | The Lighthouse Lab in Milton Keynes and Alex Alderton, Roberto Amato, Jeffrey Barrett, Sonia Goncalves, Ewan Harrison, David K. Jackson, Ian Johnston, Dominic Kwiatkowski, Cordelia Langford, John Sillitoe on behalf of the Wellcome Sanger Institute COVID-19 Surveillance Team |
| EPI_ISL_2914959 | Berkshire and Surrey Pathology Services Lighthouse Laboratory | Wellcome Sanger Institute for the COVID-19 Genomics UK (COG-UK) Consortium | Berkshire and Surrey Pathology Services Lighthouse Laboratory and Alex Alderton, Roberto Amato, Jeffrey Barrett, Sonia Goncalves, Ewan Harrison, David K. Jackson, Ian Johnston, Dominic Kwiatkowski, Cordelia Langford, John Sillitoe on behalf of the Wellcome Sanger Institute COVID-19 Surveillance Team |
| EPI_ISL_2914960 | Lighthouse Lab in Milton Keynes | Wellcome Sanger Institute for the COVID-19 Genomics UK (COG-UK) Consortium | The Lighthouse Lab in Milton Keynes and Alex Alderton, Roberto Amato, Jeffrey Barrett, Sonia Goncalves, Ewan Harrison, David K. Jackson, Ian Johnston, Dominic Kwiatkowski, Cordelia Langford, John Sillitoe on behalf of the Wellcome Sanger Institute COVID-19 Surveillance Team |
| EPI_ISL_2914961 | Berkshire and Surrey Pathology Services Lighthouse Laboratory | Wellcome Sanger Institute for the COVID-19 Genomics UK (COG-UK) Consortium | Berkshire and Surrey Pathology Services Lighthouse Laboratory and Alex Alderton, Roberto Amato, Jeffrey Barrett, Sonia Goncalves, Ewan Harrison, David K. Jackson, Ian Johnston, Dominic Kwiatkowski, Cordelia Langford, John Sillitoe on behalf of the Wellcome Sanger Institute COVID-19 Surveillance Team |
| EPI_ISL_2914962 | Lighthouse Lab in Milton Keynes | Wellcome Sanger Institute for the COVID-19 Genomics UK (COG-UK) Consortium | The Lighthouse Lab in Milton Keynes and Alex Alderton, Roberto Amato, Jeffrey Barrett, Sonia Goncalves, Ewan Harrison, David K. Jackson, Ian Johnston, Dominic Kwiatkowski, Cordelia Langford, John Sillitoe on behalf of the Wellcome Sanger Institute COVID-19 Surveillance Team |
| EPI_ISL_2914963 | Lighthouse Lab in Glasgow | Wellcome Sanger Institute for the COVID-19 Genomics UK (COG-UK) Consortium | Harper VanSteenhouse, Yumi Kasai, David Gray, Carol Clugston, Anna Dominiczak and Alex Alderton, Roberto Amato, Jeffrey Barrett, Sonia Goncalves, Ewan Harrison, David K. Jackson, Ian Johnston, Dominic Kwiatkowski, Cordelia Langford, John Sillitoe on behalf of the Wellcome Sanger Institute COVID-19 Surveillance Team |
| EPI_ISL_2914964 | Berkshire and Surrey Pathology Services Lighthouse Laboratory | Wellcome Sanger Institute for the COVID-19 Genomics UK (COG-UK) Consortium | Berkshire and Surrey Pathology Services Lighthouse Laboratory and Alex Alderton, Roberto Amato, Jeffrey Barrett, Sonia Goncalves, Ewan Harrison, David K. Jackson, Ian Johnston, Dominic Kwiatkowski, Cordelia Langford, John Sillitoe on behalf of the Wellcome Sanger Institute COVID-19 Surveillance Team |
| EPI_ISL_2914965, EPI_ISL_2914966 | Lighthouse Lab in Milton Keynes | Wellcome Sanger Institute for the COVID-19 Genomics UK (COG-UK) Consortium | The Lighthouse Lab in Milton Keynes and Alex Alderton, Roberto Amato, Jeffrey Barrett, Sonia Goncalves, Ewan Harrison, David K. Jackson, Ian Johnston, Dominic Kwiatkowski, Cordelia Langford, John Sillitoe on behalf of the Wellcome Sanger Institute COVID-19 Surveillance Team |
| EPI_ISL_2914967, EPI_ISL_2914968, EPI_ISL_2914969, EPI_ISL_2914970, EPI_ISL_2914971, EPI_ISL_2914972, EPI_ISL_2914973, EPI_ISL_2914974, EPI_ISL_2914975, EPI_ISL_2914976, EPI_ISL_2914977, EPI_ISL_2914978, EPI_ISL_2914979, EPI_ISL_2914980, EPI_ISL_2914981, EPI_ISL_2914982, EPI_ISL_2914983, EPI_ISL_2914984, EPI_ISL_2914985, EPI_ISL_2914986, EPI_ISL_2914987, EPI_ISL_2914988, EPI_ISL_2914989, EPI_ISL_2914990, EPI_ISL_2914991, EPI_ISL_2914992, EPI_ISL_2914993, EPI_ISL_2914994, EPI_ISL_2914995, EPI_ISL_2914996, EPI_ISL_2914997, EPI_ISL_2914998, EPI_ISL_2914999, EPI_ISL_2915000, EPI_ISL_2915001, EPI_ISL_2915002, EPI_ISL_2915003, EPI_ISL_2915004, EPI_ISL_2915005, EPI_ISL_2915006, EPI_ISL_2915007, EPI_ISL_2915008, EPI_ISL_2915009, EPI_ISL_2915010, EPI_ISL_2915011, EPI_ISL_2915012, EPI_ISL_2915013, EPI_ISL_2915014, EPI_ISL_2915015, EPI_ISL_2915016, EPI_ISL_2915017, EPI_ISL_2915018, EPI_ISL_2915019, EPI_ISL_2915020, EPI_ISL_2915021, EPI_ISL_2915022, EPI_ISL_2915023, EPI_ISL_2915024, EPI_ISL_2915025, EPI_ISL_2915026, EPI_ISL_2915027, EPI_ISL_2915028, EPI_ISL_2915029, EPI_ISL_2915030, EPI_ISL_2915031, EPI_ISL_2915032, EPI_ISL_2915033, EPI_ISL_2915035, EPI_ISL_2915036, EPI_ISL_2915037, EPI_ISL_2915038, EPI_ISL_2915039, EPI_ISL_2915040, EPI_ISL_2915041, EPI_ISL_2915042, EPI_ISL_2915043, EPI_ISL_2915044, EPI_ISL_2915045, EPI_ISL_2915046, EPI_ISL_2915047, EPI_ISL_2915048, EPI_ISL_2915049, EPI_ISL_2915050, EPI_ISL_2915051, EPI_ISL_2915052, EPI_ISL_2915053, EPI_ISL_2915054, EPI_ISL_2915055, EPI_ISL_2915056, EPI_ISL_2915057, EPI_ISL_2915058, EPI_ISL_2915059, EPI_ISL_2915060, EPI_ISL_2915061, EPI_ISL_2915062, EPI_ISL_2915063, EPI_ISL_2915064, EPI_ISL_2915065, EPI_ISL_2915066, EPI_ISL_2915067, EPI_ISL_2915068, EPI_ISL_2915069, EPI_ISL_2915070, EPI_ISL_2915071, EPI_ISL_2915072, EPI_ISL_2915073, EPI_ISL_2915074, EPI_ISL_2915075, EPI_ISL_2915076, EPI_ISL_2915077, EPI_ISL_2915078, EPI_ISL_2915079, EPI_ISL_2915080, EPI_ISL_2915081, EPI_ISL_2915082, EPI_ISL_2915083, EPI_ISL_2915084, EPI_ISL_2915085, EPI_ISL_2915086, EPI_ISL_2915087, EPI_ISL_2915088, EPI_ISL_2915089, EPI_ISL_2915090, EPI_ISL_2915091, EPI_ISL_2915092, EPI_ISL_2915093, EPI_ISL_2915095, EPI_ISL_2915096, EPI_ISL_2915097, EPI_ISL_2915098, EPI_ISL_2915099, EPI_ISL_2915100, EPI_ISL_2915101, EPI_ISL_2915102, EPI_ISL_2915103, EPI_ISL_2915104, EPI_ISL_2915105, EPI_ISL_2915106, EPI_ISL_2915108, EPI_ISL_2915109, EPI_ISL_2915110, EPI_ISL_2915111, EPI_ISL_2915112, EPI_ISL_2915113, EPI_ISL_2915114, EPI_ISL_2915115, EPI_ISL_2915116, EPI_ISL_2915117, EPI_ISL_2915118, EPI_ISL_2915119, EPI_ISL_2915120, EPI_ISL_2915121, EPI_ISL_2915122, EPI_ISL_2915123, EPI_ISL_2915124, EPI_ISL_2915125, EPI_ISL_2915126, EPI_ISL_2915127, EPI_ISL_2915128, EPI_ISL_2915129, EPI_ISL_2915130, EPI_ISL_2915131, EPI_ISL_2915132, EPI_ISL_2915133, EPI_ISL_2915134, EPI_ISL_2915135, EPI_ISL_2915136, EPI_ISL_2915137, EPI_ISL_2915138, EPI_ISL_2915139, EPI_ISL_2915140, EPI_ISL_2915141, EPI_ISL_2915142, EPI_ISL_2915143, EPI_ISL_2915144, EPI_ISL_2915145, EPI_ISL_2915146, EPI_ISL_2915147, EPI_ISL_2915148, EPI_ISL_2915149, EPI_ISL_2915150, EPI_ISL_2915151, EPI_ISL_2915152, EPI_ISL_2915153, EPI_ISL_2915154, EPI_ISL_2915155, EPI_ISL_2915156, EPI_ISL_2915157, EPI_ISL_2915158, EPI_ISL_2915159, EPI_ISL_2915160, EPI_ISL_2915161, EPI_ISL_2915162, EPI_ISL_2915163, EPI_ISL_2915164, EPI_ISL_2915165, EPI_ISL_2915166, EPI_ISL_2915167, EPI_ISL_2915168, EPI_ISL_2915169, EPI_ISL_2915170, EPI_ISL_2915171, EPI_ISL_2915172, EPI_ISL_2915173, EPI_ISL_2915174, EPI_ISL_2915175, EPI_ISL_2915177, EPI_ISL_2915178, EPI_ISL_2915179, EPI_ISL_2915180, EPI_ISL_2915181, EPI_ISL_2915182, EPI_ISL_2915183, EPI_ISL_2915184, EPI_ISL_2915185, EPI_ISL_2915186, EPI_ISL_2915187, EPI_ISL_2915188, EPI_ISL_2915189, EPI_ISL_2915190, EPI_ISL_2915191, EPI_ISL_2915192, EPI_ISL_2915193, EPI_ISL_2915194, EPI_ISL_2915195, EPI_ISL_2915196, EPI_ISL_2915197, EPI_ISL_2915198, EPI_ISL_2915199, EPI_ISL_2915200, EPI_ISL_2915201, EPI_ISL_2915202, EPI_ISL_2915203, EPI_ISL_2915204, EPI_ISL_2915205, EPI_ISL_2915206, EPI_ISL_2915207, EPI_ISL_2915208, EPI_ISL_2915209, EPI_ISL_2915210, EPI_ISL_2915211, EPI_ISL_2915212, EPI_ISL_2915213, EPI_ISL_2915214, EPI_ISL_2915215, EPI_ISL_2915216, EPI_ISL_2915217, EPI_ISL_2915218, EPI_ISL_2915219, EPI_ISL_2915220, EPI_ISL_2915221, EPI_ISL_2915222, EPI_ISL_2915223, EPI_ISL_2915224, EPI_ISL_2915225, EPI_ISL_2915226, EPI_ISL_2915227, EPI_ISL_2915228, EPI_ISL_2915229, EPI_ISL_2915230, EPI_ISL_2915231, EPI_ISL_2915232, EPI_ISL_2915233, EPI_ISL_2915234, EPI_ISL_2915235, EPI_ISL_2915236, EPI_ISL_2915237, EPI_ISL_2915238, EPI_ISL_2915239, EPI_ISL_2915240, EPI_ISL_2915241, EPI_ISL_2915242, EPI_ISL_2915243, EPI_ISL_2915244, EPI_ISL_2915245, EPI_ISL_2915246, EPI_ISL_2915247, EPI_ISL_2915248, EPI_ISL_2915249, EPI_ISL_2915250, EPI_ISL_2915251, EPI_ISL_2915252, EPI_ISL_2915253, EPI_ISL_2915254, EPI_ISL_2915255, EPI_ISL_2915256, EPI_ISL_2915257, EPI_ISL_2915258, EPI_ISL_2915259, EPI_ISL_2915260, EPI_ISL_2915261, EPI_ISL_2915262, EPI_ISL_2915263, EPI_ISL_2915264, EPI_ISL_2915265, EPI_ISL_2915266, EPI_ISL_2915267, EPI_ISL_2915268, EPI_ISL_2915269, EPI_ISL_2915270, EPI_ISL_2915271, EPI_ISL_2915272, EPI_ISL_2915273, EPI_ISL_2915274, EPI_ISL_2915275, EPI_ISL_2915276, EPI_ISL_2915277, EPI_ISL_2915278, EPI_ISL_2915279, EPI_ISL_2915280, EPI_ISL_2915281, EPI_ISL_2915282, EPI_ISL_2915283, EPI_ISL_2915284, EPI_ISL_2915285, EPI_ISL_2915286, EPI_ISL_2915287, EPI_ISL_2915288, EPI_ISL_2915289, EPI_ISL_2915290, EPI_ISL_2915291, EPI_ISL_2915292, EPI_ISL_2915293, EPI_ISL_2915295, EPI_ISL_2915296, EPI_ISL_2915297, EPI_ISL_2915298, EPI_ISL_2915299, EPI_ISL_2915300, EPI_ISL_2915301, EPI_ISL_2915302, EPI_ISL_2915303, EPI_ISL_2915304, EPI_ISL_2915305, EPI_ISL_2915306, EPI_ISL_2915307, EPI_ISL_2915308, EPI_ISL_2915309, EPI_ISL_2915310, EPI_ISL_2915311, EPI_ISL_2915312, EPI_ISL_2915313, EPI_ISL_2915314, EPI_ISL_2915315, EPI_ISL_2915316, EPI_ISL_2915317, EPI_ISL_2915318, EPI_ISL_2915319, EPI_ISL_2915320, EPI_ISL_2915321, EPI_ISL_2915322, EPI_ISL_2915323, EPI_ISL_2915324, EPI_ISL_2915325, EPI_ISL_2915326, EPI_ISL_2915327, EPI_ISL_2915328, EPI_ISL_2915329, EPI_ISL_2915330 | | | |
| see above | Lighthouse Lab in Glasgow | Wellcome Sanger Institute for the COVID-19 Genomics UK (COG-UK) Consortium | Harper VanSteenhouse, Yumi Kasai, David Gray, Carol Clugston, Anna Dominiczak and Alex Alderton, Roberto Amato, Jeffrey Barrett, Sonia Goncalves, Ewan Harrison, David K. Jackson, Ian Johnston, Dominic Kwiatkowski, Cordelia Langford, John Sillitoe on behalf of the Wellcome Sanger Institute COVID-19 Surveillance Team |
| EPI_ISL_2915331 | Lighthouse Laboratory Plymouth | Wellcome Sanger Institute for the COVID-19 Genomics UK (COG-UK) | Lighthouse Laboratory Plymouth and Alex Alderton, Roberto Amato, Jeffrey Barrett, Sonia Goncalves, Ewan Harrison, David K. Jackson, Ian Johnston, Dominic Kwiatkowski, Cordelia Langford, John Sillitoe on behalf of the Wellcome Sanger |

| | | | |
|---|---|---|---|
| | | Consortium | Institute COVID-19 Surveillance Team |
| EPI_ISL_2915332 | Berkshire and Surrey Pathology Services Lighthouse Laboratory | Wellcome Sanger Institute for the COVID-19 Genomics UK (COG-UK) Consortium | Berkshire and Surrey Pathology Services Lighthouse Laboratory and Alex Alderton, Roberto Amato, Jeffrey Barrett, Sonia Goncalves, Ewan Harrison, David K. Jackson, Ian Johnston, Dominic Kwiatkowski, Cordelia Langford, John Sillitoe on behalf of the Wellcome Sanger Institute COVID-19 Surveillance Team |
| EPI_ISL_2915333, EPI_ISL_2915334 | Lighthouse Lab in Alderley Park | Wellcome Sanger Institute for the COVID-19 Genomics UK (COG-UK) Consortium | Jacquelyn Wynn, Mairead Hyland, The Lighthouse Lab in Alderley Park and Alex Alderton, Roberto Amato, Jeffrey Barrett, Sonia Goncalves, Ewan Harrison, David K. Jackson, Ian Johnston, Dominic Kwiatkowski, Cordelia Langford, John Sillitoe on behalf of the Wellcome Sanger Institute COVID-19 Surveillance Team |
| EPI_ISL_2915335 | Berkshire and Surrey Pathology Services Lighthouse Laboratory | Wellcome Sanger Institute for the COVID-19 Genomics UK (COG-UK) Consortium | Berkshire and Surrey Pathology Services Lighthouse Laboratory and Alex Alderton, Roberto Amato, Jeffrey Barrett, Sonia Goncalves, Ewan Harrison, David K. Jackson, Ian Johnston, Dominic Kwiatkowski, Cordelia Langford, John Sillitoe on behalf of the Wellcome Sanger Institute COVID-19 Surveillance Team |
| EPI_ISL_2915336 | Lighthouse Lab in Glasgow | Wellcome Sanger Institute for the COVID-19 Genomics UK (COG-UK) Consortium | Harper VanSteenhouse, Yumi Kasai, David Gray, Carol Clugston, Anna Dominiczak and Alex Alderton, Roberto Amato, Jeffrey Barrett, Sonia Goncalves, Ewan Harrison, David K. Jackson, Ian Johnston, Dominic Kwiatkowski, Cordelia Langford, John Sillitoe on behalf of the Wellcome Sanger Institute COVID-19 Surveillance Team |
| EPI_ISL_2915337 | Berkshire and Surrey Pathology Services Lighthouse Laboratory | Wellcome Sanger Institute for the COVID-19 Genomics UK (COG-UK) Consortium | Berkshire and Surrey Pathology Services Lighthouse Laboratory and Alex Alderton, Roberto Amato, Jeffrey Barrett, Sonia Goncalves, Ewan Harrison, David K. Jackson, Ian Johnston, Dominic Kwiatkowski, Cordelia Langford, John Sillitoe on behalf of the Wellcome Sanger Institute COVID-19 Surveillance Team |
| EPI_ISL_2915338 | Lighthouse Laboratory Plymouth | Wellcome Sanger Institute for the COVID-19 Genomics UK (COG-UK) Consortium | Lighthouse Laboratory Plymouth and Alex Alderton, Roberto Amato, Jeffrey Barrett, Sonia Goncalves, Ewan Harrison, David K. Jackson, Ian Johnston, Dominic Kwiatkowski, Cordelia Langford, John Sillitoe on behalf of the Wellcome Sanger Institute COVID-19 Surveillance Team |
| EPI_ISL_2915339 | Berkshire and Surrey Pathology Services Lighthouse Laboratory | Wellcome Sanger Institute for the COVID-19 Genomics UK (COG-UK) Consortium | Berkshire and Surrey Pathology Services Lighthouse Laboratory and Alex Alderton, Roberto Amato, Jeffrey Barrett, Sonia Goncalves, Ewan Harrison, David K. Jackson, Ian Johnston, Dominic Kwiatkowski, Cordelia Langford, John Sillitoe on behalf of the Wellcome Sanger Institute COVID-19 Surveillance Team |
| EPI_ISL_2915340 | Lighthouse Lab in Glasgow | Wellcome Sanger Institute for the COVID-19 Genomics UK (COG-UK) Consortium | Harper VanSteenhouse, Yumi Kasai, David Gray, Carol Clugston, Anna Dominiczak and Alex Alderton, Roberto Amato, Jeffrey Barrett, Sonia Goncalves, Ewan Harrison, David K. Jackson, Ian Johnston, Dominic Kwiatkowski, Cordelia Langford, John Sillitoe on behalf of the Wellcome Sanger Institute COVID-19 Surveillance Team |
| EPI_ISL_2915341 | Lighthouse Lab in Alderley Park | Wellcome Sanger Institute for the COVID-19 Genomics UK (COG-UK) Consortium | Jacquelyn Wynn, Mairead Hyland, The Lighthouse Lab in Alderley Park and Alex Alderton, Roberto Amato, Jeffrey Barrett, Sonia Goncalves, Ewan Harrison, David K. Jackson, Ian Johnston, Dominic Kwiatkowski, Cordelia Langford, John Sillitoe on behalf of the Wellcome Sanger Institute COVID-19 Surveillance Team |
| EPI_ISL_2915342 | Lighthouse Lab in Glasgow | Wellcome Sanger Institute for the COVID-19 Genomics UK (COG-UK) Consortium | Harper VanSteenhouse, Yumi Kasai, David Gray, Carol Clugston, Anna Dominiczak and Alex Alderton, Roberto Amato, Jeffrey Barrett, Sonia Goncalves, Ewan Harrison, David K. Jackson, Ian Johnston, Dominic Kwiatkowski, Cordelia Langford, John Sillitoe on behalf of the Wellcome Sanger Institute COVID-19 Surveillance Team |
| EPI_ISL_2915343 | Berkshire and Surrey Pathology Services Lighthouse Laboratory | Wellcome Sanger Institute for the COVID-19 Genomics UK (COG-UK) Consortium | Berkshire and Surrey Pathology Services Lighthouse Laboratory and Alex Alderton, Roberto Amato, Jeffrey Barrett, Sonia Goncalves, Ewan Harrison, David K. Jackson, Ian Johnston, Dominic Kwiatkowski, Cordelia Langford, John Sillitoe on behalf of the Wellcome Sanger Institute COVID-19 Surveillance Team |
| EPI_ISL_2915344 | Lighthouse Lab in Glasgow | Wellcome Sanger Institute for the COVID-19 Genomics UK (COG-UK) Consortium | Harper VanSteenhouse, Yumi Kasai, David Gray, Carol Clugston, Anna Dominiczak and Alex Alderton, Roberto Amato, Jeffrey Barrett, Sonia Goncalves, Ewan Harrison, David K. Jackson, Ian Johnston, Dominic Kwiatkowski, Cordelia Langford, John Sillitoe on behalf of the Wellcome Sanger Institute COVID-19 Surveillance Team |
| EPI_ISL_2915345, EPI_ISL_2915346 | Lighthouse Lab in Alderley Park | Wellcome Sanger Institute for the COVID-19 Genomics UK (COG-UK) Consortium | Jacquelyn Wynn, Mairead Hyland, The Lighthouse Lab in Alderley Park and Alex Alderton, Roberto Amato, Jeffrey Barrett, Sonia Goncalves, Ewan Harrison, David K. Jackson, Ian Johnston, Dominic Kwiatkowski, Cordelia Langford, John Sillitoe on behalf of the Wellcome Sanger Institute COVID-19 Surveillance Team |
| EPI_ISL_2915347 | Berkshire and Surrey Pathology Services Lighthouse Laboratory | Wellcome Sanger Institute for the COVID-19 Genomics UK (COG-UK) Consortium | Berkshire and Surrey Pathology Services Lighthouse Laboratory and Alex Alderton, Roberto Amato, Jeffrey Barrett, Sonia Goncalves, Ewan Harrison, David K. Jackson, Ian Johnston, Dominic Kwiatkowski, Cordelia Langford, John Sillitoe on behalf of the Wellcome Sanger Institute COVID-19 Surveillance Team |
| EPI_ISL_2915348 | Lighthouse Laboratory Plymouth | Wellcome Sanger Institute for the COVID-19 Genomics UK (COG-UK) Consortium | Lighthouse Laboratory Plymouth and Alex Alderton, Roberto Amato, Jeffrey Barrett, Sonia Goncalves, Ewan Harrison, David K. Jackson, Ian Johnston, Dominic Kwiatkowski, Cordelia Langford, John Sillitoe on behalf of the Wellcome Sanger Institute COVID-19 Surveillance Team |
| EPI_ISL_2915349 | Lighthouse Lab in Glasgow | Wellcome Sanger Institute for the COVID-19 Genomics UK (COG-UK) Consortium | Harper VanSteenhouse, Yumi Kasai, David Gray, Carol Clugston, Anna Dominiczak and Alex Alderton, Roberto Amato, Jeffrey Barrett, Sonia Goncalves, Ewan Harrison, David K. Jackson, Ian Johnston, Dominic Kwiatkowski, Cordelia Langford, John Sillitoe on behalf of the Wellcome Sanger Institute COVID-19 Surveillance Team |
| EPI_ISL_2915350 | Berkshire and Surrey Pathology Services Lighthouse Laboratory | Wellcome Sanger Institute for the COVID-19 Genomics UK (COG-UK) Consortium | Berkshire and Surrey Pathology Services Lighthouse Laboratory and Alex Alderton, Roberto Amato, Jeffrey Barrett, Sonia Goncalves, Ewan Harrison, David K. Jackson, Ian Johnston, Dominic Kwiatkowski, Cordelia Langford, John Sillitoe on behalf of the Wellcome Sanger Institute COVID-19 Surveillance Team |
| EPI_ISL_2915351 | Lighthouse Lab in Alderley Park | Wellcome Sanger Institute for the COVID-19 Genomics UK (COG-UK) Consortium | Jacquelyn Wynn, Mairead Hyland, The Lighthouse Lab in Alderley Park and Alex Alderton, Roberto Amato, Jeffrey Barrett, Sonia Goncalves, Ewan Harrison, David K. Jackson, Ian Johnston, Dominic Kwiatkowski, Cordelia Langford, John Sillitoe on behalf of the Wellcome Sanger Institute COVID-19 Surveillance Team |
| EPI_ISL_2915352 | Lighthouse Lab in Glasgow | Wellcome Sanger Institute for the COVID-19 Genomics UK (COG-UK) Consortium | Harper VanSteenhouse, Yumi Kasai, David Gray, Carol Clugston, Anna Dominiczak and Alex Alderton, Roberto Amato, Jeffrey Barrett, Sonia Goncalves, Ewan Harrison, David K. Jackson, Ian Johnston, Dominic Kwiatkowski, Cordelia Langford, John Sillitoe on behalf of the Wellcome Sanger Institute COVID-19 Surveillance Team |
| EPI_ISL_2915353, EPI_ISL_2915354 | Lighthouse Lab in Alderley Park | Wellcome Sanger Institute for the COVID-19 Genomics UK (COG-UK) Consortium | Jacquelyn Wynn, Mairead Hyland, The Lighthouse Lab in Alderley Park and Alex Alderton, Roberto Amato, Jeffrey Barrett, Sonia Goncalves, Ewan Harrison, David K. Jackson, Ian Johnston, Dominic Kwiatkowski, Cordelia Langford, John Sillitoe on behalf of the Wellcome Sanger Institute COVID-19 Surveillance Team |
| EPI_ISL_2915355 | Lighthouse Lab in Glasgow | Wellcome Sanger Institute for the COVID-19 Genomics UK (COG-UK) Consortium | Harper VanSteenhouse, Yumi Kasai, David Gray, Carol Clugston, Anna Dominiczak and Alex Alderton, Roberto Amato, Jeffrey Barrett, Sonia Goncalves, Ewan Harrison, David K. Jackson, Ian Johnston, Dominic Kwiatkowski, Cordelia Langford, John Sillitoe on behalf of the Wellcome Sanger Institute COVID-19 Surveillance Team |
| EPI_ISL_2915356 | Lighthouse Lab in Alderley Park | Wellcome Sanger Institute for the COVID-19 Genomics UK (COG-UK) Consortium | Jacquelyn Wynn, Mairead Hyland, The Lighthouse Lab in Alderley Park and Alex Alderton, Roberto Amato, Jeffrey Barrett, Sonia Goncalves, Ewan Harrison, David K. Jackson, Ian Johnston, Dominic Kwiatkowski, Cordelia Langford, John Sillitoe on behalf of the Wellcome Sanger Institute COVID-19 Surveillance Team |
| EPI_ISL_2915357, EPI_ISL_2915358, EPI_ISL_2915359 | Lighthouse Laboratory Plymouth | Wellcome Sanger Institute for the COVID-19 Genomics UK (COG-UK) Consortium | Lighthouse Laboratory Plymouth and Alex Alderton, Roberto Amato, Jeffrey Barrett, Sonia Goncalves, Ewan Harrison, David K. Jackson, Ian Johnston, Dominic Kwiatkowski, Cordelia Langford, John Sillitoe on behalf of the Wellcome Sanger Institute COVID-19 Surveillance Team |
| EPI_ISL_2915360 | Lighthouse Lab in Alderley Park | Wellcome Sanger Institute for the COVID-19 Genomics UK (COG-UK) Consortium | Jacquelyn Wynn, Mairead Hyland, The Lighthouse Lab in Alderley Park and Alex Alderton, Roberto Amato, Jeffrey Barrett, Sonia Goncalves, Ewan Harrison, David K. Jackson, Ian Johnston, Dominic Kwiatkowski, Cordelia Langford, John Sillitoe on behalf of the Wellcome Sanger Institute COVID-19 Surveillance Team |
| EPI_ISL_2915361 | Berkshire and Surrey Pathology Services Lighthouse Laboratory | Wellcome Sanger Institute for the COVID-19 Genomics UK (COG-UK) Consortium | Berkshire and Surrey Pathology Services Lighthouse Laboratory and Alex Alderton, Roberto Amato, Jeffrey Barrett, Sonia Goncalves, Ewan Harrison, David K. Jackson, Ian Johnston, Dominic Kwiatkowski, Cordelia Langford, John Sillitoe on behalf of the Wellcome Sanger Institute COVID-19 Surveillance Team |
| EPI_ISL_2915362 | Lighthouse Laboratory Plymouth | Wellcome Sanger Institute for the COVID-19 Genomics UK (COG-UK) Consortium | Lighthouse Laboratory Plymouth and Alex Alderton, Roberto Amato, Jeffrey Barrett, Sonia Goncalves, Ewan Harrison, David K. Jackson, Ian Johnston, Dominic Kwiatkowski, Cordelia Langford, John Sillitoe on behalf of the Wellcome Sanger Institute COVID-19 Surveillance Team |
| EPI_ISL_2915363 | Berkshire and Surrey Pathology Services Lighthouse Laboratory | Wellcome Sanger Institute for the COVID-19 Genomics UK (COG-UK) Consortium | Berkshire and Surrey Pathology Services Lighthouse Laboratory and Alex Alderton, Roberto Amato, Jeffrey Barrett, Sonia Goncalves, Ewan Harrison, David K. Jackson, Ian Johnston, Dominic Kwiatkowski, Cordelia Langford, John Sillitoe on behalf of the Wellcome Sanger Institute COVID-19 Surveillance Team |
| EPI_ISL_2915364 | Lighthouse Lab in Glasgow | Wellcome Sanger Institute for the COVID-19 Genomics UK (COG-UK) Consortium | Harper VanSteenhouse, Yumi Kasai, David Gray, Carol Clugston, Anna Dominiczak and Alex Alderton, Roberto Amato, Jeffrey Barrett, Sonia Goncalves, Ewan Harrison, David K. Jackson, Ian Johnston, Dominic Kwiatkowski, Cordelia Langford, John Sillitoe on behalf of the Wellcome Sanger Institute COVID-19 Surveillance Team |
| EPI_ISL_2915365 | Lighthouse Lab in Alderley Park | Wellcome Sanger Institute for the COVID-19 Genomics UK (COG-UK) Consortium | Jacquelyn Wynn, Mairead Hyland, The Lighthouse Lab in Alderley Park and Alex Alderton, Roberto Amato, Jeffrey Barrett, Sonia Goncalves, Ewan Harrison, David K. Jackson, Ian Johnston, Dominic Kwiatkowski, Cordelia Langford, John Sillitoe on behalf of the Wellcome Sanger Institute COVID-19 Surveillance Team |
| EPI_ISL_2915366, EPI_ISL_2915367 | Berkshire and Surrey Pathology Services Lighthouse Laboratory | Wellcome Sanger Institute for the COVID-19 Genomics UK (COG-UK) Consortium | Berkshire and Surrey Pathology Services Lighthouse Laboratory and Alex Alderton, Roberto Amato, Jeffrey Barrett, Sonia Goncalves, Ewan Harrison, David K. Jackson, Ian Johnston, Dominic Kwiatkowski, Cordelia Langford, John Sillitoe on behalf of the Wellcome Sanger Institute COVID-19 Surveillance Team |
| EPI_ISL_2915368 | Lighthouse Laboratory Plymouth | Wellcome Sanger Institute for the COVID-19 Genomics UK (COG-UK) Consortium | Lighthouse Laboratory Plymouth and Alex Alderton, Roberto Amato, Jeffrey Barrett, Sonia Goncalves, Ewan Harrison, David K. Jackson, Ian Johnston, Dominic Kwiatkowski, Cordelia Langford, John Sillitoe on behalf of the Wellcome Sanger Institute COVID-19 Surveillance Team |
| EPI_ISL_2915369 | Lighthouse Lab in Alderley Park | Wellcome Sanger Institute for the COVID-19 Genomics UK (COG-UK) Consortium | Jacquelyn Wynn, Mairead Hyland, The Lighthouse Lab in Alderley Park and Alex Alderton, Roberto Amato, Jeffrey Barrett, Sonia Goncalves, Ewan Harrison, David K. Jackson, Ian Johnston, Dominic Kwiatkowski, Cordelia Langford, John Sillitoe on behalf of the Wellcome Sanger Institute COVID-19 Surveillance Team |
| EPI_ISL_2915370, EPI_ISL_2915371 | Lighthouse Laboratory Plymouth | Wellcome Sanger Institute for the COVID-19 Genomics UK (COG-UK) Consortium | Lighthouse Laboratory Plymouth and Alex Alderton, Roberto Amato, Jeffrey Barrett, Sonia Goncalves, Ewan Harrison, David K. Jackson, Ian Johnston, Dominic Kwiatkowski, Cordelia Langford, John Sillitoe on behalf of the Wellcome Sanger Institute COVID-19 Surveillance Team |
| EPI_ISL_2915372 | Lighthouse Lab in Alderley Park | Wellcome Sanger Institute for the COVID-19 Genomics UK (COG-UK) Consortium | Jacquelyn Wynn, Mairead Hyland, The Lighthouse Lab in Alderley Park and Alex Alderton, Roberto Amato, Jeffrey Barrett, Sonia Goncalves, Ewan Harrison, David K. Jackson, Ian Johnston, Dominic Kwiatkowski, Cordelia Langford, John Sillitoe on behalf of the Wellcome Sanger Institute COVID-19 Surveillance Team |
| EPI_ISL_2915373 | Lighthouse Laboratory Plymouth | Wellcome Sanger Institute for the COVID-19 Genomics UK (COG-UK) Consortium | Lighthouse Laboratory Plymouth and Alex Alderton, Roberto Amato, Jeffrey Barrett, Sonia Goncalves, Ewan Harrison, David K. Jackson, Ian Johnston, Dominic Kwiatkowski, Cordelia Langford, John Sillitoe on behalf of the Wellcome Sanger Institute COVID-19 Surveillance Team |
| EPI_ISL_2915374 | Lighthouse Lab in Alderley Park | Wellcome Sanger Institute for the COVID-19 Genomics UK (COG-UK) Consortium | Jacquelyn Wynn, Mairead Hyland, The Lighthouse Lab in Alderley Park and Alex Alderton, Roberto Amato, Jeffrey Barrett, Sonia Goncalves, Ewan Harrison, David K. Jackson, Ian Johnston, Dominic Kwiatkowski, Cordelia Langford, John Sillitoe on behalf of the Wellcome Sanger Institute COVID-19 Surveillance Team |
| EPI_ISL_2915375 | Berkshire and Surrey Pathology Services Lighthouse Laboratory | Wellcome Sanger Institute for the COVID-19 Genomics UK (COG-UK) Consortium | Berkshire and Surrey Pathology Services Lighthouse Laboratory and Alex Alderton, Roberto Amato, Jeffrey Barrett, Sonia Goncalves, Ewan Harrison, David K. Jackson, Ian Johnston, Dominic Kwiatkowski, Cordelia Langford, John Sillitoe on behalf of the Wellcome Sanger Institute COVID-19 Surveillance Team |
| EPI_ISL_2915376 | Lighthouse Lab in Glasgow | Wellcome Sanger Institute for the COVID-19 Genomics UK (COG-UK) Consortium | Harper VanSteenhouse, Yumi Kasai, David Gray, Carol Clugston, Anna Dominiczak and Alex Alderton, Roberto Amato, Jeffrey Barrett, Sonia Goncalves, Ewan Harrison, David K. Jackson, Ian Johnston, Dominic Kwiatkowski, Cordelia Langford, John Sillitoe on behalf of the Wellcome Sanger Institute COVID-19 Surveillance Team |
| EPI_ISL_2915377, EPI_ISL_2915378 | Berkshire and Surrey Pathology Services Lighthouse Laboratory | Wellcome Sanger Institute for the COVID-19 Genomics UK (COG-UK) Consortium | Berkshire and Surrey Pathology Services Lighthouse Laboratory and Alex Alderton, Roberto Amato, Jeffrey Barrett, Sonia Goncalves, Ewan Harrison, David K. Jackson, Ian Johnston, Dominic Kwiatkowski, Cordelia Langford, John Sillitoe on behalf of the Wellcome Sanger Institute COVID-19 Surveillance Team |
| EPI_ISL_2915379 | Lighthouse Lab in Alderley Park | Wellcome Sanger Institute for the COVID-19 Genomics UK (COG-UK) Consortium | Jacquelyn Wynn, Mairead Hyland, The Lighthouse Lab in Alderley Park and Alex Alderton, Roberto Amato, Jeffrey Barrett, Sonia Goncalves, Ewan Harrison, David K. Jackson, Ian Johnston, Dominic Kwiatkowski, Cordelia Langford, John Sillitoe on behalf of the Wellcome Sanger Institute COVID-19 Surveillance Team |
| EPI_ISL_2915380 | Lighthouse Laboratory Plymouth | Wellcome Sanger Institute for the COVID-19 Genomics UK (COG-UK) Consortium | Lighthouse Laboratory Plymouth and Alex Alderton, Roberto Amato, Jeffrey Barrett, Sonia Goncalves, Ewan Harrison, David K. Jackson, Ian Johnston, Dominic Kwiatkowski, Cordelia Langford, John Sillitoe on behalf of the Wellcome Sanger Institute COVID-19 Surveillance Team |
| EPI_ISL_2915381, EPI_ISL_2915382 | Berkshire and Surrey Pathology Services Lighthouse Laboratory | Wellcome Sanger Institute for the COVID-19 Genomics UK (COG-UK) Consortium | Berkshire and Surrey Pathology Services Lighthouse Laboratory and Alex Alderton, Roberto Amato, Jeffrey Barrett, Sonia Goncalves, Ewan Harrison, David K. Jackson, Ian Johnston, Dominic Kwiatkowski, Cordelia Langford, John Sillitoe on behalf of the Wellcome Sanger Institute COVID-19 Surveillance Team |
| EPI_ISL_2915383, EPI_ISL_2915384 | Lighthouse Lab in Alderley Park | Wellcome Sanger Institute for the COVID-19 Genomics UK (COG-UK) Consortium | Jacquelyn Wynn, Mairead Hyland, The Lighthouse Lab in Alderley Park and Alex Alderton, Roberto Amato, Jeffrey Barrett, Sonia Goncalves, Ewan Harrison, David K. Jackson, Ian Johnston, Dominic Kwiatkowski, Cordelia Langford, John Sillitoe on behalf of the Wellcome Sanger Institute COVID-19 Surveillance Team |
| EPI_ISL_2915385 | Lighthouse Lab in Glasgow | Wellcome Sanger Institute for the COVID-19 Genomics UK (COG-UK) Consortium | Harper VanSteenhouse, Yumi Kasai, David Gray, Carol Clugston, Anna Dominiczak and Alex Alderton, Roberto Amato, Jeffrey Barrett, Sonia Goncalves, Ewan Harrison, David K. Jackson, Ian Johnston, Dominic Kwiatkowski, Cordelia Langford, John Sillitoe on behalf of the Wellcome Sanger Institute COVID-19 Surveillance Team |

| | | | |
|---|---|---|---|
| EPI_ISL_2915386 | Lighthouse Lab in Alderley Park | Wellcome Sanger Institute for the COVID-19 Genomics UK (COG-UK) Consortium | Jacquelyn Wynn, Mairead Hyland, The Lighthouse Lab in Alderley Park and Alex Alderton, Roberto Amato, Jeffrey Barrett, Sonia Goncalves, Ewan Harrison, David K. Jackson, Ian Johnston, Dominic Kwiatkowski, Cordelia Langford, John Sillitoe on behalf of the Wellcome Sanger Institute COVID-19 Surveillance Team |
| EPI_ISL_2915387 | Lighthouse Laboratory Plymouth | Wellcome Sanger Institute for the COVID-19 Genomics UK (COG-UK) Consortium | Lighthouse Laboratory Plymouth and Alex Alderton, Roberto Amato, Jeffrey Barrett, Sonia Goncalves, Ewan Harrison, David K. Jackson, Ian Johnston, Dominic Kwiatkowski, Cordelia Langford, John Sillitoe on behalf of the Wellcome Sanger Institute COVID-19 Surveillance Team |
| EPI_ISL_2915388 | Berkshire and Surrey Pathology Services Lighthouse Laboratory | Wellcome Sanger Institute for the COVID-19 Genomics UK (COG-UK) Consortium | Berkshire and Surrey Pathology Services Lighthouse Laboratory and Alex Alderton, Roberto Amato, Jeffrey Barrett, Sonia Goncalves, Ewan Harrison, David K. Jackson, Ian Johnston, Dominic Kwiatkowski, Cordelia Langford, John Sillitoe on behalf of the Wellcome Sanger Institute COVID-19 Surveillance Team |
| EPI_ISL_2915389 | Lighthouse Lab in Alderley Park | Wellcome Sanger Institute for the COVID-19 Genomics UK (COG-UK) Consortium | Jacquelyn Wynn, Mairead Hyland, The Lighthouse Lab in Alderley Park and Alex Alderton, Roberto Amato, Jeffrey Barrett, Sonia Goncalves, Ewan Harrison, David K. Jackson, Ian Johnston, Dominic Kwiatkowski, Cordelia Langford, John Sillitoe on behalf of the Wellcome Sanger Institute COVID-19 Surveillance Team |
| EPI_ISL_2915390 | Berkshire and Surrey Pathology Services Lighthouse Laboratory | Wellcome Sanger Institute for the COVID-19 Genomics UK (COG-UK) Consortium | Berkshire and Surrey Pathology Services Lighthouse Laboratory and Alex Alderton, Roberto Amato, Jeffrey Barrett, Sonia Goncalves, Ewan Harrison, David K. Jackson, Ian Johnston, Dominic Kwiatkowski, Cordelia Langford, John Sillitoe on behalf of the Wellcome Sanger Institute COVID-19 Surveillance Team |
| EPI_ISL_2915391 | Lighthouse Laboratory Plymouth | Wellcome Sanger Institute for the COVID-19 Genomics UK (COG-UK) Consortium | Lighthouse Laboratory Plymouth and Alex Alderton, Roberto Amato, Jeffrey Barrett, Sonia Goncalves, Ewan Harrison, David K. Jackson, Ian Johnston, Dominic Kwiatkowski, Cordelia Langford, John Sillitoe on behalf of the Wellcome Sanger Institute COVID-19 Surveillance Team |
| EPI_ISL_2915392, EPI_ISL_2915393 | Lighthouse Lab in Glasgow | Wellcome Sanger Institute for the COVID-19 Genomics UK (COG-UK) Consortium | Harper VanSteenhouse, Yumi Kasai, David Gray, Carol Clugston, Anna Dominiczak and Alex Alderton, Roberto Amato, Jeffrey Barrett, Sonia Goncalves, Ewan Harrison, David K. Jackson, Ian Johnston, Dominic Kwiatkowski, Cordelia Langford, John Sillitoe on behalf of the Wellcome Sanger Institute COVID-19 Surveillance Team |
| EPI_ISL_2915394 | Lighthouse Lab in Alderley Park | Wellcome Sanger Institute for the COVID-19 Genomics UK (COG-UK) Consortium | Jacquelyn Wynn, Mairead Hyland, The Lighthouse Lab in Alderley Park and Alex Alderton, Roberto Amato, Jeffrey Barrett, Sonia Goncalves, Ewan Harrison, David K. Jackson, Ian Johnston, Dominic Kwiatkowski, Cordelia Langford, John Sillitoe on behalf of the Wellcome Sanger Institute COVID-19 Surveillance Team |
| EPI_ISL_2915395 | Lighthouse Lab in Glasgow | Wellcome Sanger Institute for the COVID-19 Genomics UK (COG-UK) Consortium | Harper VanSteenhouse, Yumi Kasai, David Gray, Carol Clugston, Anna Dominiczak and Alex Alderton, Roberto Amato, Jeffrey Barrett, Sonia Goncalves, Ewan Harrison, David K. Jackson, Ian Johnston, Dominic Kwiatkowski, Cordelia Langford, John Sillitoe on behalf of the Wellcome Sanger Institute COVID-19 Surveillance Team |
| EPI_ISL_2915396 | Lighthouse Lab in Alderley Park | Wellcome Sanger Institute for the COVID-19 Genomics UK (COG-UK) Consortium | Jacquelyn Wynn, Mairead Hyland, The Lighthouse Lab in Alderley Park and Alex Alderton, Roberto Amato, Jeffrey Barrett, Sonia Goncalves, Ewan Harrison, David K. Jackson, Ian Johnston, Dominic Kwiatkowski, Cordelia Langford, John Sillitoe on behalf of the Wellcome Sanger Institute COVID-19 Surveillance Team |
| EPI_ISL_2915397 | Berkshire and Surrey Pathology Services Lighthouse Laboratory | Wellcome Sanger Institute for the COVID-19 Genomics UK (COG-UK) Consortium | Berkshire and Surrey Pathology Services Lighthouse Laboratory and Alex Alderton, Roberto Amato, Jeffrey Barrett, Sonia Goncalves, Ewan Harrison, David K. Jackson, Ian Johnston, Dominic Kwiatkowski, Cordelia Langford, John Sillitoe on behalf of the Wellcome Sanger Institute COVID-19 Surveillance Team |
| EPI_ISL_2915398 | Lighthouse Lab in Alderley Park | Wellcome Sanger Institute for the COVID-19 Genomics UK (COG-UK) Consortium | Jacquelyn Wynn, Mairead Hyland, The Lighthouse Lab in Alderley Park and Alex Alderton, Roberto Amato, Jeffrey Barrett, Sonia Goncalves, Ewan Harrison, David K. Jackson, Ian Johnston, Dominic Kwiatkowski, Cordelia Langford, John Sillitoe on behalf of the Wellcome Sanger Institute COVID-19 Surveillance Team |
| EPI_ISL_2915399 | Lighthouse Lab in Glasgow | Wellcome Sanger Institute for the COVID-19 Genomics UK (COG-UK) Consortium | Harper VanSteenhouse, Yumi Kasai, David Gray, Carol Clugston, Anna Dominiczak and Alex Alderton, Roberto Amato, Jeffrey Barrett, Sonia Goncalves, Ewan Harrison, David K. Jackson, Ian Johnston, Dominic Kwiatkowski, Cordelia Langford, John Sillitoe on behalf of the Wellcome Sanger Institute COVID-19 Surveillance Team |
| EPI_ISL_2915400 | Lighthouse Laboratory Plymouth | Wellcome Sanger Institute for the COVID-19 Genomics UK (COG-UK) Consortium | Lighthouse Laboratory Plymouth and Alex Alderton, Roberto Amato, Jeffrey Barrett, Sonia Goncalves, Ewan Harrison, David K. Jackson, Ian Johnston, Dominic Kwiatkowski, Cordelia Langford, John Sillitoe on behalf of the Wellcome Sanger Institute COVID-19 Surveillance Team |
| EPI_ISL_2915401 | Lighthouse Lab in Glasgow | Wellcome Sanger Institute for the COVID-19 Genomics UK (COG-UK) Consortium | Harper VanSteenhouse, Yumi Kasai, David Gray, Carol Clugston, Anna Dominiczak and Alex Alderton, Roberto Amato, Jeffrey Barrett, Sonia Goncalves, Ewan Harrison, David K. Jackson, Ian Johnston, Dominic Kwiatkowski, Cordelia Langford, John Sillitoe on behalf of the Wellcome Sanger Institute COVID-19 Surveillance Team |
| EPI_ISL_2915402, EPI_ISL_2915403 | Berkshire and Surrey Pathology Services Lighthouse Laboratory | Wellcome Sanger Institute for the COVID-19 Genomics UK (COG-UK) Consortium | Berkshire and Surrey Pathology Services Lighthouse Laboratory and Alex Alderton, Roberto Amato, Jeffrey Barrett, Sonia Goncalves, Ewan Harrison, David K. Jackson, Ian Johnston, Dominic Kwiatkowski, Cordelia Langford, John Sillitoe on behalf of the Wellcome Sanger Institute COVID-19 Surveillance Team |
| EPI_ISL_2915404 | Lighthouse Lab in Alderley Park | Wellcome Sanger Institute for the COVID-19 Genomics UK (COG-UK) Consortium | Jacquelyn Wynn, Mairead Hyland, The Lighthouse Lab in Alderley Park and Alex Alderton, Roberto Amato, Jeffrey Barrett, Sonia Goncalves, Ewan Harrison, David K. Jackson, Ian Johnston, Dominic Kwiatkowski, Cordelia Langford, John Sillitoe on behalf of the Wellcome Sanger Institute COVID-19 Surveillance Team |
| EPI_ISL_2915405 | Berkshire and Surrey Pathology Services Lighthouse Laboratory | Wellcome Sanger Institute for the COVID-19 Genomics UK (COG-UK) Consortium | Berkshire and Surrey Pathology Services Lighthouse Laboratory and Alex Alderton, Roberto Amato, Jeffrey Barrett, Sonia Goncalves, Ewan Harrison, David K. Jackson, Ian Johnston, Dominic Kwiatkowski, Cordelia Langford, John Sillitoe on behalf of the Wellcome Sanger Institute COVID-19 Surveillance Team |
| EPI_ISL_2915406 | Lighthouse Lab in Alderley Park | Wellcome Sanger Institute for the COVID-19 Genomics UK (COG-UK) Consortium | Jacquelyn Wynn, Mairead Hyland, The Lighthouse Lab in Alderley Park and Alex Alderton, Roberto Amato, Jeffrey Barrett, Sonia Goncalves, Ewan Harrison, David K. Jackson, Ian Johnston, Dominic Kwiatkowski, Cordelia Langford, John Sillitoe on behalf of the Wellcome Sanger Institute COVID-19 Surveillance Team |
| EPI_ISL_2915407, EPI_ISL_2915408, EPI_ISL_2915409 | Lighthouse Laboratory Plymouth | Wellcome Sanger Institute for the COVID-19 Genomics UK (COG-UK) Consortium | Lighthouse Laboratory Plymouth and Alex Alderton, Roberto Amato, Jeffrey Barrett, Sonia Goncalves, Ewan Harrison, David K. Jackson, Ian Johnston, Dominic Kwiatkowski, Cordelia Langford, John Sillitoe on behalf of the Wellcome Sanger Institute COVID-19 Surveillance Team |
| EPI_ISL_2915410, EPI_ISL_2915411 | Lighthouse Lab in Alderley Park | Wellcome Sanger Institute for the COVID-19 Genomics UK (COG-UK) Consortium | Jacquelyn Wynn, Mairead Hyland, The Lighthouse Lab in Alderley Park and Alex Alderton, Roberto Amato, Jeffrey Barrett, Sonia Goncalves, Ewan Harrison, David K. Jackson, Ian Johnston, Dominic Kwiatkowski, Cordelia Langford, John Sillitoe on behalf of the Wellcome Sanger Institute COVID-19 Surveillance Team |
| EPI_ISL_2915412 | Berkshire and Surrey Pathology Services Lighthouse Laboratory | Wellcome Sanger Institute for the COVID-19 Genomics UK (COG-UK) Consortium | Berkshire and Surrey Pathology Services Lighthouse Laboratory and Alex Alderton, Roberto Amato, Jeffrey Barrett, Sonia Goncalves, Ewan Harrison, David K. Jackson, Ian Johnston, Dominic Kwiatkowski, Cordelia Langford, John Sillitoe on behalf of the Wellcome Sanger Institute COVID-19 Surveillance Team |
| EPI_ISL_2915413 | Lighthouse Lab in Alderley Park | Wellcome Sanger Institute for the COVID-19 Genomics UK (COG-UK) Consortium | Jacquelyn Wynn, Mairead Hyland, The Lighthouse Lab in Alderley Park and Alex Alderton, Roberto Amato, Jeffrey Barrett, Sonia Goncalves, Ewan Harrison, David K. Jackson, Ian Johnston, Dominic Kwiatkowski, Cordelia Langford, John Sillitoe on behalf of the Wellcome Sanger Institute COVID-19 Surveillance Team |
| EPI_ISL_2915414 | Lighthouse Lab in Glasgow | Wellcome Sanger Institute for the COVID-19 Genomics UK (COG-UK) Consortium | Harper VanSteenhouse, Yumi Kasai, David Gray, Carol Clugston, Anna Dominiczak and Alex Alderton, Roberto Amato, Jeffrey Barrett, Sonia Goncalves, Ewan Harrison, David K. Jackson, Ian Johnston, Dominic Kwiatkowski, Cordelia Langford, John Sillitoe on behalf of the Wellcome Sanger Institute COVID-19 Surveillance Team |
| EPI_ISL_2915415 | Lighthouse Laboratory Plymouth | Wellcome Sanger Institute for the COVID-19 Genomics UK (COG-UK) Consortium | Lighthouse Laboratory Plymouth and Alex Alderton, Roberto Amato, Jeffrey Barrett, Sonia Goncalves, Ewan Harrison, David K. Jackson, Ian Johnston, Dominic Kwiatkowski, Cordelia Langford, John Sillitoe on behalf of the Wellcome Sanger Institute COVID-19 Surveillance Team |
| EPI_ISL_2915416 | Berkshire and Surrey Pathology Services Lighthouse Laboratory | Wellcome Sanger Institute for the COVID-19 Genomics UK (COG-UK) Consortium | Berkshire and Surrey Pathology Services Lighthouse Laboratory and Alex Alderton, Roberto Amato, Jeffrey Barrett, Sonia Goncalves, Ewan Harrison, David K. Jackson, Ian Johnston, Dominic Kwiatkowski, Cordelia Langford, John Sillitoe on behalf of the Wellcome Sanger Institute COVID-19 Surveillance Team |
| EPI_ISL_2915417, EPI_ISL_2915418 | Lighthouse Lab in Alderley Park | Wellcome Sanger Institute for the COVID-19 Genomics UK (COG-UK) Consortium | Jacquelyn Wynn, Mairead Hyland, The Lighthouse Lab in Alderley Park and Alex Alderton, Roberto Amato, Jeffrey Barrett, Sonia Goncalves, Ewan Harrison, David K. Jackson, Ian Johnston, Dominic Kwiatkowski, Cordelia Langford, John Sillitoe on behalf of the Wellcome Sanger Institute COVID-19 Surveillance Team |
| EPI_ISL_2915419 | Lighthouse Lab in Glasgow | Wellcome Sanger Institute for the COVID-19 Genomics UK (COG-UK) Consortium | Harper VanSteenhouse, Yumi Kasai, David Gray, Carol Clugston, Anna Dominiczak and Alex Alderton, Roberto Amato, Jeffrey Barrett, Sonia Goncalves, Ewan Harrison, David K. Jackson, Ian Johnston, Dominic Kwiatkowski, Cordelia Langford, John Sillitoe on behalf of the Wellcome Sanger Institute COVID-19 Surveillance Team |
| EPI_ISL_2915420 | Lighthouse Laboratory Plymouth | Wellcome Sanger Institute for the COVID-19 Genomics UK (COG-UK) Consortium | Lighthouse Laboratory Plymouth and Alex Alderton, Roberto Amato, Jeffrey Barrett, Sonia Goncalves, Ewan Harrison, David K. Jackson, Ian Johnston, Dominic Kwiatkowski, Cordelia Langford, John Sillitoe on behalf of the Wellcome Sanger Institute COVID-19 Surveillance Team |
| EPI_ISL_2915421 | Lighthouse Lab in Glasgow | Wellcome Sanger Institute for the COVID-19 Genomics UK (COG-UK) Consortium | Harper VanSteenhouse, Yumi Kasai, David Gray, Carol Clugston, Anna Dominiczak and Alex Alderton, Roberto Amato, Jeffrey Barrett, Sonia Goncalves, Ewan Harrison, David K. Jackson, Ian Johnston, Dominic Kwiatkowski, Cordelia Langford, John Sillitoe on behalf of the Wellcome Sanger Institute COVID-19 Surveillance Team |
| EPI_ISL_2915422, EPI_ISL_2915423, EPI_ISL_2915424 | Lighthouse Laboratory Plymouth | Wellcome Sanger Institute for the COVID-19 Genomics UK (COG-UK) Consortium | Lighthouse Laboratory Plymouth and Alex Alderton, Roberto Amato, Jeffrey Barrett, Sonia Goncalves, Ewan Harrison, David K. Jackson, Ian Johnston, Dominic Kwiatkowski, Cordelia Langford, John Sillitoe on behalf of the Wellcome Sanger Institute COVID-19 Surveillance Team |
| EPI_ISL_2915425, EPI_ISL_2915426 | Lighthouse Lab in Alderley Park | Wellcome Sanger Institute for the COVID-19 Genomics UK (COG-UK) Consortium | Jacquelyn Wynn, Mairead Hyland, The Lighthouse Lab in Alderley Park and Alex Alderton, Roberto Amato, Jeffrey Barrett, Sonia Goncalves, Ewan Harrison, David K. Jackson, Ian Johnston, Dominic Kwiatkowski, Cordelia Langford, John Sillitoe on behalf of the Wellcome Sanger Institute COVID-19 Surveillance Team |
| EPI_ISL_2915427, EPI_ISL_2915428 | Berkshire and Surrey Pathology Services Lighthouse Laboratory | Wellcome Sanger Institute for the COVID-19 Genomics UK (COG-UK) Consortium | Berkshire and Surrey Pathology Services Lighthouse Laboratory and Alex Alderton, Roberto Amato, Jeffrey Barrett, Sonia Goncalves, Ewan Harrison, David K. Jackson, Ian Johnston, Dominic Kwiatkowski, Cordelia Langford, John Sillitoe on behalf of the Wellcome Sanger Institute COVID-19 Surveillance Team |
| EPI_ISL_2915429 | Lighthouse Lab in Glasgow | Wellcome Sanger Institute for the COVID-19 Genomics UK (COG-UK) Consortium | Harper VanSteenhouse, Yumi Kasai, David Gray, Carol Clugston, Anna Dominiczak and Alex Alderton, Roberto Amato, Jeffrey Barrett, Sonia Goncalves, Ewan Harrison, David K. Jackson, Ian Johnston, Dominic Kwiatkowski, Cordelia Langford, John Sillitoe on behalf of the Wellcome Sanger Institute COVID-19 Surveillance Team |
| EPI_ISL_2915430 | Berkshire and Surrey Pathology Services Lighthouse Laboratory | Wellcome Sanger Institute for the COVID-19 Genomics UK (COG-UK) Consortium | Berkshire and Surrey Pathology Services Lighthouse Laboratory and Alex Alderton, Roberto Amato, Jeffrey Barrett, Sonia Goncalves, Ewan Harrison, David K. Jackson, Ian Johnston, Dominic Kwiatkowski, Cordelia Langford, John Sillitoe on behalf of the Wellcome Sanger Institute COVID-19 Surveillance Team |
| EPI_ISL_2915431 | Lighthouse Laboratory Plymouth | Wellcome Sanger Institute for the COVID-19 Genomics UK (COG-UK) Consortium | Lighthouse Laboratory Plymouth and Alex Alderton, Roberto Amato, Jeffrey Barrett, Sonia Goncalves, Ewan Harrison, David K. Jackson, Ian Johnston, Dominic Kwiatkowski, Cordelia Langford, John Sillitoe on behalf of the Wellcome Sanger Institute COVID-19 Surveillance Team |
| EPI_ISL_2915432 | Berkshire and Surrey Pathology Services Lighthouse Laboratory | Wellcome Sanger Institute for the COVID-19 Genomics UK (COG-UK) Consortium | Berkshire and Surrey Pathology Services Lighthouse Laboratory and Alex Alderton, Roberto Amato, Jeffrey Barrett, Sonia Goncalves, Ewan Harrison, David K. Jackson, Ian Johnston, Dominic Kwiatkowski, Cordelia Langford, John Sillitoe on behalf of the Wellcome Sanger Institute COVID-19 Surveillance Team |
| EPI_ISL_2915433 | Lighthouse Lab in Alderley Park | Wellcome Sanger Institute for the COVID-19 Genomics UK (COG-UK) Consortium | Jacquelyn Wynn, Mairead Hyland, The Lighthouse Lab in Alderley Park and Alex Alderton, Roberto Amato, Jeffrey Barrett, Sonia Goncalves, Ewan Harrison, David K. Jackson, Ian Johnston, Dominic Kwiatkowski, Cordelia Langford, John Sillitoe on behalf of the Wellcome Sanger Institute COVID-19 Surveillance Team |
| EPI_ISL_2915434 | Lighthouse Lab in Glasgow | Wellcome Sanger Institute for the COVID-19 Genomics UK (COG-UK) Consortium | Harper VanSteenhouse, Yumi Kasai, David Gray, Carol Clugston, Anna Dominiczak and Alex Alderton, Roberto Amato, Jeffrey Barrett, Sonia Goncalves, Ewan Harrison, David K. Jackson, Ian Johnston, Dominic Kwiatkowski, Cordelia Langford, John Sillitoe on behalf of the Wellcome Sanger Institute COVID-19 Surveillance Team |
| EPI_ISL_2915435 | Berkshire and Surrey Pathology Services Lighthouse Laboratory | Wellcome Sanger Institute for the COVID-19 Genomics UK (COG-UK) Consortium | Berkshire and Surrey Pathology Services Lighthouse Laboratory and Alex Alderton, Roberto Amato, Jeffrey Barrett, Sonia Goncalves, Ewan Harrison, David K. Jackson, Ian Johnston, Dominic Kwiatkowski, Cordelia Langford, John Sillitoe on behalf of the Wellcome Sanger Institute COVID-19 Surveillance Team |
| EPI_ISL_2915436 | Lighthouse Lab in Glasgow | Wellcome Sanger Institute for the COVID-19 Genomics UK (COG-UK) Consortium | Harper VanSteenhouse, Yumi Kasai, David Gray, Carol Clugston, Anna Dominiczak and Alex Alderton, Roberto Amato, Jeffrey Barrett, Sonia Goncalves, Ewan Harrison, David K. Jackson, Ian Johnston, Dominic Kwiatkowski, Cordelia Langford, John Sillitoe on behalf of the Wellcome Sanger Institute COVID-19 Surveillance Team |
| EPI_ISL_2915437, EPI_ISL_2915438 | Lighthouse Laboratory Plymouth | Wellcome Sanger Institute for the COVID-19 Genomics UK (COG-UK) Consortium | Lighthouse Laboratory Plymouth and Alex Alderton, Roberto Amato, Jeffrey Barrett, Sonia Goncalves, Ewan Harrison, David K. Jackson, Ian Johnston, Dominic Kwiatkowski, Cordelia Langford, John Sillitoe on behalf of the Wellcome Sanger Institute COVID-19 Surveillance Team |
| EPI_ISL_2915439, EPI_ISL_2915440 | Lighthouse Lab in Alderley Park | Wellcome Sanger Institute for the COVID-19 Genomics UK (COG-UK) Consortium | Jacquelyn Wynn, Mairead Hyland, The Lighthouse Lab in Alderley Park and Alex Alderton, Roberto Amato, Jeffrey Barrett, Sonia Goncalves, Ewan Harrison, David K. Jackson, Ian Johnston, Dominic Kwiatkowski, Cordelia Langford, John Sillitoe on behalf of the Wellcome Sanger Institute COVID-19 Surveillance Team |
| EPI_ISL_2915441 | Berkshire and Surrey Pathology Services Lighthouse Laboratory | Wellcome Sanger Institute for the COVID-19 Genomics UK (COG-UK) Consortium | Berkshire and Surrey Pathology Services Lighthouse Laboratory and Alex Alderton, Roberto Amato, Jeffrey Barrett, Sonia Goncalves, Ewan Harrison, David K. Jackson, Ian Johnston, Dominic Kwiatkowski, Cordelia Langford, John Sillitoe on behalf of the Wellcome Sanger Institute COVID-19 Surveillance Team |

| | | | |
|---|---|---|---|
| EPI_ISL_2915442 | Lighthouse Laboratory Plymouth | Wellcome Sanger Institute for the COVID-19 Genomics UK (COG-UK) Consortium | Lighthouse Laboratory Plymouth and Alex Alderton, Roberto Amato, Jeffrey Barrett, Sonia Goncalves, Ewan Harrison, David K. Jackson, Ian Johnston, Dominic Kwiatkowski, Cordelia Langford, John Sillitoe on behalf of the Wellcome Sanger Institute COVID-19 Surveillance Team |
| EPI_ISL_2915443 | Lighthouse Lab in Glasgow | Wellcome Sanger Institute for the COVID-19 Genomics UK (COG-UK) Consortium | Harper VanSteenhouse, Yumi Kasai, David Gray, Carol Clugston, Anna Dominiczak and Alex Alderton, Roberto Amato, Jeffrey Barrett, Sonia Goncalves, Ewan Harrison, David K. Jackson, Ian Johnston, Dominic Kwiatkowski, Cordelia Langford, John Sillitoe on behalf of the Wellcome Sanger Institute COVID-19 Surveillance Team |
| EPI_ISL_2915444 | Lighthouse Lab in Alderley Park | Wellcome Sanger Institute for the COVID-19 Genomics UK (COG-UK) Consortium | Jacquelyn Wynn, Mairead Hyland, The Lighthouse Lab in Alderley Park and Alex Alderton, Roberto Amato, Jeffrey Barrett, Sonia Goncalves, Ewan Harrison, David K. Jackson, Ian Johnston, Dominic Kwiatkowski, Cordelia Langford, John Sillitoe on behalf of the Wellcome Sanger Institute COVID-19 Surveillance Team |
| EPI_ISL_2915445 | Lighthouse Laboratory Plymouth | Wellcome Sanger Institute for the COVID-19 Genomics UK (COG-UK) Consortium | Lighthouse Laboratory Plymouth and Alex Alderton, Roberto Amato, Jeffrey Barrett, Sonia Goncalves, Ewan Harrison, David K. Jackson, Ian Johnston, Dominic Kwiatkowski, Cordelia Langford, John Sillitoe on behalf of the Wellcome Sanger Institute COVID-19 Surveillance Team |
| EPI_ISL_2915446 | Berkshire and Surrey Pathology Services Lighthouse Laboratory | Wellcome Sanger Institute for the COVID-19 Genomics UK (COG-UK) Consortium | Berkshire and Surrey Pathology Services Lighthouse Laboratory and Alex Alderton, Roberto Amato, Jeffrey Barrett, Sonia Goncalves, Ewan Harrison, David K. Jackson, Ian Johnston, Dominic Kwiatkowski, Cordelia Langford, John Sillitoe on behalf of the Wellcome Sanger Institute COVID-19 Surveillance Team |
| EPI_ISL_2915447 | Lighthouse Lab in Alderley Park | Wellcome Sanger Institute for the COVID-19 Genomics UK (COG-UK) Consortium | Jacquelyn Wynn, Mairead Hyland, The Lighthouse Lab in Alderley Park and Alex Alderton, Roberto Amato, Jeffrey Barrett, Sonia Goncalves, Ewan Harrison, David K. Jackson, Ian Johnston, Dominic Kwiatkowski, Cordelia Langford, John Sillitoe on behalf of the Wellcome Sanger Institute COVID-19 Surveillance Team |
| EPI_ISL_2915448, EPI_ISL_2915449 | Lighthouse Laboratory Plymouth | Wellcome Sanger Institute for the COVID-19 Genomics UK (COG-UK) Consortium | Lighthouse Laboratory Plymouth and Alex Alderton, Roberto Amato, Jeffrey Barrett, Sonia Goncalves, Ewan Harrison, David K. Jackson, Ian Johnston, Dominic Kwiatkowski, Cordelia Langford, John Sillitoe on behalf of the Wellcome Sanger Institute COVID-19 Surveillance Team |
| EPI_ISL_2915450, EPI_ISL_2915451 | Lighthouse Lab in Alderley Park | Wellcome Sanger Institute for the COVID-19 Genomics UK (COG-UK) Consortium | Jacquelyn Wynn, Mairead Hyland, The Lighthouse Lab in Alderley Park and Alex Alderton, Roberto Amato, Jeffrey Barrett, Sonia Goncalves, Ewan Harrison, David K. Jackson, Ian Johnston, Dominic Kwiatkowski, Cordelia Langford, John Sillitoe on behalf of the Wellcome Sanger Institute COVID-19 Surveillance Team |
| EPI_ISL_2915452, EPI_ISL_2915453 | Berkshire and Surrey Pathology Services Lighthouse Laboratory | Wellcome Sanger Institute for the COVID-19 Genomics UK (COG-UK) Consortium | Berkshire and Surrey Pathology Services Lighthouse Laboratory and Alex Alderton, Roberto Amato, Jeffrey Barrett, Sonia Goncalves, Ewan Harrison, David K. Jackson, Ian Johnston, Dominic Kwiatkowski, Cordelia Langford, John Sillitoe on behalf of the Wellcome Sanger Institute COVID-19 Surveillance Team |
| EPI_ISL_2915454 | Lighthouse Lab in Alderley Park | Wellcome Sanger Institute for the COVID-19 Genomics UK (COG-UK) Consortium | Jacquelyn Wynn, Mairead Hyland, The Lighthouse Lab in Alderley Park and Alex Alderton, Roberto Amato, Jeffrey Barrett, Sonia Goncalves, Ewan Harrison, David K. Jackson, Ian Johnston, Dominic Kwiatkowski, Cordelia Langford, John Sillitoe on behalf of the Wellcome Sanger Institute COVID-19 Surveillance Team |
| EPI_ISL_2915455 | Lighthouse Laboratory Plymouth | Wellcome Sanger Institute for the COVID-19 Genomics UK (COG-UK) Consortium | Lighthouse Laboratory Plymouth and Alex Alderton, Roberto Amato, Jeffrey Barrett, Sonia Goncalves, Ewan Harrison, David K. Jackson, Ian Johnston, Dominic Kwiatkowski, Cordelia Langford, John Sillitoe on behalf of the Wellcome Sanger Institute COVID-19 Surveillance Team |
| EPI_ISL_2915456 | Lighthouse Lab in Glasgow | Wellcome Sanger Institute for the COVID-19 Genomics UK (COG-UK) Consortium | Harper VanSteenhouse, Yumi Kasai, David Gray, Carol Clugston, Anna Dominiczak and Alex Alderton, Roberto Amato, Jeffrey Barrett, Sonia Goncalves, Ewan Harrison, David K. Jackson, Ian Johnston, Dominic Kwiatkowski, Cordelia Langford, John Sillitoe on behalf of the Wellcome Sanger Institute COVID-19 Surveillance Team |
| EPI_ISL_2915457 | Berkshire and Surrey Pathology Services Lighthouse Laboratory | Wellcome Sanger Institute for the COVID-19 Genomics UK (COG-UK) Consortium | Berkshire and Surrey Pathology Services Lighthouse Laboratory and Alex Alderton, Roberto Amato, Jeffrey Barrett, Sonia Goncalves, Ewan Harrison, David K. Jackson, Ian Johnston, Dominic Kwiatkowski, Cordelia Langford, John Sillitoe on behalf of the Wellcome Sanger Institute COVID-19 Surveillance Team |
| EPI_ISL_2915458, EPI_ISL_2915459 | Lighthouse Laboratory Plymouth | Wellcome Sanger Institute for the COVID-19 Genomics UK (COG-UK) Consortium | Lighthouse Laboratory Plymouth and Alex Alderton, Roberto Amato, Jeffrey Barrett, Sonia Goncalves, Ewan Harrison, David K. Jackson, Ian Johnston, Dominic Kwiatkowski, Cordelia Langford, John Sillitoe on behalf of the Wellcome Sanger Institute COVID-19 Surveillance Team |
| EPI_ISL_2915460 | Lighthouse Lab in Glasgow | Wellcome Sanger Institute for the COVID-19 Genomics UK (COG-UK) Consortium | Harper VanSteenhouse, Yumi Kasai, David Gray, Carol Clugston, Anna Dominiczak and Alex Alderton, Roberto Amato, Jeffrey Barrett, Sonia Goncalves, Ewan Harrison, David K. Jackson, Ian Johnston, Dominic Kwiatkowski, Cordelia Langford, John Sillitoe on behalf of the Wellcome Sanger Institute COVID-19 Surveillance Team |
| EPI_ISL_2915461 | Berkshire and Surrey Pathology Services Lighthouse Laboratory | Wellcome Sanger Institute for the COVID-19 Genomics UK (COG-UK) Consortium | Berkshire and Surrey Pathology Services Lighthouse Laboratory and Alex Alderton, Roberto Amato, Jeffrey Barrett, Sonia Goncalves, Ewan Harrison, David K. Jackson, Ian Johnston, Dominic Kwiatkowski, Cordelia Langford, John Sillitoe on behalf of the Wellcome Sanger Institute COVID-19 Surveillance Team |
| EPI_ISL_2915462 | Lighthouse Lab in Alderley Park | Wellcome Sanger Institute for the COVID-19 Genomics UK (COG-UK) Consortium | Jacquelyn Wynn, Mairead Hyland, The Lighthouse Lab in Alderley Park and Alex Alderton, Roberto Amato, Jeffrey Barrett, Sonia Goncalves, Ewan Harrison, David K. Jackson, Ian Johnston, Dominic Kwiatkowski, Cordelia Langford, John Sillitoe on behalf of the Wellcome Sanger Institute COVID-19 Surveillance Team |
| EPI_ISL_2915463 | Lighthouse Laboratory Plymouth | Wellcome Sanger Institute for the COVID-19 Genomics UK (COG-UK) Consortium | Lighthouse Laboratory Plymouth and Alex Alderton, Roberto Amato, Jeffrey Barrett, Sonia Goncalves, Ewan Harrison, David K. Jackson, Ian Johnston, Dominic Kwiatkowski, Cordelia Langford, John Sillitoe on behalf of the Wellcome Sanger Institute COVID-19 Surveillance Team |
| EPI_ISL_2915464 | Berkshire and Surrey Pathology Services Lighthouse Laboratory | Wellcome Sanger Institute for the COVID-19 Genomics UK (COG-UK) Consortium | Berkshire and Surrey Pathology Services Lighthouse Laboratory and Alex Alderton, Roberto Amato, Jeffrey Barrett, Sonia Goncalves, Ewan Harrison, David K. Jackson, Ian Johnston, Dominic Kwiatkowski, Cordelia Langford, John Sillitoe on behalf of the Wellcome Sanger Institute COVID-19 Surveillance Team |
| EPI_ISL_2915465 | Lighthouse Laboratory Plymouth | Wellcome Sanger Institute for the COVID-19 Genomics UK (COG-UK) Consortium | Lighthouse Laboratory Plymouth and Alex Alderton, Roberto Amato, Jeffrey Barrett, Sonia Goncalves, Ewan Harrison, David K. Jackson, Ian Johnston, Dominic Kwiatkowski, Cordelia Langford, John Sillitoe on behalf of the Wellcome Sanger Institute COVID-19 Surveillance Team |
| EPI_ISL_2915466 | Lighthouse Lab in Glasgow | Wellcome Sanger Institute for the COVID-19 Genomics UK (COG-UK) Consortium | Harper VanSteenhouse, Yumi Kasai, David Gray, Carol Clugston, Anna Dominiczak and Alex Alderton, Roberto Amato, Jeffrey Barrett, Sonia Goncalves, Ewan Harrison, David K. Jackson, Ian Johnston, Dominic Kwiatkowski, Cordelia Langford, John Sillitoe on behalf of the Wellcome Sanger Institute COVID-19 Surveillance Team |
| EPI_ISL_2915467 | Lighthouse Lab in Alderley Park | Wellcome Sanger Institute for the COVID-19 Genomics UK (COG-UK) Consortium | Jacquelyn Wynn, Mairead Hyland, The Lighthouse Lab in Alderley Park and Alex Alderton, Roberto Amato, Jeffrey Barrett, Sonia Goncalves, Ewan Harrison, David K. Jackson, Ian Johnston, Dominic Kwiatkowski, Cordelia Langford, John Sillitoe on behalf of the Wellcome Sanger Institute COVID-19 Surveillance Team |
| EPI_ISL_2915468, EPI_ISL_2915469, EPI_ISL_2915470 | Lighthouse Lab in Glasgow | Wellcome Sanger Institute for the COVID-19 Genomics UK (COG-UK) Consortium | Harper VanSteenhouse, Yumi Kasai, David Gray, Carol Clugston, Anna Dominiczak and Alex Alderton, Roberto Amato, Jeffrey Barrett, Sonia Goncalves, Ewan Harrison, David K. Jackson, Ian Johnston, Dominic Kwiatkowski, Cordelia Langford, John Sillitoe on behalf of the Wellcome Sanger Institute COVID-19 Surveillance Team |
| EPI_ISL_2915471 | Lighthouse Lab in Alderley Park | Wellcome Sanger Institute for the COVID-19 Genomics UK (COG-UK) Consortium | Jacquelyn Wynn, Mairead Hyland, The Lighthouse Lab in Alderley Park and Alex Alderton, Roberto Amato, Jeffrey Barrett, Sonia Goncalves, Ewan Harrison, David K. Jackson, Ian Johnston, Dominic Kwiatkowski, Cordelia Langford, John Sillitoe on behalf of the Wellcome Sanger Institute COVID-19 Surveillance Team |
| EPI_ISL_2915472, EPI_ISL_2915473 | Lighthouse Laboratory Plymouth | Wellcome Sanger Institute for the COVID-19 Genomics UK (COG-UK) Consortium | Lighthouse Laboratory Plymouth and Alex Alderton, Roberto Amato, Jeffrey Barrett, Sonia Goncalves, Ewan Harrison, David K. Jackson, Ian Johnston, Dominic Kwiatkowski, Cordelia Langford, John Sillitoe on behalf of the Wellcome Sanger Institute COVID-19 Surveillance Team |
| EPI_ISL_2915474 | Lighthouse Lab in Alderley Park | Wellcome Sanger Institute for the COVID-19 Genomics UK (COG-UK) Consortium | Jacquelyn Wynn, Mairead Hyland, The Lighthouse Lab in Alderley Park and Alex Alderton, Roberto Amato, Jeffrey Barrett, Sonia Goncalves, Ewan Harrison, David K. Jackson, Ian Johnston, Dominic Kwiatkowski, Cordelia Langford, John Sillitoe on behalf of the Wellcome Sanger Institute COVID-19 Surveillance Team |
| EPI_ISL_2915475 | Berkshire and Surrey Pathology Services Lighthouse Laboratory | Wellcome Sanger Institute for the COVID-19 Genomics UK (COG-UK) Consortium | Berkshire and Surrey Pathology Services Lighthouse Laboratory and Alex Alderton, Roberto Amato, Jeffrey Barrett, Sonia Goncalves, Ewan Harrison, David K. Jackson, Ian Johnston, Dominic Kwiatkowski, Cordelia Langford, John Sillitoe on behalf of the Wellcome Sanger Institute COVID-19 Surveillance Team |
| EPI_ISL_2915476 | Lighthouse Lab in Glasgow | Wellcome Sanger Institute for the COVID-19 Genomics UK (COG-UK) Consortium | Harper VanSteenhouse, Yumi Kasai, David Gray, Carol Clugston, Anna Dominiczak and Alex Alderton, Roberto Amato, Jeffrey Barrett, Sonia Goncalves, Ewan Harrison, David K. Jackson, Ian Johnston, Dominic Kwiatkowski, Cordelia Langford, John Sillitoe on behalf of the Wellcome Sanger Institute COVID-19 Surveillance Team |
| EPI_ISL_2915477 | Lighthouse Laboratory Plymouth | Wellcome Sanger Institute for the COVID-19 Genomics UK (COG-UK) Consortium | Lighthouse Laboratory Plymouth and Alex Alderton, Roberto Amato, Jeffrey Barrett, Sonia Goncalves, Ewan Harrison, David K. Jackson, Ian Johnston, Dominic Kwiatkowski, Cordelia Langford, John Sillitoe on behalf of the Wellcome Sanger Institute COVID-19 Surveillance Team |
| EPI_ISL_2915478 | Lighthouse Lab in Alderley Park | Wellcome Sanger Institute for the COVID-19 Genomics UK (COG-UK) Consortium | Jacquelyn Wynn, Mairead Hyland, The Lighthouse Lab in Alderley Park and Alex Alderton, Roberto Amato, Jeffrey Barrett, Sonia Goncalves, Ewan Harrison, David K. Jackson, Ian Johnston, Dominic Kwiatkowski, Cordelia Langford, John Sillitoe on behalf of the Wellcome Sanger Institute COVID-19 Surveillance Team |
| EPI_ISL_2915479 | Lighthouse Lab in Glasgow | Wellcome Sanger Institute for the COVID-19 Genomics UK (COG-UK) Consortium | Harper VanSteenhouse, Yumi Kasai, David Gray, Carol Clugston, Anna Dominiczak and Alex Alderton, Roberto Amato, Jeffrey Barrett, Sonia Goncalves, Ewan Harrison, David K. Jackson, Ian Johnston, Dominic Kwiatkowski, Cordelia Langford, John Sillitoe on behalf of the Wellcome Sanger Institute COVID-19 Surveillance Team |
| EPI_ISL_2915480 | Lighthouse Lab in Alderley Park | Wellcome Sanger Institute for the COVID-19 Genomics UK (COG-UK) Consortium | Jacquelyn Wynn, Mairead Hyland, The Lighthouse Lab in Alderley Park and Alex Alderton, Roberto Amato, Jeffrey Barrett, Sonia Goncalves, Ewan Harrison, David K. Jackson, Ian Johnston, Dominic Kwiatkowski, Cordelia Langford, John Sillitoe on behalf of the Wellcome Sanger Institute COVID-19 Surveillance Team |
| EPI_ISL_2915481 | Berkshire and Surrey Pathology Services Lighthouse Laboratory | Wellcome Sanger Institute for the COVID-19 Genomics UK (COG-UK) Consortium | Berkshire and Surrey Pathology Services Lighthouse Laboratory and Alex Alderton, Roberto Amato, Jeffrey Barrett, Sonia Goncalves, Ewan Harrison, David K. Jackson, Ian Johnston, Dominic Kwiatkowski, Cordelia Langford, John Sillitoe on behalf of the Wellcome Sanger Institute COVID-19 Surveillance Team |
| EPI_ISL_2915482 | Lighthouse Lab in Glasgow | Wellcome Sanger Institute for the COVID-19 Genomics UK (COG-UK) Consortium | Harper VanSteenhouse, Yumi Kasai, David Gray, Carol Clugston, Anna Dominiczak and Alex Alderton, Roberto Amato, Jeffrey Barrett, Sonia Goncalves, Ewan Harrison, David K. Jackson, Ian Johnston, Dominic Kwiatkowski, Cordelia Langford, John Sillitoe on behalf of the Wellcome Sanger Institute COVID-19 Surveillance Team |
| EPI_ISL_2915483 | Berkshire and Surrey Pathology Services Lighthouse Laboratory | Wellcome Sanger Institute for the COVID-19 Genomics UK (COG-UK) Consortium | Berkshire and Surrey Pathology Services Lighthouse Laboratory and Alex Alderton, Roberto Amato, Jeffrey Barrett, Sonia Goncalves, Ewan Harrison, David K. Jackson, Ian Johnston, Dominic Kwiatkowski, Cordelia Langford, John Sillitoe on behalf of the Wellcome Sanger Institute COVID-19 Surveillance Team |
| EPI_ISL_2915484, EPI_ISL_2915485 | Lighthouse Laboratory Plymouth | Wellcome Sanger Institute for the COVID-19 Genomics UK (COG-UK) Consortium | Lighthouse Laboratory Plymouth and Alex Alderton, Roberto Amato, Jeffrey Barrett, Sonia Goncalves, Ewan Harrison, David K. Jackson, Ian Johnston, Dominic Kwiatkowski, Cordelia Langford, John Sillitoe on behalf of the Wellcome Sanger Institute COVID-19 Surveillance Team |
| EPI_ISL_2915486 | Berkshire and Surrey Pathology Services Lighthouse Laboratory | Wellcome Sanger Institute for the COVID-19 Genomics UK (COG-UK) Consortium | Berkshire and Surrey Pathology Services Lighthouse Laboratory and Alex Alderton, Roberto Amato, Jeffrey Barrett, Sonia Goncalves, Ewan Harrison, David K. Jackson, Ian Johnston, Dominic Kwiatkowski, Cordelia Langford, John Sillitoe on behalf of the Wellcome Sanger Institute COVID-19 Surveillance Team |
| EPI_ISL_2915487 | Lighthouse Lab in Alderley Park | Wellcome Sanger Institute for the COVID-19 Genomics UK (COG-UK) Consortium | Jacquelyn Wynn, Mairead Hyland, The Lighthouse Lab in Alderley Park and Alex Alderton, Roberto Amato, Jeffrey Barrett, Sonia Goncalves, Ewan Harrison, David K. Jackson, Ian Johnston, Dominic Kwiatkowski, Cordelia Langford, John Sillitoe on behalf of the Wellcome Sanger Institute COVID-19 Surveillance Team |
| EPI_ISL_2915488 | Berkshire and Surrey Pathology Services Lighthouse Laboratory | Wellcome Sanger Institute for the COVID-19 Genomics UK (COG-UK) Consortium | Berkshire and Surrey Pathology Services Lighthouse Laboratory and Alex Alderton, Roberto Amato, Jeffrey Barrett, Sonia Goncalves, Ewan Harrison, David K. Jackson, Ian Johnston, Dominic Kwiatkowski, Cordelia Langford, John Sillitoe on behalf of the Wellcome Sanger Institute COVID-19 Surveillance Team |
| EPI_ISL_2915489, EPI_ISL_2915490 | Lighthouse Laboratory Plymouth | Wellcome Sanger Institute for the COVID-19 Genomics UK (COG-UK) Consortium | Lighthouse Laboratory Plymouth and Alex Alderton, Roberto Amato, Jeffrey Barrett, Sonia Goncalves, Ewan Harrison, David K. Jackson, Ian Johnston, Dominic Kwiatkowski, Cordelia Langford, John Sillitoe on behalf of the Wellcome Sanger Institute COVID-19 Surveillance Team |
| EPI_ISL_2915491, EPI_ISL_2915492 | Lighthouse Lab in Glasgow | Wellcome Sanger Institute for the COVID-19 Genomics UK (COG-UK) Consortium | Harper VanSteenhouse, Yumi Kasai, David Gray, Carol Clugston, Anna Dominiczak and Alex Alderton, Roberto Amato, Jeffrey Barrett, Sonia Goncalves, Ewan Harrison, David K. Jackson, Ian Johnston, Dominic Kwiatkowski, Cordelia Langford, John Sillitoe on behalf of the Wellcome Sanger Institute COVID-19 Surveillance Team |
| EPI_ISL_2915493 | Berkshire and Surrey Pathology Services Lighthouse Laboratory | Wellcome Sanger Institute for the COVID-19 Genomics UK (COG-UK) Consortium | Berkshire and Surrey Pathology Services Lighthouse Laboratory and Alex Alderton, Roberto Amato, Jeffrey Barrett, Sonia Goncalves, Ewan Harrison, David K. Jackson, Ian Johnston, Dominic Kwiatkowski, Cordelia Langford, John Sillitoe on behalf of the Wellcome Sanger Institute COVID-19 Surveillance Team |
| EPI_ISL_2915494 | Lighthouse Lab in Alderley Park | Wellcome Sanger Institute for the COVID-19 Genomics UK (COG-UK) Consortium | Jacquelyn Wynn, Mairead Hyland, The Lighthouse Lab in Alderley Park and Alex Alderton, Roberto Amato, Jeffrey Barrett, Sonia Goncalves, Ewan Harrison, David K. Jackson, Ian Johnston, Dominic Kwiatkowski, Cordelia Langford, John Sillitoe on behalf of the Wellcome Sanger Institute COVID-19 Surveillance Team |
| EPI_ISL_2915495 | Lighthouse Laboratory Plymouth | Wellcome Sanger Institute for the COVID-19 Genomics UK (COG-UK) Consortium | Lighthouse Laboratory Plymouth and Alex Alderton, Roberto Amato, Jeffrey Barrett, Sonia Goncalves, Ewan Harrison, David K. Jackson, Ian Johnston, Dominic Kwiatkowski, Cordelia Langford, John Sillitoe on behalf of the Wellcome Sanger Institute COVID-19 Surveillance Team |
| EPI_ISL_2915496, EPI_ISL_2915497 | Lighthouse Lab in Glasgow | Wellcome Sanger Institute for the COVID-19 Genomics UK (COG-UK) Consortium | Harper VanSteenhouse, Yumi Kasai, David Gray, Carol Clugston, Anna Dominiczak and Alex Alderton, Roberto Amato, Jeffrey Barrett, Sonia Goncalves, Ewan Harrison, David K. Jackson, Ian Johnston, Dominic Kwiatkowski, Cordelia Langford, John Sillitoe on behalf of the Wellcome Sanger Institute COVID-19 Surveillance Team |

| | | | |
|---|---|---|---|
| EPI_ISL_2915498 | Lighthouse Lab in Alderley Park | Wellcome Sanger Institute for the COVID-19 Genomics UK (COG-UK) Consortium | Jacquelyn Wynn, Mairead Hyland, The Lighthouse Lab in Alderley Park and Alex Alderton, Roberto Amato, Jeffrey Barrett, Sonia Goncalves, Ewan Harrison, David K. Jackson, Ian Johnston, Dominic Kwiatkowski, Cordelia Langford, John Sillitoe on behalf of the Wellcome Sanger Institute COVID-19 Surveillance Team |
| EPI_ISL_2915499 | Lighthouse Lab in Glasgow | Wellcome Sanger Institute for the COVID-19 Genomics UK (COG-UK) Consortium | Harper VanSteenhouse, Yumi Kasai, David Gray, Carol Clugston, Anna Dominiczak and Alex Alderton, Roberto Amato, Jeffrey Barrett, Sonia Goncalves, Ewan Harrison, David K. Jackson, Ian Johnston, Dominic Kwiatkowski, Cordelia Langford, John Sillitoe on behalf of the Wellcome Sanger Institute COVID-19 Surveillance Team |
| EPI_ISL_2915500 | Berkshire and Surrey Pathology Services Lighthouse Laboratory | Wellcome Sanger Institute for the COVID-19 Genomics UK (COG-UK) Consortium | Berkshire and Surrey Pathology Services Lighthouse Laboratory and Alex Alderton, Roberto Amato, Jeffrey Barrett, Sonia Goncalves, Ewan Harrison, David K. Jackson, Ian Johnston, Dominic Kwiatkowski, Cordelia Langford, John Sillitoe on behalf of the Wellcome Sanger Institute COVID-19 Surveillance Team |
| EPI_ISL_2915501 | Lighthouse Laboratory Plymouth | Wellcome Sanger Institute for the COVID-19 Genomics UK (COG-UK) Consortium | Lighthouse Laboratory Plymouth and Alex Alderton, Roberto Amato, Jeffrey Barrett, Sonia Goncalves, Ewan Harrison, David K. Jackson, Ian Johnston, Dominic Kwiatkowski, Cordelia Langford, John Sillitoe on behalf of the Wellcome Sanger Institute COVID-19 Surveillance Team |
| EPI_ISL_2915502 | Lighthouse Lab in Glasgow | Wellcome Sanger Institute for the COVID-19 Genomics UK (COG-UK) Consortium | Harper VanSteenhouse, Yumi Kasai, David Gray, Carol Clugston, Anna Dominiczak and Alex Alderton, Roberto Amato, Jeffrey Barrett, Sonia Goncalves, Ewan Harrison, David K. Jackson, Ian Johnston, Dominic Kwiatkowski, Cordelia Langford, John Sillitoe on behalf of the Wellcome Sanger Institute COVID-19 Surveillance Team |
| EPI_ISL_2915503 | Lighthouse Lab in Alderley Park | Wellcome Sanger Institute for the COVID-19 Genomics UK (COG-UK) Consortium | Jacquelyn Wynn, Mairead Hyland, The Lighthouse Lab in Alderley Park and Alex Alderton, Roberto Amato, Jeffrey Barrett, Sonia Goncalves, Ewan Harrison, David K. Jackson, Ian Johnston, Dominic Kwiatkowski, Cordelia Langford, John Sillitoe on behalf of the Wellcome Sanger Institute COVID-19 Surveillance Team |
| EPI_ISL_2915504 | Berkshire and Surrey Pathology Services Lighthouse Laboratory | Wellcome Sanger Institute for the COVID-19 Genomics UK (COG-UK) Consortium | Berkshire and Surrey Pathology Services Lighthouse Laboratory and Alex Alderton, Roberto Amato, Jeffrey Barrett, Sonia Goncalves, Ewan Harrison, David K. Jackson, Ian Johnston, Dominic Kwiatkowski, Cordelia Langford, John Sillitoe on behalf of the Wellcome Sanger Institute COVID-19 Surveillance Team |
| EPI_ISL_2915505 | Lighthouse Lab in Glasgow | Wellcome Sanger Institute for the COVID-19 Genomics UK (COG-UK) Consortium | Harper VanSteenhouse, Yumi Kasai, David Gray, Carol Clugston, Anna Dominiczak and Alex Alderton, Roberto Amato, Jeffrey Barrett, Sonia Goncalves, Ewan Harrison, David K. Jackson, Ian Johnston, Dominic Kwiatkowski, Cordelia Langford, John Sillitoe on behalf of the Wellcome Sanger Institute COVID-19 Surveillance Team |
| EPI_ISL_2915506, EPI_ISL_2915507 | Lighthouse Laboratory Plymouth | Wellcome Sanger Institute for the COVID-19 Genomics UK (COG-UK) Consortium | Lighthouse Laboratory Plymouth and Alex Alderton, Roberto Amato, Jeffrey Barrett, Sonia Goncalves, Ewan Harrison, David K. Jackson, Ian Johnston, Dominic Kwiatkowski, Cordelia Langford, John Sillitoe on behalf of the Wellcome Sanger Institute COVID-19 Surveillance Team |
| EPI_ISL_2915508 | Lighthouse Lab in Glasgow | Wellcome Sanger Institute for the COVID-19 Genomics UK (COG-UK) Consortium | Harper VanSteenhouse, Yumi Kasai, David Gray, Carol Clugston, Anna Dominiczak and Alex Alderton, Roberto Amato, Jeffrey Barrett, Sonia Goncalves, Ewan Harrison, David K. Jackson, Ian Johnston, Dominic Kwiatkowski, Cordelia Langford, John Sillitoe on behalf of the Wellcome Sanger Institute COVID-19 Surveillance Team |
| EPI_ISL_2915509, EPI_ISL_2915510 | Lighthouse Lab in Alderley Park | Wellcome Sanger Institute for the COVID-19 Genomics UK (COG-UK) Consortium | Jacquelyn Wynn, Mairead Hyland, The Lighthouse Lab in Alderley Park and Alex Alderton, Roberto Amato, Jeffrey Barrett, Sonia Goncalves, Ewan Harrison, David K. Jackson, Ian Johnston, Dominic Kwiatkowski, Cordelia Langford, John Sillitoe on behalf of the Wellcome Sanger Institute COVID-19 Surveillance Team |
| EPI_ISL_2915511 | Lighthouse Laboratory Plymouth | Wellcome Sanger Institute for the COVID-19 Genomics UK (COG-UK) Consortium | Lighthouse Laboratory Plymouth and Alex Alderton, Roberto Amato, Jeffrey Barrett, Sonia Goncalves, Ewan Harrison, David K. Jackson, Ian Johnston, Dominic Kwiatkowski, Cordelia Langford, John Sillitoe on behalf of the Wellcome Sanger Institute COVID-19 Surveillance Team |
| EPI_ISL_2915512 | Berkshire and Surrey Pathology Services Lighthouse Laboratory | Wellcome Sanger Institute for the COVID-19 Genomics UK (COG-UK) Consortium | Berkshire and Surrey Pathology Services Lighthouse Laboratory and Alex Alderton, Roberto Amato, Jeffrey Barrett, Sonia Goncalves, Ewan Harrison, David K. Jackson, Ian Johnston, Dominic Kwiatkowski, Cordelia Langford, John Sillitoe on behalf of the Wellcome Sanger Institute COVID-19 Surveillance Team |
| EPI_ISL_2915513 | Lighthouse Laboratory Plymouth | Wellcome Sanger Institute for the COVID-19 Genomics UK (COG-UK) Consortium | Lighthouse Laboratory Plymouth and Alex Alderton, Roberto Amato, Jeffrey Barrett, Sonia Goncalves, Ewan Harrison, David K. Jackson, Ian Johnston, Dominic Kwiatkowski, Cordelia Langford, John Sillitoe on behalf of the Wellcome Sanger Institute COVID-19 Surveillance Team |
| EPI_ISL_2915514 | Lighthouse Lab in Glasgow | Wellcome Sanger Institute for the COVID-19 Genomics UK (COG-UK) Consortium | Harper VanSteenhouse, Yumi Kasai, David Gray, Carol Clugston, Anna Dominiczak and Alex Alderton, Roberto Amato, Jeffrey Barrett, Sonia Goncalves, Ewan Harrison, David K. Jackson, Ian Johnston, Dominic Kwiatkowski, Cordelia Langford, John Sillitoe on behalf of the Wellcome Sanger Institute COVID-19 Surveillance Team |
| EPI_ISL_2915515 | Lighthouse Laboratory Plymouth | Wellcome Sanger Institute for the COVID-19 Genomics UK (COG-UK) Consortium | Lighthouse Laboratory Plymouth and Alex Alderton, Roberto Amato, Jeffrey Barrett, Sonia Goncalves, Ewan Harrison, David K. Jackson, Ian Johnston, Dominic Kwiatkowski, Cordelia Langford, John Sillitoe on behalf of the Wellcome Sanger Institute COVID-19 Surveillance Team |
| EPI_ISL_2915516 | Lighthouse Lab in Alderley Park | Wellcome Sanger Institute for the COVID-19 Genomics UK (COG-UK) Consortium | Jacquelyn Wynn, Mairead Hyland, The Lighthouse Lab in Alderley Park and Alex Alderton, Roberto Amato, Jeffrey Barrett, Sonia Goncalves, Ewan Harrison, David K. Jackson, Ian Johnston, Dominic Kwiatkowski, Cordelia Langford, John Sillitoe on behalf of the Wellcome Sanger Institute COVID-19 Surveillance Team |
| EPI_ISL_2915517 | Lighthouse Lab in Glasgow | Wellcome Sanger Institute for the COVID-19 Genomics UK (COG-UK) Consortium | Harper VanSteenhouse, Yumi Kasai, David Gray, Carol Clugston, Anna Dominiczak and Alex Alderton, Roberto Amato, Jeffrey Barrett, Sonia Goncalves, Ewan Harrison, David K. Jackson, Ian Johnston, Dominic Kwiatkowski, Cordelia Langford, John Sillitoe on behalf of the Wellcome Sanger Institute COVID-19 Surveillance Team |
| EPI_ISL_2915518 | Berkshire and Surrey Pathology Services Lighthouse Laboratory | Wellcome Sanger Institute for the COVID-19 Genomics UK (COG-UK) Consortium | Berkshire and Surrey Pathology Services Lighthouse Laboratory and Alex Alderton, Roberto Amato, Jeffrey Barrett, Sonia Goncalves, Ewan Harrison, David K. Jackson, Ian Johnston, Dominic Kwiatkowski, Cordelia Langford, John Sillitoe on behalf of the Wellcome Sanger Institute COVID-19 Surveillance Team |
| EPI_ISL_2915519 | Lighthouse Laboratory Plymouth | Wellcome Sanger Institute for the COVID-19 Genomics UK (COG-UK) Consortium | Lighthouse Laboratory Plymouth and Alex Alderton, Roberto Amato, Jeffrey Barrett, Sonia Goncalves, Ewan Harrison, David K. Jackson, Ian Johnston, Dominic Kwiatkowski, Cordelia Langford, John Sillitoe on behalf of the Wellcome Sanger Institute COVID-19 Surveillance Team |
| EPI_ISL_2915520 | Berkshire and Surrey Pathology Services Lighthouse Laboratory | Wellcome Sanger Institute for the COVID-19 Genomics UK (COG-UK) Consortium | Berkshire and Surrey Pathology Services Lighthouse Laboratory and Alex Alderton, Roberto Amato, Jeffrey Barrett, Sonia Goncalves, Ewan Harrison, David K. Jackson, Ian Johnston, Dominic Kwiatkowski, Cordelia Langford, John Sillitoe on behalf of the Wellcome Sanger Institute COVID-19 Surveillance Team |
| EPI_ISL_2915521 | Lighthouse Lab in Glasgow | Wellcome Sanger Institute for the COVID-19 Genomics UK (COG-UK) Consortium | Harper VanSteenhouse, Yumi Kasai, David Gray, Carol Clugston, Anna Dominiczak and Alex Alderton, Roberto Amato, Jeffrey Barrett, Sonia Goncalves, Ewan Harrison, David K. Jackson, Ian Johnston, Dominic Kwiatkowski, Cordelia Langford, John Sillitoe on behalf of the Wellcome Sanger Institute COVID-19 Surveillance Team |
| EPI_ISL_2915522 | Lighthouse Laboratory Plymouth | Wellcome Sanger Institute for the COVID-19 Genomics UK (COG-UK) Consortium | Lighthouse Laboratory Plymouth and Alex Alderton, Roberto Amato, Jeffrey Barrett, Sonia Goncalves, Ewan Harrison, David K. Jackson, Ian Johnston, Dominic Kwiatkowski, Cordelia Langford, John Sillitoe on behalf of the Wellcome Sanger Institute COVID-19 Surveillance Team |
| EPI_ISL_2915523, EPI_ISL_2915524, EPI_ISL_2915525, EPI_ISL_2915526 | Berkshire and Surrey Pathology Services Lighthouse Laboratory | Wellcome Sanger Institute for the COVID-19 Genomics UK (COG-UK) Consortium | Berkshire and Surrey Pathology Services Lighthouse Laboratory and Alex Alderton, Roberto Amato, Jeffrey Barrett, Sonia Goncalves, Ewan Harrison, David K. Jackson, Ian Johnston, Dominic Kwiatkowski, Cordelia Langford, John Sillitoe on behalf of the Wellcome Sanger Institute COVID-19 Surveillance Team |
| EPI_ISL_2915527 | Lighthouse Lab in Alderley Park | Wellcome Sanger Institute for the COVID-19 Genomics UK (COG-UK) Consortium | Jacquelyn Wynn, Mairead Hyland, The Lighthouse Lab in Alderley Park and Alex Alderton, Roberto Amato, Jeffrey Barrett, Sonia Goncalves, Ewan Harrison, David K. Jackson, Ian Johnston, Dominic Kwiatkowski, Cordelia Langford, John Sillitoe on behalf of the Wellcome Sanger Institute COVID-19 Surveillance Team |
| EPI_ISL_2915528 | Lighthouse Lab in Glasgow | Wellcome Sanger Institute for the COVID-19 Genomics UK (COG-UK) Consortium | Harper VanSteenhouse, Yumi Kasai, David Gray, Carol Clugston, Anna Dominiczak and Alex Alderton, Roberto Amato, Jeffrey Barrett, Sonia Goncalves, Ewan Harrison, David K. Jackson, Ian Johnston, Dominic Kwiatkowski, Cordelia Langford, John Sillitoe on behalf of the Wellcome Sanger Institute COVID-19 Surveillance Team |
| EPI_ISL_2915529 | Berkshire and Surrey Pathology Services Lighthouse Laboratory | Wellcome Sanger Institute for the COVID-19 Genomics UK (COG-UK) Consortium | Berkshire and Surrey Pathology Services Lighthouse Laboratory and Alex Alderton, Roberto Amato, Jeffrey Barrett, Sonia Goncalves, Ewan Harrison, David K. Jackson, Ian Johnston, Dominic Kwiatkowski, Cordelia Langford, John Sillitoe on behalf of the Wellcome Sanger Institute COVID-19 Surveillance Team |
| EPI_ISL_2915530 | Lighthouse Lab in Glasgow | Wellcome Sanger Institute for the COVID-19 Genomics UK (COG-UK) Consortium | Harper VanSteenhouse, Yumi Kasai, David Gray, Carol Clugston, Anna Dominiczak and Alex Alderton, Roberto Amato, Jeffrey Barrett, Sonia Goncalves, Ewan Harrison, David K. Jackson, Ian Johnston, Dominic Kwiatkowski, Cordelia Langford, John Sillitoe on behalf of the Wellcome Sanger Institute COVID-19 Surveillance Team |
| EPI_ISL_2915531 | Berkshire and Surrey Pathology Services Lighthouse Laboratory | Wellcome Sanger Institute for the COVID-19 Genomics UK (COG-UK) Consortium | Berkshire and Surrey Pathology Services Lighthouse Laboratory and Alex Alderton, Roberto Amato, Jeffrey Barrett, Sonia Goncalves, Ewan Harrison, David K. Jackson, Ian Johnston, Dominic Kwiatkowski, Cordelia Langford, John Sillitoe on behalf of the Wellcome Sanger Institute COVID-19 Surveillance Team |
| EPI_ISL_2915532 | Lighthouse Laboratory Plymouth | Wellcome Sanger Institute for the COVID-19 Genomics UK (COG-UK) Consortium | Lighthouse Laboratory Plymouth and Alex Alderton, Roberto Amato, Jeffrey Barrett, Sonia Goncalves, Ewan Harrison, David K. Jackson, Ian Johnston, Dominic Kwiatkowski, Cordelia Langford, John Sillitoe on behalf of the Wellcome Sanger Institute COVID-19 Surveillance Team |
| EPI_ISL_2915533, EPI_ISL_2915534 | Berkshire and Surrey Pathology Services Lighthouse Laboratory | Wellcome Sanger Institute for the COVID-19 Genomics UK (COG-UK) Consortium | Berkshire and Surrey Pathology Services Lighthouse Laboratory and Alex Alderton, Roberto Amato, Jeffrey Barrett, Sonia Goncalves, Ewan Harrison, David K. Jackson, Ian Johnston, Dominic Kwiatkowski, Cordelia Langford, John Sillitoe on behalf of the Wellcome Sanger Institute COVID-19 Surveillance Team |
| EPI_ISL_2915535, EPI_ISL_2915536 | Lighthouse Lab in Alderley Park | Wellcome Sanger Institute for the COVID-19 Genomics UK (COG-UK) Consortium | Jacquelyn Wynn, Mairead Hyland, The Lighthouse Lab in Alderley Park and Alex Alderton, Roberto Amato, Jeffrey Barrett, Sonia Goncalves, Ewan Harrison, David K. Jackson, Ian Johnston, Dominic Kwiatkowski, Cordelia Langford, John Sillitoe on behalf of the Wellcome Sanger Institute COVID-19 Surveillance Team |
| EPI_ISL_2915537 | Berkshire and Surrey Pathology Services Lighthouse Laboratory | Wellcome Sanger Institute for the COVID-19 Genomics UK (COG-UK) Consortium | Berkshire and Surrey Pathology Services Lighthouse Laboratory and Alex Alderton, Roberto Amato, Jeffrey Barrett, Sonia Goncalves, Ewan Harrison, David K. Jackson, Ian Johnston, Dominic Kwiatkowski, Cordelia Langford, John Sillitoe on behalf of the Wellcome Sanger Institute COVID-19 Surveillance Team |
| EPI_ISL_2915538, EPI_ISL_2915539, EPI_ISL_2915540 | Lighthouse Lab in Alderley Park | Wellcome Sanger Institute for the COVID-19 Genomics UK (COG-UK) Consortium | Jacquelyn Wynn, Mairead Hyland, The Lighthouse Lab in Alderley Park and Alex Alderton, Roberto Amato, Jeffrey Barrett, Sonia Goncalves, Ewan Harrison, David K. Jackson, Ian Johnston, Dominic Kwiatkowski, Cordelia Langford, John Sillitoe on behalf of the Wellcome Sanger Institute COVID-19 Surveillance Team |
| EPI_ISL_2915541 | Lighthouse Lab in Glasgow | Wellcome Sanger Institute for the COVID-19 Genomics UK (COG-UK) Consortium | Harper VanSteenhouse, Yumi Kasai, David Gray, Carol Clugston, Anna Dominiczak and Alex Alderton, Roberto Amato, Jeffrey Barrett, Sonia Goncalves, Ewan Harrison, David K. Jackson, Ian Johnston, Dominic Kwiatkowski, Cordelia Langford, John Sillitoe on behalf of the Wellcome Sanger Institute COVID-19 Surveillance Team |
| EPI_ISL_2915542 | Berkshire and Surrey Pathology Services Lighthouse Laboratory | Wellcome Sanger Institute for the COVID-19 Genomics UK (COG-UK) Consortium | Berkshire and Surrey Pathology Services Lighthouse Laboratory and Alex Alderton, Roberto Amato, Jeffrey Barrett, Sonia Goncalves, Ewan Harrison, David K. Jackson, Ian Johnston, Dominic Kwiatkowski, Cordelia Langford, John Sillitoe on behalf of the Wellcome Sanger Institute COVID-19 Surveillance Team |
| EPI_ISL_2915543 | Lighthouse Laboratory Plymouth | Wellcome Sanger Institute for the COVID-19 Genomics UK (COG-UK) Consortium | Lighthouse Laboratory Plymouth and Alex Alderton, Roberto Amato, Jeffrey Barrett, Sonia Goncalves, Ewan Harrison, David K. Jackson, Ian Johnston, Dominic Kwiatkowski, Cordelia Langford, John Sillitoe on behalf of the Wellcome Sanger Institute COVID-19 Surveillance Team |
| EPI_ISL_2915544 | Berkshire and Surrey Pathology Services Lighthouse Laboratory | Wellcome Sanger Institute for the COVID-19 Genomics UK (COG-UK) Consortium | Berkshire and Surrey Pathology Services Lighthouse Laboratory and Alex Alderton, Roberto Amato, Jeffrey Barrett, Sonia Goncalves, Ewan Harrison, David K. Jackson, Ian Johnston, Dominic Kwiatkowski, Cordelia Langford, John Sillitoe on behalf of the Wellcome Sanger Institute COVID-19 Surveillance Team |
| EPI_ISL_2915545 | Lighthouse Lab in Alderley Park | Wellcome Sanger Institute for the COVID-19 Genomics UK (COG-UK) Consortium | Jacquelyn Wynn, Mairead Hyland, The Lighthouse Lab in Alderley Park and Alex Alderton, Roberto Amato, Jeffrey Barrett, Sonia Goncalves, Ewan Harrison, David K. Jackson, Ian Johnston, Dominic Kwiatkowski, Cordelia Langford, John Sillitoe on behalf of the Wellcome Sanger Institute COVID-19 Surveillance Team |
| EPI_ISL_2915546 | Lighthouse Lab in Glasgow | Wellcome Sanger Institute for the COVID-19 Genomics UK (COG-UK) Consortium | Harper VanSteenhouse, Yumi Kasai, David Gray, Carol Clugston, Anna Dominiczak and Alex Alderton, Roberto Amato, Jeffrey Barrett, Sonia Goncalves, Ewan Harrison, David K. Jackson, Ian Johnston, Dominic Kwiatkowski, Cordelia Langford, John Sillitoe on behalf of the Wellcome Sanger Institute COVID-19 Surveillance Team |
| EPI_ISL_2915547, EPI_ISL_2915548 | Lighthouse Lab in Alderley Park | Wellcome Sanger Institute for the COVID-19 Genomics UK (COG-UK) Consortium | Jacquelyn Wynn, Mairead Hyland, The Lighthouse Lab in Alderley Park and Alex Alderton, Roberto Amato, Jeffrey Barrett, Sonia Goncalves, Ewan Harrison, David K. Jackson, Ian Johnston, Dominic Kwiatkowski, Cordelia Langford, John Sillitoe on behalf of the Wellcome Sanger Institute COVID-19 Surveillance Team |
| EPI_ISL_2915549 | Lighthouse Lab in Glasgow | Wellcome Sanger Institute for the COVID-19 Genomics UK (COG-UK) Consortium | Harper VanSteenhouse, Yumi Kasai, David Gray, Carol Clugston, Anna Dominiczak and Alex Alderton, Roberto Amato, Jeffrey Barrett, Sonia Goncalves, Ewan Harrison, David K. Jackson, Ian Johnston, Dominic Kwiatkowski, Cordelia Langford, John Sillitoe on behalf of the Wellcome Sanger Institute COVID-19 Surveillance Team |
| EPI_ISL_2915550 | Lighthouse Laboratory Plymouth | Wellcome Sanger Institute for the COVID-19 Genomics UK (COG-UK) Consortium | Lighthouse Laboratory Plymouth and Alex Alderton, Roberto Amato, Jeffrey Barrett, Sonia Goncalves, Ewan Harrison, David K. Jackson, Ian Johnston, Dominic Kwiatkowski, Cordelia Langford, John Sillitoe on behalf of the Wellcome Sanger Institute COVID-19 Surveillance Team |
| EPI_ISL_2915551 | Lighthouse Lab in Glasgow | Wellcome Sanger Institute for the COVID-19 Genomics UK (COG-UK) Consortium | Harper VanSteenhouse, Yumi Kasai, David Gray, Carol Clugston, Anna Dominiczak and Alex Alderton, Roberto Amato, Jeffrey Barrett, Sonia Goncalves, Ewan Harrison, David K. Jackson, Ian Johnston, Dominic Kwiatkowski, Cordelia Langford, John Sillitoe on behalf of the Wellcome Sanger Institute COVID-19 Surveillance Team |

| | | | |
|---|---|---|---|
| EPI_ISL_2915552 | Berkshire and Surrey Pathology Services Lighthouse Laboratory | Wellcome Sanger Institute for the COVID-19 Genomics UK (COG-UK) Consortium | Berkshire and Surrey Pathology Services Lighthouse Laboratory and Alex Alderton, Roberto Amato, Jeffrey Barrett, Sonia Goncalves, Ewan Harrison, David K. Jackson, Ian Johnston, Dominic Kwiatkowski, Cordelia Langford, John Sillitoe on behalf of the Wellcome Sanger Institute COVID-19 Surveillance Team |
| EPI_ISL_2915553 | Lighthouse Lab in Alderley Park | Wellcome Sanger Institute for the COVID-19 Genomics UK (COG-UK) Consortium | Jacquelyn Wynn, Mairead Hyland, The Lighthouse Lab in Alderley Park and Alex Alderton, Roberto Amato, Jeffrey Barrett, Sonia Goncalves, Ewan Harrison, David K. Jackson, Ian Johnston, Dominic Kwiatkowski, Cordelia Langford, John Sillitoe on behalf of the Wellcome Sanger Institute COVID-19 Surveillance Team |
| EPI_ISL_2915554 | Lighthouse Lab in Glasgow | Wellcome Sanger Institute for the COVID-19 Genomics UK (COG-UK) Consortium | Harper VanSteenhouse, Yumi Kasai, David Gray, Carol Clugston, Anna Dominiczak and Alex Alderton, Roberto Amato, Jeffrey Barrett, Sonia Goncalves, Ewan Harrison, David K. Jackson, Ian Johnston, Dominic Kwiatkowski, Cordelia Langford, John Sillitoe on behalf of the Wellcome Sanger Institute COVID-19 Surveillance Team |
| EPI_ISL_2915555 | Lighthouse Lab in Alderley Park | Wellcome Sanger Institute for the COVID-19 Genomics UK (COG-UK) Consortium | Jacquelyn Wynn, Mairead Hyland, The Lighthouse Lab in Alderley Park and Alex Alderton, Roberto Amato, Jeffrey Barrett, Sonia Goncalves, Ewan Harrison, David K. Jackson, Ian Johnston, Dominic Kwiatkowski, Cordelia Langford, John Sillitoe on behalf of the Wellcome Sanger Institute COVID-19 Surveillance Team |
| EPI_ISL_2915556 | Berkshire and Surrey Pathology Services Lighthouse Laboratory | Wellcome Sanger Institute for the COVID-19 Genomics UK (COG-UK) Consortium | Berkshire and Surrey Pathology Services Lighthouse Laboratory and Alex Alderton, Roberto Amato, Jeffrey Barrett, Sonia Goncalves, Ewan Harrison, David K. Jackson, Ian Johnston, Dominic Kwiatkowski, Cordelia Langford, John Sillitoe on behalf of the Wellcome Sanger Institute COVID-19 Surveillance Team |
| EPI_ISL_2915557 | Lighthouse Lab in Alderley Park | Wellcome Sanger Institute for the COVID-19 Genomics UK (COG-UK) Consortium | Jacquelyn Wynn, Mairead Hyland, The Lighthouse Lab in Alderley Park and Alex Alderton, Roberto Amato, Jeffrey Barrett, Sonia Goncalves, Ewan Harrison, David K. Jackson, Ian Johnston, Dominic Kwiatkowski, Cordelia Langford, John Sillitoe on behalf of the Wellcome Sanger Institute COVID-19 Surveillance Team |
| EPI_ISL_2915558 | Lighthouse Laboratory Plymouth | Wellcome Sanger Institute for the COVID-19 Genomics UK (COG-UK) Consortium | Lighthouse Laboratory Plymouth and Alex Alderton, Roberto Amato, Jeffrey Barrett, Sonia Goncalves, Ewan Harrison, David K. Jackson, Ian Johnston, Dominic Kwiatkowski, Cordelia Langford, John Sillitoe on behalf of the Wellcome Sanger Institute COVID-19 Surveillance Team |
| EPI_ISL_2915559 | Lighthouse Lab in Alderley Park | Wellcome Sanger Institute for the COVID-19 Genomics UK (COG-UK) Consortium | Jacquelyn Wynn, Mairead Hyland, The Lighthouse Lab in Alderley Park and Alex Alderton, Roberto Amato, Jeffrey Barrett, Sonia Goncalves, Ewan Harrison, David K. Jackson, Ian Johnston, Dominic Kwiatkowski, Cordelia Langford, John Sillitoe on behalf of the Wellcome Sanger Institute COVID-19 Surveillance Team |
| EPI_ISL_2915560 | Lighthouse Laboratory Plymouth | Wellcome Sanger Institute for the COVID-19 Genomics UK (COG-UK) Consortium | Lighthouse Laboratory Plymouth and Alex Alderton, Roberto Amato, Jeffrey Barrett, Sonia Goncalves, Ewan Harrison, David K. Jackson, Ian Johnston, Dominic Kwiatkowski, Cordelia Langford, John Sillitoe on behalf of the Wellcome Sanger Institute COVID-19 Surveillance Team |
| EPI_ISL_2915561 | Lighthouse Lab in Glasgow | Wellcome Sanger Institute for the COVID-19 Genomics UK (COG-UK) Consortium | Harper VanSteenhouse, Yumi Kasai, David Gray, Carol Clugston, Anna Dominiczak and Alex Alderton, Roberto Amato, Jeffrey Barrett, Sonia Goncalves, Ewan Harrison, David K. Jackson, Ian Johnston, Dominic Kwiatkowski, Cordelia Langford, John Sillitoe on behalf of the Wellcome Sanger Institute COVID-19 Surveillance Team |
| EPI_ISL_2915562 | Lighthouse Lab in Alderley Park | Wellcome Sanger Institute for the COVID-19 Genomics UK (COG-UK) Consortium | Jacquelyn Wynn, Mairead Hyland, The Lighthouse Lab in Alderley Park and Alex Alderton, Roberto Amato, Jeffrey Barrett, Sonia Goncalves, Ewan Harrison, David K. Jackson, Ian Johnston, Dominic Kwiatkowski, Cordelia Langford, John Sillitoe on behalf of the Wellcome Sanger Institute COVID-19 Surveillance Team |
| EPI_ISL_2915563 | Lighthouse Laboratory Plymouth | Wellcome Sanger Institute for the COVID-19 Genomics UK (COG-UK) Consortium | Lighthouse Laboratory Plymouth and Alex Alderton, Roberto Amato, Jeffrey Barrett, Sonia Goncalves, Ewan Harrison, David K. Jackson, Ian Johnston, Dominic Kwiatkowski, Cordelia Langford, John Sillitoe on behalf of the Wellcome Sanger Institute COVID-19 Surveillance Team |
| EPI_ISL_2915564 | Lighthouse Lab in Glasgow | Wellcome Sanger Institute for the COVID-19 Genomics UK (COG-UK) Consortium | Harper VanSteenhouse, Yumi Kasai, David Gray, Carol Clugston, Anna Dominiczak and Alex Alderton, Roberto Amato, Jeffrey Barrett, Sonia Goncalves, Ewan Harrison, David K. Jackson, Ian Johnston, Dominic Kwiatkowski, Cordelia Langford, John Sillitoe on behalf of the Wellcome Sanger Institute COVID-19 Surveillance Team |
| EPI_ISL_2915565 | Lighthouse Lab in Alderley Park | Wellcome Sanger Institute for the COVID-19 Genomics UK (COG-UK) Consortium | Jacquelyn Wynn, Mairead Hyland, The Lighthouse Lab in Alderley Park and Alex Alderton, Roberto Amato, Jeffrey Barrett, Sonia Goncalves, Ewan Harrison, David K. Jackson, Ian Johnston, Dominic Kwiatkowski, Cordelia Langford, John Sillitoe on behalf of the Wellcome Sanger Institute COVID-19 Surveillance Team |
| EPI_ISL_2915566, EPI_ISL_2915567 | Lighthouse Lab in Glasgow | Wellcome Sanger Institute for the COVID-19 Genomics UK (COG-UK) Consortium | Harper VanSteenhouse, Yumi Kasai, David Gray, Carol Clugston, Anna Dominiczak and Alex Alderton, Roberto Amato, Jeffrey Barrett, Sonia Goncalves, Ewan Harrison, David K. Jackson, Ian Johnston, Dominic Kwiatkowski, Cordelia Langford, John Sillitoe on behalf of the Wellcome Sanger Institute COVID-19 Surveillance Team |
| EPI_ISL_2915568, EPI_ISL_2915569 | Lighthouse Lab in Alderley Park | Wellcome Sanger Institute for the COVID-19 Genomics UK (COG-UK) Consortium | Jacquelyn Wynn, Mairead Hyland, The Lighthouse Lab in Alderley Park and Alex Alderton, Roberto Amato, Jeffrey Barrett, Sonia Goncalves, Ewan Harrison, David K. Jackson, Ian Johnston, Dominic Kwiatkowski, Cordelia Langford, John Sillitoe on behalf of the Wellcome Sanger Institute COVID-19 Surveillance Team |
| EPI_ISL_2915570 | Lighthouse Laboratory Plymouth | Wellcome Sanger Institute for the COVID-19 Genomics UK (COG-UK) Consortium | Lighthouse Laboratory Plymouth and Alex Alderton, Roberto Amato, Jeffrey Barrett, Sonia Goncalves, Ewan Harrison, David K. Jackson, Ian Johnston, Dominic Kwiatkowski, Cordelia Langford, John Sillitoe on behalf of the Wellcome Sanger Institute COVID-19 Surveillance Team |
| EPI_ISL_2915571 | Lighthouse Lab in Alderley Park | Wellcome Sanger Institute for the COVID-19 Genomics UK (COG-UK) Consortium | Jacquelyn Wynn, Mairead Hyland, The Lighthouse Lab in Alderley Park and Alex Alderton, Roberto Amato, Jeffrey Barrett, Sonia Goncalves, Ewan Harrison, David K. Jackson, Ian Johnston, Dominic Kwiatkowski, Cordelia Langford, John Sillitoe on behalf of the Wellcome Sanger Institute COVID-19 Surveillance Team |
| EPI_ISL_2915572 | Lighthouse Lab in Glasgow | Wellcome Sanger Institute for the COVID-19 Genomics UK (COG-UK) Consortium | Harper VanSteenhouse, Yumi Kasai, David Gray, Carol Clugston, Anna Dominiczak and Alex Alderton, Roberto Amato, Jeffrey Barrett, Sonia Goncalves, Ewan Harrison, David K. Jackson, Ian Johnston, Dominic Kwiatkowski, Cordelia Langford, John Sillitoe on behalf of the Wellcome Sanger Institute COVID-19 Surveillance Team |
| EPI_ISL_2915573, EPI_ISL_2915574 | Berkshire and Surrey Pathology Services Lighthouse Laboratory | Wellcome Sanger Institute for the COVID-19 Genomics UK (COG-UK) Consortium | Berkshire and Surrey Pathology Services Lighthouse Laboratory and Alex Alderton, Roberto Amato, Jeffrey Barrett, Sonia Goncalves, Ewan Harrison, David K. Jackson, Ian Johnston, Dominic Kwiatkowski, Cordelia Langford, John Sillitoe on behalf of the Wellcome Sanger Institute COVID-19 Surveillance Team |
| EPI_ISL_2915575 | Lighthouse Laboratory Plymouth | Wellcome Sanger Institute for the COVID-19 Genomics UK (COG-UK) Consortium | Lighthouse Laboratory Plymouth and Alex Alderton, Roberto Amato, Jeffrey Barrett, Sonia Goncalves, Ewan Harrison, David K. Jackson, Ian Johnston, Dominic Kwiatkowski, Cordelia Langford, John Sillitoe on behalf of the Wellcome Sanger Institute COVID-19 Surveillance Team |
| EPI_ISL_2915576 | Lighthouse Lab in Alderley Park | Wellcome Sanger Institute for the COVID-19 Genomics UK (COG-UK) Consortium | Jacquelyn Wynn, Mairead Hyland, The Lighthouse Lab in Alderley Park and Alex Alderton, Roberto Amato, Jeffrey Barrett, Sonia Goncalves, Ewan Harrison, David K. Jackson, Ian Johnston, Dominic Kwiatkowski, Cordelia Langford, John Sillitoe on behalf of the Wellcome Sanger Institute COVID-19 Surveillance Team |
| EPI_ISL_2915577 | Berkshire and Surrey Pathology Services Lighthouse Laboratory | Wellcome Sanger Institute for the COVID-19 Genomics UK (COG-UK) Consortium | Berkshire and Surrey Pathology Services Lighthouse Laboratory and Alex Alderton, Roberto Amato, Jeffrey Barrett, Sonia Goncalves, Ewan Harrison, David K. Jackson, Ian Johnston, Dominic Kwiatkowski, Cordelia Langford, John Sillitoe on behalf of the Wellcome Sanger Institute COVID-19 Surveillance Team |
| EPI_ISL_2915578, EPI_ISL_2915579, EPI_ISL_2915580, EPI_ISL_2915581 | Lighthouse Lab in Glasgow | Wellcome Sanger Institute for the COVID-19 Genomics UK (COG-UK) Consortium | Harper VanSteenhouse, Yumi Kasai, David Gray, Carol Clugston, Anna Dominiczak and Alex Alderton, Roberto Amato, Jeffrey Barrett, Sonia Goncalves, Ewan Harrison, David K. Jackson, Ian Johnston, Dominic Kwiatkowski, Cordelia Langford, John Sillitoe on behalf of the Wellcome Sanger Institute COVID-19 Surveillance Team |
| EPI_ISL_2915582 | Berkshire and Surrey Pathology Services Lighthouse Laboratory | Wellcome Sanger Institute for the COVID-19 Genomics UK (COG-UK) Consortium | Berkshire and Surrey Pathology Services Lighthouse Laboratory and Alex Alderton, Roberto Amato, Jeffrey Barrett, Sonia Goncalves, Ewan Harrison, David K. Jackson, Ian Johnston, Dominic Kwiatkowski, Cordelia Langford, John Sillitoe on behalf of the Wellcome Sanger Institute COVID-19 Surveillance Team |
| EPI_ISL_2915583 | Lighthouse Lab in Glasgow | Wellcome Sanger Institute for the COVID-19 Genomics UK (COG-UK) Consortium | Harper VanSteenhouse, Yumi Kasai, David Gray, Carol Clugston, Anna Dominiczak and Alex Alderton, Roberto Amato, Jeffrey Barrett, Sonia Goncalves, Ewan Harrison, David K. Jackson, Ian Johnston, Dominic Kwiatkowski, Cordelia Langford, John Sillitoe on behalf of the Wellcome Sanger Institute COVID-19 Surveillance Team |
| EPI_ISL_2915584, EPI_ISL_2915585 | Lighthouse Laboratory Plymouth | Wellcome Sanger Institute for the COVID-19 Genomics UK (COG-UK) Consortium | Lighthouse Laboratory Plymouth and Alex Alderton, Roberto Amato, Jeffrey Barrett, Sonia Goncalves, Ewan Harrison, David K. Jackson, Ian Johnston, Dominic Kwiatkowski, Cordelia Langford, John Sillitoe on behalf of the Wellcome Sanger Institute COVID-19 Surveillance Team |
| EPI_ISL_2915586 | Berkshire and Surrey Pathology Services Lighthouse Laboratory | Wellcome Sanger Institute for the COVID-19 Genomics UK (COG-UK) Consortium | Berkshire and Surrey Pathology Services Lighthouse Laboratory and Alex Alderton, Roberto Amato, Jeffrey Barrett, Sonia Goncalves, Ewan Harrison, David K. Jackson, Ian Johnston, Dominic Kwiatkowski, Cordelia Langford, John Sillitoe on behalf of the Wellcome Sanger Institute COVID-19 Surveillance Team |
| EPI_ISL_2915587, EPI_ISL_2915588, EPI_ISL_2915589, EPI_ISL_2915590 | Lighthouse Lab in Glasgow | Wellcome Sanger Institute for the COVID-19 Genomics UK (COG-UK) Consortium | Harper VanSteenhouse, Yumi Kasai, David Gray, Carol Clugston, Anna Dominiczak and Alex Alderton, Roberto Amato, Jeffrey Barrett, Sonia Goncalves, Ewan Harrison, David K. Jackson, Ian Johnston, Dominic Kwiatkowski, Cordelia Langford, John Sillitoe on behalf of the Wellcome Sanger Institute COVID-19 Surveillance Team |
| EPI_ISL_2915591 | Lighthouse Lab in Alderley Park | Wellcome Sanger Institute for the COVID-19 Genomics UK (COG-UK) Consortium | Jacquelyn Wynn, Mairead Hyland, The Lighthouse Lab in Alderley Park and Alex Alderton, Roberto Amato, Jeffrey Barrett, Sonia Goncalves, Ewan Harrison, David K. Jackson, Ian Johnston, Dominic Kwiatkowski, Cordelia Langford, John Sillitoe on behalf of the Wellcome Sanger Institute COVID-19 Surveillance Team |
| EPI_ISL_2915592 | Lighthouse Lab in Glasgow | Wellcome Sanger Institute for the COVID-19 Genomics UK (COG-UK) Consortium | Harper VanSteenhouse, Yumi Kasai, David Gray, Carol Clugston, Anna Dominiczak and Alex Alderton, Roberto Amato, Jeffrey Barrett, Sonia Goncalves, Ewan Harrison, David K. Jackson, Ian Johnston, Dominic Kwiatkowski, Cordelia Langford, John Sillitoe on behalf of the Wellcome Sanger Institute COVID-19 Surveillance Team |
| EPI_ISL_2915593 | Berkshire and Surrey Pathology Services Lighthouse Laboratory | Wellcome Sanger Institute for the COVID-19 Genomics UK (COG-UK) Consortium | Berkshire and Surrey Pathology Services Lighthouse Laboratory and Alex Alderton, Roberto Amato, Jeffrey Barrett, Sonia Goncalves, Ewan Harrison, David K. Jackson, Ian Johnston, Dominic Kwiatkowski, Cordelia Langford, John Sillitoe on behalf of the Wellcome Sanger Institute COVID-19 Surveillance Team |
| EPI_ISL_2915594 | Lighthouse Lab in Glasgow | Wellcome Sanger Institute for the COVID-19 Genomics UK (COG-UK) Consortium | Harper VanSteenhouse, Yumi Kasai, David Gray, Carol Clugston, Anna Dominiczak and Alex Alderton, Roberto Amato, Jeffrey Barrett, Sonia Goncalves, Ewan Harrison, David K. Jackson, Ian Johnston, Dominic Kwiatkowski, Cordelia Langford, John Sillitoe on behalf of the Wellcome Sanger Institute COVID-19 Surveillance Team |
| EPI_ISL_2915595 | Lighthouse Lab in Alderley Park | Wellcome Sanger Institute for the COVID-19 Genomics UK (COG-UK) Consortium | Jacquelyn Wynn, Mairead Hyland, The Lighthouse Lab in Alderley Park and Alex Alderton, Roberto Amato, Jeffrey Barrett, Sonia Goncalves, Ewan Harrison, David K. Jackson, Ian Johnston, Dominic Kwiatkowski, Cordelia Langford, John Sillitoe on behalf of the Wellcome Sanger Institute COVID-19 Surveillance Team |
| EPI_ISL_2915596, EPI_ISL_2915597 | Berkshire and Surrey Pathology Services Lighthouse Laboratory | Wellcome Sanger Institute for the COVID-19 Genomics UK (COG-UK) Consortium | Berkshire and Surrey Pathology Services Lighthouse Laboratory and Alex Alderton, Roberto Amato, Jeffrey Barrett, Sonia Goncalves, Ewan Harrison, David K. Jackson, Ian Johnston, Dominic Kwiatkowski, Cordelia Langford, John Sillitoe on behalf of the Wellcome Sanger Institute COVID-19 Surveillance Team |
| EPI_ISL_2915598 | Lighthouse Lab in Alderley Park | Wellcome Sanger Institute for the COVID-19 Genomics UK (COG-UK) Consortium | Jacquelyn Wynn, Mairead Hyland, The Lighthouse Lab in Alderley Park and Alex Alderton, Roberto Amato, Jeffrey Barrett, Sonia Goncalves, Ewan Harrison, David K. Jackson, Ian Johnston, Dominic Kwiatkowski, Cordelia Langford, John Sillitoe on behalf of the Wellcome Sanger Institute COVID-19 Surveillance Team |
| EPI_ISL_2915599 | Lighthouse Lab in Glasgow | Wellcome Sanger Institute for the COVID-19 Genomics UK (COG-UK) Consortium | Harper VanSteenhouse, Yumi Kasai, David Gray, Carol Clugston, Anna Dominiczak and Alex Alderton, Roberto Amato, Jeffrey Barrett, Sonia Goncalves, Ewan Harrison, David K. Jackson, Ian Johnston, Dominic Kwiatkowski, Cordelia Langford, John Sillitoe on behalf of the Wellcome Sanger Institute COVID-19 Surveillance Team |
| EPI_ISL_2915600 | Lighthouse Laboratory Plymouth | Wellcome Sanger Institute for the COVID-19 Genomics UK (COG-UK) Consortium | Lighthouse Laboratory Plymouth and Alex Alderton, Roberto Amato, Jeffrey Barrett, Sonia Goncalves, Ewan Harrison, David K. Jackson, Ian Johnston, Dominic Kwiatkowski, Cordelia Langford, John Sillitoe on behalf of the Wellcome Sanger Institute COVID-19 Surveillance Team |
| EPI_ISL_2915601 | Lighthouse Lab in Alderley Park | Wellcome Sanger Institute for the COVID-19 Genomics UK (COG-UK) Consortium | Jacquelyn Wynn, Mairead Hyland, The Lighthouse Lab in Alderley Park and Alex Alderton, Roberto Amato, Jeffrey Barrett, Sonia Goncalves, Ewan Harrison, David K. Jackson, Ian Johnston, Dominic Kwiatkowski, Cordelia Langford, John Sillitoe on behalf of the Wellcome Sanger Institute COVID-19 Surveillance Team |
| EPI_ISL_2915602, EPI_ISL_2915603, EPI_ISL_2915604 | Lighthouse Laboratory Plymouth | Wellcome Sanger Institute for the COVID-19 Genomics UK (COG-UK) Consortium | Lighthouse Laboratory Plymouth and Alex Alderton, Roberto Amato, Jeffrey Barrett, Sonia Goncalves, Ewan Harrison, David K. Jackson, Ian Johnston, Dominic Kwiatkowski, Cordelia Langford, John Sillitoe on behalf of the Wellcome Sanger Institute COVID-19 Surveillance Team |
| EPI_ISL_2915605 | Berkshire and Surrey Pathology Services Lighthouse Laboratory | Wellcome Sanger Institute for the COVID-19 Genomics UK (COG-UK) Consortium | Berkshire and Surrey Pathology Services Lighthouse Laboratory and Alex Alderton, Roberto Amato, Jeffrey Barrett, Sonia Goncalves, Ewan Harrison, David K. Jackson, Ian Johnston, Dominic Kwiatkowski, Cordelia Langford, John Sillitoe on behalf of the Wellcome Sanger Institute COVID-19 Surveillance Team |
| EPI_ISL_2915606 | Lighthouse Laboratory Plymouth | Wellcome Sanger Institute for the COVID-19 Genomics UK (COG-UK) Consortium | Lighthouse Laboratory Plymouth and Alex Alderton, Roberto Amato, Jeffrey Barrett, Sonia Goncalves, Ewan Harrison, David K. Jackson, Ian Johnston, Dominic Kwiatkowski, Cordelia Langford, John Sillitoe on behalf of the Wellcome Sanger Institute COVID-19 Surveillance Team |
| EPI_ISL_2915607, EPI_ISL_2915608, | Lighthouse Lab in Glasgow | Wellcome Sanger Institute for the COVID-19 Genomics UK (COG-UK) Consortium | Harper VanSteenhouse, Yumi Kasai, David Gray, Carol Clugston, Anna Dominiczak and Alex Alderton, Roberto Amato, Jeffrey Barrett, Sonia Goncalves, Ewan Harrison, David K. Jackson, Ian Johnston, Dominic Kwiatkowski, Cordelia Langford, John Sillitoe on behalf of the Wellcome Sanger Institute COVID-19 Surveillance Team |

| | | | |
|---|---|---|---|
| EPI_ISL_2915609 | | | |
| EPI_ISL_2915610 | Lighthouse Lab in Alderley Park | Wellcome Sanger Institute for the COVID-19 Genomics UK (COG-UK) Consortium | Jacquelyn Wynn, Mairead Hyland, The Lighthouse Lab in Alderley Park and Alex Alderton, Roberto Amato, Jeffrey Barrett, Sonia Goncalves, Ewan Harrison, David K. Jackson, Ian Johnston, Dominic Kwiatkowski, Cordelia Langford, John Sillitoe on behalf of the Wellcome Sanger Institute COVID-19 Surveillance Team |
| EPI_ISL_2915611 | Berkshire and Surrey Pathology Services Lighthouse Laboratory | Wellcome Sanger Institute for the COVID-19 Genomics UK (COG-UK) Consortium | Berkshire and Surrey Pathology Services Lighthouse Laboratory and Alex Alderton, Roberto Amato, Jeffrey Barrett, Sonia Goncalves, Ewan Harrison, David K. Jackson, Ian Johnston, Dominic Kwiatkowski, Cordelia Langford, John Sillitoe on behalf of the Wellcome Sanger Institute COVID-19 Surveillance Team |
| EPI_ISL_2915612 | Lighthouse Laboratory Plymouth | Wellcome Sanger Institute for the COVID-19 Genomics UK (COG-UK) Consortium | Lighthouse Laboratory Plymouth and Alex Alderton, Roberto Amato, Jeffrey Barrett, Sonia Goncalves, Ewan Harrison, David K. Jackson, Ian Johnston, Dominic Kwiatkowski, Cordelia Langford, John Sillitoe on behalf of the Wellcome Sanger Institute COVID-19 Surveillance Team |
| EPI_ISL_2915613, EPI_ISL_2915614, EPI_ISL_2915615 | Berkshire and Surrey Pathology Services Lighthouse Laboratory | Wellcome Sanger Institute for the COVID-19 Genomics UK (COG-UK) Consortium | Berkshire and Surrey Pathology Services Lighthouse Laboratory and Alex Alderton, Roberto Amato, Jeffrey Barrett, Sonia Goncalves, Ewan Harrison, David K. Jackson, Ian Johnston, Dominic Kwiatkowski, Cordelia Langford, John Sillitoe on behalf of the Wellcome Sanger Institute COVID-19 Surveillance Team |
| EPI_ISL_2915616 | Lighthouse Lab in Glasgow | Wellcome Sanger Institute for the COVID-19 Genomics UK (COG-UK) Consortium | Harper VanSteenhouse, Yumi Kasai, David Gray, Carol Clugston, Anna Dominiczak and Alex Alderton, Roberto Amato, Jeffrey Barrett, Sonia Goncalves, Ewan Harrison, David K. Jackson, Ian Johnston, Dominic Kwiatkowski, Cordelia Langford, John Sillitoe on behalf of the Wellcome Sanger Institute COVID-19 Surveillance Team |
| EPI_ISL_2915617 | Berkshire and Surrey Pathology Services Lighthouse Laboratory | Wellcome Sanger Institute for the COVID-19 Genomics UK (COG-UK) Consortium | Berkshire and Surrey Pathology Services Lighthouse Laboratory and Alex Alderton, Roberto Amato, Jeffrey Barrett, Sonia Goncalves, Ewan Harrison, David K. Jackson, Ian Johnston, Dominic Kwiatkowski, Cordelia Langford, John Sillitoe on behalf of the Wellcome Sanger Institute COVID-19 Surveillance Team |
| EPI_ISL_2915618 | Lighthouse Laboratory Plymouth | Wellcome Sanger Institute for the COVID-19 Genomics UK (COG-UK) Consortium | Lighthouse Laboratory Plymouth and Alex Alderton, Roberto Amato, Jeffrey Barrett, Sonia Goncalves, Ewan Harrison, David K. Jackson, Ian Johnston, Dominic Kwiatkowski, Cordelia Langford, John Sillitoe on behalf of the Wellcome Sanger Institute COVID-19 Surveillance Team |
| EPI_ISL_2915619, EPI_ISL_2915620, EPI_ISL_2915621 | Lighthouse Lab in Glasgow | Wellcome Sanger Institute for the COVID-19 Genomics UK (COG-UK) Consortium | Harper VanSteenhouse, Yumi Kasai, David Gray, Carol Clugston, Anna Dominiczak and Alex Alderton, Roberto Amato, Jeffrey Barrett, Sonia Goncalves, Ewan Harrison, David K. Jackson, Ian Johnston, Dominic Kwiatkowski, Cordelia Langford, John Sillitoe on behalf of the Wellcome Sanger Institute COVID-19 Surveillance Team |
| EPI_ISL_2915622 | Lighthouse Lab in Alderley Park | Wellcome Sanger Institute for the COVID-19 Genomics UK (COG-UK) Consortium | Jacquelyn Wynn, Mairead Hyland, The Lighthouse Lab in Alderley Park and Alex Alderton, Roberto Amato, Jeffrey Barrett, Sonia Goncalves, Ewan Harrison, David K. Jackson, Ian Johnston, Dominic Kwiatkowski, Cordelia Langford, John Sillitoe on behalf of the Wellcome Sanger Institute COVID-19 Surveillance Team |
| EPI_ISL_2915623, EPI_ISL_2915624, EPI_ISL_2915625, EPI_ISL_2915626 | Lighthouse Laboratory Plymouth | Wellcome Sanger Institute for the COVID-19 Genomics UK (COG-UK) Consortium | Lighthouse Laboratory Plymouth and Alex Alderton, Roberto Amato, Jeffrey Barrett, Sonia Goncalves, Ewan Harrison, David K. Jackson, Ian Johnston, Dominic Kwiatkowski, Cordelia Langford, John Sillitoe on behalf of the Wellcome Sanger Institute COVID-19 Surveillance Team |
| EPI_ISL_2915627 | Lighthouse Lab in Glasgow | Wellcome Sanger Institute for the COVID-19 Genomics UK (COG-UK) Consortium | Harper VanSteenhouse, Yumi Kasai, David Gray, Carol Clugston, Anna Dominiczak and Alex Alderton, Roberto Amato, Jeffrey Barrett, Sonia Goncalves, Ewan Harrison, David K. Jackson, Ian Johnston, Dominic Kwiatkowski, Cordelia Langford, John Sillitoe on behalf of the Wellcome Sanger Institute COVID-19 Surveillance Team |
| EPI_ISL_2915628 | Lighthouse Lab in Alderley Park | Wellcome Sanger Institute for the COVID-19 Genomics UK (COG-UK) Consortium | Jacquelyn Wynn, Mairead Hyland, The Lighthouse Lab in Alderley Park and Alex Alderton, Roberto Amato, Jeffrey Barrett, Sonia Goncalves, Ewan Harrison, David K. Jackson, Ian Johnston, Dominic Kwiatkowski, Cordelia Langford, John Sillitoe on behalf of the Wellcome Sanger Institute COVID-19 Surveillance Team |
| EPI_ISL_2915629 | Lighthouse Lab in Glasgow | Wellcome Sanger Institute for the COVID-19 Genomics UK (COG-UK) Consortium | Harper VanSteenhouse, Yumi Kasai, David Gray, Carol Clugston, Anna Dominiczak and Alex Alderton, Roberto Amato, Jeffrey Barrett, Sonia Goncalves, Ewan Harrison, David K. Jackson, Ian Johnston, Dominic Kwiatkowski, Cordelia Langford, John Sillitoe on behalf of the Wellcome Sanger Institute COVID-19 Surveillance Team |
| EPI_ISL_2915630, EPI_ISL_2915631 | Lighthouse Laboratory Plymouth | Wellcome Sanger Institute for the COVID-19 Genomics UK (COG-UK) Consortium | Lighthouse Laboratory Plymouth and Alex Alderton, Roberto Amato, Jeffrey Barrett, Sonia Goncalves, Ewan Harrison, David K. Jackson, Ian Johnston, Dominic Kwiatkowski, Cordelia Langford, John Sillitoe on behalf of the Wellcome Sanger Institute COVID-19 Surveillance Team |
| EPI_ISL_2915632, EPI_ISL_2915633, EPI_ISL_2915634 | Lighthouse Lab in Alderley Park | Wellcome Sanger Institute for the COVID-19 Genomics UK (COG-UK) Consortium | Jacquelyn Wynn, Mairead Hyland, The Lighthouse Lab in Alderley Park and Alex Alderton, Roberto Amato, Jeffrey Barrett, Sonia Goncalves, Ewan Harrison, David K. Jackson, Ian Johnston, Dominic Kwiatkowski, Cordelia Langford, John Sillitoe on behalf of the Wellcome Sanger Institute COVID-19 Surveillance Team |
| EPI_ISL_2915635 | Lighthouse Laboratory Plymouth | Wellcome Sanger Institute for the COVID-19 Genomics UK (COG-UK) Consortium | Lighthouse Laboratory Plymouth and Alex Alderton, Roberto Amato, Jeffrey Barrett, Sonia Goncalves, Ewan Harrison, David K. Jackson, Ian Johnston, Dominic Kwiatkowski, Cordelia Langford, John Sillitoe on behalf of the Wellcome Sanger Institute COVID-19 Surveillance Team |
| EPI_ISL_2915636 | Berkshire and Surrey Pathology Services Lighthouse Laboratory | Wellcome Sanger Institute for the COVID-19 Genomics UK (COG-UK) Consortium | Berkshire and Surrey Pathology Services Lighthouse Laboratory and Alex Alderton, Roberto Amato, Jeffrey Barrett, Sonia Goncalves, Ewan Harrison, David K. Jackson, Ian Johnston, Dominic Kwiatkowski, Cordelia Langford, John Sillitoe on behalf of the Wellcome Sanger Institute COVID-19 Surveillance Team |
| EPI_ISL_2915637 | Lighthouse Laboratory Plymouth | Wellcome Sanger Institute for the COVID-19 Genomics UK (COG-UK) Consortium | Lighthouse Laboratory Plymouth and Alex Alderton, Roberto Amato, Jeffrey Barrett, Sonia Goncalves, Ewan Harrison, David K. Jackson, Ian Johnston, Dominic Kwiatkowski, Cordelia Langford, John Sillitoe on behalf of the Wellcome Sanger Institute COVID-19 Surveillance Team |
| EPI_ISL_2915638 | Lighthouse Lab in Alderley Park | Wellcome Sanger Institute for the COVID-19 Genomics UK (COG-UK) Consortium | Jacquelyn Wynn, Mairead Hyland, The Lighthouse Lab in Alderley Park and Alex Alderton, Roberto Amato, Jeffrey Barrett, Sonia Goncalves, Ewan Harrison, David K. Jackson, Ian Johnston, Dominic Kwiatkowski, Cordelia Langford, John Sillitoe on behalf of the Wellcome Sanger Institute COVID-19 Surveillance Team |
| EPI_ISL_2915639 | Lighthouse Laboratory Plymouth | Wellcome Sanger Institute for the COVID-19 Genomics UK (COG-UK) Consortium | Lighthouse Laboratory Plymouth and Alex Alderton, Roberto Amato, Jeffrey Barrett, Sonia Goncalves, Ewan Harrison, David K. Jackson, Ian Johnston, Dominic Kwiatkowski, Cordelia Langford, John Sillitoe on behalf of the Wellcome Sanger Institute COVID-19 Surveillance Team |
| EPI_ISL_2915640 | Lighthouse Lab in Glasgow | Wellcome Sanger Institute for the COVID-19 Genomics UK (COG-UK) Consortium | Harper VanSteenhouse, Yumi Kasai, David Gray, Carol Clugston, Anna Dominiczak and Alex Alderton, Roberto Amato, Jeffrey Barrett, Sonia Goncalves, Ewan Harrison, David K. Jackson, Ian Johnston, Dominic Kwiatkowski, Cordelia Langford, John Sillitoe on behalf of the Wellcome Sanger Institute COVID-19 Surveillance Team |
| EPI_ISL_2915641 | Lighthouse Laboratory Plymouth | Wellcome Sanger Institute for the COVID-19 Genomics UK (COG-UK) Consortium | Lighthouse Laboratory Plymouth and Alex Alderton, Roberto Amato, Jeffrey Barrett, Sonia Goncalves, Ewan Harrison, David K. Jackson, Ian Johnston, Dominic Kwiatkowski, Cordelia Langford, John Sillitoe on behalf of the Wellcome Sanger Institute COVID-19 Surveillance Team |
| EPI_ISL_2915642, EPI_ISL_2915643 | Lighthouse Lab in Glasgow | Wellcome Sanger Institute for the COVID-19 Genomics UK (COG-UK) Consortium | Harper VanSteenhouse, Yumi Kasai, David Gray, Carol Clugston, Anna Dominiczak and Alex Alderton, Roberto Amato, Jeffrey Barrett, Sonia Goncalves, Ewan Harrison, David K. Jackson, Ian Johnston, Dominic Kwiatkowski, Cordelia Langford, John Sillitoe on behalf of the Wellcome Sanger Institute COVID-19 Surveillance Team |
| EPI_ISL_2915644 | Lighthouse Laboratory Plymouth | Wellcome Sanger Institute for the COVID-19 Genomics UK (COG-UK) Consortium | Lighthouse Laboratory Plymouth and Alex Alderton, Roberto Amato, Jeffrey Barrett, Sonia Goncalves, Ewan Harrison, David K. Jackson, Ian Johnston, Dominic Kwiatkowski, Cordelia Langford, John Sillitoe on behalf of the Wellcome Sanger Institute COVID-19 Surveillance Team |
| EPI_ISL_2915645 | Lighthouse Lab in Glasgow | Wellcome Sanger Institute for the COVID-19 Genomics UK (COG-UK) Consortium | Harper VanSteenhouse, Yumi Kasai, David Gray, Carol Clugston, Anna Dominiczak and Alex Alderton, Roberto Amato, Jeffrey Barrett, Sonia Goncalves, Ewan Harrison, David K. Jackson, Ian Johnston, Dominic Kwiatkowski, Cordelia Langford, John Sillitoe on behalf of the Wellcome Sanger Institute COVID-19 Surveillance Team |
| EPI_ISL_2915646 | Berkshire and Surrey Pathology Services Lighthouse Laboratory | Wellcome Sanger Institute for the COVID-19 Genomics UK (COG-UK) Consortium | Berkshire and Surrey Pathology Services Lighthouse Laboratory and Alex Alderton, Roberto Amato, Jeffrey Barrett, Sonia Goncalves, Ewan Harrison, David K. Jackson, Ian Johnston, Dominic Kwiatkowski, Cordelia Langford, John Sillitoe on behalf of the Wellcome Sanger Institute COVID-19 Surveillance Team |
| EPI_ISL_2915647 | Lighthouse Lab in Glasgow | Wellcome Sanger Institute for the COVID-19 Genomics UK (COG-UK) Consortium | Harper VanSteenhouse, Yumi Kasai, David Gray, Carol Clugston, Anna Dominiczak and Alex Alderton, Roberto Amato, Jeffrey Barrett, Sonia Goncalves, Ewan Harrison, David K. Jackson, Ian Johnston, Dominic Kwiatkowski, Cordelia Langford, John Sillitoe on behalf of the Wellcome Sanger Institute COVID-19 Surveillance Team |
| EPI_ISL_2915648 | Berkshire and Surrey Pathology Services Lighthouse Laboratory | Wellcome Sanger Institute for the COVID-19 Genomics UK (COG-UK) Consortium | Berkshire and Surrey Pathology Services Lighthouse Laboratory and Alex Alderton, Roberto Amato, Jeffrey Barrett, Sonia Goncalves, Ewan Harrison, David K. Jackson, Ian Johnston, Dominic Kwiatkowski, Cordelia Langford, John Sillitoe on behalf of the Wellcome Sanger Institute COVID-19 Surveillance Team |
| EPI_ISL_2915649, EPI_ISL_2915650 | Lighthouse Lab in Alderley Park | Wellcome Sanger Institute for the COVID-19 Genomics UK (COG-UK) Consortium | Jacquelyn Wynn, Mairead Hyland, The Lighthouse Lab in Alderley Park and Alex Alderton, Roberto Amato, Jeffrey Barrett, Sonia Goncalves, Ewan Harrison, David K. Jackson, Ian Johnston, Dominic Kwiatkowski, Cordelia Langford, John Sillitoe on behalf of the Wellcome Sanger Institute COVID-19 Surveillance Team |
| EPI_ISL_2915651 | Lighthouse Laboratory Plymouth | Wellcome Sanger Institute for the COVID-19 Genomics UK (COG-UK) Consortium | Lighthouse Laboratory Plymouth and Alex Alderton, Roberto Amato, Jeffrey Barrett, Sonia Goncalves, Ewan Harrison, David K. Jackson, Ian Johnston, Dominic Kwiatkowski, Cordelia Langford, John Sillitoe on behalf of the Wellcome Sanger Institute COVID-19 Surveillance Team |
| EPI_ISL_2915652 | Lighthouse Lab in Glasgow | Wellcome Sanger Institute for the COVID-19 Genomics UK (COG-UK) Consortium | Harper VanSteenhouse, Yumi Kasai, David Gray, Carol Clugston, Anna Dominiczak and Alex Alderton, Roberto Amato, Jeffrey Barrett, Sonia Goncalves, Ewan Harrison, David K. Jackson, Ian Johnston, Dominic Kwiatkowski, Cordelia Langford, John Sillitoe on behalf of the Wellcome Sanger Institute COVID-19 Surveillance Team |
| EPI_ISL_2915653 | Lighthouse Laboratory Plymouth | Wellcome Sanger Institute for the COVID-19 Genomics UK (COG-UK) Consortium | Lighthouse Laboratory Plymouth and Alex Alderton, Roberto Amato, Jeffrey Barrett, Sonia Goncalves, Ewan Harrison, David K. Jackson, Ian Johnston, Dominic Kwiatkowski, Cordelia Langford, John Sillitoe on behalf of the Wellcome Sanger Institute COVID-19 Surveillance Team |
| EPI_ISL_2915654 | Lighthouse Lab in Glasgow | Wellcome Sanger Institute for the COVID-19 Genomics UK (COG-UK) Consortium | Harper VanSteenhouse, Yumi Kasai, David Gray, Carol Clugston, Anna Dominiczak and Alex Alderton, Roberto Amato, Jeffrey Barrett, Sonia Goncalves, Ewan Harrison, David K. Jackson, Ian Johnston, Dominic Kwiatkowski, Cordelia Langford, John Sillitoe on behalf of the Wellcome Sanger Institute COVID-19 Surveillance Team |
| EPI_ISL_2915655 | Berkshire and Surrey Pathology Services Lighthouse Laboratory | Wellcome Sanger Institute for the COVID-19 Genomics UK (COG-UK) Consortium | Berkshire and Surrey Pathology Services Lighthouse Laboratory and Alex Alderton, Roberto Amato, Jeffrey Barrett, Sonia Goncalves, Ewan Harrison, David K. Jackson, Ian Johnston, Dominic Kwiatkowski, Cordelia Langford, John Sillitoe on behalf of the Wellcome Sanger Institute COVID-19 Surveillance Team |
| EPI_ISL_2915656 | Lighthouse Lab in Glasgow | Wellcome Sanger Institute for the COVID-19 Genomics UK (COG-UK) Consortium | Harper VanSteenhouse, Yumi Kasai, David Gray, Carol Clugston, Anna Dominiczak and Alex Alderton, Roberto Amato, Jeffrey Barrett, Sonia Goncalves, Ewan Harrison, David K. Jackson, Ian Johnston, Dominic Kwiatkowski, Cordelia Langford, John Sillitoe on behalf of the Wellcome Sanger Institute COVID-19 Surveillance Team |
| EPI_ISL_2915657 | Berkshire and Surrey Pathology Services Lighthouse Laboratory | Wellcome Sanger Institute for the COVID-19 Genomics UK (COG-UK) Consortium | Berkshire and Surrey Pathology Services Lighthouse Laboratory and Alex Alderton, Roberto Amato, Jeffrey Barrett, Sonia Goncalves, Ewan Harrison, David K. Jackson, Ian Johnston, Dominic Kwiatkowski, Cordelia Langford, John Sillitoe on behalf of the Wellcome Sanger Institute COVID-19 Surveillance Team |
| EPI_ISL_2915658 | Lighthouse Lab in Alderley Park | Wellcome Sanger Institute for the COVID-19 Genomics UK (COG-UK) Consortium | Jacquelyn Wynn, Mairead Hyland, The Lighthouse Lab in Alderley Park and Alex Alderton, Roberto Amato, Jeffrey Barrett, Sonia Goncalves, Ewan Harrison, David K. Jackson, Ian Johnston, Dominic Kwiatkowski, Cordelia Langford, John Sillitoe on behalf of the Wellcome Sanger Institute COVID-19 Surveillance Team |
| EPI_ISL_2915659 | Berkshire and Surrey Pathology Services Lighthouse Laboratory | Wellcome Sanger Institute for the COVID-19 Genomics UK (COG-UK) Consortium | Berkshire and Surrey Pathology Services Lighthouse Laboratory and Alex Alderton, Roberto Amato, Jeffrey Barrett, Sonia Goncalves, Ewan Harrison, David K. Jackson, Ian Johnston, Dominic Kwiatkowski, Cordelia Langford, John Sillitoe on behalf of the Wellcome Sanger Institute COVID-19 Surveillance Team |
| EPI_ISL_2915660 | Lighthouse Laboratory Plymouth | Wellcome Sanger Institute for the COVID-19 Genomics UK (COG-UK) Consortium | Lighthouse Laboratory Plymouth and Alex Alderton, Roberto Amato, Jeffrey Barrett, Sonia Goncalves, Ewan Harrison, David K. Jackson, Ian Johnston, Dominic Kwiatkowski, Cordelia Langford, John Sillitoe on behalf of the Wellcome Sanger Institute COVID-19 Surveillance Team |
| EPI_ISL_2915661 | Lighthouse Lab in Alderley Park | Wellcome Sanger Institute for the COVID-19 Genomics UK (COG-UK) Consortium | Jacquelyn Wynn, Mairead Hyland, The Lighthouse Lab in Alderley Park and Alex Alderton, Roberto Amato, Jeffrey Barrett, Sonia Goncalves, Ewan Harrison, David K. Jackson, Ian Johnston, Dominic Kwiatkowski, Cordelia Langford, John Sillitoe on behalf of the Wellcome Sanger Institute COVID-19 Surveillance Team |
| EPI_ISL_2915662 | Berkshire and Surrey Pathology Services Lighthouse Laboratory | Wellcome Sanger Institute for the COVID-19 Genomics UK (COG-UK) Consortium | Berkshire and Surrey Pathology Services Lighthouse Laboratory and Alex Alderton, Roberto Amato, Jeffrey Barrett, Sonia Goncalves, Ewan Harrison, David K. Jackson, Ian Johnston, Dominic Kwiatkowski, Cordelia Langford, John Sillitoe on behalf of the Wellcome Sanger Institute COVID-19 Surveillance Team |
| EPI_ISL_2915663 | Lighthouse Lab in Alderley Park | Wellcome Sanger Institute for the COVID-19 Genomics UK (COG-UK) Consortium | Jacquelyn Wynn, Mairead Hyland, The Lighthouse Lab in Alderley Park and Alex Alderton, Roberto Amato, Jeffrey Barrett, Sonia Goncalves, Ewan Harrison, David K. Jackson, Ian Johnston, Dominic Kwiatkowski, Cordelia Langford, John Sillitoe on behalf of the Wellcome Sanger Institute COVID-19 Surveillance Team |
| EPI_ISL_2915664 | Lighthouse Laboratory Plymouth | Wellcome Sanger Institute for the COVID-19 Genomics UK (COG-UK) | Lighthouse Laboratory Plymouth and Alex Alderton, Roberto Amato, Jeffrey Barrett, Sonia Goncalves, Ewan Harrison, David K. Jackson, Ian Johnston, Dominic Kwiatkowski, Cordelia Langford, John Sillitoe on behalf of the Wellcome Sanger |

| | | | |
|---|---|---|---|
| | | Consortium | Institute COVID-19 Surveillance Team |
| EPI_ISL_2915665 | Berkshire and Surrey Pathology Services Lighthouse Laboratory | Wellcome Sanger Institute for the COVID-19 Genomics UK (COG-UK) Consortium | Berkshire and Surrey Pathology Services Lighthouse Laboratory and Alex Alderton, Roberto Amato, Jeffrey Barrett, Sonia Goncalves, Ewan Harrison, David K. Jackson, Ian Johnston, Dominic Kwiatkowski, Cordelia Langford, John Sillitoe on behalf of the Wellcome Sanger Institute COVID-19 Surveillance Team |
| EPI_ISL_2915666 | Lighthouse Lab in Alderley Park | Wellcome Sanger Institute for the COVID-19 Genomics UK (COG-UK) Consortium | Jacquelyn Wynn, Mairead Hyland, The Lighthouse Lab in Alderley Park and Alex Alderton, Roberto Amato, Jeffrey Barrett, Sonia Goncalves, Ewan Harrison, David K. Jackson, Ian Johnston, Dominic Kwiatkowski, Cordelia Langford, John Sillitoe on behalf of the Wellcome Sanger Institute COVID-19 Surveillance Team |
| EPI_ISL_2915667 | Lighthouse Lab in Glasgow | Wellcome Sanger Institute for the COVID-19 Genomics UK (COG-UK) Consortium | Harper VanSteenhouse, Yumi Kasai, David Gray, Carol Clugston, Anna Dominiczak and Alex Alderton, Roberto Amato, Jeffrey Barrett, Sonia Goncalves, Ewan Harrison, David K. Jackson, Ian Johnston, Dominic Kwiatkowski, Cordelia Langford, John Sillitoe on behalf of the Wellcome Sanger Institute COVID-19 Surveillance Team |
| EPI_ISL_2915668 | Lighthouse Laboratory Plymouth | Wellcome Sanger Institute for the COVID-19 Genomics UK (COG-UK) Consortium | Lighthouse Laboratory Plymouth and Alex Alderton, Roberto Amato, Jeffrey Barrett, Sonia Goncalves, Ewan Harrison, David K. Jackson, Ian Johnston, Dominic Kwiatkowski, Cordelia Langford, John Sillitoe on behalf of the Wellcome Sanger Institute COVID-19 Surveillance Team |
| EPI_ISL_2915669 | Lighthouse Lab in Alderley Park | Wellcome Sanger Institute for the COVID-19 Genomics UK (COG-UK) Consortium | Jacquelyn Wynn, Mairead Hyland, The Lighthouse Lab in Alderley Park and Alex Alderton, Roberto Amato, Jeffrey Barrett, Sonia Goncalves, Ewan Harrison, David K. Jackson, Ian Johnston, Dominic Kwiatkowski, Cordelia Langford, John Sillitoe on behalf of the Wellcome Sanger Institute COVID-19 Surveillance Team |
| EPI_ISL_2915670, EPI_ISL_2915671 | Lighthouse Laboratory Plymouth | Wellcome Sanger Institute for the COVID-19 Genomics UK (COG-UK) Consortium | Lighthouse Laboratory Plymouth and Alex Alderton, Roberto Amato, Jeffrey Barrett, Sonia Goncalves, Ewan Harrison, David K. Jackson, Ian Johnston, Dominic Kwiatkowski, Cordelia Langford, John Sillitoe on behalf of the Wellcome Sanger Institute COVID-19 Surveillance Team |
| EPI_ISL_2915672, EPI_ISL_2915673 | Berkshire and Surrey Pathology Services Lighthouse Laboratory | Wellcome Sanger Institute for the COVID-19 Genomics UK (COG-UK) Consortium | Berkshire and Surrey Pathology Services Lighthouse Laboratory and Alex Alderton, Roberto Amato, Jeffrey Barrett, Sonia Goncalves, Ewan Harrison, David K. Jackson, Ian Johnston, Dominic Kwiatkowski, Cordelia Langford, John Sillitoe on behalf of the Wellcome Sanger Institute COVID-19 Surveillance Team |
| EPI_ISL_2915674 | Lighthouse Lab in Alderley Park | Wellcome Sanger Institute for the COVID-19 Genomics UK (COG-UK) Consortium | Jacquelyn Wynn, Mairead Hyland, The Lighthouse Lab in Alderley Park and Alex Alderton, Roberto Amato, Jeffrey Barrett, Sonia Goncalves, Ewan Harrison, David K. Jackson, Ian Johnston, Dominic Kwiatkowski, Cordelia Langford, John Sillitoe on behalf of the Wellcome Sanger Institute COVID-19 Surveillance Team |
| EPI_ISL_2915675 | Lighthouse Lab in Glasgow | Wellcome Sanger Institute for the COVID-19 Genomics UK (COG-UK) Consortium | Harper VanSteenhouse, Yumi Kasai, David Gray, Carol Clugston, Anna Dominiczak and Alex Alderton, Roberto Amato, Jeffrey Barrett, Sonia Goncalves, Ewan Harrison, David K. Jackson, Ian Johnston, Dominic Kwiatkowski, Cordelia Langford, John Sillitoe on behalf of the Wellcome Sanger Institute COVID-19 Surveillance Team |
| EPI_ISL_2915676 | Lighthouse Laboratory Plymouth | Wellcome Sanger Institute for the COVID-19 Genomics UK (COG-UK) Consortium | Lighthouse Laboratory Plymouth and Alex Alderton, Roberto Amato, Jeffrey Barrett, Sonia Goncalves, Ewan Harrison, David K. Jackson, Ian Johnston, Dominic Kwiatkowski, Cordelia Langford, John Sillitoe on behalf of the Wellcome Sanger Institute COVID-19 Surveillance Team |
| EPI_ISL_2915677, EPI_ISL_2915678 | Lighthouse Lab in Alderley Park | Wellcome Sanger Institute for the COVID-19 Genomics UK (COG-UK) Consortium | Jacquelyn Wynn, Mairead Hyland, The Lighthouse Lab in Alderley Park and Alex Alderton, Roberto Amato, Jeffrey Barrett, Sonia Goncalves, Ewan Harrison, David K. Jackson, Ian Johnston, Dominic Kwiatkowski, Cordelia Langford, John Sillitoe on behalf of the Wellcome Sanger Institute COVID-19 Surveillance Team |
| EPI_ISL_2915679, EPI_ISL_2915680 | Lighthouse Lab in Glasgow | Wellcome Sanger Institute for the COVID-19 Genomics UK (COG-UK) Consortium | Harper VanSteenhouse, Yumi Kasai, David Gray, Carol Clugston, Anna Dominiczak and Alex Alderton, Roberto Amato, Jeffrey Barrett, Sonia Goncalves, Ewan Harrison, David K. Jackson, Ian Johnston, Dominic Kwiatkowski, Cordelia Langford, John Sillitoe on behalf of the Wellcome Sanger Institute COVID-19 Surveillance Team |
| EPI_ISL_2915681, EPI_ISL_2915682, EPI_ISL_2915683 | Lighthouse Laboratory Plymouth | Wellcome Sanger Institute for the COVID-19 Genomics UK (COG-UK) Consortium | Lighthouse Laboratory Plymouth and Alex Alderton, Roberto Amato, Jeffrey Barrett, Sonia Goncalves, Ewan Harrison, David K. Jackson, Ian Johnston, Dominic Kwiatkowski, Cordelia Langford, John Sillitoe on behalf of the Wellcome Sanger Institute COVID-19 Surveillance Team |
| EPI_ISL_2915684 | Lighthouse Lab in Alderley Park | Wellcome Sanger Institute for the COVID-19 Genomics UK (COG-UK) Consortium | Jacquelyn Wynn, Mairead Hyland, The Lighthouse Lab in Alderley Park and Alex Alderton, Roberto Amato, Jeffrey Barrett, Sonia Goncalves, Ewan Harrison, David K. Jackson, Ian Johnston, Dominic Kwiatkowski, Cordelia Langford, John Sillitoe on behalf of the Wellcome Sanger Institute COVID-19 Surveillance Team |
| EPI_ISL_2915685 | Lighthouse Lab in Glasgow | Wellcome Sanger Institute for the COVID-19 Genomics UK (COG-UK) Consortium | Harper VanSteenhouse, Yumi Kasai, David Gray, Carol Clugston, Anna Dominiczak and Alex Alderton, Roberto Amato, Jeffrey Barrett, Sonia Goncalves, Ewan Harrison, David K. Jackson, Ian Johnston, Dominic Kwiatkowski, Cordelia Langford, John Sillitoe on behalf of the Wellcome Sanger Institute COVID-19 Surveillance Team |
| EPI_ISL_2915686 | Lighthouse Lab in Alderley Park | Wellcome Sanger Institute for the COVID-19 Genomics UK (COG-UK) Consortium | Jacquelyn Wynn, Mairead Hyland, The Lighthouse Lab in Alderley Park and Alex Alderton, Roberto Amato, Jeffrey Barrett, Sonia Goncalves, Ewan Harrison, David K. Jackson, Ian Johnston, Dominic Kwiatkowski, Cordelia Langford, John Sillitoe on behalf of the Wellcome Sanger Institute COVID-19 Surveillance Team |
| EPI_ISL_2915687, EPI_ISL_2915688, EPI_ISL_2915689, EPI_ISL_2915690, EPI_ISL_2915691 | Berkshire and Surrey Pathology Services Lighthouse Laboratory | Wellcome Sanger Institute for the COVID-19 Genomics UK (COG-UK) Consortium | Berkshire and Surrey Pathology Services Lighthouse Laboratory and Alex Alderton, Roberto Amato, Jeffrey Barrett, Sonia Goncalves, Ewan Harrison, David K. Jackson, Ian Johnston, Dominic Kwiatkowski, Cordelia Langford, John Sillitoe on behalf of the Wellcome Sanger Institute COVID-19 Surveillance Team |
| EPI_ISL_2915692 | Lighthouse Lab in Alderley Park | Wellcome Sanger Institute for the COVID-19 Genomics UK (COG-UK) Consortium | Jacquelyn Wynn, Mairead Hyland, The Lighthouse Lab in Alderley Park and Alex Alderton, Roberto Amato, Jeffrey Barrett, Sonia Goncalves, Ewan Harrison, David K. Jackson, Ian Johnston, Dominic Kwiatkowski, Cordelia Langford, John Sillitoe on behalf of the Wellcome Sanger Institute COVID-19 Surveillance Team |
| EPI_ISL_2915693, EPI_ISL_2915694, EPI_ISL_2915695, EPI_ISL_2915696, EPI_ISL_2915697 | Berkshire and Surrey Pathology Services Lighthouse Laboratory | Wellcome Sanger Institute for the COVID-19 Genomics UK (COG-UK) Consortium | Berkshire and Surrey Pathology Services Lighthouse Laboratory and Alex Alderton, Roberto Amato, Jeffrey Barrett, Sonia Goncalves, Ewan Harrison, David K. Jackson, Ian Johnston, Dominic Kwiatkowski, Cordelia Langford, John Sillitoe on behalf of the Wellcome Sanger Institute COVID-19 Surveillance Team |
| EPI_ISL_2915698, EPI_ISL_2915699 | Lighthouse Lab in Alderley Park | Wellcome Sanger Institute for the COVID-19 Genomics UK (COG-UK) Consortium | Jacquelyn Wynn, Mairead Hyland, The Lighthouse Lab in Alderley Park and Alex Alderton, Roberto Amato, Jeffrey Barrett, Sonia Goncalves, Ewan Harrison, David K. Jackson, Ian Johnston, Dominic Kwiatkowski, Cordelia Langford, John Sillitoe on behalf of the Wellcome Sanger Institute COVID-19 Surveillance Team |
| EPI_ISL_2915700, EPI_ISL_2915701 | Berkshire and Surrey Pathology Services Lighthouse Laboratory | Wellcome Sanger Institute for the COVID-19 Genomics UK (COG-UK) Consortium | Berkshire and Surrey Pathology Services Lighthouse Laboratory and Alex Alderton, Roberto Amato, Jeffrey Barrett, Sonia Goncalves, Ewan Harrison, David K. Jackson, Ian Johnston, Dominic Kwiatkowski, Cordelia Langford, John Sillitoe on behalf of the Wellcome Sanger Institute COVID-19 Surveillance Team |
| EPI_ISL_2915702 | Lighthouse Lab in Alderley Park | Wellcome Sanger Institute for the COVID-19 Genomics UK (COG-UK) Consortium | Jacquelyn Wynn, Mairead Hyland, The Lighthouse Lab in Alderley Park and Alex Alderton, Roberto Amato, Jeffrey Barrett, Sonia Goncalves, Ewan Harrison, David K. Jackson, Ian Johnston, Dominic Kwiatkowski, Cordelia Langford, John Sillitoe on behalf of the Wellcome Sanger Institute COVID-19 Surveillance Team |
| EPI_ISL_2915703, EPI_ISL_2915704, EPI_ISL_2915705, EPI_ISL_2915706 | Berkshire and Surrey Pathology Services Lighthouse Laboratory | Wellcome Sanger Institute for the COVID-19 Genomics UK (COG-UK) Consortium | Berkshire and Surrey Pathology Services Lighthouse Laboratory and Alex Alderton, Roberto Amato, Jeffrey Barrett, Sonia Goncalves, Ewan Harrison, David K. Jackson, Ian Johnston, Dominic Kwiatkowski, Cordelia Langford, John Sillitoe on behalf of the Wellcome Sanger Institute COVID-19 Surveillance Team |
| EPI_ISL_2915707, EPI_ISL_2915708, EPI_ISL_2915709 | Lighthouse Lab in Alderley Park | Wellcome Sanger Institute for the COVID-19 Genomics UK (COG-UK) Consortium | Jacquelyn Wynn, Mairead Hyland, The Lighthouse Lab in Alderley Park and Alex Alderton, Roberto Amato, Jeffrey Barrett, Sonia Goncalves, Ewan Harrison, David K. Jackson, Ian Johnston, Dominic Kwiatkowski, Cordelia Langford, John Sillitoe on behalf of the Wellcome Sanger Institute COVID-19 Surveillance Team |
| EPI_ISL_2915710, EPI_ISL_2915711, EPI_ISL_2915712 | Berkshire and Surrey Pathology Services Lighthouse Laboratory | Wellcome Sanger Institute for the COVID-19 Genomics UK (COG-UK) Consortium | Berkshire and Surrey Pathology Services Lighthouse Laboratory and Alex Alderton, Roberto Amato, Jeffrey Barrett, Sonia Goncalves, Ewan Harrison, David K. Jackson, Ian Johnston, Dominic Kwiatkowski, Cordelia Langford, John Sillitoe on behalf of the Wellcome Sanger Institute COVID-19 Surveillance Team |
| EPI_ISL_2915713, EPI_ISL_2915714 | Lighthouse Lab in Alderley Park | Wellcome Sanger Institute for the COVID-19 Genomics UK (COG-UK) Consortium | Jacquelyn Wynn, Mairead Hyland, The Lighthouse Lab in Alderley Park and Alex Alderton, Roberto Amato, Jeffrey Barrett, Sonia Goncalves, Ewan Harrison, David K. Jackson, Ian Johnston, Dominic Kwiatkowski, Cordelia Langford, John Sillitoe on behalf of the Wellcome Sanger Institute COVID-19 Surveillance Team |
| EPI_ISL_2915715, EPI_ISL_2915716, EPI_ISL_2915717, EPI_ISL_2915718, EPI_ISL_2915719, EPI_ISL_2915720 | Berkshire and Surrey Pathology Services Lighthouse Laboratory | Wellcome Sanger Institute for the COVID-19 Genomics UK (COG-UK) Consortium | Berkshire and Surrey Pathology Services Lighthouse Laboratory and Alex Alderton, Roberto Amato, Jeffrey Barrett, Sonia Goncalves, Ewan Harrison, David K. Jackson, Ian Johnston, Dominic Kwiatkowski, Cordelia Langford, John Sillitoe on behalf of the Wellcome Sanger Institute COVID-19 Surveillance Team |
| EPI_ISL_2915721, EPI_ISL_2915722 | Lighthouse Lab in Alderley Park | Wellcome Sanger Institute for the COVID-19 Genomics UK (COG-UK) Consortium | Jacquelyn Wynn, Mairead Hyland, The Lighthouse Lab in Alderley Park and Alex Alderton, Roberto Amato, Jeffrey Barrett, Sonia Goncalves, Ewan Harrison, David K. Jackson, Ian Johnston, Dominic Kwiatkowski, Cordelia Langford, John Sillitoe on behalf of the Wellcome Sanger Institute COVID-19 Surveillance Team |
| EPI_ISL_2915723 | Berkshire and Surrey Pathology Services Lighthouse Laboratory | Wellcome Sanger Institute for the COVID-19 Genomics UK (COG-UK) Consortium | Berkshire and Surrey Pathology Services Lighthouse Laboratory and Alex Alderton, Roberto Amato, Jeffrey Barrett, Sonia Goncalves, Ewan Harrison, David K. Jackson, Ian Johnston, Dominic Kwiatkowski, Cordelia Langford, John Sillitoe on behalf of the Wellcome Sanger Institute COVID-19 Surveillance Team |
| EPI_ISL_2915724 | Lighthouse Lab in Alderley Park | Wellcome Sanger Institute for the COVID-19 Genomics UK (COG-UK) Consortium | Jacquelyn Wynn, Mairead Hyland, The Lighthouse Lab in Alderley Park and Alex Alderton, Roberto Amato, Jeffrey Barrett, Sonia Goncalves, Ewan Harrison, David K. Jackson, Ian Johnston, Dominic Kwiatkowski, Cordelia Langford, John Sillitoe on behalf of the Wellcome Sanger Institute COVID-19 Surveillance Team |
| EPI_ISL_2915725 | Berkshire and Surrey Pathology Services Lighthouse Laboratory | Wellcome Sanger Institute for the COVID-19 Genomics UK (COG-UK) Consortium | Berkshire and Surrey Pathology Services Lighthouse Laboratory and Alex Alderton, Roberto Amato, Jeffrey Barrett, Sonia Goncalves, Ewan Harrison, David K. Jackson, Ian Johnston, Dominic Kwiatkowski, Cordelia Langford, John Sillitoe on behalf of the Wellcome Sanger Institute COVID-19 Surveillance Team |
| EPI_ISL_2915726 | Lighthouse Lab in Alderley Park | Wellcome Sanger Institute for the COVID-19 Genomics UK (COG-UK) Consortium | Jacquelyn Wynn, Mairead Hyland, The Lighthouse Lab in Alderley Park and Alex Alderton, Roberto Amato, Jeffrey Barrett, Sonia Goncalves, Ewan Harrison, David K. Jackson, Ian Johnston, Dominic Kwiatkowski, Cordelia Langford, John Sillitoe on behalf of the Wellcome Sanger Institute COVID-19 Surveillance Team |
| EPI_ISL_2915727 | Berkshire and Surrey Pathology Services Lighthouse Laboratory | Wellcome Sanger Institute for the COVID-19 Genomics UK (COG-UK) Consortium | Berkshire and Surrey Pathology Services Lighthouse Laboratory and Alex Alderton, Roberto Amato, Jeffrey Barrett, Sonia Goncalves, Ewan Harrison, David K. Jackson, Ian Johnston, Dominic Kwiatkowski, Cordelia Langford, John Sillitoe on behalf of the Wellcome Sanger Institute COVID-19 Surveillance Team |
| EPI_ISL_2915728 | Lighthouse Lab in Alderley Park | Wellcome Sanger Institute for the COVID-19 Genomics UK (COG-UK) Consortium | Jacquelyn Wynn, Mairead Hyland, The Lighthouse Lab in Alderley Park and Alex Alderton, Roberto Amato, Jeffrey Barrett, Sonia Goncalves, Ewan Harrison, David K. Jackson, Ian Johnston, Dominic Kwiatkowski, Cordelia Langford, John Sillitoe on behalf of the Wellcome Sanger Institute COVID-19 Surveillance Team |
| EPI_ISL_2915729 | Berkshire and Surrey Pathology Services Lighthouse Laboratory | Wellcome Sanger Institute for the COVID-19 Genomics UK (COG-UK) Consortium | Berkshire and Surrey Pathology Services Lighthouse Laboratory and Alex Alderton, Roberto Amato, Jeffrey Barrett, Sonia Goncalves, Ewan Harrison, David K. Jackson, Ian Johnston, Dominic Kwiatkowski, Cordelia Langford, John Sillitoe on behalf of the Wellcome Sanger Institute COVID-19 Surveillance Team |
| EPI_ISL_2915730, EPI_ISL_2915731 | Lighthouse Lab in Alderley Park | Wellcome Sanger Institute for the COVID-19 Genomics UK (COG-UK) Consortium | Jacquelyn Wynn, Mairead Hyland, The Lighthouse Lab in Alderley Park and Alex Alderton, Roberto Amato, Jeffrey Barrett, Sonia Goncalves, Ewan Harrison, David K. Jackson, Ian Johnston, Dominic Kwiatkowski, Cordelia Langford, John Sillitoe on behalf of the Wellcome Sanger Institute COVID-19 Surveillance Team |
| EPI_ISL_2915732 | Berkshire and Surrey Pathology Services Lighthouse Laboratory | Wellcome Sanger Institute for the COVID-19 Genomics UK (COG-UK) Consortium | Berkshire and Surrey Pathology Services Lighthouse Laboratory and Alex Alderton, Roberto Amato, Jeffrey Barrett, Sonia Goncalves, Ewan Harrison, David K. Jackson, Ian Johnston, Dominic Kwiatkowski, Cordelia Langford, John Sillitoe on behalf of the Wellcome Sanger Institute COVID-19 Surveillance Team |
| EPI_ISL_2915733 | Lighthouse Lab in Alderley Park | Wellcome Sanger Institute for the COVID-19 Genomics UK (COG-UK) Consortium | Jacquelyn Wynn, Mairead Hyland, The Lighthouse Lab in Alderley Park and Alex Alderton, Roberto Amato, Jeffrey Barrett, Sonia Goncalves, Ewan Harrison, David K. Jackson, Ian Johnston, Dominic Kwiatkowski, Cordelia Langford, John Sillitoe on behalf of the Wellcome Sanger Institute COVID-19 Surveillance Team |

| | | | |
|---|---|---|---|
| EPI_ISL_2915734, EPI_ISL_2915735 | Berkshire and Surrey Pathology Services Lighthouse Laboratory | Wellcome Sanger Institute for the COVID-19 Genomics UK (COG-UK) Consortium | Berkshire and Surrey Pathology Services Lighthouse Laboratory and Alex Alderton, Roberto Amato, Jeffrey Barrett, Sonia Goncalves, Ewan Harrison, David K. Jackson, Ian Johnston, Dominic Kwiatkowski, Cordelia Langford, John Sillitoe on behalf of the Wellcome Sanger Institute COVID-19 Surveillance Team |
| EPI_ISL_2915736 | Lighthouse Lab in Alderley Park | Wellcome Sanger Institute for the COVID-19 Genomics UK (COG-UK) Consortium | Jacquelyn Wynn, Mairead Hyland, The Lighthouse Lab in Alderley Park and Alex Alderton, Roberto Amato, Jeffrey Barrett, Sonia Goncalves, Ewan Harrison, David K. Jackson, Ian Johnston, Dominic Kwiatkowski, Cordelia Langford, John Sillitoe on behalf of the Wellcome Sanger Institute COVID-19 Surveillance Team |
| EPI_ISL_2915737, EPI_ISL_2915738, EPI_ISL_2915739 | Berkshire and Surrey Pathology Services Lighthouse Laboratory | Wellcome Sanger Institute for the COVID-19 Genomics UK (COG-UK) Consortium | Berkshire and Surrey Pathology Services Lighthouse Laboratory and Alex Alderton, Roberto Amato, Jeffrey Barrett, Sonia Goncalves, Ewan Harrison, David K. Jackson, Ian Johnston, Dominic Kwiatkowski, Cordelia Langford, John Sillitoe on behalf of the Wellcome Sanger Institute COVID-19 Surveillance Team |
| EPI_ISL_2915740, EPI_ISL_2915741 | Lighthouse Lab in Alderley Park | Wellcome Sanger Institute for the COVID-19 Genomics UK (COG-UK) Consortium | Jacquelyn Wynn, Mairead Hyland, The Lighthouse Lab in Alderley Park and Alex Alderton, Roberto Amato, Jeffrey Barrett, Sonia Goncalves, Ewan Harrison, David K. Jackson, Ian Johnston, Dominic Kwiatkowski, Cordelia Langford, John Sillitoe on behalf of the Wellcome Sanger Institute COVID-19 Surveillance Team |
| EPI_ISL_2915742, EPI_ISL_2915743, EPI_ISL_2915744 | Berkshire and Surrey Pathology Services Lighthouse Laboratory | Wellcome Sanger Institute for the COVID-19 Genomics UK (COG-UK) Consortium | Berkshire and Surrey Pathology Services Lighthouse Laboratory and Alex Alderton, Roberto Amato, Jeffrey Barrett, Sonia Goncalves, Ewan Harrison, David K. Jackson, Ian Johnston, Dominic Kwiatkowski, Cordelia Langford, John Sillitoe on behalf of the Wellcome Sanger Institute COVID-19 Surveillance Team |
| EPI_ISL_2915745 | Lighthouse Lab in Alderley Park | Wellcome Sanger Institute for the COVID-19 Genomics UK (COG-UK) Consortium | Jacquelyn Wynn, Mairead Hyland, The Lighthouse Lab in Alderley Park and Alex Alderton, Roberto Amato, Jeffrey Barrett, Sonia Goncalves, Ewan Harrison, David K. Jackson, Ian Johnston, Dominic Kwiatkowski, Cordelia Langford, John Sillitoe on behalf of the Wellcome Sanger Institute COVID-19 Surveillance Team |
| EPI_ISL_2915746 | Berkshire and Surrey Pathology Services Lighthouse Laboratory | Wellcome Sanger Institute for the COVID-19 Genomics UK (COG-UK) Consortium | Berkshire and Surrey Pathology Services Lighthouse Laboratory and Alex Alderton, Roberto Amato, Jeffrey Barrett, Sonia Goncalves, Ewan Harrison, David K. Jackson, Ian Johnston, Dominic Kwiatkowski, Cordelia Langford, John Sillitoe on behalf of the Wellcome Sanger Institute COVID-19 Surveillance Team |
| EPI_ISL_2915747, EPI_ISL_2915748, EPI_ISL_2915749, EPI_ISL_2915750 | Lighthouse Lab in Alderley Park | Wellcome Sanger Institute for the COVID-19 Genomics UK (COG-UK) Consortium | Jacquelyn Wynn, Mairead Hyland, The Lighthouse Lab in Alderley Park and Alex Alderton, Roberto Amato, Jeffrey Barrett, Sonia Goncalves, Ewan Harrison, David K. Jackson, Ian Johnston, Dominic Kwiatkowski, Cordelia Langford, John Sillitoe on behalf of the Wellcome Sanger Institute COVID-19 Surveillance Team |
| EPI_ISL_2915751, EPI_ISL_2915752 | Berkshire and Surrey Pathology Services Lighthouse Laboratory | Wellcome Sanger Institute for the COVID-19 Genomics UK (COG-UK) Consortium | Berkshire and Surrey Pathology Services Lighthouse Laboratory and Alex Alderton, Roberto Amato, Jeffrey Barrett, Sonia Goncalves, Ewan Harrison, David K. Jackson, Ian Johnston, Dominic Kwiatkowski, Cordelia Langford, John Sillitoe on behalf of the Wellcome Sanger Institute COVID-19 Surveillance Team |
| EPI_ISL_2915753, EPI_ISL_2915754, EPI_ISL_2915755 | Lighthouse Lab in Alderley Park | Wellcome Sanger Institute for the COVID-19 Genomics UK (COG-UK) Consortium | Jacquelyn Wynn, Mairead Hyland, The Lighthouse Lab in Alderley Park and Alex Alderton, Roberto Amato, Jeffrey Barrett, Sonia Goncalves, Ewan Harrison, David K. Jackson, Ian Johnston, Dominic Kwiatkowski, Cordelia Langford, John Sillitoe on behalf of the Wellcome Sanger Institute COVID-19 Surveillance Team |
| EPI_ISL_2915756 | Berkshire and Surrey Pathology Services Lighthouse Laboratory | Wellcome Sanger Institute for the COVID-19 Genomics UK (COG-UK) Consortium | Berkshire and Surrey Pathology Services Lighthouse Laboratory and Alex Alderton, Roberto Amato, Jeffrey Barrett, Sonia Goncalves, Ewan Harrison, David K. Jackson, Ian Johnston, Dominic Kwiatkowski, Cordelia Langford, John Sillitoe on behalf of the Wellcome Sanger Institute COVID-19 Surveillance Team |
| EPI_ISL_2915757, EPI_ISL_2915758, EPI_ISL_2915759 | Lighthouse Lab in Alderley Park | Wellcome Sanger Institute for the COVID-19 Genomics UK (COG-UK) Consortium | Jacquelyn Wynn, Mairead Hyland, The Lighthouse Lab in Alderley Park and Alex Alderton, Roberto Amato, Jeffrey Barrett, Sonia Goncalves, Ewan Harrison, David K. Jackson, Ian Johnston, Dominic Kwiatkowski, Cordelia Langford, John Sillitoe on behalf of the Wellcome Sanger Institute COVID-19 Surveillance Team |
| EPI_ISL_2915760, EPI_ISL_2915761, EPI_ISL_2915762, EPI_ISL_2915763 | Berkshire and Surrey Pathology Services Lighthouse Laboratory | Wellcome Sanger Institute for the COVID-19 Genomics UK (COG-UK) Consortium | Berkshire and Surrey Pathology Services Lighthouse Laboratory and Alex Alderton, Roberto Amato, Jeffrey Barrett, Sonia Goncalves, Ewan Harrison, David K. Jackson, Ian Johnston, Dominic Kwiatkowski, Cordelia Langford, John Sillitoe on behalf of the Wellcome Sanger Institute COVID-19 Surveillance Team |
| EPI_ISL_2915764 | Lighthouse Lab in Alderley Park | Wellcome Sanger Institute for the COVID-19 Genomics UK (COG-UK) Consortium | Jacquelyn Wynn, Mairead Hyland, The Lighthouse Lab in Alderley Park and Alex Alderton, Roberto Amato, Jeffrey Barrett, Sonia Goncalves, Ewan Harrison, David K. Jackson, Ian Johnston, Dominic Kwiatkowski, Cordelia Langford, John Sillitoe on behalf of the Wellcome Sanger Institute COVID-19 Surveillance Team |
| EPI_ISL_2915765, EPI_ISL_2915766 | Berkshire and Surrey Pathology Services Lighthouse Laboratory | Wellcome Sanger Institute for the COVID-19 Genomics UK (COG-UK) Consortium | Berkshire and Surrey Pathology Services Lighthouse Laboratory and Alex Alderton, Roberto Amato, Jeffrey Barrett, Sonia Goncalves, Ewan Harrison, David K. Jackson, Ian Johnston, Dominic Kwiatkowski, Cordelia Langford, John Sillitoe on behalf of the Wellcome Sanger Institute COVID-19 Surveillance Team |
| EPI_ISL_2915767, EPI_ISL_2915768, EPI_ISL_2915769 | Lighthouse Lab in Alderley Park | Wellcome Sanger Institute for the COVID-19 Genomics UK (COG-UK) Consortium | Jacquelyn Wynn, Mairead Hyland, The Lighthouse Lab in Alderley Park and Alex Alderton, Roberto Amato, Jeffrey Barrett, Sonia Goncalves, Ewan Harrison, David K. Jackson, Ian Johnston, Dominic Kwiatkowski, Cordelia Langford, John Sillitoe on behalf of the Wellcome Sanger Institute COVID-19 Surveillance Team |
| EPI_ISL_2915770, EPI_ISL_2915771, EPI_ISL_2915772, EPI_ISL_2915773, EPI_ISL_2915774, EPI_ISL_2915775 | Berkshire and Surrey Pathology Services Lighthouse Laboratory | Wellcome Sanger Institute for the COVID-19 Genomics UK (COG-UK) Consortium | Berkshire and Surrey Pathology Services Lighthouse Laboratory and Alex Alderton, Roberto Amato, Jeffrey Barrett, Sonia Goncalves, Ewan Harrison, David K. Jackson, Ian Johnston, Dominic Kwiatkowski, Cordelia Langford, John Sillitoe on behalf of the Wellcome Sanger Institute COVID-19 Surveillance Team |
| EPI_ISL_2915776 | Lighthouse Lab in Alderley Park | Wellcome Sanger Institute for the COVID-19 Genomics UK (COG-UK) Consortium | Jacquelyn Wynn, Mairead Hyland, The Lighthouse Lab in Alderley Park and Alex Alderton, Roberto Amato, Jeffrey Barrett, Sonia Goncalves, Ewan Harrison, David K. Jackson, Ian Johnston, Dominic Kwiatkowski, Cordelia Langford, John Sillitoe on behalf of the Wellcome Sanger Institute COVID-19 Surveillance Team |
| EPI_ISL_2915777 | Berkshire and Surrey Pathology Services Lighthouse Laboratory | Wellcome Sanger Institute for the COVID-19 Genomics UK (COG-UK) Consortium | Berkshire and Surrey Pathology Services Lighthouse Laboratory and Alex Alderton, Roberto Amato, Jeffrey Barrett, Sonia Goncalves, Ewan Harrison, David K. Jackson, Ian Johnston, Dominic Kwiatkowski, Cordelia Langford, John Sillitoe on behalf of the Wellcome Sanger Institute COVID-19 Surveillance Team |
| EPI_ISL_2915778 | Lighthouse Lab in Alderley Park | Wellcome Sanger Institute for the COVID-19 Genomics UK (COG-UK) Consortium | Jacquelyn Wynn, Mairead Hyland, The Lighthouse Lab in Alderley Park and Alex Alderton, Roberto Amato, Jeffrey Barrett, Sonia Goncalves, Ewan Harrison, David K. Jackson, Ian Johnston, Dominic Kwiatkowski, Cordelia Langford, John Sillitoe on behalf of the Wellcome Sanger Institute COVID-19 Surveillance Team |
| EPI_ISL_2915779, EPI_ISL_2915780 | Berkshire and Surrey Pathology Services Lighthouse Laboratory | Wellcome Sanger Institute for the COVID-19 Genomics UK (COG-UK) Consortium | Berkshire and Surrey Pathology Services Lighthouse Laboratory and Alex Alderton, Roberto Amato, Jeffrey Barrett, Sonia Goncalves, Ewan Harrison, David K. Jackson, Ian Johnston, Dominic Kwiatkowski, Cordelia Langford, John Sillitoe on behalf of the Wellcome Sanger Institute COVID-19 Surveillance Team |
| EPI_ISL_2915781 | Lighthouse Lab in Alderley Park | Wellcome Sanger Institute for the COVID-19 Genomics UK (COG-UK) Consortium | Jacquelyn Wynn, Mairead Hyland, The Lighthouse Lab in Alderley Park and Alex Alderton, Roberto Amato, Jeffrey Barrett, Sonia Goncalves, Ewan Harrison, David K. Jackson, Ian Johnston, Dominic Kwiatkowski, Cordelia Langford, John Sillitoe on behalf of the Wellcome Sanger Institute COVID-19 Surveillance Team |
| EPI_ISL_2915782 | Berkshire and Surrey Pathology Services Lighthouse Laboratory | Wellcome Sanger Institute for the COVID-19 Genomics UK (COG-UK) Consortium | Berkshire and Surrey Pathology Services Lighthouse Laboratory and Alex Alderton, Roberto Amato, Jeffrey Barrett, Sonia Goncalves, Ewan Harrison, David K. Jackson, Ian Johnston, Dominic Kwiatkowski, Cordelia Langford, John Sillitoe on behalf of the Wellcome Sanger Institute COVID-19 Surveillance Team |
| EPI_ISL_2915783 | Lighthouse Lab in Alderley Park | Wellcome Sanger Institute for the COVID-19 Genomics UK (COG-UK) Consortium | Jacquelyn Wynn, Mairead Hyland, The Lighthouse Lab in Alderley Park and Alex Alderton, Roberto Amato, Jeffrey Barrett, Sonia Goncalves, Ewan Harrison, David K. Jackson, Ian Johnston, Dominic Kwiatkowski, Cordelia Langford, John Sillitoe on behalf of the Wellcome Sanger Institute COVID-19 Surveillance Team |
| EPI_ISL_2915784 | Berkshire and Surrey Pathology Services Lighthouse Laboratory | Wellcome Sanger Institute for the COVID-19 Genomics UK (COG-UK) Consortium | Berkshire and Surrey Pathology Services Lighthouse Laboratory and Alex Alderton, Roberto Amato, Jeffrey Barrett, Sonia Goncalves, Ewan Harrison, David K. Jackson, Ian Johnston, Dominic Kwiatkowski, Cordelia Langford, John Sillitoe on behalf of the Wellcome Sanger Institute COVID-19 Surveillance Team |
| EPI_ISL_2915785, EPI_ISL_2915786, EPI_ISL_2915787 | Lighthouse Lab in Alderley Park | Wellcome Sanger Institute for the COVID-19 Genomics UK (COG-UK) Consortium | Jacquelyn Wynn, Mairead Hyland, The Lighthouse Lab in Alderley Park and Alex Alderton, Roberto Amato, Jeffrey Barrett, Sonia Goncalves, Ewan Harrison, David K. Jackson, Ian Johnston, Dominic Kwiatkowski, Cordelia Langford, John Sillitoe on behalf of the Wellcome Sanger Institute COVID-19 Surveillance Team |
| EPI_ISL_2915788, EPI_ISL_2915789, EPI_ISL_2915790, EPI_ISL_2915791, EPI_ISL_2915792 | Berkshire and Surrey Pathology Services Lighthouse Laboratory | Wellcome Sanger Institute for the COVID-19 Genomics UK (COG-UK) Consortium | Berkshire and Surrey Pathology Services Lighthouse Laboratory and Alex Alderton, Roberto Amato, Jeffrey Barrett, Sonia Goncalves, Ewan Harrison, David K. Jackson, Ian Johnston, Dominic Kwiatkowski, Cordelia Langford, John Sillitoe on behalf of the Wellcome Sanger Institute COVID-19 Surveillance Team |
| EPI_ISL_2915793 | Lighthouse Lab in Alderley Park | Wellcome Sanger Institute for the COVID-19 Genomics UK (COG-UK) Consortium | Jacquelyn Wynn, Mairead Hyland, The Lighthouse Lab in Alderley Park and Alex Alderton, Roberto Amato, Jeffrey Barrett, Sonia Goncalves, Ewan Harrison, David K. Jackson, Ian Johnston, Dominic Kwiatkowski, Cordelia Langford, John Sillitoe on behalf of the Wellcome Sanger Institute COVID-19 Surveillance Team |
| EPI_ISL_2915794 | Berkshire and Surrey Pathology Services Lighthouse Laboratory | Wellcome Sanger Institute for the COVID-19 Genomics UK (COG-UK) Consortium | Berkshire and Surrey Pathology Services Lighthouse Laboratory and Alex Alderton, Roberto Amato, Jeffrey Barrett, Sonia Goncalves, Ewan Harrison, David K. Jackson, Ian Johnston, Dominic Kwiatkowski, Cordelia Langford, John Sillitoe on behalf of the Wellcome Sanger Institute COVID-19 Surveillance Team |
| EPI_ISL_2915795, EPI_ISL_2915796, EPI_ISL_2915797, EPI_ISL_2915798, EPI_ISL_2915799, EPI_ISL_2915800 | Lighthouse Lab in Alderley Park | Wellcome Sanger Institute for the COVID-19 Genomics UK (COG-UK) Consortium | Jacquelyn Wynn, Mairead Hyland, The Lighthouse Lab in Alderley Park and Alex Alderton, Roberto Amato, Jeffrey Barrett, Sonia Goncalves, Ewan Harrison, David K. Jackson, Ian Johnston, Dominic Kwiatkowski, Cordelia Langford, John Sillitoe on behalf of the Wellcome Sanger Institute COVID-19 Surveillance Team |
| EPI_ISL_2915801, EPI_ISL_2915802 | Berkshire and Surrey Pathology Services Lighthouse Laboratory | Wellcome Sanger Institute for the COVID-19 Genomics UK (COG-UK) Consortium | Berkshire and Surrey Pathology Services Lighthouse Laboratory and Alex Alderton, Roberto Amato, Jeffrey Barrett, Sonia Goncalves, Ewan Harrison, David K. Jackson, Ian Johnston, Dominic Kwiatkowski, Cordelia Langford, John Sillitoe on behalf of the Wellcome Sanger Institute COVID-19 Surveillance Team |
| EPI_ISL_2915803 | Lighthouse Lab in Alderley Park | Wellcome Sanger Institute for the COVID-19 Genomics UK (COG-UK) Consortium | Jacquelyn Wynn, Mairead Hyland, The Lighthouse Lab in Alderley Park and Alex Alderton, Roberto Amato, Jeffrey Barrett, Sonia Goncalves, Ewan Harrison, David K. Jackson, Ian Johnston, Dominic Kwiatkowski, Cordelia Langford, John Sillitoe on behalf of the Wellcome Sanger Institute COVID-19 Surveillance Team |
| EPI_ISL_2915804, EPI_ISL_2915805 | Berkshire and Surrey Pathology Services Lighthouse Laboratory | Wellcome Sanger Institute for the COVID-19 Genomics UK (COG-UK) Consortium | Berkshire and Surrey Pathology Services Lighthouse Laboratory and Alex Alderton, Roberto Amato, Jeffrey Barrett, Sonia Goncalves, Ewan Harrison, David K. Jackson, Ian Johnston, Dominic Kwiatkowski, Cordelia Langford, John Sillitoe on behalf of the Wellcome Sanger Institute COVID-19 Surveillance Team |
| EPI_ISL_2915806, EPI_ISL_2915807, EPI_ISL_2915808 | Lighthouse Lab in Alderley Park | Wellcome Sanger Institute for the COVID-19 Genomics UK (COG-UK) Consortium | Jacquelyn Wynn, Mairead Hyland, The Lighthouse Lab in Alderley Park and Alex Alderton, Roberto Amato, Jeffrey Barrett, Sonia Goncalves, Ewan Harrison, David K. Jackson, Ian Johnston, Dominic Kwiatkowski, Cordelia Langford, John Sillitoe on behalf of the Wellcome Sanger Institute COVID-19 Surveillance Team |
| EPI_ISL_2915809 | Berkshire and Surrey Pathology Services Lighthouse Laboratory | Wellcome Sanger Institute for the COVID-19 Genomics UK (COG-UK) Consortium | Berkshire and Surrey Pathology Services Lighthouse Laboratory and Alex Alderton, Roberto Amato, Jeffrey Barrett, Sonia Goncalves, Ewan Harrison, David K. Jackson, Ian Johnston, Dominic Kwiatkowski, Cordelia Langford, John Sillitoe on behalf of the Wellcome Sanger Institute COVID-19 Surveillance Team |
| EPI_ISL_2915810, EPI_ISL_2915811, EPI_ISL_2915812, | Lighthouse Lab in Alderley Park | Wellcome Sanger Institute for the COVID-19 Genomics UK (COG-UK) Consortium | Jacquelyn Wynn, Mairead Hyland, The Lighthouse Lab in Alderley Park and Alex Alderton, Roberto Amato, Jeffrey Barrett, Sonia Goncalves, Ewan Harrison, David K. Jackson, Ian Johnston, Dominic Kwiatkowski, Cordelia Langford, John Sillitoe on behalf of the Wellcome Sanger Institute COVID-19 Surveillance Team |

| | | | |
|---|---|---|---|
| EPI_ISL_2915813 | | | |
| EPI_ISL_2915814 | Berkshire and Surrey Pathology Services Lighthouse Laboratory | Wellcome Sanger Institute for the COVID-19 Genomics UK (COG-UK) Consortium | Berkshire and Surrey Pathology Services Lighthouse Laboratory and Alex Alderton, Roberto Amato, Jeffrey Barrett, Sonia Goncalves, Ewan Harrison, David K. Jackson, Ian Johnston, Dominic Kwiatkowski, Cordelia Langford, John Sillitoe on behalf of the Wellcome Sanger Institute COVID-19 Surveillance Team |
| EPI_ISL_2915815, EPI_ISL_2915816, EPI_ISL_2915817, EPI_ISL_2915818, EPI_ISL_2915819, EPI_ISL_2915820 | Lighthouse Lab in Alderley Park | Wellcome Sanger Institute for the COVID-19 Genomics UK (COG-UK) Consortium | Jacquelyn Wynn, Mairead Hyland, The Lighthouse Lab in Alderley Park and Alex Alderton, Roberto Amato, Jeffrey Barrett, Sonia Goncalves, Ewan Harrison, David K. Jackson, Ian Johnston, Dominic Kwiatkowski, Cordelia Langford, John Sillitoe on behalf of the Wellcome Sanger Institute COVID-19 Surveillance Team |
| EPI_ISL_2915821 | Berkshire and Surrey Pathology Services Lighthouse Laboratory | Wellcome Sanger Institute for the COVID-19 Genomics UK (COG-UK) Consortium | Berkshire and Surrey Pathology Services Lighthouse Laboratory and Alex Alderton, Roberto Amato, Jeffrey Barrett, Sonia Goncalves, Ewan Harrison, David K. Jackson, Ian Johnston, Dominic Kwiatkowski, Cordelia Langford, John Sillitoe on behalf of the Wellcome Sanger Institute COVID-19 Surveillance Team |
| EPI_ISL_2915822 | Lighthouse Lab in Alderley Park | Wellcome Sanger Institute for the COVID-19 Genomics UK (COG-UK) Consortium | Jacquelyn Wynn, Mairead Hyland, The Lighthouse Lab in Alderley Park and Alex Alderton, Roberto Amato, Jeffrey Barrett, Sonia Goncalves, Ewan Harrison, David K. Jackson, Ian Johnston, Dominic Kwiatkowski, Cordelia Langford, John Sillitoe on behalf of the Wellcome Sanger Institute COVID-19 Surveillance Team |
| EPI_ISL_2915823 | Berkshire and Surrey Pathology Services Lighthouse Laboratory | Wellcome Sanger Institute for the COVID-19 Genomics UK (COG-UK) Consortium | Berkshire and Surrey Pathology Services Lighthouse Laboratory and Alex Alderton, Roberto Amato, Jeffrey Barrett, Sonia Goncalves, Ewan Harrison, David K. Jackson, Ian Johnston, Dominic Kwiatkowski, Cordelia Langford, John Sillitoe on behalf of the Wellcome Sanger Institute COVID-19 Surveillance Team |
| EPI_ISL_2915824 | Lighthouse Lab in Alderley Park | Wellcome Sanger Institute for the COVID-19 Genomics UK (COG-UK) Consortium | Jacquelyn Wynn, Mairead Hyland, The Lighthouse Lab in Alderley Park and Alex Alderton, Roberto Amato, Jeffrey Barrett, Sonia Goncalves, Ewan Harrison, David K. Jackson, Ian Johnston, Dominic Kwiatkowski, Cordelia Langford, John Sillitoe on behalf of the Wellcome Sanger Institute COVID-19 Surveillance Team |
| EPI_ISL_2915825, EPI_ISL_2915826 | Berkshire and Surrey Pathology Services Lighthouse Laboratory | Wellcome Sanger Institute for the COVID-19 Genomics UK (COG-UK) Consortium | Berkshire and Surrey Pathology Services Lighthouse Laboratory and Alex Alderton, Roberto Amato, Jeffrey Barrett, Sonia Goncalves, Ewan Harrison, David K. Jackson, Ian Johnston, Dominic Kwiatkowski, Cordelia Langford, John Sillitoe on behalf of the Wellcome Sanger Institute COVID-19 Surveillance Team |
| EPI_ISL_2915827 | Lighthouse Lab in Alderley Park | Wellcome Sanger Institute for the COVID-19 Genomics UK (COG-UK) Consortium | Jacquelyn Wynn, Mairead Hyland, The Lighthouse Lab in Alderley Park and Alex Alderton, Roberto Amato, Jeffrey Barrett, Sonia Goncalves, Ewan Harrison, David K. Jackson, Ian Johnston, Dominic Kwiatkowski, Cordelia Langford, John Sillitoe on behalf of the Wellcome Sanger Institute COVID-19 Surveillance Team |
| EPI_ISL_2915828, EPI_ISL_2915829 | Berkshire and Surrey Pathology Services Lighthouse Laboratory | Wellcome Sanger Institute for the COVID-19 Genomics UK (COG-UK) Consortium | Berkshire and Surrey Pathology Services Lighthouse Laboratory and Alex Alderton, Roberto Amato, Jeffrey Barrett, Sonia Goncalves, Ewan Harrison, David K. Jackson, Ian Johnston, Dominic Kwiatkowski, Cordelia Langford, John Sillitoe on behalf of the Wellcome Sanger Institute COVID-19 Surveillance Team |
| EPI_ISL_2915830 | Lighthouse Lab in Alderley Park | Wellcome Sanger Institute for the COVID-19 Genomics UK (COG-UK) Consortium | Jacquelyn Wynn, Mairead Hyland, The Lighthouse Lab in Alderley Park and Alex Alderton, Roberto Amato, Jeffrey Barrett, Sonia Goncalves, Ewan Harrison, David K. Jackson, Ian Johnston, Dominic Kwiatkowski, Cordelia Langford, John Sillitoe on behalf of the Wellcome Sanger Institute COVID-19 Surveillance Team |
| EPI_ISL_2915831 | Berkshire and Surrey Pathology Services Lighthouse Laboratory | Wellcome Sanger Institute for the COVID-19 Genomics UK (COG-UK) Consortium | Berkshire and Surrey Pathology Services Lighthouse Laboratory and Alex Alderton, Roberto Amato, Jeffrey Barrett, Sonia Goncalves, Ewan Harrison, David K. Jackson, Ian Johnston, Dominic Kwiatkowski, Cordelia Langford, John Sillitoe on behalf of the Wellcome Sanger Institute COVID-19 Surveillance Team |
| EPI_ISL_2915832 | Lighthouse Lab in Alderley Park | Wellcome Sanger Institute for the COVID-19 Genomics UK (COG-UK) Consortium | Jacquelyn Wynn, Mairead Hyland, The Lighthouse Lab in Alderley Park and Alex Alderton, Roberto Amato, Jeffrey Barrett, Sonia Goncalves, Ewan Harrison, David K. Jackson, Ian Johnston, Dominic Kwiatkowski, Cordelia Langford, John Sillitoe on behalf of the Wellcome Sanger Institute COVID-19 Surveillance Team |
| EPI_ISL_2915833, EPI_ISL_2915834 | Berkshire and Surrey Pathology Services Lighthouse Laboratory | Wellcome Sanger Institute for the COVID-19 Genomics UK (COG-UK) Consortium | Berkshire and Surrey Pathology Services Lighthouse Laboratory and Alex Alderton, Roberto Amato, Jeffrey Barrett, Sonia Goncalves, Ewan Harrison, David K. Jackson, Ian Johnston, Dominic Kwiatkowski, Cordelia Langford, John Sillitoe on behalf of the Wellcome Sanger Institute COVID-19 Surveillance Team |
| EPI_ISL_2915835 | Lighthouse Lab in Alderley Park | Wellcome Sanger Institute for the COVID-19 Genomics UK (COG-UK) Consortium | Jacquelyn Wynn, Mairead Hyland, The Lighthouse Lab in Alderley Park and Alex Alderton, Roberto Amato, Jeffrey Barrett, Sonia Goncalves, Ewan Harrison, David K. Jackson, Ian Johnston, Dominic Kwiatkowski, Cordelia Langford, John Sillitoe on behalf of the Wellcome Sanger Institute COVID-19 Surveillance Team |
| EPI_ISL_2915836, EPI_ISL_2915837 | Berkshire and Surrey Pathology Services Lighthouse Laboratory | Wellcome Sanger Institute for the COVID-19 Genomics UK (COG-UK) Consortium | Berkshire and Surrey Pathology Services Lighthouse Laboratory and Alex Alderton, Roberto Amato, Jeffrey Barrett, Sonia Goncalves, Ewan Harrison, David K. Jackson, Ian Johnston, Dominic Kwiatkowski, Cordelia Langford, John Sillitoe on behalf of the Wellcome Sanger Institute COVID-19 Surveillance Team |
| EPI_ISL_2915838 | Lighthouse Lab in Alderley Park | Wellcome Sanger Institute for the COVID-19 Genomics UK (COG-UK) Consortium | Jacquelyn Wynn, Mairead Hyland, The Lighthouse Lab in Alderley Park and Alex Alderton, Roberto Amato, Jeffrey Barrett, Sonia Goncalves, Ewan Harrison, David K. Jackson, Ian Johnston, Dominic Kwiatkowski, Cordelia Langford, John Sillitoe on behalf of the Wellcome Sanger Institute COVID-19 Surveillance Team |
| EPI_ISL_2915839, EPI_ISL_2915840 | Berkshire and Surrey Pathology Services Lighthouse Laboratory | Wellcome Sanger Institute for the COVID-19 Genomics UK (COG-UK) Consortium | Berkshire and Surrey Pathology Services Lighthouse Laboratory and Alex Alderton, Roberto Amato, Jeffrey Barrett, Sonia Goncalves, Ewan Harrison, David K. Jackson, Ian Johnston, Dominic Kwiatkowski, Cordelia Langford, John Sillitoe on behalf of the Wellcome Sanger Institute COVID-19 Surveillance Team |
| EPI_ISL_2915841, EPI_ISL_2915842 | Lighthouse Lab in Alderley Park | Wellcome Sanger Institute for the COVID-19 Genomics UK (COG-UK) Consortium | Jacquelyn Wynn, Mairead Hyland, The Lighthouse Lab in Alderley Park and Alex Alderton, Roberto Amato, Jeffrey Barrett, Sonia Goncalves, Ewan Harrison, David K. Jackson, Ian Johnston, Dominic Kwiatkowski, Cordelia Langford, John Sillitoe on behalf of the Wellcome Sanger Institute COVID-19 Surveillance Team |
| EPI_ISL_2915843 | Berkshire and Surrey Pathology Services Lighthouse Laboratory | Wellcome Sanger Institute for the COVID-19 Genomics UK (COG-UK) Consortium | Berkshire and Surrey Pathology Services Lighthouse Laboratory and Alex Alderton, Roberto Amato, Jeffrey Barrett, Sonia Goncalves, Ewan Harrison, David K. Jackson, Ian Johnston, Dominic Kwiatkowski, Cordelia Langford, John Sillitoe on behalf of the Wellcome Sanger Institute COVID-19 Surveillance Team |
| EPI_ISL_2915844 | Lighthouse Lab in Alderley Park | Wellcome Sanger Institute for the COVID-19 Genomics UK (COG-UK) Consortium | Jacquelyn Wynn, Mairead Hyland, The Lighthouse Lab in Alderley Park and Alex Alderton, Roberto Amato, Jeffrey Barrett, Sonia Goncalves, Ewan Harrison, David K. Jackson, Ian Johnston, Dominic Kwiatkowski, Cordelia Langford, John Sillitoe on behalf of the Wellcome Sanger Institute COVID-19 Surveillance Team |
| EPI_ISL_2915845, EPI_ISL_2915846, EPI_ISL_2915847 | Berkshire and Surrey Pathology Services Lighthouse Laboratory | Wellcome Sanger Institute for the COVID-19 Genomics UK (COG-UK) Consortium | Berkshire and Surrey Pathology Services Lighthouse Laboratory and Alex Alderton, Roberto Amato, Jeffrey Barrett, Sonia Goncalves, Ewan Harrison, David K. Jackson, Ian Johnston, Dominic Kwiatkowski, Cordelia Langford, John Sillitoe on behalf of the Wellcome Sanger Institute COVID-19 Surveillance Team |
| EPI_ISL_2915848, EPI_ISL_2915849, EPI_ISL_2915850, EPI_ISL_2915851 | Lighthouse Lab in Alderley Park | Wellcome Sanger Institute for the COVID-19 Genomics UK (COG-UK) Consortium | Jacquelyn Wynn, Mairead Hyland, The Lighthouse Lab in Alderley Park and Alex Alderton, Roberto Amato, Jeffrey Barrett, Sonia Goncalves, Ewan Harrison, David K. Jackson, Ian Johnston, Dominic Kwiatkowski, Cordelia Langford, John Sillitoe on behalf of the Wellcome Sanger Institute COVID-19 Surveillance Team |
| EPI_ISL_2915852, EPI_ISL_2915853, EPI_ISL_2915854, EPI_ISL_2915855 | Berkshire and Surrey Pathology Services Lighthouse Laboratory | Wellcome Sanger Institute for the COVID-19 Genomics UK (COG-UK) Consortium | Berkshire and Surrey Pathology Services Lighthouse Laboratory and Alex Alderton, Roberto Amato, Jeffrey Barrett, Sonia Goncalves, Ewan Harrison, David K. Jackson, Ian Johnston, Dominic Kwiatkowski, Cordelia Langford, John Sillitoe on behalf of the Wellcome Sanger Institute COVID-19 Surveillance Team |
| EPI_ISL_2915856, EPI_ISL_2915857 | Lighthouse Lab in Alderley Park | Wellcome Sanger Institute for the COVID-19 Genomics UK (COG-UK) Consortium | Jacquelyn Wynn, Mairead Hyland, The Lighthouse Lab in Alderley Park and Alex Alderton, Roberto Amato, Jeffrey Barrett, Sonia Goncalves, Ewan Harrison, David K. Jackson, Ian Johnston, Dominic Kwiatkowski, Cordelia Langford, John Sillitoe on behalf of the Wellcome Sanger Institute COVID-19 Surveillance Team |
| EPI_ISL_2915858, EPI_ISL_2915859, EPI_ISL_2915860 | Berkshire and Surrey Pathology Services Lighthouse Laboratory | Wellcome Sanger Institute for the COVID-19 Genomics UK (COG-UK) Consortium | Berkshire and Surrey Pathology Services Lighthouse Laboratory and Alex Alderton, Roberto Amato, Jeffrey Barrett, Sonia Goncalves, Ewan Harrison, David K. Jackson, Ian Johnston, Dominic Kwiatkowski, Cordelia Langford, John Sillitoe on behalf of the Wellcome Sanger Institute COVID-19 Surveillance Team |
| EPI_ISL_2915861 | Lighthouse Lab in Alderley Park | Wellcome Sanger Institute for the COVID-19 Genomics UK (COG-UK) Consortium | Jacquelyn Wynn, Mairead Hyland, The Lighthouse Lab in Alderley Park and Alex Alderton, Roberto Amato, Jeffrey Barrett, Sonia Goncalves, Ewan Harrison, David K. Jackson, Ian Johnston, Dominic Kwiatkowski, Cordelia Langford, John Sillitoe on behalf of the Wellcome Sanger Institute COVID-19 Surveillance Team |
| EPI_ISL_2915862 | Berkshire and Surrey Pathology Services Lighthouse Laboratory | Wellcome Sanger Institute for the COVID-19 Genomics UK (COG-UK) Consortium | Berkshire and Surrey Pathology Services Lighthouse Laboratory and Alex Alderton, Roberto Amato, Jeffrey Barrett, Sonia Goncalves, Ewan Harrison, David K. Jackson, Ian Johnston, Dominic Kwiatkowski, Cordelia Langford, John Sillitoe on behalf of the Wellcome Sanger Institute COVID-19 Surveillance Team |
| EPI_ISL_2915863, EPI_ISL_2915864, EPI_ISL_2915865, EPI_ISL_2915866, EPI_ISL_2915867, EPI_ISL_2915868 | Lighthouse Lab in Alderley Park | Wellcome Sanger Institute for the COVID-19 Genomics UK (COG-UK) Consortium | Jacquelyn Wynn, Mairead Hyland, The Lighthouse Lab in Alderley Park and Alex Alderton, Roberto Amato, Jeffrey Barrett, Sonia Goncalves, Ewan Harrison, David K. Jackson, Ian Johnston, Dominic Kwiatkowski, Cordelia Langford, John Sillitoe on behalf of the Wellcome Sanger Institute COVID-19 Surveillance Team |
| EPI_ISL_2915869 | Berkshire and Surrey Pathology Services Lighthouse Laboratory | Wellcome Sanger Institute for the COVID-19 Genomics UK (COG-UK) Consortium | Berkshire and Surrey Pathology Services Lighthouse Laboratory and Alex Alderton, Roberto Amato, Jeffrey Barrett, Sonia Goncalves, Ewan Harrison, David K. Jackson, Ian Johnston, Dominic Kwiatkowski, Cordelia Langford, John Sillitoe on behalf of the Wellcome Sanger Institute COVID-19 Surveillance Team |
| EPI_ISL_2915870 | Lighthouse Lab in Alderley Park | Wellcome Sanger Institute for the COVID-19 Genomics UK (COG-UK) Consortium | Jacquelyn Wynn, Mairead Hyland, The Lighthouse Lab in Alderley Park and Alex Alderton, Roberto Amato, Jeffrey Barrett, Sonia Goncalves, Ewan Harrison, David K. Jackson, Ian Johnston, Dominic Kwiatkowski, Cordelia Langford, John Sillitoe on behalf of the Wellcome Sanger Institute COVID-19 Surveillance Team |
| EPI_ISL_2915871 | Berkshire and Surrey Pathology Services Lighthouse Laboratory | Wellcome Sanger Institute for the COVID-19 Genomics UK (COG-UK) Consortium | Berkshire and Surrey Pathology Services Lighthouse Laboratory and Alex Alderton, Roberto Amato, Jeffrey Barrett, Sonia Goncalves, Ewan Harrison, David K. Jackson, Ian Johnston, Dominic Kwiatkowski, Cordelia Langford, John Sillitoe on behalf of the Wellcome Sanger Institute COVID-19 Surveillance Team |
| EPI_ISL_2915872, EPI_ISL_2915873 | Lighthouse Lab in Alderley Park | Wellcome Sanger Institute for the COVID-19 Genomics UK (COG-UK) Consortium | Jacquelyn Wynn, Mairead Hyland, The Lighthouse Lab in Alderley Park and Alex Alderton, Roberto Amato, Jeffrey Barrett, Sonia Goncalves, Ewan Harrison, David K. Jackson, Ian Johnston, Dominic Kwiatkowski, Cordelia Langford, John Sillitoe on behalf of the Wellcome Sanger Institute COVID-19 Surveillance Team |
| EPI_ISL_2915874 | Berkshire and Surrey Pathology Services Lighthouse Laboratory | Wellcome Sanger Institute for the COVID-19 Genomics UK (COG-UK) Consortium | Berkshire and Surrey Pathology Services Lighthouse Laboratory and Alex Alderton, Roberto Amato, Jeffrey Barrett, Sonia Goncalves, Ewan Harrison, David K. Jackson, Ian Johnston, Dominic Kwiatkowski, Cordelia Langford, John Sillitoe on behalf of the Wellcome Sanger Institute COVID-19 Surveillance Team |
| EPI_ISL_2915875, EPI_ISL_2915876 | Lighthouse Lab in Alderley Park | Wellcome Sanger Institute for the COVID-19 Genomics UK (COG-UK) Consortium | Jacquelyn Wynn, Mairead Hyland, The Lighthouse Lab in Alderley Park and Alex Alderton, Roberto Amato, Jeffrey Barrett, Sonia Goncalves, Ewan Harrison, David K. Jackson, Ian Johnston, Dominic Kwiatkowski, Cordelia Langford, John Sillitoe on behalf of the Wellcome Sanger Institute COVID-19 Surveillance Team |
| EPI_ISL_2915877 | Berkshire and Surrey Pathology Services Lighthouse Laboratory | Wellcome Sanger Institute for the COVID-19 Genomics UK (COG-UK) Consortium | Berkshire and Surrey Pathology Services Lighthouse Laboratory and Alex Alderton, Roberto Amato, Jeffrey Barrett, Sonia Goncalves, Ewan Harrison, David K. Jackson, Ian Johnston, Dominic Kwiatkowski, Cordelia Langford, John Sillitoe on behalf of the Wellcome Sanger Institute COVID-19 Surveillance Team |
| EPI_ISL_2915878, EPI_ISL_2915879, EPI_ISL_2915880 | Lighthouse Lab in Alderley Park | Wellcome Sanger Institute for the COVID-19 Genomics UK (COG-UK) Consortium | Jacquelyn Wynn, Mairead Hyland, The Lighthouse Lab in Alderley Park and Alex Alderton, Roberto Amato, Jeffrey Barrett, Sonia Goncalves, Ewan Harrison, David K. Jackson, Ian Johnston, Dominic Kwiatkowski, Cordelia Langford, John Sillitoe on behalf of the Wellcome Sanger Institute COVID-19 Surveillance Team |
| EPI_ISL_2915881 | Berkshire and Surrey Pathology Services Lighthouse Laboratory | Wellcome Sanger Institute for the COVID-19 Genomics UK (COG-UK) Consortium | Berkshire and Surrey Pathology Services Lighthouse Laboratory and Alex Alderton, Roberto Amato, Jeffrey Barrett, Sonia Goncalves, Ewan Harrison, David K. Jackson, Ian Johnston, Dominic Kwiatkowski, Cordelia Langford, John Sillitoe on behalf of the Wellcome Sanger Institute COVID-19 Surveillance Team |
| EPI_ISL_2915882, EPI_ISL_2915883, EPI_ISL_2915884, | Lighthouse Lab in Alderley Park | Wellcome Sanger Institute for the COVID-19 Genomics UK (COG-UK) Consortium | Jacquelyn Wynn, Mairead Hyland, The Lighthouse Lab in Alderley Park and Alex Alderton, Roberto Amato, Jeffrey Barrett, Sonia Goncalves, Ewan Harrison, David K. Jackson, Ian Johnston, Dominic Kwiatkowski, Cordelia Langford, John Sillitoe on behalf of the Wellcome Sanger Institute COVID-19 Surveillance Team |

| | | | |
|---|---|---|---|
| EPI_ISL_2915885 | | | |
| EPI_ISL_2915886 | Berkshire and Surrey Pathology Services Lighthouse Laboratory | Wellcome Sanger Institute for the COVID-19 Genomics UK (COG-UK) Consortium | Berkshire and Surrey Pathology Services Lighthouse Laboratory and Alex Alderton, Roberto Amato, Jeffrey Barrett, Sonia Goncalves, Ewan Harrison, David K. Jackson, Ian Johnston, Dominic Kwiatkowski, Cordelia Langford, John Sillitoe on behalf of the Wellcome Sanger Institute COVID-19 Surveillance Team |
| EPI_ISL_2915887, EPI_ISL_2915888 | Lighthouse Lab in Alderley Park | Wellcome Sanger Institute for the COVID-19 Genomics UK (COG-UK) Consortium | Jacquelyn Wynn, Mairead Hyland, The Lighthouse Lab in Alderley Park and Alex Alderton, Roberto Amato, Jeffrey Barrett, Sonia Goncalves, Ewan Harrison, David K. Jackson, Ian Johnston, Dominic Kwiatkowski, Cordelia Langford, John Sillitoe on behalf of the Wellcome Sanger Institute COVID-19 Surveillance Team |
| EPI_ISL_2915889, EPI_ISL_2915890 | Berkshire and Surrey Pathology Services Lighthouse Laboratory | Wellcome Sanger Institute for the COVID-19 Genomics UK (COG-UK) Consortium | Berkshire and Surrey Pathology Services Lighthouse Laboratory and Alex Alderton, Roberto Amato, Jeffrey Barrett, Sonia Goncalves, Ewan Harrison, David K. Jackson, Ian Johnston, Dominic Kwiatkowski, Cordelia Langford, John Sillitoe on behalf of the Wellcome Sanger Institute COVID-19 Surveillance Team |
| EPI_ISL_2915891, EPI_ISL_2915892, EPI_ISL_2915893, EPI_ISL_2915894, EPI_ISL_2915895 | Lighthouse Lab in Alderley Park | Wellcome Sanger Institute for the COVID-19 Genomics UK (COG-UK) Consortium | Jacquelyn Wynn, Mairead Hyland, The Lighthouse Lab in Alderley Park and Alex Alderton, Roberto Amato, Jeffrey Barrett, Sonia Goncalves, Ewan Harrison, David K. Jackson, Ian Johnston, Dominic Kwiatkowski, Cordelia Langford, John Sillitoe on behalf of the Wellcome Sanger Institute COVID-19 Surveillance Team |
| EPI_ISL_2915896 | Berkshire and Surrey Pathology Services Lighthouse Laboratory | Wellcome Sanger Institute for the COVID-19 Genomics UK (COG-UK) Consortium | Berkshire and Surrey Pathology Services Lighthouse Laboratory and Alex Alderton, Roberto Amato, Jeffrey Barrett, Sonia Goncalves, Ewan Harrison, David K. Jackson, Ian Johnston, Dominic Kwiatkowski, Cordelia Langford, John Sillitoe on behalf of the Wellcome Sanger Institute COVID-19 Surveillance Team |
| EPI_ISL_2915897, EPI_ISL_2915898, EPI_ISL_2915899 | Lighthouse Lab in Alderley Park | Wellcome Sanger Institute for the COVID-19 Genomics UK (COG-UK) Consortium | Jacquelyn Wynn, Mairead Hyland, The Lighthouse Lab in Alderley Park and Alex Alderton, Roberto Amato, Jeffrey Barrett, Sonia Goncalves, Ewan Harrison, David K. Jackson, Ian Johnston, Dominic Kwiatkowski, Cordelia Langford, John Sillitoe on behalf of the Wellcome Sanger Institute COVID-19 Surveillance Team |
| EPI_ISL_2915900 | Berkshire and Surrey Pathology Services Lighthouse Laboratory | Wellcome Sanger Institute for the COVID-19 Genomics UK (COG-UK) Consortium | Berkshire and Surrey Pathology Services Lighthouse Laboratory and Alex Alderton, Roberto Amato, Jeffrey Barrett, Sonia Goncalves, Ewan Harrison, David K. Jackson, Ian Johnston, Dominic Kwiatkowski, Cordelia Langford, John Sillitoe on behalf of the Wellcome Sanger Institute COVID-19 Surveillance Team |
| EPI_ISL_2915901 | Lighthouse Lab in Alderley Park | Wellcome Sanger Institute for the COVID-19 Genomics UK (COG-UK) Consortium | Jacquelyn Wynn, Mairead Hyland, The Lighthouse Lab in Alderley Park and Alex Alderton, Roberto Amato, Jeffrey Barrett, Sonia Goncalves, Ewan Harrison, David K. Jackson, Ian Johnston, Dominic Kwiatkowski, Cordelia Langford, John Sillitoe on behalf of the Wellcome Sanger Institute COVID-19 Surveillance Team |
| EPI_ISL_2915902, EPI_ISL_2915903 | Berkshire and Surrey Pathology Services Lighthouse Laboratory | Wellcome Sanger Institute for the COVID-19 Genomics UK (COG-UK) Consortium | Berkshire and Surrey Pathology Services Lighthouse Laboratory and Alex Alderton, Roberto Amato, Jeffrey Barrett, Sonia Goncalves, Ewan Harrison, David K. Jackson, Ian Johnston, Dominic Kwiatkowski, Cordelia Langford, John Sillitoe on behalf of the Wellcome Sanger Institute COVID-19 Surveillance Team |
| EPI_ISL_2915904, EPI_ISL_2915905 | Lighthouse Lab in Alderley Park | Wellcome Sanger Institute for the COVID-19 Genomics UK (COG-UK) Consortium | Jacquelyn Wynn, Mairead Hyland, The Lighthouse Lab in Alderley Park and Alex Alderton, Roberto Amato, Jeffrey Barrett, Sonia Goncalves, Ewan Harrison, David K. Jackson, Ian Johnston, Dominic Kwiatkowski, Cordelia Langford, John Sillitoe on behalf of the Wellcome Sanger Institute COVID-19 Surveillance Team |
| EPI_ISL_2915906, EPI_ISL_2915907, EPI_ISL_2915908 | Berkshire and Surrey Pathology Services Lighthouse Laboratory | Wellcome Sanger Institute for the COVID-19 Genomics UK (COG-UK) Consortium | Berkshire and Surrey Pathology Services Lighthouse Laboratory and Alex Alderton, Roberto Amato, Jeffrey Barrett, Sonia Goncalves, Ewan Harrison, David K. Jackson, Ian Johnston, Dominic Kwiatkowski, Cordelia Langford, John Sillitoe on behalf of the Wellcome Sanger Institute COVID-19 Surveillance Team |
| EPI_ISL_2915909, EPI_ISL_2915910, EPI_ISL_2915911 | Lighthouse Lab in Alderley Park | Wellcome Sanger Institute for the COVID-19 Genomics UK (COG-UK) Consortium | Jacquelyn Wynn, Mairead Hyland, The Lighthouse Lab in Alderley Park and Alex Alderton, Roberto Amato, Jeffrey Barrett, Sonia Goncalves, Ewan Harrison, David K. Jackson, Ian Johnston, Dominic Kwiatkowski, Cordelia Langford, John Sillitoe on behalf of the Wellcome Sanger Institute COVID-19 Surveillance Team |
| EPI_ISL_2915912 | Berkshire and Surrey Pathology Services Lighthouse Laboratory | Wellcome Sanger Institute for the COVID-19 Genomics UK (COG-UK) Consortium | Berkshire and Surrey Pathology Services Lighthouse Laboratory and Alex Alderton, Roberto Amato, Jeffrey Barrett, Sonia Goncalves, Ewan Harrison, David K. Jackson, Ian Johnston, Dominic Kwiatkowski, Cordelia Langford, John Sillitoe on behalf of the Wellcome Sanger Institute COVID-19 Surveillance Team |
| EPI_ISL_2915913 | Lighthouse Lab in Alderley Park | Wellcome Sanger Institute for the COVID-19 Genomics UK (COG-UK) Consortium | Jacquelyn Wynn, Mairead Hyland, The Lighthouse Lab in Alderley Park and Alex Alderton, Roberto Amato, Jeffrey Barrett, Sonia Goncalves, Ewan Harrison, David K. Jackson, Ian Johnston, Dominic Kwiatkowski, Cordelia Langford, John Sillitoe on behalf of the Wellcome Sanger Institute COVID-19 Surveillance Team |
| EPI_ISL_2915914 | Berkshire and Surrey Pathology Services Lighthouse Laboratory | Wellcome Sanger Institute for the COVID-19 Genomics UK (COG-UK) Consortium | Berkshire and Surrey Pathology Services Lighthouse Laboratory and Alex Alderton, Roberto Amato, Jeffrey Barrett, Sonia Goncalves, Ewan Harrison, David K. Jackson, Ian Johnston, Dominic Kwiatkowski, Cordelia Langford, John Sillitoe on behalf of the Wellcome Sanger Institute COVID-19 Surveillance Team |
| EPI_ISL_2915915, EPI_ISL_2915916, EPI_ISL_2915917 | Lighthouse Lab in Alderley Park | Wellcome Sanger Institute for the COVID-19 Genomics UK (COG-UK) Consortium | Jacquelyn Wynn, Mairead Hyland, The Lighthouse Lab in Alderley Park and Alex Alderton, Roberto Amato, Jeffrey Barrett, Sonia Goncalves, Ewan Harrison, David K. Jackson, Ian Johnston, Dominic Kwiatkowski, Cordelia Langford, John Sillitoe on behalf of the Wellcome Sanger Institute COVID-19 Surveillance Team |
| EPI_ISL_2915918 | Berkshire and Surrey Pathology Services Lighthouse Laboratory | Wellcome Sanger Institute for the COVID-19 Genomics UK (COG-UK) Consortium | Berkshire and Surrey Pathology Services Lighthouse Laboratory and Alex Alderton, Roberto Amato, Jeffrey Barrett, Sonia Goncalves, Ewan Harrison, David K. Jackson, Ian Johnston, Dominic Kwiatkowski, Cordelia Langford, John Sillitoe on behalf of the Wellcome Sanger Institute COVID-19 Surveillance Team |
| EPI_ISL_2915919 | Lighthouse Lab in Alderley Park | Wellcome Sanger Institute for the COVID-19 Genomics UK (COG-UK) Consortium | Jacquelyn Wynn, Mairead Hyland, The Lighthouse Lab in Alderley Park and Alex Alderton, Roberto Amato, Jeffrey Barrett, Sonia Goncalves, Ewan Harrison, David K. Jackson, Ian Johnston, Dominic Kwiatkowski, Cordelia Langford, John Sillitoe on behalf of the Wellcome Sanger Institute COVID-19 Surveillance Team |
| EPI_ISL_2915920, EPI_ISL_2915921 | Berkshire and Surrey Pathology Services Lighthouse Laboratory | Wellcome Sanger Institute for the COVID-19 Genomics UK (COG-UK) Consortium | Berkshire and Surrey Pathology Services Lighthouse Laboratory and Alex Alderton, Roberto Amato, Jeffrey Barrett, Sonia Goncalves, Ewan Harrison, David K. Jackson, Ian Johnston, Dominic Kwiatkowski, Cordelia Langford, John Sillitoe on behalf of the Wellcome Sanger Institute COVID-19 Surveillance Team |
| EPI_ISL_2915922, EPI_ISL_2915923 | Lighthouse Lab in Alderley Park | Wellcome Sanger Institute for the COVID-19 Genomics UK (COG-UK) Consortium | Jacquelyn Wynn, Mairead Hyland, The Lighthouse Lab in Alderley Park and Alex Alderton, Roberto Amato, Jeffrey Barrett, Sonia Goncalves, Ewan Harrison, David K. Jackson, Ian Johnston, Dominic Kwiatkowski, Cordelia Langford, John Sillitoe on behalf of the Wellcome Sanger Institute COVID-19 Surveillance Team |
| EPI_ISL_2915924 | Berkshire and Surrey Pathology Services Lighthouse Laboratory | Wellcome Sanger Institute for the COVID-19 Genomics UK (COG-UK) Consortium | Berkshire and Surrey Pathology Services Lighthouse Laboratory and Alex Alderton, Roberto Amato, Jeffrey Barrett, Sonia Goncalves, Ewan Harrison, David K. Jackson, Ian Johnston, Dominic Kwiatkowski, Cordelia Langford, John Sillitoe on behalf of the Wellcome Sanger Institute COVID-19 Surveillance Team |
| EPI_ISL_2915925, EPI_ISL_2915926 | Lighthouse Lab in Alderley Park | Wellcome Sanger Institute for the COVID-19 Genomics UK (COG-UK) Consortium | Jacquelyn Wynn, Mairead Hyland, The Lighthouse Lab in Alderley Park and Alex Alderton, Roberto Amato, Jeffrey Barrett, Sonia Goncalves, Ewan Harrison, David K. Jackson, Ian Johnston, Dominic Kwiatkowski, Cordelia Langford, John Sillitoe on behalf of the Wellcome Sanger Institute COVID-19 Surveillance Team |
| EPI_ISL_2915927 | Berkshire and Surrey Pathology Services Lighthouse Laboratory | Wellcome Sanger Institute for the COVID-19 Genomics UK (COG-UK) Consortium | Berkshire and Surrey Pathology Services Lighthouse Laboratory and Alex Alderton, Roberto Amato, Jeffrey Barrett, Sonia Goncalves, Ewan Harrison, David K. Jackson, Ian Johnston, Dominic Kwiatkowski, Cordelia Langford, John Sillitoe on behalf of the Wellcome Sanger Institute COVID-19 Surveillance Team |
| EPI_ISL_2915928, EPI_ISL_2915929 | Lighthouse Lab in Alderley Park | Wellcome Sanger Institute for the COVID-19 Genomics UK (COG-UK) Consortium | Jacquelyn Wynn, Mairead Hyland, The Lighthouse Lab in Alderley Park and Alex Alderton, Roberto Amato, Jeffrey Barrett, Sonia Goncalves, Ewan Harrison, David K. Jackson, Ian Johnston, Dominic Kwiatkowski, Cordelia Langford, John Sillitoe on behalf of the Wellcome Sanger Institute COVID-19 Surveillance Team |
| EPI_ISL_2915930 | Berkshire and Surrey Pathology Services Lighthouse Laboratory | Wellcome Sanger Institute for the COVID-19 Genomics UK (COG-UK) Consortium | Berkshire and Surrey Pathology Services Lighthouse Laboratory and Alex Alderton, Roberto Amato, Jeffrey Barrett, Sonia Goncalves, Ewan Harrison, David K. Jackson, Ian Johnston, Dominic Kwiatkowski, Cordelia Langford, John Sillitoe on behalf of the Wellcome Sanger Institute COVID-19 Surveillance Team |
| EPI_ISL_2915931 | Lighthouse Lab in Alderley Park | Wellcome Sanger Institute for the COVID-19 Genomics UK (COG-UK) Consortium | Jacquelyn Wynn, Mairead Hyland, The Lighthouse Lab in Alderley Park and Alex Alderton, Roberto Amato, Jeffrey Barrett, Sonia Goncalves, Ewan Harrison, David K. Jackson, Ian Johnston, Dominic Kwiatkowski, Cordelia Langford, John Sillitoe on behalf of the Wellcome Sanger Institute COVID-19 Surveillance Team |
| EPI_ISL_2915932 | Berkshire and Surrey Pathology Services Lighthouse Laboratory | Wellcome Sanger Institute for the COVID-19 Genomics UK (COG-UK) Consortium | Berkshire and Surrey Pathology Services Lighthouse Laboratory and Alex Alderton, Roberto Amato, Jeffrey Barrett, Sonia Goncalves, Ewan Harrison, David K. Jackson, Ian Johnston, Dominic Kwiatkowski, Cordelia Langford, John Sillitoe on behalf of the Wellcome Sanger Institute COVID-19 Surveillance Team |
| EPI_ISL_2915933, EPI_ISL_2915934 | Lighthouse Lab in Alderley Park | Wellcome Sanger Institute for the COVID-19 Genomics UK (COG-UK) Consortium | Jacquelyn Wynn, Mairead Hyland, The Lighthouse Lab in Alderley Park and Alex Alderton, Roberto Amato, Jeffrey Barrett, Sonia Goncalves, Ewan Harrison, David K. Jackson, Ian Johnston, Dominic Kwiatkowski, Cordelia Langford, John Sillitoe on behalf of the Wellcome Sanger Institute COVID-19 Surveillance Team |
| EPI_ISL_2915935, EPI_ISL_2915936 | Berkshire and Surrey Pathology Services Lighthouse Laboratory | Wellcome Sanger Institute for the COVID-19 Genomics UK (COG-UK) Consortium | Berkshire and Surrey Pathology Services Lighthouse Laboratory and Alex Alderton, Roberto Amato, Jeffrey Barrett, Sonia Goncalves, Ewan Harrison, David K. Jackson, Ian Johnston, Dominic Kwiatkowski, Cordelia Langford, John Sillitoe on behalf of the Wellcome Sanger Institute COVID-19 Surveillance Team |
| EPI_ISL_2915937, EPI_ISL_2915938, EPI_ISL_2915939, EPI_ISL_2915940 | Lighthouse Lab in Alderley Park | Wellcome Sanger Institute for the COVID-19 Genomics UK (COG-UK) Consortium | Jacquelyn Wynn, Mairead Hyland, The Lighthouse Lab in Alderley Park and Alex Alderton, Roberto Amato, Jeffrey Barrett, Sonia Goncalves, Ewan Harrison, David K. Jackson, Ian Johnston, Dominic Kwiatkowski, Cordelia Langford, John Sillitoe on behalf of the Wellcome Sanger Institute COVID-19 Surveillance Team |
| EPI_ISL_2915941 | Berkshire and Surrey Pathology Services Lighthouse Laboratory | Wellcome Sanger Institute for the COVID-19 Genomics UK (COG-UK) Consortium | Berkshire and Surrey Pathology Services Lighthouse Laboratory and Alex Alderton, Roberto Amato, Jeffrey Barrett, Sonia Goncalves, Ewan Harrison, David K. Jackson, Ian Johnston, Dominic Kwiatkowski, Cordelia Langford, John Sillitoe on behalf of the Wellcome Sanger Institute COVID-19 Surveillance Team |
| EPI_ISL_2915942, EPI_ISL_2915943, EPI_ISL_2915944 | Lighthouse Lab in Alderley Park | Wellcome Sanger Institute for the COVID-19 Genomics UK (COG-UK) Consortium | Jacquelyn Wynn, Mairead Hyland, The Lighthouse Lab in Alderley Park and Alex Alderton, Roberto Amato, Jeffrey Barrett, Sonia Goncalves, Ewan Harrison, David K. Jackson, Ian Johnston, Dominic Kwiatkowski, Cordelia Langford, John Sillitoe on behalf of the Wellcome Sanger Institute COVID-19 Surveillance Team |
| EPI_ISL_2915945 | Berkshire and Surrey Pathology Services Lighthouse Laboratory | Wellcome Sanger Institute for the COVID-19 Genomics UK (COG-UK) Consortium | Berkshire and Surrey Pathology Services Lighthouse Laboratory and Alex Alderton, Roberto Amato, Jeffrey Barrett, Sonia Goncalves, Ewan Harrison, David K. Jackson, Ian Johnston, Dominic Kwiatkowski, Cordelia Langford, John Sillitoe on behalf of the Wellcome Sanger Institute COVID-19 Surveillance Team |
| EPI_ISL_2915946 | Lighthouse Lab in Alderley Park | Wellcome Sanger Institute for the COVID-19 Genomics UK (COG-UK) Consortium | Jacquelyn Wynn, Mairead Hyland, The Lighthouse Lab in Alderley Park and Alex Alderton, Roberto Amato, Jeffrey Barrett, Sonia Goncalves, Ewan Harrison, David K. Jackson, Ian Johnston, Dominic Kwiatkowski, Cordelia Langford, John Sillitoe on behalf of the Wellcome Sanger Institute COVID-19 Surveillance Team |
| EPI_ISL_2915947 | Berkshire and Surrey Pathology Services Lighthouse Laboratory | Wellcome Sanger Institute for the COVID-19 Genomics UK (COG-UK) Consortium | Berkshire and Surrey Pathology Services Lighthouse Laboratory and Alex Alderton, Roberto Amato, Jeffrey Barrett, Sonia Goncalves, Ewan Harrison, David K. Jackson, Ian Johnston, Dominic Kwiatkowski, Cordelia Langford, John Sillitoe on behalf of the Wellcome Sanger Institute COVID-19 Surveillance Team |
| EPI_ISL_2915948 | Lighthouse Lab in Alderley Park | Wellcome Sanger Institute for the COVID-19 Genomics UK (COG-UK) Consortium | Jacquelyn Wynn, Mairead Hyland, The Lighthouse Lab in Alderley Park and Alex Alderton, Roberto Amato, Jeffrey Barrett, Sonia Goncalves, Ewan Harrison, David K. Jackson, Ian Johnston, Dominic Kwiatkowski, Cordelia Langford, John Sillitoe on behalf of the Wellcome Sanger Institute COVID-19 Surveillance Team |
| EPI_ISL_2915949 | Berkshire and Surrey Pathology Services Lighthouse Laboratory | Wellcome Sanger Institute for the COVID-19 Genomics UK (COG-UK) Consortium | Berkshire and Surrey Pathology Services Lighthouse Laboratory and Alex Alderton, Roberto Amato, Jeffrey Barrett, Sonia Goncalves, Ewan Harrison, David K. Jackson, Ian Johnston, Dominic Kwiatkowski, Cordelia Langford, John Sillitoe on behalf of the Wellcome Sanger Institute COVID-19 Surveillance Team |
| EPI_ISL_2915950 | Lighthouse Lab in Alderley Park | Wellcome Sanger Institute for the COVID-19 Genomics UK (COG-UK) Consortium | Jacquelyn Wynn, Mairead Hyland, The Lighthouse Lab in Alderley Park and Alex Alderton, Roberto Amato, Jeffrey Barrett, Sonia Goncalves, Ewan Harrison, David K. Jackson, Ian Johnston, Dominic Kwiatkowski, Cordelia Langford, John Sillitoe on behalf of the Wellcome Sanger Institute COVID-19 Surveillance Team |
| EPI_ISL_2915951 | Berkshire and Surrey Pathology Services Lighthouse Laboratory | Wellcome Sanger Institute for the COVID-19 Genomics UK (COG-UK) Consortium | Berkshire and Surrey Pathology Services Lighthouse Laboratory and Alex Alderton, Roberto Amato, Jeffrey Barrett, Sonia Goncalves, Ewan Harrison, David K. Jackson, Ian Johnston, Dominic Kwiatkowski, Cordelia Langford, John Sillitoe on behalf of the Wellcome Sanger Institute COVID-19 Surveillance Team |
| EPI_ISL_2915952 | Lighthouse Lab in Alderley Park | Wellcome Sanger Institute for the COVID-19 Genomics UK (COG-UK) Consortium | Jacquelyn Wynn, Mairead Hyland, The Lighthouse Lab in Alderley Park and Alex Alderton, Roberto Amato, Jeffrey Barrett, Sonia Goncalves, Ewan Harrison, David K. Jackson, Ian Johnston, Dominic Kwiatkowski, Cordelia Langford, John Sillitoe on behalf of the Wellcome Sanger Institute COVID-19 Surveillance Team |

| | | | |
|---|---|---|---|
| EPI_ISL_2915953, EPI_ISL_2915954 | Berkshire and Surrey Pathology Services Lighthouse Laboratory | Wellcome Sanger Institute for the COVID-19 Genomics UK (COG-UK) Consortium | Berkshire and Surrey Pathology Services Lighthouse Laboratory and Alex Alderton, Roberto Amato, Jeffrey Barrett, Sonia Goncalves, Ewan Harrison, David K. Jackson, Ian Johnston, Dominic Kwiatkowski, Cordelia Langford, John Sillitoe on behalf of the Wellcome Sanger Institute COVID-19 Surveillance Team |
| EPI_ISL_2915955 | Lighthouse Lab in Alderley Park | Wellcome Sanger Institute for the COVID-19 Genomics UK (COG-UK) Consortium | Jacquelyn Wynn, Mairead Hyland, The Lighthouse Lab in Alderley Park and Alex Alderton, Roberto Amato, Jeffrey Barrett, Sonia Goncalves, Ewan Harrison, David K. Jackson, Ian Johnston, Dominic Kwiatkowski, Cordelia Langford, John Sillitoe on behalf of the Wellcome Sanger Institute COVID-19 Surveillance Team |
| EPI_ISL_2915956 | Berkshire and Surrey Pathology Services Lighthouse Laboratory | Wellcome Sanger Institute for the COVID-19 Genomics UK (COG-UK) Consortium | Berkshire and Surrey Pathology Services Lighthouse Laboratory and Alex Alderton, Roberto Amato, Jeffrey Barrett, Sonia Goncalves, Ewan Harrison, David K. Jackson, Ian Johnston, Dominic Kwiatkowski, Cordelia Langford, John Sillitoe on behalf of the Wellcome Sanger Institute COVID-19 Surveillance Team |
| EPI_ISL_2915957, EPI_ISL_2915958, EPI_ISL_2915959 | Lighthouse Lab in Alderley Park | Wellcome Sanger Institute for the COVID-19 Genomics UK (COG-UK) Consortium | Jacquelyn Wynn, Mairead Hyland, The Lighthouse Lab in Alderley Park and Alex Alderton, Roberto Amato, Jeffrey Barrett, Sonia Goncalves, Ewan Harrison, David K. Jackson, Ian Johnston, Dominic Kwiatkowski, Cordelia Langford, John Sillitoe on behalf of the Wellcome Sanger Institute COVID-19 Surveillance Team |
| EPI_ISL_2915960, EPI_ISL_2915961 | Berkshire and Surrey Pathology Services Lighthouse Laboratory | Wellcome Sanger Institute for the COVID-19 Genomics UK (COG-UK) Consortium | Berkshire and Surrey Pathology Services Lighthouse Laboratory and Alex Alderton, Roberto Amato, Jeffrey Barrett, Sonia Goncalves, Ewan Harrison, David K. Jackson, Ian Johnston, Dominic Kwiatkowski, Cordelia Langford, John Sillitoe on behalf of the Wellcome Sanger Institute COVID-19 Surveillance Team |
| EPI_ISL_2915962, EPI_ISL_2915963 | Lighthouse Lab in Alderley Park | Wellcome Sanger Institute for the COVID-19 Genomics UK (COG-UK) Consortium | Jacquelyn Wynn, Mairead Hyland, The Lighthouse Lab in Alderley Park and Alex Alderton, Roberto Amato, Jeffrey Barrett, Sonia Goncalves, Ewan Harrison, David K. Jackson, Ian Johnston, Dominic Kwiatkowski, Cordelia Langford, John Sillitoe on behalf of the Wellcome Sanger Institute COVID-19 Surveillance Team |
| EPI_ISL_2915964 | Berkshire and Surrey Pathology Services Lighthouse Laboratory | Wellcome Sanger Institute for the COVID-19 Genomics UK (COG-UK) Consortium | Berkshire and Surrey Pathology Services Lighthouse Laboratory and Alex Alderton, Roberto Amato, Jeffrey Barrett, Sonia Goncalves, Ewan Harrison, David K. Jackson, Ian Johnston, Dominic Kwiatkowski, Cordelia Langford, John Sillitoe on behalf of the Wellcome Sanger Institute COVID-19 Surveillance Team |
| EPI_ISL_2915965 | Lighthouse Lab in Alderley Park | Wellcome Sanger Institute for the COVID-19 Genomics UK (COG-UK) Consortium | Jacquelyn Wynn, Mairead Hyland, The Lighthouse Lab in Alderley Park and Alex Alderton, Roberto Amato, Jeffrey Barrett, Sonia Goncalves, Ewan Harrison, David K. Jackson, Ian Johnston, Dominic Kwiatkowski, Cordelia Langford, John Sillitoe on behalf of the Wellcome Sanger Institute COVID-19 Surveillance Team |
| EPI_ISL_2915966 | Berkshire and Surrey Pathology Services Lighthouse Laboratory | Wellcome Sanger Institute for the COVID-19 Genomics UK (COG-UK) Consortium | Berkshire and Surrey Pathology Services Lighthouse Laboratory and Alex Alderton, Roberto Amato, Jeffrey Barrett, Sonia Goncalves, Ewan Harrison, David K. Jackson, Ian Johnston, Dominic Kwiatkowski, Cordelia Langford, John Sillitoe on behalf of the Wellcome Sanger Institute COVID-19 Surveillance Team |
| EPI_ISL_2915967 | Lighthouse Lab in Alderley Park | Wellcome Sanger Institute for the COVID-19 Genomics UK (COG-UK) Consortium | Jacquelyn Wynn, Mairead Hyland, The Lighthouse Lab in Alderley Park and Alex Alderton, Roberto Amato, Jeffrey Barrett, Sonia Goncalves, Ewan Harrison, David K. Jackson, Ian Johnston, Dominic Kwiatkowski, Cordelia Langford, John Sillitoe on behalf of the Wellcome Sanger Institute COVID-19 Surveillance Team |
| EPI_ISL_2915968 | Berkshire and Surrey Pathology Services Lighthouse Laboratory | Wellcome Sanger Institute for the COVID-19 Genomics UK (COG-UK) Consortium | Berkshire and Surrey Pathology Services Lighthouse Laboratory and Alex Alderton, Roberto Amato, Jeffrey Barrett, Sonia Goncalves, Ewan Harrison, David K. Jackson, Ian Johnston, Dominic Kwiatkowski, Cordelia Langford, John Sillitoe on behalf of the Wellcome Sanger Institute COVID-19 Surveillance Team |
| EPI_ISL_2915970, EPI_ISL_2915971 | Lighthouse Lab in Alderley Park | Wellcome Sanger Institute for the COVID-19 Genomics UK (COG-UK) Consortium | Jacquelyn Wynn, Mairead Hyland, The Lighthouse Lab in Alderley Park and Alex Alderton, Roberto Amato, Jeffrey Barrett, Sonia Goncalves, Ewan Harrison, David K. Jackson, Ian Johnston, Dominic Kwiatkowski, Cordelia Langford, John Sillitoe on behalf of the Wellcome Sanger Institute COVID-19 Surveillance Team |
| EPI_ISL_2915972 | Berkshire and Surrey Pathology Services Lighthouse Laboratory | Wellcome Sanger Institute for the COVID-19 Genomics UK (COG-UK) Consortium | Berkshire and Surrey Pathology Services Lighthouse Laboratory and Alex Alderton, Roberto Amato, Jeffrey Barrett, Sonia Goncalves, Ewan Harrison, David K. Jackson, Ian Johnston, Dominic Kwiatkowski, Cordelia Langford, John Sillitoe on behalf of the Wellcome Sanger Institute COVID-19 Surveillance Team |
| EPI_ISL_2915973 | Lighthouse Lab in Alderley Park | Wellcome Sanger Institute for the COVID-19 Genomics UK (COG-UK) Consortium | Jacquelyn Wynn, Mairead Hyland, The Lighthouse Lab in Alderley Park and Alex Alderton, Roberto Amato, Jeffrey Barrett, Sonia Goncalves, Ewan Harrison, David K. Jackson, Ian Johnston, Dominic Kwiatkowski, Cordelia Langford, John Sillitoe on behalf of the Wellcome Sanger Institute COVID-19 Surveillance Team |
| EPI_ISL_2915974, EPI_ISL_2915975 | Berkshire and Surrey Pathology Services Lighthouse Laboratory | Wellcome Sanger Institute for the COVID-19 Genomics UK (COG-UK) Consortium | Berkshire and Surrey Pathology Services Lighthouse Laboratory and Alex Alderton, Roberto Amato, Jeffrey Barrett, Sonia Goncalves, Ewan Harrison, David K. Jackson, Ian Johnston, Dominic Kwiatkowski, Cordelia Langford, John Sillitoe on behalf of the Wellcome Sanger Institute COVID-19 Surveillance Team |
| EPI_ISL_2915976, EPI_ISL_2915977 | Lighthouse Lab in Alderley Park | Wellcome Sanger Institute for the COVID-19 Genomics UK (COG-UK) Consortium | Jacquelyn Wynn, Mairead Hyland, The Lighthouse Lab in Alderley Park and Alex Alderton, Roberto Amato, Jeffrey Barrett, Sonia Goncalves, Ewan Harrison, David K. Jackson, Ian Johnston, Dominic Kwiatkowski, Cordelia Langford, John Sillitoe on behalf of the Wellcome Sanger Institute COVID-19 Surveillance Team |
| EPI_ISL_2915978, EPI_ISL_2915979 | Berkshire and Surrey Pathology Services Lighthouse Laboratory | Wellcome Sanger Institute for the COVID-19 Genomics UK (COG-UK) Consortium | Berkshire and Surrey Pathology Services Lighthouse Laboratory and Alex Alderton, Roberto Amato, Jeffrey Barrett, Sonia Goncalves, Ewan Harrison, David K. Jackson, Ian Johnston, Dominic Kwiatkowski, Cordelia Langford, John Sillitoe on behalf of the Wellcome Sanger Institute COVID-19 Surveillance Team |
| EPI_ISL_2915980 | Lighthouse Lab in Alderley Park | Wellcome Sanger Institute for the COVID-19 Genomics UK (COG-UK) Consortium | Jacquelyn Wynn, Mairead Hyland, The Lighthouse Lab in Alderley Park and Alex Alderton, Roberto Amato, Jeffrey Barrett, Sonia Goncalves, Ewan Harrison, David K. Jackson, Ian Johnston, Dominic Kwiatkowski, Cordelia Langford, John Sillitoe on behalf of the Wellcome Sanger Institute COVID-19 Surveillance Team |
| EPI_ISL_2915981 | Berkshire and Surrey Pathology Services Lighthouse Laboratory | Wellcome Sanger Institute for the COVID-19 Genomics UK (COG-UK) Consortium | Berkshire and Surrey Pathology Services Lighthouse Laboratory and Alex Alderton, Roberto Amato, Jeffrey Barrett, Sonia Goncalves, Ewan Harrison, David K. Jackson, Ian Johnston, Dominic Kwiatkowski, Cordelia Langford, John Sillitoe on behalf of the Wellcome Sanger Institute COVID-19 Surveillance Team |
| EPI_ISL_2915982, EPI_ISL_2915983, EPI_ISL_2915984, EPI_ISL_2915985, EPI_ISL_2915986, EPI_ISL_2915987 | Lighthouse Lab in Alderley Park | Wellcome Sanger Institute for the COVID-19 Genomics UK (COG-UK) Consortium | Jacquelyn Wynn, Mairead Hyland, The Lighthouse Lab in Alderley Park and Alex Alderton, Roberto Amato, Jeffrey Barrett, Sonia Goncalves, Ewan Harrison, David K. Jackson, Ian Johnston, Dominic Kwiatkowski, Cordelia Langford, John Sillitoe on behalf of the Wellcome Sanger Institute COVID-19 Surveillance Team |
| EPI_ISL_2915988, EPI_ISL_2915989 | Berkshire and Surrey Pathology Services Lighthouse Laboratory | Wellcome Sanger Institute for the COVID-19 Genomics UK (COG-UK) Consortium | Berkshire and Surrey Pathology Services Lighthouse Laboratory and Alex Alderton, Roberto Amato, Jeffrey Barrett, Sonia Goncalves, Ewan Harrison, David K. Jackson, Ian Johnston, Dominic Kwiatkowski, Cordelia Langford, John Sillitoe on behalf of the Wellcome Sanger Institute COVID-19 Surveillance Team |
| EPI_ISL_2915990, EPI_ISL_2915991, EPI_ISL_2915992, EPI_ISL_2915993 | Lighthouse Lab in Alderley Park | Wellcome Sanger Institute for the COVID-19 Genomics UK (COG-UK) Consortium | Jacquelyn Wynn, Mairead Hyland, The Lighthouse Lab in Alderley Park and Alex Alderton, Roberto Amato, Jeffrey Barrett, Sonia Goncalves, Ewan Harrison, David K. Jackson, Ian Johnston, Dominic Kwiatkowski, Cordelia Langford, John Sillitoe on behalf of the Wellcome Sanger Institute COVID-19 Surveillance Team |
| EPI_ISL_2915994 | Berkshire and Surrey Pathology Services Lighthouse Laboratory | Wellcome Sanger Institute for the COVID-19 Genomics UK (COG-UK) Consortium | Berkshire and Surrey Pathology Services Lighthouse Laboratory and Alex Alderton, Roberto Amato, Jeffrey Barrett, Sonia Goncalves, Ewan Harrison, David K. Jackson, Ian Johnston, Dominic Kwiatkowski, Cordelia Langford, John Sillitoe on behalf of the Wellcome Sanger Institute COVID-19 Surveillance Team |
| EPI_ISL_2915995, EPI_ISL_2915996, EPI_ISL_2915997 | Lighthouse Lab in Alderley Park | Wellcome Sanger Institute for the COVID-19 Genomics UK (COG-UK) Consortium | Jacquelyn Wynn, Mairead Hyland, The Lighthouse Lab in Alderley Park and Alex Alderton, Roberto Amato, Jeffrey Barrett, Sonia Goncalves, Ewan Harrison, David K. Jackson, Ian Johnston, Dominic Kwiatkowski, Cordelia Langford, John Sillitoe on behalf of the Wellcome Sanger Institute COVID-19 Surveillance Team |
| EPI_ISL_2915998, EPI_ISL_2915999 | Berkshire and Surrey Pathology Services Lighthouse Laboratory | Wellcome Sanger Institute for the COVID-19 Genomics UK (COG-UK) Consortium | Berkshire and Surrey Pathology Services Lighthouse Laboratory and Alex Alderton, Roberto Amato, Jeffrey Barrett, Sonia Goncalves, Ewan Harrison, David K. Jackson, Ian Johnston, Dominic Kwiatkowski, Cordelia Langford, John Sillitoe on behalf of the Wellcome Sanger Institute COVID-19 Surveillance Team |
| EPI_ISL_2916000, EPI_ISL_2916001, EPI_ISL_2916002 | Lighthouse Lab in Alderley Park | Wellcome Sanger Institute for the COVID-19 Genomics UK (COG-UK) Consortium | Jacquelyn Wynn, Mairead Hyland, The Lighthouse Lab in Alderley Park and Alex Alderton, Roberto Amato, Jeffrey Barrett, Sonia Goncalves, Ewan Harrison, David K. Jackson, Ian Johnston, Dominic Kwiatkowski, Cordelia Langford, John Sillitoe on behalf of the Wellcome Sanger Institute COVID-19 Surveillance Team |
| EPI_ISL_2916003 | Berkshire and Surrey Pathology Services Lighthouse Laboratory | Wellcome Sanger Institute for the COVID-19 Genomics UK (COG-UK) Consortium | Berkshire and Surrey Pathology Services Lighthouse Laboratory and Alex Alderton, Roberto Amato, Jeffrey Barrett, Sonia Goncalves, Ewan Harrison, David K. Jackson, Ian Johnston, Dominic Kwiatkowski, Cordelia Langford, John Sillitoe on behalf of the Wellcome Sanger Institute COVID-19 Surveillance Team |
| EPI_ISL_2916004 | Lighthouse Lab in Alderley Park | Wellcome Sanger Institute for the COVID-19 Genomics UK (COG-UK) Consortium | Jacquelyn Wynn, Mairead Hyland, The Lighthouse Lab in Alderley Park and Alex Alderton, Roberto Amato, Jeffrey Barrett, Sonia Goncalves, Ewan Harrison, David K. Jackson, Ian Johnston, Dominic Kwiatkowski, Cordelia Langford, John Sillitoe on behalf of the Wellcome Sanger Institute COVID-19 Surveillance Team |
| EPI_ISL_2916005, EPI_ISL_2916006, EPI_ISL_2916007 | Berkshire and Surrey Pathology Services Lighthouse Laboratory | Wellcome Sanger Institute for the COVID-19 Genomics UK (COG-UK) Consortium | Berkshire and Surrey Pathology Services Lighthouse Laboratory and Alex Alderton, Roberto Amato, Jeffrey Barrett, Sonia Goncalves, Ewan Harrison, David K. Jackson, Ian Johnston, Dominic Kwiatkowski, Cordelia Langford, John Sillitoe on behalf of the Wellcome Sanger Institute COVID-19 Surveillance Team |
| EPI_ISL_2916008, EPI_ISL_2916009 | Lighthouse Lab in Alderley Park | Wellcome Sanger Institute for the COVID-19 Genomics UK (COG-UK) Consortium | Jacquelyn Wynn, Mairead Hyland, The Lighthouse Lab in Alderley Park and Alex Alderton, Roberto Amato, Jeffrey Barrett, Sonia Goncalves, Ewan Harrison, David K. Jackson, Ian Johnston, Dominic Kwiatkowski, Cordelia Langford, John Sillitoe on behalf of the Wellcome Sanger Institute COVID-19 Surveillance Team |
| EPI_ISL_2916010, EPI_ISL_2916011, EPI_ISL_2916012, EPI_ISL_2916013 | Berkshire and Surrey Pathology Services Lighthouse Laboratory | Wellcome Sanger Institute for the COVID-19 Genomics UK (COG-UK) Consortium | Berkshire and Surrey Pathology Services Lighthouse Laboratory and Alex Alderton, Roberto Amato, Jeffrey Barrett, Sonia Goncalves, Ewan Harrison, David K. Jackson, Ian Johnston, Dominic Kwiatkowski, Cordelia Langford, John Sillitoe on behalf of the Wellcome Sanger Institute COVID-19 Surveillance Team |
| EPI_ISL_2916014 | Lighthouse Lab in Alderley Park | Wellcome Sanger Institute for the COVID-19 Genomics UK (COG-UK) Consortium | Jacquelyn Wynn, Mairead Hyland, The Lighthouse Lab in Alderley Park and Alex Alderton, Roberto Amato, Jeffrey Barrett, Sonia Goncalves, Ewan Harrison, David K. Jackson, Ian Johnston, Dominic Kwiatkowski, Cordelia Langford, John Sillitoe on behalf of the Wellcome Sanger Institute COVID-19 Surveillance Team |
| EPI_ISL_2916015, EPI_ISL_2916016 | Berkshire and Surrey Pathology Services Lighthouse Laboratory | Wellcome Sanger Institute for the COVID-19 Genomics UK (COG-UK) Consortium | Berkshire and Surrey Pathology Services Lighthouse Laboratory and Alex Alderton, Roberto Amato, Jeffrey Barrett, Sonia Goncalves, Ewan Harrison, David K. Jackson, Ian Johnston, Dominic Kwiatkowski, Cordelia Langford, John Sillitoe on behalf of the Wellcome Sanger Institute COVID-19 Surveillance Team |
| EPI_ISL_2916017 | Lighthouse Lab in Alderley Park | Wellcome Sanger Institute for the COVID-19 Genomics UK (COG-UK) Consortium | Jacquelyn Wynn, Mairead Hyland, The Lighthouse Lab in Alderley Park and Alex Alderton, Roberto Amato, Jeffrey Barrett, Sonia Goncalves, Ewan Harrison, David K. Jackson, Ian Johnston, Dominic Kwiatkowski, Cordelia Langford, John Sillitoe on behalf of the Wellcome Sanger Institute COVID-19 Surveillance Team |
| EPI_ISL_2916018, EPI_ISL_2916019 | Berkshire and Surrey Pathology Services Lighthouse Laboratory | Wellcome Sanger Institute for the COVID-19 Genomics UK (COG-UK) Consortium | Berkshire and Surrey Pathology Services Lighthouse Laboratory and Alex Alderton, Roberto Amato, Jeffrey Barrett, Sonia Goncalves, Ewan Harrison, David K. Jackson, Ian Johnston, Dominic Kwiatkowski, Cordelia Langford, John Sillitoe on behalf of the Wellcome Sanger Institute COVID-19 Surveillance Team |
| EPI_ISL_2916020 | Lighthouse Lab in Alderley Park | Wellcome Sanger Institute for the COVID-19 Genomics UK (COG-UK) Consortium | Jacquelyn Wynn, Mairead Hyland, The Lighthouse Lab in Alderley Park and Alex Alderton, Roberto Amato, Jeffrey Barrett, Sonia Goncalves, Ewan Harrison, David K. Jackson, Ian Johnston, Dominic Kwiatkowski, Cordelia Langford, John Sillitoe on behalf of the Wellcome Sanger Institute COVID-19 Surveillance Team |
| EPI_ISL_2916021 | Berkshire and Surrey Pathology Services Lighthouse Laboratory | Wellcome Sanger Institute for the COVID-19 Genomics UK (COG-UK) Consortium | Berkshire and Surrey Pathology Services Lighthouse Laboratory and Alex Alderton, Roberto Amato, Jeffrey Barrett, Sonia Goncalves, Ewan Harrison, David K. Jackson, Ian Johnston, Dominic Kwiatkowski, Cordelia Langford, John Sillitoe on behalf of the Wellcome Sanger Institute COVID-19 Surveillance Team |
| EPI_ISL_2916022 | Lighthouse Lab in Alderley Park | Wellcome Sanger Institute for the COVID-19 Genomics UK (COG-UK) Consortium | Jacquelyn Wynn, Mairead Hyland, The Lighthouse Lab in Alderley Park and Alex Alderton, Roberto Amato, Jeffrey Barrett, Sonia Goncalves, Ewan Harrison, David K. Jackson, Ian Johnston, Dominic Kwiatkowski, Cordelia Langford, John Sillitoe on behalf of the Wellcome Sanger Institute COVID-19 Surveillance Team |
| EPI_ISL_2916023 | Berkshire and Surrey Pathology Services Lighthouse Laboratory | Wellcome Sanger Institute for the COVID-19 Genomics UK (COG-UK) Consortium | Berkshire and Surrey Pathology Services Lighthouse Laboratory and Alex Alderton, Roberto Amato, Jeffrey Barrett, Sonia Goncalves, Ewan Harrison, David K. Jackson, Ian Johnston, Dominic Kwiatkowski, Cordelia Langford, John Sillitoe on behalf of the Wellcome Sanger Institute COVID-19 Surveillance Team |
| EPI_ISL_2916024, EPI_ISL_2916025 | Lighthouse Lab in Alderley Park | Wellcome Sanger Institute for the COVID-19 Genomics UK (COG-UK) Consortium | Jacquelyn Wynn, Mairead Hyland, The Lighthouse Lab in Alderley Park and Alex Alderton, Roberto Amato, Jeffrey Barrett, Sonia Goncalves, Ewan Harrison, David K. Jackson, Ian Johnston, Dominic Kwiatkowski, Cordelia Langford, John Sillitoe on behalf of the Wellcome Sanger Institute COVID-19 Surveillance Team |

| | | | |
|---|---|---|---|
| EPI_ISL_2916026, EPI_ISL_2916027 | Berkshire and Surrey Pathology Services Lighthouse Laboratory | Wellcome Sanger Institute for the COVID-19 Genomics UK (COG-UK) Consortium | Berkshire and Surrey Pathology Services Lighthouse Laboratory and Alex Alderton, Roberto Amato, Jeffrey Barrett, Sonia Goncalves, Ewan Harrison, David K. Jackson, Ian Johnston, Dominic Kwiatkowski, Cordelia Langford, John Sillitoe on behalf of the Wellcome Sanger Institute COVID-19 Surveillance Team |
| EPI_ISL_2916028 | Lighthouse Lab in Alderley Park | Wellcome Sanger Institute for the COVID-19 Genomics UK (COG-UK) Consortium | Jacquelyn Wynn, Mairead Hyland, The Lighthouse Lab in Alderley Park and Alex Alderton, Roberto Amato, Jeffrey Barrett, Sonia Goncalves, Ewan Harrison, David K. Jackson, Ian Johnston, Dominic Kwiatkowski, Cordelia Langford, John Sillitoe on behalf of the Wellcome Sanger Institute COVID-19 Surveillance Team |
| EPI_ISL_2916029, EPI_ISL_2916030 | Berkshire and Surrey Pathology Services Lighthouse Laboratory | Wellcome Sanger Institute for the COVID-19 Genomics UK (COG-UK) Consortium | Berkshire and Surrey Pathology Services Lighthouse Laboratory and Alex Alderton, Roberto Amato, Jeffrey Barrett, Sonia Goncalves, Ewan Harrison, David K. Jackson, Ian Johnston, Dominic Kwiatkowski, Cordelia Langford, John Sillitoe on behalf of the Wellcome Sanger Institute COVID-19 Surveillance Team |
| EPI_ISL_2916031 | Lighthouse Lab in Alderley Park | Wellcome Sanger Institute for the COVID-19 Genomics UK (COG-UK) Consortium | Jacquelyn Wynn, Mairead Hyland, The Lighthouse Lab in Alderley Park and Alex Alderton, Roberto Amato, Jeffrey Barrett, Sonia Goncalves, Ewan Harrison, David K. Jackson, Ian Johnston, Dominic Kwiatkowski, Cordelia Langford, John Sillitoe on behalf of the Wellcome Sanger Institute COVID-19 Surveillance Team |
| EPI_ISL_2916032 | Lighthouse Lab in Milton Keynes | Wellcome Sanger Institute for the COVID-19 Genomics UK (COG-UK) Consortium | The Lighthouse Lab in Milton Keynes and Alex Alderton, Roberto Amato, Jeffrey Barrett, Sonia Goncalves, Ewan Harrison, David K. Jackson, Ian Johnston, Dominic Kwiatkowski, Cordelia Langford, John Sillitoe on behalf of the Wellcome Sanger Institute COVID-19 Surveillance Team |
| EPI_ISL_2916033 | Lighthouse Lab in Alderley Park | Wellcome Sanger Institute for the COVID-19 Genomics UK (COG-UK) Consortium | Jacquelyn Wynn, Mairead Hyland, The Lighthouse Lab in Alderley Park and Alex Alderton, Roberto Amato, Jeffrey Barrett, Sonia Goncalves, Ewan Harrison, David K. Jackson, Ian Johnston, Dominic Kwiatkowski, Cordelia Langford, John Sillitoe on behalf of the Wellcome Sanger Institute COVID-19 Surveillance Team |
| EPI_ISL_2916034, EPI_ISL_2916035 | Lighthouse Lab in Milton Keynes | Wellcome Sanger Institute for the COVID-19 Genomics UK (COG-UK) Consortium | The Lighthouse Lab in Milton Keynes and Alex Alderton, Roberto Amato, Jeffrey Barrett, Sonia Goncalves, Ewan Harrison, David K. Jackson, Ian Johnston, Dominic Kwiatkowski, Cordelia Langford, John Sillitoe on behalf of the Wellcome Sanger Institute COVID-19 Surveillance Team |
| EPI_ISL_2916036, EPI_ISL_2916037 | Berkshire and Surrey Pathology Services Lighthouse Laboratory | Wellcome Sanger Institute for the COVID-19 Genomics UK (COG-UK) Consortium | Berkshire and Surrey Pathology Services Lighthouse Laboratory and Alex Alderton, Roberto Amato, Jeffrey Barrett, Sonia Goncalves, Ewan Harrison, David K. Jackson, Ian Johnston, Dominic Kwiatkowski, Cordelia Langford, John Sillitoe on behalf of the Wellcome Sanger Institute COVID-19 Surveillance Team |
| EPI_ISL_2916038 | Lighthouse Lab in Alderley Park | Wellcome Sanger Institute for the COVID-19 Genomics UK (COG-UK) Consortium | Jacquelyn Wynn, Mairead Hyland, The Lighthouse Lab in Alderley Park and Alex Alderton, Roberto Amato, Jeffrey Barrett, Sonia Goncalves, Ewan Harrison, David K. Jackson, Ian Johnston, Dominic Kwiatkowski, Cordelia Langford, John Sillitoe on behalf of the Wellcome Sanger Institute COVID-19 Surveillance Team |
| EPI_ISL_2916039 | Lighthouse Lab in Milton Keynes | Wellcome Sanger Institute for the COVID-19 Genomics UK (COG-UK) Consortium | The Lighthouse Lab in Milton Keynes and Alex Alderton, Roberto Amato, Jeffrey Barrett, Sonia Goncalves, Ewan Harrison, David K. Jackson, Ian Johnston, Dominic Kwiatkowski, Cordelia Langford, John Sillitoe on behalf of the Wellcome Sanger Institute COVID-19 Surveillance Team |
| EPI_ISL_2916040 | Berkshire and Surrey Pathology Services Lighthouse Laboratory | Wellcome Sanger Institute for the COVID-19 Genomics UK (COG-UK) Consortium | Berkshire and Surrey Pathology Services Lighthouse Laboratory and Alex Alderton, Roberto Amato, Jeffrey Barrett, Sonia Goncalves, Ewan Harrison, David K. Jackson, Ian Johnston, Dominic Kwiatkowski, Cordelia Langford, John Sillitoe on behalf of the Wellcome Sanger Institute COVID-19 Surveillance Team |
| EPI_ISL_2916041, EPI_ISL_2916042 | Lighthouse Lab in Milton Keynes | Wellcome Sanger Institute for the COVID-19 Genomics UK (COG-UK) Consortium | The Lighthouse Lab in Milton Keynes and Alex Alderton, Roberto Amato, Jeffrey Barrett, Sonia Goncalves, Ewan Harrison, David K. Jackson, Ian Johnston, Dominic Kwiatkowski, Cordelia Langford, John Sillitoe on behalf of the Wellcome Sanger Institute COVID-19 Surveillance Team |
| EPI_ISL_2916043 | Lighthouse Lab in Alderley Park | Wellcome Sanger Institute for the COVID-19 Genomics UK (COG-UK) Consortium | Jacquelyn Wynn, Mairead Hyland, The Lighthouse Lab in Alderley Park and Alex Alderton, Roberto Amato, Jeffrey Barrett, Sonia Goncalves, Ewan Harrison, David K. Jackson, Ian Johnston, Dominic Kwiatkowski, Cordelia Langford, John Sillitoe on behalf of the Wellcome Sanger Institute COVID-19 Surveillance Team |
| EPI_ISL_2916044 | Lighthouse Lab in Milton Keynes | Wellcome Sanger Institute for the COVID-19 Genomics UK (COG-UK) Consortium | The Lighthouse Lab in Milton Keynes and Alex Alderton, Roberto Amato, Jeffrey Barrett, Sonia Goncalves, Ewan Harrison, David K. Jackson, Ian Johnston, Dominic Kwiatkowski, Cordelia Langford, John Sillitoe on behalf of the Wellcome Sanger Institute COVID-19 Surveillance Team |
| EPI_ISL_2916045, EPI_ISL_2916046 | Lighthouse Lab in Alderley Park | Wellcome Sanger Institute for the COVID-19 Genomics UK (COG-UK) Consortium | Jacquelyn Wynn, Mairead Hyland, The Lighthouse Lab in Alderley Park and Alex Alderton, Roberto Amato, Jeffrey Barrett, Sonia Goncalves, Ewan Harrison, David K. Jackson, Ian Johnston, Dominic Kwiatkowski, Cordelia Langford, John Sillitoe on behalf of the Wellcome Sanger Institute COVID-19 Surveillance Team |
| EPI_ISL_2916047 | Berkshire and Surrey Pathology Services Lighthouse Laboratory | Wellcome Sanger Institute for the COVID-19 Genomics UK (COG-UK) Consortium | Berkshire and Surrey Pathology Services Lighthouse Laboratory and Alex Alderton, Roberto Amato, Jeffrey Barrett, Sonia Goncalves, Ewan Harrison, David K. Jackson, Ian Johnston, Dominic Kwiatkowski, Cordelia Langford, John Sillitoe on behalf of the Wellcome Sanger Institute COVID-19 Surveillance Team |
| EPI_ISL_2916048 | Lighthouse Lab in Milton Keynes | Wellcome Sanger Institute for the COVID-19 Genomics UK (COG-UK) Consortium | The Lighthouse Lab in Milton Keynes and Alex Alderton, Roberto Amato, Jeffrey Barrett, Sonia Goncalves, Ewan Harrison, David K. Jackson, Ian Johnston, Dominic Kwiatkowski, Cordelia Langford, John Sillitoe on behalf of the Wellcome Sanger Institute COVID-19 Surveillance Team |
| EPI_ISL_2916049 | Lighthouse Lab in Alderley Park | Wellcome Sanger Institute for the COVID-19 Genomics UK (COG-UK) Consortium | Jacquelyn Wynn, Mairead Hyland, The Lighthouse Lab in Alderley Park and Alex Alderton, Roberto Amato, Jeffrey Barrett, Sonia Goncalves, Ewan Harrison, David K. Jackson, Ian Johnston, Dominic Kwiatkowski, Cordelia Langford, John Sillitoe on behalf of the Wellcome Sanger Institute COVID-19 Surveillance Team |
| EPI_ISL_2916050, EPI_ISL_2916051 | Berkshire and Surrey Pathology Services Lighthouse Laboratory | Wellcome Sanger Institute for the COVID-19 Genomics UK (COG-UK) Consortium | Berkshire and Surrey Pathology Services Lighthouse Laboratory and Alex Alderton, Roberto Amato, Jeffrey Barrett, Sonia Goncalves, Ewan Harrison, David K. Jackson, Ian Johnston, Dominic Kwiatkowski, Cordelia Langford, John Sillitoe on behalf of the Wellcome Sanger Institute COVID-19 Surveillance Team |
| EPI_ISL_2916052 | Lighthouse Lab in Milton Keynes | Wellcome Sanger Institute for the COVID-19 Genomics UK (COG-UK) Consortium | The Lighthouse Lab in Milton Keynes and Alex Alderton, Roberto Amato, Jeffrey Barrett, Sonia Goncalves, Ewan Harrison, David K. Jackson, Ian Johnston, Dominic Kwiatkowski, Cordelia Langford, John Sillitoe on behalf of the Wellcome Sanger Institute COVID-19 Surveillance Team |
| EPI_ISL_2916053 | Lighthouse Lab in Alderley Park | Wellcome Sanger Institute for the COVID-19 Genomics UK (COG-UK) Consortium | Jacquelyn Wynn, Mairead Hyland, The Lighthouse Lab in Alderley Park and Alex Alderton, Roberto Amato, Jeffrey Barrett, Sonia Goncalves, Ewan Harrison, David K. Jackson, Ian Johnston, Dominic Kwiatkowski, Cordelia Langford, John Sillitoe on behalf of the Wellcome Sanger Institute COVID-19 Surveillance Team |
| EPI_ISL_2916054, EPI_ISL_2916055 | Lighthouse Lab in Milton Keynes | Wellcome Sanger Institute for the COVID-19 Genomics UK (COG-UK) Consortium | The Lighthouse Lab in Milton Keynes and Alex Alderton, Roberto Amato, Jeffrey Barrett, Sonia Goncalves, Ewan Harrison, David K. Jackson, Ian Johnston, Dominic Kwiatkowski, Cordelia Langford, John Sillitoe on behalf of the Wellcome Sanger Institute COVID-19 Surveillance Team |
| EPI_ISL_2916056 | Berkshire and Surrey Pathology Services Lighthouse Laboratory | Wellcome Sanger Institute for the COVID-19 Genomics UK (COG-UK) Consortium | Berkshire and Surrey Pathology Services Lighthouse Laboratory and Alex Alderton, Roberto Amato, Jeffrey Barrett, Sonia Goncalves, Ewan Harrison, David K. Jackson, Ian Johnston, Dominic Kwiatkowski, Cordelia Langford, John Sillitoe on behalf of the Wellcome Sanger Institute COVID-19 Surveillance Team |
| EPI_ISL_2916057, EPI_ISL_2916058 | Lighthouse Lab in Milton Keynes | Wellcome Sanger Institute for the COVID-19 Genomics UK (COG-UK) Consortium | The Lighthouse Lab in Milton Keynes and Alex Alderton, Roberto Amato, Jeffrey Barrett, Sonia Goncalves, Ewan Harrison, David K. Jackson, Ian Johnston, Dominic Kwiatkowski, Cordelia Langford, John Sillitoe on behalf of the Wellcome Sanger Institute COVID-19 Surveillance Team |
| EPI_ISL_2916059 | Lighthouse Lab in Alderley Park | Wellcome Sanger Institute for the COVID-19 Genomics UK (COG-UK) Consortium | Jacquelyn Wynn, Mairead Hyland, The Lighthouse Lab in Alderley Park and Alex Alderton, Roberto Amato, Jeffrey Barrett, Sonia Goncalves, Ewan Harrison, David K. Jackson, Ian Johnston, Dominic Kwiatkowski, Cordelia Langford, John Sillitoe on behalf of the Wellcome Sanger Institute COVID-19 Surveillance Team |
| EPI_ISL_2916060, EPI_ISL_2916061 | Berkshire and Surrey Pathology Services Lighthouse Laboratory | Wellcome Sanger Institute for the COVID-19 Genomics UK (COG-UK) Consortium | Berkshire and Surrey Pathology Services Lighthouse Laboratory and Alex Alderton, Roberto Amato, Jeffrey Barrett, Sonia Goncalves, Ewan Harrison, David K. Jackson, Ian Johnston, Dominic Kwiatkowski, Cordelia Langford, John Sillitoe on behalf of the Wellcome Sanger Institute COVID-19 Surveillance Team |
| EPI_ISL_2916062 | Lighthouse Lab in Milton Keynes | Wellcome Sanger Institute for the COVID-19 Genomics UK (COG-UK) Consortium | The Lighthouse Lab in Milton Keynes and Alex Alderton, Roberto Amato, Jeffrey Barrett, Sonia Goncalves, Ewan Harrison, David K. Jackson, Ian Johnston, Dominic Kwiatkowski, Cordelia Langford, John Sillitoe on behalf of the Wellcome Sanger Institute COVID-19 Surveillance Team |
| EPI_ISL_2916063 | Lighthouse Lab in Alderley Park | Wellcome Sanger Institute for the COVID-19 Genomics UK (COG-UK) Consortium | Jacquelyn Wynn, Mairead Hyland, The Lighthouse Lab in Alderley Park and Alex Alderton, Roberto Amato, Jeffrey Barrett, Sonia Goncalves, Ewan Harrison, David K. Jackson, Ian Johnston, Dominic Kwiatkowski, Cordelia Langford, John Sillitoe on behalf of the Wellcome Sanger Institute COVID-19 Surveillance Team |
| EPI_ISL_2916064 | Berkshire and Surrey Pathology Services Lighthouse Laboratory | Wellcome Sanger Institute for the COVID-19 Genomics UK (COG-UK) Consortium | Berkshire and Surrey Pathology Services Lighthouse Laboratory and Alex Alderton, Roberto Amato, Jeffrey Barrett, Sonia Goncalves, Ewan Harrison, David K. Jackson, Ian Johnston, Dominic Kwiatkowski, Cordelia Langford, John Sillitoe on behalf of the Wellcome Sanger Institute COVID-19 Surveillance Team |
| EPI_ISL_2916065 | Lighthouse Lab in Milton Keynes | Wellcome Sanger Institute for the COVID-19 Genomics UK (COG-UK) Consortium | The Lighthouse Lab in Milton Keynes and Alex Alderton, Roberto Amato, Jeffrey Barrett, Sonia Goncalves, Ewan Harrison, David K. Jackson, Ian Johnston, Dominic Kwiatkowski, Cordelia Langford, John Sillitoe on behalf of the Wellcome Sanger Institute COVID-19 Surveillance Team |
| EPI_ISL_2916066 | Berkshire and Surrey Pathology Services Lighthouse Laboratory | Wellcome Sanger Institute for the COVID-19 Genomics UK (COG-UK) Consortium | Berkshire and Surrey Pathology Services Lighthouse Laboratory and Alex Alderton, Roberto Amato, Jeffrey Barrett, Sonia Goncalves, Ewan Harrison, David K. Jackson, Ian Johnston, Dominic Kwiatkowski, Cordelia Langford, John Sillitoe on behalf of the Wellcome Sanger Institute COVID-19 Surveillance Team |
| EPI_ISL_2916067 | Lighthouse Lab in Milton Keynes | Wellcome Sanger Institute for the COVID-19 Genomics UK (COG-UK) Consortium | The Lighthouse Lab in Milton Keynes and Alex Alderton, Roberto Amato, Jeffrey Barrett, Sonia Goncalves, Ewan Harrison, David K. Jackson, Ian Johnston, Dominic Kwiatkowski, Cordelia Langford, John Sillitoe on behalf of the Wellcome Sanger Institute COVID-19 Surveillance Team |
| EPI_ISL_2916068, EPI_ISL_2916069, EPI_ISL_2916070, EPI_ISL_2916071 | Lighthouse Lab in Alderley Park | Wellcome Sanger Institute for the COVID-19 Genomics UK (COG-UK) Consortium | Jacquelyn Wynn, Mairead Hyland, The Lighthouse Lab in Alderley Park and Alex Alderton, Roberto Amato, Jeffrey Barrett, Sonia Goncalves, Ewan Harrison, David K. Jackson, Ian Johnston, Dominic Kwiatkowski, Cordelia Langford, John Sillitoe on behalf of the Wellcome Sanger Institute COVID-19 Surveillance Team |
| EPI_ISL_2916072 | Berkshire and Surrey Pathology Services Lighthouse Laboratory | Wellcome Sanger Institute for the COVID-19 Genomics UK (COG-UK) Consortium | Berkshire and Surrey Pathology Services Lighthouse Laboratory and Alex Alderton, Roberto Amato, Jeffrey Barrett, Sonia Goncalves, Ewan Harrison, David K. Jackson, Ian Johnston, Dominic Kwiatkowski, Cordelia Langford, John Sillitoe on behalf of the Wellcome Sanger Institute COVID-19 Surveillance Team |
| EPI_ISL_2916073, EPI_ISL_2916074 | Lighthouse Lab in Alderley Park | Wellcome Sanger Institute for the COVID-19 Genomics UK (COG-UK) Consortium | Jacquelyn Wynn, Mairead Hyland, The Lighthouse Lab in Alderley Park and Alex Alderton, Roberto Amato, Jeffrey Barrett, Sonia Goncalves, Ewan Harrison, David K. Jackson, Ian Johnston, Dominic Kwiatkowski, Cordelia Langford, John Sillitoe on behalf of the Wellcome Sanger Institute COVID-19 Surveillance Team |
| EPI_ISL_2916075 | Berkshire and Surrey Pathology Services Lighthouse Laboratory | Wellcome Sanger Institute for the COVID-19 Genomics UK (COG-UK) Consortium | Berkshire and Surrey Pathology Services Lighthouse Laboratory and Alex Alderton, Roberto Amato, Jeffrey Barrett, Sonia Goncalves, Ewan Harrison, David K. Jackson, Ian Johnston, Dominic Kwiatkowski, Cordelia Langford, John Sillitoe on behalf of the Wellcome Sanger Institute COVID-19 Surveillance Team |
| EPI_ISL_2916076, EPI_ISL_2916077 | Lighthouse Lab in Alderley Park | Wellcome Sanger Institute for the COVID-19 Genomics UK (COG-UK) Consortium | Jacquelyn Wynn, Mairead Hyland, The Lighthouse Lab in Alderley Park and Alex Alderton, Roberto Amato, Jeffrey Barrett, Sonia Goncalves, Ewan Harrison, David K. Jackson, Ian Johnston, Dominic Kwiatkowski, Cordelia Langford, John Sillitoe on behalf of the Wellcome Sanger Institute COVID-19 Surveillance Team |
| EPI_ISL_2916078, EPI_ISL_2916079 | Berkshire and Surrey Pathology Services Lighthouse Laboratory | Wellcome Sanger Institute for the COVID-19 Genomics UK (COG-UK) Consortium | Berkshire and Surrey Pathology Services Lighthouse Laboratory and Alex Alderton, Roberto Amato, Jeffrey Barrett, Sonia Goncalves, Ewan Harrison, David K. Jackson, Ian Johnston, Dominic Kwiatkowski, Cordelia Langford, John Sillitoe on behalf of the Wellcome Sanger Institute COVID-19 Surveillance Team |
| EPI_ISL_2916080 | Lighthouse Lab in Alderley Park | Wellcome Sanger Institute for the COVID-19 Genomics UK (COG-UK) Consortium | Jacquelyn Wynn, Mairead Hyland, The Lighthouse Lab in Alderley Park and Alex Alderton, Roberto Amato, Jeffrey Barrett, Sonia Goncalves, Ewan Harrison, David K. Jackson, Ian Johnston, Dominic Kwiatkowski, Cordelia Langford, John Sillitoe on behalf of the Wellcome Sanger Institute COVID-19 Surveillance Team |
| EPI_ISL_2916081 | Lighthouse Lab in Milton Keynes | Wellcome Sanger Institute for the COVID-19 Genomics UK (COG-UK) Consortium | The Lighthouse Lab in Milton Keynes and Alex Alderton, Roberto Amato, Jeffrey Barrett, Sonia Goncalves, Ewan Harrison, David K. Jackson, Ian Johnston, Dominic Kwiatkowski, Cordelia Langford, John Sillitoe on behalf of the Wellcome Sanger Institute COVID-19 Surveillance Team |
| EPI_ISL_2916082 | Berkshire and Surrey Pathology Services Lighthouse Laboratory | Wellcome Sanger Institute for the COVID-19 Genomics UK (COG-UK) Consortium | Berkshire and Surrey Pathology Services Lighthouse Laboratory and Alex Alderton, Roberto Amato, Jeffrey Barrett, Sonia Goncalves, Ewan Harrison, David K. Jackson, Ian Johnston, Dominic Kwiatkowski, Cordelia Langford, John Sillitoe on behalf of the Wellcome Sanger Institute COVID-19 Surveillance Team |
| EPI_ISL_2916083 | Lighthouse Lab in Milton Keynes | Wellcome Sanger Institute for the COVID-19 Genomics UK (COG-UK) Consortium | The Lighthouse Lab in Milton Keynes and Alex Alderton, Roberto Amato, Jeffrey Barrett, Sonia Goncalves, Ewan Harrison, David K. Jackson, Ian Johnston, Dominic Kwiatkowski, Cordelia Langford, John Sillitoe on behalf of the Wellcome Sanger Institute COVID-19 Surveillance Team |
| EPI_ISL_2916084, EPI_ISL_2916085, EPI_ISL_2916086 | Berkshire and Surrey Pathology Services Lighthouse Laboratory | Wellcome Sanger Institute for the COVID-19 Genomics UK (COG-UK) Consortium | Berkshire and Surrey Pathology Services Lighthouse Laboratory and Alex Alderton, Roberto Amato, Jeffrey Barrett, Sonia Goncalves, Ewan Harrison, David K. Jackson, Ian Johnston, Dominic Kwiatkowski, Cordelia Langford, John Sillitoe on behalf of the Wellcome Sanger Institute COVID-19 Surveillance Team |
| EPI_ISL_2916087, EPI_ISL_2916088 | Lighthouse Lab in Milton Keynes | Wellcome Sanger Institute for the COVID-19 Genomics UK (COG-UK) Consortium | The Lighthouse Lab in Milton Keynes and Alex Alderton, Roberto Amato, Jeffrey Barrett, Sonia Goncalves, Ewan Harrison, David K. Jackson, Ian Johnston, Dominic Kwiatkowski, Cordelia Langford, John Sillitoe on behalf of the Wellcome Sanger Institute COVID-19 Surveillance Team |

| | | | |
|---|---|---|---|
| EPI_ISL_2916089, EPI_ISL_2916090 | Berkshire and Surrey Pathology Services Lighthouse Laboratory | Wellcome Sanger Institute for the COVID-19 Genomics UK (COG-UK) Consortium | Berkshire and Surrey Pathology Services Lighthouse Laboratory and Alex Alderton, Roberto Amato, Jeffrey Barrett, Sonia Goncalves, Ewan Harrison, David K. Jackson, Ian Johnston, Dominic Kwiatkowski, Cordelia Langford, John Sillitoe on behalf of the Wellcome Sanger Institute COVID-19 Surveillance Team |
| EPI_ISL_2916091 | Lighthouse Lab in Alderley Park | Wellcome Sanger Institute for the COVID-19 Genomics UK (COG-UK) Consortium | Jacquelyn Wynn, Mairead Hyland, The Lighthouse Lab in Alderley Park and Alex Alderton, Roberto Amato, Jeffrey Barrett, Sonia Goncalves, Ewan Harrison, David K. Jackson, Ian Johnston, Dominic Kwiatkowski, Cordelia Langford, John Sillitoe on behalf of the Wellcome Sanger Institute COVID-19 Surveillance Team |
| EPI_ISL_2916092 | Lighthouse Lab in Milton Keynes | Wellcome Sanger Institute for the COVID-19 Genomics UK (COG-UK) Consortium | The Lighthouse Lab in Milton Keynes and Alex Alderton, Roberto Amato, Jeffrey Barrett, Sonia Goncalves, Ewan Harrison, David K. Jackson, Ian Johnston, Dominic Kwiatkowski, Cordelia Langford, John Sillitoe on behalf of the Wellcome Sanger Institute COVID-19 Surveillance Team |
| EPI_ISL_2916093, EPI_ISL_2916094 | Berkshire and Surrey Pathology Services Lighthouse Laboratory | Wellcome Sanger Institute for the COVID-19 Genomics UK (COG-UK) Consortium | Berkshire and Surrey Pathology Services Lighthouse Laboratory and Alex Alderton, Roberto Amato, Jeffrey Barrett, Sonia Goncalves, Ewan Harrison, David K. Jackson, Ian Johnston, Dominic Kwiatkowski, Cordelia Langford, John Sillitoe on behalf of the Wellcome Sanger Institute COVID-19 Surveillance Team |
| EPI_ISL_2916095 | Lighthouse Lab in Alderley Park | Wellcome Sanger Institute for the COVID-19 Genomics UK (COG-UK) Consortium | Jacquelyn Wynn, Mairead Hyland, The Lighthouse Lab in Alderley Park and Alex Alderton, Roberto Amato, Jeffrey Barrett, Sonia Goncalves, Ewan Harrison, David K. Jackson, Ian Johnston, Dominic Kwiatkowski, Cordelia Langford, John Sillitoe on behalf of the Wellcome Sanger Institute COVID-19 Surveillance Team |
| EPI_ISL_2916096 | Lighthouse Lab in Milton Keynes | Wellcome Sanger Institute for the COVID-19 Genomics UK (COG-UK) Consortium | The Lighthouse Lab in Milton Keynes and Alex Alderton, Roberto Amato, Jeffrey Barrett, Sonia Goncalves, Ewan Harrison, David K. Jackson, Ian Johnston, Dominic Kwiatkowski, Cordelia Langford, John Sillitoe on behalf of the Wellcome Sanger Institute COVID-19 Surveillance Team |
| EPI_ISL_2916097, EPI_ISL_2916098 | Berkshire and Surrey Pathology Services Lighthouse Laboratory | Wellcome Sanger Institute for the COVID-19 Genomics UK (COG-UK) Consortium | Berkshire and Surrey Pathology Services Lighthouse Laboratory and Alex Alderton, Roberto Amato, Jeffrey Barrett, Sonia Goncalves, Ewan Harrison, David K. Jackson, Ian Johnston, Dominic Kwiatkowski, Cordelia Langford, John Sillitoe on behalf of the Wellcome Sanger Institute COVID-19 Surveillance Team |
| EPI_ISL_2916099 | Lighthouse Lab in Alderley Park | Wellcome Sanger Institute for the COVID-19 Genomics UK (COG-UK) Consortium | Jacquelyn Wynn, Mairead Hyland, The Lighthouse Lab in Alderley Park and Alex Alderton, Roberto Amato, Jeffrey Barrett, Sonia Goncalves, Ewan Harrison, David K. Jackson, Ian Johnston, Dominic Kwiatkowski, Cordelia Langford, John Sillitoe on behalf of the Wellcome Sanger Institute COVID-19 Surveillance Team |
| EPI_ISL_2916100 | Lighthouse Lab in Milton Keynes | Wellcome Sanger Institute for the COVID-19 Genomics UK (COG-UK) Consortium | The Lighthouse Lab in Milton Keynes and Alex Alderton, Roberto Amato, Jeffrey Barrett, Sonia Goncalves, Ewan Harrison, David K. Jackson, Ian Johnston, Dominic Kwiatkowski, Cordelia Langford, John Sillitoe on behalf of the Wellcome Sanger Institute COVID-19 Surveillance Team |
| EPI_ISL_2916101 | Berkshire and Surrey Pathology Services Lighthouse Laboratory | Wellcome Sanger Institute for the COVID-19 Genomics UK (COG-UK) Consortium | Berkshire and Surrey Pathology Services Lighthouse Laboratory and Alex Alderton, Roberto Amato, Jeffrey Barrett, Sonia Goncalves, Ewan Harrison, David K. Jackson, Ian Johnston, Dominic Kwiatkowski, Cordelia Langford, John Sillitoe on behalf of the Wellcome Sanger Institute COVID-19 Surveillance Team |
| EPI_ISL_2916102 | Lighthouse Lab in Milton Keynes | Wellcome Sanger Institute for the COVID-19 Genomics UK (COG-UK) Consortium | The Lighthouse Lab in Milton Keynes and Alex Alderton, Roberto Amato, Jeffrey Barrett, Sonia Goncalves, Ewan Harrison, David K. Jackson, Ian Johnston, Dominic Kwiatkowski, Cordelia Langford, John Sillitoe on behalf of the Wellcome Sanger Institute COVID-19 Surveillance Team |
| EPI_ISL_2916103 | Berkshire and Surrey Pathology Services Lighthouse Laboratory | Wellcome Sanger Institute for the COVID-19 Genomics UK (COG-UK) Consortium | Berkshire and Surrey Pathology Services Lighthouse Laboratory and Alex Alderton, Roberto Amato, Jeffrey Barrett, Sonia Goncalves, Ewan Harrison, David K. Jackson, Ian Johnston, Dominic Kwiatkowski, Cordelia Langford, John Sillitoe on behalf of the Wellcome Sanger Institute COVID-19 Surveillance Team |
| EPI_ISL_2916104, EPI_ISL_2916105 | Lighthouse Lab in Milton Keynes | Wellcome Sanger Institute for the COVID-19 Genomics UK (COG-UK) Consortium | The Lighthouse Lab in Milton Keynes and Alex Alderton, Roberto Amato, Jeffrey Barrett, Sonia Goncalves, Ewan Harrison, David K. Jackson, Ian Johnston, Dominic Kwiatkowski, Cordelia Langford, John Sillitoe on behalf of the Wellcome Sanger Institute COVID-19 Surveillance Team |
| EPI_ISL_2916107 | Berkshire and Surrey Pathology Services Lighthouse Laboratory | Wellcome Sanger Institute for the COVID-19 Genomics UK (COG-UK) Consortium | Berkshire and Surrey Pathology Services Lighthouse Laboratory and Alex Alderton, Roberto Amato, Jeffrey Barrett, Sonia Goncalves, Ewan Harrison, David K. Jackson, Ian Johnston, Dominic Kwiatkowski, Cordelia Langford, John Sillitoe on behalf of the Wellcome Sanger Institute COVID-19 Surveillance Team |
| EPI_ISL_2916108, EPI_ISL_2916109 | Lighthouse Lab in Milton Keynes | Wellcome Sanger Institute for the COVID-19 Genomics UK (COG-UK) Consortium | The Lighthouse Lab in Milton Keynes and Alex Alderton, Roberto Amato, Jeffrey Barrett, Sonia Goncalves, Ewan Harrison, David K. Jackson, Ian Johnston, Dominic Kwiatkowski, Cordelia Langford, John Sillitoe on behalf of the Wellcome Sanger Institute COVID-19 Surveillance Team |
| EPI_ISL_2916110 | Berkshire and Surrey Pathology Services Lighthouse Laboratory | Wellcome Sanger Institute for the COVID-19 Genomics UK (COG-UK) Consortium | Berkshire and Surrey Pathology Services Lighthouse Laboratory and Alex Alderton, Roberto Amato, Jeffrey Barrett, Sonia Goncalves, Ewan Harrison, David K. Jackson, Ian Johnston, Dominic Kwiatkowski, Cordelia Langford, John Sillitoe on behalf of the Wellcome Sanger Institute COVID-19 Surveillance Team |
| EPI_ISL_2916111 | Lighthouse Lab in Milton Keynes | Wellcome Sanger Institute for the COVID-19 Genomics UK (COG-UK) Consortium | The Lighthouse Lab in Milton Keynes and Alex Alderton, Roberto Amato, Jeffrey Barrett, Sonia Goncalves, Ewan Harrison, David K. Jackson, Ian Johnston, Dominic Kwiatkowski, Cordelia Langford, John Sillitoe on behalf of the Wellcome Sanger Institute COVID-19 Surveillance Team |
| EPI_ISL_2916112 | Lighthouse Lab in Alderley Park | Wellcome Sanger Institute for the COVID-19 Genomics UK (COG-UK) Consortium | Jacquelyn Wynn, Mairead Hyland, The Lighthouse Lab in Alderley Park and Alex Alderton, Roberto Amato, Jeffrey Barrett, Sonia Goncalves, Ewan Harrison, David K. Jackson, Ian Johnston, Dominic Kwiatkowski, Cordelia Langford, John Sillitoe on behalf of the Wellcome Sanger Institute COVID-19 Surveillance Team |
| EPI_ISL_2916113, EPI_ISL_2916114, EPI_ISL_2916115 | Berkshire and Surrey Pathology Services Lighthouse Laboratory | Wellcome Sanger Institute for the COVID-19 Genomics UK (COG-UK) Consortium | Berkshire and Surrey Pathology Services Lighthouse Laboratory and Alex Alderton, Roberto Amato, Jeffrey Barrett, Sonia Goncalves, Ewan Harrison, David K. Jackson, Ian Johnston, Dominic Kwiatkowski, Cordelia Langford, John Sillitoe on behalf of the Wellcome Sanger Institute COVID-19 Surveillance Team |
| EPI_ISL_2916116 | Lighthouse Lab in Alderley Park | Wellcome Sanger Institute for the COVID-19 Genomics UK (COG-UK) Consortium | Jacquelyn Wynn, Mairead Hyland, The Lighthouse Lab in Alderley Park and Alex Alderton, Roberto Amato, Jeffrey Barrett, Sonia Goncalves, Ewan Harrison, David K. Jackson, Ian Johnston, Dominic Kwiatkowski, Cordelia Langford, John Sillitoe on behalf of the Wellcome Sanger Institute COVID-19 Surveillance Team |
| EPI_ISL_2916117, EPI_ISL_2916118, EPI_ISL_2916119 | Berkshire and Surrey Pathology Services Lighthouse Laboratory | Wellcome Sanger Institute for the COVID-19 Genomics UK (COG-UK) Consortium | Berkshire and Surrey Pathology Services Lighthouse Laboratory and Alex Alderton, Roberto Amato, Jeffrey Barrett, Sonia Goncalves, Ewan Harrison, David K. Jackson, Ian Johnston, Dominic Kwiatkowski, Cordelia Langford, John Sillitoe on behalf of the Wellcome Sanger Institute COVID-19 Surveillance Team |
| EPI_ISL_2916120 | Lighthouse Lab in Alderley Park | Wellcome Sanger Institute for the COVID-19 Genomics UK (COG-UK) Consortium | Jacquelyn Wynn, Mairead Hyland, The Lighthouse Lab in Alderley Park and Alex Alderton, Roberto Amato, Jeffrey Barrett, Sonia Goncalves, Ewan Harrison, David K. Jackson, Ian Johnston, Dominic Kwiatkowski, Cordelia Langford, John Sillitoe on behalf of the Wellcome Sanger Institute COVID-19 Surveillance Team |
| EPI_ISL_2916121 | Lighthouse Lab in Milton Keynes | Wellcome Sanger Institute for the COVID-19 Genomics UK (COG-UK) Consortium | The Lighthouse Lab in Milton Keynes and Alex Alderton, Roberto Amato, Jeffrey Barrett, Sonia Goncalves, Ewan Harrison, David K. Jackson, Ian Johnston, Dominic Kwiatkowski, Cordelia Langford, John Sillitoe on behalf of the Wellcome Sanger Institute COVID-19 Surveillance Team |
| EPI_ISL_2916122, EPI_ISL_2916123 | Berkshire and Surrey Pathology Services Lighthouse Laboratory | Wellcome Sanger Institute for the COVID-19 Genomics UK (COG-UK) Consortium | Berkshire and Surrey Pathology Services Lighthouse Laboratory and Alex Alderton, Roberto Amato, Jeffrey Barrett, Sonia Goncalves, Ewan Harrison, David K. Jackson, Ian Johnston, Dominic Kwiatkowski, Cordelia Langford, John Sillitoe on behalf of the Wellcome Sanger Institute COVID-19 Surveillance Team |
| EPI_ISL_2916124 | Lighthouse Lab in Milton Keynes | Wellcome Sanger Institute for the COVID-19 Genomics UK (COG-UK) Consortium | The Lighthouse Lab in Milton Keynes and Alex Alderton, Roberto Amato, Jeffrey Barrett, Sonia Goncalves, Ewan Harrison, David K. Jackson, Ian Johnston, Dominic Kwiatkowski, Cordelia Langford, John Sillitoe on behalf of the Wellcome Sanger Institute COVID-19 Surveillance Team |
| EPI_ISL_2916125, EPI_ISL_2916126, EPI_ISL_2916127, EPI_ISL_2916128 | Lighthouse Lab in Alderley Park | Wellcome Sanger Institute for the COVID-19 Genomics UK (COG-UK) Consortium | Jacquelyn Wynn, Mairead Hyland, The Lighthouse Lab in Alderley Park and Alex Alderton, Roberto Amato, Jeffrey Barrett, Sonia Goncalves, Ewan Harrison, David K. Jackson, Ian Johnston, Dominic Kwiatkowski, Cordelia Langford, John Sillitoe on behalf of the Wellcome Sanger Institute COVID-19 Surveillance Team |
| EPI_ISL_2916129 | Berkshire and Surrey Pathology Services Lighthouse Laboratory | Wellcome Sanger Institute for the COVID-19 Genomics UK (COG-UK) Consortium | Berkshire and Surrey Pathology Services Lighthouse Laboratory and Alex Alderton, Roberto Amato, Jeffrey Barrett, Sonia Goncalves, Ewan Harrison, David K. Jackson, Ian Johnston, Dominic Kwiatkowski, Cordelia Langford, John Sillitoe on behalf of the Wellcome Sanger Institute COVID-19 Surveillance Team |
| EPI_ISL_2916130, EPI_ISL_2916131 | Lighthouse Lab in Alderley Park | Wellcome Sanger Institute for the COVID-19 Genomics UK (COG-UK) Consortium | Jacquelyn Wynn, Mairead Hyland, The Lighthouse Lab in Alderley Park and Alex Alderton, Roberto Amato, Jeffrey Barrett, Sonia Goncalves, Ewan Harrison, David K. Jackson, Ian Johnston, Dominic Kwiatkowski, Cordelia Langford, John Sillitoe on behalf of the Wellcome Sanger Institute COVID-19 Surveillance Team |
| EPI_ISL_2916132 | Berkshire and Surrey Pathology Services Lighthouse Laboratory | Wellcome Sanger Institute for the COVID-19 Genomics UK (COG-UK) Consortium | Berkshire and Surrey Pathology Services Lighthouse Laboratory and Alex Alderton, Roberto Amato, Jeffrey Barrett, Sonia Goncalves, Ewan Harrison, David K. Jackson, Ian Johnston, Dominic Kwiatkowski, Cordelia Langford, John Sillitoe on behalf of the Wellcome Sanger Institute COVID-19 Surveillance Team |
| EPI_ISL_2916133 | Lighthouse Lab in Milton Keynes | Wellcome Sanger Institute for the COVID-19 Genomics UK (COG-UK) Consortium | The Lighthouse Lab in Milton Keynes and Alex Alderton, Roberto Amato, Jeffrey Barrett, Sonia Goncalves, Ewan Harrison, David K. Jackson, Ian Johnston, Dominic Kwiatkowski, Cordelia Langford, John Sillitoe on behalf of the Wellcome Sanger Institute COVID-19 Surveillance Team |
| EPI_ISL_2916134, EPI_ISL_2916135, EPI_ISL_2916136 | Berkshire and Surrey Pathology Services Lighthouse Laboratory | Wellcome Sanger Institute for the COVID-19 Genomics UK (COG-UK) Consortium | Berkshire and Surrey Pathology Services Lighthouse Laboratory and Alex Alderton, Roberto Amato, Jeffrey Barrett, Sonia Goncalves, Ewan Harrison, David K. Jackson, Ian Johnston, Dominic Kwiatkowski, Cordelia Langford, John Sillitoe on behalf of the Wellcome Sanger Institute COVID-19 Surveillance Team |
| EPI_ISL_2916137 | Lighthouse Lab in Alderley Park | Wellcome Sanger Institute for the COVID-19 Genomics UK (COG-UK) Consortium | Jacquelyn Wynn, Mairead Hyland, The Lighthouse Lab in Alderley Park and Alex Alderton, Roberto Amato, Jeffrey Barrett, Sonia Goncalves, Ewan Harrison, David K. Jackson, Ian Johnston, Dominic Kwiatkowski, Cordelia Langford, John Sillitoe on behalf of the Wellcome Sanger Institute COVID-19 Surveillance Team |
| EPI_ISL_2916138 | Berkshire and Surrey Pathology Services Lighthouse Laboratory | Wellcome Sanger Institute for the COVID-19 Genomics UK (COG-UK) Consortium | Berkshire and Surrey Pathology Services Lighthouse Laboratory and Alex Alderton, Roberto Amato, Jeffrey Barrett, Sonia Goncalves, Ewan Harrison, David K. Jackson, Ian Johnston, Dominic Kwiatkowski, Cordelia Langford, John Sillitoe on behalf of the Wellcome Sanger Institute COVID-19 Surveillance Team |
| EPI_ISL_2916139 | Lighthouse Lab in Alderley Park | Wellcome Sanger Institute for the COVID-19 Genomics UK (COG-UK) Consortium | Jacquelyn Wynn, Mairead Hyland, The Lighthouse Lab in Alderley Park and Alex Alderton, Roberto Amato, Jeffrey Barrett, Sonia Goncalves, Ewan Harrison, David K. Jackson, Ian Johnston, Dominic Kwiatkowski, Cordelia Langford, John Sillitoe on behalf of the Wellcome Sanger Institute COVID-19 Surveillance Team |
| EPI_ISL_2916140 | Lighthouse Lab in Milton Keynes | Wellcome Sanger Institute for the COVID-19 Genomics UK (COG-UK) Consortium | The Lighthouse Lab in Milton Keynes and Alex Alderton, Roberto Amato, Jeffrey Barrett, Sonia Goncalves, Ewan Harrison, David K. Jackson, Ian Johnston, Dominic Kwiatkowski, Cordelia Langford, John Sillitoe on behalf of the Wellcome Sanger Institute COVID-19 Surveillance Team |
| EPI_ISL_2916141 | Berkshire and Surrey Pathology Services Lighthouse Laboratory | Wellcome Sanger Institute for the COVID-19 Genomics UK (COG-UK) Consortium | Berkshire and Surrey Pathology Services Lighthouse Laboratory and Alex Alderton, Roberto Amato, Jeffrey Barrett, Sonia Goncalves, Ewan Harrison, David K. Jackson, Ian Johnston, Dominic Kwiatkowski, Cordelia Langford, John Sillitoe on behalf of the Wellcome Sanger Institute COVID-19 Surveillance Team |
| EPI_ISL_2916142, EPI_ISL_2916143, EPI_ISL_2916144 | Lighthouse Lab in Alderley Park | Wellcome Sanger Institute for the COVID-19 Genomics UK (COG-UK) Consortium | Jacquelyn Wynn, Mairead Hyland, The Lighthouse Lab in Alderley Park and Alex Alderton, Roberto Amato, Jeffrey Barrett, Sonia Goncalves, Ewan Harrison, David K. Jackson, Ian Johnston, Dominic Kwiatkowski, Cordelia Langford, John Sillitoe on behalf of the Wellcome Sanger Institute COVID-19 Surveillance Team |
| EPI_ISL_2916145, EPI_ISL_2916146 | Berkshire and Surrey Pathology Services Lighthouse Laboratory | Wellcome Sanger Institute for the COVID-19 Genomics UK (COG-UK) Consortium | Berkshire and Surrey Pathology Services Lighthouse Laboratory and Alex Alderton, Roberto Amato, Jeffrey Barrett, Sonia Goncalves, Ewan Harrison, David K. Jackson, Ian Johnston, Dominic Kwiatkowski, Cordelia Langford, John Sillitoe on behalf of the Wellcome Sanger Institute COVID-19 Surveillance Team |
| EPI_ISL_2916147 | Lighthouse Lab in Alderley Park | Wellcome Sanger Institute for the COVID-19 Genomics UK (COG-UK) Consortium | Jacquelyn Wynn, Mairead Hyland, The Lighthouse Lab in Alderley Park and Alex Alderton, Roberto Amato, Jeffrey Barrett, Sonia Goncalves, Ewan Harrison, David K. Jackson, Ian Johnston, Dominic Kwiatkowski, Cordelia Langford, John Sillitoe on behalf of the Wellcome Sanger Institute COVID-19 Surveillance Team |
| EPI_ISL_2916148 | Lighthouse Lab in Milton Keynes | Wellcome Sanger Institute for the COVID-19 Genomics UK (COG-UK) Consortium | The Lighthouse Lab in Milton Keynes and Alex Alderton, Roberto Amato, Jeffrey Barrett, Sonia Goncalves, Ewan Harrison, David K. Jackson, Ian Johnston, Dominic Kwiatkowski, Cordelia Langford, John Sillitoe on behalf of the Wellcome Sanger Institute COVID-19 Surveillance Team |
| EPI_ISL_2916149 | Berkshire and Surrey Pathology Services Lighthouse Laboratory | Wellcome Sanger Institute for the COVID-19 Genomics UK (COG-UK) Consortium | Berkshire and Surrey Pathology Services Lighthouse Laboratory and Alex Alderton, Roberto Amato, Jeffrey Barrett, Sonia Goncalves, Ewan Harrison, David K. Jackson, Ian Johnston, Dominic Kwiatkowski, Cordelia Langford, John Sillitoe on behalf of the Wellcome Sanger Institute COVID-19 Surveillance Team |
| EPI_ISL_2916150 | Lighthouse Lab in Alderley Park | Wellcome Sanger Institute for the COVID-19 Genomics UK (COG-UK) Consortium | Jacquelyn Wynn, Mairead Hyland, The Lighthouse Lab in Alderley Park and Alex Alderton, Roberto Amato, Jeffrey Barrett, Sonia Goncalves, Ewan Harrison, David K. Jackson, Ian Johnston, Dominic Kwiatkowski, Cordelia Langford, John Sillitoe on behalf of the Wellcome Sanger Institute COVID-19 Surveillance Team |
| EPI_ISL_2916151 | Berkshire and Surrey Pathology Services Lighthouse Laboratory | Wellcome Sanger Institute for the COVID-19 Genomics UK (COG-UK) | Berkshire and Surrey Pathology Services Lighthouse Laboratory and Alex Alderton, Roberto Amato, Jeffrey Barrett, Sonia Goncalves, Ewan Harrison, David K. Jackson, Ian Johnston, Dominic Kwiatkowski, Cordelia Langford, John Sillitoe on behalf |

| | | | |
|---|---|---|---|
| | | Consortium | of the Wellcome Sanger Institute COVID-19 Surveillance Team |
| EPI_ISL_2916152 | Lighthouse Lab in Milton Keynes | Wellcome Sanger Institute for the COVID-19 Genomics UK (COG-UK) Consortium | The Lighthouse Lab in Milton Keynes and Alex Alderton, Roberto Amato, Jeffrey Barrett, Sonia Goncalves, Ewan Harrison, David K. Jackson, Ian Johnston, Dominic Kwiatkowski, Cordelia Langford, John Sillitoe on behalf of the Wellcome Sanger Institute COVID-19 Surveillance Team |
| EPI_ISL_2916153 | Lighthouse Lab in Alderley Park | Wellcome Sanger Institute for the COVID-19 Genomics UK (COG-UK) Consortium | Jacquelyn Wynn, Mairead Hyland, The Lighthouse Lab in Alderley Park and Alex Alderton, Roberto Amato, Jeffrey Barrett, Sonia Goncalves, Ewan Harrison, David K. Jackson, Ian Johnston, Dominic Kwiatkowski, Cordelia Langford, John Sillitoe on behalf of the Wellcome Sanger Institute COVID-19 Surveillance Team |
| EPI_ISL_2916154 | Berkshire and Surrey Pathology Services Lighthouse Laboratory | Wellcome Sanger Institute for the COVID-19 Genomics UK (COG-UK) Consortium | Berkshire and Surrey Pathology Services Lighthouse Laboratory and Alex Alderton, Roberto Amato, Jeffrey Barrett, Sonia Goncalves, Ewan Harrison, David K. Jackson, Ian Johnston, Dominic Kwiatkowski, Cordelia Langford, John Sillitoe on behalf of the Wellcome Sanger Institute COVID-19 Surveillance Team |
| EPI_ISL_2916155 | Lighthouse Lab in Milton Keynes | Wellcome Sanger Institute for the COVID-19 Genomics UK (COG-UK) Consortium | The Lighthouse Lab in Milton Keynes and Alex Alderton, Roberto Amato, Jeffrey Barrett, Sonia Goncalves, Ewan Harrison, David K. Jackson, Ian Johnston, Dominic Kwiatkowski, Cordelia Langford, John Sillitoe on behalf of the Wellcome Sanger Institute COVID-19 Surveillance Team |
| EPI_ISL_2916156, EPI_ISL_2916157, EPI_ISL_2916158, EPI_ISL_2916159 | Berkshire and Surrey Pathology Services Lighthouse Laboratory | Wellcome Sanger Institute for the COVID-19 Genomics UK (COG-UK) Consortium | Berkshire and Surrey Pathology Services Lighthouse Laboratory and Alex Alderton, Roberto Amato, Jeffrey Barrett, Sonia Goncalves, Ewan Harrison, David K. Jackson, Ian Johnston, Dominic Kwiatkowski, Cordelia Langford, John Sillitoe on behalf of the Wellcome Sanger Institute COVID-19 Surveillance Team |
| EPI_ISL_2916160 | Lighthouse Lab in Alderley Park | Wellcome Sanger Institute for the COVID-19 Genomics UK (COG-UK) Consortium | Jacquelyn Wynn, Mairead Hyland, The Lighthouse Lab in Alderley Park and Alex Alderton, Roberto Amato, Jeffrey Barrett, Sonia Goncalves, Ewan Harrison, David K. Jackson, Ian Johnston, Dominic Kwiatkowski, Cordelia Langford, John Sillitoe on behalf of the Wellcome Sanger Institute COVID-19 Surveillance Team |
| EPI_ISL_2916161 | Berkshire and Surrey Pathology Services Lighthouse Laboratory | Wellcome Sanger Institute for the COVID-19 Genomics UK (COG-UK) Consortium | Berkshire and Surrey Pathology Services Lighthouse Laboratory and Alex Alderton, Roberto Amato, Jeffrey Barrett, Sonia Goncalves, Ewan Harrison, David K. Jackson, Ian Johnston, Dominic Kwiatkowski, Cordelia Langford, John Sillitoe on behalf of the Wellcome Sanger Institute COVID-19 Surveillance Team |
| EPI_ISL_2916162 | Lighthouse Lab in Alderley Park | Wellcome Sanger Institute for the COVID-19 Genomics UK (COG-UK) Consortium | Jacquelyn Wynn, Mairead Hyland, The Lighthouse Lab in Alderley Park and Alex Alderton, Roberto Amato, Jeffrey Barrett, Sonia Goncalves, Ewan Harrison, David K. Jackson, Ian Johnston, Dominic Kwiatkowski, Cordelia Langford, John Sillitoe on behalf of the Wellcome Sanger Institute COVID-19 Surveillance Team |
| EPI_ISL_2916163 | Berkshire and Surrey Pathology Services Lighthouse Laboratory | Wellcome Sanger Institute for the COVID-19 Genomics UK (COG-UK) Consortium | Berkshire and Surrey Pathology Services Lighthouse Laboratory and Alex Alderton, Roberto Amato, Jeffrey Barrett, Sonia Goncalves, Ewan Harrison, David K. Jackson, Ian Johnston, Dominic Kwiatkowski, Cordelia Langford, John Sillitoe on behalf of the Wellcome Sanger Institute COVID-19 Surveillance Team |
| EPI_ISL_2916164, EPI_ISL_2916165 | Lighthouse Lab in Alderley Park | Wellcome Sanger Institute for the COVID-19 Genomics UK (COG-UK) Consortium | Jacquelyn Wynn, Mairead Hyland, The Lighthouse Lab in Alderley Park and Alex Alderton, Roberto Amato, Jeffrey Barrett, Sonia Goncalves, Ewan Harrison, David K. Jackson, Ian Johnston, Dominic Kwiatkowski, Cordelia Langford, John Sillitoe on behalf of the Wellcome Sanger Institute COVID-19 Surveillance Team |
| EPI_ISL_2916166, EPI_ISL_2916167, EPI_ISL_2916168 | Lighthouse Lab in Milton Keynes | Wellcome Sanger Institute for the COVID-19 Genomics UK (COG-UK) Consortium | The Lighthouse Lab in Milton Keynes and Alex Alderton, Roberto Amato, Jeffrey Barrett, Sonia Goncalves, Ewan Harrison, David K. Jackson, Ian Johnston, Dominic Kwiatkowski, Cordelia Langford, John Sillitoe on behalf of the Wellcome Sanger Institute COVID-19 Surveillance Team |
| EPI_ISL_2916169 | Lighthouse Lab in Alderley Park | Wellcome Sanger Institute for the COVID-19 Genomics UK (COG-UK) Consortium | Jacquelyn Wynn, Mairead Hyland, The Lighthouse Lab in Alderley Park and Alex Alderton, Roberto Amato, Jeffrey Barrett, Sonia Goncalves, Ewan Harrison, David K. Jackson, Ian Johnston, Dominic Kwiatkowski, Cordelia Langford, John Sillitoe on behalf of the Wellcome Sanger Institute COVID-19 Surveillance Team |
| EPI_ISL_2916170 | Lighthouse Lab in Milton Keynes | Wellcome Sanger Institute for the COVID-19 Genomics UK (COG-UK) Consortium | The Lighthouse Lab in Milton Keynes and Alex Alderton, Roberto Amato, Jeffrey Barrett, Sonia Goncalves, Ewan Harrison, David K. Jackson, Ian Johnston, Dominic Kwiatkowski, Cordelia Langford, John Sillitoe on behalf of the Wellcome Sanger Institute COVID-19 Surveillance Team |
| EPI_ISL_2916171 | Berkshire and Surrey Pathology Services Lighthouse Laboratory | Wellcome Sanger Institute for the COVID-19 Genomics UK (COG-UK) Consortium | Berkshire and Surrey Pathology Services Lighthouse Laboratory and Alex Alderton, Roberto Amato, Jeffrey Barrett, Sonia Goncalves, Ewan Harrison, David K. Jackson, Ian Johnston, Dominic Kwiatkowski, Cordelia Langford, John Sillitoe on behalf of the Wellcome Sanger Institute COVID-19 Surveillance Team |
| EPI_ISL_2916172, EPI_ISL_2916173 | Lighthouse Lab in Alderley Park | Wellcome Sanger Institute for the COVID-19 Genomics UK (COG-UK) Consortium | Jacquelyn Wynn, Mairead Hyland, The Lighthouse Lab in Alderley Park and Alex Alderton, Roberto Amato, Jeffrey Barrett, Sonia Goncalves, Ewan Harrison, David K. Jackson, Ian Johnston, Dominic Kwiatkowski, Cordelia Langford, John Sillitoe on behalf of the Wellcome Sanger Institute COVID-19 Surveillance Team |
| EPI_ISL_2916174, EPI_ISL_2916175 | Berkshire and Surrey Pathology Services Lighthouse Laboratory | Wellcome Sanger Institute for the COVID-19 Genomics UK (COG-UK) Consortium | Berkshire and Surrey Pathology Services Lighthouse Laboratory and Alex Alderton, Roberto Amato, Jeffrey Barrett, Sonia Goncalves, Ewan Harrison, David K. Jackson, Ian Johnston, Dominic Kwiatkowski, Cordelia Langford, John Sillitoe on behalf of the Wellcome Sanger Institute COVID-19 Surveillance Team |
| EPI_ISL_2916176 | Lighthouse Lab in Milton Keynes | Wellcome Sanger Institute for the COVID-19 Genomics UK (COG-UK) Consortium | The Lighthouse Lab in Milton Keynes and Alex Alderton, Roberto Amato, Jeffrey Barrett, Sonia Goncalves, Ewan Harrison, David K. Jackson, Ian Johnston, Dominic Kwiatkowski, Cordelia Langford, John Sillitoe on behalf of the Wellcome Sanger Institute COVID-19 Surveillance Team |
| EPI_ISL_2916177 | Lighthouse Lab in Alderley Park | Wellcome Sanger Institute for the COVID-19 Genomics UK (COG-UK) Consortium | Jacquelyn Wynn, Mairead Hyland, The Lighthouse Lab in Alderley Park and Alex Alderton, Roberto Amato, Jeffrey Barrett, Sonia Goncalves, Ewan Harrison, David K. Jackson, Ian Johnston, Dominic Kwiatkowski, Cordelia Langford, John Sillitoe on behalf of the Wellcome Sanger Institute COVID-19 Surveillance Team |
| EPI_ISL_2916178, EPI_ISL_2916179 | Lighthouse Lab in Milton Keynes | Wellcome Sanger Institute for the COVID-19 Genomics UK (COG-UK) Consortium | The Lighthouse Lab in Milton Keynes and Alex Alderton, Roberto Amato, Jeffrey Barrett, Sonia Goncalves, Ewan Harrison, David K. Jackson, Ian Johnston, Dominic Kwiatkowski, Cordelia Langford, John Sillitoe on behalf of the Wellcome Sanger Institute COVID-19 Surveillance Team |
| EPI_ISL_2916180, EPI_ISL_2916181 | Lighthouse Lab in Alderley Park | Wellcome Sanger Institute for the COVID-19 Genomics UK (COG-UK) Consortium | Jacquelyn Wynn, Mairead Hyland, The Lighthouse Lab in Alderley Park and Alex Alderton, Roberto Amato, Jeffrey Barrett, Sonia Goncalves, Ewan Harrison, David K. Jackson, Ian Johnston, Dominic Kwiatkowski, Cordelia Langford, John Sillitoe on behalf of the Wellcome Sanger Institute COVID-19 Surveillance Team |
| EPI_ISL_2916182 | Lighthouse Lab in Milton Keynes | Wellcome Sanger Institute for the COVID-19 Genomics UK (COG-UK) Consortium | The Lighthouse Lab in Milton Keynes and Alex Alderton, Roberto Amato, Jeffrey Barrett, Sonia Goncalves, Ewan Harrison, David K. Jackson, Ian Johnston, Dominic Kwiatkowski, Cordelia Langford, John Sillitoe on behalf of the Wellcome Sanger Institute COVID-19 Surveillance Team |
| EPI_ISL_2916183, EPI_ISL_2916184, EPI_ISL_2916185 | Lighthouse Lab in Alderley Park | Wellcome Sanger Institute for the COVID-19 Genomics UK (COG-UK) Consortium | Jacquelyn Wynn, Mairead Hyland, The Lighthouse Lab in Alderley Park and Alex Alderton, Roberto Amato, Jeffrey Barrett, Sonia Goncalves, Ewan Harrison, David K. Jackson, Ian Johnston, Dominic Kwiatkowski, Cordelia Langford, John Sillitoe on behalf of the Wellcome Sanger Institute COVID-19 Surveillance Team |
| EPI_ISL_2916186, EPI_ISL_2916187, EPI_ISL_2916188 | Lighthouse Lab in Milton Keynes | Wellcome Sanger Institute for the COVID-19 Genomics UK (COG-UK) Consortium | The Lighthouse Lab in Milton Keynes and Alex Alderton, Roberto Amato, Jeffrey Barrett, Sonia Goncalves, Ewan Harrison, David K. Jackson, Ian Johnston, Dominic Kwiatkowski, Cordelia Langford, John Sillitoe on behalf of the Wellcome Sanger Institute COVID-19 Surveillance Team |
| EPI_ISL_2916189, EPI_ISL_2916190 | Lighthouse Lab in Alderley Park | Wellcome Sanger Institute for the COVID-19 Genomics UK (COG-UK) Consortium | Jacquelyn Wynn, Mairead Hyland, The Lighthouse Lab in Alderley Park and Alex Alderton, Roberto Amato, Jeffrey Barrett, Sonia Goncalves, Ewan Harrison, David K. Jackson, Ian Johnston, Dominic Kwiatkowski, Cordelia Langford, John Sillitoe on behalf of the Wellcome Sanger Institute COVID-19 Surveillance Team |
| EPI_ISL_2916191, EPI_ISL_2916192 | Lighthouse Lab in Milton Keynes | Wellcome Sanger Institute for the COVID-19 Genomics UK (COG-UK) Consortium | The Lighthouse Lab in Milton Keynes and Alex Alderton, Roberto Amato, Jeffrey Barrett, Sonia Goncalves, Ewan Harrison, David K. Jackson, Ian Johnston, Dominic Kwiatkowski, Cordelia Langford, John Sillitoe on behalf of the Wellcome Sanger Institute COVID-19 Surveillance Team |
| EPI_ISL_2916193, EPI_ISL_2916194 | Lighthouse Lab in Alderley Park | Wellcome Sanger Institute for the COVID-19 Genomics UK (COG-UK) Consortium | Jacquelyn Wynn, Mairead Hyland, The Lighthouse Lab in Alderley Park and Alex Alderton, Roberto Amato, Jeffrey Barrett, Sonia Goncalves, Ewan Harrison, David K. Jackson, Ian Johnston, Dominic Kwiatkowski, Cordelia Langford, John Sillitoe on behalf of the Wellcome Sanger Institute COVID-19 Surveillance Team |
| EPI_ISL_2916195 | Berkshire and Surrey Pathology Services Lighthouse Laboratory | Wellcome Sanger Institute for the COVID-19 Genomics UK (COG-UK) Consortium | Berkshire and Surrey Pathology Services Lighthouse Laboratory and Alex Alderton, Roberto Amato, Jeffrey Barrett, Sonia Goncalves, Ewan Harrison, David K. Jackson, Ian Johnston, Dominic Kwiatkowski, Cordelia Langford, John Sillitoe on behalf of the Wellcome Sanger Institute COVID-19 Surveillance Team |
| EPI_ISL_2916196 | Lighthouse Lab in Milton Keynes | Wellcome Sanger Institute for the COVID-19 Genomics UK (COG-UK) Consortium | The Lighthouse Lab in Milton Keynes and Alex Alderton, Roberto Amato, Jeffrey Barrett, Sonia Goncalves, Ewan Harrison, David K. Jackson, Ian Johnston, Dominic Kwiatkowski, Cordelia Langford, John Sillitoe on behalf of the Wellcome Sanger Institute COVID-19 Surveillance Team |
| EPI_ISL_2916197 | Lighthouse Lab in Alderley Park | Wellcome Sanger Institute for the COVID-19 Genomics UK (COG-UK) Consortium | Jacquelyn Wynn, Mairead Hyland, The Lighthouse Lab in Alderley Park and Alex Alderton, Roberto Amato, Jeffrey Barrett, Sonia Goncalves, Ewan Harrison, David K. Jackson, Ian Johnston, Dominic Kwiatkowski, Cordelia Langford, John Sillitoe on behalf of the Wellcome Sanger Institute COVID-19 Surveillance Team |
| EPI_ISL_2916198 | Lighthouse Lab in Milton Keynes | Wellcome Sanger Institute for the COVID-19 Genomics UK (COG-UK) Consortium | The Lighthouse Lab in Milton Keynes and Alex Alderton, Roberto Amato, Jeffrey Barrett, Sonia Goncalves, Ewan Harrison, David K. Jackson, Ian Johnston, Dominic Kwiatkowski, Cordelia Langford, John Sillitoe on behalf of the Wellcome Sanger Institute COVID-19 Surveillance Team |
| EPI_ISL_2916199 | Lighthouse Lab in Alderley Park | Wellcome Sanger Institute for the COVID-19 Genomics UK (COG-UK) Consortium | Jacquelyn Wynn, Mairead Hyland, The Lighthouse Lab in Alderley Park and Alex Alderton, Roberto Amato, Jeffrey Barrett, Sonia Goncalves, Ewan Harrison, David K. Jackson, Ian Johnston, Dominic Kwiatkowski, Cordelia Langford, John Sillitoe on behalf of the Wellcome Sanger Institute COVID-19 Surveillance Team |
| EPI_ISL_2916200 | Berkshire and Surrey Pathology Services Lighthouse Laboratory | Wellcome Sanger Institute for the COVID-19 Genomics UK (COG-UK) Consortium | Berkshire and Surrey Pathology Services Lighthouse Laboratory and Alex Alderton, Roberto Amato, Jeffrey Barrett, Sonia Goncalves, Ewan Harrison, David K. Jackson, Ian Johnston, Dominic Kwiatkowski, Cordelia Langford, John Sillitoe on behalf of the Wellcome Sanger Institute COVID-19 Surveillance Team |
| EPI_ISL_2916201 | Lighthouse Lab in Alderley Park | Wellcome Sanger Institute for the COVID-19 Genomics UK (COG-UK) Consortium | Jacquelyn Wynn, Mairead Hyland, The Lighthouse Lab in Alderley Park and Alex Alderton, Roberto Amato, Jeffrey Barrett, Sonia Goncalves, Ewan Harrison, David K. Jackson, Ian Johnston, Dominic Kwiatkowski, Cordelia Langford, John Sillitoe on behalf of the Wellcome Sanger Institute COVID-19 Surveillance Team |
| EPI_ISL_2916202 | Lighthouse Lab in Milton Keynes | Wellcome Sanger Institute for the COVID-19 Genomics UK (COG-UK) Consortium | The Lighthouse Lab in Milton Keynes and Alex Alderton, Roberto Amato, Jeffrey Barrett, Sonia Goncalves, Ewan Harrison, David K. Jackson, Ian Johnston, Dominic Kwiatkowski, Cordelia Langford, John Sillitoe on behalf of the Wellcome Sanger Institute COVID-19 Surveillance Team |
| EPI_ISL_2916203 | Berkshire and Surrey Pathology Services Lighthouse Laboratory | Wellcome Sanger Institute for the COVID-19 Genomics UK (COG-UK) Consortium | Berkshire and Surrey Pathology Services Lighthouse Laboratory and Alex Alderton, Roberto Amato, Jeffrey Barrett, Sonia Goncalves, Ewan Harrison, David K. Jackson, Ian Johnston, Dominic Kwiatkowski, Cordelia Langford, John Sillitoe on behalf of the Wellcome Sanger Institute COVID-19 Surveillance Team |
| EPI_ISL_2916204, EPI_ISL_2916205 | Lighthouse Lab in Milton Keynes | Wellcome Sanger Institute for the COVID-19 Genomics UK (COG-UK) Consortium | The Lighthouse Lab in Milton Keynes and Alex Alderton, Roberto Amato, Jeffrey Barrett, Sonia Goncalves, Ewan Harrison, David K. Jackson, Ian Johnston, Dominic Kwiatkowski, Cordelia Langford, John Sillitoe on behalf of the Wellcome Sanger Institute COVID-19 Surveillance Team |
| EPI_ISL_2916206, EPI_ISL_2916207 | Lighthouse Lab in Alderley Park | Wellcome Sanger Institute for the COVID-19 Genomics UK (COG-UK) Consortium | Jacquelyn Wynn, Mairead Hyland, The Lighthouse Lab in Alderley Park and Alex Alderton, Roberto Amato, Jeffrey Barrett, Sonia Goncalves, Ewan Harrison, David K. Jackson, Ian Johnston, Dominic Kwiatkowski, Cordelia Langford, John Sillitoe on behalf of the Wellcome Sanger Institute COVID-19 Surveillance Team |
| EPI_ISL_2916208, EPI_ISL_2916209 | Lighthouse Lab in Milton Keynes | Wellcome Sanger Institute for the COVID-19 Genomics UK (COG-UK) Consortium | The Lighthouse Lab in Milton Keynes and Alex Alderton, Roberto Amato, Jeffrey Barrett, Sonia Goncalves, Ewan Harrison, David K. Jackson, Ian Johnston, Dominic Kwiatkowski, Cordelia Langford, John Sillitoe on behalf of the Wellcome Sanger Institute COVID-19 Surveillance Team |
| EPI_ISL_2916210 | Lighthouse Lab in Alderley Park | Wellcome Sanger Institute for the COVID-19 Genomics UK (COG-UK) Consortium | Jacquelyn Wynn, Mairead Hyland, The Lighthouse Lab in Alderley Park and Alex Alderton, Roberto Amato, Jeffrey Barrett, Sonia Goncalves, Ewan Harrison, David K. Jackson, Ian Johnston, Dominic Kwiatkowski, Cordelia Langford, John Sillitoe on behalf of the Wellcome Sanger Institute COVID-19 Surveillance Team |
| EPI_ISL_2916211 | Berkshire and Surrey Pathology Services Lighthouse Laboratory | Wellcome Sanger Institute for the COVID-19 Genomics UK (COG-UK) Consortium | Berkshire and Surrey Pathology Services Lighthouse Laboratory and Alex Alderton, Roberto Amato, Jeffrey Barrett, Sonia Goncalves, Ewan Harrison, David K. Jackson, Ian Johnston, Dominic Kwiatkowski, Cordelia Langford, John Sillitoe on behalf of the Wellcome Sanger Institute COVID-19 Surveillance Team |
| EPI_ISL_2916212 | Lighthouse Lab in Milton Keynes | Wellcome Sanger Institute for the COVID-19 Genomics UK (COG-UK) Consortium | The Lighthouse Lab in Milton Keynes and Alex Alderton, Roberto Amato, Jeffrey Barrett, Sonia Goncalves, Ewan Harrison, David K. Jackson, Ian Johnston, Dominic Kwiatkowski, Cordelia Langford, John Sillitoe on behalf of the Wellcome Sanger Institute COVID-19 Surveillance Team |
| EPI_ISL_2916213, EPI_ISL_2916214 | Lighthouse Lab in Alderley Park | Wellcome Sanger Institute for the COVID-19 Genomics UK (COG-UK) Consortium | Jacquelyn Wynn, Mairead Hyland, The Lighthouse Lab in Alderley Park and Alex Alderton, Roberto Amato, Jeffrey Barrett, Sonia Goncalves, Ewan Harrison, David K. Jackson, Ian Johnston, Dominic Kwiatkowski, Cordelia Langford, John Sillitoe on behalf of the Wellcome Sanger Institute COVID-19 Surveillance Team |
| EPI_ISL_2916215 | Lighthouse Lab in Milton Keynes | Wellcome Sanger Institute for the COVID-19 Genomics UK (COG-UK) | The Lighthouse Lab in Milton Keynes and Alex Alderton, Roberto Amato, Jeffrey Barrett, Sonia Goncalves, Ewan Harrison, David K. Jackson, Ian Johnston, Dominic Kwiatkowski, Cordelia Langford, John Sillitoe on behalf of the Wellcome Sanger |

| | | | |
|---|---|---|---|
| | | Consortium | Institute COVID-19 Surveillance Team |
| EPI_ISL_2916216, EPI_ISL_2916217 | Berkshire and Surrey Pathology Services Lighthouse Laboratory | Wellcome Sanger Institute for the COVID-19 Genomics UK (COG-UK) Consortium | Berkshire and Surrey Pathology Services Lighthouse Laboratory and Alex Alderton, Roberto Amato, Jeffrey Barrett, Sonia Goncalves, Ewan Harrison, David K. Jackson, Ian Johnston, Dominic Kwiatkowski, Cordelia Langford, John Sillitoe on behalf of the Wellcome Sanger Institute COVID-19 Surveillance Team |
| EPI_ISL_2916218 | Lighthouse Lab in Milton Keynes | Wellcome Sanger Institute for the COVID-19 Genomics UK (COG-UK) Consortium | The Lighthouse Lab in Milton Keynes and Alex Alderton, Roberto Amato, Jeffrey Barrett, Sonia Goncalves, Ewan Harrison, David K. Jackson, Ian Johnston, Dominic Kwiatkowski, Cordelia Langford, John Sillitoe on behalf of the Wellcome Sanger Institute COVID-19 Surveillance Team |
| EPI_ISL_2916219, EPI_ISL_2916220 | Berkshire and Surrey Pathology Services Lighthouse Laboratory | Wellcome Sanger Institute for the COVID-19 Genomics UK (COG-UK) Consortium | Berkshire and Surrey Pathology Services Lighthouse Laboratory and Alex Alderton, Roberto Amato, Jeffrey Barrett, Sonia Goncalves, Ewan Harrison, David K. Jackson, Ian Johnston, Dominic Kwiatkowski, Cordelia Langford, John Sillitoe on behalf of the Wellcome Sanger Institute COVID-19 Surveillance Team |
| EPI_ISL_2916221, EPI_ISL_2916222, EPI_ISL_2916223 | Lighthouse Lab in Alderley Park | Wellcome Sanger Institute for the COVID-19 Genomics UK (COG-UK) Consortium | Jacquelyn Wynn, Mairead Hyland, The Lighthouse Lab in Alderley Park and Alex Alderton, Roberto Amato, Jeffrey Barrett, Sonia Goncalves, Ewan Harrison, David K. Jackson, Ian Johnston, Dominic Kwiatkowski, Cordelia Langford, John Sillitoe on behalf of the Wellcome Sanger Institute COVID-19 Surveillance Team |
| EPI_ISL_2916224 | Berkshire and Surrey Pathology Services Lighthouse Laboratory | Wellcome Sanger Institute for the COVID-19 Genomics UK (COG-UK) Consortium | Berkshire and Surrey Pathology Services Lighthouse Laboratory and Alex Alderton, Roberto Amato, Jeffrey Barrett, Sonia Goncalves, Ewan Harrison, David K. Jackson, Ian Johnston, Dominic Kwiatkowski, Cordelia Langford, John Sillitoe on behalf of the Wellcome Sanger Institute COVID-19 Surveillance Team |
| EPI_ISL_2916225 | Lighthouse Lab in Milton Keynes | Wellcome Sanger Institute for the COVID-19 Genomics UK (COG-UK) Consortium | The Lighthouse Lab in Milton Keynes and Alex Alderton, Roberto Amato, Jeffrey Barrett, Sonia Goncalves, Ewan Harrison, David K. Jackson, Ian Johnston, Dominic Kwiatkowski, Cordelia Langford, John Sillitoe on behalf of the Wellcome Sanger Institute COVID-19 Surveillance Team |
| EPI_ISL_2916226, EPI_ISL_2916227 | Berkshire and Surrey Pathology Services Lighthouse Laboratory | Wellcome Sanger Institute for the COVID-19 Genomics UK (COG-UK) Consortium | Berkshire and Surrey Pathology Services Lighthouse Laboratory and Alex Alderton, Roberto Amato, Jeffrey Barrett, Sonia Goncalves, Ewan Harrison, David K. Jackson, Ian Johnston, Dominic Kwiatkowski, Cordelia Langford, John Sillitoe on behalf of the Wellcome Sanger Institute COVID-19 Surveillance Team |
| EPI_ISL_2916228, EPI_ISL_2916229 | Lighthouse Lab in Milton Keynes | Wellcome Sanger Institute for the COVID-19 Genomics UK (COG-UK) Consortium | The Lighthouse Lab in Milton Keynes and Alex Alderton, Roberto Amato, Jeffrey Barrett, Sonia Goncalves, Ewan Harrison, David K. Jackson, Ian Johnston, Dominic Kwiatkowski, Cordelia Langford, John Sillitoe on behalf of the Wellcome Sanger Institute COVID-19 Surveillance Team |
| EPI_ISL_2916230 | Lighthouse Lab in Alderley Park | Wellcome Sanger Institute for the COVID-19 Genomics UK (COG-UK) Consortium | Jacquelyn Wynn, Mairead Hyland, The Lighthouse Lab in Alderley Park and Alex Alderton, Roberto Amato, Jeffrey Barrett, Sonia Goncalves, Ewan Harrison, David K. Jackson, Ian Johnston, Dominic Kwiatkowski, Cordelia Langford, John Sillitoe on behalf of the Wellcome Sanger Institute COVID-19 Surveillance Team |
| EPI_ISL_2916231 | Berkshire and Surrey Pathology Services Lighthouse Laboratory | Wellcome Sanger Institute for the COVID-19 Genomics UK (COG-UK) Consortium | Berkshire and Surrey Pathology Services Lighthouse Laboratory and Alex Alderton, Roberto Amato, Jeffrey Barrett, Sonia Goncalves, Ewan Harrison, David K. Jackson, Ian Johnston, Dominic Kwiatkowski, Cordelia Langford, John Sillitoe on behalf of the Wellcome Sanger Institute COVID-19 Surveillance Team |
| EPI_ISL_2916232 | Lighthouse Lab in Alderley Park | Wellcome Sanger Institute for the COVID-19 Genomics UK (COG-UK) Consortium | Jacquelyn Wynn, Mairead Hyland, The Lighthouse Lab in Alderley Park and Alex Alderton, Roberto Amato, Jeffrey Barrett, Sonia Goncalves, Ewan Harrison, David K. Jackson, Ian Johnston, Dominic Kwiatkowski, Cordelia Langford, John Sillitoe on behalf of the Wellcome Sanger Institute COVID-19 Surveillance Team |
| EPI_ISL_2916233, EPI_ISL_2916234 | Lighthouse Lab in Milton Keynes | Wellcome Sanger Institute for the COVID-19 Genomics UK (COG-UK) Consortium | The Lighthouse Lab in Milton Keynes and Alex Alderton, Roberto Amato, Jeffrey Barrett, Sonia Goncalves, Ewan Harrison, David K. Jackson, Ian Johnston, Dominic Kwiatkowski, Cordelia Langford, John Sillitoe on behalf of the Wellcome Sanger Institute COVID-19 Surveillance Team |
| EPI_ISL_2916235, EPI_ISL_2916236 | Berkshire and Surrey Pathology Services Lighthouse Laboratory | Wellcome Sanger Institute for the COVID-19 Genomics UK (COG-UK) Consortium | Berkshire and Surrey Pathology Services Lighthouse Laboratory and Alex Alderton, Roberto Amato, Jeffrey Barrett, Sonia Goncalves, Ewan Harrison, David K. Jackson, Ian Johnston, Dominic Kwiatkowski, Cordelia Langford, John Sillitoe on behalf of the Wellcome Sanger Institute COVID-19 Surveillance Team |
| EPI_ISL_2916237 | Lighthouse Lab in Alderley Park | Wellcome Sanger Institute for the COVID-19 Genomics UK (COG-UK) Consortium | Jacquelyn Wynn, Mairead Hyland, The Lighthouse Lab in Alderley Park and Alex Alderton, Roberto Amato, Jeffrey Barrett, Sonia Goncalves, Ewan Harrison, David K. Jackson, Ian Johnston, Dominic Kwiatkowski, Cordelia Langford, John Sillitoe on behalf of the Wellcome Sanger Institute COVID-19 Surveillance Team |
| EPI_ISL_2916238, EPI_ISL_2916239, EPI_ISL_2916240 | Lighthouse Lab in Milton Keynes | Wellcome Sanger Institute for the COVID-19 Genomics UK (COG-UK) Consortium | The Lighthouse Lab in Milton Keynes and Alex Alderton, Roberto Amato, Jeffrey Barrett, Sonia Goncalves, Ewan Harrison, David K. Jackson, Ian Johnston, Dominic Kwiatkowski, Cordelia Langford, John Sillitoe on behalf of the Wellcome Sanger Institute COVID-19 Surveillance Team |
| EPI_ISL_2916241 | Lighthouse Lab in Alderley Park | Wellcome Sanger Institute for the COVID-19 Genomics UK (COG-UK) Consortium | Jacquelyn Wynn, Mairead Hyland, The Lighthouse Lab in Alderley Park and Alex Alderton, Roberto Amato, Jeffrey Barrett, Sonia Goncalves, Ewan Harrison, David K. Jackson, Ian Johnston, Dominic Kwiatkowski, Cordelia Langford, John Sillitoe on behalf of the Wellcome Sanger Institute COVID-19 Surveillance Team |
| EPI_ISL_2916242, EPI_ISL_2916243 | Lighthouse Lab in Milton Keynes | Wellcome Sanger Institute for the COVID-19 Genomics UK (COG-UK) Consortium | The Lighthouse Lab in Milton Keynes and Alex Alderton, Roberto Amato, Jeffrey Barrett, Sonia Goncalves, Ewan Harrison, David K. Jackson, Ian Johnston, Dominic Kwiatkowski, Cordelia Langford, John Sillitoe on behalf of the Wellcome Sanger Institute COVID-19 Surveillance Team |
| EPI_ISL_2916244 | Berkshire and Surrey Pathology Services Lighthouse Laboratory | Wellcome Sanger Institute for the COVID-19 Genomics UK (COG-UK) Consortium | Berkshire and Surrey Pathology Services Lighthouse Laboratory and Alex Alderton, Roberto Amato, Jeffrey Barrett, Sonia Goncalves, Ewan Harrison, David K. Jackson, Ian Johnston, Dominic Kwiatkowski, Cordelia Langford, John Sillitoe on behalf of the Wellcome Sanger Institute COVID-19 Surveillance Team |
| EPI_ISL_2916245 | Lighthouse Lab in Alderley Park | Wellcome Sanger Institute for the COVID-19 Genomics UK (COG-UK) Consortium | Jacquelyn Wynn, Mairead Hyland, The Lighthouse Lab in Alderley Park and Alex Alderton, Roberto Amato, Jeffrey Barrett, Sonia Goncalves, Ewan Harrison, David K. Jackson, Ian Johnston, Dominic Kwiatkowski, Cordelia Langford, John Sillitoe on behalf of the Wellcome Sanger Institute COVID-19 Surveillance Team |
| EPI_ISL_2916246, EPI_ISL_2916247 | Lighthouse Lab in Milton Keynes | Wellcome Sanger Institute for the COVID-19 Genomics UK (COG-UK) Consortium | The Lighthouse Lab in Milton Keynes and Alex Alderton, Roberto Amato, Jeffrey Barrett, Sonia Goncalves, Ewan Harrison, David K. Jackson, Ian Johnston, Dominic Kwiatkowski, Cordelia Langford, John Sillitoe on behalf of the Wellcome Sanger Institute COVID-19 Surveillance Team |
| EPI_ISL_2916248 | Lighthouse Lab in Alderley Park | Wellcome Sanger Institute for the COVID-19 Genomics UK (COG-UK) Consortium | Jacquelyn Wynn, Mairead Hyland, The Lighthouse Lab in Alderley Park and Alex Alderton, Roberto Amato, Jeffrey Barrett, Sonia Goncalves, Ewan Harrison, David K. Jackson, Ian Johnston, Dominic Kwiatkowski, Cordelia Langford, John Sillitoe on behalf of the Wellcome Sanger Institute COVID-19 Surveillance Team |
| EPI_ISL_2916249 | Lighthouse Lab in Milton Keynes | Wellcome Sanger Institute for the COVID-19 Genomics UK (COG-UK) Consortium | The Lighthouse Lab in Milton Keynes and Alex Alderton, Roberto Amato, Jeffrey Barrett, Sonia Goncalves, Ewan Harrison, David K. Jackson, Ian Johnston, Dominic Kwiatkowski, Cordelia Langford, John Sillitoe on behalf of the Wellcome Sanger Institute COVID-19 Surveillance Team |
| EPI_ISL_2916250 | Lighthouse Lab in Alderley Park | Wellcome Sanger Institute for the COVID-19 Genomics UK (COG-UK) Consortium | Jacquelyn Wynn, Mairead Hyland, The Lighthouse Lab in Alderley Park and Alex Alderton, Roberto Amato, Jeffrey Barrett, Sonia Goncalves, Ewan Harrison, David K. Jackson, Ian Johnston, Dominic Kwiatkowski, Cordelia Langford, John Sillitoe on behalf of the Wellcome Sanger Institute COVID-19 Surveillance Team |
| EPI_ISL_2916251, EPI_ISL_2916252 | Lighthouse Lab in Milton Keynes | Wellcome Sanger Institute for the COVID-19 Genomics UK (COG-UK) Consortium | The Lighthouse Lab in Milton Keynes and Alex Alderton, Roberto Amato, Jeffrey Barrett, Sonia Goncalves, Ewan Harrison, David K. Jackson, Ian Johnston, Dominic Kwiatkowski, Cordelia Langford, John Sillitoe on behalf of the Wellcome Sanger Institute COVID-19 Surveillance Team |
| EPI_ISL_2916253 | Lighthouse Lab in Alderley Park | Wellcome Sanger Institute for the COVID-19 Genomics UK (COG-UK) Consortium | Jacquelyn Wynn, Mairead Hyland, The Lighthouse Lab in Alderley Park and Alex Alderton, Roberto Amato, Jeffrey Barrett, Sonia Goncalves, Ewan Harrison, David K. Jackson, Ian Johnston, Dominic Kwiatkowski, Cordelia Langford, John Sillitoe on behalf of the Wellcome Sanger Institute COVID-19 Surveillance Team |
| EPI_ISL_2916254 | Berkshire and Surrey Pathology Services Lighthouse Laboratory | Wellcome Sanger Institute for the COVID-19 Genomics UK (COG-UK) Consortium | Berkshire and Surrey Pathology Services Lighthouse Laboratory and Alex Alderton, Roberto Amato, Jeffrey Barrett, Sonia Goncalves, Ewan Harrison, David K. Jackson, Ian Johnston, Dominic Kwiatkowski, Cordelia Langford, John Sillitoe on behalf of the Wellcome Sanger Institute COVID-19 Surveillance Team |
| EPI_ISL_2916255, EPI_ISL_2916256 | Lighthouse Lab in Milton Keynes | Wellcome Sanger Institute for the COVID-19 Genomics UK (COG-UK) Consortium | The Lighthouse Lab in Milton Keynes and Alex Alderton, Roberto Amato, Jeffrey Barrett, Sonia Goncalves, Ewan Harrison, David K. Jackson, Ian Johnston, Dominic Kwiatkowski, Cordelia Langford, John Sillitoe on behalf of the Wellcome Sanger Institute COVID-19 Surveillance Team |
| EPI_ISL_2916257 | Lighthouse Lab in Alderley Park | Wellcome Sanger Institute for the COVID-19 Genomics UK (COG-UK) Consortium | Jacquelyn Wynn, Mairead Hyland, The Lighthouse Lab in Alderley Park and Alex Alderton, Roberto Amato, Jeffrey Barrett, Sonia Goncalves, Ewan Harrison, David K. Jackson, Ian Johnston, Dominic Kwiatkowski, Cordelia Langford, John Sillitoe on behalf of the Wellcome Sanger Institute COVID-19 Surveillance Team |
| EPI_ISL_2916258 | Lighthouse Lab in Milton Keynes | Wellcome Sanger Institute for the COVID-19 Genomics UK (COG-UK) Consortium | The Lighthouse Lab in Milton Keynes and Alex Alderton, Roberto Amato, Jeffrey Barrett, Sonia Goncalves, Ewan Harrison, David K. Jackson, Ian Johnston, Dominic Kwiatkowski, Cordelia Langford, John Sillitoe on behalf of the Wellcome Sanger Institute COVID-19 Surveillance Team |
| EPI_ISL_2916259 | Berkshire and Surrey Pathology Services Lighthouse Laboratory | Wellcome Sanger Institute for the COVID-19 Genomics UK (COG-UK) Consortium | Berkshire and Surrey Pathology Services Lighthouse Laboratory and Alex Alderton, Roberto Amato, Jeffrey Barrett, Sonia Goncalves, Ewan Harrison, David K. Jackson, Ian Johnston, Dominic Kwiatkowski, Cordelia Langford, John Sillitoe on behalf of the Wellcome Sanger Institute COVID-19 Surveillance Team |
| EPI_ISL_2916260 | Lighthouse Lab in Milton Keynes | Wellcome Sanger Institute for the COVID-19 Genomics UK (COG-UK) Consortium | The Lighthouse Lab in Milton Keynes and Alex Alderton, Roberto Amato, Jeffrey Barrett, Sonia Goncalves, Ewan Harrison, David K. Jackson, Ian Johnston, Dominic Kwiatkowski, Cordelia Langford, John Sillitoe on behalf of the Wellcome Sanger Institute COVID-19 Surveillance Team |
| EPI_ISL_2916261, EPI_ISL_2916262 | Lighthouse Lab in Alderley Park | Wellcome Sanger Institute for the COVID-19 Genomics UK (COG-UK) Consortium | Jacquelyn Wynn, Mairead Hyland, The Lighthouse Lab in Alderley Park and Alex Alderton, Roberto Amato, Jeffrey Barrett, Sonia Goncalves, Ewan Harrison, David K. Jackson, Ian Johnston, Dominic Kwiatkowski, Cordelia Langford, John Sillitoe on behalf of the Wellcome Sanger Institute COVID-19 Surveillance Team |
| EPI_ISL_2916263 | Berkshire and Surrey Pathology Services Lighthouse Laboratory | Wellcome Sanger Institute for the COVID-19 Genomics UK (COG-UK) Consortium | Berkshire and Surrey Pathology Services Lighthouse Laboratory and Alex Alderton, Roberto Amato, Jeffrey Barrett, Sonia Goncalves, Ewan Harrison, David K. Jackson, Ian Johnston, Dominic Kwiatkowski, Cordelia Langford, John Sillitoe on behalf of the Wellcome Sanger Institute COVID-19 Surveillance Team |
| EPI_ISL_2916264, EPI_ISL_2916265 | Lighthouse Lab in Alderley Park | Wellcome Sanger Institute for the COVID-19 Genomics UK (COG-UK) Consortium | Jacquelyn Wynn, Mairead Hyland, The Lighthouse Lab in Alderley Park and Alex Alderton, Roberto Amato, Jeffrey Barrett, Sonia Goncalves, Ewan Harrison, David K. Jackson, Ian Johnston, Dominic Kwiatkowski, Cordelia Langford, John Sillitoe on behalf of the Wellcome Sanger Institute COVID-19 Surveillance Team |
| EPI_ISL_2916266 | Lighthouse Lab in Milton Keynes | Wellcome Sanger Institute for the COVID-19 Genomics UK (COG-UK) Consortium | The Lighthouse Lab in Milton Keynes and Alex Alderton, Roberto Amato, Jeffrey Barrett, Sonia Goncalves, Ewan Harrison, David K. Jackson, Ian Johnston, Dominic Kwiatkowski, Cordelia Langford, John Sillitoe on behalf of the Wellcome Sanger Institute COVID-19 Surveillance Team |
| EPI_ISL_2916267 | Lighthouse Lab in Alderley Park | Wellcome Sanger Institute for the COVID-19 Genomics UK (COG-UK) Consortium | Jacquelyn Wynn, Mairead Hyland, The Lighthouse Lab in Alderley Park and Alex Alderton, Roberto Amato, Jeffrey Barrett, Sonia Goncalves, Ewan Harrison, David K. Jackson, Ian Johnston, Dominic Kwiatkowski, Cordelia Langford, John Sillitoe on behalf of the Wellcome Sanger Institute COVID-19 Surveillance Team |
| EPI_ISL_2916268 | Berkshire and Surrey Pathology Services Lighthouse Laboratory | Wellcome Sanger Institute for the COVID-19 Genomics UK (COG-UK) Consortium | Berkshire and Surrey Pathology Services Lighthouse Laboratory and Alex Alderton, Roberto Amato, Jeffrey Barrett, Sonia Goncalves, Ewan Harrison, David K. Jackson, Ian Johnston, Dominic Kwiatkowski, Cordelia Langford, John Sillitoe on behalf of the Wellcome Sanger Institute COVID-19 Surveillance Team |
| EPI_ISL_2916269, EPI_ISL_2916270 | Lighthouse Lab in Alderley Park | Wellcome Sanger Institute for the COVID-19 Genomics UK (COG-UK) Consortium | Jacquelyn Wynn, Mairead Hyland, The Lighthouse Lab in Alderley Park and Alex Alderton, Roberto Amato, Jeffrey Barrett, Sonia Goncalves, Ewan Harrison, David K. Jackson, Ian Johnston, Dominic Kwiatkowski, Cordelia Langford, John Sillitoe on behalf of the Wellcome Sanger Institute COVID-19 Surveillance Team |
| EPI_ISL_2916271, EPI_ISL_2916272 | Berkshire and Surrey Pathology Services Lighthouse Laboratory | Wellcome Sanger Institute for the COVID-19 Genomics UK (COG-UK) Consortium | Berkshire and Surrey Pathology Services Lighthouse Laboratory and Alex Alderton, Roberto Amato, Jeffrey Barrett, Sonia Goncalves, Ewan Harrison, David K. Jackson, Ian Johnston, Dominic Kwiatkowski, Cordelia Langford, John Sillitoe on behalf of the Wellcome Sanger Institute COVID-19 Surveillance Team |
| EPI_ISL_2916273, EPI_ISL_2916274 | Lighthouse Lab in Alderley Park | Wellcome Sanger Institute for the COVID-19 Genomics UK (COG-UK) Consortium | Jacquelyn Wynn, Mairead Hyland, The Lighthouse Lab in Alderley Park and Alex Alderton, Roberto Amato, Jeffrey Barrett, Sonia Goncalves, Ewan Harrison, David K. Jackson, Ian Johnston, Dominic Kwiatkowski, Cordelia Langford, John Sillitoe on behalf of the Wellcome Sanger Institute COVID-19 Surveillance Team |
| EPI_ISL_2916275, EPI_ISL_2916276 | Lighthouse Lab in Milton Keynes | Wellcome Sanger Institute for the COVID-19 Genomics UK (COG-UK) Consortium | The Lighthouse Lab in Milton Keynes and Alex Alderton, Roberto Amato, Jeffrey Barrett, Sonia Goncalves, Ewan Harrison, David K. Jackson, Ian Johnston, Dominic Kwiatkowski, Cordelia Langford, John Sillitoe on behalf of the Wellcome Sanger Institute COVID-19 Surveillance Team |
| EPI_ISL_2916277 | Lighthouse Lab in Alderley Park | Wellcome Sanger Institute for the COVID-19 Genomics UK (COG-UK) Consortium | Jacquelyn Wynn, Mairead Hyland, The Lighthouse Lab in Alderley Park and Alex Alderton, Roberto Amato, Jeffrey Barrett, Sonia Goncalves, Ewan Harrison, David K. Jackson, Ian Johnston, Dominic Kwiatkowski, Cordelia Langford, John Sillitoe on behalf of the Wellcome Sanger Institute COVID-19 Surveillance Team |
| EPI_ISL_2916278 | Berkshire and Surrey Pathology Services Lighthouse Laboratory | Wellcome Sanger Institute for the COVID-19 Genomics UK (COG-UK) Consortium | Berkshire and Surrey Pathology Services Lighthouse Laboratory and Alex Alderton, Roberto Amato, Jeffrey Barrett, Sonia Goncalves, Ewan Harrison, David K. Jackson, Ian Johnston, Dominic Kwiatkowski, Cordelia Langford, John Sillitoe on behalf of the Wellcome Sanger Institute COVID-19 Surveillance Team |
| EPI_ISL_2916279 | Lighthouse Lab in Milton Keynes | Wellcome Sanger Institute for the COVID-19 Genomics UK (COG-UK) Consortium | The Lighthouse Lab in Milton Keynes and Alex Alderton, Roberto Amato, Jeffrey Barrett, Sonia Goncalves, Ewan Harrison, David K. Jackson, Ian Johnston, Dominic Kwiatkowski, Cordelia Langford, John Sillitoe on behalf of the Wellcome Sanger Institute COVID-19 Surveillance Team |

| | | | |
|---|---|---|---|
| EPI_ISL_2916280 | Lighthouse Lab in Alderley Park | Wellcome Sanger Institute for the COVID-19 Genomics UK (COG-UK) Consortium | Jacquelyn Wynn, Mairead Hyland, The Lighthouse Lab in Alderley Park and Alex Alderton, Roberto Amato, Jeffrey Barrett, Sonia Goncalves, Ewan Harrison, David K. Jackson, Ian Johnston, Dominic Kwiatkowski, Cordelia Langford, John Sillitoe on behalf of the Wellcome Sanger Institute COVID-19 Surveillance Team |
| EPI_ISL_2916281 | Lighthouse Lab in Milton Keynes | Wellcome Sanger Institute for the COVID-19 Genomics UK (COG-UK) Consortium | The Lighthouse Lab in Milton Keynes and Alex Alderton, Roberto Amato, Jeffrey Barrett, Sonia Goncalves, Ewan Harrison, David K. Jackson, Ian Johnston, Dominic Kwiatkowski, Cordelia Langford, John Sillitoe on behalf of the Wellcome Sanger Institute COVID-19 Surveillance Team |
| EPI_ISL_2916282, EPI_ISL_2916283 | Lighthouse Lab in Alderley Park | Wellcome Sanger Institute for the COVID-19 Genomics UK (COG-UK) Consortium | Jacquelyn Wynn, Mairead Hyland, The Lighthouse Lab in Alderley Park and Alex Alderton, Roberto Amato, Jeffrey Barrett, Sonia Goncalves, Ewan Harrison, David K. Jackson, Ian Johnston, Dominic Kwiatkowski, Cordelia Langford, John Sillitoe on behalf of the Wellcome Sanger Institute COVID-19 Surveillance Team |
| EPI_ISL_2916284 | Berkshire and Surrey Pathology Services Lighthouse Laboratory | Wellcome Sanger Institute for the COVID-19 Genomics UK (COG-UK) Consortium | Berkshire and Surrey Pathology Services Lighthouse Laboratory and Alex Alderton, Roberto Amato, Jeffrey Barrett, Sonia Goncalves, Ewan Harrison, David K. Jackson, Ian Johnston, Dominic Kwiatkowski, Cordelia Langford, John Sillitoe on behalf of the Wellcome Sanger Institute COVID-19 Surveillance Team |
| EPI_ISL_2916285 | Lighthouse Lab in Milton Keynes | Wellcome Sanger Institute for the COVID-19 Genomics UK (COG-UK) Consortium | The Lighthouse Lab in Milton Keynes and Alex Alderton, Roberto Amato, Jeffrey Barrett, Sonia Goncalves, Ewan Harrison, David K. Jackson, Ian Johnston, Dominic Kwiatkowski, Cordelia Langford, John Sillitoe on behalf of the Wellcome Sanger Institute COVID-19 Surveillance Team |
| EPI_ISL_2916286 | Lighthouse Lab in Alderley Park | Wellcome Sanger Institute for the COVID-19 Genomics UK (COG-UK) Consortium | Jacquelyn Wynn, Mairead Hyland, The Lighthouse Lab in Alderley Park and Alex Alderton, Roberto Amato, Jeffrey Barrett, Sonia Goncalves, Ewan Harrison, David K. Jackson, Ian Johnston, Dominic Kwiatkowski, Cordelia Langford, John Sillitoe on behalf of the Wellcome Sanger Institute COVID-19 Surveillance Team |
| EPI_ISL_2916287 | Lighthouse Lab in Milton Keynes | Wellcome Sanger Institute for the COVID-19 Genomics UK (COG-UK) Consortium | The Lighthouse Lab in Milton Keynes and Alex Alderton, Roberto Amato, Jeffrey Barrett, Sonia Goncalves, Ewan Harrison, David K. Jackson, Ian Johnston, Dominic Kwiatkowski, Cordelia Langford, John Sillitoe on behalf of the Wellcome Sanger Institute COVID-19 Surveillance Team |
| EPI_ISL_2916288 | Berkshire and Surrey Pathology Services Lighthouse Laboratory | Wellcome Sanger Institute for the COVID-19 Genomics UK (COG-UK) Consortium | Berkshire and Surrey Pathology Services Lighthouse Laboratory and Alex Alderton, Roberto Amato, Jeffrey Barrett, Sonia Goncalves, Ewan Harrison, David K. Jackson, Ian Johnston, Dominic Kwiatkowski, Cordelia Langford, John Sillitoe on behalf of the Wellcome Sanger Institute COVID-19 Surveillance Team |
| EPI_ISL_2916289 | Lighthouse Lab in Milton Keynes | Wellcome Sanger Institute for the COVID-19 Genomics UK (COG-UK) Consortium | The Lighthouse Lab in Milton Keynes and Alex Alderton, Roberto Amato, Jeffrey Barrett, Sonia Goncalves, Ewan Harrison, David K. Jackson, Ian Johnston, Dominic Kwiatkowski, Cordelia Langford, John Sillitoe on behalf of the Wellcome Sanger Institute COVID-19 Surveillance Team |
| EPI_ISL_2916290 | Berkshire and Surrey Pathology Services Lighthouse Laboratory | Wellcome Sanger Institute for the COVID-19 Genomics UK (COG-UK) Consortium | Berkshire and Surrey Pathology Services Lighthouse Laboratory and Alex Alderton, Roberto Amato, Jeffrey Barrett, Sonia Goncalves, Ewan Harrison, David K. Jackson, Ian Johnston, Dominic Kwiatkowski, Cordelia Langford, John Sillitoe on behalf of the Wellcome Sanger Institute COVID-19 Surveillance Team |
| EPI_ISL_2916291 | Lighthouse Lab in Milton Keynes | Wellcome Sanger Institute for the COVID-19 Genomics UK (COG-UK) Consortium | The Lighthouse Lab in Milton Keynes and Alex Alderton, Roberto Amato, Jeffrey Barrett, Sonia Goncalves, Ewan Harrison, David K. Jackson, Ian Johnston, Dominic Kwiatkowski, Cordelia Langford, John Sillitoe on behalf of the Wellcome Sanger Institute COVID-19 Surveillance Team |
| EPI_ISL_2916292 | Lighthouse Lab in Alderley Park | Wellcome Sanger Institute for the COVID-19 Genomics UK (COG-UK) Consortium | Jacquelyn Wynn, Mairead Hyland, The Lighthouse Lab in Alderley Park and Alex Alderton, Roberto Amato, Jeffrey Barrett, Sonia Goncalves, Ewan Harrison, David K. Jackson, Ian Johnston, Dominic Kwiatkowski, Cordelia Langford, John Sillitoe on behalf of the Wellcome Sanger Institute COVID-19 Surveillance Team |
| EPI_ISL_2916293 | Berkshire and Surrey Pathology Services Lighthouse Laboratory | Wellcome Sanger Institute for the COVID-19 Genomics UK (COG-UK) Consortium | Berkshire and Surrey Pathology Services Lighthouse Laboratory and Alex Alderton, Roberto Amato, Jeffrey Barrett, Sonia Goncalves, Ewan Harrison, David K. Jackson, Ian Johnston, Dominic Kwiatkowski, Cordelia Langford, John Sillitoe on behalf of the Wellcome Sanger Institute COVID-19 Surveillance Team |
| EPI_ISL_2916294 | Lighthouse Lab in Milton Keynes | Wellcome Sanger Institute for the COVID-19 Genomics UK (COG-UK) Consortium | The Lighthouse Lab in Milton Keynes and Alex Alderton, Roberto Amato, Jeffrey Barrett, Sonia Goncalves, Ewan Harrison, David K. Jackson, Ian Johnston, Dominic Kwiatkowski, Cordelia Langford, John Sillitoe on behalf of the Wellcome Sanger Institute COVID-19 Surveillance Team |
| EPI_ISL_2916295, EPI_ISL_2916296 | Berkshire and Surrey Pathology Services Lighthouse Laboratory | Wellcome Sanger Institute for the COVID-19 Genomics UK (COG-UK) Consortium | Berkshire and Surrey Pathology Services Lighthouse Laboratory and Alex Alderton, Roberto Amato, Jeffrey Barrett, Sonia Goncalves, Ewan Harrison, David K. Jackson, Ian Johnston, Dominic Kwiatkowski, Cordelia Langford, John Sillitoe on behalf of the Wellcome Sanger Institute COVID-19 Surveillance Team |
| EPI_ISL_2916297 | Lighthouse Lab in Milton Keynes | Wellcome Sanger Institute for the COVID-19 Genomics UK (COG-UK) Consortium | The Lighthouse Lab in Milton Keynes and Alex Alderton, Roberto Amato, Jeffrey Barrett, Sonia Goncalves, Ewan Harrison, David K. Jackson, Ian Johnston, Dominic Kwiatkowski, Cordelia Langford, John Sillitoe on behalf of the Wellcome Sanger Institute COVID-19 Surveillance Team |
| EPI_ISL_2916298 | Lighthouse Lab in Alderley Park | Wellcome Sanger Institute for the COVID-19 Genomics UK (COG-UK) Consortium | Jacquelyn Wynn, Mairead Hyland, The Lighthouse Lab in Alderley Park and Alex Alderton, Roberto Amato, Jeffrey Barrett, Sonia Goncalves, Ewan Harrison, David K. Jackson, Ian Johnston, Dominic Kwiatkowski, Cordelia Langford, John Sillitoe on behalf of the Wellcome Sanger Institute COVID-19 Surveillance Team |
| EPI_ISL_2916299, EPI_ISL_2916300, EPI_ISL_2916301, EPI_ISL_2916302 | Lighthouse Lab in Milton Keynes | Wellcome Sanger Institute for the COVID-19 Genomics UK (COG-UK) Consortium | The Lighthouse Lab in Milton Keynes and Alex Alderton, Roberto Amato, Jeffrey Barrett, Sonia Goncalves, Ewan Harrison, David K. Jackson, Ian Johnston, Dominic Kwiatkowski, Cordelia Langford, John Sillitoe on behalf of the Wellcome Sanger Institute COVID-19 Surveillance Team |
| EPI_ISL_2916303 | Lighthouse Lab in Alderley Park | Wellcome Sanger Institute for the COVID-19 Genomics UK (COG-UK) Consortium | Jacquelyn Wynn, Mairead Hyland, The Lighthouse Lab in Alderley Park and Alex Alderton, Roberto Amato, Jeffrey Barrett, Sonia Goncalves, Ewan Harrison, David K. Jackson, Ian Johnston, Dominic Kwiatkowski, Cordelia Langford, John Sillitoe on behalf of the Wellcome Sanger Institute COVID-19 Surveillance Team |
| EPI_ISL_2916304 | Lighthouse Lab in Glasgow | Wellcome Sanger Institute for the COVID-19 Genomics UK (COG-UK) Consortium | Harper VanSteenhouse, Yumi Kasai, David Gray, Carol Clugston, Anna Dominiczak and Alex Alderton, Roberto Amato, Jeffrey Barrett, Sonia Goncalves, Ewan Harrison, David K. Jackson, Ian Johnston, Dominic Kwiatkowski, Cordelia Langford, John Sillitoe on behalf of the Wellcome Sanger Institute COVID-19 Surveillance Team |
| EPI_ISL_2916305, EPI_ISL_2916306, EPI_ISL_2916307, EPI_ISL_2916308 | Lighthouse Lab in Milton Keynes | Wellcome Sanger Institute for the COVID-19 Genomics UK (COG-UK) Consortium | The Lighthouse Lab in Milton Keynes and Alex Alderton, Roberto Amato, Jeffrey Barrett, Sonia Goncalves, Ewan Harrison, David K. Jackson, Ian Johnston, Dominic Kwiatkowski, Cordelia Langford, John Sillitoe on behalf of the Wellcome Sanger Institute COVID-19 Surveillance Team |
| EPI_ISL_2916309, EPI_ISL_2916310 | Lighthouse Lab in Glasgow | Wellcome Sanger Institute for the COVID-19 Genomics UK (COG-UK) Consortium | Harper VanSteenhouse, Yumi Kasai, David Gray, Carol Clugston, Anna Dominiczak and Alex Alderton, Roberto Amato, Jeffrey Barrett, Sonia Goncalves, Ewan Harrison, David K. Jackson, Ian Johnston, Dominic Kwiatkowski, Cordelia Langford, John Sillitoe on behalf of the Wellcome Sanger Institute COVID-19 Surveillance Team |
| EPI_ISL_2916311, EPI_ISL_2916312 | Lighthouse Lab in Alderley Park | Wellcome Sanger Institute for the COVID-19 Genomics UK (COG-UK) Consortium | Jacquelyn Wynn, Mairead Hyland, The Lighthouse Lab in Alderley Park and Alex Alderton, Roberto Amato, Jeffrey Barrett, Sonia Goncalves, Ewan Harrison, David K. Jackson, Ian Johnston, Dominic Kwiatkowski, Cordelia Langford, John Sillitoe on behalf of the Wellcome Sanger Institute COVID-19 Surveillance Team |
| EPI_ISL_2916313 | Lighthouse Lab in Milton Keynes | Wellcome Sanger Institute for the COVID-19 Genomics UK (COG-UK) Consortium | The Lighthouse Lab in Milton Keynes and Alex Alderton, Roberto Amato, Jeffrey Barrett, Sonia Goncalves, Ewan Harrison, David K. Jackson, Ian Johnston, Dominic Kwiatkowski, Cordelia Langford, John Sillitoe on behalf of the Wellcome Sanger Institute COVID-19 Surveillance Team |
| EPI_ISL_2916314, EPI_ISL_2916315 | Lighthouse Lab in Alderley Park | Wellcome Sanger Institute for the COVID-19 Genomics UK (COG-UK) Consortium | Jacquelyn Wynn, Mairead Hyland, The Lighthouse Lab in Alderley Park and Alex Alderton, Roberto Amato, Jeffrey Barrett, Sonia Goncalves, Ewan Harrison, David K. Jackson, Ian Johnston, Dominic Kwiatkowski, Cordelia Langford, John Sillitoe on behalf of the Wellcome Sanger Institute COVID-19 Surveillance Team |
| EPI_ISL_2916317, EPI_ISL_2916318 | Lighthouse Lab in Milton Keynes | Wellcome Sanger Institute for the COVID-19 Genomics UK (COG-UK) Consortium | The Lighthouse Lab in Milton Keynes and Alex Alderton, Roberto Amato, Jeffrey Barrett, Sonia Goncalves, Ewan Harrison, David K. Jackson, Ian Johnston, Dominic Kwiatkowski, Cordelia Langford, John Sillitoe on behalf of the Wellcome Sanger Institute COVID-19 Surveillance Team |
| EPI_ISL_2916319 | Lighthouse Lab in Alderley Park | Wellcome Sanger Institute for the COVID-19 Genomics UK (COG-UK) Consortium | Jacquelyn Wynn, Mairead Hyland, The Lighthouse Lab in Alderley Park and Alex Alderton, Roberto Amato, Jeffrey Barrett, Sonia Goncalves, Ewan Harrison, David K. Jackson, Ian Johnston, Dominic Kwiatkowski, Cordelia Langford, John Sillitoe on behalf of the Wellcome Sanger Institute COVID-19 Surveillance Team |
| EPI_ISL_2916320 | Lighthouse Lab in Milton Keynes | Wellcome Sanger Institute for the COVID-19 Genomics UK (COG-UK) Consortium | The Lighthouse Lab in Milton Keynes and Alex Alderton, Roberto Amato, Jeffrey Barrett, Sonia Goncalves, Ewan Harrison, David K. Jackson, Ian Johnston, Dominic Kwiatkowski, Cordelia Langford, John Sillitoe on behalf of the Wellcome Sanger Institute COVID-19 Surveillance Team |
| EPI_ISL_2916321 | Lighthouse Lab in Glasgow | Wellcome Sanger Institute for the COVID-19 Genomics UK (COG-UK) Consortium | Harper VanSteenhouse, Yumi Kasai, David Gray, Carol Clugston, Anna Dominiczak and Alex Alderton, Roberto Amato, Jeffrey Barrett, Sonia Goncalves, Ewan Harrison, David K. Jackson, Ian Johnston, Dominic Kwiatkowski, Cordelia Langford, John Sillitoe on behalf of the Wellcome Sanger Institute COVID-19 Surveillance Team |
| EPI_ISL_2916322 | Lighthouse Lab in Milton Keynes | Wellcome Sanger Institute for the COVID-19 Genomics UK (COG-UK) Consortium | The Lighthouse Lab in Milton Keynes and Alex Alderton, Roberto Amato, Jeffrey Barrett, Sonia Goncalves, Ewan Harrison, David K. Jackson, Ian Johnston, Dominic Kwiatkowski, Cordelia Langford, John Sillitoe on behalf of the Wellcome Sanger Institute COVID-19 Surveillance Team |
| EPI_ISL_2916323 | Lighthouse Lab in Glasgow | Wellcome Sanger Institute for the COVID-19 Genomics UK (COG-UK) Consortium | Harper VanSteenhouse, Yumi Kasai, David Gray, Carol Clugston, Anna Dominiczak and Alex Alderton, Roberto Amato, Jeffrey Barrett, Sonia Goncalves, Ewan Harrison, David K. Jackson, Ian Johnston, Dominic Kwiatkowski, Cordelia Langford, John Sillitoe on behalf of the Wellcome Sanger Institute COVID-19 Surveillance Team |
| EPI_ISL_2916324 | Lighthouse Lab in Alderley Park | Wellcome Sanger Institute for the COVID-19 Genomics UK (COG-UK) Consortium | Jacquelyn Wynn, Mairead Hyland, The Lighthouse Lab in Alderley Park and Alex Alderton, Roberto Amato, Jeffrey Barrett, Sonia Goncalves, Ewan Harrison, David K. Jackson, Ian Johnston, Dominic Kwiatkowski, Cordelia Langford, John Sillitoe on behalf of the Wellcome Sanger Institute COVID-19 Surveillance Team |
| EPI_ISL_2916325, EPI_ISL_2916326 | Lighthouse Lab in Milton Keynes | Wellcome Sanger Institute for the COVID-19 Genomics UK (COG-UK) Consortium | The Lighthouse Lab in Milton Keynes and Alex Alderton, Roberto Amato, Jeffrey Barrett, Sonia Goncalves, Ewan Harrison, David K. Jackson, Ian Johnston, Dominic Kwiatkowski, Cordelia Langford, John Sillitoe on behalf of the Wellcome Sanger Institute COVID-19 Surveillance Team |
| EPI_ISL_2916327 | Lighthouse Lab in Alderley Park | Wellcome Sanger Institute for the COVID-19 Genomics UK (COG-UK) Consortium | Jacquelyn Wynn, Mairead Hyland, The Lighthouse Lab in Alderley Park and Alex Alderton, Roberto Amato, Jeffrey Barrett, Sonia Goncalves, Ewan Harrison, David K. Jackson, Ian Johnston, Dominic Kwiatkowski, Cordelia Langford, John Sillitoe on behalf of the Wellcome Sanger Institute COVID-19 Surveillance Team |
| EPI_ISL_2916328 | Lighthouse Lab in Milton Keynes | Wellcome Sanger Institute for the COVID-19 Genomics UK (COG-UK) Consortium | The Lighthouse Lab in Milton Keynes and Alex Alderton, Roberto Amato, Jeffrey Barrett, Sonia Goncalves, Ewan Harrison, David K. Jackson, Ian Johnston, Dominic Kwiatkowski, Cordelia Langford, John Sillitoe on behalf of the Wellcome Sanger Institute COVID-19 Surveillance Team |
| EPI_ISL_2916329 | Lighthouse Lab in Glasgow | Wellcome Sanger Institute for the COVID-19 Genomics UK (COG-UK) Consortium | Harper VanSteenhouse, Yumi Kasai, David Gray, Carol Clugston, Anna Dominiczak and Alex Alderton, Roberto Amato, Jeffrey Barrett, Sonia Goncalves, Ewan Harrison, David K. Jackson, Ian Johnston, Dominic Kwiatkowski, Cordelia Langford, John Sillitoe on behalf of the Wellcome Sanger Institute COVID-19 Surveillance Team |
| EPI_ISL_2916330, EPI_ISL_2916331 | Lighthouse Lab in Milton Keynes | Wellcome Sanger Institute for the COVID-19 Genomics UK (COG-UK) Consortium | The Lighthouse Lab in Milton Keynes and Alex Alderton, Roberto Amato, Jeffrey Barrett, Sonia Goncalves, Ewan Harrison, David K. Jackson, Ian Johnston, Dominic Kwiatkowski, Cordelia Langford, John Sillitoe on behalf of the Wellcome Sanger Institute COVID-19 Surveillance Team |
| EPI_ISL_2916332 | Lighthouse Lab in Glasgow | Wellcome Sanger Institute for the COVID-19 Genomics UK (COG-UK) Consortium | Harper VanSteenhouse, Yumi Kasai, David Gray, Carol Clugston, Anna Dominiczak and Alex Alderton, Roberto Amato, Jeffrey Barrett, Sonia Goncalves, Ewan Harrison, David K. Jackson, Ian Johnston, Dominic Kwiatkowski, Cordelia Langford, John Sillitoe on behalf of the Wellcome Sanger Institute COVID-19 Surveillance Team |
| EPI_ISL_2916333 | Lighthouse Lab in Milton Keynes | Wellcome Sanger Institute for the COVID-19 Genomics UK (COG-UK) Consortium | The Lighthouse Lab in Milton Keynes and Alex Alderton, Roberto Amato, Jeffrey Barrett, Sonia Goncalves, Ewan Harrison, David K. Jackson, Ian Johnston, Dominic Kwiatkowski, Cordelia Langford, John Sillitoe on behalf of the Wellcome Sanger Institute COVID-19 Surveillance Team |
| EPI_ISL_2916334 | Lighthouse Lab in Glasgow | Wellcome Sanger Institute for the COVID-19 Genomics UK (COG-UK) Consortium | Harper VanSteenhouse, Yumi Kasai, David Gray, Carol Clugston, Anna Dominiczak and Alex Alderton, Roberto Amato, Jeffrey Barrett, Sonia Goncalves, Ewan Harrison, David K. Jackson, Ian Johnston, Dominic Kwiatkowski, Cordelia Langford, John Sillitoe on behalf of the Wellcome Sanger Institute COVID-19 Surveillance Team |
| EPI_ISL_2916335 | Lighthouse Lab in Milton Keynes | Wellcome Sanger Institute for the COVID-19 Genomics UK (COG-UK) Consortium | The Lighthouse Lab in Milton Keynes and Alex Alderton, Roberto Amato, Jeffrey Barrett, Sonia Goncalves, Ewan Harrison, David K. Jackson, Ian Johnston, Dominic Kwiatkowski, Cordelia Langford, John Sillitoe on behalf of the Wellcome Sanger Institute COVID-19 Surveillance Team |
| EPI_ISL_2916336, EPI_ISL_2916337, EPI_ISL_2916338, EPI_ISL_2916339 | Lighthouse Lab in Alderley Park | Wellcome Sanger Institute for the COVID-19 Genomics UK (COG-UK) Consortium | Jacquelyn Wynn, Mairead Hyland, The Lighthouse Lab in Alderley Park and Alex Alderton, Roberto Amato, Jeffrey Barrett, Sonia Goncalves, Ewan Harrison, David K. Jackson, Ian Johnston, Dominic Kwiatkowski, Cordelia Langford, John Sillitoe on behalf of the Wellcome Sanger Institute COVID-19 Surveillance Team |

| | | | |
|---|---|---|---|
| EPI_ISL_2916340 | Lighthouse Lab in Milton Keynes | Wellcome Sanger Institute for the COVID-19 Genomics UK (COG-UK) Consortium | The Lighthouse Lab in Milton Keynes and Alex Alderton, Roberto Amato, Jeffrey Barrett, Sonia Goncalves, Ewan Harrison, David K. Jackson, Ian Johnston, Dominic Kwiatkowski, Cordelia Langford, John Sillitoe on behalf of the Wellcome Sanger Institute COVID-19 Surveillance Team |
| EPI_ISL_2916341, EPI_ISL_2916342 | Lighthouse Lab in Alderley Park | Wellcome Sanger Institute for the COVID-19 Genomics UK (COG-UK) Consortium | Jacquelyn Wynn, Mairead Hyland, The Lighthouse Lab in Alderley Park and Alex Alderton, Roberto Amato, Jeffrey Barrett, Sonia Goncalves, Ewan Harrison, David K. Jackson, Ian Johnston, Dominic Kwiatkowski, Cordelia Langford, John Sillitoe on behalf of the Wellcome Sanger Institute COVID-19 Surveillance Team |
| EPI_ISL_2916344 | Lighthouse Lab in Glasgow | Wellcome Sanger Institute for the COVID-19 Genomics UK (COG-UK) Consortium | Harper VanSteenhouse, Yumi Kasai, David Gray, Carol Clugston, Anna Dominiczak and Alex Alderton, Roberto Amato, Jeffrey Barrett, Sonia Goncalves, Ewan Harrison, David K. Jackson, Ian Johnston, Dominic Kwiatkowski, Cordelia Langford, John Sillitoe on behalf of the Wellcome Sanger Institute COVID-19 Surveillance Team |
| EPI_ISL_2916345, EPI_ISL_2916346, EPI_ISL_2916347, EPI_ISL_2916348, EPI_ISL_2916349 | Lighthouse Lab in Milton Keynes | Wellcome Sanger Institute for the COVID-19 Genomics UK (COG-UK) Consortium | The Lighthouse Lab in Milton Keynes and Alex Alderton, Roberto Amato, Jeffrey Barrett, Sonia Goncalves, Ewan Harrison, David K. Jackson, Ian Johnston, Dominic Kwiatkowski, Cordelia Langford, John Sillitoe on behalf of the Wellcome Sanger Institute COVID-19 Surveillance Team |
| EPI_ISL_2916350 | Lighthouse Lab in Glasgow | Wellcome Sanger Institute for the COVID-19 Genomics UK (COG-UK) Consortium | Harper VanSteenhouse, Yumi Kasai, David Gray, Carol Clugston, Anna Dominiczak and Alex Alderton, Roberto Amato, Jeffrey Barrett, Sonia Goncalves, Ewan Harrison, David K. Jackson, Ian Johnston, Dominic Kwiatkowski, Cordelia Langford, John Sillitoe on behalf of the Wellcome Sanger Institute COVID-19 Surveillance Team |
| EPI_ISL_2916351, EPI_ISL_2916352 | Lighthouse Lab in Milton Keynes | Wellcome Sanger Institute for the COVID-19 Genomics UK (COG-UK) Consortium | The Lighthouse Lab in Milton Keynes and Alex Alderton, Roberto Amato, Jeffrey Barrett, Sonia Goncalves, Ewan Harrison, David K. Jackson, Ian Johnston, Dominic Kwiatkowski, Cordelia Langford, John Sillitoe on behalf of the Wellcome Sanger Institute COVID-19 Surveillance Team |
| EPI_ISL_2916353 | Lighthouse Lab in Glasgow | Wellcome Sanger Institute for the COVID-19 Genomics UK (COG-UK) Consortium | Harper VanSteenhouse, Yumi Kasai, David Gray, Carol Clugston, Anna Dominiczak and Alex Alderton, Roberto Amato, Jeffrey Barrett, Sonia Goncalves, Ewan Harrison, David K. Jackson, Ian Johnston, Dominic Kwiatkowski, Cordelia Langford, John Sillitoe on behalf of the Wellcome Sanger Institute COVID-19 Surveillance Team |
| EPI_ISL_2916354 | Lighthouse Lab in Milton Keynes | Wellcome Sanger Institute for the COVID-19 Genomics UK (COG-UK) Consortium | The Lighthouse Lab in Milton Keynes and Alex Alderton, Roberto Amato, Jeffrey Barrett, Sonia Goncalves, Ewan Harrison, David K. Jackson, Ian Johnston, Dominic Kwiatkowski, Cordelia Langford, John Sillitoe on behalf of the Wellcome Sanger Institute COVID-19 Surveillance Team |
| EPI_ISL_2916355, EPI_ISL_2916356 | Lighthouse Lab in Alderley Park | Wellcome Sanger Institute for the COVID-19 Genomics UK (COG-UK) Consortium | Jacquelyn Wynn, Mairead Hyland, The Lighthouse Lab in Alderley Park and Alex Alderton, Roberto Amato, Jeffrey Barrett, Sonia Goncalves, Ewan Harrison, David K. Jackson, Ian Johnston, Dominic Kwiatkowski, Cordelia Langford, John Sillitoe on behalf of the Wellcome Sanger Institute COVID-19 Surveillance Team |
| EPI_ISL_2916357 | Lighthouse Lab in Milton Keynes | Wellcome Sanger Institute for the COVID-19 Genomics UK (COG-UK) Consortium | The Lighthouse Lab in Milton Keynes and Alex Alderton, Roberto Amato, Jeffrey Barrett, Sonia Goncalves, Ewan Harrison, David K. Jackson, Ian Johnston, Dominic Kwiatkowski, Cordelia Langford, John Sillitoe on behalf of the Wellcome Sanger Institute COVID-19 Surveillance Team |
| EPI_ISL_2916358, EPI_ISL_2916359 | Lighthouse Lab in Glasgow | Wellcome Sanger Institute for the COVID-19 Genomics UK (COG-UK) Consortium | Harper VanSteenhouse, Yumi Kasai, David Gray, Carol Clugston, Anna Dominiczak and Alex Alderton, Roberto Amato, Jeffrey Barrett, Sonia Goncalves, Ewan Harrison, David K. Jackson, Ian Johnston, Dominic Kwiatkowski, Cordelia Langford, John Sillitoe on behalf of the Wellcome Sanger Institute COVID-19 Surveillance Team |
| EPI_ISL_2916360 | Lighthouse Lab in Alderley Park | Wellcome Sanger Institute for the COVID-19 Genomics UK (COG-UK) Consortium | Jacquelyn Wynn, Mairead Hyland, The Lighthouse Lab in Alderley Park and Alex Alderton, Roberto Amato, Jeffrey Barrett, Sonia Goncalves, Ewan Harrison, David K. Jackson, Ian Johnston, Dominic Kwiatkowski, Cordelia Langford, John Sillitoe on behalf of the Wellcome Sanger Institute COVID-19 Surveillance Team |
| EPI_ISL_2916361, EPI_ISL_2916362, EPI_ISL_2916363, EPI_ISL_2916364 | Lighthouse Lab in Milton Keynes | Wellcome Sanger Institute for the COVID-19 Genomics UK (COG-UK) Consortium | The Lighthouse Lab in Milton Keynes and Alex Alderton, Roberto Amato, Jeffrey Barrett, Sonia Goncalves, Ewan Harrison, David K. Jackson, Ian Johnston, Dominic Kwiatkowski, Cordelia Langford, John Sillitoe on behalf of the Wellcome Sanger Institute COVID-19 Surveillance Team |
| EPI_ISL_2916365 | Lighthouse Lab in Alderley Park | Wellcome Sanger Institute for the COVID-19 Genomics UK (COG-UK) Consortium | Jacquelyn Wynn, Mairead Hyland, The Lighthouse Lab in Alderley Park and Alex Alderton, Roberto Amato, Jeffrey Barrett, Sonia Goncalves, Ewan Harrison, David K. Jackson, Ian Johnston, Dominic Kwiatkowski, Cordelia Langford, John Sillitoe on behalf of the Wellcome Sanger Institute COVID-19 Surveillance Team |
| EPI_ISL_2916366, EPI_ISL_2916367, EPI_ISL_2916368 | Lighthouse Lab in Milton Keynes | Wellcome Sanger Institute for the COVID-19 Genomics UK (COG-UK) Consortium | The Lighthouse Lab in Milton Keynes and Alex Alderton, Roberto Amato, Jeffrey Barrett, Sonia Goncalves, Ewan Harrison, David K. Jackson, Ian Johnston, Dominic Kwiatkowski, Cordelia Langford, John Sillitoe on behalf of the Wellcome Sanger Institute COVID-19 Surveillance Team |
| EPI_ISL_2916369, EPI_ISL_2916370 | Lighthouse Lab in Glasgow | Wellcome Sanger Institute for the COVID-19 Genomics UK (COG-UK) Consortium | Harper VanSteenhouse, Yumi Kasai, David Gray, Carol Clugston, Anna Dominiczak and Alex Alderton, Roberto Amato, Jeffrey Barrett, Sonia Goncalves, Ewan Harrison, David K. Jackson, Ian Johnston, Dominic Kwiatkowski, Cordelia Langford, John Sillitoe on behalf of the Wellcome Sanger Institute COVID-19 Surveillance Team |
| EPI_ISL_2916371, EPI_ISL_2916372 | Lighthouse Lab in Alderley Park | Wellcome Sanger Institute for the COVID-19 Genomics UK (COG-UK) Consortium | Jacquelyn Wynn, Mairead Hyland, The Lighthouse Lab in Alderley Park and Alex Alderton, Roberto Amato, Jeffrey Barrett, Sonia Goncalves, Ewan Harrison, David K. Jackson, Ian Johnston, Dominic Kwiatkowski, Cordelia Langford, John Sillitoe on behalf of the Wellcome Sanger Institute COVID-19 Surveillance Team |
| EPI_ISL_2916373, EPI_ISL_2916374 | Lighthouse Lab in Milton Keynes | Wellcome Sanger Institute for the COVID-19 Genomics UK (COG-UK) Consortium | The Lighthouse Lab in Milton Keynes and Alex Alderton, Roberto Amato, Jeffrey Barrett, Sonia Goncalves, Ewan Harrison, David K. Jackson, Ian Johnston, Dominic Kwiatkowski, Cordelia Langford, John Sillitoe on behalf of the Wellcome Sanger Institute COVID-19 Surveillance Team |
| EPI_ISL_2916375, EPI_ISL_2916376 | Lighthouse Lab in Alderley Park | Wellcome Sanger Institute for the COVID-19 Genomics UK (COG-UK) Consortium | Jacquelyn Wynn, Mairead Hyland, The Lighthouse Lab in Alderley Park and Alex Alderton, Roberto Amato, Jeffrey Barrett, Sonia Goncalves, Ewan Harrison, David K. Jackson, Ian Johnston, Dominic Kwiatkowski, Cordelia Langford, John Sillitoe on behalf of the Wellcome Sanger Institute COVID-19 Surveillance Team |
| EPI_ISL_2916377 | Lighthouse Lab in Glasgow | Wellcome Sanger Institute for the COVID-19 Genomics UK (COG-UK) Consortium | Harper VanSteenhouse, Yumi Kasai, David Gray, Carol Clugston, Anna Dominiczak and Alex Alderton, Roberto Amato, Jeffrey Barrett, Sonia Goncalves, Ewan Harrison, David K. Jackson, Ian Johnston, Dominic Kwiatkowski, Cordelia Langford, John Sillitoe on behalf of the Wellcome Sanger Institute COVID-19 Surveillance Team |
| EPI_ISL_2916379, EPI_ISL_2916380 | Lighthouse Lab in Alderley Park | Wellcome Sanger Institute for the COVID-19 Genomics UK (COG-UK) Consortium | Jacquelyn Wynn, Mairead Hyland, The Lighthouse Lab in Alderley Park and Alex Alderton, Roberto Amato, Jeffrey Barrett, Sonia Goncalves, Ewan Harrison, David K. Jackson, Ian Johnston, Dominic Kwiatkowski, Cordelia Langford, John Sillitoe on behalf of the Wellcome Sanger Institute COVID-19 Surveillance Team |
| EPI_ISL_2916381, EPI_ISL_2916382 | Lighthouse Lab in Glasgow | Wellcome Sanger Institute for the COVID-19 Genomics UK (COG-UK) Consortium | Harper VanSteenhouse, Yumi Kasai, David Gray, Carol Clugston, Anna Dominiczak and Alex Alderton, Roberto Amato, Jeffrey Barrett, Sonia Goncalves, Ewan Harrison, David K. Jackson, Ian Johnston, Dominic Kwiatkowski, Cordelia Langford, John Sillitoe on behalf of the Wellcome Sanger Institute COVID-19 Surveillance Team |
| EPI_ISL_2916383, EPI_ISL_2916384 | Lighthouse Lab in Milton Keynes | Wellcome Sanger Institute for the COVID-19 Genomics UK (COG-UK) Consortium | The Lighthouse Lab in Milton Keynes and Alex Alderton, Roberto Amato, Jeffrey Barrett, Sonia Goncalves, Ewan Harrison, David K. Jackson, Ian Johnston, Dominic Kwiatkowski, Cordelia Langford, John Sillitoe on behalf of the Wellcome Sanger Institute COVID-19 Surveillance Team |
| EPI_ISL_2916385, EPI_ISL_2916386, EPI_ISL_2916387, EPI_ISL_2916388, EPI_ISL_2916389 | Lighthouse Lab in Glasgow | Wellcome Sanger Institute for the COVID-19 Genomics UK (COG-UK) Consortium | Harper VanSteenhouse, Yumi Kasai, David Gray, Carol Clugston, Anna Dominiczak and Alex Alderton, Roberto Amato, Jeffrey Barrett, Sonia Goncalves, Ewan Harrison, David K. Jackson, Ian Johnston, Dominic Kwiatkowski, Cordelia Langford, John Sillitoe on behalf of the Wellcome Sanger Institute COVID-19 Surveillance Team |
| EPI_ISL_2916390 | Lighthouse Lab in Milton Keynes | Wellcome Sanger Institute for the COVID-19 Genomics UK (COG-UK) Consortium | The Lighthouse Lab in Milton Keynes and Alex Alderton, Roberto Amato, Jeffrey Barrett, Sonia Goncalves, Ewan Harrison, David K. Jackson, Ian Johnston, Dominic Kwiatkowski, Cordelia Langford, John Sillitoe on behalf of the Wellcome Sanger Institute COVID-19 Surveillance Team |
| EPI_ISL_2916391 | Lighthouse Lab in Alderley Park | Wellcome Sanger Institute for the COVID-19 Genomics UK (COG-UK) Consortium | Jacquelyn Wynn, Mairead Hyland, The Lighthouse Lab in Alderley Park and Alex Alderton, Roberto Amato, Jeffrey Barrett, Sonia Goncalves, Ewan Harrison, David K. Jackson, Ian Johnston, Dominic Kwiatkowski, Cordelia Langford, John Sillitoe on behalf of the Wellcome Sanger Institute COVID-19 Surveillance Team |
| EPI_ISL_2916392, EPI_ISL_2916393 | Lighthouse Lab in Milton Keynes | Wellcome Sanger Institute for the COVID-19 Genomics UK (COG-UK) Consortium | The Lighthouse Lab in Milton Keynes and Alex Alderton, Roberto Amato, Jeffrey Barrett, Sonia Goncalves, Ewan Harrison, David K. Jackson, Ian Johnston, Dominic Kwiatkowski, Cordelia Langford, John Sillitoe on behalf of the Wellcome Sanger Institute COVID-19 Surveillance Team |
| EPI_ISL_2916394 | Lighthouse Lab in Alderley Park | Wellcome Sanger Institute for the COVID-19 Genomics UK (COG-UK) Consortium | Jacquelyn Wynn, Mairead Hyland, The Lighthouse Lab in Alderley Park and Alex Alderton, Roberto Amato, Jeffrey Barrett, Sonia Goncalves, Ewan Harrison, David K. Jackson, Ian Johnston, Dominic Kwiatkowski, Cordelia Langford, John Sillitoe on behalf of the Wellcome Sanger Institute COVID-19 Surveillance Team |
| EPI_ISL_2916395 | Lighthouse Lab in Milton Keynes | Wellcome Sanger Institute for the COVID-19 Genomics UK (COG-UK) Consortium | The Lighthouse Lab in Milton Keynes and Alex Alderton, Roberto Amato, Jeffrey Barrett, Sonia Goncalves, Ewan Harrison, David K. Jackson, Ian Johnston, Dominic Kwiatkowski, Cordelia Langford, John Sillitoe on behalf of the Wellcome Sanger Institute COVID-19 Surveillance Team |
| EPI_ISL_2916396 | Lighthouse Lab in Glasgow | Wellcome Sanger Institute for the COVID-19 Genomics UK (COG-UK) Consortium | Harper VanSteenhouse, Yumi Kasai, David Gray, Carol Clugston, Anna Dominiczak and Alex Alderton, Roberto Amato, Jeffrey Barrett, Sonia Goncalves, Ewan Harrison, David K. Jackson, Ian Johnston, Dominic Kwiatkowski, Cordelia Langford, John Sillitoe on behalf of the Wellcome Sanger Institute COVID-19 Surveillance Team |
| EPI_ISL_2916397 | Lighthouse Lab in Milton Keynes | Wellcome Sanger Institute for the COVID-19 Genomics UK (COG-UK) Consortium | The Lighthouse Lab in Milton Keynes and Alex Alderton, Roberto Amato, Jeffrey Barrett, Sonia Goncalves, Ewan Harrison, David K. Jackson, Ian Johnston, Dominic Kwiatkowski, Cordelia Langford, John Sillitoe on behalf of the Wellcome Sanger Institute COVID-19 Surveillance Team |
| EPI_ISL_2916398 | Lighthouse Lab in Glasgow | Wellcome Sanger Institute for the COVID-19 Genomics UK (COG-UK) Consortium | Harper VanSteenhouse, Yumi Kasai, David Gray, Carol Clugston, Anna Dominiczak and Alex Alderton, Roberto Amato, Jeffrey Barrett, Sonia Goncalves, Ewan Harrison, David K. Jackson, Ian Johnston, Dominic Kwiatkowski, Cordelia Langford, John Sillitoe on behalf of the Wellcome Sanger Institute COVID-19 Surveillance Team |
| EPI_ISL_2916399, EPI_ISL_2916400, EPI_ISL_2916401 | Lighthouse Lab in Alderley Park | Wellcome Sanger Institute for the COVID-19 Genomics UK (COG-UK) Consortium | Jacquelyn Wynn, Mairead Hyland, The Lighthouse Lab in Alderley Park and Alex Alderton, Roberto Amato, Jeffrey Barrett, Sonia Goncalves, Ewan Harrison, David K. Jackson, Ian Johnston, Dominic Kwiatkowski, Cordelia Langford, John Sillitoe on behalf of the Wellcome Sanger Institute COVID-19 Surveillance Team |
| EPI_ISL_2916402 | Lighthouse Lab in Glasgow | Wellcome Sanger Institute for the COVID-19 Genomics UK (COG-UK) Consortium | Harper VanSteenhouse, Yumi Kasai, David Gray, Carol Clugston, Anna Dominiczak and Alex Alderton, Roberto Amato, Jeffrey Barrett, Sonia Goncalves, Ewan Harrison, David K. Jackson, Ian Johnston, Dominic Kwiatkowski, Cordelia Langford, John Sillitoe on behalf of the Wellcome Sanger Institute COVID-19 Surveillance Team |
| EPI_ISL_2916403, EPI_ISL_2916404, EPI_ISL_2916405, EPI_ISL_2916406, EPI_ISL_2916407, EPI_ISL_2916408 | Lighthouse Lab in Milton Keynes | Wellcome Sanger Institute for the COVID-19 Genomics UK (COG-UK) Consortium | The Lighthouse Lab in Milton Keynes and Alex Alderton, Roberto Amato, Jeffrey Barrett, Sonia Goncalves, Ewan Harrison, David K. Jackson, Ian Johnston, Dominic Kwiatkowski, Cordelia Langford, John Sillitoe on behalf of the Wellcome Sanger Institute COVID-19 Surveillance Team |
| EPI_ISL_2916409 | Lighthouse Lab in Alderley Park | Wellcome Sanger Institute for the COVID-19 Genomics UK (COG-UK) Consortium | Jacquelyn Wynn, Mairead Hyland, The Lighthouse Lab in Alderley Park and Alex Alderton, Roberto Amato, Jeffrey Barrett, Sonia Goncalves, Ewan Harrison, David K. Jackson, Ian Johnston, Dominic Kwiatkowski, Cordelia Langford, John Sillitoe on behalf of the Wellcome Sanger Institute COVID-19 Surveillance Team |
| EPI_ISL_2916410 | Lighthouse Lab in Glasgow | Wellcome Sanger Institute for the COVID-19 Genomics UK (COG-UK) Consortium | Harper VanSteenhouse, Yumi Kasai, David Gray, Carol Clugston, Anna Dominiczak and Alex Alderton, Roberto Amato, Jeffrey Barrett, Sonia Goncalves, Ewan Harrison, David K. Jackson, Ian Johnston, Dominic Kwiatkowski, Cordelia Langford, John Sillitoe on behalf of the Wellcome Sanger Institute COVID-19 Surveillance Team |
| EPI_ISL_2916411, EPI_ISL_2916412, EPI_ISL_2916413, EPI_ISL_2916414 | Lighthouse Lab in Milton Keynes | Wellcome Sanger Institute for the COVID-19 Genomics UK (COG-UK) Consortium | The Lighthouse Lab in Milton Keynes and Alex Alderton, Roberto Amato, Jeffrey Barrett, Sonia Goncalves, Ewan Harrison, David K. Jackson, Ian Johnston, Dominic Kwiatkowski, Cordelia Langford, John Sillitoe on behalf of the Wellcome Sanger Institute COVID-19 Surveillance Team |
| EPI_ISL_2916415 | Lighthouse Lab in Glasgow | Wellcome Sanger Institute for the COVID-19 Genomics UK (COG-UK) | Harper VanSteenhouse, Yumi Kasai, David Gray, Carol Clugston, Anna Dominiczak and Alex Alderton, Roberto Amato, Jeffrey Barrett, Sonia Goncalves, Ewan Harrison, David K. Jackson, Ian Johnston, Dominic Kwiatkowski, Cordelia Langford, |

| | | | |
|---|---|---|---|
| | | Consortium | John Sillitoe on behalf of the Wellcome Sanger Institute COVID-19 Surveillance Team |
| EPI_ISL_2916416 | Lighthouse Lab in Milton Keynes | Wellcome Sanger Institute for the COVID-19 Genomics UK (COG-UK) Consortium | The Lighthouse Lab in Milton Keynes and Alex Alderton, Roberto Amato, Jeffrey Barrett, Sonia Goncalves, Ewan Harrison, David K. Jackson, Ian Johnston, Dominic Kwiatkowski, Cordelia Langford, John Sillitoe on behalf of the Wellcome Sanger Institute COVID-19 Surveillance Team |
| EPI_ISL_2916417, EPI_ISL_2916418 | Lighthouse Lab in Alderley Park | Wellcome Sanger Institute for the COVID-19 Genomics UK (COG-UK) Consortium | Jacquelyn Wynn, Mairead Hyland, The Lighthouse Lab in Alderley Park and Alex Alderton, Roberto Amato, Jeffrey Barrett, Sonia Goncalves, Ewan Harrison, David K. Jackson, Ian Johnston, Dominic Kwiatkowski, Cordelia Langford, John Sillitoe on behalf of the Wellcome Sanger Institute COVID-19 Surveillance Team |
| EPI_ISL_2916419, EPI_ISL_2916420, EPI_ISL_2916421 | Lighthouse Lab in Milton Keynes | Wellcome Sanger Institute for the COVID-19 Genomics UK (COG-UK) Consortium | The Lighthouse Lab in Milton Keynes and Alex Alderton, Roberto Amato, Jeffrey Barrett, Sonia Goncalves, Ewan Harrison, David K. Jackson, Ian Johnston, Dominic Kwiatkowski, Cordelia Langford, John Sillitoe on behalf of the Wellcome Sanger Institute COVID-19 Surveillance Team |
| EPI_ISL_2916422 | Lighthouse Lab in Glasgow | Wellcome Sanger Institute for the COVID-19 Genomics UK (COG-UK) Consortium | Harper VanSteenhouse, Yumi Kasai, David Gray, Carol Clugston, Anna Dominiczak and Alex Alderton, Roberto Amato, Jeffrey Barrett, Sonia Goncalves, Ewan Harrison, David K. Jackson, Ian Johnston, Dominic Kwiatkowski, Cordelia Langford, John Sillitoe on behalf of the Wellcome Sanger Institute COVID-19 Surveillance Team |
| EPI_ISL_2916423, EPI_ISL_2916424, EPI_ISL_2916425, EPI_ISL_2916426 | Lighthouse Lab in Milton Keynes | Wellcome Sanger Institute for the COVID-19 Genomics UK (COG-UK) Consortium | The Lighthouse Lab in Milton Keynes and Alex Alderton, Roberto Amato, Jeffrey Barrett, Sonia Goncalves, Ewan Harrison, David K. Jackson, Ian Johnston, Dominic Kwiatkowski, Cordelia Langford, John Sillitoe on behalf of the Wellcome Sanger Institute COVID-19 Surveillance Team |
| EPI_ISL_2916427 | Lighthouse Lab in Alderley Park | Wellcome Sanger Institute for the COVID-19 Genomics UK (COG-UK) Consortium | Jacquelyn Wynn, Mairead Hyland, The Lighthouse Lab in Alderley Park and Alex Alderton, Roberto Amato, Jeffrey Barrett, Sonia Goncalves, Ewan Harrison, David K. Jackson, Ian Johnston, Dominic Kwiatkowski, Cordelia Langford, John Sillitoe on behalf of the Wellcome Sanger Institute COVID-19 Surveillance Team |
| EPI_ISL_2916428 | Lighthouse Lab in Milton Keynes | Wellcome Sanger Institute for the COVID-19 Genomics UK (COG-UK) Consortium | The Lighthouse Lab in Milton Keynes and Alex Alderton, Roberto Amato, Jeffrey Barrett, Sonia Goncalves, Ewan Harrison, David K. Jackson, Ian Johnston, Dominic Kwiatkowski, Cordelia Langford, John Sillitoe on behalf of the Wellcome Sanger Institute COVID-19 Surveillance Team |
| EPI_ISL_2916429, EPI_ISL_2916430 | Lighthouse Lab in Glasgow | Wellcome Sanger Institute for the COVID-19 Genomics UK (COG-UK) Consortium | Harper VanSteenhouse, Yumi Kasai, David Gray, Carol Clugston, Anna Dominiczak and Alex Alderton, Roberto Amato, Jeffrey Barrett, Sonia Goncalves, Ewan Harrison, David K. Jackson, Ian Johnston, Dominic Kwiatkowski, Cordelia Langford, John Sillitoe on behalf of the Wellcome Sanger Institute COVID-19 Surveillance Team |
| EPI_ISL_2916431 | Lighthouse Lab in Milton Keynes | Wellcome Sanger Institute for the COVID-19 Genomics UK (COG-UK) Consortium | The Lighthouse Lab in Milton Keynes and Alex Alderton, Roberto Amato, Jeffrey Barrett, Sonia Goncalves, Ewan Harrison, David K. Jackson, Ian Johnston, Dominic Kwiatkowski, Cordelia Langford, John Sillitoe on behalf of the Wellcome Sanger Institute COVID-19 Surveillance Team |
| EPI_ISL_2916432 | Lighthouse Lab in Alderley Park | Wellcome Sanger Institute for the COVID-19 Genomics UK (COG-UK) Consortium | Jacquelyn Wynn, Mairead Hyland, The Lighthouse Lab in Alderley Park and Alex Alderton, Roberto Amato, Jeffrey Barrett, Sonia Goncalves, Ewan Harrison, David K. Jackson, Ian Johnston, Dominic Kwiatkowski, Cordelia Langford, John Sillitoe on behalf of the Wellcome Sanger Institute COVID-19 Surveillance Team |
| EPI_ISL_2916433 | Lighthouse Lab in Glasgow | Wellcome Sanger Institute for the COVID-19 Genomics UK (COG-UK) Consortium | Harper VanSteenhouse, Yumi Kasai, David Gray, Carol Clugston, Anna Dominiczak and Alex Alderton, Roberto Amato, Jeffrey Barrett, Sonia Goncalves, Ewan Harrison, David K. Jackson, Ian Johnston, Dominic Kwiatkowski, Cordelia Langford, John Sillitoe on behalf of the Wellcome Sanger Institute COVID-19 Surveillance Team |
| EPI_ISL_2916434 | Lighthouse Lab in Milton Keynes | Wellcome Sanger Institute for the COVID-19 Genomics UK (COG-UK) Consortium | The Lighthouse Lab in Milton Keynes and Alex Alderton, Roberto Amato, Jeffrey Barrett, Sonia Goncalves, Ewan Harrison, David K. Jackson, Ian Johnston, Dominic Kwiatkowski, Cordelia Langford, John Sillitoe on behalf of the Wellcome Sanger Institute COVID-19 Surveillance Team |
| EPI_ISL_2916435, EPI_ISL_2916436, EPI_ISL_2916437 | Lighthouse Lab in Glasgow | Wellcome Sanger Institute for the COVID-19 Genomics UK (COG-UK) Consortium | Harper VanSteenhouse, Yumi Kasai, David Gray, Carol Clugston, Anna Dominiczak and Alex Alderton, Roberto Amato, Jeffrey Barrett, Sonia Goncalves, Ewan Harrison, David K. Jackson, Ian Johnston, Dominic Kwiatkowski, Cordelia Langford, John Sillitoe on behalf of the Wellcome Sanger Institute COVID-19 Surveillance Team |
| EPI_ISL_2916438 | Lighthouse Lab in Alderley Park | Wellcome Sanger Institute for the COVID-19 Genomics UK (COG-UK) Consortium | Jacquelyn Wynn, Mairead Hyland, The Lighthouse Lab in Alderley Park and Alex Alderton, Roberto Amato, Jeffrey Barrett, Sonia Goncalves, Ewan Harrison, David K. Jackson, Ian Johnston, Dominic Kwiatkowski, Cordelia Langford, John Sillitoe on behalf of the Wellcome Sanger Institute COVID-19 Surveillance Team |
| EPI_ISL_2916439 | Lighthouse Lab in Glasgow | Wellcome Sanger Institute for the COVID-19 Genomics UK (COG-UK) Consortium | Harper VanSteenhouse, Yumi Kasai, David Gray, Carol Clugston, Anna Dominiczak and Alex Alderton, Roberto Amato, Jeffrey Barrett, Sonia Goncalves, Ewan Harrison, David K. Jackson, Ian Johnston, Dominic Kwiatkowski, Cordelia Langford, John Sillitoe on behalf of the Wellcome Sanger Institute COVID-19 Surveillance Team |
| EPI_ISL_2916440, EPI_ISL_2916441, EPI_ISL_2916442, EPI_ISL_2916443, EPI_ISL_2916444 | Lighthouse Lab in Milton Keynes | Wellcome Sanger Institute for the COVID-19 Genomics UK (COG-UK) Consortium | The Lighthouse Lab in Milton Keynes and Alex Alderton, Roberto Amato, Jeffrey Barrett, Sonia Goncalves, Ewan Harrison, David K. Jackson, Ian Johnston, Dominic Kwiatkowski, Cordelia Langford, John Sillitoe on behalf of the Wellcome Sanger Institute COVID-19 Surveillance Team |
| EPI_ISL_2916445 | Lighthouse Lab in Glasgow | Wellcome Sanger Institute for the COVID-19 Genomics UK (COG-UK) Consortium | Harper VanSteenhouse, Yumi Kasai, David Gray, Carol Clugston, Anna Dominiczak and Alex Alderton, Roberto Amato, Jeffrey Barrett, Sonia Goncalves, Ewan Harrison, David K. Jackson, Ian Johnston, Dominic Kwiatkowski, Cordelia Langford, John Sillitoe on behalf of the Wellcome Sanger Institute COVID-19 Surveillance Team |
| EPI_ISL_2916446 | Lighthouse Lab in Alderley Park | Wellcome Sanger Institute for the COVID-19 Genomics UK (COG-UK) Consortium | Jacquelyn Wynn, Mairead Hyland, The Lighthouse Lab in Alderley Park and Alex Alderton, Roberto Amato, Jeffrey Barrett, Sonia Goncalves, Ewan Harrison, David K. Jackson, Ian Johnston, Dominic Kwiatkowski, Cordelia Langford, John Sillitoe on behalf of the Wellcome Sanger Institute COVID-19 Surveillance Team |
| EPI_ISL_2916447 | Lighthouse Lab in Glasgow | Wellcome Sanger Institute for the COVID-19 Genomics UK (COG-UK) Consortium | Harper VanSteenhouse, Yumi Kasai, David Gray, Carol Clugston, Anna Dominiczak and Alex Alderton, Roberto Amato, Jeffrey Barrett, Sonia Goncalves, Ewan Harrison, David K. Jackson, Ian Johnston, Dominic Kwiatkowski, Cordelia Langford, John Sillitoe on behalf of the Wellcome Sanger Institute COVID-19 Surveillance Team |
| EPI_ISL_2916448, EPI_ISL_2916449, EPI_ISL_2916450, EPI_ISL_2916451, EPI_ISL_2916452 | Lighthouse Lab in Milton Keynes | Wellcome Sanger Institute for the COVID-19 Genomics UK (COG-UK) Consortium | The Lighthouse Lab in Milton Keynes and Alex Alderton, Roberto Amato, Jeffrey Barrett, Sonia Goncalves, Ewan Harrison, David K. Jackson, Ian Johnston, Dominic Kwiatkowski, Cordelia Langford, John Sillitoe on behalf of the Wellcome Sanger Institute COVID-19 Surveillance Team |
| EPI_ISL_2916453 | Lighthouse Lab in Alderley Park | Wellcome Sanger Institute for the COVID-19 Genomics UK (COG-UK) Consortium | Jacquelyn Wynn, Mairead Hyland, The Lighthouse Lab in Alderley Park and Alex Alderton, Roberto Amato, Jeffrey Barrett, Sonia Goncalves, Ewan Harrison, David K. Jackson, Ian Johnston, Dominic Kwiatkowski, Cordelia Langford, John Sillitoe on behalf of the Wellcome Sanger Institute COVID-19 Surveillance Team |
| EPI_ISL_2916454, EPI_ISL_2916455 | Lighthouse Lab in Glasgow | Wellcome Sanger Institute for the COVID-19 Genomics UK (COG-UK) Consortium | Harper VanSteenhouse, Yumi Kasai, David Gray, Carol Clugston, Anna Dominiczak and Alex Alderton, Roberto Amato, Jeffrey Barrett, Sonia Goncalves, Ewan Harrison, David K. Jackson, Ian Johnston, Dominic Kwiatkowski, Cordelia Langford, John Sillitoe on behalf of the Wellcome Sanger Institute COVID-19 Surveillance Team |
| EPI_ISL_2916456 | Lighthouse Lab in Alderley Park | Wellcome Sanger Institute for the COVID-19 Genomics UK (COG-UK) Consortium | Jacquelyn Wynn, Mairead Hyland, The Lighthouse Lab in Alderley Park and Alex Alderton, Roberto Amato, Jeffrey Barrett, Sonia Goncalves, Ewan Harrison, David K. Jackson, Ian Johnston, Dominic Kwiatkowski, Cordelia Langford, John Sillitoe on behalf of the Wellcome Sanger Institute COVID-19 Surveillance Team |
| EPI_ISL_2916457, EPI_ISL_2916458 | Lighthouse Lab in Milton Keynes | Wellcome Sanger Institute for the COVID-19 Genomics UK (COG-UK) Consortium | The Lighthouse Lab in Milton Keynes and Alex Alderton, Roberto Amato, Jeffrey Barrett, Sonia Goncalves, Ewan Harrison, David K. Jackson, Ian Johnston, Dominic Kwiatkowski, Cordelia Langford, John Sillitoe on behalf of the Wellcome Sanger Institute COVID-19 Surveillance Team |
| EPI_ISL_2916459, EPI_ISL_2916460 | Lighthouse Lab in Glasgow | Wellcome Sanger Institute for the COVID-19 Genomics UK (COG-UK) Consortium | Harper VanSteenhouse, Yumi Kasai, David Gray, Carol Clugston, Anna Dominiczak and Alex Alderton, Roberto Amato, Jeffrey Barrett, Sonia Goncalves, Ewan Harrison, David K. Jackson, Ian Johnston, Dominic Kwiatkowski, Cordelia Langford, John Sillitoe on behalf of the Wellcome Sanger Institute COVID-19 Surveillance Team |
| EPI_ISL_2916461, EPI_ISL_2916462 | Lighthouse Lab in Alderley Park | Wellcome Sanger Institute for the COVID-19 Genomics UK (COG-UK) Consortium | Jacquelyn Wynn, Mairead Hyland, The Lighthouse Lab in Alderley Park and Alex Alderton, Roberto Amato, Jeffrey Barrett, Sonia Goncalves, Ewan Harrison, David K. Jackson, Ian Johnston, Dominic Kwiatkowski, Cordelia Langford, John Sillitoe on behalf of the Wellcome Sanger Institute COVID-19 Surveillance Team |
| EPI_ISL_2916463 | Lighthouse Lab in Milton Keynes | Wellcome Sanger Institute for the COVID-19 Genomics UK (COG-UK) Consortium | The Lighthouse Lab in Milton Keynes and Alex Alderton, Roberto Amato, Jeffrey Barrett, Sonia Goncalves, Ewan Harrison, David K. Jackson, Ian Johnston, Dominic Kwiatkowski, Cordelia Langford, John Sillitoe on behalf of the Wellcome Sanger Institute COVID-19 Surveillance Team |
| EPI_ISL_2916464, EPI_ISL_2916465 | Lighthouse Lab in Alderley Park | Wellcome Sanger Institute for the COVID-19 Genomics UK (COG-UK) Consortium | Jacquelyn Wynn, Mairead Hyland, The Lighthouse Lab in Alderley Park and Alex Alderton, Roberto Amato, Jeffrey Barrett, Sonia Goncalves, Ewan Harrison, David K. Jackson, Ian Johnston, Dominic Kwiatkowski, Cordelia Langford, John Sillitoe on behalf of the Wellcome Sanger Institute COVID-19 Surveillance Team |
| EPI_ISL_2916466, EPI_ISL_2916467 | Lighthouse Lab in Glasgow | Wellcome Sanger Institute for the COVID-19 Genomics UK (COG-UK) Consortium | Harper VanSteenhouse, Yumi Kasai, David Gray, Carol Clugston, Anna Dominiczak and Alex Alderton, Roberto Amato, Jeffrey Barrett, Sonia Goncalves, Ewan Harrison, David K. Jackson, Ian Johnston, Dominic Kwiatkowski, Cordelia Langford, John Sillitoe on behalf of the Wellcome Sanger Institute COVID-19 Surveillance Team |
| EPI_ISL_2916468, EPI_ISL_2916469, EPI_ISL_2916470, EPI_ISL_2916471 | Lighthouse Lab in Milton Keynes | Wellcome Sanger Institute for the COVID-19 Genomics UK (COG-UK) Consortium | The Lighthouse Lab in Milton Keynes and Alex Alderton, Roberto Amato, Jeffrey Barrett, Sonia Goncalves, Ewan Harrison, David K. Jackson, Ian Johnston, Dominic Kwiatkowski, Cordelia Langford, John Sillitoe on behalf of the Wellcome Sanger Institute COVID-19 Surveillance Team |
| EPI_ISL_2916472 | Lighthouse Lab in Glasgow | Wellcome Sanger Institute for the COVID-19 Genomics UK (COG-UK) Consortium | Harper VanSteenhouse, Yumi Kasai, David Gray, Carol Clugston, Anna Dominiczak and Alex Alderton, Roberto Amato, Jeffrey Barrett, Sonia Goncalves, Ewan Harrison, David K. Jackson, Ian Johnston, Dominic Kwiatkowski, Cordelia Langford, John Sillitoe on behalf of the Wellcome Sanger Institute COVID-19 Surveillance Team |
| EPI_ISL_2916473 | Lighthouse Lab in Alderley Park | Wellcome Sanger Institute for the COVID-19 Genomics UK (COG-UK) Consortium | Jacquelyn Wynn, Mairead Hyland, The Lighthouse Lab in Alderley Park and Alex Alderton, Roberto Amato, Jeffrey Barrett, Sonia Goncalves, Ewan Harrison, David K. Jackson, Ian Johnston, Dominic Kwiatkowski, Cordelia Langford, John Sillitoe on behalf of the Wellcome Sanger Institute COVID-19 Surveillance Team |
| EPI_ISL_2916474, EPI_ISL_2916475, EPI_ISL_2916476 | Lighthouse Lab in Milton Keynes | Wellcome Sanger Institute for the COVID-19 Genomics UK (COG-UK) Consortium | The Lighthouse Lab in Milton Keynes and Alex Alderton, Roberto Amato, Jeffrey Barrett, Sonia Goncalves, Ewan Harrison, David K. Jackson, Ian Johnston, Dominic Kwiatkowski, Cordelia Langford, John Sillitoe on behalf of the Wellcome Sanger Institute COVID-19 Surveillance Team |
| EPI_ISL_2916477 | Lighthouse Lab in Alderley Park | Wellcome Sanger Institute for the COVID-19 Genomics UK (COG-UK) Consortium | Jacquelyn Wynn, Mairead Hyland, The Lighthouse Lab in Alderley Park and Alex Alderton, Roberto Amato, Jeffrey Barrett, Sonia Goncalves, Ewan Harrison, David K. Jackson, Ian Johnston, Dominic Kwiatkowski, Cordelia Langford, John Sillitoe on behalf of the Wellcome Sanger Institute COVID-19 Surveillance Team |
| EPI_ISL_2916478, EPI_ISL_2916479 | Lighthouse Lab in Milton Keynes | Wellcome Sanger Institute for the COVID-19 Genomics UK (COG-UK) Consortium | The Lighthouse Lab in Milton Keynes and Alex Alderton, Roberto Amato, Jeffrey Barrett, Sonia Goncalves, Ewan Harrison, David K. Jackson, Ian Johnston, Dominic Kwiatkowski, Cordelia Langford, John Sillitoe on behalf of the Wellcome Sanger Institute COVID-19 Surveillance Team |
| EPI_ISL_2916480 | Lighthouse Lab in Glasgow | Wellcome Sanger Institute for the COVID-19 Genomics UK (COG-UK) Consortium | Harper VanSteenhouse, Yumi Kasai, David Gray, Carol Clugston, Anna Dominiczak and Alex Alderton, Roberto Amato, Jeffrey Barrett, Sonia Goncalves, Ewan Harrison, David K. Jackson, Ian Johnston, Dominic Kwiatkowski, Cordelia Langford, John Sillitoe on behalf of the Wellcome Sanger Institute COVID-19 Surveillance Team |
| EPI_ISL_2916481 | Lighthouse Lab in Milton Keynes | Wellcome Sanger Institute for the COVID-19 Genomics UK (COG-UK) Consortium | The Lighthouse Lab in Milton Keynes and Alex Alderton, Roberto Amato, Jeffrey Barrett, Sonia Goncalves, Ewan Harrison, David K. Jackson, Ian Johnston, Dominic Kwiatkowski, Cordelia Langford, John Sillitoe on behalf of the Wellcome Sanger Institute COVID-19 Surveillance Team |
| EPI_ISL_2916482 | Lighthouse Lab in Alderley Park | Wellcome Sanger Institute for the COVID-19 Genomics UK (COG-UK) Consortium | Jacquelyn Wynn, Mairead Hyland, The Lighthouse Lab in Alderley Park and Alex Alderton, Roberto Amato, Jeffrey Barrett, Sonia Goncalves, Ewan Harrison, David K. Jackson, Ian Johnston, Dominic Kwiatkowski, Cordelia Langford, John Sillitoe on behalf of the Wellcome Sanger Institute COVID-19 Surveillance Team |
| EPI_ISL_2916483 | Lighthouse Lab in Milton Keynes | Wellcome Sanger Institute for the COVID-19 Genomics UK (COG-UK) Consortium | The Lighthouse Lab in Milton Keynes and Alex Alderton, Roberto Amato, Jeffrey Barrett, Sonia Goncalves, Ewan Harrison, David K. Jackson, Ian Johnston, Dominic Kwiatkowski, Cordelia Langford, John Sillitoe on behalf of the Wellcome Sanger Institute COVID-19 Surveillance Team |
| EPI_ISL_2916484, | Lighthouse Lab in Alderley Park | Wellcome Sanger Institute for the COVID-19 Genomics UK (COG-UK) | Jacquelyn Wynn, Mairead Hyland, The Lighthouse Lab in Alderley Park and Alex Alderton, Roberto Amato, Jeffrey Barrett, Sonia Goncalves, Ewan Harrison, David K. Jackson, Ian Johnston, Dominic Kwiatkowski, Cordelia Langford, John Sillitoe on |

| | | | |
|---|---|---|---|
| EPI_ISL_2916485, EPI_ISL_2916486 | | Consortium | behalf of the Wellcome Sanger Institute COVID-19 Surveillance Team |
| EPI_ISL_2916487, EPI_ISL_2916488, EPI_ISL_2916489, EPI_ISL_2916490 | Lighthouse Lab in Milton Keynes | Wellcome Sanger Institute for the COVID-19 Genomics UK (COG-UK) Consortium | The Lighthouse Lab in Milton Keynes and Alex Alderton, Roberto Amato, Jeffrey Barrett, Sonia Goncalves, Ewan Harrison, David K. Jackson, Ian Johnston, Dominic Kwiatkowski, Cordelia Langford, John Sillitoe on behalf of the Wellcome Sanger Institute COVID-19 Surveillance Team |
| EPI_ISL_2916491 | Lighthouse Lab in Alderley Park | Wellcome Sanger Institute for the COVID-19 Genomics UK (COG-UK) Consortium | Jacquelyn Wynn, Mairead Hyland, The Lighthouse Lab in Alderley Park and Alex Alderton, Roberto Amato, Jeffrey Barrett, Sonia Goncalves, Ewan Harrison, David K. Jackson, Ian Johnston, Dominic Kwiatkowski, Cordelia Langford, John Sillitoe on behalf of the Wellcome Sanger Institute COVID-19 Surveillance Team |
| EPI_ISL_2916492 | Lighthouse Lab in Milton Keynes | Wellcome Sanger Institute for the COVID-19 Genomics UK (COG-UK) Consortium | The Lighthouse Lab in Milton Keynes and Alex Alderton, Roberto Amato, Jeffrey Barrett, Sonia Goncalves, Ewan Harrison, David K. Jackson, Ian Johnston, Dominic Kwiatkowski, Cordelia Langford, John Sillitoe on behalf of the Wellcome Sanger Institute COVID-19 Surveillance Team |
| EPI_ISL_2916493 | Lighthouse Lab in Alderley Park | Wellcome Sanger Institute for the COVID-19 Genomics UK (COG-UK) Consortium | Jacquelyn Wynn, Mairead Hyland, The Lighthouse Lab in Alderley Park and Alex Alderton, Roberto Amato, Jeffrey Barrett, Sonia Goncalves, Ewan Harrison, David K. Jackson, Ian Johnston, Dominic Kwiatkowski, Cordelia Langford, John Sillitoe on behalf of the Wellcome Sanger Institute COVID-19 Surveillance Team |
| EPI_ISL_2916494, EPI_ISL_2916495 | Lighthouse Lab in Milton Keynes | Wellcome Sanger Institute for the COVID-19 Genomics UK (COG-UK) Consortium | The Lighthouse Lab in Milton Keynes and Alex Alderton, Roberto Amato, Jeffrey Barrett, Sonia Goncalves, Ewan Harrison, David K. Jackson, Ian Johnston, Dominic Kwiatkowski, Cordelia Langford, John Sillitoe on behalf of the Wellcome Sanger Institute COVID-19 Surveillance Team |
| EPI_ISL_2916496 | Lighthouse Lab in Alderley Park | Wellcome Sanger Institute for the COVID-19 Genomics UK (COG-UK) Consortium | Jacquelyn Wynn, Mairead Hyland, The Lighthouse Lab in Alderley Park and Alex Alderton, Roberto Amato, Jeffrey Barrett, Sonia Goncalves, Ewan Harrison, David K. Jackson, Ian Johnston, Dominic Kwiatkowski, Cordelia Langford, John Sillitoe on behalf of the Wellcome Sanger Institute COVID-19 Surveillance Team |
| EPI_ISL_2916497, EPI_ISL_2916498, EPI_ISL_2916499, EPI_ISL_2916500 | Lighthouse Lab in Milton Keynes | Wellcome Sanger Institute for the COVID-19 Genomics UK (COG-UK) Consortium | The Lighthouse Lab in Milton Keynes and Alex Alderton, Roberto Amato, Jeffrey Barrett, Sonia Goncalves, Ewan Harrison, David K. Jackson, Ian Johnston, Dominic Kwiatkowski, Cordelia Langford, John Sillitoe on behalf of the Wellcome Sanger Institute COVID-19 Surveillance Team |
| EPI_ISL_2916501 | Lighthouse Lab in Alderley Park | Wellcome Sanger Institute for the COVID-19 Genomics UK (COG-UK) Consortium | Jacquelyn Wynn, Mairead Hyland, The Lighthouse Lab in Alderley Park and Alex Alderton, Roberto Amato, Jeffrey Barrett, Sonia Goncalves, Ewan Harrison, David K. Jackson, Ian Johnston, Dominic Kwiatkowski, Cordelia Langford, John Sillitoe on behalf of the Wellcome Sanger Institute COVID-19 Surveillance Team |
| EPI_ISL_2916502 | Lighthouse Lab in Milton Keynes | Wellcome Sanger Institute for the COVID-19 Genomics UK (COG-UK) Consortium | The Lighthouse Lab in Milton Keynes and Alex Alderton, Roberto Amato, Jeffrey Barrett, Sonia Goncalves, Ewan Harrison, David K. Jackson, Ian Johnston, Dominic Kwiatkowski, Cordelia Langford, John Sillitoe on behalf of the Wellcome Sanger Institute COVID-19 Surveillance Team |
| EPI_ISL_2916503 | Lighthouse Lab in Alderley Park | Wellcome Sanger Institute for the COVID-19 Genomics UK (COG-UK) Consortium | Jacquelyn Wynn, Mairead Hyland, The Lighthouse Lab in Alderley Park and Alex Alderton, Roberto Amato, Jeffrey Barrett, Sonia Goncalves, Ewan Harrison, David K. Jackson, Ian Johnston, Dominic Kwiatkowski, Cordelia Langford, John Sillitoe on behalf of the Wellcome Sanger Institute COVID-19 Surveillance Team |
| EPI_ISL_2916504, EPI_ISL_2916505, EPI_ISL_2916506 | Lighthouse Lab in Milton Keynes | Wellcome Sanger Institute for the COVID-19 Genomics UK (COG-UK) Consortium | The Lighthouse Lab in Milton Keynes and Alex Alderton, Roberto Amato, Jeffrey Barrett, Sonia Goncalves, Ewan Harrison, David K. Jackson, Ian Johnston, Dominic Kwiatkowski, Cordelia Langford, John Sillitoe on behalf of the Wellcome Sanger Institute COVID-19 Surveillance Team |
| EPI_ISL_2916507 | Lighthouse Lab in Glasgow | Wellcome Sanger Institute for the COVID-19 Genomics UK (COG-UK) Consortium | Harper VanSteenhouse, Yumi Kasai, David Gray, Carol Clugston, Anna Dominiczak and Alex Alderton, Roberto Amato, Jeffrey Barrett, Sonia Goncalves, Ewan Harrison, David K. Jackson, Ian Johnston, Dominic Kwiatkowski, Cordelia Langford, John Sillitoe on behalf of the Wellcome Sanger Institute COVID-19 Surveillance Team |
| EPI_ISL_2916508 | Lighthouse Lab in Milton Keynes | Wellcome Sanger Institute for the COVID-19 Genomics UK (COG-UK) Consortium | The Lighthouse Lab in Milton Keynes and Alex Alderton, Roberto Amato, Jeffrey Barrett, Sonia Goncalves, Ewan Harrison, David K. Jackson, Ian Johnston, Dominic Kwiatkowski, Cordelia Langford, John Sillitoe on behalf of the Wellcome Sanger Institute COVID-19 Surveillance Team |
| EPI_ISL_2916509, EPI_ISL_2916510 | Lighthouse Lab in Alderley Park | Wellcome Sanger Institute for the COVID-19 Genomics UK (COG-UK) Consortium | Jacquelyn Wynn, Mairead Hyland, The Lighthouse Lab in Alderley Park and Alex Alderton, Roberto Amato, Jeffrey Barrett, Sonia Goncalves, Ewan Harrison, David K. Jackson, Ian Johnston, Dominic Kwiatkowski, Cordelia Langford, John Sillitoe on behalf of the Wellcome Sanger Institute COVID-19 Surveillance Team |
| EPI_ISL_2916511 | Lighthouse Lab in Milton Keynes | Wellcome Sanger Institute for the COVID-19 Genomics UK (COG-UK) Consortium | The Lighthouse Lab in Milton Keynes and Alex Alderton, Roberto Amato, Jeffrey Barrett, Sonia Goncalves, Ewan Harrison, David K. Jackson, Ian Johnston, Dominic Kwiatkowski, Cordelia Langford, John Sillitoe on behalf of the Wellcome Sanger Institute COVID-19 Surveillance Team |
| EPI_ISL_2916512, EPI_ISL_2916513 | Lighthouse Lab in Alderley Park | Wellcome Sanger Institute for the COVID-19 Genomics UK (COG-UK) Consortium | Jacquelyn Wynn, Mairead Hyland, The Lighthouse Lab in Alderley Park and Alex Alderton, Roberto Amato, Jeffrey Barrett, Sonia Goncalves, Ewan Harrison, David K. Jackson, Ian Johnston, Dominic Kwiatkowski, Cordelia Langford, John Sillitoe on behalf of the Wellcome Sanger Institute COVID-19 Surveillance Team |
| EPI_ISL_2916514, EPI_ISL_2916515, EPI_ISL_2916516, EPI_ISL_2916517, EPI_ISL_2916518, EPI_ISL_2916519 | Lighthouse Lab in Milton Keynes | Wellcome Sanger Institute for the COVID-19 Genomics UK (COG-UK) Consortium | The Lighthouse Lab in Milton Keynes and Alex Alderton, Roberto Amato, Jeffrey Barrett, Sonia Goncalves, Ewan Harrison, David K. Jackson, Ian Johnston, Dominic Kwiatkowski, Cordelia Langford, John Sillitoe on behalf of the Wellcome Sanger Institute COVID-19 Surveillance Team |
| EPI_ISL_2916520 | Lighthouse Lab in Glasgow | Wellcome Sanger Institute for the COVID-19 Genomics UK (COG-UK) Consortium | Harper VanSteenhouse, Yumi Kasai, David Gray, Carol Clugston, Anna Dominiczak and Alex Alderton, Roberto Amato, Jeffrey Barrett, Sonia Goncalves, Ewan Harrison, David K. Jackson, Ian Johnston, Dominic Kwiatkowski, Cordelia Langford, John Sillitoe on behalf of the Wellcome Sanger Institute COVID-19 Surveillance Team |
| EPI_ISL_2916521, EPI_ISL_2916522, EPI_ISL_2916523 | Lighthouse Lab in Milton Keynes | Wellcome Sanger Institute for the COVID-19 Genomics UK (COG-UK) Consortium | The Lighthouse Lab in Milton Keynes and Alex Alderton, Roberto Amato, Jeffrey Barrett, Sonia Goncalves, Ewan Harrison, David K. Jackson, Ian Johnston, Dominic Kwiatkowski, Cordelia Langford, John Sillitoe on behalf of the Wellcome Sanger Institute COVID-19 Surveillance Team |
| EPI_ISL_2916524 | Lighthouse Lab in Glasgow | Wellcome Sanger Institute for the COVID-19 Genomics UK (COG-UK) Consortium | Harper VanSteenhouse, Yumi Kasai, David Gray, Carol Clugston, Anna Dominiczak and Alex Alderton, Roberto Amato, Jeffrey Barrett, Sonia Goncalves, Ewan Harrison, David K. Jackson, Ian Johnston, Dominic Kwiatkowski, Cordelia Langford, John Sillitoe on behalf of the Wellcome Sanger Institute COVID-19 Surveillance Team |
| EPI_ISL_2916525, EPI_ISL_2916526 | Lighthouse Lab in Milton Keynes | Wellcome Sanger Institute for the COVID-19 Genomics UK (COG-UK) Consortium | The Lighthouse Lab in Milton Keynes and Alex Alderton, Roberto Amato, Jeffrey Barrett, Sonia Goncalves, Ewan Harrison, David K. Jackson, Ian Johnston, Dominic Kwiatkowski, Cordelia Langford, John Sillitoe on behalf of the Wellcome Sanger Institute COVID-19 Surveillance Team |
| EPI_ISL_2916527 | Lighthouse Lab in Alderley Park | Wellcome Sanger Institute for the COVID-19 Genomics UK (COG-UK) Consortium | Jacquelyn Wynn, Mairead Hyland, The Lighthouse Lab in Alderley Park and Alex Alderton, Roberto Amato, Jeffrey Barrett, Sonia Goncalves, Ewan Harrison, David K. Jackson, Ian Johnston, Dominic Kwiatkowski, Cordelia Langford, John Sillitoe on behalf of the Wellcome Sanger Institute COVID-19 Surveillance Team |
| EPI_ISL_2916528 | Lighthouse Lab in Glasgow | Wellcome Sanger Institute for the COVID-19 Genomics UK (COG-UK) Consortium | Harper VanSteenhouse, Yumi Kasai, David Gray, Carol Clugston, Anna Dominiczak and Alex Alderton, Roberto Amato, Jeffrey Barrett, Sonia Goncalves, Ewan Harrison, David K. Jackson, Ian Johnston, Dominic Kwiatkowski, Cordelia Langford, John Sillitoe on behalf of the Wellcome Sanger Institute COVID-19 Surveillance Team |
| EPI_ISL_2916529, EPI_ISL_2916530 | Lighthouse Lab in Milton Keynes | Wellcome Sanger Institute for the COVID-19 Genomics UK (COG-UK) Consortium | The Lighthouse Lab in Milton Keynes and Alex Alderton, Roberto Amato, Jeffrey Barrett, Sonia Goncalves, Ewan Harrison, David K. Jackson, Ian Johnston, Dominic Kwiatkowski, Cordelia Langford, John Sillitoe on behalf of the Wellcome Sanger Institute COVID-19 Surveillance Team |
| EPI_ISL_2916531 | Lighthouse Lab in Glasgow | Wellcome Sanger Institute for the COVID-19 Genomics UK (COG-UK) Consortium | Harper VanSteenhouse, Yumi Kasai, David Gray, Carol Clugston, Anna Dominiczak and Alex Alderton, Roberto Amato, Jeffrey Barrett, Sonia Goncalves, Ewan Harrison, David K. Jackson, Ian Johnston, Dominic Kwiatkowski, Cordelia Langford, John Sillitoe on behalf of the Wellcome Sanger Institute COVID-19 Surveillance Team |
| EPI_ISL_2916532 | Lighthouse Lab in Alderley Park | Wellcome Sanger Institute for the COVID-19 Genomics UK (COG-UK) Consortium | Jacquelyn Wynn, Mairead Hyland, The Lighthouse Lab in Alderley Park and Alex Alderton, Roberto Amato, Jeffrey Barrett, Sonia Goncalves, Ewan Harrison, David K. Jackson, Ian Johnston, Dominic Kwiatkowski, Cordelia Langford, John Sillitoe on behalf of the Wellcome Sanger Institute COVID-19 Surveillance Team |
| EPI_ISL_2916533 | Lighthouse Lab in Milton Keynes | Wellcome Sanger Institute for the COVID-19 Genomics UK (COG-UK) Consortium | The Lighthouse Lab in Milton Keynes and Alex Alderton, Roberto Amato, Jeffrey Barrett, Sonia Goncalves, Ewan Harrison, David K. Jackson, Ian Johnston, Dominic Kwiatkowski, Cordelia Langford, John Sillitoe on behalf of the Wellcome Sanger Institute COVID-19 Surveillance Team |
| EPI_ISL_2916534 | Lighthouse Lab in Glasgow | Wellcome Sanger Institute for the COVID-19 Genomics UK (COG-UK) Consortium | Harper VanSteenhouse, Yumi Kasai, David Gray, Carol Clugston, Anna Dominiczak and Alex Alderton, Roberto Amato, Jeffrey Barrett, Sonia Goncalves, Ewan Harrison, David K. Jackson, Ian Johnston, Dominic Kwiatkowski, Cordelia Langford, John Sillitoe on behalf of the Wellcome Sanger Institute COVID-19 Surveillance Team |
| EPI_ISL_2916535 | Lighthouse Lab in Milton Keynes | Wellcome Sanger Institute for the COVID-19 Genomics UK (COG-UK) Consortium | The Lighthouse Lab in Milton Keynes and Alex Alderton, Roberto Amato, Jeffrey Barrett, Sonia Goncalves, Ewan Harrison, David K. Jackson, Ian Johnston, Dominic Kwiatkowski, Cordelia Langford, John Sillitoe on behalf of the Wellcome Sanger Institute COVID-19 Surveillance Team |
| EPI_ISL_2916536 | Lighthouse Lab in Alderley Park | Wellcome Sanger Institute for the COVID-19 Genomics UK (COG-UK) Consortium | Jacquelyn Wynn, Mairead Hyland, The Lighthouse Lab in Alderley Park and Alex Alderton, Roberto Amato, Jeffrey Barrett, Sonia Goncalves, Ewan Harrison, David K. Jackson, Ian Johnston, Dominic Kwiatkowski, Cordelia Langford, John Sillitoe on behalf of the Wellcome Sanger Institute COVID-19 Surveillance Team |
| EPI_ISL_2916537, EPI_ISL_2916538, EPI_ISL_2916539 | Lighthouse Lab in Glasgow | Wellcome Sanger Institute for the COVID-19 Genomics UK (COG-UK) Consortium | Harper VanSteenhouse, Yumi Kasai, David Gray, Carol Clugston, Anna Dominiczak and Alex Alderton, Roberto Amato, Jeffrey Barrett, Sonia Goncalves, Ewan Harrison, David K. Jackson, Ian Johnston, Dominic Kwiatkowski, Cordelia Langford, John Sillitoe on behalf of the Wellcome Sanger Institute COVID-19 Surveillance Team |
| EPI_ISL_2916540, EPI_ISL_2916541, EPI_ISL_2916542, EPI_ISL_2916543 | Lighthouse Lab in Milton Keynes | Wellcome Sanger Institute for the COVID-19 Genomics UK (COG-UK) Consortium | The Lighthouse Lab in Milton Keynes and Alex Alderton, Roberto Amato, Jeffrey Barrett, Sonia Goncalves, Ewan Harrison, David K. Jackson, Ian Johnston, Dominic Kwiatkowski, Cordelia Langford, John Sillitoe on behalf of the Wellcome Sanger Institute COVID-19 Surveillance Team |
| EPI_ISL_2916544 | Lighthouse Lab in Glasgow | Wellcome Sanger Institute for the COVID-19 Genomics UK (COG-UK) Consortium | Harper VanSteenhouse, Yumi Kasai, David Gray, Carol Clugston, Anna Dominiczak and Alex Alderton, Roberto Amato, Jeffrey Barrett, Sonia Goncalves, Ewan Harrison, David K. Jackson, Ian Johnston, Dominic Kwiatkowski, Cordelia Langford, John Sillitoe on behalf of the Wellcome Sanger Institute COVID-19 Surveillance Team |
| EPI_ISL_2916545, EPI_ISL_2916546 | Lighthouse Lab in Milton Keynes | Wellcome Sanger Institute for the COVID-19 Genomics UK (COG-UK) Consortium | The Lighthouse Lab in Milton Keynes and Alex Alderton, Roberto Amato, Jeffrey Barrett, Sonia Goncalves, Ewan Harrison, David K. Jackson, Ian Johnston, Dominic Kwiatkowski, Cordelia Langford, John Sillitoe on behalf of the Wellcome Sanger Institute COVID-19 Surveillance Team |
| EPI_ISL_2916547 | Lighthouse Lab in Glasgow | Wellcome Sanger Institute for the COVID-19 Genomics UK (COG-UK) Consortium | Harper VanSteenhouse, Yumi Kasai, David Gray, Carol Clugston, Anna Dominiczak and Alex Alderton, Roberto Amato, Jeffrey Barrett, Sonia Goncalves, Ewan Harrison, David K. Jackson, Ian Johnston, Dominic Kwiatkowski, Cordelia Langford, John Sillitoe on behalf of the Wellcome Sanger Institute COVID-19 Surveillance Team |
| EPI_ISL_2916548 | Lighthouse Lab in Alderley Park | Wellcome Sanger Institute for the COVID-19 Genomics UK (COG-UK) Consortium | Jacquelyn Wynn, Mairead Hyland, The Lighthouse Lab in Alderley Park and Alex Alderton, Roberto Amato, Jeffrey Barrett, Sonia Goncalves, Ewan Harrison, David K. Jackson, Ian Johnston, Dominic Kwiatkowski, Cordelia Langford, John Sillitoe on behalf of the Wellcome Sanger Institute COVID-19 Surveillance Team |
| EPI_ISL_2916549 | Lighthouse Lab in Milton Keynes | Wellcome Sanger Institute for the COVID-19 Genomics UK (COG-UK) Consortium | The Lighthouse Lab in Milton Keynes and Alex Alderton, Roberto Amato, Jeffrey Barrett, Sonia Goncalves, Ewan Harrison, David K. Jackson, Ian Johnston, Dominic Kwiatkowski, Cordelia Langford, John Sillitoe on behalf of the Wellcome Sanger Institute COVID-19 Surveillance Team |
| EPI_ISL_2916550, EPI_ISL_2916551 | Lighthouse Lab in Alderley Park | Wellcome Sanger Institute for the COVID-19 Genomics UK (COG-UK) Consortium | Jacquelyn Wynn, Mairead Hyland, The Lighthouse Lab in Alderley Park and Alex Alderton, Roberto Amato, Jeffrey Barrett, Sonia Goncalves, Ewan Harrison, David K. Jackson, Ian Johnston, Dominic Kwiatkowski, Cordelia Langford, John Sillitoe on behalf of the Wellcome Sanger Institute COVID-19 Surveillance Team |
| EPI_ISL_2916552 | Lighthouse Lab in Milton Keynes | Wellcome Sanger Institute for the COVID-19 Genomics UK (COG-UK) Consortium | The Lighthouse Lab in Milton Keynes and Alex Alderton, Roberto Amato, Jeffrey Barrett, Sonia Goncalves, Ewan Harrison, David K. Jackson, Ian Johnston, Dominic Kwiatkowski, Cordelia Langford, John Sillitoe on behalf of the Wellcome Sanger Institute COVID-19 Surveillance Team |

| | | | |
|---|---|---|---|
| EPI_ISL_2916553 | Lighthouse Lab in Glasgow | Wellcome Sanger Institute for the COVID-19 Genomics UK (COG-UK) Consortium | Harper VanSteenhouse, Yumi Kasai, David Gray, Carol Clugston, Anna Dominiczak and Alex Alderton, Roberto Amato, Jeffrey Barrett, Sonia Goncalves, Ewan Harrison, David K. Jackson, Ian Johnston, Dominic Kwiatkowski, Cordelia Langford, John Sillitoe on behalf of the Wellcome Sanger Institute COVID-19 Surveillance Team |
| EPI_ISL_2916554 | Lighthouse Lab in Milton Keynes | Wellcome Sanger Institute for the COVID-19 Genomics UK (COG-UK) Consortium | The Lighthouse Lab in Milton Keynes and Alex Alderton, Roberto Amato, Jeffrey Barrett, Sonia Goncalves, Ewan Harrison, David K. Jackson, Ian Johnston, Dominic Kwiatkowski, Cordelia Langford, John Sillitoe on behalf of the Wellcome Sanger Institute COVID-19 Surveillance Team |
| EPI_ISL_2916555 | Lighthouse Lab in Glasgow | Wellcome Sanger Institute for the COVID-19 Genomics UK (COG-UK) Consortium | Harper VanSteenhouse, Yumi Kasai, David Gray, Carol Clugston, Anna Dominiczak and Alex Alderton, Roberto Amato, Jeffrey Barrett, Sonia Goncalves, Ewan Harrison, David K. Jackson, Ian Johnston, Dominic Kwiatkowski, Cordelia Langford, John Sillitoe on behalf of the Wellcome Sanger Institute COVID-19 Surveillance Team |
| EPI_ISL_2916556 | Lighthouse Lab in Alderley Park | Wellcome Sanger Institute for the COVID-19 Genomics UK (COG-UK) Consortium | Jacquelyn Wynn, Mairead Hyland, The Lighthouse Lab in Alderley Park and Alex Alderton, Roberto Amato, Jeffrey Barrett, Sonia Goncalves, Ewan Harrison, David K. Jackson, Ian Johnston, Dominic Kwiatkowski, Cordelia Langford, John Sillitoe on behalf of the Wellcome Sanger Institute COVID-19 Surveillance Team |
| EPI_ISL_2916557, EPI_ISL_2916558 | Lighthouse Lab in Milton Keynes | Wellcome Sanger Institute for the COVID-19 Genomics UK (COG-UK) Consortium | The Lighthouse Lab in Milton Keynes and Alex Alderton, Roberto Amato, Jeffrey Barrett, Sonia Goncalves, Ewan Harrison, David K. Jackson, Ian Johnston, Dominic Kwiatkowski, Cordelia Langford, John Sillitoe on behalf of the Wellcome Sanger Institute COVID-19 Surveillance Team |
| EPI_ISL_2916559, EPI_ISL_2916560 | Lighthouse Lab in Glasgow | Wellcome Sanger Institute for the COVID-19 Genomics UK (COG-UK) Consortium | Harper VanSteenhouse, Yumi Kasai, David Gray, Carol Clugston, Anna Dominiczak and Alex Alderton, Roberto Amato, Jeffrey Barrett, Sonia Goncalves, Ewan Harrison, David K. Jackson, Ian Johnston, Dominic Kwiatkowski, Cordelia Langford, John Sillitoe on behalf of the Wellcome Sanger Institute COVID-19 Surveillance Team |
| EPI_ISL_2916561 | Lighthouse Lab in Milton Keynes | Wellcome Sanger Institute for the COVID-19 Genomics UK (COG-UK) Consortium | The Lighthouse Lab in Milton Keynes and Alex Alderton, Roberto Amato, Jeffrey Barrett, Sonia Goncalves, Ewan Harrison, David K. Jackson, Ian Johnston, Dominic Kwiatkowski, Cordelia Langford, John Sillitoe on behalf of the Wellcome Sanger Institute COVID-19 Surveillance Team |
| EPI_ISL_2916562 | Lighthouse Lab in Glasgow | Wellcome Sanger Institute for the COVID-19 Genomics UK (COG-UK) Consortium | Harper VanSteenhouse, Yumi Kasai, David Gray, Carol Clugston, Anna Dominiczak and Alex Alderton, Roberto Amato, Jeffrey Barrett, Sonia Goncalves, Ewan Harrison, David K. Jackson, Ian Johnston, Dominic Kwiatkowski, Cordelia Langford, John Sillitoe on behalf of the Wellcome Sanger Institute COVID-19 Surveillance Team |
| EPI_ISL_2916563 | Lighthouse Lab in Alderley Park | Wellcome Sanger Institute for the COVID-19 Genomics UK (COG-UK) Consortium | Jacquelyn Wynn, Mairead Hyland, The Lighthouse Lab in Alderley Park and Alex Alderton, Roberto Amato, Jeffrey Barrett, Sonia Goncalves, Ewan Harrison, David K. Jackson, Ian Johnston, Dominic Kwiatkowski, Cordelia Langford, John Sillitoe on behalf of the Wellcome Sanger Institute COVID-19 Surveillance Team |
| EPI_ISL_2916564 | Lighthouse Lab in Glasgow | Wellcome Sanger Institute for the COVID-19 Genomics UK (COG-UK) Consortium | Harper VanSteenhouse, Yumi Kasai, David Gray, Carol Clugston, Anna Dominiczak and Alex Alderton, Roberto Amato, Jeffrey Barrett, Sonia Goncalves, Ewan Harrison, David K. Jackson, Ian Johnston, Dominic Kwiatkowski, Cordelia Langford, John Sillitoe on behalf of the Wellcome Sanger Institute COVID-19 Surveillance Team |
| EPI_ISL_2916565 | Lighthouse Lab in Milton Keynes | Wellcome Sanger Institute for the COVID-19 Genomics UK (COG-UK) Consortium | The Lighthouse Lab in Milton Keynes and Alex Alderton, Roberto Amato, Jeffrey Barrett, Sonia Goncalves, Ewan Harrison, David K. Jackson, Ian Johnston, Dominic Kwiatkowski, Cordelia Langford, John Sillitoe on behalf of the Wellcome Sanger Institute COVID-19 Surveillance Team |
| EPI_ISL_2916566 | Lighthouse Lab in Glasgow | Wellcome Sanger Institute for the COVID-19 Genomics UK (COG-UK) Consortium | Harper VanSteenhouse, Yumi Kasai, David Gray, Carol Clugston, Anna Dominiczak and Alex Alderton, Roberto Amato, Jeffrey Barrett, Sonia Goncalves, Ewan Harrison, David K. Jackson, Ian Johnston, Dominic Kwiatkowski, Cordelia Langford, John Sillitoe on behalf of the Wellcome Sanger Institute COVID-19 Surveillance Team |
| EPI_ISL_2916567 | Lighthouse Lab in Milton Keynes | Wellcome Sanger Institute for the COVID-19 Genomics UK (COG-UK) Consortium | The Lighthouse Lab in Milton Keynes and Alex Alderton, Roberto Amato, Jeffrey Barrett, Sonia Goncalves, Ewan Harrison, David K. Jackson, Ian Johnston, Dominic Kwiatkowski, Cordelia Langford, John Sillitoe on behalf of the Wellcome Sanger Institute COVID-19 Surveillance Team |
| EPI_ISL_2916568 | Lighthouse Lab in Alderley Park | Wellcome Sanger Institute for the COVID-19 Genomics UK (COG-UK) Consortium | Jacquelyn Wynn, Mairead Hyland, The Lighthouse Lab in Alderley Park and Alex Alderton, Roberto Amato, Jeffrey Barrett, Sonia Goncalves, Ewan Harrison, David K. Jackson, Ian Johnston, Dominic Kwiatkowski, Cordelia Langford, John Sillitoe on behalf of the Wellcome Sanger Institute COVID-19 Surveillance Team |
| EPI_ISL_2916569 | Lighthouse Lab in Glasgow | Wellcome Sanger Institute for the COVID-19 Genomics UK (COG-UK) Consortium | Harper VanSteenhouse, Yumi Kasai, David Gray, Carol Clugston, Anna Dominiczak and Alex Alderton, Roberto Amato, Jeffrey Barrett, Sonia Goncalves, Ewan Harrison, David K. Jackson, Ian Johnston, Dominic Kwiatkowski, Cordelia Langford, John Sillitoe on behalf of the Wellcome Sanger Institute COVID-19 Surveillance Team |
| EPI_ISL_2916570 | Lighthouse Lab in Alderley Park | Wellcome Sanger Institute for the COVID-19 Genomics UK (COG-UK) Consortium | Jacquelyn Wynn, Mairead Hyland, The Lighthouse Lab in Alderley Park and Alex Alderton, Roberto Amato, Jeffrey Barrett, Sonia Goncalves, Ewan Harrison, David K. Jackson, Ian Johnston, Dominic Kwiatkowski, Cordelia Langford, John Sillitoe on behalf of the Wellcome Sanger Institute COVID-19 Surveillance Team |
| EPI_ISL_2916571 | Lighthouse Lab in Milton Keynes | Wellcome Sanger Institute for the COVID-19 Genomics UK (COG-UK) Consortium | The Lighthouse Lab in Milton Keynes and Alex Alderton, Roberto Amato, Jeffrey Barrett, Sonia Goncalves, Ewan Harrison, David K. Jackson, Ian Johnston, Dominic Kwiatkowski, Cordelia Langford, John Sillitoe on behalf of the Wellcome Sanger Institute COVID-19 Surveillance Team |
| EPI_ISL_2916572 | Lighthouse Lab in Glasgow | Wellcome Sanger Institute for the COVID-19 Genomics UK (COG-UK) Consortium | Harper VanSteenhouse, Yumi Kasai, David Gray, Carol Clugston, Anna Dominiczak and Alex Alderton, Roberto Amato, Jeffrey Barrett, Sonia Goncalves, Ewan Harrison, David K. Jackson, Ian Johnston, Dominic Kwiatkowski, Cordelia Langford, John Sillitoe on behalf of the Wellcome Sanger Institute COVID-19 Surveillance Team |
| EPI_ISL_2916573 | Lighthouse Lab in Milton Keynes | Wellcome Sanger Institute for the COVID-19 Genomics UK (COG-UK) Consortium | The Lighthouse Lab in Milton Keynes and Alex Alderton, Roberto Amato, Jeffrey Barrett, Sonia Goncalves, Ewan Harrison, David K. Jackson, Ian Johnston, Dominic Kwiatkowski, Cordelia Langford, John Sillitoe on behalf of the Wellcome Sanger Institute COVID-19 Surveillance Team |
| EPI_ISL_2916574 | Lighthouse Lab in Alderley Park | Wellcome Sanger Institute for the COVID-19 Genomics UK (COG-UK) Consortium | Jacquelyn Wynn, Mairead Hyland, The Lighthouse Lab in Alderley Park and Alex Alderton, Roberto Amato, Jeffrey Barrett, Sonia Goncalves, Ewan Harrison, David K. Jackson, Ian Johnston, Dominic Kwiatkowski, Cordelia Langford, John Sillitoe on behalf of the Wellcome Sanger Institute COVID-19 Surveillance Team |
| EPI_ISL_2916575 | Lighthouse Lab in Glasgow | Wellcome Sanger Institute for the COVID-19 Genomics UK (COG-UK) Consortium | Harper VanSteenhouse, Yumi Kasai, David Gray, Carol Clugston, Anna Dominiczak and Alex Alderton, Roberto Amato, Jeffrey Barrett, Sonia Goncalves, Ewan Harrison, David K. Jackson, Ian Johnston, Dominic Kwiatkowski, Cordelia Langford, John Sillitoe on behalf of the Wellcome Sanger Institute COVID-19 Surveillance Team |
| EPI_ISL_2916576, EPI_ISL_2916577 | Lighthouse Lab in Alderley Park | Wellcome Sanger Institute for the COVID-19 Genomics UK (COG-UK) Consortium | Jacquelyn Wynn, Mairead Hyland, The Lighthouse Lab in Alderley Park and Alex Alderton, Roberto Amato, Jeffrey Barrett, Sonia Goncalves, Ewan Harrison, David K. Jackson, Ian Johnston, Dominic Kwiatkowski, Cordelia Langford, John Sillitoe on behalf of the Wellcome Sanger Institute COVID-19 Surveillance Team |
| EPI_ISL_2916578 | Lighthouse Lab in Milton Keynes | Wellcome Sanger Institute for the COVID-19 Genomics UK (COG-UK) Consortium | The Lighthouse Lab in Milton Keynes and Alex Alderton, Roberto Amato, Jeffrey Barrett, Sonia Goncalves, Ewan Harrison, David K. Jackson, Ian Johnston, Dominic Kwiatkowski, Cordelia Langford, John Sillitoe on behalf of the Wellcome Sanger Institute COVID-19 Surveillance Team |
| EPI_ISL_2916579 | Lighthouse Lab in Glasgow | Wellcome Sanger Institute for the COVID-19 Genomics UK (COG-UK) Consortium | Harper VanSteenhouse, Yumi Kasai, David Gray, Carol Clugston, Anna Dominiczak and Alex Alderton, Roberto Amato, Jeffrey Barrett, Sonia Goncalves, Ewan Harrison, David K. Jackson, Ian Johnston, Dominic Kwiatkowski, Cordelia Langford, John Sillitoe on behalf of the Wellcome Sanger Institute COVID-19 Surveillance Team |
| EPI_ISL_2916580 | Lighthouse Lab in Alderley Park | Wellcome Sanger Institute for the COVID-19 Genomics UK (COG-UK) Consortium | Jacquelyn Wynn, Mairead Hyland, The Lighthouse Lab in Alderley Park and Alex Alderton, Roberto Amato, Jeffrey Barrett, Sonia Goncalves, Ewan Harrison, David K. Jackson, Ian Johnston, Dominic Kwiatkowski, Cordelia Langford, John Sillitoe on behalf of the Wellcome Sanger Institute COVID-19 Surveillance Team |
| EPI_ISL_2916581 | Lighthouse Lab in Milton Keynes | Wellcome Sanger Institute for the COVID-19 Genomics UK (COG-UK) Consortium | The Lighthouse Lab in Milton Keynes and Alex Alderton, Roberto Amato, Jeffrey Barrett, Sonia Goncalves, Ewan Harrison, David K. Jackson, Ian Johnston, Dominic Kwiatkowski, Cordelia Langford, John Sillitoe on behalf of the Wellcome Sanger Institute COVID-19 Surveillance Team |
| EPI_ISL_2916582 | Lighthouse Lab in Glasgow | Wellcome Sanger Institute for the COVID-19 Genomics UK (COG-UK) Consortium | Harper VanSteenhouse, Yumi Kasai, David Gray, Carol Clugston, Anna Dominiczak and Alex Alderton, Roberto Amato, Jeffrey Barrett, Sonia Goncalves, Ewan Harrison, David K. Jackson, Ian Johnston, Dominic Kwiatkowski, Cordelia Langford, John Sillitoe on behalf of the Wellcome Sanger Institute COVID-19 Surveillance Team |
| EPI_ISL_2916583, EPI_ISL_2916584, EPI_ISL_2916585, EPI_ISL_2916586, EPI_ISL_2916587, EPI_ISL_2916588 | Lighthouse Lab in Milton Keynes | Wellcome Sanger Institute for the COVID-19 Genomics UK (COG-UK) Consortium | The Lighthouse Lab in Milton Keynes and Alex Alderton, Roberto Amato, Jeffrey Barrett, Sonia Goncalves, Ewan Harrison, David K. Jackson, Ian Johnston, Dominic Kwiatkowski, Cordelia Langford, John Sillitoe on behalf of the Wellcome Sanger Institute COVID-19 Surveillance Team |
| EPI_ISL_2916589 | Lighthouse Lab in Glasgow | Wellcome Sanger Institute for the COVID-19 Genomics UK (COG-UK) Consortium | Harper VanSteenhouse, Yumi Kasai, David Gray, Carol Clugston, Anna Dominiczak and Alex Alderton, Roberto Amato, Jeffrey Barrett, Sonia Goncalves, Ewan Harrison, David K. Jackson, Ian Johnston, Dominic Kwiatkowski, Cordelia Langford, John Sillitoe on behalf of the Wellcome Sanger Institute COVID-19 Surveillance Team |
| EPI_ISL_2916590 | Lighthouse Lab in Alderley Park | Wellcome Sanger Institute for the COVID-19 Genomics UK (COG-UK) Consortium | Jacquelyn Wynn, Mairead Hyland, The Lighthouse Lab in Alderley Park and Alex Alderton, Roberto Amato, Jeffrey Barrett, Sonia Goncalves, Ewan Harrison, David K. Jackson, Ian Johnston, Dominic Kwiatkowski, Cordelia Langford, John Sillitoe on behalf of the Wellcome Sanger Institute COVID-19 Surveillance Team |
| EPI_ISL_2916591, EPI_ISL_2916592, EPI_ISL_2916593 | Lighthouse Lab in Milton Keynes | Wellcome Sanger Institute for the COVID-19 Genomics UK (COG-UK) Consortium | The Lighthouse Lab in Milton Keynes and Alex Alderton, Roberto Amato, Jeffrey Barrett, Sonia Goncalves, Ewan Harrison, David K. Jackson, Ian Johnston, Dominic Kwiatkowski, Cordelia Langford, John Sillitoe on behalf of the Wellcome Sanger Institute COVID-19 Surveillance Team |
| EPI_ISL_2916594, EPI_ISL_2916595, EPI_ISL_2916596, EPI_ISL_2916597, EPI_ISL_2916598 | Lighthouse Lab in Glasgow | Wellcome Sanger Institute for the COVID-19 Genomics UK (COG-UK) Consortium | Harper VanSteenhouse, Yumi Kasai, David Gray, Carol Clugston, Anna Dominiczak and Alex Alderton, Roberto Amato, Jeffrey Barrett, Sonia Goncalves, Ewan Harrison, David K. Jackson, Ian Johnston, Dominic Kwiatkowski, Cordelia Langford, John Sillitoe on behalf of the Wellcome Sanger Institute COVID-19 Surveillance Team |
| EPI_ISL_2916599, EPI_ISL_2916600 | Lighthouse Lab in Milton Keynes | Wellcome Sanger Institute for the COVID-19 Genomics UK (COG-UK) Consortium | The Lighthouse Lab in Milton Keynes and Alex Alderton, Roberto Amato, Jeffrey Barrett, Sonia Goncalves, Ewan Harrison, David K. Jackson, Ian Johnston, Dominic Kwiatkowski, Cordelia Langford, John Sillitoe on behalf of the Wellcome Sanger Institute COVID-19 Surveillance Team |
| EPI_ISL_2916601 | Lighthouse Lab in Glasgow | Wellcome Sanger Institute for the COVID-19 Genomics UK (COG-UK) Consortium | Harper VanSteenhouse, Yumi Kasai, David Gray, Carol Clugston, Anna Dominiczak and Alex Alderton, Roberto Amato, Jeffrey Barrett, Sonia Goncalves, Ewan Harrison, David K. Jackson, Ian Johnston, Dominic Kwiatkowski, Cordelia Langford, John Sillitoe on behalf of the Wellcome Sanger Institute COVID-19 Surveillance Team |
| EPI_ISL_2916602 | Lighthouse Lab in Milton Keynes | Wellcome Sanger Institute for the COVID-19 Genomics UK (COG-UK) Consortium | The Lighthouse Lab in Milton Keynes and Alex Alderton, Roberto Amato, Jeffrey Barrett, Sonia Goncalves, Ewan Harrison, David K. Jackson, Ian Johnston, Dominic Kwiatkowski, Cordelia Langford, John Sillitoe on behalf of the Wellcome Sanger Institute COVID-19 Surveillance Team |
| EPI_ISL_2916603 | Lighthouse Lab in Alderley Park | Wellcome Sanger Institute for the COVID-19 Genomics UK (COG-UK) Consortium | Jacquelyn Wynn, Mairead Hyland, The Lighthouse Lab in Alderley Park and Alex Alderton, Roberto Amato, Jeffrey Barrett, Sonia Goncalves, Ewan Harrison, David K. Jackson, Ian Johnston, Dominic Kwiatkowski, Cordelia Langford, John Sillitoe on behalf of the Wellcome Sanger Institute COVID-19 Surveillance Team |
| EPI_ISL_2916604 | Lighthouse Lab in Glasgow | Wellcome Sanger Institute for the COVID-19 Genomics UK (COG-UK) Consortium | Harper VanSteenhouse, Yumi Kasai, David Gray, Carol Clugston, Anna Dominiczak and Alex Alderton, Roberto Amato, Jeffrey Barrett, Sonia Goncalves, Ewan Harrison, David K. Jackson, Ian Johnston, Dominic Kwiatkowski, Cordelia Langford, John Sillitoe on behalf of the Wellcome Sanger Institute COVID-19 Surveillance Team |
| EPI_ISL_2916605 | Lighthouse Lab in Milton Keynes | Wellcome Sanger Institute for the COVID-19 Genomics UK (COG-UK) Consortium | The Lighthouse Lab in Milton Keynes and Alex Alderton, Roberto Amato, Jeffrey Barrett, Sonia Goncalves, Ewan Harrison, David K. Jackson, Ian Johnston, Dominic Kwiatkowski, Cordelia Langford, John Sillitoe on behalf of the Wellcome Sanger Institute COVID-19 Surveillance Team |
| EPI_ISL_2916606 | Lighthouse Lab in Alderley Park | Wellcome Sanger Institute for the COVID-19 Genomics UK (COG-UK) Consortium | Jacquelyn Wynn, Mairead Hyland, The Lighthouse Lab in Alderley Park and Alex Alderton, Roberto Amato, Jeffrey Barrett, Sonia Goncalves, Ewan Harrison, David K. Jackson, Ian Johnston, Dominic Kwiatkowski, Cordelia Langford, John Sillitoe on behalf of the Wellcome Sanger Institute COVID-19 Surveillance Team |
| EPI_ISL_2916607 | Lighthouse Lab in Milton Keynes | Wellcome Sanger Institute for the COVID-19 Genomics UK (COG-UK) Consortium | The Lighthouse Lab in Milton Keynes and Alex Alderton, Roberto Amato, Jeffrey Barrett, Sonia Goncalves, Ewan Harrison, David K. Jackson, Ian Johnston, Dominic Kwiatkowski, Cordelia Langford, John Sillitoe on behalf of the Wellcome Sanger Institute COVID-19 Surveillance Team |
| EPI_ISL_2916608 | Lighthouse Lab in Alderley Park | Wellcome Sanger Institute for the COVID-19 Genomics UK (COG-UK) Consortium | Jacquelyn Wynn, Mairead Hyland, The Lighthouse Lab in Alderley Park and Alex Alderton, Roberto Amato, Jeffrey Barrett, Sonia Goncalves, Ewan Harrison, David K. Jackson, Ian Johnston, Dominic Kwiatkowski, Cordelia Langford, John Sillitoe on behalf of the Wellcome Sanger Institute COVID-19 Surveillance Team |
| EPI_ISL_2916609 | Lighthouse Lab in Milton Keynes | Wellcome Sanger Institute for the COVID-19 Genomics UK (COG-UK) Consortium | The Lighthouse Lab in Milton Keynes and Alex Alderton, Roberto Amato, Jeffrey Barrett, Sonia Goncalves, Ewan Harrison, David K. Jackson, Ian Johnston, Dominic Kwiatkowski, Cordelia Langford, John Sillitoe on behalf of the Wellcome Sanger Institute COVID-19 Surveillance Team |

| | | | |
|---|---|---|---|
| EPI_ISL_2916610 | Lighthouse Lab in Alderley Park | Wellcome Sanger Institute for the COVID-19 Genomics UK (COG-UK) Consortium | Jacquelyn Wynn, Mairead Hyland, The Lighthouse Lab in Alderley Park and Alex Alderton, Roberto Amato, Jeffrey Barrett, Sonia Goncalves, Ewan Harrison, David K. Jackson, Ian Johnston, Dominic Kwiatkowski, Cordelia Langford, John Sillitoe on behalf of the Wellcome Sanger Institute COVID-19 Surveillance Team |
| EPI_ISL_2916611 | Lighthouse Lab in Milton Keynes | Wellcome Sanger Institute for the COVID-19 Genomics UK (COG-UK) Consortium | The Lighthouse Lab in Milton Keynes and Alex Alderton, Roberto Amato, Jeffrey Barrett, Sonia Goncalves, Ewan Harrison, David K. Jackson, Ian Johnston, Dominic Kwiatkowski, Cordelia Langford, John Sillitoe on behalf of the Wellcome Sanger Institute COVID-19 Surveillance Team |
| EPI_ISL_2916612 | Lighthouse Lab in Glasgow | Wellcome Sanger Institute for the COVID-19 Genomics UK (COG-UK) Consortium | Harper VanSteenhouse, Yumi Kasai, David Gray, Carol Clugston, Anna Dominiczak and Alex Alderton, Roberto Amato, Jeffrey Barrett, Sonia Goncalves, Ewan Harrison, David K. Jackson, Ian Johnston, Dominic Kwiatkowski, Cordelia Langford, John Sillitoe on behalf of the Wellcome Sanger Institute COVID-19 Surveillance Team |
| EPI_ISL_2916613, EPI_ISL_2916614 | Lighthouse Lab in Milton Keynes | Wellcome Sanger Institute for the COVID-19 Genomics UK (COG-UK) Consortium | The Lighthouse Lab in Milton Keynes and Alex Alderton, Roberto Amato, Jeffrey Barrett, Sonia Goncalves, Ewan Harrison, David K. Jackson, Ian Johnston, Dominic Kwiatkowski, Cordelia Langford, John Sillitoe on behalf of the Wellcome Sanger Institute COVID-19 Surveillance Team |
| EPI_ISL_2916615 | Lighthouse Lab in Alderley Park | Wellcome Sanger Institute for the COVID-19 Genomics UK (COG-UK) Consortium | Jacquelyn Wynn, Mairead Hyland, The Lighthouse Lab in Alderley Park and Alex Alderton, Roberto Amato, Jeffrey Barrett, Sonia Goncalves, Ewan Harrison, David K. Jackson, Ian Johnston, Dominic Kwiatkowski, Cordelia Langford, John Sillitoe on behalf of the Wellcome Sanger Institute COVID-19 Surveillance Team |
| EPI_ISL_2916616 | Lighthouse Lab in Glasgow | Wellcome Sanger Institute for the COVID-19 Genomics UK (COG-UK) Consortium | Harper VanSteenhouse, Yumi Kasai, David Gray, Carol Clugston, Anna Dominiczak and Alex Alderton, Roberto Amato, Jeffrey Barrett, Sonia Goncalves, Ewan Harrison, David K. Jackson, Ian Johnston, Dominic Kwiatkowski, Cordelia Langford, John Sillitoe on behalf of the Wellcome Sanger Institute COVID-19 Surveillance Team |
| EPI_ISL_2916617, EPI_ISL_2916618, EPI_ISL_2916619 | Lighthouse Lab in Milton Keynes | Wellcome Sanger Institute for the COVID-19 Genomics UK (COG-UK) Consortium | The Lighthouse Lab in Milton Keynes and Alex Alderton, Roberto Amato, Jeffrey Barrett, Sonia Goncalves, Ewan Harrison, David K. Jackson, Ian Johnston, Dominic Kwiatkowski, Cordelia Langford, John Sillitoe on behalf of the Wellcome Sanger Institute COVID-19 Surveillance Team |
| EPI_ISL_2916620 | Lighthouse Lab in Alderley Park | Wellcome Sanger Institute for the COVID-19 Genomics UK (COG-UK) Consortium | Jacquelyn Wynn, Mairead Hyland, The Lighthouse Lab in Alderley Park and Alex Alderton, Roberto Amato, Jeffrey Barrett, Sonia Goncalves, Ewan Harrison, David K. Jackson, Ian Johnston, Dominic Kwiatkowski, Cordelia Langford, John Sillitoe on behalf of the Wellcome Sanger Institute COVID-19 Surveillance Team |
| EPI_ISL_2916621 | Lighthouse Lab in Glasgow | Wellcome Sanger Institute for the COVID-19 Genomics UK (COG-UK) Consortium | Harper VanSteenhouse, Yumi Kasai, David Gray, Carol Clugston, Anna Dominiczak and Alex Alderton, Roberto Amato, Jeffrey Barrett, Sonia Goncalves, Ewan Harrison, David K. Jackson, Ian Johnston, Dominic Kwiatkowski, Cordelia Langford, John Sillitoe on behalf of the Wellcome Sanger Institute COVID-19 Surveillance Team |
| EPI_ISL_2916622 | Lighthouse Lab in Alderley Park | Wellcome Sanger Institute for the COVID-19 Genomics UK (COG-UK) Consortium | Jacquelyn Wynn, Mairead Hyland, The Lighthouse Lab in Alderley Park and Alex Alderton, Roberto Amato, Jeffrey Barrett, Sonia Goncalves, Ewan Harrison, David K. Jackson, Ian Johnston, Dominic Kwiatkowski, Cordelia Langford, John Sillitoe on behalf of the Wellcome Sanger Institute COVID-19 Surveillance Team |
| EPI_ISL_2916623 | Lighthouse Lab in Glasgow | Wellcome Sanger Institute for the COVID-19 Genomics UK (COG-UK) Consortium | Harper VanSteenhouse, Yumi Kasai, David Gray, Carol Clugston, Anna Dominiczak and Alex Alderton, Roberto Amato, Jeffrey Barrett, Sonia Goncalves, Ewan Harrison, David K. Jackson, Ian Johnston, Dominic Kwiatkowski, Cordelia Langford, John Sillitoe on behalf of the Wellcome Sanger Institute COVID-19 Surveillance Team |
| EPI_ISL_2916624, EPI_ISL_2916625 | Lighthouse Lab in Milton Keynes | Wellcome Sanger Institute for the COVID-19 Genomics UK (COG-UK) Consortium | The Lighthouse Lab in Milton Keynes and Alex Alderton, Roberto Amato, Jeffrey Barrett, Sonia Goncalves, Ewan Harrison, David K. Jackson, Ian Johnston, Dominic Kwiatkowski, Cordelia Langford, John Sillitoe on behalf of the Wellcome Sanger Institute COVID-19 Surveillance Team |
| EPI_ISL_2916626, EPI_ISL_2916627 | Lighthouse Lab in Alderley Park | Wellcome Sanger Institute for the COVID-19 Genomics UK (COG-UK) Consortium | Jacquelyn Wynn, Mairead Hyland, The Lighthouse Lab in Alderley Park and Alex Alderton, Roberto Amato, Jeffrey Barrett, Sonia Goncalves, Ewan Harrison, David K. Jackson, Ian Johnston, Dominic Kwiatkowski, Cordelia Langford, John Sillitoe on behalf of the Wellcome Sanger Institute COVID-19 Surveillance Team |
| EPI_ISL_2916628, EPI_ISL_2916629 | Lighthouse Lab in Milton Keynes | Wellcome Sanger Institute for the COVID-19 Genomics UK (COG-UK) Consortium | The Lighthouse Lab in Milton Keynes and Alex Alderton, Roberto Amato, Jeffrey Barrett, Sonia Goncalves, Ewan Harrison, David K. Jackson, Ian Johnston, Dominic Kwiatkowski, Cordelia Langford, John Sillitoe on behalf of the Wellcome Sanger Institute COVID-19 Surveillance Team |
| EPI_ISL_2916630 | Lighthouse Lab in Glasgow | Wellcome Sanger Institute for the COVID-19 Genomics UK (COG-UK) Consortium | Harper VanSteenhouse, Yumi Kasai, David Gray, Carol Clugston, Anna Dominiczak and Alex Alderton, Roberto Amato, Jeffrey Barrett, Sonia Goncalves, Ewan Harrison, David K. Jackson, Ian Johnston, Dominic Kwiatkowski, Cordelia Langford, John Sillitoe on behalf of the Wellcome Sanger Institute COVID-19 Surveillance Team |
| EPI_ISL_2916631 | Lighthouse Lab in Alderley Park | Wellcome Sanger Institute for the COVID-19 Genomics UK (COG-UK) Consortium | Jacquelyn Wynn, Mairead Hyland, The Lighthouse Lab in Alderley Park and Alex Alderton, Roberto Amato, Jeffrey Barrett, Sonia Goncalves, Ewan Harrison, David K. Jackson, Ian Johnston, Dominic Kwiatkowski, Cordelia Langford, John Sillitoe on behalf of the Wellcome Sanger Institute COVID-19 Surveillance Team |
| EPI_ISL_2916632, EPI_ISL_2916633 | Lighthouse Lab in Milton Keynes | Wellcome Sanger Institute for the COVID-19 Genomics UK (COG-UK) Consortium | The Lighthouse Lab in Milton Keynes and Alex Alderton, Roberto Amato, Jeffrey Barrett, Sonia Goncalves, Ewan Harrison, David K. Jackson, Ian Johnston, Dominic Kwiatkowski, Cordelia Langford, John Sillitoe on behalf of the Wellcome Sanger Institute COVID-19 Surveillance Team |
| EPI_ISL_2916634 | Lighthouse Lab in Alderley Park | Wellcome Sanger Institute for the COVID-19 Genomics UK (COG-UK) Consortium | Jacquelyn Wynn, Mairead Hyland, The Lighthouse Lab in Alderley Park and Alex Alderton, Roberto Amato, Jeffrey Barrett, Sonia Goncalves, Ewan Harrison, David K. Jackson, Ian Johnston, Dominic Kwiatkowski, Cordelia Langford, John Sillitoe on behalf of the Wellcome Sanger Institute COVID-19 Surveillance Team |
| EPI_ISL_2916635, EPI_ISL_2916636, EPI_ISL_2916637, EPI_ISL_2916638 | Lighthouse Lab in Milton Keynes | Wellcome Sanger Institute for the COVID-19 Genomics UK (COG-UK) Consortium | The Lighthouse Lab in Milton Keynes and Alex Alderton, Roberto Amato, Jeffrey Barrett, Sonia Goncalves, Ewan Harrison, David K. Jackson, Ian Johnston, Dominic Kwiatkowski, Cordelia Langford, John Sillitoe on behalf of the Wellcome Sanger Institute COVID-19 Surveillance Team |
| EPI_ISL_2916639 | Lighthouse Lab in Glasgow | Wellcome Sanger Institute for the COVID-19 Genomics UK (COG-UK) Consortium | Harper VanSteenhouse, Yumi Kasai, David Gray, Carol Clugston, Anna Dominiczak and Alex Alderton, Roberto Amato, Jeffrey Barrett, Sonia Goncalves, Ewan Harrison, David K. Jackson, Ian Johnston, Dominic Kwiatkowski, Cordelia Langford, John Sillitoe on behalf of the Wellcome Sanger Institute COVID-19 Surveillance Team |
| EPI_ISL_2916640 | Lighthouse Lab in Alderley Park | Wellcome Sanger Institute for the COVID-19 Genomics UK (COG-UK) Consortium | Jacquelyn Wynn, Mairead Hyland, The Lighthouse Lab in Alderley Park and Alex Alderton, Roberto Amato, Jeffrey Barrett, Sonia Goncalves, Ewan Harrison, David K. Jackson, Ian Johnston, Dominic Kwiatkowski, Cordelia Langford, John Sillitoe on behalf of the Wellcome Sanger Institute COVID-19 Surveillance Team |
| EPI_ISL_2916641 | Lighthouse Lab in Milton Keynes | Wellcome Sanger Institute for the COVID-19 Genomics UK (COG-UK) Consortium | The Lighthouse Lab in Milton Keynes and Alex Alderton, Roberto Amato, Jeffrey Barrett, Sonia Goncalves, Ewan Harrison, David K. Jackson, Ian Johnston, Dominic Kwiatkowski, Cordelia Langford, John Sillitoe on behalf of the Wellcome Sanger Institute COVID-19 Surveillance Team |
| EPI_ISL_2916642 | Lighthouse Lab in Glasgow | Wellcome Sanger Institute for the COVID-19 Genomics UK (COG-UK) Consortium | Harper VanSteenhouse, Yumi Kasai, David Gray, Carol Clugston, Anna Dominiczak and Alex Alderton, Roberto Amato, Jeffrey Barrett, Sonia Goncalves, Ewan Harrison, David K. Jackson, Ian Johnston, Dominic Kwiatkowski, Cordelia Langford, John Sillitoe on behalf of the Wellcome Sanger Institute COVID-19 Surveillance Team |
| EPI_ISL_2916643 | Lighthouse Lab in Alderley Park | Wellcome Sanger Institute for the COVID-19 Genomics UK (COG-UK) Consortium | Jacquelyn Wynn, Mairead Hyland, The Lighthouse Lab in Alderley Park and Alex Alderton, Roberto Amato, Jeffrey Barrett, Sonia Goncalves, Ewan Harrison, David K. Jackson, Ian Johnston, Dominic Kwiatkowski, Cordelia Langford, John Sillitoe on behalf of the Wellcome Sanger Institute COVID-19 Surveillance Team |
| EPI_ISL_2916644, EPI_ISL_2916645 | Lighthouse Lab in Glasgow | Wellcome Sanger Institute for the COVID-19 Genomics UK (COG-UK) Consortium | Harper VanSteenhouse, Yumi Kasai, David Gray, Carol Clugston, Anna Dominiczak and Alex Alderton, Roberto Amato, Jeffrey Barrett, Sonia Goncalves, Ewan Harrison, David K. Jackson, Ian Johnston, Dominic Kwiatkowski, Cordelia Langford, John Sillitoe on behalf of the Wellcome Sanger Institute COVID-19 Surveillance Team |
| EPI_ISL_2916646 | Lighthouse Lab in Milton Keynes | Wellcome Sanger Institute for the COVID-19 Genomics UK (COG-UK) Consortium | The Lighthouse Lab in Milton Keynes and Alex Alderton, Roberto Amato, Jeffrey Barrett, Sonia Goncalves, Ewan Harrison, David K. Jackson, Ian Johnston, Dominic Kwiatkowski, Cordelia Langford, John Sillitoe on behalf of the Wellcome Sanger Institute COVID-19 Surveillance Team |
| EPI_ISL_2916647 | Lighthouse Lab in Glasgow | Wellcome Sanger Institute for the COVID-19 Genomics UK (COG-UK) Consortium | Harper VanSteenhouse, Yumi Kasai, David Gray, Carol Clugston, Anna Dominiczak and Alex Alderton, Roberto Amato, Jeffrey Barrett, Sonia Goncalves, Ewan Harrison, David K. Jackson, Ian Johnston, Dominic Kwiatkowski, Cordelia Langford, John Sillitoe on behalf of the Wellcome Sanger Institute COVID-19 Surveillance Team |
| EPI_ISL_2916648, EPI_ISL_2916649 | Lighthouse Lab in Milton Keynes | Wellcome Sanger Institute for the COVID-19 Genomics UK (COG-UK) Consortium | The Lighthouse Lab in Milton Keynes and Alex Alderton, Roberto Amato, Jeffrey Barrett, Sonia Goncalves, Ewan Harrison, David K. Jackson, Ian Johnston, Dominic Kwiatkowski, Cordelia Langford, John Sillitoe on behalf of the Wellcome Sanger Institute COVID-19 Surveillance Team |
| EPI_ISL_2916650 | Lighthouse Lab in Alderley Park | Wellcome Sanger Institute for the COVID-19 Genomics UK (COG-UK) Consortium | Jacquelyn Wynn, Mairead Hyland, The Lighthouse Lab in Alderley Park and Alex Alderton, Roberto Amato, Jeffrey Barrett, Sonia Goncalves, Ewan Harrison, David K. Jackson, Ian Johnston, Dominic Kwiatkowski, Cordelia Langford, John Sillitoe on behalf of the Wellcome Sanger Institute COVID-19 Surveillance Team |
| EPI_ISL_2916651 | Lighthouse Lab in Milton Keynes | Wellcome Sanger Institute for the COVID-19 Genomics UK (COG-UK) Consortium | The Lighthouse Lab in Milton Keynes and Alex Alderton, Roberto Amato, Jeffrey Barrett, Sonia Goncalves, Ewan Harrison, David K. Jackson, Ian Johnston, Dominic Kwiatkowski, Cordelia Langford, John Sillitoe on behalf of the Wellcome Sanger Institute COVID-19 Surveillance Team |
| EPI_ISL_2916652 | Lighthouse Lab in Alderley Park | Wellcome Sanger Institute for the COVID-19 Genomics UK (COG-UK) Consortium | Jacquelyn Wynn, Mairead Hyland, The Lighthouse Lab in Alderley Park and Alex Alderton, Roberto Amato, Jeffrey Barrett, Sonia Goncalves, Ewan Harrison, David K. Jackson, Ian Johnston, Dominic Kwiatkowski, Cordelia Langford, John Sillitoe on behalf of the Wellcome Sanger Institute COVID-19 Surveillance Team |
| EPI_ISL_2916653, EPI_ISL_2916654 | Lighthouse Lab in Milton Keynes | Wellcome Sanger Institute for the COVID-19 Genomics UK (COG-UK) Consortium | The Lighthouse Lab in Milton Keynes and Alex Alderton, Roberto Amato, Jeffrey Barrett, Sonia Goncalves, Ewan Harrison, David K. Jackson, Ian Johnston, Dominic Kwiatkowski, Cordelia Langford, John Sillitoe on behalf of the Wellcome Sanger Institute COVID-19 Surveillance Team |
| EPI_ISL_2916655 | Lighthouse Lab in Glasgow | Wellcome Sanger Institute for the COVID-19 Genomics UK (COG-UK) Consortium | Harper VanSteenhouse, Yumi Kasai, David Gray, Carol Clugston, Anna Dominiczak and Alex Alderton, Roberto Amato, Jeffrey Barrett, Sonia Goncalves, Ewan Harrison, David K. Jackson, Ian Johnston, Dominic Kwiatkowski, Cordelia Langford, John Sillitoe on behalf of the Wellcome Sanger Institute COVID-19 Surveillance Team |
| EPI_ISL_2916657 | Lighthouse Lab in Alderley Park | Wellcome Sanger Institute for the COVID-19 Genomics UK (COG-UK) Consortium | Jacquelyn Wynn, Mairead Hyland, The Lighthouse Lab in Alderley Park and Alex Alderton, Roberto Amato, Jeffrey Barrett, Sonia Goncalves, Ewan Harrison, David K. Jackson, Ian Johnston, Dominic Kwiatkowski, Cordelia Langford, John Sillitoe on behalf of the Wellcome Sanger Institute COVID-19 Surveillance Team |
| EPI_ISL_2916658, EPI_ISL_2916659 | Lighthouse Lab in Glasgow | Wellcome Sanger Institute for the COVID-19 Genomics UK (COG-UK) Consortium | Harper VanSteenhouse, Yumi Kasai, David Gray, Carol Clugston, Anna Dominiczak and Alex Alderton, Roberto Amato, Jeffrey Barrett, Sonia Goncalves, Ewan Harrison, David K. Jackson, Ian Johnston, Dominic Kwiatkowski, Cordelia Langford, John Sillitoe on behalf of the Wellcome Sanger Institute COVID-19 Surveillance Team |
| EPI_ISL_2916660 | Lighthouse Lab in Milton Keynes | Wellcome Sanger Institute for the COVID-19 Genomics UK (COG-UK) Consortium | The Lighthouse Lab in Milton Keynes and Alex Alderton, Roberto Amato, Jeffrey Barrett, Sonia Goncalves, Ewan Harrison, David K. Jackson, Ian Johnston, Dominic Kwiatkowski, Cordelia Langford, John Sillitoe on behalf of the Wellcome Sanger Institute COVID-19 Surveillance Team |
| EPI_ISL_2916661 | Lighthouse Lab in Alderley Park | Wellcome Sanger Institute for the COVID-19 Genomics UK (COG-UK) Consortium | Jacquelyn Wynn, Mairead Hyland, The Lighthouse Lab in Alderley Park and Alex Alderton, Roberto Amato, Jeffrey Barrett, Sonia Goncalves, Ewan Harrison, David K. Jackson, Ian Johnston, Dominic Kwiatkowski, Cordelia Langford, John Sillitoe on behalf of the Wellcome Sanger Institute COVID-19 Surveillance Team |
| EPI_ISL_2916662, EPI_ISL_2916663, EPI_ISL_2916664, EPI_ISL_2916665, EPI_ISL_2916666, EPI_ISL_2916667, EPI_ISL_2916668, EPI_ISL_2916669, EPI_ISL_2916670, EPI_ISL_2916671 | Lighthouse Lab in Glasgow | Wellcome Sanger Institute for the COVID-19 Genomics UK (COG-UK) Consortium | Harper VanSteenhouse, Yumi Kasai, David Gray, Carol Clugston, Anna Dominiczak and Alex Alderton, Roberto Amato, Jeffrey Barrett, Sonia Goncalves, Ewan Harrison, David K. Jackson, Ian Johnston, Dominic Kwiatkowski, Cordelia Langford, John Sillitoe on behalf of the Wellcome Sanger Institute COVID-19 Surveillance Team |
| EPI_ISL_2916672 | Berkshire and Surrey Pathology Services Lighthouse Laboratory | Wellcome Sanger Institute for the COVID-19 Genomics UK (COG-UK) Consortium | Berkshire and Surrey Pathology Services Lighthouse Laboratory and Alex Alderton, Roberto Amato, Jeffrey Barrett, Sonia Goncalves, Ewan Harrison, David K. Jackson, Ian Johnston, Dominic Kwiatkowski, Cordelia Langford, John Sillitoe on behalf of the Wellcome Sanger Institute COVID-19 Surveillance Team |
| EPI_ISL_2916673 | Lighthouse Lab in Glasgow | Wellcome Sanger Institute for the COVID-19 Genomics UK (COG-UK) Consortium | Harper VanSteenhouse, Yumi Kasai, David Gray, Carol Clugston, Anna Dominiczak and Alex Alderton, Roberto Amato, Jeffrey Barrett, Sonia Goncalves, Ewan Harrison, David K. Jackson, Ian Johnston, Dominic Kwiatkowski, Cordelia Langford, John Sillitoe on behalf of the Wellcome Sanger Institute COVID-19 Surveillance Team |
| EPI_ISL_2916674 | Berkshire and Surrey Pathology Services Lighthouse Laboratory | Wellcome Sanger Institute for the COVID-19 Genomics UK (COG-UK) | Berkshire and Surrey Pathology Services Lighthouse Laboratory and Alex Alderton, Roberto Amato, Jeffrey Barrett, Sonia Goncalves, Ewan Harrison, David K. Jackson, Ian Johnston, Dominic Kwiatkowski, Cordelia Langford, John Sillitoe on behalf |

| | | | |
|---|---|---|---|
| | | Consortium | of the Wellcome Sanger Institute COVID-19 Surveillance Team |
| EPI_ISL_2916675 | Lighthouse Lab in Glasgow | Wellcome Sanger Institute for the COVID-19 Genomics UK (COG-UK) Consortium | Harper VanSteenhouse, Yumi Kasai, David Gray, Carol Clugston, Anna Dominiczak and Alex Alderton, Roberto Amato, Jeffrey Barrett, Sonia Goncalves, Ewan Harrison, David K. Jackson, Ian Johnston, Dominic Kwiatkowski, Cordelia Langford, John Sillitoe on behalf of the Wellcome Sanger Institute COVID-19 Surveillance Team |
| EPI_ISL_2916676, EPI_ISL_2916677, EPI_ISL_2916678 | Berkshire and Surrey Pathology Services Lighthouse Laboratory | Wellcome Sanger Institute for the COVID-19 Genomics UK (COG-UK) Consortium | Berkshire and Surrey Pathology Services Lighthouse Laboratory and Alex Alderton, Roberto Amato, Jeffrey Barrett, Sonia Goncalves, Ewan Harrison, David K. Jackson, Ian Johnston, Dominic Kwiatkowski, Cordelia Langford, John Sillitoe on behalf of the Wellcome Sanger Institute COVID-19 Surveillance Team |
| EPI_ISL_2916679, EPI_ISL_2916680, EPI_ISL_2916681 | Lighthouse Lab in Glasgow | Wellcome Sanger Institute for the COVID-19 Genomics UK (COG-UK) Consortium | Harper VanSteenhouse, Yumi Kasai, David Gray, Carol Clugston, Anna Dominiczak and Alex Alderton, Roberto Amato, Jeffrey Barrett, Sonia Goncalves, Ewan Harrison, David K. Jackson, Ian Johnston, Dominic Kwiatkowski, Cordelia Langford, John Sillitoe on behalf of the Wellcome Sanger Institute COVID-19 Surveillance Team |
| EPI_ISL_2916682, EPI_ISL_2916683 | Berkshire and Surrey Pathology Services Lighthouse Laboratory | Wellcome Sanger Institute for the COVID-19 Genomics UK (COG-UK) Consortium | Berkshire and Surrey Pathology Services Lighthouse Laboratory and Alex Alderton, Roberto Amato, Jeffrey Barrett, Sonia Goncalves, Ewan Harrison, David K. Jackson, Ian Johnston, Dominic Kwiatkowski, Cordelia Langford, John Sillitoe on behalf of the Wellcome Sanger Institute COVID-19 Surveillance Team |
| EPI_ISL_2916684, EPI_ISL_2916685, EPI_ISL_2916686 | Lighthouse Lab in Glasgow | Wellcome Sanger Institute for the COVID-19 Genomics UK (COG-UK) Consortium | Harper VanSteenhouse, Yumi Kasai, David Gray, Carol Clugston, Anna Dominiczak and Alex Alderton, Roberto Amato, Jeffrey Barrett, Sonia Goncalves, Ewan Harrison, David K. Jackson, Ian Johnston, Dominic Kwiatkowski, Cordelia Langford, John Sillitoe on behalf of the Wellcome Sanger Institute COVID-19 Surveillance Team |
| EPI_ISL_2916687 | Berkshire and Surrey Pathology Services Lighthouse Laboratory | Wellcome Sanger Institute for the COVID-19 Genomics UK (COG-UK) Consortium | Berkshire and Surrey Pathology Services Lighthouse Laboratory and Alex Alderton, Roberto Amato, Jeffrey Barrett, Sonia Goncalves, Ewan Harrison, David K. Jackson, Ian Johnston, Dominic Kwiatkowski, Cordelia Langford, John Sillitoe on behalf of the Wellcome Sanger Institute COVID-19 Surveillance Team |
| EPI_ISL_2916688 | Lighthouse Lab in Glasgow | Wellcome Sanger Institute for the COVID-19 Genomics UK (COG-UK) Consortium | Harper VanSteenhouse, Yumi Kasai, David Gray, Carol Clugston, Anna Dominiczak and Alex Alderton, Roberto Amato, Jeffrey Barrett, Sonia Goncalves, Ewan Harrison, David K. Jackson, Ian Johnston, Dominic Kwiatkowski, Cordelia Langford, John Sillitoe on behalf of the Wellcome Sanger Institute COVID-19 Surveillance Team |
| EPI_ISL_2916689, EPI_ISL_2916690, EPI_ISL_2916691, EPI_ISL_2916692 | Berkshire and Surrey Pathology Services Lighthouse Laboratory | Wellcome Sanger Institute for the COVID-19 Genomics UK (COG-UK) Consortium | Berkshire and Surrey Pathology Services Lighthouse Laboratory and Alex Alderton, Roberto Amato, Jeffrey Barrett, Sonia Goncalves, Ewan Harrison, David K. Jackson, Ian Johnston, Dominic Kwiatkowski, Cordelia Langford, John Sillitoe on behalf of the Wellcome Sanger Institute COVID-19 Surveillance Team |
| EPI_ISL_2916693 | Lighthouse Lab in Glasgow | Wellcome Sanger Institute for the COVID-19 Genomics UK (COG-UK) Consortium | Harper VanSteenhouse, Yumi Kasai, David Gray, Carol Clugston, Anna Dominiczak and Alex Alderton, Roberto Amato, Jeffrey Barrett, Sonia Goncalves, Ewan Harrison, David K. Jackson, Ian Johnston, Dominic Kwiatkowski, Cordelia Langford, John Sillitoe on behalf of the Wellcome Sanger Institute COVID-19 Surveillance Team |
| EPI_ISL_2916694, EPI_ISL_2916695 | Berkshire and Surrey Pathology Services Lighthouse Laboratory | Wellcome Sanger Institute for the COVID-19 Genomics UK (COG-UK) Consortium | Berkshire and Surrey Pathology Services Lighthouse Laboratory and Alex Alderton, Roberto Amato, Jeffrey Barrett, Sonia Goncalves, Ewan Harrison, David K. Jackson, Ian Johnston, Dominic Kwiatkowski, Cordelia Langford, John Sillitoe on behalf of the Wellcome Sanger Institute COVID-19 Surveillance Team |
| EPI_ISL_2916696 | Lighthouse Lab in Glasgow | Wellcome Sanger Institute for the COVID-19 Genomics UK (COG-UK) Consortium | Harper VanSteenhouse, Yumi Kasai, David Gray, Carol Clugston, Anna Dominiczak and Alex Alderton, Roberto Amato, Jeffrey Barrett, Sonia Goncalves, Ewan Harrison, David K. Jackson, Ian Johnston, Dominic Kwiatkowski, Cordelia Langford, John Sillitoe on behalf of the Wellcome Sanger Institute COVID-19 Surveillance Team |
| EPI_ISL_2916697, EPI_ISL_2916698, EPI_ISL_2916699 | Berkshire and Surrey Pathology Services Lighthouse Laboratory | Wellcome Sanger Institute for the COVID-19 Genomics UK (COG-UK) Consortium | Berkshire and Surrey Pathology Services Lighthouse Laboratory and Alex Alderton, Roberto Amato, Jeffrey Barrett, Sonia Goncalves, Ewan Harrison, David K. Jackson, Ian Johnston, Dominic Kwiatkowski, Cordelia Langford, John Sillitoe on behalf of the Wellcome Sanger Institute COVID-19 Surveillance Team |
| EPI_ISL_2916700, EPI_ISL_2916701 | Lighthouse Lab in Glasgow | Wellcome Sanger Institute for the COVID-19 Genomics UK (COG-UK) Consortium | Harper VanSteenhouse, Yumi Kasai, David Gray, Carol Clugston, Anna Dominiczak and Alex Alderton, Roberto Amato, Jeffrey Barrett, Sonia Goncalves, Ewan Harrison, David K. Jackson, Ian Johnston, Dominic Kwiatkowski, Cordelia Langford, John Sillitoe on behalf of the Wellcome Sanger Institute COVID-19 Surveillance Team |
| EPI_ISL_2916702 | Berkshire and Surrey Pathology Services Lighthouse Laboratory | Wellcome Sanger Institute for the COVID-19 Genomics UK (COG-UK) Consortium | Berkshire and Surrey Pathology Services Lighthouse Laboratory and Alex Alderton, Roberto Amato, Jeffrey Barrett, Sonia Goncalves, Ewan Harrison, David K. Jackson, Ian Johnston, Dominic Kwiatkowski, Cordelia Langford, John Sillitoe on behalf of the Wellcome Sanger Institute COVID-19 Surveillance Team |
| EPI_ISL_2916703 | Lighthouse Lab in Glasgow | Wellcome Sanger Institute for the COVID-19 Genomics UK (COG-UK) Consortium | Harper VanSteenhouse, Yumi Kasai, David Gray, Carol Clugston, Anna Dominiczak and Alex Alderton, Roberto Amato, Jeffrey Barrett, Sonia Goncalves, Ewan Harrison, David K. Jackson, Ian Johnston, Dominic Kwiatkowski, Cordelia Langford, John Sillitoe on behalf of the Wellcome Sanger Institute COVID-19 Surveillance Team |
| EPI_ISL_2916704 | Berkshire and Surrey Pathology Services Lighthouse Laboratory | Wellcome Sanger Institute for the COVID-19 Genomics UK (COG-UK) Consortium | Berkshire and Surrey Pathology Services Lighthouse Laboratory and Alex Alderton, Roberto Amato, Jeffrey Barrett, Sonia Goncalves, Ewan Harrison, David K. Jackson, Ian Johnston, Dominic Kwiatkowski, Cordelia Langford, John Sillitoe on behalf of the Wellcome Sanger Institute COVID-19 Surveillance Team |
| EPI_ISL_2916705, EPI_ISL_2916706 | Lighthouse Lab in Glasgow | Wellcome Sanger Institute for the COVID-19 Genomics UK (COG-UK) Consortium | Harper VanSteenhouse, Yumi Kasai, David Gray, Carol Clugston, Anna Dominiczak and Alex Alderton, Roberto Amato, Jeffrey Barrett, Sonia Goncalves, Ewan Harrison, David K. Jackson, Ian Johnston, Dominic Kwiatkowski, Cordelia Langford, John Sillitoe on behalf of the Wellcome Sanger Institute COVID-19 Surveillance Team |
| EPI_ISL_2916707, EPI_ISL_2916708 | Berkshire and Surrey Pathology Services Lighthouse Laboratory | Wellcome Sanger Institute for the COVID-19 Genomics UK (COG-UK) Consortium | Berkshire and Surrey Pathology Services Lighthouse Laboratory and Alex Alderton, Roberto Amato, Jeffrey Barrett, Sonia Goncalves, Ewan Harrison, David K. Jackson, Ian Johnston, Dominic Kwiatkowski, Cordelia Langford, John Sillitoe on behalf of the Wellcome Sanger Institute COVID-19 Surveillance Team |
| EPI_ISL_2916709, EPI_ISL_2916710 | Lighthouse Lab in Glasgow | Wellcome Sanger Institute for the COVID-19 Genomics UK (COG-UK) Consortium | Harper VanSteenhouse, Yumi Kasai, David Gray, Carol Clugston, Anna Dominiczak and Alex Alderton, Roberto Amato, Jeffrey Barrett, Sonia Goncalves, Ewan Harrison, David K. Jackson, Ian Johnston, Dominic Kwiatkowski, Cordelia Langford, John Sillitoe on behalf of the Wellcome Sanger Institute COVID-19 Surveillance Team |
| EPI_ISL_2916711 | Berkshire and Surrey Pathology Services Lighthouse Laboratory | Wellcome Sanger Institute for the COVID-19 Genomics UK (COG-UK) Consortium | Berkshire and Surrey Pathology Services Lighthouse Laboratory and Alex Alderton, Roberto Amato, Jeffrey Barrett, Sonia Goncalves, Ewan Harrison, David K. Jackson, Ian Johnston, Dominic Kwiatkowski, Cordelia Langford, John Sillitoe on behalf of the Wellcome Sanger Institute COVID-19 Surveillance Team |
| EPI_ISL_2916712, EPI_ISL_2916713, EPI_ISL_2916714, EPI_ISL_2916715, EPI_ISL_2916716 | Lighthouse Lab in Glasgow | Wellcome Sanger Institute for the COVID-19 Genomics UK (COG-UK) Consortium | Harper VanSteenhouse, Yumi Kasai, David Gray, Carol Clugston, Anna Dominiczak and Alex Alderton, Roberto Amato, Jeffrey Barrett, Sonia Goncalves, Ewan Harrison, David K. Jackson, Ian Johnston, Dominic Kwiatkowski, Cordelia Langford, John Sillitoe on behalf of the Wellcome Sanger Institute COVID-19 Surveillance Team |
| EPI_ISL_2916717, EPI_ISL_2916718 | Berkshire and Surrey Pathology Services Lighthouse Laboratory | Wellcome Sanger Institute for the COVID-19 Genomics UK (COG-UK) Consortium | Berkshire and Surrey Pathology Services Lighthouse Laboratory and Alex Alderton, Roberto Amato, Jeffrey Barrett, Sonia Goncalves, Ewan Harrison, David K. Jackson, Ian Johnston, Dominic Kwiatkowski, Cordelia Langford, John Sillitoe on behalf of the Wellcome Sanger Institute COVID-19 Surveillance Team |
| EPI_ISL_2916719 | Lighthouse Lab in Glasgow | Wellcome Sanger Institute for the COVID-19 Genomics UK (COG-UK) Consortium | Harper VanSteenhouse, Yumi Kasai, David Gray, Carol Clugston, Anna Dominiczak and Alex Alderton, Roberto Amato, Jeffrey Barrett, Sonia Goncalves, Ewan Harrison, David K. Jackson, Ian Johnston, Dominic Kwiatkowski, Cordelia Langford, John Sillitoe on behalf of the Wellcome Sanger Institute COVID-19 Surveillance Team |
| EPI_ISL_2916720 | Berkshire and Surrey Pathology Services Lighthouse Laboratory | Wellcome Sanger Institute for the COVID-19 Genomics UK (COG-UK) Consortium | Berkshire and Surrey Pathology Services Lighthouse Laboratory and Alex Alderton, Roberto Amato, Jeffrey Barrett, Sonia Goncalves, Ewan Harrison, David K. Jackson, Ian Johnston, Dominic Kwiatkowski, Cordelia Langford, John Sillitoe on behalf of the Wellcome Sanger Institute COVID-19 Surveillance Team |
| EPI_ISL_2916721 | Lighthouse Lab in Glasgow | Wellcome Sanger Institute for the COVID-19 Genomics UK (COG-UK) Consortium | Harper VanSteenhouse, Yumi Kasai, David Gray, Carol Clugston, Anna Dominiczak and Alex Alderton, Roberto Amato, Jeffrey Barrett, Sonia Goncalves, Ewan Harrison, David K. Jackson, Ian Johnston, Dominic Kwiatkowski, Cordelia Langford, John Sillitoe on behalf of the Wellcome Sanger Institute COVID-19 Surveillance Team |
| EPI_ISL_2916722 | Berkshire and Surrey Pathology Services Lighthouse Laboratory | Wellcome Sanger Institute for the COVID-19 Genomics UK (COG-UK) Consortium | Berkshire and Surrey Pathology Services Lighthouse Laboratory and Alex Alderton, Roberto Amato, Jeffrey Barrett, Sonia Goncalves, Ewan Harrison, David K. Jackson, Ian Johnston, Dominic Kwiatkowski, Cordelia Langford, John Sillitoe on behalf of the Wellcome Sanger Institute COVID-19 Surveillance Team |
| EPI_ISL_2916723 | Lighthouse Lab in Glasgow | Wellcome Sanger Institute for the COVID-19 Genomics UK (COG-UK) Consortium | Harper VanSteenhouse, Yumi Kasai, David Gray, Carol Clugston, Anna Dominiczak and Alex Alderton, Roberto Amato, Jeffrey Barrett, Sonia Goncalves, Ewan Harrison, David K. Jackson, Ian Johnston, Dominic Kwiatkowski, Cordelia Langford, John Sillitoe on behalf of the Wellcome Sanger Institute COVID-19 Surveillance Team |
| EPI_ISL_2916724, EPI_ISL_2916725 | Berkshire and Surrey Pathology Services Lighthouse Laboratory | Wellcome Sanger Institute for the COVID-19 Genomics UK (COG-UK) Consortium | Berkshire and Surrey Pathology Services Lighthouse Laboratory and Alex Alderton, Roberto Amato, Jeffrey Barrett, Sonia Goncalves, Ewan Harrison, David K. Jackson, Ian Johnston, Dominic Kwiatkowski, Cordelia Langford, John Sillitoe on behalf of the Wellcome Sanger Institute COVID-19 Surveillance Team |
| EPI_ISL_2916726 | Lighthouse Lab in Glasgow | Wellcome Sanger Institute for the COVID-19 Genomics UK (COG-UK) Consortium | Harper VanSteenhouse, Yumi Kasai, David Gray, Carol Clugston, Anna Dominiczak and Alex Alderton, Roberto Amato, Jeffrey Barrett, Sonia Goncalves, Ewan Harrison, David K. Jackson, Ian Johnston, Dominic Kwiatkowski, Cordelia Langford, John Sillitoe on behalf of the Wellcome Sanger Institute COVID-19 Surveillance Team |
| EPI_ISL_2916727 | Berkshire and Surrey Pathology Services Lighthouse Laboratory | Wellcome Sanger Institute for the COVID-19 Genomics UK (COG-UK) Consortium | Berkshire and Surrey Pathology Services Lighthouse Laboratory and Alex Alderton, Roberto Amato, Jeffrey Barrett, Sonia Goncalves, Ewan Harrison, David K. Jackson, Ian Johnston, Dominic Kwiatkowski, Cordelia Langford, John Sillitoe on behalf of the Wellcome Sanger Institute COVID-19 Surveillance Team |
| EPI_ISL_2916728 | Lighthouse Lab in Glasgow | Wellcome Sanger Institute for the COVID-19 Genomics UK (COG-UK) Consortium | Harper VanSteenhouse, Yumi Kasai, David Gray, Carol Clugston, Anna Dominiczak and Alex Alderton, Roberto Amato, Jeffrey Barrett, Sonia Goncalves, Ewan Harrison, David K. Jackson, Ian Johnston, Dominic Kwiatkowski, Cordelia Langford, John Sillitoe on behalf of the Wellcome Sanger Institute COVID-19 Surveillance Team |
| EPI_ISL_2916729 | Berkshire and Surrey Pathology Services Lighthouse Laboratory | Wellcome Sanger Institute for the COVID-19 Genomics UK (COG-UK) Consortium | Berkshire and Surrey Pathology Services Lighthouse Laboratory and Alex Alderton, Roberto Amato, Jeffrey Barrett, Sonia Goncalves, Ewan Harrison, David K. Jackson, Ian Johnston, Dominic Kwiatkowski, Cordelia Langford, John Sillitoe on behalf of the Wellcome Sanger Institute COVID-19 Surveillance Team |
| EPI_ISL_2916730, EPI_ISL_2916731 | Lighthouse Lab in Glasgow | Wellcome Sanger Institute for the COVID-19 Genomics UK (COG-UK) Consortium | Harper VanSteenhouse, Yumi Kasai, David Gray, Carol Clugston, Anna Dominiczak and Alex Alderton, Roberto Amato, Jeffrey Barrett, Sonia Goncalves, Ewan Harrison, David K. Jackson, Ian Johnston, Dominic Kwiatkowski, Cordelia Langford, John Sillitoe on behalf of the Wellcome Sanger Institute COVID-19 Surveillance Team |
| EPI_ISL_2916732 | Berkshire and Surrey Pathology Services Lighthouse Laboratory | Wellcome Sanger Institute for the COVID-19 Genomics UK (COG-UK) Consortium | Berkshire and Surrey Pathology Services Lighthouse Laboratory and Alex Alderton, Roberto Amato, Jeffrey Barrett, Sonia Goncalves, Ewan Harrison, David K. Jackson, Ian Johnston, Dominic Kwiatkowski, Cordelia Langford, John Sillitoe on behalf of the Wellcome Sanger Institute COVID-19 Surveillance Team |
| EPI_ISL_2916733, EPI_ISL_2916734 | Lighthouse Lab in Glasgow | Wellcome Sanger Institute for the COVID-19 Genomics UK (COG-UK) Consortium | Harper VanSteenhouse, Yumi Kasai, David Gray, Carol Clugston, Anna Dominiczak and Alex Alderton, Roberto Amato, Jeffrey Barrett, Sonia Goncalves, Ewan Harrison, David K. Jackson, Ian Johnston, Dominic Kwiatkowski, Cordelia Langford, John Sillitoe on behalf of the Wellcome Sanger Institute COVID-19 Surveillance Team |
| EPI_ISL_2916735, EPI_ISL_2916736, EPI_ISL_2916737 | Berkshire and Surrey Pathology Services Lighthouse Laboratory | Wellcome Sanger Institute for the COVID-19 Genomics UK (COG-UK) Consortium | Berkshire and Surrey Pathology Services Lighthouse Laboratory and Alex Alderton, Roberto Amato, Jeffrey Barrett, Sonia Goncalves, Ewan Harrison, David K. Jackson, Ian Johnston, Dominic Kwiatkowski, Cordelia Langford, John Sillitoe on behalf of the Wellcome Sanger Institute COVID-19 Surveillance Team |
| EPI_ISL_2916739, EPI_ISL_2916740 | Lighthouse Lab in Glasgow | Wellcome Sanger Institute for the COVID-19 Genomics UK (COG-UK) Consortium | Harper VanSteenhouse, Yumi Kasai, David Gray, Carol Clugston, Anna Dominiczak and Alex Alderton, Roberto Amato, Jeffrey Barrett, Sonia Goncalves, Ewan Harrison, David K. Jackson, Ian Johnston, Dominic Kwiatkowski, Cordelia Langford, John Sillitoe on behalf of the Wellcome Sanger Institute COVID-19 Surveillance Team |
| EPI_ISL_2916741 | Berkshire and Surrey Pathology Services Lighthouse Laboratory | Wellcome Sanger Institute for the COVID-19 Genomics UK (COG-UK) Consortium | Berkshire and Surrey Pathology Services Lighthouse Laboratory and Alex Alderton, Roberto Amato, Jeffrey Barrett, Sonia Goncalves, Ewan Harrison, David K. Jackson, Ian Johnston, Dominic Kwiatkowski, Cordelia Langford, John Sillitoe on behalf of the Wellcome Sanger Institute COVID-19 Surveillance Team |
| EPI_ISL_2916742, EPI_ISL_2916743, EPI_ISL_2916744, EPI_ISL_2916745, EPI_ISL_2916746, | Lighthouse Lab in Glasgow | Wellcome Sanger Institute for the COVID-19 Genomics UK (COG-UK) Consortium | Harper VanSteenhouse, Yumi Kasai, David Gray, Carol Clugston, Anna Dominiczak and Alex Alderton, Roberto Amato, Jeffrey Barrett, Sonia Goncalves, Ewan Harrison, David K. Jackson, Ian Johnston, Dominic Kwiatkowski, Cordelia Langford, John Sillitoe on behalf of the Wellcome Sanger Institute COVID-19 Surveillance Team |

| | | | |
|---|---|---|---|
| EPI_ISL_2916747 | | | |
| EPI_ISL_2916748, EPI_ISL_2916749 | Berkshire and Surrey Pathology Services Lighthouse Laboratory | Wellcome Sanger Institute for the COVID-19 Genomics UK (COG-UK) Consortium | Berkshire and Surrey Pathology Services Lighthouse Laboratory and Alex Alderton, Roberto Amato, Jeffrey Barrett, Sonia Goncalves, Ewan Harrison, David K. Jackson, Ian Johnston, Dominic Kwiatkowski, Cordelia Langford, John Sillitoe on behalf of the Wellcome Sanger Institute COVID-19 Surveillance Team |
| EPI_ISL_2916750, EPI_ISL_2916751, EPI_ISL_2916752, EPI_ISL_2916753 | Lighthouse Lab in Glasgow | Wellcome Sanger Institute for the COVID-19 Genomics UK (COG-UK) Consortium | Harper VanSteenhouse, Yumi Kasai, David Gray, Carol Clugston, Anna Dominiczak and Alex Alderton, Roberto Amato, Jeffrey Barrett, Sonia Goncalves, Ewan Harrison, David K. Jackson, Ian Johnston, Dominic Kwiatkowski, Cordelia Langford, John Sillitoe on behalf of the Wellcome Sanger Institute COVID-19 Surveillance Team |
| EPI_ISL_2916754, EPI_ISL_2916755 | Berkshire and Surrey Pathology Services Lighthouse Laboratory | Wellcome Sanger Institute for the COVID-19 Genomics UK (COG-UK) Consortium | Berkshire and Surrey Pathology Services Lighthouse Laboratory and Alex Alderton, Roberto Amato, Jeffrey Barrett, Sonia Goncalves, Ewan Harrison, David K. Jackson, Ian Johnston, Dominic Kwiatkowski, Cordelia Langford, John Sillitoe on behalf of the Wellcome Sanger Institute COVID-19 Surveillance Team |
| EPI_ISL_2916756, EPI_ISL_2916757, EPI_ISL_2916758, EPI_ISL_2916759 | Lighthouse Lab in Glasgow | Wellcome Sanger Institute for the COVID-19 Genomics UK (COG-UK) Consortium | Harper VanSteenhouse, Yumi Kasai, David Gray, Carol Clugston, Anna Dominiczak and Alex Alderton, Roberto Amato, Jeffrey Barrett, Sonia Goncalves, Ewan Harrison, David K. Jackson, Ian Johnston, Dominic Kwiatkowski, Cordelia Langford, John Sillitoe on behalf of the Wellcome Sanger Institute COVID-19 Surveillance Team |
| EPI_ISL_2916760, EPI_ISL_2916761 | Berkshire and Surrey Pathology Services Lighthouse Laboratory | Wellcome Sanger Institute for the COVID-19 Genomics UK (COG-UK) Consortium | Berkshire and Surrey Pathology Services Lighthouse Laboratory and Alex Alderton, Roberto Amato, Jeffrey Barrett, Sonia Goncalves, Ewan Harrison, David K. Jackson, Ian Johnston, Dominic Kwiatkowski, Cordelia Langford, John Sillitoe on behalf of the Wellcome Sanger Institute COVID-19 Surveillance Team |
| EPI_ISL_2916762 | Lighthouse Lab in Glasgow | Wellcome Sanger Institute for the COVID-19 Genomics UK (COG-UK) Consortium | Harper VanSteenhouse, Yumi Kasai, David Gray, Carol Clugston, Anna Dominiczak and Alex Alderton, Roberto Amato, Jeffrey Barrett, Sonia Goncalves, Ewan Harrison, David K. Jackson, Ian Johnston, Dominic Kwiatkowski, Cordelia Langford, John Sillitoe on behalf of the Wellcome Sanger Institute COVID-19 Surveillance Team |
| EPI_ISL_2916763 | Berkshire and Surrey Pathology Services Lighthouse Laboratory | Wellcome Sanger Institute for the COVID-19 Genomics UK (COG-UK) Consortium | Berkshire and Surrey Pathology Services Lighthouse Laboratory and Alex Alderton, Roberto Amato, Jeffrey Barrett, Sonia Goncalves, Ewan Harrison, David K. Jackson, Ian Johnston, Dominic Kwiatkowski, Cordelia Langford, John Sillitoe on behalf of the Wellcome Sanger Institute COVID-19 Surveillance Team |
| EPI_ISL_2916764, EPI_ISL_2916766 | Lighthouse Lab in Glasgow | Wellcome Sanger Institute for the COVID-19 Genomics UK (COG-UK) Consortium | Harper VanSteenhouse, Yumi Kasai, David Gray, Carol Clugston, Anna Dominiczak and Alex Alderton, Roberto Amato, Jeffrey Barrett, Sonia Goncalves, Ewan Harrison, David K. Jackson, Ian Johnston, Dominic Kwiatkowski, Cordelia Langford, John Sillitoe on behalf of the Wellcome Sanger Institute COVID-19 Surveillance Team |
| EPI_ISL_2916767 | Berkshire and Surrey Pathology Services Lighthouse Laboratory | Wellcome Sanger Institute for the COVID-19 Genomics UK (COG-UK) Consortium | Berkshire and Surrey Pathology Services Lighthouse Laboratory and Alex Alderton, Roberto Amato, Jeffrey Barrett, Sonia Goncalves, Ewan Harrison, David K. Jackson, Ian Johnston, Dominic Kwiatkowski, Cordelia Langford, John Sillitoe on behalf of the Wellcome Sanger Institute COVID-19 Surveillance Team |
| EPI_ISL_2916768, EPI_ISL_2916769, EPI_ISL_2916770 | Lighthouse Lab in Glasgow | Wellcome Sanger Institute for the COVID-19 Genomics UK (COG-UK) Consortium | Harper VanSteenhouse, Yumi Kasai, David Gray, Carol Clugston, Anna Dominiczak and Alex Alderton, Roberto Amato, Jeffrey Barrett, Sonia Goncalves, Ewan Harrison, David K. Jackson, Ian Johnston, Dominic Kwiatkowski, Cordelia Langford, John Sillitoe on behalf of the Wellcome Sanger Institute COVID-19 Surveillance Team |
| EPI_ISL_2916771 | Berkshire and Surrey Pathology Services Lighthouse Laboratory | Wellcome Sanger Institute for the COVID-19 Genomics UK (COG-UK) Consortium | Berkshire and Surrey Pathology Services Lighthouse Laboratory and Alex Alderton, Roberto Amato, Jeffrey Barrett, Sonia Goncalves, Ewan Harrison, David K. Jackson, Ian Johnston, Dominic Kwiatkowski, Cordelia Langford, John Sillitoe on behalf of the Wellcome Sanger Institute COVID-19 Surveillance Team |
| EPI_ISL_2916772, EPI_ISL_2916773 | Lighthouse Lab in Glasgow | Wellcome Sanger Institute for the COVID-19 Genomics UK (COG-UK) Consortium | Harper VanSteenhouse, Yumi Kasai, David Gray, Carol Clugston, Anna Dominiczak and Alex Alderton, Roberto Amato, Jeffrey Barrett, Sonia Goncalves, Ewan Harrison, David K. Jackson, Ian Johnston, Dominic Kwiatkowski, Cordelia Langford, John Sillitoe on behalf of the Wellcome Sanger Institute COVID-19 Surveillance Team |
| EPI_ISL_2916774 | Berkshire and Surrey Pathology Services Lighthouse Laboratory | Wellcome Sanger Institute for the COVID-19 Genomics UK (COG-UK) Consortium | Berkshire and Surrey Pathology Services Lighthouse Laboratory and Alex Alderton, Roberto Amato, Jeffrey Barrett, Sonia Goncalves, Ewan Harrison, David K. Jackson, Ian Johnston, Dominic Kwiatkowski, Cordelia Langford, John Sillitoe on behalf of the Wellcome Sanger Institute COVID-19 Surveillance Team |
| EPI_ISL_2916775, EPI_ISL_2916776, EPI_ISL_2916777 | Lighthouse Lab in Glasgow | Wellcome Sanger Institute for the COVID-19 Genomics UK (COG-UK) Consortium | Harper VanSteenhouse, Yumi Kasai, David Gray, Carol Clugston, Anna Dominiczak and Alex Alderton, Roberto Amato, Jeffrey Barrett, Sonia Goncalves, Ewan Harrison, David K. Jackson, Ian Johnston, Dominic Kwiatkowski, Cordelia Langford, John Sillitoe on behalf of the Wellcome Sanger Institute COVID-19 Surveillance Team |
| EPI_ISL_2916778 | Berkshire and Surrey Pathology Services Lighthouse Laboratory | Wellcome Sanger Institute for the COVID-19 Genomics UK (COG-UK) Consortium | Berkshire and Surrey Pathology Services Lighthouse Laboratory and Alex Alderton, Roberto Amato, Jeffrey Barrett, Sonia Goncalves, Ewan Harrison, David K. Jackson, Ian Johnston, Dominic Kwiatkowski, Cordelia Langford, John Sillitoe on behalf of the Wellcome Sanger Institute COVID-19 Surveillance Team |
| EPI_ISL_2916779 | Lighthouse Lab in Glasgow | Wellcome Sanger Institute for the COVID-19 Genomics UK (COG-UK) Consortium | Harper VanSteenhouse, Yumi Kasai, David Gray, Carol Clugston, Anna Dominiczak and Alex Alderton, Roberto Amato, Jeffrey Barrett, Sonia Goncalves, Ewan Harrison, David K. Jackson, Ian Johnston, Dominic Kwiatkowski, Cordelia Langford, John Sillitoe on behalf of the Wellcome Sanger Institute COVID-19 Surveillance Team |
| EPI_ISL_2916780, EPI_ISL_2916781, EPI_ISL_2916782 | Berkshire and Surrey Pathology Services Lighthouse Laboratory | Wellcome Sanger Institute for the COVID-19 Genomics UK (COG-UK) Consortium | Berkshire and Surrey Pathology Services Lighthouse Laboratory and Alex Alderton, Roberto Amato, Jeffrey Barrett, Sonia Goncalves, Ewan Harrison, David K. Jackson, Ian Johnston, Dominic Kwiatkowski, Cordelia Langford, John Sillitoe on behalf of the Wellcome Sanger Institute COVID-19 Surveillance Team |
| EPI_ISL_2916783 | Lighthouse Lab in Glasgow | Wellcome Sanger Institute for the COVID-19 Genomics UK (COG-UK) Consortium | Harper VanSteenhouse, Yumi Kasai, David Gray, Carol Clugston, Anna Dominiczak and Alex Alderton, Roberto Amato, Jeffrey Barrett, Sonia Goncalves, Ewan Harrison, David K. Jackson, Ian Johnston, Dominic Kwiatkowski, Cordelia Langford, John Sillitoe on behalf of the Wellcome Sanger Institute COVID-19 Surveillance Team |
| EPI_ISL_2916784, EPI_ISL_2916785, EPI_ISL_2916786 | Berkshire and Surrey Pathology Services Lighthouse Laboratory | Wellcome Sanger Institute for the COVID-19 Genomics UK (COG-UK) Consortium | Berkshire and Surrey Pathology Services Lighthouse Laboratory and Alex Alderton, Roberto Amato, Jeffrey Barrett, Sonia Goncalves, Ewan Harrison, David K. Jackson, Ian Johnston, Dominic Kwiatkowski, Cordelia Langford, John Sillitoe on behalf of the Wellcome Sanger Institute COVID-19 Surveillance Team |
| EPI_ISL_2916787 | Lighthouse Lab in Glasgow | Wellcome Sanger Institute for the COVID-19 Genomics UK (COG-UK) Consortium | Harper VanSteenhouse, Yumi Kasai, David Gray, Carol Clugston, Anna Dominiczak and Alex Alderton, Roberto Amato, Jeffrey Barrett, Sonia Goncalves, Ewan Harrison, David K. Jackson, Ian Johnston, Dominic Kwiatkowski, Cordelia Langford, John Sillitoe on behalf of the Wellcome Sanger Institute COVID-19 Surveillance Team |
| EPI_ISL_2916788 | Berkshire and Surrey Pathology Services Lighthouse Laboratory | Wellcome Sanger Institute for the COVID-19 Genomics UK (COG-UK) Consortium | Berkshire and Surrey Pathology Services Lighthouse Laboratory and Alex Alderton, Roberto Amato, Jeffrey Barrett, Sonia Goncalves, Ewan Harrison, David K. Jackson, Ian Johnston, Dominic Kwiatkowski, Cordelia Langford, John Sillitoe on behalf of the Wellcome Sanger Institute COVID-19 Surveillance Team |
| EPI_ISL_2916789, EPI_ISL_2916790 | Lighthouse Lab in Glasgow | Wellcome Sanger Institute for the COVID-19 Genomics UK (COG-UK) Consortium | Harper VanSteenhouse, Yumi Kasai, David Gray, Carol Clugston, Anna Dominiczak and Alex Alderton, Roberto Amato, Jeffrey Barrett, Sonia Goncalves, Ewan Harrison, David K. Jackson, Ian Johnston, Dominic Kwiatkowski, Cordelia Langford, John Sillitoe on behalf of the Wellcome Sanger Institute COVID-19 Surveillance Team |
| EPI_ISL_2916791 | Berkshire and Surrey Pathology Services Lighthouse Laboratory | Wellcome Sanger Institute for the COVID-19 Genomics UK (COG-UK) Consortium | Berkshire and Surrey Pathology Services Lighthouse Laboratory and Alex Alderton, Roberto Amato, Jeffrey Barrett, Sonia Goncalves, Ewan Harrison, David K. Jackson, Ian Johnston, Dominic Kwiatkowski, Cordelia Langford, John Sillitoe on behalf of the Wellcome Sanger Institute COVID-19 Surveillance Team |
| EPI_ISL_2916792 | Lighthouse Lab in Glasgow | Wellcome Sanger Institute for the COVID-19 Genomics UK (COG-UK) Consortium | Harper VanSteenhouse, Yumi Kasai, David Gray, Carol Clugston, Anna Dominiczak and Alex Alderton, Roberto Amato, Jeffrey Barrett, Sonia Goncalves, Ewan Harrison, David K. Jackson, Ian Johnston, Dominic Kwiatkowski, Cordelia Langford, John Sillitoe on behalf of the Wellcome Sanger Institute COVID-19 Surveillance Team |
| EPI_ISL_2916793 | Berkshire and Surrey Pathology Services Lighthouse Laboratory | Wellcome Sanger Institute for the COVID-19 Genomics UK (COG-UK) Consortium | Berkshire and Surrey Pathology Services Lighthouse Laboratory and Alex Alderton, Roberto Amato, Jeffrey Barrett, Sonia Goncalves, Ewan Harrison, David K. Jackson, Ian Johnston, Dominic Kwiatkowski, Cordelia Langford, John Sillitoe on behalf of the Wellcome Sanger Institute COVID-19 Surveillance Team |
| EPI_ISL_2916794 | Lighthouse Lab in Glasgow | Wellcome Sanger Institute for the COVID-19 Genomics UK (COG-UK) Consortium | Harper VanSteenhouse, Yumi Kasai, David Gray, Carol Clugston, Anna Dominiczak and Alex Alderton, Roberto Amato, Jeffrey Barrett, Sonia Goncalves, Ewan Harrison, David K. Jackson, Ian Johnston, Dominic Kwiatkowski, Cordelia Langford, John Sillitoe on behalf of the Wellcome Sanger Institute COVID-19 Surveillance Team |
| EPI_ISL_2916795 | Berkshire and Surrey Pathology Services Lighthouse Laboratory | Wellcome Sanger Institute for the COVID-19 Genomics UK (COG-UK) Consortium | Berkshire and Surrey Pathology Services Lighthouse Laboratory and Alex Alderton, Roberto Amato, Jeffrey Barrett, Sonia Goncalves, Ewan Harrison, David K. Jackson, Ian Johnston, Dominic Kwiatkowski, Cordelia Langford, John Sillitoe on behalf of the Wellcome Sanger Institute COVID-19 Surveillance Team |
| EPI_ISL_2916796 | Lighthouse Lab in Glasgow | Wellcome Sanger Institute for the COVID-19 Genomics UK (COG-UK) Consortium | Harper VanSteenhouse, Yumi Kasai, David Gray, Carol Clugston, Anna Dominiczak and Alex Alderton, Roberto Amato, Jeffrey Barrett, Sonia Goncalves, Ewan Harrison, David K. Jackson, Ian Johnston, Dominic Kwiatkowski, Cordelia Langford, John Sillitoe on behalf of the Wellcome Sanger Institute COVID-19 Surveillance Team |
| EPI_ISL_2916797 | Berkshire and Surrey Pathology Services Lighthouse Laboratory | Wellcome Sanger Institute for the COVID-19 Genomics UK (COG-UK) Consortium | Berkshire and Surrey Pathology Services Lighthouse Laboratory and Alex Alderton, Roberto Amato, Jeffrey Barrett, Sonia Goncalves, Ewan Harrison, David K. Jackson, Ian Johnston, Dominic Kwiatkowski, Cordelia Langford, John Sillitoe on behalf of the Wellcome Sanger Institute COVID-19 Surveillance Team |
| EPI_ISL_2916798 | Lighthouse Lab in Glasgow | Wellcome Sanger Institute for the COVID-19 Genomics UK (COG-UK) Consortium | Harper VanSteenhouse, Yumi Kasai, David Gray, Carol Clugston, Anna Dominiczak and Alex Alderton, Roberto Amato, Jeffrey Barrett, Sonia Goncalves, Ewan Harrison, David K. Jackson, Ian Johnston, Dominic Kwiatkowski, Cordelia Langford, John Sillitoe on behalf of the Wellcome Sanger Institute COVID-19 Surveillance Team |
| EPI_ISL_2916799, EPI_ISL_2916800, EPI_ISL_2916801, EPI_ISL_2916802 | Berkshire and Surrey Pathology Services Lighthouse Laboratory | Wellcome Sanger Institute for the COVID-19 Genomics UK (COG-UK) Consortium | Berkshire and Surrey Pathology Services Lighthouse Laboratory and Alex Alderton, Roberto Amato, Jeffrey Barrett, Sonia Goncalves, Ewan Harrison, David K. Jackson, Ian Johnston, Dominic Kwiatkowski, Cordelia Langford, John Sillitoe on behalf of the Wellcome Sanger Institute COVID-19 Surveillance Team |
| EPI_ISL_2916803, EPI_ISL_2916804, EPI_ISL_2916805, EPI_ISL_2916806 | Lighthouse Lab in Glasgow | Wellcome Sanger Institute for the COVID-19 Genomics UK (COG-UK) Consortium | Harper VanSteenhouse, Yumi Kasai, David Gray, Carol Clugston, Anna Dominiczak and Alex Alderton, Roberto Amato, Jeffrey Barrett, Sonia Goncalves, Ewan Harrison, David K. Jackson, Ian Johnston, Dominic Kwiatkowski, Cordelia Langford, John Sillitoe on behalf of the Wellcome Sanger Institute COVID-19 Surveillance Team |
| EPI_ISL_2916807 | Berkshire and Surrey Pathology Services Lighthouse Laboratory | Wellcome Sanger Institute for the COVID-19 Genomics UK (COG-UK) Consortium | Berkshire and Surrey Pathology Services Lighthouse Laboratory and Alex Alderton, Roberto Amato, Jeffrey Barrett, Sonia Goncalves, Ewan Harrison, David K. Jackson, Ian Johnston, Dominic Kwiatkowski, Cordelia Langford, John Sillitoe on behalf of the Wellcome Sanger Institute COVID-19 Surveillance Team |
| EPI_ISL_2916808 | Lighthouse Lab in Glasgow | Wellcome Sanger Institute for the COVID-19 Genomics UK (COG-UK) Consortium | Harper VanSteenhouse, Yumi Kasai, David Gray, Carol Clugston, Anna Dominiczak and Alex Alderton, Roberto Amato, Jeffrey Barrett, Sonia Goncalves, Ewan Harrison, David K. Jackson, Ian Johnston, Dominic Kwiatkowski, Cordelia Langford, John Sillitoe on behalf of the Wellcome Sanger Institute COVID-19 Surveillance Team |
| EPI_ISL_2916809, EPI_ISL_2916810 | Berkshire and Surrey Pathology Services Lighthouse Laboratory | Wellcome Sanger Institute for the COVID-19 Genomics UK (COG-UK) Consortium | Berkshire and Surrey Pathology Services Lighthouse Laboratory and Alex Alderton, Roberto Amato, Jeffrey Barrett, Sonia Goncalves, Ewan Harrison, David K. Jackson, Ian Johnston, Dominic Kwiatkowski, Cordelia Langford, John Sillitoe on behalf of the Wellcome Sanger Institute COVID-19 Surveillance Team |
| EPI_ISL_2916811, EPI_ISL_2916812, EPI_ISL_2916813 | Lighthouse Lab in Glasgow | Wellcome Sanger Institute for the COVID-19 Genomics UK (COG-UK) Consortium | Harper VanSteenhouse, Yumi Kasai, David Gray, Carol Clugston, Anna Dominiczak and Alex Alderton, Roberto Amato, Jeffrey Barrett, Sonia Goncalves, Ewan Harrison, David K. Jackson, Ian Johnston, Dominic Kwiatkowski, Cordelia Langford, John Sillitoe on behalf of the Wellcome Sanger Institute COVID-19 Surveillance Team |
| EPI_ISL_2916814, EPI_ISL_2916815 | Berkshire and Surrey Pathology Services Lighthouse Laboratory | Wellcome Sanger Institute for the COVID-19 Genomics UK (COG-UK) Consortium | Berkshire and Surrey Pathology Services Lighthouse Laboratory and Alex Alderton, Roberto Amato, Jeffrey Barrett, Sonia Goncalves, Ewan Harrison, David K. Jackson, Ian Johnston, Dominic Kwiatkowski, Cordelia Langford, John Sillitoe on behalf of the Wellcome Sanger Institute COVID-19 Surveillance Team |
| EPI_ISL_2916816 | Lighthouse Lab in Glasgow | Wellcome Sanger Institute for the COVID-19 Genomics UK (COG-UK) Consortium | Harper VanSteenhouse, Yumi Kasai, David Gray, Carol Clugston, Anna Dominiczak and Alex Alderton, Roberto Amato, Jeffrey Barrett, Sonia Goncalves, Ewan Harrison, David K. Jackson, Ian Johnston, Dominic Kwiatkowski, Cordelia Langford, John Sillitoe on behalf of the Wellcome Sanger Institute COVID-19 Surveillance Team |
| EPI_ISL_2916817 | Berkshire and Surrey Pathology Services Lighthouse Laboratory | Wellcome Sanger Institute for the COVID-19 Genomics UK (COG-UK) Consortium | Berkshire and Surrey Pathology Services Lighthouse Laboratory and Alex Alderton, Roberto Amato, Jeffrey Barrett, Sonia Goncalves, Ewan Harrison, David K. Jackson, Ian Johnston, Dominic Kwiatkowski, Cordelia Langford, John Sillitoe on behalf of the Wellcome Sanger Institute COVID-19 Surveillance Team |
| EPI_ISL_2916818, | Lighthouse Lab in Glasgow | Wellcome Sanger Institute for the COVID-19 Genomics UK (COG-UK) | Harper VanSteenhouse, Yumi Kasai, David Gray, Carol Clugston, Anna Dominiczak and Alex Alderton, Roberto Amato, Jeffrey Barrett, Sonia Goncalves, Ewan Harrison, David K. Jackson, Ian Johnston, Dominic Kwiatkowski, Cordelia Langford, |

| | | | |
|---|---|---|---|
| EPI_ISL_2916819 | | Consortium | John Sillitoe on behalf of the Wellcome Sanger Institute COVID-19 Surveillance Team |
| EPI_ISL_2916820 | Berkshire and Surrey Pathology Services Lighthouse Laboratory | Wellcome Sanger Institute for the COVID-19 Genomics UK (COG-UK) Consortium | Berkshire and Surrey Pathology Services Lighthouse Laboratory and Alex Alderton, Roberto Amato, Jeffrey Barrett, Sonia Goncalves, Ewan Harrison, David K. Jackson, Ian Johnston, Dominic Kwiatkowski, Cordelia Langford, John Sillitoe on behalf of the Wellcome Sanger Institute COVID-19 Surveillance Team |
| EPI_ISL_2916821, EPI_ISL_2916822, EPI_ISL_2916823 | Lighthouse Lab in Glasgow | Wellcome Sanger Institute for the COVID-19 Genomics UK (COG-UK) Consortium | Harper VanSteenhouse, Yumi Kasai, David Gray, Carol Clugston, Anna Dominiczak and Alex Alderton, Roberto Amato, Jeffrey Barrett, Sonia Goncalves, Ewan Harrison, David K. Jackson, Ian Johnston, Dominic Kwiatkowski, Cordelia Langford, John Sillitoe on behalf of the Wellcome Sanger Institute COVID-19 Surveillance Team |
| EPI_ISL_2916824, EPI_ISL_2916825 | Berkshire and Surrey Pathology Services Lighthouse Laboratory | Wellcome Sanger Institute for the COVID-19 Genomics UK (COG-UK) Consortium | Berkshire and Surrey Pathology Services Lighthouse Laboratory and Alex Alderton, Roberto Amato, Jeffrey Barrett, Sonia Goncalves, Ewan Harrison, David K. Jackson, Ian Johnston, Dominic Kwiatkowski, Cordelia Langford, John Sillitoe on behalf of the Wellcome Sanger Institute COVID-19 Surveillance Team |
| EPI_ISL_2916826, EPI_ISL_2916827 | Lighthouse Lab in Glasgow | Wellcome Sanger Institute for the COVID-19 Genomics UK (COG-UK) Consortium | Harper VanSteenhouse, Yumi Kasai, David Gray, Carol Clugston, Anna Dominiczak and Alex Alderton, Roberto Amato, Jeffrey Barrett, Sonia Goncalves, Ewan Harrison, David K. Jackson, Ian Johnston, Dominic Kwiatkowski, Cordelia Langford, John Sillitoe on behalf of the Wellcome Sanger Institute COVID-19 Surveillance Team |
| EPI_ISL_2916828 | Berkshire and Surrey Pathology Services Lighthouse Laboratory | Wellcome Sanger Institute for the COVID-19 Genomics UK (COG-UK) Consortium | Berkshire and Surrey Pathology Services Lighthouse Laboratory and Alex Alderton, Roberto Amato, Jeffrey Barrett, Sonia Goncalves, Ewan Harrison, David K. Jackson, Ian Johnston, Dominic Kwiatkowski, Cordelia Langford, John Sillitoe on behalf of the Wellcome Sanger Institute COVID-19 Surveillance Team |
| EPI_ISL_2916829 | Lighthouse Lab in Glasgow | Wellcome Sanger Institute for the COVID-19 Genomics UK (COG-UK) Consortium | Harper VanSteenhouse, Yumi Kasai, David Gray, Carol Clugston, Anna Dominiczak and Alex Alderton, Roberto Amato, Jeffrey Barrett, Sonia Goncalves, Ewan Harrison, David K. Jackson, Ian Johnston, Dominic Kwiatkowski, Cordelia Langford, John Sillitoe on behalf of the Wellcome Sanger Institute COVID-19 Surveillance Team |
| EPI_ISL_2916830, EPI_ISL_2916831, EPI_ISL_2916832 | Berkshire and Surrey Pathology Services Lighthouse Laboratory | Wellcome Sanger Institute for the COVID-19 Genomics UK (COG-UK) Consortium | Berkshire and Surrey Pathology Services Lighthouse Laboratory and Alex Alderton, Roberto Amato, Jeffrey Barrett, Sonia Goncalves, Ewan Harrison, David K. Jackson, Ian Johnston, Dominic Kwiatkowski, Cordelia Langford, John Sillitoe on behalf of the Wellcome Sanger Institute COVID-19 Surveillance Team |
| EPI_ISL_2916833, EPI_ISL_2916834, EPI_ISL_2916835, EPI_ISL_2916836 | Lighthouse Lab in Glasgow | Wellcome Sanger Institute for the COVID-19 Genomics UK (COG-UK) Consortium | Harper VanSteenhouse, Yumi Kasai, David Gray, Carol Clugston, Anna Dominiczak and Alex Alderton, Roberto Amato, Jeffrey Barrett, Sonia Goncalves, Ewan Harrison, David K. Jackson, Ian Johnston, Dominic Kwiatkowski, Cordelia Langford, John Sillitoe on behalf of the Wellcome Sanger Institute COVID-19 Surveillance Team |
| EPI_ISL_2916837 | Berkshire and Surrey Pathology Services Lighthouse Laboratory | Wellcome Sanger Institute for the COVID-19 Genomics UK (COG-UK) Consortium | Berkshire and Surrey Pathology Services Lighthouse Laboratory and Alex Alderton, Roberto Amato, Jeffrey Barrett, Sonia Goncalves, Ewan Harrison, David K. Jackson, Ian Johnston, Dominic Kwiatkowski, Cordelia Langford, John Sillitoe on behalf of the Wellcome Sanger Institute COVID-19 Surveillance Team |
| EPI_ISL_2916838, EPI_ISL_2916839, EPI_ISL_2916840, EPI_ISL_2916841, EPI_ISL_2916842 | Lighthouse Lab in Glasgow | Wellcome Sanger Institute for the COVID-19 Genomics UK (COG-UK) Consortium | Harper VanSteenhouse, Yumi Kasai, David Gray, Carol Clugston, Anna Dominiczak and Alex Alderton, Roberto Amato, Jeffrey Barrett, Sonia Goncalves, Ewan Harrison, David K. Jackson, Ian Johnston, Dominic Kwiatkowski, Cordelia Langford, John Sillitoe on behalf of the Wellcome Sanger Institute COVID-19 Surveillance Team |
| EPI_ISL_2916843, EPI_ISL_2916844 | Berkshire and Surrey Pathology Services Lighthouse Laboratory | Wellcome Sanger Institute for the COVID-19 Genomics UK (COG-UK) Consortium | Berkshire and Surrey Pathology Services Lighthouse Laboratory and Alex Alderton, Roberto Amato, Jeffrey Barrett, Sonia Goncalves, Ewan Harrison, David K. Jackson, Ian Johnston, Dominic Kwiatkowski, Cordelia Langford, John Sillitoe on behalf of the Wellcome Sanger Institute COVID-19 Surveillance Team |
| EPI_ISL_2916845 | Lighthouse Lab in Glasgow | Wellcome Sanger Institute for the COVID-19 Genomics UK (COG-UK) Consortium | Harper VanSteenhouse, Yumi Kasai, David Gray, Carol Clugston, Anna Dominiczak and Alex Alderton, Roberto Amato, Jeffrey Barrett, Sonia Goncalves, Ewan Harrison, David K. Jackson, Ian Johnston, Dominic Kwiatkowski, Cordelia Langford, John Sillitoe on behalf of the Wellcome Sanger Institute COVID-19 Surveillance Team |
| EPI_ISL_2916846 | Berkshire and Surrey Pathology Services Lighthouse Laboratory | Wellcome Sanger Institute for the COVID-19 Genomics UK (COG-UK) Consortium | Berkshire and Surrey Pathology Services Lighthouse Laboratory and Alex Alderton, Roberto Amato, Jeffrey Barrett, Sonia Goncalves, Ewan Harrison, David K. Jackson, Ian Johnston, Dominic Kwiatkowski, Cordelia Langford, John Sillitoe on behalf of the Wellcome Sanger Institute COVID-19 Surveillance Team |
| EPI_ISL_2916847 | Lighthouse Lab in Glasgow | Wellcome Sanger Institute for the COVID-19 Genomics UK (COG-UK) Consortium | Harper VanSteenhouse, Yumi Kasai, David Gray, Carol Clugston, Anna Dominiczak and Alex Alderton, Roberto Amato, Jeffrey Barrett, Sonia Goncalves, Ewan Harrison, David K. Jackson, Ian Johnston, Dominic Kwiatkowski, Cordelia Langford, John Sillitoe on behalf of the Wellcome Sanger Institute COVID-19 Surveillance Team |
| EPI_ISL_2916848, EPI_ISL_2916849, EPI_ISL_2916850 | Berkshire and Surrey Pathology Services Lighthouse Laboratory | Wellcome Sanger Institute for the COVID-19 Genomics UK (COG-UK) Consortium | Berkshire and Surrey Pathology Services Lighthouse Laboratory and Alex Alderton, Roberto Amato, Jeffrey Barrett, Sonia Goncalves, Ewan Harrison, David K. Jackson, Ian Johnston, Dominic Kwiatkowski, Cordelia Langford, John Sillitoe on behalf of the Wellcome Sanger Institute COVID-19 Surveillance Team |
| EPI_ISL_2916851, EPI_ISL_2916852, EPI_ISL_2916853, EPI_ISL_2916854, EPI_ISL_2916855 | Lighthouse Lab in Glasgow | Wellcome Sanger Institute for the COVID-19 Genomics UK (COG-UK) Consortium | Harper VanSteenhouse, Yumi Kasai, David Gray, Carol Clugston, Anna Dominiczak and Alex Alderton, Roberto Amato, Jeffrey Barrett, Sonia Goncalves, Ewan Harrison, David K. Jackson, Ian Johnston, Dominic Kwiatkowski, Cordelia Langford, John Sillitoe on behalf of the Wellcome Sanger Institute COVID-19 Surveillance Team |
| EPI_ISL_2916856, EPI_ISL_2916857 | Berkshire and Surrey Pathology Services Lighthouse Laboratory | Wellcome Sanger Institute for the COVID-19 Genomics UK (COG-UK) Consortium | Berkshire and Surrey Pathology Services Lighthouse Laboratory and Alex Alderton, Roberto Amato, Jeffrey Barrett, Sonia Goncalves, Ewan Harrison, David K. Jackson, Ian Johnston, Dominic Kwiatkowski, Cordelia Langford, John Sillitoe on behalf of the Wellcome Sanger Institute COVID-19 Surveillance Team |
| EPI_ISL_2916858 | Lighthouse Lab in Glasgow | Wellcome Sanger Institute for the COVID-19 Genomics UK (COG-UK) Consortium | Harper VanSteenhouse, Yumi Kasai, David Gray, Carol Clugston, Anna Dominiczak and Alex Alderton, Roberto Amato, Jeffrey Barrett, Sonia Goncalves, Ewan Harrison, David K. Jackson, Ian Johnston, Dominic Kwiatkowski, Cordelia Langford, John Sillitoe on behalf of the Wellcome Sanger Institute COVID-19 Surveillance Team |
| EPI_ISL_2916859 | Berkshire and Surrey Pathology Services Lighthouse Laboratory | Wellcome Sanger Institute for the COVID-19 Genomics UK (COG-UK) Consortium | Berkshire and Surrey Pathology Services Lighthouse Laboratory and Alex Alderton, Roberto Amato, Jeffrey Barrett, Sonia Goncalves, Ewan Harrison, David K. Jackson, Ian Johnston, Dominic Kwiatkowski, Cordelia Langford, John Sillitoe on behalf of the Wellcome Sanger Institute COVID-19 Surveillance Team |
| EPI_ISL_2916860, EPI_ISL_2916861, EPI_ISL_2916862 | Lighthouse Lab in Glasgow | Wellcome Sanger Institute for the COVID-19 Genomics UK (COG-UK) Consortium | Harper VanSteenhouse, Yumi Kasai, David Gray, Carol Clugston, Anna Dominiczak and Alex Alderton, Roberto Amato, Jeffrey Barrett, Sonia Goncalves, Ewan Harrison, David K. Jackson, Ian Johnston, Dominic Kwiatkowski, Cordelia Langford, John Sillitoe on behalf of the Wellcome Sanger Institute COVID-19 Surveillance Team |
| EPI_ISL_2916863 | Berkshire and Surrey Pathology Services Lighthouse Laboratory | Wellcome Sanger Institute for the COVID-19 Genomics UK (COG-UK) Consortium | Berkshire and Surrey Pathology Services Lighthouse Laboratory and Alex Alderton, Roberto Amato, Jeffrey Barrett, Sonia Goncalves, Ewan Harrison, David K. Jackson, Ian Johnston, Dominic Kwiatkowski, Cordelia Langford, John Sillitoe on behalf of the Wellcome Sanger Institute COVID-19 Surveillance Team |
| EPI_ISL_2916864, EPI_ISL_2916865 | Lighthouse Lab in Glasgow | Wellcome Sanger Institute for the COVID-19 Genomics UK (COG-UK) Consortium | Harper VanSteenhouse, Yumi Kasai, David Gray, Carol Clugston, Anna Dominiczak and Alex Alderton, Roberto Amato, Jeffrey Barrett, Sonia Goncalves, Ewan Harrison, David K. Jackson, Ian Johnston, Dominic Kwiatkowski, Cordelia Langford, John Sillitoe on behalf of the Wellcome Sanger Institute COVID-19 Surveillance Team |
| EPI_ISL_2916866 | Berkshire and Surrey Pathology Services Lighthouse Laboratory | Wellcome Sanger Institute for the COVID-19 Genomics UK (COG-UK) Consortium | Berkshire and Surrey Pathology Services Lighthouse Laboratory and Alex Alderton, Roberto Amato, Jeffrey Barrett, Sonia Goncalves, Ewan Harrison, David K. Jackson, Ian Johnston, Dominic Kwiatkowski, Cordelia Langford, John Sillitoe on behalf of the Wellcome Sanger Institute COVID-19 Surveillance Team |
| EPI_ISL_2916867 | Lighthouse Lab in Glasgow | Wellcome Sanger Institute for the COVID-19 Genomics UK (COG-UK) Consortium | Harper VanSteenhouse, Yumi Kasai, David Gray, Carol Clugston, Anna Dominiczak and Alex Alderton, Roberto Amato, Jeffrey Barrett, Sonia Goncalves, Ewan Harrison, David K. Jackson, Ian Johnston, Dominic Kwiatkowski, Cordelia Langford, John Sillitoe on behalf of the Wellcome Sanger Institute COVID-19 Surveillance Team |
| EPI_ISL_2916868, EPI_ISL_2916869 | Berkshire and Surrey Pathology Services Lighthouse Laboratory | Wellcome Sanger Institute for the COVID-19 Genomics UK (COG-UK) Consortium | Berkshire and Surrey Pathology Services Lighthouse Laboratory and Alex Alderton, Roberto Amato, Jeffrey Barrett, Sonia Goncalves, Ewan Harrison, David K. Jackson, Ian Johnston, Dominic Kwiatkowski, Cordelia Langford, John Sillitoe on behalf of the Wellcome Sanger Institute COVID-19 Surveillance Team |
| EPI_ISL_2916870 | Lighthouse Lab in Glasgow | Wellcome Sanger Institute for the COVID-19 Genomics UK (COG-UK) Consortium | Harper VanSteenhouse, Yumi Kasai, David Gray, Carol Clugston, Anna Dominiczak and Alex Alderton, Roberto Amato, Jeffrey Barrett, Sonia Goncalves, Ewan Harrison, David K. Jackson, Ian Johnston, Dominic Kwiatkowski, Cordelia Langford, John Sillitoe on behalf of the Wellcome Sanger Institute COVID-19 Surveillance Team |
| EPI_ISL_2916871 | Berkshire and Surrey Pathology Services Lighthouse Laboratory | Wellcome Sanger Institute for the COVID-19 Genomics UK (COG-UK) Consortium | Berkshire and Surrey Pathology Services Lighthouse Laboratory and Alex Alderton, Roberto Amato, Jeffrey Barrett, Sonia Goncalves, Ewan Harrison, David K. Jackson, Ian Johnston, Dominic Kwiatkowski, Cordelia Langford, John Sillitoe on behalf of the Wellcome Sanger Institute COVID-19 Surveillance Team |
| EPI_ISL_2916872 | Lighthouse Lab in Glasgow | Wellcome Sanger Institute for the COVID-19 Genomics UK (COG-UK) Consortium | Harper VanSteenhouse, Yumi Kasai, David Gray, Carol Clugston, Anna Dominiczak and Alex Alderton, Roberto Amato, Jeffrey Barrett, Sonia Goncalves, Ewan Harrison, David K. Jackson, Ian Johnston, Dominic Kwiatkowski, Cordelia Langford, John Sillitoe on behalf of the Wellcome Sanger Institute COVID-19 Surveillance Team |
| EPI_ISL_2916873 | Berkshire and Surrey Pathology Services Lighthouse Laboratory | Wellcome Sanger Institute for the COVID-19 Genomics UK (COG-UK) Consortium | Berkshire and Surrey Pathology Services Lighthouse Laboratory and Alex Alderton, Roberto Amato, Jeffrey Barrett, Sonia Goncalves, Ewan Harrison, David K. Jackson, Ian Johnston, Dominic Kwiatkowski, Cordelia Langford, John Sillitoe on behalf of the Wellcome Sanger Institute COVID-19 Surveillance Team |
| EPI_ISL_2916874, EPI_ISL_2916875 | Lighthouse Lab in Glasgow | Wellcome Sanger Institute for the COVID-19 Genomics UK (COG-UK) Consortium | Harper VanSteenhouse, Yumi Kasai, David Gray, Carol Clugston, Anna Dominiczak and Alex Alderton, Roberto Amato, Jeffrey Barrett, Sonia Goncalves, Ewan Harrison, David K. Jackson, Ian Johnston, Dominic Kwiatkowski, Cordelia Langford, John Sillitoe on behalf of the Wellcome Sanger Institute COVID-19 Surveillance Team |
| EPI_ISL_2916876, EPI_ISL_2916877, EPI_ISL_2916878, EPI_ISL_2916879, EPI_ISL_2916880, EPI_ISL_2916881, EPI_ISL_2916882 | Berkshire and Surrey Pathology Services Lighthouse Laboratory | Wellcome Sanger Institute for the COVID-19 Genomics UK (COG-UK) Consortium | Berkshire and Surrey Pathology Services Lighthouse Laboratory and Alex Alderton, Roberto Amato, Jeffrey Barrett, Sonia Goncalves, Ewan Harrison, David K. Jackson, Ian Johnston, Dominic Kwiatkowski, Cordelia Langford, John Sillitoe on behalf of the Wellcome Sanger Institute COVID-19 Surveillance Team |
| EPI_ISL_2916883, EPI_ISL_2916884, EPI_ISL_2916885 | Lighthouse Lab in Glasgow | Wellcome Sanger Institute for the COVID-19 Genomics UK (COG-UK) Consortium | Harper VanSteenhouse, Yumi Kasai, David Gray, Carol Clugston, Anna Dominiczak and Alex Alderton, Roberto Amato, Jeffrey Barrett, Sonia Goncalves, Ewan Harrison, David K. Jackson, Ian Johnston, Dominic Kwiatkowski, Cordelia Langford, John Sillitoe on behalf of the Wellcome Sanger Institute COVID-19 Surveillance Team |
| EPI_ISL_2916886, EPI_ISL_2916887, EPI_ISL_2916888 | Berkshire and Surrey Pathology Services Lighthouse Laboratory | Wellcome Sanger Institute for the COVID-19 Genomics UK (COG-UK) Consortium | Berkshire and Surrey Pathology Services Lighthouse Laboratory and Alex Alderton, Roberto Amato, Jeffrey Barrett, Sonia Goncalves, Ewan Harrison, David K. Jackson, Ian Johnston, Dominic Kwiatkowski, Cordelia Langford, John Sillitoe on behalf of the Wellcome Sanger Institute COVID-19 Surveillance Team |
| EPI_ISL_2916889 | Lighthouse Lab in Glasgow | Wellcome Sanger Institute for the COVID-19 Genomics UK (COG-UK) Consortium | Harper VanSteenhouse, Yumi Kasai, David Gray, Carol Clugston, Anna Dominiczak and Alex Alderton, Roberto Amato, Jeffrey Barrett, Sonia Goncalves, Ewan Harrison, David K. Jackson, Ian Johnston, Dominic Kwiatkowski, Cordelia Langford, John Sillitoe on behalf of the Wellcome Sanger Institute COVID-19 Surveillance Team |
| EPI_ISL_2916890, EPI_ISL_2916891, EPI_ISL_2916892 | Berkshire and Surrey Pathology Services Lighthouse Laboratory | Wellcome Sanger Institute for the COVID-19 Genomics UK (COG-UK) Consortium | Berkshire and Surrey Pathology Services Lighthouse Laboratory and Alex Alderton, Roberto Amato, Jeffrey Barrett, Sonia Goncalves, Ewan Harrison, David K. Jackson, Ian Johnston, Dominic Kwiatkowski, Cordelia Langford, John Sillitoe on behalf of the Wellcome Sanger Institute COVID-19 Surveillance Team |
| EPI_ISL_2916893 | Lighthouse Lab in Glasgow | Wellcome Sanger Institute for the COVID-19 Genomics UK (COG-UK) Consortium | Harper VanSteenhouse, Yumi Kasai, David Gray, Carol Clugston, Anna Dominiczak and Alex Alderton, Roberto Amato, Jeffrey Barrett, Sonia Goncalves, Ewan Harrison, David K. Jackson, Ian Johnston, Dominic Kwiatkowski, Cordelia Langford, John Sillitoe on behalf of the Wellcome Sanger Institute COVID-19 Surveillance Team |

| | | | |
|---|---|---|---|
| EPI_ISL_2916894, EPI_ISL_2916895 | Berkshire and Surrey Pathology Services Lighthouse Laboratory | Wellcome Sanger Institute for the COVID-19 Genomics UK (COG-UK) Consortium | Berkshire and Surrey Pathology Services Lighthouse Laboratory and Alex Alderton, Roberto Amato, Jeffrey Barrett, Sonia Goncalves, Ewan Harrison, David K. Jackson, Ian Johnston, Dominic Kwiatkowski, Cordelia Langford, John Sillitoe on behalf of the Wellcome Sanger Institute COVID-19 Surveillance Team |
| EPI_ISL_2916896, EPI_ISL_2916897, EPI_ISL_2916898, EPI_ISL_2916899, EPI_ISL_2916900 | Lighthouse Lab in Glasgow | Wellcome Sanger Institute for the COVID-19 Genomics UK (COG-UK) Consortium | Harper VanSteenhouse, Yumi Kasai, David Gray, Carol Clugston, Anna Dominiczak and Alex Alderton, Roberto Amato, Jeffrey Barrett, Sonia Goncalves, Ewan Harrison, David K. Jackson, Ian Johnston, Dominic Kwiatkowski, Cordelia Langford, John Sillitoe on behalf of the Wellcome Sanger Institute COVID-19 Surveillance Team |
| EPI_ISL_2916901, EPI_ISL_2916902, EPI_ISL_2916903, EPI_ISL_2916904 | Berkshire and Surrey Pathology Services Lighthouse Laboratory | Wellcome Sanger Institute for the COVID-19 Genomics UK (COG-UK) Consortium | Berkshire and Surrey Pathology Services Lighthouse Laboratory and Alex Alderton, Roberto Amato, Jeffrey Barrett, Sonia Goncalves, Ewan Harrison, David K. Jackson, Ian Johnston, Dominic Kwiatkowski, Cordelia Langford, John Sillitoe on behalf of the Wellcome Sanger Institute COVID-19 Surveillance Team |
| EPI_ISL_2916905, EPI_ISL_2916906, EPI_ISL_2916907 | Lighthouse Lab in Glasgow | Wellcome Sanger Institute for the COVID-19 Genomics UK (COG-UK) Consortium | Harper VanSteenhouse, Yumi Kasai, David Gray, Carol Clugston, Anna Dominiczak and Alex Alderton, Roberto Amato, Jeffrey Barrett, Sonia Goncalves, Ewan Harrison, David K. Jackson, Ian Johnston, Dominic Kwiatkowski, Cordelia Langford, John Sillitoe on behalf of the Wellcome Sanger Institute COVID-19 Surveillance Team |
| EPI_ISL_2916908 | Berkshire and Surrey Pathology Services Lighthouse Laboratory | Wellcome Sanger Institute for the COVID-19 Genomics UK (COG-UK) Consortium | Berkshire and Surrey Pathology Services Lighthouse Laboratory and Alex Alderton, Roberto Amato, Jeffrey Barrett, Sonia Goncalves, Ewan Harrison, David K. Jackson, Ian Johnston, Dominic Kwiatkowski, Cordelia Langford, John Sillitoe on behalf of the Wellcome Sanger Institute COVID-19 Surveillance Team |
| EPI_ISL_2916909 | Lighthouse Lab in Glasgow | Wellcome Sanger Institute for the COVID-19 Genomics UK (COG-UK) Consortium | Harper VanSteenhouse, Yumi Kasai, David Gray, Carol Clugston, Anna Dominiczak and Alex Alderton, Roberto Amato, Jeffrey Barrett, Sonia Goncalves, Ewan Harrison, David K. Jackson, Ian Johnston, Dominic Kwiatkowski, Cordelia Langford, John Sillitoe on behalf of the Wellcome Sanger Institute COVID-19 Surveillance Team |
| EPI_ISL_2916910 | Berkshire and Surrey Pathology Services Lighthouse Laboratory | Wellcome Sanger Institute for the COVID-19 Genomics UK (COG-UK) Consortium | Berkshire and Surrey Pathology Services Lighthouse Laboratory and Alex Alderton, Roberto Amato, Jeffrey Barrett, Sonia Goncalves, Ewan Harrison, David K. Jackson, Ian Johnston, Dominic Kwiatkowski, Cordelia Langford, John Sillitoe on behalf of the Wellcome Sanger Institute COVID-19 Surveillance Team |
| EPI_ISL_2916911, EPI_ISL_2916912 | Lighthouse Lab in Glasgow | Wellcome Sanger Institute for the COVID-19 Genomics UK (COG-UK) Consortium | Harper VanSteenhouse, Yumi Kasai, David Gray, Carol Clugston, Anna Dominiczak and Alex Alderton, Roberto Amato, Jeffrey Barrett, Sonia Goncalves, Ewan Harrison, David K. Jackson, Ian Johnston, Dominic Kwiatkowski, Cordelia Langford, John Sillitoe on behalf of the Wellcome Sanger Institute COVID-19 Surveillance Team |
| EPI_ISL_2916913, EPI_ISL_2916914 | Berkshire and Surrey Pathology Services Lighthouse Laboratory | Wellcome Sanger Institute for the COVID-19 Genomics UK (COG-UK) Consortium | Berkshire and Surrey Pathology Services Lighthouse Laboratory and Alex Alderton, Roberto Amato, Jeffrey Barrett, Sonia Goncalves, Ewan Harrison, David K. Jackson, Ian Johnston, Dominic Kwiatkowski, Cordelia Langford, John Sillitoe on behalf of the Wellcome Sanger Institute COVID-19 Surveillance Team |
| EPI_ISL_2916915, EPI_ISL_2916916, EPI_ISL_2916917, EPI_ISL_2916918, EPI_ISL_2916919 | Lighthouse Lab in Glasgow | Wellcome Sanger Institute for the COVID-19 Genomics UK (COG-UK) Consortium | Harper VanSteenhouse, Yumi Kasai, David Gray, Carol Clugston, Anna Dominiczak and Alex Alderton, Roberto Amato, Jeffrey Barrett, Sonia Goncalves, Ewan Harrison, David K. Jackson, Ian Johnston, Dominic Kwiatkowski, Cordelia Langford, John Sillitoe on behalf of the Wellcome Sanger Institute COVID-19 Surveillance Team |
| EPI_ISL_2916920, EPI_ISL_2916921 | Berkshire and Surrey Pathology Services Lighthouse Laboratory | Wellcome Sanger Institute for the COVID-19 Genomics UK (COG-UK) Consortium | Berkshire and Surrey Pathology Services Lighthouse Laboratory and Alex Alderton, Roberto Amato, Jeffrey Barrett, Sonia Goncalves, Ewan Harrison, David K. Jackson, Ian Johnston, Dominic Kwiatkowski, Cordelia Langford, John Sillitoe on behalf of the Wellcome Sanger Institute COVID-19 Surveillance Team |
| EPI_ISL_2916922, EPI_ISL_2916923 | Lighthouse Lab in Glasgow | Wellcome Sanger Institute for the COVID-19 Genomics UK (COG-UK) Consortium | Harper VanSteenhouse, Yumi Kasai, David Gray, Carol Clugston, Anna Dominiczak and Alex Alderton, Roberto Amato, Jeffrey Barrett, Sonia Goncalves, Ewan Harrison, David K. Jackson, Ian Johnston, Dominic Kwiatkowski, Cordelia Langford, John Sillitoe on behalf of the Wellcome Sanger Institute COVID-19 Surveillance Team |
| EPI_ISL_2916924, EPI_ISL_2916925, EPI_ISL_2916926, EPI_ISL_2916927 | Berkshire and Surrey Pathology Services Lighthouse Laboratory | Wellcome Sanger Institute for the COVID-19 Genomics UK (COG-UK) Consortium | Berkshire and Surrey Pathology Services Lighthouse Laboratory and Alex Alderton, Roberto Amato, Jeffrey Barrett, Sonia Goncalves, Ewan Harrison, David K. Jackson, Ian Johnston, Dominic Kwiatkowski, Cordelia Langford, John Sillitoe on behalf of the Wellcome Sanger Institute COVID-19 Surveillance Team |
| EPI_ISL_2916928, EPI_ISL_2916929 | Lighthouse Lab in Glasgow | Wellcome Sanger Institute for the COVID-19 Genomics UK (COG-UK) Consortium | Harper VanSteenhouse, Yumi Kasai, David Gray, Carol Clugston, Anna Dominiczak and Alex Alderton, Roberto Amato, Jeffrey Barrett, Sonia Goncalves, Ewan Harrison, David K. Jackson, Ian Johnston, Dominic Kwiatkowski, Cordelia Langford, John Sillitoe on behalf of the Wellcome Sanger Institute COVID-19 Surveillance Team |
| EPI_ISL_2916930, EPI_ISL_2916931 | Berkshire and Surrey Pathology Services Lighthouse Laboratory | Wellcome Sanger Institute for the COVID-19 Genomics UK (COG-UK) Consortium | Berkshire and Surrey Pathology Services Lighthouse Laboratory and Alex Alderton, Roberto Amato, Jeffrey Barrett, Sonia Goncalves, Ewan Harrison, David K. Jackson, Ian Johnston, Dominic Kwiatkowski, Cordelia Langford, John Sillitoe on behalf of the Wellcome Sanger Institute COVID-19 Surveillance Team |
| EPI_ISL_2916932 | Lighthouse Lab in Glasgow | Wellcome Sanger Institute for the COVID-19 Genomics UK (COG-UK) Consortium | Harper VanSteenhouse, Yumi Kasai, David Gray, Carol Clugston, Anna Dominiczak and Alex Alderton, Roberto Amato, Jeffrey Barrett, Sonia Goncalves, Ewan Harrison, David K. Jackson, Ian Johnston, Dominic Kwiatkowski, Cordelia Langford, John Sillitoe on behalf of the Wellcome Sanger Institute COVID-19 Surveillance Team |
| EPI_ISL_2916933, EPI_ISL_2916934 | Berkshire and Surrey Pathology Services Lighthouse Laboratory | Wellcome Sanger Institute for the COVID-19 Genomics UK (COG-UK) Consortium | Berkshire and Surrey Pathology Services Lighthouse Laboratory and Alex Alderton, Roberto Amato, Jeffrey Barrett, Sonia Goncalves, Ewan Harrison, David K. Jackson, Ian Johnston, Dominic Kwiatkowski, Cordelia Langford, John Sillitoe on behalf of the Wellcome Sanger Institute COVID-19 Surveillance Team |
| EPI_ISL_2916935, EPI_ISL_2916936, EPI_ISL_2916937, EPI_ISL_2916938 | Lighthouse Lab in Glasgow | Wellcome Sanger Institute for the COVID-19 Genomics UK (COG-UK) Consortium | Harper VanSteenhouse, Yumi Kasai, David Gray, Carol Clugston, Anna Dominiczak and Alex Alderton, Roberto Amato, Jeffrey Barrett, Sonia Goncalves, Ewan Harrison, David K. Jackson, Ian Johnston, Dominic Kwiatkowski, Cordelia Langford, John Sillitoe on behalf of the Wellcome Sanger Institute COVID-19 Surveillance Team |
| EPI_ISL_2916939, EPI_ISL_2916940, EPI_ISL_2916941 | Berkshire and Surrey Pathology Services Lighthouse Laboratory | Wellcome Sanger Institute for the COVID-19 Genomics UK (COG-UK) Consortium | Berkshire and Surrey Pathology Services Lighthouse Laboratory and Alex Alderton, Roberto Amato, Jeffrey Barrett, Sonia Goncalves, Ewan Harrison, David K. Jackson, Ian Johnston, Dominic Kwiatkowski, Cordelia Langford, John Sillitoe on behalf of the Wellcome Sanger Institute COVID-19 Surveillance Team |
| EPI_ISL_2916942, EPI_ISL_2916943 | Lighthouse Lab in Glasgow | Wellcome Sanger Institute for the COVID-19 Genomics UK (COG-UK) Consortium | Harper VanSteenhouse, Yumi Kasai, David Gray, Carol Clugston, Anna Dominiczak and Alex Alderton, Roberto Amato, Jeffrey Barrett, Sonia Goncalves, Ewan Harrison, David K. Jackson, Ian Johnston, Dominic Kwiatkowski, Cordelia Langford, John Sillitoe on behalf of the Wellcome Sanger Institute COVID-19 Surveillance Team |
| EPI_ISL_2916944, EPI_ISL_2916945 | Berkshire and Surrey Pathology Services Lighthouse Laboratory | Wellcome Sanger Institute for the COVID-19 Genomics UK (COG-UK) Consortium | Berkshire and Surrey Pathology Services Lighthouse Laboratory and Alex Alderton, Roberto Amato, Jeffrey Barrett, Sonia Goncalves, Ewan Harrison, David K. Jackson, Ian Johnston, Dominic Kwiatkowski, Cordelia Langford, John Sillitoe on behalf of the Wellcome Sanger Institute COVID-19 Surveillance Team |
| EPI_ISL_2916946, EPI_ISL_2916947 | Lighthouse Lab in Glasgow | Wellcome Sanger Institute for the COVID-19 Genomics UK (COG-UK) Consortium | Harper VanSteenhouse, Yumi Kasai, David Gray, Carol Clugston, Anna Dominiczak and Alex Alderton, Roberto Amato, Jeffrey Barrett, Sonia Goncalves, Ewan Harrison, David K. Jackson, Ian Johnston, Dominic Kwiatkowski, Cordelia Langford, John Sillitoe on behalf of the Wellcome Sanger Institute COVID-19 Surveillance Team |
| EPI_ISL_2916948, EPI_ISL_2916949 | Berkshire and Surrey Pathology Services Lighthouse Laboratory | Wellcome Sanger Institute for the COVID-19 Genomics UK (COG-UK) Consortium | Berkshire and Surrey Pathology Services Lighthouse Laboratory and Alex Alderton, Roberto Amato, Jeffrey Barrett, Sonia Goncalves, Ewan Harrison, David K. Jackson, Ian Johnston, Dominic Kwiatkowski, Cordelia Langford, John Sillitoe on behalf of the Wellcome Sanger Institute COVID-19 Surveillance Team |
| EPI_ISL_2916950, EPI_ISL_2916951 | Lighthouse Lab in Glasgow | Wellcome Sanger Institute for the COVID-19 Genomics UK (COG-UK) Consortium | Harper VanSteenhouse, Yumi Kasai, David Gray, Carol Clugston, Anna Dominiczak and Alex Alderton, Roberto Amato, Jeffrey Barrett, Sonia Goncalves, Ewan Harrison, David K. Jackson, Ian Johnston, Dominic Kwiatkowski, Cordelia Langford, John Sillitoe on behalf of the Wellcome Sanger Institute COVID-19 Surveillance Team |
| EPI_ISL_2916952, EPI_ISL_2916953, EPI_ISL_2916954 | Berkshire and Surrey Pathology Services Lighthouse Laboratory | Wellcome Sanger Institute for the COVID-19 Genomics UK (COG-UK) Consortium | Berkshire and Surrey Pathology Services Lighthouse Laboratory and Alex Alderton, Roberto Amato, Jeffrey Barrett, Sonia Goncalves, Ewan Harrison, David K. Jackson, Ian Johnston, Dominic Kwiatkowski, Cordelia Langford, John Sillitoe on behalf of the Wellcome Sanger Institute COVID-19 Surveillance Team |
| EPI_ISL_2916955, EPI_ISL_2916956 | Lighthouse Lab in Glasgow | Wellcome Sanger Institute for the COVID-19 Genomics UK (COG-UK) Consortium | Harper VanSteenhouse, Yumi Kasai, David Gray, Carol Clugston, Anna Dominiczak and Alex Alderton, Roberto Amato, Jeffrey Barrett, Sonia Goncalves, Ewan Harrison, David K. Jackson, Ian Johnston, Dominic Kwiatkowski, Cordelia Langford, John Sillitoe on behalf of the Wellcome Sanger Institute COVID-19 Surveillance Team |
| EPI_ISL_2916957, EPI_ISL_2916958, EPI_ISL_2916959, EPI_ISL_2916960, EPI_ISL_2916961 | Berkshire and Surrey Pathology Services Lighthouse Laboratory | Wellcome Sanger Institute for the COVID-19 Genomics UK (COG-UK) Consortium | Berkshire and Surrey Pathology Services Lighthouse Laboratory and Alex Alderton, Roberto Amato, Jeffrey Barrett, Sonia Goncalves, Ewan Harrison, David K. Jackson, Ian Johnston, Dominic Kwiatkowski, Cordelia Langford, John Sillitoe on behalf of the Wellcome Sanger Institute COVID-19 Surveillance Team |
| EPI_ISL_2916962, EPI_ISL_2916963, EPI_ISL_2916964, EPI_ISL_2916965, EPI_ISL_2916966 | Lighthouse Lab in Glasgow | Wellcome Sanger Institute for the COVID-19 Genomics UK (COG-UK) Consortium | Harper VanSteenhouse, Yumi Kasai, David Gray, Carol Clugston, Anna Dominiczak and Alex Alderton, Roberto Amato, Jeffrey Barrett, Sonia Goncalves, Ewan Harrison, David K. Jackson, Ian Johnston, Dominic Kwiatkowski, Cordelia Langford, John Sillitoe on behalf of the Wellcome Sanger Institute COVID-19 Surveillance Team |
| EPI_ISL_2916967, EPI_ISL_2916968, EPI_ISL_2916969, EPI_ISL_2916970 | Berkshire and Surrey Pathology Services Lighthouse Laboratory | Wellcome Sanger Institute for the COVID-19 Genomics UK (COG-UK) Consortium | Berkshire and Surrey Pathology Services Lighthouse Laboratory and Alex Alderton, Roberto Amato, Jeffrey Barrett, Sonia Goncalves, Ewan Harrison, David K. Jackson, Ian Johnston, Dominic Kwiatkowski, Cordelia Langford, John Sillitoe on behalf of the Wellcome Sanger Institute COVID-19 Surveillance Team |
| EPI_ISL_2916971, EPI_ISL_2916972 | Lighthouse Lab in Glasgow | Wellcome Sanger Institute for the COVID-19 Genomics UK (COG-UK) Consortium | Harper VanSteenhouse, Yumi Kasai, David Gray, Carol Clugston, Anna Dominiczak and Alex Alderton, Roberto Amato, Jeffrey Barrett, Sonia Goncalves, Ewan Harrison, David K. Jackson, Ian Johnston, Dominic Kwiatkowski, Cordelia Langford, John Sillitoe on behalf of the Wellcome Sanger Institute COVID-19 Surveillance Team |
| EPI_ISL_2916973 | Berkshire and Surrey Pathology Services Lighthouse Laboratory | Wellcome Sanger Institute for the COVID-19 Genomics UK (COG-UK) Consortium | Berkshire and Surrey Pathology Services Lighthouse Laboratory and Alex Alderton, Roberto Amato, Jeffrey Barrett, Sonia Goncalves, Ewan Harrison, David K. Jackson, Ian Johnston, Dominic Kwiatkowski, Cordelia Langford, John Sillitoe on behalf of the Wellcome Sanger Institute COVID-19 Surveillance Team |
| EPI_ISL_2916974, EPI_ISL_2916975 | Lighthouse Lab in Glasgow | Wellcome Sanger Institute for the COVID-19 Genomics UK (COG-UK) Consortium | Harper VanSteenhouse, Yumi Kasai, David Gray, Carol Clugston, Anna Dominiczak and Alex Alderton, Roberto Amato, Jeffrey Barrett, Sonia Goncalves, Ewan Harrison, David K. Jackson, Ian Johnston, Dominic Kwiatkowski, Cordelia Langford, John Sillitoe on behalf of the Wellcome Sanger Institute COVID-19 Surveillance Team |
| EPI_ISL_2916976 | Berkshire and Surrey Pathology Services Lighthouse Laboratory | Wellcome Sanger Institute for the COVID-19 Genomics UK (COG-UK) Consortium | Berkshire and Surrey Pathology Services Lighthouse Laboratory and Alex Alderton, Roberto Amato, Jeffrey Barrett, Sonia Goncalves, Ewan Harrison, David K. Jackson, Ian Johnston, Dominic Kwiatkowski, Cordelia Langford, John Sillitoe on behalf of the Wellcome Sanger Institute COVID-19 Surveillance Team |
| EPI_ISL_2916977, EPI_ISL_2916978 | Lighthouse Lab in Glasgow | Wellcome Sanger Institute for the COVID-19 Genomics UK (COG-UK) Consortium | Harper VanSteenhouse, Yumi Kasai, David Gray, Carol Clugston, Anna Dominiczak and Alex Alderton, Roberto Amato, Jeffrey Barrett, Sonia Goncalves, Ewan Harrison, David K. Jackson, Ian Johnston, Dominic Kwiatkowski, Cordelia Langford, John Sillitoe on behalf of the Wellcome Sanger Institute COVID-19 Surveillance Team |
| EPI_ISL_2916979 | Berkshire and Surrey Pathology Services Lighthouse Laboratory | Wellcome Sanger Institute for the COVID-19 Genomics UK (COG-UK) Consortium | Berkshire and Surrey Pathology Services Lighthouse Laboratory and Alex Alderton, Roberto Amato, Jeffrey Barrett, Sonia Goncalves, Ewan Harrison, David K. Jackson, Ian Johnston, Dominic Kwiatkowski, Cordelia Langford, John Sillitoe on behalf of the Wellcome Sanger Institute COVID-19 Surveillance Team |
| EPI_ISL_2916980 | Lighthouse Lab in Glasgow | Wellcome Sanger Institute for the COVID-19 Genomics UK (COG-UK) Consortium | Harper VanSteenhouse, Yumi Kasai, David Gray, Carol Clugston, Anna Dominiczak and Alex Alderton, Roberto Amato, Jeffrey Barrett, Sonia Goncalves, Ewan Harrison, David K. Jackson, Ian Johnston, Dominic Kwiatkowski, Cordelia Langford, John Sillitoe on behalf of the Wellcome Sanger Institute COVID-19 Surveillance Team |

| | | | |
|---|---|---|---|
| EPI_ISL_2916981 | Berkshire and Surrey Pathology Services Lighthouse Laboratory | Wellcome Sanger Institute for the COVID-19 Genomics UK (COG-UK) Consortium | Berkshire and Surrey Pathology Services Lighthouse Laboratory and Alex Alderton, Roberto Amato, Jeffrey Barrett, Sonia Goncalves, Ewan Harrison, David K. Jackson, Ian Johnston, Dominic Kwiatkowski, Cordelia Langford, John Sillitoe on behalf of the Wellcome Sanger Institute COVID-19 Surveillance Team |
| EPI_ISL_2916982 | Lighthouse Lab in Glasgow | Wellcome Sanger Institute for the COVID-19 Genomics UK (COG-UK) Consortium | Harper VanSteenhouse, Yumi Kasai, David Gray, Carol Clugston, Anna Dominiczak and Alex Alderton, Roberto Amato, Jeffrey Barrett, Sonia Goncalves, Ewan Harrison, David K. Jackson, Ian Johnston, Dominic Kwiatkowski, Cordelia Langford, John Sillitoe on behalf of the Wellcome Sanger Institute COVID-19 Surveillance Team |
| EPI_ISL_2916983, EPI_ISL_2916984, EPI_ISL_2916985, EPI_ISL_2916986 | Berkshire and Surrey Pathology Services Lighthouse Laboratory | Wellcome Sanger Institute for the COVID-19 Genomics UK (COG-UK) Consortium | Berkshire and Surrey Pathology Services Lighthouse Laboratory and Alex Alderton, Roberto Amato, Jeffrey Barrett, Sonia Goncalves, Ewan Harrison, David K. Jackson, Ian Johnston, Dominic Kwiatkowski, Cordelia Langford, John Sillitoe on behalf of the Wellcome Sanger Institute COVID-19 Surveillance Team |
| EPI_ISL_2916987 | Lighthouse Lab in Glasgow | Wellcome Sanger Institute for the COVID-19 Genomics UK (COG-UK) Consortium | Harper VanSteenhouse, Yumi Kasai, David Gray, Carol Clugston, Anna Dominiczak and Alex Alderton, Roberto Amato, Jeffrey Barrett, Sonia Goncalves, Ewan Harrison, David K. Jackson, Ian Johnston, Dominic Kwiatkowski, Cordelia Langford, John Sillitoe on behalf of the Wellcome Sanger Institute COVID-19 Surveillance Team |
| EPI_ISL_2916988, EPI_ISL_2916989, EPI_ISL_2916990, EPI_ISL_2916991, EPI_ISL_2916992 | Berkshire and Surrey Pathology Services Lighthouse Laboratory | Wellcome Sanger Institute for the COVID-19 Genomics UK (COG-UK) Consortium | Berkshire and Surrey Pathology Services Lighthouse Laboratory and Alex Alderton, Roberto Amato, Jeffrey Barrett, Sonia Goncalves, Ewan Harrison, David K. Jackson, Ian Johnston, Dominic Kwiatkowski, Cordelia Langford, John Sillitoe on behalf of the Wellcome Sanger Institute COVID-19 Surveillance Team |
| EPI_ISL_2916993 | Lighthouse Lab in Glasgow | Wellcome Sanger Institute for the COVID-19 Genomics UK (COG-UK) Consortium | Harper VanSteenhouse, Yumi Kasai, David Gray, Carol Clugston, Anna Dominiczak and Alex Alderton, Roberto Amato, Jeffrey Barrett, Sonia Goncalves, Ewan Harrison, David K. Jackson, Ian Johnston, Dominic Kwiatkowski, Cordelia Langford, John Sillitoe on behalf of the Wellcome Sanger Institute COVID-19 Surveillance Team |
| EPI_ISL_2916994 | Berkshire and Surrey Pathology Services Lighthouse Laboratory | Wellcome Sanger Institute for the COVID-19 Genomics UK (COG-UK) Consortium | Berkshire and Surrey Pathology Services Lighthouse Laboratory and Alex Alderton, Roberto Amato, Jeffrey Barrett, Sonia Goncalves, Ewan Harrison, David K. Jackson, Ian Johnston, Dominic Kwiatkowski, Cordelia Langford, John Sillitoe on behalf of the Wellcome Sanger Institute COVID-19 Surveillance Team |
| EPI_ISL_2916995, EPI_ISL_2916996, EPI_ISL_2916997 | Lighthouse Lab in Glasgow | Wellcome Sanger Institute for the COVID-19 Genomics UK (COG-UK) Consortium | Harper VanSteenhouse, Yumi Kasai, David Gray, Carol Clugston, Anna Dominiczak and Alex Alderton, Roberto Amato, Jeffrey Barrett, Sonia Goncalves, Ewan Harrison, David K. Jackson, Ian Johnston, Dominic Kwiatkowski, Cordelia Langford, John Sillitoe on behalf of the Wellcome Sanger Institute COVID-19 Surveillance Team |
| EPI_ISL_2916998, EPI_ISL_2916999, EPI_ISL_2917000, EPI_ISL_2917001, EPI_ISL_2917002, EPI_ISL_2917003, EPI_ISL_2917004 | Berkshire and Surrey Pathology Services Lighthouse Laboratory | Wellcome Sanger Institute for the COVID-19 Genomics UK (COG-UK) Consortium | Berkshire and Surrey Pathology Services Lighthouse Laboratory and Alex Alderton, Roberto Amato, Jeffrey Barrett, Sonia Goncalves, Ewan Harrison, David K. Jackson, Ian Johnston, Dominic Kwiatkowski, Cordelia Langford, John Sillitoe on behalf of the Wellcome Sanger Institute COVID-19 Surveillance Team |
| EPI_ISL_2917005, EPI_ISL_2917006 | Lighthouse Lab in Glasgow | Wellcome Sanger Institute for the COVID-19 Genomics UK (COG-UK) Consortium | Harper VanSteenhouse, Yumi Kasai, David Gray, Carol Clugston, Anna Dominiczak and Alex Alderton, Roberto Amato, Jeffrey Barrett, Sonia Goncalves, Ewan Harrison, David K. Jackson, Ian Johnston, Dominic Kwiatkowski, Cordelia Langford, John Sillitoe on behalf of the Wellcome Sanger Institute COVID-19 Surveillance Team |
| EPI_ISL_2917007 | Berkshire and Surrey Pathology Services Lighthouse Laboratory | Wellcome Sanger Institute for the COVID-19 Genomics UK (COG-UK) Consortium | Berkshire and Surrey Pathology Services Lighthouse Laboratory and Alex Alderton, Roberto Amato, Jeffrey Barrett, Sonia Goncalves, Ewan Harrison, David K. Jackson, Ian Johnston, Dominic Kwiatkowski, Cordelia Langford, John Sillitoe on behalf of the Wellcome Sanger Institute COVID-19 Surveillance Team |
| EPI_ISL_2917008 | Lighthouse Lab in Glasgow | Wellcome Sanger Institute for the COVID-19 Genomics UK (COG-UK) Consortium | Harper VanSteenhouse, Yumi Kasai, David Gray, Carol Clugston, Anna Dominiczak and Alex Alderton, Roberto Amato, Jeffrey Barrett, Sonia Goncalves, Ewan Harrison, David K. Jackson, Ian Johnston, Dominic Kwiatkowski, Cordelia Langford, John Sillitoe on behalf of the Wellcome Sanger Institute COVID-19 Surveillance Team |
| EPI_ISL_2917009 | Berkshire and Surrey Pathology Services Lighthouse Laboratory | Wellcome Sanger Institute for the COVID-19 Genomics UK (COG-UK) Consortium | Berkshire and Surrey Pathology Services Lighthouse Laboratory and Alex Alderton, Roberto Amato, Jeffrey Barrett, Sonia Goncalves, Ewan Harrison, David K. Jackson, Ian Johnston, Dominic Kwiatkowski, Cordelia Langford, John Sillitoe on behalf of the Wellcome Sanger Institute COVID-19 Surveillance Team |
| EPI_ISL_2917010 | Lighthouse Lab in Glasgow | Wellcome Sanger Institute for the COVID-19 Genomics UK (COG-UK) Consortium | Harper VanSteenhouse, Yumi Kasai, David Gray, Carol Clugston, Anna Dominiczak and Alex Alderton, Roberto Amato, Jeffrey Barrett, Sonia Goncalves, Ewan Harrison, David K. Jackson, Ian Johnston, Dominic Kwiatkowski, Cordelia Langford, John Sillitoe on behalf of the Wellcome Sanger Institute COVID-19 Surveillance Team |
| EPI_ISL_2917011, EPI_ISL_2917012, EPI_ISL_2917013 | Berkshire and Surrey Pathology Services Lighthouse Laboratory | Wellcome Sanger Institute for the COVID-19 Genomics UK (COG-UK) Consortium | Berkshire and Surrey Pathology Services Lighthouse Laboratory and Alex Alderton, Roberto Amato, Jeffrey Barrett, Sonia Goncalves, Ewan Harrison, David K. Jackson, Ian Johnston, Dominic Kwiatkowski, Cordelia Langford, John Sillitoe on behalf of the Wellcome Sanger Institute COVID-19 Surveillance Team |
| EPI_ISL_2917014 | Lighthouse Lab in Glasgow | Wellcome Sanger Institute for the COVID-19 Genomics UK (COG-UK) Consortium | Harper VanSteenhouse, Yumi Kasai, David Gray, Carol Clugston, Anna Dominiczak and Alex Alderton, Roberto Amato, Jeffrey Barrett, Sonia Goncalves, Ewan Harrison, David K. Jackson, Ian Johnston, Dominic Kwiatkowski, Cordelia Langford, John Sillitoe on behalf of the Wellcome Sanger Institute COVID-19 Surveillance Team |
| EPI_ISL_2917015, EPI_ISL_2917016 | Berkshire and Surrey Pathology Services Lighthouse Laboratory | Wellcome Sanger Institute for the COVID-19 Genomics UK (COG-UK) Consortium | Berkshire and Surrey Pathology Services Lighthouse Laboratory and Alex Alderton, Roberto Amato, Jeffrey Barrett, Sonia Goncalves, Ewan Harrison, David K. Jackson, Ian Johnston, Dominic Kwiatkowski, Cordelia Langford, John Sillitoe on behalf of the Wellcome Sanger Institute COVID-19 Surveillance Team |
| EPI_ISL_2917017 | Lighthouse Lab in Glasgow | Wellcome Sanger Institute for the COVID-19 Genomics UK (COG-UK) Consortium | Harper VanSteenhouse, Yumi Kasai, David Gray, Carol Clugston, Anna Dominiczak and Alex Alderton, Roberto Amato, Jeffrey Barrett, Sonia Goncalves, Ewan Harrison, David K. Jackson, Ian Johnston, Dominic Kwiatkowski, Cordelia Langford, John Sillitoe on behalf of the Wellcome Sanger Institute COVID-19 Surveillance Team |
| EPI_ISL_2917018 | Berkshire and Surrey Pathology Services Lighthouse Laboratory | Wellcome Sanger Institute for the COVID-19 Genomics UK (COG-UK) Consortium | Berkshire and Surrey Pathology Services Lighthouse Laboratory and Alex Alderton, Roberto Amato, Jeffrey Barrett, Sonia Goncalves, Ewan Harrison, David K. Jackson, Ian Johnston, Dominic Kwiatkowski, Cordelia Langford, John Sillitoe on behalf of the Wellcome Sanger Institute COVID-19 Surveillance Team |
| EPI_ISL_2917019, EPI_ISL_2917020 | Lighthouse Lab in Glasgow | Wellcome Sanger Institute for the COVID-19 Genomics UK (COG-UK) Consortium | Harper VanSteenhouse, Yumi Kasai, David Gray, Carol Clugston, Anna Dominiczak and Alex Alderton, Roberto Amato, Jeffrey Barrett, Sonia Goncalves, Ewan Harrison, David K. Jackson, Ian Johnston, Dominic Kwiatkowski, Cordelia Langford, John Sillitoe on behalf of the Wellcome Sanger Institute COVID-19 Surveillance Team |
| EPI_ISL_2917021, EPI_ISL_2917022, EPI_ISL_2917023, EPI_ISL_2917024, EPI_ISL_2917025 | Berkshire and Surrey Pathology Services Lighthouse Laboratory | Wellcome Sanger Institute for the COVID-19 Genomics UK (COG-UK) Consortium | Berkshire and Surrey Pathology Services Lighthouse Laboratory and Alex Alderton, Roberto Amato, Jeffrey Barrett, Sonia Goncalves, Ewan Harrison, David K. Jackson, Ian Johnston, Dominic Kwiatkowski, Cordelia Langford, John Sillitoe on behalf of the Wellcome Sanger Institute COVID-19 Surveillance Team |
| EPI_ISL_2917026, EPI_ISL_2917027 | Lighthouse Lab in Glasgow | Wellcome Sanger Institute for the COVID-19 Genomics UK (COG-UK) Consortium | Harper VanSteenhouse, Yumi Kasai, David Gray, Carol Clugston, Anna Dominiczak and Alex Alderton, Roberto Amato, Jeffrey Barrett, Sonia Goncalves, Ewan Harrison, David K. Jackson, Ian Johnston, Dominic Kwiatkowski, Cordelia Langford, John Sillitoe on behalf of the Wellcome Sanger Institute COVID-19 Surveillance Team |
| EPI_ISL_2917028 | Berkshire and Surrey Pathology Services Lighthouse Laboratory | Wellcome Sanger Institute for the COVID-19 Genomics UK (COG-UK) Consortium | Berkshire and Surrey Pathology Services Lighthouse Laboratory and Alex Alderton, Roberto Amato, Jeffrey Barrett, Sonia Goncalves, Ewan Harrison, David K. Jackson, Ian Johnston, Dominic Kwiatkowski, Cordelia Langford, John Sillitoe on behalf of the Wellcome Sanger Institute COVID-19 Surveillance Team |
| EPI_ISL_2917029 | Lighthouse Lab in Glasgow | Wellcome Sanger Institute for the COVID-19 Genomics UK (COG-UK) Consortium | Harper VanSteenhouse, Yumi Kasai, David Gray, Carol Clugston, Anna Dominiczak and Alex Alderton, Roberto Amato, Jeffrey Barrett, Sonia Goncalves, Ewan Harrison, David K. Jackson, Ian Johnston, Dominic Kwiatkowski, Cordelia Langford, John Sillitoe on behalf of the Wellcome Sanger Institute COVID-19 Surveillance Team |
| EPI_ISL_2917030 | Berkshire and Surrey Pathology Services Lighthouse Laboratory | Wellcome Sanger Institute for the COVID-19 Genomics UK (COG-UK) Consortium | Berkshire and Surrey Pathology Services Lighthouse Laboratory and Alex Alderton, Roberto Amato, Jeffrey Barrett, Sonia Goncalves, Ewan Harrison, David K. Jackson, Ian Johnston, Dominic Kwiatkowski, Cordelia Langford, John Sillitoe on behalf of the Wellcome Sanger Institute COVID-19 Surveillance Team |
| EPI_ISL_2917031, EPI_ISL_2917032 | Lighthouse Lab in Glasgow | Wellcome Sanger Institute for the COVID-19 Genomics UK (COG-UK) Consortium | Harper VanSteenhouse, Yumi Kasai, David Gray, Carol Clugston, Anna Dominiczak and Alex Alderton, Roberto Amato, Jeffrey Barrett, Sonia Goncalves, Ewan Harrison, David K. Jackson, Ian Johnston, Dominic Kwiatkowski, Cordelia Langford, John Sillitoe on behalf of the Wellcome Sanger Institute COVID-19 Surveillance Team |
| EPI_ISL_2917033 | Berkshire and Surrey Pathology Services Lighthouse Laboratory | Wellcome Sanger Institute for the COVID-19 Genomics UK (COG-UK) Consortium | Berkshire and Surrey Pathology Services Lighthouse Laboratory and Alex Alderton, Roberto Amato, Jeffrey Barrett, Sonia Goncalves, Ewan Harrison, David K. Jackson, Ian Johnston, Dominic Kwiatkowski, Cordelia Langford, John Sillitoe on behalf of the Wellcome Sanger Institute COVID-19 Surveillance Team |
| EPI_ISL_2917034 | Lighthouse Lab in Glasgow | Wellcome Sanger Institute for the COVID-19 Genomics UK (COG-UK) Consortium | Harper VanSteenhouse, Yumi Kasai, David Gray, Carol Clugston, Anna Dominiczak and Alex Alderton, Roberto Amato, Jeffrey Barrett, Sonia Goncalves, Ewan Harrison, David K. Jackson, Ian Johnston, Dominic Kwiatkowski, Cordelia Langford, John Sillitoe on behalf of the Wellcome Sanger Institute COVID-19 Surveillance Team |
| EPI_ISL_2917035 | Berkshire and Surrey Pathology Services Lighthouse Laboratory | Wellcome Sanger Institute for the COVID-19 Genomics UK (COG-UK) Consortium | Berkshire and Surrey Pathology Services Lighthouse Laboratory and Alex Alderton, Roberto Amato, Jeffrey Barrett, Sonia Goncalves, Ewan Harrison, David K. Jackson, Ian Johnston, Dominic Kwiatkowski, Cordelia Langford, John Sillitoe on behalf of the Wellcome Sanger Institute COVID-19 Surveillance Team |
| EPI_ISL_2917036, EPI_ISL_2917037, EPI_ISL_2917038, EPI_ISL_2917039, EPI_ISL_2917040 | Lighthouse Lab in Glasgow | Wellcome Sanger Institute for the COVID-19 Genomics UK (COG-UK) Consortium | Harper VanSteenhouse, Yumi Kasai, David Gray, Carol Clugston, Anna Dominiczak and Alex Alderton, Roberto Amato, Jeffrey Barrett, Sonia Goncalves, Ewan Harrison, David K. Jackson, Ian Johnston, Dominic Kwiatkowski, Cordelia Langford, John Sillitoe on behalf of the Wellcome Sanger Institute COVID-19 Surveillance Team |
| EPI_ISL_2917041 | Berkshire and Surrey Pathology Services Lighthouse Laboratory | Wellcome Sanger Institute for the COVID-19 Genomics UK (COG-UK) Consortium | Berkshire and Surrey Pathology Services Lighthouse Laboratory and Alex Alderton, Roberto Amato, Jeffrey Barrett, Sonia Goncalves, Ewan Harrison, David K. Jackson, Ian Johnston, Dominic Kwiatkowski, Cordelia Langford, John Sillitoe on behalf of the Wellcome Sanger Institute COVID-19 Surveillance Team |
| EPI_ISL_2917042, EPI_ISL_2917043, EPI_ISL_2917044 | Lighthouse Lab in Glasgow | Wellcome Sanger Institute for the COVID-19 Genomics UK (COG-UK) Consortium | Harper VanSteenhouse, Yumi Kasai, David Gray, Carol Clugston, Anna Dominiczak and Alex Alderton, Roberto Amato, Jeffrey Barrett, Sonia Goncalves, Ewan Harrison, David K. Jackson, Ian Johnston, Dominic Kwiatkowski, Cordelia Langford, John Sillitoe on behalf of the Wellcome Sanger Institute COVID-19 Surveillance Team |
| EPI_ISL_2917045, EPI_ISL_2917046, EPI_ISL_2917047 | Berkshire and Surrey Pathology Services Lighthouse Laboratory | Wellcome Sanger Institute for the COVID-19 Genomics UK (COG-UK) Consortium | Berkshire and Surrey Pathology Services Lighthouse Laboratory and Alex Alderton, Roberto Amato, Jeffrey Barrett, Sonia Goncalves, Ewan Harrison, David K. Jackson, Ian Johnston, Dominic Kwiatkowski, Cordelia Langford, John Sillitoe on behalf of the Wellcome Sanger Institute COVID-19 Surveillance Team |
| EPI_ISL_2917048, EPI_ISL_2917049 | Lighthouse Lab in Glasgow | Wellcome Sanger Institute for the COVID-19 Genomics UK (COG-UK) Consortium | Harper VanSteenhouse, Yumi Kasai, David Gray, Carol Clugston, Anna Dominiczak and Alex Alderton, Roberto Amato, Jeffrey Barrett, Sonia Goncalves, Ewan Harrison, David K. Jackson, Ian Johnston, Dominic Kwiatkowski, Cordelia Langford, John Sillitoe on behalf of the Wellcome Sanger Institute COVID-19 Surveillance Team |
| EPI_ISL_2917050 | Berkshire and Surrey Pathology Services Lighthouse Laboratory | Wellcome Sanger Institute for the COVID-19 Genomics UK (COG-UK) Consortium | Berkshire and Surrey Pathology Services Lighthouse Laboratory and Alex Alderton, Roberto Amato, Jeffrey Barrett, Sonia Goncalves, Ewan Harrison, David K. Jackson, Ian Johnston, Dominic Kwiatkowski, Cordelia Langford, John Sillitoe on behalf of the Wellcome Sanger Institute COVID-19 Surveillance Team |
| EPI_ISL_2917051 | Lighthouse Lab in Glasgow | Wellcome Sanger Institute for the COVID-19 Genomics UK (COG-UK) Consortium | Harper VanSteenhouse, Yumi Kasai, David Gray, Carol Clugston, Anna Dominiczak and Alex Alderton, Roberto Amato, Jeffrey Barrett, Sonia Goncalves, Ewan Harrison, David K. Jackson, Ian Johnston, Dominic Kwiatkowski, Cordelia Langford, John Sillitoe on behalf of the Wellcome Sanger Institute COVID-19 Surveillance Team |
| EPI_ISL_2917052 | Berkshire and Surrey Pathology Services Lighthouse Laboratory | Wellcome Sanger Institute for the COVID-19 Genomics UK (COG-UK) Consortium | Berkshire and Surrey Pathology Services Lighthouse Laboratory and Alex Alderton, Roberto Amato, Jeffrey Barrett, Sonia Goncalves, Ewan Harrison, David K. Jackson, Ian Johnston, Dominic Kwiatkowski, Cordelia Langford, John Sillitoe on behalf of the Wellcome Sanger Institute COVID-19 Surveillance Team |

| | | | |
|---|---|---|---|
| EPI_ISL_2917053 | Lighthouse Lab in Glasgow | Wellcome Sanger Institute for the COVID-19 Genomics UK (COG-UK) Consortium | Harper VanSteenhouse, Yumi Kasai, David Gray, Carol Clugston, Anna Dominiczak and Alex Alderton, Roberto Amato, Jeffrey Barrett, Sonia Goncalves, Ewan Harrison, David K. Jackson, Ian Johnston, Dominic Kwiatkowski, Cordelia Langford, John Sillitoe on behalf of the Wellcome Sanger Institute COVID-19 Surveillance Team |
| EPI_ISL_2917054 | Berkshire and Surrey Pathology Services Lighthouse Laboratory | Wellcome Sanger Institute for the COVID-19 Genomics UK (COG-UK) Consortium | Berkshire and Surrey Pathology Services Lighthouse Laboratory and Alex Alderton, Roberto Amato, Jeffrey Barrett, Sonia Goncalves, Ewan Harrison, David K. Jackson, Ian Johnston, Dominic Kwiatkowski, Cordelia Langford, John Sillitoe on behalf of the Wellcome Sanger Institute COVID-19 Surveillance Team |
| EPI_ISL_2917055 | Lighthouse Lab in Glasgow | Wellcome Sanger Institute for the COVID-19 Genomics UK (COG-UK) Consortium | Harper VanSteenhouse, Yumi Kasai, David Gray, Carol Clugston, Anna Dominiczak and Alex Alderton, Roberto Amato, Jeffrey Barrett, Sonia Goncalves, Ewan Harrison, David K. Jackson, Ian Johnston, Dominic Kwiatkowski, Cordelia Langford, John Sillitoe on behalf of the Wellcome Sanger Institute COVID-19 Surveillance Team |
| EPI_ISL_2917056 | Berkshire and Surrey Pathology Services Lighthouse Laboratory | Wellcome Sanger Institute for the COVID-19 Genomics UK (COG-UK) Consortium | Berkshire and Surrey Pathology Services Lighthouse Laboratory and Alex Alderton, Roberto Amato, Jeffrey Barrett, Sonia Goncalves, Ewan Harrison, David K. Jackson, Ian Johnston, Dominic Kwiatkowski, Cordelia Langford, John Sillitoe on behalf of the Wellcome Sanger Institute COVID-19 Surveillance Team |
| EPI_ISL_2917057, EPI_ISL_2917058, EPI_ISL_2917059 | Lighthouse Lab in Glasgow | Wellcome Sanger Institute for the COVID-19 Genomics UK (COG-UK) Consortium | Harper VanSteenhouse, Yumi Kasai, David Gray, Carol Clugston, Anna Dominiczak and Alex Alderton, Roberto Amato, Jeffrey Barrett, Sonia Goncalves, Ewan Harrison, David K. Jackson, Ian Johnston, Dominic Kwiatkowski, Cordelia Langford, John Sillitoe on behalf of the Wellcome Sanger Institute COVID-19 Surveillance Team |
| EPI_ISL_2917060 | Berkshire and Surrey Pathology Services Lighthouse Laboratory | Wellcome Sanger Institute for the COVID-19 Genomics UK (COG-UK) Consortium | Berkshire and Surrey Pathology Services Lighthouse Laboratory and Alex Alderton, Roberto Amato, Jeffrey Barrett, Sonia Goncalves, Ewan Harrison, David K. Jackson, Ian Johnston, Dominic Kwiatkowski, Cordelia Langford, John Sillitoe on behalf of the Wellcome Sanger Institute COVID-19 Surveillance Team |
| EPI_ISL_2917061 | Lighthouse Lab in Glasgow | Wellcome Sanger Institute for the COVID-19 Genomics UK (COG-UK) Consortium | Harper VanSteenhouse, Yumi Kasai, David Gray, Carol Clugston, Anna Dominiczak and Alex Alderton, Roberto Amato, Jeffrey Barrett, Sonia Goncalves, Ewan Harrison, David K. Jackson, Ian Johnston, Dominic Kwiatkowski, Cordelia Langford, John Sillitoe on behalf of the Wellcome Sanger Institute COVID-19 Surveillance Team |
| EPI_ISL_2917062, EPI_ISL_2917063, EPI_ISL_2917064 | Berkshire and Surrey Pathology Services Lighthouse Laboratory | Wellcome Sanger Institute for the COVID-19 Genomics UK (COG-UK) Consortium | Berkshire and Surrey Pathology Services Lighthouse Laboratory and Alex Alderton, Roberto Amato, Jeffrey Barrett, Sonia Goncalves, Ewan Harrison, David K. Jackson, Ian Johnston, Dominic Kwiatkowski, Cordelia Langford, John Sillitoe on behalf of the Wellcome Sanger Institute COVID-19 Surveillance Team |
| EPI_ISL_2917065 | Lighthouse Lab in Glasgow | Wellcome Sanger Institute for the COVID-19 Genomics UK (COG-UK) Consortium | Harper VanSteenhouse, Yumi Kasai, David Gray, Carol Clugston, Anna Dominiczak and Alex Alderton, Roberto Amato, Jeffrey Barrett, Sonia Goncalves, Ewan Harrison, David K. Jackson, Ian Johnston, Dominic Kwiatkowski, Cordelia Langford, John Sillitoe on behalf of the Wellcome Sanger Institute COVID-19 Surveillance Team |
| EPI_ISL_2917066, EPI_ISL_2917067, EPI_ISL_2917068, EPI_ISL_2917069 | Berkshire and Surrey Pathology Services Lighthouse Laboratory | Wellcome Sanger Institute for the COVID-19 Genomics UK (COG-UK) Consortium | Berkshire and Surrey Pathology Services Lighthouse Laboratory and Alex Alderton, Roberto Amato, Jeffrey Barrett, Sonia Goncalves, Ewan Harrison, David K. Jackson, Ian Johnston, Dominic Kwiatkowski, Cordelia Langford, John Sillitoe on behalf of the Wellcome Sanger Institute COVID-19 Surveillance Team |
| EPI_ISL_2917070 | Lighthouse Lab in Glasgow | Wellcome Sanger Institute for the COVID-19 Genomics UK (COG-UK) Consortium | Harper VanSteenhouse, Yumi Kasai, David Gray, Carol Clugston, Anna Dominiczak and Alex Alderton, Roberto Amato, Jeffrey Barrett, Sonia Goncalves, Ewan Harrison, David K. Jackson, Ian Johnston, Dominic Kwiatkowski, Cordelia Langford, John Sillitoe on behalf of the Wellcome Sanger Institute COVID-19 Surveillance Team |
| EPI_ISL_2917071 | Berkshire and Surrey Pathology Services Lighthouse Laboratory | Wellcome Sanger Institute for the COVID-19 Genomics UK (COG-UK) Consortium | Berkshire and Surrey Pathology Services Lighthouse Laboratory and Alex Alderton, Roberto Amato, Jeffrey Barrett, Sonia Goncalves, Ewan Harrison, David K. Jackson, Ian Johnston, Dominic Kwiatkowski, Cordelia Langford, John Sillitoe on behalf of the Wellcome Sanger Institute COVID-19 Surveillance Team |
| EPI_ISL_2917072 | Lighthouse Lab in Glasgow | Wellcome Sanger Institute for the COVID-19 Genomics UK (COG-UK) Consortium | Harper VanSteenhouse, Yumi Kasai, David Gray, Carol Clugston, Anna Dominiczak and Alex Alderton, Roberto Amato, Jeffrey Barrett, Sonia Goncalves, Ewan Harrison, David K. Jackson, Ian Johnston, Dominic Kwiatkowski, Cordelia Langford, John Sillitoe on behalf of the Wellcome Sanger Institute COVID-19 Surveillance Team |
| EPI_ISL_2917073, EPI_ISL_2917074, EPI_ISL_2917075 | Berkshire and Surrey Pathology Services Lighthouse Laboratory | Wellcome Sanger Institute for the COVID-19 Genomics UK (COG-UK) Consortium | Berkshire and Surrey Pathology Services Lighthouse Laboratory and Alex Alderton, Roberto Amato, Jeffrey Barrett, Sonia Goncalves, Ewan Harrison, David K. Jackson, Ian Johnston, Dominic Kwiatkowski, Cordelia Langford, John Sillitoe on behalf of the Wellcome Sanger Institute COVID-19 Surveillance Team |
| EPI_ISL_2917076 | Lighthouse Lab in Glasgow | Wellcome Sanger Institute for the COVID-19 Genomics UK (COG-UK) Consortium | Harper VanSteenhouse, Yumi Kasai, David Gray, Carol Clugston, Anna Dominiczak and Alex Alderton, Roberto Amato, Jeffrey Barrett, Sonia Goncalves, Ewan Harrison, David K. Jackson, Ian Johnston, Dominic Kwiatkowski, Cordelia Langford, John Sillitoe on behalf of the Wellcome Sanger Institute COVID-19 Surveillance Team |
| EPI_ISL_2917078, EPI_ISL_2917079, EPI_ISL_2917080 | Berkshire and Surrey Pathology Services Lighthouse Laboratory | Wellcome Sanger Institute for the COVID-19 Genomics UK (COG-UK) Consortium | Berkshire and Surrey Pathology Services Lighthouse Laboratory and Alex Alderton, Roberto Amato, Jeffrey Barrett, Sonia Goncalves, Ewan Harrison, David K. Jackson, Ian Johnston, Dominic Kwiatkowski, Cordelia Langford, John Sillitoe on behalf of the Wellcome Sanger Institute COVID-19 Surveillance Team |
| EPI_ISL_2917081 | Lighthouse Lab in Glasgow | Wellcome Sanger Institute for the COVID-19 Genomics UK (COG-UK) Consortium | Harper VanSteenhouse, Yumi Kasai, David Gray, Carol Clugston, Anna Dominiczak and Alex Alderton, Roberto Amato, Jeffrey Barrett, Sonia Goncalves, Ewan Harrison, David K. Jackson, Ian Johnston, Dominic Kwiatkowski, Cordelia Langford, John Sillitoe on behalf of the Wellcome Sanger Institute COVID-19 Surveillance Team |
| EPI_ISL_2917082, EPI_ISL_2917083 | Berkshire and Surrey Pathology Services Lighthouse Laboratory | Wellcome Sanger Institute for the COVID-19 Genomics UK (COG-UK) Consortium | Berkshire and Surrey Pathology Services Lighthouse Laboratory and Alex Alderton, Roberto Amato, Jeffrey Barrett, Sonia Goncalves, Ewan Harrison, David K. Jackson, Ian Johnston, Dominic Kwiatkowski, Cordelia Langford, John Sillitoe on behalf of the Wellcome Sanger Institute COVID-19 Surveillance Team |
| EPI_ISL_2917084, EPI_ISL_2917085 | Lighthouse Lab in Glasgow | Wellcome Sanger Institute for the COVID-19 Genomics UK (COG-UK) Consortium | Harper VanSteenhouse, Yumi Kasai, David Gray, Carol Clugston, Anna Dominiczak and Alex Alderton, Roberto Amato, Jeffrey Barrett, Sonia Goncalves, Ewan Harrison, David K. Jackson, Ian Johnston, Dominic Kwiatkowski, Cordelia Langford, John Sillitoe on behalf of the Wellcome Sanger Institute COVID-19 Surveillance Team |
| EPI_ISL_2917086 | Berkshire and Surrey Pathology Services Lighthouse Laboratory | Wellcome Sanger Institute for the COVID-19 Genomics UK (COG-UK) Consortium | Berkshire and Surrey Pathology Services Lighthouse Laboratory and Alex Alderton, Roberto Amato, Jeffrey Barrett, Sonia Goncalves, Ewan Harrison, David K. Jackson, Ian Johnston, Dominic Kwiatkowski, Cordelia Langford, John Sillitoe on behalf of the Wellcome Sanger Institute COVID-19 Surveillance Team |
| EPI_ISL_2917087 | Lighthouse Lab in Glasgow | Wellcome Sanger Institute for the COVID-19 Genomics UK (COG-UK) Consortium | Harper VanSteenhouse, Yumi Kasai, David Gray, Carol Clugston, Anna Dominiczak and Alex Alderton, Roberto Amato, Jeffrey Barrett, Sonia Goncalves, Ewan Harrison, David K. Jackson, Ian Johnston, Dominic Kwiatkowski, Cordelia Langford, John Sillitoe on behalf of the Wellcome Sanger Institute COVID-19 Surveillance Team |
| EPI_ISL_2917088 | Berkshire and Surrey Pathology Services Lighthouse Laboratory | Wellcome Sanger Institute for the COVID-19 Genomics UK (COG-UK) Consortium | Berkshire and Surrey Pathology Services Lighthouse Laboratory and Alex Alderton, Roberto Amato, Jeffrey Barrett, Sonia Goncalves, Ewan Harrison, David K. Jackson, Ian Johnston, Dominic Kwiatkowski, Cordelia Langford, John Sillitoe on behalf of the Wellcome Sanger Institute COVID-19 Surveillance Team |
| EPI_ISL_2917089, EPI_ISL_2917090 | Lighthouse Lab in Glasgow | Wellcome Sanger Institute for the COVID-19 Genomics UK (COG-UK) Consortium | Harper VanSteenhouse, Yumi Kasai, David Gray, Carol Clugston, Anna Dominiczak and Alex Alderton, Roberto Amato, Jeffrey Barrett, Sonia Goncalves, Ewan Harrison, David K. Jackson, Ian Johnston, Dominic Kwiatkowski, Cordelia Langford, John Sillitoe on behalf of the Wellcome Sanger Institute COVID-19 Surveillance Team |
| EPI_ISL_2917091, EPI_ISL_2917092, EPI_ISL_2917093, EPI_ISL_2917094 | Berkshire and Surrey Pathology Services Lighthouse Laboratory | Wellcome Sanger Institute for the COVID-19 Genomics UK (COG-UK) Consortium | Berkshire and Surrey Pathology Services Lighthouse Laboratory and Alex Alderton, Roberto Amato, Jeffrey Barrett, Sonia Goncalves, Ewan Harrison, David K. Jackson, Ian Johnston, Dominic Kwiatkowski, Cordelia Langford, John Sillitoe on behalf of the Wellcome Sanger Institute COVID-19 Surveillance Team |
| EPI_ISL_2917095 | Lighthouse Lab in Glasgow | Wellcome Sanger Institute for the COVID-19 Genomics UK (COG-UK) Consortium | Harper VanSteenhouse, Yumi Kasai, David Gray, Carol Clugston, Anna Dominiczak and Alex Alderton, Roberto Amato, Jeffrey Barrett, Sonia Goncalves, Ewan Harrison, David K. Jackson, Ian Johnston, Dominic Kwiatkowski, Cordelia Langford, John Sillitoe on behalf of the Wellcome Sanger Institute COVID-19 Surveillance Team |
| EPI_ISL_2917096 | Berkshire and Surrey Pathology Services Lighthouse Laboratory | Wellcome Sanger Institute for the COVID-19 Genomics UK (COG-UK) Consortium | Berkshire and Surrey Pathology Services Lighthouse Laboratory and Alex Alderton, Roberto Amato, Jeffrey Barrett, Sonia Goncalves, Ewan Harrison, David K. Jackson, Ian Johnston, Dominic Kwiatkowski, Cordelia Langford, John Sillitoe on behalf of the Wellcome Sanger Institute COVID-19 Surveillance Team |
| EPI_ISL_2917097, EPI_ISL_2917098, EPI_ISL_2917099, EPI_ISL_2917100, EPI_ISL_2917101, EPI_ISL_2917102 | Lighthouse Lab in Glasgow | Wellcome Sanger Institute for the COVID-19 Genomics UK (COG-UK) Consortium | Harper VanSteenhouse, Yumi Kasai, David Gray, Carol Clugston, Anna Dominiczak and Alex Alderton, Roberto Amato, Jeffrey Barrett, Sonia Goncalves, Ewan Harrison, David K. Jackson, Ian Johnston, Dominic Kwiatkowski, Cordelia Langford, John Sillitoe on behalf of the Wellcome Sanger Institute COVID-19 Surveillance Team |
| EPI_ISL_2917103 | Berkshire and Surrey Pathology Services Lighthouse Laboratory | Wellcome Sanger Institute for the COVID-19 Genomics UK (COG-UK) Consortium | Berkshire and Surrey Pathology Services Lighthouse Laboratory and Alex Alderton, Roberto Amato, Jeffrey Barrett, Sonia Goncalves, Ewan Harrison, David K. Jackson, Ian Johnston, Dominic Kwiatkowski, Cordelia Langford, John Sillitoe on behalf of the Wellcome Sanger Institute COVID-19 Surveillance Team |
| EPI_ISL_2917104 | Lighthouse Lab in Glasgow | Wellcome Sanger Institute for the COVID-19 Genomics UK (COG-UK) Consortium | Harper VanSteenhouse, Yumi Kasai, David Gray, Carol Clugston, Anna Dominiczak and Alex Alderton, Roberto Amato, Jeffrey Barrett, Sonia Goncalves, Ewan Harrison, David K. Jackson, Ian Johnston, Dominic Kwiatkowski, Cordelia Langford, John Sillitoe on behalf of the Wellcome Sanger Institute COVID-19 Surveillance Team |
| EPI_ISL_2917105, EPI_ISL_2917106 | Berkshire and Surrey Pathology Services Lighthouse Laboratory | Wellcome Sanger Institute for the COVID-19 Genomics UK (COG-UK) Consortium | Berkshire and Surrey Pathology Services Lighthouse Laboratory and Alex Alderton, Roberto Amato, Jeffrey Barrett, Sonia Goncalves, Ewan Harrison, David K. Jackson, Ian Johnston, Dominic Kwiatkowski, Cordelia Langford, John Sillitoe on behalf of the Wellcome Sanger Institute COVID-19 Surveillance Team |
| EPI_ISL_2917107, EPI_ISL_2917108 | Lighthouse Lab in Glasgow | Wellcome Sanger Institute for the COVID-19 Genomics UK (COG-UK) Consortium | Harper VanSteenhouse, Yumi Kasai, David Gray, Carol Clugston, Anna Dominiczak and Alex Alderton, Roberto Amato, Jeffrey Barrett, Sonia Goncalves, Ewan Harrison, David K. Jackson, Ian Johnston, Dominic Kwiatkowski, Cordelia Langford, John Sillitoe on behalf of the Wellcome Sanger Institute COVID-19 Surveillance Team |
| EPI_ISL_2917109 | Berkshire and Surrey Pathology Services Lighthouse Laboratory | Wellcome Sanger Institute for the COVID-19 Genomics UK (COG-UK) Consortium | Berkshire and Surrey Pathology Services Lighthouse Laboratory and Alex Alderton, Roberto Amato, Jeffrey Barrett, Sonia Goncalves, Ewan Harrison, David K. Jackson, Ian Johnston, Dominic Kwiatkowski, Cordelia Langford, John Sillitoe on behalf of the Wellcome Sanger Institute COVID-19 Surveillance Team |
| EPI_ISL_2917110 | Lighthouse Lab in Glasgow | Wellcome Sanger Institute for the COVID-19 Genomics UK (COG-UK) Consortium | Harper VanSteenhouse, Yumi Kasai, David Gray, Carol Clugston, Anna Dominiczak and Alex Alderton, Roberto Amato, Jeffrey Barrett, Sonia Goncalves, Ewan Harrison, David K. Jackson, Ian Johnston, Dominic Kwiatkowski, Cordelia Langford, John Sillitoe on behalf of the Wellcome Sanger Institute COVID-19 Surveillance Team |
| EPI_ISL_2917111, EPI_ISL_2917112 | Berkshire and Surrey Pathology Services Lighthouse Laboratory | Wellcome Sanger Institute for the COVID-19 Genomics UK (COG-UK) Consortium | Berkshire and Surrey Pathology Services Lighthouse Laboratory and Alex Alderton, Roberto Amato, Jeffrey Barrett, Sonia Goncalves, Ewan Harrison, David K. Jackson, Ian Johnston, Dominic Kwiatkowski, Cordelia Langford, John Sillitoe on behalf of the Wellcome Sanger Institute COVID-19 Surveillance Team |
| EPI_ISL_2917113, EPI_ISL_2917114, EPI_ISL_2917115 | Lighthouse Lab in Glasgow | Wellcome Sanger Institute for the COVID-19 Genomics UK (COG-UK) Consortium | Harper VanSteenhouse, Yumi Kasai, David Gray, Carol Clugston, Anna Dominiczak and Alex Alderton, Roberto Amato, Jeffrey Barrett, Sonia Goncalves, Ewan Harrison, David K. Jackson, Ian Johnston, Dominic Kwiatkowski, Cordelia Langford, John Sillitoe on behalf of the Wellcome Sanger Institute COVID-19 Surveillance Team |
| EPI_ISL_2917116 | Berkshire and Surrey Pathology Services Lighthouse Laboratory | Wellcome Sanger Institute for the COVID-19 Genomics UK (COG-UK) Consortium | Berkshire and Surrey Pathology Services Lighthouse Laboratory and Alex Alderton, Roberto Amato, Jeffrey Barrett, Sonia Goncalves, Ewan Harrison, David K. Jackson, Ian Johnston, Dominic Kwiatkowski, Cordelia Langford, John Sillitoe on behalf of the Wellcome Sanger Institute COVID-19 Surveillance Team |
| EPI_ISL_2917117 | Lighthouse Lab in Glasgow | Wellcome Sanger Institute for the COVID-19 Genomics UK (COG-UK) Consortium | Harper VanSteenhouse, Yumi Kasai, David Gray, Carol Clugston, Anna Dominiczak and Alex Alderton, Roberto Amato, Jeffrey Barrett, Sonia Goncalves, Ewan Harrison, David K. Jackson, Ian Johnston, Dominic Kwiatkowski, Cordelia Langford, John Sillitoe on behalf of the Wellcome Sanger Institute COVID-19 Surveillance Team |
| EPI_ISL_2917118 | Berkshire and Surrey Pathology Services Lighthouse Laboratory | Wellcome Sanger Institute for the COVID-19 Genomics UK (COG-UK) Consortium | Berkshire and Surrey Pathology Services Lighthouse Laboratory and Alex Alderton, Roberto Amato, Jeffrey Barrett, Sonia Goncalves, Ewan Harrison, David K. Jackson, Ian Johnston, Dominic Kwiatkowski, Cordelia Langford, John Sillitoe on behalf of the Wellcome Sanger Institute COVID-19 Surveillance Team |
| EPI_ISL_2917119, EPI_ISL_2917120 | Lighthouse Lab in Glasgow | Wellcome Sanger Institute for the COVID-19 Genomics UK (COG-UK) Consortium | Harper VanSteenhouse, Yumi Kasai, David Gray, Carol Clugston, Anna Dominiczak and Alex Alderton, Roberto Amato, Jeffrey Barrett, Sonia Goncalves, Ewan Harrison, David K. Jackson, Ian Johnston, Dominic Kwiatkowski, Cordelia Langford, John Sillitoe on behalf of the Wellcome Sanger Institute COVID-19 Surveillance Team |
| EPI_ISL_2917121 | Berkshire and Surrey Pathology Services Lighthouse Laboratory | Wellcome Sanger Institute for the COVID-19 Genomics UK (COG-UK) Consortium | Berkshire and Surrey Pathology Services Lighthouse Laboratory and Alex Alderton, Roberto Amato, Jeffrey Barrett, Sonia Goncalves, Ewan Harrison, David K. Jackson, Ian Johnston, Dominic Kwiatkowski, Cordelia Langford, John Sillitoe on behalf of the Wellcome Sanger Institute COVID-19 Surveillance Team |

| | | | |
|---|---|---|---|
| EPI_ISL_2917122 | Lighthouse Lab in Glasgow | Wellcome Sanger Institute for the COVID-19 Genomics UK (COG-UK) Consortium | Harper VanSteenhouse, Yumi Kasai, David Gray, Carol Clugston, Anna Dominiczak and Alex Alderton, Roberto Amato, Jeffrey Barrett, Sonia Goncalves, Ewan Harrison, David K. Jackson, Ian Johnston, Dominic Kwiatkowski, Cordelia Langford, John Sillitoe on behalf of the Wellcome Sanger Institute COVID-19 Surveillance Team |
| EPI_ISL_2917123 | Berkshire and Surrey Pathology Services Lighthouse Laboratory | Wellcome Sanger Institute for the COVID-19 Genomics UK (COG-UK) Consortium | Berkshire and Surrey Pathology Services Lighthouse Laboratory and Alex Alderton, Roberto Amato, Jeffrey Barrett, Sonia Goncalves, Ewan Harrison, David K. Jackson, Ian Johnston, Dominic Kwiatkowski, Cordelia Langford, John Sillitoe on behalf of the Wellcome Sanger Institute COVID-19 Surveillance Team |
| EPI_ISL_2917124, EPI_ISL_2917125, EPI_ISL_2917126, EPI_ISL_2917127, EPI_ISL_2917128 | Lighthouse Lab in Glasgow | Wellcome Sanger Institute for the COVID-19 Genomics UK (COG-UK) Consortium | Harper VanSteenhouse, Yumi Kasai, David Gray, Carol Clugston, Anna Dominiczak and Alex Alderton, Roberto Amato, Jeffrey Barrett, Sonia Goncalves, Ewan Harrison, David K. Jackson, Ian Johnston, Dominic Kwiatkowski, Cordelia Langford, John Sillitoe on behalf of the Wellcome Sanger Institute COVID-19 Surveillance Team |
| EPI_ISL_2917129 | Berkshire and Surrey Pathology Services Lighthouse Laboratory | Wellcome Sanger Institute for the COVID-19 Genomics UK (COG-UK) Consortium | Berkshire and Surrey Pathology Services Lighthouse Laboratory and Alex Alderton, Roberto Amato, Jeffrey Barrett, Sonia Goncalves, Ewan Harrison, David K. Jackson, Ian Johnston, Dominic Kwiatkowski, Cordelia Langford, John Sillitoe on behalf of the Wellcome Sanger Institute COVID-19 Surveillance Team |
| EPI_ISL_2917130, EPI_ISL_2917131, EPI_ISL_2917132 | Lighthouse Lab in Glasgow | Wellcome Sanger Institute for the COVID-19 Genomics UK (COG-UK) Consortium | Harper VanSteenhouse, Yumi Kasai, David Gray, Carol Clugston, Anna Dominiczak and Alex Alderton, Roberto Amato, Jeffrey Barrett, Sonia Goncalves, Ewan Harrison, David K. Jackson, Ian Johnston, Dominic Kwiatkowski, Cordelia Langford, John Sillitoe on behalf of the Wellcome Sanger Institute COVID-19 Surveillance Team |
| EPI_ISL_2917133, EPI_ISL_2917134 | Berkshire and Surrey Pathology Services Lighthouse Laboratory | Wellcome Sanger Institute for the COVID-19 Genomics UK (COG-UK) Consortium | Berkshire and Surrey Pathology Services Lighthouse Laboratory and Alex Alderton, Roberto Amato, Jeffrey Barrett, Sonia Goncalves, Ewan Harrison, David K. Jackson, Ian Johnston, Dominic Kwiatkowski, Cordelia Langford, John Sillitoe on behalf of the Wellcome Sanger Institute COVID-19 Surveillance Team |
| EPI_ISL_2917135, EPI_ISL_2917136, EPI_ISL_2917137 | Lighthouse Lab in Glasgow | Wellcome Sanger Institute for the COVID-19 Genomics UK (COG-UK) Consortium | Harper VanSteenhouse, Yumi Kasai, David Gray, Carol Clugston, Anna Dominiczak and Alex Alderton, Roberto Amato, Jeffrey Barrett, Sonia Goncalves, Ewan Harrison, David K. Jackson, Ian Johnston, Dominic Kwiatkowski, Cordelia Langford, John Sillitoe on behalf of the Wellcome Sanger Institute COVID-19 Surveillance Team |
| EPI_ISL_2917138, EPI_ISL_2917139 | Berkshire and Surrey Pathology Services Lighthouse Laboratory | Wellcome Sanger Institute for the COVID-19 Genomics UK (COG-UK) Consortium | Berkshire and Surrey Pathology Services Lighthouse Laboratory and Alex Alderton, Roberto Amato, Jeffrey Barrett, Sonia Goncalves, Ewan Harrison, David K. Jackson, Ian Johnston, Dominic Kwiatkowski, Cordelia Langford, John Sillitoe on behalf of the Wellcome Sanger Institute COVID-19 Surveillance Team |
| EPI_ISL_2917140 | Lighthouse Lab in Glasgow | Wellcome Sanger Institute for the COVID-19 Genomics UK (COG-UK) Consortium | Harper VanSteenhouse, Yumi Kasai, David Gray, Carol Clugston, Anna Dominiczak and Alex Alderton, Roberto Amato, Jeffrey Barrett, Sonia Goncalves, Ewan Harrison, David K. Jackson, Ian Johnston, Dominic Kwiatkowski, Cordelia Langford, John Sillitoe on behalf of the Wellcome Sanger Institute COVID-19 Surveillance Team |
| EPI_ISL_2917141 | Berkshire and Surrey Pathology Services Lighthouse Laboratory | Wellcome Sanger Institute for the COVID-19 Genomics UK (COG-UK) Consortium | Berkshire and Surrey Pathology Services Lighthouse Laboratory and Alex Alderton, Roberto Amato, Jeffrey Barrett, Sonia Goncalves, Ewan Harrison, David K. Jackson, Ian Johnston, Dominic Kwiatkowski, Cordelia Langford, John Sillitoe on behalf of the Wellcome Sanger Institute COVID-19 Surveillance Team |
| EPI_ISL_2917142 | Lighthouse Lab in Glasgow | Wellcome Sanger Institute for the COVID-19 Genomics UK (COG-UK) Consortium | Harper VanSteenhouse, Yumi Kasai, David Gray, Carol Clugston, Anna Dominiczak and Alex Alderton, Roberto Amato, Jeffrey Barrett, Sonia Goncalves, Ewan Harrison, David K. Jackson, Ian Johnston, Dominic Kwiatkowski, Cordelia Langford, John Sillitoe on behalf of the Wellcome Sanger Institute COVID-19 Surveillance Team |
| EPI_ISL_2917143 | Berkshire and Surrey Pathology Services Lighthouse Laboratory | Wellcome Sanger Institute for the COVID-19 Genomics UK (COG-UK) Consortium | Berkshire and Surrey Pathology Services Lighthouse Laboratory and Alex Alderton, Roberto Amato, Jeffrey Barrett, Sonia Goncalves, Ewan Harrison, David K. Jackson, Ian Johnston, Dominic Kwiatkowski, Cordelia Langford, John Sillitoe on behalf of the Wellcome Sanger Institute COVID-19 Surveillance Team |
| EPI_ISL_2917144 | Lighthouse Lab in Glasgow | Wellcome Sanger Institute for the COVID-19 Genomics UK (COG-UK) Consortium | Harper VanSteenhouse, Yumi Kasai, David Gray, Carol Clugston, Anna Dominiczak and Alex Alderton, Roberto Amato, Jeffrey Barrett, Sonia Goncalves, Ewan Harrison, David K. Jackson, Ian Johnston, Dominic Kwiatkowski, Cordelia Langford, John Sillitoe on behalf of the Wellcome Sanger Institute COVID-19 Surveillance Team |
| EPI_ISL_2917145, EPI_ISL_2917146, EPI_ISL_2917147, EPI_ISL_2917148 | Berkshire and Surrey Pathology Services Lighthouse Laboratory | Wellcome Sanger Institute for the COVID-19 Genomics UK (COG-UK) Consortium | Berkshire and Surrey Pathology Services Lighthouse Laboratory and Alex Alderton, Roberto Amato, Jeffrey Barrett, Sonia Goncalves, Ewan Harrison, David K. Jackson, Ian Johnston, Dominic Kwiatkowski, Cordelia Langford, John Sillitoe on behalf of the Wellcome Sanger Institute COVID-19 Surveillance Team |
| EPI_ISL_2917149, EPI_ISL_2917150 | Lighthouse Lab in Glasgow | Wellcome Sanger Institute for the COVID-19 Genomics UK (COG-UK) Consortium | Harper VanSteenhouse, Yumi Kasai, David Gray, Carol Clugston, Anna Dominiczak and Alex Alderton, Roberto Amato, Jeffrey Barrett, Sonia Goncalves, Ewan Harrison, David K. Jackson, Ian Johnston, Dominic Kwiatkowski, Cordelia Langford, John Sillitoe on behalf of the Wellcome Sanger Institute COVID-19 Surveillance Team |
| EPI_ISL_2917151, EPI_ISL_2917153, EPI_ISL_2917154, EPI_ISL_2917155, EPI_ISL_2917156 | Berkshire and Surrey Pathology Services Lighthouse Laboratory | Wellcome Sanger Institute for the COVID-19 Genomics UK (COG-UK) Consortium | Berkshire and Surrey Pathology Services Lighthouse Laboratory and Alex Alderton, Roberto Amato, Jeffrey Barrett, Sonia Goncalves, Ewan Harrison, David K. Jackson, Ian Johnston, Dominic Kwiatkowski, Cordelia Langford, John Sillitoe on behalf of the Wellcome Sanger Institute COVID-19 Surveillance Team |
| EPI_ISL_2917157 | Lighthouse Lab in Glasgow | Wellcome Sanger Institute for the COVID-19 Genomics UK (COG-UK) Consortium | Harper VanSteenhouse, Yumi Kasai, David Gray, Carol Clugston, Anna Dominiczak and Alex Alderton, Roberto Amato, Jeffrey Barrett, Sonia Goncalves, Ewan Harrison, David K. Jackson, Ian Johnston, Dominic Kwiatkowski, Cordelia Langford, John Sillitoe on behalf of the Wellcome Sanger Institute COVID-19 Surveillance Team |
| EPI_ISL_2917158, EPI_ISL_2917159 | Berkshire and Surrey Pathology Services Lighthouse Laboratory | Wellcome Sanger Institute for the COVID-19 Genomics UK (COG-UK) Consortium | Berkshire and Surrey Pathology Services Lighthouse Laboratory and Alex Alderton, Roberto Amato, Jeffrey Barrett, Sonia Goncalves, Ewan Harrison, David K. Jackson, Ian Johnston, Dominic Kwiatkowski, Cordelia Langford, John Sillitoe on behalf of the Wellcome Sanger Institute COVID-19 Surveillance Team |
| EPI_ISL_2917160 | Lighthouse Lab in Glasgow | Wellcome Sanger Institute for the COVID-19 Genomics UK (COG-UK) Consortium | Harper VanSteenhouse, Yumi Kasai, David Gray, Carol Clugston, Anna Dominiczak and Alex Alderton, Roberto Amato, Jeffrey Barrett, Sonia Goncalves, Ewan Harrison, David K. Jackson, Ian Johnston, Dominic Kwiatkowski, Cordelia Langford, John Sillitoe on behalf of the Wellcome Sanger Institute COVID-19 Surveillance Team |
| EPI_ISL_2917161, EPI_ISL_2917162, EPI_ISL_2917163, EPI_ISL_2917164 | Berkshire and Surrey Pathology Services Lighthouse Laboratory | Wellcome Sanger Institute for the COVID-19 Genomics UK (COG-UK) Consortium | Berkshire and Surrey Pathology Services Lighthouse Laboratory and Alex Alderton, Roberto Amato, Jeffrey Barrett, Sonia Goncalves, Ewan Harrison, David K. Jackson, Ian Johnston, Dominic Kwiatkowski, Cordelia Langford, John Sillitoe on behalf of the Wellcome Sanger Institute COVID-19 Surveillance Team |
| EPI_ISL_2917165, EPI_ISL_2917166 | Lighthouse Lab in Glasgow | Wellcome Sanger Institute for the COVID-19 Genomics UK (COG-UK) Consortium | Harper VanSteenhouse, Yumi Kasai, David Gray, Carol Clugston, Anna Dominiczak and Alex Alderton, Roberto Amato, Jeffrey Barrett, Sonia Goncalves, Ewan Harrison, David K. Jackson, Ian Johnston, Dominic Kwiatkowski, Cordelia Langford, John Sillitoe on behalf of the Wellcome Sanger Institute COVID-19 Surveillance Team |
| EPI_ISL_2917167, EPI_ISL_2917168 | Berkshire and Surrey Pathology Services Lighthouse Laboratory | Wellcome Sanger Institute for the COVID-19 Genomics UK (COG-UK) Consortium | Berkshire and Surrey Pathology Services Lighthouse Laboratory and Alex Alderton, Roberto Amato, Jeffrey Barrett, Sonia Goncalves, Ewan Harrison, David K. Jackson, Ian Johnston, Dominic Kwiatkowski, Cordelia Langford, John Sillitoe on behalf of the Wellcome Sanger Institute COVID-19 Surveillance Team |
| EPI_ISL_2917169 | Lighthouse Lab in Glasgow | Wellcome Sanger Institute for the COVID-19 Genomics UK (COG-UK) Consortium | Harper VanSteenhouse, Yumi Kasai, David Gray, Carol Clugston, Anna Dominiczak and Alex Alderton, Roberto Amato, Jeffrey Barrett, Sonia Goncalves, Ewan Harrison, David K. Jackson, Ian Johnston, Dominic Kwiatkowski, Cordelia Langford, John Sillitoe on behalf of the Wellcome Sanger Institute COVID-19 Surveillance Team |
| EPI_ISL_2917170 | Berkshire and Surrey Pathology Services Lighthouse Laboratory | Wellcome Sanger Institute for the COVID-19 Genomics UK (COG-UK) Consortium | Berkshire and Surrey Pathology Services Lighthouse Laboratory and Alex Alderton, Roberto Amato, Jeffrey Barrett, Sonia Goncalves, Ewan Harrison, David K. Jackson, Ian Johnston, Dominic Kwiatkowski, Cordelia Langford, John Sillitoe on behalf of the Wellcome Sanger Institute COVID-19 Surveillance Team |
| EPI_ISL_2917171 | Lighthouse Lab in Glasgow | Wellcome Sanger Institute for the COVID-19 Genomics UK (COG-UK) Consortium | Harper VanSteenhouse, Yumi Kasai, David Gray, Carol Clugston, Anna Dominiczak and Alex Alderton, Roberto Amato, Jeffrey Barrett, Sonia Goncalves, Ewan Harrison, David K. Jackson, Ian Johnston, Dominic Kwiatkowski, Cordelia Langford, John Sillitoe on behalf of the Wellcome Sanger Institute COVID-19 Surveillance Team |
| EPI_ISL_2917172, EPI_ISL_2917173 | Berkshire and Surrey Pathology Services Lighthouse Laboratory | Wellcome Sanger Institute for the COVID-19 Genomics UK (COG-UK) Consortium | Berkshire and Surrey Pathology Services Lighthouse Laboratory and Alex Alderton, Roberto Amato, Jeffrey Barrett, Sonia Goncalves, Ewan Harrison, David K. Jackson, Ian Johnston, Dominic Kwiatkowski, Cordelia Langford, John Sillitoe on behalf of the Wellcome Sanger Institute COVID-19 Surveillance Team |
| EPI_ISL_2917174, EPI_ISL_2917175, EPI_ISL_2917176, EPI_ISL_2917177, EPI_ISL_2917178, EPI_ISL_2917179 | Lighthouse Lab in Glasgow | Wellcome Sanger Institute for the COVID-19 Genomics UK (COG-UK) Consortium | Harper VanSteenhouse, Yumi Kasai, David Gray, Carol Clugston, Anna Dominiczak and Alex Alderton, Roberto Amato, Jeffrey Barrett, Sonia Goncalves, Ewan Harrison, David K. Jackson, Ian Johnston, Dominic Kwiatkowski, Cordelia Langford, John Sillitoe on behalf of the Wellcome Sanger Institute COVID-19 Surveillance Team |
| EPI_ISL_2917180, EPI_ISL_2917181, EPI_ISL_2917182, EPI_ISL_2917183, EPI_ISL_2917184 | Berkshire and Surrey Pathology Services Lighthouse Laboratory | Wellcome Sanger Institute for the COVID-19 Genomics UK (COG-UK) Consortium | Berkshire and Surrey Pathology Services Lighthouse Laboratory and Alex Alderton, Roberto Amato, Jeffrey Barrett, Sonia Goncalves, Ewan Harrison, David K. Jackson, Ian Johnston, Dominic Kwiatkowski, Cordelia Langford, John Sillitoe on behalf of the Wellcome Sanger Institute COVID-19 Surveillance Team |
| EPI_ISL_2917185, EPI_ISL_2917186, EPI_ISL_2917187 | Lighthouse Lab in Glasgow | Wellcome Sanger Institute for the COVID-19 Genomics UK (COG-UK) Consortium | Harper VanSteenhouse, Yumi Kasai, David Gray, Carol Clugston, Anna Dominiczak and Alex Alderton, Roberto Amato, Jeffrey Barrett, Sonia Goncalves, Ewan Harrison, David K. Jackson, Ian Johnston, Dominic Kwiatkowski, Cordelia Langford, John Sillitoe on behalf of the Wellcome Sanger Institute COVID-19 Surveillance Team |
| EPI_ISL_2917188, EPI_ISL_2917189 | Berkshire and Surrey Pathology Services Lighthouse Laboratory | Wellcome Sanger Institute for the COVID-19 Genomics UK (COG-UK) Consortium | Berkshire and Surrey Pathology Services Lighthouse Laboratory and Alex Alderton, Roberto Amato, Jeffrey Barrett, Sonia Goncalves, Ewan Harrison, David K. Jackson, Ian Johnston, Dominic Kwiatkowski, Cordelia Langford, John Sillitoe on behalf of the Wellcome Sanger Institute COVID-19 Surveillance Team |
| EPI_ISL_2917190 | Lighthouse Lab in Glasgow | Wellcome Sanger Institute for the COVID-19 Genomics UK (COG-UK) Consortium | Harper VanSteenhouse, Yumi Kasai, David Gray, Carol Clugston, Anna Dominiczak and Alex Alderton, Roberto Amato, Jeffrey Barrett, Sonia Goncalves, Ewan Harrison, David K. Jackson, Ian Johnston, Dominic Kwiatkowski, Cordelia Langford, John Sillitoe on behalf of the Wellcome Sanger Institute COVID-19 Surveillance Team |
| EPI_ISL_2917191 | Berkshire and Surrey Pathology Services Lighthouse Laboratory | Wellcome Sanger Institute for the COVID-19 Genomics UK (COG-UK) Consortium | Berkshire and Surrey Pathology Services Lighthouse Laboratory and Alex Alderton, Roberto Amato, Jeffrey Barrett, Sonia Goncalves, Ewan Harrison, David K. Jackson, Ian Johnston, Dominic Kwiatkowski, Cordelia Langford, John Sillitoe on behalf of the Wellcome Sanger Institute COVID-19 Surveillance Team |
| EPI_ISL_2917192 | Lighthouse Lab in Glasgow | Wellcome Sanger Institute for the COVID-19 Genomics UK (COG-UK) Consortium | Harper VanSteenhouse, Yumi Kasai, David Gray, Carol Clugston, Anna Dominiczak and Alex Alderton, Roberto Amato, Jeffrey Barrett, Sonia Goncalves, Ewan Harrison, David K. Jackson, Ian Johnston, Dominic Kwiatkowski, Cordelia Langford, John Sillitoe on behalf of the Wellcome Sanger Institute COVID-19 Surveillance Team |
| EPI_ISL_2917193, EPI_ISL_2917194, EPI_ISL_2917195, EPI_ISL_2917196 | Berkshire and Surrey Pathology Services Lighthouse Laboratory | Wellcome Sanger Institute for the COVID-19 Genomics UK (COG-UK) Consortium | Berkshire and Surrey Pathology Services Lighthouse Laboratory and Alex Alderton, Roberto Amato, Jeffrey Barrett, Sonia Goncalves, Ewan Harrison, David K. Jackson, Ian Johnston, Dominic Kwiatkowski, Cordelia Langford, John Sillitoe on behalf of the Wellcome Sanger Institute COVID-19 Surveillance Team |
| EPI_ISL_2917197 | Lighthouse Lab in Glasgow | Wellcome Sanger Institute for the COVID-19 Genomics UK (COG-UK) Consortium | Harper VanSteenhouse, Yumi Kasai, David Gray, Carol Clugston, Anna Dominiczak and Alex Alderton, Roberto Amato, Jeffrey Barrett, Sonia Goncalves, Ewan Harrison, David K. Jackson, Ian Johnston, Dominic Kwiatkowski, Cordelia Langford, John Sillitoe on behalf of the Wellcome Sanger Institute COVID-19 Surveillance Team |
| EPI_ISL_2917198 | Berkshire and Surrey Pathology Services Lighthouse Laboratory | Wellcome Sanger Institute for the COVID-19 Genomics UK (COG-UK) Consortium | Berkshire and Surrey Pathology Services Lighthouse Laboratory and Alex Alderton, Roberto Amato, Jeffrey Barrett, Sonia Goncalves, Ewan Harrison, David K. Jackson, Ian Johnston, Dominic Kwiatkowski, Cordelia Langford, John Sillitoe on behalf of the Wellcome Sanger Institute COVID-19 Surveillance Team |

| | | | |
|---|---|---|---|
| EPI_ISL_2917199, EPI_ISL_2917200, EPI_ISL_2917201 | Lighthouse Lab in Glasgow | Wellcome Sanger Institute for the COVID-19 Genomics UK (COG-UK) Consortium | Harper VanSteenhouse, Yumi Kasai, David Gray, Carol Clugston, Anna Dominiczak and Alex Alderton, Roberto Amato, Jeffrey Barrett, Sonia Goncalves, Ewan Harrison, David K. Jackson, Ian Johnston, Dominic Kwiatkowski, Cordelia Langford, John Sillitoe on behalf of the Wellcome Sanger Institute COVID-19 Surveillance Team |
| EPI_ISL_2917202, EPI_ISL_2917203, EPI_ISL_2917204 | Berkshire and Surrey Pathology Services Lighthouse Laboratory | Wellcome Sanger Institute for the COVID-19 Genomics UK (COG-UK) Consortium | Berkshire and Surrey Pathology Services Lighthouse Laboratory and Alex Alderton, Roberto Amato, Jeffrey Barrett, Sonia Goncalves, Ewan Harrison, David K. Jackson, Ian Johnston, Dominic Kwiatkowski, Cordelia Langford, John Sillitoe on behalf of the Wellcome Sanger Institute COVID-19 Surveillance Team |
| EPI_ISL_2917205, EPI_ISL_2917206, EPI_ISL_2917207, EPI_ISL_2917208, EPI_ISL_2917209 | Lighthouse Lab in Glasgow | Wellcome Sanger Institute for the COVID-19 Genomics UK (COG-UK) Consortium | Harper VanSteenhouse, Yumi Kasai, David Gray, Carol Clugston, Anna Dominiczak and Alex Alderton, Roberto Amato, Jeffrey Barrett, Sonia Goncalves, Ewan Harrison, David K. Jackson, Ian Johnston, Dominic Kwiatkowski, Cordelia Langford, John Sillitoe on behalf of the Wellcome Sanger Institute COVID-19 Surveillance Team |
| EPI_ISL_2917210, EPI_ISL_2917211 | Berkshire and Surrey Pathology Services Lighthouse Laboratory | Wellcome Sanger Institute for the COVID-19 Genomics UK (COG-UK) Consortium | Berkshire and Surrey Pathology Services Lighthouse Laboratory and Alex Alderton, Roberto Amato, Jeffrey Barrett, Sonia Goncalves, Ewan Harrison, David K. Jackson, Ian Johnston, Dominic Kwiatkowski, Cordelia Langford, John Sillitoe on behalf of the Wellcome Sanger Institute COVID-19 Surveillance Team |
| EPI_ISL_2917212 | Lighthouse Lab in Glasgow | Wellcome Sanger Institute for the COVID-19 Genomics UK (COG-UK) Consortium | Harper VanSteenhouse, Yumi Kasai, David Gray, Carol Clugston, Anna Dominiczak and Alex Alderton, Roberto Amato, Jeffrey Barrett, Sonia Goncalves, Ewan Harrison, David K. Jackson, Ian Johnston, Dominic Kwiatkowski, Cordelia Langford, John Sillitoe on behalf of the Wellcome Sanger Institute COVID-19 Surveillance Team |
| EPI_ISL_2917213 | Berkshire and Surrey Pathology Services Lighthouse Laboratory | Wellcome Sanger Institute for the COVID-19 Genomics UK (COG-UK) Consortium | Berkshire and Surrey Pathology Services Lighthouse Laboratory and Alex Alderton, Roberto Amato, Jeffrey Barrett, Sonia Goncalves, Ewan Harrison, David K. Jackson, Ian Johnston, Dominic Kwiatkowski, Cordelia Langford, John Sillitoe on behalf of the Wellcome Sanger Institute COVID-19 Surveillance Team |
| EPI_ISL_2917214, EPI_ISL_2917215, EPI_ISL_2917216, EPI_ISL_2917217 | Lighthouse Lab in Glasgow | Wellcome Sanger Institute for the COVID-19 Genomics UK (COG-UK) Consortium | Harper VanSteenhouse, Yumi Kasai, David Gray, Carol Clugston, Anna Dominiczak and Alex Alderton, Roberto Amato, Jeffrey Barrett, Sonia Goncalves, Ewan Harrison, David K. Jackson, Ian Johnston, Dominic Kwiatkowski, Cordelia Langford, John Sillitoe on behalf of the Wellcome Sanger Institute COVID-19 Surveillance Team |
| EPI_ISL_2917218 | Berkshire and Surrey Pathology Services Lighthouse Laboratory | Wellcome Sanger Institute for the COVID-19 Genomics UK (COG-UK) Consortium | Berkshire and Surrey Pathology Services Lighthouse Laboratory and Alex Alderton, Roberto Amato, Jeffrey Barrett, Sonia Goncalves, Ewan Harrison, David K. Jackson, Ian Johnston, Dominic Kwiatkowski, Cordelia Langford, John Sillitoe on behalf of the Wellcome Sanger Institute COVID-19 Surveillance Team |
| EPI_ISL_2917219, EPI_ISL_2917220 | Lighthouse Lab in Glasgow | Wellcome Sanger Institute for the COVID-19 Genomics UK (COG-UK) Consortium | Harper VanSteenhouse, Yumi Kasai, David Gray, Carol Clugston, Anna Dominiczak and Alex Alderton, Roberto Amato, Jeffrey Barrett, Sonia Goncalves, Ewan Harrison, David K. Jackson, Ian Johnston, Dominic Kwiatkowski, Cordelia Langford, John Sillitoe on behalf of the Wellcome Sanger Institute COVID-19 Surveillance Team |
| EPI_ISL_2917221, EPI_ISL_2917222 | Berkshire and Surrey Pathology Services Lighthouse Laboratory | Wellcome Sanger Institute for the COVID-19 Genomics UK (COG-UK) Consortium | Berkshire and Surrey Pathology Services Lighthouse Laboratory and Alex Alderton, Roberto Amato, Jeffrey Barrett, Sonia Goncalves, Ewan Harrison, David K. Jackson, Ian Johnston, Dominic Kwiatkowski, Cordelia Langford, John Sillitoe on behalf of the Wellcome Sanger Institute COVID-19 Surveillance Team |
| EPI_ISL_2917223, EPI_ISL_2917224 | Lighthouse Lab in Glasgow | Wellcome Sanger Institute for the COVID-19 Genomics UK (COG-UK) Consortium | Harper VanSteenhouse, Yumi Kasai, David Gray, Carol Clugston, Anna Dominiczak and Alex Alderton, Roberto Amato, Jeffrey Barrett, Sonia Goncalves, Ewan Harrison, David K. Jackson, Ian Johnston, Dominic Kwiatkowski, Cordelia Langford, John Sillitoe on behalf of the Wellcome Sanger Institute COVID-19 Surveillance Team |
| EPI_ISL_2917225 | Berkshire and Surrey Pathology Services Lighthouse Laboratory | Wellcome Sanger Institute for the COVID-19 Genomics UK (COG-UK) Consortium | Berkshire and Surrey Pathology Services Lighthouse Laboratory and Alex Alderton, Roberto Amato, Jeffrey Barrett, Sonia Goncalves, Ewan Harrison, David K. Jackson, Ian Johnston, Dominic Kwiatkowski, Cordelia Langford, John Sillitoe on behalf of the Wellcome Sanger Institute COVID-19 Surveillance Team |
| EPI_ISL_2917226, EPI_ISL_2917227 | Lighthouse Lab in Glasgow | Wellcome Sanger Institute for the COVID-19 Genomics UK (COG-UK) Consortium | Harper VanSteenhouse, Yumi Kasai, David Gray, Carol Clugston, Anna Dominiczak and Alex Alderton, Roberto Amato, Jeffrey Barrett, Sonia Goncalves, Ewan Harrison, David K. Jackson, Ian Johnston, Dominic Kwiatkowski, Cordelia Langford, John Sillitoe on behalf of the Wellcome Sanger Institute COVID-19 Surveillance Team |
| EPI_ISL_2917228, EPI_ISL_2917229 | Berkshire and Surrey Pathology Services Lighthouse Laboratory | Wellcome Sanger Institute for the COVID-19 Genomics UK (COG-UK) Consortium | Berkshire and Surrey Pathology Services Lighthouse Laboratory and Alex Alderton, Roberto Amato, Jeffrey Barrett, Sonia Goncalves, Ewan Harrison, David K. Jackson, Ian Johnston, Dominic Kwiatkowski, Cordelia Langford, John Sillitoe on behalf of the Wellcome Sanger Institute COVID-19 Surveillance Team |
| EPI_ISL_2917230, EPI_ISL_2917231 | Lighthouse Lab in Glasgow | Wellcome Sanger Institute for the COVID-19 Genomics UK (COG-UK) Consortium | Harper VanSteenhouse, Yumi Kasai, David Gray, Carol Clugston, Anna Dominiczak and Alex Alderton, Roberto Amato, Jeffrey Barrett, Sonia Goncalves, Ewan Harrison, David K. Jackson, Ian Johnston, Dominic Kwiatkowski, Cordelia Langford, John Sillitoe on behalf of the Wellcome Sanger Institute COVID-19 Surveillance Team |
| EPI_ISL_2917232 | Berkshire and Surrey Pathology Services Lighthouse Laboratory | Wellcome Sanger Institute for the COVID-19 Genomics UK (COG-UK) Consortium | Berkshire and Surrey Pathology Services Lighthouse Laboratory and Alex Alderton, Roberto Amato, Jeffrey Barrett, Sonia Goncalves, Ewan Harrison, David K. Jackson, Ian Johnston, Dominic Kwiatkowski, Cordelia Langford, John Sillitoe on behalf of the Wellcome Sanger Institute COVID-19 Surveillance Team |
| EPI_ISL_2917233 | Lighthouse Lab in Glasgow | Wellcome Sanger Institute for the COVID-19 Genomics UK (COG-UK) Consortium | Harper VanSteenhouse, Yumi Kasai, David Gray, Carol Clugston, Anna Dominiczak and Alex Alderton, Roberto Amato, Jeffrey Barrett, Sonia Goncalves, Ewan Harrison, David K. Jackson, Ian Johnston, Dominic Kwiatkowski, Cordelia Langford, John Sillitoe on behalf of the Wellcome Sanger Institute COVID-19 Surveillance Team |
| EPI_ISL_2917234 | Berkshire and Surrey Pathology Services Lighthouse Laboratory | Wellcome Sanger Institute for the COVID-19 Genomics UK (COG-UK) Consortium | Berkshire and Surrey Pathology Services Lighthouse Laboratory and Alex Alderton, Roberto Amato, Jeffrey Barrett, Sonia Goncalves, Ewan Harrison, David K. Jackson, Ian Johnston, Dominic Kwiatkowski, Cordelia Langford, John Sillitoe on behalf of the Wellcome Sanger Institute COVID-19 Surveillance Team |
| EPI_ISL_2917235 | Lighthouse Lab in Glasgow | Wellcome Sanger Institute for the COVID-19 Genomics UK (COG-UK) Consortium | Harper VanSteenhouse, Yumi Kasai, David Gray, Carol Clugston, Anna Dominiczak and Alex Alderton, Roberto Amato, Jeffrey Barrett, Sonia Goncalves, Ewan Harrison, David K. Jackson, Ian Johnston, Dominic Kwiatkowski, Cordelia Langford, John Sillitoe on behalf of the Wellcome Sanger Institute COVID-19 Surveillance Team |
| EPI_ISL_2917236 | Berkshire and Surrey Pathology Services Lighthouse Laboratory | Wellcome Sanger Institute for the COVID-19 Genomics UK (COG-UK) Consortium | Berkshire and Surrey Pathology Services Lighthouse Laboratory and Alex Alderton, Roberto Amato, Jeffrey Barrett, Sonia Goncalves, Ewan Harrison, David K. Jackson, Ian Johnston, Dominic Kwiatkowski, Cordelia Langford, John Sillitoe on behalf of the Wellcome Sanger Institute COVID-19 Surveillance Team |
| EPI_ISL_2917237, EPI_ISL_2917238 | Lighthouse Lab in Glasgow | Wellcome Sanger Institute for the COVID-19 Genomics UK (COG-UK) Consortium | Harper VanSteenhouse, Yumi Kasai, David Gray, Carol Clugston, Anna Dominiczak and Alex Alderton, Roberto Amato, Jeffrey Barrett, Sonia Goncalves, Ewan Harrison, David K. Jackson, Ian Johnston, Dominic Kwiatkowski, Cordelia Langford, John Sillitoe on behalf of the Wellcome Sanger Institute COVID-19 Surveillance Team |
| EPI_ISL_2917239 | Berkshire and Surrey Pathology Services Lighthouse Laboratory | Wellcome Sanger Institute for the COVID-19 Genomics UK (COG-UK) Consortium | Berkshire and Surrey Pathology Services Lighthouse Laboratory and Alex Alderton, Roberto Amato, Jeffrey Barrett, Sonia Goncalves, Ewan Harrison, David K. Jackson, Ian Johnston, Dominic Kwiatkowski, Cordelia Langford, John Sillitoe on behalf of the Wellcome Sanger Institute COVID-19 Surveillance Team |
| EPI_ISL_2917240, EPI_ISL_2917241 | Lighthouse Lab in Glasgow | Wellcome Sanger Institute for the COVID-19 Genomics UK (COG-UK) Consortium | Harper VanSteenhouse, Yumi Kasai, David Gray, Carol Clugston, Anna Dominiczak and Alex Alderton, Roberto Amato, Jeffrey Barrett, Sonia Goncalves, Ewan Harrison, David K. Jackson, Ian Johnston, Dominic Kwiatkowski, Cordelia Langford, John Sillitoe on behalf of the Wellcome Sanger Institute COVID-19 Surveillance Team |
| EPI_ISL_2917242, EPI_ISL_2917243, EPI_ISL_2917244, EPI_ISL_2917245, EPI_ISL_2917246, EPI_ISL_2917247, EPI_ISL_2917248 | Berkshire and Surrey Pathology Services Lighthouse Laboratory | Wellcome Sanger Institute for the COVID-19 Genomics UK (COG-UK) Consortium | Berkshire and Surrey Pathology Services Lighthouse Laboratory and Alex Alderton, Roberto Amato, Jeffrey Barrett, Sonia Goncalves, Ewan Harrison, David K. Jackson, Ian Johnston, Dominic Kwiatkowski, Cordelia Langford, John Sillitoe on behalf of the Wellcome Sanger Institute COVID-19 Surveillance Team |
| EPI_ISL_2917249, EPI_ISL_2917250, EPI_ISL_2917251, EPI_ISL_2917252, EPI_ISL_2917253, EPI_ISL_2917254, EPI_ISL_2917255 | Lighthouse Lab in Glasgow | Wellcome Sanger Institute for the COVID-19 Genomics UK (COG-UK) Consortium | Harper VanSteenhouse, Yumi Kasai, David Gray, Carol Clugston, Anna Dominiczak and Alex Alderton, Roberto Amato, Jeffrey Barrett, Sonia Goncalves, Ewan Harrison, David K. Jackson, Ian Johnston, Dominic Kwiatkowski, Cordelia Langford, John Sillitoe on behalf of the Wellcome Sanger Institute COVID-19 Surveillance Team |
| EPI_ISL_2917256 | Berkshire and Surrey Pathology Services Lighthouse Laboratory | Wellcome Sanger Institute for the COVID-19 Genomics UK (COG-UK) Consortium | Berkshire and Surrey Pathology Services Lighthouse Laboratory and Alex Alderton, Roberto Amato, Jeffrey Barrett, Sonia Goncalves, Ewan Harrison, David K. Jackson, Ian Johnston, Dominic Kwiatkowski, Cordelia Langford, John Sillitoe on behalf of the Wellcome Sanger Institute COVID-19 Surveillance Team |
| EPI_ISL_2917257 | Lighthouse Lab in Glasgow | Wellcome Sanger Institute for the COVID-19 Genomics UK (COG-UK) Consortium | Harper VanSteenhouse, Yumi Kasai, David Gray, Carol Clugston, Anna Dominiczak and Alex Alderton, Roberto Amato, Jeffrey Barrett, Sonia Goncalves, Ewan Harrison, David K. Jackson, Ian Johnston, Dominic Kwiatkowski, Cordelia Langford, John Sillitoe on behalf of the Wellcome Sanger Institute COVID-19 Surveillance Team |
| EPI_ISL_2917258, EPI_ISL_2917259, EPI_ISL_2917260, EPI_ISL_2917261 | Berkshire and Surrey Pathology Services Lighthouse Laboratory | Wellcome Sanger Institute for the COVID-19 Genomics UK (COG-UK) Consortium | Berkshire and Surrey Pathology Services Lighthouse Laboratory and Alex Alderton, Roberto Amato, Jeffrey Barrett, Sonia Goncalves, Ewan Harrison, David K. Jackson, Ian Johnston, Dominic Kwiatkowski, Cordelia Langford, John Sillitoe on behalf of the Wellcome Sanger Institute COVID-19 Surveillance Team |
| EPI_ISL_2917262, EPI_ISL_2917263 | Lighthouse Lab in Glasgow | Wellcome Sanger Institute for the COVID-19 Genomics UK (COG-UK) Consortium | Harper VanSteenhouse, Yumi Kasai, David Gray, Carol Clugston, Anna Dominiczak and Alex Alderton, Roberto Amato, Jeffrey Barrett, Sonia Goncalves, Ewan Harrison, David K. Jackson, Ian Johnston, Dominic Kwiatkowski, Cordelia Langford, John Sillitoe on behalf of the Wellcome Sanger Institute COVID-19 Surveillance Team |
| EPI_ISL_2917264, EPI_ISL_2917265 | Berkshire and Surrey Pathology Services Lighthouse Laboratory | Wellcome Sanger Institute for the COVID-19 Genomics UK (COG-UK) Consortium | Berkshire and Surrey Pathology Services Lighthouse Laboratory and Alex Alderton, Roberto Amato, Jeffrey Barrett, Sonia Goncalves, Ewan Harrison, David K. Jackson, Ian Johnston, Dominic Kwiatkowski, Cordelia Langford, John Sillitoe on behalf of the Wellcome Sanger Institute COVID-19 Surveillance Team |
| EPI_ISL_2917266, EPI_ISL_2917267, EPI_ISL_2917268, EPI_ISL_2917269 | Lighthouse Lab in Glasgow | Wellcome Sanger Institute for the COVID-19 Genomics UK (COG-UK) Consortium | Harper VanSteenhouse, Yumi Kasai, David Gray, Carol Clugston, Anna Dominiczak and Alex Alderton, Roberto Amato, Jeffrey Barrett, Sonia Goncalves, Ewan Harrison, David K. Jackson, Ian Johnston, Dominic Kwiatkowski, Cordelia Langford, John Sillitoe on behalf of the Wellcome Sanger Institute COVID-19 Surveillance Team |
| EPI_ISL_2917270 | Berkshire and Surrey Pathology Services Lighthouse Laboratory | Wellcome Sanger Institute for the COVID-19 Genomics UK (COG-UK) Consortium | Berkshire and Surrey Pathology Services Lighthouse Laboratory and Alex Alderton, Roberto Amato, Jeffrey Barrett, Sonia Goncalves, Ewan Harrison, David K. Jackson, Ian Johnston, Dominic Kwiatkowski, Cordelia Langford, John Sillitoe on behalf of the Wellcome Sanger Institute COVID-19 Surveillance Team |
| EPI_ISL_2917271 | Lighthouse Lab in Glasgow | Wellcome Sanger Institute for the COVID-19 Genomics UK (COG-UK) Consortium | Harper VanSteenhouse, Yumi Kasai, David Gray, Carol Clugston, Anna Dominiczak and Alex Alderton, Roberto Amato, Jeffrey Barrett, Sonia Goncalves, Ewan Harrison, David K. Jackson, Ian Johnston, Dominic Kwiatkowski, Cordelia Langford, John Sillitoe on behalf of the Wellcome Sanger Institute COVID-19 Surveillance Team |
| EPI_ISL_2917272 | Berkshire and Surrey Pathology Services Lighthouse Laboratory | Wellcome Sanger Institute for the COVID-19 Genomics UK (COG-UK) Consortium | Berkshire and Surrey Pathology Services Lighthouse Laboratory and Alex Alderton, Roberto Amato, Jeffrey Barrett, Sonia Goncalves, Ewan Harrison, David K. Jackson, Ian Johnston, Dominic Kwiatkowski, Cordelia Langford, John Sillitoe on behalf of the Wellcome Sanger Institute COVID-19 Surveillance Team |
| EPI_ISL_2917273 | Lighthouse Lab in Glasgow | Wellcome Sanger Institute for the COVID-19 Genomics UK (COG-UK) Consortium | Harper VanSteenhouse, Yumi Kasai, David Gray, Carol Clugston, Anna Dominiczak and Alex Alderton, Roberto Amato, Jeffrey Barrett, Sonia Goncalves, Ewan Harrison, David K. Jackson, Ian Johnston, Dominic Kwiatkowski, Cordelia Langford, John Sillitoe on behalf of the Wellcome Sanger Institute COVID-19 Surveillance Team |
| EPI_ISL_2917274 | Berkshire and Surrey Pathology Services Lighthouse Laboratory | Wellcome Sanger Institute for the COVID-19 Genomics UK (COG-UK) Consortium | Berkshire and Surrey Pathology Services Lighthouse Laboratory and Alex Alderton, Roberto Amato, Jeffrey Barrett, Sonia Goncalves, Ewan Harrison, David K. Jackson, Ian Johnston, Dominic Kwiatkowski, Cordelia Langford, John Sillitoe on behalf of the Wellcome Sanger Institute COVID-19 Surveillance Team |
| EPI_ISL_2917275, | Lighthouse Lab in Glasgow | Wellcome Sanger Institute for the COVID-19 Genomics UK (COG-UK) | Harper VanSteenhouse, Yumi Kasai, David Gray, Carol Clugston, Anna Dominiczak and Alex Alderton, Roberto Amato, Jeffrey Barrett, Sonia Goncalves, Ewan Harrison, David K. Jackson, Ian Johnston, Dominic Kwiatkowski, Cordelia Langford, |

| | | | |
|---|---|---|---|
| EPI_ISL_2917276, EPI_ISL_2917277, EPI_ISL_2917278 | | Consortium | John Sillitoe on behalf of the Wellcome Sanger Institute COVID-19 Surveillance Team |
| EPI_ISL_2917279, EPI_ISL_2917280, EPI_ISL_2917281, EPI_ISL_2917282 | Berkshire and Surrey Pathology Services Lighthouse Laboratory | Wellcome Sanger Institute for the COVID-19 Genomics UK (COG-UK) Consortium | Berkshire and Surrey Pathology Services Lighthouse Laboratory and Alex Alderton, Roberto Amato, Jeffrey Barrett, Sonia Goncalves, Ewan Harrison, David K. Jackson, Ian Johnston, Dominic Kwiatkowski, Cordelia Langford, John Sillitoe on behalf of the Wellcome Sanger Institute COVID-19 Surveillance Team |
| EPI_ISL_2917283 | Lighthouse Lab in Glasgow | Wellcome Sanger Institute for the COVID-19 Genomics UK (COG-UK) Consortium | Harper VanSteenhouse, Yumi Kasai, David Gray, Carol Clugston, Anna Dominiczak and Alex Alderton, Roberto Amato, Jeffrey Barrett, Sonia Goncalves, Ewan Harrison, David K. Jackson, Ian Johnston, Dominic Kwiatkowski, Cordelia Langford, John Sillitoe on behalf of the Wellcome Sanger Institute COVID-19 Surveillance Team |
| EPI_ISL_2917284 | Berkshire and Surrey Pathology Services Lighthouse Laboratory | Wellcome Sanger Institute for the COVID-19 Genomics UK (COG-UK) Consortium | Berkshire and Surrey Pathology Services Lighthouse Laboratory and Alex Alderton, Roberto Amato, Jeffrey Barrett, Sonia Goncalves, Ewan Harrison, David K. Jackson, Ian Johnston, Dominic Kwiatkowski, Cordelia Langford, John Sillitoe on behalf of the Wellcome Sanger Institute COVID-19 Surveillance Team |
| EPI_ISL_2917285, EPI_ISL_2917286, EPI_ISL_2917287, EPI_ISL_2917288, EPI_ISL_2917289, EPI_ISL_2917290, EPI_ISL_2917291, EPI_ISL_2917292 | Lighthouse Lab in Glasgow | Wellcome Sanger Institute for the COVID-19 Genomics UK (COG-UK) Consortium | Harper VanSteenhouse, Yumi Kasai, David Gray, Carol Clugston, Anna Dominiczak and Alex Alderton, Roberto Amato, Jeffrey Barrett, Sonia Goncalves, Ewan Harrison, David K. Jackson, Ian Johnston, Dominic Kwiatkowski, Cordelia Langford, John Sillitoe on behalf of the Wellcome Sanger Institute COVID-19 Surveillance Team |
| EPI_ISL_2917293, EPI_ISL_2917294, EPI_ISL_2917295, EPI_ISL_2917296, EPI_ISL_2917297, EPI_ISL_2917298 | Berkshire and Surrey Pathology Services Lighthouse Laboratory | Wellcome Sanger Institute for the COVID-19 Genomics UK (COG-UK) Consortium | Berkshire and Surrey Pathology Services Lighthouse Laboratory and Alex Alderton, Roberto Amato, Jeffrey Barrett, Sonia Goncalves, Ewan Harrison, David K. Jackson, Ian Johnston, Dominic Kwiatkowski, Cordelia Langford, John Sillitoe on behalf of the Wellcome Sanger Institute COVID-19 Surveillance Team |
| EPI_ISL_2917299, EPI_ISL_2917300 | Lighthouse Lab in Glasgow | Wellcome Sanger Institute for the COVID-19 Genomics UK (COG-UK) Consortium | Harper VanSteenhouse, Yumi Kasai, David Gray, Carol Clugston, Anna Dominiczak and Alex Alderton, Roberto Amato, Jeffrey Barrett, Sonia Goncalves, Ewan Harrison, David K. Jackson, Ian Johnston, Dominic Kwiatkowski, Cordelia Langford, John Sillitoe on behalf of the Wellcome Sanger Institute COVID-19 Surveillance Team |
| EPI_ISL_2917301 | Berkshire and Surrey Pathology Services Lighthouse Laboratory | Wellcome Sanger Institute for the COVID-19 Genomics UK (COG-UK) Consortium | Berkshire and Surrey Pathology Services Lighthouse Laboratory and Alex Alderton, Roberto Amato, Jeffrey Barrett, Sonia Goncalves, Ewan Harrison, David K. Jackson, Ian Johnston, Dominic Kwiatkowski, Cordelia Langford, John Sillitoe on behalf of the Wellcome Sanger Institute COVID-19 Surveillance Team |
| EPI_ISL_2917302 | Lighthouse Lab in Glasgow | Wellcome Sanger Institute for the COVID-19 Genomics UK (COG-UK) Consortium | Harper VanSteenhouse, Yumi Kasai, David Gray, Carol Clugston, Anna Dominiczak and Alex Alderton, Roberto Amato, Jeffrey Barrett, Sonia Goncalves, Ewan Harrison, David K. Jackson, Ian Johnston, Dominic Kwiatkowski, Cordelia Langford, John Sillitoe on behalf of the Wellcome Sanger Institute COVID-19 Surveillance Team |
| EPI_ISL_2917303 | Berkshire and Surrey Pathology Services Lighthouse Laboratory | Wellcome Sanger Institute for the COVID-19 Genomics UK (COG-UK) Consortium | Berkshire and Surrey Pathology Services Lighthouse Laboratory and Alex Alderton, Roberto Amato, Jeffrey Barrett, Sonia Goncalves, Ewan Harrison, David K. Jackson, Ian Johnston, Dominic Kwiatkowski, Cordelia Langford, John Sillitoe on behalf of the Wellcome Sanger Institute COVID-19 Surveillance Team |
| EPI_ISL_2917304, EPI_ISL_2917305, EPI_ISL_2917306 | Lighthouse Lab in Glasgow | Wellcome Sanger Institute for the COVID-19 Genomics UK (COG-UK) Consortium | Harper VanSteenhouse, Yumi Kasai, David Gray, Carol Clugston, Anna Dominiczak and Alex Alderton, Roberto Amato, Jeffrey Barrett, Sonia Goncalves, Ewan Harrison, David K. Jackson, Ian Johnston, Dominic Kwiatkowski, Cordelia Langford, John Sillitoe on behalf of the Wellcome Sanger Institute COVID-19 Surveillance Team |
| EPI_ISL_2917307 | Berkshire and Surrey Pathology Services Lighthouse Laboratory | Wellcome Sanger Institute for the COVID-19 Genomics UK (COG-UK) Consortium | Berkshire and Surrey Pathology Services Lighthouse Laboratory and Alex Alderton, Roberto Amato, Jeffrey Barrett, Sonia Goncalves, Ewan Harrison, David K. Jackson, Ian Johnston, Dominic Kwiatkowski, Cordelia Langford, John Sillitoe on behalf of the Wellcome Sanger Institute COVID-19 Surveillance Team |
| EPI_ISL_2917308, EPI_ISL_2917309, EPI_ISL_2917310 | Lighthouse Lab in Glasgow | Wellcome Sanger Institute for the COVID-19 Genomics UK (COG-UK) Consortium | Harper VanSteenhouse, Yumi Kasai, David Gray, Carol Clugston, Anna Dominiczak and Alex Alderton, Roberto Amato, Jeffrey Barrett, Sonia Goncalves, Ewan Harrison, David K. Jackson, Ian Johnston, Dominic Kwiatkowski, Cordelia Langford, John Sillitoe on behalf of the Wellcome Sanger Institute COVID-19 Surveillance Team |
| EPI_ISL_2917311 | Berkshire and Surrey Pathology Services Lighthouse Laboratory | Wellcome Sanger Institute for the COVID-19 Genomics UK (COG-UK) Consortium | Berkshire and Surrey Pathology Services Lighthouse Laboratory and Alex Alderton, Roberto Amato, Jeffrey Barrett, Sonia Goncalves, Ewan Harrison, David K. Jackson, Ian Johnston, Dominic Kwiatkowski, Cordelia Langford, John Sillitoe on behalf of the Wellcome Sanger Institute COVID-19 Surveillance Team |
| EPI_ISL_2917312 | Lighthouse Lab in Glasgow | Wellcome Sanger Institute for the COVID-19 Genomics UK (COG-UK) Consortium | Harper VanSteenhouse, Yumi Kasai, David Gray, Carol Clugston, Anna Dominiczak and Alex Alderton, Roberto Amato, Jeffrey Barrett, Sonia Goncalves, Ewan Harrison, David K. Jackson, Ian Johnston, Dominic Kwiatkowski, Cordelia Langford, John Sillitoe on behalf of the Wellcome Sanger Institute COVID-19 Surveillance Team |
| EPI_ISL_2917313 | Berkshire and Surrey Pathology Services Lighthouse Laboratory | Wellcome Sanger Institute for the COVID-19 Genomics UK (COG-UK) Consortium | Berkshire and Surrey Pathology Services Lighthouse Laboratory and Alex Alderton, Roberto Amato, Jeffrey Barrett, Sonia Goncalves, Ewan Harrison, David K. Jackson, Ian Johnston, Dominic Kwiatkowski, Cordelia Langford, John Sillitoe on behalf of the Wellcome Sanger Institute COVID-19 Surveillance Team |
| EPI_ISL_2917314 | Lighthouse Lab in Glasgow | Wellcome Sanger Institute for the COVID-19 Genomics UK (COG-UK) Consortium | Harper VanSteenhouse, Yumi Kasai, David Gray, Carol Clugston, Anna Dominiczak and Alex Alderton, Roberto Amato, Jeffrey Barrett, Sonia Goncalves, Ewan Harrison, David K. Jackson, Ian Johnston, Dominic Kwiatkowski, Cordelia Langford, John Sillitoe on behalf of the Wellcome Sanger Institute COVID-19 Surveillance Team |
| EPI_ISL_2917315 | Berkshire and Surrey Pathology Services Lighthouse Laboratory | Wellcome Sanger Institute for the COVID-19 Genomics UK (COG-UK) Consortium | Berkshire and Surrey Pathology Services Lighthouse Laboratory and Alex Alderton, Roberto Amato, Jeffrey Barrett, Sonia Goncalves, Ewan Harrison, David K. Jackson, Ian Johnston, Dominic Kwiatkowski, Cordelia Langford, John Sillitoe on behalf of the Wellcome Sanger Institute COVID-19 Surveillance Team |
| EPI_ISL_2917316, EPI_ISL_2917317, EPI_ISL_2917318 | Lighthouse Lab in Glasgow | Wellcome Sanger Institute for the COVID-19 Genomics UK (COG-UK) Consortium | Harper VanSteenhouse, Yumi Kasai, David Gray, Carol Clugston, Anna Dominiczak and Alex Alderton, Roberto Amato, Jeffrey Barrett, Sonia Goncalves, Ewan Harrison, David K. Jackson, Ian Johnston, Dominic Kwiatkowski, Cordelia Langford, John Sillitoe on behalf of the Wellcome Sanger Institute COVID-19 Surveillance Team |
| EPI_ISL_2917319 | Berkshire and Surrey Pathology Services Lighthouse Laboratory | Wellcome Sanger Institute for the COVID-19 Genomics UK (COG-UK) Consortium | Berkshire and Surrey Pathology Services Lighthouse Laboratory and Alex Alderton, Roberto Amato, Jeffrey Barrett, Sonia Goncalves, Ewan Harrison, David K. Jackson, Ian Johnston, Dominic Kwiatkowski, Cordelia Langford, John Sillitoe on behalf of the Wellcome Sanger Institute COVID-19 Surveillance Team |
| EPI_ISL_2917320 | Lighthouse Lab in Glasgow | Wellcome Sanger Institute for the COVID-19 Genomics UK (COG-UK) Consortium | Harper VanSteenhouse, Yumi Kasai, David Gray, Carol Clugston, Anna Dominiczak and Alex Alderton, Roberto Amato, Jeffrey Barrett, Sonia Goncalves, Ewan Harrison, David K. Jackson, Ian Johnston, Dominic Kwiatkowski, Cordelia Langford, John Sillitoe on behalf of the Wellcome Sanger Institute COVID-19 Surveillance Team |
| EPI_ISL_2917321 | Berkshire and Surrey Pathology Services Lighthouse Laboratory | Wellcome Sanger Institute for the COVID-19 Genomics UK (COG-UK) Consortium | Berkshire and Surrey Pathology Services Lighthouse Laboratory and Alex Alderton, Roberto Amato, Jeffrey Barrett, Sonia Goncalves, Ewan Harrison, David K. Jackson, Ian Johnston, Dominic Kwiatkowski, Cordelia Langford, John Sillitoe on behalf of the Wellcome Sanger Institute COVID-19 Surveillance Team |
| EPI_ISL_2917322, EPI_ISL_2917323 | Lighthouse Lab in Glasgow | Wellcome Sanger Institute for the COVID-19 Genomics UK (COG-UK) Consortium | Harper VanSteenhouse, Yumi Kasai, David Gray, Carol Clugston, Anna Dominiczak and Alex Alderton, Roberto Amato, Jeffrey Barrett, Sonia Goncalves, Ewan Harrison, David K. Jackson, Ian Johnston, Dominic Kwiatkowski, Cordelia Langford, John Sillitoe on behalf of the Wellcome Sanger Institute COVID-19 Surveillance Team |
| EPI_ISL_2917324, EPI_ISL_2917325 | Berkshire and Surrey Pathology Services Lighthouse Laboratory | Wellcome Sanger Institute for the COVID-19 Genomics UK (COG-UK) Consortium | Berkshire and Surrey Pathology Services Lighthouse Laboratory and Alex Alderton, Roberto Amato, Jeffrey Barrett, Sonia Goncalves, Ewan Harrison, David K. Jackson, Ian Johnston, Dominic Kwiatkowski, Cordelia Langford, John Sillitoe on behalf of the Wellcome Sanger Institute COVID-19 Surveillance Team |
| EPI_ISL_2917326 | Lighthouse Lab in Glasgow | Wellcome Sanger Institute for the COVID-19 Genomics UK (COG-UK) Consortium | Harper VanSteenhouse, Yumi Kasai, David Gray, Carol Clugston, Anna Dominiczak and Alex Alderton, Roberto Amato, Jeffrey Barrett, Sonia Goncalves, Ewan Harrison, David K. Jackson, Ian Johnston, Dominic Kwiatkowski, Cordelia Langford, John Sillitoe on behalf of the Wellcome Sanger Institute COVID-19 Surveillance Team |
| EPI_ISL_2917327 | Berkshire and Surrey Pathology Services Lighthouse Laboratory | Wellcome Sanger Institute for the COVID-19 Genomics UK (COG-UK) Consortium | Berkshire and Surrey Pathology Services Lighthouse Laboratory and Alex Alderton, Roberto Amato, Jeffrey Barrett, Sonia Goncalves, Ewan Harrison, David K. Jackson, Ian Johnston, Dominic Kwiatkowski, Cordelia Langford, John Sillitoe on behalf of the Wellcome Sanger Institute COVID-19 Surveillance Team |
| EPI_ISL_2917328 | Lighthouse Lab in Glasgow | Wellcome Sanger Institute for the COVID-19 Genomics UK (COG-UK) Consortium | Harper VanSteenhouse, Yumi Kasai, David Gray, Carol Clugston, Anna Dominiczak and Alex Alderton, Roberto Amato, Jeffrey Barrett, Sonia Goncalves, Ewan Harrison, David K. Jackson, Ian Johnston, Dominic Kwiatkowski, Cordelia Langford, John Sillitoe on behalf of the Wellcome Sanger Institute COVID-19 Surveillance Team |
| EPI_ISL_2917329 | Berkshire and Surrey Pathology Services Lighthouse Laboratory | Wellcome Sanger Institute for the COVID-19 Genomics UK (COG-UK) Consortium | Berkshire and Surrey Pathology Services Lighthouse Laboratory and Alex Alderton, Roberto Amato, Jeffrey Barrett, Sonia Goncalves, Ewan Harrison, David K. Jackson, Ian Johnston, Dominic Kwiatkowski, Cordelia Langford, John Sillitoe on behalf of the Wellcome Sanger Institute COVID-19 Surveillance Team |
| EPI_ISL_2917330 | Lighthouse Lab in Glasgow | Wellcome Sanger Institute for the COVID-19 Genomics UK (COG-UK) Consortium | Harper VanSteenhouse, Yumi Kasai, David Gray, Carol Clugston, Anna Dominiczak and Alex Alderton, Roberto Amato, Jeffrey Barrett, Sonia Goncalves, Ewan Harrison, David K. Jackson, Ian Johnston, Dominic Kwiatkowski, Cordelia Langford, John Sillitoe on behalf of the Wellcome Sanger Institute COVID-19 Surveillance Team |
| EPI_ISL_2917331, EPI_ISL_2917332, EPI_ISL_2917333 | Berkshire and Surrey Pathology Services Lighthouse Laboratory | Wellcome Sanger Institute for the COVID-19 Genomics UK (COG-UK) Consortium | Berkshire and Surrey Pathology Services Lighthouse Laboratory and Alex Alderton, Roberto Amato, Jeffrey Barrett, Sonia Goncalves, Ewan Harrison, David K. Jackson, Ian Johnston, Dominic Kwiatkowski, Cordelia Langford, John Sillitoe on behalf of the Wellcome Sanger Institute COVID-19 Surveillance Team |
| EPI_ISL_2917334 | Lighthouse Lab in Glasgow | Wellcome Sanger Institute for the COVID-19 Genomics UK (COG-UK) Consortium | Harper VanSteenhouse, Yumi Kasai, David Gray, Carol Clugston, Anna Dominiczak and Alex Alderton, Roberto Amato, Jeffrey Barrett, Sonia Goncalves, Ewan Harrison, David K. Jackson, Ian Johnston, Dominic Kwiatkowski, Cordelia Langford, John Sillitoe on behalf of the Wellcome Sanger Institute COVID-19 Surveillance Team |
| EPI_ISL_2917335 | Berkshire and Surrey Pathology Services Lighthouse Laboratory | Wellcome Sanger Institute for the COVID-19 Genomics UK (COG-UK) Consortium | Berkshire and Surrey Pathology Services Lighthouse Laboratory and Alex Alderton, Roberto Amato, Jeffrey Barrett, Sonia Goncalves, Ewan Harrison, David K. Jackson, Ian Johnston, Dominic Kwiatkowski, Cordelia Langford, John Sillitoe on behalf of the Wellcome Sanger Institute COVID-19 Surveillance Team |
| EPI_ISL_2917336 | Lighthouse Lab in Glasgow | Wellcome Sanger Institute for the COVID-19 Genomics UK (COG-UK) Consortium | Harper VanSteenhouse, Yumi Kasai, David Gray, Carol Clugston, Anna Dominiczak and Alex Alderton, Roberto Amato, Jeffrey Barrett, Sonia Goncalves, Ewan Harrison, David K. Jackson, Ian Johnston, Dominic Kwiatkowski, Cordelia Langford, John Sillitoe on behalf of the Wellcome Sanger Institute COVID-19 Surveillance Team |
| EPI_ISL_2917337 | Berkshire and Surrey Pathology Services Lighthouse Laboratory | Wellcome Sanger Institute for the COVID-19 Genomics UK (COG-UK) Consortium | Berkshire and Surrey Pathology Services Lighthouse Laboratory and Alex Alderton, Roberto Amato, Jeffrey Barrett, Sonia Goncalves, Ewan Harrison, David K. Jackson, Ian Johnston, Dominic Kwiatkowski, Cordelia Langford, John Sillitoe on behalf of the Wellcome Sanger Institute COVID-19 Surveillance Team |
| EPI_ISL_2917338, EPI_ISL_2917339, EPI_ISL_2917340, EPI_ISL_2917341 | Lighthouse Lab in Glasgow | Wellcome Sanger Institute for the COVID-19 Genomics UK (COG-UK) Consortium | Harper VanSteenhouse, Yumi Kasai, David Gray, Carol Clugston, Anna Dominiczak and Alex Alderton, Roberto Amato, Jeffrey Barrett, Sonia Goncalves, Ewan Harrison, David K. Jackson, Ian Johnston, Dominic Kwiatkowski, Cordelia Langford, John Sillitoe on behalf of the Wellcome Sanger Institute COVID-19 Surveillance Team |
| EPI_ISL_2917342 | Berkshire and Surrey Pathology Services Lighthouse Laboratory | Wellcome Sanger Institute for the COVID-19 Genomics UK (COG-UK) Consortium | Berkshire and Surrey Pathology Services Lighthouse Laboratory and Alex Alderton, Roberto Amato, Jeffrey Barrett, Sonia Goncalves, Ewan Harrison, David K. Jackson, Ian Johnston, Dominic Kwiatkowski, Cordelia Langford, John Sillitoe on behalf of the Wellcome Sanger Institute COVID-19 Surveillance Team |
| EPI_ISL_2917343 | Lighthouse Lab in Glasgow | Wellcome Sanger Institute for the COVID-19 Genomics UK (COG-UK) Consortium | Harper VanSteenhouse, Yumi Kasai, David Gray, Carol Clugston, Anna Dominiczak and Alex Alderton, Roberto Amato, Jeffrey Barrett, Sonia Goncalves, Ewan Harrison, David K. Jackson, Ian Johnston, Dominic Kwiatkowski, Cordelia Langford, John Sillitoe on behalf of the Wellcome Sanger Institute COVID-19 Surveillance Team |
| EPI_ISL_2917344 | Berkshire and Surrey Pathology Services Lighthouse Laboratory | Wellcome Sanger Institute for the COVID-19 Genomics UK (COG-UK) | Berkshire and Surrey Pathology Services Lighthouse Laboratory and Alex Alderton, Roberto Amato, Jeffrey Barrett, Sonia Goncalves, Ewan Harrison, David K. Jackson, Ian Johnston, Dominic Kwiatkowski, Cordelia Langford, John Sillitoe on behalf |

| | | | |
|---|---|---|---|
| | | Consortium | of the Wellcome Sanger Institute COVID-19 Surveillance Team |
| EPI_ISL_2917345 | Lighthouse Lab in Glasgow | Wellcome Sanger Institute for the COVID-19 Genomics UK (COG-UK) Consortium | Harper VanSteenhouse, Yumi Kasai, David Gray, Carol Clugston, Anna Dominiczak and Alex Alderton, Roberto Amato, Jeffrey Barrett, Sonia Goncalves, Ewan Harrison, David K. Jackson, Ian Johnston, Dominic Kwiatkowski, Cordelia Langford, John Sillitoe on behalf of the Wellcome Sanger Institute COVID-19 Surveillance Team |
| EPI_ISL_2917346, EPI_ISL_2917347 | Lighthouse Laboratory Plymouth | Wellcome Sanger Institute for the COVID-19 Genomics UK (COG-UK) Consortium | Lighthouse Laboratory Plymouth and Alex Alderton, Roberto Amato, Jeffrey Barrett, Sonia Goncalves, Ewan Harrison, David K. Jackson, Ian Johnston, Dominic Kwiatkowski, Cordelia Langford, John Sillitoe on behalf of the Wellcome Sanger Institute COVID-19 Surveillance Team |
| EPI_ISL_2917348 | Berkshire and Surrey Pathology Services Lighthouse Laboratory | Wellcome Sanger Institute for the COVID-19 Genomics UK (COG-UK) Consortium | Berkshire and Surrey Pathology Services Lighthouse Laboratory and Alex Alderton, Roberto Amato, Jeffrey Barrett, Sonia Goncalves, Ewan Harrison, David K. Jackson, Ian Johnston, Dominic Kwiatkowski, Cordelia Langford, John Sillitoe on behalf of the Wellcome Sanger Institute COVID-19 Surveillance Team |
| EPI_ISL_2917349, EPI_ISL_2917350 | Lighthouse Laboratory Plymouth | Wellcome Sanger Institute for the COVID-19 Genomics UK (COG-UK) Consortium | Lighthouse Laboratory Plymouth and Alex Alderton, Roberto Amato, Jeffrey Barrett, Sonia Goncalves, Ewan Harrison, David K. Jackson, Ian Johnston, Dominic Kwiatkowski, Cordelia Langford, John Sillitoe on behalf of the Wellcome Sanger Institute COVID-19 Surveillance Team |
| EPI_ISL_2917351 | Lighthouse Lab in Milton Keynes | Wellcome Sanger Institute for the COVID-19 Genomics UK (COG-UK) Consortium | The Lighthouse Lab in Milton Keynes and Alex Alderton, Roberto Amato, Jeffrey Barrett, Sonia Goncalves, Ewan Harrison, David K. Jackson, Ian Johnston, Dominic Kwiatkowski, Cordelia Langford, John Sillitoe on behalf of the Wellcome Sanger Institute COVID-19 Surveillance Team |
| EPI_ISL_2917352 | Lighthouse Laboratory Plymouth | Wellcome Sanger Institute for the COVID-19 Genomics UK (COG-UK) Consortium | Lighthouse Laboratory Plymouth and Alex Alderton, Roberto Amato, Jeffrey Barrett, Sonia Goncalves, Ewan Harrison, David K. Jackson, Ian Johnston, Dominic Kwiatkowski, Cordelia Langford, John Sillitoe on behalf of the Wellcome Sanger Institute COVID-19 Surveillance Team |
| EPI_ISL_2917353 | Berkshire and Surrey Pathology Services Lighthouse Laboratory | Wellcome Sanger Institute for the COVID-19 Genomics UK (COG-UK) Consortium | Berkshire and Surrey Pathology Services Lighthouse Laboratory and Alex Alderton, Roberto Amato, Jeffrey Barrett, Sonia Goncalves, Ewan Harrison, David K. Jackson, Ian Johnston, Dominic Kwiatkowski, Cordelia Langford, John Sillitoe on behalf of the Wellcome Sanger Institute COVID-19 Surveillance Team |
| EPI_ISL_2917354, EPI_ISL_2917355 | Lighthouse Lab in Milton Keynes | Wellcome Sanger Institute for the COVID-19 Genomics UK (COG-UK) Consortium | The Lighthouse Lab in Milton Keynes and Alex Alderton, Roberto Amato, Jeffrey Barrett, Sonia Goncalves, Ewan Harrison, David K. Jackson, Ian Johnston, Dominic Kwiatkowski, Cordelia Langford, John Sillitoe on behalf of the Wellcome Sanger Institute COVID-19 Surveillance Team |
| EPI_ISL_2917356 | Berkshire and Surrey Pathology Services Lighthouse Laboratory | Wellcome Sanger Institute for the COVID-19 Genomics UK (COG-UK) Consortium | Berkshire and Surrey Pathology Services Lighthouse Laboratory and Alex Alderton, Roberto Amato, Jeffrey Barrett, Sonia Goncalves, Ewan Harrison, David K. Jackson, Ian Johnston, Dominic Kwiatkowski, Cordelia Langford, John Sillitoe on behalf of the Wellcome Sanger Institute COVID-19 Surveillance Team |
| EPI_ISL_2917357 | Lighthouse Lab in Milton Keynes | Wellcome Sanger Institute for the COVID-19 Genomics UK (COG-UK) Consortium | The Lighthouse Lab in Milton Keynes and Alex Alderton, Roberto Amato, Jeffrey Barrett, Sonia Goncalves, Ewan Harrison, David K. Jackson, Ian Johnston, Dominic Kwiatkowski, Cordelia Langford, John Sillitoe on behalf of the Wellcome Sanger Institute COVID-19 Surveillance Team |
| EPI_ISL_2917358 | Lighthouse Laboratory Plymouth | Wellcome Sanger Institute for the COVID-19 Genomics UK (COG-UK) Consortium | Lighthouse Laboratory Plymouth and Alex Alderton, Roberto Amato, Jeffrey Barrett, Sonia Goncalves, Ewan Harrison, David K. Jackson, Ian Johnston, Dominic Kwiatkowski, Cordelia Langford, John Sillitoe on behalf of the Wellcome Sanger Institute COVID-19 Surveillance Team |
| EPI_ISL_2917359, EPI_ISL_2917360, EPI_ISL_2917361 | Lighthouse Lab in Milton Keynes | Wellcome Sanger Institute for the COVID-19 Genomics UK (COG-UK) Consortium | The Lighthouse Lab in Milton Keynes and Alex Alderton, Roberto Amato, Jeffrey Barrett, Sonia Goncalves, Ewan Harrison, David K. Jackson, Ian Johnston, Dominic Kwiatkowski, Cordelia Langford, John Sillitoe on behalf of the Wellcome Sanger Institute COVID-19 Surveillance Team |
| EPI_ISL_2917362 | Lighthouse Laboratory Plymouth | Wellcome Sanger Institute for the COVID-19 Genomics UK (COG-UK) Consortium | Lighthouse Laboratory Plymouth and Alex Alderton, Roberto Amato, Jeffrey Barrett, Sonia Goncalves, Ewan Harrison, David K. Jackson, Ian Johnston, Dominic Kwiatkowski, Cordelia Langford, John Sillitoe on behalf of the Wellcome Sanger Institute COVID-19 Surveillance Team |
| EPI_ISL_2917363 | Berkshire and Surrey Pathology Services Lighthouse Laboratory | Wellcome Sanger Institute for the COVID-19 Genomics UK (COG-UK) Consortium | Berkshire and Surrey Pathology Services Lighthouse Laboratory and Alex Alderton, Roberto Amato, Jeffrey Barrett, Sonia Goncalves, Ewan Harrison, David K. Jackson, Ian Johnston, Dominic Kwiatkowski, Cordelia Langford, John Sillitoe on behalf of the Wellcome Sanger Institute COVID-19 Surveillance Team |
| EPI_ISL_2917364 | Lighthouse Lab in Milton Keynes | Wellcome Sanger Institute for the COVID-19 Genomics UK (COG-UK) Consortium | The Lighthouse Lab in Milton Keynes and Alex Alderton, Roberto Amato, Jeffrey Barrett, Sonia Goncalves, Ewan Harrison, David K. Jackson, Ian Johnston, Dominic Kwiatkowski, Cordelia Langford, John Sillitoe on behalf of the Wellcome Sanger Institute COVID-19 Surveillance Team |
| EPI_ISL_2917365 | Berkshire and Surrey Pathology Services Lighthouse Laboratory | Wellcome Sanger Institute for the COVID-19 Genomics UK (COG-UK) Consortium | Berkshire and Surrey Pathology Services Lighthouse Laboratory and Alex Alderton, Roberto Amato, Jeffrey Barrett, Sonia Goncalves, Ewan Harrison, David K. Jackson, Ian Johnston, Dominic Kwiatkowski, Cordelia Langford, John Sillitoe on behalf of the Wellcome Sanger Institute COVID-19 Surveillance Team |
| EPI_ISL_2917366 | Lighthouse Lab in Milton Keynes | Wellcome Sanger Institute for the COVID-19 Genomics UK (COG-UK) Consortium | The Lighthouse Lab in Milton Keynes and Alex Alderton, Roberto Amato, Jeffrey Barrett, Sonia Goncalves, Ewan Harrison, David K. Jackson, Ian Johnston, Dominic Kwiatkowski, Cordelia Langford, John Sillitoe on behalf of the Wellcome Sanger Institute COVID-19 Surveillance Team |
| EPI_ISL_2917367, EPI_ISL_2917368 | Lighthouse Laboratory Plymouth | Wellcome Sanger Institute for the COVID-19 Genomics UK (COG-UK) Consortium | Lighthouse Laboratory Plymouth and Alex Alderton, Roberto Amato, Jeffrey Barrett, Sonia Goncalves, Ewan Harrison, David K. Jackson, Ian Johnston, Dominic Kwiatkowski, Cordelia Langford, John Sillitoe on behalf of the Wellcome Sanger Institute COVID-19 Surveillance Team |
| EPI_ISL_2917369, EPI_ISL_2917370, EPI_ISL_2917371 | Berkshire and Surrey Pathology Services Lighthouse Laboratory | Wellcome Sanger Institute for the COVID-19 Genomics UK (COG-UK) Consortium | Berkshire and Surrey Pathology Services Lighthouse Laboratory and Alex Alderton, Roberto Amato, Jeffrey Barrett, Sonia Goncalves, Ewan Harrison, David K. Jackson, Ian Johnston, Dominic Kwiatkowski, Cordelia Langford, John Sillitoe on behalf of the Wellcome Sanger Institute COVID-19 Surveillance Team |
| EPI_ISL_2917372 | Lighthouse Lab in Milton Keynes | Wellcome Sanger Institute for the COVID-19 Genomics UK (COG-UK) Consortium | The Lighthouse Lab in Milton Keynes and Alex Alderton, Roberto Amato, Jeffrey Barrett, Sonia Goncalves, Ewan Harrison, David K. Jackson, Ian Johnston, Dominic Kwiatkowski, Cordelia Langford, John Sillitoe on behalf of the Wellcome Sanger Institute COVID-19 Surveillance Team |
| EPI_ISL_2917373 | Lighthouse Laboratory Plymouth | Wellcome Sanger Institute for the COVID-19 Genomics UK (COG-UK) Consortium | Lighthouse Laboratory Plymouth and Alex Alderton, Roberto Amato, Jeffrey Barrett, Sonia Goncalves, Ewan Harrison, David K. Jackson, Ian Johnston, Dominic Kwiatkowski, Cordelia Langford, John Sillitoe on behalf of the Wellcome Sanger Institute COVID-19 Surveillance Team |
| EPI_ISL_2917374 | Lighthouse Lab in Milton Keynes | Wellcome Sanger Institute for the COVID-19 Genomics UK (COG-UK) Consortium | The Lighthouse Lab in Milton Keynes and Alex Alderton, Roberto Amato, Jeffrey Barrett, Sonia Goncalves, Ewan Harrison, David K. Jackson, Ian Johnston, Dominic Kwiatkowski, Cordelia Langford, John Sillitoe on behalf of the Wellcome Sanger Institute COVID-19 Surveillance Team |
| EPI_ISL_2917375 | Berkshire and Surrey Pathology Services Lighthouse Laboratory | Wellcome Sanger Institute for the COVID-19 Genomics UK (COG-UK) Consortium | Berkshire and Surrey Pathology Services Lighthouse Laboratory and Alex Alderton, Roberto Amato, Jeffrey Barrett, Sonia Goncalves, Ewan Harrison, David K. Jackson, Ian Johnston, Dominic Kwiatkowski, Cordelia Langford, John Sillitoe on behalf of the Wellcome Sanger Institute COVID-19 Surveillance Team |
| EPI_ISL_2917376 | Lighthouse Laboratory Plymouth | Wellcome Sanger Institute for the COVID-19 Genomics UK (COG-UK) Consortium | Lighthouse Laboratory Plymouth and Alex Alderton, Roberto Amato, Jeffrey Barrett, Sonia Goncalves, Ewan Harrison, David K. Jackson, Ian Johnston, Dominic Kwiatkowski, Cordelia Langford, John Sillitoe on behalf of the Wellcome Sanger Institute COVID-19 Surveillance Team |
| EPI_ISL_2917377 | Berkshire and Surrey Pathology Services Lighthouse Laboratory | Wellcome Sanger Institute for the COVID-19 Genomics UK (COG-UK) Consortium | Berkshire and Surrey Pathology Services Lighthouse Laboratory and Alex Alderton, Roberto Amato, Jeffrey Barrett, Sonia Goncalves, Ewan Harrison, David K. Jackson, Ian Johnston, Dominic Kwiatkowski, Cordelia Langford, John Sillitoe on behalf of the Wellcome Sanger Institute COVID-19 Surveillance Team |
| EPI_ISL_2917378 | Lighthouse Lab in Milton Keynes | Wellcome Sanger Institute for the COVID-19 Genomics UK (COG-UK) Consortium | The Lighthouse Lab in Milton Keynes and Alex Alderton, Roberto Amato, Jeffrey Barrett, Sonia Goncalves, Ewan Harrison, David K. Jackson, Ian Johnston, Dominic Kwiatkowski, Cordelia Langford, John Sillitoe on behalf of the Wellcome Sanger Institute COVID-19 Surveillance Team |
| EPI_ISL_2917379 | Lighthouse Laboratory Plymouth | Wellcome Sanger Institute for the COVID-19 Genomics UK (COG-UK) Consortium | Lighthouse Laboratory Plymouth and Alex Alderton, Roberto Amato, Jeffrey Barrett, Sonia Goncalves, Ewan Harrison, David K. Jackson, Ian Johnston, Dominic Kwiatkowski, Cordelia Langford, John Sillitoe on behalf of the Wellcome Sanger Institute COVID-19 Surveillance Team |
| EPI_ISL_2917380 | Berkshire and Surrey Pathology Services Lighthouse Laboratory | Wellcome Sanger Institute for the COVID-19 Genomics UK (COG-UK) Consortium | Berkshire and Surrey Pathology Services Lighthouse Laboratory and Alex Alderton, Roberto Amato, Jeffrey Barrett, Sonia Goncalves, Ewan Harrison, David K. Jackson, Ian Johnston, Dominic Kwiatkowski, Cordelia Langford, John Sillitoe on behalf of the Wellcome Sanger Institute COVID-19 Surveillance Team |
| EPI_ISL_2917381 | Lighthouse Lab in Milton Keynes | Wellcome Sanger Institute for the COVID-19 Genomics UK (COG-UK) Consortium | The Lighthouse Lab in Milton Keynes and Alex Alderton, Roberto Amato, Jeffrey Barrett, Sonia Goncalves, Ewan Harrison, David K. Jackson, Ian Johnston, Dominic Kwiatkowski, Cordelia Langford, John Sillitoe on behalf of the Wellcome Sanger Institute COVID-19 Surveillance Team |
| EPI_ISL_2917382 | Berkshire and Surrey Pathology Services Lighthouse Laboratory | Wellcome Sanger Institute for the COVID-19 Genomics UK (COG-UK) Consortium | Berkshire and Surrey Pathology Services Lighthouse Laboratory and Alex Alderton, Roberto Amato, Jeffrey Barrett, Sonia Goncalves, Ewan Harrison, David K. Jackson, Ian Johnston, Dominic Kwiatkowski, Cordelia Langford, John Sillitoe on behalf of the Wellcome Sanger Institute COVID-19 Surveillance Team |
| EPI_ISL_2917383 | Lighthouse Laboratory Plymouth | Wellcome Sanger Institute for the COVID-19 Genomics UK (COG-UK) Consortium | Lighthouse Laboratory Plymouth and Alex Alderton, Roberto Amato, Jeffrey Barrett, Sonia Goncalves, Ewan Harrison, David K. Jackson, Ian Johnston, Dominic Kwiatkowski, Cordelia Langford, John Sillitoe on behalf of the Wellcome Sanger Institute COVID-19 Surveillance Team |
| EPI_ISL_2917384, EPI_ISL_2917385, EPI_ISL_2917386 | Berkshire and Surrey Pathology Services Lighthouse Laboratory | Wellcome Sanger Institute for the COVID-19 Genomics UK (COG-UK) Consortium | Berkshire and Surrey Pathology Services Lighthouse Laboratory and Alex Alderton, Roberto Amato, Jeffrey Barrett, Sonia Goncalves, Ewan Harrison, David K. Jackson, Ian Johnston, Dominic Kwiatkowski, Cordelia Langford, John Sillitoe on behalf of the Wellcome Sanger Institute COVID-19 Surveillance Team |
| EPI_ISL_2917387, EPI_ISL_2917388 | Lighthouse Lab in Milton Keynes | Wellcome Sanger Institute for the COVID-19 Genomics UK (COG-UK) Consortium | The Lighthouse Lab in Milton Keynes and Alex Alderton, Roberto Amato, Jeffrey Barrett, Sonia Goncalves, Ewan Harrison, David K. Jackson, Ian Johnston, Dominic Kwiatkowski, Cordelia Langford, John Sillitoe on behalf of the Wellcome Sanger Institute COVID-19 Surveillance Team |
| EPI_ISL_2917389 | Lighthouse Laboratory Plymouth | Wellcome Sanger Institute for the COVID-19 Genomics UK (COG-UK) Consortium | Lighthouse Laboratory Plymouth and Alex Alderton, Roberto Amato, Jeffrey Barrett, Sonia Goncalves, Ewan Harrison, David K. Jackson, Ian Johnston, Dominic Kwiatkowski, Cordelia Langford, John Sillitoe on behalf of the Wellcome Sanger Institute COVID-19 Surveillance Team |
| EPI_ISL_2917390, EPI_ISL_2917391, EPI_ISL_2917392 | Berkshire and Surrey Pathology Services Lighthouse Laboratory | Wellcome Sanger Institute for the COVID-19 Genomics UK (COG-UK) Consortium | Berkshire and Surrey Pathology Services Lighthouse Laboratory and Alex Alderton, Roberto Amato, Jeffrey Barrett, Sonia Goncalves, Ewan Harrison, David K. Jackson, Ian Johnston, Dominic Kwiatkowski, Cordelia Langford, John Sillitoe on behalf of the Wellcome Sanger Institute COVID-19 Surveillance Team |
| EPI_ISL_2917393 | Lighthouse Lab in Milton Keynes | Wellcome Sanger Institute for the COVID-19 Genomics UK (COG-UK) Consortium | The Lighthouse Lab in Milton Keynes and Alex Alderton, Roberto Amato, Jeffrey Barrett, Sonia Goncalves, Ewan Harrison, David K. Jackson, Ian Johnston, Dominic Kwiatkowski, Cordelia Langford, John Sillitoe on behalf of the Wellcome Sanger Institute COVID-19 Surveillance Team |
| EPI_ISL_2917394 | Lighthouse Laboratory Plymouth | Wellcome Sanger Institute for the COVID-19 Genomics UK (COG-UK) Consortium | Lighthouse Laboratory Plymouth and Alex Alderton, Roberto Amato, Jeffrey Barrett, Sonia Goncalves, Ewan Harrison, David K. Jackson, Ian Johnston, Dominic Kwiatkowski, Cordelia Langford, John Sillitoe on behalf of the Wellcome Sanger Institute COVID-19 Surveillance Team |
| EPI_ISL_2917395 | Lighthouse Lab in Milton Keynes | Wellcome Sanger Institute for the COVID-19 Genomics UK (COG-UK) Consortium | The Lighthouse Lab in Milton Keynes and Alex Alderton, Roberto Amato, Jeffrey Barrett, Sonia Goncalves, Ewan Harrison, David K. Jackson, Ian Johnston, Dominic Kwiatkowski, Cordelia Langford, John Sillitoe on behalf of the Wellcome Sanger Institute COVID-19 Surveillance Team |
| EPI_ISL_2917396, EPI_ISL_2917397 | Berkshire and Surrey Pathology Services Lighthouse Laboratory | Wellcome Sanger Institute for the COVID-19 Genomics UK (COG-UK) Consortium | Berkshire and Surrey Pathology Services Lighthouse Laboratory and Alex Alderton, Roberto Amato, Jeffrey Barrett, Sonia Goncalves, Ewan Harrison, David K. Jackson, Ian Johnston, Dominic Kwiatkowski, Cordelia Langford, John Sillitoe on behalf of the Wellcome Sanger Institute COVID-19 Surveillance Team |
| EPI_ISL_2917398 | Lighthouse Laboratory Plymouth | Wellcome Sanger Institute for the COVID-19 Genomics UK (COG-UK) Consortium | Lighthouse Laboratory Plymouth and Alex Alderton, Roberto Amato, Jeffrey Barrett, Sonia Goncalves, Ewan Harrison, David K. Jackson, Ian Johnston, Dominic Kwiatkowski, Cordelia Langford, John Sillitoe on behalf of the Wellcome Sanger Institute COVID-19 Surveillance Team |
| EPI_ISL_2917399 | Lighthouse Lab in Milton Keynes | Wellcome Sanger Institute for the COVID-19 Genomics UK (COG-UK) Consortium | The Lighthouse Lab in Milton Keynes and Alex Alderton, Roberto Amato, Jeffrey Barrett, Sonia Goncalves, Ewan Harrison, David K. Jackson, Ian Johnston, Dominic Kwiatkowski, Cordelia Langford, John Sillitoe on behalf of the Wellcome Sanger Institute COVID-19 Surveillance Team |
| EPI_ISL_2917400, EPI_ISL_2917401 | Berkshire and Surrey Pathology Services Lighthouse Laboratory | Wellcome Sanger Institute for the COVID-19 Genomics UK (COG-UK) Consortium | Berkshire and Surrey Pathology Services Lighthouse Laboratory and Alex Alderton, Roberto Amato, Jeffrey Barrett, Sonia Goncalves, Ewan Harrison, David K. Jackson, Ian Johnston, Dominic Kwiatkowski, Cordelia Langford, John Sillitoe on behalf of the Wellcome Sanger Institute COVID-19 Surveillance Team |
| EPI_ISL_2917402 | Lighthouse Laboratory Plymouth | Wellcome Sanger Institute for the COVID-19 Genomics UK (COG-UK) Consortium | Lighthouse Laboratory Plymouth and Alex Alderton, Roberto Amato, Jeffrey Barrett, Sonia Goncalves, Ewan Harrison, David K. Jackson, Ian Johnston, Dominic Kwiatkowski, Cordelia Langford, John Sillitoe on behalf of the Wellcome Sanger Institute COVID-19 Surveillance Team |

| | | | |
|---|---|---|---|
| EPI_ISL_2917403 | Berkshire and Surrey Pathology Services Lighthouse Laboratory | Wellcome Sanger Institute for the COVID-19 Genomics UK (COG-UK) Consortium | Berkshire and Surrey Pathology Services Lighthouse Laboratory and Alex Alderton, Roberto Amato, Jeffrey Barrett, Sonia Goncalves, Ewan Harrison, David K. Jackson, Ian Johnston, Dominic Kwiatkowski, Cordelia Langford, John Sillitoe on behalf of the Wellcome Sanger Institute COVID-19 Surveillance Team |
| EPI_ISL_2917404, EPI_ISL_2917405 | Lighthouse Lab in Milton Keynes | Wellcome Sanger Institute for the COVID-19 Genomics UK (COG-UK) Consortium | The Lighthouse Lab in Milton Keynes and Alex Alderton, Roberto Amato, Jeffrey Barrett, Sonia Goncalves, Ewan Harrison, David K. Jackson, Ian Johnston, Dominic Kwiatkowski, Cordelia Langford, John Sillitoe on behalf of the Wellcome Sanger Institute COVID-19 Surveillance Team |
| EPI_ISL_2917406 | Lighthouse Laboratory Plymouth | Wellcome Sanger Institute for the COVID-19 Genomics UK (COG-UK) Consortium | Lighthouse Laboratory Plymouth and Alex Alderton, Roberto Amato, Jeffrey Barrett, Sonia Goncalves, Ewan Harrison, David K. Jackson, Ian Johnston, Dominic Kwiatkowski, Cordelia Langford, John Sillitoe on behalf of the Wellcome Sanger Institute COVID-19 Surveillance Team |
| EPI_ISL_2917407, EPI_ISL_2917408 | Lighthouse Lab in Milton Keynes | Wellcome Sanger Institute for the COVID-19 Genomics UK (COG-UK) Consortium | The Lighthouse Lab in Milton Keynes and Alex Alderton, Roberto Amato, Jeffrey Barrett, Sonia Goncalves, Ewan Harrison, David K. Jackson, Ian Johnston, Dominic Kwiatkowski, Cordelia Langford, John Sillitoe on behalf of the Wellcome Sanger Institute COVID-19 Surveillance Team |
| EPI_ISL_2917409 | Berkshire and Surrey Pathology Services Lighthouse Laboratory | Wellcome Sanger Institute for the COVID-19 Genomics UK (COG-UK) Consortium | Berkshire and Surrey Pathology Services Lighthouse Laboratory and Alex Alderton, Roberto Amato, Jeffrey Barrett, Sonia Goncalves, Ewan Harrison, David K. Jackson, Ian Johnston, Dominic Kwiatkowski, Cordelia Langford, John Sillitoe on behalf of the Wellcome Sanger Institute COVID-19 Surveillance Team |
| EPI_ISL_2917410, EPI_ISL_2917411 | Lighthouse Laboratory Plymouth | Wellcome Sanger Institute for the COVID-19 Genomics UK (COG-UK) Consortium | Lighthouse Laboratory Plymouth and Alex Alderton, Roberto Amato, Jeffrey Barrett, Sonia Goncalves, Ewan Harrison, David K. Jackson, Ian Johnston, Dominic Kwiatkowski, Cordelia Langford, John Sillitoe on behalf of the Wellcome Sanger Institute COVID-19 Surveillance Team |
| EPI_ISL_2917412 | Berkshire and Surrey Pathology Services Lighthouse Laboratory | Wellcome Sanger Institute for the COVID-19 Genomics UK (COG-UK) Consortium | Berkshire and Surrey Pathology Services Lighthouse Laboratory and Alex Alderton, Roberto Amato, Jeffrey Barrett, Sonia Goncalves, Ewan Harrison, David K. Jackson, Ian Johnston, Dominic Kwiatkowski, Cordelia Langford, John Sillitoe on behalf of the Wellcome Sanger Institute COVID-19 Surveillance Team |
| EPI_ISL_2917413 | Lighthouse Lab in Milton Keynes | Wellcome Sanger Institute for the COVID-19 Genomics UK (COG-UK) Consortium | The Lighthouse Lab in Milton Keynes and Alex Alderton, Roberto Amato, Jeffrey Barrett, Sonia Goncalves, Ewan Harrison, David K. Jackson, Ian Johnston, Dominic Kwiatkowski, Cordelia Langford, John Sillitoe on behalf of the Wellcome Sanger Institute COVID-19 Surveillance Team |
| EPI_ISL_2917414 | Berkshire and Surrey Pathology Services Lighthouse Laboratory | Wellcome Sanger Institute for the COVID-19 Genomics UK (COG-UK) Consortium | Berkshire and Surrey Pathology Services Lighthouse Laboratory and Alex Alderton, Roberto Amato, Jeffrey Barrett, Sonia Goncalves, Ewan Harrison, David K. Jackson, Ian Johnston, Dominic Kwiatkowski, Cordelia Langford, John Sillitoe on behalf of the Wellcome Sanger Institute COVID-19 Surveillance Team |
| EPI_ISL_2917415 | Lighthouse Laboratory Plymouth | Wellcome Sanger Institute for the COVID-19 Genomics UK (COG-UK) Consortium | Lighthouse Laboratory Plymouth and Alex Alderton, Roberto Amato, Jeffrey Barrett, Sonia Goncalves, Ewan Harrison, David K. Jackson, Ian Johnston, Dominic Kwiatkowski, Cordelia Langford, John Sillitoe on behalf of the Wellcome Sanger Institute COVID-19 Surveillance Team |
| EPI_ISL_2917416 | Lighthouse Lab in Milton Keynes | Wellcome Sanger Institute for the COVID-19 Genomics UK (COG-UK) Consortium | The Lighthouse Lab in Milton Keynes and Alex Alderton, Roberto Amato, Jeffrey Barrett, Sonia Goncalves, Ewan Harrison, David K. Jackson, Ian Johnston, Dominic Kwiatkowski, Cordelia Langford, John Sillitoe on behalf of the Wellcome Sanger Institute COVID-19 Surveillance Team |
| EPI_ISL_2917417 | Lighthouse Laboratory Plymouth | Wellcome Sanger Institute for the COVID-19 Genomics UK (COG-UK) Consortium | Lighthouse Laboratory Plymouth and Alex Alderton, Roberto Amato, Jeffrey Barrett, Sonia Goncalves, Ewan Harrison, David K. Jackson, Ian Johnston, Dominic Kwiatkowski, Cordelia Langford, John Sillitoe on behalf of the Wellcome Sanger Institute COVID-19 Surveillance Team |
| EPI_ISL_2917418 | Berkshire and Surrey Pathology Services Lighthouse Laboratory | Wellcome Sanger Institute for the COVID-19 Genomics UK (COG-UK) Consortium | Berkshire and Surrey Pathology Services Lighthouse Laboratory and Alex Alderton, Roberto Amato, Jeffrey Barrett, Sonia Goncalves, Ewan Harrison, David K. Jackson, Ian Johnston, Dominic Kwiatkowski, Cordelia Langford, John Sillitoe on behalf of the Wellcome Sanger Institute COVID-19 Surveillance Team |
| EPI_ISL_2917419, EPI_ISL_2917420 | Lighthouse Lab in Milton Keynes | Wellcome Sanger Institute for the COVID-19 Genomics UK (COG-UK) Consortium | The Lighthouse Lab in Milton Keynes and Alex Alderton, Roberto Amato, Jeffrey Barrett, Sonia Goncalves, Ewan Harrison, David K. Jackson, Ian Johnston, Dominic Kwiatkowski, Cordelia Langford, John Sillitoe on behalf of the Wellcome Sanger Institute COVID-19 Surveillance Team |
| EPI_ISL_2917421 | Lighthouse Laboratory Plymouth | Wellcome Sanger Institute for the COVID-19 Genomics UK (COG-UK) Consortium | Lighthouse Laboratory Plymouth and Alex Alderton, Roberto Amato, Jeffrey Barrett, Sonia Goncalves, Ewan Harrison, David K. Jackson, Ian Johnston, Dominic Kwiatkowski, Cordelia Langford, John Sillitoe on behalf of the Wellcome Sanger Institute COVID-19 Surveillance Team |
| EPI_ISL_2917422, EPI_ISL_2917423, EPI_ISL_2917424, EPI_ISL_2917425, EPI_ISL_2917426, EPI_ISL_2917427 | Berkshire and Surrey Pathology Services Lighthouse Laboratory | Wellcome Sanger Institute for the COVID-19 Genomics UK (COG-UK) Consortium | Berkshire and Surrey Pathology Services Lighthouse Laboratory and Alex Alderton, Roberto Amato, Jeffrey Barrett, Sonia Goncalves, Ewan Harrison, David K. Jackson, Ian Johnston, Dominic Kwiatkowski, Cordelia Langford, John Sillitoe on behalf of the Wellcome Sanger Institute COVID-19 Surveillance Team |
| EPI_ISL_2917428 | Lighthouse Lab in Milton Keynes | Wellcome Sanger Institute for the COVID-19 Genomics UK (COG-UK) Consortium | The Lighthouse Lab in Milton Keynes and Alex Alderton, Roberto Amato, Jeffrey Barrett, Sonia Goncalves, Ewan Harrison, David K. Jackson, Ian Johnston, Dominic Kwiatkowski, Cordelia Langford, John Sillitoe on behalf of the Wellcome Sanger Institute COVID-19 Surveillance Team |
| EPI_ISL_2917429, EPI_ISL_2917430 | Lighthouse Laboratory Plymouth | Wellcome Sanger Institute for the COVID-19 Genomics UK (COG-UK) Consortium | Lighthouse Laboratory Plymouth and Alex Alderton, Roberto Amato, Jeffrey Barrett, Sonia Goncalves, Ewan Harrison, David K. Jackson, Ian Johnston, Dominic Kwiatkowski, Cordelia Langford, John Sillitoe on behalf of the Wellcome Sanger Institute COVID-19 Surveillance Team |
| EPI_ISL_2917431 | Berkshire and Surrey Pathology Services Lighthouse Laboratory | Wellcome Sanger Institute for the COVID-19 Genomics UK (COG-UK) Consortium | Berkshire and Surrey Pathology Services Lighthouse Laboratory and Alex Alderton, Roberto Amato, Jeffrey Barrett, Sonia Goncalves, Ewan Harrison, David K. Jackson, Ian Johnston, Dominic Kwiatkowski, Cordelia Langford, John Sillitoe on behalf of the Wellcome Sanger Institute COVID-19 Surveillance Team |
| EPI_ISL_2917432 | Lighthouse Laboratory Plymouth | Wellcome Sanger Institute for the COVID-19 Genomics UK (COG-UK) Consortium | Lighthouse Laboratory Plymouth and Alex Alderton, Roberto Amato, Jeffrey Barrett, Sonia Goncalves, Ewan Harrison, David K. Jackson, Ian Johnston, Dominic Kwiatkowski, Cordelia Langford, John Sillitoe on behalf of the Wellcome Sanger Institute COVID-19 Surveillance Team |
| EPI_ISL_2917433 | Berkshire and Surrey Pathology Services Lighthouse Laboratory | Wellcome Sanger Institute for the COVID-19 Genomics UK (COG-UK) Consortium | Berkshire and Surrey Pathology Services Lighthouse Laboratory and Alex Alderton, Roberto Amato, Jeffrey Barrett, Sonia Goncalves, Ewan Harrison, David K. Jackson, Ian Johnston, Dominic Kwiatkowski, Cordelia Langford, John Sillitoe on behalf of the Wellcome Sanger Institute COVID-19 Surveillance Team |
| EPI_ISL_2917434 | Lighthouse Lab in Milton Keynes | Wellcome Sanger Institute for the COVID-19 Genomics UK (COG-UK) Consortium | The Lighthouse Lab in Milton Keynes and Alex Alderton, Roberto Amato, Jeffrey Barrett, Sonia Goncalves, Ewan Harrison, David K. Jackson, Ian Johnston, Dominic Kwiatkowski, Cordelia Langford, John Sillitoe on behalf of the Wellcome Sanger Institute COVID-19 Surveillance Team |
| EPI_ISL_2917435, EPI_ISL_2917436 | Lighthouse Laboratory Plymouth | Wellcome Sanger Institute for the COVID-19 Genomics UK (COG-UK) Consortium | Lighthouse Laboratory Plymouth and Alex Alderton, Roberto Amato, Jeffrey Barrett, Sonia Goncalves, Ewan Harrison, David K. Jackson, Ian Johnston, Dominic Kwiatkowski, Cordelia Langford, John Sillitoe on behalf of the Wellcome Sanger Institute COVID-19 Surveillance Team |
| EPI_ISL_2917437 | Lighthouse Lab in Milton Keynes | Wellcome Sanger Institute for the COVID-19 Genomics UK (COG-UK) Consortium | The Lighthouse Lab in Milton Keynes and Alex Alderton, Roberto Amato, Jeffrey Barrett, Sonia Goncalves, Ewan Harrison, David K. Jackson, Ian Johnston, Dominic Kwiatkowski, Cordelia Langford, John Sillitoe on behalf of the Wellcome Sanger Institute COVID-19 Surveillance Team |
| EPI_ISL_2917438 | Lighthouse Laboratory Plymouth | Wellcome Sanger Institute for the COVID-19 Genomics UK (COG-UK) Consortium | Lighthouse Laboratory Plymouth and Alex Alderton, Roberto Amato, Jeffrey Barrett, Sonia Goncalves, Ewan Harrison, David K. Jackson, Ian Johnston, Dominic Kwiatkowski, Cordelia Langford, John Sillitoe on behalf of the Wellcome Sanger Institute COVID-19 Surveillance Team |
| EPI_ISL_2917439, EPI_ISL_2917440 | Lighthouse Lab in Milton Keynes | Wellcome Sanger Institute for the COVID-19 Genomics UK (COG-UK) Consortium | The Lighthouse Lab in Milton Keynes and Alex Alderton, Roberto Amato, Jeffrey Barrett, Sonia Goncalves, Ewan Harrison, David K. Jackson, Ian Johnston, Dominic Kwiatkowski, Cordelia Langford, John Sillitoe on behalf of the Wellcome Sanger Institute COVID-19 Surveillance Team |
| EPI_ISL_2917441 | Berkshire and Surrey Pathology Services Lighthouse Laboratory | Wellcome Sanger Institute for the COVID-19 Genomics UK (COG-UK) Consortium | Berkshire and Surrey Pathology Services Lighthouse Laboratory and Alex Alderton, Roberto Amato, Jeffrey Barrett, Sonia Goncalves, Ewan Harrison, David K. Jackson, Ian Johnston, Dominic Kwiatkowski, Cordelia Langford, John Sillitoe on behalf of the Wellcome Sanger Institute COVID-19 Surveillance Team |
| EPI_ISL_2917442 | Lighthouse Lab in Milton Keynes | Wellcome Sanger Institute for the COVID-19 Genomics UK (COG-UK) Consortium | The Lighthouse Lab in Milton Keynes and Alex Alderton, Roberto Amato, Jeffrey Barrett, Sonia Goncalves, Ewan Harrison, David K. Jackson, Ian Johnston, Dominic Kwiatkowski, Cordelia Langford, John Sillitoe on behalf of the Wellcome Sanger Institute COVID-19 Surveillance Team |
| EPI_ISL_2917443 | Berkshire and Surrey Pathology Services Lighthouse Laboratory | Wellcome Sanger Institute for the COVID-19 Genomics UK (COG-UK) Consortium | Berkshire and Surrey Pathology Services Lighthouse Laboratory and Alex Alderton, Roberto Amato, Jeffrey Barrett, Sonia Goncalves, Ewan Harrison, David K. Jackson, Ian Johnston, Dominic Kwiatkowski, Cordelia Langford, John Sillitoe on behalf of the Wellcome Sanger Institute COVID-19 Surveillance Team |
| EPI_ISL_2917444 | Lighthouse Lab in Milton Keynes | Wellcome Sanger Institute for the COVID-19 Genomics UK (COG-UK) Consortium | The Lighthouse Lab in Milton Keynes and Alex Alderton, Roberto Amato, Jeffrey Barrett, Sonia Goncalves, Ewan Harrison, David K. Jackson, Ian Johnston, Dominic Kwiatkowski, Cordelia Langford, John Sillitoe on behalf of the Wellcome Sanger Institute COVID-19 Surveillance Team |
| EPI_ISL_2917445 | Berkshire and Surrey Pathology Services Lighthouse Laboratory | Wellcome Sanger Institute for the COVID-19 Genomics UK (COG-UK) Consortium | Berkshire and Surrey Pathology Services Lighthouse Laboratory and Alex Alderton, Roberto Amato, Jeffrey Barrett, Sonia Goncalves, Ewan Harrison, David K. Jackson, Ian Johnston, Dominic Kwiatkowski, Cordelia Langford, John Sillitoe on behalf of the Wellcome Sanger Institute COVID-19 Surveillance Team |
| EPI_ISL_2917446 | Lighthouse Laboratory Plymouth | Wellcome Sanger Institute for the COVID-19 Genomics UK (COG-UK) Consortium | Lighthouse Laboratory Plymouth and Alex Alderton, Roberto Amato, Jeffrey Barrett, Sonia Goncalves, Ewan Harrison, David K. Jackson, Ian Johnston, Dominic Kwiatkowski, Cordelia Langford, John Sillitoe on behalf of the Wellcome Sanger Institute COVID-19 Surveillance Team |
| EPI_ISL_2917447, EPI_ISL_2917448, EPI_ISL_2917449 | Lighthouse Lab in Milton Keynes | Wellcome Sanger Institute for the COVID-19 Genomics UK (COG-UK) Consortium | The Lighthouse Lab in Milton Keynes and Alex Alderton, Roberto Amato, Jeffrey Barrett, Sonia Goncalves, Ewan Harrison, David K. Jackson, Ian Johnston, Dominic Kwiatkowski, Cordelia Langford, John Sillitoe on behalf of the Wellcome Sanger Institute COVID-19 Surveillance Team |
| EPI_ISL_2917450 | Lighthouse Laboratory Plymouth | Wellcome Sanger Institute for the COVID-19 Genomics UK (COG-UK) Consortium | Lighthouse Laboratory Plymouth and Alex Alderton, Roberto Amato, Jeffrey Barrett, Sonia Goncalves, Ewan Harrison, David K. Jackson, Ian Johnston, Dominic Kwiatkowski, Cordelia Langford, John Sillitoe on behalf of the Wellcome Sanger Institute COVID-19 Surveillance Team |
| EPI_ISL_2917451 | Lighthouse Lab in Milton Keynes | Wellcome Sanger Institute for the COVID-19 Genomics UK (COG-UK) Consortium | The Lighthouse Lab in Milton Keynes and Alex Alderton, Roberto Amato, Jeffrey Barrett, Sonia Goncalves, Ewan Harrison, David K. Jackson, Ian Johnston, Dominic Kwiatkowski, Cordelia Langford, John Sillitoe on behalf of the Wellcome Sanger Institute COVID-19 Surveillance Team |
| EPI_ISL_2917452, EPI_ISL_2917453, EPI_ISL_2917454 | Lighthouse Laboratory Plymouth | Wellcome Sanger Institute for the COVID-19 Genomics UK (COG-UK) Consortium | Lighthouse Laboratory Plymouth and Alex Alderton, Roberto Amato, Jeffrey Barrett, Sonia Goncalves, Ewan Harrison, David K. Jackson, Ian Johnston, Dominic Kwiatkowski, Cordelia Langford, John Sillitoe on behalf of the Wellcome Sanger Institute COVID-19 Surveillance Team |
| EPI_ISL_2917455, EPI_ISL_2917456 | Lighthouse Lab in Milton Keynes | Wellcome Sanger Institute for the COVID-19 Genomics UK (COG-UK) Consortium | The Lighthouse Lab in Milton Keynes and Alex Alderton, Roberto Amato, Jeffrey Barrett, Sonia Goncalves, Ewan Harrison, David K. Jackson, Ian Johnston, Dominic Kwiatkowski, Cordelia Langford, John Sillitoe on behalf of the Wellcome Sanger Institute COVID-19 Surveillance Team |
| EPI_ISL_2917457 | Berkshire and Surrey Pathology Services Lighthouse Laboratory | Wellcome Sanger Institute for the COVID-19 Genomics UK (COG-UK) Consortium | Berkshire and Surrey Pathology Services Lighthouse Laboratory and Alex Alderton, Roberto Amato, Jeffrey Barrett, Sonia Goncalves, Ewan Harrison, David K. Jackson, Ian Johnston, Dominic Kwiatkowski, Cordelia Langford, John Sillitoe on behalf of the Wellcome Sanger Institute COVID-19 Surveillance Team |
| EPI_ISL_2917458 | Lighthouse Lab in Milton Keynes | Wellcome Sanger Institute for the COVID-19 Genomics UK (COG-UK) Consortium | The Lighthouse Lab in Milton Keynes and Alex Alderton, Roberto Amato, Jeffrey Barrett, Sonia Goncalves, Ewan Harrison, David K. Jackson, Ian Johnston, Dominic Kwiatkowski, Cordelia Langford, John Sillitoe on behalf of the Wellcome Sanger Institute COVID-19 Surveillance Team |
| EPI_ISL_2917459, EPI_ISL_2917460 | Berkshire and Surrey Pathology Services Lighthouse Laboratory | Wellcome Sanger Institute for the COVID-19 Genomics UK (COG-UK) Consortium | Berkshire and Surrey Pathology Services Lighthouse Laboratory and Alex Alderton, Roberto Amato, Jeffrey Barrett, Sonia Goncalves, Ewan Harrison, David K. Jackson, Ian Johnston, Dominic Kwiatkowski, Cordelia Langford, John Sillitoe on behalf of the Wellcome Sanger Institute COVID-19 Surveillance Team |
| EPI_ISL_2917461, EPI_ISL_2917462 | Lighthouse Laboratory Plymouth | Wellcome Sanger Institute for the COVID-19 Genomics UK (COG-UK) Consortium | Lighthouse Laboratory Plymouth and Alex Alderton, Roberto Amato, Jeffrey Barrett, Sonia Goncalves, Ewan Harrison, David K. Jackson, Ian Johnston, Dominic Kwiatkowski, Cordelia Langford, John Sillitoe on behalf of the Wellcome Sanger Institute COVID-19 Surveillance Team |
| EPI_ISL_2917463 | Berkshire and Surrey Pathology Services Lighthouse Laboratory | Wellcome Sanger Institute for the COVID-19 Genomics UK (COG-UK) Consortium | Berkshire and Surrey Pathology Services Lighthouse Laboratory and Alex Alderton, Roberto Amato, Jeffrey Barrett, Sonia Goncalves, Ewan Harrison, David K. Jackson, Ian Johnston, Dominic Kwiatkowski, Cordelia Langford, John Sillitoe on behalf of the Wellcome Sanger Institute COVID-19 Surveillance Team |
| EPI_ISL_2917464, | Lighthouse Laboratory Plymouth | Wellcome Sanger Institute for the COVID-19 Genomics UK (COG-UK) | Lighthouse Laboratory Plymouth and Alex Alderton, Roberto Amato, Jeffrey Barrett, Sonia Goncalves, Ewan Harrison, David K. Jackson, Ian Johnston, Dominic Kwiatkowski, Cordelia Langford, John Sillitoe on behalf of the Wellcome Sanger |

| | | | |
|---|---|---|---|
| EPI_ISL_2917465 | | Consortium | Institute COVID-19 Surveillance Team |
| EPI_ISL_2917466 | Berkshire and Surrey Pathology Services Lighthouse Laboratory | Wellcome Sanger Institute for the COVID-19 Genomics UK (COG-UK) Consortium | Berkshire and Surrey Pathology Services Lighthouse Laboratory and Alex Alderton, Roberto Amato, Jeffrey Barrett, Sonia Goncalves, Ewan Harrison, David K. Jackson, Ian Johnston, Dominic Kwiatkowski, Cordelia Langford, John Sillitoe on behalf of the Wellcome Sanger Institute COVID-19 Surveillance Team |
| EPI_ISL_2917467 | Lighthouse Lab in Milton Keynes | Wellcome Sanger Institute for the COVID-19 Genomics UK (COG-UK) Consortium | The Lighthouse Lab in Milton Keynes and Alex Alderton, Roberto Amato, Jeffrey Barrett, Sonia Goncalves, Ewan Harrison, David K. Jackson, Ian Johnston, Dominic Kwiatkowski, Cordelia Langford, John Sillitoe on behalf of the Wellcome Sanger Institute COVID-19 Surveillance Team |
| EPI_ISL_2917468, EPI_ISL_2917469 | Lighthouse Laboratory Plymouth | Wellcome Sanger Institute for the COVID-19 Genomics UK (COG-UK) Consortium | Lighthouse Laboratory Plymouth and Alex Alderton, Roberto Amato, Jeffrey Barrett, Sonia Goncalves, Ewan Harrison, David K. Jackson, Ian Johnston, Dominic Kwiatkowski, Cordelia Langford, John Sillitoe on behalf of the Wellcome Sanger Institute COVID-19 Surveillance Team |
| EPI_ISL_2917470 | Lighthouse Lab in Milton Keynes | Wellcome Sanger Institute for the COVID-19 Genomics UK (COG-UK) Consortium | The Lighthouse Lab in Milton Keynes and Alex Alderton, Roberto Amato, Jeffrey Barrett, Sonia Goncalves, Ewan Harrison, David K. Jackson, Ian Johnston, Dominic Kwiatkowski, Cordelia Langford, John Sillitoe on behalf of the Wellcome Sanger Institute COVID-19 Surveillance Team |
| EPI_ISL_2917471 | Lighthouse Laboratory Plymouth | Wellcome Sanger Institute for the COVID-19 Genomics UK (COG-UK) Consortium | Lighthouse Laboratory Plymouth and Alex Alderton, Roberto Amato, Jeffrey Barrett, Sonia Goncalves, Ewan Harrison, David K. Jackson, Ian Johnston, Dominic Kwiatkowski, Cordelia Langford, John Sillitoe on behalf of the Wellcome Sanger Institute COVID-19 Surveillance Team |
| EPI_ISL_2917472 | Lighthouse Lab in Milton Keynes | Wellcome Sanger Institute for the COVID-19 Genomics UK (COG-UK) Consortium | The Lighthouse Lab in Milton Keynes and Alex Alderton, Roberto Amato, Jeffrey Barrett, Sonia Goncalves, Ewan Harrison, David K. Jackson, Ian Johnston, Dominic Kwiatkowski, Cordelia Langford, John Sillitoe on behalf of the Wellcome Sanger Institute COVID-19 Surveillance Team |
| EPI_ISL_2917473 | Berkshire and Surrey Pathology Services Lighthouse Laboratory | Wellcome Sanger Institute for the COVID-19 Genomics UK (COG-UK) Consortium | Berkshire and Surrey Pathology Services Lighthouse Laboratory and Alex Alderton, Roberto Amato, Jeffrey Barrett, Sonia Goncalves, Ewan Harrison, David K. Jackson, Ian Johnston, Dominic Kwiatkowski, Cordelia Langford, John Sillitoe on behalf of the Wellcome Sanger Institute COVID-19 Surveillance Team |
| EPI_ISL_2917474 | Lighthouse Lab in Milton Keynes | Wellcome Sanger Institute for the COVID-19 Genomics UK (COG-UK) Consortium | The Lighthouse Lab in Milton Keynes and Alex Alderton, Roberto Amato, Jeffrey Barrett, Sonia Goncalves, Ewan Harrison, David K. Jackson, Ian Johnston, Dominic Kwiatkowski, Cordelia Langford, John Sillitoe on behalf of the Wellcome Sanger Institute COVID-19 Surveillance Team |
| EPI_ISL_2917475, EPI_ISL_2917476 | Lighthouse Laboratory Plymouth | Wellcome Sanger Institute for the COVID-19 Genomics UK (COG-UK) Consortium | Lighthouse Laboratory Plymouth and Alex Alderton, Roberto Amato, Jeffrey Barrett, Sonia Goncalves, Ewan Harrison, David K. Jackson, Ian Johnston, Dominic Kwiatkowski, Cordelia Langford, John Sillitoe on behalf of the Wellcome Sanger Institute COVID-19 Surveillance Team |
| EPI_ISL_2917477 | Lighthouse Lab in Milton Keynes | Wellcome Sanger Institute for the COVID-19 Genomics UK (COG-UK) Consortium | The Lighthouse Lab in Milton Keynes and Alex Alderton, Roberto Amato, Jeffrey Barrett, Sonia Goncalves, Ewan Harrison, David K. Jackson, Ian Johnston, Dominic Kwiatkowski, Cordelia Langford, John Sillitoe on behalf of the Wellcome Sanger Institute COVID-19 Surveillance Team |
| EPI_ISL_2917478 | Berkshire and Surrey Pathology Services Lighthouse Laboratory | Wellcome Sanger Institute for the COVID-19 Genomics UK (COG-UK) Consortium | Berkshire and Surrey Pathology Services Lighthouse Laboratory and Alex Alderton, Roberto Amato, Jeffrey Barrett, Sonia Goncalves, Ewan Harrison, David K. Jackson, Ian Johnston, Dominic Kwiatkowski, Cordelia Langford, John Sillitoe on behalf of the Wellcome Sanger Institute COVID-19 Surveillance Team |
| EPI_ISL_2917479 | Lighthouse Laboratory Plymouth | Wellcome Sanger Institute for the COVID-19 Genomics UK (COG-UK) Consortium | Lighthouse Laboratory Plymouth and Alex Alderton, Roberto Amato, Jeffrey Barrett, Sonia Goncalves, Ewan Harrison, David K. Jackson, Ian Johnston, Dominic Kwiatkowski, Cordelia Langford, John Sillitoe on behalf of the Wellcome Sanger Institute COVID-19 Surveillance Team |
| EPI_ISL_2917480 | Berkshire and Surrey Pathology Services Lighthouse Laboratory | Wellcome Sanger Institute for the COVID-19 Genomics UK (COG-UK) Consortium | Berkshire and Surrey Pathology Services Lighthouse Laboratory and Alex Alderton, Roberto Amato, Jeffrey Barrett, Sonia Goncalves, Ewan Harrison, David K. Jackson, Ian Johnston, Dominic Kwiatkowski, Cordelia Langford, John Sillitoe on behalf of the Wellcome Sanger Institute COVID-19 Surveillance Team |
| EPI_ISL_2917481 | Lighthouse Laboratory Plymouth | Wellcome Sanger Institute for the COVID-19 Genomics UK (COG-UK) Consortium | Lighthouse Laboratory Plymouth and Alex Alderton, Roberto Amato, Jeffrey Barrett, Sonia Goncalves, Ewan Harrison, David K. Jackson, Ian Johnston, Dominic Kwiatkowski, Cordelia Langford, John Sillitoe on behalf of the Wellcome Sanger Institute COVID-19 Surveillance Team |
| EPI_ISL_2917482 | Lighthouse Lab in Milton Keynes | Wellcome Sanger Institute for the COVID-19 Genomics UK (COG-UK) Consortium | The Lighthouse Lab in Milton Keynes and Alex Alderton, Roberto Amato, Jeffrey Barrett, Sonia Goncalves, Ewan Harrison, David K. Jackson, Ian Johnston, Dominic Kwiatkowski, Cordelia Langford, John Sillitoe on behalf of the Wellcome Sanger Institute COVID-19 Surveillance Team |
| EPI_ISL_2917483 | Berkshire and Surrey Pathology Services Lighthouse Laboratory | Wellcome Sanger Institute for the COVID-19 Genomics UK (COG-UK) Consortium | Berkshire and Surrey Pathology Services Lighthouse Laboratory and Alex Alderton, Roberto Amato, Jeffrey Barrett, Sonia Goncalves, Ewan Harrison, David K. Jackson, Ian Johnston, Dominic Kwiatkowski, Cordelia Langford, John Sillitoe on behalf of the Wellcome Sanger Institute COVID-19 Surveillance Team |
| EPI_ISL_2917484, EPI_ISL_2917485 | Lighthouse Lab in Milton Keynes | Wellcome Sanger Institute for the COVID-19 Genomics UK (COG-UK) Consortium | The Lighthouse Lab in Milton Keynes and Alex Alderton, Roberto Amato, Jeffrey Barrett, Sonia Goncalves, Ewan Harrison, David K. Jackson, Ian Johnston, Dominic Kwiatkowski, Cordelia Langford, John Sillitoe on behalf of the Wellcome Sanger Institute COVID-19 Surveillance Team |
| EPI_ISL_2917486 | Lighthouse Laboratory Plymouth | Wellcome Sanger Institute for the COVID-19 Genomics UK (COG-UK) Consortium | Lighthouse Laboratory Plymouth and Alex Alderton, Roberto Amato, Jeffrey Barrett, Sonia Goncalves, Ewan Harrison, David K. Jackson, Ian Johnston, Dominic Kwiatkowski, Cordelia Langford, John Sillitoe on behalf of the Wellcome Sanger Institute COVID-19 Surveillance Team |
| EPI_ISL_2917487 | Lighthouse Lab in Milton Keynes | Wellcome Sanger Institute for the COVID-19 Genomics UK (COG-UK) Consortium | The Lighthouse Lab in Milton Keynes and Alex Alderton, Roberto Amato, Jeffrey Barrett, Sonia Goncalves, Ewan Harrison, David K. Jackson, Ian Johnston, Dominic Kwiatkowski, Cordelia Langford, John Sillitoe on behalf of the Wellcome Sanger Institute COVID-19 Surveillance Team |
| EPI_ISL_2917488 | Lighthouse Laboratory Plymouth | Wellcome Sanger Institute for the COVID-19 Genomics UK (COG-UK) Consortium | Lighthouse Laboratory Plymouth and Alex Alderton, Roberto Amato, Jeffrey Barrett, Sonia Goncalves, Ewan Harrison, David K. Jackson, Ian Johnston, Dominic Kwiatkowski, Cordelia Langford, John Sillitoe on behalf of the Wellcome Sanger Institute COVID-19 Surveillance Team |
| EPI_ISL_2917489 | Berkshire and Surrey Pathology Services Lighthouse Laboratory | Wellcome Sanger Institute for the COVID-19 Genomics UK (COG-UK) Consortium | Berkshire and Surrey Pathology Services Lighthouse Laboratory and Alex Alderton, Roberto Amato, Jeffrey Barrett, Sonia Goncalves, Ewan Harrison, David K. Jackson, Ian Johnston, Dominic Kwiatkowski, Cordelia Langford, John Sillitoe on behalf of the Wellcome Sanger Institute COVID-19 Surveillance Team |
| EPI_ISL_2917490 | Lighthouse Laboratory Plymouth | Wellcome Sanger Institute for the COVID-19 Genomics UK (COG-UK) Consortium | Lighthouse Laboratory Plymouth and Alex Alderton, Roberto Amato, Jeffrey Barrett, Sonia Goncalves, Ewan Harrison, David K. Jackson, Ian Johnston, Dominic Kwiatkowski, Cordelia Langford, John Sillitoe on behalf of the Wellcome Sanger Institute COVID-19 Surveillance Team |
| EPI_ISL_2917491, EPI_ISL_2917492 | Berkshire and Surrey Pathology Services Lighthouse Laboratory | Wellcome Sanger Institute for the COVID-19 Genomics UK (COG-UK) Consortium | Berkshire and Surrey Pathology Services Lighthouse Laboratory and Alex Alderton, Roberto Amato, Jeffrey Barrett, Sonia Goncalves, Ewan Harrison, David K. Jackson, Ian Johnston, Dominic Kwiatkowski, Cordelia Langford, John Sillitoe on behalf of the Wellcome Sanger Institute COVID-19 Surveillance Team |
| EPI_ISL_2917493 | Lighthouse Laboratory Plymouth | Wellcome Sanger Institute for the COVID-19 Genomics UK (COG-UK) Consortium | Lighthouse Laboratory Plymouth and Alex Alderton, Roberto Amato, Jeffrey Barrett, Sonia Goncalves, Ewan Harrison, David K. Jackson, Ian Johnston, Dominic Kwiatkowski, Cordelia Langford, John Sillitoe on behalf of the Wellcome Sanger Institute COVID-19 Surveillance Team |
| EPI_ISL_2917494 | Lighthouse Lab in Milton Keynes | Wellcome Sanger Institute for the COVID-19 Genomics UK (COG-UK) Consortium | The Lighthouse Lab in Milton Keynes and Alex Alderton, Roberto Amato, Jeffrey Barrett, Sonia Goncalves, Ewan Harrison, David K. Jackson, Ian Johnston, Dominic Kwiatkowski, Cordelia Langford, John Sillitoe on behalf of the Wellcome Sanger Institute COVID-19 Surveillance Team |
| EPI_ISL_2917495 | Lighthouse Laboratory Plymouth | Wellcome Sanger Institute for the COVID-19 Genomics UK (COG-UK) Consortium | Lighthouse Laboratory Plymouth and Alex Alderton, Roberto Amato, Jeffrey Barrett, Sonia Goncalves, Ewan Harrison, David K. Jackson, Ian Johnston, Dominic Kwiatkowski, Cordelia Langford, John Sillitoe on behalf of the Wellcome Sanger Institute COVID-19 Surveillance Team |
| EPI_ISL_2917496, EPI_ISL_2917497 | Berkshire and Surrey Pathology Services Lighthouse Laboratory | Wellcome Sanger Institute for the COVID-19 Genomics UK (COG-UK) Consortium | Berkshire and Surrey Pathology Services Lighthouse Laboratory and Alex Alderton, Roberto Amato, Jeffrey Barrett, Sonia Goncalves, Ewan Harrison, David K. Jackson, Ian Johnston, Dominic Kwiatkowski, Cordelia Langford, John Sillitoe on behalf of the Wellcome Sanger Institute COVID-19 Surveillance Team |
| EPI_ISL_2917498, EPI_ISL_2917499, EPI_ISL_2917500, EPI_ISL_2917501, EPI_ISL_2917502 | Lighthouse Laboratory Plymouth | Wellcome Sanger Institute for the COVID-19 Genomics UK (COG-UK) Consortium | Lighthouse Laboratory Plymouth and Alex Alderton, Roberto Amato, Jeffrey Barrett, Sonia Goncalves, Ewan Harrison, David K. Jackson, Ian Johnston, Dominic Kwiatkowski, Cordelia Langford, John Sillitoe on behalf of the Wellcome Sanger Institute COVID-19 Surveillance Team |
| EPI_ISL_2917503, EPI_ISL_2917504, EPI_ISL_2917505 | Lighthouse Lab in Milton Keynes | Wellcome Sanger Institute for the COVID-19 Genomics UK (COG-UK) Consortium | The Lighthouse Lab in Milton Keynes and Alex Alderton, Roberto Amato, Jeffrey Barrett, Sonia Goncalves, Ewan Harrison, David K. Jackson, Ian Johnston, Dominic Kwiatkowski, Cordelia Langford, John Sillitoe on behalf of the Wellcome Sanger Institute COVID-19 Surveillance Team |
| EPI_ISL_2917506, EPI_ISL_2917507 | Berkshire and Surrey Pathology Services Lighthouse Laboratory | Wellcome Sanger Institute for the COVID-19 Genomics UK (COG-UK) Consortium | Berkshire and Surrey Pathology Services Lighthouse Laboratory and Alex Alderton, Roberto Amato, Jeffrey Barrett, Sonia Goncalves, Ewan Harrison, David K. Jackson, Ian Johnston, Dominic Kwiatkowski, Cordelia Langford, John Sillitoe on behalf of the Wellcome Sanger Institute COVID-19 Surveillance Team |
| EPI_ISL_2917508 | Lighthouse Laboratory Plymouth | Wellcome Sanger Institute for the COVID-19 Genomics UK (COG-UK) Consortium | Lighthouse Laboratory Plymouth and Alex Alderton, Roberto Amato, Jeffrey Barrett, Sonia Goncalves, Ewan Harrison, David K. Jackson, Ian Johnston, Dominic Kwiatkowski, Cordelia Langford, John Sillitoe on behalf of the Wellcome Sanger Institute COVID-19 Surveillance Team |
| EPI_ISL_2917509 | Berkshire and Surrey Pathology Services Lighthouse Laboratory | Wellcome Sanger Institute for the COVID-19 Genomics UK (COG-UK) Consortium | Berkshire and Surrey Pathology Services Lighthouse Laboratory and Alex Alderton, Roberto Amato, Jeffrey Barrett, Sonia Goncalves, Ewan Harrison, David K. Jackson, Ian Johnston, Dominic Kwiatkowski, Cordelia Langford, John Sillitoe on behalf of the Wellcome Sanger Institute COVID-19 Surveillance Team |
| EPI_ISL_2917510 | Lighthouse Lab in Milton Keynes | Wellcome Sanger Institute for the COVID-19 Genomics UK (COG-UK) Consortium | The Lighthouse Lab in Milton Keynes and Alex Alderton, Roberto Amato, Jeffrey Barrett, Sonia Goncalves, Ewan Harrison, David K. Jackson, Ian Johnston, Dominic Kwiatkowski, Cordelia Langford, John Sillitoe on behalf of the Wellcome Sanger Institute COVID-19 Surveillance Team |
| EPI_ISL_2917511, EPI_ISL_2917512, EPI_ISL_2917513, EPI_ISL_2917514 | Lighthouse Laboratory Plymouth | Wellcome Sanger Institute for the COVID-19 Genomics UK (COG-UK) Consortium | Lighthouse Laboratory Plymouth and Alex Alderton, Roberto Amato, Jeffrey Barrett, Sonia Goncalves, Ewan Harrison, David K. Jackson, Ian Johnston, Dominic Kwiatkowski, Cordelia Langford, John Sillitoe on behalf of the Wellcome Sanger Institute COVID-19 Surveillance Team |
| EPI_ISL_2917515, EPI_ISL_2917516 | Lighthouse Lab in Milton Keynes | Wellcome Sanger Institute for the COVID-19 Genomics UK (COG-UK) Consortium | The Lighthouse Lab in Milton Keynes and Alex Alderton, Roberto Amato, Jeffrey Barrett, Sonia Goncalves, Ewan Harrison, David K. Jackson, Ian Johnston, Dominic Kwiatkowski, Cordelia Langford, John Sillitoe on behalf of the Wellcome Sanger Institute COVID-19 Surveillance Team |
| EPI_ISL_2917517, EPI_ISL_2917518, EPI_ISL_2917519 | Lighthouse Laboratory Plymouth | Wellcome Sanger Institute for the COVID-19 Genomics UK (COG-UK) Consortium | Lighthouse Laboratory Plymouth and Alex Alderton, Roberto Amato, Jeffrey Barrett, Sonia Goncalves, Ewan Harrison, David K. Jackson, Ian Johnston, Dominic Kwiatkowski, Cordelia Langford, John Sillitoe on behalf of the Wellcome Sanger Institute COVID-19 Surveillance Team |
| EPI_ISL_2917520 | Berkshire and Surrey Pathology Services Lighthouse Laboratory | Wellcome Sanger Institute for the COVID-19 Genomics UK (COG-UK) Consortium | Berkshire and Surrey Pathology Services Lighthouse Laboratory and Alex Alderton, Roberto Amato, Jeffrey Barrett, Sonia Goncalves, Ewan Harrison, David K. Jackson, Ian Johnston, Dominic Kwiatkowski, Cordelia Langford, John Sillitoe on behalf of the Wellcome Sanger Institute COVID-19 Surveillance Team |
| EPI_ISL_2917522 | Lighthouse Laboratory Plymouth | Wellcome Sanger Institute for the COVID-19 Genomics UK (COG-UK) Consortium | Lighthouse Laboratory Plymouth and Alex Alderton, Roberto Amato, Jeffrey Barrett, Sonia Goncalves, Ewan Harrison, David K. Jackson, Ian Johnston, Dominic Kwiatkowski, Cordelia Langford, John Sillitoe on behalf of the Wellcome Sanger Institute COVID-19 Surveillance Team |
| EPI_ISL_2917523 | Berkshire and Surrey Pathology Services Lighthouse Laboratory | Wellcome Sanger Institute for the COVID-19 Genomics UK (COG-UK) Consortium | Berkshire and Surrey Pathology Services Lighthouse Laboratory and Alex Alderton, Roberto Amato, Jeffrey Barrett, Sonia Goncalves, Ewan Harrison, David K. Jackson, Ian Johnston, Dominic Kwiatkowski, Cordelia Langford, John Sillitoe on behalf of the Wellcome Sanger Institute COVID-19 Surveillance Team |
| EPI_ISL_2917524 | Lighthouse Laboratory Plymouth | Wellcome Sanger Institute for the COVID-19 Genomics UK (COG-UK) Consortium | Lighthouse Laboratory Plymouth and Alex Alderton, Roberto Amato, Jeffrey Barrett, Sonia Goncalves, Ewan Harrison, David K. Jackson, Ian Johnston, Dominic Kwiatkowski, Cordelia Langford, John Sillitoe on behalf of the Wellcome Sanger Institute COVID-19 Surveillance Team |
| EPI_ISL_2917525 | Lighthouse Lab in Milton Keynes | Wellcome Sanger Institute for the COVID-19 Genomics UK (COG-UK) Consortium | The Lighthouse Lab in Milton Keynes and Alex Alderton, Roberto Amato, Jeffrey Barrett, Sonia Goncalves, Ewan Harrison, David K. Jackson, Ian Johnston, Dominic Kwiatkowski, Cordelia Langford, John Sillitoe on behalf of the Wellcome Sanger Institute COVID-19 Surveillance Team |
| EPI_ISL_2917526 | Berkshire and Surrey Pathology Services Lighthouse Laboratory | Wellcome Sanger Institute for the COVID-19 Genomics UK (COG-UK) Consortium | Berkshire and Surrey Pathology Services Lighthouse Laboratory and Alex Alderton, Roberto Amato, Jeffrey Barrett, Sonia Goncalves, Ewan Harrison, David K. Jackson, Ian Johnston, Dominic Kwiatkowski, Cordelia Langford, John Sillitoe on behalf of the Wellcome Sanger Institute COVID-19 Surveillance Team |

| | | | |
|---|---|---|---|
| EPI_ISL_2917527 | Lighthouse Laboratory Plymouth | Wellcome Sanger Institute for the COVID-19 Genomics UK (COG-UK) Consortium | Lighthouse Laboratory Plymouth and Alex Alderton, Roberto Amato, Jeffrey Barrett, Sonia Goncalves, Ewan Harrison, David K. Jackson, Ian Johnston, Dominic Kwiatkowski, Cordelia Langford, John Sillitoe on behalf of the Wellcome Sanger Institute COVID-19 Surveillance Team |
| EPI_ISL_2917528, EPI_ISL_2917529 | Lighthouse Lab in Milton Keynes | Wellcome Sanger Institute for the COVID-19 Genomics UK (COG-UK) Consortium | The Lighthouse Lab in Milton Keynes and Alex Alderton, Roberto Amato, Jeffrey Barrett, Sonia Goncalves, Ewan Harrison, David K. Jackson, Ian Johnston, Dominic Kwiatkowski, Cordelia Langford, John Sillitoe on behalf of the Wellcome Sanger Institute COVID-19 Surveillance Team |
| EPI_ISL_2917530 | Berkshire and Surrey Pathology Services Lighthouse Laboratory | Wellcome Sanger Institute for the COVID-19 Genomics UK (COG-UK) Consortium | Berkshire and Surrey Pathology Services Lighthouse Laboratory and Alex Alderton, Roberto Amato, Jeffrey Barrett, Sonia Goncalves, Ewan Harrison, David K. Jackson, Ian Johnston, Dominic Kwiatkowski, Cordelia Langford, John Sillitoe on behalf of the Wellcome Sanger Institute COVID-19 Surveillance Team |
| EPI_ISL_2917531 | Lighthouse Laboratory Plymouth | Wellcome Sanger Institute for the COVID-19 Genomics UK (COG-UK) Consortium | Lighthouse Laboratory Plymouth and Alex Alderton, Roberto Amato, Jeffrey Barrett, Sonia Goncalves, Ewan Harrison, David K. Jackson, Ian Johnston, Dominic Kwiatkowski, Cordelia Langford, John Sillitoe on behalf of the Wellcome Sanger Institute COVID-19 Surveillance Team |
| EPI_ISL_2917532 | Berkshire and Surrey Pathology Services Lighthouse Laboratory | Wellcome Sanger Institute for the COVID-19 Genomics UK (COG-UK) Consortium | Berkshire and Surrey Pathology Services Lighthouse Laboratory and Alex Alderton, Roberto Amato, Jeffrey Barrett, Sonia Goncalves, Ewan Harrison, David K. Jackson, Ian Johnston, Dominic Kwiatkowski, Cordelia Langford, John Sillitoe on behalf of the Wellcome Sanger Institute COVID-19 Surveillance Team |
| EPI_ISL_2917533 | Lighthouse Laboratory Plymouth | Wellcome Sanger Institute for the COVID-19 Genomics UK (COG-UK) Consortium | Lighthouse Laboratory Plymouth and Alex Alderton, Roberto Amato, Jeffrey Barrett, Sonia Goncalves, Ewan Harrison, David K. Jackson, Ian Johnston, Dominic Kwiatkowski, Cordelia Langford, John Sillitoe on behalf of the Wellcome Sanger Institute COVID-19 Surveillance Team |
| EPI_ISL_2917534 | Lighthouse Lab in Milton Keynes | Wellcome Sanger Institute for the COVID-19 Genomics UK (COG-UK) Consortium | The Lighthouse Lab in Milton Keynes and Alex Alderton, Roberto Amato, Jeffrey Barrett, Sonia Goncalves, Ewan Harrison, David K. Jackson, Ian Johnston, Dominic Kwiatkowski, Cordelia Langford, John Sillitoe on behalf of the Wellcome Sanger Institute COVID-19 Surveillance Team |
| EPI_ISL_2917535 | Berkshire and Surrey Pathology Services Lighthouse Laboratory | Wellcome Sanger Institute for the COVID-19 Genomics UK (COG-UK) Consortium | Berkshire and Surrey Pathology Services Lighthouse Laboratory and Alex Alderton, Roberto Amato, Jeffrey Barrett, Sonia Goncalves, Ewan Harrison, David K. Jackson, Ian Johnston, Dominic Kwiatkowski, Cordelia Langford, John Sillitoe on behalf of the Wellcome Sanger Institute COVID-19 Surveillance Team |
| EPI_ISL_2917536 | Lighthouse Laboratory Plymouth | Wellcome Sanger Institute for the COVID-19 Genomics UK (COG-UK) Consortium | Lighthouse Laboratory Plymouth and Alex Alderton, Roberto Amato, Jeffrey Barrett, Sonia Goncalves, Ewan Harrison, David K. Jackson, Ian Johnston, Dominic Kwiatkowski, Cordelia Langford, John Sillitoe on behalf of the Wellcome Sanger Institute COVID-19 Surveillance Team |
| EPI_ISL_2917537 | Berkshire and Surrey Pathology Services Lighthouse Laboratory | Wellcome Sanger Institute for the COVID-19 Genomics UK (COG-UK) Consortium | Berkshire and Surrey Pathology Services Lighthouse Laboratory and Alex Alderton, Roberto Amato, Jeffrey Barrett, Sonia Goncalves, Ewan Harrison, David K. Jackson, Ian Johnston, Dominic Kwiatkowski, Cordelia Langford, John Sillitoe on behalf of the Wellcome Sanger Institute COVID-19 Surveillance Team |
| EPI_ISL_2917538 | Lighthouse Lab in Milton Keynes | Wellcome Sanger Institute for the COVID-19 Genomics UK (COG-UK) Consortium | The Lighthouse Lab in Milton Keynes and Alex Alderton, Roberto Amato, Jeffrey Barrett, Sonia Goncalves, Ewan Harrison, David K. Jackson, Ian Johnston, Dominic Kwiatkowski, Cordelia Langford, John Sillitoe on behalf of the Wellcome Sanger Institute COVID-19 Surveillance Team |
| EPI_ISL_2917539 | Lighthouse Laboratory Plymouth | Wellcome Sanger Institute for the COVID-19 Genomics UK (COG-UK) Consortium | Lighthouse Laboratory Plymouth and Alex Alderton, Roberto Amato, Jeffrey Barrett, Sonia Goncalves, Ewan Harrison, David K. Jackson, Ian Johnston, Dominic Kwiatkowski, Cordelia Langford, John Sillitoe on behalf of the Wellcome Sanger Institute COVID-19 Surveillance Team |
| EPI_ISL_2917540 | Berkshire and Surrey Pathology Services Lighthouse Laboratory | Wellcome Sanger Institute for the COVID-19 Genomics UK (COG-UK) Consortium | Berkshire and Surrey Pathology Services Lighthouse Laboratory and Alex Alderton, Roberto Amato, Jeffrey Barrett, Sonia Goncalves, Ewan Harrison, David K. Jackson, Ian Johnston, Dominic Kwiatkowski, Cordelia Langford, John Sillitoe on behalf of the Wellcome Sanger Institute COVID-19 Surveillance Team |
| EPI_ISL_2917541 | Lighthouse Laboratory Plymouth | Wellcome Sanger Institute for the COVID-19 Genomics UK (COG-UK) Consortium | Lighthouse Laboratory Plymouth and Alex Alderton, Roberto Amato, Jeffrey Barrett, Sonia Goncalves, Ewan Harrison, David K. Jackson, Ian Johnston, Dominic Kwiatkowski, Cordelia Langford, John Sillitoe on behalf of the Wellcome Sanger Institute COVID-19 Surveillance Team |
| EPI_ISL_2917542, EPI_ISL_2917543 | Berkshire and Surrey Pathology Services Lighthouse Laboratory | Wellcome Sanger Institute for the COVID-19 Genomics UK (COG-UK) Consortium | Berkshire and Surrey Pathology Services Lighthouse Laboratory and Alex Alderton, Roberto Amato, Jeffrey Barrett, Sonia Goncalves, Ewan Harrison, David K. Jackson, Ian Johnston, Dominic Kwiatkowski, Cordelia Langford, John Sillitoe on behalf of the Wellcome Sanger Institute COVID-19 Surveillance Team |
| EPI_ISL_2917544 | Lighthouse Laboratory Plymouth | Wellcome Sanger Institute for the COVID-19 Genomics UK (COG-UK) Consortium | Lighthouse Laboratory Plymouth and Alex Alderton, Roberto Amato, Jeffrey Barrett, Sonia Goncalves, Ewan Harrison, David K. Jackson, Ian Johnston, Dominic Kwiatkowski, Cordelia Langford, John Sillitoe on behalf of the Wellcome Sanger Institute COVID-19 Surveillance Team |
| EPI_ISL_2917545 | Lighthouse Lab in Milton Keynes | Wellcome Sanger Institute for the COVID-19 Genomics UK (COG-UK) Consortium | The Lighthouse Lab in Milton Keynes and Alex Alderton, Roberto Amato, Jeffrey Barrett, Sonia Goncalves, Ewan Harrison, David K. Jackson, Ian Johnston, Dominic Kwiatkowski, Cordelia Langford, John Sillitoe on behalf of the Wellcome Sanger Institute COVID-19 Surveillance Team |
| EPI_ISL_2917546 | Berkshire and Surrey Pathology Services Lighthouse Laboratory | Wellcome Sanger Institute for the COVID-19 Genomics UK (COG-UK) Consortium | Berkshire and Surrey Pathology Services Lighthouse Laboratory and Alex Alderton, Roberto Amato, Jeffrey Barrett, Sonia Goncalves, Ewan Harrison, David K. Jackson, Ian Johnston, Dominic Kwiatkowski, Cordelia Langford, John Sillitoe on behalf of the Wellcome Sanger Institute COVID-19 Surveillance Team |
| EPI_ISL_2917547 | Lighthouse Laboratory Plymouth | Wellcome Sanger Institute for the COVID-19 Genomics UK (COG-UK) Consortium | Lighthouse Laboratory Plymouth and Alex Alderton, Roberto Amato, Jeffrey Barrett, Sonia Goncalves, Ewan Harrison, David K. Jackson, Ian Johnston, Dominic Kwiatkowski, Cordelia Langford, John Sillitoe on behalf of the Wellcome Sanger Institute COVID-19 Surveillance Team |
| EPI_ISL_2917548, EPI_ISL_2917549 | Lighthouse Lab in Milton Keynes | Wellcome Sanger Institute for the COVID-19 Genomics UK (COG-UK) Consortium | The Lighthouse Lab in Milton Keynes and Alex Alderton, Roberto Amato, Jeffrey Barrett, Sonia Goncalves, Ewan Harrison, David K. Jackson, Ian Johnston, Dominic Kwiatkowski, Cordelia Langford, John Sillitoe on behalf of the Wellcome Sanger Institute COVID-19 Surveillance Team |
| EPI_ISL_2917550 | Lighthouse Laboratory Plymouth | Wellcome Sanger Institute for the COVID-19 Genomics UK (COG-UK) Consortium | Lighthouse Laboratory Plymouth and Alex Alderton, Roberto Amato, Jeffrey Barrett, Sonia Goncalves, Ewan Harrison, David K. Jackson, Ian Johnston, Dominic Kwiatkowski, Cordelia Langford, John Sillitoe on behalf of the Wellcome Sanger Institute COVID-19 Surveillance Team |
| EPI_ISL_2917551 | Lighthouse Lab in Milton Keynes | Wellcome Sanger Institute for the COVID-19 Genomics UK (COG-UK) Consortium | The Lighthouse Lab in Milton Keynes and Alex Alderton, Roberto Amato, Jeffrey Barrett, Sonia Goncalves, Ewan Harrison, David K. Jackson, Ian Johnston, Dominic Kwiatkowski, Cordelia Langford, John Sillitoe on behalf of the Wellcome Sanger Institute COVID-19 Surveillance Team |
| EPI_ISL_2917552 | Lighthouse Laboratory Plymouth | Wellcome Sanger Institute for the COVID-19 Genomics UK (COG-UK) Consortium | Lighthouse Laboratory Plymouth and Alex Alderton, Roberto Amato, Jeffrey Barrett, Sonia Goncalves, Ewan Harrison, David K. Jackson, Ian Johnston, Dominic Kwiatkowski, Cordelia Langford, John Sillitoe on behalf of the Wellcome Sanger Institute COVID-19 Surveillance Team |
| EPI_ISL_2917553, EPI_ISL_2917554 | Berkshire and Surrey Pathology Services Lighthouse Laboratory | Wellcome Sanger Institute for the COVID-19 Genomics UK (COG-UK) Consortium | Berkshire and Surrey Pathology Services Lighthouse Laboratory and Alex Alderton, Roberto Amato, Jeffrey Barrett, Sonia Goncalves, Ewan Harrison, David K. Jackson, Ian Johnston, Dominic Kwiatkowski, Cordelia Langford, John Sillitoe on behalf of the Wellcome Sanger Institute COVID-19 Surveillance Team |
| EPI_ISL_2917555 | Lighthouse Laboratory Plymouth | Wellcome Sanger Institute for the COVID-19 Genomics UK (COG-UK) Consortium | Lighthouse Laboratory Plymouth and Alex Alderton, Roberto Amato, Jeffrey Barrett, Sonia Goncalves, Ewan Harrison, David K. Jackson, Ian Johnston, Dominic Kwiatkowski, Cordelia Langford, John Sillitoe on behalf of the Wellcome Sanger Institute COVID-19 Surveillance Team |
| EPI_ISL_2917556 | Berkshire and Surrey Pathology Services Lighthouse Laboratory | Wellcome Sanger Institute for the COVID-19 Genomics UK (COG-UK) Consortium | Berkshire and Surrey Pathology Services Lighthouse Laboratory and Alex Alderton, Roberto Amato, Jeffrey Barrett, Sonia Goncalves, Ewan Harrison, David K. Jackson, Ian Johnston, Dominic Kwiatkowski, Cordelia Langford, John Sillitoe on behalf of the Wellcome Sanger Institute COVID-19 Surveillance Team |
| EPI_ISL_2917557 | Lighthouse Lab in Milton Keynes | Wellcome Sanger Institute for the COVID-19 Genomics UK (COG-UK) Consortium | The Lighthouse Lab in Milton Keynes and Alex Alderton, Roberto Amato, Jeffrey Barrett, Sonia Goncalves, Ewan Harrison, David K. Jackson, Ian Johnston, Dominic Kwiatkowski, Cordelia Langford, John Sillitoe on behalf of the Wellcome Sanger Institute COVID-19 Surveillance Team |
| EPI_ISL_2917558 | Berkshire and Surrey Pathology Services Lighthouse Laboratory | Wellcome Sanger Institute for the COVID-19 Genomics UK (COG-UK) Consortium | Berkshire and Surrey Pathology Services Lighthouse Laboratory and Alex Alderton, Roberto Amato, Jeffrey Barrett, Sonia Goncalves, Ewan Harrison, David K. Jackson, Ian Johnston, Dominic Kwiatkowski, Cordelia Langford, John Sillitoe on behalf of the Wellcome Sanger Institute COVID-19 Surveillance Team |
| EPI_ISL_2917559, EPI_ISL_2917560 | Lighthouse Lab in Milton Keynes | Wellcome Sanger Institute for the COVID-19 Genomics UK (COG-UK) Consortium | The Lighthouse Lab in Milton Keynes and Alex Alderton, Roberto Amato, Jeffrey Barrett, Sonia Goncalves, Ewan Harrison, David K. Jackson, Ian Johnston, Dominic Kwiatkowski, Cordelia Langford, John Sillitoe on behalf of the Wellcome Sanger Institute COVID-19 Surveillance Team |
| EPI_ISL_2917561 | Lighthouse Laboratory Plymouth | Wellcome Sanger Institute for the COVID-19 Genomics UK (COG-UK) Consortium | Lighthouse Laboratory Plymouth and Alex Alderton, Roberto Amato, Jeffrey Barrett, Sonia Goncalves, Ewan Harrison, David K. Jackson, Ian Johnston, Dominic Kwiatkowski, Cordelia Langford, John Sillitoe on behalf of the Wellcome Sanger Institute COVID-19 Surveillance Team |
| EPI_ISL_2917562 | Berkshire and Surrey Pathology Services Lighthouse Laboratory | Wellcome Sanger Institute for the COVID-19 Genomics UK (COG-UK) Consortium | Berkshire and Surrey Pathology Services Lighthouse Laboratory and Alex Alderton, Roberto Amato, Jeffrey Barrett, Sonia Goncalves, Ewan Harrison, David K. Jackson, Ian Johnston, Dominic Kwiatkowski, Cordelia Langford, John Sillitoe on behalf of the Wellcome Sanger Institute COVID-19 Surveillance Team |
| EPI_ISL_2917563 | Lighthouse Lab in Milton Keynes | Wellcome Sanger Institute for the COVID-19 Genomics UK (COG-UK) Consortium | The Lighthouse Lab in Milton Keynes and Alex Alderton, Roberto Amato, Jeffrey Barrett, Sonia Goncalves, Ewan Harrison, David K. Jackson, Ian Johnston, Dominic Kwiatkowski, Cordelia Langford, John Sillitoe on behalf of the Wellcome Sanger Institute COVID-19 Surveillance Team |
| EPI_ISL_2917564, EPI_ISL_2917565 | Lighthouse Laboratory Plymouth | Wellcome Sanger Institute for the COVID-19 Genomics UK (COG-UK) Consortium | Lighthouse Laboratory Plymouth and Alex Alderton, Roberto Amato, Jeffrey Barrett, Sonia Goncalves, Ewan Harrison, David K. Jackson, Ian Johnston, Dominic Kwiatkowski, Cordelia Langford, John Sillitoe on behalf of the Wellcome Sanger Institute COVID-19 Surveillance Team |
| EPI_ISL_2917566 | Lighthouse Lab in Milton Keynes | Wellcome Sanger Institute for the COVID-19 Genomics UK (COG-UK) Consortium | The Lighthouse Lab in Milton Keynes and Alex Alderton, Roberto Amato, Jeffrey Barrett, Sonia Goncalves, Ewan Harrison, David K. Jackson, Ian Johnston, Dominic Kwiatkowski, Cordelia Langford, John Sillitoe on behalf of the Wellcome Sanger Institute COVID-19 Surveillance Team |
| EPI_ISL_2917567, EPI_ISL_2917568, EPI_ISL_2917569 | Lighthouse Laboratory Plymouth | Wellcome Sanger Institute for the COVID-19 Genomics UK (COG-UK) Consortium | Lighthouse Laboratory Plymouth and Alex Alderton, Roberto Amato, Jeffrey Barrett, Sonia Goncalves, Ewan Harrison, David K. Jackson, Ian Johnston, Dominic Kwiatkowski, Cordelia Langford, John Sillitoe on behalf of the Wellcome Sanger Institute COVID-19 Surveillance Team |
| EPI_ISL_2917570 | Lighthouse Lab in Milton Keynes | Wellcome Sanger Institute for the COVID-19 Genomics UK (COG-UK) Consortium | The Lighthouse Lab in Milton Keynes and Alex Alderton, Roberto Amato, Jeffrey Barrett, Sonia Goncalves, Ewan Harrison, David K. Jackson, Ian Johnston, Dominic Kwiatkowski, Cordelia Langford, John Sillitoe on behalf of the Wellcome Sanger Institute COVID-19 Surveillance Team |
| EPI_ISL_2917571 | Berkshire and Surrey Pathology Services Lighthouse Laboratory | Wellcome Sanger Institute for the COVID-19 Genomics UK (COG-UK) Consortium | Berkshire and Surrey Pathology Services Lighthouse Laboratory and Alex Alderton, Roberto Amato, Jeffrey Barrett, Sonia Goncalves, Ewan Harrison, David K. Jackson, Ian Johnston, Dominic Kwiatkowski, Cordelia Langford, John Sillitoe on behalf of the Wellcome Sanger Institute COVID-19 Surveillance Team |
| EPI_ISL_2917572 | Lighthouse Lab in Milton Keynes | Wellcome Sanger Institute for the COVID-19 Genomics UK (COG-UK) Consortium | The Lighthouse Lab in Milton Keynes and Alex Alderton, Roberto Amato, Jeffrey Barrett, Sonia Goncalves, Ewan Harrison, David K. Jackson, Ian Johnston, Dominic Kwiatkowski, Cordelia Langford, John Sillitoe on behalf of the Wellcome Sanger Institute COVID-19 Surveillance Team |
| EPI_ISL_2917573 | Lighthouse Laboratory Plymouth | Wellcome Sanger Institute for the COVID-19 Genomics UK (COG-UK) Consortium | Lighthouse Laboratory Plymouth and Alex Alderton, Roberto Amato, Jeffrey Barrett, Sonia Goncalves, Ewan Harrison, David K. Jackson, Ian Johnston, Dominic Kwiatkowski, Cordelia Langford, John Sillitoe on behalf of the Wellcome Sanger Institute COVID-19 Surveillance Team |
| EPI_ISL_2917574, EPI_ISL_2917575 | Berkshire and Surrey Pathology Services Lighthouse Laboratory | Wellcome Sanger Institute for the COVID-19 Genomics UK (COG-UK) Consortium | Berkshire and Surrey Pathology Services Lighthouse Laboratory and Alex Alderton, Roberto Amato, Jeffrey Barrett, Sonia Goncalves, Ewan Harrison, David K. Jackson, Ian Johnston, Dominic Kwiatkowski, Cordelia Langford, John Sillitoe on behalf of the Wellcome Sanger Institute COVID-19 Surveillance Team |
| EPI_ISL_2917576 | Lighthouse Lab in Milton Keynes | Wellcome Sanger Institute for the COVID-19 Genomics UK (COG-UK) Consortium | The Lighthouse Lab in Milton Keynes and Alex Alderton, Roberto Amato, Jeffrey Barrett, Sonia Goncalves, Ewan Harrison, David K. Jackson, Ian Johnston, Dominic Kwiatkowski, Cordelia Langford, John Sillitoe on behalf of the Wellcome Sanger Institute COVID-19 Surveillance Team |
| EPI_ISL_2917577 | Berkshire and Surrey Pathology Services Lighthouse Laboratory | Wellcome Sanger Institute for the COVID-19 Genomics UK (COG-UK) Consortium | Berkshire and Surrey Pathology Services Lighthouse Laboratory and Alex Alderton, Roberto Amato, Jeffrey Barrett, Sonia Goncalves, Ewan Harrison, David K. Jackson, Ian Johnston, Dominic Kwiatkowski, Cordelia Langford, John Sillitoe on behalf of the Wellcome Sanger Institute COVID-19 Surveillance Team |
| EPI_ISL_2917578 | Lighthouse Lab in Milton Keynes | Wellcome Sanger Institute for the COVID-19 Genomics UK (COG-UK) Consortium | The Lighthouse Lab in Milton Keynes and Alex Alderton, Roberto Amato, Jeffrey Barrett, Sonia Goncalves, Ewan Harrison, David K. Jackson, Ian Johnston, Dominic Kwiatkowski, Cordelia Langford, John Sillitoe on behalf of the Wellcome Sanger Institute COVID-19 Surveillance Team |
| EPI_ISL_2917579 | Lighthouse Laboratory Plymouth | Wellcome Sanger Institute for the COVID-19 Genomics UK (COG-UK) Consortium | Lighthouse Laboratory Plymouth and Alex Alderton, Roberto Amato, Jeffrey Barrett, Sonia Goncalves, Ewan Harrison, David K. Jackson, Ian Johnston, Dominic Kwiatkowski, Cordelia Langford, John Sillitoe on behalf of the Wellcome Sanger Institute COVID-19 Surveillance Team |
| EPI_ISL_2917580 | Lighthouse Lab in Milton Keynes | Wellcome Sanger Institute for the COVID-19 Genomics UK (COG-UK) | The Lighthouse Lab in Milton Keynes and Alex Alderton, Roberto Amato, Jeffrey Barrett, Sonia Goncalves, Ewan Harrison, David K. Jackson, Ian Johnston, Dominic Kwiatkowski, Cordelia Langford, John Sillitoe on behalf of the Wellcome Sanger |

| | | | |
|---|---|---|---|
| | | Consortium | Institute COVID-19 Surveillance Team |
| EPI_ISL_2917581 | Berkshire and Surrey Pathology Services Lighthouse Laboratory | Wellcome Sanger Institute for the COVID-19 Genomics UK (COG-UK) Consortium | Berkshire and Surrey Pathology Services Lighthouse Laboratory and Alex Alderton, Roberto Amato, Jeffrey Barrett, Sonia Goncalves, Ewan Harrison, David K. Jackson, Ian Johnston, Dominic Kwiatkowski, Cordelia Langford, John Sillitoe on behalf of the Wellcome Sanger Institute COVID-19 Surveillance Team |
| EPI_ISL_2917582 | Lighthouse Lab in Milton Keynes | Wellcome Sanger Institute for the COVID-19 Genomics UK (COG-UK) Consortium | The Lighthouse Lab in Milton Keynes and Alex Alderton, Roberto Amato, Jeffrey Barrett, Sonia Goncalves, Ewan Harrison, David K. Jackson, Ian Johnston, Dominic Kwiatkowski, Cordelia Langford, John Sillitoe on behalf of the Wellcome Sanger Institute COVID-19 Surveillance Team |
| EPI_ISL_2917583 | Berkshire and Surrey Pathology Services Lighthouse Laboratory | Wellcome Sanger Institute for the COVID-19 Genomics UK (COG-UK) Consortium | Berkshire and Surrey Pathology Services Lighthouse Laboratory and Alex Alderton, Roberto Amato, Jeffrey Barrett, Sonia Goncalves, Ewan Harrison, David K. Jackson, Ian Johnston, Dominic Kwiatkowski, Cordelia Langford, John Sillitoe on behalf of the Wellcome Sanger Institute COVID-19 Surveillance Team |
| EPI_ISL_2917584 | Lighthouse Lab in Milton Keynes | Wellcome Sanger Institute for the COVID-19 Genomics UK (COG-UK) Consortium | The Lighthouse Lab in Milton Keynes and Alex Alderton, Roberto Amato, Jeffrey Barrett, Sonia Goncalves, Ewan Harrison, David K. Jackson, Ian Johnston, Dominic Kwiatkowski, Cordelia Langford, John Sillitoe on behalf of the Wellcome Sanger Institute COVID-19 Surveillance Team |
| EPI_ISL_2917585 | Lighthouse Laboratory Plymouth | Wellcome Sanger Institute for the COVID-19 Genomics UK (COG-UK) Consortium | Lighthouse Laboratory Plymouth and Alex Alderton, Roberto Amato, Jeffrey Barrett, Sonia Goncalves, Ewan Harrison, David K. Jackson, Ian Johnston, Dominic Kwiatkowski, Cordelia Langford, John Sillitoe on behalf of the Wellcome Sanger Institute COVID-19 Surveillance Team |
| EPI_ISL_2917586 | Berkshire and Surrey Pathology Services Lighthouse Laboratory | Wellcome Sanger Institute for the COVID-19 Genomics UK (COG-UK) Consortium | Berkshire and Surrey Pathology Services Lighthouse Laboratory and Alex Alderton, Roberto Amato, Jeffrey Barrett, Sonia Goncalves, Ewan Harrison, David K. Jackson, Ian Johnston, Dominic Kwiatkowski, Cordelia Langford, John Sillitoe on behalf of the Wellcome Sanger Institute COVID-19 Surveillance Team |
| EPI_ISL_2917587, EPI_ISL_2917588 | Lighthouse Laboratory Plymouth | Wellcome Sanger Institute for the COVID-19 Genomics UK (COG-UK) Consortium | Lighthouse Laboratory Plymouth and Alex Alderton, Roberto Amato, Jeffrey Barrett, Sonia Goncalves, Ewan Harrison, David K. Jackson, Ian Johnston, Dominic Kwiatkowski, Cordelia Langford, John Sillitoe on behalf of the Wellcome Sanger Institute COVID-19 Surveillance Team |
| EPI_ISL_2917589 | Berkshire and Surrey Pathology Services Lighthouse Laboratory | Wellcome Sanger Institute for the COVID-19 Genomics UK (COG-UK) Consortium | Berkshire and Surrey Pathology Services Lighthouse Laboratory and Alex Alderton, Roberto Amato, Jeffrey Barrett, Sonia Goncalves, Ewan Harrison, David K. Jackson, Ian Johnston, Dominic Kwiatkowski, Cordelia Langford, John Sillitoe on behalf of the Wellcome Sanger Institute COVID-19 Surveillance Team |
| EPI_ISL_2917590, EPI_ISL_2917591, EPI_ISL_2917592, EPI_ISL_2917593, EPI_ISL_2917594 | Lighthouse Lab in Milton Keynes | Wellcome Sanger Institute for the COVID-19 Genomics UK (COG-UK) Consortium | The Lighthouse Lab in Milton Keynes and Alex Alderton, Roberto Amato, Jeffrey Barrett, Sonia Goncalves, Ewan Harrison, David K. Jackson, Ian Johnston, Dominic Kwiatkowski, Cordelia Langford, John Sillitoe on behalf of the Wellcome Sanger Institute COVID-19 Surveillance Team |
| EPI_ISL_2917595, EPI_ISL_2917596 | Berkshire and Surrey Pathology Services Lighthouse Laboratory | Wellcome Sanger Institute for the COVID-19 Genomics UK (COG-UK) Consortium | Berkshire and Surrey Pathology Services Lighthouse Laboratory and Alex Alderton, Roberto Amato, Jeffrey Barrett, Sonia Goncalves, Ewan Harrison, David K. Jackson, Ian Johnston, Dominic Kwiatkowski, Cordelia Langford, John Sillitoe on behalf of the Wellcome Sanger Institute COVID-19 Surveillance Team |
| EPI_ISL_2917597, EPI_ISL_2917598 | Lighthouse Lab in Milton Keynes | Wellcome Sanger Institute for the COVID-19 Genomics UK (COG-UK) Consortium | The Lighthouse Lab in Milton Keynes and Alex Alderton, Roberto Amato, Jeffrey Barrett, Sonia Goncalves, Ewan Harrison, David K. Jackson, Ian Johnston, Dominic Kwiatkowski, Cordelia Langford, John Sillitoe on behalf of the Wellcome Sanger Institute COVID-19 Surveillance Team |
| EPI_ISL_2917599, EPI_ISL_2917600, EPI_ISL_2917601, EPI_ISL_2917602 | Lighthouse Lab in Glasgow | Wellcome Sanger Institute for the COVID-19 Genomics UK (COG-UK) Consortium | Harper VanSteenhouse, Yumi Kasai, David Gray, Carol Clugston, Anna Dominiczak and Alex Alderton, Roberto Amato, Jeffrey Barrett, Sonia Goncalves, Ewan Harrison, David K. Jackson, Ian Johnston, Dominic Kwiatkowski, Cordelia Langford, John Sillitoe on behalf of the Wellcome Sanger Institute COVID-19 Surveillance Team |
| EPI_ISL_2917603, EPI_ISL_2917604 | Berkshire and Surrey Pathology Services Lighthouse Laboratory | Wellcome Sanger Institute for the COVID-19 Genomics UK (COG-UK) Consortium | Berkshire and Surrey Pathology Services Lighthouse Laboratory and Alex Alderton, Roberto Amato, Jeffrey Barrett, Sonia Goncalves, Ewan Harrison, David K. Jackson, Ian Johnston, Dominic Kwiatkowski, Cordelia Langford, John Sillitoe on behalf of the Wellcome Sanger Institute COVID-19 Surveillance Team |
| EPI_ISL_2917605, EPI_ISL_2917606, EPI_ISL_2917607, EPI_ISL_2917608, EPI_ISL_2917609, EPI_ISL_2917610, EPI_ISL_2917611, EPI_ISL_2917612 | Lighthouse Lab in Glasgow | Wellcome Sanger Institute for the COVID-19 Genomics UK (COG-UK) Consortium | Harper VanSteenhouse, Yumi Kasai, David Gray, Carol Clugston, Anna Dominiczak and Alex Alderton, Roberto Amato, Jeffrey Barrett, Sonia Goncalves, Ewan Harrison, David K. Jackson, Ian Johnston, Dominic Kwiatkowski, Cordelia Langford, John Sillitoe on behalf of the Wellcome Sanger Institute COVID-19 Surveillance Team |
| EPI_ISL_2917613 | Berkshire and Surrey Pathology Services Lighthouse Laboratory | Wellcome Sanger Institute for the COVID-19 Genomics UK (COG-UK) Consortium | Berkshire and Surrey Pathology Services Lighthouse Laboratory and Alex Alderton, Roberto Amato, Jeffrey Barrett, Sonia Goncalves, Ewan Harrison, David K. Jackson, Ian Johnston, Dominic Kwiatkowski, Cordelia Langford, John Sillitoe on behalf of the Wellcome Sanger Institute COVID-19 Surveillance Team |
| EPI_ISL_2917614, EPI_ISL_2917615 | Lighthouse Lab in Glasgow | Wellcome Sanger Institute for the COVID-19 Genomics UK (COG-UK) Consortium | Harper VanSteenhouse, Yumi Kasai, David Gray, Carol Clugston, Anna Dominiczak and Alex Alderton, Roberto Amato, Jeffrey Barrett, Sonia Goncalves, Ewan Harrison, David K. Jackson, Ian Johnston, Dominic Kwiatkowski, Cordelia Langford, John Sillitoe on behalf of the Wellcome Sanger Institute COVID-19 Surveillance Team |
| EPI_ISL_2917616 | Berkshire and Surrey Pathology Services Lighthouse Laboratory | Wellcome Sanger Institute for the COVID-19 Genomics UK (COG-UK) Consortium | Berkshire and Surrey Pathology Services Lighthouse Laboratory and Alex Alderton, Roberto Amato, Jeffrey Barrett, Sonia Goncalves, Ewan Harrison, David K. Jackson, Ian Johnston, Dominic Kwiatkowski, Cordelia Langford, John Sillitoe on behalf of the Wellcome Sanger Institute COVID-19 Surveillance Team |
| EPI_ISL_2917617, EPI_ISL_2917618 | Lighthouse Lab in Glasgow | Wellcome Sanger Institute for the COVID-19 Genomics UK (COG-UK) Consortium | Harper VanSteenhouse, Yumi Kasai, David Gray, Carol Clugston, Anna Dominiczak and Alex Alderton, Roberto Amato, Jeffrey Barrett, Sonia Goncalves, Ewan Harrison, David K. Jackson, Ian Johnston, Dominic Kwiatkowski, Cordelia Langford, John Sillitoe on behalf of the Wellcome Sanger Institute COVID-19 Surveillance Team |
| EPI_ISL_2917619, EPI_ISL_2917620 | Berkshire and Surrey Pathology Services Lighthouse Laboratory | Wellcome Sanger Institute for the COVID-19 Genomics UK (COG-UK) Consortium | Berkshire and Surrey Pathology Services Lighthouse Laboratory and Alex Alderton, Roberto Amato, Jeffrey Barrett, Sonia Goncalves, Ewan Harrison, David K. Jackson, Ian Johnston, Dominic Kwiatkowski, Cordelia Langford, John Sillitoe on behalf of the Wellcome Sanger Institute COVID-19 Surveillance Team |
| EPI_ISL_2917621, EPI_ISL_2917622 | Lighthouse Lab in Glasgow | Wellcome Sanger Institute for the COVID-19 Genomics UK (COG-UK) Consortium | Harper VanSteenhouse, Yumi Kasai, David Gray, Carol Clugston, Anna Dominiczak and Alex Alderton, Roberto Amato, Jeffrey Barrett, Sonia Goncalves, Ewan Harrison, David K. Jackson, Ian Johnston, Dominic Kwiatkowski, Cordelia Langford, John Sillitoe on behalf of the Wellcome Sanger Institute COVID-19 Surveillance Team |
| EPI_ISL_2917623 | Berkshire and Surrey Pathology Services Lighthouse Laboratory | Wellcome Sanger Institute for the COVID-19 Genomics UK (COG-UK) Consortium | Berkshire and Surrey Pathology Services Lighthouse Laboratory and Alex Alderton, Roberto Amato, Jeffrey Barrett, Sonia Goncalves, Ewan Harrison, David K. Jackson, Ian Johnston, Dominic Kwiatkowski, Cordelia Langford, John Sillitoe on behalf of the Wellcome Sanger Institute COVID-19 Surveillance Team |
| EPI_ISL_2917624, EPI_ISL_2917625, EPI_ISL_2917626, EPI_ISL_2917627, EPI_ISL_2917628, EPI_ISL_2917629, EPI_ISL_2917630, EPI_ISL_2917631, EPI_ISL_2917632, EPI_ISL_2917633 | Lighthouse Lab in Glasgow | Wellcome Sanger Institute for the COVID-19 Genomics UK (COG-UK) Consortium | Harper VanSteenhouse, Yumi Kasai, David Gray, Carol Clugston, Anna Dominiczak and Alex Alderton, Roberto Amato, Jeffrey Barrett, Sonia Goncalves, Ewan Harrison, David K. Jackson, Ian Johnston, Dominic Kwiatkowski, Cordelia Langford, John Sillitoe on behalf of the Wellcome Sanger Institute COVID-19 Surveillance Team |
| EPI_ISL_2917634 | Berkshire and Surrey Pathology Services Lighthouse Laboratory | Wellcome Sanger Institute for the COVID-19 Genomics UK (COG-UK) Consortium | Berkshire and Surrey Pathology Services Lighthouse Laboratory and Alex Alderton, Roberto Amato, Jeffrey Barrett, Sonia Goncalves, Ewan Harrison, David K. Jackson, Ian Johnston, Dominic Kwiatkowski, Cordelia Langford, John Sillitoe on behalf of the Wellcome Sanger Institute COVID-19 Surveillance Team |
| EPI_ISL_2917635 | Lighthouse Lab in Glasgow | Wellcome Sanger Institute for the COVID-19 Genomics UK (COG-UK) Consortium | Harper VanSteenhouse, Yumi Kasai, David Gray, Carol Clugston, Anna Dominiczak and Alex Alderton, Roberto Amato, Jeffrey Barrett, Sonia Goncalves, Ewan Harrison, David K. Jackson, Ian Johnston, Dominic Kwiatkowski, Cordelia Langford, John Sillitoe on behalf of the Wellcome Sanger Institute COVID-19 Surveillance Team |
| EPI_ISL_2917636 | Berkshire and Surrey Pathology Services Lighthouse Laboratory | Wellcome Sanger Institute for the COVID-19 Genomics UK (COG-UK) Consortium | Berkshire and Surrey Pathology Services Lighthouse Laboratory and Alex Alderton, Roberto Amato, Jeffrey Barrett, Sonia Goncalves, Ewan Harrison, David K. Jackson, Ian Johnston, Dominic Kwiatkowski, Cordelia Langford, John Sillitoe on behalf of the Wellcome Sanger Institute COVID-19 Surveillance Team |
| EPI_ISL_2917637, EPI_ISL_2917638, EPI_ISL_2917639, EPI_ISL_2917640 | Lighthouse Lab in Glasgow | Wellcome Sanger Institute for the COVID-19 Genomics UK (COG-UK) Consortium | Harper VanSteenhouse, Yumi Kasai, David Gray, Carol Clugston, Anna Dominiczak and Alex Alderton, Roberto Amato, Jeffrey Barrett, Sonia Goncalves, Ewan Harrison, David K. Jackson, Ian Johnston, Dominic Kwiatkowski, Cordelia Langford, John Sillitoe on behalf of the Wellcome Sanger Institute COVID-19 Surveillance Team |
| EPI_ISL_2917641 | Berkshire and Surrey Pathology Services Lighthouse Laboratory | Wellcome Sanger Institute for the COVID-19 Genomics UK (COG-UK) Consortium | Berkshire and Surrey Pathology Services Lighthouse Laboratory and Alex Alderton, Roberto Amato, Jeffrey Barrett, Sonia Goncalves, Ewan Harrison, David K. Jackson, Ian Johnston, Dominic Kwiatkowski, Cordelia Langford, John Sillitoe on behalf of the Wellcome Sanger Institute COVID-19 Surveillance Team |
| EPI_ISL_2917642 | Lighthouse Lab in Glasgow | Wellcome Sanger Institute for the COVID-19 Genomics UK (COG-UK) Consortium | Harper VanSteenhouse, Yumi Kasai, David Gray, Carol Clugston, Anna Dominiczak and Alex Alderton, Roberto Amato, Jeffrey Barrett, Sonia Goncalves, Ewan Harrison, David K. Jackson, Ian Johnston, Dominic Kwiatkowski, Cordelia Langford, John Sillitoe on behalf of the Wellcome Sanger Institute COVID-19 Surveillance Team |
| EPI_ISL_2917643 | Berkshire and Surrey Pathology Services Lighthouse Laboratory | Wellcome Sanger Institute for the COVID-19 Genomics UK (COG-UK) Consortium | Berkshire and Surrey Pathology Services Lighthouse Laboratory and Alex Alderton, Roberto Amato, Jeffrey Barrett, Sonia Goncalves, Ewan Harrison, David K. Jackson, Ian Johnston, Dominic Kwiatkowski, Cordelia Langford, John Sillitoe on behalf of the Wellcome Sanger Institute COVID-19 Surveillance Team |
| EPI_ISL_2917644, EPI_ISL_2917645 | Lighthouse Lab in Glasgow | Wellcome Sanger Institute for the COVID-19 Genomics UK (COG-UK) Consortium | Harper VanSteenhouse, Yumi Kasai, David Gray, Carol Clugston, Anna Dominiczak and Alex Alderton, Roberto Amato, Jeffrey Barrett, Sonia Goncalves, Ewan Harrison, David K. Jackson, Ian Johnston, Dominic Kwiatkowski, Cordelia Langford, John Sillitoe on behalf of the Wellcome Sanger Institute COVID-19 Surveillance Team |
| EPI_ISL_2917646 | Berkshire and Surrey Pathology Services Lighthouse Laboratory | Wellcome Sanger Institute for the COVID-19 Genomics UK (COG-UK) Consortium | Berkshire and Surrey Pathology Services Lighthouse Laboratory and Alex Alderton, Roberto Amato, Jeffrey Barrett, Sonia Goncalves, Ewan Harrison, David K. Jackson, Ian Johnston, Dominic Kwiatkowski, Cordelia Langford, John Sillitoe on behalf of the Wellcome Sanger Institute COVID-19 Surveillance Team |
| EPI_ISL_2917647 | Lighthouse Lab in Glasgow | Wellcome Sanger Institute for the COVID-19 Genomics UK (COG-UK) Consortium | Harper VanSteenhouse, Yumi Kasai, David Gray, Carol Clugston, Anna Dominiczak and Alex Alderton, Roberto Amato, Jeffrey Barrett, Sonia Goncalves, Ewan Harrison, David K. Jackson, Ian Johnston, Dominic Kwiatkowski, Cordelia Langford, John Sillitoe on behalf of the Wellcome Sanger Institute COVID-19 Surveillance Team |
| EPI_ISL_2917648, EPI_ISL_2917649 | Berkshire and Surrey Pathology Services Lighthouse Laboratory | Wellcome Sanger Institute for the COVID-19 Genomics UK (COG-UK) Consortium | Berkshire and Surrey Pathology Services Lighthouse Laboratory and Alex Alderton, Roberto Amato, Jeffrey Barrett, Sonia Goncalves, Ewan Harrison, David K. Jackson, Ian Johnston, Dominic Kwiatkowski, Cordelia Langford, John Sillitoe on behalf of the Wellcome Sanger Institute COVID-19 Surveillance Team |
| EPI_ISL_2917650, EPI_ISL_2917651 | Lighthouse Lab in Glasgow | Wellcome Sanger Institute for the COVID-19 Genomics UK (COG-UK) Consortium | Harper VanSteenhouse, Yumi Kasai, David Gray, Carol Clugston, Anna Dominiczak and Alex Alderton, Roberto Amato, Jeffrey Barrett, Sonia Goncalves, Ewan Harrison, David K. Jackson, Ian Johnston, Dominic Kwiatkowski, Cordelia Langford, John Sillitoe on behalf of the Wellcome Sanger Institute COVID-19 Surveillance Team |
| EPI_ISL_2917652 | Berkshire and Surrey Pathology Services Lighthouse Laboratory | Wellcome Sanger Institute for the COVID-19 Genomics UK (COG-UK) Consortium | Berkshire and Surrey Pathology Services Lighthouse Laboratory and Alex Alderton, Roberto Amato, Jeffrey Barrett, Sonia Goncalves, Ewan Harrison, David K. Jackson, Ian Johnston, Dominic Kwiatkowski, Cordelia Langford, John Sillitoe on behalf of the Wellcome Sanger Institute COVID-19 Surveillance Team |
| EPI_ISL_2917653, EPI_ISL_2917654, | Lighthouse Lab in Glasgow | Wellcome Sanger Institute for the COVID-19 Genomics UK (COG-UK) Consortium | Harper VanSteenhouse, Yumi Kasai, David Gray, Carol Clugston, Anna Dominiczak and Alex Alderton, Roberto Amato, Jeffrey Barrett, Sonia Goncalves, Ewan Harrison, David K. Jackson, Ian Johnston, Dominic Kwiatkowski, Cordelia Langford, John Sillitoe on behalf of the Wellcome Sanger Institute COVID-19 Surveillance Team |

| | | | |
|---|---|---|---|
| EPI_ISL_2917655, EPI_ISL_2917656 | | | |
| EPI_ISL_2917657, EPI_ISL_2917658 | Berkshire and Surrey Pathology Services Lighthouse Laboratory | Wellcome Sanger Institute for the COVID-19 Genomics UK (COG-UK) Consortium | Berkshire and Surrey Pathology Services Lighthouse Laboratory and Alex Alderton, Roberto Amato, Jeffrey Barrett, Sonia Goncalves, Ewan Harrison, David K. Jackson, Ian Johnston, Dominic Kwiatkowski, Cordelia Langford, John Sillitoe on behalf of the Wellcome Sanger Institute COVID-19 Surveillance Team |
| EPI_ISL_2917659, EPI_ISL_2917660, EPI_ISL_2917661, EPI_ISL_2917662, EPI_ISL_2917663, EPI_ISL_2917664, EPI_ISL_2917665, EPI_ISL_2917666, EPI_ISL_2917667, EPI_ISL_2917668, EPI_ISL_2917669, EPI_ISL_2917670 | | | |
| see above | Lighthouse Lab in Glasgow | Wellcome Sanger Institute for the COVID-19 Genomics UK (COG-UK) Consortium | Harper VanSteenhouse, Yumi Kasai, David Gray, Carol Clugston, Anna Dominiczak and Alex Alderton, Roberto Amato, Jeffrey Barrett, Sonia Goncalves, Ewan Harrison, David K. Jackson, Ian Johnston, Dominic Kwiatkowski, Cordelia Langford, John Sillitoe on behalf of the Wellcome Sanger Institute COVID-19 Surveillance Team |
| EPI_ISL_2917671 | Berkshire and Surrey Pathology Services Lighthouse Laboratory | Wellcome Sanger Institute for the COVID-19 Genomics UK (COG-UK) Consortium | Berkshire and Surrey Pathology Services Lighthouse Laboratory and Alex Alderton, Roberto Amato, Jeffrey Barrett, Sonia Goncalves, Ewan Harrison, David K. Jackson, Ian Johnston, Dominic Kwiatkowski, Cordelia Langford, John Sillitoe on behalf of the Wellcome Sanger Institute COVID-19 Surveillance Team |
| EPI_ISL_2917672, EPI_ISL_2917673, EPI_ISL_2917674, EPI_ISL_2917675, EPI_ISL_2917676, EPI_ISL_2917677, EPI_ISL_2917678, EPI_ISL_2917679, EPI_ISL_2917680, EPI_ISL_2917681 | Lighthouse Lab in Glasgow | Wellcome Sanger Institute for the COVID-19 Genomics UK (COG-UK) Consortium | Harper VanSteenhouse, Yumi Kasai, David Gray, Carol Clugston, Anna Dominiczak and Alex Alderton, Roberto Amato, Jeffrey Barrett, Sonia Goncalves, Ewan Harrison, David K. Jackson, Ian Johnston, Dominic Kwiatkowski, Cordelia Langford, John Sillitoe on behalf of the Wellcome Sanger Institute COVID-19 Surveillance Team |
| EPI_ISL_2917682 | Berkshire and Surrey Pathology Services Lighthouse Laboratory | Wellcome Sanger Institute for the COVID-19 Genomics UK (COG-UK) Consortium | Berkshire and Surrey Pathology Services Lighthouse Laboratory and Alex Alderton, Roberto Amato, Jeffrey Barrett, Sonia Goncalves, Ewan Harrison, David K. Jackson, Ian Johnston, Dominic Kwiatkowski, Cordelia Langford, John Sillitoe on behalf of the Wellcome Sanger Institute COVID-19 Surveillance Team |
| EPI_ISL_2917683 | Lighthouse Lab in Glasgow | Wellcome Sanger Institute for the COVID-19 Genomics UK (COG-UK) Consortium | Harper VanSteenhouse, Yumi Kasai, David Gray, Carol Clugston, Anna Dominiczak and Alex Alderton, Roberto Amato, Jeffrey Barrett, Sonia Goncalves, Ewan Harrison, David K. Jackson, Ian Johnston, Dominic Kwiatkowski, Cordelia Langford, John Sillitoe on behalf of the Wellcome Sanger Institute COVID-19 Surveillance Team |
| EPI_ISL_2917684 | Berkshire and Surrey Pathology Services Lighthouse Laboratory | Wellcome Sanger Institute for the COVID-19 Genomics UK (COG-UK) Consortium | Berkshire and Surrey Pathology Services Lighthouse Laboratory and Alex Alderton, Roberto Amato, Jeffrey Barrett, Sonia Goncalves, Ewan Harrison, David K. Jackson, Ian Johnston, Dominic Kwiatkowski, Cordelia Langford, John Sillitoe on behalf of the Wellcome Sanger Institute COVID-19 Surveillance Team |
| EPI_ISL_2917685, EPI_ISL_2917686 | Lighthouse Lab in Glasgow | Wellcome Sanger Institute for the COVID-19 Genomics UK (COG-UK) Consortium | Harper VanSteenhouse, Yumi Kasai, David Gray, Carol Clugston, Anna Dominiczak and Alex Alderton, Roberto Amato, Jeffrey Barrett, Sonia Goncalves, Ewan Harrison, David K. Jackson, Ian Johnston, Dominic Kwiatkowski, Cordelia Langford, John Sillitoe on behalf of the Wellcome Sanger Institute COVID-19 Surveillance Team |
| EPI_ISL_2917687 | Berkshire and Surrey Pathology Services Lighthouse Laboratory | Wellcome Sanger Institute for the COVID-19 Genomics UK (COG-UK) Consortium | Berkshire and Surrey Pathology Services Lighthouse Laboratory and Alex Alderton, Roberto Amato, Jeffrey Barrett, Sonia Goncalves, Ewan Harrison, David K. Jackson, Ian Johnston, Dominic Kwiatkowski, Cordelia Langford, John Sillitoe on behalf of the Wellcome Sanger Institute COVID-19 Surveillance Team |
| EPI_ISL_2917688, EPI_ISL_2917689, EPI_ISL_2917690 | Lighthouse Lab in Glasgow | Wellcome Sanger Institute for the COVID-19 Genomics UK (COG-UK) Consortium | Harper VanSteenhouse, Yumi Kasai, David Gray, Carol Clugston, Anna Dominiczak and Alex Alderton, Roberto Amato, Jeffrey Barrett, Sonia Goncalves, Ewan Harrison, David K. Jackson, Ian Johnston, Dominic Kwiatkowski, Cordelia Langford, John Sillitoe on behalf of the Wellcome Sanger Institute COVID-19 Surveillance Team |
| EPI_ISL_2917691 | Berkshire and Surrey Pathology Services Lighthouse Laboratory | Wellcome Sanger Institute for the COVID-19 Genomics UK (COG-UK) Consortium | Berkshire and Surrey Pathology Services Lighthouse Laboratory and Alex Alderton, Roberto Amato, Jeffrey Barrett, Sonia Goncalves, Ewan Harrison, David K. Jackson, Ian Johnston, Dominic Kwiatkowski, Cordelia Langford, John Sillitoe on behalf of the Wellcome Sanger Institute COVID-19 Surveillance Team |
| EPI_ISL_2917692 | Lighthouse Lab in Glasgow | Wellcome Sanger Institute for the COVID-19 Genomics UK (COG-UK) Consortium | Harper VanSteenhouse, Yumi Kasai, David Gray, Carol Clugston, Anna Dominiczak and Alex Alderton, Roberto Amato, Jeffrey Barrett, Sonia Goncalves, Ewan Harrison, David K. Jackson, Ian Johnston, Dominic Kwiatkowski, Cordelia Langford, John Sillitoe on behalf of the Wellcome Sanger Institute COVID-19 Surveillance Team |
| EPI_ISL_2917693 | Berkshire and Surrey Pathology Services Lighthouse Laboratory | Wellcome Sanger Institute for the COVID-19 Genomics UK (COG-UK) Consortium | Berkshire and Surrey Pathology Services Lighthouse Laboratory and Alex Alderton, Roberto Amato, Jeffrey Barrett, Sonia Goncalves, Ewan Harrison, David K. Jackson, Ian Johnston, Dominic Kwiatkowski, Cordelia Langford, John Sillitoe on behalf of the Wellcome Sanger Institute COVID-19 Surveillance Team |
| EPI_ISL_2917694 | Lighthouse Lab in Glasgow | Wellcome Sanger Institute for the COVID-19 Genomics UK (COG-UK) Consortium | Harper VanSteenhouse, Yumi Kasai, David Gray, Carol Clugston, Anna Dominiczak and Alex Alderton, Roberto Amato, Jeffrey Barrett, Sonia Goncalves, Ewan Harrison, David K. Jackson, Ian Johnston, Dominic Kwiatkowski, Cordelia Langford, John Sillitoe on behalf of the Wellcome Sanger Institute COVID-19 Surveillance Team |
| EPI_ISL_2917695, EPI_ISL_2917696 | Berkshire and Surrey Pathology Services Lighthouse Laboratory | Wellcome Sanger Institute for the COVID-19 Genomics UK (COG-UK) Consortium | Berkshire and Surrey Pathology Services Lighthouse Laboratory and Alex Alderton, Roberto Amato, Jeffrey Barrett, Sonia Goncalves, Ewan Harrison, David K. Jackson, Ian Johnston, Dominic Kwiatkowski, Cordelia Langford, John Sillitoe on behalf of the Wellcome Sanger Institute COVID-19 Surveillance Team |
| EPI_ISL_2917697, EPI_ISL_2917698, EPI_ISL_2917699, EPI_ISL_2917700 | Lighthouse Lab in Glasgow | Wellcome Sanger Institute for the COVID-19 Genomics UK (COG-UK) Consortium | Harper VanSteenhouse, Yumi Kasai, David Gray, Carol Clugston, Anna Dominiczak and Alex Alderton, Roberto Amato, Jeffrey Barrett, Sonia Goncalves, Ewan Harrison, David K. Jackson, Ian Johnston, Dominic Kwiatkowski, Cordelia Langford, John Sillitoe on behalf of the Wellcome Sanger Institute COVID-19 Surveillance Team |
| EPI_ISL_2917701, EPI_ISL_2917702 | Berkshire and Surrey Pathology Services Lighthouse Laboratory | Wellcome Sanger Institute for the COVID-19 Genomics UK (COG-UK) Consortium | Berkshire and Surrey Pathology Services Lighthouse Laboratory and Alex Alderton, Roberto Amato, Jeffrey Barrett, Sonia Goncalves, Ewan Harrison, David K. Jackson, Ian Johnston, Dominic Kwiatkowski, Cordelia Langford, John Sillitoe on behalf of the Wellcome Sanger Institute COVID-19 Surveillance Team |
| EPI_ISL_2917703, EPI_ISL_2917704, EPI_ISL_2917705, EPI_ISL_2917706, EPI_ISL_2917707, EPI_ISL_2917708, EPI_ISL_2917709, EPI_ISL_2917710, EPI_ISL_2917711, EPI_ISL_2917712, EPI_ISL_2917713 | | | |
| see above | Lighthouse Lab in Glasgow | Wellcome Sanger Institute for the COVID-19 Genomics UK (COG-UK) Consortium | Harper VanSteenhouse, Yumi Kasai, David Gray, Carol Clugston, Anna Dominiczak and Alex Alderton, Roberto Amato, Jeffrey Barrett, Sonia Goncalves, Ewan Harrison, David K. Jackson, Ian Johnston, Dominic Kwiatkowski, Cordelia Langford, John Sillitoe on behalf of the Wellcome Sanger Institute COVID-19 Surveillance Team |
| EPI_ISL_2917714 | Berkshire and Surrey Pathology Services Lighthouse Laboratory | Wellcome Sanger Institute for the COVID-19 Genomics UK (COG-UK) Consortium | Berkshire and Surrey Pathology Services Lighthouse Laboratory and Alex Alderton, Roberto Amato, Jeffrey Barrett, Sonia Goncalves, Ewan Harrison, David K. Jackson, Ian Johnston, Dominic Kwiatkowski, Cordelia Langford, John Sillitoe on behalf of the Wellcome Sanger Institute COVID-19 Surveillance Team |
| EPI_ISL_2917715, EPI_ISL_2917716 | Lighthouse Lab in Glasgow | Wellcome Sanger Institute for the COVID-19 Genomics UK (COG-UK) Consortium | Harper VanSteenhouse, Yumi Kasai, David Gray, Carol Clugston, Anna Dominiczak and Alex Alderton, Roberto Amato, Jeffrey Barrett, Sonia Goncalves, Ewan Harrison, David K. Jackson, Ian Johnston, Dominic Kwiatkowski, Cordelia Langford, John Sillitoe on behalf of the Wellcome Sanger Institute COVID-19 Surveillance Team |
| EPI_ISL_2917717 | Berkshire and Surrey Pathology Services Lighthouse Laboratory | Wellcome Sanger Institute for the COVID-19 Genomics UK (COG-UK) Consortium | Berkshire and Surrey Pathology Services Lighthouse Laboratory and Alex Alderton, Roberto Amato, Jeffrey Barrett, Sonia Goncalves, Ewan Harrison, David K. Jackson, Ian Johnston, Dominic Kwiatkowski, Cordelia Langford, John Sillitoe on behalf of the Wellcome Sanger Institute COVID-19 Surveillance Team |
| EPI_ISL_2917718 | Lighthouse Lab in Glasgow | Wellcome Sanger Institute for the COVID-19 Genomics UK (COG-UK) Consortium | Harper VanSteenhouse, Yumi Kasai, David Gray, Carol Clugston, Anna Dominiczak and Alex Alderton, Roberto Amato, Jeffrey Barrett, Sonia Goncalves, Ewan Harrison, David K. Jackson, Ian Johnston, Dominic Kwiatkowski, Cordelia Langford, John Sillitoe on behalf of the Wellcome Sanger Institute COVID-19 Surveillance Team |
| EPI_ISL_2917719, EPI_ISL_2917720, EPI_ISL_2917721 | Berkshire and Surrey Pathology Services Lighthouse Laboratory | Wellcome Sanger Institute for the COVID-19 Genomics UK (COG-UK) Consortium | Berkshire and Surrey Pathology Services Lighthouse Laboratory and Alex Alderton, Roberto Amato, Jeffrey Barrett, Sonia Goncalves, Ewan Harrison, David K. Jackson, Ian Johnston, Dominic Kwiatkowski, Cordelia Langford, John Sillitoe on behalf of the Wellcome Sanger Institute COVID-19 Surveillance Team |
| EPI_ISL_2917722, EPI_ISL_2917723, EPI_ISL_2917724, EPI_ISL_2917725, EPI_ISL_2917726, EPI_ISL_2917727, EPI_ISL_2917728, EPI_ISL_2917729, EPI_ISL_2917730 | Lighthouse Lab in Glasgow | Wellcome Sanger Institute for the COVID-19 Genomics UK (COG-UK) Consortium | Harper VanSteenhouse, Yumi Kasai, David Gray, Carol Clugston, Anna Dominiczak and Alex Alderton, Roberto Amato, Jeffrey Barrett, Sonia Goncalves, Ewan Harrison, David K. Jackson, Ian Johnston, Dominic Kwiatkowski, Cordelia Langford, John Sillitoe on behalf of the Wellcome Sanger Institute COVID-19 Surveillance Team |
| EPI_ISL_2917731 | Berkshire and Surrey Pathology Services Lighthouse Laboratory | Wellcome Sanger Institute for the COVID-19 Genomics UK (COG-UK) Consortium | Berkshire and Surrey Pathology Services Lighthouse Laboratory and Alex Alderton, Roberto Amato, Jeffrey Barrett, Sonia Goncalves, Ewan Harrison, David K. Jackson, Ian Johnston, Dominic Kwiatkowski, Cordelia Langford, John Sillitoe on behalf of the Wellcome Sanger Institute COVID-19 Surveillance Team |
| EPI_ISL_2917732, EPI_ISL_2917733, EPI_ISL_2917734 | Lighthouse Lab in Glasgow | Wellcome Sanger Institute for the COVID-19 Genomics UK (COG-UK) Consortium | Harper VanSteenhouse, Yumi Kasai, David Gray, Carol Clugston, Anna Dominiczak and Alex Alderton, Roberto Amato, Jeffrey Barrett, Sonia Goncalves, Ewan Harrison, David K. Jackson, Ian Johnston, Dominic Kwiatkowski, Cordelia Langford, John Sillitoe on behalf of the Wellcome Sanger Institute COVID-19 Surveillance Team |
| EPI_ISL_2917735 | Berkshire and Surrey Pathology Services Lighthouse Laboratory | Wellcome Sanger Institute for the COVID-19 Genomics UK (COG-UK) Consortium | Berkshire and Surrey Pathology Services Lighthouse Laboratory and Alex Alderton, Roberto Amato, Jeffrey Barrett, Sonia Goncalves, Ewan Harrison, David K. Jackson, Ian Johnston, Dominic Kwiatkowski, Cordelia Langford, John Sillitoe on behalf of the Wellcome Sanger Institute COVID-19 Surveillance Team |
| EPI_ISL_2917736, EPI_ISL_2917737, EPI_ISL_2917738, EPI_ISL_2917739, EPI_ISL_2917740, EPI_ISL_2917741, EPI_ISL_2917742, EPI_ISL_2917743, EPI_ISL_2917744 | Lighthouse Lab in Glasgow | Wellcome Sanger Institute for the COVID-19 Genomics UK (COG-UK) Consortium | Harper VanSteenhouse, Yumi Kasai, David Gray, Carol Clugston, Anna Dominiczak and Alex Alderton, Roberto Amato, Jeffrey Barrett, Sonia Goncalves, Ewan Harrison, David K. Jackson, Ian Johnston, Dominic Kwiatkowski, Cordelia Langford, John Sillitoe on behalf of the Wellcome Sanger Institute COVID-19 Surveillance Team |
| EPI_ISL_2917745 | Berkshire and Surrey Pathology Services Lighthouse Laboratory | Wellcome Sanger Institute for the COVID-19 Genomics UK (COG-UK) Consortium | Berkshire and Surrey Pathology Services Lighthouse Laboratory and Alex Alderton, Roberto Amato, Jeffrey Barrett, Sonia Goncalves, Ewan Harrison, David K. Jackson, Ian Johnston, Dominic Kwiatkowski, Cordelia Langford, John Sillitoe on behalf of the Wellcome Sanger Institute COVID-19 Surveillance Team |
| EPI_ISL_2917746, EPI_ISL_2917747, EPI_ISL_2917748, EPI_ISL_2917749 | Lighthouse Lab in Glasgow | Wellcome Sanger Institute for the COVID-19 Genomics UK (COG-UK) Consortium | Harper VanSteenhouse, Yumi Kasai, David Gray, Carol Clugston, Anna Dominiczak and Alex Alderton, Roberto Amato, Jeffrey Barrett, Sonia Goncalves, Ewan Harrison, David K. Jackson, Ian Johnston, Dominic Kwiatkowski, Cordelia Langford, John Sillitoe on behalf of the Wellcome Sanger Institute COVID-19 Surveillance Team |
| EPI_ISL_2917750 | Berkshire and Surrey Pathology Services Lighthouse Laboratory | Wellcome Sanger Institute for the COVID-19 Genomics UK (COG-UK) Consortium | Berkshire and Surrey Pathology Services Lighthouse Laboratory and Alex Alderton, Roberto Amato, Jeffrey Barrett, Sonia Goncalves, Ewan Harrison, David K. Jackson, Ian Johnston, Dominic Kwiatkowski, Cordelia Langford, John Sillitoe on behalf of the Wellcome Sanger Institute COVID-19 Surveillance Team |

| | | | |
|---|---|---|---|
| EPI_ISL_2917751, EPI_ISL_2917752 | Lighthouse Lab in Glasgow | Wellcome Sanger Institute for the COVID-19 Genomics UK (COG-UK) Consortium | Harper VanSteenhouse, Yumi Kasai, David Gray, Carol Clugston, Anna Dominiczak and Alex Alderton, Roberto Amato, Jeffrey Barrett, Sonia Goncalves, Ewan Harrison, David K. Jackson, Ian Johnston, Dominic Kwiatkowski, Cordelia Langford, John Sillitoe on behalf of the Wellcome Sanger Institute COVID-19 Surveillance Team |
| EPI_ISL_2917753 | Berkshire and Surrey Pathology Services Lighthouse Laboratory | Wellcome Sanger Institute for the COVID-19 Genomics UK (COG-UK) Consortium | Berkshire and Surrey Pathology Services Lighthouse Laboratory and Alex Alderton, Roberto Amato, Jeffrey Barrett, Sonia Goncalves, Ewan Harrison, David K. Jackson, Ian Johnston, Dominic Kwiatkowski, Cordelia Langford, John Sillitoe on behalf of the Wellcome Sanger Institute COVID-19 Surveillance Team |
| EPI_ISL_2917754 | Lighthouse Lab in Glasgow | Wellcome Sanger Institute for the COVID-19 Genomics UK (COG-UK) Consortium | Harper VanSteenhouse, Yumi Kasai, David Gray, Carol Clugston, Anna Dominiczak and Alex Alderton, Roberto Amato, Jeffrey Barrett, Sonia Goncalves, Ewan Harrison, David K. Jackson, Ian Johnston, Dominic Kwiatkowski, Cordelia Langford, John Sillitoe on behalf of the Wellcome Sanger Institute COVID-19 Surveillance Team |
| EPI_ISL_2917755 | Berkshire and Surrey Pathology Services Lighthouse Laboratory | Wellcome Sanger Institute for the COVID-19 Genomics UK (COG-UK) Consortium | Berkshire and Surrey Pathology Services Lighthouse Laboratory and Alex Alderton, Roberto Amato, Jeffrey Barrett, Sonia Goncalves, Ewan Harrison, David K. Jackson, Ian Johnston, Dominic Kwiatkowski, Cordelia Langford, John Sillitoe on behalf of the Wellcome Sanger Institute COVID-19 Surveillance Team |
| EPI_ISL_2917756, EPI_ISL_2917757 | Lighthouse Lab in Glasgow | Wellcome Sanger Institute for the COVID-19 Genomics UK (COG-UK) Consortium | Harper VanSteenhouse, Yumi Kasai, David Gray, Carol Clugston, Anna Dominiczak and Alex Alderton, Roberto Amato, Jeffrey Barrett, Sonia Goncalves, Ewan Harrison, David K. Jackson, Ian Johnston, Dominic Kwiatkowski, Cordelia Langford, John Sillitoe on behalf of the Wellcome Sanger Institute COVID-19 Surveillance Team |
| EPI_ISL_2917758 | Berkshire and Surrey Pathology Services Lighthouse Laboratory | Wellcome Sanger Institute for the COVID-19 Genomics UK (COG-UK) Consortium | Berkshire and Surrey Pathology Services Lighthouse Laboratory and Alex Alderton, Roberto Amato, Jeffrey Barrett, Sonia Goncalves, Ewan Harrison, David K. Jackson, Ian Johnston, Dominic Kwiatkowski, Cordelia Langford, John Sillitoe on behalf of the Wellcome Sanger Institute COVID-19 Surveillance Team |
| EPI_ISL_2917759, EPI_ISL_2917760, EPI_ISL_2917761, EPI_ISL_2917762, EPI_ISL_2917763, EPI_ISL_2917764, EPI_ISL_2917765, EPI_ISL_2917766, EPI_ISL_2917767 | Lighthouse Lab in Glasgow | Wellcome Sanger Institute for the COVID-19 Genomics UK (COG-UK) Consortium | Harper VanSteenhouse, Yumi Kasai, David Gray, Carol Clugston, Anna Dominiczak and Alex Alderton, Roberto Amato, Jeffrey Barrett, Sonia Goncalves, Ewan Harrison, David K. Jackson, Ian Johnston, Dominic Kwiatkowski, Cordelia Langford, John Sillitoe on behalf of the Wellcome Sanger Institute COVID-19 Surveillance Team |
| EPI_ISL_2917768, EPI_ISL_2917769 | Berkshire and Surrey Pathology Services Lighthouse Laboratory | Wellcome Sanger Institute for the COVID-19 Genomics UK (COG-UK) Consortium | Berkshire and Surrey Pathology Services Lighthouse Laboratory and Alex Alderton, Roberto Amato, Jeffrey Barrett, Sonia Goncalves, Ewan Harrison, David K. Jackson, Ian Johnston, Dominic Kwiatkowski, Cordelia Langford, John Sillitoe on behalf of the Wellcome Sanger Institute COVID-19 Surveillance Team |
| EPI_ISL_2917770 | Lighthouse Lab in Glasgow | Wellcome Sanger Institute for the COVID-19 Genomics UK (COG-UK) Consortium | Harper VanSteenhouse, Yumi Kasai, David Gray, Carol Clugston, Anna Dominiczak and Alex Alderton, Roberto Amato, Jeffrey Barrett, Sonia Goncalves, Ewan Harrison, David K. Jackson, Ian Johnston, Dominic Kwiatkowski, Cordelia Langford, John Sillitoe on behalf of the Wellcome Sanger Institute COVID-19 Surveillance Team |
| EPI_ISL_2917771 | Berkshire and Surrey Pathology Services Lighthouse Laboratory | Wellcome Sanger Institute for the COVID-19 Genomics UK (COG-UK) Consortium | Berkshire and Surrey Pathology Services Lighthouse Laboratory and Alex Alderton, Roberto Amato, Jeffrey Barrett, Sonia Goncalves, Ewan Harrison, David K. Jackson, Ian Johnston, Dominic Kwiatkowski, Cordelia Langford, John Sillitoe on behalf of the Wellcome Sanger Institute COVID-19 Surveillance Team |
| EPI_ISL_2917772, EPI_ISL_2917773 | Lighthouse Lab in Glasgow | Wellcome Sanger Institute for the COVID-19 Genomics UK (COG-UK) Consortium | Harper VanSteenhouse, Yumi Kasai, David Gray, Carol Clugston, Anna Dominiczak and Alex Alderton, Roberto Amato, Jeffrey Barrett, Sonia Goncalves, Ewan Harrison, David K. Jackson, Ian Johnston, Dominic Kwiatkowski, Cordelia Langford, John Sillitoe on behalf of the Wellcome Sanger Institute COVID-19 Surveillance Team |
| EPI_ISL_2917774 | Berkshire and Surrey Pathology Services Lighthouse Laboratory | Wellcome Sanger Institute for the COVID-19 Genomics UK (COG-UK) Consortium | Berkshire and Surrey Pathology Services Lighthouse Laboratory and Alex Alderton, Roberto Amato, Jeffrey Barrett, Sonia Goncalves, Ewan Harrison, David K. Jackson, Ian Johnston, Dominic Kwiatkowski, Cordelia Langford, John Sillitoe on behalf of the Wellcome Sanger Institute COVID-19 Surveillance Team |
| EPI_ISL_2917775, EPI_ISL_2917776, EPI_ISL_2917777, EPI_ISL_2917778, EPI_ISL_2917779, EPI_ISL_2917780, EPI_ISL_2917781, EPI_ISL_2917782, EPI_ISL_2917783, EPI_ISL_2917784, EPI_ISL_2917785, EPI_ISL_2917786, EPI_ISL_2917787, EPI_ISL_2917788, EPI_ISL_2917789, EPI_ISL_2917790 | | | |
| see above | Lighthouse Lab in Glasgow | Wellcome Sanger Institute for the COVID-19 Genomics UK (COG-UK) Consortium | Harper VanSteenhouse, Yumi Kasai, David Gray, Carol Clugston, Anna Dominiczak and Alex Alderton, Roberto Amato, Jeffrey Barrett, Sonia Goncalves, Ewan Harrison, David K. Jackson, Ian Johnston, Dominic Kwiatkowski, Cordelia Langford, John Sillitoe on behalf of the Wellcome Sanger Institute COVID-19 Surveillance Team |
| EPI_ISL_2917791, EPI_ISL_2917792 | Berkshire and Surrey Pathology Services Lighthouse Laboratory | Wellcome Sanger Institute for the COVID-19 Genomics UK (COG-UK) Consortium | Berkshire and Surrey Pathology Services Lighthouse Laboratory and Alex Alderton, Roberto Amato, Jeffrey Barrett, Sonia Goncalves, Ewan Harrison, David K. Jackson, Ian Johnston, Dominic Kwiatkowski, Cordelia Langford, John Sillitoe on behalf of the Wellcome Sanger Institute COVID-19 Surveillance Team |
| EPI_ISL_2917793 | Lighthouse Lab in Glasgow | Wellcome Sanger Institute for the COVID-19 Genomics UK (COG-UK) Consortium | Harper VanSteenhouse, Yumi Kasai, David Gray, Carol Clugston, Anna Dominiczak and Alex Alderton, Roberto Amato, Jeffrey Barrett, Sonia Goncalves, Ewan Harrison, David K. Jackson, Ian Johnston, Dominic Kwiatkowski, Cordelia Langford, John Sillitoe on behalf of the Wellcome Sanger Institute COVID-19 Surveillance Team |
| EPI_ISL_2917794 | Berkshire and Surrey Pathology Services Lighthouse Laboratory | Wellcome Sanger Institute for the COVID-19 Genomics UK (COG-UK) Consortium | Berkshire and Surrey Pathology Services Lighthouse Laboratory and Alex Alderton, Roberto Amato, Jeffrey Barrett, Sonia Goncalves, Ewan Harrison, David K. Jackson, Ian Johnston, Dominic Kwiatkowski, Cordelia Langford, John Sillitoe on behalf of the Wellcome Sanger Institute COVID-19 Surveillance Team |
| EPI_ISL_2917795, EPI_ISL_2917796 | Lighthouse Lab in Glasgow | Wellcome Sanger Institute for the COVID-19 Genomics UK (COG-UK) Consortium | Harper VanSteenhouse, Yumi Kasai, David Gray, Carol Clugston, Anna Dominiczak and Alex Alderton, Roberto Amato, Jeffrey Barrett, Sonia Goncalves, Ewan Harrison, David K. Jackson, Ian Johnston, Dominic Kwiatkowski, Cordelia Langford, John Sillitoe on behalf of the Wellcome Sanger Institute COVID-19 Surveillance Team |
| EPI_ISL_2917797, EPI_ISL_2917798 | Berkshire and Surrey Pathology Services Lighthouse Laboratory | Wellcome Sanger Institute for the COVID-19 Genomics UK (COG-UK) Consortium | Berkshire and Surrey Pathology Services Lighthouse Laboratory and Alex Alderton, Roberto Amato, Jeffrey Barrett, Sonia Goncalves, Ewan Harrison, David K. Jackson, Ian Johnston, Dominic Kwiatkowski, Cordelia Langford, John Sillitoe on behalf of the Wellcome Sanger Institute COVID-19 Surveillance Team |
| EPI_ISL_2917799, EPI_ISL_2917800, EPI_ISL_2917801, EPI_ISL_2917802, EPI_ISL_2917803, EPI_ISL_2917804, EPI_ISL_2917805, EPI_ISL_2917806, EPI_ISL_2917807, EPI_ISL_2917808, EPI_ISL_2917809, EPI_ISL_2917810, EPI_ISL_2917811 | | | |
| see above | Lighthouse Lab in Glasgow | Wellcome Sanger Institute for the COVID-19 Genomics UK (COG-UK) Consortium | Harper VanSteenhouse, Yumi Kasai, David Gray, Carol Clugston, Anna Dominiczak and Alex Alderton, Roberto Amato, Jeffrey Barrett, Sonia Goncalves, Ewan Harrison, David K. Jackson, Ian Johnston, Dominic Kwiatkowski, Cordelia Langford, John Sillitoe on behalf of the Wellcome Sanger Institute COVID-19 Surveillance Team |
| EPI_ISL_2917812 | Berkshire and Surrey Pathology Services Lighthouse Laboratory | Wellcome Sanger Institute for the COVID-19 Genomics UK (COG-UK) Consortium | Berkshire and Surrey Pathology Services Lighthouse Laboratory and Alex Alderton, Roberto Amato, Jeffrey Barrett, Sonia Goncalves, Ewan Harrison, David K. Jackson, Ian Johnston, Dominic Kwiatkowski, Cordelia Langford, John Sillitoe on behalf of the Wellcome Sanger Institute COVID-19 Surveillance Team |
| EPI_ISL_2917813, EPI_ISL_2917814 | Lighthouse Lab in Glasgow | Wellcome Sanger Institute for the COVID-19 Genomics UK (COG-UK) Consortium | Harper VanSteenhouse, Yumi Kasai, David Gray, Carol Clugston, Anna Dominiczak and Alex Alderton, Roberto Amato, Jeffrey Barrett, Sonia Goncalves, Ewan Harrison, David K. Jackson, Ian Johnston, Dominic Kwiatkowski, Cordelia Langford, John Sillitoe on behalf of the Wellcome Sanger Institute COVID-19 Surveillance Team |
| EPI_ISL_2917815 | Berkshire and Surrey Pathology Services Lighthouse Laboratory | Wellcome Sanger Institute for the COVID-19 Genomics UK (COG-UK) Consortium | Berkshire and Surrey Pathology Services Lighthouse Laboratory and Alex Alderton, Roberto Amato, Jeffrey Barrett, Sonia Goncalves, Ewan Harrison, David K. Jackson, Ian Johnston, Dominic Kwiatkowski, Cordelia Langford, John Sillitoe on behalf of the Wellcome Sanger Institute COVID-19 Surveillance Team |
| EPI_ISL_2917816, EPI_ISL_2917817, EPI_ISL_2917818, EPI_ISL_2917819, EPI_ISL_2917820 | Lighthouse Lab in Glasgow | Wellcome Sanger Institute for the COVID-19 Genomics UK (COG-UK) Consortium | Harper VanSteenhouse, Yumi Kasai, David Gray, Carol Clugston, Anna Dominiczak and Alex Alderton, Roberto Amato, Jeffrey Barrett, Sonia Goncalves, Ewan Harrison, David K. Jackson, Ian Johnston, Dominic Kwiatkowski, Cordelia Langford, John Sillitoe on behalf of the Wellcome Sanger Institute COVID-19 Surveillance Team |
| EPI_ISL_2917821 | Berkshire and Surrey Pathology Services Lighthouse Laboratory | Wellcome Sanger Institute for the COVID-19 Genomics UK (COG-UK) Consortium | Berkshire and Surrey Pathology Services Lighthouse Laboratory and Alex Alderton, Roberto Amato, Jeffrey Barrett, Sonia Goncalves, Ewan Harrison, David K. Jackson, Ian Johnston, Dominic Kwiatkowski, Cordelia Langford, John Sillitoe on behalf of the Wellcome Sanger Institute COVID-19 Surveillance Team |
| EPI_ISL_2917822, EPI_ISL_2917823 | Lighthouse Lab in Glasgow | Wellcome Sanger Institute for the COVID-19 Genomics UK (COG-UK) Consortium | Harper VanSteenhouse, Yumi Kasai, David Gray, Carol Clugston, Anna Dominiczak and Alex Alderton, Roberto Amato, Jeffrey Barrett, Sonia Goncalves, Ewan Harrison, David K. Jackson, Ian Johnston, Dominic Kwiatkowski, Cordelia Langford, John Sillitoe on behalf of the Wellcome Sanger Institute COVID-19 Surveillance Team |
| EPI_ISL_2917824 | Berkshire and Surrey Pathology Services Lighthouse Laboratory | Wellcome Sanger Institute for the COVID-19 Genomics UK (COG-UK) Consortium | Berkshire and Surrey Pathology Services Lighthouse Laboratory and Alex Alderton, Roberto Amato, Jeffrey Barrett, Sonia Goncalves, Ewan Harrison, David K. Jackson, Ian Johnston, Dominic Kwiatkowski, Cordelia Langford, John Sillitoe on behalf of the Wellcome Sanger Institute COVID-19 Surveillance Team |
| EPI_ISL_2917825, EPI_ISL_2917826, EPI_ISL_2917827, EPI_ISL_2917828, EPI_ISL_2917829, EPI_ISL_2917830 | Lighthouse Lab in Glasgow | Wellcome Sanger Institute for the COVID-19 Genomics UK (COG-UK) Consortium | Harper VanSteenhouse, Yumi Kasai, David Gray, Carol Clugston, Anna Dominiczak and Alex Alderton, Roberto Amato, Jeffrey Barrett, Sonia Goncalves, Ewan Harrison, David K. Jackson, Ian Johnston, Dominic Kwiatkowski, Cordelia Langford, John Sillitoe on behalf of the Wellcome Sanger Institute COVID-19 Surveillance Team |
| EPI_ISL_2917831 | Berkshire and Surrey Pathology Services Lighthouse Laboratory | Wellcome Sanger Institute for the COVID-19 Genomics UK (COG-UK) Consortium | Berkshire and Surrey Pathology Services Lighthouse Laboratory and Alex Alderton, Roberto Amato, Jeffrey Barrett, Sonia Goncalves, Ewan Harrison, David K. Jackson, Ian Johnston, Dominic Kwiatkowski, Cordelia Langford, John Sillitoe on behalf of the Wellcome Sanger Institute COVID-19 Surveillance Team |
| EPI_ISL_2917832, EPI_ISL_2917833 | Lighthouse Lab in Glasgow | Wellcome Sanger Institute for the COVID-19 Genomics UK (COG-UK) Consortium | Harper VanSteenhouse, Yumi Kasai, David Gray, Carol Clugston, Anna Dominiczak and Alex Alderton, Roberto Amato, Jeffrey Barrett, Sonia Goncalves, Ewan Harrison, David K. Jackson, Ian Johnston, Dominic Kwiatkowski, Cordelia Langford, John Sillitoe on behalf of the Wellcome Sanger Institute COVID-19 Surveillance Team |
| EPI_ISL_2917834 | Berkshire and Surrey Pathology Services Lighthouse Laboratory | Wellcome Sanger Institute for the COVID-19 Genomics UK (COG-UK) Consortium | Berkshire and Surrey Pathology Services Lighthouse Laboratory and Alex Alderton, Roberto Amato, Jeffrey Barrett, Sonia Goncalves, Ewan Harrison, David K. Jackson, Ian Johnston, Dominic Kwiatkowski, Cordelia Langford, John Sillitoe on behalf of the Wellcome Sanger Institute COVID-19 Surveillance Team |
| EPI_ISL_2917835, EPI_ISL_2917836, EPI_ISL_2917837, EPI_ISL_2917838, EPI_ISL_2917839, EPI_ISL_2917840, EPI_ISL_2917841, EPI_ISL_2917842, EPI_ISL_2917843, EPI_ISL_2917844, EPI_ISL_2917845, EPI_ISL_2917846, EPI_ISL_2917847, EPI_ISL_2917848, EPI_ISL_2917849, EPI_ISL_2917850, EPI_ISL_2917851, EPI_ISL_2917852, EPI_ISL_2917853 | | | |
| see above | Lighthouse Lab in Glasgow | Wellcome Sanger Institute for the COVID-19 Genomics UK (COG-UK) Consortium | Harper VanSteenhouse, Yumi Kasai, David Gray, Carol Clugston, Anna Dominiczak and Alex Alderton, Roberto Amato, Jeffrey Barrett, Sonia Goncalves, Ewan Harrison, David K. Jackson, Ian Johnston, Dominic Kwiatkowski, Cordelia Langford, John Sillitoe on behalf of the Wellcome Sanger Institute COVID-19 Surveillance Team |
| EPI_ISL_2917854 | Berkshire and Surrey Pathology Services Lighthouse Laboratory | Wellcome Sanger Institute for the COVID-19 Genomics UK (COG-UK) Consortium | Berkshire and Surrey Pathology Services Lighthouse Laboratory and Alex Alderton, Roberto Amato, Jeffrey Barrett, Sonia Goncalves, Ewan Harrison, David K. Jackson, Ian Johnston, Dominic Kwiatkowski, Cordelia Langford, John Sillitoe on behalf of the Wellcome Sanger Institute COVID-19 Surveillance Team |
| EPI_ISL_2917855, EPI_ISL_2917856 | Lighthouse Lab in Glasgow | Wellcome Sanger Institute for the COVID-19 Genomics UK (COG-UK) Consortium | Harper VanSteenhouse, Yumi Kasai, David Gray, Carol Clugston, Anna Dominiczak and Alex Alderton, Roberto Amato, Jeffrey Barrett, Sonia Goncalves, Ewan Harrison, David K. Jackson, Ian Johnston, Dominic Kwiatkowski, Cordelia Langford, John Sillitoe on behalf of the Wellcome Sanger Institute COVID-19 Surveillance Team |
| EPI_ISL_2917857 | Berkshire and Surrey Pathology Services Lighthouse Laboratory | Wellcome Sanger Institute for the COVID-19 Genomics UK (COG-UK) Consortium | Berkshire and Surrey Pathology Services Lighthouse Laboratory and Alex Alderton, Roberto Amato, Jeffrey Barrett, Sonia Goncalves, Ewan Harrison, David K. Jackson, Ian Johnston, Dominic Kwiatkowski, Cordelia Langford, John Sillitoe on behalf of the Wellcome Sanger Institute COVID-19 Surveillance Team |
| EPI_ISL_2917858 | Lighthouse Lab in Glasgow | Wellcome Sanger Institute for the COVID-19 Genomics UK (COG-UK) Consortium | Harper VanSteenhouse, Yumi Kasai, David Gray, Carol Clugston, Anna Dominiczak and Alex Alderton, Roberto Amato, Jeffrey Barrett, Sonia Goncalves, Ewan Harrison, David K. Jackson, Ian Johnston, Dominic Kwiatkowski, Cordelia Langford, John Sillitoe on behalf of the Wellcome Sanger Institute COVID-19 Surveillance Team |
| EPI_ISL_2917859 | Berkshire and Surrey Pathology Services Lighthouse Laboratory | Wellcome Sanger Institute for the COVID-19 Genomics UK (COG-UK) Consortium | Berkshire and Surrey Pathology Services Lighthouse Laboratory and Alex Alderton, Roberto Amato, Jeffrey Barrett, Sonia Goncalves, Ewan Harrison, David K. Jackson, Ian Johnston, Dominic Kwiatkowski, Cordelia Langford, John Sillitoe on behalf of the Wellcome Sanger Institute COVID-19 Surveillance Team |
| EPI_ISL_2917860 | Lighthouse Lab in Glasgow | Wellcome Sanger Institute for the COVID-19 Genomics UK (COG-UK) Consortium | Harper VanSteenhouse, Yumi Kasai, David Gray, Carol Clugston, Anna Dominiczak and Alex Alderton, Roberto Amato, Jeffrey Barrett, Sonia Goncalves, Ewan Harrison, David K. Jackson, Ian Johnston, Dominic Kwiatkowski, Cordelia Langford, John Sillitoe on behalf of the Wellcome Sanger Institute COVID-19 Surveillance Team |
| EPI_ISL_2917861, EPI_ISL_2917862 | Berkshire and Surrey Pathology Services Lighthouse Laboratory | Wellcome Sanger Institute for the COVID-19 Genomics UK (COG-UK) Consortium | Berkshire and Surrey Pathology Services Lighthouse Laboratory and Alex Alderton, Roberto Amato, Jeffrey Barrett, Sonia Goncalves, Ewan Harrison, David K. Jackson, Ian Johnston, Dominic Kwiatkowski, Cordelia Langford, John Sillitoe on behalf of the Wellcome Sanger Institute COVID-19 Surveillance Team |

| | | | |
|---|---|---|---|
| EPI_ISL_2917863, EPI_ISL_2917864 | Lighthouse Lab in Glasgow | Wellcome Sanger Institute for the COVID-19 Genomics UK (COG-UK) Consortium | Harper VanSteenhouse, Yumi Kasai, David Gray, Carol Clugston, Anna Dominiczak and Alex Alderton, Roberto Amato, Jeffrey Barrett, Sonia Goncalves, Ewan Harrison, David K. Jackson, Ian Johnston, Dominic Kwiatkowski, Cordelia Langford, John Sillitoe on behalf of the Wellcome Sanger Institute COVID-19 Surveillance Team |
| EPI_ISL_2917865 | Berkshire and Surrey Pathology Services Lighthouse Laboratory | Wellcome Sanger Institute for the COVID-19 Genomics UK (COG-UK) Consortium | Berkshire and Surrey Pathology Services Lighthouse Laboratory and Alex Alderton, Roberto Amato, Jeffrey Barrett, Sonia Goncalves, Ewan Harrison, David K. Jackson, Ian Johnston, Dominic Kwiatkowski, Cordelia Langford, John Sillitoe on behalf of the Wellcome Sanger Institute COVID-19 Surveillance Team |
| EPI_ISL_2917866 | Lighthouse Lab in Glasgow | Wellcome Sanger Institute for the COVID-19 Genomics UK (COG-UK) Consortium | Harper VanSteenhouse, Yumi Kasai, David Gray, Carol Clugston, Anna Dominiczak and Alex Alderton, Roberto Amato, Jeffrey Barrett, Sonia Goncalves, Ewan Harrison, David K. Jackson, Ian Johnston, Dominic Kwiatkowski, Cordelia Langford, John Sillitoe on behalf of the Wellcome Sanger Institute COVID-19 Surveillance Team |
| EPI_ISL_2917867 | Berkshire and Surrey Pathology Services Lighthouse Laboratory | Wellcome Sanger Institute for the COVID-19 Genomics UK (COG-UK) Consortium | Berkshire and Surrey Pathology Services Lighthouse Laboratory and Alex Alderton, Roberto Amato, Jeffrey Barrett, Sonia Goncalves, Ewan Harrison, David K. Jackson, Ian Johnston, Dominic Kwiatkowski, Cordelia Langford, John Sillitoe on behalf of the Wellcome Sanger Institute COVID-19 Surveillance Team |
| EPI_ISL_2917868, EPI_ISL_2917869, EPI_ISL_2917870, EPI_ISL_2917871 | Lighthouse Lab in Glasgow | Wellcome Sanger Institute for the COVID-19 Genomics UK (COG-UK) Consortium | Harper VanSteenhouse, Yumi Kasai, David Gray, Carol Clugston, Anna Dominiczak and Alex Alderton, Roberto Amato, Jeffrey Barrett, Sonia Goncalves, Ewan Harrison, David K. Jackson, Ian Johnston, Dominic Kwiatkowski, Cordelia Langford, John Sillitoe on behalf of the Wellcome Sanger Institute COVID-19 Surveillance Team |
| EPI_ISL_2917872 | Berkshire and Surrey Pathology Services Lighthouse Laboratory | Wellcome Sanger Institute for the COVID-19 Genomics UK (COG-UK) Consortium | Berkshire and Surrey Pathology Services Lighthouse Laboratory and Alex Alderton, Roberto Amato, Jeffrey Barrett, Sonia Goncalves, Ewan Harrison, David K. Jackson, Ian Johnston, Dominic Kwiatkowski, Cordelia Langford, John Sillitoe on behalf of the Wellcome Sanger Institute COVID-19 Surveillance Team |
| EPI_ISL_2917873, EPI_ISL_2917874 | Lighthouse Lab in Glasgow | Wellcome Sanger Institute for the COVID-19 Genomics UK (COG-UK) Consortium | Harper VanSteenhouse, Yumi Kasai, David Gray, Carol Clugston, Anna Dominiczak and Alex Alderton, Roberto Amato, Jeffrey Barrett, Sonia Goncalves, Ewan Harrison, David K. Jackson, Ian Johnston, Dominic Kwiatkowski, Cordelia Langford, John Sillitoe on behalf of the Wellcome Sanger Institute COVID-19 Surveillance Team |
| EPI_ISL_2917875 | Berkshire and Surrey Pathology Services Lighthouse Laboratory | Wellcome Sanger Institute for the COVID-19 Genomics UK (COG-UK) Consortium | Berkshire and Surrey Pathology Services Lighthouse Laboratory and Alex Alderton, Roberto Amato, Jeffrey Barrett, Sonia Goncalves, Ewan Harrison, David K. Jackson, Ian Johnston, Dominic Kwiatkowski, Cordelia Langford, John Sillitoe on behalf of the Wellcome Sanger Institute COVID-19 Surveillance Team |
| EPI_ISL_2917876, EPI_ISL_2917877 | Lighthouse Lab in Glasgow | Wellcome Sanger Institute for the COVID-19 Genomics UK (COG-UK) Consortium | Harper VanSteenhouse, Yumi Kasai, David Gray, Carol Clugston, Anna Dominiczak and Alex Alderton, Roberto Amato, Jeffrey Barrett, Sonia Goncalves, Ewan Harrison, David K. Jackson, Ian Johnston, Dominic Kwiatkowski, Cordelia Langford, John Sillitoe on behalf of the Wellcome Sanger Institute COVID-19 Surveillance Team |
| EPI_ISL_2917878 | Berkshire and Surrey Pathology Services Lighthouse Laboratory | Wellcome Sanger Institute for the COVID-19 Genomics UK (COG-UK) Consortium | Berkshire and Surrey Pathology Services Lighthouse Laboratory and Alex Alderton, Roberto Amato, Jeffrey Barrett, Sonia Goncalves, Ewan Harrison, David K. Jackson, Ian Johnston, Dominic Kwiatkowski, Cordelia Langford, John Sillitoe on behalf of the Wellcome Sanger Institute COVID-19 Surveillance Team |
| EPI_ISL_2917879, EPI_ISL_2917880, EPI_ISL_2917881, EPI_ISL_2917882, EPI_ISL_2917883, EPI_ISL_2917884, EPI_ISL_2917885, EPI_ISL_2917886 | Lighthouse Lab in Glasgow | Wellcome Sanger Institute for the COVID-19 Genomics UK (COG-UK) Consortium | Harper VanSteenhouse, Yumi Kasai, David Gray, Carol Clugston, Anna Dominiczak and Alex Alderton, Roberto Amato, Jeffrey Barrett, Sonia Goncalves, Ewan Harrison, David K. Jackson, Ian Johnston, Dominic Kwiatkowski, Cordelia Langford, John Sillitoe on behalf of the Wellcome Sanger Institute COVID-19 Surveillance Team |
| EPI_ISL_2917887 | Berkshire and Surrey Pathology Services Lighthouse Laboratory | Wellcome Sanger Institute for the COVID-19 Genomics UK (COG-UK) Consortium | Berkshire and Surrey Pathology Services Lighthouse Laboratory and Alex Alderton, Roberto Amato, Jeffrey Barrett, Sonia Goncalves, Ewan Harrison, David K. Jackson, Ian Johnston, Dominic Kwiatkowski, Cordelia Langford, John Sillitoe on behalf of the Wellcome Sanger Institute COVID-19 Surveillance Team |
| EPI_ISL_2917888, EPI_ISL_2917889 | Lighthouse Lab in Glasgow | Wellcome Sanger Institute for the COVID-19 Genomics UK (COG-UK) Consortium | Harper VanSteenhouse, Yumi Kasai, David Gray, Carol Clugston, Anna Dominiczak and Alex Alderton, Roberto Amato, Jeffrey Barrett, Sonia Goncalves, Ewan Harrison, David K. Jackson, Ian Johnston, Dominic Kwiatkowski, Cordelia Langford, John Sillitoe on behalf of the Wellcome Sanger Institute COVID-19 Surveillance Team |
| EPI_ISL_2917890, EPI_ISL_2917891 | Berkshire and Surrey Pathology Services Lighthouse Laboratory | Wellcome Sanger Institute for the COVID-19 Genomics UK (COG-UK) Consortium | Berkshire and Surrey Pathology Services Lighthouse Laboratory and Alex Alderton, Roberto Amato, Jeffrey Barrett, Sonia Goncalves, Ewan Harrison, David K. Jackson, Ian Johnston, Dominic Kwiatkowski, Cordelia Langford, John Sillitoe on behalf of the Wellcome Sanger Institute COVID-19 Surveillance Team |
| EPI_ISL_2917892, EPI_ISL_2917893 | Lighthouse Lab in Glasgow | Wellcome Sanger Institute for the COVID-19 Genomics UK (COG-UK) Consortium | Harper VanSteenhouse, Yumi Kasai, David Gray, Carol Clugston, Anna Dominiczak and Alex Alderton, Roberto Amato, Jeffrey Barrett, Sonia Goncalves, Ewan Harrison, David K. Jackson, Ian Johnston, Dominic Kwiatkowski, Cordelia Langford, John Sillitoe on behalf of the Wellcome Sanger Institute COVID-19 Surveillance Team |
| EPI_ISL_2917894 | Berkshire and Surrey Pathology Services Lighthouse Laboratory | Wellcome Sanger Institute for the COVID-19 Genomics UK (COG-UK) Consortium | Berkshire and Surrey Pathology Services Lighthouse Laboratory and Alex Alderton, Roberto Amato, Jeffrey Barrett, Sonia Goncalves, Ewan Harrison, David K. Jackson, Ian Johnston, Dominic Kwiatkowski, Cordelia Langford, John Sillitoe on behalf of the Wellcome Sanger Institute COVID-19 Surveillance Team |
| EPI_ISL_2917895 | Lighthouse Lab in Glasgow | Wellcome Sanger Institute for the COVID-19 Genomics UK (COG-UK) Consortium | Harper VanSteenhouse, Yumi Kasai, David Gray, Carol Clugston, Anna Dominiczak and Alex Alderton, Roberto Amato, Jeffrey Barrett, Sonia Goncalves, Ewan Harrison, David K. Jackson, Ian Johnston, Dominic Kwiatkowski, Cordelia Langford, John Sillitoe on behalf of the Wellcome Sanger Institute COVID-19 Surveillance Team |
| EPI_ISL_2917896 | Berkshire and Surrey Pathology Services Lighthouse Laboratory | Wellcome Sanger Institute for the COVID-19 Genomics UK (COG-UK) Consortium | Berkshire and Surrey Pathology Services Lighthouse Laboratory and Alex Alderton, Roberto Amato, Jeffrey Barrett, Sonia Goncalves, Ewan Harrison, David K. Jackson, Ian Johnston, Dominic Kwiatkowski, Cordelia Langford, John Sillitoe on behalf of the Wellcome Sanger Institute COVID-19 Surveillance Team |
| EPI_ISL_2917897, EPI_ISL_2917898, EPI_ISL_2917899, EPI_ISL_2917900, EPI_ISL_2917901 | Lighthouse Lab in Glasgow | Wellcome Sanger Institute for the COVID-19 Genomics UK (COG-UK) Consortium | Harper VanSteenhouse, Yumi Kasai, David Gray, Carol Clugston, Anna Dominiczak and Alex Alderton, Roberto Amato, Jeffrey Barrett, Sonia Goncalves, Ewan Harrison, David K. Jackson, Ian Johnston, Dominic Kwiatkowski, Cordelia Langford, John Sillitoe on behalf of the Wellcome Sanger Institute COVID-19 Surveillance Team |
| EPI_ISL_2917902, EPI_ISL_2917903 | Berkshire and Surrey Pathology Services Lighthouse Laboratory | Wellcome Sanger Institute for the COVID-19 Genomics UK (COG-UK) Consortium | Berkshire and Surrey Pathology Services Lighthouse Laboratory and Alex Alderton, Roberto Amato, Jeffrey Barrett, Sonia Goncalves, Ewan Harrison, David K. Jackson, Ian Johnston, Dominic Kwiatkowski, Cordelia Langford, John Sillitoe on behalf of the Wellcome Sanger Institute COVID-19 Surveillance Team |
| EPI_ISL_2917904, EPI_ISL_2917905, EPI_ISL_2917906, EPI_ISL_2917907, EPI_ISL_2917908, EPI_ISL_2917909, EPI_ISL_2917910, EPI_ISL_2917911, EPI_ISL_2917912 | Lighthouse Lab in Glasgow | Wellcome Sanger Institute for the COVID-19 Genomics UK (COG-UK) Consortium | Harper VanSteenhouse, Yumi Kasai, David Gray, Carol Clugston, Anna Dominiczak and Alex Alderton, Roberto Amato, Jeffrey Barrett, Sonia Goncalves, Ewan Harrison, David K. Jackson, Ian Johnston, Dominic Kwiatkowski, Cordelia Langford, John Sillitoe on behalf of the Wellcome Sanger Institute COVID-19 Surveillance Team |
| EPI_ISL_2917913, EPI_ISL_2917914 | Berkshire and Surrey Pathology Services Lighthouse Laboratory | Wellcome Sanger Institute for the COVID-19 Genomics UK (COG-UK) Consortium | Berkshire and Surrey Pathology Services Lighthouse Laboratory and Alex Alderton, Roberto Amato, Jeffrey Barrett, Sonia Goncalves, Ewan Harrison, David K. Jackson, Ian Johnston, Dominic Kwiatkowski, Cordelia Langford, John Sillitoe on behalf of the Wellcome Sanger Institute COVID-19 Surveillance Team |
| EPI_ISL_2917915 | Lighthouse Lab in Glasgow | Wellcome Sanger Institute for the COVID-19 Genomics UK (COG-UK) Consortium | Harper VanSteenhouse, Yumi Kasai, David Gray, Carol Clugston, Anna Dominiczak and Alex Alderton, Roberto Amato, Jeffrey Barrett, Sonia Goncalves, Ewan Harrison, David K. Jackson, Ian Johnston, Dominic Kwiatkowski, Cordelia Langford, John Sillitoe on behalf of the Wellcome Sanger Institute COVID-19 Surveillance Team |
| EPI_ISL_2917916 | Berkshire and Surrey Pathology Services Lighthouse Laboratory | Wellcome Sanger Institute for the COVID-19 Genomics UK (COG-UK) Consortium | Berkshire and Surrey Pathology Services Lighthouse Laboratory and Alex Alderton, Roberto Amato, Jeffrey Barrett, Sonia Goncalves, Ewan Harrison, David K. Jackson, Ian Johnston, Dominic Kwiatkowski, Cordelia Langford, John Sillitoe on behalf of the Wellcome Sanger Institute COVID-19 Surveillance Team |
| EPI_ISL_2917917, EPI_ISL_2917918, EPI_ISL_2917919 | Lighthouse Lab in Glasgow | Wellcome Sanger Institute for the COVID-19 Genomics UK (COG-UK) Consortium | Harper VanSteenhouse, Yumi Kasai, David Gray, Carol Clugston, Anna Dominiczak and Alex Alderton, Roberto Amato, Jeffrey Barrett, Sonia Goncalves, Ewan Harrison, David K. Jackson, Ian Johnston, Dominic Kwiatkowski, Cordelia Langford, John Sillitoe on behalf of the Wellcome Sanger Institute COVID-19 Surveillance Team |
| EPI_ISL_2917920, EPI_ISL_2917921 | Berkshire and Surrey Pathology Services Lighthouse Laboratory | Wellcome Sanger Institute for the COVID-19 Genomics UK (COG-UK) Consortium | Berkshire and Surrey Pathology Services Lighthouse Laboratory and Alex Alderton, Roberto Amato, Jeffrey Barrett, Sonia Goncalves, Ewan Harrison, David K. Jackson, Ian Johnston, Dominic Kwiatkowski, Cordelia Langford, John Sillitoe on behalf of the Wellcome Sanger Institute COVID-19 Surveillance Team |
| EPI_ISL_2917922, EPI_ISL_2917923, EPI_ISL_2917924, EPI_ISL_2917925, EPI_ISL_2917926 | Lighthouse Lab in Glasgow | Wellcome Sanger Institute for the COVID-19 Genomics UK (COG-UK) Consortium | Harper VanSteenhouse, Yumi Kasai, David Gray, Carol Clugston, Anna Dominiczak and Alex Alderton, Roberto Amato, Jeffrey Barrett, Sonia Goncalves, Ewan Harrison, David K. Jackson, Ian Johnston, Dominic Kwiatkowski, Cordelia Langford, John Sillitoe on behalf of the Wellcome Sanger Institute COVID-19 Surveillance Team |
| EPI_ISL_2917927 | Berkshire and Surrey Pathology Services Lighthouse Laboratory | Wellcome Sanger Institute for the COVID-19 Genomics UK (COG-UK) Consortium | Berkshire and Surrey Pathology Services Lighthouse Laboratory and Alex Alderton, Roberto Amato, Jeffrey Barrett, Sonia Goncalves, Ewan Harrison, David K. Jackson, Ian Johnston, Dominic Kwiatkowski, Cordelia Langford, John Sillitoe on behalf of the Wellcome Sanger Institute COVID-19 Surveillance Team |
| EPI_ISL_2917928, EPI_ISL_2917929, EPI_ISL_2917930, EPI_ISL_2917931, EPI_ISL_2917932 | Lighthouse Lab in Glasgow | Wellcome Sanger Institute for the COVID-19 Genomics UK (COG-UK) Consortium | Harper VanSteenhouse, Yumi Kasai, David Gray, Carol Clugston, Anna Dominiczak and Alex Alderton, Roberto Amato, Jeffrey Barrett, Sonia Goncalves, Ewan Harrison, David K. Jackson, Ian Johnston, Dominic Kwiatkowski, Cordelia Langford, John Sillitoe on behalf of the Wellcome Sanger Institute COVID-19 Surveillance Team |
| EPI_ISL_2917933 | Berkshire and Surrey Pathology Services Lighthouse Laboratory | Wellcome Sanger Institute for the COVID-19 Genomics UK (COG-UK) Consortium | Berkshire and Surrey Pathology Services Lighthouse Laboratory and Alex Alderton, Roberto Amato, Jeffrey Barrett, Sonia Goncalves, Ewan Harrison, David K. Jackson, Ian Johnston, Dominic Kwiatkowski, Cordelia Langford, John Sillitoe on behalf of the Wellcome Sanger Institute COVID-19 Surveillance Team |
| EPI_ISL_2917934, EPI_ISL_2917935, EPI_ISL_2917936, EPI_ISL_2917937 | Lighthouse Lab in Glasgow | Wellcome Sanger Institute for the COVID-19 Genomics UK (COG-UK) Consortium | Harper VanSteenhouse, Yumi Kasai, David Gray, Carol Clugston, Anna Dominiczak and Alex Alderton, Roberto Amato, Jeffrey Barrett, Sonia Goncalves, Ewan Harrison, David K. Jackson, Ian Johnston, Dominic Kwiatkowski, Cordelia Langford, John Sillitoe on behalf of the Wellcome Sanger Institute COVID-19 Surveillance Team |
| EPI_ISL_2917938 | Berkshire and Surrey Pathology Services Lighthouse Laboratory | Wellcome Sanger Institute for the COVID-19 Genomics UK (COG-UK) Consortium | Berkshire and Surrey Pathology Services Lighthouse Laboratory and Alex Alderton, Roberto Amato, Jeffrey Barrett, Sonia Goncalves, Ewan Harrison, David K. Jackson, Ian Johnston, Dominic Kwiatkowski, Cordelia Langford, John Sillitoe on behalf of the Wellcome Sanger Institute COVID-19 Surveillance Team |
| EPI_ISL_2917939, EPI_ISL_2917940, EPI_ISL_2917941, EPI_ISL_2917942 | Lighthouse Lab in Glasgow | Wellcome Sanger Institute for the COVID-19 Genomics UK (COG-UK) Consortium | Harper VanSteenhouse, Yumi Kasai, David Gray, Carol Clugston, Anna Dominiczak and Alex Alderton, Roberto Amato, Jeffrey Barrett, Sonia Goncalves, Ewan Harrison, David K. Jackson, Ian Johnston, Dominic Kwiatkowski, Cordelia Langford, John Sillitoe on behalf of the Wellcome Sanger Institute COVID-19 Surveillance Team |

| | | | |
|---|---|---|---|
| EPI_ISL_2917943, EPI_ISL_2917944, EPI_ISL_2917945 | Berkshire and Surrey Pathology Services Lighthouse Laboratory | Wellcome Sanger Institute for the COVID-19 Genomics UK (COG-UK) Consortium | Berkshire and Surrey Pathology Services Lighthouse Laboratory and Alex Alderton, Roberto Amato, Jeffrey Barrett, Sonia Goncalves, Ewan Harrison, David K. Jackson, Ian Johnston, Dominic Kwiatkowski, Cordelia Langford, John Sillitoe on behalf of the Wellcome Sanger Institute COVID-19 Surveillance Team |
| EPI_ISL_2917946 | Lighthouse Lab in Glasgow | Wellcome Sanger Institute for the COVID-19 Genomics UK (COG-UK) Consortium | Harper VanSteenhouse, Yumi Kasai, David Gray, Carol Clugston, Anna Dominiczak and Alex Alderton, Roberto Amato, Jeffrey Barrett, Sonia Goncalves, Ewan Harrison, David K. Jackson, Ian Johnston, Dominic Kwiatkowski, Cordelia Langford, John Sillitoe on behalf of the Wellcome Sanger Institute COVID-19 Surveillance Team |
| EPI_ISL_2917947 | Berkshire and Surrey Pathology Services Lighthouse Laboratory | Wellcome Sanger Institute for the COVID-19 Genomics UK (COG-UK) Consortium | Berkshire and Surrey Pathology Services Lighthouse Laboratory and Alex Alderton, Roberto Amato, Jeffrey Barrett, Sonia Goncalves, Ewan Harrison, David K. Jackson, Ian Johnston, Dominic Kwiatkowski, Cordelia Langford, John Sillitoe on behalf of the Wellcome Sanger Institute COVID-19 Surveillance Team |
| EPI_ISL_2917948 | Lighthouse Lab in Glasgow | Wellcome Sanger Institute for the COVID-19 Genomics UK (COG-UK) Consortium | Harper VanSteenhouse, Yumi Kasai, David Gray, Carol Clugston, Anna Dominiczak and Alex Alderton, Roberto Amato, Jeffrey Barrett, Sonia Goncalves, Ewan Harrison, David K. Jackson, Ian Johnston, Dominic Kwiatkowski, Cordelia Langford, John Sillitoe on behalf of the Wellcome Sanger Institute COVID-19 Surveillance Team |
| EPI_ISL_2917949 | Berkshire and Surrey Pathology Services Lighthouse Laboratory | Wellcome Sanger Institute for the COVID-19 Genomics UK (COG-UK) Consortium | Berkshire and Surrey Pathology Services Lighthouse Laboratory and Alex Alderton, Roberto Amato, Jeffrey Barrett, Sonia Goncalves, Ewan Harrison, David K. Jackson, Ian Johnston, Dominic Kwiatkowski, Cordelia Langford, John Sillitoe on behalf of the Wellcome Sanger Institute COVID-19 Surveillance Team |
| EPI_ISL_2917950, EPI_ISL_2917952, EPI_ISL_2917953, EPI_ISL_2917954, EPI_ISL_2917955, EPI_ISL_2917956, EPI_ISL_2917957, EPI_ISL_2917958, EPI_ISL_2917959 | Lighthouse Lab in Glasgow | Wellcome Sanger Institute for the COVID-19 Genomics UK (COG-UK) Consortium | Harper VanSteenhouse, Yumi Kasai, David Gray, Carol Clugston, Anna Dominiczak and Alex Alderton, Roberto Amato, Jeffrey Barrett, Sonia Goncalves, Ewan Harrison, David K. Jackson, Ian Johnston, Dominic Kwiatkowski, Cordelia Langford, John Sillitoe on behalf of the Wellcome Sanger Institute COVID-19 Surveillance Team |
| EPI_ISL_2917960 | Berkshire and Surrey Pathology Services Lighthouse Laboratory | Wellcome Sanger Institute for the COVID-19 Genomics UK (COG-UK) Consortium | Berkshire and Surrey Pathology Services Lighthouse Laboratory and Alex Alderton, Roberto Amato, Jeffrey Barrett, Sonia Goncalves, Ewan Harrison, David K. Jackson, Ian Johnston, Dominic Kwiatkowski, Cordelia Langford, John Sillitoe on behalf of the Wellcome Sanger Institute COVID-19 Surveillance Team |
| EPI_ISL_2917961, EPI_ISL_2917962, EPI_ISL_2917963, EPI_ISL_2917964, EPI_ISL_2917965, EPI_ISL_2917966, EPI_ISL_2917967, EPI_ISL_2917968, EPI_ISL_2917969, EPI_ISL_2917970, EPI_ISL_2917971, EPI_ISL_2917972, EPI_ISL_2917973, EPI_ISL_2917974, EPI_ISL_2917975, EPI_ISL_2917976, EPI_ISL_2917977, EPI_ISL_2917978, EPI_ISL_2917979, EPI_ISL_2917980, EPI_ISL_2917981, EPI_ISL_2917982, EPI_ISL_2917983, EPI_ISL_2917984, EPI_ISL_2917985, EPI_ISL_2917986, EPI_ISL_2917987, EPI_ISL_2917988, EPI_ISL_2917989, EPI_ISL_2917990, EPI_ISL_2917991, EPI_ISL_2917992, EPI_ISL_2917993, EPI_ISL_2917994, EPI_ISL_2917995, EPI_ISL_2917996, EPI_ISL_2917997, EPI_ISL_2917998, EPI_ISL_2917999, EPI_ISL_2918000, EPI_ISL_2918001, EPI_ISL_2918002, EPI_ISL_2918003, EPI_ISL_2918004, EPI_ISL_2918005, EPI_ISL_2918006, EPI_ISL_2918007, EPI_ISL_2918008, EPI_ISL_2918009, EPI_ISL_2918010, EPI_ISL_2918011, EPI_ISL_2918012, EPI_ISL_2918013, EPI_ISL_2918014, EPI_ISL_2918015, EPI_ISL_2918016, EPI_ISL_2918017, EPI_ISL_2918018, EPI_ISL_2918019, EPI_ISL_2918020, EPI_ISL_2918021, EPI_ISL_2918022, EPI_ISL_2918023, EPI_ISL_2918024, EPI_ISL_2918025, EPI_ISL_2918026, EPI_ISL_2918027, EPI_ISL_2918028, EPI_ISL_2918029, EPI_ISL_2918030, EPI_ISL_2918031, EPI_ISL_2918032, EPI_ISL_2918033, EPI_ISL_2918034, EPI_ISL_2918035, EPI_ISL_2918036, EPI_ISL_2918037, EPI_ISL_2918038, EPI_ISL_2918039, EPI_ISL_2918040, EPI_ISL_2918041, EPI_ISL_2918042, EPI_ISL_2918043, EPI_ISL_2918044, EPI_ISL_2918045, EPI_ISL_2918046, EPI_ISL_2918047, EPI_ISL_2918048, EPI_ISL_2918049, EPI_ISL_2918050, EPI_ISL_2918051, EPI_ISL_2918052, EPI_ISL_2918053, EPI_ISL_2918054, EPI_ISL_2918055, EPI_ISL_2918056, EPI_ISL_2918057, EPI_ISL_2918058, EPI_ISL_2918059, EPI_ISL_2918060, EPI_ISL_2918061, EPI_ISL_2918062, EPI_ISL_2918063, EPI_ISL_2918064 | | | |
| see above | Lighthouse Lab in Glasgow | Wellcome Sanger Institute for the COVID-19 Genomics UK (COG-UK) Consortium | Harper VanSteenhouse, Yumi Kasai, David Gray, Carol Clugston, Anna Dominiczak and Alex Alderton, Roberto Amato, Jeffrey Barrett, Sonia Goncalves, Ewan Harrison, David K. Jackson, Ian Johnston, Dominic Kwiatkowski, Cordelia Langford, John Sillitoe on behalf of the Wellcome Sanger Institute COVID-19 Surveillance Team |
| EPI_ISL_2918065 | Health Services Laboratories | Wellcome Sanger Institute for the COVID-19 Genomics UK (COG-UK) Consortium | Health Services Laboratories and Alex Alderton, Roberto Amato, Jeffrey Barrett, Sonia Goncalves, Ewan Harrison, David K. Jackson, Ian Johnston, Dominic Kwiatkowski, Cordelia Langford, John Sillitoe on behalf of the Wellcome Sanger Institute COVID-19 Surveillance Team |
| EPI_ISL_2918066, EPI_ISL_2918067 | Lighthouse Lab in Milton Keynes | Wellcome Sanger Institute for the COVID-19 Genomics UK (COG-UK) Consortium | The Lighthouse Lab in Milton Keynes and Alex Alderton, Roberto Amato, Jeffrey Barrett, Sonia Goncalves, Ewan Harrison, David K. Jackson, Ian Johnston, Dominic Kwiatkowski, Cordelia Langford, John Sillitoe on behalf of the Wellcome Sanger Institute COVID-19 Surveillance Team |
| EPI_ISL_2918068, EPI_ISL_2918069 | Lighthouse Lab in Alderley Park | Wellcome Sanger Institute for the COVID-19 Genomics UK (COG-UK) Consortium | Jacquelyn Wynn, Mairead Hyland, The Lighthouse Lab in Alderley Park and Alex Alderton, Roberto Amato, Jeffrey Barrett, Sonia Goncalves, Ewan Harrison, David K. Jackson, Ian Johnston, Dominic Kwiatkowski, Cordelia Langford, John Sillitoe on behalf of the Wellcome Sanger Institute COVID-19 Surveillance Team |
| EPI_ISL_2918070 | Berkshire and Surrey Pathology Services Lighthouse Laboratory | Wellcome Sanger Institute for the COVID-19 Genomics UK (COG-UK) Consortium | Berkshire and Surrey Pathology Services Lighthouse Laboratory and Alex Alderton, Roberto Amato, Jeffrey Barrett, Sonia Goncalves, Ewan Harrison, David K. Jackson, Ian Johnston, Dominic Kwiatkowski, Cordelia Langford, John Sillitoe on behalf of the Wellcome Sanger Institute COVID-19 Surveillance Team |
| EPI_ISL_2918071, EPI_ISL_2918072 | Lighthouse Lab in Milton Keynes | Wellcome Sanger Institute for the COVID-19 Genomics UK (COG-UK) Consortium | The Lighthouse Lab in Milton Keynes and Alex Alderton, Roberto Amato, Jeffrey Barrett, Sonia Goncalves, Ewan Harrison, David K. Jackson, Ian Johnston, Dominic Kwiatkowski, Cordelia Langford, John Sillitoe on behalf of the Wellcome Sanger Institute COVID-19 Surveillance Team |
| EPI_ISL_2918073, EPI_ISL_2918074 | Lighthouse Lab in Alderley Park | Wellcome Sanger Institute for the COVID-19 Genomics UK (COG-UK) Consortium | Jacquelyn Wynn, Mairead Hyland, The Lighthouse Lab in Alderley Park and Alex Alderton, Roberto Amato, Jeffrey Barrett, Sonia Goncalves, Ewan Harrison, David K. Jackson, Ian Johnston, Dominic Kwiatkowski, Cordelia Langford, John Sillitoe on behalf of the Wellcome Sanger Institute COVID-19 Surveillance Team |
| EPI_ISL_2918075 | Berkshire and Surrey Pathology Services Lighthouse Laboratory | Wellcome Sanger Institute for the COVID-19 Genomics UK (COG-UK) Consortium | Berkshire and Surrey Pathology Services Lighthouse Laboratory and Alex Alderton, Roberto Amato, Jeffrey Barrett, Sonia Goncalves, Ewan Harrison, David K. Jackson, Ian Johnston, Dominic Kwiatkowski, Cordelia Langford, John Sillitoe on behalf of the Wellcome Sanger Institute COVID-19 Surveillance Team |
| EPI_ISL_2918076 | Lighthouse Lab in Milton Keynes | Wellcome Sanger Institute for the COVID-19 Genomics UK (COG-UK) Consortium | The Lighthouse Lab in Milton Keynes and Alex Alderton, Roberto Amato, Jeffrey Barrett, Sonia Goncalves, Ewan Harrison, David K. Jackson, Ian Johnston, Dominic Kwiatkowski, Cordelia Langford, John Sillitoe on behalf of the Wellcome Sanger Institute COVID-19 Surveillance Team |
| EPI_ISL_2918077 | Health Services Laboratories | Wellcome Sanger Institute for the COVID-19 Genomics UK (COG-UK) Consortium | Health Services Laboratories and Alex Alderton, Roberto Amato, Jeffrey Barrett, Sonia Goncalves, Ewan Harrison, David K. Jackson, Ian Johnston, Dominic Kwiatkowski, Cordelia Langford, John Sillitoe on behalf of the Wellcome Sanger Institute COVID-19 Surveillance Team |
| EPI_ISL_2918078 | Lighthouse Lab in Alderley Park | Wellcome Sanger Institute for the COVID-19 Genomics UK (COG-UK) Consortium | Jacquelyn Wynn, Mairead Hyland, The Lighthouse Lab in Alderley Park and Alex Alderton, Roberto Amato, Jeffrey Barrett, Sonia Goncalves, Ewan Harrison, David K. Jackson, Ian Johnston, Dominic Kwiatkowski, Cordelia Langford, John Sillitoe on behalf of the Wellcome Sanger Institute COVID-19 Surveillance Team |
| EPI_ISL_2918079 | Berkshire and Surrey Pathology Services Lighthouse Laboratory | Wellcome Sanger Institute for the COVID-19 Genomics UK (COG-UK) Consortium | Berkshire and Surrey Pathology Services Lighthouse Laboratory and Alex Alderton, Roberto Amato, Jeffrey Barrett, Sonia Goncalves, Ewan Harrison, David K. Jackson, Ian Johnston, Dominic Kwiatkowski, Cordelia Langford, John Sillitoe on behalf of the Wellcome Sanger Institute COVID-19 Surveillance Team |
| EPI_ISL_2918080 | Health Services Laboratories | Wellcome Sanger Institute for the COVID-19 Genomics UK (COG-UK) Consortium | Health Services Laboratories and Alex Alderton, Roberto Amato, Jeffrey Barrett, Sonia Goncalves, Ewan Harrison, David K. Jackson, Ian Johnston, Dominic Kwiatkowski, Cordelia Langford, John Sillitoe on behalf of the Wellcome Sanger Institute COVID-19 Surveillance Team |
| EPI_ISL_2918081, EPI_ISL_2918082 | Lighthouse Lab in Alderley Park | Wellcome Sanger Institute for the COVID-19 Genomics UK (COG-UK) Consortium | Jacquelyn Wynn, Mairead Hyland, The Lighthouse Lab in Alderley Park and Alex Alderton, Roberto Amato, Jeffrey Barrett, Sonia Goncalves, Ewan Harrison, David K. Jackson, Ian Johnston, Dominic Kwiatkowski, Cordelia Langford, John Sillitoe on behalf of the Wellcome Sanger Institute COVID-19 Surveillance Team |
| EPI_ISL_2918083 | Lighthouse Lab in Milton Keynes | Wellcome Sanger Institute for the COVID-19 Genomics UK (COG-UK) Consortium | The Lighthouse Lab in Milton Keynes and Alex Alderton, Roberto Amato, Jeffrey Barrett, Sonia Goncalves, Ewan Harrison, David K. Jackson, Ian Johnston, Dominic Kwiatkowski, Cordelia Langford, John Sillitoe on behalf of the Wellcome Sanger Institute COVID-19 Surveillance Team |
| EPI_ISL_2918084 | Berkshire and Surrey Pathology Services Lighthouse Laboratory | Wellcome Sanger Institute for the COVID-19 Genomics UK (COG-UK) Consortium | Berkshire and Surrey Pathology Services Lighthouse Laboratory and Alex Alderton, Roberto Amato, Jeffrey Barrett, Sonia Goncalves, Ewan Harrison, David K. Jackson, Ian Johnston, Dominic Kwiatkowski, Cordelia Langford, John Sillitoe on behalf of the Wellcome Sanger Institute COVID-19 Surveillance Team |
| EPI_ISL_2918085 | Health Services Laboratories | Wellcome Sanger Institute for the COVID-19 Genomics UK (COG-UK) Consortium | Health Services Laboratories and Alex Alderton, Roberto Amato, Jeffrey Barrett, Sonia Goncalves, Ewan Harrison, David K. Jackson, Ian Johnston, Dominic Kwiatkowski, Cordelia Langford, John Sillitoe on behalf of the Wellcome Sanger Institute COVID-19 Surveillance Team |
| EPI_ISL_2918086, EPI_ISL_2918087 | Lighthouse Lab in Alderley Park | Wellcome Sanger Institute for the COVID-19 Genomics UK (COG-UK) Consortium | Jacquelyn Wynn, Mairead Hyland, The Lighthouse Lab in Alderley Park and Alex Alderton, Roberto Amato, Jeffrey Barrett, Sonia Goncalves, Ewan Harrison, David K. Jackson, Ian Johnston, Dominic Kwiatkowski, Cordelia Langford, John Sillitoe on behalf of the Wellcome Sanger Institute COVID-19 Surveillance Team |
| EPI_ISL_2918088 | Berkshire and Surrey Pathology Services Lighthouse Laboratory | Wellcome Sanger Institute for the COVID-19 Genomics UK (COG-UK) Consortium | Berkshire and Surrey Pathology Services Lighthouse Laboratory and Alex Alderton, Roberto Amato, Jeffrey Barrett, Sonia Goncalves, Ewan Harrison, David K. Jackson, Ian Johnston, Dominic Kwiatkowski, Cordelia Langford, John Sillitoe on behalf of the Wellcome Sanger Institute COVID-19 Surveillance Team |
| EPI_ISL_2918089 | Lighthouse Lab in Alderley Park | Wellcome Sanger Institute for the COVID-19 Genomics UK (COG-UK) Consortium | Jacquelyn Wynn, Mairead Hyland, The Lighthouse Lab in Alderley Park and Alex Alderton, Roberto Amato, Jeffrey Barrett, Sonia Goncalves, Ewan Harrison, David K. Jackson, Ian Johnston, Dominic Kwiatkowski, Cordelia Langford, John Sillitoe on behalf of the Wellcome Sanger Institute COVID-19 Surveillance Team |
| EPI_ISL_2918090 | Health Services Laboratories | Wellcome Sanger Institute for the COVID-19 Genomics UK (COG-UK) Consortium | Health Services Laboratories and Alex Alderton, Roberto Amato, Jeffrey Barrett, Sonia Goncalves, Ewan Harrison, David K. Jackson, Ian Johnston, Dominic Kwiatkowski, Cordelia Langford, John Sillitoe on behalf of the Wellcome Sanger Institute COVID-19 Surveillance Team |
| EPI_ISL_2918091 | Randox Laboratories | Wellcome Sanger Institute for the COVID-19 Genomics UK (COG-UK) Consortium | Randox Laboratories and Alex Alderton, Roberto Amato, Jeffrey Barrett, Sonia Goncalves, Ewan Harrison, David K. Jackson, Ian Johnston, Dominic Kwiatkowski, Cordelia Langford, John Sillitoe on behalf of the Wellcome Sanger Institute COVID-19 Surveillance Team |
| EPI_ISL_2918092 | Lighthouse Lab in Alderley Park | Wellcome Sanger Institute for the COVID-19 Genomics UK (COG-UK) Consortium | Jacquelyn Wynn, Mairead Hyland, The Lighthouse Lab in Alderley Park and Alex Alderton, Roberto Amato, Jeffrey Barrett, Sonia Goncalves, Ewan Harrison, David K. Jackson, Ian Johnston, Dominic Kwiatkowski, Cordelia Langford, John Sillitoe on behalf of the Wellcome Sanger Institute COVID-19 Surveillance Team |
| EPI_ISL_2918093 | Berkshire and Surrey Pathology Services Lighthouse Laboratory | Wellcome Sanger Institute for the COVID-19 Genomics UK (COG-UK) Consortium | Berkshire and Surrey Pathology Services Lighthouse Laboratory and Alex Alderton, Roberto Amato, Jeffrey Barrett, Sonia Goncalves, Ewan Harrison, David K. Jackson, Ian Johnston, Dominic Kwiatkowski, Cordelia Langford, John Sillitoe on behalf of the Wellcome Sanger Institute COVID-19 Surveillance Team |
| EPI_ISL_2918094 | Lighthouse Lab in Alderley Park | Wellcome Sanger Institute for the COVID-19 Genomics UK (COG-UK) Consortium | Jacquelyn Wynn, Mairead Hyland, The Lighthouse Lab in Alderley Park and Alex Alderton, Roberto Amato, Jeffrey Barrett, Sonia Goncalves, Ewan Harrison, David K. Jackson, Ian Johnston, Dominic Kwiatkowski, Cordelia Langford, John Sillitoe on behalf of the Wellcome Sanger Institute COVID-19 Surveillance Team |
| EPI_ISL_2918095 | Lighthouse Lab in Milton Keynes | Wellcome Sanger Institute for the COVID-19 Genomics UK (COG-UK) Consortium | The Lighthouse Lab in Milton Keynes and Alex Alderton, Roberto Amato, Jeffrey Barrett, Sonia Goncalves, Ewan Harrison, David K. Jackson, Ian Johnston, Dominic Kwiatkowski, Cordelia Langford, John Sillitoe on behalf of the Wellcome Sanger Institute COVID-19 Surveillance Team |
| EPI_ISL_2918096 | Health Services Laboratories | Wellcome Sanger Institute for the COVID-19 Genomics UK (COG-UK) Consortium | Health Services Laboratories and Alex Alderton, Roberto Amato, Jeffrey Barrett, Sonia Goncalves, Ewan Harrison, David K. Jackson, Ian Johnston, Dominic Kwiatkowski, Cordelia Langford, John Sillitoe on behalf of the Wellcome Sanger Institute COVID-19 Surveillance Team |
| EPI_ISL_2918097 | Berkshire and Surrey Pathology Services Lighthouse Laboratory | Wellcome Sanger Institute for the COVID-19 Genomics UK (COG-UK) Consortium | Berkshire and Surrey Pathology Services Lighthouse Laboratory and Alex Alderton, Roberto Amato, Jeffrey Barrett, Sonia Goncalves, Ewan Harrison, David K. Jackson, Ian Johnston, Dominic Kwiatkowski, Cordelia Langford, John Sillitoe on behalf of the Wellcome Sanger Institute COVID-19 Surveillance Team |
| EPI_ISL_2918098 | Lighthouse Lab in Alderley Park | Wellcome Sanger Institute for the COVID-19 Genomics UK (COG-UK) Consortium | Jacquelyn Wynn, Mairead Hyland, The Lighthouse Lab in Alderley Park and Alex Alderton, Roberto Amato, Jeffrey Barrett, Sonia Goncalves, Ewan Harrison, David K. Jackson, Ian Johnston, Dominic Kwiatkowski, Cordelia Langford, John Sillitoe on behalf of the Wellcome Sanger Institute COVID-19 Surveillance Team |
| EPI_ISL_2918099 | Lighthouse Lab in Milton Keynes | Wellcome Sanger Institute for the COVID-19 Genomics UK (COG-UK) Consortium | The Lighthouse Lab in Milton Keynes and Alex Alderton, Roberto Amato, Jeffrey Barrett, Sonia Goncalves, Ewan Harrison, David K. Jackson, Ian Johnston, Dominic Kwiatkowski, Cordelia Langford, John Sillitoe on behalf of the Wellcome Sanger Institute COVID-19 Surveillance Team |
| EPI_ISL_2918100 | Health Services Laboratories | Wellcome Sanger Institute for the COVID-19 Genomics UK (COG-UK) Consortium | Health Services Laboratories and Alex Alderton, Roberto Amato, Jeffrey Barrett, Sonia Goncalves, Ewan Harrison, David K. Jackson, Ian Johnston, Dominic Kwiatkowski, Cordelia Langford, John Sillitoe on behalf of the Wellcome Sanger Institute COVID-19 Surveillance Team |
| EPI_ISL_2918101, EPI_ISL_2918102, EPI_ISL_2918103 | Lighthouse Lab in Alderley Park | Wellcome Sanger Institute for the COVID-19 Genomics UK (COG-UK) Consortium | Jacquelyn Wynn, Mairead Hyland, The Lighthouse Lab in Alderley Park and Alex Alderton, Roberto Amato, Jeffrey Barrett, Sonia Goncalves, Ewan Harrison, David K. Jackson, Ian Johnston, Dominic Kwiatkowski, Cordelia Langford, John Sillitoe on behalf of the Wellcome Sanger Institute COVID-19 Surveillance Team |

| | | | |
|---|---|---|---|
| EPI_ISL_2918104, EPI_ISL_2918105 | Lighthouse Lab in Milton Keynes | Wellcome Sanger Institute for the COVID-19 Genomics UK (COG-UK) Consortium | The Lighthouse Lab in Milton Keynes and Alex Alderton, Roberto Amato, Jeffrey Barrett, Sonia Goncalves, Ewan Harrison, David K. Jackson, Ian Johnston, Dominic Kwiatkowski, Cordelia Langford, John Sillitoe on behalf of the Wellcome Sanger Institute COVID-19 Surveillance Team |
| EPI_ISL_2918106 | Berkshire and Surrey Pathology Services Lighthouse Laboratory | Wellcome Sanger Institute for the COVID-19 Genomics UK (COG-UK) Consortium | Berkshire and Surrey Pathology Services Lighthouse Laboratory and Alex Alderton, Roberto Amato, Jeffrey Barrett, Sonia Goncalves, Ewan Harrison, David K. Jackson, Ian Johnston, Dominic Kwiatkowski, Cordelia Langford, John Sillitoe on behalf of the Wellcome Sanger Institute COVID-19 Surveillance Team |
| EPI_ISL_2918107, EPI_ISL_2918108 | Lighthouse Lab in Alderley Park | Wellcome Sanger Institute for the COVID-19 Genomics UK (COG-UK) Consortium | Jacquelyn Wynn, Mairead Hyland, The Lighthouse Lab in Alderley Park and Alex Alderton, Roberto Amato, Jeffrey Barrett, Sonia Goncalves, Ewan Harrison, David K. Jackson, Ian Johnston, Dominic Kwiatkowski, Cordelia Langford, John Sillitoe on behalf of the Wellcome Sanger Institute COVID-19 Surveillance Team |
| EPI_ISL_2918109 | Berkshire and Surrey Pathology Services Lighthouse Laboratory | Wellcome Sanger Institute for the COVID-19 Genomics UK (COG-UK) Consortium | Berkshire and Surrey Pathology Services Lighthouse Laboratory and Alex Alderton, Roberto Amato, Jeffrey Barrett, Sonia Goncalves, Ewan Harrison, David K. Jackson, Ian Johnston, Dominic Kwiatkowski, Cordelia Langford, John Sillitoe on behalf of the Wellcome Sanger Institute COVID-19 Surveillance Team |
| EPI_ISL_2918110 | Lighthouse Lab in Milton Keynes | Wellcome Sanger Institute for the COVID-19 Genomics UK (COG-UK) Consortium | The Lighthouse Lab in Milton Keynes and Alex Alderton, Roberto Amato, Jeffrey Barrett, Sonia Goncalves, Ewan Harrison, David K. Jackson, Ian Johnston, Dominic Kwiatkowski, Cordelia Langford, John Sillitoe on behalf of the Wellcome Sanger Institute COVID-19 Surveillance Team |
| EPI_ISL_2918111 | Randox Laboratories | Wellcome Sanger Institute for the COVID-19 Genomics UK (COG-UK) Consortium | Randox Laboratories and Alex Alderton, Roberto Amato, Jeffrey Barrett, Sonia Goncalves, Ewan Harrison, David K. Jackson, Ian Johnston, Dominic Kwiatkowski, Cordelia Langford, John Sillitoe on behalf of the Wellcome Sanger Institute COVID-19 Surveillance Team |
| EPI_ISL_2918112, EPI_ISL_2918113 | Lighthouse Lab in Alderley Park | Wellcome Sanger Institute for the COVID-19 Genomics UK (COG-UK) Consortium | Jacquelyn Wynn, Mairead Hyland, The Lighthouse Lab in Alderley Park and Alex Alderton, Roberto Amato, Jeffrey Barrett, Sonia Goncalves, Ewan Harrison, David K. Jackson, Ian Johnston, Dominic Kwiatkowski, Cordelia Langford, John Sillitoe on behalf of the Wellcome Sanger Institute COVID-19 Surveillance Team |
| EPI_ISL_2918114, EPI_ISL_2918115 | Berkshire and Surrey Pathology Services Lighthouse Laboratory | Wellcome Sanger Institute for the COVID-19 Genomics UK (COG-UK) Consortium | Berkshire and Surrey Pathology Services Lighthouse Laboratory and Alex Alderton, Roberto Amato, Jeffrey Barrett, Sonia Goncalves, Ewan Harrison, David K. Jackson, Ian Johnston, Dominic Kwiatkowski, Cordelia Langford, John Sillitoe on behalf of the Wellcome Sanger Institute COVID-19 Surveillance Team |
| EPI_ISL_2918116 | Lighthouse Lab in Alderley Park | Wellcome Sanger Institute for the COVID-19 Genomics UK (COG-UK) Consortium | Jacquelyn Wynn, Mairead Hyland, The Lighthouse Lab in Alderley Park and Alex Alderton, Roberto Amato, Jeffrey Barrett, Sonia Goncalves, Ewan Harrison, David K. Jackson, Ian Johnston, Dominic Kwiatkowski, Cordelia Langford, John Sillitoe on behalf of the Wellcome Sanger Institute COVID-19 Surveillance Team |
| EPI_ISL_2918117 | Health Services Laboratories | Wellcome Sanger Institute for the COVID-19 Genomics UK (COG-UK) Consortium | Health Services Laboratories and Alex Alderton, Roberto Amato, Jeffrey Barrett, Sonia Goncalves, Ewan Harrison, David K. Jackson, Ian Johnston, Dominic Kwiatkowski, Cordelia Langford, John Sillitoe on behalf of the Wellcome Sanger Institute COVID-19 Surveillance Team |
| EPI_ISL_2918118, EPI_ISL_2918119 | Lighthouse Lab in Alderley Park | Wellcome Sanger Institute for the COVID-19 Genomics UK (COG-UK) Consortium | Jacquelyn Wynn, Mairead Hyland, The Lighthouse Lab in Alderley Park and Alex Alderton, Roberto Amato, Jeffrey Barrett, Sonia Goncalves, Ewan Harrison, David K. Jackson, Ian Johnston, Dominic Kwiatkowski, Cordelia Langford, John Sillitoe on behalf of the Wellcome Sanger Institute COVID-19 Surveillance Team |
| EPI_ISL_2918120, EPI_ISL_2918121 | Lighthouse Lab in Milton Keynes | Wellcome Sanger Institute for the COVID-19 Genomics UK (COG-UK) Consortium | The Lighthouse Lab in Milton Keynes and Alex Alderton, Roberto Amato, Jeffrey Barrett, Sonia Goncalves, Ewan Harrison, David K. Jackson, Ian Johnston, Dominic Kwiatkowski, Cordelia Langford, John Sillitoe on behalf of the Wellcome Sanger Institute COVID-19 Surveillance Team |
| EPI_ISL_2918122 | Berkshire and Surrey Pathology Services Lighthouse Laboratory | Wellcome Sanger Institute for the COVID-19 Genomics UK (COG-UK) Consortium | Berkshire and Surrey Pathology Services Lighthouse Laboratory and Alex Alderton, Roberto Amato, Jeffrey Barrett, Sonia Goncalves, Ewan Harrison, David K. Jackson, Ian Johnston, Dominic Kwiatkowski, Cordelia Langford, John Sillitoe on behalf of the Wellcome Sanger Institute COVID-19 Surveillance Team |
| EPI_ISL_2918123, EPI_ISL_2918124 | Health Services Laboratories | Wellcome Sanger Institute for the COVID-19 Genomics UK (COG-UK) Consortium | Health Services Laboratories and Alex Alderton, Roberto Amato, Jeffrey Barrett, Sonia Goncalves, Ewan Harrison, David K. Jackson, Ian Johnston, Dominic Kwiatkowski, Cordelia Langford, John Sillitoe on behalf of the Wellcome Sanger Institute COVID-19 Surveillance Team |
| EPI_ISL_2918125 | Lighthouse Lab in Milton Keynes | Wellcome Sanger Institute for the COVID-19 Genomics UK (COG-UK) Consortium | The Lighthouse Lab in Milton Keynes and Alex Alderton, Roberto Amato, Jeffrey Barrett, Sonia Goncalves, Ewan Harrison, David K. Jackson, Ian Johnston, Dominic Kwiatkowski, Cordelia Langford, John Sillitoe on behalf of the Wellcome Sanger Institute COVID-19 Surveillance Team |
| EPI_ISL_2918126 | Health Services Laboratories | Wellcome Sanger Institute for the COVID-19 Genomics UK (COG-UK) Consortium | Health Services Laboratories and Alex Alderton, Roberto Amato, Jeffrey Barrett, Sonia Goncalves, Ewan Harrison, David K. Jackson, Ian Johnston, Dominic Kwiatkowski, Cordelia Langford, John Sillitoe on behalf of the Wellcome Sanger Institute COVID-19 Surveillance Team |
| EPI_ISL_2918127, EPI_ISL_2918128 | Lighthouse Lab in Alderley Park | Wellcome Sanger Institute for the COVID-19 Genomics UK (COG-UK) Consortium | Jacquelyn Wynn, Mairead Hyland, The Lighthouse Lab in Alderley Park and Alex Alderton, Roberto Amato, Jeffrey Barrett, Sonia Goncalves, Ewan Harrison, David K. Jackson, Ian Johnston, Dominic Kwiatkowski, Cordelia Langford, John Sillitoe on behalf of the Wellcome Sanger Institute COVID-19 Surveillance Team |
| EPI_ISL_2918129 | Health Services Laboratories | Wellcome Sanger Institute for the COVID-19 Genomics UK (COG-UK) Consortium | Health Services Laboratories and Alex Alderton, Roberto Amato, Jeffrey Barrett, Sonia Goncalves, Ewan Harrison, David K. Jackson, Ian Johnston, Dominic Kwiatkowski, Cordelia Langford, John Sillitoe on behalf of the Wellcome Sanger Institute COVID-19 Surveillance Team |
| EPI_ISL_2918130 | Lighthouse Lab in Alderley Park | Wellcome Sanger Institute for the COVID-19 Genomics UK (COG-UK) Consortium | Jacquelyn Wynn, Mairead Hyland, The Lighthouse Lab in Alderley Park and Alex Alderton, Roberto Amato, Jeffrey Barrett, Sonia Goncalves, Ewan Harrison, David K. Jackson, Ian Johnston, Dominic Kwiatkowski, Cordelia Langford, John Sillitoe on behalf of the Wellcome Sanger Institute COVID-19 Surveillance Team |
| EPI_ISL_2918131 | Lighthouse Lab in Milton Keynes | Wellcome Sanger Institute for the COVID-19 Genomics UK (COG-UK) Consortium | The Lighthouse Lab in Milton Keynes and Alex Alderton, Roberto Amato, Jeffrey Barrett, Sonia Goncalves, Ewan Harrison, David K. Jackson, Ian Johnston, Dominic Kwiatkowski, Cordelia Langford, John Sillitoe on behalf of the Wellcome Sanger Institute COVID-19 Surveillance Team |
| EPI_ISL_2918132, EPI_ISL_2918133, EPI_ISL_2918134, EPI_ISL_2918135, EPI_ISL_2918136 | Lighthouse Lab in Alderley Park | Wellcome Sanger Institute for the COVID-19 Genomics UK (COG-UK) Consortium | Jacquelyn Wynn, Mairead Hyland, The Lighthouse Lab in Alderley Park and Alex Alderton, Roberto Amato, Jeffrey Barrett, Sonia Goncalves, Ewan Harrison, David K. Jackson, Ian Johnston, Dominic Kwiatkowski, Cordelia Langford, John Sillitoe on behalf of the Wellcome Sanger Institute COVID-19 Surveillance Team |
| EPI_ISL_2918137 | Lighthouse Lab in Milton Keynes | Wellcome Sanger Institute for the COVID-19 Genomics UK (COG-UK) Consortium | The Lighthouse Lab in Milton Keynes and Alex Alderton, Roberto Amato, Jeffrey Barrett, Sonia Goncalves, Ewan Harrison, David K. Jackson, Ian Johnston, Dominic Kwiatkowski, Cordelia Langford, John Sillitoe on behalf of the Wellcome Sanger Institute COVID-19 Surveillance Team |
| EPI_ISL_2918138 | Health Services Laboratories | Wellcome Sanger Institute for the COVID-19 Genomics UK (COG-UK) Consortium | Health Services Laboratories and Alex Alderton, Roberto Amato, Jeffrey Barrett, Sonia Goncalves, Ewan Harrison, David K. Jackson, Ian Johnston, Dominic Kwiatkowski, Cordelia Langford, John Sillitoe on behalf of the Wellcome Sanger Institute COVID-19 Surveillance Team |
| EPI_ISL_2918139 | Lighthouse Lab in Milton Keynes | Wellcome Sanger Institute for the COVID-19 Genomics UK (COG-UK) Consortium | The Lighthouse Lab in Milton Keynes and Alex Alderton, Roberto Amato, Jeffrey Barrett, Sonia Goncalves, Ewan Harrison, David K. Jackson, Ian Johnston, Dominic Kwiatkowski, Cordelia Langford, John Sillitoe on behalf of the Wellcome Sanger Institute COVID-19 Surveillance Team |
| EPI_ISL_2918140 | Lighthouse Lab in Alderley Park | Wellcome Sanger Institute for the COVID-19 Genomics UK (COG-UK) Consortium | Jacquelyn Wynn, Mairead Hyland, The Lighthouse Lab in Alderley Park and Alex Alderton, Roberto Amato, Jeffrey Barrett, Sonia Goncalves, Ewan Harrison, David K. Jackson, Ian Johnston, Dominic Kwiatkowski, Cordelia Langford, John Sillitoe on behalf of the Wellcome Sanger Institute COVID-19 Surveillance Team |
| EPI_ISL_2918141 | Berkshire and Surrey Pathology Services Lighthouse Laboratory | Wellcome Sanger Institute for the COVID-19 Genomics UK (COG-UK) Consortium | Berkshire and Surrey Pathology Services Lighthouse Laboratory and Alex Alderton, Roberto Amato, Jeffrey Barrett, Sonia Goncalves, Ewan Harrison, David K. Jackson, Ian Johnston, Dominic Kwiatkowski, Cordelia Langford, John Sillitoe on behalf of the Wellcome Sanger Institute COVID-19 Surveillance Team |
| EPI_ISL_2918142 | Lighthouse Lab in Alderley Park | Wellcome Sanger Institute for the COVID-19 Genomics UK (COG-UK) Consortium | Jacquelyn Wynn, Mairead Hyland, The Lighthouse Lab in Alderley Park and Alex Alderton, Roberto Amato, Jeffrey Barrett, Sonia Goncalves, Ewan Harrison, David K. Jackson, Ian Johnston, Dominic Kwiatkowski, Cordelia Langford, John Sillitoe on behalf of the Wellcome Sanger Institute COVID-19 Surveillance Team |
| EPI_ISL_2918143, EPI_ISL_2918144 | Lighthouse Lab in Milton Keynes | Wellcome Sanger Institute for the COVID-19 Genomics UK (COG-UK) Consortium | The Lighthouse Lab in Milton Keynes and Alex Alderton, Roberto Amato, Jeffrey Barrett, Sonia Goncalves, Ewan Harrison, David K. Jackson, Ian Johnston, Dominic Kwiatkowski, Cordelia Langford, John Sillitoe on behalf of the Wellcome Sanger Institute COVID-19 Surveillance Team |
| EPI_ISL_2918145, EPI_ISL_2918146 | Berkshire and Surrey Pathology Services Lighthouse Laboratory | Wellcome Sanger Institute for the COVID-19 Genomics UK (COG-UK) Consortium | Berkshire and Surrey Pathology Services Lighthouse Laboratory and Alex Alderton, Roberto Amato, Jeffrey Barrett, Sonia Goncalves, Ewan Harrison, David K. Jackson, Ian Johnston, Dominic Kwiatkowski, Cordelia Langford, John Sillitoe on behalf of the Wellcome Sanger Institute COVID-19 Surveillance Team |
| EPI_ISL_2918147 | Lighthouse Lab in Milton Keynes | Wellcome Sanger Institute for the COVID-19 Genomics UK (COG-UK) Consortium | The Lighthouse Lab in Milton Keynes and Alex Alderton, Roberto Amato, Jeffrey Barrett, Sonia Goncalves, Ewan Harrison, David K. Jackson, Ian Johnston, Dominic Kwiatkowski, Cordelia Langford, John Sillitoe on behalf of the Wellcome Sanger Institute COVID-19 Surveillance Team |
| EPI_ISL_2918148, EPI_ISL_2918149 | Berkshire and Surrey Pathology Services Lighthouse Laboratory | Wellcome Sanger Institute for the COVID-19 Genomics UK (COG-UK) Consortium | Berkshire and Surrey Pathology Services Lighthouse Laboratory and Alex Alderton, Roberto Amato, Jeffrey Barrett, Sonia Goncalves, Ewan Harrison, David K. Jackson, Ian Johnston, Dominic Kwiatkowski, Cordelia Langford, John Sillitoe on behalf of the Wellcome Sanger Institute COVID-19 Surveillance Team |
| EPI_ISL_2918150 | Health Services Laboratories | Wellcome Sanger Institute for the COVID-19 Genomics UK (COG-UK) Consortium | Health Services Laboratories and Alex Alderton, Roberto Amato, Jeffrey Barrett, Sonia Goncalves, Ewan Harrison, David K. Jackson, Ian Johnston, Dominic Kwiatkowski, Cordelia Langford, John Sillitoe on behalf of the Wellcome Sanger Institute COVID-19 Surveillance Team |
| EPI_ISL_2918151 | Lighthouse Lab in Milton Keynes | Wellcome Sanger Institute for the COVID-19 Genomics UK (COG-UK) Consortium | The Lighthouse Lab in Milton Keynes and Alex Alderton, Roberto Amato, Jeffrey Barrett, Sonia Goncalves, Ewan Harrison, David K. Jackson, Ian Johnston, Dominic Kwiatkowski, Cordelia Langford, John Sillitoe on behalf of the Wellcome Sanger Institute COVID-19 Surveillance Team |
| EPI_ISL_2918152 | Berkshire and Surrey Pathology Services Lighthouse Laboratory | Wellcome Sanger Institute for the COVID-19 Genomics UK (COG-UK) Consortium | Berkshire and Surrey Pathology Services Lighthouse Laboratory and Alex Alderton, Roberto Amato, Jeffrey Barrett, Sonia Goncalves, Ewan Harrison, David K. Jackson, Ian Johnston, Dominic Kwiatkowski, Cordelia Langford, John Sillitoe on behalf of the Wellcome Sanger Institute COVID-19 Surveillance Team |
| EPI_ISL_2918153 | Health Services Laboratories | Wellcome Sanger Institute for the COVID-19 Genomics UK (COG-UK) Consortium | Health Services Laboratories and Alex Alderton, Roberto Amato, Jeffrey Barrett, Sonia Goncalves, Ewan Harrison, David K. Jackson, Ian Johnston, Dominic Kwiatkowski, Cordelia Langford, John Sillitoe on behalf of the Wellcome Sanger Institute COVID-19 Surveillance Team |
| EPI_ISL_2918154 | Lighthouse Lab in Milton Keynes | Wellcome Sanger Institute for the COVID-19 Genomics UK (COG-UK) Consortium | The Lighthouse Lab in Milton Keynes and Alex Alderton, Roberto Amato, Jeffrey Barrett, Sonia Goncalves, Ewan Harrison, David K. Jackson, Ian Johnston, Dominic Kwiatkowski, Cordelia Langford, John Sillitoe on behalf of the Wellcome Sanger Institute COVID-19 Surveillance Team |
| EPI_ISL_2918155 | Health Services Laboratories | Wellcome Sanger Institute for the COVID-19 Genomics UK (COG-UK) Consortium | Health Services Laboratories and Alex Alderton, Roberto Amato, Jeffrey Barrett, Sonia Goncalves, Ewan Harrison, David K. Jackson, Ian Johnston, Dominic Kwiatkowski, Cordelia Langford, John Sillitoe on behalf of the Wellcome Sanger Institute COVID-19 Surveillance Team |
| EPI_ISL_2918156 | Berkshire and Surrey Pathology Services Lighthouse Laboratory | Wellcome Sanger Institute for the COVID-19 Genomics UK (COG-UK) Consortium | Berkshire and Surrey Pathology Services Lighthouse Laboratory and Alex Alderton, Roberto Amato, Jeffrey Barrett, Sonia Goncalves, Ewan Harrison, David K. Jackson, Ian Johnston, Dominic Kwiatkowski, Cordelia Langford, John Sillitoe on behalf of the Wellcome Sanger Institute COVID-19 Surveillance Team |
| EPI_ISL_2918157, EPI_ISL_2918158, EPI_ISL_2918159 | Lighthouse Lab in Milton Keynes | Wellcome Sanger Institute for the COVID-19 Genomics UK (COG-UK) Consortium | The Lighthouse Lab in Milton Keynes and Alex Alderton, Roberto Amato, Jeffrey Barrett, Sonia Goncalves, Ewan Harrison, David K. Jackson, Ian Johnston, Dominic Kwiatkowski, Cordelia Langford, John Sillitoe on behalf of the Wellcome Sanger Institute COVID-19 Surveillance Team |
| EPI_ISL_2918160, EPI_ISL_2918161 | Health Services Laboratories | Wellcome Sanger Institute for the COVID-19 Genomics UK (COG-UK) Consortium | Health Services Laboratories and Alex Alderton, Roberto Amato, Jeffrey Barrett, Sonia Goncalves, Ewan Harrison, David K. Jackson, Ian Johnston, Dominic Kwiatkowski, Cordelia Langford, John Sillitoe on behalf of the Wellcome Sanger Institute COVID-19 Surveillance Team |
| EPI_ISL_2918162 | Berkshire and Surrey Pathology Services Lighthouse Laboratory | Wellcome Sanger Institute for the COVID-19 Genomics UK (COG-UK) Consortium | Berkshire and Surrey Pathology Services Lighthouse Laboratory and Alex Alderton, Roberto Amato, Jeffrey Barrett, Sonia Goncalves, Ewan Harrison, David K. Jackson, Ian Johnston, Dominic Kwiatkowski, Cordelia Langford, John Sillitoe on behalf of the Wellcome Sanger Institute COVID-19 Surveillance Team |
| EPI_ISL_2918163, EPI_ISL_2918164 | Health Services Laboratories | Wellcome Sanger Institute for the COVID-19 Genomics UK (COG-UK) Consortium | Health Services Laboratories and Alex Alderton, Roberto Amato, Jeffrey Barrett, Sonia Goncalves, Ewan Harrison, David K. Jackson, Ian Johnston, Dominic Kwiatkowski, Cordelia Langford, John Sillitoe on behalf of the Wellcome Sanger Institute COVID-19 Surveillance Team |
| EPI_ISL_2918165 | Lighthouse Lab in Alderley Park | Wellcome Sanger Institute for the COVID-19 Genomics UK (COG-UK) Consortium | Jacquelyn Wynn, Mairead Hyland, The Lighthouse Lab in Alderley Park and Alex Alderton, Roberto Amato, Jeffrey Barrett, Sonia Goncalves, Ewan Harrison, David K. Jackson, Ian Johnston, Dominic Kwiatkowski, Cordelia Langford, John Sillitoe on behalf of the Wellcome Sanger Institute COVID-19 Surveillance Team |

| | | | |
|---|---|---|---|
| EPI_ISL_2918166 | Lighthouse Lab in Milton Keynes | Wellcome Sanger Institute for the COVID-19 Genomics UK (COG-UK) Consortium | The Lighthouse Lab in Milton Keynes and Alex Alderton, Roberto Amato, Jeffrey Barrett, Sonia Goncalves, Ewan Harrison, David K. Jackson, Ian Johnston, Dominic Kwiatkowski, Cordelia Langford, John Sillitoe on behalf of the Wellcome Sanger Institute COVID-19 Surveillance Team |
| EPI_ISL_2918167 | Health Services Laboratories | Wellcome Sanger Institute for the COVID-19 Genomics UK (COG-UK) Consortium | Health Services Laboratories and Alex Alderton, Roberto Amato, Jeffrey Barrett, Sonia Goncalves, Ewan Harrison, David K. Jackson, Ian Johnston, Dominic Kwiatkowski, Cordelia Langford, John Sillitoe on behalf of the Wellcome Sanger Institute COVID-19 Surveillance Team |
| EPI_ISL_2918168 | Lighthouse Lab in Alderley Park | Wellcome Sanger Institute for the COVID-19 Genomics UK (COG-UK) Consortium | Jacquelyn Wynn, Mairead Hyland, The Lighthouse Lab in Alderley Park and Alex Alderton, Roberto Amato, Jeffrey Barrett, Sonia Goncalves, Ewan Harrison, David K. Jackson, Ian Johnston, Dominic Kwiatkowski, Cordelia Langford, John Sillitoe on behalf of the Wellcome Sanger Institute COVID-19 Surveillance Team |
| EPI_ISL_2918169 | Berkshire and Surrey Pathology Services Lighthouse Laboratory | Wellcome Sanger Institute for the COVID-19 Genomics UK (COG-UK) Consortium | Berkshire and Surrey Pathology Services Lighthouse Laboratory and Alex Alderton, Roberto Amato, Jeffrey Barrett, Sonia Goncalves, Ewan Harrison, David K. Jackson, Ian Johnston, Dominic Kwiatkowski, Cordelia Langford, John Sillitoe on behalf of the Wellcome Sanger Institute COVID-19 Surveillance Team |
| EPI_ISL_2918170 | Lighthouse Lab in Alderley Park | Wellcome Sanger Institute for the COVID-19 Genomics UK (COG-UK) Consortium | Jacquelyn Wynn, Mairead Hyland, The Lighthouse Lab in Alderley Park and Alex Alderton, Roberto Amato, Jeffrey Barrett, Sonia Goncalves, Ewan Harrison, David K. Jackson, Ian Johnston, Dominic Kwiatkowski, Cordelia Langford, John Sillitoe on behalf of the Wellcome Sanger Institute COVID-19 Surveillance Team |
| EPI_ISL_2918171 | Lighthouse Lab in Milton Keynes | Wellcome Sanger Institute for the COVID-19 Genomics UK (COG-UK) Consortium | The Lighthouse Lab in Milton Keynes and Alex Alderton, Roberto Amato, Jeffrey Barrett, Sonia Goncalves, Ewan Harrison, David K. Jackson, Ian Johnston, Dominic Kwiatkowski, Cordelia Langford, John Sillitoe on behalf of the Wellcome Sanger Institute COVID-19 Surveillance Team |
| EPI_ISL_2918172, EPI_ISL_2918173, EPI_ISL_2918174, EPI_ISL_2918175 | Lighthouse Lab in Alderley Park | Wellcome Sanger Institute for the COVID-19 Genomics UK (COG-UK) Consortium | Jacquelyn Wynn, Mairead Hyland, The Lighthouse Lab in Alderley Park and Alex Alderton, Roberto Amato, Jeffrey Barrett, Sonia Goncalves, Ewan Harrison, David K. Jackson, Ian Johnston, Dominic Kwiatkowski, Cordelia Langford, John Sillitoe on behalf of the Wellcome Sanger Institute COVID-19 Surveillance Team |
| EPI_ISL_2918176 | Lighthouse Lab in Milton Keynes | Wellcome Sanger Institute for the COVID-19 Genomics UK (COG-UK) Consortium | The Lighthouse Lab in Milton Keynes and Alex Alderton, Roberto Amato, Jeffrey Barrett, Sonia Goncalves, Ewan Harrison, David K. Jackson, Ian Johnston, Dominic Kwiatkowski, Cordelia Langford, John Sillitoe on behalf of the Wellcome Sanger Institute COVID-19 Surveillance Team |
| EPI_ISL_2918177 | Lighthouse Lab in Alderley Park | Wellcome Sanger Institute for the COVID-19 Genomics UK (COG-UK) Consortium | Jacquelyn Wynn, Mairead Hyland, The Lighthouse Lab in Alderley Park and Alex Alderton, Roberto Amato, Jeffrey Barrett, Sonia Goncalves, Ewan Harrison, David K. Jackson, Ian Johnston, Dominic Kwiatkowski, Cordelia Langford, John Sillitoe on behalf of the Wellcome Sanger Institute COVID-19 Surveillance Team |
| EPI_ISL_2918178, EPI_ISL_2918179 | Lighthouse Lab in Milton Keynes | Wellcome Sanger Institute for the COVID-19 Genomics UK (COG-UK) Consortium | The Lighthouse Lab in Milton Keynes and Alex Alderton, Roberto Amato, Jeffrey Barrett, Sonia Goncalves, Ewan Harrison, David K. Jackson, Ian Johnston, Dominic Kwiatkowski, Cordelia Langford, John Sillitoe on behalf of the Wellcome Sanger Institute COVID-19 Surveillance Team |
| EPI_ISL_2918180 | Berkshire and Surrey Pathology Services Lighthouse Laboratory | Wellcome Sanger Institute for the COVID-19 Genomics UK (COG-UK) Consortium | Berkshire and Surrey Pathology Services Lighthouse Laboratory and Alex Alderton, Roberto Amato, Jeffrey Barrett, Sonia Goncalves, Ewan Harrison, David K. Jackson, Ian Johnston, Dominic Kwiatkowski, Cordelia Langford, John Sillitoe on behalf of the Wellcome Sanger Institute COVID-19 Surveillance Team |
| EPI_ISL_2918181, EPI_ISL_2918182 | Lighthouse Lab in Milton Keynes | Wellcome Sanger Institute for the COVID-19 Genomics UK (COG-UK) Consortium | The Lighthouse Lab in Milton Keynes and Alex Alderton, Roberto Amato, Jeffrey Barrett, Sonia Goncalves, Ewan Harrison, David K. Jackson, Ian Johnston, Dominic Kwiatkowski, Cordelia Langford, John Sillitoe on behalf of the Wellcome Sanger Institute COVID-19 Surveillance Team |
| EPI_ISL_2918183 | Health Services Laboratories | Wellcome Sanger Institute for the COVID-19 Genomics UK (COG-UK) Consortium | Health Services Laboratories and Alex Alderton, Roberto Amato, Jeffrey Barrett, Sonia Goncalves, Ewan Harrison, David K. Jackson, Ian Johnston, Dominic Kwiatkowski, Cordelia Langford, John Sillitoe on behalf of the Wellcome Sanger Institute COVID-19 Surveillance Team |
| EPI_ISL_2918184 | Lighthouse Lab in Milton Keynes | Wellcome Sanger Institute for the COVID-19 Genomics UK (COG-UK) Consortium | The Lighthouse Lab in Milton Keynes and Alex Alderton, Roberto Amato, Jeffrey Barrett, Sonia Goncalves, Ewan Harrison, David K. Jackson, Ian Johnston, Dominic Kwiatkowski, Cordelia Langford, John Sillitoe on behalf of the Wellcome Sanger Institute COVID-19 Surveillance Team |
| EPI_ISL_2918185 | Health Services Laboratories | Wellcome Sanger Institute for the COVID-19 Genomics UK (COG-UK) Consortium | Health Services Laboratories and Alex Alderton, Roberto Amato, Jeffrey Barrett, Sonia Goncalves, Ewan Harrison, David K. Jackson, Ian Johnston, Dominic Kwiatkowski, Cordelia Langford, John Sillitoe on behalf of the Wellcome Sanger Institute COVID-19 Surveillance Team |
| EPI_ISL_2918186 | Berkshire and Surrey Pathology Services Lighthouse Laboratory | Wellcome Sanger Institute for the COVID-19 Genomics UK (COG-UK) Consortium | Berkshire and Surrey Pathology Services Lighthouse Laboratory and Alex Alderton, Roberto Amato, Jeffrey Barrett, Sonia Goncalves, Ewan Harrison, David K. Jackson, Ian Johnston, Dominic Kwiatkowski, Cordelia Langford, John Sillitoe on behalf of the Wellcome Sanger Institute COVID-19 Surveillance Team |
| EPI_ISL_2918187 | Lighthouse Lab in Milton Keynes | Wellcome Sanger Institute for the COVID-19 Genomics UK (COG-UK) Consortium | The Lighthouse Lab in Milton Keynes and Alex Alderton, Roberto Amato, Jeffrey Barrett, Sonia Goncalves, Ewan Harrison, David K. Jackson, Ian Johnston, Dominic Kwiatkowski, Cordelia Langford, John Sillitoe on behalf of the Wellcome Sanger Institute COVID-19 Surveillance Team |
| EPI_ISL_2918188 | Lighthouse Lab in Alderley Park | Wellcome Sanger Institute for the COVID-19 Genomics UK (COG-UK) Consortium | Jacquelyn Wynn, Mairead Hyland, The Lighthouse Lab in Alderley Park and Alex Alderton, Roberto Amato, Jeffrey Barrett, Sonia Goncalves, Ewan Harrison, David K. Jackson, Ian Johnston, Dominic Kwiatkowski, Cordelia Langford, John Sillitoe on behalf of the Wellcome Sanger Institute COVID-19 Surveillance Team |
| EPI_ISL_2918189, EPI_ISL_2918190, EPI_ISL_2918191 | Lighthouse Lab in Milton Keynes | Wellcome Sanger Institute for the COVID-19 Genomics UK (COG-UK) Consortium | The Lighthouse Lab in Milton Keynes and Alex Alderton, Roberto Amato, Jeffrey Barrett, Sonia Goncalves, Ewan Harrison, David K. Jackson, Ian Johnston, Dominic Kwiatkowski, Cordelia Langford, John Sillitoe on behalf of the Wellcome Sanger Institute COVID-19 Surveillance Team |
| EPI_ISL_2918192 | Berkshire and Surrey Pathology Services Lighthouse Laboratory | Wellcome Sanger Institute for the COVID-19 Genomics UK (COG-UK) Consortium | Berkshire and Surrey Pathology Services Lighthouse Laboratory and Alex Alderton, Roberto Amato, Jeffrey Barrett, Sonia Goncalves, Ewan Harrison, David K. Jackson, Ian Johnston, Dominic Kwiatkowski, Cordelia Langford, John Sillitoe on behalf of the Wellcome Sanger Institute COVID-19 Surveillance Team |
| EPI_ISL_2918193 | Lighthouse Lab in Alderley Park | Wellcome Sanger Institute for the COVID-19 Genomics UK (COG-UK) Consortium | Jacquelyn Wynn, Mairead Hyland, The Lighthouse Lab in Alderley Park and Alex Alderton, Roberto Amato, Jeffrey Barrett, Sonia Goncalves, Ewan Harrison, David K. Jackson, Ian Johnston, Dominic Kwiatkowski, Cordelia Langford, John Sillitoe on behalf of the Wellcome Sanger Institute COVID-19 Surveillance Team |
| EPI_ISL_2918194 | Lighthouse Lab in Milton Keynes | Wellcome Sanger Institute for the COVID-19 Genomics UK (COG-UK) Consortium | The Lighthouse Lab in Milton Keynes and Alex Alderton, Roberto Amato, Jeffrey Barrett, Sonia Goncalves, Ewan Harrison, David K. Jackson, Ian Johnston, Dominic Kwiatkowski, Cordelia Langford, John Sillitoe on behalf of the Wellcome Sanger Institute COVID-19 Surveillance Team |
| EPI_ISL_2918195 | Health Services Laboratories | Wellcome Sanger Institute for the COVID-19 Genomics UK (COG-UK) Consortium | Health Services Laboratories and Alex Alderton, Roberto Amato, Jeffrey Barrett, Sonia Goncalves, Ewan Harrison, David K. Jackson, Ian Johnston, Dominic Kwiatkowski, Cordelia Langford, John Sillitoe on behalf of the Wellcome Sanger Institute COVID-19 Surveillance Team |
| EPI_ISL_2918196 | Lighthouse Lab in Milton Keynes | Wellcome Sanger Institute for the COVID-19 Genomics UK (COG-UK) Consortium | The Lighthouse Lab in Milton Keynes and Alex Alderton, Roberto Amato, Jeffrey Barrett, Sonia Goncalves, Ewan Harrison, David K. Jackson, Ian Johnston, Dominic Kwiatkowski, Cordelia Langford, John Sillitoe on behalf of the Wellcome Sanger Institute COVID-19 Surveillance Team |
| EPI_ISL_2918197 | Lighthouse Lab in Alderley Park | Wellcome Sanger Institute for the COVID-19 Genomics UK (COG-UK) Consortium | Jacquelyn Wynn, Mairead Hyland, The Lighthouse Lab in Alderley Park and Alex Alderton, Roberto Amato, Jeffrey Barrett, Sonia Goncalves, Ewan Harrison, David K. Jackson, Ian Johnston, Dominic Kwiatkowski, Cordelia Langford, John Sillitoe on behalf of the Wellcome Sanger Institute COVID-19 Surveillance Team |
| EPI_ISL_2918198, EPI_ISL_2918199 | Lighthouse Lab in Milton Keynes | Wellcome Sanger Institute for the COVID-19 Genomics UK (COG-UK) Consortium | The Lighthouse Lab in Milton Keynes and Alex Alderton, Roberto Amato, Jeffrey Barrett, Sonia Goncalves, Ewan Harrison, David K. Jackson, Ian Johnston, Dominic Kwiatkowski, Cordelia Langford, John Sillitoe on behalf of the Wellcome Sanger Institute COVID-19 Surveillance Team |
| EPI_ISL_2918200, EPI_ISL_2918201 | Berkshire and Surrey Pathology Services Lighthouse Laboratory | Wellcome Sanger Institute for the COVID-19 Genomics UK (COG-UK) Consortium | Berkshire and Surrey Pathology Services Lighthouse Laboratory and Alex Alderton, Roberto Amato, Jeffrey Barrett, Sonia Goncalves, Ewan Harrison, David K. Jackson, Ian Johnston, Dominic Kwiatkowski, Cordelia Langford, John Sillitoe on behalf of the Wellcome Sanger Institute COVID-19 Surveillance Team |
| EPI_ISL_2918202 | Lighthouse Lab in Milton Keynes | Wellcome Sanger Institute for the COVID-19 Genomics UK (COG-UK) Consortium | The Lighthouse Lab in Milton Keynes and Alex Alderton, Roberto Amato, Jeffrey Barrett, Sonia Goncalves, Ewan Harrison, David K. Jackson, Ian Johnston, Dominic Kwiatkowski, Cordelia Langford, John Sillitoe on behalf of the Wellcome Sanger Institute COVID-19 Surveillance Team |
| EPI_ISL_2918203 | Berkshire and Surrey Pathology Services Lighthouse Laboratory | Wellcome Sanger Institute for the COVID-19 Genomics UK (COG-UK) Consortium | Berkshire and Surrey Pathology Services Lighthouse Laboratory and Alex Alderton, Roberto Amato, Jeffrey Barrett, Sonia Goncalves, Ewan Harrison, David K. Jackson, Ian Johnston, Dominic Kwiatkowski, Cordelia Langford, John Sillitoe on behalf of the Wellcome Sanger Institute COVID-19 Surveillance Team |
| EPI_ISL_2918204 | Health Services Laboratories | Wellcome Sanger Institute for the COVID-19 Genomics UK (COG-UK) Consortium | Health Services Laboratories and Alex Alderton, Roberto Amato, Jeffrey Barrett, Sonia Goncalves, Ewan Harrison, David K. Jackson, Ian Johnston, Dominic Kwiatkowski, Cordelia Langford, John Sillitoe on behalf of the Wellcome Sanger Institute COVID-19 Surveillance Team |
| EPI_ISL_2918205, EPI_ISL_2918206 | Berkshire and Surrey Pathology Services Lighthouse Laboratory | Wellcome Sanger Institute for the COVID-19 Genomics UK (COG-UK) Consortium | Berkshire and Surrey Pathology Services Lighthouse Laboratory and Alex Alderton, Roberto Amato, Jeffrey Barrett, Sonia Goncalves, Ewan Harrison, David K. Jackson, Ian Johnston, Dominic Kwiatkowski, Cordelia Langford, John Sillitoe on behalf of the Wellcome Sanger Institute COVID-19 Surveillance Team |
| EPI_ISL_2918207 | Lighthouse Lab in Alderley Park | Wellcome Sanger Institute for the COVID-19 Genomics UK (COG-UK) Consortium | Jacquelyn Wynn, Mairead Hyland, The Lighthouse Lab in Alderley Park and Alex Alderton, Roberto Amato, Jeffrey Barrett, Sonia Goncalves, Ewan Harrison, David K. Jackson, Ian Johnston, Dominic Kwiatkowski, Cordelia Langford, John Sillitoe on behalf of the Wellcome Sanger Institute COVID-19 Surveillance Team |
| EPI_ISL_2918208 | Lighthouse Lab in Milton Keynes | Wellcome Sanger Institute for the COVID-19 Genomics UK (COG-UK) Consortium | The Lighthouse Lab in Milton Keynes and Alex Alderton, Roberto Amato, Jeffrey Barrett, Sonia Goncalves, Ewan Harrison, David K. Jackson, Ian Johnston, Dominic Kwiatkowski, Cordelia Langford, John Sillitoe on behalf of the Wellcome Sanger Institute COVID-19 Surveillance Team |
| EPI_ISL_2918209 | Health Services Laboratories | Wellcome Sanger Institute for the COVID-19 Genomics UK (COG-UK) Consortium | Health Services Laboratories and Alex Alderton, Roberto Amato, Jeffrey Barrett, Sonia Goncalves, Ewan Harrison, David K. Jackson, Ian Johnston, Dominic Kwiatkowski, Cordelia Langford, John Sillitoe on behalf of the Wellcome Sanger Institute COVID-19 Surveillance Team |
| EPI_ISL_2918210 | Lighthouse Lab in Alderley Park | Wellcome Sanger Institute for the COVID-19 Genomics UK (COG-UK) Consortium | Jacquelyn Wynn, Mairead Hyland, The Lighthouse Lab in Alderley Park and Alex Alderton, Roberto Amato, Jeffrey Barrett, Sonia Goncalves, Ewan Harrison, David K. Jackson, Ian Johnston, Dominic Kwiatkowski, Cordelia Langford, John Sillitoe on behalf of the Wellcome Sanger Institute COVID-19 Surveillance Team |
| EPI_ISL_2918211 | Health Services Laboratories | Wellcome Sanger Institute for the COVID-19 Genomics UK (COG-UK) Consortium | Health Services Laboratories and Alex Alderton, Roberto Amato, Jeffrey Barrett, Sonia Goncalves, Ewan Harrison, David K. Jackson, Ian Johnston, Dominic Kwiatkowski, Cordelia Langford, John Sillitoe on behalf of the Wellcome Sanger Institute COVID-19 Surveillance Team |
| EPI_ISL_2918212 | Lighthouse Lab in Milton Keynes | Wellcome Sanger Institute for the COVID-19 Genomics UK (COG-UK) Consortium | The Lighthouse Lab in Milton Keynes and Alex Alderton, Roberto Amato, Jeffrey Barrett, Sonia Goncalves, Ewan Harrison, David K. Jackson, Ian Johnston, Dominic Kwiatkowski, Cordelia Langford, John Sillitoe on behalf of the Wellcome Sanger Institute COVID-19 Surveillance Team |
| EPI_ISL_2918213, EPI_ISL_2918214 | Berkshire and Surrey Pathology Services Lighthouse Laboratory | Wellcome Sanger Institute for the COVID-19 Genomics UK (COG-UK) Consortium | Berkshire and Surrey Pathology Services Lighthouse Laboratory and Alex Alderton, Roberto Amato, Jeffrey Barrett, Sonia Goncalves, Ewan Harrison, David K. Jackson, Ian Johnston, Dominic Kwiatkowski, Cordelia Langford, John Sillitoe on behalf of the Wellcome Sanger Institute COVID-19 Surveillance Team |
| EPI_ISL_2918215 | Lighthouse Lab in Alderley Park | Wellcome Sanger Institute for the COVID-19 Genomics UK (COG-UK) Consortium | Jacquelyn Wynn, Mairead Hyland, The Lighthouse Lab in Alderley Park and Alex Alderton, Roberto Amato, Jeffrey Barrett, Sonia Goncalves, Ewan Harrison, David K. Jackson, Ian Johnston, Dominic Kwiatkowski, Cordelia Langford, John Sillitoe on behalf of the Wellcome Sanger Institute COVID-19 Surveillance Team |
| EPI_ISL_2918216 | Berkshire and Surrey Pathology Services Lighthouse Laboratory | Wellcome Sanger Institute for the COVID-19 Genomics UK (COG-UK) Consortium | Berkshire and Surrey Pathology Services Lighthouse Laboratory and Alex Alderton, Roberto Amato, Jeffrey Barrett, Sonia Goncalves, Ewan Harrison, David K. Jackson, Ian Johnston, Dominic Kwiatkowski, Cordelia Langford, John Sillitoe on behalf of the Wellcome Sanger Institute COVID-19 Surveillance Team |
| EPI_ISL_2918217 | Lighthouse Lab in Milton Keynes | Wellcome Sanger Institute for the COVID-19 Genomics UK (COG-UK) Consortium | The Lighthouse Lab in Milton Keynes and Alex Alderton, Roberto Amato, Jeffrey Barrett, Sonia Goncalves, Ewan Harrison, David K. Jackson, Ian Johnston, Dominic Kwiatkowski, Cordelia Langford, John Sillitoe on behalf of the Wellcome Sanger Institute COVID-19 Surveillance Team |
| EPI_ISL_2918218 | Berkshire and Surrey Pathology Services Lighthouse Laboratory | Wellcome Sanger Institute for the COVID-19 Genomics UK (COG-UK) Consortium | Berkshire and Surrey Pathology Services Lighthouse Laboratory and Alex Alderton, Roberto Amato, Jeffrey Barrett, Sonia Goncalves, Ewan Harrison, David K. Jackson, Ian Johnston, Dominic Kwiatkowski, Cordelia Langford, John Sillitoe on behalf of the Wellcome Sanger Institute COVID-19 Surveillance Team |
| EPI_ISL_2918219 | Lighthouse Lab in Milton Keynes | Wellcome Sanger Institute for the COVID-19 Genomics UK (COG-UK) Consortium | The Lighthouse Lab in Milton Keynes and Alex Alderton, Roberto Amato, Jeffrey Barrett, Sonia Goncalves, Ewan Harrison, David K. Jackson, Ian Johnston, Dominic Kwiatkowski, Cordelia Langford, John Sillitoe on behalf of the Wellcome Sanger Institute COVID-19 Surveillance Team |
| EPI_ISL_2918220 | Health Services Laboratories | Wellcome Sanger Institute for the COVID-19 Genomics UK (COG-UK) Consortium | Health Services Laboratories and Alex Alderton, Roberto Amato, Jeffrey Barrett, Sonia Goncalves, Ewan Harrison, David K. Jackson, Ian Johnston, Dominic Kwiatkowski, Cordelia Langford, John Sillitoe on behalf of the Wellcome Sanger Institute COVID-19 Surveillance Team |

| | | | |
|---|---|---|---|
| EPI_ISL_2918221 | Lighthouse Lab in Alderley Park | Wellcome Sanger Institute for the COVID-19 Genomics UK (COG-UK) Consortium | Jacquelyn Wynn, Mairead Hyland, The Lighthouse Lab in Alderley Park and Alex Alderton, Roberto Amato, Jeffrey Barrett, Sonia Goncalves, Ewan Harrison, David K. Jackson, Ian Johnston, Dominic Kwiatkowski, Cordelia Langford, John Sillitoe on behalf of the Wellcome Sanger Institute COVID-19 Surveillance Team |
| EPI_ISL_2918222, EPI_ISL_2918223 | Health Services Laboratories | Wellcome Sanger Institute for the COVID-19 Genomics UK (COG-UK) Consortium | Health Services Laboratories and Alex Alderton, Roberto Amato, Jeffrey Barrett, Sonia Goncalves, Ewan Harrison, David K. Jackson, Ian Johnston, Dominic Kwiatkowski, Cordelia Langford, John Sillitoe on behalf of the Wellcome Sanger Institute COVID-19 Surveillance Team |
| EPI_ISL_2918224 | Lighthouse Lab in Alderley Park | Wellcome Sanger Institute for the COVID-19 Genomics UK (COG-UK) Consortium | Jacquelyn Wynn, Mairead Hyland, The Lighthouse Lab in Alderley Park and Alex Alderton, Roberto Amato, Jeffrey Barrett, Sonia Goncalves, Ewan Harrison, David K. Jackson, Ian Johnston, Dominic Kwiatkowski, Cordelia Langford, John Sillitoe on behalf of the Wellcome Sanger Institute COVID-19 Surveillance Team |
| EPI_ISL_2918225 | Lighthouse Lab in Milton Keynes | Wellcome Sanger Institute for the COVID-19 Genomics UK (COG-UK) Consortium | The Lighthouse Lab in Milton Keynes and Alex Alderton, Roberto Amato, Jeffrey Barrett, Sonia Goncalves, Ewan Harrison, David K. Jackson, Ian Johnston, Dominic Kwiatkowski, Cordelia Langford, John Sillitoe on behalf of the Wellcome Sanger Institute COVID-19 Surveillance Team |
| EPI_ISL_2918226 | Health Services Laboratories | Wellcome Sanger Institute for the COVID-19 Genomics UK (COG-UK) Consortium | Health Services Laboratories and Alex Alderton, Roberto Amato, Jeffrey Barrett, Sonia Goncalves, Ewan Harrison, David K. Jackson, Ian Johnston, Dominic Kwiatkowski, Cordelia Langford, John Sillitoe on behalf of the Wellcome Sanger Institute COVID-19 Surveillance Team |
| EPI_ISL_2918227 | Berkshire and Surrey Pathology Services Lighthouse Laboratory | Wellcome Sanger Institute for the COVID-19 Genomics UK (COG-UK) Consortium | Berkshire and Surrey Pathology Services Lighthouse Laboratory and Alex Alderton, Roberto Amato, Jeffrey Barrett, Sonia Goncalves, Ewan Harrison, David K. Jackson, Ian Johnston, Dominic Kwiatkowski, Cordelia Langford, John Sillitoe on behalf of the Wellcome Sanger Institute COVID-19 Surveillance Team |
| EPI_ISL_2918228 | Lighthouse Lab in Milton Keynes | Wellcome Sanger Institute for the COVID-19 Genomics UK (COG-UK) Consortium | The Lighthouse Lab in Milton Keynes and Alex Alderton, Roberto Amato, Jeffrey Barrett, Sonia Goncalves, Ewan Harrison, David K. Jackson, Ian Johnston, Dominic Kwiatkowski, Cordelia Langford, John Sillitoe on behalf of the Wellcome Sanger Institute COVID-19 Surveillance Team |
| EPI_ISL_2918229 | Lighthouse Lab in Alderley Park | Wellcome Sanger Institute for the COVID-19 Genomics UK (COG-UK) Consortium | Jacquelyn Wynn, Mairead Hyland, The Lighthouse Lab in Alderley Park and Alex Alderton, Roberto Amato, Jeffrey Barrett, Sonia Goncalves, Ewan Harrison, David K. Jackson, Ian Johnston, Dominic Kwiatkowski, Cordelia Langford, John Sillitoe on behalf of the Wellcome Sanger Institute COVID-19 Surveillance Team |
| EPI_ISL_2918230 | Health Services Laboratories | Wellcome Sanger Institute for the COVID-19 Genomics UK (COG-UK) Consortium | Health Services Laboratories and Alex Alderton, Roberto Amato, Jeffrey Barrett, Sonia Goncalves, Ewan Harrison, David K. Jackson, Ian Johnston, Dominic Kwiatkowski, Cordelia Langford, John Sillitoe on behalf of the Wellcome Sanger Institute COVID-19 Surveillance Team |
| EPI_ISL_2918231, EPI_ISL_2918232 | Lighthouse Lab in Milton Keynes | Wellcome Sanger Institute for the COVID-19 Genomics UK (COG-UK) Consortium | The Lighthouse Lab in Milton Keynes and Alex Alderton, Roberto Amato, Jeffrey Barrett, Sonia Goncalves, Ewan Harrison, David K. Jackson, Ian Johnston, Dominic Kwiatkowski, Cordelia Langford, John Sillitoe on behalf of the Wellcome Sanger Institute COVID-19 Surveillance Team |
| EPI_ISL_2918233 | Lighthouse Lab in Alderley Park | Wellcome Sanger Institute for the COVID-19 Genomics UK (COG-UK) Consortium | Jacquelyn Wynn, Mairead Hyland, The Lighthouse Lab in Alderley Park and Alex Alderton, Roberto Amato, Jeffrey Barrett, Sonia Goncalves, Ewan Harrison, David K. Jackson, Ian Johnston, Dominic Kwiatkowski, Cordelia Langford, John Sillitoe on behalf of the Wellcome Sanger Institute COVID-19 Surveillance Team |
| EPI_ISL_2918234 | Berkshire and Surrey Pathology Services Lighthouse Laboratory | Wellcome Sanger Institute for the COVID-19 Genomics UK (COG-UK) Consortium | Berkshire and Surrey Pathology Services Lighthouse Laboratory and Alex Alderton, Roberto Amato, Jeffrey Barrett, Sonia Goncalves, Ewan Harrison, David K. Jackson, Ian Johnston, Dominic Kwiatkowski, Cordelia Langford, John Sillitoe on behalf of the Wellcome Sanger Institute COVID-19 Surveillance Team |
| EPI_ISL_2918235 | Lighthouse Lab in Alderley Park | Wellcome Sanger Institute for the COVID-19 Genomics UK (COG-UK) Consortium | Jacquelyn Wynn, Mairead Hyland, The Lighthouse Lab in Alderley Park and Alex Alderton, Roberto Amato, Jeffrey Barrett, Sonia Goncalves, Ewan Harrison, David K. Jackson, Ian Johnston, Dominic Kwiatkowski, Cordelia Langford, John Sillitoe on behalf of the Wellcome Sanger Institute COVID-19 Surveillance Team |
| EPI_ISL_2918236 | Berkshire and Surrey Pathology Services Lighthouse Laboratory | Wellcome Sanger Institute for the COVID-19 Genomics UK (COG-UK) Consortium | Berkshire and Surrey Pathology Services Lighthouse Laboratory and Alex Alderton, Roberto Amato, Jeffrey Barrett, Sonia Goncalves, Ewan Harrison, David K. Jackson, Ian Johnston, Dominic Kwiatkowski, Cordelia Langford, John Sillitoe on behalf of the Wellcome Sanger Institute COVID-19 Surveillance Team |
| EPI_ISL_2918237 | Health Services Laboratories | Wellcome Sanger Institute for the COVID-19 Genomics UK (COG-UK) Consortium | Health Services Laboratories and Alex Alderton, Roberto Amato, Jeffrey Barrett, Sonia Goncalves, Ewan Harrison, David K. Jackson, Ian Johnston, Dominic Kwiatkowski, Cordelia Langford, John Sillitoe on behalf of the Wellcome Sanger Institute COVID-19 Surveillance Team |
| EPI_ISL_2918238 | Berkshire and Surrey Pathology Services Lighthouse Laboratory | Wellcome Sanger Institute for the COVID-19 Genomics UK (COG-UK) Consortium | Berkshire and Surrey Pathology Services Lighthouse Laboratory and Alex Alderton, Roberto Amato, Jeffrey Barrett, Sonia Goncalves, Ewan Harrison, David K. Jackson, Ian Johnston, Dominic Kwiatkowski, Cordelia Langford, John Sillitoe on behalf of the Wellcome Sanger Institute COVID-19 Surveillance Team |
| EPI_ISL_2918239, EPI_ISL_2918240 | Lighthouse Lab in Milton Keynes | Wellcome Sanger Institute for the COVID-19 Genomics UK (COG-UK) Consortium | The Lighthouse Lab in Milton Keynes and Alex Alderton, Roberto Amato, Jeffrey Barrett, Sonia Goncalves, Ewan Harrison, David K. Jackson, Ian Johnston, Dominic Kwiatkowski, Cordelia Langford, John Sillitoe on behalf of the Wellcome Sanger Institute COVID-19 Surveillance Team |
| EPI_ISL_2918241 | Lighthouse Lab in Alderley Park | Wellcome Sanger Institute for the COVID-19 Genomics UK (COG-UK) Consortium | Jacquelyn Wynn, Mairead Hyland, The Lighthouse Lab in Alderley Park and Alex Alderton, Roberto Amato, Jeffrey Barrett, Sonia Goncalves, Ewan Harrison, David K. Jackson, Ian Johnston, Dominic Kwiatkowski, Cordelia Langford, John Sillitoe on behalf of the Wellcome Sanger Institute COVID-19 Surveillance Team |
| EPI_ISL_2918242, EPI_ISL_2918243 | Health Services Laboratories | Wellcome Sanger Institute for the COVID-19 Genomics UK (COG-UK) Consortium | Health Services Laboratories and Alex Alderton, Roberto Amato, Jeffrey Barrett, Sonia Goncalves, Ewan Harrison, David K. Jackson, Ian Johnston, Dominic Kwiatkowski, Cordelia Langford, John Sillitoe on behalf of the Wellcome Sanger Institute COVID-19 Surveillance Team |
| EPI_ISL_2918244 | Berkshire and Surrey Pathology Services Lighthouse Laboratory | Wellcome Sanger Institute for the COVID-19 Genomics UK (COG-UK) Consortium | Berkshire and Surrey Pathology Services Lighthouse Laboratory and Alex Alderton, Roberto Amato, Jeffrey Barrett, Sonia Goncalves, Ewan Harrison, David K. Jackson, Ian Johnston, Dominic Kwiatkowski, Cordelia Langford, John Sillitoe on behalf of the Wellcome Sanger Institute COVID-19 Surveillance Team |
| EPI_ISL_2918245, EPI_ISL_2918246 | Lighthouse Lab in Alderley Park | Wellcome Sanger Institute for the COVID-19 Genomics UK (COG-UK) Consortium | Jacquelyn Wynn, Mairead Hyland, The Lighthouse Lab in Alderley Park and Alex Alderton, Roberto Amato, Jeffrey Barrett, Sonia Goncalves, Ewan Harrison, David K. Jackson, Ian Johnston, Dominic Kwiatkowski, Cordelia Langford, John Sillitoe on behalf of the Wellcome Sanger Institute COVID-19 Surveillance Team |
| EPI_ISL_2918247 | Lighthouse Lab in Milton Keynes | Wellcome Sanger Institute for the COVID-19 Genomics UK (COG-UK) Consortium | The Lighthouse Lab in Milton Keynes and Alex Alderton, Roberto Amato, Jeffrey Barrett, Sonia Goncalves, Ewan Harrison, David K. Jackson, Ian Johnston, Dominic Kwiatkowski, Cordelia Langford, John Sillitoe on behalf of the Wellcome Sanger Institute COVID-19 Surveillance Team |
| EPI_ISL_2918248 | Berkshire and Surrey Pathology Services Lighthouse Laboratory | Wellcome Sanger Institute for the COVID-19 Genomics UK (COG-UK) Consortium | Berkshire and Surrey Pathology Services Lighthouse Laboratory and Alex Alderton, Roberto Amato, Jeffrey Barrett, Sonia Goncalves, Ewan Harrison, David K. Jackson, Ian Johnston, Dominic Kwiatkowski, Cordelia Langford, John Sillitoe on behalf of the Wellcome Sanger Institute COVID-19 Surveillance Team |
| EPI_ISL_2918249 | Health Services Laboratories | Wellcome Sanger Institute for the COVID-19 Genomics UK (COG-UK) Consortium | Health Services Laboratories and Alex Alderton, Roberto Amato, Jeffrey Barrett, Sonia Goncalves, Ewan Harrison, David K. Jackson, Ian Johnston, Dominic Kwiatkowski, Cordelia Langford, John Sillitoe on behalf of the Wellcome Sanger Institute COVID-19 Surveillance Team |
| EPI_ISL_2918250 | Lighthouse Lab in Milton Keynes | Wellcome Sanger Institute for the COVID-19 Genomics UK (COG-UK) Consortium | The Lighthouse Lab in Milton Keynes and Alex Alderton, Roberto Amato, Jeffrey Barrett, Sonia Goncalves, Ewan Harrison, David K. Jackson, Ian Johnston, Dominic Kwiatkowski, Cordelia Langford, John Sillitoe on behalf of the Wellcome Sanger Institute COVID-19 Surveillance Team |
| EPI_ISL_2918251 | Berkshire and Surrey Pathology Services Lighthouse Laboratory | Wellcome Sanger Institute for the COVID-19 Genomics UK (COG-UK) Consortium | Berkshire and Surrey Pathology Services Lighthouse Laboratory and Alex Alderton, Roberto Amato, Jeffrey Barrett, Sonia Goncalves, Ewan Harrison, David K. Jackson, Ian Johnston, Dominic Kwiatkowski, Cordelia Langford, John Sillitoe on behalf of the Wellcome Sanger Institute COVID-19 Surveillance Team |
| EPI_ISL_2918252, EPI_ISL_2918253 | Health Services Laboratories | Wellcome Sanger Institute for the COVID-19 Genomics UK (COG-UK) Consortium | Health Services Laboratories and Alex Alderton, Roberto Amato, Jeffrey Barrett, Sonia Goncalves, Ewan Harrison, David K. Jackson, Ian Johnston, Dominic Kwiatkowski, Cordelia Langford, John Sillitoe on behalf of the Wellcome Sanger Institute COVID-19 Surveillance Team |
| EPI_ISL_2918254 | Lighthouse Lab in Alderley Park | Wellcome Sanger Institute for the COVID-19 Genomics UK (COG-UK) Consortium | Jacquelyn Wynn, Mairead Hyland, The Lighthouse Lab in Alderley Park and Alex Alderton, Roberto Amato, Jeffrey Barrett, Sonia Goncalves, Ewan Harrison, David K. Jackson, Ian Johnston, Dominic Kwiatkowski, Cordelia Langford, John Sillitoe on behalf of the Wellcome Sanger Institute COVID-19 Surveillance Team |
| EPI_ISL_2918255, EPI_ISL_2918256 | Lighthouse Lab in Milton Keynes | Wellcome Sanger Institute for the COVID-19 Genomics UK (COG-UK) Consortium | The Lighthouse Lab in Milton Keynes and Alex Alderton, Roberto Amato, Jeffrey Barrett, Sonia Goncalves, Ewan Harrison, David K. Jackson, Ian Johnston, Dominic Kwiatkowski, Cordelia Langford, John Sillitoe on behalf of the Wellcome Sanger Institute COVID-19 Surveillance Team |
| EPI_ISL_2918257 | Health Services Laboratories | Wellcome Sanger Institute for the COVID-19 Genomics UK (COG-UK) Consortium | Health Services Laboratories and Alex Alderton, Roberto Amato, Jeffrey Barrett, Sonia Goncalves, Ewan Harrison, David K. Jackson, Ian Johnston, Dominic Kwiatkowski, Cordelia Langford, John Sillitoe on behalf of the Wellcome Sanger Institute COVID-19 Surveillance Team |
| EPI_ISL_2918258, EPI_ISL_2918259 | Lighthouse Lab in Alderley Park | Wellcome Sanger Institute for the COVID-19 Genomics UK (COG-UK) Consortium | Jacquelyn Wynn, Mairead Hyland, The Lighthouse Lab in Alderley Park and Alex Alderton, Roberto Amato, Jeffrey Barrett, Sonia Goncalves, Ewan Harrison, David K. Jackson, Ian Johnston, Dominic Kwiatkowski, Cordelia Langford, John Sillitoe on behalf of the Wellcome Sanger Institute COVID-19 Surveillance Team |
| EPI_ISL_2918260 | Health Services Laboratories | Wellcome Sanger Institute for the COVID-19 Genomics UK (COG-UK) Consortium | Health Services Laboratories and Alex Alderton, Roberto Amato, Jeffrey Barrett, Sonia Goncalves, Ewan Harrison, David K. Jackson, Ian Johnston, Dominic Kwiatkowski, Cordelia Langford, John Sillitoe on behalf of the Wellcome Sanger Institute COVID-19 Surveillance Team |
| EPI_ISL_2918261 | Berkshire and Surrey Pathology Services Lighthouse Laboratory | Wellcome Sanger Institute for the COVID-19 Genomics UK (COG-UK) Consortium | Berkshire and Surrey Pathology Services Lighthouse Laboratory and Alex Alderton, Roberto Amato, Jeffrey Barrett, Sonia Goncalves, Ewan Harrison, David K. Jackson, Ian Johnston, Dominic Kwiatkowski, Cordelia Langford, John Sillitoe on behalf of the Wellcome Sanger Institute COVID-19 Surveillance Team |
| EPI_ISL_2918262 | Lighthouse Lab in Alderley Park | Wellcome Sanger Institute for the COVID-19 Genomics UK (COG-UK) Consortium | Jacquelyn Wynn, Mairead Hyland, The Lighthouse Lab in Alderley Park and Alex Alderton, Roberto Amato, Jeffrey Barrett, Sonia Goncalves, Ewan Harrison, David K. Jackson, Ian Johnston, Dominic Kwiatkowski, Cordelia Langford, John Sillitoe on behalf of the Wellcome Sanger Institute COVID-19 Surveillance Team |
| EPI_ISL_2918263 | Lighthouse Lab in Milton Keynes | Wellcome Sanger Institute for the COVID-19 Genomics UK (COG-UK) Consortium | The Lighthouse Lab in Milton Keynes and Alex Alderton, Roberto Amato, Jeffrey Barrett, Sonia Goncalves, Ewan Harrison, David K. Jackson, Ian Johnston, Dominic Kwiatkowski, Cordelia Langford, John Sillitoe on behalf of the Wellcome Sanger Institute COVID-19 Surveillance Team |
| EPI_ISL_2918264 | Berkshire and Surrey Pathology Services Lighthouse Laboratory | Wellcome Sanger Institute for the COVID-19 Genomics UK (COG-UK) Consortium | Berkshire and Surrey Pathology Services Lighthouse Laboratory and Alex Alderton, Roberto Amato, Jeffrey Barrett, Sonia Goncalves, Ewan Harrison, David K. Jackson, Ian Johnston, Dominic Kwiatkowski, Cordelia Langford, John Sillitoe on behalf of the Wellcome Sanger Institute COVID-19 Surveillance Team |
| EPI_ISL_2918265 | Lighthouse Lab in Milton Keynes | Wellcome Sanger Institute for the COVID-19 Genomics UK (COG-UK) Consortium | The Lighthouse Lab in Milton Keynes and Alex Alderton, Roberto Amato, Jeffrey Barrett, Sonia Goncalves, Ewan Harrison, David K. Jackson, Ian Johnston, Dominic Kwiatkowski, Cordelia Langford, John Sillitoe on behalf of the Wellcome Sanger Institute COVID-19 Surveillance Team |
| EPI_ISL_2918266 | Lighthouse Lab in Alderley Park | Wellcome Sanger Institute for the COVID-19 Genomics UK (COG-UK) Consortium | Jacquelyn Wynn, Mairead Hyland, The Lighthouse Lab in Alderley Park and Alex Alderton, Roberto Amato, Jeffrey Barrett, Sonia Goncalves, Ewan Harrison, David K. Jackson, Ian Johnston, Dominic Kwiatkowski, Cordelia Langford, John Sillitoe on behalf of the Wellcome Sanger Institute COVID-19 Surveillance Team |
| EPI_ISL_2918267 | Lighthouse Lab in Milton Keynes | Wellcome Sanger Institute for the COVID-19 Genomics UK (COG-UK) Consortium | The Lighthouse Lab in Milton Keynes and Alex Alderton, Roberto Amato, Jeffrey Barrett, Sonia Goncalves, Ewan Harrison, David K. Jackson, Ian Johnston, Dominic Kwiatkowski, Cordelia Langford, John Sillitoe on behalf of the Wellcome Sanger Institute COVID-19 Surveillance Team |
| EPI_ISL_2918268, EPI_ISL_2918269, EPI_ISL_2918270 | Health Services Laboratories | Wellcome Sanger Institute for the COVID-19 Genomics UK (COG-UK) Consortium | Health Services Laboratories and Alex Alderton, Roberto Amato, Jeffrey Barrett, Sonia Goncalves, Ewan Harrison, David K. Jackson, Ian Johnston, Dominic Kwiatkowski, Cordelia Langford, John Sillitoe on behalf of the Wellcome Sanger Institute COVID-19 Surveillance Team |
| EPI_ISL_2918271 | Lighthouse Lab in Milton Keynes | Wellcome Sanger Institute for the COVID-19 Genomics UK (COG-UK) Consortium | The Lighthouse Lab in Milton Keynes and Alex Alderton, Roberto Amato, Jeffrey Barrett, Sonia Goncalves, Ewan Harrison, David K. Jackson, Ian Johnston, Dominic Kwiatkowski, Cordelia Langford, John Sillitoe on behalf of the Wellcome Sanger Institute COVID-19 Surveillance Team |
| EPI_ISL_2918272 | Randox Laboratories | Wellcome Sanger Institute for the COVID-19 Genomics UK (COG-UK) Consortium | Randox Laboratories and Alex Alderton, Roberto Amato, Jeffrey Barrett, Sonia Goncalves, Ewan Harrison, David K. Jackson, Ian Johnston, Dominic Kwiatkowski, Cordelia Langford, John Sillitoe on behalf of the Wellcome Sanger Institute COVID-19 Surveillance Team |
| EPI_ISL_2918273 | Health Services Laboratories | Wellcome Sanger Institute for the COVID-19 Genomics UK (COG-UK) Consortium | Health Services Laboratories and Alex Alderton, Roberto Amato, Jeffrey Barrett, Sonia Goncalves, Ewan Harrison, David K. Jackson, Ian Johnston, Dominic Kwiatkowski, Cordelia Langford, John Sillitoe on behalf of the Wellcome Sanger Institute COVID-19 Surveillance Team |
| EPI_ISL_2918274 | Lighthouse Lab in Alderley Park | Wellcome Sanger Institute for the COVID-19 Genomics UK (COG-UK) Consortium | Jacquelyn Wynn, Mairead Hyland, The Lighthouse Lab in Alderley Park and Alex Alderton, Roberto Amato, Jeffrey Barrett, Sonia Goncalves, Ewan Harrison, David K. Jackson, Ian Johnston, Dominic Kwiatkowski, Cordelia Langford, John Sillitoe on behalf of the Wellcome Sanger Institute COVID-19 Surveillance Team |
| EPI_ISL_2918275 | Health Services Laboratories | Wellcome Sanger Institute for the COVID-19 Genomics UK (COG-UK) | Health Services Laboratories and Alex Alderton, Roberto Amato, Jeffrey Barrett, Sonia Goncalves, Ewan Harrison, David K. Jackson, Ian Johnston, Dominic Kwiatkowski, Cordelia Langford, John Sillitoe on behalf of the Wellcome Sanger Institute |

| | | | |
|---|---|---|---|
| | | Consortium | COVID-19 Surveillance Team |
| EPI_ISL_2918276 | Lighthouse Lab in Alderley Park | Wellcome Sanger Institute for the COVID-19 Genomics UK (COG-UK) Consortium | Jacquelyn Wynn, Mairead Hyland, The Lighthouse Lab in Alderley Park and Alex Alderton, Roberto Amato, Jeffrey Barrett, Sonia Goncalves, Ewan Harrison, David K. Jackson, Ian Johnston, Dominic Kwiatkowski, Cordelia Langford, John Sillitoe on behalf of the Wellcome Sanger Institute COVID-19 Surveillance Team |
| EPI_ISL_2918277, EPI_ISL_2918278 | Health Services Laboratories | Wellcome Sanger Institute for the COVID-19 Genomics UK (COG-UK) Consortium | Health Services Laboratories and Alex Alderton, Roberto Amato, Jeffrey Barrett, Sonia Goncalves, Ewan Harrison, David K. Jackson, Ian Johnston, Dominic Kwiatkowski, Cordelia Langford, John Sillitoe on behalf of the Wellcome Sanger Institute COVID-19 Surveillance Team |
| EPI_ISL_2918279, EPI_ISL_2918280 | Lighthouse Lab in Alderley Park | Wellcome Sanger Institute for the COVID-19 Genomics UK (COG-UK) Consortium | Jacquelyn Wynn, Mairead Hyland, The Lighthouse Lab in Alderley Park and Alex Alderton, Roberto Amato, Jeffrey Barrett, Sonia Goncalves, Ewan Harrison, David K. Jackson, Ian Johnston, Dominic Kwiatkowski, Cordelia Langford, John Sillitoe on behalf of the Wellcome Sanger Institute COVID-19 Surveillance Team |
| EPI_ISL_2918281 | Health Services Laboratories | Wellcome Sanger Institute for the COVID-19 Genomics UK (COG-UK) Consortium | Health Services Laboratories and Alex Alderton, Roberto Amato, Jeffrey Barrett, Sonia Goncalves, Ewan Harrison, David K. Jackson, Ian Johnston, Dominic Kwiatkowski, Cordelia Langford, John Sillitoe on behalf of the Wellcome Sanger Institute COVID-19 Surveillance Team |
| EPI_ISL_2918282 | Lighthouse Lab in Milton Keynes | Wellcome Sanger Institute for the COVID-19 Genomics UK (COG-UK) Consortium | The Lighthouse Lab in Milton Keynes and Alex Alderton, Roberto Amato, Jeffrey Barrett, Sonia Goncalves, Ewan Harrison, David K. Jackson, Ian Johnston, Dominic Kwiatkowski, Cordelia Langford, John Sillitoe on behalf of the Wellcome Sanger Institute COVID-19 Surveillance Team |
| EPI_ISL_2918283, EPI_ISL_2918284, EPI_ISL_2918285 | Berkshire and Surrey Pathology Services Lighthouse Laboratory | Wellcome Sanger Institute for the COVID-19 Genomics UK (COG-UK) Consortium | Berkshire and Surrey Pathology Services Lighthouse Laboratory and Alex Alderton, Roberto Amato, Jeffrey Barrett, Sonia Goncalves, Ewan Harrison, David K. Jackson, Ian Johnston, Dominic Kwiatkowski, Cordelia Langford, John Sillitoe on behalf of the Wellcome Sanger Institute COVID-19 Surveillance Team |
| EPI_ISL_2918286 | Lighthouse Lab in Alderley Park | Wellcome Sanger Institute for the COVID-19 Genomics UK (COG-UK) Consortium | Jacquelyn Wynn, Mairead Hyland, The Lighthouse Lab in Alderley Park and Alex Alderton, Roberto Amato, Jeffrey Barrett, Sonia Goncalves, Ewan Harrison, David K. Jackson, Ian Johnston, Dominic Kwiatkowski, Cordelia Langford, John Sillitoe on behalf of the Wellcome Sanger Institute COVID-19 Surveillance Team |
| EPI_ISL_2918287 | Lighthouse Lab in Milton Keynes | Wellcome Sanger Institute for the COVID-19 Genomics UK (COG-UK) Consortium | The Lighthouse Lab in Milton Keynes and Alex Alderton, Roberto Amato, Jeffrey Barrett, Sonia Goncalves, Ewan Harrison, David K. Jackson, Ian Johnston, Dominic Kwiatkowski, Cordelia Langford, John Sillitoe on behalf of the Wellcome Sanger Institute COVID-19 Surveillance Team |
| EPI_ISL_2918288 | Lighthouse Lab in Alderley Park | Wellcome Sanger Institute for the COVID-19 Genomics UK (COG-UK) Consortium | Jacquelyn Wynn, Mairead Hyland, The Lighthouse Lab in Alderley Park and Alex Alderton, Roberto Amato, Jeffrey Barrett, Sonia Goncalves, Ewan Harrison, David K. Jackson, Ian Johnston, Dominic Kwiatkowski, Cordelia Langford, John Sillitoe on behalf of the Wellcome Sanger Institute COVID-19 Surveillance Team |
| EPI_ISL_2918289 | Health Services Laboratories | Wellcome Sanger Institute for the COVID-19 Genomics UK (COG-UK) Consortium | Health Services Laboratories and Alex Alderton, Roberto Amato, Jeffrey Barrett, Sonia Goncalves, Ewan Harrison, David K. Jackson, Ian Johnston, Dominic Kwiatkowski, Cordelia Langford, John Sillitoe on behalf of the Wellcome Sanger Institute COVID-19 Surveillance Team |
| EPI_ISL_2918290 | Berkshire and Surrey Pathology Services Lighthouse Laboratory | Wellcome Sanger Institute for the COVID-19 Genomics UK (COG-UK) Consortium | Berkshire and Surrey Pathology Services Lighthouse Laboratory and Alex Alderton, Roberto Amato, Jeffrey Barrett, Sonia Goncalves, Ewan Harrison, David K. Jackson, Ian Johnston, Dominic Kwiatkowski, Cordelia Langford, John Sillitoe on behalf of the Wellcome Sanger Institute COVID-19 Surveillance Team |
| EPI_ISL_2918291 | Health Services Laboratories | Wellcome Sanger Institute for the COVID-19 Genomics UK (COG-UK) Consortium | Health Services Laboratories and Alex Alderton, Roberto Amato, Jeffrey Barrett, Sonia Goncalves, Ewan Harrison, David K. Jackson, Ian Johnston, Dominic Kwiatkowski, Cordelia Langford, John Sillitoe on behalf of the Wellcome Sanger Institute COVID-19 Surveillance Team |
| EPI_ISL_2918292 | Lighthouse Lab in Milton Keynes | Wellcome Sanger Institute for the COVID-19 Genomics UK (COG-UK) Consortium | The Lighthouse Lab in Milton Keynes and Alex Alderton, Roberto Amato, Jeffrey Barrett, Sonia Goncalves, Ewan Harrison, David K. Jackson, Ian Johnston, Dominic Kwiatkowski, Cordelia Langford, John Sillitoe on behalf of the Wellcome Sanger Institute COVID-19 Surveillance Team |
| EPI_ISL_2918293 | Lighthouse Lab in Alderley Park | Wellcome Sanger Institute for the COVID-19 Genomics UK (COG-UK) Consortium | Jacquelyn Wynn, Mairead Hyland, The Lighthouse Lab in Alderley Park and Alex Alderton, Roberto Amato, Jeffrey Barrett, Sonia Goncalves, Ewan Harrison, David K. Jackson, Ian Johnston, Dominic Kwiatkowski, Cordelia Langford, John Sillitoe on behalf of the Wellcome Sanger Institute COVID-19 Surveillance Team |
| EPI_ISL_2918294 | Randox Laboratories | Wellcome Sanger Institute for the COVID-19 Genomics UK (COG-UK) Consortium | Randox Laboratories and Alex Alderton, Roberto Amato, Jeffrey Barrett, Sonia Goncalves, Ewan Harrison, David K. Jackson, Ian Johnston, Dominic Kwiatkowski, Cordelia Langford, John Sillitoe on behalf of the Wellcome Sanger Institute COVID-19 Surveillance Team |
| EPI_ISL_2918295 | Berkshire and Surrey Pathology Services Lighthouse Laboratory | Wellcome Sanger Institute for the COVID-19 Genomics UK (COG-UK) Consortium | Berkshire and Surrey Pathology Services Lighthouse Laboratory and Alex Alderton, Roberto Amato, Jeffrey Barrett, Sonia Goncalves, Ewan Harrison, David K. Jackson, Ian Johnston, Dominic Kwiatkowski, Cordelia Langford, John Sillitoe on behalf of the Wellcome Sanger Institute COVID-19 Surveillance Team |
| EPI_ISL_2918296 | Lighthouse Lab in Milton Keynes | Wellcome Sanger Institute for the COVID-19 Genomics UK (COG-UK) Consortium | The Lighthouse Lab in Milton Keynes and Alex Alderton, Roberto Amato, Jeffrey Barrett, Sonia Goncalves, Ewan Harrison, David K. Jackson, Ian Johnston, Dominic Kwiatkowski, Cordelia Langford, John Sillitoe on behalf of the Wellcome Sanger Institute COVID-19 Surveillance Team |
| EPI_ISL_2918297 | Berkshire and Surrey Pathology Services Lighthouse Laboratory | Wellcome Sanger Institute for the COVID-19 Genomics UK (COG-UK) Consortium | Berkshire and Surrey Pathology Services Lighthouse Laboratory and Alex Alderton, Roberto Amato, Jeffrey Barrett, Sonia Goncalves, Ewan Harrison, David K. Jackson, Ian Johnston, Dominic Kwiatkowski, Cordelia Langford, John Sillitoe on behalf of the Wellcome Sanger Institute COVID-19 Surveillance Team |
| EPI_ISL_2918298 | Randox Laboratories | Wellcome Sanger Institute for the COVID-19 Genomics UK (COG-UK) Consortium | Randox Laboratories and Alex Alderton, Roberto Amato, Jeffrey Barrett, Sonia Goncalves, Ewan Harrison, David K. Jackson, Ian Johnston, Dominic Kwiatkowski, Cordelia Langford, John Sillitoe on behalf of the Wellcome Sanger Institute COVID-19 Surveillance Team |
| EPI_ISL_2918299, EPI_ISL_2918300 | Berkshire and Surrey Pathology Services Lighthouse Laboratory | Wellcome Sanger Institute for the COVID-19 Genomics UK (COG-UK) Consortium | Berkshire and Surrey Pathology Services Lighthouse Laboratory and Alex Alderton, Roberto Amato, Jeffrey Barrett, Sonia Goncalves, Ewan Harrison, David K. Jackson, Ian Johnston, Dominic Kwiatkowski, Cordelia Langford, John Sillitoe on behalf of the Wellcome Sanger Institute COVID-19 Surveillance Team |
| EPI_ISL_2918301 | Health Services Laboratories | Wellcome Sanger Institute for the COVID-19 Genomics UK (COG-UK) Consortium | Health Services Laboratories and Alex Alderton, Roberto Amato, Jeffrey Barrett, Sonia Goncalves, Ewan Harrison, David K. Jackson, Ian Johnston, Dominic Kwiatkowski, Cordelia Langford, John Sillitoe on behalf of the Wellcome Sanger Institute COVID-19 Surveillance Team |
| EPI_ISL_2918302 | Berkshire and Surrey Pathology Services Lighthouse Laboratory | Wellcome Sanger Institute for the COVID-19 Genomics UK (COG-UK) Consortium | Berkshire and Surrey Pathology Services Lighthouse Laboratory and Alex Alderton, Roberto Amato, Jeffrey Barrett, Sonia Goncalves, Ewan Harrison, David K. Jackson, Ian Johnston, Dominic Kwiatkowski, Cordelia Langford, John Sillitoe on behalf of the Wellcome Sanger Institute COVID-19 Surveillance Team |
| EPI_ISL_2918303 | Randox Laboratories | Wellcome Sanger Institute for the COVID-19 Genomics UK (COG-UK) Consortium | Randox Laboratories and Alex Alderton, Roberto Amato, Jeffrey Barrett, Sonia Goncalves, Ewan Harrison, David K. Jackson, Ian Johnston, Dominic Kwiatkowski, Cordelia Langford, John Sillitoe on behalf of the Wellcome Sanger Institute COVID-19 Surveillance Team |
| EPI_ISL_2918304 | Health Services Laboratories | Wellcome Sanger Institute for the COVID-19 Genomics UK (COG-UK) Consortium | Health Services Laboratories and Alex Alderton, Roberto Amato, Jeffrey Barrett, Sonia Goncalves, Ewan Harrison, David K. Jackson, Ian Johnston, Dominic Kwiatkowski, Cordelia Langford, John Sillitoe on behalf of the Wellcome Sanger Institute COVID-19 Surveillance Team |
| EPI_ISL_2918305 | Lighthouse Lab in Alderley Park | Wellcome Sanger Institute for the COVID-19 Genomics UK (COG-UK) Consortium | Jacquelyn Wynn, Mairead Hyland, The Lighthouse Lab in Alderley Park and Alex Alderton, Roberto Amato, Jeffrey Barrett, Sonia Goncalves, Ewan Harrison, David K. Jackson, Ian Johnston, Dominic Kwiatkowski, Cordelia Langford, John Sillitoe on behalf of the Wellcome Sanger Institute COVID-19 Surveillance Team |
| EPI_ISL_2918306, EPI_ISL_2918307 | Lighthouse Lab in Milton Keynes | Wellcome Sanger Institute for the COVID-19 Genomics UK (COG-UK) Consortium | The Lighthouse Lab in Milton Keynes and Alex Alderton, Roberto Amato, Jeffrey Barrett, Sonia Goncalves, Ewan Harrison, David K. Jackson, Ian Johnston, Dominic Kwiatkowski, Cordelia Langford, John Sillitoe on behalf of the Wellcome Sanger Institute COVID-19 Surveillance Team |
| EPI_ISL_2918308 | Health Services Laboratories | Wellcome Sanger Institute for the COVID-19 Genomics UK (COG-UK) Consortium | Health Services Laboratories and Alex Alderton, Roberto Amato, Jeffrey Barrett, Sonia Goncalves, Ewan Harrison, David K. Jackson, Ian Johnston, Dominic Kwiatkowski, Cordelia Langford, John Sillitoe on behalf of the Wellcome Sanger Institute COVID-19 Surveillance Team |
| EPI_ISL_2918309 | Lighthouse Lab in Milton Keynes | Wellcome Sanger Institute for the COVID-19 Genomics UK (COG-UK) Consortium | The Lighthouse Lab in Milton Keynes and Alex Alderton, Roberto Amato, Jeffrey Barrett, Sonia Goncalves, Ewan Harrison, David K. Jackson, Ian Johnston, Dominic Kwiatkowski, Cordelia Langford, John Sillitoe on behalf of the Wellcome Sanger Institute COVID-19 Surveillance Team |
| EPI_ISL_2918310, EPI_ISL_2918311 | Berkshire and Surrey Pathology Services Lighthouse Laboratory | Wellcome Sanger Institute for the COVID-19 Genomics UK (COG-UK) Consortium | Berkshire and Surrey Pathology Services Lighthouse Laboratory and Alex Alderton, Roberto Amato, Jeffrey Barrett, Sonia Goncalves, Ewan Harrison, David K. Jackson, Ian Johnston, Dominic Kwiatkowski, Cordelia Langford, John Sillitoe on behalf of the Wellcome Sanger Institute COVID-19 Surveillance Team |
| EPI_ISL_2918312 | Lighthouse Lab in Alderley Park | Wellcome Sanger Institute for the COVID-19 Genomics UK (COG-UK) Consortium | Jacquelyn Wynn, Mairead Hyland, The Lighthouse Lab in Alderley Park and Alex Alderton, Roberto Amato, Jeffrey Barrett, Sonia Goncalves, Ewan Harrison, David K. Jackson, Ian Johnston, Dominic Kwiatkowski, Cordelia Langford, John Sillitoe on behalf of the Wellcome Sanger Institute COVID-19 Surveillance Team |
| EPI_ISL_2918313, EPI_ISL_2918314 | Berkshire and Surrey Pathology Services Lighthouse Laboratory | Wellcome Sanger Institute for the COVID-19 Genomics UK (COG-UK) Consortium | Berkshire and Surrey Pathology Services Lighthouse Laboratory and Alex Alderton, Roberto Amato, Jeffrey Barrett, Sonia Goncalves, Ewan Harrison, David K. Jackson, Ian Johnston, Dominic Kwiatkowski, Cordelia Langford, John Sillitoe on behalf of the Wellcome Sanger Institute COVID-19 Surveillance Team |
| EPI_ISL_2918315, EPI_ISL_2918316 | Health Services Laboratories | Wellcome Sanger Institute for the COVID-19 Genomics UK (COG-UK) Consortium | Health Services Laboratories and Alex Alderton, Roberto Amato, Jeffrey Barrett, Sonia Goncalves, Ewan Harrison, David K. Jackson, Ian Johnston, Dominic Kwiatkowski, Cordelia Langford, John Sillitoe on behalf of the Wellcome Sanger Institute COVID-19 Surveillance Team |
| EPI_ISL_2918317 | Berkshire and Surrey Pathology Services Lighthouse Laboratory | Wellcome Sanger Institute for the COVID-19 Genomics UK (COG-UK) Consortium | Berkshire and Surrey Pathology Services Lighthouse Laboratory and Alex Alderton, Roberto Amato, Jeffrey Barrett, Sonia Goncalves, Ewan Harrison, David K. Jackson, Ian Johnston, Dominic Kwiatkowski, Cordelia Langford, John Sillitoe on behalf of the Wellcome Sanger Institute COVID-19 Surveillance Team |
| EPI_ISL_2918318 | Health Services Laboratories | Wellcome Sanger Institute for the COVID-19 Genomics UK (COG-UK) Consortium | Health Services Laboratories and Alex Alderton, Roberto Amato, Jeffrey Barrett, Sonia Goncalves, Ewan Harrison, David K. Jackson, Ian Johnston, Dominic Kwiatkowski, Cordelia Langford, John Sillitoe on behalf of the Wellcome Sanger Institute COVID-19 Surveillance Team |
| EPI_ISL_2918319 | Berkshire and Surrey Pathology Services Lighthouse Laboratory | Wellcome Sanger Institute for the COVID-19 Genomics UK (COG-UK) Consortium | Berkshire and Surrey Pathology Services Lighthouse Laboratory and Alex Alderton, Roberto Amato, Jeffrey Barrett, Sonia Goncalves, Ewan Harrison, David K. Jackson, Ian Johnston, Dominic Kwiatkowski, Cordelia Langford, John Sillitoe on behalf of the Wellcome Sanger Institute COVID-19 Surveillance Team |
| EPI_ISL_2918320, EPI_ISL_2918321 | Health Services Laboratories | Wellcome Sanger Institute for the COVID-19 Genomics UK (COG-UK) Consortium | Health Services Laboratories and Alex Alderton, Roberto Amato, Jeffrey Barrett, Sonia Goncalves, Ewan Harrison, David K. Jackson, Ian Johnston, Dominic Kwiatkowski, Cordelia Langford, John Sillitoe on behalf of the Wellcome Sanger Institute COVID-19 Surveillance Team |
| EPI_ISL_2918322 | Berkshire and Surrey Pathology Services Lighthouse Laboratory | Wellcome Sanger Institute for the COVID-19 Genomics UK (COG-UK) Consortium | Berkshire and Surrey Pathology Services Lighthouse Laboratory and Alex Alderton, Roberto Amato, Jeffrey Barrett, Sonia Goncalves, Ewan Harrison, David K. Jackson, Ian Johnston, Dominic Kwiatkowski, Cordelia Langford, John Sillitoe on behalf of the Wellcome Sanger Institute COVID-19 Surveillance Team |
| EPI_ISL_2918323, EPI_ISL_2918324, EPI_ISL_2918325 | Lighthouse Lab in Milton Keynes | Wellcome Sanger Institute for the COVID-19 Genomics UK (COG-UK) Consortium | The Lighthouse Lab in Milton Keynes and Alex Alderton, Roberto Amato, Jeffrey Barrett, Sonia Goncalves, Ewan Harrison, David K. Jackson, Ian Johnston, Dominic Kwiatkowski, Cordelia Langford, John Sillitoe on behalf of the Wellcome Sanger Institute COVID-19 Surveillance Team |
| EPI_ISL_2918326 | Berkshire and Surrey Pathology Services Lighthouse Laboratory | Wellcome Sanger Institute for the COVID-19 Genomics UK (COG-UK) Consortium | Berkshire and Surrey Pathology Services Lighthouse Laboratory and Alex Alderton, Roberto Amato, Jeffrey Barrett, Sonia Goncalves, Ewan Harrison, David K. Jackson, Ian Johnston, Dominic Kwiatkowski, Cordelia Langford, John Sillitoe on behalf of the Wellcome Sanger Institute COVID-19 Surveillance Team |
| EPI_ISL_2918327 | Health Services Laboratories | Wellcome Sanger Institute for the COVID-19 Genomics UK (COG-UK) Consortium | Health Services Laboratories and Alex Alderton, Roberto Amato, Jeffrey Barrett, Sonia Goncalves, Ewan Harrison, David K. Jackson, Ian Johnston, Dominic Kwiatkowski, Cordelia Langford, John Sillitoe on behalf of the Wellcome Sanger Institute COVID-19 Surveillance Team |
| EPI_ISL_2918328, EPI_ISL_2918329, EPI_ISL_2918330 | Lighthouse Lab in Milton Keynes | Wellcome Sanger Institute for the COVID-19 Genomics UK (COG-UK) Consortium | The Lighthouse Lab in Milton Keynes and Alex Alderton, Roberto Amato, Jeffrey Barrett, Sonia Goncalves, Ewan Harrison, David K. Jackson, Ian Johnston, Dominic Kwiatkowski, Cordelia Langford, John Sillitoe on behalf of the Wellcome Sanger Institute COVID-19 Surveillance Team |
| EPI_ISL_2918331 | Berkshire and Surrey Pathology Services Lighthouse Laboratory | Wellcome Sanger Institute for the COVID-19 Genomics UK (COG-UK) Consortium | Berkshire and Surrey Pathology Services Lighthouse Laboratory and Alex Alderton, Roberto Amato, Jeffrey Barrett, Sonia Goncalves, Ewan Harrison, David K. Jackson, Ian Johnston, Dominic Kwiatkowski, Cordelia Langford, John Sillitoe on behalf of the Wellcome Sanger Institute COVID-19 Surveillance Team |
| EPI_ISL_2918332, EPI_ISL_2918333, | Lighthouse Lab in Alderley Park | Wellcome Sanger Institute for the COVID-19 Genomics UK (COG-UK) Consortium | Jacquelyn Wynn, Mairead Hyland, The Lighthouse Lab in Alderley Park and Alex Alderton, Roberto Amato, Jeffrey Barrett, Sonia Goncalves, Ewan Harrison, David K. Jackson, Ian Johnston, Dominic Kwiatkowski, Cordelia Langford, John Sillitoe on behalf of the Wellcome Sanger Institute COVID-19 Surveillance Team |

| | | | |
|---|---|---|---|
| EPI_ISL_2918334 | | | |
| EPI_ISL_2918335 | Lighthouse Lab in Milton Keynes | Wellcome Sanger Institute for the COVID-19 Genomics UK (COG-UK) Consortium | The Lighthouse Lab in Milton Keynes and Alex Alderton, Roberto Amato, Jeffrey Barrett, Sonia Goncalves, Ewan Harrison, David K. Jackson, Ian Johnston, Dominic Kwiatkowski, Cordelia Langford, John Sillitoe on behalf of the Wellcome Sanger Institute COVID-19 Surveillance Team |
| EPI_ISL_2918336, EPI_ISL_2918337 | Health Services Laboratories | Wellcome Sanger Institute for the COVID-19 Genomics UK (COG-UK) Consortium | Health Services Laboratories and Alex Alderton, Roberto Amato, Jeffrey Barrett, Sonia Goncalves, Ewan Harrison, David K. Jackson, Ian Johnston, Dominic Kwiatkowski, Cordelia Langford, John Sillitoe on behalf of the Wellcome Sanger Institute COVID-19 Surveillance Team |
| EPI_ISL_2918338, EPI_ISL_2918339 | Lighthouse Lab in Alderley Park | Wellcome Sanger Institute for the COVID-19 Genomics UK (COG-UK) Consortium | Jacquelyn Wynn, Mairead Hyland, The Lighthouse Lab in Alderley Park and Alex Alderton, Roberto Amato, Jeffrey Barrett, Sonia Goncalves, Ewan Harrison, David K. Jackson, Ian Johnston, Dominic Kwiatkowski, Cordelia Langford, John Sillitoe on behalf of the Wellcome Sanger Institute COVID-19 Surveillance Team |
| EPI_ISL_2918340 | Lighthouse Lab in Milton Keynes | Wellcome Sanger Institute for the COVID-19 Genomics UK (COG-UK) Consortium | The Lighthouse Lab in Milton Keynes and Alex Alderton, Roberto Amato, Jeffrey Barrett, Sonia Goncalves, Ewan Harrison, David K. Jackson, Ian Johnston, Dominic Kwiatkowski, Cordelia Langford, John Sillitoe on behalf of the Wellcome Sanger Institute COVID-19 Surveillance Team |
| EPI_ISL_2918341 | Lighthouse Lab in Alderley Park | Wellcome Sanger Institute for the COVID-19 Genomics UK (COG-UK) Consortium | Jacquelyn Wynn, Mairead Hyland, The Lighthouse Lab in Alderley Park and Alex Alderton, Roberto Amato, Jeffrey Barrett, Sonia Goncalves, Ewan Harrison, David K. Jackson, Ian Johnston, Dominic Kwiatkowski, Cordelia Langford, John Sillitoe on behalf of the Wellcome Sanger Institute COVID-19 Surveillance Team |
| EPI_ISL_2918342, EPI_ISL_2918343 | Health Services Laboratories | Wellcome Sanger Institute for the COVID-19 Genomics UK (COG-UK) Consortium | Health Services Laboratories and Alex Alderton, Roberto Amato, Jeffrey Barrett, Sonia Goncalves, Ewan Harrison, David K. Jackson, Ian Johnston, Dominic Kwiatkowski, Cordelia Langford, John Sillitoe on behalf of the Wellcome Sanger Institute COVID-19 Surveillance Team |
| EPI_ISL_2918344, EPI_ISL_2918345, EPI_ISL_2918346 | Berkshire and Surrey Pathology Services Lighthouse Laboratory | Wellcome Sanger Institute for the COVID-19 Genomics UK (COG-UK) Consortium | Berkshire and Surrey Pathology Services Lighthouse Laboratory and Alex Alderton, Roberto Amato, Jeffrey Barrett, Sonia Goncalves, Ewan Harrison, David K. Jackson, Ian Johnston, Dominic Kwiatkowski, Cordelia Langford, John Sillitoe on behalf of the Wellcome Sanger Institute COVID-19 Surveillance Team |
| EPI_ISL_2918347, EPI_ISL_2918348 | Health Services Laboratories | Wellcome Sanger Institute for the COVID-19 Genomics UK (COG-UK) Consortium | Health Services Laboratories and Alex Alderton, Roberto Amato, Jeffrey Barrett, Sonia Goncalves, Ewan Harrison, David K. Jackson, Ian Johnston, Dominic Kwiatkowski, Cordelia Langford, John Sillitoe on behalf of the Wellcome Sanger Institute COVID-19 Surveillance Team |
| EPI_ISL_2918349 | Lighthouse Lab in Milton Keynes | Wellcome Sanger Institute for the COVID-19 Genomics UK (COG-UK) Consortium | The Lighthouse Lab in Milton Keynes and Alex Alderton, Roberto Amato, Jeffrey Barrett, Sonia Goncalves, Ewan Harrison, David K. Jackson, Ian Johnston, Dominic Kwiatkowski, Cordelia Langford, John Sillitoe on behalf of the Wellcome Sanger Institute COVID-19 Surveillance Team |
| EPI_ISL_2918350 | Berkshire and Surrey Pathology Services Lighthouse Laboratory | Wellcome Sanger Institute for the COVID-19 Genomics UK (COG-UK) Consortium | Berkshire and Surrey Pathology Services Lighthouse Laboratory and Alex Alderton, Roberto Amato, Jeffrey Barrett, Sonia Goncalves, Ewan Harrison, David K. Jackson, Ian Johnston, Dominic Kwiatkowski, Cordelia Langford, John Sillitoe on behalf of the Wellcome Sanger Institute COVID-19 Surveillance Team |
| EPI_ISL_2918351 | Lighthouse Lab in Alderley Park | Wellcome Sanger Institute for the COVID-19 Genomics UK (COG-UK) Consortium | Jacquelyn Wynn, Mairead Hyland, The Lighthouse Lab in Alderley Park and Alex Alderton, Roberto Amato, Jeffrey Barrett, Sonia Goncalves, Ewan Harrison, David K. Jackson, Ian Johnston, Dominic Kwiatkowski, Cordelia Langford, John Sillitoe on behalf of the Wellcome Sanger Institute COVID-19 Surveillance Team |
| EPI_ISL_2918352, EPI_ISL_2918353 | Health Services Laboratories | Wellcome Sanger Institute for the COVID-19 Genomics UK (COG-UK) Consortium | Health Services Laboratories and Alex Alderton, Roberto Amato, Jeffrey Barrett, Sonia Goncalves, Ewan Harrison, David K. Jackson, Ian Johnston, Dominic Kwiatkowski, Cordelia Langford, John Sillitoe on behalf of the Wellcome Sanger Institute COVID-19 Surveillance Team |
| EPI_ISL_2918354 | Berkshire and Surrey Pathology Services Lighthouse Laboratory | Wellcome Sanger Institute for the COVID-19 Genomics UK (COG-UK) Consortium | Berkshire and Surrey Pathology Services Lighthouse Laboratory and Alex Alderton, Roberto Amato, Jeffrey Barrett, Sonia Goncalves, Ewan Harrison, David K. Jackson, Ian Johnston, Dominic Kwiatkowski, Cordelia Langford, John Sillitoe on behalf of the Wellcome Sanger Institute COVID-19 Surveillance Team |
| EPI_ISL_2918355, EPI_ISL_2918356 | Lighthouse Lab in Alderley Park | Wellcome Sanger Institute for the COVID-19 Genomics UK (COG-UK) Consortium | Jacquelyn Wynn, Mairead Hyland, The Lighthouse Lab in Alderley Park and Alex Alderton, Roberto Amato, Jeffrey Barrett, Sonia Goncalves, Ewan Harrison, David K. Jackson, Ian Johnston, Dominic Kwiatkowski, Cordelia Langford, John Sillitoe on behalf of the Wellcome Sanger Institute COVID-19 Surveillance Team |
| EPI_ISL_2918357 | Health Services Laboratories | Wellcome Sanger Institute for the COVID-19 Genomics UK (COG-UK) Consortium | Health Services Laboratories and Alex Alderton, Roberto Amato, Jeffrey Barrett, Sonia Goncalves, Ewan Harrison, David K. Jackson, Ian Johnston, Dominic Kwiatkowski, Cordelia Langford, John Sillitoe on behalf of the Wellcome Sanger Institute COVID-19 Surveillance Team |
| EPI_ISL_2918358 | Lighthouse Lab in Milton Keynes | Wellcome Sanger Institute for the COVID-19 Genomics UK (COG-UK) Consortium | The Lighthouse Lab in Milton Keynes and Alex Alderton, Roberto Amato, Jeffrey Barrett, Sonia Goncalves, Ewan Harrison, David K. Jackson, Ian Johnston, Dominic Kwiatkowski, Cordelia Langford, John Sillitoe on behalf of the Wellcome Sanger Institute COVID-19 Surveillance Team |
| EPI_ISL_2918359 | Lighthouse Lab in Alderley Park | Wellcome Sanger Institute for the COVID-19 Genomics UK (COG-UK) Consortium | Jacquelyn Wynn, Mairead Hyland, The Lighthouse Lab in Alderley Park and Alex Alderton, Roberto Amato, Jeffrey Barrett, Sonia Goncalves, Ewan Harrison, David K. Jackson, Ian Johnston, Dominic Kwiatkowski, Cordelia Langford, John Sillitoe on behalf of the Wellcome Sanger Institute COVID-19 Surveillance Team |
| EPI_ISL_2918360 | Health Services Laboratories | Wellcome Sanger Institute for the COVID-19 Genomics UK (COG-UK) Consortium | Health Services Laboratories and Alex Alderton, Roberto Amato, Jeffrey Barrett, Sonia Goncalves, Ewan Harrison, David K. Jackson, Ian Johnston, Dominic Kwiatkowski, Cordelia Langford, John Sillitoe on behalf of the Wellcome Sanger Institute COVID-19 Surveillance Team |
| EPI_ISL_2918361 | Lighthouse Lab in Alderley Park | Wellcome Sanger Institute for the COVID-19 Genomics UK (COG-UK) Consortium | Jacquelyn Wynn, Mairead Hyland, The Lighthouse Lab in Alderley Park and Alex Alderton, Roberto Amato, Jeffrey Barrett, Sonia Goncalves, Ewan Harrison, David K. Jackson, Ian Johnston, Dominic Kwiatkowski, Cordelia Langford, John Sillitoe on behalf of the Wellcome Sanger Institute COVID-19 Surveillance Team |
| EPI_ISL_2918362 | Health Services Laboratories | Wellcome Sanger Institute for the COVID-19 Genomics UK (COG-UK) Consortium | Health Services Laboratories and Alex Alderton, Roberto Amato, Jeffrey Barrett, Sonia Goncalves, Ewan Harrison, David K. Jackson, Ian Johnston, Dominic Kwiatkowski, Cordelia Langford, John Sillitoe on behalf of the Wellcome Sanger Institute COVID-19 Surveillance Team |
| EPI_ISL_2918363 | Lighthouse Lab in Alderley Park | Wellcome Sanger Institute for the COVID-19 Genomics UK (COG-UK) Consortium | Jacquelyn Wynn, Mairead Hyland, The Lighthouse Lab in Alderley Park and Alex Alderton, Roberto Amato, Jeffrey Barrett, Sonia Goncalves, Ewan Harrison, David K. Jackson, Ian Johnston, Dominic Kwiatkowski, Cordelia Langford, John Sillitoe on behalf of the Wellcome Sanger Institute COVID-19 Surveillance Team |
| EPI_ISL_2918364, EPI_ISL_2918365 | Lighthouse Lab in Milton Keynes | Wellcome Sanger Institute for the COVID-19 Genomics UK (COG-UK) Consortium | The Lighthouse Lab in Milton Keynes and Alex Alderton, Roberto Amato, Jeffrey Barrett, Sonia Goncalves, Ewan Harrison, David K. Jackson, Ian Johnston, Dominic Kwiatkowski, Cordelia Langford, John Sillitoe on behalf of the Wellcome Sanger Institute COVID-19 Surveillance Team |
| EPI_ISL_2918366 | Health Services Laboratories | Wellcome Sanger Institute for the COVID-19 Genomics UK (COG-UK) Consortium | Health Services Laboratories and Alex Alderton, Roberto Amato, Jeffrey Barrett, Sonia Goncalves, Ewan Harrison, David K. Jackson, Ian Johnston, Dominic Kwiatkowski, Cordelia Langford, John Sillitoe on behalf of the Wellcome Sanger Institute COVID-19 Surveillance Team |
| EPI_ISL_2918367 | Lighthouse Lab in Milton Keynes | Wellcome Sanger Institute for the COVID-19 Genomics UK (COG-UK) Consortium | The Lighthouse Lab in Milton Keynes and Alex Alderton, Roberto Amato, Jeffrey Barrett, Sonia Goncalves, Ewan Harrison, David K. Jackson, Ian Johnston, Dominic Kwiatkowski, Cordelia Langford, John Sillitoe on behalf of the Wellcome Sanger Institute COVID-19 Surveillance Team |
| EPI_ISL_2918368 | Berkshire and Surrey Pathology Services Lighthouse Laboratory | Wellcome Sanger Institute for the COVID-19 Genomics UK (COG-UK) Consortium | Berkshire and Surrey Pathology Services Lighthouse Laboratory and Alex Alderton, Roberto Amato, Jeffrey Barrett, Sonia Goncalves, Ewan Harrison, David K. Jackson, Ian Johnston, Dominic Kwiatkowski, Cordelia Langford, John Sillitoe on behalf of the Wellcome Sanger Institute COVID-19 Surveillance Team |
| EPI_ISL_2918369 | Health Services Laboratories | Wellcome Sanger Institute for the COVID-19 Genomics UK (COG-UK) Consortium | Health Services Laboratories and Alex Alderton, Roberto Amato, Jeffrey Barrett, Sonia Goncalves, Ewan Harrison, David K. Jackson, Ian Johnston, Dominic Kwiatkowski, Cordelia Langford, John Sillitoe on behalf of the Wellcome Sanger Institute COVID-19 Surveillance Team |
| EPI_ISL_2918370, EPI_ISL_2918371 | Berkshire and Surrey Pathology Services Lighthouse Laboratory | Wellcome Sanger Institute for the COVID-19 Genomics UK (COG-UK) Consortium | Berkshire and Surrey Pathology Services Lighthouse Laboratory and Alex Alderton, Roberto Amato, Jeffrey Barrett, Sonia Goncalves, Ewan Harrison, David K. Jackson, Ian Johnston, Dominic Kwiatkowski, Cordelia Langford, John Sillitoe on behalf of the Wellcome Sanger Institute COVID-19 Surveillance Team |
| EPI_ISL_2918372 | Lighthouse Lab in Milton Keynes | Wellcome Sanger Institute for the COVID-19 Genomics UK (COG-UK) Consortium | The Lighthouse Lab in Milton Keynes and Alex Alderton, Roberto Amato, Jeffrey Barrett, Sonia Goncalves, Ewan Harrison, David K. Jackson, Ian Johnston, Dominic Kwiatkowski, Cordelia Langford, John Sillitoe on behalf of the Wellcome Sanger Institute COVID-19 Surveillance Team |
| EPI_ISL_2918373 | Berkshire and Surrey Pathology Services Lighthouse Laboratory | Wellcome Sanger Institute for the COVID-19 Genomics UK (COG-UK) Consortium | Berkshire and Surrey Pathology Services Lighthouse Laboratory and Alex Alderton, Roberto Amato, Jeffrey Barrett, Sonia Goncalves, Ewan Harrison, David K. Jackson, Ian Johnston, Dominic Kwiatkowski, Cordelia Langford, John Sillitoe on behalf of the Wellcome Sanger Institute COVID-19 Surveillance Team |
| EPI_ISL_2918374, EPI_ISL_2918375 | Lighthouse Lab in Alderley Park | Wellcome Sanger Institute for the COVID-19 Genomics UK (COG-UK) Consortium | Jacquelyn Wynn, Mairead Hyland, The Lighthouse Lab in Alderley Park and Alex Alderton, Roberto Amato, Jeffrey Barrett, Sonia Goncalves, Ewan Harrison, David K. Jackson, Ian Johnston, Dominic Kwiatkowski, Cordelia Langford, John Sillitoe on behalf of the Wellcome Sanger Institute COVID-19 Surveillance Team |
| EPI_ISL_2918376, EPI_ISL_2918377 | Lighthouse Lab in Milton Keynes | Wellcome Sanger Institute for the COVID-19 Genomics UK (COG-UK) Consortium | The Lighthouse Lab in Milton Keynes and Alex Alderton, Roberto Amato, Jeffrey Barrett, Sonia Goncalves, Ewan Harrison, David K. Jackson, Ian Johnston, Dominic Kwiatkowski, Cordelia Langford, John Sillitoe on behalf of the Wellcome Sanger Institute COVID-19 Surveillance Team |
| EPI_ISL_2918378 | Berkshire and Surrey Pathology Services Lighthouse Laboratory | Wellcome Sanger Institute for the COVID-19 Genomics UK (COG-UK) Consortium | Berkshire and Surrey Pathology Services Lighthouse Laboratory and Alex Alderton, Roberto Amato, Jeffrey Barrett, Sonia Goncalves, Ewan Harrison, David K. Jackson, Ian Johnston, Dominic Kwiatkowski, Cordelia Langford, John Sillitoe on behalf of the Wellcome Sanger Institute COVID-19 Surveillance Team |
| EPI_ISL_2918379, EPI_ISL_2918380 | Lighthouse Lab in Alderley Park | Wellcome Sanger Institute for the COVID-19 Genomics UK (COG-UK) Consortium | Jacquelyn Wynn, Mairead Hyland, The Lighthouse Lab in Alderley Park and Alex Alderton, Roberto Amato, Jeffrey Barrett, Sonia Goncalves, Ewan Harrison, David K. Jackson, Ian Johnston, Dominic Kwiatkowski, Cordelia Langford, John Sillitoe on behalf of the Wellcome Sanger Institute COVID-19 Surveillance Team |
| EPI_ISL_2918381 | Berkshire and Surrey Pathology Services Lighthouse Laboratory | Wellcome Sanger Institute for the COVID-19 Genomics UK (COG-UK) Consortium | Berkshire and Surrey Pathology Services Lighthouse Laboratory and Alex Alderton, Roberto Amato, Jeffrey Barrett, Sonia Goncalves, Ewan Harrison, David K. Jackson, Ian Johnston, Dominic Kwiatkowski, Cordelia Langford, John Sillitoe on behalf of the Wellcome Sanger Institute COVID-19 Surveillance Team |
| EPI_ISL_2918382 | Lighthouse Lab in Alderley Park | Wellcome Sanger Institute for the COVID-19 Genomics UK (COG-UK) Consortium | Jacquelyn Wynn, Mairead Hyland, The Lighthouse Lab in Alderley Park and Alex Alderton, Roberto Amato, Jeffrey Barrett, Sonia Goncalves, Ewan Harrison, David K. Jackson, Ian Johnston, Dominic Kwiatkowski, Cordelia Langford, John Sillitoe on behalf of the Wellcome Sanger Institute COVID-19 Surveillance Team |
| EPI_ISL_2918383 | Lighthouse Lab in Milton Keynes | Wellcome Sanger Institute for the COVID-19 Genomics UK (COG-UK) Consortium | The Lighthouse Lab in Milton Keynes and Alex Alderton, Roberto Amato, Jeffrey Barrett, Sonia Goncalves, Ewan Harrison, David K. Jackson, Ian Johnston, Dominic Kwiatkowski, Cordelia Langford, John Sillitoe on behalf of the Wellcome Sanger Institute COVID-19 Surveillance Team |
| EPI_ISL_2918384 | Lighthouse Lab in Alderley Park | Wellcome Sanger Institute for the COVID-19 Genomics UK (COG-UK) Consortium | Jacquelyn Wynn, Mairead Hyland, The Lighthouse Lab in Alderley Park and Alex Alderton, Roberto Amato, Jeffrey Barrett, Sonia Goncalves, Ewan Harrison, David K. Jackson, Ian Johnston, Dominic Kwiatkowski, Cordelia Langford, John Sillitoe on behalf of the Wellcome Sanger Institute COVID-19 Surveillance Team |
| EPI_ISL_2918385 | Lighthouse Lab in Milton Keynes | Wellcome Sanger Institute for the COVID-19 Genomics UK (COG-UK) Consortium | The Lighthouse Lab in Milton Keynes and Alex Alderton, Roberto Amato, Jeffrey Barrett, Sonia Goncalves, Ewan Harrison, David K. Jackson, Ian Johnston, Dominic Kwiatkowski, Cordelia Langford, John Sillitoe on behalf of the Wellcome Sanger Institute COVID-19 Surveillance Team |
| EPI_ISL_2918386 | Health Services Laboratories | Wellcome Sanger Institute for the COVID-19 Genomics UK (COG-UK) Consortium | Health Services Laboratories and Alex Alderton, Roberto Amato, Jeffrey Barrett, Sonia Goncalves, Ewan Harrison, David K. Jackson, Ian Johnston, Dominic Kwiatkowski, Cordelia Langford, John Sillitoe on behalf of the Wellcome Sanger Institute COVID-19 Surveillance Team |
| EPI_ISL_2918387 | Lighthouse Lab in Milton Keynes | Wellcome Sanger Institute for the COVID-19 Genomics UK (COG-UK) Consortium | The Lighthouse Lab in Milton Keynes and Alex Alderton, Roberto Amato, Jeffrey Barrett, Sonia Goncalves, Ewan Harrison, David K. Jackson, Ian Johnston, Dominic Kwiatkowski, Cordelia Langford, John Sillitoe on behalf of the Wellcome Sanger Institute COVID-19 Surveillance Team |
| EPI_ISL_2918388, EPI_ISL_2918389 | Lighthouse Lab in Alderley Park | Wellcome Sanger Institute for the COVID-19 Genomics UK (COG-UK) Consortium | Jacquelyn Wynn, Mairead Hyland, The Lighthouse Lab in Alderley Park and Alex Alderton, Roberto Amato, Jeffrey Barrett, Sonia Goncalves, Ewan Harrison, David K. Jackson, Ian Johnston, Dominic Kwiatkowski, Cordelia Langford, John Sillitoe on behalf of the Wellcome Sanger Institute COVID-19 Surveillance Team |
| EPI_ISL_2918390 | Lighthouse Lab in Milton Keynes | Wellcome Sanger Institute for the COVID-19 Genomics UK (COG-UK) Consortium | The Lighthouse Lab in Milton Keynes and Alex Alderton, Roberto Amato, Jeffrey Barrett, Sonia Goncalves, Ewan Harrison, David K. Jackson, Ian Johnston, Dominic Kwiatkowski, Cordelia Langford, John Sillitoe on behalf of the Wellcome Sanger Institute COVID-19 Surveillance Team |
| EPI_ISL_2918391, EPI_ISL_2918392 | Lighthouse Lab in Alderley Park | Wellcome Sanger Institute for the COVID-19 Genomics UK (COG-UK) Consortium | Jacquelyn Wynn, Mairead Hyland, The Lighthouse Lab in Alderley Park and Alex Alderton, Roberto Amato, Jeffrey Barrett, Sonia Goncalves, Ewan Harrison, David K. Jackson, Ian Johnston, Dominic Kwiatkowski, Cordelia Langford, John Sillitoe on behalf of the Wellcome Sanger Institute COVID-19 Surveillance Team |
| EPI_ISL_2918393 | Health Services Laboratories | Wellcome Sanger Institute for the COVID-19 Genomics UK (COG-UK) Consortium | Health Services Laboratories and Alex Alderton, Roberto Amato, Jeffrey Barrett, Sonia Goncalves, Ewan Harrison, David K. Jackson, Ian Johnston, Dominic Kwiatkowski, Cordelia Langford, John Sillitoe on behalf of the Wellcome Sanger Institute COVID-19 Surveillance Team |

| | | | |
|---|---|---|---|
| EPI_ISL_2918394 | Lighthouse Lab in Milton Keynes | Wellcome Sanger Institute for the COVID-19 Genomics UK (COG-UK) Consortium | The Lighthouse Lab in Milton Keynes and Alex Alderton, Roberto Amato, Jeffrey Barrett, Sonia Goncalves, Ewan Harrison, David K. Jackson, Ian Johnston, Dominic Kwiatkowski, Cordelia Langford, John Sillitoe on behalf of the Wellcome Sanger Institute COVID-19 Surveillance Team |
| EPI_ISL_2918395 | Health Services Laboratories | Wellcome Sanger Institute for the COVID-19 Genomics UK (COG-UK) Consortium | Health Services Laboratories and Alex Alderton, Roberto Amato, Jeffrey Barrett, Sonia Goncalves, Ewan Harrison, David K. Jackson, Ian Johnston, Dominic Kwiatkowski, Cordelia Langford, John Sillitoe on behalf of the Wellcome Sanger Institute COVID-19 Surveillance Team |
| EPI_ISL_2918396, EPI_ISL_2918397 | Lighthouse Lab in Glasgow | Wellcome Sanger Institute for the COVID-19 Genomics UK (COG-UK) Consortium | Harper VanSteenhouse, Yumi Kasai, David Gray, Carol Clugston, Anna Dominiczak and Alex Alderton, Roberto Amato, Jeffrey Barrett, Sonia Goncalves, Ewan Harrison, David K. Jackson, Ian Johnston, Dominic Kwiatkowski, Cordelia Langford, John Sillitoe on behalf of the Wellcome Sanger Institute COVID-19 Surveillance Team |
| EPI_ISL_2918398, EPI_ISL_2918399 | Berkshire and Surrey Pathology Services Lighthouse Laboratory | Wellcome Sanger Institute for the COVID-19 Genomics UK (COG-UK) Consortium | Berkshire and Surrey Pathology Services Lighthouse Laboratory and Alex Alderton, Roberto Amato, Jeffrey Barrett, Sonia Goncalves, Ewan Harrison, David K. Jackson, Ian Johnston, Dominic Kwiatkowski, Cordelia Langford, John Sillitoe on behalf of the Wellcome Sanger Institute COVID-19 Surveillance Team |
| EPI_ISL_2918400 | Lighthouse Lab in Glasgow | Wellcome Sanger Institute for the COVID-19 Genomics UK (COG-UK) Consortium | Harper VanSteenhouse, Yumi Kasai, David Gray, Carol Clugston, Anna Dominiczak and Alex Alderton, Roberto Amato, Jeffrey Barrett, Sonia Goncalves, Ewan Harrison, David K. Jackson, Ian Johnston, Dominic Kwiatkowski, Cordelia Langford, John Sillitoe on behalf of the Wellcome Sanger Institute COVID-19 Surveillance Team |
| EPI_ISL_2918401 | Berkshire and Surrey Pathology Services Lighthouse Laboratory | Wellcome Sanger Institute for the COVID-19 Genomics UK (COG-UK) Consortium | Berkshire and Surrey Pathology Services Lighthouse Laboratory and Alex Alderton, Roberto Amato, Jeffrey Barrett, Sonia Goncalves, Ewan Harrison, David K. Jackson, Ian Johnston, Dominic Kwiatkowski, Cordelia Langford, John Sillitoe on behalf of the Wellcome Sanger Institute COVID-19 Surveillance Team |
| EPI_ISL_2918402, EPI_ISL_2918403 | Lighthouse Lab in Glasgow | Wellcome Sanger Institute for the COVID-19 Genomics UK (COG-UK) Consortium | Harper VanSteenhouse, Yumi Kasai, David Gray, Carol Clugston, Anna Dominiczak and Alex Alderton, Roberto Amato, Jeffrey Barrett, Sonia Goncalves, Ewan Harrison, David K. Jackson, Ian Johnston, Dominic Kwiatkowski, Cordelia Langford, John Sillitoe on behalf of the Wellcome Sanger Institute COVID-19 Surveillance Team |
| EPI_ISL_2918404 | Berkshire and Surrey Pathology Services Lighthouse Laboratory | Wellcome Sanger Institute for the COVID-19 Genomics UK (COG-UK) Consortium | Berkshire and Surrey Pathology Services Lighthouse Laboratory and Alex Alderton, Roberto Amato, Jeffrey Barrett, Sonia Goncalves, Ewan Harrison, David K. Jackson, Ian Johnston, Dominic Kwiatkowski, Cordelia Langford, John Sillitoe on behalf of the Wellcome Sanger Institute COVID-19 Surveillance Team |
| EPI_ISL_2918405, EPI_ISL_2918406, EPI_ISL_2918407, EPI_ISL_2918408 | Lighthouse Lab in Glasgow | Wellcome Sanger Institute for the COVID-19 Genomics UK (COG-UK) Consortium | Harper VanSteenhouse, Yumi Kasai, David Gray, Carol Clugston, Anna Dominiczak and Alex Alderton, Roberto Amato, Jeffrey Barrett, Sonia Goncalves, Ewan Harrison, David K. Jackson, Ian Johnston, Dominic Kwiatkowski, Cordelia Langford, John Sillitoe on behalf of the Wellcome Sanger Institute COVID-19 Surveillance Team |
| EPI_ISL_2918409 | Berkshire and Surrey Pathology Services Lighthouse Laboratory | Wellcome Sanger Institute for the COVID-19 Genomics UK (COG-UK) Consortium | Berkshire and Surrey Pathology Services Lighthouse Laboratory and Alex Alderton, Roberto Amato, Jeffrey Barrett, Sonia Goncalves, Ewan Harrison, David K. Jackson, Ian Johnston, Dominic Kwiatkowski, Cordelia Langford, John Sillitoe on behalf of the Wellcome Sanger Institute COVID-19 Surveillance Team |
| EPI_ISL_2918410 | Randox Laboratories | Wellcome Sanger Institute for the COVID-19 Genomics UK (COG-UK) Consortium | Randox Laboratories and Alex Alderton, Roberto Amato, Jeffrey Barrett, Sonia Goncalves, Ewan Harrison, David K. Jackson, Ian Johnston, Dominic Kwiatkowski, Cordelia Langford, John Sillitoe on behalf of the Wellcome Sanger Institute COVID-19 Surveillance Team |
| EPI_ISL_2918411 | Berkshire and Surrey Pathology Services Lighthouse Laboratory | Wellcome Sanger Institute for the COVID-19 Genomics UK (COG-UK) Consortium | Berkshire and Surrey Pathology Services Lighthouse Laboratory and Alex Alderton, Roberto Amato, Jeffrey Barrett, Sonia Goncalves, Ewan Harrison, David K. Jackson, Ian Johnston, Dominic Kwiatkowski, Cordelia Langford, John Sillitoe on behalf of the Wellcome Sanger Institute COVID-19 Surveillance Team |
| EPI_ISL_2918412, EPI_ISL_2918413, EPI_ISL_2918414, EPI_ISL_2918415 | Lighthouse Lab in Glasgow | Wellcome Sanger Institute for the COVID-19 Genomics UK (COG-UK) Consortium | Harper VanSteenhouse, Yumi Kasai, David Gray, Carol Clugston, Anna Dominiczak and Alex Alderton, Roberto Amato, Jeffrey Barrett, Sonia Goncalves, Ewan Harrison, David K. Jackson, Ian Johnston, Dominic Kwiatkowski, Cordelia Langford, John Sillitoe on behalf of the Wellcome Sanger Institute COVID-19 Surveillance Team |
| EPI_ISL_2918416, EPI_ISL_2918417, EPI_ISL_2918418, EPI_ISL_2918419 | Berkshire and Surrey Pathology Services Lighthouse Laboratory | Wellcome Sanger Institute for the COVID-19 Genomics UK (COG-UK) Consortium | Berkshire and Surrey Pathology Services Lighthouse Laboratory and Alex Alderton, Roberto Amato, Jeffrey Barrett, Sonia Goncalves, Ewan Harrison, David K. Jackson, Ian Johnston, Dominic Kwiatkowski, Cordelia Langford, John Sillitoe on behalf of the Wellcome Sanger Institute COVID-19 Surveillance Team |
| EPI_ISL_2918420 | Randox Laboratories | Wellcome Sanger Institute for the COVID-19 Genomics UK (COG-UK) Consortium | Randox Laboratories and Alex Alderton, Roberto Amato, Jeffrey Barrett, Sonia Goncalves, Ewan Harrison, David K. Jackson, Ian Johnston, Dominic Kwiatkowski, Cordelia Langford, John Sillitoe on behalf of the Wellcome Sanger Institute COVID-19 Surveillance Team |
| EPI_ISL_2918421 | Berkshire and Surrey Pathology Services Lighthouse Laboratory | Wellcome Sanger Institute for the COVID-19 Genomics UK (COG-UK) Consortium | Berkshire and Surrey Pathology Services Lighthouse Laboratory and Alex Alderton, Roberto Amato, Jeffrey Barrett, Sonia Goncalves, Ewan Harrison, David K. Jackson, Ian Johnston, Dominic Kwiatkowski, Cordelia Langford, John Sillitoe on behalf of the Wellcome Sanger Institute COVID-19 Surveillance Team |
| EPI_ISL_2918422 | Randox Laboratories | Wellcome Sanger Institute for the COVID-19 Genomics UK (COG-UK) Consortium | Randox Laboratories and Alex Alderton, Roberto Amato, Jeffrey Barrett, Sonia Goncalves, Ewan Harrison, David K. Jackson, Ian Johnston, Dominic Kwiatkowski, Cordelia Langford, John Sillitoe on behalf of the Wellcome Sanger Institute COVID-19 Surveillance Team |
| EPI_ISL_2918423, EPI_ISL_2918424 | Lighthouse Lab in Glasgow | Wellcome Sanger Institute for the COVID-19 Genomics UK (COG-UK) Consortium | Harper VanSteenhouse, Yumi Kasai, David Gray, Carol Clugston, Anna Dominiczak and Alex Alderton, Roberto Amato, Jeffrey Barrett, Sonia Goncalves, Ewan Harrison, David K. Jackson, Ian Johnston, Dominic Kwiatkowski, Cordelia Langford, John Sillitoe on behalf of the Wellcome Sanger Institute COVID-19 Surveillance Team |
| EPI_ISL_2918425 | Berkshire and Surrey Pathology Services Lighthouse Laboratory | Wellcome Sanger Institute for the COVID-19 Genomics UK (COG-UK) Consortium | Berkshire and Surrey Pathology Services Lighthouse Laboratory and Alex Alderton, Roberto Amato, Jeffrey Barrett, Sonia Goncalves, Ewan Harrison, David K. Jackson, Ian Johnston, Dominic Kwiatkowski, Cordelia Langford, John Sillitoe on behalf of the Wellcome Sanger Institute COVID-19 Surveillance Team |
| EPI_ISL_2918426 | Randox Laboratories | Wellcome Sanger Institute for the COVID-19 Genomics UK (COG-UK) Consortium | Randox Laboratories and Alex Alderton, Roberto Amato, Jeffrey Barrett, Sonia Goncalves, Ewan Harrison, David K. Jackson, Ian Johnston, Dominic Kwiatkowski, Cordelia Langford, John Sillitoe on behalf of the Wellcome Sanger Institute COVID-19 Surveillance Team |
| EPI_ISL_2918427 | Lighthouse Lab in Glasgow | Wellcome Sanger Institute for the COVID-19 Genomics UK (COG-UK) Consortium | Harper VanSteenhouse, Yumi Kasai, David Gray, Carol Clugston, Anna Dominiczak and Alex Alderton, Roberto Amato, Jeffrey Barrett, Sonia Goncalves, Ewan Harrison, David K. Jackson, Ian Johnston, Dominic Kwiatkowski, Cordelia Langford, John Sillitoe on behalf of the Wellcome Sanger Institute COVID-19 Surveillance Team |
| EPI_ISL_2918428 | Randox Laboratories | Wellcome Sanger Institute for the COVID-19 Genomics UK (COG-UK) Consortium | Randox Laboratories and Alex Alderton, Roberto Amato, Jeffrey Barrett, Sonia Goncalves, Ewan Harrison, David K. Jackson, Ian Johnston, Dominic Kwiatkowski, Cordelia Langford, John Sillitoe on behalf of the Wellcome Sanger Institute COVID-19 Surveillance Team |
| EPI_ISL_2918429 | Berkshire and Surrey Pathology Services Lighthouse Laboratory | Wellcome Sanger Institute for the COVID-19 Genomics UK (COG-UK) Consortium | Berkshire and Surrey Pathology Services Lighthouse Laboratory and Alex Alderton, Roberto Amato, Jeffrey Barrett, Sonia Goncalves, Ewan Harrison, David K. Jackson, Ian Johnston, Dominic Kwiatkowski, Cordelia Langford, John Sillitoe on behalf of the Wellcome Sanger Institute COVID-19 Surveillance Team |
| EPI_ISL_2918430, EPI_ISL_2918431 | Lighthouse Lab in Glasgow | Wellcome Sanger Institute for the COVID-19 Genomics UK (COG-UK) Consortium | Harper VanSteenhouse, Yumi Kasai, David Gray, Carol Clugston, Anna Dominiczak and Alex Alderton, Roberto Amato, Jeffrey Barrett, Sonia Goncalves, Ewan Harrison, David K. Jackson, Ian Johnston, Dominic Kwiatkowski, Cordelia Langford, John Sillitoe on behalf of the Wellcome Sanger Institute COVID-19 Surveillance Team |
| EPI_ISL_2918432, EPI_ISL_2918433, EPI_ISL_2918434 | Berkshire and Surrey Pathology Services Lighthouse Laboratory | Wellcome Sanger Institute for the COVID-19 Genomics UK (COG-UK) Consortium | Berkshire and Surrey Pathology Services Lighthouse Laboratory and Alex Alderton, Roberto Amato, Jeffrey Barrett, Sonia Goncalves, Ewan Harrison, David K. Jackson, Ian Johnston, Dominic Kwiatkowski, Cordelia Langford, John Sillitoe on behalf of the Wellcome Sanger Institute COVID-19 Surveillance Team |
| EPI_ISL_2918435 | Randox Laboratories | Wellcome Sanger Institute for the COVID-19 Genomics UK (COG-UK) Consortium | Randox Laboratories and Alex Alderton, Roberto Amato, Jeffrey Barrett, Sonia Goncalves, Ewan Harrison, David K. Jackson, Ian Johnston, Dominic Kwiatkowski, Cordelia Langford, John Sillitoe on behalf of the Wellcome Sanger Institute COVID-19 Surveillance Team |
| EPI_ISL_2918436, EPI_ISL_2918437, EPI_ISL_2918438 | Lighthouse Lab in Glasgow | Wellcome Sanger Institute for the COVID-19 Genomics UK (COG-UK) Consortium | Harper VanSteenhouse, Yumi Kasai, David Gray, Carol Clugston, Anna Dominiczak and Alex Alderton, Roberto Amato, Jeffrey Barrett, Sonia Goncalves, Ewan Harrison, David K. Jackson, Ian Johnston, Dominic Kwiatkowski, Cordelia Langford, John Sillitoe on behalf of the Wellcome Sanger Institute COVID-19 Surveillance Team |
| EPI_ISL_2918439 | Berkshire and Surrey Pathology Services Lighthouse Laboratory | Wellcome Sanger Institute for the COVID-19 Genomics UK (COG-UK) Consortium | Berkshire and Surrey Pathology Services Lighthouse Laboratory and Alex Alderton, Roberto Amato, Jeffrey Barrett, Sonia Goncalves, Ewan Harrison, David K. Jackson, Ian Johnston, Dominic Kwiatkowski, Cordelia Langford, John Sillitoe on behalf of the Wellcome Sanger Institute COVID-19 Surveillance Team |
| EPI_ISL_2918440 | Lighthouse Lab in Glasgow | Wellcome Sanger Institute for the COVID-19 Genomics UK (COG-UK) Consortium | Harper VanSteenhouse, Yumi Kasai, David Gray, Carol Clugston, Anna Dominiczak and Alex Alderton, Roberto Amato, Jeffrey Barrett, Sonia Goncalves, Ewan Harrison, David K. Jackson, Ian Johnston, Dominic Kwiatkowski, Cordelia Langford, John Sillitoe on behalf of the Wellcome Sanger Institute COVID-19 Surveillance Team |
| EPI_ISL_2918441 | Berkshire and Surrey Pathology Services Lighthouse Laboratory | Wellcome Sanger Institute for the COVID-19 Genomics UK (COG-UK) Consortium | Berkshire and Surrey Pathology Services Lighthouse Laboratory and Alex Alderton, Roberto Amato, Jeffrey Barrett, Sonia Goncalves, Ewan Harrison, David K. Jackson, Ian Johnston, Dominic Kwiatkowski, Cordelia Langford, John Sillitoe on behalf of the Wellcome Sanger Institute COVID-19 Surveillance Team |
| EPI_ISL_2918442, EPI_ISL_2918443, EPI_ISL_2918444 | Lighthouse Lab in Glasgow | Wellcome Sanger Institute for the COVID-19 Genomics UK (COG-UK) Consortium | Harper VanSteenhouse, Yumi Kasai, David Gray, Carol Clugston, Anna Dominiczak and Alex Alderton, Roberto Amato, Jeffrey Barrett, Sonia Goncalves, Ewan Harrison, David K. Jackson, Ian Johnston, Dominic Kwiatkowski, Cordelia Langford, John Sillitoe on behalf of the Wellcome Sanger Institute COVID-19 Surveillance Team |
| EPI_ISL_2918445, EPI_ISL_2918446, EPI_ISL_2918447, EPI_ISL_2918448 | Berkshire and Surrey Pathology Services Lighthouse Laboratory | Wellcome Sanger Institute for the COVID-19 Genomics UK (COG-UK) Consortium | Berkshire and Surrey Pathology Services Lighthouse Laboratory and Alex Alderton, Roberto Amato, Jeffrey Barrett, Sonia Goncalves, Ewan Harrison, David K. Jackson, Ian Johnston, Dominic Kwiatkowski, Cordelia Langford, John Sillitoe on behalf of the Wellcome Sanger Institute COVID-19 Surveillance Team |
| EPI_ISL_2918449 | Lighthouse Lab in Glasgow | Wellcome Sanger Institute for the COVID-19 Genomics UK (COG-UK) Consortium | Harper VanSteenhouse, Yumi Kasai, David Gray, Carol Clugston, Anna Dominiczak and Alex Alderton, Roberto Amato, Jeffrey Barrett, Sonia Goncalves, Ewan Harrison, David K. Jackson, Ian Johnston, Dominic Kwiatkowski, Cordelia Langford, John Sillitoe on behalf of the Wellcome Sanger Institute COVID-19 Surveillance Team |
| EPI_ISL_2918450 | Berkshire and Surrey Pathology Services Lighthouse Laboratory | Wellcome Sanger Institute for the COVID-19 Genomics UK (COG-UK) Consortium | Berkshire and Surrey Pathology Services Lighthouse Laboratory and Alex Alderton, Roberto Amato, Jeffrey Barrett, Sonia Goncalves, Ewan Harrison, David K. Jackson, Ian Johnston, Dominic Kwiatkowski, Cordelia Langford, John Sillitoe on behalf of the Wellcome Sanger Institute COVID-19 Surveillance Team |
| EPI_ISL_2918451, EPI_ISL_2918452, EPI_ISL_2918453 | Lighthouse Lab in Glasgow | Wellcome Sanger Institute for the COVID-19 Genomics UK (COG-UK) Consortium | Harper VanSteenhouse, Yumi Kasai, David Gray, Carol Clugston, Anna Dominiczak and Alex Alderton, Roberto Amato, Jeffrey Barrett, Sonia Goncalves, Ewan Harrison, David K. Jackson, Ian Johnston, Dominic Kwiatkowski, Cordelia Langford, John Sillitoe on behalf of the Wellcome Sanger Institute COVID-19 Surveillance Team |
| EPI_ISL_2918454 | Berkshire and Surrey Pathology Services Lighthouse Laboratory | Wellcome Sanger Institute for the COVID-19 Genomics UK (COG-UK) Consortium | Berkshire and Surrey Pathology Services Lighthouse Laboratory and Alex Alderton, Roberto Amato, Jeffrey Barrett, Sonia Goncalves, Ewan Harrison, David K. Jackson, Ian Johnston, Dominic Kwiatkowski, Cordelia Langford, John Sillitoe on behalf of the Wellcome Sanger Institute COVID-19 Surveillance Team |
| EPI_ISL_2918455 | Lighthouse Lab in Glasgow | Wellcome Sanger Institute for the COVID-19 Genomics UK (COG-UK) Consortium | Harper VanSteenhouse, Yumi Kasai, David Gray, Carol Clugston, Anna Dominiczak and Alex Alderton, Roberto Amato, Jeffrey Barrett, Sonia Goncalves, Ewan Harrison, David K. Jackson, Ian Johnston, Dominic Kwiatkowski, Cordelia Langford, John Sillitoe on behalf of the Wellcome Sanger Institute COVID-19 Surveillance Team |
| EPI_ISL_2918456 | Berkshire and Surrey Pathology Services Lighthouse Laboratory | Wellcome Sanger Institute for the COVID-19 Genomics UK (COG-UK) Consortium | Berkshire and Surrey Pathology Services Lighthouse Laboratory and Alex Alderton, Roberto Amato, Jeffrey Barrett, Sonia Goncalves, Ewan Harrison, David K. Jackson, Ian Johnston, Dominic Kwiatkowski, Cordelia Langford, John Sillitoe on behalf of the Wellcome Sanger Institute COVID-19 Surveillance Team |
| EPI_ISL_2918457, EPI_ISL_2918458 | Randox Laboratories | Wellcome Sanger Institute for the COVID-19 Genomics UK (COG-UK) Consortium | Randox Laboratories and Alex Alderton, Roberto Amato, Jeffrey Barrett, Sonia Goncalves, Ewan Harrison, David K. Jackson, Ian Johnston, Dominic Kwiatkowski, Cordelia Langford, John Sillitoe on behalf of the Wellcome Sanger Institute COVID-19 Surveillance Team |
| EPI_ISL_2918459 | Berkshire and Surrey Pathology Services Lighthouse Laboratory | Wellcome Sanger Institute for the COVID-19 Genomics UK (COG-UK) Consortium | Berkshire and Surrey Pathology Services Lighthouse Laboratory and Alex Alderton, Roberto Amato, Jeffrey Barrett, Sonia Goncalves, Ewan Harrison, David K. Jackson, Ian Johnston, Dominic Kwiatkowski, Cordelia Langford, John Sillitoe on behalf of the Wellcome Sanger Institute COVID-19 Surveillance Team |

| | | | |
|---|---|---|---|
| EPI_ISL_2918460 | Lighthouse Lab in Glasgow | Wellcome Sanger Institute for the COVID-19 Genomics UK (COG-UK) Consortium | Harper VanSteenhouse, Yumi Kasai, David Gray, Carol Clugston, Anna Dominiczak and Alex Alderton, Roberto Amato, Jeffrey Barrett, Sonia Goncalves, Ewan Harrison, David K. Jackson, Ian Johnston, Dominic Kwiatkowski, Cordelia Langford, John Sillitoe on behalf of the Wellcome Sanger Institute COVID-19 Surveillance Team |
| EPI_ISL_2918461, EPI_ISL_2918462, EPI_ISL_2918463, EPI_ISL_2918464 | Berkshire and Surrey Pathology Services Lighthouse Laboratory | Wellcome Sanger Institute for the COVID-19 Genomics UK (COG-UK) Consortium | Berkshire and Surrey Pathology Services Lighthouse Laboratory and Alex Alderton, Roberto Amato, Jeffrey Barrett, Sonia Goncalves, Ewan Harrison, David K. Jackson, Ian Johnston, Dominic Kwiatkowski, Cordelia Langford, John Sillitoe on behalf of the Wellcome Sanger Institute COVID-19 Surveillance Team |
| EPI_ISL_2918465 | Lighthouse Lab in Glasgow | Wellcome Sanger Institute for the COVID-19 Genomics UK (COG-UK) Consortium | Harper VanSteenhouse, Yumi Kasai, David Gray, Carol Clugston, Anna Dominiczak and Alex Alderton, Roberto Amato, Jeffrey Barrett, Sonia Goncalves, Ewan Harrison, David K. Jackson, Ian Johnston, Dominic Kwiatkowski, Cordelia Langford, John Sillitoe on behalf of the Wellcome Sanger Institute COVID-19 Surveillance Team |
| EPI_ISL_2918466 | Berkshire and Surrey Pathology Services Lighthouse Laboratory | Wellcome Sanger Institute for the COVID-19 Genomics UK (COG-UK) Consortium | Berkshire and Surrey Pathology Services Lighthouse Laboratory and Alex Alderton, Roberto Amato, Jeffrey Barrett, Sonia Goncalves, Ewan Harrison, David K. Jackson, Ian Johnston, Dominic Kwiatkowski, Cordelia Langford, John Sillitoe on behalf of the Wellcome Sanger Institute COVID-19 Surveillance Team |
| EPI_ISL_2918467 | Lighthouse Lab in Glasgow | Wellcome Sanger Institute for the COVID-19 Genomics UK (COG-UK) Consortium | Harper VanSteenhouse, Yumi Kasai, David Gray, Carol Clugston, Anna Dominiczak and Alex Alderton, Roberto Amato, Jeffrey Barrett, Sonia Goncalves, Ewan Harrison, David K. Jackson, Ian Johnston, Dominic Kwiatkowski, Cordelia Langford, John Sillitoe on behalf of the Wellcome Sanger Institute COVID-19 Surveillance Team |
| EPI_ISL_2918468 | Randox Laboratories | Wellcome Sanger Institute for the COVID-19 Genomics UK (COG-UK) Consortium | Randox Laboratories and Alex Alderton, Roberto Amato, Jeffrey Barrett, Sonia Goncalves, Ewan Harrison, David K. Jackson, Ian Johnston, Dominic Kwiatkowski, Cordelia Langford, John Sillitoe on behalf of the Wellcome Sanger Institute COVID-19 Surveillance Team |
| EPI_ISL_2918469, EPI_ISL_2918470 | Lighthouse Lab in Glasgow | Wellcome Sanger Institute for the COVID-19 Genomics UK (COG-UK) Consortium | Harper VanSteenhouse, Yumi Kasai, David Gray, Carol Clugston, Anna Dominiczak and Alex Alderton, Roberto Amato, Jeffrey Barrett, Sonia Goncalves, Ewan Harrison, David K. Jackson, Ian Johnston, Dominic Kwiatkowski, Cordelia Langford, John Sillitoe on behalf of the Wellcome Sanger Institute COVID-19 Surveillance Team |
| EPI_ISL_2918471 | Berkshire and Surrey Pathology Services Lighthouse Laboratory | Wellcome Sanger Institute for the COVID-19 Genomics UK (COG-UK) Consortium | Berkshire and Surrey Pathology Services Lighthouse Laboratory and Alex Alderton, Roberto Amato, Jeffrey Barrett, Sonia Goncalves, Ewan Harrison, David K. Jackson, Ian Johnston, Dominic Kwiatkowski, Cordelia Langford, John Sillitoe on behalf of the Wellcome Sanger Institute COVID-19 Surveillance Team |
| EPI_ISL_2918472 | Lighthouse Lab in Glasgow | Wellcome Sanger Institute for the COVID-19 Genomics UK (COG-UK) Consortium | Harper VanSteenhouse, Yumi Kasai, David Gray, Carol Clugston, Anna Dominiczak and Alex Alderton, Roberto Amato, Jeffrey Barrett, Sonia Goncalves, Ewan Harrison, David K. Jackson, Ian Johnston, Dominic Kwiatkowski, Cordelia Langford, John Sillitoe on behalf of the Wellcome Sanger Institute COVID-19 Surveillance Team |
| EPI_ISL_2918473 | Randox Laboratories | Wellcome Sanger Institute for the COVID-19 Genomics UK (COG-UK) Consortium | Randox Laboratories and Alex Alderton, Roberto Amato, Jeffrey Barrett, Sonia Goncalves, Ewan Harrison, David K. Jackson, Ian Johnston, Dominic Kwiatkowski, Cordelia Langford, John Sillitoe on behalf of the Wellcome Sanger Institute COVID-19 Surveillance Team |
| EPI_ISL_2918474 | Lighthouse Lab in Glasgow | Wellcome Sanger Institute for the COVID-19 Genomics UK (COG-UK) Consortium | Harper VanSteenhouse, Yumi Kasai, David Gray, Carol Clugston, Anna Dominiczak and Alex Alderton, Roberto Amato, Jeffrey Barrett, Sonia Goncalves, Ewan Harrison, David K. Jackson, Ian Johnston, Dominic Kwiatkowski, Cordelia Langford, John Sillitoe on behalf of the Wellcome Sanger Institute COVID-19 Surveillance Team |
| EPI_ISL_2918475, EPI_ISL_2918476 | Berkshire and Surrey Pathology Services Lighthouse Laboratory | Wellcome Sanger Institute for the COVID-19 Genomics UK (COG-UK) Consortium | Berkshire and Surrey Pathology Services Lighthouse Laboratory and Alex Alderton, Roberto Amato, Jeffrey Barrett, Sonia Goncalves, Ewan Harrison, David K. Jackson, Ian Johnston, Dominic Kwiatkowski, Cordelia Langford, John Sillitoe on behalf of the Wellcome Sanger Institute COVID-19 Surveillance Team |
| EPI_ISL_2918477 | Lighthouse Lab in Glasgow | Wellcome Sanger Institute for the COVID-19 Genomics UK (COG-UK) Consortium | Harper VanSteenhouse, Yumi Kasai, David Gray, Carol Clugston, Anna Dominiczak and Alex Alderton, Roberto Amato, Jeffrey Barrett, Sonia Goncalves, Ewan Harrison, David K. Jackson, Ian Johnston, Dominic Kwiatkowski, Cordelia Langford, John Sillitoe on behalf of the Wellcome Sanger Institute COVID-19 Surveillance Team |
| EPI_ISL_2918478 | Randox Laboratories | Wellcome Sanger Institute for the COVID-19 Genomics UK (COG-UK) Consortium | Randox Laboratories and Alex Alderton, Roberto Amato, Jeffrey Barrett, Sonia Goncalves, Ewan Harrison, David K. Jackson, Ian Johnston, Dominic Kwiatkowski, Cordelia Langford, John Sillitoe on behalf of the Wellcome Sanger Institute COVID-19 Surveillance Team |
| EPI_ISL_2918479, EPI_ISL_2918480, EPI_ISL_2918481 | Lighthouse Lab in Glasgow | Wellcome Sanger Institute for the COVID-19 Genomics UK (COG-UK) Consortium | Harper VanSteenhouse, Yumi Kasai, David Gray, Carol Clugston, Anna Dominiczak and Alex Alderton, Roberto Amato, Jeffrey Barrett, Sonia Goncalves, Ewan Harrison, David K. Jackson, Ian Johnston, Dominic Kwiatkowski, Cordelia Langford, John Sillitoe on behalf of the Wellcome Sanger Institute COVID-19 Surveillance Team |
| EPI_ISL_2918482, EPI_ISL_2918483, EPI_ISL_2918484, EPI_ISL_2918485 | Berkshire and Surrey Pathology Services Lighthouse Laboratory | Wellcome Sanger Institute for the COVID-19 Genomics UK (COG-UK) Consortium | Berkshire and Surrey Pathology Services Lighthouse Laboratory and Alex Alderton, Roberto Amato, Jeffrey Barrett, Sonia Goncalves, Ewan Harrison, David K. Jackson, Ian Johnston, Dominic Kwiatkowski, Cordelia Langford, John Sillitoe on behalf of the Wellcome Sanger Institute COVID-19 Surveillance Team |
| EPI_ISL_2918486 | Lighthouse Lab in Glasgow | Wellcome Sanger Institute for the COVID-19 Genomics UK (COG-UK) Consortium | Harper VanSteenhouse, Yumi Kasai, David Gray, Carol Clugston, Anna Dominiczak and Alex Alderton, Roberto Amato, Jeffrey Barrett, Sonia Goncalves, Ewan Harrison, David K. Jackson, Ian Johnston, Dominic Kwiatkowski, Cordelia Langford, John Sillitoe on behalf of the Wellcome Sanger Institute COVID-19 Surveillance Team |
| EPI_ISL_2918487 | Randox Laboratories | Wellcome Sanger Institute for the COVID-19 Genomics UK (COG-UK) Consortium | Randox Laboratories and Alex Alderton, Roberto Amato, Jeffrey Barrett, Sonia Goncalves, Ewan Harrison, David K. Jackson, Ian Johnston, Dominic Kwiatkowski, Cordelia Langford, John Sillitoe on behalf of the Wellcome Sanger Institute COVID-19 Surveillance Team |
| EPI_ISL_2918488, EPI_ISL_2918489, EPI_ISL_2918490 | Lighthouse Lab in Glasgow | Wellcome Sanger Institute for the COVID-19 Genomics UK (COG-UK) Consortium | Harper VanSteenhouse, Yumi Kasai, David Gray, Carol Clugston, Anna Dominiczak and Alex Alderton, Roberto Amato, Jeffrey Barrett, Sonia Goncalves, Ewan Harrison, David K. Jackson, Ian Johnston, Dominic Kwiatkowski, Cordelia Langford, John Sillitoe on behalf of the Wellcome Sanger Institute COVID-19 Surveillance Team |
| EPI_ISL_2918491, EPI_ISL_2918492 | Berkshire and Surrey Pathology Services Lighthouse Laboratory | Wellcome Sanger Institute for the COVID-19 Genomics UK (COG-UK) Consortium | Berkshire and Surrey Pathology Services Lighthouse Laboratory and Alex Alderton, Roberto Amato, Jeffrey Barrett, Sonia Goncalves, Ewan Harrison, David K. Jackson, Ian Johnston, Dominic Kwiatkowski, Cordelia Langford, John Sillitoe on behalf of the Wellcome Sanger Institute COVID-19 Surveillance Team |
| EPI_ISL_2918493, EPI_ISL_2918494 | Lighthouse Lab in Glasgow | Wellcome Sanger Institute for the COVID-19 Genomics UK (COG-UK) Consortium | Harper VanSteenhouse, Yumi Kasai, David Gray, Carol Clugston, Anna Dominiczak and Alex Alderton, Roberto Amato, Jeffrey Barrett, Sonia Goncalves, Ewan Harrison, David K. Jackson, Ian Johnston, Dominic Kwiatkowski, Cordelia Langford, John Sillitoe on behalf of the Wellcome Sanger Institute COVID-19 Surveillance Team |
| EPI_ISL_2918495 | Randox Laboratories | Wellcome Sanger Institute for the COVID-19 Genomics UK (COG-UK) Consortium | Randox Laboratories and Alex Alderton, Roberto Amato, Jeffrey Barrett, Sonia Goncalves, Ewan Harrison, David K. Jackson, Ian Johnston, Dominic Kwiatkowski, Cordelia Langford, John Sillitoe on behalf of the Wellcome Sanger Institute COVID-19 Surveillance Team |
| EPI_ISL_2918496 | Berkshire and Surrey Pathology Services Lighthouse Laboratory | Wellcome Sanger Institute for the COVID-19 Genomics UK (COG-UK) Consortium | Berkshire and Surrey Pathology Services Lighthouse Laboratory and Alex Alderton, Roberto Amato, Jeffrey Barrett, Sonia Goncalves, Ewan Harrison, David K. Jackson, Ian Johnston, Dominic Kwiatkowski, Cordelia Langford, John Sillitoe on behalf of the Wellcome Sanger Institute COVID-19 Surveillance Team |
| EPI_ISL_2918497 | Randox Laboratories | Wellcome Sanger Institute for the COVID-19 Genomics UK (COG-UK) Consortium | Randox Laboratories and Alex Alderton, Roberto Amato, Jeffrey Barrett, Sonia Goncalves, Ewan Harrison, David K. Jackson, Ian Johnston, Dominic Kwiatkowski, Cordelia Langford, John Sillitoe on behalf of the Wellcome Sanger Institute COVID-19 Surveillance Team |
| EPI_ISL_2918498, EPI_ISL_2918499, EPI_ISL_2918500 | Berkshire and Surrey Pathology Services Lighthouse Laboratory | Wellcome Sanger Institute for the COVID-19 Genomics UK (COG-UK) Consortium | Berkshire and Surrey Pathology Services Lighthouse Laboratory and Alex Alderton, Roberto Amato, Jeffrey Barrett, Sonia Goncalves, Ewan Harrison, David K. Jackson, Ian Johnston, Dominic Kwiatkowski, Cordelia Langford, John Sillitoe on behalf of the Wellcome Sanger Institute COVID-19 Surveillance Team |
| EPI_ISL_2918501 | Lighthouse Lab in Glasgow | Wellcome Sanger Institute for the COVID-19 Genomics UK (COG-UK) Consortium | Harper VanSteenhouse, Yumi Kasai, David Gray, Carol Clugston, Anna Dominiczak and Alex Alderton, Roberto Amato, Jeffrey Barrett, Sonia Goncalves, Ewan Harrison, David K. Jackson, Ian Johnston, Dominic Kwiatkowski, Cordelia Langford, John Sillitoe on behalf of the Wellcome Sanger Institute COVID-19 Surveillance Team |
| EPI_ISL_2918502 | Berkshire and Surrey Pathology Services Lighthouse Laboratory | Wellcome Sanger Institute for the COVID-19 Genomics UK (COG-UK) Consortium | Berkshire and Surrey Pathology Services Lighthouse Laboratory and Alex Alderton, Roberto Amato, Jeffrey Barrett, Sonia Goncalves, Ewan Harrison, David K. Jackson, Ian Johnston, Dominic Kwiatkowski, Cordelia Langford, John Sillitoe on behalf of the Wellcome Sanger Institute COVID-19 Surveillance Team |
| EPI_ISL_2918503, EPI_ISL_2918504, EPI_ISL_2918505 | Lighthouse Lab in Glasgow | Wellcome Sanger Institute for the COVID-19 Genomics UK (COG-UK) Consortium | Harper VanSteenhouse, Yumi Kasai, David Gray, Carol Clugston, Anna Dominiczak and Alex Alderton, Roberto Amato, Jeffrey Barrett, Sonia Goncalves, Ewan Harrison, David K. Jackson, Ian Johnston, Dominic Kwiatkowski, Cordelia Langford, John Sillitoe on behalf of the Wellcome Sanger Institute COVID-19 Surveillance Team |
| EPI_ISL_2918506 | Berkshire and Surrey Pathology Services Lighthouse Laboratory | Wellcome Sanger Institute for the COVID-19 Genomics UK (COG-UK) Consortium | Berkshire and Surrey Pathology Services Lighthouse Laboratory and Alex Alderton, Roberto Amato, Jeffrey Barrett, Sonia Goncalves, Ewan Harrison, David K. Jackson, Ian Johnston, Dominic Kwiatkowski, Cordelia Langford, John Sillitoe on behalf of the Wellcome Sanger Institute COVID-19 Surveillance Team |
| EPI_ISL_2918507 | Lighthouse Lab in Glasgow | Wellcome Sanger Institute for the COVID-19 Genomics UK (COG-UK) Consortium | Harper VanSteenhouse, Yumi Kasai, David Gray, Carol Clugston, Anna Dominiczak and Alex Alderton, Roberto Amato, Jeffrey Barrett, Sonia Goncalves, Ewan Harrison, David K. Jackson, Ian Johnston, Dominic Kwiatkowski, Cordelia Langford, John Sillitoe on behalf of the Wellcome Sanger Institute COVID-19 Surveillance Team |
| EPI_ISL_2918508, EPI_ISL_2918509 | Berkshire and Surrey Pathology Services Lighthouse Laboratory | Wellcome Sanger Institute for the COVID-19 Genomics UK (COG-UK) Consortium | Berkshire and Surrey Pathology Services Lighthouse Laboratory and Alex Alderton, Roberto Amato, Jeffrey Barrett, Sonia Goncalves, Ewan Harrison, David K. Jackson, Ian Johnston, Dominic Kwiatkowski, Cordelia Langford, John Sillitoe on behalf of the Wellcome Sanger Institute COVID-19 Surveillance Team |
| EPI_ISL_2918510, EPI_ISL_2918511 | Lighthouse Lab in Glasgow | Wellcome Sanger Institute for the COVID-19 Genomics UK (COG-UK) Consortium | Harper VanSteenhouse, Yumi Kasai, David Gray, Carol Clugston, Anna Dominiczak and Alex Alderton, Roberto Amato, Jeffrey Barrett, Sonia Goncalves, Ewan Harrison, David K. Jackson, Ian Johnston, Dominic Kwiatkowski, Cordelia Langford, John Sillitoe on behalf of the Wellcome Sanger Institute COVID-19 Surveillance Team |
| EPI_ISL_2918512 | Randox Laboratories | Wellcome Sanger Institute for the COVID-19 Genomics UK (COG-UK) Consortium | Randox Laboratories and Alex Alderton, Roberto Amato, Jeffrey Barrett, Sonia Goncalves, Ewan Harrison, David K. Jackson, Ian Johnston, Dominic Kwiatkowski, Cordelia Langford, John Sillitoe on behalf of the Wellcome Sanger Institute COVID-19 Surveillance Team |
| EPI_ISL_2918513 | Lighthouse Lab in Glasgow | Wellcome Sanger Institute for the COVID-19 Genomics UK (COG-UK) Consortium | Harper VanSteenhouse, Yumi Kasai, David Gray, Carol Clugston, Anna Dominiczak and Alex Alderton, Roberto Amato, Jeffrey Barrett, Sonia Goncalves, Ewan Harrison, David K. Jackson, Ian Johnston, Dominic Kwiatkowski, Cordelia Langford, John Sillitoe on behalf of the Wellcome Sanger Institute COVID-19 Surveillance Team |
| EPI_ISL_2918514 | Berkshire and Surrey Pathology Services Lighthouse Laboratory | Wellcome Sanger Institute for the COVID-19 Genomics UK (COG-UK) Consortium | Berkshire and Surrey Pathology Services Lighthouse Laboratory and Alex Alderton, Roberto Amato, Jeffrey Barrett, Sonia Goncalves, Ewan Harrison, David K. Jackson, Ian Johnston, Dominic Kwiatkowski, Cordelia Langford, John Sillitoe on behalf of the Wellcome Sanger Institute COVID-19 Surveillance Team |
| EPI_ISL_2918515 | Lighthouse Lab in Glasgow | Wellcome Sanger Institute for the COVID-19 Genomics UK (COG-UK) Consortium | Harper VanSteenhouse, Yumi Kasai, David Gray, Carol Clugston, Anna Dominiczak and Alex Alderton, Roberto Amato, Jeffrey Barrett, Sonia Goncalves, Ewan Harrison, David K. Jackson, Ian Johnston, Dominic Kwiatkowski, Cordelia Langford, John Sillitoe on behalf of the Wellcome Sanger Institute COVID-19 Surveillance Team |
| EPI_ISL_2918516 | Berkshire and Surrey Pathology Services Lighthouse Laboratory | Wellcome Sanger Institute for the COVID-19 Genomics UK (COG-UK) Consortium | Berkshire and Surrey Pathology Services Lighthouse Laboratory and Alex Alderton, Roberto Amato, Jeffrey Barrett, Sonia Goncalves, Ewan Harrison, David K. Jackson, Ian Johnston, Dominic Kwiatkowski, Cordelia Langford, John Sillitoe on behalf of the Wellcome Sanger Institute COVID-19 Surveillance Team |
| EPI_ISL_2918517 | Lighthouse Lab in Glasgow | Wellcome Sanger Institute for the COVID-19 Genomics UK (COG-UK) Consortium | Harper VanSteenhouse, Yumi Kasai, David Gray, Carol Clugston, Anna Dominiczak and Alex Alderton, Roberto Amato, Jeffrey Barrett, Sonia Goncalves, Ewan Harrison, David K. Jackson, Ian Johnston, Dominic Kwiatkowski, Cordelia Langford, John Sillitoe on behalf of the Wellcome Sanger Institute COVID-19 Surveillance Team |
| EPI_ISL_2918518 | Randox Laboratories | Wellcome Sanger Institute for the COVID-19 Genomics UK (COG-UK) Consortium | Randox Laboratories and Alex Alderton, Roberto Amato, Jeffrey Barrett, Sonia Goncalves, Ewan Harrison, David K. Jackson, Ian Johnston, Dominic Kwiatkowski, Cordelia Langford, John Sillitoe on behalf of the Wellcome Sanger Institute COVID-19 Surveillance Team |
| EPI_ISL_2918519, EPI_ISL_2918520 | Berkshire and Surrey Pathology Services Lighthouse Laboratory | Wellcome Sanger Institute for the COVID-19 Genomics UK (COG-UK) Consortium | Berkshire and Surrey Pathology Services Lighthouse Laboratory and Alex Alderton, Roberto Amato, Jeffrey Barrett, Sonia Goncalves, Ewan Harrison, David K. Jackson, Ian Johnston, Dominic Kwiatkowski, Cordelia Langford, John Sillitoe on behalf of the Wellcome Sanger Institute COVID-19 Surveillance Team |
| EPI_ISL_2918521 | Randox Laboratories | Wellcome Sanger Institute for the COVID-19 Genomics UK (COG-UK) Consortium | Randox Laboratories and Alex Alderton, Roberto Amato, Jeffrey Barrett, Sonia Goncalves, Ewan Harrison, David K. Jackson, Ian Johnston, Dominic Kwiatkowski, Cordelia Langford, John Sillitoe on behalf of the Wellcome Sanger Institute COVID-19 Surveillance Team |
| EPI_ISL_2918522 | Berkshire and Surrey Pathology Services Lighthouse Laboratory | Wellcome Sanger Institute for the COVID-19 Genomics UK (COG-UK) Consortium | Berkshire and Surrey Pathology Services Lighthouse Laboratory and Alex Alderton, Roberto Amato, Jeffrey Barrett, Sonia Goncalves, Ewan Harrison, David K. Jackson, Ian Johnston, Dominic Kwiatkowski, Cordelia Langford, John Sillitoe on behalf of the Wellcome Sanger Institute COVID-19 Surveillance Team |

| | | | |
|---|---|---|---|
| EPI_ISL_2918523 | Lighthouse Lab in Glasgow | Wellcome Sanger Institute for the COVID-19 Genomics UK (COG-UK) Consortium | Harper VanSteenhouse, Yumi Kasai, David Gray, Carol Clugston, Anna Dominiczak and Alex Alderton, Roberto Amato, Jeffrey Barrett, Sonia Goncalves, Ewan Harrison, David K. Jackson, Ian Johnston, Dominic Kwiatkowski, Cordelia Langford, John Sillitoe on behalf of the Wellcome Sanger Institute COVID-19 Surveillance Team |
| EPI_ISL_2918524, EPI_ISL_2918525 | Berkshire and Surrey Pathology Services Lighthouse Laboratory | Wellcome Sanger Institute for the COVID-19 Genomics UK (COG-UK) Consortium | Berkshire and Surrey Pathology Services Lighthouse Laboratory and Alex Alderton, Roberto Amato, Jeffrey Barrett, Sonia Goncalves, Ewan Harrison, David K. Jackson, Ian Johnston, Dominic Kwiatkowski, Cordelia Langford, John Sillitoe on behalf of the Wellcome Sanger Institute COVID-19 Surveillance Team |
| EPI_ISL_2918526 | Lighthouse Lab in Glasgow | Wellcome Sanger Institute for the COVID-19 Genomics UK (COG-UK) Consortium | Harper VanSteenhouse, Yumi Kasai, David Gray, Carol Clugston, Anna Dominiczak and Alex Alderton, Roberto Amato, Jeffrey Barrett, Sonia Goncalves, Ewan Harrison, David K. Jackson, Ian Johnston, Dominic Kwiatkowski, Cordelia Langford, John Sillitoe on behalf of the Wellcome Sanger Institute COVID-19 Surveillance Team |
| EPI_ISL_2918527 | Randox Laboratories | Wellcome Sanger Institute for the COVID-19 Genomics UK (COG-UK) Consortium | Randox Laboratories and Alex Alderton, Roberto Amato, Jeffrey Barrett, Sonia Goncalves, Ewan Harrison, David K. Jackson, Ian Johnston, Dominic Kwiatkowski, Cordelia Langford, John Sillitoe on behalf of the Wellcome Sanger Institute COVID-19 Surveillance Team |
| EPI_ISL_2918528 | Berkshire and Surrey Pathology Services Lighthouse Laboratory | Wellcome Sanger Institute for the COVID-19 Genomics UK (COG-UK) Consortium | Berkshire and Surrey Pathology Services Lighthouse Laboratory and Alex Alderton, Roberto Amato, Jeffrey Barrett, Sonia Goncalves, Ewan Harrison, David K. Jackson, Ian Johnston, Dominic Kwiatkowski, Cordelia Langford, John Sillitoe on behalf of the Wellcome Sanger Institute COVID-19 Surveillance Team |
| EPI_ISL_2918529 | Lighthouse Lab in Glasgow | Wellcome Sanger Institute for the COVID-19 Genomics UK (COG-UK) Consortium | Harper VanSteenhouse, Yumi Kasai, David Gray, Carol Clugston, Anna Dominiczak and Alex Alderton, Roberto Amato, Jeffrey Barrett, Sonia Goncalves, Ewan Harrison, David K. Jackson, Ian Johnston, Dominic Kwiatkowski, Cordelia Langford, John Sillitoe on behalf of the Wellcome Sanger Institute COVID-19 Surveillance Team |
| EPI_ISL_2918530 | Randox Laboratories | Wellcome Sanger Institute for the COVID-19 Genomics UK (COG-UK) Consortium | Randox Laboratories and Alex Alderton, Roberto Amato, Jeffrey Barrett, Sonia Goncalves, Ewan Harrison, David K. Jackson, Ian Johnston, Dominic Kwiatkowski, Cordelia Langford, John Sillitoe on behalf of the Wellcome Sanger Institute COVID-19 Surveillance Team |
| EPI_ISL_2918531, EPI_ISL_2918532 | Berkshire and Surrey Pathology Services Lighthouse Laboratory | Wellcome Sanger Institute for the COVID-19 Genomics UK (COG-UK) Consortium | Berkshire and Surrey Pathology Services Lighthouse Laboratory and Alex Alderton, Roberto Amato, Jeffrey Barrett, Sonia Goncalves, Ewan Harrison, David K. Jackson, Ian Johnston, Dominic Kwiatkowski, Cordelia Langford, John Sillitoe on behalf of the Wellcome Sanger Institute COVID-19 Surveillance Team |
| EPI_ISL_2918533 | Lighthouse Lab in Glasgow | Wellcome Sanger Institute for the COVID-19 Genomics UK (COG-UK) Consortium | Harper VanSteenhouse, Yumi Kasai, David Gray, Carol Clugston, Anna Dominiczak and Alex Alderton, Roberto Amato, Jeffrey Barrett, Sonia Goncalves, Ewan Harrison, David K. Jackson, Ian Johnston, Dominic Kwiatkowski, Cordelia Langford, John Sillitoe on behalf of the Wellcome Sanger Institute COVID-19 Surveillance Team |
| EPI_ISL_2918534 | Randox Laboratories | Wellcome Sanger Institute for the COVID-19 Genomics UK (COG-UK) Consortium | Randox Laboratories and Alex Alderton, Roberto Amato, Jeffrey Barrett, Sonia Goncalves, Ewan Harrison, David K. Jackson, Ian Johnston, Dominic Kwiatkowski, Cordelia Langford, John Sillitoe on behalf of the Wellcome Sanger Institute COVID-19 Surveillance Team |
| EPI_ISL_2918535 | Berkshire and Surrey Pathology Services Lighthouse Laboratory | Wellcome Sanger Institute for the COVID-19 Genomics UK (COG-UK) Consortium | Berkshire and Surrey Pathology Services Lighthouse Laboratory and Alex Alderton, Roberto Amato, Jeffrey Barrett, Sonia Goncalves, Ewan Harrison, David K. Jackson, Ian Johnston, Dominic Kwiatkowski, Cordelia Langford, John Sillitoe on behalf of the Wellcome Sanger Institute COVID-19 Surveillance Team |
| EPI_ISL_2918536 | Randox Laboratories | Wellcome Sanger Institute for the COVID-19 Genomics UK (COG-UK) Consortium | Randox Laboratories and Alex Alderton, Roberto Amato, Jeffrey Barrett, Sonia Goncalves, Ewan Harrison, David K. Jackson, Ian Johnston, Dominic Kwiatkowski, Cordelia Langford, John Sillitoe on behalf of the Wellcome Sanger Institute COVID-19 Surveillance Team |
| EPI_ISL_2918537 | Berkshire and Surrey Pathology Services Lighthouse Laboratory | Wellcome Sanger Institute for the COVID-19 Genomics UK (COG-UK) Consortium | Berkshire and Surrey Pathology Services Lighthouse Laboratory and Alex Alderton, Roberto Amato, Jeffrey Barrett, Sonia Goncalves, Ewan Harrison, David K. Jackson, Ian Johnston, Dominic Kwiatkowski, Cordelia Langford, John Sillitoe on behalf of the Wellcome Sanger Institute COVID-19 Surveillance Team |
| EPI_ISL_2918538 | Lighthouse Lab in Glasgow | Wellcome Sanger Institute for the COVID-19 Genomics UK (COG-UK) Consortium | Harper VanSteenhouse, Yumi Kasai, David Gray, Carol Clugston, Anna Dominiczak and Alex Alderton, Roberto Amato, Jeffrey Barrett, Sonia Goncalves, Ewan Harrison, David K. Jackson, Ian Johnston, Dominic Kwiatkowski, Cordelia Langford, John Sillitoe on behalf of the Wellcome Sanger Institute COVID-19 Surveillance Team |
| EPI_ISL_2918539 | Randox Laboratories | Wellcome Sanger Institute for the COVID-19 Genomics UK (COG-UK) Consortium | Randox Laboratories and Alex Alderton, Roberto Amato, Jeffrey Barrett, Sonia Goncalves, Ewan Harrison, David K. Jackson, Ian Johnston, Dominic Kwiatkowski, Cordelia Langford, John Sillitoe on behalf of the Wellcome Sanger Institute COVID-19 Surveillance Team |
| EPI_ISL_2918540 | Berkshire and Surrey Pathology Services Lighthouse Laboratory | Wellcome Sanger Institute for the COVID-19 Genomics UK (COG-UK) Consortium | Berkshire and Surrey Pathology Services Lighthouse Laboratory and Alex Alderton, Roberto Amato, Jeffrey Barrett, Sonia Goncalves, Ewan Harrison, David K. Jackson, Ian Johnston, Dominic Kwiatkowski, Cordelia Langford, John Sillitoe on behalf of the Wellcome Sanger Institute COVID-19 Surveillance Team |
| EPI_ISL_2918541 | Lighthouse Lab in Glasgow | Wellcome Sanger Institute for the COVID-19 Genomics UK (COG-UK) Consortium | Harper VanSteenhouse, Yumi Kasai, David Gray, Carol Clugston, Anna Dominiczak and Alex Alderton, Roberto Amato, Jeffrey Barrett, Sonia Goncalves, Ewan Harrison, David K. Jackson, Ian Johnston, Dominic Kwiatkowski, Cordelia Langford, John Sillitoe on behalf of the Wellcome Sanger Institute COVID-19 Surveillance Team |
| EPI_ISL_2918542 | Berkshire and Surrey Pathology Services Lighthouse Laboratory | Wellcome Sanger Institute for the COVID-19 Genomics UK (COG-UK) Consortium | Berkshire and Surrey Pathology Services Lighthouse Laboratory and Alex Alderton, Roberto Amato, Jeffrey Barrett, Sonia Goncalves, Ewan Harrison, David K. Jackson, Ian Johnston, Dominic Kwiatkowski, Cordelia Langford, John Sillitoe on behalf of the Wellcome Sanger Institute COVID-19 Surveillance Team |
| EPI_ISL_2918543 | Lighthouse Lab in Glasgow | Wellcome Sanger Institute for the COVID-19 Genomics UK (COG-UK) Consortium | Harper VanSteenhouse, Yumi Kasai, David Gray, Carol Clugston, Anna Dominiczak and Alex Alderton, Roberto Amato, Jeffrey Barrett, Sonia Goncalves, Ewan Harrison, David K. Jackson, Ian Johnston, Dominic Kwiatkowski, Cordelia Langford, John Sillitoe on behalf of the Wellcome Sanger Institute COVID-19 Surveillance Team |
| EPI_ISL_2918544 | Randox Laboratories | Wellcome Sanger Institute for the COVID-19 Genomics UK (COG-UK) Consortium | Randox Laboratories and Alex Alderton, Roberto Amato, Jeffrey Barrett, Sonia Goncalves, Ewan Harrison, David K. Jackson, Ian Johnston, Dominic Kwiatkowski, Cordelia Langford, John Sillitoe on behalf of the Wellcome Sanger Institute COVID-19 Surveillance Team |
| EPI_ISL_2918545 | Lighthouse Lab in Glasgow | Wellcome Sanger Institute for the COVID-19 Genomics UK (COG-UK) Consortium | Harper VanSteenhouse, Yumi Kasai, David Gray, Carol Clugston, Anna Dominiczak and Alex Alderton, Roberto Amato, Jeffrey Barrett, Sonia Goncalves, Ewan Harrison, David K. Jackson, Ian Johnston, Dominic Kwiatkowski, Cordelia Langford, John Sillitoe on behalf of the Wellcome Sanger Institute COVID-19 Surveillance Team |
| EPI_ISL_2918546, EPI_ISL_2918547 | Berkshire and Surrey Pathology Services Lighthouse Laboratory | Wellcome Sanger Institute for the COVID-19 Genomics UK (COG-UK) Consortium | Berkshire and Surrey Pathology Services Lighthouse Laboratory and Alex Alderton, Roberto Amato, Jeffrey Barrett, Sonia Goncalves, Ewan Harrison, David K. Jackson, Ian Johnston, Dominic Kwiatkowski, Cordelia Langford, John Sillitoe on behalf of the Wellcome Sanger Institute COVID-19 Surveillance Team |
| EPI_ISL_2918548, EPI_ISL_2918549 | Lighthouse Lab in Glasgow | Wellcome Sanger Institute for the COVID-19 Genomics UK (COG-UK) Consortium | Harper VanSteenhouse, Yumi Kasai, David Gray, Carol Clugston, Anna Dominiczak and Alex Alderton, Roberto Amato, Jeffrey Barrett, Sonia Goncalves, Ewan Harrison, David K. Jackson, Ian Johnston, Dominic Kwiatkowski, Cordelia Langford, John Sillitoe on behalf of the Wellcome Sanger Institute COVID-19 Surveillance Team |
| EPI_ISL_2918550 | Randox Laboratories | Wellcome Sanger Institute for the COVID-19 Genomics UK (COG-UK) Consortium | Randox Laboratories and Alex Alderton, Roberto Amato, Jeffrey Barrett, Sonia Goncalves, Ewan Harrison, David K. Jackson, Ian Johnston, Dominic Kwiatkowski, Cordelia Langford, John Sillitoe on behalf of the Wellcome Sanger Institute COVID-19 Surveillance Team |
| EPI_ISL_2918551 | Berkshire and Surrey Pathology Services Lighthouse Laboratory | Wellcome Sanger Institute for the COVID-19 Genomics UK (COG-UK) Consortium | Berkshire and Surrey Pathology Services Lighthouse Laboratory and Alex Alderton, Roberto Amato, Jeffrey Barrett, Sonia Goncalves, Ewan Harrison, David K. Jackson, Ian Johnston, Dominic Kwiatkowski, Cordelia Langford, John Sillitoe on behalf of the Wellcome Sanger Institute COVID-19 Surveillance Team |
| EPI_ISL_2918552, EPI_ISL_2918553, EPI_ISL_2918554 | Lighthouse Lab in Glasgow | Wellcome Sanger Institute for the COVID-19 Genomics UK (COG-UK) Consortium | Harper VanSteenhouse, Yumi Kasai, David Gray, Carol Clugston, Anna Dominiczak and Alex Alderton, Roberto Amato, Jeffrey Barrett, Sonia Goncalves, Ewan Harrison, David K. Jackson, Ian Johnston, Dominic Kwiatkowski, Cordelia Langford, John Sillitoe on behalf of the Wellcome Sanger Institute COVID-19 Surveillance Team |
| EPI_ISL_2918555, EPI_ISL_2918556 | Berkshire and Surrey Pathology Services Lighthouse Laboratory | Wellcome Sanger Institute for the COVID-19 Genomics UK (COG-UK) Consortium | Berkshire and Surrey Pathology Services Lighthouse Laboratory and Alex Alderton, Roberto Amato, Jeffrey Barrett, Sonia Goncalves, Ewan Harrison, David K. Jackson, Ian Johnston, Dominic Kwiatkowski, Cordelia Langford, John Sillitoe on behalf of the Wellcome Sanger Institute COVID-19 Surveillance Team |
| EPI_ISL_2918557, EPI_ISL_2918558 | Lighthouse Lab in Glasgow | Wellcome Sanger Institute for the COVID-19 Genomics UK (COG-UK) Consortium | Harper VanSteenhouse, Yumi Kasai, David Gray, Carol Clugston, Anna Dominiczak and Alex Alderton, Roberto Amato, Jeffrey Barrett, Sonia Goncalves, Ewan Harrison, David K. Jackson, Ian Johnston, Dominic Kwiatkowski, Cordelia Langford, John Sillitoe on behalf of the Wellcome Sanger Institute COVID-19 Surveillance Team |
| EPI_ISL_2918559 | Berkshire and Surrey Pathology Services Lighthouse Laboratory | Wellcome Sanger Institute for the COVID-19 Genomics UK (COG-UK) Consortium | Berkshire and Surrey Pathology Services Lighthouse Laboratory and Alex Alderton, Roberto Amato, Jeffrey Barrett, Sonia Goncalves, Ewan Harrison, David K. Jackson, Ian Johnston, Dominic Kwiatkowski, Cordelia Langford, John Sillitoe on behalf of the Wellcome Sanger Institute COVID-19 Surveillance Team |
| EPI_ISL_2918560 | Randox Laboratories | Wellcome Sanger Institute for the COVID-19 Genomics UK (COG-UK) Consortium | Randox Laboratories and Alex Alderton, Roberto Amato, Jeffrey Barrett, Sonia Goncalves, Ewan Harrison, David K. Jackson, Ian Johnston, Dominic Kwiatkowski, Cordelia Langford, John Sillitoe on behalf of the Wellcome Sanger Institute COVID-19 Surveillance Team |
| EPI_ISL_2918561, EPI_ISL_2918562, EPI_ISL_2918563, EPI_ISL_2918564, EPI_ISL_2918565 | Lighthouse Lab in Glasgow | Wellcome Sanger Institute for the COVID-19 Genomics UK (COG-UK) Consortium | Harper VanSteenhouse, Yumi Kasai, David Gray, Carol Clugston, Anna Dominiczak and Alex Alderton, Roberto Amato, Jeffrey Barrett, Sonia Goncalves, Ewan Harrison, David K. Jackson, Ian Johnston, Dominic Kwiatkowski, Cordelia Langford, John Sillitoe on behalf of the Wellcome Sanger Institute COVID-19 Surveillance Team |
| EPI_ISL_2918566 | Berkshire and Surrey Pathology Services Lighthouse Laboratory | Wellcome Sanger Institute for the COVID-19 Genomics UK (COG-UK) Consortium | Berkshire and Surrey Pathology Services Lighthouse Laboratory and Alex Alderton, Roberto Amato, Jeffrey Barrett, Sonia Goncalves, Ewan Harrison, David K. Jackson, Ian Johnston, Dominic Kwiatkowski, Cordelia Langford, John Sillitoe on behalf of the Wellcome Sanger Institute COVID-19 Surveillance Team |
| EPI_ISL_2918567 | Randox Laboratories | Wellcome Sanger Institute for the COVID-19 Genomics UK (COG-UK) Consortium | Randox Laboratories and Alex Alderton, Roberto Amato, Jeffrey Barrett, Sonia Goncalves, Ewan Harrison, David K. Jackson, Ian Johnston, Dominic Kwiatkowski, Cordelia Langford, John Sillitoe on behalf of the Wellcome Sanger Institute COVID-19 Surveillance Team |
| EPI_ISL_2918568 | Berkshire and Surrey Pathology Services Lighthouse Laboratory | Wellcome Sanger Institute for the COVID-19 Genomics UK (COG-UK) Consortium | Berkshire and Surrey Pathology Services Lighthouse Laboratory and Alex Alderton, Roberto Amato, Jeffrey Barrett, Sonia Goncalves, Ewan Harrison, David K. Jackson, Ian Johnston, Dominic Kwiatkowski, Cordelia Langford, John Sillitoe on behalf of the Wellcome Sanger Institute COVID-19 Surveillance Team |
| EPI_ISL_2918569 | Lighthouse Lab in Glasgow | Wellcome Sanger Institute for the COVID-19 Genomics UK (COG-UK) Consortium | Harper VanSteenhouse, Yumi Kasai, David Gray, Carol Clugston, Anna Dominiczak and Alex Alderton, Roberto Amato, Jeffrey Barrett, Sonia Goncalves, Ewan Harrison, David K. Jackson, Ian Johnston, Dominic Kwiatkowski, Cordelia Langford, John Sillitoe on behalf of the Wellcome Sanger Institute COVID-19 Surveillance Team |
| EPI_ISL_2918570 | Berkshire and Surrey Pathology Services Lighthouse Laboratory | Wellcome Sanger Institute for the COVID-19 Genomics UK (COG-UK) Consortium | Berkshire and Surrey Pathology Services Lighthouse Laboratory and Alex Alderton, Roberto Amato, Jeffrey Barrett, Sonia Goncalves, Ewan Harrison, David K. Jackson, Ian Johnston, Dominic Kwiatkowski, Cordelia Langford, John Sillitoe on behalf of the Wellcome Sanger Institute COVID-19 Surveillance Team |
| EPI_ISL_2918571 | Lighthouse Lab in Glasgow | Wellcome Sanger Institute for the COVID-19 Genomics UK (COG-UK) Consortium | Harper VanSteenhouse, Yumi Kasai, David Gray, Carol Clugston, Anna Dominiczak and Alex Alderton, Roberto Amato, Jeffrey Barrett, Sonia Goncalves, Ewan Harrison, David K. Jackson, Ian Johnston, Dominic Kwiatkowski, Cordelia Langford, John Sillitoe on behalf of the Wellcome Sanger Institute COVID-19 Surveillance Team |
| EPI_ISL_2918572 | Berkshire and Surrey Pathology Services Lighthouse Laboratory | Wellcome Sanger Institute for the COVID-19 Genomics UK (COG-UK) Consortium | Berkshire and Surrey Pathology Services Lighthouse Laboratory and Alex Alderton, Roberto Amato, Jeffrey Barrett, Sonia Goncalves, Ewan Harrison, David K. Jackson, Ian Johnston, Dominic Kwiatkowski, Cordelia Langford, John Sillitoe on behalf of the Wellcome Sanger Institute COVID-19 Surveillance Team |
| EPI_ISL_2918573, EPI_ISL_2918574 | Lighthouse Lab in Glasgow | Wellcome Sanger Institute for the COVID-19 Genomics UK (COG-UK) Consortium | Harper VanSteenhouse, Yumi Kasai, David Gray, Carol Clugston, Anna Dominiczak and Alex Alderton, Roberto Amato, Jeffrey Barrett, Sonia Goncalves, Ewan Harrison, David K. Jackson, Ian Johnston, Dominic Kwiatkowski, Cordelia Langford, John Sillitoe on behalf of the Wellcome Sanger Institute COVID-19 Surveillance Team |
| EPI_ISL_2918575 | Randox Laboratories | Wellcome Sanger Institute for the COVID-19 Genomics UK (COG-UK) Consortium | Randox Laboratories and Alex Alderton, Roberto Amato, Jeffrey Barrett, Sonia Goncalves, Ewan Harrison, David K. Jackson, Ian Johnston, Dominic Kwiatkowski, Cordelia Langford, John Sillitoe on behalf of the Wellcome Sanger Institute COVID-19 Surveillance Team |
| EPI_ISL_2918576, EPI_ISL_2918577 | Berkshire and Surrey Pathology Services Lighthouse Laboratory | Wellcome Sanger Institute for the COVID-19 Genomics UK (COG-UK) Consortium | Berkshire and Surrey Pathology Services Lighthouse Laboratory and Alex Alderton, Roberto Amato, Jeffrey Barrett, Sonia Goncalves, Ewan Harrison, David K. Jackson, Ian Johnston, Dominic Kwiatkowski, Cordelia Langford, John Sillitoe on behalf of the Wellcome Sanger Institute COVID-19 Surveillance Team |
| EPI_ISL_2918578, EPI_ISL_2918579 | Lighthouse Lab in Glasgow | Wellcome Sanger Institute for the COVID-19 Genomics UK (COG-UK) Consortium | Harper VanSteenhouse, Yumi Kasai, David Gray, Carol Clugston, Anna Dominiczak and Alex Alderton, Roberto Amato, Jeffrey Barrett, Sonia Goncalves, Ewan Harrison, David K. Jackson, Ian Johnston, Dominic Kwiatkowski, Cordelia Langford, John Sillitoe on behalf of the Wellcome Sanger Institute COVID-19 Surveillance Team |
| EPI_ISL_2918580 | Berkshire and Surrey Pathology Services Lighthouse Laboratory | Wellcome Sanger Institute for the COVID-19 Genomics UK (COG-UK) Consortium | Berkshire and Surrey Pathology Services Lighthouse Laboratory and Alex Alderton, Roberto Amato, Jeffrey Barrett, Sonia Goncalves, Ewan Harrison, David K. Jackson, Ian Johnston, Dominic Kwiatkowski, Cordelia Langford, John Sillitoe on behalf of the Wellcome Sanger Institute COVID-19 Surveillance Team |
| EPI_ISL_2918581, | Lighthouse Lab in Glasgow | Wellcome Sanger Institute for the COVID-19 Genomics UK (COG-UK) | Harper VanSteenhouse, Yumi Kasai, David Gray, Carol Clugston, Anna Dominiczak and Alex Alderton, Roberto Amato, Jeffrey Barrett, Sonia Goncalves, Ewan Harrison, David K. Jackson, Ian Johnston, Dominic Kwiatkowski, Cordelia Langford, |

| | | | |
|---|---|---|---|
| EPI_ISL_2918582, EPI_ISL_2918583 | | Consortium | John Sillitoe on behalf of the Wellcome Sanger Institute COVID-19 Surveillance Team |
| EPI_ISL_2918584 | Berkshire and Surrey Pathology Services Lighthouse Laboratory | Wellcome Sanger Institute for the COVID-19 Genomics UK (COG-UK) Consortium | Berkshire and Surrey Pathology Services Lighthouse Laboratory and Alex Alderton, Roberto Amato, Jeffrey Barrett, Sonia Goncalves, Ewan Harrison, David K. Jackson, Ian Johnston, Dominic Kwiatkowski, Cordelia Langford, John Sillitoe on behalf of the Wellcome Sanger Institute COVID-19 Surveillance Team |
| EPI_ISL_2918585, EPI_ISL_2918586 | Randox Laboratories | Wellcome Sanger Institute for the COVID-19 Genomics UK (COG-UK) Consortium | Randox Laboratories and Alex Alderton, Roberto Amato, Jeffrey Barrett, Sonia Goncalves, Ewan Harrison, David K. Jackson, Ian Johnston, Dominic Kwiatkowski, Cordelia Langford, John Sillitoe on behalf of the Wellcome Sanger Institute COVID-19 Surveillance Team |
| EPI_ISL_2918587 | Berkshire and Surrey Pathology Services Lighthouse Laboratory | Wellcome Sanger Institute for the COVID-19 Genomics UK (COG-UK) Consortium | Berkshire and Surrey Pathology Services Lighthouse Laboratory and Alex Alderton, Roberto Amato, Jeffrey Barrett, Sonia Goncalves, Ewan Harrison, David K. Jackson, Ian Johnston, Dominic Kwiatkowski, Cordelia Langford, John Sillitoe on behalf of the Wellcome Sanger Institute COVID-19 Surveillance Team |
| EPI_ISL_2918588, EPI_ISL_2918589, EPI_ISL_2918590 | Lighthouse Lab in Glasgow | Wellcome Sanger Institute for the COVID-19 Genomics UK (COG-UK) Consortium | Harper VanSteenhouse, Yumi Kasai, David Gray, Carol Clugston, Anna Dominiczak and Alex Alderton, Roberto Amato, Jeffrey Barrett, Sonia Goncalves, Ewan Harrison, David K. Jackson, Ian Johnston, Dominic Kwiatkowski, Cordelia Langford, John Sillitoe on behalf of the Wellcome Sanger Institute COVID-19 Surveillance Team |
| EPI_ISL_2918591, EPI_ISL_2918592, EPI_ISL_2918593, EPI_ISL_2918594, EPI_ISL_2918595 | Berkshire and Surrey Pathology Services Lighthouse Laboratory | Wellcome Sanger Institute for the COVID-19 Genomics UK (COG-UK) Consortium | Berkshire and Surrey Pathology Services Lighthouse Laboratory and Alex Alderton, Roberto Amato, Jeffrey Barrett, Sonia Goncalves, Ewan Harrison, David K. Jackson, Ian Johnston, Dominic Kwiatkowski, Cordelia Langford, John Sillitoe on behalf of the Wellcome Sanger Institute COVID-19 Surveillance Team |
| EPI_ISL_2918596 | Lighthouse Lab in Glasgow | Wellcome Sanger Institute for the COVID-19 Genomics UK (COG-UK) Consortium | Harper VanSteenhouse, Yumi Kasai, David Gray, Carol Clugston, Anna Dominiczak and Alex Alderton, Roberto Amato, Jeffrey Barrett, Sonia Goncalves, Ewan Harrison, David K. Jackson, Ian Johnston, Dominic Kwiatkowski, Cordelia Langford, John Sillitoe on behalf of the Wellcome Sanger Institute COVID-19 Surveillance Team |
| EPI_ISL_2918597, EPI_ISL_2918598, EPI_ISL_2918599 | Berkshire and Surrey Pathology Services Lighthouse Laboratory | Wellcome Sanger Institute for the COVID-19 Genomics UK (COG-UK) Consortium | Berkshire and Surrey Pathology Services Lighthouse Laboratory and Alex Alderton, Roberto Amato, Jeffrey Barrett, Sonia Goncalves, Ewan Harrison, David K. Jackson, Ian Johnston, Dominic Kwiatkowski, Cordelia Langford, John Sillitoe on behalf of the Wellcome Sanger Institute COVID-19 Surveillance Team |
| EPI_ISL_2918600 | Lighthouse Lab in Glasgow | Wellcome Sanger Institute for the COVID-19 Genomics UK (COG-UK) Consortium | Harper VanSteenhouse, Yumi Kasai, David Gray, Carol Clugston, Anna Dominiczak and Alex Alderton, Roberto Amato, Jeffrey Barrett, Sonia Goncalves, Ewan Harrison, David K. Jackson, Ian Johnston, Dominic Kwiatkowski, Cordelia Langford, John Sillitoe on behalf of the Wellcome Sanger Institute COVID-19 Surveillance Team |
| EPI_ISL_2918601, EPI_ISL_2918602 | Lighthouse Laboratory Plymouth | Wellcome Sanger Institute for the COVID-19 Genomics UK (COG-UK) Consortium | Lighthouse Laboratory Plymouth and Alex Alderton, Roberto Amato, Jeffrey Barrett, Sonia Goncalves, Ewan Harrison, David K. Jackson, Ian Johnston, Dominic Kwiatkowski, Cordelia Langford, John Sillitoe on behalf of the Wellcome Sanger Institute COVID-19 Surveillance Team |
| EPI_ISL_2918603 | Berkshire and Surrey Pathology Services Lighthouse Laboratory | Wellcome Sanger Institute for the COVID-19 Genomics UK (COG-UK) Consortium | Berkshire and Surrey Pathology Services Lighthouse Laboratory and Alex Alderton, Roberto Amato, Jeffrey Barrett, Sonia Goncalves, Ewan Harrison, David K. Jackson, Ian Johnston, Dominic Kwiatkowski, Cordelia Langford, John Sillitoe on behalf of the Wellcome Sanger Institute COVID-19 Surveillance Team |
| EPI_ISL_2918604 | RS Pondok Indah Pondok Indah | Eijkman Institute for Molecular Biology, National Research and Innovation Agency | Lidwina Priliani, Edison Johar, Frilasita A Yudhaputri, Muhammad Rezki Rasyak, Willy Agustine, Hidayat Trimarsanto, Lydia V. Panggalo, Iskandar Adnan, Sukma Oktavianthi, Safarina G Malik, Khin Saw Myint, Amin Soebandrio |
| EPI_ISL_2918605 | RS Pondok Indah Pondok Indah | Eijkman Institute for Molecular Biology, National Research and Innovation Agency | Edison Johar, Frilasita A Yudhaputri, Muhammad Rezki Rasyak, Willy Agustine, Hidayat Trimarsanto, Lydia V. Panggalo, Iskandar Adnan, Sukma Oktavianthi, Lidwina Priliani, Safarina G Malik, Khin Saw Myint, Amin Soebandrio |
| EPI_ISL_2918606 | RS Pondok Indah Pondok Indah | Eijkman Institute for Molecular Biology, National Research and Innovation Agency | Muhammad Rezki Rasyak, Willy Agustine, Hidayat Trimarsanto, Lydia V. Panggalo, Iskandar Adnan, Sukma Oktavianthi, Lidwina Priliani, Edison Johar, Frilasita A Yudhaputri, Safarina G Malik, Khin Saw Myint, Amin Soebandrio |
| EPI_ISL_2918607 | RS Pondok Indah Pondok Indah | Eijkman Institute for Molecular Biology, National Research and Innovation Agency | Willy Agustine, Hidayat Trimarsanto, Lydia V. Panggalo, Iskandar Adnan, Sukma Oktavianthi, Lidwina Priliani, Edison Johar, Frilasita A Yudhaputri, Muhammad Rezki Rasyak, Safarina G Malik, Khin Saw Myint, Amin Soebandrio |
| EPI_ISL_2918608 | RSCM | Eijkman Institute for Molecular Biology, National Research and Innovation Agency | Hidayat Trimarsanto, Lydia V. Panggalo, Iskandar Adnan, Sukma Oktavianthi, Lidwina Priliani, Edison Johar, Frilasita A Yudhaputri, Muhammad Rezki Rasyak, Willy Agustine, Safarina G Malik, Khin Saw Myint, Amin Soebandrio |
| EPI_ISL_2918609 | RS Pondok Indah Pondok Indah | Eijkman Institute for Molecular Biology, National Research and Innovation Agency | Lydia V. Panggalo, Iskandar Adnan, Sukma Oktavianthi, Lidwina Priliani, Edison Johar, Frilasita A Yudhaputri, Muhammad Rezki Rasyak, Willy Agustine, Hidayat Trimarsanto, Safarina G Malik, Khin Saw Myint, Amin Soebandrio |
| EPI_ISL_2918610 | RS Pondok Indah Pondok Indah | Eijkman Institute for Molecular Biology, National Research and Innovation Agency | Iskandar Adnan, Sukma Oktavianthi, Lidwina Priliani, Edison Johar, Frilasita A Yudhaputri, Muhammad Rezki Rasyak, Willy Agustine, Hidayat Trimarsanto, Lydia V. Panggalo, Safarina G Malik, Khin Saw Myint, Amin Soebandrio |
| EPI_ISL_2918611 | RS Pondok Indah Pondok Indah | Eijkman Institute for Molecular Biology, National Research and Innovation Agency | Sukma Oktavianthi, Lidwina Priliani, Edison Johar, Frilasita A Yudhaputri, Muhammad Rezki Rasyak, Willy Agustine, Hidayat Trimarsanto, Lydia V. Panggalo, Iskandar Adnan, Safarina G Malik, Khin Saw Myint, Amin Soebandrio |
| EPI_ISL_2918612 | RS Pondok Indah Pondok Indah | Eijkman Institute for Molecular Biology, National Research and Innovation Agency | Lidwina Priliani, Edison Johar, Frilasita A Yudhaputri, Muhammad Rezki Rasyak, Willy Agustine, Hidayat Trimarsanto, Lydia V. Panggalo, Iskandar Adnan, Sukma Oktavianthi, Safarina G Malik, Khin Saw Myint, Amin Soebandrio |
| EPI_ISL_2918613 | RS Medistra | Eijkman Institute for Molecular Biology, National Research and Innovation Agency | Edison Johar, Frilasita A Yudhaputri, Muhammad Rezki Rasyak, Willy Agustine, Hidayat Trimarsanto, Lydia V. Panggalo, Iskandar Adnan, Sukma Oktavianthi, Lidwina Priliani, Safarina G Malik, Khin Saw Myint, Amin Soebandrio |
| EPI_ISL_2918614 | RSUP Dr. Kariadi | Eijkman Institute for Molecular Biology, National Research and Innovation Agency | Frilasita A Yudhaputri, Muhammad Rezki Rasyak, Willy Agustine, Hidayat Trimarsanto, Lydia V. Panggalo, Iskandar Adnan, Sukma Oktavianthi, Lidwina Priliani, Edison Johar, Safarina G Malik, Khin Saw Myint, Amin Soebandrio |
| EPI_ISL_2918615 | RSUP Dr. Kariadi | Eijkman Institute for Molecular Biology, National Research and Innovation Agency | Muhammad Rezki Rasyak, Willy Agustine, Hidayat Trimarsanto, Lydia V. Panggalo, Iskandar Adnan, Sukma Oktavianthi, Lidwina Priliani, Edison Johar, Frilasita A Yudhaputri, Safarina G Malik, Khin Saw Myint, Amin Soebandrio |
| EPI_ISL_2918616 | RSUP Dr. Kariadi | Eijkman Institute for Molecular Biology, National Research and Innovation Agency | Willy Agustine, Hidayat Trimarsanto, Lydia V. Panggalo, Iskandar Adnan, Sukma Oktavianthi, Lidwina Priliani, Edison Johar, Frilasita A Yudhaputri, Muhammad Rezki Rasyak, Safarina G Malik, Khin Saw Myint, Amin Soebandrio |
| EPI_ISL_2918617 | RSUP Dr. Kariadi | Eijkman Institute for Molecular Biology, National Research and Innovation Agency | Hidayat Trimarsanto, Lydia V. Panggalo, Iskandar Adnan, Sukma Oktavianthi, Lidwina Priliani, Edison Johar, Frilasita A Yudhaputri, Muhammad Rezki Rasyak, Willy Agustine, Safarina G Malik, Khin Saw Myint, Amin Soebandrio |
| EPI_ISL_2918618 | RSUP Dr. Kariadi | Eijkman Institute for Molecular Biology, National Research and Innovation Agency | Lydia V. Panggalo, Iskandar Adnan, Sukma Oktavianthi, Lidwina Priliani, Edison Johar, Frilasita A Yudhaputri, Muhammad Rezki Rasyak, Willy Agustine, Hidayat Trimarsanto, Safarina G Malik, Khin Saw Myint, Amin Soebandrio |
| EPI_ISL_2918619 | RSUP Dr. Kariadi | Eijkman Institute for Molecular Biology, National Research and Innovation Agency | Iskandar Adnan, Sukma Oktavianthi, Lidwina Priliani, Edison Johar, Frilasita A Yudhaputri, Muhammad Rezki Rasyak, Willy Agustine, Hidayat Trimarsanto, Lydia V. Panggalo, Safarina G Malik, Khin Saw Myint, Amin Soebandrio |
| EPI_ISL_2918620 | RSUP Dr. Kariadi | Eijkman Institute for Molecular Biology, National Research and Innovation Agency | Sukma Oktavianthi, Lidwina Priliani, Edison Johar, Frilasita A Yudhaputri, Muhammad Rezki Rasyak, Willy Agustine, Hidayat Trimarsanto, Lydia V. Panggalo, Iskandar Adnan, Safarina G Malik, Khin Saw Myint, Amin Soebandrio |
| EPI_ISL_2918621 | RSUP Dr. Kariadi | Eijkman Institute for Molecular Biology, National Research and Innovation Agency | Lidwina Priliani, Edison Johar, Frilasita A Yudhaputri, Muhammad Rezki Rasyak, Willy Agustine, Hidayat Trimarsanto, Lydia V. Panggalo, Iskandar Adnan, Sukma Oktavianthi, Safarina G Malik, Khin Saw Myint, Amin Soebandrio |
| EPI_ISL_2918622 | RSUP Dr. Kariadi | Eijkman Institute for Molecular Biology, National Research and Innovation Agency | Edison Johar, Frilasita A Yudhaputri, Muhammad Rezki Rasyak, Willy Agustine, Hidayat Trimarsanto, Lydia V. Panggalo, Iskandar Adnan, Sukma Oktavianthi, Lidwina Priliani, Safarina G Malik, Khin Saw Myint, Amin Soebandrio |
| EPI_ISL_2918623 | RSUP Dr. Kariadi | Eijkman Institute for Molecular Biology, National Research and Innovation Agency | Frilasita A Yudhaputri, Muhammad Rezki Rasyak, Willy Agustine, Hidayat Trimarsanto, Lydia V. Panggalo, Iskandar Adnan, Sukma Oktavianthi, Lidwina Priliani, Edison Johar, Safarina G Malik, Khin Saw Myint, Amin Soebandrio |
| EPI_ISL_2918626 | RSUP Dr. Kariadi | Eijkman Institute for Molecular Biology, National Research and Innovation Agency | Hidayat Trimarsanto, Lydia V. Panggalo, Iskandar Adnan, Sukma Oktavianthi, Lidwina Priliani, Edison Johar, Frilasita A Yudhaputri, Muhammad Rezki Rasyak, Willy Agustine, Safarina G Malik, Khin Saw Myint, Amin Soebandrio |
| EPI_ISL_2918627 | Puskesmas Sugihmukti | Eijkman Institute for Molecular Biology, National Research and Innovation Agency; West Java Health Laboratory | Hidayat Trimarsanto, Lydia V. Panggalo, Iskandar Adnan, Sukma Oktavianthi, Lidwina Priliani, Edison Johar, Frilasita A Yudhaputri, Muhammad Rezki Rasyak, Willy Agustine, Azzania Fibriani, Ema Rahmawati, Ryan Bayusantika Ristandi, Rifky Waluyajati Rachman, Cut Nur Cinthia Alamanda, Safarina G Malik, Khin Saw Myint, Amin Soebandrio |
| EPI_ISL_2918628 | Puskesmas Pameungpeuk | Eijkman Institute for Molecular Biology, National Research and Innovation Agency; West Java Health Laboratory | Lydia V. Panggalo, Iskandar Adnan, Sukma Oktavianthi, Lidwina Priliani, Edison Johar, Frilasita A Yudhaputri, Muhammad Rezki Rasyak, Willy Agustine, Hidayat Trimarsanto, Azzania Fibriani, Ema Rahmawati, Ryan Bayusantika Ristandi, Rifky Waluyajati Rachman, Cut Nur Cinthia Alamanda, Safarina G Malik, Khin Saw Myint, Amin Soebandrio |
| EPI_ISL_2918629 | Puskesmas Sangkanhurip | Eijkman Institute for Molecular Biology, National Research and Innovation Agency; West Java Health Laboratory | Iskandar Adnan, Sukma Oktavianthi, Lidwina Priliani, Edison Johar, Frilasita A Yudhaputri, Muhammad Rezki Rasyak, Willy Agustine, Hidayat Trimarsanto, Lydia V. Panggalo, Azzania Fibriani, Ema Rahmawati, Ryan Bayusantika Ristandi, Rifky Waluyajati Rachman, Cut Nur Cinthia Alamanda, Safarina G Malik, Khin Saw Myint, Amin Soebandrio |
| EPI_ISL_2918630 | Puskesmas Pangalengan | Eijkman Institute for Molecular Biology, National Research and Innovation Agency; West Java Health Laboratory | Sukma Oktavianthi, Lidwina Priliani, Edison Johar, Frilasita A Yudhaputri, Muhammad Rezki Rasyak, Willy Agustine, Hidayat Trimarsanto, Lydia V. Panggalo, Iskandar Adnan, Azzania Fibriani, Ema Rahmawati, Ryan Bayusantika Ristandi, Rifky Waluyajati Rachman, Cut Nur Cinthia Alamanda, Safarina G Malik, Khin Saw Myint, Amin Soebandrio |
| EPI_ISL_2918631 | Puskesmas Kramatmulya | Eijkman Institute for Molecular Biology, National Research and Innovation Agency; West Java Health Laboratory | Lidwina Priliani, Edison Johar, Frilasita A Yudhaputri, Muhammad Rezki Rasyak, Willy Agustine, Hidayat Trimarsanto, Lydia V. Panggalo, Iskandar Adnan, Sukma Oktavianthi, Azzania Fibriani, Ema Rahmawati, Ryan Bayusantika Ristandi, Rifky Waluyajati Rachman, Cut Nur Cinthia Alamanda, Safarina G Malik, Khin Saw Myint, Amin Soebandrio |
| EPI_ISL_2918632 | Puskesmas Margajaya | Eijkman Institute for Molecular Biology, National Research and Innovation Agency; West Java Health Laboratory | Edison Johar, Frilasita A Yudhaputri, Muhammad Rezki Rasyak, Willy Agustine, Hidayat Trimarsanto, Lydia V. Panggalo, Iskandar Adnan, Sukma Oktavianthi, Lidwina Priliani, Azzania Fibriani, Ema Rahmawati, Ryan Bayusantika Ristandi, Rifky Waluyajati Rachman, Cut Nur Cinthia Alamanda, Safarina G Malik, Khin Saw Myint, Amin Soebandrio |
| EPI_ISL_2918634 | Dinas Kesehatan Kabupaten Indramayu | Eijkman Institute for Molecular Biology, National Research and Innovation Agency; West Java Health Laboratory | Muhammad Rezki Rasyak, Willy Agustine, Hidayat Trimarsanto, Lydia V. Panggalo, Iskandar Adnan, Sukma Oktavianthi, Lidwina Priliani, Edison Johar, Frilasita A Yudhaputri, Azzania Fibriani, Ema Rahmawati, Ryan Bayusantika Ristandi, Rifky Waluyajati Rachman, Cut Nur Cinthia Alamanda, Safarina G Malik, Khin Saw Myint, Amin Soebandrio |
| EPI_ISL_2918635 | Puskesmas Rancakalong | Eijkman Institute for Molecular Biology, National Research and Innovation Agency; West Java Health Laboratory | Willy Agustine, Hidayat Trimarsanto, Lydia V. Panggalo, Iskandar Adnan, Sukma Oktavianthi, Lidwina Priliani, Edison Johar, Frilasita A Yudhaputri, Muhammad Rezki Rasyak, Azzania Fibriani, Ema Rahmawati, Ryan Bayusantika Ristandi, Rifky Waluyajati Rachman, Cut Nur Cinthia Alamanda, Safarina G Malik, Khin Saw Myint, Amin Soebandrio |
| EPI_ISL_2918636 | Puskesmas Margajaya | Eijkman Institute for Molecular Biology, National Research and Innovation Agency; West Java Health Laboratory | Hidayat Trimarsanto, Lydia V. Panggalo, Iskandar Adnan, Sukma Oktavianthi, Lidwina Priliani, Edison Johar, Frilasita A Yudhaputri, Muhammad Rezki Rasyak, Willy Agustine, Azzania Fibriani, Ema Rahmawati, Ryan Bayusantika Ristandi, Rifky Waluyajati Rachman, Cut Nur Cinthia Alamanda, Safarina G Malik, Khin Saw Myint, Amin Soebandrio |
| EPI_ISL_2918637 | Puskesmas Pangalengan | Eijkman Institute for Molecular Biology, National Research and Innovation Agency; West Java Health Laboratory | Lydia V. Panggalo, Iskandar Adnan, Sukma Oktavianthi, Lidwina Priliani, Edison Johar, Frilasita A Yudhaputri, Muhammad Rezki Rasyak, Willy Agustine, Hidayat Trimarsanto, Azzania Fibriani, Ema Rahmawati, Ryan Bayusantika Ristandi, Rifky Waluyajati Rachman, Cut Nur Cinthia Alamanda, Safarina G Malik, Khin Saw Myint, Amin Soebandrio |
| EPI_ISL_2918638 | Puskesmas Pakutandang | Eijkman Institute for Molecular Biology, National Research and Innovation Agency; West Java Health Laboratory | Iskandar Adnan, Sukma Oktavianthi, Lidwina Priliani, Edison Johar, Frilasita A Yudhaputri, Muhammad Rezki Rasyak, Willy Agustine, Hidayat Trimarsanto, Lydia V. Panggalo, Azzania Fibriani, Ema Rahmawati, Ryan Bayusantika Ristandi, Rifky Waluyajati Rachman, Cut Nur Cinthia Alamanda, Safarina G Malik, Khin Saw Myint, Amin Soebandrio |

| | | | |
|---|---|---|---|
| EPI_ISL_2918639 | Puskesmas Banjaran Kota | Eijkman Institute for Molecular Biology, National Research and Innovation Agency; West Java Health Laboratory | Sukma Oktavianthi, Lidwina Priliani, Edison Johar, Frilasita A Yudhaputri, Muhammad Rezki Rasyak, Willy Agustine, Hidayat Trimarsanto, Lydia V. Panggalo, Iskandar Adnan, Azzania Fibriani, Ema Rahmawati, Ryan Bayusantika Ristandi, Rifky Waluyajati Rachman, Cut Nur Cinthia Alamanda, Safarina G Malik, Khin Saw Myint, Amin Soebandrio |
| EPI_ISL_2918640 | Puskesmas Kiangroke | Eijkman Institute for Molecular Biology, National Research and Innovation Agency; West Java Health Laboratory | Lidwina Priliani, Edison Johar, Frilasita A Yudhaputri, Muhammad Rezki Rasyak, Willy Agustine, Hidayat Trimarsanto, Lydia V. Panggalo, Iskandar Adnan, Sukma Oktavianthi, Azzania Fibriani, Ema Rahmawati, Ryan Bayusantika Ristandi, Rifky Waluyajati Rachman, Cut Nur Cinthia Alamanda, Safarina G Malik, Khin Saw Myint, Amin Soebandrio |
| EPI_ISL_2918641 | Puskesmas Rancamanyar | Eijkman Institute for Molecular Biology, National Research and Innovation Agency; West Java Health Laboratory | Edison Johar, Frilasita A Yudhaputri, Muhammad Rezki Rasyak, Willy Agustine, Hidayat Trimarsanto, Lydia V. Panggalo, Iskandar Adnan, Sukma Oktavianthi, Lidwina Priliani, Azzania Fibriani, Ema Rahmawati, Ryan Bayusantika Ristandi, Rifky Waluyajati Rachman, Cut Nur Cinthia Alamanda, Safarina G Malik, Khin Saw Myint, Amin Soebandrio |
| EPI_ISL_2918642 | Puskesmas Rancamanyar | Eijkman Institute for Molecular Biology, National Research and Innovation Agency; West Java Health Laboratory | Frilasita A Yudhaputri, Muhammad Rezki Rasyak, Willy Agustine, Hidayat Trimarsanto, Lydia V. Panggalo, Iskandar Adnan, Sukma Oktavianthi, Lidwina Priliani, Edison Johar, Azzania Fibriani, Ema Rahmawati, Ryan Bayusantika Ristandi, Rifky Waluyajati Rachman, Cut Nur Cinthia Alamanda, Safarina G Malik, Khin Saw Myint, Amin Soebandrio |
| EPI_ISL_2918643 | Puskesmas Rancamanyar | Eijkman Institute for Molecular Biology, National Research and Innovation Agency; West Java Health Laboratory | Muhammad Rezki Rasyak, Willy Agustine, Hidayat Trimarsanto, Lydia V. Panggalo, Iskandar Adnan, Sukma Oktavianthi, Lidwina Priliani, Edison Johar, Frilasita A Yudhaputri, Azzania Fibriani, Ema Rahmawati, Ryan Bayusantika Ristandi, Rifky Waluyajati Rachman, Cut Nur Cinthia Alamanda, Safarina G Malik, Khin Saw Myint, Amin Soebandrio |
| EPI_ISL_2918644 | Puskesmas Rancamanyar | Eijkman Institute for Molecular Biology, National Research and Innovation Agency; West Java Health Laboratory | Willy Agustine, Hidayat Trimarsanto, Lydia V. Panggalo, Iskandar Adnan, Sukma Oktavianthi, Lidwina Priliani, Edison Johar, Frilasita A Yudhaputri, Muhammad Rezki Rasyak, Azzania Fibriani, Ema Rahmawati, Ryan Bayusantika Ristandi, Rifky Waluyajati Rachman, Cut Nur Cinthia Alamanda, Safarina G Malik, Khin Saw Myint, Amin Soebandrio |
| EPI_ISL_2918645 | Puskesmas Rancamanyar | Eijkman Institute for Molecular Biology, National Research and Innovation Agency; West Java Health Laboratory | Hidayat Trimarsanto, Lydia V. Panggalo, Iskandar Adnan, Sukma Oktavianthi, Lidwina Priliani, Edison Johar, Frilasita A Yudhaputri, Muhammad Rezki Rasyak, Willy Agustine, Azzania Fibriani, Ema Rahmawati, Ryan Bayusantika Ristandi, Rifky Waluyajati Rachman, Cut Nur Cinthia Alamanda, Safarina G Malik, Khin Saw Myint, Amin Soebandrio |
| EPI_ISL_2918646 | Puskesmas Baleendah | Eijkman Institute for Molecular Biology, National Research and Innovation Agency; West Java Health Laboratory | Lydia V. Panggalo, Iskandar Adnan, Sukma Oktavianthi, Lidwina Priliani, Edison Johar, Frilasita A Yudhaputri, Muhammad Rezki Rasyak, Willy Agustine, Hidayat Trimarsanto, Azzania Fibriani, Ema Rahmawati, Ryan Bayusantika Ristandi, Rifky Waluyajati Rachman, Cut Nur Cinthia Alamanda, Safarina G Malik, Khin Saw Myint, Amin Soebandrio |
| EPI_ISL_2918647 | Puskesmas Baleendah | Eijkman Institute for Molecular Biology, National Research and Innovation Agency; West Java Health Laboratory | Iskandar Adnan, Sukma Oktavianthi, Lidwina Priliani, Edison Johar, Frilasita A Yudhaputri, Muhammad Rezki Rasyak, Willy Agustine, Hidayat Trimarsanto, Lydia V. Panggalo, Azzania Fibriani, Ema Rahmawati, Ryan Bayusantika Ristandi, Rifky Waluyajati Rachman, Cut Nur Cinthia Alamanda, Safarina G Malik, Khin Saw Myint, Amin Soebandrio |
| EPI_ISL_2918648 | Puskesmas Baleendah | Eijkman Institute for Molecular Biology, National Research and Innovation Agency; West Java Health Laboratory | Sukma Oktavianthi, Lidwina Priliani, Edison Johar, Frilasita A Yudhaputri, Muhammad Rezki Rasyak, Willy Agustine, Hidayat Trimarsanto, Lydia V. Panggalo, Iskandar Adnan, Azzania Fibriani, Ema Rahmawati, Ryan Bayusantika Ristandi, Rifky Waluyajati Rachman, Cut Nur Cinthia Alamanda, Safarina G Malik, Khin Saw Myint, Amin Soebandrio |
| EPI_ISL_2918649 | Puskesmas Baleendah | Eijkman Institute for Molecular Biology, National Research and Innovation Agency; West Java Health Laboratory | Lidwina Priliani, Edison Johar, Frilasita A Yudhaputri, Muhammad Rezki Rasyak, Willy Agustine, Hidayat Trimarsanto, Lydia V. Panggalo, Iskandar Adnan, Sukma Oktavianthi, Azzania Fibriani, Ema Rahmawati, Ryan Bayusantika Ristandi, Rifky Waluyajati Rachman, Cut Nur Cinthia Alamanda, Safarina G Malik, Khin Saw Myint, Amin Soebandrio |
| EPI_ISL_2918754 | New Mexico Department of Health Scientific Laboratory | New Mexico Department of Health Scientific Laboratory | Ellie Johnson, D'eldra Malone, Jennifer Benoit, Ratheesh Rajan, Linda Salazar, Keila Gutierrez, Anastacia Griego-Fisher |
| EPI_ISL_2918756, EPI_ISL_2918758, EPI_ISL_2918759 | Labormedizinisches Zentrum Dr Risch | Clinical Bacteriology | Tim Roloff, Fanny Wegner, Helena MB Seth-Smith, Alfredo Mari, Karoline Leuzinger, Julia Bielicki, Nadia Wohlwend,Martin Risch, Lorenz Risch, Manuel Battegay, Hans Hirsch, Adrian Egli |
| EPI_ISL_2918762 | SMS, Medical College, Jaipur | SMS, Medical College, Jaipur | Bharti Malhotra, Sudhher Bhandari, Swati Gatuam, Pratibha Mishra, Dinesh parsoya, Himanshu Sharma, Neha Bhomia |
| EPI_ISL_2918763, EPI_ISL_2918764, EPI_ISL_2918767 | Center for Laboratory Medicine | Center for Laboratory Medicine | Yannick Gerth |
| EPI_ISL_2918791 | Spital Männedorf AG | Institute of Medical Virology | Daniel Ehrsam, Isabel Stürmer, Catharine Aquino, Joel Wirz, Weihong Qi, Hubert Rehrauer, Verena Kufner, Gabriela Ziltener, Maryam Zaheri, Stefan Schmutz, Annette Audigé, Maria Grünberg, Kevin Steiner, Jon Huder, Cyril Shah, Riccarda Capaul, Guido Bloemberg, Jürg Böni, Michael Huber, Alexandra Trkola |
| EPI_ISL_2918835 | NCSLPH | NCSLPH | Chase K, Miller MC, Greene S, Glover W |
| EPI_ISL_2918838 | Kantonsspital Schaffhausen | Institute of Medical Virology | Verena Kufner, Gabriela Ziltener, Maryam Zaheri, Stefan Schmutz, Annette Audigé, Maria Grünberg, Kevin Steiner, Jon Huder, Cyril Shah, Riccarda Capaul, Guido Bloemberg, Jürg Böni, Michael Huber, Alexandra Trkola |
| EPI_ISL_2918839 | UniversitätsSpital Zürich 009 | Institute of Medical Virology | Verena Kufner, Gabriela Ziltener, Maryam Zaheri, Stefan Schmutz, Annette Audigé, Maria Grünberg, Kevin Steiner, Jon Huder, Cyril Shah, Riccarda Capaul, Guido Bloemberg, Jürg Böni, Michael Huber, Alexandra Trkola |
| EPI_ISL_2918840 | UniversitätsSpital Zürich | Institute of Medical Virology | Verena Kufner, Gabriela Ziltener, Maryam Zaheri, Stefan Schmutz, Annette Audigé, Maria Grünberg, Kevin Steiner, Jon Huder, Cyril Shah, Riccarda Capaul, Guido Bloemberg, Jürg Böni, Michael Huber, Alexandra Trkola |
| EPI_ISL_2918841 | Spital Limmattal | Institute of Medical Virology | Verena Kufner, Gabriela Ziltener, Maryam Zaheri, Stefan Schmutz, Annette Audigé, Maria Grünberg, Kevin Steiner, Jon Huder, Cyril Shah, Riccarda Capaul, Guido Bloemberg, Jürg Böni, Michael Huber, Alexandra Trkola |
| EPI_ISL_2918842, EPI_ISL_2918843, EPI_ISL_2918844 | Stadtspital Triemli | Institute of Medical Virology | Verena Kufner, Gabriela Ziltener, Maryam Zaheri, Stefan Schmutz, Annette Audigé, Maria Grünberg, Kevin Steiner, Jon Huder, Cyril Shah, Riccarda Capaul, Guido Bloemberg, Jürg Böni, Michael Huber, Alexandra Trkola |
| EPI_ISL_2918847 | UniversitätsSpital Zürich | Institute of Medical Virology | Verena Kufner, Gabriela Ziltener, Maryam Zaheri, Stefan Schmutz, Annette Audigé, Maria Grünberg, Kevin Steiner, Jon Huder, Cyril Shah, Riccarda Capaul, Guido Bloemberg, Jürg Böni, Michael Huber, Alexandra Trkola |
| EPI_ISL_2918849, EPI_ISL_2918850, EPI_ISL_2918851 | Stadtspital Triemli | Institute of Medical Virology | Verena Kufner, Gabriela Ziltener, Maryam Zaheri, Stefan Schmutz, Annette Audigé, Maria Grünberg, Kevin Steiner, Jon Huder, Cyril Shah, Riccarda Capaul, Guido Bloemberg, Jürg Böni, Michael Huber, Alexandra Trkola |
| EPI_ISL_2918852 | Spital Männedorf AG | Institute of Medical Virology | Verena Kufner, Gabriela Ziltener, Maryam Zaheri, Stefan Schmutz, Annette Audigé, Maria Grünberg, Kevin Steiner, Jon Huder, Cyril Shah, Riccarda Capaul, Guido Bloemberg, Jürg Böni, Michael Huber, Alexandra Trkola |
| EPI_ISL_2918854, EPI_ISL_2918855, EPI_ISL_2918856, EPI_ISL_2918857 | UniversitätsSpital Zürich | Institute of Medical Virology | Verena Kufner, Gabriela Ziltener, Maryam Zaheri, Stefan Schmutz, Annette Audigé, Maria Grünberg, Kevin Steiner, Jon Huder, Cyril Shah, Riccarda Capaul, Guido Bloemberg, Jürg Böni, Michael Huber, Alexandra Trkola |
| EPI_ISL_2918858 | Spital Limmattal | Institute of Medical Virology | Verena Kufner, Gabriela Ziltener, Maryam Zaheri, Stefan Schmutz, Annette Audigé, Maria Grünberg, Kevin Steiner, Jon Huder, Cyril Shah, Riccarda Capaul, Guido Bloemberg, Jürg Böni, Michael Huber, Alexandra Trkola |
| EPI_ISL_2918860, EPI_ISL_2918861, EPI_ISL_2918862 | UniversitätsSpital Zürich | Institute of Medical Virology | Verena Kufner, Gabriela Ziltener, Maryam Zaheri, Stefan Schmutz, Annette Audigé, Maria Grünberg, Kevin Steiner, Jon Huder, Cyril Shah, Riccarda Capaul, Guido Bloemberg, Jürg Böni, Michael Huber, Alexandra Trkola |
| EPI_ISL_2918864, EPI_ISL_2918866 | Kantonsspital Baden AG | Institute of Medical Virology | Verena Kufner, Gabriela Ziltener, Maryam Zaheri, Stefan Schmutz, Annette Audigé, Maria Grünberg, Kevin Steiner, Jon Huder, Cyril Shah, Riccarda Capaul, Guido Bloemberg, Jürg Böni, Michael Huber, Alexandra Trkola |
| EPI_ISL_2918868, EPI_ISL_2918869 | Spital Männedorf AG | Institute of Medical Virology | Verena Kufner, Gabriela Ziltener, Maryam Zaheri, Stefan Schmutz, Annette Audigé, Maria Grünberg, Kevin Steiner, Jon Huder, Cyril Shah, Riccarda Capaul, Guido Bloemberg, Jürg Böni, Michael Huber, Alexandra Trkola |
| EPI_ISL_2918870 | Spital Limmattal | Institute of Medical Virology | Verena Kufner, Gabriela Ziltener, Maryam Zaheri, Stefan Schmutz, Annette Audigé, Maria Grünberg, Kevin Steiner, Jon Huder, Cyril Shah, Riccarda Capaul, Guido Bloemberg, Jürg Böni, Michael Huber, Alexandra Trkola |
| EPI_ISL_2918871, EPI_ISL_2918873 | UniversitätsSpital Zürich | Institute of Medical Virology | Verena Kufner, Gabriela Ziltener, Maryam Zaheri, Stefan Schmutz, Annette Audigé, Maria Grünberg, Kevin Steiner, Jon Huder, Cyril Shah, Riccarda Capaul, Guido Bloemberg, Jürg Böni, Michael Huber, Alexandra Trkola |
| EPI_ISL_2918875, EPI_ISL_2918877 | Kantonsspital Winterthur | Institute of Medical Virology | Verena Kufner, Gabriela Ziltener, Maryam Zaheri, Stefan Schmutz, Annette Audigé, Maria Grünberg, Kevin Steiner, Jon Huder, Cyril Shah, Riccarda Capaul, Guido Bloemberg, Jürg Böni, Michael Huber, Alexandra Trkola |
| EPI_ISL_2918880, EPI_ISL_2918881, EPI_ISL_2918883, EPI_ISL_2918884, EPI_ISL_2918887, EPI_ISL_2918888, EPI_ISL_2918889, EPI_ISL_2918891, EPI_ISL_2918892, EPI_ISL_2918893, EPI_ISL_2918894, EPI_ISL_2918895, EPI_ISL_2918896, EPI_ISL_2918897, EPI_ISL_2918898, EPI_ISL_2918899, EPI_ISL_2918900, EPI_ISL_2918901, EPI_ISL_2918902, EPI_ISL_2918903, EPI_ISL_2918904, EPI_ISL_2918905, EPI_ISL_2918907, EPI_ISL_2918910 | | | |
| see above | Laboratorio di Microbiologia | Laboratorio di Microbiologia | Martinetti Lucchini Gladys, Valeria Spina |
| EPI_ISL_2918912, EPI_ISL_2918915 | HOSPITAL UNIVERSITARIO SON ESPASES | HOSPITAL UNIVERSITARIO SON ESPASES | Servicio de Microbiología, Hospital Universitario Son Espases |
| EPI_ISL_2918916 | HOSPITAL CAN MISSES | HOSPITAL UNIVERSITARIO SON ESPASES | Servicio de Microbiología, Hospital Universitario Son Espases |
| EPI_ISL_2918917, EPI_ISL_2918918 | HOSPITAL UNIVERSITARIO SON ESPASES | HOSPITAL UNIVERSITARIO SON ESPASES | Servicio de Microbiología, Hospital Universitario Son Espases |
| EPI_ISL_2918919 | HOSPITAL UNIVERSITARIO SON LLATZER | HOSPITAL UNIVERSITARIO SON ESPASES | Servicio de Microbiología, Hospital Universitario Son Espases |
| EPI_ISL_2918921, EPI_ISL_2918922 | HOSPITAL UNIVERSITARIO SON ESPASES | HOSPITAL UNIVERSITARIO SON ESPASES | Servicio de Microbiología, Hospital Universitario Son Espases |
| EPI_ISL_2918923 | HOSPITAL CAN MISSES | HOSPITAL UNIVERSITARIO SON ESPASES | Servicio de Microbiología, Hospital Universitario Son Espases |
| EPI_ISL_2918925 | HOSPITAL UNIVERSITARIO SON LLATZER | HOSPITAL UNIVERSITARIO SON ESPASES | Servicio de Microbiología, Hospital Universitario Son Espases |
| EPI_ISL_2918926 | HOSPITAL UNIVERSITARIO SON ESPASES | HOSPITAL UNIVERSITARIO SON ESPASES | Servicio de Microbiología, Hospital Universitario Son Espases |
| EPI_ISL_2918927 | HOSPITAL MATEU ORFILA | HOSPITAL UNIVERSITARIO SON ESPASES | Servicio de Microbiología, Hospital Universitario Son Espases |
| EPI_ISL_2918928 | HOSPITAL CAN MISSES | HOSPITAL UNIVERSITARIO SON ESPASES | Servicio de Microbiología, Hospital Universitario Son Espases |
| EPI_ISL_2918929 | HOSPITAL UNIVERSITARIO SON ESPASES | HOSPITAL UNIVERSITARIO SON ESPASES | Servicio de Microbiología, Hospital Universitario Son Espases |
| EPI_ISL_2918930 | HOSPITAL MANACOR | HOSPITAL UNIVERSITARIO SON ESPASES | Servicio de Microbiología, Hospital Universitario Son Espases |
| EPI_ISL_2918931, EPI_ISL_2918932, EPI_ISL_2918934, EPI_ISL_2918935 | HOSPITAL UNIVERSITARIO SON ESPASES | HOSPITAL UNIVERSITARIO SON ESPASES | Servicio de Microbiología, Hospital Universitario Son Espases |
| EPI_ISL_2918936 | HOSPITAL CAN MISSES | HOSPITAL UNIVERSITARIO SON ESPASES | Servicio de Microbiología, Hospital Universitario Son Espases |
| EPI_ISL_2918938, EPI_ISL_2918939 | HOSPITAL UNIVERSITARIO SON ESPASES | HOSPITAL UNIVERSITARIO SON ESPASES | Servicio de Microbiología, Hospital Universitario Son Espases |

| | | | |
|---|---|---|---|
| EPI_ISL_2918941, EPI_ISL_2918942, EPI_ISL_2918943, EPI_ISL_2918944, EPI_ISL_2918945, EPI_ISL_2918946, EPI_ISL_2918947, EPI_ISL_2918948, EPI_ISL_2918949, EPI_ISL_2918950, EPI_ISL_2918951, EPI_ISL_2918952, EPI_ISL_2918953, EPI_ISL_2918954, EPI_ISL_2918955, EPI_ISL_2918956, EPI_ISL_2918957, EPI_ISL_2918958, EPI_ISL_2918959 | | | |
| see above | Babina Diagnostics | CDFD | Murali Bashyam, Asmita Gupta, Reelina Basu, Vinay Donipadi, Divya Vashisht, Ashwin Dalal |
| EPI_ISL_2919042, EPI_ISL_2919043, EPI_ISL_2919044, EPI_ISL_2919045, EPI_ISL_2919046, EPI_ISL_2919047, EPI_ISL_2919048, EPI_ISL_2919049, EPI_ISL_2919050, EPI_ISL_2919051, EPI_ISL_2919052 | | | |
| see above | Kansas Health and Environmental Lab | Kansas Health and Environmental Lab | Mike Grose, Katherine Wiggins, Jonathan Barnell, Ben Olsen, and Phil Adam |
| EPI_ISL_2919053, EPI_ISL_2919054, EPI_ISL_2919057, EPI_ISL_2919059, EPI_ISL_2919060, EPI_ISL_2919061, EPI_ISL_2919062, EPI_ISL_2919063, EPI_ISL_2919064, EPI_ISL_2919065 | Clinical Reference Laboratory | Kansas Health and Environmental Lab | Mike Grose, Katherine Wiggins, Jonathan Barnell, Ben Olsen, and Phil Adam |
| EPI_ISL_2919066, EPI_ISL_2919067, EPI_ISL_2919068, EPI_ISL_2919069 | Wichita State University - Molecular Diagnostics Lab | Kansas Health and Environmental Lab | Mike Grose, Katherine Wiggins, Jonathan Barnell, Ben Olsen, and Phil Adam |
| EPI_ISL_2919070, EPI_ISL_2919077, EPI_ISL_2919085, EPI_ISL_2919086, EPI_ISL_2919087, EPI_ISL_2919088, EPI_ISL_2919090, EPI_ISL_2919091, EPI_ISL_2919094, EPI_ISL_2919095, EPI_ISL_2919098, EPI_ISL_2919099, EPI_ISL_2919100, EPI_ISL_2919102, EPI_ISL_2919103 | | | |
| see above | Arizona State Public Health Laboratory | Arizona State Public Health Laboratory | Trung Huynh, Jessica Escobar, Katherine Fullerton, Nobuko Fukushima, Matthew Contursi, Stacy White, Linda Getsinger, Victor Waddell |
| EPI_ISL_2919105, EPI_ISL_2919106, EPI_ISL_2919112, EPI_ISL_2919114, EPI_ISL_2919115 | Università degli Studi di Perugia | Istituto Zooprofilattico Sperimentale dell'Abruzzo e Molise "G. Caporale" | Mencacci A, Camilloni B, Biagetti M, Giammarioli M, Lorusso A, Marcacci M, Di Domenico M, Ancora M, Curini V, Mangone I, Rinaldi A, Delli Compagni E, Scialabba S, Di Pasquale A, Cammà C, Puglia I, Calistri P, Savini G |
| EPI_ISL_2919118, EPI_ISL_2919119 | Laboratorio Analisi Osp. Città di Castello - Azienda USL Umbria1 | Istituto Zooprofilattico Sperimentale dell'Abruzzo e Molise "G. Caporale" | Malagigi V, Tacconi P,Lorusso A, Marcacci M, Di Domenico M, Ancora M, Curini V, Mangone I, Rinaldi A, Delli Compagni E, Scialabba S, Di Pasquale A, Cammà C, Puglia I, Calistri P, Savini G |
| EPI_ISL_2919120 | Università degli Studi di Perugia | Istituto Zooprofilattico Sperimentale dell'Abruzzo e Molise "G. Caporale" | Mencacci A, Camilloni B, Biagetti M, Giammarioli M, Lorusso A, Marcacci M, Di Domenico M, Ancora M, Curini V, Mangone I, Rinaldi A, Delli Compagni E, Scialabba S, Di Pasquale A, Cammà C, Puglia I, Calistri P, Savini G |
| EPI_ISL_2919124 | Azienda USL Umbria 2 | Istituto Zooprofilattico Sperimentale dell'Abruzzo e Molise "G. Caporale" | Proietti A, Pistoni E, Lorusso A, Marcacci M, Di Domenico M, Ancora M, Curini V, Mangone I, Rinaldi A, Delli Compagni E, Scialabba S, Di Pasquale A, Cammà C, Puglia I, Calistri P, Savini G |
| EPI_ISL_2919137, EPI_ISL_2919138, EPI_ISL_2919139, EPI_ISL_2919140 | Dianalabs SA | Genesupport | Tanguy ARAUD, Geraldine Jost, Katia Jaton, Nadia Liassine |
| EPI_ISL_2919142, EPI_ISL_2919143, EPI_ISL_2919145, EPI_ISL_2919146, EPI_ISL_2919149, EPI_ISL_2919150, EPI_ISL_2919152, EPI_ISL_2919154, EPI_ISL_2919155, EPI_ISL_2919161 | Fakultní nemocnice Hradec Králové | Fakultní nemocnice Hradec Králové | Marketa Gancarcikova, Helena Kovarikova |
| EPI_ISL_2919163, EPI_ISL_2919164, EPI_ISL_2919165, EPI_ISL_2919166, EPI_ISL_2919167, EPI_ISL_2919168, EPI_ISL_2919169, EPI_ISL_2919170, EPI_ISL_2919171, EPI_ISL_2919172, EPI_ISL_2919173, EPI_ISL_2919174, EPI_ISL_2919175, EPI_ISL_2919176, EPI_ISL_2919177, EPI_ISL_2919178, EPI_ISL_2919179, EPI_ISL_2919180, EPI_ISL_2919181, EPI_ISL_2919182, EPI_ISL_2919183, EPI_ISL_2919184, EPI_ISL_2919185, EPI_ISL_2919186, EPI_ISL_2919187, EPI_ISL_2919188, EPI_ISL_2919189 | | | |
| see above | Kansas Health and Environmental Lab | Kansas Health and Environmental Lab | Mike Grose, Katherine Wiggins, Jonathan Barnell, Ben Olsen, and Phil Adam |
| EPI_ISL_2919318, EPI_ISL_2919401, EPI_ISL_2919405, EPI_ISL_2919448, EPI_ISL_2919517, EPI_ISL_2919554, EPI_ISL_2919586, EPI_ISL_2919611, EPI_ISL_2919627, EPI_ISL_2919717, EPI_ISL_2919774, EPI_ISL_2919784, EPI_ISL_2919803, EPI_ISL_2919824, EPI_ISL_2919836, EPI_ISL_2919871 | | | |
| see above | TGen North | TGen North | "Jolene Bowers, Heather Centner, Brett Van Tassel, Chris French, Hayley Yaglom, Darrin Lemmer, Dave Engelthaler" |
| EPI_ISL_2919995, EPI_ISL_2919998, EPI_ISL_2920000, EPI_ISL_2920001, EPI_ISL_2920004 | County Hospital of Liberec, Dept. of genetic and molecular diagnostic | Institute of Medical Microbiology and Virology, University Hospital Carl Gustav Carus, TU Dresden | Marlena Stadtmüller, Alexa Laubner, Johanna Beil, Fabian Rost, Sylvia Klemroth, Grit Mehnert, Montserrat Palau de Miquel, Susanne Reinhardt, Leo Büttner, Anett Zabzinski, Sylke Winkler, Alexander Dalpke, Iva Dolinová |
| EPI_ISL_2920014, EPI_ISL_2920056 | Oddělení klinické mikrobiologie | Institute of Medical Microbiology and Virology, University Hospital Carl Gustav Carus, TU Dresden | Marlena Stadtmüller, Alexa Laubner, Johanna Beil, Fabian Rost, Sylvia Klemroth, Grit Mehnert, Montserrat Palau de Miquel, Susanne Reinhardt, Leo Büttner, Anett Zabzinski, Sylke Winkler, Alexander Dalpke, Eva Patrasová, Lenka Šimůnková, Lenka Dvořáková |
| EPI_ISL_2920079, EPI_ISL_2920080 | County Hospital of Liberec, Dept. of genetic and molecular diagnostic | Institute of Medical Microbiology and Virology, University Hospital Carl Gustav Carus, TU Dresden | Marlena Stadtmüller, Alexa Laubner, Johanna Beil, Fabian Rost, Sylvia Klemroth, Grit Mehnert, Montserrat Palau de Miquel, Susanne Reinhardt, Leo Büttner, Anett Zabzinski, Sylke Winkler, Alexander Dalpke, Iva Dolinová |
| EPI_ISL_2920088, EPI_ISL_2920089 | University Hospital Brno, OKMI | University Hospital Brno, CMBG | Matej Bezdicek, Jan Svaton, Martina Lengerova, Monika Dolejska |
| EPI_ISL_2920091 | OKMI | CMBG | Matej Bezdicek, Jan Svaton, Martina Lengerova, Monika Dolejska |
| EPI_ISL_2920092, EPI_ISL_2920093 | FNUSA | CMBG | Matej Bezdicek, Jan Svaton, Martina Lengerova, Monika Dolejska |
| EPI_ISL_2920095, EPI_ISL_2920096 | OKMI | CMBG | Matej Bezdicek, Jan Svaton, Martina Lengerova, Monika Dolejska |
| EPI_ISL_2920217, EPI_ISL_2920219, EPI_ISL_2920220, EPI_ISL_2920225, EPI_ISL_2920226, EPI_ISL_2920228, EPI_ISL_2920231, EPI_ISL_2920235, EPI_ISL_2920239, EPI_ISL_2920240, EPI_ISL_2920241, EPI_ISL_2920242, EPI_ISL_2920244, EPI_ISL_2920245 | | | |
| see above | URMC LABS | URMC LABS | Dwight Hardy, Andrew Cameron, Stepan Voljehn, Kelly DeLary |
| EPI_ISL_2920257 | Massachusetts State Public Health Laboratory | Massachusetts State Public Health Laboratory | Andrew Lang, Timelia Fink, Glen Gallagher, Sandra Smole |
| EPI_ISL_2920281 | City of Milwaukee Health Department Laboratory | City of Milwaukee Health Department Laboratory | Manjeet Khubbar, Amy Bauer, Jennifer Lentz, Samantha Scott, Sanjib Bhattacharyya |
| EPI_ISL_2920432, EPI_ISL_2920433, EPI_ISL_2920434, EPI_ISL_2920436, EPI_ISL_2920437, EPI_ISL_2920438, EPI_ISL_2920439, EPI_ISL_2920441, EPI_ISL_2920442, EPI_ISL_2920443 | Sharp HealthCare Laboratory | Andersen lab at Scripps Research | SEARCH Alliance San Diego with Aaron Harding, Jacquelyn Berumen, Cathy Woerle, Liam McGinnis, Art Mendoza, Omid Bakhtar |
| EPI_ISL_2920460, EPI_ISL_2920488, EPI_ISL_2920510, EPI_ISL_2920511, EPI_ISL_2920519, EPI_ISL_2920557, EPI_ISL_2920578, EPI_ISL_2920596, EPI_ISL_2920600, EPI_ISL_2920609, EPI_ISL_2920618, EPI_ISL_2920649, EPI_ISL_2920653, EPI_ISL_2920655, EPI_ISL_2920656, EPI_ISL_2920657, EPI_ISL_2920658, EPI_ISL_2920664, EPI_ISL_2920667, EPI_ISL_2920669, EPI_ISL_2920671, EPI_ISL_2920672, EPI_ISL_2920674, EPI_ISL_2920677, EPI_ISL_2920679, EPI_ISL_2920680 | | | |
| see above | Florida Bureau of Public Health Laboratories | Florida Bureau of Public Health Laboratories | Sarah Schmedes, Jason Blanton |
| EPI_ISL_2920792, EPI_ISL_2920795, EPI_ISL_2920801, EPI_ISL_2920804, EPI_ISL_2920806, EPI_ISL_2920808, EPI_ISL_2920809, EPI_ISL_2920814, EPI_ISL_2920815, EPI_ISL_2920816, EPI_ISL_2920817, EPI_ISL_2920818, EPI_ISL_2920819, EPI_ISL_2920820, EPI_ISL_2920821, EPI_ISL_2920822, EPI_ISL_2920823 | | | |
| see above | Virginia Division of Consolidated Laboratory Services | Virginia Division of Consolidated Laboratory Services | Virginia Division of Consolidated Laboratory Services |
| EPI_ISL_2920824, EPI_ISL_2920825, EPI_ISL_2920826, EPI_ISL_2920827, EPI_ISL_2920828, EPI_ISL_2920829, EPI_ISL_2920830, EPI_ISL_2920831, EPI_ISL_2920833, EPI_ISL_2920834, EPI_ISL_2920835, EPI_ISL_2920836, EPI_ISL_2920837, EPI_ISL_2920838, EPI_ISL_2920839, EPI_ISL_2920840, EPI_ISL_2920841, EPI_ISL_2920842, EPI_ISL_2920843, EPI_ISL_2920844, EPI_ISL_2920845, EPI_ISL_2920846, EPI_ISL_2920847, EPI_ISL_2920848, EPI_ISL_2920849, EPI_ISL_2920850 | | | |
| see above | University of Mississippi Medical Center, Department of Pathology | University of Mississippi Medical Center, Molecular and Genomics Core Facility | Ashley C. Johnson, Wenjie Wu, Ithiel J. Frame, Krishna K. Ayyalasomayajula, Michael R. Garrett, D. Ashley Robinson |
| EPI_ISL_2920851, EPI_ISL_2920852, EPI_ISL_2920853, EPI_ISL_2920854, EPI_ISL_2920855, EPI_ISL_2920856, EPI_ISL_2920857, EPI_ISL_2920858, EPI_ISL_2920859, EPI_ISL_2920860, EPI_ISL_2920861, EPI_ISL_2920864, EPI_ISL_2920866, EPI_ISL_2920867, EPI_ISL_2920868, EPI_ISL_2920872, EPI_ISL_2920874, EPI_ISL_2920875, EPI_ISL_2920876, EPI_ISL_2920877, EPI_ISL_2920878, EPI_ISL_2920879, EPI_ISL_2920880, EPI_ISL_2920883, EPI_ISL_2920885, EPI_ISL_2920886, EPI_ISL_2920887, EPI_ISL_2920888, EPI_ISL_2920889, EPI_ISL_2920890, EPI_ISL_2920891, EPI_ISL_2920892, EPI_ISL_2920893, EPI_ISL_2920895, EPI_ISL_2920896, EPI_ISL_2920897, EPI_ISL_2920899, EPI_ISL_2920900, EPI_ISL_2920902, EPI_ISL_2920903, EPI_ISL_2920904, EPI_ISL_2920905, EPI_ISL_2920906, EPI_ISL_2920907, EPI_ISL_2920908, EPI_ISL_2920909, EPI_ISL_2920911, EPI_ISL_2920912, EPI_ISL_2920913, EPI_ISL_2920914, EPI_ISL_2920915, EPI_ISL_2920916, EPI_ISL_2920917, EPI_ISL_2920918, EPI_ISL_2920919, EPI_ISL_2920920, EPI_ISL_2920921, EPI_ISL_2920922, EPI_ISL_2920923, EPI_ISL_2920924 | | | |
| see above | MS Public Health Laboratory | University of Mississippi Medical Center, Molecular and Genomics Core Facility | Ashley C. Johnson, Wenjie Wu, Ithiel J. Frame, Krishna K. Ayyalasomayajula, Michael R. Garrett, D. Ashley Robinson |

| | | | |
|---|---|---|---|
| EPI_ISL_2920932, EPI_ISL_2920936, EPI_ISL_2920941, EPI_ISL_2920942, EPI_ISL_2920943, EPI_ISL_2920954, EPI_ISL_2920956, EPI_ISL_2920958, EPI_ISL_2920964, EPI_ISL_2920971, EPI_ISL_2920972, EPI_ISL_2920973, EPI_ISL_2920974, EPI_ISL_2920975, EPI_ISL_2920976, EPI_ISL_2920977, EPI_ISL_2920978, EPI_ISL_2920979, EPI_ISL_2920980, EPI_ISL_2920982, EPI_ISL_2920983, EPI_ISL_2920984, EPI_ISL_2920985, EPI_ISL_2920987, EPI_ISL_2920988, EPI_ISL_2920990, EPI_ISL_2920992, EPI_ISL_2920993, EPI_ISL_2920994, EPI_ISL_2920995, EPI_ISL_2921000 | | | |
| see above | Victorian Infectious Diseases Reference Laboratory (VIDRL) | VIDRL and MDU-PHL | Caly L., Seemann T., Sait, M.L., Druce J., Sherry, N.L. |
| EPI_ISL_2921027 | Shenzhen Center for Disease Control and Prevention | National Institute for Viral Disease Control and Prevention, China CDC | Long Chen, Ying Sun,Xiaoliang Xiao,Renli Zhang,Kai Nie,Weihua Wu,Xinyi Wei,Yue Li,Shaoyu Deng and Yaqing He |
| EPI_ISL_2921028, EPI_ISL_2921029 | County of San Luis Obispo Public Health Laboratory | County of San Luis Obispo Public Health Laboratory | Katrina Erwin, Frances Sidhu |
| EPI_ISL_2921031, EPI_ISL_2921032, EPI_ISL_2921033, EPI_ISL_2921034, EPI_ISL_2921035, EPI_ISL_2921036, EPI_ISL_2921037, EPI_ISL_2921038, EPI_ISL_2921039, EPI_ISL_2921040, EPI_ISL_2921041, EPI_ISL_2921042, EPI_ISL_2921043, EPI_ISL_2921044 | | | |
| see above | Servicio Microbiología Hospital La Paz | Servicio Microbiología Hospital La Paz | Fernando Lázaro, Rubén Cáceres, Jesús Mingorance Cruz, Elie Dahdouh |
| EPI_ISL_2921141, EPI_ISL_2921145, EPI_ISL_2921146, EPI_ISL_2921147, EPI_ISL_2921148, EPI_ISL_2921150, EPI_ISL_2921151, EPI_ISL_2921156, EPI_ISL_2921158, EPI_ISL_2921161, EPI_ISL_2921162, EPI_ISL_2921163, EPI_ISL_2921165, EPI_ISL_2921166 | | | |
| see above | AZDelta | AZ Delta Medical Laboratories in Roeselare, Belgium | Geert Martens, Dieter De Smet, Merijn Vanhee, on behalf of AZ Delta COVID-19 Genomics core (member of Genomic surveillance of SARS-CoV-2 in Belgium network) |
| EPI_ISL_2921181, EPI_ISL_2921183, EPI_ISL_2921184, EPI_ISL_2921185 | Oman-NIC | Oman-National Influenza Center | Laila Al Balushi, Aisha Al-Busaidi, Samiha Al Kharusi, Amina Al Jardani, Hanan Al Kindi, Intisar Al-Shukri, Fahad Al Balushi, Bilal Hussain. |
| EPI_ISL_2921200 | Oman-NIC | Oman-National Influenza Center | Samiha Al Kharusi, Laila Al Balushi, Aisha Al-Busaidi, Amina Al Jardani, Hanan Al Kindi, Intisar Al-Shukri. |
| EPI_ISL_2921208 | OSPEDALE SAN SALVATORE - MEDICINA DI LABORATORIO | Istituto Zooprofilattico Sperimentale dell'Abruzzo e Molise "G. Caporale" | Lorusso A, Marcacci M, Di Domenico M, Ancora M, Curini V, Di Lollo Valeria, Mangone I, Rinaldi A, Delli Compagni E, Scialabba S, Caporale M, Di Pasquale A, Cammà C, Puglia I, Calistri P, Savini G |
| EPI_ISL_2921216, EPI_ISL_2921218 | PO AVEZZANO - U.O.C. Laboratorio Analisi | Istituto Zooprofilattico Sperimentale dell'Abruzzo e Molise "G. Caporale" | Lorusso A, Marcacci M, Di Domenico M, Ancora M, Curini V, Di Lollo Valeria, Mangone I, Rinaldi A, Delli Compagni E, Scialabba S, Caporale M, Di Pasquale A, Cammà C, Puglia I, Calistri P, Savini G |
| EPI_ISL_2921226, EPI_ISL_2921227, EPI_ISL_2921229, EPI_ISL_2921230, EPI_ISL_2921233, EPI_ISL_2921234, EPI_ISL_2921235, EPI_ISL_2921236, EPI_ISL_2921237, EPI_ISL_2921238, EPI_ISL_2921239, EPI_ISL_2921240, EPI_ISL_2921241, EPI_ISL_2921242 | | | |
| see above | OSPEDALE CIVILE TERAMO - CENTRO TRASFUSIONALE | Istituto Zooprofilattico Sperimentale dell'Abruzzo e Molise "G. Caporale" | Lorusso A, Marcacci M, Di Domenico M, Ancora M, Curini V, Di Lollo Valeria, Mangone I, Rinaldi A, Delli Compagni E, Scialabba S, Caporale M, Di Pasquale A, Cammà C, Puglia I, Calistri P, Savini G |
| EPI_ISL_2921255 | PO AVEZZANO - U.O.C. Laboratorio Analisi | Istituto Zooprofilattico Sperimentale dell'Abruzzo e Molise "G. Caporale" | Lorusso A, Marcacci M, Di Domenico M, Ancora M, Curini V, Di Lollo Valeria, Mangone I, Rinaldi A, Delli Compagni E, Scialabba S, Caporale M, Di Pasquale A, Cammà C, Puglia I, Calistri P, Savini G |
| EPI_ISL_2921262 | Pantai Hospital Kuala Lumpur | Institute for Medical Research, Infectious Disease Research Centre, National Institutes of Health, Ministry of Health Malaysia | Suppiah J, Kamel K, Mohd Zawawi Z, Azizan MA, Ramly N, Robert F, Anasir MI, Thayan R |
| EPI_ISL_2921263, EPI_ISL_2921264, EPI_ISL_2921265, EPI_ISL_2921266 | Pahang State Health Department | Institute for Medical Research, Infectious Disease Research Centre, National Institutes of Health, Ministry of Health Malaysia | Suppiah J, Kamel K, Mohd Zawawi Z, Azizan MA, Ramly N, Robert F, Anasir MI, Thayan R |
| EPI_ISL_2921611 | Labuan Hospital | Institute for Medical Research, Infectious Disease Research Centre, National Institutes of Health, Ministry of Health Malaysia | Suppiah J, Kamel K, Mohd Zawawi Z, Azizan MA, Ramly N, Robert F, Anasir MI, Thayan R |
| EPI_ISL_2921612 | Sultan Haji Ahmad Shah Hospital | Institute for Medical Research, Infectious Disease Research Centre, National Institutes of Health, Ministry of Health Malaysia | Suppiah J, Kamel K, Mohd Zawawi Z, Azizan MA, Ramly N, Robert F, Anasir MI, Thayan R |
| EPI_ISL_2921613, EPI_ISL_2921614, EPI_ISL_2921615, EPI_ISL_2921616, EPI_ISL_2921617, EPI_ISL_2921618, EPI_ISL_2921619, EPI_ISL_2921620, EPI_ISL_2921621, EPI_ISL_2921622, EPI_ISL_2921843, EPI_ISL_2921844, EPI_ISL_2921845 | | | |
| see above | New South Wales Health Pathology Royal Prince Alfred Hospital | Microbiology RPAH | Foster, C.; Au, J.; Ruiz Silva, M.; Deveson, I.; Bull, R.; Van Hal, S.; Rawlinson, W. |
| EPI_ISL_2921996 | Melaka Hospital | Institute for Medical Research, Infectious Disease Research Centre, National Institutes of Health, Ministry of Health Malaysia | Suppiah J, Kamel K, Mohd Zawawi Z, Azizan MA, Ramly N, Robert F, Anasir MI, Thayan R |
| EPI_ISL_2921999 | SMS, Medical college | SMS, Medical college | Bharti Malhotra, Sudheer Bhandari, Swati Gautam, Pratibha Mishra, Dinesh Parsoya, Himanshu Sharma, Neha Bhomia |
| EPI_ISL_2922000 | SMS Medical College | SMS Medical College Jaipur | Bharti Malhotra, Sudheer Bhandari, Swati Gautam, Pratibha Sharma, Dinesh Parsoya, Himanshu Sharma, Neha Bhomia |
| EPI_ISL_2922001 | SMS Medical College, Jaipur | SMS Medical College | Bharti Malhotra, Sudheer Bhandari, Swati Gautam, Pratibha Mishra, Dinesh Parsoya, Himanshu Sharma, Neha Bhomia |
| EPI_ISL_2922021, EPI_ISL_2922086, EPI_ISL_2922122, EPI_ISL_2922128, EPI_ISL_2922194, EPI_ISL_2922209, EPI_ISL_2922262 | CDPH VBL | California Department of Public Health | CDPH-COVIDNet |
| EPI_ISL_2922327, EPI_ISL_2922360, EPI_ISL_2922361, EPI_ISL_2922372, EPI_ISL_2922374, EPI_ISL_2922384, EPI_ISL_2922394, EPI_ISL_2922395, EPI_ISL_2922396, EPI_ISL_2922397, EPI_ISL_2922399, EPI_ISL_2922400, EPI_ISL_2922402, EPI_ISL_2922404, EPI_ISL_2922405, EPI_ISL_2922408, EPI_ISL_2922409, EPI_ISL_2922410, EPI_ISL_2922411, EPI_ISL_2922412, EPI_ISL_2922414, EPI_ISL_2922416, EPI_ISL_2922417, EPI_ISL_2922418, EPI_ISL_2922434, EPI_ISL_2922450, EPI_ISL_2922451, EPI_ISL_2922452, EPI_ISL_2922453, EPI_ISL_2922454, EPI_ISL_2922457, EPI_ISL_2922460, EPI_ISL_2922461, EPI_ISL_2922463, EPI_ISL_2922467, EPI_ISL_2922468, EPI_ISL_2922469, EPI_ISL_2922470, EPI_ISL_2922472, EPI_ISL_2922473, EPI_ISL_2922474, EPI_ISL_2922475, EPI_ISL_2922476, EPI_ISL_2922477, EPI_ISL_2922478, EPI_ISL_2922479, EPI_ISL_2922480, EPI_ISL_2922482, EPI_ISL_2922483, EPI_ISL_2922484, EPI_ISL_2922485, EPI_ISL_2922486, EPI_ISL_2922487, EPI_ISL_2922488, EPI_ISL_2922489, EPI_ISL_2922490, EPI_ISL_2922491, EPI_ISL_2922492, EPI_ISL_2922493, EPI_ISL_2922494, EPI_ISL_2922495, EPI_ISL_2922496, EPI_ISL_2922497, EPI_ISL_2922509, EPI_ISL_2922510, EPI_ISL_2922511, EPI_ISL_2922512, EPI_ISL_2922516, EPI_ISL_2922517, EPI_ISL_2922522, EPI_ISL_2922537, EPI_ISL_2922540, EPI_ISL_2922549, EPI_ISL_2922552, EPI_ISL_2922554, EPI_ISL_2922556, EPI_ISL_2922557, EPI_ISL_2922589, EPI_ISL_2922600, EPI_ISL_2922601, EPI_ISL_2922602, EPI_ISL_2922603, EPI_ISL_2922604, EPI_ISL_2922605, EPI_ISL_2922606, EPI_ISL_2922607, EPI_ISL_2922608, EPI_ISL_2922611, EPI_ISL_2922612, EPI_ISL_2922613, EPI_ISL_2922615, EPI_ISL_2922617, EPI_ISL_2922622, EPI_ISL_2922623, EPI_ISL_2922624, EPI_ISL_2922625, EPI_ISL_2922626, EPI_ISL_2922627, EPI_ISL_2922628, EPI_ISL_2922629, EPI_ISL_2922630, EPI_ISL_2922632, EPI_ISL_2922633, EPI_ISL_2922634, EPI_ISL_2922635, EPI_ISL_2922636, EPI_ISL_2922637, EPI_ISL_2922638, EPI_ISL_2922639, EPI_ISL_2922640, EPI_ISL_2922641, EPI_ISL_2922642, EPI_ISL_2922643, EPI_ISL_2922644, EPI_ISL_2922645, EPI_ISL_2922646, EPI_ISL_2922647, EPI_ISL_2922648, EPI_ISL_2922649, EPI_ISL_2922650, EPI_ISL_2922651, EPI_ISL_2922652, EPI_ISL_2922653, EPI_ISL_2922654, EPI_ISL_2922655, EPI_ISL_2922656, EPI_ISL_2922657, EPI_ISL_2922658, EPI_ISL_2922659, EPI_ISL_2922660, EPI_ISL_2922661, EPI_ISL_2922662, EPI_ISL_2922663, EPI_ISL_2922664, EPI_ISL_2922665, EPI_ISL_2922666, EPI_ISL_2922667, EPI_ISL_2922668, EPI_ISL_2922669, EPI_ISL_2922670, EPI_ISL_2922671, EPI_ISL_2922672, EPI_ISL_2922673, EPI_ISL_2922674, EPI_ISL_2922675, EPI_ISL_2922676, EPI_ISL_2922677, EPI_ISL_2922678, EPI_ISL_2922679, EPI_ISL_2922680, EPI_ISL_2922681, EPI_ISL_2922686, EPI_ISL_2922690, EPI_ISL_2922691, EPI_ISL_2922692, EPI_ISL_2922693, EPI_ISL_2922696, EPI_ISL_2922697, EPI_ISL_2922698, EPI_ISL_2922699, EPI_ISL_2922700, EPI_ISL_2922703, EPI_ISL_2922704, EPI_ISL_2922705, EPI_ISL_2922709, EPI_ISL_2922710, EPI_ISL_2922712, EPI_ISL_2922714, EPI_ISL_2922715, EPI_ISL_2922716, EPI_ISL_2922717, EPI_ISL_2922718, EPI_ISL_2922719, EPI_ISL_2922720, EPI_ISL_2922721, EPI_ISL_2922722, EPI_ISL_2922723, EPI_ISL_2922724, EPI_ISL_2922725, EPI_ISL_2922726, EPI_ISL_2922727, EPI_ISL_2922728, EPI_ISL_2922734, EPI_ISL_2922735, EPI_ISL_2922736, EPI_ISL_2922738, EPI_ISL_2922739, EPI_ISL_2922740, EPI_ISL_2922741, EPI_ISL_2922742, EPI_ISL_2922743, EPI_ISL_2922744, EPI_ISL_2922745, EPI_ISL_2922746, EPI_ISL_2922748, EPI_ISL_2922750, EPI_ISL_2922763, EPI_ISL_2922764, EPI_ISL_2922765, EPI_ISL_2922766, EPI_ISL_2922768, EPI_ISL_2922770, EPI_ISL_2922774, EPI_ISL_2922776, EPI_ISL_2922779, EPI_ISL_2922780, EPI_ISL_2922781, EPI_ISL_2922782, EPI_ISL_2922786, EPI_ISL_2922787, EPI_ISL_2922794, EPI_ISL_2922795, EPI_ISL_2922796, EPI_ISL_2922797, EPI_ISL_2922803, EPI_ISL_2922804, EPI_ISL_2922805, EPI_ISL_2922806, EPI_ISL_2922807, EPI_ISL_2922808, EPI_ISL_2922809, EPI_ISL_2922810, EPI_ISL_2922811, EPI_ISL_2922812, EPI_ISL_2922813, EPI_ISL_2922814, EPI_ISL_2922815, EPI_ISL_2922816, EPI_ISL_2922820, EPI_ISL_2922821, EPI_ISL_2922826, EPI_ISL_2922828, EPI_ISL_2922830, EPI_ISL_2922832, EPI_ISL_2922833, EPI_ISL_2922836, EPI_ISL_2922848, EPI_ISL_2922849, EPI_ISL_2922850, EPI_ISL_2922851, EPI_ISL_2922853, EPI_ISL_2922856, EPI_ISL_2922857, EPI_ISL_2922858, EPI_ISL_2922859, EPI_ISL_2922860, EPI_ISL_2922861, EPI_ISL_2922862, EPI_ISL_2922863, EPI_ISL_2922864, EPI_ISL_2922865, EPI_ISL_2922866, EPI_ISL_2922867, EPI_ISL_2922870, EPI_ISL_2922871, EPI_ISL_2922872, EPI_ISL_2922873, EPI_ISL_2922874, EPI_ISL_2922878, EPI_ISL_2922881, EPI_ISL_2922882, EPI_ISL_2922885, EPI_ISL_2922886, EPI_ISL_2922887, EPI_ISL_2922888, EPI_ISL_2922889, EPI_ISL_2922890, EPI_ISL_2922891, EPI_ISL_2922892, EPI_ISL_2922893, EPI_ISL_2922894, EPI_ISL_2922895, EPI_ISL_2922896, EPI_ISL_2922897, EPI_ISL_2922898, EPI_ISL_2922899, EPI_ISL_2922900, EPI_ISL_2922901, EPI_ISL_2922902, EPI_ISL_2922903, EPI_ISL_2922904, EPI_ISL_2922905, EPI_ISL_2922906, EPI_ISL_2922907, EPI_ISL_2922908, EPI_ISL_2922909, EPI_ISL_2922910, EPI_ISL_2922911, EPI_ISL_2922912, EPI_ISL_2922913, EPI_ISL_2922914, EPI_ISL_2922915, EPI_ISL_2922916, EPI_ISL_2922917, EPI_ISL_2922918, EPI_ISL_2922919, EPI_ISL_2922920, EPI_ISL_2922921, EPI_ISL_2922922, EPI_ISL_2922923, EPI_ISL_2922924, EPI_ISL_2922925, EPI_ISL_2922926, EPI_ISL_2922927, EPI_ISL_2922928, EPI_ISL_2922929, EPI_ISL_2922930, EPI_ISL_2922931, EPI_ISL_2922932, EPI_ISL_2922933, EPI_ISL_2922937, EPI_ISL_2922938, EPI_ISL_2922939, EPI_ISL_2922940, EPI_ISL_2922942, EPI_ISL_2922943, EPI_ISL_2922944, EPI_ISL_2922945, EPI_ISL_2922946, EPI_ISL_2922947, EPI_ISL_2922948, EPI_ISL_2922949, EPI_ISL_2922950, EPI_ISL_2922951, EPI_ISL_2922954, EPI_ISL_2922955, EPI_ISL_2922956, EPI_ISL_2922958, EPI_ISL_2922959, EPI_ISL_2922960, EPI_ISL_2922961, EPI_ISL_2922962, EPI_ISL_2922963, EPI_ISL_2922965, EPI_ISL_2922966, EPI_ISL_2922967, EPI_ISL_2922971, EPI_ISL_2922972, EPI_ISL_2922973, EPI_ISL_2922974, EPI_ISL_2922976, EPI_ISL_2922977, EPI_ISL_2922978, EPI_ISL_2922979, EPI_ISL_2922980, EPI_ISL_2922981, EPI_ISL_2922983, EPI_ISL_2922984, EPI_ISL_2922985, EPI_ISL_2922986, EPI_ISL_2922987, EPI_ISL_2922988, EPI_ISL_2922989, EPI_ISL_2922990, EPI_ISL_2922992, EPI_ISL_2922993 | | | |
| see above | Public Health Ontario Laboratory | Public Health Ontario Laboratory | Vanessa G Allen, Philip Banh, Yao Chen, Richard de Borja, Alireza Eshaghi, Nahuel Fittipaldi, Christine Frantz, Jonathan B Gubbay, Jennifer L Guthrie, Lawrence Heisler, Esha Joshi, Michael Laszloffy, Aimin Li, Michael CY Li, Dean Maxwell, Sandeep Nagra, Samir N Patel, Jared Simpson, Karthikeyan Sivaraman, Ashleigh Sullivan, Yogi Sundaravadanam, Sarah Teatero, Andre Villegas, Matthew Watson, Sandra Zittermann |
| EPI_ISL_2922997 | IRCCS San Gallicano Dermatological Institute | IRCCS Regina Elena National Cancer Institute | Francesca De Nicola, Giulia Orlandi, Eleonora Sperandio, Elisabetta Trento, Fabrizio Ensoli, Fulvia Pimpinelli, Maurizio Fanciulli, Giovanni Blandino, Aldo Morrone, Gennaro Ciliberto |
| EPI_ISL_2922998 | IRCCS San Gallicano Dermatological Institute | IRCCS Regina Elena National Cancer Institute | Ludovica Ciuffreda, Sara Donzelli, Alice Massacci, Arianna Mastrofrancesco, Fabrizio Ensoli, Fulvia Pimpinelli, Maurizio Fanciulli, Giovanni Blandino, Aldo Morrone, Gennaro Ciliberto |
| EPI_ISL_2922999 | IRCCS San Gallicano Dermatological Institute | IRCCS Regina Elena National Cancer Institute | Frauke Goeman, Giulia Orlandi, Matteo Pallocca, Giovanna D'agosto, Fabrizio Ensoli, Fulvia Pimpinelli, Maurizio Fanciulli, Giovanni Blandino, Aldo Morrone, Gennaro Ciliberto |
| EPI_ISL_2923001 | IRCCS San Gallicano Dermatological Institute | IRCCS Regina Elena National Cancer Institute | Francesca De Nicola, Giulia Orlandi, Alice Massacci, Martina Pontone, Fabrizio Ensoli, Fulvia Pimpinelli, Maurizio Fanciulli, Giovanni Blandino, Aldo Morrone, Gennaro Ciliberto |
| EPI_ISL_2923002 | IRCCS San Gallicano Dermatological Institute | IRCCS Regina Elena National Cancer Institute | Ludovica Ciuffreda, Sara Donzelli, Matteo Pallocca, di Domenico Enea Gino, Fabrizio Ensoli, Fulvia Pimpinelli, Maurizio Fanciulli, Giovanni Blandino, Aldo Morrone, Gennaro Ciliberto |
| EPI_ISL_2923004 | IRCCS San Gallicano Dermatological Institute | IRCCS Regina Elena National Cancer Institute | Sabrina Strano, Sara Donzelli, Alice Massacci, Francesca Sivori, Fabrizio Ensoli, Fulvia Pimpinelli, Maurizio Fanciulli, Giovanni Blandino, Aldo Morrone, Gennaro Ciliberto |
| EPI_ISL_2923005 | IRCCS San Gallicano Dermatological Institute | IRCCS Regina Elena National Cancer Institute | Francesca De Nicola, Giulia Orlandi, Matteo Pallocca, Andrea Cazzani, Fabrizio Ensoli, Fulvia Pimpinelli, Maurizio Fanciulli, Giovanni Blandino, Aldo Morrone, Gennaro Ciliberto |
| EPI_ISL_2923006 | IRCCS San Gallicano Dermatological Institute | IRCCS Regina Elena National Cancer Institute | Ludovica Ciuffreda, Sara Donzelli, Eleonora Sperandio, Ilaria Celesti, Fabrizio Ensoli, Fulvia Pimpinelli, Maurizio Fanciulli, Giovanni Blandino, Aldo Morrone, Gennaro Ciliberto |
| EPI_ISL_2923007 | IRCCS San Gallicano Dermatological Institute | IRCCS Regina Elena National Cancer Institute | Frauke Goeman, Giulia Orlandi, Alice Massacci, Martina Diano, Fabrizio Ensoli, Fulvia Pimpinelli, Maurizio Fanciulli, Giovanni Blandino, Aldo Morrone, Gennaro Ciliberto |
| EPI_ISL_2923008 | IRCCS San Gallicano Dermatological Institute | IRCCS Regina Elena National Cancer Institute | Francesca De Nicola, Giulia Orlandi, Eleonora Sperandio, Fulvia Fraticelli, Fabrizio Ensoli, Fulvia Pimpinelli, Maurizio Fanciulli, Giovanni Blandino, Aldo Morrone, Gennaro Ciliberto |
| EPI_ISL_2923009 | IRCCS San Gallicano Dermatological Institute | IRCCS Regina Elena National Cancer Institute | Ludovica Ciuffreda, Sara Donzelli, Alice Massacci, Lorenzo Furzi, Fabrizio Ensoli, Fulvia Pimpinelli, Maurizio Fanciulli, Giovanni Blandino, Aldo Morrone, Gennaro Ciliberto |
| EPI_ISL_2923010 | IRCCS San Gallicano Dermatological Institute | IRCCS Regina Elena National Cancer Institute | Frauke Goeman, Giulia Orlandi, Matteo Pallocca, Alessia Lauretti, Fabrizio Ensoli, Fulvia Pimpinelli, Maurizio Fanciulli, Giovanni Blandino, Aldo Morrone, Gennaro Ciliberto |
| EPI_ISL_2923013 | IRCCS San Gallicano Dermatological Institute | IRCCS Regina Elena National Cancer Institute | Ludovica Ciuffreda, Sara Donzelli, Matteo Pallocca, Silvia Paluzzi, Fabrizio Ensoli, Fulvia Pimpinelli, Maurizio Fanciulli, Giovanni Blandino, Aldo Morrone, Gennaro Ciliberto |
| EPI_ISL_2923014 | IRCCS San Gallicano Dermatological Institute | IRCCS Regina Elena National Cancer Institute | Frauke Goeman, Giulia Orlandi, Eleonora Sperandio, Sara Petrolo, Fabrizio Ensoli, Fulvia Pimpinelli, Maurizio Fanciulli, Giovanni Blandino, Aldo Morrone, Gennaro Ciliberto |
| EPI_ISL_2923015 | IRCCS San Gallicano Dermatological Institute | IRCCS Regina Elena National Cancer Institute | Sabrina Strano, Sara Donzelli, Alice Massacci, Valentina Ricca, Fabrizio Ensoli, Fulvia Pimpinelli, Maurizio Fanciulli, Giovanni Blandino, Aldo Morrone, Gennaro Ciliberto |
| EPI_ISL_2923017 | IRCCS San Gallicano Dermatological Institute | IRCCS Regina Elena National Cancer Institute | Ludovica Ciuffreda, Sara Donzelli, Eleonora Sperandio, Grazia Prignano, Fabrizio Ensoli, Fulvia Pimpinelli, Maurizio Fanciulli, Giovanni Blandino, Aldo Morrone, Gennaro Ciliberto |
| EPI_ISL_2923018 | IRCCS San Gallicano Dermatological Institute | IRCCS Regina Elena National Cancer Institute | Frauke Goeman, Giulia Orlandi, Alice Massacci, Elisabetta Trento, Fabrizio Ensoli, Fulvia Pimpinelli, Maurizio Fanciulli, Giovanni Blandino, Aldo Morrone, Gennaro Ciliberto |
| EPI_ISL_2923019 | IRCCS San Gallicano Dermatological Institute | IRCCS Regina Elena National Cancer Institute | Sabrina Strano, Sara Donzelli, Matteo Pallocca, Arianna Mastrofrancesco, Fabrizio Ensoli, Fulvia Pimpinelli, Maurizio Fanciulli, Giovanni Blandino, Aldo Morrone, Gennaro Ciliberto |
| EPI_ISL_2923021 | IRCCS San Gallicano Dermatological Institute | IRCCS Regina Elena National Cancer Institute | Ludovica Ciuffreda, Sara Donzelli, Alice Massacci, Carla Mottini, Fabrizio Ensoli, Fulvia Pimpinelli, Maurizio Fanciulli, Giovanni Blandino, Aldo Morrone, Gennaro Ciliberto |
| EPI_ISL_2923026 | IRCCS San Gallicano Dermatological Institute | IRCCS Regina Elena National Cancer Institute | Frauke Goeman, Giulia Orlandi, Eleonora Sperandio, Andrea Cazzani, Fabrizio Ensoli, Fulvia Pimpinelli, Maurizio Fanciulli, Giovanni Blandino, Aldo Morrone, Gennaro Ciliberto |
| EPI_ISL_2923027 | IRCCS San Gallicano Dermatological Institute | IRCCS Regina Elena National Cancer Institute | Francesca De Nicola, Giulia Orlandi, Matteo Pallocca, Martina Diano, Fabrizio Ensoli, Fulvia Pimpinelli, Maurizio Fanciulli, Giovanni Blandino, Aldo Morrone, Gennaro Ciliberto |

| | | | |
|---|---|---|---|
| EPI_ISL_2923030 | IRCCS San Gallicano Dermatological Institute | IRCCS Regina Elena National Cancer Institute | Sabrina Strano, Sara Donzelli, Matteo Pallocca, Lorenzo Furzi, Fabrizio Ensoli, Fulvia Pimpinelli, Maurizio Fanciulli, Giovanni Blandino, Aldo Morrone, Gennaro Ciliberto |
| EPI_ISL_2923031 | IRCCS San Gallicano Dermatological Institute | IRCCS Regina Elena National Cancer Institute | Francesca De Nicola, Giulia Orlandi, Eleonora Sperandio, Alessia Lauretti, Fabrizio Ensoli, Fulvia Pimpinelli, Maurizio Fanciulli, Giovanni Blandino, Aldo Morrone, Gennaro Ciliberto |
| EPI_ISL_2923033 | IRCCS San Gallicano Dermatological Institute | IRCCS Regina Elena National Cancer Institute | Frauke Goeman, Giulia Orlandi, Matteo Pallocca, Francisco Obregon, Fabrizio Ensoli, Fulvia Pimpinelli, Maurizio Fanciulli, Giovanni Blandino, Aldo Morrone, Gennaro Ciliberto |
| EPI_ISL_2923034 | IRCCS San Gallicano Dermatological Institute | IRCCS Regina Elena National Cancer Institute | Sabrina Strano, Sara Donzelli, Eleonora Sperandio, Silvia Paluzzi, Fabrizio Ensoli, Fulvia Pimpinelli, Maurizio Fanciulli, Giovanni Blandino, Aldo Morrone, Gennaro Ciliberto |
| EPI_ISL_2923035 | IRCCS San Gallicano Dermatological Institute | IRCCS Regina Elena National Cancer Institute | Francesca De Nicola, Giulia Orlandi, Alice Massacci, Sara Petrolo, Fabrizio Ensoli, Fulvia Pimpinelli, Maurizio Fanciulli, Giovanni Blandino, Aldo Morrone, Gennaro Ciliberto |
| EPI_ISL_2923036 | IRCCS San Gallicano Dermatological Institute | IRCCS Regina Elena National Cancer Institute | Ludovica Ciuffreda, Sara Donzelli, Matteo Pallocca, Valentina Ricca, Fabrizio Ensoli, Fulvia Pimpinelli, Maurizio Fanciulli, Giovanni Blandino, Aldo Morrone, Gennaro Ciliberto |
| EPI_ISL_2923037 | IRCCS San Gallicano Dermatological Institute | IRCCS Regina Elena National Cancer Institute | Frauke Goeman, Giulia Orlandi, Eleonora Sperandio, Serena Salvo, Fabrizio Ensoli, Fulvia Pimpinelli, Maurizio Fanciulli, Giovanni Blandino, Aldo Morrone, Gennaro Ciliberto |
| EPI_ISL_2923038 | IRCCS San Gallicano Dermatological Institute | IRCCS Regina Elena National Cancer Institute | Sabrina Strano, Sara Donzelli, Alice Massacci, Grazia Prignano, Fabrizio Ensoli, Fulvia Pimpinelli, Maurizio Fanciulli, Giovanni Blandino, Aldo Morrone, Gennaro Ciliberto |
| EPI_ISL_2923040 | IRCCS San Gallicano Dermatological Institute | IRCCS Regina Elena National Cancer Institute | Ludovica Ciuffreda, Sara Donzelli, Eleonora Sperandio, Arianna Mastrofrancesco, Fabrizio Ensoli, Fulvia Pimpinelli, Maurizio Fanciulli, Giovanni Blandino, Aldo Morrone, Gennaro Ciliberto |
| EPI_ISL_2923041 | IRCCS San Gallicano Dermatological Institute | IRCCS Regina Elena National Cancer Institute | Frauke Goeman, Giulia Orlandi, Alice Massacci, Giovanna D'agosto, Fabrizio Ensoli, Fulvia Pimpinelli, Maurizio Fanciulli, Giovanni Blandino, Aldo Morrone, Gennaro Ciliberto |
| EPI_ISL_2923042 | IRCCS San Gallicano Dermatological Institute | IRCCS Regina Elena National Cancer Institute | Sabrina Strano, Sara Donzelli, Matteo Pallocca, Carla Mottini, Fabrizio Ensoli, Fulvia Pimpinelli, Maurizio Fanciulli, Giovanni Blandino, Aldo Morrone, Gennaro Ciliberto |
| EPI_ISL_2923043 | IRCCS San Gallicano Dermatological Institute | IRCCS Regina Elena National Cancer Institute | Francesca De Nicola, Giulia Orlandi, Eleonora Sperandio, Martina Pontone, Fabrizio Ensoli, Fulvia Pimpinelli, Maurizio Fanciulli, Giovanni Blandino, Aldo Morrone, Gennaro Ciliberto |
| EPI_ISL_2923044 | IRCCS San Gallicano Dermatological Institute | IRCCS Regina Elena National Cancer Institute | Ludovica Ciuffreda, Sara Donzelli, Alice Massacci, di Domenico Enea Gino, Fabrizio Ensoli, Fulvia Pimpinelli, Maurizio Fanciulli, Giovanni Blandino, Aldo Morrone, Gennaro Ciliberto |
| EPI_ISL_2923046 | IRCCS San Gallicano Dermatological Institute | IRCCS Regina Elena National Cancer Institute | Sabrina Strano, Sara Donzelli, Eleonora Sperandio, Francesca Sivori, Fabrizio Ensoli, Fulvia Pimpinelli, Maurizio Fanciulli, Giovanni Blandino, Aldo Morrone, Gennaro Ciliberto |
| EPI_ISL_2923048 | IRCCS San Gallicano Dermatological Institute | IRCCS Regina Elena National Cancer Institute | Ludovica Ciuffreda, Sara Donzelli, Matteo Pallocca, Ilaria Celesti, Fabrizio Ensoli, Fulvia Pimpinelli, Maurizio Fanciulli, Giovanni Blandino, Aldo Morrone, Gennaro Ciliberto |
| EPI_ISL_2923049 | IRCCS San Gallicano Dermatological Institute | IRCCS Regina Elena National Cancer Institute | Frauke Goeman, Giulia Orlandi, Eleonora Sperandio, Martina Diano, Fabrizio Ensoli, Fulvia Pimpinelli, Maurizio Fanciulli, Giovanni Blandino, Aldo Morrone, Gennaro Ciliberto |
| EPI_ISL_2923050 | IRCCS San Gallicano Dermatological Institute | IRCCS Regina Elena National Cancer Institute | Sabrina Strano, Sara Donzelli, Alice Massacci, Antonio Federico, Fabrizio Ensoli, Fulvia Pimpinelli, Maurizio Fanciulli, Giovanni Blandino, Aldo Morrone, Gennaro Ciliberto |
| EPI_ISL_2923051 | IRCCS San Gallicano Dermatological Institute | IRCCS Regina Elena National Cancer Institute | Francesca De Nicola, Giulia Orlandi, Matteo Pallocca, Fulvia Fraticelli, Fabrizio Ensoli, Fulvia Pimpinelli, Maurizio Fanciulli, Giovanni Blandino, Aldo Morrone, Gennaro Ciliberto |
| EPI_ISL_2923052 | IRCCS San Gallicano Dermatological Institute | IRCCS Regina Elena National Cancer Institute | Ludovica Ciuffreda, Sara Donzelli, Eleonora Sperandio, Lorenzo Furzi, Fabrizio Ensoli, Fulvia Pimpinelli, Maurizio Fanciulli, Giovanni Blandino, Aldo Morrone, Gennaro Ciliberto |
| EPI_ISL_2923053 | IRCCS San Gallicano Dermatological Institute | IRCCS Regina Elena National Cancer Institute | Frauke Goeman, Giulia Orlandi, Alice Massacci, Alessia Lauretti, Fabrizio Ensoli, Fulvia Pimpinelli, Maurizio Fanciulli, Giovanni Blandino, Aldo Morrone, Gennaro Ciliberto |
| EPI_ISL_2923054 | IRCCS San Gallicano Dermatological Institute | IRCCS Regina Elena National Cancer Institute | Sabrina Strano, Sara Donzelli, Matteo Pallocca, Francesca Maione, Fabrizio Ensoli, Fulvia Pimpinelli, Maurizio Fanciulli, Giovanni Blandino, Aldo Morrone, Gennaro Ciliberto |
| EPI_ISL_2923055 | IRCCS San Gallicano Dermatological Institute | IRCCS Regina Elena National Cancer Institute | Francesca De Nicola, Giulia Orlandi, Eleonora Sperandio, Francisco Obregon, Fabrizio Ensoli, Fulvia Pimpinelli, Maurizio Fanciulli, Giovanni Blandino, Aldo Morrone, Gennaro Ciliberto |
| EPI_ISL_2923056 | IRCCS San Gallicano Dermatological Institute | IRCCS Regina Elena National Cancer Institute | Ludovica Ciuffreda, Sara Donzelli, Alice Massacci, Silvia Paluzzi, Fabrizio Ensoli, Fulvia Pimpinelli, Maurizio Fanciulli, Giovanni Blandino, Aldo Morrone, Gennaro Ciliberto |
| EPI_ISL_2923057 | IRCCS San Gallicano Dermatological Institute | IRCCS Regina Elena National Cancer Institute | Frauke Goeman, Giulia Orlandi, Matteo Pallocca, Sara Petrolo, Fabrizio Ensoli, Fulvia Pimpinelli, Maurizio Fanciulli, Giovanni Blandino, Aldo Morrone, Gennaro Ciliberto |
| EPI_ISL_2923059 | IRCCS San Gallicano Dermatological Institute | IRCCS Regina Elena National Cancer Institute | Francesca De Nicola, Giulia Orlandi, Alice Massacci, Serena Salvo, Fabrizio Ensoli, Fulvia Pimpinelli, Maurizio Fanciulli, Giovanni Blandino, Aldo Morrone, Gennaro Ciliberto |
| EPI_ISL_2923060 | IRCCS San Gallicano Dermatological Institute | IRCCS Regina Elena National Cancer Institute | Ludovica Ciuffreda, Sara Donzelli, Matteo Pallocca, Grazia Prignano, Fabrizio Ensoli, Fulvia Pimpinelli, Maurizio Fanciulli, Giovanni Blandino, Aldo Morrone, Gennaro Ciliberto |
| EPI_ISL_2923061 | IRCCS San Gallicano Dermatological Institute | IRCCS Regina Elena National Cancer Institute | Frauke Goeman, Giulia Orlandi, Eleonora Sperandio, Elisabetta Trento, Fabrizio Ensoli, Fulvia Pimpinelli, Maurizio Fanciulli, Giovanni Blandino, Aldo Morrone, Gennaro Ciliberto |
| EPI_ISL_2923062 | IRCCS San Gallicano Dermatological Institute | IRCCS Regina Elena National Cancer Institute | Sabrina Strano, Sara Donzelli, Alice Massacci, Arianna Mastrofrancesco, Fabrizio Ensoli, Fulvia Pimpinelli, Maurizio Fanciulli, Giovanni Blandino, Aldo Morrone, Gennaro Ciliberto |
| EPI_ISL_2923064 | IRCCS San Gallicano Dermatological Institute | IRCCS Regina Elena National Cancer Institute | Ludovica Ciuffreda, Sara Donzelli, Eleonora Sperandio, Carla Mottini, Fabrizio Ensoli, Fulvia Pimpinelli, Maurizio Fanciulli, Giovanni Blandino, Aldo Morrone, Gennaro Ciliberto |
| EPI_ISL_2923065 | IRCCS San Gallicano Dermatological Institute | IRCCS Regina Elena National Cancer Institute | Frauke Goeman, Giulia Orlandi, Alice Massacci, Martina Pontone, Fabrizio Ensoli, Fulvia Pimpinelli, Maurizio Fanciulli, Giovanni Blandino, Aldo Morrone, Gennaro Ciliberto |
| EPI_ISL_2923068 | IRCCS San Gallicano Dermatological Institute | IRCCS Regina Elena National Cancer Institute | Ludovica Ciuffreda, Sara Donzelli, Alice Massacci, Francesca Sivori, Fabrizio Ensoli, Fulvia Pimpinelli, Maurizio Fanciulli, Giovanni Blandino, Aldo Morrone, Gennaro Ciliberto |
| EPI_ISL_2923069 | IRCCS San Gallicano Dermatological Institute | IRCCS Regina Elena National Cancer Institute | Frauke Goeman, Giulia Orlandi, Matteo Pallocca, Andrea Cazzani, Fabrizio Ensoli, Fulvia Pimpinelli, Maurizio Fanciulli, Giovanni Blandino, Aldo Morrone, Gennaro Ciliberto |
| EPI_ISL_2923071 | IRCCS San Gallicano Dermatological Institute | IRCCS Regina Elena National Cancer Institute | Francesca De Nicola, Giulia Orlandi, Alice Massacci, Martina Diano, Fabrizio Ensoli, Fulvia Pimpinelli, Maurizio Fanciulli, Giovanni Blandino, Aldo Morrone, Gennaro Ciliberto |
| EPI_ISL_2923072 | IRCCS San Gallicano Dermatological Institute | IRCCS Regina Elena National Cancer Institute | Ludovica Ciuffreda, Sara Donzelli, Matteo Pallocca, Antonio Federico, Fabrizio Ensoli, Fulvia Pimpinelli, Maurizio Fanciulli, Giovanni Blandino, Aldo Morrone, Gennaro Ciliberto |
| EPI_ISL_2923073 | IRCCS San Gallicano Dermatological Institute | IRCCS Regina Elena National Cancer Institute | Frauke Goeman, Giulia Orlandi, Eleonora Sperandio, Fulvia Fraticelli, Fabrizio Ensoli, Fulvia Pimpinelli, Maurizio Fanciulli, Giovanni Blandino, Aldo Morrone, Gennaro Ciliberto |
| EPI_ISL_2923074 | IRCCS San Gallicano Dermatological Institute | IRCCS Regina Elena National Cancer Institute | Sabrina Strano, Sara Donzelli, Alice Massacci, Lorenzo Furzi, Fabrizio Ensoli, Fulvia Pimpinelli, Maurizio Fanciulli, Giovanni Blandino, Aldo Morrone, Gennaro Ciliberto |
| EPI_ISL_2923076 | IRCCS San Gallicano Dermatological Institute | IRCCS Regina Elena National Cancer Institute | Ludovica Ciuffreda, Sara Donzelli, Eleonora Sperandio, Francesca Maione, Fabrizio Ensoli, Fulvia Pimpinelli, Maurizio Fanciulli, Giovanni Blandino, Aldo Morrone, Gennaro Ciliberto |
| EPI_ISL_2923077 | IRCCS San Gallicano Dermatological Institute | IRCCS Regina Elena National Cancer Institute | Frauke Goeman, Giulia Orlandi, Alice Massacci, Francisco Obregon, Fabrizio Ensoli, Fulvia Pimpinelli, Maurizio Fanciulli, Giovanni Blandino, Aldo Morrone, Gennaro Ciliberto |
| EPI_ISL_2923079 | IRCCS San Gallicano Dermatological Institute | IRCCS Regina Elena National Cancer Institute | Francesca De Nicola, Giulia Orlandi, Eleonora Sperandio, Sara Petrolo, Fabrizio Ensoli, Fulvia Pimpinelli, Maurizio Fanciulli, Giovanni Blandino, Aldo Morrone, Gennaro Ciliberto |
| EPI_ISL_2923080 | IRCCS San Gallicano Dermatological Institute | IRCCS Regina Elena National Cancer Institute | Ludovica Ciuffreda, Sara Donzelli, Alice Massacci, Valentina Ricca, Fabrizio Ensoli, Fulvia Pimpinelli, Maurizio Fanciulli, Giovanni Blandino, Aldo Morrone, Gennaro Ciliberto |
| EPI_ISL_2923082 | IRCCS San Gallicano Dermatological Institute | IRCCS Regina Elena National Cancer Institute | Sabrina Strano, Sara Donzelli, Eleonora Sperandio, Grazia Prignano, Fabrizio Ensoli, Fulvia Pimpinelli, Maurizio Fanciulli, Giovanni Blandino, Aldo Morrone, Gennaro Ciliberto |
| EPI_ISL_2923084 | IRCCS San Gallicano Dermatological Institute | IRCCS Regina Elena National Cancer Institute | Ludovica Ciuffreda, Sara Donzelli, Matteo Pallocca, Arianna Mastrofrancesco, Fabrizio Ensoli, Fulvia Pimpinelli, Maurizio Fanciulli, Giovanni Blandino, Aldo Morrone, Gennaro Ciliberto |
| EPI_ISL_2923086 | IRCCS San Gallicano Dermatological Institute | IRCCS Regina Elena National Cancer Institute | Sabrina Strano, Sara Donzelli, Alice Massacci, Carla Mottini, Fabrizio Ensoli, Fulvia Pimpinelli, Maurizio Fanciulli, Giovanni Blandino, Aldo Morrone, Gennaro Ciliberto |
| EPI_ISL_2923087 | IRCCS San Gallicano Dermatological Institute | IRCCS Regina Elena National Cancer Institute | Francesca De Nicola, Giulia Orlandi, Matteo Pallocca, Martina Pontone, Fabrizio Ensoli, Fulvia Pimpinelli, Maurizio Fanciulli, Giovanni Blandino, Aldo Morrone, Gennaro Ciliberto |
| EPI_ISL_2923088 | IRCCS San Gallicano Dermatological Institute | IRCCS Regina Elena National Cancer Institute | Ludovica Ciuffreda, Sara Donzelli, Eleonora Sperandio, di Domenico Enea Gino, Fabrizio Ensoli, Fulvia Pimpinelli, Maurizio Fanciulli, Giovanni Blandino, Aldo Morrone, Gennaro Ciliberto |
| EPI_ISL_2923089 | IRCCS San Gallicano Dermatological Institute | IRCCS Regina Elena National Cancer Institute | Frauke Goeman, Giulia Orlandi, Alice Massacci, Ilaria Cavallo, Fabrizio Ensoli, Fulvia Pimpinelli, Maurizio Fanciulli, Giovanni Blandino, Aldo Morrone, Gennaro Ciliberto |
| EPI_ISL_2923091 | IRCCS San Gallicano Dermatological Institute | IRCCS Regina Elena National Cancer Institute | Francesca De Nicola, Giulia Orlandi, Eleonora Sperandio, Andrea Cazzani, Fabrizio Ensoli, Fulvia Pimpinelli, Maurizio Fanciulli, Giovanni Blandino, Aldo Morrone, Gennaro Ciliberto |
| EPI_ISL_2923092 | IRCCS San Gallicano Dermatological Institute | IRCCS Regina Elena National Cancer Institute | Ludovica Ciuffreda, Sara Donzelli, Alice Massacci, Ilaria Celesti, Fabrizio Ensoli, Fulvia Pimpinelli, Maurizio Fanciulli, Giovanni Blandino, Aldo Morrone, Gennaro Ciliberto |
| EPI_ISL_2923094 | IRCCS San Gallicano Dermatological Institute | IRCCS Regina Elena National Cancer Institute | Sabrina Strano, Sara Donzelli, Eleonora Sperandio, Antonio Federico, Fabrizio Ensoli, Fulvia Pimpinelli, Maurizio Fanciulli, Giovanni Blandino, Aldo Morrone, Gennaro Ciliberto |
| EPI_ISL_2923095 | IRCCS San Gallicano Dermatological Institute | IRCCS Regina Elena National Cancer Institute | Francesca De Nicola, Giulia Orlandi, Alice Massacci, Fulvia Fraticelli, Fabrizio Ensoli, Fulvia Pimpinelli, Maurizio Fanciulli, Giovanni Blandino, Aldo Morrone, Gennaro Ciliberto |
| EPI_ISL_2923096 | IRCCS San Gallicano Dermatological Institute | IRCCS Regina Elena National Cancer Institute | Ludovica Ciuffreda, Sara Donzelli, Matteo Pallocca, Lorenzo Furzi, Fabrizio Ensoli, Fulvia Pimpinelli, Maurizio Fanciulli, Giovanni Blandino, Aldo Morrone, Gennaro Ciliberto |
| EPI_ISL_2923097 | IRCCS San Gallicano Dermatological Institute | IRCCS Regina Elena National Cancer Institute | Frauke Goeman, Giulia Orlandi, Eleonora Sperandio, Alessia Lauretti, Fabrizio Ensoli, Fulvia Pimpinelli, Maurizio Fanciulli, Giovanni Blandino, Aldo Morrone, Gennaro Ciliberto |
| EPI_ISL_2923098 | IRCCS San Gallicano Dermatological Institute | IRCCS Regina Elena National Cancer Institute | Sabrina Strano, Sara Donzelli, Alice Massacci, Francesca Maione, Fabrizio Ensoli, Fulvia Pimpinelli, Maurizio Fanciulli, Giovanni Blandino, Aldo Morrone, Gennaro Ciliberto |
| EPI_ISL_2923099 | IRCCS San Gallicano Dermatological Institute | IRCCS Regina Elena National Cancer Institute | Ludovica Ciuffreda, Sara Donzelli, Eleonora Sperandio, Silvia Paluzzi, Fabrizio Ensoli, Fulvia Pimpinelli, Maurizio Fanciulli, Giovanni Blandino, Aldo Morrone, Gennaro Ciliberto |
| EPI_ISL_2923100 | IRCCS San Gallicano Dermatological Institute | IRCCS Regina Elena National Cancer Institute | Frauke Goeman, Giulia Orlandi, Alice Massacci, Sara Petrolo, Fabrizio Ensoli, Fulvia Pimpinelli, Maurizio Fanciulli, Giovanni Blandino, Aldo Morrone, Gennaro Ciliberto |
| EPI_ISL_2923101 | IRCCS San Gallicano Dermatological Institute | IRCCS Regina Elena National Cancer Institute | Sabrina Strano, Sara Donzelli, Matteo Pallocca, Valentina Ricca, Fabrizio Ensoli, Fulvia Pimpinelli, Maurizio Fanciulli, Giovanni Blandino, Aldo Morrone, Gennaro Ciliberto |
| EPI_ISL_2923102 | IRCCS San Gallicano Dermatological Institute | IRCCS Regina Elena National Cancer Institute | Francesca De Nicola, Giulia Orlandi, Eleonora Sperandio, Serena Salvo, Fabrizio Ensoli, Fulvia Pimpinelli, Maurizio Fanciulli, Giovanni Blandino, Aldo Morrone, Gennaro Ciliberto |
| EPI_ISL_2923103 | IRCCS San Gallicano Dermatological Institute | IRCCS Regina Elena National Cancer Institute | Ludovica Ciuffreda, Sara Donzelli, Alice Massacci, Grazia Prignano, Fabrizio Ensoli, Fulvia Pimpinelli, Maurizio Fanciulli, Giovanni Blandino, Aldo Morrone, Gennaro Ciliberto |
| EPI_ISL_2923104 | IRCCS San Gallicano Dermatological Institute | IRCCS Regina Elena National Cancer Institute | Frauke Goeman, Giulia Orlandi, Matteo Pallocca, Elisabetta Trento, Fabrizio Ensoli, Fulvia Pimpinelli, Maurizio Fanciulli, Giovanni Blandino, Aldo Morrone, Gennaro Ciliberto |
| EPI_ISL_2923105 | IRCCS San Gallicano Dermatological Institute | IRCCS Regina Elena National Cancer Institute | Sabrina Strano, Sara Donzelli, Eleonora Sperandio, Arianna Mastrofrancesco, Fabrizio Ensoli, Fulvia Pimpinelli, Maurizio Fanciulli, Giovanni Blandino, Aldo Morrone, Gennaro Ciliberto |
| EPI_ISL_2923106 | IRCCS San Gallicano Dermatological Institute | IRCCS Regina Elena National Cancer Institute | Ludovica Ciuffreda, Sara Donzelli, Matteo Pallocca, Carla Mottini, Fabrizio Ensoli, Fulvia Pimpinelli, Maurizio Fanciulli, Giovanni Blandino, Aldo Morrone, Gennaro Ciliberto |
| EPI_ISL_2923108 | IRCCS San Gallicano Dermatological Institute | IRCCS Regina Elena National Cancer Institute | Sabrina Strano, Sara Donzelli, Alice Massacci, di Domenico Enea Gino, Fabrizio Ensoli, Fulvia Pimpinelli, Maurizio Fanciulli, Giovanni Blandino, Aldo Morrone, Gennaro Ciliberto |
| EPI_ISL_2923109 | IRCCS San Gallicano Dermatological Institute | IRCCS Regina Elena National Cancer Institute | Francesca De Nicola, Giulia Orlandi, Matteo Pallocca, Ilaria Cavallo, Fabrizio Ensoli, Fulvia Pimpinelli, Maurizio Fanciulli, Giovanni Blandino, Aldo Morrone, Gennaro Ciliberto |
| EPI_ISL_2923110 | IRCCS San Gallicano Dermatological Institute | IRCCS Regina Elena National Cancer Institute | Ludovica Ciuffreda, Sara Donzelli, Eleonora Sperandio, Francesca Sivori, Fabrizio Ensoli, Fulvia Pimpinelli, Maurizio Fanciulli, Giovanni Blandino, Aldo Morrone, Gennaro Ciliberto |
| EPI_ISL_2923111 | IRCCS San Gallicano Dermatological Institute | IRCCS Regina Elena National Cancer Institute | Frauke Goeman, Giulia Orlandi, Alice Massacci, Andrea Cazzani, Fabrizio Ensoli, Fulvia Pimpinelli, Maurizio Fanciulli, Giovanni Blandino, Aldo Morrone, Gennaro Ciliberto |
| EPI_ISL_2923115 | Hospital | National Reference Center for Viruses of Respiratory Infections, Institut Pasteur, Paris | Marion Barbet, Sylvie Behillil, Méline Bizard, Angela Brisebarre, Camille Capel, Vincent Enouf, Louise Lefrançois, Frédéric Lemoine, Christophe Malabat, Corinne Maufrais, Hub Bioinformatique Biostatistiques, Etienne Simon-Lorière, Maud Vanpeene, Sylvie Van der Werf ,FARFOUR Eric |
| EPI_ISL_2923118 | Labo Analyses Med | National Reference Center for Viruses of Respiratory Infections, Institut Pasteur, Paris | Marion Barbet, Sylvie Behillil, Méline Bizard, Angela Brisebarre, Camille Capel, Vincent Enouf, Louise Lefrançois, Frédéric Lemoine, Christophe Malabat, Corinne Maufrais, Damien Mornico, Etienne Simon-Lorière, Maud Vanpeene, Sylvie Van der Werf ,TROMBERT Sabine |
| EPI_ISL_2923119 | Labo Analyses Med | National Reference Center for Viruses of Respiratory Infections, Institut Pasteur, Paris | Marion Barbet, Sylvie Behillil, Méline Bizard, Angela Brisebarre, Camille Capel, Vincent Enouf, Louise Lefrançois, Frédéric Lemoine, Christophe Malabat, Corinne Maufrais, Hub Bioinformatique Biostatistiques, Etienne Simon-Lorière, Maud Vanpeene, Sylvie Van der Werf ,TROMBERT Sabine |
| EPI_ISL_2923120 | Hospital | National Reference Center for Viruses of Respiratory Infections, Institut Pasteur, Paris | Marion Barbet, Sylvie Behillil, Méline Bizard, Angela Brisebarre, Camille Capel, Vincent Enouf, Louise Lefrançois, Frédéric Lemoine, Christophe Malabat, Corinne Maufrais, Damien Mornico, Etienne Simon-Lorière, Maud Vanpeene, Sylvie Van der Werf ,RAMIREZ José-Manuel |
| EPI_ISL_2923121 | Labo Analyses Med | National Reference Center for Viruses of Respiratory Infections, Institut Pasteur, Paris | Marion Barbet, Sylvie Behillil, Méline Bizard, Angela Brisebarre, Camille Capel, Vincent Enouf, Louise Lefrançois, Frédéric Lemoine, Christophe Malabat, Corinne Maufrais, Damien Mornico, Etienne Simon-Lorière, Maud Vanpeene, Sylvie Van der Werf ,LABORATOIRE Palissy |
| EPI_ISL_2923122 | Labo Analyses Med | National Reference Center for Viruses of Respiratory Infections, Institut Pasteur, Paris | Marion Barbet, Sylvie Behillil, Méline Bizard, Angela Brisebarre, Camille Capel, Vincent Enouf, Louise Lefrançois, Frédéric Lemoine, Christophe Malabat, Corinne Maufrais, Victoire Baillet, Etienne Simon-Lorière, Maud Vanpeene, Sylvie Van der Werf ,LABORATOIRE Palissy |
| EPI_ISL_2923123, EPI_ISL_2923124 | Labo Analyses Med | National Reference Center for Viruses of Respiratory Infections, Institut Pasteur, Paris | Marion Barbet, Sylvie Behillil, Méline Bizard, Angela Brisebarre, Camille Capel, Vincent Enouf, Louise Lefrançois, Frédéric Lemoine, Christophe Malabat, Corinne Maufrais, Damien Mornico, Etienne Simon-Lorière, Maud Vanpeene, Sylvie Van der Werf ,LABORATOIRE Palissy |

| | | | |
|---|---|---|---|
| EPI_ISL_2923125, EPI_ISL_2923126 | Hospital | National Reference Center for Viruses of Respiratory Infections, Institut Pasteur, Paris | Marion Barbet, Sylvie Behillil, Méline Bizard, Angela Brisebarre, Camille Capel, Vincent Enouf, Louise Lefrançois, Frédéric Lemoine, Christophe Malabat, Corinne Maufrais, Hub Bioinformatique Biostatistiques, Etienne Simon-Lorière, Maud Vanpeene, Sylvie Van der Werf ,HOLSTEIN Anne |
| EPI_ISL_2923127 | Hospital | National Reference Center for Viruses of Respiratory Infections, Institut Pasteur, Paris | Marion Barbet, Sylvie Behillil, Méline Bizard, Angela Brisebarre, Camille Capel, Vincent Enouf, Louise Lefrançois, Frédéric Lemoine, Christophe Malabat, Corinne Maufrais, Victoire Baillet, Etienne Simon-Lorière, Maud Vanpeene, Sylvie Van der Werf ,HOLSTEIN Anne |
| EPI_ISL_2923128 | Hospital | National Reference Center for Viruses of Respiratory Infections, Institut Pasteur, Paris | Marion Barbet, Sylvie Behillil, Méline Bizard, Angela Brisebarre, Camille Capel, Vincent Enouf, Louise Lefrançois, Frédéric Lemoine, Christophe Malabat, Corinne Maufrais, Victoire Baillet, Etienne Simon-Lorière, Maud Vanpeene, Sylvie Van der Werf ,SERAZIN Valérie |
| EPI_ISL_2923130 | Labo Analyses Med | National Reference Center for Viruses of Respiratory Infections, Institut Pasteur, Paris | Marion Barbet, Sylvie Behillil, Méline Bizard, Angela Brisebarre, Camille Capel, Vincent Enouf, Louise Lefrançois, Frédéric Lemoine, Christophe Malabat, Corinne Maufrais, Victoire Baillet, Etienne Simon-Lorière, Maud Vanpeene, Sylvie Van der Werf ,TROMBERT Sabine |
| EPI_ISL_2923131, EPI_ISL_2923132, EPI_ISL_2923133, EPI_ISL_2923137 | Labo Analyses Med | National Reference Center for Viruses of Respiratory Infections, Institut Pasteur, Paris | Marion Barbet, Sylvie Behillil, Méline Bizard, Angela Brisebarre, Camille Capel, Vincent Enouf, Louise Lefrançois, Frédéric Lemoine, Christophe Malabat, Corinne Maufrais, Hub Bioinformatique Biostatistiques, Etienne Simon-Lorière, Maud Vanpeene, Sylvie Van der Werf ,TROMBERT Sabine |
| EPI_ISL_2923138 | Hospital | National Reference Center for Viruses of Respiratory Infections, Institut Pasteur, Paris | Marion Barbet, Sylvie Behillil, Méline Bizard, Angela Brisebarre, Camille Capel, Vincent Enouf, Louise Lefrançois, Frédéric Lemoine, Christophe Malabat, Corinne Maufrais, Hub Bioinformatique Biostatistiques, Etienne Simon-Lorière, Maud Vanpeene, Sylvie Van der Werf ,MARTRES Pascale |
| EPI_ISL_2923140 | Labo Analyses Med | National Reference Center for Viruses of Respiratory Infections, Institut Pasteur, Paris | Marion Barbet, Sylvie Behillil, Méline Bizard, Angela Brisebarre, Camille Capel, Vincent Enouf, Louise Lefrançois, Frédéric Lemoine, Christophe Malabat, Corinne Maufrais, Amaury Vaysse, Etienne Simon-Lorière, Maud Vanpeene, Sylvie Van der Werf ,POTIRON Grégoire |
| EPI_ISL_2923143 | Labo Analyses Med | National Reference Center for Viruses of Respiratory Infections, Institut Pasteur, Paris | Marion Barbet, Sylvie Behillil, Méline Bizard, Angela Brisebarre, Camille Capel, Vincent Enouf, Louise Lefrançois, Frédéric Lemoine, Christophe Malabat, Corinne Maufrais, Hub Bioinformatique Biostatistiques, Etienne Simon-Lorière, Maud Vanpeene, Sylvie Van der Werf ,LEONARD Pierre-Yves |
| EPI_ISL_2923144, EPI_ISL_2923145, EPI_ISL_2923146, EPI_ISL_2923147 | Labo Analyses Med | National Reference Center for Viruses of Respiratory Infections, Institut Pasteur, Paris | Marion Barbet, Sylvie Behillil, Méline Bizard, Angela Brisebarre, Camille Capel, Vincent Enouf, Louise Lefrançois, Frédéric Lemoine, Christophe Malabat, Corinne Maufrais, Hub Bioinformatique Biostatistiques, Etienne Simon-Lorière, Maud Vanpeene, Sylvie Van der Werf ,HAGUENOER Eve |
| EPI_ISL_2923148, EPI_ISL_2923149, EPI_ISL_2923150, EPI_ISL_2923151, EPI_ISL_2923152, EPI_ISL_2923153, EPI_ISL_2923154, EPI_ISL_2923155, EPI_ISL_2923156 | Labo Analyses Med | National Reference Center for Viruses of Respiratory Infections, Institut Pasteur, Paris | Marion Barbet, Sylvie Behillil, Méline Bizard, Angela Brisebarre, Camille Capel, Vincent Enouf, Louise Lefrançois, Frédéric Lemoine, Christophe Malabat, Corinne Maufrais, Hub Bioinformatique Biostatistiques, Etienne Simon-Lorière, Maud Vanpeene, Sylvie Van der Werf ,DURIVAULT Jérôme |
| EPI_ISL_2923157 | Labo Analyses Med | National Reference Center for Viruses of Respiratory Infections, Institut Pasteur, Paris | Marion Barbet, Sylvie Behillil, Méline Bizard, Angela Brisebarre, Camille Capel, Vincent Enouf, Louise Lefrançois, Frédéric Lemoine, Christophe Malabat, Corinne Maufrais,Hub Bioinformatique Biostatistiques, Etienne Simon-Lorière, Maud Vanpeene, Sylvie Van der Werf ,DURIVAULT Jérôme |
| EPI_ISL_2923158 | Labo Analyses Med | National Reference Center for Viruses of Respiratory Infections, Institut Pasteur, Paris | Marion Barbet, Sylvie Behillil, Méline Bizard, Angela Brisebarre, Camille Capel, Vincent Enouf, Louise Lefrançois, Frédéric Lemoine, Christophe Malabat, Corinne Maufrais, Hub Bioinformatique Biostatistiques, Etienne Simon-Lorière, Maud Vanpeene, Sylvie Van der Werf ,DURIVAULT Jérôme |
| EPI_ISL_2923159 | Labo Analyses Med | National Reference Center for Viruses of Respiratory Infections, Institut Pasteur, Paris | Marion Barbet, Sylvie Behillil, Méline Bizard, Angela Brisebarre, Camille Capel, Vincent Enouf, Louise Lefrançois, Frédéric Lemoine, Christophe Malabat, Corinne Maufrais,Hub Bioinformatique Biostatistiques, Etienne Simon-Lorière, Maud Vanpeene, Sylvie Van der Werf ,DURIVAULT Jérôme |
| EPI_ISL_2923160, EPI_ISL_2923161 | Labo Analyses Med | National Reference Center for Viruses of Respiratory Infections, Institut Pasteur, Paris | Marion Barbet, Sylvie Behillil, Méline Bizard, Angela Brisebarre, Camille Capel, Vincent Enouf, Louise Lefrançois, Frédéric Lemoine, Christophe Malabat, Corinne Maufrais, Amaury Vaysse, Etienne Simon-Lorière, Maud Vanpeene, Sylvie Van der Werf ,DURIVAULT Jérôme |
| EPI_ISL_2923162, EPI_ISL_2923163 | Hospital | National Reference Center for Viruses of Respiratory Infections, Institut Pasteur, Paris | Marion Barbet, Sylvie Behillil, Méline Bizard, Angela Brisebarre, Camille Capel, Vincent Enouf, Louise Lefrançois, Frédéric Lemoine, Christophe Malabat, Corinne Maufrais, Hub Bioinformatique Biostatistiques, Etienne Simon-Lorière, Maud Vanpeene, Sylvie Van der Werf ,DJAMDJIAN Laura |
| EPI_ISL_2923164 | Hospital | National Reference Center for Viruses of Respiratory Infections, Institut Pasteur, Paris | Marion Barbet, Sylvie Behillil, Méline Bizard, Angela Brisebarre, Camille Capel, Vincent Enouf, Louise Lefrançois, Frédéric Lemoine, Christophe Malabat, Corinne Maufrais, Amaury Vaysse, Etienne Simon-Lorière, Maud Vanpeene, Sylvie Van der Werf ,GUINARD Jérome |
| EPI_ISL_2923165 | Hospital | National Reference Center for Viruses of Respiratory Infections, Institut Pasteur, Paris | Marion Barbet, Sylvie Behillil, Méline Bizard, Angela Brisebarre, Camille Capel, Vincent Enouf, Louise Lefrançois, Frédéric Lemoine, Christophe Malabat, Corinne Maufrais, Hub Bioinformatique Biostatistiques, Etienne Simon-Lorière, Maud Vanpeene, Sylvie Van der Werf ,GUINARD Jérome |
| EPI_ISL_2923167 | Hospital | National Reference Center for Viruses of Respiratory Infections, Institut Pasteur, Paris | Marion Barbet, Sylvie Behillil, Méline Bizard, Angela Brisebarre, Camille Capel, Vincent Enouf, Louise Lefrançois, Frédéric Lemoine, Christophe Malabat, Corinne Maufrais, Amaury Vaysse, Etienne Simon-Lorière, Maud Vanpeene, Sylvie Van der Werf ,GALEAZZI GUY |
| EPI_ISL_2923169 | Labo Analyses Med | National Reference Center for Viruses of Respiratory Infections, Institut Pasteur, Paris | Marion Barbet, Sylvie Behillil, Méline Bizard, Angela Brisebarre, Camille Capel, Vincent Enouf, Louise Lefrançois, Frédéric Lemoine, Christophe Malabat, Corinne Maufrais, Hub Bioinformatique Biostatistiques, Etienne Simon-Lorière, Maud Vanpeene, Sylvie Van der Werf ,MIARA Philippe |
| EPI_ISL_2923175 | Labo Analyses Med | National Reference Center for Viruses of Respiratory Infections, Institut Pasteur, Paris | Marion Barbet, Sylvie Behillil, Méline Bizard, Angela Brisebarre, Camille Capel, Vincent Enouf, Louise Lefrançois, Frédéric Lemoine, Christophe Malabat, Corinne Maufrais, Hub Bioinformatique Biostatistiques, Etienne Simon-Lorière, Maud Vanpeene, Sylvie Van der Werf ,TROMBERT Sabine |
| EPI_ISL_2923182 | Labo Analyses Med | National Reference Center for Viruses of Respiratory Infections, Institut Pasteur, Paris | Marion Barbet, Sylvie Behillil, Méline Bizard, Angela Brisebarre, Camille Capel, Vincent Enouf, Louise Lefrançois, Frédéric Lemoine, Christophe Malabat, Corinne Maufrais, Amaury Vaysse, Etienne Simon-Lorière, Maud Vanpeene, Sylvie Van der Werf ,TROMBERT Sabine |
| EPI_ISL_2923183, EPI_ISL_2923185 | Labo Analyses Med | National Reference Center for Viruses of Respiratory Infections, Institut Pasteur, Paris | Marion Barbet, Sylvie Behillil, Méline Bizard, Angela Brisebarre, Camille Capel, Vincent Enouf, Louise Lefrançois, Frédéric Lemoine, Christophe Malabat, Corinne Maufrais, Hub Bioinformatique Biostatistiques, Etienne Simon-Lorière, Maud Vanpeene, Sylvie Van der Werf ,TROMBERT Sabine |
| EPI_ISL_2923190 | Labo Analyses Med | National Reference Center for Viruses of Respiratory Infections, Institut Pasteur, Paris | Marion Barbet, Sylvie Behillil, Méline Bizard, Angela Brisebarre, Camille Capel, Vincent Enouf, Louise Lefrançois, Frédéric Lemoine, Christophe Malabat, Corinne Maufrais, Amaury Vaysse, Etienne Simon-Lorière, Maud Vanpeene, Sylvie Van der Werf ,TROMBERT Sabine |
| EPI_ISL_2923192 | Labo Analyses Med | National Reference Center for Viruses of Respiratory Infections, Institut Pasteur, Paris | Marion Barbet, Sylvie Behillil, Méline Bizard, Angela Brisebarre, Camille Capel, Vincent Enouf, Louise Lefrançois, Frédéric Lemoine, Christophe Malabat, Corinne Maufrais, Hub Bioinformatique Biostatistiques, Etienne Simon-Lorière, Maud Vanpeene, Sylvie Van der Werf ,TROMBERT Sabine |
| EPI_ISL_2923194 | Labo Analyses Med | National Reference Center for Viruses of Respiratory Infections, Institut Pasteur, Paris | Marion Barbet, Sylvie Behillil, Méline Bizard, Angela Brisebarre, Camille Capel, Vincent Enouf, Louise Lefrançois, Frédéric Lemoine, Christophe Malabat, Corinne Maufrais, Hub Bioinformatique Biostatistiques, Etienne Simon-Lorière, Maud Vanpeene, Sylvie Van der Werf ,VIEILLEFOND VINCENT |
| EPI_ISL_2923195 | Labo Analyses Med | National Reference Center for Viruses of Respiratory Infections, Institut Pasteur, Paris | Marion Barbet, Sylvie Behillil, Méline Bizard, Angela Brisebarre, Camille Capel, Vincent Enouf, Louise Lefrançois, Frédéric Lemoine, Christophe Malabat, Corinne Maufrais, Amaury Vaysse, Etienne Simon-Lorière, Maud Vanpeene, Sylvie Van der Werf ,VIEILLEFOND VINCENT |
| EPI_ISL_2923199 | Labo Analyses Med | National Reference Center for Viruses of Respiratory Infections, Institut Pasteur, Paris | Marion Barbet, Sylvie Behillil, Méline Bizard, Angela Brisebarre, Camille Capel, Vincent Enouf, Louise Lefrançois, Frédéric Lemoine, Christophe Malabat, Corinne Maufrais, Hub Bioinformatique Biostatistiques, Etienne Simon-Lorière, Maud Vanpeene, Sylvie Van der Werf ,VIEILLEFOND VINCENT |
| EPI_ISL_2923200, EPI_ISL_2923201 | Labo Analyses Med | National Reference Center for Viruses of Respiratory Infections, Institut Pasteur, Paris | Marion Barbet, Sylvie Behillil, Méline Bizard, Angela Brisebarre, Camille Capel, Vincent Enouf, Louise Lefrançois, Frédéric Lemoine, Christophe Malabat, Corinne Maufrais, Amaury Vaysse, Etienne Simon-Lorière, Maud Vanpeene, Sylvie Van der Werf ,VIEILLEFOND VINCENT |
| EPI_ISL_2923202, EPI_ISL_2923203, EPI_ISL_2923204 | Labo Analyses Med | National Reference Center for Viruses of Respiratory Infections, Institut Pasteur, Paris | Marion Barbet, Sylvie Behillil, Méline Bizard, Angela Brisebarre, Camille Capel, Vincent Enouf, Louise Lefrançois, Frédéric Lemoine, Christophe Malabat, Corinne Maufrais, Hub Bioinformatique Biostatistiques, Etienne Simon-Lorière, Maud Vanpeene, Sylvie Van der Werf ,VIEILLEFOND VINCENT |
| EPI_ISL_2923206 | Labo Analyses Med | National Reference Center for Viruses of Respiratory Infections, Institut Pasteur, Paris | Marion Barbet, Sylvie Behillil, Méline Bizard, Angela Brisebarre, Camille Capel, Vincent Enouf, Louise Lefrançois, Frédéric Lemoine, Christophe Malabat, Corinne Maufrais, Pierre Lechat, Etienne Simon-Lorière, Maud Vanpeene, Sylvie Van der Werf ,VIEILLEFOND VINCENT |
| EPI_ISL_2923207 | Labo Analyses Med | National Reference Center for Viruses of Respiratory Infections, Institut Pasteur, Paris | Marion Barbet, Sylvie Behillil, Méline Bizard, Angela Brisebarre, Camille Capel, Vincent Enouf, Louise Lefrançois, Frédéric Lemoine, Christophe Malabat, Corinne Maufrais, Hub Bioinformatique Biostatistiques, Etienne Simon-Lorière, Maud Vanpeene, Sylvie Van der Werf ,VIEILLEFOND VINCENT |
| EPI_ISL_2923210 | Labo Analyses Med | National Reference Center for Viruses of Respiratory Infections, Institut Pasteur, Paris | Marion Barbet, Sylvie Behillil, Méline Bizard, Angela Brisebarre, Camille Capel, Vincent Enouf, Louise Lefrançois, Frédéric Lemoine, Christophe Malabat, Corinne Maufrais, Pierre Lechat, Etienne Simon-Lorière, Maud Vanpeene, Sylvie Van der Werf ,VIEILLEFOND VINCENT |
| EPI_ISL_2923213 | Labo Analyses Med | National Reference Center for Viruses of Respiratory Infections, Institut Pasteur, Paris | Marion Barbet, Sylvie Behillil, Méline Bizard, Angela Brisebarre, Camille Capel, Vincent Enouf, Louise Lefrançois, Frédéric Lemoine, Christophe Malabat, Corinne Maufrais, Hub Bioinformatique Biostatistiques, Etienne Simon-Lorière, Maud Vanpeene, Sylvie Van der Werf ,FOULQUIER Jean-Baptiste |
| EPI_ISL_2923215 | Labo Analyses Med | National Reference Center for Viruses of Respiratory Infections, Institut Pasteur, Paris | Marion Barbet, Sylvie Behillil, Méline Bizard, Angela Brisebarre, Camille Capel, Vincent Enouf, Louise Lefrançois, Frédéric Lemoine, Christophe Malabat, Corinne Maufrais, Gael Millot, Etienne Simon-Lorière, Maud Vanpeene, Sylvie Van der Werf ,PASQUIER Emmanuel |
| EPI_ISL_2923216, EPI_ISL_2923217 | Labo Analyses Med | National Reference Center for Viruses of Respiratory Infections, Institut Pasteur, Paris | Marion Barbet, Sylvie Behillil, Méline Bizard, Angela Brisebarre, Camille Capel, Vincent Enouf, Louise Lefrançois, Frédéric Lemoine, Christophe Malabat, Corinne Maufrais, Hub Bioinformatique Biostatistiques, Etienne Simon-Lorière, Maud Vanpeene, Sylvie Van der Werf ,PASQUIER Emmanuel |
| EPI_ISL_2923218, EPI_ISL_2923220 | Labo Analyses Med | National Reference Center for Viruses of Respiratory Infections, Institut Pasteur, Paris | Marion Barbet, Sylvie Behillil, Méline Bizard, Angela Brisebarre, Camille Capel, Vincent Enouf, Louise Lefrançois, Frédéric Lemoine, Christophe Malabat, Corinne Maufrais, Hub Bioinformatique Biostatistiques, Etienne Simon-Lorière, Maud Vanpeene, Sylvie Van der Werf ,FELLONI Claire |
| EPI_ISL_2923221 | Labo Analyses Med | National Reference Center for Viruses of Respiratory Infections, Institut Pasteur, Paris | Marion Barbet, Sylvie Behillil, Méline Bizard, Angela Brisebarre, Camille Capel, Vincent Enouf, Louise Lefrançois, Frédéric Lemoine, Christophe Malabat, Corinne Maufrais, Gael Millot, Etienne Simon-Lorière, Maud Vanpeene, Sylvie Van der Werf ,HOLSTEIN Anne |
| EPI_ISL_2923222, EPI_ISL_2923224 | Labo Analyses Med | National Reference Center for Viruses of Respiratory Infections, Institut Pasteur, Paris | Marion Barbet, Sylvie Behillil, Méline Bizard, Angela Brisebarre, Camille Capel, Vincent Enouf, Louise Lefrançois, Frédéric Lemoine, Christophe Malabat, Corinne Maufrais, Hub Bioinformatique Biostatistiques, Etienne Simon-Lorière, Maud Vanpeene, Sylvie Van der Werf ,HOLSTEIN Anne |
| EPI_ISL_2923226 | Labo Analyses Med | National Reference Center for Viruses of Respiratory Infections, Institut Pasteur, Paris | Marion Barbet, Sylvie Behillil, Méline Bizard, Angela Brisebarre, Camille Capel, Vincent Enouf, Louise Lefrançois, Frédéric Lemoine, Christophe Malabat, Corinne Maufrais, Gael Millot, Etienne Simon-Lorière, Maud Vanpeene, Sylvie Van der Werf ,HOLSTEIN Anne |
| EPI_ISL_2923227 | Labo Analyses Med | National Reference Center for Viruses of Respiratory Infections, Institut Pasteur, Paris | Marion Barbet, Sylvie Behillil, Méline Bizard, Angela Brisebarre, Camille Capel, Vincent Enouf, Louise Lefrançois, Frédéric Lemoine, Christophe Malabat, Corinne Maufrais, Hub Bioinformatique Biostatistiques, Etienne Simon-Lorière, Maud Vanpeene, Sylvie Van der Werf ,HOLSTEIN Anne |

| | | | |
|---|---|---|---|
| EPI_ISL_2923231 | Labo Analyses Med | National Reference Center for Viruses of Respiratory Infections, Institut Pasteur, Paris | Marion Barbet, Sylvie Behillil, Méline Bizard, Angela Brisebarre, Camille Capel, Vincent Enouf, Louise Lefrançois, Frédéric Lemoine, Christophe Malabat, Corinne Maufrais, Gael Millot, Etienne Simon-Lorière, Maud Vanpeene, Sylvie Van der Werf ,GESTIN Brieuc |
| EPI_ISL_2923232, EPI_ISL_2923235 | Labo Analyses Med | National Reference Center for Viruses of Respiratory Infections, Institut Pasteur, Paris | Marion Barbet, Sylvie Behillil, Méline Bizard, Angela Brisebarre, Camille Capel, Vincent Enouf, Louise Lefrançois, Frédéric Lemoine, Christophe Malabat, Corinne Maufrais, Hub Bioinformatique Biostatistiques, Etienne Simon-Lorière, Maud Vanpeene, Sylvie Van der Werf ,GESTIN Brieuc |
| EPI_ISL_2923241 | Labo Analyses Med | National Reference Center for Viruses of Respiratory Infections, Institut Pasteur, Paris | Marion Barbet, Sylvie Behillil, Méline Bizard, Angela Brisebarre, Camille Capel, Vincent Enouf, Louise Lefrançois, Frédéric Lemoine, Christophe Malabat, Corinne Maufrais, Hub Bioinformatique Biostatistiques, Etienne Simon-Lorière, Maud Vanpeene, Sylvie Van der Werf ,CERBALLIANCE Ile de France sud |
| EPI_ISL_2923242 | Labo Analyses Med | National Reference Center for Viruses of Respiratory Infections, Institut Pasteur, Paris | Marion Barbet, Sylvie Behillil, Méline Bizard, Angela Brisebarre, Camille Capel, Vincent Enouf, Louise Lefrançois, Frédéric Lemoine, Christophe Malabat, Corinne Maufrais, Pierre Lechat, Etienne Simon-Lorière, Maud Vanpeene, Sylvie Van der Werf ,CERBALLIANCE Ile de France sud |
| EPI_ISL_2923243, EPI_ISL_2923245 | Labo Analyses Med | National Reference Center for Viruses of Respiratory Infections, Institut Pasteur, Paris | Marion Barbet, Sylvie Behillil, Méline Bizard, Angela Brisebarre, Camille Capel, Vincent Enouf, Louise Lefrançois, Frédéric Lemoine, Christophe Malabat, Corinne Maufrais, Gael Millot, Etienne Simon-Lorière, Maud Vanpeene, Sylvie Van der Werf ,BESSON Jérôme |
| EPI_ISL_2923246 | Labo Analyses Med | National Reference Center for Viruses of Respiratory Infections, Institut Pasteur, Paris | Marion Barbet, Sylvie Behillil, Méline Bizard, Angela Brisebarre, Camille Capel, Vincent Enouf, Louise Lefrançois, Frédéric Lemoine, Christophe Malabat, Corinne Maufrais, Pierre Lechat, Etienne Simon-Lorière, Maud Vanpeene, Sylvie Van der Werf ,BESSON Jérôme |
| EPI_ISL_2923247 | Labo Analyses Med | National Reference Center for Viruses of Respiratory Infections, Institut Pasteur, Paris | Marion Barbet, Sylvie Behillil, Méline Bizard, Angela Brisebarre, Camille Capel, Vincent Enouf, Louise Lefrançois, Frédéric Lemoine, Christophe Malabat, Corinne Maufrais, Hub Bioinformatique Biostatistiques, Etienne Simon-Lorière, Maud Vanpeene, Sylvie Van der Werf ,BESSON Jérôme |
| EPI_ISL_2923249, EPI_ISL_2923250, EPI_ISL_2923253, EPI_ISL_2923256 | Labo Analyses Med | National Reference Center for Viruses of Respiratory Infections, Institut Pasteur, Paris | Marion Barbet, Sylvie Behillil, Méline Bizard, Angela Brisebarre, Camille Capel, Vincent Enouf, Louise Lefrançois, Frédéric Lemoine, Christophe Malabat, Corinne Maufrais, Hub Bioinformatique Biostatistiques, Etienne Simon-Lorière, Maud Vanpeene, Sylvie Van der Werf ,COCCO ARDITI |
| EPI_ISL_2923258 | Labo Analyses Med | National Reference Center for Viruses of Respiratory Infections, Institut Pasteur, Paris | Marion Barbet, Sylvie Behillil, Méline Bizard, Angela Brisebarre, Camille Capel, Vincent Enouf, Louise Lefrançois, Frédéric Lemoine, Christophe Malabat, Corinne Maufrais, Gael Millot, Etienne Simon-Lorière, Maud Vanpeene, Sylvie Van der Werf ,COCCO ARDITI |
| EPI_ISL_2923260 | Labo Analyses Med | National Reference Center for Viruses of Respiratory Infections, Institut Pasteur, Paris | Marion Barbet, Sylvie Behillil, Méline Bizard, Angela Brisebarre, Camille Capel, Vincent Enouf, Louise Lefrançois, Frédéric Lemoine, Christophe Malabat, Corinne Maufrais, Hub Bioinformatique Biostatistiques, Etienne Simon-Lorière, Maud Vanpeene, Sylvie Van der Werf ,COCCO ARDITI |
| EPI_ISL_2923262 | Labo Analyses Med | National Reference Center for Viruses of Respiratory Infections, Institut Pasteur, Paris | Marion Barbet, Sylvie Behillil, Méline Bizard, Angela Brisebarre, Camille Capel, Vincent Enouf, Louise Lefrançois, Frédéric Lemoine, Christophe Malabat, Corinne Maufrais, Gael Millot, Etienne Simon-Lorière, Maud Vanpeene, Sylvie Van der Werf ,COMES Jean-François |
| EPI_ISL_2923265, EPI_ISL_2923266 | Labo Analyses Med | National Reference Center for Viruses of Respiratory Infections, Institut Pasteur, Paris | Marion Barbet, Sylvie Behillil, Méline Bizard, Angela Brisebarre, Camille Capel, Vincent Enouf, Louise Lefrançois, Frédéric Lemoine, Christophe Malabat, Corinne Maufrais, Hub Bioinformatique Biostatistiques, Etienne Simon-Lorière, Maud Vanpeene, Sylvie Van der Werf ,COMES Jean-François |
| EPI_ISL_2923268 | Hospital | National Reference Center for Viruses of Respiratory Infections, Institut Pasteur, Paris | Marion Barbet, Sylvie Behillil, Méline Bizard, Angela Brisebarre, Camille Capel, Vincent Enouf, Louise Lefrançois, Frédéric Lemoine, Christophe Malabat, Corinne Maufrais, Gael Millot, Etienne Simon-Lorière, Maud Vanpeene, Sylvie Van der Werf ,GAUDY GRAFFIN Catherine |
| EPI_ISL_2923269 | Hospital | National Reference Center for Viruses of Respiratory Infections, Institut Pasteur, Paris | Marion Barbet, Sylvie Behillil, Méline Bizard, Angela Brisebarre, Camille Capel, Vincent Enouf, Louise Lefrançois, Frédéric Lemoine, Christophe Malabat, Corinne Maufrais, Hub Bioinformatique Biostatistiques, Etienne Simon-Lorière, Maud Vanpeene, Sylvie Van der Werf ,GAUDY GRAFFIN Catherine |
| EPI_ISL_2923271 | Labo Analyses Med | National Reference Center for Viruses of Respiratory Infections, Institut Pasteur, Paris | Marion Barbet, Sylvie Behillil, Méline Bizard, Angela Brisebarre, Camille Capel, Vincent Enouf, Louise Lefrançois, Frédéric Lemoine, Christophe Malabat, Corinne Maufrais, Hub Bioinformatique Biostatistiques, Etienne Simon-Lorière, Maud Vanpeene, Sylvie Van der Werf ,WEBER Valérie |
| EPI_ISL_2923272 | Labo Analyses Med | National Reference Center for Viruses of Respiratory Infections, Institut Pasteur, Paris | Marion Barbet, Sylvie Behillil, Méline Bizard, Angela Brisebarre, Camille Capel, Vincent Enouf, Louise Lefrançois, Frédéric Lemoine, Christophe Malabat, Corinne Maufrais, Gael Millot, Etienne Simon-Lorière, Maud Vanpeene, Sylvie Van der Werf ,WEBER Valérie |
| EPI_ISL_2923273, EPI_ISL_2923274 | Labo Analyses Med | National Reference Center for Viruses of Respiratory Infections, Institut Pasteur, Paris | Marion Barbet, Sylvie Behillil, Méline Bizard, Angela Brisebarre, Camille Capel, Vincent Enouf, Louise Lefrançois, Frédéric Lemoine, Christophe Malabat, Corinne Maufrais, Hub Bioinformatique Biostatistiques, Etienne Simon-Lorière, Maud Vanpeene, Sylvie Van der Werf ,TROMBERT Sabine |
| EPI_ISL_2923275 | Labo Analyses Med | National Reference Center for Viruses of Respiratory Infections, Institut Pasteur, Paris | Marion Barbet, Sylvie Behillil, Méline Bizard, Angela Brisebarre, Camille Capel, Vincent Enouf, Louise Lefrançois, Frédéric Lemoine, Christophe Malabat, Corinne Maufrais, Gael Millot, Etienne Simon-Lorière, Maud Vanpeene, Sylvie Van der Werf ,TROMBERT Sabine |
| EPI_ISL_2923276 | Labo Analyses Med | National Reference Center for Viruses of Respiratory Infections, Institut Pasteur, Paris | Marion Barbet, Sylvie Behillil, Méline Bizard, Angela Brisebarre, Camille Capel, Vincent Enouf, Louise Lefrançois, Frédéric Lemoine, Christophe Malabat, Corinne Maufrais, Hub Bioinformatique Biostatistiques, Etienne Simon-Lorière, Maud Vanpeene, Sylvie Van der Werf ,TROMBERT Sabine |
| EPI_ISL_2923279, EPI_ISL_2923280 | Labo Analyses Med | National Reference Center for Viruses of Respiratory Infections, Institut Pasteur, Paris | Marion Barbet, Sylvie Behillil, Méline Bizard, Angela Brisebarre, Camille Capel, Vincent Enouf, Louise Lefrançois, Frédéric Lemoine, Christophe Malabat, Corinne Maufrais, Gael Millot, Etienne Simon-Lorière, Maud Vanpeene, Sylvie Van der Werf ,MICHEZ Karine |
| EPI_ISL_2923281 | Labo Analyses Med | National Reference Center for Viruses of Respiratory Infections, Institut Pasteur, Paris | Marion Barbet, Sylvie Behillil, Méline Bizard, Angela Brisebarre, Camille Capel, Vincent Enouf, Louise Lefrançois, Frédéric Lemoine, Christophe Malabat, Corinne Maufrais, Hub Bioinformatique Biostatistiques, Etienne Simon-Lorière, Maud Vanpeene, Sylvie Van der Werf ,MICHEZ Karine |
| EPI_ISL_2923282 | Labo Analyses Med | National Reference Center for Viruses of Respiratory Infections, Institut Pasteur, Paris | Marion Barbet, Sylvie Behillil, Méline Bizard, Angela Brisebarre, Camille Capel, Vincent Enouf, Louise Lefrançois, Frédéric Lemoine, Christophe Malabat, Corinne Maufrais, Hub Bioinformatique Biostatistiques, Etienne Simon-Lorière, Maud Vanpeene, Sylvie Van der Werf ,POTIRON Grégoire |
| EPI_ISL_2923293, EPI_ISL_2923295, EPI_ISL_2923303, EPI_ISL_2923307, EPI_ISL_2923323, EPI_ISL_2923327, EPI_ISL_2923328, EPI_ISL_2923340, EPI_ISL_2923373, EPI_ISL_2923402, EPI_ISL_2923408, EPI_ISL_2923411, EPI_ISL_2923455, EPI_ISL_2923476, EPI_ISL_2923477, EPI_ISL_2923511, EPI_ISL_2923515, EPI_ISL_2923541, EPI_ISL_2923558, EPI_ISL_2923560, EPI_ISL_2923561, EPI_ISL_2923568, EPI_ISL_2923572, EPI_ISL_2923573, EPI_ISL_2923574, EPI_ISL_2923575, EPI_ISL_2923576, EPI_ISL_2923577, EPI_ISL_2923578, EPI_ISL_2923579, EPI_ISL_2923623, EPI_ISL_2923664, EPI_ISL_2923672, EPI_ISL_2923688, EPI_ISL_2923689, EPI_ISL_2923690, EPI_ISL_2923710, EPI_ISL_2923711, EPI_ISL_2923713, EPI_ISL_2923730 | | | |
| see above | CDPH VBL | California Department of Public Health | CDPH-COVIDNet |
| EPI_ISL_2923908, EPI_ISL_2923909, EPI_ISL_2923910, EPI_ISL_2923911, EPI_ISL_2923913, EPI_ISL_2923916, EPI_ISL_2923917, EPI_ISL_2923919, EPI_ISL_2923920, EPI_ISL_2923921, EPI_ISL_2923922, EPI_ISL_2923924, EPI_ISL_2923925, EPI_ISL_2923926, EPI_ISL_2923927, EPI_ISL_2923929, EPI_ISL_2923930, EPI_ISL_2923932, EPI_ISL_2923933, EPI_ISL_2923934, EPI_ISL_2923935, EPI_ISL_2923936, EPI_ISL_2923937, EPI_ISL_2923938, EPI_ISL_2923939, EPI_ISL_2923940, EPI_ISL_2923941, EPI_ISL_2923944, EPI_ISL_2923946, EPI_ISL_2923947, EPI_ISL_2923948, EPI_ISL_2923949 | | | |
| see above | Pandemic Response Lab - NYC | Pandemic Response Lab, R&D | Henry Lee, Michael Hammerling, Melissa Hopkins, Cybill del Castillo, Shinyoung Clair Kang, William Ward, Pradeep Bugga, Sol Rey, Dylan Law, Isabel Fernandez Escapa, Haiping Hao, Jon Laurent |
| EPI_ISL_2924057 | Labuan Hospital | Institute for Medical Research, Infectious Disease Research Centre, National Institutes of Health, Ministry of Health Malaysia | Suppiah J, Kamel K, Mohd Zawawi Z, Azizan MA, Ramly N, Robert F, Anasir MI, Thayan R |
| EPI_ISL_2924137 | Sultan Haji Ahmad Shah Hospital | Institute for Medical Research, Infectious Disease Research Centre, National Institutes of Health, Ministry of Health Malaysia | Suppiah J, Kamel K, Mohd Zawawi Z, Azizan MA, Ramly N, Robert F, Anasir MI, Thayan R |
| EPI_ISL_2924138 | AOU Polclinico "G Rodolico-S.Marco" PO Rodolico Catania | AOU Polclinico "G Rodolico-S.Marco" PO Rodolico and BIOMETEC UNICT CATANIA | STEFANI AND SCALIA |
| EPI_ISL_2924139 | Sultan Haji Ahmad Shah Hospital | Institute for Medical Research, Infectious Disease Research Centre, National Institutes of Health, Ministry of Health Malaysia | Suppiah J, Kamel K, Mohd Zawawi Z, Azizan MA, Ramly N, Robert F, Anasir MI, Thayan R |
| EPI_ISL_2924140 | AOU Polclinico "G Rodolico-S.Marco" PO Rodolico CATANIA | AOU Polclinico "G Rodolico-S.Marco" PO Rodolico and BIOMETEC UNICT CATANIA | STEFANI AND SCALIA |
| EPI_ISL_2924142, EPI_ISL_2924143, EPI_ISL_2924145, EPI_ISL_2924149 | National Platform bis UMONS/Jolimont | National Platform bis UMONS/Jolimont | François Dufrasne, Guillaume Bayon-Vicente, Florian Juszczak, Eric Tarantino, Gautier Detry, Ruddy Wattiez |
| EPI_ISL_2924152, EPI_ISL_2924153, EPI_ISL_2924154, EPI_ISL_2924155, EPI_ISL_2924156, EPI_ISL_2924157, EPI_ISL_2924158, EPI_ISL_2924159, EPI_ISL_2924160, EPI_ISL_2924161, EPI_ISL_2924162, EPI_ISL_2924163, EPI_ISL_2924164, EPI_ISL_2924165, EPI_ISL_2924166, EPI_ISL_2924167, EPI_ISL_2924168, EPI_ISL_2924169, EPI_ISL_2924170 | | | |
| see above | Gandhi Hospital | CDFD | Murali Bashyam, Asmita Gupta, Reelina Basu, Vinay Donipadi, Nagamani Kammili, Divya Vashisht, Ashwin Dalal |
| EPI_ISL_2924188, EPI_ISL_2924190, EPI_ISL_2924191, EPI_ISL_2924192, EPI_ISL_2924193, EPI_ISL_2924194, EPI_ISL_2924196, EPI_ISL_2924197, EPI_ISL_2924198, EPI_ISL_2924199, EPI_ISL_2924200, EPI_ISL_2924201, EPI_ISL_2924202 | | | |
| see above | Institute of Preventive Medicine | CDFD | Murali Bashyam, Asmita Gupta, Reelina Basu, Vinay Donipadi, Nagamani Kammili, Divya Vashisht, Ashwin Dalal |
| EPI_ISL_2924203, EPI_ISL_2924204, EPI_ISL_2924205, EPI_ISL_2924206, EPI_ISL_2924207, EPI_ISL_2924208, EPI_ISL_2924209, EPI_ISL_2924210, EPI_ISL_2924211, EPI_ISL_2924212, EPI_ISL_2924213, EPI_ISL_2924214, EPI_ISL_2924215 | | | |
| see above | Government Fever Hospital | CDFD | Murali Bashyam, Asmita Gupta, Reelina Basu, Vinay Donipadi, Nagamani Kammili, Divya Vashisht, Ashwin Dalal |
| EPI_ISL_2924216, EPI_ISL_2924218, EPI_ISL_2924219, EPI_ISL_2924220, EPI_ISL_2924221, EPI_ISL_2924222, EPI_ISL_2924223, EPI_ISL_2924224, EPI_ISL_2924225, EPI_ISL_2924226, EPI_ISL_2924227, EPI_ISL_2924228, EPI_ISL_2924229, EPI_ISL_2924230, EPI_ISL_2924231, EPI_ISL_2924232, EPI_ISL_2924234, EPI_ISL_2924235 | | | |
| see above | Gandhi Hospital | CDFD | Murali Bashyam, Asmita Gupta, Reelina Basu, Vinay Donipadi, Nagamani Kammili, Divya Vashisht, Ashwin Dalal |
| EPI_ISL_2924236, EPI_ISL_2924237, EPI_ISL_2924238, EPI_ISL_2924239, EPI_ISL_2924240, EPI_ISL_2924241, EPI_ISL_2924242, EPI_ISL_2924243, EPI_ISL_2924244, EPI_ISL_2924245, EPI_ISL_2924246, EPI_ISL_2924247, EPI_ISL_2924248, EPI_ISL_2924249 | | | |
| see above | Institute of Preventive Medicine | CDFD | Murali Bashyam, Asmita Gupta, Reelina Basu, Vinay Donipadi, Nagamani Kammili, Divya Vashisht, Ashwin Dalal |
| EPI_ISL_2924250, EPI_ISL_2924251, EPI_ISL_2924252, EPI_ISL_2924253, EPI_ISL_2924254, EPI_ISL_2924255, EPI_ISL_2924256, EPI_ISL_2924257, EPI_ISL_2924258, EPI_ISL_2924259, EPI_ISL_2924260, EPI_ISL_2924261, EPI_ISL_2924262, EPI_ISL_2924263 | | | |
| see above | Osmania Medical College | CDFD | Murali Bashyam, Asmita Gupta, Reelina Basu, Vinay Donipadi, Nagamani Kammili, Divya Vashisht, Ashwin Dalal |
| EPI_ISL_2924264, EPI_ISL_2924265, EPI_ISL_2924266, EPI_ISL_2924267 | Gandhi Hospital | CDFD | Murali Bashyam, Asmita Gupta, Reelina Basu, Vinay Donipadi, Nagamani Kammili, Divya Vashisht, Ashwin Dalal |
| EPI_ISL_2924268 | National Platform bis UMONS/Jolimont | National Platform bis UMONS/Jolimont | François Dufrasne, Guillaume Bayon-Vicente, Florian Juszczak, Eric Tarantino, Gautier Detry, Ruddy Wattiez |
| EPI_ISL_2924269 | Lighthouse Lab in Alderley Park | Wellcome Sanger Institute for the COVID-19 Genomics UK (COG-UK) Consortium | Jacquelyn Wynn, Mairead Hyland, The Lighthouse Lab in Alderley Park and Alex Alderton, Roberto Amato, Jeffrey Barrett, Sonia Goncalves, Ewan Harrison, David K. Jackson, Ian Johnston, Dominic Kwiatkowski, Cordelia Langford, John Sillitoe on behalf of the Wellcome Sanger Institute COVID-19 Surveillance Team |
| EPI_ISL_2924270 | Lighthouse Lab in Glasgow | Wellcome Sanger Institute for the COVID-19 Genomics UK (COG-UK) Consortium | Harper VanSteenhouse, Yumi Kasai, David Gray, Carol Clugston, Anna Dominiczak and Alex Alderton, Roberto Amato, Jeffrey Barrett, Sonia Goncalves, Ewan Harrison, David K. Jackson, Ian Johnston, Dominic Kwiatkowski, Cordelia Langford, John Sillitoe on behalf of the Wellcome Sanger Institute COVID-19 Surveillance Team |
| EPI_ISL_2924271, | Lighthouse Lab in Alderley Park | Wellcome Sanger Institute for the COVID-19 Genomics UK (COG-UK) | Jacquelyn Wynn, Mairead Hyland, The Lighthouse Lab in Alderley Park and Alex Alderton, Roberto Amato, Jeffrey Barrett, Sonia Goncalves, Ewan Harrison, David K. Jackson, Ian Johnston, Dominic Kwiatkowski, Cordelia Langford, John Sillitoe on |

| | | | |
|---|---|---|---|
| EPI_ISL_2924272 | | Consortium | behalf of the Wellcome Sanger Institute COVID-19 Surveillance Team |
| EPI_ISL_2924273, EPI_ISL_2924274 | Lighthouse Lab in Glasgow | Wellcome Sanger Institute for the COVID-19 Genomics UK (COG-UK) Consortium | Harper VanSteenhouse, Yumi Kasai, David Gray, Carol Clugston, Anna Dominiczak and Alex Alderton, Roberto Amato, Jeffrey Barrett, Sonia Goncalves, Ewan Harrison, David K. Jackson, Ian Johnston, Dominic Kwiatkowski, Cordelia Langford, John Sillitoe on behalf of the Wellcome Sanger Institute COVID-19 Surveillance Team |
| EPI_ISL_2924275, EPI_ISL_2924276, EPI_ISL_2924277, EPI_ISL_2924278, EPI_ISL_2924279, EPI_ISL_2924280, EPI_ISL_2924281, EPI_ISL_2924282, EPI_ISL_2924283, EPI_ISL_2924284, EPI_ISL_2924285, EPI_ISL_2924286, EPI_ISL_2924288, EPI_ISL_2924289, EPI_ISL_2924290, EPI_ISL_2924291, EPI_ISL_2924292, EPI_ISL_2924293, EPI_ISL_2924294 | | | |
| see above | Lighthouse Lab in Alderley Park | Wellcome Sanger Institute for the COVID-19 Genomics UK (COG-UK) Consortium | Jacquelyn Wynn, Mairead Hyland, The Lighthouse Lab in Alderley Park and Alex Alderton, Roberto Amato, Jeffrey Barrett, Sonia Goncalves, Ewan Harrison, David K. Jackson, Ian Johnston, Dominic Kwiatkowski, Cordelia Langford, John Sillitoe on behalf of the Wellcome Sanger Institute COVID-19 Surveillance Team |
| EPI_ISL_2924295 | Lighthouse Lab in Glasgow | Wellcome Sanger Institute for the COVID-19 Genomics UK (COG-UK) Consortium | Harper VanSteenhouse, Yumi Kasai, David Gray, Carol Clugston, Anna Dominiczak and Alex Alderton, Roberto Amato, Jeffrey Barrett, Sonia Goncalves, Ewan Harrison, David K. Jackson, Ian Johnston, Dominic Kwiatkowski, Cordelia Langford, John Sillitoe on behalf of the Wellcome Sanger Institute COVID-19 Surveillance Team |
| EPI_ISL_2924296, EPI_ISL_2924297, EPI_ISL_2924298, EPI_ISL_2924299, EPI_ISL_2924300, EPI_ISL_2924301, EPI_ISL_2924302, EPI_ISL_2924303, EPI_ISL_2924304, EPI_ISL_2924305 | Lighthouse Lab in Alderley Park | Wellcome Sanger Institute for the COVID-19 Genomics UK (COG-UK) Consortium | Jacquelyn Wynn, Mairead Hyland, The Lighthouse Lab in Alderley Park and Alex Alderton, Roberto Amato, Jeffrey Barrett, Sonia Goncalves, Ewan Harrison, David K. Jackson, Ian Johnston, Dominic Kwiatkowski, Cordelia Langford, John Sillitoe on behalf of the Wellcome Sanger Institute COVID-19 Surveillance Team |
| EPI_ISL_2924306 | Lighthouse Lab in Glasgow | Wellcome Sanger Institute for the COVID-19 Genomics UK (COG-UK) Consortium | Harper VanSteenhouse, Yumi Kasai, David Gray, Carol Clugston, Anna Dominiczak and Alex Alderton, Roberto Amato, Jeffrey Barrett, Sonia Goncalves, Ewan Harrison, David K. Jackson, Ian Johnston, Dominic Kwiatkowski, Cordelia Langford, John Sillitoe on behalf of the Wellcome Sanger Institute COVID-19 Surveillance Team |
| EPI_ISL_2924307 | Lighthouse Lab in Alderley Park | Wellcome Sanger Institute for the COVID-19 Genomics UK (COG-UK) Consortium | Jacquelyn Wynn, Mairead Hyland, The Lighthouse Lab in Alderley Park and Alex Alderton, Roberto Amato, Jeffrey Barrett, Sonia Goncalves, Ewan Harrison, David K. Jackson, Ian Johnston, Dominic Kwiatkowski, Cordelia Langford, John Sillitoe on behalf of the Wellcome Sanger Institute COVID-19 Surveillance Team |
| EPI_ISL_2924308, EPI_ISL_2924309 | Lighthouse Lab in Glasgow | Wellcome Sanger Institute for the COVID-19 Genomics UK (COG-UK) Consortium | Harper VanSteenhouse, Yumi Kasai, David Gray, Carol Clugston, Anna Dominiczak and Alex Alderton, Roberto Amato, Jeffrey Barrett, Sonia Goncalves, Ewan Harrison, David K. Jackson, Ian Johnston, Dominic Kwiatkowski, Cordelia Langford, John Sillitoe on behalf of the Wellcome Sanger Institute COVID-19 Surveillance Team |
| EPI_ISL_2924310, EPI_ISL_2924311, EPI_ISL_2924312, EPI_ISL_2924313, EPI_ISL_2924314, EPI_ISL_2924315 | Lighthouse Lab in Alderley Park | Wellcome Sanger Institute for the COVID-19 Genomics UK (COG-UK) Consortium | Jacquelyn Wynn, Mairead Hyland, The Lighthouse Lab in Alderley Park and Alex Alderton, Roberto Amato, Jeffrey Barrett, Sonia Goncalves, Ewan Harrison, David K. Jackson, Ian Johnston, Dominic Kwiatkowski, Cordelia Langford, John Sillitoe on behalf of the Wellcome Sanger Institute COVID-19 Surveillance Team |
| EPI_ISL_2924316 | Lighthouse Lab in Glasgow | Wellcome Sanger Institute for the COVID-19 Genomics UK (COG-UK) Consortium | Harper VanSteenhouse, Yumi Kasai, David Gray, Carol Clugston, Anna Dominiczak and Alex Alderton, Roberto Amato, Jeffrey Barrett, Sonia Goncalves, Ewan Harrison, David K. Jackson, Ian Johnston, Dominic Kwiatkowski, Cordelia Langford, John Sillitoe on behalf of the Wellcome Sanger Institute COVID-19 Surveillance Team |
| EPI_ISL_2924317, EPI_ISL_2924318, EPI_ISL_2924319, EPI_ISL_2924320, EPI_ISL_2924321, EPI_ISL_2924322, EPI_ISL_2924323, EPI_ISL_2924324, EPI_ISL_2924325, EPI_ISL_2924326, EPI_ISL_2924327 | | | |
| see above | Lighthouse Lab in Alderley Park | Wellcome Sanger Institute for the COVID-19 Genomics UK (COG-UK) Consortium | Jacquelyn Wynn, Mairead Hyland, The Lighthouse Lab in Alderley Park and Alex Alderton, Roberto Amato, Jeffrey Barrett, Sonia Goncalves, Ewan Harrison, David K. Jackson, Ian Johnston, Dominic Kwiatkowski, Cordelia Langford, John Sillitoe on behalf of the Wellcome Sanger Institute COVID-19 Surveillance Team |
| EPI_ISL_2924328 | Lighthouse Lab in Glasgow | Wellcome Sanger Institute for the COVID-19 Genomics UK (COG-UK) Consortium | Harper VanSteenhouse, Yumi Kasai, David Gray, Carol Clugston, Anna Dominiczak and Alex Alderton, Roberto Amato, Jeffrey Barrett, Sonia Goncalves, Ewan Harrison, David K. Jackson, Ian Johnston, Dominic Kwiatkowski, Cordelia Langford, John Sillitoe on behalf of the Wellcome Sanger Institute COVID-19 Surveillance Team |
| EPI_ISL_2924329, EPI_ISL_2924330, EPI_ISL_2924331, EPI_ISL_2924332, EPI_ISL_2924333, EPI_ISL_2924334, EPI_ISL_2924335, EPI_ISL_2924336, EPI_ISL_2924337 | Lighthouse Lab in Alderley Park | Wellcome Sanger Institute for the COVID-19 Genomics UK (COG-UK) Consortium | Jacquelyn Wynn, Mairead Hyland, The Lighthouse Lab in Alderley Park and Alex Alderton, Roberto Amato, Jeffrey Barrett, Sonia Goncalves, Ewan Harrison, David K. Jackson, Ian Johnston, Dominic Kwiatkowski, Cordelia Langford, John Sillitoe on behalf of the Wellcome Sanger Institute COVID-19 Surveillance Team |
| EPI_ISL_2924338 | Lighthouse Lab in Glasgow | Wellcome Sanger Institute for the COVID-19 Genomics UK (COG-UK) Consortium | Harper VanSteenhouse, Yumi Kasai, David Gray, Carol Clugston, Anna Dominiczak and Alex Alderton, Roberto Amato, Jeffrey Barrett, Sonia Goncalves, Ewan Harrison, David K. Jackson, Ian Johnston, Dominic Kwiatkowski, Cordelia Langford, John Sillitoe on behalf of the Wellcome Sanger Institute COVID-19 Surveillance Team |
| EPI_ISL_2924339, EPI_ISL_2924340, EPI_ISL_2924341, EPI_ISL_2924342 | Lighthouse Lab in Alderley Park | Wellcome Sanger Institute for the COVID-19 Genomics UK (COG-UK) Consortium | Jacquelyn Wynn, Mairead Hyland, The Lighthouse Lab in Alderley Park and Alex Alderton, Roberto Amato, Jeffrey Barrett, Sonia Goncalves, Ewan Harrison, David K. Jackson, Ian Johnston, Dominic Kwiatkowski, Cordelia Langford, John Sillitoe on behalf of the Wellcome Sanger Institute COVID-19 Surveillance Team |
| EPI_ISL_2924343 | Lighthouse Lab in Glasgow | Wellcome Sanger Institute for the COVID-19 Genomics UK (COG-UK) Consortium | Harper VanSteenhouse, Yumi Kasai, David Gray, Carol Clugston, Anna Dominiczak and Alex Alderton, Roberto Amato, Jeffrey Barrett, Sonia Goncalves, Ewan Harrison, David K. Jackson, Ian Johnston, Dominic Kwiatkowski, Cordelia Langford, John Sillitoe on behalf of the Wellcome Sanger Institute COVID-19 Surveillance Team |
| EPI_ISL_2924344 | Lighthouse Lab in Alderley Park | Wellcome Sanger Institute for the COVID-19 Genomics UK (COG-UK) Consortium | Jacquelyn Wynn, Mairead Hyland, The Lighthouse Lab in Alderley Park and Alex Alderton, Roberto Amato, Jeffrey Barrett, Sonia Goncalves, Ewan Harrison, David K. Jackson, Ian Johnston, Dominic Kwiatkowski, Cordelia Langford, John Sillitoe on behalf of the Wellcome Sanger Institute COVID-19 Surveillance Team |
| EPI_ISL_2924345, EPI_ISL_2924346 | Lighthouse Lab in Glasgow | Wellcome Sanger Institute for the COVID-19 Genomics UK (COG-UK) Consortium | Harper VanSteenhouse, Yumi Kasai, David Gray, Carol Clugston, Anna Dominiczak and Alex Alderton, Roberto Amato, Jeffrey Barrett, Sonia Goncalves, Ewan Harrison, David K. Jackson, Ian Johnston, Dominic Kwiatkowski, Cordelia Langford, John Sillitoe on behalf of the Wellcome Sanger Institute COVID-19 Surveillance Team |
| EPI_ISL_2924347, EPI_ISL_2924348, EPI_ISL_2924349, EPI_ISL_2924350, EPI_ISL_2924351, EPI_ISL_2924352, EPI_ISL_2924353, EPI_ISL_2924354, EPI_ISL_2924355, EPI_ISL_2924356, EPI_ISL_2924357 | | | |
| see above | Lighthouse Lab in Alderley Park | Wellcome Sanger Institute for the COVID-19 Genomics UK (COG-UK) Consortium | Jacquelyn Wynn, Mairead Hyland, The Lighthouse Lab in Alderley Park and Alex Alderton, Roberto Amato, Jeffrey Barrett, Sonia Goncalves, Ewan Harrison, David K. Jackson, Ian Johnston, Dominic Kwiatkowski, Cordelia Langford, John Sillitoe on behalf of the Wellcome Sanger Institute COVID-19 Surveillance Team |
| EPI_ISL_2924358 | Lighthouse Lab in Glasgow | Wellcome Sanger Institute for the COVID-19 Genomics UK (COG-UK) Consortium | Harper VanSteenhouse, Yumi Kasai, David Gray, Carol Clugston, Anna Dominiczak and Alex Alderton, Roberto Amato, Jeffrey Barrett, Sonia Goncalves, Ewan Harrison, David K. Jackson, Ian Johnston, Dominic Kwiatkowski, Cordelia Langford, John Sillitoe on behalf of the Wellcome Sanger Institute COVID-19 Surveillance Team |
| EPI_ISL_2924359, EPI_ISL_2924360 | Lighthouse Lab in Alderley Park | Wellcome Sanger Institute for the COVID-19 Genomics UK (COG-UK) Consortium | Jacquelyn Wynn, Mairead Hyland, The Lighthouse Lab in Alderley Park and Alex Alderton, Roberto Amato, Jeffrey Barrett, Sonia Goncalves, Ewan Harrison, David K. Jackson, Ian Johnston, Dominic Kwiatkowski, Cordelia Langford, John Sillitoe on behalf of the Wellcome Sanger Institute COVID-19 Surveillance Team |
| EPI_ISL_2924361 | Lighthouse Lab in Glasgow | Wellcome Sanger Institute for the COVID-19 Genomics UK (COG-UK) Consortium | Harper VanSteenhouse, Yumi Kasai, David Gray, Carol Clugston, Anna Dominiczak and Alex Alderton, Roberto Amato, Jeffrey Barrett, Sonia Goncalves, Ewan Harrison, David K. Jackson, Ian Johnston, Dominic Kwiatkowski, Cordelia Langford, John Sillitoe on behalf of the Wellcome Sanger Institute COVID-19 Surveillance Team |
| EPI_ISL_2924362 | Lighthouse Lab in Alderley Park | Wellcome Sanger Institute for the COVID-19 Genomics UK (COG-UK) Consortium | Jacquelyn Wynn, Mairead Hyland, The Lighthouse Lab in Alderley Park and Alex Alderton, Roberto Amato, Jeffrey Barrett, Sonia Goncalves, Ewan Harrison, David K. Jackson, Ian Johnston, Dominic Kwiatkowski, Cordelia Langford, John Sillitoe on behalf of the Wellcome Sanger Institute COVID-19 Surveillance Team |
| EPI_ISL_2924363 | Lighthouse Lab in Glasgow | Wellcome Sanger Institute for the COVID-19 Genomics UK (COG-UK) Consortium | Harper VanSteenhouse, Yumi Kasai, David Gray, Carol Clugston, Anna Dominiczak and Alex Alderton, Roberto Amato, Jeffrey Barrett, Sonia Goncalves, Ewan Harrison, David K. Jackson, Ian Johnston, Dominic Kwiatkowski, Cordelia Langford, John Sillitoe on behalf of the Wellcome Sanger Institute COVID-19 Surveillance Team |
| EPI_ISL_2924364, EPI_ISL_2924365, EPI_ISL_2924366, EPI_ISL_2924367, EPI_ISL_2924368, EPI_ISL_2924369 | Lighthouse Lab in Alderley Park | Wellcome Sanger Institute for the COVID-19 Genomics UK (COG-UK) Consortium | Jacquelyn Wynn, Mairead Hyland, The Lighthouse Lab in Alderley Park and Alex Alderton, Roberto Amato, Jeffrey Barrett, Sonia Goncalves, Ewan Harrison, David K. Jackson, Ian Johnston, Dominic Kwiatkowski, Cordelia Langford, John Sillitoe on behalf of the Wellcome Sanger Institute COVID-19 Surveillance Team |
| EPI_ISL_2924370 | Lighthouse Lab in Glasgow | Wellcome Sanger Institute for the COVID-19 Genomics UK (COG-UK) Consortium | Harper VanSteenhouse, Yumi Kasai, David Gray, Carol Clugston, Anna Dominiczak and Alex Alderton, Roberto Amato, Jeffrey Barrett, Sonia Goncalves, Ewan Harrison, David K. Jackson, Ian Johnston, Dominic Kwiatkowski, Cordelia Langford, John Sillitoe on behalf of the Wellcome Sanger Institute COVID-19 Surveillance Team |
| EPI_ISL_2924371 | Lighthouse Lab in Alderley Park | Wellcome Sanger Institute for the COVID-19 Genomics UK (COG-UK) Consortium | Jacquelyn Wynn, Mairead Hyland, The Lighthouse Lab in Alderley Park and Alex Alderton, Roberto Amato, Jeffrey Barrett, Sonia Goncalves, Ewan Harrison, David K. Jackson, Ian Johnston, Dominic Kwiatkowski, Cordelia Langford, John Sillitoe on behalf of the Wellcome Sanger Institute COVID-19 Surveillance Team |
| EPI_ISL_2924372 | Lighthouse Lab in Glasgow | Wellcome Sanger Institute for the COVID-19 Genomics UK (COG-UK) Consortium | Harper VanSteenhouse, Yumi Kasai, David Gray, Carol Clugston, Anna Dominiczak and Alex Alderton, Roberto Amato, Jeffrey Barrett, Sonia Goncalves, Ewan Harrison, David K. Jackson, Ian Johnston, Dominic Kwiatkowski, Cordelia Langford, John Sillitoe on behalf of the Wellcome Sanger Institute COVID-19 Surveillance Team |
| EPI_ISL_2924373, EPI_ISL_2924374, EPI_ISL_2924375, EPI_ISL_2924376, EPI_ISL_2924377, EPI_ISL_2924378, EPI_ISL_2924379, EPI_ISL_2924380, EPI_ISL_2924381, EPI_ISL_2924382 | Lighthouse Lab in Alderley Park | Wellcome Sanger Institute for the COVID-19 Genomics UK (COG-UK) Consortium | Jacquelyn Wynn, Mairead Hyland, The Lighthouse Lab in Alderley Park and Alex Alderton, Roberto Amato, Jeffrey Barrett, Sonia Goncalves, Ewan Harrison, David K. Jackson, Ian Johnston, Dominic Kwiatkowski, Cordelia Langford, John Sillitoe on behalf of the Wellcome Sanger Institute COVID-19 Surveillance Team |
| EPI_ISL_2924383 | Lighthouse Lab in Glasgow | Wellcome Sanger Institute for the COVID-19 Genomics UK (COG-UK) Consortium | Harper VanSteenhouse, Yumi Kasai, David Gray, Carol Clugston, Anna Dominiczak and Alex Alderton, Roberto Amato, Jeffrey Barrett, Sonia Goncalves, Ewan Harrison, David K. Jackson, Ian Johnston, Dominic Kwiatkowski, Cordelia Langford, John Sillitoe on behalf of the Wellcome Sanger Institute COVID-19 Surveillance Team |
| EPI_ISL_2924384, EPI_ISL_2924385, EPI_ISL_2924386, EPI_ISL_2924387, EPI_ISL_2924388, EPI_ISL_2924389, EPI_ISL_2924390, EPI_ISL_2924391, EPI_ISL_2924392, EPI_ISL_2924393, EPI_ISL_2924394, EPI_ISL_2924395, EPI_ISL_2924396, EPI_ISL_2924397, EPI_ISL_2924398, EPI_ISL_2924399, EPI_ISL_2924400, EPI_ISL_2924401, EPI_ISL_2924402, EPI_ISL_2924403 | | | |

| | | | |
|---|---|---|---|
| see above | Lighthouse Lab in Alderley Park | Wellcome Sanger Institute for the COVID-19 Genomics UK (COG-UK) Consortium | Jacquelyn Wynn, Mairead Hyland, The Lighthouse Lab in Alderley Park and Alex Alderton, Roberto Amato, Jeffrey Barrett, Sonia Goncalves, Ewan Harrison, David K. Jackson, Ian Johnston, Dominic Kwiatkowski, Cordelia Langford, John Sillitoe on behalf of the Wellcome Sanger Institute COVID-19 Surveillance Team |
| EPI_ISL_2924404 | Lighthouse Lab in Glasgow | Wellcome Sanger Institute for the COVID-19 Genomics UK (COG-UK) Consortium | Harper VanSteenhouse, Yumi Kasai, David Gray, Carol Clugston, Anna Dominiczak and Alex Alderton, Roberto Amato, Jeffrey Barrett, Sonia Goncalves, Ewan Harrison, David K. Jackson, Ian Johnston, Dominic Kwiatkowski, Cordelia Langford, John Sillitoe on behalf of the Wellcome Sanger Institute COVID-19 Surveillance Team |
| EPI_ISL_2924405, EPI_ISL_2924406, EPI_ISL_2924407, EPI_ISL_2924408, EPI_ISL_2924409, EPI_ISL_2924410 | Lighthouse Lab in Alderley Park | Wellcome Sanger Institute for the COVID-19 Genomics UK (COG-UK) Consortium | Jacquelyn Wynn, Mairead Hyland, The Lighthouse Lab in Alderley Park and Alex Alderton, Roberto Amato, Jeffrey Barrett, Sonia Goncalves, Ewan Harrison, David K. Jackson, Ian Johnston, Dominic Kwiatkowski, Cordelia Langford, John Sillitoe on behalf of the Wellcome Sanger Institute COVID-19 Surveillance Team |
| EPI_ISL_2924411 | Lighthouse Lab in Glasgow | Wellcome Sanger Institute for the COVID-19 Genomics UK (COG-UK) Consortium | Harper VanSteenhouse, Yumi Kasai, David Gray, Carol Clugston, Anna Dominiczak and Alex Alderton, Roberto Amato, Jeffrey Barrett, Sonia Goncalves, Ewan Harrison, David K. Jackson, Ian Johnston, Dominic Kwiatkowski, Cordelia Langford, John Sillitoe on behalf of the Wellcome Sanger Institute COVID-19 Surveillance Team |
| EPI_ISL_2924412, EPI_ISL_2924413, EPI_ISL_2924414, EPI_ISL_2924415, EPI_ISL_2924416, EPI_ISL_2924417 | Lighthouse Lab in Alderley Park | Wellcome Sanger Institute for the COVID-19 Genomics UK (COG-UK) Consortium | Jacquelyn Wynn, Mairead Hyland, The Lighthouse Lab in Alderley Park and Alex Alderton, Roberto Amato, Jeffrey Barrett, Sonia Goncalves, Ewan Harrison, David K. Jackson, Ian Johnston, Dominic Kwiatkowski, Cordelia Langford, John Sillitoe on behalf of the Wellcome Sanger Institute COVID-19 Surveillance Team |
| EPI_ISL_2924418 | Lighthouse Lab in Glasgow | Wellcome Sanger Institute for the COVID-19 Genomics UK (COG-UK) Consortium | Harper VanSteenhouse, Yumi Kasai, David Gray, Carol Clugston, Anna Dominiczak and Alex Alderton, Roberto Amato, Jeffrey Barrett, Sonia Goncalves, Ewan Harrison, David K. Jackson, Ian Johnston, Dominic Kwiatkowski, Cordelia Langford, John Sillitoe on behalf of the Wellcome Sanger Institute COVID-19 Surveillance Team |
| EPI_ISL_2924419, EPI_ISL_2924420, EPI_ISL_2924421, EPI_ISL_2924422, EPI_ISL_2924423, EPI_ISL_2924424, EPI_ISL_2924425 | Lighthouse Lab in Alderley Park | Wellcome Sanger Institute for the COVID-19 Genomics UK (COG-UK) Consortium | Jacquelyn Wynn, Mairead Hyland, The Lighthouse Lab in Alderley Park and Alex Alderton, Roberto Amato, Jeffrey Barrett, Sonia Goncalves, Ewan Harrison, David K. Jackson, Ian Johnston, Dominic Kwiatkowski, Cordelia Langford, John Sillitoe on behalf of the Wellcome Sanger Institute COVID-19 Surveillance Team |
| EPI_ISL_2924426, EPI_ISL_2924427, EPI_ISL_2924428 | Lighthouse Lab in Glasgow | Wellcome Sanger Institute for the COVID-19 Genomics UK (COG-UK) Consortium | Harper VanSteenhouse, Yumi Kasai, David Gray, Carol Clugston, Anna Dominiczak and Alex Alderton, Roberto Amato, Jeffrey Barrett, Sonia Goncalves, Ewan Harrison, David K. Jackson, Ian Johnston, Dominic Kwiatkowski, Cordelia Langford, John Sillitoe on behalf of the Wellcome Sanger Institute COVID-19 Surveillance Team |
| EPI_ISL_2924429, EPI_ISL_2924430, EPI_ISL_2924431, EPI_ISL_2924432, EPI_ISL_2924433, EPI_ISL_2924434, EPI_ISL_2924435, EPI_ISL_2924436 | Lighthouse Lab in Alderley Park | Wellcome Sanger Institute for the COVID-19 Genomics UK (COG-UK) Consortium | Jacquelyn Wynn, Mairead Hyland, The Lighthouse Lab in Alderley Park and Alex Alderton, Roberto Amato, Jeffrey Barrett, Sonia Goncalves, Ewan Harrison, David K. Jackson, Ian Johnston, Dominic Kwiatkowski, Cordelia Langford, John Sillitoe on behalf of the Wellcome Sanger Institute COVID-19 Surveillance Team |
| EPI_ISL_2924437, EPI_ISL_2924438, EPI_ISL_2924439 | Lighthouse Lab in Glasgow | Wellcome Sanger Institute for the COVID-19 Genomics UK (COG-UK) Consortium | Harper VanSteenhouse, Yumi Kasai, David Gray, Carol Clugston, Anna Dominiczak and Alex Alderton, Roberto Amato, Jeffrey Barrett, Sonia Goncalves, Ewan Harrison, David K. Jackson, Ian Johnston, Dominic Kwiatkowski, Cordelia Langford, John Sillitoe on behalf of the Wellcome Sanger Institute COVID-19 Surveillance Team |
| EPI_ISL_2924440, EPI_ISL_2924441, EPI_ISL_2924442, EPI_ISL_2924443 | Lighthouse Lab in Alderley Park | Wellcome Sanger Institute for the COVID-19 Genomics UK (COG-UK) Consortium | Jacquelyn Wynn, Mairead Hyland, The Lighthouse Lab in Alderley Park and Alex Alderton, Roberto Amato, Jeffrey Barrett, Sonia Goncalves, Ewan Harrison, David K. Jackson, Ian Johnston, Dominic Kwiatkowski, Cordelia Langford, John Sillitoe on behalf of the Wellcome Sanger Institute COVID-19 Surveillance Team |
| EPI_ISL_2924444 | Lighthouse Lab in Glasgow | Wellcome Sanger Institute for the COVID-19 Genomics UK (COG-UK) Consortium | Harper VanSteenhouse, Yumi Kasai, David Gray, Carol Clugston, Anna Dominiczak and Alex Alderton, Roberto Amato, Jeffrey Barrett, Sonia Goncalves, Ewan Harrison, David K. Jackson, Ian Johnston, Dominic Kwiatkowski, Cordelia Langford, John Sillitoe on behalf of the Wellcome Sanger Institute COVID-19 Surveillance Team |
| EPI_ISL_2924445 | Lighthouse Lab in Alderley Park | Wellcome Sanger Institute for the COVID-19 Genomics UK (COG-UK) Consortium | Jacquelyn Wynn, Mairead Hyland, The Lighthouse Lab in Alderley Park and Alex Alderton, Roberto Amato, Jeffrey Barrett, Sonia Goncalves, Ewan Harrison, David K. Jackson, Ian Johnston, Dominic Kwiatkowski, Cordelia Langford, John Sillitoe on behalf of the Wellcome Sanger Institute COVID-19 Surveillance Team |
| EPI_ISL_2924446 | Lighthouse Lab in Glasgow | Wellcome Sanger Institute for the COVID-19 Genomics UK (COG-UK) Consortium | Harper VanSteenhouse, Yumi Kasai, David Gray, Carol Clugston, Anna Dominiczak and Alex Alderton, Roberto Amato, Jeffrey Barrett, Sonia Goncalves, Ewan Harrison, David K. Jackson, Ian Johnston, Dominic Kwiatkowski, Cordelia Langford, John Sillitoe on behalf of the Wellcome Sanger Institute COVID-19 Surveillance Team |
| EPI_ISL_2924447 | Lighthouse Lab in Alderley Park | Wellcome Sanger Institute for the COVID-19 Genomics UK (COG-UK) Consortium | Jacquelyn Wynn, Mairead Hyland, The Lighthouse Lab in Alderley Park and Alex Alderton, Roberto Amato, Jeffrey Barrett, Sonia Goncalves, Ewan Harrison, David K. Jackson, Ian Johnston, Dominic Kwiatkowski, Cordelia Langford, John Sillitoe on behalf of the Wellcome Sanger Institute COVID-19 Surveillance Team |
| EPI_ISL_2924448 | Lighthouse Lab in Glasgow | Wellcome Sanger Institute for the COVID-19 Genomics UK (COG-UK) Consortium | Harper VanSteenhouse, Yumi Kasai, David Gray, Carol Clugston, Anna Dominiczak and Alex Alderton, Roberto Amato, Jeffrey Barrett, Sonia Goncalves, Ewan Harrison, David K. Jackson, Ian Johnston, Dominic Kwiatkowski, Cordelia Langford, John Sillitoe on behalf of the Wellcome Sanger Institute COVID-19 Surveillance Team |
| EPI_ISL_2924449, EPI_ISL_2924450 | Lighthouse Lab in Alderley Park | Wellcome Sanger Institute for the COVID-19 Genomics UK (COG-UK) Consortium | Jacquelyn Wynn, Mairead Hyland, The Lighthouse Lab in Alderley Park and Alex Alderton, Roberto Amato, Jeffrey Barrett, Sonia Goncalves, Ewan Harrison, David K. Jackson, Ian Johnston, Dominic Kwiatkowski, Cordelia Langford, John Sillitoe on behalf of the Wellcome Sanger Institute COVID-19 Surveillance Team |
| EPI_ISL_2924451 | Lighthouse Lab in Glasgow | Wellcome Sanger Institute for the COVID-19 Genomics UK (COG-UK) Consortium | Harper VanSteenhouse, Yumi Kasai, David Gray, Carol Clugston, Anna Dominiczak and Alex Alderton, Roberto Amato, Jeffrey Barrett, Sonia Goncalves, Ewan Harrison, David K. Jackson, Ian Johnston, Dominic Kwiatkowski, Cordelia Langford, John Sillitoe on behalf of the Wellcome Sanger Institute COVID-19 Surveillance Team |
| EPI_ISL_2924452, EPI_ISL_2924453, EPI_ISL_2924454, EPI_ISL_2924455 | Lighthouse Lab in Alderley Park | Wellcome Sanger Institute for the COVID-19 Genomics UK (COG-UK) Consortium | Jacquelyn Wynn, Mairead Hyland, The Lighthouse Lab in Alderley Park and Alex Alderton, Roberto Amato, Jeffrey Barrett, Sonia Goncalves, Ewan Harrison, David K. Jackson, Ian Johnston, Dominic Kwiatkowski, Cordelia Langford, John Sillitoe on behalf of the Wellcome Sanger Institute COVID-19 Surveillance Team |
| EPI_ISL_2924456, EPI_ISL_2924457 | Lighthouse Lab in Glasgow | Wellcome Sanger Institute for the COVID-19 Genomics UK (COG-UK) Consortium | Harper VanSteenhouse, Yumi Kasai, David Gray, Carol Clugston, Anna Dominiczak and Alex Alderton, Roberto Amato, Jeffrey Barrett, Sonia Goncalves, Ewan Harrison, David K. Jackson, Ian Johnston, Dominic Kwiatkowski, Cordelia Langford, John Sillitoe on behalf of the Wellcome Sanger Institute COVID-19 Surveillance Team |
| EPI_ISL_2924458, EPI_ISL_2924459, EPI_ISL_2924460, EPI_ISL_2924461 | Lighthouse Lab in Alderley Park | Wellcome Sanger Institute for the COVID-19 Genomics UK (COG-UK) Consortium | Jacquelyn Wynn, Mairead Hyland, The Lighthouse Lab in Alderley Park and Alex Alderton, Roberto Amato, Jeffrey Barrett, Sonia Goncalves, Ewan Harrison, David K. Jackson, Ian Johnston, Dominic Kwiatkowski, Cordelia Langford, John Sillitoe on behalf of the Wellcome Sanger Institute COVID-19 Surveillance Team |
| EPI_ISL_2924462 | Lighthouse Lab in Glasgow | Wellcome Sanger Institute for the COVID-19 Genomics UK (COG-UK) Consortium | Harper VanSteenhouse, Yumi Kasai, David Gray, Carol Clugston, Anna Dominiczak and Alex Alderton, Roberto Amato, Jeffrey Barrett, Sonia Goncalves, Ewan Harrison, David K. Jackson, Ian Johnston, Dominic Kwiatkowski, Cordelia Langford, John Sillitoe on behalf of the Wellcome Sanger Institute COVID-19 Surveillance Team |
| EPI_ISL_2924463, EPI_ISL_2924464, EPI_ISL_2924465, EPI_ISL_2924466, EPI_ISL_2924467 | Lighthouse Lab in Alderley Park | Wellcome Sanger Institute for the COVID-19 Genomics UK (COG-UK) Consortium | Jacquelyn Wynn, Mairead Hyland, The Lighthouse Lab in Alderley Park and Alex Alderton, Roberto Amato, Jeffrey Barrett, Sonia Goncalves, Ewan Harrison, David K. Jackson, Ian Johnston, Dominic Kwiatkowski, Cordelia Langford, John Sillitoe on behalf of the Wellcome Sanger Institute COVID-19 Surveillance Team |
| EPI_ISL_2924468 | Lighthouse Lab in Glasgow | Wellcome Sanger Institute for the COVID-19 Genomics UK (COG-UK) Consortium | Harper VanSteenhouse, Yumi Kasai, David Gray, Carol Clugston, Anna Dominiczak and Alex Alderton, Roberto Amato, Jeffrey Barrett, Sonia Goncalves, Ewan Harrison, David K. Jackson, Ian Johnston, Dominic Kwiatkowski, Cordelia Langford, John Sillitoe on behalf of the Wellcome Sanger Institute COVID-19 Surveillance Team |
| EPI_ISL_2924469 | Lighthouse Lab in Alderley Park | Wellcome Sanger Institute for the COVID-19 Genomics UK (COG-UK) Consortium | Jacquelyn Wynn, Mairead Hyland, The Lighthouse Lab in Alderley Park and Alex Alderton, Roberto Amato, Jeffrey Barrett, Sonia Goncalves, Ewan Harrison, David K. Jackson, Ian Johnston, Dominic Kwiatkowski, Cordelia Langford, John Sillitoe on behalf of the Wellcome Sanger Institute COVID-19 Surveillance Team |
| EPI_ISL_2924470 | Lighthouse Lab in Glasgow | Wellcome Sanger Institute for the COVID-19 Genomics UK (COG-UK) Consortium | Harper VanSteenhouse, Yumi Kasai, David Gray, Carol Clugston, Anna Dominiczak and Alex Alderton, Roberto Amato, Jeffrey Barrett, Sonia Goncalves, Ewan Harrison, David K. Jackson, Ian Johnston, Dominic Kwiatkowski, Cordelia Langford, John Sillitoe on behalf of the Wellcome Sanger Institute COVID-19 Surveillance Team |
| EPI_ISL_2924471, EPI_ISL_2924472, EPI_ISL_2924473, EPI_ISL_2924474, EPI_ISL_2924475, EPI_ISL_2924476, EPI_ISL_2924477, EPI_ISL_2924478, EPI_ISL_2924479, EPI_ISL_2924480 | Lighthouse Lab in Alderley Park | Wellcome Sanger Institute for the COVID-19 Genomics UK (COG-UK) Consortium | Jacquelyn Wynn, Mairead Hyland, The Lighthouse Lab in Alderley Park and Alex Alderton, Roberto Amato, Jeffrey Barrett, Sonia Goncalves, Ewan Harrison, David K. Jackson, Ian Johnston, Dominic Kwiatkowski, Cordelia Langford, John Sillitoe on behalf of the Wellcome Sanger Institute COVID-19 Surveillance Team |
| EPI_ISL_2924481 | Lighthouse Lab in Glasgow | Wellcome Sanger Institute for the COVID-19 Genomics UK (COG-UK) Consortium | Harper VanSteenhouse, Yumi Kasai, David Gray, Carol Clugston, Anna Dominiczak and Alex Alderton, Roberto Amato, Jeffrey Barrett, Sonia Goncalves, Ewan Harrison, David K. Jackson, Ian Johnston, Dominic Kwiatkowski, Cordelia Langford, John Sillitoe on behalf of the Wellcome Sanger Institute COVID-19 Surveillance Team |
| EPI_ISL_2924482, EPI_ISL_2924483, | Lighthouse Lab in Alderley Park | Wellcome Sanger Institute for the COVID-19 Genomics UK (COG-UK) Consortium | Jacquelyn Wynn, Mairead Hyland, The Lighthouse Lab in Alderley Park and Alex Alderton, Roberto Amato, Jeffrey Barrett, Sonia Goncalves, Ewan Harrison, David K. Jackson, Ian Johnston, Dominic Kwiatkowski, Cordelia Langford, John Sillitoe on behalf of the Wellcome Sanger Institute COVID-19 Surveillance Team |

| | | | |
|---|---|---|---|
| EPI_ISL_2924484, EPI_ISL_2924485, EPI_ISL_2924486, EPI_ISL_2924487 | | | |
| EPI_ISL_2924488 | Lighthouse Lab in Glasgow | Wellcome Sanger Institute for the COVID-19 Genomics UK (COG-UK) Consortium | Harper VanSteenhouse, Yumi Kasai, David Gray, Carol Clugston, Anna Dominiczak and Alex Alderton, Roberto Amato, Jeffrey Barrett, Sonia Goncalves, Ewan Harrison, David K. Jackson, Ian Johnston, Dominic Kwiatkowski, Cordelia Langford, John Sillitoe on behalf of the Wellcome Sanger Institute COVID-19 Surveillance Team |
| EPI_ISL_2924489, EPI_ISL_2924490, EPI_ISL_2924491 | Lighthouse Lab in Alderley Park | Wellcome Sanger Institute for the COVID-19 Genomics UK (COG-UK) Consortium | Jacquelyn Wynn, Mairead Hyland, The Lighthouse Lab in Alderley Park and Alex Alderton, Roberto Amato, Jeffrey Barrett, Sonia Goncalves, Ewan Harrison, David K. Jackson, Ian Johnston, Dominic Kwiatkowski, Cordelia Langford, John Sillitoe on behalf of the Wellcome Sanger Institute COVID-19 Surveillance Team |
| EPI_ISL_2924492 | Lighthouse Lab in Glasgow | Wellcome Sanger Institute for the COVID-19 Genomics UK (COG-UK) Consortium | Harper VanSteenhouse, Yumi Kasai, David Gray, Carol Clugston, Anna Dominiczak and Alex Alderton, Roberto Amato, Jeffrey Barrett, Sonia Goncalves, Ewan Harrison, David K. Jackson, Ian Johnston, Dominic Kwiatkowski, Cordelia Langford, John Sillitoe on behalf of the Wellcome Sanger Institute COVID-19 Surveillance Team |
| EPI_ISL_2924493 | Lighthouse Lab in Alderley Park | Wellcome Sanger Institute for the COVID-19 Genomics UK (COG-UK) Consortium | Jacquelyn Wynn, Mairead Hyland, The Lighthouse Lab in Alderley Park and Alex Alderton, Roberto Amato, Jeffrey Barrett, Sonia Goncalves, Ewan Harrison, David K. Jackson, Ian Johnston, Dominic Kwiatkowski, Cordelia Langford, John Sillitoe on behalf of the Wellcome Sanger Institute COVID-19 Surveillance Team |
| EPI_ISL_2924494, EPI_ISL_2924495 | Lighthouse Lab in Glasgow | Wellcome Sanger Institute for the COVID-19 Genomics UK (COG-UK) Consortium | Harper VanSteenhouse, Yumi Kasai, David Gray, Carol Clugston, Anna Dominiczak and Alex Alderton, Roberto Amato, Jeffrey Barrett, Sonia Goncalves, Ewan Harrison, David K. Jackson, Ian Johnston, Dominic Kwiatkowski, Cordelia Langford, John Sillitoe on behalf of the Wellcome Sanger Institute COVID-19 Surveillance Team |
| EPI_ISL_2924496, EPI_ISL_2924497, EPI_ISL_2924498, EPI_ISL_2924499, EPI_ISL_2924500, EPI_ISL_2924501, EPI_ISL_2924502, EPI_ISL_2924503, EPI_ISL_2924504, EPI_ISL_2924505, EPI_ISL_2924506, EPI_ISL_2924507, EPI_ISL_2924508, EPI_ISL_2924509, EPI_ISL_2924510, EPI_ISL_2924511, EPI_ISL_2924512, EPI_ISL_2924513, EPI_ISL_2924514 | | | |
| see above | Lighthouse Lab in Alderley Park | Wellcome Sanger Institute for the COVID-19 Genomics UK (COG-UK) Consortium | Jacquelyn Wynn, Mairead Hyland, The Lighthouse Lab in Alderley Park and Alex Alderton, Roberto Amato, Jeffrey Barrett, Sonia Goncalves, Ewan Harrison, David K. Jackson, Ian Johnston, Dominic Kwiatkowski, Cordelia Langford, John Sillitoe on behalf of the Wellcome Sanger Institute COVID-19 Surveillance Team |
| EPI_ISL_2924515 | Lighthouse Lab in Glasgow | Wellcome Sanger Institute for the COVID-19 Genomics UK (COG-UK) Consortium | Harper VanSteenhouse, Yumi Kasai, David Gray, Carol Clugston, Anna Dominiczak and Alex Alderton, Roberto Amato, Jeffrey Barrett, Sonia Goncalves, Ewan Harrison, David K. Jackson, Ian Johnston, Dominic Kwiatkowski, Cordelia Langford, John Sillitoe on behalf of the Wellcome Sanger Institute COVID-19 Surveillance Team |
| EPI_ISL_2924516, EPI_ISL_2924517, EPI_ISL_2924518, EPI_ISL_2924519, EPI_ISL_2924520 | Lighthouse Lab in Alderley Park | Wellcome Sanger Institute for the COVID-19 Genomics UK (COG-UK) Consortium | Jacquelyn Wynn, Mairead Hyland, The Lighthouse Lab in Alderley Park and Alex Alderton, Roberto Amato, Jeffrey Barrett, Sonia Goncalves, Ewan Harrison, David K. Jackson, Ian Johnston, Dominic Kwiatkowski, Cordelia Langford, John Sillitoe on behalf of the Wellcome Sanger Institute COVID-19 Surveillance Team |
| EPI_ISL_2924521 | Lighthouse Lab in Glasgow | Wellcome Sanger Institute for the COVID-19 Genomics UK (COG-UK) Consortium | Harper VanSteenhouse, Yumi Kasai, David Gray, Carol Clugston, Anna Dominiczak and Alex Alderton, Roberto Amato, Jeffrey Barrett, Sonia Goncalves, Ewan Harrison, David K. Jackson, Ian Johnston, Dominic Kwiatkowski, Cordelia Langford, John Sillitoe on behalf of the Wellcome Sanger Institute COVID-19 Surveillance Team |
| EPI_ISL_2924522, EPI_ISL_2924523 | Lighthouse Lab in Alderley Park | Wellcome Sanger Institute for the COVID-19 Genomics UK (COG-UK) Consortium | Jacquelyn Wynn, Mairead Hyland, The Lighthouse Lab in Alderley Park and Alex Alderton, Roberto Amato, Jeffrey Barrett, Sonia Goncalves, Ewan Harrison, David K. Jackson, Ian Johnston, Dominic Kwiatkowski, Cordelia Langford, John Sillitoe on behalf of the Wellcome Sanger Institute COVID-19 Surveillance Team |
| EPI_ISL_2924524 | Lighthouse Lab in Glasgow | Wellcome Sanger Institute for the COVID-19 Genomics UK (COG-UK) Consortium | Harper VanSteenhouse, Yumi Kasai, David Gray, Carol Clugston, Anna Dominiczak and Alex Alderton, Roberto Amato, Jeffrey Barrett, Sonia Goncalves, Ewan Harrison, David K. Jackson, Ian Johnston, Dominic Kwiatkowski, Cordelia Langford, John Sillitoe on behalf of the Wellcome Sanger Institute COVID-19 Surveillance Team |
| EPI_ISL_2924525, EPI_ISL_2924526, EPI_ISL_2924527, EPI_ISL_2924528, EPI_ISL_2924529, EPI_ISL_2924530, EPI_ISL_2924531 | Lighthouse Lab in Alderley Park | Wellcome Sanger Institute for the COVID-19 Genomics UK (COG-UK) Consortium | Jacquelyn Wynn, Mairead Hyland, The Lighthouse Lab in Alderley Park and Alex Alderton, Roberto Amato, Jeffrey Barrett, Sonia Goncalves, Ewan Harrison, David K. Jackson, Ian Johnston, Dominic Kwiatkowski, Cordelia Langford, John Sillitoe on behalf of the Wellcome Sanger Institute COVID-19 Surveillance Team |
| EPI_ISL_2924532 | Lighthouse Lab in Glasgow | Wellcome Sanger Institute for the COVID-19 Genomics UK (COG-UK) Consortium | Harper VanSteenhouse, Yumi Kasai, David Gray, Carol Clugston, Anna Dominiczak and Alex Alderton, Roberto Amato, Jeffrey Barrett, Sonia Goncalves, Ewan Harrison, David K. Jackson, Ian Johnston, Dominic Kwiatkowski, Cordelia Langford, John Sillitoe on behalf of the Wellcome Sanger Institute COVID-19 Surveillance Team |
| EPI_ISL_2924533, EPI_ISL_2924534, EPI_ISL_2924535, EPI_ISL_2924536, EPI_ISL_2924537 | Lighthouse Lab in Alderley Park | Wellcome Sanger Institute for the COVID-19 Genomics UK (COG-UK) Consortium | Jacquelyn Wynn, Mairead Hyland, The Lighthouse Lab in Alderley Park and Alex Alderton, Roberto Amato, Jeffrey Barrett, Sonia Goncalves, Ewan Harrison, David K. Jackson, Ian Johnston, Dominic Kwiatkowski, Cordelia Langford, John Sillitoe on behalf of the Wellcome Sanger Institute COVID-19 Surveillance Team |
| EPI_ISL_2924538 | Lighthouse Lab in Glasgow | Wellcome Sanger Institute for the COVID-19 Genomics UK (COG-UK) Consortium | Harper VanSteenhouse, Yumi Kasai, David Gray, Carol Clugston, Anna Dominiczak and Alex Alderton, Roberto Amato, Jeffrey Barrett, Sonia Goncalves, Ewan Harrison, David K. Jackson, Ian Johnston, Dominic Kwiatkowski, Cordelia Langford, John Sillitoe on behalf of the Wellcome Sanger Institute COVID-19 Surveillance Team |
| EPI_ISL_2924539, EPI_ISL_2924540, EPI_ISL_2924541 | Lighthouse Lab in Alderley Park | Wellcome Sanger Institute for the COVID-19 Genomics UK (COG-UK) Consortium | Jacquelyn Wynn, Mairead Hyland, The Lighthouse Lab in Alderley Park and Alex Alderton, Roberto Amato, Jeffrey Barrett, Sonia Goncalves, Ewan Harrison, David K. Jackson, Ian Johnston, Dominic Kwiatkowski, Cordelia Langford, John Sillitoe on behalf of the Wellcome Sanger Institute COVID-19 Surveillance Team |
| EPI_ISL_2924542, EPI_ISL_2924543 | Lighthouse Lab in Glasgow | Wellcome Sanger Institute for the COVID-19 Genomics UK (COG-UK) Consortium | Harper VanSteenhouse, Yumi Kasai, David Gray, Carol Clugston, Anna Dominiczak and Alex Alderton, Roberto Amato, Jeffrey Barrett, Sonia Goncalves, Ewan Harrison, David K. Jackson, Ian Johnston, Dominic Kwiatkowski, Cordelia Langford, John Sillitoe on behalf of the Wellcome Sanger Institute COVID-19 Surveillance Team |
| EPI_ISL_2924544, EPI_ISL_2924545 | Lighthouse Lab in Alderley Park | Wellcome Sanger Institute for the COVID-19 Genomics UK (COG-UK) Consortium | Jacquelyn Wynn, Mairead Hyland, The Lighthouse Lab in Alderley Park and Alex Alderton, Roberto Amato, Jeffrey Barrett, Sonia Goncalves, Ewan Harrison, David K. Jackson, Ian Johnston, Dominic Kwiatkowski, Cordelia Langford, John Sillitoe on behalf of the Wellcome Sanger Institute COVID-19 Surveillance Team |
| EPI_ISL_2924546 | Lighthouse Lab in Glasgow | Wellcome Sanger Institute for the COVID-19 Genomics UK (COG-UK) Consortium | Harper VanSteenhouse, Yumi Kasai, David Gray, Carol Clugston, Anna Dominiczak and Alex Alderton, Roberto Amato, Jeffrey Barrett, Sonia Goncalves, Ewan Harrison, David K. Jackson, Ian Johnston, Dominic Kwiatkowski, Cordelia Langford, John Sillitoe on behalf of the Wellcome Sanger Institute COVID-19 Surveillance Team |
| EPI_ISL_2924548, EPI_ISL_2924549, EPI_ISL_2924550, EPI_ISL_2924551, EPI_ISL_2924552, EPI_ISL_2924553, EPI_ISL_2924554 | Lighthouse Lab in Alderley Park | Wellcome Sanger Institute for the COVID-19 Genomics UK (COG-UK) Consortium | Jacquelyn Wynn, Mairead Hyland, The Lighthouse Lab in Alderley Park and Alex Alderton, Roberto Amato, Jeffrey Barrett, Sonia Goncalves, Ewan Harrison, David K. Jackson, Ian Johnston, Dominic Kwiatkowski, Cordelia Langford, John Sillitoe on behalf of the Wellcome Sanger Institute COVID-19 Surveillance Team |
| EPI_ISL_2924555 | Lighthouse Lab in Glasgow | Wellcome Sanger Institute for the COVID-19 Genomics UK (COG-UK) Consortium | Harper VanSteenhouse, Yumi Kasai, David Gray, Carol Clugston, Anna Dominiczak and Alex Alderton, Roberto Amato, Jeffrey Barrett, Sonia Goncalves, Ewan Harrison, David K. Jackson, Ian Johnston, Dominic Kwiatkowski, Cordelia Langford, John Sillitoe on behalf of the Wellcome Sanger Institute COVID-19 Surveillance Team |
| EPI_ISL_2924556, EPI_ISL_2924557, EPI_ISL_2924558, EPI_ISL_2924559, EPI_ISL_2924560 | Lighthouse Lab in Alderley Park | Wellcome Sanger Institute for the COVID-19 Genomics UK (COG-UK) Consortium | Jacquelyn Wynn, Mairead Hyland, The Lighthouse Lab in Alderley Park and Alex Alderton, Roberto Amato, Jeffrey Barrett, Sonia Goncalves, Ewan Harrison, David K. Jackson, Ian Johnston, Dominic Kwiatkowski, Cordelia Langford, John Sillitoe on behalf of the Wellcome Sanger Institute COVID-19 Surveillance Team |
| EPI_ISL_2924561 | Lighthouse Lab in Glasgow | Wellcome Sanger Institute for the COVID-19 Genomics UK (COG-UK) Consortium | Harper VanSteenhouse, Yumi Kasai, David Gray, Carol Clugston, Anna Dominiczak and Alex Alderton, Roberto Amato, Jeffrey Barrett, Sonia Goncalves, Ewan Harrison, David K. Jackson, Ian Johnston, Dominic Kwiatkowski, Cordelia Langford, John Sillitoe on behalf of the Wellcome Sanger Institute COVID-19 Surveillance Team |
| EPI_ISL_2924562, EPI_ISL_2924563, EPI_ISL_2924564, EPI_ISL_2924565, EPI_ISL_2924566, EPI_ISL_2924567, EPI_ISL_2924568 | Lighthouse Lab in Alderley Park | Wellcome Sanger Institute for the COVID-19 Genomics UK (COG-UK) Consortium | Jacquelyn Wynn, Mairead Hyland, The Lighthouse Lab in Alderley Park and Alex Alderton, Roberto Amato, Jeffrey Barrett, Sonia Goncalves, Ewan Harrison, David K. Jackson, Ian Johnston, Dominic Kwiatkowski, Cordelia Langford, John Sillitoe on behalf of the Wellcome Sanger Institute COVID-19 Surveillance Team |
| EPI_ISL_2924569 | Lighthouse Lab in Glasgow | Wellcome Sanger Institute for the COVID-19 Genomics UK (COG-UK) Consortium | Harper VanSteenhouse, Yumi Kasai, David Gray, Carol Clugston, Anna Dominiczak and Alex Alderton, Roberto Amato, Jeffrey Barrett, Sonia Goncalves, Ewan Harrison, David K. Jackson, Ian Johnston, Dominic Kwiatkowski, Cordelia Langford, John Sillitoe on behalf of the Wellcome Sanger Institute COVID-19 Surveillance Team |
| EPI_ISL_2924570, EPI_ISL_2924571, EPI_ISL_2924572, EPI_ISL_2924573, EPI_ISL_2924574, EPI_ISL_2924575, EPI_ISL_2924576, EPI_ISL_2924577, EPI_ISL_2924578, EPI_ISL_2924579 | Lighthouse Lab in Alderley Park | Wellcome Sanger Institute for the COVID-19 Genomics UK (COG-UK) Consortium | Jacquelyn Wynn, Mairead Hyland, The Lighthouse Lab in Alderley Park and Alex Alderton, Roberto Amato, Jeffrey Barrett, Sonia Goncalves, Ewan Harrison, David K. Jackson, Ian Johnston, Dominic Kwiatkowski, Cordelia Langford, John Sillitoe on behalf of the Wellcome Sanger Institute COVID-19 Surveillance Team |
| EPI_ISL_2924580, EPI_ISL_2924581 | Lighthouse Lab in Glasgow | Wellcome Sanger Institute for the COVID-19 Genomics UK (COG-UK) Consortium | Harper VanSteenhouse, Yumi Kasai, David Gray, Carol Clugston, Anna Dominiczak and Alex Alderton, Roberto Amato, Jeffrey Barrett, Sonia Goncalves, Ewan Harrison, David K. Jackson, Ian Johnston, Dominic Kwiatkowski, Cordelia Langford, John Sillitoe on behalf of the Wellcome Sanger Institute COVID-19 Surveillance Team |
| EPI_ISL_2924582 | Lighthouse Lab in Alderley Park | Wellcome Sanger Institute for the COVID-19 Genomics UK (COG-UK) Consortium | Jacquelyn Wynn, Mairead Hyland, The Lighthouse Lab in Alderley Park and Alex Alderton, Roberto Amato, Jeffrey Barrett, Sonia Goncalves, Ewan Harrison, David K. Jackson, Ian Johnston, Dominic Kwiatkowski, Cordelia Langford, John Sillitoe on behalf of the Wellcome Sanger Institute COVID-19 Surveillance Team |
| EPI_ISL_2924583 | Lighthouse Lab in Glasgow | Wellcome Sanger Institute for the COVID-19 Genomics UK (COG-UK) Consortium | Harper VanSteenhouse, Yumi Kasai, David Gray, Carol Clugston, Anna Dominiczak and Alex Alderton, Roberto Amato, Jeffrey Barrett, Sonia Goncalves, Ewan Harrison, David K. Jackson, Ian Johnston, Dominic Kwiatkowski, Cordelia Langford, John Sillitoe on behalf of the Wellcome Sanger Institute COVID-19 Surveillance Team |

| | | | |
|---|---|---|---|
| EPI_ISL_2924584 | Lighthouse Lab in Alderley Park | Wellcome Sanger Institute for the COVID-19 Genomics UK (COG-UK) Consortium | Jacquelyn Wynn, Mairead Hyland, The Lighthouse Lab in Alderley Park and Alex Alderton, Roberto Amato, Jeffrey Barrett, Sonia Goncalves, Ewan Harrison, David K. Jackson, Ian Johnston, Dominic Kwiatkowski, Cordelia Langford, John Sillitoe on behalf of the Wellcome Sanger Institute COVID-19 Surveillance Team |
| EPI_ISL_2924585 | Lighthouse Lab in Glasgow | Wellcome Sanger Institute for the COVID-19 Genomics UK (COG-UK) Consortium | Harper VanSteenhouse, Yumi Kasai, David Gray, Carol Clugston, Anna Dominiczak and Alex Alderton, Roberto Amato, Jeffrey Barrett, Sonia Goncalves, Ewan Harrison, David K. Jackson, Ian Johnston, Dominic Kwiatkowski, Cordelia Langford, John Sillitoe on behalf of the Wellcome Sanger Institute COVID-19 Surveillance Team |
| EPI_ISL_2924586, EPI_ISL_2924587, EPI_ISL_2924588, EPI_ISL_2924589 | Lighthouse Lab in Alderley Park | Wellcome Sanger Institute for the COVID-19 Genomics UK (COG-UK) Consortium | Jacquelyn Wynn, Mairead Hyland, The Lighthouse Lab in Alderley Park and Alex Alderton, Roberto Amato, Jeffrey Barrett, Sonia Goncalves, Ewan Harrison, David K. Jackson, Ian Johnston, Dominic Kwiatkowski, Cordelia Langford, John Sillitoe on behalf of the Wellcome Sanger Institute COVID-19 Surveillance Team |
| EPI_ISL_2924590 | Lighthouse Lab in Glasgow | Wellcome Sanger Institute for the COVID-19 Genomics UK (COG-UK) Consortium | Harper VanSteenhouse, Yumi Kasai, David Gray, Carol Clugston, Anna Dominiczak and Alex Alderton, Roberto Amato, Jeffrey Barrett, Sonia Goncalves, Ewan Harrison, David K. Jackson, Ian Johnston, Dominic Kwiatkowski, Cordelia Langford, John Sillitoe on behalf of the Wellcome Sanger Institute COVID-19 Surveillance Team |
| EPI_ISL_2924591, EPI_ISL_2924592, EPI_ISL_2924593, EPI_ISL_2924594 | Lighthouse Lab in Alderley Park | Wellcome Sanger Institute for the COVID-19 Genomics UK (COG-UK) Consortium | Jacquelyn Wynn, Mairead Hyland, The Lighthouse Lab in Alderley Park and Alex Alderton, Roberto Amato, Jeffrey Barrett, Sonia Goncalves, Ewan Harrison, David K. Jackson, Ian Johnston, Dominic Kwiatkowski, Cordelia Langford, John Sillitoe on behalf of the Wellcome Sanger Institute COVID-19 Surveillance Team |
| EPI_ISL_2924595 | Lighthouse Lab in Glasgow | Wellcome Sanger Institute for the COVID-19 Genomics UK (COG-UK) Consortium | Harper VanSteenhouse, Yumi Kasai, David Gray, Carol Clugston, Anna Dominiczak and Alex Alderton, Roberto Amato, Jeffrey Barrett, Sonia Goncalves, Ewan Harrison, David K. Jackson, Ian Johnston, Dominic Kwiatkowski, Cordelia Langford, John Sillitoe on behalf of the Wellcome Sanger Institute COVID-19 Surveillance Team |
| EPI_ISL_2924596, EPI_ISL_2924597, EPI_ISL_2924598, EPI_ISL_2924600 | Lighthouse Lab in Alderley Park | Wellcome Sanger Institute for the COVID-19 Genomics UK (COG-UK) Consortium | Jacquelyn Wynn, Mairead Hyland, The Lighthouse Lab in Alderley Park and Alex Alderton, Roberto Amato, Jeffrey Barrett, Sonia Goncalves, Ewan Harrison, David K. Jackson, Ian Johnston, Dominic Kwiatkowski, Cordelia Langford, John Sillitoe on behalf of the Wellcome Sanger Institute COVID-19 Surveillance Team |
| EPI_ISL_2924601, EPI_ISL_2924602 | Lighthouse Lab in Glasgow | Wellcome Sanger Institute for the COVID-19 Genomics UK (COG-UK) Consortium | Harper VanSteenhouse, Yumi Kasai, David Gray, Carol Clugston, Anna Dominiczak and Alex Alderton, Roberto Amato, Jeffrey Barrett, Sonia Goncalves, Ewan Harrison, David K. Jackson, Ian Johnston, Dominic Kwiatkowski, Cordelia Langford, John Sillitoe on behalf of the Wellcome Sanger Institute COVID-19 Surveillance Team |
| EPI_ISL_2924603 | Lighthouse Lab in Alderley Park | Wellcome Sanger Institute for the COVID-19 Genomics UK (COG-UK) Consortium | Jacquelyn Wynn, Mairead Hyland, The Lighthouse Lab in Alderley Park and Alex Alderton, Roberto Amato, Jeffrey Barrett, Sonia Goncalves, Ewan Harrison, David K. Jackson, Ian Johnston, Dominic Kwiatkowski, Cordelia Langford, John Sillitoe on behalf of the Wellcome Sanger Institute COVID-19 Surveillance Team |
| EPI_ISL_2924604, EPI_ISL_2924605, EPI_ISL_2924606 | Lighthouse Lab in Glasgow | Wellcome Sanger Institute for the COVID-19 Genomics UK (COG-UK) Consortium | Harper VanSteenhouse, Yumi Kasai, David Gray, Carol Clugston, Anna Dominiczak and Alex Alderton, Roberto Amato, Jeffrey Barrett, Sonia Goncalves, Ewan Harrison, David K. Jackson, Ian Johnston, Dominic Kwiatkowski, Cordelia Langford, John Sillitoe on behalf of the Wellcome Sanger Institute COVID-19 Surveillance Team |
| EPI_ISL_2924607 | Lighthouse Lab in Alderley Park | Wellcome Sanger Institute for the COVID-19 Genomics UK (COG-UK) Consortium | Jacquelyn Wynn, Mairead Hyland, The Lighthouse Lab in Alderley Park and Alex Alderton, Roberto Amato, Jeffrey Barrett, Sonia Goncalves, Ewan Harrison, David K. Jackson, Ian Johnston, Dominic Kwiatkowski, Cordelia Langford, John Sillitoe on behalf of the Wellcome Sanger Institute COVID-19 Surveillance Team |
| EPI_ISL_2924608 | Lighthouse Lab in Glasgow | Wellcome Sanger Institute for the COVID-19 Genomics UK (COG-UK) Consortium | Harper VanSteenhouse, Yumi Kasai, David Gray, Carol Clugston, Anna Dominiczak and Alex Alderton, Roberto Amato, Jeffrey Barrett, Sonia Goncalves, Ewan Harrison, David K. Jackson, Ian Johnston, Dominic Kwiatkowski, Cordelia Langford, John Sillitoe on behalf of the Wellcome Sanger Institute COVID-19 Surveillance Team |
| EPI_ISL_2924609, EPI_ISL_2924610, EPI_ISL_2924611 | Lighthouse Lab in Alderley Park | Wellcome Sanger Institute for the COVID-19 Genomics UK (COG-UK) Consortium | Jacquelyn Wynn, Mairead Hyland, The Lighthouse Lab in Alderley Park and Alex Alderton, Roberto Amato, Jeffrey Barrett, Sonia Goncalves, Ewan Harrison, David K. Jackson, Ian Johnston, Dominic Kwiatkowski, Cordelia Langford, John Sillitoe on behalf of the Wellcome Sanger Institute COVID-19 Surveillance Team |
| EPI_ISL_2924612 | Lighthouse Lab in Glasgow | Wellcome Sanger Institute for the COVID-19 Genomics UK (COG-UK) Consortium | Harper VanSteenhouse, Yumi Kasai, David Gray, Carol Clugston, Anna Dominiczak and Alex Alderton, Roberto Amato, Jeffrey Barrett, Sonia Goncalves, Ewan Harrison, David K. Jackson, Ian Johnston, Dominic Kwiatkowski, Cordelia Langford, John Sillitoe on behalf of the Wellcome Sanger Institute COVID-19 Surveillance Team |
| EPI_ISL_2924613, EPI_ISL_2924614, EPI_ISL_2924615 | Lighthouse Lab in Alderley Park | Wellcome Sanger Institute for the COVID-19 Genomics UK (COG-UK) Consortium | Jacquelyn Wynn, Mairead Hyland, The Lighthouse Lab in Alderley Park and Alex Alderton, Roberto Amato, Jeffrey Barrett, Sonia Goncalves, Ewan Harrison, David K. Jackson, Ian Johnston, Dominic Kwiatkowski, Cordelia Langford, John Sillitoe on behalf of the Wellcome Sanger Institute COVID-19 Surveillance Team |
| EPI_ISL_2924616, EPI_ISL_2924617, EPI_ISL_2924618 | Lighthouse Lab in Glasgow | Wellcome Sanger Institute for the COVID-19 Genomics UK (COG-UK) Consortium | Harper VanSteenhouse, Yumi Kasai, David Gray, Carol Clugston, Anna Dominiczak and Alex Alderton, Roberto Amato, Jeffrey Barrett, Sonia Goncalves, Ewan Harrison, David K. Jackson, Ian Johnston, Dominic Kwiatkowski, Cordelia Langford, John Sillitoe on behalf of the Wellcome Sanger Institute COVID-19 Surveillance Team |
| EPI_ISL_2924619 | Lighthouse Lab in Alderley Park | Wellcome Sanger Institute for the COVID-19 Genomics UK (COG-UK) Consortium | Jacquelyn Wynn, Mairead Hyland, The Lighthouse Lab in Alderley Park and Alex Alderton, Roberto Amato, Jeffrey Barrett, Sonia Goncalves, Ewan Harrison, David K. Jackson, Ian Johnston, Dominic Kwiatkowski, Cordelia Langford, John Sillitoe on behalf of the Wellcome Sanger Institute COVID-19 Surveillance Team |
| EPI_ISL_2924620 | Lighthouse Lab in Glasgow | Wellcome Sanger Institute for the COVID-19 Genomics UK (COG-UK) Consortium | Harper VanSteenhouse, Yumi Kasai, David Gray, Carol Clugston, Anna Dominiczak and Alex Alderton, Roberto Amato, Jeffrey Barrett, Sonia Goncalves, Ewan Harrison, David K. Jackson, Ian Johnston, Dominic Kwiatkowski, Cordelia Langford, John Sillitoe on behalf of the Wellcome Sanger Institute COVID-19 Surveillance Team |
| EPI_ISL_2924621, EPI_ISL_2924622, EPI_ISL_2924623, EPI_ISL_2924624, EPI_ISL_2924625, EPI_ISL_2924626, EPI_ISL_2924627, EPI_ISL_2924628, EPI_ISL_2924629, EPI_ISL_2924630, EPI_ISL_2924631, EPI_ISL_2924632, EPI_ISL_2924633, EPI_ISL_2924634, EPI_ISL_2924635, EPI_ISL_2924636 | | | |
| see above | Lighthouse Lab in Alderley Park | Wellcome Sanger Institute for the COVID-19 Genomics UK (COG-UK) Consortium | Jacquelyn Wynn, Mairead Hyland, The Lighthouse Lab in Alderley Park and Alex Alderton, Roberto Amato, Jeffrey Barrett, Sonia Goncalves, Ewan Harrison, David K. Jackson, Ian Johnston, Dominic Kwiatkowski, Cordelia Langford, John Sillitoe on behalf of the Wellcome Sanger Institute COVID-19 Surveillance Team |
| EPI_ISL_2924637 | Lighthouse Lab in Glasgow | Wellcome Sanger Institute for the COVID-19 Genomics UK (COG-UK) Consortium | Harper VanSteenhouse, Yumi Kasai, David Gray, Carol Clugston, Anna Dominiczak and Alex Alderton, Roberto Amato, Jeffrey Barrett, Sonia Goncalves, Ewan Harrison, David K. Jackson, Ian Johnston, Dominic Kwiatkowski, Cordelia Langford, John Sillitoe on behalf of the Wellcome Sanger Institute COVID-19 Surveillance Team |
| EPI_ISL_2924638 | Lighthouse Lab in Alderley Park | Wellcome Sanger Institute for the COVID-19 Genomics UK (COG-UK) Consortium | Jacquelyn Wynn, Mairead Hyland, The Lighthouse Lab in Alderley Park and Alex Alderton, Roberto Amato, Jeffrey Barrett, Sonia Goncalves, Ewan Harrison, David K. Jackson, Ian Johnston, Dominic Kwiatkowski, Cordelia Langford, John Sillitoe on behalf of the Wellcome Sanger Institute COVID-19 Surveillance Team |
| EPI_ISL_2924639, EPI_ISL_2924640, EPI_ISL_2924641 | Lighthouse Lab in Glasgow | Wellcome Sanger Institute for the COVID-19 Genomics UK (COG-UK) Consortium | Harper VanSteenhouse, Yumi Kasai, David Gray, Carol Clugston, Anna Dominiczak and Alex Alderton, Roberto Amato, Jeffrey Barrett, Sonia Goncalves, Ewan Harrison, David K. Jackson, Ian Johnston, Dominic Kwiatkowski, Cordelia Langford, John Sillitoe on behalf of the Wellcome Sanger Institute COVID-19 Surveillance Team |
| EPI_ISL_2924642, EPI_ISL_2924643, EPI_ISL_2924644, EPI_ISL_2924645, EPI_ISL_2924646, EPI_ISL_2924647, EPI_ISL_2924648, EPI_ISL_2924649, EPI_ISL_2924650, EPI_ISL_2924651, EPI_ISL_2924652, EPI_ISL_2924653 | | | |
| see above | Lighthouse Lab in Alderley Park | Wellcome Sanger Institute for the COVID-19 Genomics UK (COG-UK) Consortium | Jacquelyn Wynn, Mairead Hyland, The Lighthouse Lab in Alderley Park and Alex Alderton, Roberto Amato, Jeffrey Barrett, Sonia Goncalves, Ewan Harrison, David K. Jackson, Ian Johnston, Dominic Kwiatkowski, Cordelia Langford, John Sillitoe on behalf of the Wellcome Sanger Institute COVID-19 Surveillance Team |
| EPI_ISL_2924654 | Lighthouse Lab in Glasgow | Wellcome Sanger Institute for the COVID-19 Genomics UK (COG-UK) Consortium | Harper VanSteenhouse, Yumi Kasai, David Gray, Carol Clugston, Anna Dominiczak and Alex Alderton, Roberto Amato, Jeffrey Barrett, Sonia Goncalves, Ewan Harrison, David K. Jackson, Ian Johnston, Dominic Kwiatkowski, Cordelia Langford, John Sillitoe on behalf of the Wellcome Sanger Institute COVID-19 Surveillance Team |
| EPI_ISL_2924655, EPI_ISL_2924656, EPI_ISL_2924657, EPI_ISL_2924658, EPI_ISL_2924659, EPI_ISL_2924660 | Lighthouse Lab in Alderley Park | Wellcome Sanger Institute for the COVID-19 Genomics UK (COG-UK) Consortium | Jacquelyn Wynn, Mairead Hyland, The Lighthouse Lab in Alderley Park and Alex Alderton, Roberto Amato, Jeffrey Barrett, Sonia Goncalves, Ewan Harrison, David K. Jackson, Ian Johnston, Dominic Kwiatkowski, Cordelia Langford, John Sillitoe on behalf of the Wellcome Sanger Institute COVID-19 Surveillance Team |
| EPI_ISL_2924661 | Lighthouse Lab in Glasgow | Wellcome Sanger Institute for the COVID-19 Genomics UK (COG-UK) Consortium | Harper VanSteenhouse, Yumi Kasai, David Gray, Carol Clugston, Anna Dominiczak and Alex Alderton, Roberto Amato, Jeffrey Barrett, Sonia Goncalves, Ewan Harrison, David K. Jackson, Ian Johnston, Dominic Kwiatkowski, Cordelia Langford, John Sillitoe on behalf of the Wellcome Sanger Institute COVID-19 Surveillance Team |
| EPI_ISL_2924662, EPI_ISL_2924663, EPI_ISL_2924664, EPI_ISL_2924665, EPI_ISL_2924666, EPI_ISL_2924667, EPI_ISL_2924668, EPI_ISL_2924669, EPI_ISL_2924670 | Lighthouse Lab in Alderley Park | Wellcome Sanger Institute for the COVID-19 Genomics UK (COG-UK) Consortium | Jacquelyn Wynn, Mairead Hyland, The Lighthouse Lab in Alderley Park and Alex Alderton, Roberto Amato, Jeffrey Barrett, Sonia Goncalves, Ewan Harrison, David K. Jackson, Ian Johnston, Dominic Kwiatkowski, Cordelia Langford, John Sillitoe on behalf of the Wellcome Sanger Institute COVID-19 Surveillance Team |
| EPI_ISL_2924671 | Lighthouse Lab in Glasgow | Wellcome Sanger Institute for the COVID-19 Genomics UK (COG-UK) Consortium | Harper VanSteenhouse, Yumi Kasai, David Gray, Carol Clugston, Anna Dominiczak and Alex Alderton, Roberto Amato, Jeffrey Barrett, Sonia Goncalves, Ewan Harrison, David K. Jackson, Ian Johnston, Dominic Kwiatkowski, Cordelia Langford, John Sillitoe on behalf of the Wellcome Sanger Institute COVID-19 Surveillance Team |
| EPI_ISL_2924672, EPI_ISL_2924673, EPI_ISL_2924674, EPI_ISL_2924675, EPI_ISL_2924676, EPI_ISL_2924677 | Lighthouse Lab in Alderley Park | Wellcome Sanger Institute for the COVID-19 Genomics UK (COG-UK) Consortium | Jacquelyn Wynn, Mairead Hyland, The Lighthouse Lab in Alderley Park and Alex Alderton, Roberto Amato, Jeffrey Barrett, Sonia Goncalves, Ewan Harrison, David K. Jackson, Ian Johnston, Dominic Kwiatkowski, Cordelia Langford, John Sillitoe on behalf of the Wellcome Sanger Institute COVID-19 Surveillance Team |
| EPI_ISL_2924678 | Lighthouse Lab in Glasgow | Wellcome Sanger Institute for the COVID-19 Genomics UK (COG-UK) Consortium | Harper VanSteenhouse, Yumi Kasai, David Gray, Carol Clugston, Anna Dominiczak and Alex Alderton, Roberto Amato, Jeffrey Barrett, Sonia Goncalves, Ewan Harrison, David K. Jackson, Ian Johnston, Dominic Kwiatkowski, Cordelia Langford, John Sillitoe on behalf of the Wellcome Sanger Institute COVID-19 Surveillance Team |
| EPI_ISL_2924679, EPI_ISL_2924680, EPI_ISL_2924681, EPI_ISL_2924682 | Lighthouse Lab in Alderley Park | Wellcome Sanger Institute for the COVID-19 Genomics UK (COG-UK) Consortium | Jacquelyn Wynn, Mairead Hyland, The Lighthouse Lab in Alderley Park and Alex Alderton, Roberto Amato, Jeffrey Barrett, Sonia Goncalves, Ewan Harrison, David K. Jackson, Ian Johnston, Dominic Kwiatkowski, Cordelia Langford, John Sillitoe on behalf of the Wellcome Sanger Institute COVID-19 Surveillance Team |
| EPI_ISL_2924683 | Lighthouse Lab in Glasgow | Wellcome Sanger Institute for the COVID-19 Genomics UK (COG-UK) Consortium | Harper VanSteenhouse, Yumi Kasai, David Gray, Carol Clugston, Anna Dominiczak and Alex Alderton, Roberto Amato, Jeffrey Barrett, Sonia Goncalves, Ewan Harrison, David K. Jackson, Ian Johnston, Dominic Kwiatkowski, Cordelia Langford, John Sillitoe on behalf of the Wellcome Sanger Institute COVID-19 Surveillance Team |

| | | | |
|---|---|---|---|
| EPI_ISL_2924684, EPI_ISL_2924685 | Lighthouse Lab in Alderley Park | Wellcome Sanger Institute for the COVID-19 Genomics UK (COG-UK) Consortium | Jacquelyn Wynn, Mairead Hyland, The Lighthouse Lab in Alderley Park and Alex Alderton, Roberto Amato, Jeffrey Barrett, Sonia Goncalves, Ewan Harrison, David K. Jackson, Ian Johnston, Dominic Kwiatkowski, Cordelia Langford, John Sillitoe on behalf of the Wellcome Sanger Institute COVID-19 Surveillance Team |
| EPI_ISL_2924686 | Lighthouse Lab in Glasgow | Wellcome Sanger Institute for the COVID-19 Genomics UK (COG-UK) Consortium | Harper VanSteenhouse, Yumi Kasai, David Gray, Carol Clugston, Anna Dominiczak and Alex Alderton, Roberto Amato, Jeffrey Barrett, Sonia Goncalves, Ewan Harrison, David K. Jackson, Ian Johnston, Dominic Kwiatkowski, Cordelia Langford, John Sillitoe on behalf of the Wellcome Sanger Institute COVID-19 Surveillance Team |
| EPI_ISL_2924687 | Lighthouse Lab in Alderley Park | Wellcome Sanger Institute for the COVID-19 Genomics UK (COG-UK) Consortium | Jacquelyn Wynn, Mairead Hyland, The Lighthouse Lab in Alderley Park and Alex Alderton, Roberto Amato, Jeffrey Barrett, Sonia Goncalves, Ewan Harrison, David K. Jackson, Ian Johnston, Dominic Kwiatkowski, Cordelia Langford, John Sillitoe on behalf of the Wellcome Sanger Institute COVID-19 Surveillance Team |
| EPI_ISL_2924688 | Lighthouse Lab in Glasgow | Wellcome Sanger Institute for the COVID-19 Genomics UK (COG-UK) Consortium | Harper VanSteenhouse, Yumi Kasai, David Gray, Carol Clugston, Anna Dominiczak and Alex Alderton, Roberto Amato, Jeffrey Barrett, Sonia Goncalves, Ewan Harrison, David K. Jackson, Ian Johnston, Dominic Kwiatkowski, Cordelia Langford, John Sillitoe on behalf of the Wellcome Sanger Institute COVID-19 Surveillance Team |
| EPI_ISL_2924689 | Lighthouse Lab in Alderley Park | Wellcome Sanger Institute for the COVID-19 Genomics UK (COG-UK) Consortium | Jacquelyn Wynn, Mairead Hyland, The Lighthouse Lab in Alderley Park and Alex Alderton, Roberto Amato, Jeffrey Barrett, Sonia Goncalves, Ewan Harrison, David K. Jackson, Ian Johnston, Dominic Kwiatkowski, Cordelia Langford, John Sillitoe on behalf of the Wellcome Sanger Institute COVID-19 Surveillance Team |
| EPI_ISL_2924690 | Lighthouse Lab in Glasgow | Wellcome Sanger Institute for the COVID-19 Genomics UK (COG-UK) Consortium | Harper VanSteenhouse, Yumi Kasai, David Gray, Carol Clugston, Anna Dominiczak and Alex Alderton, Roberto Amato, Jeffrey Barrett, Sonia Goncalves, Ewan Harrison, David K. Jackson, Ian Johnston, Dominic Kwiatkowski, Cordelia Langford, John Sillitoe on behalf of the Wellcome Sanger Institute COVID-19 Surveillance Team |
| EPI_ISL_2924691, EPI_ISL_2924692, EPI_ISL_2924693, EPI_ISL_2924694, EPI_ISL_2924695, EPI_ISL_2924696, EPI_ISL_2924697, EPI_ISL_2924698, EPI_ISL_2924699, EPI_ISL_2924700, EPI_ISL_2924701 | | | |
| see above | Lighthouse Lab in Alderley Park | Wellcome Sanger Institute for the COVID-19 Genomics UK (COG-UK) Consortium | Jacquelyn Wynn, Mairead Hyland, The Lighthouse Lab in Alderley Park and Alex Alderton, Roberto Amato, Jeffrey Barrett, Sonia Goncalves, Ewan Harrison, David K. Jackson, Ian Johnston, Dominic Kwiatkowski, Cordelia Langford, John Sillitoe on behalf of the Wellcome Sanger Institute COVID-19 Surveillance Team |
| EPI_ISL_2924702 | Lighthouse Lab in Glasgow | Wellcome Sanger Institute for the COVID-19 Genomics UK (COG-UK) Consortium | Harper VanSteenhouse, Yumi Kasai, David Gray, Carol Clugston, Anna Dominiczak and Alex Alderton, Roberto Amato, Jeffrey Barrett, Sonia Goncalves, Ewan Harrison, David K. Jackson, Ian Johnston, Dominic Kwiatkowski, Cordelia Langford, John Sillitoe on behalf of the Wellcome Sanger Institute COVID-19 Surveillance Team |
| EPI_ISL_2924703, EPI_ISL_2924704, EPI_ISL_2924705, EPI_ISL_2924706, EPI_ISL_2924707, EPI_ISL_2924708, EPI_ISL_2924709 | Lighthouse Lab in Alderley Park | Wellcome Sanger Institute for the COVID-19 Genomics UK (COG-UK) Consortium | Jacquelyn Wynn, Mairead Hyland, The Lighthouse Lab in Alderley Park and Alex Alderton, Roberto Amato, Jeffrey Barrett, Sonia Goncalves, Ewan Harrison, David K. Jackson, Ian Johnston, Dominic Kwiatkowski, Cordelia Langford, John Sillitoe on behalf of the Wellcome Sanger Institute COVID-19 Surveillance Team |
| EPI_ISL_2924710 | Lighthouse Lab in Glasgow | Wellcome Sanger Institute for the COVID-19 Genomics UK (COG-UK) Consortium | Harper VanSteenhouse, Yumi Kasai, David Gray, Carol Clugston, Anna Dominiczak and Alex Alderton, Roberto Amato, Jeffrey Barrett, Sonia Goncalves, Ewan Harrison, David K. Jackson, Ian Johnston, Dominic Kwiatkowski, Cordelia Langford, John Sillitoe on behalf of the Wellcome Sanger Institute COVID-19 Surveillance Team |
| EPI_ISL_2924712, EPI_ISL_2924713 | Lighthouse Lab in Alderley Park | Wellcome Sanger Institute for the COVID-19 Genomics UK (COG-UK) Consortium | Jacquelyn Wynn, Mairead Hyland, The Lighthouse Lab in Alderley Park and Alex Alderton, Roberto Amato, Jeffrey Barrett, Sonia Goncalves, Ewan Harrison, David K. Jackson, Ian Johnston, Dominic Kwiatkowski, Cordelia Langford, John Sillitoe on behalf of the Wellcome Sanger Institute COVID-19 Surveillance Team |
| EPI_ISL_2924714 | Lighthouse Lab in Glasgow | Wellcome Sanger Institute for the COVID-19 Genomics UK (COG-UK) Consortium | Harper VanSteenhouse, Yumi Kasai, David Gray, Carol Clugston, Anna Dominiczak and Alex Alderton, Roberto Amato, Jeffrey Barrett, Sonia Goncalves, Ewan Harrison, David K. Jackson, Ian Johnston, Dominic Kwiatkowski, Cordelia Langford, John Sillitoe on behalf of the Wellcome Sanger Institute COVID-19 Surveillance Team |
| EPI_ISL_2924715, EPI_ISL_2924716, EPI_ISL_2924717, EPI_ISL_2924718, EPI_ISL_2924719, EPI_ISL_2924720, EPI_ISL_2924722, EPI_ISL_2924723 | Lighthouse Lab in Alderley Park | Wellcome Sanger Institute for the COVID-19 Genomics UK (COG-UK) Consortium | Jacquelyn Wynn, Mairead Hyland, The Lighthouse Lab in Alderley Park and Alex Alderton, Roberto Amato, Jeffrey Barrett, Sonia Goncalves, Ewan Harrison, David K. Jackson, Ian Johnston, Dominic Kwiatkowski, Cordelia Langford, John Sillitoe on behalf of the Wellcome Sanger Institute COVID-19 Surveillance Team |
| EPI_ISL_2924724 | Lighthouse Lab in Glasgow | Wellcome Sanger Institute for the COVID-19 Genomics UK (COG-UK) Consortium | Harper VanSteenhouse, Yumi Kasai, David Gray, Carol Clugston, Anna Dominiczak and Alex Alderton, Roberto Amato, Jeffrey Barrett, Sonia Goncalves, Ewan Harrison, David K. Jackson, Ian Johnston, Dominic Kwiatkowski, Cordelia Langford, John Sillitoe on behalf of the Wellcome Sanger Institute COVID-19 Surveillance Team |
| EPI_ISL_2924725, EPI_ISL_2924726, EPI_ISL_2924727, EPI_ISL_2924728, EPI_ISL_2924729, EPI_ISL_2924730, EPI_ISL_2924731, EPI_ISL_2924732 | Lighthouse Lab in Alderley Park | Wellcome Sanger Institute for the COVID-19 Genomics UK (COG-UK) Consortium | Jacquelyn Wynn, Mairead Hyland, The Lighthouse Lab in Alderley Park and Alex Alderton, Roberto Amato, Jeffrey Barrett, Sonia Goncalves, Ewan Harrison, David K. Jackson, Ian Johnston, Dominic Kwiatkowski, Cordelia Langford, John Sillitoe on behalf of the Wellcome Sanger Institute COVID-19 Surveillance Team |
| EPI_ISL_2924733 | Lighthouse Lab in Glasgow | Wellcome Sanger Institute for the COVID-19 Genomics UK (COG-UK) Consortium | Harper VanSteenhouse, Yumi Kasai, David Gray, Carol Clugston, Anna Dominiczak and Alex Alderton, Roberto Amato, Jeffrey Barrett, Sonia Goncalves, Ewan Harrison, David K. Jackson, Ian Johnston, Dominic Kwiatkowski, Cordelia Langford, John Sillitoe on behalf of the Wellcome Sanger Institute COVID-19 Surveillance Team |
| EPI_ISL_2924734 | Lighthouse Lab in Alderley Park | Wellcome Sanger Institute for the COVID-19 Genomics UK (COG-UK) Consortium | Jacquelyn Wynn, Mairead Hyland, The Lighthouse Lab in Alderley Park and Alex Alderton, Roberto Amato, Jeffrey Barrett, Sonia Goncalves, Ewan Harrison, David K. Jackson, Ian Johnston, Dominic Kwiatkowski, Cordelia Langford, John Sillitoe on behalf of the Wellcome Sanger Institute COVID-19 Surveillance Team |
| EPI_ISL_2924735 | Lighthouse Lab in Glasgow | Wellcome Sanger Institute for the COVID-19 Genomics UK (COG-UK) Consortium | Harper VanSteenhouse, Yumi Kasai, David Gray, Carol Clugston, Anna Dominiczak and Alex Alderton, Roberto Amato, Jeffrey Barrett, Sonia Goncalves, Ewan Harrison, David K. Jackson, Ian Johnston, Dominic Kwiatkowski, Cordelia Langford, John Sillitoe on behalf of the Wellcome Sanger Institute COVID-19 Surveillance Team |
| EPI_ISL_2924736, EPI_ISL_2924737 | Lighthouse Lab in Alderley Park | Wellcome Sanger Institute for the COVID-19 Genomics UK (COG-UK) Consortium | Jacquelyn Wynn, Mairead Hyland, The Lighthouse Lab in Alderley Park and Alex Alderton, Roberto Amato, Jeffrey Barrett, Sonia Goncalves, Ewan Harrison, David K. Jackson, Ian Johnston, Dominic Kwiatkowski, Cordelia Langford, John Sillitoe on behalf of the Wellcome Sanger Institute COVID-19 Surveillance Team |
| EPI_ISL_2924738 | Lighthouse Lab in Glasgow | Wellcome Sanger Institute for the COVID-19 Genomics UK (COG-UK) Consortium | Harper VanSteenhouse, Yumi Kasai, David Gray, Carol Clugston, Anna Dominiczak and Alex Alderton, Roberto Amato, Jeffrey Barrett, Sonia Goncalves, Ewan Harrison, David K. Jackson, Ian Johnston, Dominic Kwiatkowski, Cordelia Langford, John Sillitoe on behalf of the Wellcome Sanger Institute COVID-19 Surveillance Team |
| EPI_ISL_2924739, EPI_ISL_2924740, EPI_ISL_2924741, EPI_ISL_2924742, EPI_ISL_2924743, EPI_ISL_2924744 | Lighthouse Lab in Alderley Park | Wellcome Sanger Institute for the COVID-19 Genomics UK (COG-UK) Consortium | Jacquelyn Wynn, Mairead Hyland, The Lighthouse Lab in Alderley Park and Alex Alderton, Roberto Amato, Jeffrey Barrett, Sonia Goncalves, Ewan Harrison, David K. Jackson, Ian Johnston, Dominic Kwiatkowski, Cordelia Langford, John Sillitoe on behalf of the Wellcome Sanger Institute COVID-19 Surveillance Team |
| EPI_ISL_2924745 | Lighthouse Lab in Glasgow | Wellcome Sanger Institute for the COVID-19 Genomics UK (COG-UK) Consortium | Harper VanSteenhouse, Yumi Kasai, David Gray, Carol Clugston, Anna Dominiczak and Alex Alderton, Roberto Amato, Jeffrey Barrett, Sonia Goncalves, Ewan Harrison, David K. Jackson, Ian Johnston, Dominic Kwiatkowski, Cordelia Langford, John Sillitoe on behalf of the Wellcome Sanger Institute COVID-19 Surveillance Team |
| EPI_ISL_2924746 | Lighthouse Lab in Alderley Park | Wellcome Sanger Institute for the COVID-19 Genomics UK (COG-UK) Consortium | Jacquelyn Wynn, Mairead Hyland, The Lighthouse Lab in Alderley Park and Alex Alderton, Roberto Amato, Jeffrey Barrett, Sonia Goncalves, Ewan Harrison, David K. Jackson, Ian Johnston, Dominic Kwiatkowski, Cordelia Langford, John Sillitoe on behalf of the Wellcome Sanger Institute COVID-19 Surveillance Team |
| EPI_ISL_2924747 | Lighthouse Lab in Glasgow | Wellcome Sanger Institute for the COVID-19 Genomics UK (COG-UK) Consortium | Harper VanSteenhouse, Yumi Kasai, David Gray, Carol Clugston, Anna Dominiczak and Alex Alderton, Roberto Amato, Jeffrey Barrett, Sonia Goncalves, Ewan Harrison, David K. Jackson, Ian Johnston, Dominic Kwiatkowski, Cordelia Langford, John Sillitoe on behalf of the Wellcome Sanger Institute COVID-19 Surveillance Team |
| EPI_ISL_2924748, EPI_ISL_2924749, EPI_ISL_2924750 | Lighthouse Lab in Alderley Park | Wellcome Sanger Institute for the COVID-19 Genomics UK (COG-UK) Consortium | Jacquelyn Wynn, Mairead Hyland, The Lighthouse Lab in Alderley Park and Alex Alderton, Roberto Amato, Jeffrey Barrett, Sonia Goncalves, Ewan Harrison, David K. Jackson, Ian Johnston, Dominic Kwiatkowski, Cordelia Langford, John Sillitoe on behalf of the Wellcome Sanger Institute COVID-19 Surveillance Team |
| EPI_ISL_2924751 | Lighthouse Lab in Glasgow | Wellcome Sanger Institute for the COVID-19 Genomics UK (COG-UK) Consortium | Harper VanSteenhouse, Yumi Kasai, David Gray, Carol Clugston, Anna Dominiczak and Alex Alderton, Roberto Amato, Jeffrey Barrett, Sonia Goncalves, Ewan Harrison, David K. Jackson, Ian Johnston, Dominic Kwiatkowski, Cordelia Langford, John Sillitoe on behalf of the Wellcome Sanger Institute COVID-19 Surveillance Team |
| EPI_ISL_2924752, EPI_ISL_2924753, EPI_ISL_2924754, EPI_ISL_2924755, EPI_ISL_2924756, EPI_ISL_2924757, EPI_ISL_2924758 | Lighthouse Lab in Alderley Park | Wellcome Sanger Institute for the COVID-19 Genomics UK (COG-UK) Consortium | Jacquelyn Wynn, Mairead Hyland, The Lighthouse Lab in Alderley Park and Alex Alderton, Roberto Amato, Jeffrey Barrett, Sonia Goncalves, Ewan Harrison, David K. Jackson, Ian Johnston, Dominic Kwiatkowski, Cordelia Langford, John Sillitoe on behalf of the Wellcome Sanger Institute COVID-19 Surveillance Team |
| EPI_ISL_2924759 | Lighthouse Lab in Glasgow | Wellcome Sanger Institute for the COVID-19 Genomics UK (COG-UK) Consortium | Harper VanSteenhouse, Yumi Kasai, David Gray, Carol Clugston, Anna Dominiczak and Alex Alderton, Roberto Amato, Jeffrey Barrett, Sonia Goncalves, Ewan Harrison, David K. Jackson, Ian Johnston, Dominic Kwiatkowski, Cordelia Langford, John Sillitoe on behalf of the Wellcome Sanger Institute COVID-19 Surveillance Team |
| EPI_ISL_2924760, EPI_ISL_2924761 | Lighthouse Lab in Alderley Park | Wellcome Sanger Institute for the COVID-19 Genomics UK (COG-UK) Consortium | Jacquelyn Wynn, Mairead Hyland, The Lighthouse Lab in Alderley Park and Alex Alderton, Roberto Amato, Jeffrey Barrett, Sonia Goncalves, Ewan Harrison, David K. Jackson, Ian Johnston, Dominic Kwiatkowski, Cordelia Langford, John Sillitoe on behalf of the Wellcome Sanger Institute COVID-19 Surveillance Team |
| EPI_ISL_2924762 | Lighthouse Lab in Glasgow | Wellcome Sanger Institute for the COVID-19 Genomics UK (COG-UK) Consortium | Harper VanSteenhouse, Yumi Kasai, David Gray, Carol Clugston, Anna Dominiczak and Alex Alderton, Roberto Amato, Jeffrey Barrett, Sonia Goncalves, Ewan Harrison, David K. Jackson, Ian Johnston, Dominic Kwiatkowski, Cordelia Langford, John Sillitoe on behalf of the Wellcome Sanger Institute COVID-19 Surveillance Team |
| EPI_ISL_2924763 | Lighthouse Lab in Alderley Park | Wellcome Sanger Institute for the COVID-19 Genomics UK (COG-UK) Consortium | Jacquelyn Wynn, Mairead Hyland, The Lighthouse Lab in Alderley Park and Alex Alderton, Roberto Amato, Jeffrey Barrett, Sonia Goncalves, Ewan Harrison, David K. Jackson, Ian Johnston, Dominic Kwiatkowski, Cordelia Langford, John Sillitoe on behalf of the Wellcome Sanger Institute COVID-19 Surveillance Team |
| EPI_ISL_2924764 | Lighthouse Lab in Glasgow | Wellcome Sanger Institute for the COVID-19 Genomics UK (COG-UK) Consortium | Harper VanSteenhouse, Yumi Kasai, David Gray, Carol Clugston, Anna Dominiczak and Alex Alderton, Roberto Amato, Jeffrey Barrett, Sonia Goncalves, Ewan Harrison, David K. Jackson, Ian Johnston, Dominic Kwiatkowski, Cordelia Langford, John Sillitoe on behalf of the Wellcome Sanger Institute COVID-19 Surveillance Team |
| EPI_ISL_2924765, EPI_ISL_2924766, | Lighthouse Lab in Alderley Park | Wellcome Sanger Institute for the COVID-19 Genomics UK (COG-UK) Consortium | Jacquelyn Wynn, Mairead Hyland, The Lighthouse Lab in Alderley Park and Alex Alderton, Roberto Amato, Jeffrey Barrett, Sonia Goncalves, Ewan Harrison, David K. Jackson, Ian Johnston, Dominic Kwiatkowski, Cordelia Langford, John Sillitoe on behalf of the Wellcome Sanger Institute COVID-19 Surveillance Team |

| | | | |
|---|---|---|---|
| EPI_ISL_2924767, EPI_ISL_2924768, EPI_ISL_2924769 | | | |
| EPI_ISL_2924770 | Lighthouse Lab in Glasgow | Wellcome Sanger Institute for the COVID-19 Genomics UK (COG-UK) Consortium | Harper VanSteenhouse, Yumi Kasai, David Gray, Carol Clugston, Anna Dominiczak and Alex Alderton, Roberto Amato, Jeffrey Barrett, Sonia Goncalves, Ewan Harrison, David K. Jackson, Ian Johnston, Dominic Kwiatkowski, Cordelia Langford, John Sillitoe on behalf of the Wellcome Sanger Institute COVID-19 Surveillance Team |
| EPI_ISL_2924771, EPI_ISL_2924772 | Lighthouse Lab in Alderley Park | Wellcome Sanger Institute for the COVID-19 Genomics UK (COG-UK) Consortium | Jacquelyn Wynn, Mairead Hyland, The Lighthouse Lab in Alderley Park and Alex Alderton, Roberto Amato, Jeffrey Barrett, Sonia Goncalves, Ewan Harrison, David K. Jackson, Ian Johnston, Dominic Kwiatkowski, Cordelia Langford, John Sillitoe on behalf of the Wellcome Sanger Institute COVID-19 Surveillance Team |
| EPI_ISL_2924773 | Lighthouse Lab in Glasgow | Wellcome Sanger Institute for the COVID-19 Genomics UK (COG-UK) Consortium | Harper VanSteenhouse, Yumi Kasai, David Gray, Carol Clugston, Anna Dominiczak and Alex Alderton, Roberto Amato, Jeffrey Barrett, Sonia Goncalves, Ewan Harrison, David K. Jackson, Ian Johnston, Dominic Kwiatkowski, Cordelia Langford, John Sillitoe on behalf of the Wellcome Sanger Institute COVID-19 Surveillance Team |
| EPI_ISL_2924774 | Lighthouse Lab in Alderley Park | Wellcome Sanger Institute for the COVID-19 Genomics UK (COG-UK) Consortium | Jacquelyn Wynn, Mairead Hyland, The Lighthouse Lab in Alderley Park and Alex Alderton, Roberto Amato, Jeffrey Barrett, Sonia Goncalves, Ewan Harrison, David K. Jackson, Ian Johnston, Dominic Kwiatkowski, Cordelia Langford, John Sillitoe on behalf of the Wellcome Sanger Institute COVID-19 Surveillance Team |
| EPI_ISL_2924775, EPI_ISL_2924776 | Lighthouse Lab in Glasgow | Wellcome Sanger Institute for the COVID-19 Genomics UK (COG-UK) Consortium | Harper VanSteenhouse, Yumi Kasai, David Gray, Carol Clugston, Anna Dominiczak and Alex Alderton, Roberto Amato, Jeffrey Barrett, Sonia Goncalves, Ewan Harrison, David K. Jackson, Ian Johnston, Dominic Kwiatkowski, Cordelia Langford, John Sillitoe on behalf of the Wellcome Sanger Institute COVID-19 Surveillance Team |
| EPI_ISL_2924777 | Lighthouse Lab in Alderley Park | Wellcome Sanger Institute for the COVID-19 Genomics UK (COG-UK) Consortium | Jacquelyn Wynn, Mairead Hyland, The Lighthouse Lab in Alderley Park and Alex Alderton, Roberto Amato, Jeffrey Barrett, Sonia Goncalves, Ewan Harrison, David K. Jackson, Ian Johnston, Dominic Kwiatkowski, Cordelia Langford, John Sillitoe on behalf of the Wellcome Sanger Institute COVID-19 Surveillance Team |
| EPI_ISL_2924778 | Lighthouse Lab in Glasgow | Wellcome Sanger Institute for the COVID-19 Genomics UK (COG-UK) Consortium | Harper VanSteenhouse, Yumi Kasai, David Gray, Carol Clugston, Anna Dominiczak and Alex Alderton, Roberto Amato, Jeffrey Barrett, Sonia Goncalves, Ewan Harrison, David K. Jackson, Ian Johnston, Dominic Kwiatkowski, Cordelia Langford, John Sillitoe on behalf of the Wellcome Sanger Institute COVID-19 Surveillance Team |
| EPI_ISL_2924779, EPI_ISL_2924780 | Lighthouse Lab in Alderley Park | Wellcome Sanger Institute for the COVID-19 Genomics UK (COG-UK) Consortium | Jacquelyn Wynn, Mairead Hyland, The Lighthouse Lab in Alderley Park and Alex Alderton, Roberto Amato, Jeffrey Barrett, Sonia Goncalves, Ewan Harrison, David K. Jackson, Ian Johnston, Dominic Kwiatkowski, Cordelia Langford, John Sillitoe on behalf of the Wellcome Sanger Institute COVID-19 Surveillance Team |
| EPI_ISL_2924781 | Lighthouse Lab in Glasgow | Wellcome Sanger Institute for the COVID-19 Genomics UK (COG-UK) Consortium | Harper VanSteenhouse, Yumi Kasai, David Gray, Carol Clugston, Anna Dominiczak and Alex Alderton, Roberto Amato, Jeffrey Barrett, Sonia Goncalves, Ewan Harrison, David K. Jackson, Ian Johnston, Dominic Kwiatkowski, Cordelia Langford, John Sillitoe on behalf of the Wellcome Sanger Institute COVID-19 Surveillance Team |
| EPI_ISL_2924782 | Lighthouse Lab in Alderley Park | Wellcome Sanger Institute for the COVID-19 Genomics UK (COG-UK) Consortium | Jacquelyn Wynn, Mairead Hyland, The Lighthouse Lab in Alderley Park and Alex Alderton, Roberto Amato, Jeffrey Barrett, Sonia Goncalves, Ewan Harrison, David K. Jackson, Ian Johnston, Dominic Kwiatkowski, Cordelia Langford, John Sillitoe on behalf of the Wellcome Sanger Institute COVID-19 Surveillance Team |
| EPI_ISL_2924783 | Lighthouse Lab in Glasgow | Wellcome Sanger Institute for the COVID-19 Genomics UK (COG-UK) Consortium | Harper VanSteenhouse, Yumi Kasai, David Gray, Carol Clugston, Anna Dominiczak and Alex Alderton, Roberto Amato, Jeffrey Barrett, Sonia Goncalves, Ewan Harrison, David K. Jackson, Ian Johnston, Dominic Kwiatkowski, Cordelia Langford, John Sillitoe on behalf of the Wellcome Sanger Institute COVID-19 Surveillance Team |
| EPI_ISL_2924784, EPI_ISL_2924785, EPI_ISL_2924786, EPI_ISL_2924787, EPI_ISL_2924788, EPI_ISL_2924789 | Lighthouse Lab in Alderley Park | Wellcome Sanger Institute for the COVID-19 Genomics UK (COG-UK) Consortium | Jacquelyn Wynn, Mairead Hyland, The Lighthouse Lab in Alderley Park and Alex Alderton, Roberto Amato, Jeffrey Barrett, Sonia Goncalves, Ewan Harrison, David K. Jackson, Ian Johnston, Dominic Kwiatkowski, Cordelia Langford, John Sillitoe on behalf of the Wellcome Sanger Institute COVID-19 Surveillance Team |
| EPI_ISL_2924790, EPI_ISL_2924791 | Lighthouse Lab in Glasgow | Wellcome Sanger Institute for the COVID-19 Genomics UK (COG-UK) Consortium | Harper VanSteenhouse, Yumi Kasai, David Gray, Carol Clugston, Anna Dominiczak and Alex Alderton, Roberto Amato, Jeffrey Barrett, Sonia Goncalves, Ewan Harrison, David K. Jackson, Ian Johnston, Dominic Kwiatkowski, Cordelia Langford, John Sillitoe on behalf of the Wellcome Sanger Institute COVID-19 Surveillance Team |
| EPI_ISL_2924792, EPI_ISL_2924793, EPI_ISL_2924794, EPI_ISL_2924795, EPI_ISL_2924796, EPI_ISL_2924797, EPI_ISL_2924798 | Lighthouse Lab in Alderley Park | Wellcome Sanger Institute for the COVID-19 Genomics UK (COG-UK) Consortium | Jacquelyn Wynn, Mairead Hyland, The Lighthouse Lab in Alderley Park and Alex Alderton, Roberto Amato, Jeffrey Barrett, Sonia Goncalves, Ewan Harrison, David K. Jackson, Ian Johnston, Dominic Kwiatkowski, Cordelia Langford, John Sillitoe on behalf of the Wellcome Sanger Institute COVID-19 Surveillance Team |
| EPI_ISL_2924799 | Lighthouse Lab in Glasgow | Wellcome Sanger Institute for the COVID-19 Genomics UK (COG-UK) Consortium | Harper VanSteenhouse, Yumi Kasai, David Gray, Carol Clugston, Anna Dominiczak and Alex Alderton, Roberto Amato, Jeffrey Barrett, Sonia Goncalves, Ewan Harrison, David K. Jackson, Ian Johnston, Dominic Kwiatkowski, Cordelia Langford, John Sillitoe on behalf of the Wellcome Sanger Institute COVID-19 Surveillance Team |
| EPI_ISL_2924800, EPI_ISL_2924801 | Lighthouse Lab in Alderley Park | Wellcome Sanger Institute for the COVID-19 Genomics UK (COG-UK) Consortium | Jacquelyn Wynn, Mairead Hyland, The Lighthouse Lab in Alderley Park and Alex Alderton, Roberto Amato, Jeffrey Barrett, Sonia Goncalves, Ewan Harrison, David K. Jackson, Ian Johnston, Dominic Kwiatkowski, Cordelia Langford, John Sillitoe on behalf of the Wellcome Sanger Institute COVID-19 Surveillance Team |
| EPI_ISL_2924802 | Lighthouse Lab in Glasgow | Wellcome Sanger Institute for the COVID-19 Genomics UK (COG-UK) Consortium | Harper VanSteenhouse, Yumi Kasai, David Gray, Carol Clugston, Anna Dominiczak and Alex Alderton, Roberto Amato, Jeffrey Barrett, Sonia Goncalves, Ewan Harrison, David K. Jackson, Ian Johnston, Dominic Kwiatkowski, Cordelia Langford, John Sillitoe on behalf of the Wellcome Sanger Institute COVID-19 Surveillance Team |
| EPI_ISL_2924803, EPI_ISL_2924804, EPI_ISL_2924805, EPI_ISL_2924806, EPI_ISL_2924807, EPI_ISL_2924808, EPI_ISL_2924809 | Lighthouse Lab in Alderley Park | Wellcome Sanger Institute for the COVID-19 Genomics UK (COG-UK) Consortium | Jacquelyn Wynn, Mairead Hyland, The Lighthouse Lab in Alderley Park and Alex Alderton, Roberto Amato, Jeffrey Barrett, Sonia Goncalves, Ewan Harrison, David K. Jackson, Ian Johnston, Dominic Kwiatkowski, Cordelia Langford, John Sillitoe on behalf of the Wellcome Sanger Institute COVID-19 Surveillance Team |
| EPI_ISL_2924810 | Lighthouse Lab in Glasgow | Wellcome Sanger Institute for the COVID-19 Genomics UK (COG-UK) Consortium | Harper VanSteenhouse, Yumi Kasai, David Gray, Carol Clugston, Anna Dominiczak and Alex Alderton, Roberto Amato, Jeffrey Barrett, Sonia Goncalves, Ewan Harrison, David K. Jackson, Ian Johnston, Dominic Kwiatkowski, Cordelia Langford, John Sillitoe on behalf of the Wellcome Sanger Institute COVID-19 Surveillance Team |
| EPI_ISL_2924811, EPI_ISL_2924812, EPI_ISL_2924813, EPI_ISL_2924814, EPI_ISL_2924815, EPI_ISL_2924816, EPI_ISL_2924817, EPI_ISL_2924818, EPI_ISL_2924819, EPI_ISL_2924820, EPI_ISL_2924821 | | | |
| see above | Lighthouse Lab in Alderley Park | Wellcome Sanger Institute for the COVID-19 Genomics UK (COG-UK) Consortium | Jacquelyn Wynn, Mairead Hyland, The Lighthouse Lab in Alderley Park and Alex Alderton, Roberto Amato, Jeffrey Barrett, Sonia Goncalves, Ewan Harrison, David K. Jackson, Ian Johnston, Dominic Kwiatkowski, Cordelia Langford, John Sillitoe on behalf of the Wellcome Sanger Institute COVID-19 Surveillance Team |
| EPI_ISL_2924822, EPI_ISL_2924823, EPI_ISL_2924824 | Lighthouse Lab in Glasgow | Wellcome Sanger Institute for the COVID-19 Genomics UK (COG-UK) Consortium | Harper VanSteenhouse, Yumi Kasai, David Gray, Carol Clugston, Anna Dominiczak and Alex Alderton, Roberto Amato, Jeffrey Barrett, Sonia Goncalves, Ewan Harrison, David K. Jackson, Ian Johnston, Dominic Kwiatkowski, Cordelia Langford, John Sillitoe on behalf of the Wellcome Sanger Institute COVID-19 Surveillance Team |
| EPI_ISL_2924825, EPI_ISL_2924826, EPI_ISL_2924827 | Lighthouse Lab in Alderley Park | Wellcome Sanger Institute for the COVID-19 Genomics UK (COG-UK) Consortium | Jacquelyn Wynn, Mairead Hyland, The Lighthouse Lab in Alderley Park and Alex Alderton, Roberto Amato, Jeffrey Barrett, Sonia Goncalves, Ewan Harrison, David K. Jackson, Ian Johnston, Dominic Kwiatkowski, Cordelia Langford, John Sillitoe on behalf of the Wellcome Sanger Institute COVID-19 Surveillance Team |
| EPI_ISL_2924828, EPI_ISL_2924829 | Lighthouse Lab in Glasgow | Wellcome Sanger Institute for the COVID-19 Genomics UK (COG-UK) Consortium | Harper VanSteenhouse, Yumi Kasai, David Gray, Carol Clugston, Anna Dominiczak and Alex Alderton, Roberto Amato, Jeffrey Barrett, Sonia Goncalves, Ewan Harrison, David K. Jackson, Ian Johnston, Dominic Kwiatkowski, Cordelia Langford, John Sillitoe on behalf of the Wellcome Sanger Institute COVID-19 Surveillance Team |
| EPI_ISL_2924830 | Lighthouse Lab in Alderley Park | Wellcome Sanger Institute for the COVID-19 Genomics UK (COG-UK) Consortium | Jacquelyn Wynn, Mairead Hyland, The Lighthouse Lab in Alderley Park and Alex Alderton, Roberto Amato, Jeffrey Barrett, Sonia Goncalves, Ewan Harrison, David K. Jackson, Ian Johnston, Dominic Kwiatkowski, Cordelia Langford, John Sillitoe on behalf of the Wellcome Sanger Institute COVID-19 Surveillance Team |
| EPI_ISL_2924831 | Lighthouse Lab in Glasgow | Wellcome Sanger Institute for the COVID-19 Genomics UK (COG-UK) Consortium | Harper VanSteenhouse, Yumi Kasai, David Gray, Carol Clugston, Anna Dominiczak and Alex Alderton, Roberto Amato, Jeffrey Barrett, Sonia Goncalves, Ewan Harrison, David K. Jackson, Ian Johnston, Dominic Kwiatkowski, Cordelia Langford, John Sillitoe on behalf of the Wellcome Sanger Institute COVID-19 Surveillance Team |
| EPI_ISL_2924832 | Lighthouse Lab in Alderley Park | Wellcome Sanger Institute for the COVID-19 Genomics UK (COG-UK) Consortium | Jacquelyn Wynn, Mairead Hyland, The Lighthouse Lab in Alderley Park and Alex Alderton, Roberto Amato, Jeffrey Barrett, Sonia Goncalves, Ewan Harrison, David K. Jackson, Ian Johnston, Dominic Kwiatkowski, Cordelia Langford, John Sillitoe on behalf of the Wellcome Sanger Institute COVID-19 Surveillance Team |
| EPI_ISL_2924833 | Lighthouse Lab in Glasgow | Wellcome Sanger Institute for the COVID-19 Genomics UK (COG-UK) Consortium | Harper VanSteenhouse, Yumi Kasai, David Gray, Carol Clugston, Anna Dominiczak and Alex Alderton, Roberto Amato, Jeffrey Barrett, Sonia Goncalves, Ewan Harrison, David K. Jackson, Ian Johnston, Dominic Kwiatkowski, Cordelia Langford, John Sillitoe on behalf of the Wellcome Sanger Institute COVID-19 Surveillance Team |
| EPI_ISL_2924834, EPI_ISL_2924835, EPI_ISL_2924836, EPI_ISL_2924837, EPI_ISL_2924838, EPI_ISL_2924839, EPI_ISL_2924840 | Lighthouse Lab in Alderley Park | Wellcome Sanger Institute for the COVID-19 Genomics UK (COG-UK) Consortium | Jacquelyn Wynn, Mairead Hyland, The Lighthouse Lab in Alderley Park and Alex Alderton, Roberto Amato, Jeffrey Barrett, Sonia Goncalves, Ewan Harrison, David K. Jackson, Ian Johnston, Dominic Kwiatkowski, Cordelia Langford, John Sillitoe on behalf of the Wellcome Sanger Institute COVID-19 Surveillance Team |
| EPI_ISL_2924841 | Lighthouse Lab in Glasgow | Wellcome Sanger Institute for the COVID-19 Genomics UK (COG-UK) Consortium | Harper VanSteenhouse, Yumi Kasai, David Gray, Carol Clugston, Anna Dominiczak and Alex Alderton, Roberto Amato, Jeffrey Barrett, Sonia Goncalves, Ewan Harrison, David K. Jackson, Ian Johnston, Dominic Kwiatkowski, Cordelia Langford, John Sillitoe on behalf of the Wellcome Sanger Institute COVID-19 Surveillance Team |
| EPI_ISL_2924842 | Lighthouse Lab in Milton Keynes | Wellcome Sanger Institute for the COVID-19 Genomics UK (COG-UK) Consortium | The Lighthouse Lab in Milton Keynes and Alex Alderton, Roberto Amato, Jeffrey Barrett, Sonia Goncalves, Ewan Harrison, David K. Jackson, Ian Johnston, Dominic Kwiatkowski, Cordelia Langford, John Sillitoe on behalf of the Wellcome Sanger Institute COVID-19 Surveillance Team |
| EPI_ISL_2924843 | Lighthouse Lab in Glasgow | Wellcome Sanger Institute for the COVID-19 Genomics UK (COG-UK) Consortium | Harper VanSteenhouse, Yumi Kasai, David Gray, Carol Clugston, Anna Dominiczak and Alex Alderton, Roberto Amato, Jeffrey Barrett, Sonia Goncalves, Ewan Harrison, David K. Jackson, Ian Johnston, Dominic Kwiatkowski, Cordelia Langford, John Sillitoe on behalf of the Wellcome Sanger Institute COVID-19 Surveillance Team |
| EPI_ISL_2924844 | Lighthouse Lab in Alderley Park | Wellcome Sanger Institute for the COVID-19 Genomics UK (COG-UK) Consortium | Jacquelyn Wynn, Mairead Hyland, The Lighthouse Lab in Alderley Park and Alex Alderton, Roberto Amato, Jeffrey Barrett, Sonia Goncalves, Ewan Harrison, David K. Jackson, Ian Johnston, Dominic Kwiatkowski, Cordelia Langford, John Sillitoe on behalf of the Wellcome Sanger Institute COVID-19 Surveillance Team |
| EPI_ISL_2924845 | Lighthouse Lab in Glasgow | Wellcome Sanger Institute for the COVID-19 Genomics UK (COG-UK) Consortium | Harper VanSteenhouse, Yumi Kasai, David Gray, Carol Clugston, Anna Dominiczak and Alex Alderton, Roberto Amato, Jeffrey Barrett, Sonia Goncalves, Ewan Harrison, David K. Jackson, Ian Johnston, Dominic Kwiatkowski, Cordelia Langford, John Sillitoe on behalf of the Wellcome Sanger Institute COVID-19 Surveillance Team |
| EPI_ISL_2924846 | Lighthouse Lab in Milton Keynes | Wellcome Sanger Institute for the COVID-19 Genomics UK (COG-UK) Consortium | The Lighthouse Lab in Milton Keynes and Alex Alderton, Roberto Amato, Jeffrey Barrett, Sonia Goncalves, Ewan Harrison, David K. Jackson, Ian Johnston, Dominic Kwiatkowski, Cordelia Langford, John Sillitoe on behalf of the Wellcome Sanger Institute COVID-19 Surveillance Team |
| EPI_ISL_2924847 | Lighthouse Lab in Alderley Park | Wellcome Sanger Institute for the COVID-19 Genomics UK (COG-UK) | Jacquelyn Wynn, Mairead Hyland, The Lighthouse Lab in Alderley Park and Alex Alderton, Roberto Amato, Jeffrey Barrett, Sonia Goncalves, Ewan Harrison, David K. Jackson, Ian Johnston, Dominic Kwiatkowski, Cordelia Langford, John Sillitoe on |

| | | | |
|---|---|---|---|
| | | Consortium | behalf of the Wellcome Sanger Institute COVID-19 Surveillance Team |
| EPI_ISL_2924848 | Lighthouse Lab in Glasgow | Wellcome Sanger Institute for the COVID-19 Genomics UK (COG-UK) Consortium | Harper VanSteenhouse, Yumi Kasai, David Gray, Carol Clugston, Anna Dominiczak and Alex Alderton, Roberto Amato, Jeffrey Barrett, Sonia Goncalves, Ewan Harrison, David K. Jackson, Ian Johnston, Dominic Kwiatkowski, Cordelia Langford, John Sillitoe on behalf of the Wellcome Sanger Institute COVID-19 Surveillance Team |
| EPI_ISL_2924849, EPI_ISL_2924850 | Lighthouse Lab in Alderley Park | Wellcome Sanger Institute for the COVID-19 Genomics UK (COG-UK) Consortium | Jacquelyn Wynn, Mairead Hyland, The Lighthouse Lab in Alderley Park and Alex Alderton, Roberto Amato, Jeffrey Barrett, Sonia Goncalves, Ewan Harrison, David K. Jackson, Ian Johnston, Dominic Kwiatkowski, Cordelia Langford, John Sillitoe on behalf of the Wellcome Sanger Institute COVID-19 Surveillance Team |
| EPI_ISL_2924851, EPI_ISL_2924852 | Lighthouse Lab in Milton Keynes | Wellcome Sanger Institute for the COVID-19 Genomics UK (COG-UK) Consortium | The Lighthouse Lab in Milton Keynes and Alex Alderton, Roberto Amato, Jeffrey Barrett, Sonia Goncalves, Ewan Harrison, David K. Jackson, Ian Johnston, Dominic Kwiatkowski, Cordelia Langford, John Sillitoe on behalf of the Wellcome Sanger Institute COVID-19 Surveillance Team |
| EPI_ISL_2924853 | Lighthouse Lab in Glasgow | Wellcome Sanger Institute for the COVID-19 Genomics UK (COG-UK) Consortium | Harper VanSteenhouse, Yumi Kasai, David Gray, Carol Clugston, Anna Dominiczak and Alex Alderton, Roberto Amato, Jeffrey Barrett, Sonia Goncalves, Ewan Harrison, David K. Jackson, Ian Johnston, Dominic Kwiatkowski, Cordelia Langford, John Sillitoe on behalf of the Wellcome Sanger Institute COVID-19 Surveillance Team |
| EPI_ISL_2924854 | Lighthouse Lab in Milton Keynes | Wellcome Sanger Institute for the COVID-19 Genomics UK (COG-UK) Consortium | The Lighthouse Lab in Milton Keynes and Alex Alderton, Roberto Amato, Jeffrey Barrett, Sonia Goncalves, Ewan Harrison, David K. Jackson, Ian Johnston, Dominic Kwiatkowski, Cordelia Langford, John Sillitoe on behalf of the Wellcome Sanger Institute COVID-19 Surveillance Team |
| EPI_ISL_2924855 | Lighthouse Lab in Glasgow | Wellcome Sanger Institute for the COVID-19 Genomics UK (COG-UK) Consortium | Harper VanSteenhouse, Yumi Kasai, David Gray, Carol Clugston, Anna Dominiczak and Alex Alderton, Roberto Amato, Jeffrey Barrett, Sonia Goncalves, Ewan Harrison, David K. Jackson, Ian Johnston, Dominic Kwiatkowski, Cordelia Langford, John Sillitoe on behalf of the Wellcome Sanger Institute COVID-19 Surveillance Team |
| EPI_ISL_2924856 | Lighthouse Lab in Alderley Park | Wellcome Sanger Institute for the COVID-19 Genomics UK (COG-UK) Consortium | Jacquelyn Wynn, Mairead Hyland, The Lighthouse Lab in Alderley Park and Alex Alderton, Roberto Amato, Jeffrey Barrett, Sonia Goncalves, Ewan Harrison, David K. Jackson, Ian Johnston, Dominic Kwiatkowski, Cordelia Langford, John Sillitoe on behalf of the Wellcome Sanger Institute COVID-19 Surveillance Team |
| EPI_ISL_2924857, EPI_ISL_2924858, EPI_ISL_2924859 | Lighthouse Lab in Glasgow | Wellcome Sanger Institute for the COVID-19 Genomics UK (COG-UK) Consortium | Harper VanSteenhouse, Yumi Kasai, David Gray, Carol Clugston, Anna Dominiczak and Alex Alderton, Roberto Amato, Jeffrey Barrett, Sonia Goncalves, Ewan Harrison, David K. Jackson, Ian Johnston, Dominic Kwiatkowski, Cordelia Langford, John Sillitoe on behalf of the Wellcome Sanger Institute COVID-19 Surveillance Team |
| EPI_ISL_2924860 | Lighthouse Lab in Milton Keynes | Wellcome Sanger Institute for the COVID-19 Genomics UK (COG-UK) Consortium | The Lighthouse Lab in Milton Keynes and Alex Alderton, Roberto Amato, Jeffrey Barrett, Sonia Goncalves, Ewan Harrison, David K. Jackson, Ian Johnston, Dominic Kwiatkowski, Cordelia Langford, John Sillitoe on behalf of the Wellcome Sanger Institute COVID-19 Surveillance Team |
| EPI_ISL_2924861 | Lighthouse Lab in Glasgow | Wellcome Sanger Institute for the COVID-19 Genomics UK (COG-UK) Consortium | Harper VanSteenhouse, Yumi Kasai, David Gray, Carol Clugston, Anna Dominiczak and Alex Alderton, Roberto Amato, Jeffrey Barrett, Sonia Goncalves, Ewan Harrison, David K. Jackson, Ian Johnston, Dominic Kwiatkowski, Cordelia Langford, John Sillitoe on behalf of the Wellcome Sanger Institute COVID-19 Surveillance Team |
| EPI_ISL_2924862 | Lighthouse Lab in Milton Keynes | Wellcome Sanger Institute for the COVID-19 Genomics UK (COG-UK) Consortium | The Lighthouse Lab in Milton Keynes and Alex Alderton, Roberto Amato, Jeffrey Barrett, Sonia Goncalves, Ewan Harrison, David K. Jackson, Ian Johnston, Dominic Kwiatkowski, Cordelia Langford, John Sillitoe on behalf of the Wellcome Sanger Institute COVID-19 Surveillance Team |
| EPI_ISL_2924863 | Lighthouse Lab in Alderley Park | Wellcome Sanger Institute for the COVID-19 Genomics UK (COG-UK) Consortium | Jacquelyn Wynn, Mairead Hyland, The Lighthouse Lab in Alderley Park and Alex Alderton, Roberto Amato, Jeffrey Barrett, Sonia Goncalves, Ewan Harrison, David K. Jackson, Ian Johnston, Dominic Kwiatkowski, Cordelia Langford, John Sillitoe on behalf of the Wellcome Sanger Institute COVID-19 Surveillance Team |
| EPI_ISL_2924864 | Lighthouse Lab in Glasgow | Wellcome Sanger Institute for the COVID-19 Genomics UK (COG-UK) Consortium | Harper VanSteenhouse, Yumi Kasai, David Gray, Carol Clugston, Anna Dominiczak and Alex Alderton, Roberto Amato, Jeffrey Barrett, Sonia Goncalves, Ewan Harrison, David K. Jackson, Ian Johnston, Dominic Kwiatkowski, Cordelia Langford, John Sillitoe on behalf of the Wellcome Sanger Institute COVID-19 Surveillance Team |
| EPI_ISL_2924865 | Lighthouse Lab in Milton Keynes | Wellcome Sanger Institute for the COVID-19 Genomics UK (COG-UK) Consortium | The Lighthouse Lab in Milton Keynes and Alex Alderton, Roberto Amato, Jeffrey Barrett, Sonia Goncalves, Ewan Harrison, David K. Jackson, Ian Johnston, Dominic Kwiatkowski, Cordelia Langford, John Sillitoe on behalf of the Wellcome Sanger Institute COVID-19 Surveillance Team |
| EPI_ISL_2924866, EPI_ISL_2924867 | Lighthouse Lab in Glasgow | Wellcome Sanger Institute for the COVID-19 Genomics UK (COG-UK) Consortium | Harper VanSteenhouse, Yumi Kasai, David Gray, Carol Clugston, Anna Dominiczak and Alex Alderton, Roberto Amato, Jeffrey Barrett, Sonia Goncalves, Ewan Harrison, David K. Jackson, Ian Johnston, Dominic Kwiatkowski, Cordelia Langford, John Sillitoe on behalf of the Wellcome Sanger Institute COVID-19 Surveillance Team |
| EPI_ISL_2924868 | Lighthouse Lab in Alderley Park | Wellcome Sanger Institute for the COVID-19 Genomics UK (COG-UK) Consortium | Jacquelyn Wynn, Mairead Hyland, The Lighthouse Lab in Alderley Park and Alex Alderton, Roberto Amato, Jeffrey Barrett, Sonia Goncalves, Ewan Harrison, David K. Jackson, Ian Johnston, Dominic Kwiatkowski, Cordelia Langford, John Sillitoe on behalf of the Wellcome Sanger Institute COVID-19 Surveillance Team |
| EPI_ISL_2924869 | Lighthouse Lab in Milton Keynes | Wellcome Sanger Institute for the COVID-19 Genomics UK (COG-UK) Consortium | The Lighthouse Lab in Milton Keynes and Alex Alderton, Roberto Amato, Jeffrey Barrett, Sonia Goncalves, Ewan Harrison, David K. Jackson, Ian Johnston, Dominic Kwiatkowski, Cordelia Langford, John Sillitoe on behalf of the Wellcome Sanger Institute COVID-19 Surveillance Team |
| EPI_ISL_2924870 | Lighthouse Lab in Glasgow | Wellcome Sanger Institute for the COVID-19 Genomics UK (COG-UK) Consortium | Harper VanSteenhouse, Yumi Kasai, David Gray, Carol Clugston, Anna Dominiczak and Alex Alderton, Roberto Amato, Jeffrey Barrett, Sonia Goncalves, Ewan Harrison, David K. Jackson, Ian Johnston, Dominic Kwiatkowski, Cordelia Langford, John Sillitoe on behalf of the Wellcome Sanger Institute COVID-19 Surveillance Team |
| EPI_ISL_2924871, EPI_ISL_2924872 | Lighthouse Lab in Milton Keynes | Wellcome Sanger Institute for the COVID-19 Genomics UK (COG-UK) Consortium | The Lighthouse Lab in Milton Keynes and Alex Alderton, Roberto Amato, Jeffrey Barrett, Sonia Goncalves, Ewan Harrison, David K. Jackson, Ian Johnston, Dominic Kwiatkowski, Cordelia Langford, John Sillitoe on behalf of the Wellcome Sanger Institute COVID-19 Surveillance Team |
| EPI_ISL_2924873 | Lighthouse Lab in Glasgow | Wellcome Sanger Institute for the COVID-19 Genomics UK (COG-UK) Consortium | Harper VanSteenhouse, Yumi Kasai, David Gray, Carol Clugston, Anna Dominiczak and Alex Alderton, Roberto Amato, Jeffrey Barrett, Sonia Goncalves, Ewan Harrison, David K. Jackson, Ian Johnston, Dominic Kwiatkowski, Cordelia Langford, John Sillitoe on behalf of the Wellcome Sanger Institute COVID-19 Surveillance Team |
| EPI_ISL_2924874 | Lighthouse Lab in Milton Keynes | Wellcome Sanger Institute for the COVID-19 Genomics UK (COG-UK) Consortium | The Lighthouse Lab in Milton Keynes and Alex Alderton, Roberto Amato, Jeffrey Barrett, Sonia Goncalves, Ewan Harrison, David K. Jackson, Ian Johnston, Dominic Kwiatkowski, Cordelia Langford, John Sillitoe on behalf of the Wellcome Sanger Institute COVID-19 Surveillance Team |
| EPI_ISL_2924875 | Lighthouse Lab in Glasgow | Wellcome Sanger Institute for the COVID-19 Genomics UK (COG-UK) Consortium | Harper VanSteenhouse, Yumi Kasai, David Gray, Carol Clugston, Anna Dominiczak and Alex Alderton, Roberto Amato, Jeffrey Barrett, Sonia Goncalves, Ewan Harrison, David K. Jackson, Ian Johnston, Dominic Kwiatkowski, Cordelia Langford, John Sillitoe on behalf of the Wellcome Sanger Institute COVID-19 Surveillance Team |
| EPI_ISL_2924876, EPI_ISL_2924877 | Lighthouse Lab in Milton Keynes | Wellcome Sanger Institute for the COVID-19 Genomics UK (COG-UK) Consortium | The Lighthouse Lab in Milton Keynes and Alex Alderton, Roberto Amato, Jeffrey Barrett, Sonia Goncalves, Ewan Harrison, David K. Jackson, Ian Johnston, Dominic Kwiatkowski, Cordelia Langford, John Sillitoe on behalf of the Wellcome Sanger Institute COVID-19 Surveillance Team |
| EPI_ISL_2924878 | Lighthouse Lab in Glasgow | Wellcome Sanger Institute for the COVID-19 Genomics UK (COG-UK) Consortium | Harper VanSteenhouse, Yumi Kasai, David Gray, Carol Clugston, Anna Dominiczak and Alex Alderton, Roberto Amato, Jeffrey Barrett, Sonia Goncalves, Ewan Harrison, David K. Jackson, Ian Johnston, Dominic Kwiatkowski, Cordelia Langford, John Sillitoe on behalf of the Wellcome Sanger Institute COVID-19 Surveillance Team |
| EPI_ISL_2924879 | Lighthouse Lab in Alderley Park | Wellcome Sanger Institute for the COVID-19 Genomics UK (COG-UK) Consortium | Jacquelyn Wynn, Mairead Hyland, The Lighthouse Lab in Alderley Park and Alex Alderton, Roberto Amato, Jeffrey Barrett, Sonia Goncalves, Ewan Harrison, David K. Jackson, Ian Johnston, Dominic Kwiatkowski, Cordelia Langford, John Sillitoe on behalf of the Wellcome Sanger Institute COVID-19 Surveillance Team |
| EPI_ISL_2924880 | Lighthouse Lab in Milton Keynes | Wellcome Sanger Institute for the COVID-19 Genomics UK (COG-UK) Consortium | The Lighthouse Lab in Milton Keynes and Alex Alderton, Roberto Amato, Jeffrey Barrett, Sonia Goncalves, Ewan Harrison, David K. Jackson, Ian Johnston, Dominic Kwiatkowski, Cordelia Langford, John Sillitoe on behalf of the Wellcome Sanger Institute COVID-19 Surveillance Team |
| EPI_ISL_2924881 | Lighthouse Lab in Glasgow | Wellcome Sanger Institute for the COVID-19 Genomics UK (COG-UK) Consortium | Harper VanSteenhouse, Yumi Kasai, David Gray, Carol Clugston, Anna Dominiczak and Alex Alderton, Roberto Amato, Jeffrey Barrett, Sonia Goncalves, Ewan Harrison, David K. Jackson, Ian Johnston, Dominic Kwiatkowski, Cordelia Langford, John Sillitoe on behalf of the Wellcome Sanger Institute COVID-19 Surveillance Team |
| EPI_ISL_2924882 | Lighthouse Lab in Alderley Park | Wellcome Sanger Institute for the COVID-19 Genomics UK (COG-UK) Consortium | Jacquelyn Wynn, Mairead Hyland, The Lighthouse Lab in Alderley Park and Alex Alderton, Roberto Amato, Jeffrey Barrett, Sonia Goncalves, Ewan Harrison, David K. Jackson, Ian Johnston, Dominic Kwiatkowski, Cordelia Langford, John Sillitoe on behalf of the Wellcome Sanger Institute COVID-19 Surveillance Team |
| EPI_ISL_2924883 | Lighthouse Lab in Milton Keynes | Wellcome Sanger Institute for the COVID-19 Genomics UK (COG-UK) Consortium | The Lighthouse Lab in Milton Keynes and Alex Alderton, Roberto Amato, Jeffrey Barrett, Sonia Goncalves, Ewan Harrison, David K. Jackson, Ian Johnston, Dominic Kwiatkowski, Cordelia Langford, John Sillitoe on behalf of the Wellcome Sanger Institute COVID-19 Surveillance Team |
| EPI_ISL_2924884 | Lighthouse Lab in Alderley Park | Wellcome Sanger Institute for the COVID-19 Genomics UK (COG-UK) Consortium | Jacquelyn Wynn, Mairead Hyland, The Lighthouse Lab in Alderley Park and Alex Alderton, Roberto Amato, Jeffrey Barrett, Sonia Goncalves, Ewan Harrison, David K. Jackson, Ian Johnston, Dominic Kwiatkowski, Cordelia Langford, John Sillitoe on behalf of the Wellcome Sanger Institute COVID-19 Surveillance Team |
| EPI_ISL_2924885 | Lighthouse Lab in Milton Keynes | Wellcome Sanger Institute for the COVID-19 Genomics UK (COG-UK) Consortium | The Lighthouse Lab in Milton Keynes and Alex Alderton, Roberto Amato, Jeffrey Barrett, Sonia Goncalves, Ewan Harrison, David K. Jackson, Ian Johnston, Dominic Kwiatkowski, Cordelia Langford, John Sillitoe on behalf of the Wellcome Sanger Institute COVID-19 Surveillance Team |
| EPI_ISL_2924886, EPI_ISL_2924887 | Lighthouse Lab in Alderley Park | Wellcome Sanger Institute for the COVID-19 Genomics UK (COG-UK) Consortium | Jacquelyn Wynn, Mairead Hyland, The Lighthouse Lab in Alderley Park and Alex Alderton, Roberto Amato, Jeffrey Barrett, Sonia Goncalves, Ewan Harrison, David K. Jackson, Ian Johnston, Dominic Kwiatkowski, Cordelia Langford, John Sillitoe on behalf of the Wellcome Sanger Institute COVID-19 Surveillance Team |
| EPI_ISL_2924888 | Lighthouse Lab in Glasgow | Wellcome Sanger Institute for the COVID-19 Genomics UK (COG-UK) Consortium | Harper VanSteenhouse, Yumi Kasai, David Gray, Carol Clugston, Anna Dominiczak and Alex Alderton, Roberto Amato, Jeffrey Barrett, Sonia Goncalves, Ewan Harrison, David K. Jackson, Ian Johnston, Dominic Kwiatkowski, Cordelia Langford, John Sillitoe on behalf of the Wellcome Sanger Institute COVID-19 Surveillance Team |
| EPI_ISL_2924889 | Lighthouse Lab in Milton Keynes | Wellcome Sanger Institute for the COVID-19 Genomics UK (COG-UK) Consortium | The Lighthouse Lab in Milton Keynes and Alex Alderton, Roberto Amato, Jeffrey Barrett, Sonia Goncalves, Ewan Harrison, David K. Jackson, Ian Johnston, Dominic Kwiatkowski, Cordelia Langford, John Sillitoe on behalf of the Wellcome Sanger Institute COVID-19 Surveillance Team |
| EPI_ISL_2924890 | Lighthouse Lab in Alderley Park | Wellcome Sanger Institute for the COVID-19 Genomics UK (COG-UK) Consortium | Jacquelyn Wynn, Mairead Hyland, The Lighthouse Lab in Alderley Park and Alex Alderton, Roberto Amato, Jeffrey Barrett, Sonia Goncalves, Ewan Harrison, David K. Jackson, Ian Johnston, Dominic Kwiatkowski, Cordelia Langford, John Sillitoe on behalf of the Wellcome Sanger Institute COVID-19 Surveillance Team |
| EPI_ISL_2924891 | Lighthouse Lab in Milton Keynes | Wellcome Sanger Institute for the COVID-19 Genomics UK (COG-UK) Consortium | The Lighthouse Lab in Milton Keynes and Alex Alderton, Roberto Amato, Jeffrey Barrett, Sonia Goncalves, Ewan Harrison, David K. Jackson, Ian Johnston, Dominic Kwiatkowski, Cordelia Langford, John Sillitoe on behalf of the Wellcome Sanger Institute COVID-19 Surveillance Team |
| EPI_ISL_2924892, EPI_ISL_2924893, EPI_ISL_2924894, EPI_ISL_2924895, EPI_ISL_2924896 | Lighthouse Lab in Alderley Park | Wellcome Sanger Institute for the COVID-19 Genomics UK (COG-UK) Consortium | Jacquelyn Wynn, Mairead Hyland, The Lighthouse Lab in Alderley Park and Alex Alderton, Roberto Amato, Jeffrey Barrett, Sonia Goncalves, Ewan Harrison, David K. Jackson, Ian Johnston, Dominic Kwiatkowski, Cordelia Langford, John Sillitoe on behalf of the Wellcome Sanger Institute COVID-19 Surveillance Team |
| EPI_ISL_2924897 | Lighthouse Lab in Glasgow | Wellcome Sanger Institute for the COVID-19 Genomics UK (COG-UK) Consortium | Harper VanSteenhouse, Yumi Kasai, David Gray, Carol Clugston, Anna Dominiczak and Alex Alderton, Roberto Amato, Jeffrey Barrett, Sonia Goncalves, Ewan Harrison, David K. Jackson, Ian Johnston, Dominic Kwiatkowski, Cordelia Langford, John Sillitoe on behalf of the Wellcome Sanger Institute COVID-19 Surveillance Team |
| EPI_ISL_2924898 | Lighthouse Lab in Milton Keynes | Wellcome Sanger Institute for the COVID-19 Genomics UK (COG-UK) Consortium | The Lighthouse Lab in Milton Keynes and Alex Alderton, Roberto Amato, Jeffrey Barrett, Sonia Goncalves, Ewan Harrison, David K. Jackson, Ian Johnston, Dominic Kwiatkowski, Cordelia Langford, John Sillitoe on behalf of the Wellcome Sanger Institute COVID-19 Surveillance Team |
| EPI_ISL_2924899, EPI_ISL_2924900 | Lighthouse Lab in Glasgow | Wellcome Sanger Institute for the COVID-19 Genomics UK (COG-UK) Consortium | Harper VanSteenhouse, Yumi Kasai, David Gray, Carol Clugston, Anna Dominiczak and Alex Alderton, Roberto Amato, Jeffrey Barrett, Sonia Goncalves, Ewan Harrison, David K. Jackson, Ian Johnston, Dominic Kwiatkowski, Cordelia Langford, John Sillitoe on behalf of the Wellcome Sanger Institute COVID-19 Surveillance Team |
| EPI_ISL_2924901, EPI_ISL_2924902 | Lighthouse Lab in Milton Keynes | Wellcome Sanger Institute for the COVID-19 Genomics UK (COG-UK) Consortium | The Lighthouse Lab in Milton Keynes and Alex Alderton, Roberto Amato, Jeffrey Barrett, Sonia Goncalves, Ewan Harrison, David K. Jackson, Ian Johnston, Dominic Kwiatkowski, Cordelia Langford, John Sillitoe on behalf of the Wellcome Sanger Institute COVID-19 Surveillance Team |
| EPI_ISL_2924903 | Lighthouse Lab in Alderley Park | Wellcome Sanger Institute for the COVID-19 Genomics UK (COG-UK) Consortium | Jacquelyn Wynn, Mairead Hyland, The Lighthouse Lab in Alderley Park and Alex Alderton, Roberto Amato, Jeffrey Barrett, Sonia Goncalves, Ewan Harrison, David K. Jackson, Ian Johnston, Dominic Kwiatkowski, Cordelia Langford, John Sillitoe on behalf of the Wellcome Sanger Institute COVID-19 Surveillance Team |
| EPI_ISL_2924904 | Lighthouse Lab in Milton Keynes | Wellcome Sanger Institute for the COVID-19 Genomics UK (COG-UK) Consortium | The Lighthouse Lab in Milton Keynes and Alex Alderton, Roberto Amato, Jeffrey Barrett, Sonia Goncalves, Ewan Harrison, David K. Jackson, Ian Johnston, Dominic Kwiatkowski, Cordelia Langford, John Sillitoe on behalf of the Wellcome Sanger Institute COVID-19 Surveillance Team |

| | | | |
|---|---|---|---|
| EPI_ISL_2924905, EPI_ISL_2924906 | Lighthouse Lab in Glasgow | Wellcome Sanger Institute for the COVID-19 Genomics UK (COG-UK) Consortium | Harper VanSteenhouse, Yumi Kasai, David Gray, Carol Clugston, Anna Dominiczak and Alex Alderton, Roberto Amato, Jeffrey Barrett, Sonia Goncalves, Ewan Harrison, David K. Jackson, Ian Johnston, Dominic Kwiatkowski, Cordelia Langford, John Sillitoe on behalf of the Wellcome Sanger Institute COVID-19 Surveillance Team |
| EPI_ISL_2924907 | Lighthouse Lab in Milton Keynes | Wellcome Sanger Institute for the COVID-19 Genomics UK (COG-UK) Consortium | The Lighthouse Lab in Milton Keynes and Alex Alderton, Roberto Amato, Jeffrey Barrett, Sonia Goncalves, Ewan Harrison, David K. Jackson, Ian Johnston, Dominic Kwiatkowski, Cordelia Langford, John Sillitoe on behalf of the Wellcome Sanger Institute COVID-19 Surveillance Team |
| EPI_ISL_2924908, EPI_ISL_2924909, EPI_ISL_2924910 | Lighthouse Lab in Glasgow | Wellcome Sanger Institute for the COVID-19 Genomics UK (COG-UK) Consortium | Harper VanSteenhouse, Yumi Kasai, David Gray, Carol Clugston, Anna Dominiczak and Alex Alderton, Roberto Amato, Jeffrey Barrett, Sonia Goncalves, Ewan Harrison, David K. Jackson, Ian Johnston, Dominic Kwiatkowski, Cordelia Langford, John Sillitoe on behalf of the Wellcome Sanger Institute COVID-19 Surveillance Team |
| EPI_ISL_2924911, EPI_ISL_2924912 | Lighthouse Lab in Alderley Park | Wellcome Sanger Institute for the COVID-19 Genomics UK (COG-UK) Consortium | Jacquelyn Wynn, Mairead Hyland, The Lighthouse Lab in Alderley Park and Alex Alderton, Roberto Amato, Jeffrey Barrett, Sonia Goncalves, Ewan Harrison, David K. Jackson, Ian Johnston, Dominic Kwiatkowski, Cordelia Langford, John Sillitoe on behalf of the Wellcome Sanger Institute COVID-19 Surveillance Team |
| EPI_ISL_2924913 | Lighthouse Lab in Glasgow | Wellcome Sanger Institute for the COVID-19 Genomics UK (COG-UK) Consortium | Harper VanSteenhouse, Yumi Kasai, David Gray, Carol Clugston, Anna Dominiczak and Alex Alderton, Roberto Amato, Jeffrey Barrett, Sonia Goncalves, Ewan Harrison, David K. Jackson, Ian Johnston, Dominic Kwiatkowski, Cordelia Langford, John Sillitoe on behalf of the Wellcome Sanger Institute COVID-19 Surveillance Team |
| EPI_ISL_2924914 | Lighthouse Lab in Milton Keynes | Wellcome Sanger Institute for the COVID-19 Genomics UK (COG-UK) Consortium | The Lighthouse Lab in Milton Keynes and Alex Alderton, Roberto Amato, Jeffrey Barrett, Sonia Goncalves, Ewan Harrison, David K. Jackson, Ian Johnston, Dominic Kwiatkowski, Cordelia Langford, John Sillitoe on behalf of the Wellcome Sanger Institute COVID-19 Surveillance Team |
| EPI_ISL_2924915, EPI_ISL_2924916 | Lighthouse Lab in Glasgow | Wellcome Sanger Institute for the COVID-19 Genomics UK (COG-UK) Consortium | Harper VanSteenhouse, Yumi Kasai, David Gray, Carol Clugston, Anna Dominiczak and Alex Alderton, Roberto Amato, Jeffrey Barrett, Sonia Goncalves, Ewan Harrison, David K. Jackson, Ian Johnston, Dominic Kwiatkowski, Cordelia Langford, John Sillitoe on behalf of the Wellcome Sanger Institute COVID-19 Surveillance Team |
| EPI_ISL_2924917 | Lighthouse Lab in Alderley Park | Wellcome Sanger Institute for the COVID-19 Genomics UK (COG-UK) Consortium | Jacquelyn Wynn, Mairead Hyland, The Lighthouse Lab in Alderley Park and Alex Alderton, Roberto Amato, Jeffrey Barrett, Sonia Goncalves, Ewan Harrison, David K. Jackson, Ian Johnston, Dominic Kwiatkowski, Cordelia Langford, John Sillitoe on behalf of the Wellcome Sanger Institute COVID-19 Surveillance Team |
| EPI_ISL_2924918 | Lighthouse Lab in Milton Keynes | Wellcome Sanger Institute for the COVID-19 Genomics UK (COG-UK) Consortium | The Lighthouse Lab in Milton Keynes and Alex Alderton, Roberto Amato, Jeffrey Barrett, Sonia Goncalves, Ewan Harrison, David K. Jackson, Ian Johnston, Dominic Kwiatkowski, Cordelia Langford, John Sillitoe on behalf of the Wellcome Sanger Institute COVID-19 Surveillance Team |
| EPI_ISL_2924919 | Lighthouse Lab in Glasgow | Wellcome Sanger Institute for the COVID-19 Genomics UK (COG-UK) Consortium | Harper VanSteenhouse, Yumi Kasai, David Gray, Carol Clugston, Anna Dominiczak and Alex Alderton, Roberto Amato, Jeffrey Barrett, Sonia Goncalves, Ewan Harrison, David K. Jackson, Ian Johnston, Dominic Kwiatkowski, Cordelia Langford, John Sillitoe on behalf of the Wellcome Sanger Institute COVID-19 Surveillance Team |
| EPI_ISL_2924920 | Lighthouse Lab in Alderley Park | Wellcome Sanger Institute for the COVID-19 Genomics UK (COG-UK) Consortium | Jacquelyn Wynn, Mairead Hyland, The Lighthouse Lab in Alderley Park and Alex Alderton, Roberto Amato, Jeffrey Barrett, Sonia Goncalves, Ewan Harrison, David K. Jackson, Ian Johnston, Dominic Kwiatkowski, Cordelia Langford, John Sillitoe on behalf of the Wellcome Sanger Institute COVID-19 Surveillance Team |
| EPI_ISL_2924921, EPI_ISL_2924922 | Lighthouse Lab in Glasgow | Wellcome Sanger Institute for the COVID-19 Genomics UK (COG-UK) Consortium | Harper VanSteenhouse, Yumi Kasai, David Gray, Carol Clugston, Anna Dominiczak and Alex Alderton, Roberto Amato, Jeffrey Barrett, Sonia Goncalves, Ewan Harrison, David K. Jackson, Ian Johnston, Dominic Kwiatkowski, Cordelia Langford, John Sillitoe on behalf of the Wellcome Sanger Institute COVID-19 Surveillance Team |
| EPI_ISL_2924923 | Lighthouse Lab in Alderley Park | Wellcome Sanger Institute for the COVID-19 Genomics UK (COG-UK) Consortium | Jacquelyn Wynn, Mairead Hyland, The Lighthouse Lab in Alderley Park and Alex Alderton, Roberto Amato, Jeffrey Barrett, Sonia Goncalves, Ewan Harrison, David K. Jackson, Ian Johnston, Dominic Kwiatkowski, Cordelia Langford, John Sillitoe on behalf of the Wellcome Sanger Institute COVID-19 Surveillance Team |
| EPI_ISL_2924924 | Lighthouse Lab in Glasgow | Wellcome Sanger Institute for the COVID-19 Genomics UK (COG-UK) Consortium | Harper VanSteenhouse, Yumi Kasai, David Gray, Carol Clugston, Anna Dominiczak and Alex Alderton, Roberto Amato, Jeffrey Barrett, Sonia Goncalves, Ewan Harrison, David K. Jackson, Ian Johnston, Dominic Kwiatkowski, Cordelia Langford, John Sillitoe on behalf of the Wellcome Sanger Institute COVID-19 Surveillance Team |
| EPI_ISL_2924925, EPI_ISL_2924926 | Lighthouse Lab in Milton Keynes | Wellcome Sanger Institute for the COVID-19 Genomics UK (COG-UK) Consortium | The Lighthouse Lab in Milton Keynes and Alex Alderton, Roberto Amato, Jeffrey Barrett, Sonia Goncalves, Ewan Harrison, David K. Jackson, Ian Johnston, Dominic Kwiatkowski, Cordelia Langford, John Sillitoe on behalf of the Wellcome Sanger Institute COVID-19 Surveillance Team |
| EPI_ISL_2924927 | Lighthouse Lab in Alderley Park | Wellcome Sanger Institute for the COVID-19 Genomics UK (COG-UK) Consortium | Jacquelyn Wynn, Mairead Hyland, The Lighthouse Lab in Alderley Park and Alex Alderton, Roberto Amato, Jeffrey Barrett, Sonia Goncalves, Ewan Harrison, David K. Jackson, Ian Johnston, Dominic Kwiatkowski, Cordelia Langford, John Sillitoe on behalf of the Wellcome Sanger Institute COVID-19 Surveillance Team |
| EPI_ISL_2924928 | Lighthouse Lab in Milton Keynes | Wellcome Sanger Institute for the COVID-19 Genomics UK (COG-UK) Consortium | The Lighthouse Lab in Milton Keynes and Alex Alderton, Roberto Amato, Jeffrey Barrett, Sonia Goncalves, Ewan Harrison, David K. Jackson, Ian Johnston, Dominic Kwiatkowski, Cordelia Langford, John Sillitoe on behalf of the Wellcome Sanger Institute COVID-19 Surveillance Team |
| EPI_ISL_2924929, EPI_ISL_2924930 | Lighthouse Lab in Glasgow | Wellcome Sanger Institute for the COVID-19 Genomics UK (COG-UK) Consortium | Harper VanSteenhouse, Yumi Kasai, David Gray, Carol Clugston, Anna Dominiczak and Alex Alderton, Roberto Amato, Jeffrey Barrett, Sonia Goncalves, Ewan Harrison, David K. Jackson, Ian Johnston, Dominic Kwiatkowski, Cordelia Langford, John Sillitoe on behalf of the Wellcome Sanger Institute COVID-19 Surveillance Team |
| EPI_ISL_2924931, EPI_ISL_2924932, EPI_ISL_2924933, EPI_ISL_2924934 | Lighthouse Lab in Milton Keynes | Wellcome Sanger Institute for the COVID-19 Genomics UK (COG-UK) Consortium | The Lighthouse Lab in Milton Keynes and Alex Alderton, Roberto Amato, Jeffrey Barrett, Sonia Goncalves, Ewan Harrison, David K. Jackson, Ian Johnston, Dominic Kwiatkowski, Cordelia Langford, John Sillitoe on behalf of the Wellcome Sanger Institute COVID-19 Surveillance Team |
| EPI_ISL_2924935 | Lighthouse Lab in Glasgow | Wellcome Sanger Institute for the COVID-19 Genomics UK (COG-UK) Consortium | Harper VanSteenhouse, Yumi Kasai, David Gray, Carol Clugston, Anna Dominiczak and Alex Alderton, Roberto Amato, Jeffrey Barrett, Sonia Goncalves, Ewan Harrison, David K. Jackson, Ian Johnston, Dominic Kwiatkowski, Cordelia Langford, John Sillitoe on behalf of the Wellcome Sanger Institute COVID-19 Surveillance Team |
| EPI_ISL_2924936 | Lighthouse Lab in Milton Keynes | Wellcome Sanger Institute for the COVID-19 Genomics UK (COG-UK) Consortium | The Lighthouse Lab in Milton Keynes and Alex Alderton, Roberto Amato, Jeffrey Barrett, Sonia Goncalves, Ewan Harrison, David K. Jackson, Ian Johnston, Dominic Kwiatkowski, Cordelia Langford, John Sillitoe on behalf of the Wellcome Sanger Institute COVID-19 Surveillance Team |
| EPI_ISL_2924937 | Lighthouse Lab in Alderley Park | Wellcome Sanger Institute for the COVID-19 Genomics UK (COG-UK) Consortium | Jacquelyn Wynn, Mairead Hyland, The Lighthouse Lab in Alderley Park and Alex Alderton, Roberto Amato, Jeffrey Barrett, Sonia Goncalves, Ewan Harrison, David K. Jackson, Ian Johnston, Dominic Kwiatkowski, Cordelia Langford, John Sillitoe on behalf of the Wellcome Sanger Institute COVID-19 Surveillance Team |
| EPI_ISL_2924938, EPI_ISL_2924939 | Lighthouse Lab in Glasgow | Wellcome Sanger Institute for the COVID-19 Genomics UK (COG-UK) Consortium | Harper VanSteenhouse, Yumi Kasai, David Gray, Carol Clugston, Anna Dominiczak and Alex Alderton, Roberto Amato, Jeffrey Barrett, Sonia Goncalves, Ewan Harrison, David K. Jackson, Ian Johnston, Dominic Kwiatkowski, Cordelia Langford, John Sillitoe on behalf of the Wellcome Sanger Institute COVID-19 Surveillance Team |
| EPI_ISL_2924940 | Lighthouse Lab in Alderley Park | Wellcome Sanger Institute for the COVID-19 Genomics UK (COG-UK) Consortium | Jacquelyn Wynn, Mairead Hyland, The Lighthouse Lab in Alderley Park and Alex Alderton, Roberto Amato, Jeffrey Barrett, Sonia Goncalves, Ewan Harrison, David K. Jackson, Ian Johnston, Dominic Kwiatkowski, Cordelia Langford, John Sillitoe on behalf of the Wellcome Sanger Institute COVID-19 Surveillance Team |
| EPI_ISL_2924941 | Lighthouse Lab in Glasgow | Wellcome Sanger Institute for the COVID-19 Genomics UK (COG-UK) Consortium | Harper VanSteenhouse, Yumi Kasai, David Gray, Carol Clugston, Anna Dominiczak and Alex Alderton, Roberto Amato, Jeffrey Barrett, Sonia Goncalves, Ewan Harrison, David K. Jackson, Ian Johnston, Dominic Kwiatkowski, Cordelia Langford, John Sillitoe on behalf of the Wellcome Sanger Institute COVID-19 Surveillance Team |
| EPI_ISL_2924942 | Lighthouse Lab in Alderley Park | Wellcome Sanger Institute for the COVID-19 Genomics UK (COG-UK) Consortium | Jacquelyn Wynn, Mairead Hyland, The Lighthouse Lab in Alderley Park and Alex Alderton, Roberto Amato, Jeffrey Barrett, Sonia Goncalves, Ewan Harrison, David K. Jackson, Ian Johnston, Dominic Kwiatkowski, Cordelia Langford, John Sillitoe on behalf of the Wellcome Sanger Institute COVID-19 Surveillance Team |
| EPI_ISL_2924943 | Lighthouse Lab in Glasgow | Wellcome Sanger Institute for the COVID-19 Genomics UK (COG-UK) Consortium | Harper VanSteenhouse, Yumi Kasai, David Gray, Carol Clugston, Anna Dominiczak and Alex Alderton, Roberto Amato, Jeffrey Barrett, Sonia Goncalves, Ewan Harrison, David K. Jackson, Ian Johnston, Dominic Kwiatkowski, Cordelia Langford, John Sillitoe on behalf of the Wellcome Sanger Institute COVID-19 Surveillance Team |
| EPI_ISL_2924944, EPI_ISL_2924945 | Lighthouse Lab in Alderley Park | Wellcome Sanger Institute for the COVID-19 Genomics UK (COG-UK) Consortium | Jacquelyn Wynn, Mairead Hyland, The Lighthouse Lab in Alderley Park and Alex Alderton, Roberto Amato, Jeffrey Barrett, Sonia Goncalves, Ewan Harrison, David K. Jackson, Ian Johnston, Dominic Kwiatkowski, Cordelia Langford, John Sillitoe on behalf of the Wellcome Sanger Institute COVID-19 Surveillance Team |
| EPI_ISL_2924946, EPI_ISL_2924947, EPI_ISL_2924948 | Lighthouse Lab in Milton Keynes | Wellcome Sanger Institute for the COVID-19 Genomics UK (COG-UK) Consortium | The Lighthouse Lab in Milton Keynes and Alex Alderton, Roberto Amato, Jeffrey Barrett, Sonia Goncalves, Ewan Harrison, David K. Jackson, Ian Johnston, Dominic Kwiatkowski, Cordelia Langford, John Sillitoe on behalf of the Wellcome Sanger Institute COVID-19 Surveillance Team |
| EPI_ISL_2924949 | Lighthouse Lab in Alderley Park | Wellcome Sanger Institute for the COVID-19 Genomics UK (COG-UK) Consortium | Jacquelyn Wynn, Mairead Hyland, The Lighthouse Lab in Alderley Park and Alex Alderton, Roberto Amato, Jeffrey Barrett, Sonia Goncalves, Ewan Harrison, David K. Jackson, Ian Johnston, Dominic Kwiatkowski, Cordelia Langford, John Sillitoe on behalf of the Wellcome Sanger Institute COVID-19 Surveillance Team |
| EPI_ISL_2924950 | Lighthouse Lab in Glasgow | Wellcome Sanger Institute for the COVID-19 Genomics UK (COG-UK) Consortium | Harper VanSteenhouse, Yumi Kasai, David Gray, Carol Clugston, Anna Dominiczak and Alex Alderton, Roberto Amato, Jeffrey Barrett, Sonia Goncalves, Ewan Harrison, David K. Jackson, Ian Johnston, Dominic Kwiatkowski, Cordelia Langford, John Sillitoe on behalf of the Wellcome Sanger Institute COVID-19 Surveillance Team |
| EPI_ISL_2924951 | Lighthouse Lab in Milton Keynes | Wellcome Sanger Institute for the COVID-19 Genomics UK (COG-UK) Consortium | The Lighthouse Lab in Milton Keynes and Alex Alderton, Roberto Amato, Jeffrey Barrett, Sonia Goncalves, Ewan Harrison, David K. Jackson, Ian Johnston, Dominic Kwiatkowski, Cordelia Langford, John Sillitoe on behalf of the Wellcome Sanger Institute COVID-19 Surveillance Team |
| EPI_ISL_2924952, EPI_ISL_2924953, EPI_ISL_2924954, EPI_ISL_2924955 | Lighthouse Lab in Glasgow | Wellcome Sanger Institute for the COVID-19 Genomics UK (COG-UK) Consortium | Harper VanSteenhouse, Yumi Kasai, David Gray, Carol Clugston, Anna Dominiczak and Alex Alderton, Roberto Amato, Jeffrey Barrett, Sonia Goncalves, Ewan Harrison, David K. Jackson, Ian Johnston, Dominic Kwiatkowski, Cordelia Langford, John Sillitoe on behalf of the Wellcome Sanger Institute COVID-19 Surveillance Team |
| EPI_ISL_2924956 | Lighthouse Lab in Alderley Park | Wellcome Sanger Institute for the COVID-19 Genomics UK (COG-UK) Consortium | Jacquelyn Wynn, Mairead Hyland, The Lighthouse Lab in Alderley Park and Alex Alderton, Roberto Amato, Jeffrey Barrett, Sonia Goncalves, Ewan Harrison, David K. Jackson, Ian Johnston, Dominic Kwiatkowski, Cordelia Langford, John Sillitoe on behalf of the Wellcome Sanger Institute COVID-19 Surveillance Team |
| EPI_ISL_2924957 | Lighthouse Lab in Milton Keynes | Wellcome Sanger Institute for the COVID-19 Genomics UK (COG-UK) Consortium | The Lighthouse Lab in Milton Keynes and Alex Alderton, Roberto Amato, Jeffrey Barrett, Sonia Goncalves, Ewan Harrison, David K. Jackson, Ian Johnston, Dominic Kwiatkowski, Cordelia Langford, John Sillitoe on behalf of the Wellcome Sanger Institute COVID-19 Surveillance Team |
| EPI_ISL_2924958, EPI_ISL_2924959 | Lighthouse Lab in Glasgow | Wellcome Sanger Institute for the COVID-19 Genomics UK (COG-UK) Consortium | Harper VanSteenhouse, Yumi Kasai, David Gray, Carol Clugston, Anna Dominiczak and Alex Alderton, Roberto Amato, Jeffrey Barrett, Sonia Goncalves, Ewan Harrison, David K. Jackson, Ian Johnston, Dominic Kwiatkowski, Cordelia Langford, John Sillitoe on behalf of the Wellcome Sanger Institute COVID-19 Surveillance Team |
| EPI_ISL_2924960 | Lighthouse Lab in Alderley Park | Wellcome Sanger Institute for the COVID-19 Genomics UK (COG-UK) Consortium | Jacquelyn Wynn, Mairead Hyland, The Lighthouse Lab in Alderley Park and Alex Alderton, Roberto Amato, Jeffrey Barrett, Sonia Goncalves, Ewan Harrison, David K. Jackson, Ian Johnston, Dominic Kwiatkowski, Cordelia Langford, John Sillitoe on behalf of the Wellcome Sanger Institute COVID-19 Surveillance Team |
| EPI_ISL_2924961 | Lighthouse Lab in Glasgow | Wellcome Sanger Institute for the COVID-19 Genomics UK (COG-UK) Consortium | Harper VanSteenhouse, Yumi Kasai, David Gray, Carol Clugston, Anna Dominiczak and Alex Alderton, Roberto Amato, Jeffrey Barrett, Sonia Goncalves, Ewan Harrison, David K. Jackson, Ian Johnston, Dominic Kwiatkowski, Cordelia Langford, John Sillitoe on behalf of the Wellcome Sanger Institute COVID-19 Surveillance Team |
| EPI_ISL_2924962 | Lighthouse Lab in Alderley Park | Wellcome Sanger Institute for the COVID-19 Genomics UK (COG-UK) Consortium | Jacquelyn Wynn, Mairead Hyland, The Lighthouse Lab in Alderley Park and Alex Alderton, Roberto Amato, Jeffrey Barrett, Sonia Goncalves, Ewan Harrison, David K. Jackson, Ian Johnston, Dominic Kwiatkowski, Cordelia Langford, John Sillitoe on behalf of the Wellcome Sanger Institute COVID-19 Surveillance Team |
| EPI_ISL_2924963 | Lighthouse Lab in Glasgow | Wellcome Sanger Institute for the COVID-19 Genomics UK (COG-UK) Consortium | Harper VanSteenhouse, Yumi Kasai, David Gray, Carol Clugston, Anna Dominiczak and Alex Alderton, Roberto Amato, Jeffrey Barrett, Sonia Goncalves, Ewan Harrison, David K. Jackson, Ian Johnston, Dominic Kwiatkowski, Cordelia Langford, John Sillitoe on behalf of the Wellcome Sanger Institute COVID-19 Surveillance Team |
| EPI_ISL_2924964 | Lighthouse Lab in Alderley Park | Wellcome Sanger Institute for the COVID-19 Genomics UK (COG-UK) Consortium | Jacquelyn Wynn, Mairead Hyland, The Lighthouse Lab in Alderley Park and Alex Alderton, Roberto Amato, Jeffrey Barrett, Sonia Goncalves, Ewan Harrison, David K. Jackson, Ian Johnston, Dominic Kwiatkowski, Cordelia Langford, John Sillitoe on behalf of the Wellcome Sanger Institute COVID-19 Surveillance Team |
| EPI_ISL_2924965, EPI_ISL_2924966 | Lighthouse Lab in Milton Keynes | Wellcome Sanger Institute for the COVID-19 Genomics UK (COG-UK) Consortium | The Lighthouse Lab in Milton Keynes and Alex Alderton, Roberto Amato, Jeffrey Barrett, Sonia Goncalves, Ewan Harrison, David K. Jackson, Ian Johnston, Dominic Kwiatkowski, Cordelia Langford, John Sillitoe on behalf of the Wellcome Sanger Institute COVID-19 Surveillance Team |
| EPI_ISL_2924967, | Lighthouse Lab in Glasgow | Wellcome Sanger Institute for the COVID-19 Genomics UK (COG-UK) | Harper VanSteenhouse, Yumi Kasai, David Gray, Carol Clugston, Anna Dominiczak and Alex Alderton, Roberto Amato, Jeffrey Barrett, Sonia Goncalves, Ewan Harrison, David K. Jackson, Ian Johnston, Dominic Kwiatkowski, Cordelia Langford, |

| | | | |
|---|---|---|---|
| EPI_ISL_2924968 | | Consortium | John Sillitoe on behalf of the Wellcome Sanger Institute COVID-19 Surveillance Team |
| EPI_ISL_2924969 | Lighthouse Lab in Milton Keynes | Wellcome Sanger Institute for the COVID-19 Genomics UK (COG-UK) Consortium | The Lighthouse Lab in Milton Keynes and Alex Alderton, Roberto Amato, Jeffrey Barrett, Sonia Goncalves, Ewan Harrison, David K. Jackson, Ian Johnston, Dominic Kwiatkowski, Cordelia Langford, John Sillitoe on behalf of the Wellcome Sanger Institute COVID-19 Surveillance Team |
| EPI_ISL_2924970 | Lighthouse Lab in Glasgow | Wellcome Sanger Institute for the COVID-19 Genomics UK (COG-UK) Consortium | Harper VanSteenhouse, Yumi Kasai, David Gray, Carol Clugston, Anna Dominiczak and Alex Alderton, Roberto Amato, Jeffrey Barrett, Sonia Goncalves, Ewan Harrison, David K. Jackson, Ian Johnston, Dominic Kwiatkowski, Cordelia Langford, John Sillitoe on behalf of the Wellcome Sanger Institute COVID-19 Surveillance Team |
| EPI_ISL_2924971 | Lighthouse Lab in Alderley Park | Wellcome Sanger Institute for the COVID-19 Genomics UK (COG-UK) Consortium | Jacquelyn Wynn, Mairead Hyland, The Lighthouse Lab in Alderley Park and Alex Alderton, Roberto Amato, Jeffrey Barrett, Sonia Goncalves, Ewan Harrison, David K. Jackson, Ian Johnston, Dominic Kwiatkowski, Cordelia Langford, John Sillitoe on behalf of the Wellcome Sanger Institute COVID-19 Surveillance Team |
| EPI_ISL_2924972, EPI_ISL_2924973, EPI_ISL_2924974 | Lighthouse Lab in Milton Keynes | Wellcome Sanger Institute for the COVID-19 Genomics UK (COG-UK) Consortium | The Lighthouse Lab in Milton Keynes and Alex Alderton, Roberto Amato, Jeffrey Barrett, Sonia Goncalves, Ewan Harrison, David K. Jackson, Ian Johnston, Dominic Kwiatkowski, Cordelia Langford, John Sillitoe on behalf of the Wellcome Sanger Institute COVID-19 Surveillance Team |
| EPI_ISL_2924975, EPI_ISL_2924976 | Lighthouse Lab in Glasgow | Wellcome Sanger Institute for the COVID-19 Genomics UK (COG-UK) Consortium | Harper VanSteenhouse, Yumi Kasai, David Gray, Carol Clugston, Anna Dominiczak and Alex Alderton, Roberto Amato, Jeffrey Barrett, Sonia Goncalves, Ewan Harrison, David K. Jackson, Ian Johnston, Dominic Kwiatkowski, Cordelia Langford, John Sillitoe on behalf of the Wellcome Sanger Institute COVID-19 Surveillance Team |
| EPI_ISL_2924977 | Lighthouse Lab in Milton Keynes | Wellcome Sanger Institute for the COVID-19 Genomics UK (COG-UK) Consortium | The Lighthouse Lab in Milton Keynes and Alex Alderton, Roberto Amato, Jeffrey Barrett, Sonia Goncalves, Ewan Harrison, David K. Jackson, Ian Johnston, Dominic Kwiatkowski, Cordelia Langford, John Sillitoe on behalf of the Wellcome Sanger Institute COVID-19 Surveillance Team |
| EPI_ISL_2924978, EPI_ISL_2924979 | Lighthouse Lab in Alderley Park | Wellcome Sanger Institute for the COVID-19 Genomics UK (COG-UK) Consortium | Jacquelyn Wynn, Mairead Hyland, The Lighthouse Lab in Alderley Park and Alex Alderton, Roberto Amato, Jeffrey Barrett, Sonia Goncalves, Ewan Harrison, David K. Jackson, Ian Johnston, Dominic Kwiatkowski, Cordelia Langford, John Sillitoe on behalf of the Wellcome Sanger Institute COVID-19 Surveillance Team |
| EPI_ISL_2924980, EPI_ISL_2924981, EPI_ISL_2924982 | Lighthouse Lab in Glasgow | Wellcome Sanger Institute for the COVID-19 Genomics UK (COG-UK) Consortium | Harper VanSteenhouse, Yumi Kasai, David Gray, Carol Clugston, Anna Dominiczak and Alex Alderton, Roberto Amato, Jeffrey Barrett, Sonia Goncalves, Ewan Harrison, David K. Jackson, Ian Johnston, Dominic Kwiatkowski, Cordelia Langford, John Sillitoe on behalf of the Wellcome Sanger Institute COVID-19 Surveillance Team |
| EPI_ISL_2924983, EPI_ISL_2924984, EPI_ISL_2924985 | Lighthouse Lab in Alderley Park | Wellcome Sanger Institute for the COVID-19 Genomics UK (COG-UK) Consortium | Jacquelyn Wynn, Mairead Hyland, The Lighthouse Lab in Alderley Park and Alex Alderton, Roberto Amato, Jeffrey Barrett, Sonia Goncalves, Ewan Harrison, David K. Jackson, Ian Johnston, Dominic Kwiatkowski, Cordelia Langford, John Sillitoe on behalf of the Wellcome Sanger Institute COVID-19 Surveillance Team |
| EPI_ISL_2924986, EPI_ISL_2924987 | Lighthouse Lab in Glasgow | Wellcome Sanger Institute for the COVID-19 Genomics UK (COG-UK) Consortium | Harper VanSteenhouse, Yumi Kasai, David Gray, Carol Clugston, Anna Dominiczak and Alex Alderton, Roberto Amato, Jeffrey Barrett, Sonia Goncalves, Ewan Harrison, David K. Jackson, Ian Johnston, Dominic Kwiatkowski, Cordelia Langford, John Sillitoe on behalf of the Wellcome Sanger Institute COVID-19 Surveillance Team |
| EPI_ISL_2924988, EPI_ISL_2924989, EPI_ISL_2924990, EPI_ISL_2924991, EPI_ISL_2924992 | Lighthouse Lab in Alderley Park | Wellcome Sanger Institute for the COVID-19 Genomics UK (COG-UK) Consortium | Jacquelyn Wynn, Mairead Hyland, The Lighthouse Lab in Alderley Park and Alex Alderton, Roberto Amato, Jeffrey Barrett, Sonia Goncalves, Ewan Harrison, David K. Jackson, Ian Johnston, Dominic Kwiatkowski, Cordelia Langford, John Sillitoe on behalf of the Wellcome Sanger Institute COVID-19 Surveillance Team |
| EPI_ISL_2924993 | Lighthouse Lab in Milton Keynes | Wellcome Sanger Institute for the COVID-19 Genomics UK (COG-UK) Consortium | The Lighthouse Lab in Milton Keynes and Alex Alderton, Roberto Amato, Jeffrey Barrett, Sonia Goncalves, Ewan Harrison, David K. Jackson, Ian Johnston, Dominic Kwiatkowski, Cordelia Langford, John Sillitoe on behalf of the Wellcome Sanger Institute COVID-19 Surveillance Team |
| EPI_ISL_2924994 | Lighthouse Lab in Alderley Park | Wellcome Sanger Institute for the COVID-19 Genomics UK (COG-UK) Consortium | Jacquelyn Wynn, Mairead Hyland, The Lighthouse Lab in Alderley Park and Alex Alderton, Roberto Amato, Jeffrey Barrett, Sonia Goncalves, Ewan Harrison, David K. Jackson, Ian Johnston, Dominic Kwiatkowski, Cordelia Langford, John Sillitoe on behalf of the Wellcome Sanger Institute COVID-19 Surveillance Team |
| EPI_ISL_2924995 | Lighthouse Lab in Glasgow | Wellcome Sanger Institute for the COVID-19 Genomics UK (COG-UK) Consortium | Harper VanSteenhouse, Yumi Kasai, David Gray, Carol Clugston, Anna Dominiczak and Alex Alderton, Roberto Amato, Jeffrey Barrett, Sonia Goncalves, Ewan Harrison, David K. Jackson, Ian Johnston, Dominic Kwiatkowski, Cordelia Langford, John Sillitoe on behalf of the Wellcome Sanger Institute COVID-19 Surveillance Team |
| EPI_ISL_2924996 | Lighthouse Lab in Alderley Park | Wellcome Sanger Institute for the COVID-19 Genomics UK (COG-UK) Consortium | Jacquelyn Wynn, Mairead Hyland, The Lighthouse Lab in Alderley Park and Alex Alderton, Roberto Amato, Jeffrey Barrett, Sonia Goncalves, Ewan Harrison, David K. Jackson, Ian Johnston, Dominic Kwiatkowski, Cordelia Langford, John Sillitoe on behalf of the Wellcome Sanger Institute COVID-19 Surveillance Team |
| EPI_ISL_2924997 | Lighthouse Lab in Glasgow | Wellcome Sanger Institute for the COVID-19 Genomics UK (COG-UK) Consortium | Harper VanSteenhouse, Yumi Kasai, David Gray, Carol Clugston, Anna Dominiczak and Alex Alderton, Roberto Amato, Jeffrey Barrett, Sonia Goncalves, Ewan Harrison, David K. Jackson, Ian Johnston, Dominic Kwiatkowski, Cordelia Langford, John Sillitoe on behalf of the Wellcome Sanger Institute COVID-19 Surveillance Team |
| EPI_ISL_2924998, EPI_ISL_2924999 | Lighthouse Lab in Milton Keynes | Wellcome Sanger Institute for the COVID-19 Genomics UK (COG-UK) Consortium | The Lighthouse Lab in Milton Keynes and Alex Alderton, Roberto Amato, Jeffrey Barrett, Sonia Goncalves, Ewan Harrison, David K. Jackson, Ian Johnston, Dominic Kwiatkowski, Cordelia Langford, John Sillitoe on behalf of the Wellcome Sanger Institute COVID-19 Surveillance Team |
| EPI_ISL_2925000 | Lighthouse Lab in Glasgow | Wellcome Sanger Institute for the COVID-19 Genomics UK (COG-UK) Consortium | Harper VanSteenhouse, Yumi Kasai, David Gray, Carol Clugston, Anna Dominiczak and Alex Alderton, Roberto Amato, Jeffrey Barrett, Sonia Goncalves, Ewan Harrison, David K. Jackson, Ian Johnston, Dominic Kwiatkowski, Cordelia Langford, John Sillitoe on behalf of the Wellcome Sanger Institute COVID-19 Surveillance Team |
| EPI_ISL_2925001, EPI_ISL_2925002 | Lighthouse Lab in Alderley Park | Wellcome Sanger Institute for the COVID-19 Genomics UK (COG-UK) Consortium | Jacquelyn Wynn, Mairead Hyland, The Lighthouse Lab in Alderley Park and Alex Alderton, Roberto Amato, Jeffrey Barrett, Sonia Goncalves, Ewan Harrison, David K. Jackson, Ian Johnston, Dominic Kwiatkowski, Cordelia Langford, John Sillitoe on behalf of the Wellcome Sanger Institute COVID-19 Surveillance Team |
| EPI_ISL_2925003 | Lighthouse Lab in Glasgow | Wellcome Sanger Institute for the COVID-19 Genomics UK (COG-UK) Consortium | Harper VanSteenhouse, Yumi Kasai, David Gray, Carol Clugston, Anna Dominiczak and Alex Alderton, Roberto Amato, Jeffrey Barrett, Sonia Goncalves, Ewan Harrison, David K. Jackson, Ian Johnston, Dominic Kwiatkowski, Cordelia Langford, John Sillitoe on behalf of the Wellcome Sanger Institute COVID-19 Surveillance Team |
| EPI_ISL_2925004 | Lighthouse Lab in Alderley Park | Wellcome Sanger Institute for the COVID-19 Genomics UK (COG-UK) Consortium | Jacquelyn Wynn, Mairead Hyland, The Lighthouse Lab in Alderley Park and Alex Alderton, Roberto Amato, Jeffrey Barrett, Sonia Goncalves, Ewan Harrison, David K. Jackson, Ian Johnston, Dominic Kwiatkowski, Cordelia Langford, John Sillitoe on behalf of the Wellcome Sanger Institute COVID-19 Surveillance Team |
| EPI_ISL_2925005, EPI_ISL_2925006 | Lighthouse Lab in Glasgow | Wellcome Sanger Institute for the COVID-19 Genomics UK (COG-UK) Consortium | Harper VanSteenhouse, Yumi Kasai, David Gray, Carol Clugston, Anna Dominiczak and Alex Alderton, Roberto Amato, Jeffrey Barrett, Sonia Goncalves, Ewan Harrison, David K. Jackson, Ian Johnston, Dominic Kwiatkowski, Cordelia Langford, John Sillitoe on behalf of the Wellcome Sanger Institute COVID-19 Surveillance Team |
| EPI_ISL_2925007 | Lighthouse Lab in Milton Keynes | Wellcome Sanger Institute for the COVID-19 Genomics UK (COG-UK) Consortium | The Lighthouse Lab in Milton Keynes and Alex Alderton, Roberto Amato, Jeffrey Barrett, Sonia Goncalves, Ewan Harrison, David K. Jackson, Ian Johnston, Dominic Kwiatkowski, Cordelia Langford, John Sillitoe on behalf of the Wellcome Sanger Institute COVID-19 Surveillance Team |
| EPI_ISL_2925008, EPI_ISL_2925009 | Lighthouse Lab in Glasgow | Wellcome Sanger Institute for the COVID-19 Genomics UK (COG-UK) Consortium | Harper VanSteenhouse, Yumi Kasai, David Gray, Carol Clugston, Anna Dominiczak and Alex Alderton, Roberto Amato, Jeffrey Barrett, Sonia Goncalves, Ewan Harrison, David K. Jackson, Ian Johnston, Dominic Kwiatkowski, Cordelia Langford, John Sillitoe on behalf of the Wellcome Sanger Institute COVID-19 Surveillance Team |
| EPI_ISL_2925010 | Lighthouse Lab in Milton Keynes | Wellcome Sanger Institute for the COVID-19 Genomics UK (COG-UK) Consortium | The Lighthouse Lab in Milton Keynes and Alex Alderton, Roberto Amato, Jeffrey Barrett, Sonia Goncalves, Ewan Harrison, David K. Jackson, Ian Johnston, Dominic Kwiatkowski, Cordelia Langford, John Sillitoe on behalf of the Wellcome Sanger Institute COVID-19 Surveillance Team |
| EPI_ISL_2925011 | Lighthouse Lab in Alderley Park | Wellcome Sanger Institute for the COVID-19 Genomics UK (COG-UK) Consortium | Jacquelyn Wynn, Mairead Hyland, The Lighthouse Lab in Alderley Park and Alex Alderton, Roberto Amato, Jeffrey Barrett, Sonia Goncalves, Ewan Harrison, David K. Jackson, Ian Johnston, Dominic Kwiatkowski, Cordelia Langford, John Sillitoe on behalf of the Wellcome Sanger Institute COVID-19 Surveillance Team |
| EPI_ISL_2925012 | Lighthouse Lab in Milton Keynes | Wellcome Sanger Institute for the COVID-19 Genomics UK (COG-UK) Consortium | The Lighthouse Lab in Milton Keynes and Alex Alderton, Roberto Amato, Jeffrey Barrett, Sonia Goncalves, Ewan Harrison, David K. Jackson, Ian Johnston, Dominic Kwiatkowski, Cordelia Langford, John Sillitoe on behalf of the Wellcome Sanger Institute COVID-19 Surveillance Team |
| EPI_ISL_2925013 | Lighthouse Lab in Glasgow | Wellcome Sanger Institute for the COVID-19 Genomics UK (COG-UK) Consortium | Harper VanSteenhouse, Yumi Kasai, David Gray, Carol Clugston, Anna Dominiczak and Alex Alderton, Roberto Amato, Jeffrey Barrett, Sonia Goncalves, Ewan Harrison, David K. Jackson, Ian Johnston, Dominic Kwiatkowski, Cordelia Langford, John Sillitoe on behalf of the Wellcome Sanger Institute COVID-19 Surveillance Team |
| EPI_ISL_2925014, EPI_ISL_2925015 | Lighthouse Lab in Alderley Park | Wellcome Sanger Institute for the COVID-19 Genomics UK (COG-UK) Consortium | Jacquelyn Wynn, Mairead Hyland, The Lighthouse Lab in Alderley Park and Alex Alderton, Roberto Amato, Jeffrey Barrett, Sonia Goncalves, Ewan Harrison, David K. Jackson, Ian Johnston, Dominic Kwiatkowski, Cordelia Langford, John Sillitoe on behalf of the Wellcome Sanger Institute COVID-19 Surveillance Team |
| EPI_ISL_2925016 | Lighthouse Lab in Glasgow | Wellcome Sanger Institute for the COVID-19 Genomics UK (COG-UK) Consortium | Harper VanSteenhouse, Yumi Kasai, David Gray, Carol Clugston, Anna Dominiczak and Alex Alderton, Roberto Amato, Jeffrey Barrett, Sonia Goncalves, Ewan Harrison, David K. Jackson, Ian Johnston, Dominic Kwiatkowski, Cordelia Langford, John Sillitoe on behalf of the Wellcome Sanger Institute COVID-19 Surveillance Team |
| EPI_ISL_2925017 | Lighthouse Lab in Alderley Park | Wellcome Sanger Institute for the COVID-19 Genomics UK (COG-UK) Consortium | Jacquelyn Wynn, Mairead Hyland, The Lighthouse Lab in Alderley Park and Alex Alderton, Roberto Amato, Jeffrey Barrett, Sonia Goncalves, Ewan Harrison, David K. Jackson, Ian Johnston, Dominic Kwiatkowski, Cordelia Langford, John Sillitoe on behalf of the Wellcome Sanger Institute COVID-19 Surveillance Team |
| EPI_ISL_2925018 | Lighthouse Lab in Milton Keynes | Wellcome Sanger Institute for the COVID-19 Genomics UK (COG-UK) Consortium | The Lighthouse Lab in Milton Keynes and Alex Alderton, Roberto Amato, Jeffrey Barrett, Sonia Goncalves, Ewan Harrison, David K. Jackson, Ian Johnston, Dominic Kwiatkowski, Cordelia Langford, John Sillitoe on behalf of the Wellcome Sanger Institute COVID-19 Surveillance Team |
| EPI_ISL_2925019 | Lighthouse Lab in Alderley Park | Wellcome Sanger Institute for the COVID-19 Genomics UK (COG-UK) Consortium | Jacquelyn Wynn, Mairead Hyland, The Lighthouse Lab in Alderley Park and Alex Alderton, Roberto Amato, Jeffrey Barrett, Sonia Goncalves, Ewan Harrison, David K. Jackson, Ian Johnston, Dominic Kwiatkowski, Cordelia Langford, John Sillitoe on behalf of the Wellcome Sanger Institute COVID-19 Surveillance Team |
| EPI_ISL_2925020 | Lighthouse Lab in Milton Keynes | Wellcome Sanger Institute for the COVID-19 Genomics UK (COG-UK) Consortium | The Lighthouse Lab in Milton Keynes and Alex Alderton, Roberto Amato, Jeffrey Barrett, Sonia Goncalves, Ewan Harrison, David K. Jackson, Ian Johnston, Dominic Kwiatkowski, Cordelia Langford, John Sillitoe on behalf of the Wellcome Sanger Institute COVID-19 Surveillance Team |
| EPI_ISL_2925021 | Lighthouse Lab in Glasgow | Wellcome Sanger Institute for the COVID-19 Genomics UK (COG-UK) Consortium | Harper VanSteenhouse, Yumi Kasai, David Gray, Carol Clugston, Anna Dominiczak and Alex Alderton, Roberto Amato, Jeffrey Barrett, Sonia Goncalves, Ewan Harrison, David K. Jackson, Ian Johnston, Dominic Kwiatkowski, Cordelia Langford, John Sillitoe on behalf of the Wellcome Sanger Institute COVID-19 Surveillance Team |
| EPI_ISL_2925022 | Lighthouse Lab in Milton Keynes | Wellcome Sanger Institute for the COVID-19 Genomics UK (COG-UK) Consortium | The Lighthouse Lab in Milton Keynes and Alex Alderton, Roberto Amato, Jeffrey Barrett, Sonia Goncalves, Ewan Harrison, David K. Jackson, Ian Johnston, Dominic Kwiatkowski, Cordelia Langford, John Sillitoe on behalf of the Wellcome Sanger Institute COVID-19 Surveillance Team |
| EPI_ISL_2925023 | Lighthouse Lab in Alderley Park | Wellcome Sanger Institute for the COVID-19 Genomics UK (COG-UK) Consortium | Jacquelyn Wynn, Mairead Hyland, The Lighthouse Lab in Alderley Park and Alex Alderton, Roberto Amato, Jeffrey Barrett, Sonia Goncalves, Ewan Harrison, David K. Jackson, Ian Johnston, Dominic Kwiatkowski, Cordelia Langford, John Sillitoe on behalf of the Wellcome Sanger Institute COVID-19 Surveillance Team |
| EPI_ISL_2925024, EPI_ISL_2925025, EPI_ISL_2925026 | Lighthouse Lab in Milton Keynes | Wellcome Sanger Institute for the COVID-19 Genomics UK (COG-UK) Consortium | The Lighthouse Lab in Milton Keynes and Alex Alderton, Roberto Amato, Jeffrey Barrett, Sonia Goncalves, Ewan Harrison, David K. Jackson, Ian Johnston, Dominic Kwiatkowski, Cordelia Langford, John Sillitoe on behalf of the Wellcome Sanger Institute COVID-19 Surveillance Team |
| EPI_ISL_2925027 | Lighthouse Lab in Alderley Park | Wellcome Sanger Institute for the COVID-19 Genomics UK (COG-UK) Consortium | Jacquelyn Wynn, Mairead Hyland, The Lighthouse Lab in Alderley Park and Alex Alderton, Roberto Amato, Jeffrey Barrett, Sonia Goncalves, Ewan Harrison, David K. Jackson, Ian Johnston, Dominic Kwiatkowski, Cordelia Langford, John Sillitoe on behalf of the Wellcome Sanger Institute COVID-19 Surveillance Team |
| EPI_ISL_2925028 | Lighthouse Lab in Glasgow | Wellcome Sanger Institute for the COVID-19 Genomics UK (COG-UK) Consortium | Harper VanSteenhouse, Yumi Kasai, David Gray, Carol Clugston, Anna Dominiczak and Alex Alderton, Roberto Amato, Jeffrey Barrett, Sonia Goncalves, Ewan Harrison, David K. Jackson, Ian Johnston, Dominic Kwiatkowski, Cordelia Langford, John Sillitoe on behalf of the Wellcome Sanger Institute COVID-19 Surveillance Team |
| EPI_ISL_2925029, EPI_ISL_2925030 | Lighthouse Lab in Milton Keynes | Wellcome Sanger Institute for the COVID-19 Genomics UK (COG-UK) Consortium | The Lighthouse Lab in Milton Keynes and Alex Alderton, Roberto Amato, Jeffrey Barrett, Sonia Goncalves, Ewan Harrison, David K. Jackson, Ian Johnston, Dominic Kwiatkowski, Cordelia Langford, John Sillitoe on behalf of the Wellcome Sanger Institute COVID-19 Surveillance Team |
| EPI_ISL_2925031 | Lighthouse Lab in Alderley Park | Wellcome Sanger Institute for the COVID-19 Genomics UK (COG-UK) | Jacquelyn Wynn, Mairead Hyland, The Lighthouse Lab in Alderley Park and Alex Alderton, Roberto Amato, Jeffrey Barrett, Sonia Goncalves, Ewan Harrison, David K. Jackson, Ian Johnston, Dominic Kwiatkowski, Cordelia Langford, John Sillitoe on |

| | | | |
|---|---|---|---|
| | | Consortium | behalf of the Wellcome Sanger Institute COVID-19 Surveillance Team |
| EPI_ISL_2925032 | Lighthouse Lab in Glasgow | Wellcome Sanger Institute for the COVID-19 Genomics UK (COG-UK) Consortium | Harper VanSteenhouse, Yumi Kasai, David Gray, Carol Clugston, Anna Dominiczak and Alex Alderton, Roberto Amato, Jeffrey Barrett, Sonia Goncalves, Ewan Harrison, David K. Jackson, Ian Johnston, Dominic Kwiatkowski, Cordelia Langford, John Sillitoe on behalf of the Wellcome Sanger Institute COVID-19 Surveillance Team |
| EPI_ISL_2925033 | Lighthouse Lab in Alderley Park | Wellcome Sanger Institute for the COVID-19 Genomics UK (COG-UK) Consortium | Jacquelyn Wynn, Mairead Hyland, The Lighthouse Lab in Alderley Park and Alex Alderton, Roberto Amato, Jeffrey Barrett, Sonia Goncalves, Ewan Harrison, David K. Jackson, Ian Johnston, Dominic Kwiatkowski, Cordelia Langford, John Sillitoe on behalf of the Wellcome Sanger Institute COVID-19 Surveillance Team |
| EPI_ISL_2925034 | Lighthouse Lab in Glasgow | Wellcome Sanger Institute for the COVID-19 Genomics UK (COG-UK) Consortium | Harper VanSteenhouse, Yumi Kasai, David Gray, Carol Clugston, Anna Dominiczak and Alex Alderton, Roberto Amato, Jeffrey Barrett, Sonia Goncalves, Ewan Harrison, David K. Jackson, Ian Johnston, Dominic Kwiatkowski, Cordelia Langford, John Sillitoe on behalf of the Wellcome Sanger Institute COVID-19 Surveillance Team |
| EPI_ISL_2925035, EPI_ISL_2925036, EPI_ISL_2925037, EPI_ISL_2925038 | Lighthouse Lab in Alderley Park | Wellcome Sanger Institute for the COVID-19 Genomics UK (COG-UK) Consortium | Jacquelyn Wynn, Mairead Hyland, The Lighthouse Lab in Alderley Park and Alex Alderton, Roberto Amato, Jeffrey Barrett, Sonia Goncalves, Ewan Harrison, David K. Jackson, Ian Johnston, Dominic Kwiatkowski, Cordelia Langford, John Sillitoe on behalf of the Wellcome Sanger Institute COVID-19 Surveillance Team |
| EPI_ISL_2925039, EPI_ISL_2925040 | Lighthouse Lab in Milton Keynes | Wellcome Sanger Institute for the COVID-19 Genomics UK (COG-UK) Consortium | The Lighthouse Lab in Milton Keynes and Alex Alderton, Roberto Amato, Jeffrey Barrett, Sonia Goncalves, Ewan Harrison, David K. Jackson, Ian Johnston, Dominic Kwiatkowski, Cordelia Langford, John Sillitoe on behalf of the Wellcome Sanger Institute COVID-19 Surveillance Team |
| EPI_ISL_2925041 | Lighthouse Lab in Alderley Park | Wellcome Sanger Institute for the COVID-19 Genomics UK (COG-UK) Consortium | Jacquelyn Wynn, Mairead Hyland, The Lighthouse Lab in Alderley Park and Alex Alderton, Roberto Amato, Jeffrey Barrett, Sonia Goncalves, Ewan Harrison, David K. Jackson, Ian Johnston, Dominic Kwiatkowski, Cordelia Langford, John Sillitoe on behalf of the Wellcome Sanger Institute COVID-19 Surveillance Team |
| EPI_ISL_2925042, EPI_ISL_2925043, EPI_ISL_2925044 | Lighthouse Lab in Milton Keynes | Wellcome Sanger Institute for the COVID-19 Genomics UK (COG-UK) Consortium | The Lighthouse Lab in Milton Keynes and Alex Alderton, Roberto Amato, Jeffrey Barrett, Sonia Goncalves, Ewan Harrison, David K. Jackson, Ian Johnston, Dominic Kwiatkowski, Cordelia Langford, John Sillitoe on behalf of the Wellcome Sanger Institute COVID-19 Surveillance Team |
| EPI_ISL_2925045 | Lighthouse Lab in Glasgow | Wellcome Sanger Institute for the COVID-19 Genomics UK (COG-UK) Consortium | Harper VanSteenhouse, Yumi Kasai, David Gray, Carol Clugston, Anna Dominiczak and Alex Alderton, Roberto Amato, Jeffrey Barrett, Sonia Goncalves, Ewan Harrison, David K. Jackson, Ian Johnston, Dominic Kwiatkowski, Cordelia Langford, John Sillitoe on behalf of the Wellcome Sanger Institute COVID-19 Surveillance Team |
| EPI_ISL_2925046 | Lighthouse Lab in Milton Keynes | Wellcome Sanger Institute for the COVID-19 Genomics UK (COG-UK) Consortium | The Lighthouse Lab in Milton Keynes and Alex Alderton, Roberto Amato, Jeffrey Barrett, Sonia Goncalves, Ewan Harrison, David K. Jackson, Ian Johnston, Dominic Kwiatkowski, Cordelia Langford, John Sillitoe on behalf of the Wellcome Sanger Institute COVID-19 Surveillance Team |
| EPI_ISL_2925047, EPI_ISL_2925048, EPI_ISL_2925049 | Lighthouse Lab in Alderley Park | Wellcome Sanger Institute for the COVID-19 Genomics UK (COG-UK) Consortium | Jacquelyn Wynn, Mairead Hyland, The Lighthouse Lab in Alderley Park and Alex Alderton, Roberto Amato, Jeffrey Barrett, Sonia Goncalves, Ewan Harrison, David K. Jackson, Ian Johnston, Dominic Kwiatkowski, Cordelia Langford, John Sillitoe on behalf of the Wellcome Sanger Institute COVID-19 Surveillance Team |
| EPI_ISL_2925050 | Lighthouse Lab in Milton Keynes | Wellcome Sanger Institute for the COVID-19 Genomics UK (COG-UK) Consortium | The Lighthouse Lab in Milton Keynes and Alex Alderton, Roberto Amato, Jeffrey Barrett, Sonia Goncalves, Ewan Harrison, David K. Jackson, Ian Johnston, Dominic Kwiatkowski, Cordelia Langford, John Sillitoe on behalf of the Wellcome Sanger Institute COVID-19 Surveillance Team |
| EPI_ISL_2925051, EPI_ISL_2925052, EPI_ISL_2925053 | Lighthouse Lab in Alderley Park | Wellcome Sanger Institute for the COVID-19 Genomics UK (COG-UK) Consortium | Jacquelyn Wynn, Mairead Hyland, The Lighthouse Lab in Alderley Park and Alex Alderton, Roberto Amato, Jeffrey Barrett, Sonia Goncalves, Ewan Harrison, David K. Jackson, Ian Johnston, Dominic Kwiatkowski, Cordelia Langford, John Sillitoe on behalf of the Wellcome Sanger Institute COVID-19 Surveillance Team |
| EPI_ISL_2925054 | Lighthouse Lab in Glasgow | Wellcome Sanger Institute for the COVID-19 Genomics UK (COG-UK) Consortium | Harper VanSteenhouse, Yumi Kasai, David Gray, Carol Clugston, Anna Dominiczak and Alex Alderton, Roberto Amato, Jeffrey Barrett, Sonia Goncalves, Ewan Harrison, David K. Jackson, Ian Johnston, Dominic Kwiatkowski, Cordelia Langford, John Sillitoe on behalf of the Wellcome Sanger Institute COVID-19 Surveillance Team |
| EPI_ISL_2925055 | Lighthouse Lab in Milton Keynes | Wellcome Sanger Institute for the COVID-19 Genomics UK (COG-UK) Consortium | The Lighthouse Lab in Milton Keynes and Alex Alderton, Roberto Amato, Jeffrey Barrett, Sonia Goncalves, Ewan Harrison, David K. Jackson, Ian Johnston, Dominic Kwiatkowski, Cordelia Langford, John Sillitoe on behalf of the Wellcome Sanger Institute COVID-19 Surveillance Team |
| EPI_ISL_2925056 | Lighthouse Lab in Alderley Park | Wellcome Sanger Institute for the COVID-19 Genomics UK (COG-UK) Consortium | Jacquelyn Wynn, Mairead Hyland, The Lighthouse Lab in Alderley Park and Alex Alderton, Roberto Amato, Jeffrey Barrett, Sonia Goncalves, Ewan Harrison, David K. Jackson, Ian Johnston, Dominic Kwiatkowski, Cordelia Langford, John Sillitoe on behalf of the Wellcome Sanger Institute COVID-19 Surveillance Team |
| EPI_ISL_2925057 | Lighthouse Lab in Milton Keynes | Wellcome Sanger Institute for the COVID-19 Genomics UK (COG-UK) Consortium | The Lighthouse Lab in Milton Keynes and Alex Alderton, Roberto Amato, Jeffrey Barrett, Sonia Goncalves, Ewan Harrison, David K. Jackson, Ian Johnston, Dominic Kwiatkowski, Cordelia Langford, John Sillitoe on behalf of the Wellcome Sanger Institute COVID-19 Surveillance Team |
| EPI_ISL_2925058 | Lighthouse Lab in Glasgow | Wellcome Sanger Institute for the COVID-19 Genomics UK (COG-UK) Consortium | Harper VanSteenhouse, Yumi Kasai, David Gray, Carol Clugston, Anna Dominiczak and Alex Alderton, Roberto Amato, Jeffrey Barrett, Sonia Goncalves, Ewan Harrison, David K. Jackson, Ian Johnston, Dominic Kwiatkowski, Cordelia Langford, John Sillitoe on behalf of the Wellcome Sanger Institute COVID-19 Surveillance Team |
| EPI_ISL_2925059 | Lighthouse Lab in Milton Keynes | Wellcome Sanger Institute for the COVID-19 Genomics UK (COG-UK) Consortium | The Lighthouse Lab in Milton Keynes and Alex Alderton, Roberto Amato, Jeffrey Barrett, Sonia Goncalves, Ewan Harrison, David K. Jackson, Ian Johnston, Dominic Kwiatkowski, Cordelia Langford, John Sillitoe on behalf of the Wellcome Sanger Institute COVID-19 Surveillance Team |
| EPI_ISL_2925060, EPI_ISL_2925061 | Lighthouse Lab in Glasgow | Wellcome Sanger Institute for the COVID-19 Genomics UK (COG-UK) Consortium | Harper VanSteenhouse, Yumi Kasai, David Gray, Carol Clugston, Anna Dominiczak and Alex Alderton, Roberto Amato, Jeffrey Barrett, Sonia Goncalves, Ewan Harrison, David K. Jackson, Ian Johnston, Dominic Kwiatkowski, Cordelia Langford, John Sillitoe on behalf of the Wellcome Sanger Institute COVID-19 Surveillance Team |
| EPI_ISL_2925062 | Lighthouse Lab in Alderley Park | Wellcome Sanger Institute for the COVID-19 Genomics UK (COG-UK) Consortium | Jacquelyn Wynn, Mairead Hyland, The Lighthouse Lab in Alderley Park and Alex Alderton, Roberto Amato, Jeffrey Barrett, Sonia Goncalves, Ewan Harrison, David K. Jackson, Ian Johnston, Dominic Kwiatkowski, Cordelia Langford, John Sillitoe on behalf of the Wellcome Sanger Institute COVID-19 Surveillance Team |
| EPI_ISL_2925063 | Lighthouse Lab in Milton Keynes | Wellcome Sanger Institute for the COVID-19 Genomics UK (COG-UK) Consortium | The Lighthouse Lab in Milton Keynes and Alex Alderton, Roberto Amato, Jeffrey Barrett, Sonia Goncalves, Ewan Harrison, David K. Jackson, Ian Johnston, Dominic Kwiatkowski, Cordelia Langford, John Sillitoe on behalf of the Wellcome Sanger Institute COVID-19 Surveillance Team |
| EPI_ISL_2925064, EPI_ISL_2925065, EPI_ISL_2925066 | Lighthouse Lab in Glasgow | Wellcome Sanger Institute for the COVID-19 Genomics UK (COG-UK) Consortium | Harper VanSteenhouse, Yumi Kasai, David Gray, Carol Clugston, Anna Dominiczak and Alex Alderton, Roberto Amato, Jeffrey Barrett, Sonia Goncalves, Ewan Harrison, David K. Jackson, Ian Johnston, Dominic Kwiatkowski, Cordelia Langford, John Sillitoe on behalf of the Wellcome Sanger Institute COVID-19 Surveillance Team |
| EPI_ISL_2925067 | Lighthouse Lab in Milton Keynes | Wellcome Sanger Institute for the COVID-19 Genomics UK (COG-UK) Consortium | The Lighthouse Lab in Milton Keynes and Alex Alderton, Roberto Amato, Jeffrey Barrett, Sonia Goncalves, Ewan Harrison, David K. Jackson, Ian Johnston, Dominic Kwiatkowski, Cordelia Langford, John Sillitoe on behalf of the Wellcome Sanger Institute COVID-19 Surveillance Team |
| EPI_ISL_2925068, EPI_ISL_2925069, EPI_ISL_2925070, EPI_ISL_2925071 | Lighthouse Lab in Glasgow | Wellcome Sanger Institute for the COVID-19 Genomics UK (COG-UK) Consortium | Harper VanSteenhouse, Yumi Kasai, David Gray, Carol Clugston, Anna Dominiczak and Alex Alderton, Roberto Amato, Jeffrey Barrett, Sonia Goncalves, Ewan Harrison, David K. Jackson, Ian Johnston, Dominic Kwiatkowski, Cordelia Langford, John Sillitoe on behalf of the Wellcome Sanger Institute COVID-19 Surveillance Team |
| EPI_ISL_2925072 | Lighthouse Lab in Alderley Park | Wellcome Sanger Institute for the COVID-19 Genomics UK (COG-UK) Consortium | Jacquelyn Wynn, Mairead Hyland, The Lighthouse Lab in Alderley Park and Alex Alderton, Roberto Amato, Jeffrey Barrett, Sonia Goncalves, Ewan Harrison, David K. Jackson, Ian Johnston, Dominic Kwiatkowski, Cordelia Langford, John Sillitoe on behalf of the Wellcome Sanger Institute COVID-19 Surveillance Team |
| EPI_ISL_2925073, EPI_ISL_2925074 | Lighthouse Lab in Glasgow | Wellcome Sanger Institute for the COVID-19 Genomics UK (COG-UK) Consortium | Harper VanSteenhouse, Yumi Kasai, David Gray, Carol Clugston, Anna Dominiczak and Alex Alderton, Roberto Amato, Jeffrey Barrett, Sonia Goncalves, Ewan Harrison, David K. Jackson, Ian Johnston, Dominic Kwiatkowski, Cordelia Langford, John Sillitoe on behalf of the Wellcome Sanger Institute COVID-19 Surveillance Team |
| EPI_ISL_2925075, EPI_ISL_2925076, EPI_ISL_2925077, EPI_ISL_2925078 | Lighthouse Lab in Alderley Park | Wellcome Sanger Institute for the COVID-19 Genomics UK (COG-UK) Consortium | Jacquelyn Wynn, Mairead Hyland, The Lighthouse Lab in Alderley Park and Alex Alderton, Roberto Amato, Jeffrey Barrett, Sonia Goncalves, Ewan Harrison, David K. Jackson, Ian Johnston, Dominic Kwiatkowski, Cordelia Langford, John Sillitoe on behalf of the Wellcome Sanger Institute COVID-19 Surveillance Team |
| EPI_ISL_2925079 | Lighthouse Lab in Milton Keynes | Wellcome Sanger Institute for the COVID-19 Genomics UK (COG-UK) Consortium | The Lighthouse Lab in Milton Keynes and Alex Alderton, Roberto Amato, Jeffrey Barrett, Sonia Goncalves, Ewan Harrison, David K. Jackson, Ian Johnston, Dominic Kwiatkowski, Cordelia Langford, John Sillitoe on behalf of the Wellcome Sanger Institute COVID-19 Surveillance Team |
| EPI_ISL_2925080 | Lighthouse Lab in Alderley Park | Wellcome Sanger Institute for the COVID-19 Genomics UK (COG-UK) Consortium | Jacquelyn Wynn, Mairead Hyland, The Lighthouse Lab in Alderley Park and Alex Alderton, Roberto Amato, Jeffrey Barrett, Sonia Goncalves, Ewan Harrison, David K. Jackson, Ian Johnston, Dominic Kwiatkowski, Cordelia Langford, John Sillitoe on behalf of the Wellcome Sanger Institute COVID-19 Surveillance Team |
| EPI_ISL_2925081 | Lighthouse Lab in Milton Keynes | Wellcome Sanger Institute for the COVID-19 Genomics UK (COG-UK) Consortium | The Lighthouse Lab in Milton Keynes and Alex Alderton, Roberto Amato, Jeffrey Barrett, Sonia Goncalves, Ewan Harrison, David K. Jackson, Ian Johnston, Dominic Kwiatkowski, Cordelia Langford, John Sillitoe on behalf of the Wellcome Sanger Institute COVID-19 Surveillance Team |
| EPI_ISL_2925082, EPI_ISL_2925083, EPI_ISL_2925084 | Lighthouse Lab in Glasgow | Wellcome Sanger Institute for the COVID-19 Genomics UK (COG-UK) Consortium | Harper VanSteenhouse, Yumi Kasai, David Gray, Carol Clugston, Anna Dominiczak and Alex Alderton, Roberto Amato, Jeffrey Barrett, Sonia Goncalves, Ewan Harrison, David K. Jackson, Ian Johnston, Dominic Kwiatkowski, Cordelia Langford, John Sillitoe on behalf of the Wellcome Sanger Institute COVID-19 Surveillance Team |
| EPI_ISL_2925085, EPI_ISL_2925086, EPI_ISL_2925087 | Lighthouse Lab in Milton Keynes | Wellcome Sanger Institute for the COVID-19 Genomics UK (COG-UK) Consortium | The Lighthouse Lab in Milton Keynes and Alex Alderton, Roberto Amato, Jeffrey Barrett, Sonia Goncalves, Ewan Harrison, David K. Jackson, Ian Johnston, Dominic Kwiatkowski, Cordelia Langford, John Sillitoe on behalf of the Wellcome Sanger Institute COVID-19 Surveillance Team |
| EPI_ISL_2925088, EPI_ISL_2925089, EPI_ISL_2925090 | Lighthouse Lab in Glasgow | Wellcome Sanger Institute for the COVID-19 Genomics UK (COG-UK) Consortium | Harper VanSteenhouse, Yumi Kasai, David Gray, Carol Clugston, Anna Dominiczak and Alex Alderton, Roberto Amato, Jeffrey Barrett, Sonia Goncalves, Ewan Harrison, David K. Jackson, Ian Johnston, Dominic Kwiatkowski, Cordelia Langford, John Sillitoe on behalf of the Wellcome Sanger Institute COVID-19 Surveillance Team |
| EPI_ISL_2925091, EPI_ISL_2925092 | Lighthouse Lab in Milton Keynes | Wellcome Sanger Institute for the COVID-19 Genomics UK (COG-UK) Consortium | The Lighthouse Lab in Milton Keynes and Alex Alderton, Roberto Amato, Jeffrey Barrett, Sonia Goncalves, Ewan Harrison, David K. Jackson, Ian Johnston, Dominic Kwiatkowski, Cordelia Langford, John Sillitoe on behalf of the Wellcome Sanger Institute COVID-19 Surveillance Team |
| EPI_ISL_2925093 | Lighthouse Lab in Alderley Park | Wellcome Sanger Institute for the COVID-19 Genomics UK (COG-UK) Consortium | Jacquelyn Wynn, Mairead Hyland, The Lighthouse Lab in Alderley Park and Alex Alderton, Roberto Amato, Jeffrey Barrett, Sonia Goncalves, Ewan Harrison, David K. Jackson, Ian Johnston, Dominic Kwiatkowski, Cordelia Langford, John Sillitoe on behalf of the Wellcome Sanger Institute COVID-19 Surveillance Team |
| EPI_ISL_2925094 | Lighthouse Lab in Glasgow | Wellcome Sanger Institute for the COVID-19 Genomics UK (COG-UK) Consortium | Harper VanSteenhouse, Yumi Kasai, David Gray, Carol Clugston, Anna Dominiczak and Alex Alderton, Roberto Amato, Jeffrey Barrett, Sonia Goncalves, Ewan Harrison, David K. Jackson, Ian Johnston, Dominic Kwiatkowski, Cordelia Langford, John Sillitoe on behalf of the Wellcome Sanger Institute COVID-19 Surveillance Team |
| EPI_ISL_2925095, EPI_ISL_2925096 | Lighthouse Lab in Alderley Park | Wellcome Sanger Institute for the COVID-19 Genomics UK (COG-UK) Consortium | Jacquelyn Wynn, Mairead Hyland, The Lighthouse Lab in Alderley Park and Alex Alderton, Roberto Amato, Jeffrey Barrett, Sonia Goncalves, Ewan Harrison, David K. Jackson, Ian Johnston, Dominic Kwiatkowski, Cordelia Langford, John Sillitoe on behalf of the Wellcome Sanger Institute COVID-19 Surveillance Team |
| EPI_ISL_2925097 | Lighthouse Lab in Glasgow | Wellcome Sanger Institute for the COVID-19 Genomics UK (COG-UK) Consortium | Harper VanSteenhouse, Yumi Kasai, David Gray, Carol Clugston, Anna Dominiczak and Alex Alderton, Roberto Amato, Jeffrey Barrett, Sonia Goncalves, Ewan Harrison, David K. Jackson, Ian Johnston, Dominic Kwiatkowski, Cordelia Langford, John Sillitoe on behalf of the Wellcome Sanger Institute COVID-19 Surveillance Team |
| EPI_ISL_2925098 | Lighthouse Lab in Milton Keynes | Wellcome Sanger Institute for the COVID-19 Genomics UK (COG-UK) Consortium | The Lighthouse Lab in Milton Keynes and Alex Alderton, Roberto Amato, Jeffrey Barrett, Sonia Goncalves, Ewan Harrison, David K. Jackson, Ian Johnston, Dominic Kwiatkowski, Cordelia Langford, John Sillitoe on behalf of the Wellcome Sanger Institute COVID-19 Surveillance Team |

| | | | |
|---|---|---|---|
| EPI_ISL_2925099 | Lighthouse Lab in Glasgow | Wellcome Sanger Institute for the COVID-19 Genomics UK (COG-UK) Consortium | Harper VanSteenhouse, Yumi Kasai, David Gray, Carol Clugston, Anna Dominiczak and Alex Alderton, Roberto Amato, Jeffrey Barrett, Sonia Goncalves, Ewan Harrison, David K. Jackson, Ian Johnston, Dominic Kwiatkowski, Cordelia Langford, John Sillitoe on behalf of the Wellcome Sanger Institute COVID-19 Surveillance Team |
| EPI_ISL_2925100 | Lighthouse Lab in Alderley Park | Wellcome Sanger Institute for the COVID-19 Genomics UK (COG-UK) Consortium | Jacquelyn Wynn, Mairead Hyland, The Lighthouse Lab in Alderley Park and Alex Alderton, Roberto Amato, Jeffrey Barrett, Sonia Goncalves, Ewan Harrison, David K. Jackson, Ian Johnston, Dominic Kwiatkowski, Cordelia Langford, John Sillitoe on behalf of the Wellcome Sanger Institute COVID-19 Surveillance Team |
| EPI_ISL_2925101 | Lighthouse Lab in Glasgow | Wellcome Sanger Institute for the COVID-19 Genomics UK (COG-UK) Consortium | Harper VanSteenhouse, Yumi Kasai, David Gray, Carol Clugston, Anna Dominiczak and Alex Alderton, Roberto Amato, Jeffrey Barrett, Sonia Goncalves, Ewan Harrison, David K. Jackson, Ian Johnston, Dominic Kwiatkowski, Cordelia Langford, John Sillitoe on behalf of the Wellcome Sanger Institute COVID-19 Surveillance Team |
| EPI_ISL_2925102 | Lighthouse Lab in Alderley Park | Wellcome Sanger Institute for the COVID-19 Genomics UK (COG-UK) Consortium | Jacquelyn Wynn, Mairead Hyland, The Lighthouse Lab in Alderley Park and Alex Alderton, Roberto Amato, Jeffrey Barrett, Sonia Goncalves, Ewan Harrison, David K. Jackson, Ian Johnston, Dominic Kwiatkowski, Cordelia Langford, John Sillitoe on behalf of the Wellcome Sanger Institute COVID-19 Surveillance Team |
| EPI_ISL_2925103, EPI_ISL_2925104, EPI_ISL_2925105 | Lighthouse Lab in Glasgow | Wellcome Sanger Institute for the COVID-19 Genomics UK (COG-UK) Consortium | Harper VanSteenhouse, Yumi Kasai, David Gray, Carol Clugston, Anna Dominiczak and Alex Alderton, Roberto Amato, Jeffrey Barrett, Sonia Goncalves, Ewan Harrison, David K. Jackson, Ian Johnston, Dominic Kwiatkowski, Cordelia Langford, John Sillitoe on behalf of the Wellcome Sanger Institute COVID-19 Surveillance Team |
| EPI_ISL_2925106 | Lighthouse Lab in Milton Keynes | Wellcome Sanger Institute for the COVID-19 Genomics UK (COG-UK) Consortium | The Lighthouse Lab in Milton Keynes and Alex Alderton, Roberto Amato, Jeffrey Barrett, Sonia Goncalves, Ewan Harrison, David K. Jackson, Ian Johnston, Dominic Kwiatkowski, Cordelia Langford, John Sillitoe on behalf of the Wellcome Sanger Institute COVID-19 Surveillance Team |
| EPI_ISL_2925107, EPI_ISL_2925108, EPI_ISL_2925109 | Lighthouse Lab in Alderley Park | Wellcome Sanger Institute for the COVID-19 Genomics UK (COG-UK) Consortium | Jacquelyn Wynn, Mairead Hyland, The Lighthouse Lab in Alderley Park and Alex Alderton, Roberto Amato, Jeffrey Barrett, Sonia Goncalves, Ewan Harrison, David K. Jackson, Ian Johnston, Dominic Kwiatkowski, Cordelia Langford, John Sillitoe on behalf of the Wellcome Sanger Institute COVID-19 Surveillance Team |
| EPI_ISL_2925110, EPI_ISL_2925111 | Lighthouse Lab in Glasgow | Wellcome Sanger Institute for the COVID-19 Genomics UK (COG-UK) Consortium | Harper VanSteenhouse, Yumi Kasai, David Gray, Carol Clugston, Anna Dominiczak and Alex Alderton, Roberto Amato, Jeffrey Barrett, Sonia Goncalves, Ewan Harrison, David K. Jackson, Ian Johnston, Dominic Kwiatkowski, Cordelia Langford, John Sillitoe on behalf of the Wellcome Sanger Institute COVID-19 Surveillance Team |
| EPI_ISL_2925112 | Lighthouse Lab in Alderley Park | Wellcome Sanger Institute for the COVID-19 Genomics UK (COG-UK) Consortium | Jacquelyn Wynn, Mairead Hyland, The Lighthouse Lab in Alderley Park and Alex Alderton, Roberto Amato, Jeffrey Barrett, Sonia Goncalves, Ewan Harrison, David K. Jackson, Ian Johnston, Dominic Kwiatkowski, Cordelia Langford, John Sillitoe on behalf of the Wellcome Sanger Institute COVID-19 Surveillance Team |
| EPI_ISL_2925113 | Lighthouse Lab in Milton Keynes | Wellcome Sanger Institute for the COVID-19 Genomics UK (COG-UK) Consortium | The Lighthouse Lab in Milton Keynes and Alex Alderton, Roberto Amato, Jeffrey Barrett, Sonia Goncalves, Ewan Harrison, David K. Jackson, Ian Johnston, Dominic Kwiatkowski, Cordelia Langford, John Sillitoe on behalf of the Wellcome Sanger Institute COVID-19 Surveillance Team |
| EPI_ISL_2925114 | Lighthouse Lab in Alderley Park | Wellcome Sanger Institute for the COVID-19 Genomics UK (COG-UK) Consortium | Jacquelyn Wynn, Mairead Hyland, The Lighthouse Lab in Alderley Park and Alex Alderton, Roberto Amato, Jeffrey Barrett, Sonia Goncalves, Ewan Harrison, David K. Jackson, Ian Johnston, Dominic Kwiatkowski, Cordelia Langford, John Sillitoe on behalf of the Wellcome Sanger Institute COVID-19 Surveillance Team |
| EPI_ISL_2925115 | Lighthouse Lab in Milton Keynes | Wellcome Sanger Institute for the COVID-19 Genomics UK (COG-UK) Consortium | The Lighthouse Lab in Milton Keynes and Alex Alderton, Roberto Amato, Jeffrey Barrett, Sonia Goncalves, Ewan Harrison, David K. Jackson, Ian Johnston, Dominic Kwiatkowski, Cordelia Langford, John Sillitoe on behalf of the Wellcome Sanger Institute COVID-19 Surveillance Team |
| EPI_ISL_2925116 | Lighthouse Lab in Alderley Park | Wellcome Sanger Institute for the COVID-19 Genomics UK (COG-UK) Consortium | Jacquelyn Wynn, Mairead Hyland, The Lighthouse Lab in Alderley Park and Alex Alderton, Roberto Amato, Jeffrey Barrett, Sonia Goncalves, Ewan Harrison, David K. Jackson, Ian Johnston, Dominic Kwiatkowski, Cordelia Langford, John Sillitoe on behalf of the Wellcome Sanger Institute COVID-19 Surveillance Team |
| EPI_ISL_2925117 | Lighthouse Lab in Milton Keynes | Wellcome Sanger Institute for the COVID-19 Genomics UK (COG-UK) Consortium | The Lighthouse Lab in Milton Keynes and Alex Alderton, Roberto Amato, Jeffrey Barrett, Sonia Goncalves, Ewan Harrison, David K. Jackson, Ian Johnston, Dominic Kwiatkowski, Cordelia Langford, John Sillitoe on behalf of the Wellcome Sanger Institute COVID-19 Surveillance Team |
| EPI_ISL_2925118 | Lighthouse Lab in Alderley Park | Wellcome Sanger Institute for the COVID-19 Genomics UK (COG-UK) Consortium | Jacquelyn Wynn, Mairead Hyland, The Lighthouse Lab in Alderley Park and Alex Alderton, Roberto Amato, Jeffrey Barrett, Sonia Goncalves, Ewan Harrison, David K. Jackson, Ian Johnston, Dominic Kwiatkowski, Cordelia Langford, John Sillitoe on behalf of the Wellcome Sanger Institute COVID-19 Surveillance Team |
| EPI_ISL_2925119 | Lighthouse Lab in Glasgow | Wellcome Sanger Institute for the COVID-19 Genomics UK (COG-UK) Consortium | Harper VanSteenhouse, Yumi Kasai, David Gray, Carol Clugston, Anna Dominiczak and Alex Alderton, Roberto Amato, Jeffrey Barrett, Sonia Goncalves, Ewan Harrison, David K. Jackson, Ian Johnston, Dominic Kwiatkowski, Cordelia Langford, John Sillitoe on behalf of the Wellcome Sanger Institute COVID-19 Surveillance Team |
| EPI_ISL_2925120 | Lighthouse Lab in Milton Keynes | Wellcome Sanger Institute for the COVID-19 Genomics UK (COG-UK) Consortium | The Lighthouse Lab in Milton Keynes and Alex Alderton, Roberto Amato, Jeffrey Barrett, Sonia Goncalves, Ewan Harrison, David K. Jackson, Ian Johnston, Dominic Kwiatkowski, Cordelia Langford, John Sillitoe on behalf of the Wellcome Sanger Institute COVID-19 Surveillance Team |
| EPI_ISL_2925121, EPI_ISL_2925122 | Lighthouse Lab in Glasgow | Wellcome Sanger Institute for the COVID-19 Genomics UK (COG-UK) Consortium | Harper VanSteenhouse, Yumi Kasai, David Gray, Carol Clugston, Anna Dominiczak and Alex Alderton, Roberto Amato, Jeffrey Barrett, Sonia Goncalves, Ewan Harrison, David K. Jackson, Ian Johnston, Dominic Kwiatkowski, Cordelia Langford, John Sillitoe on behalf of the Wellcome Sanger Institute COVID-19 Surveillance Team |
| EPI_ISL_2925123 | Lighthouse Lab in Alderley Park | Wellcome Sanger Institute for the COVID-19 Genomics UK (COG-UK) Consortium | Jacquelyn Wynn, Mairead Hyland, The Lighthouse Lab in Alderley Park and Alex Alderton, Roberto Amato, Jeffrey Barrett, Sonia Goncalves, Ewan Harrison, David K. Jackson, Ian Johnston, Dominic Kwiatkowski, Cordelia Langford, John Sillitoe on behalf of the Wellcome Sanger Institute COVID-19 Surveillance Team |
| EPI_ISL_2925124 | Lighthouse Lab in Milton Keynes | Wellcome Sanger Institute for the COVID-19 Genomics UK (COG-UK) Consortium | The Lighthouse Lab in Milton Keynes and Alex Alderton, Roberto Amato, Jeffrey Barrett, Sonia Goncalves, Ewan Harrison, David K. Jackson, Ian Johnston, Dominic Kwiatkowski, Cordelia Langford, John Sillitoe on behalf of the Wellcome Sanger Institute COVID-19 Surveillance Team |
| EPI_ISL_2925125 | Lighthouse Lab in Alderley Park | Wellcome Sanger Institute for the COVID-19 Genomics UK (COG-UK) Consortium | Jacquelyn Wynn, Mairead Hyland, The Lighthouse Lab in Alderley Park and Alex Alderton, Roberto Amato, Jeffrey Barrett, Sonia Goncalves, Ewan Harrison, David K. Jackson, Ian Johnston, Dominic Kwiatkowski, Cordelia Langford, John Sillitoe on behalf of the Wellcome Sanger Institute COVID-19 Surveillance Team |
| EPI_ISL_2925126 | Lighthouse Lab in Milton Keynes | Wellcome Sanger Institute for the COVID-19 Genomics UK (COG-UK) Consortium | The Lighthouse Lab in Milton Keynes and Alex Alderton, Roberto Amato, Jeffrey Barrett, Sonia Goncalves, Ewan Harrison, David K. Jackson, Ian Johnston, Dominic Kwiatkowski, Cordelia Langford, John Sillitoe on behalf of the Wellcome Sanger Institute COVID-19 Surveillance Team |
| EPI_ISL_2925127, EPI_ISL_2925128, EPI_ISL_2925129 | Lighthouse Lab in Alderley Park | Wellcome Sanger Institute for the COVID-19 Genomics UK (COG-UK) Consortium | Jacquelyn Wynn, Mairead Hyland, The Lighthouse Lab in Alderley Park and Alex Alderton, Roberto Amato, Jeffrey Barrett, Sonia Goncalves, Ewan Harrison, David K. Jackson, Ian Johnston, Dominic Kwiatkowski, Cordelia Langford, John Sillitoe on behalf of the Wellcome Sanger Institute COVID-19 Surveillance Team |
| EPI_ISL_2925130 | Lighthouse Lab in Milton Keynes | Wellcome Sanger Institute for the COVID-19 Genomics UK (COG-UK) Consortium | The Lighthouse Lab in Milton Keynes and Alex Alderton, Roberto Amato, Jeffrey Barrett, Sonia Goncalves, Ewan Harrison, David K. Jackson, Ian Johnston, Dominic Kwiatkowski, Cordelia Langford, John Sillitoe on behalf of the Wellcome Sanger Institute COVID-19 Surveillance Team |
| EPI_ISL_2925131 | Lighthouse Lab in Alderley Park | Wellcome Sanger Institute for the COVID-19 Genomics UK (COG-UK) Consortium | Jacquelyn Wynn, Mairead Hyland, The Lighthouse Lab in Alderley Park and Alex Alderton, Roberto Amato, Jeffrey Barrett, Sonia Goncalves, Ewan Harrison, David K. Jackson, Ian Johnston, Dominic Kwiatkowski, Cordelia Langford, John Sillitoe on behalf of the Wellcome Sanger Institute COVID-19 Surveillance Team |
| EPI_ISL_2925132 | Lighthouse Lab in Milton Keynes | Wellcome Sanger Institute for the COVID-19 Genomics UK (COG-UK) Consortium | The Lighthouse Lab in Milton Keynes and Alex Alderton, Roberto Amato, Jeffrey Barrett, Sonia Goncalves, Ewan Harrison, David K. Jackson, Ian Johnston, Dominic Kwiatkowski, Cordelia Langford, John Sillitoe on behalf of the Wellcome Sanger Institute COVID-19 Surveillance Team |
| EPI_ISL_2925133, EPI_ISL_2925134, EPI_ISL_2925135, EPI_ISL_2925136, EPI_ISL_2925137 | Lighthouse Lab in Alderley Park | Wellcome Sanger Institute for the COVID-19 Genomics UK (COG-UK) Consortium | Jacquelyn Wynn, Mairead Hyland, The Lighthouse Lab in Alderley Park and Alex Alderton, Roberto Amato, Jeffrey Barrett, Sonia Goncalves, Ewan Harrison, David K. Jackson, Ian Johnston, Dominic Kwiatkowski, Cordelia Langford, John Sillitoe on behalf of the Wellcome Sanger Institute COVID-19 Surveillance Team |
| EPI_ISL_2925138 | Lighthouse Lab in Milton Keynes | Wellcome Sanger Institute for the COVID-19 Genomics UK (COG-UK) Consortium | The Lighthouse Lab in Milton Keynes and Alex Alderton, Roberto Amato, Jeffrey Barrett, Sonia Goncalves, Ewan Harrison, David K. Jackson, Ian Johnston, Dominic Kwiatkowski, Cordelia Langford, John Sillitoe on behalf of the Wellcome Sanger Institute COVID-19 Surveillance Team |
| EPI_ISL_2925139 | Lighthouse Lab in Alderley Park | Wellcome Sanger Institute for the COVID-19 Genomics UK (COG-UK) Consortium | Jacquelyn Wynn, Mairead Hyland, The Lighthouse Lab in Alderley Park and Alex Alderton, Roberto Amato, Jeffrey Barrett, Sonia Goncalves, Ewan Harrison, David K. Jackson, Ian Johnston, Dominic Kwiatkowski, Cordelia Langford, John Sillitoe on behalf of the Wellcome Sanger Institute COVID-19 Surveillance Team |
| EPI_ISL_2925140 | Lighthouse Lab in Milton Keynes | Wellcome Sanger Institute for the COVID-19 Genomics UK (COG-UK) Consortium | The Lighthouse Lab in Milton Keynes and Alex Alderton, Roberto Amato, Jeffrey Barrett, Sonia Goncalves, Ewan Harrison, David K. Jackson, Ian Johnston, Dominic Kwiatkowski, Cordelia Langford, John Sillitoe on behalf of the Wellcome Sanger Institute COVID-19 Surveillance Team |
| EPI_ISL_2925141, EPI_ISL_2925142 | Lighthouse Lab in Alderley Park | Wellcome Sanger Institute for the COVID-19 Genomics UK (COG-UK) Consortium | Jacquelyn Wynn, Mairead Hyland, The Lighthouse Lab in Alderley Park and Alex Alderton, Roberto Amato, Jeffrey Barrett, Sonia Goncalves, Ewan Harrison, David K. Jackson, Ian Johnston, Dominic Kwiatkowski, Cordelia Langford, John Sillitoe on behalf of the Wellcome Sanger Institute COVID-19 Surveillance Team |
| EPI_ISL_2925143 | Lighthouse Lab in Milton Keynes | Wellcome Sanger Institute for the COVID-19 Genomics UK (COG-UK) Consortium | The Lighthouse Lab in Milton Keynes and Alex Alderton, Roberto Amato, Jeffrey Barrett, Sonia Goncalves, Ewan Harrison, David K. Jackson, Ian Johnston, Dominic Kwiatkowski, Cordelia Langford, John Sillitoe on behalf of the Wellcome Sanger Institute COVID-19 Surveillance Team |
| EPI_ISL_2925144, EPI_ISL_2925145, EPI_ISL_2925146 | Lighthouse Lab in Alderley Park | Wellcome Sanger Institute for the COVID-19 Genomics UK (COG-UK) Consortium | Jacquelyn Wynn, Mairead Hyland, The Lighthouse Lab in Alderley Park and Alex Alderton, Roberto Amato, Jeffrey Barrett, Sonia Goncalves, Ewan Harrison, David K. Jackson, Ian Johnston, Dominic Kwiatkowski, Cordelia Langford, John Sillitoe on behalf of the Wellcome Sanger Institute COVID-19 Surveillance Team |
| EPI_ISL_2925147 | Lighthouse Lab in Milton Keynes | Wellcome Sanger Institute for the COVID-19 Genomics UK (COG-UK) Consortium | The Lighthouse Lab in Milton Keynes and Alex Alderton, Roberto Amato, Jeffrey Barrett, Sonia Goncalves, Ewan Harrison, David K. Jackson, Ian Johnston, Dominic Kwiatkowski, Cordelia Langford, John Sillitoe on behalf of the Wellcome Sanger Institute COVID-19 Surveillance Team |
| EPI_ISL_2925148, EPI_ISL_2925149, EPI_ISL_2925150, EPI_ISL_2925151, EPI_ISL_2925152 | Lighthouse Lab in Alderley Park | Wellcome Sanger Institute for the COVID-19 Genomics UK (COG-UK) Consortium | Jacquelyn Wynn, Mairead Hyland, The Lighthouse Lab in Alderley Park and Alex Alderton, Roberto Amato, Jeffrey Barrett, Sonia Goncalves, Ewan Harrison, David K. Jackson, Ian Johnston, Dominic Kwiatkowski, Cordelia Langford, John Sillitoe on behalf of the Wellcome Sanger Institute COVID-19 Surveillance Team |
| EPI_ISL_2925153 | Lighthouse Lab in Milton Keynes | Wellcome Sanger Institute for the COVID-19 Genomics UK (COG-UK) Consortium | The Lighthouse Lab in Milton Keynes and Alex Alderton, Roberto Amato, Jeffrey Barrett, Sonia Goncalves, Ewan Harrison, David K. Jackson, Ian Johnston, Dominic Kwiatkowski, Cordelia Langford, John Sillitoe on behalf of the Wellcome Sanger Institute COVID-19 Surveillance Team |
| EPI_ISL_2925154 | Lighthouse Lab in Alderley Park | Wellcome Sanger Institute for the COVID-19 Genomics UK (COG-UK) Consortium | Jacquelyn Wynn, Mairead Hyland, The Lighthouse Lab in Alderley Park and Alex Alderton, Roberto Amato, Jeffrey Barrett, Sonia Goncalves, Ewan Harrison, David K. Jackson, Ian Johnston, Dominic Kwiatkowski, Cordelia Langford, John Sillitoe on behalf of the Wellcome Sanger Institute COVID-19 Surveillance Team |
| EPI_ISL_2925155 | Lighthouse Lab in Milton Keynes | Wellcome Sanger Institute for the COVID-19 Genomics UK (COG-UK) Consortium | The Lighthouse Lab in Milton Keynes and Alex Alderton, Roberto Amato, Jeffrey Barrett, Sonia Goncalves, Ewan Harrison, David K. Jackson, Ian Johnston, Dominic Kwiatkowski, Cordelia Langford, John Sillitoe on behalf of the Wellcome Sanger Institute COVID-19 Surveillance Team |
| EPI_ISL_2925156, EPI_ISL_2925157, EPI_ISL_2925158 | Lighthouse Lab in Alderley Park | Wellcome Sanger Institute for the COVID-19 Genomics UK (COG-UK) Consortium | Jacquelyn Wynn, Mairead Hyland, The Lighthouse Lab in Alderley Park and Alex Alderton, Roberto Amato, Jeffrey Barrett, Sonia Goncalves, Ewan Harrison, David K. Jackson, Ian Johnston, Dominic Kwiatkowski, Cordelia Langford, John Sillitoe on behalf of the Wellcome Sanger Institute COVID-19 Surveillance Team |
| EPI_ISL_2925159 | Lighthouse Lab in Milton Keynes | Wellcome Sanger Institute for the COVID-19 Genomics UK (COG-UK) Consortium | The Lighthouse Lab in Milton Keynes and Alex Alderton, Roberto Amato, Jeffrey Barrett, Sonia Goncalves, Ewan Harrison, David K. Jackson, Ian Johnston, Dominic Kwiatkowski, Cordelia Langford, John Sillitoe on behalf of the Wellcome Sanger Institute COVID-19 Surveillance Team |
| EPI_ISL_2925160, EPI_ISL_2925161, | Lighthouse Lab in Alderley Park | Wellcome Sanger Institute for the COVID-19 Genomics UK (COG-UK) Consortium | Jacquelyn Wynn, Mairead Hyland, The Lighthouse Lab in Alderley Park and Alex Alderton, Roberto Amato, Jeffrey Barrett, Sonia Goncalves, Ewan Harrison, David K. Jackson, Ian Johnston, Dominic Kwiatkowski, Cordelia Langford, John Sillitoe on behalf of the Wellcome Sanger Institute COVID-19 Surveillance Team |

| | | | |
|---|---|---|---|
| EPI_ISL_2925162 | | | |
| EPI_ISL_2925163, EPI_ISL_2925164 | Lighthouse Lab in Milton Keynes | Wellcome Sanger Institute for the COVID-19 Genomics UK (COG-UK) Consortium | The Lighthouse Lab in Milton Keynes and Alex Alderton, Roberto Amato, Jeffrey Barrett, Sonia Goncalves, Ewan Harrison, David K. Jackson, Ian Johnston, Dominic Kwiatkowski, Cordelia Langford, John Sillitoe on behalf of the Wellcome Sanger Institute COVID-19 Surveillance Team |
| EPI_ISL_2925165, EPI_ISL_2925166, EPI_ISL_2925167, EPI_ISL_2925168, EPI_ISL_2925169, EPI_ISL_2925170, EPI_ISL_2925171, EPI_ISL_2925172 | Lighthouse Lab in Alderley Park | Wellcome Sanger Institute for the COVID-19 Genomics UK (COG-UK) Consortium | Jacquelyn Wynn, Mairead Hyland, The Lighthouse Lab in Alderley Park and Alex Alderton, Roberto Amato, Jeffrey Barrett, Sonia Goncalves, Ewan Harrison, David K. Jackson, Ian Johnston, Dominic Kwiatkowski, Cordelia Langford, John Sillitoe on behalf of the Wellcome Sanger Institute COVID-19 Surveillance Team |
| EPI_ISL_2925173 | Lighthouse Lab in Milton Keynes | Wellcome Sanger Institute for the COVID-19 Genomics UK (COG-UK) Consortium | The Lighthouse Lab in Milton Keynes and Alex Alderton, Roberto Amato, Jeffrey Barrett, Sonia Goncalves, Ewan Harrison, David K. Jackson, Ian Johnston, Dominic Kwiatkowski, Cordelia Langford, John Sillitoe on behalf of the Wellcome Sanger Institute COVID-19 Surveillance Team |
| EPI_ISL_2925174 | National Platform bis UMONS/Jolimont | National Platform bis UMONS/Jolimont | François Dufrasne, Guillaume Bayon-Vicente, Florian Juszczak, Eric Tarantino, Gautier Detry, Ruddy Wattiez |
| EPI_ISL_2925175, EPI_ISL_2925176, EPI_ISL_2925177, EPI_ISL_2925178, EPI_ISL_2925179 | Lighthouse Lab in Alderley Park | Wellcome Sanger Institute for the COVID-19 Genomics UK (COG-UK) Consortium | Jacquelyn Wynn, Mairead Hyland, The Lighthouse Lab in Alderley Park and Alex Alderton, Roberto Amato, Jeffrey Barrett, Sonia Goncalves, Ewan Harrison, David K. Jackson, Ian Johnston, Dominic Kwiatkowski, Cordelia Langford, John Sillitoe on behalf of the Wellcome Sanger Institute COVID-19 Surveillance Team |
| EPI_ISL_2925180 | Lighthouse Lab in Milton Keynes | Wellcome Sanger Institute for the COVID-19 Genomics UK (COG-UK) Consortium | The Lighthouse Lab in Milton Keynes and Alex Alderton, Roberto Amato, Jeffrey Barrett, Sonia Goncalves, Ewan Harrison, David K. Jackson, Ian Johnston, Dominic Kwiatkowski, Cordelia Langford, John Sillitoe on behalf of the Wellcome Sanger Institute COVID-19 Surveillance Team |
| EPI_ISL_2925181 | Lighthouse Lab in Alderley Park | Wellcome Sanger Institute for the COVID-19 Genomics UK (COG-UK) Consortium | Jacquelyn Wynn, Mairead Hyland, The Lighthouse Lab in Alderley Park and Alex Alderton, Roberto Amato, Jeffrey Barrett, Sonia Goncalves, Ewan Harrison, David K. Jackson, Ian Johnston, Dominic Kwiatkowski, Cordelia Langford, John Sillitoe on behalf of the Wellcome Sanger Institute COVID-19 Surveillance Team |
| EPI_ISL_2925182 | Lighthouse Lab in Milton Keynes | Wellcome Sanger Institute for the COVID-19 Genomics UK (COG-UK) Consortium | The Lighthouse Lab in Milton Keynes and Alex Alderton, Roberto Amato, Jeffrey Barrett, Sonia Goncalves, Ewan Harrison, David K. Jackson, Ian Johnston, Dominic Kwiatkowski, Cordelia Langford, John Sillitoe on behalf of the Wellcome Sanger Institute COVID-19 Surveillance Team |
| EPI_ISL_2925183 | Lighthouse Lab in Alderley Park | Wellcome Sanger Institute for the COVID-19 Genomics UK (COG-UK) Consortium | Jacquelyn Wynn, Mairead Hyland, The Lighthouse Lab in Alderley Park and Alex Alderton, Roberto Amato, Jeffrey Barrett, Sonia Goncalves, Ewan Harrison, David K. Jackson, Ian Johnston, Dominic Kwiatkowski, Cordelia Langford, John Sillitoe on behalf of the Wellcome Sanger Institute COVID-19 Surveillance Team |
| EPI_ISL_2925184 | Lighthouse Lab in Milton Keynes | Wellcome Sanger Institute for the COVID-19 Genomics UK (COG-UK) Consortium | The Lighthouse Lab in Milton Keynes and Alex Alderton, Roberto Amato, Jeffrey Barrett, Sonia Goncalves, Ewan Harrison, David K. Jackson, Ian Johnston, Dominic Kwiatkowski, Cordelia Langford, John Sillitoe on behalf of the Wellcome Sanger Institute COVID-19 Surveillance Team |
| EPI_ISL_2925185 | Lighthouse Lab in Alderley Park | Wellcome Sanger Institute for the COVID-19 Genomics UK (COG-UK) Consortium | Jacquelyn Wynn, Mairead Hyland, The Lighthouse Lab in Alderley Park and Alex Alderton, Roberto Amato, Jeffrey Barrett, Sonia Goncalves, Ewan Harrison, David K. Jackson, Ian Johnston, Dominic Kwiatkowski, Cordelia Langford, John Sillitoe on behalf of the Wellcome Sanger Institute COVID-19 Surveillance Team |
| EPI_ISL_2925186 | Lighthouse Lab in Milton Keynes | Wellcome Sanger Institute for the COVID-19 Genomics UK (COG-UK) Consortium | The Lighthouse Lab in Milton Keynes and Alex Alderton, Roberto Amato, Jeffrey Barrett, Sonia Goncalves, Ewan Harrison, David K. Jackson, Ian Johnston, Dominic Kwiatkowski, Cordelia Langford, John Sillitoe on behalf of the Wellcome Sanger Institute COVID-19 Surveillance Team |
| EPI_ISL_2925187, EPI_ISL_2925188, EPI_ISL_2925189, EPI_ISL_2925190, EPI_ISL_2925191 | Lighthouse Lab in Alderley Park | Wellcome Sanger Institute for the COVID-19 Genomics UK (COG-UK) Consortium | Jacquelyn Wynn, Mairead Hyland, The Lighthouse Lab in Alderley Park and Alex Alderton, Roberto Amato, Jeffrey Barrett, Sonia Goncalves, Ewan Harrison, David K. Jackson, Ian Johnston, Dominic Kwiatkowski, Cordelia Langford, John Sillitoe on behalf of the Wellcome Sanger Institute COVID-19 Surveillance Team |
| EPI_ISL_2925192 | Lighthouse Lab in Milton Keynes | Wellcome Sanger Institute for the COVID-19 Genomics UK (COG-UK) Consortium | The Lighthouse Lab in Milton Keynes and Alex Alderton, Roberto Amato, Jeffrey Barrett, Sonia Goncalves, Ewan Harrison, David K. Jackson, Ian Johnston, Dominic Kwiatkowski, Cordelia Langford, John Sillitoe on behalf of the Wellcome Sanger Institute COVID-19 Surveillance Team |
| EPI_ISL_2925193, EPI_ISL_2925194, EPI_ISL_2925195, EPI_ISL_2925196 | Lighthouse Lab in Alderley Park | Wellcome Sanger Institute for the COVID-19 Genomics UK (COG-UK) Consortium | Jacquelyn Wynn, Mairead Hyland, The Lighthouse Lab in Alderley Park and Alex Alderton, Roberto Amato, Jeffrey Barrett, Sonia Goncalves, Ewan Harrison, David K. Jackson, Ian Johnston, Dominic Kwiatkowski, Cordelia Langford, John Sillitoe on behalf of the Wellcome Sanger Institute COVID-19 Surveillance Team |
| EPI_ISL_2925197 | Lighthouse Lab in Milton Keynes | Wellcome Sanger Institute for the COVID-19 Genomics UK (COG-UK) Consortium | The Lighthouse Lab in Milton Keynes and Alex Alderton, Roberto Amato, Jeffrey Barrett, Sonia Goncalves, Ewan Harrison, David K. Jackson, Ian Johnston, Dominic Kwiatkowski, Cordelia Langford, John Sillitoe on behalf of the Wellcome Sanger Institute COVID-19 Surveillance Team |
| EPI_ISL_2925199, EPI_ISL_2925200 | Lighthouse Lab in Alderley Park | Wellcome Sanger Institute for the COVID-19 Genomics UK (COG-UK) Consortium | Jacquelyn Wynn, Mairead Hyland, The Lighthouse Lab in Alderley Park and Alex Alderton, Roberto Amato, Jeffrey Barrett, Sonia Goncalves, Ewan Harrison, David K. Jackson, Ian Johnston, Dominic Kwiatkowski, Cordelia Langford, John Sillitoe on behalf of the Wellcome Sanger Institute COVID-19 Surveillance Team |
| EPI_ISL_2925201, EPI_ISL_2925202 | Lighthouse Lab in Milton Keynes | Wellcome Sanger Institute for the COVID-19 Genomics UK (COG-UK) Consortium | The Lighthouse Lab in Milton Keynes and Alex Alderton, Roberto Amato, Jeffrey Barrett, Sonia Goncalves, Ewan Harrison, David K. Jackson, Ian Johnston, Dominic Kwiatkowski, Cordelia Langford, John Sillitoe on behalf of the Wellcome Sanger Institute COVID-19 Surveillance Team |
| EPI_ISL_2925203 | Lighthouse Lab in Alderley Park | Wellcome Sanger Institute for the COVID-19 Genomics UK (COG-UK) Consortium | Jacquelyn Wynn, Mairead Hyland, The Lighthouse Lab in Alderley Park and Alex Alderton, Roberto Amato, Jeffrey Barrett, Sonia Goncalves, Ewan Harrison, David K. Jackson, Ian Johnston, Dominic Kwiatkowski, Cordelia Langford, John Sillitoe on behalf of the Wellcome Sanger Institute COVID-19 Surveillance Team |
| EPI_ISL_2925204, EPI_ISL_2925205 | Lighthouse Lab in Milton Keynes | Wellcome Sanger Institute for the COVID-19 Genomics UK (COG-UK) Consortium | The Lighthouse Lab in Milton Keynes and Alex Alderton, Roberto Amato, Jeffrey Barrett, Sonia Goncalves, Ewan Harrison, David K. Jackson, Ian Johnston, Dominic Kwiatkowski, Cordelia Langford, John Sillitoe on behalf of the Wellcome Sanger Institute COVID-19 Surveillance Team |
| EPI_ISL_2925206 | Lighthouse Lab in Alderley Park | Wellcome Sanger Institute for the COVID-19 Genomics UK (COG-UK) Consortium | Jacquelyn Wynn, Mairead Hyland, The Lighthouse Lab in Alderley Park and Alex Alderton, Roberto Amato, Jeffrey Barrett, Sonia Goncalves, Ewan Harrison, David K. Jackson, Ian Johnston, Dominic Kwiatkowski, Cordelia Langford, John Sillitoe on behalf of the Wellcome Sanger Institute COVID-19 Surveillance Team |
| EPI_ISL_2925207 | Lighthouse Lab in Milton Keynes | Wellcome Sanger Institute for the COVID-19 Genomics UK (COG-UK) Consortium | The Lighthouse Lab in Milton Keynes and Alex Alderton, Roberto Amato, Jeffrey Barrett, Sonia Goncalves, Ewan Harrison, David K. Jackson, Ian Johnston, Dominic Kwiatkowski, Cordelia Langford, John Sillitoe on behalf of the Wellcome Sanger Institute COVID-19 Surveillance Team |
| EPI_ISL_2925208, EPI_ISL_2925209, EPI_ISL_2925210 | Lighthouse Lab in Alderley Park | Wellcome Sanger Institute for the COVID-19 Genomics UK (COG-UK) Consortium | Jacquelyn Wynn, Mairead Hyland, The Lighthouse Lab in Alderley Park and Alex Alderton, Roberto Amato, Jeffrey Barrett, Sonia Goncalves, Ewan Harrison, David K. Jackson, Ian Johnston, Dominic Kwiatkowski, Cordelia Langford, John Sillitoe on behalf of the Wellcome Sanger Institute COVID-19 Surveillance Team |
| EPI_ISL_2925211 | Lighthouse Lab in Milton Keynes | Wellcome Sanger Institute for the COVID-19 Genomics UK (COG-UK) Consortium | The Lighthouse Lab in Milton Keynes and Alex Alderton, Roberto Amato, Jeffrey Barrett, Sonia Goncalves, Ewan Harrison, David K. Jackson, Ian Johnston, Dominic Kwiatkowski, Cordelia Langford, John Sillitoe on behalf of the Wellcome Sanger Institute COVID-19 Surveillance Team |
| EPI_ISL_2925212 | Lighthouse Lab in Alderley Park | Wellcome Sanger Institute for the COVID-19 Genomics UK (COG-UK) Consortium | Jacquelyn Wynn, Mairead Hyland, The Lighthouse Lab in Alderley Park and Alex Alderton, Roberto Amato, Jeffrey Barrett, Sonia Goncalves, Ewan Harrison, David K. Jackson, Ian Johnston, Dominic Kwiatkowski, Cordelia Langford, John Sillitoe on behalf of the Wellcome Sanger Institute COVID-19 Surveillance Team |
| EPI_ISL_2925213 | Lighthouse Lab in Milton Keynes | Wellcome Sanger Institute for the COVID-19 Genomics UK (COG-UK) Consortium | The Lighthouse Lab in Milton Keynes and Alex Alderton, Roberto Amato, Jeffrey Barrett, Sonia Goncalves, Ewan Harrison, David K. Jackson, Ian Johnston, Dominic Kwiatkowski, Cordelia Langford, John Sillitoe on behalf of the Wellcome Sanger Institute COVID-19 Surveillance Team |
| EPI_ISL_2925214, EPI_ISL_2925215 | Lighthouse Lab in Alderley Park | Wellcome Sanger Institute for the COVID-19 Genomics UK (COG-UK) Consortium | Jacquelyn Wynn, Mairead Hyland, The Lighthouse Lab in Alderley Park and Alex Alderton, Roberto Amato, Jeffrey Barrett, Sonia Goncalves, Ewan Harrison, David K. Jackson, Ian Johnston, Dominic Kwiatkowski, Cordelia Langford, John Sillitoe on behalf of the Wellcome Sanger Institute COVID-19 Surveillance Team |
| EPI_ISL_2925216 | Lighthouse Lab in Milton Keynes | Wellcome Sanger Institute for the COVID-19 Genomics UK (COG-UK) Consortium | The Lighthouse Lab in Milton Keynes and Alex Alderton, Roberto Amato, Jeffrey Barrett, Sonia Goncalves, Ewan Harrison, David K. Jackson, Ian Johnston, Dominic Kwiatkowski, Cordelia Langford, John Sillitoe on behalf of the Wellcome Sanger Institute COVID-19 Surveillance Team |
| EPI_ISL_2925217, EPI_ISL_2925218, EPI_ISL_2925219, EPI_ISL_2925220, EPI_ISL_2925221, EPI_ISL_2925222, EPI_ISL_2925223, EPI_ISL_2925224, EPI_ISL_2925225 | Lighthouse Lab in Alderley Park | Wellcome Sanger Institute for the COVID-19 Genomics UK (COG-UK) Consortium | Jacquelyn Wynn, Mairead Hyland, The Lighthouse Lab in Alderley Park and Alex Alderton, Roberto Amato, Jeffrey Barrett, Sonia Goncalves, Ewan Harrison, David K. Jackson, Ian Johnston, Dominic Kwiatkowski, Cordelia Langford, John Sillitoe on behalf of the Wellcome Sanger Institute COVID-19 Surveillance Team |
| EPI_ISL_2925226 | Lighthouse Lab in Milton Keynes | Wellcome Sanger Institute for the COVID-19 Genomics UK (COG-UK) Consortium | The Lighthouse Lab in Milton Keynes and Alex Alderton, Roberto Amato, Jeffrey Barrett, Sonia Goncalves, Ewan Harrison, David K. Jackson, Ian Johnston, Dominic Kwiatkowski, Cordelia Langford, John Sillitoe on behalf of the Wellcome Sanger Institute COVID-19 Surveillance Team |
| EPI_ISL_2925227, EPI_ISL_2925228, EPI_ISL_2925229, EPI_ISL_2925230, EPI_ISL_2925231 | Lighthouse Lab in Alderley Park | Wellcome Sanger Institute for the COVID-19 Genomics UK (COG-UK) Consortium | Jacquelyn Wynn, Mairead Hyland, The Lighthouse Lab in Alderley Park and Alex Alderton, Roberto Amato, Jeffrey Barrett, Sonia Goncalves, Ewan Harrison, David K. Jackson, Ian Johnston, Dominic Kwiatkowski, Cordelia Langford, John Sillitoe on behalf of the Wellcome Sanger Institute COVID-19 Surveillance Team |
| EPI_ISL_2925232 | Lighthouse Lab in Milton Keynes | Wellcome Sanger Institute for the COVID-19 Genomics UK (COG-UK) Consortium | The Lighthouse Lab in Milton Keynes and Alex Alderton, Roberto Amato, Jeffrey Barrett, Sonia Goncalves, Ewan Harrison, David K. Jackson, Ian Johnston, Dominic Kwiatkowski, Cordelia Langford, John Sillitoe on behalf of the Wellcome Sanger Institute COVID-19 Surveillance Team |
| EPI_ISL_2925233, EPI_ISL_2925234, EPI_ISL_2925235, EPI_ISL_2925236, EPI_ISL_2925237 | Lighthouse Lab in Alderley Park | Wellcome Sanger Institute for the COVID-19 Genomics UK (COG-UK) Consortium | Jacquelyn Wynn, Mairead Hyland, The Lighthouse Lab in Alderley Park and Alex Alderton, Roberto Amato, Jeffrey Barrett, Sonia Goncalves, Ewan Harrison, David K. Jackson, Ian Johnston, Dominic Kwiatkowski, Cordelia Langford, John Sillitoe on behalf of the Wellcome Sanger Institute COVID-19 Surveillance Team |

| | | | |
|---|---|---|---|
| EPI_ISL_2925238 | Lighthouse Lab in Milton Keynes | Wellcome Sanger Institute for the COVID-19 Genomics UK (COG-UK) Consortium | The Lighthouse Lab in Milton Keynes and Alex Alderton, Roberto Amato, Jeffrey Barrett, Sonia Goncalves, Ewan Harrison, David K. Jackson, Ian Johnston, Dominic Kwiatkowski, Cordelia Langford, John Sillitoe on behalf of the Wellcome Sanger Institute COVID-19 Surveillance Team |
| EPI_ISL_2925239, EPI_ISL_2925240, EPI_ISL_2925241 | Lighthouse Lab in Alderley Park | Wellcome Sanger Institute for the COVID-19 Genomics UK (COG-UK) Consortium | Jacquelyn Wynn, Mairead Hyland, The Lighthouse Lab in Alderley Park and Alex Alderton, Roberto Amato, Jeffrey Barrett, Sonia Goncalves, Ewan Harrison, David K. Jackson, Ian Johnston, Dominic Kwiatkowski, Cordelia Langford, John Sillitoe on behalf of the Wellcome Sanger Institute COVID-19 Surveillance Team |
| EPI_ISL_2925242 | Lighthouse Lab in Milton Keynes | Wellcome Sanger Institute for the COVID-19 Genomics UK (COG-UK) Consortium | The Lighthouse Lab in Milton Keynes and Alex Alderton, Roberto Amato, Jeffrey Barrett, Sonia Goncalves, Ewan Harrison, David K. Jackson, Ian Johnston, Dominic Kwiatkowski, Cordelia Langford, John Sillitoe on behalf of the Wellcome Sanger Institute COVID-19 Surveillance Team |
| EPI_ISL_2925243, EPI_ISL_2925244, EPI_ISL_2925245, EPI_ISL_2925246, EPI_ISL_2925247, EPI_ISL_2925248 | Lighthouse Lab in Alderley Park | Wellcome Sanger Institute for the COVID-19 Genomics UK (COG-UK) Consortium | Jacquelyn Wynn, Mairead Hyland, The Lighthouse Lab in Alderley Park and Alex Alderton, Roberto Amato, Jeffrey Barrett, Sonia Goncalves, Ewan Harrison, David K. Jackson, Ian Johnston, Dominic Kwiatkowski, Cordelia Langford, John Sillitoe on behalf of the Wellcome Sanger Institute COVID-19 Surveillance Team |
| EPI_ISL_2925249 | Lighthouse Lab in Milton Keynes | Wellcome Sanger Institute for the COVID-19 Genomics UK (COG-UK) Consortium | The Lighthouse Lab in Milton Keynes and Alex Alderton, Roberto Amato, Jeffrey Barrett, Sonia Goncalves, Ewan Harrison, David K. Jackson, Ian Johnston, Dominic Kwiatkowski, Cordelia Langford, John Sillitoe on behalf of the Wellcome Sanger Institute COVID-19 Surveillance Team |
| EPI_ISL_2925250 | Lighthouse Lab in Alderley Park | Wellcome Sanger Institute for the COVID-19 Genomics UK (COG-UK) Consortium | Jacquelyn Wynn, Mairead Hyland, The Lighthouse Lab in Alderley Park and Alex Alderton, Roberto Amato, Jeffrey Barrett, Sonia Goncalves, Ewan Harrison, David K. Jackson, Ian Johnston, Dominic Kwiatkowski, Cordelia Langford, John Sillitoe on behalf of the Wellcome Sanger Institute COVID-19 Surveillance Team |
| EPI_ISL_2925251 | Lighthouse Lab in Milton Keynes | Wellcome Sanger Institute for the COVID-19 Genomics UK (COG-UK) Consortium | The Lighthouse Lab in Milton Keynes and Alex Alderton, Roberto Amato, Jeffrey Barrett, Sonia Goncalves, Ewan Harrison, David K. Jackson, Ian Johnston, Dominic Kwiatkowski, Cordelia Langford, John Sillitoe on behalf of the Wellcome Sanger Institute COVID-19 Surveillance Team |
| EPI_ISL_2925252, EPI_ISL_2925253 | Lighthouse Lab in Alderley Park | Wellcome Sanger Institute for the COVID-19 Genomics UK (COG-UK) Consortium | Jacquelyn Wynn, Mairead Hyland, The Lighthouse Lab in Alderley Park and Alex Alderton, Roberto Amato, Jeffrey Barrett, Sonia Goncalves, Ewan Harrison, David K. Jackson, Ian Johnston, Dominic Kwiatkowski, Cordelia Langford, John Sillitoe on behalf of the Wellcome Sanger Institute COVID-19 Surveillance Team |
| EPI_ISL_2925254 | Lighthouse Lab in Milton Keynes | Wellcome Sanger Institute for the COVID-19 Genomics UK (COG-UK) Consortium | The Lighthouse Lab in Milton Keynes and Alex Alderton, Roberto Amato, Jeffrey Barrett, Sonia Goncalves, Ewan Harrison, David K. Jackson, Ian Johnston, Dominic Kwiatkowski, Cordelia Langford, John Sillitoe on behalf of the Wellcome Sanger Institute COVID-19 Surveillance Team |
| EPI_ISL_2925255 | Lighthouse Lab in Alderley Park | Wellcome Sanger Institute for the COVID-19 Genomics UK (COG-UK) Consortium | Jacquelyn Wynn, Mairead Hyland, The Lighthouse Lab in Alderley Park and Alex Alderton, Roberto Amato, Jeffrey Barrett, Sonia Goncalves, Ewan Harrison, David K. Jackson, Ian Johnston, Dominic Kwiatkowski, Cordelia Langford, John Sillitoe on behalf of the Wellcome Sanger Institute COVID-19 Surveillance Team |
| EPI_ISL_2925256 | Lighthouse Lab in Milton Keynes | Wellcome Sanger Institute for the COVID-19 Genomics UK (COG-UK) Consortium | The Lighthouse Lab in Milton Keynes and Alex Alderton, Roberto Amato, Jeffrey Barrett, Sonia Goncalves, Ewan Harrison, David K. Jackson, Ian Johnston, Dominic Kwiatkowski, Cordelia Langford, John Sillitoe on behalf of the Wellcome Sanger Institute COVID-19 Surveillance Team |
| EPI_ISL_2925257, EPI_ISL_2925258 | Lighthouse Lab in Alderley Park | Wellcome Sanger Institute for the COVID-19 Genomics UK (COG-UK) Consortium | Jacquelyn Wynn, Mairead Hyland, The Lighthouse Lab in Alderley Park and Alex Alderton, Roberto Amato, Jeffrey Barrett, Sonia Goncalves, Ewan Harrison, David K. Jackson, Ian Johnston, Dominic Kwiatkowski, Cordelia Langford, John Sillitoe on behalf of the Wellcome Sanger Institute COVID-19 Surveillance Team |
| EPI_ISL_2925259 | Lighthouse Lab in Milton Keynes | Wellcome Sanger Institute for the COVID-19 Genomics UK (COG-UK) Consortium | The Lighthouse Lab in Milton Keynes and Alex Alderton, Roberto Amato, Jeffrey Barrett, Sonia Goncalves, Ewan Harrison, David K. Jackson, Ian Johnston, Dominic Kwiatkowski, Cordelia Langford, John Sillitoe on behalf of the Wellcome Sanger Institute COVID-19 Surveillance Team |
| EPI_ISL_2925260, EPI_ISL_2925261, EPI_ISL_2925262, EPI_ISL_2925263, EPI_ISL_2925264, EPI_ISL_2925265 | Lighthouse Lab in Alderley Park | Wellcome Sanger Institute for the COVID-19 Genomics UK (COG-UK) Consortium | Jacquelyn Wynn, Mairead Hyland, The Lighthouse Lab in Alderley Park and Alex Alderton, Roberto Amato, Jeffrey Barrett, Sonia Goncalves, Ewan Harrison, David K. Jackson, Ian Johnston, Dominic Kwiatkowski, Cordelia Langford, John Sillitoe on behalf of the Wellcome Sanger Institute COVID-19 Surveillance Team |
| EPI_ISL_2925266 | Lighthouse Lab in Milton Keynes | Wellcome Sanger Institute for the COVID-19 Genomics UK (COG-UK) Consortium | The Lighthouse Lab in Milton Keynes and Alex Alderton, Roberto Amato, Jeffrey Barrett, Sonia Goncalves, Ewan Harrison, David K. Jackson, Ian Johnston, Dominic Kwiatkowski, Cordelia Langford, John Sillitoe on behalf of the Wellcome Sanger Institute COVID-19 Surveillance Team |
| EPI_ISL_2925267 | Lighthouse Lab in Alderley Park | Wellcome Sanger Institute for the COVID-19 Genomics UK (COG-UK) Consortium | Jacquelyn Wynn, Mairead Hyland, The Lighthouse Lab in Alderley Park and Alex Alderton, Roberto Amato, Jeffrey Barrett, Sonia Goncalves, Ewan Harrison, David K. Jackson, Ian Johnston, Dominic Kwiatkowski, Cordelia Langford, John Sillitoe on behalf of the Wellcome Sanger Institute COVID-19 Surveillance Team |
| EPI_ISL_2925268, EPI_ISL_2925269 | Lighthouse Lab in Milton Keynes | Wellcome Sanger Institute for the COVID-19 Genomics UK (COG-UK) Consortium | The Lighthouse Lab in Milton Keynes and Alex Alderton, Roberto Amato, Jeffrey Barrett, Sonia Goncalves, Ewan Harrison, David K. Jackson, Ian Johnston, Dominic Kwiatkowski, Cordelia Langford, John Sillitoe on behalf of the Wellcome Sanger Institute COVID-19 Surveillance Team |
| EPI_ISL_2925270, EPI_ISL_2925271, EPI_ISL_2925272, EPI_ISL_2925273 | Lighthouse Lab in Alderley Park | Wellcome Sanger Institute for the COVID-19 Genomics UK (COG-UK) Consortium | Jacquelyn Wynn, Mairead Hyland, The Lighthouse Lab in Alderley Park and Alex Alderton, Roberto Amato, Jeffrey Barrett, Sonia Goncalves, Ewan Harrison, David K. Jackson, Ian Johnston, Dominic Kwiatkowski, Cordelia Langford, John Sillitoe on behalf of the Wellcome Sanger Institute COVID-19 Surveillance Team |
| EPI_ISL_2925274 | Lighthouse Lab in Milton Keynes | Wellcome Sanger Institute for the COVID-19 Genomics UK (COG-UK) Consortium | The Lighthouse Lab in Milton Keynes and Alex Alderton, Roberto Amato, Jeffrey Barrett, Sonia Goncalves, Ewan Harrison, David K. Jackson, Ian Johnston, Dominic Kwiatkowski, Cordelia Langford, John Sillitoe on behalf of the Wellcome Sanger Institute COVID-19 Surveillance Team |
| EPI_ISL_2925275 | Lighthouse Lab in Alderley Park | Wellcome Sanger Institute for the COVID-19 Genomics UK (COG-UK) Consortium | Jacquelyn Wynn, Mairead Hyland, The Lighthouse Lab in Alderley Park and Alex Alderton, Roberto Amato, Jeffrey Barrett, Sonia Goncalves, Ewan Harrison, David K. Jackson, Ian Johnston, Dominic Kwiatkowski, Cordelia Langford, John Sillitoe on behalf of the Wellcome Sanger Institute COVID-19 Surveillance Team |
| EPI_ISL_2925276 | Lighthouse Lab in Milton Keynes | Wellcome Sanger Institute for the COVID-19 Genomics UK (COG-UK) Consortium | The Lighthouse Lab in Milton Keynes and Alex Alderton, Roberto Amato, Jeffrey Barrett, Sonia Goncalves, Ewan Harrison, David K. Jackson, Ian Johnston, Dominic Kwiatkowski, Cordelia Langford, John Sillitoe on behalf of the Wellcome Sanger Institute COVID-19 Surveillance Team |
| EPI_ISL_2925277, EPI_ISL_2925278, EPI_ISL_2925279, EPI_ISL_2925280 | Lighthouse Lab in Alderley Park | Wellcome Sanger Institute for the COVID-19 Genomics UK (COG-UK) Consortium | Jacquelyn Wynn, Mairead Hyland, The Lighthouse Lab in Alderley Park and Alex Alderton, Roberto Amato, Jeffrey Barrett, Sonia Goncalves, Ewan Harrison, David K. Jackson, Ian Johnston, Dominic Kwiatkowski, Cordelia Langford, John Sillitoe on behalf of the Wellcome Sanger Institute COVID-19 Surveillance Team |
| EPI_ISL_2925281 | Lighthouse Lab in Milton Keynes | Wellcome Sanger Institute for the COVID-19 Genomics UK (COG-UK) Consortium | The Lighthouse Lab in Milton Keynes and Alex Alderton, Roberto Amato, Jeffrey Barrett, Sonia Goncalves, Ewan Harrison, David K. Jackson, Ian Johnston, Dominic Kwiatkowski, Cordelia Langford, John Sillitoe on behalf of the Wellcome Sanger Institute COVID-19 Surveillance Team |
| EPI_ISL_2925282, EPI_ISL_2925283 | Lighthouse Lab in Alderley Park | Wellcome Sanger Institute for the COVID-19 Genomics UK (COG-UK) Consortium | Jacquelyn Wynn, Mairead Hyland, The Lighthouse Lab in Alderley Park and Alex Alderton, Roberto Amato, Jeffrey Barrett, Sonia Goncalves, Ewan Harrison, David K. Jackson, Ian Johnston, Dominic Kwiatkowski, Cordelia Langford, John Sillitoe on behalf of the Wellcome Sanger Institute COVID-19 Surveillance Team |
| EPI_ISL_2925284 | Lighthouse Lab in Milton Keynes | Wellcome Sanger Institute for the COVID-19 Genomics UK (COG-UK) Consortium | The Lighthouse Lab in Milton Keynes and Alex Alderton, Roberto Amato, Jeffrey Barrett, Sonia Goncalves, Ewan Harrison, David K. Jackson, Ian Johnston, Dominic Kwiatkowski, Cordelia Langford, John Sillitoe on behalf of the Wellcome Sanger Institute COVID-19 Surveillance Team |
| EPI_ISL_2925285, EPI_ISL_2925286, EPI_ISL_2925287, EPI_ISL_2925288 | Lighthouse Lab in Alderley Park | Wellcome Sanger Institute for the COVID-19 Genomics UK (COG-UK) Consortium | Jacquelyn Wynn, Mairead Hyland, The Lighthouse Lab in Alderley Park and Alex Alderton, Roberto Amato, Jeffrey Barrett, Sonia Goncalves, Ewan Harrison, David K. Jackson, Ian Johnston, Dominic Kwiatkowski, Cordelia Langford, John Sillitoe on behalf of the Wellcome Sanger Institute COVID-19 Surveillance Team |
| EPI_ISL_2925289 | Lighthouse Lab in Milton Keynes | Wellcome Sanger Institute for the COVID-19 Genomics UK (COG-UK) Consortium | The Lighthouse Lab in Milton Keynes and Alex Alderton, Roberto Amato, Jeffrey Barrett, Sonia Goncalves, Ewan Harrison, David K. Jackson, Ian Johnston, Dominic Kwiatkowski, Cordelia Langford, John Sillitoe on behalf of the Wellcome Sanger Institute COVID-19 Surveillance Team |
| EPI_ISL_2925290, EPI_ISL_2925291, EPI_ISL_2925292, EPI_ISL_2925293, EPI_ISL_2925294 | Lighthouse Lab in Alderley Park | Wellcome Sanger Institute for the COVID-19 Genomics UK (COG-UK) Consortium | Jacquelyn Wynn, Mairead Hyland, The Lighthouse Lab in Alderley Park and Alex Alderton, Roberto Amato, Jeffrey Barrett, Sonia Goncalves, Ewan Harrison, David K. Jackson, Ian Johnston, Dominic Kwiatkowski, Cordelia Langford, John Sillitoe on behalf of the Wellcome Sanger Institute COVID-19 Surveillance Team |
| EPI_ISL_2925295 | Lighthouse Lab in Milton Keynes | Wellcome Sanger Institute for the COVID-19 Genomics UK (COG-UK) Consortium | The Lighthouse Lab in Milton Keynes and Alex Alderton, Roberto Amato, Jeffrey Barrett, Sonia Goncalves, Ewan Harrison, David K. Jackson, Ian Johnston, Dominic Kwiatkowski, Cordelia Langford, John Sillitoe on behalf of the Wellcome Sanger Institute COVID-19 Surveillance Team |
| EPI_ISL_2925297, EPI_ISL_2925298, EPI_ISL_2925299, EPI_ISL_2925300, EPI_ISL_2925301, EPI_ISL_2925302 | Lighthouse Lab in Alderley Park | Wellcome Sanger Institute for the COVID-19 Genomics UK (COG-UK) Consortium | Jacquelyn Wynn, Mairead Hyland, The Lighthouse Lab in Alderley Park and Alex Alderton, Roberto Amato, Jeffrey Barrett, Sonia Goncalves, Ewan Harrison, David K. Jackson, Ian Johnston, Dominic Kwiatkowski, Cordelia Langford, John Sillitoe on behalf of the Wellcome Sanger Institute COVID-19 Surveillance Team |
| EPI_ISL_2925303 | Lighthouse Lab in Milton Keynes | Wellcome Sanger Institute for the COVID-19 Genomics UK (COG-UK) Consortium | The Lighthouse Lab in Milton Keynes and Alex Alderton, Roberto Amato, Jeffrey Barrett, Sonia Goncalves, Ewan Harrison, David K. Jackson, Ian Johnston, Dominic Kwiatkowski, Cordelia Langford, John Sillitoe on behalf of the Wellcome Sanger Institute COVID-19 Surveillance Team |
| EPI_ISL_2925304 | Lighthouse Lab in Alderley Park | Wellcome Sanger Institute for the COVID-19 Genomics UK (COG-UK) Consortium | Jacquelyn Wynn, Mairead Hyland, The Lighthouse Lab in Alderley Park and Alex Alderton, Roberto Amato, Jeffrey Barrett, Sonia Goncalves, Ewan Harrison, David K. Jackson, Ian Johnston, Dominic Kwiatkowski, Cordelia Langford, John Sillitoe on behalf of the Wellcome Sanger Institute COVID-19 Surveillance Team |
| EPI_ISL_2925305 | Lighthouse Lab in Milton Keynes | Wellcome Sanger Institute for the COVID-19 Genomics UK (COG-UK) Consortium | The Lighthouse Lab in Milton Keynes and Alex Alderton, Roberto Amato, Jeffrey Barrett, Sonia Goncalves, Ewan Harrison, David K. Jackson, Ian Johnston, Dominic Kwiatkowski, Cordelia Langford, John Sillitoe on behalf of the Wellcome Sanger Institute COVID-19 Surveillance Team |
| EPI_ISL_2925306, EPI_ISL_2925307, EPI_ISL_2925308, EPI_ISL_2925309, EPI_ISL_2925310 | Lighthouse Lab in Alderley Park | Wellcome Sanger Institute for the COVID-19 Genomics UK (COG-UK) Consortium | Jacquelyn Wynn, Mairead Hyland, The Lighthouse Lab in Alderley Park and Alex Alderton, Roberto Amato, Jeffrey Barrett, Sonia Goncalves, Ewan Harrison, David K. Jackson, Ian Johnston, Dominic Kwiatkowski, Cordelia Langford, John Sillitoe on behalf of the Wellcome Sanger Institute COVID-19 Surveillance Team |
| EPI_ISL_2925311, EPI_ISL_2925312, | Lighthouse Lab in Milton Keynes | Wellcome Sanger Institute for the COVID-19 Genomics UK (COG-UK) Consortium | The Lighthouse Lab in Milton Keynes and Alex Alderton, Roberto Amato, Jeffrey Barrett, Sonia Goncalves, Ewan Harrison, David K. Jackson, Ian Johnston, Dominic Kwiatkowski, Cordelia Langford, John Sillitoe on behalf of the Wellcome Sanger Institute COVID-19 Surveillance Team |

| | | | |
|---|---|---|---|
| EPI_ISL_2925313, EPI_ISL_2925314 | | | |
| EPI_ISL_2925315, EPI_ISL_2925316, EPI_ISL_2925317 | Lighthouse Lab in Alderley Park | Wellcome Sanger Institute for the COVID-19 Genomics UK (COG-UK) Consortium | Jacquelyn Wynn, Mairead Hyland, The Lighthouse Lab in Alderley Park and Alex Alderton, Roberto Amato, Jeffrey Barrett, Sonia Goncalves, Ewan Harrison, David K. Jackson, Ian Johnston, Dominic Kwiatkowski, Cordelia Langford, John Sillitoe on behalf of the Wellcome Sanger Institute COVID-19 Surveillance Team |
| EPI_ISL_2925318 | Lighthouse Lab in Milton Keynes | Wellcome Sanger Institute for the COVID-19 Genomics UK (COG-UK) Consortium | The Lighthouse Lab in Milton Keynes and Alex Alderton, Roberto Amato, Jeffrey Barrett, Sonia Goncalves, Ewan Harrison, David K. Jackson, Ian Johnston, Dominic Kwiatkowski, Cordelia Langford, John Sillitoe on behalf of the Wellcome Sanger Institute COVID-19 Surveillance Team |
| EPI_ISL_2925319, EPI_ISL_2925320 | Lighthouse Lab in Alderley Park | Wellcome Sanger Institute for the COVID-19 Genomics UK (COG-UK) Consortium | Jacquelyn Wynn, Mairead Hyland, The Lighthouse Lab in Alderley Park and Alex Alderton, Roberto Amato, Jeffrey Barrett, Sonia Goncalves, Ewan Harrison, David K. Jackson, Ian Johnston, Dominic Kwiatkowski, Cordelia Langford, John Sillitoe on behalf of the Wellcome Sanger Institute COVID-19 Surveillance Team |
| EPI_ISL_2925321 | Lighthouse Lab in Milton Keynes | Wellcome Sanger Institute for the COVID-19 Genomics UK (COG-UK) Consortium | The Lighthouse Lab in Milton Keynes and Alex Alderton, Roberto Amato, Jeffrey Barrett, Sonia Goncalves, Ewan Harrison, David K. Jackson, Ian Johnston, Dominic Kwiatkowski, Cordelia Langford, John Sillitoe on behalf of the Wellcome Sanger Institute COVID-19 Surveillance Team |
| EPI_ISL_2925322 | Lighthouse Lab in Alderley Park | Wellcome Sanger Institute for the COVID-19 Genomics UK (COG-UK) Consortium | Jacquelyn Wynn, Mairead Hyland, The Lighthouse Lab in Alderley Park and Alex Alderton, Roberto Amato, Jeffrey Barrett, Sonia Goncalves, Ewan Harrison, David K. Jackson, Ian Johnston, Dominic Kwiatkowski, Cordelia Langford, John Sillitoe on behalf of the Wellcome Sanger Institute COVID-19 Surveillance Team |
| EPI_ISL_2925323 | Lighthouse Lab in Milton Keynes | Wellcome Sanger Institute for the COVID-19 Genomics UK (COG-UK) Consortium | The Lighthouse Lab in Milton Keynes and Alex Alderton, Roberto Amato, Jeffrey Barrett, Sonia Goncalves, Ewan Harrison, David K. Jackson, Ian Johnston, Dominic Kwiatkowski, Cordelia Langford, John Sillitoe on behalf of the Wellcome Sanger Institute COVID-19 Surveillance Team |
| EPI_ISL_2925324, EPI_ISL_2925325 | Lighthouse Lab in Alderley Park | Wellcome Sanger Institute for the COVID-19 Genomics UK (COG-UK) Consortium | Jacquelyn Wynn, Mairead Hyland, The Lighthouse Lab in Alderley Park and Alex Alderton, Roberto Amato, Jeffrey Barrett, Sonia Goncalves, Ewan Harrison, David K. Jackson, Ian Johnston, Dominic Kwiatkowski, Cordelia Langford, John Sillitoe on behalf of the Wellcome Sanger Institute COVID-19 Surveillance Team |
| EPI_ISL_2925326 | Lighthouse Lab in Milton Keynes | Wellcome Sanger Institute for the COVID-19 Genomics UK (COG-UK) Consortium | The Lighthouse Lab in Milton Keynes and Alex Alderton, Roberto Amato, Jeffrey Barrett, Sonia Goncalves, Ewan Harrison, David K. Jackson, Ian Johnston, Dominic Kwiatkowski, Cordelia Langford, John Sillitoe on behalf of the Wellcome Sanger Institute COVID-19 Surveillance Team |
| EPI_ISL_2925327 | Lighthouse Lab in Alderley Park | Wellcome Sanger Institute for the COVID-19 Genomics UK (COG-UK) Consortium | Jacquelyn Wynn, Mairead Hyland, The Lighthouse Lab in Alderley Park and Alex Alderton, Roberto Amato, Jeffrey Barrett, Sonia Goncalves, Ewan Harrison, David K. Jackson, Ian Johnston, Dominic Kwiatkowski, Cordelia Langford, John Sillitoe on behalf of the Wellcome Sanger Institute COVID-19 Surveillance Team |
| EPI_ISL_2925328 | Lighthouse Lab in Milton Keynes | Wellcome Sanger Institute for the COVID-19 Genomics UK (COG-UK) Consortium | The Lighthouse Lab in Milton Keynes and Alex Alderton, Roberto Amato, Jeffrey Barrett, Sonia Goncalves, Ewan Harrison, David K. Jackson, Ian Johnston, Dominic Kwiatkowski, Cordelia Langford, John Sillitoe on behalf of the Wellcome Sanger Institute COVID-19 Surveillance Team |
| EPI_ISL_2925329, EPI_ISL_2925330 | Lighthouse Lab in Alderley Park | Wellcome Sanger Institute for the COVID-19 Genomics UK (COG-UK) Consortium | Jacquelyn Wynn, Mairead Hyland, The Lighthouse Lab in Alderley Park and Alex Alderton, Roberto Amato, Jeffrey Barrett, Sonia Goncalves, Ewan Harrison, David K. Jackson, Ian Johnston, Dominic Kwiatkowski, Cordelia Langford, John Sillitoe on behalf of the Wellcome Sanger Institute COVID-19 Surveillance Team |
| EPI_ISL_2925331, EPI_ISL_2925332, EPI_ISL_2925333 | Lighthouse Lab in Milton Keynes | Wellcome Sanger Institute for the COVID-19 Genomics UK (COG-UK) Consortium | The Lighthouse Lab in Milton Keynes and Alex Alderton, Roberto Amato, Jeffrey Barrett, Sonia Goncalves, Ewan Harrison, David K. Jackson, Ian Johnston, Dominic Kwiatkowski, Cordelia Langford, John Sillitoe on behalf of the Wellcome Sanger Institute COVID-19 Surveillance Team |
| EPI_ISL_2925334 | Lighthouse Lab in Alderley Park | Wellcome Sanger Institute for the COVID-19 Genomics UK (COG-UK) Consortium | Jacquelyn Wynn, Mairead Hyland, The Lighthouse Lab in Alderley Park and Alex Alderton, Roberto Amato, Jeffrey Barrett, Sonia Goncalves, Ewan Harrison, David K. Jackson, Ian Johnston, Dominic Kwiatkowski, Cordelia Langford, John Sillitoe on behalf of the Wellcome Sanger Institute COVID-19 Surveillance Team |
| EPI_ISL_2925335, EPI_ISL_2925336 | Lighthouse Lab in Milton Keynes | Wellcome Sanger Institute for the COVID-19 Genomics UK (COG-UK) Consortium | The Lighthouse Lab in Milton Keynes and Alex Alderton, Roberto Amato, Jeffrey Barrett, Sonia Goncalves, Ewan Harrison, David K. Jackson, Ian Johnston, Dominic Kwiatkowski, Cordelia Langford, John Sillitoe on behalf of the Wellcome Sanger Institute COVID-19 Surveillance Team |
| EPI_ISL_2925337, EPI_ISL_2925338, EPI_ISL_2925339, EPI_ISL_2925340 | Lighthouse Lab in Alderley Park | Wellcome Sanger Institute for the COVID-19 Genomics UK (COG-UK) Consortium | Jacquelyn Wynn, Mairead Hyland, The Lighthouse Lab in Alderley Park and Alex Alderton, Roberto Amato, Jeffrey Barrett, Sonia Goncalves, Ewan Harrison, David K. Jackson, Ian Johnston, Dominic Kwiatkowski, Cordelia Langford, John Sillitoe on behalf of the Wellcome Sanger Institute COVID-19 Surveillance Team |
| EPI_ISL_2925341 | Lighthouse Lab in Milton Keynes | Wellcome Sanger Institute for the COVID-19 Genomics UK (COG-UK) Consortium | The Lighthouse Lab in Milton Keynes and Alex Alderton, Roberto Amato, Jeffrey Barrett, Sonia Goncalves, Ewan Harrison, David K. Jackson, Ian Johnston, Dominic Kwiatkowski, Cordelia Langford, John Sillitoe on behalf of the Wellcome Sanger Institute COVID-19 Surveillance Team |
| EPI_ISL_2925342 | Lighthouse Lab in Alderley Park | Wellcome Sanger Institute for the COVID-19 Genomics UK (COG-UK) Consortium | Jacquelyn Wynn, Mairead Hyland, The Lighthouse Lab in Alderley Park and Alex Alderton, Roberto Amato, Jeffrey Barrett, Sonia Goncalves, Ewan Harrison, David K. Jackson, Ian Johnston, Dominic Kwiatkowski, Cordelia Langford, John Sillitoe on behalf of the Wellcome Sanger Institute COVID-19 Surveillance Team |
| EPI_ISL_2925343 | Lighthouse Lab in Milton Keynes | Wellcome Sanger Institute for the COVID-19 Genomics UK (COG-UK) Consortium | The Lighthouse Lab in Milton Keynes and Alex Alderton, Roberto Amato, Jeffrey Barrett, Sonia Goncalves, Ewan Harrison, David K. Jackson, Ian Johnston, Dominic Kwiatkowski, Cordelia Langford, John Sillitoe on behalf of the Wellcome Sanger Institute COVID-19 Surveillance Team |
| EPI_ISL_2925344, EPI_ISL_2925345, EPI_ISL_2925346, EPI_ISL_2925347, EPI_ISL_2925348, EPI_ISL_2925349, EPI_ISL_2925350, EPI_ISL_2925351 | Lighthouse Lab in Alderley Park | Wellcome Sanger Institute for the COVID-19 Genomics UK (COG-UK) Consortium | Jacquelyn Wynn, Mairead Hyland, The Lighthouse Lab in Alderley Park and Alex Alderton, Roberto Amato, Jeffrey Barrett, Sonia Goncalves, Ewan Harrison, David K. Jackson, Ian Johnston, Dominic Kwiatkowski, Cordelia Langford, John Sillitoe on behalf of the Wellcome Sanger Institute COVID-19 Surveillance Team |
| EPI_ISL_2925352, EPI_ISL_2925353 | Lighthouse Lab in Milton Keynes | Wellcome Sanger Institute for the COVID-19 Genomics UK (COG-UK) Consortium | The Lighthouse Lab in Milton Keynes and Alex Alderton, Roberto Amato, Jeffrey Barrett, Sonia Goncalves, Ewan Harrison, David K. Jackson, Ian Johnston, Dominic Kwiatkowski, Cordelia Langford, John Sillitoe on behalf of the Wellcome Sanger Institute COVID-19 Surveillance Team |
| EPI_ISL_2925354, EPI_ISL_2925355, EPI_ISL_2925356, EPI_ISL_2925357, EPI_ISL_2925358, EPI_ISL_2925359, EPI_ISL_2925360, EPI_ISL_2925361, EPI_ISL_2925362, EPI_ISL_2925363, EPI_ISL_2925364 | | | |
| see above | Lighthouse Lab in Alderley Park | Wellcome Sanger Institute for the COVID-19 Genomics UK (COG-UK) Consortium | Jacquelyn Wynn, Mairead Hyland, The Lighthouse Lab in Alderley Park and Alex Alderton, Roberto Amato, Jeffrey Barrett, Sonia Goncalves, Ewan Harrison, David K. Jackson, Ian Johnston, Dominic Kwiatkowski, Cordelia Langford, John Sillitoe on behalf of the Wellcome Sanger Institute COVID-19 Surveillance Team |
| EPI_ISL_2925365, EPI_ISL_2925366 | Lighthouse Lab in Milton Keynes | Wellcome Sanger Institute for the COVID-19 Genomics UK (COG-UK) Consortium | The Lighthouse Lab in Milton Keynes and Alex Alderton, Roberto Amato, Jeffrey Barrett, Sonia Goncalves, Ewan Harrison, David K. Jackson, Ian Johnston, Dominic Kwiatkowski, Cordelia Langford, John Sillitoe on behalf of the Wellcome Sanger Institute COVID-19 Surveillance Team |
| EPI_ISL_2925367, EPI_ISL_2925368 | Lighthouse Lab in Alderley Park | Wellcome Sanger Institute for the COVID-19 Genomics UK (COG-UK) Consortium | Jacquelyn Wynn, Mairead Hyland, The Lighthouse Lab in Alderley Park and Alex Alderton, Roberto Amato, Jeffrey Barrett, Sonia Goncalves, Ewan Harrison, David K. Jackson, Ian Johnston, Dominic Kwiatkowski, Cordelia Langford, John Sillitoe on behalf of the Wellcome Sanger Institute COVID-19 Surveillance Team |
| EPI_ISL_2925369, EPI_ISL_2925370 | Lighthouse Lab in Milton Keynes | Wellcome Sanger Institute for the COVID-19 Genomics UK (COG-UK) Consortium | The Lighthouse Lab in Milton Keynes and Alex Alderton, Roberto Amato, Jeffrey Barrett, Sonia Goncalves, Ewan Harrison, David K. Jackson, Ian Johnston, Dominic Kwiatkowski, Cordelia Langford, John Sillitoe on behalf of the Wellcome Sanger Institute COVID-19 Surveillance Team |
| EPI_ISL_2925371, EPI_ISL_2925372, EPI_ISL_2925373, EPI_ISL_2925374, EPI_ISL_2925375, EPI_ISL_2925376, EPI_ISL_2925377, EPI_ISL_2925378 | Lighthouse Lab in Alderley Park | Wellcome Sanger Institute for the COVID-19 Genomics UK (COG-UK) Consortium | Jacquelyn Wynn, Mairead Hyland, The Lighthouse Lab in Alderley Park and Alex Alderton, Roberto Amato, Jeffrey Barrett, Sonia Goncalves, Ewan Harrison, David K. Jackson, Ian Johnston, Dominic Kwiatkowski, Cordelia Langford, John Sillitoe on behalf of the Wellcome Sanger Institute COVID-19 Surveillance Team |
| EPI_ISL_2925379 | Lighthouse Lab in Milton Keynes | Wellcome Sanger Institute for the COVID-19 Genomics UK (COG-UK) Consortium | The Lighthouse Lab in Milton Keynes and Alex Alderton, Roberto Amato, Jeffrey Barrett, Sonia Goncalves, Ewan Harrison, David K. Jackson, Ian Johnston, Dominic Kwiatkowski, Cordelia Langford, John Sillitoe on behalf of the Wellcome Sanger Institute COVID-19 Surveillance Team |
| EPI_ISL_2925380 | Lighthouse Lab in Alderley Park | Wellcome Sanger Institute for the COVID-19 Genomics UK (COG-UK) Consortium | Jacquelyn Wynn, Mairead Hyland, The Lighthouse Lab in Alderley Park and Alex Alderton, Roberto Amato, Jeffrey Barrett, Sonia Goncalves, Ewan Harrison, David K. Jackson, Ian Johnston, Dominic Kwiatkowski, Cordelia Langford, John Sillitoe on behalf of the Wellcome Sanger Institute COVID-19 Surveillance Team |
| EPI_ISL_2925381, EPI_ISL_2925382 | Lighthouse Lab in Milton Keynes | Wellcome Sanger Institute for the COVID-19 Genomics UK (COG-UK) Consortium | The Lighthouse Lab in Milton Keynes and Alex Alderton, Roberto Amato, Jeffrey Barrett, Sonia Goncalves, Ewan Harrison, David K. Jackson, Ian Johnston, Dominic Kwiatkowski, Cordelia Langford, John Sillitoe on behalf of the Wellcome Sanger Institute COVID-19 Surveillance Team |
| EPI_ISL_2925383, EPI_ISL_2925384, EPI_ISL_2925385 | Lighthouse Lab in Alderley Park | Wellcome Sanger Institute for the COVID-19 Genomics UK (COG-UK) Consortium | Jacquelyn Wynn, Mairead Hyland, The Lighthouse Lab in Alderley Park and Alex Alderton, Roberto Amato, Jeffrey Barrett, Sonia Goncalves, Ewan Harrison, David K. Jackson, Ian Johnston, Dominic Kwiatkowski, Cordelia Langford, John Sillitoe on behalf of the Wellcome Sanger Institute COVID-19 Surveillance Team |
| EPI_ISL_2925386, EPI_ISL_2925387 | Lighthouse Lab in Milton Keynes | Wellcome Sanger Institute for the COVID-19 Genomics UK (COG-UK) Consortium | The Lighthouse Lab in Milton Keynes and Alex Alderton, Roberto Amato, Jeffrey Barrett, Sonia Goncalves, Ewan Harrison, David K. Jackson, Ian Johnston, Dominic Kwiatkowski, Cordelia Langford, John Sillitoe on behalf of the Wellcome Sanger Institute COVID-19 Surveillance Team |
| EPI_ISL_2925388, EPI_ISL_2925389, EPI_ISL_2925390, EPI_ISL_2925391, EPI_ISL_2925392, EPI_ISL_2925393, EPI_ISL_2925394, EPI_ISL_2925395, EPI_ISL_2925396, EPI_ISL_2925397, EPI_ISL_2925398, EPI_ISL_2925399, EPI_ISL_2925400, EPI_ISL_2925401, EPI_ISL_2925402, EPI_ISL_2925403 | | | |
| see above | Lighthouse Lab in Alderley Park | Wellcome Sanger Institute for the COVID-19 Genomics UK (COG-UK) Consortium | Jacquelyn Wynn, Mairead Hyland, The Lighthouse Lab in Alderley Park and Alex Alderton, Roberto Amato, Jeffrey Barrett, Sonia Goncalves, Ewan Harrison, David K. Jackson, Ian Johnston, Dominic Kwiatkowski, Cordelia Langford, John Sillitoe on behalf of the Wellcome Sanger Institute COVID-19 Surveillance Team |
| EPI_ISL_2925404 | Lighthouse Lab in Milton Keynes | Wellcome Sanger Institute for the COVID-19 Genomics UK (COG-UK) Consortium | The Lighthouse Lab in Milton Keynes and Alex Alderton, Roberto Amato, Jeffrey Barrett, Sonia Goncalves, Ewan Harrison, David K. Jackson, Ian Johnston, Dominic Kwiatkowski, Cordelia Langford, John Sillitoe on behalf of the Wellcome Sanger Institute COVID-19 Surveillance Team |
| EPI_ISL_2925405, EPI_ISL_2925406, EPI_ISL_2925407 | Lighthouse Lab in Alderley Park | Wellcome Sanger Institute for the COVID-19 Genomics UK (COG-UK) Consortium | Jacquelyn Wynn, Mairead Hyland, The Lighthouse Lab in Alderley Park and Alex Alderton, Roberto Amato, Jeffrey Barrett, Sonia Goncalves, Ewan Harrison, David K. Jackson, Ian Johnston, Dominic Kwiatkowski, Cordelia Langford, John Sillitoe on behalf of the Wellcome Sanger Institute COVID-19 Surveillance Team |
| EPI_ISL_2925408 | Lighthouse Lab in Milton Keynes | Wellcome Sanger Institute for the COVID-19 Genomics UK (COG-UK) Consortium | The Lighthouse Lab in Milton Keynes and Alex Alderton, Roberto Amato, Jeffrey Barrett, Sonia Goncalves, Ewan Harrison, David K. Jackson, Ian Johnston, Dominic Kwiatkowski, Cordelia Langford, John Sillitoe on behalf of the Wellcome Sanger Institute COVID-19 Surveillance Team |
| EPI_ISL_2925409, EPI_ISL_2925410, EPI_ISL_2925411, EPI_ISL_2925412, EPI_ISL_2925413, EPI_ISL_2925414, EPI_ISL_2925415, EPI_ISL_2925416, EPI_ISL_2925417, EPI_ISL_2925418, EPI_ISL_2925419, EPI_ISL_2925420, EPI_ISL_2925421, EPI_ISL_2925422, EPI_ISL_2925423, EPI_ISL_2925424 | | | |
| see above | Lighthouse Lab in Alderley Park | Wellcome Sanger Institute for the COVID-19 Genomics UK (COG-UK) Consortium | Jacquelyn Wynn, Mairead Hyland, The Lighthouse Lab in Alderley Park and Alex Alderton, Roberto Amato, Jeffrey Barrett, Sonia Goncalves, Ewan Harrison, David K. Jackson, Ian Johnston, Dominic Kwiatkowski, Cordelia Langford, John Sillitoe on behalf of the Wellcome Sanger Institute COVID-19 Surveillance Team |

| | | | |
|---|---|---|---|
| EPI_ISL_2925425 | Lighthouse Lab in Milton Keynes | Wellcome Sanger Institute for the COVID-19 Genomics UK (COG-UK) Consortium | The Lighthouse Lab in Milton Keynes and Alex Alderton, Roberto Amato, Jeffrey Barrett, Sonia Goncalves, Ewan Harrison, David K. Jackson, Ian Johnston, Dominic Kwiatkowski, Cordelia Langford, John Sillitoe on behalf of the Wellcome Sanger Institute COVID-19 Surveillance Team |
| EPI_ISL_2925426, EPI_ISL_2925427 | Lighthouse Lab in Alderley Park | Wellcome Sanger Institute for the COVID-19 Genomics UK (COG-UK) Consortium | Jacquelyn Wynn, Mairead Hyland, The Lighthouse Lab in Alderley Park and Alex Alderton, Roberto Amato, Jeffrey Barrett, Sonia Goncalves, Ewan Harrison, David K. Jackson, Ian Johnston, Dominic Kwiatkowski, Cordelia Langford, John Sillitoe on behalf of the Wellcome Sanger Institute COVID-19 Surveillance Team |
| EPI_ISL_2925428 | Lighthouse Lab in Milton Keynes | Wellcome Sanger Institute for the COVID-19 Genomics UK (COG-UK) Consortium | The Lighthouse Lab in Milton Keynes and Alex Alderton, Roberto Amato, Jeffrey Barrett, Sonia Goncalves, Ewan Harrison, David K. Jackson, Ian Johnston, Dominic Kwiatkowski, Cordelia Langford, John Sillitoe on behalf of the Wellcome Sanger Institute COVID-19 Surveillance Team |
| EPI_ISL_2925429 | Lighthouse Lab in Alderley Park | Wellcome Sanger Institute for the COVID-19 Genomics UK (COG-UK) Consortium | Jacquelyn Wynn, Mairead Hyland, The Lighthouse Lab in Alderley Park and Alex Alderton, Roberto Amato, Jeffrey Barrett, Sonia Goncalves, Ewan Harrison, David K. Jackson, Ian Johnston, Dominic Kwiatkowski, Cordelia Langford, John Sillitoe on behalf of the Wellcome Sanger Institute COVID-19 Surveillance Team |
| EPI_ISL_2925430, EPI_ISL_2925431 | Lighthouse Lab in Milton Keynes | Wellcome Sanger Institute for the COVID-19 Genomics UK (COG-UK) Consortium | The Lighthouse Lab in Milton Keynes and Alex Alderton, Roberto Amato, Jeffrey Barrett, Sonia Goncalves, Ewan Harrison, David K. Jackson, Ian Johnston, Dominic Kwiatkowski, Cordelia Langford, John Sillitoe on behalf of the Wellcome Sanger Institute COVID-19 Surveillance Team |
| EPI_ISL_2925432, EPI_ISL_2925433, EPI_ISL_2925434, EPI_ISL_2925435, EPI_ISL_2925436, EPI_ISL_2925437, EPI_ISL_2925438, EPI_ISL_2925439, EPI_ISL_2925440, EPI_ISL_2925441, EPI_ISL_2925442, EPI_ISL_2925443, EPI_ISL_2925444, EPI_ISL_2925445, EPI_ISL_2925446, EPI_ISL_2925447, EPI_ISL_2925448, EPI_ISL_2925449, EPI_ISL_2925450, EPI_ISL_2925451, EPI_ISL_2925452, EPI_ISL_2925453, EPI_ISL_2925454, EPI_ISL_2925455 | | | |
| see above | Lighthouse Lab in Alderley Park | Wellcome Sanger Institute for the COVID-19 Genomics UK (COG-UK) Consortium | Jacquelyn Wynn, Mairead Hyland, The Lighthouse Lab in Alderley Park and Alex Alderton, Roberto Amato, Jeffrey Barrett, Sonia Goncalves, Ewan Harrison, David K. Jackson, Ian Johnston, Dominic Kwiatkowski, Cordelia Langford, John Sillitoe on behalf of the Wellcome Sanger Institute COVID-19 Surveillance Team |
| EPI_ISL_2925456 | Lighthouse Lab in Milton Keynes | Wellcome Sanger Institute for the COVID-19 Genomics UK (COG-UK) Consortium | The Lighthouse Lab in Milton Keynes and Alex Alderton, Roberto Amato, Jeffrey Barrett, Sonia Goncalves, Ewan Harrison, David K. Jackson, Ian Johnston, Dominic Kwiatkowski, Cordelia Langford, John Sillitoe on behalf of the Wellcome Sanger Institute COVID-19 Surveillance Team |
| EPI_ISL_2925457, EPI_ISL_2925458, EPI_ISL_2925459 | Lighthouse Lab in Alderley Park | Wellcome Sanger Institute for the COVID-19 Genomics UK (COG-UK) Consortium | Jacquelyn Wynn, Mairead Hyland, The Lighthouse Lab in Alderley Park and Alex Alderton, Roberto Amato, Jeffrey Barrett, Sonia Goncalves, Ewan Harrison, David K. Jackson, Ian Johnston, Dominic Kwiatkowski, Cordelia Langford, John Sillitoe on behalf of the Wellcome Sanger Institute COVID-19 Surveillance Team |
| EPI_ISL_2925460, EPI_ISL_2925461 | Lighthouse Lab in Milton Keynes | Wellcome Sanger Institute for the COVID-19 Genomics UK (COG-UK) Consortium | The Lighthouse Lab in Milton Keynes and Alex Alderton, Roberto Amato, Jeffrey Barrett, Sonia Goncalves, Ewan Harrison, David K. Jackson, Ian Johnston, Dominic Kwiatkowski, Cordelia Langford, John Sillitoe on behalf of the Wellcome Sanger Institute COVID-19 Surveillance Team |
| EPI_ISL_2925462, EPI_ISL_2925463 | Lighthouse Lab in Alderley Park | Wellcome Sanger Institute for the COVID-19 Genomics UK (COG-UK) Consortium | Jacquelyn Wynn, Mairead Hyland, The Lighthouse Lab in Alderley Park and Alex Alderton, Roberto Amato, Jeffrey Barrett, Sonia Goncalves, Ewan Harrison, David K. Jackson, Ian Johnston, Dominic Kwiatkowski, Cordelia Langford, John Sillitoe on behalf of the Wellcome Sanger Institute COVID-19 Surveillance Team |
| EPI_ISL_2925464 | Lighthouse Lab in Milton Keynes | Wellcome Sanger Institute for the COVID-19 Genomics UK (COG-UK) Consortium | The Lighthouse Lab in Milton Keynes and Alex Alderton, Roberto Amato, Jeffrey Barrett, Sonia Goncalves, Ewan Harrison, David K. Jackson, Ian Johnston, Dominic Kwiatkowski, Cordelia Langford, John Sillitoe on behalf of the Wellcome Sanger Institute COVID-19 Surveillance Team |
| EPI_ISL_2925465 | Lighthouse Lab in Alderley Park | Wellcome Sanger Institute for the COVID-19 Genomics UK (COG-UK) Consortium | Jacquelyn Wynn, Mairead Hyland, The Lighthouse Lab in Alderley Park and Alex Alderton, Roberto Amato, Jeffrey Barrett, Sonia Goncalves, Ewan Harrison, David K. Jackson, Ian Johnston, Dominic Kwiatkowski, Cordelia Langford, John Sillitoe on behalf of the Wellcome Sanger Institute COVID-19 Surveillance Team |
| EPI_ISL_2925466 | Lighthouse Lab in Glasgow | Wellcome Sanger Institute for the COVID-19 Genomics UK (COG-UK) Consortium | Harper VanSteenhouse, Yumi Kasai, David Gray, Carol Clugston, Anna Dominiczak and Alex Alderton, Roberto Amato, Jeffrey Barrett, Sonia Goncalves, Ewan Harrison, David K. Jackson, Ian Johnston, Dominic Kwiatkowski, Cordelia Langford, John Sillitoe on behalf of the Wellcome Sanger Institute COVID-19 Surveillance Team |
| EPI_ISL_2925467 | Lighthouse Lab in Milton Keynes | Wellcome Sanger Institute for the COVID-19 Genomics UK (COG-UK) Consortium | The Lighthouse Lab in Milton Keynes and Alex Alderton, Roberto Amato, Jeffrey Barrett, Sonia Goncalves, Ewan Harrison, David K. Jackson, Ian Johnston, Dominic Kwiatkowski, Cordelia Langford, John Sillitoe on behalf of the Wellcome Sanger Institute COVID-19 Surveillance Team |
| EPI_ISL_2925468 | Lighthouse Lab in Alderley Park | Wellcome Sanger Institute for the COVID-19 Genomics UK (COG-UK) Consortium | Jacquelyn Wynn, Mairead Hyland, The Lighthouse Lab in Alderley Park and Alex Alderton, Roberto Amato, Jeffrey Barrett, Sonia Goncalves, Ewan Harrison, David K. Jackson, Ian Johnston, Dominic Kwiatkowski, Cordelia Langford, John Sillitoe on behalf of the Wellcome Sanger Institute COVID-19 Surveillance Team |
| EPI_ISL_2925469 | Lighthouse Lab in Glasgow | Wellcome Sanger Institute for the COVID-19 Genomics UK (COG-UK) Consortium | Harper VanSteenhouse, Yumi Kasai, David Gray, Carol Clugston, Anna Dominiczak and Alex Alderton, Roberto Amato, Jeffrey Barrett, Sonia Goncalves, Ewan Harrison, David K. Jackson, Ian Johnston, Dominic Kwiatkowski, Cordelia Langford, John Sillitoe on behalf of the Wellcome Sanger Institute COVID-19 Surveillance Team |
| EPI_ISL_2925470 | Lighthouse Lab in Milton Keynes | Wellcome Sanger Institute for the COVID-19 Genomics UK (COG-UK) Consortium | The Lighthouse Lab in Milton Keynes and Alex Alderton, Roberto Amato, Jeffrey Barrett, Sonia Goncalves, Ewan Harrison, David K. Jackson, Ian Johnston, Dominic Kwiatkowski, Cordelia Langford, John Sillitoe on behalf of the Wellcome Sanger Institute COVID-19 Surveillance Team |
| EPI_ISL_2925471, EPI_ISL_2925472 | Lighthouse Lab in Alderley Park | Wellcome Sanger Institute for the COVID-19 Genomics UK (COG-UK) Consortium | Jacquelyn Wynn, Mairead Hyland, The Lighthouse Lab in Alderley Park and Alex Alderton, Roberto Amato, Jeffrey Barrett, Sonia Goncalves, Ewan Harrison, David K. Jackson, Ian Johnston, Dominic Kwiatkowski, Cordelia Langford, John Sillitoe on behalf of the Wellcome Sanger Institute COVID-19 Surveillance Team |
| EPI_ISL_2925473 | Lighthouse Lab in Milton Keynes | Wellcome Sanger Institute for the COVID-19 Genomics UK (COG-UK) Consortium | The Lighthouse Lab in Milton Keynes and Alex Alderton, Roberto Amato, Jeffrey Barrett, Sonia Goncalves, Ewan Harrison, David K. Jackson, Ian Johnston, Dominic Kwiatkowski, Cordelia Langford, John Sillitoe on behalf of the Wellcome Sanger Institute COVID-19 Surveillance Team |
| EPI_ISL_2925482 | INMI | National Institute for Infectious Diseases (INMI) L. Spallanzani I.R.C.C.S | E Giombini, B Bartolini, M Rueca, G Bonfiglio, O Butera, CEM Gruber, F Santini, F Messina, A Di Caro, MR Capobianchi |
| EPI_ISL_2925483, EPI_ISL_2925485, EPI_ISL_2925486, EPI_ISL_2925487, EPI_ISL_2925489, EPI_ISL_2925490 | Policlinico Tor Vergata | National Institute for Infectious Diseases (INMI) L. Spallanzani I.R.C.C.S | CEM Gruber, F Messina, F Santini, M Rueca, G Bonfiglio, O Butera, B Bartolini, E Giombini, MR Capobianchi, A Di Caro |
| EPI_ISL_2925491 | Spaziani | National Institute for Infectious Diseases (INMI) L. Spallanzani I.R.C.C.S | CEM Gruber, F Messina, F Santini, M Rueca, G Bonfiglio, O Butera, B Bartolini, E Giombini, MR Capobianchi, A Di Caro |
| EPI_ISL_2925493, EPI_ISL_2925495 | S. M. Goretti | National Institute for Infectious Diseases (INMI) L. Spallanzani I.R.C.C.S | CEM Gruber, F Messina, F Santini, M Rueca, G Bonfiglio, O Butera, B Bartolini, E Giombini, MR Capobianchi, A Di Caro |
| EPI_ISL_2925496, EPI_ISL_2925497 | S. Camillo De Lellis | National Institute for Infectious Diseases (INMI) L. Spallanzani I.R.C.C.S | CEM Gruber, F Messina, F Santini, M Rueca, G Bonfiglio, O Butera, B Bartolini, E Giombini, MR Capobianchi, A Di Caro |
| EPI_ISL_2925498 | Ospedale S. M. Goretti | National Institute for Infectious Diseases (INMI) L. Spallanzani I.R.C.C.S | CEM Gruber, F Messina, F Santini, M Rueca, G Bonfiglio, O Butera, B Bartolini, E Giombini, MR Capobianchi, A Di Caro |
| EPI_ISL_2925499 | Campus Biomedico | National Institute for Infectious Diseases (INMI) L. Spallanzani I.R.C.C.S | CEM Gruber, F Messina, F Santini, M Rueca, G Bonfiglio, O Butera, B Bartolini, E Giombini, MR Capobianchi, A Di Caro |
| EPI_ISL_2925500 | I.F.O. | National Institute for Infectious Diseases (INMI) L. Spallanzani I.R.C.C.S | CEM Gruber, F Messina, F Santini, M Rueca, G Bonfiglio, O Butera, B Bartolini, E Giombini, MR Capobianchi, A Di Caro |
| EPI_ISL_2925502, EPI_ISL_2925503 | Ospedale di Genzano | National Institute for Infectious Diseases (INMI) L. Spallanzani I.R.C.C.S | CEM Gruber, F Messina, F Santini, M Rueca, G Bonfiglio, O Butera, B Bartolini, E Giombini, MR Capobianchi, A Di Caro |
| EPI_ISL_2925504 | RM6 | National Institute for Infectious Diseases (INMI) L. Spallanzani I.R.C.C.S | CEM Gruber, F Messina, F Santini, M Rueca, G Bonfiglio, O Butera, B Bartolini, E Giombini, MR Capobianchi, A Di Caro |
| EPI_ISL_2925505, EPI_ISL_2925506, EPI_ISL_2925508, EPI_ISL_2925510, EPI_ISL_2925511, EPI_ISL_2925512, EPI_ISL_2925513 | I.F.O. | National Institute for Infectious Diseases (INMI) L. Spallanzani I.R.C.C.S | CEM Gruber, F Messina, F Santini, M Rueca, G Bonfiglio, O Butera, B Bartolini, E Giombini, MR Capobianchi, A Di Caro |
| EPI_ISL_2925516, EPI_ISL_2925518 | A.S.L. ALESSANDRIA | Fondazione del Piemonte per l'Oncologia IRCCS | Antonino Sottile, Silvia Brossa, Paola Marino, Giorgio Giardina |
| EPI_ISL_2925522 | A.S.L. ASTI | Fondazione del Piemonte per l'Oncologia IRCCS | Antonino Sottile, Silvia Brossa, Paola Marino, Giorgio Giardina |
| EPI_ISL_2925524, EPI_ISL_2925525, EPI_ISL_2925526, EPI_ISL_2925529 | A.S.L. NOVARA-BORGOMANERO | Fondazione del Piemonte per l'Oncologia IRCCS | Antonino Sottile, Silvia Brossa, Paola Marino, Giorgio Giardina |
| EPI_ISL_2925532 | A.S.L. VCO | Fondazione del Piemonte per l'Oncologia IRCCS | Antonino Sottile, Silvia Brossa, Paola Marino, Giorgio Giardina |
| EPI_ISL_2925536 | A.S.L. ALESSANDRIA | Fondazione del Piemonte per l'Oncologia IRCCS | Antonino Sottile, Silvia Brossa, Paola Marino, Giorgio Giardina |
| EPI_ISL_2925541 | A.S.L. VCO | Fondazione del Piemonte per l'Oncologia IRCCS | Antonino Sottile, Silvia Brossa, Paola Marino, Giorgio Giardina |
| EPI_ISL_2925543 | A.S.L. TO4 | Fondazione del Piemonte per l'Oncologia IRCCS | Antonino Sottile, Silvia Brossa, Paola Marino, Giorgio Giardina |
| EPI_ISL_2925544, EPI_ISL_2925545, EPI_ISL_2925547, EPI_ISL_2925548 | A.S.L.VERCELLI | Fondazione del Piemonte per l'Oncologia IRCCS | Antonino Sottile, Silvia Brossa, Paola Marino, Giorgio Giardina |
| EPI_ISL_2925550, EPI_ISL_2925551, EPI_ISL_2925553, EPI_ISL_2925554, EPI_ISL_2925555 | Policlinico Tor Vergata | National Institute for Infectious Diseases (INMI) L. Spallanzani I.R.C.C.S | E Giombini, B Bartolini, M Rueca, G Bonfiglio, O Butera, CEM Gruber, F Santini, F Messina, A Di Caro, MR Capobianchi |

| | | | |
|---|---|---|---|
| EPI_ISL_2925557, EPI_ISL_2925560 | Pertini | National Institute for Infectious Diseases (INMI) L. Spallanzani I.R.C.C.S | E Giombini, B Bartolini, M Rueca, G Bonfiglio, O Butera, CEM Gruber, F Santini, F Messina, A Di Caro, MR Capobianchi |
| EPI_ISL_2925563, EPI_ISL_2925564, EPI_ISL_2925565, EPI_ISL_2925566, EPI_ISL_2925567, EPI_ISL_2925568 | Policlinico Tor Vergata | National Institute for Infectious Diseases (INMI) L. Spallanzani I.R.C.C.S | E Giombini, B Bartolini, M Rueca, G Bonfiglio, O Butera, CEM Gruber, F Santini, F Messina, A Di Caro, MR Capobianchi |
| EPI_ISL_2925569 | Bambino Gesù | National Institute for Infectious Diseases (INMI) L. Spallanzani I.R.C.C.S | E Giombini, B Bartolini, M Rueca, G Bonfiglio, O Butera, CEM Gruber, F Santini, F Messina, A Di Caro, MR Capobianchi |
| EPI_ISL_2925572, EPI_ISL_2925573 | INMI "L.Spallanzani" | National Institute for Infectious Diseases (INMI) L. Spallanzani I.R.C.C.S | E Giombini, B Bartolini, M Rueca, G Bonfiglio, O Butera, CEM Gruber, F Santini, F Messina, A Di Caro, MR Capobianchi |
| EPI_ISL_2925577, EPI_ISL_2925578, EPI_ISL_2925584, EPI_ISL_2925593, EPI_ISL_2925606 | Microvida | Microvida | Suzan D. Pas, Jaco J. Verweij, Joep J. J. M. Stohr |
| EPI_ISL_2925621, EPI_ISL_2925624, EPI_ISL_2925625 | Oman-NIC | Oman-National Influenza Center | Laila Al Balushi, Aisha Al-Busaidi, Samiha Al Kharusi, Amina Al Jardani, Hanan Al Kindi, Intisar Al-Shukri, Fahad Al Balushi, Bilal Hussain. |
| EPI_ISL_2925673 | Hospital del Oriente | Laboratorio de Virología HUCA | Sandoval M, Castelló C, Gómez de Oña J, Boga JA, Rojo S, Alvarez-Arguelles ME, Abreu F, Costales I, Perez-Martínez Z, Martín-Rodríguez G, Coto E, Melón S |
| EPI_ISL_2925692 | Laboratorio de Virología HUCA | Laboratorio de Virología HUCA | Sandoval M, Castelló C, Gómez de Oña J, Boga JA, Rojo S, Alvarez-Arguelles ME, Abreu F, Costales I, Perez-Martínez Z, Martín-Rodríguez G, Coto E, Melón S |
| EPI_ISL_2925693 | Osp. S.Giovanni Evangelista | National Institute for Infectious Diseases (INMI) L. Spallanzani I.R.C.C.S | E Giombini, B Bartolini, M Rueca, G Bonfiglio, O Butera, CEM Gruber, F Santini, F Messina, A Di Caro, MR Capobianchi |
| EPI_ISL_2925694 | PTV | National Institute for Infectious Diseases (INMI) L. Spallanzani I.R.C.C.S | E Giombini, B Bartolini, M Rueca, G Bonfiglio, O Butera, CEM Gruber, F Santini, F Messina, A Di Caro, MR Capobianchi |
| EPI_ISL_2925697 | Pertini | National Institute for Infectious Diseases (INMI) L. Spallanzani I.R.C.C.S | E Giombini, B Bartolini, M Rueca, G Bonfiglio, O Butera, CEM Gruber, F Santini, F Messina, A Di Caro, MR Capobianchi |
| EPI_ISL_2925701 | S. Camillo | National Institute for Infectious Diseases (INMI) L. Spallanzani I.R.C.C.S | E Giombini, B Bartolini, M Rueca, G Bonfiglio, O Butera, CEM Gruber, F Santini, F Messina, A Di Caro, MR Capobianchi |
| EPI_ISL_2925702 | Ospedale Sandro Pertini ASL Roma 2 | National Institute for Infectious Diseases (INMI) L. Spallanzani I.R.C.C.S | E Giombini, B Bartolini, M Rueca, G Bonfiglio, O Butera, CEM Gruber, F Santini, F Messina, A Di Caro, MR Capobianchi |
| EPI_ISL_2925703 | INMI Lazzaro Spallanzani IRCCS | National Institute for Infectious Diseases (INMI) L. Spallanzani I.R.C.C.S | E Giombini, B Bartolini, M Rueca, G Bonfiglio, O Butera, CEM Gruber, F Santini, F Messina, A Di Caro, MR Capobianchi |
| EPI_ISL_2925705, EPI_ISL_2925706 | Ospedale Sandro Pertini ASL Roma 2 | National Institute for Infectious Diseases (INMI) L. Spallanzani I.R.C.C.S | E Giombini, B Bartolini, M Rueca, G Bonfiglio, O Butera, CEM Gruber, F Santini, F Messina, A Di Caro, MR Capobianchi |
| EPI_ISL_2925708 | Policlinico Tor Vergata | National Institute for Infectious Diseases (INMI) L. Spallanzani I.R.C.C.S | E Giombini, B Bartolini, M Rueca, G Bonfiglio, O Butera, CEM Gruber, F Santini, F Messina, A Di Caro, MR Capobianchi |
| EPI_ISL_2925709, EPI_ISL_2925710 | Ospedale Sandro Pertini ASL Roma 2 | National Institute for Infectious Diseases (INMI) L. Spallanzani I.R.C.C.S | E Giombini, B Bartolini, M Rueca, G Bonfiglio, O Butera, CEM Gruber, F Santini, F Messina, A Di Caro, MR Capobianchi |
| EPI_ISL_2925713 | Policlinico Tor Vergata | National Institute for Infectious Diseases (INMI) L. Spallanzani I.R.C.C.S | E Giombini, B Bartolini, M Rueca, G Bonfiglio, O Butera, CEM Gruber, F Santini, F Messina, A Di Caro, MR Capobianchi |
| EPI_ISL_2925714, EPI_ISL_2925715 | P.O.U. Rieti | National Institute for Infectious Diseases (INMI) L. Spallanzani I.R.C.C.S | E Giombini, B Bartolini, M Rueca, G Bonfiglio, O Butera, CEM Gruber, F Santini, F Messina, A Di Caro, MR Capobianchi |
| EPI_ISL_2925716 | Policlinico Tor Vergata | National Institute for Infectious Diseases (INMI) L. Spallanzani I.R.C.C.S | E Giombini, B Bartolini, M Rueca, G Bonfiglio, O Butera, CEM Gruber, F Santini, F Messina, A Di Caro, MR Capobianchi |
| EPI_ISL_2925720, EPI_ISL_2925721 | Bambino Gesù | National Institute for Infectious Diseases (INMI) L. Spallanzani I.R.C.C.S | E Giombini, B Bartolini, M Rueca, G Bonfiglio, O Butera, CEM Gruber, F Santini, F Messina, A Di Caro, MR Capobianchi |
| EPI_ISL_2925723 | INMI | National Institute for Infectious Diseases (INMI) L. Spallanzani I.R.C.C.S | E Giombini, B Bartolini, M Rueca, G Bonfiglio, O Butera, CEM Gruber, F Santini, F Messina, A Di Caro, MR Capobianchi |
| EPI_ISL_2925726 | Osp. S.Giovanni Evengelista | National Institute for Infectious Diseases (INMI) L. Spallanzani I.R.C.C.S | E Giombini, B Bartolini, M Rueca, G Bonfiglio, O Butera, CEM Gruber, F Santini, F Messina, A Di Caro, MR Capobianchi |
| EPI_ISL_2925728 | Sant'Andrea | National Institute for Infectious Diseases (INMI) L. Spallanzani I.R.C.C.S | E Giombini, B Bartolini, M Rueca, G Bonfiglio, O Butera, CEM Gruber, F Santini, F Messina, A Di Caro, MR Capobianchi |
| EPI_ISL_2925729 | INMI "L.Spallanzani" | National Institute for Infectious Diseases (INMI) L. Spallanzani I.R.C.C.S | E Giombini, B Bartolini, M Rueca, G Bonfiglio, O Butera, CEM Gruber, F Santini, F Messina, A Di Caro, MR Capobianchi |
| EPI_ISL_2925730 | Bambino Gesù | National Institute for Infectious Diseases (INMI) L. Spallanzani I.R.C.C.S | E Giombini, B Bartolini, M Rueca, G Bonfiglio, O Butera, CEM Gruber, F Santini, F Messina, A Di Caro, MR Capobianchi |
| EPI_ISL_2925731, EPI_ISL_2925732 | Sant'Eugenio | National Institute for Infectious Diseases (INMI) L. Spallanzani I.R.C.C.S | E Giombini, B Bartolini, M Rueca, G Bonfiglio, O Butera, CEM Gruber, F Santini, F Messina, A Di Caro, MR Capobianchi |
| EPI_ISL_2925735, EPI_ISL_2925737, EPI_ISL_2925738 | Hospital del Oriente | Laboratorio de Virología HUCA | Sandoval M, Castelló C, Gómez de Oña J, Boga JA, Rojo S, Alvarez-Arguelles ME, Abreu F, Costales I, Perez-Martínez Z, Martín-Rodríguez G, Coto E, Melón S |
| EPI_ISL_2925742 | Laboratorio de Virología HUCA | Laboratorio de Virología HUCA | Sandoval M, Castelló C, Gómez de Oña J, Boga JA, Rojo S, Alvarez-Arguelles ME, Abreu F, Costales I, Perez-Martínez Z, Martín-Rodríguez G, Coto E, Melón S |
| EPI_ISL_2925763, EPI_ISL_2925764, EPI_ISL_2925768, EPI_ISL_2925770, EPI_ISL_2925771, EPI_ISL_2925772 | Furst Medical Laboratory | Norwegian Institute of Public Health, Department of Virology | Kathrine Stene-Johansen, Kamilla Heddeland Instefjord, Hilde Elshaug, Garcia Llorente Ignacio, Jon Bråte, Engebretsen Serina Beate,Pedersen Benedikte Nevjen, Line Victoria Moen, Debech Nadia, Atiya R Ali,Marie Paulsen Madsen, Rasmus Riis Kopperud, Hilde Vollan, Karoline Bragstad, Olav Hungnes |
| EPI_ISL_2925773 | Ostfold Hospital Trust - Kalnes, Centre for Laboratory Medicine, Section for gene technology and infection serology | Norwegian Institute of Public Health, Department of Virology | Kathrine Stene-Johansen, Kamilla Heddeland Instefjord, Hilde Elshaug, Garcia Llorente Ignacio, Jon Bråte, Engebretsen Serina Beate,Pedersen Benedikte Nevjen, Line Victoria Moen, Debech Nadia, Atiya R Ali,Marie Paulsen Madsen, Rasmus Riis Kopperud, Hilde Vollan, Karoline Bragstad, Olav Hungnes |
| EPI_ISL_2925775, EPI_ISL_2925776, EPI_ISL_2925777, EPI_ISL_2925780, EPI_ISL_2925785, EPI_ISL_2925790, EPI_ISL_2925791, EPI_ISL_2925792, EPI_ISL_2925795, EPI_ISL_2925796, EPI_ISL_2925797, EPI_ISL_2925801, EPI_ISL_2925804, EPI_ISL_2925807, EPI_ISL_2925808, EPI_ISL_2925809 | | | |
| see above | Furst Medical Laboratory | Norwegian Institute of Public Health, Department of Virology | Kathrine Stene-Johansen, Kamilla Heddeland Instefjord, Hilde Elshaug, Garcia Llorente Ignacio, Jon Bråte, Engebretsen Serina Beate,Pedersen Benedikte Nevjen, Line Victoria Moen, Debech Nadia, Atiya R Ali,Marie Paulsen Madsen, Rasmus Riis Kopperud, Hilde Vollan, Karoline Bragstad, Olav Hungnes |
| EPI_ISL_2925811 | Ostfold Hospital Trust - Kalnes, Centre for Laboratory Medicine, Section for gene technology and infection serology | Norwegian Institute of Public Health, Department of Virology | Kathrine Stene-Johansen, Kamilla Heddeland Instefjord, Hilde Elshaug, Garcia Llorente Ignacio, Jon Bråte, Engebretsen Serina Beate,Pedersen Benedikte Nevjen, Line Victoria Moen, Debech Nadia, Atiya R Ali,Marie Paulsen Madsen, Rasmus Riis Kopperud, Hilde Vollan, Karoline Bragstad, Olav Hungnes |
| EPI_ISL_2925812, EPI_ISL_2925813 | Furst Medical Laboratory | Norwegian Institute of Public Health, Department of Virology | Kathrine Stene-Johansen, Kamilla Heddeland Instefjord, Hilde Elshaug, Garcia Llorente Ignacio, Jon Bråte, Engebretsen Serina Beate,Pedersen Benedikte Nevjen, Line Victoria Moen, Debech Nadia, Atiya R Ali,Marie Paulsen Madsen, Rasmus Riis Kopperud, Hilde Vollan, Karoline Bragstad, Olav Hungnes |
| EPI_ISL_2925823 | Clinical Microbiology, Nationwide Children's Hospital | James Molecular Laboratory | Huanyu Wang, Pam Snyder, Zachary Rudy, Dan Jones, Amy Leber |
| EPI_ISL_2925832, EPI_ISL_2925833, EPI_ISL_2925835, EPI_ISL_2925836, EPI_ISL_2925837, EPI_ISL_2925839, EPI_ISL_2925840, EPI_ISL_2925841, EPI_ISL_2925842, EPI_ISL_2925843, EPI_ISL_2925844, EPI_ISL_2925845, EPI_ISL_2925846, EPI_ISL_2925847, EPI_ISL_2925848, EPI_ISL_2925849, EPI_ISL_2925850, EPI_ISL_2925851, EPI_ISL_2925853, EPI_ISL_2925854, EPI_ISL_2925856, EPI_ISL_2925858 | | | |
| see above | AULSS 2 Marca Trevigiana | Istituto Zooprofilattico Sperimentale delle Venezie | Adelaide Milani, Alessia Schivo, Annalisa Salviato, Elisa Palumbo, Erika Giorgia Quaranta, Luca Tassoni, Ambra Pastori, Edoardo Giussani, Alice Fusaro, Isabella Monne, Calogero Terregino, Antonia Ricci |
| EPI_ISL_2925874, EPI_ISL_2925875, EPI_ISL_2925876, EPI_ISL_2925877, EPI_ISL_2925878, EPI_ISL_2925879, EPI_ISL_2925880, EPI_ISL_2925881, EPI_ISL_2925882, EPI_ISL_2925883, EPI_ISL_2925885, EPI_ISL_2925886, EPI_ISL_2925887, EPI_ISL_2925888, EPI_ISL_2925889, EPI_ISL_2925890, EPI_ISL_2925891, EPI_ISL_2925892, EPI_ISL_2925893, EPI_ISL_2925894, EPI_ISL_2925895, EPI_ISL_2925896, EPI_ISL_2925897, EPI_ISL_2925898, EPI_ISL_2925900, EPI_ISL_2925912, EPI_ISL_2925914, EPI_ISL_2925915, EPI_ISL_2925916, EPI_ISL_2925917, EPI_ISL_2925921, EPI_ISL_2925922, EPI_ISL_2925923, EPI_ISL_2925926, EPI_ISL_2925928, EPI_ISL_2925929, EPI_ISL_2925930, EPI_ISL_2925931, EPI_ISL_2925932, EPI_ISL_2925936, EPI_ISL_2925937, EPI_ISL_2925938, EPI_ISL_2925944, EPI_ISL_2925945, EPI_ISL_2925946, EPI_ISL_2925947, EPI_ISL_2925949, EPI_ISL_2925950, EPI_ISL_2925955, EPI_ISL_2925964, EPI_ISL_2925966, EPI_ISL_2925967, EPI_ISL_2925976, EPI_ISL_2926001, EPI_ISL_2926019, EPI_ISL_2926030, EPI_ISL_2926032, EPI_ISL_2926055, EPI_ISL_2926056, EPI_ISL_2926057, EPI_ISL_2926060, EPI_ISL_2926061, EPI_ISL_2926067 | | | |
| see above | Laboratorio CQRC | CQRC_QUALITY CONTROL CHEMICAL BIOLOGICAL RISK_AOOR Villa Sofia Cervello Palermo | Di Gaudio F., Contino F., Brunacci G. |
| EPI_ISL_2926092, EPI_ISL_2926093, EPI_ISL_2926094, EPI_ISL_2926095, EPI_ISL_2926097, EPI_ISL_2926098, EPI_ISL_2926099, EPI_ISL_2926100, EPI_ISL_2926101, EPI_ISL_2926102, EPI_ISL_2926103, EPI_ISL_2926104, EPI_ISL_2926105, EPI_ISL_2926106, EPI_ISL_2926108, EPI_ISL_2926110, EPI_ISL_2926111, EPI_ISL_2926112, EPI_ISL_2926113, EPI_ISL_2926115, EPI_ISL_2926116, EPI_ISL_2926117, EPI_ISL_2926118, EPI_ISL_2926120, EPI_ISL_2926121, EPI_ISL_2926122, EPI_ISL_2926123, EPI_ISL_2926125, EPI_ISL_2926126, EPI_ISL_2926127, EPI_ISL_2926128, EPI_ISL_2926129, EPI_ISL_2926130, EPI_ISL_2926131, EPI_ISL_2926133, EPI_ISL_2926134, EPI_ISL_2926135, EPI_ISL_2926139, EPI_ISL_2926142, EPI_ISL_2926143, EPI_ISL_2926144, EPI_ISL_2926145, EPI_ISL_2926146, EPI_ISL_2926147, EPI_ISL_2926148, EPI_ISL_2926149, EPI_ISL_2926150, EPI_ISL_2926151, EPI_ISL_2926155, EPI_ISL_2926157, EPI_ISL_2926158, EPI_ISL_2926160, EPI_ISL_2926162, EPI_ISL_2926163, EPI_ISL_2926164, EPI_ISL_2926165, EPI_ISL_2926166, EPI_ISL_2926167, EPI_ISL_2926169, EPI_ISL_2926170, EPI_ISL_2926171, EPI_ISL_2926174, EPI_ISL_2926175, EPI_ISL_2926176, EPI_ISL_2926177, EPI_ISL_2926178, EPI_ISL_2926180, EPI_ISL_2926181, EPI_ISL_2926182, EPI_ISL_2926183, EPI_ISL_2926186, EPI_ISL_2926187, EPI_ISL_2926188, EPI_ISL_2926189, EPI_ISL_2926190, EPI_ISL_2926191, EPI_ISL_2926192, EPI_ISL_2926193, EPI_ISL_2926194, EPI_ISL_2926195, EPI_ISL_2926198, EPI_ISL_2926199, EPI_ISL_2926200, EPI_ISL_2926202, EPI_ISL_2926204, EPI_ISL_2926205, EPI_ISL_2926206, EPI_ISL_2926207 | | | |
| see above | Microbiology Department, Laboratori Clínic Metropolitana Nord. | Can Ruti SARS-CoV-2 Sequencing Hub (HUGTiP/IrsiCaixa/IGTP) | Marc Noguera-Julian, Pilar Armengol, Ignacio Blanco, Antoni E Bordoy, Francesc Catala-Moll, Pere-Joan Cardona, Maria Casadellà, Cristina Casañ, Gemma Clara, Bonaventura Clotet, Montserrat Giménez, Mercedes Guerrero, Anna Not, David |

| | | | |
|---|---|---|---|
| | Hospital Universitari Germans Trias i Pujol | | Panisello, Roger Paredes, Mariona Parera, Alexia París, Verónica Saludes, Alba Sánchez, Laia Soler, Lauro Sumoy, and Elisa Martró on behalf of the Can Ruti SARS-CoV-2 Sequencing Hub |
| EPI_ISL_2926210, EPI_ISL_2926219, EPI_ISL_2926220, EPI_ISL_2926224, EPI_ISL_2926228, EPI_ISL_2926230, EPI_ISL_2926231, EPI_ISL_2926232, EPI_ISL_2926233, EPI_ISL_2926235, EPI_ISL_2926236, EPI_ISL_2926238, EPI_ISL_2926239, EPI_ISL_2926241, EPI_ISL_2926243, EPI_ISL_2926247, EPI_ISL_2926251, EPI_ISL_2926252, EPI_ISL_2926253, EPI_ISL_2926258, EPI_ISL_2926260, EPI_ISL_2926262, EPI_ISL_2926263, EPI_ISL_2926265, EPI_ISL_2926266, EPI_ISL_2926268, EPI_ISL_2926270, EPI_ISL_2926273, EPI_ISL_2926274, EPI_ISL_2926275, EPI_ISL_2926276, EPI_ISL_2926278, EPI_ISL_2926280, EPI_ISL_2926283, EPI_ISL_2926284, EPI_ISL_2926286, EPI_ISL_2926287, EPI_ISL_2926288, EPI_ISL_2926289, EPI_ISL_2926290, EPI_ISL_2926291, EPI_ISL_2926292, EPI_ISL_2926293, EPI_ISL_2926294, EPI_ISL_2926295, EPI_ISL_2926301, EPI_ISL_2926302, EPI_ISL_2926306, EPI_ISL_2926307, EPI_ISL_2926311, EPI_ISL_2926312, EPI_ISL_2926313, EPI_ISL_2926314, EPI_ISL_2926315, EPI_ISL_2926316, EPI_ISL_2926317, EPI_ISL_2926318, EPI_ISL_2926320, EPI_ISL_2926322, EPI_ISL_2926325, EPI_ISL_2926327, EPI_ISL_2926329, EPI_ISL_2926330, EPI_ISL_2926331, EPI_ISL_2926333, EPI_ISL_2926334, EPI_ISL_2926335, EPI_ISL_2926337, EPI_ISL_2926338, EPI_ISL_2926340, EPI_ISL_2926344, EPI_ISL_2926345, EPI_ISL_2926348, EPI_ISL_2926349, EPI_ISL_2926350, EPI_ISL_2926351, EPI_ISL_2926352, EPI_ISL_2926355, EPI_ISL_2926357, EPI_ISL_2926358, EPI_ISL_2926359, EPI_ISL_2926361, EPI_ISL_2926364, EPI_ISL_2926365, EPI_ISL_2926367, EPI_ISL_2926370, EPI_ISL_2926371, EPI_ISL_2926373, EPI_ISL_2926374, EPI_ISL_2926375, EPI_ISL_2926377, EPI_ISL_2926381, EPI_ISL_2926382, EPI_ISL_2926383, EPI_ISL_2926384, EPI_ISL_2926386, EPI_ISL_2926388, EPI_ISL_2926389, EPI_ISL_2926390, EPI_ISL_2926391, EPI_ISL_2926393, EPI_ISL_2926397, EPI_ISL_2926400, EPI_ISL_2926401, EPI_ISL_2926404, EPI_ISL_2926406, EPI_ISL_2926408, EPI_ISL_2926409, EPI_ISL_2926411, EPI_ISL_2926412, EPI_ISL_2926413, EPI_ISL_2926416, EPI_ISL_2926417, EPI_ISL_2926418, EPI_ISL_2926419, EPI_ISL_2926421, EPI_ISL_2926424, EPI_ISL_2926425, EPI_ISL_2926426, EPI_ISL_2926427, EPI_ISL_2926428, EPI_ISL_2926430, EPI_ISL_2926431, EPI_ISL_2926432, EPI_ISL_2926433, EPI_ISL_2926439, EPI_ISL_2926441, EPI_ISL_2926443, EPI_ISL_2926444, EPI_ISL_2926445, EPI_ISL_2926446, EPI_ISL_2926447, EPI_ISL_2926448, EPI_ISL_2926449, EPI_ISL_2926450, EPI_ISL_2926452, EPI_ISL_2926454, EPI_ISL_2926457, EPI_ISL_2926458, EPI_ISL_2926459, EPI_ISL_2926460, EPI_ISL_2926461, EPI_ISL_2926462, EPI_ISL_2926464, EPI_ISL_2926465, EPI_ISL_2926466, EPI_ISL_2926467, EPI_ISL_2926468, EPI_ISL_2926469, EPI_ISL_2926470, EPI_ISL_2926471, EPI_ISL_2926472, EPI_ISL_2926473, EPI_ISL_2926474, EPI_ISL_2926475, EPI_ISL_2926476, EPI_ISL_2926477, EPI_ISL_2926478, EPI_ISL_2926479, EPI_ISL_2926481, EPI_ISL_2926482, EPI_ISL_2926483, EPI_ISL_2926486, EPI_ISL_2926487, EPI_ISL_2926488, EPI_ISL_2926491, EPI_ISL_2926493, EPI_ISL_2926494, EPI_ISL_2926496, EPI_ISL_2926497, EPI_ISL_2926499, EPI_ISL_2926501, EPI_ISL_2926502, EPI_ISL_2926503, EPI_ISL_2926505, EPI_ISL_2926506, EPI_ISL_2926509, EPI_ISL_2926510, EPI_ISL_2926511, EPI_ISL_2926513, EPI_ISL_2926518, EPI_ISL_2926519, EPI_ISL_2926522, EPI_ISL_2926524, EPI_ISL_2926525, EPI_ISL_2926526, EPI_ISL_2926527, EPI_ISL_2926533, EPI_ISL_2926535, EPI_ISL_2926536, EPI_ISL_2926538, EPI_ISL_2926539, EPI_ISL_2926540, EPI_ISL_2926541, EPI_ISL_2926542, EPI_ISL_2926543, EPI_ISL_2926544, EPI_ISL_2926545, EPI_ISL_2926546, EPI_ISL_2926547, EPI_ISL_2926548, EPI_ISL_2926549, EPI_ISL_2926550, EPI_ISL_2926551, EPI_ISL_2926554, EPI_ISL_2926555, EPI_ISL_2926557, EPI_ISL_2926558, EPI_ISL_2926562, EPI_ISL_2926563, EPI_ISL_2926564, EPI_ISL_2926567, EPI_ISL_2926570, EPI_ISL_2926571, EPI_ISL_2926572, EPI_ISL_2926573, EPI_ISL_2926574, EPI_ISL_2926575, EPI_ISL_2926576, EPI_ISL_2926577, EPI_ISL_2926578, EPI_ISL_2926579, EPI_ISL_2926580, EPI_ISL_2926581, EPI_ISL_2926582, EPI_ISL_2926583, EPI_ISL_2926584, EPI_ISL_2926585, EPI_ISL_2926587, EPI_ISL_2926588, EPI_ISL_2926589, EPI_ISL_2926591, EPI_ISL_2926592, EPI_ISL_2926593, EPI_ISL_2926594, EPI_ISL_2926595, EPI_ISL_2926596, EPI_ISL_2926597, EPI_ISL_2926598, EPI_ISL_2926599, EPI_ISL_2926601, EPI_ISL_2926602, EPI_ISL_2926603, EPI_ISL_2926604, EPI_ISL_2926605, EPI_ISL_2926606, EPI_ISL_2926608, EPI_ISL_2926614, EPI_ISL_2926615, EPI_ISL_2926619, EPI_ISL_2926622, EPI_ISL_2926624, EPI_ISL_2926625, EPI_ISL_2926627, EPI_ISL_2926629, EPI_ISL_2926630, EPI_ISL_2926633, EPI_ISL_2926634, EPI_ISL_2926635, EPI_ISL_2926636, EPI_ISL_2926637, EPI_ISL_2926638, EPI_ISL_2926639, EPI_ISL_2926642, EPI_ISL_2926643, EPI_ISL_2926644, EPI_ISL_2926645, EPI_ISL_2926646, EPI_ISL_2926647, EPI_ISL_2926648, EPI_ISL_2926649, EPI_ISL_2926650, EPI_ISL_2926651, EPI_ISL_2926652, EPI_ISL_2926653, EPI_ISL_2926657, EPI_ISL_2926658, EPI_ISL_2926659, EPI_ISL_2926662, EPI_ISL_2926668, EPI_ISL_2926669, EPI_ISL_2926671, EPI_ISL_2926672, EPI_ISL_2926674, EPI_ISL_2926677, EPI_ISL_2926678, EPI_ISL_2926679, EPI_ISL_2926680, EPI_ISL_2926681, EPI_ISL_2926682, EPI_ISL_2926684, EPI_ISL_2926685, EPI_ISL_2926686, EPI_ISL_2926687, EPI_ISL_2926689, EPI_ISL_2926690, EPI_ISL_2926691, EPI_ISL_2926694, EPI_ISL_2926695, EPI_ISL_2926697, EPI_ISL_2926700, EPI_ISL_2926702, EPI_ISL_2926705, EPI_ISL_2926706, EPI_ISL_2926707, EPI_ISL_2926709, EPI_ISL_2926711, EPI_ISL_2926712, EPI_ISL_2926713, EPI_ISL_2926714, EPI_ISL_2926716, EPI_ISL_2926718, EPI_ISL_2926719, EPI_ISL_2926720, EPI_ISL_2926721, EPI_ISL_2926722, EPI_ISL_2926723, EPI_ISL_2926725, EPI_ISL_2926726, EPI_ISL_2926727, EPI_ISL_2926728, EPI_ISL_2926729, EPI_ISL_2926730, EPI_ISL_2926734, EPI_ISL_2926735, EPI_ISL_2926736, EPI_ISL_2926737, EPI_ISL_2926739, EPI_ISL_2926741, EPI_ISL_2926742, EPI_ISL_2926743, EPI_ISL_2926744, EPI_ISL_2926747, EPI_ISL_2926748, EPI_ISL_2926749, EPI_ISL_2926750, EPI_ISL_2926752, EPI_ISL_2926753, EPI_ISL_2926754, EPI_ISL_2926755, EPI_ISL_2926756, EPI_ISL_2926757, EPI_ISL_2926758, EPI_ISL_2926760, EPI_ISL_2926761, EPI_ISL_2926762, EPI_ISL_2926763, EPI_ISL_2926764, EPI_ISL_2926765, EPI_ISL_2926767, EPI_ISL_2926770, EPI_ISL_2926774, EPI_ISL_2926775, EPI_ISL_2926777, EPI_ISL_2926779, EPI_ISL_2926780, EPI_ISL_2926781, EPI_ISL_2926782, EPI_ISL_2926784, EPI_ISL_2926786, EPI_ISL_2926787, EPI_ISL_2926789, EPI_ISL_2926791, EPI_ISL_2926792, EPI_ISL_2926793, EPI_ISL_2926797, EPI_ISL_2926798, EPI_ISL_2926799, EPI_ISL_2926800, EPI_ISL_2926801, EPI_ISL_2926802, EPI_ISL_2926803, EPI_ISL_2926804, EPI_ISL_2926805, EPI_ISL_2926806, EPI_ISL_2926807, EPI_ISL_2926808, EPI_ISL_2926809, EPI_ISL_2926810, EPI_ISL_2926811, EPI_ISL_2926812, EPI_ISL_2926814, EPI_ISL_2926815, EPI_ISL_2926816, EPI_ISL_2926817, EPI_ISL_2926818, EPI_ISL_2926819, EPI_ISL_2926820, EPI_ISL_2926821, EPI_ISL_2926822, EPI_ISL_2926823, EPI_ISL_2926824, EPI_ISL_2926825, EPI_ISL_2926826, EPI_ISL_2926827, EPI_ISL_2926828, EPI_ISL_2926829, EPI_ISL_2926830, EPI_ISL_2926832, EPI_ISL_2926833, EPI_ISL_2926834, EPI_ISL_2926835, EPI_ISL_2926836, EPI_ISL_2926837, EPI_ISL_2926838, EPI_ISL_2926839, EPI_ISL_2926840, EPI_ISL_2926841, EPI_ISL_2926843, EPI_ISL_2926845, EPI_ISL_2926846, EPI_ISL_2926848, EPI_ISL_2926849, EPI_ISL_2926850, EPI_ISL_2926852, EPI_ISL_2926854, EPI_ISL_2926855, EPI_ISL_2926861, EPI_ISL_2926863, EPI_ISL_2926865, EPI_ISL_2926866, EPI_ISL_2926867, EPI_ISL_2926869, EPI_ISL_2926872, EPI_ISL_2926876, EPI_ISL_2926877, EPI_ISL_2926878, EPI_ISL_2926879, EPI_ISL_2926880, EPI_ISL_2926881, EPI_ISL_2926882, EPI_ISL_2926883, EPI_ISL_2926884, EPI_ISL_2926885, EPI_ISL_2926886, EPI_ISL_2926887, EPI_ISL_2926888, EPI_ISL_2926889, EPI_ISL_2926890, EPI_ISL_2926892, EPI_ISL_2926893, EPI_ISL_2926894, EPI_ISL_2926895, EPI_ISL_2926896, EPI_ISL_2926899, EPI_ISL_2926900, EPI_ISL_2926902, EPI_ISL_2926904, EPI_ISL_2926905, EPI_ISL_2926907, EPI_ISL_2926908, EPI_ISL_2926909, EPI_ISL_2926910, EPI_ISL_2926912, EPI_ISL_2926914, EPI_ISL_2926915, EPI_ISL_2926916, EPI_ISL_2926918, EPI_ISL_2926922, EPI_ISL_2926923, EPI_ISL_2926925, EPI_ISL_2926927, EPI_ISL_2926928, EPI_ISL_2926930, EPI_ISL_2926932, EPI_ISL_2926933, EPI_ISL_2926936, EPI_ISL_2926938, EPI_ISL_2926943, EPI_ISL_2926947, EPI_ISL_2926949, EPI_ISL_2926951, EPI_ISL_2926952, EPI_ISL_2926953, EPI_ISL_2926954, EPI_ISL_2926956, EPI_ISL_2926957, EPI_ISL_2926958, EPI_ISL_2926962, EPI_ISL_2926964, EPI_ISL_2926967, EPI_ISL_2926968, EPI_ISL_2926970, EPI_ISL_2926971, EPI_ISL_2926972, EPI_ISL_2926974, EPI_ISL_2926975, EPI_ISL_2926976, EPI_ISL_2926977, EPI_ISL_2926980, EPI_ISL_2926982, EPI_ISL_2926984, EPI_ISL_2926985, EPI_ISL_2926986, EPI_ISL_2926987, EPI_ISL_2926989, EPI_ISL_2926990, EPI_ISL_2926992, EPI_ISL_2926993, EPI_ISL_2926994, EPI_ISL_2926995, EPI_ISL_2926996, EPI_ISL_2926997, EPI_ISL_2926998, EPI_ISL_2926999, EPI_ISL_2927000, EPI_ISL_2927002, EPI_ISL_2927003, EPI_ISL_2927004, EPI_ISL_2927005, EPI_ISL_2927008, EPI_ISL_2927009, EPI_ISL_2927010, EPI_ISL_2927011, EPI_ISL_2927012, EPI_ISL_2927014, EPI_ISL_2927017, EPI_ISL_2927018, EPI_ISL_2927019, EPI_ISL_2927020, EPI_ISL_2927023, EPI_ISL_2927026, EPI_ISL_2927027, EPI_ISL_2927028, EPI_ISL_2927029, EPI_ISL_2927030, EPI_ISL_2927032, EPI_ISL_2927033, EPI_ISL_2927034, EPI_ISL_2927035, EPI_ISL_2927036, EPI_ISL_2927037, EPI_ISL_2927038, EPI_ISL_2927039, EPI_ISL_2927040, EPI_ISL_2927042, EPI_ISL_2927043, EPI_ISL_2927044, EPI_ISL_2927045, EPI_ISL_2927048, EPI_ISL_2927049, EPI_ISL_2927050, EPI_ISL_2927052, EPI_ISL_2927053, EPI_ISL_2927054, EPI_ISL_2927055, EPI_ISL_2927056, EPI_ISL_2927057, EPI_ISL_2927059, EPI_ISL_2927061, EPI_ISL_2927063, EPI_ISL_2927064, EPI_ISL_2927065, EPI_ISL_2927066, EPI_ISL_2927067, EPI_ISL_2927068, EPI_ISL_2927069, EPI_ISL_2927070, EPI_ISL_2927071, EPI_ISL_2927073, EPI_ISL_2927074, EPI_ISL_2927075, EPI_ISL_2927077, EPI_ISL_2927078, EPI_ISL_2927079, EPI_ISL_2927080, EPI_ISL_2927081, EPI_ISL_2927082, EPI_ISL_2927085, EPI_ISL_2927086, EPI_ISL_2927087, EPI_ISL_2927088, EPI_ISL_2927089, EPI_ISL_2927090, EPI_ISL_2927091, EPI_ISL_2927092, EPI_ISL_2927093, EPI_ISL_2927095, EPI_ISL_2927096, EPI_ISL_2927097, EPI_ISL_2927098, EPI_ISL_2927099, EPI_ISL_2927100, EPI_ISL_2927101, EPI_ISL_2927102, EPI_ISL_2927103, EPI_ISL_2927104, EPI_ISL_2927105, EPI_ISL_2927106, EPI_ISL_2927107, EPI_ISL_2927108, EPI_ISL_2927109, EPI_ISL_2927110, EPI_ISL_2927111, EPI_ISL_2927112, EPI_ISL_2927116, EPI_ISL_2927117, EPI_ISL_2927118, EPI_ISL_2927119, EPI_ISL_2927120, EPI_ISL_2927121, EPI_ISL_2927122, EPI_ISL_2927125, EPI_ISL_2927126, EPI_ISL_2927133, EPI_ISL_2927134, EPI_ISL_2927142, EPI_ISL_2927147, EPI_ISL_2927152, EPI_ISL_2927153, EPI_ISL_2927154, EPI_ISL_2927158, EPI_ISL_2927161, EPI_ISL_2927162, EPI_ISL_2927164, EPI_ISL_2927170, EPI_ISL_2927171, EPI_ISL_2927172, EPI_ISL_2927173, EPI_ISL_2927174, EPI_ISL_2927176, EPI_ISL_2927180, EPI_ISL_2927181, EPI_ISL_2927182, EPI_ISL_2927186, EPI_ISL_2927189, EPI_ISL_2927190, EPI_ISL_2927191, EPI_ISL_2927192, EPI_ISL_2927193, EPI_ISL_2927194, EPI_ISL_2927200, EPI_ISL_2927201, EPI_ISL_2927202, EPI_ISL_2927203, EPI_ISL_2927204, EPI_ISL_2927208, EPI_ISL_2927209, EPI_ISL_2927210, EPI_ISL_2927211, EPI_ISL_2927215, EPI_ISL_2927217, EPI_ISL_2927222, EPI_ISL_2927223, EPI_ISL_2927224, EPI_ISL_2927226, EPI_ISL_2927228, EPI_ISL_2927231, EPI_ISL_2927234, EPI_ISL_2927235, EPI_ISL_2927236, EPI_ISL_2927237, EPI_ISL_2927238, EPI_ISL_2927239, EPI_ISL_2927243, EPI_ISL_2927248, EPI_ISL_2927249, EPI_ISL_2927250, EPI_ISL_2927251, EPI_ISL_2927252, EPI_ISL_2927254, EPI_ISL_2927256, EPI_ISL_2927257, EPI_ISL_2927260, EPI_ISL_2927261, EPI_ISL_2927262, EPI_ISL_2927263, EPI_ISL_2927264, EPI_ISL_2927266, EPI_ISL_2927268, EPI_ISL_2927271, EPI_ISL_2927272, EPI_ISL_2927273, EPI_ISL_2927274, EPI_ISL_2927275, EPI_ISL_2927278, EPI_ISL_2927279, EPI_ISL_2927280, EPI_ISL_2927287, EPI_ISL_2927288, EPI_ISL_2927290, EPI_ISL_2927291, EPI_ISL_2927292, EPI_ISL_2927294, EPI_ISL_2927295, EPI_ISL_2927296, EPI_ISL_2927299, EPI_ISL_2927301, EPI_ISL_2927302, EPI_ISL_2927303, EPI_ISL_2927305, EPI_ISL_2927308, EPI_ISL_2927310, EPI_ISL_2927311, EPI_ISL_2927313, EPI_ISL_2927316, EPI_ISL_2927317, EPI_ISL_2927318, EPI_ISL_2927322, EPI_ISL_2927323, EPI_ISL_2927324, EPI_ISL_2927325, EPI_ISL_2927329, EPI_ISL_2927331, EPI_ISL_2927336, EPI_ISL_2927338, EPI_ISL_2927339, EPI_ISL_2927341, EPI_ISL_2927342, EPI_ISL_2927343, EPI_ISL_2927344, EPI_ISL_2927345, EPI_ISL_2927346, EPI_ISL_2927347, EPI_ISL_2927349, EPI_ISL_2927351, EPI_ISL_2927353, EPI_ISL_2927354, EPI_ISL_2927355, EPI_ISL_2927356, EPI_ISL_2927357, EPI_ISL_2927358, EPI_ISL_2927362, EPI_ISL_2927363, EPI_ISL_2927365, EPI_ISL_2927366, EPI_ISL_2927368, EPI_ISL_2927369, EPI_ISL_2927370, EPI_ISL_2927371, EPI_ISL_2927372, EPI_ISL_2927373, EPI_ISL_2927374, EPI_ISL_2927375, EPI_ISL_2927376, EPI_ISL_2927380, EPI_ISL_2927382, EPI_ISL_2927383, EPI_ISL_2927384, EPI_ISL_2927385, EPI_ISL_2927386, EPI_ISL_2927389, EPI_ISL_2927390, EPI_ISL_2927391, EPI_ISL_2927394, EPI_ISL_2927396, EPI_ISL_2927397, EPI_ISL_2927399, EPI_ISL_2927400, EPI_ISL_2927405, EPI_ISL_2927406, EPI_ISL_2927407, EPI_ISL_2927411, EPI_ISL_2927412, EPI_ISL_2927414, EPI_ISL_2927415, EPI_ISL_2927416, EPI_ISL_2927417, EPI_ISL_2927418, EPI_ISL_2927419, EPI_ISL_2927420, EPI_ISL_2927421, EPI_ISL_2927422, EPI_ISL_2927423, EPI_ISL_2927425, EPI_ISL_2927426, EPI_ISL_2927428, EPI_ISL_2927429, EPI_ISL_2927430, EPI_ISL_2927431, EPI_ISL_2927432, EPI_ISL_2927433, EPI_ISL_2927434, EPI_ISL_2927438, EPI_ISL_2927440, EPI_ISL_2927442, EPI_ISL_2927446, EPI_ISL_2927448, EPI_ISL_2927451, EPI_ISL_2927453, EPI_ISL_2927454, EPI_ISL_2927455, EPI_ISL_2927457, EPI_ISL_2927458, EPI_ISL_2927459, EPI_ISL_2927462, EPI_ISL_2927463, EPI_ISL_2927464, EPI_ISL_2927465, EPI_ISL_2927466, EPI_ISL_2927468, EPI_ISL_2927469, EPI_ISL_2927470, EPI_ISL_2927471, EPI_ISL_2927473, EPI_ISL_2927474, EPI_ISL_2927477, EPI_ISL_2927478, EPI_ISL_2927480, EPI_ISL_2927481, EPI_ISL_2927482, EPI_ISL_2927484, EPI_ISL_2927491, EPI_ISL_2927492, EPI_ISL_2927493, EPI_ISL_2927494, EPI_ISL_2927495, EPI_ISL_2927496, EPI_ISL_2927498, EPI_ISL_2927499, EPI_ISL_2927500, EPI_ISL_2927503, EPI_ISL_2927507, EPI_ISL_2927508, EPI_ISL_2927510, EPI_ISL_2927511, EPI_ISL_2927512, EPI_ISL_2927514, EPI_ISL_2927515, EPI_ISL_2927518, EPI_ISL_2927520, EPI_ISL_2927521, EPI_ISL_2927524, EPI_ISL_2927525, EPI_ISL_2927526, EPI_ISL_2927527, EPI_ISL_2927528, EPI_ISL_2927529, EPI_ISL_2927530, EPI_ISL_2927533, EPI_ISL_2927534, EPI_ISL_2927535, EPI_ISL_2927538, EPI_ISL_2927539, EPI_ISL_2927546, EPI_ISL_2927551, EPI_ISL_2927554, EPI_ISL_2927555, EPI_ISL_2927558, EPI_ISL_2927559, EPI_ISL_2927561, EPI_ISL_2927563, EPI_ISL_2927564, EPI_ISL_2927565, EPI_ISL_2927566, EPI_ISL_2927567, EPI_ISL_2927568, EPI_ISL_2927569, EPI_ISL_2927570, EPI_ISL_2927573, EPI_ISL_2927574, EPI_ISL_2927575, EPI_ISL_2927576, EPI_ISL_2927577, EPI_ISL_2927578, EPI_ISL_2927579, EPI_ISL_2927580, EPI_ISL_2927581, EPI_ISL_2927582, EPI_ISL_2927583, EPI_ISL_2927584, EPI_ISL_2927585, EPI_ISL_2927589, EPI_ISL_2927590, EPI_ISL_2927592, EPI_ISL_2927593, EPI_ISL_2927594, EPI_ISL_2927596, EPI_ISL_2927598, EPI_ISL_2927600, EPI_ISL_2927601, EPI_ISL_2927603, EPI_ISL_2927605, EPI_ISL_2927606, EPI_ISL_2927607, EPI_ISL_2927608, EPI_ISL_2927609, EPI_ISL_2927610, EPI_ISL_2927612, EPI_ISL_2927613, EPI_ISL_2927614, EPI_ISL_2927615, EPI_ISL_2927618, EPI_ISL_2927620, EPI_ISL_2927621, EPI_ISL_2927624, EPI_ISL_2927625, EPI_ISL_2927628, EPI_ISL_2927629, EPI_ISL_2927631, EPI_ISL_2927632, EPI_ISL_2927634, EPI_ISL_2927635, EPI_ISL_2927636, EPI_ISL_2927637, EPI_ISL_2927639, EPI_ISL_2927640, EPI_ISL_2927641, EPI_ISL_2927642, EPI_ISL_2927643, EPI_ISL_2927644, EPI_ISL_2927645, EPI_ISL_2927647, EPI_ISL_2927648, EPI_ISL_2927649, EPI_ISL_2927650, EPI_ISL_2927651, EPI_ISL_2927652, EPI_ISL_2927655, EPI_ISL_2927656, EPI_ISL_2927657, EPI_ISL_2927658, EPI_ISL_2927663, EPI_ISL_2927664, EPI_ISL_2927665, EPI_ISL_2927666, EPI_ISL_2927667, EPI_ISL_2927668, EPI_ISL_2927669, EPI_ISL_2927670, EPI_ISL_2927671, EPI_ISL_2927673, EPI_ISL_2927675, EPI_ISL_2927677, EPI_ISL_2927678, EPI_ISL_2927679, EPI_ISL_2927681, EPI_ISL_2927683, EPI_ISL_2927687, EPI_ISL_2927689, EPI_ISL_2927690, EPI_ISL_2927692, EPI_ISL_2927693, EPI_ISL_2927696, EPI_ISL_2927697, EPI_ISL_2927698, EPI_ISL_2927699, EPI_ISL_2927700, EPI_ISL_2927702, EPI_ISL_2927703, EPI_ISL_2927705, EPI_ISL_2927706, EPI_ISL_2927710, EPI_ISL_2927713, EPI_ISL_2927714, EPI_ISL_2927716, EPI_ISL_2927717, EPI_ISL_2927721, EPI_ISL_2927722, EPI_ISL_2927724, EPI_ISL_2927725, EPI_ISL_2927727, EPI_ISL_2927728, EPI_ISL_2927730, EPI_ISL_2927731, EPI_ISL_2927732, EPI_ISL_2927733, EPI_ISL_2927734, EPI_ISL_2927735, EPI_ISL_2927736, EPI_ISL_2927737, EPI_ISL_2927740, EPI_ISL_2927741 | | | |
| see above | Laboratory Corporation of America | Centers for Disease Control and Prevention Division of Viral Diseases, Pathogen Discovery | Dakota Howard, Dhwani Batra, Peter W. Cook, Kara Moser, Adrian Paskey, Jason Caravas, Benjamin Rambo-Martin, Shatavia Morrison, Christopher Gulvick, Scott Sammons, Yvette Unoarumhi, Darlene Wagner, Matthew Schmerer, Minoo Agarwal, Eyad Almasri, Debbie Boles, Ayla Burns, Nuthawin Charoensri, Oren Cohen, Susan Countryman, Mary Ann Cristobal, Bobbi Croy, Suzanne Dale, Hrushikesh Deshmukh, Amanda Douglas, Vincent Drouillon, Marcia Eisenberg, Howard Engler, Rama Ghatti, Prashant Gupta, Susan Hicks, Jake Humphrey, Lax Iyer, Lisa Pfefferle, Manoj Jain, Matthew Robinson, Mohan Kolli, Brian Krueger, Tim Kuphal, Stanley Letovsky, Michael Levandoski, Craig Lukasik, Jonathan Meltzer, Brian Norvell, Mindy Nye, Scott Parker, Christos Petropoulos, John Pruitt, Steven Ragan, Scott Ryan, Mike Sapeta, Jana Schroth, Suresh Babu Selvaraju, Goran Stevovic, Amanda Suchanek, Andrea Throop, Lyndon Tilson, Thomas Urban, Joe Voshell, Kimberly Wagner, Jonathan Williams, Mary Williamson, Qian Zeng, Tricia Zwiefelhofer, Clinton R. Paden, Duncan MacCannell |
| EPI_ISL_2927798 | Furst Medical Laboratory | Norwegian Institute of Public Health, Department of Virology | Kathrine Stene-Johansen, Kamilla Heddeland Instefjord, Hilde Elshaug, Garcia Llorente Ignacio, Jon Bråte, Engebretsen Serina Beate,Pedersen Benedikte Nevjen, Line Victoria Moen, Debech Nadia, Atiya R Ali,Marie Paulsen Madsen, Rasmus Riis Kopperud, Hilde Vollan, Karoline Bragstad, Olav Hungnes |
| EPI_ISL_2927801, EPI_ISL_2927803 | Medical Microbiology Unit, Department for Laboratory Medicine, Drammen Hospital, Vestre Viken Health Trust, | Norwegian Institute of Public Health, Department of Virology | Kathrine Stene-Johansen, Kamilla Heddeland Instefjord, Hilde Elshaug, Garcia Llorente Ignacio, Jon Bråte, Engebretsen Serina Beate,Pedersen Benedikte Nevjen, Line Victoria Moen, Debech Nadia, Atiya R Ali,Marie Paulsen Madsen, Rasmus Riis Kopperud, Hilde Vollan, Karoline Bragstad, Olav Hungnes |
| EPI_ISL_2927804, EPI_ISL_2927806, EPI_ISL_2927808, EPI_ISL_2927810, EPI_ISL_2927812, EPI_ISL_2927814, EPI_ISL_2927817, EPI_ISL_2927818, EPI_ISL_2927820, EPI_ISL_2927821, EPI_ISL_2927822, EPI_ISL_2927825, EPI_ISL_2927829, EPI_ISL_2927830, EPI_ISL_2927834, EPI_ISL_2927835, EPI_ISL_2927836, EPI_ISL_2927840, EPI_ISL_2927843, EPI_ISL_2927844, EPI_ISL_2927847, EPI_ISL_2927850, EPI_ISL_2927854, EPI_ISL_2927857, EPI_ISL_2927862, EPI_ISL_2927866, EPI_ISL_2927870, EPI_ISL_2927873, EPI_ISL_2927878 | | | |
| see above | Furst Medical Laboratory | Norwegian Institute of Public Health, Department of Virology | Kathrine Stene-Johansen, Kamilla Heddeland Instefjord, Hilde Elshaug, Garcia Llorente Ignacio, Jon Bråte, Engebretsen Serina Beate,Pedersen Benedikte Nevjen, Line Victoria Moen, Debech Nadia, Atiya R Ali,Marie Paulsen Madsen, Rasmus Riis Kopperud, Hilde Vollan, Karoline Bragstad, Olav Hungnes |
| EPI_ISL_2927889 | Medical Microbiology Unit, Department for Laboratory Medicine, Drammen Hospital, Vestre Viken Health Trust, | Norwegian Institute of Public Health, Department of Virology | Kathrine Stene-Johansen, Kamilla Heddeland Instefjord, Hilde Elshaug, Garcia Llorente Ignacio, Jon Bråte, Engebretsen Serina Beate,Pedersen Benedikte Nevjen, Line Victoria Moen, Debech Nadia, Atiya R Ali,Marie Paulsen Madsen, Rasmus Riis Kopperud, Hilde Vollan, Karoline Bragstad, Olav Hungnes |
| EPI_ISL_2927903 | Furst Medical Laboratory | Norwegian Institute of Public Health, Department of Virology | Kathrine Stene-Johansen, Kamilla Heddeland Instefjord, Hilde Elshaug, Garcia Llorente Ignacio, Jon Bråte, Engebretsen Serina Beate,Pedersen Benedikte Nevjen, Line Victoria Moen, Debech Nadia, Atiya R Ali,Marie Paulsen Madsen, Rasmus Riis Kopperud, Hilde Vollan, Karoline Bragstad, Olav Hungnes |
| EPI_ISL_2927914, EPI_ISL_2927916, EPI_ISL_2927917, EPI_ISL_2927921 | Medical Microbiology Unit, Department for Laboratory Medicine, Drammen Hospital, Vestre Viken Health Trust, | Norwegian Institute of Public Health, Department of Virology | Kathrine Stene-Johansen, Kamilla Heddeland Instefjord, Hilde Elshaug, Garcia Llorente Ignacio, Jon Bråte, Engebretsen Serina Beate,Pedersen Benedikte Nevjen, Line Victoria Moen, Debech Nadia, Atiya R Ali,Marie Paulsen Madsen, Rasmus Riis Kopperud, Hilde Vollan, Karoline Bragstad, Olav Hungnes |
| EPI_ISL_2927945 | Department of Medical Microbiology - section Molde, Molde Hospital | Norwegian Institute of Public Health, Department of Virology | Kathrine Stene-Johansen, Kamilla Heddeland Instefjord, Hilde Elshaug, Garcia Llorente Ignacio, Jon Bråte, Engebretsen Serina Beate,Pedersen Benedikte Nevjen, Line Victoria Moen, Debech Nadia, Atiya R Ali,Marie Paulsen Madsen, Rasmus Riis Kopperud, Hilde Vollan, Karoline Bragstad, Olav Hungnes |
| EPI_ISL_2927948 | Medical Microbiology Unit, Department for Laboratory Medicine, Drammen Hospital, Vestre Viken Health Trust, | Norwegian Institute of Public Health, Department of Virology | Kathrine Stene-Johansen, Kamilla Heddeland Instefjord, Hilde Elshaug, Garcia Llorente Ignacio, Jon Bråte, Engebretsen Serina Beate,Pedersen Benedikte Nevjen, Line Victoria Moen, Debech Nadia, Atiya R Ali,Marie Paulsen Madsen, Rasmus Riis Kopperud, Hilde Vollan, Karoline Bragstad, Olav Hungnes |
| EPI_ISL_2927952 | Department of Medical Microbiology, St. Olavs hospital | Norwegian Institute of Public Health, Department of Virology | Kathrine Stene-Johansen, Kamilla Heddeland Instefjord, Hilde Elshaug, Garcia Llorente Ignacio, Jon Bråte, Engebretsen Serina Beate,Pedersen Benedikte Nevjen, Line Victoria Moen, Debech Nadia, Atiya R Ali,Marie Paulsen Madsen, Rasmus Riis Kopperud, Hilde Vollan, Karoline Bragstad, Olav Hungnes |
| EPI_ISL_2927958, EPI_ISL_2927959, EPI_ISL_2927960 | Medical Microbiology Unit, Department for Laboratory Medicine, Drammen Hospital, Vestre Viken Health Trust, | Norwegian Institute of Public Health, Department of Virology | Kathrine Stene-Johansen, Kamilla Heddeland Instefjord, Hilde Elshaug, Garcia Llorente Ignacio, Jon Bråte, Engebretsen Serina Beate,Pedersen Benedikte Nevjen, Line Victoria Moen, Debech Nadia, Atiya R Ali,Marie Paulsen Madsen, Rasmus Riis Kopperud, Hilde Vollan, Karoline Bragstad, Olav Hungnes |
| EPI_ISL_2927969 | Laboratorios Delia Barraza | Laboratorio nacional-LANIIA-CIAD | Consorcio Mexicano de Vigilancia Genómica (CoViGen-Mex). Authors (in alphabetical order): Julio Elias Alvarado-Yaah, Carlos F. Arias, Santiago Ávila-Ríos, Eduardo Becerril-Vargas, Víctor Hugo Borja-Aburto, Celia Boukadida, Cristóbal Cháidez-Quiróz, Juan Bautista Chale-Dzul, Ricardo Ciria Merce, Andreu Comas-García, Célida Duque Molina, Julissa Enciso-Ibarra, José Antonio Enciso-Moreno, Gloria Elena Espinosa-Ayala, Fernando Fontove-Herrera, Daniel Fregoso-Rueda, Victor Eduardo García-Arias, Alejandra García-Gasca, Bruno Gómez-Gil, Jean Pierre González, Irvin González-López, Concepción Grajales-Muñiz, Ricardo Grande, Rosa María Gutiérrez Rios, Jesús Hernández, Alejandra Hernández-Terán, Alfredo Herrera-Estrella, Carla Ivón Herrera-Najera, Pavel Isa, Verónica Mata-Haro, Daniel Lira Morales, Susana Lopez, Antonio Loza Román, Brenda Irasema Maldonado-Meza, Bernardo Martínez-Miguel, José Arturo Martínez-Orozco, Célida Martínez-Rodríguez, Margarita Matías-Florentino, Fidencio Mejía-Nepomuceno, María Guadalupe de Jesús Mireles-Rivera, Gloria María Molina-Salinas, Hector Montoya-Fuentes, , Mario Mújica-Sánchez, José Esteban Muñoz-Medina, José de Jesús Nuñez-Contreras, Alicia Ocaña-Mondragón, Luis Alberto Ochoa-Carrera, Hector Esteban Paz-Juárez, Marissa Perez-Garcia, Francisco Pulido, Helen Haydee Fernanda Ramírez-Plascencia, Jorge Salas-Hernández, Angel Gustavo Salas-Lais, Alejandro Sánchez-Flores, Clara Esperanza Santacruz-Tinoco, María Guadalupe Santiago-Mauricio, Selene Zárate, Nelly Sélem-Mojica, Blanca Taboada, Gloria Vazquez, Joel Armando Vázquez-Pérez. Laboratorios Delia Barraza: María Delia Barraza Sámano, Claudia Soto Félix, Melissa García Angulo. |
| EPI_ISL_2927970 | Servicio Diagnóstico Clínico | Laboratorio nacional-LANIIA-CIAD | Consorcio Mexicano de Vigilancia Genómica (CoViGen-Mex). Authors (in alphabetical order): Julio Elias Alvarado-Yaah, Carlos F. Arias, Santiago Ávila-Ríos, Eduardo Becerril-Vargas, Víctor Hugo Borja-Aburto, Celia Boukadida, Cristóbal Cháidez-Quiróz, Juan Bautista Chale-Dzul, Ricardo Ciria Merce, Andreu Comas-García, Célida Duque Molina, Julissa Enciso-Ibarra, José Antonio Enciso-Moreno, Gloria Elena Espinosa-Ayala, Fernando Fontove-Herrera, Daniel Fregoso-Rueda, Victor Eduardo García-Arias, Alejandra García-Gasca, Bruno Gómez-Gil, Jean Pierre González, Irvin González-López, Concepción Grajales-Muñiz, Ricardo Grande, Rosa María Gutiérrez Rios, Jesús Hernández, Alejandra Hernández-Terán, Alfredo Herrera-Estrella, Carla Ivón Herrera-Najera, Pavel Isa, Verónica Mata-Haro, Daniel Lira Morales, Susana Lopez, Antonio Loza Román, Brenda Irasema Maldonado-Meza, Bernardo Martínez-Miguel, José Arturo Martínez-Orozco, Célida Martínez-Rodríguez, Margarita Matías-Florentino, Fidencio Mejía-Nepomuceno, María Guadalupe de Jesús Mireles-Rivera, Gloria María Molina-Salinas, Hector Montoya-Fuentes, , Mario Mújica-Sánchez, José Esteban Muñoz-Medina, José de Jesús Nuñez-Contreras, Alicia Ocaña-Mondragón, Luis Alberto Ochoa-Carrera, Hector Esteban Paz-Juárez, Marissa Perez-Garcia, Francisco Pulido, Helen Haydee Fernanda Ramírez-Plascencia, Jorge Salas-Hernández, Angel Gustavo Salas-Lais, Alejandro Sánchez-Flores, Clara Esperanza Santacruz-Tinoco, María Guadalupe Santiago-Mauricio, Selene Zárate, Nelly Sélem-Mojica, Blanca Taboada, Gloria Vazquez, Joel Armando Vázquez-Pérez. |
| EPI_ISL_2927982 | AOUIVR | Istituto Zooprofilattico Sperimentale delle Venezie | Adelaide Milani, Alessia Schivo, Annalisa Salviato, Elisa Palumbo, Erika Giorgia Quaranta, Luca Tassoni, Ambra Pastori, Edoardo Giussani, Alice Fusaro, Isabella Monne, Calogero Terregino, Antonia Ricci |
| EPI_ISL_2927992, EPI_ISL_2927996 | AULSS 9 Scaligera | Istituto Zooprofilattico Sperimentale delle Venezie | Adelaide Milani, Alessia Schivo, Annalisa Salviato, Elisa Palumbo, Erika Giorgia Quaranta, Luca Tassoni, Ambra Pastori, Edoardo Giussani, Alice Fusaro, Isabella Monne, Calogero Terregino, Antonia Ricci |
| EPI_ISL_2928006, EPI_ISL_2928008, EPI_ISL_2928011 | National Institute of Public Health, Bujumbura,Burundi, MRC/UVRI & LSHTM Uganda Research Unit | MRC/UVRI & LSHTM Uganda Research Unit, National Institute of Public Health | Jerome Nkurunziza , Alexis Niyomwungere, Léopold Ouedraogo, Anatole Nkeshimana, Dan Lule Bugembe, Deogratius Ssemwanga, Pontiano Kaleebu, Matthew Cotten, My V.T. Phan |
| EPI_ISL_2928016, EPI_ISL_2928023, EPI_ISL_2928024 | South Sudan Ministry of Health, WHO South Sudan, MRC/UVRI & LSHTM Uganda Research Unit | MRC/UVRI & LSHTM Uganda Research Unit, South Sudan Ministry of Health, WHO South Sudan | Joseph Francis Wamala, Abe G. Abias, Dennis Kenyi Lodiongo, Sudhir Bunga, James Ayei, Lul Lojok Deng, Richard Lino Loro Lako, John Rumunu, Juma John HM, Dan Lule Bugembe, Pontiano Kaleebu, Matthew Cotten, My V.T. Phan |
| EPI_ISL_2928041, EPI_ISL_2928042, | New Mexico Department of Health Scientific Laboratory | New Mexico Department of Health Scientific Laboratory | Ellie Johnson, D'eldra Malone, Jennifer Benoit, Ratheesh Rajan, Linda Salazar, Keila Gutierrez, Anastacia Griego-Fisher |

| | | | |
|---|---|---|---|
| EPI_ISL_2928043, EPI_ISL_2928044 | | | |
| EPI_ISL_2928110, EPI_ISL_2928115 | Curative Labs | New Mexico Department of Health Scientific Laboratory | Ellie Johnson, D'eldra Malone, Jennifer Benoit, Ratheesh Rajan, Linda Salazar, Keila Gutierrez, Anastacia Griego-Fisher |
| EPI_ISL_2928119, EPI_ISL_2928120, EPI_ISL_2928122, EPI_ISL_2928127 | New Mexico Department of Health Scientific Laboratory | New Mexico Department of Health Scientific Laboratory | Ellie Johnson, D'eldra Malone, Jennifer Benoit, Ratheesh Rajan, Linda Salazar, Keila Gutierrez, Anastacia Griego-Fisher |
| EPI_ISL_2928155, EPI_ISL_2928157, EPI_ISL_2928160, EPI_ISL_2928165, EPI_ISL_2928166, EPI_ISL_2928177, EPI_ISL_2928181, EPI_ISL_2928182, EPI_ISL_2928184, EPI_ISL_2928185, EPI_ISL_2928186, EPI_ISL_2928187, EPI_ISL_2928188, EPI_ISL_2928189, EPI_ISL_2928199, EPI_ISL_2928200, EPI_ISL_2928201, EPI_ISL_2928202, EPI_ISL_2928206, EPI_ISL_2928209, EPI_ISL_2928210, EPI_ISL_2928211, EPI_ISL_2928212, EPI_ISL_2928213, EPI_ISL_2928214, EPI_ISL_2928217, EPI_ISL_2928218, EPI_ISL_2928219, EPI_ISL_2928220 | | | |
| see above | Orange County Public Health Laboratory | Orange County Public Health Laboratory | Angelica Torres, Julia Wolfe |
| EPI_ISL_2928226, EPI_ISL_2928227, EPI_ISL_2928228, EPI_ISL_2928229, EPI_ISL_2928230, EPI_ISL_2928231, EPI_ISL_2928232, EPI_ISL_2928233, EPI_ISL_2928234, EPI_ISL_2928235, EPI_ISL_2928236, EPI_ISL_2928237, EPI_ISL_2928238, EPI_ISL_2928239, EPI_ISL_2928240, EPI_ISL_2928241, EPI_ISL_2928243, EPI_ISL_2928244, EPI_ISL_2928246, EPI_ISL_2928248, EPI_ISL_2928249, EPI_ISL_2928250, EPI_ISL_2928251, EPI_ISL_2928252, EPI_ISL_2928253, EPI_ISL_2928255, EPI_ISL_2928258, EPI_ISL_2928260, EPI_ISL_2928261, EPI_ISL_2928262, EPI_ISL_2928263 | | | |
| see above | Baylor Scott & White-Temple | Baylor Scott & White-Temple | Ari Rao, Linden Morales, Kimberly Walker, Marcus Volz, Shelby Johnson |
| EPI_ISL_2928264 | Baylor Scott & White-Waxahachie | Baylor Scott & White-Temple | Ari Rao, Linden Morales, Kimberly Walker, Marcus Volz, Shelby Johnson |
| EPI_ISL_2928265, EPI_ISL_2928266, EPI_ISL_2928267, EPI_ISL_2928268, EPI_ISL_2928269, EPI_ISL_2928270, EPI_ISL_2928271, EPI_ISL_2928272, EPI_ISL_2928273, EPI_ISL_2928274, EPI_ISL_2928276, EPI_ISL_2928277, EPI_ISL_2928278, EPI_ISL_2928279, EPI_ISL_2928282, EPI_ISL_2928283, EPI_ISL_2928284, EPI_ISL_2928285, EPI_ISL_2928286, EPI_ISL_2928287, EPI_ISL_2928288, EPI_ISL_2928289, EPI_ISL_2928290, EPI_ISL_2928291, EPI_ISL_2928292, EPI_ISL_2928293, EPI_ISL_2928294, EPI_ISL_2928295, EPI_ISL_2928296 | | | |
| see above | Baylor Scott & White-Temple | Baylor Scott & White-Temple | Ari Rao, Linden Morales, Kimberly Walker, Marcus Volz, Shelby Johnson |
| EPI_ISL_2928345, EPI_ISL_2928346, EPI_ISL_2928347 | Mobile County Health Department | Synergy Laboratories | Megan Cornwell |
| EPI_ISL_2928348, EPI_ISL_2928349, EPI_ISL_2928350 | USA Health Care Authority | Synergy Laboratories | Megan Cornwell |
| EPI_ISL_2928351, EPI_ISL_2928352, EPI_ISL_2928353, EPI_ISL_2928354, EPI_ISL_2928355, EPI_ISL_2928356, EPI_ISL_2928357, EPI_ISL_2928358, EPI_ISL_2928360 | Mobile County Health Department | Synergy Laboratories | Megan Cornwell |
| EPI_ISL_2928361, EPI_ISL_2928362 | Mobile Infirmary Medical Center | Synergy Laboratories | Megan Cornwell |
| EPI_ISL_2928363, EPI_ISL_2928364, EPI_ISL_2928365, EPI_ISL_2928366, EPI_ISL_2928369, EPI_ISL_2928370, EPI_ISL_2928371, EPI_ISL_2928372, EPI_ISL_2928373, EPI_ISL_2928374, EPI_ISL_2928375, EPI_ISL_2928376 | | | |
| see above | Mobile County Health Department | Synergy Laboratories | Megan Cornwell |
| EPI_ISL_2928379 | Synergy Laboratories | Synergy Laboratories | Megan Cornwell |
| EPI_ISL_2928381, EPI_ISL_2928383 | Minnesota Department of Health, Public Health Laboratory | Minnesota Department of Health, Public Health Laboratory | Alexandra Lorentz, Jacob Garfin, Matt Plumb, and Xiong Wang |
| EPI_ISL_2928385 | Regions Hospital | Minnesota Department of Health, Public Health Laboratory | Alexandra Lorentz, Jacob Garfin, Matt Plumb, and Xiong Wang |
| EPI_ISL_2928386 | Hennepin County Medical Center | Minnesota Department of Health, Public Health Laboratory | Alexandra Lorentz, Jacob Garfin, Matt Plumb, and Xiong Wang |
| EPI_ISL_2928388 | Centracare Laboratory Services | Minnesota Department of Health, Public Health Laboratory | Alexandra Lorentz, Jacob Garfin, Matt Plumb, and Xiong Wang |
| EPI_ISL_2928391, EPI_ISL_2928393, EPI_ISL_2928394, EPI_ISL_2928395, EPI_ISL_2928396, EPI_ISL_2928397, EPI_ISL_2928398, EPI_ISL_2928399, EPI_ISL_2928400, EPI_ISL_2928401 | Minnesota Department of Health, Public Health Laboratory | Minnesota Department of Health, Public Health Laboratory | Alexandra Lorentz, Jacob Garfin, Matt Plumb, and Xiong Wang |
| EPI_ISL_2928405, EPI_ISL_2928408 | Medical Laboratories Duesseldorf | Center of Medical Microbiology, Virology, and Hospital Hygiene, University of Duesseldorf | Maximilian Damagnez;Alexander Dilthey;Angelika Helmer;Torsten Houwaart;Lisanna Hülse;Christian Lange;Malte Kohns Vasconcelos;Nadine Lübke;Jessica Nicolai;Klaus Pfeffer;Daniel Strelow;Jörg Timm;Andreas Walker;Tobias Wienemann |
| EPI_ISL_2928424, EPI_ISL_2928427, EPI_ISL_2928429, EPI_ISL_2928430, EPI_ISL_2928434, EPI_ISL_2928437 | Humboldt County Public Health Laboratory | Humboldt County Public Health Laboratory | Jeremy Corrigan |
| EPI_ISL_2928525, EPI_ISL_2928526, EPI_ISL_2928527, EPI_ISL_2928528, EPI_ISL_2928530, EPI_ISL_2928533, EPI_ISL_2928534, EPI_ISL_2928535, EPI_ISL_2928538, EPI_ISL_2928540 | Ramathibodi Hospital | COVID-19 Network Investigations (CONI) Alliance | Elizabeth Batty, Wasun Chantratita, Thanat Chookajorn, Stefan Fernandez, Angkana Huang, Anthony R. Jones, Khajohn Joonlasak, Chonticha Klungtong, Theerarat Kochakarn, Namfon Kotanan, Krittikorn Kumpornsin, Duangkamon Loesbanluechai, Wudtichai Manasatienkij, Bhakbhoom Panthan, Ekawat Pasomsub, Kingkan Rakmanee, Insee Sensorn, Janjira Thaipadungpanit, Arporn Wangwiwatsin, Treewat Watthanachockchai |
| EPI_ISL_2928542, EPI_ISL_2928543, EPI_ISL_2928544, EPI_ISL_2928545, EPI_ISL_2928546 | Institute for Urban Disease Control and Prevention | COVID-19 Network Investigations (CONI) Alliance | Elizabeth Batty, Wasun Chantratita, Thanat Chookajorn, Stefan Fernandez, Angkana Huang, Anthony R. Jones, Khajohn Joonlasak, Chonticha Klungtong, Theerarat Kochakarn, Namfon Kotanan, Krittikorn Kumpornsin, Duangkamon Loesbanluechai, Wudtichai Manasatienkij, Bhakbhoom Panthan, Ekawat Pasomsub, Kingkan Rakmanee, Insee Sensorn, Janjira Thaipadungpanit, Arporn Wangwiwatsin, Treewat Watthanachockchai, Kamolthip Atsawawaranunt, Prayuth Kaewmalang, Amornmas Kongklieng, Pukkaporn Parnwijitkul, Vichan Pawun |
| EPI_ISL_2928553, EPI_ISL_2928555, EPI_ISL_2928556, EPI_ISL_2928557, EPI_ISL_2928559, EPI_ISL_2928560, EPI_ISL_2928561, EPI_ISL_2928562, EPI_ISL_2928564, EPI_ISL_2928565, EPI_ISL_2928566, EPI_ISL_2928568, EPI_ISL_2928569, EPI_ISL_2928570, EPI_ISL_2928571, EPI_ISL_2928573, EPI_ISL_2928574, EPI_ISL_2928575, EPI_ISL_2928576, EPI_ISL_2928577, EPI_ISL_2928578, EPI_ISL_2928579, EPI_ISL_2928580, EPI_ISL_2928581, EPI_ISL_2928582, EPI_ISL_2928583, EPI_ISL_2928584, EPI_ISL_2928585, EPI_ISL_2928586, EPI_ISL_2928587, EPI_ISL_2928588, EPI_ISL_2928589, EPI_ISL_2928590, EPI_ISL_2928591, EPI_ISL_2928592, EPI_ISL_2928593, EPI_ISL_2928594, EPI_ISL_2928596, EPI_ISL_2928597, EPI_ISL_2928599, EPI_ISL_2928600, EPI_ISL_2928601, EPI_ISL_2928602, EPI_ISL_2928603, EPI_ISL_2928604, EPI_ISL_2928605, EPI_ISL_2928606, EPI_ISL_2928607, EPI_ISL_2928610, EPI_ISL_2928611, EPI_ISL_2928613, EPI_ISL_2928614, EPI_ISL_2928615, EPI_ISL_2928616, EPI_ISL_2928617, EPI_ISL_2928618, EPI_ISL_2928619, EPI_ISL_2928620, EPI_ISL_2928621, EPI_ISL_2928622, EPI_ISL_2928623, EPI_ISL_2928625, EPI_ISL_2928626, EPI_ISL_2928627, EPI_ISL_2928628, EPI_ISL_2928629, EPI_ISL_2928630, EPI_ISL_2928631, EPI_ISL_2928632, EPI_ISL_2928635, EPI_ISL_2928636, EPI_ISL_2928637, EPI_ISL_2928638, EPI_ISL_2928641, EPI_ISL_2928642, EPI_ISL_2928643, EPI_ISL_2928644, EPI_ISL_2928646, EPI_ISL_2928648, EPI_ISL_2928650, EPI_ISL_2928652, EPI_ISL_2928654, EPI_ISL_2928655, EPI_ISL_2928658, EPI_ISL_2928660, EPI_ISL_2928661, EPI_ISL_2928663, EPI_ISL_2928665, EPI_ISL_2928666, EPI_ISL_2928668, EPI_ISL_2928669, EPI_ISL_2928670, EPI_ISL_2928671, EPI_ISL_2928673, EPI_ISL_2928674, EPI_ISL_2928675, EPI_ISL_2928676, EPI_ISL_2928677, EPI_ISL_2928679, EPI_ISL_2928680, EPI_ISL_2928683, EPI_ISL_2928684, EPI_ISL_2928685, EPI_ISL_2928686, EPI_ISL_2928687, EPI_ISL_2928688, EPI_ISL_2928690, EPI_ISL_2928691, EPI_ISL_2928692, EPI_ISL_2928693, EPI_ISL_2928694, EPI_ISL_2928695, EPI_ISL_2928696, EPI_ISL_2928698, EPI_ISL_2928699, EPI_ISL_2928700, EPI_ISL_2928701, EPI_ISL_2928703, EPI_ISL_2928705, EPI_ISL_2928706, EPI_ISL_2928707, EPI_ISL_2928708, EPI_ISL_2928709, EPI_ISL_2928710, EPI_ISL_2928711, EPI_ISL_2928712, EPI_ISL_2928714, EPI_ISL_2928715, EPI_ISL_2928716, EPI_ISL_2928718, EPI_ISL_2928719, EPI_ISL_2928720, EPI_ISL_2928722, EPI_ISL_2928723, EPI_ISL_2928724, EPI_ISL_2928726, EPI_ISL_2928727, EPI_ISL_2928728, EPI_ISL_2928730, EPI_ISL_2928731, EPI_ISL_2928732, EPI_ISL_2928733, EPI_ISL_2928734, EPI_ISL_2928735, EPI_ISL_2928736, EPI_ISL_2928737, EPI_ISL_2928738, EPI_ISL_2928739, EPI_ISL_2928740, EPI_ISL_2928741, EPI_ISL_2928742, EPI_ISL_2928743, EPI_ISL_2928745, EPI_ISL_2928746, EPI_ISL_2928747, EPI_ISL_2928748, EPI_ISL_2928749, EPI_ISL_2928750, EPI_ISL_2928751, EPI_ISL_2928752, EPI_ISL_2928753, EPI_ISL_2928754, EPI_ISL_2928755, EPI_ISL_2928756, EPI_ISL_2928757, EPI_ISL_2928758, EPI_ISL_2928759, EPI_ISL_2928760, EPI_ISL_2928761, EPI_ISL_2928762, EPI_ISL_2928763, EPI_ISL_2928766, EPI_ISL_2928767, EPI_ISL_2928768, EPI_ISL_2928769, EPI_ISL_2928770, EPI_ISL_2928771, EPI_ISL_2928772, EPI_ISL_2928773, EPI_ISL_2928775, EPI_ISL_2928779, EPI_ISL_2928780, EPI_ISL_2928781, EPI_ISL_2928782, EPI_ISL_2928783, EPI_ISL_2928785, EPI_ISL_2928786, EPI_ISL_2928787, EPI_ISL_2928788, EPI_ISL_2928789, EPI_ISL_2928790, EPI_ISL_2928791, EPI_ISL_2928792, EPI_ISL_2928795, EPI_ISL_2928796, EPI_ISL_2928797, EPI_ISL_2928798, EPI_ISL_2928799, EPI_ISL_2928800, EPI_ISL_2928801, EPI_ISL_2928804, EPI_ISL_2928805, EPI_ISL_2928806, EPI_ISL_2928807, EPI_ISL_2928810, EPI_ISL_2928811, EPI_ISL_2928812, EPI_ISL_2928813, EPI_ISL_2928814, EPI_ISL_2928815, EPI_ISL_2928816, EPI_ISL_2928817, EPI_ISL_2928818, EPI_ISL_2928819, EPI_ISL_2928821, EPI_ISL_2928822, EPI_ISL_2928823, EPI_ISL_2928825, EPI_ISL_2928826, EPI_ISL_2928827, EPI_ISL_2928829, EPI_ISL_2928830, EPI_ISL_2928831, EPI_ISL_2928834, EPI_ISL_2928835, EPI_ISL_2928836, EPI_ISL_2928837, EPI_ISL_2928839, EPI_ISL_2928840, EPI_ISL_2928841, EPI_ISL_2928842, EPI_ISL_2928843, EPI_ISL_2928845, EPI_ISL_2928846, EPI_ISL_2928847, EPI_ISL_2928848, EPI_ISL_2928849, EPI_ISL_2928850, EPI_ISL_2928851, EPI_ISL_2928852, EPI_ISL_2928853, EPI_ISL_2928854, EPI_ISL_2928858, EPI_ISL_2928859, EPI_ISL_2928860, EPI_ISL_2928861, EPI_ISL_2928862, EPI_ISL_2928864, EPI_ISL_2928865, EPI_ISL_2928866, EPI_ISL_2928867, EPI_ISL_2928868, EPI_ISL_2928869, EPI_ISL_2928870, EPI_ISL_2928871, EPI_ISL_2928872, EPI_ISL_2928873, EPI_ISL_2928874, EPI_ISL_2928875, EPI_ISL_2928877, EPI_ISL_2928878, EPI_ISL_2928880, EPI_ISL_2928881, EPI_ISL_2928882, EPI_ISL_2928883, EPI_ISL_2928884, EPI_ISL_2928885, EPI_ISL_2928887, EPI_ISL_2928888, EPI_ISL_2928889, EPI_ISL_2928890, EPI_ISL_2928892, EPI_ISL_2928894, EPI_ISL_2928895, EPI_ISL_2928896, EPI_ISL_2928897, EPI_ISL_2928898, EPI_ISL_2928899, EPI_ISL_2928900, EPI_ISL_2928901, EPI_ISL_2928902, EPI_ISL_2928903, EPI_ISL_2928906, EPI_ISL_2928908 | | | |
| see above | Fulgent Genetics | Centers for Disease Control and Prevention Division of Viral Diseases, Pathogen Discovery | Dakota Howard, Dhwani Batra, Peter W. Cook, Kara Moser, Adrian Paskey, Jason Caravas, Benjamin Rambo-Martin, Shatavia Morrison, Christopher Gulvick, Scott Sammons, Yvette Unoarumhi, Darlene Wagner, Matthew Schmerer, Harry Gao, Mickey Li, John Gao, Joseph Fierro, Benafsh Sapra, Becky Tsai, Yan Meng, Doreen Ng, James Xie, Clinton R. Paden, Duncan MacCannell |
| EPI_ISL_2928909, EPI_ISL_2928910 | Oregon State Public Health Laboratory | Oregon State Public Health Laboratory | Rafia Razzaque, Eugene Yeboah, Vanda Makris, Laura Tsaknaridis, John Fontana and Shane Sevey |
| EPI_ISL_2928911 | Fulgent Genetics | Centers for Disease Control and Prevention Division of Viral Diseases, Pathogen Discovery | Dakota Howard, Dhwani Batra, Peter W. Cook, Kara Moser, Adrian Paskey, Jason Caravas, Benjamin Rambo-Martin, Shatavia Morrison, Christopher Gulvick, Scott Sammons, Yvette Unoarumhi, Darlene Wagner, Matthew Schmerer, Harry Gao, Mickey Li, John Gao, Joseph Fierro, Benafsh Sapra, Becky Tsai, Yan Meng, Doreen Ng, James Xie, Clinton R. Paden, Duncan MacCannell |
| EPI_ISL_2928912, | Oregon State Public Health Laboratory | Oregon State Public Health Laboratory | Rafia Razzaque, Eugene Yeboah, Vanda Makris, Laura Tsaknaridis, John Fontana and Shane Sevey |

| | | | |
|---|---|---|---|
| EPI_ISL_2928913 | | | |
| EPI_ISL_2928914 | Fulgent Genetics | Centers for Disease Control and Prevention Division of Viral Diseases, Pathogen Discovery | Dakota Howard, Dhwani Batra, Peter W. Cook, Kara Moser, Adrian Paskey, Jason Caravas, Benjamin Rambo-Martin, Shatavia Morrison, Christopher Gulvick, Scott Sammons, Yvette Unoarumhi, Darlene Wagner, Matthew Schmerer, Harry Gao, Mickey Li, John Gao, Joseph Fierro, Benafsh Sapra, Becky Tsai, Yan Meng, Doreen Ng, James Xie, Clinton R. Paden, Duncan MacCannell |
| EPI_ISL_2928916 | Oregon State Public Health Laboratory | Oregon State Public Health Laboratory | Rafia Razzaque, Eugene Yeboah, Vanda Makris, Laura Tsaknaridis, John Fontana and Shane Sevey |
| EPI_ISL_2928917 | Fulgent Genetics | Centers for Disease Control and Prevention Division of Viral Diseases, Pathogen Discovery | Dakota Howard, Dhwani Batra, Peter W. Cook, Kara Moser, Adrian Paskey, Jason Caravas, Benjamin Rambo-Martin, Shatavia Morrison, Christopher Gulvick, Scott Sammons, Yvette Unoarumhi, Darlene Wagner, Matthew Schmerer, Harry Gao, Mickey Li, John Gao, Joseph Fierro, Benafsh Sapra, Becky Tsai, Yan Meng, Doreen Ng, James Xie, Clinton R. Paden, Duncan MacCannell |
| EPI_ISL_2928918 | Oregon State Public Health Laboratory | Oregon State Public Health Laboratory | Rafia Razzaque, Eugene Yeboah, Vanda Makris, Laura Tsaknaridis, John Fontana and Shane Sevey |
| EPI_ISL_2928920, EPI_ISL_2928921, EPI_ISL_2928922, EPI_ISL_2928923, EPI_ISL_2928924, EPI_ISL_2928925, EPI_ISL_2928926, EPI_ISL_2928927, EPI_ISL_2928928, EPI_ISL_2928929, EPI_ISL_2928932, EPI_ISL_2928933, EPI_ISL_2928934, EPI_ISL_2928936, EPI_ISL_2928937, EPI_ISL_2928938, EPI_ISL_2928939, EPI_ISL_2928940, EPI_ISL_2928941, EPI_ISL_2928942, EPI_ISL_2928943, EPI_ISL_2928945, EPI_ISL_2928946, EPI_ISL_2928947, EPI_ISL_2928948, EPI_ISL_2928949, EPI_ISL_2928950, EPI_ISL_2928952, EPI_ISL_2928953, EPI_ISL_2928954, EPI_ISL_2928955, EPI_ISL_2928956, EPI_ISL_2928957, EPI_ISL_2928958, EPI_ISL_2928959, EPI_ISL_2928960, EPI_ISL_2928961, EPI_ISL_2928962, EPI_ISL_2928963, EPI_ISL_2928964, EPI_ISL_2928965, EPI_ISL_2928966, EPI_ISL_2928967, EPI_ISL_2928968, EPI_ISL_2928969, EPI_ISL_2928970, EPI_ISL_2928971, EPI_ISL_2928972, EPI_ISL_2928973, EPI_ISL_2928974, EPI_ISL_2928975, EPI_ISL_2928976, EPI_ISL_2928977, EPI_ISL_2928978, EPI_ISL_2928979, EPI_ISL_2928980, EPI_ISL_2928981, EPI_ISL_2928983, EPI_ISL_2928984, EPI_ISL_2928986, EPI_ISL_2928988, EPI_ISL_2928989, EPI_ISL_2928990, EPI_ISL_2928991, EPI_ISL_2928994, EPI_ISL_2928995, EPI_ISL_2928997, EPI_ISL_2928998, EPI_ISL_2928999, EPI_ISL_2929000, EPI_ISL_2929001, EPI_ISL_2929002, EPI_ISL_2929003, EPI_ISL_2929005, EPI_ISL_2929006, EPI_ISL_2929007, EPI_ISL_2929008, EPI_ISL_2929009, EPI_ISL_2929011, EPI_ISL_2929012, EPI_ISL_2929013, EPI_ISL_2929014, EPI_ISL_2929018, EPI_ISL_2929019, EPI_ISL_2929023, EPI_ISL_2929024, EPI_ISL_2929025, EPI_ISL_2929026, EPI_ISL_2929027, EPI_ISL_2929028, EPI_ISL_2929029, EPI_ISL_2929030, EPI_ISL_2929031, EPI_ISL_2929032, EPI_ISL_2929033, EPI_ISL_2929034, EPI_ISL_2929035, EPI_ISL_2929036, EPI_ISL_2929037, EPI_ISL_2929038, EPI_ISL_2929039, EPI_ISL_2929040, EPI_ISL_2929041, EPI_ISL_2929042, EPI_ISL_2929043, EPI_ISL_2929044, EPI_ISL_2929045, EPI_ISL_2929046, EPI_ISL_2929047, EPI_ISL_2929049, EPI_ISL_2929050, EPI_ISL_2929053, EPI_ISL_2929054, EPI_ISL_2929055, EPI_ISL_2929056, EPI_ISL_2929057, EPI_ISL_2929059, EPI_ISL_2929061, EPI_ISL_2929062, EPI_ISL_2929064, EPI_ISL_2929065, EPI_ISL_2929066, EPI_ISL_2929067, EPI_ISL_2929069, EPI_ISL_2929070, EPI_ISL_2929071, EPI_ISL_2929072, EPI_ISL_2929074, EPI_ISL_2929075, EPI_ISL_2929076, EPI_ISL_2929079, EPI_ISL_2929081, EPI_ISL_2929082, EPI_ISL_2929084, EPI_ISL_2929087, EPI_ISL_2929088, EPI_ISL_2929089, EPI_ISL_2929090, EPI_ISL_2929091, EPI_ISL_2929092, EPI_ISL_2929094, EPI_ISL_2929095, EPI_ISL_2929096, EPI_ISL_2929097, EPI_ISL_2929098, EPI_ISL_2929099, EPI_ISL_2929100, EPI_ISL_2929101, EPI_ISL_2929102, EPI_ISL_2929104, EPI_ISL_2929105, EPI_ISL_2929106, EPI_ISL_2929107, EPI_ISL_2929108, EPI_ISL_2929109, EPI_ISL_2929110, EPI_ISL_2929111, EPI_ISL_2929112, EPI_ISL_2929113, EPI_ISL_2929114, EPI_ISL_2929115, EPI_ISL_2929116, EPI_ISL_2929117, EPI_ISL_2929118, EPI_ISL_2929119, EPI_ISL_2929120, EPI_ISL_2929121, EPI_ISL_2929122, EPI_ISL_2929123, EPI_ISL_2929124, EPI_ISL_2929126, EPI_ISL_2929127, EPI_ISL_2929129, EPI_ISL_2929130, EPI_ISL_2929131, EPI_ISL_2929132, EPI_ISL_2929133, EPI_ISL_2929134, EPI_ISL_2929135, EPI_ISL_2929140, EPI_ISL_2929145, EPI_ISL_2929147, EPI_ISL_2929148, EPI_ISL_2929151, EPI_ISL_2929154, EPI_ISL_2929155, EPI_ISL_2929157, EPI_ISL_2929159 | | | |
| see above | Fulgent Genetics | Centers for Disease Control and Prevention Division of Viral Diseases, Pathogen Discovery | Dakota Howard, Dhwani Batra, Peter W. Cook, Kara Moser, Adrian Paskey, Jason Caravas, Benjamin Rambo-Martin, Shatavia Morrison, Christopher Gulvick, Scott Sammons, Yvette Unoarumhi, Darlene Wagner, Matthew Schmerer, Harry Gao, Mickey Li, John Gao, Joseph Fierro, Benafsh Sapra, Becky Tsai, Yan Meng, Doreen Ng, James Xie, Clinton R. Paden, Duncan MacCannell |
| EPI_ISL_2929163, EPI_ISL_2929164, EPI_ISL_2929165, EPI_ISL_2929166, EPI_ISL_2929167, EPI_ISL_2929168, EPI_ISL_2929169, EPI_ISL_2929170, EPI_ISL_2929171, EPI_ISL_2929172, EPI_ISL_2929173, EPI_ISL_2929174, EPI_ISL_2929175, EPI_ISL_2929176, EPI_ISL_2929177, EPI_ISL_2929178, EPI_ISL_2929179, EPI_ISL_2929180, EPI_ISL_2929181, EPI_ISL_2929182, EPI_ISL_2929183, EPI_ISL_2929184, EPI_ISL_2929185, EPI_ISL_2929186, EPI_ISL_2929187, EPI_ISL_2929188, EPI_ISL_2929189, EPI_ISL_2929190, EPI_ISL_2929191, EPI_ISL_2929192, EPI_ISL_2929193, EPI_ISL_2929194, EPI_ISL_2929195, EPI_ISL_2929196, EPI_ISL_2929197, EPI_ISL_2929198, EPI_ISL_2929199, EPI_ISL_2929200, EPI_ISL_2929201, EPI_ISL_2929202, EPI_ISL_2929203, EPI_ISL_2929204, EPI_ISL_2929205, EPI_ISL_2929206, EPI_ISL_2929207, EPI_ISL_2929208, EPI_ISL_2929209, EPI_ISL_2929210, EPI_ISL_2929211, EPI_ISL_2929212, EPI_ISL_2929213, EPI_ISL_2929214, EPI_ISL_2929216, EPI_ISL_2929218, EPI_ISL_2929219, EPI_ISL_2929220, EPI_ISL_2929221, EPI_ISL_2929222, EPI_ISL_2929225, EPI_ISL_2929226, EPI_ISL_2929229, EPI_ISL_2929230, EPI_ISL_2929231, EPI_ISL_2929232, EPI_ISL_2929233, EPI_ISL_2929236, EPI_ISL_2929237, EPI_ISL_2929240, EPI_ISL_2929241, EPI_ISL_2929242, EPI_ISL_2929243, EPI_ISL_2929244, EPI_ISL_2929246, EPI_ISL_2929247, EPI_ISL_2929248, EPI_ISL_2929249, EPI_ISL_2929251, EPI_ISL_2929252, EPI_ISL_2929253, EPI_ISL_2929254, EPI_ISL_2929255, EPI_ISL_2929256, EPI_ISL_2929259, EPI_ISL_2929260, EPI_ISL_2929261, EPI_ISL_2929262, EPI_ISL_2929263, EPI_ISL_2929264, EPI_ISL_2929268, EPI_ISL_2929269, EPI_ISL_2929270, EPI_ISL_2929271, EPI_ISL_2929272, EPI_ISL_2929273, EPI_ISL_2929274, EPI_ISL_2929275, EPI_ISL_2929276, EPI_ISL_2929278, EPI_ISL_2929279, EPI_ISL_2929280, EPI_ISL_2929281, EPI_ISL_2929282, EPI_ISL_2929283, EPI_ISL_2929284, EPI_ISL_2929285, EPI_ISL_2929286, EPI_ISL_2929287, EPI_ISL_2929288, EPI_ISL_2929289, EPI_ISL_2929291, EPI_ISL_2929292, EPI_ISL_2929293, EPI_ISL_2929294, EPI_ISL_2929296, EPI_ISL_2929298, EPI_ISL_2929299, EPI_ISL_2929301, EPI_ISL_2929302, EPI_ISL_2929303, EPI_ISL_2929304, EPI_ISL_2929305, EPI_ISL_2929306, EPI_ISL_2929307, EPI_ISL_2929308, EPI_ISL_2929310, EPI_ISL_2929311, EPI_ISL_2929314, EPI_ISL_2929317, EPI_ISL_2929318, EPI_ISL_2929319, EPI_ISL_2929320, EPI_ISL_2929321, EPI_ISL_2929322, EPI_ISL_2929324, EPI_ISL_2929328, EPI_ISL_2929329, EPI_ISL_2929331, EPI_ISL_2929334, EPI_ISL_2929335, EPI_ISL_2929336, EPI_ISL_2929337, EPI_ISL_2929338, EPI_ISL_2929340, EPI_ISL_2929341, EPI_ISL_2929342, EPI_ISL_2929346, EPI_ISL_2929347, EPI_ISL_2929348, EPI_ISL_2929349, EPI_ISL_2929350, EPI_ISL_2929351, EPI_ISL_2929352, EPI_ISL_2929355, EPI_ISL_2929356, EPI_ISL_2929357, EPI_ISL_2929358, EPI_ISL_2929360, EPI_ISL_2929362, EPI_ISL_2929363, EPI_ISL_2929364, EPI_ISL_2929366, EPI_ISL_2929367, EPI_ISL_2929368, EPI_ISL_2929369, EPI_ISL_2929370, EPI_ISL_2929372, EPI_ISL_2929375, EPI_ISL_2929377, EPI_ISL_2929379, EPI_ISL_2929380, EPI_ISL_2929381, EPI_ISL_2929382, EPI_ISL_2929383, EPI_ISL_2929384, EPI_ISL_2929385, EPI_ISL_2929387, EPI_ISL_2929388, EPI_ISL_2929390, EPI_ISL_2929391, EPI_ISL_2929392, EPI_ISL_2929393, EPI_ISL_2929394, EPI_ISL_2929395, EPI_ISL_2929396, EPI_ISL_2929397, EPI_ISL_2929398, EPI_ISL_2929399, EPI_ISL_2929400, EPI_ISL_2929403, EPI_ISL_2929404, EPI_ISL_2929406, EPI_ISL_2929408, EPI_ISL_2929409, EPI_ISL_2929410, EPI_ISL_2929411, EPI_ISL_2929412, EPI_ISL_2929413, EPI_ISL_2929414, EPI_ISL_2929418, EPI_ISL_2929419, EPI_ISL_2929420, EPI_ISL_2929422, EPI_ISL_2929424, EPI_ISL_2929425, EPI_ISL_2929426, EPI_ISL_2929427, EPI_ISL_2929429, EPI_ISL_2929431, EPI_ISL_2929433, EPI_ISL_2929434, EPI_ISL_2929435, EPI_ISL_2929436, EPI_ISL_2929437, EPI_ISL_2929438, EPI_ISL_2929439, EPI_ISL_2929440, EPI_ISL_2929444, EPI_ISL_2929445, EPI_ISL_2929446, EPI_ISL_2929447, EPI_ISL_2929448, EPI_ISL_2929449, EPI_ISL_2929451, EPI_ISL_2929452, EPI_ISL_2929453, EPI_ISL_2929454, EPI_ISL_2929455, EPI_ISL_2929456, EPI_ISL_2929458, EPI_ISL_2929459, EPI_ISL_2929460, EPI_ISL_2929461, EPI_ISL_2929462, EPI_ISL_2929463, EPI_ISL_2929464, EPI_ISL_2929465, EPI_ISL_2929466, EPI_ISL_2929467, EPI_ISL_2929468, EPI_ISL_2929469, EPI_ISL_2929470, EPI_ISL_2929471, EPI_ISL_2929472, EPI_ISL_2929473, EPI_ISL_2929474, EPI_ISL_2929475, EPI_ISL_2929476, EPI_ISL_2929477, EPI_ISL_2929478, EPI_ISL_2929483, EPI_ISL_2929484, EPI_ISL_2929485, EPI_ISL_2929486, EPI_ISL_2929487, EPI_ISL_2929488, EPI_ISL_2929490, EPI_ISL_2929491, EPI_ISL_2929492, EPI_ISL_2929493, EPI_ISL_2929494, EPI_ISL_2929495, EPI_ISL_2929496, EPI_ISL_2929497, EPI_ISL_2929498, EPI_ISL_2929499, EPI_ISL_2929500, EPI_ISL_2929501, EPI_ISL_2929503, EPI_ISL_2929504, EPI_ISL_2929505, EPI_ISL_2929508, EPI_ISL_2929509, EPI_ISL_2929510, EPI_ISL_2929511, EPI_ISL_2929512, EPI_ISL_2929513, EPI_ISL_2929514, EPI_ISL_2929516, EPI_ISL_2929517, EPI_ISL_2929518, EPI_ISL_2929522, EPI_ISL_2929523, EPI_ISL_2929527, EPI_ISL_2929528, EPI_ISL_2929529, EPI_ISL_2929530, EPI_ISL_2929531, EPI_ISL_2929532, EPI_ISL_2929533, EPI_ISL_2929534, EPI_ISL_2929535, EPI_ISL_2929536, EPI_ISL_2929537, EPI_ISL_2929538, EPI_ISL_2929540, EPI_ISL_2929541, EPI_ISL_2929542, EPI_ISL_2929543, EPI_ISL_2929544, EPI_ISL_2929545, EPI_ISL_2929546, EPI_ISL_2929547, EPI_ISL_2929548, EPI_ISL_2929549, EPI_ISL_2929551, EPI_ISL_2929552, EPI_ISL_2929554, EPI_ISL_2929557, EPI_ISL_2929558, EPI_ISL_2929560, EPI_ISL_2929561, EPI_ISL_2929562, EPI_ISL_2929563, EPI_ISL_2929564, EPI_ISL_2929565, EPI_ISL_2929566, EPI_ISL_2929570, EPI_ISL_2929571, EPI_ISL_2929574, EPI_ISL_2929575, EPI_ISL_2929576, EPI_ISL_2929578, EPI_ISL_2929579, EPI_ISL_2929580, EPI_ISL_2929581, EPI_ISL_2929582, EPI_ISL_2929584, EPI_ISL_2929586, EPI_ISL_2929588, EPI_ISL_2929591, EPI_ISL_2929592, EPI_ISL_2929594, EPI_ISL_2929595, EPI_ISL_2929598, EPI_ISL_2929599, EPI_ISL_2929600, EPI_ISL_2929601, EPI_ISL_2929603, EPI_ISL_2929604, EPI_ISL_2929605, EPI_ISL_2929607, EPI_ISL_2929609, EPI_ISL_2929610, EPI_ISL_2929611, EPI_ISL_2929614, EPI_ISL_2929615, EPI_ISL_2929616, EPI_ISL_2929617, EPI_ISL_2929618, EPI_ISL_2929622, EPI_ISL_2929623, EPI_ISL_2929624, EPI_ISL_2929625, EPI_ISL_2929627, EPI_ISL_2929628, EPI_ISL_2929630, EPI_ISL_2929632, EPI_ISL_2929633, EPI_ISL_2929634, EPI_ISL_2929635, EPI_ISL_2929636, EPI_ISL_2929637, EPI_ISL_2929638, EPI_ISL_2929639, EPI_ISL_2929640, EPI_ISL_2929642, EPI_ISL_2929643, EPI_ISL_2929645, EPI_ISL_2929646, EPI_ISL_2929647, EPI_ISL_2929648, EPI_ISL_2929651, EPI_ISL_2929652, EPI_ISL_2929654, EPI_ISL_2929655, EPI_ISL_2929656, EPI_ISL_2929658, EPI_ISL_2929659, EPI_ISL_2929660, EPI_ISL_2929661, EPI_ISL_2929662, EPI_ISL_2929663, EPI_ISL_2929664, EPI_ISL_2929665, EPI_ISL_2929666, EPI_ISL_2929668, EPI_ISL_2929669, EPI_ISL_2929670, EPI_ISL_2929671, EPI_ISL_2929673, EPI_ISL_2929674, EPI_ISL_2929675, EPI_ISL_2929677, EPI_ISL_2929678, EPI_ISL_2929680, EPI_ISL_2929681, EPI_ISL_2929682, EPI_ISL_2929683, EPI_ISL_2929684, EPI_ISL_2929685, EPI_ISL_2929686, EPI_ISL_2929687, EPI_ISL_2929688, EPI_ISL_2929691, EPI_ISL_2929692, EPI_ISL_2929693, EPI_ISL_2929694, EPI_ISL_2929695, EPI_ISL_2929696, EPI_ISL_2929698, EPI_ISL_2929699, EPI_ISL_2929700, EPI_ISL_2929704, EPI_ISL_2929705, EPI_ISL_2929706, EPI_ISL_2929709, EPI_ISL_2929710, EPI_ISL_2929711, EPI_ISL_2929712, EPI_ISL_2929713, EPI_ISL_2929714, EPI_ISL_2929715, EPI_ISL_2929716, EPI_ISL_2929717, EPI_ISL_2929719, EPI_ISL_2929720, EPI_ISL_2929722, EPI_ISL_2929723, EPI_ISL_2929724, EPI_ISL_2929725, EPI_ISL_2929726, EPI_ISL_2929727, EPI_ISL_2929728, EPI_ISL_2929729, EPI_ISL_2929730, EPI_ISL_2929733, EPI_ISL_2929734, EPI_ISL_2929735, EPI_ISL_2929736, EPI_ISL_2929738, EPI_ISL_2929740, EPI_ISL_2929741, EPI_ISL_2929742, EPI_ISL_2929743, EPI_ISL_2929744, EPI_ISL_2929745, EPI_ISL_2929746, EPI_ISL_2929749, EPI_ISL_2929751, EPI_ISL_2929752, EPI_ISL_2929753, EPI_ISL_2929754, EPI_ISL_2929755, EPI_ISL_2929758, EPI_ISL_2929759, EPI_ISL_2929760, EPI_ISL_2929761, EPI_ISL_2929762, EPI_ISL_2929763, EPI_ISL_2929764, EPI_ISL_2929765, EPI_ISL_2929766, EPI_ISL_2929767, EPI_ISL_2929768, EPI_ISL_2929771, EPI_ISL_2929772, EPI_ISL_2929773, EPI_ISL_2929775, EPI_ISL_2929776, EPI_ISL_2929778, EPI_ISL_2929779, EPI_ISL_2929780, EPI_ISL_2929782, EPI_ISL_2929783, EPI_ISL_2929784, EPI_ISL_2929785, EPI_ISL_2929786, EPI_ISL_2929788, EPI_ISL_2929790, EPI_ISL_2929791, EPI_ISL_2929792, EPI_ISL_2929793, EPI_ISL_2929794, EPI_ISL_2929797, EPI_ISL_2929798, EPI_ISL_2929799, EPI_ISL_2929800, EPI_ISL_2929801, EPI_ISL_2929802, EPI_ISL_2929803, EPI_ISL_2929804, EPI_ISL_2929805, EPI_ISL_2929808, EPI_ISL_2929809, EPI_ISL_2929810, EPI_ISL_2929811, EPI_ISL_2929812, EPI_ISL_2929814, EPI_ISL_2929815, EPI_ISL_2929816, EPI_ISL_2929818, EPI_ISL_2929819, EPI_ISL_2929820, EPI_ISL_2929821, EPI_ISL_2929822, EPI_ISL_2929823, EPI_ISL_2929824, EPI_ISL_2929825, EPI_ISL_2929826, EPI_ISL_2929827, EPI_ISL_2929828, EPI_ISL_2929829, EPI_ISL_2929830, EPI_ISL_2929831, EPI_ISL_2929832, EPI_ISL_2929833, EPI_ISL_2929834, EPI_ISL_2929836, EPI_ISL_2929837, EPI_ISL_2929839, EPI_ISL_2929840, EPI_ISL_2929841, EPI_ISL_2929843, EPI_ISL_2929844, EPI_ISL_2929845, EPI_ISL_2929846, EPI_ISL_2929849, EPI_ISL_2929850, EPI_ISL_2929851, EPI_ISL_2929852, EPI_ISL_2929854, EPI_ISL_2929855, EPI_ISL_2929858, EPI_ISL_2929859, EPI_ISL_2929860, EPI_ISL_2929861, EPI_ISL_2929862, EPI_ISL_2929864, EPI_ISL_2929865, EPI_ISL_2929866, EPI_ISL_2929870, EPI_ISL_2929871, EPI_ISL_2929872, EPI_ISL_2929875, EPI_ISL_2929876, EPI_ISL_2929877, EPI_ISL_2929878, EPI_ISL_2929879, EPI_ISL_2929880, EPI_ISL_2929881, EPI_ISL_2929882, EPI_ISL_2929883, EPI_ISL_2929884, EPI_ISL_2929885, EPI_ISL_2929886, EPI_ISL_2929887, EPI_ISL_2929888, EPI_ISL_2929889, EPI_ISL_2929891, EPI_ISL_2929893, EPI_ISL_2929894, EPI_ISL_2929895, EPI_ISL_2929896, EPI_ISL_2929897, EPI_ISL_2929899, EPI_ISL_2929900, EPI_ISL_2929901, EPI_ISL_2929902, EPI_ISL_2929904, EPI_ISL_2929905, EPI_ISL_2929907, EPI_ISL_2929908, EPI_ISL_2929909, EPI_ISL_2929910, EPI_ISL_2929911, EPI_ISL_2929912, EPI_ISL_2929914, EPI_ISL_2929915, EPI_ISL_2929916, EPI_ISL_2929917, EPI_ISL_2929918, EPI_ISL_2929919, EPI_ISL_2929921, EPI_ISL_2929922, EPI_ISL_2929923, EPI_ISL_2929924, EPI_ISL_2929925, EPI_ISL_2929926, EPI_ISL_2929927, EPI_ISL_2929929, EPI_ISL_2929930, EPI_ISL_2929931, EPI_ISL_2929933, EPI_ISL_2929935, EPI_ISL_2929936, EPI_ISL_2929937, EPI_ISL_2929938, EPI_ISL_2929939, EPI_ISL_2929940, EPI_ISL_2929941, EPI_ISL_2929942, EPI_ISL_2929944, EPI_ISL_2929946, EPI_ISL_2929949, EPI_ISL_2929951, EPI_ISL_2929952, EPI_ISL_2929953, EPI_ISL_2929955, EPI_ISL_2929956, EPI_ISL_2929957, EPI_ISL_2929958, EPI_ISL_2929959, EPI_ISL_2929960, EPI_ISL_2929962, EPI_ISL_2929963, EPI_ISL_2929964, EPI_ISL_2929965, EPI_ISL_2929966, EPI_ISL_2929967, EPI_ISL_2929968, EPI_ISL_2929969, EPI_ISL_2929970, EPI_ISL_2929972, EPI_ISL_2929974, EPI_ISL_2929975, EPI_ISL_2929976, EPI_ISL_2929977, EPI_ISL_2929978, EPI_ISL_2929979, EPI_ISL_2929980, EPI_ISL_2929981, EPI_ISL_2929982, EPI_ISL_2929983, EPI_ISL_2929984, EPI_ISL_2929985, EPI_ISL_2929986, EPI_ISL_2929987, EPI_ISL_2929988, EPI_ISL_2929989, EPI_ISL_2929990, EPI_ISL_2929991, EPI_ISL_2929992, EPI_ISL_2929993, EPI_ISL_2929994, EPI_ISL_2929995, EPI_ISL_2929998, EPI_ISL_2929999, EPI_ISL_2930000, EPI_ISL_2930001, EPI_ISL_2930004, EPI_ISL_2930005, EPI_ISL_2930007, EPI_ISL_2930008, EPI_ISL_2930009, EPI_ISL_2930010, EPI_ISL_2930011, EPI_ISL_2930012, EPI_ISL_2930013, EPI_ISL_2930014, EPI_ISL_2930015, EPI_ISL_2930016, EPI_ISL_2930017, EPI_ISL_2930018, EPI_ISL_2930019, EPI_ISL_2930020, EPI_ISL_2930021, EPI_ISL_2930022, EPI_ISL_2930024, EPI_ISL_2930026, EPI_ISL_2930027, EPI_ISL_2930028, EPI_ISL_2930029, EPI_ISL_2930030, EPI_ISL_2930031, EPI_ISL_2930032, EPI_ISL_2930033, EPI_ISL_2930034, EPI_ISL_2930035, EPI_ISL_2930036, EPI_ISL_2930037, EPI_ISL_2930038, EPI_ISL_2930039, EPI_ISL_2930040, EPI_ISL_2930041, EPI_ISL_2930042, EPI_ISL_2930043, EPI_ISL_2930044, EPI_ISL_2930045, EPI_ISL_2930046, EPI_ISL_2930047, EPI_ISL_2930048, EPI_ISL_2930049, EPI_ISL_2930050, EPI_ISL_2930051, EPI_ISL_2930052, EPI_ISL_2930053, EPI_ISL_2930054, EPI_ISL_2930055, EPI_ISL_2930056, EPI_ISL_2930057, EPI_ISL_2930058, EPI_ISL_2930061, EPI_ISL_2930062, EPI_ISL_2930063, EPI_ISL_2930065, EPI_ISL_2930066, EPI_ISL_2930068, EPI_ISL_2930069, EPI_ISL_2930070, EPI_ISL_2930073, EPI_ISL_2930074, EPI_ISL_2930075, EPI_ISL_2930077, EPI_ISL_2930078, EPI_ISL_2930079, EPI_ISL_2930082, EPI_ISL_2930083, EPI_ISL_2930086, EPI_ISL_2930087, EPI_ISL_2930088, EPI_ISL_2930089, EPI_ISL_2930090, EPI_ISL_2930091, EPI_ISL_2930092, EPI_ISL_2930093, EPI_ISL_2930096, EPI_ISL_2930097, EPI_ISL_2930098, EPI_ISL_2930099, EPI_ISL_2930100, EPI_ISL_2930101, EPI_ISL_2930102, EPI_ISL_2930103, EPI_ISL_2930105, EPI_ISL_2930106, EPI_ISL_2930108, EPI_ISL_2930109, EPI_ISL_2930110, EPI_ISL_2930111, EPI_ISL_2930112, EPI_ISL_2930116, EPI_ISL_2930117, EPI_ISL_2930118, EPI_ISL_2930119, EPI_ISL_2930120, EPI_ISL_2930121, EPI_ISL_2930122, EPI_ISL_2930123, EPI_ISL_2930124, EPI_ISL_2930125, EPI_ISL_2930126, EPI_ISL_2930127, EPI_ISL_2930129, EPI_ISL_2930130, EPI_ISL_2930131, EPI_ISL_2930132, EPI_ISL_2930133, EPI_ISL_2930137, EPI_ISL_2930140, EPI_ISL_2930143, EPI_ISL_2930144, EPI_ISL_2930145, EPI_ISL_2930147, EPI_ISL_2930148, EPI_ISL_2930150, EPI_ISL_2930152, EPI_ISL_2930153, EPI_ISL_2930154, EPI_ISL_2930155, EPI_ISL_2930156, EPI_ISL_2930157, EPI_ISL_2930158, EPI_ISL_2930159, EPI_ISL_2930160, EPI_ISL_2930161, EPI_ISL_2930164, EPI_ISL_2930165, EPI_ISL_2930166, EPI_ISL_2930167, EPI_ISL_2930168, EPI_ISL_2930170, EPI_ISL_2930172, EPI_ISL_2930174, EPI_ISL_2930175, EPI_ISL_2930176, EPI_ISL_2930177, EPI_ISL_2930178, EPI_ISL_2930179, EPI_ISL_2930181, EPI_ISL_2930183, EPI_ISL_2930184, EPI_ISL_2930185, EPI_ISL_2930186, EPI_ISL_2930187, EPI_ISL_2930188, EPI_ISL_2930190, EPI_ISL_2930191, EPI_ISL_2930192, EPI_ISL_2930193, EPI_ISL_2930194, EPI_ISL_2930195, EPI_ISL_2930196, EPI_ISL_2930199, EPI_ISL_2930200, EPI_ISL_2930201, EPI_ISL_2930203, EPI_ISL_2930204, EPI_ISL_2930206, EPI_ISL_2930207, EPI_ISL_2930208, EPI_ISL_2930210, EPI_ISL_2930211, EPI_ISL_2930214, EPI_ISL_2930217, EPI_ISL_2930218, EPI_ISL_2930219, EPI_ISL_2930220, EPI_ISL_2930222, EPI_ISL_2930224, EPI_ISL_2930225, EPI_ISL_2930226, EPI_ISL_2930228, EPI_ISL_2930230, EPI_ISL_2930231, EPI_ISL_2930233, EPI_ISL_2930234, EPI_ISL_2930236, EPI_ISL_2930237, EPI_ISL_2930238, EPI_ISL_2930239, EPI_ISL_2930240, EPI_ISL_2930241, EPI_ISL_2930242, EPI_ISL_2930243, EPI_ISL_2930244, EPI_ISL_2930246, EPI_ISL_2930247, EPI_ISL_2930248, EPI_ISL_2930250, EPI_ISL_2930251, EPI_ISL_2930253, EPI_ISL_2930254, EPI_ISL_2930255, EPI_ISL_2930256, EPI_ISL_2930257, EPI_ISL_2930258, EPI_ISL_2930259, EPI_ISL_2930261, EPI_ISL_2930262, EPI_ISL_2930263, EPI_ISL_2930265, EPI_ISL_2930266, EPI_ISL_2930267, EPI_ISL_2930268, EPI_ISL_2930269, EPI_ISL_2930270, EPI_ISL_2930271, EPI_ISL_2930274, EPI_ISL_2930275, EPI_ISL_2930276, EPI_ISL_2930277, EPI_ISL_2930278, EPI_ISL_2930279, EPI_ISL_2930280, EPI_ISL_2930281, EPI_ISL_2930282, EPI_ISL_2930283, EPI_ISL_2930284, EPI_ISL_2930285, EPI_ISL_2930286, EPI_ISL_2930287, EPI_ISL_2930289, EPI_ISL_2930290, EPI_ISL_2930291, EPI_ISL_2930292, EPI_ISL_2930293, EPI_ISL_2930294, EPI_ISL_2930295, EPI_ISL_2930296, EPI_ISL_2930297, EPI_ISL_2930298, EPI_ISL_2930299, EPI_ISL_2930301, EPI_ISL_2930302, EPI_ISL_2930304, EPI_ISL_2930309, EPI_ISL_2930310, EPI_ISL_2930311, EPI_ISL_2930312, EPI_ISL_2930315, EPI_ISL_2930316, EPI_ISL_2930317, EPI_ISL_2930318, EPI_ISL_2930319, EPI_ISL_2930320, EPI_ISL_2930321 | | | |
| see above | Quest Diagnostics Incorporated | Centers for Disease Control and Prevention Division of Viral Diseases, Pathogen Discovery | Dakota Howard, Dhwani Batra, Peter W. Cook, Kara Moser, Adrian Paskey, Jason Caravas, Benjamin Rambo-Martin, Shatavia Morrison, Christopher Gulvick, Scott Sammons, Yvette Unoarumhi, Darlene Wagner, Matthew Schmerer, S. H. Rosenthal, A. Gerasimova, R. M. Kagan, B. Anderson, M. Hua, Y. Liu, L.E. Bernstein, K.E. Livingston, A. Perez, I. A. Shlyakhter, R. V. Rolando, R. Owen, P. Tanpaiboon, F. Lacbawan, Clinton R. Paden, Duncan MacCannell |
| EPI_ISL_2930323, EPI_ISL_2930326, EPI_ISL_2930328, EPI_ISL_2930329, EPI_ISL_2930332, EPI_ISL_2930333, EPI_ISL_2930334, EPI_ISL_2930336, EPI_ISL_2930337, EPI_ISL_2930339, EPI_ISL_2930340, EPI_ISL_2930345, EPI_ISL_2930346, EPI_ISL_2930348, EPI_ISL_2930349, EPI_ISL_2930350, EPI_ISL_2930351, EPI_ISL_2930352, EPI_ISL_2930353, EPI_ISL_2930355, EPI_ISL_2930356, EPI_ISL_2930357, EPI_ISL_2930358, EPI_ISL_2930359, EPI_ISL_2930360, EPI_ISL_2930361, EPI_ISL_2930365, EPI_ISL_2930366, EPI_ISL_2930367, EPI_ISL_2930368, EPI_ISL_2930369, EPI_ISL_2930370, EPI_ISL_2930371, EPI_ISL_2930372, EPI_ISL_2930373, EPI_ISL_2930374, EPI_ISL_2930375, EPI_ISL_2930376, EPI_ISL_2930377, EPI_ISL_2930378, EPI_ISL_2930379, EPI_ISL_2930380, EPI_ISL_2930381, EPI_ISL_2930382, EPI_ISL_2930383, EPI_ISL_2930384, EPI_ISL_2930385, EPI_ISL_2930386, EPI_ISL_2930387, EPI_ISL_2930388, EPI_ISL_2930389, EPI_ISL_2930391, EPI_ISL_2930392, EPI_ISL_2930393, EPI_ISL_2930394, EPI_ISL_2930395, EPI_ISL_2930396, EPI_ISL_2930397, EPI_ISL_2930399, EPI_ISL_2930400, EPI_ISL_2930401, EPI_ISL_2930405, EPI_ISL_2930409, EPI_ISL_2930411, EPI_ISL_2930413, EPI_ISL_2930415, EPI_ISL_2930416, EPI_ISL_2930418, EPI_ISL_2930419, EPI_ISL_2930421, EPI_ISL_2930422, EPI_ISL_2930423, EPI_ISL_2930424, EPI_ISL_2930425, EPI_ISL_2930427, EPI_ISL_2930428, EPI_ISL_2930431 | | | |
| see above | Cotugno | TIGEM | Antonio Grimaldi Patrizia Annunziata Francesco Panariello Biancamaria Pierri Claudia Tiberio Teresa Giuliano Valentina Bouche Chiara Colantuono Maria Concetta Cuomo Denise Di Concilio Lucio Di Filippo Anna Manfredi Marcello Salvi Antonio Limone Luigi Atripaldi Pellegrino Cerino Andrea Ballabio Davide Cacchiarelli |
| EPI_ISL_2930476, EPI_ISL_2930480 | Tampa General Hospital Esoteric Lab | Tampa General Hospital Esoteric Research & Development Lab | Jon Faughn, Deanna Becker, Dominic Uy, Jorge Mecias-Frias, Elaine Vendrone, Vicki Healer, Amorce Lima, Suzane Silbert |
| EPI_ISL_2930562, EPI_ISL_2930563, EPI_ISL_2930564, EPI_ISL_2930566, EPI_ISL_2930567, EPI_ISL_2930568, EPI_ISL_2930569, EPI_ISL_2930570, EPI_ISL_2930572, EPI_ISL_2930573, EPI_ISL_2930574, EPI_ISL_2930575, EPI_ISL_2930576, EPI_ISL_2930578, EPI_ISL_2930580, EPI_ISL_2930582, EPI_ISL_2930583, EPI_ISL_2930584, EPI_ISL_2930585, EPI_ISL_2930587, EPI_ISL_2930588, EPI_ISL_2930589, EPI_ISL_2930590, EPI_ISL_2930591, EPI_ISL_2930592, EPI_ISL_2930593, EPI_ISL_2930594, EPI_ISL_2930595, EPI_ISL_2930596, EPI_ISL_2930597, EPI_ISL_2930598, EPI_ISL_2930599, EPI_ISL_2930600, EPI_ISL_2930601, EPI_ISL_2930602, EPI_ISL_2930603, EPI_ISL_2930604, EPI_ISL_2930605, EPI_ISL_2930606, EPI_ISL_2930607, EPI_ISL_2930608, EPI_ISL_2930609, EPI_ISL_2930610, EPI_ISL_2930612, EPI_ISL_2930613, EPI_ISL_2930614, EPI_ISL_2930615, EPI_ISL_2930616, EPI_ISL_2930617, EPI_ISL_2930618, EPI_ISL_2930619, EPI_ISL_2930620, EPI_ISL_2930621, EPI_ISL_2930622, EPI_ISL_2930623, EPI_ISL_2930624, EPI_ISL_2930625, EPI_ISL_2930626, EPI_ISL_2930627, EPI_ISL_2930628 | | | |
| see above | SARS-CoV-2 testing team, National Institute of Infectious Diseases | Pathogen Genomics Center, National Institute of Infectious Diseases | Tsuyoshi Sekizuka, Kentaro Itokawa, Rina Tanaka, Masanori Hashino, Nozomu Hanaoka, Masumichi Saito, Naomi Nojiri, Hazuka Y Furihata, Hiromizu Takahashi, Tsuguto Fujimoto, Makoto Kuroda |
| EPI_ISL_2930636, EPI_ISL_2930637, EPI_ISL_2930641, EPI_ISL_2930644 | SYNLAB | GIGA Medical Genomics | Keith Durkin, Maria Artesi, Bouchra Boujemla, Nathalie Renotte, Nadine Cambisano, Sébastien Bontems, Cécile Meex, Claire Gourzonès, Olivier Ek, Laurent Gillet, Marie-Pierre Hayette, Vincent Bours |
| EPI_ISL_2930655, EPI_ISL_2930656, EPI_ISL_2930658, EPI_ISL_2930661, EPI_ISL_2930666, EPI_ISL_2930671, EPI_ISL_2930672, EPI_ISL_2930673, EPI_ISL_2930675, EPI_ISL_2930676, EPI_ISL_2930678, EPI_ISL_2930679, EPI_ISL_2930681, EPI_ISL_2930682, EPI_ISL_2930683, EPI_ISL_2930686, EPI_ISL_2930689 | | | |
| see above | Department of Clinical Microbiology | GIGA Medical Genomics | Keith Durkin, Maria Artesi, Bouchra Boujemla, Nathalie Renotte, Nadine Cambisano, Sébastien Bontems, Cécile Meex, Claire Gourzonès, Olivier Ek, Laurent Gillet, Marie-Pierre Hayette, Vincent Bours |
| EPI_ISL_2930691, EPI_ISL_2930694 | SYNLAB | GIGA Medical Genomics | Keith Durkin, Maria Artesi, Bouchra Boujemla, Nathalie Renotte, Nadine Cambisano, Sébastien Bontems, Cécile Meex, Claire Gourzonès, Olivier Ek, Laurent Gillet, Marie-Pierre Hayette, Vincent Bours |
| EPI_ISL_2930696 | C.H.C MontLégia | GIGA Medical Genomics | Keith Durkin, Maria Artesi, Bouchra Boujemla, Nathalie Renotte, Nadine Cambisano, Sébastien Bontems, Cécile Meex, Claire Gourzonès, Olivier Ek, Laurent Gillet, Marie-Pierre Hayette, Vincent Bours |
| EPI_ISL_2930698, EPI_ISL_2930701, EPI_ISL_2930702 | SYNLAB | GIGA Medical Genomics | Keith Durkin, Maria Artesi, Bouchra Boujemla, Nathalie Renotte, Nadine Cambisano, Sébastien Bontems, Cécile Meex, Claire Gourzonès, Olivier Ek, Laurent Gillet, Marie-Pierre Hayette, Vincent Bours |
| EPI_ISL_2930705 | C.H.C MontLégia | GIGA Medical Genomics | Keith Durkin, Maria Artesi, Bouchra Boujemla, Nathalie Renotte, Nadine Cambisano, Sébastien Bontems, Cécile Meex, Claire Gourzonès, Olivier Ek, Laurent Gillet, Marie-Pierre Hayette, Vincent Bours |
| EPI_ISL_2930706 | SYNLAB | GIGA Medical Genomics | Keith Durkin, Maria Artesi, Bouchra Boujemla, Nathalie Renotte, Nadine Cambisano, Sébastien Bontems, Cécile Meex, Claire Gourzonès, Olivier Ek, Laurent Gillet, Marie-Pierre Hayette, Vincent Bours |
| EPI_ISL_2930707, EPI_ISL_2930708, EPI_ISL_2930709, EPI_ISL_2930710, EPI_ISL_2930711, EPI_ISL_2930712, EPI_ISL_2930714, EPI_ISL_2930717, EPI_ISL_2930718, EPI_ISL_2930719, EPI_ISL_2930720, EPI_ISL_2930723, EPI_ISL_2930724, EPI_ISL_2930725, EPI_ISL_2930728, EPI_ISL_2930729, EPI_ISL_2930730, EPI_ISL_2930731, EPI_ISL_2930732, EPI_ISL_2930735, EPI_ISL_2930737 | | | |
| see above | Department of Clinical Microbiology | GIGA Medical Genomics | Keith Durkin, Maria Artesi, Bouchra Boujemla, Nathalie Renotte, Nadine Cambisano, Sébastien Bontems, Cécile Meex, Claire Gourzonès, Olivier Ek, Laurent Gillet, Marie-Pierre Hayette, Vincent Bours |
| EPI_ISL_2930741, EPI_ISL_2930744, EPI_ISL_2930745, EPI_ISL_2930746, EPI_ISL_2930747 | SYNLAB | GIGA Medical Genomics | Keith Durkin, Maria Artesi, Bouchra Boujemla, Nathalie Renotte, Nadine Cambisano, Sébastien Bontems, Cécile Meex, Claire Gourzonès, Olivier Ek, Laurent Gillet, Marie-Pierre Hayette, Vincent Bours |
| EPI_ISL_2930750, EPI_ISL_2930751, EPI_ISL_2930752, EPI_ISL_2930753, | Department of Clinical Microbiology | GIGA Medical Genomics | Keith Durkin, Maria Artesi, Bouchra Boujemla, Nathalie Renotte, Nadine Cambisano, Sébastien Bontems, Cécile Meex, Claire Gourzonès, Olivier Ek, Laurent Gillet, Marie-Pierre Hayette, Vincent Bours |

| | | | |
|---|---|---|---|
| EPI_ISL_2930754, EPI_ISL_2930755, EPI_ISL_2930756, EPI_ISL_2930757, EPI_ISL_2930758 | | | |
| EPI_ISL_2930759, EPI_ISL_2930760, EPI_ISL_2930761 | SYNLAB | GIGA Medical Genomics | Keith Durkin, Maria Artesi, Bouchra Boujemla, Nathalie Renotte, Nadine Cambisano, Sébastien Bontems, Cécile Meex, Claire Gourzonès, Olivier Ek, Laurent Gillet, Marie-Pierre Hayette, Vincent Bours |
| EPI_ISL_2930769, EPI_ISL_2930778, EPI_ISL_2930779, EPI_ISL_2930780, EPI_ISL_2930782 | Department of Clinical Microbiology | GIGA Medical Genomics | Keith Durkin, Maria Artesi, Bouchra Boujemla, Nathalie Renotte, Nadine Cambisano, Sébastien Bontems, Cécile Meex, Claire Gourzonès, Olivier Ek, Laurent Gillet, Marie-Pierre Hayette, Vincent Bours |
| EPI_ISL_2930785, EPI_ISL_2930787, EPI_ISL_2930790, EPI_ISL_2930792, EPI_ISL_2930793, EPI_ISL_2930795, EPI_ISL_2930796, EPI_ISL_2930797, EPI_ISL_2930798, EPI_ISL_2930800, EPI_ISL_2930801 | | | |
| see above | SYNLAB | GIGA Medical Genomics | Keith Durkin, Maria Artesi, Bouchra Boujemla, Nathalie Renotte, Nadine Cambisano, Sébastien Bontems, Cécile Meex, Claire Gourzonès, Olivier Ek, Laurent Gillet, Marie-Pierre Hayette, Vincent Bours |
| EPI_ISL_2930803 | Department of Clinical Microbiology | GIGA Medical Genomics | Keith Durkin, Maria Artesi, Bouchra Boujemla, Nathalie Renotte, Nadine Cambisano, Sébastien Bontems, Cécile Meex, Claire Gourzonès, Olivier Ek, Laurent Gillet, Marie-Pierre Hayette, Vincent Bours |
| EPI_ISL_2930806 | SYNLAB | GIGA Medical Genomics | Keith Durkin, Maria Artesi, Bouchra Boujemla, Nathalie Renotte, Nadine Cambisano, Sébastien Bontems, Cécile Meex, Claire Gourzonès, Olivier Ek, Laurent Gillet, Marie-Pierre Hayette, Vincent Bours |
| EPI_ISL_2930807 | Department of Clinical Microbiology | GIGA Medical Genomics | Keith Durkin, Maria Artesi, Bouchra Boujemla, Nathalie Renotte, Nadine Cambisano, Sébastien Bontems, Cécile Meex, Claire Gourzonès, Olivier Ek, Laurent Gillet, Marie-Pierre Hayette, Vincent Bours |
| EPI_ISL_2930808 | SYNLAB | GIGA Medical Genomics | Keith Durkin, Maria Artesi, Bouchra Boujemla, Nathalie Renotte, Nadine Cambisano, Sébastien Bontems, Cécile Meex, Claire Gourzonès, Olivier Ek, Laurent Gillet, Marie-Pierre Hayette, Vincent Bours |
| EPI_ISL_2930819 | Department of Clinical Microbiology | GIGA Medical Genomics | Keith Durkin, Maria Artesi, Bouchra Boujemla, Nathalie Renotte, Nadine Cambisano, Sébastien Bontems, Cécile Meex, Claire Gourzonès, Olivier Ek, Laurent Gillet, Marie-Pierre Hayette, Vincent Bours |
| EPI_ISL_2930821, EPI_ISL_2930822, EPI_ISL_2930823, EPI_ISL_2930824, EPI_ISL_2930825, EPI_ISL_2930826, EPI_ISL_2930827, EPI_ISL_2930828, EPI_ISL_2930833, EPI_ISL_2930834, EPI_ISL_2930837, EPI_ISL_2930838, EPI_ISL_2930840, EPI_ISL_2930841, EPI_ISL_2930842 | | | |
| see above | SYNLAB | GIGA Medical Genomics | Keith Durkin, Maria Artesi, Bouchra Boujemla, Nathalie Renotte, Nadine Cambisano, Sébastien Bontems, Cécile Meex, Claire Gourzonès, Olivier Ek, Laurent Gillet, Marie-Pierre Hayette, Vincent Bours |
| EPI_ISL_2930843, EPI_ISL_2930845, EPI_ISL_2930846, EPI_ISL_2930849 | Department of Clinical Microbiology | GIGA Medical Genomics | Keith Durkin, Maria Artesi, Bouchra Boujemla, Nathalie Renotte, Nadine Cambisano, Sébastien Bontems, Cécile Meex, Claire Gourzonès, Olivier Ek, Laurent Gillet, Marie-Pierre Hayette, Vincent Bours |
| EPI_ISL_2930852, EPI_ISL_2930853, EPI_ISL_2930855, EPI_ISL_2930856 | SYNLAB | GIGA Medical Genomics | Keith Durkin, Maria Artesi, Bouchra Boujemla, Nathalie Renotte, Nadine Cambisano, Sébastien Bontems, Cécile Meex, Claire Gourzonès, Olivier Ek, Laurent Gillet, Marie-Pierre Hayette, Vincent Bours |
| EPI_ISL_2930857, EPI_ISL_2930858, EPI_ISL_2930859, EPI_ISL_2930860, EPI_ISL_2930861, EPI_ISL_2930862, EPI_ISL_2930863, EPI_ISL_2930864, EPI_ISL_2930865, EPI_ISL_2930866, EPI_ISL_2930867, EPI_ISL_2930868, EPI_ISL_2930869, EPI_ISL_2930870, EPI_ISL_2930871, EPI_ISL_2930875, EPI_ISL_2930877, EPI_ISL_2930878, EPI_ISL_2930879, EPI_ISL_2930880, EPI_ISL_2930881, EPI_ISL_2930882, EPI_ISL_2930883, EPI_ISL_2930884, EPI_ISL_2930885, EPI_ISL_2930886, EPI_ISL_2930888, EPI_ISL_2930890, EPI_ISL_2930892, EPI_ISL_2930893, EPI_ISL_2930894, EPI_ISL_2930895, EPI_ISL_2930896, EPI_ISL_2930902, EPI_ISL_2930903, EPI_ISL_2930907, EPI_ISL_2930908, EPI_ISL_2930909, EPI_ISL_2930911, EPI_ISL_2930912, EPI_ISL_2930913, EPI_ISL_2930914, EPI_ISL_2930915, EPI_ISL_2930916, EPI_ISL_2930917, EPI_ISL_2930918, EPI_ISL_2930919, EPI_ISL_2930920, EPI_ISL_2930921, EPI_ISL_2930922, EPI_ISL_2930923, EPI_ISL_2930924, EPI_ISL_2930925, EPI_ISL_2930926, EPI_ISL_2930927, EPI_ISL_2930928, EPI_ISL_2930929, EPI_ISL_2930930, EPI_ISL_2930931, EPI_ISL_2930932, EPI_ISL_2930933, EPI_ISL_2930934, EPI_ISL_2930935, EPI_ISL_2930936, EPI_ISL_2930937, EPI_ISL_2930938, EPI_ISL_2930939, EPI_ISL_2930940, EPI_ISL_2930941, EPI_ISL_2930942, EPI_ISL_2930943, EPI_ISL_2930944, EPI_ISL_2930945, EPI_ISL_2930946, EPI_ISL_2930947, EPI_ISL_2930948, EPI_ISL_2930949, EPI_ISL_2930950, EPI_ISL_2930951, EPI_ISL_2930952, EPI_ISL_2930953, EPI_ISL_2930954, EPI_ISL_2930956, EPI_ISL_2930960, EPI_ISL_2930961, EPI_ISL_2930962, EPI_ISL_2930963, EPI_ISL_2930964, EPI_ISL_2930970, EPI_ISL_2930971, EPI_ISL_2930973, EPI_ISL_2930974, EPI_ISL_2930975 | | | |
| see above | Department of Clinical Microbiology | GIGA Medical Genomics | Keith Durkin, Maria Artesi, Bouchra Boujemla, Nathalie Renotte, Nadine Cambisano, Sébastien Bontems, Cécile Meex, Claire Gourzonès, Olivier Ek, Laurent Gillet, Marie-Pierre Hayette, Vincent Bours |
| EPI_ISL_2930977 | University of Liège COVID-19 testing center | GIGA Medical Genomics | Keith Durkin, Maria Artesi, Bouchra Boujemla, Nathalie Renotte, Nadine Cambisano, Sébastien Bontems, Cécile Meex, Claire Gourzonès, Olivier Ek, Laurent Gillet, Marie-Pierre Hayette, Vincent Bours |
| EPI_ISL_2930981 | SYNLAB | GIGA Medical Genomics | Keith Durkin, Maria Artesi, Bouchra Boujemla, Nathalie Renotte, Nadine Cambisano, Sébastien Bontems, Cécile Meex, Claire Gourzonès, Olivier Ek, Laurent Gillet, Marie-Pierre Hayette, Vincent Bours |
| EPI_ISL_2930983 | Department of Clinical Microbiology | GIGA Medical Genomics | Keith Durkin, Maria Artesi, Sébastien Bontems, Raphaël Boreux, Bouchra Boujemla, Nathalie Renotte, Cécile Meex, Pierrette Melin, Marie-Pierre Hayette, Vincent Bours |
| EPI_ISL_2931006, EPI_ISL_2931012 | University of Arkansas for Medical Sciences | Center for Global Health, University of New Mexico Health Sciences Center | Daryl Domman, Kurt Schwalm, Valerie Morley, Susan Harley, Jennifer Laudadio, Joshua L. Kennedy, Catherine Kirkpatrick, Uday Chalwadi, David Ussery, Darrell Dinwiddie |
| EPI_ISL_2931022 | Arkansas Department of Health | Center for Global Health, University of New Mexico Health Sciences Center | Daryl Domman, Kurt Schwalm, Valerie Morley, Atul Kothari, Joshua L. Kennedy, Darrell Dinwiddie |
| EPI_ISL_2931026, EPI_ISL_2931029, EPI_ISL_2931031, EPI_ISL_2931036, EPI_ISL_2931041, EPI_ISL_2931043, EPI_ISL_2931044 | Arkansas Children's Hospital | Center for Global Health, University of New Mexico Health Sciences Center | Daryl Domman, Kurt Schwalm, Valerie Morley, Amanda Novack, Dirk Haselow, Joshua L. Kennedy, Darrell Dinwiddie |
| EPI_ISL_2931099 | Rush University Medical Center | RIPHL at Rush University Medical Center | Stefan Green, Kevin Kunstman, Max Kolton, Marieta Hyde, Laura Furtado, Felix Araujo Perez, Mary Hayden, Joyce Houlihan, Diane Springer |
| EPI_ISL_2931125 | John H Stroger Jr Hospital of Cook County | RIPHL at Rush University Medical Center | Stefan Green, Kevin Kunstman, Max Kolton, Marieta Hyde, Laura Furtado, Felix Araujo Perez, Ah-Young Sung, Sharon Welbel |
| EPI_ISL_2931131, EPI_ISL_2931138 | Biology Department, College of Science, Al Muthanna University and Public Health Laboratory, Al-Muthanna Health Directorate | Department of Virology, Faculty of Medicine, University of Helsinki, Helsinki, Finland | Nihad Al-Rashedi, Hussein Alburkat, Murad Munahi, Ali Jasim, Batool Kadham Salman, Hussein Riadh Kitab, Olli Vapalahti, Tarja Sironen,Teemu Smura |
| EPI_ISL_2931139, EPI_ISL_2931140, EPI_ISL_2931141, EPI_ISL_2931142, EPI_ISL_2931143, EPI_ISL_2931144, EPI_ISL_2931145, EPI_ISL_2931146, EPI_ISL_2931147, EPI_ISL_2931148, EPI_ISL_2931149, EPI_ISL_2931150, EPI_ISL_2931151, EPI_ISL_2931152, EPI_ISL_2931153, EPI_ISL_2931154, EPI_ISL_2931155, EPI_ISL_2931156 | | | |
| see above | Kansas Health and Environmental Lab | Kansas Health and Environmental Lab | Mike Grose, Katherine Wiggins, Jonathan Barnell, Ben Olsen, and Phil Adam |
| EPI_ISL_2931161, EPI_ISL_2931167, EPI_ISL_2931168, EPI_ISL_2931171, EPI_ISL_2931175 | Baptist Health Medical Center | Center for Global Health, University of New Mexico Health Sciences Center | Daryl Domman, Kurt Schwalm, Valerie Morley, Amanda Novack, Dirk Haselow, Joshua L. Kennedy, Darrell Dinwiddie |
| EPI_ISL_2931184, EPI_ISL_2931185, EPI_ISL_2931186, EPI_ISL_2931192, EPI_ISL_2931193, EPI_ISL_2931194, EPI_ISL_2931198, EPI_ISL_2931199, EPI_ISL_2931204, EPI_ISL_2931210, EPI_ISL_2931214 | | | |
| see above | Arkansas Children's Hospital | Center for Global Health, University of New Mexico Health Sciences Center | Daryl Domman, Kurt Schwalm, Valerie Morley, Ashton Ingold, Bobby Boyanton, Joshua L. Kennedy, Darrell Dinwiddie |
| EPI_ISL_2931252, EPI_ISL_2931254 | Wyoming Public Health Laboratory | Wyoming Public Health Laboratory | Jim Mildenberger, Taylor Fearing, Channing Weber, Ashley Norberg, Chayse Rowley, Marley Goetz, Brian Dominguez, Elliot Thomasson, Sam Britz, Cari Sloma, Robert Petit, and Rob Christensen |
| EPI_ISL_2931263 | Hospital | National Reference Center for Viruses of Respiratory Infections, Institut Pasteur, Paris | Marion Barbet, Sylvie Behillil, Méline Bizard, Angela Brisebarre, Camille Capel, Vincent Enouf, Louise Lefrançois, Frédéric Lemoine, Christophe Malabat, Corinne Maufrais, Victoire Baillet, Etienne Simon-Lorière, Maud Vanpeene, Sylvie Van der Werf ,DUCANCELLE Alexandra |
| EPI_ISL_2931267 | Labo Analyses Med | National Reference Center for Viruses of Respiratory Infections, Institut Pasteur, Paris | Marion Barbet, Sylvie Behillil, Méline Bizard, Angela Brisebarre, Camille Capel, Vincent Enouf, Louise Lefrançois, Frédéric Lemoine, Christophe Malabat, Corinne Maufrais, Amaury Vaysse, Etienne Simon-Lorière, Maud Vanpeene, Sylvie Van der Werf ,TROMBERT Sabine |
| EPI_ISL_2931268 | Labo Analyses Med | National Reference Center for Viruses of Respiratory Infections, Institut Pasteur, Paris | Marion Barbet, Sylvie Behillil, Méline Bizard, Angela Brisebarre, Camille Capel, Vincent Enouf, Louise Lefrançois, Frédéric Lemoine, Christophe Malabat, Corinne Maufrais, Hub Bioinformatique Biostatistiques, Etienne Simon-Lorière, Maud Vanpeene, Sylvie Van der Werf ,TROMBERT Sabine |
| EPI_ISL_2931271 | Labo Analyses Med | National Reference Center for Viruses of Respiratory Infections, Institut Pasteur, Paris | Marion Barbet, Sylvie Behillil, Méline Bizard, Angela Brisebarre, Camille Capel, Vincent Enouf, Louise Lefrançois, Frédéric Lemoine, Christophe Malabat, Corinne Maufrais, Pierre Lechat, Etienne Simon-Lorière, Maud Vanpeene, Sylvie Van der Werf ,CERBALLIANCE Ile de France sud |
| EPI_ISL_2931272 | Shenzhen Center for Disease Control and Prevention | National Institute for Viral Disease Control and Prevention, China CDC | Long Chen, Xinyi Wei, Shimin Li, Renli Zhang,Kai Nie, Yenan Feng, Chunli Wu, Can Zhu, Dana Huang,and Yaqing He |
| EPI_ISL_2931273 | Shenzhen Center for Disease Control and Prevention | National Institute for Viral Disease Control and Prevention, China CDC | Long Chen, Hang Zhang, Ying Sun, Renli Zhang,Kai Nie, Yenan Feng, Chunli Wu, Can Zhu, Dana Huang,and Yaqing He |
| EPI_ISL_2931274 | Shenzhen Center for Disease Control and Prevention | National Institute for Viral Disease Control and Prevention, China CDC | Long Chen, Xinyi Wei, Shimin Li, Renli Zhang,Peihua Niu, Kai Nie, Shisong Fang,Xiaoliang Xiao,Fan Yang,Qingju Lu,and Yaqing He |
| EPI_ISL_2931275 | Shenzhen Center for Disease Control and Prevention | National Institute for Viral Disease Control and Prevention, China CDC | Long Chen, Xinyi Wei, Can Zhu, Renli Zhang,Peihua Niu, Kai Nie, Bo Peng,Shimin Li,Yijun Tang,Zidan Lin and Yaqing He |
| EPI_ISL_2931276 | Shenzhen Center for Disease Control and Prevention | National Institute for Viral Disease Control and Prevention, China CDC | Long Chen,Can Zhu,Jiancheng Chen,Renli Zhang,Peihua Niu, Yenan Feng, Bo Peng,Shimin Li,Yijun Tang,Zidan Lin and Yaqin He |
| EPI_ISL_2931277 | Shenzhen Center for Disease Control and Prevention | National Institute for Viral Disease Control and Prevention, China CDC | Long Chen,Can Zhu,Jiancheng Chen,Renli Zhang, Kai Nie, Peihua Niu,Bo Peng,Shimin Li,Yijun Tang,Zidan Lin and Yaqin He |
| EPI_ISL_2931279 | Shenzhen Center for Disease Control and Prevention | National Institute for Viral Disease Control and Prevention, China CDC | Long Chen, Shimin Li,Jiancheng Chen,Renli Zhang,Yenan Feng, Bo Peng,Yijun Tang,Zidan Lin and Yaqing He |
| EPI_ISL_2931280 | Shenzhen Center for Disease Control and Prevention | National Institute for Viral Disease Control and Prevention, China CDC | Long Chen, Can Zhu,Xinyi Wei,Renli Zhang, Peihua Niu,Weihua Wu,Yue Li,Shaoyu Deng and Yaqing He |
| EPI_ISL_2931282 | Shenzhen Center for Disease Control and Prevention | National Institute for Viral Disease Control and Prevention, China | Long Chen, Can Zhu,Xinyi Wei,Renli Zhang,Kai Nie, Peihua Niu,Weihua Wu,Yue Li,Shaoyu Deng and Yaqing He |

| | | | |
|---|---|---|---|
| | | CDC | |
| EPI_ISL_2931283 | Shenzhen Center for Disease Control and Prevention | National Institute for Viral Disease Control and Prevention, China CDC | Long Chen, Shimin Li, Jiancheng Chen,Renli Zhang, Yenan Feng, Shisong Fang, Xiaoliang Xiao,Qingju Lu,Fan Yang and Yaqing He |
| EPI_ISL_2931284 | Shenzhen Center for Disease Control and Prevention | National Institute for Viral Disease Control and Prevention, China CDC | Long Chen,Xiaoliang Xiao,Qingju Lu,Renli Zhang,Peihua Niu,Weihua Wu,Xinyi Wei,Yue Li,Shaoyu Deng and Yaqing He |
| EPI_ISL_2931287 | Nemocnice Vyskov | CMBG | Matej Bezdicek, Jan Svaton, Martina Lengerova, Monika Dolejska |
| EPI_ISL_2931288 | Nemocnice Trebic | CMBG | Matej Bezdicek, Jan Svaton, Martina Lengerova, Monika Dolejska |
| EPI_ISL_2931289 | OKMI | CMBG | Matej Bezdicek, Jan Svaton, Martina Lengerova, Monika Dolejska |
| EPI_ISL_2931296, EPI_ISL_2931297, EPI_ISL_2931299 | FNUSA | CMBG | Matej Bezdicek, Jan Svaton, Martina Lengerova, Monika Dolejska |
| EPI_ISL_2931301, EPI_ISL_2931302 | St. Anne´s University Hospital Brno | University Hospital Brno, CMBG | Matej Bezdicek, Jan Svaton, Martina Lengerova, Monika Dolejska |
| EPI_ISL_2931330, EPI_ISL_2931331, EPI_ISL_2931332, EPI_ISL_2931333, EPI_ISL_2931334, EPI_ISL_2931335 | Temporary Specimen Collection Centre at the AsiaWorld-Expo | Hong Kong Department of Health | Alan K.L. Tsang, Peter C.W. Yip, Ken H.L. Ng, Edman T.K. Lam, Rickjason C.W. Chan |
| EPI_ISL_2931336 | Home Quarantine Taskforce | Hong Kong Department of Health | Alan K.L. Tsang, Peter C.W. Yip, Ken H.L. Ng, Edman T.K. Lam, Rickjason C.W. Chan |
| EPI_ISL_2931337, EPI_ISL_2931338 | Temporary Specimen Collection Centre at the AsiaWorld-Expo | Hong Kong Department of Health | Alan K.L. Tsang, Peter C.W. Yip, Ken H.L. Ng, Edman T.K. Lam, Rickjason C.W. Chan |
| EPI_ISL_2931339 | Queen Elizabeth Hospital | Hong Kong Department of Health | Alan K.L. Tsang, Peter C.W. Yip, Ken H.L. Ng, Edman T.K. Lam, Rickjason C.W. Chan |
| EPI_ISL_2931340, EPI_ISL_2931341 | Temporary Specimen Collection Centre at the AsiaWorld-Expo | Hong Kong Department of Health | Alan K.L. Tsang, Peter C.W. Yip, Ken H.L. Ng, Edman T.K. Lam, Rickjason C.W. Chan |
| EPI_ISL_2931342, EPI_ISL_2931343, EPI_ISL_2931344, EPI_ISL_2931345 | Home Quarantine Taskforce | Hong Kong Department of Health | Alan K.L. Tsang, Peter C.W. Yip, Ken H.L. Ng, Edman T.K. Lam, Rickjason C.W. Chan |
| EPI_ISL_2931346 | Queen Elizabeth Hospital | Hong Kong Department of Health | Alan K.L. Tsang, Peter C.W. Yip, Ken H.L. Ng, Edman T.K. Lam, Rickjason C.W. Chan |
| EPI_ISL_2931347, EPI_ISL_2931348, EPI_ISL_2931349 | Home Quarantine Taskforce | Hong Kong Department of Health | Alan K.L. Tsang, Peter C.W. Yip, Ken H.L. Ng, Edman T.K. Lam, Rickjason C.W. Chan |
| EPI_ISL_2931350 | Temporary Specimen Collection Centre at the AsiaWorld-Expo | Hong Kong Department of Health | Alan K.L. Tsang, Peter C.W. Yip, Ken H.L. Ng, Edman T.K. Lam, Rickjason C.W. Chan |
| EPI_ISL_2931351 | Home Quarantine Taskforce | Hong Kong Department of Health | Alan K.L. Tsang, Peter C.W. Yip, Ken H.L. Ng, Edman T.K. Lam, Rickjason C.W. Chan |
| EPI_ISL_2931352 | Temporary Specimen Collection Centre at the AsiaWorld-Expo | Hong Kong Department of Health | Alan K.L. Tsang, Peter C.W. Yip, Ken H.L. Ng, Edman T.K. Lam, Rickjason C.W. Chan |
| EPI_ISL_2931353, EPI_ISL_2931354, EPI_ISL_2931355, EPI_ISL_2931356, EPI_ISL_2931357, EPI_ISL_2931358, EPI_ISL_2931359, EPI_ISL_2931360 | Home Quarantine Taskforce | Hong Kong Department of Health | Alan K.L. Tsang, Peter C.W. Yip, Ken H.L. Ng, Edman T.K. Lam, Rickjason C.W. Chan |
| EPI_ISL_2931361 | Temporary Specimen Collection Centre at the AsiaWorld-Expo | Hong Kong Department of Health | Alan K.L. Tsang, Peter C.W. Yip, Ken H.L. Ng, Edman T.K. Lam, Rickjason C.W. Chan |
| EPI_ISL_2931362 | Home Quarantine Taskforce | Hong Kong Department of Health | Alan K.L. Tsang, Peter C.W. Yip, Ken H.L. Ng, Edman T.K. Lam, Rickjason C.W. Chan |
| EPI_ISL_2931363, EPI_ISL_2931364 | Temporary Specimen Collection Centre at the AsiaWorld-Expo | Hong Kong Department of Health | Alan K.L. Tsang, Peter C.W. Yip, Ken H.L. Ng, Edman T.K. Lam, Rickjason C.W. Chan |
| EPI_ISL_2931365 | Home Quarantine Taskforce | Hong Kong Department of Health | Alan K.L. Tsang, Peter C.W. Yip, Ken H.L. Ng, Edman T.K. Lam, Rickjason C.W. Chan |
| EPI_ISL_2931366 | Quarantine Centre Silka Tsuen Wan | Hong Kong Department of Health | Alan K.L. Tsang, Peter C.W. Yip, Ken H.L. Ng, Edman T.K. Lam, Rickjason C.W. Chan |
| EPI_ISL_2931367, EPI_ISL_2931368, EPI_ISL_2931369 | North Lantau Hospital | Hong Kong Department of Health | Alan K.L. Tsang, Peter C.W. Yip, Ken H.L. Ng, Edman T.K. Lam, Rickjason C.W. Chan |
| EPI_ISL_2931385 | Laboratori de Referencia de Catalunya | Laboratori de Referencia de Catalunya | Ramirez A., Bellosillo B., Padilla E., Vilas A., Hernandez JJ., Canal M. |
| EPI_ISL_2931388 | Hospital Universitari i Politècnic La Fe de València | SeqCOVID-SPAIN consortium/IBV(CSIC) | María Dolores Gómez Ruiz, Eva González Barberá, Ana Gil Brusola, Salvador Giner Almaraz, José Luis López Hontangas and SeqCOVID-SPAIN consortium |
| EPI_ISL_2931389, EPI_ISL_2931390, EPI_ISL_2931391, EPI_ISL_2931392, EPI_ISL_2931393, EPI_ISL_2931394, EPI_ISL_2931395, EPI_ISL_2931398, EPI_ISL_2931399, EPI_ISL_2931400, EPI_ISL_2931401, EPI_ISL_2931403, EPI_ISL_2931405, EPI_ISL_2931407, EPI_ISL_2931408, EPI_ISL_2931409, EPI_ISL_2931410, EPI_ISL_2931412, EPI_ISL_2931413, EPI_ISL_2931419, EPI_ISL_2931420, EPI_ISL_2931421, EPI_ISL_2931422, EPI_ISL_2931423, EPI_ISL_2931424 | | | |
| see above | Servicio de Microbiología. Hospital Clínico Universitario de Valencia | SeqCOVID-SPAIN consortium/IBV(CSIC) | David Navarro Ortega, Eliseo Albert Vicent, Ignacio Torres and SeqCOVID-SPAIN consortium |
| EPI_ISL_2931426, EPI_ISL_2931427, EPI_ISL_2931428, EPI_ISL_2931430, EPI_ISL_2931431, EPI_ISL_2931432, EPI_ISL_2931433, EPI_ISL_2931435, EPI_ISL_2931436, EPI_ISL_2931437, EPI_ISL_2931438, EPI_ISL_2931439, EPI_ISL_2931440, EPI_ISL_2931442, EPI_ISL_2931443, EPI_ISL_2931445, EPI_ISL_2931447, EPI_ISL_2931448, EPI_ISL_2931451, EPI_ISL_2931452, EPI_ISL_2931453, EPI_ISL_2931455, EPI_ISL_2931456, EPI_ISL_2931457, EPI_ISL_2931458, EPI_ISL_2931459, EPI_ISL_2931460, EPI_ISL_2931461, EPI_ISL_2931462, EPI_ISL_2931463, EPI_ISL_2931464, EPI_ISL_2931465, EPI_ISL_2931466 | | | |
| see above | Hospital Universitari i Politècnic La Fe de València | SeqCOVID-SPAIN consortium/IBV(CSIC) | María Dolores Gómez Ruiz, Eva González Barberá, Ana Gil Brusola, Salvador Giner Almaraz, José Luis López Hontangas and SeqCOVID-SPAIN consortium |
| EPI_ISL_2931468, EPI_ISL_2931469, EPI_ISL_2931470, EPI_ISL_2931471, EPI_ISL_2931472, EPI_ISL_2931473, EPI_ISL_2931474, EPI_ISL_2931475, EPI_ISL_2931476, EPI_ISL_2931477, EPI_ISL_2931478, EPI_ISL_2931479, EPI_ISL_2931481, EPI_ISL_2931482, EPI_ISL_2931483, EPI_ISL_2931484, EPI_ISL_2931485, EPI_ISL_2931486, EPI_ISL_2931487, EPI_ISL_2931488, EPI_ISL_2931489, EPI_ISL_2931490, EPI_ISL_2931491, EPI_ISL_2931492, EPI_ISL_2931493, EPI_ISL_2931494, EPI_ISL_2931495, EPI_ISL_2931496, EPI_ISL_2931497, EPI_ISL_2931498, EPI_ISL_2931499, EPI_ISL_2931500, EPI_ISL_2931501, EPI_ISL_2931502, EPI_ISL_2931503, EPI_ISL_2931504, EPI_ISL_2931506, EPI_ISL_2931507, EPI_ISL_2931508, EPI_ISL_2931510, EPI_ISL_2931511, EPI_ISL_2931512, EPI_ISL_2931513, EPI_ISL_2931514, EPI_ISL_2931515, EPI_ISL_2931516, EPI_ISL_2931517, EPI_ISL_2931518, EPI_ISL_2931520, EPI_ISL_2931521, EPI_ISL_2931522, EPI_ISL_2931523, EPI_ISL_2931524, EPI_ISL_2931525, EPI_ISL_2931526, EPI_ISL_2931527, EPI_ISL_2931528, EPI_ISL_2931529, EPI_ISL_2931530, EPI_ISL_2931532, EPI_ISL_2931533, EPI_ISL_2931534, EPI_ISL_2931535, EPI_ISL_2931536, EPI_ISL_2931538, EPI_ISL_2931539, EPI_ISL_2931540, EPI_ISL_2931541, EPI_ISL_2931542, EPI_ISL_2931544, EPI_ISL_2931545, EPI_ISL_2931546, EPI_ISL_2931548, EPI_ISL_2931549, EPI_ISL_2931550, EPI_ISL_2931553, EPI_ISL_2931554, EPI_ISL_2931555, EPI_ISL_2931556, EPI_ISL_2931557, EPI_ISL_2931558, EPI_ISL_2931559, EPI_ISL_2931560, EPI_ISL_2931561, EPI_ISL_2931562, EPI_ISL_2931563, EPI_ISL_2931564, EPI_ISL_2931565, EPI_ISL_2931566, EPI_ISL_2931567, EPI_ISL_2931568, EPI_ISL_2931569, EPI_ISL_2931570, EPI_ISL_2931571, EPI_ISL_2931572, EPI_ISL_2931573, EPI_ISL_2931574, EPI_ISL_2931575, EPI_ISL_2931577, EPI_ISL_2931578, EPI_ISL_2931579, EPI_ISL_2931580, EPI_ISL_2931582, EPI_ISL_2931583, EPI_ISL_2931584, EPI_ISL_2931585, EPI_ISL_2931586, EPI_ISL_2931587 | | | |
| see above | Arkansas Public Health Laboratory, Arkansas Department of Health | University of Minnesota Genomics Center | Daryl M. Gohl, John Garbe, Jaquelyn Kuriger-Laber, Corbin Dirkx |
| EPI_ISL_2931690, EPI_ISL_2931691 | I.R.C.C.S. "S. De Bellis" - Ente Ospedaliero | Istituto Zooprofilattico Sperimentale della Puglia e della Basilicata | Parisi A., Bianco A., Capozzi L., Del Sambro L., Simone D., Difato L., Notarnicola M., Lippolis A. |
| EPI_ISL_2931692, EPI_ISL_2931693, EPI_ISL_2931694, EPI_ISL_2931696, EPI_ISL_2931697, EPI_ISL_2931698, EPI_ISL_2931699, EPI_ISL_2931700, EPI_ISL_2931701 | Ospedale Santa Caterina Novella | Istituto Zooprofilattico Sperimentale della Puglia e della Basilicata | Parisi A., Bianco A., Capozzi L., Del Sambro L., Simone D., Difato L., Bruno A. R. |
| EPI_ISL_2931702, EPI_ISL_2931703, EPI_ISL_2931704 | Ospedale Vito Fazzi | Istituto Zooprofilattico Sperimentale della Puglia e della Basilicata | Parisi A., Bianco A., Capozzi L., Del Sambro L., Simone D., Difato L., Lobreglio G. |
| EPI_ISL_2931708, EPI_ISL_2931709, EPI_ISL_2931710, EPI_ISL_2931712 | Ospedale Santissima Annunziata | Istituto Zooprofilattico Sperimentale della Puglia e della Basilicata | Parisi A., Bianco A., Capozzi L., Del Sambro L., Simone D., Difato L., Ridolfi D., Giannico A, Tampoia M. |
| EPI_ISL_2931716, EPI_ISL_2931721, EPI_ISL_2931722, EPI_ISL_2931724 | Presidio di Brindisi Di Summa - Perrino | Istituto Zooprofilattico Sperimentale della Puglia e della Basilicata | Parisi A., Bianco A., Capozzi L., Del Sambro L., Simone D., Difato L., Santoro A., Ridolfi D., Giannico A. |
| EPI_ISL_2931727 | RSKIA Kota Bandung | National Institute of Health Research and Development | Vivi Setiawaty, Hana Apsari Pawestri, Subangkit, Kartika Dewi Puspa, Arie Ardiansyah Nugraha, Hartanti Dian Ikawati, Krisna Pangesti, Triyani Soekarso, Nelly Puspandari |

| | | | |
|---|---|---|---|
| EPI_ISL_2931737 | RSUD dr Soediran Mangun Sumarso, Wonogiri | National Institute of Health Research and Development | Vivi Setiawaty, Hana Apsari Pawestri, Subangkit, Kartika Dewi Puspa, Arie Ardiansyah Nugraha, Hartanti Dian Ikawati, Krisna Pangesti, Triyani Soekarso, Nelly Puspandari |
| EPI_ISL_2931743 | BTKL PP Kelas I Batam | National Institute of Health Research and Development | Vivi Setiawaty, Hana Apsari Pawestri, Subangkit, Kartika Dewi Puspa, Arie Ardiansyah Nugraha, Hartanti Dian Ikawati, Krisna Pangesti, Triyani Soekarso, Nelly Puspandari |
| EPI_ISL_2931745 | Dinkes Musi Banyu Asin | National Institute of Health Research and Development | Hana Apsari Pawestri, Subangkit, Kartika Dewi Puspa, Arie Ardiansyah Nugraha, Hartanti Dian Ikawati, Krisna Pangesti, Triyani Soekarso, Nelly Puspandari, Vivi Setiawaty |
| EPI_ISL_2931746 | RSUD dr Soediran Mangun Sumarso, Wonogiri | National Institute of Health Research and Development | Hana Apsari Pawestri, Subangkit, Kartika Dewi Puspa, Arie Ardiansyah Nugraha, Hartanti Dian Ikawati, Krisna Pangesti, Triyani Soekarso, Nelly Puspandari, Vivi Setiawaty |
| EPI_ISL_2931747 | PKM Abiansemal III, Bali | National Institute of Health Research and Development | Hana Apsari Pawestri, Subangkit, Kartika Dewi Puspa, Arie Ardiansyah Nugraha, Hartanti Dian Ikawati, Krisna Pangesti, Triyani Soekarso, Nelly Puspandari, Vivi Setiawaty |
| EPI_ISL_2931751 | Dinkes Provinsi Lampung | National Institute of Health Research and Development | Hana Apsari Pawestri, Subangkit, Kartika Dewi Puspa, Arie Ardiansyah Nugraha, Hartanti Dian Ikawati, Krisna Pangesti, Triyani Soekarso, Nelly Puspandari, Vivi Setiawaty |
| EPI_ISL_2931755 | Dinkes Musi Banyu Asin | National Institute of Health Research and Development | Hana Apsari Pawestri, Subangkit, Kartika Dewi Puspa, Arie Ardiansyah Nugraha, Hartanti Dian Ikawati, Krisna Pangesti, Triyani Soekarso, Nelly Puspandari, Vivi Setiawaty |
| EPI_ISL_2931756, EPI_ISL_2931757 | RSUD dr Soediran Mangun Sumarso, Wonogiri | National Institute of Health Research and Development | Hana Apsari Pawestri, Subangkit, Kartika Dewi Puspa, Arie Ardiansyah Nugraha, Hartanti Dian Ikawati, Krisna Pangesti, Triyani Soekarso, Nelly Puspandari, Vivi Setiawaty |
| EPI_ISL_2931761 | Labkesda Depok | National Institute of Health Research and Development | Hana Apsari Pawestri, Subangkit, Kartika Dewi Puspa, Arie Ardiansyah Nugraha, Hartanti Dian Ikawati, Krisna Pangesti, Triyani Soekarso, Nelly Puspandari, Vivi Setiawaty |
| EPI_ISL_2931763 | RSKIA Kota Bandung | National Institute of Health Research and Development | Hana Apsari Pawestri, Subangkit, Kartika Dewi Puspa, Arie Ardiansyah Nugraha, Hartanti Dian Ikawati, Krisna Pangesti, Triyani Soekarso, Nelly Puspandari, Vivi Setiawaty |
| EPI_ISL_2931765 | RSUD dr Soediran Mangun Sumarso, Wonogiri | National Institute of Health Research and Development | Hana Apsari Pawestri, Subangkit, Kartika Dewi Puspa, Arie Ardiansyah Nugraha, Hartanti Dian Ikawati, Krisna Pangesti, Triyani Soekarso, Nelly Puspandari, Vivi Setiawaty |
| EPI_ISL_2931771 | DINKES KOTA BOGOR | National Institute of Health Research and Development | Hana Apsari Pawestri, Subangkit, Kartika Dewi Puspa, Arie Ardiansyah Nugraha, Hartanti Dian Ikawati, Krisna Pangesti, Triyani Soekarso, Nelly Puspandari, Vivi Setiawaty |
| EPI_ISL_2931772 | SmartCO Lab | National Institute of Health Research and Development | Subangkit, Hana Apsari Pawestri, , Kartika Dewi Puspa, Arie Ardiansyah Nugraha, Hartanti Dian Ikawati, Krisna Pangesti, Triyani Soekarso, Nelly Puspandari, Vivi Setiawaty |
| EPI_ISL_2931774 | RSKIA Kota Bandung | National Institute of Health Research and Development | Subangkit, Hana Apsari Pawestri, , Kartika Dewi Puspa, Arie Ardiansyah Nugraha, Hartanti Dian Ikawati, Krisna Pangesti, Triyani Soekarso, Nelly Puspandari, Vivi Setiawaty |
| EPI_ISL_2931775 | Dinkes Musi Banyu Asin | National Institute of Health Research and Development | Subangkit, Hana Apsari Pawestri, , Kartika Dewi Puspa, Arie Ardiansyah Nugraha, Hartanti Dian Ikawati, Krisna Pangesti, Triyani Soekarso, Nelly Puspandari, Vivi Setiawaty |
| EPI_ISL_2931776 | RSUD dr Soediran Mangun Sumarso, Wonogiri | National Institute of Health Research and Development | Subangkit, Hana Apsari Pawestri, , Kartika Dewi Puspa, Arie Ardiansyah Nugraha, Hartanti Dian Ikawati, Krisna Pangesti, Triyani Soekarso, Nelly Puspandari, Vivi Setiawaty |
| EPI_ISL_2931780 | DINKES KOTA BOGOR | National Institute of Health Research and Development | Subangkit, Hana Apsari Pawestri, , Kartika Dewi Puspa, Arie Ardiansyah Nugraha, Hartanti Dian Ikawati, Krisna Pangesti, Triyani Soekarso, Nelly Puspandari, Vivi Setiawaty |
| EPI_ISL_2931781 | RSKIA Kota Bandung | National Institute of Health Research and Development | Subangkit, Hana Apsari Pawestri, , Kartika Dewi Puspa, Arie Ardiansyah Nugraha, Hartanti Dian Ikawati, Krisna Pangesti, Triyani Soekarso, Nelly Puspandari, Vivi Setiawaty |
| EPI_ISL_2931787 | DINKES KOTA BOGOR | National Institute of Health Research and Development | Subangkit, Hana Apsari Pawestri, , Kartika Dewi Puspa, Arie Ardiansyah Nugraha, Hartanti Dian Ikawati, Krisna Pangesti, Triyani Soekarso, Nelly Puspandari, Vivi Setiawaty |
| EPI_ISL_2931790 | Dinkes Musi Banyu Asin | National Institute of Health Research and Development | Subangkit, Hana Apsari Pawestri, , Kartika Dewi Puspa, Arie Ardiansyah Nugraha, Hartanti Dian Ikawati, Krisna Pangesti, Triyani Soekarso, Nelly Puspandari, Vivi Setiawaty |
| EPI_ISL_2931791 | RSUD dr Soediran Mangun Sumarso, Wonogiri | National Institute of Health Research and Development | Subangkit, Hana Apsari Pawestri, , Kartika Dewi Puspa, Arie Ardiansyah Nugraha, Hartanti Dian Ikawati, Krisna Pangesti, Triyani Soekarso, Nelly Puspandari, Vivi Setiawaty |
| EPI_ISL_2931796 | RS Advent Bandar Lampung | National Institute of Health Research and Development | Subangkit, Hana Apsari Pawestri, , Kartika Dewi Puspa, Arie Ardiansyah Nugraha, Hartanti Dian Ikawati, Krisna Pangesti, Triyani Soekarso, Nelly Puspandari, Vivi Setiawaty |
| EPI_ISL_2931797 | CIPAKU KOTA BOGOR | National Institute of Health Research and Development | Subangkit, Hana Apsari Pawestri, , Kartika Dewi Puspa, Arie Ardiansyah Nugraha, Hartanti Dian Ikawati, Krisna Pangesti, Triyani Soekarso, Nelly Puspandari, Vivi Setiawaty |
| EPI_ISL_2931837, EPI_ISL_2931838, EPI_ISL_2931839, EPI_ISL_2931840, EPI_ISL_2931841, EPI_ISL_2931842, EPI_ISL_2931843, EPI_ISL_2931844, EPI_ISL_2931845, EPI_ISL_2931846, EPI_ISL_2931847, EPI_ISL_2931848, EPI_ISL_2931849, EPI_ISL_2931850, EPI_ISL_2931851, EPI_ISL_2931852, EPI_ISL_2931866 | | | |
| see above | Microbiology Department. Complexo Hospitalario Universitario de Vigo | Microbiology Department. Complexo Hospitalario Universitario de Vigo | Alfaya N, Alonso I, Alvarez M, Cabrera JJ, Carballo R, Cores O, Cortizo S, del-Campo V, Martinez L, Mediero G, Perez S, Davina C, Potel C, Regueiro B, Rey S, Vasallo FJ |
| EPI_ISL_2931920 | Sultanah Fatimah Specialist Hospital, Muar | Institute for Medical Research, Infectious Disease Research Centre, National Institutes of Health, Ministry of Health Malaysia | Suppiah J, Kamel K, Mohd Zawawi Z, Azizan MA, Ramly N, Robert F, Anasir MI, Thayan R |
| EPI_ISL_2931921 | Labuan Hospital | Institute for Medical Research, Infectious Disease Research Centre, National Institutes of Health, Ministry of Health Malaysia | Suppiah J, Kamel K, Mohd Zawawi Z, Azizan MA, Ramly N, Robert F, Anasir MI, Thayan R |
| EPI_ISL_2931922 | Tuanku Fauziah Hospital | Institute for Medical Research, Infectious Disease Research Centre, National Institutes of Health, Ministry of Health Malaysia | Suppiah J, Kamel K, Mohd Zawawi Z, Azizan MA, Ramly N, Robert F, Anasir MI, Thayan R |
| EPI_ISL_2931923 | Sultanah Fatimah Specialist Hospital, Muar | Institute for Medical Research, Infectious Disease Research Centre, National Institutes of Health, Ministry of Health Malaysia | Suppiah J, Kamel K, Mohd Zawawi Z, Azizan MA, Ramly N, Robert F, Anasir MI, Thayan R |
| EPI_ISL_2931924 | Queen Elizabeth Hospital, Kota Kinabalu | Institute for Medical Research, Infectious Disease Research Centre, National Institutes of Health, Ministry of Health Malaysia | Suppiah J, Kamel K, Mohd Zawawi Z, Azizan MA, Ramly N, Robert F, Anasir MI, Thayan R |
| EPI_ISL_2931925, EPI_ISL_2931926, EPI_ISL_2931927, EPI_ISL_2931928, EPI_ISL_2931929, EPI_ISL_2931930, EPI_ISL_2931931, EPI_ISL_2931932, EPI_ISL_2931933, EPI_ISL_2931934, EPI_ISL_2931935, EPI_ISL_2931936, EPI_ISL_2931937, EPI_ISL_2931938, EPI_ISL_2931939, EPI_ISL_2931940, EPI_ISL_2931941, EPI_ISL_2931942, EPI_ISL_2931943, EPI_ISL_2931944, EPI_ISL_2931945, EPI_ISL_2931946, EPI_ISL_2931947, EPI_ISL_2931948, EPI_ISL_2931949, EPI_ISL_2931950, EPI_ISL_2931951, EPI_ISL_2931952, EPI_ISL_2931953, EPI_ISL_2931954, EPI_ISL_2931955, EPI_ISL_2931956, EPI_ISL_2931957, EPI_ISL_2931958, EPI_ISL_2931959, EPI_ISL_2931960, EPI_ISL_2931961, EPI_ISL_2931962, EPI_ISL_2931963, EPI_ISL_2931964, EPI_ISL_2931965, EPI_ISL_2931966, EPI_ISL_2931967, EPI_ISL_2931968, EPI_ISL_2931969, EPI_ISL_2931970, EPI_ISL_2931971, EPI_ISL_2931972, EPI_ISL_2931973, EPI_ISL_2931974, EPI_ISL_2931975, EPI_ISL_2931976, EPI_ISL_2931977, EPI_ISL_2931978, EPI_ISL_2931979, EPI_ISL_2931980, EPI_ISL_2931981, EPI_ISL_2931982, EPI_ISL_2931983, EPI_ISL_2931984, EPI_ISL_2931985, EPI_ISL_2931986, EPI_ISL_2931987, EPI_ISL_2931988, EPI_ISL_2931989, EPI_ISL_2931990, EPI_ISL_2931991, EPI_ISL_2931992, EPI_ISL_2931993, EPI_ISL_2931994, EPI_ISL_2931995, EPI_ISL_2931996, EPI_ISL_2931997, EPI_ISL_2931998, EPI_ISL_2931999, EPI_ISL_2932000, EPI_ISL_2932001, EPI_ISL_2932002, EPI_ISL_2932003, EPI_ISL_2932004, EPI_ISL_2932005, EPI_ISL_2932006, EPI_ISL_2932007, EPI_ISL_2932008, EPI_ISL_2932009, EPI_ISL_2932010, EPI_ISL_2932012, EPI_ISL_2932013, EPI_ISL_2932014, EPI_ISL_2932015, EPI_ISL_2932016, EPI_ISL_2932017, EPI_ISL_2932018, EPI_ISL_2932019, EPI_ISL_2932020, EPI_ISL_2932021, EPI_ISL_2932022, EPI_ISL_2932023, EPI_ISL_2932024, EPI_ISL_2932025, EPI_ISL_2932026, EPI_ISL_2932027, EPI_ISL_2932028, EPI_ISL_2932029, EPI_ISL_2932030, EPI_ISL_2932031, EPI_ISL_2932032, EPI_ISL_2932033, EPI_ISL_2932034, EPI_ISL_2932035, EPI_ISL_2932036, EPI_ISL_2932037, EPI_ISL_2932038, EPI_ISL_2932039, EPI_ISL_2932040, EPI_ISL_2932042, EPI_ISL_2932043, EPI_ISL_2932046, EPI_ISL_2932047, EPI_ISL_2932048, EPI_ISL_2932049, EPI_ISL_2932056, EPI_ISL_2932057, EPI_ISL_2932070, EPI_ISL_2932072, EPI_ISL_2932073, EPI_ISL_2932085, EPI_ISL_2932086, EPI_ISL_2932087, EPI_ISL_2932091, EPI_ISL_2932092, EPI_ISL_2932103, EPI_ISL_2932123, EPI_ISL_2932127, EPI_ISL_2932128, EPI_ISL_2932129, EPI_ISL_2932135, EPI_ISL_2932136, EPI_ISL_2932138, EPI_ISL_2932142, EPI_ISL_2932143, EPI_ISL_2932144, EPI_ISL_2932145, EPI_ISL_2932147, EPI_ISL_2932149, EPI_ISL_2932150, EPI_ISL_2932154, EPI_ISL_2932155, EPI_ISL_2932157, EPI_ISL_2932158, EPI_ISL_2932159, EPI_ISL_2932160, EPI_ISL_2932161, EPI_ISL_2932162, EPI_ISL_2932163, EPI_ISL_2932164, EPI_ISL_2932166, EPI_ISL_2932167, EPI_ISL_2932168, EPI_ISL_2932169, EPI_ISL_2932170, EPI_ISL_2932171, EPI_ISL_2932173, EPI_ISL_2932174, EPI_ISL_2932177, EPI_ISL_2932178, EPI_ISL_2932179, EPI_ISL_2932180, EPI_ISL_2932183, EPI_ISL_2932184, EPI_ISL_2932185, EPI_ISL_2932186, EPI_ISL_2932190, EPI_ISL_2932196, EPI_ISL_2932199, EPI_ISL_2932201, EPI_ISL_2932202, EPI_ISL_2932204, EPI_ISL_2932205, EPI_ISL_2932207, EPI_ISL_2932208, EPI_ISL_2932211, EPI_ISL_2932212, EPI_ISL_2932213, EPI_ISL_2932214, EPI_ISL_2932215, EPI_ISL_2932216, EPI_ISL_2932217, EPI_ISL_2932218, EPI_ISL_2932219, EPI_ISL_2932221, EPI_ISL_2932222, EPI_ISL_2932224, EPI_ISL_2932225, EPI_ISL_2932226, EPI_ISL_2932227, EPI_ISL_2932230, EPI_ISL_2932232, EPI_ISL_2932235, EPI_ISL_2932236, EPI_ISL_2932237, EPI_ISL_2932238, EPI_ISL_2932241, EPI_ISL_2932242, EPI_ISL_2932243, EPI_ISL_2932244, EPI_ISL_2932245, EPI_ISL_2932246, EPI_ISL_2932247, EPI_ISL_2932248, EPI_ISL_2932249, EPI_ISL_2932255, EPI_ISL_2932256, EPI_ISL_2932257, EPI_ISL_2932258 | | | |
| see above | National Virus Reference Laboratory | National Virus Reference Laboratory | Zoe Yandle, Charlene Bennett, Gabriel Gonzalez, Michael Carr, Jonathan Dean, Cillian F De Gascun |
| EPI_ISL_2932263 | PathWest Laboratory Medicine WA | PathWest Laboratory Medicine WA | PathWest Laboratory Medicine WA Microbial Surveillance Unit |
| EPI_ISL_2932266, EPI_ISL_2932270, EPI_ISL_2932271, EPI_ISL_2932273, EPI_ISL_2932274, EPI_ISL_2932300, EPI_ISL_2932301, EPI_ISL_2932302, EPI_ISL_2932303 | Pathogen Genomics Center, National Institute of Infectious Diseases | Pathogen Genomics Center, National Institute of Infectious Diseases | Tsuyoshi Sekizuka, Kentaro Itokawa, Rina Tanaka, Masanori Hashino, Makoto Kuroda |
| EPI_ISL_2932318, EPI_ISL_2932319, EPI_ISL_2932320, EPI_ISL_2932322, EPI_ISL_2932323, EPI_ISL_2932324, EPI_ISL_2932325, EPI_ISL_2932326, EPI_ISL_2932327, EPI_ISL_2932328, EPI_ISL_2932329, EPI_ISL_2932330, EPI_ISL_2932331, EPI_ISL_2932332, EPI_ISL_2932333, EPI_ISL_2932334 | | | |
| see above | Platform BIS UZA/UAntwerpen | Labo Klinische Biologie, UZA | Marie Le Mercier, Jasmine Coppens, Kathleen Holemans, Ines Verbesselt, Basil Britto Xavier, Christine Lammens, Veerle Matheeussen, Herman Goossens |
| EPI_ISL_2932336 | CENTRUM VOOR MEDISCHE ANALYSE | Labo Klinische Biologie, UZA | Marie Le Mercier, Jasmine Coppens, Kathleen Holemans, Ines Verbesselt, Basil Britto Xavier, Christine Lammens, Veerle Matheeussen, Herman Goossens |
| EPI_ISL_2932337, EPI_ISL_2932338, EPI_ISL_2932339, EPI_ISL_2932340, EPI_ISL_2932341, EPI_ISL_2932342, EPI_ISL_2932343, EPI_ISL_2932344, EPI_ISL_2932345, EPI_ISL_2932346, EPI_ISL_2932347, EPI_ISL_2932348, EPI_ISL_2932349, EPI_ISL_2932350, EPI_ISL_2932352, EPI_ISL_2932353, EPI_ISL_2932354, EPI_ISL_2932355, EPI_ISL_2932356, EPI_ISL_2932357, EPI_ISL_2932358, EPI_ISL_2932359, EPI_ISL_2932360, EPI_ISL_2932361, EPI_ISL_2932363, EPI_ISL_2932365, EPI_ISL_2932366, EPI_ISL_2932367, EPI_ISL_2932369, EPI_ISL_2932370, EPI_ISL_2932375, EPI_ISL_2932376, EPI_ISL_2932377, EPI_ISL_2932379, EPI_ISL_2932380 | | | |
| see above | Platform BIS UZA/UAntwerpen | Labo Klinische Biologie, UZA | Marie Le Mercier, Jasmine Coppens, Kathleen Holemans, Ines Verbesselt, Basil Britto Xavier, Christine Lammens, Veerle Matheeussen, Herman Goossens |
| EPI_ISL_2932381, EPI_ISL_2932382, EPI_ISL_2932385, EPI_ISL_2932386, EPI_ISL_2932389, EPI_ISL_2932390, EPI_ISL_2932391, EPI_ISL_2932392, EPI_ISL_2932393, EPI_ISL_2932394, EPI_ISL_2932395, EPI_ISL_2932396, EPI_ISL_2932397, EPI_ISL_2932398 | | | |
| see above | Algemeen Medisch Labo | Labo Klinische Biologie, UZA | Marie Le Mercier, Jasmine Coppens, Kathleen Holemans, Ines Verbesselt, Basil Britto Xavier, Christine Lammens, Veerle Matheeussen, Herman Goossens |
| EPI_ISL_2932503 | National Institute of Infectious Diseases-Prof. Dr. Matei Bals Molecular Diagnostics Laboratory | National Institute of Infectious Diseases-Prof. Dr. Matei Bals Molecular Diagnostics Laboratory | Corina Casangiu, Leontina Banica, Marius Surleac, Petre Milu, Robert Hohan, Simona Paraschiv, Dan Otelea |
| EPI_ISL_2932506 | Department of Public Health Bucharest | National Institute of Infectious Diseases-Prof. Dr. Matei Bals Molecular Diagnostics Laboratory | Corina Casangiu, Leontina Banica, Marius Surleac, Petre Milu, Robert Hohan, Simona Paraschiv, Dan Otelea |
| EPI_ISL_2932518 | Clinical Hospital of Infectious Diseases „Sfanta Parascheva" Iasi | National Institute of Infectious Diseases-Prof. Dr. Matei Bals Molecular Diagnostics Laboratory | Corina Casangiu, Leontina Banica, Marius Surleac, Petre Milu, Robert Hohan, Simona Paraschiv, Dan Otelea |
| EPI_ISL_2932519 | National Institute of Infectious Diseases-Prof. Dr. Matei Bals Molecular Diagnostics Laboratory | National Institute of Infectious Diseases-Prof. Dr. Matei Bals Molecular Diagnostics Laboratory | Corina Casangiu, Leontina Banica, Marius Surleac, Petre Milu, Robert Hohan, Simona Paraschiv, Dan Otelea |
| EPI_ISL_2932520, EPI_ISL_2932522, EPI_ISL_2932523, EPI_ISL_2932525 | Department of Public Health Bucharest | National Institute of Infectious Diseases-Prof. Dr. Matei Bals Molecular Diagnostics Laboratory | Corina Casangiu, Leontina Banica, Marius Surleac, Petre Milu, Robert Hohan, Simona Paraschiv, Dan Otelea |
| EPI_ISL_2932526 | National Institute of Infectious Diseases-Prof. Dr. Matei Bals Molecular Diagnostics Laboratory | National Institute of Infectious Diseases-Prof. Dr. Matei Bals Molecular Diagnostics Laboratory | Corina Casangiu, Leontina Banica, Marius Surleac, Petre Milu, Robert Hohan, Simona Paraschiv, Dan Otelea |
| EPI_ISL_2932529 | Department of Public Health Bucharest | National Institute of Infectious Diseases-Prof. Dr. Matei Bals Molecular Diagnostics Laboratory | Corina Casangiu, Leontina Banica, Marius Surleac, Petre Milu, Robert Hohan, Simona Paraschiv, Dan Otelea |
| EPI_ISL_2932530, EPI_ISL_2932531 | Institute of Public Health Kraljevo | Veterinary Specialized Institute Kraljevo | Vidanovic,D., Tesovic,B., Knezevic,A., Jovanovic,T., Jankovic,M., Sekler,M., Djukic V,., Banovic Djeri,B., Petrovic,T., Volkening,J., Afonso,C. |
| EPI_ISL_2932594 | Laboratorio de Virología HUCA | Laboratorio de Virología HUCA | Sandoval M, Castelló C, Gómez de Oña J, Boga JA, Rojo S, Alvarez-Arguelles ME, Abreu F, Costales I, Perez-Martínez Z, Martín-Rodríguez G, Coto E, Melón S |
| EPI_ISL_2932610 | KOTA Yogya DKK Gondokusuman II | Genetics Working Group (Pokja Genetik) Faculty of Medicine, Public Health and Nursing Universitas Gadjah Mada (FK-KMK UGM); Disease Investigation Center Wates Ministry of Agriculture | Gunadi, Hendra Wibawa, Marcellus, Mohamad S. Hakim, Edwin W. Daniwijaya, Ludhang P. Rizki, Endah Supriyati, Eggi Arguni, Titik Nuryastuti, Tri Wibawa, Dwi AA Nugrahaningsih, Afiahayati, Siswanto,Kristy Iskandar,Nungki Anggorowati, Pramesti G Dewi, Dwi Indaryati,Indarto Sulistiyono, Khanza a Vujira, Alvina A Setiawan |

| | | | |
|---|---|---|---|
| | | Indonesia; Department of Microbiology FK-KMK UGM; Laboratorium Diagnostik Yayasan Tahija World Mosquito Program (WMP) Yogyakarta Center for | |
| EPI_ISL_2932611 | Jordan University of Science Technology/ Irbid filed Hospital/ MOH | Princess Haya Biotechnology Center/ Jordan University of Science Technology | Saied Jaradat, Hazem Haddad, Areej Alquran, Maha Karam, Shereen Issa,Suha Hasan, Emad Abu Yaqeen |
| EPI_ISL_2932613 | DKK Sleman/Puskesmas Depok 1 | Genetics Working Group (Pokja Genetik) Faculty of Medicine, Public Health and Nursing Universitas Gadjah Mada (FK-KMK UGM); Disease Investigation Center Wates Ministry of Agriculture Indonesia; Department of Microbiology FK-KMK UGM; Laboratorium Diagnostik Yayasan Tahija World Mosquito Program (WMP) Yogyakarta Center for Tropical Medicine FK-KMK UGM; Integrated Research Center FK-KMK UGM; Department of Computer Science and Electronics FMIPA UGM; RSUP Dr. Sardjito | Gunadi, Hendra Wibawa, Marcellus, Mohamad S. Hakim, Edwin W. Daniwijaya, Ludhang P. Rizki, Endah Supriyati, Eggi Arguni, Titik Nuryastuti, Tri Wibawa, Dwi AA Nugrahaningsih, Afiahayati, Siswanto,Kristy Iskandar,Nungki Anggorowati, Pramesti G Dewi, Dwi Indaryati,Indarto Sulistiyono, Irene Tania, Hana F Hanifin |
| EPI_ISL_2932615, EPI_ISL_2932618 | Jordan University of Science Technology/ Irbid filed Hospital/ MOH | Princess Haya Biotechnology Center/ Jordan University of Science Technology | Saied Jaradat, Hazem Haddad, Areej Alquran, Maha Karam, Shereen Issa,Suha Hasan, Emad Abu Yaqeen |
| EPI_ISL_2932620, EPI_ISL_2932621, EPI_ISL_2932622, EPI_ISL_2932623, EPI_ISL_2932624, EPI_ISL_2932625, EPI_ISL_2932626, EPI_ISL_2932627, EPI_ISL_2932628, EPI_ISL_2932629, EPI_ISL_2932630, EPI_ISL_2932631, EPI_ISL_2932632, EPI_ISL_2932633, EPI_ISL_2932634, EPI_ISL_2932635, EPI_ISL_2932636, EPI_ISL_2932637, EPI_ISL_2932639, EPI_ISL_2932640, EPI_ISL_2932641, EPI_ISL_2932642, EPI_ISL_2932643, EPI_ISL_2932644, EPI_ISL_2932645, EPI_ISL_2932646, EPI_ISL_2932647, EPI_ISL_2932648, EPI_ISL_2932649, EPI_ISL_2932650, EPI_ISL_2932651, EPI_ISL_2932652, EPI_ISL_2932653, EPI_ISL_2932654, EPI_ISL_2932656, EPI_ISL_2932657, EPI_ISL_2932660, EPI_ISL_2932661, EPI_ISL_2932662, EPI_ISL_2932663, EPI_ISL_2932664, EPI_ISL_2932665, EPI_ISL_2932666, EPI_ISL_2932669, EPI_ISL_2932670, EPI_ISL_2932671, EPI_ISL_2932672, EPI_ISL_2932673, EPI_ISL_2932674, EPI_ISL_2932675, EPI_ISL_2932676, EPI_ISL_2932677, EPI_ISL_2932678, EPI_ISL_2932679, EPI_ISL_2932680 | | | |
| see above | Communicable Disease Laboratory, Public Health Directorate | Communicable Disease Laboratory, Public Health Directorate | Alwasti,H., AlHujairi,Z., AlAbbas,Z., Marhoon,A., Almoamen,G. |
| EPI_ISL_2932684 | Arlon | Plateforme de testing Namuroise | Lesly Nyinkeu Kemamen; Nicolas Gilliard; Demars Aurore ; Maschietto Céline ; Nobis Chloé ; Otto Gaetan ; Denis Olivier ; Degosserie Jonathan ; Mullier François |
| EPI_ISL_2932685, EPI_ISL_2932687, EPI_ISL_2932689, EPI_ISL_2932690, EPI_ISL_2932692, EPI_ISL_2932693, EPI_ISL_2932695 | Plateforme de testing Namuroise | Plateforme de testing Namuroise | Lesly Nyinkeu Kemamen; Nicolas Gilliard; Demars Aurore ; Maschietto Céline ; Nobis Chloé ; Otto Gaetan ; Denis Olivier ; Degosserie Jonathan ; Mullier François |
| EPI_ISL_2932696 | Arlon | Plateforme de testing Namuroise | Lesly Nyinkeu Kemamen; Nicolas Gilliard; Demars Aurore ; Maschietto Céline ; Nobis Chloé ; Otto Gaetan ; Denis Olivier ; Degosserie Jonathan ; Mullier François |
| EPI_ISL_2932697, EPI_ISL_2932698, EPI_ISL_2932699, EPI_ISL_2932701, EPI_ISL_2932702, EPI_ISL_2932704 | Plateforme de testing Namuroise | Plateforme de testing Namuroise | Lesly Nyinkeu Kemamen; Nicolas Gilliard; Demars Aurore ; Maschietto Céline ; Nobis Chloé ; Otto Gaetan ; Denis Olivier ; Degosserie Jonathan ; Mullier François |
| EPI_ISL_2932708 | Cliniques universitaires Saint-Luc | UCLouvain/IREC/MBLG | Jean Ruelle, Bertrand Bearzatto, Benoit Kabamba Mukadi |
| EPI_ISL_2932742 | National Hematology and Transfusiology Center, Department of Medical Genetics | National Hematology and Transfusiology Center, Department of Medical Genetics | Agha Rza Aghayev |
| EPI_ISL_2932744 | Molekylær Medicinsk Afdeling, Aarhus University Hospital, Aarhus, Denmark | Statens Serum Institut Bioinformatics and Microbial Genomics | Danish Covid-19 Genome Consortium |
| EPI_ISL_2932745, EPI_ISL_2932746, EPI_ISL_2932747, EPI_ISL_2932748, EPI_ISL_2932749, EPI_ISL_2932750, EPI_ISL_2932751, EPI_ISL_2932754, EPI_ISL_2932755, EPI_ISL_2932756, EPI_ISL_2932757, EPI_ISL_2932758, EPI_ISL_2932759, EPI_ISL_2932760, EPI_ISL_2932761, EPI_ISL_2932762, EPI_ISL_2932763, EPI_ISL_2932764, EPI_ISL_2932765, EPI_ISL_2932766, EPI_ISL_2932767, EPI_ISL_2932768, EPI_ISL_2932769, EPI_ISL_2932770, EPI_ISL_2932771, EPI_ISL_2932773, EPI_ISL_2932774, EPI_ISL_2932775, EPI_ISL_2932776, EPI_ISL_2932777, EPI_ISL_2932778, EPI_ISL_2932779, EPI_ISL_2932780, EPI_ISL_2932781, EPI_ISL_2932782, EPI_ISL_2932783, EPI_ISL_2932784, EPI_ISL_2932786, EPI_ISL_2932787, EPI_ISL_2932788, EPI_ISL_2932789, EPI_ISL_2932790, EPI_ISL_2932791, EPI_ISL_2932792, EPI_ISL_2932793, EPI_ISL_2932794, EPI_ISL_2932795, EPI_ISL_2932796, EPI_ISL_2932799, EPI_ISL_2932800, EPI_ISL_2932801, EPI_ISL_2932802, EPI_ISL_2932804, EPI_ISL_2932805, EPI_ISL_2932806, EPI_ISL_2932807, EPI_ISL_2932808, EPI_ISL_2932809, EPI_ISL_2932810, EPI_ISL_2932811, EPI_ISL_2932812, EPI_ISL_2932813, EPI_ISL_2932814, EPI_ISL_2932815, EPI_ISL_2932816, EPI_ISL_2932817, EPI_ISL_2932818, EPI_ISL_2932819, EPI_ISL_2932820, EPI_ISL_2932821, EPI_ISL_2932822, EPI_ISL_2932823 | | | |
| see above | Department of Bacteria, Parasites and Fungi, Statens Serum Institut, Copenhagen, Denmark | Statens Serum Institut Bioinformatics and Microbial Genomics | Danish Covid-19 Genome Consortium |
| EPI_ISL_2932824 | Department of Clinical Microbiology, Odense University Hospital, Odense, Denmark | Statens Serum Institut Bioinformatics and Microbial Genomics | Danish Covid-19 Genome Consortium |
| EPI_ISL_2932826, EPI_ISL_2932827, EPI_ISL_2932828, EPI_ISL_2932829, EPI_ISL_2932830, EPI_ISL_2932831, EPI_ISL_2932832, EPI_ISL_2932833, EPI_ISL_2932834, EPI_ISL_2932835, EPI_ISL_2932836, EPI_ISL_2932837, EPI_ISL_2932838, EPI_ISL_2932839, EPI_ISL_2932840, EPI_ISL_2932841, EPI_ISL_2932842, EPI_ISL_2932843, EPI_ISL_2932844, EPI_ISL_2932845, EPI_ISL_2932846, EPI_ISL_2932847, EPI_ISL_2932848, EPI_ISL_2932849, EPI_ISL_2932850, EPI_ISL_2932851, EPI_ISL_2932852, EPI_ISL_2932854, EPI_ISL_2932855, EPI_ISL_2932856, EPI_ISL_2932857, EPI_ISL_2932858, EPI_ISL_2932859, EPI_ISL_2932860, EPI_ISL_2932861, EPI_ISL_2932862, EPI_ISL_2932863, EPI_ISL_2932864, EPI_ISL_2932865, EPI_ISL_2932866, EPI_ISL_2932867, EPI_ISL_2932868, EPI_ISL_2932869, EPI_ISL_2932870, EPI_ISL_2932871, EPI_ISL_2932872, EPI_ISL_2932873, EPI_ISL_2932874, EPI_ISL_2932875, EPI_ISL_2932876, EPI_ISL_2932877, EPI_ISL_2932878, EPI_ISL_2932880, EPI_ISL_2932881, EPI_ISL_2932882, EPI_ISL_2932883, EPI_ISL_2932884, EPI_ISL_2932885, EPI_ISL_2932886, EPI_ISL_2932887, EPI_ISL_2932888, EPI_ISL_2932889, EPI_ISL_2932890, EPI_ISL_2932891, EPI_ISL_2932892, EPI_ISL_2932893, EPI_ISL_2932894, EPI_ISL_2932895, EPI_ISL_2932896, EPI_ISL_2932897, EPI_ISL_2932899, EPI_ISL_2932900, EPI_ISL_2932901, EPI_ISL_2932902, EPI_ISL_2932903, EPI_ISL_2932904, EPI_ISL_2932905, EPI_ISL_2932906, EPI_ISL_2932907, EPI_ISL_2932908, EPI_ISL_2932909, EPI_ISL_2932910, EPI_ISL_2932911, EPI_ISL_2932912, EPI_ISL_2932913 | | | |
| see above | Department of Bacteria, Parasites and Fungi, Statens Serum Institut, Copenhagen, Denmark | Statens Serum Institut Bioinformatics and Microbial Genomics | Danish Covid-19 Genome Consortium |
| EPI_ISL_2932914 | Molekylær Medicinsk Afdeling, Aarhus University Hospital, Aarhus, Denmark | Statens Serum Institut Bioinformatics and Microbial Genomics | Danish Covid-19 Genome Consortium |
| EPI_ISL_2932915, EPI_ISL_2932916, EPI_ISL_2932917, EPI_ISL_2932918, EPI_ISL_2932919, EPI_ISL_2932920, EPI_ISL_2932921, EPI_ISL_2932922, EPI_ISL_2932923, EPI_ISL_2932924, EPI_ISL_2932925, EPI_ISL_2932926, EPI_ISL_2932927, EPI_ISL_2932928, EPI_ISL_2932929, EPI_ISL_2932930, EPI_ISL_2932931, EPI_ISL_2932932, EPI_ISL_2932933, EPI_ISL_2932934, EPI_ISL_2932935, EPI_ISL_2932936, EPI_ISL_2932937, EPI_ISL_2932938, EPI_ISL_2932939, EPI_ISL_2932940, EPI_ISL_2932941, EPI_ISL_2932942, EPI_ISL_2932943, EPI_ISL_2932944, EPI_ISL_2932945, EPI_ISL_2932946, EPI_ISL_2932947, EPI_ISL_2932948, EPI_ISL_2932949, EPI_ISL_2932950, EPI_ISL_2932951, EPI_ISL_2932952, EPI_ISL_2932953, EPI_ISL_2932954, EPI_ISL_2932955, EPI_ISL_2932956, EPI_ISL_2932957, EPI_ISL_2932958, EPI_ISL_2932959, EPI_ISL_2932961, EPI_ISL_2932962, EPI_ISL_2932963, EPI_ISL_2932964, EPI_ISL_2932965, EPI_ISL_2932966, EPI_ISL_2932967, EPI_ISL_2932968, EPI_ISL_2932969, EPI_ISL_2932970, EPI_ISL_2932971, EPI_ISL_2932972, EPI_ISL_2932973, EPI_ISL_2932974, EPI_ISL_2932975, EPI_ISL_2932976, EPI_ISL_2932977, EPI_ISL_2932978, EPI_ISL_2932979, EPI_ISL_2932980, EPI_ISL_2932981, EPI_ISL_2932982, EPI_ISL_2932983, EPI_ISL_2932984, EPI_ISL_2932986, EPI_ISL_2932987, EPI_ISL_2932988, EPI_ISL_2932989, EPI_ISL_2932990, EPI_ISL_2932991, EPI_ISL_2932992, EPI_ISL_2932993, EPI_ISL_2932994, EPI_ISL_2932995, EPI_ISL_2932996, EPI_ISL_2932997, EPI_ISL_2932998, EPI_ISL_2932999, EPI_ISL_2933000, EPI_ISL_2933001, EPI_ISL_2933002, EPI_ISL_2933003, EPI_ISL_2933004, EPI_ISL_2933005, EPI_ISL_2933006, EPI_ISL_2933007, EPI_ISL_2933008, EPI_ISL_2933009, EPI_ISL_2933010, EPI_ISL_2933011, EPI_ISL_2933012, EPI_ISL_2933013, EPI_ISL_2933014, EPI_ISL_2933015, EPI_ISL_2933016, EPI_ISL_2933017, EPI_ISL_2933018, EPI_ISL_2933019, EPI_ISL_2933020, EPI_ISL_2933021, EPI_ISL_2933022, EPI_ISL_2933023, EPI_ISL_2933024, EPI_ISL_2933025, EPI_ISL_2933026, EPI_ISL_2933027, EPI_ISL_2933028, EPI_ISL_2933029, EPI_ISL_2933030, EPI_ISL_2933031, EPI_ISL_2933032, EPI_ISL_2933033, EPI_ISL_2933034, EPI_ISL_2933035, EPI_ISL_2933036, EPI_ISL_2933037, EPI_ISL_2933038, EPI_ISL_2933041, EPI_ISL_2933042, EPI_ISL_2933043, EPI_ISL_2933044, EPI_ISL_2933046, EPI_ISL_2933047, EPI_ISL_2933048, EPI_ISL_2933049, EPI_ISL_2933051, EPI_ISL_2933052, EPI_ISL_2933053, EPI_ISL_2933054, EPI_ISL_2933055, EPI_ISL_2933056, EPI_ISL_2933057, EPI_ISL_2933058, EPI_ISL_2933059, EPI_ISL_2933060, EPI_ISL_2933061, EPI_ISL_2933062, EPI_ISL_2933063, EPI_ISL_2933065, EPI_ISL_2933066, EPI_ISL_2933067, EPI_ISL_2933068, EPI_ISL_2933069, EPI_ISL_2933070, EPI_ISL_2933072, EPI_ISL_2933073, EPI_ISL_2933075, EPI_ISL_2933076, EPI_ISL_2933077, EPI_ISL_2933078, EPI_ISL_2933079, EPI_ISL_2933080, EPI_ISL_2933081, EPI_ISL_2933082, EPI_ISL_2933083, EPI_ISL_2933084, EPI_ISL_2933085 | | | |
| see above | Department of Bacteria, Parasites and Fungi, Statens Serum Institut, Copenhagen, Denmark | Statens Serum Institut Bioinformatics and Microbial Genomics | Danish Covid-19 Genome Consortium |
| EPI_ISL_2933086 | Molekylær Medicinsk Afdeling, Aarhus University Hospital, Aarhus, Denmark | Statens Serum Institut Bioinformatics and Microbial Genomics | Danish Covid-19 Genome Consortium |
| EPI_ISL_2933087, EPI_ISL_2933088, EPI_ISL_2933089, EPI_ISL_2933090, EPI_ISL_2933091, EPI_ISL_2933092, EPI_ISL_2933093, EPI_ISL_2933094, EPI_ISL_2933095, EPI_ISL_2933096, EPI_ISL_2933097, EPI_ISL_2933098 | | | |
| see above | Department of Bacteria, Parasites and Fungi, Statens Serum Institut, Copenhagen, Denmark | Statens Serum Institut Bioinformatics and Microbial Genomics | Danish Covid-19 Genome Consortium |
| EPI_ISL_2933099 | Molekylær Medicinsk Afdeling, Aarhus University Hospital, Aarhus, Denmark | Statens Serum Institut Bioinformatics and Microbial Genomics | Danish Covid-19 Genome Consortium |
| EPI_ISL_2933100, EPI_ISL_2933101 | Department of Bacteria, Parasites and Fungi, Statens Serum Institut, Copenhagen, Denmark | Statens Serum Institut Bioinformatics and Microbial Genomics | Danish Covid-19 Genome Consortium |
| EPI_ISL_2933109 | DKK Sleman/Puskesmas Depok 1 | Genetics Working Group (Pokja Genetik) Faculty of Medicine, Public Health and Nursing Universitas Gadjah Mada (FK-KMK UGM); Disease Investigation Center Wates Ministry of Agriculture Indonesia; Department of Microbiology FK-KMK UGM; Laboratorium Diagnostik Yayasan Tahija World Mosquito Program (WMP) Yogyakarta Center for Tropical Medicine FK-KMK UGM; Integrated Research Center FK-KMK UGM; Department of Computer Science and Electronics FMIPA UGM; RSUP Dr. Sardjito | Gunadi, Hendra Wibawa, Marcellus, Mohamad S. Hakim, Edwin W. Daniwijaya, Ludhang P. Rizki, Endah Supriyati, Eggi Arguni, Titik Nuryastuti, Tri Wibawa, Dwi AA Nugrahaningsih, Afiahayati, Siswanto,Kristy Iskandar,Nungki Anggorowati, Pramesti G Dewi, Dwi Indaryati,Indarto Sulistiyono, Susan Simanjaya, Dyah A Puspitarani |
| EPI_ISL_2933110 | DKK Sleman/Puskesmas Depok 1 | Genetics Working Group (Pokja Genetik) Faculty of Medicine, Public Health and Nursing Universitas Gadjah Mada (FK-KMK UGM); Disease Investigation Center Wates Ministry of Agriculture Indonesia; Department of Microbiology FK-KMK UGM; Laboratorium Diagnostik Yayasan Tahija World Mosquito Program (WMP) Yogyakarta Center for Tropical Medicine FK-KMK UGM; Integrated Research Center FK-KMK UGM; Department of Computer Science and | Gunadi, Hendra Wibawa, Marcellus, Mohamad S. Hakim, Edwin W. Daniwijaya, Ludhang P. Rizki, Endah Supriyati, Eggi Arguni, Titik Nuryastuti, Tri Wibawa, Dwi AA Nugrahaningsih, Afiahayati, Siswanto,Kristy Iskandar,Nungki Anggorowati, Pramesti G Dewi, Dwi Indaryati,Indarto Sulistiyono, Khanza a Vujira, Alvina A Setiawan |
| EPI_ISL_2933111, EPI_ISL_2933112, EPI_ISL_2933114, EPI_ISL_2933115, EPI_ISL_2933116, EPI_ISL_2933117, EPI_ISL_2933118, | National Platform bis UMONS/Jolimont | National Platform bis UMONS/Jolimont | François Dufrasne, Guillaume Bayon-Vicente, Florian Juszczak, Eric Tarantino, Gautier Detry, Ruddy Wattiez |

| | | | |
|---|---|---|---|
| EPI_ISL_2933119, EPI_ISL_2933123 | | | |
| EPI_ISL_2933129 | AX BIO OCEAN | CNR Virus des Infections Respiratoires - France SUD | Antonin Bal, Gregory Destras, Gwendolyne Burfin, Hadrien Regue, Quentin Semanas, Martine Valette, Bruno Lina, Laurence Josset |
| EPI_ISL_2933131 | MIRIALIS CLUSES BECHET | CNR Virus des Infections Respiratoires - France SUD | Antonin Bal, Gregory Destras, Gwendolyne Burfin, Hadrien Regue, Quentin Semanas, Martine Valette, Bruno Lina, Laurence Josset |
| EPI_ISL_2933134 | BIOMNIS LYON | CNR Virus des Infections Respiratoires - France SUD | Antonin Bal, Gregory Destras, Gwendolyne Burfin, Hadrien Regue, Quentin Semanas, Martine Valette, Bruno Lina, Laurence Josset |
| EPI_ISL_2933140 | AX BIO OCEAN | CNR Virus des Infections Respiratoires - France SUD | Antonin Bal, Gregory Destras, Gwendolyne Burfin, Hadrien Regue, Quentin Semanas, Martine Valette, Bruno Lina, Laurence Josset |
| EPI_ISL_2933142 | BIOMNIS LYON | CNR Virus des Infections Respiratoires - France SUD | Antonin Bal, Gregory Destras, Gwendolyne Burfin, Hadrien Regue, Quentin Semanas, Martine Valette, Bruno Lina, Laurence Josset |
| EPI_ISL_2933148 | AX BIO OCEAN | CNR Virus des Infections Respiratoires - France SUD | Antonin Bal, Gregory Destras, Gwendolyne Burfin, Hadrien Regue, Quentin Semanas, Martine Valette, Bruno Lina, Laurence Josset |
| EPI_ISL_2933150 | ORIAPOLE | CNR Virus des Infections Respiratoires - France SUD | Antonin Bal, Gregory Destras, Gwendolyne Burfin, Hadrien Regue, Quentin Semanas, Martine Valette, Bruno Lina, Laurence Josset |
| EPI_ISL_2933151 | BIOMNIS LYON | CNR Virus des Infections Respiratoires - France SUD | Antonin Bal, Gregory Destras, Gwendolyne Burfin, Hadrien Regue, Quentin Semanas, Martine Valette, Bruno Lina, Laurence Josset |
| EPI_ISL_2933156 | DYOMEDEA-LABORATOIRE DE LA SAUVEGARDE | CNR Virus des Infections Respiratoires - France SUD | Antonin Bal, Gregory Destras, Gwendolyne Burfin, Hadrien Regue, Quentin Semanas, Martine Valette, Bruno Lina, Laurence Josset |
| EPI_ISL_2933161, EPI_ISL_2933165 | BIOMNIS LYON | CNR Virus des Infections Respiratoires - France SUD | Antonin Bal, Gregory Destras, Gwendolyne Burfin, Hadrien Regue, Quentin Semanas, Martine Valette, Bruno Lina, Laurence Josset |
| EPI_ISL_2933166 | CNR Virus des Infections Respiratoires - France SUD | CNR Virus des Infections Respiratoires - France SUD | Antonin Bal, Gregory Destras, Gwendolyne Burfin, Hadrien Regue, Quentin Semanas, Martine Valette, Bruno Lina, Laurence Josset |
| EPI_ISL_2933167 | SCM B12 | CNR Virus des Infections Respiratoires - France SUD | Antonin Bal, Gregory Destras, Gwendolyne Burfin, Hadrien Regue, Quentin Semanas, Martine Valette, Bruno Lina, Laurence Josset |
| EPI_ISL_2933173 | LABORATOIRE CBM DE MURET | CNR Virus des Infections Respiratoires - France SUD | Antonin Bal, Gregory Destras, Gwendolyne Burfin, Hadrien Regue, Quentin Semanas, Martine Valette, Bruno Lina, Laurence Josset |
| EPI_ISL_2933179 | LABORATOIRE CERBALLIANCE PLT VILLON | CNR Virus des Infections Respiratoires - France SUD | Antonin Bal, Gregory Destras, Gwendolyne Burfin, Hadrien Regue, Quentin Semanas, Martine Valette, Bruno Lina, Laurence Josset |
| EPI_ISL_2933181 | DYOMEDEA-LABORATOIRE DE LA SAUVEGARDE | CNR Virus des Infections Respiratoires - France SUD | Antonin Bal, Gregory Destras, Gwendolyne Burfin, Hadrien Regue, Quentin Semanas, Martine Valette, Bruno Lina, Laurence Josset |
| EPI_ISL_2933188 | LABORATOIRE CERBALLIANCE PLT VILLON | CNR Virus des Infections Respiratoires - France SUD | Antonin Bal, Gregory Destras, Gwendolyne Burfin, Hadrien Regue, Quentin Semanas, Martine Valette, Bruno Lina, Laurence Josset |
| EPI_ISL_2933193 | MIRIALIS CLUSES BECHET | CNR Virus des Infections Respiratoires - France SUD | Antonin Bal, Gregory Destras, Gwendolyne Burfin, Hadrien Regue, Quentin Semanas, Martine Valette, Bruno Lina, Laurence Josset |
| EPI_ISL_2933196 | BIOMNIS LYON | CNR Virus des Infections Respiratoires - France SUD | Antonin Bal, Gregory Destras, Gwendolyne Burfin, Hadrien Regue, Quentin Semanas, Martine Valette, Bruno Lina, Laurence Josset |
| EPI_ISL_2933197 | MIRIALIS CLUSES BECHET | CNR Virus des Infections Respiratoires - France SUD | Antonin Bal, Gregory Destras, Gwendolyne Burfin, Hadrien Regue, Quentin Semanas, Martine Valette, Bruno Lina, Laurence Josset |
| EPI_ISL_2933202 | BIOMNIS LYON | CNR Virus des Infections Respiratoires - France SUD | Antonin Bal, Gregory Destras, Gwendolyne Burfin, Hadrien Regue, Quentin Semanas, Martine Valette, Bruno Lina, Laurence Josset |
| EPI_ISL_2933204 | SCM B12 | CNR Virus des Infections Respiratoires - France SUD | Antonin Bal, Gregory Destras, Gwendolyne Burfin, Hadrien Regue, Quentin Semanas, Martine Valette, Bruno Lina, Laurence Josset |
| EPI_ISL_2933343 | BIOMNIS LYON | CNR Virus des Infections Respiratoires - France SUD | Antonin Bal, Gregory Destras, Gwendolyne Burfin, Hadrien Regue, Quentin Semanas, Martine Valette, Bruno Lina, Laurence Josset |
| EPI_ISL_2933346 | MIRIALIS CLUSES BECHET | CNR Virus des Infections Respiratoires - France SUD | Antonin Bal, Gregory Destras, Gwendolyne Burfin, Hadrien Regue, Quentin Semanas, Martine Valette, Bruno Lina, Laurence Josset |
| EPI_ISL_2933348, EPI_ISL_2933349 | CNR Virus des Infections Respiratoires - France SUD | CNR Virus des Infections Respiratoires - France SUD | Antonin Bal, Gregory Destras, Gwendolyne Burfin, Hadrien Regue, Quentin Semanas, Martine Valette, Bruno Lina, Laurence Josset |
| EPI_ISL_2933443, EPI_ISL_2933447, EPI_ISL_2933450, EPI_ISL_2933456, EPI_ISL_2933457 | BIOMNIS LYON | CNR Virus des Infections Respiratoires - France SUD | Antonin Bal, Gregory Destras, Gwendolyne Burfin, Hadrien Regue, Quentin Semanas, Martine Valette, Bruno Lina, Laurence Josset |
| EPI_ISL_2933458, EPI_ISL_2933459, EPI_ISL_2933460 | MIRIALIS CLUSES BECHET | CNR Virus des Infections Respiratoires - France SUD | Antonin Bal, Gregory Destras, Gwendolyne Burfin, Hadrien Regue, Quentin Semanas, Martine Valette, Bruno Lina, Laurence Josset |
| EPI_ISL_2933461, EPI_ISL_2933462 | BIOMNIS LYON | CNR Virus des Infections Respiratoires - France SUD | Antonin Bal, Gregory Destras, Gwendolyne Burfin, Hadrien Regue, Quentin Semanas, Martine Valette, Bruno Lina, Laurence Josset |
| EPI_ISL_2933492 | LABORATOIRE CERBALLIANCE PLT VILLON | CNR Virus des Infections Respiratoires - France SUD | Antonin Bal, Gregory Destras, Gwendolyne Burfin, Hadrien Regue, Quentin Semanas, Martine Valette, Bruno Lina, Laurence Josset |
| EPI_ISL_2933493, EPI_ISL_2933494, EPI_ISL_2933495 | BIOMNIS LYON | CNR Virus des Infections Respiratoires - France SUD | Antonin Bal, Gregory Destras, Gwendolyne Burfin, Hadrien Regue, Quentin Semanas, Martine Valette, Bruno Lina, Laurence Josset |
| EPI_ISL_2933496 | LABORATOIRE BIOLAB AVENIR | CNR Virus des Infections Respiratoires - France SUD | Antonin Bal, Gregory Destras, Gwendolyne Burfin, Hadrien Regue, Quentin Semanas, Martine Valette, Bruno Lina, Laurence Josset |
| EPI_ISL_2933497, EPI_ISL_2933498, EPI_ISL_2933499, EPI_ISL_2933500, EPI_ISL_2933501, EPI_ISL_2933502 | BIOMNIS LYON | CNR Virus des Infections Respiratoires - France SUD | Antonin Bal, Gregory Destras, Gwendolyne Burfin, Hadrien Regue, Quentin Semanas, Martine Valette, Bruno Lina, Laurence Josset |
| EPI_ISL_2933503 | LABORATOIRE DES CEDRES | CNR Virus des Infections Respiratoires - France SUD | Antonin Bal, Gregory Destras, Gwendolyne Burfin, Hadrien Regue, Quentin Semanas, Martine Valette, Bruno Lina, Laurence Josset |
| EPI_ISL_2933504, EPI_ISL_2933505, EPI_ISL_2933506 | BIOMNIS LYON | CNR Virus des Infections Respiratoires - France SUD | Antonin Bal, Gregory Destras, Gwendolyne Burfin, Hadrien Regue, Quentin Semanas, Martine Valette, Bruno Lina, Laurence Josset |
| EPI_ISL_2933507 | ASTRALAB TASSIGNY | CNR Virus des Infections Respiratoires - France SUD | Antonin Bal, Gregory Destras, Gwendolyne Burfin, Hadrien Regue, Quentin Semanas, Martine Valette, Bruno Lina, Laurence Josset |
| EPI_ISL_2933508 | CNR Virus des Infections Respiratoires - France SUD | CNR Virus des Infections Respiratoires - France SUD | Antonin Bal, Gregory Destras, Gwendolyne Burfin, Hadrien Regue, Quentin Semanas, Martine Valette, Bruno Lina, Laurence Josset |
| EPI_ISL_2933509 | NEOLAB LIMAS VILLEFRANCHE | CNR Virus des Infections Respiratoires - France SUD | Antonin Bal, Gregory Destras, Gwendolyne Burfin, Hadrien Regue, Quentin Semanas, Martine Valette, Bruno Lina, Laurence Josset |
| EPI_ISL_2933510, EPI_ISL_2933511 | LABORATOIRE CBM DE MURET | CNR Virus des Infections Respiratoires - France SUD | Antonin Bal, Gregory Destras, Gwendolyne Burfin, Hadrien Regue, Quentin Semanas, Martine Valette, Bruno Lina, Laurence Josset |
| EPI_ISL_2933512, EPI_ISL_2933513, EPI_ISL_2933514, EPI_ISL_2933515 | AX BIO OCEAN | CNR Virus des Infections Respiratoires - France SUD | Antonin Bal, Gregory Destras, Gwendolyne Burfin, Hadrien Regue, Quentin Semanas, Martine Valette, Bruno Lina, Laurence Josset |
| EPI_ISL_2933516 | SCM B12 | CNR Virus des Infections Respiratoires - France SUD | Antonin Bal, Gregory Destras, Gwendolyne Burfin, Hadrien Regue, Quentin Semanas, Martine Valette, Bruno Lina, Laurence Josset |
| EPI_ISL_2933517, EPI_ISL_2933518, EPI_ISL_2933519, EPI_ISL_2933520 | AX BIO OCEAN | CNR Virus des Infections Respiratoires - France SUD | Antonin Bal, Gregory Destras, Gwendolyne Burfin, Hadrien Regue, Quentin Semanas, Martine Valette, Bruno Lina, Laurence Josset |
| EPI_ISL_2933521 | ORIAPOLE | CNR Virus des Infections Respiratoires - France SUD | Antonin Bal, Gregory Destras, Gwendolyne Burfin, Hadrien Regue, Quentin Semanas, Martine Valette, Bruno Lina, Laurence Josset |
| EPI_ISL_2933522, EPI_ISL_2933523, EPI_ISL_2933524, EPI_ISL_2933525, EPI_ISL_2933526, EPI_ISL_2933527 | LABORATOIRE FORTE BIO ST PAUL LES DAX | CNR Virus des Infections Respiratoires - France SUD | Antonin Bal, Gregory Destras, Gwendolyne Burfin, Hadrien Regue, Quentin Semanas, Martine Valette, Bruno Lina, Laurence Josset |
| EPI_ISL_2933528 | CH METROPOLE SAVOIE | CNR Virus des Infections Respiratoires - France SUD | Antonin Bal, Gregory Destras, Gwendolyne Burfin, Hadrien Regue, Quentin Semanas, Martine Valette, Bruno Lina, Laurence Josset |
| EPI_ISL_2933529 | LABORATOIRE ACCOLAB SUD OUEST | CNR Virus des Infections Respiratoires - France SUD | Antonin Bal, Gregory Destras, Gwendolyne Burfin, Hadrien Regue, Quentin Semanas, Martine Valette, Bruno Lina, Laurence Josset |
| EPI_ISL_2933530 | LAM CERBALLIANCE | CNR Virus des Infections Respiratoires - France SUD | Antonin Bal, Gregory Destras, Gwendolyne Burfin, Hadrien Regue, Quentin Semanas, Martine Valette, Bruno Lina, Laurence Josset |
| EPI_ISL_2933531 | CERBALLIANCE MIDI-PYRENEES | CNR Virus des Infections Respiratoires - France SUD | Antonin Bal, Gregory Destras, Gwendolyne Burfin, Hadrien Regue, Quentin Semanas, Martine Valette, Bruno Lina, Laurence Josset |
| EPI_ISL_2933532 | LABORATOIRE BIO-VAL | CNR Virus des Infections Respiratoires - France SUD | Antonin Bal, Gregory Destras, Gwendolyne Burfin, Hadrien Regue, Quentin Semanas, Martine Valette, Bruno Lina, Laurence Josset |
| EPI_ISL_2933533 | ASTRALAB TASSIGNY | CNR Virus des Infections Respiratoires - France SUD | Antonin Bal, Gregory Destras, Gwendolyne Burfin, Hadrien Regue, Quentin Semanas, Martine Valette, Bruno Lina, Laurence Josset |
| EPI_ISL_2933534, EPI_ISL_2933535 | ORIAPOLE | CNR Virus des Infections Respiratoires - France SUD | Antonin Bal, Gregory Destras, Gwendolyne Burfin, Hadrien Regue, Quentin Semanas, Martine Valette, Bruno Lina, Laurence Josset |
| EPI_ISL_2933536 | AX BIO OCEAN | CNR Virus des Infections Respiratoires - France SUD | Antonin Bal, Gregory Destras, Gwendolyne Burfin, Hadrien Regue, Quentin Semanas, Martine Valette, Bruno Lina, Laurence Josset |
| EPI_ISL_2933537, EPI_ISL_2933538, EPI_ISL_2933539 | ORIAPOLE | CNR Virus des Infections Respiratoires - France SUD | Antonin Bal, Gregory Destras, Gwendolyne Burfin, Hadrien Regue, Quentin Semanas, Martine Valette, Bruno Lina, Laurence Josset |
| EPI_ISL_2933540 | AX BIO OCEAN | CNR Virus des Infections Respiratoires - France SUD | Antonin Bal, Gregory Destras, Gwendolyne Burfin, Hadrien Regue, Quentin Semanas, Martine Valette, Bruno Lina, Laurence Josset |

| | | | |
|---|---|---|---|
| EPI_ISL_2933541, EPI_ISL_2933542, EPI_ISL_2933543, EPI_ISL_2933544 | ORIAPOLE | CNR Virus des Infections Respiratoires - France SUD | Antonin Bal, Gregory Destras, Gwendolyne Burfin, Hadrien Regue, Quentin Semanas, Martine Valette, Bruno Lina, Laurence Josset |
| EPI_ISL_2933545, EPI_ISL_2933546 | LABORATOIRE CBM DE MURET | CNR Virus des Infections Respiratoires - France SUD | Antonin Bal, Gregory Destras, Gwendolyne Burfin, Hadrien Regue, Quentin Semanas, Martine Valette, Bruno Lina, Laurence Josset |
| EPI_ISL_2933547 | CENTRE HOSPITALIER DU PAYS D'AIX | CNR Virus des Infections Respiratoires - France SUD | Antonin Bal, Gregory Destras, Gwendolyne Burfin, Hadrien Regue, Quentin Semanas, Martine Valette, Bruno Lina, Laurence Josset |
| EPI_ISL_2933548, EPI_ISL_2933549 | LABORATOIRE MAYMAT | CNR Virus des Infections Respiratoires - France SUD | Antonin Bal, Gregory Destras, Gwendolyne Burfin, Hadrien Regue, Quentin Semanas, Martine Valette, Bruno Lina, Laurence Josset |
| EPI_ISL_2933550, EPI_ISL_2933551, EPI_ISL_2933552 | LABORATOIRE CBM DE MURET | CNR Virus des Infections Respiratoires - France SUD | Antonin Bal, Gregory Destras, Gwendolyne Burfin, Hadrien Regue, Quentin Semanas, Martine Valette, Bruno Lina, Laurence Josset |
| EPI_ISL_2933553, EPI_ISL_2933554 | SCM B12 | CNR Virus des Infections Respiratoires - France SUD | Antonin Bal, Gregory Destras, Gwendolyne Burfin, Hadrien Regue, Quentin Semanas, Martine Valette, Bruno Lina, Laurence Josset |
| EPI_ISL_2933555 | LABORATOIRE CBM DE MURET | CNR Virus des Infections Respiratoires - France SUD | Antonin Bal, Gregory Destras, Gwendolyne Burfin, Hadrien Regue, Quentin Semanas, Martine Valette, Bruno Lina, Laurence Josset |
| EPI_ISL_2933556 | SCM B12 | CNR Virus des Infections Respiratoires - France SUD | Antonin Bal, Gregory Destras, Gwendolyne Burfin, Hadrien Regue, Quentin Semanas, Martine Valette, Bruno Lina, Laurence Josset |
| EPI_ISL_2933557, EPI_ISL_2933558, EPI_ISL_2933559 | LABORATOIRE CBM DE MURET | CNR Virus des Infections Respiratoires - France SUD | Antonin Bal, Gregory Destras, Gwendolyne Burfin, Hadrien Regue, Quentin Semanas, Martine Valette, Bruno Lina, Laurence Josset |
| EPI_ISL_2933560 | CENTRE HOSPITALIER PIERRE OUDOT | CNR Virus des Infections Respiratoires - France SUD | Antonin Bal, Gregory Destras, Gwendolyne Burfin, Hadrien Regue, Quentin Semanas, Martine Valette, Bruno Lina, Laurence Josset |
| EPI_ISL_2933561, EPI_ISL_2933562, EPI_ISL_2933563 | LABORATOIRE CERBALLIANCE PLT VILLON | CNR Virus des Infections Respiratoires - France SUD | Antonin Bal, Gregory Destras, Gwendolyne Burfin, Hadrien Regue, Quentin Semanas, Martine Valette, Bruno Lina, Laurence Josset |
| EPI_ISL_2933564, EPI_ISL_2933565 | BIOMNIS LYON | CNR Virus des Infections Respiratoires - France SUD | Antonin Bal, Gregory Destras, Gwendolyne Burfin, Hadrien Regue, Quentin Semanas, Martine Valette, Bruno Lina, Laurence Josset |
| EPI_ISL_2933566 | BIOMNIS PARIS | CNR Virus des Infections Respiratoires - France SUD | Antonin Bal, Gregory Destras, Gwendolyne Burfin, Hadrien Regue, Quentin Semanas, Martine Valette, Bruno Lina, Laurence Josset |
| EPI_ISL_2933567, EPI_ISL_2933568, EPI_ISL_2933569, EPI_ISL_2933570, EPI_ISL_2933571, EPI_ISL_2933572, EPI_ISL_2933573, EPI_ISL_2933574, EPI_ISL_2933575, EPI_ISL_2933576, EPI_ISL_2933577, EPI_ISL_2933578, EPI_ISL_2933579, EPI_ISL_2933580, EPI_ISL_2933581, EPI_ISL_2933582 | | | |
| see above | BIOMNIS LYON | CNR Virus des Infections Respiratoires - France SUD | Antonin Bal, Gregory Destras, Gwendolyne Burfin, Hadrien Regue, Quentin Semanas, Martine Valette, Bruno Lina, Laurence Josset |
| EPI_ISL_2933583 | CABINET MEDICAL DU CARREAU | CNR Virus des Infections Respiratoires - France SUD | Antonin Bal, Gregory Destras, Gwendolyne Burfin, Hadrien Regue, Quentin Semanas, Martine Valette, Bruno Lina, Laurence Josset |
| EPI_ISL_2933584, EPI_ISL_2933585, EPI_ISL_2933586, EPI_ISL_2933587, EPI_ISL_2933588, EPI_ISL_2933589, EPI_ISL_2933590, EPI_ISL_2933591 | BIOMNIS LYON | CNR Virus des Infections Respiratoires - France SUD | Antonin Bal, Gregory Destras, Gwendolyne Burfin, Hadrien Regue, Quentin Semanas, Martine Valette, Bruno Lina, Laurence Josset |
| EPI_ISL_2933592, EPI_ISL_2933593, EPI_ISL_2933594, EPI_ISL_2933595, EPI_ISL_2933596 | LABORATOIRE CBM DE MURET | CNR Virus des Infections Respiratoires - France SUD | Antonin Bal, Gregory Destras, Gwendolyne Burfin, Hadrien Regue, Quentin Semanas, Martine Valette, Bruno Lina, Laurence Josset |
| EPI_ISL_2933597 | LABORATOIRE BOUVIER | CNR Virus des Infections Respiratoires - France SUD | Antonin Bal, Gregory Destras, Gwendolyne Burfin, Hadrien Regue, Quentin Semanas, Martine Valette, Bruno Lina, Laurence Josset |
| EPI_ISL_2933598, EPI_ISL_2933599, EPI_ISL_2933600, EPI_ISL_2933601 | MIRIALIS CLUSES BECHET | CNR Virus des Infections Respiratoires - France SUD | Antonin Bal, Gregory Destras, Gwendolyne Burfin, Hadrien Regue, Quentin Semanas, Martine Valette, Bruno Lina, Laurence Josset |
| EPI_ISL_2933602 | AX BIO OCEAN | CNR Virus des Infections Respiratoires - France SUD | Antonin Bal, Gregory Destras, Gwendolyne Burfin, Hadrien Regue, Quentin Semanas, Martine Valette, Bruno Lina, Laurence Josset |
| EPI_ISL_2933603 | MIRIALIS CLUSES BECHET | CNR Virus des Infections Respiratoires - France SUD | Antonin Bal, Gregory Destras, Gwendolyne Burfin, Hadrien Regue, Quentin Semanas, Martine Valette, Bruno Lina, Laurence Josset |
| EPI_ISL_2933604, EPI_ISL_2933605, EPI_ISL_2933606 | BIOMNIS LYON | CNR Virus des Infections Respiratoires - France SUD | Antonin Bal, Gregory Destras, Gwendolyne Burfin, Hadrien Regue, Quentin Semanas, Martine Valette, Bruno Lina, Laurence Josset |
| EPI_ISL_2933607 | MIRIALIS CLUSES BECHET | CNR Virus des Infections Respiratoires - France SUD | Antonin Bal, Gregory Destras, Gwendolyne Burfin, Hadrien Regue, Quentin Semanas, Martine Valette, Bruno Lina, Laurence Josset |
| EPI_ISL_2933608 | BIOMNIS LYON | CNR Virus des Infections Respiratoires - France SUD | Antonin Bal, Gregory Destras, Gwendolyne Burfin, Hadrien Regue, Quentin Semanas, Martine Valette, Bruno Lina, Laurence Josset |
| EPI_ISL_2933609 | AX BIO OCEAN | CNR Virus des Infections Respiratoires - France SUD | Antonin Bal, Gregory Destras, Gwendolyne Burfin, Hadrien Regue, Quentin Semanas, Martine Valette, Bruno Lina, Laurence Josset |
| EPI_ISL_2933610 | LABORATOIRE CERBA | CNR Virus des Infections Respiratoires - France SUD | Antonin Bal, Gregory Destras, Gwendolyne Burfin, Hadrien Regue, Quentin Semanas, Martine Valette, Bruno Lina, Laurence Josset |
| EPI_ISL_2933611, EPI_ISL_2933612, EPI_ISL_2933613, EPI_ISL_2933614 | AX BIO OCEAN | CNR Virus des Infections Respiratoires - France SUD | Antonin Bal, Gregory Destras, Gwendolyne Burfin, Hadrien Regue, Quentin Semanas, Martine Valette, Bruno Lina, Laurence Josset |
| EPI_ISL_2933615 | BIOMNIS LYON | CNR Virus des Infections Respiratoires - France SUD | Antonin Bal, Gregory Destras, Gwendolyne Burfin, Hadrien Regue, Quentin Semanas, Martine Valette, Bruno Lina, Laurence Josset |
| EPI_ISL_2933616, EPI_ISL_2933617 | AX BIO OCEAN | CNR Virus des Infections Respiratoires - France SUD | Antonin Bal, Gregory Destras, Gwendolyne Burfin, Hadrien Regue, Quentin Semanas, Martine Valette, Bruno Lina, Laurence Josset |
| EPI_ISL_2933618 | MIRIALIS CLUSES BECHET | CNR Virus des Infections Respiratoires - France SUD | Antonin Bal, Gregory Destras, Gwendolyne Burfin, Hadrien Regue, Quentin Semanas, Martine Valette, Bruno Lina, Laurence Josset |
| EPI_ISL_2933619, EPI_ISL_2933620 | AX BIO OCEAN | CNR Virus des Infections Respiratoires - France SUD | Antonin Bal, Gregory Destras, Gwendolyne Burfin, Hadrien Regue, Quentin Semanas, Martine Valette, Bruno Lina, Laurence Josset |
| EPI_ISL_2933621 | MIRIALIS CLUSES BECHET | CNR Virus des Infections Respiratoires - France SUD | Antonin Bal, Gregory Destras, Gwendolyne Burfin, Hadrien Regue, Quentin Semanas, Martine Valette, Bruno Lina, Laurence Josset |
| EPI_ISL_2933622 | AX BIO OCEAN | CNR Virus des Infections Respiratoires - France SUD | Antonin Bal, Gregory Destras, Gwendolyne Burfin, Hadrien Regue, Quentin Semanas, Martine Valette, Bruno Lina, Laurence Josset |
| EPI_ISL_2933623 | MIRIALIS CLUSES BECHET | CNR Virus des Infections Respiratoires - France SUD | Antonin Bal, Gregory Destras, Gwendolyne Burfin, Hadrien Regue, Quentin Semanas, Martine Valette, Bruno Lina, Laurence Josset |
| EPI_ISL_2933624 | AX BIO OCEAN | CNR Virus des Infections Respiratoires - France SUD | Antonin Bal, Gregory Destras, Gwendolyne Burfin, Hadrien Regue, Quentin Semanas, Martine Valette, Bruno Lina, Laurence Josset |
| EPI_ISL_2933625 | MIRIALIS CLUSES BECHET | CNR Virus des Infections Respiratoires - France SUD | Antonin Bal, Gregory Destras, Gwendolyne Burfin, Hadrien Regue, Quentin Semanas, Martine Valette, Bruno Lina, Laurence Josset |
| EPI_ISL_2933626 | AX BIO OCEAN | CNR Virus des Infections Respiratoires - France SUD | Antonin Bal, Gregory Destras, Gwendolyne Burfin, Hadrien Regue, Quentin Semanas, Martine Valette, Bruno Lina, Laurence Josset |
| EPI_ISL_2933627 | MIRIALIS CLUSES BECHET | CNR Virus des Infections Respiratoires - France SUD | Antonin Bal, Gregory Destras, Gwendolyne Burfin, Hadrien Regue, Quentin Semanas, Martine Valette, Bruno Lina, Laurence Josset |
| EPI_ISL_2933628 | AX BIO OCEAN | CNR Virus des Infections Respiratoires - France SUD | Antonin Bal, Gregory Destras, Gwendolyne Burfin, Hadrien Regue, Quentin Semanas, Martine Valette, Bruno Lina, Laurence Josset |
| EPI_ISL_2933629 | MIRIALIS CLUSES BECHET | CNR Virus des Infections Respiratoires - France SUD | Antonin Bal, Gregory Destras, Gwendolyne Burfin, Hadrien Regue, Quentin Semanas, Martine Valette, Bruno Lina, Laurence Josset |
| EPI_ISL_2933630 | AX BIO OCEAN | CNR Virus des Infections Respiratoires - France SUD | Antonin Bal, Gregory Destras, Gwendolyne Burfin, Hadrien Regue, Quentin Semanas, Martine Valette, Bruno Lina, Laurence Josset |
| EPI_ISL_2933631 | MIRIALIS CLUSES BECHET | CNR Virus des Infections Respiratoires - France SUD | Antonin Bal, Gregory Destras, Gwendolyne Burfin, Hadrien Regue, Quentin Semanas, Martine Valette, Bruno Lina, Laurence Josset |
| EPI_ISL_2933632 | AX BIO OCEAN | CNR Virus des Infections Respiratoires - France SUD | Antonin Bal, Gregory Destras, Gwendolyne Burfin, Hadrien Regue, Quentin Semanas, Martine Valette, Bruno Lina, Laurence Josset |
| EPI_ISL_2933633 | BIOMNIS LYON | CNR Virus des Infections Respiratoires - France SUD | Antonin Bal, Gregory Destras, Gwendolyne Burfin, Hadrien Regue, Quentin Semanas, Martine Valette, Bruno Lina, Laurence Josset |
| EPI_ISL_2933634 | LABORATOIRE CERBA | CNR Virus des Infections Respiratoires - France SUD | Antonin Bal, Gregory Destras, Gwendolyne Burfin, Hadrien Regue, Quentin Semanas, Martine Valette, Bruno Lina, Laurence Josset |
| EPI_ISL_2933635 | MIRIALIS CLUSES BECHET | CNR Virus des Infections Respiratoires - France SUD | Antonin Bal, Gregory Destras, Gwendolyne Burfin, Hadrien Regue, Quentin Semanas, Martine Valette, Bruno Lina, Laurence Josset |
| EPI_ISL_2933636 | AX BIO OCEAN | CNR Virus des Infections Respiratoires - France SUD | Antonin Bal, Gregory Destras, Gwendolyne Burfin, Hadrien Regue, Quentin Semanas, Martine Valette, Bruno Lina, Laurence Josset |
| EPI_ISL_2933637 | MIRIALIS CLUSES BECHET | CNR Virus des Infections Respiratoires - France SUD | Antonin Bal, Gregory Destras, Gwendolyne Burfin, Hadrien Regue, Quentin Semanas, Martine Valette, Bruno Lina, Laurence Josset |
| EPI_ISL_2933638 | AX BIO OCEAN | CNR Virus des Infections Respiratoires - France SUD | Antonin Bal, Gregory Destras, Gwendolyne Burfin, Hadrien Regue, Quentin Semanas, Martine Valette, Bruno Lina, Laurence Josset |
| EPI_ISL_2933639, EPI_ISL_2933640, EPI_ISL_2933641, EPI_ISL_2933642, | MIRIALIS CLUSES BECHET | CNR Virus des Infections Respiratoires - France SUD | Antonin Bal, Gregory Destras, Gwendolyne Burfin, Hadrien Regue, Quentin Semanas, Martine Valette, Bruno Lina, Laurence Josset |

| | | | |
|---|---|---|---|
| EPI_ISL_2933643, EPI_ISL_2933644, EPI_ISL_2933645 | | | |
| EPI_ISL_2933646 | BIOMNIS LYON | CNR Virus des Infections Respiratoires - France SUD | Antonin Bal, Gregory Destras, Gwendolyne Burfin, Hadrien Regue, Quentin Semanas, Martine Valette, Bruno Lina, Laurence Josset |
| EPI_ISL_2933647 | LABORATOIRE CERBALLIANCE PLT VILLON | CNR Virus des Infections Respiratoires - France SUD | Antonin Bal, Gregory Destras, Gwendolyne Burfin, Hadrien Regue, Quentin Semanas, Martine Valette, Bruno Lina, Laurence Josset |
| EPI_ISL_2933648, EPI_ISL_2933649 | LABORATOIRE DE L'OSPEDALE - 2A 2B | CNR Virus des Infections Respiratoires - France SUD | Antonin Bal, Gregory Destras, Gwendolyne Burfin, Hadrien Regue, Quentin Semanas, Martine Valette, Bruno Lina, Laurence Josset |
| EPI_ISL_2933650, EPI_ISL_2933651, EPI_ISL_2933652, EPI_ISL_2933653 | LABORATOIRE BIOFUSION | CNR Virus des Infections Respiratoires - France SUD | Antonin Bal, Gregory Destras, Gwendolyne Burfin, Hadrien Regue, Quentin Semanas, Martine Valette, Bruno Lina, Laurence Josset |
| EPI_ISL_2933654, EPI_ISL_2933655 | LABORATOIRE LABAZUR | CNR Virus des Infections Respiratoires - France SUD | Antonin Bal, Gregory Destras, Gwendolyne Burfin, Hadrien Regue, Quentin Semanas, Martine Valette, Bruno Lina, Laurence Josset |
| EPI_ISL_2933656 | LABORATOIRE NOVELAB | CNR Virus des Infections Respiratoires - France SUD | Antonin Bal, Gregory Destras, Gwendolyne Burfin, Hadrien Regue, Quentin Semanas, Martine Valette, Bruno Lina, Laurence Josset |
| EPI_ISL_2933657, EPI_ISL_2933658 | CERBALLIANCE MIDI-PYRENEES | CNR Virus des Infections Respiratoires - France SUD | Antonin Bal, Gregory Destras, Gwendolyne Burfin, Hadrien Regue, Quentin Semanas, Martine Valette, Bruno Lina, Laurence Josset |
| EPI_ISL_2933659, EPI_ISL_2933660 | LABORATOIRE UNILIANS DECINES | CNR Virus des Infections Respiratoires - France SUD | Antonin Bal, Gregory Destras, Gwendolyne Burfin, Hadrien Regue, Quentin Semanas, Martine Valette, Bruno Lina, Laurence Josset |
| EPI_ISL_2933661, EPI_ISL_2933662 | LABORATOIRE CBM DE MURET | CNR Virus des Infections Respiratoires - France SUD | Antonin Bal, Gregory Destras, Gwendolyne Burfin, Hadrien Regue, Quentin Semanas, Martine Valette, Bruno Lina, Laurence Josset |
| EPI_ISL_2933663, EPI_ISL_2933664 | LABORATOIRE UNILIANS DECINES | CNR Virus des Infections Respiratoires - France SUD | Antonin Bal, Gregory Destras, Gwendolyne Burfin, Hadrien Regue, Quentin Semanas, Martine Valette, Bruno Lina, Laurence Josset |
| EPI_ISL_2933665, EPI_ISL_2933666, EPI_ISL_2933667, EPI_ISL_2933668, EPI_ISL_2933669 | LABORATOIRE BIOESTEREL | CNR Virus des Infections Respiratoires - France SUD | Antonin Bal, Gregory Destras, Gwendolyne Burfin, Hadrien Regue, Quentin Semanas, Martine Valette, Bruno Lina, Laurence Josset |
| EPI_ISL_2933670, EPI_ISL_2933671, EPI_ISL_2933672, EPI_ISL_2933673, EPI_ISL_2933674, EPI_ISL_2933675, EPI_ISL_2933676, EPI_ISL_2933677, EPI_ISL_2933678, EPI_ISL_2933679, EPI_ISL_2933680, EPI_ISL_2933681, EPI_ISL_2933682, EPI_ISL_2933683, EPI_ISL_2933684, EPI_ISL_2933685, EPI_ISL_2933686, EPI_ISL_2933687, EPI_ISL_2933688, EPI_ISL_2933689, EPI_ISL_2933690, EPI_ISL_2933691, EPI_ISL_2933692 | | | |
| see above | BIOMNIS LYON | CNR Virus des Infections Respiratoires - France SUD | Antonin Bal, Gregory Destras, Gwendolyne Burfin, Hadrien Regue, Quentin Semanas, Martine Valette, Bruno Lina, Laurence Josset |
| EPI_ISL_2933693, EPI_ISL_2933694, EPI_ISL_2933695 | AX BIO OCEAN | CNR Virus des Infections Respiratoires - France SUD | Antonin Bal, Gregory Destras, Gwendolyne Burfin, Hadrien Regue, Quentin Semanas, Martine Valette, Bruno Lina, Laurence Josset |
| EPI_ISL_2933696, EPI_ISL_2933697 | BIOMNIS LYON | CNR Virus des Infections Respiratoires - France SUD | Antonin Bal, Gregory Destras, Gwendolyne Burfin, Hadrien Regue, Quentin Semanas, Martine Valette, Bruno Lina, Laurence Josset |
| EPI_ISL_2933698 | AX BIO OCEAN | CNR Virus des Infections Respiratoires - France SUD | Antonin Bal, Gregory Destras, Gwendolyne Burfin, Hadrien Regue, Quentin Semanas, Martine Valette, Bruno Lina, Laurence Josset |
| EPI_ISL_2933699, EPI_ISL_2933700, EPI_ISL_2933701, EPI_ISL_2933702, EPI_ISL_2933703 | LABORATOIRE CBM DE MURET | CNR Virus des Infections Respiratoires - France SUD | Antonin Bal, Gregory Destras, Gwendolyne Burfin, Hadrien Regue, Quentin Semanas, Martine Valette, Bruno Lina, Laurence Josset |
| EPI_ISL_2933704, EPI_ISL_2933705 | AX BIO OCEAN | CNR Virus des Infections Respiratoires - France SUD | Antonin Bal, Gregory Destras, Gwendolyne Burfin, Hadrien Regue, Quentin Semanas, Martine Valette, Bruno Lina, Laurence Josset |
| EPI_ISL_2933706, EPI_ISL_2933707 | LABORATOIRE DES CEDRES | CNR Virus des Infections Respiratoires - France SUD | Antonin Bal, Gregory Destras, Gwendolyne Burfin, Hadrien Regue, Quentin Semanas, Martine Valette, Bruno Lina, Laurence Josset |
| EPI_ISL_2933708, EPI_ISL_2933709, EPI_ISL_2933710, EPI_ISL_2933711, EPI_ISL_2933712, EPI_ISL_2933713, EPI_ISL_2933714, EPI_ISL_2933715 | AX BIO OCEAN | CNR Virus des Infections Respiratoires - France SUD | Antonin Bal, Gregory Destras, Gwendolyne Burfin, Hadrien Regue, Quentin Semanas, Martine Valette, Bruno Lina, Laurence Josset |
| EPI_ISL_2933716 | BIOMNIS LYON | CNR Virus des Infections Respiratoires - France SUD | Antonin Bal, Gregory Destras, Gwendolyne Burfin, Hadrien Regue, Quentin Semanas, Martine Valette, Bruno Lina, Laurence Josset |
| EPI_ISL_2933717 | AX BIO OCEAN | CNR Virus des Infections Respiratoires - France SUD | Antonin Bal, Gregory Destras, Gwendolyne Burfin, Hadrien Regue, Quentin Semanas, Martine Valette, Bruno Lina, Laurence Josset |
| EPI_ISL_2933718, EPI_ISL_2933719, EPI_ISL_2933720 | BIOMNIS LYON | CNR Virus des Infections Respiratoires - France SUD | Antonin Bal, Gregory Destras, Gwendolyne Burfin, Hadrien Regue, Quentin Semanas, Martine Valette, Bruno Lina, Laurence Josset |
| EPI_ISL_2933721, EPI_ISL_2933722 | ORIAPOLE | CNR Virus des Infections Respiratoires - France SUD | Antonin Bal, Gregory Destras, Gwendolyne Burfin, Hadrien Regue, Quentin Semanas, Martine Valette, Bruno Lina, Laurence Josset |
| EPI_ISL_2933723 | BIOMNIS LYON | CNR Virus des Infections Respiratoires - France SUD | Antonin Bal, Gregory Destras, Gwendolyne Burfin, Hadrien Regue, Quentin Semanas, Martine Valette, Bruno Lina, Laurence Josset |
| EPI_ISL_2933724, EPI_ISL_2933725, EPI_ISL_2933726, EPI_ISL_2933727, EPI_ISL_2933728 | ORIAPOLE | CNR Virus des Infections Respiratoires - France SUD | Antonin Bal, Gregory Destras, Gwendolyne Burfin, Hadrien Regue, Quentin Semanas, Martine Valette, Bruno Lina, Laurence Josset |
| EPI_ISL_2933729 | BIOMNIS LYON | CNR Virus des Infections Respiratoires - France SUD | Antonin Bal, Gregory Destras, Gwendolyne Burfin, Hadrien Regue, Quentin Semanas, Martine Valette, Bruno Lina, Laurence Josset |
| EPI_ISL_2933730, EPI_ISL_2933731, EPI_ISL_2933732 | ORIAPOLE | CNR Virus des Infections Respiratoires - France SUD | Antonin Bal, Gregory Destras, Gwendolyne Burfin, Hadrien Regue, Quentin Semanas, Martine Valette, Bruno Lina, Laurence Josset |
| EPI_ISL_2933733, EPI_ISL_2933734 | BIOMNIS LYON | CNR Virus des Infections Respiratoires - France SUD | Antonin Bal, Gregory Destras, Gwendolyne Burfin, Hadrien Regue, Quentin Semanas, Martine Valette, Bruno Lina, Laurence Josset |
| EPI_ISL_2933735 | ORIAPOLE | CNR Virus des Infections Respiratoires - France SUD | Antonin Bal, Gregory Destras, Gwendolyne Burfin, Hadrien Regue, Quentin Semanas, Martine Valette, Bruno Lina, Laurence Josset |
| EPI_ISL_2933736 | BIOMNIS LYON | CNR Virus des Infections Respiratoires - France SUD | Antonin Bal, Gregory Destras, Gwendolyne Burfin, Hadrien Regue, Quentin Semanas, Martine Valette, Bruno Lina, Laurence Josset |
| EPI_ISL_2933737, EPI_ISL_2933738, EPI_ISL_2933739, EPI_ISL_2933740 | MIRIALIS CLUSES BECHET | CNR Virus des Infections Respiratoires - France SUD | Antonin Bal, Gregory Destras, Gwendolyne Burfin, Hadrien Regue, Quentin Semanas, Martine Valette, Bruno Lina, Laurence Josset |
| EPI_ISL_2933741 | BIOMNIS LYON | CNR Virus des Infections Respiratoires - France SUD | Antonin Bal, Gregory Destras, Gwendolyne Burfin, Hadrien Regue, Quentin Semanas, Martine Valette, Bruno Lina, Laurence Josset |
| EPI_ISL_2933742 | LABORATOIRE DES CEDRES | CNR Virus des Infections Respiratoires - France SUD | Antonin Bal, Gregory Destras, Gwendolyne Burfin, Hadrien Regue, Quentin Semanas, Martine Valette, Bruno Lina, Laurence Josset |
| EPI_ISL_2933743, EPI_ISL_2933744, EPI_ISL_2933745, EPI_ISL_2933746 | MIRIALIS CLUSES BECHET | CNR Virus des Infections Respiratoires - France SUD | Antonin Bal, Gregory Destras, Gwendolyne Burfin, Hadrien Regue, Quentin Semanas, Martine Valette, Bruno Lina, Laurence Josset |
| EPI_ISL_2933747 | CNR Virus des Infections Respiratoires - France SUD | CNR Virus des Infections Respiratoires - France SUD | Antonin Bal, Gregory Destras, Gwendolyne Burfin, Hadrien Regue, Quentin Semanas, Martine Valette, Bruno Lina, Laurence Josset |
| EPI_ISL_2933748 | LAM GEN-BIO GRAVANCHES | CNR Virus des Infections Respiratoires - France SUD | Antonin Bal, Gregory Destras, Gwendolyne Burfin, Hadrien Regue, Quentin Semanas, Martine Valette, Bruno Lina, Laurence Josset |
| EPI_ISL_2933749 | LABORATOIRE BIOLAB AVENIR | CNR Virus des Infections Respiratoires - France SUD | Antonin Bal, Gregory Destras, Gwendolyne Burfin, Hadrien Regue, Quentin Semanas, Martine Valette, Bruno Lina, Laurence Josset |
| EPI_ISL_2933750, EPI_ISL_2933751, EPI_ISL_2933752 | BIOMNIS LYON | CNR Virus des Infections Respiratoires - France SUD | Antonin Bal, Gregory Destras, Gwendolyne Burfin, Hadrien Regue, Quentin Semanas, Martine Valette, Bruno Lina, Laurence Josset |
| EPI_ISL_2933753 | BIOLITTORAL BIOGROUP PLATEAU TECH | CNR Virus des Infections Respiratoires - France SUD | Antonin Bal, Gregory Destras, Gwendolyne Burfin, Hadrien Regue, Quentin Semanas, Martine Valette, Bruno Lina, Laurence Josset |

| | | | |
|---|---|---|---|
| EPI_ISL_2933754 | BIOMNIS LYON | CNR Virus des Infections Respiratoires - France SUD | Antonin Bal, Gregory Destras, Gwendolyne Burfin, Hadrien Regue, Quentin Semanas, Martine Valette, Bruno Lina, Laurence Josset |
| EPI_ISL_2933755 | BIOLITTORAL BIOGROUP PLATEAU TECH | CNR Virus des Infections Respiratoires - France SUD | Antonin Bal, Gregory Destras, Gwendolyne Burfin, Hadrien Regue, Quentin Semanas, Martine Valette, Bruno Lina, Laurence Josset |
| EPI_ISL_2933756, EPI_ISL_2933757, EPI_ISL_2933758, EPI_ISL_2933759, EPI_ISL_2933760, EPI_ISL_2933761, EPI_ISL_2933762, EPI_ISL_2933763, EPI_ISL_2933764, EPI_ISL_2933765, EPI_ISL_2933766, EPI_ISL_2933767 | | | |
| see above | BIOMNIS LYON | CNR Virus des Infections Respiratoires - France SUD | Antonin Bal, Gregory Destras, Gwendolyne Burfin, Hadrien Regue, Quentin Semanas, Martine Valette, Bruno Lina, Laurence Josset |
| EPI_ISL_2933768 | LABORATOIRE BIOESTEREL | CNR Virus des Infections Respiratoires - France SUD | Antonin Bal, Gregory Destras, Gwendolyne Burfin, Hadrien Regue, Quentin Semanas, Martine Valette, Bruno Lina, Laurence Josset |
| EPI_ISL_2933769 | BIOMNIS LYON | CNR Virus des Infections Respiratoires - France SUD | Antonin Bal, Gregory Destras, Gwendolyne Burfin, Hadrien Regue, Quentin Semanas, Martine Valette, Bruno Lina, Laurence Josset |
| EPI_ISL_2933770 | LABORATOIRE BIO-VAL | CNR Virus des Infections Respiratoires - France SUD | Antonin Bal, Gregory Destras, Gwendolyne Burfin, Hadrien Regue, Quentin Semanas, Martine Valette, Bruno Lina, Laurence Josset |
| EPI_ISL_2933771 | BIOLITTORAL SANARY ROSE | CNR Virus des Infections Respiratoires - France SUD | Antonin Bal, Gregory Destras, Gwendolyne Burfin, Hadrien Regue, Quentin Semanas, Martine Valette, Bruno Lina, Laurence Josset |
| EPI_ISL_2933772 | LABORATOIRE BIO-VAL | CNR Virus des Infections Respiratoires - France SUD | Antonin Bal, Gregory Destras, Gwendolyne Burfin, Hadrien Regue, Quentin Semanas, Martine Valette, Bruno Lina, Laurence Josset |
| EPI_ISL_2933773 | LABORATOIRE d'ANALYSES de BIOLOGIE MEDICALES | CNR Virus des Infections Respiratoires - France SUD | Antonin Bal, Gregory Destras, Gwendolyne Burfin, Hadrien Regue, Quentin Semanas, Martine Valette, Bruno Lina, Laurence Josset |
| EPI_ISL_2933774 | BIOMNIS LYON | CNR Virus des Infections Respiratoires - France SUD | Antonin Bal, Gregory Destras, Gwendolyne Burfin, Hadrien Regue, Quentin Semanas, Martine Valette, Bruno Lina, Laurence Josset |
| EPI_ISL_2933775 | LABORATOIRE BIO-VAL | CNR Virus des Infections Respiratoires - France SUD | Antonin Bal, Gregory Destras, Gwendolyne Burfin, Hadrien Regue, Quentin Semanas, Martine Valette, Bruno Lina, Laurence Josset |
| EPI_ISL_2933776 | LABORATOIRE d'ANALYSES de BIOLOGIE MEDICALES | CNR Virus des Infections Respiratoires - France SUD | Antonin Bal, Gregory Destras, Gwendolyne Burfin, Hadrien Regue, Quentin Semanas, Martine Valette, Bruno Lina, Laurence Josset |
| EPI_ISL_2933777 | BIOMNIS LYON | CNR Virus des Infections Respiratoires - France SUD | Antonin Bal, Gregory Destras, Gwendolyne Burfin, Hadrien Regue, Quentin Semanas, Martine Valette, Bruno Lina, Laurence Josset |
| EPI_ISL_2933778, EPI_ISL_2933779 | LABORATOIRE d'ANALYSES de BIOLOGIE MEDICALES | CNR Virus des Infections Respiratoires - France SUD | Antonin Bal, Gregory Destras, Gwendolyne Burfin, Hadrien Regue, Quentin Semanas, Martine Valette, Bruno Lina, Laurence Josset |
| EPI_ISL_2933780, EPI_ISL_2933781 | ASTRALAB TASSIGNY | CNR Virus des Infections Respiratoires - France SUD | Antonin Bal, Gregory Destras, Gwendolyne Burfin, Hadrien Regue, Quentin Semanas, Martine Valette, Bruno Lina, Laurence Josset |
| EPI_ISL_2933782 | BIOMNIS LYON | CNR Virus des Infections Respiratoires - France SUD | Antonin Bal, Gregory Destras, Gwendolyne Burfin, Hadrien Regue, Quentin Semanas, Martine Valette, Bruno Lina, Laurence Josset |
| EPI_ISL_2933783, EPI_ISL_2933784 | ASTRALAB TASSIGNY | CNR Virus des Infections Respiratoires - France SUD | Antonin Bal, Gregory Destras, Gwendolyne Burfin, Hadrien Regue, Quentin Semanas, Martine Valette, Bruno Lina, Laurence Josset |
| EPI_ISL_2933785 | BIOMNIS LYON | CNR Virus des Infections Respiratoires - France SUD | Antonin Bal, Gregory Destras, Gwendolyne Burfin, Hadrien Regue, Quentin Semanas, Martine Valette, Bruno Lina, Laurence Josset |
| EPI_ISL_2933786, EPI_ISL_2933787 | CERBALLIANCE CHARENTES SAINTES | CNR Virus des Infections Respiratoires - France SUD | Antonin Bal, Gregory Destras, Gwendolyne Burfin, Hadrien Regue, Quentin Semanas, Martine Valette, Bruno Lina, Laurence Josset |
| EPI_ISL_2933788, EPI_ISL_2933789, EPI_ISL_2933790 | LAM BIOPOLE ORTHEZ | CNR Virus des Infections Respiratoires - France SUD | Antonin Bal, Gregory Destras, Gwendolyne Burfin, Hadrien Regue, Quentin Semanas, Martine Valette, Bruno Lina, Laurence Josset |
| EPI_ISL_2933791 | LABORATOIRE CERBA VAL DE CAUSSELS | CNR Virus des Infections Respiratoires - France SUD | Antonin Bal, Gregory Destras, Gwendolyne Burfin, Hadrien Regue, Quentin Semanas, Martine Valette, Bruno Lina, Laurence Josset |
| EPI_ISL_2933792 | BIOMNIS LYON | CNR Virus des Infections Respiratoires - France SUD | Antonin Bal, Gregory Destras, Gwendolyne Burfin, Hadrien Regue, Quentin Semanas, Martine Valette, Bruno Lina, Laurence Josset |
| EPI_ISL_2933793, EPI_ISL_2933794 | DYOMEDEA-LABORATOIRE DE LA SAUVEGARDE | CNR Virus des Infections Respiratoires - France SUD | Antonin Bal, Gregory Destras, Gwendolyne Burfin, Hadrien Regue, Quentin Semanas, Martine Valette, Bruno Lina, Laurence Josset |
| EPI_ISL_2933795 | BIOMNIS LYON | CNR Virus des Infections Respiratoires - France SUD | Antonin Bal, Gregory Destras, Gwendolyne Burfin, Hadrien Regue, Quentin Semanas, Martine Valette, Bruno Lina, Laurence Josset |
| EPI_ISL_2933796, EPI_ISL_2933797, EPI_ISL_2933798 | DYOMEDEA-LABORATOIRE DE LA SAUVEGARDE | CNR Virus des Infections Respiratoires - France SUD | Antonin Bal, Gregory Destras, Gwendolyne Burfin, Hadrien Regue, Quentin Semanas, Martine Valette, Bruno Lina, Laurence Josset |
| EPI_ISL_2933799, EPI_ISL_2933800, EPI_ISL_2933801, EPI_ISL_2933802, EPI_ISL_2933803, EPI_ISL_2933804 | BIOMNIS LYON | CNR Virus des Infections Respiratoires - France SUD | Antonin Bal, Gregory Destras, Gwendolyne Burfin, Hadrien Regue, Quentin Semanas, Martine Valette, Bruno Lina, Laurence Josset |
| EPI_ISL_2933805, EPI_ISL_2933806, EPI_ISL_2933807, EPI_ISL_2933808 | LABORATOIRE CERBALLIANCE PLT VILLON | CNR Virus des Infections Respiratoires - France SUD | Antonin Bal, Gregory Destras, Gwendolyne Burfin, Hadrien Regue, Quentin Semanas, Martine Valette, Bruno Lina, Laurence Josset |
| EPI_ISL_2933809 | BIOMNIS LYON | CNR Virus des Infections Respiratoires - France SUD | Antonin Bal, Gregory Destras, Gwendolyne Burfin, Hadrien Regue, Quentin Semanas, Martine Valette, Bruno Lina, Laurence Josset |
| EPI_ISL_2933810 | CNR Virus des Infections Respiratoires - France SUD | CNR Virus des Infections Respiratoires - France SUD | Antonin Bal, Gregory Destras, Gwendolyne Burfin, Hadrien Regue, Quentin Semanas, Martine Valette, Bruno Lina, Laurence Josset |
| EPI_ISL_2933814 | SARS-CoV-2 testing team, National Institute of Infectious Diseases | Pathogen Genomics Center, National Institute of Infectious Diseases | Tsuyoshi Sekizuka, Kentaro Itokawa, Rina Tanaka, Masanori Hashino, Nozomu Hanaoka, Masumichi Saito, Naomi Nojiri, Hazuka Y Furihata, Hiromizu Takahashi, Tsuguto Fujimoto, Makoto Kuroda |
| EPI_ISL_2933888, EPI_ISL_2933892, EPI_ISL_2933894, EPI_ISL_2933895, EPI_ISL_2933899, EPI_ISL_2933909, EPI_ISL_2933912, EPI_ISL_2933924 | National Virus Reference Laboratory | National Virus Reference Laboratory | Guerrino Macori, Gabriel Gonzalez, Michael Carr, Zoe Yandle, Charlene Bennett, Jonathan Dean, Seamus Fanning, Cillian F De Gascun |
| EPI_ISL_2933963, EPI_ISL_2933969, EPI_ISL_2933970, EPI_ISL_2933979 | Royal GD | Royal GD | Remco Dijkman |
| EPI_ISL_2933987 | Santa Clara Valley Medical Center | Santa Clara County Public Health Laboratory | Santa Clara County Public Health Department |
| EPI_ISL_2933988 | Kaiser Regional | Santa Clara County Public Health Laboratory | Santa Clara County Public Health Department |
| EPI_ISL_2933989 | Santa Clara Valley Medical Center | Santa Clara County Public Health Laboratory | Santa Clara County Public Health Department |
| EPI_ISL_2933992 | Kaiser Regional | Santa Clara County Public Health Laboratory | Santa Clara County Public Health Department |
| EPI_ISL_2933997, EPI_ISL_2933998 | Santa Clara Valley Medical Center | Santa Clara County Public Health Laboratory | Santa Clara County Public Health Department |
| EPI_ISL_2933999 | Kaiser Regional | Santa Clara County Public Health Laboratory | Santa Clara County Public Health Department |
| EPI_ISL_2934000 | Santa Clara Valley Medical Center | Santa Clara County Public Health Laboratory | Santa Clara County Public Health Department |
| EPI_ISL_2934001 | Good Samaritan Hospital | Santa Clara County Public Health Laboratory | Santa Clara County Public Health Department |
| EPI_ISL_2934003 | Kaiser Regional | Santa Clara County Public Health Laboratory | Santa Clara County Public Health Department |
| EPI_ISL_2934004 | Kaiser San Jose | Santa Clara County Public Health Laboratory | Santa Clara County Public Health Department |
| EPI_ISL_2934005, EPI_ISL_2934011, EPI_ISL_2934012 | Kaiser Regional | Santa Clara County Public Health Laboratory | Santa Clara County Public Health Department |
| EPI_ISL_2934035, EPI_ISL_2934036 | Santa Clara Valley Medical Center | Santa Clara County Public Health Laboratory | Santa Clara County Public Health Department |
| EPI_ISL_2934038 | PAMF Palo Alto | Santa Clara County Public Health Laboratory | Santa Clara County Public Health Department |
| EPI_ISL_2934041 | Santa Clara Valley Medical Center | Santa Clara County Public Health Laboratory | Santa Clara County Public Health Department |
| EPI_ISL_2934044 | Kaiser Regional | Santa Clara County Public Health Laboratory | Santa Clara County Public Health Department |
| EPI_ISL_2934046, EPI_ISL_2934047 | Santa Clara Valley Medical Center | Santa Clara County Public Health Laboratory | Santa Clara County Public Health Department |
| EPI_ISL_2934048 | PAMF Palo Alto | Santa Clara County Public Health Laboratory | Santa Clara County Public Health Department |
| EPI_ISL_2934051 | Santa Clara Valley Medical Center | Santa Clara County Public Health Laboratory | Santa Clara County Public Health Department |
| EPI_ISL_2934052 | Regional Medical Center | Santa Clara County Public Health Laboratory | Santa Clara County Public Health Department |
| EPI_ISL_2934053 | Good Samaritan Hospital | Santa Clara County Public Health Laboratory | Santa Clara County Public Health Department |
| EPI_ISL_2934056 | Santa Clara Valley Medical Center | Santa Clara County Public Health Laboratory | Santa Clara County Public Health Department |

| | | | |
|---|---|---|---|
| EPI_ISL_2934058 | El Camino Hospital | Santa Clara County Public Health Laboratory | Santa Clara County Public Health Department |
| EPI_ISL_2934059 | Santa Clara Valley Medical Center | Santa Clara County Public Health Laboratory | Santa Clara County Public Health Department |
| EPI_ISL_2934061 | Kaiser San Jose | Santa Clara County Public Health Laboratory | Santa Clara County Public Health Department |
| EPI_ISL_2934063, EPI_ISL_2934064, EPI_ISL_2934072 | Santa Clara Valley Medical Center | Santa Clara County Public Health Laboratory | Santa Clara County Public Health Department |
| EPI_ISL_2934075, EPI_ISL_2934076, EPI_ISL_2934078, EPI_ISL_2934080 | Kaiser Regional | Santa Clara County Public Health Laboratory | Santa Clara County Public Health Department |
| EPI_ISL_2934081, EPI_ISL_2934082, EPI_ISL_2934083 | Santa Clara Valley Medical Center | Santa Clara County Public Health Laboratory | Santa Clara County Public Health Department |
| EPI_ISL_2934569, EPI_ISL_2934574 | SECP MHU | The Institute of Molecular Biology and Genetics of NASU | M.Tukalo et al. |
| EPI_ISL_2934591, EPI_ISL_2934592, EPI_ISL_2934593, EPI_ISL_2934595, EPI_ISL_2934604, EPI_ISL_2934605, EPI_ISL_2934606, EPI_ISL_2934607, EPI_ISL_2934608, EPI_ISL_2934609, EPI_ISL_2934610, EPI_ISL_2934611, EPI_ISL_2934612, EPI_ISL_2934613, EPI_ISL_2934614, EPI_ISL_2934615, EPI_ISL_2934616, EPI_ISL_2934636, EPI_ISL_2934637, EPI_ISL_2934641, EPI_ISL_2934643, EPI_ISL_2934644, EPI_ISL_2934691, EPI_ISL_2934692, EPI_ISL_2934693, EPI_ISL_2934694, EPI_ISL_2934695, EPI_ISL_2934696, EPI_ISL_2934697, EPI_ISL_2934698, EPI_ISL_2934701, EPI_ISL_2934702, EPI_ISL_2934738, EPI_ISL_2934739, EPI_ISL_2934740, EPI_ISL_2934741, EPI_ISL_2934742, EPI_ISL_2934743, EPI_ISL_2934744, EPI_ISL_2934745, EPI_ISL_2934746, EPI_ISL_2934747, EPI_ISL_2934838, EPI_ISL_2934839, EPI_ISL_2934840, EPI_ISL_2934841, EPI_ISL_2934842, EPI_ISL_2934843, EPI_ISL_2934856, EPI_ISL_2934857, EPI_ISL_2934858, EPI_ISL_2934859, EPI_ISL_2934860, EPI_ISL_2934861, EPI_ISL_2934862, EPI_ISL_2934864, EPI_ISL_2934865, EPI_ISL_2934866, EPI_ISL_2934867, EPI_ISL_2934868, EPI_ISL_2934869, EPI_ISL_2934871, EPI_ISL_2934872, EPI_ISL_2934873, EPI_ISL_2934874, EPI_ISL_2934875, EPI_ISL_2934876, EPI_ISL_2934877, EPI_ISL_2934878, EPI_ISL_2934879, EPI_ISL_2934880, EPI_ISL_2934881, EPI_ISL_2934882, EPI_ISL_2934883, EPI_ISL_2934885, EPI_ISL_2934886, EPI_ISL_2934887, EPI_ISL_2934888, EPI_ISL_2934889, EPI_ISL_2934894, EPI_ISL_2934895, EPI_ISL_2934900, EPI_ISL_2934901, EPI_ISL_2934902, EPI_ISL_2934903, EPI_ISL_2934904, EPI_ISL_2934907, EPI_ISL_2934909, EPI_ISL_2934911, EPI_ISL_2934912, EPI_ISL_2934914, EPI_ISL_2934915, EPI_ISL_2934916, EPI_ISL_2934920, EPI_ISL_2934921, EPI_ISL_2934923 | | | |
| see above | Servicio de Microbiología Clínica (Complejo Hospitalario de Navarra, Pamplona) | Centro de Secuenciación NASERTIC | Carmen Ezpeleta Baquedano, Ana Navascués, Ana Miqueleiz |
| EPI_ISL_2934990 | Lighthouse Lab in Milton Keynes | Wellcome Sanger Institute for the COVID-19 Genomics UK (COG-UK) Consortium | The Lighthouse Lab in Milton Keynes and Alex Alderton, Roberto Amato, Jeffrey Barrett, Sonia Goncalves, Ewan Harrison, David K. Jackson, Ian Johnston, Dominic Kwiatkowski, Cordelia Langford, John Sillitoe on behalf of the Wellcome Sanger Institute COVID-19 Surveillance Team |
| EPI_ISL_2934991 | Health Services Laboratories | Wellcome Sanger Institute for the COVID-19 Genomics UK (COG-UK) Consortium | Health Services Laboratories and Alex Alderton, Roberto Amato, Jeffrey Barrett, Sonia Goncalves, Ewan Harrison, David K. Jackson, Ian Johnston, Dominic Kwiatkowski, Cordelia Langford, John Sillitoe on behalf of the Wellcome Sanger Institute COVID-19 Surveillance Team |
| EPI_ISL_2934992 | Lighthouse Laboratory Plymouth | Wellcome Sanger Institute for the COVID-19 Genomics UK (COG-UK) Consortium | Lighthouse Laboratory Plymouth and Alex Alderton, Roberto Amato, Jeffrey Barrett, Sonia Goncalves, Ewan Harrison, David K. Jackson, Ian Johnston, Dominic Kwiatkowski, Cordelia Langford, John Sillitoe on behalf of the Wellcome Sanger Institute COVID-19 Surveillance Team |
| EPI_ISL_2934993, EPI_ISL_2934994 | Lighthouse Lab in Milton Keynes | Wellcome Sanger Institute for the COVID-19 Genomics UK (COG-UK) Consortium | The Lighthouse Lab in Milton Keynes and Alex Alderton, Roberto Amato, Jeffrey Barrett, Sonia Goncalves, Ewan Harrison, David K. Jackson, Ian Johnston, Dominic Kwiatkowski, Cordelia Langford, John Sillitoe on behalf of the Wellcome Sanger Institute COVID-19 Surveillance Team |
| EPI_ISL_2934995, EPI_ISL_2934996 | Lighthouse Lab in Glasgow | Wellcome Sanger Institute for the COVID-19 Genomics UK (COG-UK) Consortium | Harper VanSteenhouse, Yumi Kasai, David Gray, Carol Clugston, Anna Dominiczak and Alex Alderton, Roberto Amato, Jeffrey Barrett, Sonia Goncalves, Ewan Harrison, David K. Jackson, Ian Johnston, Dominic Kwiatkowski, Cordelia Langford, John Sillitoe on behalf of the Wellcome Sanger Institute COVID-19 Surveillance Team |
| EPI_ISL_2934997 | Lighthouse Lab in Milton Keynes | Wellcome Sanger Institute for the COVID-19 Genomics UK (COG-UK) Consortium | The Lighthouse Lab in Milton Keynes and Alex Alderton, Roberto Amato, Jeffrey Barrett, Sonia Goncalves, Ewan Harrison, David K. Jackson, Ian Johnston, Dominic Kwiatkowski, Cordelia Langford, John Sillitoe on behalf of the Wellcome Sanger Institute COVID-19 Surveillance Team |
| EPI_ISL_2934998 | Berkshire and Surrey Pathology Services Lighthouse Laboratory | Wellcome Sanger Institute for the COVID-19 Genomics UK (COG-UK) Consortium | Berkshire and Surrey Pathology Services Lighthouse Laboratory and Alex Alderton, Roberto Amato, Jeffrey Barrett, Sonia Goncalves, Ewan Harrison, David K. Jackson, Ian Johnston, Dominic Kwiatkowski, Cordelia Langford, John Sillitoe on behalf of the Wellcome Sanger Institute COVID-19 Surveillance Team |
| EPI_ISL_2934999, EPI_ISL_2935000, EPI_ISL_2935001 | Lighthouse Lab in Glasgow | Wellcome Sanger Institute for the COVID-19 Genomics UK (COG-UK) Consortium | Harper VanSteenhouse, Yumi Kasai, David Gray, Carol Clugston, Anna Dominiczak and Alex Alderton, Roberto Amato, Jeffrey Barrett, Sonia Goncalves, Ewan Harrison, David K. Jackson, Ian Johnston, Dominic Kwiatkowski, Cordelia Langford, John Sillitoe on behalf of the Wellcome Sanger Institute COVID-19 Surveillance Team |
| EPI_ISL_2935002 | Lighthouse Laboratory Plymouth | Wellcome Sanger Institute for the COVID-19 Genomics UK (COG-UK) Consortium | Lighthouse Laboratory Plymouth and Alex Alderton, Roberto Amato, Jeffrey Barrett, Sonia Goncalves, Ewan Harrison, David K. Jackson, Ian Johnston, Dominic Kwiatkowski, Cordelia Langford, John Sillitoe on behalf of the Wellcome Sanger Institute COVID-19 Surveillance Team |
| EPI_ISL_2935003 | Lighthouse Lab in Milton Keynes | Wellcome Sanger Institute for the COVID-19 Genomics UK (COG-UK) Consortium | The Lighthouse Lab in Milton Keynes and Alex Alderton, Roberto Amato, Jeffrey Barrett, Sonia Goncalves, Ewan Harrison, David K. Jackson, Ian Johnston, Dominic Kwiatkowski, Cordelia Langford, John Sillitoe on behalf of the Wellcome Sanger Institute COVID-19 Surveillance Team |
| EPI_ISL_2935004 | Berkshire and Surrey Pathology Services Lighthouse Laboratory | Wellcome Sanger Institute for the COVID-19 Genomics UK (COG-UK) Consortium | Berkshire and Surrey Pathology Services Lighthouse Laboratory and Alex Alderton, Roberto Amato, Jeffrey Barrett, Sonia Goncalves, Ewan Harrison, David K. Jackson, Ian Johnston, Dominic Kwiatkowski, Cordelia Langford, John Sillitoe on behalf of the Wellcome Sanger Institute COVID-19 Surveillance Team |
| EPI_ISL_2935005, EPI_ISL_2935006 | Lighthouse Lab in Milton Keynes | Wellcome Sanger Institute for the COVID-19 Genomics UK (COG-UK) Consortium | The Lighthouse Lab in Milton Keynes and Alex Alderton, Roberto Amato, Jeffrey Barrett, Sonia Goncalves, Ewan Harrison, David K. Jackson, Ian Johnston, Dominic Kwiatkowski, Cordelia Langford, John Sillitoe on behalf of the Wellcome Sanger Institute COVID-19 Surveillance Team |
| EPI_ISL_2935007 | Lighthouse Lab in Glasgow | Wellcome Sanger Institute for the COVID-19 Genomics UK (COG-UK) Consortium | Harper VanSteenhouse, Yumi Kasai, David Gray, Carol Clugston, Anna Dominiczak and Alex Alderton, Roberto Amato, Jeffrey Barrett, Sonia Goncalves, Ewan Harrison, David K. Jackson, Ian Johnston, Dominic Kwiatkowski, Cordelia Langford, John Sillitoe on behalf of the Wellcome Sanger Institute COVID-19 Surveillance Team |
| EPI_ISL_2935008 | Berkshire and Surrey Pathology Services Lighthouse Laboratory | Wellcome Sanger Institute for the COVID-19 Genomics UK (COG-UK) Consortium | Berkshire and Surrey Pathology Services Lighthouse Laboratory and Alex Alderton, Roberto Amato, Jeffrey Barrett, Sonia Goncalves, Ewan Harrison, David K. Jackson, Ian Johnston, Dominic Kwiatkowski, Cordelia Langford, John Sillitoe on behalf of the Wellcome Sanger Institute COVID-19 Surveillance Team |
| EPI_ISL_2935009 | Lighthouse Lab in Milton Keynes | Wellcome Sanger Institute for the COVID-19 Genomics UK (COG-UK) Consortium | The Lighthouse Lab in Milton Keynes and Alex Alderton, Roberto Amato, Jeffrey Barrett, Sonia Goncalves, Ewan Harrison, David K. Jackson, Ian Johnston, Dominic Kwiatkowski, Cordelia Langford, John Sillitoe on behalf of the Wellcome Sanger Institute COVID-19 Surveillance Team |
| EPI_ISL_2935010 | Berkshire and Surrey Pathology Services Lighthouse Laboratory | Wellcome Sanger Institute for the COVID-19 Genomics UK (COG-UK) Consortium | Berkshire and Surrey Pathology Services Lighthouse Laboratory and Alex Alderton, Roberto Amato, Jeffrey Barrett, Sonia Goncalves, Ewan Harrison, David K. Jackson, Ian Johnston, Dominic Kwiatkowski, Cordelia Langford, John Sillitoe on behalf of the Wellcome Sanger Institute COVID-19 Surveillance Team |
| EPI_ISL_2935011, EPI_ISL_2935012, EPI_ISL_2935013 | Lighthouse Laboratory Plymouth | Wellcome Sanger Institute for the COVID-19 Genomics UK (COG-UK) Consortium | Lighthouse Laboratory Plymouth and Alex Alderton, Roberto Amato, Jeffrey Barrett, Sonia Goncalves, Ewan Harrison, David K. Jackson, Ian Johnston, Dominic Kwiatkowski, Cordelia Langford, John Sillitoe on behalf of the Wellcome Sanger Institute COVID-19 Surveillance Team |
| EPI_ISL_2935014, EPI_ISL_2935015 | Berkshire and Surrey Pathology Services Lighthouse Laboratory | Wellcome Sanger Institute for the COVID-19 Genomics UK (COG-UK) Consortium | Berkshire and Surrey Pathology Services Lighthouse Laboratory and Alex Alderton, Roberto Amato, Jeffrey Barrett, Sonia Goncalves, Ewan Harrison, David K. Jackson, Ian Johnston, Dominic Kwiatkowski, Cordelia Langford, John Sillitoe on behalf of the Wellcome Sanger Institute COVID-19 Surveillance Team |
| EPI_ISL_2935016, EPI_ISL_2935017 | Lighthouse Lab in Glasgow | Wellcome Sanger Institute for the COVID-19 Genomics UK (COG-UK) Consortium | Harper VanSteenhouse, Yumi Kasai, David Gray, Carol Clugston, Anna Dominiczak and Alex Alderton, Roberto Amato, Jeffrey Barrett, Sonia Goncalves, Ewan Harrison, David K. Jackson, Ian Johnston, Dominic Kwiatkowski, Cordelia Langford, John Sillitoe on behalf of the Wellcome Sanger Institute COVID-19 Surveillance Team |
| EPI_ISL_2935018, EPI_ISL_2935019, EPI_ISL_2935020 | Lighthouse Lab in Milton Keynes | Wellcome Sanger Institute for the COVID-19 Genomics UK (COG-UK) Consortium | The Lighthouse Lab in Milton Keynes and Alex Alderton, Roberto Amato, Jeffrey Barrett, Sonia Goncalves, Ewan Harrison, David K. Jackson, Ian Johnston, Dominic Kwiatkowski, Cordelia Langford, John Sillitoe on behalf of the Wellcome Sanger Institute COVID-19 Surveillance Team |
| EPI_ISL_2935021 | Lighthouse Laboratory Plymouth | Wellcome Sanger Institute for the COVID-19 Genomics UK (COG-UK) Consortium | Lighthouse Laboratory Plymouth and Alex Alderton, Roberto Amato, Jeffrey Barrett, Sonia Goncalves, Ewan Harrison, David K. Jackson, Ian Johnston, Dominic Kwiatkowski, Cordelia Langford, John Sillitoe on behalf of the Wellcome Sanger Institute COVID-19 Surveillance Team |
| EPI_ISL_2935022, EPI_ISL_2935023 | Berkshire and Surrey Pathology Services Lighthouse Laboratory | Wellcome Sanger Institute for the COVID-19 Genomics UK (COG-UK) Consortium | Berkshire and Surrey Pathology Services Lighthouse Laboratory and Alex Alderton, Roberto Amato, Jeffrey Barrett, Sonia Goncalves, Ewan Harrison, David K. Jackson, Ian Johnston, Dominic Kwiatkowski, Cordelia Langford, John Sillitoe on behalf of the Wellcome Sanger Institute COVID-19 Surveillance Team |
| EPI_ISL_2935024 | Lighthouse Laboratory Plymouth | Wellcome Sanger Institute for the COVID-19 Genomics UK (COG-UK) Consortium | Lighthouse Laboratory Plymouth and Alex Alderton, Roberto Amato, Jeffrey Barrett, Sonia Goncalves, Ewan Harrison, David K. Jackson, Ian Johnston, Dominic Kwiatkowski, Cordelia Langford, John Sillitoe on behalf of the Wellcome Sanger Institute COVID-19 Surveillance Team |
| EPI_ISL_2935025, EPI_ISL_2935026 | Lighthouse Lab in Milton Keynes | Wellcome Sanger Institute for the COVID-19 Genomics UK (COG-UK) Consortium | The Lighthouse Lab in Milton Keynes and Alex Alderton, Roberto Amato, Jeffrey Barrett, Sonia Goncalves, Ewan Harrison, David K. Jackson, Ian Johnston, Dominic Kwiatkowski, Cordelia Langford, John Sillitoe on behalf of the Wellcome Sanger Institute COVID-19 Surveillance Team |
| EPI_ISL_2935027 | Lighthouse Laboratory Plymouth | Wellcome Sanger Institute for the COVID-19 Genomics UK (COG-UK) Consortium | Lighthouse Laboratory Plymouth and Alex Alderton, Roberto Amato, Jeffrey Barrett, Sonia Goncalves, Ewan Harrison, David K. Jackson, Ian Johnston, Dominic Kwiatkowski, Cordelia Langford, John Sillitoe on behalf of the Wellcome Sanger Institute COVID-19 Surveillance Team |
| EPI_ISL_2935028 | Lighthouse Lab in Milton Keynes | Wellcome Sanger Institute for the COVID-19 Genomics UK (COG-UK) Consortium | The Lighthouse Lab in Milton Keynes and Alex Alderton, Roberto Amato, Jeffrey Barrett, Sonia Goncalves, Ewan Harrison, David K. Jackson, Ian Johnston, Dominic Kwiatkowski, Cordelia Langford, John Sillitoe on behalf of the Wellcome Sanger Institute COVID-19 Surveillance Team |
| EPI_ISL_2935029 | Berkshire and Surrey Pathology Services Lighthouse Laboratory | Wellcome Sanger Institute for the COVID-19 Genomics UK (COG-UK) Consortium | Berkshire and Surrey Pathology Services Lighthouse Laboratory and Alex Alderton, Roberto Amato, Jeffrey Barrett, Sonia Goncalves, Ewan Harrison, David K. Jackson, Ian Johnston, Dominic Kwiatkowski, Cordelia Langford, John Sillitoe on behalf of the Wellcome Sanger Institute COVID-19 Surveillance Team |
| EPI_ISL_2935030 | Lighthouse Lab in Milton Keynes | Wellcome Sanger Institute for the COVID-19 Genomics UK (COG-UK) Consortium | The Lighthouse Lab in Milton Keynes and Alex Alderton, Roberto Amato, Jeffrey Barrett, Sonia Goncalves, Ewan Harrison, David K. Jackson, Ian Johnston, Dominic Kwiatkowski, Cordelia Langford, John Sillitoe on behalf of the Wellcome Sanger Institute COVID-19 Surveillance Team |
| EPI_ISL_2935031 | Berkshire and Surrey Pathology Services Lighthouse Laboratory | Wellcome Sanger Institute for the COVID-19 Genomics UK (COG-UK) Consortium | Berkshire and Surrey Pathology Services Lighthouse Laboratory and Alex Alderton, Roberto Amato, Jeffrey Barrett, Sonia Goncalves, Ewan Harrison, David K. Jackson, Ian Johnston, Dominic Kwiatkowski, Cordelia Langford, John Sillitoe on behalf of the Wellcome Sanger Institute COVID-19 Surveillance Team |
| EPI_ISL_2935032 | Lighthouse Lab in Glasgow | Wellcome Sanger Institute for the COVID-19 Genomics UK (COG-UK) Consortium | Harper VanSteenhouse, Yumi Kasai, David Gray, Carol Clugston, Anna Dominiczak and Alex Alderton, Roberto Amato, Jeffrey Barrett, Sonia Goncalves, Ewan Harrison, David K. Jackson, Ian Johnston, Dominic Kwiatkowski, Cordelia Langford, John Sillitoe on behalf of the Wellcome Sanger Institute COVID-19 Surveillance Team |
| EPI_ISL_2935033, EPI_ISL_2935034 | Lighthouse Laboratory Plymouth | Wellcome Sanger Institute for the COVID-19 Genomics UK (COG-UK) Consortium | Lighthouse Laboratory Plymouth and Alex Alderton, Roberto Amato, Jeffrey Barrett, Sonia Goncalves, Ewan Harrison, David K. Jackson, Ian Johnston, Dominic Kwiatkowski, Cordelia Langford, John Sillitoe on behalf of the Wellcome Sanger Institute COVID-19 Surveillance Team |
| EPI_ISL_2935035, EPI_ISL_2935036, EPI_ISL_2935037 | Berkshire and Surrey Pathology Services Lighthouse Laboratory | Wellcome Sanger Institute for the COVID-19 Genomics UK (COG-UK) Consortium | Berkshire and Surrey Pathology Services Lighthouse Laboratory and Alex Alderton, Roberto Amato, Jeffrey Barrett, Sonia Goncalves, Ewan Harrison, David K. Jackson, Ian Johnston, Dominic Kwiatkowski, Cordelia Langford, John Sillitoe on behalf of the Wellcome Sanger Institute COVID-19 Surveillance Team |
| EPI_ISL_2935038 | Lighthouse Laboratory Plymouth | Wellcome Sanger Institute for the COVID-19 Genomics UK (COG-UK) Consortium | Lighthouse Laboratory Plymouth and Alex Alderton, Roberto Amato, Jeffrey Barrett, Sonia Goncalves, Ewan Harrison, David K. Jackson, Ian Johnston, Dominic Kwiatkowski, Cordelia Langford, John Sillitoe on behalf of the Wellcome Sanger Institute COVID-19 Surveillance Team |

| | | | |
|---|---|---|---|
| EPI_ISL_2935039 | Berkshire and Surrey Pathology Services Lighthouse Laboratory | Wellcome Sanger Institute for the COVID-19 Genomics UK (COG-UK) Consortium | Berkshire and Surrey Pathology Services Lighthouse Laboratory and Alex Alderton, Roberto Amato, Jeffrey Barrett, Sonia Goncalves, Ewan Harrison, David K. Jackson, Ian Johnston, Dominic Kwiatkowski, Cordelia Langford, John Sillitoe on behalf of the Wellcome Sanger Institute COVID-19 Surveillance Team |
| EPI_ISL_2935040 | Lighthouse Lab in Glasgow | Wellcome Sanger Institute for the COVID-19 Genomics UK (COG-UK) Consortium | Harper VanSteenhouse, Yumi Kasai, David Gray, Carol Clugston, Anna Dominiczak and Alex Alderton, Roberto Amato, Jeffrey Barrett, Sonia Goncalves, Ewan Harrison, David K. Jackson, Ian Johnston, Dominic Kwiatkowski, Cordelia Langford, John Sillitoe on behalf of the Wellcome Sanger Institute COVID-19 Surveillance Team |
| EPI_ISL_2935041 | Berkshire and Surrey Pathology Services Lighthouse Laboratory | Wellcome Sanger Institute for the COVID-19 Genomics UK (COG-UK) Consortium | Berkshire and Surrey Pathology Services Lighthouse Laboratory and Alex Alderton, Roberto Amato, Jeffrey Barrett, Sonia Goncalves, Ewan Harrison, David K. Jackson, Ian Johnston, Dominic Kwiatkowski, Cordelia Langford, John Sillitoe on behalf of the Wellcome Sanger Institute COVID-19 Surveillance Team |
| EPI_ISL_2935042 | Lighthouse Lab in Glasgow | Wellcome Sanger Institute for the COVID-19 Genomics UK (COG-UK) Consortium | Harper VanSteenhouse, Yumi Kasai, David Gray, Carol Clugston, Anna Dominiczak and Alex Alderton, Roberto Amato, Jeffrey Barrett, Sonia Goncalves, Ewan Harrison, David K. Jackson, Ian Johnston, Dominic Kwiatkowski, Cordelia Langford, John Sillitoe on behalf of the Wellcome Sanger Institute COVID-19 Surveillance Team |
| EPI_ISL_2935043, EPI_ISL_2935044, EPI_ISL_2935045 | Lighthouse Laboratory Plymouth | Wellcome Sanger Institute for the COVID-19 Genomics UK (COG-UK) Consortium | Lighthouse Laboratory Plymouth and Alex Alderton, Roberto Amato, Jeffrey Barrett, Sonia Goncalves, Ewan Harrison, David K. Jackson, Ian Johnston, Dominic Kwiatkowski, Cordelia Langford, John Sillitoe on behalf of the Wellcome Sanger Institute COVID-19 Surveillance Team |
| EPI_ISL_2935046 | Lighthouse Lab in Glasgow | Wellcome Sanger Institute for the COVID-19 Genomics UK (COG-UK) Consortium | Harper VanSteenhouse, Yumi Kasai, David Gray, Carol Clugston, Anna Dominiczak and Alex Alderton, Roberto Amato, Jeffrey Barrett, Sonia Goncalves, Ewan Harrison, David K. Jackson, Ian Johnston, Dominic Kwiatkowski, Cordelia Langford, John Sillitoe on behalf of the Wellcome Sanger Institute COVID-19 Surveillance Team |
| EPI_ISL_2935047 | Berkshire and Surrey Pathology Services Lighthouse Laboratory | Wellcome Sanger Institute for the COVID-19 Genomics UK (COG-UK) Consortium | Berkshire and Surrey Pathology Services Lighthouse Laboratory and Alex Alderton, Roberto Amato, Jeffrey Barrett, Sonia Goncalves, Ewan Harrison, David K. Jackson, Ian Johnston, Dominic Kwiatkowski, Cordelia Langford, John Sillitoe on behalf of the Wellcome Sanger Institute COVID-19 Surveillance Team |
| EPI_ISL_2935048 | Lighthouse Lab in Milton Keynes | Wellcome Sanger Institute for the COVID-19 Genomics UK (COG-UK) Consortium | The Lighthouse Lab in Milton Keynes and Alex Alderton, Roberto Amato, Jeffrey Barrett, Sonia Goncalves, Ewan Harrison, David K. Jackson, Ian Johnston, Dominic Kwiatkowski, Cordelia Langford, John Sillitoe on behalf of the Wellcome Sanger Institute COVID-19 Surveillance Team |
| EPI_ISL_2935049, EPI_ISL_2935050 | Berkshire and Surrey Pathology Services Lighthouse Laboratory | Wellcome Sanger Institute for the COVID-19 Genomics UK (COG-UK) Consortium | Berkshire and Surrey Pathology Services Lighthouse Laboratory and Alex Alderton, Roberto Amato, Jeffrey Barrett, Sonia Goncalves, Ewan Harrison, David K. Jackson, Ian Johnston, Dominic Kwiatkowski, Cordelia Langford, John Sillitoe on behalf of the Wellcome Sanger Institute COVID-19 Surveillance Team |
| EPI_ISL_2935051, EPI_ISL_2935052 | Lighthouse Lab in Milton Keynes | Wellcome Sanger Institute for the COVID-19 Genomics UK (COG-UK) Consortium | The Lighthouse Lab in Milton Keynes and Alex Alderton, Roberto Amato, Jeffrey Barrett, Sonia Goncalves, Ewan Harrison, David K. Jackson, Ian Johnston, Dominic Kwiatkowski, Cordelia Langford, John Sillitoe on behalf of the Wellcome Sanger Institute COVID-19 Surveillance Team |
| EPI_ISL_2935053 | Lighthouse Laboratory Plymouth | Wellcome Sanger Institute for the COVID-19 Genomics UK (COG-UK) Consortium | Lighthouse Laboratory Plymouth and Alex Alderton, Roberto Amato, Jeffrey Barrett, Sonia Goncalves, Ewan Harrison, David K. Jackson, Ian Johnston, Dominic Kwiatkowski, Cordelia Langford, John Sillitoe on behalf of the Wellcome Sanger Institute COVID-19 Surveillance Team |
| EPI_ISL_2935054 | Lighthouse Lab in Glasgow | Wellcome Sanger Institute for the COVID-19 Genomics UK (COG-UK) Consortium | Harper VanSteenhouse, Yumi Kasai, David Gray, Carol Clugston, Anna Dominiczak and Alex Alderton, Roberto Amato, Jeffrey Barrett, Sonia Goncalves, Ewan Harrison, David K. Jackson, Ian Johnston, Dominic Kwiatkowski, Cordelia Langford, John Sillitoe on behalf of the Wellcome Sanger Institute COVID-19 Surveillance Team |
| EPI_ISL_2935055 | Lighthouse Laboratory Plymouth | Wellcome Sanger Institute for the COVID-19 Genomics UK (COG-UK) Consortium | Lighthouse Laboratory Plymouth and Alex Alderton, Roberto Amato, Jeffrey Barrett, Sonia Goncalves, Ewan Harrison, David K. Jackson, Ian Johnston, Dominic Kwiatkowski, Cordelia Langford, John Sillitoe on behalf of the Wellcome Sanger Institute COVID-19 Surveillance Team |
| EPI_ISL_2935056 | Berkshire and Surrey Pathology Services Lighthouse Laboratory | Wellcome Sanger Institute for the COVID-19 Genomics UK (COG-UK) Consortium | Berkshire and Surrey Pathology Services Lighthouse Laboratory and Alex Alderton, Roberto Amato, Jeffrey Barrett, Sonia Goncalves, Ewan Harrison, David K. Jackson, Ian Johnston, Dominic Kwiatkowski, Cordelia Langford, John Sillitoe on behalf of the Wellcome Sanger Institute COVID-19 Surveillance Team |
| EPI_ISL_2935057 | Lighthouse Laboratory Plymouth | Wellcome Sanger Institute for the COVID-19 Genomics UK (COG-UK) Consortium | Lighthouse Laboratory Plymouth and Alex Alderton, Roberto Amato, Jeffrey Barrett, Sonia Goncalves, Ewan Harrison, David K. Jackson, Ian Johnston, Dominic Kwiatkowski, Cordelia Langford, John Sillitoe on behalf of the Wellcome Sanger Institute COVID-19 Surveillance Team |
| EPI_ISL_2935058, EPI_ISL_2935059 | Lighthouse Lab in Glasgow | Wellcome Sanger Institute for the COVID-19 Genomics UK (COG-UK) Consortium | Harper VanSteenhouse, Yumi Kasai, David Gray, Carol Clugston, Anna Dominiczak and Alex Alderton, Roberto Amato, Jeffrey Barrett, Sonia Goncalves, Ewan Harrison, David K. Jackson, Ian Johnston, Dominic Kwiatkowski, Cordelia Langford, John Sillitoe on behalf of the Wellcome Sanger Institute COVID-19 Surveillance Team |
| EPI_ISL_2935060 | Lighthouse Lab in Milton Keynes | Wellcome Sanger Institute for the COVID-19 Genomics UK (COG-UK) Consortium | The Lighthouse Lab in Milton Keynes and Alex Alderton, Roberto Amato, Jeffrey Barrett, Sonia Goncalves, Ewan Harrison, David K. Jackson, Ian Johnston, Dominic Kwiatkowski, Cordelia Langford, John Sillitoe on behalf of the Wellcome Sanger Institute COVID-19 Surveillance Team |
| EPI_ISL_2935061 | Berkshire and Surrey Pathology Services Lighthouse Laboratory | Wellcome Sanger Institute for the COVID-19 Genomics UK (COG-UK) Consortium | Berkshire and Surrey Pathology Services Lighthouse Laboratory and Alex Alderton, Roberto Amato, Jeffrey Barrett, Sonia Goncalves, Ewan Harrison, David K. Jackson, Ian Johnston, Dominic Kwiatkowski, Cordelia Langford, John Sillitoe on behalf of the Wellcome Sanger Institute COVID-19 Surveillance Team |
| EPI_ISL_2935062 | Lighthouse Lab in Glasgow | Wellcome Sanger Institute for the COVID-19 Genomics UK (COG-UK) Consortium | Harper VanSteenhouse, Yumi Kasai, David Gray, Carol Clugston, Anna Dominiczak and Alex Alderton, Roberto Amato, Jeffrey Barrett, Sonia Goncalves, Ewan Harrison, David K. Jackson, Ian Johnston, Dominic Kwiatkowski, Cordelia Langford, John Sillitoe on behalf of the Wellcome Sanger Institute COVID-19 Surveillance Team |
| EPI_ISL_2935063, EPI_ISL_2935064 | Lighthouse Laboratory Plymouth | Wellcome Sanger Institute for the COVID-19 Genomics UK (COG-UK) Consortium | Lighthouse Laboratory Plymouth and Alex Alderton, Roberto Amato, Jeffrey Barrett, Sonia Goncalves, Ewan Harrison, David K. Jackson, Ian Johnston, Dominic Kwiatkowski, Cordelia Langford, John Sillitoe on behalf of the Wellcome Sanger Institute COVID-19 Surveillance Team |
| EPI_ISL_2935065 | Berkshire and Surrey Pathology Services Lighthouse Laboratory | Wellcome Sanger Institute for the COVID-19 Genomics UK (COG-UK) Consortium | Berkshire and Surrey Pathology Services Lighthouse Laboratory and Alex Alderton, Roberto Amato, Jeffrey Barrett, Sonia Goncalves, Ewan Harrison, David K. Jackson, Ian Johnston, Dominic Kwiatkowski, Cordelia Langford, John Sillitoe on behalf of the Wellcome Sanger Institute COVID-19 Surveillance Team |
| EPI_ISL_2935066 | Lighthouse Laboratory Plymouth | Wellcome Sanger Institute for the COVID-19 Genomics UK (COG-UK) Consortium | Lighthouse Laboratory Plymouth and Alex Alderton, Roberto Amato, Jeffrey Barrett, Sonia Goncalves, Ewan Harrison, David K. Jackson, Ian Johnston, Dominic Kwiatkowski, Cordelia Langford, John Sillitoe on behalf of the Wellcome Sanger Institute COVID-19 Surveillance Team |
| EPI_ISL_2935067 | Lighthouse Lab in Milton Keynes | Wellcome Sanger Institute for the COVID-19 Genomics UK (COG-UK) Consortium | The Lighthouse Lab in Milton Keynes and Alex Alderton, Roberto Amato, Jeffrey Barrett, Sonia Goncalves, Ewan Harrison, David K. Jackson, Ian Johnston, Dominic Kwiatkowski, Cordelia Langford, John Sillitoe on behalf of the Wellcome Sanger Institute COVID-19 Surveillance Team |
| EPI_ISL_2935068, EPI_ISL_2935069, EPI_ISL_2935070 | Lighthouse Lab in Glasgow | Wellcome Sanger Institute for the COVID-19 Genomics UK (COG-UK) Consortium | Harper VanSteenhouse, Yumi Kasai, David Gray, Carol Clugston, Anna Dominiczak and Alex Alderton, Roberto Amato, Jeffrey Barrett, Sonia Goncalves, Ewan Harrison, David K. Jackson, Ian Johnston, Dominic Kwiatkowski, Cordelia Langford, John Sillitoe on behalf of the Wellcome Sanger Institute COVID-19 Surveillance Team |
| EPI_ISL_2935071 | Berkshire and Surrey Pathology Services Lighthouse Laboratory | Wellcome Sanger Institute for the COVID-19 Genomics UK (COG-UK) Consortium | Berkshire and Surrey Pathology Services Lighthouse Laboratory and Alex Alderton, Roberto Amato, Jeffrey Barrett, Sonia Goncalves, Ewan Harrison, David K. Jackson, Ian Johnston, Dominic Kwiatkowski, Cordelia Langford, John Sillitoe on behalf of the Wellcome Sanger Institute COVID-19 Surveillance Team |
| EPI_ISL_2935072, EPI_ISL_2935073 | Lighthouse Laboratory Plymouth | Wellcome Sanger Institute for the COVID-19 Genomics UK (COG-UK) Consortium | Lighthouse Laboratory Plymouth and Alex Alderton, Roberto Amato, Jeffrey Barrett, Sonia Goncalves, Ewan Harrison, David K. Jackson, Ian Johnston, Dominic Kwiatkowski, Cordelia Langford, John Sillitoe on behalf of the Wellcome Sanger Institute COVID-19 Surveillance Team |
| EPI_ISL_2935074, EPI_ISL_2935075 | Berkshire and Surrey Pathology Services Lighthouse Laboratory | Wellcome Sanger Institute for the COVID-19 Genomics UK (COG-UK) Consortium | Berkshire and Surrey Pathology Services Lighthouse Laboratory and Alex Alderton, Roberto Amato, Jeffrey Barrett, Sonia Goncalves, Ewan Harrison, David K. Jackson, Ian Johnston, Dominic Kwiatkowski, Cordelia Langford, John Sillitoe on behalf of the Wellcome Sanger Institute COVID-19 Surveillance Team |
| EPI_ISL_2935076 | Lighthouse Lab in Glasgow | Wellcome Sanger Institute for the COVID-19 Genomics UK (COG-UK) Consortium | Harper VanSteenhouse, Yumi Kasai, David Gray, Carol Clugston, Anna Dominiczak and Alex Alderton, Roberto Amato, Jeffrey Barrett, Sonia Goncalves, Ewan Harrison, David K. Jackson, Ian Johnston, Dominic Kwiatkowski, Cordelia Langford, John Sillitoe on behalf of the Wellcome Sanger Institute COVID-19 Surveillance Team |
| EPI_ISL_2935077 | Lighthouse Lab in Milton Keynes | Wellcome Sanger Institute for the COVID-19 Genomics UK (COG-UK) Consortium | The Lighthouse Lab in Milton Keynes and Alex Alderton, Roberto Amato, Jeffrey Barrett, Sonia Goncalves, Ewan Harrison, David K. Jackson, Ian Johnston, Dominic Kwiatkowski, Cordelia Langford, John Sillitoe on behalf of the Wellcome Sanger Institute COVID-19 Surveillance Team |
| EPI_ISL_2935078 | Lighthouse Lab in Glasgow | Wellcome Sanger Institute for the COVID-19 Genomics UK (COG-UK) Consortium | Harper VanSteenhouse, Yumi Kasai, David Gray, Carol Clugston, Anna Dominiczak and Alex Alderton, Roberto Amato, Jeffrey Barrett, Sonia Goncalves, Ewan Harrison, David K. Jackson, Ian Johnston, Dominic Kwiatkowski, Cordelia Langford, John Sillitoe on behalf of the Wellcome Sanger Institute COVID-19 Surveillance Team |
| EPI_ISL_2935079 | Berkshire and Surrey Pathology Services Lighthouse Laboratory | Wellcome Sanger Institute for the COVID-19 Genomics UK (COG-UK) Consortium | Berkshire and Surrey Pathology Services Lighthouse Laboratory and Alex Alderton, Roberto Amato, Jeffrey Barrett, Sonia Goncalves, Ewan Harrison, David K. Jackson, Ian Johnston, Dominic Kwiatkowski, Cordelia Langford, John Sillitoe on behalf of the Wellcome Sanger Institute COVID-19 Surveillance Team |
| EPI_ISL_2935080 | Lighthouse Laboratory Plymouth | Wellcome Sanger Institute for the COVID-19 Genomics UK (COG-UK) Consortium | Lighthouse Laboratory Plymouth and Alex Alderton, Roberto Amato, Jeffrey Barrett, Sonia Goncalves, Ewan Harrison, David K. Jackson, Ian Johnston, Dominic Kwiatkowski, Cordelia Langford, John Sillitoe on behalf of the Wellcome Sanger Institute COVID-19 Surveillance Team |
| EPI_ISL_2935081 | Lighthouse Lab in Glasgow | Wellcome Sanger Institute for the COVID-19 Genomics UK (COG-UK) Consortium | Harper VanSteenhouse, Yumi Kasai, David Gray, Carol Clugston, Anna Dominiczak and Alex Alderton, Roberto Amato, Jeffrey Barrett, Sonia Goncalves, Ewan Harrison, David K. Jackson, Ian Johnston, Dominic Kwiatkowski, Cordelia Langford, John Sillitoe on behalf of the Wellcome Sanger Institute COVID-19 Surveillance Team |
| EPI_ISL_2935082, EPI_ISL_2935083 | Lighthouse Lab in Milton Keynes | Wellcome Sanger Institute for the COVID-19 Genomics UK (COG-UK) Consortium | The Lighthouse Lab in Milton Keynes and Alex Alderton, Roberto Amato, Jeffrey Barrett, Sonia Goncalves, Ewan Harrison, David K. Jackson, Ian Johnston, Dominic Kwiatkowski, Cordelia Langford, John Sillitoe on behalf of the Wellcome Sanger Institute COVID-19 Surveillance Team |
| EPI_ISL_2935084 | Lighthouse Laboratory Plymouth | Wellcome Sanger Institute for the COVID-19 Genomics UK (COG-UK) Consortium | Lighthouse Laboratory Plymouth and Alex Alderton, Roberto Amato, Jeffrey Barrett, Sonia Goncalves, Ewan Harrison, David K. Jackson, Ian Johnston, Dominic Kwiatkowski, Cordelia Langford, John Sillitoe on behalf of the Wellcome Sanger Institute COVID-19 Surveillance Team |
| EPI_ISL_2935085, EPI_ISL_2935086, EPI_ISL_2935087 | Berkshire and Surrey Pathology Services Lighthouse Laboratory | Wellcome Sanger Institute for the COVID-19 Genomics UK (COG-UK) Consortium | Berkshire and Surrey Pathology Services Lighthouse Laboratory and Alex Alderton, Roberto Amato, Jeffrey Barrett, Sonia Goncalves, Ewan Harrison, David K. Jackson, Ian Johnston, Dominic Kwiatkowski, Cordelia Langford, John Sillitoe on behalf of the Wellcome Sanger Institute COVID-19 Surveillance Team |
| EPI_ISL_2935088 | Lighthouse Lab in Glasgow | Wellcome Sanger Institute for the COVID-19 Genomics UK (COG-UK) Consortium | Harper VanSteenhouse, Yumi Kasai, David Gray, Carol Clugston, Anna Dominiczak and Alex Alderton, Roberto Amato, Jeffrey Barrett, Sonia Goncalves, Ewan Harrison, David K. Jackson, Ian Johnston, Dominic Kwiatkowski, Cordelia Langford, John Sillitoe on behalf of the Wellcome Sanger Institute COVID-19 Surveillance Team |
| EPI_ISL_2935089, EPI_ISL_2935090 | Lighthouse Laboratory Plymouth | Wellcome Sanger Institute for the COVID-19 Genomics UK (COG-UK) Consortium | Lighthouse Laboratory Plymouth and Alex Alderton, Roberto Amato, Jeffrey Barrett, Sonia Goncalves, Ewan Harrison, David K. Jackson, Ian Johnston, Dominic Kwiatkowski, Cordelia Langford, John Sillitoe on behalf of the Wellcome Sanger Institute COVID-19 Surveillance Team |
| EPI_ISL_2935091 | Berkshire and Surrey Pathology Services Lighthouse Laboratory | Wellcome Sanger Institute for the COVID-19 Genomics UK (COG-UK) Consortium | Berkshire and Surrey Pathology Services Lighthouse Laboratory and Alex Alderton, Roberto Amato, Jeffrey Barrett, Sonia Goncalves, Ewan Harrison, David K. Jackson, Ian Johnston, Dominic Kwiatkowski, Cordelia Langford, John Sillitoe on behalf of the Wellcome Sanger Institute COVID-19 Surveillance Team |
| EPI_ISL_2935092 | Lighthouse Lab in Glasgow | Wellcome Sanger Institute for the COVID-19 Genomics UK (COG-UK) Consortium | Harper VanSteenhouse, Yumi Kasai, David Gray, Carol Clugston, Anna Dominiczak and Alex Alderton, Roberto Amato, Jeffrey Barrett, Sonia Goncalves, Ewan Harrison, David K. Jackson, Ian Johnston, Dominic Kwiatkowski, Cordelia Langford, John Sillitoe on behalf of the Wellcome Sanger Institute COVID-19 Surveillance Team |
| EPI_ISL_2935093 | Berkshire and Surrey Pathology Services Lighthouse Laboratory | Wellcome Sanger Institute for the COVID-19 Genomics UK (COG-UK) Consortium | Berkshire and Surrey Pathology Services Lighthouse Laboratory and Alex Alderton, Roberto Amato, Jeffrey Barrett, Sonia Goncalves, Ewan Harrison, David K. Jackson, Ian Johnston, Dominic Kwiatkowski, Cordelia Langford, John Sillitoe on behalf of the Wellcome Sanger Institute COVID-19 Surveillance Team |
| EPI_ISL_2935094 | Lighthouse Lab in Glasgow | Wellcome Sanger Institute for the COVID-19 Genomics UK (COG-UK) Consortium | Harper VanSteenhouse, Yumi Kasai, David Gray, Carol Clugston, Anna Dominiczak and Alex Alderton, Roberto Amato, Jeffrey Barrett, Sonia Goncalves, Ewan Harrison, David K. Jackson, Ian Johnston, Dominic Kwiatkowski, Cordelia Langford, John Sillitoe on behalf of the Wellcome Sanger Institute COVID-19 Surveillance Team |
| EPI_ISL_2935095 | Lighthouse Lab in Milton Keynes | Wellcome Sanger Institute for the COVID-19 Genomics UK (COG-UK) Consortium | The Lighthouse Lab in Milton Keynes and Alex Alderton, Roberto Amato, Jeffrey Barrett, Sonia Goncalves, Ewan Harrison, David K. Jackson, Ian Johnston, Dominic Kwiatkowski, Cordelia Langford, John Sillitoe on behalf of the Wellcome Sanger Institute COVID-19 Surveillance Team |
| EPI_ISL_2935096, | Lighthouse Laboratory Plymouth | Wellcome Sanger Institute for the COVID-19 Genomics UK (COG-UK) | Lighthouse Laboratory Plymouth and Alex Alderton, Roberto Amato, Jeffrey Barrett, Sonia Goncalves, Ewan Harrison, David K. Jackson, Ian Johnston, Dominic Kwiatkowski, Cordelia Langford, John Sillitoe on behalf of the Wellcome Sanger |

| | | | |
|---|---|---|---|
| EPI_ISL_2935097, EPI_ISL_2935098 | | Consortium | Institute COVID-19 Surveillance Team |
| EPI_ISL_2935099 | Health Services Laboratories | Wellcome Sanger Institute for the COVID-19 Genomics UK (COG-UK) Consortium | Health Services Laboratories and Alex Alderton, Roberto Amato, Jeffrey Barrett, Sonia Goncalves, Ewan Harrison, David K. Jackson, Ian Johnston, Dominic Kwiatkowski, Cordelia Langford, John Sillitoe on behalf of the Wellcome Sanger Institute COVID-19 Surveillance Team |
| EPI_ISL_2935100 | Lighthouse Lab in Glasgow | Wellcome Sanger Institute for the COVID-19 Genomics UK (COG-UK) Consortium | Harper VanSteenhouse, Yumi Kasai, David Gray, Carol Clugston, Anna Dominiczak and Alex Alderton, Roberto Amato, Jeffrey Barrett, Sonia Goncalves, Ewan Harrison, David K. Jackson, Ian Johnston, Dominic Kwiatkowski, Cordelia Langford, John Sillitoe on behalf of the Wellcome Sanger Institute COVID-19 Surveillance Team |
| EPI_ISL_2935101 | Lighthouse Laboratory Plymouth | Wellcome Sanger Institute for the COVID-19 Genomics UK (COG-UK) Consortium | Lighthouse Laboratory Plymouth and Alex Alderton, Roberto Amato, Jeffrey Barrett, Sonia Goncalves, Ewan Harrison, David K. Jackson, Ian Johnston, Dominic Kwiatkowski, Cordelia Langford, John Sillitoe on behalf of the Wellcome Sanger Institute COVID-19 Surveillance Team |
| EPI_ISL_2935102 | Lighthouse Lab in Milton Keynes | Wellcome Sanger Institute for the COVID-19 Genomics UK (COG-UK) Consortium | The Lighthouse Lab in Milton Keynes and Alex Alderton, Roberto Amato, Jeffrey Barrett, Sonia Goncalves, Ewan Harrison, David K. Jackson, Ian Johnston, Dominic Kwiatkowski, Cordelia Langford, John Sillitoe on behalf of the Wellcome Sanger Institute COVID-19 Surveillance Team |
| EPI_ISL_2935103 | Lighthouse Lab in Glasgow | Wellcome Sanger Institute for the COVID-19 Genomics UK (COG-UK) Consortium | Harper VanSteenhouse, Yumi Kasai, David Gray, Carol Clugston, Anna Dominiczak and Alex Alderton, Roberto Amato, Jeffrey Barrett, Sonia Goncalves, Ewan Harrison, David K. Jackson, Ian Johnston, Dominic Kwiatkowski, Cordelia Langford, John Sillitoe on behalf of the Wellcome Sanger Institute COVID-19 Surveillance Team |
| EPI_ISL_2935104 | Lighthouse Lab in Milton Keynes | Wellcome Sanger Institute for the COVID-19 Genomics UK (COG-UK) Consortium | The Lighthouse Lab in Milton Keynes and Alex Alderton, Roberto Amato, Jeffrey Barrett, Sonia Goncalves, Ewan Harrison, David K. Jackson, Ian Johnston, Dominic Kwiatkowski, Cordelia Langford, John Sillitoe on behalf of the Wellcome Sanger Institute COVID-19 Surveillance Team |
| EPI_ISL_2935105, EPI_ISL_2935106 | Lighthouse Laboratory Plymouth | Wellcome Sanger Institute for the COVID-19 Genomics UK (COG-UK) Consortium | Lighthouse Laboratory Plymouth and Alex Alderton, Roberto Amato, Jeffrey Barrett, Sonia Goncalves, Ewan Harrison, David K. Jackson, Ian Johnston, Dominic Kwiatkowski, Cordelia Langford, John Sillitoe on behalf of the Wellcome Sanger Institute COVID-19 Surveillance Team |
| EPI_ISL_2935107 | Lighthouse Lab in Glasgow | Wellcome Sanger Institute for the COVID-19 Genomics UK (COG-UK) Consortium | Harper VanSteenhouse, Yumi Kasai, David Gray, Carol Clugston, Anna Dominiczak and Alex Alderton, Roberto Amato, Jeffrey Barrett, Sonia Goncalves, Ewan Harrison, David K. Jackson, Ian Johnston, Dominic Kwiatkowski, Cordelia Langford, John Sillitoe on behalf of the Wellcome Sanger Institute COVID-19 Surveillance Team |
| EPI_ISL_2935108, EPI_ISL_2935109 | Lighthouse Lab in Milton Keynes | Wellcome Sanger Institute for the COVID-19 Genomics UK (COG-UK) Consortium | The Lighthouse Lab in Milton Keynes and Alex Alderton, Roberto Amato, Jeffrey Barrett, Sonia Goncalves, Ewan Harrison, David K. Jackson, Ian Johnston, Dominic Kwiatkowski, Cordelia Langford, John Sillitoe on behalf of the Wellcome Sanger Institute COVID-19 Surveillance Team |
| EPI_ISL_2935110 | Lighthouse Lab in Glasgow | Wellcome Sanger Institute for the COVID-19 Genomics UK (COG-UK) Consortium | Harper VanSteenhouse, Yumi Kasai, David Gray, Carol Clugston, Anna Dominiczak and Alex Alderton, Roberto Amato, Jeffrey Barrett, Sonia Goncalves, Ewan Harrison, David K. Jackson, Ian Johnston, Dominic Kwiatkowski, Cordelia Langford, John Sillitoe on behalf of the Wellcome Sanger Institute COVID-19 Surveillance Team |
| EPI_ISL_2935111 | Lighthouse Laboratory Plymouth | Wellcome Sanger Institute for the COVID-19 Genomics UK (COG-UK) Consortium | Lighthouse Laboratory Plymouth and Alex Alderton, Roberto Amato, Jeffrey Barrett, Sonia Goncalves, Ewan Harrison, David K. Jackson, Ian Johnston, Dominic Kwiatkowski, Cordelia Langford, John Sillitoe on behalf of the Wellcome Sanger Institute COVID-19 Surveillance Team |
| EPI_ISL_2935112 | Lighthouse Lab in Milton Keynes | Wellcome Sanger Institute for the COVID-19 Genomics UK (COG-UK) Consortium | The Lighthouse Lab in Milton Keynes and Alex Alderton, Roberto Amato, Jeffrey Barrett, Sonia Goncalves, Ewan Harrison, David K. Jackson, Ian Johnston, Dominic Kwiatkowski, Cordelia Langford, John Sillitoe on behalf of the Wellcome Sanger Institute COVID-19 Surveillance Team |
| EPI_ISL_2935113, EPI_ISL_2935114 | Berkshire and Surrey Pathology Services Lighthouse Laboratory | Wellcome Sanger Institute for the COVID-19 Genomics UK (COG-UK) Consortium | Berkshire and Surrey Pathology Services Lighthouse Laboratory and Alex Alderton, Roberto Amato, Jeffrey Barrett, Sonia Goncalves, Ewan Harrison, David K. Jackson, Ian Johnston, Dominic Kwiatkowski, Cordelia Langford, John Sillitoe on behalf of the Wellcome Sanger Institute COVID-19 Surveillance Team |
| EPI_ISL_2935115, EPI_ISL_2935116, EPI_ISL_2935117 | Lighthouse Lab in Glasgow | Wellcome Sanger Institute for the COVID-19 Genomics UK (COG-UK) Consortium | Harper VanSteenhouse, Yumi Kasai, David Gray, Carol Clugston, Anna Dominiczak and Alex Alderton, Roberto Amato, Jeffrey Barrett, Sonia Goncalves, Ewan Harrison, David K. Jackson, Ian Johnston, Dominic Kwiatkowski, Cordelia Langford, John Sillitoe on behalf of the Wellcome Sanger Institute COVID-19 Surveillance Team |
| EPI_ISL_2935118, EPI_ISL_2935119 | Lighthouse Lab in Milton Keynes | Wellcome Sanger Institute for the COVID-19 Genomics UK (COG-UK) Consortium | The Lighthouse Lab in Milton Keynes and Alex Alderton, Roberto Amato, Jeffrey Barrett, Sonia Goncalves, Ewan Harrison, David K. Jackson, Ian Johnston, Dominic Kwiatkowski, Cordelia Langford, John Sillitoe on behalf of the Wellcome Sanger Institute COVID-19 Surveillance Team |
| EPI_ISL_2935120 | Lighthouse Lab in Glasgow | Wellcome Sanger Institute for the COVID-19 Genomics UK (COG-UK) Consortium | Harper VanSteenhouse, Yumi Kasai, David Gray, Carol Clugston, Anna Dominiczak and Alex Alderton, Roberto Amato, Jeffrey Barrett, Sonia Goncalves, Ewan Harrison, David K. Jackson, Ian Johnston, Dominic Kwiatkowski, Cordelia Langford, John Sillitoe on behalf of the Wellcome Sanger Institute COVID-19 Surveillance Team |
| EPI_ISL_2935121 | Berkshire and Surrey Pathology Services Lighthouse Laboratory | Wellcome Sanger Institute for the COVID-19 Genomics UK (COG-UK) Consortium | Berkshire and Surrey Pathology Services Lighthouse Laboratory and Alex Alderton, Roberto Amato, Jeffrey Barrett, Sonia Goncalves, Ewan Harrison, David K. Jackson, Ian Johnston, Dominic Kwiatkowski, Cordelia Langford, John Sillitoe on behalf of the Wellcome Sanger Institute COVID-19 Surveillance Team |
| EPI_ISL_2935122 | Lighthouse Lab in Milton Keynes | Wellcome Sanger Institute for the COVID-19 Genomics UK (COG-UK) Consortium | The Lighthouse Lab in Milton Keynes and Alex Alderton, Roberto Amato, Jeffrey Barrett, Sonia Goncalves, Ewan Harrison, David K. Jackson, Ian Johnston, Dominic Kwiatkowski, Cordelia Langford, John Sillitoe on behalf of the Wellcome Sanger Institute COVID-19 Surveillance Team |
| EPI_ISL_2935123 | Lighthouse Laboratory Plymouth | Wellcome Sanger Institute for the COVID-19 Genomics UK (COG-UK) Consortium | Lighthouse Laboratory Plymouth and Alex Alderton, Roberto Amato, Jeffrey Barrett, Sonia Goncalves, Ewan Harrison, David K. Jackson, Ian Johnston, Dominic Kwiatkowski, Cordelia Langford, John Sillitoe on behalf of the Wellcome Sanger Institute COVID-19 Surveillance Team |
| EPI_ISL_2935124 | Lighthouse Lab in Milton Keynes | Wellcome Sanger Institute for the COVID-19 Genomics UK (COG-UK) Consortium | The Lighthouse Lab in Milton Keynes and Alex Alderton, Roberto Amato, Jeffrey Barrett, Sonia Goncalves, Ewan Harrison, David K. Jackson, Ian Johnston, Dominic Kwiatkowski, Cordelia Langford, John Sillitoe on behalf of the Wellcome Sanger Institute COVID-19 Surveillance Team |
| EPI_ISL_2935125, EPI_ISL_2935126 | Lighthouse Lab in Glasgow | Wellcome Sanger Institute for the COVID-19 Genomics UK (COG-UK) Consortium | Harper VanSteenhouse, Yumi Kasai, David Gray, Carol Clugston, Anna Dominiczak and Alex Alderton, Roberto Amato, Jeffrey Barrett, Sonia Goncalves, Ewan Harrison, David K. Jackson, Ian Johnston, Dominic Kwiatkowski, Cordelia Langford, John Sillitoe on behalf of the Wellcome Sanger Institute COVID-19 Surveillance Team |
| EPI_ISL_2935127, EPI_ISL_2935128 | Berkshire and Surrey Pathology Services Lighthouse Laboratory | Wellcome Sanger Institute for the COVID-19 Genomics UK (COG-UK) Consortium | Berkshire and Surrey Pathology Services Lighthouse Laboratory and Alex Alderton, Roberto Amato, Jeffrey Barrett, Sonia Goncalves, Ewan Harrison, David K. Jackson, Ian Johnston, Dominic Kwiatkowski, Cordelia Langford, John Sillitoe on behalf of the Wellcome Sanger Institute COVID-19 Surveillance Team |
| EPI_ISL_2935129 | Lighthouse Laboratory Plymouth | Wellcome Sanger Institute for the COVID-19 Genomics UK (COG-UK) Consortium | Lighthouse Laboratory Plymouth and Alex Alderton, Roberto Amato, Jeffrey Barrett, Sonia Goncalves, Ewan Harrison, David K. Jackson, Ian Johnston, Dominic Kwiatkowski, Cordelia Langford, John Sillitoe on behalf of the Wellcome Sanger Institute COVID-19 Surveillance Team |
| EPI_ISL_2935130 | Lighthouse Lab in Milton Keynes | Wellcome Sanger Institute for the COVID-19 Genomics UK (COG-UK) Consortium | The Lighthouse Lab in Milton Keynes and Alex Alderton, Roberto Amato, Jeffrey Barrett, Sonia Goncalves, Ewan Harrison, David K. Jackson, Ian Johnston, Dominic Kwiatkowski, Cordelia Langford, John Sillitoe on behalf of the Wellcome Sanger Institute COVID-19 Surveillance Team |
| EPI_ISL_2935131 | Lighthouse Lab in Glasgow | Wellcome Sanger Institute for the COVID-19 Genomics UK (COG-UK) Consortium | Harper VanSteenhouse, Yumi Kasai, David Gray, Carol Clugston, Anna Dominiczak and Alex Alderton, Roberto Amato, Jeffrey Barrett, Sonia Goncalves, Ewan Harrison, David K. Jackson, Ian Johnston, Dominic Kwiatkowski, Cordelia Langford, John Sillitoe on behalf of the Wellcome Sanger Institute COVID-19 Surveillance Team |
| EPI_ISL_2935132, EPI_ISL_2935133 | Lighthouse Laboratory Plymouth | Wellcome Sanger Institute for the COVID-19 Genomics UK (COG-UK) Consortium | Lighthouse Laboratory Plymouth and Alex Alderton, Roberto Amato, Jeffrey Barrett, Sonia Goncalves, Ewan Harrison, David K. Jackson, Ian Johnston, Dominic Kwiatkowski, Cordelia Langford, John Sillitoe on behalf of the Wellcome Sanger Institute COVID-19 Surveillance Team |
| EPI_ISL_2935134 | Lighthouse Lab in Milton Keynes | Wellcome Sanger Institute for the COVID-19 Genomics UK (COG-UK) Consortium | The Lighthouse Lab in Milton Keynes and Alex Alderton, Roberto Amato, Jeffrey Barrett, Sonia Goncalves, Ewan Harrison, David K. Jackson, Ian Johnston, Dominic Kwiatkowski, Cordelia Langford, John Sillitoe on behalf of the Wellcome Sanger Institute COVID-19 Surveillance Team |
| EPI_ISL_2935135 | Berkshire and Surrey Pathology Services Lighthouse Laboratory | Wellcome Sanger Institute for the COVID-19 Genomics UK (COG-UK) Consortium | Berkshire and Surrey Pathology Services Lighthouse Laboratory and Alex Alderton, Roberto Amato, Jeffrey Barrett, Sonia Goncalves, Ewan Harrison, David K. Jackson, Ian Johnston, Dominic Kwiatkowski, Cordelia Langford, John Sillitoe on behalf of the Wellcome Sanger Institute COVID-19 Surveillance Team |
| EPI_ISL_2935136 | Lighthouse Laboratory Plymouth | Wellcome Sanger Institute for the COVID-19 Genomics UK (COG-UK) Consortium | Lighthouse Laboratory Plymouth and Alex Alderton, Roberto Amato, Jeffrey Barrett, Sonia Goncalves, Ewan Harrison, David K. Jackson, Ian Johnston, Dominic Kwiatkowski, Cordelia Langford, John Sillitoe on behalf of the Wellcome Sanger Institute COVID-19 Surveillance Team |
| EPI_ISL_2935137 | Lighthouse Lab in Glasgow | Wellcome Sanger Institute for the COVID-19 Genomics UK (COG-UK) Consortium | Harper VanSteenhouse, Yumi Kasai, David Gray, Carol Clugston, Anna Dominiczak and Alex Alderton, Roberto Amato, Jeffrey Barrett, Sonia Goncalves, Ewan Harrison, David K. Jackson, Ian Johnston, Dominic Kwiatkowski, Cordelia Langford, John Sillitoe on behalf of the Wellcome Sanger Institute COVID-19 Surveillance Team |
| EPI_ISL_2935138 | Berkshire and Surrey Pathology Services Lighthouse Laboratory | Wellcome Sanger Institute for the COVID-19 Genomics UK (COG-UK) Consortium | Berkshire and Surrey Pathology Services Lighthouse Laboratory and Alex Alderton, Roberto Amato, Jeffrey Barrett, Sonia Goncalves, Ewan Harrison, David K. Jackson, Ian Johnston, Dominic Kwiatkowski, Cordelia Langford, John Sillitoe on behalf of the Wellcome Sanger Institute COVID-19 Surveillance Team |
| EPI_ISL_2935139, EPI_ISL_2935140 | Lighthouse Lab in Milton Keynes | Wellcome Sanger Institute for the COVID-19 Genomics UK (COG-UK) Consortium | The Lighthouse Lab in Milton Keynes and Alex Alderton, Roberto Amato, Jeffrey Barrett, Sonia Goncalves, Ewan Harrison, David K. Jackson, Ian Johnston, Dominic Kwiatkowski, Cordelia Langford, John Sillitoe on behalf of the Wellcome Sanger Institute COVID-19 Surveillance Team |
| EPI_ISL_2935141, EPI_ISL_2935142 | Lighthouse Laboratory Plymouth | Wellcome Sanger Institute for the COVID-19 Genomics UK (COG-UK) Consortium | Lighthouse Laboratory Plymouth and Alex Alderton, Roberto Amato, Jeffrey Barrett, Sonia Goncalves, Ewan Harrison, David K. Jackson, Ian Johnston, Dominic Kwiatkowski, Cordelia Langford, John Sillitoe on behalf of the Wellcome Sanger Institute COVID-19 Surveillance Team |
| EPI_ISL_2935143 | Lighthouse Lab in Glasgow | Wellcome Sanger Institute for the COVID-19 Genomics UK (COG-UK) Consortium | Harper VanSteenhouse, Yumi Kasai, David Gray, Carol Clugston, Anna Dominiczak and Alex Alderton, Roberto Amato, Jeffrey Barrett, Sonia Goncalves, Ewan Harrison, David K. Jackson, Ian Johnston, Dominic Kwiatkowski, Cordelia Langford, John Sillitoe on behalf of the Wellcome Sanger Institute COVID-19 Surveillance Team |
| EPI_ISL_2935144 | Berkshire and Surrey Pathology Services Lighthouse Laboratory | Wellcome Sanger Institute for the COVID-19 Genomics UK (COG-UK) Consortium | Berkshire and Surrey Pathology Services Lighthouse Laboratory and Alex Alderton, Roberto Amato, Jeffrey Barrett, Sonia Goncalves, Ewan Harrison, David K. Jackson, Ian Johnston, Dominic Kwiatkowski, Cordelia Langford, John Sillitoe on behalf of the Wellcome Sanger Institute COVID-19 Surveillance Team |
| EPI_ISL_2935145 | Lighthouse Laboratory Plymouth | Wellcome Sanger Institute for the COVID-19 Genomics UK (COG-UK) Consortium | Lighthouse Laboratory Plymouth and Alex Alderton, Roberto Amato, Jeffrey Barrett, Sonia Goncalves, Ewan Harrison, David K. Jackson, Ian Johnston, Dominic Kwiatkowski, Cordelia Langford, John Sillitoe on behalf of the Wellcome Sanger Institute COVID-19 Surveillance Team |
| EPI_ISL_2935146, EPI_ISL_2935147, EPI_ISL_2935148 | Lighthouse Lab in Milton Keynes | Wellcome Sanger Institute for the COVID-19 Genomics UK (COG-UK) Consortium | The Lighthouse Lab in Milton Keynes and Alex Alderton, Roberto Amato, Jeffrey Barrett, Sonia Goncalves, Ewan Harrison, David K. Jackson, Ian Johnston, Dominic Kwiatkowski, Cordelia Langford, John Sillitoe on behalf of the Wellcome Sanger Institute COVID-19 Surveillance Team |
| EPI_ISL_2935149 | Lighthouse Lab in Glasgow | Wellcome Sanger Institute for the COVID-19 Genomics UK (COG-UK) Consortium | Harper VanSteenhouse, Yumi Kasai, David Gray, Carol Clugston, Anna Dominiczak and Alex Alderton, Roberto Amato, Jeffrey Barrett, Sonia Goncalves, Ewan Harrison, David K. Jackson, Ian Johnston, Dominic Kwiatkowski, Cordelia Langford, John Sillitoe on behalf of the Wellcome Sanger Institute COVID-19 Surveillance Team |
| EPI_ISL_2935150, EPI_ISL_2935151 | Lighthouse Lab in Milton Keynes | Wellcome Sanger Institute for the COVID-19 Genomics UK (COG-UK) Consortium | The Lighthouse Lab in Milton Keynes and Alex Alderton, Roberto Amato, Jeffrey Barrett, Sonia Goncalves, Ewan Harrison, David K. Jackson, Ian Johnston, Dominic Kwiatkowski, Cordelia Langford, John Sillitoe on behalf of the Wellcome Sanger Institute COVID-19 Surveillance Team |
| EPI_ISL_2935152 | Lighthouse Laboratory Plymouth | Wellcome Sanger Institute for the COVID-19 Genomics UK (COG-UK) Consortium | Lighthouse Laboratory Plymouth and Alex Alderton, Roberto Amato, Jeffrey Barrett, Sonia Goncalves, Ewan Harrison, David K. Jackson, Ian Johnston, Dominic Kwiatkowski, Cordelia Langford, John Sillitoe on behalf of the Wellcome Sanger Institute COVID-19 Surveillance Team |
| EPI_ISL_2935153 | Lighthouse Lab in Milton Keynes | Wellcome Sanger Institute for the COVID-19 Genomics UK (COG-UK) Consortium | The Lighthouse Lab in Milton Keynes and Alex Alderton, Roberto Amato, Jeffrey Barrett, Sonia Goncalves, Ewan Harrison, David K. Jackson, Ian Johnston, Dominic Kwiatkowski, Cordelia Langford, John Sillitoe on behalf of the Wellcome Sanger Institute COVID-19 Surveillance Team |
| EPI_ISL_2935154, EPI_ISL_2935155 | Lighthouse Laboratory Plymouth | Wellcome Sanger Institute for the COVID-19 Genomics UK (COG-UK) Consortium | Lighthouse Laboratory Plymouth and Alex Alderton, Roberto Amato, Jeffrey Barrett, Sonia Goncalves, Ewan Harrison, David K. Jackson, Ian Johnston, Dominic Kwiatkowski, Cordelia Langford, John Sillitoe on behalf of the Wellcome Sanger Institute COVID-19 Surveillance Team |
| EPI_ISL_2935156 | Lighthouse Lab in Glasgow | Wellcome Sanger Institute for the COVID-19 Genomics UK (COG-UK) Consortium | Harper VanSteenhouse, Yumi Kasai, David Gray, Carol Clugston, Anna Dominiczak and Alex Alderton, Roberto Amato, Jeffrey Barrett, Sonia Goncalves, Ewan Harrison, David K. Jackson, Ian Johnston, Dominic Kwiatkowski, Cordelia Langford, John Sillitoe on behalf of the Wellcome Sanger Institute COVID-19 Surveillance Team |
| EPI_ISL_2935157 | Health Services Laboratories | Wellcome Sanger Institute for the COVID-19 Genomics UK (COG-UK) | Health Services Laboratories and Alex Alderton, Roberto Amato, Jeffrey Barrett, Sonia Goncalves, Ewan Harrison, David K. Jackson, Ian Johnston, Dominic Kwiatkowski, Cordelia Langford, John Sillitoe on behalf of the Wellcome Sanger Institute |

| | | | |
|---|---|---|---|
| | | Consortium | COVID-19 Surveillance Team |
| EPI_ISL_2935158, EPI_ISL_2935159 | Lighthouse Laboratory Plymouth | Wellcome Sanger Institute for the COVID-19 Genomics UK (COG-UK) Consortium | Lighthouse Laboratory Plymouth and Alex Alderton, Roberto Amato, Jeffrey Barrett, Sonia Goncalves, Ewan Harrison, David K. Jackson, Ian Johnston, Dominic Kwiatkowski, Cordelia Langford, John Sillitoe on behalf of the Wellcome Sanger Institute COVID-19 Surveillance Team |
| EPI_ISL_2935160 | Lighthouse Lab in Milton Keynes | Wellcome Sanger Institute for the COVID-19 Genomics UK (COG-UK) Consortium | The Lighthouse Lab in Milton Keynes and Alex Alderton, Roberto Amato, Jeffrey Barrett, Sonia Goncalves, Ewan Harrison, David K. Jackson, Ian Johnston, Dominic Kwiatkowski, Cordelia Langford, John Sillitoe on behalf of the Wellcome Sanger Institute COVID-19 Surveillance Team |
| EPI_ISL_2935161 | Berkshire and Surrey Pathology Services Lighthouse Laboratory | Wellcome Sanger Institute for the COVID-19 Genomics UK (COG-UK) Consortium | Berkshire and Surrey Pathology Services Lighthouse Laboratory and Alex Alderton, Roberto Amato, Jeffrey Barrett, Sonia Goncalves, Ewan Harrison, David K. Jackson, Ian Johnston, Dominic Kwiatkowski, Cordelia Langford, John Sillitoe on behalf of the Wellcome Sanger Institute COVID-19 Surveillance Team |
| EPI_ISL_2935162, EPI_ISL_2935163 | Lighthouse Laboratory Plymouth | Wellcome Sanger Institute for the COVID-19 Genomics UK (COG-UK) Consortium | Lighthouse Laboratory Plymouth and Alex Alderton, Roberto Amato, Jeffrey Barrett, Sonia Goncalves, Ewan Harrison, David K. Jackson, Ian Johnston, Dominic Kwiatkowski, Cordelia Langford, John Sillitoe on behalf of the Wellcome Sanger Institute COVID-19 Surveillance Team |
| EPI_ISL_2935164 | Berkshire and Surrey Pathology Services Lighthouse Laboratory | Wellcome Sanger Institute for the COVID-19 Genomics UK (COG-UK) Consortium | Berkshire and Surrey Pathology Services Lighthouse Laboratory and Alex Alderton, Roberto Amato, Jeffrey Barrett, Sonia Goncalves, Ewan Harrison, David K. Jackson, Ian Johnston, Dominic Kwiatkowski, Cordelia Langford, John Sillitoe on behalf of the Wellcome Sanger Institute COVID-19 Surveillance Team |
| EPI_ISL_2935165 | Lighthouse Laboratory Plymouth | Wellcome Sanger Institute for the COVID-19 Genomics UK (COG-UK) Consortium | Lighthouse Laboratory Plymouth and Alex Alderton, Roberto Amato, Jeffrey Barrett, Sonia Goncalves, Ewan Harrison, David K. Jackson, Ian Johnston, Dominic Kwiatkowski, Cordelia Langford, John Sillitoe on behalf of the Wellcome Sanger Institute COVID-19 Surveillance Team |
| EPI_ISL_2935166 | Lighthouse Lab in Glasgow | Wellcome Sanger Institute for the COVID-19 Genomics UK (COG-UK) Consortium | Harper VanSteenhouse, Yumi Kasai, David Gray, Carol Clugston, Anna Dominiczak and Alex Alderton, Roberto Amato, Jeffrey Barrett, Sonia Goncalves, Ewan Harrison, David K. Jackson, Ian Johnston, Dominic Kwiatkowski, Cordelia Langford, John Sillitoe on behalf of the Wellcome Sanger Institute COVID-19 Surveillance Team |
| EPI_ISL_2935167 | Lighthouse Lab in Milton Keynes | Wellcome Sanger Institute for the COVID-19 Genomics UK (COG-UK) Consortium | The Lighthouse Lab in Milton Keynes and Alex Alderton, Roberto Amato, Jeffrey Barrett, Sonia Goncalves, Ewan Harrison, David K. Jackson, Ian Johnston, Dominic Kwiatkowski, Cordelia Langford, John Sillitoe on behalf of the Wellcome Sanger Institute COVID-19 Surveillance Team |
| EPI_ISL_2935168 | Lighthouse Lab in Glasgow | Wellcome Sanger Institute for the COVID-19 Genomics UK (COG-UK) Consortium | Harper VanSteenhouse, Yumi Kasai, David Gray, Carol Clugston, Anna Dominiczak and Alex Alderton, Roberto Amato, Jeffrey Barrett, Sonia Goncalves, Ewan Harrison, David K. Jackson, Ian Johnston, Dominic Kwiatkowski, Cordelia Langford, John Sillitoe on behalf of the Wellcome Sanger Institute COVID-19 Surveillance Team |
| EPI_ISL_2935169 | Berkshire and Surrey Pathology Services Lighthouse Laboratory | Wellcome Sanger Institute for the COVID-19 Genomics UK (COG-UK) Consortium | Berkshire and Surrey Pathology Services Lighthouse Laboratory and Alex Alderton, Roberto Amato, Jeffrey Barrett, Sonia Goncalves, Ewan Harrison, David K. Jackson, Ian Johnston, Dominic Kwiatkowski, Cordelia Langford, John Sillitoe on behalf of the Wellcome Sanger Institute COVID-19 Surveillance Team |
| EPI_ISL_2935170 | Lighthouse Lab in Glasgow | Wellcome Sanger Institute for the COVID-19 Genomics UK (COG-UK) Consortium | Harper VanSteenhouse, Yumi Kasai, David Gray, Carol Clugston, Anna Dominiczak and Alex Alderton, Roberto Amato, Jeffrey Barrett, Sonia Goncalves, Ewan Harrison, David K. Jackson, Ian Johnston, Dominic Kwiatkowski, Cordelia Langford, John Sillitoe on behalf of the Wellcome Sanger Institute COVID-19 Surveillance Team |
| EPI_ISL_2935171, EPI_ISL_2935172, EPI_ISL_2935173 | Lighthouse Laboratory Plymouth | Wellcome Sanger Institute for the COVID-19 Genomics UK (COG-UK) Consortium | Lighthouse Laboratory Plymouth and Alex Alderton, Roberto Amato, Jeffrey Barrett, Sonia Goncalves, Ewan Harrison, David K. Jackson, Ian Johnston, Dominic Kwiatkowski, Cordelia Langford, John Sillitoe on behalf of the Wellcome Sanger Institute COVID-19 Surveillance Team |
| EPI_ISL_2935174 | Lighthouse Lab in Milton Keynes | Wellcome Sanger Institute for the COVID-19 Genomics UK (COG-UK) Consortium | The Lighthouse Lab in Milton Keynes and Alex Alderton, Roberto Amato, Jeffrey Barrett, Sonia Goncalves, Ewan Harrison, David K. Jackson, Ian Johnston, Dominic Kwiatkowski, Cordelia Langford, John Sillitoe on behalf of the Wellcome Sanger Institute COVID-19 Surveillance Team |
| EPI_ISL_2935175, EPI_ISL_2935176 | Berkshire and Surrey Pathology Services Lighthouse Laboratory | Wellcome Sanger Institute for the COVID-19 Genomics UK (COG-UK) Consortium | Berkshire and Surrey Pathology Services Lighthouse Laboratory and Alex Alderton, Roberto Amato, Jeffrey Barrett, Sonia Goncalves, Ewan Harrison, David K. Jackson, Ian Johnston, Dominic Kwiatkowski, Cordelia Langford, John Sillitoe on behalf of the Wellcome Sanger Institute COVID-19 Surveillance Team |
| EPI_ISL_2935177 | Lighthouse Laboratory Plymouth | Wellcome Sanger Institute for the COVID-19 Genomics UK (COG-UK) Consortium | Lighthouse Laboratory Plymouth and Alex Alderton, Roberto Amato, Jeffrey Barrett, Sonia Goncalves, Ewan Harrison, David K. Jackson, Ian Johnston, Dominic Kwiatkowski, Cordelia Langford, John Sillitoe on behalf of the Wellcome Sanger Institute COVID-19 Surveillance Team |
| EPI_ISL_2935178 | Lighthouse Lab in Glasgow | Wellcome Sanger Institute for the COVID-19 Genomics UK (COG-UK) Consortium | Harper VanSteenhouse, Yumi Kasai, David Gray, Carol Clugston, Anna Dominiczak and Alex Alderton, Roberto Amato, Jeffrey Barrett, Sonia Goncalves, Ewan Harrison, David K. Jackson, Ian Johnston, Dominic Kwiatkowski, Cordelia Langford, John Sillitoe on behalf of the Wellcome Sanger Institute COVID-19 Surveillance Team |
| EPI_ISL_2935179 | Lighthouse Laboratory Plymouth | Wellcome Sanger Institute for the COVID-19 Genomics UK (COG-UK) Consortium | Lighthouse Laboratory Plymouth and Alex Alderton, Roberto Amato, Jeffrey Barrett, Sonia Goncalves, Ewan Harrison, David K. Jackson, Ian Johnston, Dominic Kwiatkowski, Cordelia Langford, John Sillitoe on behalf of the Wellcome Sanger Institute COVID-19 Surveillance Team |
| EPI_ISL_2935180, EPI_ISL_2935181 | Berkshire and Surrey Pathology Services Lighthouse Laboratory | Wellcome Sanger Institute for the COVID-19 Genomics UK (COG-UK) Consortium | Berkshire and Surrey Pathology Services Lighthouse Laboratory and Alex Alderton, Roberto Amato, Jeffrey Barrett, Sonia Goncalves, Ewan Harrison, David K. Jackson, Ian Johnston, Dominic Kwiatkowski, Cordelia Langford, John Sillitoe on behalf of the Wellcome Sanger Institute COVID-19 Surveillance Team |
| EPI_ISL_2935182 | Lighthouse Laboratory Plymouth | Wellcome Sanger Institute for the COVID-19 Genomics UK (COG-UK) Consortium | Lighthouse Laboratory Plymouth and Alex Alderton, Roberto Amato, Jeffrey Barrett, Sonia Goncalves, Ewan Harrison, David K. Jackson, Ian Johnston, Dominic Kwiatkowski, Cordelia Langford, John Sillitoe on behalf of the Wellcome Sanger Institute COVID-19 Surveillance Team |
| EPI_ISL_2935183 | Berkshire and Surrey Pathology Services Lighthouse Laboratory | Wellcome Sanger Institute for the COVID-19 Genomics UK (COG-UK) Consortium | Berkshire and Surrey Pathology Services Lighthouse Laboratory and Alex Alderton, Roberto Amato, Jeffrey Barrett, Sonia Goncalves, Ewan Harrison, David K. Jackson, Ian Johnston, Dominic Kwiatkowski, Cordelia Langford, John Sillitoe on behalf of the Wellcome Sanger Institute COVID-19 Surveillance Team |
| EPI_ISL_2935184 | Lighthouse Lab in Glasgow | Wellcome Sanger Institute for the COVID-19 Genomics UK (COG-UK) Consortium | Harper VanSteenhouse, Yumi Kasai, David Gray, Carol Clugston, Anna Dominiczak and Alex Alderton, Roberto Amato, Jeffrey Barrett, Sonia Goncalves, Ewan Harrison, David K. Jackson, Ian Johnston, Dominic Kwiatkowski, Cordelia Langford, John Sillitoe on behalf of the Wellcome Sanger Institute COVID-19 Surveillance Team |
| EPI_ISL_2935185, EPI_ISL_2935186 | Berkshire and Surrey Pathology Services Lighthouse Laboratory | Wellcome Sanger Institute for the COVID-19 Genomics UK (COG-UK) Consortium | Berkshire and Surrey Pathology Services Lighthouse Laboratory and Alex Alderton, Roberto Amato, Jeffrey Barrett, Sonia Goncalves, Ewan Harrison, David K. Jackson, Ian Johnston, Dominic Kwiatkowski, Cordelia Langford, John Sillitoe on behalf of the Wellcome Sanger Institute COVID-19 Surveillance Team |
| EPI_ISL_2935187, EPI_ISL_2935188 | Lighthouse Lab in Glasgow | Wellcome Sanger Institute for the COVID-19 Genomics UK (COG-UK) Consortium | Harper VanSteenhouse, Yumi Kasai, David Gray, Carol Clugston, Anna Dominiczak and Alex Alderton, Roberto Amato, Jeffrey Barrett, Sonia Goncalves, Ewan Harrison, David K. Jackson, Ian Johnston, Dominic Kwiatkowski, Cordelia Langford, John Sillitoe on behalf of the Wellcome Sanger Institute COVID-19 Surveillance Team |
| EPI_ISL_2935189 | Health Services Laboratories | Wellcome Sanger Institute for the COVID-19 Genomics UK (COG-UK) Consortium | Health Services Laboratories and Alex Alderton, Roberto Amato, Jeffrey Barrett, Sonia Goncalves, Ewan Harrison, David K. Jackson, Ian Johnston, Dominic Kwiatkowski, Cordelia Langford, John Sillitoe on behalf of the Wellcome Sanger Institute COVID-19 Surveillance Team |
| EPI_ISL_2935190 | Lighthouse Lab in Glasgow | Wellcome Sanger Institute for the COVID-19 Genomics UK (COG-UK) Consortium | Harper VanSteenhouse, Yumi Kasai, David Gray, Carol Clugston, Anna Dominiczak and Alex Alderton, Roberto Amato, Jeffrey Barrett, Sonia Goncalves, Ewan Harrison, David K. Jackson, Ian Johnston, Dominic Kwiatkowski, Cordelia Langford, John Sillitoe on behalf of the Wellcome Sanger Institute COVID-19 Surveillance Team |
| EPI_ISL_2935191 | Lighthouse Laboratory Plymouth | Wellcome Sanger Institute for the COVID-19 Genomics UK (COG-UK) Consortium | Lighthouse Laboratory Plymouth and Alex Alderton, Roberto Amato, Jeffrey Barrett, Sonia Goncalves, Ewan Harrison, David K. Jackson, Ian Johnston, Dominic Kwiatkowski, Cordelia Langford, John Sillitoe on behalf of the Wellcome Sanger Institute COVID-19 Surveillance Team |
| EPI_ISL_2935192 | Lighthouse Lab in Milton Keynes | Wellcome Sanger Institute for the COVID-19 Genomics UK (COG-UK) Consortium | The Lighthouse Lab in Milton Keynes and Alex Alderton, Roberto Amato, Jeffrey Barrett, Sonia Goncalves, Ewan Harrison, David K. Jackson, Ian Johnston, Dominic Kwiatkowski, Cordelia Langford, John Sillitoe on behalf of the Wellcome Sanger Institute COVID-19 Surveillance Team |
| EPI_ISL_2935193, EPI_ISL_2935194 | Lighthouse Lab in Glasgow | Wellcome Sanger Institute for the COVID-19 Genomics UK (COG-UK) Consortium | Harper VanSteenhouse, Yumi Kasai, David Gray, Carol Clugston, Anna Dominiczak and Alex Alderton, Roberto Amato, Jeffrey Barrett, Sonia Goncalves, Ewan Harrison, David K. Jackson, Ian Johnston, Dominic Kwiatkowski, Cordelia Langford, John Sillitoe on behalf of the Wellcome Sanger Institute COVID-19 Surveillance Team |
| EPI_ISL_2935195 | Lighthouse Lab in Milton Keynes | Wellcome Sanger Institute for the COVID-19 Genomics UK (COG-UK) Consortium | The Lighthouse Lab in Milton Keynes and Alex Alderton, Roberto Amato, Jeffrey Barrett, Sonia Goncalves, Ewan Harrison, David K. Jackson, Ian Johnston, Dominic Kwiatkowski, Cordelia Langford, John Sillitoe on behalf of the Wellcome Sanger Institute COVID-19 Surveillance Team |
| EPI_ISL_2935196 | Lighthouse Lab in Glasgow | Wellcome Sanger Institute for the COVID-19 Genomics UK (COG-UK) Consortium | Harper VanSteenhouse, Yumi Kasai, David Gray, Carol Clugston, Anna Dominiczak and Alex Alderton, Roberto Amato, Jeffrey Barrett, Sonia Goncalves, Ewan Harrison, David K. Jackson, Ian Johnston, Dominic Kwiatkowski, Cordelia Langford, John Sillitoe on behalf of the Wellcome Sanger Institute COVID-19 Surveillance Team |
| EPI_ISL_2935197 | Berkshire and Surrey Pathology Services Lighthouse Laboratory | Wellcome Sanger Institute for the COVID-19 Genomics UK (COG-UK) Consortium | Berkshire and Surrey Pathology Services Lighthouse Laboratory and Alex Alderton, Roberto Amato, Jeffrey Barrett, Sonia Goncalves, Ewan Harrison, David K. Jackson, Ian Johnston, Dominic Kwiatkowski, Cordelia Langford, John Sillitoe on behalf of the Wellcome Sanger Institute COVID-19 Surveillance Team |
| EPI_ISL_2935198 | Lighthouse Laboratory Plymouth | Wellcome Sanger Institute for the COVID-19 Genomics UK (COG-UK) Consortium | Lighthouse Laboratory Plymouth and Alex Alderton, Roberto Amato, Jeffrey Barrett, Sonia Goncalves, Ewan Harrison, David K. Jackson, Ian Johnston, Dominic Kwiatkowski, Cordelia Langford, John Sillitoe on behalf of the Wellcome Sanger Institute COVID-19 Surveillance Team |
| EPI_ISL_2935199, EPI_ISL_2935200 | Lighthouse Lab in Milton Keynes | Wellcome Sanger Institute for the COVID-19 Genomics UK (COG-UK) Consortium | The Lighthouse Lab in Milton Keynes and Alex Alderton, Roberto Amato, Jeffrey Barrett, Sonia Goncalves, Ewan Harrison, David K. Jackson, Ian Johnston, Dominic Kwiatkowski, Cordelia Langford, John Sillitoe on behalf of the Wellcome Sanger Institute COVID-19 Surveillance Team |
| EPI_ISL_2935201 | Berkshire and Surrey Pathology Services Lighthouse Laboratory | Wellcome Sanger Institute for the COVID-19 Genomics UK (COG-UK) Consortium | Berkshire and Surrey Pathology Services Lighthouse Laboratory and Alex Alderton, Roberto Amato, Jeffrey Barrett, Sonia Goncalves, Ewan Harrison, David K. Jackson, Ian Johnston, Dominic Kwiatkowski, Cordelia Langford, John Sillitoe on behalf of the Wellcome Sanger Institute COVID-19 Surveillance Team |
| EPI_ISL_2935202 | Lighthouse Lab in Glasgow | Wellcome Sanger Institute for the COVID-19 Genomics UK (COG-UK) Consortium | Harper VanSteenhouse, Yumi Kasai, David Gray, Carol Clugston, Anna Dominiczak and Alex Alderton, Roberto Amato, Jeffrey Barrett, Sonia Goncalves, Ewan Harrison, David K. Jackson, Ian Johnston, Dominic Kwiatkowski, Cordelia Langford, John Sillitoe on behalf of the Wellcome Sanger Institute COVID-19 Surveillance Team |
| EPI_ISL_2935203 | Berkshire and Surrey Pathology Services Lighthouse Laboratory | Wellcome Sanger Institute for the COVID-19 Genomics UK (COG-UK) Consortium | Berkshire and Surrey Pathology Services Lighthouse Laboratory and Alex Alderton, Roberto Amato, Jeffrey Barrett, Sonia Goncalves, Ewan Harrison, David K. Jackson, Ian Johnston, Dominic Kwiatkowski, Cordelia Langford, John Sillitoe on behalf of the Wellcome Sanger Institute COVID-19 Surveillance Team |
| EPI_ISL_2935204, EPI_ISL_2935205 | Lighthouse Lab in Glasgow | Wellcome Sanger Institute for the COVID-19 Genomics UK (COG-UK) Consortium | Harper VanSteenhouse, Yumi Kasai, David Gray, Carol Clugston, Anna Dominiczak and Alex Alderton, Roberto Amato, Jeffrey Barrett, Sonia Goncalves, Ewan Harrison, David K. Jackson, Ian Johnston, Dominic Kwiatkowski, Cordelia Langford, John Sillitoe on behalf of the Wellcome Sanger Institute COVID-19 Surveillance Team |
| EPI_ISL_2935206 | Berkshire and Surrey Pathology Services Lighthouse Laboratory | Wellcome Sanger Institute for the COVID-19 Genomics UK (COG-UK) Consortium | Berkshire and Surrey Pathology Services Lighthouse Laboratory and Alex Alderton, Roberto Amato, Jeffrey Barrett, Sonia Goncalves, Ewan Harrison, David K. Jackson, Ian Johnston, Dominic Kwiatkowski, Cordelia Langford, John Sillitoe on behalf of the Wellcome Sanger Institute COVID-19 Surveillance Team |
| EPI_ISL_2935207 | Lighthouse Lab in Milton Keynes | Wellcome Sanger Institute for the COVID-19 Genomics UK (COG-UK) Consortium | The Lighthouse Lab in Milton Keynes and Alex Alderton, Roberto Amato, Jeffrey Barrett, Sonia Goncalves, Ewan Harrison, David K. Jackson, Ian Johnston, Dominic Kwiatkowski, Cordelia Langford, John Sillitoe on behalf of the Wellcome Sanger Institute COVID-19 Surveillance Team |
| EPI_ISL_2935208 | Health Services Laboratories | Wellcome Sanger Institute for the COVID-19 Genomics UK (COG-UK) Consortium | Health Services Laboratories and Alex Alderton, Roberto Amato, Jeffrey Barrett, Sonia Goncalves, Ewan Harrison, David K. Jackson, Ian Johnston, Dominic Kwiatkowski, Cordelia Langford, John Sillitoe on behalf of the Wellcome Sanger Institute COVID-19 Surveillance Team |
| EPI_ISL_2935209, EPI_ISL_2935210 | Berkshire and Surrey Pathology Services Lighthouse Laboratory | Wellcome Sanger Institute for the COVID-19 Genomics UK (COG-UK) Consortium | Berkshire and Surrey Pathology Services Lighthouse Laboratory and Alex Alderton, Roberto Amato, Jeffrey Barrett, Sonia Goncalves, Ewan Harrison, David K. Jackson, Ian Johnston, Dominic Kwiatkowski, Cordelia Langford, John Sillitoe on behalf of the Wellcome Sanger Institute COVID-19 Surveillance Team |
| EPI_ISL_2935211, EPI_ISL_2935212 | Lighthouse Lab in Glasgow | Wellcome Sanger Institute for the COVID-19 Genomics UK (COG-UK) Consortium | Harper VanSteenhouse, Yumi Kasai, David Gray, Carol Clugston, Anna Dominiczak and Alex Alderton, Roberto Amato, Jeffrey Barrett, Sonia Goncalves, Ewan Harrison, David K. Jackson, Ian Johnston, Dominic Kwiatkowski, Cordelia Langford, John Sillitoe on behalf of the Wellcome Sanger Institute COVID-19 Surveillance Team |
| EPI_ISL_2935213, EPI_ISL_2935214 | Lighthouse Lab in Milton Keynes | Wellcome Sanger Institute for the COVID-19 Genomics UK (COG-UK) Consortium | The Lighthouse Lab in Milton Keynes and Alex Alderton, Roberto Amato, Jeffrey Barrett, Sonia Goncalves, Ewan Harrison, David K. Jackson, Ian Johnston, Dominic Kwiatkowski, Cordelia Langford, John Sillitoe on behalf of the Wellcome Sanger Institute COVID-19 Surveillance Team |
| EPI_ISL_2935215 | Lighthouse Lab in Glasgow | Wellcome Sanger Institute for the COVID-19 Genomics UK (COG-UK) Consortium | Harper VanSteenhouse, Yumi Kasai, David Gray, Carol Clugston, Anna Dominiczak and Alex Alderton, Roberto Amato, Jeffrey Barrett, Sonia Goncalves, Ewan Harrison, David K. Jackson, Ian Johnston, Dominic Kwiatkowski, Cordelia Langford, John Sillitoe on behalf of the Wellcome Sanger Institute COVID-19 Surveillance Team |

| | | | |
|---|---|---|---|
| EPI_ISL_2935216 | Berkshire and Surrey Pathology Services Lighthouse Laboratory | Wellcome Sanger Institute for the COVID-19 Genomics UK (COG-UK) Consortium | Berkshire and Surrey Pathology Services Lighthouse Laboratory and Alex Alderton, Roberto Amato, Jeffrey Barrett, Sonia Goncalves, Ewan Harrison, David K. Jackson, Ian Johnston, Dominic Kwiatkowski, Cordelia Langford, John Sillitoe on behalf of the Wellcome Sanger Institute COVID-19 Surveillance Team |
| EPI_ISL_2935217 | Lighthouse Lab in Milton Keynes | Wellcome Sanger Institute for the COVID-19 Genomics UK (COG-UK) Consortium | The Lighthouse Lab in Milton Keynes and Alex Alderton, Roberto Amato, Jeffrey Barrett, Sonia Goncalves, Ewan Harrison, David K. Jackson, Ian Johnston, Dominic Kwiatkowski, Cordelia Langford, John Sillitoe on behalf of the Wellcome Sanger Institute COVID-19 Surveillance Team |
| EPI_ISL_2935218, EPI_ISL_2935219 | Lighthouse Laboratory Plymouth | Wellcome Sanger Institute for the COVID-19 Genomics UK (COG-UK) Consortium | Lighthouse Laboratory Plymouth and Alex Alderton, Roberto Amato, Jeffrey Barrett, Sonia Goncalves, Ewan Harrison, David K. Jackson, Ian Johnston, Dominic Kwiatkowski, Cordelia Langford, John Sillitoe on behalf of the Wellcome Sanger Institute COVID-19 Surveillance Team |
| EPI_ISL_2935220 | Lighthouse Lab in Glasgow | Wellcome Sanger Institute for the COVID-19 Genomics UK (COG-UK) Consortium | Harper VanSteenhouse, Yumi Kasai, David Gray, Carol Clugston, Anna Dominiczak and Alex Alderton, Roberto Amato, Jeffrey Barrett, Sonia Goncalves, Ewan Harrison, David K. Jackson, Ian Johnston, Dominic Kwiatkowski, Cordelia Langford, John Sillitoe on behalf of the Wellcome Sanger Institute COVID-19 Surveillance Team |
| EPI_ISL_2935221, EPI_ISL_2935222 | Lighthouse Laboratory Plymouth | Wellcome Sanger Institute for the COVID-19 Genomics UK (COG-UK) Consortium | Lighthouse Laboratory Plymouth and Alex Alderton, Roberto Amato, Jeffrey Barrett, Sonia Goncalves, Ewan Harrison, David K. Jackson, Ian Johnston, Dominic Kwiatkowski, Cordelia Langford, John Sillitoe on behalf of the Wellcome Sanger Institute COVID-19 Surveillance Team |
| EPI_ISL_2935223 | Lighthouse Lab in Glasgow | Wellcome Sanger Institute for the COVID-19 Genomics UK (COG-UK) Consortium | Harper VanSteenhouse, Yumi Kasai, David Gray, Carol Clugston, Anna Dominiczak and Alex Alderton, Roberto Amato, Jeffrey Barrett, Sonia Goncalves, Ewan Harrison, David K. Jackson, Ian Johnston, Dominic Kwiatkowski, Cordelia Langford, John Sillitoe on behalf of the Wellcome Sanger Institute COVID-19 Surveillance Team |
| EPI_ISL_2935224 | Lighthouse Laboratory Plymouth | Wellcome Sanger Institute for the COVID-19 Genomics UK (COG-UK) Consortium | Lighthouse Laboratory Plymouth and Alex Alderton, Roberto Amato, Jeffrey Barrett, Sonia Goncalves, Ewan Harrison, David K. Jackson, Ian Johnston, Dominic Kwiatkowski, Cordelia Langford, John Sillitoe on behalf of the Wellcome Sanger Institute COVID-19 Surveillance Team |
| EPI_ISL_2935225 | Lighthouse Lab in Glasgow | Wellcome Sanger Institute for the COVID-19 Genomics UK (COG-UK) Consortium | Harper VanSteenhouse, Yumi Kasai, David Gray, Carol Clugston, Anna Dominiczak and Alex Alderton, Roberto Amato, Jeffrey Barrett, Sonia Goncalves, Ewan Harrison, David K. Jackson, Ian Johnston, Dominic Kwiatkowski, Cordelia Langford, John Sillitoe on behalf of the Wellcome Sanger Institute COVID-19 Surveillance Team |
| EPI_ISL_2935226, EPI_ISL_2935227 | Berkshire and Surrey Pathology Services Lighthouse Laboratory | Wellcome Sanger Institute for the COVID-19 Genomics UK (COG-UK) Consortium | Berkshire and Surrey Pathology Services Lighthouse Laboratory and Alex Alderton, Roberto Amato, Jeffrey Barrett, Sonia Goncalves, Ewan Harrison, David K. Jackson, Ian Johnston, Dominic Kwiatkowski, Cordelia Langford, John Sillitoe on behalf of the Wellcome Sanger Institute COVID-19 Surveillance Team |
| EPI_ISL_2935228 | Lighthouse Lab in Glasgow | Wellcome Sanger Institute for the COVID-19 Genomics UK (COG-UK) Consortium | Harper VanSteenhouse, Yumi Kasai, David Gray, Carol Clugston, Anna Dominiczak and Alex Alderton, Roberto Amato, Jeffrey Barrett, Sonia Goncalves, Ewan Harrison, David K. Jackson, Ian Johnston, Dominic Kwiatkowski, Cordelia Langford, John Sillitoe on behalf of the Wellcome Sanger Institute COVID-19 Surveillance Team |
| EPI_ISL_2935229 | Berkshire and Surrey Pathology Services Lighthouse Laboratory | Wellcome Sanger Institute for the COVID-19 Genomics UK (COG-UK) Consortium | Berkshire and Surrey Pathology Services Lighthouse Laboratory and Alex Alderton, Roberto Amato, Jeffrey Barrett, Sonia Goncalves, Ewan Harrison, David K. Jackson, Ian Johnston, Dominic Kwiatkowski, Cordelia Langford, John Sillitoe on behalf of the Wellcome Sanger Institute COVID-19 Surveillance Team |
| EPI_ISL_2935230 | Lighthouse Lab in Glasgow | Wellcome Sanger Institute for the COVID-19 Genomics UK (COG-UK) Consortium | Harper VanSteenhouse, Yumi Kasai, David Gray, Carol Clugston, Anna Dominiczak and Alex Alderton, Roberto Amato, Jeffrey Barrett, Sonia Goncalves, Ewan Harrison, David K. Jackson, Ian Johnston, Dominic Kwiatkowski, Cordelia Langford, John Sillitoe on behalf of the Wellcome Sanger Institute COVID-19 Surveillance Team |
| EPI_ISL_2935231 | Berkshire and Surrey Pathology Services Lighthouse Laboratory | Wellcome Sanger Institute for the COVID-19 Genomics UK (COG-UK) Consortium | Berkshire and Surrey Pathology Services Lighthouse Laboratory and Alex Alderton, Roberto Amato, Jeffrey Barrett, Sonia Goncalves, Ewan Harrison, David K. Jackson, Ian Johnston, Dominic Kwiatkowski, Cordelia Langford, John Sillitoe on behalf of the Wellcome Sanger Institute COVID-19 Surveillance Team |
| EPI_ISL_2935232 | Lighthouse Lab in Milton Keynes | Wellcome Sanger Institute for the COVID-19 Genomics UK (COG-UK) Consortium | The Lighthouse Lab in Milton Keynes and Alex Alderton, Roberto Amato, Jeffrey Barrett, Sonia Goncalves, Ewan Harrison, David K. Jackson, Ian Johnston, Dominic Kwiatkowski, Cordelia Langford, John Sillitoe on behalf of the Wellcome Sanger Institute COVID-19 Surveillance Team |
| EPI_ISL_2935233 | Lighthouse Lab in Glasgow | Wellcome Sanger Institute for the COVID-19 Genomics UK (COG-UK) Consortium | Harper VanSteenhouse, Yumi Kasai, David Gray, Carol Clugston, Anna Dominiczak and Alex Alderton, Roberto Amato, Jeffrey Barrett, Sonia Goncalves, Ewan Harrison, David K. Jackson, Ian Johnston, Dominic Kwiatkowski, Cordelia Langford, John Sillitoe on behalf of the Wellcome Sanger Institute COVID-19 Surveillance Team |
| EPI_ISL_2935234 | Lighthouse Lab in Milton Keynes | Wellcome Sanger Institute for the COVID-19 Genomics UK (COG-UK) Consortium | The Lighthouse Lab in Milton Keynes and Alex Alderton, Roberto Amato, Jeffrey Barrett, Sonia Goncalves, Ewan Harrison, David K. Jackson, Ian Johnston, Dominic Kwiatkowski, Cordelia Langford, John Sillitoe on behalf of the Wellcome Sanger Institute COVID-19 Surveillance Team |
| EPI_ISL_2935235 | Lighthouse Lab in Glasgow | Wellcome Sanger Institute for the COVID-19 Genomics UK (COG-UK) Consortium | Harper VanSteenhouse, Yumi Kasai, David Gray, Carol Clugston, Anna Dominiczak and Alex Alderton, Roberto Amato, Jeffrey Barrett, Sonia Goncalves, Ewan Harrison, David K. Jackson, Ian Johnston, Dominic Kwiatkowski, Cordelia Langford, John Sillitoe on behalf of the Wellcome Sanger Institute COVID-19 Surveillance Team |
| EPI_ISL_2935236 | Lighthouse Lab in Milton Keynes | Wellcome Sanger Institute for the COVID-19 Genomics UK (COG-UK) Consortium | The Lighthouse Lab in Milton Keynes and Alex Alderton, Roberto Amato, Jeffrey Barrett, Sonia Goncalves, Ewan Harrison, David K. Jackson, Ian Johnston, Dominic Kwiatkowski, Cordelia Langford, John Sillitoe on behalf of the Wellcome Sanger Institute COVID-19 Surveillance Team |
| EPI_ISL_2935237 | Health Services Laboratories | Wellcome Sanger Institute for the COVID-19 Genomics UK (COG-UK) Consortium | Health Services Laboratories and Alex Alderton, Roberto Amato, Jeffrey Barrett, Sonia Goncalves, Ewan Harrison, David K. Jackson, Ian Johnston, Dominic Kwiatkowski, Cordelia Langford, John Sillitoe on behalf of the Wellcome Sanger Institute COVID-19 Surveillance Team |
| EPI_ISL_2935238 | Berkshire and Surrey Pathology Services Lighthouse Laboratory | Wellcome Sanger Institute for the COVID-19 Genomics UK (COG-UK) Consortium | Berkshire and Surrey Pathology Services Lighthouse Laboratory and Alex Alderton, Roberto Amato, Jeffrey Barrett, Sonia Goncalves, Ewan Harrison, David K. Jackson, Ian Johnston, Dominic Kwiatkowski, Cordelia Langford, John Sillitoe on behalf of the Wellcome Sanger Institute COVID-19 Surveillance Team |
| EPI_ISL_2935239, EPI_ISL_2935240, EPI_ISL_2935241 | Lighthouse Laboratory Plymouth | Wellcome Sanger Institute for the COVID-19 Genomics UK (COG-UK) Consortium | Lighthouse Laboratory Plymouth and Alex Alderton, Roberto Amato, Jeffrey Barrett, Sonia Goncalves, Ewan Harrison, David K. Jackson, Ian Johnston, Dominic Kwiatkowski, Cordelia Langford, John Sillitoe on behalf of the Wellcome Sanger Institute COVID-19 Surveillance Team |
| EPI_ISL_2935242, EPI_ISL_2935243 | Lighthouse Lab in Glasgow | Wellcome Sanger Institute for the COVID-19 Genomics UK (COG-UK) Consortium | Harper VanSteenhouse, Yumi Kasai, David Gray, Carol Clugston, Anna Dominiczak and Alex Alderton, Roberto Amato, Jeffrey Barrett, Sonia Goncalves, Ewan Harrison, David K. Jackson, Ian Johnston, Dominic Kwiatkowski, Cordelia Langford, John Sillitoe on behalf of the Wellcome Sanger Institute COVID-19 Surveillance Team |
| EPI_ISL_2935244 | Lighthouse Lab in Milton Keynes | Wellcome Sanger Institute for the COVID-19 Genomics UK (COG-UK) Consortium | The Lighthouse Lab in Milton Keynes and Alex Alderton, Roberto Amato, Jeffrey Barrett, Sonia Goncalves, Ewan Harrison, David K. Jackson, Ian Johnston, Dominic Kwiatkowski, Cordelia Langford, John Sillitoe on behalf of the Wellcome Sanger Institute COVID-19 Surveillance Team |
| EPI_ISL_2935245 | Berkshire and Surrey Pathology Services Lighthouse Laboratory | Wellcome Sanger Institute for the COVID-19 Genomics UK (COG-UK) Consortium | Berkshire and Surrey Pathology Services Lighthouse Laboratory and Alex Alderton, Roberto Amato, Jeffrey Barrett, Sonia Goncalves, Ewan Harrison, David K. Jackson, Ian Johnston, Dominic Kwiatkowski, Cordelia Langford, John Sillitoe on behalf of the Wellcome Sanger Institute COVID-19 Surveillance Team |
| EPI_ISL_2935246 | Lighthouse Lab in Glasgow | Wellcome Sanger Institute for the COVID-19 Genomics UK (COG-UK) Consortium | Harper VanSteenhouse, Yumi Kasai, David Gray, Carol Clugston, Anna Dominiczak and Alex Alderton, Roberto Amato, Jeffrey Barrett, Sonia Goncalves, Ewan Harrison, David K. Jackson, Ian Johnston, Dominic Kwiatkowski, Cordelia Langford, John Sillitoe on behalf of the Wellcome Sanger Institute COVID-19 Surveillance Team |
| EPI_ISL_2935247 | Berkshire and Surrey Pathology Services Lighthouse Laboratory | Wellcome Sanger Institute for the COVID-19 Genomics UK (COG-UK) Consortium | Berkshire and Surrey Pathology Services Lighthouse Laboratory and Alex Alderton, Roberto Amato, Jeffrey Barrett, Sonia Goncalves, Ewan Harrison, David K. Jackson, Ian Johnston, Dominic Kwiatkowski, Cordelia Langford, John Sillitoe on behalf of the Wellcome Sanger Institute COVID-19 Surveillance Team |
| EPI_ISL_2935248, EPI_ISL_2935249 | Lighthouse Laboratory Plymouth | Wellcome Sanger Institute for the COVID-19 Genomics UK (COG-UK) Consortium | Lighthouse Laboratory Plymouth and Alex Alderton, Roberto Amato, Jeffrey Barrett, Sonia Goncalves, Ewan Harrison, David K. Jackson, Ian Johnston, Dominic Kwiatkowski, Cordelia Langford, John Sillitoe on behalf of the Wellcome Sanger Institute COVID-19 Surveillance Team |
| EPI_ISL_2935250 | Lighthouse Lab in Glasgow | Wellcome Sanger Institute for the COVID-19 Genomics UK (COG-UK) Consortium | Harper VanSteenhouse, Yumi Kasai, David Gray, Carol Clugston, Anna Dominiczak and Alex Alderton, Roberto Amato, Jeffrey Barrett, Sonia Goncalves, Ewan Harrison, David K. Jackson, Ian Johnston, Dominic Kwiatkowski, Cordelia Langford, John Sillitoe on behalf of the Wellcome Sanger Institute COVID-19 Surveillance Team |
| EPI_ISL_2935251 | Lighthouse Laboratory Plymouth | Wellcome Sanger Institute for the COVID-19 Genomics UK (COG-UK) Consortium | Lighthouse Laboratory Plymouth and Alex Alderton, Roberto Amato, Jeffrey Barrett, Sonia Goncalves, Ewan Harrison, David K. Jackson, Ian Johnston, Dominic Kwiatkowski, Cordelia Langford, John Sillitoe on behalf of the Wellcome Sanger Institute COVID-19 Surveillance Team |
| EPI_ISL_2935252, EPI_ISL_2935253, EPI_ISL_2935254 | Berkshire and Surrey Pathology Services Lighthouse Laboratory | Wellcome Sanger Institute for the COVID-19 Genomics UK (COG-UK) Consortium | Berkshire and Surrey Pathology Services Lighthouse Laboratory and Alex Alderton, Roberto Amato, Jeffrey Barrett, Sonia Goncalves, Ewan Harrison, David K. Jackson, Ian Johnston, Dominic Kwiatkowski, Cordelia Langford, John Sillitoe on behalf of the Wellcome Sanger Institute COVID-19 Surveillance Team |
| EPI_ISL_2935255, EPI_ISL_2935256 | Lighthouse Lab in Glasgow | Wellcome Sanger Institute for the COVID-19 Genomics UK (COG-UK) Consortium | Harper VanSteenhouse, Yumi Kasai, David Gray, Carol Clugston, Anna Dominiczak and Alex Alderton, Roberto Amato, Jeffrey Barrett, Sonia Goncalves, Ewan Harrison, David K. Jackson, Ian Johnston, Dominic Kwiatkowski, Cordelia Langford, John Sillitoe on behalf of the Wellcome Sanger Institute COVID-19 Surveillance Team |
| EPI_ISL_2935257 | Lighthouse Lab in Milton Keynes | Wellcome Sanger Institute for the COVID-19 Genomics UK (COG-UK) Consortium | The Lighthouse Lab in Milton Keynes and Alex Alderton, Roberto Amato, Jeffrey Barrett, Sonia Goncalves, Ewan Harrison, David K. Jackson, Ian Johnston, Dominic Kwiatkowski, Cordelia Langford, John Sillitoe on behalf of the Wellcome Sanger Institute COVID-19 Surveillance Team |
| EPI_ISL_2935258, EPI_ISL_2935259 | Lighthouse Laboratory Plymouth | Wellcome Sanger Institute for the COVID-19 Genomics UK (COG-UK) Consortium | Lighthouse Laboratory Plymouth and Alex Alderton, Roberto Amato, Jeffrey Barrett, Sonia Goncalves, Ewan Harrison, David K. Jackson, Ian Johnston, Dominic Kwiatkowski, Cordelia Langford, John Sillitoe on behalf of the Wellcome Sanger Institute COVID-19 Surveillance Team |
| EPI_ISL_2935260 | Lighthouse Lab in Milton Keynes | Wellcome Sanger Institute for the COVID-19 Genomics UK (COG-UK) Consortium | The Lighthouse Lab in Milton Keynes and Alex Alderton, Roberto Amato, Jeffrey Barrett, Sonia Goncalves, Ewan Harrison, David K. Jackson, Ian Johnston, Dominic Kwiatkowski, Cordelia Langford, John Sillitoe on behalf of the Wellcome Sanger Institute COVID-19 Surveillance Team |
| EPI_ISL_2935261 | Lighthouse Laboratory Plymouth | Wellcome Sanger Institute for the COVID-19 Genomics UK (COG-UK) Consortium | Lighthouse Laboratory Plymouth and Alex Alderton, Roberto Amato, Jeffrey Barrett, Sonia Goncalves, Ewan Harrison, David K. Jackson, Ian Johnston, Dominic Kwiatkowski, Cordelia Langford, John Sillitoe on behalf of the Wellcome Sanger Institute COVID-19 Surveillance Team |
| EPI_ISL_2935262 | Lighthouse Lab in Glasgow | Wellcome Sanger Institute for the COVID-19 Genomics UK (COG-UK) Consortium | Harper VanSteenhouse, Yumi Kasai, David Gray, Carol Clugston, Anna Dominiczak and Alex Alderton, Roberto Amato, Jeffrey Barrett, Sonia Goncalves, Ewan Harrison, David K. Jackson, Ian Johnston, Dominic Kwiatkowski, Cordelia Langford, John Sillitoe on behalf of the Wellcome Sanger Institute COVID-19 Surveillance Team |
| EPI_ISL_2935264 | Berkshire and Surrey Pathology Services Lighthouse Laboratory | Wellcome Sanger Institute for the COVID-19 Genomics UK (COG-UK) Consortium | Berkshire and Surrey Pathology Services Lighthouse Laboratory and Alex Alderton, Roberto Amato, Jeffrey Barrett, Sonia Goncalves, Ewan Harrison, David K. Jackson, Ian Johnston, Dominic Kwiatkowski, Cordelia Langford, John Sillitoe on behalf of the Wellcome Sanger Institute COVID-19 Surveillance Team |
| EPI_ISL_2935265 | Lighthouse Lab in Glasgow | Wellcome Sanger Institute for the COVID-19 Genomics UK (COG-UK) Consortium | Harper VanSteenhouse, Yumi Kasai, David Gray, Carol Clugston, Anna Dominiczak and Alex Alderton, Roberto Amato, Jeffrey Barrett, Sonia Goncalves, Ewan Harrison, David K. Jackson, Ian Johnston, Dominic Kwiatkowski, Cordelia Langford, John Sillitoe on behalf of the Wellcome Sanger Institute COVID-19 Surveillance Team |
| EPI_ISL_2935266 | Lighthouse Laboratory Plymouth | Wellcome Sanger Institute for the COVID-19 Genomics UK (COG-UK) Consortium | Lighthouse Laboratory Plymouth and Alex Alderton, Roberto Amato, Jeffrey Barrett, Sonia Goncalves, Ewan Harrison, David K. Jackson, Ian Johnston, Dominic Kwiatkowski, Cordelia Langford, John Sillitoe on behalf of the Wellcome Sanger Institute COVID-19 Surveillance Team |
| EPI_ISL_2935267 | Lighthouse Lab in Glasgow | Wellcome Sanger Institute for the COVID-19 Genomics UK (COG-UK) Consortium | Harper VanSteenhouse, Yumi Kasai, David Gray, Carol Clugston, Anna Dominiczak and Alex Alderton, Roberto Amato, Jeffrey Barrett, Sonia Goncalves, Ewan Harrison, David K. Jackson, Ian Johnston, Dominic Kwiatkowski, Cordelia Langford, John Sillitoe on behalf of the Wellcome Sanger Institute COVID-19 Surveillance Team |
| EPI_ISL_2935268 | Lighthouse Laboratory Plymouth | Wellcome Sanger Institute for the COVID-19 Genomics UK (COG-UK) Consortium | Lighthouse Laboratory Plymouth and Alex Alderton, Roberto Amato, Jeffrey Barrett, Sonia Goncalves, Ewan Harrison, David K. Jackson, Ian Johnston, Dominic Kwiatkowski, Cordelia Langford, John Sillitoe on behalf of the Wellcome Sanger Institute COVID-19 Surveillance Team |
| EPI_ISL_2935269, EPI_ISL_2935270, EPI_ISL_2935271, EPI_ISL_2935273 | Berkshire and Surrey Pathology Services Lighthouse Laboratory | Wellcome Sanger Institute for the COVID-19 Genomics UK (COG-UK) Consortium | Berkshire and Surrey Pathology Services Lighthouse Laboratory and Alex Alderton, Roberto Amato, Jeffrey Barrett, Sonia Goncalves, Ewan Harrison, David K. Jackson, Ian Johnston, Dominic Kwiatkowski, Cordelia Langford, John Sillitoe on behalf of the Wellcome Sanger Institute COVID-19 Surveillance Team |
| EPI_ISL_2935274 | Lighthouse Lab in Glasgow | Wellcome Sanger Institute for the COVID-19 Genomics UK (COG-UK) Consortium | Harper VanSteenhouse, Yumi Kasai, David Gray, Carol Clugston, Anna Dominiczak and Alex Alderton, Roberto Amato, Jeffrey Barrett, Sonia Goncalves, Ewan Harrison, David K. Jackson, Ian Johnston, Dominic Kwiatkowski, Cordelia Langford, John Sillitoe on behalf of the Wellcome Sanger Institute COVID-19 Surveillance Team |
| EPI_ISL_2935275 | Lighthouse Lab in Milton Keynes | Wellcome Sanger Institute for the COVID-19 Genomics UK (COG-UK) | The Lighthouse Lab in Milton Keynes and Alex Alderton, Roberto Amato, Jeffrey Barrett, Sonia Goncalves, Ewan Harrison, David K. Jackson, Ian Johnston, Dominic Kwiatkowski, Cordelia Langford, John Sillitoe on behalf of the Wellcome Sanger |

| | | | |
|---|---|---|---|
| | | Consortium | Institute COVID-19 Surveillance Team |
| EPI_ISL_2935276 | Lighthouse Laboratory Plymouth | Wellcome Sanger Institute for the COVID-19 Genomics UK (COG-UK) Consortium | Lighthouse Laboratory Plymouth and Alex Alderton, Roberto Amato, Jeffrey Barrett, Sonia Goncalves, Ewan Harrison, David K. Jackson, Ian Johnston, Dominic Kwiatkowski, Cordelia Langford, John Sillitoe on behalf of the Wellcome Sanger Institute COVID-19 Surveillance Team |
| EPI_ISL_2935277 | Lighthouse Lab in Glasgow | Wellcome Sanger Institute for the COVID-19 Genomics UK (COG-UK) Consortium | Harper VanSteenhouse, Yumi Kasai, David Gray, Carol Clugston, Anna Dominiczak and Alex Alderton, Roberto Amato, Jeffrey Barrett, Sonia Goncalves, Ewan Harrison, David K. Jackson, Ian Johnston, Dominic Kwiatkowski, Cordelia Langford, John Sillitoe on behalf of the Wellcome Sanger Institute COVID-19 Surveillance Team |
| EPI_ISL_2935278 | Lighthouse Lab in Milton Keynes | Wellcome Sanger Institute for the COVID-19 Genomics UK (COG-UK) Consortium | The Lighthouse Lab in Milton Keynes and Alex Alderton, Roberto Amato, Jeffrey Barrett, Sonia Goncalves, Ewan Harrison, David K. Jackson, Ian Johnston, Dominic Kwiatkowski, Cordelia Langford, John Sillitoe on behalf of the Wellcome Sanger Institute COVID-19 Surveillance Team |
| EPI_ISL_2935279 | Lighthouse Laboratory Plymouth | Wellcome Sanger Institute for the COVID-19 Genomics UK (COG-UK) Consortium | Lighthouse Laboratory Plymouth and Alex Alderton, Roberto Amato, Jeffrey Barrett, Sonia Goncalves, Ewan Harrison, David K. Jackson, Ian Johnston, Dominic Kwiatkowski, Cordelia Langford, John Sillitoe on behalf of the Wellcome Sanger Institute COVID-19 Surveillance Team |
| EPI_ISL_2935280 | Lighthouse Lab in Milton Keynes | Wellcome Sanger Institute for the COVID-19 Genomics UK (COG-UK) Consortium | The Lighthouse Lab in Milton Keynes and Alex Alderton, Roberto Amato, Jeffrey Barrett, Sonia Goncalves, Ewan Harrison, David K. Jackson, Ian Johnston, Dominic Kwiatkowski, Cordelia Langford, John Sillitoe on behalf of the Wellcome Sanger Institute COVID-19 Surveillance Team |
| EPI_ISL_2935281, EPI_ISL_2935282, EPI_ISL_2935283 | Lighthouse Lab in Glasgow | Wellcome Sanger Institute for the COVID-19 Genomics UK (COG-UK) Consortium | Harper VanSteenhouse, Yumi Kasai, David Gray, Carol Clugston, Anna Dominiczak and Alex Alderton, Roberto Amato, Jeffrey Barrett, Sonia Goncalves, Ewan Harrison, David K. Jackson, Ian Johnston, Dominic Kwiatkowski, Cordelia Langford, John Sillitoe on behalf of the Wellcome Sanger Institute COVID-19 Surveillance Team |
| EPI_ISL_2935284 | Lighthouse Laboratory Plymouth | Wellcome Sanger Institute for the COVID-19 Genomics UK (COG-UK) Consortium | Lighthouse Laboratory Plymouth and Alex Alderton, Roberto Amato, Jeffrey Barrett, Sonia Goncalves, Ewan Harrison, David K. Jackson, Ian Johnston, Dominic Kwiatkowski, Cordelia Langford, John Sillitoe on behalf of the Wellcome Sanger Institute COVID-19 Surveillance Team |
| EPI_ISL_2935285 | Lighthouse Lab in Glasgow | Wellcome Sanger Institute for the COVID-19 Genomics UK (COG-UK) Consortium | Harper VanSteenhouse, Yumi Kasai, David Gray, Carol Clugston, Anna Dominiczak and Alex Alderton, Roberto Amato, Jeffrey Barrett, Sonia Goncalves, Ewan Harrison, David K. Jackson, Ian Johnston, Dominic Kwiatkowski, Cordelia Langford, John Sillitoe on behalf of the Wellcome Sanger Institute COVID-19 Surveillance Team |
| EPI_ISL_2935286, EPI_ISL_2935287, EPI_ISL_2935288 | Lighthouse Lab in Milton Keynes | Wellcome Sanger Institute for the COVID-19 Genomics UK (COG-UK) Consortium | The Lighthouse Lab in Milton Keynes and Alex Alderton, Roberto Amato, Jeffrey Barrett, Sonia Goncalves, Ewan Harrison, David K. Jackson, Ian Johnston, Dominic Kwiatkowski, Cordelia Langford, John Sillitoe on behalf of the Wellcome Sanger Institute COVID-19 Surveillance Team |
| EPI_ISL_2935289 | Lighthouse Laboratory Plymouth | Wellcome Sanger Institute for the COVID-19 Genomics UK (COG-UK) Consortium | Lighthouse Laboratory Plymouth and Alex Alderton, Roberto Amato, Jeffrey Barrett, Sonia Goncalves, Ewan Harrison, David K. Jackson, Ian Johnston, Dominic Kwiatkowski, Cordelia Langford, John Sillitoe on behalf of the Wellcome Sanger Institute COVID-19 Surveillance Team |
| EPI_ISL_2935290, EPI_ISL_2935291 | Lighthouse Lab in Glasgow | Wellcome Sanger Institute for the COVID-19 Genomics UK (COG-UK) Consortium | Harper VanSteenhouse, Yumi Kasai, David Gray, Carol Clugston, Anna Dominiczak and Alex Alderton, Roberto Amato, Jeffrey Barrett, Sonia Goncalves, Ewan Harrison, David K. Jackson, Ian Johnston, Dominic Kwiatkowski, Cordelia Langford, John Sillitoe on behalf of the Wellcome Sanger Institute COVID-19 Surveillance Team |
| EPI_ISL_2935292, EPI_ISL_2935293 | Lighthouse Laboratory Plymouth | Wellcome Sanger Institute for the COVID-19 Genomics UK (COG-UK) Consortium | Lighthouse Laboratory Plymouth and Alex Alderton, Roberto Amato, Jeffrey Barrett, Sonia Goncalves, Ewan Harrison, David K. Jackson, Ian Johnston, Dominic Kwiatkowski, Cordelia Langford, John Sillitoe on behalf of the Wellcome Sanger Institute COVID-19 Surveillance Team |
| EPI_ISL_2935294, EPI_ISL_2935295 | Berkshire and Surrey Pathology Services Lighthouse Laboratory | Wellcome Sanger Institute for the COVID-19 Genomics UK (COG-UK) Consortium | Berkshire and Surrey Pathology Services Lighthouse Laboratory and Alex Alderton, Roberto Amato, Jeffrey Barrett, Sonia Goncalves, Ewan Harrison, David K. Jackson, Ian Johnston, Dominic Kwiatkowski, Cordelia Langford, John Sillitoe on behalf of the Wellcome Sanger Institute COVID-19 Surveillance Team |
| EPI_ISL_2935296 | Lighthouse Laboratory Plymouth | Wellcome Sanger Institute for the COVID-19 Genomics UK (COG-UK) Consortium | Lighthouse Laboratory Plymouth and Alex Alderton, Roberto Amato, Jeffrey Barrett, Sonia Goncalves, Ewan Harrison, David K. Jackson, Ian Johnston, Dominic Kwiatkowski, Cordelia Langford, John Sillitoe on behalf of the Wellcome Sanger Institute COVID-19 Surveillance Team |
| EPI_ISL_2935297 | Lighthouse Lab in Milton Keynes | Wellcome Sanger Institute for the COVID-19 Genomics UK (COG-UK) Consortium | The Lighthouse Lab in Milton Keynes and Alex Alderton, Roberto Amato, Jeffrey Barrett, Sonia Goncalves, Ewan Harrison, David K. Jackson, Ian Johnston, Dominic Kwiatkowski, Cordelia Langford, John Sillitoe on behalf of the Wellcome Sanger Institute COVID-19 Surveillance Team |
| EPI_ISL_2935298 | Berkshire and Surrey Pathology Services Lighthouse Laboratory | Wellcome Sanger Institute for the COVID-19 Genomics UK (COG-UK) Consortium | Berkshire and Surrey Pathology Services Lighthouse Laboratory and Alex Alderton, Roberto Amato, Jeffrey Barrett, Sonia Goncalves, Ewan Harrison, David K. Jackson, Ian Johnston, Dominic Kwiatkowski, Cordelia Langford, John Sillitoe on behalf of the Wellcome Sanger Institute COVID-19 Surveillance Team |
| EPI_ISL_2935299 | Lighthouse Lab in Milton Keynes | Wellcome Sanger Institute for the COVID-19 Genomics UK (COG-UK) Consortium | The Lighthouse Lab in Milton Keynes and Alex Alderton, Roberto Amato, Jeffrey Barrett, Sonia Goncalves, Ewan Harrison, David K. Jackson, Ian Johnston, Dominic Kwiatkowski, Cordelia Langford, John Sillitoe on behalf of the Wellcome Sanger Institute COVID-19 Surveillance Team |
| EPI_ISL_2935300, EPI_ISL_2935301 | Berkshire and Surrey Pathology Services Lighthouse Laboratory | Wellcome Sanger Institute for the COVID-19 Genomics UK (COG-UK) Consortium | Berkshire and Surrey Pathology Services Lighthouse Laboratory and Alex Alderton, Roberto Amato, Jeffrey Barrett, Sonia Goncalves, Ewan Harrison, David K. Jackson, Ian Johnston, Dominic Kwiatkowski, Cordelia Langford, John Sillitoe on behalf of the Wellcome Sanger Institute COVID-19 Surveillance Team |
| EPI_ISL_2935302, EPI_ISL_2935303 | Lighthouse Lab in Glasgow | Wellcome Sanger Institute for the COVID-19 Genomics UK (COG-UK) Consortium | Harper VanSteenhouse, Yumi Kasai, David Gray, Carol Clugston, Anna Dominiczak and Alex Alderton, Roberto Amato, Jeffrey Barrett, Sonia Goncalves, Ewan Harrison, David K. Jackson, Ian Johnston, Dominic Kwiatkowski, Cordelia Langford, John Sillitoe on behalf of the Wellcome Sanger Institute COVID-19 Surveillance Team |
| EPI_ISL_2935304 | Lighthouse Lab in Milton Keynes | Wellcome Sanger Institute for the COVID-19 Genomics UK (COG-UK) Consortium | The Lighthouse Lab in Milton Keynes and Alex Alderton, Roberto Amato, Jeffrey Barrett, Sonia Goncalves, Ewan Harrison, David K. Jackson, Ian Johnston, Dominic Kwiatkowski, Cordelia Langford, John Sillitoe on behalf of the Wellcome Sanger Institute COVID-19 Surveillance Team |
| EPI_ISL_2935305 | Lighthouse Lab in Glasgow | Wellcome Sanger Institute for the COVID-19 Genomics UK (COG-UK) Consortium | Harper VanSteenhouse, Yumi Kasai, David Gray, Carol Clugston, Anna Dominiczak and Alex Alderton, Roberto Amato, Jeffrey Barrett, Sonia Goncalves, Ewan Harrison, David K. Jackson, Ian Johnston, Dominic Kwiatkowski, Cordelia Langford, John Sillitoe on behalf of the Wellcome Sanger Institute COVID-19 Surveillance Team |
| EPI_ISL_2935306 | Lighthouse Lab in Milton Keynes | Wellcome Sanger Institute for the COVID-19 Genomics UK (COG-UK) Consortium | The Lighthouse Lab in Milton Keynes and Alex Alderton, Roberto Amato, Jeffrey Barrett, Sonia Goncalves, Ewan Harrison, David K. Jackson, Ian Johnston, Dominic Kwiatkowski, Cordelia Langford, John Sillitoe on behalf of the Wellcome Sanger Institute COVID-19 Surveillance Team |
| EPI_ISL_2935307 | Lighthouse Lab in Glasgow | Wellcome Sanger Institute for the COVID-19 Genomics UK (COG-UK) Consortium | Harper VanSteenhouse, Yumi Kasai, David Gray, Carol Clugston, Anna Dominiczak and Alex Alderton, Roberto Amato, Jeffrey Barrett, Sonia Goncalves, Ewan Harrison, David K. Jackson, Ian Johnston, Dominic Kwiatkowski, Cordelia Langford, John Sillitoe on behalf of the Wellcome Sanger Institute COVID-19 Surveillance Team |
| EPI_ISL_2935308 | Lighthouse Lab in Milton Keynes | Wellcome Sanger Institute for the COVID-19 Genomics UK (COG-UK) Consortium | The Lighthouse Lab in Milton Keynes and Alex Alderton, Roberto Amato, Jeffrey Barrett, Sonia Goncalves, Ewan Harrison, David K. Jackson, Ian Johnston, Dominic Kwiatkowski, Cordelia Langford, John Sillitoe on behalf of the Wellcome Sanger Institute COVID-19 Surveillance Team |
| EPI_ISL_2935309 | Berkshire and Surrey Pathology Services Lighthouse Laboratory | Wellcome Sanger Institute for the COVID-19 Genomics UK (COG-UK) Consortium | Berkshire and Surrey Pathology Services Lighthouse Laboratory and Alex Alderton, Roberto Amato, Jeffrey Barrett, Sonia Goncalves, Ewan Harrison, David K. Jackson, Ian Johnston, Dominic Kwiatkowski, Cordelia Langford, John Sillitoe on behalf of the Wellcome Sanger Institute COVID-19 Surveillance Team |
| EPI_ISL_2935310 | Lighthouse Lab in Milton Keynes | Wellcome Sanger Institute for the COVID-19 Genomics UK (COG-UK) Consortium | The Lighthouse Lab in Milton Keynes and Alex Alderton, Roberto Amato, Jeffrey Barrett, Sonia Goncalves, Ewan Harrison, David K. Jackson, Ian Johnston, Dominic Kwiatkowski, Cordelia Langford, John Sillitoe on behalf of the Wellcome Sanger Institute COVID-19 Surveillance Team |
| EPI_ISL_2935311, EPI_ISL_2935312 | Lighthouse Laboratory Plymouth | Wellcome Sanger Institute for the COVID-19 Genomics UK (COG-UK) Consortium | Lighthouse Laboratory Plymouth and Alex Alderton, Roberto Amato, Jeffrey Barrett, Sonia Goncalves, Ewan Harrison, David K. Jackson, Ian Johnston, Dominic Kwiatkowski, Cordelia Langford, John Sillitoe on behalf of the Wellcome Sanger Institute COVID-19 Surveillance Team |
| EPI_ISL_2935313 | Berkshire and Surrey Pathology Services Lighthouse Laboratory | Wellcome Sanger Institute for the COVID-19 Genomics UK (COG-UK) Consortium | Berkshire and Surrey Pathology Services Lighthouse Laboratory and Alex Alderton, Roberto Amato, Jeffrey Barrett, Sonia Goncalves, Ewan Harrison, David K. Jackson, Ian Johnston, Dominic Kwiatkowski, Cordelia Langford, John Sillitoe on behalf of the Wellcome Sanger Institute COVID-19 Surveillance Team |
| EPI_ISL_2935314 | Lighthouse Lab in Glasgow | Wellcome Sanger Institute for the COVID-19 Genomics UK (COG-UK) Consortium | Harper VanSteenhouse, Yumi Kasai, David Gray, Carol Clugston, Anna Dominiczak and Alex Alderton, Roberto Amato, Jeffrey Barrett, Sonia Goncalves, Ewan Harrison, David K. Jackson, Ian Johnston, Dominic Kwiatkowski, Cordelia Langford, John Sillitoe on behalf of the Wellcome Sanger Institute COVID-19 Surveillance Team |
| EPI_ISL_2935315 | Berkshire and Surrey Pathology Services Lighthouse Laboratory | Wellcome Sanger Institute for the COVID-19 Genomics UK (COG-UK) Consortium | Berkshire and Surrey Pathology Services Lighthouse Laboratory and Alex Alderton, Roberto Amato, Jeffrey Barrett, Sonia Goncalves, Ewan Harrison, David K. Jackson, Ian Johnston, Dominic Kwiatkowski, Cordelia Langford, John Sillitoe on behalf of the Wellcome Sanger Institute COVID-19 Surveillance Team |
| EPI_ISL_2935316 | Lighthouse Lab in Milton Keynes | Wellcome Sanger Institute for the COVID-19 Genomics UK (COG-UK) Consortium | The Lighthouse Lab in Milton Keynes and Alex Alderton, Roberto Amato, Jeffrey Barrett, Sonia Goncalves, Ewan Harrison, David K. Jackson, Ian Johnston, Dominic Kwiatkowski, Cordelia Langford, John Sillitoe on behalf of the Wellcome Sanger Institute COVID-19 Surveillance Team |
| EPI_ISL_2935317 | Lighthouse Laboratory Plymouth | Wellcome Sanger Institute for the COVID-19 Genomics UK (COG-UK) Consortium | Lighthouse Laboratory Plymouth and Alex Alderton, Roberto Amato, Jeffrey Barrett, Sonia Goncalves, Ewan Harrison, David K. Jackson, Ian Johnston, Dominic Kwiatkowski, Cordelia Langford, John Sillitoe on behalf of the Wellcome Sanger Institute COVID-19 Surveillance Team |
| EPI_ISL_2935318 | Berkshire and Surrey Pathology Services Lighthouse Laboratory | Wellcome Sanger Institute for the COVID-19 Genomics UK (COG-UK) Consortium | Berkshire and Surrey Pathology Services Lighthouse Laboratory and Alex Alderton, Roberto Amato, Jeffrey Barrett, Sonia Goncalves, Ewan Harrison, David K. Jackson, Ian Johnston, Dominic Kwiatkowski, Cordelia Langford, John Sillitoe on behalf of the Wellcome Sanger Institute COVID-19 Surveillance Team |
| EPI_ISL_2935319 | Lighthouse Laboratory Plymouth | Wellcome Sanger Institute for the COVID-19 Genomics UK (COG-UK) Consortium | Lighthouse Laboratory Plymouth and Alex Alderton, Roberto Amato, Jeffrey Barrett, Sonia Goncalves, Ewan Harrison, David K. Jackson, Ian Johnston, Dominic Kwiatkowski, Cordelia Langford, John Sillitoe on behalf of the Wellcome Sanger Institute COVID-19 Surveillance Team |
| EPI_ISL_2935320 | Lighthouse Lab in Glasgow | Wellcome Sanger Institute for the COVID-19 Genomics UK (COG-UK) Consortium | Harper VanSteenhouse, Yumi Kasai, David Gray, Carol Clugston, Anna Dominiczak and Alex Alderton, Roberto Amato, Jeffrey Barrett, Sonia Goncalves, Ewan Harrison, David K. Jackson, Ian Johnston, Dominic Kwiatkowski, Cordelia Langford, John Sillitoe on behalf of the Wellcome Sanger Institute COVID-19 Surveillance Team |
| EPI_ISL_2935322 | Lighthouse Lab in Milton Keynes | Wellcome Sanger Institute for the COVID-19 Genomics UK (COG-UK) Consortium | The Lighthouse Lab in Milton Keynes and Alex Alderton, Roberto Amato, Jeffrey Barrett, Sonia Goncalves, Ewan Harrison, David K. Jackson, Ian Johnston, Dominic Kwiatkowski, Cordelia Langford, John Sillitoe on behalf of the Wellcome Sanger Institute COVID-19 Surveillance Team |
| EPI_ISL_2935323 | Lighthouse Lab in Glasgow | Wellcome Sanger Institute for the COVID-19 Genomics UK (COG-UK) Consortium | Harper VanSteenhouse, Yumi Kasai, David Gray, Carol Clugston, Anna Dominiczak and Alex Alderton, Roberto Amato, Jeffrey Barrett, Sonia Goncalves, Ewan Harrison, David K. Jackson, Ian Johnston, Dominic Kwiatkowski, Cordelia Langford, John Sillitoe on behalf of the Wellcome Sanger Institute COVID-19 Surveillance Team |
| EPI_ISL_2935324 | Lighthouse Laboratory Plymouth | Wellcome Sanger Institute for the COVID-19 Genomics UK (COG-UK) Consortium | Lighthouse Laboratory Plymouth and Alex Alderton, Roberto Amato, Jeffrey Barrett, Sonia Goncalves, Ewan Harrison, David K. Jackson, Ian Johnston, Dominic Kwiatkowski, Cordelia Langford, John Sillitoe on behalf of the Wellcome Sanger Institute COVID-19 Surveillance Team |
| EPI_ISL_2935325 | Lighthouse Lab in Alderley Park | Wellcome Sanger Institute for the COVID-19 Genomics UK (COG-UK) Consortium | Jacquelyn Wynn, Mairead Hyland, The Lighthouse Lab in Alderley Park and Alex Alderton, Roberto Amato, Jeffrey Barrett, Sonia Goncalves, Ewan Harrison, David K. Jackson, Ian Johnston, Dominic Kwiatkowski, Cordelia Langford, John Sillitoe on behalf of the Wellcome Sanger Institute COVID-19 Surveillance Team |
| EPI_ISL_2935326, EPI_ISL_2935327, EPI_ISL_2935328, EPI_ISL_2935329 | Lighthouse Laboratory Plymouth | Wellcome Sanger Institute for the COVID-19 Genomics UK (COG-UK) Consortium | Lighthouse Laboratory Plymouth and Alex Alderton, Roberto Amato, Jeffrey Barrett, Sonia Goncalves, Ewan Harrison, David K. Jackson, Ian Johnston, Dominic Kwiatkowski, Cordelia Langford, John Sillitoe on behalf of the Wellcome Sanger Institute COVID-19 Surveillance Team |
| EPI_ISL_2935330, EPI_ISL_2935331 | Lighthouse Lab in Milton Keynes | Wellcome Sanger Institute for the COVID-19 Genomics UK (COG-UK) Consortium | The Lighthouse Lab in Milton Keynes and Alex Alderton, Roberto Amato, Jeffrey Barrett, Sonia Goncalves, Ewan Harrison, David K. Jackson, Ian Johnston, Dominic Kwiatkowski, Cordelia Langford, John Sillitoe on behalf of the Wellcome Sanger Institute COVID-19 Surveillance Team |
| EPI_ISL_2935332, EPI_ISL_2935333, EPI_ISL_2935334, EPI_ISL_2935335, | Lighthouse Laboratory Plymouth | Wellcome Sanger Institute for the COVID-19 Genomics UK (COG-UK) Consortium | Lighthouse Laboratory Plymouth and Alex Alderton, Roberto Amato, Jeffrey Barrett, Sonia Goncalves, Ewan Harrison, David K. Jackson, Ian Johnston, Dominic Kwiatkowski, Cordelia Langford, John Sillitoe on behalf of the Wellcome Sanger Institute COVID-19 Surveillance Team |

| | | | |
|---|---|---|---|
| EPI_ISL_2935336 | | | |
| EPI_ISL_2935337 | Lighthouse Lab in Alderley Park | Wellcome Sanger Institute for the COVID-19 Genomics UK (COG-UK) Consortium | Jacquelyn Wynn, Mairead Hyland, The Lighthouse Lab in Alderley Park and Alex Alderton, Roberto Amato, Jeffrey Barrett, Sonia Goncalves, Ewan Harrison, David K. Jackson, Ian Johnston, Dominic Kwiatkowski, Cordelia Langford, John Sillitoe on behalf of the Wellcome Sanger Institute COVID-19 Surveillance Team |
| EPI_ISL_2935338 | Lighthouse Laboratory Plymouth | Wellcome Sanger Institute for the COVID-19 Genomics UK (COG-UK) Consortium | Lighthouse Laboratory Plymouth and Alex Alderton, Roberto Amato, Jeffrey Barrett, Sonia Goncalves, Ewan Harrison, David K. Jackson, Ian Johnston, Dominic Kwiatkowski, Cordelia Langford, John Sillitoe on behalf of the Wellcome Sanger Institute COVID-19 Surveillance Team |
| EPI_ISL_2935339, EPI_ISL_2935340 | Lighthouse Lab in Alderley Park | Wellcome Sanger Institute for the COVID-19 Genomics UK (COG-UK) Consortium | Jacquelyn Wynn, Mairead Hyland, The Lighthouse Lab in Alderley Park and Alex Alderton, Roberto Amato, Jeffrey Barrett, Sonia Goncalves, Ewan Harrison, David K. Jackson, Ian Johnston, Dominic Kwiatkowski, Cordelia Langford, John Sillitoe on behalf of the Wellcome Sanger Institute COVID-19 Surveillance Team |
| EPI_ISL_2935341 | Lighthouse Lab in Milton Keynes | Wellcome Sanger Institute for the COVID-19 Genomics UK (COG-UK) Consortium | The Lighthouse Lab in Milton Keynes and Alex Alderton, Roberto Amato, Jeffrey Barrett, Sonia Goncalves, Ewan Harrison, David K. Jackson, Ian Johnston, Dominic Kwiatkowski, Cordelia Langford, John Sillitoe on behalf of the Wellcome Sanger Institute COVID-19 Surveillance Team |
| EPI_ISL_2935342 | Lighthouse Lab in Alderley Park | Wellcome Sanger Institute for the COVID-19 Genomics UK (COG-UK) Consortium | Jacquelyn Wynn, Mairead Hyland, The Lighthouse Lab in Alderley Park and Alex Alderton, Roberto Amato, Jeffrey Barrett, Sonia Goncalves, Ewan Harrison, David K. Jackson, Ian Johnston, Dominic Kwiatkowski, Cordelia Langford, John Sillitoe on behalf of the Wellcome Sanger Institute COVID-19 Surveillance Team |
| EPI_ISL_2935343 | Lighthouse Laboratory Plymouth | Wellcome Sanger Institute for the COVID-19 Genomics UK (COG-UK) Consortium | Lighthouse Laboratory Plymouth and Alex Alderton, Roberto Amato, Jeffrey Barrett, Sonia Goncalves, Ewan Harrison, David K. Jackson, Ian Johnston, Dominic Kwiatkowski, Cordelia Langford, John Sillitoe on behalf of the Wellcome Sanger Institute COVID-19 Surveillance Team |
| EPI_ISL_2935344, EPI_ISL_2935345, EPI_ISL_2935346, EPI_ISL_2935347 | Lighthouse Lab in Milton Keynes | Wellcome Sanger Institute for the COVID-19 Genomics UK (COG-UK) Consortium | The Lighthouse Lab in Milton Keynes and Alex Alderton, Roberto Amato, Jeffrey Barrett, Sonia Goncalves, Ewan Harrison, David K. Jackson, Ian Johnston, Dominic Kwiatkowski, Cordelia Langford, John Sillitoe on behalf of the Wellcome Sanger Institute COVID-19 Surveillance Team |
| EPI_ISL_2935348, EPI_ISL_2935349, EPI_ISL_2935350, EPI_ISL_2935351 | Lighthouse Laboratory Plymouth | Wellcome Sanger Institute for the COVID-19 Genomics UK (COG-UK) Consortium | Lighthouse Laboratory Plymouth and Alex Alderton, Roberto Amato, Jeffrey Barrett, Sonia Goncalves, Ewan Harrison, David K. Jackson, Ian Johnston, Dominic Kwiatkowski, Cordelia Langford, John Sillitoe on behalf of the Wellcome Sanger Institute COVID-19 Surveillance Team |
| EPI_ISL_2935352 | Lighthouse Lab in Alderley Park | Wellcome Sanger Institute for the COVID-19 Genomics UK (COG-UK) Consortium | Jacquelyn Wynn, Mairead Hyland, The Lighthouse Lab in Alderley Park and Alex Alderton, Roberto Amato, Jeffrey Barrett, Sonia Goncalves, Ewan Harrison, David K. Jackson, Ian Johnston, Dominic Kwiatkowski, Cordelia Langford, John Sillitoe on behalf of the Wellcome Sanger Institute COVID-19 Surveillance Team |
| EPI_ISL_2935353 | Lighthouse Lab in Milton Keynes | Wellcome Sanger Institute for the COVID-19 Genomics UK (COG-UK) Consortium | The Lighthouse Lab in Milton Keynes and Alex Alderton, Roberto Amato, Jeffrey Barrett, Sonia Goncalves, Ewan Harrison, David K. Jackson, Ian Johnston, Dominic Kwiatkowski, Cordelia Langford, John Sillitoe on behalf of the Wellcome Sanger Institute COVID-19 Surveillance Team |
| EPI_ISL_2935354 | Lighthouse Lab in Alderley Park | Wellcome Sanger Institute for the COVID-19 Genomics UK (COG-UK) Consortium | Jacquelyn Wynn, Mairead Hyland, The Lighthouse Lab in Alderley Park and Alex Alderton, Roberto Amato, Jeffrey Barrett, Sonia Goncalves, Ewan Harrison, David K. Jackson, Ian Johnston, Dominic Kwiatkowski, Cordelia Langford, John Sillitoe on behalf of the Wellcome Sanger Institute COVID-19 Surveillance Team |
| EPI_ISL_2935355, EPI_ISL_2935356 | Lighthouse Laboratory Plymouth | Wellcome Sanger Institute for the COVID-19 Genomics UK (COG-UK) Consortium | Lighthouse Laboratory Plymouth and Alex Alderton, Roberto Amato, Jeffrey Barrett, Sonia Goncalves, Ewan Harrison, David K. Jackson, Ian Johnston, Dominic Kwiatkowski, Cordelia Langford, John Sillitoe on behalf of the Wellcome Sanger Institute COVID-19 Surveillance Team |
| EPI_ISL_2935357 | Lighthouse Lab in Alderley Park | Wellcome Sanger Institute for the COVID-19 Genomics UK (COG-UK) Consortium | Jacquelyn Wynn, Mairead Hyland, The Lighthouse Lab in Alderley Park and Alex Alderton, Roberto Amato, Jeffrey Barrett, Sonia Goncalves, Ewan Harrison, David K. Jackson, Ian Johnston, Dominic Kwiatkowski, Cordelia Langford, John Sillitoe on behalf of the Wellcome Sanger Institute COVID-19 Surveillance Team |
| EPI_ISL_2935358 | Lighthouse Lab in Milton Keynes | Wellcome Sanger Institute for the COVID-19 Genomics UK (COG-UK) Consortium | The Lighthouse Lab in Milton Keynes and Alex Alderton, Roberto Amato, Jeffrey Barrett, Sonia Goncalves, Ewan Harrison, David K. Jackson, Ian Johnston, Dominic Kwiatkowski, Cordelia Langford, John Sillitoe on behalf of the Wellcome Sanger Institute COVID-19 Surveillance Team |
| EPI_ISL_2935359, EPI_ISL_2935360, EPI_ISL_2935361, EPI_ISL_2935362 | Lighthouse Lab in Alderley Park | Wellcome Sanger Institute for the COVID-19 Genomics UK (COG-UK) Consortium | Jacquelyn Wynn, Mairead Hyland, The Lighthouse Lab in Alderley Park and Alex Alderton, Roberto Amato, Jeffrey Barrett, Sonia Goncalves, Ewan Harrison, David K. Jackson, Ian Johnston, Dominic Kwiatkowski, Cordelia Langford, John Sillitoe on behalf of the Wellcome Sanger Institute COVID-19 Surveillance Team |
| EPI_ISL_2935363 | Lighthouse Laboratory Plymouth | Wellcome Sanger Institute for the COVID-19 Genomics UK (COG-UK) Consortium | Lighthouse Laboratory Plymouth and Alex Alderton, Roberto Amato, Jeffrey Barrett, Sonia Goncalves, Ewan Harrison, David K. Jackson, Ian Johnston, Dominic Kwiatkowski, Cordelia Langford, John Sillitoe on behalf of the Wellcome Sanger Institute COVID-19 Surveillance Team |
| EPI_ISL_2935364 | Lighthouse Lab in Milton Keynes | Wellcome Sanger Institute for the COVID-19 Genomics UK (COG-UK) Consortium | The Lighthouse Lab in Milton Keynes and Alex Alderton, Roberto Amato, Jeffrey Barrett, Sonia Goncalves, Ewan Harrison, David K. Jackson, Ian Johnston, Dominic Kwiatkowski, Cordelia Langford, John Sillitoe on behalf of the Wellcome Sanger Institute COVID-19 Surveillance Team |
| EPI_ISL_2935365 | Lighthouse Laboratory Plymouth | Wellcome Sanger Institute for the COVID-19 Genomics UK (COG-UK) Consortium | Lighthouse Laboratory Plymouth and Alex Alderton, Roberto Amato, Jeffrey Barrett, Sonia Goncalves, Ewan Harrison, David K. Jackson, Ian Johnston, Dominic Kwiatkowski, Cordelia Langford, John Sillitoe on behalf of the Wellcome Sanger Institute COVID-19 Surveillance Team |
| EPI_ISL_2935366, EPI_ISL_2935367, EPI_ISL_2935368, EPI_ISL_2935369 | Lighthouse Lab in Milton Keynes | Wellcome Sanger Institute for the COVID-19 Genomics UK (COG-UK) Consortium | The Lighthouse Lab in Milton Keynes and Alex Alderton, Roberto Amato, Jeffrey Barrett, Sonia Goncalves, Ewan Harrison, David K. Jackson, Ian Johnston, Dominic Kwiatkowski, Cordelia Langford, John Sillitoe on behalf of the Wellcome Sanger Institute COVID-19 Surveillance Team |
| EPI_ISL_2935370 | Lighthouse Lab in Alderley Park | Wellcome Sanger Institute for the COVID-19 Genomics UK (COG-UK) Consortium | Jacquelyn Wynn, Mairead Hyland, The Lighthouse Lab in Alderley Park and Alex Alderton, Roberto Amato, Jeffrey Barrett, Sonia Goncalves, Ewan Harrison, David K. Jackson, Ian Johnston, Dominic Kwiatkowski, Cordelia Langford, John Sillitoe on behalf of the Wellcome Sanger Institute COVID-19 Surveillance Team |
| EPI_ISL_2935371, EPI_ISL_2935372, EPI_ISL_2935373, EPI_ISL_2935374, EPI_ISL_2935375, EPI_ISL_2935376 | Lighthouse Laboratory Plymouth | Wellcome Sanger Institute for the COVID-19 Genomics UK (COG-UK) Consortium | Lighthouse Laboratory Plymouth and Alex Alderton, Roberto Amato, Jeffrey Barrett, Sonia Goncalves, Ewan Harrison, David K. Jackson, Ian Johnston, Dominic Kwiatkowski, Cordelia Langford, John Sillitoe on behalf of the Wellcome Sanger Institute COVID-19 Surveillance Team |
| EPI_ISL_2935377 | Lighthouse Lab in Milton Keynes | Wellcome Sanger Institute for the COVID-19 Genomics UK (COG-UK) Consortium | The Lighthouse Lab in Milton Keynes and Alex Alderton, Roberto Amato, Jeffrey Barrett, Sonia Goncalves, Ewan Harrison, David K. Jackson, Ian Johnston, Dominic Kwiatkowski, Cordelia Langford, John Sillitoe on behalf of the Wellcome Sanger Institute COVID-19 Surveillance Team |
| EPI_ISL_2935378 | Lighthouse Laboratory Plymouth | Wellcome Sanger Institute for the COVID-19 Genomics UK (COG-UK) Consortium | Lighthouse Laboratory Plymouth and Alex Alderton, Roberto Amato, Jeffrey Barrett, Sonia Goncalves, Ewan Harrison, David K. Jackson, Ian Johnston, Dominic Kwiatkowski, Cordelia Langford, John Sillitoe on behalf of the Wellcome Sanger Institute COVID-19 Surveillance Team |
| EPI_ISL_2935379, EPI_ISL_2935380, EPI_ISL_2935381 | Lighthouse Lab in Alderley Park | Wellcome Sanger Institute for the COVID-19 Genomics UK (COG-UK) Consortium | Jacquelyn Wynn, Mairead Hyland, The Lighthouse Lab in Alderley Park and Alex Alderton, Roberto Amato, Jeffrey Barrett, Sonia Goncalves, Ewan Harrison, David K. Jackson, Ian Johnston, Dominic Kwiatkowski, Cordelia Langford, John Sillitoe on behalf of the Wellcome Sanger Institute COVID-19 Surveillance Team |
| EPI_ISL_2935382 | Lighthouse Laboratory Plymouth | Wellcome Sanger Institute for the COVID-19 Genomics UK (COG-UK) Consortium | Lighthouse Laboratory Plymouth and Alex Alderton, Roberto Amato, Jeffrey Barrett, Sonia Goncalves, Ewan Harrison, David K. Jackson, Ian Johnston, Dominic Kwiatkowski, Cordelia Langford, John Sillitoe on behalf of the Wellcome Sanger Institute COVID-19 Surveillance Team |
| EPI_ISL_2935383, EPI_ISL_2935384, EPI_ISL_2935385, EPI_ISL_2935386 | Lighthouse Lab in Alderley Park | Wellcome Sanger Institute for the COVID-19 Genomics UK (COG-UK) Consortium | Jacquelyn Wynn, Mairead Hyland, The Lighthouse Lab in Alderley Park and Alex Alderton, Roberto Amato, Jeffrey Barrett, Sonia Goncalves, Ewan Harrison, David K. Jackson, Ian Johnston, Dominic Kwiatkowski, Cordelia Langford, John Sillitoe on behalf of the Wellcome Sanger Institute COVID-19 Surveillance Team |
| EPI_ISL_2935387 | Lighthouse Lab in Milton Keynes | Wellcome Sanger Institute for the COVID-19 Genomics UK (COG-UK) Consortium | The Lighthouse Lab in Milton Keynes and Alex Alderton, Roberto Amato, Jeffrey Barrett, Sonia Goncalves, Ewan Harrison, David K. Jackson, Ian Johnston, Dominic Kwiatkowski, Cordelia Langford, John Sillitoe on behalf of the Wellcome Sanger Institute COVID-19 Surveillance Team |
| EPI_ISL_2935388, EPI_ISL_2935389 | Lighthouse Laboratory Plymouth | Wellcome Sanger Institute for the COVID-19 Genomics UK (COG-UK) Consortium | Lighthouse Laboratory Plymouth and Alex Alderton, Roberto Amato, Jeffrey Barrett, Sonia Goncalves, Ewan Harrison, David K. Jackson, Ian Johnston, Dominic Kwiatkowski, Cordelia Langford, John Sillitoe on behalf of the Wellcome Sanger Institute COVID-19 Surveillance Team |
| EPI_ISL_2935390 | Lighthouse Lab in Alderley Park | Wellcome Sanger Institute for the COVID-19 Genomics UK (COG-UK) Consortium | Jacquelyn Wynn, Mairead Hyland, The Lighthouse Lab in Alderley Park and Alex Alderton, Roberto Amato, Jeffrey Barrett, Sonia Goncalves, Ewan Harrison, David K. Jackson, Ian Johnston, Dominic Kwiatkowski, Cordelia Langford, John Sillitoe on behalf of the Wellcome Sanger Institute COVID-19 Surveillance Team |
| EPI_ISL_2935391 | Lighthouse Lab in Milton Keynes | Wellcome Sanger Institute for the COVID-19 Genomics UK (COG-UK) Consortium | The Lighthouse Lab in Milton Keynes and Alex Alderton, Roberto Amato, Jeffrey Barrett, Sonia Goncalves, Ewan Harrison, David K. Jackson, Ian Johnston, Dominic Kwiatkowski, Cordelia Langford, John Sillitoe on behalf of the Wellcome Sanger Institute COVID-19 Surveillance Team |
| EPI_ISL_2935392, EPI_ISL_2935393 | Lighthouse Lab in Alderley Park | Wellcome Sanger Institute for the COVID-19 Genomics UK (COG-UK) Consortium | Jacquelyn Wynn, Mairead Hyland, The Lighthouse Lab in Alderley Park and Alex Alderton, Roberto Amato, Jeffrey Barrett, Sonia Goncalves, Ewan Harrison, David K. Jackson, Ian Johnston, Dominic Kwiatkowski, Cordelia Langford, John Sillitoe on behalf of the Wellcome Sanger Institute COVID-19 Surveillance Team |
| EPI_ISL_2935394, EPI_ISL_2935395 | Lighthouse Lab in Milton Keynes | Wellcome Sanger Institute for the COVID-19 Genomics UK (COG-UK) Consortium | The Lighthouse Lab in Milton Keynes and Alex Alderton, Roberto Amato, Jeffrey Barrett, Sonia Goncalves, Ewan Harrison, David K. Jackson, Ian Johnston, Dominic Kwiatkowski, Cordelia Langford, John Sillitoe on behalf of the Wellcome Sanger Institute COVID-19 Surveillance Team |
| EPI_ISL_2935396, EPI_ISL_2935397, EPI_ISL_2935398, EPI_ISL_2935399 | Lighthouse Lab in Alderley Park | Wellcome Sanger Institute for the COVID-19 Genomics UK (COG-UK) Consortium | Jacquelyn Wynn, Mairead Hyland, The Lighthouse Lab in Alderley Park and Alex Alderton, Roberto Amato, Jeffrey Barrett, Sonia Goncalves, Ewan Harrison, David K. Jackson, Ian Johnston, Dominic Kwiatkowski, Cordelia Langford, John Sillitoe on behalf of the Wellcome Sanger Institute COVID-19 Surveillance Team |
| EPI_ISL_2935400 | Lighthouse Lab in Milton Keynes | Wellcome Sanger Institute for the COVID-19 Genomics UK (COG-UK) Consortium | The Lighthouse Lab in Milton Keynes and Alex Alderton, Roberto Amato, Jeffrey Barrett, Sonia Goncalves, Ewan Harrison, David K. Jackson, Ian Johnston, Dominic Kwiatkowski, Cordelia Langford, John Sillitoe on behalf of the Wellcome Sanger Institute COVID-19 Surveillance Team |
| EPI_ISL_2935401 | Lighthouse Laboratory Plymouth | Wellcome Sanger Institute for the COVID-19 Genomics UK (COG-UK) Consortium | Lighthouse Laboratory Plymouth and Alex Alderton, Roberto Amato, Jeffrey Barrett, Sonia Goncalves, Ewan Harrison, David K. Jackson, Ian Johnston, Dominic Kwiatkowski, Cordelia Langford, John Sillitoe on behalf of the Wellcome Sanger Institute COVID-19 Surveillance Team |
| EPI_ISL_2935402 | Lighthouse Lab in Milton Keynes | Wellcome Sanger Institute for the COVID-19 Genomics UK (COG-UK) Consortium | The Lighthouse Lab in Milton Keynes and Alex Alderton, Roberto Amato, Jeffrey Barrett, Sonia Goncalves, Ewan Harrison, David K. Jackson, Ian Johnston, Dominic Kwiatkowski, Cordelia Langford, John Sillitoe on behalf of the Wellcome Sanger Institute COVID-19 Surveillance Team |
| EPI_ISL_2935403, EPI_ISL_2935404 | Lighthouse Lab in Alderley Park | Wellcome Sanger Institute for the COVID-19 Genomics UK (COG-UK) Consortium | Jacquelyn Wynn, Mairead Hyland, The Lighthouse Lab in Alderley Park and Alex Alderton, Roberto Amato, Jeffrey Barrett, Sonia Goncalves, Ewan Harrison, David K. Jackson, Ian Johnston, Dominic Kwiatkowski, Cordelia Langford, John Sillitoe on behalf of the Wellcome Sanger Institute COVID-19 Surveillance Team |
| EPI_ISL_2935405 | Lighthouse Laboratory Plymouth | Wellcome Sanger Institute for the COVID-19 Genomics UK (COG-UK) | Lighthouse Laboratory Plymouth and Alex Alderton, Roberto Amato, Jeffrey Barrett, Sonia Goncalves, Ewan Harrison, David K. Jackson, Ian Johnston, Dominic Kwiatkowski, Cordelia Langford, John Sillitoe on behalf of the Wellcome Sanger |

| | | | |
|---|---|---|---|
| | | Consortium | Institute COVID-19 Surveillance Team |
| EPI_ISL_2935406 | Lighthouse Lab in Alderley Park | Wellcome Sanger Institute for the COVID-19 Genomics UK (COG-UK) Consortium | Jacquelyn Wynn, Mairead Hyland, The Lighthouse Lab in Alderley Park and Alex Alderton, Roberto Amato, Jeffrey Barrett, Sonia Goncalves, Ewan Harrison, David K. Jackson, Ian Johnston, Dominic Kwiatkowski, Cordelia Langford, John Sillitoe on behalf of the Wellcome Sanger Institute COVID-19 Surveillance Team |
| EPI_ISL_2935407, EPI_ISL_2935408, EPI_ISL_2935409 | Lighthouse Laboratory Plymouth | Wellcome Sanger Institute for the COVID-19 Genomics UK (COG-UK) Consortium | Lighthouse Laboratory Plymouth and Alex Alderton, Roberto Amato, Jeffrey Barrett, Sonia Goncalves, Ewan Harrison, David K. Jackson, Ian Johnston, Dominic Kwiatkowski, Cordelia Langford, John Sillitoe on behalf of the Wellcome Sanger Institute COVID-19 Surveillance Team |
| EPI_ISL_2935410 | Lighthouse Lab in Alderley Park | Wellcome Sanger Institute for the COVID-19 Genomics UK (COG-UK) Consortium | Jacquelyn Wynn, Mairead Hyland, The Lighthouse Lab in Alderley Park and Alex Alderton, Roberto Amato, Jeffrey Barrett, Sonia Goncalves, Ewan Harrison, David K. Jackson, Ian Johnston, Dominic Kwiatkowski, Cordelia Langford, John Sillitoe on behalf of the Wellcome Sanger Institute COVID-19 Surveillance Team |
| EPI_ISL_2935411, EPI_ISL_2935412 | Lighthouse Laboratory Plymouth | Wellcome Sanger Institute for the COVID-19 Genomics UK (COG-UK) Consortium | Lighthouse Laboratory Plymouth and Alex Alderton, Roberto Amato, Jeffrey Barrett, Sonia Goncalves, Ewan Harrison, David K. Jackson, Ian Johnston, Dominic Kwiatkowski, Cordelia Langford, John Sillitoe on behalf of the Wellcome Sanger Institute COVID-19 Surveillance Team |
| EPI_ISL_2935413 | Lighthouse Lab in Alderley Park | Wellcome Sanger Institute for the COVID-19 Genomics UK (COG-UK) Consortium | Jacquelyn Wynn, Mairead Hyland, The Lighthouse Lab in Alderley Park and Alex Alderton, Roberto Amato, Jeffrey Barrett, Sonia Goncalves, Ewan Harrison, David K. Jackson, Ian Johnston, Dominic Kwiatkowski, Cordelia Langford, John Sillitoe on behalf of the Wellcome Sanger Institute COVID-19 Surveillance Team |
| EPI_ISL_2935414 | Lighthouse Laboratory Plymouth | Wellcome Sanger Institute for the COVID-19 Genomics UK (COG-UK) Consortium | Lighthouse Laboratory Plymouth and Alex Alderton, Roberto Amato, Jeffrey Barrett, Sonia Goncalves, Ewan Harrison, David K. Jackson, Ian Johnston, Dominic Kwiatkowski, Cordelia Langford, John Sillitoe on behalf of the Wellcome Sanger Institute COVID-19 Surveillance Team |
| EPI_ISL_2935415 | Lighthouse Lab in Alderley Park | Wellcome Sanger Institute for the COVID-19 Genomics UK (COG-UK) Consortium | Jacquelyn Wynn, Mairead Hyland, The Lighthouse Lab in Alderley Park and Alex Alderton, Roberto Amato, Jeffrey Barrett, Sonia Goncalves, Ewan Harrison, David K. Jackson, Ian Johnston, Dominic Kwiatkowski, Cordelia Langford, John Sillitoe on behalf of the Wellcome Sanger Institute COVID-19 Surveillance Team |
| EPI_ISL_2935416 | Lighthouse Lab in Milton Keynes | Wellcome Sanger Institute for the COVID-19 Genomics UK (COG-UK) Consortium | The Lighthouse Lab in Milton Keynes and Alex Alderton, Roberto Amato, Jeffrey Barrett, Sonia Goncalves, Ewan Harrison, David K. Jackson, Ian Johnston, Dominic Kwiatkowski, Cordelia Langford, John Sillitoe on behalf of the Wellcome Sanger Institute COVID-19 Surveillance Team |
| EPI_ISL_2935417, EPI_ISL_2935418 | Lighthouse Lab in Alderley Park | Wellcome Sanger Institute for the COVID-19 Genomics UK (COG-UK) Consortium | Jacquelyn Wynn, Mairead Hyland, The Lighthouse Lab in Alderley Park and Alex Alderton, Roberto Amato, Jeffrey Barrett, Sonia Goncalves, Ewan Harrison, David K. Jackson, Ian Johnston, Dominic Kwiatkowski, Cordelia Langford, John Sillitoe on behalf of the Wellcome Sanger Institute COVID-19 Surveillance Team |
| EPI_ISL_2935419 | Lighthouse Lab in Milton Keynes | Wellcome Sanger Institute for the COVID-19 Genomics UK (COG-UK) Consortium | The Lighthouse Lab in Milton Keynes and Alex Alderton, Roberto Amato, Jeffrey Barrett, Sonia Goncalves, Ewan Harrison, David K. Jackson, Ian Johnston, Dominic Kwiatkowski, Cordelia Langford, John Sillitoe on behalf of the Wellcome Sanger Institute COVID-19 Surveillance Team |
| EPI_ISL_2935420, EPI_ISL_2935421 | Lighthouse Laboratory Plymouth | Wellcome Sanger Institute for the COVID-19 Genomics UK (COG-UK) Consortium | Lighthouse Laboratory Plymouth and Alex Alderton, Roberto Amato, Jeffrey Barrett, Sonia Goncalves, Ewan Harrison, David K. Jackson, Ian Johnston, Dominic Kwiatkowski, Cordelia Langford, John Sillitoe on behalf of the Wellcome Sanger Institute COVID-19 Surveillance Team |
| EPI_ISL_2935422 | Lighthouse Lab in Alderley Park | Wellcome Sanger Institute for the COVID-19 Genomics UK (COG-UK) Consortium | Jacquelyn Wynn, Mairead Hyland, The Lighthouse Lab in Alderley Park and Alex Alderton, Roberto Amato, Jeffrey Barrett, Sonia Goncalves, Ewan Harrison, David K. Jackson, Ian Johnston, Dominic Kwiatkowski, Cordelia Langford, John Sillitoe on behalf of the Wellcome Sanger Institute COVID-19 Surveillance Team |
| EPI_ISL_2935423, EPI_ISL_2935424, EPI_ISL_2935425 | Lighthouse Laboratory Plymouth | Wellcome Sanger Institute for the COVID-19 Genomics UK (COG-UK) Consortium | Lighthouse Laboratory Plymouth and Alex Alderton, Roberto Amato, Jeffrey Barrett, Sonia Goncalves, Ewan Harrison, David K. Jackson, Ian Johnston, Dominic Kwiatkowski, Cordelia Langford, John Sillitoe on behalf of the Wellcome Sanger Institute COVID-19 Surveillance Team |
| EPI_ISL_2935426 | Lighthouse Lab in Alderley Park | Wellcome Sanger Institute for the COVID-19 Genomics UK (COG-UK) Consortium | Jacquelyn Wynn, Mairead Hyland, The Lighthouse Lab in Alderley Park and Alex Alderton, Roberto Amato, Jeffrey Barrett, Sonia Goncalves, Ewan Harrison, David K. Jackson, Ian Johnston, Dominic Kwiatkowski, Cordelia Langford, John Sillitoe on behalf of the Wellcome Sanger Institute COVID-19 Surveillance Team |
| EPI_ISL_2935427 | Lighthouse Lab in Milton Keynes | Wellcome Sanger Institute for the COVID-19 Genomics UK (COG-UK) Consortium | The Lighthouse Lab in Milton Keynes and Alex Alderton, Roberto Amato, Jeffrey Barrett, Sonia Goncalves, Ewan Harrison, David K. Jackson, Ian Johnston, Dominic Kwiatkowski, Cordelia Langford, John Sillitoe on behalf of the Wellcome Sanger Institute COVID-19 Surveillance Team |
| EPI_ISL_2935428, EPI_ISL_2935429, EPI_ISL_2935430 | Lighthouse Lab in Alderley Park | Wellcome Sanger Institute for the COVID-19 Genomics UK (COG-UK) Consortium | Jacquelyn Wynn, Mairead Hyland, The Lighthouse Lab in Alderley Park and Alex Alderton, Roberto Amato, Jeffrey Barrett, Sonia Goncalves, Ewan Harrison, David K. Jackson, Ian Johnston, Dominic Kwiatkowski, Cordelia Langford, John Sillitoe on behalf of the Wellcome Sanger Institute COVID-19 Surveillance Team |
| EPI_ISL_2935431, EPI_ISL_2935432, EPI_ISL_2935433 | Lighthouse Laboratory Plymouth | Wellcome Sanger Institute for the COVID-19 Genomics UK (COG-UK) Consortium | Lighthouse Laboratory Plymouth and Alex Alderton, Roberto Amato, Jeffrey Barrett, Sonia Goncalves, Ewan Harrison, David K. Jackson, Ian Johnston, Dominic Kwiatkowski, Cordelia Langford, John Sillitoe on behalf of the Wellcome Sanger Institute COVID-19 Surveillance Team |
| EPI_ISL_2935434 | Lighthouse Lab in Milton Keynes | Wellcome Sanger Institute for the COVID-19 Genomics UK (COG-UK) Consortium | The Lighthouse Lab in Milton Keynes and Alex Alderton, Roberto Amato, Jeffrey Barrett, Sonia Goncalves, Ewan Harrison, David K. Jackson, Ian Johnston, Dominic Kwiatkowski, Cordelia Langford, John Sillitoe on behalf of the Wellcome Sanger Institute COVID-19 Surveillance Team |
| EPI_ISL_2935435 | Lighthouse Laboratory Plymouth | Wellcome Sanger Institute for the COVID-19 Genomics UK (COG-UK) Consortium | Lighthouse Laboratory Plymouth and Alex Alderton, Roberto Amato, Jeffrey Barrett, Sonia Goncalves, Ewan Harrison, David K. Jackson, Ian Johnston, Dominic Kwiatkowski, Cordelia Langford, John Sillitoe on behalf of the Wellcome Sanger Institute COVID-19 Surveillance Team |
| EPI_ISL_2935436, EPI_ISL_2935437 | Lighthouse Lab in Milton Keynes | Wellcome Sanger Institute for the COVID-19 Genomics UK (COG-UK) Consortium | The Lighthouse Lab in Milton Keynes and Alex Alderton, Roberto Amato, Jeffrey Barrett, Sonia Goncalves, Ewan Harrison, David K. Jackson, Ian Johnston, Dominic Kwiatkowski, Cordelia Langford, John Sillitoe on behalf of the Wellcome Sanger Institute COVID-19 Surveillance Team |
| EPI_ISL_2935438 | Lighthouse Lab in Alderley Park | Wellcome Sanger Institute for the COVID-19 Genomics UK (COG-UK) Consortium | Jacquelyn Wynn, Mairead Hyland, The Lighthouse Lab in Alderley Park and Alex Alderton, Roberto Amato, Jeffrey Barrett, Sonia Goncalves, Ewan Harrison, David K. Jackson, Ian Johnston, Dominic Kwiatkowski, Cordelia Langford, John Sillitoe on behalf of the Wellcome Sanger Institute COVID-19 Surveillance Team |
| EPI_ISL_2935439 | Lighthouse Laboratory Plymouth | Wellcome Sanger Institute for the COVID-19 Genomics UK (COG-UK) Consortium | Lighthouse Laboratory Plymouth and Alex Alderton, Roberto Amato, Jeffrey Barrett, Sonia Goncalves, Ewan Harrison, David K. Jackson, Ian Johnston, Dominic Kwiatkowski, Cordelia Langford, John Sillitoe on behalf of the Wellcome Sanger Institute COVID-19 Surveillance Team |
| EPI_ISL_2935440 | Lighthouse Lab in Milton Keynes | Wellcome Sanger Institute for the COVID-19 Genomics UK (COG-UK) Consortium | The Lighthouse Lab in Milton Keynes and Alex Alderton, Roberto Amato, Jeffrey Barrett, Sonia Goncalves, Ewan Harrison, David K. Jackson, Ian Johnston, Dominic Kwiatkowski, Cordelia Langford, John Sillitoe on behalf of the Wellcome Sanger Institute COVID-19 Surveillance Team |
| EPI_ISL_2935441 | Lighthouse Lab in Alderley Park | Wellcome Sanger Institute for the COVID-19 Genomics UK (COG-UK) Consortium | Jacquelyn Wynn, Mairead Hyland, The Lighthouse Lab in Alderley Park and Alex Alderton, Roberto Amato, Jeffrey Barrett, Sonia Goncalves, Ewan Harrison, David K. Jackson, Ian Johnston, Dominic Kwiatkowski, Cordelia Langford, John Sillitoe on behalf of the Wellcome Sanger Institute COVID-19 Surveillance Team |
| EPI_ISL_2935442 | Lighthouse Lab in Milton Keynes | Wellcome Sanger Institute for the COVID-19 Genomics UK (COG-UK) Consortium | The Lighthouse Lab in Milton Keynes and Alex Alderton, Roberto Amato, Jeffrey Barrett, Sonia Goncalves, Ewan Harrison, David K. Jackson, Ian Johnston, Dominic Kwiatkowski, Cordelia Langford, John Sillitoe on behalf of the Wellcome Sanger Institute COVID-19 Surveillance Team |
| EPI_ISL_2935443 | Lighthouse Laboratory Plymouth | Wellcome Sanger Institute for the COVID-19 Genomics UK (COG-UK) Consortium | Lighthouse Laboratory Plymouth and Alex Alderton, Roberto Amato, Jeffrey Barrett, Sonia Goncalves, Ewan Harrison, David K. Jackson, Ian Johnston, Dominic Kwiatkowski, Cordelia Langford, John Sillitoe on behalf of the Wellcome Sanger Institute COVID-19 Surveillance Team |
| EPI_ISL_2935444 | Lighthouse Lab in Alderley Park | Wellcome Sanger Institute for the COVID-19 Genomics UK (COG-UK) Consortium | Jacquelyn Wynn, Mairead Hyland, The Lighthouse Lab in Alderley Park and Alex Alderton, Roberto Amato, Jeffrey Barrett, Sonia Goncalves, Ewan Harrison, David K. Jackson, Ian Johnston, Dominic Kwiatkowski, Cordelia Langford, John Sillitoe on behalf of the Wellcome Sanger Institute COVID-19 Surveillance Team |
| EPI_ISL_2935445 | Lighthouse Laboratory Plymouth | Wellcome Sanger Institute for the COVID-19 Genomics UK (COG-UK) Consortium | Lighthouse Laboratory Plymouth and Alex Alderton, Roberto Amato, Jeffrey Barrett, Sonia Goncalves, Ewan Harrison, David K. Jackson, Ian Johnston, Dominic Kwiatkowski, Cordelia Langford, John Sillitoe on behalf of the Wellcome Sanger Institute COVID-19 Surveillance Team |
| EPI_ISL_2935446 | Lighthouse Lab in Alderley Park | Wellcome Sanger Institute for the COVID-19 Genomics UK (COG-UK) Consortium | Jacquelyn Wynn, Mairead Hyland, The Lighthouse Lab in Alderley Park and Alex Alderton, Roberto Amato, Jeffrey Barrett, Sonia Goncalves, Ewan Harrison, David K. Jackson, Ian Johnston, Dominic Kwiatkowski, Cordelia Langford, John Sillitoe on behalf of the Wellcome Sanger Institute COVID-19 Surveillance Team |
| EPI_ISL_2935447, EPI_ISL_2935448 | Lighthouse Laboratory Plymouth | Wellcome Sanger Institute for the COVID-19 Genomics UK (COG-UK) Consortium | Lighthouse Laboratory Plymouth and Alex Alderton, Roberto Amato, Jeffrey Barrett, Sonia Goncalves, Ewan Harrison, David K. Jackson, Ian Johnston, Dominic Kwiatkowski, Cordelia Langford, John Sillitoe on behalf of the Wellcome Sanger Institute COVID-19 Surveillance Team |
| EPI_ISL_2935449 | Lighthouse Lab in Milton Keynes | Wellcome Sanger Institute for the COVID-19 Genomics UK (COG-UK) Consortium | The Lighthouse Lab in Milton Keynes and Alex Alderton, Roberto Amato, Jeffrey Barrett, Sonia Goncalves, Ewan Harrison, David K. Jackson, Ian Johnston, Dominic Kwiatkowski, Cordelia Langford, John Sillitoe on behalf of the Wellcome Sanger Institute COVID-19 Surveillance Team |
| EPI_ISL_2935450 | Lighthouse Laboratory Plymouth | Wellcome Sanger Institute for the COVID-19 Genomics UK (COG-UK) Consortium | Lighthouse Laboratory Plymouth and Alex Alderton, Roberto Amato, Jeffrey Barrett, Sonia Goncalves, Ewan Harrison, David K. Jackson, Ian Johnston, Dominic Kwiatkowski, Cordelia Langford, John Sillitoe on behalf of the Wellcome Sanger Institute COVID-19 Surveillance Team |
| EPI_ISL_2935451 | Lighthouse Lab in Milton Keynes | Wellcome Sanger Institute for the COVID-19 Genomics UK (COG-UK) Consortium | The Lighthouse Lab in Milton Keynes and Alex Alderton, Roberto Amato, Jeffrey Barrett, Sonia Goncalves, Ewan Harrison, David K. Jackson, Ian Johnston, Dominic Kwiatkowski, Cordelia Langford, John Sillitoe on behalf of the Wellcome Sanger Institute COVID-19 Surveillance Team |
| EPI_ISL_2935452 | Lighthouse Lab in Alderley Park | Wellcome Sanger Institute for the COVID-19 Genomics UK (COG-UK) Consortium | Jacquelyn Wynn, Mairead Hyland, The Lighthouse Lab in Alderley Park and Alex Alderton, Roberto Amato, Jeffrey Barrett, Sonia Goncalves, Ewan Harrison, David K. Jackson, Ian Johnston, Dominic Kwiatkowski, Cordelia Langford, John Sillitoe on behalf of the Wellcome Sanger Institute COVID-19 Surveillance Team |
| EPI_ISL_2935453 | Lighthouse Lab in Milton Keynes | Wellcome Sanger Institute for the COVID-19 Genomics UK (COG-UK) Consortium | The Lighthouse Lab in Milton Keynes and Alex Alderton, Roberto Amato, Jeffrey Barrett, Sonia Goncalves, Ewan Harrison, David K. Jackson, Ian Johnston, Dominic Kwiatkowski, Cordelia Langford, John Sillitoe on behalf of the Wellcome Sanger Institute COVID-19 Surveillance Team |
| EPI_ISL_2935454 | Lighthouse Laboratory Plymouth | Wellcome Sanger Institute for the COVID-19 Genomics UK (COG-UK) Consortium | Lighthouse Laboratory Plymouth and Alex Alderton, Roberto Amato, Jeffrey Barrett, Sonia Goncalves, Ewan Harrison, David K. Jackson, Ian Johnston, Dominic Kwiatkowski, Cordelia Langford, John Sillitoe on behalf of the Wellcome Sanger Institute COVID-19 Surveillance Team |
| EPI_ISL_2935455 | Lighthouse Lab in Milton Keynes | Wellcome Sanger Institute for the COVID-19 Genomics UK (COG-UK) Consortium | The Lighthouse Lab in Milton Keynes and Alex Alderton, Roberto Amato, Jeffrey Barrett, Sonia Goncalves, Ewan Harrison, David K. Jackson, Ian Johnston, Dominic Kwiatkowski, Cordelia Langford, John Sillitoe on behalf of the Wellcome Sanger Institute COVID-19 Surveillance Team |
| EPI_ISL_2935456 | Lighthouse Lab in Alderley Park | Wellcome Sanger Institute for the COVID-19 Genomics UK (COG-UK) Consortium | Jacquelyn Wynn, Mairead Hyland, The Lighthouse Lab in Alderley Park and Alex Alderton, Roberto Amato, Jeffrey Barrett, Sonia Goncalves, Ewan Harrison, David K. Jackson, Ian Johnston, Dominic Kwiatkowski, Cordelia Langford, John Sillitoe on behalf of the Wellcome Sanger Institute COVID-19 Surveillance Team |
| EPI_ISL_2935457 | Lighthouse Lab in Milton Keynes | Wellcome Sanger Institute for the COVID-19 Genomics UK (COG-UK) Consortium | The Lighthouse Lab in Milton Keynes and Alex Alderton, Roberto Amato, Jeffrey Barrett, Sonia Goncalves, Ewan Harrison, David K. Jackson, Ian Johnston, Dominic Kwiatkowski, Cordelia Langford, John Sillitoe on behalf of the Wellcome Sanger Institute COVID-19 Surveillance Team |
| EPI_ISL_2935458 | Lighthouse Lab in Alderley Park | Wellcome Sanger Institute for the COVID-19 Genomics UK (COG-UK) Consortium | Jacquelyn Wynn, Mairead Hyland, The Lighthouse Lab in Alderley Park and Alex Alderton, Roberto Amato, Jeffrey Barrett, Sonia Goncalves, Ewan Harrison, David K. Jackson, Ian Johnston, Dominic Kwiatkowski, Cordelia Langford, John Sillitoe on behalf of the Wellcome Sanger Institute COVID-19 Surveillance Team |
| EPI_ISL_2935459, EPI_ISL_2935460, EPI_ISL_2935461, EPI_ISL_2935462 | Lighthouse Laboratory Plymouth | Wellcome Sanger Institute for the COVID-19 Genomics UK (COG-UK) Consortium | Lighthouse Laboratory Plymouth and Alex Alderton, Roberto Amato, Jeffrey Barrett, Sonia Goncalves, Ewan Harrison, David K. Jackson, Ian Johnston, Dominic Kwiatkowski, Cordelia Langford, John Sillitoe on behalf of the Wellcome Sanger Institute COVID-19 Surveillance Team |
| EPI_ISL_2935463 | Lighthouse Lab in Alderley Park | Wellcome Sanger Institute for the COVID-19 Genomics UK (COG-UK) Consortium | Jacquelyn Wynn, Mairead Hyland, The Lighthouse Lab in Alderley Park and Alex Alderton, Roberto Amato, Jeffrey Barrett, Sonia Goncalves, Ewan Harrison, David K. Jackson, Ian Johnston, Dominic Kwiatkowski, Cordelia Langford, John Sillitoe on behalf of the Wellcome Sanger Institute COVID-19 Surveillance Team |

| | | | |
|---|---|---|---|
| EPI_ISL_2935464, EPI_ISL_2935465 | Lighthouse Laboratory Plymouth | Wellcome Sanger Institute for the COVID-19 Genomics UK (COG-UK) Consortium | Lighthouse Laboratory Plymouth and Alex Alderton, Roberto Amato, Jeffrey Barrett, Sonia Goncalves, Ewan Harrison, David K. Jackson, Ian Johnston, Dominic Kwiatkowski, Cordelia Langford, John Sillitoe on behalf of the Wellcome Sanger Institute COVID-19 Surveillance Team |
| EPI_ISL_2935466 | Lighthouse Lab in Alderley Park | Wellcome Sanger Institute for the COVID-19 Genomics UK (COG-UK) Consortium | Jacquelyn Wynn, Mairead Hyland, The Lighthouse Lab in Alderley Park and Alex Alderton, Roberto Amato, Jeffrey Barrett, Sonia Goncalves, Ewan Harrison, David K. Jackson, Ian Johnston, Dominic Kwiatkowski, Cordelia Langford, John Sillitoe on behalf of the Wellcome Sanger Institute COVID-19 Surveillance Team |
| EPI_ISL_2935467 | Lighthouse Laboratory Plymouth | Wellcome Sanger Institute for the COVID-19 Genomics UK (COG-UK) Consortium | Lighthouse Laboratory Plymouth and Alex Alderton, Roberto Amato, Jeffrey Barrett, Sonia Goncalves, Ewan Harrison, David K. Jackson, Ian Johnston, Dominic Kwiatkowski, Cordelia Langford, John Sillitoe on behalf of the Wellcome Sanger Institute COVID-19 Surveillance Team |
| EPI_ISL_2935468 | Lighthouse Lab in Milton Keynes | Wellcome Sanger Institute for the COVID-19 Genomics UK (COG-UK) Consortium | The Lighthouse Lab in Milton Keynes and Alex Alderton, Roberto Amato, Jeffrey Barrett, Sonia Goncalves, Ewan Harrison, David K. Jackson, Ian Johnston, Dominic Kwiatkowski, Cordelia Langford, John Sillitoe on behalf of the Wellcome Sanger Institute COVID-19 Surveillance Team |
| EPI_ISL_2935469, EPI_ISL_2935470, EPI_ISL_2935471, EPI_ISL_2935472, EPI_ISL_2935473, EPI_ISL_2935474 | Lighthouse Laboratory Plymouth | Wellcome Sanger Institute for the COVID-19 Genomics UK (COG-UK) Consortium | Lighthouse Laboratory Plymouth and Alex Alderton, Roberto Amato, Jeffrey Barrett, Sonia Goncalves, Ewan Harrison, David K. Jackson, Ian Johnston, Dominic Kwiatkowski, Cordelia Langford, John Sillitoe on behalf of the Wellcome Sanger Institute COVID-19 Surveillance Team |
| EPI_ISL_2935475 | Lighthouse Lab in Milton Keynes | Wellcome Sanger Institute for the COVID-19 Genomics UK (COG-UK) Consortium | The Lighthouse Lab in Milton Keynes and Alex Alderton, Roberto Amato, Jeffrey Barrett, Sonia Goncalves, Ewan Harrison, David K. Jackson, Ian Johnston, Dominic Kwiatkowski, Cordelia Langford, John Sillitoe on behalf of the Wellcome Sanger Institute COVID-19 Surveillance Team |
| EPI_ISL_2935476 | Lighthouse Lab in Alderley Park | Wellcome Sanger Institute for the COVID-19 Genomics UK (COG-UK) Consortium | Jacquelyn Wynn, Mairead Hyland, The Lighthouse Lab in Alderley Park and Alex Alderton, Roberto Amato, Jeffrey Barrett, Sonia Goncalves, Ewan Harrison, David K. Jackson, Ian Johnston, Dominic Kwiatkowski, Cordelia Langford, John Sillitoe on behalf of the Wellcome Sanger Institute COVID-19 Surveillance Team |
| EPI_ISL_2935477, EPI_ISL_2935478, EPI_ISL_2935479 | Lighthouse Laboratory Plymouth | Wellcome Sanger Institute for the COVID-19 Genomics UK (COG-UK) Consortium | Lighthouse Laboratory Plymouth and Alex Alderton, Roberto Amato, Jeffrey Barrett, Sonia Goncalves, Ewan Harrison, David K. Jackson, Ian Johnston, Dominic Kwiatkowski, Cordelia Langford, John Sillitoe on behalf of the Wellcome Sanger Institute COVID-19 Surveillance Team |
| EPI_ISL_2935480 | Lighthouse Lab in Milton Keynes | Wellcome Sanger Institute for the COVID-19 Genomics UK (COG-UK) Consortium | The Lighthouse Lab in Milton Keynes and Alex Alderton, Roberto Amato, Jeffrey Barrett, Sonia Goncalves, Ewan Harrison, David K. Jackson, Ian Johnston, Dominic Kwiatkowski, Cordelia Langford, John Sillitoe on behalf of the Wellcome Sanger Institute COVID-19 Surveillance Team |
| EPI_ISL_2935481 | Lighthouse Lab in Alderley Park | Wellcome Sanger Institute for the COVID-19 Genomics UK (COG-UK) Consortium | Jacquelyn Wynn, Mairead Hyland, The Lighthouse Lab in Alderley Park and Alex Alderton, Roberto Amato, Jeffrey Barrett, Sonia Goncalves, Ewan Harrison, David K. Jackson, Ian Johnston, Dominic Kwiatkowski, Cordelia Langford, John Sillitoe on behalf of the Wellcome Sanger Institute COVID-19 Surveillance Team |
| EPI_ISL_2935482 | Lighthouse Laboratory Plymouth | Wellcome Sanger Institute for the COVID-19 Genomics UK (COG-UK) Consortium | Lighthouse Laboratory Plymouth and Alex Alderton, Roberto Amato, Jeffrey Barrett, Sonia Goncalves, Ewan Harrison, David K. Jackson, Ian Johnston, Dominic Kwiatkowski, Cordelia Langford, John Sillitoe on behalf of the Wellcome Sanger Institute COVID-19 Surveillance Team |
| EPI_ISL_2935483 | Lighthouse Lab in Milton Keynes | Wellcome Sanger Institute for the COVID-19 Genomics UK (COG-UK) Consortium | The Lighthouse Lab in Milton Keynes and Alex Alderton, Roberto Amato, Jeffrey Barrett, Sonia Goncalves, Ewan Harrison, David K. Jackson, Ian Johnston, Dominic Kwiatkowski, Cordelia Langford, John Sillitoe on behalf of the Wellcome Sanger Institute COVID-19 Surveillance Team |
| EPI_ISL_2935484, EPI_ISL_2935485 | Lighthouse Laboratory Plymouth | Wellcome Sanger Institute for the COVID-19 Genomics UK (COG-UK) Consortium | Lighthouse Laboratory Plymouth and Alex Alderton, Roberto Amato, Jeffrey Barrett, Sonia Goncalves, Ewan Harrison, David K. Jackson, Ian Johnston, Dominic Kwiatkowski, Cordelia Langford, John Sillitoe on behalf of the Wellcome Sanger Institute COVID-19 Surveillance Team |
| EPI_ISL_2935486 | Lighthouse Lab in Milton Keynes | Wellcome Sanger Institute for the COVID-19 Genomics UK (COG-UK) Consortium | The Lighthouse Lab in Milton Keynes and Alex Alderton, Roberto Amato, Jeffrey Barrett, Sonia Goncalves, Ewan Harrison, David K. Jackson, Ian Johnston, Dominic Kwiatkowski, Cordelia Langford, John Sillitoe on behalf of the Wellcome Sanger Institute COVID-19 Surveillance Team |
| EPI_ISL_2935487, EPI_ISL_2935488 | Lighthouse Lab in Alderley Park | Wellcome Sanger Institute for the COVID-19 Genomics UK (COG-UK) Consortium | Jacquelyn Wynn, Mairead Hyland, The Lighthouse Lab in Alderley Park and Alex Alderton, Roberto Amato, Jeffrey Barrett, Sonia Goncalves, Ewan Harrison, David K. Jackson, Ian Johnston, Dominic Kwiatkowski, Cordelia Langford, John Sillitoe on behalf of the Wellcome Sanger Institute COVID-19 Surveillance Team |
| EPI_ISL_2935489 | Lighthouse Laboratory Plymouth | Wellcome Sanger Institute for the COVID-19 Genomics UK (COG-UK) Consortium | Lighthouse Laboratory Plymouth and Alex Alderton, Roberto Amato, Jeffrey Barrett, Sonia Goncalves, Ewan Harrison, David K. Jackson, Ian Johnston, Dominic Kwiatkowski, Cordelia Langford, John Sillitoe on behalf of the Wellcome Sanger Institute COVID-19 Surveillance Team |
| EPI_ISL_2935490 | Lighthouse Lab in Alderley Park | Wellcome Sanger Institute for the COVID-19 Genomics UK (COG-UK) Consortium | Jacquelyn Wynn, Mairead Hyland, The Lighthouse Lab in Alderley Park and Alex Alderton, Roberto Amato, Jeffrey Barrett, Sonia Goncalves, Ewan Harrison, David K. Jackson, Ian Johnston, Dominic Kwiatkowski, Cordelia Langford, John Sillitoe on behalf of the Wellcome Sanger Institute COVID-19 Surveillance Team |
| EPI_ISL_2935491 | Lighthouse Laboratory Plymouth | Wellcome Sanger Institute for the COVID-19 Genomics UK (COG-UK) Consortium | Lighthouse Laboratory Plymouth and Alex Alderton, Roberto Amato, Jeffrey Barrett, Sonia Goncalves, Ewan Harrison, David K. Jackson, Ian Johnston, Dominic Kwiatkowski, Cordelia Langford, John Sillitoe on behalf of the Wellcome Sanger Institute COVID-19 Surveillance Team |
| EPI_ISL_2935492, EPI_ISL_2935493 | Lighthouse Lab in Alderley Park | Wellcome Sanger Institute for the COVID-19 Genomics UK (COG-UK) Consortium | Jacquelyn Wynn, Mairead Hyland, The Lighthouse Lab in Alderley Park and Alex Alderton, Roberto Amato, Jeffrey Barrett, Sonia Goncalves, Ewan Harrison, David K. Jackson, Ian Johnston, Dominic Kwiatkowski, Cordelia Langford, John Sillitoe on behalf of the Wellcome Sanger Institute COVID-19 Surveillance Team |
| EPI_ISL_2935494, EPI_ISL_2935495 | Lighthouse Laboratory Plymouth | Wellcome Sanger Institute for the COVID-19 Genomics UK (COG-UK) Consortium | Lighthouse Laboratory Plymouth and Alex Alderton, Roberto Amato, Jeffrey Barrett, Sonia Goncalves, Ewan Harrison, David K. Jackson, Ian Johnston, Dominic Kwiatkowski, Cordelia Langford, John Sillitoe on behalf of the Wellcome Sanger Institute COVID-19 Surveillance Team |
| EPI_ISL_2935496, EPI_ISL_2935497 | Lighthouse Lab in Milton Keynes | Wellcome Sanger Institute for the COVID-19 Genomics UK (COG-UK) Consortium | The Lighthouse Lab in Milton Keynes and Alex Alderton, Roberto Amato, Jeffrey Barrett, Sonia Goncalves, Ewan Harrison, David K. Jackson, Ian Johnston, Dominic Kwiatkowski, Cordelia Langford, John Sillitoe on behalf of the Wellcome Sanger Institute COVID-19 Surveillance Team |
| EPI_ISL_2935498, EPI_ISL_2935499, EPI_ISL_2935500, EPI_ISL_2935501, EPI_ISL_2935502, EPI_ISL_2935503, EPI_ISL_2935504, EPI_ISL_2935505 | Lighthouse Laboratory Plymouth | Wellcome Sanger Institute for the COVID-19 Genomics UK (COG-UK) Consortium | Lighthouse Laboratory Plymouth and Alex Alderton, Roberto Amato, Jeffrey Barrett, Sonia Goncalves, Ewan Harrison, David K. Jackson, Ian Johnston, Dominic Kwiatkowski, Cordelia Langford, John Sillitoe on behalf of the Wellcome Sanger Institute COVID-19 Surveillance Team |
| EPI_ISL_2935506 | Lighthouse Lab in Alderley Park | Wellcome Sanger Institute for the COVID-19 Genomics UK (COG-UK) Consortium | Jacquelyn Wynn, Mairead Hyland, The Lighthouse Lab in Alderley Park and Alex Alderton, Roberto Amato, Jeffrey Barrett, Sonia Goncalves, Ewan Harrison, David K. Jackson, Ian Johnston, Dominic Kwiatkowski, Cordelia Langford, John Sillitoe on behalf of the Wellcome Sanger Institute COVID-19 Surveillance Team |
| EPI_ISL_2935507 | Lighthouse Lab in Milton Keynes | Wellcome Sanger Institute for the COVID-19 Genomics UK (COG-UK) Consortium | The Lighthouse Lab in Milton Keynes and Alex Alderton, Roberto Amato, Jeffrey Barrett, Sonia Goncalves, Ewan Harrison, David K. Jackson, Ian Johnston, Dominic Kwiatkowski, Cordelia Langford, John Sillitoe on behalf of the Wellcome Sanger Institute COVID-19 Surveillance Team |
| EPI_ISL_2935508 | Lighthouse Laboratory Plymouth | Wellcome Sanger Institute for the COVID-19 Genomics UK (COG-UK) Consortium | Lighthouse Laboratory Plymouth and Alex Alderton, Roberto Amato, Jeffrey Barrett, Sonia Goncalves, Ewan Harrison, David K. Jackson, Ian Johnston, Dominic Kwiatkowski, Cordelia Langford, John Sillitoe on behalf of the Wellcome Sanger Institute COVID-19 Surveillance Team |
| EPI_ISL_2935509 | Lighthouse Lab in Milton Keynes | Wellcome Sanger Institute for the COVID-19 Genomics UK (COG-UK) Consortium | The Lighthouse Lab in Milton Keynes and Alex Alderton, Roberto Amato, Jeffrey Barrett, Sonia Goncalves, Ewan Harrison, David K. Jackson, Ian Johnston, Dominic Kwiatkowski, Cordelia Langford, John Sillitoe on behalf of the Wellcome Sanger Institute COVID-19 Surveillance Team |
| EPI_ISL_2935510, EPI_ISL_2935511, EPI_ISL_2935512 | Lighthouse Laboratory Plymouth | Wellcome Sanger Institute for the COVID-19 Genomics UK (COG-UK) Consortium | Lighthouse Laboratory Plymouth and Alex Alderton, Roberto Amato, Jeffrey Barrett, Sonia Goncalves, Ewan Harrison, David K. Jackson, Ian Johnston, Dominic Kwiatkowski, Cordelia Langford, John Sillitoe on behalf of the Wellcome Sanger Institute COVID-19 Surveillance Team |
| EPI_ISL_2935513 | Lighthouse Lab in Milton Keynes | Wellcome Sanger Institute for the COVID-19 Genomics UK (COG-UK) Consortium | The Lighthouse Lab in Milton Keynes and Alex Alderton, Roberto Amato, Jeffrey Barrett, Sonia Goncalves, Ewan Harrison, David K. Jackson, Ian Johnston, Dominic Kwiatkowski, Cordelia Langford, John Sillitoe on behalf of the Wellcome Sanger Institute COVID-19 Surveillance Team |
| EPI_ISL_2935514 | Lighthouse Lab in Alderley Park | Wellcome Sanger Institute for the COVID-19 Genomics UK (COG-UK) Consortium | Jacquelyn Wynn, Mairead Hyland, The Lighthouse Lab in Alderley Park and Alex Alderton, Roberto Amato, Jeffrey Barrett, Sonia Goncalves, Ewan Harrison, David K. Jackson, Ian Johnston, Dominic Kwiatkowski, Cordelia Langford, John Sillitoe on behalf of the Wellcome Sanger Institute COVID-19 Surveillance Team |
| EPI_ISL_2935515, EPI_ISL_2935516, EPI_ISL_2935517, EPI_ISL_2935518 | Lighthouse Laboratory Plymouth | Wellcome Sanger Institute for the COVID-19 Genomics UK (COG-UK) Consortium | Lighthouse Laboratory Plymouth and Alex Alderton, Roberto Amato, Jeffrey Barrett, Sonia Goncalves, Ewan Harrison, David K. Jackson, Ian Johnston, Dominic Kwiatkowski, Cordelia Langford, John Sillitoe on behalf of the Wellcome Sanger Institute COVID-19 Surveillance Team |
| EPI_ISL_2935519 | Lighthouse Lab in Alderley Park | Wellcome Sanger Institute for the COVID-19 Genomics UK (COG-UK) Consortium | Jacquelyn Wynn, Mairead Hyland, The Lighthouse Lab in Alderley Park and Alex Alderton, Roberto Amato, Jeffrey Barrett, Sonia Goncalves, Ewan Harrison, David K. Jackson, Ian Johnston, Dominic Kwiatkowski, Cordelia Langford, John Sillitoe on behalf of the Wellcome Sanger Institute COVID-19 Surveillance Team |
| EPI_ISL_2935520 | Lighthouse Lab in Milton Keynes | Wellcome Sanger Institute for the COVID-19 Genomics UK (COG-UK) Consortium | The Lighthouse Lab in Milton Keynes and Alex Alderton, Roberto Amato, Jeffrey Barrett, Sonia Goncalves, Ewan Harrison, David K. Jackson, Ian Johnston, Dominic Kwiatkowski, Cordelia Langford, John Sillitoe on behalf of the Wellcome Sanger Institute COVID-19 Surveillance Team |
| EPI_ISL_2935521, EPI_ISL_2935522, EPI_ISL_2935523 | Lighthouse Laboratory Plymouth | Wellcome Sanger Institute for the COVID-19 Genomics UK (COG-UK) Consortium | Lighthouse Laboratory Plymouth and Alex Alderton, Roberto Amato, Jeffrey Barrett, Sonia Goncalves, Ewan Harrison, David K. Jackson, Ian Johnston, Dominic Kwiatkowski, Cordelia Langford, John Sillitoe on behalf of the Wellcome Sanger Institute COVID-19 Surveillance Team |
| EPI_ISL_2935524, EPI_ISL_2935525 | Lighthouse Lab in Milton Keynes | Wellcome Sanger Institute for the COVID-19 Genomics UK (COG-UK) Consortium | The Lighthouse Lab in Milton Keynes and Alex Alderton, Roberto Amato, Jeffrey Barrett, Sonia Goncalves, Ewan Harrison, David K. Jackson, Ian Johnston, Dominic Kwiatkowski, Cordelia Langford, John Sillitoe on behalf of the Wellcome Sanger Institute COVID-19 Surveillance Team |
| EPI_ISL_2935526, EPI_ISL_2935527 | Lighthouse Laboratory Plymouth | Wellcome Sanger Institute for the COVID-19 Genomics UK (COG-UK) Consortium | Lighthouse Laboratory Plymouth and Alex Alderton, Roberto Amato, Jeffrey Barrett, Sonia Goncalves, Ewan Harrison, David K. Jackson, Ian Johnston, Dominic Kwiatkowski, Cordelia Langford, John Sillitoe on behalf of the Wellcome Sanger Institute COVID-19 Surveillance Team |
| EPI_ISL_2935528 | Lighthouse Lab in Milton Keynes | Wellcome Sanger Institute for the COVID-19 Genomics UK (COG-UK) Consortium | The Lighthouse Lab in Milton Keynes and Alex Alderton, Roberto Amato, Jeffrey Barrett, Sonia Goncalves, Ewan Harrison, David K. Jackson, Ian Johnston, Dominic Kwiatkowski, Cordelia Langford, John Sillitoe on behalf of the Wellcome Sanger Institute COVID-19 Surveillance Team |
| EPI_ISL_2935529 | Lighthouse Laboratory Plymouth | Wellcome Sanger Institute for the COVID-19 Genomics UK (COG-UK) Consortium | Lighthouse Laboratory Plymouth and Alex Alderton, Roberto Amato, Jeffrey Barrett, Sonia Goncalves, Ewan Harrison, David K. Jackson, Ian Johnston, Dominic Kwiatkowski, Cordelia Langford, John Sillitoe on behalf of the Wellcome Sanger Institute COVID-19 Surveillance Team |
| EPI_ISL_2935530 | Lighthouse Lab in Milton Keynes | Wellcome Sanger Institute for the COVID-19 Genomics UK (COG-UK) Consortium | The Lighthouse Lab in Milton Keynes and Alex Alderton, Roberto Amato, Jeffrey Barrett, Sonia Goncalves, Ewan Harrison, David K. Jackson, Ian Johnston, Dominic Kwiatkowski, Cordelia Langford, John Sillitoe on behalf of the Wellcome Sanger Institute COVID-19 Surveillance Team |
| EPI_ISL_2935531 | Lighthouse Lab in Alderley Park | Wellcome Sanger Institute for the COVID-19 Genomics UK (COG-UK) Consortium | Jacquelyn Wynn, Mairead Hyland, The Lighthouse Lab in Alderley Park and Alex Alderton, Roberto Amato, Jeffrey Barrett, Sonia Goncalves, Ewan Harrison, David K. Jackson, Ian Johnston, Dominic Kwiatkowski, Cordelia Langford, John Sillitoe on behalf of the Wellcome Sanger Institute COVID-19 Surveillance Team |

| | | | |
|---|---|---|---|
| EPI_ISL_2935532 | Lighthouse Lab in Milton Keynes | Wellcome Sanger Institute for the COVID-19 Genomics UK (COG-UK) Consortium | The Lighthouse Lab in Milton Keynes and Alex Alderton, Roberto Amato, Jeffrey Barrett, Sonia Goncalves, Ewan Harrison, David K. Jackson, Ian Johnston, Dominic Kwiatkowski, Cordelia Langford, John Sillitoe on behalf of the Wellcome Sanger Institute COVID-19 Surveillance Team |
| EPI_ISL_2935533 | Lighthouse Laboratory Plymouth | Wellcome Sanger Institute for the COVID-19 Genomics UK (COG-UK) Consortium | Lighthouse Laboratory Plymouth and Alex Alderton, Roberto Amato, Jeffrey Barrett, Sonia Goncalves, Ewan Harrison, David K. Jackson, Ian Johnston, Dominic Kwiatkowski, Cordelia Langford, John Sillitoe on behalf of the Wellcome Sanger Institute COVID-19 Surveillance Team |
| EPI_ISL_2935534, EPI_ISL_2935535 | Lighthouse Lab in Alderley Park | Wellcome Sanger Institute for the COVID-19 Genomics UK (COG-UK) Consortium | Jacquelyn Wynn, Mairead Hyland, The Lighthouse Lab in Alderley Park and Alex Alderton, Roberto Amato, Jeffrey Barrett, Sonia Goncalves, Ewan Harrison, David K. Jackson, Ian Johnston, Dominic Kwiatkowski, Cordelia Langford, John Sillitoe on behalf of the Wellcome Sanger Institute COVID-19 Surveillance Team |
| EPI_ISL_2935536 | Lighthouse Lab in Milton Keynes | Wellcome Sanger Institute for the COVID-19 Genomics UK (COG-UK) Consortium | The Lighthouse Lab in Milton Keynes and Alex Alderton, Roberto Amato, Jeffrey Barrett, Sonia Goncalves, Ewan Harrison, David K. Jackson, Ian Johnston, Dominic Kwiatkowski, Cordelia Langford, John Sillitoe on behalf of the Wellcome Sanger Institute COVID-19 Surveillance Team |
| EPI_ISL_2935537, EPI_ISL_2935538 | Lighthouse Laboratory Plymouth | Wellcome Sanger Institute for the COVID-19 Genomics UK (COG-UK) Consortium | Lighthouse Laboratory Plymouth and Alex Alderton, Roberto Amato, Jeffrey Barrett, Sonia Goncalves, Ewan Harrison, David K. Jackson, Ian Johnston, Dominic Kwiatkowski, Cordelia Langford, John Sillitoe on behalf of the Wellcome Sanger Institute COVID-19 Surveillance Team |
| EPI_ISL_2935539 | Lighthouse Lab in Milton Keynes | Wellcome Sanger Institute for the COVID-19 Genomics UK (COG-UK) Consortium | The Lighthouse Lab in Milton Keynes and Alex Alderton, Roberto Amato, Jeffrey Barrett, Sonia Goncalves, Ewan Harrison, David K. Jackson, Ian Johnston, Dominic Kwiatkowski, Cordelia Langford, John Sillitoe on behalf of the Wellcome Sanger Institute COVID-19 Surveillance Team |
| EPI_ISL_2935540 | Lighthouse Laboratory Plymouth | Wellcome Sanger Institute for the COVID-19 Genomics UK (COG-UK) Consortium | Lighthouse Laboratory Plymouth and Alex Alderton, Roberto Amato, Jeffrey Barrett, Sonia Goncalves, Ewan Harrison, David K. Jackson, Ian Johnston, Dominic Kwiatkowski, Cordelia Langford, John Sillitoe on behalf of the Wellcome Sanger Institute COVID-19 Surveillance Team |
| EPI_ISL_2935541, EPI_ISL_2935542 | Lighthouse Lab in Milton Keynes | Wellcome Sanger Institute for the COVID-19 Genomics UK (COG-UK) Consortium | The Lighthouse Lab in Milton Keynes and Alex Alderton, Roberto Amato, Jeffrey Barrett, Sonia Goncalves, Ewan Harrison, David K. Jackson, Ian Johnston, Dominic Kwiatkowski, Cordelia Langford, John Sillitoe on behalf of the Wellcome Sanger Institute COVID-19 Surveillance Team |
| EPI_ISL_2935543 | Lighthouse Lab in Alderley Park | Wellcome Sanger Institute for the COVID-19 Genomics UK (COG-UK) Consortium | Jacquelyn Wynn, Mairead Hyland, The Lighthouse Lab in Alderley Park and Alex Alderton, Roberto Amato, Jeffrey Barrett, Sonia Goncalves, Ewan Harrison, David K. Jackson, Ian Johnston, Dominic Kwiatkowski, Cordelia Langford, John Sillitoe on behalf of the Wellcome Sanger Institute COVID-19 Surveillance Team |
| EPI_ISL_2935544, EPI_ISL_2935545, EPI_ISL_2935546, EPI_ISL_2935547 | Lighthouse Laboratory Plymouth | Wellcome Sanger Institute for the COVID-19 Genomics UK (COG-UK) Consortium | Lighthouse Laboratory Plymouth and Alex Alderton, Roberto Amato, Jeffrey Barrett, Sonia Goncalves, Ewan Harrison, David K. Jackson, Ian Johnston, Dominic Kwiatkowski, Cordelia Langford, John Sillitoe on behalf of the Wellcome Sanger Institute COVID-19 Surveillance Team |
| EPI_ISL_2935548 | Lighthouse Lab in Alderley Park | Wellcome Sanger Institute for the COVID-19 Genomics UK (COG-UK) Consortium | Jacquelyn Wynn, Mairead Hyland, The Lighthouse Lab in Alderley Park and Alex Alderton, Roberto Amato, Jeffrey Barrett, Sonia Goncalves, Ewan Harrison, David K. Jackson, Ian Johnston, Dominic Kwiatkowski, Cordelia Langford, John Sillitoe on behalf of the Wellcome Sanger Institute COVID-19 Surveillance Team |
| EPI_ISL_2935549, EPI_ISL_2935550, EPI_ISL_2935551, EPI_ISL_2935552, EPI_ISL_2935553, EPI_ISL_2935554, EPI_ISL_2935555, EPI_ISL_2935556 | Lighthouse Laboratory Plymouth | Wellcome Sanger Institute for the COVID-19 Genomics UK (COG-UK) Consortium | Lighthouse Laboratory Plymouth and Alex Alderton, Roberto Amato, Jeffrey Barrett, Sonia Goncalves, Ewan Harrison, David K. Jackson, Ian Johnston, Dominic Kwiatkowski, Cordelia Langford, John Sillitoe on behalf of the Wellcome Sanger Institute COVID-19 Surveillance Team |
| EPI_ISL_2935557 | Lighthouse Lab in Milton Keynes | Wellcome Sanger Institute for the COVID-19 Genomics UK (COG-UK) Consortium | The Lighthouse Lab in Milton Keynes and Alex Alderton, Roberto Amato, Jeffrey Barrett, Sonia Goncalves, Ewan Harrison, David K. Jackson, Ian Johnston, Dominic Kwiatkowski, Cordelia Langford, John Sillitoe on behalf of the Wellcome Sanger Institute COVID-19 Surveillance Team |
| EPI_ISL_2935558 | Lighthouse Laboratory Plymouth | Wellcome Sanger Institute for the COVID-19 Genomics UK (COG-UK) Consortium | Lighthouse Laboratory Plymouth and Alex Alderton, Roberto Amato, Jeffrey Barrett, Sonia Goncalves, Ewan Harrison, David K. Jackson, Ian Johnston, Dominic Kwiatkowski, Cordelia Langford, John Sillitoe on behalf of the Wellcome Sanger Institute COVID-19 Surveillance Team |
| EPI_ISL_2935559 | Lighthouse Lab in Milton Keynes | Wellcome Sanger Institute for the COVID-19 Genomics UK (COG-UK) Consortium | The Lighthouse Lab in Milton Keynes and Alex Alderton, Roberto Amato, Jeffrey Barrett, Sonia Goncalves, Ewan Harrison, David K. Jackson, Ian Johnston, Dominic Kwiatkowski, Cordelia Langford, John Sillitoe on behalf of the Wellcome Sanger Institute COVID-19 Surveillance Team |
| EPI_ISL_2935560 | Lighthouse Lab in Alderley Park | Wellcome Sanger Institute for the COVID-19 Genomics UK (COG-UK) Consortium | Jacquelyn Wynn, Mairead Hyland, The Lighthouse Lab in Alderley Park and Alex Alderton, Roberto Amato, Jeffrey Barrett, Sonia Goncalves, Ewan Harrison, David K. Jackson, Ian Johnston, Dominic Kwiatkowski, Cordelia Langford, John Sillitoe on behalf of the Wellcome Sanger Institute COVID-19 Surveillance Team |
| EPI_ISL_2935561 | Lighthouse Lab in Milton Keynes | Wellcome Sanger Institute for the COVID-19 Genomics UK (COG-UK) Consortium | The Lighthouse Lab in Milton Keynes and Alex Alderton, Roberto Amato, Jeffrey Barrett, Sonia Goncalves, Ewan Harrison, David K. Jackson, Ian Johnston, Dominic Kwiatkowski, Cordelia Langford, John Sillitoe on behalf of the Wellcome Sanger Institute COVID-19 Surveillance Team |
| EPI_ISL_2935562, EPI_ISL_2935563 | Lighthouse Laboratory Plymouth | Wellcome Sanger Institute for the COVID-19 Genomics UK (COG-UK) Consortium | Lighthouse Laboratory Plymouth and Alex Alderton, Roberto Amato, Jeffrey Barrett, Sonia Goncalves, Ewan Harrison, David K. Jackson, Ian Johnston, Dominic Kwiatkowski, Cordelia Langford, John Sillitoe on behalf of the Wellcome Sanger Institute COVID-19 Surveillance Team |
| EPI_ISL_2935564 | Lighthouse Lab in Milton Keynes | Wellcome Sanger Institute for the COVID-19 Genomics UK (COG-UK) Consortium | The Lighthouse Lab in Milton Keynes and Alex Alderton, Roberto Amato, Jeffrey Barrett, Sonia Goncalves, Ewan Harrison, David K. Jackson, Ian Johnston, Dominic Kwiatkowski, Cordelia Langford, John Sillitoe on behalf of the Wellcome Sanger Institute COVID-19 Surveillance Team |
| EPI_ISL_2935565 | Lighthouse Laboratory Plymouth | Wellcome Sanger Institute for the COVID-19 Genomics UK (COG-UK) Consortium | Lighthouse Laboratory Plymouth and Alex Alderton, Roberto Amato, Jeffrey Barrett, Sonia Goncalves, Ewan Harrison, David K. Jackson, Ian Johnston, Dominic Kwiatkowski, Cordelia Langford, John Sillitoe on behalf of the Wellcome Sanger Institute COVID-19 Surveillance Team |
| EPI_ISL_2935566 | Lighthouse Lab in Alderley Park | Wellcome Sanger Institute for the COVID-19 Genomics UK (COG-UK) Consortium | Jacquelyn Wynn, Mairead Hyland, The Lighthouse Lab in Alderley Park and Alex Alderton, Roberto Amato, Jeffrey Barrett, Sonia Goncalves, Ewan Harrison, David K. Jackson, Ian Johnston, Dominic Kwiatkowski, Cordelia Langford, John Sillitoe on behalf of the Wellcome Sanger Institute COVID-19 Surveillance Team |
| EPI_ISL_2935567, EPI_ISL_2935568, EPI_ISL_2935569 | Lighthouse Laboratory Plymouth | Wellcome Sanger Institute for the COVID-19 Genomics UK (COG-UK) Consortium | Lighthouse Laboratory Plymouth and Alex Alderton, Roberto Amato, Jeffrey Barrett, Sonia Goncalves, Ewan Harrison, David K. Jackson, Ian Johnston, Dominic Kwiatkowski, Cordelia Langford, John Sillitoe on behalf of the Wellcome Sanger Institute COVID-19 Surveillance Team |
| EPI_ISL_2935570, EPI_ISL_2935571 | Lighthouse Lab in Milton Keynes | Wellcome Sanger Institute for the COVID-19 Genomics UK (COG-UK) Consortium | The Lighthouse Lab in Milton Keynes and Alex Alderton, Roberto Amato, Jeffrey Barrett, Sonia Goncalves, Ewan Harrison, David K. Jackson, Ian Johnston, Dominic Kwiatkowski, Cordelia Langford, John Sillitoe on behalf of the Wellcome Sanger Institute COVID-19 Surveillance Team |
| EPI_ISL_2935572 | Lighthouse Lab in Alderley Park | Wellcome Sanger Institute for the COVID-19 Genomics UK (COG-UK) Consortium | Jacquelyn Wynn, Mairead Hyland, The Lighthouse Lab in Alderley Park and Alex Alderton, Roberto Amato, Jeffrey Barrett, Sonia Goncalves, Ewan Harrison, David K. Jackson, Ian Johnston, Dominic Kwiatkowski, Cordelia Langford, John Sillitoe on behalf of the Wellcome Sanger Institute COVID-19 Surveillance Team |
| EPI_ISL_2935573, EPI_ISL_2935574, EPI_ISL_2935575 | Lighthouse Laboratory Plymouth | Wellcome Sanger Institute for the COVID-19 Genomics UK (COG-UK) Consortium | Lighthouse Laboratory Plymouth and Alex Alderton, Roberto Amato, Jeffrey Barrett, Sonia Goncalves, Ewan Harrison, David K. Jackson, Ian Johnston, Dominic Kwiatkowski, Cordelia Langford, John Sillitoe on behalf of the Wellcome Sanger Institute COVID-19 Surveillance Team |
| EPI_ISL_2935576, EPI_ISL_2935577, EPI_ISL_2935578 | Lighthouse Lab in Alderley Park | Wellcome Sanger Institute for the COVID-19 Genomics UK (COG-UK) Consortium | Jacquelyn Wynn, Mairead Hyland, The Lighthouse Lab in Alderley Park and Alex Alderton, Roberto Amato, Jeffrey Barrett, Sonia Goncalves, Ewan Harrison, David K. Jackson, Ian Johnston, Dominic Kwiatkowski, Cordelia Langford, John Sillitoe on behalf of the Wellcome Sanger Institute COVID-19 Surveillance Team |
| EPI_ISL_2935579, EPI_ISL_2935580 | Lighthouse Laboratory Plymouth | Wellcome Sanger Institute for the COVID-19 Genomics UK (COG-UK) Consortium | Lighthouse Laboratory Plymouth and Alex Alderton, Roberto Amato, Jeffrey Barrett, Sonia Goncalves, Ewan Harrison, David K. Jackson, Ian Johnston, Dominic Kwiatkowski, Cordelia Langford, John Sillitoe on behalf of the Wellcome Sanger Institute COVID-19 Surveillance Team |
| EPI_ISL_2935581 | Lighthouse Lab in Milton Keynes | Wellcome Sanger Institute for the COVID-19 Genomics UK (COG-UK) Consortium | The Lighthouse Lab in Milton Keynes and Alex Alderton, Roberto Amato, Jeffrey Barrett, Sonia Goncalves, Ewan Harrison, David K. Jackson, Ian Johnston, Dominic Kwiatkowski, Cordelia Langford, John Sillitoe on behalf of the Wellcome Sanger Institute COVID-19 Surveillance Team |
| EPI_ISL_2935582, EPI_ISL_2935583 | Lighthouse Laboratory Plymouth | Wellcome Sanger Institute for the COVID-19 Genomics UK (COG-UK) Consortium | Lighthouse Laboratory Plymouth and Alex Alderton, Roberto Amato, Jeffrey Barrett, Sonia Goncalves, Ewan Harrison, David K. Jackson, Ian Johnston, Dominic Kwiatkowski, Cordelia Langford, John Sillitoe on behalf of the Wellcome Sanger Institute COVID-19 Surveillance Team |
| EPI_ISL_2935584, EPI_ISL_2935585 | Lighthouse Lab in Milton Keynes | Wellcome Sanger Institute for the COVID-19 Genomics UK (COG-UK) Consortium | The Lighthouse Lab in Milton Keynes and Alex Alderton, Roberto Amato, Jeffrey Barrett, Sonia Goncalves, Ewan Harrison, David K. Jackson, Ian Johnston, Dominic Kwiatkowski, Cordelia Langford, John Sillitoe on behalf of the Wellcome Sanger Institute COVID-19 Surveillance Team |
| EPI_ISL_2935586, EPI_ISL_2935587, EPI_ISL_2935588 | Lighthouse Lab in Alderley Park | Wellcome Sanger Institute for the COVID-19 Genomics UK (COG-UK) Consortium | Jacquelyn Wynn, Mairead Hyland, The Lighthouse Lab in Alderley Park and Alex Alderton, Roberto Amato, Jeffrey Barrett, Sonia Goncalves, Ewan Harrison, David K. Jackson, Ian Johnston, Dominic Kwiatkowski, Cordelia Langford, John Sillitoe on behalf of the Wellcome Sanger Institute COVID-19 Surveillance Team |
| EPI_ISL_2935589 | Lighthouse Laboratory Plymouth | Wellcome Sanger Institute for the COVID-19 Genomics UK (COG-UK) Consortium | Lighthouse Laboratory Plymouth and Alex Alderton, Roberto Amato, Jeffrey Barrett, Sonia Goncalves, Ewan Harrison, David K. Jackson, Ian Johnston, Dominic Kwiatkowski, Cordelia Langford, John Sillitoe on behalf of the Wellcome Sanger Institute COVID-19 Surveillance Team |
| EPI_ISL_2935590 | Lighthouse Lab in Milton Keynes | Wellcome Sanger Institute for the COVID-19 Genomics UK (COG-UK) Consortium | The Lighthouse Lab in Milton Keynes and Alex Alderton, Roberto Amato, Jeffrey Barrett, Sonia Goncalves, Ewan Harrison, David K. Jackson, Ian Johnston, Dominic Kwiatkowski, Cordelia Langford, John Sillitoe on behalf of the Wellcome Sanger Institute COVID-19 Surveillance Team |
| EPI_ISL_2935591 | Lighthouse Laboratory Plymouth | Wellcome Sanger Institute for the COVID-19 Genomics UK (COG-UK) Consortium | Lighthouse Laboratory Plymouth and Alex Alderton, Roberto Amato, Jeffrey Barrett, Sonia Goncalves, Ewan Harrison, David K. Jackson, Ian Johnston, Dominic Kwiatkowski, Cordelia Langford, John Sillitoe on behalf of the Wellcome Sanger Institute COVID-19 Surveillance Team |
| EPI_ISL_2935592 | Lighthouse Lab in Alderley Park | Wellcome Sanger Institute for the COVID-19 Genomics UK (COG-UK) Consortium | Jacquelyn Wynn, Mairead Hyland, The Lighthouse Lab in Alderley Park and Alex Alderton, Roberto Amato, Jeffrey Barrett, Sonia Goncalves, Ewan Harrison, David K. Jackson, Ian Johnston, Dominic Kwiatkowski, Cordelia Langford, John Sillitoe on behalf of the Wellcome Sanger Institute COVID-19 Surveillance Team |
| EPI_ISL_2935593 | Lighthouse Laboratory Plymouth | Wellcome Sanger Institute for the COVID-19 Genomics UK (COG-UK) Consortium | Lighthouse Laboratory Plymouth and Alex Alderton, Roberto Amato, Jeffrey Barrett, Sonia Goncalves, Ewan Harrison, David K. Jackson, Ian Johnston, Dominic Kwiatkowski, Cordelia Langford, John Sillitoe on behalf of the Wellcome Sanger Institute COVID-19 Surveillance Team |
| EPI_ISL_2935594, EPI_ISL_2935595, EPI_ISL_2935596 | Lighthouse Lab in Milton Keynes | Wellcome Sanger Institute for the COVID-19 Genomics UK (COG-UK) Consortium | The Lighthouse Lab in Milton Keynes and Alex Alderton, Roberto Amato, Jeffrey Barrett, Sonia Goncalves, Ewan Harrison, David K. Jackson, Ian Johnston, Dominic Kwiatkowski, Cordelia Langford, John Sillitoe on behalf of the Wellcome Sanger Institute COVID-19 Surveillance Team |
| EPI_ISL_2935597, EPI_ISL_2935598, EPI_ISL_2935599, EPI_ISL_2935600 | Lighthouse Laboratory Plymouth | Wellcome Sanger Institute for the COVID-19 Genomics UK (COG-UK) Consortium | Lighthouse Laboratory Plymouth and Alex Alderton, Roberto Amato, Jeffrey Barrett, Sonia Goncalves, Ewan Harrison, David K. Jackson, Ian Johnston, Dominic Kwiatkowski, Cordelia Langford, John Sillitoe on behalf of the Wellcome Sanger Institute COVID-19 Surveillance Team |
| EPI_ISL_2935601 | Lighthouse Lab in Milton Keynes | Wellcome Sanger Institute for the COVID-19 Genomics UK (COG-UK) Consortium | The Lighthouse Lab in Milton Keynes and Alex Alderton, Roberto Amato, Jeffrey Barrett, Sonia Goncalves, Ewan Harrison, David K. Jackson, Ian Johnston, Dominic Kwiatkowski, Cordelia Langford, John Sillitoe on behalf of the Wellcome Sanger Institute COVID-19 Surveillance Team |

| | | | |
|---|---|---|---|
| EPI_ISL_2935602, EPI_ISL_2935603, EPI_ISL_2935604 | Lighthouse Lab in Alderley Park | Wellcome Sanger Institute for the COVID-19 Genomics UK (COG-UK) Consortium | Jacquelyn Wynn, Mairead Hyland, The Lighthouse Lab in Alderley Park and Alex Alderton, Roberto Amato, Jeffrey Barrett, Sonia Goncalves, Ewan Harrison, David K. Jackson, Ian Johnston, Dominic Kwiatkowski, Cordelia Langford, John Sillitoe on behalf of the Wellcome Sanger Institute COVID-19 Surveillance Team |
| EPI_ISL_2935605 | Lighthouse Laboratory Plymouth | Wellcome Sanger Institute for the COVID-19 Genomics UK (COG-UK) Consortium | Lighthouse Laboratory Plymouth and Alex Alderton, Roberto Amato, Jeffrey Barrett, Sonia Goncalves, Ewan Harrison, David K. Jackson, Ian Johnston, Dominic Kwiatkowski, Cordelia Langford, John Sillitoe on behalf of the Wellcome Sanger Institute COVID-19 Surveillance Team |
| EPI_ISL_2935606 | Lighthouse Lab in Milton Keynes | Wellcome Sanger Institute for the COVID-19 Genomics UK (COG-UK) Consortium | The Lighthouse Lab in Milton Keynes and Alex Alderton, Roberto Amato, Jeffrey Barrett, Sonia Goncalves, Ewan Harrison, David K. Jackson, Ian Johnston, Dominic Kwiatkowski, Cordelia Langford, John Sillitoe on behalf of the Wellcome Sanger Institute COVID-19 Surveillance Team |
| EPI_ISL_2935607 | Lighthouse Laboratory Plymouth | Wellcome Sanger Institute for the COVID-19 Genomics UK (COG-UK) Consortium | Lighthouse Laboratory Plymouth and Alex Alderton, Roberto Amato, Jeffrey Barrett, Sonia Goncalves, Ewan Harrison, David K. Jackson, Ian Johnston, Dominic Kwiatkowski, Cordelia Langford, John Sillitoe on behalf of the Wellcome Sanger Institute COVID-19 Surveillance Team |
| EPI_ISL_2935608 | Lighthouse Lab in Milton Keynes | Wellcome Sanger Institute for the COVID-19 Genomics UK (COG-UK) Consortium | The Lighthouse Lab in Milton Keynes and Alex Alderton, Roberto Amato, Jeffrey Barrett, Sonia Goncalves, Ewan Harrison, David K. Jackson, Ian Johnston, Dominic Kwiatkowski, Cordelia Langford, John Sillitoe on behalf of the Wellcome Sanger Institute COVID-19 Surveillance Team |
| EPI_ISL_2935609, EPI_ISL_2935610 | Lighthouse Lab in Alderley Park | Wellcome Sanger Institute for the COVID-19 Genomics UK (COG-UK) Consortium | Jacquelyn Wynn, Mairead Hyland, The Lighthouse Lab in Alderley Park and Alex Alderton, Roberto Amato, Jeffrey Barrett, Sonia Goncalves, Ewan Harrison, David K. Jackson, Ian Johnston, Dominic Kwiatkowski, Cordelia Langford, John Sillitoe on behalf of the Wellcome Sanger Institute COVID-19 Surveillance Team |
| EPI_ISL_2935611 | Lighthouse Lab in Milton Keynes | Wellcome Sanger Institute for the COVID-19 Genomics UK (COG-UK) Consortium | The Lighthouse Lab in Milton Keynes and Alex Alderton, Roberto Amato, Jeffrey Barrett, Sonia Goncalves, Ewan Harrison, David K. Jackson, Ian Johnston, Dominic Kwiatkowski, Cordelia Langford, John Sillitoe on behalf of the Wellcome Sanger Institute COVID-19 Surveillance Team |
| EPI_ISL_2935612, EPI_ISL_2935613 | Lighthouse Laboratory Plymouth | Wellcome Sanger Institute for the COVID-19 Genomics UK (COG-UK) Consortium | Lighthouse Laboratory Plymouth and Alex Alderton, Roberto Amato, Jeffrey Barrett, Sonia Goncalves, Ewan Harrison, David K. Jackson, Ian Johnston, Dominic Kwiatkowski, Cordelia Langford, John Sillitoe on behalf of the Wellcome Sanger Institute COVID-19 Surveillance Team |
| EPI_ISL_2935614, EPI_ISL_2935615, EPI_ISL_2935616 | Lighthouse Lab in Alderley Park | Wellcome Sanger Institute for the COVID-19 Genomics UK (COG-UK) Consortium | Jacquelyn Wynn, Mairead Hyland, The Lighthouse Lab in Alderley Park and Alex Alderton, Roberto Amato, Jeffrey Barrett, Sonia Goncalves, Ewan Harrison, David K. Jackson, Ian Johnston, Dominic Kwiatkowski, Cordelia Langford, John Sillitoe on behalf of the Wellcome Sanger Institute COVID-19 Surveillance Team |
| EPI_ISL_2935618, EPI_ISL_2935619 | Lighthouse Laboratory Plymouth | Wellcome Sanger Institute for the COVID-19 Genomics UK (COG-UK) Consortium | Lighthouse Laboratory Plymouth and Alex Alderton, Roberto Amato, Jeffrey Barrett, Sonia Goncalves, Ewan Harrison, David K. Jackson, Ian Johnston, Dominic Kwiatkowski, Cordelia Langford, John Sillitoe on behalf of the Wellcome Sanger Institute COVID-19 Surveillance Team |
| EPI_ISL_2935620 | Lighthouse Lab in Milton Keynes | Wellcome Sanger Institute for the COVID-19 Genomics UK (COG-UK) Consortium | The Lighthouse Lab in Milton Keynes and Alex Alderton, Roberto Amato, Jeffrey Barrett, Sonia Goncalves, Ewan Harrison, David K. Jackson, Ian Johnston, Dominic Kwiatkowski, Cordelia Langford, John Sillitoe on behalf of the Wellcome Sanger Institute COVID-19 Surveillance Team |
| EPI_ISL_2935621 | Lighthouse Laboratory Plymouth | Wellcome Sanger Institute for the COVID-19 Genomics UK (COG-UK) Consortium | Lighthouse Laboratory Plymouth and Alex Alderton, Roberto Amato, Jeffrey Barrett, Sonia Goncalves, Ewan Harrison, David K. Jackson, Ian Johnston, Dominic Kwiatkowski, Cordelia Langford, John Sillitoe on behalf of the Wellcome Sanger Institute COVID-19 Surveillance Team |
| EPI_ISL_2935622, EPI_ISL_2935623, EPI_ISL_2935624, EPI_ISL_2935625, EPI_ISL_2935626 | Lighthouse Lab in Milton Keynes | Wellcome Sanger Institute for the COVID-19 Genomics UK (COG-UK) Consortium | The Lighthouse Lab in Milton Keynes and Alex Alderton, Roberto Amato, Jeffrey Barrett, Sonia Goncalves, Ewan Harrison, David K. Jackson, Ian Johnston, Dominic Kwiatkowski, Cordelia Langford, John Sillitoe on behalf of the Wellcome Sanger Institute COVID-19 Surveillance Team |
| EPI_ISL_2935627 | Lighthouse Laboratory Plymouth | Wellcome Sanger Institute for the COVID-19 Genomics UK (COG-UK) Consortium | Lighthouse Laboratory Plymouth and Alex Alderton, Roberto Amato, Jeffrey Barrett, Sonia Goncalves, Ewan Harrison, David K. Jackson, Ian Johnston, Dominic Kwiatkowski, Cordelia Langford, John Sillitoe on behalf of the Wellcome Sanger Institute COVID-19 Surveillance Team |
| EPI_ISL_2935628, EPI_ISL_2935629 | Lighthouse Lab in Milton Keynes | Wellcome Sanger Institute for the COVID-19 Genomics UK (COG-UK) Consortium | The Lighthouse Lab in Milton Keynes and Alex Alderton, Roberto Amato, Jeffrey Barrett, Sonia Goncalves, Ewan Harrison, David K. Jackson, Ian Johnston, Dominic Kwiatkowski, Cordelia Langford, John Sillitoe on behalf of the Wellcome Sanger Institute COVID-19 Surveillance Team |
| EPI_ISL_2935630, EPI_ISL_2935631, EPI_ISL_2935632 | Lighthouse Laboratory Plymouth | Wellcome Sanger Institute for the COVID-19 Genomics UK (COG-UK) Consortium | Lighthouse Laboratory Plymouth and Alex Alderton, Roberto Amato, Jeffrey Barrett, Sonia Goncalves, Ewan Harrison, David K. Jackson, Ian Johnston, Dominic Kwiatkowski, Cordelia Langford, John Sillitoe on behalf of the Wellcome Sanger Institute COVID-19 Surveillance Team |
| EPI_ISL_2935633 | Lighthouse Lab in Milton Keynes | Wellcome Sanger Institute for the COVID-19 Genomics UK (COG-UK) Consortium | The Lighthouse Lab in Milton Keynes and Alex Alderton, Roberto Amato, Jeffrey Barrett, Sonia Goncalves, Ewan Harrison, David K. Jackson, Ian Johnston, Dominic Kwiatkowski, Cordelia Langford, John Sillitoe on behalf of the Wellcome Sanger Institute COVID-19 Surveillance Team |
| EPI_ISL_2935634 | Lighthouse Laboratory Plymouth | Wellcome Sanger Institute for the COVID-19 Genomics UK (COG-UK) Consortium | Lighthouse Laboratory Plymouth and Alex Alderton, Roberto Amato, Jeffrey Barrett, Sonia Goncalves, Ewan Harrison, David K. Jackson, Ian Johnston, Dominic Kwiatkowski, Cordelia Langford, John Sillitoe on behalf of the Wellcome Sanger Institute COVID-19 Surveillance Team |
| EPI_ISL_2935635 | Lighthouse Lab in Milton Keynes | Wellcome Sanger Institute for the COVID-19 Genomics UK (COG-UK) Consortium | The Lighthouse Lab in Milton Keynes and Alex Alderton, Roberto Amato, Jeffrey Barrett, Sonia Goncalves, Ewan Harrison, David K. Jackson, Ian Johnston, Dominic Kwiatkowski, Cordelia Langford, John Sillitoe on behalf of the Wellcome Sanger Institute COVID-19 Surveillance Team |
| EPI_ISL_2935636 | Lighthouse Lab in Alderley Park | Wellcome Sanger Institute for the COVID-19 Genomics UK (COG-UK) Consortium | Jacquelyn Wynn, Mairead Hyland, The Lighthouse Lab in Alderley Park and Alex Alderton, Roberto Amato, Jeffrey Barrett, Sonia Goncalves, Ewan Harrison, David K. Jackson, Ian Johnston, Dominic Kwiatkowski, Cordelia Langford, John Sillitoe on behalf of the Wellcome Sanger Institute COVID-19 Surveillance Team |
| EPI_ISL_2935637, EPI_ISL_2935638 | Lighthouse Lab in Milton Keynes | Wellcome Sanger Institute for the COVID-19 Genomics UK (COG-UK) Consortium | The Lighthouse Lab in Milton Keynes and Alex Alderton, Roberto Amato, Jeffrey Barrett, Sonia Goncalves, Ewan Harrison, David K. Jackson, Ian Johnston, Dominic Kwiatkowski, Cordelia Langford, John Sillitoe on behalf of the Wellcome Sanger Institute COVID-19 Surveillance Team |
| EPI_ISL_2935639, EPI_ISL_2935640 | Lighthouse Laboratory Plymouth | Wellcome Sanger Institute for the COVID-19 Genomics UK (COG-UK) Consortium | Lighthouse Laboratory Plymouth and Alex Alderton, Roberto Amato, Jeffrey Barrett, Sonia Goncalves, Ewan Harrison, David K. Jackson, Ian Johnston, Dominic Kwiatkowski, Cordelia Langford, John Sillitoe on behalf of the Wellcome Sanger Institute COVID-19 Surveillance Team |
| EPI_ISL_2935641 | Lighthouse Lab in Alderley Park | Wellcome Sanger Institute for the COVID-19 Genomics UK (COG-UK) Consortium | Jacquelyn Wynn, Mairead Hyland, The Lighthouse Lab in Alderley Park and Alex Alderton, Roberto Amato, Jeffrey Barrett, Sonia Goncalves, Ewan Harrison, David K. Jackson, Ian Johnston, Dominic Kwiatkowski, Cordelia Langford, John Sillitoe on behalf of the Wellcome Sanger Institute COVID-19 Surveillance Team |
| EPI_ISL_2935642, EPI_ISL_2935643 | Lighthouse Laboratory Plymouth | Wellcome Sanger Institute for the COVID-19 Genomics UK (COG-UK) Consortium | Lighthouse Laboratory Plymouth and Alex Alderton, Roberto Amato, Jeffrey Barrett, Sonia Goncalves, Ewan Harrison, David K. Jackson, Ian Johnston, Dominic Kwiatkowski, Cordelia Langford, John Sillitoe on behalf of the Wellcome Sanger Institute COVID-19 Surveillance Team |
| EPI_ISL_2935644 | Lighthouse Lab in Alderley Park | Wellcome Sanger Institute for the COVID-19 Genomics UK (COG-UK) Consortium | Jacquelyn Wynn, Mairead Hyland, The Lighthouse Lab in Alderley Park and Alex Alderton, Roberto Amato, Jeffrey Barrett, Sonia Goncalves, Ewan Harrison, David K. Jackson, Ian Johnston, Dominic Kwiatkowski, Cordelia Langford, John Sillitoe on behalf of the Wellcome Sanger Institute COVID-19 Surveillance Team |
| EPI_ISL_2935645 | Lighthouse Lab in Milton Keynes | Wellcome Sanger Institute for the COVID-19 Genomics UK (COG-UK) Consortium | The Lighthouse Lab in Milton Keynes and Alex Alderton, Roberto Amato, Jeffrey Barrett, Sonia Goncalves, Ewan Harrison, David K. Jackson, Ian Johnston, Dominic Kwiatkowski, Cordelia Langford, John Sillitoe on behalf of the Wellcome Sanger Institute COVID-19 Surveillance Team |
| EPI_ISL_2935646, EPI_ISL_2935647, EPI_ISL_2935648, EPI_ISL_2935649 | Lighthouse Laboratory Plymouth | Wellcome Sanger Institute for the COVID-19 Genomics UK (COG-UK) Consortium | Lighthouse Laboratory Plymouth and Alex Alderton, Roberto Amato, Jeffrey Barrett, Sonia Goncalves, Ewan Harrison, David K. Jackson, Ian Johnston, Dominic Kwiatkowski, Cordelia Langford, John Sillitoe on behalf of the Wellcome Sanger Institute COVID-19 Surveillance Team |
| EPI_ISL_2935650 | Lighthouse Lab in Milton Keynes | Wellcome Sanger Institute for the COVID-19 Genomics UK (COG-UK) Consortium | The Lighthouse Lab in Milton Keynes and Alex Alderton, Roberto Amato, Jeffrey Barrett, Sonia Goncalves, Ewan Harrison, David K. Jackson, Ian Johnston, Dominic Kwiatkowski, Cordelia Langford, John Sillitoe on behalf of the Wellcome Sanger Institute COVID-19 Surveillance Team |
| EPI_ISL_2935651, EPI_ISL_2935652 | Lighthouse Laboratory Plymouth | Wellcome Sanger Institute for the COVID-19 Genomics UK (COG-UK) Consortium | Lighthouse Laboratory Plymouth and Alex Alderton, Roberto Amato, Jeffrey Barrett, Sonia Goncalves, Ewan Harrison, David K. Jackson, Ian Johnston, Dominic Kwiatkowski, Cordelia Langford, John Sillitoe on behalf of the Wellcome Sanger Institute COVID-19 Surveillance Team |
| EPI_ISL_2935653 | Lighthouse Lab in Alderley Park | Wellcome Sanger Institute for the COVID-19 Genomics UK (COG-UK) Consortium | Jacquelyn Wynn, Mairead Hyland, The Lighthouse Lab in Alderley Park and Alex Alderton, Roberto Amato, Jeffrey Barrett, Sonia Goncalves, Ewan Harrison, David K. Jackson, Ian Johnston, Dominic Kwiatkowski, Cordelia Langford, John Sillitoe on behalf of the Wellcome Sanger Institute COVID-19 Surveillance Team |
| EPI_ISL_2935654 | Lighthouse Lab in Milton Keynes | Wellcome Sanger Institute for the COVID-19 Genomics UK (COG-UK) Consortium | The Lighthouse Lab in Milton Keynes and Alex Alderton, Roberto Amato, Jeffrey Barrett, Sonia Goncalves, Ewan Harrison, David K. Jackson, Ian Johnston, Dominic Kwiatkowski, Cordelia Langford, John Sillitoe on behalf of the Wellcome Sanger Institute COVID-19 Surveillance Team |
| EPI_ISL_2935655, EPI_ISL_2935656, EPI_ISL_2935657 | Lighthouse Lab in Alderley Park | Wellcome Sanger Institute for the COVID-19 Genomics UK (COG-UK) Consortium | Jacquelyn Wynn, Mairead Hyland, The Lighthouse Lab in Alderley Park and Alex Alderton, Roberto Amato, Jeffrey Barrett, Sonia Goncalves, Ewan Harrison, David K. Jackson, Ian Johnston, Dominic Kwiatkowski, Cordelia Langford, John Sillitoe on behalf of the Wellcome Sanger Institute COVID-19 Surveillance Team |
| EPI_ISL_2935658, EPI_ISL_2935659, EPI_ISL_2935660, EPI_ISL_2935661 | Lighthouse Laboratory Plymouth | Wellcome Sanger Institute for the COVID-19 Genomics UK (COG-UK) Consortium | Lighthouse Laboratory Plymouth and Alex Alderton, Roberto Amato, Jeffrey Barrett, Sonia Goncalves, Ewan Harrison, David K. Jackson, Ian Johnston, Dominic Kwiatkowski, Cordelia Langford, John Sillitoe on behalf of the Wellcome Sanger Institute COVID-19 Surveillance Team |
| EPI_ISL_2935662 | Lighthouse Lab in Alderley Park | Wellcome Sanger Institute for the COVID-19 Genomics UK (COG-UK) Consortium | Jacquelyn Wynn, Mairead Hyland, The Lighthouse Lab in Alderley Park and Alex Alderton, Roberto Amato, Jeffrey Barrett, Sonia Goncalves, Ewan Harrison, David K. Jackson, Ian Johnston, Dominic Kwiatkowski, Cordelia Langford, John Sillitoe on behalf of the Wellcome Sanger Institute COVID-19 Surveillance Team |
| EPI_ISL_2935663 | Lighthouse Laboratory Plymouth | Wellcome Sanger Institute for the COVID-19 Genomics UK (COG-UK) Consortium | Lighthouse Laboratory Plymouth and Alex Alderton, Roberto Amato, Jeffrey Barrett, Sonia Goncalves, Ewan Harrison, David K. Jackson, Ian Johnston, Dominic Kwiatkowski, Cordelia Langford, John Sillitoe on behalf of the Wellcome Sanger Institute COVID-19 Surveillance Team |
| EPI_ISL_2935664 | Lighthouse Lab in Alderley Park | Wellcome Sanger Institute for the COVID-19 Genomics UK (COG-UK) Consortium | Jacquelyn Wynn, Mairead Hyland, The Lighthouse Lab in Alderley Park and Alex Alderton, Roberto Amato, Jeffrey Barrett, Sonia Goncalves, Ewan Harrison, David K. Jackson, Ian Johnston, Dominic Kwiatkowski, Cordelia Langford, John Sillitoe on behalf of the Wellcome Sanger Institute COVID-19 Surveillance Team |
| EPI_ISL_2935665 | Lighthouse Laboratory Plymouth | Wellcome Sanger Institute for the COVID-19 Genomics UK (COG-UK) Consortium | Lighthouse Laboratory Plymouth and Alex Alderton, Roberto Amato, Jeffrey Barrett, Sonia Goncalves, Ewan Harrison, David K. Jackson, Ian Johnston, Dominic Kwiatkowski, Cordelia Langford, John Sillitoe on behalf of the Wellcome Sanger Institute COVID-19 Surveillance Team |
| EPI_ISL_2935666 | Lighthouse Lab in Milton Keynes | Wellcome Sanger Institute for the COVID-19 Genomics UK (COG-UK) Consortium | The Lighthouse Lab in Milton Keynes and Alex Alderton, Roberto Amato, Jeffrey Barrett, Sonia Goncalves, Ewan Harrison, David K. Jackson, Ian Johnston, Dominic Kwiatkowski, Cordelia Langford, John Sillitoe on behalf of the Wellcome Sanger Institute COVID-19 Surveillance Team |
| EPI_ISL_2935667 | Lighthouse Lab in Alderley Park | Wellcome Sanger Institute for the COVID-19 Genomics UK (COG-UK) Consortium | Jacquelyn Wynn, Mairead Hyland, The Lighthouse Lab in Alderley Park and Alex Alderton, Roberto Amato, Jeffrey Barrett, Sonia Goncalves, Ewan Harrison, David K. Jackson, Ian Johnston, Dominic Kwiatkowski, Cordelia Langford, John Sillitoe on behalf of the Wellcome Sanger Institute COVID-19 Surveillance Team |
| EPI_ISL_2935668 | Lighthouse Laboratory Plymouth | Wellcome Sanger Institute for the COVID-19 Genomics UK (COG-UK) Consortium | Lighthouse Laboratory Plymouth and Alex Alderton, Roberto Amato, Jeffrey Barrett, Sonia Goncalves, Ewan Harrison, David K. Jackson, Ian Johnston, Dominic Kwiatkowski, Cordelia Langford, John Sillitoe on behalf of the Wellcome Sanger Institute COVID-19 Surveillance Team |
| EPI_ISL_2935669 | Lighthouse Lab in Alderley Park | Wellcome Sanger Institute for the COVID-19 Genomics UK (COG-UK) | Jacquelyn Wynn, Mairead Hyland, The Lighthouse Lab in Alderley Park and Alex Alderton, Roberto Amato, Jeffrey Barrett, Sonia Goncalves, Ewan Harrison, David K. Jackson, Ian Johnston, Dominic Kwiatkowski, Cordelia Langford, John Sillitoe on |

| | | | |
|---|---|---|---|
| | | Consortium | behalf of the Wellcome Sanger Institute COVID-19 Surveillance Team |
| EPI_ISL_2935670 | Lighthouse Laboratory Plymouth | Wellcome Sanger Institute for the COVID-19 Genomics UK (COG-UK) Consortium | Lighthouse Laboratory Plymouth and Alex Alderton, Roberto Amato, Jeffrey Barrett, Sonia Goncalves, Ewan Harrison, David K. Jackson, Ian Johnston, Dominic Kwiatkowski, Cordelia Langford, John Sillitoe on behalf of the Wellcome Sanger Institute COVID-19 Surveillance Team |
| EPI_ISL_2935671, EPI_ISL_2935672 | Lighthouse Lab in Milton Keynes | Wellcome Sanger Institute for the COVID-19 Genomics UK (COG-UK) Consortium | The Lighthouse Lab in Milton Keynes and Alex Alderton, Roberto Amato, Jeffrey Barrett, Sonia Goncalves, Ewan Harrison, David K. Jackson, Ian Johnston, Dominic Kwiatkowski, Cordelia Langford, John Sillitoe on behalf of the Wellcome Sanger Institute COVID-19 Surveillance Team |
| EPI_ISL_2935673, EPI_ISL_2935674, EPI_ISL_2935675, EPI_ISL_2935676 | Lighthouse Laboratory Plymouth | Wellcome Sanger Institute for the COVID-19 Genomics UK (COG-UK) Consortium | Lighthouse Laboratory Plymouth and Alex Alderton, Roberto Amato, Jeffrey Barrett, Sonia Goncalves, Ewan Harrison, David K. Jackson, Ian Johnston, Dominic Kwiatkowski, Cordelia Langford, John Sillitoe on behalf of the Wellcome Sanger Institute COVID-19 Surveillance Team |
| EPI_ISL_2935677, EPI_ISL_2935678 | Lighthouse Lab in Milton Keynes | Wellcome Sanger Institute for the COVID-19 Genomics UK (COG-UK) Consortium | The Lighthouse Lab in Milton Keynes and Alex Alderton, Roberto Amato, Jeffrey Barrett, Sonia Goncalves, Ewan Harrison, David K. Jackson, Ian Johnston, Dominic Kwiatkowski, Cordelia Langford, John Sillitoe on behalf of the Wellcome Sanger Institute COVID-19 Surveillance Team |
| EPI_ISL_2935679, EPI_ISL_2935680 | Lighthouse Laboratory Plymouth | Wellcome Sanger Institute for the COVID-19 Genomics UK (COG-UK) Consortium | Lighthouse Laboratory Plymouth and Alex Alderton, Roberto Amato, Jeffrey Barrett, Sonia Goncalves, Ewan Harrison, David K. Jackson, Ian Johnston, Dominic Kwiatkowski, Cordelia Langford, John Sillitoe on behalf of the Wellcome Sanger Institute COVID-19 Surveillance Team |
| EPI_ISL_2935681 | Lighthouse Lab in Milton Keynes | Wellcome Sanger Institute for the COVID-19 Genomics UK (COG-UK) Consortium | The Lighthouse Lab in Milton Keynes and Alex Alderton, Roberto Amato, Jeffrey Barrett, Sonia Goncalves, Ewan Harrison, David K. Jackson, Ian Johnston, Dominic Kwiatkowski, Cordelia Langford, John Sillitoe on behalf of the Wellcome Sanger Institute COVID-19 Surveillance Team |
| EPI_ISL_2935682 | Lighthouse Laboratory Plymouth | Wellcome Sanger Institute for the COVID-19 Genomics UK (COG-UK) Consortium | Lighthouse Laboratory Plymouth and Alex Alderton, Roberto Amato, Jeffrey Barrett, Sonia Goncalves, Ewan Harrison, David K. Jackson, Ian Johnston, Dominic Kwiatkowski, Cordelia Langford, John Sillitoe on behalf of the Wellcome Sanger Institute COVID-19 Surveillance Team |
| EPI_ISL_2935683, EPI_ISL_2935684, EPI_ISL_2935685 | Lighthouse Lab in Glasgow | Wellcome Sanger Institute for the COVID-19 Genomics UK (COG-UK) Consortium | Harper VanSteenhouse, Yumi Kasai, David Gray, Carol Clugston, Anna Dominiczak and Alex Alderton, Roberto Amato, Jeffrey Barrett, Sonia Goncalves, Ewan Harrison, David K. Jackson, Ian Johnston, Dominic Kwiatkowski, Cordelia Langford, John Sillitoe on behalf of the Wellcome Sanger Institute COVID-19 Surveillance Team |
| EPI_ISL_2935686, EPI_ISL_2935687, EPI_ISL_2935688, EPI_ISL_2935689 | Lighthouse Lab in Milton Keynes | Wellcome Sanger Institute for the COVID-19 Genomics UK (COG-UK) Consortium | The Lighthouse Lab in Milton Keynes and Alex Alderton, Roberto Amato, Jeffrey Barrett, Sonia Goncalves, Ewan Harrison, David K. Jackson, Ian Johnston, Dominic Kwiatkowski, Cordelia Langford, John Sillitoe on behalf of the Wellcome Sanger Institute COVID-19 Surveillance Team |
| EPI_ISL_2935690 | Lighthouse Lab in Glasgow | Wellcome Sanger Institute for the COVID-19 Genomics UK (COG-UK) Consortium | Harper VanSteenhouse, Yumi Kasai, David Gray, Carol Clugston, Anna Dominiczak and Alex Alderton, Roberto Amato, Jeffrey Barrett, Sonia Goncalves, Ewan Harrison, David K. Jackson, Ian Johnston, Dominic Kwiatkowski, Cordelia Langford, John Sillitoe on behalf of the Wellcome Sanger Institute COVID-19 Surveillance Team |
| EPI_ISL_2935691 | Lighthouse Lab in Milton Keynes | Wellcome Sanger Institute for the COVID-19 Genomics UK (COG-UK) Consortium | The Lighthouse Lab in Milton Keynes and Alex Alderton, Roberto Amato, Jeffrey Barrett, Sonia Goncalves, Ewan Harrison, David K. Jackson, Ian Johnston, Dominic Kwiatkowski, Cordelia Langford, John Sillitoe on behalf of the Wellcome Sanger Institute COVID-19 Surveillance Team |
| EPI_ISL_2935692, EPI_ISL_2935693, EPI_ISL_2935694, EPI_ISL_2935695 | Lighthouse Lab in Glasgow | Wellcome Sanger Institute for the COVID-19 Genomics UK (COG-UK) Consortium | Harper VanSteenhouse, Yumi Kasai, David Gray, Carol Clugston, Anna Dominiczak and Alex Alderton, Roberto Amato, Jeffrey Barrett, Sonia Goncalves, Ewan Harrison, David K. Jackson, Ian Johnston, Dominic Kwiatkowski, Cordelia Langford, John Sillitoe on behalf of the Wellcome Sanger Institute COVID-19 Surveillance Team |
| EPI_ISL_2935696, EPI_ISL_2935697 | Lighthouse Lab in Milton Keynes | Wellcome Sanger Institute for the COVID-19 Genomics UK (COG-UK) Consortium | The Lighthouse Lab in Milton Keynes and Alex Alderton, Roberto Amato, Jeffrey Barrett, Sonia Goncalves, Ewan Harrison, David K. Jackson, Ian Johnston, Dominic Kwiatkowski, Cordelia Langford, John Sillitoe on behalf of the Wellcome Sanger Institute COVID-19 Surveillance Team |
| EPI_ISL_2935698, EPI_ISL_2935699, EPI_ISL_2935700, EPI_ISL_2935701, EPI_ISL_2935702, EPI_ISL_2935703 | Lighthouse Lab in Glasgow | Wellcome Sanger Institute for the COVID-19 Genomics UK (COG-UK) Consortium | Harper VanSteenhouse, Yumi Kasai, David Gray, Carol Clugston, Anna Dominiczak and Alex Alderton, Roberto Amato, Jeffrey Barrett, Sonia Goncalves, Ewan Harrison, David K. Jackson, Ian Johnston, Dominic Kwiatkowski, Cordelia Langford, John Sillitoe on behalf of the Wellcome Sanger Institute COVID-19 Surveillance Team |
| EPI_ISL_2935704 | Lighthouse Lab in Milton Keynes | Wellcome Sanger Institute for the COVID-19 Genomics UK (COG-UK) Consortium | The Lighthouse Lab in Milton Keynes and Alex Alderton, Roberto Amato, Jeffrey Barrett, Sonia Goncalves, Ewan Harrison, David K. Jackson, Ian Johnston, Dominic Kwiatkowski, Cordelia Langford, John Sillitoe on behalf of the Wellcome Sanger Institute COVID-19 Surveillance Team |
| EPI_ISL_2935705, EPI_ISL_2935706, EPI_ISL_2935707, EPI_ISL_2935708 | Lighthouse Lab in Glasgow | Wellcome Sanger Institute for the COVID-19 Genomics UK (COG-UK) Consortium | Harper VanSteenhouse, Yumi Kasai, David Gray, Carol Clugston, Anna Dominiczak and Alex Alderton, Roberto Amato, Jeffrey Barrett, Sonia Goncalves, Ewan Harrison, David K. Jackson, Ian Johnston, Dominic Kwiatkowski, Cordelia Langford, John Sillitoe on behalf of the Wellcome Sanger Institute COVID-19 Surveillance Team |
| EPI_ISL_2935709 | Lighthouse Lab in Milton Keynes | Wellcome Sanger Institute for the COVID-19 Genomics UK (COG-UK) Consortium | The Lighthouse Lab in Milton Keynes and Alex Alderton, Roberto Amato, Jeffrey Barrett, Sonia Goncalves, Ewan Harrison, David K. Jackson, Ian Johnston, Dominic Kwiatkowski, Cordelia Langford, John Sillitoe on behalf of the Wellcome Sanger Institute COVID-19 Surveillance Team |
| EPI_ISL_2935710, EPI_ISL_2935711, EPI_ISL_2935712 | Lighthouse Lab in Glasgow | Wellcome Sanger Institute for the COVID-19 Genomics UK (COG-UK) Consortium | Harper VanSteenhouse, Yumi Kasai, David Gray, Carol Clugston, Anna Dominiczak and Alex Alderton, Roberto Amato, Jeffrey Barrett, Sonia Goncalves, Ewan Harrison, David K. Jackson, Ian Johnston, Dominic Kwiatkowski, Cordelia Langford, John Sillitoe on behalf of the Wellcome Sanger Institute COVID-19 Surveillance Team |
| EPI_ISL_2935713, EPI_ISL_2935714 | Lighthouse Lab in Milton Keynes | Wellcome Sanger Institute for the COVID-19 Genomics UK (COG-UK) Consortium | The Lighthouse Lab in Milton Keynes and Alex Alderton, Roberto Amato, Jeffrey Barrett, Sonia Goncalves, Ewan Harrison, David K. Jackson, Ian Johnston, Dominic Kwiatkowski, Cordelia Langford, John Sillitoe on behalf of the Wellcome Sanger Institute COVID-19 Surveillance Team |
| EPI_ISL_2935715, EPI_ISL_2935716 | Lighthouse Lab in Glasgow | Wellcome Sanger Institute for the COVID-19 Genomics UK (COG-UK) Consortium | Harper VanSteenhouse, Yumi Kasai, David Gray, Carol Clugston, Anna Dominiczak and Alex Alderton, Roberto Amato, Jeffrey Barrett, Sonia Goncalves, Ewan Harrison, David K. Jackson, Ian Johnston, Dominic Kwiatkowski, Cordelia Langford, John Sillitoe on behalf of the Wellcome Sanger Institute COVID-19 Surveillance Team |
| EPI_ISL_2935717, EPI_ISL_2935718 | Lighthouse Lab in Milton Keynes | Wellcome Sanger Institute for the COVID-19 Genomics UK (COG-UK) Consortium | The Lighthouse Lab in Milton Keynes and Alex Alderton, Roberto Amato, Jeffrey Barrett, Sonia Goncalves, Ewan Harrison, David K. Jackson, Ian Johnston, Dominic Kwiatkowski, Cordelia Langford, John Sillitoe on behalf of the Wellcome Sanger Institute COVID-19 Surveillance Team |
| EPI_ISL_2935719 | Lighthouse Lab in Glasgow | Wellcome Sanger Institute for the COVID-19 Genomics UK (COG-UK) Consortium | Harper VanSteenhouse, Yumi Kasai, David Gray, Carol Clugston, Anna Dominiczak and Alex Alderton, Roberto Amato, Jeffrey Barrett, Sonia Goncalves, Ewan Harrison, David K. Jackson, Ian Johnston, Dominic Kwiatkowski, Cordelia Langford, John Sillitoe on behalf of the Wellcome Sanger Institute COVID-19 Surveillance Team |
| EPI_ISL_2935720 | Lighthouse Lab in Milton Keynes | Wellcome Sanger Institute for the COVID-19 Genomics UK (COG-UK) Consortium | The Lighthouse Lab in Milton Keynes and Alex Alderton, Roberto Amato, Jeffrey Barrett, Sonia Goncalves, Ewan Harrison, David K. Jackson, Ian Johnston, Dominic Kwiatkowski, Cordelia Langford, John Sillitoe on behalf of the Wellcome Sanger Institute COVID-19 Surveillance Team |
| EPI_ISL_2935721 | Lighthouse Lab in Glasgow | Wellcome Sanger Institute for the COVID-19 Genomics UK (COG-UK) Consortium | Harper VanSteenhouse, Yumi Kasai, David Gray, Carol Clugston, Anna Dominiczak and Alex Alderton, Roberto Amato, Jeffrey Barrett, Sonia Goncalves, Ewan Harrison, David K. Jackson, Ian Johnston, Dominic Kwiatkowski, Cordelia Langford, John Sillitoe on behalf of the Wellcome Sanger Institute COVID-19 Surveillance Team |
| EPI_ISL_2935722 | Lighthouse Lab in Milton Keynes | Wellcome Sanger Institute for the COVID-19 Genomics UK (COG-UK) Consortium | The Lighthouse Lab in Milton Keynes and Alex Alderton, Roberto Amato, Jeffrey Barrett, Sonia Goncalves, Ewan Harrison, David K. Jackson, Ian Johnston, Dominic Kwiatkowski, Cordelia Langford, John Sillitoe on behalf of the Wellcome Sanger Institute COVID-19 Surveillance Team |
| EPI_ISL_2935723, EPI_ISL_2935724, EPI_ISL_2935725 | Lighthouse Lab in Glasgow | Wellcome Sanger Institute for the COVID-19 Genomics UK (COG-UK) Consortium | Harper VanSteenhouse, Yumi Kasai, David Gray, Carol Clugston, Anna Dominiczak and Alex Alderton, Roberto Amato, Jeffrey Barrett, Sonia Goncalves, Ewan Harrison, David K. Jackson, Ian Johnston, Dominic Kwiatkowski, Cordelia Langford, John Sillitoe on behalf of the Wellcome Sanger Institute COVID-19 Surveillance Team |
| EPI_ISL_2935726 | Lighthouse Lab in Milton Keynes | Wellcome Sanger Institute for the COVID-19 Genomics UK (COG-UK) Consortium | The Lighthouse Lab in Milton Keynes and Alex Alderton, Roberto Amato, Jeffrey Barrett, Sonia Goncalves, Ewan Harrison, David K. Jackson, Ian Johnston, Dominic Kwiatkowski, Cordelia Langford, John Sillitoe on behalf of the Wellcome Sanger Institute COVID-19 Surveillance Team |
| EPI_ISL_2935727, EPI_ISL_2935728, EPI_ISL_2935729, EPI_ISL_2935730, EPI_ISL_2935731, EPI_ISL_2935732, EPI_ISL_2935733, EPI_ISL_2935734 | Lighthouse Lab in Glasgow | Wellcome Sanger Institute for the COVID-19 Genomics UK (COG-UK) Consortium | Harper VanSteenhouse, Yumi Kasai, David Gray, Carol Clugston, Anna Dominiczak and Alex Alderton, Roberto Amato, Jeffrey Barrett, Sonia Goncalves, Ewan Harrison, David K. Jackson, Ian Johnston, Dominic Kwiatkowski, Cordelia Langford, John Sillitoe on behalf of the Wellcome Sanger Institute COVID-19 Surveillance Team |
| EPI_ISL_2935735, EPI_ISL_2935736 | Lighthouse Lab in Milton Keynes | Wellcome Sanger Institute for the COVID-19 Genomics UK (COG-UK) Consortium | The Lighthouse Lab in Milton Keynes and Alex Alderton, Roberto Amato, Jeffrey Barrett, Sonia Goncalves, Ewan Harrison, David K. Jackson, Ian Johnston, Dominic Kwiatkowski, Cordelia Langford, John Sillitoe on behalf of the Wellcome Sanger Institute COVID-19 Surveillance Team |
| EPI_ISL_2935737, EPI_ISL_2935738, EPI_ISL_2935739 | Lighthouse Lab in Glasgow | Wellcome Sanger Institute for the COVID-19 Genomics UK (COG-UK) Consortium | Harper VanSteenhouse, Yumi Kasai, David Gray, Carol Clugston, Anna Dominiczak and Alex Alderton, Roberto Amato, Jeffrey Barrett, Sonia Goncalves, Ewan Harrison, David K. Jackson, Ian Johnston, Dominic Kwiatkowski, Cordelia Langford, John Sillitoe on behalf of the Wellcome Sanger Institute COVID-19 Surveillance Team |
| EPI_ISL_2935740 | Lighthouse Lab in Milton Keynes | Wellcome Sanger Institute for the COVID-19 Genomics UK (COG-UK) Consortium | The Lighthouse Lab in Milton Keynes and Alex Alderton, Roberto Amato, Jeffrey Barrett, Sonia Goncalves, Ewan Harrison, David K. Jackson, Ian Johnston, Dominic Kwiatkowski, Cordelia Langford, John Sillitoe on behalf of the Wellcome Sanger Institute COVID-19 Surveillance Team |
| EPI_ISL_2935741, EPI_ISL_2935742, EPI_ISL_2935743 | Lighthouse Lab in Glasgow | Wellcome Sanger Institute for the COVID-19 Genomics UK (COG-UK) Consortium | Harper VanSteenhouse, Yumi Kasai, David Gray, Carol Clugston, Anna Dominiczak and Alex Alderton, Roberto Amato, Jeffrey Barrett, Sonia Goncalves, Ewan Harrison, David K. Jackson, Ian Johnston, Dominic Kwiatkowski, Cordelia Langford, John Sillitoe on behalf of the Wellcome Sanger Institute COVID-19 Surveillance Team |
| EPI_ISL_2935744, EPI_ISL_2935745 | Lighthouse Lab in Milton Keynes | Wellcome Sanger Institute for the COVID-19 Genomics UK (COG-UK) Consortium | The Lighthouse Lab in Milton Keynes and Alex Alderton, Roberto Amato, Jeffrey Barrett, Sonia Goncalves, Ewan Harrison, David K. Jackson, Ian Johnston, Dominic Kwiatkowski, Cordelia Langford, John Sillitoe on behalf of the Wellcome Sanger Institute COVID-19 Surveillance Team |
| EPI_ISL_2935746, EPI_ISL_2935747, EPI_ISL_2935748 | Lighthouse Lab in Glasgow | Wellcome Sanger Institute for the COVID-19 Genomics UK (COG-UK) Consortium | Harper VanSteenhouse, Yumi Kasai, David Gray, Carol Clugston, Anna Dominiczak and Alex Alderton, Roberto Amato, Jeffrey Barrett, Sonia Goncalves, Ewan Harrison, David K. Jackson, Ian Johnston, Dominic Kwiatkowski, Cordelia Langford, John Sillitoe on behalf of the Wellcome Sanger Institute COVID-19 Surveillance Team |
| EPI_ISL_2935749, EPI_ISL_2935750 | Lighthouse Lab in Milton Keynes | Wellcome Sanger Institute for the COVID-19 Genomics UK (COG-UK) Consortium | The Lighthouse Lab in Milton Keynes and Alex Alderton, Roberto Amato, Jeffrey Barrett, Sonia Goncalves, Ewan Harrison, David K. Jackson, Ian Johnston, Dominic Kwiatkowski, Cordelia Langford, John Sillitoe on behalf of the Wellcome Sanger Institute COVID-19 Surveillance Team |

| | | | |
|---|---|---|---|
| EPI_ISL_2935751 | Lighthouse Lab in Glasgow | Wellcome Sanger Institute for the COVID-19 Genomics UK (COG-UK) Consortium | Harper VanSteenhouse, Yumi Kasai, David Gray, Carol Clugston, Anna Dominiczak and Alex Alderton, Roberto Amato, Jeffrey Barrett, Sonia Goncalves, Ewan Harrison, David K. Jackson, Ian Johnston, Dominic Kwiatkowski, Cordelia Langford, John Sillitoe on behalf of the Wellcome Sanger Institute COVID-19 Surveillance Team |
| EPI_ISL_2935752 | Lighthouse Lab in Milton Keynes | Wellcome Sanger Institute for the COVID-19 Genomics UK (COG-UK) Consortium | The Lighthouse Lab in Milton Keynes and Alex Alderton, Roberto Amato, Jeffrey Barrett, Sonia Goncalves, Ewan Harrison, David K. Jackson, Ian Johnston, Dominic Kwiatkowski, Cordelia Langford, John Sillitoe on behalf of the Wellcome Sanger Institute COVID-19 Surveillance Team |
| EPI_ISL_2935753, EPI_ISL_2935754 | Lighthouse Lab in Glasgow | Wellcome Sanger Institute for the COVID-19 Genomics UK (COG-UK) Consortium | Harper VanSteenhouse, Yumi Kasai, David Gray, Carol Clugston, Anna Dominiczak and Alex Alderton, Roberto Amato, Jeffrey Barrett, Sonia Goncalves, Ewan Harrison, David K. Jackson, Ian Johnston, Dominic Kwiatkowski, Cordelia Langford, John Sillitoe on behalf of the Wellcome Sanger Institute COVID-19 Surveillance Team |
| EPI_ISL_2935755, EPI_ISL_2935756 | Lighthouse Lab in Milton Keynes | Wellcome Sanger Institute for the COVID-19 Genomics UK (COG-UK) Consortium | The Lighthouse Lab in Milton Keynes and Alex Alderton, Roberto Amato, Jeffrey Barrett, Sonia Goncalves, Ewan Harrison, David K. Jackson, Ian Johnston, Dominic Kwiatkowski, Cordelia Langford, John Sillitoe on behalf of the Wellcome Sanger Institute COVID-19 Surveillance Team |
| EPI_ISL_2935757, EPI_ISL_2935758, EPI_ISL_2935759 | Lighthouse Lab in Glasgow | Wellcome Sanger Institute for the COVID-19 Genomics UK (COG-UK) Consortium | Harper VanSteenhouse, Yumi Kasai, David Gray, Carol Clugston, Anna Dominiczak and Alex Alderton, Roberto Amato, Jeffrey Barrett, Sonia Goncalves, Ewan Harrison, David K. Jackson, Ian Johnston, Dominic Kwiatkowski, Cordelia Langford, John Sillitoe on behalf of the Wellcome Sanger Institute COVID-19 Surveillance Team |
| EPI_ISL_2935760, EPI_ISL_2935761 | Lighthouse Lab in Milton Keynes | Wellcome Sanger Institute for the COVID-19 Genomics UK (COG-UK) Consortium | The Lighthouse Lab in Milton Keynes and Alex Alderton, Roberto Amato, Jeffrey Barrett, Sonia Goncalves, Ewan Harrison, David K. Jackson, Ian Johnston, Dominic Kwiatkowski, Cordelia Langford, John Sillitoe on behalf of the Wellcome Sanger Institute COVID-19 Surveillance Team |
| EPI_ISL_2935762 | Lighthouse Lab in Glasgow | Wellcome Sanger Institute for the COVID-19 Genomics UK (COG-UK) Consortium | Harper VanSteenhouse, Yumi Kasai, David Gray, Carol Clugston, Anna Dominiczak and Alex Alderton, Roberto Amato, Jeffrey Barrett, Sonia Goncalves, Ewan Harrison, David K. Jackson, Ian Johnston, Dominic Kwiatkowski, Cordelia Langford, John Sillitoe on behalf of the Wellcome Sanger Institute COVID-19 Surveillance Team |
| EPI_ISL_2935763 | Lighthouse Lab in Milton Keynes | Wellcome Sanger Institute for the COVID-19 Genomics UK (COG-UK) Consortium | The Lighthouse Lab in Milton Keynes and Alex Alderton, Roberto Amato, Jeffrey Barrett, Sonia Goncalves, Ewan Harrison, David K. Jackson, Ian Johnston, Dominic Kwiatkowski, Cordelia Langford, John Sillitoe on behalf of the Wellcome Sanger Institute COVID-19 Surveillance Team |
| EPI_ISL_2935764, EPI_ISL_2935765, EPI_ISL_2935766, EPI_ISL_2935767 | Lighthouse Lab in Glasgow | Wellcome Sanger Institute for the COVID-19 Genomics UK (COG-UK) Consortium | Harper VanSteenhouse, Yumi Kasai, David Gray, Carol Clugston, Anna Dominiczak and Alex Alderton, Roberto Amato, Jeffrey Barrett, Sonia Goncalves, Ewan Harrison, David K. Jackson, Ian Johnston, Dominic Kwiatkowski, Cordelia Langford, John Sillitoe on behalf of the Wellcome Sanger Institute COVID-19 Surveillance Team |
| EPI_ISL_2935768, EPI_ISL_2935769 | Lighthouse Lab in Milton Keynes | Wellcome Sanger Institute for the COVID-19 Genomics UK (COG-UK) Consortium | The Lighthouse Lab in Milton Keynes and Alex Alderton, Roberto Amato, Jeffrey Barrett, Sonia Goncalves, Ewan Harrison, David K. Jackson, Ian Johnston, Dominic Kwiatkowski, Cordelia Langford, John Sillitoe on behalf of the Wellcome Sanger Institute COVID-19 Surveillance Team |
| EPI_ISL_2935770, EPI_ISL_2935771, EPI_ISL_2935772 | Lighthouse Lab in Glasgow | Wellcome Sanger Institute for the COVID-19 Genomics UK (COG-UK) Consortium | Harper VanSteenhouse, Yumi Kasai, David Gray, Carol Clugston, Anna Dominiczak and Alex Alderton, Roberto Amato, Jeffrey Barrett, Sonia Goncalves, Ewan Harrison, David K. Jackson, Ian Johnston, Dominic Kwiatkowski, Cordelia Langford, John Sillitoe on behalf of the Wellcome Sanger Institute COVID-19 Surveillance Team |
| EPI_ISL_2935773, EPI_ISL_2935775, EPI_ISL_2935776, EPI_ISL_2935777 | Lighthouse Lab in Milton Keynes | Wellcome Sanger Institute for the COVID-19 Genomics UK (COG-UK) Consortium | The Lighthouse Lab in Milton Keynes and Alex Alderton, Roberto Amato, Jeffrey Barrett, Sonia Goncalves, Ewan Harrison, David K. Jackson, Ian Johnston, Dominic Kwiatkowski, Cordelia Langford, John Sillitoe on behalf of the Wellcome Sanger Institute COVID-19 Surveillance Team |
| EPI_ISL_2935778 | Lighthouse Lab in Glasgow | Wellcome Sanger Institute for the COVID-19 Genomics UK (COG-UK) Consortium | Harper VanSteenhouse, Yumi Kasai, David Gray, Carol Clugston, Anna Dominiczak and Alex Alderton, Roberto Amato, Jeffrey Barrett, Sonia Goncalves, Ewan Harrison, David K. Jackson, Ian Johnston, Dominic Kwiatkowski, Cordelia Langford, John Sillitoe on behalf of the Wellcome Sanger Institute COVID-19 Surveillance Team |
| EPI_ISL_2935779, EPI_ISL_2935780 | Lighthouse Lab in Milton Keynes | Wellcome Sanger Institute for the COVID-19 Genomics UK (COG-UK) Consortium | The Lighthouse Lab in Milton Keynes and Alex Alderton, Roberto Amato, Jeffrey Barrett, Sonia Goncalves, Ewan Harrison, David K. Jackson, Ian Johnston, Dominic Kwiatkowski, Cordelia Langford, John Sillitoe on behalf of the Wellcome Sanger Institute COVID-19 Surveillance Team |
| EPI_ISL_2935781, EPI_ISL_2935782, EPI_ISL_2935783, EPI_ISL_2935784 | Lighthouse Lab in Glasgow | Wellcome Sanger Institute for the COVID-19 Genomics UK (COG-UK) Consortium | Harper VanSteenhouse, Yumi Kasai, David Gray, Carol Clugston, Anna Dominiczak and Alex Alderton, Roberto Amato, Jeffrey Barrett, Sonia Goncalves, Ewan Harrison, David K. Jackson, Ian Johnston, Dominic Kwiatkowski, Cordelia Langford, John Sillitoe on behalf of the Wellcome Sanger Institute COVID-19 Surveillance Team |
| EPI_ISL_2935785, EPI_ISL_2935786, EPI_ISL_2935787, EPI_ISL_2935788 | Lighthouse Lab in Milton Keynes | Wellcome Sanger Institute for the COVID-19 Genomics UK (COG-UK) Consortium | The Lighthouse Lab in Milton Keynes and Alex Alderton, Roberto Amato, Jeffrey Barrett, Sonia Goncalves, Ewan Harrison, David K. Jackson, Ian Johnston, Dominic Kwiatkowski, Cordelia Langford, John Sillitoe on behalf of the Wellcome Sanger Institute COVID-19 Surveillance Team |
| EPI_ISL_2935789, EPI_ISL_2935790, EPI_ISL_2935791 | Lighthouse Lab in Glasgow | Wellcome Sanger Institute for the COVID-19 Genomics UK (COG-UK) Consortium | Harper VanSteenhouse, Yumi Kasai, David Gray, Carol Clugston, Anna Dominiczak and Alex Alderton, Roberto Amato, Jeffrey Barrett, Sonia Goncalves, Ewan Harrison, David K. Jackson, Ian Johnston, Dominic Kwiatkowski, Cordelia Langford, John Sillitoe on behalf of the Wellcome Sanger Institute COVID-19 Surveillance Team |
| EPI_ISL_2935792 | Lighthouse Lab in Milton Keynes | Wellcome Sanger Institute for the COVID-19 Genomics UK (COG-UK) Consortium | The Lighthouse Lab in Milton Keynes and Alex Alderton, Roberto Amato, Jeffrey Barrett, Sonia Goncalves, Ewan Harrison, David K. Jackson, Ian Johnston, Dominic Kwiatkowski, Cordelia Langford, John Sillitoe on behalf of the Wellcome Sanger Institute COVID-19 Surveillance Team |
| EPI_ISL_2935793, EPI_ISL_2935794, EPI_ISL_2935795, EPI_ISL_2935796, EPI_ISL_2935797, EPI_ISL_2935798 | Lighthouse Lab in Glasgow | Wellcome Sanger Institute for the COVID-19 Genomics UK (COG-UK) Consortium | Harper VanSteenhouse, Yumi Kasai, David Gray, Carol Clugston, Anna Dominiczak and Alex Alderton, Roberto Amato, Jeffrey Barrett, Sonia Goncalves, Ewan Harrison, David K. Jackson, Ian Johnston, Dominic Kwiatkowski, Cordelia Langford, John Sillitoe on behalf of the Wellcome Sanger Institute COVID-19 Surveillance Team |
| EPI_ISL_2935799 | Lighthouse Lab in Milton Keynes | Wellcome Sanger Institute for the COVID-19 Genomics UK (COG-UK) Consortium | The Lighthouse Lab in Milton Keynes and Alex Alderton, Roberto Amato, Jeffrey Barrett, Sonia Goncalves, Ewan Harrison, David K. Jackson, Ian Johnston, Dominic Kwiatkowski, Cordelia Langford, John Sillitoe on behalf of the Wellcome Sanger Institute COVID-19 Surveillance Team |
| EPI_ISL_2935800 | Lighthouse Lab in Glasgow | Wellcome Sanger Institute for the COVID-19 Genomics UK (COG-UK) Consortium | Harper VanSteenhouse, Yumi Kasai, David Gray, Carol Clugston, Anna Dominiczak and Alex Alderton, Roberto Amato, Jeffrey Barrett, Sonia Goncalves, Ewan Harrison, David K. Jackson, Ian Johnston, Dominic Kwiatkowski, Cordelia Langford, John Sillitoe on behalf of the Wellcome Sanger Institute COVID-19 Surveillance Team |
| EPI_ISL_2935801 | Lighthouse Lab in Milton Keynes | Wellcome Sanger Institute for the COVID-19 Genomics UK (COG-UK) Consortium | The Lighthouse Lab in Milton Keynes and Alex Alderton, Roberto Amato, Jeffrey Barrett, Sonia Goncalves, Ewan Harrison, David K. Jackson, Ian Johnston, Dominic Kwiatkowski, Cordelia Langford, John Sillitoe on behalf of the Wellcome Sanger Institute COVID-19 Surveillance Team |
| EPI_ISL_2935802, EPI_ISL_2935803, EPI_ISL_2935804, EPI_ISL_2935805, EPI_ISL_2935806 | Lighthouse Lab in Glasgow | Wellcome Sanger Institute for the COVID-19 Genomics UK (COG-UK) Consortium | Harper VanSteenhouse, Yumi Kasai, David Gray, Carol Clugston, Anna Dominiczak and Alex Alderton, Roberto Amato, Jeffrey Barrett, Sonia Goncalves, Ewan Harrison, David K. Jackson, Ian Johnston, Dominic Kwiatkowski, Cordelia Langford, John Sillitoe on behalf of the Wellcome Sanger Institute COVID-19 Surveillance Team |
| EPI_ISL_2935807 | Lighthouse Lab in Milton Keynes | Wellcome Sanger Institute for the COVID-19 Genomics UK (COG-UK) Consortium | The Lighthouse Lab in Milton Keynes and Alex Alderton, Roberto Amato, Jeffrey Barrett, Sonia Goncalves, Ewan Harrison, David K. Jackson, Ian Johnston, Dominic Kwiatkowski, Cordelia Langford, John Sillitoe on behalf of the Wellcome Sanger Institute COVID-19 Surveillance Team |
| EPI_ISL_2935808 | Lighthouse Lab in Glasgow | Wellcome Sanger Institute for the COVID-19 Genomics UK (COG-UK) Consortium | Harper VanSteenhouse, Yumi Kasai, David Gray, Carol Clugston, Anna Dominiczak and Alex Alderton, Roberto Amato, Jeffrey Barrett, Sonia Goncalves, Ewan Harrison, David K. Jackson, Ian Johnston, Dominic Kwiatkowski, Cordelia Langford, John Sillitoe on behalf of the Wellcome Sanger Institute COVID-19 Surveillance Team |
| EPI_ISL_2935809 | Lighthouse Lab in Milton Keynes | Wellcome Sanger Institute for the COVID-19 Genomics UK (COG-UK) Consortium | The Lighthouse Lab in Milton Keynes and Alex Alderton, Roberto Amato, Jeffrey Barrett, Sonia Goncalves, Ewan Harrison, David K. Jackson, Ian Johnston, Dominic Kwiatkowski, Cordelia Langford, John Sillitoe on behalf of the Wellcome Sanger Institute COVID-19 Surveillance Team |
| EPI_ISL_2935810, EPI_ISL_2935811, EPI_ISL_2935812, EPI_ISL_2935813, EPI_ISL_2935814, EPI_ISL_2935815, EPI_ISL_2935816, EPI_ISL_2935817, EPI_ISL_2935818 | Lighthouse Lab in Glasgow | Wellcome Sanger Institute for the COVID-19 Genomics UK (COG-UK) Consortium | Harper VanSteenhouse, Yumi Kasai, David Gray, Carol Clugston, Anna Dominiczak and Alex Alderton, Roberto Amato, Jeffrey Barrett, Sonia Goncalves, Ewan Harrison, David K. Jackson, Ian Johnston, Dominic Kwiatkowski, Cordelia Langford, John Sillitoe on behalf of the Wellcome Sanger Institute COVID-19 Surveillance Team |
| EPI_ISL_2935819 | Lighthouse Lab in Milton Keynes | Wellcome Sanger Institute for the COVID-19 Genomics UK (COG-UK) Consortium | The Lighthouse Lab in Milton Keynes and Alex Alderton, Roberto Amato, Jeffrey Barrett, Sonia Goncalves, Ewan Harrison, David K. Jackson, Ian Johnston, Dominic Kwiatkowski, Cordelia Langford, John Sillitoe on behalf of the Wellcome Sanger Institute COVID-19 Surveillance Team |
| EPI_ISL_2935820, EPI_ISL_2935821, EPI_ISL_2935822, EPI_ISL_2935823 | Lighthouse Lab in Glasgow | Wellcome Sanger Institute for the COVID-19 Genomics UK (COG-UK) Consortium | Harper VanSteenhouse, Yumi Kasai, David Gray, Carol Clugston, Anna Dominiczak and Alex Alderton, Roberto Amato, Jeffrey Barrett, Sonia Goncalves, Ewan Harrison, David K. Jackson, Ian Johnston, Dominic Kwiatkowski, Cordelia Langford, John Sillitoe on behalf of the Wellcome Sanger Institute COVID-19 Surveillance Team |
| EPI_ISL_2935824, EPI_ISL_2935825 | Lighthouse Lab in Milton Keynes | Wellcome Sanger Institute for the COVID-19 Genomics UK (COG-UK) Consortium | The Lighthouse Lab in Milton Keynes and Alex Alderton, Roberto Amato, Jeffrey Barrett, Sonia Goncalves, Ewan Harrison, David K. Jackson, Ian Johnston, Dominic Kwiatkowski, Cordelia Langford, John Sillitoe on behalf of the Wellcome Sanger Institute COVID-19 Surveillance Team |
| EPI_ISL_2935826, EPI_ISL_2935827, EPI_ISL_2935828, EPI_ISL_2935829, EPI_ISL_2935830, EPI_ISL_2935831 | Lighthouse Lab in Glasgow | Wellcome Sanger Institute for the COVID-19 Genomics UK (COG-UK) Consortium | Harper VanSteenhouse, Yumi Kasai, David Gray, Carol Clugston, Anna Dominiczak and Alex Alderton, Roberto Amato, Jeffrey Barrett, Sonia Goncalves, Ewan Harrison, David K. Jackson, Ian Johnston, Dominic Kwiatkowski, Cordelia Langford, John Sillitoe on behalf of the Wellcome Sanger Institute COVID-19 Surveillance Team |
| EPI_ISL_2935832 | Lighthouse Lab in Milton Keynes | Wellcome Sanger Institute for the COVID-19 Genomics UK (COG-UK) Consortium | The Lighthouse Lab in Milton Keynes and Alex Alderton, Roberto Amato, Jeffrey Barrett, Sonia Goncalves, Ewan Harrison, David K. Jackson, Ian Johnston, Dominic Kwiatkowski, Cordelia Langford, John Sillitoe on behalf of the Wellcome Sanger Institute COVID-19 Surveillance Team |

| | | | |
|---|---|---|---|
| EPI_ISL_2935833, EPI_ISL_2935834 | Lighthouse Lab in Glasgow | Wellcome Sanger Institute for the COVID-19 Genomics UK (COG-UK) Consortium | Harper VanSteenhouse, Yumi Kasai, David Gray, Carol Clugston, Anna Dominiczak and Alex Alderton, Roberto Amato, Jeffrey Barrett, Sonia Goncalves, Ewan Harrison, David K. Jackson, Ian Johnston, Dominic Kwiatkowski, Cordelia Langford, John Sillitoe on behalf of the Wellcome Sanger Institute COVID-19 Surveillance Team |
| EPI_ISL_2935835, EPI_ISL_2935836, EPI_ISL_2935837, EPI_ISL_2935838, EPI_ISL_2935839 | Lighthouse Lab in Milton Keynes | Wellcome Sanger Institute for the COVID-19 Genomics UK (COG-UK) Consortium | The Lighthouse Lab in Milton Keynes and Alex Alderton, Roberto Amato, Jeffrey Barrett, Sonia Goncalves, Ewan Harrison, David K. Jackson, Ian Johnston, Dominic Kwiatkowski, Cordelia Langford, John Sillitoe on behalf of the Wellcome Sanger Institute COVID-19 Surveillance Team |
| EPI_ISL_2935840, EPI_ISL_2935841, EPI_ISL_2935842, EPI_ISL_2935843 | Lighthouse Lab in Glasgow | Wellcome Sanger Institute for the COVID-19 Genomics UK (COG-UK) Consortium | Harper VanSteenhouse, Yumi Kasai, David Gray, Carol Clugston, Anna Dominiczak and Alex Alderton, Roberto Amato, Jeffrey Barrett, Sonia Goncalves, Ewan Harrison, David K. Jackson, Ian Johnston, Dominic Kwiatkowski, Cordelia Langford, John Sillitoe on behalf of the Wellcome Sanger Institute COVID-19 Surveillance Team |
| EPI_ISL_2935844 | Lighthouse Lab in Milton Keynes | Wellcome Sanger Institute for the COVID-19 Genomics UK (COG-UK) Consortium | The Lighthouse Lab in Milton Keynes and Alex Alderton, Roberto Amato, Jeffrey Barrett, Sonia Goncalves, Ewan Harrison, David K. Jackson, Ian Johnston, Dominic Kwiatkowski, Cordelia Langford, John Sillitoe on behalf of the Wellcome Sanger Institute COVID-19 Surveillance Team |
| EPI_ISL_2935845, EPI_ISL_2935846, EPI_ISL_2935847 | Lighthouse Lab in Glasgow | Wellcome Sanger Institute for the COVID-19 Genomics UK (COG-UK) Consortium | Harper VanSteenhouse, Yumi Kasai, David Gray, Carol Clugston, Anna Dominiczak and Alex Alderton, Roberto Amato, Jeffrey Barrett, Sonia Goncalves, Ewan Harrison, David K. Jackson, Ian Johnston, Dominic Kwiatkowski, Cordelia Langford, John Sillitoe on behalf of the Wellcome Sanger Institute COVID-19 Surveillance Team |
| EPI_ISL_2935848, EPI_ISL_2935849 | Lighthouse Lab in Milton Keynes | Wellcome Sanger Institute for the COVID-19 Genomics UK (COG-UK) Consortium | The Lighthouse Lab in Milton Keynes and Alex Alderton, Roberto Amato, Jeffrey Barrett, Sonia Goncalves, Ewan Harrison, David K. Jackson, Ian Johnston, Dominic Kwiatkowski, Cordelia Langford, John Sillitoe on behalf of the Wellcome Sanger Institute COVID-19 Surveillance Team |
| EPI_ISL_2935850, EPI_ISL_2935851, EPI_ISL_2935852 | Lighthouse Lab in Glasgow | Wellcome Sanger Institute for the COVID-19 Genomics UK (COG-UK) Consortium | Harper VanSteenhouse, Yumi Kasai, David Gray, Carol Clugston, Anna Dominiczak and Alex Alderton, Roberto Amato, Jeffrey Barrett, Sonia Goncalves, Ewan Harrison, David K. Jackson, Ian Johnston, Dominic Kwiatkowski, Cordelia Langford, John Sillitoe on behalf of the Wellcome Sanger Institute COVID-19 Surveillance Team |
| EPI_ISL_2935853 | Lighthouse Lab in Milton Keynes | Wellcome Sanger Institute for the COVID-19 Genomics UK (COG-UK) Consortium | The Lighthouse Lab in Milton Keynes and Alex Alderton, Roberto Amato, Jeffrey Barrett, Sonia Goncalves, Ewan Harrison, David K. Jackson, Ian Johnston, Dominic Kwiatkowski, Cordelia Langford, John Sillitoe on behalf of the Wellcome Sanger Institute COVID-19 Surveillance Team |
| EPI_ISL_2935854, EPI_ISL_2935855 | Lighthouse Lab in Glasgow | Wellcome Sanger Institute for the COVID-19 Genomics UK (COG-UK) Consortium | Harper VanSteenhouse, Yumi Kasai, David Gray, Carol Clugston, Anna Dominiczak and Alex Alderton, Roberto Amato, Jeffrey Barrett, Sonia Goncalves, Ewan Harrison, David K. Jackson, Ian Johnston, Dominic Kwiatkowski, Cordelia Langford, John Sillitoe on behalf of the Wellcome Sanger Institute COVID-19 Surveillance Team |
| EPI_ISL_2935856 | Lighthouse Lab in Milton Keynes | Wellcome Sanger Institute for the COVID-19 Genomics UK (COG-UK) Consortium | The Lighthouse Lab in Milton Keynes and Alex Alderton, Roberto Amato, Jeffrey Barrett, Sonia Goncalves, Ewan Harrison, David K. Jackson, Ian Johnston, Dominic Kwiatkowski, Cordelia Langford, John Sillitoe on behalf of the Wellcome Sanger Institute COVID-19 Surveillance Team |
| EPI_ISL_2935857, EPI_ISL_2935858 | Lighthouse Lab in Glasgow | Wellcome Sanger Institute for the COVID-19 Genomics UK (COG-UK) Consortium | Harper VanSteenhouse, Yumi Kasai, David Gray, Carol Clugston, Anna Dominiczak and Alex Alderton, Roberto Amato, Jeffrey Barrett, Sonia Goncalves, Ewan Harrison, David K. Jackson, Ian Johnston, Dominic Kwiatkowski, Cordelia Langford, John Sillitoe on behalf of the Wellcome Sanger Institute COVID-19 Surveillance Team |
| EPI_ISL_2935859, EPI_ISL_2935860, EPI_ISL_2935861 | Lighthouse Lab in Milton Keynes | Wellcome Sanger Institute for the COVID-19 Genomics UK (COG-UK) Consortium | The Lighthouse Lab in Milton Keynes and Alex Alderton, Roberto Amato, Jeffrey Barrett, Sonia Goncalves, Ewan Harrison, David K. Jackson, Ian Johnston, Dominic Kwiatkowski, Cordelia Langford, John Sillitoe on behalf of the Wellcome Sanger Institute COVID-19 Surveillance Team |
| EPI_ISL_2935862, EPI_ISL_2935863 | Lighthouse Lab in Glasgow | Wellcome Sanger Institute for the COVID-19 Genomics UK (COG-UK) Consortium | Harper VanSteenhouse, Yumi Kasai, David Gray, Carol Clugston, Anna Dominiczak and Alex Alderton, Roberto Amato, Jeffrey Barrett, Sonia Goncalves, Ewan Harrison, David K. Jackson, Ian Johnston, Dominic Kwiatkowski, Cordelia Langford, John Sillitoe on behalf of the Wellcome Sanger Institute COVID-19 Surveillance Team |
| EPI_ISL_2935864, EPI_ISL_2935865 | Lighthouse Lab in Milton Keynes | Wellcome Sanger Institute for the COVID-19 Genomics UK (COG-UK) Consortium | The Lighthouse Lab in Milton Keynes and Alex Alderton, Roberto Amato, Jeffrey Barrett, Sonia Goncalves, Ewan Harrison, David K. Jackson, Ian Johnston, Dominic Kwiatkowski, Cordelia Langford, John Sillitoe on behalf of the Wellcome Sanger Institute COVID-19 Surveillance Team |
| EPI_ISL_2935866, EPI_ISL_2935867 | Lighthouse Lab in Glasgow | Wellcome Sanger Institute for the COVID-19 Genomics UK (COG-UK) Consortium | Harper VanSteenhouse, Yumi Kasai, David Gray, Carol Clugston, Anna Dominiczak and Alex Alderton, Roberto Amato, Jeffrey Barrett, Sonia Goncalves, Ewan Harrison, David K. Jackson, Ian Johnston, Dominic Kwiatkowski, Cordelia Langford, John Sillitoe on behalf of the Wellcome Sanger Institute COVID-19 Surveillance Team |
| EPI_ISL_2935868, EPI_ISL_2935869 | Lighthouse Lab in Milton Keynes | Wellcome Sanger Institute for the COVID-19 Genomics UK (COG-UK) Consortium | The Lighthouse Lab in Milton Keynes and Alex Alderton, Roberto Amato, Jeffrey Barrett, Sonia Goncalves, Ewan Harrison, David K. Jackson, Ian Johnston, Dominic Kwiatkowski, Cordelia Langford, John Sillitoe on behalf of the Wellcome Sanger Institute COVID-19 Surveillance Team |
| EPI_ISL_2935870 | Lighthouse Lab in Glasgow | Wellcome Sanger Institute for the COVID-19 Genomics UK (COG-UK) Consortium | Harper VanSteenhouse, Yumi Kasai, David Gray, Carol Clugston, Anna Dominiczak and Alex Alderton, Roberto Amato, Jeffrey Barrett, Sonia Goncalves, Ewan Harrison, David K. Jackson, Ian Johnston, Dominic Kwiatkowski, Cordelia Langford, John Sillitoe on behalf of the Wellcome Sanger Institute COVID-19 Surveillance Team |
| EPI_ISL_2935871 | Lighthouse Lab in Milton Keynes | Wellcome Sanger Institute for the COVID-19 Genomics UK (COG-UK) Consortium | The Lighthouse Lab in Milton Keynes and Alex Alderton, Roberto Amato, Jeffrey Barrett, Sonia Goncalves, Ewan Harrison, David K. Jackson, Ian Johnston, Dominic Kwiatkowski, Cordelia Langford, John Sillitoe on behalf of the Wellcome Sanger Institute COVID-19 Surveillance Team |
| EPI_ISL_2935872, EPI_ISL_2935873, EPI_ISL_2935874, EPI_ISL_2935875, EPI_ISL_2935876 | Lighthouse Lab in Glasgow | Wellcome Sanger Institute for the COVID-19 Genomics UK (COG-UK) Consortium | Harper VanSteenhouse, Yumi Kasai, David Gray, Carol Clugston, Anna Dominiczak and Alex Alderton, Roberto Amato, Jeffrey Barrett, Sonia Goncalves, Ewan Harrison, David K. Jackson, Ian Johnston, Dominic Kwiatkowski, Cordelia Langford, John Sillitoe on behalf of the Wellcome Sanger Institute COVID-19 Surveillance Team |
| EPI_ISL_2935877 | Lighthouse Lab in Milton Keynes | Wellcome Sanger Institute for the COVID-19 Genomics UK (COG-UK) Consortium | The Lighthouse Lab in Milton Keynes and Alex Alderton, Roberto Amato, Jeffrey Barrett, Sonia Goncalves, Ewan Harrison, David K. Jackson, Ian Johnston, Dominic Kwiatkowski, Cordelia Langford, John Sillitoe on behalf of the Wellcome Sanger Institute COVID-19 Surveillance Team |
| EPI_ISL_2935878 | Lighthouse Lab in Glasgow | Wellcome Sanger Institute for the COVID-19 Genomics UK (COG-UK) Consortium | Harper VanSteenhouse, Yumi Kasai, David Gray, Carol Clugston, Anna Dominiczak and Alex Alderton, Roberto Amato, Jeffrey Barrett, Sonia Goncalves, Ewan Harrison, David K. Jackson, Ian Johnston, Dominic Kwiatkowski, Cordelia Langford, John Sillitoe on behalf of the Wellcome Sanger Institute COVID-19 Surveillance Team |
| EPI_ISL_2935879 | Lighthouse Lab in Milton Keynes | Wellcome Sanger Institute for the COVID-19 Genomics UK (COG-UK) Consortium | The Lighthouse Lab in Milton Keynes and Alex Alderton, Roberto Amato, Jeffrey Barrett, Sonia Goncalves, Ewan Harrison, David K. Jackson, Ian Johnston, Dominic Kwiatkowski, Cordelia Langford, John Sillitoe on behalf of the Wellcome Sanger Institute COVID-19 Surveillance Team |
| EPI_ISL_2935880, EPI_ISL_2935881 | Lighthouse Lab in Glasgow | Wellcome Sanger Institute for the COVID-19 Genomics UK (COG-UK) Consortium | Harper VanSteenhouse, Yumi Kasai, David Gray, Carol Clugston, Anna Dominiczak and Alex Alderton, Roberto Amato, Jeffrey Barrett, Sonia Goncalves, Ewan Harrison, David K. Jackson, Ian Johnston, Dominic Kwiatkowski, Cordelia Langford, John Sillitoe on behalf of the Wellcome Sanger Institute COVID-19 Surveillance Team |
| EPI_ISL_2935882 | Lighthouse Lab in Milton Keynes | Wellcome Sanger Institute for the COVID-19 Genomics UK (COG-UK) Consortium | The Lighthouse Lab in Milton Keynes and Alex Alderton, Roberto Amato, Jeffrey Barrett, Sonia Goncalves, Ewan Harrison, David K. Jackson, Ian Johnston, Dominic Kwiatkowski, Cordelia Langford, John Sillitoe on behalf of the Wellcome Sanger Institute COVID-19 Surveillance Team |
| EPI_ISL_2935883 | Lighthouse Lab in Glasgow | Wellcome Sanger Institute for the COVID-19 Genomics UK (COG-UK) Consortium | Harper VanSteenhouse, Yumi Kasai, David Gray, Carol Clugston, Anna Dominiczak and Alex Alderton, Roberto Amato, Jeffrey Barrett, Sonia Goncalves, Ewan Harrison, David K. Jackson, Ian Johnston, Dominic Kwiatkowski, Cordelia Langford, John Sillitoe on behalf of the Wellcome Sanger Institute COVID-19 Surveillance Team |
| EPI_ISL_2935885 | Lighthouse Lab in Milton Keynes | Wellcome Sanger Institute for the COVID-19 Genomics UK (COG-UK) Consortium | The Lighthouse Lab in Milton Keynes and Alex Alderton, Roberto Amato, Jeffrey Barrett, Sonia Goncalves, Ewan Harrison, David K. Jackson, Ian Johnston, Dominic Kwiatkowski, Cordelia Langford, John Sillitoe on behalf of the Wellcome Sanger Institute COVID-19 Surveillance Team |
| EPI_ISL_2935886 | Lighthouse Lab in Glasgow | Wellcome Sanger Institute for the COVID-19 Genomics UK (COG-UK) Consortium | Harper VanSteenhouse, Yumi Kasai, David Gray, Carol Clugston, Anna Dominiczak and Alex Alderton, Roberto Amato, Jeffrey Barrett, Sonia Goncalves, Ewan Harrison, David K. Jackson, Ian Johnston, Dominic Kwiatkowski, Cordelia Langford, John Sillitoe on behalf of the Wellcome Sanger Institute COVID-19 Surveillance Team |
| EPI_ISL_2935887 | Lighthouse Lab in Milton Keynes | Wellcome Sanger Institute for the COVID-19 Genomics UK (COG-UK) Consortium | The Lighthouse Lab in Milton Keynes and Alex Alderton, Roberto Amato, Jeffrey Barrett, Sonia Goncalves, Ewan Harrison, David K. Jackson, Ian Johnston, Dominic Kwiatkowski, Cordelia Langford, John Sillitoe on behalf of the Wellcome Sanger Institute COVID-19 Surveillance Team |
| EPI_ISL_2935888, EPI_ISL_2935889, EPI_ISL_2935890, EPI_ISL_2935891, EPI_ISL_2935892 | Lighthouse Lab in Glasgow | Wellcome Sanger Institute for the COVID-19 Genomics UK (COG-UK) Consortium | Harper VanSteenhouse, Yumi Kasai, David Gray, Carol Clugston, Anna Dominiczak and Alex Alderton, Roberto Amato, Jeffrey Barrett, Sonia Goncalves, Ewan Harrison, David K. Jackson, Ian Johnston, Dominic Kwiatkowski, Cordelia Langford, John Sillitoe on behalf of the Wellcome Sanger Institute COVID-19 Surveillance Team |
| EPI_ISL_2935893, EPI_ISL_2935894, EPI_ISL_2935895 | Lighthouse Lab in Milton Keynes | Wellcome Sanger Institute for the COVID-19 Genomics UK (COG-UK) Consortium | The Lighthouse Lab in Milton Keynes and Alex Alderton, Roberto Amato, Jeffrey Barrett, Sonia Goncalves, Ewan Harrison, David K. Jackson, Ian Johnston, Dominic Kwiatkowski, Cordelia Langford, John Sillitoe on behalf of the Wellcome Sanger Institute COVID-19 Surveillance Team |
| EPI_ISL_2935896 | Lighthouse Lab in Glasgow | Wellcome Sanger Institute for the COVID-19 Genomics UK (COG-UK) Consortium | Harper VanSteenhouse, Yumi Kasai, David Gray, Carol Clugston, Anna Dominiczak and Alex Alderton, Roberto Amato, Jeffrey Barrett, Sonia Goncalves, Ewan Harrison, David K. Jackson, Ian Johnston, Dominic Kwiatkowski, Cordelia Langford, John Sillitoe on behalf of the Wellcome Sanger Institute COVID-19 Surveillance Team |
| EPI_ISL_2935897, EPI_ISL_2935898 | Lighthouse Lab in Milton Keynes | Wellcome Sanger Institute for the COVID-19 Genomics UK (COG-UK) Consortium | The Lighthouse Lab in Milton Keynes and Alex Alderton, Roberto Amato, Jeffrey Barrett, Sonia Goncalves, Ewan Harrison, David K. Jackson, Ian Johnston, Dominic Kwiatkowski, Cordelia Langford, John Sillitoe on behalf of the Wellcome Sanger Institute COVID-19 Surveillance Team |
| EPI_ISL_2935899, EPI_ISL_2935900 | Lighthouse Lab in Glasgow | Wellcome Sanger Institute for the COVID-19 Genomics UK (COG-UK) Consortium | Harper VanSteenhouse, Yumi Kasai, David Gray, Carol Clugston, Anna Dominiczak and Alex Alderton, Roberto Amato, Jeffrey Barrett, Sonia Goncalves, Ewan Harrison, David K. Jackson, Ian Johnston, Dominic Kwiatkowski, Cordelia Langford, John Sillitoe on behalf of the Wellcome Sanger Institute COVID-19 Surveillance Team |
| EPI_ISL_2935901, EPI_ISL_2935902 | Lighthouse Lab in Milton Keynes | Wellcome Sanger Institute for the COVID-19 Genomics UK (COG-UK) Consortium | The Lighthouse Lab in Milton Keynes and Alex Alderton, Roberto Amato, Jeffrey Barrett, Sonia Goncalves, Ewan Harrison, David K. Jackson, Ian Johnston, Dominic Kwiatkowski, Cordelia Langford, John Sillitoe on behalf of the Wellcome Sanger Institute COVID-19 Surveillance Team |
| EPI_ISL_2935903, EPI_ISL_2935904 | Lighthouse Lab in Glasgow | Wellcome Sanger Institute for the COVID-19 Genomics UK (COG-UK) Consortium | Harper VanSteenhouse, Yumi Kasai, David Gray, Carol Clugston, Anna Dominiczak and Alex Alderton, Roberto Amato, Jeffrey Barrett, Sonia Goncalves, Ewan Harrison, David K. Jackson, Ian Johnston, Dominic Kwiatkowski, Cordelia Langford, John Sillitoe on behalf of the Wellcome Sanger Institute COVID-19 Surveillance Team |
| EPI_ISL_2935905 | Lighthouse Lab in Milton Keynes | Wellcome Sanger Institute for the COVID-19 Genomics UK (COG-UK) Consortium | The Lighthouse Lab in Milton Keynes and Alex Alderton, Roberto Amato, Jeffrey Barrett, Sonia Goncalves, Ewan Harrison, David K. Jackson, Ian Johnston, Dominic Kwiatkowski, Cordelia Langford, John Sillitoe on behalf of the Wellcome Sanger Institute COVID-19 Surveillance Team |
| EPI_ISL_2935907 | Lighthouse Lab in Glasgow | Wellcome Sanger Institute for the COVID-19 Genomics UK (COG-UK) Consortium | Harper VanSteenhouse, Yumi Kasai, David Gray, Carol Clugston, Anna Dominiczak and Alex Alderton, Roberto Amato, Jeffrey Barrett, Sonia Goncalves, Ewan Harrison, David K. Jackson, Ian Johnston, Dominic Kwiatkowski, Cordelia Langford, John Sillitoe on behalf of the Wellcome Sanger Institute COVID-19 Surveillance Team |
| EPI_ISL_2935908, EPI_ISL_2935909 | Lighthouse Lab in Milton Keynes | Wellcome Sanger Institute for the COVID-19 Genomics UK (COG-UK) Consortium | The Lighthouse Lab in Milton Keynes and Alex Alderton, Roberto Amato, Jeffrey Barrett, Sonia Goncalves, Ewan Harrison, David K. Jackson, Ian Johnston, Dominic Kwiatkowski, Cordelia Langford, John Sillitoe on behalf of the Wellcome Sanger Institute COVID-19 Surveillance Team |
| EPI_ISL_2935910 | Lighthouse Lab in Glasgow | Wellcome Sanger Institute for the COVID-19 Genomics UK (COG-UK) Consortium | Harper VanSteenhouse, Yumi Kasai, David Gray, Carol Clugston, Anna Dominiczak and Alex Alderton, Roberto Amato, Jeffrey Barrett, Sonia Goncalves, Ewan Harrison, David K. Jackson, Ian Johnston, Dominic Kwiatkowski, Cordelia Langford, John Sillitoe on behalf of the Wellcome Sanger Institute COVID-19 Surveillance Team |

| | | | |
|---|---|---|---|
| EPI_ISL_2935911 | Lighthouse Lab in Milton Keynes | Wellcome Sanger Institute for the COVID-19 Genomics UK (COG-UK) Consortium | The Lighthouse Lab in Milton Keynes and Alex Alderton, Roberto Amato, Jeffrey Barrett, Sonia Goncalves, Ewan Harrison, David K. Jackson, Ian Johnston, Dominic Kwiatkowski, Cordelia Langford, John Sillitoe on behalf of the Wellcome Sanger Institute COVID-19 Surveillance Team |
| EPI_ISL_2935913 | Lighthouse Lab in Glasgow | Wellcome Sanger Institute for the COVID-19 Genomics UK (COG-UK) Consortium | Harper VanSteenhouse, Yumi Kasai, David Gray, Carol Clugston, Anna Dominiczak and Alex Alderton, Roberto Amato, Jeffrey Barrett, Sonia Goncalves, Ewan Harrison, David K. Jackson, Ian Johnston, Dominic Kwiatkowski, Cordelia Langford, John Sillitoe on behalf of the Wellcome Sanger Institute COVID-19 Surveillance Team |
| EPI_ISL_2935914, EPI_ISL_2935915, EPI_ISL_2935916, EPI_ISL_2935917, EPI_ISL_2935918 | Lighthouse Lab in Milton Keynes | Wellcome Sanger Institute for the COVID-19 Genomics UK (COG-UK) Consortium | The Lighthouse Lab in Milton Keynes and Alex Alderton, Roberto Amato, Jeffrey Barrett, Sonia Goncalves, Ewan Harrison, David K. Jackson, Ian Johnston, Dominic Kwiatkowski, Cordelia Langford, John Sillitoe on behalf of the Wellcome Sanger Institute COVID-19 Surveillance Team |
| EPI_ISL_2935919 | Lighthouse Lab in Glasgow | Wellcome Sanger Institute for the COVID-19 Genomics UK (COG-UK) Consortium | Harper VanSteenhouse, Yumi Kasai, David Gray, Carol Clugston, Anna Dominiczak and Alex Alderton, Roberto Amato, Jeffrey Barrett, Sonia Goncalves, Ewan Harrison, David K. Jackson, Ian Johnston, Dominic Kwiatkowski, Cordelia Langford, John Sillitoe on behalf of the Wellcome Sanger Institute COVID-19 Surveillance Team |
| EPI_ISL_2935920, EPI_ISL_2935921 | Lighthouse Lab in Milton Keynes | Wellcome Sanger Institute for the COVID-19 Genomics UK (COG-UK) Consortium | The Lighthouse Lab in Milton Keynes and Alex Alderton, Roberto Amato, Jeffrey Barrett, Sonia Goncalves, Ewan Harrison, David K. Jackson, Ian Johnston, Dominic Kwiatkowski, Cordelia Langford, John Sillitoe on behalf of the Wellcome Sanger Institute COVID-19 Surveillance Team |
| EPI_ISL_2935922, EPI_ISL_2935923 | Lighthouse Lab in Glasgow | Wellcome Sanger Institute for the COVID-19 Genomics UK (COG-UK) Consortium | Harper VanSteenhouse, Yumi Kasai, David Gray, Carol Clugston, Anna Dominiczak and Alex Alderton, Roberto Amato, Jeffrey Barrett, Sonia Goncalves, Ewan Harrison, David K. Jackson, Ian Johnston, Dominic Kwiatkowski, Cordelia Langford, John Sillitoe on behalf of the Wellcome Sanger Institute COVID-19 Surveillance Team |
| EPI_ISL_2935924, EPI_ISL_2935925, EPI_ISL_2935926, EPI_ISL_2935927 | Lighthouse Lab in Milton Keynes | Wellcome Sanger Institute for the COVID-19 Genomics UK (COG-UK) Consortium | The Lighthouse Lab in Milton Keynes and Alex Alderton, Roberto Amato, Jeffrey Barrett, Sonia Goncalves, Ewan Harrison, David K. Jackson, Ian Johnston, Dominic Kwiatkowski, Cordelia Langford, John Sillitoe on behalf of the Wellcome Sanger Institute COVID-19 Surveillance Team |
| EPI_ISL_2935928 | Lighthouse Lab in Glasgow | Wellcome Sanger Institute for the COVID-19 Genomics UK (COG-UK) Consortium | Harper VanSteenhouse, Yumi Kasai, David Gray, Carol Clugston, Anna Dominiczak and Alex Alderton, Roberto Amato, Jeffrey Barrett, Sonia Goncalves, Ewan Harrison, David K. Jackson, Ian Johnston, Dominic Kwiatkowski, Cordelia Langford, John Sillitoe on behalf of the Wellcome Sanger Institute COVID-19 Surveillance Team |
| EPI_ISL_2935929 | Lighthouse Lab in Milton Keynes | Wellcome Sanger Institute for the COVID-19 Genomics UK (COG-UK) Consortium | The Lighthouse Lab in Milton Keynes and Alex Alderton, Roberto Amato, Jeffrey Barrett, Sonia Goncalves, Ewan Harrison, David K. Jackson, Ian Johnston, Dominic Kwiatkowski, Cordelia Langford, John Sillitoe on behalf of the Wellcome Sanger Institute COVID-19 Surveillance Team |
| EPI_ISL_2935930, EPI_ISL_2935931, EPI_ISL_2935932, EPI_ISL_2935933 | Lighthouse Lab in Glasgow | Wellcome Sanger Institute for the COVID-19 Genomics UK (COG-UK) Consortium | Harper VanSteenhouse, Yumi Kasai, David Gray, Carol Clugston, Anna Dominiczak and Alex Alderton, Roberto Amato, Jeffrey Barrett, Sonia Goncalves, Ewan Harrison, David K. Jackson, Ian Johnston, Dominic Kwiatkowski, Cordelia Langford, John Sillitoe on behalf of the Wellcome Sanger Institute COVID-19 Surveillance Team |
| EPI_ISL_2935934, EPI_ISL_2935935 | Lighthouse Lab in Milton Keynes | Wellcome Sanger Institute for the COVID-19 Genomics UK (COG-UK) Consortium | The Lighthouse Lab in Milton Keynes and Alex Alderton, Roberto Amato, Jeffrey Barrett, Sonia Goncalves, Ewan Harrison, David K. Jackson, Ian Johnston, Dominic Kwiatkowski, Cordelia Langford, John Sillitoe on behalf of the Wellcome Sanger Institute COVID-19 Surveillance Team |
| EPI_ISL_2935936 | Lighthouse Lab in Glasgow | Wellcome Sanger Institute for the COVID-19 Genomics UK (COG-UK) Consortium | Harper VanSteenhouse, Yumi Kasai, David Gray, Carol Clugston, Anna Dominiczak and Alex Alderton, Roberto Amato, Jeffrey Barrett, Sonia Goncalves, Ewan Harrison, David K. Jackson, Ian Johnston, Dominic Kwiatkowski, Cordelia Langford, John Sillitoe on behalf of the Wellcome Sanger Institute COVID-19 Surveillance Team |
| EPI_ISL_2935937, EPI_ISL_2935938, EPI_ISL_2935939, EPI_ISL_2935940 | Lighthouse Lab in Milton Keynes | Wellcome Sanger Institute for the COVID-19 Genomics UK (COG-UK) Consortium | The Lighthouse Lab in Milton Keynes and Alex Alderton, Roberto Amato, Jeffrey Barrett, Sonia Goncalves, Ewan Harrison, David K. Jackson, Ian Johnston, Dominic Kwiatkowski, Cordelia Langford, John Sillitoe on behalf of the Wellcome Sanger Institute COVID-19 Surveillance Team |
| EPI_ISL_2935941 | Lighthouse Lab in Glasgow | Wellcome Sanger Institute for the COVID-19 Genomics UK (COG-UK) Consortium | Harper VanSteenhouse, Yumi Kasai, David Gray, Carol Clugston, Anna Dominiczak and Alex Alderton, Roberto Amato, Jeffrey Barrett, Sonia Goncalves, Ewan Harrison, David K. Jackson, Ian Johnston, Dominic Kwiatkowski, Cordelia Langford, John Sillitoe on behalf of the Wellcome Sanger Institute COVID-19 Surveillance Team |
| EPI_ISL_2935942, EPI_ISL_2935943 | Lighthouse Lab in Milton Keynes | Wellcome Sanger Institute for the COVID-19 Genomics UK (COG-UK) Consortium | The Lighthouse Lab in Milton Keynes and Alex Alderton, Roberto Amato, Jeffrey Barrett, Sonia Goncalves, Ewan Harrison, David K. Jackson, Ian Johnston, Dominic Kwiatkowski, Cordelia Langford, John Sillitoe on behalf of the Wellcome Sanger Institute COVID-19 Surveillance Team |
| EPI_ISL_2935944, EPI_ISL_2935945 | Lighthouse Lab in Glasgow | Wellcome Sanger Institute for the COVID-19 Genomics UK (COG-UK) Consortium | Harper VanSteenhouse, Yumi Kasai, David Gray, Carol Clugston, Anna Dominiczak and Alex Alderton, Roberto Amato, Jeffrey Barrett, Sonia Goncalves, Ewan Harrison, David K. Jackson, Ian Johnston, Dominic Kwiatkowski, Cordelia Langford, John Sillitoe on behalf of the Wellcome Sanger Institute COVID-19 Surveillance Team |
| EPI_ISL_2935946, EPI_ISL_2935947, EPI_ISL_2935948, EPI_ISL_2935949 | Lighthouse Lab in Milton Keynes | Wellcome Sanger Institute for the COVID-19 Genomics UK (COG-UK) Consortium | The Lighthouse Lab in Milton Keynes and Alex Alderton, Roberto Amato, Jeffrey Barrett, Sonia Goncalves, Ewan Harrison, David K. Jackson, Ian Johnston, Dominic Kwiatkowski, Cordelia Langford, John Sillitoe on behalf of the Wellcome Sanger Institute COVID-19 Surveillance Team |
| EPI_ISL_2935950 | Lighthouse Lab in Glasgow | Wellcome Sanger Institute for the COVID-19 Genomics UK (COG-UK) Consortium | Harper VanSteenhouse, Yumi Kasai, David Gray, Carol Clugston, Anna Dominiczak and Alex Alderton, Roberto Amato, Jeffrey Barrett, Sonia Goncalves, Ewan Harrison, David K. Jackson, Ian Johnston, Dominic Kwiatkowski, Cordelia Langford, John Sillitoe on behalf of the Wellcome Sanger Institute COVID-19 Surveillance Team |
| EPI_ISL_2935951, EPI_ISL_2935952 | Lighthouse Lab in Milton Keynes | Wellcome Sanger Institute for the COVID-19 Genomics UK (COG-UK) Consortium | The Lighthouse Lab in Milton Keynes and Alex Alderton, Roberto Amato, Jeffrey Barrett, Sonia Goncalves, Ewan Harrison, David K. Jackson, Ian Johnston, Dominic Kwiatkowski, Cordelia Langford, John Sillitoe on behalf of the Wellcome Sanger Institute COVID-19 Surveillance Team |
| EPI_ISL_2935953 | Lighthouse Lab in Glasgow | Wellcome Sanger Institute for the COVID-19 Genomics UK (COG-UK) Consortium | Harper VanSteenhouse, Yumi Kasai, David Gray, Carol Clugston, Anna Dominiczak and Alex Alderton, Roberto Amato, Jeffrey Barrett, Sonia Goncalves, Ewan Harrison, David K. Jackson, Ian Johnston, Dominic Kwiatkowski, Cordelia Langford, John Sillitoe on behalf of the Wellcome Sanger Institute COVID-19 Surveillance Team |
| EPI_ISL_2935954, EPI_ISL_2935955, EPI_ISL_2935956 | Lighthouse Lab in Milton Keynes | Wellcome Sanger Institute for the COVID-19 Genomics UK (COG-UK) Consortium | The Lighthouse Lab in Milton Keynes and Alex Alderton, Roberto Amato, Jeffrey Barrett, Sonia Goncalves, Ewan Harrison, David K. Jackson, Ian Johnston, Dominic Kwiatkowski, Cordelia Langford, John Sillitoe on behalf of the Wellcome Sanger Institute COVID-19 Surveillance Team |
| EPI_ISL_2935957 | Lighthouse Lab in Glasgow | Wellcome Sanger Institute for the COVID-19 Genomics UK (COG-UK) Consortium | Harper VanSteenhouse, Yumi Kasai, David Gray, Carol Clugston, Anna Dominiczak and Alex Alderton, Roberto Amato, Jeffrey Barrett, Sonia Goncalves, Ewan Harrison, David K. Jackson, Ian Johnston, Dominic Kwiatkowski, Cordelia Langford, John Sillitoe on behalf of the Wellcome Sanger Institute COVID-19 Surveillance Team |
| EPI_ISL_2935958, EPI_ISL_2935959 | Lighthouse Lab in Milton Keynes | Wellcome Sanger Institute for the COVID-19 Genomics UK (COG-UK) Consortium | The Lighthouse Lab in Milton Keynes and Alex Alderton, Roberto Amato, Jeffrey Barrett, Sonia Goncalves, Ewan Harrison, David K. Jackson, Ian Johnston, Dominic Kwiatkowski, Cordelia Langford, John Sillitoe on behalf of the Wellcome Sanger Institute COVID-19 Surveillance Team |
| EPI_ISL_2935960 | Lighthouse Lab in Glasgow | Wellcome Sanger Institute for the COVID-19 Genomics UK (COG-UK) Consortium | Harper VanSteenhouse, Yumi Kasai, David Gray, Carol Clugston, Anna Dominiczak and Alex Alderton, Roberto Amato, Jeffrey Barrett, Sonia Goncalves, Ewan Harrison, David K. Jackson, Ian Johnston, Dominic Kwiatkowski, Cordelia Langford, John Sillitoe on behalf of the Wellcome Sanger Institute COVID-19 Surveillance Team |
| EPI_ISL_2935961 | Lighthouse Lab in Milton Keynes | Wellcome Sanger Institute for the COVID-19 Genomics UK (COG-UK) Consortium | The Lighthouse Lab in Milton Keynes and Alex Alderton, Roberto Amato, Jeffrey Barrett, Sonia Goncalves, Ewan Harrison, David K. Jackson, Ian Johnston, Dominic Kwiatkowski, Cordelia Langford, John Sillitoe on behalf of the Wellcome Sanger Institute COVID-19 Surveillance Team |
| EPI_ISL_2935962 | Lighthouse Lab in Glasgow | Wellcome Sanger Institute for the COVID-19 Genomics UK (COG-UK) Consortium | Harper VanSteenhouse, Yumi Kasai, David Gray, Carol Clugston, Anna Dominiczak and Alex Alderton, Roberto Amato, Jeffrey Barrett, Sonia Goncalves, Ewan Harrison, David K. Jackson, Ian Johnston, Dominic Kwiatkowski, Cordelia Langford, John Sillitoe on behalf of the Wellcome Sanger Institute COVID-19 Surveillance Team |
| EPI_ISL_2935963 | Lighthouse Lab in Milton Keynes | Wellcome Sanger Institute for the COVID-19 Genomics UK (COG-UK) Consortium | The Lighthouse Lab in Milton Keynes and Alex Alderton, Roberto Amato, Jeffrey Barrett, Sonia Goncalves, Ewan Harrison, David K. Jackson, Ian Johnston, Dominic Kwiatkowski, Cordelia Langford, John Sillitoe on behalf of the Wellcome Sanger Institute COVID-19 Surveillance Team |
| EPI_ISL_2935964 | Lighthouse Lab in Glasgow | Wellcome Sanger Institute for the COVID-19 Genomics UK (COG-UK) Consortium | Harper VanSteenhouse, Yumi Kasai, David Gray, Carol Clugston, Anna Dominiczak and Alex Alderton, Roberto Amato, Jeffrey Barrett, Sonia Goncalves, Ewan Harrison, David K. Jackson, Ian Johnston, Dominic Kwiatkowski, Cordelia Langford, John Sillitoe on behalf of the Wellcome Sanger Institute COVID-19 Surveillance Team |
| EPI_ISL_2935965, EPI_ISL_2935966, EPI_ISL_2935967, EPI_ISL_2935968, EPI_ISL_2935969, EPI_ISL_2935970, EPI_ISL_2935971, EPI_ISL_2935972, EPI_ISL_2935973, EPI_ISL_2935974 | Lighthouse Lab in Milton Keynes | Wellcome Sanger Institute for the COVID-19 Genomics UK (COG-UK) Consortium | The Lighthouse Lab in Milton Keynes and Alex Alderton, Roberto Amato, Jeffrey Barrett, Sonia Goncalves, Ewan Harrison, David K. Jackson, Ian Johnston, Dominic Kwiatkowski, Cordelia Langford, John Sillitoe on behalf of the Wellcome Sanger Institute COVID-19 Surveillance Team |
| EPI_ISL_2935975 | Lighthouse Lab in Glasgow | Wellcome Sanger Institute for the COVID-19 Genomics UK (COG-UK) Consortium | Harper VanSteenhouse, Yumi Kasai, David Gray, Carol Clugston, Anna Dominiczak and Alex Alderton, Roberto Amato, Jeffrey Barrett, Sonia Goncalves, Ewan Harrison, David K. Jackson, Ian Johnston, Dominic Kwiatkowski, Cordelia Langford, John Sillitoe on behalf of the Wellcome Sanger Institute COVID-19 Surveillance Team |
| EPI_ISL_2935977 | Lighthouse Lab in Milton Keynes | Wellcome Sanger Institute for the COVID-19 Genomics UK (COG-UK) Consortium | The Lighthouse Lab in Milton Keynes and Alex Alderton, Roberto Amato, Jeffrey Barrett, Sonia Goncalves, Ewan Harrison, David K. Jackson, Ian Johnston, Dominic Kwiatkowski, Cordelia Langford, John Sillitoe on behalf of the Wellcome Sanger Institute COVID-19 Surveillance Team |
| EPI_ISL_2935978, EPI_ISL_2935979, EPI_ISL_2935980 | Lighthouse Lab in Glasgow | Wellcome Sanger Institute for the COVID-19 Genomics UK (COG-UK) Consortium | Harper VanSteenhouse, Yumi Kasai, David Gray, Carol Clugston, Anna Dominiczak and Alex Alderton, Roberto Amato, Jeffrey Barrett, Sonia Goncalves, Ewan Harrison, David K. Jackson, Ian Johnston, Dominic Kwiatkowski, Cordelia Langford, John Sillitoe on behalf of the Wellcome Sanger Institute COVID-19 Surveillance Team |
| EPI_ISL_2935981 | Lighthouse Lab in Milton Keynes | Wellcome Sanger Institute for the COVID-19 Genomics UK (COG-UK) Consortium | The Lighthouse Lab in Milton Keynes and Alex Alderton, Roberto Amato, Jeffrey Barrett, Sonia Goncalves, Ewan Harrison, David K. Jackson, Ian Johnston, Dominic Kwiatkowski, Cordelia Langford, John Sillitoe on behalf of the Wellcome Sanger Institute COVID-19 Surveillance Team |
| EPI_ISL_2935982, EPI_ISL_2935983, EPI_ISL_2935984 | Lighthouse Lab in Glasgow | Wellcome Sanger Institute for the COVID-19 Genomics UK (COG-UK) Consortium | Harper VanSteenhouse, Yumi Kasai, David Gray, Carol Clugston, Anna Dominiczak and Alex Alderton, Roberto Amato, Jeffrey Barrett, Sonia Goncalves, Ewan Harrison, David K. Jackson, Ian Johnston, Dominic Kwiatkowski, Cordelia Langford, John Sillitoe on behalf of the Wellcome Sanger Institute COVID-19 Surveillance Team |
| EPI_ISL_2935985 | Lighthouse Lab in Milton Keynes | Wellcome Sanger Institute for the COVID-19 Genomics UK (COG-UK) Consortium | The Lighthouse Lab in Milton Keynes and Alex Alderton, Roberto Amato, Jeffrey Barrett, Sonia Goncalves, Ewan Harrison, David K. Jackson, Ian Johnston, Dominic Kwiatkowski, Cordelia Langford, John Sillitoe on behalf of the Wellcome Sanger Institute COVID-19 Surveillance Team |
| EPI_ISL_2935986 | Lighthouse Lab in Glasgow | Wellcome Sanger Institute for the COVID-19 Genomics UK (COG-UK) Consortium | Harper VanSteenhouse, Yumi Kasai, David Gray, Carol Clugston, Anna Dominiczak and Alex Alderton, Roberto Amato, Jeffrey Barrett, Sonia Goncalves, Ewan Harrison, David K. Jackson, Ian Johnston, Dominic Kwiatkowski, Cordelia Langford, John Sillitoe on behalf of the Wellcome Sanger Institute COVID-19 Surveillance Team |

| | | | |
|---|---|---|---|
| EPI_ISL_2935987 | Lighthouse Lab in Milton Keynes | Wellcome Sanger Institute for the COVID-19 Genomics UK (COG-UK) Consortium | The Lighthouse Lab in Milton Keynes and Alex Alderton, Roberto Amato, Jeffrey Barrett, Sonia Goncalves, Ewan Harrison, David K. Jackson, Ian Johnston, Dominic Kwiatkowski, Cordelia Langford, John Sillitoe on behalf of the Wellcome Sanger Institute COVID-19 Surveillance Team |
| EPI_ISL_2935988, EPI_ISL_2935989, EPI_ISL_2935990 | Lighthouse Lab in Glasgow | Wellcome Sanger Institute for the COVID-19 Genomics UK (COG-UK) Consortium | Harper VanSteenhouse, Yumi Kasai, David Gray, Carol Clugston, Anna Dominiczak and Alex Alderton, Roberto Amato, Jeffrey Barrett, Sonia Goncalves, Ewan Harrison, David K. Jackson, Ian Johnston, Dominic Kwiatkowski, Cordelia Langford, John Sillitoe on behalf of the Wellcome Sanger Institute COVID-19 Surveillance Team |
| EPI_ISL_2935991, EPI_ISL_2935992, EPI_ISL_2935993 | Lighthouse Lab in Milton Keynes | Wellcome Sanger Institute for the COVID-19 Genomics UK (COG-UK) Consortium | The Lighthouse Lab in Milton Keynes and Alex Alderton, Roberto Amato, Jeffrey Barrett, Sonia Goncalves, Ewan Harrison, David K. Jackson, Ian Johnston, Dominic Kwiatkowski, Cordelia Langford, John Sillitoe on behalf of the Wellcome Sanger Institute COVID-19 Surveillance Team |
| EPI_ISL_2935994 | Lighthouse Lab in Glasgow | Wellcome Sanger Institute for the COVID-19 Genomics UK (COG-UK) Consortium | Harper VanSteenhouse, Yumi Kasai, David Gray, Carol Clugston, Anna Dominiczak and Alex Alderton, Roberto Amato, Jeffrey Barrett, Sonia Goncalves, Ewan Harrison, David K. Jackson, Ian Johnston, Dominic Kwiatkowski, Cordelia Langford, John Sillitoe on behalf of the Wellcome Sanger Institute COVID-19 Surveillance Team |
| EPI_ISL_2935995, EPI_ISL_2935996, EPI_ISL_2935997, EPI_ISL_2935998, EPI_ISL_2935999, EPI_ISL_2936000 | Lighthouse Lab in Milton Keynes | Wellcome Sanger Institute for the COVID-19 Genomics UK (COG-UK) Consortium | The Lighthouse Lab in Milton Keynes and Alex Alderton, Roberto Amato, Jeffrey Barrett, Sonia Goncalves, Ewan Harrison, David K. Jackson, Ian Johnston, Dominic Kwiatkowski, Cordelia Langford, John Sillitoe on behalf of the Wellcome Sanger Institute COVID-19 Surveillance Team |
| EPI_ISL_2936001, EPI_ISL_2936002 | Lighthouse Lab in Glasgow | Wellcome Sanger Institute for the COVID-19 Genomics UK (COG-UK) Consortium | Harper VanSteenhouse, Yumi Kasai, David Gray, Carol Clugston, Anna Dominiczak and Alex Alderton, Roberto Amato, Jeffrey Barrett, Sonia Goncalves, Ewan Harrison, David K. Jackson, Ian Johnston, Dominic Kwiatkowski, Cordelia Langford, John Sillitoe on behalf of the Wellcome Sanger Institute COVID-19 Surveillance Team |
| EPI_ISL_2936003 | Lighthouse Lab in Milton Keynes | Wellcome Sanger Institute for the COVID-19 Genomics UK (COG-UK) Consortium | The Lighthouse Lab in Milton Keynes and Alex Alderton, Roberto Amato, Jeffrey Barrett, Sonia Goncalves, Ewan Harrison, David K. Jackson, Ian Johnston, Dominic Kwiatkowski, Cordelia Langford, John Sillitoe on behalf of the Wellcome Sanger Institute COVID-19 Surveillance Team |
| EPI_ISL_2936004 | Lighthouse Lab in Glasgow | Wellcome Sanger Institute for the COVID-19 Genomics UK (COG-UK) Consortium | Harper VanSteenhouse, Yumi Kasai, David Gray, Carol Clugston, Anna Dominiczak and Alex Alderton, Roberto Amato, Jeffrey Barrett, Sonia Goncalves, Ewan Harrison, David K. Jackson, Ian Johnston, Dominic Kwiatkowski, Cordelia Langford, John Sillitoe on behalf of the Wellcome Sanger Institute COVID-19 Surveillance Team |
| EPI_ISL_2936005, EPI_ISL_2936006 | Lighthouse Lab in Milton Keynes | Wellcome Sanger Institute for the COVID-19 Genomics UK (COG-UK) Consortium | The Lighthouse Lab in Milton Keynes and Alex Alderton, Roberto Amato, Jeffrey Barrett, Sonia Goncalves, Ewan Harrison, David K. Jackson, Ian Johnston, Dominic Kwiatkowski, Cordelia Langford, John Sillitoe on behalf of the Wellcome Sanger Institute COVID-19 Surveillance Team |
| EPI_ISL_2936007, EPI_ISL_2936008 | Lighthouse Lab in Glasgow | Wellcome Sanger Institute for the COVID-19 Genomics UK (COG-UK) Consortium | Harper VanSteenhouse, Yumi Kasai, David Gray, Carol Clugston, Anna Dominiczak and Alex Alderton, Roberto Amato, Jeffrey Barrett, Sonia Goncalves, Ewan Harrison, David K. Jackson, Ian Johnston, Dominic Kwiatkowski, Cordelia Langford, John Sillitoe on behalf of the Wellcome Sanger Institute COVID-19 Surveillance Team |
| EPI_ISL_2936009, EPI_ISL_2936010 | Lighthouse Lab in Milton Keynes | Wellcome Sanger Institute for the COVID-19 Genomics UK (COG-UK) Consortium | The Lighthouse Lab in Milton Keynes and Alex Alderton, Roberto Amato, Jeffrey Barrett, Sonia Goncalves, Ewan Harrison, David K. Jackson, Ian Johnston, Dominic Kwiatkowski, Cordelia Langford, John Sillitoe on behalf of the Wellcome Sanger Institute COVID-19 Surveillance Team |
| EPI_ISL_2936011 | Lighthouse Lab in Glasgow | Wellcome Sanger Institute for the COVID-19 Genomics UK (COG-UK) Consortium | Harper VanSteenhouse, Yumi Kasai, David Gray, Carol Clugston, Anna Dominiczak and Alex Alderton, Roberto Amato, Jeffrey Barrett, Sonia Goncalves, Ewan Harrison, David K. Jackson, Ian Johnston, Dominic Kwiatkowski, Cordelia Langford, John Sillitoe on behalf of the Wellcome Sanger Institute COVID-19 Surveillance Team |
| EPI_ISL_2936012 | Lighthouse Lab in Milton Keynes | Wellcome Sanger Institute for the COVID-19 Genomics UK (COG-UK) Consortium | The Lighthouse Lab in Milton Keynes and Alex Alderton, Roberto Amato, Jeffrey Barrett, Sonia Goncalves, Ewan Harrison, David K. Jackson, Ian Johnston, Dominic Kwiatkowski, Cordelia Langford, John Sillitoe on behalf of the Wellcome Sanger Institute COVID-19 Surveillance Team |
| EPI_ISL_2936013, EPI_ISL_2936014 | Lighthouse Lab in Glasgow | Wellcome Sanger Institute for the COVID-19 Genomics UK (COG-UK) Consortium | Harper VanSteenhouse, Yumi Kasai, David Gray, Carol Clugston, Anna Dominiczak and Alex Alderton, Roberto Amato, Jeffrey Barrett, Sonia Goncalves, Ewan Harrison, David K. Jackson, Ian Johnston, Dominic Kwiatkowski, Cordelia Langford, John Sillitoe on behalf of the Wellcome Sanger Institute COVID-19 Surveillance Team |
| EPI_ISL_2936015 | Lighthouse Lab in Milton Keynes | Wellcome Sanger Institute for the COVID-19 Genomics UK (COG-UK) Consortium | The Lighthouse Lab in Milton Keynes and Alex Alderton, Roberto Amato, Jeffrey Barrett, Sonia Goncalves, Ewan Harrison, David K. Jackson, Ian Johnston, Dominic Kwiatkowski, Cordelia Langford, John Sillitoe on behalf of the Wellcome Sanger Institute COVID-19 Surveillance Team |
| EPI_ISL_2936016, EPI_ISL_2936017 | Lighthouse Lab in Glasgow | Wellcome Sanger Institute for the COVID-19 Genomics UK (COG-UK) Consortium | Harper VanSteenhouse, Yumi Kasai, David Gray, Carol Clugston, Anna Dominiczak and Alex Alderton, Roberto Amato, Jeffrey Barrett, Sonia Goncalves, Ewan Harrison, David K. Jackson, Ian Johnston, Dominic Kwiatkowski, Cordelia Langford, John Sillitoe on behalf of the Wellcome Sanger Institute COVID-19 Surveillance Team |
| EPI_ISL_2936018 | Lighthouse Lab in Milton Keynes | Wellcome Sanger Institute for the COVID-19 Genomics UK (COG-UK) Consortium | The Lighthouse Lab in Milton Keynes and Alex Alderton, Roberto Amato, Jeffrey Barrett, Sonia Goncalves, Ewan Harrison, David K. Jackson, Ian Johnston, Dominic Kwiatkowski, Cordelia Langford, John Sillitoe on behalf of the Wellcome Sanger Institute COVID-19 Surveillance Team |
| EPI_ISL_2936019 | Lighthouse Lab in Glasgow | Wellcome Sanger Institute for the COVID-19 Genomics UK (COG-UK) Consortium | Harper VanSteenhouse, Yumi Kasai, David Gray, Carol Clugston, Anna Dominiczak and Alex Alderton, Roberto Amato, Jeffrey Barrett, Sonia Goncalves, Ewan Harrison, David K. Jackson, Ian Johnston, Dominic Kwiatkowski, Cordelia Langford, John Sillitoe on behalf of the Wellcome Sanger Institute COVID-19 Surveillance Team |
| EPI_ISL_2936020, EPI_ISL_2936021, EPI_ISL_2936022, EPI_ISL_2936023, EPI_ISL_2936024 | Lighthouse Lab in Milton Keynes | Wellcome Sanger Institute for the COVID-19 Genomics UK (COG-UK) Consortium | The Lighthouse Lab in Milton Keynes and Alex Alderton, Roberto Amato, Jeffrey Barrett, Sonia Goncalves, Ewan Harrison, David K. Jackson, Ian Johnston, Dominic Kwiatkowski, Cordelia Langford, John Sillitoe on behalf of the Wellcome Sanger Institute COVID-19 Surveillance Team |
| EPI_ISL_2936025 | Lighthouse Lab in Glasgow | Wellcome Sanger Institute for the COVID-19 Genomics UK (COG-UK) Consortium | Harper VanSteenhouse, Yumi Kasai, David Gray, Carol Clugston, Anna Dominiczak and Alex Alderton, Roberto Amato, Jeffrey Barrett, Sonia Goncalves, Ewan Harrison, David K. Jackson, Ian Johnston, Dominic Kwiatkowski, Cordelia Langford, John Sillitoe on behalf of the Wellcome Sanger Institute COVID-19 Surveillance Team |
| EPI_ISL_2936026 | Lighthouse Lab in Milton Keynes | Wellcome Sanger Institute for the COVID-19 Genomics UK (COG-UK) Consortium | The Lighthouse Lab in Milton Keynes and Alex Alderton, Roberto Amato, Jeffrey Barrett, Sonia Goncalves, Ewan Harrison, David K. Jackson, Ian Johnston, Dominic Kwiatkowski, Cordelia Langford, John Sillitoe on behalf of the Wellcome Sanger Institute COVID-19 Surveillance Team |
| EPI_ISL_2936027 | Lighthouse Lab in Glasgow | Wellcome Sanger Institute for the COVID-19 Genomics UK (COG-UK) Consortium | Harper VanSteenhouse, Yumi Kasai, David Gray, Carol Clugston, Anna Dominiczak and Alex Alderton, Roberto Amato, Jeffrey Barrett, Sonia Goncalves, Ewan Harrison, David K. Jackson, Ian Johnston, Dominic Kwiatkowski, Cordelia Langford, John Sillitoe on behalf of the Wellcome Sanger Institute COVID-19 Surveillance Team |
| EPI_ISL_2936028 | Lighthouse Lab in Milton Keynes | Wellcome Sanger Institute for the COVID-19 Genomics UK (COG-UK) Consortium | The Lighthouse Lab in Milton Keynes and Alex Alderton, Roberto Amato, Jeffrey Barrett, Sonia Goncalves, Ewan Harrison, David K. Jackson, Ian Johnston, Dominic Kwiatkowski, Cordelia Langford, John Sillitoe on behalf of the Wellcome Sanger Institute COVID-19 Surveillance Team |
| EPI_ISL_2936029 | Lighthouse Lab in Glasgow | Wellcome Sanger Institute for the COVID-19 Genomics UK (COG-UK) Consortium | Harper VanSteenhouse, Yumi Kasai, David Gray, Carol Clugston, Anna Dominiczak and Alex Alderton, Roberto Amato, Jeffrey Barrett, Sonia Goncalves, Ewan Harrison, David K. Jackson, Ian Johnston, Dominic Kwiatkowski, Cordelia Langford, John Sillitoe on behalf of the Wellcome Sanger Institute COVID-19 Surveillance Team |
| EPI_ISL_2936030, EPI_ISL_2936031 | Lighthouse Lab in Milton Keynes | Wellcome Sanger Institute for the COVID-19 Genomics UK (COG-UK) Consortium | The Lighthouse Lab in Milton Keynes and Alex Alderton, Roberto Amato, Jeffrey Barrett, Sonia Goncalves, Ewan Harrison, David K. Jackson, Ian Johnston, Dominic Kwiatkowski, Cordelia Langford, John Sillitoe on behalf of the Wellcome Sanger Institute COVID-19 Surveillance Team |
| EPI_ISL_2936032, EPI_ISL_2936033, EPI_ISL_2936034 | Lighthouse Lab in Glasgow | Wellcome Sanger Institute for the COVID-19 Genomics UK (COG-UK) Consortium | Harper VanSteenhouse, Yumi Kasai, David Gray, Carol Clugston, Anna Dominiczak and Alex Alderton, Roberto Amato, Jeffrey Barrett, Sonia Goncalves, Ewan Harrison, David K. Jackson, Ian Johnston, Dominic Kwiatkowski, Cordelia Langford, John Sillitoe on behalf of the Wellcome Sanger Institute COVID-19 Surveillance Team |
| EPI_ISL_2936035 | Lighthouse Lab in Milton Keynes | Wellcome Sanger Institute for the COVID-19 Genomics UK (COG-UK) Consortium | The Lighthouse Lab in Milton Keynes and Alex Alderton, Roberto Amato, Jeffrey Barrett, Sonia Goncalves, Ewan Harrison, David K. Jackson, Ian Johnston, Dominic Kwiatkowski, Cordelia Langford, John Sillitoe on behalf of the Wellcome Sanger Institute COVID-19 Surveillance Team |
| EPI_ISL_2936036 | Lighthouse Lab in Glasgow | Wellcome Sanger Institute for the COVID-19 Genomics UK (COG-UK) Consortium | Harper VanSteenhouse, Yumi Kasai, David Gray, Carol Clugston, Anna Dominiczak and Alex Alderton, Roberto Amato, Jeffrey Barrett, Sonia Goncalves, Ewan Harrison, David K. Jackson, Ian Johnston, Dominic Kwiatkowski, Cordelia Langford, John Sillitoe on behalf of the Wellcome Sanger Institute COVID-19 Surveillance Team |
| EPI_ISL_2936037, EPI_ISL_2936038, EPI_ISL_2936039, EPI_ISL_2936040, EPI_ISL_2936041, EPI_ISL_2936042 | Lighthouse Lab in Milton Keynes | Wellcome Sanger Institute for the COVID-19 Genomics UK (COG-UK) Consortium | The Lighthouse Lab in Milton Keynes and Alex Alderton, Roberto Amato, Jeffrey Barrett, Sonia Goncalves, Ewan Harrison, David K. Jackson, Ian Johnston, Dominic Kwiatkowski, Cordelia Langford, John Sillitoe on behalf of the Wellcome Sanger Institute COVID-19 Surveillance Team |
| EPI_ISL_2936043, EPI_ISL_2936044 | Lighthouse Lab in Glasgow | Wellcome Sanger Institute for the COVID-19 Genomics UK (COG-UK) Consortium | Harper VanSteenhouse, Yumi Kasai, David Gray, Carol Clugston, Anna Dominiczak and Alex Alderton, Roberto Amato, Jeffrey Barrett, Sonia Goncalves, Ewan Harrison, David K. Jackson, Ian Johnston, Dominic Kwiatkowski, Cordelia Langford, John Sillitoe on behalf of the Wellcome Sanger Institute COVID-19 Surveillance Team |
| EPI_ISL_2936045, EPI_ISL_2936046, EPI_ISL_2936047, EPI_ISL_2936048 | Lighthouse Lab in Milton Keynes | Wellcome Sanger Institute for the COVID-19 Genomics UK (COG-UK) Consortium | The Lighthouse Lab in Milton Keynes and Alex Alderton, Roberto Amato, Jeffrey Barrett, Sonia Goncalves, Ewan Harrison, David K. Jackson, Ian Johnston, Dominic Kwiatkowski, Cordelia Langford, John Sillitoe on behalf of the Wellcome Sanger Institute COVID-19 Surveillance Team |
| EPI_ISL_2936049 | Lighthouse Lab in Glasgow | Wellcome Sanger Institute for the COVID-19 Genomics UK (COG-UK) Consortium | Harper VanSteenhouse, Yumi Kasai, David Gray, Carol Clugston, Anna Dominiczak and Alex Alderton, Roberto Amato, Jeffrey Barrett, Sonia Goncalves, Ewan Harrison, David K. Jackson, Ian Johnston, Dominic Kwiatkowski, Cordelia Langford, John Sillitoe on behalf of the Wellcome Sanger Institute COVID-19 Surveillance Team |
| EPI_ISL_2936050, EPI_ISL_2936051, EPI_ISL_2936052, EPI_ISL_2936053, EPI_ISL_2936054 | Lighthouse Lab in Milton Keynes | Wellcome Sanger Institute for the COVID-19 Genomics UK (COG-UK) Consortium | The Lighthouse Lab in Milton Keynes and Alex Alderton, Roberto Amato, Jeffrey Barrett, Sonia Goncalves, Ewan Harrison, David K. Jackson, Ian Johnston, Dominic Kwiatkowski, Cordelia Langford, John Sillitoe on behalf of the Wellcome Sanger Institute COVID-19 Surveillance Team |
| EPI_ISL_2936055 | Lighthouse Lab in Glasgow | Wellcome Sanger Institute for the COVID-19 Genomics UK (COG-UK) Consortium | Harper VanSteenhouse, Yumi Kasai, David Gray, Carol Clugston, Anna Dominiczak and Alex Alderton, Roberto Amato, Jeffrey Barrett, Sonia Goncalves, Ewan Harrison, David K. Jackson, Ian Johnston, Dominic Kwiatkowski, Cordelia Langford, John Sillitoe on behalf of the Wellcome Sanger Institute COVID-19 Surveillance Team |
| EPI_ISL_2936056 | Lighthouse Lab in Milton Keynes | Wellcome Sanger Institute for the COVID-19 Genomics UK (COG-UK) Consortium | The Lighthouse Lab in Milton Keynes and Alex Alderton, Roberto Amato, Jeffrey Barrett, Sonia Goncalves, Ewan Harrison, David K. Jackson, Ian Johnston, Dominic Kwiatkowski, Cordelia Langford, John Sillitoe on behalf of the Wellcome Sanger Institute COVID-19 Surveillance Team |
| EPI_ISL_2936057 | Lighthouse Lab in Glasgow | Wellcome Sanger Institute for the COVID-19 Genomics UK (COG-UK) Consortium | Harper VanSteenhouse, Yumi Kasai, David Gray, Carol Clugston, Anna Dominiczak and Alex Alderton, Roberto Amato, Jeffrey Barrett, Sonia Goncalves, Ewan Harrison, David K. Jackson, Ian Johnston, Dominic Kwiatkowski, Cordelia Langford, John Sillitoe on behalf of the Wellcome Sanger Institute COVID-19 Surveillance Team |
| EPI_ISL_2936058 | Lighthouse Lab in Milton Keynes | Wellcome Sanger Institute for the COVID-19 Genomics UK (COG-UK) Consortium | The Lighthouse Lab in Milton Keynes and Alex Alderton, Roberto Amato, Jeffrey Barrett, Sonia Goncalves, Ewan Harrison, David K. Jackson, Ian Johnston, Dominic Kwiatkowski, Cordelia Langford, John Sillitoe on behalf of the Wellcome Sanger Institute COVID-19 Surveillance Team |
| EPI_ISL_2936059, | Lighthouse Lab in Glasgow | Wellcome Sanger Institute for the COVID-19 Genomics UK (COG-UK) | Harper VanSteenhouse, Yumi Kasai, David Gray, Carol Clugston, Anna Dominiczak and Alex Alderton, Roberto Amato, Jeffrey Barrett, Sonia Goncalves, Ewan Harrison, David K. Jackson, Ian Johnston, Dominic Kwiatkowski, Cordelia Langford, |

| | | | |
|---|---|---|---|
| EPI_ISL_2936060 | | Consortium | John Sillitoe on behalf of the Wellcome Sanger Institute COVID-19 Surveillance Team |
| EPI_ISL_2936061, EPI_ISL_2936062, EPI_ISL_2936063, EPI_ISL_2936064, EPI_ISL_2936065 | Lighthouse Lab in Milton Keynes | Wellcome Sanger Institute for the COVID-19 Genomics UK (COG-UK) Consortium | The Lighthouse Lab in Milton Keynes and Alex Alderton, Roberto Amato, Jeffrey Barrett, Sonia Goncalves, Ewan Harrison, David K. Jackson, Ian Johnston, Dominic Kwiatkowski, Cordelia Langford, John Sillitoe on behalf of the Wellcome Sanger Institute COVID-19 Surveillance Team |
| EPI_ISL_2936066 | Lighthouse Lab in Glasgow | Wellcome Sanger Institute for the COVID-19 Genomics UK (COG-UK) Consortium | Harper VanSteenhouse, Yumi Kasai, David Gray, Carol Clugston, Anna Dominiczak and Alex Alderton, Roberto Amato, Jeffrey Barrett, Sonia Goncalves, Ewan Harrison, David K. Jackson, Ian Johnston, Dominic Kwiatkowski, Cordelia Langford, John Sillitoe on behalf of the Wellcome Sanger Institute COVID-19 Surveillance Team |
| EPI_ISL_2936067, EPI_ISL_2936068 | Lighthouse Lab in Milton Keynes | Wellcome Sanger Institute for the COVID-19 Genomics UK (COG-UK) Consortium | The Lighthouse Lab in Milton Keynes and Alex Alderton, Roberto Amato, Jeffrey Barrett, Sonia Goncalves, Ewan Harrison, David K. Jackson, Ian Johnston, Dominic Kwiatkowski, Cordelia Langford, John Sillitoe on behalf of the Wellcome Sanger Institute COVID-19 Surveillance Team |
| EPI_ISL_2936069 | Lighthouse Lab in Glasgow | Wellcome Sanger Institute for the COVID-19 Genomics UK (COG-UK) Consortium | Harper VanSteenhouse, Yumi Kasai, David Gray, Carol Clugston, Anna Dominiczak and Alex Alderton, Roberto Amato, Jeffrey Barrett, Sonia Goncalves, Ewan Harrison, David K. Jackson, Ian Johnston, Dominic Kwiatkowski, Cordelia Langford, John Sillitoe on behalf of the Wellcome Sanger Institute COVID-19 Surveillance Team |
| EPI_ISL_2936070 | Lighthouse Lab in Milton Keynes | Wellcome Sanger Institute for the COVID-19 Genomics UK (COG-UK) Consortium | The Lighthouse Lab in Milton Keynes and Alex Alderton, Roberto Amato, Jeffrey Barrett, Sonia Goncalves, Ewan Harrison, David K. Jackson, Ian Johnston, Dominic Kwiatkowski, Cordelia Langford, John Sillitoe on behalf of the Wellcome Sanger Institute COVID-19 Surveillance Team |
| EPI_ISL_2936071 | Lighthouse Lab in Glasgow | Wellcome Sanger Institute for the COVID-19 Genomics UK (COG-UK) Consortium | Harper VanSteenhouse, Yumi Kasai, David Gray, Carol Clugston, Anna Dominiczak and Alex Alderton, Roberto Amato, Jeffrey Barrett, Sonia Goncalves, Ewan Harrison, David K. Jackson, Ian Johnston, Dominic Kwiatkowski, Cordelia Langford, John Sillitoe on behalf of the Wellcome Sanger Institute COVID-19 Surveillance Team |
| EPI_ISL_2936072, EPI_ISL_2936073, EPI_ISL_2936074 | Lighthouse Lab in Milton Keynes | Wellcome Sanger Institute for the COVID-19 Genomics UK (COG-UK) Consortium | The Lighthouse Lab in Milton Keynes and Alex Alderton, Roberto Amato, Jeffrey Barrett, Sonia Goncalves, Ewan Harrison, David K. Jackson, Ian Johnston, Dominic Kwiatkowski, Cordelia Langford, John Sillitoe on behalf of the Wellcome Sanger Institute COVID-19 Surveillance Team |
| EPI_ISL_2936075 | Lighthouse Lab in Glasgow | Wellcome Sanger Institute for the COVID-19 Genomics UK (COG-UK) Consortium | Harper VanSteenhouse, Yumi Kasai, David Gray, Carol Clugston, Anna Dominiczak and Alex Alderton, Roberto Amato, Jeffrey Barrett, Sonia Goncalves, Ewan Harrison, David K. Jackson, Ian Johnston, Dominic Kwiatkowski, Cordelia Langford, John Sillitoe on behalf of the Wellcome Sanger Institute COVID-19 Surveillance Team |
| EPI_ISL_2936076, EPI_ISL_2936077 | Lighthouse Lab in Milton Keynes | Wellcome Sanger Institute for the COVID-19 Genomics UK (COG-UK) Consortium | The Lighthouse Lab in Milton Keynes and Alex Alderton, Roberto Amato, Jeffrey Barrett, Sonia Goncalves, Ewan Harrison, David K. Jackson, Ian Johnston, Dominic Kwiatkowski, Cordelia Langford, John Sillitoe on behalf of the Wellcome Sanger Institute COVID-19 Surveillance Team |
| EPI_ISL_2936078 | Lighthouse Lab in Glasgow | Wellcome Sanger Institute for the COVID-19 Genomics UK (COG-UK) Consortium | Harper VanSteenhouse, Yumi Kasai, David Gray, Carol Clugston, Anna Dominiczak and Alex Alderton, Roberto Amato, Jeffrey Barrett, Sonia Goncalves, Ewan Harrison, David K. Jackson, Ian Johnston, Dominic Kwiatkowski, Cordelia Langford, John Sillitoe on behalf of the Wellcome Sanger Institute COVID-19 Surveillance Team |
| EPI_ISL_2936079, EPI_ISL_2936080, EPI_ISL_2936081 | Lighthouse Lab in Milton Keynes | Wellcome Sanger Institute for the COVID-19 Genomics UK (COG-UK) Consortium | The Lighthouse Lab in Milton Keynes and Alex Alderton, Roberto Amato, Jeffrey Barrett, Sonia Goncalves, Ewan Harrison, David K. Jackson, Ian Johnston, Dominic Kwiatkowski, Cordelia Langford, John Sillitoe on behalf of the Wellcome Sanger Institute COVID-19 Surveillance Team |
| EPI_ISL_2936082 | Lighthouse Lab in Glasgow | Wellcome Sanger Institute for the COVID-19 Genomics UK (COG-UK) Consortium | Harper VanSteenhouse, Yumi Kasai, David Gray, Carol Clugston, Anna Dominiczak and Alex Alderton, Roberto Amato, Jeffrey Barrett, Sonia Goncalves, Ewan Harrison, David K. Jackson, Ian Johnston, Dominic Kwiatkowski, Cordelia Langford, John Sillitoe on behalf of the Wellcome Sanger Institute COVID-19 Surveillance Team |
| EPI_ISL_2936083 | Lighthouse Lab in Milton Keynes | Wellcome Sanger Institute for the COVID-19 Genomics UK (COG-UK) Consortium | The Lighthouse Lab in Milton Keynes and Alex Alderton, Roberto Amato, Jeffrey Barrett, Sonia Goncalves, Ewan Harrison, David K. Jackson, Ian Johnston, Dominic Kwiatkowski, Cordelia Langford, John Sillitoe on behalf of the Wellcome Sanger Institute COVID-19 Surveillance Team |
| EPI_ISL_2936084 | Lighthouse Lab in Glasgow | Wellcome Sanger Institute for the COVID-19 Genomics UK (COG-UK) Consortium | Harper VanSteenhouse, Yumi Kasai, David Gray, Carol Clugston, Anna Dominiczak and Alex Alderton, Roberto Amato, Jeffrey Barrett, Sonia Goncalves, Ewan Harrison, David K. Jackson, Ian Johnston, Dominic Kwiatkowski, Cordelia Langford, John Sillitoe on behalf of the Wellcome Sanger Institute COVID-19 Surveillance Team |
| EPI_ISL_2936085, EPI_ISL_2936086 | Lighthouse Lab in Milton Keynes | Wellcome Sanger Institute for the COVID-19 Genomics UK (COG-UK) Consortium | The Lighthouse Lab in Milton Keynes and Alex Alderton, Roberto Amato, Jeffrey Barrett, Sonia Goncalves, Ewan Harrison, David K. Jackson, Ian Johnston, Dominic Kwiatkowski, Cordelia Langford, John Sillitoe on behalf of the Wellcome Sanger Institute COVID-19 Surveillance Team |
| EPI_ISL_2936087, EPI_ISL_2936088 | Lighthouse Lab in Glasgow | Wellcome Sanger Institute for the COVID-19 Genomics UK (COG-UK) Consortium | Harper VanSteenhouse, Yumi Kasai, David Gray, Carol Clugston, Anna Dominiczak and Alex Alderton, Roberto Amato, Jeffrey Barrett, Sonia Goncalves, Ewan Harrison, David K. Jackson, Ian Johnston, Dominic Kwiatkowski, Cordelia Langford, John Sillitoe on behalf of the Wellcome Sanger Institute COVID-19 Surveillance Team |
| EPI_ISL_2936089 | Lighthouse Lab in Milton Keynes | Wellcome Sanger Institute for the COVID-19 Genomics UK (COG-UK) Consortium | The Lighthouse Lab in Milton Keynes and Alex Alderton, Roberto Amato, Jeffrey Barrett, Sonia Goncalves, Ewan Harrison, David K. Jackson, Ian Johnston, Dominic Kwiatkowski, Cordelia Langford, John Sillitoe on behalf of the Wellcome Sanger Institute COVID-19 Surveillance Team |
| EPI_ISL_2936090, EPI_ISL_2936091, EPI_ISL_2936092, EPI_ISL_2936093 | Lighthouse Lab in Glasgow | Wellcome Sanger Institute for the COVID-19 Genomics UK (COG-UK) Consortium | Harper VanSteenhouse, Yumi Kasai, David Gray, Carol Clugston, Anna Dominiczak and Alex Alderton, Roberto Amato, Jeffrey Barrett, Sonia Goncalves, Ewan Harrison, David K. Jackson, Ian Johnston, Dominic Kwiatkowski, Cordelia Langford, John Sillitoe on behalf of the Wellcome Sanger Institute COVID-19 Surveillance Team |
| EPI_ISL_2936094, EPI_ISL_2936095, EPI_ISL_2936096, EPI_ISL_2936097, EPI_ISL_2936098, EPI_ISL_2936099, EPI_ISL_2936100, EPI_ISL_2936101, EPI_ISL_2936102 | Lighthouse Lab in Milton Keynes | Wellcome Sanger Institute for the COVID-19 Genomics UK (COG-UK) Consortium | The Lighthouse Lab in Milton Keynes and Alex Alderton, Roberto Amato, Jeffrey Barrett, Sonia Goncalves, Ewan Harrison, David K. Jackson, Ian Johnston, Dominic Kwiatkowski, Cordelia Langford, John Sillitoe on behalf of the Wellcome Sanger Institute COVID-19 Surveillance Team |
| EPI_ISL_2936103, EPI_ISL_2936104 | Lighthouse Lab in Glasgow | Wellcome Sanger Institute for the COVID-19 Genomics UK (COG-UK) Consortium | Harper VanSteenhouse, Yumi Kasai, David Gray, Carol Clugston, Anna Dominiczak and Alex Alderton, Roberto Amato, Jeffrey Barrett, Sonia Goncalves, Ewan Harrison, David K. Jackson, Ian Johnston, Dominic Kwiatkowski, Cordelia Langford, John Sillitoe on behalf of the Wellcome Sanger Institute COVID-19 Surveillance Team |
| EPI_ISL_2936105 | Lighthouse Lab in Milton Keynes | Wellcome Sanger Institute for the COVID-19 Genomics UK (COG-UK) Consortium | The Lighthouse Lab in Milton Keynes and Alex Alderton, Roberto Amato, Jeffrey Barrett, Sonia Goncalves, Ewan Harrison, David K. Jackson, Ian Johnston, Dominic Kwiatkowski, Cordelia Langford, John Sillitoe on behalf of the Wellcome Sanger Institute COVID-19 Surveillance Team |
| EPI_ISL_2936106, EPI_ISL_2936107 | Lighthouse Lab in Glasgow | Wellcome Sanger Institute for the COVID-19 Genomics UK (COG-UK) Consortium | Harper VanSteenhouse, Yumi Kasai, David Gray, Carol Clugston, Anna Dominiczak and Alex Alderton, Roberto Amato, Jeffrey Barrett, Sonia Goncalves, Ewan Harrison, David K. Jackson, Ian Johnston, Dominic Kwiatkowski, Cordelia Langford, John Sillitoe on behalf of the Wellcome Sanger Institute COVID-19 Surveillance Team |
| EPI_ISL_2936108 | Lighthouse Lab in Milton Keynes | Wellcome Sanger Institute for the COVID-19 Genomics UK (COG-UK) Consortium | The Lighthouse Lab in Milton Keynes and Alex Alderton, Roberto Amato, Jeffrey Barrett, Sonia Goncalves, Ewan Harrison, David K. Jackson, Ian Johnston, Dominic Kwiatkowski, Cordelia Langford, John Sillitoe on behalf of the Wellcome Sanger Institute COVID-19 Surveillance Team |
| EPI_ISL_2936109 | Lighthouse Lab in Glasgow | Wellcome Sanger Institute for the COVID-19 Genomics UK (COG-UK) Consortium | Harper VanSteenhouse, Yumi Kasai, David Gray, Carol Clugston, Anna Dominiczak and Alex Alderton, Roberto Amato, Jeffrey Barrett, Sonia Goncalves, Ewan Harrison, David K. Jackson, Ian Johnston, Dominic Kwiatkowski, Cordelia Langford, John Sillitoe on behalf of the Wellcome Sanger Institute COVID-19 Surveillance Team |
| EPI_ISL_2936110 | Lighthouse Lab in Milton Keynes | Wellcome Sanger Institute for the COVID-19 Genomics UK (COG-UK) Consortium | The Lighthouse Lab in Milton Keynes and Alex Alderton, Roberto Amato, Jeffrey Barrett, Sonia Goncalves, Ewan Harrison, David K. Jackson, Ian Johnston, Dominic Kwiatkowski, Cordelia Langford, John Sillitoe on behalf of the Wellcome Sanger Institute COVID-19 Surveillance Team |
| EPI_ISL_2936111 | Lighthouse Lab in Glasgow | Wellcome Sanger Institute for the COVID-19 Genomics UK (COG-UK) Consortium | Harper VanSteenhouse, Yumi Kasai, David Gray, Carol Clugston, Anna Dominiczak and Alex Alderton, Roberto Amato, Jeffrey Barrett, Sonia Goncalves, Ewan Harrison, David K. Jackson, Ian Johnston, Dominic Kwiatkowski, Cordelia Langford, John Sillitoe on behalf of the Wellcome Sanger Institute COVID-19 Surveillance Team |
| EPI_ISL_2936112, EPI_ISL_2936113 | Lighthouse Lab in Milton Keynes | Wellcome Sanger Institute for the COVID-19 Genomics UK (COG-UK) Consortium | The Lighthouse Lab in Milton Keynes and Alex Alderton, Roberto Amato, Jeffrey Barrett, Sonia Goncalves, Ewan Harrison, David K. Jackson, Ian Johnston, Dominic Kwiatkowski, Cordelia Langford, John Sillitoe on behalf of the Wellcome Sanger Institute COVID-19 Surveillance Team |
| EPI_ISL_2936114 | Lighthouse Lab in Glasgow | Wellcome Sanger Institute for the COVID-19 Genomics UK (COG-UK) Consortium | Harper VanSteenhouse, Yumi Kasai, David Gray, Carol Clugston, Anna Dominiczak and Alex Alderton, Roberto Amato, Jeffrey Barrett, Sonia Goncalves, Ewan Harrison, David K. Jackson, Ian Johnston, Dominic Kwiatkowski, Cordelia Langford, John Sillitoe on behalf of the Wellcome Sanger Institute COVID-19 Surveillance Team |
| EPI_ISL_2936115, EPI_ISL_2936116 | Lighthouse Lab in Milton Keynes | Wellcome Sanger Institute for the COVID-19 Genomics UK (COG-UK) Consortium | The Lighthouse Lab in Milton Keynes and Alex Alderton, Roberto Amato, Jeffrey Barrett, Sonia Goncalves, Ewan Harrison, David K. Jackson, Ian Johnston, Dominic Kwiatkowski, Cordelia Langford, John Sillitoe on behalf of the Wellcome Sanger Institute COVID-19 Surveillance Team |
| EPI_ISL_2936117 | Lighthouse Lab in Glasgow | Wellcome Sanger Institute for the COVID-19 Genomics UK (COG-UK) Consortium | Harper VanSteenhouse, Yumi Kasai, David Gray, Carol Clugston, Anna Dominiczak and Alex Alderton, Roberto Amato, Jeffrey Barrett, Sonia Goncalves, Ewan Harrison, David K. Jackson, Ian Johnston, Dominic Kwiatkowski, Cordelia Langford, John Sillitoe on behalf of the Wellcome Sanger Institute COVID-19 Surveillance Team |
| EPI_ISL_2936118 | Lighthouse Lab in Milton Keynes | Wellcome Sanger Institute for the COVID-19 Genomics UK (COG-UK) Consortium | The Lighthouse Lab in Milton Keynes and Alex Alderton, Roberto Amato, Jeffrey Barrett, Sonia Goncalves, Ewan Harrison, David K. Jackson, Ian Johnston, Dominic Kwiatkowski, Cordelia Langford, John Sillitoe on behalf of the Wellcome Sanger Institute COVID-19 Surveillance Team |
| EPI_ISL_2936119 | Lighthouse Lab in Glasgow | Wellcome Sanger Institute for the COVID-19 Genomics UK (COG-UK) Consortium | Harper VanSteenhouse, Yumi Kasai, David Gray, Carol Clugston, Anna Dominiczak and Alex Alderton, Roberto Amato, Jeffrey Barrett, Sonia Goncalves, Ewan Harrison, David K. Jackson, Ian Johnston, Dominic Kwiatkowski, Cordelia Langford, John Sillitoe on behalf of the Wellcome Sanger Institute COVID-19 Surveillance Team |
| EPI_ISL_2936120, EPI_ISL_2936121, EPI_ISL_2936123, EPI_ISL_2936124 | Lighthouse Lab in Milton Keynes | Wellcome Sanger Institute for the COVID-19 Genomics UK (COG-UK) Consortium | The Lighthouse Lab in Milton Keynes and Alex Alderton, Roberto Amato, Jeffrey Barrett, Sonia Goncalves, Ewan Harrison, David K. Jackson, Ian Johnston, Dominic Kwiatkowski, Cordelia Langford, John Sillitoe on behalf of the Wellcome Sanger Institute COVID-19 Surveillance Team |
| EPI_ISL_2936126 | Lighthouse Lab in Glasgow | Wellcome Sanger Institute for the COVID-19 Genomics UK (COG-UK) Consortium | Harper VanSteenhouse, Yumi Kasai, David Gray, Carol Clugston, Anna Dominiczak and Alex Alderton, Roberto Amato, Jeffrey Barrett, Sonia Goncalves, Ewan Harrison, David K. Jackson, Ian Johnston, Dominic Kwiatkowski, Cordelia Langford, John Sillitoe on behalf of the Wellcome Sanger Institute COVID-19 Surveillance Team |
| EPI_ISL_2936127 | Lighthouse Lab in Milton Keynes | Wellcome Sanger Institute for the COVID-19 Genomics UK (COG-UK) Consortium | The Lighthouse Lab in Milton Keynes and Alex Alderton, Roberto Amato, Jeffrey Barrett, Sonia Goncalves, Ewan Harrison, David K. Jackson, Ian Johnston, Dominic Kwiatkowski, Cordelia Langford, John Sillitoe on behalf of the Wellcome Sanger Institute COVID-19 Surveillance Team |
| EPI_ISL_2936129 | Lighthouse Lab in Glasgow | Wellcome Sanger Institute for the COVID-19 Genomics UK (COG-UK) Consortium | Harper VanSteenhouse, Yumi Kasai, David Gray, Carol Clugston, Anna Dominiczak and Alex Alderton, Roberto Amato, Jeffrey Barrett, Sonia Goncalves, Ewan Harrison, David K. Jackson, Ian Johnston, Dominic Kwiatkowski, Cordelia Langford, John Sillitoe on behalf of the Wellcome Sanger Institute COVID-19 Surveillance Team |
| EPI_ISL_2936131 | Lighthouse Lab in Milton Keynes | Wellcome Sanger Institute for the COVID-19 Genomics UK (COG-UK) Consortium | The Lighthouse Lab in Milton Keynes and Alex Alderton, Roberto Amato, Jeffrey Barrett, Sonia Goncalves, Ewan Harrison, David K. Jackson, Ian Johnston, Dominic Kwiatkowski, Cordelia Langford, John Sillitoe on behalf of the Wellcome Sanger Institute COVID-19 Surveillance Team |
| EPI_ISL_2936132 | Lighthouse Lab in Glasgow | Wellcome Sanger Institute for the COVID-19 Genomics UK (COG-UK) Consortium | Harper VanSteenhouse, Yumi Kasai, David Gray, Carol Clugston, Anna Dominiczak and Alex Alderton, Roberto Amato, Jeffrey Barrett, Sonia Goncalves, Ewan Harrison, David K. Jackson, Ian Johnston, Dominic Kwiatkowski, Cordelia Langford, John Sillitoe on behalf of the Wellcome Sanger Institute COVID-19 Surveillance Team |

| | | | |
|---|---|---|---|
| EPI_ISL_2936134, EPI_ISL_2936136, EPI_ISL_2936137, EPI_ISL_2936138 | Lighthouse Lab in Milton Keynes | Wellcome Sanger Institute for the COVID-19 Genomics UK (COG-UK) Consortium | The Lighthouse Lab in Milton Keynes and Alex Alderton, Roberto Amato, Jeffrey Barrett, Sonia Goncalves, Ewan Harrison, David K. Jackson, Ian Johnston, Dominic Kwiatkowski, Cordelia Langford, John Sillitoe on behalf of the Wellcome Sanger Institute COVID-19 Surveillance Team |
| EPI_ISL_2936140 | Lighthouse Lab in Glasgow | Wellcome Sanger Institute for the COVID-19 Genomics UK (COG-UK) Consortium | Harper VanSteenhouse, Yumi Kasai, David Gray, Carol Clugston, Anna Dominiczak and Alex Alderton, Roberto Amato, Jeffrey Barrett, Sonia Goncalves, Ewan Harrison, David K. Jackson, Ian Johnston, Dominic Kwiatkowski, Cordelia Langford, John Sillitoe on behalf of the Wellcome Sanger Institute COVID-19 Surveillance Team |
| EPI_ISL_2936142 | Lighthouse Lab in Milton Keynes | Wellcome Sanger Institute for the COVID-19 Genomics UK (COG-UK) Consortium | The Lighthouse Lab in Milton Keynes and Alex Alderton, Roberto Amato, Jeffrey Barrett, Sonia Goncalves, Ewan Harrison, David K. Jackson, Ian Johnston, Dominic Kwiatkowski, Cordelia Langford, John Sillitoe on behalf of the Wellcome Sanger Institute COVID-19 Surveillance Team |
| EPI_ISL_2936143 | Lighthouse Lab in Glasgow | Wellcome Sanger Institute for the COVID-19 Genomics UK (COG-UK) Consortium | Harper VanSteenhouse, Yumi Kasai, David Gray, Carol Clugston, Anna Dominiczak and Alex Alderton, Roberto Amato, Jeffrey Barrett, Sonia Goncalves, Ewan Harrison, David K. Jackson, Ian Johnston, Dominic Kwiatkowski, Cordelia Langford, John Sillitoe on behalf of the Wellcome Sanger Institute COVID-19 Surveillance Team |
| EPI_ISL_2936145, EPI_ISL_2936147, EPI_ISL_2936148 | Lighthouse Lab in Milton Keynes | Wellcome Sanger Institute for the COVID-19 Genomics UK (COG-UK) Consortium | The Lighthouse Lab in Milton Keynes and Alex Alderton, Roberto Amato, Jeffrey Barrett, Sonia Goncalves, Ewan Harrison, David K. Jackson, Ian Johnston, Dominic Kwiatkowski, Cordelia Langford, John Sillitoe on behalf of the Wellcome Sanger Institute COVID-19 Surveillance Team |
| EPI_ISL_2936150, EPI_ISL_2936151, EPI_ISL_2936153, EPI_ISL_2936154 | Lighthouse Lab in Glasgow | Wellcome Sanger Institute for the COVID-19 Genomics UK (COG-UK) Consortium | Harper VanSteenhouse, Yumi Kasai, David Gray, Carol Clugston, Anna Dominiczak and Alex Alderton, Roberto Amato, Jeffrey Barrett, Sonia Goncalves, Ewan Harrison, David K. Jackson, Ian Johnston, Dominic Kwiatkowski, Cordelia Langford, John Sillitoe on behalf of the Wellcome Sanger Institute COVID-19 Surveillance Team |
| EPI_ISL_2936157, EPI_ISL_2936159, EPI_ISL_2936160, EPI_ISL_2936162, EPI_ISL_2936163, EPI_ISL_2936165, EPI_ISL_2936166, EPI_ISL_2936168, EPI_ISL_2936170, EPI_ISL_2936171 | Lighthouse Lab in Milton Keynes | Wellcome Sanger Institute for the COVID-19 Genomics UK (COG-UK) Consortium | The Lighthouse Lab in Milton Keynes and Alex Alderton, Roberto Amato, Jeffrey Barrett, Sonia Goncalves, Ewan Harrison, David K. Jackson, Ian Johnston, Dominic Kwiatkowski, Cordelia Langford, John Sillitoe on behalf of the Wellcome Sanger Institute COVID-19 Surveillance Team |
| EPI_ISL_2936173, EPI_ISL_2936174 | Lighthouse Lab in Glasgow | Wellcome Sanger Institute for the COVID-19 Genomics UK (COG-UK) Consortium | Harper VanSteenhouse, Yumi Kasai, David Gray, Carol Clugston, Anna Dominiczak and Alex Alderton, Roberto Amato, Jeffrey Barrett, Sonia Goncalves, Ewan Harrison, David K. Jackson, Ian Johnston, Dominic Kwiatkowski, Cordelia Langford, John Sillitoe on behalf of the Wellcome Sanger Institute COVID-19 Surveillance Team |
| EPI_ISL_2936176, EPI_ISL_2936177, EPI_ISL_2936179, EPI_ISL_2936181, EPI_ISL_2936182, EPI_ISL_2936184 | Lighthouse Lab in Milton Keynes | Wellcome Sanger Institute for the COVID-19 Genomics UK (COG-UK) Consortium | The Lighthouse Lab in Milton Keynes and Alex Alderton, Roberto Amato, Jeffrey Barrett, Sonia Goncalves, Ewan Harrison, David K. Jackson, Ian Johnston, Dominic Kwiatkowski, Cordelia Langford, John Sillitoe on behalf of the Wellcome Sanger Institute COVID-19 Surveillance Team |
| EPI_ISL_2936185 | Lighthouse Lab in Glasgow | Wellcome Sanger Institute for the COVID-19 Genomics UK (COG-UK) Consortium | Harper VanSteenhouse, Yumi Kasai, David Gray, Carol Clugston, Anna Dominiczak and Alex Alderton, Roberto Amato, Jeffrey Barrett, Sonia Goncalves, Ewan Harrison, David K. Jackson, Ian Johnston, Dominic Kwiatkowski, Cordelia Langford, John Sillitoe on behalf of the Wellcome Sanger Institute COVID-19 Surveillance Team |
| EPI_ISL_2936187 | Lighthouse Laboratory Plymouth | Wellcome Sanger Institute for the COVID-19 Genomics UK (COG-UK) Consortium | Lighthouse Laboratory Plymouth and Alex Alderton, Roberto Amato, Jeffrey Barrett, Sonia Goncalves, Ewan Harrison, David K. Jackson, Ian Johnston, Dominic Kwiatkowski, Cordelia Langford, John Sillitoe on behalf of the Wellcome Sanger Institute COVID-19 Surveillance Team |
| EPI_ISL_2936188, EPI_ISL_2936190, EPI_ISL_2936191 | Lighthouse Lab in Milton Keynes | Wellcome Sanger Institute for the COVID-19 Genomics UK (COG-UK) Consortium | The Lighthouse Lab in Milton Keynes and Alex Alderton, Roberto Amato, Jeffrey Barrett, Sonia Goncalves, Ewan Harrison, David K. Jackson, Ian Johnston, Dominic Kwiatkowski, Cordelia Langford, John Sillitoe on behalf of the Wellcome Sanger Institute COVID-19 Surveillance Team |
| EPI_ISL_2936193, EPI_ISL_2936195, EPI_ISL_2936196, EPI_ISL_2936198 | Lighthouse Laboratory Plymouth | Wellcome Sanger Institute for the COVID-19 Genomics UK (COG-UK) Consortium | Lighthouse Laboratory Plymouth and Alex Alderton, Roberto Amato, Jeffrey Barrett, Sonia Goncalves, Ewan Harrison, David K. Jackson, Ian Johnston, Dominic Kwiatkowski, Cordelia Langford, John Sillitoe on behalf of the Wellcome Sanger Institute COVID-19 Surveillance Team |
| EPI_ISL_2936199, EPI_ISL_2936201 | Lighthouse Lab in Milton Keynes | Wellcome Sanger Institute for the COVID-19 Genomics UK (COG-UK) Consortium | The Lighthouse Lab in Milton Keynes and Alex Alderton, Roberto Amato, Jeffrey Barrett, Sonia Goncalves, Ewan Harrison, David K. Jackson, Ian Johnston, Dominic Kwiatkowski, Cordelia Langford, John Sillitoe on behalf of the Wellcome Sanger Institute COVID-19 Surveillance Team |
| EPI_ISL_2936202, EPI_ISL_2936204 | Lighthouse Laboratory Plymouth | Wellcome Sanger Institute for the COVID-19 Genomics UK (COG-UK) Consortium | Lighthouse Laboratory Plymouth and Alex Alderton, Roberto Amato, Jeffrey Barrett, Sonia Goncalves, Ewan Harrison, David K. Jackson, Ian Johnston, Dominic Kwiatkowski, Cordelia Langford, John Sillitoe on behalf of the Wellcome Sanger Institute COVID-19 Surveillance Team |
| EPI_ISL_2936206 | Lighthouse Lab in Milton Keynes | Wellcome Sanger Institute for the COVID-19 Genomics UK (COG-UK) Consortium | The Lighthouse Lab in Milton Keynes and Alex Alderton, Roberto Amato, Jeffrey Barrett, Sonia Goncalves, Ewan Harrison, David K. Jackson, Ian Johnston, Dominic Kwiatkowski, Cordelia Langford, John Sillitoe on behalf of the Wellcome Sanger Institute COVID-19 Surveillance Team |
| EPI_ISL_2936207, EPI_ISL_2936209 | Lighthouse Laboratory Plymouth | Wellcome Sanger Institute for the COVID-19 Genomics UK (COG-UK) Consortium | Lighthouse Laboratory Plymouth and Alex Alderton, Roberto Amato, Jeffrey Barrett, Sonia Goncalves, Ewan Harrison, David K. Jackson, Ian Johnston, Dominic Kwiatkowski, Cordelia Langford, John Sillitoe on behalf of the Wellcome Sanger Institute COVID-19 Surveillance Team |
| EPI_ISL_2936210 | Lighthouse Lab in Milton Keynes | Wellcome Sanger Institute for the COVID-19 Genomics UK (COG-UK) Consortium | The Lighthouse Lab in Milton Keynes and Alex Alderton, Roberto Amato, Jeffrey Barrett, Sonia Goncalves, Ewan Harrison, David K. Jackson, Ian Johnston, Dominic Kwiatkowski, Cordelia Langford, John Sillitoe on behalf of the Wellcome Sanger Institute COVID-19 Surveillance Team |
| EPI_ISL_2936212, EPI_ISL_2936213, EPI_ISL_2936215, EPI_ISL_2936216, EPI_ISL_2936218 | Lighthouse Laboratory Plymouth | Wellcome Sanger Institute for the COVID-19 Genomics UK (COG-UK) Consortium | Lighthouse Laboratory Plymouth and Alex Alderton, Roberto Amato, Jeffrey Barrett, Sonia Goncalves, Ewan Harrison, David K. Jackson, Ian Johnston, Dominic Kwiatkowski, Cordelia Langford, John Sillitoe on behalf of the Wellcome Sanger Institute COVID-19 Surveillance Team |
| EPI_ISL_2936219 | Lighthouse Lab in Milton Keynes | Wellcome Sanger Institute for the COVID-19 Genomics UK (COG-UK) Consortium | The Lighthouse Lab in Milton Keynes and Alex Alderton, Roberto Amato, Jeffrey Barrett, Sonia Goncalves, Ewan Harrison, David K. Jackson, Ian Johnston, Dominic Kwiatkowski, Cordelia Langford, John Sillitoe on behalf of the Wellcome Sanger Institute COVID-19 Surveillance Team |
| EPI_ISL_2936221, EPI_ISL_2936222, EPI_ISL_2936224 | Lighthouse Laboratory Plymouth | Wellcome Sanger Institute for the COVID-19 Genomics UK (COG-UK) Consortium | Lighthouse Laboratory Plymouth and Alex Alderton, Roberto Amato, Jeffrey Barrett, Sonia Goncalves, Ewan Harrison, David K. Jackson, Ian Johnston, Dominic Kwiatkowski, Cordelia Langford, John Sillitoe on behalf of the Wellcome Sanger Institute COVID-19 Surveillance Team |
| EPI_ISL_2936226 | Lighthouse Lab in Milton Keynes | Wellcome Sanger Institute for the COVID-19 Genomics UK (COG-UK) Consortium | The Lighthouse Lab in Milton Keynes and Alex Alderton, Roberto Amato, Jeffrey Barrett, Sonia Goncalves, Ewan Harrison, David K. Jackson, Ian Johnston, Dominic Kwiatkowski, Cordelia Langford, John Sillitoe on behalf of the Wellcome Sanger Institute COVID-19 Surveillance Team |
| EPI_ISL_2936227, EPI_ISL_2936228 | Lighthouse Laboratory Plymouth | Wellcome Sanger Institute for the COVID-19 Genomics UK (COG-UK) Consortium | Lighthouse Laboratory Plymouth and Alex Alderton, Roberto Amato, Jeffrey Barrett, Sonia Goncalves, Ewan Harrison, David K. Jackson, Ian Johnston, Dominic Kwiatkowski, Cordelia Langford, John Sillitoe on behalf of the Wellcome Sanger Institute COVID-19 Surveillance Team |
| EPI_ISL_2936230 | Lighthouse Lab in Milton Keynes | Wellcome Sanger Institute for the COVID-19 Genomics UK (COG-UK) Consortium | The Lighthouse Lab in Milton Keynes and Alex Alderton, Roberto Amato, Jeffrey Barrett, Sonia Goncalves, Ewan Harrison, David K. Jackson, Ian Johnston, Dominic Kwiatkowski, Cordelia Langford, John Sillitoe on behalf of the Wellcome Sanger Institute COVID-19 Surveillance Team |
| EPI_ISL_2936232, EPI_ISL_2936233, EPI_ISL_2936235, EPI_ISL_2936236 | Lighthouse Laboratory Plymouth | Wellcome Sanger Institute for the COVID-19 Genomics UK (COG-UK) Consortium | Lighthouse Laboratory Plymouth and Alex Alderton, Roberto Amato, Jeffrey Barrett, Sonia Goncalves, Ewan Harrison, David K. Jackson, Ian Johnston, Dominic Kwiatkowski, Cordelia Langford, John Sillitoe on behalf of the Wellcome Sanger Institute COVID-19 Surveillance Team |
| EPI_ISL_2936238 | Lighthouse Lab in Milton Keynes | Wellcome Sanger Institute for the COVID-19 Genomics UK (COG-UK) Consortium | The Lighthouse Lab in Milton Keynes and Alex Alderton, Roberto Amato, Jeffrey Barrett, Sonia Goncalves, Ewan Harrison, David K. Jackson, Ian Johnston, Dominic Kwiatkowski, Cordelia Langford, John Sillitoe on behalf of the Wellcome Sanger Institute COVID-19 Surveillance Team |
| EPI_ISL_2936239, EPI_ISL_2936241, EPI_ISL_2936242, EPI_ISL_2936244, EPI_ISL_2936245 | Lighthouse Laboratory Plymouth | Wellcome Sanger Institute for the COVID-19 Genomics UK (COG-UK) Consortium | Lighthouse Laboratory Plymouth and Alex Alderton, Roberto Amato, Jeffrey Barrett, Sonia Goncalves, Ewan Harrison, David K. Jackson, Ian Johnston, Dominic Kwiatkowski, Cordelia Langford, John Sillitoe on behalf of the Wellcome Sanger Institute COVID-19 Surveillance Team |
| EPI_ISL_2936247 | Lighthouse Lab in Milton Keynes | Wellcome Sanger Institute for the COVID-19 Genomics UK (COG-UK) Consortium | The Lighthouse Lab in Milton Keynes and Alex Alderton, Roberto Amato, Jeffrey Barrett, Sonia Goncalves, Ewan Harrison, David K. Jackson, Ian Johnston, Dominic Kwiatkowski, Cordelia Langford, John Sillitoe on behalf of the Wellcome Sanger Institute COVID-19 Surveillance Team |
| EPI_ISL_2936249, EPI_ISL_2936250 | Lighthouse Laboratory Plymouth | Wellcome Sanger Institute for the COVID-19 Genomics UK (COG-UK) Consortium | Lighthouse Laboratory Plymouth and Alex Alderton, Roberto Amato, Jeffrey Barrett, Sonia Goncalves, Ewan Harrison, David K. Jackson, Ian Johnston, Dominic Kwiatkowski, Cordelia Langford, John Sillitoe on behalf of the Wellcome Sanger Institute COVID-19 Surveillance Team |
| EPI_ISL_2936252 | Lighthouse Lab in Milton Keynes | Wellcome Sanger Institute for the COVID-19 Genomics UK (COG-UK) Consortium | The Lighthouse Lab in Milton Keynes and Alex Alderton, Roberto Amato, Jeffrey Barrett, Sonia Goncalves, Ewan Harrison, David K. Jackson, Ian Johnston, Dominic Kwiatkowski, Cordelia Langford, John Sillitoe on behalf of the Wellcome Sanger Institute COVID-19 Surveillance Team |
| EPI_ISL_2936253, EPI_ISL_2936255, EPI_ISL_2936257, EPI_ISL_2936259, EPI_ISL_2936260, EPI_ISL_2936262, EPI_ISL_2936263, | Lighthouse Laboratory Plymouth | Wellcome Sanger Institute for the COVID-19 Genomics UK (COG-UK) Consortium | Lighthouse Laboratory Plymouth and Alex Alderton, Roberto Amato, Jeffrey Barrett, Sonia Goncalves, Ewan Harrison, David K. Jackson, Ian Johnston, Dominic Kwiatkowski, Cordelia Langford, John Sillitoe on behalf of the Wellcome Sanger Institute COVID-19 Surveillance Team |

| | | | |
|---|---|---|---|
| EPI_ISL_2936265, EPI_ISL_2936266 | | | |
| EPI_ISL_2936268 | Lighthouse Lab in Milton Keynes | Wellcome Sanger Institute for the COVID-19 Genomics UK (COG-UK) Consortium | The Lighthouse Lab in Milton Keynes and Alex Alderton, Roberto Amato, Jeffrey Barrett, Sonia Goncalves, Ewan Harrison, David K. Jackson, Ian Johnston, Dominic Kwiatkowski, Cordelia Langford, John Sillitoe on behalf of the Wellcome Sanger Institute COVID-19 Surveillance Team |
| EPI_ISL_2936269, EPI_ISL_2936271, EPI_ISL_2936273, EPI_ISL_2936274, EPI_ISL_2936276, EPI_ISL_2936277, EPI_ISL_2936279 | Lighthouse Laboratory Plymouth | Wellcome Sanger Institute for the COVID-19 Genomics UK (COG-UK) Consortium | Lighthouse Laboratory Plymouth and Alex Alderton, Roberto Amato, Jeffrey Barrett, Sonia Goncalves, Ewan Harrison, David K. Jackson, Ian Johnston, Dominic Kwiatkowski, Cordelia Langford, John Sillitoe on behalf of the Wellcome Sanger Institute COVID-19 Surveillance Team |
| EPI_ISL_2936281 | Lighthouse Lab in Milton Keynes | Wellcome Sanger Institute for the COVID-19 Genomics UK (COG-UK) Consortium | The Lighthouse Lab in Milton Keynes and Alex Alderton, Roberto Amato, Jeffrey Barrett, Sonia Goncalves, Ewan Harrison, David K. Jackson, Ian Johnston, Dominic Kwiatkowski, Cordelia Langford, John Sillitoe on behalf of the Wellcome Sanger Institute COVID-19 Surveillance Team |
| EPI_ISL_2936282 | Lighthouse Laboratory Plymouth | Wellcome Sanger Institute for the COVID-19 Genomics UK (COG-UK) Consortium | Lighthouse Laboratory Plymouth and Alex Alderton, Roberto Amato, Jeffrey Barrett, Sonia Goncalves, Ewan Harrison, David K. Jackson, Ian Johnston, Dominic Kwiatkowski, Cordelia Langford, John Sillitoe on behalf of the Wellcome Sanger Institute COVID-19 Surveillance Team |
| EPI_ISL_2936283, EPI_ISL_2936284 | Lighthouse Lab in Milton Keynes | Wellcome Sanger Institute for the COVID-19 Genomics UK (COG-UK) Consortium | The Lighthouse Lab in Milton Keynes and Alex Alderton, Roberto Amato, Jeffrey Barrett, Sonia Goncalves, Ewan Harrison, David K. Jackson, Ian Johnston, Dominic Kwiatkowski, Cordelia Langford, John Sillitoe on behalf of the Wellcome Sanger Institute COVID-19 Surveillance Team |
| EPI_ISL_2936285 | Lighthouse Laboratory Plymouth | Wellcome Sanger Institute for the COVID-19 Genomics UK (COG-UK) Consortium | Lighthouse Laboratory Plymouth and Alex Alderton, Roberto Amato, Jeffrey Barrett, Sonia Goncalves, Ewan Harrison, David K. Jackson, Ian Johnston, Dominic Kwiatkowski, Cordelia Langford, John Sillitoe on behalf of the Wellcome Sanger Institute COVID-19 Surveillance Team |
| EPI_ISL_2936287 | Lighthouse Lab in Milton Keynes | Wellcome Sanger Institute for the COVID-19 Genomics UK (COG-UK) Consortium | The Lighthouse Lab in Milton Keynes and Alex Alderton, Roberto Amato, Jeffrey Barrett, Sonia Goncalves, Ewan Harrison, David K. Jackson, Ian Johnston, Dominic Kwiatkowski, Cordelia Langford, John Sillitoe on behalf of the Wellcome Sanger Institute COVID-19 Surveillance Team |
| EPI_ISL_2936288, EPI_ISL_2936290, EPI_ISL_2936291, EPI_ISL_2936293, EPI_ISL_2936295 | Lighthouse Laboratory Plymouth | Wellcome Sanger Institute for the COVID-19 Genomics UK (COG-UK) Consortium | Lighthouse Laboratory Plymouth and Alex Alderton, Roberto Amato, Jeffrey Barrett, Sonia Goncalves, Ewan Harrison, David K. Jackson, Ian Johnston, Dominic Kwiatkowski, Cordelia Langford, John Sillitoe on behalf of the Wellcome Sanger Institute COVID-19 Surveillance Team |
| EPI_ISL_2936297, EPI_ISL_2936298, EPI_ISL_2936300, EPI_ISL_2936301 | Lighthouse Lab in Milton Keynes | Wellcome Sanger Institute for the COVID-19 Genomics UK (COG-UK) Consortium | The Lighthouse Lab in Milton Keynes and Alex Alderton, Roberto Amato, Jeffrey Barrett, Sonia Goncalves, Ewan Harrison, David K. Jackson, Ian Johnston, Dominic Kwiatkowski, Cordelia Langford, John Sillitoe on behalf of the Wellcome Sanger Institute COVID-19 Surveillance Team |
| EPI_ISL_2936302 | Lighthouse Laboratory Plymouth | Wellcome Sanger Institute for the COVID-19 Genomics UK (COG-UK) Consortium | Lighthouse Laboratory Plymouth and Alex Alderton, Roberto Amato, Jeffrey Barrett, Sonia Goncalves, Ewan Harrison, David K. Jackson, Ian Johnston, Dominic Kwiatkowski, Cordelia Langford, John Sillitoe on behalf of the Wellcome Sanger Institute COVID-19 Surveillance Team |
| EPI_ISL_2936304 | Randox Laboratories | Wellcome Sanger Institute for the COVID-19 Genomics UK (COG-UK) Consortium | Randox Laboratories and Alex Alderton, Roberto Amato, Jeffrey Barrett, Sonia Goncalves, Ewan Harrison, David K. Jackson, Ian Johnston, Dominic Kwiatkowski, Cordelia Langford, John Sillitoe on behalf of the Wellcome Sanger Institute COVID-19 Surveillance Team |
| EPI_ISL_2936305 | Lighthouse Lab in Milton Keynes | Wellcome Sanger Institute for the COVID-19 Genomics UK (COG-UK) Consortium | The Lighthouse Lab in Milton Keynes and Alex Alderton, Roberto Amato, Jeffrey Barrett, Sonia Goncalves, Ewan Harrison, David K. Jackson, Ian Johnston, Dominic Kwiatkowski, Cordelia Langford, John Sillitoe on behalf of the Wellcome Sanger Institute COVID-19 Surveillance Team |
| EPI_ISL_2936307, EPI_ISL_2936309, EPI_ISL_2936310, EPI_ISL_2936312, EPI_ISL_2936313 | Lighthouse Laboratory Plymouth | Wellcome Sanger Institute for the COVID-19 Genomics UK (COG-UK) Consortium | Lighthouse Laboratory Plymouth and Alex Alderton, Roberto Amato, Jeffrey Barrett, Sonia Goncalves, Ewan Harrison, David K. Jackson, Ian Johnston, Dominic Kwiatkowski, Cordelia Langford, John Sillitoe on behalf of the Wellcome Sanger Institute COVID-19 Surveillance Team |
| EPI_ISL_2936315 | Lighthouse Lab in Milton Keynes | Wellcome Sanger Institute for the COVID-19 Genomics UK (COG-UK) Consortium | The Lighthouse Lab in Milton Keynes and Alex Alderton, Roberto Amato, Jeffrey Barrett, Sonia Goncalves, Ewan Harrison, David K. Jackson, Ian Johnston, Dominic Kwiatkowski, Cordelia Langford, John Sillitoe on behalf of the Wellcome Sanger Institute COVID-19 Surveillance Team |
| EPI_ISL_2936317, EPI_ISL_2936318, EPI_ISL_2936320, EPI_ISL_2936322, EPI_ISL_2936323 | Lighthouse Laboratory Plymouth | Wellcome Sanger Institute for the COVID-19 Genomics UK (COG-UK) Consortium | Lighthouse Laboratory Plymouth and Alex Alderton, Roberto Amato, Jeffrey Barrett, Sonia Goncalves, Ewan Harrison, David K. Jackson, Ian Johnston, Dominic Kwiatkowski, Cordelia Langford, John Sillitoe on behalf of the Wellcome Sanger Institute COVID-19 Surveillance Team |
| EPI_ISL_2936325, EPI_ISL_2936327, EPI_ISL_2936328, EPI_ISL_2936330, EPI_ISL_2936331, EPI_ISL_2936333 | Lighthouse Lab in Milton Keynes | Wellcome Sanger Institute for the COVID-19 Genomics UK (COG-UK) Consortium | The Lighthouse Lab in Milton Keynes and Alex Alderton, Roberto Amato, Jeffrey Barrett, Sonia Goncalves, Ewan Harrison, David K. Jackson, Ian Johnston, Dominic Kwiatkowski, Cordelia Langford, John Sillitoe on behalf of the Wellcome Sanger Institute COVID-19 Surveillance Team |
| EPI_ISL_2936334, EPI_ISL_2936336, EPI_ISL_2936337 | Lighthouse Laboratory Plymouth | Wellcome Sanger Institute for the COVID-19 Genomics UK (COG-UK) Consortium | Lighthouse Laboratory Plymouth and Alex Alderton, Roberto Amato, Jeffrey Barrett, Sonia Goncalves, Ewan Harrison, David K. Jackson, Ian Johnston, Dominic Kwiatkowski, Cordelia Langford, John Sillitoe on behalf of the Wellcome Sanger Institute COVID-19 Surveillance Team |
| EPI_ISL_2936339 | Lighthouse Lab in Milton Keynes | Wellcome Sanger Institute for the COVID-19 Genomics UK (COG-UK) Consortium | The Lighthouse Lab in Milton Keynes and Alex Alderton, Roberto Amato, Jeffrey Barrett, Sonia Goncalves, Ewan Harrison, David K. Jackson, Ian Johnston, Dominic Kwiatkowski, Cordelia Langford, John Sillitoe on behalf of the Wellcome Sanger Institute COVID-19 Surveillance Team |
| EPI_ISL_2936341, EPI_ISL_2936342 | Lighthouse Laboratory Plymouth | Wellcome Sanger Institute for the COVID-19 Genomics UK (COG-UK) Consortium | Lighthouse Laboratory Plymouth and Alex Alderton, Roberto Amato, Jeffrey Barrett, Sonia Goncalves, Ewan Harrison, David K. Jackson, Ian Johnston, Dominic Kwiatkowski, Cordelia Langford, John Sillitoe on behalf of the Wellcome Sanger Institute COVID-19 Surveillance Team |
| EPI_ISL_2936344 | Lighthouse Lab in Milton Keynes | Wellcome Sanger Institute for the COVID-19 Genomics UK (COG-UK) Consortium | The Lighthouse Lab in Milton Keynes and Alex Alderton, Roberto Amato, Jeffrey Barrett, Sonia Goncalves, Ewan Harrison, David K. Jackson, Ian Johnston, Dominic Kwiatkowski, Cordelia Langford, John Sillitoe on behalf of the Wellcome Sanger Institute COVID-19 Surveillance Team |
| EPI_ISL_2936346, EPI_ISL_2936349, EPI_ISL_2936351, EPI_ISL_2936353, EPI_ISL_2936354, EPI_ISL_2936355 | Lighthouse Laboratory Plymouth | Wellcome Sanger Institute for the COVID-19 Genomics UK (COG-UK) Consortium | Lighthouse Laboratory Plymouth and Alex Alderton, Roberto Amato, Jeffrey Barrett, Sonia Goncalves, Ewan Harrison, David K. Jackson, Ian Johnston, Dominic Kwiatkowski, Cordelia Langford, John Sillitoe on behalf of the Wellcome Sanger Institute COVID-19 Surveillance Team |
| EPI_ISL_2936357 | Lighthouse Lab in Milton Keynes | Wellcome Sanger Institute for the COVID-19 Genomics UK (COG-UK) Consortium | The Lighthouse Lab in Milton Keynes and Alex Alderton, Roberto Amato, Jeffrey Barrett, Sonia Goncalves, Ewan Harrison, David K. Jackson, Ian Johnston, Dominic Kwiatkowski, Cordelia Langford, John Sillitoe on behalf of the Wellcome Sanger Institute COVID-19 Surveillance Team |
| EPI_ISL_2936358 | Lighthouse Laboratory Plymouth | Wellcome Sanger Institute for the COVID-19 Genomics UK (COG-UK) Consortium | Lighthouse Laboratory Plymouth and Alex Alderton, Roberto Amato, Jeffrey Barrett, Sonia Goncalves, Ewan Harrison, David K. Jackson, Ian Johnston, Dominic Kwiatkowski, Cordelia Langford, John Sillitoe on behalf of the Wellcome Sanger Institute COVID-19 Surveillance Team |
| EPI_ISL_2936360 | Lighthouse Lab in Milton Keynes | Wellcome Sanger Institute for the COVID-19 Genomics UK (COG-UK) Consortium | The Lighthouse Lab in Milton Keynes and Alex Alderton, Roberto Amato, Jeffrey Barrett, Sonia Goncalves, Ewan Harrison, David K. Jackson, Ian Johnston, Dominic Kwiatkowski, Cordelia Langford, John Sillitoe on behalf of the Wellcome Sanger Institute COVID-19 Surveillance Team |
| EPI_ISL_2936362 | Lighthouse Laboratory Plymouth | Wellcome Sanger Institute for the COVID-19 Genomics UK (COG-UK) Consortium | Lighthouse Laboratory Plymouth and Alex Alderton, Roberto Amato, Jeffrey Barrett, Sonia Goncalves, Ewan Harrison, David K. Jackson, Ian Johnston, Dominic Kwiatkowski, Cordelia Langford, John Sillitoe on behalf of the Wellcome Sanger Institute COVID-19 Surveillance Team |
| EPI_ISL_2936363 | Lighthouse Lab in Milton Keynes | Wellcome Sanger Institute for the COVID-19 Genomics UK (COG-UK) Consortium | The Lighthouse Lab in Milton Keynes and Alex Alderton, Roberto Amato, Jeffrey Barrett, Sonia Goncalves, Ewan Harrison, David K. Jackson, Ian Johnston, Dominic Kwiatkowski, Cordelia Langford, John Sillitoe on behalf of the Wellcome Sanger Institute COVID-19 Surveillance Team |
| EPI_ISL_2936365, EPI_ISL_2936367, EPI_ISL_2936368, EPI_ISL_2936370, EPI_ISL_2936372, EPI_ISL_2936373, EPI_ISL_2936375, EPI_ISL_2936377 | Lighthouse Laboratory Plymouth | Wellcome Sanger Institute for the COVID-19 Genomics UK (COG-UK) Consortium | Lighthouse Laboratory Plymouth and Alex Alderton, Roberto Amato, Jeffrey Barrett, Sonia Goncalves, Ewan Harrison, David K. Jackson, Ian Johnston, Dominic Kwiatkowski, Cordelia Langford, John Sillitoe on behalf of the Wellcome Sanger Institute COVID-19 Surveillance Team |
| EPI_ISL_2936378 | Lighthouse Lab in Milton Keynes | Wellcome Sanger Institute for the COVID-19 Genomics UK (COG-UK) Consortium | The Lighthouse Lab in Milton Keynes and Alex Alderton, Roberto Amato, Jeffrey Barrett, Sonia Goncalves, Ewan Harrison, David K. Jackson, Ian Johnston, Dominic Kwiatkowski, Cordelia Langford, John Sillitoe on behalf of the Wellcome Sanger Institute COVID-19 Surveillance Team |
| EPI_ISL_2936380, EPI_ISL_2936381, EPI_ISL_2936383, EPI_ISL_2936384, EPI_ISL_2936386 | Lighthouse Laboratory Plymouth | Wellcome Sanger Institute for the COVID-19 Genomics UK (COG-UK) Consortium | Lighthouse Laboratory Plymouth and Alex Alderton, Roberto Amato, Jeffrey Barrett, Sonia Goncalves, Ewan Harrison, David K. Jackson, Ian Johnston, Dominic Kwiatkowski, Cordelia Langford, John Sillitoe on behalf of the Wellcome Sanger Institute COVID-19 Surveillance Team |
| EPI_ISL_2936387, EPI_ISL_2936389 | Lighthouse Lab in Milton Keynes | Wellcome Sanger Institute for the COVID-19 Genomics UK (COG-UK) Consortium | The Lighthouse Lab in Milton Keynes and Alex Alderton, Roberto Amato, Jeffrey Barrett, Sonia Goncalves, Ewan Harrison, David K. Jackson, Ian Johnston, Dominic Kwiatkowski, Cordelia Langford, John Sillitoe on behalf of the Wellcome Sanger Institute COVID-19 Surveillance Team |

| | | | |
|---|---|---|---|
| EPI_ISL_2936391, EPI_ISL_2936392, EPI_ISL_2936394 | Lighthouse Laboratory Plymouth | Wellcome Sanger Institute for the COVID-19 Genomics UK (COG-UK) Consortium | Lighthouse Laboratory Plymouth and Alex Alderton, Roberto Amato, Jeffrey Barrett, Sonia Goncalves, Ewan Harrison, David K. Jackson, Ian Johnston, Dominic Kwiatkowski, Cordelia Langford, John Sillitoe on behalf of the Wellcome Sanger Institute COVID-19 Surveillance Team |
| EPI_ISL_2936395 | Lighthouse Lab in Milton Keynes | Wellcome Sanger Institute for the COVID-19 Genomics UK (COG-UK) Consortium | The Lighthouse Lab in Milton Keynes and Alex Alderton, Roberto Amato, Jeffrey Barrett, Sonia Goncalves, Ewan Harrison, David K. Jackson, Ian Johnston, Dominic Kwiatkowski, Cordelia Langford, John Sillitoe on behalf of the Wellcome Sanger Institute COVID-19 Surveillance Team |
| EPI_ISL_2936397, EPI_ISL_2936399 | Lighthouse Laboratory Plymouth | Wellcome Sanger Institute for the COVID-19 Genomics UK (COG-UK) Consortium | Lighthouse Laboratory Plymouth and Alex Alderton, Roberto Amato, Jeffrey Barrett, Sonia Goncalves, Ewan Harrison, David K. Jackson, Ian Johnston, Dominic Kwiatkowski, Cordelia Langford, John Sillitoe on behalf of the Wellcome Sanger Institute COVID-19 Surveillance Team |
| EPI_ISL_2936400 | Lighthouse Lab in Milton Keynes | Wellcome Sanger Institute for the COVID-19 Genomics UK (COG-UK) Consortium | The Lighthouse Lab in Milton Keynes and Alex Alderton, Roberto Amato, Jeffrey Barrett, Sonia Goncalves, Ewan Harrison, David K. Jackson, Ian Johnston, Dominic Kwiatkowski, Cordelia Langford, John Sillitoe on behalf of the Wellcome Sanger Institute COVID-19 Surveillance Team |
| EPI_ISL_2936402, EPI_ISL_2936404, EPI_ISL_2936405 | Lighthouse Laboratory Plymouth | Wellcome Sanger Institute for the COVID-19 Genomics UK (COG-UK) Consortium | Lighthouse Laboratory Plymouth and Alex Alderton, Roberto Amato, Jeffrey Barrett, Sonia Goncalves, Ewan Harrison, David K. Jackson, Ian Johnston, Dominic Kwiatkowski, Cordelia Langford, John Sillitoe on behalf of the Wellcome Sanger Institute COVID-19 Surveillance Team |
| EPI_ISL_2936407 | Lighthouse Lab in Milton Keynes | Wellcome Sanger Institute for the COVID-19 Genomics UK (COG-UK) Consortium | The Lighthouse Lab in Milton Keynes and Alex Alderton, Roberto Amato, Jeffrey Barrett, Sonia Goncalves, Ewan Harrison, David K. Jackson, Ian Johnston, Dominic Kwiatkowski, Cordelia Langford, John Sillitoe on behalf of the Wellcome Sanger Institute COVID-19 Surveillance Team |
| EPI_ISL_2936409, EPI_ISL_2936410, EPI_ISL_2936412, EPI_ISL_2936413, EPI_ISL_2936415 | Lighthouse Laboratory Plymouth | Wellcome Sanger Institute for the COVID-19 Genomics UK (COG-UK) Consortium | Lighthouse Laboratory Plymouth and Alex Alderton, Roberto Amato, Jeffrey Barrett, Sonia Goncalves, Ewan Harrison, David K. Jackson, Ian Johnston, Dominic Kwiatkowski, Cordelia Langford, John Sillitoe on behalf of the Wellcome Sanger Institute COVID-19 Surveillance Team |
| EPI_ISL_2936416 | Lighthouse Lab in Milton Keynes | Wellcome Sanger Institute for the COVID-19 Genomics UK (COG-UK) Consortium | The Lighthouse Lab in Milton Keynes and Alex Alderton, Roberto Amato, Jeffrey Barrett, Sonia Goncalves, Ewan Harrison, David K. Jackson, Ian Johnston, Dominic Kwiatkowski, Cordelia Langford, John Sillitoe on behalf of the Wellcome Sanger Institute COVID-19 Surveillance Team |
| EPI_ISL_2936418, EPI_ISL_2936419, EPI_ISL_2936421, EPI_ISL_2936422, EPI_ISL_2936424, EPI_ISL_2936425, EPI_ISL_2936427 | Lighthouse Laboratory Plymouth | Wellcome Sanger Institute for the COVID-19 Genomics UK (COG-UK) Consortium | Lighthouse Laboratory Plymouth and Alex Alderton, Roberto Amato, Jeffrey Barrett, Sonia Goncalves, Ewan Harrison, David K. Jackson, Ian Johnston, Dominic Kwiatkowski, Cordelia Langford, John Sillitoe on behalf of the Wellcome Sanger Institute COVID-19 Surveillance Team |
| EPI_ISL_2936429, EPI_ISL_2936430 | Lighthouse Lab in Milton Keynes | Wellcome Sanger Institute for the COVID-19 Genomics UK (COG-UK) Consortium | The Lighthouse Lab in Milton Keynes and Alex Alderton, Roberto Amato, Jeffrey Barrett, Sonia Goncalves, Ewan Harrison, David K. Jackson, Ian Johnston, Dominic Kwiatkowski, Cordelia Langford, John Sillitoe on behalf of the Wellcome Sanger Institute COVID-19 Surveillance Team |
| EPI_ISL_2936432, EPI_ISL_2936433, EPI_ISL_2936435 | Lighthouse Laboratory Plymouth | Wellcome Sanger Institute for the COVID-19 Genomics UK (COG-UK) Consortium | Lighthouse Laboratory Plymouth and Alex Alderton, Roberto Amato, Jeffrey Barrett, Sonia Goncalves, Ewan Harrison, David K. Jackson, Ian Johnston, Dominic Kwiatkowski, Cordelia Langford, John Sillitoe on behalf of the Wellcome Sanger Institute COVID-19 Surveillance Team |
| EPI_ISL_2936436, EPI_ISL_2936437 | Lighthouse Lab in Milton Keynes | Wellcome Sanger Institute for the COVID-19 Genomics UK (COG-UK) Consortium | The Lighthouse Lab in Milton Keynes and Alex Alderton, Roberto Amato, Jeffrey Barrett, Sonia Goncalves, Ewan Harrison, David K. Jackson, Ian Johnston, Dominic Kwiatkowski, Cordelia Langford, John Sillitoe on behalf of the Wellcome Sanger Institute COVID-19 Surveillance Team |
| EPI_ISL_2936438 | Ministry of Health Turkey | Ministry of Health Turkey | Fatma Bayrakdar, Yasemin Cosgun, Suleyman Yalcin, Gulay Korukluoglu |
| EPI_ISL_2936439, EPI_ISL_2936441, EPI_ISL_2936442, EPI_ISL_2936444 | Lighthouse Laboratory Plymouth | Wellcome Sanger Institute for the COVID-19 Genomics UK (COG-UK) Consortium | Lighthouse Laboratory Plymouth and Alex Alderton, Roberto Amato, Jeffrey Barrett, Sonia Goncalves, Ewan Harrison, David K. Jackson, Ian Johnston, Dominic Kwiatkowski, Cordelia Langford, John Sillitoe on behalf of the Wellcome Sanger Institute COVID-19 Surveillance Team |
| EPI_ISL_2936445, EPI_ISL_2936447 | Lighthouse Lab in Milton Keynes | Wellcome Sanger Institute for the COVID-19 Genomics UK (COG-UK) Consortium | The Lighthouse Lab in Milton Keynes and Alex Alderton, Roberto Amato, Jeffrey Barrett, Sonia Goncalves, Ewan Harrison, David K. Jackson, Ian Johnston, Dominic Kwiatkowski, Cordelia Langford, John Sillitoe on behalf of the Wellcome Sanger Institute COVID-19 Surveillance Team |
| EPI_ISL_2936448, EPI_ISL_2936450 | Lighthouse Laboratory Plymouth | Wellcome Sanger Institute for the COVID-19 Genomics UK (COG-UK) Consortium | Lighthouse Laboratory Plymouth and Alex Alderton, Roberto Amato, Jeffrey Barrett, Sonia Goncalves, Ewan Harrison, David K. Jackson, Ian Johnston, Dominic Kwiatkowski, Cordelia Langford, John Sillitoe on behalf of the Wellcome Sanger Institute COVID-19 Surveillance Team |
| EPI_ISL_2936451, EPI_ISL_2936453 | Lighthouse Lab in Milton Keynes | Wellcome Sanger Institute for the COVID-19 Genomics UK (COG-UK) Consortium | The Lighthouse Lab in Milton Keynes and Alex Alderton, Roberto Amato, Jeffrey Barrett, Sonia Goncalves, Ewan Harrison, David K. Jackson, Ian Johnston, Dominic Kwiatkowski, Cordelia Langford, John Sillitoe on behalf of the Wellcome Sanger Institute COVID-19 Surveillance Team |
| EPI_ISL_2936454, EPI_ISL_2936456, EPI_ISL_2936457 | Lighthouse Laboratory Plymouth | Wellcome Sanger Institute for the COVID-19 Genomics UK (COG-UK) Consortium | Lighthouse Laboratory Plymouth and Alex Alderton, Roberto Amato, Jeffrey Barrett, Sonia Goncalves, Ewan Harrison, David K. Jackson, Ian Johnston, Dominic Kwiatkowski, Cordelia Langford, John Sillitoe on behalf of the Wellcome Sanger Institute COVID-19 Surveillance Team |
| EPI_ISL_2936459 | Lighthouse Lab in Milton Keynes | Wellcome Sanger Institute for the COVID-19 Genomics UK (COG-UK) Consortium | The Lighthouse Lab in Milton Keynes and Alex Alderton, Roberto Amato, Jeffrey Barrett, Sonia Goncalves, Ewan Harrison, David K. Jackson, Ian Johnston, Dominic Kwiatkowski, Cordelia Langford, John Sillitoe on behalf of the Wellcome Sanger Institute COVID-19 Surveillance Team |
| EPI_ISL_2936461, EPI_ISL_2936462, EPI_ISL_2936464 | Lighthouse Laboratory Plymouth | Wellcome Sanger Institute for the COVID-19 Genomics UK (COG-UK) Consortium | Lighthouse Laboratory Plymouth and Alex Alderton, Roberto Amato, Jeffrey Barrett, Sonia Goncalves, Ewan Harrison, David K. Jackson, Ian Johnston, Dominic Kwiatkowski, Cordelia Langford, John Sillitoe on behalf of the Wellcome Sanger Institute COVID-19 Surveillance Team |
| EPI_ISL_2936465 | Lighthouse Lab in Milton Keynes | Wellcome Sanger Institute for the COVID-19 Genomics UK (COG-UK) Consortium | The Lighthouse Lab in Milton Keynes and Alex Alderton, Roberto Amato, Jeffrey Barrett, Sonia Goncalves, Ewan Harrison, David K. Jackson, Ian Johnston, Dominic Kwiatkowski, Cordelia Langford, John Sillitoe on behalf of the Wellcome Sanger Institute COVID-19 Surveillance Team |
| EPI_ISL_2936467, EPI_ISL_2936469, EPI_ISL_2936470 | Lighthouse Laboratory Plymouth | Wellcome Sanger Institute for the COVID-19 Genomics UK (COG-UK) Consortium | Lighthouse Laboratory Plymouth and Alex Alderton, Roberto Amato, Jeffrey Barrett, Sonia Goncalves, Ewan Harrison, David K. Jackson, Ian Johnston, Dominic Kwiatkowski, Cordelia Langford, John Sillitoe on behalf of the Wellcome Sanger Institute COVID-19 Surveillance Team |
| EPI_ISL_2936472 | Lighthouse Lab in Milton Keynes | Wellcome Sanger Institute for the COVID-19 Genomics UK (COG-UK) Consortium | The Lighthouse Lab in Milton Keynes and Alex Alderton, Roberto Amato, Jeffrey Barrett, Sonia Goncalves, Ewan Harrison, David K. Jackson, Ian Johnston, Dominic Kwiatkowski, Cordelia Langford, John Sillitoe on behalf of the Wellcome Sanger Institute COVID-19 Surveillance Team |
| EPI_ISL_2936474, EPI_ISL_2936475 | Lighthouse Laboratory Plymouth | Wellcome Sanger Institute for the COVID-19 Genomics UK (COG-UK) Consortium | Lighthouse Laboratory Plymouth and Alex Alderton, Roberto Amato, Jeffrey Barrett, Sonia Goncalves, Ewan Harrison, David K. Jackson, Ian Johnston, Dominic Kwiatkowski, Cordelia Langford, John Sillitoe on behalf of the Wellcome Sanger Institute COVID-19 Surveillance Team |
| EPI_ISL_2936477 | Lighthouse Lab in Milton Keynes | Wellcome Sanger Institute for the COVID-19 Genomics UK (COG-UK) Consortium | The Lighthouse Lab in Milton Keynes and Alex Alderton, Roberto Amato, Jeffrey Barrett, Sonia Goncalves, Ewan Harrison, David K. Jackson, Ian Johnston, Dominic Kwiatkowski, Cordelia Langford, John Sillitoe on behalf of the Wellcome Sanger Institute COVID-19 Surveillance Team |
| EPI_ISL_2936478, EPI_ISL_2936480 | Lighthouse Laboratory Plymouth | Wellcome Sanger Institute for the COVID-19 Genomics UK (COG-UK) Consortium | Lighthouse Laboratory Plymouth and Alex Alderton, Roberto Amato, Jeffrey Barrett, Sonia Goncalves, Ewan Harrison, David K. Jackson, Ian Johnston, Dominic Kwiatkowski, Cordelia Langford, John Sillitoe on behalf of the Wellcome Sanger Institute COVID-19 Surveillance Team |
| EPI_ISL_2936481 | Lighthouse Lab in Milton Keynes | Wellcome Sanger Institute for the COVID-19 Genomics UK (COG-UK) Consortium | The Lighthouse Lab in Milton Keynes and Alex Alderton, Roberto Amato, Jeffrey Barrett, Sonia Goncalves, Ewan Harrison, David K. Jackson, Ian Johnston, Dominic Kwiatkowski, Cordelia Langford, John Sillitoe on behalf of the Wellcome Sanger Institute COVID-19 Surveillance Team |
| EPI_ISL_2936483, EPI_ISL_2936484, EPI_ISL_2936486 | Lighthouse Laboratory Plymouth | Wellcome Sanger Institute for the COVID-19 Genomics UK (COG-UK) Consortium | Lighthouse Laboratory Plymouth and Alex Alderton, Roberto Amato, Jeffrey Barrett, Sonia Goncalves, Ewan Harrison, David K. Jackson, Ian Johnston, Dominic Kwiatkowski, Cordelia Langford, John Sillitoe on behalf of the Wellcome Sanger Institute COVID-19 Surveillance Team |
| EPI_ISL_2936487 | Lighthouse Lab in Milton Keynes | Wellcome Sanger Institute for the COVID-19 Genomics UK (COG-UK) Consortium | The Lighthouse Lab in Milton Keynes and Alex Alderton, Roberto Amato, Jeffrey Barrett, Sonia Goncalves, Ewan Harrison, David K. Jackson, Ian Johnston, Dominic Kwiatkowski, Cordelia Langford, John Sillitoe on behalf of the Wellcome Sanger Institute COVID-19 Surveillance Team |
| EPI_ISL_2936488 | Ministry of Health Turkey | Ministry of Health Turkey | Fatma Bayrakdar, Yasemin Cosgun, Suleyman Yalcin, Gulay Korukluoglu |
| EPI_ISL_2936489, EPI_ISL_2936490, EPI_ISL_2936493, EPI_ISL_2936495, EPI_ISL_2936496, EPI_ISL_2936498, EPI_ISL_2936500, EPI_ISL_2936501, EPI_ISL_2936503 | Lighthouse Laboratory Plymouth | Wellcome Sanger Institute for the COVID-19 Genomics UK (COG-UK) Consortium | Lighthouse Laboratory Plymouth and Alex Alderton, Roberto Amato, Jeffrey Barrett, Sonia Goncalves, Ewan Harrison, David K. Jackson, Ian Johnston, Dominic Kwiatkowski, Cordelia Langford, John Sillitoe on behalf of the Wellcome Sanger Institute COVID-19 Surveillance Team |
| EPI_ISL_2936504 | Lighthouse Lab in Milton Keynes | Wellcome Sanger Institute for the COVID-19 Genomics UK (COG-UK) Consortium | The Lighthouse Lab in Milton Keynes and Alex Alderton, Roberto Amato, Jeffrey Barrett, Sonia Goncalves, Ewan Harrison, David K. Jackson, Ian Johnston, Dominic Kwiatkowski, Cordelia Langford, John Sillitoe on behalf of the Wellcome Sanger Institute COVID-19 Surveillance Team |
| EPI_ISL_2936506, EPI_ISL_2936507, EPI_ISL_2936509 | Lighthouse Laboratory Plymouth | Wellcome Sanger Institute for the COVID-19 Genomics UK (COG-UK) Consortium | Lighthouse Laboratory Plymouth and Alex Alderton, Roberto Amato, Jeffrey Barrett, Sonia Goncalves, Ewan Harrison, David K. Jackson, Ian Johnston, Dominic Kwiatkowski, Cordelia Langford, John Sillitoe on behalf of the Wellcome Sanger Institute COVID-19 Surveillance Team |
| EPI_ISL_2936510, EPI_ISL_2936511 | Lighthouse Lab in Milton Keynes | Wellcome Sanger Institute for the COVID-19 Genomics UK (COG-UK) Consortium | The Lighthouse Lab in Milton Keynes and Alex Alderton, Roberto Amato, Jeffrey Barrett, Sonia Goncalves, Ewan Harrison, David K. Jackson, Ian Johnston, Dominic Kwiatkowski, Cordelia Langford, John Sillitoe on behalf of the Wellcome Sanger Institute COVID-19 Surveillance Team |
| EPI_ISL_2936513, EPI_ISL_2936514, EPI_ISL_2936516, EPI_ISL_2936517 | Lighthouse Laboratory Plymouth | Wellcome Sanger Institute for the COVID-19 Genomics UK (COG-UK) Consortium | Lighthouse Laboratory Plymouth and Alex Alderton, Roberto Amato, Jeffrey Barrett, Sonia Goncalves, Ewan Harrison, David K. Jackson, Ian Johnston, Dominic Kwiatkowski, Cordelia Langford, John Sillitoe on behalf of the Wellcome Sanger Institute COVID-19 Surveillance Team |

| | | | |
|---|---|---|---|
| EPI_ISL_2936519, EPI_ISL_2936521 | Lighthouse Lab in Milton Keynes | Wellcome Sanger Institute for the COVID-19 Genomics UK (COG-UK) Consortium | The Lighthouse Lab in Milton Keynes and Alex Alderton, Roberto Amato, Jeffrey Barrett, Sonia Goncalves, Ewan Harrison, David K. Jackson, Ian Johnston, Dominic Kwiatkowski, Cordelia Langford, John Sillitoe on behalf of the Wellcome Sanger Institute COVID-19 Surveillance Team |
| EPI_ISL_2936522, EPI_ISL_2936524 | Lighthouse Laboratory Plymouth | Wellcome Sanger Institute for the COVID-19 Genomics UK (COG-UK) Consortium | Lighthouse Laboratory Plymouth and Alex Alderton, Roberto Amato, Jeffrey Barrett, Sonia Goncalves, Ewan Harrison, David K. Jackson, Ian Johnston, Dominic Kwiatkowski, Cordelia Langford, John Sillitoe on behalf of the Wellcome Sanger Institute COVID-19 Surveillance Team |
| EPI_ISL_2936525 | Lighthouse Lab in Milton Keynes | Wellcome Sanger Institute for the COVID-19 Genomics UK (COG-UK) Consortium | The Lighthouse Lab in Milton Keynes and Alex Alderton, Roberto Amato, Jeffrey Barrett, Sonia Goncalves, Ewan Harrison, David K. Jackson, Ian Johnston, Dominic Kwiatkowski, Cordelia Langford, John Sillitoe on behalf of the Wellcome Sanger Institute COVID-19 Surveillance Team |
| EPI_ISL_2936527, EPI_ISL_2936529, EPI_ISL_2936530, EPI_ISL_2936532 | Lighthouse Laboratory Plymouth | Wellcome Sanger Institute for the COVID-19 Genomics UK (COG-UK) Consortium | Lighthouse Laboratory Plymouth and Alex Alderton, Roberto Amato, Jeffrey Barrett, Sonia Goncalves, Ewan Harrison, David K. Jackson, Ian Johnston, Dominic Kwiatkowski, Cordelia Langford, John Sillitoe on behalf of the Wellcome Sanger Institute COVID-19 Surveillance Team |
| EPI_ISL_2936534 | Lighthouse Lab in Milton Keynes | Wellcome Sanger Institute for the COVID-19 Genomics UK (COG-UK) Consortium | The Lighthouse Lab in Milton Keynes and Alex Alderton, Roberto Amato, Jeffrey Barrett, Sonia Goncalves, Ewan Harrison, David K. Jackson, Ian Johnston, Dominic Kwiatkowski, Cordelia Langford, John Sillitoe on behalf of the Wellcome Sanger Institute COVID-19 Surveillance Team |
| EPI_ISL_2936535, EPI_ISL_2936537, EPI_ISL_2936539, EPI_ISL_2936540 | Lighthouse Laboratory Plymouth | Wellcome Sanger Institute for the COVID-19 Genomics UK (COG-UK) Consortium | Lighthouse Laboratory Plymouth and Alex Alderton, Roberto Amato, Jeffrey Barrett, Sonia Goncalves, Ewan Harrison, David K. Jackson, Ian Johnston, Dominic Kwiatkowski, Cordelia Langford, John Sillitoe on behalf of the Wellcome Sanger Institute COVID-19 Surveillance Team |
| EPI_ISL_2936542, EPI_ISL_2936543 | Lighthouse Lab in Milton Keynes | Wellcome Sanger Institute for the COVID-19 Genomics UK (COG-UK) Consortium | The Lighthouse Lab in Milton Keynes and Alex Alderton, Roberto Amato, Jeffrey Barrett, Sonia Goncalves, Ewan Harrison, David K. Jackson, Ian Johnston, Dominic Kwiatkowski, Cordelia Langford, John Sillitoe on behalf of the Wellcome Sanger Institute COVID-19 Surveillance Team |
| EPI_ISL_2936545, EPI_ISL_2936547, EPI_ISL_2936548, EPI_ISL_2936550 | Lighthouse Laboratory Plymouth | Wellcome Sanger Institute for the COVID-19 Genomics UK (COG-UK) Consortium | Lighthouse Laboratory Plymouth and Alex Alderton, Roberto Amato, Jeffrey Barrett, Sonia Goncalves, Ewan Harrison, David K. Jackson, Ian Johnston, Dominic Kwiatkowski, Cordelia Langford, John Sillitoe on behalf of the Wellcome Sanger Institute COVID-19 Surveillance Team |
| EPI_ISL_2936551 | Lighthouse Lab in Milton Keynes | Wellcome Sanger Institute for the COVID-19 Genomics UK (COG-UK) Consortium | The Lighthouse Lab in Milton Keynes and Alex Alderton, Roberto Amato, Jeffrey Barrett, Sonia Goncalves, Ewan Harrison, David K. Jackson, Ian Johnston, Dominic Kwiatkowski, Cordelia Langford, John Sillitoe on behalf of the Wellcome Sanger Institute COVID-19 Surveillance Team |
| EPI_ISL_2936553, EPI_ISL_2936554 | Lighthouse Laboratory Plymouth | Wellcome Sanger Institute for the COVID-19 Genomics UK (COG-UK) Consortium | Lighthouse Laboratory Plymouth and Alex Alderton, Roberto Amato, Jeffrey Barrett, Sonia Goncalves, Ewan Harrison, David K. Jackson, Ian Johnston, Dominic Kwiatkowski, Cordelia Langford, John Sillitoe on behalf of the Wellcome Sanger Institute COVID-19 Surveillance Team |
| EPI_ISL_2936556 | Lighthouse Lab in Milton Keynes | Wellcome Sanger Institute for the COVID-19 Genomics UK (COG-UK) Consortium | The Lighthouse Lab in Milton Keynes and Alex Alderton, Roberto Amato, Jeffrey Barrett, Sonia Goncalves, Ewan Harrison, David K. Jackson, Ian Johnston, Dominic Kwiatkowski, Cordelia Langford, John Sillitoe on behalf of the Wellcome Sanger Institute COVID-19 Surveillance Team |
| EPI_ISL_2936557, EPI_ISL_2936558 | Lighthouse Laboratory Plymouth | Wellcome Sanger Institute for the COVID-19 Genomics UK (COG-UK) Consortium | Lighthouse Laboratory Plymouth and Alex Alderton, Roberto Amato, Jeffrey Barrett, Sonia Goncalves, Ewan Harrison, David K. Jackson, Ian Johnston, Dominic Kwiatkowski, Cordelia Langford, John Sillitoe on behalf of the Wellcome Sanger Institute COVID-19 Surveillance Team |
| EPI_ISL_2936560, EPI_ISL_2936561 | Lighthouse Lab in Milton Keynes | Wellcome Sanger Institute for the COVID-19 Genomics UK (COG-UK) Consortium | The Lighthouse Lab in Milton Keynes and Alex Alderton, Roberto Amato, Jeffrey Barrett, Sonia Goncalves, Ewan Harrison, David K. Jackson, Ian Johnston, Dominic Kwiatkowski, Cordelia Langford, John Sillitoe on behalf of the Wellcome Sanger Institute COVID-19 Surveillance Team |
| EPI_ISL_2936563, EPI_ISL_2936565, EPI_ISL_2936566, EPI_ISL_2936568 | Lighthouse Laboratory Plymouth | Wellcome Sanger Institute for the COVID-19 Genomics UK (COG-UK) Consortium | Lighthouse Laboratory Plymouth and Alex Alderton, Roberto Amato, Jeffrey Barrett, Sonia Goncalves, Ewan Harrison, David K. Jackson, Ian Johnston, Dominic Kwiatkowski, Cordelia Langford, John Sillitoe on behalf of the Wellcome Sanger Institute COVID-19 Surveillance Team |
| EPI_ISL_2936569 | Lighthouse Lab in Milton Keynes | Wellcome Sanger Institute for the COVID-19 Genomics UK (COG-UK) Consortium | The Lighthouse Lab in Milton Keynes and Alex Alderton, Roberto Amato, Jeffrey Barrett, Sonia Goncalves, Ewan Harrison, David K. Jackson, Ian Johnston, Dominic Kwiatkowski, Cordelia Langford, John Sillitoe on behalf of the Wellcome Sanger Institute COVID-19 Surveillance Team |
| EPI_ISL_2936571 | Lighthouse Laboratory Plymouth | Wellcome Sanger Institute for the COVID-19 Genomics UK (COG-UK) Consortium | Lighthouse Laboratory Plymouth and Alex Alderton, Roberto Amato, Jeffrey Barrett, Sonia Goncalves, Ewan Harrison, David K. Jackson, Ian Johnston, Dominic Kwiatkowski, Cordelia Langford, John Sillitoe on behalf of the Wellcome Sanger Institute COVID-19 Surveillance Team |
| EPI_ISL_2936572 | Lighthouse Lab in Milton Keynes | Wellcome Sanger Institute for the COVID-19 Genomics UK (COG-UK) Consortium | The Lighthouse Lab in Milton Keynes and Alex Alderton, Roberto Amato, Jeffrey Barrett, Sonia Goncalves, Ewan Harrison, David K. Jackson, Ian Johnston, Dominic Kwiatkowski, Cordelia Langford, John Sillitoe on behalf of the Wellcome Sanger Institute COVID-19 Surveillance Team |
| EPI_ISL_2936574, EPI_ISL_2936576, EPI_ISL_2936577, EPI_ISL_2936579, EPI_ISL_2936580 | Lighthouse Laboratory Plymouth | Wellcome Sanger Institute for the COVID-19 Genomics UK (COG-UK) Consortium | Lighthouse Laboratory Plymouth and Alex Alderton, Roberto Amato, Jeffrey Barrett, Sonia Goncalves, Ewan Harrison, David K. Jackson, Ian Johnston, Dominic Kwiatkowski, Cordelia Langford, John Sillitoe on behalf of the Wellcome Sanger Institute COVID-19 Surveillance Team |
| EPI_ISL_2936582, EPI_ISL_2936584 | Lighthouse Lab in Milton Keynes | Wellcome Sanger Institute for the COVID-19 Genomics UK (COG-UK) Consortium | The Lighthouse Lab in Milton Keynes and Alex Alderton, Roberto Amato, Jeffrey Barrett, Sonia Goncalves, Ewan Harrison, David K. Jackson, Ian Johnston, Dominic Kwiatkowski, Cordelia Langford, John Sillitoe on behalf of the Wellcome Sanger Institute COVID-19 Surveillance Team |
| EPI_ISL_2936585 | Lighthouse Laboratory Plymouth | Wellcome Sanger Institute for the COVID-19 Genomics UK (COG-UK) Consortium | Lighthouse Laboratory Plymouth and Alex Alderton, Roberto Amato, Jeffrey Barrett, Sonia Goncalves, Ewan Harrison, David K. Jackson, Ian Johnston, Dominic Kwiatkowski, Cordelia Langford, John Sillitoe on behalf of the Wellcome Sanger Institute COVID-19 Surveillance Team |
| EPI_ISL_2936587, EPI_ISL_2936589, EPI_ISL_2936590, EPI_ISL_2936592 | Lighthouse Lab in Milton Keynes | Wellcome Sanger Institute for the COVID-19 Genomics UK (COG-UK) Consortium | The Lighthouse Lab in Milton Keynes and Alex Alderton, Roberto Amato, Jeffrey Barrett, Sonia Goncalves, Ewan Harrison, David K. Jackson, Ian Johnston, Dominic Kwiatkowski, Cordelia Langford, John Sillitoe on behalf of the Wellcome Sanger Institute COVID-19 Surveillance Team |
| EPI_ISL_2936593 | Lighthouse Lab in Alderley Park | Wellcome Sanger Institute for the COVID-19 Genomics UK (COG-UK) Consortium | Jacquelyn Wynn, Mairead Hyland, The Lighthouse Lab in Alderley Park and Alex Alderton, Roberto Amato, Jeffrey Barrett, Sonia Goncalves, Ewan Harrison, David K. Jackson, Ian Johnston, Dominic Kwiatkowski, Cordelia Langford, John Sillitoe on behalf of the Wellcome Sanger Institute COVID-19 Surveillance Team |
| EPI_ISL_2936595, EPI_ISL_2936597, EPI_ISL_2936598, EPI_ISL_2936600 | Lighthouse Lab in Milton Keynes | Wellcome Sanger Institute for the COVID-19 Genomics UK (COG-UK) Consortium | The Lighthouse Lab in Milton Keynes and Alex Alderton, Roberto Amato, Jeffrey Barrett, Sonia Goncalves, Ewan Harrison, David K. Jackson, Ian Johnston, Dominic Kwiatkowski, Cordelia Langford, John Sillitoe on behalf of the Wellcome Sanger Institute COVID-19 Surveillance Team |
| EPI_ISL_2936601 | Lighthouse Lab in Alderley Park | Wellcome Sanger Institute for the COVID-19 Genomics UK (COG-UK) Consortium | Jacquelyn Wynn, Mairead Hyland, The Lighthouse Lab in Alderley Park and Alex Alderton, Roberto Amato, Jeffrey Barrett, Sonia Goncalves, Ewan Harrison, David K. Jackson, Ian Johnston, Dominic Kwiatkowski, Cordelia Langford, John Sillitoe on behalf of the Wellcome Sanger Institute COVID-19 Surveillance Team |
| EPI_ISL_2936603, EPI_ISL_2936605 | Lighthouse Lab in Milton Keynes | Wellcome Sanger Institute for the COVID-19 Genomics UK (COG-UK) Consortium | The Lighthouse Lab in Milton Keynes and Alex Alderton, Roberto Amato, Jeffrey Barrett, Sonia Goncalves, Ewan Harrison, David K. Jackson, Ian Johnston, Dominic Kwiatkowski, Cordelia Langford, John Sillitoe on behalf of the Wellcome Sanger Institute COVID-19 Surveillance Team |
| EPI_ISL_2936607 | Lighthouse Lab in Glasgow | Wellcome Sanger Institute for the COVID-19 Genomics UK (COG-UK) Consortium | Harper VanSteenhouse, Yumi Kasai, David Gray, Carol Clugston, Anna Dominiczak and Alex Alderton, Roberto Amato, Jeffrey Barrett, Sonia Goncalves, Ewan Harrison, David K. Jackson, Ian Johnston, Dominic Kwiatkowski, Cordelia Langford, John Sillitoe on behalf of the Wellcome Sanger Institute COVID-19 Surveillance Team |
| EPI_ISL_2936608, EPI_ISL_2936610, EPI_ISL_2936611, EPI_ISL_2936613 | Lighthouse Lab in Milton Keynes | Wellcome Sanger Institute for the COVID-19 Genomics UK (COG-UK) Consortium | The Lighthouse Lab in Milton Keynes and Alex Alderton, Roberto Amato, Jeffrey Barrett, Sonia Goncalves, Ewan Harrison, David K. Jackson, Ian Johnston, Dominic Kwiatkowski, Cordelia Langford, John Sillitoe on behalf of the Wellcome Sanger Institute COVID-19 Surveillance Team |
| EPI_ISL_2936614, EPI_ISL_2936616 | Lighthouse Lab in Alderley Park | Wellcome Sanger Institute for the COVID-19 Genomics UK (COG-UK) Consortium | Jacquelyn Wynn, Mairead Hyland, The Lighthouse Lab in Alderley Park and Alex Alderton, Roberto Amato, Jeffrey Barrett, Sonia Goncalves, Ewan Harrison, David K. Jackson, Ian Johnston, Dominic Kwiatkowski, Cordelia Langford, John Sillitoe on behalf of the Wellcome Sanger Institute COVID-19 Surveillance Team |
| EPI_ISL_2936617 | Lighthouse Lab in Glasgow | Wellcome Sanger Institute for the COVID-19 Genomics UK (COG-UK) Consortium | Harper VanSteenhouse, Yumi Kasai, David Gray, Carol Clugston, Anna Dominiczak and Alex Alderton, Roberto Amato, Jeffrey Barrett, Sonia Goncalves, Ewan Harrison, David K. Jackson, Ian Johnston, Dominic Kwiatkowski, Cordelia Langford, John Sillitoe on behalf of the Wellcome Sanger Institute COVID-19 Surveillance Team |
| EPI_ISL_2936619 | Lighthouse Lab in Milton Keynes | Wellcome Sanger Institute for the COVID-19 Genomics UK (COG-UK) Consortium | The Lighthouse Lab in Milton Keynes and Alex Alderton, Roberto Amato, Jeffrey Barrett, Sonia Goncalves, Ewan Harrison, David K. Jackson, Ian Johnston, Dominic Kwiatkowski, Cordelia Langford, John Sillitoe on behalf of the Wellcome Sanger Institute COVID-19 Surveillance Team |
| EPI_ISL_2936621, EPI_ISL_2936622, EPI_ISL_2936623 | Lighthouse Lab in Alderley Park | Wellcome Sanger Institute for the COVID-19 Genomics UK (COG-UK) Consortium | Jacquelyn Wynn, Mairead Hyland, The Lighthouse Lab in Alderley Park and Alex Alderton, Roberto Amato, Jeffrey Barrett, Sonia Goncalves, Ewan Harrison, David K. Jackson, Ian Johnston, Dominic Kwiatkowski, Cordelia Langford, John Sillitoe on behalf of the Wellcome Sanger Institute COVID-19 Surveillance Team |
| EPI_ISL_2936625, EPI_ISL_2936626, EPI_ISL_2936628, EPI_ISL_2936629 | Lighthouse Lab in Milton Keynes | Wellcome Sanger Institute for the COVID-19 Genomics UK (COG-UK) Consortium | The Lighthouse Lab in Milton Keynes and Alex Alderton, Roberto Amato, Jeffrey Barrett, Sonia Goncalves, Ewan Harrison, David K. Jackson, Ian Johnston, Dominic Kwiatkowski, Cordelia Langford, John Sillitoe on behalf of the Wellcome Sanger Institute COVID-19 Surveillance Team |
| EPI_ISL_2936631 | Lighthouse Lab in Glasgow | Wellcome Sanger Institute for the COVID-19 Genomics UK (COG-UK) Consortium | Harper VanSteenhouse, Yumi Kasai, David Gray, Carol Clugston, Anna Dominiczak and Alex Alderton, Roberto Amato, Jeffrey Barrett, Sonia Goncalves, Ewan Harrison, David K. Jackson, Ian Johnston, Dominic Kwiatkowski, Cordelia Langford, John Sillitoe on behalf of the Wellcome Sanger Institute COVID-19 Surveillance Team |
| EPI_ISL_2936632, EPI_ISL_2936633, EPI_ISL_2936635, EPI_ISL_2936637, EPI_ISL_2936638 | Lighthouse Lab in Milton Keynes | Wellcome Sanger Institute for the COVID-19 Genomics UK (COG-UK) Consortium | The Lighthouse Lab in Milton Keynes and Alex Alderton, Roberto Amato, Jeffrey Barrett, Sonia Goncalves, Ewan Harrison, David K. Jackson, Ian Johnston, Dominic Kwiatkowski, Cordelia Langford, John Sillitoe on behalf of the Wellcome Sanger Institute COVID-19 Surveillance Team |
| EPI_ISL_2936640 | Lighthouse Lab in Alderley Park | Wellcome Sanger Institute for the COVID-19 Genomics UK (COG-UK) Consortium | Jacquelyn Wynn, Mairead Hyland, The Lighthouse Lab in Alderley Park and Alex Alderton, Roberto Amato, Jeffrey Barrett, Sonia Goncalves, Ewan Harrison, David K. Jackson, Ian Johnston, Dominic Kwiatkowski, Cordelia Langford, John Sillitoe on behalf of the Wellcome Sanger Institute COVID-19 Surveillance Team |
| EPI_ISL_2936641, | Lighthouse Lab in Milton Keynes | Wellcome Sanger Institute for the COVID-19 Genomics UK (COG-UK) | The Lighthouse Lab in Milton Keynes and Alex Alderton, Roberto Amato, Jeffrey Barrett, Sonia Goncalves, Ewan Harrison, David K. Jackson, Ian Johnston, Dominic Kwiatkowski, Cordelia Langford, John Sillitoe on behalf of the Wellcome Sanger |

| | | | |
|---|---|---|---|
| EPI_ISL_2936643, EPI_ISL_2936644 | | Consortium | Institute COVID-19 Surveillance Team |
| EPI_ISL_2936645 | Lighthouse Lab in Alderley Park | Wellcome Sanger Institute for the COVID-19 Genomics UK (COG-UK) Consortium | Jacquelyn Wynn, Mairead Hyland, The Lighthouse Lab in Alderley Park and Alex Alderton, Roberto Amato, Jeffrey Barrett, Sonia Goncalves, Ewan Harrison, David K. Jackson, Ian Johnston, Dominic Kwiatkowski, Cordelia Langford, John Sillitoe on behalf of the Wellcome Sanger Institute COVID-19 Surveillance Team |
| EPI_ISL_2936647 | Lighthouse Lab in Milton Keynes | Wellcome Sanger Institute for the COVID-19 Genomics UK (COG-UK) Consortium | The Lighthouse Lab in Milton Keynes and Alex Alderton, Roberto Amato, Jeffrey Barrett, Sonia Goncalves, Ewan Harrison, David K. Jackson, Ian Johnston, Dominic Kwiatkowski, Cordelia Langford, John Sillitoe on behalf of the Wellcome Sanger Institute COVID-19 Surveillance Team |
| EPI_ISL_2936649 | Lighthouse Lab in Glasgow | Wellcome Sanger Institute for the COVID-19 Genomics UK (COG-UK) Consortium | Harper VanSteenhouse, Yumi Kasai, David Gray, Carol Clugston, Anna Dominiczak and Alex Alderton, Roberto Amato, Jeffrey Barrett, Sonia Goncalves, Ewan Harrison, David K. Jackson, Ian Johnston, Dominic Kwiatkowski, Cordelia Langford, John Sillitoe on behalf of the Wellcome Sanger Institute COVID-19 Surveillance Team |
| EPI_ISL_2936651, EPI_ISL_2936652 | Lighthouse Lab in Alderley Park | Wellcome Sanger Institute for the COVID-19 Genomics UK (COG-UK) Consortium | Jacquelyn Wynn, Mairead Hyland, The Lighthouse Lab in Alderley Park and Alex Alderton, Roberto Amato, Jeffrey Barrett, Sonia Goncalves, Ewan Harrison, David K. Jackson, Ian Johnston, Dominic Kwiatkowski, Cordelia Langford, John Sillitoe on behalf of the Wellcome Sanger Institute COVID-19 Surveillance Team |
| EPI_ISL_2936654, EPI_ISL_2936655, EPI_ISL_2936657, EPI_ISL_2936658, EPI_ISL_2936660, EPI_ISL_2936662, EPI_ISL_2936663 | Lighthouse Lab in Milton Keynes | Wellcome Sanger Institute for the COVID-19 Genomics UK (COG-UK) Consortium | The Lighthouse Lab in Milton Keynes and Alex Alderton, Roberto Amato, Jeffrey Barrett, Sonia Goncalves, Ewan Harrison, David K. Jackson, Ian Johnston, Dominic Kwiatkowski, Cordelia Langford, John Sillitoe on behalf of the Wellcome Sanger Institute COVID-19 Surveillance Team |
| EPI_ISL_2936665 | Lighthouse Lab in Alderley Park | Wellcome Sanger Institute for the COVID-19 Genomics UK (COG-UK) Consortium | Jacquelyn Wynn, Mairead Hyland, The Lighthouse Lab in Alderley Park and Alex Alderton, Roberto Amato, Jeffrey Barrett, Sonia Goncalves, Ewan Harrison, David K. Jackson, Ian Johnston, Dominic Kwiatkowski, Cordelia Langford, John Sillitoe on behalf of the Wellcome Sanger Institute COVID-19 Surveillance Team |
| EPI_ISL_2936666, EPI_ISL_2936668, EPI_ISL_2936670 | Lighthouse Lab in Milton Keynes | Wellcome Sanger Institute for the COVID-19 Genomics UK (COG-UK) Consortium | The Lighthouse Lab in Milton Keynes and Alex Alderton, Roberto Amato, Jeffrey Barrett, Sonia Goncalves, Ewan Harrison, David K. Jackson, Ian Johnston, Dominic Kwiatkowski, Cordelia Langford, John Sillitoe on behalf of the Wellcome Sanger Institute COVID-19 Surveillance Team |
| EPI_ISL_2936672 | Lighthouse Lab in Glasgow | Wellcome Sanger Institute for the COVID-19 Genomics UK (COG-UK) Consortium | Harper VanSteenhouse, Yumi Kasai, David Gray, Carol Clugston, Anna Dominiczak and Alex Alderton, Roberto Amato, Jeffrey Barrett, Sonia Goncalves, Ewan Harrison, David K. Jackson, Ian Johnston, Dominic Kwiatkowski, Cordelia Langford, John Sillitoe on behalf of the Wellcome Sanger Institute COVID-19 Surveillance Team |
| EPI_ISL_2936674, EPI_ISL_2936675 | Lighthouse Lab in Milton Keynes | Wellcome Sanger Institute for the COVID-19 Genomics UK (COG-UK) Consortium | The Lighthouse Lab in Milton Keynes and Alex Alderton, Roberto Amato, Jeffrey Barrett, Sonia Goncalves, Ewan Harrison, David K. Jackson, Ian Johnston, Dominic Kwiatkowski, Cordelia Langford, John Sillitoe on behalf of the Wellcome Sanger Institute COVID-19 Surveillance Team |
| EPI_ISL_2936677 | Lighthouse Lab in Glasgow | Wellcome Sanger Institute for the COVID-19 Genomics UK (COG-UK) Consortium | Harper VanSteenhouse, Yumi Kasai, David Gray, Carol Clugston, Anna Dominiczak and Alex Alderton, Roberto Amato, Jeffrey Barrett, Sonia Goncalves, Ewan Harrison, David K. Jackson, Ian Johnston, Dominic Kwiatkowski, Cordelia Langford, John Sillitoe on behalf of the Wellcome Sanger Institute COVID-19 Surveillance Team |
| EPI_ISL_2936678, EPI_ISL_2936680, EPI_ISL_2936681, EPI_ISL_2936683, EPI_ISL_2936684, EPI_ISL_2936685, EPI_ISL_2936687 | Lighthouse Lab in Milton Keynes | Wellcome Sanger Institute for the COVID-19 Genomics UK (COG-UK) Consortium | The Lighthouse Lab in Milton Keynes and Alex Alderton, Roberto Amato, Jeffrey Barrett, Sonia Goncalves, Ewan Harrison, David K. Jackson, Ian Johnston, Dominic Kwiatkowski, Cordelia Langford, John Sillitoe on behalf of the Wellcome Sanger Institute COVID-19 Surveillance Team |
| EPI_ISL_2936688 | Lighthouse Lab in Glasgow | Wellcome Sanger Institute for the COVID-19 Genomics UK (COG-UK) Consortium | Harper VanSteenhouse, Yumi Kasai, David Gray, Carol Clugston, Anna Dominiczak and Alex Alderton, Roberto Amato, Jeffrey Barrett, Sonia Goncalves, Ewan Harrison, David K. Jackson, Ian Johnston, Dominic Kwiatkowski, Cordelia Langford, John Sillitoe on behalf of the Wellcome Sanger Institute COVID-19 Surveillance Team |
| EPI_ISL_2936689, EPI_ISL_2936691 | Lighthouse Lab in Milton Keynes | Wellcome Sanger Institute for the COVID-19 Genomics UK (COG-UK) Consortium | The Lighthouse Lab in Milton Keynes and Alex Alderton, Roberto Amato, Jeffrey Barrett, Sonia Goncalves, Ewan Harrison, David K. Jackson, Ian Johnston, Dominic Kwiatkowski, Cordelia Langford, John Sillitoe on behalf of the Wellcome Sanger Institute COVID-19 Surveillance Team |
| EPI_ISL_2936693 | Lighthouse Lab in Glasgow | Wellcome Sanger Institute for the COVID-19 Genomics UK (COG-UK) Consortium | Harper VanSteenhouse, Yumi Kasai, David Gray, Carol Clugston, Anna Dominiczak and Alex Alderton, Roberto Amato, Jeffrey Barrett, Sonia Goncalves, Ewan Harrison, David K. Jackson, Ian Johnston, Dominic Kwiatkowski, Cordelia Langford, John Sillitoe on behalf of the Wellcome Sanger Institute COVID-19 Surveillance Team |
| EPI_ISL_2936694 | Lighthouse Lab in Alderley Park | Wellcome Sanger Institute for the COVID-19 Genomics UK (COG-UK) Consortium | Jacquelyn Wynn, Mairead Hyland, The Lighthouse Lab in Alderley Park and Alex Alderton, Roberto Amato, Jeffrey Barrett, Sonia Goncalves, Ewan Harrison, David K. Jackson, Ian Johnston, Dominic Kwiatkowski, Cordelia Langford, John Sillitoe on behalf of the Wellcome Sanger Institute COVID-19 Surveillance Team |
| EPI_ISL_2936696 | Lighthouse Lab in Milton Keynes | Wellcome Sanger Institute for the COVID-19 Genomics UK (COG-UK) Consortium | The Lighthouse Lab in Milton Keynes and Alex Alderton, Roberto Amato, Jeffrey Barrett, Sonia Goncalves, Ewan Harrison, David K. Jackson, Ian Johnston, Dominic Kwiatkowski, Cordelia Langford, John Sillitoe on behalf of the Wellcome Sanger Institute COVID-19 Surveillance Team |
| EPI_ISL_2936723 | Lighthouse Lab in Alderley Park | Wellcome Sanger Institute for the COVID-19 Genomics UK (COG-UK) Consortium | Jacquelyn Wynn, Mairead Hyland, The Lighthouse Lab in Alderley Park and Alex Alderton, Roberto Amato, Jeffrey Barrett, Sonia Goncalves, Ewan Harrison, David K. Jackson, Ian Johnston, Dominic Kwiatkowski, Cordelia Langford, John Sillitoe on behalf of the Wellcome Sanger Institute COVID-19 Surveillance Team |
| EPI_ISL_2936724 | Lighthouse Lab in Milton Keynes | Wellcome Sanger Institute for the COVID-19 Genomics UK (COG-UK) Consortium | The Lighthouse Lab in Milton Keynes and Alex Alderton, Roberto Amato, Jeffrey Barrett, Sonia Goncalves, Ewan Harrison, David K. Jackson, Ian Johnston, Dominic Kwiatkowski, Cordelia Langford, John Sillitoe on behalf of the Wellcome Sanger Institute COVID-19 Surveillance Team |
| EPI_ISL_2936725 | Lighthouse Lab in Alderley Park | Wellcome Sanger Institute for the COVID-19 Genomics UK (COG-UK) Consortium | Jacquelyn Wynn, Mairead Hyland, The Lighthouse Lab in Alderley Park and Alex Alderton, Roberto Amato, Jeffrey Barrett, Sonia Goncalves, Ewan Harrison, David K. Jackson, Ian Johnston, Dominic Kwiatkowski, Cordelia Langford, John Sillitoe on behalf of the Wellcome Sanger Institute COVID-19 Surveillance Team |
| EPI_ISL_2936726 | Lighthouse Lab in Milton Keynes | Wellcome Sanger Institute for the COVID-19 Genomics UK (COG-UK) Consortium | The Lighthouse Lab in Milton Keynes and Alex Alderton, Roberto Amato, Jeffrey Barrett, Sonia Goncalves, Ewan Harrison, David K. Jackson, Ian Johnston, Dominic Kwiatkowski, Cordelia Langford, John Sillitoe on behalf of the Wellcome Sanger Institute COVID-19 Surveillance Team |
| EPI_ISL_2936727, EPI_ISL_2936728 | Lighthouse Lab in Alderley Park | Wellcome Sanger Institute for the COVID-19 Genomics UK (COG-UK) Consortium | Jacquelyn Wynn, Mairead Hyland, The Lighthouse Lab in Alderley Park and Alex Alderton, Roberto Amato, Jeffrey Barrett, Sonia Goncalves, Ewan Harrison, David K. Jackson, Ian Johnston, Dominic Kwiatkowski, Cordelia Langford, John Sillitoe on behalf of the Wellcome Sanger Institute COVID-19 Surveillance Team |
| EPI_ISL_2936729 | Lighthouse Lab in Glasgow | Wellcome Sanger Institute for the COVID-19 Genomics UK (COG-UK) Consortium | Harper VanSteenhouse, Yumi Kasai, David Gray, Carol Clugston, Anna Dominiczak and Alex Alderton, Roberto Amato, Jeffrey Barrett, Sonia Goncalves, Ewan Harrison, David K. Jackson, Ian Johnston, Dominic Kwiatkowski, Cordelia Langford, John Sillitoe on behalf of the Wellcome Sanger Institute COVID-19 Surveillance Team |
| EPI_ISL_2936730 | Lighthouse Lab in Alderley Park | Wellcome Sanger Institute for the COVID-19 Genomics UK (COG-UK) Consortium | Jacquelyn Wynn, Mairead Hyland, The Lighthouse Lab in Alderley Park and Alex Alderton, Roberto Amato, Jeffrey Barrett, Sonia Goncalves, Ewan Harrison, David K. Jackson, Ian Johnston, Dominic Kwiatkowski, Cordelia Langford, John Sillitoe on behalf of the Wellcome Sanger Institute COVID-19 Surveillance Team |
| EPI_ISL_2936731 | Lighthouse Lab in Glasgow | Wellcome Sanger Institute for the COVID-19 Genomics UK (COG-UK) Consortium | Harper VanSteenhouse, Yumi Kasai, David Gray, Carol Clugston, Anna Dominiczak and Alex Alderton, Roberto Amato, Jeffrey Barrett, Sonia Goncalves, Ewan Harrison, David K. Jackson, Ian Johnston, Dominic Kwiatkowski, Cordelia Langford, John Sillitoe on behalf of the Wellcome Sanger Institute COVID-19 Surveillance Team |
| EPI_ISL_2936732, EPI_ISL_2936733 | Lighthouse Lab in Alderley Park | Wellcome Sanger Institute for the COVID-19 Genomics UK (COG-UK) Consortium | Jacquelyn Wynn, Mairead Hyland, The Lighthouse Lab in Alderley Park and Alex Alderton, Roberto Amato, Jeffrey Barrett, Sonia Goncalves, Ewan Harrison, David K. Jackson, Ian Johnston, Dominic Kwiatkowski, Cordelia Langford, John Sillitoe on behalf of the Wellcome Sanger Institute COVID-19 Surveillance Team |
| EPI_ISL_2936734 | Lighthouse Lab in Milton Keynes | Wellcome Sanger Institute for the COVID-19 Genomics UK (COG-UK) Consortium | The Lighthouse Lab in Milton Keynes and Alex Alderton, Roberto Amato, Jeffrey Barrett, Sonia Goncalves, Ewan Harrison, David K. Jackson, Ian Johnston, Dominic Kwiatkowski, Cordelia Langford, John Sillitoe on behalf of the Wellcome Sanger Institute COVID-19 Surveillance Team |
| EPI_ISL_2936735, EPI_ISL_2936736 | Lighthouse Lab in Alderley Park | Wellcome Sanger Institute for the COVID-19 Genomics UK (COG-UK) Consortium | Jacquelyn Wynn, Mairead Hyland, The Lighthouse Lab in Alderley Park and Alex Alderton, Roberto Amato, Jeffrey Barrett, Sonia Goncalves, Ewan Harrison, David K. Jackson, Ian Johnston, Dominic Kwiatkowski, Cordelia Langford, John Sillitoe on behalf of the Wellcome Sanger Institute COVID-19 Surveillance Team |
| EPI_ISL_2936737, EPI_ISL_2936738, EPI_ISL_2936739 | Lighthouse Lab in Milton Keynes | Wellcome Sanger Institute for the COVID-19 Genomics UK (COG-UK) Consortium | The Lighthouse Lab in Milton Keynes and Alex Alderton, Roberto Amato, Jeffrey Barrett, Sonia Goncalves, Ewan Harrison, David K. Jackson, Ian Johnston, Dominic Kwiatkowski, Cordelia Langford, John Sillitoe on behalf of the Wellcome Sanger Institute COVID-19 Surveillance Team |
| EPI_ISL_2936740 | Lighthouse Lab in Alderley Park | Wellcome Sanger Institute for the COVID-19 Genomics UK (COG-UK) Consortium | Jacquelyn Wynn, Mairead Hyland, The Lighthouse Lab in Alderley Park and Alex Alderton, Roberto Amato, Jeffrey Barrett, Sonia Goncalves, Ewan Harrison, David K. Jackson, Ian Johnston, Dominic Kwiatkowski, Cordelia Langford, John Sillitoe on behalf of the Wellcome Sanger Institute COVID-19 Surveillance Team |
| EPI_ISL_2936741 | Lighthouse Lab in Milton Keynes | Wellcome Sanger Institute for the COVID-19 Genomics UK (COG-UK) Consortium | The Lighthouse Lab in Milton Keynes and Alex Alderton, Roberto Amato, Jeffrey Barrett, Sonia Goncalves, Ewan Harrison, David K. Jackson, Ian Johnston, Dominic Kwiatkowski, Cordelia Langford, John Sillitoe on behalf of the Wellcome Sanger Institute COVID-19 Surveillance Team |
| EPI_ISL_2936742 | Lighthouse Lab in Alderley Park | Wellcome Sanger Institute for the COVID-19 Genomics UK (COG-UK) Consortium | Jacquelyn Wynn, Mairead Hyland, The Lighthouse Lab in Alderley Park and Alex Alderton, Roberto Amato, Jeffrey Barrett, Sonia Goncalves, Ewan Harrison, David K. Jackson, Ian Johnston, Dominic Kwiatkowski, Cordelia Langford, John Sillitoe on behalf of the Wellcome Sanger Institute COVID-19 Surveillance Team |
| EPI_ISL_2936743 | Lighthouse Lab in Milton Keynes | Wellcome Sanger Institute for the COVID-19 Genomics UK (COG-UK) Consortium | The Lighthouse Lab in Milton Keynes and Alex Alderton, Roberto Amato, Jeffrey Barrett, Sonia Goncalves, Ewan Harrison, David K. Jackson, Ian Johnston, Dominic Kwiatkowski, Cordelia Langford, John Sillitoe on behalf of the Wellcome Sanger Institute COVID-19 Surveillance Team |
| EPI_ISL_2936744, EPI_ISL_2936745 | Lighthouse Lab in Alderley Park | Wellcome Sanger Institute for the COVID-19 Genomics UK (COG-UK) Consortium | Jacquelyn Wynn, Mairead Hyland, The Lighthouse Lab in Alderley Park and Alex Alderton, Roberto Amato, Jeffrey Barrett, Sonia Goncalves, Ewan Harrison, David K. Jackson, Ian Johnston, Dominic Kwiatkowski, Cordelia Langford, John Sillitoe on behalf of the Wellcome Sanger Institute COVID-19 Surveillance Team |
| EPI_ISL_2936746, EPI_ISL_2936747 | Lighthouse Lab in Milton Keynes | Wellcome Sanger Institute for the COVID-19 Genomics UK (COG-UK) Consortium | The Lighthouse Lab in Milton Keynes and Alex Alderton, Roberto Amato, Jeffrey Barrett, Sonia Goncalves, Ewan Harrison, David K. Jackson, Ian Johnston, Dominic Kwiatkowski, Cordelia Langford, John Sillitoe on behalf of the Wellcome Sanger Institute COVID-19 Surveillance Team |
| EPI_ISL_2936748, EPI_ISL_2936749 | Lighthouse Lab in Alderley Park | Wellcome Sanger Institute for the COVID-19 Genomics UK (COG-UK) Consortium | Jacquelyn Wynn, Mairead Hyland, The Lighthouse Lab in Alderley Park and Alex Alderton, Roberto Amato, Jeffrey Barrett, Sonia Goncalves, Ewan Harrison, David K. Jackson, Ian Johnston, Dominic Kwiatkowski, Cordelia Langford, John Sillitoe on behalf of the Wellcome Sanger Institute COVID-19 Surveillance Team |
| EPI_ISL_2936750, EPI_ISL_2936751, EPI_ISL_2936752 | Lighthouse Lab in Milton Keynes | Wellcome Sanger Institute for the COVID-19 Genomics UK (COG-UK) Consortium | The Lighthouse Lab in Milton Keynes and Alex Alderton, Roberto Amato, Jeffrey Barrett, Sonia Goncalves, Ewan Harrison, David K. Jackson, Ian Johnston, Dominic Kwiatkowski, Cordelia Langford, John Sillitoe on behalf of the Wellcome Sanger Institute COVID-19 Surveillance Team |
| EPI_ISL_2936753, EPI_ISL_2936754 | Lighthouse Lab in Glasgow | Wellcome Sanger Institute for the COVID-19 Genomics UK (COG-UK) Consortium | Harper VanSteenhouse, Yumi Kasai, David Gray, Carol Clugston, Anna Dominiczak and Alex Alderton, Roberto Amato, Jeffrey Barrett, Sonia Goncalves, Ewan Harrison, David K. Jackson, Ian Johnston, Dominic Kwiatkowski, Cordelia Langford, John Sillitoe on behalf of the Wellcome Sanger Institute COVID-19 Surveillance Team |
| EPI_ISL_2936755, EPI_ISL_2936756 | Lighthouse Lab in Milton Keynes | Wellcome Sanger Institute for the COVID-19 Genomics UK (COG-UK) Consortium | The Lighthouse Lab in Milton Keynes and Alex Alderton, Roberto Amato, Jeffrey Barrett, Sonia Goncalves, Ewan Harrison, David K. Jackson, Ian Johnston, Dominic Kwiatkowski, Cordelia Langford, John Sillitoe on behalf of the Wellcome Sanger Institute COVID-19 Surveillance Team |
| EPI_ISL_2936757 | Lighthouse Lab in Alderley Park | Wellcome Sanger Institute for the COVID-19 Genomics UK (COG-UK) Consortium | Jacquelyn Wynn, Mairead Hyland, The Lighthouse Lab in Alderley Park and Alex Alderton, Roberto Amato, Jeffrey Barrett, Sonia Goncalves, Ewan Harrison, David K. Jackson, Ian Johnston, Dominic Kwiatkowski, Cordelia Langford, John Sillitoe on behalf of the Wellcome Sanger Institute COVID-19 Surveillance Team |

| | | | |
|---|---|---|---|
| EPI_ISL_2936758, EPI_ISL_2936759, EPI_ISL_2936760, EPI_ISL_2936761 | Lighthouse Lab in Milton Keynes | Wellcome Sanger Institute for the COVID-19 Genomics UK (COG-UK) Consortium | The Lighthouse Lab in Milton Keynes and Alex Alderton, Roberto Amato, Jeffrey Barrett, Sonia Goncalves, Ewan Harrison, David K. Jackson, Ian Johnston, Dominic Kwiatkowski, Cordelia Langford, John Sillitoe on behalf of the Wellcome Sanger Institute COVID-19 Surveillance Team |
| EPI_ISL_2936762, EPI_ISL_2936763 | Lighthouse Lab in Alderley Park | Wellcome Sanger Institute for the COVID-19 Genomics UK (COG-UK) Consortium | Jacquelyn Wynn, Mairead Hyland, The Lighthouse Lab in Alderley Park and Alex Alderton, Roberto Amato, Jeffrey Barrett, Sonia Goncalves, Ewan Harrison, David K. Jackson, Ian Johnston, Dominic Kwiatkowski, Cordelia Langford, John Sillitoe on behalf of the Wellcome Sanger Institute COVID-19 Surveillance Team |
| EPI_ISL_2936764, EPI_ISL_2936765 | Lighthouse Lab in Milton Keynes | Wellcome Sanger Institute for the COVID-19 Genomics UK (COG-UK) Consortium | The Lighthouse Lab in Milton Keynes and Alex Alderton, Roberto Amato, Jeffrey Barrett, Sonia Goncalves, Ewan Harrison, David K. Jackson, Ian Johnston, Dominic Kwiatkowski, Cordelia Langford, John Sillitoe on behalf of the Wellcome Sanger Institute COVID-19 Surveillance Team |
| EPI_ISL_2936766 | Lighthouse Lab in Glasgow | Wellcome Sanger Institute for the COVID-19 Genomics UK (COG-UK) Consortium | Harper VanSteenhouse, Yumi Kasai, David Gray, Carol Clugston, Anna Dominiczak and Alex Alderton, Roberto Amato, Jeffrey Barrett, Sonia Goncalves, Ewan Harrison, David K. Jackson, Ian Johnston, Dominic Kwiatkowski, Cordelia Langford, John Sillitoe on behalf of the Wellcome Sanger Institute COVID-19 Surveillance Team |
| EPI_ISL_2936767 | Lighthouse Lab in Milton Keynes | Wellcome Sanger Institute for the COVID-19 Genomics UK (COG-UK) Consortium | The Lighthouse Lab in Milton Keynes and Alex Alderton, Roberto Amato, Jeffrey Barrett, Sonia Goncalves, Ewan Harrison, David K. Jackson, Ian Johnston, Dominic Kwiatkowski, Cordelia Langford, John Sillitoe on behalf of the Wellcome Sanger Institute COVID-19 Surveillance Team |
| EPI_ISL_2936768 | Lighthouse Lab in Glasgow | Wellcome Sanger Institute for the COVID-19 Genomics UK (COG-UK) Consortium | Harper VanSteenhouse, Yumi Kasai, David Gray, Carol Clugston, Anna Dominiczak and Alex Alderton, Roberto Amato, Jeffrey Barrett, Sonia Goncalves, Ewan Harrison, David K. Jackson, Ian Johnston, Dominic Kwiatkowski, Cordelia Langford, John Sillitoe on behalf of the Wellcome Sanger Institute COVID-19 Surveillance Team |
| EPI_ISL_2936769 | Lighthouse Lab in Milton Keynes | Wellcome Sanger Institute for the COVID-19 Genomics UK (COG-UK) Consortium | The Lighthouse Lab in Milton Keynes and Alex Alderton, Roberto Amato, Jeffrey Barrett, Sonia Goncalves, Ewan Harrison, David K. Jackson, Ian Johnston, Dominic Kwiatkowski, Cordelia Langford, John Sillitoe on behalf of the Wellcome Sanger Institute COVID-19 Surveillance Team |
| EPI_ISL_2936770 | Lighthouse Lab in Alderley Park | Wellcome Sanger Institute for the COVID-19 Genomics UK (COG-UK) Consortium | Jacquelyn Wynn, Mairead Hyland, The Lighthouse Lab in Alderley Park and Alex Alderton, Roberto Amato, Jeffrey Barrett, Sonia Goncalves, Ewan Harrison, David K. Jackson, Ian Johnston, Dominic Kwiatkowski, Cordelia Langford, John Sillitoe on behalf of the Wellcome Sanger Institute COVID-19 Surveillance Team |
| EPI_ISL_2936771, EPI_ISL_2936772, EPI_ISL_2936773, EPI_ISL_2936774 | Lighthouse Lab in Milton Keynes | Wellcome Sanger Institute for the COVID-19 Genomics UK (COG-UK) Consortium | The Lighthouse Lab in Milton Keynes and Alex Alderton, Roberto Amato, Jeffrey Barrett, Sonia Goncalves, Ewan Harrison, David K. Jackson, Ian Johnston, Dominic Kwiatkowski, Cordelia Langford, John Sillitoe on behalf of the Wellcome Sanger Institute COVID-19 Surveillance Team |
| EPI_ISL_2936775 | Lighthouse Lab in Alderley Park | Wellcome Sanger Institute for the COVID-19 Genomics UK (COG-UK) Consortium | Jacquelyn Wynn, Mairead Hyland, The Lighthouse Lab in Alderley Park and Alex Alderton, Roberto Amato, Jeffrey Barrett, Sonia Goncalves, Ewan Harrison, David K. Jackson, Ian Johnston, Dominic Kwiatkowski, Cordelia Langford, John Sillitoe on behalf of the Wellcome Sanger Institute COVID-19 Surveillance Team |
| EPI_ISL_2936776, EPI_ISL_2936777, EPI_ISL_2936778 | Lighthouse Lab in Milton Keynes | Wellcome Sanger Institute for the COVID-19 Genomics UK (COG-UK) Consortium | The Lighthouse Lab in Milton Keynes and Alex Alderton, Roberto Amato, Jeffrey Barrett, Sonia Goncalves, Ewan Harrison, David K. Jackson, Ian Johnston, Dominic Kwiatkowski, Cordelia Langford, John Sillitoe on behalf of the Wellcome Sanger Institute COVID-19 Surveillance Team |
| EPI_ISL_2936779 | Lighthouse Lab in Alderley Park | Wellcome Sanger Institute for the COVID-19 Genomics UK (COG-UK) Consortium | Jacquelyn Wynn, Mairead Hyland, The Lighthouse Lab in Alderley Park and Alex Alderton, Roberto Amato, Jeffrey Barrett, Sonia Goncalves, Ewan Harrison, David K. Jackson, Ian Johnston, Dominic Kwiatkowski, Cordelia Langford, John Sillitoe on behalf of the Wellcome Sanger Institute COVID-19 Surveillance Team |
| EPI_ISL_2936780 | Lighthouse Lab in Glasgow | Wellcome Sanger Institute for the COVID-19 Genomics UK (COG-UK) Consortium | Harper VanSteenhouse, Yumi Kasai, David Gray, Carol Clugston, Anna Dominiczak and Alex Alderton, Roberto Amato, Jeffrey Barrett, Sonia Goncalves, Ewan Harrison, David K. Jackson, Ian Johnston, Dominic Kwiatkowski, Cordelia Langford, John Sillitoe on behalf of the Wellcome Sanger Institute COVID-19 Surveillance Team |
| EPI_ISL_2936781 | Lighthouse Lab in Milton Keynes | Wellcome Sanger Institute for the COVID-19 Genomics UK (COG-UK) Consortium | The Lighthouse Lab in Milton Keynes and Alex Alderton, Roberto Amato, Jeffrey Barrett, Sonia Goncalves, Ewan Harrison, David K. Jackson, Ian Johnston, Dominic Kwiatkowski, Cordelia Langford, John Sillitoe on behalf of the Wellcome Sanger Institute COVID-19 Surveillance Team |
| EPI_ISL_2936782 | Lighthouse Lab in Glasgow | Wellcome Sanger Institute for the COVID-19 Genomics UK (COG-UK) Consortium | Harper VanSteenhouse, Yumi Kasai, David Gray, Carol Clugston, Anna Dominiczak and Alex Alderton, Roberto Amato, Jeffrey Barrett, Sonia Goncalves, Ewan Harrison, David K. Jackson, Ian Johnston, Dominic Kwiatkowski, Cordelia Langford, John Sillitoe on behalf of the Wellcome Sanger Institute COVID-19 Surveillance Team |
| EPI_ISL_2936783, EPI_ISL_2936784 | Lighthouse Lab in Milton Keynes | Wellcome Sanger Institute for the COVID-19 Genomics UK (COG-UK) Consortium | The Lighthouse Lab in Milton Keynes and Alex Alderton, Roberto Amato, Jeffrey Barrett, Sonia Goncalves, Ewan Harrison, David K. Jackson, Ian Johnston, Dominic Kwiatkowski, Cordelia Langford, John Sillitoe on behalf of the Wellcome Sanger Institute COVID-19 Surveillance Team |
| EPI_ISL_2936785 | Lighthouse Lab in Glasgow | Wellcome Sanger Institute for the COVID-19 Genomics UK (COG-UK) Consortium | Harper VanSteenhouse, Yumi Kasai, David Gray, Carol Clugston, Anna Dominiczak and Alex Alderton, Roberto Amato, Jeffrey Barrett, Sonia Goncalves, Ewan Harrison, David K. Jackson, Ian Johnston, Dominic Kwiatkowski, Cordelia Langford, John Sillitoe on behalf of the Wellcome Sanger Institute COVID-19 Surveillance Team |
| EPI_ISL_2936786, EPI_ISL_2936787, EPI_ISL_2936788 | Lighthouse Lab in Milton Keynes | Wellcome Sanger Institute for the COVID-19 Genomics UK (COG-UK) Consortium | The Lighthouse Lab in Milton Keynes and Alex Alderton, Roberto Amato, Jeffrey Barrett, Sonia Goncalves, Ewan Harrison, David K. Jackson, Ian Johnston, Dominic Kwiatkowski, Cordelia Langford, John Sillitoe on behalf of the Wellcome Sanger Institute COVID-19 Surveillance Team |
| EPI_ISL_2936789 | Lighthouse Lab in Alderley Park | Wellcome Sanger Institute for the COVID-19 Genomics UK (COG-UK) Consortium | Jacquelyn Wynn, Mairead Hyland, The Lighthouse Lab in Alderley Park and Alex Alderton, Roberto Amato, Jeffrey Barrett, Sonia Goncalves, Ewan Harrison, David K. Jackson, Ian Johnston, Dominic Kwiatkowski, Cordelia Langford, John Sillitoe on behalf of the Wellcome Sanger Institute COVID-19 Surveillance Team |
| EPI_ISL_2936790, EPI_ISL_2936791, EPI_ISL_2936792 | Lighthouse Lab in Milton Keynes | Wellcome Sanger Institute for the COVID-19 Genomics UK (COG-UK) Consortium | The Lighthouse Lab in Milton Keynes and Alex Alderton, Roberto Amato, Jeffrey Barrett, Sonia Goncalves, Ewan Harrison, David K. Jackson, Ian Johnston, Dominic Kwiatkowski, Cordelia Langford, John Sillitoe on behalf of the Wellcome Sanger Institute COVID-19 Surveillance Team |
| EPI_ISL_2936793, EPI_ISL_2936794 | Lighthouse Lab in Alderley Park | Wellcome Sanger Institute for the COVID-19 Genomics UK (COG-UK) Consortium | Jacquelyn Wynn, Mairead Hyland, The Lighthouse Lab in Alderley Park and Alex Alderton, Roberto Amato, Jeffrey Barrett, Sonia Goncalves, Ewan Harrison, David K. Jackson, Ian Johnston, Dominic Kwiatkowski, Cordelia Langford, John Sillitoe on behalf of the Wellcome Sanger Institute COVID-19 Surveillance Team |
| EPI_ISL_2936795 | Lighthouse Lab in Glasgow | Wellcome Sanger Institute for the COVID-19 Genomics UK (COG-UK) Consortium | Harper VanSteenhouse, Yumi Kasai, David Gray, Carol Clugston, Anna Dominiczak and Alex Alderton, Roberto Amato, Jeffrey Barrett, Sonia Goncalves, Ewan Harrison, David K. Jackson, Ian Johnston, Dominic Kwiatkowski, Cordelia Langford, John Sillitoe on behalf of the Wellcome Sanger Institute COVID-19 Surveillance Team |
| EPI_ISL_2936796 | Lighthouse Lab in Alderley Park | Wellcome Sanger Institute for the COVID-19 Genomics UK (COG-UK) Consortium | Jacquelyn Wynn, Mairead Hyland, The Lighthouse Lab in Alderley Park and Alex Alderton, Roberto Amato, Jeffrey Barrett, Sonia Goncalves, Ewan Harrison, David K. Jackson, Ian Johnston, Dominic Kwiatkowski, Cordelia Langford, John Sillitoe on behalf of the Wellcome Sanger Institute COVID-19 Surveillance Team |
| EPI_ISL_2936797 | Lighthouse Lab in Milton Keynes | Wellcome Sanger Institute for the COVID-19 Genomics UK (COG-UK) Consortium | The Lighthouse Lab in Milton Keynes and Alex Alderton, Roberto Amato, Jeffrey Barrett, Sonia Goncalves, Ewan Harrison, David K. Jackson, Ian Johnston, Dominic Kwiatkowski, Cordelia Langford, John Sillitoe on behalf of the Wellcome Sanger Institute COVID-19 Surveillance Team |
| EPI_ISL_2936798 | Lighthouse Lab in Alderley Park | Wellcome Sanger Institute for the COVID-19 Genomics UK (COG-UK) Consortium | Jacquelyn Wynn, Mairead Hyland, The Lighthouse Lab in Alderley Park and Alex Alderton, Roberto Amato, Jeffrey Barrett, Sonia Goncalves, Ewan Harrison, David K. Jackson, Ian Johnston, Dominic Kwiatkowski, Cordelia Langford, John Sillitoe on behalf of the Wellcome Sanger Institute COVID-19 Surveillance Team |
| EPI_ISL_2936799, EPI_ISL_2936800, EPI_ISL_2936801 | Lighthouse Lab in Glasgow | Wellcome Sanger Institute for the COVID-19 Genomics UK (COG-UK) Consortium | Harper VanSteenhouse, Yumi Kasai, David Gray, Carol Clugston, Anna Dominiczak and Alex Alderton, Roberto Amato, Jeffrey Barrett, Sonia Goncalves, Ewan Harrison, David K. Jackson, Ian Johnston, Dominic Kwiatkowski, Cordelia Langford, John Sillitoe on behalf of the Wellcome Sanger Institute COVID-19 Surveillance Team |
| EPI_ISL_2936802 | Lighthouse Lab in Alderley Park | Wellcome Sanger Institute for the COVID-19 Genomics UK (COG-UK) Consortium | Jacquelyn Wynn, Mairead Hyland, The Lighthouse Lab in Alderley Park and Alex Alderton, Roberto Amato, Jeffrey Barrett, Sonia Goncalves, Ewan Harrison, David K. Jackson, Ian Johnston, Dominic Kwiatkowski, Cordelia Langford, John Sillitoe on behalf of the Wellcome Sanger Institute COVID-19 Surveillance Team |
| EPI_ISL_2936803, EPI_ISL_2936804, EPI_ISL_2936805 | Lighthouse Lab in Milton Keynes | Wellcome Sanger Institute for the COVID-19 Genomics UK (COG-UK) Consortium | The Lighthouse Lab in Milton Keynes and Alex Alderton, Roberto Amato, Jeffrey Barrett, Sonia Goncalves, Ewan Harrison, David K. Jackson, Ian Johnston, Dominic Kwiatkowski, Cordelia Langford, John Sillitoe on behalf of the Wellcome Sanger Institute COVID-19 Surveillance Team |
| EPI_ISL_2936806 | Lighthouse Lab in Glasgow | Wellcome Sanger Institute for the COVID-19 Genomics UK (COG-UK) Consortium | Harper VanSteenhouse, Yumi Kasai, David Gray, Carol Clugston, Anna Dominiczak and Alex Alderton, Roberto Amato, Jeffrey Barrett, Sonia Goncalves, Ewan Harrison, David K. Jackson, Ian Johnston, Dominic Kwiatkowski, Cordelia Langford, John Sillitoe on behalf of the Wellcome Sanger Institute COVID-19 Surveillance Team |
| EPI_ISL_2936807 | Lighthouse Lab in Alderley Park | Wellcome Sanger Institute for the COVID-19 Genomics UK (COG-UK) Consortium | Jacquelyn Wynn, Mairead Hyland, The Lighthouse Lab in Alderley Park and Alex Alderton, Roberto Amato, Jeffrey Barrett, Sonia Goncalves, Ewan Harrison, David K. Jackson, Ian Johnston, Dominic Kwiatkowski, Cordelia Langford, John Sillitoe on behalf of the Wellcome Sanger Institute COVID-19 Surveillance Team |
| EPI_ISL_2936808 | Lighthouse Lab in Milton Keynes | Wellcome Sanger Institute for the COVID-19 Genomics UK (COG-UK) Consortium | The Lighthouse Lab in Milton Keynes and Alex Alderton, Roberto Amato, Jeffrey Barrett, Sonia Goncalves, Ewan Harrison, David K. Jackson, Ian Johnston, Dominic Kwiatkowski, Cordelia Langford, John Sillitoe on behalf of the Wellcome Sanger Institute COVID-19 Surveillance Team |
| EPI_ISL_2936809 | Lighthouse Lab in Alderley Park | Wellcome Sanger Institute for the COVID-19 Genomics UK (COG-UK) Consortium | Jacquelyn Wynn, Mairead Hyland, The Lighthouse Lab in Alderley Park and Alex Alderton, Roberto Amato, Jeffrey Barrett, Sonia Goncalves, Ewan Harrison, David K. Jackson, Ian Johnston, Dominic Kwiatkowski, Cordelia Langford, John Sillitoe on behalf of the Wellcome Sanger Institute COVID-19 Surveillance Team |
| EPI_ISL_2936810 | Lighthouse Lab in Glasgow | Wellcome Sanger Institute for the COVID-19 Genomics UK (COG-UK) Consortium | Harper VanSteenhouse, Yumi Kasai, David Gray, Carol Clugston, Anna Dominiczak and Alex Alderton, Roberto Amato, Jeffrey Barrett, Sonia Goncalves, Ewan Harrison, David K. Jackson, Ian Johnston, Dominic Kwiatkowski, Cordelia Langford, John Sillitoe on behalf of the Wellcome Sanger Institute COVID-19 Surveillance Team |
| EPI_ISL_2936811, EPI_ISL_2936813 | Lighthouse Lab in Alderley Park | Wellcome Sanger Institute for the COVID-19 Genomics UK (COG-UK) Consortium | Jacquelyn Wynn, Mairead Hyland, The Lighthouse Lab in Alderley Park and Alex Alderton, Roberto Amato, Jeffrey Barrett, Sonia Goncalves, Ewan Harrison, David K. Jackson, Ian Johnston, Dominic Kwiatkowski, Cordelia Langford, John Sillitoe on behalf of the Wellcome Sanger Institute COVID-19 Surveillance Team |
| EPI_ISL_2936814, EPI_ISL_2936815, EPI_ISL_2936816 | Lighthouse Lab in Milton Keynes | Wellcome Sanger Institute for the COVID-19 Genomics UK (COG-UK) Consortium | The Lighthouse Lab in Milton Keynes and Alex Alderton, Roberto Amato, Jeffrey Barrett, Sonia Goncalves, Ewan Harrison, David K. Jackson, Ian Johnston, Dominic Kwiatkowski, Cordelia Langford, John Sillitoe on behalf of the Wellcome Sanger Institute COVID-19 Surveillance Team |
| EPI_ISL_2936817, EPI_ISL_2936818 | Lighthouse Lab in Alderley Park | Wellcome Sanger Institute for the COVID-19 Genomics UK (COG-UK) Consortium | Jacquelyn Wynn, Mairead Hyland, The Lighthouse Lab in Alderley Park and Alex Alderton, Roberto Amato, Jeffrey Barrett, Sonia Goncalves, Ewan Harrison, David K. Jackson, Ian Johnston, Dominic Kwiatkowski, Cordelia Langford, John Sillitoe on behalf of the Wellcome Sanger Institute COVID-19 Surveillance Team |
| EPI_ISL_2936819 | Lighthouse Lab in Milton Keynes | Wellcome Sanger Institute for the COVID-19 Genomics UK (COG-UK) Consortium | The Lighthouse Lab in Milton Keynes and Alex Alderton, Roberto Amato, Jeffrey Barrett, Sonia Goncalves, Ewan Harrison, David K. Jackson, Ian Johnston, Dominic Kwiatkowski, Cordelia Langford, John Sillitoe on behalf of the Wellcome Sanger Institute COVID-19 Surveillance Team |
| EPI_ISL_2936820 | Lighthouse Lab in Alderley Park | Wellcome Sanger Institute for the COVID-19 Genomics UK (COG-UK) Consortium | Jacquelyn Wynn, Mairead Hyland, The Lighthouse Lab in Alderley Park and Alex Alderton, Roberto Amato, Jeffrey Barrett, Sonia Goncalves, Ewan Harrison, David K. Jackson, Ian Johnston, Dominic Kwiatkowski, Cordelia Langford, John Sillitoe on behalf of the Wellcome Sanger Institute COVID-19 Surveillance Team |
| EPI_ISL_2936821, EPI_ISL_2936822, EPI_ISL_2936823, EPI_ISL_2936824 | Lighthouse Lab in Milton Keynes | Wellcome Sanger Institute for the COVID-19 Genomics UK (COG-UK) Consortium | The Lighthouse Lab in Milton Keynes and Alex Alderton, Roberto Amato, Jeffrey Barrett, Sonia Goncalves, Ewan Harrison, David K. Jackson, Ian Johnston, Dominic Kwiatkowski, Cordelia Langford, John Sillitoe on behalf of the Wellcome Sanger Institute COVID-19 Surveillance Team |
| EPI_ISL_2936825 | Lighthouse Lab in Glasgow | Wellcome Sanger Institute for the COVID-19 Genomics UK (COG-UK) Consortium | Harper VanSteenhouse, Yumi Kasai, David Gray, Carol Clugston, Anna Dominiczak and Alex Alderton, Roberto Amato, Jeffrey Barrett, Sonia Goncalves, Ewan Harrison, David K. Jackson, Ian Johnston, Dominic Kwiatkowski, Cordelia Langford, John Sillitoe on behalf of the Wellcome Sanger Institute COVID-19 Surveillance Team |

| | | | |
|---|---|---|---|
| EPI_ISL_2936826, EPI_ISL_2936827 | Lighthouse Lab in Milton Keynes | Wellcome Sanger Institute for the COVID-19 Genomics UK (COG-UK) Consortium | The Lighthouse Lab in Milton Keynes and Alex Alderton, Roberto Amato, Jeffrey Barrett, Sonia Goncalves, Ewan Harrison, David K. Jackson, Ian Johnston, Dominic Kwiatkowski, Cordelia Langford, John Sillitoe on behalf of the Wellcome Sanger Institute COVID-19 Surveillance Team |
| EPI_ISL_2936828 | Lighthouse Lab in Alderley Park | Wellcome Sanger Institute for the COVID-19 Genomics UK (COG-UK) Consortium | Jacquelyn Wynn, Mairead Hyland, The Lighthouse Lab in Alderley Park and Alex Alderton, Roberto Amato, Jeffrey Barrett, Sonia Goncalves, Ewan Harrison, David K. Jackson, Ian Johnston, Dominic Kwiatkowski, Cordelia Langford, John Sillitoe on behalf of the Wellcome Sanger Institute COVID-19 Surveillance Team |
| EPI_ISL_2936829 | Lighthouse Lab in Glasgow | Wellcome Sanger Institute for the COVID-19 Genomics UK (COG-UK) Consortium | Harper VanSteenhouse, Yumi Kasai, David Gray, Carol Clugston, Anna Dominiczak and Alex Alderton, Roberto Amato, Jeffrey Barrett, Sonia Goncalves, Ewan Harrison, David K. Jackson, Ian Johnston, Dominic Kwiatkowski, Cordelia Langford, John Sillitoe on behalf of the Wellcome Sanger Institute COVID-19 Surveillance Team |
| EPI_ISL_2936830, EPI_ISL_2936831, EPI_ISL_2936832, EPI_ISL_2936833, EPI_ISL_2936834 | Lighthouse Lab in Alderley Park | Wellcome Sanger Institute for the COVID-19 Genomics UK (COG-UK) Consortium | Jacquelyn Wynn, Mairead Hyland, The Lighthouse Lab in Alderley Park and Alex Alderton, Roberto Amato, Jeffrey Barrett, Sonia Goncalves, Ewan Harrison, David K. Jackson, Ian Johnston, Dominic Kwiatkowski, Cordelia Langford, John Sillitoe on behalf of the Wellcome Sanger Institute COVID-19 Surveillance Team |
| EPI_ISL_2936835 | Lighthouse Lab in Glasgow | Wellcome Sanger Institute for the COVID-19 Genomics UK (COG-UK) Consortium | Harper VanSteenhouse, Yumi Kasai, David Gray, Carol Clugston, Anna Dominiczak and Alex Alderton, Roberto Amato, Jeffrey Barrett, Sonia Goncalves, Ewan Harrison, David K. Jackson, Ian Johnston, Dominic Kwiatkowski, Cordelia Langford, John Sillitoe on behalf of the Wellcome Sanger Institute COVID-19 Surveillance Team |
| EPI_ISL_2936836 | Lighthouse Lab in Milton Keynes | Wellcome Sanger Institute for the COVID-19 Genomics UK (COG-UK) Consortium | The Lighthouse Lab in Milton Keynes and Alex Alderton, Roberto Amato, Jeffrey Barrett, Sonia Goncalves, Ewan Harrison, David K. Jackson, Ian Johnston, Dominic Kwiatkowski, Cordelia Langford, John Sillitoe on behalf of the Wellcome Sanger Institute COVID-19 Surveillance Team |
| EPI_ISL_2936837 | Lighthouse Lab in Alderley Park | Wellcome Sanger Institute for the COVID-19 Genomics UK (COG-UK) Consortium | Jacquelyn Wynn, Mairead Hyland, The Lighthouse Lab in Alderley Park and Alex Alderton, Roberto Amato, Jeffrey Barrett, Sonia Goncalves, Ewan Harrison, David K. Jackson, Ian Johnston, Dominic Kwiatkowski, Cordelia Langford, John Sillitoe on behalf of the Wellcome Sanger Institute COVID-19 Surveillance Team |
| EPI_ISL_2936838 | Lighthouse Lab in Milton Keynes | Wellcome Sanger Institute for the COVID-19 Genomics UK (COG-UK) Consortium | The Lighthouse Lab in Milton Keynes and Alex Alderton, Roberto Amato, Jeffrey Barrett, Sonia Goncalves, Ewan Harrison, David K. Jackson, Ian Johnston, Dominic Kwiatkowski, Cordelia Langford, John Sillitoe on behalf of the Wellcome Sanger Institute COVID-19 Surveillance Team |
| EPI_ISL_2936839, EPI_ISL_2936840 | Lighthouse Lab in Glasgow | Wellcome Sanger Institute for the COVID-19 Genomics UK (COG-UK) Consortium | Harper VanSteenhouse, Yumi Kasai, David Gray, Carol Clugston, Anna Dominiczak and Alex Alderton, Roberto Amato, Jeffrey Barrett, Sonia Goncalves, Ewan Harrison, David K. Jackson, Ian Johnston, Dominic Kwiatkowski, Cordelia Langford, John Sillitoe on behalf of the Wellcome Sanger Institute COVID-19 Surveillance Team |
| EPI_ISL_2936841 | Lighthouse Lab in Milton Keynes | Wellcome Sanger Institute for the COVID-19 Genomics UK (COG-UK) Consortium | The Lighthouse Lab in Milton Keynes and Alex Alderton, Roberto Amato, Jeffrey Barrett, Sonia Goncalves, Ewan Harrison, David K. Jackson, Ian Johnston, Dominic Kwiatkowski, Cordelia Langford, John Sillitoe on behalf of the Wellcome Sanger Institute COVID-19 Surveillance Team |
| EPI_ISL_2936842 | Lighthouse Lab in Alderley Park | Wellcome Sanger Institute for the COVID-19 Genomics UK (COG-UK) Consortium | Jacquelyn Wynn, Mairead Hyland, The Lighthouse Lab in Alderley Park and Alex Alderton, Roberto Amato, Jeffrey Barrett, Sonia Goncalves, Ewan Harrison, David K. Jackson, Ian Johnston, Dominic Kwiatkowski, Cordelia Langford, John Sillitoe on behalf of the Wellcome Sanger Institute COVID-19 Surveillance Team |
| EPI_ISL_2936843 | Lighthouse Lab in Milton Keynes | Wellcome Sanger Institute for the COVID-19 Genomics UK (COG-UK) Consortium | The Lighthouse Lab in Milton Keynes and Alex Alderton, Roberto Amato, Jeffrey Barrett, Sonia Goncalves, Ewan Harrison, David K. Jackson, Ian Johnston, Dominic Kwiatkowski, Cordelia Langford, John Sillitoe on behalf of the Wellcome Sanger Institute COVID-19 Surveillance Team |
| EPI_ISL_2936844, EPI_ISL_2936845 | Lighthouse Lab in Alderley Park | Wellcome Sanger Institute for the COVID-19 Genomics UK (COG-UK) Consortium | Jacquelyn Wynn, Mairead Hyland, The Lighthouse Lab in Alderley Park and Alex Alderton, Roberto Amato, Jeffrey Barrett, Sonia Goncalves, Ewan Harrison, David K. Jackson, Ian Johnston, Dominic Kwiatkowski, Cordelia Langford, John Sillitoe on behalf of the Wellcome Sanger Institute COVID-19 Surveillance Team |
| EPI_ISL_2936846, EPI_ISL_2936847 | Lighthouse Lab in Glasgow | Wellcome Sanger Institute for the COVID-19 Genomics UK (COG-UK) Consortium | Harper VanSteenhouse, Yumi Kasai, David Gray, Carol Clugston, Anna Dominiczak and Alex Alderton, Roberto Amato, Jeffrey Barrett, Sonia Goncalves, Ewan Harrison, David K. Jackson, Ian Johnston, Dominic Kwiatkowski, Cordelia Langford, John Sillitoe on behalf of the Wellcome Sanger Institute COVID-19 Surveillance Team |
| EPI_ISL_2936848, EPI_ISL_2936849, EPI_ISL_2936850 | Lighthouse Lab in Milton Keynes | Wellcome Sanger Institute for the COVID-19 Genomics UK (COG-UK) Consortium | The Lighthouse Lab in Milton Keynes and Alex Alderton, Roberto Amato, Jeffrey Barrett, Sonia Goncalves, Ewan Harrison, David K. Jackson, Ian Johnston, Dominic Kwiatkowski, Cordelia Langford, John Sillitoe on behalf of the Wellcome Sanger Institute COVID-19 Surveillance Team |
| EPI_ISL_2936851 | Lighthouse Lab in Alderley Park | Wellcome Sanger Institute for the COVID-19 Genomics UK (COG-UK) Consortium | Jacquelyn Wynn, Mairead Hyland, The Lighthouse Lab in Alderley Park and Alex Alderton, Roberto Amato, Jeffrey Barrett, Sonia Goncalves, Ewan Harrison, David K. Jackson, Ian Johnston, Dominic Kwiatkowski, Cordelia Langford, John Sillitoe on behalf of the Wellcome Sanger Institute COVID-19 Surveillance Team |
| EPI_ISL_2936852, EPI_ISL_2936853 | Lighthouse Lab in Glasgow | Wellcome Sanger Institute for the COVID-19 Genomics UK (COG-UK) Consortium | Harper VanSteenhouse, Yumi Kasai, David Gray, Carol Clugston, Anna Dominiczak and Alex Alderton, Roberto Amato, Jeffrey Barrett, Sonia Goncalves, Ewan Harrison, David K. Jackson, Ian Johnston, Dominic Kwiatkowski, Cordelia Langford, John Sillitoe on behalf of the Wellcome Sanger Institute COVID-19 Surveillance Team |
| EPI_ISL_2936854, EPI_ISL_2936855 | Lighthouse Lab in Milton Keynes | Wellcome Sanger Institute for the COVID-19 Genomics UK (COG-UK) Consortium | The Lighthouse Lab in Milton Keynes and Alex Alderton, Roberto Amato, Jeffrey Barrett, Sonia Goncalves, Ewan Harrison, David K. Jackson, Ian Johnston, Dominic Kwiatkowski, Cordelia Langford, John Sillitoe on behalf of the Wellcome Sanger Institute COVID-19 Surveillance Team |
| EPI_ISL_2936856 | Lighthouse Lab in Alderley Park | Wellcome Sanger Institute for the COVID-19 Genomics UK (COG-UK) Consortium | Jacquelyn Wynn, Mairead Hyland, The Lighthouse Lab in Alderley Park and Alex Alderton, Roberto Amato, Jeffrey Barrett, Sonia Goncalves, Ewan Harrison, David K. Jackson, Ian Johnston, Dominic Kwiatkowski, Cordelia Langford, John Sillitoe on behalf of the Wellcome Sanger Institute COVID-19 Surveillance Team |
| EPI_ISL_2936857 | Lighthouse Lab in Milton Keynes | Wellcome Sanger Institute for the COVID-19 Genomics UK (COG-UK) Consortium | The Lighthouse Lab in Milton Keynes and Alex Alderton, Roberto Amato, Jeffrey Barrett, Sonia Goncalves, Ewan Harrison, David K. Jackson, Ian Johnston, Dominic Kwiatkowski, Cordelia Langford, John Sillitoe on behalf of the Wellcome Sanger Institute COVID-19 Surveillance Team |
| EPI_ISL_2936858, EPI_ISL_2936859 | Lighthouse Lab in Glasgow | Wellcome Sanger Institute for the COVID-19 Genomics UK (COG-UK) Consortium | Harper VanSteenhouse, Yumi Kasai, David Gray, Carol Clugston, Anna Dominiczak and Alex Alderton, Roberto Amato, Jeffrey Barrett, Sonia Goncalves, Ewan Harrison, David K. Jackson, Ian Johnston, Dominic Kwiatkowski, Cordelia Langford, John Sillitoe on behalf of the Wellcome Sanger Institute COVID-19 Surveillance Team |
| EPI_ISL_2936860 | Lighthouse Lab in Alderley Park | Wellcome Sanger Institute for the COVID-19 Genomics UK (COG-UK) Consortium | Jacquelyn Wynn, Mairead Hyland, The Lighthouse Lab in Alderley Park and Alex Alderton, Roberto Amato, Jeffrey Barrett, Sonia Goncalves, Ewan Harrison, David K. Jackson, Ian Johnston, Dominic Kwiatkowski, Cordelia Langford, John Sillitoe on behalf of the Wellcome Sanger Institute COVID-19 Surveillance Team |
| EPI_ISL_2936861 | Lighthouse Lab in Milton Keynes | Wellcome Sanger Institute for the COVID-19 Genomics UK (COG-UK) Consortium | The Lighthouse Lab in Milton Keynes and Alex Alderton, Roberto Amato, Jeffrey Barrett, Sonia Goncalves, Ewan Harrison, David K. Jackson, Ian Johnston, Dominic Kwiatkowski, Cordelia Langford, John Sillitoe on behalf of the Wellcome Sanger Institute COVID-19 Surveillance Team |
| EPI_ISL_2936862 | Lighthouse Lab in Alderley Park | Wellcome Sanger Institute for the COVID-19 Genomics UK (COG-UK) Consortium | Jacquelyn Wynn, Mairead Hyland, The Lighthouse Lab in Alderley Park and Alex Alderton, Roberto Amato, Jeffrey Barrett, Sonia Goncalves, Ewan Harrison, David K. Jackson, Ian Johnston, Dominic Kwiatkowski, Cordelia Langford, John Sillitoe on behalf of the Wellcome Sanger Institute COVID-19 Surveillance Team |
| EPI_ISL_2936863, EPI_ISL_2936864, EPI_ISL_2936865, EPI_ISL_2936866, EPI_ISL_2936867, EPI_ISL_2936868, EPI_ISL_2936869 | Lighthouse Lab in Milton Keynes | Wellcome Sanger Institute for the COVID-19 Genomics UK (COG-UK) Consortium | The Lighthouse Lab in Milton Keynes and Alex Alderton, Roberto Amato, Jeffrey Barrett, Sonia Goncalves, Ewan Harrison, David K. Jackson, Ian Johnston, Dominic Kwiatkowski, Cordelia Langford, John Sillitoe on behalf of the Wellcome Sanger Institute COVID-19 Surveillance Team |
| EPI_ISL_2936870 | Lighthouse Lab in Alderley Park | Wellcome Sanger Institute for the COVID-19 Genomics UK (COG-UK) Consortium | Jacquelyn Wynn, Mairead Hyland, The Lighthouse Lab in Alderley Park and Alex Alderton, Roberto Amato, Jeffrey Barrett, Sonia Goncalves, Ewan Harrison, David K. Jackson, Ian Johnston, Dominic Kwiatkowski, Cordelia Langford, John Sillitoe on behalf of the Wellcome Sanger Institute COVID-19 Surveillance Team |
| EPI_ISL_2936871 | Lighthouse Lab in Milton Keynes | Wellcome Sanger Institute for the COVID-19 Genomics UK (COG-UK) Consortium | The Lighthouse Lab in Milton Keynes and Alex Alderton, Roberto Amato, Jeffrey Barrett, Sonia Goncalves, Ewan Harrison, David K. Jackson, Ian Johnston, Dominic Kwiatkowski, Cordelia Langford, John Sillitoe on behalf of the Wellcome Sanger Institute COVID-19 Surveillance Team |
| EPI_ISL_2936872 | Lighthouse Lab in Glasgow | Wellcome Sanger Institute for the COVID-19 Genomics UK (COG-UK) Consortium | Harper VanSteenhouse, Yumi Kasai, David Gray, Carol Clugston, Anna Dominiczak and Alex Alderton, Roberto Amato, Jeffrey Barrett, Sonia Goncalves, Ewan Harrison, David K. Jackson, Ian Johnston, Dominic Kwiatkowski, Cordelia Langford, John Sillitoe on behalf of the Wellcome Sanger Institute COVID-19 Surveillance Team |
| EPI_ISL_2936873, EPI_ISL_2936874, EPI_ISL_2936875, EPI_ISL_2936876 | Lighthouse Lab in Alderley Park | Wellcome Sanger Institute for the COVID-19 Genomics UK (COG-UK) Consortium | Jacquelyn Wynn, Mairead Hyland, The Lighthouse Lab in Alderley Park and Alex Alderton, Roberto Amato, Jeffrey Barrett, Sonia Goncalves, Ewan Harrison, David K. Jackson, Ian Johnston, Dominic Kwiatkowski, Cordelia Langford, John Sillitoe on behalf of the Wellcome Sanger Institute COVID-19 Surveillance Team |
| EPI_ISL_2936877 | Lighthouse Lab in Milton Keynes | Wellcome Sanger Institute for the COVID-19 Genomics UK (COG-UK) Consortium | The Lighthouse Lab in Milton Keynes and Alex Alderton, Roberto Amato, Jeffrey Barrett, Sonia Goncalves, Ewan Harrison, David K. Jackson, Ian Johnston, Dominic Kwiatkowski, Cordelia Langford, John Sillitoe on behalf of the Wellcome Sanger Institute COVID-19 Surveillance Team |
| EPI_ISL_2936878 | Lighthouse Lab in Alderley Park | Wellcome Sanger Institute for the COVID-19 Genomics UK (COG-UK) Consortium | Jacquelyn Wynn, Mairead Hyland, The Lighthouse Lab in Alderley Park and Alex Alderton, Roberto Amato, Jeffrey Barrett, Sonia Goncalves, Ewan Harrison, David K. Jackson, Ian Johnston, Dominic Kwiatkowski, Cordelia Langford, John Sillitoe on behalf of the Wellcome Sanger Institute COVID-19 Surveillance Team |
| EPI_ISL_2936879, EPI_ISL_2936880, EPI_ISL_2936881 | Lighthouse Lab in Milton Keynes | Wellcome Sanger Institute for the COVID-19 Genomics UK (COG-UK) Consortium | The Lighthouse Lab in Milton Keynes and Alex Alderton, Roberto Amato, Jeffrey Barrett, Sonia Goncalves, Ewan Harrison, David K. Jackson, Ian Johnston, Dominic Kwiatkowski, Cordelia Langford, John Sillitoe on behalf of the Wellcome Sanger Institute COVID-19 Surveillance Team |
| EPI_ISL_2936882 | Lighthouse Lab in Alderley Park | Wellcome Sanger Institute for the COVID-19 Genomics UK (COG-UK) Consortium | Jacquelyn Wynn, Mairead Hyland, The Lighthouse Lab in Alderley Park and Alex Alderton, Roberto Amato, Jeffrey Barrett, Sonia Goncalves, Ewan Harrison, David K. Jackson, Ian Johnston, Dominic Kwiatkowski, Cordelia Langford, John Sillitoe on behalf of the Wellcome Sanger Institute COVID-19 Surveillance Team |
| EPI_ISL_2936883, EPI_ISL_2936884 | Lighthouse Lab in Milton Keynes | Wellcome Sanger Institute for the COVID-19 Genomics UK (COG-UK) Consortium | The Lighthouse Lab in Milton Keynes and Alex Alderton, Roberto Amato, Jeffrey Barrett, Sonia Goncalves, Ewan Harrison, David K. Jackson, Ian Johnston, Dominic Kwiatkowski, Cordelia Langford, John Sillitoe on behalf of the Wellcome Sanger Institute COVID-19 Surveillance Team |
| EPI_ISL_2936885, EPI_ISL_2936886 | Lighthouse Lab in Alderley Park | Wellcome Sanger Institute for the COVID-19 Genomics UK (COG-UK) Consortium | Jacquelyn Wynn, Mairead Hyland, The Lighthouse Lab in Alderley Park and Alex Alderton, Roberto Amato, Jeffrey Barrett, Sonia Goncalves, Ewan Harrison, David K. Jackson, Ian Johnston, Dominic Kwiatkowski, Cordelia Langford, John Sillitoe on behalf of the Wellcome Sanger Institute COVID-19 Surveillance Team |
| EPI_ISL_2936887, EPI_ISL_2936888, EPI_ISL_2936889 | Lighthouse Lab in Milton Keynes | Wellcome Sanger Institute for the COVID-19 Genomics UK (COG-UK) Consortium | The Lighthouse Lab in Milton Keynes and Alex Alderton, Roberto Amato, Jeffrey Barrett, Sonia Goncalves, Ewan Harrison, David K. Jackson, Ian Johnston, Dominic Kwiatkowski, Cordelia Langford, John Sillitoe on behalf of the Wellcome Sanger Institute COVID-19 Surveillance Team |
| EPI_ISL_2936890, EPI_ISL_2936891 | Lighthouse Lab in Alderley Park | Wellcome Sanger Institute for the COVID-19 Genomics UK (COG-UK) Consortium | Jacquelyn Wynn, Mairead Hyland, The Lighthouse Lab in Alderley Park and Alex Alderton, Roberto Amato, Jeffrey Barrett, Sonia Goncalves, Ewan Harrison, David K. Jackson, Ian Johnston, Dominic Kwiatkowski, Cordelia Langford, John Sillitoe on behalf of the Wellcome Sanger Institute COVID-19 Surveillance Team |
| EPI_ISL_2936892 | Lighthouse Lab in Glasgow | Wellcome Sanger Institute for the COVID-19 Genomics UK (COG-UK) Consortium | Harper VanSteenhouse, Yumi Kasai, David Gray, Carol Clugston, Anna Dominiczak and Alex Alderton, Roberto Amato, Jeffrey Barrett, Sonia Goncalves, Ewan Harrison, David K. Jackson, Ian Johnston, Dominic Kwiatkowski, Cordelia Langford, John Sillitoe on behalf of the Wellcome Sanger Institute COVID-19 Surveillance Team |
| EPI_ISL_2936893 | Lighthouse Lab in Alderley Park | Wellcome Sanger Institute for the COVID-19 Genomics UK (COG-UK) Consortium | Jacquelyn Wynn, Mairead Hyland, The Lighthouse Lab in Alderley Park and Alex Alderton, Roberto Amato, Jeffrey Barrett, Sonia Goncalves, Ewan Harrison, David K. Jackson, Ian Johnston, Dominic Kwiatkowski, Cordelia Langford, John Sillitoe on behalf of the Wellcome Sanger Institute COVID-19 Surveillance Team |
| EPI_ISL_2936894 | Lighthouse Lab in Milton Keynes | Wellcome Sanger Institute for the COVID-19 Genomics UK (COG-UK) Consortium | The Lighthouse Lab in Milton Keynes and Alex Alderton, Roberto Amato, Jeffrey Barrett, Sonia Goncalves, Ewan Harrison, David K. Jackson, Ian Johnston, Dominic Kwiatkowski, Cordelia Langford, John Sillitoe on behalf of the Wellcome Sanger Institute COVID-19 Surveillance Team |

| | | | |
|---|---|---|---|
| EPI_ISL_2936895, EPI_ISL_2936896 | Lighthouse Lab in Alderley Park | Wellcome Sanger Institute for the COVID-19 Genomics UK (COG-UK) Consortium | Jacquelyn Wynn, Mairead Hyland, The Lighthouse Lab in Alderley Park and Alex Alderton, Roberto Amato, Jeffrey Barrett, Sonia Goncalves, Ewan Harrison, David K. Jackson, Ian Johnston, Dominic Kwiatkowski, Cordelia Langford, John Sillitoe on behalf of the Wellcome Sanger Institute COVID-19 Surveillance Team |
| EPI_ISL_2936897, EPI_ISL_2936898, EPI_ISL_2936899, EPI_ISL_2936900, EPI_ISL_2936901 | Lighthouse Lab in Milton Keynes | Wellcome Sanger Institute for the COVID-19 Genomics UK (COG-UK) Consortium | The Lighthouse Lab in Milton Keynes and Alex Alderton, Roberto Amato, Jeffrey Barrett, Sonia Goncalves, Ewan Harrison, David K. Jackson, Ian Johnston, Dominic Kwiatkowski, Cordelia Langford, John Sillitoe on behalf of the Wellcome Sanger Institute COVID-19 Surveillance Team |
| EPI_ISL_2936902, EPI_ISL_2936903 | Lighthouse Lab in Alderley Park | Wellcome Sanger Institute for the COVID-19 Genomics UK (COG-UK) Consortium | Jacquelyn Wynn, Mairead Hyland, The Lighthouse Lab in Alderley Park and Alex Alderton, Roberto Amato, Jeffrey Barrett, Sonia Goncalves, Ewan Harrison, David K. Jackson, Ian Johnston, Dominic Kwiatkowski, Cordelia Langford, John Sillitoe on behalf of the Wellcome Sanger Institute COVID-19 Surveillance Team |
| EPI_ISL_2936904 | Lighthouse Lab in Milton Keynes | Wellcome Sanger Institute for the COVID-19 Genomics UK (COG-UK) Consortium | The Lighthouse Lab in Milton Keynes and Alex Alderton, Roberto Amato, Jeffrey Barrett, Sonia Goncalves, Ewan Harrison, David K. Jackson, Ian Johnston, Dominic Kwiatkowski, Cordelia Langford, John Sillitoe on behalf of the Wellcome Sanger Institute COVID-19 Surveillance Team |
| EPI_ISL_2936905 | Lighthouse Lab in Alderley Park | Wellcome Sanger Institute for the COVID-19 Genomics UK (COG-UK) Consortium | Jacquelyn Wynn, Mairead Hyland, The Lighthouse Lab in Alderley Park and Alex Alderton, Roberto Amato, Jeffrey Barrett, Sonia Goncalves, Ewan Harrison, David K. Jackson, Ian Johnston, Dominic Kwiatkowski, Cordelia Langford, John Sillitoe on behalf of the Wellcome Sanger Institute COVID-19 Surveillance Team |
| EPI_ISL_2936906, EPI_ISL_2936907, EPI_ISL_2936908, EPI_ISL_2936909, EPI_ISL_2936910 | Lighthouse Lab in Milton Keynes | Wellcome Sanger Institute for the COVID-19 Genomics UK (COG-UK) Consortium | The Lighthouse Lab in Milton Keynes and Alex Alderton, Roberto Amato, Jeffrey Barrett, Sonia Goncalves, Ewan Harrison, David K. Jackson, Ian Johnston, Dominic Kwiatkowski, Cordelia Langford, John Sillitoe on behalf of the Wellcome Sanger Institute COVID-19 Surveillance Team |
| EPI_ISL_2936911 | Lighthouse Lab in Alderley Park | Wellcome Sanger Institute for the COVID-19 Genomics UK (COG-UK) Consortium | Jacquelyn Wynn, Mairead Hyland, The Lighthouse Lab in Alderley Park and Alex Alderton, Roberto Amato, Jeffrey Barrett, Sonia Goncalves, Ewan Harrison, David K. Jackson, Ian Johnston, Dominic Kwiatkowski, Cordelia Langford, John Sillitoe on behalf of the Wellcome Sanger Institute COVID-19 Surveillance Team |
| EPI_ISL_2936912 | Lighthouse Lab in Milton Keynes | Wellcome Sanger Institute for the COVID-19 Genomics UK (COG-UK) Consortium | The Lighthouse Lab in Milton Keynes and Alex Alderton, Roberto Amato, Jeffrey Barrett, Sonia Goncalves, Ewan Harrison, David K. Jackson, Ian Johnston, Dominic Kwiatkowski, Cordelia Langford, John Sillitoe on behalf of the Wellcome Sanger Institute COVID-19 Surveillance Team |
| EPI_ISL_2936913 | Lighthouse Lab in Alderley Park | Wellcome Sanger Institute for the COVID-19 Genomics UK (COG-UK) Consortium | Jacquelyn Wynn, Mairead Hyland, The Lighthouse Lab in Alderley Park and Alex Alderton, Roberto Amato, Jeffrey Barrett, Sonia Goncalves, Ewan Harrison, David K. Jackson, Ian Johnston, Dominic Kwiatkowski, Cordelia Langford, John Sillitoe on behalf of the Wellcome Sanger Institute COVID-19 Surveillance Team |
| EPI_ISL_2936914 | Lighthouse Lab in Milton Keynes | Wellcome Sanger Institute for the COVID-19 Genomics UK (COG-UK) Consortium | The Lighthouse Lab in Milton Keynes and Alex Alderton, Roberto Amato, Jeffrey Barrett, Sonia Goncalves, Ewan Harrison, David K. Jackson, Ian Johnston, Dominic Kwiatkowski, Cordelia Langford, John Sillitoe on behalf of the Wellcome Sanger Institute COVID-19 Surveillance Team |
| EPI_ISL_2936915, EPI_ISL_2936916, EPI_ISL_2936917, EPI_ISL_2936918 | Lighthouse Lab in Alderley Park | Wellcome Sanger Institute for the COVID-19 Genomics UK (COG-UK) Consortium | Jacquelyn Wynn, Mairead Hyland, The Lighthouse Lab in Alderley Park and Alex Alderton, Roberto Amato, Jeffrey Barrett, Sonia Goncalves, Ewan Harrison, David K. Jackson, Ian Johnston, Dominic Kwiatkowski, Cordelia Langford, John Sillitoe on behalf of the Wellcome Sanger Institute COVID-19 Surveillance Team |
| EPI_ISL_2936919, EPI_ISL_2936920, EPI_ISL_2936921, EPI_ISL_2936922, EPI_ISL_2936923, EPI_ISL_2936924, EPI_ISL_2936925, EPI_ISL_2936926, EPI_ISL_2936927, EPI_ISL_2936928, EPI_ISL_2936929, EPI_ISL_2936930, EPI_ISL_2936931, EPI_ISL_2936932, EPI_ISL_2936933, EPI_ISL_2936934, EPI_ISL_2936935, EPI_ISL_2936936, EPI_ISL_2936937, EPI_ISL_2936938, EPI_ISL_2936939, EPI_ISL_2936940, EPI_ISL_2936941, EPI_ISL_2936942, EPI_ISL_2936943, EPI_ISL_2936944, EPI_ISL_2936945, EPI_ISL_2936946, EPI_ISL_2936947, EPI_ISL_2936948, EPI_ISL_2936949, EPI_ISL_2936950, EPI_ISL_2936951, EPI_ISL_2936952, EPI_ISL_2936953, EPI_ISL_2936954, EPI_ISL_2936955, EPI_ISL_2936956, EPI_ISL_2936957, EPI_ISL_2936958, EPI_ISL_2936959, EPI_ISL_2936960, EPI_ISL_2936961, EPI_ISL_2936962, EPI_ISL_2936963, EPI_ISL_2936964, EPI_ISL_2936965, EPI_ISL_2936966, EPI_ISL_2936967, EPI_ISL_2936968, EPI_ISL_2936969, EPI_ISL_2936970, EPI_ISL_2936971, EPI_ISL_2936972, EPI_ISL_2936973, EPI_ISL_2936974, EPI_ISL_2936975, EPI_ISL_2936976, EPI_ISL_2936977, EPI_ISL_2936978, EPI_ISL_2936979 | | | |
| see above | Lighthouse Lab in Glasgow | Wellcome Sanger Institute for the COVID-19 Genomics UK (COG-UK) Consortium | Harper VanSteenhouse, Yumi Kasai, David Gray, Carol Clugston, Anna Dominiczak and Alex Alderton, Roberto Amato, Jeffrey Barrett, Sonia Goncalves, Ewan Harrison, David K. Jackson, Ian Johnston, Dominic Kwiatkowski, Cordelia Langford, John Sillitoe on behalf of the Wellcome Sanger Institute COVID-19 Surveillance Team |
| EPI_ISL_2936980 | Lighthouse Lab in Alderley Park | Wellcome Sanger Institute for the COVID-19 Genomics UK (COG-UK) Consortium | Jacquelyn Wynn, Mairead Hyland, The Lighthouse Lab in Alderley Park and Alex Alderton, Roberto Amato, Jeffrey Barrett, Sonia Goncalves, Ewan Harrison, David K. Jackson, Ian Johnston, Dominic Kwiatkowski, Cordelia Langford, John Sillitoe on behalf of the Wellcome Sanger Institute COVID-19 Surveillance Team |
| EPI_ISL_2936981, EPI_ISL_2936982, EPI_ISL_2936983, EPI_ISL_2936984, EPI_ISL_2936985, EPI_ISL_2936986, EPI_ISL_2936988, EPI_ISL_2936989, EPI_ISL_2936990, EPI_ISL_2936991, EPI_ISL_2936992 | | | |
| see above | Lighthouse Lab in Glasgow | Wellcome Sanger Institute for the COVID-19 Genomics UK (COG-UK) Consortium | Harper VanSteenhouse, Yumi Kasai, David Gray, Carol Clugston, Anna Dominiczak and Alex Alderton, Roberto Amato, Jeffrey Barrett, Sonia Goncalves, Ewan Harrison, David K. Jackson, Ian Johnston, Dominic Kwiatkowski, Cordelia Langford, John Sillitoe on behalf of the Wellcome Sanger Institute COVID-19 Surveillance Team |
| EPI_ISL_2936993, EPI_ISL_2936994, EPI_ISL_2936995, EPI_ISL_2936996, EPI_ISL_2936997 | Lighthouse Lab in Alderley Park | Wellcome Sanger Institute for the COVID-19 Genomics UK (COG-UK) Consortium | Jacquelyn Wynn, Mairead Hyland, The Lighthouse Lab in Alderley Park and Alex Alderton, Roberto Amato, Jeffrey Barrett, Sonia Goncalves, Ewan Harrison, David K. Jackson, Ian Johnston, Dominic Kwiatkowski, Cordelia Langford, John Sillitoe on behalf of the Wellcome Sanger Institute COVID-19 Surveillance Team |
| EPI_ISL_2936998 | Lighthouse Lab in Milton Keynes | Wellcome Sanger Institute for the COVID-19 Genomics UK (COG-UK) Consortium | The Lighthouse Lab in Milton Keynes and Alex Alderton, Roberto Amato, Jeffrey Barrett, Sonia Goncalves, Ewan Harrison, David K. Jackson, Ian Johnston, Dominic Kwiatkowski, Cordelia Langford, John Sillitoe on behalf of the Wellcome Sanger Institute COVID-19 Surveillance Team |
| EPI_ISL_2936999, EPI_ISL_2937000 | Lighthouse Lab in Alderley Park | Wellcome Sanger Institute for the COVID-19 Genomics UK (COG-UK) Consortium | Jacquelyn Wynn, Mairead Hyland, The Lighthouse Lab in Alderley Park and Alex Alderton, Roberto Amato, Jeffrey Barrett, Sonia Goncalves, Ewan Harrison, David K. Jackson, Ian Johnston, Dominic Kwiatkowski, Cordelia Langford, John Sillitoe on behalf of the Wellcome Sanger Institute COVID-19 Surveillance Team |
| EPI_ISL_2937001 | Lighthouse Lab in Milton Keynes | Wellcome Sanger Institute for the COVID-19 Genomics UK (COG-UK) Consortium | The Lighthouse Lab in Milton Keynes and Alex Alderton, Roberto Amato, Jeffrey Barrett, Sonia Goncalves, Ewan Harrison, David K. Jackson, Ian Johnston, Dominic Kwiatkowski, Cordelia Langford, John Sillitoe on behalf of the Wellcome Sanger Institute COVID-19 Surveillance Team |
| EPI_ISL_2937002, EPI_ISL_2937003, EPI_ISL_2937004, EPI_ISL_2937005, EPI_ISL_2937006, EPI_ISL_2937007, EPI_ISL_2937008, EPI_ISL_2937009, EPI_ISL_2937010 | Lighthouse Lab in Glasgow | Wellcome Sanger Institute for the COVID-19 Genomics UK (COG-UK) Consortium | Harper VanSteenhouse, Yumi Kasai, David Gray, Carol Clugston, Anna Dominiczak and Alex Alderton, Roberto Amato, Jeffrey Barrett, Sonia Goncalves, Ewan Harrison, David K. Jackson, Ian Johnston, Dominic Kwiatkowski, Cordelia Langford, John Sillitoe on behalf of the Wellcome Sanger Institute COVID-19 Surveillance Team |
| EPI_ISL_2937011 | Lighthouse Lab in Alderley Park | Wellcome Sanger Institute for the COVID-19 Genomics UK (COG-UK) Consortium | Jacquelyn Wynn, Mairead Hyland, The Lighthouse Lab in Alderley Park and Alex Alderton, Roberto Amato, Jeffrey Barrett, Sonia Goncalves, Ewan Harrison, David K. Jackson, Ian Johnston, Dominic Kwiatkowski, Cordelia Langford, John Sillitoe on behalf of the Wellcome Sanger Institute COVID-19 Surveillance Team |
| EPI_ISL_2937012, EPI_ISL_2937013, EPI_ISL_2937014, EPI_ISL_2937015, EPI_ISL_2937016, EPI_ISL_2937017, EPI_ISL_2937018 | Lighthouse Lab in Glasgow | Wellcome Sanger Institute for the COVID-19 Genomics UK (COG-UK) Consortium | Harper VanSteenhouse, Yumi Kasai, David Gray, Carol Clugston, Anna Dominiczak and Alex Alderton, Roberto Amato, Jeffrey Barrett, Sonia Goncalves, Ewan Harrison, David K. Jackson, Ian Johnston, Dominic Kwiatkowski, Cordelia Langford, John Sillitoe on behalf of the Wellcome Sanger Institute COVID-19 Surveillance Team |
| EPI_ISL_2937019 | Lighthouse Lab in Alderley Park | Wellcome Sanger Institute for the COVID-19 Genomics UK (COG-UK) Consortium | Jacquelyn Wynn, Mairead Hyland, The Lighthouse Lab in Alderley Park and Alex Alderton, Roberto Amato, Jeffrey Barrett, Sonia Goncalves, Ewan Harrison, David K. Jackson, Ian Johnston, Dominic Kwiatkowski, Cordelia Langford, John Sillitoe on behalf of the Wellcome Sanger Institute COVID-19 Surveillance Team |
| EPI_ISL_2937020 | Lighthouse Lab in Glasgow | Wellcome Sanger Institute for the COVID-19 Genomics UK (COG-UK) Consortium | Harper VanSteenhouse, Yumi Kasai, David Gray, Carol Clugston, Anna Dominiczak and Alex Alderton, Roberto Amato, Jeffrey Barrett, Sonia Goncalves, Ewan Harrison, David K. Jackson, Ian Johnston, Dominic Kwiatkowski, Cordelia Langford, John Sillitoe on behalf of the Wellcome Sanger Institute COVID-19 Surveillance Team |
| EPI_ISL_2937021 | Lighthouse Lab in Alderley Park | Wellcome Sanger Institute for the COVID-19 Genomics UK (COG-UK) Consortium | Jacquelyn Wynn, Mairead Hyland, The Lighthouse Lab in Alderley Park and Alex Alderton, Roberto Amato, Jeffrey Barrett, Sonia Goncalves, Ewan Harrison, David K. Jackson, Ian Johnston, Dominic Kwiatkowski, Cordelia Langford, John Sillitoe on behalf of the Wellcome Sanger Institute COVID-19 Surveillance Team |
| EPI_ISL_2937022, EPI_ISL_2937023, EPI_ISL_2937024, EPI_ISL_2937025, EPI_ISL_2937026, EPI_ISL_2937027, EPI_ISL_2937028, EPI_ISL_2937029, EPI_ISL_2937030, EPI_ISL_2937031, EPI_ISL_2937032, EPI_ISL_2937033, EPI_ISL_2937034, EPI_ISL_2937035, EPI_ISL_2937036, EPI_ISL_2937037, EPI_ISL_2937038, EPI_ISL_2937039, EPI_ISL_2937040, EPI_ISL_2937041, EPI_ISL_2937042, EPI_ISL_2937043, EPI_ISL_2937044, EPI_ISL_2937045, EPI_ISL_2937046, EPI_ISL_2937047, EPI_ISL_2937048, EPI_ISL_2937049, EPI_ISL_2937050, EPI_ISL_2937051, EPI_ISL_2937052, EPI_ISL_2937053, EPI_ISL_2937054, EPI_ISL_2937055, EPI_ISL_2937056, EPI_ISL_2937057, EPI_ISL_2937058, EPI_ISL_2937059, EPI_ISL_2937060, EPI_ISL_2937061, EPI_ISL_2937062, EPI_ISL_2937063, EPI_ISL_2937064, EPI_ISL_2937065, EPI_ISL_2937066, EPI_ISL_2937067, EPI_ISL_2937068, EPI_ISL_2937069, EPI_ISL_2937070, EPI_ISL_2937071, EPI_ISL_2937072, EPI_ISL_2937073, EPI_ISL_2937074, EPI_ISL_2937075, EPI_ISL_2937076, EPI_ISL_2937077, EPI_ISL_2937078, EPI_ISL_2937079 | | | |
| see above | Lighthouse Lab in Glasgow | Wellcome Sanger Institute for the COVID-19 Genomics UK (COG-UK) Consortium | Harper VanSteenhouse, Yumi Kasai, David Gray, Carol Clugston, Anna Dominiczak and Alex Alderton, Roberto Amato, Jeffrey Barrett, Sonia Goncalves, Ewan Harrison, David K. Jackson, Ian Johnston, Dominic Kwiatkowski, Cordelia Langford, John Sillitoe on behalf of the Wellcome Sanger Institute COVID-19 Surveillance Team |
| EPI_ISL_2937080 | Lighthouse Lab in Alderley Park | Wellcome Sanger Institute for the COVID-19 Genomics UK (COG-UK) Consortium | Jacquelyn Wynn, Mairead Hyland, The Lighthouse Lab in Alderley Park and Alex Alderton, Roberto Amato, Jeffrey Barrett, Sonia Goncalves, Ewan Harrison, David K. Jackson, Ian Johnston, Dominic Kwiatkowski, Cordelia Langford, John Sillitoe on behalf of the Wellcome Sanger Institute COVID-19 Surveillance Team |
| EPI_ISL_2937081, EPI_ISL_2937082, EPI_ISL_2937083, EPI_ISL_2937084, EPI_ISL_2937085, EPI_ISL_2937086, EPI_ISL_2937087, EPI_ISL_2937088, EPI_ISL_2937089, EPI_ISL_2937090, EPI_ISL_2937091, EPI_ISL_2937092, EPI_ISL_2937093, EPI_ISL_2937094, EPI_ISL_2937095, EPI_ISL_2937096, EPI_ISL_2937097, EPI_ISL_2937098, EPI_ISL_2937099, EPI_ISL_2937100, EPI_ISL_2937101, EPI_ISL_2937102, EPI_ISL_2937103, EPI_ISL_2937104, EPI_ISL_2937105, EPI_ISL_2937106, EPI_ISL_2937107, EPI_ISL_2937108, EPI_ISL_2937109, EPI_ISL_2937110, EPI_ISL_2937111, EPI_ISL_2937113, EPI_ISL_2937114, EPI_ISL_2937115, EPI_ISL_2937116, EPI_ISL_2937117, EPI_ISL_2937118, EPI_ISL_2937119, EPI_ISL_2937120, EPI_ISL_2937121, EPI_ISL_2937122, EPI_ISL_2937123, EPI_ISL_2937124, EPI_ISL_2937125, EPI_ISL_2937126, EPI_ISL_2937127, EPI_ISL_2937128, EPI_ISL_2937129, EPI_ISL_2937130, EPI_ISL_2937131, EPI_ISL_2937132, EPI_ISL_2937133, EPI_ISL_2937134, EPI_ISL_2937135, EPI_ISL_2937136, EPI_ISL_2937137, EPI_ISL_2937138, EPI_ISL_2937139, EPI_ISL_2937140, EPI_ISL_2937141, EPI_ISL_2937142, EPI_ISL_2937143, EPI_ISL_2937144, EPI_ISL_2937145, EPI_ISL_2937146, EPI_ISL_2937147, EPI_ISL_2937148, EPI_ISL_2937149, EPI_ISL_2937150, EPI_ISL_2937151, EPI_ISL_2937152, EPI_ISL_2937153, EPI_ISL_2937154, EPI_ISL_2937155, EPI_ISL_2937156 | | | |
| see above | Lighthouse Lab in Glasgow | Wellcome Sanger Institute for the COVID-19 Genomics UK (COG-UK) Consortium | Harper VanSteenhouse, Yumi Kasai, David Gray, Carol Clugston, Anna Dominiczak and Alex Alderton, Roberto Amato, Jeffrey Barrett, Sonia Goncalves, Ewan Harrison, David K. Jackson, Ian Johnston, Dominic Kwiatkowski, Cordelia Langford, John Sillitoe on behalf of the Wellcome Sanger Institute COVID-19 Surveillance Team |
| EPI_ISL_2937157, EPI_ISL_2937158, EPI_ISL_2937159, EPI_ISL_2937160 | Lighthouse Lab in Alderley Park | Wellcome Sanger Institute for the COVID-19 Genomics UK (COG-UK) Consortium | Jacquelyn Wynn, Mairead Hyland, The Lighthouse Lab in Alderley Park and Alex Alderton, Roberto Amato, Jeffrey Barrett, Sonia Goncalves, Ewan Harrison, David K. Jackson, Ian Johnston, Dominic Kwiatkowski, Cordelia Langford, John Sillitoe on behalf of the Wellcome Sanger Institute COVID-19 Surveillance Team |
| EPI_ISL_2937161, EPI_ISL_2937162 | Lighthouse Laboratory Plymouth | Wellcome Sanger Institute for the COVID-19 Genomics UK (COG-UK) Consortium | Lighthouse Laboratory Plymouth and Alex Alderton, Roberto Amato, Jeffrey Barrett, Sonia Goncalves, Ewan Harrison, David K. Jackson, Ian Johnston, Dominic Kwiatkowski, Cordelia Langford, John Sillitoe on behalf of the Wellcome Sanger Institute COVID-19 Surveillance Team |
| EPI_ISL_2937163, EPI_ISL_2937164, EPI_ISL_2937165, EPI_ISL_2937166, EPI_ISL_2937167, EPI_ISL_2937168, EPI_ISL_2937169, EPI_ISL_2937170, EPI_ISL_2937171, EPI_ISL_2937172, EPI_ISL_2937173, EPI_ISL_2937174, EPI_ISL_2937175, EPI_ISL_2937176, EPI_ISL_2937177, EPI_ISL_2937178, EPI_ISL_2937179, EPI_ISL_2937180, EPI_ISL_2937181, EPI_ISL_2937182, EPI_ISL_2937183, EPI_ISL_2937184, | | | |

| | | | |
|---|---|---|---|
| EPI_ISL_2937185, EPI_ISL_2937186, EPI_ISL_2937187, EPI_ISL_2937188, EPI_ISL_2937189, EPI_ISL_2937190, EPI_ISL_2937191, EPI_ISL_2937192, EPI_ISL_2937193, EPI_ISL_2937194, EPI_ISL_2937195, EPI_ISL_2937196, EPI_ISL_2937197 | | | |
| see above | Lighthouse Lab in Glasgow | Wellcome Sanger Institute for the COVID-19 Genomics UK (COG-UK) Consortium | Harper VanSteenhouse, Yumi Kasai, David Gray, Carol Clugston, Anna Dominiczak and Alex Alderton, Roberto Amato, Jeffrey Barrett, Sonia Goncalves, Ewan Harrison, David K. Jackson, Ian Johnston, Dominic Kwiatkowski, Cordelia Langford, John Sillitoe on behalf of the Wellcome Sanger Institute COVID-19 Surveillance Team |
| EPI_ISL_2937198 | Lighthouse Lab in Milton Keynes | Wellcome Sanger Institute for the COVID-19 Genomics UK (COG-UK) Consortium | The Lighthouse Lab in Milton Keynes and Alex Alderton, Roberto Amato, Jeffrey Barrett, Sonia Goncalves, Ewan Harrison, David K. Jackson, Ian Johnston, Dominic Kwiatkowski, Cordelia Langford, John Sillitoe on behalf of the Wellcome Sanger Institute COVID-19 Surveillance Team |
| EPI_ISL_2937199, EPI_ISL_2937200, EPI_ISL_2937201, EPI_ISL_2937202, EPI_ISL_2937203, EPI_ISL_2937204, EPI_ISL_2937205, EPI_ISL_2937206, EPI_ISL_2937207, EPI_ISL_2937208, EPI_ISL_2937209, EPI_ISL_2937210, EPI_ISL_2937211, EPI_ISL_2937212, EPI_ISL_2937213, EPI_ISL_2937214, EPI_ISL_2937215, EPI_ISL_2937216, EPI_ISL_2937217, EPI_ISL_2937218, EPI_ISL_2937219, EPI_ISL_2937220, EPI_ISL_2937221, EPI_ISL_2937222, EPI_ISL_2937223, EPI_ISL_2937224, EPI_ISL_2937225, EPI_ISL_2937226, EPI_ISL_2937227, EPI_ISL_2937228, EPI_ISL_2937229, EPI_ISL_2937230, EPI_ISL_2937231, EPI_ISL_2937232, EPI_ISL_2937233, EPI_ISL_2937234, EPI_ISL_2937235, EPI_ISL_2937236, EPI_ISL_2937237, EPI_ISL_2937238, EPI_ISL_2937239, EPI_ISL_2937240, EPI_ISL_2937241, EPI_ISL_2937242, EPI_ISL_2937243, EPI_ISL_2937244, EPI_ISL_2937245, EPI_ISL_2937246, EPI_ISL_2937247, EPI_ISL_2937248, EPI_ISL_2937249, EPI_ISL_2937250, EPI_ISL_2937251, EPI_ISL_2937252, EPI_ISL_2937253, EPI_ISL_2937254, EPI_ISL_2937255, EPI_ISL_2937256, EPI_ISL_2937257, EPI_ISL_2937258, EPI_ISL_2937259, EPI_ISL_2937260, EPI_ISL_2937261, EPI_ISL_2937262, EPI_ISL_2937263, EPI_ISL_2937264, EPI_ISL_2937265, EPI_ISL_2937266, EPI_ISL_2937267, EPI_ISL_2937268, EPI_ISL_2937269, EPI_ISL_2937270, EPI_ISL_2937271, EPI_ISL_2937272, EPI_ISL_2937273, EPI_ISL_2937274, EPI_ISL_2937275, EPI_ISL_2937276, EPI_ISL_2937277, EPI_ISL_2937278, EPI_ISL_2937279, EPI_ISL_2937280, EPI_ISL_2937281, EPI_ISL_2937282, EPI_ISL_2937283, EPI_ISL_2937284, EPI_ISL_2937285, EPI_ISL_2937286, EPI_ISL_2937287, EPI_ISL_2937288, EPI_ISL_2937289, EPI_ISL_2937290, EPI_ISL_2937291, EPI_ISL_2937292, EPI_ISL_2937293, EPI_ISL_2937294, EPI_ISL_2937295, EPI_ISL_2937296, EPI_ISL_2937297, EPI_ISL_2937299, EPI_ISL_2937300, EPI_ISL_2937301, EPI_ISL_2937302, EPI_ISL_2937303, EPI_ISL_2937304, EPI_ISL_2937305, EPI_ISL_2937306, EPI_ISL_2937307, EPI_ISL_2937308, EPI_ISL_2937309, EPI_ISL_2937310, EPI_ISL_2937311, EPI_ISL_2937312, EPI_ISL_2937313, EPI_ISL_2937314, EPI_ISL_2937315, EPI_ISL_2937316, EPI_ISL_2937317, EPI_ISL_2937318, EPI_ISL_2937319, EPI_ISL_2937320, EPI_ISL_2937321, EPI_ISL_2937322, EPI_ISL_2937323, EPI_ISL_2937324, EPI_ISL_2937325, EPI_ISL_2937326, EPI_ISL_2937327, EPI_ISL_2937328, EPI_ISL_2937329, EPI_ISL_2937330, EPI_ISL_2937331, EPI_ISL_2937332, EPI_ISL_2937333, EPI_ISL_2937334, EPI_ISL_2937335, EPI_ISL_2937336, EPI_ISL_2937337, EPI_ISL_2937338, EPI_ISL_2937339, EPI_ISL_2937340, EPI_ISL_2937341, EPI_ISL_2937342, EPI_ISL_2937343, EPI_ISL_2937344, EPI_ISL_2937345, EPI_ISL_2937346, EPI_ISL_2937347, EPI_ISL_2937348, EPI_ISL_2937349, EPI_ISL_2937350, EPI_ISL_2937351, EPI_ISL_2937352, EPI_ISL_2937353, EPI_ISL_2937354, EPI_ISL_2937355, EPI_ISL_2937356, EPI_ISL_2937357, EPI_ISL_2937358, EPI_ISL_2937359, EPI_ISL_2937360, EPI_ISL_2937361, EPI_ISL_2937362, EPI_ISL_2937363, EPI_ISL_2937364, EPI_ISL_2937365, EPI_ISL_2937366, EPI_ISL_2937367, EPI_ISL_2937368, EPI_ISL_2937369, EPI_ISL_2937370, EPI_ISL_2937371, EPI_ISL_2937372, EPI_ISL_2937373, EPI_ISL_2937374, EPI_ISL_2937375, EPI_ISL_2937376, EPI_ISL_2937377, EPI_ISL_2937378, EPI_ISL_2937379, EPI_ISL_2937380, EPI_ISL_2937381, EPI_ISL_2937382, EPI_ISL_2937383, EPI_ISL_2937384, EPI_ISL_2937385, EPI_ISL_2937386, EPI_ISL_2937387, EPI_ISL_2937388, EPI_ISL_2937389, EPI_ISL_2937390, EPI_ISL_2937391, EPI_ISL_2937392, EPI_ISL_2937393, EPI_ISL_2937394, EPI_ISL_2937395, EPI_ISL_2937396, EPI_ISL_2937397, EPI_ISL_2937398, EPI_ISL_2937399, EPI_ISL_2937400, EPI_ISL_2937401, EPI_ISL_2937402, EPI_ISL_2937403, EPI_ISL_2937404, EPI_ISL_2937405, EPI_ISL_2937406, EPI_ISL_2937407, EPI_ISL_2937408, EPI_ISL_2937409, EPI_ISL_2937410, EPI_ISL_2937411, EPI_ISL_2937412, EPI_ISL_2937413, EPI_ISL_2937414, EPI_ISL_2937415, EPI_ISL_2937416, EPI_ISL_2937417, EPI_ISL_2937418, EPI_ISL_2937419, EPI_ISL_2937420, EPI_ISL_2937421, EPI_ISL_2937422, EPI_ISL_2937423, EPI_ISL_2937424, EPI_ISL_2937425, EPI_ISL_2937426, EPI_ISL_2937427, EPI_ISL_2937428, EPI_ISL_2937429, EPI_ISL_2937430, EPI_ISL_2937432, EPI_ISL_2937433, EPI_ISL_2937434, EPI_ISL_2937435, EPI_ISL_2937436, EPI_ISL_2937437, EPI_ISL_2937438, EPI_ISL_2937439, EPI_ISL_2937440, EPI_ISL_2937441, EPI_ISL_2937442, EPI_ISL_2937443, EPI_ISL_2937444, EPI_ISL_2937445, EPI_ISL_2937446, EPI_ISL_2937447, EPI_ISL_2937448, EPI_ISL_2937449, EPI_ISL_2937450, EPI_ISL_2937451, EPI_ISL_2937452, EPI_ISL_2937453, EPI_ISL_2937454, EPI_ISL_2937455, EPI_ISL_2937456, EPI_ISL_2937457, EPI_ISL_2937458, EPI_ISL_2937459, EPI_ISL_2937460, EPI_ISL_2937461, EPI_ISL_2937462, EPI_ISL_2937463, EPI_ISL_2937464, EPI_ISL_2937465, EPI_ISL_2937466, EPI_ISL_2937467, EPI_ISL_2937468, EPI_ISL_2937469, EPI_ISL_2937470, EPI_ISL_2937471, EPI_ISL_2937472, EPI_ISL_2937473, EPI_ISL_2937474, EPI_ISL_2937475, EPI_ISL_2937476, EPI_ISL_2937477, EPI_ISL_2937478, EPI_ISL_2937479, EPI_ISL_2937480, EPI_ISL_2937481, EPI_ISL_2937482, EPI_ISL_2937483, EPI_ISL_2937484, EPI_ISL_2937485, EPI_ISL_2937486, EPI_ISL_2937487, EPI_ISL_2937488, EPI_ISL_2937489, EPI_ISL_2937490, EPI_ISL_2937491, EPI_ISL_2937492, EPI_ISL_2937493, EPI_ISL_2937494, EPI_ISL_2937495, EPI_ISL_2937496, EPI_ISL_2937497, EPI_ISL_2937498, EPI_ISL_2937499, EPI_ISL_2937500, EPI_ISL_2937501, EPI_ISL_2937502, EPI_ISL_2937503, EPI_ISL_2937504, EPI_ISL_2937505, EPI_ISL_2937506, EPI_ISL_2937507, EPI_ISL_2937508, EPI_ISL_2937509, EPI_ISL_2937510, EPI_ISL_2937511, EPI_ISL_2937512, EPI_ISL_2937513, EPI_ISL_2937514, EPI_ISL_2937515, EPI_ISL_2937516, EPI_ISL_2937517, EPI_ISL_2937518, EPI_ISL_2937519, EPI_ISL_2937520, EPI_ISL_2937521, EPI_ISL_2937522, EPI_ISL_2937523, EPI_ISL_2937524, EPI_ISL_2937525, EPI_ISL_2937526, EPI_ISL_2937527, EPI_ISL_2937528, EPI_ISL_2937529, EPI_ISL_2937530, EPI_ISL_2937531, EPI_ISL_2937532, EPI_ISL_2937533, EPI_ISL_2937534, EPI_ISL_2937535, EPI_ISL_2937536, EPI_ISL_2937537, EPI_ISL_2937538, EPI_ISL_2937539, EPI_ISL_2937540, EPI_ISL_2937541, EPI_ISL_2937542, EPI_ISL_2937543, EPI_ISL_2937544, EPI_ISL_2937545, EPI_ISL_2937546, EPI_ISL_2937547, EPI_ISL_2937548, EPI_ISL_2937549, EPI_ISL_2937550, EPI_ISL_2937551, EPI_ISL_2937552, EPI_ISL_2937553, EPI_ISL_2937554, EPI_ISL_2937555, EPI_ISL_2937556, EPI_ISL_2937557, EPI_ISL_2937558, EPI_ISL_2937559, EPI_ISL_2937560, EPI_ISL_2937561, EPI_ISL_2937562, EPI_ISL_2937563, EPI_ISL_2937564, EPI_ISL_2937565, EPI_ISL_2937566, EPI_ISL_2937567, EPI_ISL_2937568, EPI_ISL_2937569, EPI_ISL_2937570, EPI_ISL_2937571, EPI_ISL_2937572, EPI_ISL_2937573, EPI_ISL_2937574, EPI_ISL_2937575, EPI_ISL_2937576, EPI_ISL_2937577, EPI_ISL_2937578, EPI_ISL_2937579, EPI_ISL_2937580, EPI_ISL_2937581, EPI_ISL_2937582, EPI_ISL_2937583, EPI_ISL_2937584, EPI_ISL_2937586, EPI_ISL_2937587, EPI_ISL_2937588, EPI_ISL_2937589, EPI_ISL_2937590, EPI_ISL_2937591, EPI_ISL_2937592, EPI_ISL_2937593, EPI_ISL_2937594, EPI_ISL_2937595, EPI_ISL_2937596, EPI_ISL_2937597, EPI_ISL_2937598, EPI_ISL_2937599, EPI_ISL_2937600, EPI_ISL_2937601, EPI_ISL_2937602, EPI_ISL_2937603, EPI_ISL_2937604, EPI_ISL_2937605, EPI_ISL_2937606, EPI_ISL_2937607, EPI_ISL_2937608, EPI_ISL_2937609, EPI_ISL_2937610, EPI_ISL_2937611, EPI_ISL_2937612, EPI_ISL_2937613, EPI_ISL_2937614, EPI_ISL_2937615, EPI_ISL_2937616, EPI_ISL_2937617, EPI_ISL_2937618, EPI_ISL_2937619, EPI_ISL_2937620, EPI_ISL_2937621, EPI_ISL_2937622, EPI_ISL_2937623, EPI_ISL_2937624, EPI_ISL_2937625, EPI_ISL_2937626, EPI_ISL_2937627, EPI_ISL_2937628, EPI_ISL_2937629, EPI_ISL_2937630, EPI_ISL_2937631, EPI_ISL_2937632, EPI_ISL_2937633, EPI_ISL_2937634, EPI_ISL_2937635, EPI_ISL_2937636, EPI_ISL_2937637, EPI_ISL_2937638, EPI_ISL_2937639, EPI_ISL_2937640 | | | |
| see above | Lighthouse Lab in Glasgow | Wellcome Sanger Institute for the COVID-19 Genomics UK (COG-UK) Consortium | Harper VanSteenhouse, Yumi Kasai, David Gray, Carol Clugston, Anna Dominiczak and Alex Alderton, Roberto Amato, Jeffrey Barrett, Sonia Goncalves, Ewan Harrison, David K. Jackson, Ian Johnston, Dominic Kwiatkowski, Cordelia Langford, John Sillitoe on behalf of the Wellcome Sanger Institute COVID-19 Surveillance Team |
| EPI_ISL_2937641 | Lighthouse Lab in Alderley Park | Wellcome Sanger Institute for the COVID-19 Genomics UK (COG-UK) Consortium | Jacquelyn Wynn, Mairead Hyland, The Lighthouse Lab in Alderley Park and Alex Alderton, Roberto Amato, Jeffrey Barrett, Sonia Goncalves, Ewan Harrison, David K. Jackson, Ian Johnston, Dominic Kwiatkowski, Cordelia Langford, John Sillitoe on behalf of the Wellcome Sanger Institute COVID-19 Surveillance Team |
| EPI_ISL_2937642, EPI_ISL_2937643, EPI_ISL_2937644, EPI_ISL_2937645, EPI_ISL_2937646, EPI_ISL_2937647, EPI_ISL_2937648, EPI_ISL_2937649, EPI_ISL_2937650, EPI_ISL_2937651, EPI_ISL_2937652, EPI_ISL_2937653, EPI_ISL_2937654, EPI_ISL_2937655, EPI_ISL_2937656, EPI_ISL_2937657, EPI_ISL_2937658, EPI_ISL_2937659, EPI_ISL_2937660, EPI_ISL_2937661, EPI_ISL_2937662, EPI_ISL_2937663, EPI_ISL_2937664, EPI_ISL_2937665, EPI_ISL_2937666, EPI_ISL_2937667, EPI_ISL_2937668, EPI_ISL_2937669, EPI_ISL_2937670, EPI_ISL_2937671, EPI_ISL_2937672, EPI_ISL_2937673, EPI_ISL_2937674, EPI_ISL_2937675, EPI_ISL_2937676, EPI_ISL_2937677, EPI_ISL_2937678, EPI_ISL_2937679, EPI_ISL_2937680, EPI_ISL_2937681 | | | |
| see above | Lighthouse Lab in Glasgow | Wellcome Sanger Institute for the COVID-19 Genomics UK (COG-UK) Consortium | Harper VanSteenhouse, Yumi Kasai, David Gray, Carol Clugston, Anna Dominiczak and Alex Alderton, Roberto Amato, Jeffrey Barrett, Sonia Goncalves, Ewan Harrison, David K. Jackson, Ian Johnston, Dominic Kwiatkowski, Cordelia Langford, John Sillitoe on behalf of the Wellcome Sanger Institute COVID-19 Surveillance Team |
| EPI_ISL_2937682 | Lighthouse Lab in Milton Keynes | Wellcome Sanger Institute for the COVID-19 Genomics UK (COG-UK) Consortium | The Lighthouse Lab in Milton Keynes and Alex Alderton, Roberto Amato, Jeffrey Barrett, Sonia Goncalves, Ewan Harrison, David K. Jackson, Ian Johnston, Dominic Kwiatkowski, Cordelia Langford, John Sillitoe on behalf of the Wellcome Sanger Institute COVID-19 Surveillance Team |
| EPI_ISL_2937683, EPI_ISL_2937684, EPI_ISL_2937685, EPI_ISL_2937686, EPI_ISL_2937687, EPI_ISL_2937688, EPI_ISL_2937689, EPI_ISL_2937690, EPI_ISL_2937691, EPI_ISL_2937692, EPI_ISL_2937693, EPI_ISL_2937694, EPI_ISL_2937695, EPI_ISL_2937696, EPI_ISL_2937697, EPI_ISL_2937698, EPI_ISL_2937699, EPI_ISL_2937700, EPI_ISL_2937701, EPI_ISL_2937702, EPI_ISL_2937703, EPI_ISL_2937704, EPI_ISL_2937705, EPI_ISL_2937706, EPI_ISL_2937707, EPI_ISL_2937708, EPI_ISL_2937709, EPI_ISL_2937710, EPI_ISL_2937711, EPI_ISL_2937712, EPI_ISL_2937713 | | | |
| see above | Lighthouse Lab in Glasgow | Wellcome Sanger Institute for the COVID-19 Genomics UK (COG-UK) Consortium | Harper VanSteenhouse, Yumi Kasai, David Gray, Carol Clugston, Anna Dominiczak and Alex Alderton, Roberto Amato, Jeffrey Barrett, Sonia Goncalves, Ewan Harrison, David K. Jackson, Ian Johnston, Dominic Kwiatkowski, Cordelia Langford, John Sillitoe on behalf of the Wellcome Sanger Institute COVID-19 Surveillance Team |
| EPI_ISL_2937714 | Lighthouse Lab in Milton Keynes | Wellcome Sanger Institute for the COVID-19 Genomics UK (COG-UK) Consortium | The Lighthouse Lab in Milton Keynes and Alex Alderton, Roberto Amato, Jeffrey Barrett, Sonia Goncalves, Ewan Harrison, David K. Jackson, Ian Johnston, Dominic Kwiatkowski, Cordelia Langford, John Sillitoe on behalf of the Wellcome Sanger Institute COVID-19 Surveillance Team |
| EPI_ISL_2937715, EPI_ISL_2937716, EPI_ISL_2937717, EPI_ISL_2937718, EPI_ISL_2937719, EPI_ISL_2937720, EPI_ISL_2937721, EPI_ISL_2937722, EPI_ISL_2937723, EPI_ISL_2937724, EPI_ISL_2937725, EPI_ISL_2937726, EPI_ISL_2937727, EPI_ISL_2937728, EPI_ISL_2937729 | | | |
| see above | Lighthouse Lab in Glasgow | Wellcome Sanger Institute for the COVID-19 Genomics UK (COG-UK) Consortium | Harper VanSteenhouse, Yumi Kasai, David Gray, Carol Clugston, Anna Dominiczak and Alex Alderton, Roberto Amato, Jeffrey Barrett, Sonia Goncalves, Ewan Harrison, David K. Jackson, Ian Johnston, Dominic Kwiatkowski, Cordelia Langford, John Sillitoe on behalf of the Wellcome Sanger Institute COVID-19 Surveillance Team |
| EPI_ISL_2937730 | Lighthouse Lab in Milton Keynes | Wellcome Sanger Institute for the COVID-19 Genomics UK (COG-UK) Consortium | The Lighthouse Lab in Milton Keynes and Alex Alderton, Roberto Amato, Jeffrey Barrett, Sonia Goncalves, Ewan Harrison, David K. Jackson, Ian Johnston, Dominic Kwiatkowski, Cordelia Langford, John Sillitoe on behalf of the Wellcome Sanger Institute COVID-19 Surveillance Team |
| EPI_ISL_2937731, EPI_ISL_2937732, EPI_ISL_2937733, EPI_ISL_2937734, EPI_ISL_2937735 | Lighthouse Lab in Glasgow | Wellcome Sanger Institute for the COVID-19 Genomics UK (COG-UK) Consortium | Harper VanSteenhouse, Yumi Kasai, David Gray, Carol Clugston, Anna Dominiczak and Alex Alderton, Roberto Amato, Jeffrey Barrett, Sonia Goncalves, Ewan Harrison, David K. Jackson, Ian Johnston, Dominic Kwiatkowski, Cordelia Langford, John Sillitoe on behalf of the Wellcome Sanger Institute COVID-19 Surveillance Team |
| EPI_ISL_2937736 | Lighthouse Laboratory Plymouth | Wellcome Sanger Institute for the COVID-19 Genomics UK (COG-UK) Consortium | Lighthouse Laboratory Plymouth and Alex Alderton, Roberto Amato, Jeffrey Barrett, Sonia Goncalves, Ewan Harrison, David K. Jackson, Ian Johnston, Dominic Kwiatkowski, Cordelia Langford, John Sillitoe on behalf of the Wellcome Sanger Institute COVID-19 Surveillance Team |
| EPI_ISL_2937737 | Lighthouse Lab in Milton Keynes | Wellcome Sanger Institute for the COVID-19 Genomics UK (COG-UK) Consortium | The Lighthouse Lab in Milton Keynes and Alex Alderton, Roberto Amato, Jeffrey Barrett, Sonia Goncalves, Ewan Harrison, David K. Jackson, Ian Johnston, Dominic Kwiatkowski, Cordelia Langford, John Sillitoe on behalf of the Wellcome Sanger Institute COVID-19 Surveillance Team |
| EPI_ISL_2937738 | Lighthouse Laboratory Plymouth | Wellcome Sanger Institute for the COVID-19 Genomics UK (COG-UK) Consortium | Lighthouse Laboratory Plymouth and Alex Alderton, Roberto Amato, Jeffrey Barrett, Sonia Goncalves, Ewan Harrison, David K. Jackson, Ian Johnston, Dominic Kwiatkowski, Cordelia Langford, John Sillitoe on behalf of the Wellcome Sanger Institute COVID-19 Surveillance Team |
| EPI_ISL_2937739 | Lighthouse Lab in Milton Keynes | Wellcome Sanger Institute for the COVID-19 Genomics UK (COG-UK) Consortium | The Lighthouse Lab in Milton Keynes and Alex Alderton, Roberto Amato, Jeffrey Barrett, Sonia Goncalves, Ewan Harrison, David K. Jackson, Ian Johnston, Dominic Kwiatkowski, Cordelia Langford, John Sillitoe on behalf of the Wellcome Sanger Institute COVID-19 Surveillance Team |
| EPI_ISL_2937740 | Berkshire and Surrey Pathology Services Lighthouse Laboratory | Wellcome Sanger Institute for the COVID-19 Genomics UK (COG-UK) Consortium | Berkshire and Surrey Pathology Services Lighthouse Laboratory and Alex Alderton, Roberto Amato, Jeffrey Barrett, Sonia Goncalves, Ewan Harrison, David K. Jackson, Ian Johnston, Dominic Kwiatkowski, Cordelia Langford, John Sillitoe on behalf of the Wellcome Sanger Institute COVID-19 Surveillance Team |
| EPI_ISL_2937741 | Lighthouse Lab in Milton Keynes | Wellcome Sanger Institute for the COVID-19 Genomics UK (COG-UK) Consortium | The Lighthouse Lab in Milton Keynes and Alex Alderton, Roberto Amato, Jeffrey Barrett, Sonia Goncalves, Ewan Harrison, David K. Jackson, Ian Johnston, Dominic Kwiatkowski, Cordelia Langford, John Sillitoe on behalf of the Wellcome Sanger Institute COVID-19 Surveillance Team |
| EPI_ISL_2937742 | Berkshire and Surrey Pathology Services Lighthouse Laboratory | Wellcome Sanger Institute for the COVID-19 Genomics UK (COG-UK) Consortium | Berkshire and Surrey Pathology Services Lighthouse Laboratory and Alex Alderton, Roberto Amato, Jeffrey Barrett, Sonia Goncalves, Ewan Harrison, David K. Jackson, Ian Johnston, Dominic Kwiatkowski, Cordelia Langford, John Sillitoe on behalf of the Wellcome Sanger Institute COVID-19 Surveillance Team |
| EPI_ISL_2937743 | Lighthouse Laboratory Plymouth | Wellcome Sanger Institute for the COVID-19 Genomics UK (COG-UK) Consortium | Lighthouse Laboratory Plymouth and Alex Alderton, Roberto Amato, Jeffrey Barrett, Sonia Goncalves, Ewan Harrison, David K. Jackson, Ian Johnston, Dominic Kwiatkowski, Cordelia Langford, John Sillitoe on behalf of the Wellcome Sanger Institute COVID-19 Surveillance Team |
| EPI_ISL_2937744 | Lighthouse Lab in Glasgow | Wellcome Sanger Institute for the COVID-19 Genomics UK (COG-UK) Consortium | Harper VanSteenhouse, Yumi Kasai, David Gray, Carol Clugston, Anna Dominiczak and Alex Alderton, Roberto Amato, Jeffrey Barrett, Sonia Goncalves, Ewan Harrison, David K. Jackson, Ian Johnston, Dominic Kwiatkowski, Cordelia Langford, John Sillitoe on behalf of the Wellcome Sanger Institute COVID-19 Surveillance Team |
| EPI_ISL_2937745 | Randox Laboratories | Wellcome Sanger Institute for the COVID-19 Genomics UK (COG-UK) Consortium | Randox Laboratories and Alex Alderton, Roberto Amato, Jeffrey Barrett, Sonia Goncalves, Ewan Harrison, David K. Jackson, Ian Johnston, Dominic Kwiatkowski, Cordelia Langford, John Sillitoe on behalf of the Wellcome Sanger Institute COVID-19 Surveillance Team |
| EPI_ISL_2937746 | Lighthouse Laboratory Plymouth | Wellcome Sanger Institute for the COVID-19 Genomics UK (COG-UK) Consortium | Lighthouse Laboratory Plymouth and Alex Alderton, Roberto Amato, Jeffrey Barrett, Sonia Goncalves, Ewan Harrison, David K. Jackson, Ian Johnston, Dominic Kwiatkowski, Cordelia Langford, John Sillitoe on behalf of the Wellcome Sanger Institute COVID-19 Surveillance Team |
| EPI_ISL_2937748 | LESP Aguascalientes | Instituto de Diagnostico y Referencia Epidemiologicos (INDRE) | Claudia Wong-Arambula, Abril Rodriguez-Maldonado, Vanessa Rivero-Arredondo, Ariadna Medina-Benitez, Joaquin Quiroz-Mercado, Sergio Rangel-Guerrero, Maribel Gonzalez-Villa, Natividad Cruz-Ortiz, Tatiana Nunez-Garcia, Gisela Barrera-Badillo, Lucia Hernandez-Rivas, Irma Lopez-Martinez, Ernesto Ramirez-Gonzalez. |
| EPI_ISL_2937749 | LESP Tabasco | Instituto de Diagnostico y Referencia Epidemiologicos (INDRE) | Claudia Wong-Arambula, Abril Rodriguez-Maldonado, Vanessa Rivero-Arredondo, Ariadna Medina-Benitez, Joaquin Quiroz-Mercado, Sergio Rangel-Guerrero, Maribel Gonzalez-Villa, Natividad Cruz-Ortiz, Tatiana Nunez-Garcia, Gisela Barrera-Badillo, Lucia Hernandez-Rivas, Irma Lopez-Martinez, Ernesto Ramirez-Gonzalez. |
| EPI_ISL_2937750 | LESP Aguascalientes | Instituto de Diagnostico y Referencia Epidemiologicos (INDRE) | Claudia Wong-Arambula, Abril Rodriguez-Maldonado, Vanessa Rivero-Arredondo, Ariadna Medina-Benitez, Joaquin Quiroz-Mercado, Sergio Rangel-Guerrero, Maribel Gonzalez-Villa, Natividad Cruz-Ortiz, Tatiana Nunez-Garcia, Gisela Barrera-Badillo, Lucia Hernandez-Rivas, Irma Lopez-Martinez, Ernesto Ramirez-Gonzalez. |
| EPI_ISL_2937752, EPI_ISL_2937832, EPI_ISL_2937833, EPI_ISL_2937834, EPI_ISL_2937835 | LESP Ciudad de Mexico | Instituto de Diagnostico y Referencia Epidemiologicos (INDRE) | Claudia Wong-Arambula, Abril Rodriguez-Maldonado, Vanessa Rivero-Arredondo, Ariadna Medina-Benitez, Joaquin Quiroz-Mercado, Sergio Rangel-Guerrero, Maribel Gonzalez-Villa, Natividad Cruz-Ortiz, Tatiana Nunez-Garcia, Gisela Barrera-Badillo, Lucia Hernandez-Rivas, Irma Lopez-Martinez, Ernesto Ramirez-Gonzalez. |
| EPI_ISL_2937836 | LESP Baja California Sur | Instituto de Diagnostico y Referencia Epidemiologicos (INDRE) | Claudia Wong-Arambula, Abril Rodriguez-Maldonado, Vanessa Rivero-Arredondo, Ariadna Medina-Benitez, Joaquin Quiroz-Mercado, Sergio Rangel-Guerrero, Maribel Gonzalez-Villa, Natividad Cruz-Ortiz, Tatiana Nunez-Garcia, Gisela Barrera-Badillo, Lucia Hernandez-Rivas, Irma Lopez-Martinez, Ernesto Ramirez-Gonzalez. |
| EPI_ISL_2937837 | LESP Zacatecas | Instituto de Diagnostico y Referencia Epidemiologicos (INDRE) | Claudia Wong-Arambula, Abril Rodriguez-Maldonado, Vanessa Rivero-Arredondo, Ariadna Medina-Benitez, Joaquin Quiroz-Mercado, Sergio Rangel-Guerrero, Maribel Gonzalez-Villa, Natividad Cruz-Ortiz, Tatiana Nunez-Garcia, Gisela Barrera-Badillo, Lucia Hernandez-Rivas, Irma Lopez-Martinez, Ernesto Ramirez-Gonzalez. |
| EPI_ISL_2937838, EPI_ISL_2937839, EPI_ISL_2937840, EPI_ISL_2937841, EPI_ISL_2937842, EPI_ISL_2937843 | LESP Ciudad de Mexico | Instituto de Diagnostico y Referencia Epidemiologicos (INDRE) | Claudia Wong-Arambula, Abril Rodriguez-Maldonado, Vanessa Rivero-Arredondo, Ariadna Medina-Benitez, Joaquin Quiroz-Mercado, Sergio Rangel-Guerrero, Maribel Gonzalez-Villa, Natividad Cruz-Ortiz, Tatiana Nunez-Garcia, Gisela Barrera-Badillo, Lucia Hernandez-Rivas, Irma Lopez-Martinez, Ernesto Ramirez-Gonzalez. |
| EPI_ISL_2937844 | LESP Nuevo Leon | Instituto de Diagnostico y Referencia Epidemiologicos (INDRE) | Claudia Wong-Arambula, Abril Rodriguez-Maldonado, Vanessa Rivero-Arredondo, Ariadna Medina-Benitez, Joaquin Quiroz-Mercado, Sergio Rangel-Guerrero, Maribel Gonzalez-Villa, Natividad Cruz-Ortiz, Tatiana Nunez-Garcia, Gisela Barrera- |

| | | | |
|---|---|---|---|
| | | | Badillo, Lucia Hernandez-Rivas, Irma Lopez-Martinez, Ernesto Ramirez-Gonzalez. |
| EPI_ISL_2937845 | LESP Tlaxcala | Instituto de Diagnostico y Referencia Epidemiologicos (INDRE) | Claudia Wong-Arambula, Abril Rodriguez-Maldonado, Vanessa Rivero-Arredondo, Ariadna Medina-Benitez, Joaquin Quiroz-Mercado, Sergio Rangel-Guerrero, Maribel Gonzalez-Villa, Natividad Cruz-Ortiz, Tatiana Nunez-Garcia, Gisela Barrera-Badillo, Lucia Hernandez-Rivas, Irma Lopez-Martinez, Ernesto Ramirez-Gonzalez. |
| EPI_ISL_2937846 | LESP Michoacan | Instituto de Diagnostico y Referencia Epidemiologicos (INDRE) | Claudia Wong-Arambula, Abril Rodriguez-Maldonado, Vanessa Rivero-Arredondo, Ariadna Medina-Benitez, Joaquin Quiroz-Mercado, Sergio Rangel-Guerrero, Maribel Gonzalez-Villa, Natividad Cruz-Ortiz, Tatiana Nunez-Garcia, Gisela Barrera-Badillo, Lucia Hernandez-Rivas, Irma Lopez-Martinez, Ernesto Ramirez-Gonzalez. |
| EPI_ISL_2937847, EPI_ISL_2937848 | LESP Baja California Sur | Instituto de Diagnostico y Referencia Epidemiologicos (INDRE) | Claudia Wong-Arambula, Abril Rodriguez-Maldonado, Vanessa Rivero-Arredondo, Ariadna Medina-Benitez, Joaquin Quiroz-Mercado, Sergio Rangel-Guerrero, Maribel Gonzalez-Villa, Natividad Cruz-Ortiz, Tatiana Nunez-Garcia, Gisela Barrera-Badillo, Lucia Hernandez-Rivas, Irma Lopez-Martinez, Ernesto Ramirez-Gonzalez. |
| EPI_ISL_2937849, EPI_ISL_2937850 | LESP Veracruz | Instituto de Diagnostico y Referencia Epidemiologicos (INDRE) | Claudia Wong-Arambula, Abril Rodriguez-Maldonado, Vanessa Rivero-Arredondo, Ariadna Medina-Benitez, Joaquin Quiroz-Mercado, Sergio Rangel-Guerrero, Maribel Gonzalez-Villa, Natividad Cruz-Ortiz, Tatiana Nunez-Garcia, Gisela Barrera-Badillo, Lucia Hernandez-Rivas, Irma Lopez-Martinez, Ernesto Ramirez-Gonzalez. |
| EPI_ISL_2937851, EPI_ISL_2937852, EPI_ISL_2937853, EPI_ISL_2937854, EPI_ISL_2937855 | LESP Sonora | Instituto de Diagnostico y Referencia Epidemiologicos (INDRE) | Claudia Wong-Arambula, Abril Rodriguez-Maldonado, Vanessa Rivero-Arredondo, Ariadna Medina-Benitez, Joaquin Quiroz-Mercado, Sergio Rangel-Guerrero, Maribel Gonzalez-Villa, Natividad Cruz-Ortiz, Tatiana Nunez-Garcia, Gisela Barrera-Badillo, Lucia Hernandez-Rivas, Irma Lopez-Martinez, Ernesto Ramirez-Gonzalez. |
| EPI_ISL_2937856 | LESP Quintana Roo | Instituto de Diagnostico y Referencia Epidemiologicos (INDRE) | Claudia Wong-Arambula, Abril Rodriguez-Maldonado, Vanessa Rivero-Arredondo, Ariadna Medina-Benitez, Joaquin Quiroz-Mercado, Sergio Rangel-Guerrero, Maribel Gonzalez-Villa, Natividad Cruz-Ortiz, Tatiana Nunez-Garcia, Gisela Barrera-Badillo, Lucia Hernandez-Rivas, Irma Lopez-Martinez, Ernesto Ramirez-Gonzalez. |
| EPI_ISL_2937857, EPI_ISL_2937858, EPI_ISL_2937859, EPI_ISL_2937860 | LESP Ciudad de Mexico | Instituto de Diagnostico y Referencia Epidemiologicos (INDRE) | Claudia Wong-Arambula, Abril Rodriguez-Maldonado, Vanessa Rivero-Arredondo, Ariadna Medina-Benitez, Joaquin Quiroz-Mercado, Sergio Rangel-Guerrero, Maribel Gonzalez-Villa, Natividad Cruz-Ortiz, Tatiana Nunez-Garcia, Gisela Barrera-Badillo, Lucia Hernandez-Rivas, Irma Lopez-Martinez, Ernesto Ramirez-Gonzalez. |
| EPI_ISL_2937861 | LESP Estado de Mexico | Instituto de Diagnostico y Referencia Epidemiologicos (INDRE) | Claudia Wong-Arambula, Abril Rodriguez-Maldonado, Vanessa Rivero-Arredondo, Ariadna Medina-Benitez, Joaquin Quiroz-Mercado, Sergio Rangel-Guerrero, Maribel Gonzalez-Villa, Natividad Cruz-Ortiz, Tatiana Nunez-Garcia, Gisela Barrera-Badillo, Lucia Hernandez-Rivas, Irma Lopez-Martinez, Ernesto Ramirez-Gonzalez. |
| EPI_ISL_2937862, EPI_ISL_2937863, EPI_ISL_2937864, EPI_ISL_2937865, EPI_ISL_2937866, EPI_ISL_2937867, EPI_ISL_2937868, EPI_ISL_2937869, EPI_ISL_2937870, EPI_ISL_2937871, EPI_ISL_2937872, EPI_ISL_2937873, EPI_ISL_2937874, EPI_ISL_2937875, EPI_ISL_2937876, EPI_ISL_2937877, EPI_ISL_2937878, EPI_ISL_2937879, EPI_ISL_2937880, EPI_ISL_2937881, EPI_ISL_2937882, EPI_ISL_2937883, EPI_ISL_2937884, EPI_ISL_2937885, EPI_ISL_2937886, EPI_ISL_2937887, EPI_ISL_2937888, EPI_ISL_2937889, EPI_ISL_2937890, EPI_ISL_2937891, EPI_ISL_2937892, EPI_ISL_2937893, EPI_ISL_2937894, EPI_ISL_2937895 | | | |
| see above | LESP Ciudad de Mexico | Instituto de Diagnostico y Referencia Epidemiologicos (INDRE) | Claudia Wong-Arambula, Abril Rodriguez-Maldonado, Vanessa Rivero-Arredondo, Ariadna Medina-Benitez, Joaquin Quiroz-Mercado, Sergio Rangel-Guerrero, Maribel Gonzalez-Villa, Natividad Cruz-Ortiz, Tatiana Nunez-Garcia, Gisela Barrera-Badillo, Lucia Hernandez-Rivas, Irma Lopez-Martinez, Ernesto Ramirez-Gonzalez. |
| EPI_ISL_2937896, EPI_ISL_2937897, EPI_ISL_2937898 | LESP Jalisco | Instituto de Diagnostico y Referencia Epidemiologicos (INDRE) | Claudia Wong-Arambula, Abril Rodriguez-Maldonado, Vanessa Rivero-Arredondo, Ariadna Medina-Benitez, Joaquin Quiroz-Mercado, Sergio Rangel-Guerrero, Maribel Gonzalez-Villa, Natividad Cruz-Ortiz, Tatiana Nunez-Garcia, Gisela Barrera-Badillo, Lucia Hernandez-Rivas, Irma Lopez-Martinez, Ernesto Ramirez-Gonzalez. |
| EPI_ISL_2937899 | LESP Tabasco | Instituto de Diagnostico y Referencia Epidemiologicos (INDRE) | Claudia Wong-Arambula, Abril Rodriguez-Maldonado, Vanessa Rivero-Arredondo, Ariadna Medina-Benitez, Joaquin Quiroz-Mercado, Sergio Rangel-Guerrero, Maribel Gonzalez-Villa, Natividad Cruz-Ortiz, Tatiana Nunez-Garcia, Gisela Barrera-Badillo, Lucia Hernandez-Rivas, Irma Lopez-Martinez, Ernesto Ramirez-Gonzalez. |
| EPI_ISL_2937900, EPI_ISL_2937901, EPI_ISL_2937902, EPI_ISL_2937903, EPI_ISL_2937904, EPI_ISL_2937905 | LESP Aguascalientes | Instituto de Diagnostico y Referencia Epidemiologicos (INDRE) | Claudia Wong-Arambula, Abril Rodriguez-Maldonado, Vanessa Rivero-Arredondo, Ariadna Medina-Benitez, Joaquin Quiroz-Mercado, Sergio Rangel-Guerrero, Maribel Gonzalez-Villa, Natividad Cruz-Ortiz, Tatiana Nunez-Garcia, Gisela Barrera-Badillo, Lucia Hernandez-Rivas, Irma Lopez-Martinez, Ernesto Ramirez-Gonzalez. |
| EPI_ISL_2937908, EPI_ISL_2937909, EPI_ISL_2937910, EPI_ISL_2937911, EPI_ISL_2937912, EPI_ISL_2937913, EPI_ISL_2937914, EPI_ISL_2937915, EPI_ISL_2937916, EPI_ISL_2937917, EPI_ISL_2937918, EPI_ISL_2937919, EPI_ISL_2937920, EPI_ISL_2937921, EPI_ISL_2937922, EPI_ISL_2937923, EPI_ISL_2937924, EPI_ISL_2937925 | | | |
| see above | LESP Ciudad de Mexico | Instituto de Diagnostico y Referencia Epidemiologicos (INDRE) | Claudia Wong-Arambula, Abril Rodriguez-Maldonado, Vanessa Rivero-Arredondo, Ariadna Medina-Benitez, Joaquin Quiroz-Mercado, Sergio Rangel-Guerrero, Maribel Gonzalez-Villa, Natividad Cruz-Ortiz, Tatiana Nunez-Garcia, Gisela Barrera-Badillo, Lucia Hernandez-Rivas, Irma Lopez-Martinez, Ernesto Ramirez-Gonzalez. |
| EPI_ISL_2937926, EPI_ISL_2937927 | LESP Queretaro | Instituto de Diagnostico y Referencia Epidemiologicos (INDRE) | Claudia Wong-Arambula, Abril Rodriguez-Maldonado, Vanessa Rivero-Arredondo, Ariadna Medina-Benitez, Joaquin Quiroz-Mercado, Sergio Rangel-Guerrero, Maribel Gonzalez-Villa, Natividad Cruz-Ortiz, Tatiana Nunez-Garcia, Gisela Barrera-Badillo, Lucia Hernandez-Rivas, Irma Lopez-Martinez, Ernesto Ramirez-Gonzalez. |
| EPI_ISL_2937932, EPI_ISL_2937939 | UW Virology Lab | UW Virology Lab | Pavitra Roychoudhury, Hong Xie, Lasata Shrestha, Shah Mohamed Bakhash, Tien V. Nguyen, Noah R. Baker, Sean Ellis, Meei-Li Huang, Keith R Jerome, Alexander Greninger |
| EPI_ISL_2937940 | Policlinico Tor Vergata | INMI Lazzaro Spallanzani IRCCS | CEM Gruber, B Bartolini, E Giombini, F Messina, M Rueca, O Butera, F Santini, A Di Caro, MR Capobianchi |
| EPI_ISL_2937965, EPI_ISL_2937967, EPI_ISL_2937984, EPI_ISL_2937985, EPI_ISL_2937986, EPI_ISL_2937990, EPI_ISL_2937991, EPI_ISL_2938003, EPI_ISL_2938005, EPI_ISL_2938007, EPI_ISL_2938018 | | | |
| see above | Diagnostyka. Laboratoria Medyczne. | 1. ViroGenetics - BSL3 Laboratory of Virology, Małopolska Centre of Biotechnology, Jagiellonian University; 2. genXone SA, Research & Development Laboratory | Mazur-Panasiuk,N., Grzegorz Nowicki, Wydmanski,W., Gromowski,T., Natalia Drweska-Matelska, Jakub Grabowski, Anna Brylak, Aleksandra Gidlewicz, Karol Szeszko, Maciej Sykulski, Łukasz Krych, Kowalski,M., Szulc,P., Sylwia Januszczak, Labaj,P.P., Michał Kaszuba, Pyrc,K. |
| EPI_ISL_2938020, EPI_ISL_2938021, EPI_ISL_2938023, EPI_ISL_2938024, EPI_ISL_2938025, EPI_ISL_2938026, EPI_ISL_2938027, EPI_ISL_2938028, EPI_ISL_2938029, EPI_ISL_2938030, EPI_ISL_2938031, EPI_ISL_2938033, EPI_ISL_2938034, EPI_ISL_2938035, EPI_ISL_2938036, EPI_ISL_2938037, EPI_ISL_2938038, EPI_ISL_2938039, EPI_ISL_2938040, EPI_ISL_2938041, EPI_ISL_2938042, EPI_ISL_2938043, EPI_ISL_2938044, EPI_ISL_2938045, EPI_ISL_2938046, EPI_ISL_2938047, EPI_ISL_2938049, EPI_ISL_2938050, EPI_ISL_2938051, EPI_ISL_2938052, EPI_ISL_2938053, EPI_ISL_2938054, EPI_ISL_2938055, EPI_ISL_2938056, EPI_ISL_2938057, EPI_ISL_2938058, EPI_ISL_2938059, EPI_ISL_2938060, EPI_ISL_2938062, EPI_ISL_2938063, EPI_ISL_2938065, EPI_ISL_2938066, EPI_ISL_2938067, EPI_ISL_2938068, EPI_ISL_2938069, EPI_ISL_2938070, EPI_ISL_2938071, EPI_ISL_2938072, EPI_ISL_2938073, EPI_ISL_2938074, EPI_ISL_2938077, EPI_ISL_2938078, EPI_ISL_2938079, EPI_ISL_2938080, EPI_ISL_2938081, EPI_ISL_2938082, EPI_ISL_2938083, EPI_ISL_2938084, EPI_ISL_2938085, EPI_ISL_2938088, EPI_ISL_2938089, EPI_ISL_2938090, EPI_ISL_2938091, EPI_ISL_2938092, EPI_ISL_2938093, EPI_ISL_2938094 | | | |
| see above | Hospital Universitari Vall d'Hebron - Vall d'Hebron Institut de Recerca | Hospital Universitari Vall d'Hebron - Vall d'Hebron Institut de Recerca | Cristina Andrés, Maria Piñana, Alejandra González-Sánchez, Damir Garcia-Cehic, Ariadna Rando, Juliana Esperalba, Maria Gema Codina, Carla Castillo, Karen García, Rodrigo Vásquez, Maria Carmen Martin, Tomàs Pumarola, Josep Quer, Andrés Antón |
| EPI_ISL_2938095 | LABORATORIO DR PAULO EMILIO DALESSANDRO PINDAMONHANGABA | Instituto Butantan | Dimas Tadeu Covas, Antonio Jorge Martins, Claudia Renata dos Santos Barros, David Schlesinger, Debora Botequio Moretti, Elaine Cristina Marqueze, Elaine Vieira Santos, Evandra Strazza Rodrigues, Heidge Fukumasu, Jayme Augusto de Souza-Neto, José Salvatore Leister Patané, Luiz Alcantara, Luiz Lehmann Coutinho, Maria Carolina Elias, Maurício Lacerda Nogueira, Rafael dos Santos Bezerra, Raul Machado Neto, Rejane Maria Tommasini Grotto, Ricardo Haddad, Sandra Coccuzzo Sampaio Vessoni, Simone Kashima, Svetoslav Nanev Slavov, Vincent Louis Viala |
| EPI_ISL_2938096 | SECRETARIA MUNICIPAL DA SAUDE DE GUARATINGUETA LABORATÓRIO 01 | Instituto Butantan | Dimas Tadeu Covas, Antonio Jorge Martins, Claudia Renata dos Santos Barros, David Schlesinger, Debora Botequio Moretti, Elaine Cristina Marqueze, Elaine Vieira Santos, Evandra Strazza Rodrigues, Heidge Fukumasu, Jayme Augusto de Souza-Neto, José Salvatore Leister Patané, Luiz Alcantara, Luiz Lehmann Coutinho, Maria Carolina Elias, Maurício Lacerda Nogueira, Rafael dos Santos Bezerra, Raul Machado Neto, Rejane Maria Tommasini Grotto, Ricardo Haddad, Sandra Coccuzzo Sampaio Vessoni, Simone Kashima, Svetoslav Nanev Slavov, Vincent Louis Viala |
| EPI_ISL_2938097 | Health Center Velika Kladuša | Clinical Center, University of Sarajevo; Unit for Clinical Microbiology | Irma Salimović-Bešić, Amela Dedeić-Ljubović, Edina Zahirović, Sandra Vegar-Zubović, Sebija Izetbegović |
| EPI_ISL_2938099, EPI_ISL_2938100, EPI_ISL_2938101 | Health Center Cazin | Clinical Center, University of Sarajevo; Unit for Clinical Microbiology | Irma Salimović-Bešić, Amela Dedeić-Ljubović, Edina Zahirović, Sandra Vegar-Zubović, Sebija Izetbegović |
| EPI_ISL_2938103 | Health Center Novo Sarajevo | Clinical Center, University of Sarajevo; Unit for Clinical Microbiology | Irma Salimović-Bešić, Amela Dedeić-Ljubović, Edina Zahirović, Sandra Vegar-Zubović, Sebija Izetbegović |
| EPI_ISL_2938104 | Clinical Center, University of Sarajevo; Unit for Clinical Microbiology | Clinical Center, University of Sarajevo; Unit for Clinical Microbiology | Irma Salimović-Bešić, Amela Dedeić-Ljubović, Edina Zahirović, Sandra Vegar-Zubović, Sebija Izetbegović |
| EPI_ISL_2938109 | Health Center Sarajevo- Center | Clinical Center, University of Sarajevo; Unit for Clinical Microbiology | Irma Salimović-Bešić, Amela Dedeić-Ljubović, Edina Zahirović, Sandra Vegar-Zubović, Sebija Izetbegović |
| EPI_ISL_2938111 | Health Center Velika Kladuša | Clinical Center, University of Sarajevo; Unit for Clinical Microbiology | Irma Salimović-Bešić, Amela Dedeić-Ljubović, Edina Zahirović, Sandra Vegar-Zubović, Sebija Izetbegović |
| EPI_ISL_2938112 | Health Center Cazin | Clinical Center, University of Sarajevo; Unit for Clinical Microbiology | Irma Salimović-Bešić, Amela Dedeić-Ljubović, Edina Zahirović, Sandra Vegar-Zubović, Sebija Izetbegović |
| EPI_ISL_2938116 | Health Center Vogošća | Clinical Center, University of Sarajevo; Unit for Clinical Microbiology | Irma Salimović-Bešić, Amela Dedeić-Ljubović, Edina Zahirović, Sandra Vegar-Zubović, Sebija Izetbegović |
| EPI_ISL_2938117 | Health Center Novo Sarajevo | Clinical Center, University of Sarajevo; Unit for Clinical Microbiology | Irma Salimović-Bešić, Amela Dedeić-Ljubović, Edina Zahirović, Sandra Vegar-Zubović, Sebija Izetbegović |
| EPI_ISL_2938119 | Health Center Novi Grad Sarajevo | Clinical Center, University of Sarajevo; Unit for Clinical Microbiology | Irma Salimović-Bešić, Amela Dedeić-Ljubović, Edina Zahirović, Sandra Vegar-Zubović, Sebija Izetbegović |
| EPI_ISL_2938278, EPI_ISL_2938279, EPI_ISL_2938280, EPI_ISL_2938281, EPI_ISL_2938282, EPI_ISL_2938283, EPI_ISL_2938284, EPI_ISL_2938285, EPI_ISL_2938286, EPI_ISL_2938287, EPI_ISL_2938288, EPI_ISL_2938289, EPI_ISL_2938290, EPI_ISL_2938291, EPI_ISL_2938292, EPI_ISL_2938293 | | | |
| see above | UNAM Molecular Diagnostic Laboratory | Forschungszentrum Borstel | Emmanuel Nepolo, Lusia Mhuulu, Azaria Diergaardt, Loide Shipingana, Christian Utpatel, Ivan Barilar, Stefan Niemann, Vanessa Mohr, Tanja Niemann, Jasmin Scharnberg |
| EPI_ISL_2938306, EPI_ISL_2938309, EPI_ISL_2938310 | Mobile County Health Department | Synergy Laboratories | Megan Cornwell |
| EPI_ISL_2938311, EPI_ISL_2938312 | St James's Hospital, Dublin 8, Ireland. | ELDA biotech | Deirdre M. Broderick, Peter R. Flanagan, Brendan Crowley |
| EPI_ISL_2938331, EPI_ISL_2938333 | Synlab Suisse SA | Clinical Bacteriology | Tim Roloff, Fanny Wegner, Helena MB Seth-Smith, Alfredo Mari, Karoline Leuzinger, Julia Bielicki, Manuel Battegay, Hans Hirsch, Adrian Egli |
| EPI_ISL_2938334 | Viollier AG | Department of Biosystems Science and Engineering, ETH Zürich | Chaoran Chen, Sarah Nadeau, Catharine Aquino, Ivan Topolsky, Kim Philipp Jablonski, Lara Fuhrmann, Daniel Ehrsam, Isabel Stürmer, Andreia Cabral de Gouvea, Maria Domenica Moccia, Simon Grüter, Timothy Sykes, Lennart Opitz, Griffin White, Laura Neff, Doris Popovic, Andrea Patrignani, Jay Tracy, Ralph Schlapbach, Christiane Beckmann, Maurice Redondo, Olivier Kobel, Christoph Noppen, Niko Beerenwinkel, Tanja Stadler |
| EPI_ISL_2938338, EPI_ISL_2938339, EPI_ISL_2938341, EPI_ISL_2938345, EPI_ISL_2938351, EPI_ISL_2938352 | Viollier AG | Department of Biosystems Science and Engineering, ETH Zürich | Christian Beisel, Sarah Nadeau, Chaoran Chen, Ivan Topolsky, Kim Philipp Jablonski, Lara Fuhrmann, Rebecca Denes, Mirjam Feldkamp, Ina Nissen, Natascha Santacroce, Elodie Burcklen, Christiane Beckmann, Maurice Redondo, Olivier Kobel, Christoph Noppen, Niko Beerenwinkel, Tanja Stadler |
| EPI_ISL_2938355 | Viollier AG | Department of Biosystems Science and Engineering, ETH Zürich | Chaoran Chen, Sarah Nadeau, Catharine Aquino, Ivan Topolsky, Kim Philipp Jablonski, Lara Fuhrmann, Daniel Ehrsam, Isabel Stürmer, Andreia Cabral de Gouvea, Maria Domenica Moccia, Simon Grüter, Timothy Sykes, Lennart Opitz, Griffin |

| | | | |
|---|---|---|---|
| | | | White, Laura Neff, Doris Popovic, Andrea Patrignani, Jay Tracy, Ralph Schlapbach, Christiane Beckmann, Maurice Redondo, Olivier Kobel, Christoph Noppen, Niko Beerenwinkel, Tanja Stadler |
| EPI_ISL_2938360, EPI_ISL_2938362 | Viollier AG | Department of Biosystems Science and Engineering, ETH Zürich | Christian Beisel, Sarah Nadeau, Chaoran Chen, Ivan Topolsky, Kim Philipp Jablonski, Lara Fuhrmann, Rebecca Denes, Mirjam Feldkamp, Ina Nissen, Natascha Santacroce, Elodie Burcklen, Christiane Beckmann, Maurice Redondo, Olivier Kobel, Christoph Noppen, Niko Beerenwinkel, Tanja Stadler |
| EPI_ISL_2938363 | Viollier AG | Department of Biosystems Science and Engineering, ETH Zürich | Chaoran Chen, Sarah Nadeau, Catharine Aquino, Ivan Topolsky, Kim Philipp Jablonski, Lara Fuhrmann, Daniel Ehrsam, Isabel Stürmer, Andreia Cabral de Gouvea, Maria Domenica Moccia, Simon Grüter, Timothy Sykes, Lennart Opitz, Griffin White, Laura Neff, Doris Popovic, Andrea Patrignani, Jay Tracy, Ralph Schlapbach, Christiane Beckmann, Maurice Redondo, Olivier Kobel, Christoph Noppen, Niko Beerenwinkel, Tanja Stadler |
| EPI_ISL_2938370, EPI_ISL_2938371, EPI_ISL_2938372, EPI_ISL_2938373, EPI_ISL_2938376, EPI_ISL_2938377 | Viollier AG | Department of Biosystems Science and Engineering, ETH Zürich | Christian Beisel, Sarah Nadeau, Chaoran Chen, Ivan Topolsky, Kim Philipp Jablonski, Lara Fuhrmann, Rebecca Denes, Mirjam Feldkamp, Ina Nissen, Natascha Santacroce, Elodie Burcklen, Christiane Beckmann, Maurice Redondo, Olivier Kobel, Christoph Noppen, Niko Beerenwinkel, Tanja Stadler |
| EPI_ISL_2938379 | Viollier AG | Department of Biosystems Science and Engineering, ETH Zürich | Chaoran Chen, Sarah Nadeau, Catharine Aquino, Ivan Topolsky, Kim Philipp Jablonski, Lara Fuhrmann, Daniel Ehrsam, Isabel Stürmer, Andreia Cabral de Gouvea, Maria Domenica Moccia, Simon Grüter, Timothy Sykes, Lennart Opitz, Griffin White, Laura Neff, Doris Popovic, Andrea Patrignani, Jay Tracy, Ralph Schlapbach, Christiane Beckmann, Maurice Redondo, Olivier Kobel, Christoph Noppen, Niko Beerenwinkel, Tanja Stadler |
| EPI_ISL_2938381 | Viollier AG | Department of Biosystems Science and Engineering, ETH Zürich | Christian Beisel, Sarah Nadeau, Chaoran Chen, Ivan Topolsky, Kim Philipp Jablonski, Lara Fuhrmann, Rebecca Denes, Mirjam Feldkamp, Ina Nissen, Natascha Santacroce, Elodie Burcklen, Christiane Beckmann, Maurice Redondo, Olivier Kobel, Christoph Noppen, Niko Beerenwinkel, Tanja Stadler |
| EPI_ISL_2938383, EPI_ISL_2938384 | Viollier AG | Department of Biosystems Science and Engineering, ETH Zürich | Chaoran Chen, Sarah Nadeau, Catharine Aquino, Ivan Topolsky, Kim Philipp Jablonski, Lara Fuhrmann, Daniel Ehrsam, Isabel Stürmer, Andreia Cabral de Gouvea, Maria Domenica Moccia, Simon Grüter, Timothy Sykes, Lennart Opitz, Griffin White, Laura Neff, Doris Popovic, Andrea Patrignani, Jay Tracy, Ralph Schlapbach, Christiane Beckmann, Maurice Redondo, Olivier Kobel, Christoph Noppen, Niko Beerenwinkel, Tanja Stadler |
| EPI_ISL_2938385, EPI_ISL_2938386, EPI_ISL_2938387, EPI_ISL_2938388, EPI_ISL_2938389, EPI_ISL_2938391, EPI_ISL_2938393, EPI_ISL_2938394, EPI_ISL_2938395, EPI_ISL_2938396 | Viollier AG | Department of Biosystems Science and Engineering, ETH Zürich | Christian Beisel, Sarah Nadeau, Chaoran Chen, Ivan Topolsky, Kim Philipp Jablonski, Lara Fuhrmann, Rebecca Denes, Mirjam Feldkamp, Ina Nissen, Natascha Santacroce, Elodie Burcklen, Christiane Beckmann, Maurice Redondo, Olivier Kobel, Christoph Noppen, Niko Beerenwinkel, Tanja Stadler |
| EPI_ISL_2938397 | Viollier AG | Department of Biosystems Science and Engineering, ETH Zürich | Chaoran Chen, Sarah Nadeau, Catharine Aquino, Ivan Topolsky, Kim Philipp Jablonski, Lara Fuhrmann, Daniel Ehrsam, Isabel Stürmer, Andreia Cabral de Gouvea, Maria Domenica Moccia, Simon Grüter, Timothy Sykes, Lennart Opitz, Griffin White, Laura Neff, Doris Popovic, Andrea Patrignani, Jay Tracy, Ralph Schlapbach, Christiane Beckmann, Maurice Redondo, Olivier Kobel, Christoph Noppen, Niko Beerenwinkel, Tanja Stadler |
| EPI_ISL_2938398, EPI_ISL_2938399, EPI_ISL_2938401, EPI_ISL_2938402, EPI_ISL_2938403, EPI_ISL_2938404, EPI_ISL_2938405, EPI_ISL_2938406, EPI_ISL_2938407, EPI_ISL_2938408, EPI_ISL_2938409, EPI_ISL_2938410, EPI_ISL_2938411, EPI_ISL_2938412, EPI_ISL_2938413, EPI_ISL_2938414, EPI_ISL_2938416, EPI_ISL_2938417, EPI_ISL_2938418 | | | |
| see above | Viollier AG | Department of Biosystems Science and Engineering, ETH Zürich | Christian Beisel, Sarah Nadeau, Chaoran Chen, Ivan Topolsky, Kim Philipp Jablonski, Lara Fuhrmann, Rebecca Denes, Mirjam Feldkamp, Ina Nissen, Natascha Santacroce, Elodie Burcklen, Christiane Beckmann, Maurice Redondo, Olivier Kobel, Christoph Noppen, Niko Beerenwinkel, Tanja Stadler |
| EPI_ISL_2938419 | Viollier AG | Department of Biosystems Science and Engineering, ETH Zürich | Chaoran Chen, Sarah Nadeau, Catharine Aquino, Ivan Topolsky, Kim Philipp Jablonski, Lara Fuhrmann, Daniel Ehrsam, Isabel Stürmer, Andreia Cabral de Gouvea, Maria Domenica Moccia, Simon Grüter, Timothy Sykes, Lennart Opitz, Griffin White, Laura Neff, Doris Popovic, Andrea Patrignani, Jay Tracy, Ralph Schlapbach, Christiane Beckmann, Maurice Redondo, Olivier Kobel, Christoph Noppen, Niko Beerenwinkel, Tanja Stadler |
| EPI_ISL_2938420, EPI_ISL_2938428 | Viollier AG | Department of Biosystems Science and Engineering, ETH Zürich | Christian Beisel, Sarah Nadeau, Chaoran Chen, Ivan Topolsky, Kim Philipp Jablonski, Lara Fuhrmann, Rebecca Denes, Mirjam Feldkamp, Ina Nissen, Natascha Santacroce, Elodie Burcklen, Christiane Beckmann, Maurice Redondo, Olivier Kobel, Christoph Noppen, Niko Beerenwinkel, Tanja Stadler |
| EPI_ISL_2938429, EPI_ISL_2938430 | Viollier AG | Department of Biosystems Science and Engineering, ETH Zürich | Chaoran Chen, Sarah Nadeau, Catharine Aquino, Ivan Topolsky, Kim Philipp Jablonski, Lara Fuhrmann, Daniel Ehrsam, Isabel Stürmer, Andreia Cabral de Gouvea, Maria Domenica Moccia, Simon Grüter, Timothy Sykes, Lennart Opitz, Griffin White, Laura Neff, Doris Popovic, Andrea Patrignani, Jay Tracy, Ralph Schlapbach, Christiane Beckmann, Maurice Redondo, Olivier Kobel, Christoph Noppen, Niko Beerenwinkel, Tanja Stadler |
| EPI_ISL_2938431 | Viollier AG | Department of Biosystems Science and Engineering, ETH Zürich | Christian Beisel, Sarah Nadeau, Chaoran Chen, Ivan Topolsky, Kim Philipp Jablonski, Lara Fuhrmann, Rebecca Denes, Mirjam Feldkamp, Ina Nissen, Natascha Santacroce, Elodie Burcklen, Christiane Beckmann, Maurice Redondo, Olivier Kobel, Christoph Noppen, Niko Beerenwinkel, Tanja Stadler |
| EPI_ISL_2938438, EPI_ISL_2938439 | Viollier AG | Department of Biosystems Science and Engineering, ETH Zürich | Chaoran Chen, Sarah Nadeau, Catharine Aquino, Ivan Topolsky, Kim Philipp Jablonski, Lara Fuhrmann, Daniel Ehrsam, Isabel Stürmer, Andreia Cabral de Gouvea, Maria Domenica Moccia, Simon Grüter, Timothy Sykes, Lennart Opitz, Griffin White, Laura Neff, Doris Popovic, Andrea Patrignani, Jay Tracy, Ralph Schlapbach, Christiane Beckmann, Maurice Redondo, Olivier Kobel, Christoph Noppen, Niko Beerenwinkel, Tanja Stadler |
| EPI_ISL_2938440, EPI_ISL_2938441, EPI_ISL_2938445, EPI_ISL_2938446, EPI_ISL_2938448, EPI_ISL_2938450, EPI_ISL_2938451, EPI_ISL_2938453, EPI_ISL_2938454, EPI_ISL_2938456 | Viollier AG | Department of Biosystems Science and Engineering, ETH Zürich | Christian Beisel, Sarah Nadeau, Chaoran Chen, Ivan Topolsky, Kim Philipp Jablonski, Lara Fuhrmann, Rebecca Denes, Mirjam Feldkamp, Ina Nissen, Natascha Santacroce, Elodie Burcklen, Christiane Beckmann, Maurice Redondo, Olivier Kobel, Christoph Noppen, Niko Beerenwinkel, Tanja Stadler |
| EPI_ISL_2938457 | Viollier AG | Department of Biosystems Science and Engineering, ETH Zürich | Chaoran Chen, Sarah Nadeau, Catharine Aquino, Ivan Topolsky, Kim Philipp Jablonski, Lara Fuhrmann, Daniel Ehrsam, Isabel Stürmer, Andreia Cabral de Gouvea, Maria Domenica Moccia, Simon Grüter, Timothy Sykes, Lennart Opitz, Griffin White, Laura Neff, Doris Popovic, Andrea Patrignani, Jay Tracy, Ralph Schlapbach, Christiane Beckmann, Maurice Redondo, Olivier Kobel, Christoph Noppen, Niko Beerenwinkel, Tanja Stadler |
| EPI_ISL_2938462, EPI_ISL_2938463, EPI_ISL_2938465, EPI_ISL_2938470, EPI_ISL_2938474, EPI_ISL_2938475, EPI_ISL_2938477 | Viollier AG | Department of Biosystems Science and Engineering, ETH Zürich | Christian Beisel, Sarah Nadeau, Chaoran Chen, Ivan Topolsky, Kim Philipp Jablonski, Lara Fuhrmann, Rebecca Denes, Mirjam Feldkamp, Ina Nissen, Natascha Santacroce, Elodie Burcklen, Christiane Beckmann, Maurice Redondo, Olivier Kobel, Christoph Noppen, Niko Beerenwinkel, Tanja Stadler |
| EPI_ISL_2938619, EPI_ISL_2938622, EPI_ISL_2938623, EPI_ISL_2938624, EPI_ISL_2938628, EPI_ISL_2938629, EPI_ISL_2938630, EPI_ISL_2938635, EPI_ISL_2938638, EPI_ISL_2938639, EPI_ISL_2938640, EPI_ISL_2938643, EPI_ISL_2938644, EPI_ISL_2938645, EPI_ISL_2938650, EPI_ISL_2938651, EPI_ISL_2938655, EPI_ISL_2938662, EPI_ISL_2938972, EPI_ISL_2938973, EPI_ISL_2938974, EPI_ISL_2938976, EPI_ISL_2938980, EPI_ISL_2938982, EPI_ISL_2938994, EPI_ISL_2938995, EPI_ISL_2938997, EPI_ISL_2938999, EPI_ISL_2939000, EPI_ISL_2939002, EPI_ISL_2939003, EPI_ISL_2939004, EPI_ISL_2939005, EPI_ISL_2939006, EPI_ISL_2939007, EPI_ISL_2939008, EPI_ISL_2939009, EPI_ISL_2939010, EPI_ISL_2939011, EPI_ISL_2939012, EPI_ISL_2939013, EPI_ISL_2939019 | | | |
| see above | Edmonton Provincial Lab | Public Health Agency of Canada (PHAC) National Microbiology Laboratory | Buss, E, Croxen M, Deo A, Dieu P, Gill K, Ferrato C, Khan F, Koleva P, Li V, Lloyd C, Lynch T, Ma R, Murphy S, Pabbaraju K, Shokoples S, Tipples G, Thayer J, Whitehouse M, Wong A, Yu C, Zelyas N |
| EPI_ISL_2939023 | Baptist Health Medical Center | Center for Global Health, University of New Mexico Health Sciences Center | Daryl Domman, Kurt Schwalm, Valerie Morley, Amanda Novack, Dirk Haselow, Joshua L. Kennedy, Darrell Dinwiddie |
| EPI_ISL_2939025, EPI_ISL_2939028 | Edmonton Provincial Lab | Public Health Agency of Canada (PHAC) National Microbiology Laboratory | Buss, E, Croxen M, Deo A, Dieu P, Gill K, Ferrato C, Khan F, Koleva P, Li V, Lloyd C, Lynch T, Ma R, Murphy S, Pabbaraju K, Shokoples S, Tipples G, Thayer J, Whitehouse M, Wong A, Yu C, Zelyas N |
| EPI_ISL_2939029, EPI_ISL_2939030, EPI_ISL_2939032, EPI_ISL_2939033 | Baptist Health Medical Center | Center for Global Health, University of New Mexico Health Sciences Center | Daryl Domman, Kurt Schwalm, Valerie Morley, Amanda Novack, Dirk Haselow, Joshua L. Kennedy, Darrell Dinwiddie |
| EPI_ISL_2939037 | Edmonton Provincial Lab | Public Health Agency of Canada (PHAC) National Microbiology Laboratory | Buss, E, Croxen M, Deo A, Dieu P, Gill K, Ferrato C, Khan F, Koleva P, Li V, Lloyd C, Lynch T, Ma R, Murphy S, Pabbaraju K, Shokoples S, Tipples G, Thayer J, Whitehouse M, Wong A, Yu C, Zelyas N |
| EPI_ISL_2939039 | Arkansas Department of Health | Center for Global Health, University of New Mexico Health Sciences Center | Daryl Domman, Kurt Schwalm, Valerie Morley, Atul Kothari, Joshua L. Kennedy, Darrell Dinwiddie |
| EPI_ISL_2939040, EPI_ISL_2939043, EPI_ISL_2939045, EPI_ISL_2939059, EPI_ISL_2939062, EPI_ISL_2939065, EPI_ISL_2939070, EPI_ISL_2939073, EPI_ISL_2939080, EPI_ISL_2939082, EPI_ISL_2939083, EPI_ISL_2939087, EPI_ISL_2939089, EPI_ISL_2939094, EPI_ISL_2939095, EPI_ISL_2939097, EPI_ISL_2939099, EPI_ISL_2939101, EPI_ISL_2939110, EPI_ISL_2939127, EPI_ISL_2939130, EPI_ISL_2939137, EPI_ISL_2939138, EPI_ISL_2939164, EPI_ISL_2939167, EPI_ISL_2939168, EPI_ISL_2939169, EPI_ISL_2939171, EPI_ISL_2939180, EPI_ISL_2939192, EPI_ISL_2939197, EPI_ISL_2939199, EPI_ISL_2939201, EPI_ISL_2939202, EPI_ISL_2939205, EPI_ISL_2939209, EPI_ISL_2939210, EPI_ISL_2939352, EPI_ISL_2939353, EPI_ISL_2939354, EPI_ISL_2939355, EPI_ISL_2939356, EPI_ISL_2939357, EPI_ISL_2939358, EPI_ISL_2939359, EPI_ISL_2939360, EPI_ISL_2939361, EPI_ISL_2939362, EPI_ISL_2939363, EPI_ISL_2939364, EPI_ISL_2939365, EPI_ISL_2939366, EPI_ISL_2939367, EPI_ISL_2939368, EPI_ISL_2939369, EPI_ISL_2939370, EPI_ISL_2939371, EPI_ISL_2939373, EPI_ISL_2939374, EPI_ISL_2939375, EPI_ISL_2939376, EPI_ISL_2939447, EPI_ISL_2939456, EPI_ISL_2939462, EPI_ISL_2939464, EPI_ISL_2939482, EPI_ISL_2939484, EPI_ISL_2939525, EPI_ISL_2939526, EPI_ISL_2939530, EPI_ISL_2939534, EPI_ISL_2939535, EPI_ISL_2939538, EPI_ISL_2939546, EPI_ISL_2939549, EPI_ISL_2939560, EPI_ISL_2939652, EPI_ISL_2939656, EPI_ISL_2939657, EPI_ISL_2939661, EPI_ISL_2939692, EPI_ISL_2939694, EPI_ISL_2939696, EPI_ISL_2939697, EPI_ISL_2939700, EPI_ISL_2939703, EPI_ISL_2939719, EPI_ISL_2939724, EPI_ISL_2939726, EPI_ISL_2939727, EPI_ISL_2939784, EPI_ISL_2939786, EPI_ISL_2939787, EPI_ISL_2939788, EPI_ISL_2939789, EPI_ISL_2939790, EPI_ISL_2939791, EPI_ISL_2939792, EPI_ISL_2939793, EPI_ISL_2939794, EPI_ISL_2939795, EPI_ISL_2939796, EPI_ISL_2939797, EPI_ISL_2939798, EPI_ISL_2939799, EPI_ISL_2939840, EPI_ISL_2940054, EPI_ISL_2940061, EPI_ISL_2940065, EPI_ISL_2940070, EPI_ISL_2940094, EPI_ISL_2940105, EPI_ISL_2940108, EPI_ISL_2940110, EPI_ISL_2940113, EPI_ISL_2940114, EPI_ISL_2940115, EPI_ISL_2940191 | | | |
| see above | Edmonton Provincial Lab | Public Health Agency of Canada (PHAC) National Microbiology Laboratory | Buss, E, Croxen M, Deo A, Dieu P, Gill K, Ferrato C, Khan F, Koleva P, Li V, Lloyd C, Lynch T, Ma R, Murphy S, Pabbaraju K, Shokoples S, Tipples G, Thayer J, Whitehouse M, Wong A, Yu C, Zelyas N |
| EPI_ISL_2940309, EPI_ISL_2940314, EPI_ISL_2940316, EPI_ISL_2940319, EPI_ISL_2940322, EPI_ISL_2940326, EPI_ISL_2940328, EPI_ISL_2940331, EPI_ISL_2940333, EPI_ISL_2940336 | NJDOH, Public Health and Environmental Laboratories | NJ_PHEL | Lindsey Bodnar, Shiv K. Verma, Ryan Pachucki, Chelsea San Filippo, Jacquelyn Deverell, Dana Woell, Allison Roder, Byeong Jeong |
| EPI_ISL_2940338 | Edmonton Provincial Lab | Public Health Agency of Canada (PHAC) National Microbiology Laboratory | Buss, E, Croxen M, Deo A, Dieu P, Gill K, Ferrato C, Khan F, Koleva P, Li V, Lloyd C, Lynch T, Ma R, Murphy S, Pabbaraju K, Shokoples S, Tipples G, Thayer J, Whitehouse M, Wong A, Yu C, Zelyas N |

| | | | |
|---|---|---|---|
| EPI_ISL_2940339, EPI_ISL_2940340, EPI_ISL_2940341, EPI_ISL_2940342, EPI_ISL_2940344, EPI_ISL_2940349, EPI_ISL_2940351, EPI_ISL_2940352, EPI_ISL_2940354, EPI_ISL_2940359, EPI_ISL_2940362, EPI_ISL_2940363, EPI_ISL_2940392 | | | |
| see above | NJDOH, Public Health and Environmental Laboratories | NJ_PHEL | Lindsey Bodnar, Shiv K. Verma, Ryan Pachucki, Chelsea San Filippo, Jacquelyn Deverell, Dana Woell, Allison Roder, Byeong Jeong |
| EPI_ISL_2940397, EPI_ISL_2940401, EPI_ISL_2940403, EPI_ISL_2940406, EPI_ISL_2940420, EPI_ISL_2940422, EPI_ISL_2940428, EPI_ISL_2940431, EPI_ISL_2940433, EPI_ISL_2940445, EPI_ISL_2940447, EPI_ISL_2940450, EPI_ISL_2940453, EPI_ISL_2940459, EPI_ISL_2940461, EPI_ISL_2940467, EPI_ISL_2940477, EPI_ISL_2940480, EPI_ISL_2940481, EPI_ISL_2940487, EPI_ISL_2940491, EPI_ISL_2940493, EPI_ISL_2940508, EPI_ISL_2940510, EPI_ISL_2940513, EPI_ISL_2940519, EPI_ISL_2940522, EPI_ISL_2940526, EPI_ISL_2940528, EPI_ISL_2940578, EPI_ISL_2940590, EPI_ISL_2940596, EPI_ISL_2940604, EPI_ISL_2940606, EPI_ISL_2940609, EPI_ISL_2940611, EPI_ISL_2940612, EPI_ISL_2940613, EPI_ISL_2940614, EPI_ISL_2940615, EPI_ISL_2940616, EPI_ISL_2940617, EPI_ISL_2940619, EPI_ISL_2940620, EPI_ISL_2940622, EPI_ISL_2940623, EPI_ISL_2940631, EPI_ISL_2940636, EPI_ISL_2940637, EPI_ISL_2940643, EPI_ISL_2940644, EPI_ISL_2940681, EPI_ISL_2940683, EPI_ISL_2940686, EPI_ISL_2940688, EPI_ISL_2940697, EPI_ISL_2940700, EPI_ISL_2940709, EPI_ISL_2940717, EPI_ISL_2940735, EPI_ISL_2940737, EPI_ISL_2940738, EPI_ISL_2940753 | | | |
| see above | Edmonton Provincial Lab | Public Health Agency of Canada (PHAC) National Microbiology Laboratory | Buss, E, Croxen M, Deo A, Dieu P, Gill K, Ferrato C, Khan F, Koleva P, Li V, Lloyd C, Lynch T, Ma R, Murphy S, Pabbaraju K, Shokoples S, Tipples G, Thayer J, Whitehouse M, Wong A, Yu C, Zelyas N |
| EPI_ISL_2940757 | Laboratorium Diagnostyczne Centrum Medycznego Luxmed | 1. Tricity SARS-CoV-2 sequencing consortium: University of Gdansk, Medical University of Gdansk, Vaxican Ltd., Invicta Ltd. 2. National Institute of Public Health - National Institute of Hygiene, Warsaw, Poland | Maciej Kosinski, Celina Cybulska, Krystyna Bienkowska Szewczyk, Maciej Grzybek, Katarzyna Gackowska, Marcin Lubocki, Katarzyna Groth, Lukasz Rabalski, Katarzyna Zacharczuk, Magdalena Nowakowska, Małgorzata Sadkowska-Todys, Tomasz Wołkowicz |
| EPI_ISL_2940763, EPI_ISL_2940764, EPI_ISL_2940769, EPI_ISL_2940783, EPI_ISL_2940786, EPI_ISL_2940788, EPI_ISL_2940790, EPI_ISL_2940791, EPI_ISL_2940792, EPI_ISL_2940795, EPI_ISL_2940800, EPI_ISL_2940808, EPI_ISL_2940809, EPI_ISL_2940817, EPI_ISL_2940825, EPI_ISL_2940826, EPI_ISL_2940827, EPI_ISL_2940828, EPI_ISL_2940831, EPI_ISL_2940838, EPI_ISL_2940850, EPI_ISL_2940906, EPI_ISL_2941109, EPI_ISL_2941121, EPI_ISL_2941289 | | | |
| see above | Edmonton Provincial Lab | Public Health Agency of Canada (PHAC) National Microbiology Laboratory | Buss, E, Croxen M, Deo A, Dieu P, Gill K, Ferrato C, Khan F, Koleva P, Li V, Lloyd C, Lynch T, Ma R, Murphy S, Pabbaraju K, Shokoples S, Tipples G, Thayer J, Whitehouse M, Wong A, Yu C, Zelyas N |
| EPI_ISL_2941510, EPI_ISL_2941518 | Mobile County Health Department | Synergy Laboratories | Megan Cornwell |
| EPI_ISL_2941550, EPI_ISL_2941551 | Wyoming Public Health Laboratory | Wyoming Public Health Laboratory | Jim Mildenberger, Taylor Fearing, Channing Weber, Ashley Norberg, Chayse Rowley, Marley Goetz, Brian Dominguez, Elliot Thomasson, Sam Britz, Cari Sloma, Robert Petit, and Rob Christensen |
| EPI_ISL_2941585, EPI_ISL_2941587, EPI_ISL_2941588, EPI_ISL_2941589, EPI_ISL_2941590, EPI_ISL_2941591, EPI_ISL_2941592, EPI_ISL_2941595, EPI_ISL_2941596, EPI_ISL_2941597, EPI_ISL_2941598, EPI_ISL_2941599, EPI_ISL_2941600, EPI_ISL_2941601, EPI_ISL_2941602, EPI_ISL_2941603, EPI_ISL_2941604, EPI_ISL_2941606, EPI_ISL_2941607, EPI_ISL_2941621, EPI_ISL_2941629, EPI_ISL_2941631, EPI_ISL_2941632, EPI_ISL_2941633, EPI_ISL_2941634, EPI_ISL_2941638, EPI_ISL_2941639, EPI_ISL_2941640 | | | |
| see above | Colorado Department of Public Health and Environment | Colorado Department of Public Health and Environment | Laura Bankers, Molly C. Hetherington-Rauth, Diana Ir, Alexandria Rossheim, Michael A. Martin, Mandy Waters, Shannon R. Matzinger, Sarah Elizabeth Totten, Emily A. Travanty |
| EPI_ISL_2941645 | New Mexico Department of Health Scientific Laboratory | New Mexico Department of Health Scientific Laboratory | Ellie Johnson, D'eldra Malone, Jennifer Benoit, Ratheesh Rajan, Linda Salazar, Keila Gutierrez, Anastacia Griego-Fisher, |
| EPI_ISL_2941660, EPI_ISL_2941669, EPI_ISL_2941670, EPI_ISL_2941679 | Curative Labs | New Mexico Department of Health Scientific Laboratory | Ellie Johnson, D'eldra Malone, Jennifer Benoit, Ratheesh Rajan, Linda Salazar, Keila Gutierrez, Anastacia Griego-Fisher, |
| EPI_ISL_2941722, EPI_ISL_2941724, EPI_ISL_2941727, EPI_ISL_2941728, EPI_ISL_2941734, EPI_ISL_2941738, EPI_ISL_2941740, EPI_ISL_2941741, EPI_ISL_2941744, EPI_ISL_2941751, EPI_ISL_2941764, EPI_ISL_2941779 | | | |
| see above | Montana Public Health Laboratory | Montana Public Health Laboratory | Joy Ritter, Michelle Mozer, Carrie Biskupiak, Deborah Gibson, Michael Dills |
| EPI_ISL_2941814, EPI_ISL_2941815, EPI_ISL_2941816, EPI_ISL_2941817 | Wyoming Public Health Laboratory | Wyoming Public Health Laboratory | Jim Mildenberger, Taylor Fearing, Channing Weber, Ashley Norberg, Chayse Rowley, Marley Goetz, Brian Dominguez, Elliot Thomasson, Sam Britz, Cari Sloma, Robert Petit, and Rob Christensen |
| EPI_ISL_2941840, EPI_ISL_2941842, EPI_ISL_2941843, EPI_ISL_2941844, EPI_ISL_2941845, EPI_ISL_2941853, EPI_ISL_2941855, EPI_ISL_2941856, EPI_ISL_2941857, EPI_ISL_2941859, EPI_ISL_2941861, EPI_ISL_2941863, EPI_ISL_2941864, EPI_ISL_2941870, EPI_ISL_2941871, EPI_ISL_2941874, EPI_ISL_2941879, EPI_ISL_2941881, EPI_ISL_2941882, EPI_ISL_2941883, EPI_ISL_2941884, EPI_ISL_2941886, EPI_ISL_2941888, EPI_ISL_2941889, EPI_ISL_2941892, EPI_ISL_2941897, EPI_ISL_2941898, EPI_ISL_2941901, EPI_ISL_2941904, EPI_ISL_2941909, EPI_ISL_2941911, EPI_ISL_2941913, EPI_ISL_2941916, EPI_ISL_2941917, EPI_ISL_2941920, EPI_ISL_2941921, EPI_ISL_2941922, EPI_ISL_2941924, EPI_ISL_2941925, EPI_ISL_2941944, EPI_ISL_2941949, EPI_ISL_2941958, EPI_ISL_2941961, EPI_ISL_2941974, EPI_ISL_2941976, EPI_ISL_2941984, EPI_ISL_2941985, EPI_ISL_2941986, EPI_ISL_2941988, EPI_ISL_2941992, EPI_ISL_2941993, EPI_ISL_2941995, EPI_ISL_2941997, EPI_ISL_2941998, EPI_ISL_2941999, EPI_ISL_2942001, EPI_ISL_2942003, EPI_ISL_2942012, EPI_ISL_2942017, EPI_ISL_2942018, EPI_ISL_2942019, EPI_ISL_2942021, EPI_ISL_2942023, EPI_ISL_2942024, EPI_ISL_2942025, EPI_ISL_2942026, EPI_ISL_2942028, EPI_ISL_2942029, EPI_ISL_2942030, EPI_ISL_2942031, EPI_ISL_2942033, EPI_ISL_2942036, EPI_ISL_2942038, EPI_ISL_2942039, EPI_ISL_2942041, EPI_ISL_2942043, EPI_ISL_2942044, EPI_ISL_2942045, EPI_ISL_2942046, EPI_ISL_2942047, EPI_ISL_2942048, EPI_ISL_2942049, EPI_ISL_2942050, EPI_ISL_2942051, EPI_ISL_2942052, EPI_ISL_2942053, EPI_ISL_2942054, EPI_ISL_2942055, EPI_ISL_2942056, EPI_ISL_2942057, EPI_ISL_2942059, EPI_ISL_2942060, EPI_ISL_2942061, EPI_ISL_2942062, EPI_ISL_2942063, EPI_ISL_2942064, EPI_ISL_2942065, EPI_ISL_2942066, EPI_ISL_2942067, EPI_ISL_2942068, EPI_ISL_2942070, EPI_ISL_2942071, EPI_ISL_2942073, EPI_ISL_2942075, EPI_ISL_2942076, EPI_ISL_2942077, EPI_ISL_2942078, EPI_ISL_2942079, EPI_ISL_2942080, EPI_ISL_2942082, EPI_ISL_2942083, EPI_ISL_2942084, EPI_ISL_2942085, EPI_ISL_2942088, EPI_ISL_2942089, EPI_ISL_2942091, EPI_ISL_2942092, EPI_ISL_2942093, EPI_ISL_2942094, EPI_ISL_2942095, EPI_ISL_2942096, EPI_ISL_2942098, EPI_ISL_2942099, EPI_ISL_2942119, EPI_ISL_2942130, EPI_ISL_2942141, EPI_ISL_2942149, EPI_ISL_2942150, EPI_ISL_2942157, EPI_ISL_2942160, EPI_ISL_2942162, EPI_ISL_2942166, EPI_ISL_2942182, EPI_ISL_2942183, EPI_ISL_2942186, EPI_ISL_2942187 | | | |
| see above | Curative Labs | Curative Labs | Mikhail Hanewich-Hollatz, J.Cesar Ignacio-Espinoza, Elias L. Salfati, Nabjot Sandhu, George Way, Zhiyi Xie, Eugenia Khorosheva, Sophia Quasem, Janet Chen, Vladimir Slepnev |
| EPI_ISL_2942188, EPI_ISL_2942189, EPI_ISL_2942190, EPI_ISL_2942191, EPI_ISL_2942195, EPI_ISL_2942196, EPI_ISL_2942197, EPI_ISL_2942198, EPI_ISL_2942199, EPI_ISL_2942200, EPI_ISL_2942201, EPI_ISL_2942202, EPI_ISL_2942203, EPI_ISL_2942205, EPI_ISL_2942207, EPI_ISL_2942208, EPI_ISL_2942209, EPI_ISL_2942210, EPI_ISL_2942211, EPI_ISL_2942212, EPI_ISL_2942213, EPI_ISL_2942214, EPI_ISL_2942215, EPI_ISL_2942216, EPI_ISL_2942217, EPI_ISL_2942218, EPI_ISL_2942220, EPI_ISL_2942221, EPI_ISL_2942222, EPI_ISL_2942223, EPI_ISL_2942224, EPI_ISL_2942225, EPI_ISL_2942226, EPI_ISL_2942227, EPI_ISL_2942228, EPI_ISL_2942229, EPI_ISL_2942230, EPI_ISL_2942231, EPI_ISL_2942232, EPI_ISL_2942233, EPI_ISL_2942234, EPI_ISL_2942235, EPI_ISL_2942236, EPI_ISL_2942237, EPI_ISL_2942239, EPI_ISL_2942240, EPI_ISL_2942241, EPI_ISL_2942243, EPI_ISL_2942244, EPI_ISL_2942245, EPI_ISL_2942246, EPI_ISL_2942247, EPI_ISL_2942248, EPI_ISL_2942249, EPI_ISL_2942250, EPI_ISL_2942251, EPI_ISL_2942252, EPI_ISL_2942253, EPI_ISL_2942254, EPI_ISL_2942255, EPI_ISL_2942256, EPI_ISL_2942257, EPI_ISL_2942258, EPI_ISL_2942259, EPI_ISL_2942260, EPI_ISL_2942261, EPI_ISL_2942262, EPI_ISL_2942263, EPI_ISL_2942264, EPI_ISL_2942265, EPI_ISL_2942266, EPI_ISL_2942267, EPI_ISL_2942268, EPI_ISL_2942269, EPI_ISL_2942270, EPI_ISL_2942271, EPI_ISL_2942272, EPI_ISL_2942273, EPI_ISL_2942275, EPI_ISL_2942276, EPI_ISL_2942277, EPI_ISL_2942278 | | | |
| see above | Nevada State Public Health Laboratory | Nevada State Public Health Laboratory | Andrew Gorzalski, Mark Pandori |
| EPI_ISL_2942281 | RUSTENBURG LABORATORY | National Institute for Communicable Diseases of the National Health Laboratory Service | Amoako DG, Everatt J, Scheepers C, Mohale T, Ntuli N, Mahlangu B, Mnguni A, Ismail A, Bhiman JN |
| EPI_ISL_2942283 | KOPANONG LABORATORY | National Institute for Communicable Diseases of the National Health Laboratory Service | Amoako DG, Everatt J, Scheepers C, Mohale T, Ntuli N, Mahlangu B, Mnguni A, Ismail A, Bhiman JN |
| EPI_ISL_2942288 | PORT ELIZABETH LABORATORY | National Institute for Communicable Diseases of the National Health Laboratory Service | Amoako DG, Everatt J, Scheepers C, Mohale T, Ntuli N, Mahlangu B, Mnguni A, Ismail A, Bhiman JN |
| EPI_ISL_2942336, EPI_ISL_2942339, EPI_ISL_2942345, EPI_ISL_2942364, EPI_ISL_2942384, EPI_ISL_2942393, EPI_ISL_2942396, EPI_ISL_2942397 | Unidad de Investigacion Medica de Yucatan (UIMY) | Unidad de Genomica Avanzada | Consorcio Mexicano de Vigilancia Genomica (CoViGen-Mex). Authors (in alphabetical order): Julio Elias Alvarado-Yaah, Carlos F. Arias, Santiago avila-Rios, Eduardo Becerril-Vargas, Victor Hugo Borja-Aburto, Celia Boukadida, Cristobal Chaidez-Quiroz, Juan Bautista Chale-Dzul, Ricardo Ciria Merce, Andreu Comas-Garcia, Celida Duque Molina, Julissa Enciso-Ibarra, Jose Antonio Enciso-Moreno, Gloria Elena Espinosa-Ayala, Fernando Fontove-Herrera, Daniel Fregoso-Rueda, Victor Eduardo Garcia-Arias, Alejandra Garcia-Gasca, Bruno Gomez-Gil, Jean Pierre Gonzalez, Irvin Gonzalez-Lopez, Concepcion Grajales-Muñiz, Ricardo Grande, Rosa Maria Gutierrez Rios, Jesus Hernandez, Alejandra Hernandez-Teran, Alfredo Herrera-Estrella, Carla Ivon Herrera-Najera, Pavel Isa, Veronica Mata-Haro, Daniel Lira Morales, Susana Lopez, Antonio Loza Roman, Brenda Irasema Maldonado-Meza, Bernardo Martinez-Miguel, Jose Arturo Martinez-Orozco, Celida Martinez-Rodriguez, Margarita Matias-Florentino, Fidencio Mejia-Nepomuceno, Maria Guadalupe de Jesus Mireles-Rivera, Gloria Maria Molina-Salinas, Hector Montoya-Fuentes, , Mario Mujica-Sanchez, Jose Esteban Muñoz-Medina, Jose de Jesus Nuñez-Contreras, Alicia Ocaña-Mondragon, Luis Alberto Ochoa-Carrera, Hector Esteban Paz-Juarez, Marissa Perez-Garcia, Francisco Pulido, Helen Haydee Fernanda Ramirez-Plascencia, Jorge Salas-Hernandez, Angel Gustavo Salas-Lais, Alejandro Sanchez-Flores, Clara Esperanza Santacruz-Tinoco, Maria Guadalupe Santiago-Mauricio, Selene Zarate, Nelly Selem-Mojica, Blanca Taboada, Gloria Vazquez, Joel Armando Vazquez-Perez. |
| EPI_ISL_2942412, EPI_ISL_2942413, EPI_ISL_2942435, EPI_ISL_2942437, EPI_ISL_2942439, EPI_ISL_2942447, EPI_ISL_2942448, EPI_ISL_2942449, EPI_ISL_2942451, EPI_ISL_2942452, EPI_ISL_2942453, EPI_ISL_2942454, EPI_ISL_2942456, EPI_ISL_2942457, EPI_ISL_2942458, EPI_ISL_2942459, EPI_ISL_2942461, EPI_ISL_2942465 | | | |
| see above | Centro de Investigacion Biomedica de Occidente (CIBO) | Unidad de Genomica Avanzada | Consorcio Mexicano de Vigilancia Genomica (CoViGen-Mex). Authors (in alphabetical order): Julio Elias Alvarado-Yaah, Carlos F. Arias, Santiago avila-Rios, Eduardo Becerril-Vargas, Victor Hugo Borja-Aburto, Celia Boukadida, Cristobal Chaidez-Quiroz, Juan Bautista Chale-Dzul, Ricardo Ciria Merce, Andreu Comas-Garcia, Celida Duque Molina, Julissa Enciso-Ibarra, Jose Antonio Enciso-Moreno, Gloria Elena Espinosa-Ayala, Fernando Fontove-Herrera, Daniel Fregoso-Rueda, Victor Eduardo Garcia-Arias, Alejandra Garcia-Gasca, Bruno Gomez-Gil, Jean Pierre Gonzalez, Irvin Gonzalez-Lopez, Concepcion Grajales-Muñiz, Ricardo Grande, Rosa Maria Gutierrez Rios, Jesus Hernandez, Alejandra Hernandez-Teran, Alfredo Herrera-Estrella, Carla Ivon Herrera-Najera, Pavel Isa, Veronica Mata-Haro, Daniel Lira Morales, Susana Lopez, Antonio Loza Roman, Brenda Irasema Maldonado-Meza, Bernardo Martinez-Miguel, Jose Arturo Martinez-Orozco, Celida Martinez-Rodriguez, Margarita Matias-Florentino, Fidencio Mejia-Nepomuceno, Maria Guadalupe de Jesus Mireles-Rivera, Gloria Maria Molina-Salinas, Hector Montoya-Fuentes, , Mario Mujica-Sanchez, Jose Esteban Muñoz-Medina, Jose de Jesus Nuñez-Contreras, Alicia Ocaña-Mondragon, Luis Alberto Ochoa-Carrera, Hector Esteban Paz-Juarez, Marissa Perez-Garcia, Francisco Pulido, Helen Haydee Fernanda Ramirez-Plascencia, Jorge Salas-Hernandez, Angel Gustavo Salas-Lais, Alejandro Sanchez-Flores, Clara Esperanza Santacruz-Tinoco, Maria Guadalupe Santiago-Mauricio, Selene Zarate, Nelly Selem-Mojica, Blanca Taboada, Gloria Vazquez, Joel Armando Vazquez-Perez. |
| EPI_ISL_2942469, EPI_ISL_2942475, EPI_ISL_2942478, EPI_ISL_2942479, EPI_ISL_2942480 | Unidad de Investigacion Biomedica de Zacatecas (UIBZ) | Unidad de Genomica Avanzada | Consorcio Mexicano de Vigilancia Genomica (CoViGen-Mex). Authors (in alphabetical order): Julio Elias Alvarado-Yaah, Carlos F. Arias, Santiago avila-Rios, Eduardo Becerril-Vargas, Victor Hugo Borja-Aburto, Celia Boukadida, Cristobal Chaidez-Quiroz, Juan Bautista Chale-Dzul, Ricardo Ciria Merce, Andreu Comas-Garcia, Celida Duque Molina, Julissa Enciso-Ibarra, Jose Antonio Enciso-Moreno, Gloria Elena Espinosa-Ayala, Fernando Fontove-Herrera, Daniel Fregoso-Rueda, Victor Eduardo Garcia-Arias, Alejandra Garcia-Gasca, Bruno Gomez-Gil, Jean Pierre Gonzalez, Irvin Gonzalez-Lopez, Concepcion Grajales-Muñiz, Ricardo Grande, Rosa Maria Gutierrez Rios, Jesus Hernandez, Alejandra Hernandez-Teran, Alfredo Herrera-Estrella, Carla Ivon Herrera-Najera, Pavel Isa, Veronica Mata-Haro, Daniel Lira Morales, Susana Lopez, Antonio Loza Roman, Brenda Irasema Maldonado-Meza, Bernardo Martinez-Miguel, Jose Arturo Martinez-Orozco, Celida Martinez-Rodriguez, Margarita Matias-Florentino, Fidencio Mejia-Nepomuceno, Maria Guadalupe de Jesus Mireles-Rivera, Gloria Maria Molina-Salinas, Hector Montoya-Fuentes, , Mario Mujica-Sanchez, Jose Esteban Muñoz-Medina, Jose de Jesus Nuñez-Contreras, Alicia Ocaña-Mondragon, Luis Alberto Ochoa-Carrera, Hector Esteban Paz-Juarez, Marissa Perez-Garcia, Francisco Pulido, Helen Haydee Fernanda Ramirez-Plascencia, Jorge Salas-Hernandez, Angel Gustavo Salas-Lais, Alejandro Sanchez-Flores, Clara Esperanza Santacruz-Tinoco, Maria Guadalupe Santiago-Mauricio, Selene Zarate, Nelly Selem-Mojica, Blanca Taboada, Gloria Vazquez, Joel Armando Vazquez-Perez. |
| EPI_ISL_2942494, EPI_ISL_2942503, EPI_ISL_2942511, EPI_ISL_2942512 | Centro de Investigacion Biomedica del Noreste (CIBIN) | Unidad de Genomica Avanzada | Consorcio Mexicano de Vigilancia Genomica (CoViGen-Mex). Authors (in alphabetical order): Julio Elias Alvarado-Yaah, Carlos F. Arias, Santiago avila-Rios, Eduardo Becerril-Vargas, Victor Hugo Borja-Aburto, Celia Boukadida, Cristobal Chaidez-Quiroz, Juan Bautista Chale-Dzul, Ricardo Ciria Merce, Andreu Comas-Garcia, Celida Duque Molina, Julissa Enciso-Ibarra, Jose Antonio Enciso-Moreno, Gloria Elena Espinosa-Ayala, Fernando Fontove-Herrera, Daniel Fregoso-Rueda, Victor Eduardo Garcia-Arias, Alejandra Garcia-Gasca, Bruno Gomez-Gil, Jean Pierre Gonzalez, Irvin Gonzalez-Lopez, Concepcion Grajales-Muñiz, Ricardo Grande, Rosa Maria Gutierrez Rios, Jesus Hernandez, Alejandra Hernandez-Teran, Alfredo Herrera-Estrella, Carla Ivon Herrera-Najera, Pavel Isa, Veronica Mata-Haro, Daniel Lira Morales, Susana Lopez, Antonio Loza Roman, Brenda Irasema Maldonado-Meza, Bernardo Martinez-Miguel, Jose Arturo Martinez-Orozco, Celida Martinez-Rodriguez, Margarita Matias-Florentino, Fidencio Mejia-Nepomuceno, Maria Guadalupe de Jesus Mireles-Rivera, Gloria Maria Molina-Salinas, Hector Montoya-Fuentes, , Mario Mujica-Sanchez, Jose Esteban Muñoz-Medina, Jose de Jesus Nuñez-Contreras, Alicia Ocaña-Mondragon, Luis Alberto Ochoa-Carrera, Hector Esteban Paz-Juarez, Marissa Perez-Garcia, Francisco Pulido, Helen Haydee Fernanda Ramirez-Plascencia, Jorge Salas-Hernandez, Angel Gustavo Salas-Lais, Alejandro Sanchez-Flores, Clara Esperanza Santacruz-Tinoco, Maria Guadalupe Santiago-Mauricio, Selene Zarate, Nelly Selem-Mojica, Blanca Taboada, Gloria Vazquez, Joel Armando Vazquez-Perez. |
| EPI_ISL_2942534, EPI_ISL_2942537, EPI_ISL_2942538, EPI_ISL_2942539, EPI_ISL_2942549, EPI_ISL_2942553, EPI_ISL_2942557, EPI_ISL_2942559, EPI_ISL_2942570, EPI_ISL_2942572, EPI_ISL_2942575, EPI_ISL_2942576, EPI_ISL_2942578, EPI_ISL_2942579, EPI_ISL_2942583, EPI_ISL_2942585, EPI_ISL_2942586, EPI_ISL_2942587, EPI_ISL_2942598, EPI_ISL_2942599, EPI_ISL_2942600, EPI_ISL_2942601, EPI_ISL_2942603, EPI_ISL_2942607, EPI_ISL_2942608, EPI_ISL_2942609, EPI_ISL_2942610, EPI_ISL_2942611, EPI_ISL_2942612, EPI_ISL_2942613, EPI_ISL_2942614, EPI_ISL_2942615, EPI_ISL_2942616, EPI_ISL_2942618, EPI_ISL_2942619, EPI_ISL_2942620, EPI_ISL_2942622, EPI_ISL_2942623, EPI_ISL_2942624 | | | |
| see above | Laboratorio Central de Epidemiologia (LCE) | Unidad de Genomica Avanzada | Consorcio Mexicano de Vigilancia Genomica (CoViGen-Mex). Authors (in alphabetical order): Julio Elias Alvarado-Yaah, Carlos F. Arias, Santiago avila-Rios, Eduardo Becerril-Vargas, Victor Hugo Borja-Aburto, Celia Boukadida, Cristobal Chaidez-Quiroz, Juan Bautista Chale-Dzul, Ricardo Ciria Merce, Andreu Comas-Garcia, Celida Duque Molina, Julissa Enciso-Ibarra, Jose Antonio Enciso-Moreno, Gloria Elena Espinosa-Ayala, Fernando Fontove-Herrera, Daniel Fregoso-Rueda, Victor Eduardo Garcia-Arias, Alejandra Garcia-Gasca, Bruno Gomez-Gil, Jean Pierre Gonzalez, Irvin Gonzalez-Lopez, Concepcion Grajales-Muñiz, Ricardo Grande, Rosa Maria Gutierrez Rios, Jesus Hernandez, Alejandra Hernandez-Teran, Alfredo Herrera-Estrella, Carla Ivon Herrera-Najera, Pavel Isa, Veronica Mata-Haro, Daniel Lira Morales, Susana Lopez, Antonio Loza Roman, Brenda Irasema Maldonado-Meza, Bernardo Martinez-Miguel, Jose Arturo Martinez-Orozco, Celida Martinez-Rodriguez, Margarita Matias-Florentino, Fidencio Mejia-Nepomuceno, Maria Guadalupe de Jesus Mireles-Rivera, Gloria Maria Molina-Salinas, Hector Montoya-Fuentes, , Mario Mujica-Sanchez, Jose Esteban Muñoz-Medina, Jose de Jesus Nuñez-Contreras, Alicia Ocaña-Mondragon, Luis Alberto Ochoa-Carrera, Hector Esteban Paz-Juarez, Marissa Perez-Garcia, Francisco Pulido, Helen Haydee Fernanda Ramirez-Plascencia, Jorge Salas-Hernandez, Angel Gustavo Salas-Lais, Alejandro Sanchez-Flores, Clara Esperanza Santacruz-Tinoco, Maria Guadalupe Santiago-Mauricio, Selene Zarate, Nelly Selem-Mojica, Blanca Taboada, Gloria Vazquez, Joel Armando Vazquez-Perez. |
| EPI_ISL_2942625 | Department of Health Technology and Informatics, The Hong Kong Polytechnic University | Department of Health Technology and Informatics, The Hong Kong Polytechnic University | Gilman Kit-Hang Siu, Lam-Kwong Lee, Kenneth Siu-Sing Leung, Jake Siu-Lun Leung, Wing-Hei Lo, Timothy Ting-Leung Ng, Chloe Toi-Mei Chan, Hoi-Ching Jim, Ivan Tak-Fai Wong, Kingsley King-Gee Tam, Hiu-Yin Lao, Denise Sze-Hang Wong, Alan Ka-Lun Wu, Miranda Chong-Yee Yau, Yvette Wai-Man Lai, Kitty Sau-Chun Fung, Sandy Ka-Yee Chau, Barry Kin-Chung Wong, Wing-Kin To, Kristine Luk, Alex Yat-Man Ho, Tak-Lun Que, Kam-Tong Yip, Wing Cheong Yam, David Ho-Keung Shum, Shea Ping Yip |
| EPI_ISL_2942638, EPI_ISL_2942644, | Unidad de Investigacion Medica de Yucatan (UIMY) | Unidad de Genomica Avanzada | Consorcio Mexicano de Vigilancia Genomica (CoViGen-Mex). Authors (in alphabetical order): Julio Elias Alvarado-Yaah, Carlos F. Arias, Santiago avila-Rios, Eduardo Becerril-Vargas, Victor Hugo Borja-Aburto, Celia Boukadida, Cristobal Chaidez-Quiroz, Juan Bautista Chale-Dzul, Ricardo Ciria Merce, Andreu Comas-Garcia, Celida Duque Molina, Julissa Enciso-Ibarra, Jose Antonio Enciso-Moreno, Gloria Elena Espinosa-Ayala, Fernando Fontove-Herrera, Daniel Fregoso-Rueda, Victor |

| | | | |
|---|---|---|---|
| EPI_ISL_2942674, EPI_ISL_2942690, EPI_ISL_2942693 | | | Eduardo Garcia-Arias, Alejandra Garcia-Gasca, Bruno Gomez-Gil, Jean Pierre Gonzalez, Irvin Gonzalez-Lopez, Concepcion Grajales-Muñiz, Ricardo Grande, Rosa Maria Gutierrez Rios, Jesus Hernandez, Alejandra Hernandez-Teran, Alfredo Herrera-Estrella, Carla Ivon Herrera-Najera, Pavel Isa, Veronica Mata-Haro, Daniel Lira Morales, Susana Lopez, Antonio Loza Roman, Brenda Irasema Maldonado-Meza, Bernardo Martinez-Miguel, Jose Arturo Martinez-Orozco, Celida Martinez-Rodriguez, Margarita Matias-Florentino, Fidencio Mejia-Nepomuceno, Maria Guadalupe de Jesus Mireles-Rivera, Gloria Maria Molina-Salinas, Hector Montoya-Fuentes, , Mario Mujica-Sanchez, Jose Esteban Muñoz-Medina, Jose de Jesus Nuñez-Contreras, Alicia Ocaña-Mondragon, Luis Alberto Ochoa-Carrera, Hector Esteban Paz-Juarez, Marissa Perez-Garcia, Francisco Pulido, Helen Haydee Fernanda Ramirez-Plascencia, Jorge Salas-Hernandez, Angel Gustavo Salas-Lais, Alejandro Sanchez-Flores, Clara Esperanza Santacruz-Tinoco, Maria Guadalupe Santiago-Mauricio, Selene Zarate, Nelly Selem-Mojica, Blanca Taboada, Gloria Vazquez, Joel Armando Vazquez-Perez. |
| EPI_ISL_2942720, EPI_ISL_2942721, EPI_ISL_2942722, EPI_ISL_2942723, EPI_ISL_2942724, EPI_ISL_2942726, EPI_ISL_2942727, EPI_ISL_2942729, EPI_ISL_2942730, EPI_ISL_2942731, EPI_ISL_2942733, EPI_ISL_2942734, EPI_ISL_2942736, EPI_ISL_2942738, EPI_ISL_2942740, EPI_ISL_2942741, EPI_ISL_2942742, EPI_ISL_2942743, EPI_ISL_2942744, EPI_ISL_2942745, EPI_ISL_2942746, EPI_ISL_2942747, EPI_ISL_2942748, EPI_ISL_2942749 | | | |
| see above | Centro de Investigacion Biomedica de Occidente (CIBO) | Unidad de Genomica Avanzada | Consorcio Mexicano de Vigilancia Genomica (CoViGen-Mex). Authors (in alphabetical order): Julio Elias Alvarado-Yaah, Carlos F. Arias, Santiago avila-Rios, Eduardo Becerril-Vargas, Victor Hugo Borja-Aburto, Celia Boukadida, Cristobal Chaidez-Quiroz, Juan Bautista Chale-Dzul, Ricardo Ciria Merce, Andreu Comas-Garcia, Celida Duque Molina, Julissa Enciso-Ibarra, Jose Antonio Enciso-Moreno, Gloria Elena Espinosa-Ayala, Fernando Fontove-Herrera, Daniel Fregoso-Rueda, Victor Eduardo Garcia-Arias, Alejandra Garcia-Gasca, Bruno Gomez-Gil, Jean Pierre Gonzalez, Irvin Gonzalez-Lopez, Concepcion Grajales-Muñiz, Ricardo Grande, Rosa Maria Gutierrez Rios, Jesus Hernandez, Alejandra Hernandez-Teran, Alfredo Herrera-Estrella, Carla Ivon Herrera-Najera, Pavel Isa, Veronica Mata-Haro, Daniel Lira Morales, Susana Lopez, Antonio Loza Roman, Brenda Irasema Maldonado-Meza, Bernardo Martinez-Miguel, Jose Arturo Martinez-Orozco, Celida Martinez-Rodriguez, Margarita Matias-Florentino, Fidencio Mejia-Nepomuceno, Maria Guadalupe de Jesus Mireles-Rivera, Gloria Maria Molina-Salinas, Hector Montoya-Fuentes, , Mario Mujica-Sanchez, Jose Esteban Muñoz-Medina, Jose de Jesus Nuñez-Contreras, Alicia Ocaña-Mondragon, Luis Alberto Ochoa-Carrera, Hector Esteban Paz-Juarez, Marissa Perez-Garcia, Francisco Pulido, Helen Haydee Fernanda Ramirez-Plascencia, Jorge Salas-Hernandez, Angel Gustavo Salas-Lais, Alejandro Sanchez-Flores, Clara Esperanza Santacruz-Tinoco, Maria Guadalupe Santiago-Mauricio, Selene Zarate, Nelly Selem-Mojica, Blanca Taboada, Gloria Vazquez, Joel Armando Vazquez-Perez. |
| EPI_ISL_2942750, EPI_ISL_2942752, EPI_ISL_2942753 | Unidad de Investigacion Biomedica de Zacatecas (UIBZ) | Unidad de Genomica Avanzada | Consorcio Mexicano de Vigilancia Genomica (CoViGen-Mex). Authors (in alphabetical order): Julio Elias Alvarado-Yaah, Carlos F. Arias, Santiago avila-Rios, Eduardo Becerril-Vargas, Victor Hugo Borja-Aburto, Celia Boukadida, Cristobal Chaidez-Quiroz, Juan Bautista Chale-Dzul, Ricardo Ciria Merce, Andreu Comas-Garcia, Celida Duque Molina, Julissa Enciso-Ibarra, Jose Antonio Enciso-Moreno, Gloria Elena Espinosa-Ayala, Fernando Fontove-Herrera, Daniel Fregoso-Rueda, Victor Eduardo Garcia-Arias, Alejandra Garcia-Gasca, Bruno Gomez-Gil, Jean Pierre Gonzalez, Irvin Gonzalez-Lopez, Concepcion Grajales-Muñiz, Ricardo Grande, Rosa Maria Gutierrez Rios, Jesus Hernandez, Alejandra Hernandez-Teran, Alfredo Herrera-Estrella, Carla Ivon Herrera-Najera, Pavel Isa, Veronica Mata-Haro, Daniel Lira Morales, Susana Lopez, Antonio Loza Roman, Brenda Irasema Maldonado-Meza, Bernardo Martinez-Miguel, Jose Arturo Martinez-Orozco, Celida Martinez-Rodriguez, Margarita Matias-Florentino, Fidencio Mejia-Nepomuceno, Maria Guadalupe de Jesus Mireles-Rivera, Gloria Maria Molina-Salinas, Hector Montoya-Fuentes, , Mario Mujica-Sanchez, Jose Esteban Muñoz-Medina, Jose de Jesus Nuñez-Contreras, Alicia Ocaña-Mondragon, Luis Alberto Ochoa-Carrera, Hector Esteban Paz-Juarez, Marissa Perez-Garcia, Francisco Pulido, Helen Haydee Fernanda Ramirez-Plascencia, Jorge Salas-Hernandez, Angel Gustavo Salas-Lais, Alejandro Sanchez-Flores, Clara Esperanza Santacruz-Tinoco, Maria Guadalupe Santiago-Mauricio, Selene Zarate, Nelly Selem-Mojica, Blanca Taboada, Gloria Vazquez, Joel Armando Vazquez-Perez. |
| EPI_ISL_2942758, EPI_ISL_2942759, EPI_ISL_2942760, EPI_ISL_2942761, EPI_ISL_2942763, EPI_ISL_2942773, EPI_ISL_2942775, EPI_ISL_2942776, EPI_ISL_2942777, EPI_ISL_2942779, EPI_ISL_2942780, EPI_ISL_2942781, EPI_ISL_2942782, EPI_ISL_2942783, EPI_ISL_2942784, EPI_ISL_2942786, EPI_ISL_2942787, EPI_ISL_2942788, EPI_ISL_2942790, EPI_ISL_2942791, EPI_ISL_2942792 | | | |
| see above | Centro de Investigacion Biomedica del Noreste (CIBIN) | Unidad de Genomica Avanzada | Consorcio Mexicano de Vigilancia Genomica (CoViGen-Mex). Authors (in alphabetical order): Julio Elias Alvarado-Yaah, Carlos F. Arias, Santiago avila-Rios, Eduardo Becerril-Vargas, Victor Hugo Borja-Aburto, Celia Boukadida, Cristobal Chaidez-Quiroz, Juan Bautista Chale-Dzul, Ricardo Ciria Merce, Andreu Comas-Garcia, Celida Duque Molina, Julissa Enciso-Ibarra, Jose Antonio Enciso-Moreno, Gloria Elena Espinosa-Ayala, Fernando Fontove-Herrera, Daniel Fregoso-Rueda, Victor Eduardo Garcia-Arias, Alejandra Garcia-Gasca, Bruno Gomez-Gil, Jean Pierre Gonzalez, Irvin Gonzalez-Lopez, Concepcion Grajales-Muñiz, Ricardo Grande, Rosa Maria Gutierrez Rios, Jesus Hernandez, Alejandra Hernandez-Teran, Alfredo Herrera-Estrella, Carla Ivon Herrera-Najera, Pavel Isa, Veronica Mata-Haro, Daniel Lira Morales, Susana Lopez, Antonio Loza Roman, Brenda Irasema Maldonado-Meza, Bernardo Martinez-Miguel, Jose Arturo Martinez-Orozco, Celida Martinez-Rodriguez, Margarita Matias-Florentino, Fidencio Mejia-Nepomuceno, Maria Guadalupe de Jesus Mireles-Rivera, Gloria Maria Molina-Salinas, Hector Montoya-Fuentes, , Mario Mujica-Sanchez, Jose Esteban Muñoz-Medina, Jose de Jesus Nuñez-Contreras, Alicia Ocaña-Mondragon, Luis Alberto Ochoa-Carrera, Hector Esteban Paz-Juarez, Marissa Perez-Garcia, Francisco Pulido, Helen Haydee Fernanda Ramirez-Plascencia, Jorge Salas-Hernandez, Angel Gustavo Salas-Lais, Alejandro Sanchez-Flores, Clara Esperanza Santacruz-Tinoco, Maria Guadalupe Santiago-Mauricio, Selene Zarate, Nelly Selem-Mojica, Blanca Taboada, Gloria Vazquez, Joel Armando Vazquez-Perez. |
| EPI_ISL_2942793, EPI_ISL_2942794, EPI_ISL_2942795, EPI_ISL_2942797, EPI_ISL_2942798, EPI_ISL_2942799, EPI_ISL_2942800, EPI_ISL_2942801, EPI_ISL_2942802, EPI_ISL_2942803, EPI_ISL_2942804, EPI_ISL_2942807, EPI_ISL_2942808, EPI_ISL_2942809, EPI_ISL_2942810, EPI_ISL_2942811, EPI_ISL_2942813, EPI_ISL_2942814, EPI_ISL_2942815, EPI_ISL_2942822, EPI_ISL_2942823, EPI_ISL_2942824, EPI_ISL_2942825, EPI_ISL_2942826, EPI_ISL_2942832, EPI_ISL_2942834, EPI_ISL_2942835, EPI_ISL_2942836, EPI_ISL_2942838, EPI_ISL_2942840, EPI_ISL_2942841, EPI_ISL_2942842, EPI_ISL_2942843, EPI_ISL_2942844, EPI_ISL_2942846, EPI_ISL_2942847, EPI_ISL_2942850, EPI_ISL_2942852, EPI_ISL_2942853, EPI_ISL_2942854, EPI_ISL_2942855, EPI_ISL_2942856, EPI_ISL_2942857, EPI_ISL_2942858, EPI_ISL_2942859, EPI_ISL_2942860, EPI_ISL_2942861, EPI_ISL_2942862, EPI_ISL_2942863, EPI_ISL_2942864, EPI_ISL_2942867, EPI_ISL_2942868, EPI_ISL_2942869, EPI_ISL_2942870, EPI_ISL_2942871, EPI_ISL_2942875, EPI_ISL_2942876, EPI_ISL_2942879, EPI_ISL_2942880, EPI_ISL_2942881, EPI_ISL_2942882, EPI_ISL_2942883, EPI_ISL_2942884, EPI_ISL_2942885, EPI_ISL_2942886, EPI_ISL_2942887, EPI_ISL_2942888, EPI_ISL_2942889, EPI_ISL_2942890, EPI_ISL_2942891, EPI_ISL_2942892, EPI_ISL_2942894, EPI_ISL_2942895, EPI_ISL_2942896, EPI_ISL_2942897, EPI_ISL_2942901, EPI_ISL_2942906, EPI_ISL_2942907, EPI_ISL_2942910, EPI_ISL_2942911, EPI_ISL_2942912, EPI_ISL_2942913, EPI_ISL_2942914, EPI_ISL_2942915, EPI_ISL_2942916, EPI_ISL_2942917, EPI_ISL_2942918, EPI_ISL_2942925, EPI_ISL_2942927, EPI_ISL_2942928, EPI_ISL_2942929, EPI_ISL_2942931, EPI_ISL_2942932, EPI_ISL_2942935 | | | |
| see above | Laboratorio Central de Epidemiologia (LCE) | Unidad de Genomica Avanzada | Consorcio Mexicano de Vigilancia Genomica (CoViGen-Mex). Authors (in alphabetical order): Julio Elias Alvarado-Yaah, Carlos F. Arias, Santiago avila-Rios, Eduardo Becerril-Vargas, Victor Hugo Borja-Aburto, Celia Boukadida, Cristobal Chaidez-Quiroz, Juan Bautista Chale-Dzul, Ricardo Ciria Merce, Andreu Comas-Garcia, Celida Duque Molina, Julissa Enciso-Ibarra, Jose Antonio Enciso-Moreno, Gloria Elena Espinosa-Ayala, Fernando Fontove-Herrera, Daniel Fregoso-Rueda, Victor Eduardo Garcia-Arias, Alejandra Garcia-Gasca, Bruno Gomez-Gil, Jean Pierre Gonzalez, Irvin Gonzalez-Lopez, Concepcion Grajales-Muñiz, Ricardo Grande, Rosa Maria Gutierrez Rios, Jesus Hernandez, Alejandra Hernandez-Teran, Alfredo Herrera-Estrella, Carla Ivon Herrera-Najera, Pavel Isa, Veronica Mata-Haro, Daniel Lira Morales, Susana Lopez, Antonio Loza Roman, Brenda Irasema Maldonado-Meza, Bernardo Martinez-Miguel, Jose Arturo Martinez-Orozco, Celida Martinez-Rodriguez, Margarita Matias-Florentino, Fidencio Mejia-Nepomuceno, Maria Guadalupe de Jesus Mireles-Rivera, Gloria Maria Molina-Salinas, Hector Montoya-Fuentes, , Mario Mujica-Sanchez, Jose Esteban Muñoz-Medina, Jose de Jesus Nuñez-Contreras, Alicia Ocaña-Mondragon, Luis Alberto Ochoa-Carrera, Hector Esteban Paz-Juarez, Marissa Perez-Garcia, Francisco Pulido, Helen Haydee Fernanda Ramirez-Plascencia, Jorge Salas-Hernandez, Angel Gustavo Salas-Lais, Alejandro Sanchez-Flores, Clara Esperanza Santacruz-Tinoco, Maria Guadalupe Santiago-Mauricio, Selene Zarate, Nelly Selem-Mojica, Blanca Taboada, Gloria Vazquez, Joel Armando Vazquez-Perez. |
| EPI_ISL_2942936, EPI_ISL_2942937, EPI_ISL_2942938, EPI_ISL_2942939, EPI_ISL_2942940, EPI_ISL_2942941, EPI_ISL_2942942, EPI_ISL_2942943, EPI_ISL_2942944, EPI_ISL_2942945, EPI_ISL_2942946, EPI_ISL_2942947, EPI_ISL_2942948, EPI_ISL_2942949, EPI_ISL_2942950, EPI_ISL_2942951, EPI_ISL_2942952, EPI_ISL_2942954, EPI_ISL_2942956, EPI_ISL_2942957, EPI_ISL_2942958, EPI_ISL_2942959, EPI_ISL_2942960, EPI_ISL_2942961, EPI_ISL_2942962, EPI_ISL_2942963, EPI_ISL_2942964 | | | |
| see above | Kansas Health and Environmental Lab | Kansas Health and Environmental Lab | Mike Grose, Jonathan Barnell, Ben Olsen, and Phil Adam |
| EPI_ISL_2942965, EPI_ISL_2942966 | PathWest Laboratory Medicine WA | PathWest Laboratory Medicine WA Microbial Surveillance Unit | PathWest Laboratory Medicine WA Microbial Surveillance Unit |
| EPI_ISL_2942968, EPI_ISL_2942969, EPI_ISL_2942970, EPI_ISL_2942971, EPI_ISL_2942972, EPI_ISL_2942973, EPI_ISL_2942974, EPI_ISL_2942977, EPI_ISL_2942978, EPI_ISL_2942979, EPI_ISL_2942982, EPI_ISL_2942984, EPI_ISL_2942985, EPI_ISL_2942986, EPI_ISL_2942987, EPI_ISL_2942988, EPI_ISL_2942989, EPI_ISL_2942990, EPI_ISL_2942991, EPI_ISL_2942992, EPI_ISL_2942993, EPI_ISL_2942994, EPI_ISL_2942995, EPI_ISL_2942997, EPI_ISL_2942998, EPI_ISL_2942999, EPI_ISL_2943000 | | | |
| see above | Platform BIS UZA/UAntwerpen | Labo Klinische Biologie, UZA | Marie Le Mercier, Jasmine Coppens, Kathleen Holemans, Ines Verbesselt, Basil Britto Xavier, Christine Lammens, Veerle Matheeussen, Herman Goossens |
| EPI_ISL_2943001, EPI_ISL_2943002 | CENTRUM VOOR MEDISCHE ANALYSE | Labo Klinische Biologie, UZA | Marie Le Mercier, Jasmine Coppens, Kathleen Holemans, Ines Verbesselt, Basil Britto Xavier, Christine Lammens, Veerle Matheeussen, Herman Goossens |
| EPI_ISL_2943003, EPI_ISL_2943004, EPI_ISL_2943005, EPI_ISL_2943006, EPI_ISL_2943008, EPI_ISL_2943010, EPI_ISL_2943011, EPI_ISL_2943012, EPI_ISL_2943013, EPI_ISL_2943014, EPI_ISL_2943015, EPI_ISL_2943016, EPI_ISL_2943017 | | | |
| see above | Platform BIS UZA/UAntwerpen | Labo Klinische Biologie, UZA | Marie Le Mercier, Jasmine Coppens, Kathleen Holemans, Ines Verbesselt, Basil Britto Xavier, Christine Lammens, Veerle Matheeussen, Herman Goossens |
| EPI_ISL_2943019 | CENTRUM VOOR MEDISCHE ANALYSE | Labo Klinische Biologie, UZA | Marie Le Mercier, Jasmine Coppens, Kathleen Holemans, Ines Verbesselt, Basil Britto Xavier, Christine Lammens, Veerle Matheeussen, Herman Goossens |
| EPI_ISL_2943020, EPI_ISL_2943021 | Platform BIS UZA/UAntwerpen | Labo Klinische Biologie, UZA | Marie Le Mercier, Jasmine Coppens, Kathleen Holemans, Ines Verbesselt, Basil Britto Xavier, Christine Lammens, Veerle Matheeussen, Herman Goossens |
| EPI_ISL_2943022 | #N/A | Labo Klinische Biologie, UZA | Marie Le Mercier, Jasmine Coppens, Kathleen Holemans, Ines Verbesselt, Basil Britto Xavier, Christine Lammens, Veerle Matheeussen, Herman Goossens |
| EPI_ISL_2943023, EPI_ISL_2943024, EPI_ISL_2943025, EPI_ISL_2943026, EPI_ISL_2943027, EPI_ISL_2943028, EPI_ISL_2943029, EPI_ISL_2943031, EPI_ISL_2943032, EPI_ISL_2943033, EPI_ISL_2943034, EPI_ISL_2943035, EPI_ISL_2943036, EPI_ISL_2943037 | | | |
| see above | LHUB-ULB | Labo Klinische Biologie, UZA | Marie Le Mercier, Jasmine Coppens, Kathleen Holemans, Ines Verbesselt, Basil Britto Xavier, Christine Lammens, Veerle Matheeussen, Herman Goossens |
| EPI_ISL_2943038, EPI_ISL_2943039, EPI_ISL_2943040, EPI_ISL_2943041, EPI_ISL_2943042, EPI_ISL_2943044, EPI_ISL_2943045, EPI_ISL_2943046, EPI_ISL_2943048, EPI_ISL_2943049 | Platform BIS UZA/UAntwerpen | Labo Klinische Biologie, UZA | Marie Le Mercier, Jasmine Coppens, Kathleen Holemans, Ines Verbesselt, Basil Britto Xavier, Christine Lammens, Veerle Matheeussen, Herman Goossens |
| EPI_ISL_2943054, EPI_ISL_2943055, EPI_ISL_2943056, EPI_ISL_2943058, EPI_ISL_2943059, EPI_ISL_2943063, EPI_ISL_2943065, EPI_ISL_2943066 | Arizona State Public Health Laboratory | Arizona State Public Health Laboratory | Trung Huynh, Jessica Escobar, Katherine Fullerton, Nobuko Fukushima, Matthew Contursi, Stacy White, Linda Getsinger, Victor Waddell |
| EPI_ISL_2943075, EPI_ISL_2943078 | Department of Public Health Bucharest | National Institute of Infectious Diseases-Prof. Dr. Matei Bals Molecular Diagnostics Laboratory | Corina Casangiu, Leontina Banica, Marius Surleac, Petre Milu, Robert Hohan, Simona Paraschiv, Dan Otelea |
| EPI_ISL_2943144, EPI_ISL_2943145 | 1. Diagnostyka. Laboratoria Medyczne. 2. Wojewódzka Stacja Sanitarno-Epidemiologiczna w Krakowie (WSSE Kraków). | 1. ViroGenetics - BSL3 Laboratory of Virology, Małopolska Centre of Biotechnology, Jagiellonian University; 2. Human Genome Variation Research Group, Malopolska Centre of Biotechnology, Jagiellonian University; | Herda K., Mazur-Panasiuk N., Gromowski T., Branicki W., Kowalski M., Labaj P. P., Zielinska M., Foremny J., Starowicz A., Pyrc K. |
| EPI_ISL_2943148 | CENTRO MEDICO NAVAL SEMAR LAB BIOLOGIA MOLECULAR | CENTRO MEDICO NAVAL SEMAR LABORATORIO DE PATOLOGIA MOLECULAR | MEIXUEIRO CALDERON CLAUDIA. MORALES RAMIREZ OMAR. BALTIERRA URIBE SHANTAL. ZUNO CORONADO TEUTLI. GARCIA ALEGRIA IRMA.ALVAREZ JIMENEZ VIOLETA DEJANIRA. LOPEZ BUSTOS ABRAHAM. CABRERA GONZALEZ MOISES. ROMERO ROMAN EDGAR. LIEVANO ESQUINCA JAZMIN. FIGUEROA ANTONIO VICTOR. DIAZ PONCE MEDRANO JUAN ALBERTO. |
| EPI_ISL_2943150, EPI_ISL_2943151, EPI_ISL_2943152, EPI_ISL_2943153, EPI_ISL_2943154, EPI_ISL_2943155, EPI_ISL_2943156, EPI_ISL_2943157, EPI_ISL_2943158, EPI_ISL_2943159, EPI_ISL_2943160, EPI_ISL_2943161, EPI_ISL_2943162, EPI_ISL_2943163, EPI_ISL_2943164, EPI_ISL_2943165, EPI_ISL_2943166, EPI_ISL_2943167, EPI_ISL_2943168 | | | |
| see above | Wyoming Public Health Laboratory | Wyoming Public Health Laboratory | Jim Mildenberger, Taylor Fearing, Channing Weber, Ashley Norberg, Chayse Rowley, Marley Goetz, Brian Dominguez, Elliot Thomasson, Sam Britz, Cari Sloma, Robert Petit, and Rob Christensen |
| EPI_ISL_2943172 | CENTRO MEDICO NAVAL SEMAR LAB BIOLOGIA MOLECULAR | CENTRO MEDICO NAVAL SEMAR LAB PATOLOGIA MOLECULAR | MEIXUEIRO CALDERON CLAUDIA. MORALES RAMIREZ OMAR. BALTIERRA URIBE SHANTAL. ZUNO CORONADO TEUTLI. GARCIA ALEGRIA IRMA.ALVAREZ JIMENEZ VIOLETA DEJANIRA. LOPEZ BUSTOS ABRAHAM. CABRERA GONZALEZ MOISES. ROMERO ROMAN EDGAR. LIEVANO ESQUINCA JAZMIN. FIGUEROA ANTONIO VICTOR. DIAZ PONCE MEDRANO JUAN ALBERTO. |
| EPI_ISL_2943173 | RSUP Dr. Sardjito | Genetics Working Group (Pokja Genetik) Faculty of Medicine, Public Health and Nursing Universitas Gadjah Mada (FK-KMK UGM); Disease Investigation Center Wates Ministry of Agriculture Indonesia; Department of Microbiology FK-KMK UGM; Laboratorium Diagnostik Yayasan Tahija World Mosquito Program | Gunadi, Hendra Wibawa, . Marcellus, Mohamad S. Hakim, Edwin W. Daniwijaya, Ludhang P. Rizki, Endah Supriyati, Eggi Arguni, Titik Nuryastuti, Tri Wibawa, Dwi AA Nugrahaningsih, Afiahayati, Siswanto, Kristy Iskandar, Nungki Anggorowati, Ika Trisnawati, Sumardi, Eko Budiono, Bambang Sigit Riyanto, Heni Retnowulan, Munawar Gani, Satria Maulana, Nur Rahmi Ananda, Riat El Khair, Yunika Puspadewi, Osman Sianipar, Umi Solekhah Intansari, Elizabeth Henny Herningtiyas, Ira Puspitawati, Nur Imma Fatimah Harahap, Irene Tania, Hana F Hanifin |
| EPI_ISL_2943178, EPI_ISL_2943179, EPI_ISL_2943192, EPI_ISL_2943194, EPI_ISL_2943195, EPI_ISL_2943196, EPI_ISL_2943197, EPI_ISL_2943201, EPI_ISL_2943205, EPI_ISL_2943206, EPI_ISL_2943210, EPI_ISL_2943213, EPI_ISL_2943215, EPI_ISL_2943216, EPI_ISL_2943217, EPI_ISL_2943221, EPI_ISL_2943222, EPI_ISL_2943224, EPI_ISL_2943226, EPI_ISL_2943227, EPI_ISL_2943228, EPI_ISL_2943229 | | | |
| see above | Montana Public Health Laboratory | Montana Public Health Laboratory | Joy Ritter, Michelle Mozer, Carrie Biskupiak, Deborah Gibson, Michael Dills |

| | | | |
|---|---|---|---|
| EPI_ISL_2943235 | BBTKLPP Yogyakarta | Genetics Working Group (Pokja Genetik) Faculty of Medicine, Public Health and Nursing Universitas Gadjah Mada (FK-KMK UGM); Disease Investigation Center Wates Ministry of Agriculture Indonesia; Department of Microbiology FK-KMK UGM; Laboratorium Diagnostik Yayasan Tahija World Mosquito Program (WMP) Yogyakarta Center for Tropical Medicine FK-KMK UGM; Integrated Research Center FK-KMK UGM; Department of Computer Science and Electronics FMIPA UGM; RSUP Dr. Sardjito | Gunadi, Hendra Wibawa, Marcellus, Mohamad S. Hakim, Edwin W. Daniwijaya, Ludhang P. Rizki, Endah Supriyati, Eggi Arguni, Titik Nuryastuti, Tri Wibawa, Dwi AA Nugrahaningsih, Afiahayati, Siswanto,Kristy Iskandar,Nungki Anggorowati, Pramesti G Dewi, Dwi Indaryati,Indarto Sulistiyono, Irene Tania, Hana F Hanifin |
| EPI_ISL_2943236 | CENTRO MEDICO NAVAL SEMAR LAB BIOLOGIA MOLECULAR | CENTRO MEDICO NAVAL SEMAR LABORATORIO DE PATOLOGIA MOLECULAR | MEIXUEIRO CALDERON CLAUDIA. MORALES RAMIREZ OMAR. BALTIERRA URIBE SHANTAL. ZUNO CORONADO TEUTLI. GARCIA ALEGRIA IRMA.ALVAREZ JIMENEZ VIOLETA DEJANIRA. LOPEZ BUSTOS ABRAHAM. CABRERA GONZALEZ MOISES. ROMERO ROMAN EDGAR. LIEVANO ESQUINCA JAZMIN. FIGUEROA ANTONIO VICTOR. DIAZ PONCE MEDRANO JUAN ALBERTO. |
| EPI_ISL_2943237 | DKK Sleman/Puskesmas Depok 1 | Genetics Working Group (Pokja Genetik) Faculty of Medicine, Public Health and Nursing Universitas Gadjah Mada (FK-KMK UGM); Disease Investigation Center Wates Ministry of Agriculture Indonesia; Department of Microbiology FK-KMK UGM; Laboratorium Diagnostik Yayasan Tahija World Mosquito Program (WMP) Yogyakarta Center for Tropical Medicine FK-KMK UGM; Integrated Research Center FK-KMK UGM; Department of Computer Science and Electronics FMIPA UGM; RSUP Dr. Sardjito | Gunadi, Hendra Wibawa, Marcellus, Mohamad S. Hakim, Edwin W. Daniwijaya, Ludhang P. Rizki, Endah Supriyati, Eggi Arguni, Titik Nuryastuti, Tri Wibawa, Dwi AA Nugrahaningsih, Afiahayati, Siswanto,Kristy Iskandar,Nungki Anggorowati, Pramesti G Dewi, Dwi Indaryati,Indarto Sulistiyono, Cita S Amalia, Alvina A Setiawan |
| EPI_ISL_2954064 | RSUD Panembahan Senopati, Bantul | Genetics Working Group (Pokja Genetik) Faculty of Medicine, Public Health and Nursing Universitas Gadjah Mada (FK-KMK UGM); Disease Investigation Center Wates Ministry of Agriculture Indonesia; Department of Microbiology FK-KMK UGM; Laboratorium Diagnostik Yayasan Tahija World Mosquito Program (WMP) Yogyakarta Center for Tropical Medicine FK-KMK UGM; Integrated Research Center FK-KMK UGM; Department of Computer | Gunadi, Hendra Wibawa, Marcellus, Mohamad S. Hakim, Edwin W. Daniwijaya, Ludhang P. Rizki, Endah Supriyati, Eggi Arguni, Titik Nuryastuti, Tri Wibawa, Dwi AA Nugrahaningsih, Afiahayati, Siswanto,Kristy Iskandar,Nungki Anggorowati, Pramesti G Dewi, Dwi Indaryati,Indarto Sulistiyono, Irene Tania, Hana F Hanifin |
| EPI_ISL_2954684 | DKK Gunungkidul/ Puskesmas Girisubo | Genetics Working Group (Pokja Genetik) Faculty of Medicine, Public Health and Nursing Universitas Gadjah Mada (FK-KMK UGM); Disease Investigation Center Wates Ministry of Agriculture Indonesia; Department of Microbiology FK-KMK UGM; Laboratorium Diagnostik Yayasan Tahija World Mosquito Program (WMP) Yogyakarta Center for Tropical Medicine FK-KMK UGM; Integrated Research Center FK-KMK UGM; Department of Computer Science and Electronics FMIPA UGM; RSUP Dr. Sardjito | Gunadi, Hendra Wibawa, Marcellus, Mohamad S. Hakim, Edwin W. Daniwijaya, Ludhang P. Rizki, Endah Supriyati, Eggi Arguni, Titik Nuryastuti, Tri Wibawa, Dwi AA Nugrahaningsih, Afiahayati, Siswanto,Kristy Iskandar,Nungki Anggorowati, Pramesti G Dewi, Dwi Indaryati,Indarto Sulistiyono, Cita S Amalia, Alvina A Setiawan |
| EPI_ISL_2955330 | Labkesda Bantul | Genetics Working Group (Pokja Genetik) Faculty of Medicine, Public Health and Nursing Universitas Gadjah Mada (FK-KMK UGM); Disease Investigation Center Wates Ministry of Agriculture Indonesia; Department of Microbiology FK-KMK UGM; Laboratorium Diagnostik Yayasan Tahija World Mosquito Program (WMP) Yogyakarta Center for Tropical Medicine FK-KMK UGM; Integrated Research Center FK-KMK UGM; Department of Computer Science and Electronics FMIPA UGM; RSUP Dr. Sardjito | Gunadi, Hendra Wibawa, Marcellus, Mohamad S. Hakim, Edwin W. Daniwijaya, Ludhang P. Rizki, Endah Supriyati, Eggi Arguni, Titik Nuryastuti, Tri Wibawa, Dwi AA Nugrahaningsih, Afiahayati, Siswanto,Kristy Iskandar,Nungki Anggorowati, Pramesti G Dewi, Dwi Indaryati,Indarto Sulistiyono, Cita S Amalia, Alvina A Setiawan |
| EPI_ISL_2955331, EPI_ISL_2955334 | Nationa Laboratory of Health, Environment and Food | Department of Public Health Microbiology Ljubljana, National Laboratory for Health, Environment and Food | Tom Koritnik, Martin Bosilj, Katarina Prosenc, Natasa Berginc, Verica Mioč, Marija Trkov |
| EPI_ISL_2955335 | DKK Gunungkidul/ Puskesmas Karangmojo I | Genetics Working Group (Pokja Genetik) Faculty of Medicine, Public Health and Nursing Universitas Gadjah Mada (FK-KMK UGM); Disease Investigation Center Wates Ministry of Agriculture Indonesia; Department of Microbiology FK-KMK UGM; Laboratorium Diagnostik Yayasan Tahija World Mosquito Program (WMP) Yogyakarta Center for Tropical Medicine FK-KMK UGM; Integrated Research Center FK-KMK UGM; Department of Computer Science and Electronics FMIPA UGM; RSUP Dr. Sardjito | Gunadi, Hendra Wibawa, Marcellus, Mohamad S. Hakim, Edwin W. Daniwijaya, Ludhang P. Rizki, Endah Supriyati, Eggi Arguni, Titik Nuryastuti, Tri Wibawa, Dwi AA Nugrahaningsih, Afiahayati, Siswanto,Kristy Iskandar,Nungki Anggorowati, Pramesti G Dewi, Dwi Indaryati,Indarto Sulistiyono, Irene Tania, Hana F Hanifin |
| EPI_ISL_2955346, EPI_ISL_2955347, EPI_ISL_2955348, EPI_ISL_2955349, EPI_ISL_2955350, EPI_ISL_2955351, EPI_ISL_2955352 | Servicio Microbiología Hospital La Paz | Servicio Microbiología Hospital La Paz | Fernando Lázaro, Rubén Cáceres, Jesús Mingorance Cruz, Elie Dahdouh |
| EPI_ISL_2955364, EPI_ISL_2955365, EPI_ISL_2955366 | Monterey County Public Health Laboratory | Monterey County Public Health Laboratory | Monterey County Public Health Laboratory |
| EPI_ISL_2955406, EPI_ISL_2955408, EPI_ISL_2955409, EPI_ISL_2955410, EPI_ISL_2955411, EPI_ISL_2955413, EPI_ISL_2955415, EPI_ISL_2955416, EPI_ISL_2955418, EPI_ISL_2955419, EPI_ISL_2955420 | | | |
| see above | Oregon State Public Health Laboratory | Oregon State Public Health Laboratory | Rafia Razzaque, Eugene Yeboah, Vanda Makris, Laura Tsaknaridis, John Fontana and Shane Sevey |
| EPI_ISL_2955421, EPI_ISL_2955422, EPI_ISL_2955423, EPI_ISL_2955424, EPI_ISL_2955425, EPI_ISL_2955427, EPI_ISL_2955429, EPI_ISL_2955430, EPI_ISL_2955431, EPI_ISL_2955432, EPI_ISL_2955433, EPI_ISL_2955434, EPI_ISL_2955435, EPI_ISL_2955436, EPI_ISL_2955437, EPI_ISL_2955438, EPI_ISL_2955439, EPI_ISL_2955440, EPI_ISL_2955441, EPI_ISL_2955442, EPI_ISL_2955443, EPI_ISL_2955444, EPI_ISL_2955445, EPI_ISL_2955446, EPI_ISL_2955447, EPI_ISL_2955448, EPI_ISL_2955449, EPI_ISL_2955451, EPI_ISL_2955452 | | | |
| see above | ZARV/NHLS, Department Medical Virology, University of Pretoria | KRISP, KZn Research Innovation and Sequencing Platform | Amy Strydom, Adriano Mendes, Micheala Davids, Sim Mayaphi and Marietjie Venter, Giandhari J, Pillay S, Naidoo Y, Ramphal U, Lessells R, Emmanuel SJ, Tegally H, Wilkinson E, de Oliveira T |
| EPI_ISL_2955456, EPI_ISL_2955457, EPI_ISL_2955459, EPI_ISL_2955460, EPI_ISL_2955461, EPI_ISL_2955462, EPI_ISL_2955463, EPI_ISL_2955464, EPI_ISL_2955465, EPI_ISL_2955466, EPI_ISL_2955467, EPI_ISL_2955468, EPI_ISL_2955469, EPI_ISL_2955470, EPI_ISL_2955471, EPI_ISL_2955472, EPI_ISL_2955474, EPI_ISL_2955476, EPI_ISL_2955477, EPI_ISL_2955478, EPI_ISL_2955479, EPI_ISL_2955480, EPI_ISL_2955481, EPI_ISL_2955482, EPI_ISL_2955483, EPI_ISL_2955486, EPI_ISL_2955489, EPI_ISL_2955490, EPI_ISL_2955492, EPI_ISL_2955493, EPI_ISL_2955494, EPI_ISL_2955495, EPI_ISL_2955498, EPI_ISL_2955499 | | | |
| see above | NHLS_VIRO Eastern Cape | KRISP, KZn Research Innovation and Sequencing Platform | Laguda-Akingba O, Giandhari J, Pillay S, Lessells R, Mdlalose K, York D, Khan S, Emmanuel SJ, Tegally H, Wilkinson E, de Oliveira T |
| EPI_ISL_2955500 | Netcare | KRISP, KZn Research Innovation and Sequencing Platform | Giandhari J, Pillay S, Naidoo Y, Ramphal U, Lessells R, Emmanuel SJ, Tegally H, Wilkinson E, de Oliveira T |
| EPI_ISL_2955518, EPI_ISL_2955521, EPI_ISL_2955522, EPI_ISL_2955524, EPI_ISL_2955525, EPI_ISL_2955526, EPI_ISL_2955527, EPI_ISL_2955528, EPI_ISL_2955559, EPI_ISL_2955560, EPI_ISL_2955563, EPI_ISL_2955565, EPI_ISL_2955567, EPI_ISL_2955568, EPI_ISL_2955571, EPI_ISL_2955572, EPI_ISL_2955574, EPI_ISL_2955575, EPI_ISL_2955576, EPI_ISL_2955577, EPI_ISL_2955585 | | | |
| see above | MD PHL | MD PHL | Maryland Department of Health Laboratories Administration |
| EPI_ISL_2955586 | DKK Gunungkidul/ Puskesmas Karangmojo I | Genetics Working Group (Pokja Genetik) Faculty of Medicine, Public Health and Nursing Universitas Gadjah Mada (FK-KMK UGM); Disease Investigation Center Wates Ministry of Agriculture Indonesia; Department of Microbiology FK-KMK UGM; Laboratorium Diagnostik Yayasan Tahija World Mosquito Program (WMP) Yogyakarta Center for Tropical Medicine FK-KMK UGM; Integrated Research Center FK-KMK UGM; Department of Computer Science and Electronics FMIPA UGM; RSUP Dr. Sardjito | Gunadi, Hendra Wibawa, Marcellus, Mohamad S. Hakim, Edwin W. Daniwijaya, Ludhang P. Rizki, Endah Supriyati, Eggi Arguni, Titik Nuryastuti, Tri Wibawa, Dwi AA Nugrahaningsih, Afiahayati, Siswanto,Kristy Iskandar,Nungki Anggorowati, Pramesti G Dewi, Dwi Indaryati,Indarto Sulistiyono, Susan Simanjaya, Dyah A Puspitarani |
| EPI_ISL_2955597, EPI_ISL_2955598 | DKK Gunungkidul/ Puskesmas Karangmojo I | Genetics Working Group (Pokja Genetik) Faculty of Medicine, Public Health and Nursing Universitas Gadjah Mada (FK-KMK UGM); Disease Investigation Center Wates Ministry of Agriculture Indonesia; Department of Microbiology FK-KMK UGM; Laboratorium Diagnostik Yayasan Tahija World Mosquito Program (WMP) Yogyakarta Center for Tropical Medicine FK-KMK UGM; Integrated Research Center FK-KMK UGM; Department of Computer Science and Electronics FMIPA UGM; RSUP Dr. Sardjito | Gunadi, Hendra Wibawa, Marcellus, Mohamad S. Hakim, Edwin W. Daniwijaya, Ludhang P. Rizki, Endah Supriyati, Eggi Arguni, Titik Nuryastuti, Tri Wibawa, Dwi AA Nugrahaningsih, Afiahayati, Siswanto,Kristy Iskandar,Nungki Anggorowati, Pramesti G Dewi, Dwi Indaryati,Indarto Sulistiyono, Khanza a Vujira, Alvina A Setiawan |
| EPI_ISL_2955606 | RSUP Sardjito | Genetics Working Group (Pokja Genetik) Faculty of Medicine, Public Health and Nursing Universitas Gadjah Mada (FK-KMK UGM); Disease Investigation Center Wates Ministry of Agriculture Indonesia; Department of Microbiology FK-KMK UGM; Laboratorium Diagnostik Yayasan Tahija World Mosquito Program (WMP) Yogyakarta Center for Tropical Medicine FK-KMK UGM; Integrated Research Center FK-KMK UGM; Department of Computer Science and Electronics FMIPA UGM; RSUP Dr. Sardjito | Gunadi, Hendra Wibawa, . Marcellus, Mohamad S. Hakim, Edwin W. Daniwijaya, Ludhang P. Rizki, Endah Supriyati, Eggi Arguni, Titik Nuryastuti, Tri Wibawa, Dwi AA Nugrahaningsih, Afiahayati, Siswanto, Kristy Iskandar, Nungki Anggorowati, Ika Trisnawati, Sumardi, Eko Budiono, Bambang Sigit Riyanto, Heni Retnowulan, Munawar Gani, Satria Maulana, Nur Rahmi Ananda, Riat El Khair, Yunika Puspadewi, Osman Sianipar, Umi Solekhah Intansari, Elizabeth Henny Herningtiyas, Ira Puspitawati, Nur Imma Fatimah Harahap, Khanza a Vujira, Alvina A Setiawan |
| EPI_ISL_2955607 | RSUP Dr Sardjito | Genetics Working Group (Pokja Genetik) Faculty of Medicine, Public Health and Nursing Universitas Gadjah Mada (FK-KMK UGM); Disease Investigation Center Wates Ministry of Agriculture Indonesia; Department of Microbiology FK-KMK UGM; Laboratorium | Gunadi, Hendra Wibawa, . Marcellus, Mohamad S. Hakim, Edwin W. Daniwijaya, Ludhang P. Rizki, Endah Supriyati, Eggi Arguni, Titik Nuryastuti, Tri Wibawa, Dwi AA Nugrahaningsih, Afiahayati, Siswanto, Kristy Iskandar, Nungki Anggorowati, Ika Trisnawati, Sumardi, Eko Budiono, Bambang Sigit Riyanto, Heni Retnowulan, Munawar Gani, Satria Maulana, Nur Rahmi Ananda, Riat El Khair, Yunika Puspadewi, Osman Sianipar, Umi Solekhah Intansari, Elizabeth Henny Herningtiyas, Ira Puspitawati, Nur Imma Fatimah Harahap, Irene Tania, Hana F Hanifin |

| | | | |
|---|---|---|---|
| | | Diagnostik Yayasan Tahija World Mosquito Program (WMP) Yogyakarta Center for Tropical Medicine FK-KMK UGM; Integrated Research Center FK-KMK UGM; Department of Computer Science and Electronics FMIPA UGM; RSUP Dr. Sardjito | |
| EPI_ISL_2955608 | RSA UGM | Genetics Working Group (Pokja Genetik) Faculty of Medicine, Public Health and Nursing Universitas Gadjah Mada (FK-KMK UGM); Disease Investigation Center Wates Ministry of Agriculture Indonesia; Department of Microbiology FK-KMK UGM; Laboratorium Diagnostik Yayasan Tahija World Mosquito Program (WMP) Yogyakarta Center for Tropical Medicine FK-KMK UGM; Integrated Research Center FK-KMK UGM; Department of Computer Science and Electronics FMIPA UGM; RSUP Dr. Sardjito | Gunadi, Hendra Wibawa, Marcellus, Mohamad S. Hakim, Edwin W. Daniwijaya, Ludhang P. Rizki, Endah Supriyati, Eggi Arguni, Titik Nuryastuti, Tri Wibawa, Dwi AA Nugrahaningsih, Afiahayati, Siswanto,Kristy Iskandar,Nungki Anggorowati, Pramesti G Dewi, Dwi Indaryati,Indarto Sulistiyono, Susan Simanjaya, Dyah A Puspitarani |
| EPI_ISL_2955609 | DKK Sleman/Puskesmas Depok 1 | Genetics Working Group (Pokja Genetik) Faculty of Medicine, Public Health and Nursing Universitas Gadjah Mada (FK-KMK UGM); Disease Investigation Center Wates Ministry of Agriculture Indonesia; Department of Microbiology FK-KMK UGM; Laboratorium Diagnostik Yayasan Tahija World Mosquito Program (WMP) Yogyakarta Center for Tropical Medicine FK-KMK UGM; Integrated Research Center FK-KMK UGM; Department of Computer Science and Electronics FMIPA UGM; RSUP Dr. Sardjito | Gunadi, Hendra Wibawa, Marcellus, Mohamad S. Hakim, Edwin W. Daniwijaya, Ludhang P. Rizki, Endah Supriyati, Eggi Arguni, Titik Nuryastuti, Tri Wibawa, Dwi AA Nugrahaningsih, Afiahayati, Siswanto,Kristy Iskandar,Nungki Anggorowati, Pramesti G Dewi, Dwi Indaryati,Indarto Sulistiyono, Cita S Amalia, Alvina A Setiawan |
| EPI_ISL_2955611, EPI_ISL_2955614, EPI_ISL_2955616, EPI_ISL_2955617, EPI_ISL_2955619, EPI_ISL_2955621, EPI_ISL_2955623 | Microvida | Microvida | Suzan D. Pas, Jaco J. Verweij, Joep J. J. M. Stohr |
| EPI_ISL_2955627 | DKK Sleman/Puskesmas Ngemplak 2 | Genetics Working Group (Pokja Genetik) Faculty of Medicine, Public Health and Nursing Universitas Gadjah Mada (FK-KMK UGM); Disease Investigation Center Wates Ministry of Agriculture Indonesia; Department of Microbiology FK-KMK UGM; Laboratorium Diagnostik Yayasan Tahija World Mosquito Program (WMP) Yogyakarta Center for Tropical Medicine FK-KMK UGM; Integrated Research Center FK-KMK UGM; Department of Computer Science and Electronics FMIPA UGM; RSUP Dr. Sardjito | Gunadi, Hendra Wibawa, Marcellus, Mohamad S. Hakim, Edwin W. Daniwijaya, Ludhang P. Rizki, Endah Supriyati, Eggi Arguni, Titik Nuryastuti, Tri Wibawa, Dwi AA Nugrahaningsih, Afiahayati, Siswanto,Kristy Iskandar,Nungki Anggorowati, Pramesti G Dewi, Dwi Indaryati,Indarto Sulistiyono, Irene Tania, Hana F Hanifin |
| EPI_ISL_2955628 | DKK Sleman/Puskesmas Ngemplak 2 | Genetics Working Group (Pokja Genetik) Faculty of Medicine, Public Health and Nursing Universitas Gadjah Mada (FK-KMK UGM); Disease Investigation Center Wates Ministry of Agriculture Indonesia; Department of Microbiology FK-KMK UGM; Laboratorium Diagnostik Yayasan Tahija World Mosquito Program (WMP) Yogyakarta Center for Tropical Medicine FK-KMK UGM; Integrated Research Center FK-KMK UGM; Department of Computer Science and Electronics FMIPA UGM; RSUP Dr. Sardjito | Gunadi, Hendra Wibawa, Marcellus, Mohamad S. Hakim, Edwin W. Daniwijaya, Ludhang P. Rizki, Endah Supriyati, Eggi Arguni, Titik Nuryastuti, Tri Wibawa, Dwi AA Nugrahaningsih, Afiahayati, Siswanto,Kristy Iskandar,Nungki Anggorowati, Pramesti G Dewi, Dwi Indaryati,Indarto Sulistiyono, Susan Simanjaya, Dyah A Puspitarani |
| EPI_ISL_2955629 | LKO | Jessa | Berden et al. on behalf of the Jessa_cmdLab |
| EPI_ISL_2955631, EPI_ISL_2955632, EPI_ISL_2955633, EPI_ISL_2955634, EPI_ISL_2955635, EPI_ISL_2955636, EPI_ISL_2955637 | ZOL | Jessa | Berden et al. on behalf of the Jessa_cmdLab |
| EPI_ISL_2955638, EPI_ISL_2955639, EPI_ISL_2955640, EPI_ISL_2955641, EPI_ISL_2955642, EPI_ISL_2955643, EPI_ISL_2955644, EPI_ISL_2955645, EPI_ISL_2955646, EPI_ISL_2955647 | Jessa | Jessa | Berden et al. on behalf of the Jessa_cmdLab |
| EPI_ISL_2955666, EPI_ISL_2955669, EPI_ISL_2955670, EPI_ISL_2955674, EPI_ISL_2955676, EPI_ISL_2955681, EPI_ISL_2955682, EPI_ISL_2955683, EPI_ISL_2955686, EPI_ISL_2955690, EPI_ISL_2955691, EPI_ISL_2955693, EPI_ISL_2955694, EPI_ISL_2955695, EPI_ISL_2955696, EPI_ISL_2955700, EPI_ISL_2955702, EPI_ISL_2955703, EPI_ISL_2955704, EPI_ISL_2955706, EPI_ISL_2955710, EPI_ISL_2955711, EPI_ISL_2955712, EPI_ISL_2955713 | | | |
| see above | National Centre for Disease Control (NCDC) Biotechnology Division, Delhi | NCDC Delhi, Biotechnology Division INSACOG | Meena Datta, Priyanka Singh, Uma Sharma, Hemlata Lall, Manoj K Singh, Radhakrishnan V. S, Robin Marwal, Mahesh S Dhar, Kalaiarasan Ponnusamy, Hema Gogia, Preeti Madan, Sandhya Kabra, Sujeet K Singh, Partha Rakshit |
| EPI_ISL_2955715, EPI_ISL_2955716, EPI_ISL_2955717, EPI_ISL_2955718, EPI_ISL_2955719, EPI_ISL_2955720, EPI_ISL_2955721, EPI_ISL_2955722, EPI_ISL_2955724, EPI_ISL_2955726, EPI_ISL_2955728, EPI_ISL_2955729, EPI_ISL_2955730, EPI_ISL_2955732, EPI_ISL_2955733, EPI_ISL_2955734, EPI_ISL_2955735, EPI_ISL_2955736, EPI_ISL_2955738, EPI_ISL_2955739, EPI_ISL_2955740, EPI_ISL_2955741, EPI_ISL_2955742, EPI_ISL_2955744, EPI_ISL_2955746, EPI_ISL_2955747, EPI_ISL_2955748, EPI_ISL_2955749 | | | |
| see above | Indraprastha Apollo Hospital, New Delhi, India | NCDC Delhi, Biotechnology Division INSACOG | Meena Datta, Priyanka Singh, Uma Sharma, Hemlata Lall, Manoj K Singh, Radhakrishnan V. S, Robin Marwal, Mahesh S Dhar, Kalaiarasan Ponnusamy, Hema Gogia, Preeti Madan, Sandhya Kabra, Sujeet K Singh, Partha Rakshit |
| EPI_ISL_2955751, EPI_ISL_2955753, EPI_ISL_2955755, EPI_ISL_2955756, EPI_ISL_2955757, EPI_ISL_2955759, EPI_ISL_2955762, EPI_ISL_2955763, EPI_ISL_2955764, EPI_ISL_2955765, EPI_ISL_2955767 | | | |
| see above | Indraprastha Apollo Hospital, New Delhi, India | NCDC Delhi, Biotechnology Division INSACOG | Priyanka Singh, Uma Sharma, Hemlata Lall, Manoj K Singh, Radhakrishnan V. S, Robin Marwal, Mahesh S Dhar, Kalaiarasan Ponnusamy, Meena Datta, Hema Gogia, Preeti Madan, Sandhya Kabra, Sujeet K Singh, Partha Rakshit |
| EPI_ISL_2955769, EPI_ISL_2955770, EPI_ISL_2955772, EPI_ISL_2955773, EPI_ISL_2955774, EPI_ISL_2955775, EPI_ISL_2955777, EPI_ISL_2955778, EPI_ISL_2955780, EPI_ISL_2955781, EPI_ISL_2955782, EPI_ISL_2955783, EPI_ISL_2955784 | | | |
| see above | National Centre for Disease Control (NCDC) Biotechnology Division, Delhi | NCDC Delhi, Biotechnology Division INSACOG | Priyanka Singh, Uma Sharma, Hemlata Lall, Manoj K Singh, Radhakrishnan V. S, Robin Marwal, Mahesh S Dhar, Kalaiarasan Ponnusamy, Meena Datta, Hema Gogia, Preeti Madan, Sandhya Kabra, Sujeet K Singh, Partha Rakshit |
| EPI_ISL_2955786, EPI_ISL_2955789, EPI_ISL_2955790, EPI_ISL_2955791, EPI_ISL_2955792, EPI_ISL_2955793, EPI_ISL_2955796, EPI_ISL_2955797, EPI_ISL_2955798, EPI_ISL_2955799, EPI_ISL_2955800, EPI_ISL_2955806, EPI_ISL_2955807, EPI_ISL_2955808, EPI_ISL_2955810, EPI_ISL_2955812, EPI_ISL_2955813, EPI_ISL_2955814, EPI_ISL_2955815, EPI_ISL_2955816, EPI_ISL_2955817, EPI_ISL_2955818, EPI_ISL_2955819, EPI_ISL_2955820, EPI_ISL_2955821, EPI_ISL_2955822, EPI_ISL_2955823, EPI_ISL_2955824, EPI_ISL_2955825, EPI_ISL_2955826, EPI_ISL_2955827, EPI_ISL_2955828, EPI_ISL_2955829, EPI_ISL_2955830, EPI_ISL_2955831, EPI_ISL_2955832, EPI_ISL_2955834 | | | |
| see above | Northern Railway Central Hospital, New Delhi, India | NCDC Delhi, Biotechnology Division INSACOG | Priyanka Singh, Uma Sharma, Hemlata Lall, Manoj K Singh, Radhakrishnan V. S, Robin Marwal, Mahesh S Dhar, Kalaiarasan Ponnusamy, Meena Datta, Hema Gogia, Preeti Madan, Sandhya Kabra, Sujeet K Singh, Partha Rakshit |
| EPI_ISL_2955836, EPI_ISL_2955837, EPI_ISL_2955838, EPI_ISL_2955839, EPI_ISL_2955840, EPI_ISL_2955841, EPI_ISL_2955842, EPI_ISL_2955843, EPI_ISL_2955844, EPI_ISL_2955845, EPI_ISL_2955846, EPI_ISL_2955847, EPI_ISL_2955848, EPI_ISL_2955849, EPI_ISL_2955850, EPI_ISL_2955851, EPI_ISL_2955852, EPI_ISL_2955853, EPI_ISL_2955854, EPI_ISL_2955855, EPI_ISL_2955856, EPI_ISL_2955857, EPI_ISL_2955859, EPI_ISL_2955860, EPI_ISL_2955861, EPI_ISL_2955863, EPI_ISL_2955865, EPI_ISL_2955866, EPI_ISL_2955867, EPI_ISL_2955871, EPI_ISL_2955872, EPI_ISL_2955874, EPI_ISL_2955875, EPI_ISL_2955879, EPI_ISL_2955881, EPI_ISL_2955885, EPI_ISL_2955886, EPI_ISL_2955887, EPI_ISL_2955889, EPI_ISL_2955892, EPI_ISL_2955893, EPI_ISL_2955894, EPI_ISL_2955895, EPI_ISL_2955897, EPI_ISL_2955898, EPI_ISL_2955899, EPI_ISL_2955901, EPI_ISL_2955902, EPI_ISL_2955903, EPI_ISL_2955906, EPI_ISL_2955907, EPI_ISL_2955908, EPI_ISL_2955909, EPI_ISL_2955910, EPI_ISL_2955917, EPI_ISL_2955918, EPI_ISL_2955919, EPI_ISL_2955921, EPI_ISL_2955924, EPI_ISL_2955926, EPI_ISL_2955927, EPI_ISL_2955928, EPI_ISL_2955929, EPI_ISL_2955930, EPI_ISL_2955931 | | | |
| see above | National Centre for Disease Control (NCDC) Biotechnology Division, Delhi | NCDC Delhi, Biotechnology Division INSACOG | Priyanka Singh, Uma Sharma, Hemlata Lall, Manoj K Singh, Radhakrishnan V. S, Robin Marwal, Mahesh S Dhar, Kalaiarasan Ponnusamy, Meena Datta, Hema Gogia, Preeti Madan, Sandhya Kabra, Sujeet K Singh, Partha Rakshit |
| EPI_ISL_2955944, EPI_ISL_2955948, EPI_ISL_2955949, EPI_ISL_2955950, EPI_ISL_2955975, EPI_ISL_2955983, EPI_ISL_2955985, EPI_ISL_2955988, EPI_ISL_2955996, EPI_ISL_2955999, EPI_ISL_2956000, EPI_ISL_2956003, EPI_ISL_2956004 | | | |
| see above | Microbiology Department & Molecular Biology CORE \| CDB \| Hospital Clinic \| Barcelona | Microbiology Department & Molecular Biology CORE \| CDB \| Hospital Clinic \| Barcelona | Andrea Vergara Gomez , Jose Luis Villanueva Cañas , Miguel Julian Martínez Yoldi , Mª del Mar Mosquera Gutierrez , Mª Angeles Marcos Maeso |
| EPI_ISL_2956008, EPI_ISL_2956010, EPI_ISL_2956011, EPI_ISL_2956013, EPI_ISL_2956014 | Queensland Health Forensic and Scientific Services | Queensland Health Forensic and Scientific Services | Son Nguyen |
| EPI_ISL_2956019 | Swedish national genomic surveillance program of SARS-CoV-2 | The Public Health Agency of Sweden | Maximilian Riess, Maria Lind Karlberg, Alma Brolund, Swedish national genomic surveillance program of SARS-CoV-2 |
| EPI_ISL_2956048, EPI_ISL_2956049, EPI_ISL_2956050, EPI_ISL_2956051 | Tokyo Metropolitan Institute of Public Health | Tokyo Metropolitan Institute of Public Health | Hirofumi Miyake, Kenshirou Kuroki, Maya Isogai, Masaki Hayashi, Takako Yamazaki, Akane Negishi,Takushi Fujiwara, Mami Nagashima, Jun Suzuki, Kenji Sadamasu |
| EPI_ISL_2956069, EPI_ISL_2956070, EPI_ISL_2956072, EPI_ISL_2956073, EPI_ISL_2956074, | Institut für Labormedizin Mikrobiologie und Hygiene | Robert Koch Institute | unknown |

| | | | |
|---|---|---|---|
| EPI_ISL_2956075, EPI_ISL_2956077, EPI_ISL_2956078 | | | |
| EPI_ISL_2956079 | Medizinisches Versorgungszentrum für Labormedizin und Mikrobiologie Ruhr GmbH - mvzlm RUHR GmbH | Robert Koch Institute | unknown |
| EPI_ISL_2956080, EPI_ISL_2956082 | MVZ Dr. Eberhard & Partner Dortmund | Robert Koch Institute | unknown |
| EPI_ISL_2956084 | Laborarztpraxis Osnabrück | Robert Koch Institute | unknown |
| EPI_ISL_2956085 | MVZ für Laboratoriumsmedizin und Mikrobiologie Würzburg | Robert Koch Institute | unknown |
| EPI_ISL_2956086, EPI_ISL_2956087, EPI_ISL_2956088 | Laborarztpraxis Osnabrück | Robert Koch Institute | unknown |
| EPI_ISL_2956092, EPI_ISL_2956096, EPI_ISL_2956097, EPI_ISL_2956098 | Medizinische Laboratorien Düsseldorf | Robert Koch Institute | unknown |
| EPI_ISL_2956099, EPI_ISL_2956100, EPI_ISL_2956102 | Klinikum Ernst von Bergmann gemeinnützige GmbH - stationärer Bereich | Robert Koch Institute | unknown |
| EPI_ISL_2956103, EPI_ISL_2956104, EPI_ISL_2956105, EPI_ISL_2956106, EPI_ISL_2956107, EPI_ISL_2956109 | Labormedizin Darmstadt | Robert Koch Institute | unknown |
| EPI_ISL_2956110 | IMD Labor Frankfurt | Robert Koch Institute | unknown |
| EPI_ISL_2956111, EPI_ISL_2956113 | Labormedizin Darmstadt | Robert Koch Institute | unknown |
| EPI_ISL_2956114 | IMD Labor Frankfurt | Robert Koch Institute | unknown |
| EPI_ISL_2956115 | Labormedizin Darmstadt | Robert Koch Institute | unknown |
| EPI_ISL_2956116, EPI_ISL_2956117 | IMD Labor Frankfurt | Robert Koch Institute | unknown |
| EPI_ISL_2956118, EPI_ISL_2956119, EPI_ISL_2956121, EPI_ISL_2956124, EPI_ISL_2956125 | Labormedizin Darmstadt | Robert Koch Institute | unknown |
| EPI_ISL_2956126 | SYNLAB MVZ Leinfelden-Echterdingen | Robert Koch Institute | unknown |
| EPI_ISL_2956127 | Synlab MVZ Augsburg | Robert Koch Institute | unknown |
| EPI_ISL_2956128 | SYNLAB MVZ Weiden | Robert Koch Institute | unknown |
| EPI_ISL_2956129 | SYNLAB MVZ Leverkusen | Robert Koch Institute | unknown |
| EPI_ISL_2956131 | SYNLAB MVZ Leinfelden-Echterdingen | Robert Koch Institute | unknown |
| EPI_ISL_2956132 | SYNLAB MVZ Weiden | Robert Koch Institute | unknown |
| EPI_ISL_2956133, EPI_ISL_2956134 | SYNLAB Labor München Zentrum LMZ | Robert Koch Institute | unknown |
| EPI_ISL_2956135, EPI_ISL_2956137 | SYNLAB MVZ Leinfelden-Echterdingen | Robert Koch Institute | unknown |
| EPI_ISL_2956138 | SYNLAB MVZ Trier | Robert Koch Institute | unknown |
| EPI_ISL_2956141 | SYNLAB MVZ Leverkusen | Robert Koch Institute | unknown |
| EPI_ISL_2956143 | SYNLAB MVZ Trier | Robert Koch Institute | unknown |
| EPI_ISL_2956144 | SYNLAB MVZ Weiden | Robert Koch Institute | unknown |
| EPI_ISL_2956148 | SYNLAB Labor München Zentrum LMZ | Robert Koch Institute | unknown |
| EPI_ISL_2956150 | SYNLAB MVZ Leverkusen | Robert Koch Institute | unknown |
| EPI_ISL_2956151 | SYNLAB MVZ Trier | Robert Koch Institute | unknown |
| EPI_ISL_2956152, EPI_ISL_2956154, EPI_ISL_2956155 | SYNLAB Labor München Zentrum LMZ | Robert Koch Institute | unknown |
| EPI_ISL_2956156, EPI_ISL_2956157 | SYNLAB MVZ Weiden | Robert Koch Institute | unknown |
| EPI_ISL_2956159 | Synlab MVZ Augsburg | Robert Koch Institute | unknown |
| EPI_ISL_2956161 | SYNLAB MVZ Leinfelden-Echterdingen | Robert Koch Institute | unknown |
| EPI_ISL_2956162 | SYNLAB MVZ Weiden | Robert Koch Institute | unknown |
| EPI_ISL_2956163, EPI_ISL_2956164, EPI_ISL_2956165 | SYNLAB MVZ Leinfelden-Echterdingen | Robert Koch Institute | unknown |
| EPI_ISL_2956166, EPI_ISL_2956167 | SYNLAB MVZ Leverkusen | Robert Koch Institute | unknown |
| EPI_ISL_2956169 | Synlab MVZ Augsburg | Robert Koch Institute | unknown |
| EPI_ISL_2956170 | SYNLAB MVZ Leverkusen | Robert Koch Institute | unknown |
| EPI_ISL_2956171 | SYNLAB MVZ Dachau | Robert Koch Institute | unknown |
| EPI_ISL_2956173 | SYNLAB MVZ Leinfelden-Echterdingen | Robert Koch Institute | unknown |
| EPI_ISL_2956174 | SYNLAB MVZ Trier | Robert Koch Institute | unknown |
| EPI_ISL_2956178 | SYNLAB MVZ Leinfelden-Echterdingen | Robert Koch Institute | unknown |
| EPI_ISL_2956179 | SYNLAB MVZ Weiden | Robert Koch Institute | unknown |
| EPI_ISL_2956180 | SYNLAB MVZ Leinfelden-Echterdingen | Robert Koch Institute | unknown |
| EPI_ISL_2956181 | SYNLAB MVZ Leverkusen | Robert Koch Institute | unknown |
| EPI_ISL_2956182 | SYNLAB MVZ Weiden | Robert Koch Institute | unknown |
| EPI_ISL_2956184 | SYNLAB MVZ Leverkusen | Robert Koch Institute | unknown |
| EPI_ISL_2956185 | SYNLAB MVZ Weiden | Robert Koch Institute | unknown |
| EPI_ISL_2956186 | SYNLAB MVZ Leverkusen | Robert Koch Institute | unknown |
| EPI_ISL_2956187 | Synlab MVZ Augsburg | Robert Koch Institute | unknown |
| EPI_ISL_2956188 | SYNLAB Labor München Zentrum LMZ | Robert Koch Institute | unknown |
| EPI_ISL_2956189 | SYNLAB MVZ Leinfelden-Echterdingen | Robert Koch Institute | unknown |
| EPI_ISL_2956190 | SYNLAB MVZ Leverkusen | Robert Koch Institute | unknown |
| EPI_ISL_2956191, EPI_ISL_2956194 | SYNLAB MVZ Leinfelden-Echterdingen | Robert Koch Institute | unknown |

| | | | |
|---|---|---|---|
| EPI_ISL_2956195 | SYNLAB MVZ Heidelberg | Robert Koch Institute | unknown |
| EPI_ISL_2956196 | SYNLAB MVZ Weiden | Robert Koch Institute | unknown |
| EPI_ISL_2956197 | SYNLAB MVZ Leverkusen | Robert Koch Institute | unknown |
| EPI_ISL_2956198 | SYNLAB MVZ Weiden | Robert Koch Institute | unknown |
| EPI_ISL_2956199 | SYNLAB MVZ Leverkusen | Robert Koch Institute | unknown |
| EPI_ISL_2956201 | SYNLAB MVZ Leinfelden-Echterdingen | Robert Koch Institute | unknown |
| EPI_ISL_2956202 | SYNLAB MVZ Weiden | Robert Koch Institute | unknown |
| EPI_ISL_2956203, EPI_ISL_2956204, EPI_ISL_2956206 | SYNLAB MVZ Leverkusen | Robert Koch Institute | unknown |
| EPI_ISL_2956207 | SYNLAB MVZ Leinfelden-Echterdingen | Robert Koch Institute | unknown |
| EPI_ISL_2956208 | SYNLAB MVZ Leverkusen | Robert Koch Institute | unknown |
| EPI_ISL_2956209, EPI_ISL_2956210, EPI_ISL_2956211 | SYNLAB MVZ Leinfelden-Echterdingen | Robert Koch Institute | unknown |
| EPI_ISL_2956212 | SYNLAB MVZ Weiden | Robert Koch Institute | unknown |
| EPI_ISL_2956213 | SYNLAB MVZ Leverkusen | Robert Koch Institute | unknown |
| EPI_ISL_2956215, EPI_ISL_2956216 | SYNLAB MVZ Leinfelden-Echterdingen | Robert Koch Institute | unknown |
| EPI_ISL_2956217 | SYNLAB MVZ Weiden | Robert Koch Institute | unknown |
| EPI_ISL_2956218 | Synlab MVZ Augsburg | Robert Koch Institute | unknown |
| EPI_ISL_2956219 | SYNLAB MVZ Leverkusen | Robert Koch Institute | unknown |
| EPI_ISL_2956220 | SYNLAB MVZ Dachau | Robert Koch Institute | unknown |
| EPI_ISL_2956221 | SYNLAB MVZ Leinfelden-Echterdingen | Robert Koch Institute | unknown |
| EPI_ISL_2956223, EPI_ISL_2956224 | SYNLAB MVZ Weiden | Robert Koch Institute | unknown |
| EPI_ISL_2956225 | SYNLAB MVZ Leverkusen | Robert Koch Institute | unknown |
| EPI_ISL_2956228, EPI_ISL_2956232 | SYNLAB MVZ Weiden | Robert Koch Institute | unknown |
| EPI_ISL_2956233 | SYNLAB MVZ Leinfelden-Echterdingen | Robert Koch Institute | unknown |
| EPI_ISL_2956235, EPI_ISL_2956236 | SYNLAB MVZ Weiden | Robert Koch Institute | unknown |
| EPI_ISL_2956237 | SYNLAB MVZ Leinfelden-Echterdingen | Robert Koch Institute | unknown |
| EPI_ISL_2956238, EPI_ISL_2956240 | SYNLAB MVZ Weiden | Robert Koch Institute | unknown |
| EPI_ISL_2956241 | SYNLAB MVZ Leinfelden-Echterdingen | Robert Koch Institute | unknown |
| EPI_ISL_2956243 | SYNLAB MVZ Leverkusen | Robert Koch Institute | unknown |
| EPI_ISL_2956244, EPI_ISL_2956247 | SYNLAB MVZ Trier | Robert Koch Institute | unknown |
| EPI_ISL_2956248 | SYNLAB MVZ Leinfelden-Echterdingen | Robert Koch Institute | unknown |
| EPI_ISL_2956249, EPI_ISL_2956252, EPI_ISL_2956254, EPI_ISL_2956255 | SYNLAB MVZ Leverkusen | Robert Koch Institute | unknown |
| EPI_ISL_2956256 | SYNLAB MVZ Leinfelden-Echterdingen | Robert Koch Institute | unknown |
| EPI_ISL_2956257 | SYNLAB MVZ Leverkusen | Robert Koch Institute | unknown |
| EPI_ISL_2956259 | SYNLAB MVZ Trier | Robert Koch Institute | unknown |
| EPI_ISL_2956260, EPI_ISL_2956261 | SYNLAB MVZ Leverkusen | Robert Koch Institute | unknown |
| EPI_ISL_2956264 | SYNLAB MVZ Weiden | Robert Koch Institute | unknown |
| EPI_ISL_2956265 | SYNLAB MVZ Leinfelden-Echterdingen | Robert Koch Institute | unknown |
| EPI_ISL_2956266, EPI_ISL_2956269 | SYNLAB MVZ Weiden | Robert Koch Institute | unknown |
| EPI_ISL_2956270 | Synlab MVZ Augsburg | Robert Koch Institute | unknown |
| EPI_ISL_2956271 | SYNLAB MVZ Weiden | Robert Koch Institute | unknown |
| EPI_ISL_2956272 | SYNLAB Labor München Zentrum LMZ | Robert Koch Institute | unknown |
| EPI_ISL_2956274 | SYNLAB MVZ Leverkusen | Robert Koch Institute | unknown |
| EPI_ISL_2956275 | SYNLAB MVZ Weiden | Robert Koch Institute | unknown |
| EPI_ISL_2956276 | Synlab MVZ Augsburg | Robert Koch Institute | unknown |
| EPI_ISL_2956277 | SYNLAB MVZ Leverkusen | Robert Koch Institute | unknown |
| EPI_ISL_2956278 | Synlab MVZ Augsburg | Robert Koch Institute | unknown |
| EPI_ISL_2956279, EPI_ISL_2956280 | SYNLAB MVZ Leverkusen | Robert Koch Institute | unknown |
| EPI_ISL_2956281 | SYNLAB MVZ Weiden | Robert Koch Institute | unknown |
| EPI_ISL_2956282 | Synlab MVZ Augsburg | Robert Koch Institute | unknown |
| EPI_ISL_2956283 | SYNLAB MVZ Leverkusen | Robert Koch Institute | unknown |
| EPI_ISL_2956286 | SYNLAB MVZ Weiden | Robert Koch Institute | unknown |
| EPI_ISL_2956287 | SYNLAB MVZ Leverkusen | Robert Koch Institute | unknown |
| EPI_ISL_2956288 | SYNLAB MVZ Leinfelden-Echterdingen | Robert Koch Institute | unknown |
| EPI_ISL_2956289 | SYNLAB MVZ Leverkusen | Robert Koch Institute | unknown |
| EPI_ISL_2956290, EPI_ISL_2956292, EPI_ISL_2956293, EPI_ISL_2956295, EPI_ISL_2956296 | SYNLAB MVZ Leinfelden-Echterdingen | Robert Koch Institute | unknown |
| EPI_ISL_2956297 | SYNLAB MVZ Weiden | Robert Koch Institute | unknown |
| EPI_ISL_2956298 | SYNLAB MVZ Leverkusen | Robert Koch Institute | unknown |
| EPI_ISL_2956299, EPI_ISL_2956300 | Synlab MVZ Augsburg | Robert Koch Institute | unknown |
| EPI_ISL_2956301 | SYNLAB MVZ Weiden | Robert Koch Institute | unknown |
| EPI_ISL_2956303, EPI_ISL_2956306 | SYNLAB MVZ Leverkusen | Robert Koch Institute | unknown |
| EPI_ISL_2956307 | SYNLAB MVZ Weiden | Robert Koch Institute | unknown |

| | | | |
|---|---|---|---|
| EPI_ISL_2956308, EPI_ISL_2956309, EPI_ISL_2956310 | SYNLAB MVZ Leinfelden-Echterdingen | Robert Koch Institute | unknown |
| EPI_ISL_2956311, EPI_ISL_2956312 | Synlab MVZ Augsburg | Robert Koch Institute | unknown |
| EPI_ISL_2956313 | SYNLAB MVZ Weiden | Robert Koch Institute | unknown |
| EPI_ISL_2956314 | Synlab MVZ Augsburg | Robert Koch Institute | unknown |
| EPI_ISL_2956315 | SYNLAB MVZ Weiden | Robert Koch Institute | unknown |
| EPI_ISL_2956316 | SYNLAB MVZ Trier | Robert Koch Institute | unknown |
| EPI_ISL_2956317 | SYNLAB MVZ Leverkusen | Robert Koch Institute | unknown |
| EPI_ISL_2956319 | SYNLAB MVZ Weiden | Robert Koch Institute | unknown |
| EPI_ISL_2956320 | SYNLAB MVZ Trier | Robert Koch Institute | unknown |
| EPI_ISL_2956321, EPI_ISL_2956323 | SYNLAB MVZ Weiden | Robert Koch Institute | unknown |
| EPI_ISL_2956324 | SYNLAB MVZ Trier | Robert Koch Institute | unknown |
| EPI_ISL_2956326 | SYNLAB MVZ Weiden | Robert Koch Institute | unknown |
| EPI_ISL_2956327, EPI_ISL_2956328 | SYNLAB MVZ Leinfelden-Echterdingen | Robert Koch Institute | unknown |
| EPI_ISL_2956329 | SYNLAB MVZ Weiden | Robert Koch Institute | unknown |
| EPI_ISL_2956330 | SYNLAB MVZ Heidelberg | Robert Koch Institute | unknown |
| EPI_ISL_2956331 | SYNLAB MVZ Weiden | Robert Koch Institute | unknown |
| EPI_ISL_2956332 | SYNLAB Labor München Zentrum LMZ | Robert Koch Institute | unknown |
| EPI_ISL_2956334 | SYNLAB MVZ Trier | Robert Koch Institute | unknown |
| EPI_ISL_2956337 | SYNLAB MVZ Leverkusen | Robert Koch Institute | unknown |
| EPI_ISL_2956338 | amedes MVZ Hannover | Robert Koch Institute | unknown |
| EPI_ISL_2956339 | amedes MVZ für Laboratoriumsdiagnostik Raubling GmbH | Robert Koch Institute | unknown |
| EPI_ISL_2956340 | amedes MVZ DIAMEDIS Sennestadt | Robert Koch Institute | unknown |
| EPI_ISL_2956341 | nordlab - Partnerschaftspraxis für Laboratoriumsmedizin | Robert Koch Institute | unknown |
| EPI_ISL_2956342 | amedes MVZ für Labordiagnostik Rhein-Main | Robert Koch Institute | unknown |
| EPI_ISL_2956343 | CDFD | CDFD | Murali Bashyam, Asmita Gupta, Reelina Basu, Vinay Donipadi, Nagamani Kammili, Divya Vashisht, Ashwin Dalal |
| EPI_ISL_2956344 | amedes Medlab Arnold Analytik MVZ GmbH | Robert Koch Institute | unknown |
| EPI_ISL_2956345 | CDFD | CDFD | Murali Bashyam, Asmita Gupta, Reelina Basu, Vinay Donipadi, Nagamani Kammili, Divya Vashisht, Ashwin Dalal |
| EPI_ISL_2956346 | nordlab - Partnerschaftspraxis für Laboratoriumsmedizin | Robert Koch Institute | unknown |
| EPI_ISL_2956347 | amedes Medlab Arnold Analytik MVZ GmbH | Robert Koch Institute | unknown |
| EPI_ISL_2956349 | amedes MVZ Hannover | Robert Koch Institute | unknown |
| EPI_ISL_2956350 | amedes MVZ für Laboratoriumsdiagnostik Raubling GmbH | Robert Koch Institute | unknown |
| EPI_ISL_2956351 | amedes MVZ DIAMEDIS Sennestadt | Robert Koch Institute | unknown |
| EPI_ISL_2956353, EPI_ISL_2956354 | amedes MVZ Hannover | Robert Koch Institute | unknown |
| EPI_ISL_2956358 | amedes Göttingen | Robert Koch Institute | unknown |
| EPI_ISL_2956359 | nordlab - Partnerschaftspraxis für Laboratoriumsmedizin | Robert Koch Institute | unknown |
| EPI_ISL_2956361 | amedes MVZ Halle/Leipzig | Robert Koch Institute | unknown |
| EPI_ISL_2956363 | amedes MVZ DIAMEDIS Sennestadt | Robert Koch Institute | unknown |
| EPI_ISL_2956364, EPI_ISL_2956365 | amedes MVZ Jena | Robert Koch Institute | unknown |
| EPI_ISL_2956366 | amedes MVZ DIAMEDIS Sennestadt | Robert Koch Institute | unknown |
| EPI_ISL_2956367 | Uniklinikum Jena | Robert Koch Institute | unknown |
| EPI_ISL_2956368, EPI_ISL_2956369 | Niedersächsisches Landesgesundheitsamt (NLGA) | Robert Koch Institute | unknown |
| EPI_ISL_2956371 | Bioscientia Labor Wermsdorf | Robert Koch Institute | unknown |
| EPI_ISL_2956372 | Sonic - MVZ Medizinisches Labor Bremen GmbH | Robert Koch Institute | unknown |
| EPI_ISL_2956373 | Bioscientia Labor Wermsdorf | Robert Koch Institute | unknown |
| EPI_ISL_2956374 | Sonic - MVZ Medizinisches Labor Bremen GmbH | Robert Koch Institute | unknown |
| EPI_ISL_2956375, EPI_ISL_2956376 | Bioscientia Labor Wermsdorf | Robert Koch Institute | unknown |
| EPI_ISL_2956377 | Sonic - Bioscientia - MVZ Labor Saar GmbH | Robert Koch Institute | unknown |
| EPI_ISL_2956378, EPI_ISL_2956379, EPI_ISL_2956380 | Bioscientia Labor Wermsdorf | Robert Koch Institute | unknown |
| EPI_ISL_2956382 | Sonic - Bioscientia - MVZ Labor Saar GmbH | Robert Koch Institute | unknown |
| EPI_ISL_2956383 | Sonic - Labor Lademannbogen MVZ GmbH | Robert Koch Institute | unknown |
| EPI_ISL_2956384, EPI_ISL_2956385, EPI_ISL_2956386 | Bioscientia Labor Wermsdorf | Robert Koch Institute | unknown |
| EPI_ISL_2956387 | Sonic - MVZ Medizinisches Labor Bremen GmbH | Robert Koch Institute | unknown |
| EPI_ISL_2956388 | Bioscientia Labor Wermsdorf | Robert Koch Institute | unknown |
| EPI_ISL_2956389 | Sonic - Bioscientia - MVZ Labor Saar GmbH | Robert Koch Institute | unknown |
| EPI_ISL_2956390, EPI_ISL_2956391, EPI_ISL_2956392, EPI_ISL_2956393 | Bioscientia Labor Wermsdorf | Robert Koch Institute | unknown |
| EPI_ISL_2956395 | Sonic - MVZ Medizinisches Labor Bremen GmbH | Robert Koch Institute | unknown |
| EPI_ISL_2956396, EPI_ISL_2956397, EPI_ISL_2956398 | Bioscientia Labor Wermsdorf | Robert Koch Institute | unknown |
| EPI_ISL_2956399 | Sonic - MVZ Medizinisches Labor Bremen GmbH | Robert Koch Institute | unknown |
| EPI_ISL_2956400 | Bioscientia Labor Wermsdorf | Robert Koch Institute | unknown |
| EPI_ISL_2956401 | Sonic - Bioscientia - MVZ Labor Saar GmbH | Robert Koch Institute | unknown |
| EPI_ISL_2956402 | Bioscientia Labor Wermsdorf | Robert Koch Institute | unknown |
| EPI_ISL_2956403 | Sonic - MVZ Medizinisches Labor Bremen GmbH | Robert Koch Institute | unknown |
| EPI_ISL_2956404, EPI_ISL_2956405, EPI_ISL_2956406, EPI_ISL_2956407, EPI_ISL_2956408, EPI_ISL_2956410, EPI_ISL_2956411, EPI_ISL_2956412, EPI_ISL_2956413, EPI_ISL_2956414, EPI_ISL_2956415, EPI_ISL_2956416, EPI_ISL_2956418, EPI_ISL_2956419, EPI_ISL_2956420, EPI_ISL_2956422, EPI_ISL_2956423 | | | |
| see above | Bioscientia Labor Wermsdorf | Robert Koch Institute | unknown |

| | | | |
|---|---|---|---|
| EPI_ISL_2956424 | Sonic - Labor Dr. von Froreich GmbH | Robert Koch Institute | unknown |
| EPI_ISL_2956425, EPI_ISL_2956426 | Bioscientia Labor Wermsdorf | Robert Koch Institute | unknown |
| EPI_ISL_2956427 | LabKom - Labor Augsburg MVZ GmbH | Robert Koch Institute | unknown |
| EPI_ISL_2956429, EPI_ISL_2956431, EPI_ISL_2956432, EPI_ISL_2956433, EPI_ISL_2956434, EPI_ISL_2956435, EPI_ISL_2956436, EPI_ISL_2956437, EPI_ISL_2956439, EPI_ISL_2956440 | Bioscientia Labor Wermsdorf | Robert Koch Institute | unknown |
| EPI_ISL_2956442 | LabKom - Labor an der Salzbrücke MVZ GmbH | Robert Koch Institute | unknown |
| EPI_ISL_2956443, EPI_ISL_2956444 | Bioscientia MVZ Labor Karlsruhe GmbH | Robert Koch Institute | unknown |
| EPI_ISL_2956445, EPI_ISL_2956447 | Bioscientia Labor Wermsdorf | Robert Koch Institute | unknown |
| EPI_ISL_2956449 | Sonic - Dr. Staber & Kollegen GmbH München | Robert Koch Institute | unknown |
| EPI_ISL_2956450 | Bioscientia Labor Wermsdorf | Robert Koch Institute | unknown |
| EPI_ISL_2956451 | Sonic - MVZ Medizinisches Labor Bremen GmbH | Robert Koch Institute | unknown |
| EPI_ISL_2956452 | Sonic - Labor Staber Dresden (Klipphausen) | Robert Koch Institute | unknown |
| EPI_ISL_2956453 | Sonic - Dr. Staber & Kollegen GmbH München | Robert Koch Institute | unknown |
| EPI_ISL_2956454 | Bioscientia Labor Wermsdorf | Robert Koch Institute | unknown |
| EPI_ISL_2956455 | Sonic - MVZ Medizinisches Labor Bremen GmbH | Robert Koch Institute | unknown |
| EPI_ISL_2956456, EPI_ISL_2956457, EPI_ISL_2956458, EPI_ISL_2956459, EPI_ISL_2956460, EPI_ISL_2956462 | Bioscientia Labor Wermsdorf | Robert Koch Institute | unknown |
| EPI_ISL_2956463 | Sonic - Bioscientia - MVZ Labor Saar GmbH | Robert Koch Institute | unknown |
| EPI_ISL_2956464, EPI_ISL_2956465 | Bioscientia Labor Wermsdorf | Robert Koch Institute | unknown |
| EPI_ISL_2956468 | Sonic - Labor Dr. von Froreich GmbH | Robert Koch Institute | unknown |
| EPI_ISL_2956469, EPI_ISL_2956471, EPI_ISL_2956472, EPI_ISL_2956473, EPI_ISL_2956474, EPI_ISL_2956475 | Bioscientia Labor Wermsdorf | Robert Koch Institute | unknown |
| EPI_ISL_2956476, EPI_ISL_2956478 | Sonic - Labor Dr. von Froreich GmbH | Robert Koch Institute | unknown |
| EPI_ISL_2956479 | Sonic - MVZ Medizinisches Labor Bremen GmbH | Robert Koch Institute | unknown |
| EPI_ISL_2956480 | Bioscientia Labor Wermsdorf | Robert Koch Institute | unknown |
| EPI_ISL_2956481 | Sonic - Labor Staber Dresden (Klipphausen) | Robert Koch Institute | unknown |
| EPI_ISL_2956483, EPI_ISL_2956484 | Bioscientia Labor Wermsdorf | Robert Koch Institute | unknown |
| EPI_ISL_2956486 | Sonic - Labor Dr. von Froreich GmbH | Robert Koch Institute | unknown |
| EPI_ISL_2956487 | Bioscientia Labor Wermsdorf | Robert Koch Institute | unknown |
| EPI_ISL_2956488 | Sonic - Bioscientia - MVZ Labor Saar GmbH | Robert Koch Institute | unknown |
| EPI_ISL_2956490 | Sonic - MVZ Medizinisches Labor Bremen GmbH | Robert Koch Institute | unknown |
| EPI_ISL_2956491 | Bioscientia Labor Wermsdorf | Robert Koch Institute | unknown |
| EPI_ISL_2956492 | Sonic - Bioscientia - MVZ Labor Saar GmbH | Robert Koch Institute | unknown |
| EPI_ISL_2956493 | Bioscientia Labor Wermsdorf | Robert Koch Institute | unknown |
| EPI_ISL_2956494 | LabKom - Labor an der Salzbrücke MVZ GmbH | Robert Koch Institute | unknown |
| EPI_ISL_2956495 | Bioscientia MVZ Labor Karlsruhe GmbH | Robert Koch Institute | unknown |
| EPI_ISL_2956496 | Bioscientia Labor Wermsdorf | Robert Koch Institute | unknown |
| EPI_ISL_2956497 | Sonic - Labor Dr. von Froreich GmbH | Robert Koch Institute | unknown |
| EPI_ISL_2956498, EPI_ISL_2956499, EPI_ISL_2956500 | Bioscientia Labor Wermsdorf | Robert Koch Institute | unknown |
| EPI_ISL_2956502 | Sonic - Labor Dr. von Froreich GmbH | Robert Koch Institute | unknown |
| EPI_ISL_2956503 | Sonic - Bioscientia - MVZ Labor Saar GmbH | Robert Koch Institute | unknown |
| EPI_ISL_2956504 | Bioscientia Labor Wermsdorf | Robert Koch Institute | unknown |
| EPI_ISL_2956505 | Bioscientia MVZ Labor Karlsruhe GmbH | Robert Koch Institute | unknown |
| EPI_ISL_2956506 | Bioscientia Labor Wermsdorf | Robert Koch Institute | unknown |
| EPI_ISL_2956507 | LabKom - Labor Augsburg MVZ GmbH | Robert Koch Institute | unknown |
| EPI_ISL_2956508, EPI_ISL_2956510, EPI_ISL_2956511, EPI_ISL_2956514 | Bioscientia Labor Wermsdorf | Robert Koch Institute | unknown |
| EPI_ISL_2956515 | Sonic - Bioscientia - MVZ Labor Saar GmbH | Robert Koch Institute | unknown |
| EPI_ISL_2956516, EPI_ISL_2956518 | Bioscientia Labor Wermsdorf | Robert Koch Institute | unknown |
| EPI_ISL_2956524, EPI_ISL_2956525, EPI_ISL_2956526, EPI_ISL_2956527, EPI_ISL_2956528 | alphaomega Labor GbR (Medizinische Kooperationsgemeinschaft Dres. Ackermann/Tennert/Zaiß) | Robert Koch Institute | unknown |
| EPI_ISL_2956529, EPI_ISL_2956530, EPI_ISL_2956531, EPI_ISL_2956532, EPI_ISL_2956533, EPI_ISL_2956534, EPI_ISL_2956535, EPI_ISL_2956537, | Labor Dr. Heidrich & Kollegen MVZ GmbH Hamburg | Robert Koch Institute | unknown |

| | | | |
|---|---|---|---|
| EPI_ISL_2956538, EPI_ISL_2956540 | | | |
| EPI_ISL_2956541, EPI_ISL_2956542 | Philipps-Universität Marburg Fachbereich Medizin Institut für Virologie | Robert Koch Institute | unknown |
| EPI_ISL_2956543 | Laborarztpraxis Osnabrück | Robert Koch Institute | unknown |
| EPI_ISL_2956545 | MVZ Dr. Eberhard & Partner Dortmund | Robert Koch Institute | unknown |
| EPI_ISL_2956546 | Laborarztpraxis Osnabrück | Robert Koch Institute | unknown |
| EPI_ISL_2956547 | Medizinisches Versorgungszentrum für Labormedizin und Mikrobiologie Ruhr GmbH - mvzlm RUHR GmbH | Robert Koch Institute | unknown |
| EPI_ISL_2956549 | Procomcure Biotech Germany GmbH | Robert Koch Institute | unknown |
| EPI_ISL_2956550 | MVZ Dr. Eberhard & Partner Dortmund | Robert Koch Institute | unknown |
| EPI_ISL_2956551 | Medizinisches Versorgungszentrum für Labormedizin und Mikrobiologie Ruhr GmbH - mvzlm RUHR GmbH | Robert Koch Institute | unknown |
| EPI_ISL_2956552, EPI_ISL_2956553 | Laborarztpraxis Osnabrück | Robert Koch Institute | unknown |
| EPI_ISL_2956554 | MVZ Dr. Eberhard & Partner Dortmund | Robert Koch Institute | unknown |
| EPI_ISL_2956555, EPI_ISL_2956556 | Laborarztpraxis Osnabrück | Robert Koch Institute | unknown |
| EPI_ISL_2956557, EPI_ISL_2956558, EPI_ISL_2956559, EPI_ISL_2956560, EPI_ISL_2956561, EPI_ISL_2956562, EPI_ISL_2956564 | MVZ Labor Krone GbR | Robert Koch Institute | unknown |
| EPI_ISL_2956565 | Department of Clinical Microbiology | GIGA Medical Genomics | Keith Durkin, Maria Artesi, Bouchra Boujemla, Nathalie Renotte, Nadine Cambisano, Sébastien Bontems, Cécile Meex, Claire Gourzonès, Olivier Ek, Laurent Gillet, Marie-Pierre Hayette, Vincent Bours |
| EPI_ISL_2956566, EPI_ISL_2956569 | MVZ Labor Krone GbR | Robert Koch Institute | unknown |
| EPI_ISL_2956570 | Department of Clinical Microbiology | GIGA Medical Genomics | Keith Durkin, Maria Artesi, Bouchra Boujemla, Nathalie Renotte, Nadine Cambisano, Sébastien Bontems, Cécile Meex, Claire Gourzonès, Olivier Ek, Laurent Gillet, Marie-Pierre Hayette, Vincent Bours |
| EPI_ISL_2956571 | MVZ Labor Krone GbR | Robert Koch Institute | unknown |
| EPI_ISL_2956573 | Klinikum Stuttgart | Robert Koch Institute | unknown |
| EPI_ISL_2956574, EPI_ISL_2956575, EPI_ISL_2956577, EPI_ISL_2956579, EPI_ISL_2956581, EPI_ISL_2956582, EPI_ISL_2956584 | SYNLAB | GIGA Medical Genomics | Keith Durkin, Maria Artesi, Bouchra Boujemla, Nathalie Renotte, Nadine Cambisano, Sébastien Bontems, Cécile Meex, Claire Gourzonès, Olivier Ek, Laurent Gillet, Marie-Pierre Hayette, Vincent Bours |
| EPI_ISL_2956585 | Department of Clinical Microbiology | GIGA Medical Genomics | Keith Durkin, Maria Artesi, Bouchra Boujemla, Nathalie Renotte, Nadine Cambisano, Sébastien Bontems, Cécile Meex, Claire Gourzonès, Olivier Ek, Laurent Gillet, Marie-Pierre Hayette, Vincent Bours |
| EPI_ISL_2956586, EPI_ISL_2956587, EPI_ISL_2956588 | SYNLAB | GIGA Medical Genomics | Keith Durkin, Maria Artesi, Bouchra Boujemla, Nathalie Renotte, Nadine Cambisano, Sébastien Bontems, Cécile Meex, Claire Gourzonès, Olivier Ek, Laurent Gillet, Marie-Pierre Hayette, Vincent Bours |
| EPI_ISL_2956589, EPI_ISL_2956593 | Department of Clinical Microbiology | GIGA Medical Genomics | Keith Durkin, Maria Artesi, Bouchra Boujemla, Nathalie Renotte, Nadine Cambisano, Sébastien Bontems, Cécile Meex, Claire Gourzonès, Olivier Ek, Laurent Gillet, Marie-Pierre Hayette, Vincent Bours |
| EPI_ISL_2956594 | SYNLAB | GIGA Medical Genomics | Keith Durkin, Maria Artesi, Bouchra Boujemla, Nathalie Renotte, Nadine Cambisano, Sébastien Bontems, Cécile Meex, Claire Gourzonès, Olivier Ek, Laurent Gillet, Marie-Pierre Hayette, Vincent Bours |
| EPI_ISL_2956595, EPI_ISL_2956596 | Klinikum Stuttgart | Robert Koch Institute | unknown |
| EPI_ISL_2956597 | SYNLAB | GIGA Medical Genomics | Keith Durkin, Maria Artesi, Bouchra Boujemla, Nathalie Renotte, Nadine Cambisano, Sébastien Bontems, Cécile Meex, Claire Gourzonès, Olivier Ek, Laurent Gillet, Marie-Pierre Hayette, Vincent Bours |
| EPI_ISL_2956598 | Klinikum Stuttgart | Robert Koch Institute | unknown |
| EPI_ISL_2956600 | Department of Clinical Microbiology | GIGA Medical Genomics | Keith Durkin, Maria Artesi, Bouchra Boujemla, Nathalie Renotte, Nadine Cambisano, Sébastien Bontems, Cécile Meex, Claire Gourzonès, Olivier Ek, Laurent Gillet, Marie-Pierre Hayette, Vincent Bours |
| EPI_ISL_2956601 | LabKom - Labor Augsburg MVZ GmbH | Robert Koch Institute | unknown |
| EPI_ISL_2956603, EPI_ISL_2956607 | Bioscientia Labor Wermsdorf | Robert Koch Institute | unknown |
| EPI_ISL_2956608 | Department of Clinical Microbiology | GIGA Medical Genomics | Keith Durkin, Maria Artesi, Bouchra Boujemla, Nathalie Renotte, Nadine Cambisano, Sébastien Bontems, Cécile Meex, Claire Gourzonès, Olivier Ek, Laurent Gillet, Marie-Pierre Hayette, Vincent Bours |
| EPI_ISL_2956609, EPI_ISL_2956611 | Bioscientia Labor Wermsdorf | Robert Koch Institute | unknown |
| EPI_ISL_2956612, EPI_ISL_2956613, EPI_ISL_2956615 | LabKom - Labor Augsburg MVZ GmbH | Robert Koch Institute | unknown |
| EPI_ISL_2956617 | Labor Dr. Schumacher MVZ | Robert Koch Institute | unknown |
| EPI_ISL_2956619, EPI_ISL_2956620, EPI_ISL_2956621 | LabKom - Labor Augsburg MVZ GmbH | Robert Koch Institute | unknown |
| EPI_ISL_2956626 | Labor Dr. Schumacher MVZ | Robert Koch Institute | unknown |
| EPI_ISL_2956627, EPI_ISL_2956631 | LabKom - Labor Augsburg MVZ GmbH | Robert Koch Institute | unknown |
| EPI_ISL_2956632 | Bioscientia Labor Wermsdorf | Robert Koch Institute | unknown |
| EPI_ISL_2956633 | LabKom - Labor Augsburg MVZ GmbH | Robert Koch Institute | unknown |
| EPI_ISL_2956634 | Bioscientia Labor Wermsdorf | Robert Koch Institute | unknown |
| EPI_ISL_2956635 | LabKom - Labor Augsburg MVZ GmbH | Robert Koch Institute | unknown |
| EPI_ISL_2956636 | Bioscientia Labor Wermsdorf | Robert Koch Institute | unknown |
| EPI_ISL_2956637, EPI_ISL_2956638 | LabKom - Labor Augsburg MVZ GmbH | Robert Koch Institute | unknown |
| EPI_ISL_2956639 | Bioscientia Labor Wermsdorf | Robert Koch Institute | unknown |
| EPI_ISL_2956640 | LabKom - Labor Augsburg MVZ GmbH | Robert Koch Institute | unknown |
| EPI_ISL_2956641 | Bioscientia MVZ Labor Karlsruhe GmbH | Robert Koch Institute | unknown |
| EPI_ISL_2956642 | Labor 28 MVZ GmbH | Robert Koch Institute | unknown |
| EPI_ISL_2956643, EPI_ISL_2956644 | LabKom - Labor Augsburg MVZ GmbH | Robert Koch Institute | unknown |
| EPI_ISL_2956646 | Bioscientia Labor Wermsdorf | Robert Koch Institute | unknown |
| EPI_ISL_2956647 | Labor 28 MVZ GmbH | Robert Koch Institute | unknown |
| EPI_ISL_2956648 | LabKom - Labor Augsburg MVZ GmbH | Robert Koch Institute | unknown |
| EPI_ISL_2956649 | Bioscientia Labor Wermsdorf | Robert Koch Institute | unknown |
| EPI_ISL_2956650, EPI_ISL_2956651 | LabKom - Labor Augsburg MVZ GmbH | Robert Koch Institute | unknown |
| EPI_ISL_2956652 | Labor 28 MVZ GmbH | Robert Koch Institute | unknown |
| EPI_ISL_2956653, | LabKom - Labor Augsburg MVZ GmbH | Robert Koch Institute | unknown |

| | | | |
|---|---|---|---|
| EPI_ISL_2956655, EPI_ISL_2956656 | | | |
| EPI_ISL_2956657 | Bioscientia Labor Wermsdorf | Robert Koch Institute | unknown |
| EPI_ISL_2956658 | LabKom - Labor Augsburg MVZ GmbH | Robert Koch Institute | unknown |
| EPI_ISL_2956659 | Labor Dr. Schumacher MVZ | Robert Koch Institute | unknown |
| EPI_ISL_2956660, EPI_ISL_2956661, EPI_ISL_2956662, EPI_ISL_2956663 | LabKom - Labor Augsburg MVZ GmbH | Robert Koch Institute | unknown |
| EPI_ISL_2956664, EPI_ISL_2956665 | Bioscientia Labor Wermsdorf | Robert Koch Institute | unknown |
| EPI_ISL_2956666 | LabKom - Labor Augsburg MVZ GmbH | Robert Koch Institute | unknown |
| EPI_ISL_2956667 | Labor 28 MVZ GmbH | Robert Koch Institute | unknown |
| EPI_ISL_2956669 | LabKom - Labor Augsburg MVZ GmbH | Robert Koch Institute | unknown |
| EPI_ISL_2956670 | Bioscientia Labor Wermsdorf | Robert Koch Institute | unknown |
| EPI_ISL_2956671 | Bioscientia MVZ Labor Karlsruhe GmbH | Robert Koch Institute | unknown |
| EPI_ISL_2956672, EPI_ISL_2956673 | LabKom - Labor Augsburg MVZ GmbH | Robert Koch Institute | unknown |
| EPI_ISL_2956674 | Labor 28 MVZ GmbH | Robert Koch Institute | unknown |
| EPI_ISL_2956676, EPI_ISL_2956678, EPI_ISL_2956679, EPI_ISL_2956680, EPI_ISL_2956681, EPI_ISL_2956683, EPI_ISL_2956684, EPI_ISL_2956685 | LabKom - Labor Augsburg MVZ GmbH | Robert Koch Institute | unknown |
| EPI_ISL_2956686, EPI_ISL_2956691, EPI_ISL_2956692, EPI_ISL_2956693, EPI_ISL_2956695, EPI_ISL_2956696, EPI_ISL_2956697, EPI_ISL_2956698 | Eurofins MVZ Labor Gelsenkirchen | Robert Koch Institute | unknown |
| EPI_ISL_2956700 | MVZ Labor Dr. Reising-Ackermann und Kollegen | Robert Koch Institute | unknown |
| EPI_ISL_2956702 | MVZ für Laboratoriumsmedizin und Mikrobiologie Koblenz-Mittelrhein (Labor Koblenz) | Robert Koch Institute | unknown |
| EPI_ISL_2956703 | MVZ Labor Dr. Limbach & Kollegen GbR | Robert Koch Institute | unknown |
| EPI_ISL_2956704 | MVZ Medizinisches Labor Hannover GmbH | Robert Koch Institute | unknown |
| EPI_ISL_2956705, EPI_ISL_2956706 | MVZ Labor Dr. Limbach & Kollegen GbR | Robert Koch Institute | unknown |
| EPI_ISL_2956707 | MVZ für Laboratoriumsmedizin und Mikrobiologie Koblenz-Mittelrhein (Labor Koblenz) | Robert Koch Institute | unknown |
| EPI_ISL_2956709 | MVZ Labor Dr. Limbach & Kollegen GbR | Robert Koch Institute | unknown |
| EPI_ISL_2956710 | Limbach - MVZ Gemeinschaftslabor Suhl | Robert Koch Institute | unknown |
| EPI_ISL_2956712, EPI_ISL_2956713 | Eurofins MVZ Labor Gelsenkirchen | Robert Koch Institute | unknown |
| EPI_ISL_2956714, EPI_ISL_2956715, EPI_ISL_2956716, EPI_ISL_2956717, EPI_ISL_2956718, EPI_ISL_2956719 | MVZ Labor Dr. Limbach & Kollegen GbR | Robert Koch Institute | unknown |
| EPI_ISL_2956722 | MVZ Medizinisches Labor Hannover GmbH | Robert Koch Institute | unknown |
| EPI_ISL_2956723 | Limbach - MVZ Labor Ludwigsburg | Robert Koch Institute | unknown |
| EPI_ISL_2956724 | MVZ für Laboratoriumsmedizin und Mikrobiologie Koblenz-Mittelrhein (Labor Koblenz) | Robert Koch Institute | unknown |
| EPI_ISL_2956725 | MVZ Labor Dr. Limbach & Kollegen GbR | Robert Koch Institute | unknown |
| EPI_ISL_2956727 | Limbach - MVZ Labor Ludwigsburg | Robert Koch Institute | unknown |
| EPI_ISL_2956728, EPI_ISL_2956729 | MVZ für Laboratoriumsmedizin und Mikrobiologie Koblenz-Mittelrhein (Labor Koblenz) | Robert Koch Institute | unknown |
| EPI_ISL_2956730 | Klinikum Darmstadt GmbH | Robert Koch Institute | unknown |
| EPI_ISL_2956731 | Limbach - MVZ Labor Ludwigsburg | Robert Koch Institute | unknown |
| EPI_ISL_2956732 | MVZ für Laboratoriumsmedizin und Mikrobiologie Koblenz-Mittelrhein (Labor Koblenz) | Robert Koch Institute | unknown |
| EPI_ISL_2956734 | MVZ Labor Dr. Limbach & Kollegen GbR | Robert Koch Institute | unknown |
| EPI_ISL_2956735 | Limbach - MVZ Labor Ludwigsburg | Robert Koch Institute | unknown |
| EPI_ISL_2956736 | MVZ Medizinisches Labor Hannover GmbH | Robert Koch Institute | unknown |
| EPI_ISL_2956737 | MVZ für Laboratoriumsmedizin und Mikrobiologie Koblenz-Mittelrhein (Labor Koblenz) | Robert Koch Institute | unknown |
| EPI_ISL_2956739, EPI_ISL_2956740, EPI_ISL_2956741, EPI_ISL_2956742, EPI_ISL_2956743, EPI_ISL_2956744, EPI_ISL_2956745, EPI_ISL_2956746, EPI_ISL_2956747, EPI_ISL_2956748, EPI_ISL_2956749, EPI_ISL_2956750, EPI_ISL_2956752 | | | |
| see above | Eurofins LifeCodexx GmbH | Robert Koch Institute | unknown |
| EPI_ISL_2956753, EPI_ISL_2956754 | Amazon Dx, AUV2 Laboratory | Amazon Dx, AUV2 Laboratory | Luke Meredith |
| EPI_ISL_2956755 | Eurofins LifeCodexx GmbH | Robert Koch Institute | unknown |
| EPI_ISL_2956756 | Amazon Dx, AUV2 Laboratory | Amazon Dx, AUV2 Laboratory | Luke Meredith |
| EPI_ISL_2956757 | Eurofins LifeCodexx GmbH | Robert Koch Institute | unknown |
| EPI_ISL_2956758 | Amazon Dx, AUV2 Laboratory | Amazon Dx, AUV2 Laboratory | Luke Meredith |
| EPI_ISL_2956759, EPI_ISL_2956760 | Eurofins LifeCodexx GmbH | Robert Koch Institute | unknown |
| EPI_ISL_2956761 | Amazon Dx, AUV2 Laboratory | Amazon Dx, AUV2 Laboratory | Luke Meredith |
| EPI_ISL_2956762 | Eurofins LifeCodexx GmbH | Robert Koch Institute | unknown |
| EPI_ISL_2956763 | Amazon Dx, AUV2 Laboratory | Amazon Dx, AUV2 Laboratory | Luke Meredith |
| EPI_ISL_2956764 | Eurofins LifeCodexx GmbH | Robert Koch Institute | unknown |
| EPI_ISL_2956765 | Amazon Dx, AUV2 Laboratory | Amazon Dx, AUV2 Laboratory | Luke Meredith |
| EPI_ISL_2956766 | Eurofins LifeCodexx GmbH | Robert Koch Institute | unknown |

| | | | |
|---|---|---|---|
| EPI_ISL_2956767 | Amazon Dx, AUV2 Laboratory | Amazon Dx, AUV2 Laboratory | Luke Meredith |
| EPI_ISL_2956768 | Eurofins LifeCodexx GmbH | Robert Koch Institute | unknown |
| EPI_ISL_2956769, EPI_ISL_2956770 | Amazon Dx, AUV2 Laboratory | Amazon Dx, AUV2 Laboratory | Luke Meredith |
| EPI_ISL_2956771 | Eurofins LifeCodexx GmbH | Robert Koch Institute | unknown |
| EPI_ISL_2956772 | Amazon Dx, AUV2 Laboratory | Amazon Dx, AUV2 Laboratory | Luke Meredith |
| EPI_ISL_2956773 | Eurofins LifeCodexx GmbH | Robert Koch Institute | unknown |
| EPI_ISL_2956774 | Amazon Dx, AUV2 Laboratory | Amazon Dx, AUV2 Laboratory | Luke Meredith |
| EPI_ISL_2956775 | Eurofins LifeCodexx GmbH | Robert Koch Institute | unknown |
| EPI_ISL_2956776, EPI_ISL_2956777 | Amazon Dx, AUV2 Laboratory | Amazon Dx, AUV2 Laboratory | Luke Meredith |
| EPI_ISL_2956778 | Eurofins LifeCodexx GmbH | Robert Koch Institute | unknown |
| EPI_ISL_2956779 | Amazon Dx, AUV2 Laboratory | Amazon Dx, AUV2 Laboratory | Luke Meredith |
| EPI_ISL_2956780 | Eurofins LifeCodexx GmbH | Robert Koch Institute | unknown |
| EPI_ISL_2956781, EPI_ISL_2956782 | Amazon Dx, AUV2 Laboratory | Amazon Dx, AUV2 Laboratory | Luke Meredith |
| EPI_ISL_2956783 | Eurofins LifeCodexx GmbH | Robert Koch Institute | unknown |
| EPI_ISL_2956784 | Amazon Dx, AUV2 Laboratory | Amazon Dx, AUV2 Laboratory | Luke Meredith |
| EPI_ISL_2956785, EPI_ISL_2956786 | Eurofins LifeCodexx GmbH | Robert Koch Institute | unknown |
| EPI_ISL_2956787, EPI_ISL_2956788 | Amazon Dx, AUV2 Laboratory | Amazon Dx, AUV2 Laboratory | Luke Meredith |
| EPI_ISL_2956789 | Eurofins LifeCodexx GmbH | Robert Koch Institute | unknown |
| EPI_ISL_2956790 | Amazon Dx, AUV2 Laboratory | Amazon Dx, AUV2 Laboratory | Luke Meredith |
| EPI_ISL_2956791 | Eurofins LifeCodexx GmbH | Robert Koch Institute | unknown |
| EPI_ISL_2956792, EPI_ISL_2956793 | Amazon Dx, AUV2 Laboratory | Amazon Dx, AUV2 Laboratory | Luke Meredith |
| EPI_ISL_2956794 | Eurofins LifeCodexx GmbH | Robert Koch Institute | unknown |
| EPI_ISL_2956795 | Amazon Dx, AUV2 Laboratory | Amazon Dx, AUV2 Laboratory | Luke Meredith |
| EPI_ISL_2956796 | Eurofins LifeCodexx GmbH | Robert Koch Institute | unknown |
| EPI_ISL_2956797 | Amazon Dx, AUV2 Laboratory | Amazon Dx, AUV2 Laboratory | Luke Meredith |
| EPI_ISL_2956798, EPI_ISL_2956799 | Eurofins LifeCodexx GmbH | Robert Koch Institute | unknown |
| EPI_ISL_2956800 | Amazon Dx, AUV2 Laboratory | Amazon Dx, AUV2 Laboratory | Luke Meredith |
| EPI_ISL_2956801 | Eurofins LifeCodexx GmbH | Robert Koch Institute | unknown |
| EPI_ISL_2956802, EPI_ISL_2956803, EPI_ISL_2956804, EPI_ISL_2956805, EPI_ISL_2956806, EPI_ISL_2956807, EPI_ISL_2956809, EPI_ISL_2956810, EPI_ISL_2956811, EPI_ISL_2956812, EPI_ISL_2956813, EPI_ISL_2956814, EPI_ISL_2956815, EPI_ISL_2956816 | | | |
| see above | Amazon Dx, AUV2 Laboratory | Amazon Dx, AUV2 Laboratory | Luke Meredith |
| EPI_ISL_2956817, EPI_ISL_2956818, EPI_ISL_2956819, EPI_ISL_2956820, EPI_ISL_2956821, EPI_ISL_2956822, EPI_ISL_2956823, EPI_ISL_2956824, EPI_ISL_2956825, EPI_ISL_2956826, EPI_ISL_2956827, EPI_ISL_2956828, EPI_ISL_2956829, EPI_ISL_2956830, EPI_ISL_2956831, EPI_ISL_2956832, EPI_ISL_2956833, EPI_ISL_2956834, EPI_ISL_2956835, EPI_ISL_2956836, EPI_ISL_2956837 | | | |
| see above | Eurofins LifeCodexx GmbH | Robert Koch Institute | unknown |
| EPI_ISL_2956838 | Uniklinikum Carl Gustav Carus an der TU Dresden; Institut für Virologie | Robert Koch Institute | unknown |
| EPI_ISL_2956840 | Städtisches Klinikum Dresden Institut für Klinische Chemie und Laboratoriumsmedizin | Robert Koch Institute | unknown |
| EPI_ISL_2956842 | Landesuntersuchungsanstalt für das Gesundheits- und Veterinärwesen (LUA) Sachsen | Robert Koch Institute | unknown |
| EPI_ISL_2956843 | Uniklinikum Carl Gustav Carus an der TU Dresden; Institut für Virologie | Robert Koch Institute | unknown |
| EPI_ISL_2956847 | LABOR CHEMNITZ – Zentrum für Diagnostik am Klinikum Chemnitz | Robert Koch Institute | unknown |
| EPI_ISL_2956848 | Uniklinikum Carl Gustav Carus an der TU Dresden; Institut für Virologie | Robert Koch Institute | unknown |
| EPI_ISL_2956855, EPI_ISL_2956860, EPI_ISL_2956865, EPI_ISL_2956866, EPI_ISL_2956867, EPI_ISL_2956868, EPI_ISL_2956869, EPI_ISL_2956870, EPI_ISL_2956871, EPI_ISL_2956872, EPI_ISL_2956873, EPI_ISL_2956875, EPI_ISL_2956877, EPI_ISL_2956878, EPI_ISL_2956879, EPI_ISL_2956881, EPI_ISL_2956882, EPI_ISL_2956883, EPI_ISL_2956884, EPI_ISL_2956888, EPI_ISL_2956890, EPI_ISL_2956892, EPI_ISL_2956893 | | | |
| see above | Landesgesundheitsamt Baden-Württemberg | Robert Koch Institute | unknown |
| EPI_ISL_2956896, EPI_ISL_2956899 | Limbach - Ganzimmun Diagnostics AG Mainz | Robert Koch Institute | unknown |
| EPI_ISL_2956901, EPI_ISL_2956902, EPI_ISL_2956904, EPI_ISL_2956905, EPI_ISL_2956906, EPI_ISL_2956908, EPI_ISL_2956909, EPI_ISL_2956910, EPI_ISL_2956911, EPI_ISL_2956912, EPI_ISL_2956913, EPI_ISL_2956914, EPI_ISL_2956915, EPI_ISL_2956916, EPI_ISL_2956917, EPI_ISL_2956918, EPI_ISL_2956919, EPI_ISL_2956920 | | | |
| see above | MVZ Labor Dr. Fenner und Kollegen (Standort Hamburg) | Robert Koch Institute | unknown |
| EPI_ISL_2956922, EPI_ISL_2956923, EPI_ISL_2956924, EPI_ISL_2956926, EPI_ISL_2956927, EPI_ISL_2956928, EPI_ISL_2956929, EPI_ISL_2956930, EPI_ISL_2956931, EPI_ISL_2956932, EPI_ISL_2956933, EPI_ISL_2956934, EPI_ISL_2956935, EPI_ISL_2956936, EPI_ISL_2956937, EPI_ISL_2956939, EPI_ISL_2956940, EPI_ISL_2956941, EPI_ISL_2956942, EPI_ISL_2956943, EPI_ISL_2956944, EPI_ISL_2956945, EPI_ISL_2956946, EPI_ISL_2956947, EPI_ISL_2956948, EPI_ISL_2956949, EPI_ISL_2956950, EPI_ISL_2956951, EPI_ISL_2956952, EPI_ISL_2956953, EPI_ISL_2956954, EPI_ISL_2956955, EPI_ISL_2956956, EPI_ISL_2956957, EPI_ISL_2956958, EPI_ISL_2956959, EPI_ISL_2956960, EPI_ISL_2956961, EPI_ISL_2956962, EPI_ISL_2956963, EPI_ISL_2956964, EPI_ISL_2956965, EPI_ISL_2956966, EPI_ISL_2956968, EPI_ISL_2956969, EPI_ISL_2956970, EPI_ISL_2956971, EPI_ISL_2956973, EPI_ISL_2956974, EPI_ISL_2956975 | | | |
| see above | LADR Zentrallabor DR. Kramer & Kollegen Geesthacht | Robert Koch Institute | unknown |
| EPI_ISL_2956977 | Hospital Universitari Bellvitge | Microbiology Department | Sara Marti, Aida Gonzalez-Diaz, Laura Calatayud, Jordi Niubó, Miguel Fernandez-Huerta, Carmen Ardanuy, Jordi Camara, M Angeles Domínguez |
| EPI_ISL_2956979 | LADR Zentrallabor DR. Kramer & Kollegen Geesthacht | Robert Koch Institute | unknown |
| EPI_ISL_2956980 | CLILAB | Microbiology Department | Sara Marti, Aida Gonzalez-Diaz, Laura Calatayud, Jordi Niubó, Miguel Fernandez-Huerta, Carmen Ardanuy, Jordi Camara, M Angeles Domínguez |
| EPI_ISL_2956981 | LADR Zentrallabor DR. Kramer & Kollegen Geesthacht | Robert Koch Institute | unknown |
| EPI_ISL_2956982 | CLILAB | Microbiology Department | Sara Marti, Aida Gonzalez-Diaz, Laura Calatayud, Jordi Niubó, Miguel Fernandez-Huerta, Carmen Ardanuy, Jordi Camara, M Angeles Domínguez |
| EPI_ISL_2956983 | LADR Zentrallabor DR. Kramer & Kollegen Geesthacht | Robert Koch Institute | unknown |
| EPI_ISL_2956985 | CLILAB | Microbiology Department | Sara Marti, Aida Gonzalez-Diaz, Laura Calatayud, Jordi Niubó, Miguel Fernandez-Huerta, Carmen Ardanuy, Jordi Camara, M Angeles Domínguez |
| EPI_ISL_2956986, EPI_ISL_2956987 | LADR Zentrallabor DR. Kramer & Kollegen Geesthacht | Robert Koch Institute | unknown |
| EPI_ISL_2956988 | Hospital Universitari Bellvitge | Microbiology Department | Sara Marti, Aida Gonzalez-Diaz, Laura Calatayud, Jordi Niubó, Miguel Fernandez-Huerta, Carmen Ardanuy, Jordi Camara, M Angeles Domínguez |
| EPI_ISL_2956989, EPI_ISL_2956990 | LADR Zentrallabor DR. Kramer & Kollegen Geesthacht | Robert Koch Institute | unknown |
| EPI_ISL_2956991 | Hospital Universitari Bellvitge | Microbiology Department | Sara Marti, Aida Gonzalez-Diaz, Laura Calatayud, Jordi Niubó, Miguel Fernandez-Huerta, Carmen Ardanuy, Jordi Camara, M Angeles Domínguez |
| EPI_ISL_2956992, EPI_ISL_2956993 | LADR Zentrallabor DR. Kramer & Kollegen Geesthacht | Robert Koch Institute | unknown |
| EPI_ISL_2956994 | CLILAB | Microbiology Department | Sara Marti, Aida Gonzalez-Diaz, Laura Calatayud, Jordi Niubó, Miguel Fernandez-Huerta, Carmen Ardanuy, Jordi Camara, M Angeles Domínguez |
| EPI_ISL_2956995, EPI_ISL_2956996, EPI_ISL_2956997 | LADR Zentrallabor DR. Kramer & Kollegen Geesthacht | Robert Koch Institute | unknown |
| EPI_ISL_2956998 | Hospital Universitari Bellvitge | Microbiology Department | Sara Marti, Aida Gonzalez-Diaz, Laura Calatayud, Jordi Niubó, Miguel Fernandez-Huerta, Carmen Ardanuy, Jordi Camara, M Angeles Domínguez |
| EPI_ISL_2956999 | LADR Zentrallabor DR. Kramer & Kollegen Geesthacht | Robert Koch Institute | unknown |

| | | | |
|---|---|---|---|
| EPI_ISL_2957000 | CLILAB | Microbiology Department | Sara Marti, Aida Gonzalez-Diaz, Laura Calatayud, Jordi Niubó, Miguel Fernandez-Huerta, Carmen Ardanuy, Jordi Camara, M Angeles Domínguez |
| EPI_ISL_2957001 | LADR Zentrallabor DR. Kramer & Kollegen Geesthacht | Robert Koch Institute | unknown |
| EPI_ISL_2957003 | CLILAB | Microbiology Department | Sara Marti, Aida Gonzalez-Diaz, Laura Calatayud, Jordi Niubó, Miguel Fernandez-Huerta, Carmen Ardanuy, Jordi Camara, M Angeles Domínguez |
| EPI_ISL_2957004, EPI_ISL_2957005, EPI_ISL_2957007, EPI_ISL_2957008 | LADR Zentrallabor DR. Kramer & Kollegen Geesthacht | Robert Koch Institute | unknown |
| EPI_ISL_2957009 | CLILAB | Microbiology Department | Sara Marti, Aida Gonzalez-Diaz, Laura Calatayud, Jordi Niubó, Miguel Fernandez-Huerta, Carmen Ardanuy, Jordi Camara, M Angeles Domínguez |
| EPI_ISL_2957010, EPI_ISL_2957011 | LADR Zentrallabor DR. Kramer & Kollegen Geesthacht | Robert Koch Institute | unknown |
| EPI_ISL_2957012 | CLILAB | Microbiology Department | Sara Marti, Aida Gonzalez-Diaz, Laura Calatayud, Jordi Niubó, Miguel Fernandez-Huerta, Carmen Ardanuy, Jordi Camara, M Angeles Domínguez |
| EPI_ISL_2957013, EPI_ISL_2957014 | LADR Zentrallabor DR. Kramer & Kollegen Geesthacht | Robert Koch Institute | unknown |
| EPI_ISL_2957015 | CLILAB | Microbiology Department | Sara Marti, Aida Gonzalez-Diaz, Laura Calatayud, Jordi Niubó, Miguel Fernandez-Huerta, Carmen Ardanuy, Jordi Camara, M Angeles Domínguez |
| EPI_ISL_2957016, EPI_ISL_2957017 | LADR Zentrallabor DR. Kramer & Kollegen Geesthacht | Robert Koch Institute | unknown |
| EPI_ISL_2957018 | Hospital Universitari Bellvitge | Microbiology Department | Sara Marti, Aida Gonzalez-Diaz, Laura Calatayud, Jordi Niubó, Miguel Fernandez-Huerta, Carmen Ardanuy, Jordi Camara, M Angeles Domínguez |
| EPI_ISL_2957019, EPI_ISL_2957020 | LADR Zentrallabor DR. Kramer & Kollegen Geesthacht | Robert Koch Institute | unknown |
| EPI_ISL_2957021 | Hospital Universitari Bellvitge | Microbiology Department | Sara Marti, Aida Gonzalez-Diaz, Laura Calatayud, Jordi Niubó, Miguel Fernandez-Huerta, Carmen Ardanuy, Jordi Camara, M Angeles Domínguez |
| EPI_ISL_2957023, EPI_ISL_2957026 | LADR Zentrallabor DR. Kramer & Kollegen Geesthacht | Robert Koch Institute | unknown |
| EPI_ISL_2957027 | Hospital Universitari Bellvitge | Microbiology Department | Sara Marti, Aida Gonzalez-Diaz, Laura Calatayud, Jordi Niubó, Miguel Fernandez-Huerta, Carmen Ardanuy, Jordi Camara, M Angeles Domínguez |
| EPI_ISL_2957028 | LADR Zentrallabor DR. Kramer & Kollegen Geesthacht | Robert Koch Institute | unknown |
| EPI_ISL_2957029, EPI_ISL_2957030, EPI_ISL_2957031 | Hospital Universitari Bellvitge | Microbiology Department | Sara Marti, Aida Gonzalez-Diaz, Laura Calatayud, Jordi Niubó, Miguel Fernandez-Huerta, Carmen Ardanuy, Jordi Camara, M Angeles Domínguez |
| EPI_ISL_2957032 | LADR Zentrallabor DR. Kramer & Kollegen Geesthacht | Robert Koch Institute | unknown |
| EPI_ISL_2957033 | Hospital Universitari Bellvitge | Microbiology Department | Sara Marti, Aida Gonzalez-Diaz, Laura Calatayud, Jordi Niubó, Miguel Fernandez-Huerta, Carmen Ardanuy, Jordi Camara, M Angeles Domínguez |
| EPI_ISL_2957034 | CLILAB | Microbiology Department | Sara Marti, Aida Gonzalez-Diaz, Laura Calatayud, Jordi Niubó, Miguel Fernandez-Huerta, Carmen Ardanuy, Jordi Camara, M Angeles Domínguez |
| EPI_ISL_2957035, EPI_ISL_2957037 | LADR Zentrallabor DR. Kramer & Kollegen Geesthacht | Robert Koch Institute | unknown |
| EPI_ISL_2957039 | CLILAB | Microbiology Department | Sara Marti, Aida Gonzalez-Diaz, Laura Calatayud, Jordi Niubó, Miguel Fernandez-Huerta, Carmen Ardanuy, Jordi Camara, M Angeles Domínguez |
| EPI_ISL_2957040 | LADR Zentrallabor DR. Kramer & Kollegen Geesthacht | Robert Koch Institute | unknown |
| EPI_ISL_2957042, EPI_ISL_2957046, EPI_ISL_2957048 | CLILAB | Microbiology Department | Sara Marti, Aida Gonzalez-Diaz, Laura Calatayud, Jordi Niubó, Miguel Fernandez-Huerta, Carmen Ardanuy, Jordi Camara, M Angeles Domínguez |
| EPI_ISL_2957050 | Hospital Universitari Bellvitge | Microbiology Department | Sara Marti, Aida Gonzalez-Diaz, Laura Calatayud, Jordi Niubó, Miguel Fernandez-Huerta, Carmen Ardanuy, Jordi Camara, M Angeles Domínguez |
| EPI_ISL_2957054, EPI_ISL_2957055, EPI_ISL_2957057 | CLILAB | Microbiology Department | Sara Marti, Aida Gonzalez-Diaz, Laura Calatayud, Jordi Niubó, Miguel Fernandez-Huerta, Carmen Ardanuy, Jordi Camara, M Angeles Domínguez |
| EPI_ISL_2957059 | Hospital Universitari Bellvitge | Microbiology Department | Sara Marti, Aida Gonzalez-Diaz, Laura Calatayud, Jordi Niubó, Miguel Fernandez-Huerta, Carmen Ardanuy, Jordi Camara, M Angeles Domínguez |
| EPI_ISL_2957060 | Bayerisches Landesamt für Gesundheit und Lebensmittelsicherheit (LGL) | Robert Koch Institute | unknown |
| EPI_ISL_2957061 | Hospital Universitari Bellvitge | Microbiology Department | Sara Marti, Aida Gonzalez-Diaz, Laura Calatayud, Jordi Niubó, Miguel Fernandez-Huerta, Carmen Ardanuy, Jordi Camara, M Angeles Domínguez |
| EPI_ISL_2957062 | Bayerisches Landesamt für Gesundheit und Lebensmittelsicherheit (LGL) | Robert Koch Institute | unknown |
| EPI_ISL_2957063 | Hospital Universitari Bellvitge | Microbiology Department | Sara Marti, Aida Gonzalez-Diaz, Laura Calatayud, Jordi Niubó, Miguel Fernandez-Huerta, Carmen Ardanuy, Jordi Camara, M Angeles Domínguez |
| EPI_ISL_2957064 | Bayerisches Landesamt für Gesundheit und Lebensmittelsicherheit (LGL) | Robert Koch Institute | unknown |
| EPI_ISL_2957065, EPI_ISL_2957067, EPI_ISL_2957071 | Hospital Universitari Bellvitge | Microbiology Department | Sara Marti, Aida Gonzalez-Diaz, Laura Calatayud, Jordi Niubó, Miguel Fernandez-Huerta, Carmen Ardanuy, Jordi Camara, M Angeles Domínguez |
| EPI_ISL_2957075 | Bayerisches Landesamt für Gesundheit und Lebensmittelsicherheit (LGL) | Robert Koch Institute | unknown |
| EPI_ISL_2957076, EPI_ISL_2957078, EPI_ISL_2957080 | Hospital Universitari Bellvitge | Microbiology Department | Sara Marti, Aida Gonzalez-Diaz, Laura Calatayud, Jordi Niubó, Miguel Fernandez-Huerta, Carmen Ardanuy, Jordi Camara, M Angeles Domínguez |
| EPI_ISL_2957081 | Bayerisches Landesamt für Gesundheit und Lebensmittelsicherheit (LGL) | Robert Koch Institute | unknown |
| EPI_ISL_2957082 | Hospital Universitari Bellvitge | Microbiology Department | Sara Marti, Aida Gonzalez-Diaz, Laura Calatayud, Jordi Niubó, Miguel Fernandez-Huerta, Carmen Ardanuy, Jordi Camara, M Angeles Domínguez |
| EPI_ISL_2957083 | Bayerisches Landesamt für Gesundheit und Lebensmittelsicherheit (LGL) | Robert Koch Institute | unknown |
| EPI_ISL_2957084, EPI_ISL_2957086 | Hospital Universitari Bellvitge | Microbiology Department | Sara Marti, Aida Gonzalez-Diaz, Laura Calatayud, Jordi Niubó, Miguel Fernandez-Huerta, Carmen Ardanuy, Jordi Camara, M Angeles Domínguez |
| EPI_ISL_2957087, EPI_ISL_2957088, EPI_ISL_2957089 | Bayerisches Landesamt für Gesundheit und Lebensmittelsicherheit (LGL) | Robert Koch Institute | unknown |
| EPI_ISL_2957090, EPI_ISL_2957093, EPI_ISL_2957095 | Hospital Universitari Bellvitge | Microbiology Department | Sara Marti, Aida Gonzalez-Diaz, Laura Calatayud, Jordi Niubó, Miguel Fernandez-Huerta, Carmen Ardanuy, Jordi Camara, M Angeles Domínguez |
| EPI_ISL_2957096 | Bayerisches Landesamt für Gesundheit und Lebensmittelsicherheit (LGL) | Robert Koch Institute | unknown |
| EPI_ISL_2957097, EPI_ISL_2957099 | Hospital Universitari Bellvitge | Microbiology Department | Sara Marti, Aida Gonzalez-Diaz, Laura Calatayud, Jordi Niubó, Miguel Fernandez-Huerta, Carmen Ardanuy, Jordi Camara, M Angeles Domínguez |
| EPI_ISL_2957100 | Bayerisches Landesamt für Gesundheit und Lebensmittelsicherheit (LGL) | Robert Koch Institute | unknown |
| EPI_ISL_2957101 | Hospital Universitari Bellvitge | Microbiology Department | Sara Marti, Aida Gonzalez-Diaz, Laura Calatayud, Jordi Niubó, Miguel Fernandez-Huerta, Carmen Ardanuy, Jordi Camara, M Angeles Domínguez |
| EPI_ISL_2957102 | Bayerisches Landesamt für Gesundheit und Lebensmittelsicherheit (LGL) | Robert Koch Institute | unknown |
| EPI_ISL_2957104, EPI_ISL_2957106 | Hospital Universitari Bellvitge | Microbiology Department | Sara Marti, Aida Gonzalez-Diaz, Laura Calatayud, Jordi Niubó, Miguel Fernandez-Huerta, Carmen Ardanuy, Jordi Camara, M Angeles Domínguez |
| EPI_ISL_2957108, EPI_ISL_2957110, EPI_ISL_2957111 | Bayerisches Landesamt für Gesundheit und Lebensmittelsicherheit (LGL) | Robert Koch Institute | unknown |
| EPI_ISL_2957112, EPI_ISL_2957115 | Hospital Universitari Bellvitge | Microbiology Department | Sara Marti, Aida Gonzalez-Diaz, Laura Calatayud, Jordi Niubó, Miguel Fernandez-Huerta, Carmen Ardanuy, Jordi Camara, M Angeles Domínguez |
| EPI_ISL_2957116 | Bayerisches Landesamt für Gesundheit und | Robert Koch Institute | unknown |

| | | | |
|---|---|---|---|
| | Lebensmittelsicherheit (LGL) | | |
| EPI_ISL_2957118, EPI_ISL_2957119, EPI_ISL_2957120, EPI_ISL_2957121, EPI_ISL_2957123 | Hospital Universitari Bellvitge | Microbiology Department | Sara Marti, Aida Gonzalez-Diaz, Laura Calatayud, Jordi Niubó, Miguel Fernandez-Huerta, Carmen Ardanuy, Jordi Camara, M Angeles Domínguez |
| EPI_ISL_2957124 | Bayerisches Landesamt für Gesundheit und Lebensmittelsicherheit (LGL) | Robert Koch Institute | unknown |
| EPI_ISL_2957125 | Hospital Universitari Bellvitge | Microbiology Department | Sara Marti, Aida Gonzalez-Diaz, Laura Calatayud, Jordi Niubó, Miguel Fernandez-Huerta, Carmen Ardanuy, Jordi Camara, M Angeles Domínguez |
| EPI_ISL_2957126, EPI_ISL_2957130 | Bayerisches Landesamt für Gesundheit und Lebensmittelsicherheit (LGL) | Robert Koch Institute | unknown |
| EPI_ISL_2957133, EPI_ISL_2957134, EPI_ISL_2957136, EPI_ISL_2957138 | Hospital Universitari Bellvitge | Microbiology Department | Sara Marti, Aida Gonzalez-Diaz, Laura Calatayud, Jordi Niubó, Miguel Fernandez-Huerta, Carmen Ardanuy, Jordi Camara, M Angeles Domínguez |
| EPI_ISL_2957139 | Bayerisches Landesamt für Gesundheit und Lebensmittelsicherheit (LGL) | Robert Koch Institute | unknown |
| EPI_ISL_2957140 | Hospital Universitari Bellvitge | Microbiology Department | Sara Marti, Aida Gonzalez-Diaz, Laura Calatayud, Jordi Niubó, Miguel Fernandez-Huerta, Carmen Ardanuy, Jordi Camara, M Angeles Domínguez |
| EPI_ISL_2957141 | Bayerisches Landesamt für Gesundheit und Lebensmittelsicherheit (LGL) | Robert Koch Institute | unknown |
| EPI_ISL_2957142 | Hospital Universitari Bellvitge | Microbiology Department | Sara Marti, Aida Gonzalez-Diaz, Laura Calatayud, Jordi Niubó, Miguel Fernandez-Huerta, Carmen Ardanuy, Jordi Camara, M Angeles Domínguez |
| EPI_ISL_2957143 | Bayerisches Landesamt für Gesundheit und Lebensmittelsicherheit (LGL) | Robert Koch Institute | unknown |
| EPI_ISL_2957144 | Hospital Universitari Bellvitge | Microbiology Department | Sara Marti, Aida Gonzalez-Diaz, Laura Calatayud, Jordi Niubó, Miguel Fernandez-Huerta, Carmen Ardanuy, Jordi Camara, M Angeles Domínguez |
| EPI_ISL_2957145 | Bayerisches Landesamt für Gesundheit und Lebensmittelsicherheit (LGL) | Robert Koch Institute | unknown |
| EPI_ISL_2957146 | Hospital Universitari Bellvitge | Microbiology Department | Sara Marti, Aida Gonzalez-Diaz, Laura Calatayud, Jordi Niubó, Miguel Fernandez-Huerta, Carmen Ardanuy, Jordi Camara, M Angeles Domínguez |
| EPI_ISL_2957147 | Bayerisches Landesamt für Gesundheit und Lebensmittelsicherheit (LGL) | Robert Koch Institute | unknown |
| EPI_ISL_2957150 | Hospital Universitari Bellvitge | Microbiology Department | Sara Marti, Aida Gonzalez-Diaz, Laura Calatayud, Jordi Niubó, Miguel Fernandez-Huerta, Carmen Ardanuy, Jordi Camara, M Angeles Domínguez |
| EPI_ISL_2957151, EPI_ISL_2957152, EPI_ISL_2957153, EPI_ISL_2957154, EPI_ISL_2957156, EPI_ISL_2957160, EPI_ISL_2957161, EPI_ISL_2957163, EPI_ISL_2957165, EPI_ISL_2957167, EPI_ISL_2957171, EPI_ISL_2957172, EPI_ISL_2957173, EPI_ISL_2957176, EPI_ISL_2957177, EPI_ISL_2957179, EPI_ISL_2957180, EPI_ISL_2957181, EPI_ISL_2957183, EPI_ISL_2957184, EPI_ISL_2957185, EPI_ISL_2957186, EPI_ISL_2957189, EPI_ISL_2957192, EPI_ISL_2957195, EPI_ISL_2957198, EPI_ISL_2957201, EPI_ISL_2957202, EPI_ISL_2957203, EPI_ISL_2957204, EPI_ISL_2957208 | | | |
| see above | Bayerisches Landesamt für Gesundheit und Lebensmittelsicherheit (LGL) | Robert Koch Institute | unknown |
| EPI_ISL_2957210 | MVZ Medizinisches Labor Hannover GmbH | Robert Koch Institute | unknown |
| EPI_ISL_2957211, EPI_ISL_2957212, EPI_ISL_2957213 | MVZ für Laboratoriumsmedizin und Mikrobiologie Koblenz-Mittelrhein (Labor Koblenz) | Robert Koch Institute | unknown |
| EPI_ISL_2957214, EPI_ISL_2957215, EPI_ISL_2957216 | MDI Limbach Berlin GmbH; MVZ Labor Berlin | Robert Koch Institute | unknown |
| EPI_ISL_2957217 | MVZ Medizinisches Labor Hannover GmbH | Robert Koch Institute | unknown |
| EPI_ISL_2957218, EPI_ISL_2957219, EPI_ISL_2957220, EPI_ISL_2957221 | MVZ für Laboratoriumsmedizin und Mikrobiologie Koblenz-Mittelrhein (Labor Koblenz) | Robert Koch Institute | unknown |
| EPI_ISL_2957222 | MDI Limbach Berlin GmbH; MVZ Labor Berlin | Robert Koch Institute | unknown |
| EPI_ISL_2957224 | Kerckhoff-Klinik GmbH | Robert Koch Institute | unknown |
| EPI_ISL_2957225 | MVZ für Laboratoriumsmedizin und Mikrobiologie Koblenz-Mittelrhein (Labor Koblenz) | Robert Koch Institute | unknown |
| EPI_ISL_2957226 | MVZ Medizinisches Labor Hannover GmbH | Robert Koch Institute | unknown |
| EPI_ISL_2957227, EPI_ISL_2957229, EPI_ISL_2957230 | MDI Limbach Berlin GmbH; MVZ Labor Berlin | Robert Koch Institute | unknown |
| EPI_ISL_2957231 | MVZ für Laboratoriumsmedizin und Mikrobiologie Koblenz-Mittelrhein (Labor Koblenz) | Robert Koch Institute | unknown |
| EPI_ISL_2957232 | MDI Limbach Berlin GmbH; MVZ Labor Berlin | Robert Koch Institute | unknown |
| EPI_ISL_2957233, EPI_ISL_2957234 | MVZ Medizinisches Labor Hannover GmbH | Robert Koch Institute | unknown |
| EPI_ISL_2957235 | Limbach - MVZ Labor Ludwigsburg | Robert Koch Institute | unknown |
| EPI_ISL_2957237 | Diagnosticum - Labor Neukirchen | Robert Koch Institute | unknown |
| EPI_ISL_2957238 | Dept. of Microbiology and Infection Control, Akershus University Hospital HF | Dept. of Microbiology and Infection Control, Akershus University Hospital HF | Hege Vangstein Aamot, Alexander Hesselberg Løvestad |
| EPI_ISL_2957239 | Diagnosticum - Labor Neukirchen | Robert Koch Institute | unknown |
| EPI_ISL_2957240, EPI_ISL_2957241 | Dept. of Microbiology and Infection Control, Akershus University Hospital HF | Dept. of Microbiology and Infection Control, Akershus University Hospital HF | Hege Vangstein Aamot, Alexander Hesselberg Løvestad |
| EPI_ISL_2957242 | Diagnosticum - Labor Neukirchen | Robert Koch Institute | unknown |
| EPI_ISL_2957243, EPI_ISL_2957246, EPI_ISL_2957247, EPI_ISL_2957249, EPI_ISL_2957250, EPI_ISL_2957252, EPI_ISL_2957254 | Dept. of Microbiology and Infection Control, Akershus University Hospital HF | Dept. of Microbiology and Infection Control, Akershus University Hospital HF | Hege Vangstein Aamot, Alexander Hesselberg Løvestad |
| EPI_ISL_2957255 | Diagnosticum - Labor Neukirchen | Robert Koch Institute | unknown |
| EPI_ISL_2957256, EPI_ISL_2957258 | Dept. of Microbiology and Infection Control, Akershus University Hospital HF | Dept. of Microbiology and Infection Control, Akershus University Hospital HF | Hege Vangstein Aamot, Alexander Hesselberg Løvestad |
| EPI_ISL_2957259 | Diagnosticum - Labor Neukirchen | Robert Koch Institute | unknown |
| EPI_ISL_2957260 | Dept. of Microbiology and Infection Control, Akershus University Hospital HF | Dept. of Microbiology and Infection Control, Akershus University Hospital HF | Hege Vangstein Aamot, Alexander Hesselberg Løvestad |
| EPI_ISL_2957261 | Diagnosticum - Labor Neukirchen | Robert Koch Institute | unknown |
| EPI_ISL_2957262 | Dept. of Microbiology and Infection Control, Akershus University Hospital HF | Dept. of Microbiology and Infection Control, Akershus University Hospital HF | Hege Vangstein Aamot, Alexander Hesselberg Løvestad |
| EPI_ISL_2957263 | Diagnosticum - Labor Neukirchen | Robert Koch Institute | unknown |
| EPI_ISL_2957264, EPI_ISL_2957266 | Dept. of Microbiology and Infection Control, Akershus University Hospital HF | Dept. of Microbiology and Infection Control, Akershus University Hospital HF | Hege Vangstein Aamot, Alexander Hesselberg Løvestad |
| EPI_ISL_2957267 | Diagnosticum - Labor Neukirchen | Robert Koch Institute | unknown |
| EPI_ISL_2957268 | Dept. of Microbiology and Infection Control, Akershus University Hospital HF | Dept. of Microbiology and Infection Control, Akershus University Hospital HF | Hege Vangstein Aamot, Alexander Hesselberg Løvestad |

| | | | |
|---|---|---|---|
| EPI_ISL_2957269 | Diagnosticum - Labor Neukirchen | Robert Koch Institute | unknown |
| EPI_ISL_2957270 | Dept. of Microbiology and Infection Control, Akershus University Hospital HF | Dept. of Microbiology and Infection Control, Akershus University Hospital HF | Hege Vangstein Aamot, Alexander Hesselberg Løvestad |
| EPI_ISL_2957273 | Diagnosticum - Labor Neukirchen | Robert Koch Institute | unknown |
| EPI_ISL_2957274, EPI_ISL_2957276, EPI_ISL_2957280 | Dept. of Microbiology and Infection Control, Akershus University Hospital HF | Dept. of Microbiology and Infection Control, Akershus University Hospital HF | Hege Vangstein Aamot, Alexander Hesselberg Løvestad |
| EPI_ISL_2957281 | Diagnosticum - Labor Neukirchen | Robert Koch Institute | unknown |
| EPI_ISL_2957282, EPI_ISL_2957284, EPI_ISL_2957285 | Dept. of Microbiology and Infection Control, Akershus University Hospital HF | Dept. of Microbiology and Infection Control, Akershus University Hospital HF | Hege Vangstein Aamot, Alexander Hesselberg Løvestad |
| EPI_ISL_2957286 | Diagnosticum - Labor Neukirchen | Robert Koch Institute | unknown |
| EPI_ISL_2957287, EPI_ISL_2957289 | Dept. of Microbiology and Infection Control, Akershus University Hospital HF | Dept. of Microbiology and Infection Control, Akershus University Hospital HF | Hege Vangstein Aamot, Alexander Hesselberg Løvestad |
| EPI_ISL_2957290 | Diagnosticum - Labor Neukirchen | Robert Koch Institute | unknown |
| EPI_ISL_2957291, EPI_ISL_2957293, EPI_ISL_2957295, EPI_ISL_2957297 | Dept. of Microbiology and Infection Control, Akershus University Hospital HF | Dept. of Microbiology and Infection Control, Akershus University Hospital HF | Hege Vangstein Aamot, Alexander Hesselberg Løvestad |
| EPI_ISL_2957298 | Diagnosticum - Labor Neukirchen | Robert Koch Institute | unknown |
| EPI_ISL_2957299 | Dept. of Microbiology and Infection Control, Akershus University Hospital HF | Dept. of Microbiology and Infection Control, Akershus University Hospital HF | Hege Vangstein Aamot, Alexander Hesselberg Løvestad |
| EPI_ISL_2957300 | Diagnosticum - Labor Neukirchen | Robert Koch Institute | unknown |
| EPI_ISL_2957301, EPI_ISL_2957303 | Dept. of Microbiology and Infection Control, Akershus University Hospital HF | Dept. of Microbiology and Infection Control, Akershus University Hospital HF | Hege Vangstein Aamot, Alexander Hesselberg Løvestad |
| EPI_ISL_2957304, EPI_ISL_2957306, EPI_ISL_2957307, EPI_ISL_2957311, EPI_ISL_2957312 | Diagnosticum - Labor Neukirchen | Robert Koch Institute | unknown |
| EPI_ISL_2957313 | MVZ Labor Dr. Limbach & Kollegen GbR | Robert Koch Institute | unknown |
| EPI_ISL_2957314 | Labor Dr. Wisplinghoff - Köln | Robert Koch Institute | unknown |
| EPI_ISL_2957315 | Landesamt für Verbraucherschutz des Landes Sachsen-Anhalt | Robert Koch Institute | unknown |
| EPI_ISL_2957316 | MVZ Labor Dr. Limbach & Kollegen GbR | Robert Koch Institute | unknown |
| EPI_ISL_2957318 | Labor Mönchengladbach MVZ Dr. Stein + Kollegen GbR | Robert Koch Institute | unknown |
| EPI_ISL_2957319 | Limbach - MVZ Labor Eveld & Kollegen Essen | Robert Koch Institute | unknown |
| EPI_ISL_2957320, EPI_ISL_2957321, EPI_ISL_2957324 | MVZ Labor Dr. Limbach & Kollegen GbR | Robert Koch Institute | unknown |
| EPI_ISL_2957326 | Labor Mönchengladbach MVZ Dr. Stein + Kollegen GbR | Robert Koch Institute | unknown |
| EPI_ISL_2957327, EPI_ISL_2957328, EPI_ISL_2957329, EPI_ISL_2957332, EPI_ISL_2957334, EPI_ISL_2957335, EPI_ISL_2957336, EPI_ISL_2957337 | MVZ Labor Dr. Limbach & Kollegen GbR | Robert Koch Institute | unknown |
| EPI_ISL_2957343, EPI_ISL_2957344, EPI_ISL_2957345, EPI_ISL_2957346, EPI_ISL_2957347, EPI_ISL_2957348, EPI_ISL_2957349, EPI_ISL_2957350, EPI_ISL_2957351, EPI_ISL_2957352, EPI_ISL_2957353, EPI_ISL_2957354, EPI_ISL_2957355, EPI_ISL_2957356, EPI_ISL_2957359, EPI_ISL_2957360, EPI_ISL_2957361 | | | |
| see above | Labor Mönchengladbach MVZ Dr. Stein + Kollegen GbR | Robert Koch Institute | unknown |
| EPI_ISL_2957363 | MDI Limbach Berlin GmbH; MVZ Labor Berlin | Robert Koch Institute | unknown |
| EPI_ISL_2957364 | Limbach - MVZ Labor Eveld & Kollegen Essen | Robert Koch Institute | unknown |
| EPI_ISL_2957365, EPI_ISL_2957366 | Labor Mönchengladbach MVZ Dr. Stein + Kollegen GbR | Robert Koch Institute | unknown |
| EPI_ISL_2957368 | MDI Limbach Berlin GmbH; MVZ Labor Berlin | Robert Koch Institute | unknown |
| EPI_ISL_2957369 | MVZ Medizinisches Labor Hannover GmbH | Robert Koch Institute | unknown |
| EPI_ISL_2957370, EPI_ISL_2957371 | MDI Limbach Berlin GmbH; MVZ Labor Berlin | Robert Koch Institute | unknown |
| EPI_ISL_2957372 | MVZ für Laboratoriumsmedizin und Mikrobiologie Koblenz-Mittelrhein (Labor Koblenz) | Robert Koch Institute | unknown |
| EPI_ISL_2957373 | Labor Prof. Dr. G. Enders MVZ GbR | Robert Koch Institute | unknown |
| EPI_ISL_2957374 | MVZ Medizinisches Labor Hannover GmbH | Robert Koch Institute | unknown |
| EPI_ISL_2957375, EPI_ISL_2957376, EPI_ISL_2957377 | MVZ für Laboratoriumsmedizin und Mikrobiologie Koblenz-Mittelrhein (Labor Koblenz) | Robert Koch Institute | unknown |
| EPI_ISL_2957379 | MVZ Medizinisches Labor Hannover GmbH | Robert Koch Institute | unknown |
| EPI_ISL_2957381, EPI_ISL_2957382 | MVZ für Laboratoriumsmedizin und Mikrobiologie Koblenz-Mittelrhein (Labor Koblenz) | Robert Koch Institute | unknown |
| EPI_ISL_2957383, EPI_ISL_2957384 | MDI Limbach Berlin GmbH; MVZ Labor Berlin | Robert Koch Institute | unknown |
| EPI_ISL_2957385 | Labor Prof. Dr. G. Enders MVZ GbR | Robert Koch Institute | unknown |
| EPI_ISL_2957386 | MVZ für Laboratoriumsmedizin und Mikrobiologie Koblenz-Mittelrhein (Labor Koblenz) | Robert Koch Institute | unknown |
| EPI_ISL_2957387 | MVZ Medizinisches Labor Hannover GmbH | Robert Koch Institute | unknown |
| EPI_ISL_2957388 | MVZ Labor Dr. Limbach & Kollegen GbR | Robert Koch Institute | unknown |
| EPI_ISL_2957389 | Labor Prof. Dr. G. Enders MVZ GbR | Robert Koch Institute | unknown |
| EPI_ISL_2957390, EPI_ISL_2957391, EPI_ISL_2957392, EPI_ISL_2957393 | MVZ Medizinisches Labor Hannover GmbH | Robert Koch Institute | unknown |
| EPI_ISL_2957394 | Labor Mönchengladbach MVZ Dr. Stein + Kollegen GbR | Robert Koch Institute | unknown |
| EPI_ISL_2957395 | MDI Limbach Berlin GmbH; MVZ Labor Berlin | Robert Koch Institute | unknown |
| EPI_ISL_2957396, EPI_ISL_2957400 | Labor Prof. Dr. G. Enders MVZ GbR | Robert Koch Institute | unknown |
| EPI_ISL_2957401 | MVZ für Laboratoriumsmedizin und Mikrobiologie Koblenz-Mittelrhein (Labor Koblenz) | Robert Koch Institute | unknown |
| EPI_ISL_2957402, EPI_ISL_2957403 | MDI Limbach Berlin GmbH; MVZ Labor Berlin | Robert Koch Institute | unknown |

| | | | |
|---|---|---|---|
| EPI_ISL_2957404 | MVZ für Laboratoriumsmedizin und Mikrobiologie Koblenz-Mittelrhein (Labor Koblenz) | Robert Koch Institute | unknown |
| EPI_ISL_2957405, EPI_ISL_2957406 | MVZ Medizinisches Labor Hannover GmbH | Robert Koch Institute | unknown |
| EPI_ISL_2957407 | MVZ für Laboratoriumsmedizin und Mikrobiologie Koblenz-Mittelrhein (Labor Koblenz) | Robert Koch Institute | unknown |
| EPI_ISL_2957408, EPI_ISL_2957409, EPI_ISL_2957411 | IMD - Labor Oderland | Robert Koch Institute | unknown |
| EPI_ISL_2957412 | IMD - MVZ Labor Martinsried | Robert Koch Institute | unknown |
| EPI_ISL_2957413, EPI_ISL_2957414, EPI_ISL_2957418, EPI_ISL_2957422, EPI_ISL_2957427, EPI_ISL_2957434, EPI_ISL_2957435, EPI_ISL_2957437, EPI_ISL_2957438, EPI_ISL_2957439, EPI_ISL_2957441, EPI_ISL_2957443, EPI_ISL_2957444, EPI_ISL_2957445, EPI_ISL_2957448, EPI_ISL_2957449, EPI_ISL_2957450, EPI_ISL_2957451, EPI_ISL_2957452, EPI_ISL_2957453, EPI_ISL_2957454, EPI_ISL_2957455, EPI_ISL_2957456, EPI_ISL_2957459 | | | |
| see above | Microvida | Microvida | Suzan D. Pas, Jaco J. Verweij, Joep J. J. M. Stohr |
| EPI_ISL_2957565, EPI_ISL_2957584, EPI_ISL_2957831, EPI_ISL_2957887, EPI_ISL_2957897, EPI_ISL_2957941, EPI_ISL_2957954, EPI_ISL_2957963, EPI_ISL_2958035, EPI_ISL_2958045, EPI_ISL_2958046, EPI_ISL_2958069, EPI_ISL_2958070, EPI_ISL_2958094, EPI_ISL_2958206, EPI_ISL_2958226, EPI_ISL_2958227, EPI_ISL_2958274, EPI_ISL_2958299, EPI_ISL_2958304, EPI_ISL_2958311, EPI_ISL_2958312, EPI_ISL_2958314, EPI_ISL_2958317, EPI_ISL_2958318, EPI_ISL_2958319, EPI_ISL_2958325, EPI_ISL_2958326 | | | |
| see above | SARS-CoV-2 Sequencing Castilla y Leon-Spain Consortium | SARS-CoV-2 Sequencing Castilla y Leon-Spain Consortium | Antonio Orduña-Domingo, Marta Hernandez, David Abad, Marta Dominguez-Gil, Silvia Rojo, Laura Sánchez de Prada, Sonsoles Garcinuño Pérez, Carmen Aldea-Mansilla, Mª Fe Brezmes-Valdivieso, Gregoria Megías Lobón, María Antonia García Castro, Carmen Gimeno Crespo, Noelia Arenal Andrés, Carlos Fuster Foz, M. Isabel Fernandez-Natal, Jose María Eiros Bouza |
| EPI_ISL_2958364, EPI_ISL_2958365, EPI_ISL_2958366, EPI_ISL_2958367, EPI_ISL_2958368, EPI_ISL_2958370, EPI_ISL_2958373, EPI_ISL_2958374, EPI_ISL_2958376, EPI_ISL_2958377, EPI_ISL_2958378, EPI_ISL_2958379, EPI_ISL_2958380, EPI_ISL_2958381, EPI_ISL_2958383, EPI_ISL_2958384, EPI_ISL_2958385, EPI_ISL_2958386, EPI_ISL_2958388, EPI_ISL_2958390, EPI_ISL_2958391, EPI_ISL_2958392, EPI_ISL_2958393, EPI_ISL_2958394, EPI_ISL_2958395, EPI_ISL_2958396, EPI_ISL_2958397, EPI_ISL_2958399, EPI_ISL_2958400, EPI_ISL_2958401, EPI_ISL_2958402, EPI_ISL_2958403, EPI_ISL_2958404, EPI_ISL_2958405, EPI_ISL_2958407, EPI_ISL_2958408, EPI_ISL_2958409, EPI_ISL_2958410, EPI_ISL_2958411, EPI_ISL_2958412, EPI_ISL_2958415, EPI_ISL_2958416, EPI_ISL_2958417, EPI_ISL_2958418, EPI_ISL_2958419, EPI_ISL_2958420, EPI_ISL_2958421, EPI_ISL_2958422, EPI_ISL_2958423, EPI_ISL_2958424, EPI_ISL_2958425, EPI_ISL_2958426, EPI_ISL_2958428, EPI_ISL_2958429, EPI_ISL_2958430, EPI_ISL_2958431, EPI_ISL_2958432, EPI_ISL_2958433, EPI_ISL_2958434, EPI_ISL_2958438, EPI_ISL_2958439, EPI_ISL_2958440, EPI_ISL_2958441, EPI_ISL_2958442, EPI_ISL_2958443, EPI_ISL_2958454, EPI_ISL_2958456, EPI_ISL_2958459, EPI_ISL_2958464, EPI_ISL_2958465, EPI_ISL_2958466, EPI_ISL_2958467, EPI_ISL_2958468, EPI_ISL_2958469, EPI_ISL_2958470, EPI_ISL_2958472, EPI_ISL_2958473, EPI_ISL_2958474, EPI_ISL_2958475, EPI_ISL_2958476, EPI_ISL_2958477, EPI_ISL_2958478, EPI_ISL_2958479, EPI_ISL_2958480, EPI_ISL_2958481, EPI_ISL_2958482, EPI_ISL_2958483, EPI_ISL_2958484, EPI_ISL_2958485, EPI_ISL_2958487, EPI_ISL_2958488, EPI_ISL_2958489, EPI_ISL_2958490, EPI_ISL_2958491, EPI_ISL_2958492, EPI_ISL_2958493, EPI_ISL_2958494, EPI_ISL_2958495, EPI_ISL_2958496, EPI_ISL_2958497, EPI_ISL_2958498, EPI_ISL_2958499, EPI_ISL_2958500, EPI_ISL_2958503, EPI_ISL_2958504, EPI_ISL_2958505, EPI_ISL_2958506, EPI_ISL_2958507, EPI_ISL_2958508, EPI_ISL_2958509, EPI_ISL_2958510, EPI_ISL_2958511, EPI_ISL_2958512, EPI_ISL_2958513, EPI_ISL_2958514, EPI_ISL_2958515, EPI_ISL_2958516, EPI_ISL_2958517, EPI_ISL_2958518, EPI_ISL_2958519, EPI_ISL_2958520, EPI_ISL_2958521, EPI_ISL_2958522, EPI_ISL_2958523, EPI_ISL_2958524, EPI_ISL_2958525, EPI_ISL_2958526, EPI_ISL_2958527, EPI_ISL_2958528, EPI_ISL_2958529, EPI_ISL_2958530, EPI_ISL_2958531, EPI_ISL_2958532, EPI_ISL_2958533, EPI_ISL_2958534, EPI_ISL_2958535, EPI_ISL_2958539, EPI_ISL_2958540, EPI_ISL_2958541, EPI_ISL_2958542, EPI_ISL_2958543, EPI_ISL_2958544, EPI_ISL_2958545, EPI_ISL_2958546, EPI_ISL_2958547, EPI_ISL_2958548, EPI_ISL_2958549, EPI_ISL_2958550, EPI_ISL_2958553, EPI_ISL_2958554, EPI_ISL_2958556, EPI_ISL_2958558, EPI_ISL_2958559, EPI_ISL_2958560, EPI_ISL_2958564, EPI_ISL_2958565, EPI_ISL_2958566, EPI_ISL_2958567, EPI_ISL_2958568, EPI_ISL_2958569, EPI_ISL_2958570, EPI_ISL_2958571, EPI_ISL_2958572, EPI_ISL_2958573, EPI_ISL_2958574, EPI_ISL_2958576, EPI_ISL_2958578, EPI_ISL_2958579, EPI_ISL_2958581, EPI_ISL_2958582, EPI_ISL_2958583, EPI_ISL_2958584, EPI_ISL_2958586, EPI_ISL_2958587, EPI_ISL_2958588, EPI_ISL_2958589, EPI_ISL_2958593, EPI_ISL_2958594, EPI_ISL_2958595, EPI_ISL_2958596, EPI_ISL_2958597, EPI_ISL_2958598, EPI_ISL_2958599, EPI_ISL_2958600, EPI_ISL_2958601, EPI_ISL_2958602, EPI_ISL_2958603, EPI_ISL_2958604, EPI_ISL_2958606, EPI_ISL_2958607, EPI_ISL_2958608, EPI_ISL_2958609, EPI_ISL_2958610, EPI_ISL_2958611, EPI_ISL_2958612, EPI_ISL_2958614, EPI_ISL_2958615, EPI_ISL_2958616, EPI_ISL_2958617, EPI_ISL_2958618, EPI_ISL_2958619, EPI_ISL_2958620, EPI_ISL_2958621, EPI_ISL_2958622, EPI_ISL_2958623, EPI_ISL_2958624 | | | |
| see above | Ministry of Health Turkey | Ministry of Health Turkey | Fatma Bayrakdar, Yasemin Cosgun, Suleyman Yalcin, Gulay Korukluoglu |
| EPI_ISL_2958645, EPI_ISL_2958646 | MRCG at LSHTM Genomics lab | MRCG at LSHTM Genomics lab | Abdul Karim sesay, Abdoulie Kanteh, Jarra Manneh, Mariama Kujabi, Bakary Sanyang,Dabiri Damilari |
| EPI_ISL_2958686, EPI_ISL_2958687 | Microbiology and Virology Laboratory, "Policlinico Riuniti, Azienda Ospedaliero Universitaria, Foggia" | Microbiology and Virology Laboratory, "Policlinico Riuniti, Azienda Ospedaliero Universitaria, Foggia" | Fabio Arena, Settimia Altamura, Donatella Cedola, Valeria Delli Carri, Annamaria D'Aprile, Maria Rosaria Lipsi, Loris Micelli, Maria Pia Patrizio, Daniela Pisanelli, Tiziana Rollo, Maurizio Margaglione and Rosella De Nittis. |
| EPI_ISL_2958688, EPI_ISL_2958689, EPI_ISL_2958690, EPI_ISL_2958691, EPI_ISL_2958692, EPI_ISL_2958693, EPI_ISL_2958694, EPI_ISL_2958695, EPI_ISL_2958696, EPI_ISL_2958697, EPI_ISL_2958698 | | | |
| see above | Jaber Al Ahmad Hospital | Virology Lab, Jaber Al Ahmad Hospital | Dr. Zahrah Buhamad, Dr. Ebaa Al-Awadhi, Haroon Masih, Khubaid-ur-Rehman |
| EPI_ISL_2958753, EPI_ISL_2958756 | labo Analyses Med | National Reference Center for Viruses of Respiratory Infections, Institut Pasteur, Paris | Marion Barbet, Sylvie Behillil, Méline Bizard, Angela Brisebarre, Camille Capel, Vincent Enouf, Louise Lefrançois, Frédéric Lemoine, Christophe Malabat, Corinne Maufrais, Hub Bioinformatique Biostatistiques, Etienne Simon-Lorière, Maud Vanpeene, Sylvie Van der Werf ,LEONARD Pierre-Yves |
| EPI_ISL_2958757 | labo Analyses Med | National Reference Center for Viruses of Respiratory Infections, Institut Pasteur, Paris | Marion Barbet, Sylvie Behillil, Méline Bizard, Angela Brisebarre, Camille Capel, Vincent Enouf, Louise Lefrançois, Frédéric Lemoine, Christophe Malabat, Corinne Maufrais, Victoire Baillet, Etienne Simon-Lorière, Maud Vanpeene, Sylvie Van der Werf ,LEONARD Pierre-Yves |
| EPI_ISL_2958759, EPI_ISL_2958760 | labo Analyses Med | National Reference Center for Viruses of Respiratory Infections, Institut Pasteur, Paris | Marion Barbet, Sylvie Behillil, Méline Bizard, Angela Brisebarre, Camille Capel, Vincent Enouf, Louise Lefrançois, Frédéric Lemoine, Christophe Malabat, Corinne Maufrais, Hub Bioinformatique Biostatistiques, Etienne Simon-Lorière, Maud Vanpeene, Sylvie Van der Werf ,LEONARD Pierre-Yves |
| EPI_ISL_2958761, EPI_ISL_2958762 | Labo Analyses Med | National Reference Center for Viruses of Respiratory Infections, Institut Pasteur, Paris | Marion Barbet, Sylvie Behillil, Méline Bizard, Angela Brisebarre, Camille Capel, Vincent Enouf, Louise Lefrançois, Frédéric Lemoine, Christophe Malabat, Corinne Maufrais, Victoire Baillet, Etienne Simon-Lorière, Maud Vanpeene, Sylvie Van der Werf ,POTIRON Grégoire |
| EPI_ISL_2958764 | Labo Analyses Med | National Reference Center for Viruses of Respiratory Infections, Institut Pasteur, Paris | Marion Barbet, Sylvie Behillil, Méline Bizard, Angela Brisebarre, Camille Capel, Vincent Enouf, Louise Lefrançois, Frédéric Lemoine, Christophe Malabat, Corinne Maufrais, Hub Bioinformatique Biostatistiques, Etienne Simon-Lorière, Maud Vanpeene, Sylvie Van der Werf ,HOLSTEIN Anne |
| EPI_ISL_2958765 | Hospital | National Reference Center for Viruses of Respiratory Infections, Institut Pasteur, Paris | Marion Barbet, Sylvie Behillil, Méline Bizard, Angela Brisebarre, Camille Capel, Vincent Enouf, Louise Lefrançois, Frédéric Lemoine, Christophe Malabat, Corinne Maufrais, Victoire Baillet, Etienne Simon-Lorière, Maud Vanpeene, Sylvie Van der Werf ,FARRUGIA Cécile |
| EPI_ISL_2958766 | Hospital | National Reference Center for Viruses of Respiratory Infections, Institut Pasteur, Paris | Marion Barbet, Sylvie Behillil, Méline Bizard, Angela Brisebarre, Camille Capel, Vincent Enouf, Louise Lefrançois, Frédéric Lemoine, Christophe Malabat, Corinne Maufrais, Hub Bioinformatique Biostatistiques, Etienne Simon-Lorière, Maud Vanpeene, Sylvie Van der Werf ,FARRUGIA Cécile |
| EPI_ISL_2958767 | Hospital | National Reference Center for Viruses of Respiratory Infections, Institut Pasteur, Paris | Marion Barbet, Sylvie Behillil, Méline Bizard, Angela Brisebarre, Camille Capel, Vincent Enouf, Louise Lefrançois, Frédéric Lemoine, Christophe Malabat, Corinne Maufrais, Victoire Baillet, Etienne Simon-Lorière, Maud Vanpeene, Sylvie Van der Werf ,PILORGE Léa |
| EPI_ISL_2958770, EPI_ISL_2958771 | labo Analyses Med | National Reference Center for Viruses of Respiratory Infections, Institut Pasteur, Paris | Marion Barbet, Sylvie Behillil, Méline Bizard, Angela Brisebarre, Camille Capel, Vincent Enouf, Louise Lefrançois, Frédéric Lemoine, Christophe Malabat, Corinne Maufrais, Hub Bioinformatique Biostatistiques, Etienne Simon-Lorière, Maud Vanpeene, Sylvie Van der Werf ,GUSSOND Thierry |
| EPI_ISL_2958773 | labo Analyses Med | National Reference Center for Viruses of Respiratory Infections, Institut Pasteur, Paris | Marion Barbet, Sylvie Behillil, Méline Bizard, Angela Brisebarre, Camille Capel, Vincent Enouf, Louise Lefrançois, Frédéric Lemoine, Christophe Malabat, Corinne Maufrais, Victoire Baillet, Etienne Simon-Lorière, Maud Vanpeene, Sylvie Van der Werf ,GUSSOND Thierry |
| EPI_ISL_2958774, EPI_ISL_2958775 | labo Analyses Med | National Reference Center for Viruses of Respiratory Infections, Institut Pasteur, Paris | Marion Barbet, Sylvie Behillil, Méline Bizard, Angela Brisebarre, Camille Capel, Vincent Enouf, Louise Lefrançois, Frédéric Lemoine, Christophe Malabat, Corinne Maufrais, Hub Bioinformatique Biostatistiques, Etienne Simon-Lorière, Maud Vanpeene, Sylvie Van der Werf ,GUSSOND Thierry |
| EPI_ISL_2958777 | Labo Analyses Med | National Reference Center for Viruses of Respiratory Infections, Institut Pasteur, Paris | Marion Barbet, Sylvie Behillil, Méline Bizard, Angela Brisebarre, Camille Capel, Vincent Enouf, Louise Lefrançois, Frédéric Lemoine, Christophe Malabat, Corinne Maufrais, Emmanuelle Permal, Etienne Simon-Lorière, Maud Vanpeene, Sylvie Van der Werf ,GOUBARD Agathe |
| EPI_ISL_2958778 | Labo Analyses Med | National Reference Center for Viruses of Respiratory Infections, Institut Pasteur, Paris | Marion Barbet, Sylvie Behillil, Méline Bizard, Angela Brisebarre, Camille Capel, Vincent Enouf, Louise Lefrançois, Frédéric Lemoine, Christophe Malabat, Corinne Maufrais, Hub Bioinformatique Biostatistiques, Etienne Simon-Lorière, Maud Vanpeene, Sylvie Van der Werf ,BLANC-GALIBY M.-J. |
| EPI_ISL_2958779 | labo Analyses Med | National Reference Center for Viruses of Respiratory Infections, Institut Pasteur, Paris | Marion Barbet, Sylvie Behillil, Méline Bizard, Angela Brisebarre, Camille Capel, Vincent Enouf, Louise Lefrançois, Frédéric Lemoine, Christophe Malabat, Corinne Maufrais, Hub Bioinformatique Biostatistiques, Etienne Simon-Lorière, Maud Vanpeene, Sylvie Van der Werf ,LEONARD Pierre-Yves |
| EPI_ISL_2958781, EPI_ISL_2958782, EPI_ISL_2958783 | Labo Analyses Med | National Reference Center for Viruses of Respiratory Infections, Institut Pasteur, Paris | Marion Barbet, Sylvie Behillil, Méline Bizard, Angela Brisebarre, Camille Capel, Vincent Enouf, Louise Lefrançois, Frédéric Lemoine, Christophe Malabat, Corinne Maufrais, Hub Bioinformatique Biostatistiques, Etienne Simon-Lorière, Maud Vanpeene, Sylvie Van der Werf ,LEONARD Pierre-Yves |
| EPI_ISL_2958784, EPI_ISL_2958785 | Labo Analyses Med | National Reference Center for Viruses of Respiratory Infections, Institut Pasteur, Paris | Marion Barbet, Sylvie Behillil, Méline Bizard, Angela Brisebarre, Camille Capel, Vincent Enouf, Louise Lefrançois, Frédéric Lemoine, Christophe Malabat, Corinne Maufrais, Hub Bioinformatique Biostatistiques, Etienne Simon-Lorière, Maud Vanpeene, Sylvie Van der Werf |
| EPI_ISL_2958786 | Labo Analyses Med | National Reference Center for Viruses of Respiratory Infections, Institut Pasteur, Paris | Marion Barbet, Sylvie Behillil, Méline Bizard, Angela Brisebarre, Camille Capel, Vincent Enouf, Louise Lefrançois, Frédéric Lemoine, Christophe Malabat, Corinne Maufrais, Victoire Baillet, Etienne Simon-Lorière, Maud Vanpeene, Sylvie Van der Werf |
| EPI_ISL_2958787 | Labo Analyses Med | National Reference Center for Viruses of Respiratory Infections, Institut Pasteur, Paris | Marion Barbet, Sylvie Behillil, Méline Bizard, Angela Brisebarre, Camille Capel, Vincent Enouf, Louise Lefrançois, Frédéric Lemoine, Christophe Malabat, Corinne Maufrais, Hub Bioinformatique Biostatistiques, Etienne Simon-Lorière, Maud Vanpeene, Sylvie Van der Werf ,TROMBERT Sabine |
| EPI_ISL_2958791 | Labo Analyses Med | National Reference Center for Viruses of Respiratory Infections, Institut Pasteur, Paris | Marion Barbet, Sylvie Behillil, Méline Bizard, Angela Brisebarre, Camille Capel, Vincent Enouf, Louise Lefrançois, Frédéric Lemoine, Christophe Malabat, Corinne Maufrais, Hub Bioinformatique Biostatistiques, Etienne Simon-Lorière, Maud Vanpeene, Sylvie Van der Werf ,POTIRON Grégoire |
| EPI_ISL_2958792, EPI_ISL_2958793 | Labo Analyses Med | National Reference Center for Viruses of Respiratory Infections, Institut Pasteur, Paris | Marion Barbet, Sylvie Behillil, Méline Bizard, Angela Brisebarre, Camille Capel, Vincent Enouf, Louise Lefrançois, Frédéric Lemoine, Christophe Malabat, Corinne Maufrais, Victoire Baillet, Etienne Simon-Lorière, Maud Vanpeene, Sylvie Van der Werf ,POTIRON Grégoire |
| EPI_ISL_2958807 | Hospital | National Reference Center for Viruses of Respiratory Infections, Institut Pasteur, Paris | Marion Barbet, Sylvie Behillil, Méline Bizard, Angela Brisebarre, Camille Capel, Vincent Enouf, Louise Lefrançois, Frédéric Lemoine, Christophe Malabat, Corinne Maufrais, Hub Bioinformatique Biostatistiques, Etienne Simon-Lorière, Maud Vanpeene, Sylvie Van der Werf ,GUINARD Jérome |
| EPI_ISL_2958813, EPI_ISL_2958815, EPI_ISL_2958816, EPI_ISL_2958817, EPI_ISL_2958818 | Labo Analyses Med | National Reference Center for Viruses of Respiratory Infections, Institut Pasteur, Paris | Marion Barbet, Sylvie Behillil, Méline Bizard, Angela Brisebarre, Camille Capel, Vincent Enouf, Louise Lefrançois, Frédéric Lemoine, Christophe Malabat, Corinne Maufrais, Hub Bioinformatique Biostatistiques, Etienne Simon-Lorière, Maud Vanpeene, Sylvie Van der Werf ,POTIRON Grégoire |
| EPI_ISL_2958819 | Labo Analyses Med | National Reference Center for Viruses of Respiratory Infections, Institut Pasteur, Paris | Marion Barbet, Sylvie Behillil, Méline Bizard, Angela Brisebarre, Camille Capel, Vincent Enouf, Louise Lefrançois, Frédéric Lemoine, Christophe Malabat, Corinne Maufrais, Victoire Baillet, Etienne Simon-Lorière, Maud Vanpeene, Sylvie Van der Werf ,POTIRON Grégoire |
| EPI_ISL_2958820 | Labo Analyses Med | National Reference Center for Viruses of Respiratory Infections, Institut Pasteur, Paris | Marion Barbet, Sylvie Behillil, Méline Bizard, Angela Brisebarre, Camille Capel, Vincent Enouf, Louise Lefrançois, Frédéric Lemoine, Christophe Malabat, Corinne Maufrais, Hub Bioinformatique Biostatistiques, Etienne Simon-Lorière, Maud Vanpeene, Sylvie Van der Werf ,POTIRON Grégoire |
| EPI_ISL_2958821 | Labo Analyses Med | National Reference Center for Viruses of Respiratory Infections, Institut Pasteur, Paris | Marion Barbet, Sylvie Behillil, Méline Bizard, Angela Brisebarre, Camille Capel, Vincent Enouf, Louise Lefrançois, Frédéric Lemoine, Christophe Malabat, Corinne Maufrais, Victoire Baillet, Etienne Simon-Lorière, Maud Vanpeene, Sylvie Van der Werf ,POTIRON Grégoire |
| EPI_ISL_2958822 | Labo Analyses Med | National Reference Center for Viruses of Respiratory Infections, Institut Pasteur, Paris | Marion Barbet, Sylvie Behillil, Méline Bizard, Angela Brisebarre, Camille Capel, Vincent Enouf, Louise Lefrançois, Frédéric Lemoine, Christophe Malabat, Corinne Maufrais, Hub Bioinformatique Biostatistiques, Etienne Simon-Lorière, Maud Vanpeene, Sylvie Van der Werf ,POTIRON Grégoire |
| EPI_ISL_2958824, | GA Department of Public Health | GA Department of Public Health | Stacy Reeves, Jonathan Edwards, Cynthia Dixey, Tonia Parrott, Aliyah Fields, Taylor Smith |

| | | | |
|---|---|---|---|
| EPI_ISL_2958825, EPI_ISL_2958826, EPI_ISL_2958827, EPI_ISL_2958828, EPI_ISL_2958829, EPI_ISL_2958832 | | | |
| EPI_ISL_2958920, EPI_ISL_2958921 | Mobile County Health Department | Synergy Laboratories | Megan Cornwell |
| EPI_ISL_2958933 | Mayo Clinic & Mayo Clinic Laboratories | Minnesota Department of Health, Public Health Laboratory | Alexandra Lorentz, Jacob Garfin, Matt Plumb, and Xiong Wang |
| EPI_ISL_2958935, EPI_ISL_2958936, EPI_ISL_2958937, EPI_ISL_2958941, EPI_ISL_2958942 | Minnesota Department of Health, Public Health Laboratory | Minnesota Department of Health, Public Health Laboratory | Alexandra Lorentz, Jacob Garfin, Matt Plumb, and Xiong Wang |
| EPI_ISL_2958946, EPI_ISL_2958948, EPI_ISL_2958951, EPI_ISL_2958952, EPI_ISL_2958953, EPI_ISL_2958954, EPI_ISL_2958955, EPI_ISL_2958956, EPI_ISL_2958959, EPI_ISL_2958960, EPI_ISL_2958961, EPI_ISL_2958962, EPI_ISL_2958963, EPI_ISL_2958964, EPI_ISL_2958965, EPI_ISL_2958968 | | | |
| see above | Mayo Clinic & Mayo Clinic Laboratories | Minnesota Department of Health, Public Health Laboratory | Alexandra Lorentz, Jacob Garfin, Matt Plumb, and Xiong Wang |
| EPI_ISL_2958969, EPI_ISL_2958970, EPI_ISL_2958971, EPI_ISL_2958972, EPI_ISL_2958974, EPI_ISL_2958975 | Centracare Laboratory Services | Minnesota Department of Health, Public Health Laboratory | Alexandra Lorentz, Jacob Garfin, Matt Plumb, and Xiong Wang |
| EPI_ISL_2958978 | Premier Medical Lab | Minnesota Department of Health, Public Health Laboratory | Alexandra Lorentz, Jacob Garfin, Matt Plumb, and Xiong Wang |
| EPI_ISL_2958979 | ASL BARI | University of Bari Biomedical Sciences and Human Oncology | Maria Chironna |
| EPI_ISL_2958987, EPI_ISL_2958988, EPI_ISL_2958989, EPI_ISL_2958990, EPI_ISL_2958991, EPI_ISL_2958992, EPI_ISL_2958993, EPI_ISL_2958994, EPI_ISL_2958995, EPI_ISL_2958996, EPI_ISL_2958997, EPI_ISL_2958998, EPI_ISL_2959000, EPI_ISL_2959001, EPI_ISL_2959002, EPI_ISL_2959003, EPI_ISL_2959004, EPI_ISL_2959005, EPI_ISL_2959006, EPI_ISL_2959007, EPI_ISL_2959008, EPI_ISL_2959009, EPI_ISL_2959010, EPI_ISL_2959011, EPI_ISL_2959012 | | | |
| see above | Microbiology Department. Complexo Hospitalario Universitario de Vigo | Microbiology Department. Complexo Hospitalario Universitario de Vigo | Alfaya N, Alonso I, Alvarez M, Cabrera JJ, Carballo R, Cores O, Cortizo S, del-Campo V, Martinez L, Mediero G, Perez S, Davina C, Potel C, Regueiro B, Rey S, Vasallo FJ |
| EPI_ISL_2960038, EPI_ISL_2960039, EPI_ISL_2960040, EPI_ISL_2960041, EPI_ISL_2960044, EPI_ISL_2960046, EPI_ISL_2960047, EPI_ISL_2960049, EPI_ISL_2960050 | Yale Clinical Virology Lab | Grubaugh Lab - Yale School of Public Health | Joseph Fauver, Mallery Breban, Isabel Ott, Tara Alpert, Mary Petrone, Anderson Brito, Chantal Vogels, Annie Watkins, Chaney Kalinich, Jessica Rothman, Kendall Billig, Tobias Koch, Marie L. Landry, Nathan Grubaugh |
| EPI_ISL_2960051, EPI_ISL_2960052, EPI_ISL_2960053, EPI_ISL_2960054, EPI_ISL_2960055, EPI_ISL_2960056, EPI_ISL_2960057, EPI_ISL_2960058, EPI_ISL_2960059, EPI_ISL_2960060, EPI_ISL_2960061, EPI_ISL_2960062, EPI_ISL_2960063, EPI_ISL_2960064, EPI_ISL_2960065, EPI_ISL_2960066, EPI_ISL_2960067, EPI_ISL_2960068, EPI_ISL_2960069, EPI_ISL_2960070, EPI_ISL_2960071, EPI_ISL_2960072, EPI_ISL_2960073, EPI_ISL_2960074 | | | |
| see above | UCC (Universidad Central del Caribe) | Grubaugh Lab - Yale School of Public Health | Joseph Fauver, Mallery Breban, Isabel Ott, Tara Alpert, Mary Petrone, Anderson Brito, Chantal Vogels, Annie Watkins, Chaney Kalinich, Jessica Rothman, Kendall Billig, Tobias Koch, Fabiola Cruz-Lopez, David Rivera Aponte, Fabiola Fontanet, Natalie Machargo, Andrea Firpo, Vianca Aponte, Ernesto Vazquez, Patricia Serrano, Tania Mitwalli, Karolane Gonzalez, Nathan Grubaugh |
| EPI_ISL_2960104, EPI_ISL_2960105, EPI_ISL_2960106, EPI_ISL_2960107 | Yale Pathology Lab | Grubaugh Lab - Yale School of Public Health | Joseph Fauver, Mallery Breban, Isabel Ott, Tara Alpert, Mary Petrone, Anderson Brito, Chantal Vogels, Annie Watkins, Chaney Kalinich, Jessica Rothman, Kendall Billig, Tobias Koch, Jianhui Wang, Chen Liu, Pei Hui, Nathan Grubaugh |
| EPI_ISL_2960128, EPI_ISL_2960132, EPI_ISL_2960133, EPI_ISL_2960135, EPI_ISL_2960136 | Wyoming Public Health Laboratory | Wyoming Public Health Laboratory | Jim Mildenberger, Taylor Fearing, Channing Weber, Ashley Norberg, Chayse Rowley, Marley Goetz, Brian Dominguez, Elliot Thomasson, Sam Britz, Cari Sloma, Robert Petit, and Rob Christensen |
| EPI_ISL_2960183, EPI_ISL_2960188, EPI_ISL_2960192, EPI_ISL_2960193, EPI_ISL_2960194, EPI_ISL_2960201 | Illinois Department of Public Health | Gagnon Lab, Southern Illinois University | Keith Gagnon |
| EPI_ISL_2960208, EPI_ISL_2960209 | Simple Laboratories | Gagnon Lab, Southern Illinois University | Keith Gagnon |
| EPI_ISL_2960211, EPI_ISL_2960219, EPI_ISL_2960220, EPI_ISL_2960223, EPI_ISL_2960224, EPI_ISL_2960225, EPI_ISL_2960227, EPI_ISL_2960229, EPI_ISL_2960233, EPI_ISL_2960236, EPI_ISL_2960237, EPI_ISL_2960238, EPI_ISL_2960241, EPI_ISL_2960242, EPI_ISL_2960243, EPI_ISL_2960246, EPI_ISL_2960248, EPI_ISL_2960249, EPI_ISL_2960250, EPI_ISL_2960254, EPI_ISL_2960256 | | | |
| see above | Servicio de Microbiologia Hospital Ramon y Cajal | Servicio de Microbiologia Hospital Ramon y Cajal | Ponce M, Galan JC, Martinez L. Abreu M, y Gonzalez-Alba JM |
| EPI_ISL_2960292, EPI_ISL_2960294, EPI_ISL_2960295, EPI_ISL_2960298, EPI_ISL_2960299, EPI_ISL_2960300, EPI_ISL_2960302, EPI_ISL_2960303, EPI_ISL_2960304, EPI_ISL_2960305, EPI_ISL_2960306, EPI_ISL_2960307, EPI_ISL_2960308, EPI_ISL_2960309, EPI_ISL_2960310, EPI_ISL_2960311, EPI_ISL_2960312, EPI_ISL_2960313, EPI_ISL_2960314, EPI_ISL_2960316, EPI_ISL_2960317 | | | |
| see above | Wyoming Public Health Laboratory | Wyoming Public Health Laboratory | Jim Mildenberger, Taylor Fearing, Channing Weber, Ashley Norberg, Chayse Rowley, Marley Goetz, Brian Dominguez, Elliot Thomasson, Sam Britz, Cari Sloma, Robert Petit, and Rob Christensen |
| EPI_ISL_2960343 | Lighthouse Lab in Milton Keynes | Wellcome Sanger Institute for the COVID-19 Genomics UK (COG-UK) Consortium | The Lighthouse Lab in Milton Keynes and Alex Alderton, Roberto Amato, Jeffrey Barrett, Sonia Goncalves, Ewan Harrison, David K. Jackson, Ian Johnston, Dominic Kwiatkowski, Cordelia Langford, John Sillitoe on behalf of the Wellcome Sanger Institute COVID-19 Surveillance Team |
| EPI_ISL_2960344 | Lighthouse Lab in Glasgow | Wellcome Sanger Institute for the COVID-19 Genomics UK (COG-UK) Consortium | Harper VanSteenhouse, Yumi Kasai, David Gray, Carol Clugston, Anna Dominiczak and Alex Alderton, Roberto Amato, Jeffrey Barrett, Sonia Goncalves, Ewan Harrison, David K. Jackson, Ian Johnston, Dominic Kwiatkowski, Cordelia Langford, John Sillitoe on behalf of the Wellcome Sanger Institute COVID-19 Surveillance Team |
| EPI_ISL_2960345, EPI_ISL_2960346 | Lighthouse Lab in Milton Keynes | Wellcome Sanger Institute for the COVID-19 Genomics UK (COG-UK) Consortium | The Lighthouse Lab in Milton Keynes and Alex Alderton, Roberto Amato, Jeffrey Barrett, Sonia Goncalves, Ewan Harrison, David K. Jackson, Ian Johnston, Dominic Kwiatkowski, Cordelia Langford, John Sillitoe on behalf of the Wellcome Sanger Institute COVID-19 Surveillance Team |
| EPI_ISL_2960347, EPI_ISL_2960348 | Lighthouse Lab in Glasgow | Wellcome Sanger Institute for the COVID-19 Genomics UK (COG-UK) Consortium | Harper VanSteenhouse, Yumi Kasai, David Gray, Carol Clugston, Anna Dominiczak and Alex Alderton, Roberto Amato, Jeffrey Barrett, Sonia Goncalves, Ewan Harrison, David K. Jackson, Ian Johnston, Dominic Kwiatkowski, Cordelia Langford, John Sillitoe on behalf of the Wellcome Sanger Institute COVID-19 Surveillance Team |
| EPI_ISL_2960349 | Lighthouse Lab in Milton Keynes | Wellcome Sanger Institute for the COVID-19 Genomics UK (COG-UK) Consortium | The Lighthouse Lab in Milton Keynes and Alex Alderton, Roberto Amato, Jeffrey Barrett, Sonia Goncalves, Ewan Harrison, David K. Jackson, Ian Johnston, Dominic Kwiatkowski, Cordelia Langford, John Sillitoe on behalf of the Wellcome Sanger Institute COVID-19 Surveillance Team |
| EPI_ISL_2960350 | Lighthouse Laboratory Plymouth | Wellcome Sanger Institute for the COVID-19 Genomics UK (COG-UK) Consortium | Lighthouse Laboratory Plymouth and Alex Alderton, Roberto Amato, Jeffrey Barrett, Sonia Goncalves, Ewan Harrison, David K. Jackson, Ian Johnston, Dominic Kwiatkowski, Cordelia Langford, John Sillitoe on behalf of the Wellcome Sanger Institute COVID-19 Surveillance Team |
| EPI_ISL_2960351 | Lighthouse Lab in Milton Keynes | Wellcome Sanger Institute for the COVID-19 Genomics UK (COG-UK) Consortium | The Lighthouse Lab in Milton Keynes and Alex Alderton, Roberto Amato, Jeffrey Barrett, Sonia Goncalves, Ewan Harrison, David K. Jackson, Ian Johnston, Dominic Kwiatkowski, Cordelia Langford, John Sillitoe on behalf of the Wellcome Sanger Institute COVID-19 Surveillance Team |
| EPI_ISL_2960352 | Lighthouse Lab in Glasgow | Wellcome Sanger Institute for the COVID-19 Genomics UK (COG-UK) Consortium | Harper VanSteenhouse, Yumi Kasai, David Gray, Carol Clugston, Anna Dominiczak and Alex Alderton, Roberto Amato, Jeffrey Barrett, Sonia Goncalves, Ewan Harrison, David K. Jackson, Ian Johnston, Dominic Kwiatkowski, Cordelia Langford, John Sillitoe on behalf of the Wellcome Sanger Institute COVID-19 Surveillance Team |
| EPI_ISL_2960353 | Lighthouse Laboratory Plymouth | Wellcome Sanger Institute for the COVID-19 Genomics UK (COG-UK) Consortium | Lighthouse Laboratory Plymouth and Alex Alderton, Roberto Amato, Jeffrey Barrett, Sonia Goncalves, Ewan Harrison, David K. Jackson, Ian Johnston, Dominic Kwiatkowski, Cordelia Langford, John Sillitoe on behalf of the Wellcome Sanger Institute COVID-19 Surveillance Team |
| EPI_ISL_2960354 | Lighthouse Lab in Milton Keynes | Wellcome Sanger Institute for the COVID-19 Genomics UK (COG-UK) Consortium | The Lighthouse Lab in Milton Keynes and Alex Alderton, Roberto Amato, Jeffrey Barrett, Sonia Goncalves, Ewan Harrison, David K. Jackson, Ian Johnston, Dominic Kwiatkowski, Cordelia Langford, John Sillitoe on behalf of the Wellcome Sanger Institute COVID-19 Surveillance Team |
| EPI_ISL_2960355 | Lighthouse Laboratory Plymouth | Wellcome Sanger Institute for the COVID-19 Genomics UK (COG-UK) Consortium | Lighthouse Laboratory Plymouth and Alex Alderton, Roberto Amato, Jeffrey Barrett, Sonia Goncalves, Ewan Harrison, David K. Jackson, Ian Johnston, Dominic Kwiatkowski, Cordelia Langford, John Sillitoe on behalf of the Wellcome Sanger Institute COVID-19 Surveillance Team |
| EPI_ISL_2960356 | Lighthouse Lab in Glasgow | Wellcome Sanger Institute for the COVID-19 Genomics UK (COG-UK) Consortium | Harper VanSteenhouse, Yumi Kasai, David Gray, Carol Clugston, Anna Dominiczak and Alex Alderton, Roberto Amato, Jeffrey Barrett, Sonia Goncalves, Ewan Harrison, David K. Jackson, Ian Johnston, Dominic Kwiatkowski, Cordelia Langford, John Sillitoe on behalf of the Wellcome Sanger Institute COVID-19 Surveillance Team |
| EPI_ISL_2960357 | Lighthouse Laboratory Plymouth | Wellcome Sanger Institute for the COVID-19 Genomics UK (COG-UK) Consortium | Lighthouse Laboratory Plymouth and Alex Alderton, Roberto Amato, Jeffrey Barrett, Sonia Goncalves, Ewan Harrison, David K. Jackson, Ian Johnston, Dominic Kwiatkowski, Cordelia Langford, John Sillitoe on behalf of the Wellcome Sanger Institute COVID-19 Surveillance Team |
| EPI_ISL_2960358 | Lighthouse Lab in Glasgow | Wellcome Sanger Institute for the COVID-19 Genomics UK (COG-UK) Consortium | Harper VanSteenhouse, Yumi Kasai, David Gray, Carol Clugston, Anna Dominiczak and Alex Alderton, Roberto Amato, Jeffrey Barrett, Sonia Goncalves, Ewan Harrison, David K. Jackson, Ian Johnston, Dominic Kwiatkowski, Cordelia Langford, John Sillitoe on behalf of the Wellcome Sanger Institute COVID-19 Surveillance Team |
| EPI_ISL_2960359, EPI_ISL_2960360 | Lighthouse Laboratory Plymouth | Wellcome Sanger Institute for the COVID-19 Genomics UK (COG-UK) Consortium | Lighthouse Laboratory Plymouth and Alex Alderton, Roberto Amato, Jeffrey Barrett, Sonia Goncalves, Ewan Harrison, David K. Jackson, Ian Johnston, Dominic Kwiatkowski, Cordelia Langford, John Sillitoe on behalf of the Wellcome Sanger Institute COVID-19 Surveillance Team |
| EPI_ISL_2960361 | Lighthouse Lab in Milton Keynes | Wellcome Sanger Institute for the COVID-19 Genomics UK (COG-UK) Consortium | The Lighthouse Lab in Milton Keynes and Alex Alderton, Roberto Amato, Jeffrey Barrett, Sonia Goncalves, Ewan Harrison, David K. Jackson, Ian Johnston, Dominic Kwiatkowski, Cordelia Langford, John Sillitoe on behalf of the Wellcome Sanger Institute COVID-19 Surveillance Team |

| | | | |
|---|---|---|---|
| EPI_ISL_2960362 | Lighthouse Lab in Glasgow | Wellcome Sanger Institute for the COVID-19 Genomics UK (COG-UK) Consortium | Harper VanSteenhouse, Yumi Kasai, David Gray, Carol Clugston, Anna Dominiczak and Alex Alderton, Roberto Amato, Jeffrey Barrett, Sonia Goncalves, Ewan Harrison, David K. Jackson, Ian Johnston, Dominic Kwiatkowski, Cordelia Langford, John Sillitoe on behalf of the Wellcome Sanger Institute COVID-19 Surveillance Team |
| EPI_ISL_2960363, EPI_ISL_2960364 | Lighthouse Laboratory Plymouth | Wellcome Sanger Institute for the COVID-19 Genomics UK (COG-UK) Consortium | Lighthouse Laboratory Plymouth and Alex Alderton, Roberto Amato, Jeffrey Barrett, Sonia Goncalves, Ewan Harrison, David K. Jackson, Ian Johnston, Dominic Kwiatkowski, Cordelia Langford, John Sillitoe on behalf of the Wellcome Sanger Institute COVID-19 Surveillance Team |
| EPI_ISL_2960365, EPI_ISL_2960366 | Lighthouse Lab in Glasgow | Wellcome Sanger Institute for the COVID-19 Genomics UK (COG-UK) Consortium | Harper VanSteenhouse, Yumi Kasai, David Gray, Carol Clugston, Anna Dominiczak and Alex Alderton, Roberto Amato, Jeffrey Barrett, Sonia Goncalves, Ewan Harrison, David K. Jackson, Ian Johnston, Dominic Kwiatkowski, Cordelia Langford, John Sillitoe on behalf of the Wellcome Sanger Institute COVID-19 Surveillance Team |
| EPI_ISL_2960367 | Lighthouse Laboratory Plymouth | Wellcome Sanger Institute for the COVID-19 Genomics UK (COG-UK) Consortium | Lighthouse Laboratory Plymouth and Alex Alderton, Roberto Amato, Jeffrey Barrett, Sonia Goncalves, Ewan Harrison, David K. Jackson, Ian Johnston, Dominic Kwiatkowski, Cordelia Langford, John Sillitoe on behalf of the Wellcome Sanger Institute COVID-19 Surveillance Team |
| EPI_ISL_2960368 | Lighthouse Lab in Glasgow | Wellcome Sanger Institute for the COVID-19 Genomics UK (COG-UK) Consortium | Harper VanSteenhouse, Yumi Kasai, David Gray, Carol Clugston, Anna Dominiczak and Alex Alderton, Roberto Amato, Jeffrey Barrett, Sonia Goncalves, Ewan Harrison, David K. Jackson, Ian Johnston, Dominic Kwiatkowski, Cordelia Langford, John Sillitoe on behalf of the Wellcome Sanger Institute COVID-19 Surveillance Team |
| EPI_ISL_2960369 | Lighthouse Laboratory Plymouth | Wellcome Sanger Institute for the COVID-19 Genomics UK (COG-UK) Consortium | Lighthouse Laboratory Plymouth and Alex Alderton, Roberto Amato, Jeffrey Barrett, Sonia Goncalves, Ewan Harrison, David K. Jackson, Ian Johnston, Dominic Kwiatkowski, Cordelia Langford, John Sillitoe on behalf of the Wellcome Sanger Institute COVID-19 Surveillance Team |
| EPI_ISL_2960370, EPI_ISL_2960371 | Lighthouse Lab in Glasgow | Wellcome Sanger Institute for the COVID-19 Genomics UK (COG-UK) Consortium | Harper VanSteenhouse, Yumi Kasai, David Gray, Carol Clugston, Anna Dominiczak and Alex Alderton, Roberto Amato, Jeffrey Barrett, Sonia Goncalves, Ewan Harrison, David K. Jackson, Ian Johnston, Dominic Kwiatkowski, Cordelia Langford, John Sillitoe on behalf of the Wellcome Sanger Institute COVID-19 Surveillance Team |
| EPI_ISL_2960372, EPI_ISL_2960373 | Lighthouse Laboratory Plymouth | Wellcome Sanger Institute for the COVID-19 Genomics UK (COG-UK) Consortium | Lighthouse Laboratory Plymouth and Alex Alderton, Roberto Amato, Jeffrey Barrett, Sonia Goncalves, Ewan Harrison, David K. Jackson, Ian Johnston, Dominic Kwiatkowski, Cordelia Langford, John Sillitoe on behalf of the Wellcome Sanger Institute COVID-19 Surveillance Team |
| EPI_ISL_2960374 | Lighthouse Lab in Milton Keynes | Wellcome Sanger Institute for the COVID-19 Genomics UK (COG-UK) Consortium | The Lighthouse Lab in Milton Keynes and Alex Alderton, Roberto Amato, Jeffrey Barrett, Sonia Goncalves, Ewan Harrison, David K. Jackson, Ian Johnston, Dominic Kwiatkowski, Cordelia Langford, John Sillitoe on behalf of the Wellcome Sanger Institute COVID-19 Surveillance Team |
| EPI_ISL_2960375 | Lighthouse Lab in Glasgow | Wellcome Sanger Institute for the COVID-19 Genomics UK (COG-UK) Consortium | Harper VanSteenhouse, Yumi Kasai, David Gray, Carol Clugston, Anna Dominiczak and Alex Alderton, Roberto Amato, Jeffrey Barrett, Sonia Goncalves, Ewan Harrison, David K. Jackson, Ian Johnston, Dominic Kwiatkowski, Cordelia Langford, John Sillitoe on behalf of the Wellcome Sanger Institute COVID-19 Surveillance Team |
| EPI_ISL_2960376, EPI_ISL_2960377 | Lighthouse Laboratory Plymouth | Wellcome Sanger Institute for the COVID-19 Genomics UK (COG-UK) Consortium | Lighthouse Laboratory Plymouth and Alex Alderton, Roberto Amato, Jeffrey Barrett, Sonia Goncalves, Ewan Harrison, David K. Jackson, Ian Johnston, Dominic Kwiatkowski, Cordelia Langford, John Sillitoe on behalf of the Wellcome Sanger Institute COVID-19 Surveillance Team |
| EPI_ISL_2960378 | Lighthouse Lab in Glasgow | Wellcome Sanger Institute for the COVID-19 Genomics UK (COG-UK) Consortium | Harper VanSteenhouse, Yumi Kasai, David Gray, Carol Clugston, Anna Dominiczak and Alex Alderton, Roberto Amato, Jeffrey Barrett, Sonia Goncalves, Ewan Harrison, David K. Jackson, Ian Johnston, Dominic Kwiatkowski, Cordelia Langford, John Sillitoe on behalf of the Wellcome Sanger Institute COVID-19 Surveillance Team |
| EPI_ISL_2960379 | Lighthouse Laboratory Plymouth | Wellcome Sanger Institute for the COVID-19 Genomics UK (COG-UK) Consortium | Lighthouse Laboratory Plymouth and Alex Alderton, Roberto Amato, Jeffrey Barrett, Sonia Goncalves, Ewan Harrison, David K. Jackson, Ian Johnston, Dominic Kwiatkowski, Cordelia Langford, John Sillitoe on behalf of the Wellcome Sanger Institute COVID-19 Surveillance Team |
| EPI_ISL_2960380, EPI_ISL_2960381 | Lighthouse Lab in Glasgow | Wellcome Sanger Institute for the COVID-19 Genomics UK (COG-UK) Consortium | Harper VanSteenhouse, Yumi Kasai, David Gray, Carol Clugston, Anna Dominiczak and Alex Alderton, Roberto Amato, Jeffrey Barrett, Sonia Goncalves, Ewan Harrison, David K. Jackson, Ian Johnston, Dominic Kwiatkowski, Cordelia Langford, John Sillitoe on behalf of the Wellcome Sanger Institute COVID-19 Surveillance Team |
| EPI_ISL_2960382 | Lighthouse Laboratory Plymouth | Wellcome Sanger Institute for the COVID-19 Genomics UK (COG-UK) Consortium | Lighthouse Laboratory Plymouth and Alex Alderton, Roberto Amato, Jeffrey Barrett, Sonia Goncalves, Ewan Harrison, David K. Jackson, Ian Johnston, Dominic Kwiatkowski, Cordelia Langford, John Sillitoe on behalf of the Wellcome Sanger Institute COVID-19 Surveillance Team |
| EPI_ISL_2960383, EPI_ISL_2960384 | Lighthouse Lab in Milton Keynes | Wellcome Sanger Institute for the COVID-19 Genomics UK (COG-UK) Consortium | The Lighthouse Lab in Milton Keynes and Alex Alderton, Roberto Amato, Jeffrey Barrett, Sonia Goncalves, Ewan Harrison, David K. Jackson, Ian Johnston, Dominic Kwiatkowski, Cordelia Langford, John Sillitoe on behalf of the Wellcome Sanger Institute COVID-19 Surveillance Team |
| EPI_ISL_2960385 | Lighthouse Lab in Glasgow | Wellcome Sanger Institute for the COVID-19 Genomics UK (COG-UK) Consortium | Harper VanSteenhouse, Yumi Kasai, David Gray, Carol Clugston, Anna Dominiczak and Alex Alderton, Roberto Amato, Jeffrey Barrett, Sonia Goncalves, Ewan Harrison, David K. Jackson, Ian Johnston, Dominic Kwiatkowski, Cordelia Langford, John Sillitoe on behalf of the Wellcome Sanger Institute COVID-19 Surveillance Team |
| EPI_ISL_2960386 | Lighthouse Laboratory Plymouth | Wellcome Sanger Institute for the COVID-19 Genomics UK (COG-UK) Consortium | Lighthouse Laboratory Plymouth and Alex Alderton, Roberto Amato, Jeffrey Barrett, Sonia Goncalves, Ewan Harrison, David K. Jackson, Ian Johnston, Dominic Kwiatkowski, Cordelia Langford, John Sillitoe on behalf of the Wellcome Sanger Institute COVID-19 Surveillance Team |
| EPI_ISL_2960387 | Lighthouse Lab in Milton Keynes | Wellcome Sanger Institute for the COVID-19 Genomics UK (COG-UK) Consortium | The Lighthouse Lab in Milton Keynes and Alex Alderton, Roberto Amato, Jeffrey Barrett, Sonia Goncalves, Ewan Harrison, David K. Jackson, Ian Johnston, Dominic Kwiatkowski, Cordelia Langford, John Sillitoe on behalf of the Wellcome Sanger Institute COVID-19 Surveillance Team |
| EPI_ISL_2960388 | Lighthouse Laboratory Plymouth | Wellcome Sanger Institute for the COVID-19 Genomics UK (COG-UK) Consortium | Lighthouse Laboratory Plymouth and Alex Alderton, Roberto Amato, Jeffrey Barrett, Sonia Goncalves, Ewan Harrison, David K. Jackson, Ian Johnston, Dominic Kwiatkowski, Cordelia Langford, John Sillitoe on behalf of the Wellcome Sanger Institute COVID-19 Surveillance Team |
| EPI_ISL_2960389 | Lighthouse Lab in Glasgow | Wellcome Sanger Institute for the COVID-19 Genomics UK (COG-UK) Consortium | Harper VanSteenhouse, Yumi Kasai, David Gray, Carol Clugston, Anna Dominiczak and Alex Alderton, Roberto Amato, Jeffrey Barrett, Sonia Goncalves, Ewan Harrison, David K. Jackson, Ian Johnston, Dominic Kwiatkowski, Cordelia Langford, John Sillitoe on behalf of the Wellcome Sanger Institute COVID-19 Surveillance Team |
| EPI_ISL_2960390 | Lighthouse Laboratory Plymouth | Wellcome Sanger Institute for the COVID-19 Genomics UK (COG-UK) Consortium | Lighthouse Laboratory Plymouth and Alex Alderton, Roberto Amato, Jeffrey Barrett, Sonia Goncalves, Ewan Harrison, David K. Jackson, Ian Johnston, Dominic Kwiatkowski, Cordelia Langford, John Sillitoe on behalf of the Wellcome Sanger Institute COVID-19 Surveillance Team |
| EPI_ISL_2960391 | Lighthouse Lab in Glasgow | Wellcome Sanger Institute for the COVID-19 Genomics UK (COG-UK) Consortium | Harper VanSteenhouse, Yumi Kasai, David Gray, Carol Clugston, Anna Dominiczak and Alex Alderton, Roberto Amato, Jeffrey Barrett, Sonia Goncalves, Ewan Harrison, David K. Jackson, Ian Johnston, Dominic Kwiatkowski, Cordelia Langford, John Sillitoe on behalf of the Wellcome Sanger Institute COVID-19 Surveillance Team |
| EPI_ISL_2960392, EPI_ISL_2960393 | Lighthouse Laboratory Plymouth | Wellcome Sanger Institute for the COVID-19 Genomics UK (COG-UK) Consortium | Lighthouse Laboratory Plymouth and Alex Alderton, Roberto Amato, Jeffrey Barrett, Sonia Goncalves, Ewan Harrison, David K. Jackson, Ian Johnston, Dominic Kwiatkowski, Cordelia Langford, John Sillitoe on behalf of the Wellcome Sanger Institute COVID-19 Surveillance Team |
| EPI_ISL_2960394 | Lighthouse Lab in Glasgow | Wellcome Sanger Institute for the COVID-19 Genomics UK (COG-UK) Consortium | Harper VanSteenhouse, Yumi Kasai, David Gray, Carol Clugston, Anna Dominiczak and Alex Alderton, Roberto Amato, Jeffrey Barrett, Sonia Goncalves, Ewan Harrison, David K. Jackson, Ian Johnston, Dominic Kwiatkowski, Cordelia Langford, John Sillitoe on behalf of the Wellcome Sanger Institute COVID-19 Surveillance Team |
| EPI_ISL_2960395, EPI_ISL_2960396 | Lighthouse Lab in Milton Keynes | Wellcome Sanger Institute for the COVID-19 Genomics UK (COG-UK) Consortium | The Lighthouse Lab in Milton Keynes and Alex Alderton, Roberto Amato, Jeffrey Barrett, Sonia Goncalves, Ewan Harrison, David K. Jackson, Ian Johnston, Dominic Kwiatkowski, Cordelia Langford, John Sillitoe on behalf of the Wellcome Sanger Institute COVID-19 Surveillance Team |
| EPI_ISL_2960397 | Lighthouse Laboratory Plymouth | Wellcome Sanger Institute for the COVID-19 Genomics UK (COG-UK) Consortium | Lighthouse Laboratory Plymouth and Alex Alderton, Roberto Amato, Jeffrey Barrett, Sonia Goncalves, Ewan Harrison, David K. Jackson, Ian Johnston, Dominic Kwiatkowski, Cordelia Langford, John Sillitoe on behalf of the Wellcome Sanger Institute COVID-19 Surveillance Team |
| EPI_ISL_2960398, EPI_ISL_2960399 | Lighthouse Lab in Glasgow | Wellcome Sanger Institute for the COVID-19 Genomics UK (COG-UK) Consortium | Harper VanSteenhouse, Yumi Kasai, David Gray, Carol Clugston, Anna Dominiczak and Alex Alderton, Roberto Amato, Jeffrey Barrett, Sonia Goncalves, Ewan Harrison, David K. Jackson, Ian Johnston, Dominic Kwiatkowski, Cordelia Langford, John Sillitoe on behalf of the Wellcome Sanger Institute COVID-19 Surveillance Team |
| EPI_ISL_2960400 | Lighthouse Laboratory Plymouth | Wellcome Sanger Institute for the COVID-19 Genomics UK (COG-UK) Consortium | Lighthouse Laboratory Plymouth and Alex Alderton, Roberto Amato, Jeffrey Barrett, Sonia Goncalves, Ewan Harrison, David K. Jackson, Ian Johnston, Dominic Kwiatkowski, Cordelia Langford, John Sillitoe on behalf of the Wellcome Sanger Institute COVID-19 Surveillance Team |
| EPI_ISL_2960401 | Lighthouse Lab in Milton Keynes | Wellcome Sanger Institute for the COVID-19 Genomics UK (COG-UK) Consortium | The Lighthouse Lab in Milton Keynes and Alex Alderton, Roberto Amato, Jeffrey Barrett, Sonia Goncalves, Ewan Harrison, David K. Jackson, Ian Johnston, Dominic Kwiatkowski, Cordelia Langford, John Sillitoe on behalf of the Wellcome Sanger Institute COVID-19 Surveillance Team |
| EPI_ISL_2960402, EPI_ISL_2960403, EPI_ISL_2960404 | Lighthouse Lab in Glasgow | Wellcome Sanger Institute for the COVID-19 Genomics UK (COG-UK) Consortium | Harper VanSteenhouse, Yumi Kasai, David Gray, Carol Clugston, Anna Dominiczak and Alex Alderton, Roberto Amato, Jeffrey Barrett, Sonia Goncalves, Ewan Harrison, David K. Jackson, Ian Johnston, Dominic Kwiatkowski, Cordelia Langford, John Sillitoe on behalf of the Wellcome Sanger Institute COVID-19 Surveillance Team |
| EPI_ISL_2960405, EPI_ISL_2960406, EPI_ISL_2960407 | Lighthouse Laboratory Plymouth | Wellcome Sanger Institute for the COVID-19 Genomics UK (COG-UK) Consortium | Lighthouse Laboratory Plymouth and Alex Alderton, Roberto Amato, Jeffrey Barrett, Sonia Goncalves, Ewan Harrison, David K. Jackson, Ian Johnston, Dominic Kwiatkowski, Cordelia Langford, John Sillitoe on behalf of the Wellcome Sanger Institute COVID-19 Surveillance Team |
| EPI_ISL_2960408 | Lighthouse Lab in Glasgow | Wellcome Sanger Institute for the COVID-19 Genomics UK (COG-UK) Consortium | Harper VanSteenhouse, Yumi Kasai, David Gray, Carol Clugston, Anna Dominiczak and Alex Alderton, Roberto Amato, Jeffrey Barrett, Sonia Goncalves, Ewan Harrison, David K. Jackson, Ian Johnston, Dominic Kwiatkowski, Cordelia Langford, John Sillitoe on behalf of the Wellcome Sanger Institute COVID-19 Surveillance Team |
| EPI_ISL_2960409, EPI_ISL_2960410 | Lighthouse Laboratory Plymouth | Wellcome Sanger Institute for the COVID-19 Genomics UK (COG-UK) Consortium | Lighthouse Laboratory Plymouth and Alex Alderton, Roberto Amato, Jeffrey Barrett, Sonia Goncalves, Ewan Harrison, David K. Jackson, Ian Johnston, Dominic Kwiatkowski, Cordelia Langford, John Sillitoe on behalf of the Wellcome Sanger Institute COVID-19 Surveillance Team |
| EPI_ISL_2960411 | Lighthouse Lab in Glasgow | Wellcome Sanger Institute for the COVID-19 Genomics UK (COG-UK) Consortium | Harper VanSteenhouse, Yumi Kasai, David Gray, Carol Clugston, Anna Dominiczak and Alex Alderton, Roberto Amato, Jeffrey Barrett, Sonia Goncalves, Ewan Harrison, David K. Jackson, Ian Johnston, Dominic Kwiatkowski, Cordelia Langford, John Sillitoe on behalf of the Wellcome Sanger Institute COVID-19 Surveillance Team |
| EPI_ISL_2960412 | Lighthouse Laboratory Plymouth | Wellcome Sanger Institute for the COVID-19 Genomics UK (COG-UK) Consortium | Lighthouse Laboratory Plymouth and Alex Alderton, Roberto Amato, Jeffrey Barrett, Sonia Goncalves, Ewan Harrison, David K. Jackson, Ian Johnston, Dominic Kwiatkowski, Cordelia Langford, John Sillitoe on behalf of the Wellcome Sanger Institute COVID-19 Surveillance Team |
| EPI_ISL_2960413 | Lighthouse Lab in Glasgow | Wellcome Sanger Institute for the COVID-19 Genomics UK (COG-UK) Consortium | Harper VanSteenhouse, Yumi Kasai, David Gray, Carol Clugston, Anna Dominiczak and Alex Alderton, Roberto Amato, Jeffrey Barrett, Sonia Goncalves, Ewan Harrison, David K. Jackson, Ian Johnston, Dominic Kwiatkowski, Cordelia Langford, John Sillitoe on behalf of the Wellcome Sanger Institute COVID-19 Surveillance Team |
| EPI_ISL_2960414 | Lighthouse Laboratory Plymouth | Wellcome Sanger Institute for the COVID-19 Genomics UK (COG-UK) Consortium | Lighthouse Laboratory Plymouth and Alex Alderton, Roberto Amato, Jeffrey Barrett, Sonia Goncalves, Ewan Harrison, David K. Jackson, Ian Johnston, Dominic Kwiatkowski, Cordelia Langford, John Sillitoe on behalf of the Wellcome Sanger Institute COVID-19 Surveillance Team |
| EPI_ISL_2960415 | Lighthouse Lab in Glasgow | Wellcome Sanger Institute for the COVID-19 Genomics UK (COG-UK) Consortium | Harper VanSteenhouse, Yumi Kasai, David Gray, Carol Clugston, Anna Dominiczak and Alex Alderton, Roberto Amato, Jeffrey Barrett, Sonia Goncalves, Ewan Harrison, David K. Jackson, Ian Johnston, Dominic Kwiatkowski, Cordelia Langford, John Sillitoe on behalf of the Wellcome Sanger Institute COVID-19 Surveillance Team |
| EPI_ISL_2960416 | Lighthouse Laboratory Plymouth | Wellcome Sanger Institute for the COVID-19 Genomics UK (COG-UK) Consortium | Lighthouse Laboratory Plymouth and Alex Alderton, Roberto Amato, Jeffrey Barrett, Sonia Goncalves, Ewan Harrison, David K. Jackson, Ian Johnston, Dominic Kwiatkowski, Cordelia Langford, John Sillitoe on behalf of the Wellcome Sanger Institute COVID-19 Surveillance Team |
| EPI_ISL_2960417 | Lighthouse Lab in Glasgow | Wellcome Sanger Institute for the COVID-19 Genomics UK (COG-UK) Consortium | Harper VanSteenhouse, Yumi Kasai, David Gray, Carol Clugston, Anna Dominiczak and Alex Alderton, Roberto Amato, Jeffrey Barrett, Sonia Goncalves, Ewan Harrison, David K. Jackson, Ian Johnston, Dominic Kwiatkowski, Cordelia Langford, John Sillitoe on behalf of the Wellcome Sanger Institute COVID-19 Surveillance Team |
| EPI_ISL_2960418 | Lighthouse Lab in Milton Keynes | Wellcome Sanger Institute for the COVID-19 Genomics UK (COG-UK) Consortium | The Lighthouse Lab in Milton Keynes and Alex Alderton, Roberto Amato, Jeffrey Barrett, Sonia Goncalves, Ewan Harrison, David K. Jackson, Ian Johnston, Dominic Kwiatkowski, Cordelia Langford, John Sillitoe on behalf of the Wellcome Sanger Institute COVID-19 Surveillance Team |
| EPI_ISL_2960419, EPI_ISL_2960420 | Lighthouse Laboratory Plymouth | Wellcome Sanger Institute for the COVID-19 Genomics UK (COG-UK) Consortium | Lighthouse Laboratory Plymouth and Alex Alderton, Roberto Amato, Jeffrey Barrett, Sonia Goncalves, Ewan Harrison, David K. Jackson, Ian Johnston, Dominic Kwiatkowski, Cordelia Langford, John Sillitoe on behalf of the Wellcome Sanger Institute COVID-19 Surveillance Team |
| EPI_ISL_2960421, EPI_ISL_2960422 | Lighthouse Lab in Milton Keynes | Wellcome Sanger Institute for the COVID-19 Genomics UK (COG-UK) Consortium | The Lighthouse Lab in Milton Keynes and Alex Alderton, Roberto Amato, Jeffrey Barrett, Sonia Goncalves, Ewan Harrison, David K. Jackson, Ian Johnston, Dominic Kwiatkowski, Cordelia Langford, John Sillitoe on behalf of the Wellcome Sanger Institute COVID-19 Surveillance Team |
| EPI_ISL_2960423 | Lighthouse Lab in Glasgow | Wellcome Sanger Institute for the COVID-19 Genomics UK (COG-UK) | Harper VanSteenhouse, Yumi Kasai, David Gray, Carol Clugston, Anna Dominiczak and Alex Alderton, Roberto Amato, Jeffrey Barrett, Sonia Goncalves, Ewan Harrison, David K. Jackson, Ian Johnston, Dominic Kwiatkowski, Cordelia Langford, |

| | | | |
|---|---|---|---|
| | | Consortium | John Sillitoe on behalf of the Wellcome Sanger Institute COVID-19 Surveillance Team |
| EPI_ISL_2960424 | Lighthouse Lab in Milton Keynes | Wellcome Sanger Institute for the COVID-19 Genomics UK (COG-UK) Consortium | The Lighthouse Lab in Milton Keynes and Alex Alderton, Roberto Amato, Jeffrey Barrett, Sonia Goncalves, Ewan Harrison, David K. Jackson, Ian Johnston, Dominic Kwiatkowski, Cordelia Langford, John Sillitoe on behalf of the Wellcome Sanger Institute COVID-19 Surveillance Team |
| EPI_ISL_2960425 | Lighthouse Laboratory Plymouth | Wellcome Sanger Institute for the COVID-19 Genomics UK (COG-UK) Consortium | Lighthouse Laboratory Plymouth and Alex Alderton, Roberto Amato, Jeffrey Barrett, Sonia Goncalves, Ewan Harrison, David K. Jackson, Ian Johnston, Dominic Kwiatkowski, Cordelia Langford, John Sillitoe on behalf of the Wellcome Sanger Institute COVID-19 Surveillance Team |
| EPI_ISL_2960426, EPI_ISL_2960427, EPI_ISL_2960428 | Lighthouse Lab in Milton Keynes | Wellcome Sanger Institute for the COVID-19 Genomics UK (COG-UK) Consortium | The Lighthouse Lab in Milton Keynes and Alex Alderton, Roberto Amato, Jeffrey Barrett, Sonia Goncalves, Ewan Harrison, David K. Jackson, Ian Johnston, Dominic Kwiatkowski, Cordelia Langford, John Sillitoe on behalf of the Wellcome Sanger Institute COVID-19 Surveillance Team |
| EPI_ISL_2960429, EPI_ISL_2960430 | Lighthouse Lab in Glasgow | Wellcome Sanger Institute for the COVID-19 Genomics UK (COG-UK) Consortium | Harper VanSteenhouse, Yumi Kasai, David Gray, Carol Clugston, Anna Dominiczak and Alex Alderton, Roberto Amato, Jeffrey Barrett, Sonia Goncalves, Ewan Harrison, David K. Jackson, Ian Johnston, Dominic Kwiatkowski, Cordelia Langford, John Sillitoe on behalf of the Wellcome Sanger Institute COVID-19 Surveillance Team |
| EPI_ISL_2960431 | Lighthouse Laboratory Plymouth | Wellcome Sanger Institute for the COVID-19 Genomics UK (COG-UK) Consortium | Lighthouse Laboratory Plymouth and Alex Alderton, Roberto Amato, Jeffrey Barrett, Sonia Goncalves, Ewan Harrison, David K. Jackson, Ian Johnston, Dominic Kwiatkowski, Cordelia Langford, John Sillitoe on behalf of the Wellcome Sanger Institute COVID-19 Surveillance Team |
| EPI_ISL_2960432, EPI_ISL_2960433 | Lighthouse Lab in Glasgow | Wellcome Sanger Institute for the COVID-19 Genomics UK (COG-UK) Consortium | Harper VanSteenhouse, Yumi Kasai, David Gray, Carol Clugston, Anna Dominiczak and Alex Alderton, Roberto Amato, Jeffrey Barrett, Sonia Goncalves, Ewan Harrison, David K. Jackson, Ian Johnston, Dominic Kwiatkowski, Cordelia Langford, John Sillitoe on behalf of the Wellcome Sanger Institute COVID-19 Surveillance Team |
| EPI_ISL_2960434 | Lighthouse Laboratory Plymouth | Wellcome Sanger Institute for the COVID-19 Genomics UK (COG-UK) Consortium | Lighthouse Laboratory Plymouth and Alex Alderton, Roberto Amato, Jeffrey Barrett, Sonia Goncalves, Ewan Harrison, David K. Jackson, Ian Johnston, Dominic Kwiatkowski, Cordelia Langford, John Sillitoe on behalf of the Wellcome Sanger Institute COVID-19 Surveillance Team |
| EPI_ISL_2960435 | Lighthouse Lab in Milton Keynes | Wellcome Sanger Institute for the COVID-19 Genomics UK (COG-UK) Consortium | The Lighthouse Lab in Milton Keynes and Alex Alderton, Roberto Amato, Jeffrey Barrett, Sonia Goncalves, Ewan Harrison, David K. Jackson, Ian Johnston, Dominic Kwiatkowski, Cordelia Langford, John Sillitoe on behalf of the Wellcome Sanger Institute COVID-19 Surveillance Team |
| EPI_ISL_2960436 | Lighthouse Laboratory Plymouth | Wellcome Sanger Institute for the COVID-19 Genomics UK (COG-UK) Consortium | Lighthouse Laboratory Plymouth and Alex Alderton, Roberto Amato, Jeffrey Barrett, Sonia Goncalves, Ewan Harrison, David K. Jackson, Ian Johnston, Dominic Kwiatkowski, Cordelia Langford, John Sillitoe on behalf of the Wellcome Sanger Institute COVID-19 Surveillance Team |
| EPI_ISL_2960437 | Lighthouse Lab in Glasgow | Wellcome Sanger Institute for the COVID-19 Genomics UK (COG-UK) Consortium | Harper VanSteenhouse, Yumi Kasai, David Gray, Carol Clugston, Anna Dominiczak and Alex Alderton, Roberto Amato, Jeffrey Barrett, Sonia Goncalves, Ewan Harrison, David K. Jackson, Ian Johnston, Dominic Kwiatkowski, Cordelia Langford, John Sillitoe on behalf of the Wellcome Sanger Institute COVID-19 Surveillance Team |
| EPI_ISL_2960438 | Lighthouse Lab in Milton Keynes | Wellcome Sanger Institute for the COVID-19 Genomics UK (COG-UK) Consortium | The Lighthouse Lab in Milton Keynes and Alex Alderton, Roberto Amato, Jeffrey Barrett, Sonia Goncalves, Ewan Harrison, David K. Jackson, Ian Johnston, Dominic Kwiatkowski, Cordelia Langford, John Sillitoe on behalf of the Wellcome Sanger Institute COVID-19 Surveillance Team |
| EPI_ISL_2960439, EPI_ISL_2960440, EPI_ISL_2960441, EPI_ISL_2960442 | Lighthouse Lab in Glasgow | Wellcome Sanger Institute for the COVID-19 Genomics UK (COG-UK) Consortium | Harper VanSteenhouse, Yumi Kasai, David Gray, Carol Clugston, Anna Dominiczak and Alex Alderton, Roberto Amato, Jeffrey Barrett, Sonia Goncalves, Ewan Harrison, David K. Jackson, Ian Johnston, Dominic Kwiatkowski, Cordelia Langford, John Sillitoe on behalf of the Wellcome Sanger Institute COVID-19 Surveillance Team |
| EPI_ISL_2960443, EPI_ISL_2960444, EPI_ISL_2960445 | Lighthouse Laboratory Plymouth | Wellcome Sanger Institute for the COVID-19 Genomics UK (COG-UK) Consortium | Lighthouse Laboratory Plymouth and Alex Alderton, Roberto Amato, Jeffrey Barrett, Sonia Goncalves, Ewan Harrison, David K. Jackson, Ian Johnston, Dominic Kwiatkowski, Cordelia Langford, John Sillitoe on behalf of the Wellcome Sanger Institute COVID-19 Surveillance Team |
| EPI_ISL_2960446, EPI_ISL_2960447 | Lighthouse Lab in Glasgow | Wellcome Sanger Institute for the COVID-19 Genomics UK (COG-UK) Consortium | Harper VanSteenhouse, Yumi Kasai, David Gray, Carol Clugston, Anna Dominiczak and Alex Alderton, Roberto Amato, Jeffrey Barrett, Sonia Goncalves, Ewan Harrison, David K. Jackson, Ian Johnston, Dominic Kwiatkowski, Cordelia Langford, John Sillitoe on behalf of the Wellcome Sanger Institute COVID-19 Surveillance Team |
| EPI_ISL_2960448, EPI_ISL_2960449, EPI_ISL_2960450 | Lighthouse Laboratory Plymouth | Wellcome Sanger Institute for the COVID-19 Genomics UK (COG-UK) Consortium | Lighthouse Laboratory Plymouth and Alex Alderton, Roberto Amato, Jeffrey Barrett, Sonia Goncalves, Ewan Harrison, David K. Jackson, Ian Johnston, Dominic Kwiatkowski, Cordelia Langford, John Sillitoe on behalf of the Wellcome Sanger Institute COVID-19 Surveillance Team |
| EPI_ISL_2960451 | Lighthouse Lab in Glasgow | Wellcome Sanger Institute for the COVID-19 Genomics UK (COG-UK) Consortium | Harper VanSteenhouse, Yumi Kasai, David Gray, Carol Clugston, Anna Dominiczak and Alex Alderton, Roberto Amato, Jeffrey Barrett, Sonia Goncalves, Ewan Harrison, David K. Jackson, Ian Johnston, Dominic Kwiatkowski, Cordelia Langford, John Sillitoe on behalf of the Wellcome Sanger Institute COVID-19 Surveillance Team |
| EPI_ISL_2960452 | Lighthouse Lab in Milton Keynes | Wellcome Sanger Institute for the COVID-19 Genomics UK (COG-UK) Consortium | The Lighthouse Lab in Milton Keynes and Alex Alderton, Roberto Amato, Jeffrey Barrett, Sonia Goncalves, Ewan Harrison, David K. Jackson, Ian Johnston, Dominic Kwiatkowski, Cordelia Langford, John Sillitoe on behalf of the Wellcome Sanger Institute COVID-19 Surveillance Team |
| EPI_ISL_2960453, EPI_ISL_2960454 | Lighthouse Lab in Glasgow | Wellcome Sanger Institute for the COVID-19 Genomics UK (COG-UK) Consortium | Harper VanSteenhouse, Yumi Kasai, David Gray, Carol Clugston, Anna Dominiczak and Alex Alderton, Roberto Amato, Jeffrey Barrett, Sonia Goncalves, Ewan Harrison, David K. Jackson, Ian Johnston, Dominic Kwiatkowski, Cordelia Langford, John Sillitoe on behalf of the Wellcome Sanger Institute COVID-19 Surveillance Team |
| EPI_ISL_2960455, EPI_ISL_2960456, EPI_ISL_2960457 | Lighthouse Lab in Milton Keynes | Wellcome Sanger Institute for the COVID-19 Genomics UK (COG-UK) Consortium | The Lighthouse Lab in Milton Keynes and Alex Alderton, Roberto Amato, Jeffrey Barrett, Sonia Goncalves, Ewan Harrison, David K. Jackson, Ian Johnston, Dominic Kwiatkowski, Cordelia Langford, John Sillitoe on behalf of the Wellcome Sanger Institute COVID-19 Surveillance Team |
| EPI_ISL_2960458 | Lighthouse Lab in Glasgow | Wellcome Sanger Institute for the COVID-19 Genomics UK (COG-UK) Consortium | Harper VanSteenhouse, Yumi Kasai, David Gray, Carol Clugston, Anna Dominiczak and Alex Alderton, Roberto Amato, Jeffrey Barrett, Sonia Goncalves, Ewan Harrison, David K. Jackson, Ian Johnston, Dominic Kwiatkowski, Cordelia Langford, John Sillitoe on behalf of the Wellcome Sanger Institute COVID-19 Surveillance Team |
| EPI_ISL_2960459 | Lighthouse Laboratory Plymouth | Wellcome Sanger Institute for the COVID-19 Genomics UK (COG-UK) Consortium | Lighthouse Laboratory Plymouth and Alex Alderton, Roberto Amato, Jeffrey Barrett, Sonia Goncalves, Ewan Harrison, David K. Jackson, Ian Johnston, Dominic Kwiatkowski, Cordelia Langford, John Sillitoe on behalf of the Wellcome Sanger Institute COVID-19 Surveillance Team |
| EPI_ISL_2960460 | Lighthouse Lab in Glasgow | Wellcome Sanger Institute for the COVID-19 Genomics UK (COG-UK) Consortium | Harper VanSteenhouse, Yumi Kasai, David Gray, Carol Clugston, Anna Dominiczak and Alex Alderton, Roberto Amato, Jeffrey Barrett, Sonia Goncalves, Ewan Harrison, David K. Jackson, Ian Johnston, Dominic Kwiatkowski, Cordelia Langford, John Sillitoe on behalf of the Wellcome Sanger Institute COVID-19 Surveillance Team |
| EPI_ISL_2960461, EPI_ISL_2960462 | Lighthouse Laboratory Plymouth | Wellcome Sanger Institute for the COVID-19 Genomics UK (COG-UK) Consortium | Lighthouse Laboratory Plymouth and Alex Alderton, Roberto Amato, Jeffrey Barrett, Sonia Goncalves, Ewan Harrison, David K. Jackson, Ian Johnston, Dominic Kwiatkowski, Cordelia Langford, John Sillitoe on behalf of the Wellcome Sanger Institute COVID-19 Surveillance Team |
| EPI_ISL_2960463, EPI_ISL_2960464, EPI_ISL_2960465, EPI_ISL_2960466, EPI_ISL_2960467 | Lighthouse Lab in Milton Keynes | Wellcome Sanger Institute for the COVID-19 Genomics UK (COG-UK) Consortium | The Lighthouse Lab in Milton Keynes and Alex Alderton, Roberto Amato, Jeffrey Barrett, Sonia Goncalves, Ewan Harrison, David K. Jackson, Ian Johnston, Dominic Kwiatkowski, Cordelia Langford, John Sillitoe on behalf of the Wellcome Sanger Institute COVID-19 Surveillance Team |
| EPI_ISL_2960468 | Lighthouse Lab in Glasgow | Wellcome Sanger Institute for the COVID-19 Genomics UK (COG-UK) Consortium | Harper VanSteenhouse, Yumi Kasai, David Gray, Carol Clugston, Anna Dominiczak and Alex Alderton, Roberto Amato, Jeffrey Barrett, Sonia Goncalves, Ewan Harrison, David K. Jackson, Ian Johnston, Dominic Kwiatkowski, Cordelia Langford, John Sillitoe on behalf of the Wellcome Sanger Institute COVID-19 Surveillance Team |
| EPI_ISL_2960469 | Lighthouse Laboratory Plymouth | Wellcome Sanger Institute for the COVID-19 Genomics UK (COG-UK) Consortium | Lighthouse Laboratory Plymouth and Alex Alderton, Roberto Amato, Jeffrey Barrett, Sonia Goncalves, Ewan Harrison, David K. Jackson, Ian Johnston, Dominic Kwiatkowski, Cordelia Langford, John Sillitoe on behalf of the Wellcome Sanger Institute COVID-19 Surveillance Team |
| EPI_ISL_2960470 | Lighthouse Lab in Glasgow | Wellcome Sanger Institute for the COVID-19 Genomics UK (COG-UK) Consortium | Harper VanSteenhouse, Yumi Kasai, David Gray, Carol Clugston, Anna Dominiczak and Alex Alderton, Roberto Amato, Jeffrey Barrett, Sonia Goncalves, Ewan Harrison, David K. Jackson, Ian Johnston, Dominic Kwiatkowski, Cordelia Langford, John Sillitoe on behalf of the Wellcome Sanger Institute COVID-19 Surveillance Team |
| EPI_ISL_2960471 | Lighthouse Lab in Milton Keynes | Wellcome Sanger Institute for the COVID-19 Genomics UK (COG-UK) Consortium | The Lighthouse Lab in Milton Keynes and Alex Alderton, Roberto Amato, Jeffrey Barrett, Sonia Goncalves, Ewan Harrison, David K. Jackson, Ian Johnston, Dominic Kwiatkowski, Cordelia Langford, John Sillitoe on behalf of the Wellcome Sanger Institute COVID-19 Surveillance Team |
| EPI_ISL_2960472, EPI_ISL_2960473 | Lighthouse Lab in Glasgow | Wellcome Sanger Institute for the COVID-19 Genomics UK (COG-UK) Consortium | Harper VanSteenhouse, Yumi Kasai, David Gray, Carol Clugston, Anna Dominiczak and Alex Alderton, Roberto Amato, Jeffrey Barrett, Sonia Goncalves, Ewan Harrison, David K. Jackson, Ian Johnston, Dominic Kwiatkowski, Cordelia Langford, John Sillitoe on behalf of the Wellcome Sanger Institute COVID-19 Surveillance Team |
| EPI_ISL_2960474, EPI_ISL_2960475 | Lighthouse Lab in Milton Keynes | Wellcome Sanger Institute for the COVID-19 Genomics UK (COG-UK) Consortium | The Lighthouse Lab in Milton Keynes and Alex Alderton, Roberto Amato, Jeffrey Barrett, Sonia Goncalves, Ewan Harrison, David K. Jackson, Ian Johnston, Dominic Kwiatkowski, Cordelia Langford, John Sillitoe on behalf of the Wellcome Sanger Institute COVID-19 Surveillance Team |
| EPI_ISL_2960476, EPI_ISL_2960477, EPI_ISL_2960478, EPI_ISL_2960479 | Lighthouse Lab in Glasgow | Wellcome Sanger Institute for the COVID-19 Genomics UK (COG-UK) Consortium | Harper VanSteenhouse, Yumi Kasai, David Gray, Carol Clugston, Anna Dominiczak and Alex Alderton, Roberto Amato, Jeffrey Barrett, Sonia Goncalves, Ewan Harrison, David K. Jackson, Ian Johnston, Dominic Kwiatkowski, Cordelia Langford, John Sillitoe on behalf of the Wellcome Sanger Institute COVID-19 Surveillance Team |
| EPI_ISL_2960480 | Lighthouse Lab in Milton Keynes | Wellcome Sanger Institute for the COVID-19 Genomics UK (COG-UK) Consortium | The Lighthouse Lab in Milton Keynes and Alex Alderton, Roberto Amato, Jeffrey Barrett, Sonia Goncalves, Ewan Harrison, David K. Jackson, Ian Johnston, Dominic Kwiatkowski, Cordelia Langford, John Sillitoe on behalf of the Wellcome Sanger Institute COVID-19 Surveillance Team |
| EPI_ISL_2960481, EPI_ISL_2960482, EPI_ISL_2960483 | Lighthouse Lab in Glasgow | Wellcome Sanger Institute for the COVID-19 Genomics UK (COG-UK) Consortium | Harper VanSteenhouse, Yumi Kasai, David Gray, Carol Clugston, Anna Dominiczak and Alex Alderton, Roberto Amato, Jeffrey Barrett, Sonia Goncalves, Ewan Harrison, David K. Jackson, Ian Johnston, Dominic Kwiatkowski, Cordelia Langford, John Sillitoe on behalf of the Wellcome Sanger Institute COVID-19 Surveillance Team |
| EPI_ISL_2960484 | Lighthouse Laboratory Plymouth | Wellcome Sanger Institute for the COVID-19 Genomics UK (COG-UK) Consortium | Lighthouse Laboratory Plymouth and Alex Alderton, Roberto Amato, Jeffrey Barrett, Sonia Goncalves, Ewan Harrison, David K. Jackson, Ian Johnston, Dominic Kwiatkowski, Cordelia Langford, John Sillitoe on behalf of the Wellcome Sanger Institute COVID-19 Surveillance Team |
| EPI_ISL_2960485 | Lighthouse Lab in Glasgow | Wellcome Sanger Institute for the COVID-19 Genomics UK (COG-UK) Consortium | Harper VanSteenhouse, Yumi Kasai, David Gray, Carol Clugston, Anna Dominiczak and Alex Alderton, Roberto Amato, Jeffrey Barrett, Sonia Goncalves, Ewan Harrison, David K. Jackson, Ian Johnston, Dominic Kwiatkowski, Cordelia Langford, John Sillitoe on behalf of the Wellcome Sanger Institute COVID-19 Surveillance Team |
| EPI_ISL_2960486 | Lighthouse Lab in Milton Keynes | Wellcome Sanger Institute for the COVID-19 Genomics UK (COG-UK) Consortium | The Lighthouse Lab in Milton Keynes and Alex Alderton, Roberto Amato, Jeffrey Barrett, Sonia Goncalves, Ewan Harrison, David K. Jackson, Ian Johnston, Dominic Kwiatkowski, Cordelia Langford, John Sillitoe on behalf of the Wellcome Sanger Institute COVID-19 Surveillance Team |
| EPI_ISL_2960487, EPI_ISL_2960488 | Lighthouse Laboratory Plymouth | Wellcome Sanger Institute for the COVID-19 Genomics UK (COG-UK) Consortium | Lighthouse Laboratory Plymouth and Alex Alderton, Roberto Amato, Jeffrey Barrett, Sonia Goncalves, Ewan Harrison, David K. Jackson, Ian Johnston, Dominic Kwiatkowski, Cordelia Langford, John Sillitoe on behalf of the Wellcome Sanger Institute COVID-19 Surveillance Team |
| EPI_ISL_2960489 | Lighthouse Lab in Milton Keynes | Wellcome Sanger Institute for the COVID-19 Genomics UK (COG-UK) Consortium | The Lighthouse Lab in Milton Keynes and Alex Alderton, Roberto Amato, Jeffrey Barrett, Sonia Goncalves, Ewan Harrison, David K. Jackson, Ian Johnston, Dominic Kwiatkowski, Cordelia Langford, John Sillitoe on behalf of the Wellcome Sanger Institute COVID-19 Surveillance Team |
| EPI_ISL_2960490 | Lighthouse Laboratory Plymouth | Wellcome Sanger Institute for the COVID-19 Genomics UK (COG-UK) Consortium | Lighthouse Laboratory Plymouth and Alex Alderton, Roberto Amato, Jeffrey Barrett, Sonia Goncalves, Ewan Harrison, David K. Jackson, Ian Johnston, Dominic Kwiatkowski, Cordelia Langford, John Sillitoe on behalf of the Wellcome Sanger Institute COVID-19 Surveillance Team |
| EPI_ISL_2960491 | Lighthouse Lab in Glasgow | Wellcome Sanger Institute for the COVID-19 Genomics UK (COG-UK) Consortium | Harper VanSteenhouse, Yumi Kasai, David Gray, Carol Clugston, Anna Dominiczak and Alex Alderton, Roberto Amato, Jeffrey Barrett, Sonia Goncalves, Ewan Harrison, David K. Jackson, Ian Johnston, Dominic Kwiatkowski, Cordelia Langford, John Sillitoe on behalf of the Wellcome Sanger Institute COVID-19 Surveillance Team |

| | | | |
|---|---|---|---|
| EPI_ISL_2960492, EPI_ISL_2960493 | Lighthouse Lab in Milton Keynes | Wellcome Sanger Institute for the COVID-19 Genomics UK (COG-UK) Consortium | The Lighthouse Lab in Milton Keynes and Alex Alderton, Roberto Amato, Jeffrey Barrett, Sonia Goncalves, Ewan Harrison, David K. Jackson, Ian Johnston, Dominic Kwiatkowski, Cordelia Langford, John Sillitoe on behalf of the Wellcome Sanger Institute COVID-19 Surveillance Team |
| EPI_ISL_2960494 | Lighthouse Lab in Glasgow | Wellcome Sanger Institute for the COVID-19 Genomics UK (COG-UK) Consortium | Harper VanSteenhouse, Yumi Kasai, David Gray, Carol Clugston, Anna Dominiczak and Alex Alderton, Roberto Amato, Jeffrey Barrett, Sonia Goncalves, Ewan Harrison, David K. Jackson, Ian Johnston, Dominic Kwiatkowski, Cordelia Langford, John Sillitoe on behalf of the Wellcome Sanger Institute COVID-19 Surveillance Team |
| EPI_ISL_2960495 | Lighthouse Laboratory Plymouth | Wellcome Sanger Institute for the COVID-19 Genomics UK (COG-UK) Consortium | Lighthouse Laboratory Plymouth and Alex Alderton, Roberto Amato, Jeffrey Barrett, Sonia Goncalves, Ewan Harrison, David K. Jackson, Ian Johnston, Dominic Kwiatkowski, Cordelia Langford, John Sillitoe on behalf of the Wellcome Sanger Institute COVID-19 Surveillance Team |
| EPI_ISL_2960496 | Lighthouse Lab in Milton Keynes | Wellcome Sanger Institute for the COVID-19 Genomics UK (COG-UK) Consortium | The Lighthouse Lab in Milton Keynes and Alex Alderton, Roberto Amato, Jeffrey Barrett, Sonia Goncalves, Ewan Harrison, David K. Jackson, Ian Johnston, Dominic Kwiatkowski, Cordelia Langford, John Sillitoe on behalf of the Wellcome Sanger Institute COVID-19 Surveillance Team |
| EPI_ISL_2960497, EPI_ISL_2960498 | Lighthouse Lab in Glasgow | Wellcome Sanger Institute for the COVID-19 Genomics UK (COG-UK) Consortium | Harper VanSteenhouse, Yumi Kasai, David Gray, Carol Clugston, Anna Dominiczak and Alex Alderton, Roberto Amato, Jeffrey Barrett, Sonia Goncalves, Ewan Harrison, David K. Jackson, Ian Johnston, Dominic Kwiatkowski, Cordelia Langford, John Sillitoe on behalf of the Wellcome Sanger Institute COVID-19 Surveillance Team |
| EPI_ISL_2960499 | Lighthouse Laboratory Plymouth | Wellcome Sanger Institute for the COVID-19 Genomics UK (COG-UK) Consortium | Lighthouse Laboratory Plymouth and Alex Alderton, Roberto Amato, Jeffrey Barrett, Sonia Goncalves, Ewan Harrison, David K. Jackson, Ian Johnston, Dominic Kwiatkowski, Cordelia Langford, John Sillitoe on behalf of the Wellcome Sanger Institute COVID-19 Surveillance Team |
| EPI_ISL_2960500 | Lighthouse Lab in Milton Keynes | Wellcome Sanger Institute for the COVID-19 Genomics UK (COG-UK) Consortium | The Lighthouse Lab in Milton Keynes and Alex Alderton, Roberto Amato, Jeffrey Barrett, Sonia Goncalves, Ewan Harrison, David K. Jackson, Ian Johnston, Dominic Kwiatkowski, Cordelia Langford, John Sillitoe on behalf of the Wellcome Sanger Institute COVID-19 Surveillance Team |
| EPI_ISL_2960501 | Lighthouse Laboratory Plymouth | Wellcome Sanger Institute for the COVID-19 Genomics UK (COG-UK) Consortium | Lighthouse Laboratory Plymouth and Alex Alderton, Roberto Amato, Jeffrey Barrett, Sonia Goncalves, Ewan Harrison, David K. Jackson, Ian Johnston, Dominic Kwiatkowski, Cordelia Langford, John Sillitoe on behalf of the Wellcome Sanger Institute COVID-19 Surveillance Team |
| EPI_ISL_2960502 | Lighthouse Lab in Glasgow | Wellcome Sanger Institute for the COVID-19 Genomics UK (COG-UK) Consortium | Harper VanSteenhouse, Yumi Kasai, David Gray, Carol Clugston, Anna Dominiczak and Alex Alderton, Roberto Amato, Jeffrey Barrett, Sonia Goncalves, Ewan Harrison, David K. Jackson, Ian Johnston, Dominic Kwiatkowski, Cordelia Langford, John Sillitoe on behalf of the Wellcome Sanger Institute COVID-19 Surveillance Team |
| EPI_ISL_2960503 | Lighthouse Lab in Milton Keynes | Wellcome Sanger Institute for the COVID-19 Genomics UK (COG-UK) Consortium | The Lighthouse Lab in Milton Keynes and Alex Alderton, Roberto Amato, Jeffrey Barrett, Sonia Goncalves, Ewan Harrison, David K. Jackson, Ian Johnston, Dominic Kwiatkowski, Cordelia Langford, John Sillitoe on behalf of the Wellcome Sanger Institute COVID-19 Surveillance Team |
| EPI_ISL_2960504 | Lighthouse Lab in Glasgow | Wellcome Sanger Institute for the COVID-19 Genomics UK (COG-UK) Consortium | Harper VanSteenhouse, Yumi Kasai, David Gray, Carol Clugston, Anna Dominiczak and Alex Alderton, Roberto Amato, Jeffrey Barrett, Sonia Goncalves, Ewan Harrison, David K. Jackson, Ian Johnston, Dominic Kwiatkowski, Cordelia Langford, John Sillitoe on behalf of the Wellcome Sanger Institute COVID-19 Surveillance Team |
| EPI_ISL_2960505 | Lighthouse Lab in Milton Keynes | Wellcome Sanger Institute for the COVID-19 Genomics UK (COG-UK) Consortium | The Lighthouse Lab in Milton Keynes and Alex Alderton, Roberto Amato, Jeffrey Barrett, Sonia Goncalves, Ewan Harrison, David K. Jackson, Ian Johnston, Dominic Kwiatkowski, Cordelia Langford, John Sillitoe on behalf of the Wellcome Sanger Institute COVID-19 Surveillance Team |
| EPI_ISL_2960506 | Lighthouse Laboratory Plymouth | Wellcome Sanger Institute for the COVID-19 Genomics UK (COG-UK) Consortium | Lighthouse Laboratory Plymouth and Alex Alderton, Roberto Amato, Jeffrey Barrett, Sonia Goncalves, Ewan Harrison, David K. Jackson, Ian Johnston, Dominic Kwiatkowski, Cordelia Langford, John Sillitoe on behalf of the Wellcome Sanger Institute COVID-19 Surveillance Team |
| EPI_ISL_2960507 | Lighthouse Lab in Milton Keynes | Wellcome Sanger Institute for the COVID-19 Genomics UK (COG-UK) Consortium | The Lighthouse Lab in Milton Keynes and Alex Alderton, Roberto Amato, Jeffrey Barrett, Sonia Goncalves, Ewan Harrison, David K. Jackson, Ian Johnston, Dominic Kwiatkowski, Cordelia Langford, John Sillitoe on behalf of the Wellcome Sanger Institute COVID-19 Surveillance Team |
| EPI_ISL_2960508 | Lighthouse Lab in Glasgow | Wellcome Sanger Institute for the COVID-19 Genomics UK (COG-UK) Consortium | Harper VanSteenhouse, Yumi Kasai, David Gray, Carol Clugston, Anna Dominiczak and Alex Alderton, Roberto Amato, Jeffrey Barrett, Sonia Goncalves, Ewan Harrison, David K. Jackson, Ian Johnston, Dominic Kwiatkowski, Cordelia Langford, John Sillitoe on behalf of the Wellcome Sanger Institute COVID-19 Surveillance Team |
| EPI_ISL_2960509, EPI_ISL_2960510 | Lighthouse Lab in Milton Keynes | Wellcome Sanger Institute for the COVID-19 Genomics UK (COG-UK) Consortium | The Lighthouse Lab in Milton Keynes and Alex Alderton, Roberto Amato, Jeffrey Barrett, Sonia Goncalves, Ewan Harrison, David K. Jackson, Ian Johnston, Dominic Kwiatkowski, Cordelia Langford, John Sillitoe on behalf of the Wellcome Sanger Institute COVID-19 Surveillance Team |
| EPI_ISL_2960511 | Lighthouse Laboratory Plymouth | Wellcome Sanger Institute for the COVID-19 Genomics UK (COG-UK) Consortium | Lighthouse Laboratory Plymouth and Alex Alderton, Roberto Amato, Jeffrey Barrett, Sonia Goncalves, Ewan Harrison, David K. Jackson, Ian Johnston, Dominic Kwiatkowski, Cordelia Langford, John Sillitoe on behalf of the Wellcome Sanger Institute COVID-19 Surveillance Team |
| EPI_ISL_2960512 | Lighthouse Lab in Milton Keynes | Wellcome Sanger Institute for the COVID-19 Genomics UK (COG-UK) Consortium | The Lighthouse Lab in Milton Keynes and Alex Alderton, Roberto Amato, Jeffrey Barrett, Sonia Goncalves, Ewan Harrison, David K. Jackson, Ian Johnston, Dominic Kwiatkowski, Cordelia Langford, John Sillitoe on behalf of the Wellcome Sanger Institute COVID-19 Surveillance Team |
| EPI_ISL_2960513 | Lighthouse Lab in Glasgow | Wellcome Sanger Institute for the COVID-19 Genomics UK (COG-UK) Consortium | Harper VanSteenhouse, Yumi Kasai, David Gray, Carol Clugston, Anna Dominiczak and Alex Alderton, Roberto Amato, Jeffrey Barrett, Sonia Goncalves, Ewan Harrison, David K. Jackson, Ian Johnston, Dominic Kwiatkowski, Cordelia Langford, John Sillitoe on behalf of the Wellcome Sanger Institute COVID-19 Surveillance Team |
| EPI_ISL_2960514 | Lighthouse Laboratory Plymouth | Wellcome Sanger Institute for the COVID-19 Genomics UK (COG-UK) Consortium | Lighthouse Laboratory Plymouth and Alex Alderton, Roberto Amato, Jeffrey Barrett, Sonia Goncalves, Ewan Harrison, David K. Jackson, Ian Johnston, Dominic Kwiatkowski, Cordelia Langford, John Sillitoe on behalf of the Wellcome Sanger Institute COVID-19 Surveillance Team |
| EPI_ISL_2960515 | Lighthouse Lab in Glasgow | Wellcome Sanger Institute for the COVID-19 Genomics UK (COG-UK) Consortium | Harper VanSteenhouse, Yumi Kasai, David Gray, Carol Clugston, Anna Dominiczak and Alex Alderton, Roberto Amato, Jeffrey Barrett, Sonia Goncalves, Ewan Harrison, David K. Jackson, Ian Johnston, Dominic Kwiatkowski, Cordelia Langford, John Sillitoe on behalf of the Wellcome Sanger Institute COVID-19 Surveillance Team |
| EPI_ISL_2960516 | Lighthouse Lab in Milton Keynes | Wellcome Sanger Institute for the COVID-19 Genomics UK (COG-UK) Consortium | The Lighthouse Lab in Milton Keynes and Alex Alderton, Roberto Amato, Jeffrey Barrett, Sonia Goncalves, Ewan Harrison, David K. Jackson, Ian Johnston, Dominic Kwiatkowski, Cordelia Langford, John Sillitoe on behalf of the Wellcome Sanger Institute COVID-19 Surveillance Team |
| EPI_ISL_2960517, EPI_ISL_2960518 | Lighthouse Lab in Glasgow | Wellcome Sanger Institute for the COVID-19 Genomics UK (COG-UK) Consortium | Harper VanSteenhouse, Yumi Kasai, David Gray, Carol Clugston, Anna Dominiczak and Alex Alderton, Roberto Amato, Jeffrey Barrett, Sonia Goncalves, Ewan Harrison, David K. Jackson, Ian Johnston, Dominic Kwiatkowski, Cordelia Langford, John Sillitoe on behalf of the Wellcome Sanger Institute COVID-19 Surveillance Team |
| EPI_ISL_2960519, EPI_ISL_2960520 | Lighthouse Laboratory Plymouth | Wellcome Sanger Institute for the COVID-19 Genomics UK (COG-UK) Consortium | Lighthouse Laboratory Plymouth and Alex Alderton, Roberto Amato, Jeffrey Barrett, Sonia Goncalves, Ewan Harrison, David K. Jackson, Ian Johnston, Dominic Kwiatkowski, Cordelia Langford, John Sillitoe on behalf of the Wellcome Sanger Institute COVID-19 Surveillance Team |
| EPI_ISL_2960521 | Lighthouse Lab in Glasgow | Wellcome Sanger Institute for the COVID-19 Genomics UK (COG-UK) Consortium | Harper VanSteenhouse, Yumi Kasai, David Gray, Carol Clugston, Anna Dominiczak and Alex Alderton, Roberto Amato, Jeffrey Barrett, Sonia Goncalves, Ewan Harrison, David K. Jackson, Ian Johnston, Dominic Kwiatkowski, Cordelia Langford, John Sillitoe on behalf of the Wellcome Sanger Institute COVID-19 Surveillance Team |
| EPI_ISL_2960522 | Lighthouse Lab in Milton Keynes | Wellcome Sanger Institute for the COVID-19 Genomics UK (COG-UK) Consortium | The Lighthouse Lab in Milton Keynes and Alex Alderton, Roberto Amato, Jeffrey Barrett, Sonia Goncalves, Ewan Harrison, David K. Jackson, Ian Johnston, Dominic Kwiatkowski, Cordelia Langford, John Sillitoe on behalf of the Wellcome Sanger Institute COVID-19 Surveillance Team |
| EPI_ISL_2960523 | Lighthouse Laboratory Plymouth | Wellcome Sanger Institute for the COVID-19 Genomics UK (COG-UK) Consortium | Lighthouse Laboratory Plymouth and Alex Alderton, Roberto Amato, Jeffrey Barrett, Sonia Goncalves, Ewan Harrison, David K. Jackson, Ian Johnston, Dominic Kwiatkowski, Cordelia Langford, John Sillitoe on behalf of the Wellcome Sanger Institute COVID-19 Surveillance Team |
| EPI_ISL_2960524 | Lighthouse Lab in Milton Keynes | Wellcome Sanger Institute for the COVID-19 Genomics UK (COG-UK) Consortium | The Lighthouse Lab in Milton Keynes and Alex Alderton, Roberto Amato, Jeffrey Barrett, Sonia Goncalves, Ewan Harrison, David K. Jackson, Ian Johnston, Dominic Kwiatkowski, Cordelia Langford, John Sillitoe on behalf of the Wellcome Sanger Institute COVID-19 Surveillance Team |
| EPI_ISL_2960525 | Lighthouse Laboratory Plymouth | Wellcome Sanger Institute for the COVID-19 Genomics UK (COG-UK) Consortium | Lighthouse Laboratory Plymouth and Alex Alderton, Roberto Amato, Jeffrey Barrett, Sonia Goncalves, Ewan Harrison, David K. Jackson, Ian Johnston, Dominic Kwiatkowski, Cordelia Langford, John Sillitoe on behalf of the Wellcome Sanger Institute COVID-19 Surveillance Team |
| EPI_ISL_2960526 | Lighthouse Lab in Glasgow | Wellcome Sanger Institute for the COVID-19 Genomics UK (COG-UK) Consortium | Harper VanSteenhouse, Yumi Kasai, David Gray, Carol Clugston, Anna Dominiczak and Alex Alderton, Roberto Amato, Jeffrey Barrett, Sonia Goncalves, Ewan Harrison, David K. Jackson, Ian Johnston, Dominic Kwiatkowski, Cordelia Langford, John Sillitoe on behalf of the Wellcome Sanger Institute COVID-19 Surveillance Team |
| EPI_ISL_2960527 | Lighthouse Laboratory Plymouth | Wellcome Sanger Institute for the COVID-19 Genomics UK (COG-UK) Consortium | Lighthouse Laboratory Plymouth and Alex Alderton, Roberto Amato, Jeffrey Barrett, Sonia Goncalves, Ewan Harrison, David K. Jackson, Ian Johnston, Dominic Kwiatkowski, Cordelia Langford, John Sillitoe on behalf of the Wellcome Sanger Institute COVID-19 Surveillance Team |
| EPI_ISL_2960528 | Lighthouse Lab in Glasgow | Wellcome Sanger Institute for the COVID-19 Genomics UK (COG-UK) Consortium | Harper VanSteenhouse, Yumi Kasai, David Gray, Carol Clugston, Anna Dominiczak and Alex Alderton, Roberto Amato, Jeffrey Barrett, Sonia Goncalves, Ewan Harrison, David K. Jackson, Ian Johnston, Dominic Kwiatkowski, Cordelia Langford, John Sillitoe on behalf of the Wellcome Sanger Institute COVID-19 Surveillance Team |
| EPI_ISL_2960529 | Lighthouse Lab in Milton Keynes | Wellcome Sanger Institute for the COVID-19 Genomics UK (COG-UK) Consortium | The Lighthouse Lab in Milton Keynes and Alex Alderton, Roberto Amato, Jeffrey Barrett, Sonia Goncalves, Ewan Harrison, David K. Jackson, Ian Johnston, Dominic Kwiatkowski, Cordelia Langford, John Sillitoe on behalf of the Wellcome Sanger Institute COVID-19 Surveillance Team |
| EPI_ISL_2960530 | Lighthouse Laboratory Plymouth | Wellcome Sanger Institute for the COVID-19 Genomics UK (COG-UK) Consortium | Lighthouse Laboratory Plymouth and Alex Alderton, Roberto Amato, Jeffrey Barrett, Sonia Goncalves, Ewan Harrison, David K. Jackson, Ian Johnston, Dominic Kwiatkowski, Cordelia Langford, John Sillitoe on behalf of the Wellcome Sanger Institute COVID-19 Surveillance Team |
| EPI_ISL_2960531, EPI_ISL_2960532 | Lighthouse Lab in Milton Keynes | Wellcome Sanger Institute for the COVID-19 Genomics UK (COG-UK) Consortium | The Lighthouse Lab in Milton Keynes and Alex Alderton, Roberto Amato, Jeffrey Barrett, Sonia Goncalves, Ewan Harrison, David K. Jackson, Ian Johnston, Dominic Kwiatkowski, Cordelia Langford, John Sillitoe on behalf of the Wellcome Sanger Institute COVID-19 Surveillance Team |
| EPI_ISL_2960533, EPI_ISL_2960534, EPI_ISL_2960535 | Lighthouse Laboratory Plymouth | Wellcome Sanger Institute for the COVID-19 Genomics UK (COG-UK) Consortium | Lighthouse Laboratory Plymouth and Alex Alderton, Roberto Amato, Jeffrey Barrett, Sonia Goncalves, Ewan Harrison, David K. Jackson, Ian Johnston, Dominic Kwiatkowski, Cordelia Langford, John Sillitoe on behalf of the Wellcome Sanger Institute COVID-19 Surveillance Team |
| EPI_ISL_2960536 | Lighthouse Lab in Glasgow | Wellcome Sanger Institute for the COVID-19 Genomics UK (COG-UK) Consortium | Harper VanSteenhouse, Yumi Kasai, David Gray, Carol Clugston, Anna Dominiczak and Alex Alderton, Roberto Amato, Jeffrey Barrett, Sonia Goncalves, Ewan Harrison, David K. Jackson, Ian Johnston, Dominic Kwiatkowski, Cordelia Langford, John Sillitoe on behalf of the Wellcome Sanger Institute COVID-19 Surveillance Team |
| EPI_ISL_2960537 | Lighthouse Lab in Milton Keynes | Wellcome Sanger Institute for the COVID-19 Genomics UK (COG-UK) Consortium | The Lighthouse Lab in Milton Keynes and Alex Alderton, Roberto Amato, Jeffrey Barrett, Sonia Goncalves, Ewan Harrison, David K. Jackson, Ian Johnston, Dominic Kwiatkowski, Cordelia Langford, John Sillitoe on behalf of the Wellcome Sanger Institute COVID-19 Surveillance Team |
| EPI_ISL_2960538 | Lighthouse Laboratory Plymouth | Wellcome Sanger Institute for the COVID-19 Genomics UK (COG-UK) Consortium | Lighthouse Laboratory Plymouth and Alex Alderton, Roberto Amato, Jeffrey Barrett, Sonia Goncalves, Ewan Harrison, David K. Jackson, Ian Johnston, Dominic Kwiatkowski, Cordelia Langford, John Sillitoe on behalf of the Wellcome Sanger Institute COVID-19 Surveillance Team |
| EPI_ISL_2960539, EPI_ISL_2960540, EPI_ISL_2960541 | Lighthouse Lab in Milton Keynes | Wellcome Sanger Institute for the COVID-19 Genomics UK (COG-UK) Consortium | The Lighthouse Lab in Milton Keynes and Alex Alderton, Roberto Amato, Jeffrey Barrett, Sonia Goncalves, Ewan Harrison, David K. Jackson, Ian Johnston, Dominic Kwiatkowski, Cordelia Langford, John Sillitoe on behalf of the Wellcome Sanger Institute COVID-19 Surveillance Team |
| EPI_ISL_2960542 | Lighthouse Lab in Glasgow | Wellcome Sanger Institute for the COVID-19 Genomics UK (COG-UK) Consortium | Harper VanSteenhouse, Yumi Kasai, David Gray, Carol Clugston, Anna Dominiczak and Alex Alderton, Roberto Amato, Jeffrey Barrett, Sonia Goncalves, Ewan Harrison, David K. Jackson, Ian Johnston, Dominic Kwiatkowski, Cordelia Langford, John Sillitoe on behalf of the Wellcome Sanger Institute COVID-19 Surveillance Team |
| EPI_ISL_2960543 | Lighthouse Laboratory Plymouth | Wellcome Sanger Institute for the COVID-19 Genomics UK (COG-UK) Consortium | Lighthouse Laboratory Plymouth and Alex Alderton, Roberto Amato, Jeffrey Barrett, Sonia Goncalves, Ewan Harrison, David K. Jackson, Ian Johnston, Dominic Kwiatkowski, Cordelia Langford, John Sillitoe on behalf of the Wellcome Sanger Institute COVID-19 Surveillance Team |
| EPI_ISL_2960544 | Lighthouse Lab in Milton Keynes | Wellcome Sanger Institute for the COVID-19 Genomics UK (COG-UK) Consortium | The Lighthouse Lab in Milton Keynes and Alex Alderton, Roberto Amato, Jeffrey Barrett, Sonia Goncalves, Ewan Harrison, David K. Jackson, Ian Johnston, Dominic Kwiatkowski, Cordelia Langford, John Sillitoe on behalf of the Wellcome Sanger Institute COVID-19 Surveillance Team |
| EPI_ISL_2960545 | Lighthouse Laboratory Plymouth | Wellcome Sanger Institute for the COVID-19 Genomics UK (COG-UK) Consortium | Lighthouse Laboratory Plymouth and Alex Alderton, Roberto Amato, Jeffrey Barrett, Sonia Goncalves, Ewan Harrison, David K. Jackson, Ian Johnston, Dominic Kwiatkowski, Cordelia Langford, John Sillitoe on behalf of the Wellcome Sanger Institute COVID-19 Surveillance Team |

| | | | |
|---|---|---|---|
| EPI_ISL_2960546, EPI_ISL_2960547 | Lighthouse Lab in Glasgow | Wellcome Sanger Institute for the COVID-19 Genomics UK (COG-UK) Consortium | Harper VanSteenhouse, Yumi Kasai, David Gray, Carol Clugston, Anna Dominiczak and Alex Alderton, Roberto Amato, Jeffrey Barrett, Sonia Goncalves, Ewan Harrison, David K. Jackson, Ian Johnston, Dominic Kwiatkowski, Cordelia Langford, John Sillitoe on behalf of the Wellcome Sanger Institute COVID-19 Surveillance Team |
| EPI_ISL_2960548 | Lighthouse Laboratory Plymouth | Wellcome Sanger Institute for the COVID-19 Genomics UK (COG-UK) Consortium | Lighthouse Laboratory Plymouth and Alex Alderton, Roberto Amato, Jeffrey Barrett, Sonia Goncalves, Ewan Harrison, David K. Jackson, Ian Johnston, Dominic Kwiatkowski, Cordelia Langford, John Sillitoe on behalf of the Wellcome Sanger Institute COVID-19 Surveillance Team |
| EPI_ISL_2960549 | Lighthouse Lab in Glasgow | Wellcome Sanger Institute for the COVID-19 Genomics UK (COG-UK) Consortium | Harper VanSteenhouse, Yumi Kasai, David Gray, Carol Clugston, Anna Dominiczak and Alex Alderton, Roberto Amato, Jeffrey Barrett, Sonia Goncalves, Ewan Harrison, David K. Jackson, Ian Johnston, Dominic Kwiatkowski, Cordelia Langford, John Sillitoe on behalf of the Wellcome Sanger Institute COVID-19 Surveillance Team |
| EPI_ISL_2960550 | Lighthouse Laboratory Plymouth | Wellcome Sanger Institute for the COVID-19 Genomics UK (COG-UK) Consortium | Lighthouse Laboratory Plymouth and Alex Alderton, Roberto Amato, Jeffrey Barrett, Sonia Goncalves, Ewan Harrison, David K. Jackson, Ian Johnston, Dominic Kwiatkowski, Cordelia Langford, John Sillitoe on behalf of the Wellcome Sanger Institute COVID-19 Surveillance Team |
| EPI_ISL_2960551 | Lighthouse Lab in Milton Keynes | Wellcome Sanger Institute for the COVID-19 Genomics UK (COG-UK) Consortium | The Lighthouse Lab in Milton Keynes and Alex Alderton, Roberto Amato, Jeffrey Barrett, Sonia Goncalves, Ewan Harrison, David K. Jackson, Ian Johnston, Dominic Kwiatkowski, Cordelia Langford, John Sillitoe on behalf of the Wellcome Sanger Institute COVID-19 Surveillance Team |
| EPI_ISL_2960552 | Lighthouse Lab in Glasgow | Wellcome Sanger Institute for the COVID-19 Genomics UK (COG-UK) Consortium | Harper VanSteenhouse, Yumi Kasai, David Gray, Carol Clugston, Anna Dominiczak and Alex Alderton, Roberto Amato, Jeffrey Barrett, Sonia Goncalves, Ewan Harrison, David K. Jackson, Ian Johnston, Dominic Kwiatkowski, Cordelia Langford, John Sillitoe on behalf of the Wellcome Sanger Institute COVID-19 Surveillance Team |
| EPI_ISL_2960553 | Lighthouse Lab in Milton Keynes | Wellcome Sanger Institute for the COVID-19 Genomics UK (COG-UK) Consortium | The Lighthouse Lab in Milton Keynes and Alex Alderton, Roberto Amato, Jeffrey Barrett, Sonia Goncalves, Ewan Harrison, David K. Jackson, Ian Johnston, Dominic Kwiatkowski, Cordelia Langford, John Sillitoe on behalf of the Wellcome Sanger Institute COVID-19 Surveillance Team |
| EPI_ISL_2960554, EPI_ISL_2960555, EPI_ISL_2960556 | Lighthouse Lab in Glasgow | Wellcome Sanger Institute for the COVID-19 Genomics UK (COG-UK) Consortium | Harper VanSteenhouse, Yumi Kasai, David Gray, Carol Clugston, Anna Dominiczak and Alex Alderton, Roberto Amato, Jeffrey Barrett, Sonia Goncalves, Ewan Harrison, David K. Jackson, Ian Johnston, Dominic Kwiatkowski, Cordelia Langford, John Sillitoe on behalf of the Wellcome Sanger Institute COVID-19 Surveillance Team |
| EPI_ISL_2960557 | Lighthouse Laboratory Plymouth | Wellcome Sanger Institute for the COVID-19 Genomics UK (COG-UK) Consortium | Lighthouse Laboratory Plymouth and Alex Alderton, Roberto Amato, Jeffrey Barrett, Sonia Goncalves, Ewan Harrison, David K. Jackson, Ian Johnston, Dominic Kwiatkowski, Cordelia Langford, John Sillitoe on behalf of the Wellcome Sanger Institute COVID-19 Surveillance Team |
| EPI_ISL_2960558 | Lighthouse Lab in Glasgow | Wellcome Sanger Institute for the COVID-19 Genomics UK (COG-UK) Consortium | Harper VanSteenhouse, Yumi Kasai, David Gray, Carol Clugston, Anna Dominiczak and Alex Alderton, Roberto Amato, Jeffrey Barrett, Sonia Goncalves, Ewan Harrison, David K. Jackson, Ian Johnston, Dominic Kwiatkowski, Cordelia Langford, John Sillitoe on behalf of the Wellcome Sanger Institute COVID-19 Surveillance Team |
| EPI_ISL_2960559 | Lighthouse Lab in Milton Keynes | Wellcome Sanger Institute for the COVID-19 Genomics UK (COG-UK) Consortium | The Lighthouse Lab in Milton Keynes and Alex Alderton, Roberto Amato, Jeffrey Barrett, Sonia Goncalves, Ewan Harrison, David K. Jackson, Ian Johnston, Dominic Kwiatkowski, Cordelia Langford, John Sillitoe on behalf of the Wellcome Sanger Institute COVID-19 Surveillance Team |
| EPI_ISL_2960560 | Lighthouse Laboratory Plymouth | Wellcome Sanger Institute for the COVID-19 Genomics UK (COG-UK) Consortium | Lighthouse Laboratory Plymouth and Alex Alderton, Roberto Amato, Jeffrey Barrett, Sonia Goncalves, Ewan Harrison, David K. Jackson, Ian Johnston, Dominic Kwiatkowski, Cordelia Langford, John Sillitoe on behalf of the Wellcome Sanger Institute COVID-19 Surveillance Team |
| EPI_ISL_2960561, EPI_ISL_2960562, EPI_ISL_2960563 | Lighthouse Lab in Glasgow | Wellcome Sanger Institute for the COVID-19 Genomics UK (COG-UK) Consortium | Harper VanSteenhouse, Yumi Kasai, David Gray, Carol Clugston, Anna Dominiczak and Alex Alderton, Roberto Amato, Jeffrey Barrett, Sonia Goncalves, Ewan Harrison, David K. Jackson, Ian Johnston, Dominic Kwiatkowski, Cordelia Langford, John Sillitoe on behalf of the Wellcome Sanger Institute COVID-19 Surveillance Team |
| EPI_ISL_2960564 | Lighthouse Lab in Milton Keynes | Wellcome Sanger Institute for the COVID-19 Genomics UK (COG-UK) Consortium | The Lighthouse Lab in Milton Keynes and Alex Alderton, Roberto Amato, Jeffrey Barrett, Sonia Goncalves, Ewan Harrison, David K. Jackson, Ian Johnston, Dominic Kwiatkowski, Cordelia Langford, John Sillitoe on behalf of the Wellcome Sanger Institute COVID-19 Surveillance Team |
| EPI_ISL_2960565 | Lighthouse Laboratory Plymouth | Wellcome Sanger Institute for the COVID-19 Genomics UK (COG-UK) Consortium | Lighthouse Laboratory Plymouth and Alex Alderton, Roberto Amato, Jeffrey Barrett, Sonia Goncalves, Ewan Harrison, David K. Jackson, Ian Johnston, Dominic Kwiatkowski, Cordelia Langford, John Sillitoe on behalf of the Wellcome Sanger Institute COVID-19 Surveillance Team |
| EPI_ISL_2960566 | Lighthouse Lab in Glasgow | Wellcome Sanger Institute for the COVID-19 Genomics UK (COG-UK) Consortium | Harper VanSteenhouse, Yumi Kasai, David Gray, Carol Clugston, Anna Dominiczak and Alex Alderton, Roberto Amato, Jeffrey Barrett, Sonia Goncalves, Ewan Harrison, David K. Jackson, Ian Johnston, Dominic Kwiatkowski, Cordelia Langford, John Sillitoe on behalf of the Wellcome Sanger Institute COVID-19 Surveillance Team |
| EPI_ISL_2960567 | Lighthouse Laboratory Plymouth | Wellcome Sanger Institute for the COVID-19 Genomics UK (COG-UK) Consortium | Lighthouse Laboratory Plymouth and Alex Alderton, Roberto Amato, Jeffrey Barrett, Sonia Goncalves, Ewan Harrison, David K. Jackson, Ian Johnston, Dominic Kwiatkowski, Cordelia Langford, John Sillitoe on behalf of the Wellcome Sanger Institute COVID-19 Surveillance Team |
| EPI_ISL_2960568, EPI_ISL_2960569 | Lighthouse Lab in Glasgow | Wellcome Sanger Institute for the COVID-19 Genomics UK (COG-UK) Consortium | Harper VanSteenhouse, Yumi Kasai, David Gray, Carol Clugston, Anna Dominiczak and Alex Alderton, Roberto Amato, Jeffrey Barrett, Sonia Goncalves, Ewan Harrison, David K. Jackson, Ian Johnston, Dominic Kwiatkowski, Cordelia Langford, John Sillitoe on behalf of the Wellcome Sanger Institute COVID-19 Surveillance Team |
| EPI_ISL_2960570 | Lighthouse Lab in Milton Keynes | Wellcome Sanger Institute for the COVID-19 Genomics UK (COG-UK) Consortium | The Lighthouse Lab in Milton Keynes and Alex Alderton, Roberto Amato, Jeffrey Barrett, Sonia Goncalves, Ewan Harrison, David K. Jackson, Ian Johnston, Dominic Kwiatkowski, Cordelia Langford, John Sillitoe on behalf of the Wellcome Sanger Institute COVID-19 Surveillance Team |
| EPI_ISL_2960571, EPI_ISL_2960572 | Lighthouse Lab in Glasgow | Wellcome Sanger Institute for the COVID-19 Genomics UK (COG-UK) Consortium | Harper VanSteenhouse, Yumi Kasai, David Gray, Carol Clugston, Anna Dominiczak and Alex Alderton, Roberto Amato, Jeffrey Barrett, Sonia Goncalves, Ewan Harrison, David K. Jackson, Ian Johnston, Dominic Kwiatkowski, Cordelia Langford, John Sillitoe on behalf of the Wellcome Sanger Institute COVID-19 Surveillance Team |
| EPI_ISL_2960573 | Lighthouse Lab in Milton Keynes | Wellcome Sanger Institute for the COVID-19 Genomics UK (COG-UK) Consortium | The Lighthouse Lab in Milton Keynes and Alex Alderton, Roberto Amato, Jeffrey Barrett, Sonia Goncalves, Ewan Harrison, David K. Jackson, Ian Johnston, Dominic Kwiatkowski, Cordelia Langford, John Sillitoe on behalf of the Wellcome Sanger Institute COVID-19 Surveillance Team |
| EPI_ISL_2960574 | Lighthouse Lab in Glasgow | Wellcome Sanger Institute for the COVID-19 Genomics UK (COG-UK) Consortium | Harper VanSteenhouse, Yumi Kasai, David Gray, Carol Clugston, Anna Dominiczak and Alex Alderton, Roberto Amato, Jeffrey Barrett, Sonia Goncalves, Ewan Harrison, David K. Jackson, Ian Johnston, Dominic Kwiatkowski, Cordelia Langford, John Sillitoe on behalf of the Wellcome Sanger Institute COVID-19 Surveillance Team |
| EPI_ISL_2960575, EPI_ISL_2960576 | Lighthouse Laboratory Plymouth | Wellcome Sanger Institute for the COVID-19 Genomics UK (COG-UK) Consortium | Lighthouse Laboratory Plymouth and Alex Alderton, Roberto Amato, Jeffrey Barrett, Sonia Goncalves, Ewan Harrison, David K. Jackson, Ian Johnston, Dominic Kwiatkowski, Cordelia Langford, John Sillitoe on behalf of the Wellcome Sanger Institute COVID-19 Surveillance Team |
| EPI_ISL_2960577 | Lighthouse Lab in Glasgow | Wellcome Sanger Institute for the COVID-19 Genomics UK (COG-UK) Consortium | Harper VanSteenhouse, Yumi Kasai, David Gray, Carol Clugston, Anna Dominiczak and Alex Alderton, Roberto Amato, Jeffrey Barrett, Sonia Goncalves, Ewan Harrison, David K. Jackson, Ian Johnston, Dominic Kwiatkowski, Cordelia Langford, John Sillitoe on behalf of the Wellcome Sanger Institute COVID-19 Surveillance Team |
| EPI_ISL_2960578 | Lighthouse Lab in Milton Keynes | Wellcome Sanger Institute for the COVID-19 Genomics UK (COG-UK) Consortium | The Lighthouse Lab in Milton Keynes and Alex Alderton, Roberto Amato, Jeffrey Barrett, Sonia Goncalves, Ewan Harrison, David K. Jackson, Ian Johnston, Dominic Kwiatkowski, Cordelia Langford, John Sillitoe on behalf of the Wellcome Sanger Institute COVID-19 Surveillance Team |
| EPI_ISL_2960579 | Lighthouse Lab in Glasgow | Wellcome Sanger Institute for the COVID-19 Genomics UK (COG-UK) Consortium | Harper VanSteenhouse, Yumi Kasai, David Gray, Carol Clugston, Anna Dominiczak and Alex Alderton, Roberto Amato, Jeffrey Barrett, Sonia Goncalves, Ewan Harrison, David K. Jackson, Ian Johnston, Dominic Kwiatkowski, Cordelia Langford, John Sillitoe on behalf of the Wellcome Sanger Institute COVID-19 Surveillance Team |
| EPI_ISL_2960580 | Lighthouse Lab in Milton Keynes | Wellcome Sanger Institute for the COVID-19 Genomics UK (COG-UK) Consortium | The Lighthouse Lab in Milton Keynes and Alex Alderton, Roberto Amato, Jeffrey Barrett, Sonia Goncalves, Ewan Harrison, David K. Jackson, Ian Johnston, Dominic Kwiatkowski, Cordelia Langford, John Sillitoe on behalf of the Wellcome Sanger Institute COVID-19 Surveillance Team |
| EPI_ISL_2960581 | Lighthouse Lab in Glasgow | Wellcome Sanger Institute for the COVID-19 Genomics UK (COG-UK) Consortium | Harper VanSteenhouse, Yumi Kasai, David Gray, Carol Clugston, Anna Dominiczak and Alex Alderton, Roberto Amato, Jeffrey Barrett, Sonia Goncalves, Ewan Harrison, David K. Jackson, Ian Johnston, Dominic Kwiatkowski, Cordelia Langford, John Sillitoe on behalf of the Wellcome Sanger Institute COVID-19 Surveillance Team |
| EPI_ISL_2960582 | Lighthouse Laboratory Plymouth | Wellcome Sanger Institute for the COVID-19 Genomics UK (COG-UK) Consortium | Lighthouse Laboratory Plymouth and Alex Alderton, Roberto Amato, Jeffrey Barrett, Sonia Goncalves, Ewan Harrison, David K. Jackson, Ian Johnston, Dominic Kwiatkowski, Cordelia Langford, John Sillitoe on behalf of the Wellcome Sanger Institute COVID-19 Surveillance Team |
| EPI_ISL_2960583, EPI_ISL_2960584, EPI_ISL_2960585, EPI_ISL_2960586 | Lighthouse Lab in Glasgow | Wellcome Sanger Institute for the COVID-19 Genomics UK (COG-UK) Consortium | Harper VanSteenhouse, Yumi Kasai, David Gray, Carol Clugston, Anna Dominiczak and Alex Alderton, Roberto Amato, Jeffrey Barrett, Sonia Goncalves, Ewan Harrison, David K. Jackson, Ian Johnston, Dominic Kwiatkowski, Cordelia Langford, John Sillitoe on behalf of the Wellcome Sanger Institute COVID-19 Surveillance Team |
| EPI_ISL_2960587, EPI_ISL_2960588 | Lighthouse Laboratory Plymouth | Wellcome Sanger Institute for the COVID-19 Genomics UK (COG-UK) Consortium | Lighthouse Laboratory Plymouth and Alex Alderton, Roberto Amato, Jeffrey Barrett, Sonia Goncalves, Ewan Harrison, David K. Jackson, Ian Johnston, Dominic Kwiatkowski, Cordelia Langford, John Sillitoe on behalf of the Wellcome Sanger Institute COVID-19 Surveillance Team |
| EPI_ISL_2960589, EPI_ISL_2960590 | Lighthouse Lab in Glasgow | Wellcome Sanger Institute for the COVID-19 Genomics UK (COG-UK) Consortium | Harper VanSteenhouse, Yumi Kasai, David Gray, Carol Clugston, Anna Dominiczak and Alex Alderton, Roberto Amato, Jeffrey Barrett, Sonia Goncalves, Ewan Harrison, David K. Jackson, Ian Johnston, Dominic Kwiatkowski, Cordelia Langford, John Sillitoe on behalf of the Wellcome Sanger Institute COVID-19 Surveillance Team |
| EPI_ISL_2960591 | Lighthouse Lab in Milton Keynes | Wellcome Sanger Institute for the COVID-19 Genomics UK (COG-UK) Consortium | The Lighthouse Lab in Milton Keynes and Alex Alderton, Roberto Amato, Jeffrey Barrett, Sonia Goncalves, Ewan Harrison, David K. Jackson, Ian Johnston, Dominic Kwiatkowski, Cordelia Langford, John Sillitoe on behalf of the Wellcome Sanger Institute COVID-19 Surveillance Team |
| EPI_ISL_2960592, EPI_ISL_2960593 | Lighthouse Lab in Glasgow | Wellcome Sanger Institute for the COVID-19 Genomics UK (COG-UK) Consortium | Harper VanSteenhouse, Yumi Kasai, David Gray, Carol Clugston, Anna Dominiczak and Alex Alderton, Roberto Amato, Jeffrey Barrett, Sonia Goncalves, Ewan Harrison, David K. Jackson, Ian Johnston, Dominic Kwiatkowski, Cordelia Langford, John Sillitoe on behalf of the Wellcome Sanger Institute COVID-19 Surveillance Team |
| EPI_ISL_2960594 | Lighthouse Laboratory Plymouth | Wellcome Sanger Institute for the COVID-19 Genomics UK (COG-UK) Consortium | Lighthouse Laboratory Plymouth and Alex Alderton, Roberto Amato, Jeffrey Barrett, Sonia Goncalves, Ewan Harrison, David K. Jackson, Ian Johnston, Dominic Kwiatkowski, Cordelia Langford, John Sillitoe on behalf of the Wellcome Sanger Institute COVID-19 Surveillance Team |
| EPI_ISL_2960595 | Lighthouse Lab in Milton Keynes | Wellcome Sanger Institute for the COVID-19 Genomics UK (COG-UK) Consortium | The Lighthouse Lab in Milton Keynes and Alex Alderton, Roberto Amato, Jeffrey Barrett, Sonia Goncalves, Ewan Harrison, David K. Jackson, Ian Johnston, Dominic Kwiatkowski, Cordelia Langford, John Sillitoe on behalf of the Wellcome Sanger Institute COVID-19 Surveillance Team |
| EPI_ISL_2960596, EPI_ISL_2960597 | Lighthouse Lab in Glasgow | Wellcome Sanger Institute for the COVID-19 Genomics UK (COG-UK) Consortium | Harper VanSteenhouse, Yumi Kasai, David Gray, Carol Clugston, Anna Dominiczak and Alex Alderton, Roberto Amato, Jeffrey Barrett, Sonia Goncalves, Ewan Harrison, David K. Jackson, Ian Johnston, Dominic Kwiatkowski, Cordelia Langford, John Sillitoe on behalf of the Wellcome Sanger Institute COVID-19 Surveillance Team |
| EPI_ISL_2960598 | Lighthouse Lab in Milton Keynes | Wellcome Sanger Institute for the COVID-19 Genomics UK (COG-UK) Consortium | The Lighthouse Lab in Milton Keynes and Alex Alderton, Roberto Amato, Jeffrey Barrett, Sonia Goncalves, Ewan Harrison, David K. Jackson, Ian Johnston, Dominic Kwiatkowski, Cordelia Langford, John Sillitoe on behalf of the Wellcome Sanger Institute COVID-19 Surveillance Team |
| EPI_ISL_2960599 | Lighthouse Laboratory Plymouth | Wellcome Sanger Institute for the COVID-19 Genomics UK (COG-UK) Consortium | Lighthouse Laboratory Plymouth and Alex Alderton, Roberto Amato, Jeffrey Barrett, Sonia Goncalves, Ewan Harrison, David K. Jackson, Ian Johnston, Dominic Kwiatkowski, Cordelia Langford, John Sillitoe on behalf of the Wellcome Sanger Institute COVID-19 Surveillance Team |
| EPI_ISL_2960600 | Lighthouse Lab in Milton Keynes | Wellcome Sanger Institute for the COVID-19 Genomics UK (COG-UK) Consortium | The Lighthouse Lab in Milton Keynes and Alex Alderton, Roberto Amato, Jeffrey Barrett, Sonia Goncalves, Ewan Harrison, David K. Jackson, Ian Johnston, Dominic Kwiatkowski, Cordelia Langford, John Sillitoe on behalf of the Wellcome Sanger Institute COVID-19 Surveillance Team |
| EPI_ISL_2960601 | Lighthouse Lab in Glasgow | Wellcome Sanger Institute for the COVID-19 Genomics UK (COG-UK) Consortium | Harper VanSteenhouse, Yumi Kasai, David Gray, Carol Clugston, Anna Dominiczak and Alex Alderton, Roberto Amato, Jeffrey Barrett, Sonia Goncalves, Ewan Harrison, David K. Jackson, Ian Johnston, Dominic Kwiatkowski, Cordelia Langford, John Sillitoe on behalf of the Wellcome Sanger Institute COVID-19 Surveillance Team |
| EPI_ISL_2960602, EPI_ISL_2960603 | Lighthouse Laboratory Plymouth | Wellcome Sanger Institute for the COVID-19 Genomics UK (COG-UK) Consortium | Lighthouse Laboratory Plymouth and Alex Alderton, Roberto Amato, Jeffrey Barrett, Sonia Goncalves, Ewan Harrison, David K. Jackson, Ian Johnston, Dominic Kwiatkowski, Cordelia Langford, John Sillitoe on behalf of the Wellcome Sanger Institute COVID-19 Surveillance Team |
| EPI_ISL_2960604 | Lighthouse Lab in Glasgow | Wellcome Sanger Institute for the COVID-19 Genomics UK (COG-UK) Consortium | Harper VanSteenhouse, Yumi Kasai, David Gray, Carol Clugston, Anna Dominiczak and Alex Alderton, Roberto Amato, Jeffrey Barrett, Sonia Goncalves, Ewan Harrison, David K. Jackson, Ian Johnston, Dominic Kwiatkowski, Cordelia Langford, John Sillitoe on behalf of the Wellcome Sanger Institute COVID-19 Surveillance Team |
| EPI_ISL_2960605 | Lighthouse Laboratory Plymouth | Wellcome Sanger Institute for the COVID-19 Genomics UK (COG-UK) | Lighthouse Laboratory Plymouth and Alex Alderton, Roberto Amato, Jeffrey Barrett, Sonia Goncalves, Ewan Harrison, David K. Jackson, Ian Johnston, Dominic Kwiatkowski, Cordelia Langford, John Sillitoe on behalf of the Wellcome Sanger |

| | | | |
|---|---|---|---|
| | | Consortium | Institute COVID-19 Surveillance Team |
| EPI_ISL_2960606 | Lighthouse Lab in Glasgow | Wellcome Sanger Institute for the COVID-19 Genomics UK (COG-UK) Consortium | Harper VanSteenhouse, Yumi Kasai, David Gray, Carol Clugston, Anna Dominiczak and Alex Alderton, Roberto Amato, Jeffrey Barrett, Sonia Goncalves, Ewan Harrison, David K. Jackson, Ian Johnston, Dominic Kwiatkowski, Cordelia Langford, John Sillitoe on behalf of the Wellcome Sanger Institute COVID-19 Surveillance Team |
| EPI_ISL_2960607, EPI_ISL_2960608 | Lighthouse Lab in Milton Keynes | Wellcome Sanger Institute for the COVID-19 Genomics UK (COG-UK) Consortium | The Lighthouse Lab in Milton Keynes and Alex Alderton, Roberto Amato, Jeffrey Barrett, Sonia Goncalves, Ewan Harrison, David K. Jackson, Ian Johnston, Dominic Kwiatkowski, Cordelia Langford, John Sillitoe on behalf of the Wellcome Sanger Institute COVID-19 Surveillance Team |
| EPI_ISL_2960609 | Lighthouse Laboratory Plymouth | Wellcome Sanger Institute for the COVID-19 Genomics UK (COG-UK) Consortium | Lighthouse Laboratory Plymouth and Alex Alderton, Roberto Amato, Jeffrey Barrett, Sonia Goncalves, Ewan Harrison, David K. Jackson, Ian Johnston, Dominic Kwiatkowski, Cordelia Langford, John Sillitoe on behalf of the Wellcome Sanger Institute COVID-19 Surveillance Team |
| EPI_ISL_2960610 | Berkshire and Surrey Pathology Services Lighthouse Laboratory | Wellcome Sanger Institute for the COVID-19 Genomics UK (COG-UK) Consortium | Berkshire and Surrey Pathology Services Lighthouse Laboratory and Alex Alderton, Roberto Amato, Jeffrey Barrett, Sonia Goncalves, Ewan Harrison, David K. Jackson, Ian Johnston, Dominic Kwiatkowski, Cordelia Langford, John Sillitoe on behalf of the Wellcome Sanger Institute COVID-19 Surveillance Team |
| EPI_ISL_2960611, EPI_ISL_2960612, EPI_ISL_2960613 | Lighthouse Lab in Glasgow | Wellcome Sanger Institute for the COVID-19 Genomics UK (COG-UK) Consortium | Harper VanSteenhouse, Yumi Kasai, David Gray, Carol Clugston, Anna Dominiczak and Alex Alderton, Roberto Amato, Jeffrey Barrett, Sonia Goncalves, Ewan Harrison, David K. Jackson, Ian Johnston, Dominic Kwiatkowski, Cordelia Langford, John Sillitoe on behalf of the Wellcome Sanger Institute COVID-19 Surveillance Team |
| EPI_ISL_2960614 | Lighthouse Lab in Alderley Park | Wellcome Sanger Institute for the COVID-19 Genomics UK (COG-UK) Consortium | Jacquelyn Wynn, Mairead Hyland, The Lighthouse Lab in Alderley Park and Alex Alderton, Roberto Amato, Jeffrey Barrett, Sonia Goncalves, Ewan Harrison, David K. Jackson, Ian Johnston, Dominic Kwiatkowski, Cordelia Langford, John Sillitoe on behalf of the Wellcome Sanger Institute COVID-19 Surveillance Team |
| EPI_ISL_2960615, EPI_ISL_2960616 | Lighthouse Lab in Glasgow | Wellcome Sanger Institute for the COVID-19 Genomics UK (COG-UK) Consortium | Harper VanSteenhouse, Yumi Kasai, David Gray, Carol Clugston, Anna Dominiczak and Alex Alderton, Roberto Amato, Jeffrey Barrett, Sonia Goncalves, Ewan Harrison, David K. Jackson, Ian Johnston, Dominic Kwiatkowski, Cordelia Langford, John Sillitoe on behalf of the Wellcome Sanger Institute COVID-19 Surveillance Team |
| EPI_ISL_2960617, EPI_ISL_2960618 | Lighthouse Lab in Alderley Park | Wellcome Sanger Institute for the COVID-19 Genomics UK (COG-UK) Consortium | Jacquelyn Wynn, Mairead Hyland, The Lighthouse Lab in Alderley Park and Alex Alderton, Roberto Amato, Jeffrey Barrett, Sonia Goncalves, Ewan Harrison, David K. Jackson, Ian Johnston, Dominic Kwiatkowski, Cordelia Langford, John Sillitoe on behalf of the Wellcome Sanger Institute COVID-19 Surveillance Team |
| EPI_ISL_2960619 | Berkshire and Surrey Pathology Services Lighthouse Laboratory | Wellcome Sanger Institute for the COVID-19 Genomics UK (COG-UK) Consortium | Berkshire and Surrey Pathology Services Lighthouse Laboratory and Alex Alderton, Roberto Amato, Jeffrey Barrett, Sonia Goncalves, Ewan Harrison, David K. Jackson, Ian Johnston, Dominic Kwiatkowski, Cordelia Langford, John Sillitoe on behalf of the Wellcome Sanger Institute COVID-19 Surveillance Team |
| EPI_ISL_2960620 | Lighthouse Lab in Glasgow | Wellcome Sanger Institute for the COVID-19 Genomics UK (COG-UK) Consortium | Harper VanSteenhouse, Yumi Kasai, David Gray, Carol Clugston, Anna Dominiczak and Alex Alderton, Roberto Amato, Jeffrey Barrett, Sonia Goncalves, Ewan Harrison, David K. Jackson, Ian Johnston, Dominic Kwiatkowski, Cordelia Langford, John Sillitoe on behalf of the Wellcome Sanger Institute COVID-19 Surveillance Team |
| EPI_ISL_2960621, EPI_ISL_2960622, EPI_ISL_2960623 | Lighthouse Lab in Alderley Park | Wellcome Sanger Institute for the COVID-19 Genomics UK (COG-UK) Consortium | Jacquelyn Wynn, Mairead Hyland, The Lighthouse Lab in Alderley Park and Alex Alderton, Roberto Amato, Jeffrey Barrett, Sonia Goncalves, Ewan Harrison, David K. Jackson, Ian Johnston, Dominic Kwiatkowski, Cordelia Langford, John Sillitoe on behalf of the Wellcome Sanger Institute COVID-19 Surveillance Team |
| EPI_ISL_2960624 | Berkshire and Surrey Pathology Services Lighthouse Laboratory | Wellcome Sanger Institute for the COVID-19 Genomics UK (COG-UK) Consortium | Berkshire and Surrey Pathology Services Lighthouse Laboratory and Alex Alderton, Roberto Amato, Jeffrey Barrett, Sonia Goncalves, Ewan Harrison, David K. Jackson, Ian Johnston, Dominic Kwiatkowski, Cordelia Langford, John Sillitoe on behalf of the Wellcome Sanger Institute COVID-19 Surveillance Team |
| EPI_ISL_2960625 | Lighthouse Lab in Alderley Park | Wellcome Sanger Institute for the COVID-19 Genomics UK (COG-UK) Consortium | Jacquelyn Wynn, Mairead Hyland, The Lighthouse Lab in Alderley Park and Alex Alderton, Roberto Amato, Jeffrey Barrett, Sonia Goncalves, Ewan Harrison, David K. Jackson, Ian Johnston, Dominic Kwiatkowski, Cordelia Langford, John Sillitoe on behalf of the Wellcome Sanger Institute COVID-19 Surveillance Team |
| EPI_ISL_2960626, EPI_ISL_2960627 | Lighthouse Lab in Glasgow | Wellcome Sanger Institute for the COVID-19 Genomics UK (COG-UK) Consortium | Harper VanSteenhouse, Yumi Kasai, David Gray, Carol Clugston, Anna Dominiczak and Alex Alderton, Roberto Amato, Jeffrey Barrett, Sonia Goncalves, Ewan Harrison, David K. Jackson, Ian Johnston, Dominic Kwiatkowski, Cordelia Langford, John Sillitoe on behalf of the Wellcome Sanger Institute COVID-19 Surveillance Team |
| EPI_ISL_2960628, EPI_ISL_2960629, EPI_ISL_2960630 | Berkshire and Surrey Pathology Services Lighthouse Laboratory | Wellcome Sanger Institute for the COVID-19 Genomics UK (COG-UK) Consortium | Berkshire and Surrey Pathology Services Lighthouse Laboratory and Alex Alderton, Roberto Amato, Jeffrey Barrett, Sonia Goncalves, Ewan Harrison, David K. Jackson, Ian Johnston, Dominic Kwiatkowski, Cordelia Langford, John Sillitoe on behalf of the Wellcome Sanger Institute COVID-19 Surveillance Team |
| EPI_ISL_2960631 | Lighthouse Lab in Alderley Park | Wellcome Sanger Institute for the COVID-19 Genomics UK (COG-UK) Consortium | Jacquelyn Wynn, Mairead Hyland, The Lighthouse Lab in Alderley Park and Alex Alderton, Roberto Amato, Jeffrey Barrett, Sonia Goncalves, Ewan Harrison, David K. Jackson, Ian Johnston, Dominic Kwiatkowski, Cordelia Langford, John Sillitoe on behalf of the Wellcome Sanger Institute COVID-19 Surveillance Team |
| EPI_ISL_2960632 | Berkshire and Surrey Pathology Services Lighthouse Laboratory | Wellcome Sanger Institute for the COVID-19 Genomics UK (COG-UK) Consortium | Berkshire and Surrey Pathology Services Lighthouse Laboratory and Alex Alderton, Roberto Amato, Jeffrey Barrett, Sonia Goncalves, Ewan Harrison, David K. Jackson, Ian Johnston, Dominic Kwiatkowski, Cordelia Langford, John Sillitoe on behalf of the Wellcome Sanger Institute COVID-19 Surveillance Team |
| EPI_ISL_2960633, EPI_ISL_2960634 | Lighthouse Lab in Alderley Park | Wellcome Sanger Institute for the COVID-19 Genomics UK (COG-UK) Consortium | Jacquelyn Wynn, Mairead Hyland, The Lighthouse Lab in Alderley Park and Alex Alderton, Roberto Amato, Jeffrey Barrett, Sonia Goncalves, Ewan Harrison, David K. Jackson, Ian Johnston, Dominic Kwiatkowski, Cordelia Langford, John Sillitoe on behalf of the Wellcome Sanger Institute COVID-19 Surveillance Team |
| EPI_ISL_2960635, EPI_ISL_2960636 | Berkshire and Surrey Pathology Services Lighthouse Laboratory | Wellcome Sanger Institute for the COVID-19 Genomics UK (COG-UK) Consortium | Berkshire and Surrey Pathology Services Lighthouse Laboratory and Alex Alderton, Roberto Amato, Jeffrey Barrett, Sonia Goncalves, Ewan Harrison, David K. Jackson, Ian Johnston, Dominic Kwiatkowski, Cordelia Langford, John Sillitoe on behalf of the Wellcome Sanger Institute COVID-19 Surveillance Team |
| EPI_ISL_2960637 | Lighthouse Lab in Glasgow | Wellcome Sanger Institute for the COVID-19 Genomics UK (COG-UK) Consortium | Harper VanSteenhouse, Yumi Kasai, David Gray, Carol Clugston, Anna Dominiczak and Alex Alderton, Roberto Amato, Jeffrey Barrett, Sonia Goncalves, Ewan Harrison, David K. Jackson, Ian Johnston, Dominic Kwiatkowski, Cordelia Langford, John Sillitoe on behalf of the Wellcome Sanger Institute COVID-19 Surveillance Team |
| EPI_ISL_2960638, EPI_ISL_2960639, EPI_ISL_2960640, EPI_ISL_2960641 | Lighthouse Lab in Alderley Park | Wellcome Sanger Institute for the COVID-19 Genomics UK (COG-UK) Consortium | Jacquelyn Wynn, Mairead Hyland, The Lighthouse Lab in Alderley Park and Alex Alderton, Roberto Amato, Jeffrey Barrett, Sonia Goncalves, Ewan Harrison, David K. Jackson, Ian Johnston, Dominic Kwiatkowski, Cordelia Langford, John Sillitoe on behalf of the Wellcome Sanger Institute COVID-19 Surveillance Team |
| EPI_ISL_2960642 | Berkshire and Surrey Pathology Services Lighthouse Laboratory | Wellcome Sanger Institute for the COVID-19 Genomics UK (COG-UK) Consortium | Berkshire and Surrey Pathology Services Lighthouse Laboratory and Alex Alderton, Roberto Amato, Jeffrey Barrett, Sonia Goncalves, Ewan Harrison, David K. Jackson, Ian Johnston, Dominic Kwiatkowski, Cordelia Langford, John Sillitoe on behalf of the Wellcome Sanger Institute COVID-19 Surveillance Team |
| EPI_ISL_2960643, EPI_ISL_2960644, EPI_ISL_2960645, EPI_ISL_2960646, EPI_ISL_2960647, EPI_ISL_2960648, EPI_ISL_2960649 | Lighthouse Lab in Glasgow | Wellcome Sanger Institute for the COVID-19 Genomics UK (COG-UK) Consortium | Harper VanSteenhouse, Yumi Kasai, David Gray, Carol Clugston, Anna Dominiczak and Alex Alderton, Roberto Amato, Jeffrey Barrett, Sonia Goncalves, Ewan Harrison, David K. Jackson, Ian Johnston, Dominic Kwiatkowski, Cordelia Langford, John Sillitoe on behalf of the Wellcome Sanger Institute COVID-19 Surveillance Team |
| EPI_ISL_2960650 | Berkshire and Surrey Pathology Services Lighthouse Laboratory | Wellcome Sanger Institute for the COVID-19 Genomics UK (COG-UK) Consortium | Berkshire and Surrey Pathology Services Lighthouse Laboratory and Alex Alderton, Roberto Amato, Jeffrey Barrett, Sonia Goncalves, Ewan Harrison, David K. Jackson, Ian Johnston, Dominic Kwiatkowski, Cordelia Langford, John Sillitoe on behalf of the Wellcome Sanger Institute COVID-19 Surveillance Team |
| EPI_ISL_2960651 | Lighthouse Lab in Alderley Park | Wellcome Sanger Institute for the COVID-19 Genomics UK (COG-UK) Consortium | Jacquelyn Wynn, Mairead Hyland, The Lighthouse Lab in Alderley Park and Alex Alderton, Roberto Amato, Jeffrey Barrett, Sonia Goncalves, Ewan Harrison, David K. Jackson, Ian Johnston, Dominic Kwiatkowski, Cordelia Langford, John Sillitoe on behalf of the Wellcome Sanger Institute COVID-19 Surveillance Team |
| EPI_ISL_2960652, EPI_ISL_2960653, EPI_ISL_2960654 | Berkshire and Surrey Pathology Services Lighthouse Laboratory | Wellcome Sanger Institute for the COVID-19 Genomics UK (COG-UK) Consortium | Berkshire and Surrey Pathology Services Lighthouse Laboratory and Alex Alderton, Roberto Amato, Jeffrey Barrett, Sonia Goncalves, Ewan Harrison, David K. Jackson, Ian Johnston, Dominic Kwiatkowski, Cordelia Langford, John Sillitoe on behalf of the Wellcome Sanger Institute COVID-19 Surveillance Team |
| EPI_ISL_2960655 | Lighthouse Lab in Glasgow | Wellcome Sanger Institute for the COVID-19 Genomics UK (COG-UK) Consortium | Harper VanSteenhouse, Yumi Kasai, David Gray, Carol Clugston, Anna Dominiczak and Alex Alderton, Roberto Amato, Jeffrey Barrett, Sonia Goncalves, Ewan Harrison, David K. Jackson, Ian Johnston, Dominic Kwiatkowski, Cordelia Langford, John Sillitoe on behalf of the Wellcome Sanger Institute COVID-19 Surveillance Team |
| EPI_ISL_2960656 | Berkshire and Surrey Pathology Services Lighthouse Laboratory | Wellcome Sanger Institute for the COVID-19 Genomics UK (COG-UK) Consortium | Berkshire and Surrey Pathology Services Lighthouse Laboratory and Alex Alderton, Roberto Amato, Jeffrey Barrett, Sonia Goncalves, Ewan Harrison, David K. Jackson, Ian Johnston, Dominic Kwiatkowski, Cordelia Langford, John Sillitoe on behalf of the Wellcome Sanger Institute COVID-19 Surveillance Team |
| EPI_ISL_2960657, EPI_ISL_2960658, EPI_ISL_2960659, EPI_ISL_2960660, EPI_ISL_2960661, EPI_ISL_2960662 | Lighthouse Lab in Glasgow | Wellcome Sanger Institute for the COVID-19 Genomics UK (COG-UK) Consortium | Harper VanSteenhouse, Yumi Kasai, David Gray, Carol Clugston, Anna Dominiczak and Alex Alderton, Roberto Amato, Jeffrey Barrett, Sonia Goncalves, Ewan Harrison, David K. Jackson, Ian Johnston, Dominic Kwiatkowski, Cordelia Langford, John Sillitoe on behalf of the Wellcome Sanger Institute COVID-19 Surveillance Team |
| EPI_ISL_2960663 | Berkshire and Surrey Pathology Services Lighthouse Laboratory | Wellcome Sanger Institute for the COVID-19 Genomics UK (COG-UK) Consortium | Berkshire and Surrey Pathology Services Lighthouse Laboratory and Alex Alderton, Roberto Amato, Jeffrey Barrett, Sonia Goncalves, Ewan Harrison, David K. Jackson, Ian Johnston, Dominic Kwiatkowski, Cordelia Langford, John Sillitoe on behalf of the Wellcome Sanger Institute COVID-19 Surveillance Team |
| EPI_ISL_2960664 | Lighthouse Lab in Alderley Park | Wellcome Sanger Institute for the COVID-19 Genomics UK (COG-UK) Consortium | Jacquelyn Wynn, Mairead Hyland, The Lighthouse Lab in Alderley Park and Alex Alderton, Roberto Amato, Jeffrey Barrett, Sonia Goncalves, Ewan Harrison, David K. Jackson, Ian Johnston, Dominic Kwiatkowski, Cordelia Langford, John Sillitoe on behalf of the Wellcome Sanger Institute COVID-19 Surveillance Team |
| EPI_ISL_2960665 | Lighthouse Lab in Glasgow | Wellcome Sanger Institute for the COVID-19 Genomics UK (COG-UK) Consortium | Harper VanSteenhouse, Yumi Kasai, David Gray, Carol Clugston, Anna Dominiczak and Alex Alderton, Roberto Amato, Jeffrey Barrett, Sonia Goncalves, Ewan Harrison, David K. Jackson, Ian Johnston, Dominic Kwiatkowski, Cordelia Langford, John Sillitoe on behalf of the Wellcome Sanger Institute COVID-19 Surveillance Team |
| EPI_ISL_2960666, EPI_ISL_2960667, EPI_ISL_2960668, EPI_ISL_2960669, EPI_ISL_2960670 | Berkshire and Surrey Pathology Services Lighthouse Laboratory | Wellcome Sanger Institute for the COVID-19 Genomics UK (COG-UK) Consortium | Berkshire and Surrey Pathology Services Lighthouse Laboratory and Alex Alderton, Roberto Amato, Jeffrey Barrett, Sonia Goncalves, Ewan Harrison, David K. Jackson, Ian Johnston, Dominic Kwiatkowski, Cordelia Langford, John Sillitoe on behalf of the Wellcome Sanger Institute COVID-19 Surveillance Team |
| EPI_ISL_2960672 | Lighthouse Lab in Glasgow | Wellcome Sanger Institute for the COVID-19 Genomics UK (COG-UK) Consortium | Harper VanSteenhouse, Yumi Kasai, David Gray, Carol Clugston, Anna Dominiczak and Alex Alderton, Roberto Amato, Jeffrey Barrett, Sonia Goncalves, Ewan Harrison, David K. Jackson, Ian Johnston, Dominic Kwiatkowski, Cordelia Langford, John Sillitoe on behalf of the Wellcome Sanger Institute COVID-19 Surveillance Team |
| EPI_ISL_2960673 | Lighthouse Lab in Alderley Park | Wellcome Sanger Institute for the COVID-19 Genomics UK (COG-UK) Consortium | Jacquelyn Wynn, Mairead Hyland, The Lighthouse Lab in Alderley Park and Alex Alderton, Roberto Amato, Jeffrey Barrett, Sonia Goncalves, Ewan Harrison, David K. Jackson, Ian Johnston, Dominic Kwiatkowski, Cordelia Langford, John Sillitoe on behalf of the Wellcome Sanger Institute COVID-19 Surveillance Team |
| EPI_ISL_2960674 | Berkshire and Surrey Pathology Services Lighthouse Laboratory | Wellcome Sanger Institute for the COVID-19 Genomics UK (COG-UK) Consortium | Berkshire and Surrey Pathology Services Lighthouse Laboratory and Alex Alderton, Roberto Amato, Jeffrey Barrett, Sonia Goncalves, Ewan Harrison, David K. Jackson, Ian Johnston, Dominic Kwiatkowski, Cordelia Langford, John Sillitoe on behalf of the Wellcome Sanger Institute COVID-19 Surveillance Team |
| EPI_ISL_2960675, EPI_ISL_2960676, EPI_ISL_2960677 | Lighthouse Lab in Glasgow | Wellcome Sanger Institute for the COVID-19 Genomics UK (COG-UK) Consortium | Harper VanSteenhouse, Yumi Kasai, David Gray, Carol Clugston, Anna Dominiczak and Alex Alderton, Roberto Amato, Jeffrey Barrett, Sonia Goncalves, Ewan Harrison, David K. Jackson, Ian Johnston, Dominic Kwiatkowski, Cordelia Langford, John Sillitoe on behalf of the Wellcome Sanger Institute COVID-19 Surveillance Team |

| | | | |
|---|---|---|---|
| EPI_ISL_2960678 | Lighthouse Lab in Alderley Park | Wellcome Sanger Institute for the COVID-19 Genomics UK (COG-UK) Consortium | Jacquelyn Wynn, Mairead Hyland, The Lighthouse Lab in Alderley Park and Alex Alderton, Roberto Amato, Jeffrey Barrett, Sonia Goncalves, Ewan Harrison, David K. Jackson, Ian Johnston, Dominic Kwiatkowski, Cordelia Langford, John Sillitoe on behalf of the Wellcome Sanger Institute COVID-19 Surveillance Team |
| EPI_ISL_2960679 | Berkshire and Surrey Pathology Services Lighthouse Laboratory | Wellcome Sanger Institute for the COVID-19 Genomics UK (COG-UK) Consortium | Berkshire and Surrey Pathology Services Lighthouse Laboratory and Alex Alderton, Roberto Amato, Jeffrey Barrett, Sonia Goncalves, Ewan Harrison, David K. Jackson, Ian Johnston, Dominic Kwiatkowski, Cordelia Langford, John Sillitoe on behalf of the Wellcome Sanger Institute COVID-19 Surveillance Team |
| EPI_ISL_2960680 | Lighthouse Lab in Glasgow | Wellcome Sanger Institute for the COVID-19 Genomics UK (COG-UK) Consortium | Harper VanSteenhouse, Yumi Kasai, David Gray, Carol Clugston, Anna Dominiczak and Alex Alderton, Roberto Amato, Jeffrey Barrett, Sonia Goncalves, Ewan Harrison, David K. Jackson, Ian Johnston, Dominic Kwiatkowski, Cordelia Langford, John Sillitoe on behalf of the Wellcome Sanger Institute COVID-19 Surveillance Team |
| EPI_ISL_2960681 | Lighthouse Lab in Alderley Park | Wellcome Sanger Institute for the COVID-19 Genomics UK (COG-UK) Consortium | Jacquelyn Wynn, Mairead Hyland, The Lighthouse Lab in Alderley Park and Alex Alderton, Roberto Amato, Jeffrey Barrett, Sonia Goncalves, Ewan Harrison, David K. Jackson, Ian Johnston, Dominic Kwiatkowski, Cordelia Langford, John Sillitoe on behalf of the Wellcome Sanger Institute COVID-19 Surveillance Team |
| EPI_ISL_2960682, EPI_ISL_2960683 | Lighthouse Lab in Glasgow | Wellcome Sanger Institute for the COVID-19 Genomics UK (COG-UK) Consortium | Harper VanSteenhouse, Yumi Kasai, David Gray, Carol Clugston, Anna Dominiczak and Alex Alderton, Roberto Amato, Jeffrey Barrett, Sonia Goncalves, Ewan Harrison, David K. Jackson, Ian Johnston, Dominic Kwiatkowski, Cordelia Langford, John Sillitoe on behalf of the Wellcome Sanger Institute COVID-19 Surveillance Team |
| EPI_ISL_2960684, EPI_ISL_2960685 | Lighthouse Lab in Alderley Park | Wellcome Sanger Institute for the COVID-19 Genomics UK (COG-UK) Consortium | Jacquelyn Wynn, Mairead Hyland, The Lighthouse Lab in Alderley Park and Alex Alderton, Roberto Amato, Jeffrey Barrett, Sonia Goncalves, Ewan Harrison, David K. Jackson, Ian Johnston, Dominic Kwiatkowski, Cordelia Langford, John Sillitoe on behalf of the Wellcome Sanger Institute COVID-19 Surveillance Team |
| EPI_ISL_2960686 | Berkshire and Surrey Pathology Services Lighthouse Laboratory | Wellcome Sanger Institute for the COVID-19 Genomics UK (COG-UK) Consortium | Berkshire and Surrey Pathology Services Lighthouse Laboratory and Alex Alderton, Roberto Amato, Jeffrey Barrett, Sonia Goncalves, Ewan Harrison, David K. Jackson, Ian Johnston, Dominic Kwiatkowski, Cordelia Langford, John Sillitoe on behalf of the Wellcome Sanger Institute COVID-19 Surveillance Team |
| EPI_ISL_2960687, EPI_ISL_2960688, EPI_ISL_2960689 | Lighthouse Lab in Glasgow | Wellcome Sanger Institute for the COVID-19 Genomics UK (COG-UK) Consortium | Harper VanSteenhouse, Yumi Kasai, David Gray, Carol Clugston, Anna Dominiczak and Alex Alderton, Roberto Amato, Jeffrey Barrett, Sonia Goncalves, Ewan Harrison, David K. Jackson, Ian Johnston, Dominic Kwiatkowski, Cordelia Langford, John Sillitoe on behalf of the Wellcome Sanger Institute COVID-19 Surveillance Team |
| EPI_ISL_2960690 | Berkshire and Surrey Pathology Services Lighthouse Laboratory | Wellcome Sanger Institute for the COVID-19 Genomics UK (COG-UK) Consortium | Berkshire and Surrey Pathology Services Lighthouse Laboratory and Alex Alderton, Roberto Amato, Jeffrey Barrett, Sonia Goncalves, Ewan Harrison, David K. Jackson, Ian Johnston, Dominic Kwiatkowski, Cordelia Langford, John Sillitoe on behalf of the Wellcome Sanger Institute COVID-19 Surveillance Team |
| EPI_ISL_2960691 | Lighthouse Lab in Alderley Park | Wellcome Sanger Institute for the COVID-19 Genomics UK (COG-UK) Consortium | Jacquelyn Wynn, Mairead Hyland, The Lighthouse Lab in Alderley Park and Alex Alderton, Roberto Amato, Jeffrey Barrett, Sonia Goncalves, Ewan Harrison, David K. Jackson, Ian Johnston, Dominic Kwiatkowski, Cordelia Langford, John Sillitoe on behalf of the Wellcome Sanger Institute COVID-19 Surveillance Team |
| EPI_ISL_2960692 | Berkshire and Surrey Pathology Services Lighthouse Laboratory | Wellcome Sanger Institute for the COVID-19 Genomics UK (COG-UK) Consortium | Berkshire and Surrey Pathology Services Lighthouse Laboratory and Alex Alderton, Roberto Amato, Jeffrey Barrett, Sonia Goncalves, Ewan Harrison, David K. Jackson, Ian Johnston, Dominic Kwiatkowski, Cordelia Langford, John Sillitoe on behalf of the Wellcome Sanger Institute COVID-19 Surveillance Team |
| EPI_ISL_2960693 | Lighthouse Lab in Alderley Park | Wellcome Sanger Institute for the COVID-19 Genomics UK (COG-UK) Consortium | Jacquelyn Wynn, Mairead Hyland, The Lighthouse Lab in Alderley Park and Alex Alderton, Roberto Amato, Jeffrey Barrett, Sonia Goncalves, Ewan Harrison, David K. Jackson, Ian Johnston, Dominic Kwiatkowski, Cordelia Langford, John Sillitoe on behalf of the Wellcome Sanger Institute COVID-19 Surveillance Team |
| EPI_ISL_2960694 | Lighthouse Lab in Glasgow | Wellcome Sanger Institute for the COVID-19 Genomics UK (COG-UK) Consortium | Harper VanSteenhouse, Yumi Kasai, David Gray, Carol Clugston, Anna Dominiczak and Alex Alderton, Roberto Amato, Jeffrey Barrett, Sonia Goncalves, Ewan Harrison, David K. Jackson, Ian Johnston, Dominic Kwiatkowski, Cordelia Langford, John Sillitoe on behalf of the Wellcome Sanger Institute COVID-19 Surveillance Team |
| EPI_ISL_2960695 | Berkshire and Surrey Pathology Services Lighthouse Laboratory | Wellcome Sanger Institute for the COVID-19 Genomics UK (COG-UK) Consortium | Berkshire and Surrey Pathology Services Lighthouse Laboratory and Alex Alderton, Roberto Amato, Jeffrey Barrett, Sonia Goncalves, Ewan Harrison, David K. Jackson, Ian Johnston, Dominic Kwiatkowski, Cordelia Langford, John Sillitoe on behalf of the Wellcome Sanger Institute COVID-19 Surveillance Team |
| EPI_ISL_2960696 | Lighthouse Lab in Alderley Park | Wellcome Sanger Institute for the COVID-19 Genomics UK (COG-UK) Consortium | Jacquelyn Wynn, Mairead Hyland, The Lighthouse Lab in Alderley Park and Alex Alderton, Roberto Amato, Jeffrey Barrett, Sonia Goncalves, Ewan Harrison, David K. Jackson, Ian Johnston, Dominic Kwiatkowski, Cordelia Langford, John Sillitoe on behalf of the Wellcome Sanger Institute COVID-19 Surveillance Team |
| EPI_ISL_2960697, EPI_ISL_2960698 | Lighthouse Lab in Glasgow | Wellcome Sanger Institute for the COVID-19 Genomics UK (COG-UK) Consortium | Harper VanSteenhouse, Yumi Kasai, David Gray, Carol Clugston, Anna Dominiczak and Alex Alderton, Roberto Amato, Jeffrey Barrett, Sonia Goncalves, Ewan Harrison, David K. Jackson, Ian Johnston, Dominic Kwiatkowski, Cordelia Langford, John Sillitoe on behalf of the Wellcome Sanger Institute COVID-19 Surveillance Team |
| EPI_ISL_2960699, EPI_ISL_2960700 | Lighthouse Lab in Alderley Park | Wellcome Sanger Institute for the COVID-19 Genomics UK (COG-UK) Consortium | Jacquelyn Wynn, Mairead Hyland, The Lighthouse Lab in Alderley Park and Alex Alderton, Roberto Amato, Jeffrey Barrett, Sonia Goncalves, Ewan Harrison, David K. Jackson, Ian Johnston, Dominic Kwiatkowski, Cordelia Langford, John Sillitoe on behalf of the Wellcome Sanger Institute COVID-19 Surveillance Team |
| EPI_ISL_2960701 | Lighthouse Lab in Glasgow | Wellcome Sanger Institute for the COVID-19 Genomics UK (COG-UK) Consortium | Harper VanSteenhouse, Yumi Kasai, David Gray, Carol Clugston, Anna Dominiczak and Alex Alderton, Roberto Amato, Jeffrey Barrett, Sonia Goncalves, Ewan Harrison, David K. Jackson, Ian Johnston, Dominic Kwiatkowski, Cordelia Langford, John Sillitoe on behalf of the Wellcome Sanger Institute COVID-19 Surveillance Team |
| EPI_ISL_2960702 | Berkshire and Surrey Pathology Services Lighthouse Laboratory | Wellcome Sanger Institute for the COVID-19 Genomics UK (COG-UK) Consortium | Berkshire and Surrey Pathology Services Lighthouse Laboratory and Alex Alderton, Roberto Amato, Jeffrey Barrett, Sonia Goncalves, Ewan Harrison, David K. Jackson, Ian Johnston, Dominic Kwiatkowski, Cordelia Langford, John Sillitoe on behalf of the Wellcome Sanger Institute COVID-19 Surveillance Team |
| EPI_ISL_2960703 | Lighthouse Lab in Glasgow | Wellcome Sanger Institute for the COVID-19 Genomics UK (COG-UK) Consortium | Harper VanSteenhouse, Yumi Kasai, David Gray, Carol Clugston, Anna Dominiczak and Alex Alderton, Roberto Amato, Jeffrey Barrett, Sonia Goncalves, Ewan Harrison, David K. Jackson, Ian Johnston, Dominic Kwiatkowski, Cordelia Langford, John Sillitoe on behalf of the Wellcome Sanger Institute COVID-19 Surveillance Team |
| EPI_ISL_2960704 | Berkshire and Surrey Pathology Services Lighthouse Laboratory | Wellcome Sanger Institute for the COVID-19 Genomics UK (COG-UK) Consortium | Berkshire and Surrey Pathology Services Lighthouse Laboratory and Alex Alderton, Roberto Amato, Jeffrey Barrett, Sonia Goncalves, Ewan Harrison, David K. Jackson, Ian Johnston, Dominic Kwiatkowski, Cordelia Langford, John Sillitoe on behalf of the Wellcome Sanger Institute COVID-19 Surveillance Team |
| EPI_ISL_2960705 | Lighthouse Lab in Alderley Park | Wellcome Sanger Institute for the COVID-19 Genomics UK (COG-UK) Consortium | Jacquelyn Wynn, Mairead Hyland, The Lighthouse Lab in Alderley Park and Alex Alderton, Roberto Amato, Jeffrey Barrett, Sonia Goncalves, Ewan Harrison, David K. Jackson, Ian Johnston, Dominic Kwiatkowski, Cordelia Langford, John Sillitoe on behalf of the Wellcome Sanger Institute COVID-19 Surveillance Team |
| EPI_ISL_2960706 | Lighthouse Lab in Glasgow | Wellcome Sanger Institute for the COVID-19 Genomics UK (COG-UK) Consortium | Harper VanSteenhouse, Yumi Kasai, David Gray, Carol Clugston, Anna Dominiczak and Alex Alderton, Roberto Amato, Jeffrey Barrett, Sonia Goncalves, Ewan Harrison, David K. Jackson, Ian Johnston, Dominic Kwiatkowski, Cordelia Langford, John Sillitoe on behalf of the Wellcome Sanger Institute COVID-19 Surveillance Team |
| EPI_ISL_2960707 | Berkshire and Surrey Pathology Services Lighthouse Laboratory | Wellcome Sanger Institute for the COVID-19 Genomics UK (COG-UK) Consortium | Berkshire and Surrey Pathology Services Lighthouse Laboratory and Alex Alderton, Roberto Amato, Jeffrey Barrett, Sonia Goncalves, Ewan Harrison, David K. Jackson, Ian Johnston, Dominic Kwiatkowski, Cordelia Langford, John Sillitoe on behalf of the Wellcome Sanger Institute COVID-19 Surveillance Team |
| EPI_ISL_2960708 | Lighthouse Lab in Alderley Park | Wellcome Sanger Institute for the COVID-19 Genomics UK (COG-UK) Consortium | Jacquelyn Wynn, Mairead Hyland, The Lighthouse Lab in Alderley Park and Alex Alderton, Roberto Amato, Jeffrey Barrett, Sonia Goncalves, Ewan Harrison, David K. Jackson, Ian Johnston, Dominic Kwiatkowski, Cordelia Langford, John Sillitoe on behalf of the Wellcome Sanger Institute COVID-19 Surveillance Team |
| EPI_ISL_2960709, EPI_ISL_2960710, EPI_ISL_2960711 | Lighthouse Lab in Glasgow | Wellcome Sanger Institute for the COVID-19 Genomics UK (COG-UK) Consortium | Harper VanSteenhouse, Yumi Kasai, David Gray, Carol Clugston, Anna Dominiczak and Alex Alderton, Roberto Amato, Jeffrey Barrett, Sonia Goncalves, Ewan Harrison, David K. Jackson, Ian Johnston, Dominic Kwiatkowski, Cordelia Langford, John Sillitoe on behalf of the Wellcome Sanger Institute COVID-19 Surveillance Team |
| EPI_ISL_2960712 | Berkshire and Surrey Pathology Services Lighthouse Laboratory | Wellcome Sanger Institute for the COVID-19 Genomics UK (COG-UK) Consortium | Berkshire and Surrey Pathology Services Lighthouse Laboratory and Alex Alderton, Roberto Amato, Jeffrey Barrett, Sonia Goncalves, Ewan Harrison, David K. Jackson, Ian Johnston, Dominic Kwiatkowski, Cordelia Langford, John Sillitoe on behalf of the Wellcome Sanger Institute COVID-19 Surveillance Team |
| EPI_ISL_2960713 | Lighthouse Lab in Glasgow | Wellcome Sanger Institute for the COVID-19 Genomics UK (COG-UK) Consortium | Harper VanSteenhouse, Yumi Kasai, David Gray, Carol Clugston, Anna Dominiczak and Alex Alderton, Roberto Amato, Jeffrey Barrett, Sonia Goncalves, Ewan Harrison, David K. Jackson, Ian Johnston, Dominic Kwiatkowski, Cordelia Langford, John Sillitoe on behalf of the Wellcome Sanger Institute COVID-19 Surveillance Team |
| EPI_ISL_2960714, EPI_ISL_2960715 | Lighthouse Lab in Alderley Park | Wellcome Sanger Institute for the COVID-19 Genomics UK (COG-UK) Consortium | Jacquelyn Wynn, Mairead Hyland, The Lighthouse Lab in Alderley Park and Alex Alderton, Roberto Amato, Jeffrey Barrett, Sonia Goncalves, Ewan Harrison, David K. Jackson, Ian Johnston, Dominic Kwiatkowski, Cordelia Langford, John Sillitoe on behalf of the Wellcome Sanger Institute COVID-19 Surveillance Team |
| EPI_ISL_2960716 | Berkshire and Surrey Pathology Services Lighthouse Laboratory | Wellcome Sanger Institute for the COVID-19 Genomics UK (COG-UK) Consortium | Berkshire and Surrey Pathology Services Lighthouse Laboratory and Alex Alderton, Roberto Amato, Jeffrey Barrett, Sonia Goncalves, Ewan Harrison, David K. Jackson, Ian Johnston, Dominic Kwiatkowski, Cordelia Langford, John Sillitoe on behalf of the Wellcome Sanger Institute COVID-19 Surveillance Team |
| EPI_ISL_2960717 | Lighthouse Lab in Glasgow | Wellcome Sanger Institute for the COVID-19 Genomics UK (COG-UK) Consortium | Harper VanSteenhouse, Yumi Kasai, David Gray, Carol Clugston, Anna Dominiczak and Alex Alderton, Roberto Amato, Jeffrey Barrett, Sonia Goncalves, Ewan Harrison, David K. Jackson, Ian Johnston, Dominic Kwiatkowski, Cordelia Langford, John Sillitoe on behalf of the Wellcome Sanger Institute COVID-19 Surveillance Team |
| EPI_ISL_2960718 | Berkshire and Surrey Pathology Services Lighthouse Laboratory | Wellcome Sanger Institute for the COVID-19 Genomics UK (COG-UK) Consortium | Berkshire and Surrey Pathology Services Lighthouse Laboratory and Alex Alderton, Roberto Amato, Jeffrey Barrett, Sonia Goncalves, Ewan Harrison, David K. Jackson, Ian Johnston, Dominic Kwiatkowski, Cordelia Langford, John Sillitoe on behalf of the Wellcome Sanger Institute COVID-19 Surveillance Team |
| EPI_ISL_2960719 | Lighthouse Lab in Glasgow | Wellcome Sanger Institute for the COVID-19 Genomics UK (COG-UK) Consortium | Harper VanSteenhouse, Yumi Kasai, David Gray, Carol Clugston, Anna Dominiczak and Alex Alderton, Roberto Amato, Jeffrey Barrett, Sonia Goncalves, Ewan Harrison, David K. Jackson, Ian Johnston, Dominic Kwiatkowski, Cordelia Langford, John Sillitoe on behalf of the Wellcome Sanger Institute COVID-19 Surveillance Team |
| EPI_ISL_2960720 | Lighthouse Lab in Alderley Park | Wellcome Sanger Institute for the COVID-19 Genomics UK (COG-UK) Consortium | Jacquelyn Wynn, Mairead Hyland, The Lighthouse Lab in Alderley Park and Alex Alderton, Roberto Amato, Jeffrey Barrett, Sonia Goncalves, Ewan Harrison, David K. Jackson, Ian Johnston, Dominic Kwiatkowski, Cordelia Langford, John Sillitoe on behalf of the Wellcome Sanger Institute COVID-19 Surveillance Team |
| EPI_ISL_2960721 | Lighthouse Lab in Glasgow | Wellcome Sanger Institute for the COVID-19 Genomics UK (COG-UK) Consortium | Harper VanSteenhouse, Yumi Kasai, David Gray, Carol Clugston, Anna Dominiczak and Alex Alderton, Roberto Amato, Jeffrey Barrett, Sonia Goncalves, Ewan Harrison, David K. Jackson, Ian Johnston, Dominic Kwiatkowski, Cordelia Langford, John Sillitoe on behalf of the Wellcome Sanger Institute COVID-19 Surveillance Team |
| EPI_ISL_2960722 | Berkshire and Surrey Pathology Services Lighthouse Laboratory | Wellcome Sanger Institute for the COVID-19 Genomics UK (COG-UK) Consortium | Berkshire and Surrey Pathology Services Lighthouse Laboratory and Alex Alderton, Roberto Amato, Jeffrey Barrett, Sonia Goncalves, Ewan Harrison, David K. Jackson, Ian Johnston, Dominic Kwiatkowski, Cordelia Langford, John Sillitoe on behalf of the Wellcome Sanger Institute COVID-19 Surveillance Team |
| EPI_ISL_2960723 | Lighthouse Lab in Glasgow | Wellcome Sanger Institute for the COVID-19 Genomics UK (COG-UK) Consortium | Harper VanSteenhouse, Yumi Kasai, David Gray, Carol Clugston, Anna Dominiczak and Alex Alderton, Roberto Amato, Jeffrey Barrett, Sonia Goncalves, Ewan Harrison, David K. Jackson, Ian Johnston, Dominic Kwiatkowski, Cordelia Langford, John Sillitoe on behalf of the Wellcome Sanger Institute COVID-19 Surveillance Team |
| EPI_ISL_2960724, EPI_ISL_2960725 | Berkshire and Surrey Pathology Services Lighthouse Laboratory | Wellcome Sanger Institute for the COVID-19 Genomics UK (COG-UK) Consortium | Berkshire and Surrey Pathology Services Lighthouse Laboratory and Alex Alderton, Roberto Amato, Jeffrey Barrett, Sonia Goncalves, Ewan Harrison, David K. Jackson, Ian Johnston, Dominic Kwiatkowski, Cordelia Langford, John Sillitoe on behalf of the Wellcome Sanger Institute COVID-19 Surveillance Team |
| EPI_ISL_2960726 | Lighthouse Lab in Alderley Park | Wellcome Sanger Institute for the COVID-19 Genomics UK (COG-UK) Consortium | Jacquelyn Wynn, Mairead Hyland, The Lighthouse Lab in Alderley Park and Alex Alderton, Roberto Amato, Jeffrey Barrett, Sonia Goncalves, Ewan Harrison, David K. Jackson, Ian Johnston, Dominic Kwiatkowski, Cordelia Langford, John Sillitoe on behalf of the Wellcome Sanger Institute COVID-19 Surveillance Team |
| EPI_ISL_2960727 | Berkshire and Surrey Pathology Services Lighthouse Laboratory | Wellcome Sanger Institute for the COVID-19 Genomics UK (COG-UK) Consortium | Berkshire and Surrey Pathology Services Lighthouse Laboratory and Alex Alderton, Roberto Amato, Jeffrey Barrett, Sonia Goncalves, Ewan Harrison, David K. Jackson, Ian Johnston, Dominic Kwiatkowski, Cordelia Langford, John Sillitoe on behalf of the Wellcome Sanger Institute COVID-19 Surveillance Team |
| EPI_ISL_2960728 | Lighthouse Lab in Alderley Park | Wellcome Sanger Institute for the COVID-19 Genomics UK (COG-UK) Consortium | Jacquelyn Wynn, Mairead Hyland, The Lighthouse Lab in Alderley Park and Alex Alderton, Roberto Amato, Jeffrey Barrett, Sonia Goncalves, Ewan Harrison, David K. Jackson, Ian Johnston, Dominic Kwiatkowski, Cordelia Langford, John Sillitoe on behalf of the Wellcome Sanger Institute COVID-19 Surveillance Team |
| EPI_ISL_2960729 | Berkshire and Surrey Pathology Services Lighthouse Laboratory | Wellcome Sanger Institute for the COVID-19 Genomics UK (COG-UK) Consortium | Berkshire and Surrey Pathology Services Lighthouse Laboratory and Alex Alderton, Roberto Amato, Jeffrey Barrett, Sonia Goncalves, Ewan Harrison, David K. Jackson, Ian Johnston, Dominic Kwiatkowski, Cordelia Langford, John Sillitoe on behalf of the Wellcome Sanger Institute COVID-19 Surveillance Team |
| EPI_ISL_2960730 | Lighthouse Lab in Glasgow | Wellcome Sanger Institute for the COVID-19 Genomics UK (COG-UK) Consortium | Harper VanSteenhouse, Yumi Kasai, David Gray, Carol Clugston, Anna Dominiczak and Alex Alderton, Roberto Amato, Jeffrey Barrett, Sonia Goncalves, Ewan Harrison, David K. Jackson, Ian Johnston, Dominic Kwiatkowski, Cordelia Langford, John Sillitoe on behalf of the Wellcome Sanger Institute COVID-19 Surveillance Team |
| EPI_ISL_2960731 | Lighthouse Lab in Alderley Park | Wellcome Sanger Institute for the COVID-19 Genomics UK (COG-UK) Consortium | Jacquelyn Wynn, Mairead Hyland, The Lighthouse Lab in Alderley Park and Alex Alderton, Roberto Amato, Jeffrey Barrett, Sonia Goncalves, Ewan Harrison, David K. Jackson, Ian Johnston, Dominic Kwiatkowski, Cordelia Langford, John Sillitoe on behalf of the Wellcome Sanger Institute COVID-19 Surveillance Team |

| | | | |
|---|---|---|---|
| EPI_ISL_2960732 | Lighthouse Lab in Glasgow | Wellcome Sanger Institute for the COVID-19 Genomics UK (COG-UK) Consortium | Harper VanSteenhouse, Yumi Kasai, David Gray, Carol Clugston, Anna Dominiczak and Alex Alderton, Roberto Amato, Jeffrey Barrett, Sonia Goncalves, Ewan Harrison, David K. Jackson, Ian Johnston, Dominic Kwiatkowski, Cordelia Langford, John Sillitoe on behalf of the Wellcome Sanger Institute COVID-19 Surveillance Team |
| EPI_ISL_2960733 | Lighthouse Lab in Alderley Park | Wellcome Sanger Institute for the COVID-19 Genomics UK (COG-UK) Consortium | Jacquelyn Wynn, Mairead Hyland, The Lighthouse Lab in Alderley Park and Alex Alderton, Roberto Amato, Jeffrey Barrett, Sonia Goncalves, Ewan Harrison, David K. Jackson, Ian Johnston, Dominic Kwiatkowski, Cordelia Langford, John Sillitoe on behalf of the Wellcome Sanger Institute COVID-19 Surveillance Team |
| EPI_ISL_2960734, EPI_ISL_2960735, EPI_ISL_2960736 | Lighthouse Lab in Glasgow | Wellcome Sanger Institute for the COVID-19 Genomics UK (COG-UK) Consortium | Harper VanSteenhouse, Yumi Kasai, David Gray, Carol Clugston, Anna Dominiczak and Alex Alderton, Roberto Amato, Jeffrey Barrett, Sonia Goncalves, Ewan Harrison, David K. Jackson, Ian Johnston, Dominic Kwiatkowski, Cordelia Langford, John Sillitoe on behalf of the Wellcome Sanger Institute COVID-19 Surveillance Team |
| EPI_ISL_2960737, EPI_ISL_2960738 | Lighthouse Lab in Alderley Park | Wellcome Sanger Institute for the COVID-19 Genomics UK (COG-UK) Consortium | Jacquelyn Wynn, Mairead Hyland, The Lighthouse Lab in Alderley Park and Alex Alderton, Roberto Amato, Jeffrey Barrett, Sonia Goncalves, Ewan Harrison, David K. Jackson, Ian Johnston, Dominic Kwiatkowski, Cordelia Langford, John Sillitoe on behalf of the Wellcome Sanger Institute COVID-19 Surveillance Team |
| EPI_ISL_2960739 | Lighthouse Lab in Glasgow | Wellcome Sanger Institute for the COVID-19 Genomics UK (COG-UK) Consortium | Harper VanSteenhouse, Yumi Kasai, David Gray, Carol Clugston, Anna Dominiczak and Alex Alderton, Roberto Amato, Jeffrey Barrett, Sonia Goncalves, Ewan Harrison, David K. Jackson, Ian Johnston, Dominic Kwiatkowski, Cordelia Langford, John Sillitoe on behalf of the Wellcome Sanger Institute COVID-19 Surveillance Team |
| EPI_ISL_2960740, EPI_ISL_2960741 | Lighthouse Lab in Alderley Park | Wellcome Sanger Institute for the COVID-19 Genomics UK (COG-UK) Consortium | Jacquelyn Wynn, Mairead Hyland, The Lighthouse Lab in Alderley Park and Alex Alderton, Roberto Amato, Jeffrey Barrett, Sonia Goncalves, Ewan Harrison, David K. Jackson, Ian Johnston, Dominic Kwiatkowski, Cordelia Langford, John Sillitoe on behalf of the Wellcome Sanger Institute COVID-19 Surveillance Team |
| EPI_ISL_2960742 | Lighthouse Lab in Glasgow | Wellcome Sanger Institute for the COVID-19 Genomics UK (COG-UK) Consortium | Harper VanSteenhouse, Yumi Kasai, David Gray, Carol Clugston, Anna Dominiczak and Alex Alderton, Roberto Amato, Jeffrey Barrett, Sonia Goncalves, Ewan Harrison, David K. Jackson, Ian Johnston, Dominic Kwiatkowski, Cordelia Langford, John Sillitoe on behalf of the Wellcome Sanger Institute COVID-19 Surveillance Team |
| EPI_ISL_2960743, EPI_ISL_2960744 | Lighthouse Lab in Alderley Park | Wellcome Sanger Institute for the COVID-19 Genomics UK (COG-UK) Consortium | Jacquelyn Wynn, Mairead Hyland, The Lighthouse Lab in Alderley Park and Alex Alderton, Roberto Amato, Jeffrey Barrett, Sonia Goncalves, Ewan Harrison, David K. Jackson, Ian Johnston, Dominic Kwiatkowski, Cordelia Langford, John Sillitoe on behalf of the Wellcome Sanger Institute COVID-19 Surveillance Team |
| EPI_ISL_2960745 | Berkshire and Surrey Pathology Services Lighthouse Laboratory | Wellcome Sanger Institute for the COVID-19 Genomics UK (COG-UK) Consortium | Berkshire and Surrey Pathology Services Lighthouse Laboratory and Alex Alderton, Roberto Amato, Jeffrey Barrett, Sonia Goncalves, Ewan Harrison, David K. Jackson, Ian Johnston, Dominic Kwiatkowski, Cordelia Langford, John Sillitoe on behalf of the Wellcome Sanger Institute COVID-19 Surveillance Team |
| EPI_ISL_2960746 | Lighthouse Lab in Alderley Park | Wellcome Sanger Institute for the COVID-19 Genomics UK (COG-UK) Consortium | Jacquelyn Wynn, Mairead Hyland, The Lighthouse Lab in Alderley Park and Alex Alderton, Roberto Amato, Jeffrey Barrett, Sonia Goncalves, Ewan Harrison, David K. Jackson, Ian Johnston, Dominic Kwiatkowski, Cordelia Langford, John Sillitoe on behalf of the Wellcome Sanger Institute COVID-19 Surveillance Team |
| EPI_ISL_2960747 | Berkshire and Surrey Pathology Services Lighthouse Laboratory | Wellcome Sanger Institute for the COVID-19 Genomics UK (COG-UK) Consortium | Berkshire and Surrey Pathology Services Lighthouse Laboratory and Alex Alderton, Roberto Amato, Jeffrey Barrett, Sonia Goncalves, Ewan Harrison, David K. Jackson, Ian Johnston, Dominic Kwiatkowski, Cordelia Langford, John Sillitoe on behalf of the Wellcome Sanger Institute COVID-19 Surveillance Team |
| EPI_ISL_2960748 | Lighthouse Lab in Glasgow | Wellcome Sanger Institute for the COVID-19 Genomics UK (COG-UK) Consortium | Harper VanSteenhouse, Yumi Kasai, David Gray, Carol Clugston, Anna Dominiczak and Alex Alderton, Roberto Amato, Jeffrey Barrett, Sonia Goncalves, Ewan Harrison, David K. Jackson, Ian Johnston, Dominic Kwiatkowski, Cordelia Langford, John Sillitoe on behalf of the Wellcome Sanger Institute COVID-19 Surveillance Team |
| EPI_ISL_2960749 | Berkshire and Surrey Pathology Services Lighthouse Laboratory | Wellcome Sanger Institute for the COVID-19 Genomics UK (COG-UK) Consortium | Berkshire and Surrey Pathology Services Lighthouse Laboratory and Alex Alderton, Roberto Amato, Jeffrey Barrett, Sonia Goncalves, Ewan Harrison, David K. Jackson, Ian Johnston, Dominic Kwiatkowski, Cordelia Langford, John Sillitoe on behalf of the Wellcome Sanger Institute COVID-19 Surveillance Team |
| EPI_ISL_2960750 | Lighthouse Lab in Alderley Park | Wellcome Sanger Institute for the COVID-19 Genomics UK (COG-UK) Consortium | Jacquelyn Wynn, Mairead Hyland, The Lighthouse Lab in Alderley Park and Alex Alderton, Roberto Amato, Jeffrey Barrett, Sonia Goncalves, Ewan Harrison, David K. Jackson, Ian Johnston, Dominic Kwiatkowski, Cordelia Langford, John Sillitoe on behalf of the Wellcome Sanger Institute COVID-19 Surveillance Team |
| EPI_ISL_2960751, EPI_ISL_2960752 | Lighthouse Lab in Glasgow | Wellcome Sanger Institute for the COVID-19 Genomics UK (COG-UK) Consortium | Harper VanSteenhouse, Yumi Kasai, David Gray, Carol Clugston, Anna Dominiczak and Alex Alderton, Roberto Amato, Jeffrey Barrett, Sonia Goncalves, Ewan Harrison, David K. Jackson, Ian Johnston, Dominic Kwiatkowski, Cordelia Langford, John Sillitoe on behalf of the Wellcome Sanger Institute COVID-19 Surveillance Team |
| EPI_ISL_2960753 | Berkshire and Surrey Pathology Services Lighthouse Laboratory | Wellcome Sanger Institute for the COVID-19 Genomics UK (COG-UK) Consortium | Berkshire and Surrey Pathology Services Lighthouse Laboratory and Alex Alderton, Roberto Amato, Jeffrey Barrett, Sonia Goncalves, Ewan Harrison, David K. Jackson, Ian Johnston, Dominic Kwiatkowski, Cordelia Langford, John Sillitoe on behalf of the Wellcome Sanger Institute COVID-19 Surveillance Team |
| EPI_ISL_2960754, EPI_ISL_2960755 | Lighthouse Lab in Glasgow | Wellcome Sanger Institute for the COVID-19 Genomics UK (COG-UK) Consortium | Harper VanSteenhouse, Yumi Kasai, David Gray, Carol Clugston, Anna Dominiczak and Alex Alderton, Roberto Amato, Jeffrey Barrett, Sonia Goncalves, Ewan Harrison, David K. Jackson, Ian Johnston, Dominic Kwiatkowski, Cordelia Langford, John Sillitoe on behalf of the Wellcome Sanger Institute COVID-19 Surveillance Team |
| EPI_ISL_2960756 | Berkshire and Surrey Pathology Services Lighthouse Laboratory | Wellcome Sanger Institute for the COVID-19 Genomics UK (COG-UK) Consortium | Berkshire and Surrey Pathology Services Lighthouse Laboratory and Alex Alderton, Roberto Amato, Jeffrey Barrett, Sonia Goncalves, Ewan Harrison, David K. Jackson, Ian Johnston, Dominic Kwiatkowski, Cordelia Langford, John Sillitoe on behalf of the Wellcome Sanger Institute COVID-19 Surveillance Team |
| EPI_ISL_2960757 | Lighthouse Lab in Alderley Park | Wellcome Sanger Institute for the COVID-19 Genomics UK (COG-UK) Consortium | Jacquelyn Wynn, Mairead Hyland, The Lighthouse Lab in Alderley Park and Alex Alderton, Roberto Amato, Jeffrey Barrett, Sonia Goncalves, Ewan Harrison, David K. Jackson, Ian Johnston, Dominic Kwiatkowski, Cordelia Langford, John Sillitoe on behalf of the Wellcome Sanger Institute COVID-19 Surveillance Team |
| EPI_ISL_2960758, EPI_ISL_2960759, EPI_ISL_2960760 | Berkshire and Surrey Pathology Services Lighthouse Laboratory | Wellcome Sanger Institute for the COVID-19 Genomics UK (COG-UK) Consortium | Berkshire and Surrey Pathology Services Lighthouse Laboratory and Alex Alderton, Roberto Amato, Jeffrey Barrett, Sonia Goncalves, Ewan Harrison, David K. Jackson, Ian Johnston, Dominic Kwiatkowski, Cordelia Langford, John Sillitoe on behalf of the Wellcome Sanger Institute COVID-19 Surveillance Team |
| EPI_ISL_2960761, EPI_ISL_2960762, EPI_ISL_2960763 | Lighthouse Lab in Glasgow | Wellcome Sanger Institute for the COVID-19 Genomics UK (COG-UK) Consortium | Harper VanSteenhouse, Yumi Kasai, David Gray, Carol Clugston, Anna Dominiczak and Alex Alderton, Roberto Amato, Jeffrey Barrett, Sonia Goncalves, Ewan Harrison, David K. Jackson, Ian Johnston, Dominic Kwiatkowski, Cordelia Langford, John Sillitoe on behalf of the Wellcome Sanger Institute COVID-19 Surveillance Team |
| EPI_ISL_2960764 | Berkshire and Surrey Pathology Services Lighthouse Laboratory | Wellcome Sanger Institute for the COVID-19 Genomics UK (COG-UK) Consortium | Berkshire and Surrey Pathology Services Lighthouse Laboratory and Alex Alderton, Roberto Amato, Jeffrey Barrett, Sonia Goncalves, Ewan Harrison, David K. Jackson, Ian Johnston, Dominic Kwiatkowski, Cordelia Langford, John Sillitoe on behalf of the Wellcome Sanger Institute COVID-19 Surveillance Team |
| EPI_ISL_2960765 | Lighthouse Lab in Glasgow | Wellcome Sanger Institute for the COVID-19 Genomics UK (COG-UK) Consortium | Harper VanSteenhouse, Yumi Kasai, David Gray, Carol Clugston, Anna Dominiczak and Alex Alderton, Roberto Amato, Jeffrey Barrett, Sonia Goncalves, Ewan Harrison, David K. Jackson, Ian Johnston, Dominic Kwiatkowski, Cordelia Langford, John Sillitoe on behalf of the Wellcome Sanger Institute COVID-19 Surveillance Team |
| EPI_ISL_2960766, EPI_ISL_2960767 | Lighthouse Lab in Alderley Park | Wellcome Sanger Institute for the COVID-19 Genomics UK (COG-UK) Consortium | Jacquelyn Wynn, Mairead Hyland, The Lighthouse Lab in Alderley Park and Alex Alderton, Roberto Amato, Jeffrey Barrett, Sonia Goncalves, Ewan Harrison, David K. Jackson, Ian Johnston, Dominic Kwiatkowski, Cordelia Langford, John Sillitoe on behalf of the Wellcome Sanger Institute COVID-19 Surveillance Team |
| EPI_ISL_2960768 | Berkshire and Surrey Pathology Services Lighthouse Laboratory | Wellcome Sanger Institute for the COVID-19 Genomics UK (COG-UK) Consortium | Berkshire and Surrey Pathology Services Lighthouse Laboratory and Alex Alderton, Roberto Amato, Jeffrey Barrett, Sonia Goncalves, Ewan Harrison, David K. Jackson, Ian Johnston, Dominic Kwiatkowski, Cordelia Langford, John Sillitoe on behalf of the Wellcome Sanger Institute COVID-19 Surveillance Team |
| EPI_ISL_2960769 | Lighthouse Lab in Glasgow | Wellcome Sanger Institute for the COVID-19 Genomics UK (COG-UK) Consortium | Harper VanSteenhouse, Yumi Kasai, David Gray, Carol Clugston, Anna Dominiczak and Alex Alderton, Roberto Amato, Jeffrey Barrett, Sonia Goncalves, Ewan Harrison, David K. Jackson, Ian Johnston, Dominic Kwiatkowski, Cordelia Langford, John Sillitoe on behalf of the Wellcome Sanger Institute COVID-19 Surveillance Team |
| EPI_ISL_2960770 | Lighthouse Lab in Alderley Park | Wellcome Sanger Institute for the COVID-19 Genomics UK (COG-UK) Consortium | Jacquelyn Wynn, Mairead Hyland, The Lighthouse Lab in Alderley Park and Alex Alderton, Roberto Amato, Jeffrey Barrett, Sonia Goncalves, Ewan Harrison, David K. Jackson, Ian Johnston, Dominic Kwiatkowski, Cordelia Langford, John Sillitoe on behalf of the Wellcome Sanger Institute COVID-19 Surveillance Team |
| EPI_ISL_2960771 | Lighthouse Lab in Glasgow | Wellcome Sanger Institute for the COVID-19 Genomics UK (COG-UK) Consortium | Harper VanSteenhouse, Yumi Kasai, David Gray, Carol Clugston, Anna Dominiczak and Alex Alderton, Roberto Amato, Jeffrey Barrett, Sonia Goncalves, Ewan Harrison, David K. Jackson, Ian Johnston, Dominic Kwiatkowski, Cordelia Langford, John Sillitoe on behalf of the Wellcome Sanger Institute COVID-19 Surveillance Team |
| EPI_ISL_2960772, EPI_ISL_2960773 | Lighthouse Lab in Alderley Park | Wellcome Sanger Institute for the COVID-19 Genomics UK (COG-UK) Consortium | Jacquelyn Wynn, Mairead Hyland, The Lighthouse Lab in Alderley Park and Alex Alderton, Roberto Amato, Jeffrey Barrett, Sonia Goncalves, Ewan Harrison, David K. Jackson, Ian Johnston, Dominic Kwiatkowski, Cordelia Langford, John Sillitoe on behalf of the Wellcome Sanger Institute COVID-19 Surveillance Team |
| EPI_ISL_2960774 | Berkshire and Surrey Pathology Services Lighthouse Laboratory | Wellcome Sanger Institute for the COVID-19 Genomics UK (COG-UK) Consortium | Berkshire and Surrey Pathology Services Lighthouse Laboratory and Alex Alderton, Roberto Amato, Jeffrey Barrett, Sonia Goncalves, Ewan Harrison, David K. Jackson, Ian Johnston, Dominic Kwiatkowski, Cordelia Langford, John Sillitoe on behalf of the Wellcome Sanger Institute COVID-19 Surveillance Team |
| EPI_ISL_2960775, EPI_ISL_2960776 | Lighthouse Lab in Glasgow | Wellcome Sanger Institute for the COVID-19 Genomics UK (COG-UK) Consortium | Harper VanSteenhouse, Yumi Kasai, David Gray, Carol Clugston, Anna Dominiczak and Alex Alderton, Roberto Amato, Jeffrey Barrett, Sonia Goncalves, Ewan Harrison, David K. Jackson, Ian Johnston, Dominic Kwiatkowski, Cordelia Langford, John Sillitoe on behalf of the Wellcome Sanger Institute COVID-19 Surveillance Team |
| EPI_ISL_2960777 | Berkshire and Surrey Pathology Services Lighthouse Laboratory | Wellcome Sanger Institute for the COVID-19 Genomics UK (COG-UK) Consortium | Berkshire and Surrey Pathology Services Lighthouse Laboratory and Alex Alderton, Roberto Amato, Jeffrey Barrett, Sonia Goncalves, Ewan Harrison, David K. Jackson, Ian Johnston, Dominic Kwiatkowski, Cordelia Langford, John Sillitoe on behalf of the Wellcome Sanger Institute COVID-19 Surveillance Team |
| EPI_ISL_2960778, EPI_ISL_2960779 | Lighthouse Lab in Glasgow | Wellcome Sanger Institute for the COVID-19 Genomics UK (COG-UK) Consortium | Harper VanSteenhouse, Yumi Kasai, David Gray, Carol Clugston, Anna Dominiczak and Alex Alderton, Roberto Amato, Jeffrey Barrett, Sonia Goncalves, Ewan Harrison, David K. Jackson, Ian Johnston, Dominic Kwiatkowski, Cordelia Langford, John Sillitoe on behalf of the Wellcome Sanger Institute COVID-19 Surveillance Team |
| EPI_ISL_2960780, EPI_ISL_2960781 | Lighthouse Lab in Alderley Park | Wellcome Sanger Institute for the COVID-19 Genomics UK (COG-UK) Consortium | Jacquelyn Wynn, Mairead Hyland, The Lighthouse Lab in Alderley Park and Alex Alderton, Roberto Amato, Jeffrey Barrett, Sonia Goncalves, Ewan Harrison, David K. Jackson, Ian Johnston, Dominic Kwiatkowski, Cordelia Langford, John Sillitoe on behalf of the Wellcome Sanger Institute COVID-19 Surveillance Team |
| EPI_ISL_2960782, EPI_ISL_2960783 | Lighthouse Lab in Glasgow | Wellcome Sanger Institute for the COVID-19 Genomics UK (COG-UK) Consortium | Harper VanSteenhouse, Yumi Kasai, David Gray, Carol Clugston, Anna Dominiczak and Alex Alderton, Roberto Amato, Jeffrey Barrett, Sonia Goncalves, Ewan Harrison, David K. Jackson, Ian Johnston, Dominic Kwiatkowski, Cordelia Langford, John Sillitoe on behalf of the Wellcome Sanger Institute COVID-19 Surveillance Team |
| EPI_ISL_2960784, EPI_ISL_2960785, EPI_ISL_2960786, EPI_ISL_2960787 | Lighthouse Lab in Alderley Park | Wellcome Sanger Institute for the COVID-19 Genomics UK (COG-UK) Consortium | Jacquelyn Wynn, Mairead Hyland, The Lighthouse Lab in Alderley Park and Alex Alderton, Roberto Amato, Jeffrey Barrett, Sonia Goncalves, Ewan Harrison, David K. Jackson, Ian Johnston, Dominic Kwiatkowski, Cordelia Langford, John Sillitoe on behalf of the Wellcome Sanger Institute COVID-19 Surveillance Team |
| EPI_ISL_2960788 | Berkshire and Surrey Pathology Services Lighthouse Laboratory | Wellcome Sanger Institute for the COVID-19 Genomics UK (COG-UK) Consortium | Berkshire and Surrey Pathology Services Lighthouse Laboratory and Alex Alderton, Roberto Amato, Jeffrey Barrett, Sonia Goncalves, Ewan Harrison, David K. Jackson, Ian Johnston, Dominic Kwiatkowski, Cordelia Langford, John Sillitoe on behalf of the Wellcome Sanger Institute COVID-19 Surveillance Team |
| EPI_ISL_2960789 | Lighthouse Lab in Glasgow | Wellcome Sanger Institute for the COVID-19 Genomics UK (COG-UK) Consortium | Harper VanSteenhouse, Yumi Kasai, David Gray, Carol Clugston, Anna Dominiczak and Alex Alderton, Roberto Amato, Jeffrey Barrett, Sonia Goncalves, Ewan Harrison, David K. Jackson, Ian Johnston, Dominic Kwiatkowski, Cordelia Langford, John Sillitoe on behalf of the Wellcome Sanger Institute COVID-19 Surveillance Team |
| EPI_ISL_2960790 | Berkshire and Surrey Pathology Services Lighthouse Laboratory | Wellcome Sanger Institute for the COVID-19 Genomics UK (COG-UK) Consortium | Berkshire and Surrey Pathology Services Lighthouse Laboratory and Alex Alderton, Roberto Amato, Jeffrey Barrett, Sonia Goncalves, Ewan Harrison, David K. Jackson, Ian Johnston, Dominic Kwiatkowski, Cordelia Langford, John Sillitoe on behalf of the Wellcome Sanger Institute COVID-19 Surveillance Team |
| EPI_ISL_2960791, EPI_ISL_2960792 | Lighthouse Lab in Glasgow | Wellcome Sanger Institute for the COVID-19 Genomics UK (COG-UK) Consortium | Harper VanSteenhouse, Yumi Kasai, David Gray, Carol Clugston, Anna Dominiczak and Alex Alderton, Roberto Amato, Jeffrey Barrett, Sonia Goncalves, Ewan Harrison, David K. Jackson, Ian Johnston, Dominic Kwiatkowski, Cordelia Langford, John Sillitoe on behalf of the Wellcome Sanger Institute COVID-19 Surveillance Team |
| EPI_ISL_2960793 | Lighthouse Lab in Alderley Park | Wellcome Sanger Institute for the COVID-19 Genomics UK (COG-UK) Consortium | Jacquelyn Wynn, Mairead Hyland, The Lighthouse Lab in Alderley Park and Alex Alderton, Roberto Amato, Jeffrey Barrett, Sonia Goncalves, Ewan Harrison, David K. Jackson, Ian Johnston, Dominic Kwiatkowski, Cordelia Langford, John Sillitoe on behalf of the Wellcome Sanger Institute COVID-19 Surveillance Team |
| EPI_ISL_2960794 | Lighthouse Lab in Glasgow | Wellcome Sanger Institute for the COVID-19 Genomics UK (COG-UK) Consortium | Harper VanSteenhouse, Yumi Kasai, David Gray, Carol Clugston, Anna Dominiczak and Alex Alderton, Roberto Amato, Jeffrey Barrett, Sonia Goncalves, Ewan Harrison, David K. Jackson, Ian Johnston, Dominic Kwiatkowski, Cordelia Langford, John Sillitoe on behalf of the Wellcome Sanger Institute COVID-19 Surveillance Team |
| EPI_ISL_2960795 | Lighthouse Lab in Alderley Park | Wellcome Sanger Institute for the COVID-19 Genomics UK (COG-UK) Consortium | Jacquelyn Wynn, Mairead Hyland, The Lighthouse Lab in Alderley Park and Alex Alderton, Roberto Amato, Jeffrey Barrett, Sonia Goncalves, Ewan Harrison, David K. Jackson, Ian Johnston, Dominic Kwiatkowski, Cordelia Langford, John Sillitoe on behalf of the Wellcome Sanger Institute COVID-19 Surveillance Team |

| | | | |
|---|---|---|---|
| EPI_ISL_2960796 | Berkshire and Surrey Pathology Services Lighthouse Laboratory | Wellcome Sanger Institute for the COVID-19 Genomics UK (COG-UK) Consortium | Berkshire and Surrey Pathology Services Lighthouse Laboratory and Alex Alderton, Roberto Amato, Jeffrey Barrett, Sonia Goncalves, Ewan Harrison, David K. Jackson, Ian Johnston, Dominic Kwiatkowski, Cordelia Langford, John Sillitoe on behalf of the Wellcome Sanger Institute COVID-19 Surveillance Team |
| EPI_ISL_2960797 | Lighthouse Lab in Glasgow | Wellcome Sanger Institute for the COVID-19 Genomics UK (COG-UK) Consortium | Harper VanSteenhouse, Yumi Kasai, David Gray, Carol Clugston, Anna Dominiczak and Alex Alderton, Roberto Amato, Jeffrey Barrett, Sonia Goncalves, Ewan Harrison, David K. Jackson, Ian Johnston, Dominic Kwiatkowski, Cordelia Langford, John Sillitoe on behalf of the Wellcome Sanger Institute COVID-19 Surveillance Team |
| EPI_ISL_2960798, EPI_ISL_2960799 | Berkshire and Surrey Pathology Services Lighthouse Laboratory | Wellcome Sanger Institute for the COVID-19 Genomics UK (COG-UK) Consortium | Berkshire and Surrey Pathology Services Lighthouse Laboratory and Alex Alderton, Roberto Amato, Jeffrey Barrett, Sonia Goncalves, Ewan Harrison, David K. Jackson, Ian Johnston, Dominic Kwiatkowski, Cordelia Langford, John Sillitoe on behalf of the Wellcome Sanger Institute COVID-19 Surveillance Team |
| EPI_ISL_2960800 | Lighthouse Lab in Alderley Park | Wellcome Sanger Institute for the COVID-19 Genomics UK (COG-UK) Consortium | Jacquelyn Wynn, Mairead Hyland, The Lighthouse Lab in Alderley Park and Alex Alderton, Roberto Amato, Jeffrey Barrett, Sonia Goncalves, Ewan Harrison, David K. Jackson, Ian Johnston, Dominic Kwiatkowski, Cordelia Langford, John Sillitoe on behalf of the Wellcome Sanger Institute COVID-19 Surveillance Team |
| EPI_ISL_2960801, EPI_ISL_2960802, EPI_ISL_2960803, EPI_ISL_2960804, EPI_ISL_2960805, EPI_ISL_2960806, EPI_ISL_2960807, EPI_ISL_2960808 | Lighthouse Lab in Glasgow | Wellcome Sanger Institute for the COVID-19 Genomics UK (COG-UK) Consortium | Harper VanSteenhouse, Yumi Kasai, David Gray, Carol Clugston, Anna Dominiczak and Alex Alderton, Roberto Amato, Jeffrey Barrett, Sonia Goncalves, Ewan Harrison, David K. Jackson, Ian Johnston, Dominic Kwiatkowski, Cordelia Langford, John Sillitoe on behalf of the Wellcome Sanger Institute COVID-19 Surveillance Team |
| EPI_ISL_2960809 | Berkshire and Surrey Pathology Services Lighthouse Laboratory | Wellcome Sanger Institute for the COVID-19 Genomics UK (COG-UK) Consortium | Berkshire and Surrey Pathology Services Lighthouse Laboratory and Alex Alderton, Roberto Amato, Jeffrey Barrett, Sonia Goncalves, Ewan Harrison, David K. Jackson, Ian Johnston, Dominic Kwiatkowski, Cordelia Langford, John Sillitoe on behalf of the Wellcome Sanger Institute COVID-19 Surveillance Team |
| EPI_ISL_2960810 | Lighthouse Lab in Glasgow | Wellcome Sanger Institute for the COVID-19 Genomics UK (COG-UK) Consortium | Harper VanSteenhouse, Yumi Kasai, David Gray, Carol Clugston, Anna Dominiczak and Alex Alderton, Roberto Amato, Jeffrey Barrett, Sonia Goncalves, Ewan Harrison, David K. Jackson, Ian Johnston, Dominic Kwiatkowski, Cordelia Langford, John Sillitoe on behalf of the Wellcome Sanger Institute COVID-19 Surveillance Team |
| EPI_ISL_2960811 | Berkshire and Surrey Pathology Services Lighthouse Laboratory | Wellcome Sanger Institute for the COVID-19 Genomics UK (COG-UK) Consortium | Berkshire and Surrey Pathology Services Lighthouse Laboratory and Alex Alderton, Roberto Amato, Jeffrey Barrett, Sonia Goncalves, Ewan Harrison, David K. Jackson, Ian Johnston, Dominic Kwiatkowski, Cordelia Langford, John Sillitoe on behalf of the Wellcome Sanger Institute COVID-19 Surveillance Team |
| EPI_ISL_2960812 | Lighthouse Lab in Alderley Park | Wellcome Sanger Institute for the COVID-19 Genomics UK (COG-UK) Consortium | Jacquelyn Wynn, Mairead Hyland, The Lighthouse Lab in Alderley Park and Alex Alderton, Roberto Amato, Jeffrey Barrett, Sonia Goncalves, Ewan Harrison, David K. Jackson, Ian Johnston, Dominic Kwiatkowski, Cordelia Langford, John Sillitoe on behalf of the Wellcome Sanger Institute COVID-19 Surveillance Team |
| EPI_ISL_2960813 | Berkshire and Surrey Pathology Services Lighthouse Laboratory | Wellcome Sanger Institute for the COVID-19 Genomics UK (COG-UK) Consortium | Berkshire and Surrey Pathology Services Lighthouse Laboratory and Alex Alderton, Roberto Amato, Jeffrey Barrett, Sonia Goncalves, Ewan Harrison, David K. Jackson, Ian Johnston, Dominic Kwiatkowski, Cordelia Langford, John Sillitoe on behalf of the Wellcome Sanger Institute COVID-19 Surveillance Team |
| EPI_ISL_2960814, EPI_ISL_2960815, EPI_ISL_2960816 | Lighthouse Lab in Glasgow | Wellcome Sanger Institute for the COVID-19 Genomics UK (COG-UK) Consortium | Harper VanSteenhouse, Yumi Kasai, David Gray, Carol Clugston, Anna Dominiczak and Alex Alderton, Roberto Amato, Jeffrey Barrett, Sonia Goncalves, Ewan Harrison, David K. Jackson, Ian Johnston, Dominic Kwiatkowski, Cordelia Langford, John Sillitoe on behalf of the Wellcome Sanger Institute COVID-19 Surveillance Team |
| EPI_ISL_2960817 | Lighthouse Lab in Alderley Park | Wellcome Sanger Institute for the COVID-19 Genomics UK (COG-UK) Consortium | Jacquelyn Wynn, Mairead Hyland, The Lighthouse Lab in Alderley Park and Alex Alderton, Roberto Amato, Jeffrey Barrett, Sonia Goncalves, Ewan Harrison, David K. Jackson, Ian Johnston, Dominic Kwiatkowski, Cordelia Langford, John Sillitoe on behalf of the Wellcome Sanger Institute COVID-19 Surveillance Team |
| EPI_ISL_2960818 | Lighthouse Lab in Glasgow | Wellcome Sanger Institute for the COVID-19 Genomics UK (COG-UK) Consortium | Harper VanSteenhouse, Yumi Kasai, David Gray, Carol Clugston, Anna Dominiczak and Alex Alderton, Roberto Amato, Jeffrey Barrett, Sonia Goncalves, Ewan Harrison, David K. Jackson, Ian Johnston, Dominic Kwiatkowski, Cordelia Langford, John Sillitoe on behalf of the Wellcome Sanger Institute COVID-19 Surveillance Team |
| EPI_ISL_2960819 | Berkshire and Surrey Pathology Services Lighthouse Laboratory | Wellcome Sanger Institute for the COVID-19 Genomics UK (COG-UK) Consortium | Berkshire and Surrey Pathology Services Lighthouse Laboratory and Alex Alderton, Roberto Amato, Jeffrey Barrett, Sonia Goncalves, Ewan Harrison, David K. Jackson, Ian Johnston, Dominic Kwiatkowski, Cordelia Langford, John Sillitoe on behalf of the Wellcome Sanger Institute COVID-19 Surveillance Team |
| EPI_ISL_2960820 | Lighthouse Lab in Glasgow | Wellcome Sanger Institute for the COVID-19 Genomics UK (COG-UK) Consortium | Harper VanSteenhouse, Yumi Kasai, David Gray, Carol Clugston, Anna Dominiczak and Alex Alderton, Roberto Amato, Jeffrey Barrett, Sonia Goncalves, Ewan Harrison, David K. Jackson, Ian Johnston, Dominic Kwiatkowski, Cordelia Langford, John Sillitoe on behalf of the Wellcome Sanger Institute COVID-19 Surveillance Team |
| EPI_ISL_2960821, EPI_ISL_2960822 | Lighthouse Lab in Alderley Park | Wellcome Sanger Institute for the COVID-19 Genomics UK (COG-UK) Consortium | Jacquelyn Wynn, Mairead Hyland, The Lighthouse Lab in Alderley Park and Alex Alderton, Roberto Amato, Jeffrey Barrett, Sonia Goncalves, Ewan Harrison, David K. Jackson, Ian Johnston, Dominic Kwiatkowski, Cordelia Langford, John Sillitoe on behalf of the Wellcome Sanger Institute COVID-19 Surveillance Team |
| EPI_ISL_2960823, EPI_ISL_2960824 | Berkshire and Surrey Pathology Services Lighthouse Laboratory | Wellcome Sanger Institute for the COVID-19 Genomics UK (COG-UK) Consortium | Berkshire and Surrey Pathology Services Lighthouse Laboratory and Alex Alderton, Roberto Amato, Jeffrey Barrett, Sonia Goncalves, Ewan Harrison, David K. Jackson, Ian Johnston, Dominic Kwiatkowski, Cordelia Langford, John Sillitoe on behalf of the Wellcome Sanger Institute COVID-19 Surveillance Team |
| EPI_ISL_2960825 | Lighthouse Lab in Glasgow | Wellcome Sanger Institute for the COVID-19 Genomics UK (COG-UK) Consortium | Harper VanSteenhouse, Yumi Kasai, David Gray, Carol Clugston, Anna Dominiczak and Alex Alderton, Roberto Amato, Jeffrey Barrett, Sonia Goncalves, Ewan Harrison, David K. Jackson, Ian Johnston, Dominic Kwiatkowski, Cordelia Langford, John Sillitoe on behalf of the Wellcome Sanger Institute COVID-19 Surveillance Team |
| EPI_ISL_2960826, EPI_ISL_2960827, EPI_ISL_2960828 | Lighthouse Lab in Alderley Park | Wellcome Sanger Institute for the COVID-19 Genomics UK (COG-UK) Consortium | Jacquelyn Wynn, Mairead Hyland, The Lighthouse Lab in Alderley Park and Alex Alderton, Roberto Amato, Jeffrey Barrett, Sonia Goncalves, Ewan Harrison, David K. Jackson, Ian Johnston, Dominic Kwiatkowski, Cordelia Langford, John Sillitoe on behalf of the Wellcome Sanger Institute COVID-19 Surveillance Team |
| EPI_ISL_2960829 | Lighthouse Lab in Glasgow | Wellcome Sanger Institute for the COVID-19 Genomics UK (COG-UK) Consortium | Harper VanSteenhouse, Yumi Kasai, David Gray, Carol Clugston, Anna Dominiczak and Alex Alderton, Roberto Amato, Jeffrey Barrett, Sonia Goncalves, Ewan Harrison, David K. Jackson, Ian Johnston, Dominic Kwiatkowski, Cordelia Langford, John Sillitoe on behalf of the Wellcome Sanger Institute COVID-19 Surveillance Team |
| EPI_ISL_2960830, EPI_ISL_2960831, EPI_ISL_2960832 | Berkshire and Surrey Pathology Services Lighthouse Laboratory | Wellcome Sanger Institute for the COVID-19 Genomics UK (COG-UK) Consortium | Berkshire and Surrey Pathology Services Lighthouse Laboratory and Alex Alderton, Roberto Amato, Jeffrey Barrett, Sonia Goncalves, Ewan Harrison, David K. Jackson, Ian Johnston, Dominic Kwiatkowski, Cordelia Langford, John Sillitoe on behalf of the Wellcome Sanger Institute COVID-19 Surveillance Team |
| EPI_ISL_2960833, EPI_ISL_2960834 | Lighthouse Lab in Glasgow | Wellcome Sanger Institute for the COVID-19 Genomics UK (COG-UK) Consortium | Harper VanSteenhouse, Yumi Kasai, David Gray, Carol Clugston, Anna Dominiczak and Alex Alderton, Roberto Amato, Jeffrey Barrett, Sonia Goncalves, Ewan Harrison, David K. Jackson, Ian Johnston, Dominic Kwiatkowski, Cordelia Langford, John Sillitoe on behalf of the Wellcome Sanger Institute COVID-19 Surveillance Team |
| EPI_ISL_2960835 | Berkshire and Surrey Pathology Services Lighthouse Laboratory | Wellcome Sanger Institute for the COVID-19 Genomics UK (COG-UK) Consortium | Berkshire and Surrey Pathology Services Lighthouse Laboratory and Alex Alderton, Roberto Amato, Jeffrey Barrett, Sonia Goncalves, Ewan Harrison, David K. Jackson, Ian Johnston, Dominic Kwiatkowski, Cordelia Langford, John Sillitoe on behalf of the Wellcome Sanger Institute COVID-19 Surveillance Team |
| EPI_ISL_2960836 | Lighthouse Lab in Alderley Park | Wellcome Sanger Institute for the COVID-19 Genomics UK (COG-UK) Consortium | Jacquelyn Wynn, Mairead Hyland, The Lighthouse Lab in Alderley Park and Alex Alderton, Roberto Amato, Jeffrey Barrett, Sonia Goncalves, Ewan Harrison, David K. Jackson, Ian Johnston, Dominic Kwiatkowski, Cordelia Langford, John Sillitoe on behalf of the Wellcome Sanger Institute COVID-19 Surveillance Team |
| EPI_ISL_2960837 | Lighthouse Lab in Glasgow | Wellcome Sanger Institute for the COVID-19 Genomics UK (COG-UK) Consortium | Harper VanSteenhouse, Yumi Kasai, David Gray, Carol Clugston, Anna Dominiczak and Alex Alderton, Roberto Amato, Jeffrey Barrett, Sonia Goncalves, Ewan Harrison, David K. Jackson, Ian Johnston, Dominic Kwiatkowski, Cordelia Langford, John Sillitoe on behalf of the Wellcome Sanger Institute COVID-19 Surveillance Team |
| EPI_ISL_2960838 | Lighthouse Lab in Alderley Park | Wellcome Sanger Institute for the COVID-19 Genomics UK (COG-UK) Consortium | Jacquelyn Wynn, Mairead Hyland, The Lighthouse Lab in Alderley Park and Alex Alderton, Roberto Amato, Jeffrey Barrett, Sonia Goncalves, Ewan Harrison, David K. Jackson, Ian Johnston, Dominic Kwiatkowski, Cordelia Langford, John Sillitoe on behalf of the Wellcome Sanger Institute COVID-19 Surveillance Team |
| EPI_ISL_2960839, EPI_ISL_2960840 | Lighthouse Lab in Glasgow | Wellcome Sanger Institute for the COVID-19 Genomics UK (COG-UK) Consortium | Harper VanSteenhouse, Yumi Kasai, David Gray, Carol Clugston, Anna Dominiczak and Alex Alderton, Roberto Amato, Jeffrey Barrett, Sonia Goncalves, Ewan Harrison, David K. Jackson, Ian Johnston, Dominic Kwiatkowski, Cordelia Langford, John Sillitoe on behalf of the Wellcome Sanger Institute COVID-19 Surveillance Team |
| EPI_ISL_2960841, EPI_ISL_2960842 | Lighthouse Lab in Alderley Park | Wellcome Sanger Institute for the COVID-19 Genomics UK (COG-UK) Consortium | Jacquelyn Wynn, Mairead Hyland, The Lighthouse Lab in Alderley Park and Alex Alderton, Roberto Amato, Jeffrey Barrett, Sonia Goncalves, Ewan Harrison, David K. Jackson, Ian Johnston, Dominic Kwiatkowski, Cordelia Langford, John Sillitoe on behalf of the Wellcome Sanger Institute COVID-19 Surveillance Team |
| EPI_ISL_2960843, EPI_ISL_2960844, EPI_ISL_2960845 | Berkshire and Surrey Pathology Services Lighthouse Laboratory | Wellcome Sanger Institute for the COVID-19 Genomics UK (COG-UK) Consortium | Berkshire and Surrey Pathology Services Lighthouse Laboratory and Alex Alderton, Roberto Amato, Jeffrey Barrett, Sonia Goncalves, Ewan Harrison, David K. Jackson, Ian Johnston, Dominic Kwiatkowski, Cordelia Langford, John Sillitoe on behalf of the Wellcome Sanger Institute COVID-19 Surveillance Team |
| EPI_ISL_2960846 | Lighthouse Lab in Alderley Park | Wellcome Sanger Institute for the COVID-19 Genomics UK (COG-UK) Consortium | Jacquelyn Wynn, Mairead Hyland, The Lighthouse Lab in Alderley Park and Alex Alderton, Roberto Amato, Jeffrey Barrett, Sonia Goncalves, Ewan Harrison, David K. Jackson, Ian Johnston, Dominic Kwiatkowski, Cordelia Langford, John Sillitoe on behalf of the Wellcome Sanger Institute COVID-19 Surveillance Team |
| EPI_ISL_2960847, EPI_ISL_2960848 | Lighthouse Lab in Glasgow | Wellcome Sanger Institute for the COVID-19 Genomics UK (COG-UK) Consortium | Harper VanSteenhouse, Yumi Kasai, David Gray, Carol Clugston, Anna Dominiczak and Alex Alderton, Roberto Amato, Jeffrey Barrett, Sonia Goncalves, Ewan Harrison, David K. Jackson, Ian Johnston, Dominic Kwiatkowski, Cordelia Langford, John Sillitoe on behalf of the Wellcome Sanger Institute COVID-19 Surveillance Team |
| EPI_ISL_2960849 | Berkshire and Surrey Pathology Services Lighthouse Laboratory | Wellcome Sanger Institute for the COVID-19 Genomics UK (COG-UK) Consortium | Berkshire and Surrey Pathology Services Lighthouse Laboratory and Alex Alderton, Roberto Amato, Jeffrey Barrett, Sonia Goncalves, Ewan Harrison, David K. Jackson, Ian Johnston, Dominic Kwiatkowski, Cordelia Langford, John Sillitoe on behalf of the Wellcome Sanger Institute COVID-19 Surveillance Team |
| EPI_ISL_2960850, EPI_ISL_2960851 | Lighthouse Lab in Glasgow | Wellcome Sanger Institute for the COVID-19 Genomics UK (COG-UK) Consortium | Harper VanSteenhouse, Yumi Kasai, David Gray, Carol Clugston, Anna Dominiczak and Alex Alderton, Roberto Amato, Jeffrey Barrett, Sonia Goncalves, Ewan Harrison, David K. Jackson, Ian Johnston, Dominic Kwiatkowski, Cordelia Langford, John Sillitoe on behalf of the Wellcome Sanger Institute COVID-19 Surveillance Team |
| EPI_ISL_2960852 | Lighthouse Lab in Alderley Park | Wellcome Sanger Institute for the COVID-19 Genomics UK (COG-UK) Consortium | Jacquelyn Wynn, Mairead Hyland, The Lighthouse Lab in Alderley Park and Alex Alderton, Roberto Amato, Jeffrey Barrett, Sonia Goncalves, Ewan Harrison, David K. Jackson, Ian Johnston, Dominic Kwiatkowski, Cordelia Langford, John Sillitoe on behalf of the Wellcome Sanger Institute COVID-19 Surveillance Team |
| EPI_ISL_2960853, EPI_ISL_2960854, EPI_ISL_2960855 | Lighthouse Lab in Glasgow | Wellcome Sanger Institute for the COVID-19 Genomics UK (COG-UK) Consortium | Harper VanSteenhouse, Yumi Kasai, David Gray, Carol Clugston, Anna Dominiczak and Alex Alderton, Roberto Amato, Jeffrey Barrett, Sonia Goncalves, Ewan Harrison, David K. Jackson, Ian Johnston, Dominic Kwiatkowski, Cordelia Langford, John Sillitoe on behalf of the Wellcome Sanger Institute COVID-19 Surveillance Team |
| EPI_ISL_2960856, EPI_ISL_2960857 | Lighthouse Lab in Alderley Park | Wellcome Sanger Institute for the COVID-19 Genomics UK (COG-UK) Consortium | Jacquelyn Wynn, Mairead Hyland, The Lighthouse Lab in Alderley Park and Alex Alderton, Roberto Amato, Jeffrey Barrett, Sonia Goncalves, Ewan Harrison, David K. Jackson, Ian Johnston, Dominic Kwiatkowski, Cordelia Langford, John Sillitoe on behalf of the Wellcome Sanger Institute COVID-19 Surveillance Team |
| EPI_ISL_2960858 | Berkshire and Surrey Pathology Services Lighthouse Laboratory | Wellcome Sanger Institute for the COVID-19 Genomics UK (COG-UK) Consortium | Berkshire and Surrey Pathology Services Lighthouse Laboratory and Alex Alderton, Roberto Amato, Jeffrey Barrett, Sonia Goncalves, Ewan Harrison, David K. Jackson, Ian Johnston, Dominic Kwiatkowski, Cordelia Langford, John Sillitoe on behalf of the Wellcome Sanger Institute COVID-19 Surveillance Team |
| EPI_ISL_2960859, EPI_ISL_2960860 | Lighthouse Lab in Glasgow | Wellcome Sanger Institute for the COVID-19 Genomics UK (COG-UK) Consortium | Harper VanSteenhouse, Yumi Kasai, David Gray, Carol Clugston, Anna Dominiczak and Alex Alderton, Roberto Amato, Jeffrey Barrett, Sonia Goncalves, Ewan Harrison, David K. Jackson, Ian Johnston, Dominic Kwiatkowski, Cordelia Langford, John Sillitoe on behalf of the Wellcome Sanger Institute COVID-19 Surveillance Team |
| EPI_ISL_2960861 | Lighthouse Lab in Alderley Park | Wellcome Sanger Institute for the COVID-19 Genomics UK (COG-UK) Consortium | Jacquelyn Wynn, Mairead Hyland, The Lighthouse Lab in Alderley Park and Alex Alderton, Roberto Amato, Jeffrey Barrett, Sonia Goncalves, Ewan Harrison, David K. Jackson, Ian Johnston, Dominic Kwiatkowski, Cordelia Langford, John Sillitoe on behalf of the Wellcome Sanger Institute COVID-19 Surveillance Team |
| EPI_ISL_2960862 | Berkshire and Surrey Pathology Services Lighthouse Laboratory | Wellcome Sanger Institute for the COVID-19 Genomics UK (COG-UK) Consortium | Berkshire and Surrey Pathology Services Lighthouse Laboratory and Alex Alderton, Roberto Amato, Jeffrey Barrett, Sonia Goncalves, Ewan Harrison, David K. Jackson, Ian Johnston, Dominic Kwiatkowski, Cordelia Langford, John Sillitoe on behalf of the Wellcome Sanger Institute COVID-19 Surveillance Team |
| EPI_ISL_2960863 | Lighthouse Lab in Glasgow | Wellcome Sanger Institute for the COVID-19 Genomics UK (COG-UK) Consortium | Harper VanSteenhouse, Yumi Kasai, David Gray, Carol Clugston, Anna Dominiczak and Alex Alderton, Roberto Amato, Jeffrey Barrett, Sonia Goncalves, Ewan Harrison, David K. Jackson, Ian Johnston, Dominic Kwiatkowski, Cordelia Langford, John Sillitoe on behalf of the Wellcome Sanger Institute COVID-19 Surveillance Team |

| | | | |
|---|---|---|---|
| EPI_ISL_2960864 | Lighthouse Lab in Alderley Park | Wellcome Sanger Institute for the COVID-19 Genomics UK (COG-UK) Consortium | Jacquelyn Wynn, Mairead Hyland, The Lighthouse Lab in Alderley Park and Alex Alderton, Roberto Amato, Jeffrey Barrett, Sonia Goncalves, Ewan Harrison, David K. Jackson, Ian Johnston, Dominic Kwiatkowski, Cordelia Langford, John Sillitoe on behalf of the Wellcome Sanger Institute COVID-19 Surveillance Team |
| EPI_ISL_2960865 | Berkshire and Surrey Pathology Services Lighthouse Laboratory | Wellcome Sanger Institute for the COVID-19 Genomics UK (COG-UK) Consortium | Berkshire and Surrey Pathology Services Lighthouse Laboratory and Alex Alderton, Roberto Amato, Jeffrey Barrett, Sonia Goncalves, Ewan Harrison, David K. Jackson, Ian Johnston, Dominic Kwiatkowski, Cordelia Langford, John Sillitoe on behalf of the Wellcome Sanger Institute COVID-19 Surveillance Team |
| EPI_ISL_2960866, EPI_ISL_2960867 | Lighthouse Lab in Glasgow | Wellcome Sanger Institute for the COVID-19 Genomics UK (COG-UK) Consortium | Harper VanSteenhouse, Yumi Kasai, David Gray, Carol Clugston, Anna Dominiczak and Alex Alderton, Roberto Amato, Jeffrey Barrett, Sonia Goncalves, Ewan Harrison, David K. Jackson, Ian Johnston, Dominic Kwiatkowski, Cordelia Langford, John Sillitoe on behalf of the Wellcome Sanger Institute COVID-19 Surveillance Team |
| EPI_ISL_2960868 | Lighthouse Lab in Alderley Park | Wellcome Sanger Institute for the COVID-19 Genomics UK (COG-UK) Consortium | Jacquelyn Wynn, Mairead Hyland, The Lighthouse Lab in Alderley Park and Alex Alderton, Roberto Amato, Jeffrey Barrett, Sonia Goncalves, Ewan Harrison, David K. Jackson, Ian Johnston, Dominic Kwiatkowski, Cordelia Langford, John Sillitoe on behalf of the Wellcome Sanger Institute COVID-19 Surveillance Team |
| EPI_ISL_2960869 | Berkshire and Surrey Pathology Services Lighthouse Laboratory | Wellcome Sanger Institute for the COVID-19 Genomics UK (COG-UK) Consortium | Berkshire and Surrey Pathology Services Lighthouse Laboratory and Alex Alderton, Roberto Amato, Jeffrey Barrett, Sonia Goncalves, Ewan Harrison, David K. Jackson, Ian Johnston, Dominic Kwiatkowski, Cordelia Langford, John Sillitoe on behalf of the Wellcome Sanger Institute COVID-19 Surveillance Team |
| EPI_ISL_2960870, EPI_ISL_2960871 | Lighthouse Lab in Glasgow | Wellcome Sanger Institute for the COVID-19 Genomics UK (COG-UK) Consortium | Harper VanSteenhouse, Yumi Kasai, David Gray, Carol Clugston, Anna Dominiczak and Alex Alderton, Roberto Amato, Jeffrey Barrett, Sonia Goncalves, Ewan Harrison, David K. Jackson, Ian Johnston, Dominic Kwiatkowski, Cordelia Langford, John Sillitoe on behalf of the Wellcome Sanger Institute COVID-19 Surveillance Team |
| EPI_ISL_2960872 | Berkshire and Surrey Pathology Services Lighthouse Laboratory | Wellcome Sanger Institute for the COVID-19 Genomics UK (COG-UK) Consortium | Berkshire and Surrey Pathology Services Lighthouse Laboratory and Alex Alderton, Roberto Amato, Jeffrey Barrett, Sonia Goncalves, Ewan Harrison, David K. Jackson, Ian Johnston, Dominic Kwiatkowski, Cordelia Langford, John Sillitoe on behalf of the Wellcome Sanger Institute COVID-19 Surveillance Team |
| EPI_ISL_2960873, EPI_ISL_2960874, EPI_ISL_2960875 | Lighthouse Lab in Alderley Park | Wellcome Sanger Institute for the COVID-19 Genomics UK (COG-UK) Consortium | Jacquelyn Wynn, Mairead Hyland, The Lighthouse Lab in Alderley Park and Alex Alderton, Roberto Amato, Jeffrey Barrett, Sonia Goncalves, Ewan Harrison, David K. Jackson, Ian Johnston, Dominic Kwiatkowski, Cordelia Langford, John Sillitoe on behalf of the Wellcome Sanger Institute COVID-19 Surveillance Team |
| EPI_ISL_2960876 | Berkshire and Surrey Pathology Services Lighthouse Laboratory | Wellcome Sanger Institute for the COVID-19 Genomics UK (COG-UK) Consortium | Berkshire and Surrey Pathology Services Lighthouse Laboratory and Alex Alderton, Roberto Amato, Jeffrey Barrett, Sonia Goncalves, Ewan Harrison, David K. Jackson, Ian Johnston, Dominic Kwiatkowski, Cordelia Langford, John Sillitoe on behalf of the Wellcome Sanger Institute COVID-19 Surveillance Team |
| EPI_ISL_2960877, EPI_ISL_2960878 | Lighthouse Lab in Glasgow | Wellcome Sanger Institute for the COVID-19 Genomics UK (COG-UK) Consortium | Harper VanSteenhouse, Yumi Kasai, David Gray, Carol Clugston, Anna Dominiczak and Alex Alderton, Roberto Amato, Jeffrey Barrett, Sonia Goncalves, Ewan Harrison, David K. Jackson, Ian Johnston, Dominic Kwiatkowski, Cordelia Langford, John Sillitoe on behalf of the Wellcome Sanger Institute COVID-19 Surveillance Team |
| EPI_ISL_2960879 | Lighthouse Lab in Alderley Park | Wellcome Sanger Institute for the COVID-19 Genomics UK (COG-UK) Consortium | Jacquelyn Wynn, Mairead Hyland, The Lighthouse Lab in Alderley Park and Alex Alderton, Roberto Amato, Jeffrey Barrett, Sonia Goncalves, Ewan Harrison, David K. Jackson, Ian Johnston, Dominic Kwiatkowski, Cordelia Langford, John Sillitoe on behalf of the Wellcome Sanger Institute COVID-19 Surveillance Team |
| EPI_ISL_2960880, EPI_ISL_2960881 | Lighthouse Lab in Glasgow | Wellcome Sanger Institute for the COVID-19 Genomics UK (COG-UK) Consortium | Harper VanSteenhouse, Yumi Kasai, David Gray, Carol Clugston, Anna Dominiczak and Alex Alderton, Roberto Amato, Jeffrey Barrett, Sonia Goncalves, Ewan Harrison, David K. Jackson, Ian Johnston, Dominic Kwiatkowski, Cordelia Langford, John Sillitoe on behalf of the Wellcome Sanger Institute COVID-19 Surveillance Team |
| EPI_ISL_2960882 | Lighthouse Lab in Alderley Park | Wellcome Sanger Institute for the COVID-19 Genomics UK (COG-UK) Consortium | Jacquelyn Wynn, Mairead Hyland, The Lighthouse Lab in Alderley Park and Alex Alderton, Roberto Amato, Jeffrey Barrett, Sonia Goncalves, Ewan Harrison, David K. Jackson, Ian Johnston, Dominic Kwiatkowski, Cordelia Langford, John Sillitoe on behalf of the Wellcome Sanger Institute COVID-19 Surveillance Team |
| EPI_ISL_2960883 | Lighthouse Lab in Glasgow | Wellcome Sanger Institute for the COVID-19 Genomics UK (COG-UK) Consortium | Harper VanSteenhouse, Yumi Kasai, David Gray, Carol Clugston, Anna Dominiczak and Alex Alderton, Roberto Amato, Jeffrey Barrett, Sonia Goncalves, Ewan Harrison, David K. Jackson, Ian Johnston, Dominic Kwiatkowski, Cordelia Langford, John Sillitoe on behalf of the Wellcome Sanger Institute COVID-19 Surveillance Team |
| EPI_ISL_2960884 | Berkshire and Surrey Pathology Services Lighthouse Laboratory | Wellcome Sanger Institute for the COVID-19 Genomics UK (COG-UK) Consortium | Berkshire and Surrey Pathology Services Lighthouse Laboratory and Alex Alderton, Roberto Amato, Jeffrey Barrett, Sonia Goncalves, Ewan Harrison, David K. Jackson, Ian Johnston, Dominic Kwiatkowski, Cordelia Langford, John Sillitoe on behalf of the Wellcome Sanger Institute COVID-19 Surveillance Team |
| EPI_ISL_2960885 | Lighthouse Lab in Glasgow | Wellcome Sanger Institute for the COVID-19 Genomics UK (COG-UK) Consortium | Harper VanSteenhouse, Yumi Kasai, David Gray, Carol Clugston, Anna Dominiczak and Alex Alderton, Roberto Amato, Jeffrey Barrett, Sonia Goncalves, Ewan Harrison, David K. Jackson, Ian Johnston, Dominic Kwiatkowski, Cordelia Langford, John Sillitoe on behalf of the Wellcome Sanger Institute COVID-19 Surveillance Team |
| EPI_ISL_2960886, EPI_ISL_2960887 | Berkshire and Surrey Pathology Services Lighthouse Laboratory | Wellcome Sanger Institute for the COVID-19 Genomics UK (COG-UK) Consortium | Berkshire and Surrey Pathology Services Lighthouse Laboratory and Alex Alderton, Roberto Amato, Jeffrey Barrett, Sonia Goncalves, Ewan Harrison, David K. Jackson, Ian Johnston, Dominic Kwiatkowski, Cordelia Langford, John Sillitoe on behalf of the Wellcome Sanger Institute COVID-19 Surveillance Team |
| EPI_ISL_2960888 | Lighthouse Lab in Glasgow | Wellcome Sanger Institute for the COVID-19 Genomics UK (COG-UK) Consortium | Harper VanSteenhouse, Yumi Kasai, David Gray, Carol Clugston, Anna Dominiczak and Alex Alderton, Roberto Amato, Jeffrey Barrett, Sonia Goncalves, Ewan Harrison, David K. Jackson, Ian Johnston, Dominic Kwiatkowski, Cordelia Langford, John Sillitoe on behalf of the Wellcome Sanger Institute COVID-19 Surveillance Team |
| EPI_ISL_2960889 | Berkshire and Surrey Pathology Services Lighthouse Laboratory | Wellcome Sanger Institute for the COVID-19 Genomics UK (COG-UK) Consortium | Berkshire and Surrey Pathology Services Lighthouse Laboratory and Alex Alderton, Roberto Amato, Jeffrey Barrett, Sonia Goncalves, Ewan Harrison, David K. Jackson, Ian Johnston, Dominic Kwiatkowski, Cordelia Langford, John Sillitoe on behalf of the Wellcome Sanger Institute COVID-19 Surveillance Team |
| EPI_ISL_2960890 | Lighthouse Lab in Glasgow | Wellcome Sanger Institute for the COVID-19 Genomics UK (COG-UK) Consortium | Harper VanSteenhouse, Yumi Kasai, David Gray, Carol Clugston, Anna Dominiczak and Alex Alderton, Roberto Amato, Jeffrey Barrett, Sonia Goncalves, Ewan Harrison, David K. Jackson, Ian Johnston, Dominic Kwiatkowski, Cordelia Langford, John Sillitoe on behalf of the Wellcome Sanger Institute COVID-19 Surveillance Team |
| EPI_ISL_2960891 | Berkshire and Surrey Pathology Services Lighthouse Laboratory | Wellcome Sanger Institute for the COVID-19 Genomics UK (COG-UK) Consortium | Berkshire and Surrey Pathology Services Lighthouse Laboratory and Alex Alderton, Roberto Amato, Jeffrey Barrett, Sonia Goncalves, Ewan Harrison, David K. Jackson, Ian Johnston, Dominic Kwiatkowski, Cordelia Langford, John Sillitoe on behalf of the Wellcome Sanger Institute COVID-19 Surveillance Team |
| EPI_ISL_2960892 | Lighthouse Lab in Alderley Park | Wellcome Sanger Institute for the COVID-19 Genomics UK (COG-UK) Consortium | Jacquelyn Wynn, Mairead Hyland, The Lighthouse Lab in Alderley Park and Alex Alderton, Roberto Amato, Jeffrey Barrett, Sonia Goncalves, Ewan Harrison, David K. Jackson, Ian Johnston, Dominic Kwiatkowski, Cordelia Langford, John Sillitoe on behalf of the Wellcome Sanger Institute COVID-19 Surveillance Team |
| EPI_ISL_2960893 | Lighthouse Lab in Glasgow | Wellcome Sanger Institute for the COVID-19 Genomics UK (COG-UK) Consortium | Harper VanSteenhouse, Yumi Kasai, David Gray, Carol Clugston, Anna Dominiczak and Alex Alderton, Roberto Amato, Jeffrey Barrett, Sonia Goncalves, Ewan Harrison, David K. Jackson, Ian Johnston, Dominic Kwiatkowski, Cordelia Langford, John Sillitoe on behalf of the Wellcome Sanger Institute COVID-19 Surveillance Team |
| EPI_ISL_2960894 | Lighthouse Lab in Alderley Park | Wellcome Sanger Institute for the COVID-19 Genomics UK (COG-UK) Consortium | Jacquelyn Wynn, Mairead Hyland, The Lighthouse Lab in Alderley Park and Alex Alderton, Roberto Amato, Jeffrey Barrett, Sonia Goncalves, Ewan Harrison, David K. Jackson, Ian Johnston, Dominic Kwiatkowski, Cordelia Langford, John Sillitoe on behalf of the Wellcome Sanger Institute COVID-19 Surveillance Team |
| EPI_ISL_2960895 | Lighthouse Lab in Glasgow | Wellcome Sanger Institute for the COVID-19 Genomics UK (COG-UK) Consortium | Harper VanSteenhouse, Yumi Kasai, David Gray, Carol Clugston, Anna Dominiczak and Alex Alderton, Roberto Amato, Jeffrey Barrett, Sonia Goncalves, Ewan Harrison, David K. Jackson, Ian Johnston, Dominic Kwiatkowski, Cordelia Langford, John Sillitoe on behalf of the Wellcome Sanger Institute COVID-19 Surveillance Team |
| EPI_ISL_2960896, EPI_ISL_2960897, EPI_ISL_2960898 | Lighthouse Lab in Alderley Park | Wellcome Sanger Institute for the COVID-19 Genomics UK (COG-UK) Consortium | Jacquelyn Wynn, Mairead Hyland, The Lighthouse Lab in Alderley Park and Alex Alderton, Roberto Amato, Jeffrey Barrett, Sonia Goncalves, Ewan Harrison, David K. Jackson, Ian Johnston, Dominic Kwiatkowski, Cordelia Langford, John Sillitoe on behalf of the Wellcome Sanger Institute COVID-19 Surveillance Team |
| EPI_ISL_2960899, EPI_ISL_2960900, EPI_ISL_2960901 | Lighthouse Lab in Glasgow | Wellcome Sanger Institute for the COVID-19 Genomics UK (COG-UK) Consortium | Harper VanSteenhouse, Yumi Kasai, David Gray, Carol Clugston, Anna Dominiczak and Alex Alderton, Roberto Amato, Jeffrey Barrett, Sonia Goncalves, Ewan Harrison, David K. Jackson, Ian Johnston, Dominic Kwiatkowski, Cordelia Langford, John Sillitoe on behalf of the Wellcome Sanger Institute COVID-19 Surveillance Team |
| EPI_ISL_2960902, EPI_ISL_2960903 | Berkshire and Surrey Pathology Services Lighthouse Laboratory | Wellcome Sanger Institute for the COVID-19 Genomics UK (COG-UK) Consortium | Berkshire and Surrey Pathology Services Lighthouse Laboratory and Alex Alderton, Roberto Amato, Jeffrey Barrett, Sonia Goncalves, Ewan Harrison, David K. Jackson, Ian Johnston, Dominic Kwiatkowski, Cordelia Langford, John Sillitoe on behalf of the Wellcome Sanger Institute COVID-19 Surveillance Team |
| EPI_ISL_2960904, EPI_ISL_2960905 | Lighthouse Lab in Glasgow | Wellcome Sanger Institute for the COVID-19 Genomics UK (COG-UK) Consortium | Harper VanSteenhouse, Yumi Kasai, David Gray, Carol Clugston, Anna Dominiczak and Alex Alderton, Roberto Amato, Jeffrey Barrett, Sonia Goncalves, Ewan Harrison, David K. Jackson, Ian Johnston, Dominic Kwiatkowski, Cordelia Langford, John Sillitoe on behalf of the Wellcome Sanger Institute COVID-19 Surveillance Team |
| EPI_ISL_2960906 | Lighthouse Lab in Alderley Park | Wellcome Sanger Institute for the COVID-19 Genomics UK (COG-UK) Consortium | Jacquelyn Wynn, Mairead Hyland, The Lighthouse Lab in Alderley Park and Alex Alderton, Roberto Amato, Jeffrey Barrett, Sonia Goncalves, Ewan Harrison, David K. Jackson, Ian Johnston, Dominic Kwiatkowski, Cordelia Langford, John Sillitoe on behalf of the Wellcome Sanger Institute COVID-19 Surveillance Team |
| EPI_ISL_2960907, EPI_ISL_2960908, EPI_ISL_2960909 | Berkshire and Surrey Pathology Services Lighthouse Laboratory | Wellcome Sanger Institute for the COVID-19 Genomics UK (COG-UK) Consortium | Berkshire and Surrey Pathology Services Lighthouse Laboratory and Alex Alderton, Roberto Amato, Jeffrey Barrett, Sonia Goncalves, Ewan Harrison, David K. Jackson, Ian Johnston, Dominic Kwiatkowski, Cordelia Langford, John Sillitoe on behalf of the Wellcome Sanger Institute COVID-19 Surveillance Team |
| EPI_ISL_2960910 | Lighthouse Lab in Alderley Park | Wellcome Sanger Institute for the COVID-19 Genomics UK (COG-UK) Consortium | Jacquelyn Wynn, Mairead Hyland, The Lighthouse Lab in Alderley Park and Alex Alderton, Roberto Amato, Jeffrey Barrett, Sonia Goncalves, Ewan Harrison, David K. Jackson, Ian Johnston, Dominic Kwiatkowski, Cordelia Langford, John Sillitoe on behalf of the Wellcome Sanger Institute COVID-19 Surveillance Team |
| EPI_ISL_2960911 | Lighthouse Laboratory Plymouth | Wellcome Sanger Institute for the COVID-19 Genomics UK (COG-UK) Consortium | Lighthouse Laboratory Plymouth and Alex Alderton, Roberto Amato, Jeffrey Barrett, Sonia Goncalves, Ewan Harrison, David K. Jackson, Ian Johnston, Dominic Kwiatkowski, Cordelia Langford, John Sillitoe on behalf of the Wellcome Sanger Institute COVID-19 Surveillance Team |
| EPI_ISL_2960912, EPI_ISL_2960913, EPI_ISL_2960914 | Lighthouse Lab in Alderley Park | Wellcome Sanger Institute for the COVID-19 Genomics UK (COG-UK) Consortium | Jacquelyn Wynn, Mairead Hyland, The Lighthouse Lab in Alderley Park and Alex Alderton, Roberto Amato, Jeffrey Barrett, Sonia Goncalves, Ewan Harrison, David K. Jackson, Ian Johnston, Dominic Kwiatkowski, Cordelia Langford, John Sillitoe on behalf of the Wellcome Sanger Institute COVID-19 Surveillance Team |
| EPI_ISL_2960915 | Lighthouse Laboratory Plymouth | Wellcome Sanger Institute for the COVID-19 Genomics UK (COG-UK) Consortium | Lighthouse Laboratory Plymouth and Alex Alderton, Roberto Amato, Jeffrey Barrett, Sonia Goncalves, Ewan Harrison, David K. Jackson, Ian Johnston, Dominic Kwiatkowski, Cordelia Langford, John Sillitoe on behalf of the Wellcome Sanger Institute COVID-19 Surveillance Team |
| EPI_ISL_2960916 | Lighthouse Lab in Alderley Park | Wellcome Sanger Institute for the COVID-19 Genomics UK (COG-UK) Consortium | Jacquelyn Wynn, Mairead Hyland, The Lighthouse Lab in Alderley Park and Alex Alderton, Roberto Amato, Jeffrey Barrett, Sonia Goncalves, Ewan Harrison, David K. Jackson, Ian Johnston, Dominic Kwiatkowski, Cordelia Langford, John Sillitoe on behalf of the Wellcome Sanger Institute COVID-19 Surveillance Team |
| EPI_ISL_2960917 | Lighthouse Laboratory Plymouth | Wellcome Sanger Institute for the COVID-19 Genomics UK (COG-UK) Consortium | Lighthouse Laboratory Plymouth and Alex Alderton, Roberto Amato, Jeffrey Barrett, Sonia Goncalves, Ewan Harrison, David K. Jackson, Ian Johnston, Dominic Kwiatkowski, Cordelia Langford, John Sillitoe on behalf of the Wellcome Sanger Institute COVID-19 Surveillance Team |
| EPI_ISL_2960918 | Berkshire and Surrey Pathology Services Lighthouse Laboratory | Wellcome Sanger Institute for the COVID-19 Genomics UK (COG-UK) Consortium | Berkshire and Surrey Pathology Services Lighthouse Laboratory and Alex Alderton, Roberto Amato, Jeffrey Barrett, Sonia Goncalves, Ewan Harrison, David K. Jackson, Ian Johnston, Dominic Kwiatkowski, Cordelia Langford, John Sillitoe on behalf of the Wellcome Sanger Institute COVID-19 Surveillance Team |
| EPI_ISL_2960919 | Lighthouse Laboratory Plymouth | Wellcome Sanger Institute for the COVID-19 Genomics UK (COG-UK) Consortium | Lighthouse Laboratory Plymouth and Alex Alderton, Roberto Amato, Jeffrey Barrett, Sonia Goncalves, Ewan Harrison, David K. Jackson, Ian Johnston, Dominic Kwiatkowski, Cordelia Langford, John Sillitoe on behalf of the Wellcome Sanger Institute COVID-19 Surveillance Team |
| EPI_ISL_2960920, EPI_ISL_2960921 | Berkshire and Surrey Pathology Services Lighthouse Laboratory | Wellcome Sanger Institute for the COVID-19 Genomics UK (COG-UK) Consortium | Berkshire and Surrey Pathology Services Lighthouse Laboratory and Alex Alderton, Roberto Amato, Jeffrey Barrett, Sonia Goncalves, Ewan Harrison, David K. Jackson, Ian Johnston, Dominic Kwiatkowski, Cordelia Langford, John Sillitoe on behalf of the Wellcome Sanger Institute COVID-19 Surveillance Team |
| EPI_ISL_2960922 | Lighthouse Lab in Alderley Park | Wellcome Sanger Institute for the COVID-19 Genomics UK (COG-UK) Consortium | Jacquelyn Wynn, Mairead Hyland, The Lighthouse Lab in Alderley Park and Alex Alderton, Roberto Amato, Jeffrey Barrett, Sonia Goncalves, Ewan Harrison, David K. Jackson, Ian Johnston, Dominic Kwiatkowski, Cordelia Langford, John Sillitoe on behalf of the Wellcome Sanger Institute COVID-19 Surveillance Team |
| EPI_ISL_2960923 | Berkshire and Surrey Pathology Services Lighthouse Laboratory | Wellcome Sanger Institute for the COVID-19 Genomics UK (COG-UK) Consortium | Berkshire and Surrey Pathology Services Lighthouse Laboratory and Alex Alderton, Roberto Amato, Jeffrey Barrett, Sonia Goncalves, Ewan Harrison, David K. Jackson, Ian Johnston, Dominic Kwiatkowski, Cordelia Langford, John Sillitoe on behalf of the Wellcome Sanger Institute COVID-19 Surveillance Team |
| EPI_ISL_2960924, EPI_ISL_2960925 | Lighthouse Lab in Alderley Park | Wellcome Sanger Institute for the COVID-19 Genomics UK (COG-UK) Consortium | Jacquelyn Wynn, Mairead Hyland, The Lighthouse Lab in Alderley Park and Alex Alderton, Roberto Amato, Jeffrey Barrett, Sonia Goncalves, Ewan Harrison, David K. Jackson, Ian Johnston, Dominic Kwiatkowski, Cordelia Langford, John Sillitoe on behalf of the Wellcome Sanger Institute COVID-19 Surveillance Team |

| | | | |
|---|---|---|---|
| EPI_ISL_2960926 | Lighthouse Laboratory Plymouth | Wellcome Sanger Institute for the COVID-19 Genomics UK (COG-UK) Consortium | Lighthouse Laboratory Plymouth and Alex Alderton, Roberto Amato, Jeffrey Barrett, Sonia Goncalves, Ewan Harrison, David K. Jackson, Ian Johnston, Dominic Kwiatkowski, Cordelia Langford, John Sillitoe on behalf of the Wellcome Sanger Institute COVID-19 Surveillance Team |
| EPI_ISL_2960927, EPI_ISL_2960928 | Lighthouse Lab in Alderley Park | Wellcome Sanger Institute for the COVID-19 Genomics UK (COG-UK) Consortium | Jacquelyn Wynn, Mairead Hyland, The Lighthouse Lab in Alderley Park and Alex Alderton, Roberto Amato, Jeffrey Barrett, Sonia Goncalves, Ewan Harrison, David K. Jackson, Ian Johnston, Dominic Kwiatkowski, Cordelia Langford, John Sillitoe on behalf of the Wellcome Sanger Institute COVID-19 Surveillance Team |
| EPI_ISL_2960929 | Lighthouse Laboratory Plymouth | Wellcome Sanger Institute for the COVID-19 Genomics UK (COG-UK) Consortium | Lighthouse Laboratory Plymouth and Alex Alderton, Roberto Amato, Jeffrey Barrett, Sonia Goncalves, Ewan Harrison, David K. Jackson, Ian Johnston, Dominic Kwiatkowski, Cordelia Langford, John Sillitoe on behalf of the Wellcome Sanger Institute COVID-19 Surveillance Team |
| EPI_ISL_2960930, EPI_ISL_2960931 | Berkshire and Surrey Pathology Services Lighthouse Laboratory | Wellcome Sanger Institute for the COVID-19 Genomics UK (COG-UK) Consortium | Berkshire and Surrey Pathology Services Lighthouse Laboratory and Alex Alderton, Roberto Amato, Jeffrey Barrett, Sonia Goncalves, Ewan Harrison, David K. Jackson, Ian Johnston, Dominic Kwiatkowski, Cordelia Langford, John Sillitoe on behalf of the Wellcome Sanger Institute COVID-19 Surveillance Team |
| EPI_ISL_2960932, EPI_ISL_2960933, EPI_ISL_2960934, EPI_ISL_2960935 | Lighthouse Lab in Alderley Park | Wellcome Sanger Institute for the COVID-19 Genomics UK (COG-UK) Consortium | Jacquelyn Wynn, Mairead Hyland, The Lighthouse Lab in Alderley Park and Alex Alderton, Roberto Amato, Jeffrey Barrett, Sonia Goncalves, Ewan Harrison, David K. Jackson, Ian Johnston, Dominic Kwiatkowski, Cordelia Langford, John Sillitoe on behalf of the Wellcome Sanger Institute COVID-19 Surveillance Team |
| EPI_ISL_2960936 | Berkshire and Surrey Pathology Services Lighthouse Laboratory | Wellcome Sanger Institute for the COVID-19 Genomics UK (COG-UK) Consortium | Berkshire and Surrey Pathology Services Lighthouse Laboratory and Alex Alderton, Roberto Amato, Jeffrey Barrett, Sonia Goncalves, Ewan Harrison, David K. Jackson, Ian Johnston, Dominic Kwiatkowski, Cordelia Langford, John Sillitoe on behalf of the Wellcome Sanger Institute COVID-19 Surveillance Team |
| EPI_ISL_2960937 | Lighthouse Lab in Alderley Park | Wellcome Sanger Institute for the COVID-19 Genomics UK (COG-UK) Consortium | Jacquelyn Wynn, Mairead Hyland, The Lighthouse Lab in Alderley Park and Alex Alderton, Roberto Amato, Jeffrey Barrett, Sonia Goncalves, Ewan Harrison, David K. Jackson, Ian Johnston, Dominic Kwiatkowski, Cordelia Langford, John Sillitoe on behalf of the Wellcome Sanger Institute COVID-19 Surveillance Team |
| EPI_ISL_2960938 | Lighthouse Laboratory Plymouth | Wellcome Sanger Institute for the COVID-19 Genomics UK (COG-UK) Consortium | Lighthouse Laboratory Plymouth and Alex Alderton, Roberto Amato, Jeffrey Barrett, Sonia Goncalves, Ewan Harrison, David K. Jackson, Ian Johnston, Dominic Kwiatkowski, Cordelia Langford, John Sillitoe on behalf of the Wellcome Sanger Institute COVID-19 Surveillance Team |
| EPI_ISL_2960939 | Lighthouse Lab in Alderley Park | Wellcome Sanger Institute for the COVID-19 Genomics UK (COG-UK) Consortium | Jacquelyn Wynn, Mairead Hyland, The Lighthouse Lab in Alderley Park and Alex Alderton, Roberto Amato, Jeffrey Barrett, Sonia Goncalves, Ewan Harrison, David K. Jackson, Ian Johnston, Dominic Kwiatkowski, Cordelia Langford, John Sillitoe on behalf of the Wellcome Sanger Institute COVID-19 Surveillance Team |
| EPI_ISL_2960940 | Lighthouse Laboratory Plymouth | Wellcome Sanger Institute for the COVID-19 Genomics UK (COG-UK) Consortium | Lighthouse Laboratory Plymouth and Alex Alderton, Roberto Amato, Jeffrey Barrett, Sonia Goncalves, Ewan Harrison, David K. Jackson, Ian Johnston, Dominic Kwiatkowski, Cordelia Langford, John Sillitoe on behalf of the Wellcome Sanger Institute COVID-19 Surveillance Team |
| EPI_ISL_2960941 | Lighthouse Lab in Alderley Park | Wellcome Sanger Institute for the COVID-19 Genomics UK (COG-UK) Consortium | Jacquelyn Wynn, Mairead Hyland, The Lighthouse Lab in Alderley Park and Alex Alderton, Roberto Amato, Jeffrey Barrett, Sonia Goncalves, Ewan Harrison, David K. Jackson, Ian Johnston, Dominic Kwiatkowski, Cordelia Langford, John Sillitoe on behalf of the Wellcome Sanger Institute COVID-19 Surveillance Team |
| EPI_ISL_2960942, EPI_ISL_2960943, EPI_ISL_2960944 | Berkshire and Surrey Pathology Services Lighthouse Laboratory | Wellcome Sanger Institute for the COVID-19 Genomics UK (COG-UK) Consortium | Berkshire and Surrey Pathology Services Lighthouse Laboratory and Alex Alderton, Roberto Amato, Jeffrey Barrett, Sonia Goncalves, Ewan Harrison, David K. Jackson, Ian Johnston, Dominic Kwiatkowski, Cordelia Langford, John Sillitoe on behalf of the Wellcome Sanger Institute COVID-19 Surveillance Team |
| EPI_ISL_2960945 | Lighthouse Lab in Alderley Park | Wellcome Sanger Institute for the COVID-19 Genomics UK (COG-UK) Consortium | Jacquelyn Wynn, Mairead Hyland, The Lighthouse Lab in Alderley Park and Alex Alderton, Roberto Amato, Jeffrey Barrett, Sonia Goncalves, Ewan Harrison, David K. Jackson, Ian Johnston, Dominic Kwiatkowski, Cordelia Langford, John Sillitoe on behalf of the Wellcome Sanger Institute COVID-19 Surveillance Team |
| EPI_ISL_2960946, EPI_ISL_2960947 | Berkshire and Surrey Pathology Services Lighthouse Laboratory | Wellcome Sanger Institute for the COVID-19 Genomics UK (COG-UK) Consortium | Berkshire and Surrey Pathology Services Lighthouse Laboratory and Alex Alderton, Roberto Amato, Jeffrey Barrett, Sonia Goncalves, Ewan Harrison, David K. Jackson, Ian Johnston, Dominic Kwiatkowski, Cordelia Langford, John Sillitoe on behalf of the Wellcome Sanger Institute COVID-19 Surveillance Team |
| EPI_ISL_2960948 | Lighthouse Lab in Alderley Park | Wellcome Sanger Institute for the COVID-19 Genomics UK (COG-UK) Consortium | Jacquelyn Wynn, Mairead Hyland, The Lighthouse Lab in Alderley Park and Alex Alderton, Roberto Amato, Jeffrey Barrett, Sonia Goncalves, Ewan Harrison, David K. Jackson, Ian Johnston, Dominic Kwiatkowski, Cordelia Langford, John Sillitoe on behalf of the Wellcome Sanger Institute COVID-19 Surveillance Team |
| EPI_ISL_2960949, EPI_ISL_2960950 | Lighthouse Laboratory Plymouth | Wellcome Sanger Institute for the COVID-19 Genomics UK (COG-UK) Consortium | Lighthouse Laboratory Plymouth and Alex Alderton, Roberto Amato, Jeffrey Barrett, Sonia Goncalves, Ewan Harrison, David K. Jackson, Ian Johnston, Dominic Kwiatkowski, Cordelia Langford, John Sillitoe on behalf of the Wellcome Sanger Institute COVID-19 Surveillance Team |
| EPI_ISL_2960951, EPI_ISL_2960952, EPI_ISL_2960953, EPI_ISL_2960954, EPI_ISL_2960955 | Lighthouse Lab in Alderley Park | Wellcome Sanger Institute for the COVID-19 Genomics UK (COG-UK) Consortium | Jacquelyn Wynn, Mairead Hyland, The Lighthouse Lab in Alderley Park and Alex Alderton, Roberto Amato, Jeffrey Barrett, Sonia Goncalves, Ewan Harrison, David K. Jackson, Ian Johnston, Dominic Kwiatkowski, Cordelia Langford, John Sillitoe on behalf of the Wellcome Sanger Institute COVID-19 Surveillance Team |
| EPI_ISL_2960956 | Berkshire and Surrey Pathology Services Lighthouse Laboratory | Wellcome Sanger Institute for the COVID-19 Genomics UK (COG-UK) Consortium | Berkshire and Surrey Pathology Services Lighthouse Laboratory and Alex Alderton, Roberto Amato, Jeffrey Barrett, Sonia Goncalves, Ewan Harrison, David K. Jackson, Ian Johnston, Dominic Kwiatkowski, Cordelia Langford, John Sillitoe on behalf of the Wellcome Sanger Institute COVID-19 Surveillance Team |
| EPI_ISL_2960957, EPI_ISL_2960958, EPI_ISL_2960959 | Lighthouse Lab in Alderley Park | Wellcome Sanger Institute for the COVID-19 Genomics UK (COG-UK) Consortium | Jacquelyn Wynn, Mairead Hyland, The Lighthouse Lab in Alderley Park and Alex Alderton, Roberto Amato, Jeffrey Barrett, Sonia Goncalves, Ewan Harrison, David K. Jackson, Ian Johnston, Dominic Kwiatkowski, Cordelia Langford, John Sillitoe on behalf of the Wellcome Sanger Institute COVID-19 Surveillance Team |
| EPI_ISL_2960960 | Lighthouse Laboratory Plymouth | Wellcome Sanger Institute for the COVID-19 Genomics UK (COG-UK) Consortium | Lighthouse Laboratory Plymouth and Alex Alderton, Roberto Amato, Jeffrey Barrett, Sonia Goncalves, Ewan Harrison, David K. Jackson, Ian Johnston, Dominic Kwiatkowski, Cordelia Langford, John Sillitoe on behalf of the Wellcome Sanger Institute COVID-19 Surveillance Team |
| EPI_ISL_2960961, EPI_ISL_2960962, EPI_ISL_2960963, EPI_ISL_2960964, EPI_ISL_2960965 | Lighthouse Lab in Alderley Park | Wellcome Sanger Institute for the COVID-19 Genomics UK (COG-UK) Consortium | Jacquelyn Wynn, Mairead Hyland, The Lighthouse Lab in Alderley Park and Alex Alderton, Roberto Amato, Jeffrey Barrett, Sonia Goncalves, Ewan Harrison, David K. Jackson, Ian Johnston, Dominic Kwiatkowski, Cordelia Langford, John Sillitoe on behalf of the Wellcome Sanger Institute COVID-19 Surveillance Team |
| EPI_ISL_2960966 | Berkshire and Surrey Pathology Services Lighthouse Laboratory | Wellcome Sanger Institute for the COVID-19 Genomics UK (COG-UK) Consortium | Berkshire and Surrey Pathology Services Lighthouse Laboratory and Alex Alderton, Roberto Amato, Jeffrey Barrett, Sonia Goncalves, Ewan Harrison, David K. Jackson, Ian Johnston, Dominic Kwiatkowski, Cordelia Langford, John Sillitoe on behalf of the Wellcome Sanger Institute COVID-19 Surveillance Team |
| EPI_ISL_2960967, EPI_ISL_2960968 | Lighthouse Laboratory Plymouth | Wellcome Sanger Institute for the COVID-19 Genomics UK (COG-UK) Consortium | Lighthouse Laboratory Plymouth and Alex Alderton, Roberto Amato, Jeffrey Barrett, Sonia Goncalves, Ewan Harrison, David K. Jackson, Ian Johnston, Dominic Kwiatkowski, Cordelia Langford, John Sillitoe on behalf of the Wellcome Sanger Institute COVID-19 Surveillance Team |
| EPI_ISL_2960969 | Lighthouse Lab in Alderley Park | Wellcome Sanger Institute for the COVID-19 Genomics UK (COG-UK) Consortium | Jacquelyn Wynn, Mairead Hyland, The Lighthouse Lab in Alderley Park and Alex Alderton, Roberto Amato, Jeffrey Barrett, Sonia Goncalves, Ewan Harrison, David K. Jackson, Ian Johnston, Dominic Kwiatkowski, Cordelia Langford, John Sillitoe on behalf of the Wellcome Sanger Institute COVID-19 Surveillance Team |
| EPI_ISL_2960970 | Berkshire and Surrey Pathology Services Lighthouse Laboratory | Wellcome Sanger Institute for the COVID-19 Genomics UK (COG-UK) Consortium | Berkshire and Surrey Pathology Services Lighthouse Laboratory and Alex Alderton, Roberto Amato, Jeffrey Barrett, Sonia Goncalves, Ewan Harrison, David K. Jackson, Ian Johnston, Dominic Kwiatkowski, Cordelia Langford, John Sillitoe on behalf of the Wellcome Sanger Institute COVID-19 Surveillance Team |
| EPI_ISL_2960971, EPI_ISL_2960972, EPI_ISL_2960973, EPI_ISL_2960974 | Lighthouse Lab in Alderley Park | Wellcome Sanger Institute for the COVID-19 Genomics UK (COG-UK) Consortium | Jacquelyn Wynn, Mairead Hyland, The Lighthouse Lab in Alderley Park and Alex Alderton, Roberto Amato, Jeffrey Barrett, Sonia Goncalves, Ewan Harrison, David K. Jackson, Ian Johnston, Dominic Kwiatkowski, Cordelia Langford, John Sillitoe on behalf of the Wellcome Sanger Institute COVID-19 Surveillance Team |
| EPI_ISL_2960975 | Lighthouse Laboratory Plymouth | Wellcome Sanger Institute for the COVID-19 Genomics UK (COG-UK) Consortium | Lighthouse Laboratory Plymouth and Alex Alderton, Roberto Amato, Jeffrey Barrett, Sonia Goncalves, Ewan Harrison, David K. Jackson, Ian Johnston, Dominic Kwiatkowski, Cordelia Langford, John Sillitoe on behalf of the Wellcome Sanger Institute COVID-19 Surveillance Team |
| EPI_ISL_2960976 | Lighthouse Lab in Alderley Park | Wellcome Sanger Institute for the COVID-19 Genomics UK (COG-UK) Consortium | Jacquelyn Wynn, Mairead Hyland, The Lighthouse Lab in Alderley Park and Alex Alderton, Roberto Amato, Jeffrey Barrett, Sonia Goncalves, Ewan Harrison, David K. Jackson, Ian Johnston, Dominic Kwiatkowski, Cordelia Langford, John Sillitoe on behalf of the Wellcome Sanger Institute COVID-19 Surveillance Team |
| EPI_ISL_2960977, EPI_ISL_2960978 | Lighthouse Laboratory Plymouth | Wellcome Sanger Institute for the COVID-19 Genomics UK (COG-UK) Consortium | Lighthouse Laboratory Plymouth and Alex Alderton, Roberto Amato, Jeffrey Barrett, Sonia Goncalves, Ewan Harrison, David K. Jackson, Ian Johnston, Dominic Kwiatkowski, Cordelia Langford, John Sillitoe on behalf of the Wellcome Sanger Institute COVID-19 Surveillance Team |
| EPI_ISL_2960979 | Berkshire and Surrey Pathology Services Lighthouse Laboratory | Wellcome Sanger Institute for the COVID-19 Genomics UK (COG-UK) Consortium | Berkshire and Surrey Pathology Services Lighthouse Laboratory and Alex Alderton, Roberto Amato, Jeffrey Barrett, Sonia Goncalves, Ewan Harrison, David K. Jackson, Ian Johnston, Dominic Kwiatkowski, Cordelia Langford, John Sillitoe on behalf of the Wellcome Sanger Institute COVID-19 Surveillance Team |
| EPI_ISL_2960980, EPI_ISL_2960981 | Lighthouse Lab in Alderley Park | Wellcome Sanger Institute for the COVID-19 Genomics UK (COG-UK) Consortium | Jacquelyn Wynn, Mairead Hyland, The Lighthouse Lab in Alderley Park and Alex Alderton, Roberto Amato, Jeffrey Barrett, Sonia Goncalves, Ewan Harrison, David K. Jackson, Ian Johnston, Dominic Kwiatkowski, Cordelia Langford, John Sillitoe on behalf of the Wellcome Sanger Institute COVID-19 Surveillance Team |
| EPI_ISL_2960982 | Lighthouse Laboratory Plymouth | Wellcome Sanger Institute for the COVID-19 Genomics UK (COG-UK) Consortium | Lighthouse Laboratory Plymouth and Alex Alderton, Roberto Amato, Jeffrey Barrett, Sonia Goncalves, Ewan Harrison, David K. Jackson, Ian Johnston, Dominic Kwiatkowski, Cordelia Langford, John Sillitoe on behalf of the Wellcome Sanger Institute COVID-19 Surveillance Team |
| EPI_ISL_2960983 | Berkshire and Surrey Pathology Services Lighthouse Laboratory | Wellcome Sanger Institute for the COVID-19 Genomics UK (COG-UK) Consortium | Berkshire and Surrey Pathology Services Lighthouse Laboratory and Alex Alderton, Roberto Amato, Jeffrey Barrett, Sonia Goncalves, Ewan Harrison, David K. Jackson, Ian Johnston, Dominic Kwiatkowski, Cordelia Langford, John Sillitoe on behalf of the Wellcome Sanger Institute COVID-19 Surveillance Team |
| EPI_ISL_2960984 | Lighthouse Lab in Alderley Park | Wellcome Sanger Institute for the COVID-19 Genomics UK (COG-UK) Consortium | Jacquelyn Wynn, Mairead Hyland, The Lighthouse Lab in Alderley Park and Alex Alderton, Roberto Amato, Jeffrey Barrett, Sonia Goncalves, Ewan Harrison, David K. Jackson, Ian Johnston, Dominic Kwiatkowski, Cordelia Langford, John Sillitoe on behalf of the Wellcome Sanger Institute COVID-19 Surveillance Team |
| EPI_ISL_2960985 | Lighthouse Laboratory Plymouth | Wellcome Sanger Institute for the COVID-19 Genomics UK (COG-UK) Consortium | Lighthouse Laboratory Plymouth and Alex Alderton, Roberto Amato, Jeffrey Barrett, Sonia Goncalves, Ewan Harrison, David K. Jackson, Ian Johnston, Dominic Kwiatkowski, Cordelia Langford, John Sillitoe on behalf of the Wellcome Sanger Institute COVID-19 Surveillance Team |
| EPI_ISL_2960986, EPI_ISL_2960987 | Lighthouse Lab in Alderley Park | Wellcome Sanger Institute for the COVID-19 Genomics UK (COG-UK) Consortium | Jacquelyn Wynn, Mairead Hyland, The Lighthouse Lab in Alderley Park and Alex Alderton, Roberto Amato, Jeffrey Barrett, Sonia Goncalves, Ewan Harrison, David K. Jackson, Ian Johnston, Dominic Kwiatkowski, Cordelia Langford, John Sillitoe on behalf of the Wellcome Sanger Institute COVID-19 Surveillance Team |
| EPI_ISL_2960988 | Lighthouse Laboratory Plymouth | Wellcome Sanger Institute for the COVID-19 Genomics UK (COG-UK) Consortium | Lighthouse Laboratory Plymouth and Alex Alderton, Roberto Amato, Jeffrey Barrett, Sonia Goncalves, Ewan Harrison, David K. Jackson, Ian Johnston, Dominic Kwiatkowski, Cordelia Langford, John Sillitoe on behalf of the Wellcome Sanger Institute COVID-19 Surveillance Team |
| EPI_ISL_2960989 | Lighthouse Lab in Alderley Park | Wellcome Sanger Institute for the COVID-19 Genomics UK (COG-UK) Consortium | Jacquelyn Wynn, Mairead Hyland, The Lighthouse Lab in Alderley Park and Alex Alderton, Roberto Amato, Jeffrey Barrett, Sonia Goncalves, Ewan Harrison, David K. Jackson, Ian Johnston, Dominic Kwiatkowski, Cordelia Langford, John Sillitoe on behalf of the Wellcome Sanger Institute COVID-19 Surveillance Team |
| EPI_ISL_2960990 | Berkshire and Surrey Pathology Services Lighthouse Laboratory | Wellcome Sanger Institute for the COVID-19 Genomics UK (COG-UK) Consortium | Berkshire and Surrey Pathology Services Lighthouse Laboratory and Alex Alderton, Roberto Amato, Jeffrey Barrett, Sonia Goncalves, Ewan Harrison, David K. Jackson, Ian Johnston, Dominic Kwiatkowski, Cordelia Langford, John Sillitoe on behalf of the Wellcome Sanger Institute COVID-19 Surveillance Team |
| EPI_ISL_2960991, EPI_ISL_2960992, | Lighthouse Lab in Alderley Park | Wellcome Sanger Institute for the COVID-19 Genomics UK (COG-UK) Consortium | Jacquelyn Wynn, Mairead Hyland, The Lighthouse Lab in Alderley Park and Alex Alderton, Roberto Amato, Jeffrey Barrett, Sonia Goncalves, Ewan Harrison, David K. Jackson, Ian Johnston, Dominic Kwiatkowski, Cordelia Langford, John Sillitoe on behalf of the Wellcome Sanger Institute COVID-19 Surveillance Team |

| | | | |
|---|---|---|---|
| EPI_ISL_2960993 | | | |
| EPI_ISL_2960994, EPI_ISL_2960995 | Berkshire and Surrey Pathology Services Lighthouse Laboratory | Wellcome Sanger Institute for the COVID-19 Genomics UK (COG-UK) Consortium | Berkshire and Surrey Pathology Services Lighthouse Laboratory and Alex Alderton, Roberto Amato, Jeffrey Barrett, Sonia Goncalves, Ewan Harrison, David K. Jackson, Ian Johnston, Dominic Kwiatkowski, Cordelia Langford, John Sillitoe on behalf of the Wellcome Sanger Institute COVID-19 Surveillance Team |
| EPI_ISL_2960996, EPI_ISL_2960997 | Lighthouse Lab in Alderley Park | Wellcome Sanger Institute for the COVID-19 Genomics UK (COG-UK) Consortium | Jacquelyn Wynn, Mairead Hyland, The Lighthouse Lab in Alderley Park and Alex Alderton, Roberto Amato, Jeffrey Barrett, Sonia Goncalves, Ewan Harrison, David K. Jackson, Ian Johnston, Dominic Kwiatkowski, Cordelia Langford, John Sillitoe on behalf of the Wellcome Sanger Institute COVID-19 Surveillance Team |
| EPI_ISL_2960998 | Berkshire and Surrey Pathology Services Lighthouse Laboratory | Wellcome Sanger Institute for the COVID-19 Genomics UK (COG-UK) Consortium | Berkshire and Surrey Pathology Services Lighthouse Laboratory and Alex Alderton, Roberto Amato, Jeffrey Barrett, Sonia Goncalves, Ewan Harrison, David K. Jackson, Ian Johnston, Dominic Kwiatkowski, Cordelia Langford, John Sillitoe on behalf of the Wellcome Sanger Institute COVID-19 Surveillance Team |
| EPI_ISL_2960999 | Lighthouse Laboratory Plymouth | Wellcome Sanger Institute for the COVID-19 Genomics UK (COG-UK) Consortium | Lighthouse Laboratory Plymouth and Alex Alderton, Roberto Amato, Jeffrey Barrett, Sonia Goncalves, Ewan Harrison, David K. Jackson, Ian Johnston, Dominic Kwiatkowski, Cordelia Langford, John Sillitoe on behalf of the Wellcome Sanger Institute COVID-19 Surveillance Team |
| EPI_ISL_2961000 | Berkshire and Surrey Pathology Services Lighthouse Laboratory | Wellcome Sanger Institute for the COVID-19 Genomics UK (COG-UK) Consortium | Berkshire and Surrey Pathology Services Lighthouse Laboratory and Alex Alderton, Roberto Amato, Jeffrey Barrett, Sonia Goncalves, Ewan Harrison, David K. Jackson, Ian Johnston, Dominic Kwiatkowski, Cordelia Langford, John Sillitoe on behalf of the Wellcome Sanger Institute COVID-19 Surveillance Team |
| EPI_ISL_2961001 | Lighthouse Lab in Alderley Park | Wellcome Sanger Institute for the COVID-19 Genomics UK (COG-UK) Consortium | Jacquelyn Wynn, Mairead Hyland, The Lighthouse Lab in Alderley Park and Alex Alderton, Roberto Amato, Jeffrey Barrett, Sonia Goncalves, Ewan Harrison, David K. Jackson, Ian Johnston, Dominic Kwiatkowski, Cordelia Langford, John Sillitoe on behalf of the Wellcome Sanger Institute COVID-19 Surveillance Team |
| EPI_ISL_2961002 | Lighthouse Laboratory Plymouth | Wellcome Sanger Institute for the COVID-19 Genomics UK (COG-UK) Consortium | Lighthouse Laboratory Plymouth and Alex Alderton, Roberto Amato, Jeffrey Barrett, Sonia Goncalves, Ewan Harrison, David K. Jackson, Ian Johnston, Dominic Kwiatkowski, Cordelia Langford, John Sillitoe on behalf of the Wellcome Sanger Institute COVID-19 Surveillance Team |
| EPI_ISL_2961003, EPI_ISL_2961004 | Lighthouse Lab in Alderley Park | Wellcome Sanger Institute for the COVID-19 Genomics UK (COG-UK) Consortium | Jacquelyn Wynn, Mairead Hyland, The Lighthouse Lab in Alderley Park and Alex Alderton, Roberto Amato, Jeffrey Barrett, Sonia Goncalves, Ewan Harrison, David K. Jackson, Ian Johnston, Dominic Kwiatkowski, Cordelia Langford, John Sillitoe on behalf of the Wellcome Sanger Institute COVID-19 Surveillance Team |
| EPI_ISL_2961005 | Lighthouse Laboratory Plymouth | Wellcome Sanger Institute for the COVID-19 Genomics UK (COG-UK) Consortium | Lighthouse Laboratory Plymouth and Alex Alderton, Roberto Amato, Jeffrey Barrett, Sonia Goncalves, Ewan Harrison, David K. Jackson, Ian Johnston, Dominic Kwiatkowski, Cordelia Langford, John Sillitoe on behalf of the Wellcome Sanger Institute COVID-19 Surveillance Team |
| EPI_ISL_2961006 | Lighthouse Lab in Alderley Park | Wellcome Sanger Institute for the COVID-19 Genomics UK (COG-UK) Consortium | Jacquelyn Wynn, Mairead Hyland, The Lighthouse Lab in Alderley Park and Alex Alderton, Roberto Amato, Jeffrey Barrett, Sonia Goncalves, Ewan Harrison, David K. Jackson, Ian Johnston, Dominic Kwiatkowski, Cordelia Langford, John Sillitoe on behalf of the Wellcome Sanger Institute COVID-19 Surveillance Team |
| EPI_ISL_2961007 | Lighthouse Laboratory Plymouth | Wellcome Sanger Institute for the COVID-19 Genomics UK (COG-UK) Consortium | Lighthouse Laboratory Plymouth and Alex Alderton, Roberto Amato, Jeffrey Barrett, Sonia Goncalves, Ewan Harrison, David K. Jackson, Ian Johnston, Dominic Kwiatkowski, Cordelia Langford, John Sillitoe on behalf of the Wellcome Sanger Institute COVID-19 Surveillance Team |
| EPI_ISL_2961008 | Lighthouse Lab in Alderley Park | Wellcome Sanger Institute for the COVID-19 Genomics UK (COG-UK) Consortium | Jacquelyn Wynn, Mairead Hyland, The Lighthouse Lab in Alderley Park and Alex Alderton, Roberto Amato, Jeffrey Barrett, Sonia Goncalves, Ewan Harrison, David K. Jackson, Ian Johnston, Dominic Kwiatkowski, Cordelia Langford, John Sillitoe on behalf of the Wellcome Sanger Institute COVID-19 Surveillance Team |
| EPI_ISL_2961009, EPI_ISL_2961010 | Lighthouse Laboratory Plymouth | Wellcome Sanger Institute for the COVID-19 Genomics UK (COG-UK) Consortium | Lighthouse Laboratory Plymouth and Alex Alderton, Roberto Amato, Jeffrey Barrett, Sonia Goncalves, Ewan Harrison, David K. Jackson, Ian Johnston, Dominic Kwiatkowski, Cordelia Langford, John Sillitoe on behalf of the Wellcome Sanger Institute COVID-19 Surveillance Team |
| EPI_ISL_2961011 | Lighthouse Lab in Alderley Park | Wellcome Sanger Institute for the COVID-19 Genomics UK (COG-UK) Consortium | Jacquelyn Wynn, Mairead Hyland, The Lighthouse Lab in Alderley Park and Alex Alderton, Roberto Amato, Jeffrey Barrett, Sonia Goncalves, Ewan Harrison, David K. Jackson, Ian Johnston, Dominic Kwiatkowski, Cordelia Langford, John Sillitoe on behalf of the Wellcome Sanger Institute COVID-19 Surveillance Team |
| EPI_ISL_2961012, EPI_ISL_2961013, EPI_ISL_2961014 | Berkshire and Surrey Pathology Services Lighthouse Laboratory | Wellcome Sanger Institute for the COVID-19 Genomics UK (COG-UK) Consortium | Berkshire and Surrey Pathology Services Lighthouse Laboratory and Alex Alderton, Roberto Amato, Jeffrey Barrett, Sonia Goncalves, Ewan Harrison, David K. Jackson, Ian Johnston, Dominic Kwiatkowski, Cordelia Langford, John Sillitoe on behalf of the Wellcome Sanger Institute COVID-19 Surveillance Team |
| EPI_ISL_2961015, EPI_ISL_2961016 | Lighthouse Laboratory Plymouth | Wellcome Sanger Institute for the COVID-19 Genomics UK (COG-UK) Consortium | Lighthouse Laboratory Plymouth and Alex Alderton, Roberto Amato, Jeffrey Barrett, Sonia Goncalves, Ewan Harrison, David K. Jackson, Ian Johnston, Dominic Kwiatkowski, Cordelia Langford, John Sillitoe on behalf of the Wellcome Sanger Institute COVID-19 Surveillance Team |
| EPI_ISL_2961017 | Lighthouse Lab in Alderley Park | Wellcome Sanger Institute for the COVID-19 Genomics UK (COG-UK) Consortium | Jacquelyn Wynn, Mairead Hyland, The Lighthouse Lab in Alderley Park and Alex Alderton, Roberto Amato, Jeffrey Barrett, Sonia Goncalves, Ewan Harrison, David K. Jackson, Ian Johnston, Dominic Kwiatkowski, Cordelia Langford, John Sillitoe on behalf of the Wellcome Sanger Institute COVID-19 Surveillance Team |
| EPI_ISL_2961018, EPI_ISL_2961019, EPI_ISL_2961020 | Lighthouse Laboratory Plymouth | Wellcome Sanger Institute for the COVID-19 Genomics UK (COG-UK) Consortium | Lighthouse Laboratory Plymouth and Alex Alderton, Roberto Amato, Jeffrey Barrett, Sonia Goncalves, Ewan Harrison, David K. Jackson, Ian Johnston, Dominic Kwiatkowski, Cordelia Langford, John Sillitoe on behalf of the Wellcome Sanger Institute COVID-19 Surveillance Team |
| EPI_ISL_2961021 | Lighthouse Lab in Alderley Park | Wellcome Sanger Institute for the COVID-19 Genomics UK (COG-UK) Consortium | Jacquelyn Wynn, Mairead Hyland, The Lighthouse Lab in Alderley Park and Alex Alderton, Roberto Amato, Jeffrey Barrett, Sonia Goncalves, Ewan Harrison, David K. Jackson, Ian Johnston, Dominic Kwiatkowski, Cordelia Langford, John Sillitoe on behalf of the Wellcome Sanger Institute COVID-19 Surveillance Team |
| EPI_ISL_2961022, EPI_ISL_2961023 | Berkshire and Surrey Pathology Services Lighthouse Laboratory | Wellcome Sanger Institute for the COVID-19 Genomics UK (COG-UK) Consortium | Berkshire and Surrey Pathology Services Lighthouse Laboratory and Alex Alderton, Roberto Amato, Jeffrey Barrett, Sonia Goncalves, Ewan Harrison, David K. Jackson, Ian Johnston, Dominic Kwiatkowski, Cordelia Langford, John Sillitoe on behalf of the Wellcome Sanger Institute COVID-19 Surveillance Team |
| EPI_ISL_2961024, EPI_ISL_2961025 | Lighthouse Lab in Alderley Park | Wellcome Sanger Institute for the COVID-19 Genomics UK (COG-UK) Consortium | Jacquelyn Wynn, Mairead Hyland, The Lighthouse Lab in Alderley Park and Alex Alderton, Roberto Amato, Jeffrey Barrett, Sonia Goncalves, Ewan Harrison, David K. Jackson, Ian Johnston, Dominic Kwiatkowski, Cordelia Langford, John Sillitoe on behalf of the Wellcome Sanger Institute COVID-19 Surveillance Team |
| EPI_ISL_2961026 | Berkshire and Surrey Pathology Services Lighthouse Laboratory | Wellcome Sanger Institute for the COVID-19 Genomics UK (COG-UK) Consortium | Berkshire and Surrey Pathology Services Lighthouse Laboratory and Alex Alderton, Roberto Amato, Jeffrey Barrett, Sonia Goncalves, Ewan Harrison, David K. Jackson, Ian Johnston, Dominic Kwiatkowski, Cordelia Langford, John Sillitoe on behalf of the Wellcome Sanger Institute COVID-19 Surveillance Team |
| EPI_ISL_2961027 | Lighthouse Lab in Alderley Park | Wellcome Sanger Institute for the COVID-19 Genomics UK (COG-UK) Consortium | Jacquelyn Wynn, Mairead Hyland, The Lighthouse Lab in Alderley Park and Alex Alderton, Roberto Amato, Jeffrey Barrett, Sonia Goncalves, Ewan Harrison, David K. Jackson, Ian Johnston, Dominic Kwiatkowski, Cordelia Langford, John Sillitoe on behalf of the Wellcome Sanger Institute COVID-19 Surveillance Team |
| EPI_ISL_2961028, EPI_ISL_2961029 | Lighthouse Laboratory Plymouth | Wellcome Sanger Institute for the COVID-19 Genomics UK (COG-UK) Consortium | Lighthouse Laboratory Plymouth and Alex Alderton, Roberto Amato, Jeffrey Barrett, Sonia Goncalves, Ewan Harrison, David K. Jackson, Ian Johnston, Dominic Kwiatkowski, Cordelia Langford, John Sillitoe on behalf of the Wellcome Sanger Institute COVID-19 Surveillance Team |
| EPI_ISL_2961030 | Lighthouse Lab in Alderley Park | Wellcome Sanger Institute for the COVID-19 Genomics UK (COG-UK) Consortium | Jacquelyn Wynn, Mairead Hyland, The Lighthouse Lab in Alderley Park and Alex Alderton, Roberto Amato, Jeffrey Barrett, Sonia Goncalves, Ewan Harrison, David K. Jackson, Ian Johnston, Dominic Kwiatkowski, Cordelia Langford, John Sillitoe on behalf of the Wellcome Sanger Institute COVID-19 Surveillance Team |
| EPI_ISL_2961031 | Berkshire and Surrey Pathology Services Lighthouse Laboratory | Wellcome Sanger Institute for the COVID-19 Genomics UK (COG-UK) Consortium | Berkshire and Surrey Pathology Services Lighthouse Laboratory and Alex Alderton, Roberto Amato, Jeffrey Barrett, Sonia Goncalves, Ewan Harrison, David K. Jackson, Ian Johnston, Dominic Kwiatkowski, Cordelia Langford, John Sillitoe on behalf of the Wellcome Sanger Institute COVID-19 Surveillance Team |
| EPI_ISL_2961032 | Lighthouse Lab in Alderley Park | Wellcome Sanger Institute for the COVID-19 Genomics UK (COG-UK) Consortium | Jacquelyn Wynn, Mairead Hyland, The Lighthouse Lab in Alderley Park and Alex Alderton, Roberto Amato, Jeffrey Barrett, Sonia Goncalves, Ewan Harrison, David K. Jackson, Ian Johnston, Dominic Kwiatkowski, Cordelia Langford, John Sillitoe on behalf of the Wellcome Sanger Institute COVID-19 Surveillance Team |
| EPI_ISL_2961033 | Berkshire and Surrey Pathology Services Lighthouse Laboratory | Wellcome Sanger Institute for the COVID-19 Genomics UK (COG-UK) Consortium | Berkshire and Surrey Pathology Services Lighthouse Laboratory and Alex Alderton, Roberto Amato, Jeffrey Barrett, Sonia Goncalves, Ewan Harrison, David K. Jackson, Ian Johnston, Dominic Kwiatkowski, Cordelia Langford, John Sillitoe on behalf of the Wellcome Sanger Institute COVID-19 Surveillance Team |
| EPI_ISL_2961034 | Lighthouse Lab in Alderley Park | Wellcome Sanger Institute for the COVID-19 Genomics UK (COG-UK) Consortium | Jacquelyn Wynn, Mairead Hyland, The Lighthouse Lab in Alderley Park and Alex Alderton, Roberto Amato, Jeffrey Barrett, Sonia Goncalves, Ewan Harrison, David K. Jackson, Ian Johnston, Dominic Kwiatkowski, Cordelia Langford, John Sillitoe on behalf of the Wellcome Sanger Institute COVID-19 Surveillance Team |
| EPI_ISL_2961035 | Berkshire and Surrey Pathology Services Lighthouse Laboratory | Wellcome Sanger Institute for the COVID-19 Genomics UK (COG-UK) Consortium | Berkshire and Surrey Pathology Services Lighthouse Laboratory and Alex Alderton, Roberto Amato, Jeffrey Barrett, Sonia Goncalves, Ewan Harrison, David K. Jackson, Ian Johnston, Dominic Kwiatkowski, Cordelia Langford, John Sillitoe on behalf of the Wellcome Sanger Institute COVID-19 Surveillance Team |
| EPI_ISL_2961036 | Lighthouse Lab in Alderley Park | Wellcome Sanger Institute for the COVID-19 Genomics UK (COG-UK) Consortium | Jacquelyn Wynn, Mairead Hyland, The Lighthouse Lab in Alderley Park and Alex Alderton, Roberto Amato, Jeffrey Barrett, Sonia Goncalves, Ewan Harrison, David K. Jackson, Ian Johnston, Dominic Kwiatkowski, Cordelia Langford, John Sillitoe on behalf of the Wellcome Sanger Institute COVID-19 Surveillance Team |
| EPI_ISL_2961037 | Berkshire and Surrey Pathology Services Lighthouse Laboratory | Wellcome Sanger Institute for the COVID-19 Genomics UK (COG-UK) Consortium | Berkshire and Surrey Pathology Services Lighthouse Laboratory and Alex Alderton, Roberto Amato, Jeffrey Barrett, Sonia Goncalves, Ewan Harrison, David K. Jackson, Ian Johnston, Dominic Kwiatkowski, Cordelia Langford, John Sillitoe on behalf of the Wellcome Sanger Institute COVID-19 Surveillance Team |
| EPI_ISL_2961038, EPI_ISL_2961039 | Lighthouse Lab in Alderley Park | Wellcome Sanger Institute for the COVID-19 Genomics UK (COG-UK) Consortium | Jacquelyn Wynn, Mairead Hyland, The Lighthouse Lab in Alderley Park and Alex Alderton, Roberto Amato, Jeffrey Barrett, Sonia Goncalves, Ewan Harrison, David K. Jackson, Ian Johnston, Dominic Kwiatkowski, Cordelia Langford, John Sillitoe on behalf of the Wellcome Sanger Institute COVID-19 Surveillance Team |
| EPI_ISL_2961040 | Berkshire and Surrey Pathology Services Lighthouse Laboratory | Wellcome Sanger Institute for the COVID-19 Genomics UK (COG-UK) Consortium | Berkshire and Surrey Pathology Services Lighthouse Laboratory and Alex Alderton, Roberto Amato, Jeffrey Barrett, Sonia Goncalves, Ewan Harrison, David K. Jackson, Ian Johnston, Dominic Kwiatkowski, Cordelia Langford, John Sillitoe on behalf of the Wellcome Sanger Institute COVID-19 Surveillance Team |
| EPI_ISL_2961041, EPI_ISL_2961042, EPI_ISL_2961043 | Lighthouse Laboratory Plymouth | Wellcome Sanger Institute for the COVID-19 Genomics UK (COG-UK) Consortium | Lighthouse Laboratory Plymouth and Alex Alderton, Roberto Amato, Jeffrey Barrett, Sonia Goncalves, Ewan Harrison, David K. Jackson, Ian Johnston, Dominic Kwiatkowski, Cordelia Langford, John Sillitoe on behalf of the Wellcome Sanger Institute COVID-19 Surveillance Team |
| EPI_ISL_2961044 | Lighthouse Lab in Alderley Park | Wellcome Sanger Institute for the COVID-19 Genomics UK (COG-UK) Consortium | Jacquelyn Wynn, Mairead Hyland, The Lighthouse Lab in Alderley Park and Alex Alderton, Roberto Amato, Jeffrey Barrett, Sonia Goncalves, Ewan Harrison, David K. Jackson, Ian Johnston, Dominic Kwiatkowski, Cordelia Langford, John Sillitoe on behalf of the Wellcome Sanger Institute COVID-19 Surveillance Team |
| EPI_ISL_2961045 | Lighthouse Laboratory Plymouth | Wellcome Sanger Institute for the COVID-19 Genomics UK (COG-UK) Consortium | Lighthouse Laboratory Plymouth and Alex Alderton, Roberto Amato, Jeffrey Barrett, Sonia Goncalves, Ewan Harrison, David K. Jackson, Ian Johnston, Dominic Kwiatkowski, Cordelia Langford, John Sillitoe on behalf of the Wellcome Sanger Institute COVID-19 Surveillance Team |
| EPI_ISL_2961046 | Berkshire and Surrey Pathology Services Lighthouse Laboratory | Wellcome Sanger Institute for the COVID-19 Genomics UK (COG-UK) Consortium | Berkshire and Surrey Pathology Services Lighthouse Laboratory and Alex Alderton, Roberto Amato, Jeffrey Barrett, Sonia Goncalves, Ewan Harrison, David K. Jackson, Ian Johnston, Dominic Kwiatkowski, Cordelia Langford, John Sillitoe on behalf of the Wellcome Sanger Institute COVID-19 Surveillance Team |
| EPI_ISL_2961047, EPI_ISL_2961048 | Lighthouse Lab in Alderley Park | Wellcome Sanger Institute for the COVID-19 Genomics UK (COG-UK) Consortium | Jacquelyn Wynn, Mairead Hyland, The Lighthouse Lab in Alderley Park and Alex Alderton, Roberto Amato, Jeffrey Barrett, Sonia Goncalves, Ewan Harrison, David K. Jackson, Ian Johnston, Dominic Kwiatkowski, Cordelia Langford, John Sillitoe on behalf of the Wellcome Sanger Institute COVID-19 Surveillance Team |
| EPI_ISL_2961049 | Lighthouse Laboratory Plymouth | Wellcome Sanger Institute for the COVID-19 Genomics UK (COG-UK) Consortium | Lighthouse Laboratory Plymouth and Alex Alderton, Roberto Amato, Jeffrey Barrett, Sonia Goncalves, Ewan Harrison, David K. Jackson, Ian Johnston, Dominic Kwiatkowski, Cordelia Langford, John Sillitoe on behalf of the Wellcome Sanger Institute COVID-19 Surveillance Team |
| EPI_ISL_2961050, EPI_ISL_2961051 | Lighthouse Lab in Alderley Park | Wellcome Sanger Institute for the COVID-19 Genomics UK (COG-UK) Consortium | Jacquelyn Wynn, Mairead Hyland, The Lighthouse Lab in Alderley Park and Alex Alderton, Roberto Amato, Jeffrey Barrett, Sonia Goncalves, Ewan Harrison, David K. Jackson, Ian Johnston, Dominic Kwiatkowski, Cordelia Langford, John Sillitoe on behalf of the Wellcome Sanger Institute COVID-19 Surveillance Team |
| EPI_ISL_2961052 | Berkshire and Surrey Pathology Services Lighthouse Laboratory | Wellcome Sanger Institute for the COVID-19 Genomics UK (COG-UK) Consortium | Berkshire and Surrey Pathology Services Lighthouse Laboratory and Alex Alderton, Roberto Amato, Jeffrey Barrett, Sonia Goncalves, Ewan Harrison, David K. Jackson, Ian Johnston, Dominic Kwiatkowski, Cordelia Langford, John Sillitoe on behalf of the Wellcome Sanger Institute COVID-19 Surveillance Team |
| EPI_ISL_2961053, EPI_ISL_2961054 | Lighthouse Lab in Alderley Park | Wellcome Sanger Institute for the COVID-19 Genomics UK (COG-UK) Consortium | Jacquelyn Wynn, Mairead Hyland, The Lighthouse Lab in Alderley Park and Alex Alderton, Roberto Amato, Jeffrey Barrett, Sonia Goncalves, Ewan Harrison, David K. Jackson, Ian Johnston, Dominic Kwiatkowski, Cordelia Langford, John Sillitoe on behalf of the Wellcome Sanger Institute COVID-19 Surveillance Team |

| | | | |
|---|---|---|---|
| EPI_ISL_2961055 | Berkshire and Surrey Pathology Services Lighthouse Laboratory | Wellcome Sanger Institute for the COVID-19 Genomics UK (COG-UK) Consortium | Berkshire and Surrey Pathology Services Lighthouse Laboratory and Alex Alderton, Roberto Amato, Jeffrey Barrett, Sonia Goncalves, Ewan Harrison, David K. Jackson, Ian Johnston, Dominic Kwiatkowski, Cordelia Langford, John Sillitoe on behalf of the Wellcome Sanger Institute COVID-19 Surveillance Team |
| EPI_ISL_2961056 | Lighthouse Laboratory Plymouth | Wellcome Sanger Institute for the COVID-19 Genomics UK (COG-UK) Consortium | Lighthouse Laboratory Plymouth and Alex Alderton, Roberto Amato, Jeffrey Barrett, Sonia Goncalves, Ewan Harrison, David K. Jackson, Ian Johnston, Dominic Kwiatkowski, Cordelia Langford, John Sillitoe on behalf of the Wellcome Sanger Institute COVID-19 Surveillance Team |
| EPI_ISL_2961057 | Lighthouse Lab in Alderley Park | Wellcome Sanger Institute for the COVID-19 Genomics UK (COG-UK) Consortium | Jacquelyn Wynn, Mairead Hyland, The Lighthouse Lab in Alderley Park and Alex Alderton, Roberto Amato, Jeffrey Barrett, Sonia Goncalves, Ewan Harrison, David K. Jackson, Ian Johnston, Dominic Kwiatkowski, Cordelia Langford, John Sillitoe on behalf of the Wellcome Sanger Institute COVID-19 Surveillance Team |
| EPI_ISL_2961058 | Berkshire and Surrey Pathology Services Lighthouse Laboratory | Wellcome Sanger Institute for the COVID-19 Genomics UK (COG-UK) Consortium | Berkshire and Surrey Pathology Services Lighthouse Laboratory and Alex Alderton, Roberto Amato, Jeffrey Barrett, Sonia Goncalves, Ewan Harrison, David K. Jackson, Ian Johnston, Dominic Kwiatkowski, Cordelia Langford, John Sillitoe on behalf of the Wellcome Sanger Institute COVID-19 Surveillance Team |
| EPI_ISL_2961059 | Lighthouse Laboratory Plymouth | Wellcome Sanger Institute for the COVID-19 Genomics UK (COG-UK) Consortium | Lighthouse Laboratory Plymouth and Alex Alderton, Roberto Amato, Jeffrey Barrett, Sonia Goncalves, Ewan Harrison, David K. Jackson, Ian Johnston, Dominic Kwiatkowski, Cordelia Langford, John Sillitoe on behalf of the Wellcome Sanger Institute COVID-19 Surveillance Team |
| EPI_ISL_2961060 | Lighthouse Lab in Alderley Park | Wellcome Sanger Institute for the COVID-19 Genomics UK (COG-UK) Consortium | Jacquelyn Wynn, Mairead Hyland, The Lighthouse Lab in Alderley Park and Alex Alderton, Roberto Amato, Jeffrey Barrett, Sonia Goncalves, Ewan Harrison, David K. Jackson, Ian Johnston, Dominic Kwiatkowski, Cordelia Langford, John Sillitoe on behalf of the Wellcome Sanger Institute COVID-19 Surveillance Team |
| EPI_ISL_2961061 | Lighthouse Laboratory Plymouth | Wellcome Sanger Institute for the COVID-19 Genomics UK (COG-UK) Consortium | Lighthouse Laboratory Plymouth and Alex Alderton, Roberto Amato, Jeffrey Barrett, Sonia Goncalves, Ewan Harrison, David K. Jackson, Ian Johnston, Dominic Kwiatkowski, Cordelia Langford, John Sillitoe on behalf of the Wellcome Sanger Institute COVID-19 Surveillance Team |
| EPI_ISL_2961062, EPI_ISL_2961063 | Lighthouse Lab in Alderley Park | Wellcome Sanger Institute for the COVID-19 Genomics UK (COG-UK) Consortium | Jacquelyn Wynn, Mairead Hyland, The Lighthouse Lab in Alderley Park and Alex Alderton, Roberto Amato, Jeffrey Barrett, Sonia Goncalves, Ewan Harrison, David K. Jackson, Ian Johnston, Dominic Kwiatkowski, Cordelia Langford, John Sillitoe on behalf of the Wellcome Sanger Institute COVID-19 Surveillance Team |
| EPI_ISL_2961064 | Berkshire and Surrey Pathology Services Lighthouse Laboratory | Wellcome Sanger Institute for the COVID-19 Genomics UK (COG-UK) Consortium | Berkshire and Surrey Pathology Services Lighthouse Laboratory and Alex Alderton, Roberto Amato, Jeffrey Barrett, Sonia Goncalves, Ewan Harrison, David K. Jackson, Ian Johnston, Dominic Kwiatkowski, Cordelia Langford, John Sillitoe on behalf of the Wellcome Sanger Institute COVID-19 Surveillance Team |
| EPI_ISL_2961065, EPI_ISL_2961066 | Lighthouse Lab in Alderley Park | Wellcome Sanger Institute for the COVID-19 Genomics UK (COG-UK) Consortium | Jacquelyn Wynn, Mairead Hyland, The Lighthouse Lab in Alderley Park and Alex Alderton, Roberto Amato, Jeffrey Barrett, Sonia Goncalves, Ewan Harrison, David K. Jackson, Ian Johnston, Dominic Kwiatkowski, Cordelia Langford, John Sillitoe on behalf of the Wellcome Sanger Institute COVID-19 Surveillance Team |
| EPI_ISL_2961067 | Berkshire and Surrey Pathology Services Lighthouse Laboratory | Wellcome Sanger Institute for the COVID-19 Genomics UK (COG-UK) Consortium | Berkshire and Surrey Pathology Services Lighthouse Laboratory and Alex Alderton, Roberto Amato, Jeffrey Barrett, Sonia Goncalves, Ewan Harrison, David K. Jackson, Ian Johnston, Dominic Kwiatkowski, Cordelia Langford, John Sillitoe on behalf of the Wellcome Sanger Institute COVID-19 Surveillance Team |
| EPI_ISL_2961068 | Lighthouse Lab in Alderley Park | Wellcome Sanger Institute for the COVID-19 Genomics UK (COG-UK) Consortium | Jacquelyn Wynn, Mairead Hyland, The Lighthouse Lab in Alderley Park and Alex Alderton, Roberto Amato, Jeffrey Barrett, Sonia Goncalves, Ewan Harrison, David K. Jackson, Ian Johnston, Dominic Kwiatkowski, Cordelia Langford, John Sillitoe on behalf of the Wellcome Sanger Institute COVID-19 Surveillance Team |
| EPI_ISL_2961069, EPI_ISL_2961070 | Lighthouse Laboratory Plymouth | Wellcome Sanger Institute for the COVID-19 Genomics UK (COG-UK) Consortium | Lighthouse Laboratory Plymouth and Alex Alderton, Roberto Amato, Jeffrey Barrett, Sonia Goncalves, Ewan Harrison, David K. Jackson, Ian Johnston, Dominic Kwiatkowski, Cordelia Langford, John Sillitoe on behalf of the Wellcome Sanger Institute COVID-19 Surveillance Team |
| EPI_ISL_2961071 | Lighthouse Lab in Alderley Park | Wellcome Sanger Institute for the COVID-19 Genomics UK (COG-UK) Consortium | Jacquelyn Wynn, Mairead Hyland, The Lighthouse Lab in Alderley Park and Alex Alderton, Roberto Amato, Jeffrey Barrett, Sonia Goncalves, Ewan Harrison, David K. Jackson, Ian Johnston, Dominic Kwiatkowski, Cordelia Langford, John Sillitoe on behalf of the Wellcome Sanger Institute COVID-19 Surveillance Team |
| EPI_ISL_2961072 | Lighthouse Laboratory Plymouth | Wellcome Sanger Institute for the COVID-19 Genomics UK (COG-UK) Consortium | Lighthouse Laboratory Plymouth and Alex Alderton, Roberto Amato, Jeffrey Barrett, Sonia Goncalves, Ewan Harrison, David K. Jackson, Ian Johnston, Dominic Kwiatkowski, Cordelia Langford, John Sillitoe on behalf of the Wellcome Sanger Institute COVID-19 Surveillance Team |
| EPI_ISL_2961073 | Lighthouse Lab in Alderley Park | Wellcome Sanger Institute for the COVID-19 Genomics UK (COG-UK) Consortium | Jacquelyn Wynn, Mairead Hyland, The Lighthouse Lab in Alderley Park and Alex Alderton, Roberto Amato, Jeffrey Barrett, Sonia Goncalves, Ewan Harrison, David K. Jackson, Ian Johnston, Dominic Kwiatkowski, Cordelia Langford, John Sillitoe on behalf of the Wellcome Sanger Institute COVID-19 Surveillance Team |
| EPI_ISL_2961074 | Berkshire and Surrey Pathology Services Lighthouse Laboratory | Wellcome Sanger Institute for the COVID-19 Genomics UK (COG-UK) Consortium | Berkshire and Surrey Pathology Services Lighthouse Laboratory and Alex Alderton, Roberto Amato, Jeffrey Barrett, Sonia Goncalves, Ewan Harrison, David K. Jackson, Ian Johnston, Dominic Kwiatkowski, Cordelia Langford, John Sillitoe on behalf of the Wellcome Sanger Institute COVID-19 Surveillance Team |
| EPI_ISL_2961075 | Lighthouse Lab in Alderley Park | Wellcome Sanger Institute for the COVID-19 Genomics UK (COG-UK) Consortium | Jacquelyn Wynn, Mairead Hyland, The Lighthouse Lab in Alderley Park and Alex Alderton, Roberto Amato, Jeffrey Barrett, Sonia Goncalves, Ewan Harrison, David K. Jackson, Ian Johnston, Dominic Kwiatkowski, Cordelia Langford, John Sillitoe on behalf of the Wellcome Sanger Institute COVID-19 Surveillance Team |
| EPI_ISL_2961076, EPI_ISL_2961077 | Berkshire and Surrey Pathology Services Lighthouse Laboratory | Wellcome Sanger Institute for the COVID-19 Genomics UK (COG-UK) Consortium | Berkshire and Surrey Pathology Services Lighthouse Laboratory and Alex Alderton, Roberto Amato, Jeffrey Barrett, Sonia Goncalves, Ewan Harrison, David K. Jackson, Ian Johnston, Dominic Kwiatkowski, Cordelia Langford, John Sillitoe on behalf of the Wellcome Sanger Institute COVID-19 Surveillance Team |
| EPI_ISL_2961078, EPI_ISL_2961079 | Lighthouse Laboratory Plymouth | Wellcome Sanger Institute for the COVID-19 Genomics UK (COG-UK) Consortium | Lighthouse Laboratory Plymouth and Alex Alderton, Roberto Amato, Jeffrey Barrett, Sonia Goncalves, Ewan Harrison, David K. Jackson, Ian Johnston, Dominic Kwiatkowski, Cordelia Langford, John Sillitoe on behalf of the Wellcome Sanger Institute COVID-19 Surveillance Team |
| EPI_ISL_2961080 | Lighthouse Lab in Alderley Park | Wellcome Sanger Institute for the COVID-19 Genomics UK (COG-UK) Consortium | Jacquelyn Wynn, Mairead Hyland, The Lighthouse Lab in Alderley Park and Alex Alderton, Roberto Amato, Jeffrey Barrett, Sonia Goncalves, Ewan Harrison, David K. Jackson, Ian Johnston, Dominic Kwiatkowski, Cordelia Langford, John Sillitoe on behalf of the Wellcome Sanger Institute COVID-19 Surveillance Team |
| EPI_ISL_2961081 | Berkshire and Surrey Pathology Services Lighthouse Laboratory | Wellcome Sanger Institute for the COVID-19 Genomics UK (COG-UK) Consortium | Berkshire and Surrey Pathology Services Lighthouse Laboratory and Alex Alderton, Roberto Amato, Jeffrey Barrett, Sonia Goncalves, Ewan Harrison, David K. Jackson, Ian Johnston, Dominic Kwiatkowski, Cordelia Langford, John Sillitoe on behalf of the Wellcome Sanger Institute COVID-19 Surveillance Team |
| EPI_ISL_2961082 | Lighthouse Laboratory Plymouth | Wellcome Sanger Institute for the COVID-19 Genomics UK (COG-UK) Consortium | Lighthouse Laboratory Plymouth and Alex Alderton, Roberto Amato, Jeffrey Barrett, Sonia Goncalves, Ewan Harrison, David K. Jackson, Ian Johnston, Dominic Kwiatkowski, Cordelia Langford, John Sillitoe on behalf of the Wellcome Sanger Institute COVID-19 Surveillance Team |
| EPI_ISL_2961083, EPI_ISL_2961084 | Lighthouse Lab in Alderley Park | Wellcome Sanger Institute for the COVID-19 Genomics UK (COG-UK) Consortium | Jacquelyn Wynn, Mairead Hyland, The Lighthouse Lab in Alderley Park and Alex Alderton, Roberto Amato, Jeffrey Barrett, Sonia Goncalves, Ewan Harrison, David K. Jackson, Ian Johnston, Dominic Kwiatkowski, Cordelia Langford, John Sillitoe on behalf of the Wellcome Sanger Institute COVID-19 Surveillance Team |
| EPI_ISL_2961085 | Berkshire and Surrey Pathology Services Lighthouse Laboratory | Wellcome Sanger Institute for the COVID-19 Genomics UK (COG-UK) Consortium | Berkshire and Surrey Pathology Services Lighthouse Laboratory and Alex Alderton, Roberto Amato, Jeffrey Barrett, Sonia Goncalves, Ewan Harrison, David K. Jackson, Ian Johnston, Dominic Kwiatkowski, Cordelia Langford, John Sillitoe on behalf of the Wellcome Sanger Institute COVID-19 Surveillance Team |
| EPI_ISL_2961087, EPI_ISL_2961088, EPI_ISL_2961089, EPI_ISL_2961090 | Lighthouse Lab in Alderley Park | Wellcome Sanger Institute for the COVID-19 Genomics UK (COG-UK) Consortium | Jacquelyn Wynn, Mairead Hyland, The Lighthouse Lab in Alderley Park and Alex Alderton, Roberto Amato, Jeffrey Barrett, Sonia Goncalves, Ewan Harrison, David K. Jackson, Ian Johnston, Dominic Kwiatkowski, Cordelia Langford, John Sillitoe on behalf of the Wellcome Sanger Institute COVID-19 Surveillance Team |
| EPI_ISL_2961091, EPI_ISL_2961092, EPI_ISL_2961093 | Lighthouse Laboratory Plymouth | Wellcome Sanger Institute for the COVID-19 Genomics UK (COG-UK) Consortium | Lighthouse Laboratory Plymouth and Alex Alderton, Roberto Amato, Jeffrey Barrett, Sonia Goncalves, Ewan Harrison, David K. Jackson, Ian Johnston, Dominic Kwiatkowski, Cordelia Langford, John Sillitoe on behalf of the Wellcome Sanger Institute COVID-19 Surveillance Team |
| EPI_ISL_2961094 | Berkshire and Surrey Pathology Services Lighthouse Laboratory | Wellcome Sanger Institute for the COVID-19 Genomics UK (COG-UK) Consortium | Berkshire and Surrey Pathology Services Lighthouse Laboratory and Alex Alderton, Roberto Amato, Jeffrey Barrett, Sonia Goncalves, Ewan Harrison, David K. Jackson, Ian Johnston, Dominic Kwiatkowski, Cordelia Langford, John Sillitoe on behalf of the Wellcome Sanger Institute COVID-19 Surveillance Team |
| EPI_ISL_2961095, EPI_ISL_2961096, EPI_ISL_2961097 | Lighthouse Lab in Alderley Park | Wellcome Sanger Institute for the COVID-19 Genomics UK (COG-UK) Consortium | Jacquelyn Wynn, Mairead Hyland, The Lighthouse Lab in Alderley Park and Alex Alderton, Roberto Amato, Jeffrey Barrett, Sonia Goncalves, Ewan Harrison, David K. Jackson, Ian Johnston, Dominic Kwiatkowski, Cordelia Langford, John Sillitoe on behalf of the Wellcome Sanger Institute COVID-19 Surveillance Team |
| EPI_ISL_2961098 | Berkshire and Surrey Pathology Services Lighthouse Laboratory | Wellcome Sanger Institute for the COVID-19 Genomics UK (COG-UK) Consortium | Berkshire and Surrey Pathology Services Lighthouse Laboratory and Alex Alderton, Roberto Amato, Jeffrey Barrett, Sonia Goncalves, Ewan Harrison, David K. Jackson, Ian Johnston, Dominic Kwiatkowski, Cordelia Langford, John Sillitoe on behalf of the Wellcome Sanger Institute COVID-19 Surveillance Team |
| EPI_ISL_2961099 | Lighthouse Lab in Alderley Park | Wellcome Sanger Institute for the COVID-19 Genomics UK (COG-UK) Consortium | Jacquelyn Wynn, Mairead Hyland, The Lighthouse Lab in Alderley Park and Alex Alderton, Roberto Amato, Jeffrey Barrett, Sonia Goncalves, Ewan Harrison, David K. Jackson, Ian Johnston, Dominic Kwiatkowski, Cordelia Langford, John Sillitoe on behalf of the Wellcome Sanger Institute COVID-19 Surveillance Team |
| EPI_ISL_2961100 | Berkshire and Surrey Pathology Services Lighthouse Laboratory | Wellcome Sanger Institute for the COVID-19 Genomics UK (COG-UK) Consortium | Berkshire and Surrey Pathology Services Lighthouse Laboratory and Alex Alderton, Roberto Amato, Jeffrey Barrett, Sonia Goncalves, Ewan Harrison, David K. Jackson, Ian Johnston, Dominic Kwiatkowski, Cordelia Langford, John Sillitoe on behalf of the Wellcome Sanger Institute COVID-19 Surveillance Team |
| EPI_ISL_2961101, EPI_ISL_2961102, EPI_ISL_2961103 | Lighthouse Lab in Alderley Park | Wellcome Sanger Institute for the COVID-19 Genomics UK (COG-UK) Consortium | Jacquelyn Wynn, Mairead Hyland, The Lighthouse Lab in Alderley Park and Alex Alderton, Roberto Amato, Jeffrey Barrett, Sonia Goncalves, Ewan Harrison, David K. Jackson, Ian Johnston, Dominic Kwiatkowski, Cordelia Langford, John Sillitoe on behalf of the Wellcome Sanger Institute COVID-19 Surveillance Team |
| EPI_ISL_2961104 | Lighthouse Laboratory Plymouth | Wellcome Sanger Institute for the COVID-19 Genomics UK (COG-UK) Consortium | Lighthouse Laboratory Plymouth and Alex Alderton, Roberto Amato, Jeffrey Barrett, Sonia Goncalves, Ewan Harrison, David K. Jackson, Ian Johnston, Dominic Kwiatkowski, Cordelia Langford, John Sillitoe on behalf of the Wellcome Sanger Institute COVID-19 Surveillance Team |
| EPI_ISL_2961105 | Lighthouse Lab in Alderley Park | Wellcome Sanger Institute for the COVID-19 Genomics UK (COG-UK) Consortium | Jacquelyn Wynn, Mairead Hyland, The Lighthouse Lab in Alderley Park and Alex Alderton, Roberto Amato, Jeffrey Barrett, Sonia Goncalves, Ewan Harrison, David K. Jackson, Ian Johnston, Dominic Kwiatkowski, Cordelia Langford, John Sillitoe on behalf of the Wellcome Sanger Institute COVID-19 Surveillance Team |
| EPI_ISL_2961106 | Lighthouse Laboratory Plymouth | Wellcome Sanger Institute for the COVID-19 Genomics UK (COG-UK) Consortium | Lighthouse Laboratory Plymouth and Alex Alderton, Roberto Amato, Jeffrey Barrett, Sonia Goncalves, Ewan Harrison, David K. Jackson, Ian Johnston, Dominic Kwiatkowski, Cordelia Langford, John Sillitoe on behalf of the Wellcome Sanger Institute COVID-19 Surveillance Team |
| EPI_ISL_2961107 | Berkshire and Surrey Pathology Services Lighthouse Laboratory | Wellcome Sanger Institute for the COVID-19 Genomics UK (COG-UK) Consortium | Berkshire and Surrey Pathology Services Lighthouse Laboratory and Alex Alderton, Roberto Amato, Jeffrey Barrett, Sonia Goncalves, Ewan Harrison, David K. Jackson, Ian Johnston, Dominic Kwiatkowski, Cordelia Langford, John Sillitoe on behalf of the Wellcome Sanger Institute COVID-19 Surveillance Team |
| EPI_ISL_2961108 | Lighthouse Lab in Alderley Park | Wellcome Sanger Institute for the COVID-19 Genomics UK (COG-UK) Consortium | Jacquelyn Wynn, Mairead Hyland, The Lighthouse Lab in Alderley Park and Alex Alderton, Roberto Amato, Jeffrey Barrett, Sonia Goncalves, Ewan Harrison, David K. Jackson, Ian Johnston, Dominic Kwiatkowski, Cordelia Langford, John Sillitoe on behalf of the Wellcome Sanger Institute COVID-19 Surveillance Team |
| EPI_ISL_2961109 | Lighthouse Laboratory Plymouth | Wellcome Sanger Institute for the COVID-19 Genomics UK (COG-UK) Consortium | Lighthouse Laboratory Plymouth and Alex Alderton, Roberto Amato, Jeffrey Barrett, Sonia Goncalves, Ewan Harrison, David K. Jackson, Ian Johnston, Dominic Kwiatkowski, Cordelia Langford, John Sillitoe on behalf of the Wellcome Sanger Institute COVID-19 Surveillance Team |
| EPI_ISL_2961110 | Berkshire and Surrey Pathology Services Lighthouse Laboratory | Wellcome Sanger Institute for the COVID-19 Genomics UK (COG-UK) Consortium | Berkshire and Surrey Pathology Services Lighthouse Laboratory and Alex Alderton, Roberto Amato, Jeffrey Barrett, Sonia Goncalves, Ewan Harrison, David K. Jackson, Ian Johnston, Dominic Kwiatkowski, Cordelia Langford, John Sillitoe on behalf of the Wellcome Sanger Institute COVID-19 Surveillance Team |
| EPI_ISL_2961111, EPI_ISL_2961112 | Lighthouse Lab in Alderley Park | Wellcome Sanger Institute for the COVID-19 Genomics UK (COG-UK) Consortium | Jacquelyn Wynn, Mairead Hyland, The Lighthouse Lab in Alderley Park and Alex Alderton, Roberto Amato, Jeffrey Barrett, Sonia Goncalves, Ewan Harrison, David K. Jackson, Ian Johnston, Dominic Kwiatkowski, Cordelia Langford, John Sillitoe on behalf of the Wellcome Sanger Institute COVID-19 Surveillance Team |
| EPI_ISL_2961113 | Lighthouse Laboratory Plymouth | Wellcome Sanger Institute for the COVID-19 Genomics UK (COG-UK) Consortium | Lighthouse Laboratory Plymouth and Alex Alderton, Roberto Amato, Jeffrey Barrett, Sonia Goncalves, Ewan Harrison, David K. Jackson, Ian Johnston, Dominic Kwiatkowski, Cordelia Langford, John Sillitoe on behalf of the Wellcome Sanger Institute COVID-19 Surveillance Team |
| EPI_ISL_2961114 | Lighthouse Lab in Alderley Park | Wellcome Sanger Institute for the COVID-19 Genomics UK (COG-UK) Consortium | Jacquelyn Wynn, Mairead Hyland, The Lighthouse Lab in Alderley Park and Alex Alderton, Roberto Amato, Jeffrey Barrett, Sonia Goncalves, Ewan Harrison, David K. Jackson, Ian Johnston, Dominic Kwiatkowski, Cordelia Langford, John Sillitoe on behalf of the Wellcome Sanger Institute COVID-19 Surveillance Team |

| | | | |
|---|---|---|---|
| EPI_ISL_2961115 | Lighthouse Laboratory Plymouth | Wellcome Sanger Institute for the COVID-19 Genomics UK (COG-UK) Consortium | Lighthouse Laboratory Plymouth and Alex Alderton, Roberto Amato, Jeffrey Barrett, Sonia Goncalves, Ewan Harrison, David K. Jackson, Ian Johnston, Dominic Kwiatkowski, Cordelia Langford, John Sillitoe on behalf of the Wellcome Sanger Institute COVID-19 Surveillance Team |
| EPI_ISL_2961116 | Lighthouse Lab in Alderley Park | Wellcome Sanger Institute for the COVID-19 Genomics UK (COG-UK) Consortium | Jacquelyn Wynn, Mairead Hyland, The Lighthouse Lab in Alderley Park and Alex Alderton, Roberto Amato, Jeffrey Barrett, Sonia Goncalves, Ewan Harrison, David K. Jackson, Ian Johnston, Dominic Kwiatkowski, Cordelia Langford, John Sillitoe on behalf of the Wellcome Sanger Institute COVID-19 Surveillance Team |
| EPI_ISL_2961117, EPI_ISL_2961118 | Lighthouse Laboratory Plymouth | Wellcome Sanger Institute for the COVID-19 Genomics UK (COG-UK) Consortium | Lighthouse Laboratory Plymouth and Alex Alderton, Roberto Amato, Jeffrey Barrett, Sonia Goncalves, Ewan Harrison, David K. Jackson, Ian Johnston, Dominic Kwiatkowski, Cordelia Langford, John Sillitoe on behalf of the Wellcome Sanger Institute COVID-19 Surveillance Team |
| EPI_ISL_2961119, EPI_ISL_2961120, EPI_ISL_2961121, EPI_ISL_2961122 | Lighthouse Lab in Alderley Park | Wellcome Sanger Institute for the COVID-19 Genomics UK (COG-UK) Consortium | Jacquelyn Wynn, Mairead Hyland, The Lighthouse Lab in Alderley Park and Alex Alderton, Roberto Amato, Jeffrey Barrett, Sonia Goncalves, Ewan Harrison, David K. Jackson, Ian Johnston, Dominic Kwiatkowski, Cordelia Langford, John Sillitoe on behalf of the Wellcome Sanger Institute COVID-19 Surveillance Team |
| EPI_ISL_2961123 | Lighthouse Laboratory Plymouth | Wellcome Sanger Institute for the COVID-19 Genomics UK (COG-UK) Consortium | Lighthouse Laboratory Plymouth and Alex Alderton, Roberto Amato, Jeffrey Barrett, Sonia Goncalves, Ewan Harrison, David K. Jackson, Ian Johnston, Dominic Kwiatkowski, Cordelia Langford, John Sillitoe on behalf of the Wellcome Sanger Institute COVID-19 Surveillance Team |
| EPI_ISL_2961124 | Lighthouse Lab in Alderley Park | Wellcome Sanger Institute for the COVID-19 Genomics UK (COG-UK) Consortium | Jacquelyn Wynn, Mairead Hyland, The Lighthouse Lab in Alderley Park and Alex Alderton, Roberto Amato, Jeffrey Barrett, Sonia Goncalves, Ewan Harrison, David K. Jackson, Ian Johnston, Dominic Kwiatkowski, Cordelia Langford, John Sillitoe on behalf of the Wellcome Sanger Institute COVID-19 Surveillance Team |
| EPI_ISL_2961125, EPI_ISL_2961126 | Lighthouse Laboratory Plymouth | Wellcome Sanger Institute for the COVID-19 Genomics UK (COG-UK) Consortium | Lighthouse Laboratory Plymouth and Alex Alderton, Roberto Amato, Jeffrey Barrett, Sonia Goncalves, Ewan Harrison, David K. Jackson, Ian Johnston, Dominic Kwiatkowski, Cordelia Langford, John Sillitoe on behalf of the Wellcome Sanger Institute COVID-19 Surveillance Team |
| EPI_ISL_2961127 | Lighthouse Lab in Alderley Park | Wellcome Sanger Institute for the COVID-19 Genomics UK (COG-UK) Consortium | Jacquelyn Wynn, Mairead Hyland, The Lighthouse Lab in Alderley Park and Alex Alderton, Roberto Amato, Jeffrey Barrett, Sonia Goncalves, Ewan Harrison, David K. Jackson, Ian Johnston, Dominic Kwiatkowski, Cordelia Langford, John Sillitoe on behalf of the Wellcome Sanger Institute COVID-19 Surveillance Team |
| EPI_ISL_2961128 | Berkshire and Surrey Pathology Services Lighthouse Laboratory | Wellcome Sanger Institute for the COVID-19 Genomics UK (COG-UK) Consortium | Berkshire and Surrey Pathology Services Lighthouse Laboratory and Alex Alderton, Roberto Amato, Jeffrey Barrett, Sonia Goncalves, Ewan Harrison, David K. Jackson, Ian Johnston, Dominic Kwiatkowski, Cordelia Langford, John Sillitoe on behalf of the Wellcome Sanger Institute COVID-19 Surveillance Team |
| EPI_ISL_2961129, EPI_ISL_2961130, EPI_ISL_2961131 | Lighthouse Lab in Alderley Park | Wellcome Sanger Institute for the COVID-19 Genomics UK (COG-UK) Consortium | Jacquelyn Wynn, Mairead Hyland, The Lighthouse Lab in Alderley Park and Alex Alderton, Roberto Amato, Jeffrey Barrett, Sonia Goncalves, Ewan Harrison, David K. Jackson, Ian Johnston, Dominic Kwiatkowski, Cordelia Langford, John Sillitoe on behalf of the Wellcome Sanger Institute COVID-19 Surveillance Team |
| EPI_ISL_2961132 | Berkshire and Surrey Pathology Services Lighthouse Laboratory | Wellcome Sanger Institute for the COVID-19 Genomics UK (COG-UK) Consortium | Berkshire and Surrey Pathology Services Lighthouse Laboratory and Alex Alderton, Roberto Amato, Jeffrey Barrett, Sonia Goncalves, Ewan Harrison, David K. Jackson, Ian Johnston, Dominic Kwiatkowski, Cordelia Langford, John Sillitoe on behalf of the Wellcome Sanger Institute COVID-19 Surveillance Team |
| EPI_ISL_2961133, EPI_ISL_2961134, EPI_ISL_2961135, EPI_ISL_2961136, EPI_ISL_2961137 | Lighthouse Lab in Alderley Park | Wellcome Sanger Institute for the COVID-19 Genomics UK (COG-UK) Consortium | Jacquelyn Wynn, Mairead Hyland, The Lighthouse Lab in Alderley Park and Alex Alderton, Roberto Amato, Jeffrey Barrett, Sonia Goncalves, Ewan Harrison, David K. Jackson, Ian Johnston, Dominic Kwiatkowski, Cordelia Langford, John Sillitoe on behalf of the Wellcome Sanger Institute COVID-19 Surveillance Team |
| EPI_ISL_2961138 | Berkshire and Surrey Pathology Services Lighthouse Laboratory | Wellcome Sanger Institute for the COVID-19 Genomics UK (COG-UK) Consortium | Berkshire and Surrey Pathology Services Lighthouse Laboratory and Alex Alderton, Roberto Amato, Jeffrey Barrett, Sonia Goncalves, Ewan Harrison, David K. Jackson, Ian Johnston, Dominic Kwiatkowski, Cordelia Langford, John Sillitoe on behalf of the Wellcome Sanger Institute COVID-19 Surveillance Team |
| EPI_ISL_2961139 | Lighthouse Laboratory Plymouth | Wellcome Sanger Institute for the COVID-19 Genomics UK (COG-UK) Consortium | Lighthouse Laboratory Plymouth and Alex Alderton, Roberto Amato, Jeffrey Barrett, Sonia Goncalves, Ewan Harrison, David K. Jackson, Ian Johnston, Dominic Kwiatkowski, Cordelia Langford, John Sillitoe on behalf of the Wellcome Sanger Institute COVID-19 Surveillance Team |
| EPI_ISL_2961140, EPI_ISL_2961141 | Lighthouse Lab in Alderley Park | Wellcome Sanger Institute for the COVID-19 Genomics UK (COG-UK) Consortium | Jacquelyn Wynn, Mairead Hyland, The Lighthouse Lab in Alderley Park and Alex Alderton, Roberto Amato, Jeffrey Barrett, Sonia Goncalves, Ewan Harrison, David K. Jackson, Ian Johnston, Dominic Kwiatkowski, Cordelia Langford, John Sillitoe on behalf of the Wellcome Sanger Institute COVID-19 Surveillance Team |
| EPI_ISL_2961142 | Berkshire and Surrey Pathology Services Lighthouse Laboratory | Wellcome Sanger Institute for the COVID-19 Genomics UK (COG-UK) Consortium | Berkshire and Surrey Pathology Services Lighthouse Laboratory and Alex Alderton, Roberto Amato, Jeffrey Barrett, Sonia Goncalves, Ewan Harrison, David K. Jackson, Ian Johnston, Dominic Kwiatkowski, Cordelia Langford, John Sillitoe on behalf of the Wellcome Sanger Institute COVID-19 Surveillance Team |
| EPI_ISL_2961143 | Lighthouse Laboratory Plymouth | Wellcome Sanger Institute for the COVID-19 Genomics UK (COG-UK) Consortium | Lighthouse Laboratory Plymouth and Alex Alderton, Roberto Amato, Jeffrey Barrett, Sonia Goncalves, Ewan Harrison, David K. Jackson, Ian Johnston, Dominic Kwiatkowski, Cordelia Langford, John Sillitoe on behalf of the Wellcome Sanger Institute COVID-19 Surveillance Team |
| EPI_ISL_2961144 | Berkshire and Surrey Pathology Services Lighthouse Laboratory | Wellcome Sanger Institute for the COVID-19 Genomics UK (COG-UK) Consortium | Berkshire and Surrey Pathology Services Lighthouse Laboratory and Alex Alderton, Roberto Amato, Jeffrey Barrett, Sonia Goncalves, Ewan Harrison, David K. Jackson, Ian Johnston, Dominic Kwiatkowski, Cordelia Langford, John Sillitoe on behalf of the Wellcome Sanger Institute COVID-19 Surveillance Team |
| EPI_ISL_2961145 | Lighthouse Laboratory Plymouth | Wellcome Sanger Institute for the COVID-19 Genomics UK (COG-UK) Consortium | Lighthouse Laboratory Plymouth and Alex Alderton, Roberto Amato, Jeffrey Barrett, Sonia Goncalves, Ewan Harrison, David K. Jackson, Ian Johnston, Dominic Kwiatkowski, Cordelia Langford, John Sillitoe on behalf of the Wellcome Sanger Institute COVID-19 Surveillance Team |
| EPI_ISL_2961147, EPI_ISL_2961148 | Lighthouse Lab in Alderley Park | Wellcome Sanger Institute for the COVID-19 Genomics UK (COG-UK) Consortium | Jacquelyn Wynn, Mairead Hyland, The Lighthouse Lab in Alderley Park and Alex Alderton, Roberto Amato, Jeffrey Barrett, Sonia Goncalves, Ewan Harrison, David K. Jackson, Ian Johnston, Dominic Kwiatkowski, Cordelia Langford, John Sillitoe on behalf of the Wellcome Sanger Institute COVID-19 Surveillance Team |
| EPI_ISL_2961149 | Berkshire and Surrey Pathology Services Lighthouse Laboratory | Wellcome Sanger Institute for the COVID-19 Genomics UK (COG-UK) Consortium | Berkshire and Surrey Pathology Services Lighthouse Laboratory and Alex Alderton, Roberto Amato, Jeffrey Barrett, Sonia Goncalves, Ewan Harrison, David K. Jackson, Ian Johnston, Dominic Kwiatkowski, Cordelia Langford, John Sillitoe on behalf of the Wellcome Sanger Institute COVID-19 Surveillance Team |
| EPI_ISL_2961150 | Lighthouse Lab in Alderley Park | Wellcome Sanger Institute for the COVID-19 Genomics UK (COG-UK) Consortium | Jacquelyn Wynn, Mairead Hyland, The Lighthouse Lab in Alderley Park and Alex Alderton, Roberto Amato, Jeffrey Barrett, Sonia Goncalves, Ewan Harrison, David K. Jackson, Ian Johnston, Dominic Kwiatkowski, Cordelia Langford, John Sillitoe on behalf of the Wellcome Sanger Institute COVID-19 Surveillance Team |
| EPI_ISL_2961151 | Berkshire and Surrey Pathology Services Lighthouse Laboratory | Wellcome Sanger Institute for the COVID-19 Genomics UK (COG-UK) Consortium | Berkshire and Surrey Pathology Services Lighthouse Laboratory and Alex Alderton, Roberto Amato, Jeffrey Barrett, Sonia Goncalves, Ewan Harrison, David K. Jackson, Ian Johnston, Dominic Kwiatkowski, Cordelia Langford, John Sillitoe on behalf of the Wellcome Sanger Institute COVID-19 Surveillance Team |
| EPI_ISL_2961152 | Lighthouse Laboratory Plymouth | Wellcome Sanger Institute for the COVID-19 Genomics UK (COG-UK) Consortium | Lighthouse Laboratory Plymouth and Alex Alderton, Roberto Amato, Jeffrey Barrett, Sonia Goncalves, Ewan Harrison, David K. Jackson, Ian Johnston, Dominic Kwiatkowski, Cordelia Langford, John Sillitoe on behalf of the Wellcome Sanger Institute COVID-19 Surveillance Team |
| EPI_ISL_2961153 | Lighthouse Lab in Alderley Park | Wellcome Sanger Institute for the COVID-19 Genomics UK (COG-UK) Consortium | Jacquelyn Wynn, Mairead Hyland, The Lighthouse Lab in Alderley Park and Alex Alderton, Roberto Amato, Jeffrey Barrett, Sonia Goncalves, Ewan Harrison, David K. Jackson, Ian Johnston, Dominic Kwiatkowski, Cordelia Langford, John Sillitoe on behalf of the Wellcome Sanger Institute COVID-19 Surveillance Team |
| EPI_ISL_2961154 | Berkshire and Surrey Pathology Services Lighthouse Laboratory | Wellcome Sanger Institute for the COVID-19 Genomics UK (COG-UK) Consortium | Berkshire and Surrey Pathology Services Lighthouse Laboratory and Alex Alderton, Roberto Amato, Jeffrey Barrett, Sonia Goncalves, Ewan Harrison, David K. Jackson, Ian Johnston, Dominic Kwiatkowski, Cordelia Langford, John Sillitoe on behalf of the Wellcome Sanger Institute COVID-19 Surveillance Team |
| EPI_ISL_2961155, EPI_ISL_2961156, EPI_ISL_2961157 | Lighthouse Lab in Alderley Park | Wellcome Sanger Institute for the COVID-19 Genomics UK (COG-UK) Consortium | Jacquelyn Wynn, Mairead Hyland, The Lighthouse Lab in Alderley Park and Alex Alderton, Roberto Amato, Jeffrey Barrett, Sonia Goncalves, Ewan Harrison, David K. Jackson, Ian Johnston, Dominic Kwiatkowski, Cordelia Langford, John Sillitoe on behalf of the Wellcome Sanger Institute COVID-19 Surveillance Team |
| EPI_ISL_2961158 | Berkshire and Surrey Pathology Services Lighthouse Laboratory | Wellcome Sanger Institute for the COVID-19 Genomics UK (COG-UK) Consortium | Berkshire and Surrey Pathology Services Lighthouse Laboratory and Alex Alderton, Roberto Amato, Jeffrey Barrett, Sonia Goncalves, Ewan Harrison, David K. Jackson, Ian Johnston, Dominic Kwiatkowski, Cordelia Langford, John Sillitoe on behalf of the Wellcome Sanger Institute COVID-19 Surveillance Team |
| EPI_ISL_2961159 | Lighthouse Lab in Alderley Park | Wellcome Sanger Institute for the COVID-19 Genomics UK (COG-UK) Consortium | Jacquelyn Wynn, Mairead Hyland, The Lighthouse Lab in Alderley Park and Alex Alderton, Roberto Amato, Jeffrey Barrett, Sonia Goncalves, Ewan Harrison, David K. Jackson, Ian Johnston, Dominic Kwiatkowski, Cordelia Langford, John Sillitoe on behalf of the Wellcome Sanger Institute COVID-19 Surveillance Team |
| EPI_ISL_2961160 | Lighthouse Laboratory Plymouth | Wellcome Sanger Institute for the COVID-19 Genomics UK (COG-UK) Consortium | Lighthouse Laboratory Plymouth and Alex Alderton, Roberto Amato, Jeffrey Barrett, Sonia Goncalves, Ewan Harrison, David K. Jackson, Ian Johnston, Dominic Kwiatkowski, Cordelia Langford, John Sillitoe on behalf of the Wellcome Sanger Institute COVID-19 Surveillance Team |
| EPI_ISL_2961161 | Berkshire and Surrey Pathology Services Lighthouse Laboratory | Wellcome Sanger Institute for the COVID-19 Genomics UK (COG-UK) Consortium | Berkshire and Surrey Pathology Services Lighthouse Laboratory and Alex Alderton, Roberto Amato, Jeffrey Barrett, Sonia Goncalves, Ewan Harrison, David K. Jackson, Ian Johnston, Dominic Kwiatkowski, Cordelia Langford, John Sillitoe on behalf of the Wellcome Sanger Institute COVID-19 Surveillance Team |
| EPI_ISL_2961162 | Lighthouse Lab in Alderley Park | Wellcome Sanger Institute for the COVID-19 Genomics UK (COG-UK) Consortium | Jacquelyn Wynn, Mairead Hyland, The Lighthouse Lab in Alderley Park and Alex Alderton, Roberto Amato, Jeffrey Barrett, Sonia Goncalves, Ewan Harrison, David K. Jackson, Ian Johnston, Dominic Kwiatkowski, Cordelia Langford, John Sillitoe on behalf of the Wellcome Sanger Institute COVID-19 Surveillance Team |
| EPI_ISL_2961163, EPI_ISL_2961164 | Lighthouse Laboratory Plymouth | Wellcome Sanger Institute for the COVID-19 Genomics UK (COG-UK) Consortium | Lighthouse Laboratory Plymouth and Alex Alderton, Roberto Amato, Jeffrey Barrett, Sonia Goncalves, Ewan Harrison, David K. Jackson, Ian Johnston, Dominic Kwiatkowski, Cordelia Langford, John Sillitoe on behalf of the Wellcome Sanger Institute COVID-19 Surveillance Team |
| EPI_ISL_2961165 | Berkshire and Surrey Pathology Services Lighthouse Laboratory | Wellcome Sanger Institute for the COVID-19 Genomics UK (COG-UK) Consortium | Berkshire and Surrey Pathology Services Lighthouse Laboratory and Alex Alderton, Roberto Amato, Jeffrey Barrett, Sonia Goncalves, Ewan Harrison, David K. Jackson, Ian Johnston, Dominic Kwiatkowski, Cordelia Langford, John Sillitoe on behalf of the Wellcome Sanger Institute COVID-19 Surveillance Team |
| EPI_ISL_2961166, EPI_ISL_2961167 | Lighthouse Lab in Alderley Park | Wellcome Sanger Institute for the COVID-19 Genomics UK (COG-UK) Consortium | Jacquelyn Wynn, Mairead Hyland, The Lighthouse Lab in Alderley Park and Alex Alderton, Roberto Amato, Jeffrey Barrett, Sonia Goncalves, Ewan Harrison, David K. Jackson, Ian Johnston, Dominic Kwiatkowski, Cordelia Langford, John Sillitoe on behalf of the Wellcome Sanger Institute COVID-19 Surveillance Team |
| EPI_ISL_2961168 | Berkshire and Surrey Pathology Services Lighthouse Laboratory | Wellcome Sanger Institute for the COVID-19 Genomics UK (COG-UK) Consortium | Berkshire and Surrey Pathology Services Lighthouse Laboratory and Alex Alderton, Roberto Amato, Jeffrey Barrett, Sonia Goncalves, Ewan Harrison, David K. Jackson, Ian Johnston, Dominic Kwiatkowski, Cordelia Langford, John Sillitoe on behalf of the Wellcome Sanger Institute COVID-19 Surveillance Team |
| EPI_ISL_2961169, EPI_ISL_2961170 | Lighthouse Laboratory Plymouth | Wellcome Sanger Institute for the COVID-19 Genomics UK (COG-UK) Consortium | Lighthouse Laboratory Plymouth and Alex Alderton, Roberto Amato, Jeffrey Barrett, Sonia Goncalves, Ewan Harrison, David K. Jackson, Ian Johnston, Dominic Kwiatkowski, Cordelia Langford, John Sillitoe on behalf of the Wellcome Sanger Institute COVID-19 Surveillance Team |
| EPI_ISL_2961171 | Berkshire and Surrey Pathology Services Lighthouse Laboratory | Wellcome Sanger Institute for the COVID-19 Genomics UK (COG-UK) Consortium | Berkshire and Surrey Pathology Services Lighthouse Laboratory and Alex Alderton, Roberto Amato, Jeffrey Barrett, Sonia Goncalves, Ewan Harrison, David K. Jackson, Ian Johnston, Dominic Kwiatkowski, Cordelia Langford, John Sillitoe on behalf of the Wellcome Sanger Institute COVID-19 Surveillance Team |
| EPI_ISL_2961172 | Lighthouse Lab in Alderley Park | Wellcome Sanger Institute for the COVID-19 Genomics UK (COG-UK) Consortium | Jacquelyn Wynn, Mairead Hyland, The Lighthouse Lab in Alderley Park and Alex Alderton, Roberto Amato, Jeffrey Barrett, Sonia Goncalves, Ewan Harrison, David K. Jackson, Ian Johnston, Dominic Kwiatkowski, Cordelia Langford, John Sillitoe on behalf of the Wellcome Sanger Institute COVID-19 Surveillance Team |
| EPI_ISL_2961173 | Berkshire and Surrey Pathology Services Lighthouse Laboratory | Wellcome Sanger Institute for the COVID-19 Genomics UK (COG-UK) Consortium | Berkshire and Surrey Pathology Services Lighthouse Laboratory and Alex Alderton, Roberto Amato, Jeffrey Barrett, Sonia Goncalves, Ewan Harrison, David K. Jackson, Ian Johnston, Dominic Kwiatkowski, Cordelia Langford, John Sillitoe on behalf of the Wellcome Sanger Institute COVID-19 Surveillance Team |
| EPI_ISL_2961174 | Lighthouse Laboratory Plymouth | Wellcome Sanger Institute for the COVID-19 Genomics UK (COG-UK) Consortium | Lighthouse Laboratory Plymouth and Alex Alderton, Roberto Amato, Jeffrey Barrett, Sonia Goncalves, Ewan Harrison, David K. Jackson, Ian Johnston, Dominic Kwiatkowski, Cordelia Langford, John Sillitoe on behalf of the Wellcome Sanger Institute COVID-19 Surveillance Team |
| EPI_ISL_2961175 | Berkshire and Surrey Pathology Services Lighthouse Laboratory | Wellcome Sanger Institute for the COVID-19 Genomics UK (COG-UK) Consortium | Berkshire and Surrey Pathology Services Lighthouse Laboratory and Alex Alderton, Roberto Amato, Jeffrey Barrett, Sonia Goncalves, Ewan Harrison, David K. Jackson, Ian Johnston, Dominic Kwiatkowski, Cordelia Langford, John Sillitoe on behalf of the Wellcome Sanger Institute COVID-19 Surveillance Team |

| | | | |
|---|---|---|---|
| EPI_ISL_2961176 | Lighthouse Lab in Alderley Park | Wellcome Sanger Institute for the COVID-19 Genomics UK (COG-UK) Consortium | Jacquelyn Wynn, Mairead Hyland, The Lighthouse Lab in Alderley Park and Alex Alderton, Roberto Amato, Jeffrey Barrett, Sonia Goncalves, Ewan Harrison, David K. Jackson, Ian Johnston, Dominic Kwiatkowski, Cordelia Langford, John Sillitoe on behalf of the Wellcome Sanger Institute COVID-19 Surveillance Team |
| EPI_ISL_2961177 | Lighthouse Laboratory Plymouth | Wellcome Sanger Institute for the COVID-19 Genomics UK (COG-UK) Consortium | Lighthouse Laboratory Plymouth and Alex Alderton, Roberto Amato, Jeffrey Barrett, Sonia Goncalves, Ewan Harrison, David K. Jackson, Ian Johnston, Dominic Kwiatkowski, Cordelia Langford, John Sillitoe on behalf of the Wellcome Sanger Institute COVID-19 Surveillance Team |
| EPI_ISL_2961178 | Lighthouse Lab in Alderley Park | Wellcome Sanger Institute for the COVID-19 Genomics UK (COG-UK) Consortium | Jacquelyn Wynn, Mairead Hyland, The Lighthouse Lab in Alderley Park and Alex Alderton, Roberto Amato, Jeffrey Barrett, Sonia Goncalves, Ewan Harrison, David K. Jackson, Ian Johnston, Dominic Kwiatkowski, Cordelia Langford, John Sillitoe on behalf of the Wellcome Sanger Institute COVID-19 Surveillance Team |
| EPI_ISL_2961179 | Lighthouse Laboratory Plymouth | Wellcome Sanger Institute for the COVID-19 Genomics UK (COG-UK) Consortium | Lighthouse Laboratory Plymouth and Alex Alderton, Roberto Amato, Jeffrey Barrett, Sonia Goncalves, Ewan Harrison, David K. Jackson, Ian Johnston, Dominic Kwiatkowski, Cordelia Langford, John Sillitoe on behalf of the Wellcome Sanger Institute COVID-19 Surveillance Team |
| EPI_ISL_2961180, EPI_ISL_2961181, EPI_ISL_2961182 | Lighthouse Lab in Alderley Park | Wellcome Sanger Institute for the COVID-19 Genomics UK (COG-UK) Consortium | Jacquelyn Wynn, Mairead Hyland, The Lighthouse Lab in Alderley Park and Alex Alderton, Roberto Amato, Jeffrey Barrett, Sonia Goncalves, Ewan Harrison, David K. Jackson, Ian Johnston, Dominic Kwiatkowski, Cordelia Langford, John Sillitoe on behalf of the Wellcome Sanger Institute COVID-19 Surveillance Team |
| EPI_ISL_2961183 | Lighthouse Laboratory Plymouth | Wellcome Sanger Institute for the COVID-19 Genomics UK (COG-UK) Consortium | Lighthouse Laboratory Plymouth and Alex Alderton, Roberto Amato, Jeffrey Barrett, Sonia Goncalves, Ewan Harrison, David K. Jackson, Ian Johnston, Dominic Kwiatkowski, Cordelia Langford, John Sillitoe on behalf of the Wellcome Sanger Institute COVID-19 Surveillance Team |
| EPI_ISL_2961184 | Berkshire and Surrey Pathology Services Lighthouse Laboratory | Wellcome Sanger Institute for the COVID-19 Genomics UK (COG-UK) Consortium | Berkshire and Surrey Pathology Services Lighthouse Laboratory and Alex Alderton, Roberto Amato, Jeffrey Barrett, Sonia Goncalves, Ewan Harrison, David K. Jackson, Ian Johnston, Dominic Kwiatkowski, Cordelia Langford, John Sillitoe on behalf of the Wellcome Sanger Institute COVID-19 Surveillance Team |
| EPI_ISL_2961185 | Lighthouse Lab in Alderley Park | Wellcome Sanger Institute for the COVID-19 Genomics UK (COG-UK) Consortium | Jacquelyn Wynn, Mairead Hyland, The Lighthouse Lab in Alderley Park and Alex Alderton, Roberto Amato, Jeffrey Barrett, Sonia Goncalves, Ewan Harrison, David K. Jackson, Ian Johnston, Dominic Kwiatkowski, Cordelia Langford, John Sillitoe on behalf of the Wellcome Sanger Institute COVID-19 Surveillance Team |
| EPI_ISL_2961186 | Berkshire and Surrey Pathology Services Lighthouse Laboratory | Wellcome Sanger Institute for the COVID-19 Genomics UK (COG-UK) Consortium | Berkshire and Surrey Pathology Services Lighthouse Laboratory and Alex Alderton, Roberto Amato, Jeffrey Barrett, Sonia Goncalves, Ewan Harrison, David K. Jackson, Ian Johnston, Dominic Kwiatkowski, Cordelia Langford, John Sillitoe on behalf of the Wellcome Sanger Institute COVID-19 Surveillance Team |
| EPI_ISL_2961187 | Lighthouse Lab in Alderley Park | Wellcome Sanger Institute for the COVID-19 Genomics UK (COG-UK) Consortium | Jacquelyn Wynn, Mairead Hyland, The Lighthouse Lab in Alderley Park and Alex Alderton, Roberto Amato, Jeffrey Barrett, Sonia Goncalves, Ewan Harrison, David K. Jackson, Ian Johnston, Dominic Kwiatkowski, Cordelia Langford, John Sillitoe on behalf of the Wellcome Sanger Institute COVID-19 Surveillance Team |
| EPI_ISL_2961188 | Berkshire and Surrey Pathology Services Lighthouse Laboratory | Wellcome Sanger Institute for the COVID-19 Genomics UK (COG-UK) Consortium | Berkshire and Surrey Pathology Services Lighthouse Laboratory and Alex Alderton, Roberto Amato, Jeffrey Barrett, Sonia Goncalves, Ewan Harrison, David K. Jackson, Ian Johnston, Dominic Kwiatkowski, Cordelia Langford, John Sillitoe on behalf of the Wellcome Sanger Institute COVID-19 Surveillance Team |
| EPI_ISL_2961189, EPI_ISL_2961190, EPI_ISL_2961191 | Lighthouse Lab in Alderley Park | Wellcome Sanger Institute for the COVID-19 Genomics UK (COG-UK) Consortium | Jacquelyn Wynn, Mairead Hyland, The Lighthouse Lab in Alderley Park and Alex Alderton, Roberto Amato, Jeffrey Barrett, Sonia Goncalves, Ewan Harrison, David K. Jackson, Ian Johnston, Dominic Kwiatkowski, Cordelia Langford, John Sillitoe on behalf of the Wellcome Sanger Institute COVID-19 Surveillance Team |
| EPI_ISL_2961192, EPI_ISL_2961193 | Berkshire and Surrey Pathology Services Lighthouse Laboratory | Wellcome Sanger Institute for the COVID-19 Genomics UK (COG-UK) Consortium | Berkshire and Surrey Pathology Services Lighthouse Laboratory and Alex Alderton, Roberto Amato, Jeffrey Barrett, Sonia Goncalves, Ewan Harrison, David K. Jackson, Ian Johnston, Dominic Kwiatkowski, Cordelia Langford, John Sillitoe on behalf of the Wellcome Sanger Institute COVID-19 Surveillance Team |
| EPI_ISL_2961194, EPI_ISL_2961195, EPI_ISL_2961196 | Lighthouse Lab in Alderley Park | Wellcome Sanger Institute for the COVID-19 Genomics UK (COG-UK) Consortium | Jacquelyn Wynn, Mairead Hyland, The Lighthouse Lab in Alderley Park and Alex Alderton, Roberto Amato, Jeffrey Barrett, Sonia Goncalves, Ewan Harrison, David K. Jackson, Ian Johnston, Dominic Kwiatkowski, Cordelia Langford, John Sillitoe on behalf of the Wellcome Sanger Institute COVID-19 Surveillance Team |
| EPI_ISL_2961197 | Lighthouse Laboratory Plymouth | Wellcome Sanger Institute for the COVID-19 Genomics UK (COG-UK) Consortium | Lighthouse Laboratory Plymouth and Alex Alderton, Roberto Amato, Jeffrey Barrett, Sonia Goncalves, Ewan Harrison, David K. Jackson, Ian Johnston, Dominic Kwiatkowski, Cordelia Langford, John Sillitoe on behalf of the Wellcome Sanger Institute COVID-19 Surveillance Team |
| EPI_ISL_2961198 | Lighthouse Lab in Alderley Park | Wellcome Sanger Institute for the COVID-19 Genomics UK (COG-UK) Consortium | Jacquelyn Wynn, Mairead Hyland, The Lighthouse Lab in Alderley Park and Alex Alderton, Roberto Amato, Jeffrey Barrett, Sonia Goncalves, Ewan Harrison, David K. Jackson, Ian Johnston, Dominic Kwiatkowski, Cordelia Langford, John Sillitoe on behalf of the Wellcome Sanger Institute COVID-19 Surveillance Team |
| EPI_ISL_2961199 | Lighthouse Laboratory Plymouth | Wellcome Sanger Institute for the COVID-19 Genomics UK (COG-UK) Consortium | Lighthouse Laboratory Plymouth and Alex Alderton, Roberto Amato, Jeffrey Barrett, Sonia Goncalves, Ewan Harrison, David K. Jackson, Ian Johnston, Dominic Kwiatkowski, Cordelia Langford, John Sillitoe on behalf of the Wellcome Sanger Institute COVID-19 Surveillance Team |
| EPI_ISL_2961200 | Lighthouse Lab in Alderley Park | Wellcome Sanger Institute for the COVID-19 Genomics UK (COG-UK) Consortium | Jacquelyn Wynn, Mairead Hyland, The Lighthouse Lab in Alderley Park and Alex Alderton, Roberto Amato, Jeffrey Barrett, Sonia Goncalves, Ewan Harrison, David K. Jackson, Ian Johnston, Dominic Kwiatkowski, Cordelia Langford, John Sillitoe on behalf of the Wellcome Sanger Institute COVID-19 Surveillance Team |
| EPI_ISL_2961201, EPI_ISL_2961202, EPI_ISL_2961203 | Berkshire and Surrey Pathology Services Lighthouse Laboratory | Wellcome Sanger Institute for the COVID-19 Genomics UK (COG-UK) Consortium | Berkshire and Surrey Pathology Services Lighthouse Laboratory and Alex Alderton, Roberto Amato, Jeffrey Barrett, Sonia Goncalves, Ewan Harrison, David K. Jackson, Ian Johnston, Dominic Kwiatkowski, Cordelia Langford, John Sillitoe on behalf of the Wellcome Sanger Institute COVID-19 Surveillance Team |
| EPI_ISL_2961204 | Lighthouse Lab in Alderley Park | Wellcome Sanger Institute for the COVID-19 Genomics UK (COG-UK) Consortium | Jacquelyn Wynn, Mairead Hyland, The Lighthouse Lab in Alderley Park and Alex Alderton, Roberto Amato, Jeffrey Barrett, Sonia Goncalves, Ewan Harrison, David K. Jackson, Ian Johnston, Dominic Kwiatkowski, Cordelia Langford, John Sillitoe on behalf of the Wellcome Sanger Institute COVID-19 Surveillance Team |
| EPI_ISL_2961205 | Berkshire and Surrey Pathology Services Lighthouse Laboratory | Wellcome Sanger Institute for the COVID-19 Genomics UK (COG-UK) Consortium | Berkshire and Surrey Pathology Services Lighthouse Laboratory and Alex Alderton, Roberto Amato, Jeffrey Barrett, Sonia Goncalves, Ewan Harrison, David K. Jackson, Ian Johnston, Dominic Kwiatkowski, Cordelia Langford, John Sillitoe on behalf of the Wellcome Sanger Institute COVID-19 Surveillance Team |
| EPI_ISL_2961206 | Lighthouse Lab in Alderley Park | Wellcome Sanger Institute for the COVID-19 Genomics UK (COG-UK) Consortium | Jacquelyn Wynn, Mairead Hyland, The Lighthouse Lab in Alderley Park and Alex Alderton, Roberto Amato, Jeffrey Barrett, Sonia Goncalves, Ewan Harrison, David K. Jackson, Ian Johnston, Dominic Kwiatkowski, Cordelia Langford, John Sillitoe on behalf of the Wellcome Sanger Institute COVID-19 Surveillance Team |
| EPI_ISL_2961207, EPI_ISL_2961208, EPI_ISL_2961209 | Lighthouse Laboratory Plymouth | Wellcome Sanger Institute for the COVID-19 Genomics UK (COG-UK) Consortium | Lighthouse Laboratory Plymouth and Alex Alderton, Roberto Amato, Jeffrey Barrett, Sonia Goncalves, Ewan Harrison, David K. Jackson, Ian Johnston, Dominic Kwiatkowski, Cordelia Langford, John Sillitoe on behalf of the Wellcome Sanger Institute COVID-19 Surveillance Team |
| EPI_ISL_2961210, EPI_ISL_2961211 | Berkshire and Surrey Pathology Services Lighthouse Laboratory | Wellcome Sanger Institute for the COVID-19 Genomics UK (COG-UK) Consortium | Berkshire and Surrey Pathology Services Lighthouse Laboratory and Alex Alderton, Roberto Amato, Jeffrey Barrett, Sonia Goncalves, Ewan Harrison, David K. Jackson, Ian Johnston, Dominic Kwiatkowski, Cordelia Langford, John Sillitoe on behalf of the Wellcome Sanger Institute COVID-19 Surveillance Team |
| EPI_ISL_2961212 | Lighthouse Lab in Alderley Park | Wellcome Sanger Institute for the COVID-19 Genomics UK (COG-UK) Consortium | Jacquelyn Wynn, Mairead Hyland, The Lighthouse Lab in Alderley Park and Alex Alderton, Roberto Amato, Jeffrey Barrett, Sonia Goncalves, Ewan Harrison, David K. Jackson, Ian Johnston, Dominic Kwiatkowski, Cordelia Langford, John Sillitoe on behalf of the Wellcome Sanger Institute COVID-19 Surveillance Team |
| EPI_ISL_2961213 | Berkshire and Surrey Pathology Services Lighthouse Laboratory | Wellcome Sanger Institute for the COVID-19 Genomics UK (COG-UK) Consortium | Berkshire and Surrey Pathology Services Lighthouse Laboratory and Alex Alderton, Roberto Amato, Jeffrey Barrett, Sonia Goncalves, Ewan Harrison, David K. Jackson, Ian Johnston, Dominic Kwiatkowski, Cordelia Langford, John Sillitoe on behalf of the Wellcome Sanger Institute COVID-19 Surveillance Team |
| EPI_ISL_2961214, EPI_ISL_2961215 | Lighthouse Lab in Alderley Park | Wellcome Sanger Institute for the COVID-19 Genomics UK (COG-UK) Consortium | Jacquelyn Wynn, Mairead Hyland, The Lighthouse Lab in Alderley Park and Alex Alderton, Roberto Amato, Jeffrey Barrett, Sonia Goncalves, Ewan Harrison, David K. Jackson, Ian Johnston, Dominic Kwiatkowski, Cordelia Langford, John Sillitoe on behalf of the Wellcome Sanger Institute COVID-19 Surveillance Team |
| EPI_ISL_2961216 | Berkshire and Surrey Pathology Services Lighthouse Laboratory | Wellcome Sanger Institute for the COVID-19 Genomics UK (COG-UK) Consortium | Berkshire and Surrey Pathology Services Lighthouse Laboratory and Alex Alderton, Roberto Amato, Jeffrey Barrett, Sonia Goncalves, Ewan Harrison, David K. Jackson, Ian Johnston, Dominic Kwiatkowski, Cordelia Langford, John Sillitoe on behalf of the Wellcome Sanger Institute COVID-19 Surveillance Team |
| EPI_ISL_2961217, EPI_ISL_2961218 | Lighthouse Lab in Alderley Park | Wellcome Sanger Institute for the COVID-19 Genomics UK (COG-UK) Consortium | Jacquelyn Wynn, Mairead Hyland, The Lighthouse Lab in Alderley Park and Alex Alderton, Roberto Amato, Jeffrey Barrett, Sonia Goncalves, Ewan Harrison, David K. Jackson, Ian Johnston, Dominic Kwiatkowski, Cordelia Langford, John Sillitoe on behalf of the Wellcome Sanger Institute COVID-19 Surveillance Team |
| EPI_ISL_2961219 | Berkshire and Surrey Pathology Services Lighthouse Laboratory | Wellcome Sanger Institute for the COVID-19 Genomics UK (COG-UK) Consortium | Berkshire and Surrey Pathology Services Lighthouse Laboratory and Alex Alderton, Roberto Amato, Jeffrey Barrett, Sonia Goncalves, Ewan Harrison, David K. Jackson, Ian Johnston, Dominic Kwiatkowski, Cordelia Langford, John Sillitoe on behalf of the Wellcome Sanger Institute COVID-19 Surveillance Team |
| EPI_ISL_2961220, EPI_ISL_2961221, EPI_ISL_2961222, EPI_ISL_2961223 | Lighthouse Lab in Alderley Park | Wellcome Sanger Institute for the COVID-19 Genomics UK (COG-UK) Consortium | Jacquelyn Wynn, Mairead Hyland, The Lighthouse Lab in Alderley Park and Alex Alderton, Roberto Amato, Jeffrey Barrett, Sonia Goncalves, Ewan Harrison, David K. Jackson, Ian Johnston, Dominic Kwiatkowski, Cordelia Langford, John Sillitoe on behalf of the Wellcome Sanger Institute COVID-19 Surveillance Team |
| EPI_ISL_2961224, EPI_ISL_2961225 | Berkshire and Surrey Pathology Services Lighthouse Laboratory | Wellcome Sanger Institute for the COVID-19 Genomics UK (COG-UK) Consortium | Berkshire and Surrey Pathology Services Lighthouse Laboratory and Alex Alderton, Roberto Amato, Jeffrey Barrett, Sonia Goncalves, Ewan Harrison, David K. Jackson, Ian Johnston, Dominic Kwiatkowski, Cordelia Langford, John Sillitoe on behalf of the Wellcome Sanger Institute COVID-19 Surveillance Team |
| EPI_ISL_2961226, EPI_ISL_2961227, EPI_ISL_2961228, EPI_ISL_2961229 | Lighthouse Lab in Alderley Park | Wellcome Sanger Institute for the COVID-19 Genomics UK (COG-UK) Consortium | Jacquelyn Wynn, Mairead Hyland, The Lighthouse Lab in Alderley Park and Alex Alderton, Roberto Amato, Jeffrey Barrett, Sonia Goncalves, Ewan Harrison, David K. Jackson, Ian Johnston, Dominic Kwiatkowski, Cordelia Langford, John Sillitoe on behalf of the Wellcome Sanger Institute COVID-19 Surveillance Team |
| EPI_ISL_2961230 | Berkshire and Surrey Pathology Services Lighthouse Laboratory | Wellcome Sanger Institute for the COVID-19 Genomics UK (COG-UK) Consortium | Berkshire and Surrey Pathology Services Lighthouse Laboratory and Alex Alderton, Roberto Amato, Jeffrey Barrett, Sonia Goncalves, Ewan Harrison, David K. Jackson, Ian Johnston, Dominic Kwiatkowski, Cordelia Langford, John Sillitoe on behalf of the Wellcome Sanger Institute COVID-19 Surveillance Team |
| EPI_ISL_2961231 | Lighthouse Lab in Glasgow | Wellcome Sanger Institute for the COVID-19 Genomics UK (COG-UK) Consortium | Harper VanSteenhouse, Yumi Kasai, David Gray, Carol Clugston, Anna Dominiczak and Alex Alderton, Roberto Amato, Jeffrey Barrett, Sonia Goncalves, Ewan Harrison, David K. Jackson, Ian Johnston, Dominic Kwiatkowski, Cordelia Langford, John Sillitoe on behalf of the Wellcome Sanger Institute COVID-19 Surveillance Team |
| EPI_ISL_2961232 | Lighthouse Lab in Alderley Park | Wellcome Sanger Institute for the COVID-19 Genomics UK (COG-UK) Consortium | Jacquelyn Wynn, Mairead Hyland, The Lighthouse Lab in Alderley Park and Alex Alderton, Roberto Amato, Jeffrey Barrett, Sonia Goncalves, Ewan Harrison, David K. Jackson, Ian Johnston, Dominic Kwiatkowski, Cordelia Langford, John Sillitoe on behalf of the Wellcome Sanger Institute COVID-19 Surveillance Team |
| EPI_ISL_2961233, EPI_ISL_2961234, EPI_ISL_2961235 | Lighthouse Lab in Glasgow | Wellcome Sanger Institute for the COVID-19 Genomics UK (COG-UK) Consortium | Harper VanSteenhouse, Yumi Kasai, David Gray, Carol Clugston, Anna Dominiczak and Alex Alderton, Roberto Amato, Jeffrey Barrett, Sonia Goncalves, Ewan Harrison, David K. Jackson, Ian Johnston, Dominic Kwiatkowski, Cordelia Langford, John Sillitoe on behalf of the Wellcome Sanger Institute COVID-19 Surveillance Team |
| EPI_ISL_2961236, EPI_ISL_2961237 | Lighthouse Lab in Alderley Park | Wellcome Sanger Institute for the COVID-19 Genomics UK (COG-UK) Consortium | Jacquelyn Wynn, Mairead Hyland, The Lighthouse Lab in Alderley Park and Alex Alderton, Roberto Amato, Jeffrey Barrett, Sonia Goncalves, Ewan Harrison, David K. Jackson, Ian Johnston, Dominic Kwiatkowski, Cordelia Langford, John Sillitoe on behalf of the Wellcome Sanger Institute COVID-19 Surveillance Team |
| EPI_ISL_2961238, EPI_ISL_2961239 | Lighthouse Lab in Glasgow | Wellcome Sanger Institute for the COVID-19 Genomics UK (COG-UK) Consortium | Harper VanSteenhouse, Yumi Kasai, David Gray, Carol Clugston, Anna Dominiczak and Alex Alderton, Roberto Amato, Jeffrey Barrett, Sonia Goncalves, Ewan Harrison, David K. Jackson, Ian Johnston, Dominic Kwiatkowski, Cordelia Langford, John Sillitoe on behalf of the Wellcome Sanger Institute COVID-19 Surveillance Team |
| EPI_ISL_2961240 | Berkshire and Surrey Pathology Services Lighthouse Laboratory | Wellcome Sanger Institute for the COVID-19 Genomics UK (COG-UK) Consortium | Berkshire and Surrey Pathology Services Lighthouse Laboratory and Alex Alderton, Roberto Amato, Jeffrey Barrett, Sonia Goncalves, Ewan Harrison, David K. Jackson, Ian Johnston, Dominic Kwiatkowski, Cordelia Langford, John Sillitoe on behalf of the Wellcome Sanger Institute COVID-19 Surveillance Team |
| EPI_ISL_2961241 | Lighthouse Lab in Alderley Park | Wellcome Sanger Institute for the COVID-19 Genomics UK (COG-UK) Consortium | Jacquelyn Wynn, Mairead Hyland, The Lighthouse Lab in Alderley Park and Alex Alderton, Roberto Amato, Jeffrey Barrett, Sonia Goncalves, Ewan Harrison, David K. Jackson, Ian Johnston, Dominic Kwiatkowski, Cordelia Langford, John Sillitoe on behalf of the Wellcome Sanger Institute COVID-19 Surveillance Team |

| | | | |
|---|---|---|---|
| EPI_ISL_2961242 | Lighthouse Lab in Glasgow | Wellcome Sanger Institute for the COVID-19 Genomics UK (COG-UK) Consortium | Harper VanSteenhouse, Yumi Kasai, David Gray, Carol Clugston, Anna Dominiczak and Alex Alderton, Roberto Amato, Jeffrey Barrett, Sonia Goncalves, Ewan Harrison, David K. Jackson, Ian Johnston, Dominic Kwiatkowski, Cordelia Langford, John Sillitoe on behalf of the Wellcome Sanger Institute COVID-19 Surveillance Team |
| EPI_ISL_2961243 | Lighthouse Lab in Alderley Park | Wellcome Sanger Institute for the COVID-19 Genomics UK (COG-UK) Consortium | Jacquelyn Wynn, Mairead Hyland, The Lighthouse Lab in Alderley Park and Alex Alderton, Roberto Amato, Jeffrey Barrett, Sonia Goncalves, Ewan Harrison, David K. Jackson, Ian Johnston, Dominic Kwiatkowski, Cordelia Langford, John Sillitoe on behalf of the Wellcome Sanger Institute COVID-19 Surveillance Team |
| EPI_ISL_2961244, EPI_ISL_2961245 | Lighthouse Lab in Glasgow | Wellcome Sanger Institute for the COVID-19 Genomics UK (COG-UK) Consortium | Harper VanSteenhouse, Yumi Kasai, David Gray, Carol Clugston, Anna Dominiczak and Alex Alderton, Roberto Amato, Jeffrey Barrett, Sonia Goncalves, Ewan Harrison, David K. Jackson, Ian Johnston, Dominic Kwiatkowski, Cordelia Langford, John Sillitoe on behalf of the Wellcome Sanger Institute COVID-19 Surveillance Team |
| EPI_ISL_2961246 | Lighthouse Lab in Alderley Park | Wellcome Sanger Institute for the COVID-19 Genomics UK (COG-UK) Consortium | Jacquelyn Wynn, Mairead Hyland, The Lighthouse Lab in Alderley Park and Alex Alderton, Roberto Amato, Jeffrey Barrett, Sonia Goncalves, Ewan Harrison, David K. Jackson, Ian Johnston, Dominic Kwiatkowski, Cordelia Langford, John Sillitoe on behalf of the Wellcome Sanger Institute COVID-19 Surveillance Team |
| EPI_ISL_2961247, EPI_ISL_2961248 | Lighthouse Lab in Glasgow | Wellcome Sanger Institute for the COVID-19 Genomics UK (COG-UK) Consortium | Harper VanSteenhouse, Yumi Kasai, David Gray, Carol Clugston, Anna Dominiczak and Alex Alderton, Roberto Amato, Jeffrey Barrett, Sonia Goncalves, Ewan Harrison, David K. Jackson, Ian Johnston, Dominic Kwiatkowski, Cordelia Langford, John Sillitoe on behalf of the Wellcome Sanger Institute COVID-19 Surveillance Team |
| EPI_ISL_2961249, EPI_ISL_2961250 | Lighthouse Lab in Alderley Park | Wellcome Sanger Institute for the COVID-19 Genomics UK (COG-UK) Consortium | Jacquelyn Wynn, Mairead Hyland, The Lighthouse Lab in Alderley Park and Alex Alderton, Roberto Amato, Jeffrey Barrett, Sonia Goncalves, Ewan Harrison, David K. Jackson, Ian Johnston, Dominic Kwiatkowski, Cordelia Langford, John Sillitoe on behalf of the Wellcome Sanger Institute COVID-19 Surveillance Team |
| EPI_ISL_2961251, EPI_ISL_2961252 | Lighthouse Lab in Glasgow | Wellcome Sanger Institute for the COVID-19 Genomics UK (COG-UK) Consortium | Harper VanSteenhouse, Yumi Kasai, David Gray, Carol Clugston, Anna Dominiczak and Alex Alderton, Roberto Amato, Jeffrey Barrett, Sonia Goncalves, Ewan Harrison, David K. Jackson, Ian Johnston, Dominic Kwiatkowski, Cordelia Langford, John Sillitoe on behalf of the Wellcome Sanger Institute COVID-19 Surveillance Team |
| EPI_ISL_2961253, EPI_ISL_2961254 | Lighthouse Lab in Alderley Park | Wellcome Sanger Institute for the COVID-19 Genomics UK (COG-UK) Consortium | Jacquelyn Wynn, Mairead Hyland, The Lighthouse Lab in Alderley Park and Alex Alderton, Roberto Amato, Jeffrey Barrett, Sonia Goncalves, Ewan Harrison, David K. Jackson, Ian Johnston, Dominic Kwiatkowski, Cordelia Langford, John Sillitoe on behalf of the Wellcome Sanger Institute COVID-19 Surveillance Team |
| EPI_ISL_2961255, EPI_ISL_2961256, EPI_ISL_2961257 | Berkshire and Surrey Pathology Services Lighthouse Laboratory | Wellcome Sanger Institute for the COVID-19 Genomics UK (COG-UK) Consortium | Berkshire and Surrey Pathology Services Lighthouse Laboratory and Alex Alderton, Roberto Amato, Jeffrey Barrett, Sonia Goncalves, Ewan Harrison, David K. Jackson, Ian Johnston, Dominic Kwiatkowski, Cordelia Langford, John Sillitoe on behalf of the Wellcome Sanger Institute COVID-19 Surveillance Team |
| EPI_ISL_2961258 | Lighthouse Lab in Glasgow | Wellcome Sanger Institute for the COVID-19 Genomics UK (COG-UK) Consortium | Harper VanSteenhouse, Yumi Kasai, David Gray, Carol Clugston, Anna Dominiczak and Alex Alderton, Roberto Amato, Jeffrey Barrett, Sonia Goncalves, Ewan Harrison, David K. Jackson, Ian Johnston, Dominic Kwiatkowski, Cordelia Langford, John Sillitoe on behalf of the Wellcome Sanger Institute COVID-19 Surveillance Team |
| EPI_ISL_2961259 | Berkshire and Surrey Pathology Services Lighthouse Laboratory | Wellcome Sanger Institute for the COVID-19 Genomics UK (COG-UK) Consortium | Berkshire and Surrey Pathology Services Lighthouse Laboratory and Alex Alderton, Roberto Amato, Jeffrey Barrett, Sonia Goncalves, Ewan Harrison, David K. Jackson, Ian Johnston, Dominic Kwiatkowski, Cordelia Langford, John Sillitoe on behalf of the Wellcome Sanger Institute COVID-19 Surveillance Team |
| EPI_ISL_2961260, EPI_ISL_2961261 | Lighthouse Lab in Glasgow | Wellcome Sanger Institute for the COVID-19 Genomics UK (COG-UK) Consortium | Harper VanSteenhouse, Yumi Kasai, David Gray, Carol Clugston, Anna Dominiczak and Alex Alderton, Roberto Amato, Jeffrey Barrett, Sonia Goncalves, Ewan Harrison, David K. Jackson, Ian Johnston, Dominic Kwiatkowski, Cordelia Langford, John Sillitoe on behalf of the Wellcome Sanger Institute COVID-19 Surveillance Team |
| EPI_ISL_2961262 | Lighthouse Lab in Alderley Park | Wellcome Sanger Institute for the COVID-19 Genomics UK (COG-UK) Consortium | Jacquelyn Wynn, Mairead Hyland, The Lighthouse Lab in Alderley Park and Alex Alderton, Roberto Amato, Jeffrey Barrett, Sonia Goncalves, Ewan Harrison, David K. Jackson, Ian Johnston, Dominic Kwiatkowski, Cordelia Langford, John Sillitoe on behalf of the Wellcome Sanger Institute COVID-19 Surveillance Team |
| EPI_ISL_2961263 | Lighthouse Lab in Glasgow | Wellcome Sanger Institute for the COVID-19 Genomics UK (COG-UK) Consortium | Harper VanSteenhouse, Yumi Kasai, David Gray, Carol Clugston, Anna Dominiczak and Alex Alderton, Roberto Amato, Jeffrey Barrett, Sonia Goncalves, Ewan Harrison, David K. Jackson, Ian Johnston, Dominic Kwiatkowski, Cordelia Langford, John Sillitoe on behalf of the Wellcome Sanger Institute COVID-19 Surveillance Team |
| EPI_ISL_2961264 | Berkshire and Surrey Pathology Services Lighthouse Laboratory | Wellcome Sanger Institute for the COVID-19 Genomics UK (COG-UK) Consortium | Berkshire and Surrey Pathology Services Lighthouse Laboratory and Alex Alderton, Roberto Amato, Jeffrey Barrett, Sonia Goncalves, Ewan Harrison, David K. Jackson, Ian Johnston, Dominic Kwiatkowski, Cordelia Langford, John Sillitoe on behalf of the Wellcome Sanger Institute COVID-19 Surveillance Team |
| EPI_ISL_2961265 | Lighthouse Lab in Alderley Park | Wellcome Sanger Institute for the COVID-19 Genomics UK (COG-UK) Consortium | Jacquelyn Wynn, Mairead Hyland, The Lighthouse Lab in Alderley Park and Alex Alderton, Roberto Amato, Jeffrey Barrett, Sonia Goncalves, Ewan Harrison, David K. Jackson, Ian Johnston, Dominic Kwiatkowski, Cordelia Langford, John Sillitoe on behalf of the Wellcome Sanger Institute COVID-19 Surveillance Team |
| EPI_ISL_2961266 | Lighthouse Lab in Glasgow | Wellcome Sanger Institute for the COVID-19 Genomics UK (COG-UK) Consortium | Harper VanSteenhouse, Yumi Kasai, David Gray, Carol Clugston, Anna Dominiczak and Alex Alderton, Roberto Amato, Jeffrey Barrett, Sonia Goncalves, Ewan Harrison, David K. Jackson, Ian Johnston, Dominic Kwiatkowski, Cordelia Langford, John Sillitoe on behalf of the Wellcome Sanger Institute COVID-19 Surveillance Team |
| EPI_ISL_2961267 | Berkshire and Surrey Pathology Services Lighthouse Laboratory | Wellcome Sanger Institute for the COVID-19 Genomics UK (COG-UK) Consortium | Berkshire and Surrey Pathology Services Lighthouse Laboratory and Alex Alderton, Roberto Amato, Jeffrey Barrett, Sonia Goncalves, Ewan Harrison, David K. Jackson, Ian Johnston, Dominic Kwiatkowski, Cordelia Langford, John Sillitoe on behalf of the Wellcome Sanger Institute COVID-19 Surveillance Team |
| EPI_ISL_2961268 | Lighthouse Lab in Alderley Park | Wellcome Sanger Institute for the COVID-19 Genomics UK (COG-UK) Consortium | Jacquelyn Wynn, Mairead Hyland, The Lighthouse Lab in Alderley Park and Alex Alderton, Roberto Amato, Jeffrey Barrett, Sonia Goncalves, Ewan Harrison, David K. Jackson, Ian Johnston, Dominic Kwiatkowski, Cordelia Langford, John Sillitoe on behalf of the Wellcome Sanger Institute COVID-19 Surveillance Team |
| EPI_ISL_2961269 | Berkshire and Surrey Pathology Services Lighthouse Laboratory | Wellcome Sanger Institute for the COVID-19 Genomics UK (COG-UK) Consortium | Berkshire and Surrey Pathology Services Lighthouse Laboratory and Alex Alderton, Roberto Amato, Jeffrey Barrett, Sonia Goncalves, Ewan Harrison, David K. Jackson, Ian Johnston, Dominic Kwiatkowski, Cordelia Langford, John Sillitoe on behalf of the Wellcome Sanger Institute COVID-19 Surveillance Team |
| EPI_ISL_2961270 | Lighthouse Lab in Alderley Park | Wellcome Sanger Institute for the COVID-19 Genomics UK (COG-UK) Consortium | Jacquelyn Wynn, Mairead Hyland, The Lighthouse Lab in Alderley Park and Alex Alderton, Roberto Amato, Jeffrey Barrett, Sonia Goncalves, Ewan Harrison, David K. Jackson, Ian Johnston, Dominic Kwiatkowski, Cordelia Langford, John Sillitoe on behalf of the Wellcome Sanger Institute COVID-19 Surveillance Team |
| EPI_ISL_2961271 | Berkshire and Surrey Pathology Services Lighthouse Laboratory | Wellcome Sanger Institute for the COVID-19 Genomics UK (COG-UK) Consortium | Berkshire and Surrey Pathology Services Lighthouse Laboratory and Alex Alderton, Roberto Amato, Jeffrey Barrett, Sonia Goncalves, Ewan Harrison, David K. Jackson, Ian Johnston, Dominic Kwiatkowski, Cordelia Langford, John Sillitoe on behalf of the Wellcome Sanger Institute COVID-19 Surveillance Team |
| EPI_ISL_2961272 | Lighthouse Lab in Glasgow | Wellcome Sanger Institute for the COVID-19 Genomics UK (COG-UK) Consortium | Harper VanSteenhouse, Yumi Kasai, David Gray, Carol Clugston, Anna Dominiczak and Alex Alderton, Roberto Amato, Jeffrey Barrett, Sonia Goncalves, Ewan Harrison, David K. Jackson, Ian Johnston, Dominic Kwiatkowski, Cordelia Langford, John Sillitoe on behalf of the Wellcome Sanger Institute COVID-19 Surveillance Team |
| EPI_ISL_2961273, EPI_ISL_2961274 | Berkshire and Surrey Pathology Services Lighthouse Laboratory | Wellcome Sanger Institute for the COVID-19 Genomics UK (COG-UK) Consortium | Berkshire and Surrey Pathology Services Lighthouse Laboratory and Alex Alderton, Roberto Amato, Jeffrey Barrett, Sonia Goncalves, Ewan Harrison, David K. Jackson, Ian Johnston, Dominic Kwiatkowski, Cordelia Langford, John Sillitoe on behalf of the Wellcome Sanger Institute COVID-19 Surveillance Team |
| EPI_ISL_2961275 | Lighthouse Lab in Glasgow | Wellcome Sanger Institute for the COVID-19 Genomics UK (COG-UK) Consortium | Harper VanSteenhouse, Yumi Kasai, David Gray, Carol Clugston, Anna Dominiczak and Alex Alderton, Roberto Amato, Jeffrey Barrett, Sonia Goncalves, Ewan Harrison, David K. Jackson, Ian Johnston, Dominic Kwiatkowski, Cordelia Langford, John Sillitoe on behalf of the Wellcome Sanger Institute COVID-19 Surveillance Team |
| EPI_ISL_2961276 | Berkshire and Surrey Pathology Services Lighthouse Laboratory | Wellcome Sanger Institute for the COVID-19 Genomics UK (COG-UK) Consortium | Berkshire and Surrey Pathology Services Lighthouse Laboratory and Alex Alderton, Roberto Amato, Jeffrey Barrett, Sonia Goncalves, Ewan Harrison, David K. Jackson, Ian Johnston, Dominic Kwiatkowski, Cordelia Langford, John Sillitoe on behalf of the Wellcome Sanger Institute COVID-19 Surveillance Team |
| EPI_ISL_2961277 | Lighthouse Lab in Alderley Park | Wellcome Sanger Institute for the COVID-19 Genomics UK (COG-UK) Consortium | Jacquelyn Wynn, Mairead Hyland, The Lighthouse Lab in Alderley Park and Alex Alderton, Roberto Amato, Jeffrey Barrett, Sonia Goncalves, Ewan Harrison, David K. Jackson, Ian Johnston, Dominic Kwiatkowski, Cordelia Langford, John Sillitoe on behalf of the Wellcome Sanger Institute COVID-19 Surveillance Team |
| EPI_ISL_2961278 | Berkshire and Surrey Pathology Services Lighthouse Laboratory | Wellcome Sanger Institute for the COVID-19 Genomics UK (COG-UK) Consortium | Berkshire and Surrey Pathology Services Lighthouse Laboratory and Alex Alderton, Roberto Amato, Jeffrey Barrett, Sonia Goncalves, Ewan Harrison, David K. Jackson, Ian Johnston, Dominic Kwiatkowski, Cordelia Langford, John Sillitoe on behalf of the Wellcome Sanger Institute COVID-19 Surveillance Team |
| EPI_ISL_2961279 | Lighthouse Lab in Glasgow | Wellcome Sanger Institute for the COVID-19 Genomics UK (COG-UK) Consortium | Harper VanSteenhouse, Yumi Kasai, David Gray, Carol Clugston, Anna Dominiczak and Alex Alderton, Roberto Amato, Jeffrey Barrett, Sonia Goncalves, Ewan Harrison, David K. Jackson, Ian Johnston, Dominic Kwiatkowski, Cordelia Langford, John Sillitoe on behalf of the Wellcome Sanger Institute COVID-19 Surveillance Team |
| EPI_ISL_2961280 | Lighthouse Lab in Alderley Park | Wellcome Sanger Institute for the COVID-19 Genomics UK (COG-UK) Consortium | Jacquelyn Wynn, Mairead Hyland, The Lighthouse Lab in Alderley Park and Alex Alderton, Roberto Amato, Jeffrey Barrett, Sonia Goncalves, Ewan Harrison, David K. Jackson, Ian Johnston, Dominic Kwiatkowski, Cordelia Langford, John Sillitoe on behalf of the Wellcome Sanger Institute COVID-19 Surveillance Team |
| EPI_ISL_2961281 | Berkshire and Surrey Pathology Services Lighthouse Laboratory | Wellcome Sanger Institute for the COVID-19 Genomics UK (COG-UK) Consortium | Berkshire and Surrey Pathology Services Lighthouse Laboratory and Alex Alderton, Roberto Amato, Jeffrey Barrett, Sonia Goncalves, Ewan Harrison, David K. Jackson, Ian Johnston, Dominic Kwiatkowski, Cordelia Langford, John Sillitoe on behalf of the Wellcome Sanger Institute COVID-19 Surveillance Team |
| EPI_ISL_2961282 | Lighthouse Lab in Glasgow | Wellcome Sanger Institute for the COVID-19 Genomics UK (COG-UK) Consortium | Harper VanSteenhouse, Yumi Kasai, David Gray, Carol Clugston, Anna Dominiczak and Alex Alderton, Roberto Amato, Jeffrey Barrett, Sonia Goncalves, Ewan Harrison, David K. Jackson, Ian Johnston, Dominic Kwiatkowski, Cordelia Langford, John Sillitoe on behalf of the Wellcome Sanger Institute COVID-19 Surveillance Team |
| EPI_ISL_2961283 | Berkshire and Surrey Pathology Services Lighthouse Laboratory | Wellcome Sanger Institute for the COVID-19 Genomics UK (COG-UK) Consortium | Berkshire and Surrey Pathology Services Lighthouse Laboratory and Alex Alderton, Roberto Amato, Jeffrey Barrett, Sonia Goncalves, Ewan Harrison, David K. Jackson, Ian Johnston, Dominic Kwiatkowski, Cordelia Langford, John Sillitoe on behalf of the Wellcome Sanger Institute COVID-19 Surveillance Team |
| EPI_ISL_2961284, EPI_ISL_2961285, EPI_ISL_2961286 | Lighthouse Lab in Glasgow | Wellcome Sanger Institute for the COVID-19 Genomics UK (COG-UK) Consortium | Harper VanSteenhouse, Yumi Kasai, David Gray, Carol Clugston, Anna Dominiczak and Alex Alderton, Roberto Amato, Jeffrey Barrett, Sonia Goncalves, Ewan Harrison, David K. Jackson, Ian Johnston, Dominic Kwiatkowski, Cordelia Langford, John Sillitoe on behalf of the Wellcome Sanger Institute COVID-19 Surveillance Team |
| EPI_ISL_2961287, EPI_ISL_2961288 | Berkshire and Surrey Pathology Services Lighthouse Laboratory | Wellcome Sanger Institute for the COVID-19 Genomics UK (COG-UK) Consortium | Berkshire and Surrey Pathology Services Lighthouse Laboratory and Alex Alderton, Roberto Amato, Jeffrey Barrett, Sonia Goncalves, Ewan Harrison, David K. Jackson, Ian Johnston, Dominic Kwiatkowski, Cordelia Langford, John Sillitoe on behalf of the Wellcome Sanger Institute COVID-19 Surveillance Team |
| EPI_ISL_2961289, EPI_ISL_2961290 | Lighthouse Lab in Glasgow | Wellcome Sanger Institute for the COVID-19 Genomics UK (COG-UK) Consortium | Harper VanSteenhouse, Yumi Kasai, David Gray, Carol Clugston, Anna Dominiczak and Alex Alderton, Roberto Amato, Jeffrey Barrett, Sonia Goncalves, Ewan Harrison, David K. Jackson, Ian Johnston, Dominic Kwiatkowski, Cordelia Langford, John Sillitoe on behalf of the Wellcome Sanger Institute COVID-19 Surveillance Team |
| EPI_ISL_2961291 | Lighthouse Lab in Alderley Park | Wellcome Sanger Institute for the COVID-19 Genomics UK (COG-UK) Consortium | Jacquelyn Wynn, Mairead Hyland, The Lighthouse Lab in Alderley Park and Alex Alderton, Roberto Amato, Jeffrey Barrett, Sonia Goncalves, Ewan Harrison, David K. Jackson, Ian Johnston, Dominic Kwiatkowski, Cordelia Langford, John Sillitoe on behalf of the Wellcome Sanger Institute COVID-19 Surveillance Team |
| EPI_ISL_2961292 | Lighthouse Lab in Glasgow | Wellcome Sanger Institute for the COVID-19 Genomics UK (COG-UK) Consortium | Harper VanSteenhouse, Yumi Kasai, David Gray, Carol Clugston, Anna Dominiczak and Alex Alderton, Roberto Amato, Jeffrey Barrett, Sonia Goncalves, Ewan Harrison, David K. Jackson, Ian Johnston, Dominic Kwiatkowski, Cordelia Langford, John Sillitoe on behalf of the Wellcome Sanger Institute COVID-19 Surveillance Team |
| EPI_ISL_2961293, EPI_ISL_2961294 | Lighthouse Lab in Alderley Park | Wellcome Sanger Institute for the COVID-19 Genomics UK (COG-UK) Consortium | Jacquelyn Wynn, Mairead Hyland, The Lighthouse Lab in Alderley Park and Alex Alderton, Roberto Amato, Jeffrey Barrett, Sonia Goncalves, Ewan Harrison, David K. Jackson, Ian Johnston, Dominic Kwiatkowski, Cordelia Langford, John Sillitoe on behalf of the Wellcome Sanger Institute COVID-19 Surveillance Team |
| EPI_ISL_2961295 | Lighthouse Lab in Glasgow | Wellcome Sanger Institute for the COVID-19 Genomics UK (COG-UK) Consortium | Harper VanSteenhouse, Yumi Kasai, David Gray, Carol Clugston, Anna Dominiczak and Alex Alderton, Roberto Amato, Jeffrey Barrett, Sonia Goncalves, Ewan Harrison, David K. Jackson, Ian Johnston, Dominic Kwiatkowski, Cordelia Langford, John Sillitoe on behalf of the Wellcome Sanger Institute COVID-19 Surveillance Team |
| EPI_ISL_2961296 | Berkshire and Surrey Pathology Services Lighthouse Laboratory | Wellcome Sanger Institute for the COVID-19 Genomics UK (COG-UK) Consortium | Berkshire and Surrey Pathology Services Lighthouse Laboratory and Alex Alderton, Roberto Amato, Jeffrey Barrett, Sonia Goncalves, Ewan Harrison, David K. Jackson, Ian Johnston, Dominic Kwiatkowski, Cordelia Langford, John Sillitoe on behalf of the Wellcome Sanger Institute COVID-19 Surveillance Team |
| EPI_ISL_2961297 | Lighthouse Lab in Glasgow | Wellcome Sanger Institute for the COVID-19 Genomics UK (COG-UK) Consortium | Harper VanSteenhouse, Yumi Kasai, David Gray, Carol Clugston, Anna Dominiczak and Alex Alderton, Roberto Amato, Jeffrey Barrett, Sonia Goncalves, Ewan Harrison, David K. Jackson, Ian Johnston, Dominic Kwiatkowski, Cordelia Langford, John Sillitoe on behalf of the Wellcome Sanger Institute COVID-19 Surveillance Team |
| EPI_ISL_2961298 | Berkshire and Surrey Pathology Services Lighthouse Laboratory | Wellcome Sanger Institute for the COVID-19 Genomics UK (COG-UK) Consortium | Berkshire and Surrey Pathology Services Lighthouse Laboratory and Alex Alderton, Roberto Amato, Jeffrey Barrett, Sonia Goncalves, Ewan Harrison, David K. Jackson, Ian Johnston, Dominic Kwiatkowski, Cordelia Langford, John Sillitoe on behalf of the Wellcome Sanger Institute COVID-19 Surveillance Team |
| EPI_ISL_2961299 | Lighthouse Lab in Alderley Park | Wellcome Sanger Institute for the COVID-19 Genomics UK (COG-UK) Consortium | Jacquelyn Wynn, Mairead Hyland, The Lighthouse Lab in Alderley Park and Alex Alderton, Roberto Amato, Jeffrey Barrett, Sonia Goncalves, Ewan Harrison, David K. Jackson, Ian Johnston, Dominic Kwiatkowski, Cordelia Langford, John Sillitoe on behalf of the Wellcome Sanger Institute COVID-19 Surveillance Team |

| | | | |
|---|---|---|---|
| EPI_ISL_2961300, EPI_ISL_2961301 | Berkshire and Surrey Pathology Services Lighthouse Laboratory | Wellcome Sanger Institute for the COVID-19 Genomics UK (COG-UK) Consortium | Berkshire and Surrey Pathology Services Lighthouse Laboratory and Alex Alderton, Roberto Amato, Jeffrey Barrett, Sonia Goncalves, Ewan Harrison, David K. Jackson, Ian Johnston, Dominic Kwiatkowski, Cordelia Langford, John Sillitoe on behalf of the Wellcome Sanger Institute COVID-19 Surveillance Team |
| EPI_ISL_2961302 | Lighthouse Lab in Glasgow | Wellcome Sanger Institute for the COVID-19 Genomics UK (COG-UK) Consortium | Harper VanSteenhouse, Yumi Kasai, David Gray, Carol Clugston, Anna Dominiczak and Alex Alderton, Roberto Amato, Jeffrey Barrett, Sonia Goncalves, Ewan Harrison, David K. Jackson, Ian Johnston, Dominic Kwiatkowski, Cordelia Langford, John Sillitoe on behalf of the Wellcome Sanger Institute COVID-19 Surveillance Team |
| EPI_ISL_2961303 | Berkshire and Surrey Pathology Services Lighthouse Laboratory | Wellcome Sanger Institute for the COVID-19 Genomics UK (COG-UK) Consortium | Berkshire and Surrey Pathology Services Lighthouse Laboratory and Alex Alderton, Roberto Amato, Jeffrey Barrett, Sonia Goncalves, Ewan Harrison, David K. Jackson, Ian Johnston, Dominic Kwiatkowski, Cordelia Langford, John Sillitoe on behalf of the Wellcome Sanger Institute COVID-19 Surveillance Team |
| EPI_ISL_2961304, EPI_ISL_2961305 | Lighthouse Lab in Glasgow | Wellcome Sanger Institute for the COVID-19 Genomics UK (COG-UK) Consortium | Harper VanSteenhouse, Yumi Kasai, David Gray, Carol Clugston, Anna Dominiczak and Alex Alderton, Roberto Amato, Jeffrey Barrett, Sonia Goncalves, Ewan Harrison, David K. Jackson, Ian Johnston, Dominic Kwiatkowski, Cordelia Langford, John Sillitoe on behalf of the Wellcome Sanger Institute COVID-19 Surveillance Team |
| EPI_ISL_2961306 | Lighthouse Lab in Alderley Park | Wellcome Sanger Institute for the COVID-19 Genomics UK (COG-UK) Consortium | Jacquelyn Wynn, Mairead Hyland, The Lighthouse Lab in Alderley Park and Alex Alderton, Roberto Amato, Jeffrey Barrett, Sonia Goncalves, Ewan Harrison, David K. Jackson, Ian Johnston, Dominic Kwiatkowski, Cordelia Langford, John Sillitoe on behalf of the Wellcome Sanger Institute COVID-19 Surveillance Team |
| EPI_ISL_2961307 | Berkshire and Surrey Pathology Services Lighthouse Laboratory | Wellcome Sanger Institute for the COVID-19 Genomics UK (COG-UK) Consortium | Berkshire and Surrey Pathology Services Lighthouse Laboratory and Alex Alderton, Roberto Amato, Jeffrey Barrett, Sonia Goncalves, Ewan Harrison, David K. Jackson, Ian Johnston, Dominic Kwiatkowski, Cordelia Langford, John Sillitoe on behalf of the Wellcome Sanger Institute COVID-19 Surveillance Team |
| EPI_ISL_2961308 | Lighthouse Lab in Alderley Park | Wellcome Sanger Institute for the COVID-19 Genomics UK (COG-UK) Consortium | Jacquelyn Wynn, Mairead Hyland, The Lighthouse Lab in Alderley Park and Alex Alderton, Roberto Amato, Jeffrey Barrett, Sonia Goncalves, Ewan Harrison, David K. Jackson, Ian Johnston, Dominic Kwiatkowski, Cordelia Langford, John Sillitoe on behalf of the Wellcome Sanger Institute COVID-19 Surveillance Team |
| EPI_ISL_2961309 | Lighthouse Lab in Glasgow | Wellcome Sanger Institute for the COVID-19 Genomics UK (COG-UK) Consortium | Harper VanSteenhouse, Yumi Kasai, David Gray, Carol Clugston, Anna Dominiczak and Alex Alderton, Roberto Amato, Jeffrey Barrett, Sonia Goncalves, Ewan Harrison, David K. Jackson, Ian Johnston, Dominic Kwiatkowski, Cordelia Langford, John Sillitoe on behalf of the Wellcome Sanger Institute COVID-19 Surveillance Team |
| EPI_ISL_2961310 | Berkshire and Surrey Pathology Services Lighthouse Laboratory | Wellcome Sanger Institute for the COVID-19 Genomics UK (COG-UK) Consortium | Berkshire and Surrey Pathology Services Lighthouse Laboratory and Alex Alderton, Roberto Amato, Jeffrey Barrett, Sonia Goncalves, Ewan Harrison, David K. Jackson, Ian Johnston, Dominic Kwiatkowski, Cordelia Langford, John Sillitoe on behalf of the Wellcome Sanger Institute COVID-19 Surveillance Team |
| EPI_ISL_2961311 | Lighthouse Lab in Alderley Park | Wellcome Sanger Institute for the COVID-19 Genomics UK (COG-UK) Consortium | Jacquelyn Wynn, Mairead Hyland, The Lighthouse Lab in Alderley Park and Alex Alderton, Roberto Amato, Jeffrey Barrett, Sonia Goncalves, Ewan Harrison, David K. Jackson, Ian Johnston, Dominic Kwiatkowski, Cordelia Langford, John Sillitoe on behalf of the Wellcome Sanger Institute COVID-19 Surveillance Team |
| EPI_ISL_2961312, EPI_ISL_2961313 | Lighthouse Lab in Glasgow | Wellcome Sanger Institute for the COVID-19 Genomics UK (COG-UK) Consortium | Harper VanSteenhouse, Yumi Kasai, David Gray, Carol Clugston, Anna Dominiczak and Alex Alderton, Roberto Amato, Jeffrey Barrett, Sonia Goncalves, Ewan Harrison, David K. Jackson, Ian Johnston, Dominic Kwiatkowski, Cordelia Langford, John Sillitoe on behalf of the Wellcome Sanger Institute COVID-19 Surveillance Team |
| EPI_ISL_2961314 | Berkshire and Surrey Pathology Services Lighthouse Laboratory | Wellcome Sanger Institute for the COVID-19 Genomics UK (COG-UK) Consortium | Berkshire and Surrey Pathology Services Lighthouse Laboratory and Alex Alderton, Roberto Amato, Jeffrey Barrett, Sonia Goncalves, Ewan Harrison, David K. Jackson, Ian Johnston, Dominic Kwiatkowski, Cordelia Langford, John Sillitoe on behalf of the Wellcome Sanger Institute COVID-19 Surveillance Team |
| EPI_ISL_2961315 | Lighthouse Lab in Alderley Park | Wellcome Sanger Institute for the COVID-19 Genomics UK (COG-UK) Consortium | Jacquelyn Wynn, Mairead Hyland, The Lighthouse Lab in Alderley Park and Alex Alderton, Roberto Amato, Jeffrey Barrett, Sonia Goncalves, Ewan Harrison, David K. Jackson, Ian Johnston, Dominic Kwiatkowski, Cordelia Langford, John Sillitoe on behalf of the Wellcome Sanger Institute COVID-19 Surveillance Team |
| EPI_ISL_2961316, EPI_ISL_2961317 | Berkshire and Surrey Pathology Services Lighthouse Laboratory | Wellcome Sanger Institute for the COVID-19 Genomics UK (COG-UK) Consortium | Berkshire and Surrey Pathology Services Lighthouse Laboratory and Alex Alderton, Roberto Amato, Jeffrey Barrett, Sonia Goncalves, Ewan Harrison, David K. Jackson, Ian Johnston, Dominic Kwiatkowski, Cordelia Langford, John Sillitoe on behalf of the Wellcome Sanger Institute COVID-19 Surveillance Team |
| EPI_ISL_2961318 | Lighthouse Lab in Glasgow | Wellcome Sanger Institute for the COVID-19 Genomics UK (COG-UK) Consortium | Harper VanSteenhouse, Yumi Kasai, David Gray, Carol Clugston, Anna Dominiczak and Alex Alderton, Roberto Amato, Jeffrey Barrett, Sonia Goncalves, Ewan Harrison, David K. Jackson, Ian Johnston, Dominic Kwiatkowski, Cordelia Langford, John Sillitoe on behalf of the Wellcome Sanger Institute COVID-19 Surveillance Team |
| EPI_ISL_2961319, EPI_ISL_2961320 | Lighthouse Lab in Alderley Park | Wellcome Sanger Institute for the COVID-19 Genomics UK (COG-UK) Consortium | Jacquelyn Wynn, Mairead Hyland, The Lighthouse Lab in Alderley Park and Alex Alderton, Roberto Amato, Jeffrey Barrett, Sonia Goncalves, Ewan Harrison, David K. Jackson, Ian Johnston, Dominic Kwiatkowski, Cordelia Langford, John Sillitoe on behalf of the Wellcome Sanger Institute COVID-19 Surveillance Team |
| EPI_ISL_2961321 | Berkshire and Surrey Pathology Services Lighthouse Laboratory | Wellcome Sanger Institute for the COVID-19 Genomics UK (COG-UK) Consortium | Berkshire and Surrey Pathology Services Lighthouse Laboratory and Alex Alderton, Roberto Amato, Jeffrey Barrett, Sonia Goncalves, Ewan Harrison, David K. Jackson, Ian Johnston, Dominic Kwiatkowski, Cordelia Langford, John Sillitoe on behalf of the Wellcome Sanger Institute COVID-19 Surveillance Team |
| EPI_ISL_2961322 | Lighthouse Lab in Glasgow | Wellcome Sanger Institute for the COVID-19 Genomics UK (COG-UK) Consortium | Harper VanSteenhouse, Yumi Kasai, David Gray, Carol Clugston, Anna Dominiczak and Alex Alderton, Roberto Amato, Jeffrey Barrett, Sonia Goncalves, Ewan Harrison, David K. Jackson, Ian Johnston, Dominic Kwiatkowski, Cordelia Langford, John Sillitoe on behalf of the Wellcome Sanger Institute COVID-19 Surveillance Team |
| EPI_ISL_2961323 | Lighthouse Lab in Alderley Park | Wellcome Sanger Institute for the COVID-19 Genomics UK (COG-UK) Consortium | Jacquelyn Wynn, Mairead Hyland, The Lighthouse Lab in Alderley Park and Alex Alderton, Roberto Amato, Jeffrey Barrett, Sonia Goncalves, Ewan Harrison, David K. Jackson, Ian Johnston, Dominic Kwiatkowski, Cordelia Langford, John Sillitoe on behalf of the Wellcome Sanger Institute COVID-19 Surveillance Team |
| EPI_ISL_2961324 | Lighthouse Lab in Glasgow | Wellcome Sanger Institute for the COVID-19 Genomics UK (COG-UK) Consortium | Harper VanSteenhouse, Yumi Kasai, David Gray, Carol Clugston, Anna Dominiczak and Alex Alderton, Roberto Amato, Jeffrey Barrett, Sonia Goncalves, Ewan Harrison, David K. Jackson, Ian Johnston, Dominic Kwiatkowski, Cordelia Langford, John Sillitoe on behalf of the Wellcome Sanger Institute COVID-19 Surveillance Team |
| EPI_ISL_2961325 | Lighthouse Lab in Alderley Park | Wellcome Sanger Institute for the COVID-19 Genomics UK (COG-UK) Consortium | Jacquelyn Wynn, Mairead Hyland, The Lighthouse Lab in Alderley Park and Alex Alderton, Roberto Amato, Jeffrey Barrett, Sonia Goncalves, Ewan Harrison, David K. Jackson, Ian Johnston, Dominic Kwiatkowski, Cordelia Langford, John Sillitoe on behalf of the Wellcome Sanger Institute COVID-19 Surveillance Team |
| EPI_ISL_2961326 | Lighthouse Lab in Glasgow | Wellcome Sanger Institute for the COVID-19 Genomics UK (COG-UK) Consortium | Harper VanSteenhouse, Yumi Kasai, David Gray, Carol Clugston, Anna Dominiczak and Alex Alderton, Roberto Amato, Jeffrey Barrett, Sonia Goncalves, Ewan Harrison, David K. Jackson, Ian Johnston, Dominic Kwiatkowski, Cordelia Langford, John Sillitoe on behalf of the Wellcome Sanger Institute COVID-19 Surveillance Team |
| EPI_ISL_2961327, EPI_ISL_2961328 | Berkshire and Surrey Pathology Services Lighthouse Laboratory | Wellcome Sanger Institute for the COVID-19 Genomics UK (COG-UK) Consortium | Berkshire and Surrey Pathology Services Lighthouse Laboratory and Alex Alderton, Roberto Amato, Jeffrey Barrett, Sonia Goncalves, Ewan Harrison, David K. Jackson, Ian Johnston, Dominic Kwiatkowski, Cordelia Langford, John Sillitoe on behalf of the Wellcome Sanger Institute COVID-19 Surveillance Team |
| EPI_ISL_2961329, EPI_ISL_2961330 | Lighthouse Lab in Glasgow | Wellcome Sanger Institute for the COVID-19 Genomics UK (COG-UK) Consortium | Harper VanSteenhouse, Yumi Kasai, David Gray, Carol Clugston, Anna Dominiczak and Alex Alderton, Roberto Amato, Jeffrey Barrett, Sonia Goncalves, Ewan Harrison, David K. Jackson, Ian Johnston, Dominic Kwiatkowski, Cordelia Langford, John Sillitoe on behalf of the Wellcome Sanger Institute COVID-19 Surveillance Team |
| EPI_ISL_2961331, EPI_ISL_2961332 | Lighthouse Lab in Alderley Park | Wellcome Sanger Institute for the COVID-19 Genomics UK (COG-UK) Consortium | Jacquelyn Wynn, Mairead Hyland, The Lighthouse Lab in Alderley Park and Alex Alderton, Roberto Amato, Jeffrey Barrett, Sonia Goncalves, Ewan Harrison, David K. Jackson, Ian Johnston, Dominic Kwiatkowski, Cordelia Langford, John Sillitoe on behalf of the Wellcome Sanger Institute COVID-19 Surveillance Team |
| EPI_ISL_2961333 | Lighthouse Lab in Glasgow | Wellcome Sanger Institute for the COVID-19 Genomics UK (COG-UK) Consortium | Harper VanSteenhouse, Yumi Kasai, David Gray, Carol Clugston, Anna Dominiczak and Alex Alderton, Roberto Amato, Jeffrey Barrett, Sonia Goncalves, Ewan Harrison, David K. Jackson, Ian Johnston, Dominic Kwiatkowski, Cordelia Langford, John Sillitoe on behalf of the Wellcome Sanger Institute COVID-19 Surveillance Team |
| EPI_ISL_2961334 | Lighthouse Lab in Alderley Park | Wellcome Sanger Institute for the COVID-19 Genomics UK (COG-UK) Consortium | Jacquelyn Wynn, Mairead Hyland, The Lighthouse Lab in Alderley Park and Alex Alderton, Roberto Amato, Jeffrey Barrett, Sonia Goncalves, Ewan Harrison, David K. Jackson, Ian Johnston, Dominic Kwiatkowski, Cordelia Langford, John Sillitoe on behalf of the Wellcome Sanger Institute COVID-19 Surveillance Team |
| EPI_ISL_2961335 | Lighthouse Lab in Glasgow | Wellcome Sanger Institute for the COVID-19 Genomics UK (COG-UK) Consortium | Harper VanSteenhouse, Yumi Kasai, David Gray, Carol Clugston, Anna Dominiczak and Alex Alderton, Roberto Amato, Jeffrey Barrett, Sonia Goncalves, Ewan Harrison, David K. Jackson, Ian Johnston, Dominic Kwiatkowski, Cordelia Langford, John Sillitoe on behalf of the Wellcome Sanger Institute COVID-19 Surveillance Team |
| EPI_ISL_2961336 | Berkshire and Surrey Pathology Services Lighthouse Laboratory | Wellcome Sanger Institute for the COVID-19 Genomics UK (COG-UK) Consortium | Berkshire and Surrey Pathology Services Lighthouse Laboratory and Alex Alderton, Roberto Amato, Jeffrey Barrett, Sonia Goncalves, Ewan Harrison, David K. Jackson, Ian Johnston, Dominic Kwiatkowski, Cordelia Langford, John Sillitoe on behalf of the Wellcome Sanger Institute COVID-19 Surveillance Team |
| EPI_ISL_2961337 | Lighthouse Lab in Glasgow | Wellcome Sanger Institute for the COVID-19 Genomics UK (COG-UK) Consortium | Harper VanSteenhouse, Yumi Kasai, David Gray, Carol Clugston, Anna Dominiczak and Alex Alderton, Roberto Amato, Jeffrey Barrett, Sonia Goncalves, Ewan Harrison, David K. Jackson, Ian Johnston, Dominic Kwiatkowski, Cordelia Langford, John Sillitoe on behalf of the Wellcome Sanger Institute COVID-19 Surveillance Team |
| EPI_ISL_2961338, EPI_ISL_2961339, EPI_ISL_2961340 | Lighthouse Lab in Alderley Park | Wellcome Sanger Institute for the COVID-19 Genomics UK (COG-UK) Consortium | Jacquelyn Wynn, Mairead Hyland, The Lighthouse Lab in Alderley Park and Alex Alderton, Roberto Amato, Jeffrey Barrett, Sonia Goncalves, Ewan Harrison, David K. Jackson, Ian Johnston, Dominic Kwiatkowski, Cordelia Langford, John Sillitoe on behalf of the Wellcome Sanger Institute COVID-19 Surveillance Team |
| EPI_ISL_2961341, EPI_ISL_2961342, EPI_ISL_2961343, EPI_ISL_2961344 | Lighthouse Lab in Glasgow | Wellcome Sanger Institute for the COVID-19 Genomics UK (COG-UK) Consortium | Harper VanSteenhouse, Yumi Kasai, David Gray, Carol Clugston, Anna Dominiczak and Alex Alderton, Roberto Amato, Jeffrey Barrett, Sonia Goncalves, Ewan Harrison, David K. Jackson, Ian Johnston, Dominic Kwiatkowski, Cordelia Langford, John Sillitoe on behalf of the Wellcome Sanger Institute COVID-19 Surveillance Team |
| EPI_ISL_2961345 | Lighthouse Lab in Alderley Park | Wellcome Sanger Institute for the COVID-19 Genomics UK (COG-UK) Consortium | Jacquelyn Wynn, Mairead Hyland, The Lighthouse Lab in Alderley Park and Alex Alderton, Roberto Amato, Jeffrey Barrett, Sonia Goncalves, Ewan Harrison, David K. Jackson, Ian Johnston, Dominic Kwiatkowski, Cordelia Langford, John Sillitoe on behalf of the Wellcome Sanger Institute COVID-19 Surveillance Team |
| EPI_ISL_2961346, EPI_ISL_2961347, EPI_ISL_2961348 | Lighthouse Lab in Glasgow | Wellcome Sanger Institute for the COVID-19 Genomics UK (COG-UK) Consortium | Harper VanSteenhouse, Yumi Kasai, David Gray, Carol Clugston, Anna Dominiczak and Alex Alderton, Roberto Amato, Jeffrey Barrett, Sonia Goncalves, Ewan Harrison, David K. Jackson, Ian Johnston, Dominic Kwiatkowski, Cordelia Langford, John Sillitoe on behalf of the Wellcome Sanger Institute COVID-19 Surveillance Team |
| EPI_ISL_2961349 | Berkshire and Surrey Pathology Services Lighthouse Laboratory | Wellcome Sanger Institute for the COVID-19 Genomics UK (COG-UK) Consortium | Berkshire and Surrey Pathology Services Lighthouse Laboratory and Alex Alderton, Roberto Amato, Jeffrey Barrett, Sonia Goncalves, Ewan Harrison, David K. Jackson, Ian Johnston, Dominic Kwiatkowski, Cordelia Langford, John Sillitoe on behalf of the Wellcome Sanger Institute COVID-19 Surveillance Team |
| EPI_ISL_2961350 | Lighthouse Lab in Glasgow | Wellcome Sanger Institute for the COVID-19 Genomics UK (COG-UK) Consortium | Harper VanSteenhouse, Yumi Kasai, David Gray, Carol Clugston, Anna Dominiczak and Alex Alderton, Roberto Amato, Jeffrey Barrett, Sonia Goncalves, Ewan Harrison, David K. Jackson, Ian Johnston, Dominic Kwiatkowski, Cordelia Langford, John Sillitoe on behalf of the Wellcome Sanger Institute COVID-19 Surveillance Team |
| EPI_ISL_2961351 | Lighthouse Lab in Alderley Park | Wellcome Sanger Institute for the COVID-19 Genomics UK (COG-UK) Consortium | Jacquelyn Wynn, Mairead Hyland, The Lighthouse Lab in Alderley Park and Alex Alderton, Roberto Amato, Jeffrey Barrett, Sonia Goncalves, Ewan Harrison, David K. Jackson, Ian Johnston, Dominic Kwiatkowski, Cordelia Langford, John Sillitoe on behalf of the Wellcome Sanger Institute COVID-19 Surveillance Team |
| EPI_ISL_2961352, EPI_ISL_2961353, EPI_ISL_2961354 | Lighthouse Lab in Glasgow | Wellcome Sanger Institute for the COVID-19 Genomics UK (COG-UK) Consortium | Harper VanSteenhouse, Yumi Kasai, David Gray, Carol Clugston, Anna Dominiczak and Alex Alderton, Roberto Amato, Jeffrey Barrett, Sonia Goncalves, Ewan Harrison, David K. Jackson, Ian Johnston, Dominic Kwiatkowski, Cordelia Langford, John Sillitoe on behalf of the Wellcome Sanger Institute COVID-19 Surveillance Team |
| EPI_ISL_2961355, EPI_ISL_2961356 | Berkshire and Surrey Pathology Services Lighthouse Laboratory | Wellcome Sanger Institute for the COVID-19 Genomics UK (COG-UK) Consortium | Berkshire and Surrey Pathology Services Lighthouse Laboratory and Alex Alderton, Roberto Amato, Jeffrey Barrett, Sonia Goncalves, Ewan Harrison, David K. Jackson, Ian Johnston, Dominic Kwiatkowski, Cordelia Langford, John Sillitoe on behalf of the Wellcome Sanger Institute COVID-19 Surveillance Team |
| EPI_ISL_2961357, EPI_ISL_2961358 | Lighthouse Lab in Glasgow | Wellcome Sanger Institute for the COVID-19 Genomics UK (COG-UK) Consortium | Harper VanSteenhouse, Yumi Kasai, David Gray, Carol Clugston, Anna Dominiczak and Alex Alderton, Roberto Amato, Jeffrey Barrett, Sonia Goncalves, Ewan Harrison, David K. Jackson, Ian Johnston, Dominic Kwiatkowski, Cordelia Langford, John Sillitoe on behalf of the Wellcome Sanger Institute COVID-19 Surveillance Team |
| EPI_ISL_2961359 | Lighthouse Lab in Alderley Park | Wellcome Sanger Institute for the COVID-19 Genomics UK (COG-UK) Consortium | Jacquelyn Wynn, Mairead Hyland, The Lighthouse Lab in Alderley Park and Alex Alderton, Roberto Amato, Jeffrey Barrett, Sonia Goncalves, Ewan Harrison, David K. Jackson, Ian Johnston, Dominic Kwiatkowski, Cordelia Langford, John Sillitoe on behalf of the Wellcome Sanger Institute COVID-19 Surveillance Team |
| EPI_ISL_2961360 | Lighthouse Lab in Glasgow | Wellcome Sanger Institute for the COVID-19 Genomics UK (COG-UK) Consortium | Harper VanSteenhouse, Yumi Kasai, David Gray, Carol Clugston, Anna Dominiczak and Alex Alderton, Roberto Amato, Jeffrey Barrett, Sonia Goncalves, Ewan Harrison, David K. Jackson, Ian Johnston, Dominic Kwiatkowski, Cordelia Langford, John Sillitoe on behalf of the Wellcome Sanger Institute COVID-19 Surveillance Team |
| EPI_ISL_2961361, EPI_ISL_2961362, | Berkshire and Surrey Pathology Services Lighthouse Laboratory | Wellcome Sanger Institute for the COVID-19 Genomics UK (COG-UK) Consortium | Berkshire and Surrey Pathology Services Lighthouse Laboratory and Alex Alderton, Roberto Amato, Jeffrey Barrett, Sonia Goncalves, Ewan Harrison, David K. Jackson, Ian Johnston, Dominic Kwiatkowski, Cordelia Langford, John Sillitoe on behalf of the Wellcome Sanger Institute COVID-19 Surveillance Team |

| | | | |
|---|---|---|---|
| EPI_ISL_2961363 | | | |
| EPI_ISL_2961364 | Lighthouse Lab in Glasgow | Wellcome Sanger Institute for the COVID-19 Genomics UK (COG-UK) Consortium | Harper VanSteenhouse, Yumi Kasai, David Gray, Carol Clugston, Anna Dominiczak and Alex Alderton, Roberto Amato, Jeffrey Barrett, Sonia Goncalves, Ewan Harrison, David K. Jackson, Ian Johnston, Dominic Kwiatkowski, Cordelia Langford, John Sillitoe on behalf of the Wellcome Sanger Institute COVID-19 Surveillance Team |
| EPI_ISL_2961365 | Lighthouse Lab in Alderley Park | Wellcome Sanger Institute for the COVID-19 Genomics UK (COG-UK) Consortium | Jacquelyn Wynn, Mairead Hyland, The Lighthouse Lab in Alderley Park and Alex Alderton, Roberto Amato, Jeffrey Barrett, Sonia Goncalves, Ewan Harrison, David K. Jackson, Ian Johnston, Dominic Kwiatkowski, Cordelia Langford, John Sillitoe on behalf of the Wellcome Sanger Institute COVID-19 Surveillance Team |
| EPI_ISL_2961366 | Berkshire and Surrey Pathology Services Lighthouse Laboratory | Wellcome Sanger Institute for the COVID-19 Genomics UK (COG-UK) Consortium | Berkshire and Surrey Pathology Services Lighthouse Laboratory and Alex Alderton, Roberto Amato, Jeffrey Barrett, Sonia Goncalves, Ewan Harrison, David K. Jackson, Ian Johnston, Dominic Kwiatkowski, Cordelia Langford, John Sillitoe on behalf of the Wellcome Sanger Institute COVID-19 Surveillance Team |
| EPI_ISL_2961367 | Lighthouse Lab in Alderley Park | Wellcome Sanger Institute for the COVID-19 Genomics UK (COG-UK) Consortium | Jacquelyn Wynn, Mairead Hyland, The Lighthouse Lab in Alderley Park and Alex Alderton, Roberto Amato, Jeffrey Barrett, Sonia Goncalves, Ewan Harrison, David K. Jackson, Ian Johnston, Dominic Kwiatkowski, Cordelia Langford, John Sillitoe on behalf of the Wellcome Sanger Institute COVID-19 Surveillance Team |
| EPI_ISL_2961368, EPI_ISL_2961369 | Lighthouse Lab in Glasgow | Wellcome Sanger Institute for the COVID-19 Genomics UK (COG-UK) Consortium | Harper VanSteenhouse, Yumi Kasai, David Gray, Carol Clugston, Anna Dominiczak and Alex Alderton, Roberto Amato, Jeffrey Barrett, Sonia Goncalves, Ewan Harrison, David K. Jackson, Ian Johnston, Dominic Kwiatkowski, Cordelia Langford, John Sillitoe on behalf of the Wellcome Sanger Institute COVID-19 Surveillance Team |
| EPI_ISL_2961370 | Lighthouse Lab in Alderley Park | Wellcome Sanger Institute for the COVID-19 Genomics UK (COG-UK) Consortium | Jacquelyn Wynn, Mairead Hyland, The Lighthouse Lab in Alderley Park and Alex Alderton, Roberto Amato, Jeffrey Barrett, Sonia Goncalves, Ewan Harrison, David K. Jackson, Ian Johnston, Dominic Kwiatkowski, Cordelia Langford, John Sillitoe on behalf of the Wellcome Sanger Institute COVID-19 Surveillance Team |
| EPI_ISL_2961371, EPI_ISL_2961372 | Lighthouse Lab in Glasgow | Wellcome Sanger Institute for the COVID-19 Genomics UK (COG-UK) Consortium | Harper VanSteenhouse, Yumi Kasai, David Gray, Carol Clugston, Anna Dominiczak and Alex Alderton, Roberto Amato, Jeffrey Barrett, Sonia Goncalves, Ewan Harrison, David K. Jackson, Ian Johnston, Dominic Kwiatkowski, Cordelia Langford, John Sillitoe on behalf of the Wellcome Sanger Institute COVID-19 Surveillance Team |
| EPI_ISL_2961373 | Lighthouse Lab in Alderley Park | Wellcome Sanger Institute for the COVID-19 Genomics UK (COG-UK) Consortium | Jacquelyn Wynn, Mairead Hyland, The Lighthouse Lab in Alderley Park and Alex Alderton, Roberto Amato, Jeffrey Barrett, Sonia Goncalves, Ewan Harrison, David K. Jackson, Ian Johnston, Dominic Kwiatkowski, Cordelia Langford, John Sillitoe on behalf of the Wellcome Sanger Institute COVID-19 Surveillance Team |
| EPI_ISL_2961374, EPI_ISL_2961375 | Lighthouse Lab in Glasgow | Wellcome Sanger Institute for the COVID-19 Genomics UK (COG-UK) Consortium | Harper VanSteenhouse, Yumi Kasai, David Gray, Carol Clugston, Anna Dominiczak and Alex Alderton, Roberto Amato, Jeffrey Barrett, Sonia Goncalves, Ewan Harrison, David K. Jackson, Ian Johnston, Dominic Kwiatkowski, Cordelia Langford, John Sillitoe on behalf of the Wellcome Sanger Institute COVID-19 Surveillance Team |
| EPI_ISL_2961376 | Berkshire and Surrey Pathology Services Lighthouse Laboratory | Wellcome Sanger Institute for the COVID-19 Genomics UK (COG-UK) Consortium | Berkshire and Surrey Pathology Services Lighthouse Laboratory and Alex Alderton, Roberto Amato, Jeffrey Barrett, Sonia Goncalves, Ewan Harrison, David K. Jackson, Ian Johnston, Dominic Kwiatkowski, Cordelia Langford, John Sillitoe on behalf of the Wellcome Sanger Institute COVID-19 Surveillance Team |
| EPI_ISL_2961377 | Lighthouse Lab in Glasgow | Wellcome Sanger Institute for the COVID-19 Genomics UK (COG-UK) Consortium | Harper VanSteenhouse, Yumi Kasai, David Gray, Carol Clugston, Anna Dominiczak and Alex Alderton, Roberto Amato, Jeffrey Barrett, Sonia Goncalves, Ewan Harrison, David K. Jackson, Ian Johnston, Dominic Kwiatkowski, Cordelia Langford, John Sillitoe on behalf of the Wellcome Sanger Institute COVID-19 Surveillance Team |
| EPI_ISL_2961378, EPI_ISL_2961379 | Berkshire and Surrey Pathology Services Lighthouse Laboratory | Wellcome Sanger Institute for the COVID-19 Genomics UK (COG-UK) Consortium | Berkshire and Surrey Pathology Services Lighthouse Laboratory and Alex Alderton, Roberto Amato, Jeffrey Barrett, Sonia Goncalves, Ewan Harrison, David K. Jackson, Ian Johnston, Dominic Kwiatkowski, Cordelia Langford, John Sillitoe on behalf of the Wellcome Sanger Institute COVID-19 Surveillance Team |
| EPI_ISL_2961380 | Lighthouse Lab in Glasgow | Wellcome Sanger Institute for the COVID-19 Genomics UK (COG-UK) Consortium | Harper VanSteenhouse, Yumi Kasai, David Gray, Carol Clugston, Anna Dominiczak and Alex Alderton, Roberto Amato, Jeffrey Barrett, Sonia Goncalves, Ewan Harrison, David K. Jackson, Ian Johnston, Dominic Kwiatkowski, Cordelia Langford, John Sillitoe on behalf of the Wellcome Sanger Institute COVID-19 Surveillance Team |
| EPI_ISL_2961381 | Berkshire and Surrey Pathology Services Lighthouse Laboratory | Wellcome Sanger Institute for the COVID-19 Genomics UK (COG-UK) Consortium | Berkshire and Surrey Pathology Services Lighthouse Laboratory and Alex Alderton, Roberto Amato, Jeffrey Barrett, Sonia Goncalves, Ewan Harrison, David K. Jackson, Ian Johnston, Dominic Kwiatkowski, Cordelia Langford, John Sillitoe on behalf of the Wellcome Sanger Institute COVID-19 Surveillance Team |
| EPI_ISL_2961382, EPI_ISL_2961383, EPI_ISL_2961384 | Lighthouse Lab in Alderley Park | Wellcome Sanger Institute for the COVID-19 Genomics UK (COG-UK) Consortium | Jacquelyn Wynn, Mairead Hyland, The Lighthouse Lab in Alderley Park and Alex Alderton, Roberto Amato, Jeffrey Barrett, Sonia Goncalves, Ewan Harrison, David K. Jackson, Ian Johnston, Dominic Kwiatkowski, Cordelia Langford, John Sillitoe on behalf of the Wellcome Sanger Institute COVID-19 Surveillance Team |
| EPI_ISL_2961385 | Berkshire and Surrey Pathology Services Lighthouse Laboratory | Wellcome Sanger Institute for the COVID-19 Genomics UK (COG-UK) Consortium | Berkshire and Surrey Pathology Services Lighthouse Laboratory and Alex Alderton, Roberto Amato, Jeffrey Barrett, Sonia Goncalves, Ewan Harrison, David K. Jackson, Ian Johnston, Dominic Kwiatkowski, Cordelia Langford, John Sillitoe on behalf of the Wellcome Sanger Institute COVID-19 Surveillance Team |
| EPI_ISL_2961386 | Lighthouse Lab in Alderley Park | Wellcome Sanger Institute for the COVID-19 Genomics UK (COG-UK) Consortium | Jacquelyn Wynn, Mairead Hyland, The Lighthouse Lab in Alderley Park and Alex Alderton, Roberto Amato, Jeffrey Barrett, Sonia Goncalves, Ewan Harrison, David K. Jackson, Ian Johnston, Dominic Kwiatkowski, Cordelia Langford, John Sillitoe on behalf of the Wellcome Sanger Institute COVID-19 Surveillance Team |
| EPI_ISL_2961387, EPI_ISL_2961388 | Berkshire and Surrey Pathology Services Lighthouse Laboratory | Wellcome Sanger Institute for the COVID-19 Genomics UK (COG-UK) Consortium | Berkshire and Surrey Pathology Services Lighthouse Laboratory and Alex Alderton, Roberto Amato, Jeffrey Barrett, Sonia Goncalves, Ewan Harrison, David K. Jackson, Ian Johnston, Dominic Kwiatkowski, Cordelia Langford, John Sillitoe on behalf of the Wellcome Sanger Institute COVID-19 Surveillance Team |
| EPI_ISL_2961389 | Lighthouse Lab in Alderley Park | Wellcome Sanger Institute for the COVID-19 Genomics UK (COG-UK) Consortium | Jacquelyn Wynn, Mairead Hyland, The Lighthouse Lab in Alderley Park and Alex Alderton, Roberto Amato, Jeffrey Barrett, Sonia Goncalves, Ewan Harrison, David K. Jackson, Ian Johnston, Dominic Kwiatkowski, Cordelia Langford, John Sillitoe on behalf of the Wellcome Sanger Institute COVID-19 Surveillance Team |
| EPI_ISL_2961390 | Berkshire and Surrey Pathology Services Lighthouse Laboratory | Wellcome Sanger Institute for the COVID-19 Genomics UK (COG-UK) Consortium | Berkshire and Surrey Pathology Services Lighthouse Laboratory and Alex Alderton, Roberto Amato, Jeffrey Barrett, Sonia Goncalves, Ewan Harrison, David K. Jackson, Ian Johnston, Dominic Kwiatkowski, Cordelia Langford, John Sillitoe on behalf of the Wellcome Sanger Institute COVID-19 Surveillance Team |
| EPI_ISL_2961391 | Lighthouse Lab in Alderley Park | Wellcome Sanger Institute for the COVID-19 Genomics UK (COG-UK) Consortium | Jacquelyn Wynn, Mairead Hyland, The Lighthouse Lab in Alderley Park and Alex Alderton, Roberto Amato, Jeffrey Barrett, Sonia Goncalves, Ewan Harrison, David K. Jackson, Ian Johnston, Dominic Kwiatkowski, Cordelia Langford, John Sillitoe on behalf of the Wellcome Sanger Institute COVID-19 Surveillance Team |
| EPI_ISL_2961392, EPI_ISL_2961393 | Berkshire and Surrey Pathology Services Lighthouse Laboratory | Wellcome Sanger Institute for the COVID-19 Genomics UK (COG-UK) Consortium | Berkshire and Surrey Pathology Services Lighthouse Laboratory and Alex Alderton, Roberto Amato, Jeffrey Barrett, Sonia Goncalves, Ewan Harrison, David K. Jackson, Ian Johnston, Dominic Kwiatkowski, Cordelia Langford, John Sillitoe on behalf of the Wellcome Sanger Institute COVID-19 Surveillance Team |
| EPI_ISL_2961394 | Lighthouse Lab in Alderley Park | Wellcome Sanger Institute for the COVID-19 Genomics UK (COG-UK) Consortium | Jacquelyn Wynn, Mairead Hyland, The Lighthouse Lab in Alderley Park and Alex Alderton, Roberto Amato, Jeffrey Barrett, Sonia Goncalves, Ewan Harrison, David K. Jackson, Ian Johnston, Dominic Kwiatkowski, Cordelia Langford, John Sillitoe on behalf of the Wellcome Sanger Institute COVID-19 Surveillance Team |
| EPI_ISL_2961395 | Berkshire and Surrey Pathology Services Lighthouse Laboratory | Wellcome Sanger Institute for the COVID-19 Genomics UK (COG-UK) Consortium | Berkshire and Surrey Pathology Services Lighthouse Laboratory and Alex Alderton, Roberto Amato, Jeffrey Barrett, Sonia Goncalves, Ewan Harrison, David K. Jackson, Ian Johnston, Dominic Kwiatkowski, Cordelia Langford, John Sillitoe on behalf of the Wellcome Sanger Institute COVID-19 Surveillance Team |
| EPI_ISL_2961396 | Lighthouse Lab in Alderley Park | Wellcome Sanger Institute for the COVID-19 Genomics UK (COG-UK) Consortium | Jacquelyn Wynn, Mairead Hyland, The Lighthouse Lab in Alderley Park and Alex Alderton, Roberto Amato, Jeffrey Barrett, Sonia Goncalves, Ewan Harrison, David K. Jackson, Ian Johnston, Dominic Kwiatkowski, Cordelia Langford, John Sillitoe on behalf of the Wellcome Sanger Institute COVID-19 Surveillance Team |
| EPI_ISL_2961397, EPI_ISL_2961398 | Lighthouse Lab in Glasgow | Wellcome Sanger Institute for the COVID-19 Genomics UK (COG-UK) Consortium | Harper VanSteenhouse, Yumi Kasai, David Gray, Carol Clugston, Anna Dominiczak and Alex Alderton, Roberto Amato, Jeffrey Barrett, Sonia Goncalves, Ewan Harrison, David K. Jackson, Ian Johnston, Dominic Kwiatkowski, Cordelia Langford, John Sillitoe on behalf of the Wellcome Sanger Institute COVID-19 Surveillance Team |
| EPI_ISL_2961399 | Berkshire and Surrey Pathology Services Lighthouse Laboratory | Wellcome Sanger Institute for the COVID-19 Genomics UK (COG-UK) Consortium | Berkshire and Surrey Pathology Services Lighthouse Laboratory and Alex Alderton, Roberto Amato, Jeffrey Barrett, Sonia Goncalves, Ewan Harrison, David K. Jackson, Ian Johnston, Dominic Kwiatkowski, Cordelia Langford, John Sillitoe on behalf of the Wellcome Sanger Institute COVID-19 Surveillance Team |
| EPI_ISL_2961400, EPI_ISL_2961401 | Lighthouse Lab in Alderley Park | Wellcome Sanger Institute for the COVID-19 Genomics UK (COG-UK) Consortium | Jacquelyn Wynn, Mairead Hyland, The Lighthouse Lab in Alderley Park and Alex Alderton, Roberto Amato, Jeffrey Barrett, Sonia Goncalves, Ewan Harrison, David K. Jackson, Ian Johnston, Dominic Kwiatkowski, Cordelia Langford, John Sillitoe on behalf of the Wellcome Sanger Institute COVID-19 Surveillance Team |
| EPI_ISL_2961402 | Berkshire and Surrey Pathology Services Lighthouse Laboratory | Wellcome Sanger Institute for the COVID-19 Genomics UK (COG-UK) Consortium | Berkshire and Surrey Pathology Services Lighthouse Laboratory and Alex Alderton, Roberto Amato, Jeffrey Barrett, Sonia Goncalves, Ewan Harrison, David K. Jackson, Ian Johnston, Dominic Kwiatkowski, Cordelia Langford, John Sillitoe on behalf of the Wellcome Sanger Institute COVID-19 Surveillance Team |
| EPI_ISL_2961403 | Lighthouse Lab in Alderley Park | Wellcome Sanger Institute for the COVID-19 Genomics UK (COG-UK) Consortium | Jacquelyn Wynn, Mairead Hyland, The Lighthouse Lab in Alderley Park and Alex Alderton, Roberto Amato, Jeffrey Barrett, Sonia Goncalves, Ewan Harrison, David K. Jackson, Ian Johnston, Dominic Kwiatkowski, Cordelia Langford, John Sillitoe on behalf of the Wellcome Sanger Institute COVID-19 Surveillance Team |
| EPI_ISL_2961404 | Lighthouse Lab in Glasgow | Wellcome Sanger Institute for the COVID-19 Genomics UK (COG-UK) Consortium | Harper VanSteenhouse, Yumi Kasai, David Gray, Carol Clugston, Anna Dominiczak and Alex Alderton, Roberto Amato, Jeffrey Barrett, Sonia Goncalves, Ewan Harrison, David K. Jackson, Ian Johnston, Dominic Kwiatkowski, Cordelia Langford, John Sillitoe on behalf of the Wellcome Sanger Institute COVID-19 Surveillance Team |
| EPI_ISL_2961405, EPI_ISL_2961406 | Berkshire and Surrey Pathology Services Lighthouse Laboratory | Wellcome Sanger Institute for the COVID-19 Genomics UK (COG-UK) Consortium | Berkshire and Surrey Pathology Services Lighthouse Laboratory and Alex Alderton, Roberto Amato, Jeffrey Barrett, Sonia Goncalves, Ewan Harrison, David K. Jackson, Ian Johnston, Dominic Kwiatkowski, Cordelia Langford, John Sillitoe on behalf of the Wellcome Sanger Institute COVID-19 Surveillance Team |
| EPI_ISL_2961407, EPI_ISL_2961408 | Lighthouse Lab in Glasgow | Wellcome Sanger Institute for the COVID-19 Genomics UK (COG-UK) Consortium | Harper VanSteenhouse, Yumi Kasai, David Gray, Carol Clugston, Anna Dominiczak and Alex Alderton, Roberto Amato, Jeffrey Barrett, Sonia Goncalves, Ewan Harrison, David K. Jackson, Ian Johnston, Dominic Kwiatkowski, Cordelia Langford, John Sillitoe on behalf of the Wellcome Sanger Institute COVID-19 Surveillance Team |
| EPI_ISL_2961409 | Berkshire and Surrey Pathology Services Lighthouse Laboratory | Wellcome Sanger Institute for the COVID-19 Genomics UK (COG-UK) Consortium | Berkshire and Surrey Pathology Services Lighthouse Laboratory and Alex Alderton, Roberto Amato, Jeffrey Barrett, Sonia Goncalves, Ewan Harrison, David K. Jackson, Ian Johnston, Dominic Kwiatkowski, Cordelia Langford, John Sillitoe on behalf of the Wellcome Sanger Institute COVID-19 Surveillance Team |
| EPI_ISL_2961410, EPI_ISL_2961411 | Lighthouse Lab in Glasgow | Wellcome Sanger Institute for the COVID-19 Genomics UK (COG-UK) Consortium | Harper VanSteenhouse, Yumi Kasai, David Gray, Carol Clugston, Anna Dominiczak and Alex Alderton, Roberto Amato, Jeffrey Barrett, Sonia Goncalves, Ewan Harrison, David K. Jackson, Ian Johnston, Dominic Kwiatkowski, Cordelia Langford, John Sillitoe on behalf of the Wellcome Sanger Institute COVID-19 Surveillance Team |
| EPI_ISL_2961412 | Lighthouse Lab in Alderley Park | Wellcome Sanger Institute for the COVID-19 Genomics UK (COG-UK) Consortium | Jacquelyn Wynn, Mairead Hyland, The Lighthouse Lab in Alderley Park and Alex Alderton, Roberto Amato, Jeffrey Barrett, Sonia Goncalves, Ewan Harrison, David K. Jackson, Ian Johnston, Dominic Kwiatkowski, Cordelia Langford, John Sillitoe on behalf of the Wellcome Sanger Institute COVID-19 Surveillance Team |
| EPI_ISL_2961413, EPI_ISL_2961414, EPI_ISL_2961415 | Lighthouse Lab in Glasgow | Wellcome Sanger Institute for the COVID-19 Genomics UK (COG-UK) Consortium | Harper VanSteenhouse, Yumi Kasai, David Gray, Carol Clugston, Anna Dominiczak and Alex Alderton, Roberto Amato, Jeffrey Barrett, Sonia Goncalves, Ewan Harrison, David K. Jackson, Ian Johnston, Dominic Kwiatkowski, Cordelia Langford, John Sillitoe on behalf of the Wellcome Sanger Institute COVID-19 Surveillance Team |
| EPI_ISL_2961416, EPI_ISL_2961417 | Lighthouse Lab in Alderley Park | Wellcome Sanger Institute for the COVID-19 Genomics UK (COG-UK) Consortium | Jacquelyn Wynn, Mairead Hyland, The Lighthouse Lab in Alderley Park and Alex Alderton, Roberto Amato, Jeffrey Barrett, Sonia Goncalves, Ewan Harrison, David K. Jackson, Ian Johnston, Dominic Kwiatkowski, Cordelia Langford, John Sillitoe on behalf of the Wellcome Sanger Institute COVID-19 Surveillance Team |
| EPI_ISL_2961418 | Lighthouse Lab in Glasgow | Wellcome Sanger Institute for the COVID-19 Genomics UK (COG-UK) Consortium | Harper VanSteenhouse, Yumi Kasai, David Gray, Carol Clugston, Anna Dominiczak and Alex Alderton, Roberto Amato, Jeffrey Barrett, Sonia Goncalves, Ewan Harrison, David K. Jackson, Ian Johnston, Dominic Kwiatkowski, Cordelia Langford, John Sillitoe on behalf of the Wellcome Sanger Institute COVID-19 Surveillance Team |
| EPI_ISL_2961419 | Lighthouse Lab in Alderley Park | Wellcome Sanger Institute for the COVID-19 Genomics UK (COG-UK) Consortium | Jacquelyn Wynn, Mairead Hyland, The Lighthouse Lab in Alderley Park and Alex Alderton, Roberto Amato, Jeffrey Barrett, Sonia Goncalves, Ewan Harrison, David K. Jackson, Ian Johnston, Dominic Kwiatkowski, Cordelia Langford, John Sillitoe on behalf of the Wellcome Sanger Institute COVID-19 Surveillance Team |
| EPI_ISL_2961420 | Berkshire and Surrey Pathology Services Lighthouse Laboratory | Wellcome Sanger Institute for the COVID-19 Genomics UK (COG-UK) Consortium | Berkshire and Surrey Pathology Services Lighthouse Laboratory and Alex Alderton, Roberto Amato, Jeffrey Barrett, Sonia Goncalves, Ewan Harrison, David K. Jackson, Ian Johnston, Dominic Kwiatkowski, Cordelia Langford, John Sillitoe on behalf of the Wellcome Sanger Institute COVID-19 Surveillance Team |
| EPI_ISL_2961421 | Lighthouse Lab in Glasgow | Wellcome Sanger Institute for the COVID-19 Genomics UK (COG-UK) Consortium | Harper VanSteenhouse, Yumi Kasai, David Gray, Carol Clugston, Anna Dominiczak and Alex Alderton, Roberto Amato, Jeffrey Barrett, Sonia Goncalves, Ewan Harrison, David K. Jackson, Ian Johnston, Dominic Kwiatkowski, Cordelia Langford, John Sillitoe on behalf of the Wellcome Sanger Institute COVID-19 Surveillance Team |
| EPI_ISL_2961422 | Berkshire and Surrey Pathology Services Lighthouse Laboratory | Wellcome Sanger Institute for the COVID-19 Genomics UK (COG-UK) Consortium | Berkshire and Surrey Pathology Services Lighthouse Laboratory and Alex Alderton, Roberto Amato, Jeffrey Barrett, Sonia Goncalves, Ewan Harrison, David K. Jackson, Ian Johnston, Dominic Kwiatkowski, Cordelia Langford, John Sillitoe on behalf of the Wellcome Sanger Institute COVID-19 Surveillance Team |
| EPI_ISL_2961423 | Lighthouse Lab in Alderley Park | Wellcome Sanger Institute for the COVID-19 Genomics UK (COG-UK) Consortium | Jacquelyn Wynn, Mairead Hyland, The Lighthouse Lab in Alderley Park and Alex Alderton, Roberto Amato, Jeffrey Barrett, Sonia Goncalves, Ewan Harrison, David K. Jackson, Ian Johnston, Dominic Kwiatkowski, Cordelia Langford, John Sillitoe on behalf of the Wellcome Sanger Institute COVID-19 Surveillance Team |

| | | | |
|---|---|---|---|
| EPI_ISL_2961424, EPI_ISL_2961425 | Lighthouse Lab in Glasgow | Wellcome Sanger Institute for the COVID-19 Genomics UK (COG-UK) Consortium | Harper VanSteenhouse, Yumi Kasai, David Gray, Carol Clugston, Anna Dominiczak and Alex Alderton, Roberto Amato, Jeffrey Barrett, Sonia Goncalves, Ewan Harrison, David K. Jackson, Ian Johnston, Dominic Kwiatkowski, Cordelia Langford, John Sillitoe on behalf of the Wellcome Sanger Institute COVID-19 Surveillance Team |
| EPI_ISL_2961426 | Lighthouse Lab in Alderley Park | Wellcome Sanger Institute for the COVID-19 Genomics UK (COG-UK) Consortium | Jacquelyn Wynn, Mairead Hyland, The Lighthouse Lab in Alderley Park and Alex Alderton, Roberto Amato, Jeffrey Barrett, Sonia Goncalves, Ewan Harrison, David K. Jackson, Ian Johnston, Dominic Kwiatkowski, Cordelia Langford, John Sillitoe on behalf of the Wellcome Sanger Institute COVID-19 Surveillance Team |
| EPI_ISL_2961427 | Lighthouse Lab in Glasgow | Wellcome Sanger Institute for the COVID-19 Genomics UK (COG-UK) Consortium | Harper VanSteenhouse, Yumi Kasai, David Gray, Carol Clugston, Anna Dominiczak and Alex Alderton, Roberto Amato, Jeffrey Barrett, Sonia Goncalves, Ewan Harrison, David K. Jackson, Ian Johnston, Dominic Kwiatkowski, Cordelia Langford, John Sillitoe on behalf of the Wellcome Sanger Institute COVID-19 Surveillance Team |
| EPI_ISL_2961428 | Berkshire and Surrey Pathology Services Lighthouse Laboratory | Wellcome Sanger Institute for the COVID-19 Genomics UK (COG-UK) Consortium | Berkshire and Surrey Pathology Services Lighthouse Laboratory and Alex Alderton, Roberto Amato, Jeffrey Barrett, Sonia Goncalves, Ewan Harrison, David K. Jackson, Ian Johnston, Dominic Kwiatkowski, Cordelia Langford, John Sillitoe on behalf of the Wellcome Sanger Institute COVID-19 Surveillance Team |
| EPI_ISL_2961429 | Lighthouse Lab in Alderley Park | Wellcome Sanger Institute for the COVID-19 Genomics UK (COG-UK) Consortium | Jacquelyn Wynn, Mairead Hyland, The Lighthouse Lab in Alderley Park and Alex Alderton, Roberto Amato, Jeffrey Barrett, Sonia Goncalves, Ewan Harrison, David K. Jackson, Ian Johnston, Dominic Kwiatkowski, Cordelia Langford, John Sillitoe on behalf of the Wellcome Sanger Institute COVID-19 Surveillance Team |
| EPI_ISL_2961430 | Berkshire and Surrey Pathology Services Lighthouse Laboratory | Wellcome Sanger Institute for the COVID-19 Genomics UK (COG-UK) Consortium | Berkshire and Surrey Pathology Services Lighthouse Laboratory and Alex Alderton, Roberto Amato, Jeffrey Barrett, Sonia Goncalves, Ewan Harrison, David K. Jackson, Ian Johnston, Dominic Kwiatkowski, Cordelia Langford, John Sillitoe on behalf of the Wellcome Sanger Institute COVID-19 Surveillance Team |
| EPI_ISL_2961431 | Lighthouse Lab in Alderley Park | Wellcome Sanger Institute for the COVID-19 Genomics UK (COG-UK) Consortium | Jacquelyn Wynn, Mairead Hyland, The Lighthouse Lab in Alderley Park and Alex Alderton, Roberto Amato, Jeffrey Barrett, Sonia Goncalves, Ewan Harrison, David K. Jackson, Ian Johnston, Dominic Kwiatkowski, Cordelia Langford, John Sillitoe on behalf of the Wellcome Sanger Institute COVID-19 Surveillance Team |
| EPI_ISL_2961432, EPI_ISL_2961433, EPI_ISL_2961434, EPI_ISL_2961435 | Lighthouse Lab in Glasgow | Wellcome Sanger Institute for the COVID-19 Genomics UK (COG-UK) Consortium | Harper VanSteenhouse, Yumi Kasai, David Gray, Carol Clugston, Anna Dominiczak and Alex Alderton, Roberto Amato, Jeffrey Barrett, Sonia Goncalves, Ewan Harrison, David K. Jackson, Ian Johnston, Dominic Kwiatkowski, Cordelia Langford, John Sillitoe on behalf of the Wellcome Sanger Institute COVID-19 Surveillance Team |
| EPI_ISL_2961436 | Berkshire and Surrey Pathology Services Lighthouse Laboratory | Wellcome Sanger Institute for the COVID-19 Genomics UK (COG-UK) Consortium | Berkshire and Surrey Pathology Services Lighthouse Laboratory and Alex Alderton, Roberto Amato, Jeffrey Barrett, Sonia Goncalves, Ewan Harrison, David K. Jackson, Ian Johnston, Dominic Kwiatkowski, Cordelia Langford, John Sillitoe on behalf of the Wellcome Sanger Institute COVID-19 Surveillance Team |
| EPI_ISL_2961437 | Lighthouse Lab in Alderley Park | Wellcome Sanger Institute for the COVID-19 Genomics UK (COG-UK) Consortium | Jacquelyn Wynn, Mairead Hyland, The Lighthouse Lab in Alderley Park and Alex Alderton, Roberto Amato, Jeffrey Barrett, Sonia Goncalves, Ewan Harrison, David K. Jackson, Ian Johnston, Dominic Kwiatkowski, Cordelia Langford, John Sillitoe on behalf of the Wellcome Sanger Institute COVID-19 Surveillance Team |
| EPI_ISL_2961438 | Berkshire and Surrey Pathology Services Lighthouse Laboratory | Wellcome Sanger Institute for the COVID-19 Genomics UK (COG-UK) Consortium | Berkshire and Surrey Pathology Services Lighthouse Laboratory and Alex Alderton, Roberto Amato, Jeffrey Barrett, Sonia Goncalves, Ewan Harrison, David K. Jackson, Ian Johnston, Dominic Kwiatkowski, Cordelia Langford, John Sillitoe on behalf of the Wellcome Sanger Institute COVID-19 Surveillance Team |
| EPI_ISL_2961439, EPI_ISL_2961440 | Lighthouse Lab in Glasgow | Wellcome Sanger Institute for the COVID-19 Genomics UK (COG-UK) Consortium | Harper VanSteenhouse, Yumi Kasai, David Gray, Carol Clugston, Anna Dominiczak and Alex Alderton, Roberto Amato, Jeffrey Barrett, Sonia Goncalves, Ewan Harrison, David K. Jackson, Ian Johnston, Dominic Kwiatkowski, Cordelia Langford, John Sillitoe on behalf of the Wellcome Sanger Institute COVID-19 Surveillance Team |
| EPI_ISL_2961441 | Lighthouse Lab in Alderley Park | Wellcome Sanger Institute for the COVID-19 Genomics UK (COG-UK) Consortium | Jacquelyn Wynn, Mairead Hyland, The Lighthouse Lab in Alderley Park and Alex Alderton, Roberto Amato, Jeffrey Barrett, Sonia Goncalves, Ewan Harrison, David K. Jackson, Ian Johnston, Dominic Kwiatkowski, Cordelia Langford, John Sillitoe on behalf of the Wellcome Sanger Institute COVID-19 Surveillance Team |
| EPI_ISL_2961442 | Lighthouse Lab in Glasgow | Wellcome Sanger Institute for the COVID-19 Genomics UK (COG-UK) Consortium | Harper VanSteenhouse, Yumi Kasai, David Gray, Carol Clugston, Anna Dominiczak and Alex Alderton, Roberto Amato, Jeffrey Barrett, Sonia Goncalves, Ewan Harrison, David K. Jackson, Ian Johnston, Dominic Kwiatkowski, Cordelia Langford, John Sillitoe on behalf of the Wellcome Sanger Institute COVID-19 Surveillance Team |
| EPI_ISL_2961443 | Lighthouse Lab in Alderley Park | Wellcome Sanger Institute for the COVID-19 Genomics UK (COG-UK) Consortium | Jacquelyn Wynn, Mairead Hyland, The Lighthouse Lab in Alderley Park and Alex Alderton, Roberto Amato, Jeffrey Barrett, Sonia Goncalves, Ewan Harrison, David K. Jackson, Ian Johnston, Dominic Kwiatkowski, Cordelia Langford, John Sillitoe on behalf of the Wellcome Sanger Institute COVID-19 Surveillance Team |
| EPI_ISL_2961444 | Lighthouse Lab in Glasgow | Wellcome Sanger Institute for the COVID-19 Genomics UK (COG-UK) Consortium | Harper VanSteenhouse, Yumi Kasai, David Gray, Carol Clugston, Anna Dominiczak and Alex Alderton, Roberto Amato, Jeffrey Barrett, Sonia Goncalves, Ewan Harrison, David K. Jackson, Ian Johnston, Dominic Kwiatkowski, Cordelia Langford, John Sillitoe on behalf of the Wellcome Sanger Institute COVID-19 Surveillance Team |
| EPI_ISL_2961445 | Lighthouse Lab in Alderley Park | Wellcome Sanger Institute for the COVID-19 Genomics UK (COG-UK) Consortium | Jacquelyn Wynn, Mairead Hyland, The Lighthouse Lab in Alderley Park and Alex Alderton, Roberto Amato, Jeffrey Barrett, Sonia Goncalves, Ewan Harrison, David K. Jackson, Ian Johnston, Dominic Kwiatkowski, Cordelia Langford, John Sillitoe on behalf of the Wellcome Sanger Institute COVID-19 Surveillance Team |
| EPI_ISL_2961446, EPI_ISL_2961447 | Lighthouse Lab in Glasgow | Wellcome Sanger Institute for the COVID-19 Genomics UK (COG-UK) Consortium | Harper VanSteenhouse, Yumi Kasai, David Gray, Carol Clugston, Anna Dominiczak and Alex Alderton, Roberto Amato, Jeffrey Barrett, Sonia Goncalves, Ewan Harrison, David K. Jackson, Ian Johnston, Dominic Kwiatkowski, Cordelia Langford, John Sillitoe on behalf of the Wellcome Sanger Institute COVID-19 Surveillance Team |
| EPI_ISL_2961448 | Lighthouse Lab in Alderley Park | Wellcome Sanger Institute for the COVID-19 Genomics UK (COG-UK) Consortium | Jacquelyn Wynn, Mairead Hyland, The Lighthouse Lab in Alderley Park and Alex Alderton, Roberto Amato, Jeffrey Barrett, Sonia Goncalves, Ewan Harrison, David K. Jackson, Ian Johnston, Dominic Kwiatkowski, Cordelia Langford, John Sillitoe on behalf of the Wellcome Sanger Institute COVID-19 Surveillance Team |
| EPI_ISL_2961449, EPI_ISL_2961450 | Lighthouse Lab in Glasgow | Wellcome Sanger Institute for the COVID-19 Genomics UK (COG-UK) Consortium | Harper VanSteenhouse, Yumi Kasai, David Gray, Carol Clugston, Anna Dominiczak and Alex Alderton, Roberto Amato, Jeffrey Barrett, Sonia Goncalves, Ewan Harrison, David K. Jackson, Ian Johnston, Dominic Kwiatkowski, Cordelia Langford, John Sillitoe on behalf of the Wellcome Sanger Institute COVID-19 Surveillance Team |
| EPI_ISL_2961451 | Lighthouse Lab in Alderley Park | Wellcome Sanger Institute for the COVID-19 Genomics UK (COG-UK) Consortium | Jacquelyn Wynn, Mairead Hyland, The Lighthouse Lab in Alderley Park and Alex Alderton, Roberto Amato, Jeffrey Barrett, Sonia Goncalves, Ewan Harrison, David K. Jackson, Ian Johnston, Dominic Kwiatkowski, Cordelia Langford, John Sillitoe on behalf of the Wellcome Sanger Institute COVID-19 Surveillance Team |
| EPI_ISL_2961452, EPI_ISL_2961453 | Berkshire and Surrey Pathology Services Lighthouse Laboratory | Wellcome Sanger Institute for the COVID-19 Genomics UK (COG-UK) Consortium | Berkshire and Surrey Pathology Services Lighthouse Laboratory and Alex Alderton, Roberto Amato, Jeffrey Barrett, Sonia Goncalves, Ewan Harrison, David K. Jackson, Ian Johnston, Dominic Kwiatkowski, Cordelia Langford, John Sillitoe on behalf of the Wellcome Sanger Institute COVID-19 Surveillance Team |
| EPI_ISL_2961454, EPI_ISL_2961455, EPI_ISL_2961456 | Lighthouse Lab in Glasgow | Wellcome Sanger Institute for the COVID-19 Genomics UK (COG-UK) Consortium | Harper VanSteenhouse, Yumi Kasai, David Gray, Carol Clugston, Anna Dominiczak and Alex Alderton, Roberto Amato, Jeffrey Barrett, Sonia Goncalves, Ewan Harrison, David K. Jackson, Ian Johnston, Dominic Kwiatkowski, Cordelia Langford, John Sillitoe on behalf of the Wellcome Sanger Institute COVID-19 Surveillance Team |
| EPI_ISL_2961457 | Berkshire and Surrey Pathology Services Lighthouse Laboratory | Wellcome Sanger Institute for the COVID-19 Genomics UK (COG-UK) Consortium | Berkshire and Surrey Pathology Services Lighthouse Laboratory and Alex Alderton, Roberto Amato, Jeffrey Barrett, Sonia Goncalves, Ewan Harrison, David K. Jackson, Ian Johnston, Dominic Kwiatkowski, Cordelia Langford, John Sillitoe on behalf of the Wellcome Sanger Institute COVID-19 Surveillance Team |
| EPI_ISL_2961458, EPI_ISL_2961459, EPI_ISL_2961460, EPI_ISL_2961461 | Lighthouse Lab in Glasgow | Wellcome Sanger Institute for the COVID-19 Genomics UK (COG-UK) Consortium | Harper VanSteenhouse, Yumi Kasai, David Gray, Carol Clugston, Anna Dominiczak and Alex Alderton, Roberto Amato, Jeffrey Barrett, Sonia Goncalves, Ewan Harrison, David K. Jackson, Ian Johnston, Dominic Kwiatkowski, Cordelia Langford, John Sillitoe on behalf of the Wellcome Sanger Institute COVID-19 Surveillance Team |
| EPI_ISL_2961462 | Lighthouse Lab in Alderley Park | Wellcome Sanger Institute for the COVID-19 Genomics UK (COG-UK) Consortium | Jacquelyn Wynn, Mairead Hyland, The Lighthouse Lab in Alderley Park and Alex Alderton, Roberto Amato, Jeffrey Barrett, Sonia Goncalves, Ewan Harrison, David K. Jackson, Ian Johnston, Dominic Kwiatkowski, Cordelia Langford, John Sillitoe on behalf of the Wellcome Sanger Institute COVID-19 Surveillance Team |
| EPI_ISL_2961463 | Berkshire and Surrey Pathology Services Lighthouse Laboratory | Wellcome Sanger Institute for the COVID-19 Genomics UK (COG-UK) Consortium | Berkshire and Surrey Pathology Services Lighthouse Laboratory and Alex Alderton, Roberto Amato, Jeffrey Barrett, Sonia Goncalves, Ewan Harrison, David K. Jackson, Ian Johnston, Dominic Kwiatkowski, Cordelia Langford, John Sillitoe on behalf of the Wellcome Sanger Institute COVID-19 Surveillance Team |
| EPI_ISL_2961464, EPI_ISL_2961465 | Lighthouse Lab in Glasgow | Wellcome Sanger Institute for the COVID-19 Genomics UK (COG-UK) Consortium | Harper VanSteenhouse, Yumi Kasai, David Gray, Carol Clugston, Anna Dominiczak and Alex Alderton, Roberto Amato, Jeffrey Barrett, Sonia Goncalves, Ewan Harrison, David K. Jackson, Ian Johnston, Dominic Kwiatkowski, Cordelia Langford, John Sillitoe on behalf of the Wellcome Sanger Institute COVID-19 Surveillance Team |
| EPI_ISL_2961466 | Berkshire and Surrey Pathology Services Lighthouse Laboratory | Wellcome Sanger Institute for the COVID-19 Genomics UK (COG-UK) Consortium | Berkshire and Surrey Pathology Services Lighthouse Laboratory and Alex Alderton, Roberto Amato, Jeffrey Barrett, Sonia Goncalves, Ewan Harrison, David K. Jackson, Ian Johnston, Dominic Kwiatkowski, Cordelia Langford, John Sillitoe on behalf of the Wellcome Sanger Institute COVID-19 Surveillance Team |
| EPI_ISL_2961467 | Lighthouse Lab in Alderley Park | Wellcome Sanger Institute for the COVID-19 Genomics UK (COG-UK) Consortium | Jacquelyn Wynn, Mairead Hyland, The Lighthouse Lab in Alderley Park and Alex Alderton, Roberto Amato, Jeffrey Barrett, Sonia Goncalves, Ewan Harrison, David K. Jackson, Ian Johnston, Dominic Kwiatkowski, Cordelia Langford, John Sillitoe on behalf of the Wellcome Sanger Institute COVID-19 Surveillance Team |
| EPI_ISL_2961468 | Berkshire and Surrey Pathology Services Lighthouse Laboratory | Wellcome Sanger Institute for the COVID-19 Genomics UK (COG-UK) Consortium | Berkshire and Surrey Pathology Services Lighthouse Laboratory and Alex Alderton, Roberto Amato, Jeffrey Barrett, Sonia Goncalves, Ewan Harrison, David K. Jackson, Ian Johnston, Dominic Kwiatkowski, Cordelia Langford, John Sillitoe on behalf of the Wellcome Sanger Institute COVID-19 Surveillance Team |
| EPI_ISL_2961469, EPI_ISL_2961470 | Lighthouse Lab in Glasgow | Wellcome Sanger Institute for the COVID-19 Genomics UK (COG-UK) Consortium | Harper VanSteenhouse, Yumi Kasai, David Gray, Carol Clugston, Anna Dominiczak and Alex Alderton, Roberto Amato, Jeffrey Barrett, Sonia Goncalves, Ewan Harrison, David K. Jackson, Ian Johnston, Dominic Kwiatkowski, Cordelia Langford, John Sillitoe on behalf of the Wellcome Sanger Institute COVID-19 Surveillance Team |
| EPI_ISL_2961471 | Berkshire and Surrey Pathology Services Lighthouse Laboratory | Wellcome Sanger Institute for the COVID-19 Genomics UK (COG-UK) Consortium | Berkshire and Surrey Pathology Services Lighthouse Laboratory and Alex Alderton, Roberto Amato, Jeffrey Barrett, Sonia Goncalves, Ewan Harrison, David K. Jackson, Ian Johnston, Dominic Kwiatkowski, Cordelia Langford, John Sillitoe on behalf of the Wellcome Sanger Institute COVID-19 Surveillance Team |
| EPI_ISL_2961472 | Lighthouse Lab in Glasgow | Wellcome Sanger Institute for the COVID-19 Genomics UK (COG-UK) Consortium | Harper VanSteenhouse, Yumi Kasai, David Gray, Carol Clugston, Anna Dominiczak and Alex Alderton, Roberto Amato, Jeffrey Barrett, Sonia Goncalves, Ewan Harrison, David K. Jackson, Ian Johnston, Dominic Kwiatkowski, Cordelia Langford, John Sillitoe on behalf of the Wellcome Sanger Institute COVID-19 Surveillance Team |
| EPI_ISL_2961473 | Lighthouse Lab in Alderley Park | Wellcome Sanger Institute for the COVID-19 Genomics UK (COG-UK) Consortium | Jacquelyn Wynn, Mairead Hyland, The Lighthouse Lab in Alderley Park and Alex Alderton, Roberto Amato, Jeffrey Barrett, Sonia Goncalves, Ewan Harrison, David K. Jackson, Ian Johnston, Dominic Kwiatkowski, Cordelia Langford, John Sillitoe on behalf of the Wellcome Sanger Institute COVID-19 Surveillance Team |
| EPI_ISL_2961474, EPI_ISL_2961475 | Lighthouse Lab in Glasgow | Wellcome Sanger Institute for the COVID-19 Genomics UK (COG-UK) Consortium | Harper VanSteenhouse, Yumi Kasai, David Gray, Carol Clugston, Anna Dominiczak and Alex Alderton, Roberto Amato, Jeffrey Barrett, Sonia Goncalves, Ewan Harrison, David K. Jackson, Ian Johnston, Dominic Kwiatkowski, Cordelia Langford, John Sillitoe on behalf of the Wellcome Sanger Institute COVID-19 Surveillance Team |
| EPI_ISL_2961476 | Berkshire and Surrey Pathology Services Lighthouse Laboratory | Wellcome Sanger Institute for the COVID-19 Genomics UK (COG-UK) Consortium | Berkshire and Surrey Pathology Services Lighthouse Laboratory and Alex Alderton, Roberto Amato, Jeffrey Barrett, Sonia Goncalves, Ewan Harrison, David K. Jackson, Ian Johnston, Dominic Kwiatkowski, Cordelia Langford, John Sillitoe on behalf of the Wellcome Sanger Institute COVID-19 Surveillance Team |
| EPI_ISL_2961477 | Lighthouse Lab in Alderley Park | Wellcome Sanger Institute for the COVID-19 Genomics UK (COG-UK) Consortium | Jacquelyn Wynn, Mairead Hyland, The Lighthouse Lab in Alderley Park and Alex Alderton, Roberto Amato, Jeffrey Barrett, Sonia Goncalves, Ewan Harrison, David K. Jackson, Ian Johnston, Dominic Kwiatkowski, Cordelia Langford, John Sillitoe on behalf of the Wellcome Sanger Institute COVID-19 Surveillance Team |
| EPI_ISL_2961478 | Berkshire and Surrey Pathology Services Lighthouse Laboratory | Wellcome Sanger Institute for the COVID-19 Genomics UK (COG-UK) Consortium | Berkshire and Surrey Pathology Services Lighthouse Laboratory and Alex Alderton, Roberto Amato, Jeffrey Barrett, Sonia Goncalves, Ewan Harrison, David K. Jackson, Ian Johnston, Dominic Kwiatkowski, Cordelia Langford, John Sillitoe on behalf of the Wellcome Sanger Institute COVID-19 Surveillance Team |
| EPI_ISL_2961479 | Lighthouse Lab in Alderley Park | Wellcome Sanger Institute for the COVID-19 Genomics UK (COG-UK) Consortium | Jacquelyn Wynn, Mairead Hyland, The Lighthouse Lab in Alderley Park and Alex Alderton, Roberto Amato, Jeffrey Barrett, Sonia Goncalves, Ewan Harrison, David K. Jackson, Ian Johnston, Dominic Kwiatkowski, Cordelia Langford, John Sillitoe on behalf of the Wellcome Sanger Institute COVID-19 Surveillance Team |
| EPI_ISL_2961480 | Lighthouse Lab in Glasgow | Wellcome Sanger Institute for the COVID-19 Genomics UK (COG-UK) Consortium | Harper VanSteenhouse, Yumi Kasai, David Gray, Carol Clugston, Anna Dominiczak and Alex Alderton, Roberto Amato, Jeffrey Barrett, Sonia Goncalves, Ewan Harrison, David K. Jackson, Ian Johnston, Dominic Kwiatkowski, Cordelia Langford, John Sillitoe on behalf of the Wellcome Sanger Institute COVID-19 Surveillance Team |
| EPI_ISL_2961481 | Berkshire and Surrey Pathology Services Lighthouse Laboratory | Wellcome Sanger Institute for the COVID-19 Genomics UK (COG-UK) Consortium | Berkshire and Surrey Pathology Services Lighthouse Laboratory and Alex Alderton, Roberto Amato, Jeffrey Barrett, Sonia Goncalves, Ewan Harrison, David K. Jackson, Ian Johnston, Dominic Kwiatkowski, Cordelia Langford, John Sillitoe on behalf of the Wellcome Sanger Institute COVID-19 Surveillance Team |
| EPI_ISL_2961482, EPI_ISL_2961483 | Lighthouse Lab in Alderley Park | Wellcome Sanger Institute for the COVID-19 Genomics UK (COG-UK) Consortium | Jacquelyn Wynn, Mairead Hyland, The Lighthouse Lab in Alderley Park and Alex Alderton, Roberto Amato, Jeffrey Barrett, Sonia Goncalves, Ewan Harrison, David K. Jackson, Ian Johnston, Dominic Kwiatkowski, Cordelia Langford, John Sillitoe on behalf of the Wellcome Sanger Institute COVID-19 Surveillance Team |

| | | | |
|---|---|---|---|
| EPI_ISL_2961484 | Lighthouse Lab in Glasgow | Wellcome Sanger Institute for the COVID-19 Genomics UK (COG-UK) Consortium | Harper VanSteenhouse, Yumi Kasai, David Gray, Carol Clugston, Anna Dominiczak and Alex Alderton, Roberto Amato, Jeffrey Barrett, Sonia Goncalves, Ewan Harrison, David K. Jackson, Ian Johnston, Dominic Kwiatkowski, Cordelia Langford, John Sillitoe on behalf of the Wellcome Sanger Institute COVID-19 Surveillance Team |
| EPI_ISL_2961485, EPI_ISL_2961486 | Berkshire and Surrey Pathology Services Lighthouse Laboratory | Wellcome Sanger Institute for the COVID-19 Genomics UK (COG-UK) Consortium | Berkshire and Surrey Pathology Services Lighthouse Laboratory and Alex Alderton, Roberto Amato, Jeffrey Barrett, Sonia Goncalves, Ewan Harrison, David K. Jackson, Ian Johnston, Dominic Kwiatkowski, Cordelia Langford, John Sillitoe on behalf of the Wellcome Sanger Institute COVID-19 Surveillance Team |
| EPI_ISL_2961487 | Lighthouse Lab in Glasgow | Wellcome Sanger Institute for the COVID-19 Genomics UK (COG-UK) Consortium | Harper VanSteenhouse, Yumi Kasai, David Gray, Carol Clugston, Anna Dominiczak and Alex Alderton, Roberto Amato, Jeffrey Barrett, Sonia Goncalves, Ewan Harrison, David K. Jackson, Ian Johnston, Dominic Kwiatkowski, Cordelia Langford, John Sillitoe on behalf of the Wellcome Sanger Institute COVID-19 Surveillance Team |
| EPI_ISL_2961488 | Lighthouse Lab in Alderley Park | Wellcome Sanger Institute for the COVID-19 Genomics UK (COG-UK) Consortium | Jacquelyn Wynn, Mairead Hyland, The Lighthouse Lab in Alderley Park and Alex Alderton, Roberto Amato, Jeffrey Barrett, Sonia Goncalves, Ewan Harrison, David K. Jackson, Ian Johnston, Dominic Kwiatkowski, Cordelia Langford, John Sillitoe on behalf of the Wellcome Sanger Institute COVID-19 Surveillance Team |
| EPI_ISL_2961489 | Berkshire and Surrey Pathology Services Lighthouse Laboratory | Wellcome Sanger Institute for the COVID-19 Genomics UK (COG-UK) Consortium | Berkshire and Surrey Pathology Services Lighthouse Laboratory and Alex Alderton, Roberto Amato, Jeffrey Barrett, Sonia Goncalves, Ewan Harrison, David K. Jackson, Ian Johnston, Dominic Kwiatkowski, Cordelia Langford, John Sillitoe on behalf of the Wellcome Sanger Institute COVID-19 Surveillance Team |
| EPI_ISL_2961490 | Lighthouse Lab in Glasgow | Wellcome Sanger Institute for the COVID-19 Genomics UK (COG-UK) Consortium | Harper VanSteenhouse, Yumi Kasai, David Gray, Carol Clugston, Anna Dominiczak and Alex Alderton, Roberto Amato, Jeffrey Barrett, Sonia Goncalves, Ewan Harrison, David K. Jackson, Ian Johnston, Dominic Kwiatkowski, Cordelia Langford, John Sillitoe on behalf of the Wellcome Sanger Institute COVID-19 Surveillance Team |
| EPI_ISL_2961491 | Berkshire and Surrey Pathology Services Lighthouse Laboratory | Wellcome Sanger Institute for the COVID-19 Genomics UK (COG-UK) Consortium | Berkshire and Surrey Pathology Services Lighthouse Laboratory and Alex Alderton, Roberto Amato, Jeffrey Barrett, Sonia Goncalves, Ewan Harrison, David K. Jackson, Ian Johnston, Dominic Kwiatkowski, Cordelia Langford, John Sillitoe on behalf of the Wellcome Sanger Institute COVID-19 Surveillance Team |
| EPI_ISL_2961492 | Lighthouse Lab in Glasgow | Wellcome Sanger Institute for the COVID-19 Genomics UK (COG-UK) Consortium | Harper VanSteenhouse, Yumi Kasai, David Gray, Carol Clugston, Anna Dominiczak and Alex Alderton, Roberto Amato, Jeffrey Barrett, Sonia Goncalves, Ewan Harrison, David K. Jackson, Ian Johnston, Dominic Kwiatkowski, Cordelia Langford, John Sillitoe on behalf of the Wellcome Sanger Institute COVID-19 Surveillance Team |
| EPI_ISL_2961493 | Berkshire and Surrey Pathology Services Lighthouse Laboratory | Wellcome Sanger Institute for the COVID-19 Genomics UK (COG-UK) Consortium | Berkshire and Surrey Pathology Services Lighthouse Laboratory and Alex Alderton, Roberto Amato, Jeffrey Barrett, Sonia Goncalves, Ewan Harrison, David K. Jackson, Ian Johnston, Dominic Kwiatkowski, Cordelia Langford, John Sillitoe on behalf of the Wellcome Sanger Institute COVID-19 Surveillance Team |
| EPI_ISL_2961494 | Lighthouse Lab in Glasgow | Wellcome Sanger Institute for the COVID-19 Genomics UK (COG-UK) Consortium | Harper VanSteenhouse, Yumi Kasai, David Gray, Carol Clugston, Anna Dominiczak and Alex Alderton, Roberto Amato, Jeffrey Barrett, Sonia Goncalves, Ewan Harrison, David K. Jackson, Ian Johnston, Dominic Kwiatkowski, Cordelia Langford, John Sillitoe on behalf of the Wellcome Sanger Institute COVID-19 Surveillance Team |
| EPI_ISL_2961495 | Berkshire and Surrey Pathology Services Lighthouse Laboratory | Wellcome Sanger Institute for the COVID-19 Genomics UK (COG-UK) Consortium | Berkshire and Surrey Pathology Services Lighthouse Laboratory and Alex Alderton, Roberto Amato, Jeffrey Barrett, Sonia Goncalves, Ewan Harrison, David K. Jackson, Ian Johnston, Dominic Kwiatkowski, Cordelia Langford, John Sillitoe on behalf of the Wellcome Sanger Institute COVID-19 Surveillance Team |
| EPI_ISL_2961496, EPI_ISL_2961497 | Lighthouse Lab in Glasgow | Wellcome Sanger Institute for the COVID-19 Genomics UK (COG-UK) Consortium | Harper VanSteenhouse, Yumi Kasai, David Gray, Carol Clugston, Anna Dominiczak and Alex Alderton, Roberto Amato, Jeffrey Barrett, Sonia Goncalves, Ewan Harrison, David K. Jackson, Ian Johnston, Dominic Kwiatkowski, Cordelia Langford, John Sillitoe on behalf of the Wellcome Sanger Institute COVID-19 Surveillance Team |
| EPI_ISL_2961498 | Lighthouse Lab in Alderley Park | Wellcome Sanger Institute for the COVID-19 Genomics UK (COG-UK) Consortium | Jacquelyn Wynn, Mairead Hyland, The Lighthouse Lab in Alderley Park and Alex Alderton, Roberto Amato, Jeffrey Barrett, Sonia Goncalves, Ewan Harrison, David K. Jackson, Ian Johnston, Dominic Kwiatkowski, Cordelia Langford, John Sillitoe on behalf of the Wellcome Sanger Institute COVID-19 Surveillance Team |
| EPI_ISL_2961499, EPI_ISL_2961500, EPI_ISL_2961501 | Berkshire and Surrey Pathology Services Lighthouse Laboratory | Wellcome Sanger Institute for the COVID-19 Genomics UK (COG-UK) Consortium | Berkshire and Surrey Pathology Services Lighthouse Laboratory and Alex Alderton, Roberto Amato, Jeffrey Barrett, Sonia Goncalves, Ewan Harrison, David K. Jackson, Ian Johnston, Dominic Kwiatkowski, Cordelia Langford, John Sillitoe on behalf of the Wellcome Sanger Institute COVID-19 Surveillance Team |
| EPI_ISL_2961502, EPI_ISL_2961503 | Lighthouse Lab in Alderley Park | Wellcome Sanger Institute for the COVID-19 Genomics UK (COG-UK) Consortium | Jacquelyn Wynn, Mairead Hyland, The Lighthouse Lab in Alderley Park and Alex Alderton, Roberto Amato, Jeffrey Barrett, Sonia Goncalves, Ewan Harrison, David K. Jackson, Ian Johnston, Dominic Kwiatkowski, Cordelia Langford, John Sillitoe on behalf of the Wellcome Sanger Institute COVID-19 Surveillance Team |
| EPI_ISL_2961504 | Lighthouse Lab in Glasgow | Wellcome Sanger Institute for the COVID-19 Genomics UK (COG-UK) Consortium | Harper VanSteenhouse, Yumi Kasai, David Gray, Carol Clugston, Anna Dominiczak and Alex Alderton, Roberto Amato, Jeffrey Barrett, Sonia Goncalves, Ewan Harrison, David K. Jackson, Ian Johnston, Dominic Kwiatkowski, Cordelia Langford, John Sillitoe on behalf of the Wellcome Sanger Institute COVID-19 Surveillance Team |
| EPI_ISL_2961505, EPI_ISL_2961506 | Lighthouse Lab in Alderley Park | Wellcome Sanger Institute for the COVID-19 Genomics UK (COG-UK) Consortium | Jacquelyn Wynn, Mairead Hyland, The Lighthouse Lab in Alderley Park and Alex Alderton, Roberto Amato, Jeffrey Barrett, Sonia Goncalves, Ewan Harrison, David K. Jackson, Ian Johnston, Dominic Kwiatkowski, Cordelia Langford, John Sillitoe on behalf of the Wellcome Sanger Institute COVID-19 Surveillance Team |
| EPI_ISL_2961507, EPI_ISL_2961508, EPI_ISL_2961509, EPI_ISL_2961510, EPI_ISL_2961511 | Lighthouse Laboratory Plymouth | Wellcome Sanger Institute for the COVID-19 Genomics UK (COG-UK) Consortium | Lighthouse Laboratory Plymouth and Alex Alderton, Roberto Amato, Jeffrey Barrett, Sonia Goncalves, Ewan Harrison, David K. Jackson, Ian Johnston, Dominic Kwiatkowski, Cordelia Langford, John Sillitoe on behalf of the Wellcome Sanger Institute COVID-19 Surveillance Team |
| EPI_ISL_2961512 | Berkshire and Surrey Pathology Services Lighthouse Laboratory | Wellcome Sanger Institute for the COVID-19 Genomics UK (COG-UK) Consortium | Berkshire and Surrey Pathology Services Lighthouse Laboratory and Alex Alderton, Roberto Amato, Jeffrey Barrett, Sonia Goncalves, Ewan Harrison, David K. Jackson, Ian Johnston, Dominic Kwiatkowski, Cordelia Langford, John Sillitoe on behalf of the Wellcome Sanger Institute COVID-19 Surveillance Team |
| EPI_ISL_2961513 | Lighthouse Lab in Alderley Park | Wellcome Sanger Institute for the COVID-19 Genomics UK (COG-UK) Consortium | Jacquelyn Wynn, Mairead Hyland, The Lighthouse Lab in Alderley Park and Alex Alderton, Roberto Amato, Jeffrey Barrett, Sonia Goncalves, Ewan Harrison, David K. Jackson, Ian Johnston, Dominic Kwiatkowski, Cordelia Langford, John Sillitoe on behalf of the Wellcome Sanger Institute COVID-19 Surveillance Team |
| EPI_ISL_2961514, EPI_ISL_2961515 | Lighthouse Laboratory Plymouth | Wellcome Sanger Institute for the COVID-19 Genomics UK (COG-UK) Consortium | Lighthouse Laboratory Plymouth and Alex Alderton, Roberto Amato, Jeffrey Barrett, Sonia Goncalves, Ewan Harrison, David K. Jackson, Ian Johnston, Dominic Kwiatkowski, Cordelia Langford, John Sillitoe on behalf of the Wellcome Sanger Institute COVID-19 Surveillance Team |
| EPI_ISL_2961516, EPI_ISL_2961517, EPI_ISL_2961518 | Lighthouse Lab in Alderley Park | Wellcome Sanger Institute for the COVID-19 Genomics UK (COG-UK) Consortium | Jacquelyn Wynn, Mairead Hyland, The Lighthouse Lab in Alderley Park and Alex Alderton, Roberto Amato, Jeffrey Barrett, Sonia Goncalves, Ewan Harrison, David K. Jackson, Ian Johnston, Dominic Kwiatkowski, Cordelia Langford, John Sillitoe on behalf of the Wellcome Sanger Institute COVID-19 Surveillance Team |
| EPI_ISL_2961519 | Berkshire and Surrey Pathology Services Lighthouse Laboratory | Wellcome Sanger Institute for the COVID-19 Genomics UK (COG-UK) Consortium | Berkshire and Surrey Pathology Services Lighthouse Laboratory and Alex Alderton, Roberto Amato, Jeffrey Barrett, Sonia Goncalves, Ewan Harrison, David K. Jackson, Ian Johnston, Dominic Kwiatkowski, Cordelia Langford, John Sillitoe on behalf of the Wellcome Sanger Institute COVID-19 Surveillance Team |
| EPI_ISL_2961520, EPI_ISL_2961521, EPI_ISL_2961522, EPI_ISL_2961523 | Lighthouse Laboratory Plymouth | Wellcome Sanger Institute for the COVID-19 Genomics UK (COG-UK) Consortium | Lighthouse Laboratory Plymouth and Alex Alderton, Roberto Amato, Jeffrey Barrett, Sonia Goncalves, Ewan Harrison, David K. Jackson, Ian Johnston, Dominic Kwiatkowski, Cordelia Langford, John Sillitoe on behalf of the Wellcome Sanger Institute COVID-19 Surveillance Team |
| EPI_ISL_2961524, EPI_ISL_2961525 | Berkshire and Surrey Pathology Services Lighthouse Laboratory | Wellcome Sanger Institute for the COVID-19 Genomics UK (COG-UK) Consortium | Berkshire and Surrey Pathology Services Lighthouse Laboratory and Alex Alderton, Roberto Amato, Jeffrey Barrett, Sonia Goncalves, Ewan Harrison, David K. Jackson, Ian Johnston, Dominic Kwiatkowski, Cordelia Langford, John Sillitoe on behalf of the Wellcome Sanger Institute COVID-19 Surveillance Team |
| EPI_ISL_2961526 | Lighthouse Lab in Alderley Park | Wellcome Sanger Institute for the COVID-19 Genomics UK (COG-UK) Consortium | Jacquelyn Wynn, Mairead Hyland, The Lighthouse Lab in Alderley Park and Alex Alderton, Roberto Amato, Jeffrey Barrett, Sonia Goncalves, Ewan Harrison, David K. Jackson, Ian Johnston, Dominic Kwiatkowski, Cordelia Langford, John Sillitoe on behalf of the Wellcome Sanger Institute COVID-19 Surveillance Team |
| EPI_ISL_2961527, EPI_ISL_2961528, EPI_ISL_2961529 | Lighthouse Laboratory Plymouth | Wellcome Sanger Institute for the COVID-19 Genomics UK (COG-UK) Consortium | Lighthouse Laboratory Plymouth and Alex Alderton, Roberto Amato, Jeffrey Barrett, Sonia Goncalves, Ewan Harrison, David K. Jackson, Ian Johnston, Dominic Kwiatkowski, Cordelia Langford, John Sillitoe on behalf of the Wellcome Sanger Institute COVID-19 Surveillance Team |
| EPI_ISL_2961530 | Lighthouse Lab in Alderley Park | Wellcome Sanger Institute for the COVID-19 Genomics UK (COG-UK) Consortium | Jacquelyn Wynn, Mairead Hyland, The Lighthouse Lab in Alderley Park and Alex Alderton, Roberto Amato, Jeffrey Barrett, Sonia Goncalves, Ewan Harrison, David K. Jackson, Ian Johnston, Dominic Kwiatkowski, Cordelia Langford, John Sillitoe on behalf of the Wellcome Sanger Institute COVID-19 Surveillance Team |
| EPI_ISL_2961531 | Berkshire and Surrey Pathology Services Lighthouse Laboratory | Wellcome Sanger Institute for the COVID-19 Genomics UK (COG-UK) Consortium | Berkshire and Surrey Pathology Services Lighthouse Laboratory and Alex Alderton, Roberto Amato, Jeffrey Barrett, Sonia Goncalves, Ewan Harrison, David K. Jackson, Ian Johnston, Dominic Kwiatkowski, Cordelia Langford, John Sillitoe on behalf of the Wellcome Sanger Institute COVID-19 Surveillance Team |
| EPI_ISL_2961532 | Lighthouse Laboratory Plymouth | Wellcome Sanger Institute for the COVID-19 Genomics UK (COG-UK) Consortium | Lighthouse Laboratory Plymouth and Alex Alderton, Roberto Amato, Jeffrey Barrett, Sonia Goncalves, Ewan Harrison, David K. Jackson, Ian Johnston, Dominic Kwiatkowski, Cordelia Langford, John Sillitoe on behalf of the Wellcome Sanger Institute COVID-19 Surveillance Team |
| EPI_ISL_2961533, EPI_ISL_2961534, EPI_ISL_2961535, EPI_ISL_2961536 | Lighthouse Lab in Alderley Park | Wellcome Sanger Institute for the COVID-19 Genomics UK (COG-UK) Consortium | Jacquelyn Wynn, Mairead Hyland, The Lighthouse Lab in Alderley Park and Alex Alderton, Roberto Amato, Jeffrey Barrett, Sonia Goncalves, Ewan Harrison, David K. Jackson, Ian Johnston, Dominic Kwiatkowski, Cordelia Langford, John Sillitoe on behalf of the Wellcome Sanger Institute COVID-19 Surveillance Team |
| EPI_ISL_2961537, EPI_ISL_2961538 | Berkshire and Surrey Pathology Services Lighthouse Laboratory | Wellcome Sanger Institute for the COVID-19 Genomics UK (COG-UK) Consortium | Berkshire and Surrey Pathology Services Lighthouse Laboratory and Alex Alderton, Roberto Amato, Jeffrey Barrett, Sonia Goncalves, Ewan Harrison, David K. Jackson, Ian Johnston, Dominic Kwiatkowski, Cordelia Langford, John Sillitoe on behalf of the Wellcome Sanger Institute COVID-19 Surveillance Team |
| EPI_ISL_2961539 | Lighthouse Lab in Alderley Park | Wellcome Sanger Institute for the COVID-19 Genomics UK (COG-UK) Consortium | Jacquelyn Wynn, Mairead Hyland, The Lighthouse Lab in Alderley Park and Alex Alderton, Roberto Amato, Jeffrey Barrett, Sonia Goncalves, Ewan Harrison, David K. Jackson, Ian Johnston, Dominic Kwiatkowski, Cordelia Langford, John Sillitoe on behalf of the Wellcome Sanger Institute COVID-19 Surveillance Team |
| EPI_ISL_2961540 | Berkshire and Surrey Pathology Services Lighthouse Laboratory | Wellcome Sanger Institute for the COVID-19 Genomics UK (COG-UK) Consortium | Berkshire and Surrey Pathology Services Lighthouse Laboratory and Alex Alderton, Roberto Amato, Jeffrey Barrett, Sonia Goncalves, Ewan Harrison, David K. Jackson, Ian Johnston, Dominic Kwiatkowski, Cordelia Langford, John Sillitoe on behalf of the Wellcome Sanger Institute COVID-19 Surveillance Team |
| EPI_ISL_2961541 | Lighthouse Laboratory Plymouth | Wellcome Sanger Institute for the COVID-19 Genomics UK (COG-UK) Consortium | Lighthouse Laboratory Plymouth and Alex Alderton, Roberto Amato, Jeffrey Barrett, Sonia Goncalves, Ewan Harrison, David K. Jackson, Ian Johnston, Dominic Kwiatkowski, Cordelia Langford, John Sillitoe on behalf of the Wellcome Sanger Institute COVID-19 Surveillance Team |
| EPI_ISL_2961542 | Berkshire and Surrey Pathology Services Lighthouse Laboratory | Wellcome Sanger Institute for the COVID-19 Genomics UK (COG-UK) Consortium | Berkshire and Surrey Pathology Services Lighthouse Laboratory and Alex Alderton, Roberto Amato, Jeffrey Barrett, Sonia Goncalves, Ewan Harrison, David K. Jackson, Ian Johnston, Dominic Kwiatkowski, Cordelia Langford, John Sillitoe on behalf of the Wellcome Sanger Institute COVID-19 Surveillance Team |
| EPI_ISL_2961543 | Lighthouse Laboratory Plymouth | Wellcome Sanger Institute for the COVID-19 Genomics UK (COG-UK) Consortium | Lighthouse Laboratory Plymouth and Alex Alderton, Roberto Amato, Jeffrey Barrett, Sonia Goncalves, Ewan Harrison, David K. Jackson, Ian Johnston, Dominic Kwiatkowski, Cordelia Langford, John Sillitoe on behalf of the Wellcome Sanger Institute COVID-19 Surveillance Team |
| EPI_ISL_2961544 | Lighthouse Lab in Alderley Park | Wellcome Sanger Institute for the COVID-19 Genomics UK (COG-UK) Consortium | Jacquelyn Wynn, Mairead Hyland, The Lighthouse Lab in Alderley Park and Alex Alderton, Roberto Amato, Jeffrey Barrett, Sonia Goncalves, Ewan Harrison, David K. Jackson, Ian Johnston, Dominic Kwiatkowski, Cordelia Langford, John Sillitoe on behalf of the Wellcome Sanger Institute COVID-19 Surveillance Team |
| EPI_ISL_2961545, EPI_ISL_2961546 | Lighthouse Laboratory Plymouth | Wellcome Sanger Institute for the COVID-19 Genomics UK (COG-UK) Consortium | Lighthouse Laboratory Plymouth and Alex Alderton, Roberto Amato, Jeffrey Barrett, Sonia Goncalves, Ewan Harrison, David K. Jackson, Ian Johnston, Dominic Kwiatkowski, Cordelia Langford, John Sillitoe on behalf of the Wellcome Sanger Institute COVID-19 Surveillance Team |
| EPI_ISL_2961547 | Lighthouse Lab in Alderley Park | Wellcome Sanger Institute for the COVID-19 Genomics UK (COG-UK) Consortium | Jacquelyn Wynn, Mairead Hyland, The Lighthouse Lab in Alderley Park and Alex Alderton, Roberto Amato, Jeffrey Barrett, Sonia Goncalves, Ewan Harrison, David K. Jackson, Ian Johnston, Dominic Kwiatkowski, Cordelia Langford, John Sillitoe on behalf of the Wellcome Sanger Institute COVID-19 Surveillance Team |
| EPI_ISL_2961548 | Berkshire and Surrey Pathology Services Lighthouse Laboratory | Wellcome Sanger Institute for the COVID-19 Genomics UK (COG-UK) Consortium | Berkshire and Surrey Pathology Services Lighthouse Laboratory and Alex Alderton, Roberto Amato, Jeffrey Barrett, Sonia Goncalves, Ewan Harrison, David K. Jackson, Ian Johnston, Dominic Kwiatkowski, Cordelia Langford, John Sillitoe on behalf of the Wellcome Sanger Institute COVID-19 Surveillance Team |

| | | | |
|---|---|---|---|
| EPI_ISL_2961549, EPI_ISL_2961550, EPI_ISL_2961551 | Lighthouse Laboratory Plymouth | Wellcome Sanger Institute for the COVID-19 Genomics UK (COG-UK) Consortium | Lighthouse Laboratory Plymouth and Alex Alderton, Roberto Amato, Jeffrey Barrett, Sonia Goncalves, Ewan Harrison, David K. Jackson, Ian Johnston, Dominic Kwiatkowski, Cordelia Langford, John Sillitoe on behalf of the Wellcome Sanger Institute COVID-19 Surveillance Team |
| EPI_ISL_2961552 | Berkshire and Surrey Pathology Services Lighthouse Laboratory | Wellcome Sanger Institute for the COVID-19 Genomics UK (COG-UK) Consortium | Berkshire and Surrey Pathology Services Lighthouse Laboratory and Alex Alderton, Roberto Amato, Jeffrey Barrett, Sonia Goncalves, Ewan Harrison, David K. Jackson, Ian Johnston, Dominic Kwiatkowski, Cordelia Langford, John Sillitoe on behalf of the Wellcome Sanger Institute COVID-19 Surveillance Team |
| EPI_ISL_2961553, EPI_ISL_2961554, EPI_ISL_2961555 | Lighthouse Laboratory Plymouth | Wellcome Sanger Institute for the COVID-19 Genomics UK (COG-UK) Consortium | Lighthouse Laboratory Plymouth and Alex Alderton, Roberto Amato, Jeffrey Barrett, Sonia Goncalves, Ewan Harrison, David K. Jackson, Ian Johnston, Dominic Kwiatkowski, Cordelia Langford, John Sillitoe on behalf of the Wellcome Sanger Institute COVID-19 Surveillance Team |
| EPI_ISL_2961556 | Berkshire and Surrey Pathology Services Lighthouse Laboratory | Wellcome Sanger Institute for the COVID-19 Genomics UK (COG-UK) Consortium | Berkshire and Surrey Pathology Services Lighthouse Laboratory and Alex Alderton, Roberto Amato, Jeffrey Barrett, Sonia Goncalves, Ewan Harrison, David K. Jackson, Ian Johnston, Dominic Kwiatkowski, Cordelia Langford, John Sillitoe on behalf of the Wellcome Sanger Institute COVID-19 Surveillance Team |
| EPI_ISL_2961557, EPI_ISL_2961558 | Lighthouse Laboratory Plymouth | Wellcome Sanger Institute for the COVID-19 Genomics UK (COG-UK) Consortium | Lighthouse Laboratory Plymouth and Alex Alderton, Roberto Amato, Jeffrey Barrett, Sonia Goncalves, Ewan Harrison, David K. Jackson, Ian Johnston, Dominic Kwiatkowski, Cordelia Langford, John Sillitoe on behalf of the Wellcome Sanger Institute COVID-19 Surveillance Team |
| EPI_ISL_2961559, EPI_ISL_2961560, EPI_ISL_2961561 | Berkshire and Surrey Pathology Services Lighthouse Laboratory | Wellcome Sanger Institute for the COVID-19 Genomics UK (COG-UK) Consortium | Berkshire and Surrey Pathology Services Lighthouse Laboratory and Alex Alderton, Roberto Amato, Jeffrey Barrett, Sonia Goncalves, Ewan Harrison, David K. Jackson, Ian Johnston, Dominic Kwiatkowski, Cordelia Langford, John Sillitoe on behalf of the Wellcome Sanger Institute COVID-19 Surveillance Team |
| EPI_ISL_2961562, EPI_ISL_2961563 | Lighthouse Laboratory Plymouth | Wellcome Sanger Institute for the COVID-19 Genomics UK (COG-UK) Consortium | Lighthouse Laboratory Plymouth and Alex Alderton, Roberto Amato, Jeffrey Barrett, Sonia Goncalves, Ewan Harrison, David K. Jackson, Ian Johnston, Dominic Kwiatkowski, Cordelia Langford, John Sillitoe on behalf of the Wellcome Sanger Institute COVID-19 Surveillance Team |
| EPI_ISL_2961564 | Lighthouse Lab in Alderley Park | Wellcome Sanger Institute for the COVID-19 Genomics UK (COG-UK) Consortium | Jacquelyn Wynn, Mairead Hyland, The Lighthouse Lab in Alderley Park and Alex Alderton, Roberto Amato, Jeffrey Barrett, Sonia Goncalves, Ewan Harrison, David K. Jackson, Ian Johnston, Dominic Kwiatkowski, Cordelia Langford, John Sillitoe on behalf of the Wellcome Sanger Institute COVID-19 Surveillance Team |
| EPI_ISL_2961565, EPI_ISL_2961566 | Lighthouse Laboratory Plymouth | Wellcome Sanger Institute for the COVID-19 Genomics UK (COG-UK) Consortium | Lighthouse Laboratory Plymouth and Alex Alderton, Roberto Amato, Jeffrey Barrett, Sonia Goncalves, Ewan Harrison, David K. Jackson, Ian Johnston, Dominic Kwiatkowski, Cordelia Langford, John Sillitoe on behalf of the Wellcome Sanger Institute COVID-19 Surveillance Team |
| EPI_ISL_2961567 | Berkshire and Surrey Pathology Services Lighthouse Laboratory | Wellcome Sanger Institute for the COVID-19 Genomics UK (COG-UK) Consortium | Berkshire and Surrey Pathology Services Lighthouse Laboratory and Alex Alderton, Roberto Amato, Jeffrey Barrett, Sonia Goncalves, Ewan Harrison, David K. Jackson, Ian Johnston, Dominic Kwiatkowski, Cordelia Langford, John Sillitoe on behalf of the Wellcome Sanger Institute COVID-19 Surveillance Team |
| EPI_ISL_2961568 | Lighthouse Lab in Alderley Park | Wellcome Sanger Institute for the COVID-19 Genomics UK (COG-UK) Consortium | Jacquelyn Wynn, Mairead Hyland, The Lighthouse Lab in Alderley Park and Alex Alderton, Roberto Amato, Jeffrey Barrett, Sonia Goncalves, Ewan Harrison, David K. Jackson, Ian Johnston, Dominic Kwiatkowski, Cordelia Langford, John Sillitoe on behalf of the Wellcome Sanger Institute COVID-19 Surveillance Team |
| EPI_ISL_2961569 | Berkshire and Surrey Pathology Services Lighthouse Laboratory | Wellcome Sanger Institute for the COVID-19 Genomics UK (COG-UK) Consortium | Berkshire and Surrey Pathology Services Lighthouse Laboratory and Alex Alderton, Roberto Amato, Jeffrey Barrett, Sonia Goncalves, Ewan Harrison, David K. Jackson, Ian Johnston, Dominic Kwiatkowski, Cordelia Langford, John Sillitoe on behalf of the Wellcome Sanger Institute COVID-19 Surveillance Team |
| EPI_ISL_2961570, EPI_ISL_2961571 | Lighthouse Laboratory Plymouth | Wellcome Sanger Institute for the COVID-19 Genomics UK (COG-UK) Consortium | Lighthouse Laboratory Plymouth and Alex Alderton, Roberto Amato, Jeffrey Barrett, Sonia Goncalves, Ewan Harrison, David K. Jackson, Ian Johnston, Dominic Kwiatkowski, Cordelia Langford, John Sillitoe on behalf of the Wellcome Sanger Institute COVID-19 Surveillance Team |
| EPI_ISL_2961572, EPI_ISL_2961573 | Berkshire and Surrey Pathology Services Lighthouse Laboratory | Wellcome Sanger Institute for the COVID-19 Genomics UK (COG-UK) Consortium | Berkshire and Surrey Pathology Services Lighthouse Laboratory and Alex Alderton, Roberto Amato, Jeffrey Barrett, Sonia Goncalves, Ewan Harrison, David K. Jackson, Ian Johnston, Dominic Kwiatkowski, Cordelia Langford, John Sillitoe on behalf of the Wellcome Sanger Institute COVID-19 Surveillance Team |
| EPI_ISL_2961574 | Lighthouse Laboratory Plymouth | Wellcome Sanger Institute for the COVID-19 Genomics UK (COG-UK) Consortium | Lighthouse Laboratory Plymouth and Alex Alderton, Roberto Amato, Jeffrey Barrett, Sonia Goncalves, Ewan Harrison, David K. Jackson, Ian Johnston, Dominic Kwiatkowski, Cordelia Langford, John Sillitoe on behalf of the Wellcome Sanger Institute COVID-19 Surveillance Team |
| EPI_ISL_2961575 | Lighthouse Lab in Alderley Park | Wellcome Sanger Institute for the COVID-19 Genomics UK (COG-UK) Consortium | Jacquelyn Wynn, Mairead Hyland, The Lighthouse Lab in Alderley Park and Alex Alderton, Roberto Amato, Jeffrey Barrett, Sonia Goncalves, Ewan Harrison, David K. Jackson, Ian Johnston, Dominic Kwiatkowski, Cordelia Langford, John Sillitoe on behalf of the Wellcome Sanger Institute COVID-19 Surveillance Team |
| EPI_ISL_2961576, EPI_ISL_2961577, EPI_ISL_2961578, EPI_ISL_2961579 | Lighthouse Laboratory Plymouth | Wellcome Sanger Institute for the COVID-19 Genomics UK (COG-UK) Consortium | Lighthouse Laboratory Plymouth and Alex Alderton, Roberto Amato, Jeffrey Barrett, Sonia Goncalves, Ewan Harrison, David K. Jackson, Ian Johnston, Dominic Kwiatkowski, Cordelia Langford, John Sillitoe on behalf of the Wellcome Sanger Institute COVID-19 Surveillance Team |
| EPI_ISL_2961580 | Lighthouse Lab in Alderley Park | Wellcome Sanger Institute for the COVID-19 Genomics UK (COG-UK) Consortium | Jacquelyn Wynn, Mairead Hyland, The Lighthouse Lab in Alderley Park and Alex Alderton, Roberto Amato, Jeffrey Barrett, Sonia Goncalves, Ewan Harrison, David K. Jackson, Ian Johnston, Dominic Kwiatkowski, Cordelia Langford, John Sillitoe on behalf of the Wellcome Sanger Institute COVID-19 Surveillance Team |
| EPI_ISL_2961581, EPI_ISL_2961582 | Lighthouse Laboratory Plymouth | Wellcome Sanger Institute for the COVID-19 Genomics UK (COG-UK) Consortium | Lighthouse Laboratory Plymouth and Alex Alderton, Roberto Amato, Jeffrey Barrett, Sonia Goncalves, Ewan Harrison, David K. Jackson, Ian Johnston, Dominic Kwiatkowski, Cordelia Langford, John Sillitoe on behalf of the Wellcome Sanger Institute COVID-19 Surveillance Team |
| EPI_ISL_2961583 | Berkshire and Surrey Pathology Services Lighthouse Laboratory | Wellcome Sanger Institute for the COVID-19 Genomics UK (COG-UK) Consortium | Berkshire and Surrey Pathology Services Lighthouse Laboratory and Alex Alderton, Roberto Amato, Jeffrey Barrett, Sonia Goncalves, Ewan Harrison, David K. Jackson, Ian Johnston, Dominic Kwiatkowski, Cordelia Langford, John Sillitoe on behalf of the Wellcome Sanger Institute COVID-19 Surveillance Team |
| EPI_ISL_2961584 | Lighthouse Laboratory Plymouth | Wellcome Sanger Institute for the COVID-19 Genomics UK (COG-UK) Consortium | Lighthouse Laboratory Plymouth and Alex Alderton, Roberto Amato, Jeffrey Barrett, Sonia Goncalves, Ewan Harrison, David K. Jackson, Ian Johnston, Dominic Kwiatkowski, Cordelia Langford, John Sillitoe on behalf of the Wellcome Sanger Institute COVID-19 Surveillance Team |
| EPI_ISL_2961585, EPI_ISL_2961586 | Lighthouse Lab in Alderley Park | Wellcome Sanger Institute for the COVID-19 Genomics UK (COG-UK) Consortium | Jacquelyn Wynn, Mairead Hyland, The Lighthouse Lab in Alderley Park and Alex Alderton, Roberto Amato, Jeffrey Barrett, Sonia Goncalves, Ewan Harrison, David K. Jackson, Ian Johnston, Dominic Kwiatkowski, Cordelia Langford, John Sillitoe on behalf of the Wellcome Sanger Institute COVID-19 Surveillance Team |
| EPI_ISL_2961587, EPI_ISL_2961588, EPI_ISL_2961589, EPI_ISL_2961590 | Lighthouse Laboratory Plymouth | Wellcome Sanger Institute for the COVID-19 Genomics UK (COG-UK) Consortium | Lighthouse Laboratory Plymouth and Alex Alderton, Roberto Amato, Jeffrey Barrett, Sonia Goncalves, Ewan Harrison, David K. Jackson, Ian Johnston, Dominic Kwiatkowski, Cordelia Langford, John Sillitoe on behalf of the Wellcome Sanger Institute COVID-19 Surveillance Team |
| EPI_ISL_2961591 | Lighthouse Lab in Alderley Park | Wellcome Sanger Institute for the COVID-19 Genomics UK (COG-UK) Consortium | Jacquelyn Wynn, Mairead Hyland, The Lighthouse Lab in Alderley Park and Alex Alderton, Roberto Amato, Jeffrey Barrett, Sonia Goncalves, Ewan Harrison, David K. Jackson, Ian Johnston, Dominic Kwiatkowski, Cordelia Langford, John Sillitoe on behalf of the Wellcome Sanger Institute COVID-19 Surveillance Team |
| EPI_ISL_2961592 | Lighthouse Laboratory Plymouth | Wellcome Sanger Institute for the COVID-19 Genomics UK (COG-UK) Consortium | Lighthouse Laboratory Plymouth and Alex Alderton, Roberto Amato, Jeffrey Barrett, Sonia Goncalves, Ewan Harrison, David K. Jackson, Ian Johnston, Dominic Kwiatkowski, Cordelia Langford, John Sillitoe on behalf of the Wellcome Sanger Institute COVID-19 Surveillance Team |
| EPI_ISL_2961593 | Berkshire and Surrey Pathology Services Lighthouse Laboratory | Wellcome Sanger Institute for the COVID-19 Genomics UK (COG-UK) Consortium | Berkshire and Surrey Pathology Services Lighthouse Laboratory and Alex Alderton, Roberto Amato, Jeffrey Barrett, Sonia Goncalves, Ewan Harrison, David K. Jackson, Ian Johnston, Dominic Kwiatkowski, Cordelia Langford, John Sillitoe on behalf of the Wellcome Sanger Institute COVID-19 Surveillance Team |
| EPI_ISL_2961594 | Lighthouse Lab in Alderley Park | Wellcome Sanger Institute for the COVID-19 Genomics UK (COG-UK) Consortium | Jacquelyn Wynn, Mairead Hyland, The Lighthouse Lab in Alderley Park and Alex Alderton, Roberto Amato, Jeffrey Barrett, Sonia Goncalves, Ewan Harrison, David K. Jackson, Ian Johnston, Dominic Kwiatkowski, Cordelia Langford, John Sillitoe on behalf of the Wellcome Sanger Institute COVID-19 Surveillance Team |
| EPI_ISL_2961595, EPI_ISL_2961596 | Lighthouse Laboratory Plymouth | Wellcome Sanger Institute for the COVID-19 Genomics UK (COG-UK) Consortium | Lighthouse Laboratory Plymouth and Alex Alderton, Roberto Amato, Jeffrey Barrett, Sonia Goncalves, Ewan Harrison, David K. Jackson, Ian Johnston, Dominic Kwiatkowski, Cordelia Langford, John Sillitoe on behalf of the Wellcome Sanger Institute COVID-19 Surveillance Team |
| EPI_ISL_2961597, EPI_ISL_2961598 | Berkshire and Surrey Pathology Services Lighthouse Laboratory | Wellcome Sanger Institute for the COVID-19 Genomics UK (COG-UK) Consortium | Berkshire and Surrey Pathology Services Lighthouse Laboratory and Alex Alderton, Roberto Amato, Jeffrey Barrett, Sonia Goncalves, Ewan Harrison, David K. Jackson, Ian Johnston, Dominic Kwiatkowski, Cordelia Langford, John Sillitoe on behalf of the Wellcome Sanger Institute COVID-19 Surveillance Team |
| EPI_ISL_2961599, EPI_ISL_2961600 | Lighthouse Lab in Alderley Park | Wellcome Sanger Institute for the COVID-19 Genomics UK (COG-UK) Consortium | Jacquelyn Wynn, Mairead Hyland, The Lighthouse Lab in Alderley Park and Alex Alderton, Roberto Amato, Jeffrey Barrett, Sonia Goncalves, Ewan Harrison, David K. Jackson, Ian Johnston, Dominic Kwiatkowski, Cordelia Langford, John Sillitoe on behalf of the Wellcome Sanger Institute COVID-19 Surveillance Team |
| EPI_ISL_2961601, EPI_ISL_2961602 | Berkshire and Surrey Pathology Services Lighthouse Laboratory | Wellcome Sanger Institute for the COVID-19 Genomics UK (COG-UK) Consortium | Berkshire and Surrey Pathology Services Lighthouse Laboratory and Alex Alderton, Roberto Amato, Jeffrey Barrett, Sonia Goncalves, Ewan Harrison, David K. Jackson, Ian Johnston, Dominic Kwiatkowski, Cordelia Langford, John Sillitoe on behalf of the Wellcome Sanger Institute COVID-19 Surveillance Team |
| EPI_ISL_2961603, EPI_ISL_2961604, EPI_ISL_2961605, EPI_ISL_2961606 | Lighthouse Laboratory Plymouth | Wellcome Sanger Institute for the COVID-19 Genomics UK (COG-UK) Consortium | Lighthouse Laboratory Plymouth and Alex Alderton, Roberto Amato, Jeffrey Barrett, Sonia Goncalves, Ewan Harrison, David K. Jackson, Ian Johnston, Dominic Kwiatkowski, Cordelia Langford, John Sillitoe on behalf of the Wellcome Sanger Institute COVID-19 Surveillance Team |
| EPI_ISL_2961607, EPI_ISL_2961608 | Lighthouse Lab in Alderley Park | Wellcome Sanger Institute for the COVID-19 Genomics UK (COG-UK) Consortium | Jacquelyn Wynn, Mairead Hyland, The Lighthouse Lab in Alderley Park and Alex Alderton, Roberto Amato, Jeffrey Barrett, Sonia Goncalves, Ewan Harrison, David K. Jackson, Ian Johnston, Dominic Kwiatkowski, Cordelia Langford, John Sillitoe on behalf of the Wellcome Sanger Institute COVID-19 Surveillance Team |
| EPI_ISL_2961609 | Berkshire and Surrey Pathology Services Lighthouse Laboratory | Wellcome Sanger Institute for the COVID-19 Genomics UK (COG-UK) Consortium | Berkshire and Surrey Pathology Services Lighthouse Laboratory and Alex Alderton, Roberto Amato, Jeffrey Barrett, Sonia Goncalves, Ewan Harrison, David K. Jackson, Ian Johnston, Dominic Kwiatkowski, Cordelia Langford, John Sillitoe on behalf of the Wellcome Sanger Institute COVID-19 Surveillance Team |
| EPI_ISL_2961610, EPI_ISL_2961611 | Lighthouse Laboratory Plymouth | Wellcome Sanger Institute for the COVID-19 Genomics UK (COG-UK) Consortium | Lighthouse Laboratory Plymouth and Alex Alderton, Roberto Amato, Jeffrey Barrett, Sonia Goncalves, Ewan Harrison, David K. Jackson, Ian Johnston, Dominic Kwiatkowski, Cordelia Langford, John Sillitoe on behalf of the Wellcome Sanger Institute COVID-19 Surveillance Team |
| EPI_ISL_2961612 | Berkshire and Surrey Pathology Services Lighthouse Laboratory | Wellcome Sanger Institute for the COVID-19 Genomics UK (COG-UK) Consortium | Berkshire and Surrey Pathology Services Lighthouse Laboratory and Alex Alderton, Roberto Amato, Jeffrey Barrett, Sonia Goncalves, Ewan Harrison, David K. Jackson, Ian Johnston, Dominic Kwiatkowski, Cordelia Langford, John Sillitoe on behalf of the Wellcome Sanger Institute COVID-19 Surveillance Team |
| EPI_ISL_2961613, EPI_ISL_2961614 | Lighthouse Lab in Alderley Park | Wellcome Sanger Institute for the COVID-19 Genomics UK (COG-UK) Consortium | Jacquelyn Wynn, Mairead Hyland, The Lighthouse Lab in Alderley Park and Alex Alderton, Roberto Amato, Jeffrey Barrett, Sonia Goncalves, Ewan Harrison, David K. Jackson, Ian Johnston, Dominic Kwiatkowski, Cordelia Langford, John Sillitoe on behalf of the Wellcome Sanger Institute COVID-19 Surveillance Team |
| EPI_ISL_2961615 | Berkshire and Surrey Pathology Services Lighthouse Laboratory | Wellcome Sanger Institute for the COVID-19 Genomics UK (COG-UK) Consortium | Berkshire and Surrey Pathology Services Lighthouse Laboratory and Alex Alderton, Roberto Amato, Jeffrey Barrett, Sonia Goncalves, Ewan Harrison, David K. Jackson, Ian Johnston, Dominic Kwiatkowski, Cordelia Langford, John Sillitoe on behalf of the Wellcome Sanger Institute COVID-19 Surveillance Team |
| EPI_ISL_2961616, EPI_ISL_2961617 | Lighthouse Laboratory Plymouth | Wellcome Sanger Institute for the COVID-19 Genomics UK (COG-UK) Consortium | Lighthouse Laboratory Plymouth and Alex Alderton, Roberto Amato, Jeffrey Barrett, Sonia Goncalves, Ewan Harrison, David K. Jackson, Ian Johnston, Dominic Kwiatkowski, Cordelia Langford, John Sillitoe on behalf of the Wellcome Sanger Institute COVID-19 Surveillance Team |
| EPI_ISL_2961618 | Berkshire and Surrey Pathology Services Lighthouse Laboratory | Wellcome Sanger Institute for the COVID-19 Genomics UK (COG-UK) Consortium | Berkshire and Surrey Pathology Services Lighthouse Laboratory and Alex Alderton, Roberto Amato, Jeffrey Barrett, Sonia Goncalves, Ewan Harrison, David K. Jackson, Ian Johnston, Dominic Kwiatkowski, Cordelia Langford, John Sillitoe on behalf of the Wellcome Sanger Institute COVID-19 Surveillance Team |
| EPI_ISL_2961619 | Lighthouse Lab in Alderley Park | Wellcome Sanger Institute for the COVID-19 Genomics UK (COG-UK) Consortium | Jacquelyn Wynn, Mairead Hyland, The Lighthouse Lab in Alderley Park and Alex Alderton, Roberto Amato, Jeffrey Barrett, Sonia Goncalves, Ewan Harrison, David K. Jackson, Ian Johnston, Dominic Kwiatkowski, Cordelia Langford, John Sillitoe on behalf of the Wellcome Sanger Institute COVID-19 Surveillance Team |
| EPI_ISL_2961620, | Berkshire and Surrey Pathology Services Lighthouse Laboratory | Wellcome Sanger Institute for the COVID-19 Genomics UK (COG-UK) | Berkshire and Surrey Pathology Services Lighthouse Laboratory and Alex Alderton, Roberto Amato, Jeffrey Barrett, Sonia Goncalves, Ewan Harrison, David K. Jackson, Ian Johnston, Dominic Kwiatkowski, Cordelia Langford, John Sillitoe on behalf |

| | | | |
|---|---|---|---|
| EPI_ISL_2961621 | | Consortium | of the Wellcome Sanger Institute COVID-19 Surveillance Team |
| EPI_ISL_2961622, EPI_ISL_2961623 | Lighthouse Lab in Alderley Park | Wellcome Sanger Institute for the COVID-19 Genomics UK (COG-UK) Consortium | Jacquelyn Wynn, Mairead Hyland, The Lighthouse Lab in Alderley Park and Alex Alderton, Roberto Amato, Jeffrey Barrett, Sonia Goncalves, Ewan Harrison, David K. Jackson, Ian Johnston, Dominic Kwiatkowski, Cordelia Langford, John Sillitoe on behalf of the Wellcome Sanger Institute COVID-19 Surveillance Team |
| EPI_ISL_2961624 | Lighthouse Laboratory Plymouth | Wellcome Sanger Institute for the COVID-19 Genomics UK (COG-UK) Consortium | Lighthouse Laboratory Plymouth and Alex Alderton, Roberto Amato, Jeffrey Barrett, Sonia Goncalves, Ewan Harrison, David K. Jackson, Ian Johnston, Dominic Kwiatkowski, Cordelia Langford, John Sillitoe on behalf of the Wellcome Sanger Institute COVID-19 Surveillance Team |
| EPI_ISL_2961625 | Berkshire and Surrey Pathology Services Lighthouse Laboratory | Wellcome Sanger Institute for the COVID-19 Genomics UK (COG-UK) Consortium | Berkshire and Surrey Pathology Services Lighthouse Laboratory and Alex Alderton, Roberto Amato, Jeffrey Barrett, Sonia Goncalves, Ewan Harrison, David K. Jackson, Ian Johnston, Dominic Kwiatkowski, Cordelia Langford, John Sillitoe on behalf of the Wellcome Sanger Institute COVID-19 Surveillance Team |
| EPI_ISL_2961626 | Lighthouse Laboratory Plymouth | Wellcome Sanger Institute for the COVID-19 Genomics UK (COG-UK) Consortium | Lighthouse Laboratory Plymouth and Alex Alderton, Roberto Amato, Jeffrey Barrett, Sonia Goncalves, Ewan Harrison, David K. Jackson, Ian Johnston, Dominic Kwiatkowski, Cordelia Langford, John Sillitoe on behalf of the Wellcome Sanger Institute COVID-19 Surveillance Team |
| EPI_ISL_2961627 | Lighthouse Lab in Alderley Park | Wellcome Sanger Institute for the COVID-19 Genomics UK (COG-UK) Consortium | Jacquelyn Wynn, Mairead Hyland, The Lighthouse Lab in Alderley Park and Alex Alderton, Roberto Amato, Jeffrey Barrett, Sonia Goncalves, Ewan Harrison, David K. Jackson, Ian Johnston, Dominic Kwiatkowski, Cordelia Langford, John Sillitoe on behalf of the Wellcome Sanger Institute COVID-19 Surveillance Team |
| EPI_ISL_2961628 | Lighthouse Laboratory Plymouth | Wellcome Sanger Institute for the COVID-19 Genomics UK (COG-UK) Consortium | Lighthouse Laboratory Plymouth and Alex Alderton, Roberto Amato, Jeffrey Barrett, Sonia Goncalves, Ewan Harrison, David K. Jackson, Ian Johnston, Dominic Kwiatkowski, Cordelia Langford, John Sillitoe on behalf of the Wellcome Sanger Institute COVID-19 Surveillance Team |
| EPI_ISL_2961629 | Lighthouse Lab in Alderley Park | Wellcome Sanger Institute for the COVID-19 Genomics UK (COG-UK) Consortium | Jacquelyn Wynn, Mairead Hyland, The Lighthouse Lab in Alderley Park and Alex Alderton, Roberto Amato, Jeffrey Barrett, Sonia Goncalves, Ewan Harrison, David K. Jackson, Ian Johnston, Dominic Kwiatkowski, Cordelia Langford, John Sillitoe on behalf of the Wellcome Sanger Institute COVID-19 Surveillance Team |
| EPI_ISL_2961630 | Lighthouse Laboratory Plymouth | Wellcome Sanger Institute for the COVID-19 Genomics UK (COG-UK) Consortium | Lighthouse Laboratory Plymouth and Alex Alderton, Roberto Amato, Jeffrey Barrett, Sonia Goncalves, Ewan Harrison, David K. Jackson, Ian Johnston, Dominic Kwiatkowski, Cordelia Langford, John Sillitoe on behalf of the Wellcome Sanger Institute COVID-19 Surveillance Team |
| EPI_ISL_2961631 | Berkshire and Surrey Pathology Services Lighthouse Laboratory | Wellcome Sanger Institute for the COVID-19 Genomics UK (COG-UK) Consortium | Berkshire and Surrey Pathology Services Lighthouse Laboratory and Alex Alderton, Roberto Amato, Jeffrey Barrett, Sonia Goncalves, Ewan Harrison, David K. Jackson, Ian Johnston, Dominic Kwiatkowski, Cordelia Langford, John Sillitoe on behalf of the Wellcome Sanger Institute COVID-19 Surveillance Team |
| EPI_ISL_2961632, EPI_ISL_2961633 | Lighthouse Laboratory Plymouth | Wellcome Sanger Institute for the COVID-19 Genomics UK (COG-UK) Consortium | Lighthouse Laboratory Plymouth and Alex Alderton, Roberto Amato, Jeffrey Barrett, Sonia Goncalves, Ewan Harrison, David K. Jackson, Ian Johnston, Dominic Kwiatkowski, Cordelia Langford, John Sillitoe on behalf of the Wellcome Sanger Institute COVID-19 Surveillance Team |
| EPI_ISL_2961634, EPI_ISL_2961635 | Lighthouse Lab in Alderley Park | Wellcome Sanger Institute for the COVID-19 Genomics UK (COG-UK) Consortium | Jacquelyn Wynn, Mairead Hyland, The Lighthouse Lab in Alderley Park and Alex Alderton, Roberto Amato, Jeffrey Barrett, Sonia Goncalves, Ewan Harrison, David K. Jackson, Ian Johnston, Dominic Kwiatkowski, Cordelia Langford, John Sillitoe on behalf of the Wellcome Sanger Institute COVID-19 Surveillance Team |
| EPI_ISL_2961636, EPI_ISL_2961637 | Berkshire and Surrey Pathology Services Lighthouse Laboratory | Wellcome Sanger Institute for the COVID-19 Genomics UK (COG-UK) Consortium | Berkshire and Surrey Pathology Services Lighthouse Laboratory and Alex Alderton, Roberto Amato, Jeffrey Barrett, Sonia Goncalves, Ewan Harrison, David K. Jackson, Ian Johnston, Dominic Kwiatkowski, Cordelia Langford, John Sillitoe on behalf of the Wellcome Sanger Institute COVID-19 Surveillance Team |
| EPI_ISL_2961638 | Lighthouse Lab in Alderley Park | Wellcome Sanger Institute for the COVID-19 Genomics UK (COG-UK) Consortium | Jacquelyn Wynn, Mairead Hyland, The Lighthouse Lab in Alderley Park and Alex Alderton, Roberto Amato, Jeffrey Barrett, Sonia Goncalves, Ewan Harrison, David K. Jackson, Ian Johnston, Dominic Kwiatkowski, Cordelia Langford, John Sillitoe on behalf of the Wellcome Sanger Institute COVID-19 Surveillance Team |
| EPI_ISL_2961639, EPI_ISL_2961640 | Lighthouse Laboratory Plymouth | Wellcome Sanger Institute for the COVID-19 Genomics UK (COG-UK) Consortium | Lighthouse Laboratory Plymouth and Alex Alderton, Roberto Amato, Jeffrey Barrett, Sonia Goncalves, Ewan Harrison, David K. Jackson, Ian Johnston, Dominic Kwiatkowski, Cordelia Langford, John Sillitoe on behalf of the Wellcome Sanger Institute COVID-19 Surveillance Team |
| EPI_ISL_2961641 | Lighthouse Lab in Alderley Park | Wellcome Sanger Institute for the COVID-19 Genomics UK (COG-UK) Consortium | Jacquelyn Wynn, Mairead Hyland, The Lighthouse Lab in Alderley Park and Alex Alderton, Roberto Amato, Jeffrey Barrett, Sonia Goncalves, Ewan Harrison, David K. Jackson, Ian Johnston, Dominic Kwiatkowski, Cordelia Langford, John Sillitoe on behalf of the Wellcome Sanger Institute COVID-19 Surveillance Team |
| EPI_ISL_2961642, EPI_ISL_2961643, EPI_ISL_2961644, EPI_ISL_2961645, EPI_ISL_2961646 | Lighthouse Laboratory Plymouth | Wellcome Sanger Institute for the COVID-19 Genomics UK (COG-UK) Consortium | Lighthouse Laboratory Plymouth and Alex Alderton, Roberto Amato, Jeffrey Barrett, Sonia Goncalves, Ewan Harrison, David K. Jackson, Ian Johnston, Dominic Kwiatkowski, Cordelia Langford, John Sillitoe on behalf of the Wellcome Sanger Institute COVID-19 Surveillance Team |
| EPI_ISL_2961647 | Berkshire and Surrey Pathology Services Lighthouse Laboratory | Wellcome Sanger Institute for the COVID-19 Genomics UK (COG-UK) Consortium | Berkshire and Surrey Pathology Services Lighthouse Laboratory and Alex Alderton, Roberto Amato, Jeffrey Barrett, Sonia Goncalves, Ewan Harrison, David K. Jackson, Ian Johnston, Dominic Kwiatkowski, Cordelia Langford, John Sillitoe on behalf of the Wellcome Sanger Institute COVID-19 Surveillance Team |
| EPI_ISL_2961648, EPI_ISL_2961649 | Lighthouse Laboratory Plymouth | Wellcome Sanger Institute for the COVID-19 Genomics UK (COG-UK) Consortium | Lighthouse Laboratory Plymouth and Alex Alderton, Roberto Amato, Jeffrey Barrett, Sonia Goncalves, Ewan Harrison, David K. Jackson, Ian Johnston, Dominic Kwiatkowski, Cordelia Langford, John Sillitoe on behalf of the Wellcome Sanger Institute COVID-19 Surveillance Team |
| EPI_ISL_2961650 | Berkshire and Surrey Pathology Services Lighthouse Laboratory | Wellcome Sanger Institute for the COVID-19 Genomics UK (COG-UK) Consortium | Berkshire and Surrey Pathology Services Lighthouse Laboratory and Alex Alderton, Roberto Amato, Jeffrey Barrett, Sonia Goncalves, Ewan Harrison, David K. Jackson, Ian Johnston, Dominic Kwiatkowski, Cordelia Langford, John Sillitoe on behalf of the Wellcome Sanger Institute COVID-19 Surveillance Team |
| EPI_ISL_2961651 | Lighthouse Lab in Alderley Park | Wellcome Sanger Institute for the COVID-19 Genomics UK (COG-UK) Consortium | Jacquelyn Wynn, Mairead Hyland, The Lighthouse Lab in Alderley Park and Alex Alderton, Roberto Amato, Jeffrey Barrett, Sonia Goncalves, Ewan Harrison, David K. Jackson, Ian Johnston, Dominic Kwiatkowski, Cordelia Langford, John Sillitoe on behalf of the Wellcome Sanger Institute COVID-19 Surveillance Team |
| EPI_ISL_2961652 | Berkshire and Surrey Pathology Services Lighthouse Laboratory | Wellcome Sanger Institute for the COVID-19 Genomics UK (COG-UK) Consortium | Berkshire and Surrey Pathology Services Lighthouse Laboratory and Alex Alderton, Roberto Amato, Jeffrey Barrett, Sonia Goncalves, Ewan Harrison, David K. Jackson, Ian Johnston, Dominic Kwiatkowski, Cordelia Langford, John Sillitoe on behalf of the Wellcome Sanger Institute COVID-19 Surveillance Team |
| EPI_ISL_2961653 | Lighthouse Laboratory Plymouth | Wellcome Sanger Institute for the COVID-19 Genomics UK (COG-UK) Consortium | Lighthouse Laboratory Plymouth and Alex Alderton, Roberto Amato, Jeffrey Barrett, Sonia Goncalves, Ewan Harrison, David K. Jackson, Ian Johnston, Dominic Kwiatkowski, Cordelia Langford, John Sillitoe on behalf of the Wellcome Sanger Institute COVID-19 Surveillance Team |
| EPI_ISL_2961654 | Lighthouse Lab in Alderley Park | Wellcome Sanger Institute for the COVID-19 Genomics UK (COG-UK) Consortium | Jacquelyn Wynn, Mairead Hyland, The Lighthouse Lab in Alderley Park and Alex Alderton, Roberto Amato, Jeffrey Barrett, Sonia Goncalves, Ewan Harrison, David K. Jackson, Ian Johnston, Dominic Kwiatkowski, Cordelia Langford, John Sillitoe on behalf of the Wellcome Sanger Institute COVID-19 Surveillance Team |
| EPI_ISL_2961655, EPI_ISL_2961656 | Lighthouse Laboratory Plymouth | Wellcome Sanger Institute for the COVID-19 Genomics UK (COG-UK) Consortium | Lighthouse Laboratory Plymouth and Alex Alderton, Roberto Amato, Jeffrey Barrett, Sonia Goncalves, Ewan Harrison, David K. Jackson, Ian Johnston, Dominic Kwiatkowski, Cordelia Langford, John Sillitoe on behalf of the Wellcome Sanger Institute COVID-19 Surveillance Team |
| EPI_ISL_2961657 | Lighthouse Lab in Alderley Park | Wellcome Sanger Institute for the COVID-19 Genomics UK (COG-UK) Consortium | Jacquelyn Wynn, Mairead Hyland, The Lighthouse Lab in Alderley Park and Alex Alderton, Roberto Amato, Jeffrey Barrett, Sonia Goncalves, Ewan Harrison, David K. Jackson, Ian Johnston, Dominic Kwiatkowski, Cordelia Langford, John Sillitoe on behalf of the Wellcome Sanger Institute COVID-19 Surveillance Team |
| EPI_ISL_2961658, EPI_ISL_2961659 | Lighthouse Laboratory Plymouth | Wellcome Sanger Institute for the COVID-19 Genomics UK (COG-UK) Consortium | Lighthouse Laboratory Plymouth and Alex Alderton, Roberto Amato, Jeffrey Barrett, Sonia Goncalves, Ewan Harrison, David K. Jackson, Ian Johnston, Dominic Kwiatkowski, Cordelia Langford, John Sillitoe on behalf of the Wellcome Sanger Institute COVID-19 Surveillance Team |
| EPI_ISL_2961660 | Lighthouse Lab in Alderley Park | Wellcome Sanger Institute for the COVID-19 Genomics UK (COG-UK) Consortium | Jacquelyn Wynn, Mairead Hyland, The Lighthouse Lab in Alderley Park and Alex Alderton, Roberto Amato, Jeffrey Barrett, Sonia Goncalves, Ewan Harrison, David K. Jackson, Ian Johnston, Dominic Kwiatkowski, Cordelia Langford, John Sillitoe on behalf of the Wellcome Sanger Institute COVID-19 Surveillance Team |
| EPI_ISL_2961661 | Berkshire and Surrey Pathology Services Lighthouse Laboratory | Wellcome Sanger Institute for the COVID-19 Genomics UK (COG-UK) Consortium | Berkshire and Surrey Pathology Services Lighthouse Laboratory and Alex Alderton, Roberto Amato, Jeffrey Barrett, Sonia Goncalves, Ewan Harrison, David K. Jackson, Ian Johnston, Dominic Kwiatkowski, Cordelia Langford, John Sillitoe on behalf of the Wellcome Sanger Institute COVID-19 Surveillance Team |
| EPI_ISL_2961662 | Lighthouse Laboratory Plymouth | Wellcome Sanger Institute for the COVID-19 Genomics UK (COG-UK) Consortium | Lighthouse Laboratory Plymouth and Alex Alderton, Roberto Amato, Jeffrey Barrett, Sonia Goncalves, Ewan Harrison, David K. Jackson, Ian Johnston, Dominic Kwiatkowski, Cordelia Langford, John Sillitoe on behalf of the Wellcome Sanger Institute COVID-19 Surveillance Team |
| EPI_ISL_2961663 | Lighthouse Lab in Alderley Park | Wellcome Sanger Institute for the COVID-19 Genomics UK (COG-UK) Consortium | Jacquelyn Wynn, Mairead Hyland, The Lighthouse Lab in Alderley Park and Alex Alderton, Roberto Amato, Jeffrey Barrett, Sonia Goncalves, Ewan Harrison, David K. Jackson, Ian Johnston, Dominic Kwiatkowski, Cordelia Langford, John Sillitoe on behalf of the Wellcome Sanger Institute COVID-19 Surveillance Team |
| EPI_ISL_2961664 | Lighthouse Laboratory Plymouth | Wellcome Sanger Institute for the COVID-19 Genomics UK (COG-UK) Consortium | Lighthouse Laboratory Plymouth and Alex Alderton, Roberto Amato, Jeffrey Barrett, Sonia Goncalves, Ewan Harrison, David K. Jackson, Ian Johnston, Dominic Kwiatkowski, Cordelia Langford, John Sillitoe on behalf of the Wellcome Sanger Institute COVID-19 Surveillance Team |
| EPI_ISL_2961665 | Berkshire and Surrey Pathology Services Lighthouse Laboratory | Wellcome Sanger Institute for the COVID-19 Genomics UK (COG-UK) Consortium | Berkshire and Surrey Pathology Services Lighthouse Laboratory and Alex Alderton, Roberto Amato, Jeffrey Barrett, Sonia Goncalves, Ewan Harrison, David K. Jackson, Ian Johnston, Dominic Kwiatkowski, Cordelia Langford, John Sillitoe on behalf of the Wellcome Sanger Institute COVID-19 Surveillance Team |
| EPI_ISL_2961666, EPI_ISL_2961667 | Lighthouse Laboratory Plymouth | Wellcome Sanger Institute for the COVID-19 Genomics UK (COG-UK) Consortium | Lighthouse Laboratory Plymouth and Alex Alderton, Roberto Amato, Jeffrey Barrett, Sonia Goncalves, Ewan Harrison, David K. Jackson, Ian Johnston, Dominic Kwiatkowski, Cordelia Langford, John Sillitoe on behalf of the Wellcome Sanger Institute COVID-19 Surveillance Team |
| EPI_ISL_2961668 | Lighthouse Lab in Alderley Park | Wellcome Sanger Institute for the COVID-19 Genomics UK (COG-UK) Consortium | Jacquelyn Wynn, Mairead Hyland, The Lighthouse Lab in Alderley Park and Alex Alderton, Roberto Amato, Jeffrey Barrett, Sonia Goncalves, Ewan Harrison, David K. Jackson, Ian Johnston, Dominic Kwiatkowski, Cordelia Langford, John Sillitoe on behalf of the Wellcome Sanger Institute COVID-19 Surveillance Team |
| EPI_ISL_2961669 | Lighthouse Laboratory Plymouth | Wellcome Sanger Institute for the COVID-19 Genomics UK (COG-UK) Consortium | Lighthouse Laboratory Plymouth and Alex Alderton, Roberto Amato, Jeffrey Barrett, Sonia Goncalves, Ewan Harrison, David K. Jackson, Ian Johnston, Dominic Kwiatkowski, Cordelia Langford, John Sillitoe on behalf of the Wellcome Sanger Institute COVID-19 Surveillance Team |
| EPI_ISL_2961670, EPI_ISL_2961671 | Lighthouse Lab in Alderley Park | Wellcome Sanger Institute for the COVID-19 Genomics UK (COG-UK) Consortium | Jacquelyn Wynn, Mairead Hyland, The Lighthouse Lab in Alderley Park and Alex Alderton, Roberto Amato, Jeffrey Barrett, Sonia Goncalves, Ewan Harrison, David K. Jackson, Ian Johnston, Dominic Kwiatkowski, Cordelia Langford, John Sillitoe on behalf of the Wellcome Sanger Institute COVID-19 Surveillance Team |
| EPI_ISL_2961672 | Lighthouse Laboratory Plymouth | Wellcome Sanger Institute for the COVID-19 Genomics UK (COG-UK) Consortium | Lighthouse Laboratory Plymouth and Alex Alderton, Roberto Amato, Jeffrey Barrett, Sonia Goncalves, Ewan Harrison, David K. Jackson, Ian Johnston, Dominic Kwiatkowski, Cordelia Langford, John Sillitoe on behalf of the Wellcome Sanger Institute COVID-19 Surveillance Team |
| EPI_ISL_2961673, EPI_ISL_2961674 | Lighthouse Lab in Alderley Park | Wellcome Sanger Institute for the COVID-19 Genomics UK (COG-UK) Consortium | Jacquelyn Wynn, Mairead Hyland, The Lighthouse Lab in Alderley Park and Alex Alderton, Roberto Amato, Jeffrey Barrett, Sonia Goncalves, Ewan Harrison, David K. Jackson, Ian Johnston, Dominic Kwiatkowski, Cordelia Langford, John Sillitoe on behalf of the Wellcome Sanger Institute COVID-19 Surveillance Team |
| EPI_ISL_2961675, EPI_ISL_2961676, EPI_ISL_2961677 | Berkshire and Surrey Pathology Services Lighthouse Laboratory | Wellcome Sanger Institute for the COVID-19 Genomics UK (COG-UK) Consortium | Berkshire and Surrey Pathology Services Lighthouse Laboratory and Alex Alderton, Roberto Amato, Jeffrey Barrett, Sonia Goncalves, Ewan Harrison, David K. Jackson, Ian Johnston, Dominic Kwiatkowski, Cordelia Langford, John Sillitoe on behalf of the Wellcome Sanger Institute COVID-19 Surveillance Team |
| EPI_ISL_2961678 | Lighthouse Lab in Alderley Park | Wellcome Sanger Institute for the COVID-19 Genomics UK (COG-UK) Consortium | Jacquelyn Wynn, Mairead Hyland, The Lighthouse Lab in Alderley Park and Alex Alderton, Roberto Amato, Jeffrey Barrett, Sonia Goncalves, Ewan Harrison, David K. Jackson, Ian Johnston, Dominic Kwiatkowski, Cordelia Langford, John Sillitoe on behalf of the Wellcome Sanger Institute COVID-19 Surveillance Team |
| EPI_ISL_2961679 | Lighthouse Laboratory Plymouth | Wellcome Sanger Institute for the COVID-19 Genomics UK (COG-UK) Consortium | Lighthouse Laboratory Plymouth and Alex Alderton, Roberto Amato, Jeffrey Barrett, Sonia Goncalves, Ewan Harrison, David K. Jackson, Ian Johnston, Dominic Kwiatkowski, Cordelia Langford, John Sillitoe on behalf of the Wellcome Sanger Institute COVID-19 Surveillance Team |
| EPI_ISL_2961680 | Lighthouse Lab in Alderley Park | Wellcome Sanger Institute for the COVID-19 Genomics UK (COG-UK) Consortium | Jacquelyn Wynn, Mairead Hyland, The Lighthouse Lab in Alderley Park and Alex Alderton, Roberto Amato, Jeffrey Barrett, Sonia Goncalves, Ewan Harrison, David K. Jackson, Ian Johnston, Dominic Kwiatkowski, Cordelia Langford, John Sillitoe on behalf of the Wellcome Sanger Institute COVID-19 Surveillance Team |
| EPI_ISL_2961681, EPI_ISL_2961682, | Lighthouse Laboratory Plymouth | Wellcome Sanger Institute for the COVID-19 Genomics UK (COG-UK) Consortium | Lighthouse Laboratory Plymouth and Alex Alderton, Roberto Amato, Jeffrey Barrett, Sonia Goncalves, Ewan Harrison, David K. Jackson, Ian Johnston, Dominic Kwiatkowski, Cordelia Langford, John Sillitoe on behalf of the Wellcome Sanger Institute COVID-19 Surveillance Team |

| | | | |
|---|---|---|---|
| EPI_ISL_2961683 | | | |
| EPI_ISL_2961684 | Berkshire and Surrey Pathology Services Lighthouse Laboratory | Wellcome Sanger Institute for the COVID-19 Genomics UK (COG-UK) Consortium | Berkshire and Surrey Pathology Services Lighthouse Laboratory and Alex Alderton, Roberto Amato, Jeffrey Barrett, Sonia Goncalves, Ewan Harrison, David K. Jackson, Ian Johnston, Dominic Kwiatkowski, Cordelia Langford, John Sillitoe on behalf of the Wellcome Sanger Institute COVID-19 Surveillance Team |
| EPI_ISL_2961685, EPI_ISL_2961686, EPI_ISL_2961687 | Lighthouse Laboratory Plymouth | Wellcome Sanger Institute for the COVID-19 Genomics UK (COG-UK) Consortium | Lighthouse Laboratory Plymouth and Alex Alderton, Roberto Amato, Jeffrey Barrett, Sonia Goncalves, Ewan Harrison, David K. Jackson, Ian Johnston, Dominic Kwiatkowski, Cordelia Langford, John Sillitoe on behalf of the Wellcome Sanger Institute COVID-19 Surveillance Team |
| EPI_ISL_2961688 | Berkshire and Surrey Pathology Services Lighthouse Laboratory | Wellcome Sanger Institute for the COVID-19 Genomics UK (COG-UK) Consortium | Berkshire and Surrey Pathology Services Lighthouse Laboratory and Alex Alderton, Roberto Amato, Jeffrey Barrett, Sonia Goncalves, Ewan Harrison, David K. Jackson, Ian Johnston, Dominic Kwiatkowski, Cordelia Langford, John Sillitoe on behalf of the Wellcome Sanger Institute COVID-19 Surveillance Team |
| EPI_ISL_2961689, EPI_ISL_2961690 | Lighthouse Lab in Alderley Park | Wellcome Sanger Institute for the COVID-19 Genomics UK (COG-UK) Consortium | Jacquelyn Wynn, Mairead Hyland, The Lighthouse Lab in Alderley Park and Alex Alderton, Roberto Amato, Jeffrey Barrett, Sonia Goncalves, Ewan Harrison, David K. Jackson, Ian Johnston, Dominic Kwiatkowski, Cordelia Langford, John Sillitoe on behalf of the Wellcome Sanger Institute COVID-19 Surveillance Team |
| EPI_ISL_2961691 | Lighthouse Laboratory Plymouth | Wellcome Sanger Institute for the COVID-19 Genomics UK (COG-UK) Consortium | Lighthouse Laboratory Plymouth and Alex Alderton, Roberto Amato, Jeffrey Barrett, Sonia Goncalves, Ewan Harrison, David K. Jackson, Ian Johnston, Dominic Kwiatkowski, Cordelia Langford, John Sillitoe on behalf of the Wellcome Sanger Institute COVID-19 Surveillance Team |
| EPI_ISL_2961692 | Lighthouse Lab in Alderley Park | Wellcome Sanger Institute for the COVID-19 Genomics UK (COG-UK) Consortium | Jacquelyn Wynn, Mairead Hyland, The Lighthouse Lab in Alderley Park and Alex Alderton, Roberto Amato, Jeffrey Barrett, Sonia Goncalves, Ewan Harrison, David K. Jackson, Ian Johnston, Dominic Kwiatkowski, Cordelia Langford, John Sillitoe on behalf of the Wellcome Sanger Institute COVID-19 Surveillance Team |
| EPI_ISL_2961693 | Lighthouse Laboratory Plymouth | Wellcome Sanger Institute for the COVID-19 Genomics UK (COG-UK) Consortium | Lighthouse Laboratory Plymouth and Alex Alderton, Roberto Amato, Jeffrey Barrett, Sonia Goncalves, Ewan Harrison, David K. Jackson, Ian Johnston, Dominic Kwiatkowski, Cordelia Langford, John Sillitoe on behalf of the Wellcome Sanger Institute COVID-19 Surveillance Team |
| EPI_ISL_2961694 | Berkshire and Surrey Pathology Services Lighthouse Laboratory | Wellcome Sanger Institute for the COVID-19 Genomics UK (COG-UK) Consortium | Berkshire and Surrey Pathology Services Lighthouse Laboratory and Alex Alderton, Roberto Amato, Jeffrey Barrett, Sonia Goncalves, Ewan Harrison, David K. Jackson, Ian Johnston, Dominic Kwiatkowski, Cordelia Langford, John Sillitoe on behalf of the Wellcome Sanger Institute COVID-19 Surveillance Team |
| EPI_ISL_2961695, EPI_ISL_2961696, EPI_ISL_2961697 | Lighthouse Laboratory Plymouth | Wellcome Sanger Institute for the COVID-19 Genomics UK (COG-UK) Consortium | Lighthouse Laboratory Plymouth and Alex Alderton, Roberto Amato, Jeffrey Barrett, Sonia Goncalves, Ewan Harrison, David K. Jackson, Ian Johnston, Dominic Kwiatkowski, Cordelia Langford, John Sillitoe on behalf of the Wellcome Sanger Institute COVID-19 Surveillance Team |
| EPI_ISL_2961698 | Berkshire and Surrey Pathology Services Lighthouse Laboratory | Wellcome Sanger Institute for the COVID-19 Genomics UK (COG-UK) Consortium | Berkshire and Surrey Pathology Services Lighthouse Laboratory and Alex Alderton, Roberto Amato, Jeffrey Barrett, Sonia Goncalves, Ewan Harrison, David K. Jackson, Ian Johnston, Dominic Kwiatkowski, Cordelia Langford, John Sillitoe on behalf of the Wellcome Sanger Institute COVID-19 Surveillance Team |
| EPI_ISL_2961699, EPI_ISL_2961700, EPI_ISL_2961701 | Lighthouse Laboratory Plymouth | Wellcome Sanger Institute for the COVID-19 Genomics UK (COG-UK) Consortium | Lighthouse Laboratory Plymouth and Alex Alderton, Roberto Amato, Jeffrey Barrett, Sonia Goncalves, Ewan Harrison, David K. Jackson, Ian Johnston, Dominic Kwiatkowski, Cordelia Langford, John Sillitoe on behalf of the Wellcome Sanger Institute COVID-19 Surveillance Team |
| EPI_ISL_2961702 | Berkshire and Surrey Pathology Services Lighthouse Laboratory | Wellcome Sanger Institute for the COVID-19 Genomics UK (COG-UK) Consortium | Berkshire and Surrey Pathology Services Lighthouse Laboratory and Alex Alderton, Roberto Amato, Jeffrey Barrett, Sonia Goncalves, Ewan Harrison, David K. Jackson, Ian Johnston, Dominic Kwiatkowski, Cordelia Langford, John Sillitoe on behalf of the Wellcome Sanger Institute COVID-19 Surveillance Team |
| EPI_ISL_2961703, EPI_ISL_2961704 | Lighthouse Laboratory Plymouth | Wellcome Sanger Institute for the COVID-19 Genomics UK (COG-UK) Consortium | Lighthouse Laboratory Plymouth and Alex Alderton, Roberto Amato, Jeffrey Barrett, Sonia Goncalves, Ewan Harrison, David K. Jackson, Ian Johnston, Dominic Kwiatkowski, Cordelia Langford, John Sillitoe on behalf of the Wellcome Sanger Institute COVID-19 Surveillance Team |
| EPI_ISL_2961705, EPI_ISL_2961706 | Berkshire and Surrey Pathology Services Lighthouse Laboratory | Wellcome Sanger Institute for the COVID-19 Genomics UK (COG-UK) Consortium | Berkshire and Surrey Pathology Services Lighthouse Laboratory and Alex Alderton, Roberto Amato, Jeffrey Barrett, Sonia Goncalves, Ewan Harrison, David K. Jackson, Ian Johnston, Dominic Kwiatkowski, Cordelia Langford, John Sillitoe on behalf of the Wellcome Sanger Institute COVID-19 Surveillance Team |
| EPI_ISL_2961707, EPI_ISL_2961708 | Lighthouse Laboratory Plymouth | Wellcome Sanger Institute for the COVID-19 Genomics UK (COG-UK) Consortium | Lighthouse Laboratory Plymouth and Alex Alderton, Roberto Amato, Jeffrey Barrett, Sonia Goncalves, Ewan Harrison, David K. Jackson, Ian Johnston, Dominic Kwiatkowski, Cordelia Langford, John Sillitoe on behalf of the Wellcome Sanger Institute COVID-19 Surveillance Team |
| EPI_ISL_2961709 | Lighthouse Lab in Alderley Park | Wellcome Sanger Institute for the COVID-19 Genomics UK (COG-UK) Consortium | Jacquelyn Wynn, Mairead Hyland, The Lighthouse Lab in Alderley Park and Alex Alderton, Roberto Amato, Jeffrey Barrett, Sonia Goncalves, Ewan Harrison, David K. Jackson, Ian Johnston, Dominic Kwiatkowski, Cordelia Langford, John Sillitoe on behalf of the Wellcome Sanger Institute COVID-19 Surveillance Team |
| EPI_ISL_2961710 | Berkshire and Surrey Pathology Services Lighthouse Laboratory | Wellcome Sanger Institute for the COVID-19 Genomics UK (COG-UK) Consortium | Berkshire and Surrey Pathology Services Lighthouse Laboratory and Alex Alderton, Roberto Amato, Jeffrey Barrett, Sonia Goncalves, Ewan Harrison, David K. Jackson, Ian Johnston, Dominic Kwiatkowski, Cordelia Langford, John Sillitoe on behalf of the Wellcome Sanger Institute COVID-19 Surveillance Team |
| EPI_ISL_2961711 | Lighthouse Lab in Alderley Park | Wellcome Sanger Institute for the COVID-19 Genomics UK (COG-UK) Consortium | Jacquelyn Wynn, Mairead Hyland, The Lighthouse Lab in Alderley Park and Alex Alderton, Roberto Amato, Jeffrey Barrett, Sonia Goncalves, Ewan Harrison, David K. Jackson, Ian Johnston, Dominic Kwiatkowski, Cordelia Langford, John Sillitoe on behalf of the Wellcome Sanger Institute COVID-19 Surveillance Team |
| EPI_ISL_2961712 | Lighthouse Laboratory Plymouth | Wellcome Sanger Institute for the COVID-19 Genomics UK (COG-UK) Consortium | Lighthouse Laboratory Plymouth and Alex Alderton, Roberto Amato, Jeffrey Barrett, Sonia Goncalves, Ewan Harrison, David K. Jackson, Ian Johnston, Dominic Kwiatkowski, Cordelia Langford, John Sillitoe on behalf of the Wellcome Sanger Institute COVID-19 Surveillance Team |
| EPI_ISL_2961713 | Lighthouse Lab in Alderley Park | Wellcome Sanger Institute for the COVID-19 Genomics UK (COG-UK) Consortium | Jacquelyn Wynn, Mairead Hyland, The Lighthouse Lab in Alderley Park and Alex Alderton, Roberto Amato, Jeffrey Barrett, Sonia Goncalves, Ewan Harrison, David K. Jackson, Ian Johnston, Dominic Kwiatkowski, Cordelia Langford, John Sillitoe on behalf of the Wellcome Sanger Institute COVID-19 Surveillance Team |
| EPI_ISL_2961714 | Berkshire and Surrey Pathology Services Lighthouse Laboratory | Wellcome Sanger Institute for the COVID-19 Genomics UK (COG-UK) Consortium | Berkshire and Surrey Pathology Services Lighthouse Laboratory and Alex Alderton, Roberto Amato, Jeffrey Barrett, Sonia Goncalves, Ewan Harrison, David K. Jackson, Ian Johnston, Dominic Kwiatkowski, Cordelia Langford, John Sillitoe on behalf of the Wellcome Sanger Institute COVID-19 Surveillance Team |
| EPI_ISL_2961715 | Lighthouse Lab in Alderley Park | Wellcome Sanger Institute for the COVID-19 Genomics UK (COG-UK) Consortium | Jacquelyn Wynn, Mairead Hyland, The Lighthouse Lab in Alderley Park and Alex Alderton, Roberto Amato, Jeffrey Barrett, Sonia Goncalves, Ewan Harrison, David K. Jackson, Ian Johnston, Dominic Kwiatkowski, Cordelia Langford, John Sillitoe on behalf of the Wellcome Sanger Institute COVID-19 Surveillance Team |
| EPI_ISL_2961716, EPI_ISL_2961717, EPI_ISL_2961718 | Lighthouse Laboratory Plymouth | Wellcome Sanger Institute for the COVID-19 Genomics UK (COG-UK) Consortium | Lighthouse Laboratory Plymouth and Alex Alderton, Roberto Amato, Jeffrey Barrett, Sonia Goncalves, Ewan Harrison, David K. Jackson, Ian Johnston, Dominic Kwiatkowski, Cordelia Langford, John Sillitoe on behalf of the Wellcome Sanger Institute COVID-19 Surveillance Team |
| EPI_ISL_2961719 | Berkshire and Surrey Pathology Services Lighthouse Laboratory | Wellcome Sanger Institute for the COVID-19 Genomics UK (COG-UK) Consortium | Berkshire and Surrey Pathology Services Lighthouse Laboratory and Alex Alderton, Roberto Amato, Jeffrey Barrett, Sonia Goncalves, Ewan Harrison, David K. Jackson, Ian Johnston, Dominic Kwiatkowski, Cordelia Langford, John Sillitoe on behalf of the Wellcome Sanger Institute COVID-19 Surveillance Team |
| EPI_ISL_2961720 | Lighthouse Laboratory Plymouth | Wellcome Sanger Institute for the COVID-19 Genomics UK (COG-UK) Consortium | Lighthouse Laboratory Plymouth and Alex Alderton, Roberto Amato, Jeffrey Barrett, Sonia Goncalves, Ewan Harrison, David K. Jackson, Ian Johnston, Dominic Kwiatkowski, Cordelia Langford, John Sillitoe on behalf of the Wellcome Sanger Institute COVID-19 Surveillance Team |
| EPI_ISL_2961721 | Lighthouse Lab in Alderley Park | Wellcome Sanger Institute for the COVID-19 Genomics UK (COG-UK) Consortium | Jacquelyn Wynn, Mairead Hyland, The Lighthouse Lab in Alderley Park and Alex Alderton, Roberto Amato, Jeffrey Barrett, Sonia Goncalves, Ewan Harrison, David K. Jackson, Ian Johnston, Dominic Kwiatkowski, Cordelia Langford, John Sillitoe on behalf of the Wellcome Sanger Institute COVID-19 Surveillance Team |
| EPI_ISL_2961722 | Berkshire and Surrey Pathology Services Lighthouse Laboratory | Wellcome Sanger Institute for the COVID-19 Genomics UK (COG-UK) Consortium | Berkshire and Surrey Pathology Services Lighthouse Laboratory and Alex Alderton, Roberto Amato, Jeffrey Barrett, Sonia Goncalves, Ewan Harrison, David K. Jackson, Ian Johnston, Dominic Kwiatkowski, Cordelia Langford, John Sillitoe on behalf of the Wellcome Sanger Institute COVID-19 Surveillance Team |
| EPI_ISL_2961723 | Lighthouse Lab in Alderley Park | Wellcome Sanger Institute for the COVID-19 Genomics UK (COG-UK) Consortium | Jacquelyn Wynn, Mairead Hyland, The Lighthouse Lab in Alderley Park and Alex Alderton, Roberto Amato, Jeffrey Barrett, Sonia Goncalves, Ewan Harrison, David K. Jackson, Ian Johnston, Dominic Kwiatkowski, Cordelia Langford, John Sillitoe on behalf of the Wellcome Sanger Institute COVID-19 Surveillance Team |
| EPI_ISL_2961724, EPI_ISL_2961725, EPI_ISL_2961726, EPI_ISL_2961727 | Lighthouse Laboratory Plymouth | Wellcome Sanger Institute for the COVID-19 Genomics UK (COG-UK) Consortium | Lighthouse Laboratory Plymouth and Alex Alderton, Roberto Amato, Jeffrey Barrett, Sonia Goncalves, Ewan Harrison, David K. Jackson, Ian Johnston, Dominic Kwiatkowski, Cordelia Langford, John Sillitoe on behalf of the Wellcome Sanger Institute COVID-19 Surveillance Team |
| EPI_ISL_2961728 | Berkshire and Surrey Pathology Services Lighthouse Laboratory | Wellcome Sanger Institute for the COVID-19 Genomics UK (COG-UK) Consortium | Berkshire and Surrey Pathology Services Lighthouse Laboratory and Alex Alderton, Roberto Amato, Jeffrey Barrett, Sonia Goncalves, Ewan Harrison, David K. Jackson, Ian Johnston, Dominic Kwiatkowski, Cordelia Langford, John Sillitoe on behalf of the Wellcome Sanger Institute COVID-19 Surveillance Team |
| EPI_ISL_2961729, EPI_ISL_2961730, EPI_ISL_2961731, EPI_ISL_2961732, EPI_ISL_2961733, EPI_ISL_2961734 | Lighthouse Laboratory Plymouth | Wellcome Sanger Institute for the COVID-19 Genomics UK (COG-UK) Consortium | Lighthouse Laboratory Plymouth and Alex Alderton, Roberto Amato, Jeffrey Barrett, Sonia Goncalves, Ewan Harrison, David K. Jackson, Ian Johnston, Dominic Kwiatkowski, Cordelia Langford, John Sillitoe on behalf of the Wellcome Sanger Institute COVID-19 Surveillance Team |
| EPI_ISL_2961735 | Lighthouse Lab in Alderley Park | Wellcome Sanger Institute for the COVID-19 Genomics UK (COG-UK) Consortium | Jacquelyn Wynn, Mairead Hyland, The Lighthouse Lab in Alderley Park and Alex Alderton, Roberto Amato, Jeffrey Barrett, Sonia Goncalves, Ewan Harrison, David K. Jackson, Ian Johnston, Dominic Kwiatkowski, Cordelia Langford, John Sillitoe on behalf of the Wellcome Sanger Institute COVID-19 Surveillance Team |
| EPI_ISL_2961736, EPI_ISL_2961737 | Berkshire and Surrey Pathology Services Lighthouse Laboratory | Wellcome Sanger Institute for the COVID-19 Genomics UK (COG-UK) Consortium | Berkshire and Surrey Pathology Services Lighthouse Laboratory and Alex Alderton, Roberto Amato, Jeffrey Barrett, Sonia Goncalves, Ewan Harrison, David K. Jackson, Ian Johnston, Dominic Kwiatkowski, Cordelia Langford, John Sillitoe on behalf of the Wellcome Sanger Institute COVID-19 Surveillance Team |
| EPI_ISL_2961738 | Lighthouse Lab in Alderley Park | Wellcome Sanger Institute for the COVID-19 Genomics UK (COG-UK) Consortium | Jacquelyn Wynn, Mairead Hyland, The Lighthouse Lab in Alderley Park and Alex Alderton, Roberto Amato, Jeffrey Barrett, Sonia Goncalves, Ewan Harrison, David K. Jackson, Ian Johnston, Dominic Kwiatkowski, Cordelia Langford, John Sillitoe on behalf of the Wellcome Sanger Institute COVID-19 Surveillance Team |
| EPI_ISL_2961739, EPI_ISL_2961740, EPI_ISL_2961741, EPI_ISL_2961742, EPI_ISL_2961743 | Lighthouse Laboratory Plymouth | Wellcome Sanger Institute for the COVID-19 Genomics UK (COG-UK) Consortium | Lighthouse Laboratory Plymouth and Alex Alderton, Roberto Amato, Jeffrey Barrett, Sonia Goncalves, Ewan Harrison, David K. Jackson, Ian Johnston, Dominic Kwiatkowski, Cordelia Langford, John Sillitoe on behalf of the Wellcome Sanger Institute COVID-19 Surveillance Team |
| EPI_ISL_2961744 | Berkshire and Surrey Pathology Services Lighthouse Laboratory | Wellcome Sanger Institute for the COVID-19 Genomics UK (COG-UK) Consortium | Berkshire and Surrey Pathology Services Lighthouse Laboratory and Alex Alderton, Roberto Amato, Jeffrey Barrett, Sonia Goncalves, Ewan Harrison, David K. Jackson, Ian Johnston, Dominic Kwiatkowski, Cordelia Langford, John Sillitoe on behalf of the Wellcome Sanger Institute COVID-19 Surveillance Team |
| EPI_ISL_2961745 | Lighthouse Laboratory Plymouth | Wellcome Sanger Institute for the COVID-19 Genomics UK (COG-UK) Consortium | Lighthouse Laboratory Plymouth and Alex Alderton, Roberto Amato, Jeffrey Barrett, Sonia Goncalves, Ewan Harrison, David K. Jackson, Ian Johnston, Dominic Kwiatkowski, Cordelia Langford, John Sillitoe on behalf of the Wellcome Sanger Institute COVID-19 Surveillance Team |
| EPI_ISL_2961746 | Berkshire and Surrey Pathology Services Lighthouse Laboratory | Wellcome Sanger Institute for the COVID-19 Genomics UK (COG-UK) Consortium | Berkshire and Surrey Pathology Services Lighthouse Laboratory and Alex Alderton, Roberto Amato, Jeffrey Barrett, Sonia Goncalves, Ewan Harrison, David K. Jackson, Ian Johnston, Dominic Kwiatkowski, Cordelia Langford, John Sillitoe on behalf of the Wellcome Sanger Institute COVID-19 Surveillance Team |
| EPI_ISL_2961747 | Lighthouse Laboratory Plymouth | Wellcome Sanger Institute for the COVID-19 Genomics UK (COG-UK) Consortium | Lighthouse Laboratory Plymouth and Alex Alderton, Roberto Amato, Jeffrey Barrett, Sonia Goncalves, Ewan Harrison, David K. Jackson, Ian Johnston, Dominic Kwiatkowski, Cordelia Langford, John Sillitoe on behalf of the Wellcome Sanger Institute COVID-19 Surveillance Team |
| EPI_ISL_2961748 | Berkshire and Surrey Pathology Services Lighthouse Laboratory | Wellcome Sanger Institute for the COVID-19 Genomics UK (COG-UK) | Berkshire and Surrey Pathology Services Lighthouse Laboratory and Alex Alderton, Roberto Amato, Jeffrey Barrett, Sonia Goncalves, Ewan Harrison, David K. Jackson, Ian Johnston, Dominic Kwiatkowski, Cordelia Langford, John Sillitoe on behalf |

| | | | |
|---|---|---|---|
| | | Consortium | of the Wellcome Sanger Institute COVID-19 Surveillance Team |
| EPI_ISL_2961749, EPI_ISL_2961750 | Lighthouse Laboratory Plymouth | Wellcome Sanger Institute for the COVID-19 Genomics UK (COG-UK) Consortium | Lighthouse Laboratory Plymouth and Alex Alderton, Roberto Amato, Jeffrey Barrett, Sonia Goncalves, Ewan Harrison, David K. Jackson, Ian Johnston, Dominic Kwiatkowski, Cordelia Langford, John Sillitoe on behalf of the Wellcome Sanger Institute COVID-19 Surveillance Team |
| EPI_ISL_2961751, EPI_ISL_2961752 | Lighthouse Lab in Alderley Park | Wellcome Sanger Institute for the COVID-19 Genomics UK (COG-UK) Consortium | Jacquelyn Wynn, Mairead Hyland, The Lighthouse Lab in Alderley Park and Alex Alderton, Roberto Amato, Jeffrey Barrett, Sonia Goncalves, Ewan Harrison, David K. Jackson, Ian Johnston, Dominic Kwiatkowski, Cordelia Langford, John Sillitoe on behalf of the Wellcome Sanger Institute COVID-19 Surveillance Team |
| EPI_ISL_2961753 | Berkshire and Surrey Pathology Services Lighthouse Laboratory | Wellcome Sanger Institute for the COVID-19 Genomics UK (COG-UK) Consortium | Berkshire and Surrey Pathology Services Lighthouse Laboratory and Alex Alderton, Roberto Amato, Jeffrey Barrett, Sonia Goncalves, Ewan Harrison, David K. Jackson, Ian Johnston, Dominic Kwiatkowski, Cordelia Langford, John Sillitoe on behalf of the Wellcome Sanger Institute COVID-19 Surveillance Team |
| EPI_ISL_2961754, EPI_ISL_2961755, EPI_ISL_2961756 | Lighthouse Laboratory Plymouth | Wellcome Sanger Institute for the COVID-19 Genomics UK (COG-UK) Consortium | Lighthouse Laboratory Plymouth and Alex Alderton, Roberto Amato, Jeffrey Barrett, Sonia Goncalves, Ewan Harrison, David K. Jackson, Ian Johnston, Dominic Kwiatkowski, Cordelia Langford, John Sillitoe on behalf of the Wellcome Sanger Institute COVID-19 Surveillance Team |
| EPI_ISL_2961757 | Lighthouse Lab in Alderley Park | Wellcome Sanger Institute for the COVID-19 Genomics UK (COG-UK) Consortium | Jacquelyn Wynn, Mairead Hyland, The Lighthouse Lab in Alderley Park and Alex Alderton, Roberto Amato, Jeffrey Barrett, Sonia Goncalves, Ewan Harrison, David K. Jackson, Ian Johnston, Dominic Kwiatkowski, Cordelia Langford, John Sillitoe on behalf of the Wellcome Sanger Institute COVID-19 Surveillance Team |
| EPI_ISL_2961758 | Lighthouse Laboratory Plymouth | Wellcome Sanger Institute for the COVID-19 Genomics UK (COG-UK) Consortium | Lighthouse Laboratory Plymouth and Alex Alderton, Roberto Amato, Jeffrey Barrett, Sonia Goncalves, Ewan Harrison, David K. Jackson, Ian Johnston, Dominic Kwiatkowski, Cordelia Langford, John Sillitoe on behalf of the Wellcome Sanger Institute COVID-19 Surveillance Team |
| EPI_ISL_2961759, EPI_ISL_2961760, EPI_ISL_2961761 | Berkshire and Surrey Pathology Services Lighthouse Laboratory | Wellcome Sanger Institute for the COVID-19 Genomics UK (COG-UK) Consortium | Berkshire and Surrey Pathology Services Lighthouse Laboratory and Alex Alderton, Roberto Amato, Jeffrey Barrett, Sonia Goncalves, Ewan Harrison, David K. Jackson, Ian Johnston, Dominic Kwiatkowski, Cordelia Langford, John Sillitoe on behalf of the Wellcome Sanger Institute COVID-19 Surveillance Team |
| EPI_ISL_2961762, EPI_ISL_2961763, EPI_ISL_2961764 | Lighthouse Laboratory Plymouth | Wellcome Sanger Institute for the COVID-19 Genomics UK (COG-UK) Consortium | Lighthouse Laboratory Plymouth and Alex Alderton, Roberto Amato, Jeffrey Barrett, Sonia Goncalves, Ewan Harrison, David K. Jackson, Ian Johnston, Dominic Kwiatkowski, Cordelia Langford, John Sillitoe on behalf of the Wellcome Sanger Institute COVID-19 Surveillance Team |
| EPI_ISL_2961765 | Berkshire and Surrey Pathology Services Lighthouse Laboratory | Wellcome Sanger Institute for the COVID-19 Genomics UK (COG-UK) Consortium | Berkshire and Surrey Pathology Services Lighthouse Laboratory and Alex Alderton, Roberto Amato, Jeffrey Barrett, Sonia Goncalves, Ewan Harrison, David K. Jackson, Ian Johnston, Dominic Kwiatkowski, Cordelia Langford, John Sillitoe on behalf of the Wellcome Sanger Institute COVID-19 Surveillance Team |
| EPI_ISL_2961766, EPI_ISL_2961767 | Lighthouse Laboratory Plymouth | Wellcome Sanger Institute for the COVID-19 Genomics UK (COG-UK) Consortium | Lighthouse Laboratory Plymouth and Alex Alderton, Roberto Amato, Jeffrey Barrett, Sonia Goncalves, Ewan Harrison, David K. Jackson, Ian Johnston, Dominic Kwiatkowski, Cordelia Langford, John Sillitoe on behalf of the Wellcome Sanger Institute COVID-19 Surveillance Team |
| EPI_ISL_2961768, EPI_ISL_2961769 | Lighthouse Lab in Alderley Park | Wellcome Sanger Institute for the COVID-19 Genomics UK (COG-UK) Consortium | Jacquelyn Wynn, Mairead Hyland, The Lighthouse Lab in Alderley Park and Alex Alderton, Roberto Amato, Jeffrey Barrett, Sonia Goncalves, Ewan Harrison, David K. Jackson, Ian Johnston, Dominic Kwiatkowski, Cordelia Langford, John Sillitoe on behalf of the Wellcome Sanger Institute COVID-19 Surveillance Team |
| EPI_ISL_2961770, EPI_ISL_2961771, EPI_ISL_2961772, EPI_ISL_2961773, EPI_ISL_2961774, EPI_ISL_2961775, EPI_ISL_2961776 | Lighthouse Laboratory Plymouth | Wellcome Sanger Institute for the COVID-19 Genomics UK (COG-UK) Consortium | Lighthouse Laboratory Plymouth and Alex Alderton, Roberto Amato, Jeffrey Barrett, Sonia Goncalves, Ewan Harrison, David K. Jackson, Ian Johnston, Dominic Kwiatkowski, Cordelia Langford, John Sillitoe on behalf of the Wellcome Sanger Institute COVID-19 Surveillance Team |
| EPI_ISL_2961777 | Lighthouse Lab in Alderley Park | Wellcome Sanger Institute for the COVID-19 Genomics UK (COG-UK) Consortium | Jacquelyn Wynn, Mairead Hyland, The Lighthouse Lab in Alderley Park and Alex Alderton, Roberto Amato, Jeffrey Barrett, Sonia Goncalves, Ewan Harrison, David K. Jackson, Ian Johnston, Dominic Kwiatkowski, Cordelia Langford, John Sillitoe on behalf of the Wellcome Sanger Institute COVID-19 Surveillance Team |
| EPI_ISL_2961778 | Berkshire and Surrey Pathology Services Lighthouse Laboratory | Wellcome Sanger Institute for the COVID-19 Genomics UK (COG-UK) Consortium | Berkshire and Surrey Pathology Services Lighthouse Laboratory and Alex Alderton, Roberto Amato, Jeffrey Barrett, Sonia Goncalves, Ewan Harrison, David K. Jackson, Ian Johnston, Dominic Kwiatkowski, Cordelia Langford, John Sillitoe on behalf of the Wellcome Sanger Institute COVID-19 Surveillance Team |
| EPI_ISL_2961779, EPI_ISL_2961780, EPI_ISL_2961781, EPI_ISL_2961782, EPI_ISL_2961783 | Lighthouse Laboratory Plymouth | Wellcome Sanger Institute for the COVID-19 Genomics UK (COG-UK) Consortium | Lighthouse Laboratory Plymouth and Alex Alderton, Roberto Amato, Jeffrey Barrett, Sonia Goncalves, Ewan Harrison, David K. Jackson, Ian Johnston, Dominic Kwiatkowski, Cordelia Langford, John Sillitoe on behalf of the Wellcome Sanger Institute COVID-19 Surveillance Team |
| EPI_ISL_2961784 | Lighthouse Lab in Alderley Park | Wellcome Sanger Institute for the COVID-19 Genomics UK (COG-UK) Consortium | Jacquelyn Wynn, Mairead Hyland, The Lighthouse Lab in Alderley Park and Alex Alderton, Roberto Amato, Jeffrey Barrett, Sonia Goncalves, Ewan Harrison, David K. Jackson, Ian Johnston, Dominic Kwiatkowski, Cordelia Langford, John Sillitoe on behalf of the Wellcome Sanger Institute COVID-19 Surveillance Team |
| EPI_ISL_2961785 | Lighthouse Laboratory Plymouth | Wellcome Sanger Institute for the COVID-19 Genomics UK (COG-UK) Consortium | Lighthouse Laboratory Plymouth and Alex Alderton, Roberto Amato, Jeffrey Barrett, Sonia Goncalves, Ewan Harrison, David K. Jackson, Ian Johnston, Dominic Kwiatkowski, Cordelia Langford, John Sillitoe on behalf of the Wellcome Sanger Institute COVID-19 Surveillance Team |
| EPI_ISL_2961786 | Lighthouse Lab in Alderley Park | Wellcome Sanger Institute for the COVID-19 Genomics UK (COG-UK) Consortium | Jacquelyn Wynn, Mairead Hyland, The Lighthouse Lab in Alderley Park and Alex Alderton, Roberto Amato, Jeffrey Barrett, Sonia Goncalves, Ewan Harrison, David K. Jackson, Ian Johnston, Dominic Kwiatkowski, Cordelia Langford, John Sillitoe on behalf of the Wellcome Sanger Institute COVID-19 Surveillance Team |
| EPI_ISL_2961787 | Berkshire and Surrey Pathology Services Lighthouse Laboratory | Wellcome Sanger Institute for the COVID-19 Genomics UK (COG-UK) Consortium | Berkshire and Surrey Pathology Services Lighthouse Laboratory and Alex Alderton, Roberto Amato, Jeffrey Barrett, Sonia Goncalves, Ewan Harrison, David K. Jackson, Ian Johnston, Dominic Kwiatkowski, Cordelia Langford, John Sillitoe on behalf of the Wellcome Sanger Institute COVID-19 Surveillance Team |
| EPI_ISL_2961788, EPI_ISL_2961789 | Lighthouse Laboratory Plymouth | Wellcome Sanger Institute for the COVID-19 Genomics UK (COG-UK) Consortium | Lighthouse Laboratory Plymouth and Alex Alderton, Roberto Amato, Jeffrey Barrett, Sonia Goncalves, Ewan Harrison, David K. Jackson, Ian Johnston, Dominic Kwiatkowski, Cordelia Langford, John Sillitoe on behalf of the Wellcome Sanger Institute COVID-19 Surveillance Team |
| EPI_ISL_2961790, EPI_ISL_2961791 | Berkshire and Surrey Pathology Services Lighthouse Laboratory | Wellcome Sanger Institute for the COVID-19 Genomics UK (COG-UK) Consortium | Berkshire and Surrey Pathology Services Lighthouse Laboratory and Alex Alderton, Roberto Amato, Jeffrey Barrett, Sonia Goncalves, Ewan Harrison, David K. Jackson, Ian Johnston, Dominic Kwiatkowski, Cordelia Langford, John Sillitoe on behalf of the Wellcome Sanger Institute COVID-19 Surveillance Team |
| EPI_ISL_2961792, EPI_ISL_2961793 | Lighthouse Laboratory Plymouth | Wellcome Sanger Institute for the COVID-19 Genomics UK (COG-UK) Consortium | Lighthouse Laboratory Plymouth and Alex Alderton, Roberto Amato, Jeffrey Barrett, Sonia Goncalves, Ewan Harrison, David K. Jackson, Ian Johnston, Dominic Kwiatkowski, Cordelia Langford, John Sillitoe on behalf of the Wellcome Sanger Institute COVID-19 Surveillance Team |
| EPI_ISL_2961794 | Lighthouse Lab in Alderley Park | Wellcome Sanger Institute for the COVID-19 Genomics UK (COG-UK) Consortium | Jacquelyn Wynn, Mairead Hyland, The Lighthouse Lab in Alderley Park and Alex Alderton, Roberto Amato, Jeffrey Barrett, Sonia Goncalves, Ewan Harrison, David K. Jackson, Ian Johnston, Dominic Kwiatkowski, Cordelia Langford, John Sillitoe on behalf of the Wellcome Sanger Institute COVID-19 Surveillance Team |
| EPI_ISL_2961795 | Lighthouse Laboratory Plymouth | Wellcome Sanger Institute for the COVID-19 Genomics UK (COG-UK) Consortium | Lighthouse Laboratory Plymouth and Alex Alderton, Roberto Amato, Jeffrey Barrett, Sonia Goncalves, Ewan Harrison, David K. Jackson, Ian Johnston, Dominic Kwiatkowski, Cordelia Langford, John Sillitoe on behalf of the Wellcome Sanger Institute COVID-19 Surveillance Team |
| EPI_ISL_2961796 | Berkshire and Surrey Pathology Services Lighthouse Laboratory | Wellcome Sanger Institute for the COVID-19 Genomics UK (COG-UK) Consortium | Berkshire and Surrey Pathology Services Lighthouse Laboratory and Alex Alderton, Roberto Amato, Jeffrey Barrett, Sonia Goncalves, Ewan Harrison, David K. Jackson, Ian Johnston, Dominic Kwiatkowski, Cordelia Langford, John Sillitoe on behalf of the Wellcome Sanger Institute COVID-19 Surveillance Team |
| EPI_ISL_2961797, EPI_ISL_2961798 | Lighthouse Laboratory Plymouth | Wellcome Sanger Institute for the COVID-19 Genomics UK (COG-UK) Consortium | Lighthouse Laboratory Plymouth and Alex Alderton, Roberto Amato, Jeffrey Barrett, Sonia Goncalves, Ewan Harrison, David K. Jackson, Ian Johnston, Dominic Kwiatkowski, Cordelia Langford, John Sillitoe on behalf of the Wellcome Sanger Institute COVID-19 Surveillance Team |
| EPI_ISL_2961799, EPI_ISL_2961800 | Berkshire and Surrey Pathology Services Lighthouse Laboratory | Wellcome Sanger Institute for the COVID-19 Genomics UK (COG-UK) Consortium | Berkshire and Surrey Pathology Services Lighthouse Laboratory and Alex Alderton, Roberto Amato, Jeffrey Barrett, Sonia Goncalves, Ewan Harrison, David K. Jackson, Ian Johnston, Dominic Kwiatkowski, Cordelia Langford, John Sillitoe on behalf of the Wellcome Sanger Institute COVID-19 Surveillance Team |
| EPI_ISL_2961801 | Lighthouse Laboratory Plymouth | Wellcome Sanger Institute for the COVID-19 Genomics UK (COG-UK) Consortium | Lighthouse Laboratory Plymouth and Alex Alderton, Roberto Amato, Jeffrey Barrett, Sonia Goncalves, Ewan Harrison, David K. Jackson, Ian Johnston, Dominic Kwiatkowski, Cordelia Langford, John Sillitoe on behalf of the Wellcome Sanger Institute COVID-19 Surveillance Team |
| EPI_ISL_2961802 | Lighthouse Lab in Alderley Park | Wellcome Sanger Institute for the COVID-19 Genomics UK (COG-UK) Consortium | Jacquelyn Wynn, Mairead Hyland, The Lighthouse Lab in Alderley Park and Alex Alderton, Roberto Amato, Jeffrey Barrett, Sonia Goncalves, Ewan Harrison, David K. Jackson, Ian Johnston, Dominic Kwiatkowski, Cordelia Langford, John Sillitoe on behalf of the Wellcome Sanger Institute COVID-19 Surveillance Team |
| EPI_ISL_2961803, EPI_ISL_2961804 | Lighthouse Laboratory Plymouth | Wellcome Sanger Institute for the COVID-19 Genomics UK (COG-UK) Consortium | Lighthouse Laboratory Plymouth and Alex Alderton, Roberto Amato, Jeffrey Barrett, Sonia Goncalves, Ewan Harrison, David K. Jackson, Ian Johnston, Dominic Kwiatkowski, Cordelia Langford, John Sillitoe on behalf of the Wellcome Sanger Institute COVID-19 Surveillance Team |
| EPI_ISL_2961805 | Lighthouse Lab in Alderley Park | Wellcome Sanger Institute for the COVID-19 Genomics UK (COG-UK) Consortium | Jacquelyn Wynn, Mairead Hyland, The Lighthouse Lab in Alderley Park and Alex Alderton, Roberto Amato, Jeffrey Barrett, Sonia Goncalves, Ewan Harrison, David K. Jackson, Ian Johnston, Dominic Kwiatkowski, Cordelia Langford, John Sillitoe on behalf of the Wellcome Sanger Institute COVID-19 Surveillance Team |
| EPI_ISL_2961806, EPI_ISL_2961807 | Berkshire and Surrey Pathology Services Lighthouse Laboratory | Wellcome Sanger Institute for the COVID-19 Genomics UK (COG-UK) Consortium | Berkshire and Surrey Pathology Services Lighthouse Laboratory and Alex Alderton, Roberto Amato, Jeffrey Barrett, Sonia Goncalves, Ewan Harrison, David K. Jackson, Ian Johnston, Dominic Kwiatkowski, Cordelia Langford, John Sillitoe on behalf of the Wellcome Sanger Institute COVID-19 Surveillance Team |
| EPI_ISL_2961808, EPI_ISL_2961809, EPI_ISL_2961810 | Lighthouse Laboratory Plymouth | Wellcome Sanger Institute for the COVID-19 Genomics UK (COG-UK) Consortium | Lighthouse Laboratory Plymouth and Alex Alderton, Roberto Amato, Jeffrey Barrett, Sonia Goncalves, Ewan Harrison, David K. Jackson, Ian Johnston, Dominic Kwiatkowski, Cordelia Langford, John Sillitoe on behalf of the Wellcome Sanger Institute COVID-19 Surveillance Team |
| EPI_ISL_2961811 | Lighthouse Lab in Alderley Park | Wellcome Sanger Institute for the COVID-19 Genomics UK (COG-UK) Consortium | Jacquelyn Wynn, Mairead Hyland, The Lighthouse Lab in Alderley Park and Alex Alderton, Roberto Amato, Jeffrey Barrett, Sonia Goncalves, Ewan Harrison, David K. Jackson, Ian Johnston, Dominic Kwiatkowski, Cordelia Langford, John Sillitoe on behalf of the Wellcome Sanger Institute COVID-19 Surveillance Team |
| EPI_ISL_2961812 | Lighthouse Laboratory Plymouth | Wellcome Sanger Institute for the COVID-19 Genomics UK (COG-UK) Consortium | Lighthouse Laboratory Plymouth and Alex Alderton, Roberto Amato, Jeffrey Barrett, Sonia Goncalves, Ewan Harrison, David K. Jackson, Ian Johnston, Dominic Kwiatkowski, Cordelia Langford, John Sillitoe on behalf of the Wellcome Sanger Institute COVID-19 Surveillance Team |
| EPI_ISL_2961813 | Lighthouse Lab in Alderley Park | Wellcome Sanger Institute for the COVID-19 Genomics UK (COG-UK) Consortium | Jacquelyn Wynn, Mairead Hyland, The Lighthouse Lab in Alderley Park and Alex Alderton, Roberto Amato, Jeffrey Barrett, Sonia Goncalves, Ewan Harrison, David K. Jackson, Ian Johnston, Dominic Kwiatkowski, Cordelia Langford, John Sillitoe on behalf of the Wellcome Sanger Institute COVID-19 Surveillance Team |
| EPI_ISL_2961814 | Health Services Laboratories | Wellcome Sanger Institute for the COVID-19 Genomics UK (COG-UK) Consortium | Health Services Laboratories and Alex Alderton, Roberto Amato, Jeffrey Barrett, Sonia Goncalves, Ewan Harrison, David K. Jackson, Ian Johnston, Dominic Kwiatkowski, Cordelia Langford, John Sillitoe on behalf of the Wellcome Sanger Institute COVID-19 Surveillance Team |
| EPI_ISL_2961815, EPI_ISL_2961816 | Lighthouse Lab in Alderley Park | Wellcome Sanger Institute for the COVID-19 Genomics UK (COG-UK) Consortium | Jacquelyn Wynn, Mairead Hyland, The Lighthouse Lab in Alderley Park and Alex Alderton, Roberto Amato, Jeffrey Barrett, Sonia Goncalves, Ewan Harrison, David K. Jackson, Ian Johnston, Dominic Kwiatkowski, Cordelia Langford, John Sillitoe on behalf of the Wellcome Sanger Institute COVID-19 Surveillance Team |
| EPI_ISL_2961817 | Health Services Laboratories | Wellcome Sanger Institute for the COVID-19 Genomics UK (COG-UK) Consortium | Health Services Laboratories and Alex Alderton, Roberto Amato, Jeffrey Barrett, Sonia Goncalves, Ewan Harrison, David K. Jackson, Ian Johnston, Dominic Kwiatkowski, Cordelia Langford, John Sillitoe on behalf of the Wellcome Sanger Institute COVID-19 Surveillance Team |
| EPI_ISL_2961818 | Lighthouse Lab in Alderley Park | Wellcome Sanger Institute for the COVID-19 Genomics UK (COG-UK) Consortium | Jacquelyn Wynn, Mairead Hyland, The Lighthouse Lab in Alderley Park and Alex Alderton, Roberto Amato, Jeffrey Barrett, Sonia Goncalves, Ewan Harrison, David K. Jackson, Ian Johnston, Dominic Kwiatkowski, Cordelia Langford, John Sillitoe on behalf of the Wellcome Sanger Institute COVID-19 Surveillance Team |

| | | | |
|---|---|---|---|
| EPI_ISL_2961819 | Lighthouse Laboratory Plymouth | Wellcome Sanger Institute for the COVID-19 Genomics UK (COG-UK) Consortium | Lighthouse Laboratory Plymouth and Alex Alderton, Roberto Amato, Jeffrey Barrett, Sonia Goncalves, Ewan Harrison, David K. Jackson, Ian Johnston, Dominic Kwiatkowski, Cordelia Langford, John Sillitoe on behalf of the Wellcome Sanger Institute COVID-19 Surveillance Team |
| EPI_ISL_2961820, EPI_ISL_2961821, EPI_ISL_2961822 | Lighthouse Lab in Alderley Park | Wellcome Sanger Institute for the COVID-19 Genomics UK (COG-UK) Consortium | Jacquelyn Wynn, Mairead Hyland, The Lighthouse Lab in Alderley Park and Alex Alderton, Roberto Amato, Jeffrey Barrett, Sonia Goncalves, Ewan Harrison, David K. Jackson, Ian Johnston, Dominic Kwiatkowski, Cordelia Langford, John Sillitoe on behalf of the Wellcome Sanger Institute COVID-19 Surveillance Team |
| EPI_ISL_2961823 | Lighthouse Laboratory Plymouth | Wellcome Sanger Institute for the COVID-19 Genomics UK (COG-UK) Consortium | Lighthouse Laboratory Plymouth and Alex Alderton, Roberto Amato, Jeffrey Barrett, Sonia Goncalves, Ewan Harrison, David K. Jackson, Ian Johnston, Dominic Kwiatkowski, Cordelia Langford, John Sillitoe on behalf of the Wellcome Sanger Institute COVID-19 Surveillance Team |
| EPI_ISL_2961824 | Lighthouse Lab in Alderley Park | Wellcome Sanger Institute for the COVID-19 Genomics UK (COG-UK) Consortium | Jacquelyn Wynn, Mairead Hyland, The Lighthouse Lab in Alderley Park and Alex Alderton, Roberto Amato, Jeffrey Barrett, Sonia Goncalves, Ewan Harrison, David K. Jackson, Ian Johnston, Dominic Kwiatkowski, Cordelia Langford, John Sillitoe on behalf of the Wellcome Sanger Institute COVID-19 Surveillance Team |
| EPI_ISL_2961825 | Health Services Laboratories | Wellcome Sanger Institute for the COVID-19 Genomics UK (COG-UK) Consortium | Health Services Laboratories and Alex Alderton, Roberto Amato, Jeffrey Barrett, Sonia Goncalves, Ewan Harrison, David K. Jackson, Ian Johnston, Dominic Kwiatkowski, Cordelia Langford, John Sillitoe on behalf of the Wellcome Sanger Institute COVID-19 Surveillance Team |
| EPI_ISL_2961826 | Lighthouse Laboratory Plymouth | Wellcome Sanger Institute for the COVID-19 Genomics UK (COG-UK) Consortium | Lighthouse Laboratory Plymouth and Alex Alderton, Roberto Amato, Jeffrey Barrett, Sonia Goncalves, Ewan Harrison, David K. Jackson, Ian Johnston, Dominic Kwiatkowski, Cordelia Langford, John Sillitoe on behalf of the Wellcome Sanger Institute COVID-19 Surveillance Team |
| EPI_ISL_2961827, EPI_ISL_2961828 | Lighthouse Lab in Alderley Park | Wellcome Sanger Institute for the COVID-19 Genomics UK (COG-UK) Consortium | Jacquelyn Wynn, Mairead Hyland, The Lighthouse Lab in Alderley Park and Alex Alderton, Roberto Amato, Jeffrey Barrett, Sonia Goncalves, Ewan Harrison, David K. Jackson, Ian Johnston, Dominic Kwiatkowski, Cordelia Langford, John Sillitoe on behalf of the Wellcome Sanger Institute COVID-19 Surveillance Team |
| EPI_ISL_2961829 | Health Services Laboratories | Wellcome Sanger Institute for the COVID-19 Genomics UK (COG-UK) Consortium | Health Services Laboratories and Alex Alderton, Roberto Amato, Jeffrey Barrett, Sonia Goncalves, Ewan Harrison, David K. Jackson, Ian Johnston, Dominic Kwiatkowski, Cordelia Langford, John Sillitoe on behalf of the Wellcome Sanger Institute COVID-19 Surveillance Team |
| EPI_ISL_2961830, EPI_ISL_2961831 | Lighthouse Lab in Alderley Park | Wellcome Sanger Institute for the COVID-19 Genomics UK (COG-UK) Consortium | Jacquelyn Wynn, Mairead Hyland, The Lighthouse Lab in Alderley Park and Alex Alderton, Roberto Amato, Jeffrey Barrett, Sonia Goncalves, Ewan Harrison, David K. Jackson, Ian Johnston, Dominic Kwiatkowski, Cordelia Langford, John Sillitoe on behalf of the Wellcome Sanger Institute COVID-19 Surveillance Team |
| EPI_ISL_2961832 | Lighthouse Laboratory Plymouth | Wellcome Sanger Institute for the COVID-19 Genomics UK (COG-UK) Consortium | Lighthouse Laboratory Plymouth and Alex Alderton, Roberto Amato, Jeffrey Barrett, Sonia Goncalves, Ewan Harrison, David K. Jackson, Ian Johnston, Dominic Kwiatkowski, Cordelia Langford, John Sillitoe on behalf of the Wellcome Sanger Institute COVID-19 Surveillance Team |
| EPI_ISL_2961833, EPI_ISL_2961834 | Lighthouse Lab in Alderley Park | Wellcome Sanger Institute for the COVID-19 Genomics UK (COG-UK) Consortium | Jacquelyn Wynn, Mairead Hyland, The Lighthouse Lab in Alderley Park and Alex Alderton, Roberto Amato, Jeffrey Barrett, Sonia Goncalves, Ewan Harrison, David K. Jackson, Ian Johnston, Dominic Kwiatkowski, Cordelia Langford, John Sillitoe on behalf of the Wellcome Sanger Institute COVID-19 Surveillance Team |
| EPI_ISL_2961835 | Health Services Laboratories | Wellcome Sanger Institute for the COVID-19 Genomics UK (COG-UK) Consortium | Health Services Laboratories and Alex Alderton, Roberto Amato, Jeffrey Barrett, Sonia Goncalves, Ewan Harrison, David K. Jackson, Ian Johnston, Dominic Kwiatkowski, Cordelia Langford, John Sillitoe on behalf of the Wellcome Sanger Institute COVID-19 Surveillance Team |
| EPI_ISL_2961836, EPI_ISL_2961837, EPI_ISL_2961838 | Lighthouse Lab in Alderley Park | Wellcome Sanger Institute for the COVID-19 Genomics UK (COG-UK) Consortium | Jacquelyn Wynn, Mairead Hyland, The Lighthouse Lab in Alderley Park and Alex Alderton, Roberto Amato, Jeffrey Barrett, Sonia Goncalves, Ewan Harrison, David K. Jackson, Ian Johnston, Dominic Kwiatkowski, Cordelia Langford, John Sillitoe on behalf of the Wellcome Sanger Institute COVID-19 Surveillance Team |
| EPI_ISL_2961839 | Lighthouse Laboratory Plymouth | Wellcome Sanger Institute for the COVID-19 Genomics UK (COG-UK) Consortium | Lighthouse Laboratory Plymouth and Alex Alderton, Roberto Amato, Jeffrey Barrett, Sonia Goncalves, Ewan Harrison, David K. Jackson, Ian Johnston, Dominic Kwiatkowski, Cordelia Langford, John Sillitoe on behalf of the Wellcome Sanger Institute COVID-19 Surveillance Team |
| EPI_ISL_2961840 | Lighthouse Lab in Alderley Park | Wellcome Sanger Institute for the COVID-19 Genomics UK (COG-UK) Consortium | Jacquelyn Wynn, Mairead Hyland, The Lighthouse Lab in Alderley Park and Alex Alderton, Roberto Amato, Jeffrey Barrett, Sonia Goncalves, Ewan Harrison, David K. Jackson, Ian Johnston, Dominic Kwiatkowski, Cordelia Langford, John Sillitoe on behalf of the Wellcome Sanger Institute COVID-19 Surveillance Team |
| EPI_ISL_2961841 | Lighthouse Laboratory Plymouth | Wellcome Sanger Institute for the COVID-19 Genomics UK (COG-UK) Consortium | Lighthouse Laboratory Plymouth and Alex Alderton, Roberto Amato, Jeffrey Barrett, Sonia Goncalves, Ewan Harrison, David K. Jackson, Ian Johnston, Dominic Kwiatkowski, Cordelia Langford, John Sillitoe on behalf of the Wellcome Sanger Institute COVID-19 Surveillance Team |
| EPI_ISL_2961842, EPI_ISL_2961843 | Lighthouse Lab in Alderley Park | Wellcome Sanger Institute for the COVID-19 Genomics UK (COG-UK) Consortium | Jacquelyn Wynn, Mairead Hyland, The Lighthouse Lab in Alderley Park and Alex Alderton, Roberto Amato, Jeffrey Barrett, Sonia Goncalves, Ewan Harrison, David K. Jackson, Ian Johnston, Dominic Kwiatkowski, Cordelia Langford, John Sillitoe on behalf of the Wellcome Sanger Institute COVID-19 Surveillance Team |
| EPI_ISL_2961844, EPI_ISL_2961845 | Lighthouse Laboratory Plymouth | Wellcome Sanger Institute for the COVID-19 Genomics UK (COG-UK) Consortium | Lighthouse Laboratory Plymouth and Alex Alderton, Roberto Amato, Jeffrey Barrett, Sonia Goncalves, Ewan Harrison, David K. Jackson, Ian Johnston, Dominic Kwiatkowski, Cordelia Langford, John Sillitoe on behalf of the Wellcome Sanger Institute COVID-19 Surveillance Team |
| EPI_ISL_2961846 | Health Services Laboratories | Wellcome Sanger Institute for the COVID-19 Genomics UK (COG-UK) Consortium | Health Services Laboratories and Alex Alderton, Roberto Amato, Jeffrey Barrett, Sonia Goncalves, Ewan Harrison, David K. Jackson, Ian Johnston, Dominic Kwiatkowski, Cordelia Langford, John Sillitoe on behalf of the Wellcome Sanger Institute COVID-19 Surveillance Team |
| EPI_ISL_2961847 | Lighthouse Lab in Alderley Park | Wellcome Sanger Institute for the COVID-19 Genomics UK (COG-UK) Consortium | Jacquelyn Wynn, Mairead Hyland, The Lighthouse Lab in Alderley Park and Alex Alderton, Roberto Amato, Jeffrey Barrett, Sonia Goncalves, Ewan Harrison, David K. Jackson, Ian Johnston, Dominic Kwiatkowski, Cordelia Langford, John Sillitoe on behalf of the Wellcome Sanger Institute COVID-19 Surveillance Team |
| EPI_ISL_2961848, EPI_ISL_2961849 | Health Services Laboratories | Wellcome Sanger Institute for the COVID-19 Genomics UK (COG-UK) Consortium | Health Services Laboratories and Alex Alderton, Roberto Amato, Jeffrey Barrett, Sonia Goncalves, Ewan Harrison, David K. Jackson, Ian Johnston, Dominic Kwiatkowski, Cordelia Langford, John Sillitoe on behalf of the Wellcome Sanger Institute COVID-19 Surveillance Team |
| EPI_ISL_2961850 | Lighthouse Lab in Alderley Park | Wellcome Sanger Institute for the COVID-19 Genomics UK (COG-UK) Consortium | Jacquelyn Wynn, Mairead Hyland, The Lighthouse Lab in Alderley Park and Alex Alderton, Roberto Amato, Jeffrey Barrett, Sonia Goncalves, Ewan Harrison, David K. Jackson, Ian Johnston, Dominic Kwiatkowski, Cordelia Langford, John Sillitoe on behalf of the Wellcome Sanger Institute COVID-19 Surveillance Team |
| EPI_ISL_2961851, EPI_ISL_2961852 | Lighthouse Laboratory Plymouth | Wellcome Sanger Institute for the COVID-19 Genomics UK (COG-UK) Consortium | Lighthouse Laboratory Plymouth and Alex Alderton, Roberto Amato, Jeffrey Barrett, Sonia Goncalves, Ewan Harrison, David K. Jackson, Ian Johnston, Dominic Kwiatkowski, Cordelia Langford, John Sillitoe on behalf of the Wellcome Sanger Institute COVID-19 Surveillance Team |
| EPI_ISL_2961853, EPI_ISL_2961854 | Lighthouse Lab in Alderley Park | Wellcome Sanger Institute for the COVID-19 Genomics UK (COG-UK) Consortium | Jacquelyn Wynn, Mairead Hyland, The Lighthouse Lab in Alderley Park and Alex Alderton, Roberto Amato, Jeffrey Barrett, Sonia Goncalves, Ewan Harrison, David K. Jackson, Ian Johnston, Dominic Kwiatkowski, Cordelia Langford, John Sillitoe on behalf of the Wellcome Sanger Institute COVID-19 Surveillance Team |
| EPI_ISL_2961855 | Health Services Laboratories | Wellcome Sanger Institute for the COVID-19 Genomics UK (COG-UK) Consortium | Health Services Laboratories and Alex Alderton, Roberto Amato, Jeffrey Barrett, Sonia Goncalves, Ewan Harrison, David K. Jackson, Ian Johnston, Dominic Kwiatkowski, Cordelia Langford, John Sillitoe on behalf of the Wellcome Sanger Institute COVID-19 Surveillance Team |
| EPI_ISL_2961856 | Lighthouse Lab in Alderley Park | Wellcome Sanger Institute for the COVID-19 Genomics UK (COG-UK) Consortium | Jacquelyn Wynn, Mairead Hyland, The Lighthouse Lab in Alderley Park and Alex Alderton, Roberto Amato, Jeffrey Barrett, Sonia Goncalves, Ewan Harrison, David K. Jackson, Ian Johnston, Dominic Kwiatkowski, Cordelia Langford, John Sillitoe on behalf of the Wellcome Sanger Institute COVID-19 Surveillance Team |
| EPI_ISL_2961857 | Health Services Laboratories | Wellcome Sanger Institute for the COVID-19 Genomics UK (COG-UK) Consortium | Health Services Laboratories and Alex Alderton, Roberto Amato, Jeffrey Barrett, Sonia Goncalves, Ewan Harrison, David K. Jackson, Ian Johnston, Dominic Kwiatkowski, Cordelia Langford, John Sillitoe on behalf of the Wellcome Sanger Institute COVID-19 Surveillance Team |
| EPI_ISL_2961858 | Lighthouse Laboratory Plymouth | Wellcome Sanger Institute for the COVID-19 Genomics UK (COG-UK) Consortium | Lighthouse Laboratory Plymouth and Alex Alderton, Roberto Amato, Jeffrey Barrett, Sonia Goncalves, Ewan Harrison, David K. Jackson, Ian Johnston, Dominic Kwiatkowski, Cordelia Langford, John Sillitoe on behalf of the Wellcome Sanger Institute COVID-19 Surveillance Team |
| EPI_ISL_2961859 | Health Services Laboratories | Wellcome Sanger Institute for the COVID-19 Genomics UK (COG-UK) Consortium | Health Services Laboratories and Alex Alderton, Roberto Amato, Jeffrey Barrett, Sonia Goncalves, Ewan Harrison, David K. Jackson, Ian Johnston, Dominic Kwiatkowski, Cordelia Langford, John Sillitoe on behalf of the Wellcome Sanger Institute COVID-19 Surveillance Team |
| EPI_ISL_2961860, EPI_ISL_2961861 | Lighthouse Lab in Alderley Park | Wellcome Sanger Institute for the COVID-19 Genomics UK (COG-UK) Consortium | Jacquelyn Wynn, Mairead Hyland, The Lighthouse Lab in Alderley Park and Alex Alderton, Roberto Amato, Jeffrey Barrett, Sonia Goncalves, Ewan Harrison, David K. Jackson, Ian Johnston, Dominic Kwiatkowski, Cordelia Langford, John Sillitoe on behalf of the Wellcome Sanger Institute COVID-19 Surveillance Team |
| EPI_ISL_2961862 | Health Services Laboratories | Wellcome Sanger Institute for the COVID-19 Genomics UK (COG-UK) Consortium | Health Services Laboratories and Alex Alderton, Roberto Amato, Jeffrey Barrett, Sonia Goncalves, Ewan Harrison, David K. Jackson, Ian Johnston, Dominic Kwiatkowski, Cordelia Langford, John Sillitoe on behalf of the Wellcome Sanger Institute COVID-19 Surveillance Team |
| EPI_ISL_2961863 | Lighthouse Lab in Alderley Park | Wellcome Sanger Institute for the COVID-19 Genomics UK (COG-UK) Consortium | Jacquelyn Wynn, Mairead Hyland, The Lighthouse Lab in Alderley Park and Alex Alderton, Roberto Amato, Jeffrey Barrett, Sonia Goncalves, Ewan Harrison, David K. Jackson, Ian Johnston, Dominic Kwiatkowski, Cordelia Langford, John Sillitoe on behalf of the Wellcome Sanger Institute COVID-19 Surveillance Team |
| EPI_ISL_2961864 | Lighthouse Laboratory Plymouth | Wellcome Sanger Institute for the COVID-19 Genomics UK (COG-UK) Consortium | Lighthouse Laboratory Plymouth and Alex Alderton, Roberto Amato, Jeffrey Barrett, Sonia Goncalves, Ewan Harrison, David K. Jackson, Ian Johnston, Dominic Kwiatkowski, Cordelia Langford, John Sillitoe on behalf of the Wellcome Sanger Institute COVID-19 Surveillance Team |
| EPI_ISL_2961865 | Lighthouse Lab in Alderley Park | Wellcome Sanger Institute for the COVID-19 Genomics UK (COG-UK) Consortium | Jacquelyn Wynn, Mairead Hyland, The Lighthouse Lab in Alderley Park and Alex Alderton, Roberto Amato, Jeffrey Barrett, Sonia Goncalves, Ewan Harrison, David K. Jackson, Ian Johnston, Dominic Kwiatkowski, Cordelia Langford, John Sillitoe on behalf of the Wellcome Sanger Institute COVID-19 Surveillance Team |
| EPI_ISL_2961866 | Lighthouse Laboratory Plymouth | Wellcome Sanger Institute for the COVID-19 Genomics UK (COG-UK) Consortium | Lighthouse Laboratory Plymouth and Alex Alderton, Roberto Amato, Jeffrey Barrett, Sonia Goncalves, Ewan Harrison, David K. Jackson, Ian Johnston, Dominic Kwiatkowski, Cordelia Langford, John Sillitoe on behalf of the Wellcome Sanger Institute COVID-19 Surveillance Team |
| EPI_ISL_2961867 | Lighthouse Lab in Alderley Park | Wellcome Sanger Institute for the COVID-19 Genomics UK (COG-UK) Consortium | Jacquelyn Wynn, Mairead Hyland, The Lighthouse Lab in Alderley Park and Alex Alderton, Roberto Amato, Jeffrey Barrett, Sonia Goncalves, Ewan Harrison, David K. Jackson, Ian Johnston, Dominic Kwiatkowski, Cordelia Langford, John Sillitoe on behalf of the Wellcome Sanger Institute COVID-19 Surveillance Team |
| EPI_ISL_2961868 | Health Services Laboratories | Wellcome Sanger Institute for the COVID-19 Genomics UK (COG-UK) Consortium | Health Services Laboratories and Alex Alderton, Roberto Amato, Jeffrey Barrett, Sonia Goncalves, Ewan Harrison, David K. Jackson, Ian Johnston, Dominic Kwiatkowski, Cordelia Langford, John Sillitoe on behalf of the Wellcome Sanger Institute COVID-19 Surveillance Team |
| EPI_ISL_2961869 | Lighthouse Laboratory Plymouth | Wellcome Sanger Institute for the COVID-19 Genomics UK (COG-UK) Consortium | Lighthouse Laboratory Plymouth and Alex Alderton, Roberto Amato, Jeffrey Barrett, Sonia Goncalves, Ewan Harrison, David K. Jackson, Ian Johnston, Dominic Kwiatkowski, Cordelia Langford, John Sillitoe on behalf of the Wellcome Sanger Institute COVID-19 Surveillance Team |
| EPI_ISL_2961870, EPI_ISL_2961871, EPI_ISL_2961872 | Health Services Laboratories | Wellcome Sanger Institute for the COVID-19 Genomics UK (COG-UK) Consortium | Health Services Laboratories and Alex Alderton, Roberto Amato, Jeffrey Barrett, Sonia Goncalves, Ewan Harrison, David K. Jackson, Ian Johnston, Dominic Kwiatkowski, Cordelia Langford, John Sillitoe on behalf of the Wellcome Sanger Institute COVID-19 Surveillance Team |
| EPI_ISL_2961873 | Lighthouse Laboratory Plymouth | Wellcome Sanger Institute for the COVID-19 Genomics UK (COG-UK) Consortium | Lighthouse Laboratory Plymouth and Alex Alderton, Roberto Amato, Jeffrey Barrett, Sonia Goncalves, Ewan Harrison, David K. Jackson, Ian Johnston, Dominic Kwiatkowski, Cordelia Langford, John Sillitoe on behalf of the Wellcome Sanger Institute COVID-19 Surveillance Team |
| EPI_ISL_2961874, EPI_ISL_2961875, EPI_ISL_2961876 | Lighthouse Lab in Alderley Park | Wellcome Sanger Institute for the COVID-19 Genomics UK (COG-UK) Consortium | Jacquelyn Wynn, Mairead Hyland, The Lighthouse Lab in Alderley Park and Alex Alderton, Roberto Amato, Jeffrey Barrett, Sonia Goncalves, Ewan Harrison, David K. Jackson, Ian Johnston, Dominic Kwiatkowski, Cordelia Langford, John Sillitoe on behalf of the Wellcome Sanger Institute COVID-19 Surveillance Team |
| EPI_ISL_2961877, EPI_ISL_2961878 | Health Services Laboratories | Wellcome Sanger Institute for the COVID-19 Genomics UK (COG-UK) Consortium | Health Services Laboratories and Alex Alderton, Roberto Amato, Jeffrey Barrett, Sonia Goncalves, Ewan Harrison, David K. Jackson, Ian Johnston, Dominic Kwiatkowski, Cordelia Langford, John Sillitoe on behalf of the Wellcome Sanger Institute COVID-19 Surveillance Team |
| EPI_ISL_2961879 | Lighthouse Lab in Alderley Park | Wellcome Sanger Institute for the COVID-19 Genomics UK (COG-UK) Consortium | Jacquelyn Wynn, Mairead Hyland, The Lighthouse Lab in Alderley Park and Alex Alderton, Roberto Amato, Jeffrey Barrett, Sonia Goncalves, Ewan Harrison, David K. Jackson, Ian Johnston, Dominic Kwiatkowski, Cordelia Langford, John Sillitoe on behalf of the Wellcome Sanger Institute COVID-19 Surveillance Team |
| EPI_ISL_2961880 | Health Services Laboratories | Wellcome Sanger Institute for the COVID-19 Genomics UK (COG-UK) | Health Services Laboratories and Alex Alderton, Roberto Amato, Jeffrey Barrett, Sonia Goncalves, Ewan Harrison, David K. Jackson, Ian Johnston, Dominic Kwiatkowski, Cordelia Langford, John Sillitoe on behalf of the Wellcome Sanger Institute |

| | | | |
|---|---|---|---|
| | | Consortium | COVID-19 Surveillance Team |
| EPI_ISL_2961881 | Lighthouse Lab in Alderley Park | Wellcome Sanger Institute for the COVID-19 Genomics UK (COG-UK) Consortium | Jacquelyn Wynn, Mairead Hyland, The Lighthouse Lab in Alderley Park and Alex Alderton, Roberto Amato, Jeffrey Barrett, Sonia Goncalves, Ewan Harrison, David K. Jackson, Ian Johnston, Dominic Kwiatkowski, Cordelia Langford, John Sillitoe on behalf of the Wellcome Sanger Institute COVID-19 Surveillance Team |
| EPI_ISL_2961882 | Lighthouse Laboratory Plymouth | Wellcome Sanger Institute for the COVID-19 Genomics UK (COG-UK) Consortium | Lighthouse Laboratory Plymouth and Alex Alderton, Roberto Amato, Jeffrey Barrett, Sonia Goncalves, Ewan Harrison, David K. Jackson, Ian Johnston, Dominic Kwiatkowski, Cordelia Langford, John Sillitoe on behalf of the Wellcome Sanger Institute COVID-19 Surveillance Team |
| EPI_ISL_2961883 | Lighthouse Lab in Alderley Park | Wellcome Sanger Institute for the COVID-19 Genomics UK (COG-UK) Consortium | Jacquelyn Wynn, Mairead Hyland, The Lighthouse Lab in Alderley Park and Alex Alderton, Roberto Amato, Jeffrey Barrett, Sonia Goncalves, Ewan Harrison, David K. Jackson, Ian Johnston, Dominic Kwiatkowski, Cordelia Langford, John Sillitoe on behalf of the Wellcome Sanger Institute COVID-19 Surveillance Team |
| EPI_ISL_2961884 | Lighthouse Laboratory Plymouth | Wellcome Sanger Institute for the COVID-19 Genomics UK (COG-UK) Consortium | Lighthouse Laboratory Plymouth and Alex Alderton, Roberto Amato, Jeffrey Barrett, Sonia Goncalves, Ewan Harrison, David K. Jackson, Ian Johnston, Dominic Kwiatkowski, Cordelia Langford, John Sillitoe on behalf of the Wellcome Sanger Institute COVID-19 Surveillance Team |
| EPI_ISL_2961885, EPI_ISL_2961886 | Lighthouse Lab in Alderley Park | Wellcome Sanger Institute for the COVID-19 Genomics UK (COG-UK) Consortium | Jacquelyn Wynn, Mairead Hyland, The Lighthouse Lab in Alderley Park and Alex Alderton, Roberto Amato, Jeffrey Barrett, Sonia Goncalves, Ewan Harrison, David K. Jackson, Ian Johnston, Dominic Kwiatkowski, Cordelia Langford, John Sillitoe on behalf of the Wellcome Sanger Institute COVID-19 Surveillance Team |
| EPI_ISL_2961887 | Lighthouse Laboratory Plymouth | Wellcome Sanger Institute for the COVID-19 Genomics UK (COG-UK) Consortium | Lighthouse Laboratory Plymouth and Alex Alderton, Roberto Amato, Jeffrey Barrett, Sonia Goncalves, Ewan Harrison, David K. Jackson, Ian Johnston, Dominic Kwiatkowski, Cordelia Langford, John Sillitoe on behalf of the Wellcome Sanger Institute COVID-19 Surveillance Team |
| EPI_ISL_2961888, EPI_ISL_2961889, EPI_ISL_2961890, EPI_ISL_2961891, EPI_ISL_2961892, EPI_ISL_2961893 | Lighthouse Lab in Alderley Park | Wellcome Sanger Institute for the COVID-19 Genomics UK (COG-UK) Consortium | Jacquelyn Wynn, Mairead Hyland, The Lighthouse Lab in Alderley Park and Alex Alderton, Roberto Amato, Jeffrey Barrett, Sonia Goncalves, Ewan Harrison, David K. Jackson, Ian Johnston, Dominic Kwiatkowski, Cordelia Langford, John Sillitoe on behalf of the Wellcome Sanger Institute COVID-19 Surveillance Team |
| EPI_ISL_2961894 | Lighthouse Laboratory Plymouth | Wellcome Sanger Institute for the COVID-19 Genomics UK (COG-UK) Consortium | Lighthouse Laboratory Plymouth and Alex Alderton, Roberto Amato, Jeffrey Barrett, Sonia Goncalves, Ewan Harrison, David K. Jackson, Ian Johnston, Dominic Kwiatkowski, Cordelia Langford, John Sillitoe on behalf of the Wellcome Sanger Institute COVID-19 Surveillance Team |
| EPI_ISL_2961895 | Health Services Laboratories | Wellcome Sanger Institute for the COVID-19 Genomics UK (COG-UK) Consortium | Health Services Laboratories and Alex Alderton, Roberto Amato, Jeffrey Barrett, Sonia Goncalves, Ewan Harrison, David K. Jackson, Ian Johnston, Dominic Kwiatkowski, Cordelia Langford, John Sillitoe on behalf of the Wellcome Sanger Institute COVID-19 Surveillance Team |
| EPI_ISL_2961896 | Lighthouse Laboratory Plymouth | Wellcome Sanger Institute for the COVID-19 Genomics UK (COG-UK) Consortium | Lighthouse Laboratory Plymouth and Alex Alderton, Roberto Amato, Jeffrey Barrett, Sonia Goncalves, Ewan Harrison, David K. Jackson, Ian Johnston, Dominic Kwiatkowski, Cordelia Langford, John Sillitoe on behalf of the Wellcome Sanger Institute COVID-19 Surveillance Team |
| EPI_ISL_2961897 | Lighthouse Lab in Alderley Park | Wellcome Sanger Institute for the COVID-19 Genomics UK (COG-UK) Consortium | Jacquelyn Wynn, Mairead Hyland, The Lighthouse Lab in Alderley Park and Alex Alderton, Roberto Amato, Jeffrey Barrett, Sonia Goncalves, Ewan Harrison, David K. Jackson, Ian Johnston, Dominic Kwiatkowski, Cordelia Langford, John Sillitoe on behalf of the Wellcome Sanger Institute COVID-19 Surveillance Team |
| EPI_ISL_2961898 | Lighthouse Laboratory Plymouth | Wellcome Sanger Institute for the COVID-19 Genomics UK (COG-UK) Consortium | Lighthouse Laboratory Plymouth and Alex Alderton, Roberto Amato, Jeffrey Barrett, Sonia Goncalves, Ewan Harrison, David K. Jackson, Ian Johnston, Dominic Kwiatkowski, Cordelia Langford, John Sillitoe on behalf of the Wellcome Sanger Institute COVID-19 Surveillance Team |
| EPI_ISL_2961899 | Health Services Laboratories | Wellcome Sanger Institute for the COVID-19 Genomics UK (COG-UK) Consortium | Health Services Laboratories and Alex Alderton, Roberto Amato, Jeffrey Barrett, Sonia Goncalves, Ewan Harrison, David K. Jackson, Ian Johnston, Dominic Kwiatkowski, Cordelia Langford, John Sillitoe on behalf of the Wellcome Sanger Institute COVID-19 Surveillance Team |
| EPI_ISL_2961900 | Lighthouse Lab in Alderley Park | Wellcome Sanger Institute for the COVID-19 Genomics UK (COG-UK) Consortium | Jacquelyn Wynn, Mairead Hyland, The Lighthouse Lab in Alderley Park and Alex Alderton, Roberto Amato, Jeffrey Barrett, Sonia Goncalves, Ewan Harrison, David K. Jackson, Ian Johnston, Dominic Kwiatkowski, Cordelia Langford, John Sillitoe on behalf of the Wellcome Sanger Institute COVID-19 Surveillance Team |
| EPI_ISL_2961901 | Lighthouse Laboratory Plymouth | Wellcome Sanger Institute for the COVID-19 Genomics UK (COG-UK) Consortium | Lighthouse Laboratory Plymouth and Alex Alderton, Roberto Amato, Jeffrey Barrett, Sonia Goncalves, Ewan Harrison, David K. Jackson, Ian Johnston, Dominic Kwiatkowski, Cordelia Langford, John Sillitoe on behalf of the Wellcome Sanger Institute COVID-19 Surveillance Team |
| EPI_ISL_2961902 | Lighthouse Lab in Alderley Park | Wellcome Sanger Institute for the COVID-19 Genomics UK (COG-UK) Consortium | Jacquelyn Wynn, Mairead Hyland, The Lighthouse Lab in Alderley Park and Alex Alderton, Roberto Amato, Jeffrey Barrett, Sonia Goncalves, Ewan Harrison, David K. Jackson, Ian Johnston, Dominic Kwiatkowski, Cordelia Langford, John Sillitoe on behalf of the Wellcome Sanger Institute COVID-19 Surveillance Team |
| EPI_ISL_2961903 | Health Services Laboratories | Wellcome Sanger Institute for the COVID-19 Genomics UK (COG-UK) Consortium | Health Services Laboratories and Alex Alderton, Roberto Amato, Jeffrey Barrett, Sonia Goncalves, Ewan Harrison, David K. Jackson, Ian Johnston, Dominic Kwiatkowski, Cordelia Langford, John Sillitoe on behalf of the Wellcome Sanger Institute COVID-19 Surveillance Team |
| EPI_ISL_2961904 | Lighthouse Lab in Alderley Park | Wellcome Sanger Institute for the COVID-19 Genomics UK (COG-UK) Consortium | Jacquelyn Wynn, Mairead Hyland, The Lighthouse Lab in Alderley Park and Alex Alderton, Roberto Amato, Jeffrey Barrett, Sonia Goncalves, Ewan Harrison, David K. Jackson, Ian Johnston, Dominic Kwiatkowski, Cordelia Langford, John Sillitoe on behalf of the Wellcome Sanger Institute COVID-19 Surveillance Team |
| EPI_ISL_2961905 | Health Services Laboratories | Wellcome Sanger Institute for the COVID-19 Genomics UK (COG-UK) Consortium | Health Services Laboratories and Alex Alderton, Roberto Amato, Jeffrey Barrett, Sonia Goncalves, Ewan Harrison, David K. Jackson, Ian Johnston, Dominic Kwiatkowski, Cordelia Langford, John Sillitoe on behalf of the Wellcome Sanger Institute COVID-19 Surveillance Team |
| EPI_ISL_2961906 | Lighthouse Laboratory Plymouth | Wellcome Sanger Institute for the COVID-19 Genomics UK (COG-UK) Consortium | Lighthouse Laboratory Plymouth and Alex Alderton, Roberto Amato, Jeffrey Barrett, Sonia Goncalves, Ewan Harrison, David K. Jackson, Ian Johnston, Dominic Kwiatkowski, Cordelia Langford, John Sillitoe on behalf of the Wellcome Sanger Institute COVID-19 Surveillance Team |
| EPI_ISL_2961907, EPI_ISL_2961908, EPI_ISL_2961909 | Lighthouse Lab in Alderley Park | Wellcome Sanger Institute for the COVID-19 Genomics UK (COG-UK) Consortium | Jacquelyn Wynn, Mairead Hyland, The Lighthouse Lab in Alderley Park and Alex Alderton, Roberto Amato, Jeffrey Barrett, Sonia Goncalves, Ewan Harrison, David K. Jackson, Ian Johnston, Dominic Kwiatkowski, Cordelia Langford, John Sillitoe on behalf of the Wellcome Sanger Institute COVID-19 Surveillance Team |
| EPI_ISL_2961910 | Lighthouse Laboratory Plymouth | Wellcome Sanger Institute for the COVID-19 Genomics UK (COG-UK) Consortium | Lighthouse Laboratory Plymouth and Alex Alderton, Roberto Amato, Jeffrey Barrett, Sonia Goncalves, Ewan Harrison, David K. Jackson, Ian Johnston, Dominic Kwiatkowski, Cordelia Langford, John Sillitoe on behalf of the Wellcome Sanger Institute COVID-19 Surveillance Team |
| EPI_ISL_2961911, EPI_ISL_2961912, EPI_ISL_2961913, EPI_ISL_2961914 | Lighthouse Lab in Alderley Park | Wellcome Sanger Institute for the COVID-19 Genomics UK (COG-UK) Consortium | Jacquelyn Wynn, Mairead Hyland, The Lighthouse Lab in Alderley Park and Alex Alderton, Roberto Amato, Jeffrey Barrett, Sonia Goncalves, Ewan Harrison, David K. Jackson, Ian Johnston, Dominic Kwiatkowski, Cordelia Langford, John Sillitoe on behalf of the Wellcome Sanger Institute COVID-19 Surveillance Team |
| EPI_ISL_2961915 | Lighthouse Laboratory Plymouth | Wellcome Sanger Institute for the COVID-19 Genomics UK (COG-UK) Consortium | Lighthouse Laboratory Plymouth and Alex Alderton, Roberto Amato, Jeffrey Barrett, Sonia Goncalves, Ewan Harrison, David K. Jackson, Ian Johnston, Dominic Kwiatkowski, Cordelia Langford, John Sillitoe on behalf of the Wellcome Sanger Institute COVID-19 Surveillance Team |
| EPI_ISL_2961916 | Lighthouse Lab in Alderley Park | Wellcome Sanger Institute for the COVID-19 Genomics UK (COG-UK) Consortium | Jacquelyn Wynn, Mairead Hyland, The Lighthouse Lab in Alderley Park and Alex Alderton, Roberto Amato, Jeffrey Barrett, Sonia Goncalves, Ewan Harrison, David K. Jackson, Ian Johnston, Dominic Kwiatkowski, Cordelia Langford, John Sillitoe on behalf of the Wellcome Sanger Institute COVID-19 Surveillance Team |
| EPI_ISL_2961917 | Health Services Laboratories | Wellcome Sanger Institute for the COVID-19 Genomics UK (COG-UK) Consortium | Health Services Laboratories and Alex Alderton, Roberto Amato, Jeffrey Barrett, Sonia Goncalves, Ewan Harrison, David K. Jackson, Ian Johnston, Dominic Kwiatkowski, Cordelia Langford, John Sillitoe on behalf of the Wellcome Sanger Institute COVID-19 Surveillance Team |
| EPI_ISL_2961918, EPI_ISL_2961919, EPI_ISL_2961920, EPI_ISL_2961921 | Lighthouse Lab in Alderley Park | Wellcome Sanger Institute for the COVID-19 Genomics UK (COG-UK) Consortium | Jacquelyn Wynn, Mairead Hyland, The Lighthouse Lab in Alderley Park and Alex Alderton, Roberto Amato, Jeffrey Barrett, Sonia Goncalves, Ewan Harrison, David K. Jackson, Ian Johnston, Dominic Kwiatkowski, Cordelia Langford, John Sillitoe on behalf of the Wellcome Sanger Institute COVID-19 Surveillance Team |
| EPI_ISL_2961922 | Health Services Laboratories | Wellcome Sanger Institute for the COVID-19 Genomics UK (COG-UK) Consortium | Health Services Laboratories and Alex Alderton, Roberto Amato, Jeffrey Barrett, Sonia Goncalves, Ewan Harrison, David K. Jackson, Ian Johnston, Dominic Kwiatkowski, Cordelia Langford, John Sillitoe on behalf of the Wellcome Sanger Institute COVID-19 Surveillance Team |
| EPI_ISL_2961923 | Lighthouse Laboratory Plymouth | Wellcome Sanger Institute for the COVID-19 Genomics UK (COG-UK) Consortium | Lighthouse Laboratory Plymouth and Alex Alderton, Roberto Amato, Jeffrey Barrett, Sonia Goncalves, Ewan Harrison, David K. Jackson, Ian Johnston, Dominic Kwiatkowski, Cordelia Langford, John Sillitoe on behalf of the Wellcome Sanger Institute COVID-19 Surveillance Team |
| EPI_ISL_2961924, EPI_ISL_2961925, EPI_ISL_2961926 | Health Services Laboratories | Wellcome Sanger Institute for the COVID-19 Genomics UK (COG-UK) Consortium | Health Services Laboratories and Alex Alderton, Roberto Amato, Jeffrey Barrett, Sonia Goncalves, Ewan Harrison, David K. Jackson, Ian Johnston, Dominic Kwiatkowski, Cordelia Langford, John Sillitoe on behalf of the Wellcome Sanger Institute COVID-19 Surveillance Team |
| EPI_ISL_2961927 | Lighthouse Lab in Alderley Park | Wellcome Sanger Institute for the COVID-19 Genomics UK (COG-UK) Consortium | Jacquelyn Wynn, Mairead Hyland, The Lighthouse Lab in Alderley Park and Alex Alderton, Roberto Amato, Jeffrey Barrett, Sonia Goncalves, Ewan Harrison, David K. Jackson, Ian Johnston, Dominic Kwiatkowski, Cordelia Langford, John Sillitoe on behalf of the Wellcome Sanger Institute COVID-19 Surveillance Team |
| EPI_ISL_2961928 | Health Services Laboratories | Wellcome Sanger Institute for the COVID-19 Genomics UK (COG-UK) Consortium | Health Services Laboratories and Alex Alderton, Roberto Amato, Jeffrey Barrett, Sonia Goncalves, Ewan Harrison, David K. Jackson, Ian Johnston, Dominic Kwiatkowski, Cordelia Langford, John Sillitoe on behalf of the Wellcome Sanger Institute COVID-19 Surveillance Team |
| EPI_ISL_2961929 | Lighthouse Lab in Alderley Park | Wellcome Sanger Institute for the COVID-19 Genomics UK (COG-UK) Consortium | Jacquelyn Wynn, Mairead Hyland, The Lighthouse Lab in Alderley Park and Alex Alderton, Roberto Amato, Jeffrey Barrett, Sonia Goncalves, Ewan Harrison, David K. Jackson, Ian Johnston, Dominic Kwiatkowski, Cordelia Langford, John Sillitoe on behalf of the Wellcome Sanger Institute COVID-19 Surveillance Team |
| EPI_ISL_2961930, EPI_ISL_2961931 | Health Services Laboratories | Wellcome Sanger Institute for the COVID-19 Genomics UK (COG-UK) Consortium | Health Services Laboratories and Alex Alderton, Roberto Amato, Jeffrey Barrett, Sonia Goncalves, Ewan Harrison, David K. Jackson, Ian Johnston, Dominic Kwiatkowski, Cordelia Langford, John Sillitoe on behalf of the Wellcome Sanger Institute COVID-19 Surveillance Team |
| EPI_ISL_2961932 | Lighthouse Laboratory Plymouth | Wellcome Sanger Institute for the COVID-19 Genomics UK (COG-UK) Consortium | Lighthouse Laboratory Plymouth and Alex Alderton, Roberto Amato, Jeffrey Barrett, Sonia Goncalves, Ewan Harrison, David K. Jackson, Ian Johnston, Dominic Kwiatkowski, Cordelia Langford, John Sillitoe on behalf of the Wellcome Sanger Institute COVID-19 Surveillance Team |
| EPI_ISL_2961933 | Lighthouse Lab in Alderley Park | Wellcome Sanger Institute for the COVID-19 Genomics UK (COG-UK) Consortium | Jacquelyn Wynn, Mairead Hyland, The Lighthouse Lab in Alderley Park and Alex Alderton, Roberto Amato, Jeffrey Barrett, Sonia Goncalves, Ewan Harrison, David K. Jackson, Ian Johnston, Dominic Kwiatkowski, Cordelia Langford, John Sillitoe on behalf of the Wellcome Sanger Institute COVID-19 Surveillance Team |
| EPI_ISL_2961934 | Lighthouse Laboratory Plymouth | Wellcome Sanger Institute for the COVID-19 Genomics UK (COG-UK) Consortium | Lighthouse Laboratory Plymouth and Alex Alderton, Roberto Amato, Jeffrey Barrett, Sonia Goncalves, Ewan Harrison, David K. Jackson, Ian Johnston, Dominic Kwiatkowski, Cordelia Langford, John Sillitoe on behalf of the Wellcome Sanger Institute COVID-19 Surveillance Team |
| EPI_ISL_2961935 | Health Services Laboratories | Wellcome Sanger Institute for the COVID-19 Genomics UK (COG-UK) Consortium | Health Services Laboratories and Alex Alderton, Roberto Amato, Jeffrey Barrett, Sonia Goncalves, Ewan Harrison, David K. Jackson, Ian Johnston, Dominic Kwiatkowski, Cordelia Langford, John Sillitoe on behalf of the Wellcome Sanger Institute COVID-19 Surveillance Team |
| EPI_ISL_2961936, EPI_ISL_2961937 | Lighthouse Lab in Alderley Park | Wellcome Sanger Institute for the COVID-19 Genomics UK (COG-UK) Consortium | Jacquelyn Wynn, Mairead Hyland, The Lighthouse Lab in Alderley Park and Alex Alderton, Roberto Amato, Jeffrey Barrett, Sonia Goncalves, Ewan Harrison, David K. Jackson, Ian Johnston, Dominic Kwiatkowski, Cordelia Langford, John Sillitoe on behalf of the Wellcome Sanger Institute COVID-19 Surveillance Team |
| EPI_ISL_2961938 | Health Services Laboratories | Wellcome Sanger Institute for the COVID-19 Genomics UK (COG-UK) Consortium | Health Services Laboratories and Alex Alderton, Roberto Amato, Jeffrey Barrett, Sonia Goncalves, Ewan Harrison, David K. Jackson, Ian Johnston, Dominic Kwiatkowski, Cordelia Langford, John Sillitoe on behalf of the Wellcome Sanger Institute COVID-19 Surveillance Team |
| EPI_ISL_2961939 | Lighthouse Laboratory Plymouth | Wellcome Sanger Institute for the COVID-19 Genomics UK (COG-UK) | Lighthouse Laboratory Plymouth and Alex Alderton, Roberto Amato, Jeffrey Barrett, Sonia Goncalves, Ewan Harrison, David K. Jackson, Ian Johnston, Dominic Kwiatkowski, Cordelia Langford, John Sillitoe on behalf of the Wellcome Sanger |

| | | | |
|---|---|---|---|
| | | Consortium | Institute COVID-19 Surveillance Team |
| EPI_ISL_2961940 | Lighthouse Lab in Alderley Park | Wellcome Sanger Institute for the COVID-19 Genomics UK (COG-UK) Consortium | Jacquelyn Wynn, Mairead Hyland, The Lighthouse Lab in Alderley Park and Alex Alderton, Roberto Amato, Jeffrey Barrett, Sonia Goncalves, Ewan Harrison, David K. Jackson, Ian Johnston, Dominic Kwiatkowski, Cordelia Langford, John Sillitoe on behalf of the Wellcome Sanger Institute COVID-19 Surveillance Team |
| EPI_ISL_2961941 | Lighthouse Laboratory Plymouth | Wellcome Sanger Institute for the COVID-19 Genomics UK (COG-UK) Consortium | Lighthouse Laboratory Plymouth and Alex Alderton, Roberto Amato, Jeffrey Barrett, Sonia Goncalves, Ewan Harrison, David K. Jackson, Ian Johnston, Dominic Kwiatkowski, Cordelia Langford, John Sillitoe on behalf of the Wellcome Sanger Institute COVID-19 Surveillance Team |
| EPI_ISL_2961942 | Lighthouse Lab in Alderley Park | Wellcome Sanger Institute for the COVID-19 Genomics UK (COG-UK) Consortium | Jacquelyn Wynn, Mairead Hyland, The Lighthouse Lab in Alderley Park and Alex Alderton, Roberto Amato, Jeffrey Barrett, Sonia Goncalves, Ewan Harrison, David K. Jackson, Ian Johnston, Dominic Kwiatkowski, Cordelia Langford, John Sillitoe on behalf of the Wellcome Sanger Institute COVID-19 Surveillance Team |
| EPI_ISL_2961943 | Health Services Laboratories | Wellcome Sanger Institute for the COVID-19 Genomics UK (COG-UK) Consortium | Health Services Laboratories and Alex Alderton, Roberto Amato, Jeffrey Barrett, Sonia Goncalves, Ewan Harrison, David K. Jackson, Ian Johnston, Dominic Kwiatkowski, Cordelia Langford, John Sillitoe on behalf of the Wellcome Sanger Institute COVID-19 Surveillance Team |
| EPI_ISL_2961944 | Lighthouse Laboratory Plymouth | Wellcome Sanger Institute for the COVID-19 Genomics UK (COG-UK) Consortium | Lighthouse Laboratory Plymouth and Alex Alderton, Roberto Amato, Jeffrey Barrett, Sonia Goncalves, Ewan Harrison, David K. Jackson, Ian Johnston, Dominic Kwiatkowski, Cordelia Langford, John Sillitoe on behalf of the Wellcome Sanger Institute COVID-19 Surveillance Team |
| EPI_ISL_2961945, EPI_ISL_2961946 | Health Services Laboratories | Wellcome Sanger Institute for the COVID-19 Genomics UK (COG-UK) Consortium | Health Services Laboratories and Alex Alderton, Roberto Amato, Jeffrey Barrett, Sonia Goncalves, Ewan Harrison, David K. Jackson, Ian Johnston, Dominic Kwiatkowski, Cordelia Langford, John Sillitoe on behalf of the Wellcome Sanger Institute COVID-19 Surveillance Team |
| EPI_ISL_2961947, EPI_ISL_2961948, EPI_ISL_2961949 | Lighthouse Laboratory Plymouth | Wellcome Sanger Institute for the COVID-19 Genomics UK (COG-UK) Consortium | Lighthouse Laboratory Plymouth and Alex Alderton, Roberto Amato, Jeffrey Barrett, Sonia Goncalves, Ewan Harrison, David K. Jackson, Ian Johnston, Dominic Kwiatkowski, Cordelia Langford, John Sillitoe on behalf of the Wellcome Sanger Institute COVID-19 Surveillance Team |
| EPI_ISL_2961950 | Lighthouse Lab in Alderley Park | Wellcome Sanger Institute for the COVID-19 Genomics UK (COG-UK) Consortium | Jacquelyn Wynn, Mairead Hyland, The Lighthouse Lab in Alderley Park and Alex Alderton, Roberto Amato, Jeffrey Barrett, Sonia Goncalves, Ewan Harrison, David K. Jackson, Ian Johnston, Dominic Kwiatkowski, Cordelia Langford, John Sillitoe on behalf of the Wellcome Sanger Institute COVID-19 Surveillance Team |
| EPI_ISL_2961951 | Lighthouse Laboratory Plymouth | Wellcome Sanger Institute for the COVID-19 Genomics UK (COG-UK) Consortium | Lighthouse Laboratory Plymouth and Alex Alderton, Roberto Amato, Jeffrey Barrett, Sonia Goncalves, Ewan Harrison, David K. Jackson, Ian Johnston, Dominic Kwiatkowski, Cordelia Langford, John Sillitoe on behalf of the Wellcome Sanger Institute COVID-19 Surveillance Team |
| EPI_ISL_2961952, EPI_ISL_2961953 | Health Services Laboratories | Wellcome Sanger Institute for the COVID-19 Genomics UK (COG-UK) Consortium | Health Services Laboratories and Alex Alderton, Roberto Amato, Jeffrey Barrett, Sonia Goncalves, Ewan Harrison, David K. Jackson, Ian Johnston, Dominic Kwiatkowski, Cordelia Langford, John Sillitoe on behalf of the Wellcome Sanger Institute COVID-19 Surveillance Team |
| EPI_ISL_2961954 | Lighthouse Laboratory Plymouth | Wellcome Sanger Institute for the COVID-19 Genomics UK (COG-UK) Consortium | Lighthouse Laboratory Plymouth and Alex Alderton, Roberto Amato, Jeffrey Barrett, Sonia Goncalves, Ewan Harrison, David K. Jackson, Ian Johnston, Dominic Kwiatkowski, Cordelia Langford, John Sillitoe on behalf of the Wellcome Sanger Institute COVID-19 Surveillance Team |
| EPI_ISL_2961955, EPI_ISL_2961956 | Health Services Laboratories | Wellcome Sanger Institute for the COVID-19 Genomics UK (COG-UK) Consortium | Health Services Laboratories and Alex Alderton, Roberto Amato, Jeffrey Barrett, Sonia Goncalves, Ewan Harrison, David K. Jackson, Ian Johnston, Dominic Kwiatkowski, Cordelia Langford, John Sillitoe on behalf of the Wellcome Sanger Institute COVID-19 Surveillance Team |
| EPI_ISL_2961957, EPI_ISL_2961958 | Lighthouse Lab in Alderley Park | Wellcome Sanger Institute for the COVID-19 Genomics UK (COG-UK) Consortium | Jacquelyn Wynn, Mairead Hyland, The Lighthouse Lab in Alderley Park and Alex Alderton, Roberto Amato, Jeffrey Barrett, Sonia Goncalves, Ewan Harrison, David K. Jackson, Ian Johnston, Dominic Kwiatkowski, Cordelia Langford, John Sillitoe on behalf of the Wellcome Sanger Institute COVID-19 Surveillance Team |
| EPI_ISL_2961959 | Health Services Laboratories | Wellcome Sanger Institute for the COVID-19 Genomics UK (COG-UK) Consortium | Health Services Laboratories and Alex Alderton, Roberto Amato, Jeffrey Barrett, Sonia Goncalves, Ewan Harrison, David K. Jackson, Ian Johnston, Dominic Kwiatkowski, Cordelia Langford, John Sillitoe on behalf of the Wellcome Sanger Institute COVID-19 Surveillance Team |
| EPI_ISL_2961960, EPI_ISL_2961961, EPI_ISL_2961962, EPI_ISL_2961963, EPI_ISL_2961964, EPI_ISL_2961965 | Lighthouse Lab in Alderley Park | Wellcome Sanger Institute for the COVID-19 Genomics UK (COG-UK) Consortium | Jacquelyn Wynn, Mairead Hyland, The Lighthouse Lab in Alderley Park and Alex Alderton, Roberto Amato, Jeffrey Barrett, Sonia Goncalves, Ewan Harrison, David K. Jackson, Ian Johnston, Dominic Kwiatkowski, Cordelia Langford, John Sillitoe on behalf of the Wellcome Sanger Institute COVID-19 Surveillance Team |
| EPI_ISL_2961966 | Lighthouse Laboratory Plymouth | Wellcome Sanger Institute for the COVID-19 Genomics UK (COG-UK) Consortium | Lighthouse Laboratory Plymouth and Alex Alderton, Roberto Amato, Jeffrey Barrett, Sonia Goncalves, Ewan Harrison, David K. Jackson, Ian Johnston, Dominic Kwiatkowski, Cordelia Langford, John Sillitoe on behalf of the Wellcome Sanger Institute COVID-19 Surveillance Team |
| EPI_ISL_2961967 | Lighthouse Lab in Alderley Park | Wellcome Sanger Institute for the COVID-19 Genomics UK (COG-UK) Consortium | Jacquelyn Wynn, Mairead Hyland, The Lighthouse Lab in Alderley Park and Alex Alderton, Roberto Amato, Jeffrey Barrett, Sonia Goncalves, Ewan Harrison, David K. Jackson, Ian Johnston, Dominic Kwiatkowski, Cordelia Langford, John Sillitoe on behalf of the Wellcome Sanger Institute COVID-19 Surveillance Team |
| EPI_ISL_2961968 | Lighthouse Laboratory Plymouth | Wellcome Sanger Institute for the COVID-19 Genomics UK (COG-UK) Consortium | Lighthouse Laboratory Plymouth and Alex Alderton, Roberto Amato, Jeffrey Barrett, Sonia Goncalves, Ewan Harrison, David K. Jackson, Ian Johnston, Dominic Kwiatkowski, Cordelia Langford, John Sillitoe on behalf of the Wellcome Sanger Institute COVID-19 Surveillance Team |
| EPI_ISL_2961969 | Lighthouse Lab in Alderley Park | Wellcome Sanger Institute for the COVID-19 Genomics UK (COG-UK) Consortium | Jacquelyn Wynn, Mairead Hyland, The Lighthouse Lab in Alderley Park and Alex Alderton, Roberto Amato, Jeffrey Barrett, Sonia Goncalves, Ewan Harrison, David K. Jackson, Ian Johnston, Dominic Kwiatkowski, Cordelia Langford, John Sillitoe on behalf of the Wellcome Sanger Institute COVID-19 Surveillance Team |
| EPI_ISL_2961970 | Lighthouse Laboratory Plymouth | Wellcome Sanger Institute for the COVID-19 Genomics UK (COG-UK) Consortium | Lighthouse Laboratory Plymouth and Alex Alderton, Roberto Amato, Jeffrey Barrett, Sonia Goncalves, Ewan Harrison, David K. Jackson, Ian Johnston, Dominic Kwiatkowski, Cordelia Langford, John Sillitoe on behalf of the Wellcome Sanger Institute COVID-19 Surveillance Team |
| EPI_ISL_2961971, EPI_ISL_2961972, EPI_ISL_2961973, EPI_ISL_2961974 | Health Services Laboratories | Wellcome Sanger Institute for the COVID-19 Genomics UK (COG-UK) Consortium | Health Services Laboratories and Alex Alderton, Roberto Amato, Jeffrey Barrett, Sonia Goncalves, Ewan Harrison, David K. Jackson, Ian Johnston, Dominic Kwiatkowski, Cordelia Langford, John Sillitoe on behalf of the Wellcome Sanger Institute COVID-19 Surveillance Team |
| EPI_ISL_2961975 | Lighthouse Lab in Alderley Park | Wellcome Sanger Institute for the COVID-19 Genomics UK (COG-UK) Consortium | Jacquelyn Wynn, Mairead Hyland, The Lighthouse Lab in Alderley Park and Alex Alderton, Roberto Amato, Jeffrey Barrett, Sonia Goncalves, Ewan Harrison, David K. Jackson, Ian Johnston, Dominic Kwiatkowski, Cordelia Langford, John Sillitoe on behalf of the Wellcome Sanger Institute COVID-19 Surveillance Team |
| EPI_ISL_2961976, EPI_ISL_2961977 | Lighthouse Laboratory Plymouth | Wellcome Sanger Institute for the COVID-19 Genomics UK (COG-UK) Consortium | Lighthouse Laboratory Plymouth and Alex Alderton, Roberto Amato, Jeffrey Barrett, Sonia Goncalves, Ewan Harrison, David K. Jackson, Ian Johnston, Dominic Kwiatkowski, Cordelia Langford, John Sillitoe on behalf of the Wellcome Sanger Institute COVID-19 Surveillance Team |
| EPI_ISL_2961978, EPI_ISL_2961979 | Lighthouse Lab in Alderley Park | Wellcome Sanger Institute for the COVID-19 Genomics UK (COG-UK) Consortium | Jacquelyn Wynn, Mairead Hyland, The Lighthouse Lab in Alderley Park and Alex Alderton, Roberto Amato, Jeffrey Barrett, Sonia Goncalves, Ewan Harrison, David K. Jackson, Ian Johnston, Dominic Kwiatkowski, Cordelia Langford, John Sillitoe on behalf of the Wellcome Sanger Institute COVID-19 Surveillance Team |
| EPI_ISL_2961980, EPI_ISL_2961981 | Lighthouse Laboratory Plymouth | Wellcome Sanger Institute for the COVID-19 Genomics UK (COG-UK) Consortium | Lighthouse Laboratory Plymouth and Alex Alderton, Roberto Amato, Jeffrey Barrett, Sonia Goncalves, Ewan Harrison, David K. Jackson, Ian Johnston, Dominic Kwiatkowski, Cordelia Langford, John Sillitoe on behalf of the Wellcome Sanger Institute COVID-19 Surveillance Team |
| EPI_ISL_2961982, EPI_ISL_2961983, EPI_ISL_2961984 | Lighthouse Lab in Alderley Park | Wellcome Sanger Institute for the COVID-19 Genomics UK (COG-UK) Consortium | Jacquelyn Wynn, Mairead Hyland, The Lighthouse Lab in Alderley Park and Alex Alderton, Roberto Amato, Jeffrey Barrett, Sonia Goncalves, Ewan Harrison, David K. Jackson, Ian Johnston, Dominic Kwiatkowski, Cordelia Langford, John Sillitoe on behalf of the Wellcome Sanger Institute COVID-19 Surveillance Team |
| EPI_ISL_2961985 | Health Services Laboratories | Wellcome Sanger Institute for the COVID-19 Genomics UK (COG-UK) Consortium | Health Services Laboratories and Alex Alderton, Roberto Amato, Jeffrey Barrett, Sonia Goncalves, Ewan Harrison, David K. Jackson, Ian Johnston, Dominic Kwiatkowski, Cordelia Langford, John Sillitoe on behalf of the Wellcome Sanger Institute COVID-19 Surveillance Team |
| EPI_ISL_2961986, EPI_ISL_2961987, EPI_ISL_2961988, EPI_ISL_2961989, EPI_ISL_2961990 | Lighthouse Lab in Alderley Park | Wellcome Sanger Institute for the COVID-19 Genomics UK (COG-UK) Consortium | Jacquelyn Wynn, Mairead Hyland, The Lighthouse Lab in Alderley Park and Alex Alderton, Roberto Amato, Jeffrey Barrett, Sonia Goncalves, Ewan Harrison, David K. Jackson, Ian Johnston, Dominic Kwiatkowski, Cordelia Langford, John Sillitoe on behalf of the Wellcome Sanger Institute COVID-19 Surveillance Team |
| EPI_ISL_2961991, EPI_ISL_2961992 | Health Services Laboratories | Wellcome Sanger Institute for the COVID-19 Genomics UK (COG-UK) Consortium | Health Services Laboratories and Alex Alderton, Roberto Amato, Jeffrey Barrett, Sonia Goncalves, Ewan Harrison, David K. Jackson, Ian Johnston, Dominic Kwiatkowski, Cordelia Langford, John Sillitoe on behalf of the Wellcome Sanger Institute COVID-19 Surveillance Team |
| EPI_ISL_2961993 | Lighthouse Lab in Alderley Park | Wellcome Sanger Institute for the COVID-19 Genomics UK (COG-UK) Consortium | Jacquelyn Wynn, Mairead Hyland, The Lighthouse Lab in Alderley Park and Alex Alderton, Roberto Amato, Jeffrey Barrett, Sonia Goncalves, Ewan Harrison, David K. Jackson, Ian Johnston, Dominic Kwiatkowski, Cordelia Langford, John Sillitoe on behalf of the Wellcome Sanger Institute COVID-19 Surveillance Team |
| EPI_ISL_2961994 | Health Services Laboratories | Wellcome Sanger Institute for the COVID-19 Genomics UK (COG-UK) Consortium | Health Services Laboratories and Alex Alderton, Roberto Amato, Jeffrey Barrett, Sonia Goncalves, Ewan Harrison, David K. Jackson, Ian Johnston, Dominic Kwiatkowski, Cordelia Langford, John Sillitoe on behalf of the Wellcome Sanger Institute COVID-19 Surveillance Team |
| EPI_ISL_2961995 | Lighthouse Laboratory Plymouth | Wellcome Sanger Institute for the COVID-19 Genomics UK (COG-UK) Consortium | Lighthouse Laboratory Plymouth and Alex Alderton, Roberto Amato, Jeffrey Barrett, Sonia Goncalves, Ewan Harrison, David K. Jackson, Ian Johnston, Dominic Kwiatkowski, Cordelia Langford, John Sillitoe on behalf of the Wellcome Sanger Institute COVID-19 Surveillance Team |
| EPI_ISL_2961996 | Health Services Laboratories | Wellcome Sanger Institute for the COVID-19 Genomics UK (COG-UK) Consortium | Health Services Laboratories and Alex Alderton, Roberto Amato, Jeffrey Barrett, Sonia Goncalves, Ewan Harrison, David K. Jackson, Ian Johnston, Dominic Kwiatkowski, Cordelia Langford, John Sillitoe on behalf of the Wellcome Sanger Institute COVID-19 Surveillance Team |
| EPI_ISL_2961997 | Lighthouse Lab in Alderley Park | Wellcome Sanger Institute for the COVID-19 Genomics UK (COG-UK) Consortium | Jacquelyn Wynn, Mairead Hyland, The Lighthouse Lab in Alderley Park and Alex Alderton, Roberto Amato, Jeffrey Barrett, Sonia Goncalves, Ewan Harrison, David K. Jackson, Ian Johnston, Dominic Kwiatkowski, Cordelia Langford, John Sillitoe on behalf of the Wellcome Sanger Institute COVID-19 Surveillance Team |
| EPI_ISL_2961998 | Lighthouse Laboratory Plymouth | Wellcome Sanger Institute for the COVID-19 Genomics UK (COG-UK) Consortium | Lighthouse Laboratory Plymouth and Alex Alderton, Roberto Amato, Jeffrey Barrett, Sonia Goncalves, Ewan Harrison, David K. Jackson, Ian Johnston, Dominic Kwiatkowski, Cordelia Langford, John Sillitoe on behalf of the Wellcome Sanger Institute COVID-19 Surveillance Team |
| EPI_ISL_2961999 | Lighthouse Lab in Alderley Park | Wellcome Sanger Institute for the COVID-19 Genomics UK (COG-UK) Consortium | Jacquelyn Wynn, Mairead Hyland, The Lighthouse Lab in Alderley Park and Alex Alderton, Roberto Amato, Jeffrey Barrett, Sonia Goncalves, Ewan Harrison, David K. Jackson, Ian Johnston, Dominic Kwiatkowski, Cordelia Langford, John Sillitoe on behalf of the Wellcome Sanger Institute COVID-19 Surveillance Team |
| EPI_ISL_2962000, EPI_ISL_2962001, EPI_ISL_2962002 | Lighthouse Laboratory Plymouth | Wellcome Sanger Institute for the COVID-19 Genomics UK (COG-UK) Consortium | Lighthouse Laboratory Plymouth and Alex Alderton, Roberto Amato, Jeffrey Barrett, Sonia Goncalves, Ewan Harrison, David K. Jackson, Ian Johnston, Dominic Kwiatkowski, Cordelia Langford, John Sillitoe on behalf of the Wellcome Sanger Institute COVID-19 Surveillance Team |
| EPI_ISL_2962003, EPI_ISL_2962004, EPI_ISL_2962005, EPI_ISL_2962006 | Lighthouse Lab in Alderley Park | Wellcome Sanger Institute for the COVID-19 Genomics UK (COG-UK) Consortium | Jacquelyn Wynn, Mairead Hyland, The Lighthouse Lab in Alderley Park and Alex Alderton, Roberto Amato, Jeffrey Barrett, Sonia Goncalves, Ewan Harrison, David K. Jackson, Ian Johnston, Dominic Kwiatkowski, Cordelia Langford, John Sillitoe on behalf of the Wellcome Sanger Institute COVID-19 Surveillance Team |
| EPI_ISL_2962007 | Lighthouse Laboratory Plymouth | Wellcome Sanger Institute for the COVID-19 Genomics UK (COG-UK) Consortium | Lighthouse Laboratory Plymouth and Alex Alderton, Roberto Amato, Jeffrey Barrett, Sonia Goncalves, Ewan Harrison, David K. Jackson, Ian Johnston, Dominic Kwiatkowski, Cordelia Langford, John Sillitoe on behalf of the Wellcome Sanger Institute COVID-19 Surveillance Team |

| | | | |
|---|---|---|---|
| EPI_ISL_2962008 | Lighthouse Lab in Alderley Park | Wellcome Sanger Institute for the COVID-19 Genomics UK (COG-UK) Consortium | Jacquelyn Wynn, Mairead Hyland, The Lighthouse Lab in Alderley Park and Alex Alderton, Roberto Amato, Jeffrey Barrett, Sonia Goncalves, Ewan Harrison, David K. Jackson, Ian Johnston, Dominic Kwiatkowski, Cordelia Langford, John Sillitoe on behalf of the Wellcome Sanger Institute COVID-19 Surveillance Team |
| EPI_ISL_2962009 | Health Services Laboratories | Wellcome Sanger Institute for the COVID-19 Genomics UK (COG-UK) Consortium | Health Services Laboratories and Alex Alderton, Roberto Amato, Jeffrey Barrett, Sonia Goncalves, Ewan Harrison, David K. Jackson, Ian Johnston, Dominic Kwiatkowski, Cordelia Langford, John Sillitoe on behalf of the Wellcome Sanger Institute COVID-19 Surveillance Team |
| EPI_ISL_2962010 | Lighthouse Laboratory Plymouth | Wellcome Sanger Institute for the COVID-19 Genomics UK (COG-UK) Consortium | Lighthouse Laboratory Plymouth and Alex Alderton, Roberto Amato, Jeffrey Barrett, Sonia Goncalves, Ewan Harrison, David K. Jackson, Ian Johnston, Dominic Kwiatkowski, Cordelia Langford, John Sillitoe on behalf of the Wellcome Sanger Institute COVID-19 Surveillance Team |
| EPI_ISL_2962011, EPI_ISL_2962012, EPI_ISL_2962013 | Lighthouse Lab in Alderley Park | Wellcome Sanger Institute for the COVID-19 Genomics UK (COG-UK) Consortium | Jacquelyn Wynn, Mairead Hyland, The Lighthouse Lab in Alderley Park and Alex Alderton, Roberto Amato, Jeffrey Barrett, Sonia Goncalves, Ewan Harrison, David K. Jackson, Ian Johnston, Dominic Kwiatkowski, Cordelia Langford, John Sillitoe on behalf of the Wellcome Sanger Institute COVID-19 Surveillance Team |
| EPI_ISL_2962014, EPI_ISL_2962015 | Health Services Laboratories | Wellcome Sanger Institute for the COVID-19 Genomics UK (COG-UK) Consortium | Health Services Laboratories and Alex Alderton, Roberto Amato, Jeffrey Barrett, Sonia Goncalves, Ewan Harrison, David K. Jackson, Ian Johnston, Dominic Kwiatkowski, Cordelia Langford, John Sillitoe on behalf of the Wellcome Sanger Institute COVID-19 Surveillance Team |
| EPI_ISL_2962016 | Lighthouse Laboratory Plymouth | Wellcome Sanger Institute for the COVID-19 Genomics UK (COG-UK) Consortium | Lighthouse Laboratory Plymouth and Alex Alderton, Roberto Amato, Jeffrey Barrett, Sonia Goncalves, Ewan Harrison, David K. Jackson, Ian Johnston, Dominic Kwiatkowski, Cordelia Langford, John Sillitoe on behalf of the Wellcome Sanger Institute COVID-19 Surveillance Team |
| EPI_ISL_2962017, EPI_ISL_2962018 | Lighthouse Lab in Alderley Park | Wellcome Sanger Institute for the COVID-19 Genomics UK (COG-UK) Consortium | Jacquelyn Wynn, Mairead Hyland, The Lighthouse Lab in Alderley Park and Alex Alderton, Roberto Amato, Jeffrey Barrett, Sonia Goncalves, Ewan Harrison, David K. Jackson, Ian Johnston, Dominic Kwiatkowski, Cordelia Langford, John Sillitoe on behalf of the Wellcome Sanger Institute COVID-19 Surveillance Team |
| EPI_ISL_2962019 | Lighthouse Laboratory Plymouth | Wellcome Sanger Institute for the COVID-19 Genomics UK (COG-UK) Consortium | Lighthouse Laboratory Plymouth and Alex Alderton, Roberto Amato, Jeffrey Barrett, Sonia Goncalves, Ewan Harrison, David K. Jackson, Ian Johnston, Dominic Kwiatkowski, Cordelia Langford, John Sillitoe on behalf of the Wellcome Sanger Institute COVID-19 Surveillance Team |
| EPI_ISL_2962020, EPI_ISL_2962021 | Lighthouse Lab in Alderley Park | Wellcome Sanger Institute for the COVID-19 Genomics UK (COG-UK) Consortium | Jacquelyn Wynn, Mairead Hyland, The Lighthouse Lab in Alderley Park and Alex Alderton, Roberto Amato, Jeffrey Barrett, Sonia Goncalves, Ewan Harrison, David K. Jackson, Ian Johnston, Dominic Kwiatkowski, Cordelia Langford, John Sillitoe on behalf of the Wellcome Sanger Institute COVID-19 Surveillance Team |
| EPI_ISL_2962022, EPI_ISL_2962023 | Lighthouse Laboratory Plymouth | Wellcome Sanger Institute for the COVID-19 Genomics UK (COG-UK) Consortium | Lighthouse Laboratory Plymouth and Alex Alderton, Roberto Amato, Jeffrey Barrett, Sonia Goncalves, Ewan Harrison, David K. Jackson, Ian Johnston, Dominic Kwiatkowski, Cordelia Langford, John Sillitoe on behalf of the Wellcome Sanger Institute COVID-19 Surveillance Team |
| EPI_ISL_2962024 | Health Services Laboratories | Wellcome Sanger Institute for the COVID-19 Genomics UK (COG-UK) Consortium | Health Services Laboratories and Alex Alderton, Roberto Amato, Jeffrey Barrett, Sonia Goncalves, Ewan Harrison, David K. Jackson, Ian Johnston, Dominic Kwiatkowski, Cordelia Langford, John Sillitoe on behalf of the Wellcome Sanger Institute COVID-19 Surveillance Team |
| EPI_ISL_2962025, EPI_ISL_2962026, EPI_ISL_2962027 | Lighthouse Lab in Alderley Park | Wellcome Sanger Institute for the COVID-19 Genomics UK (COG-UK) Consortium | Jacquelyn Wynn, Mairead Hyland, The Lighthouse Lab in Alderley Park and Alex Alderton, Roberto Amato, Jeffrey Barrett, Sonia Goncalves, Ewan Harrison, David K. Jackson, Ian Johnston, Dominic Kwiatkowski, Cordelia Langford, John Sillitoe on behalf of the Wellcome Sanger Institute COVID-19 Surveillance Team |
| EPI_ISL_2962028 | Health Services Laboratories | Wellcome Sanger Institute for the COVID-19 Genomics UK (COG-UK) Consortium | Health Services Laboratories and Alex Alderton, Roberto Amato, Jeffrey Barrett, Sonia Goncalves, Ewan Harrison, David K. Jackson, Ian Johnston, Dominic Kwiatkowski, Cordelia Langford, John Sillitoe on behalf of the Wellcome Sanger Institute COVID-19 Surveillance Team |
| EPI_ISL_2962029 | Lighthouse Laboratory Plymouth | Wellcome Sanger Institute for the COVID-19 Genomics UK (COG-UK) Consortium | Lighthouse Laboratory Plymouth and Alex Alderton, Roberto Amato, Jeffrey Barrett, Sonia Goncalves, Ewan Harrison, David K. Jackson, Ian Johnston, Dominic Kwiatkowski, Cordelia Langford, John Sillitoe on behalf of the Wellcome Sanger Institute COVID-19 Surveillance Team |
| EPI_ISL_2962030 | Lighthouse Lab in Alderley Park | Wellcome Sanger Institute for the COVID-19 Genomics UK (COG-UK) Consortium | Jacquelyn Wynn, Mairead Hyland, The Lighthouse Lab in Alderley Park and Alex Alderton, Roberto Amato, Jeffrey Barrett, Sonia Goncalves, Ewan Harrison, David K. Jackson, Ian Johnston, Dominic Kwiatkowski, Cordelia Langford, John Sillitoe on behalf of the Wellcome Sanger Institute COVID-19 Surveillance Team |
| EPI_ISL_2962031 | Lighthouse Laboratory Plymouth | Wellcome Sanger Institute for the COVID-19 Genomics UK (COG-UK) Consortium | Lighthouse Laboratory Plymouth and Alex Alderton, Roberto Amato, Jeffrey Barrett, Sonia Goncalves, Ewan Harrison, David K. Jackson, Ian Johnston, Dominic Kwiatkowski, Cordelia Langford, John Sillitoe on behalf of the Wellcome Sanger Institute COVID-19 Surveillance Team |
| EPI_ISL_2962032 | Lighthouse Lab in Alderley Park | Wellcome Sanger Institute for the COVID-19 Genomics UK (COG-UK) Consortium | Jacquelyn Wynn, Mairead Hyland, The Lighthouse Lab in Alderley Park and Alex Alderton, Roberto Amato, Jeffrey Barrett, Sonia Goncalves, Ewan Harrison, David K. Jackson, Ian Johnston, Dominic Kwiatkowski, Cordelia Langford, John Sillitoe on behalf of the Wellcome Sanger Institute COVID-19 Surveillance Team |
| EPI_ISL_2962033 | Health Services Laboratories | Wellcome Sanger Institute for the COVID-19 Genomics UK (COG-UK) Consortium | Health Services Laboratories and Alex Alderton, Roberto Amato, Jeffrey Barrett, Sonia Goncalves, Ewan Harrison, David K. Jackson, Ian Johnston, Dominic Kwiatkowski, Cordelia Langford, John Sillitoe on behalf of the Wellcome Sanger Institute COVID-19 Surveillance Team |
| EPI_ISL_2962034, EPI_ISL_2962035 | Lighthouse Lab in Alderley Park | Wellcome Sanger Institute for the COVID-19 Genomics UK (COG-UK) Consortium | Jacquelyn Wynn, Mairead Hyland, The Lighthouse Lab in Alderley Park and Alex Alderton, Roberto Amato, Jeffrey Barrett, Sonia Goncalves, Ewan Harrison, David K. Jackson, Ian Johnston, Dominic Kwiatkowski, Cordelia Langford, John Sillitoe on behalf of the Wellcome Sanger Institute COVID-19 Surveillance Team |
| EPI_ISL_2962036, EPI_ISL_2962037 | Health Services Laboratories | Wellcome Sanger Institute for the COVID-19 Genomics UK (COG-UK) Consortium | Health Services Laboratories and Alex Alderton, Roberto Amato, Jeffrey Barrett, Sonia Goncalves, Ewan Harrison, David K. Jackson, Ian Johnston, Dominic Kwiatkowski, Cordelia Langford, John Sillitoe on behalf of the Wellcome Sanger Institute COVID-19 Surveillance Team |
| EPI_ISL_2962038, EPI_ISL_2962039 | Lighthouse Lab in Alderley Park | Wellcome Sanger Institute for the COVID-19 Genomics UK (COG-UK) Consortium | Jacquelyn Wynn, Mairead Hyland, The Lighthouse Lab in Alderley Park and Alex Alderton, Roberto Amato, Jeffrey Barrett, Sonia Goncalves, Ewan Harrison, David K. Jackson, Ian Johnston, Dominic Kwiatkowski, Cordelia Langford, John Sillitoe on behalf of the Wellcome Sanger Institute COVID-19 Surveillance Team |
| EPI_ISL_2962040 | Lighthouse Laboratory Plymouth | Wellcome Sanger Institute for the COVID-19 Genomics UK (COG-UK) Consortium | Lighthouse Laboratory Plymouth and Alex Alderton, Roberto Amato, Jeffrey Barrett, Sonia Goncalves, Ewan Harrison, David K. Jackson, Ian Johnston, Dominic Kwiatkowski, Cordelia Langford, John Sillitoe on behalf of the Wellcome Sanger Institute COVID-19 Surveillance Team |
| EPI_ISL_2962041 | Lighthouse Lab in Alderley Park | Wellcome Sanger Institute for the COVID-19 Genomics UK (COG-UK) Consortium | Jacquelyn Wynn, Mairead Hyland, The Lighthouse Lab in Alderley Park and Alex Alderton, Roberto Amato, Jeffrey Barrett, Sonia Goncalves, Ewan Harrison, David K. Jackson, Ian Johnston, Dominic Kwiatkowski, Cordelia Langford, John Sillitoe on behalf of the Wellcome Sanger Institute COVID-19 Surveillance Team |
| EPI_ISL_2962042 | Lighthouse Laboratory Plymouth | Wellcome Sanger Institute for the COVID-19 Genomics UK (COG-UK) Consortium | Lighthouse Laboratory Plymouth and Alex Alderton, Roberto Amato, Jeffrey Barrett, Sonia Goncalves, Ewan Harrison, David K. Jackson, Ian Johnston, Dominic Kwiatkowski, Cordelia Langford, John Sillitoe on behalf of the Wellcome Sanger Institute COVID-19 Surveillance Team |
| EPI_ISL_2962043 | Health Services Laboratories | Wellcome Sanger Institute for the COVID-19 Genomics UK (COG-UK) Consortium | Health Services Laboratories and Alex Alderton, Roberto Amato, Jeffrey Barrett, Sonia Goncalves, Ewan Harrison, David K. Jackson, Ian Johnston, Dominic Kwiatkowski, Cordelia Langford, John Sillitoe on behalf of the Wellcome Sanger Institute COVID-19 Surveillance Team |
| EPI_ISL_2962044 | Lighthouse Lab in Alderley Park | Wellcome Sanger Institute for the COVID-19 Genomics UK (COG-UK) Consortium | Jacquelyn Wynn, Mairead Hyland, The Lighthouse Lab in Alderley Park and Alex Alderton, Roberto Amato, Jeffrey Barrett, Sonia Goncalves, Ewan Harrison, David K. Jackson, Ian Johnston, Dominic Kwiatkowski, Cordelia Langford, John Sillitoe on behalf of the Wellcome Sanger Institute COVID-19 Surveillance Team |
| EPI_ISL_2962045 | Health Services Laboratories | Wellcome Sanger Institute for the COVID-19 Genomics UK (COG-UK) Consortium | Health Services Laboratories and Alex Alderton, Roberto Amato, Jeffrey Barrett, Sonia Goncalves, Ewan Harrison, David K. Jackson, Ian Johnston, Dominic Kwiatkowski, Cordelia Langford, John Sillitoe on behalf of the Wellcome Sanger Institute COVID-19 Surveillance Team |
| EPI_ISL_2962046, EPI_ISL_2962047, EPI_ISL_2962048 | Lighthouse Lab in Alderley Park | Wellcome Sanger Institute for the COVID-19 Genomics UK (COG-UK) Consortium | Jacquelyn Wynn, Mairead Hyland, The Lighthouse Lab in Alderley Park and Alex Alderton, Roberto Amato, Jeffrey Barrett, Sonia Goncalves, Ewan Harrison, David K. Jackson, Ian Johnston, Dominic Kwiatkowski, Cordelia Langford, John Sillitoe on behalf of the Wellcome Sanger Institute COVID-19 Surveillance Team |
| EPI_ISL_2962049 | Health Services Laboratories | Wellcome Sanger Institute for the COVID-19 Genomics UK (COG-UK) Consortium | Health Services Laboratories and Alex Alderton, Roberto Amato, Jeffrey Barrett, Sonia Goncalves, Ewan Harrison, David K. Jackson, Ian Johnston, Dominic Kwiatkowski, Cordelia Langford, John Sillitoe on behalf of the Wellcome Sanger Institute COVID-19 Surveillance Team |
| EPI_ISL_2962050, EPI_ISL_2962051 | Lighthouse Lab in Alderley Park | Wellcome Sanger Institute for the COVID-19 Genomics UK (COG-UK) Consortium | Jacquelyn Wynn, Mairead Hyland, The Lighthouse Lab in Alderley Park and Alex Alderton, Roberto Amato, Jeffrey Barrett, Sonia Goncalves, Ewan Harrison, David K. Jackson, Ian Johnston, Dominic Kwiatkowski, Cordelia Langford, John Sillitoe on behalf of the Wellcome Sanger Institute COVID-19 Surveillance Team |
| EPI_ISL_2962052, EPI_ISL_2962053, EPI_ISL_2962054 | Lighthouse Laboratory Plymouth | Wellcome Sanger Institute for the COVID-19 Genomics UK (COG-UK) Consortium | Lighthouse Laboratory Plymouth and Alex Alderton, Roberto Amato, Jeffrey Barrett, Sonia Goncalves, Ewan Harrison, David K. Jackson, Ian Johnston, Dominic Kwiatkowski, Cordelia Langford, John Sillitoe on behalf of the Wellcome Sanger Institute COVID-19 Surveillance Team |
| EPI_ISL_2962055 | Lighthouse Lab in Alderley Park | Wellcome Sanger Institute for the COVID-19 Genomics UK (COG-UK) Consortium | Jacquelyn Wynn, Mairead Hyland, The Lighthouse Lab in Alderley Park and Alex Alderton, Roberto Amato, Jeffrey Barrett, Sonia Goncalves, Ewan Harrison, David K. Jackson, Ian Johnston, Dominic Kwiatkowski, Cordelia Langford, John Sillitoe on behalf of the Wellcome Sanger Institute COVID-19 Surveillance Team |
| EPI_ISL_2962056 | Lighthouse Laboratory Plymouth | Wellcome Sanger Institute for the COVID-19 Genomics UK (COG-UK) Consortium | Lighthouse Laboratory Plymouth and Alex Alderton, Roberto Amato, Jeffrey Barrett, Sonia Goncalves, Ewan Harrison, David K. Jackson, Ian Johnston, Dominic Kwiatkowski, Cordelia Langford, John Sillitoe on behalf of the Wellcome Sanger Institute COVID-19 Surveillance Team |
| EPI_ISL_2962057 | Lighthouse Lab in Alderley Park | Wellcome Sanger Institute for the COVID-19 Genomics UK (COG-UK) Consortium | Jacquelyn Wynn, Mairead Hyland, The Lighthouse Lab in Alderley Park and Alex Alderton, Roberto Amato, Jeffrey Barrett, Sonia Goncalves, Ewan Harrison, David K. Jackson, Ian Johnston, Dominic Kwiatkowski, Cordelia Langford, John Sillitoe on behalf of the Wellcome Sanger Institute COVID-19 Surveillance Team |
| EPI_ISL_2962058 | Health Services Laboratories | Wellcome Sanger Institute for the COVID-19 Genomics UK (COG-UK) Consortium | Health Services Laboratories and Alex Alderton, Roberto Amato, Jeffrey Barrett, Sonia Goncalves, Ewan Harrison, David K. Jackson, Ian Johnston, Dominic Kwiatkowski, Cordelia Langford, John Sillitoe on behalf of the Wellcome Sanger Institute COVID-19 Surveillance Team |
| EPI_ISL_2962059 | Lighthouse Lab in Alderley Park | Wellcome Sanger Institute for the COVID-19 Genomics UK (COG-UK) Consortium | Jacquelyn Wynn, Mairead Hyland, The Lighthouse Lab in Alderley Park and Alex Alderton, Roberto Amato, Jeffrey Barrett, Sonia Goncalves, Ewan Harrison, David K. Jackson, Ian Johnston, Dominic Kwiatkowski, Cordelia Langford, John Sillitoe on behalf of the Wellcome Sanger Institute COVID-19 Surveillance Team |
| EPI_ISL_2962060 | Health Services Laboratories | Wellcome Sanger Institute for the COVID-19 Genomics UK (COG-UK) Consortium | Health Services Laboratories and Alex Alderton, Roberto Amato, Jeffrey Barrett, Sonia Goncalves, Ewan Harrison, David K. Jackson, Ian Johnston, Dominic Kwiatkowski, Cordelia Langford, John Sillitoe on behalf of the Wellcome Sanger Institute COVID-19 Surveillance Team |
| EPI_ISL_2962061 | Lighthouse Lab in Alderley Park | Wellcome Sanger Institute for the COVID-19 Genomics UK (COG-UK) Consortium | Jacquelyn Wynn, Mairead Hyland, The Lighthouse Lab in Alderley Park and Alex Alderton, Roberto Amato, Jeffrey Barrett, Sonia Goncalves, Ewan Harrison, David K. Jackson, Ian Johnston, Dominic Kwiatkowski, Cordelia Langford, John Sillitoe on behalf of the Wellcome Sanger Institute COVID-19 Surveillance Team |
| EPI_ISL_2962062, EPI_ISL_2962063 | Lighthouse Laboratory Plymouth | Wellcome Sanger Institute for the COVID-19 Genomics UK (COG-UK) Consortium | Lighthouse Laboratory Plymouth and Alex Alderton, Roberto Amato, Jeffrey Barrett, Sonia Goncalves, Ewan Harrison, David K. Jackson, Ian Johnston, Dominic Kwiatkowski, Cordelia Langford, John Sillitoe on behalf of the Wellcome Sanger Institute COVID-19 Surveillance Team |
| EPI_ISL_2962064 | Lighthouse Lab in Alderley Park | Wellcome Sanger Institute for the COVID-19 Genomics UK (COG-UK) Consortium | Jacquelyn Wynn, Mairead Hyland, The Lighthouse Lab in Alderley Park and Alex Alderton, Roberto Amato, Jeffrey Barrett, Sonia Goncalves, Ewan Harrison, David K. Jackson, Ian Johnston, Dominic Kwiatkowski, Cordelia Langford, John Sillitoe on behalf of the Wellcome Sanger Institute COVID-19 Surveillance Team |
| EPI_ISL_2962065 | Lighthouse Laboratory Plymouth | Wellcome Sanger Institute for the COVID-19 Genomics UK (COG-UK) Consortium | Lighthouse Laboratory Plymouth and Alex Alderton, Roberto Amato, Jeffrey Barrett, Sonia Goncalves, Ewan Harrison, David K. Jackson, Ian Johnston, Dominic Kwiatkowski, Cordelia Langford, John Sillitoe on behalf of the Wellcome Sanger Institute COVID-19 Surveillance Team |
| EPI_ISL_2962066, EPI_ISL_2962067, EPI_ISL_2962068, EPI_ISL_2962069, EPI_ISL_2962070, | Lighthouse Lab in Alderley Park | Wellcome Sanger Institute for the COVID-19 Genomics UK (COG-UK) Consortium | Jacquelyn Wynn, Mairead Hyland, The Lighthouse Lab in Alderley Park and Alex Alderton, Roberto Amato, Jeffrey Barrett, Sonia Goncalves, Ewan Harrison, David K. Jackson, Ian Johnston, Dominic Kwiatkowski, Cordelia Langford, John Sillitoe on behalf of the Wellcome Sanger Institute COVID-19 Surveillance Team |

| | | | |
|---|---|---|---|
| EPI_ISL_2962071 | | | |
| EPI_ISL_2962072 | Lighthouse Laboratory Plymouth | Wellcome Sanger Institute for the COVID-19 Genomics UK (COG-UK) Consortium | Lighthouse Laboratory Plymouth and Alex Alderton, Roberto Amato, Jeffrey Barrett, Sonia Goncalves, Ewan Harrison, David K. Jackson, Ian Johnston, Dominic Kwiatkowski, Cordelia Langford, John Sillitoe on behalf of the Wellcome Sanger Institute COVID-19 Surveillance Team |
| EPI_ISL_2962073 | Lighthouse Lab in Alderley Park | Wellcome Sanger Institute for the COVID-19 Genomics UK (COG-UK) Consortium | Jacquelyn Wynn, Mairead Hyland, The Lighthouse Lab in Alderley Park and Alex Alderton, Roberto Amato, Jeffrey Barrett, Sonia Goncalves, Ewan Harrison, David K. Jackson, Ian Johnston, Dominic Kwiatkowski, Cordelia Langford, John Sillitoe on behalf of the Wellcome Sanger Institute COVID-19 Surveillance Team |
| EPI_ISL_2962074 | Lighthouse Laboratory Plymouth | Wellcome Sanger Institute for the COVID-19 Genomics UK (COG-UK) Consortium | Lighthouse Laboratory Plymouth and Alex Alderton, Roberto Amato, Jeffrey Barrett, Sonia Goncalves, Ewan Harrison, David K. Jackson, Ian Johnston, Dominic Kwiatkowski, Cordelia Langford, John Sillitoe on behalf of the Wellcome Sanger Institute COVID-19 Surveillance Team |
| EPI_ISL_2962075 | Health Services Laboratories | Wellcome Sanger Institute for the COVID-19 Genomics UK (COG-UK) Consortium | Health Services Laboratories and Alex Alderton, Roberto Amato, Jeffrey Barrett, Sonia Goncalves, Ewan Harrison, David K. Jackson, Ian Johnston, Dominic Kwiatkowski, Cordelia Langford, John Sillitoe on behalf of the Wellcome Sanger Institute COVID-19 Surveillance Team |
| EPI_ISL_2962076, EPI_ISL_2962077 | Lighthouse Lab in Alderley Park | Wellcome Sanger Institute for the COVID-19 Genomics UK (COG-UK) Consortium | Jacquelyn Wynn, Mairead Hyland, The Lighthouse Lab in Alderley Park and Alex Alderton, Roberto Amato, Jeffrey Barrett, Sonia Goncalves, Ewan Harrison, David K. Jackson, Ian Johnston, Dominic Kwiatkowski, Cordelia Langford, John Sillitoe on behalf of the Wellcome Sanger Institute COVID-19 Surveillance Team |
| EPI_ISL_2962078, EPI_ISL_2962079 | Health Services Laboratories | Wellcome Sanger Institute for the COVID-19 Genomics UK (COG-UK) Consortium | Health Services Laboratories and Alex Alderton, Roberto Amato, Jeffrey Barrett, Sonia Goncalves, Ewan Harrison, David K. Jackson, Ian Johnston, Dominic Kwiatkowski, Cordelia Langford, John Sillitoe on behalf of the Wellcome Sanger Institute COVID-19 Surveillance Team |
| EPI_ISL_2962080 | Lighthouse Laboratory Plymouth | Wellcome Sanger Institute for the COVID-19 Genomics UK (COG-UK) Consortium | Lighthouse Laboratory Plymouth and Alex Alderton, Roberto Amato, Jeffrey Barrett, Sonia Goncalves, Ewan Harrison, David K. Jackson, Ian Johnston, Dominic Kwiatkowski, Cordelia Langford, John Sillitoe on behalf of the Wellcome Sanger Institute COVID-19 Surveillance Team |
| EPI_ISL_2962081 | Health Services Laboratories | Wellcome Sanger Institute for the COVID-19 Genomics UK (COG-UK) Consortium | Health Services Laboratories and Alex Alderton, Roberto Amato, Jeffrey Barrett, Sonia Goncalves, Ewan Harrison, David K. Jackson, Ian Johnston, Dominic Kwiatkowski, Cordelia Langford, John Sillitoe on behalf of the Wellcome Sanger Institute COVID-19 Surveillance Team |
| EPI_ISL_2962082, EPI_ISL_2962083, EPI_ISL_2962084, EPI_ISL_2962085 | Lighthouse Lab in Alderley Park | Wellcome Sanger Institute for the COVID-19 Genomics UK (COG-UK) Consortium | Jacquelyn Wynn, Mairead Hyland, The Lighthouse Lab in Alderley Park and Alex Alderton, Roberto Amato, Jeffrey Barrett, Sonia Goncalves, Ewan Harrison, David K. Jackson, Ian Johnston, Dominic Kwiatkowski, Cordelia Langford, John Sillitoe on behalf of the Wellcome Sanger Institute COVID-19 Surveillance Team |
| EPI_ISL_2962086 | Health Services Laboratories | Wellcome Sanger Institute for the COVID-19 Genomics UK (COG-UK) Consortium | Health Services Laboratories and Alex Alderton, Roberto Amato, Jeffrey Barrett, Sonia Goncalves, Ewan Harrison, David K. Jackson, Ian Johnston, Dominic Kwiatkowski, Cordelia Langford, John Sillitoe on behalf of the Wellcome Sanger Institute COVID-19 Surveillance Team |
| EPI_ISL_2962087, EPI_ISL_2962088, EPI_ISL_2962089 | Lighthouse Lab in Alderley Park | Wellcome Sanger Institute for the COVID-19 Genomics UK (COG-UK) Consortium | Jacquelyn Wynn, Mairead Hyland, The Lighthouse Lab in Alderley Park and Alex Alderton, Roberto Amato, Jeffrey Barrett, Sonia Goncalves, Ewan Harrison, David K. Jackson, Ian Johnston, Dominic Kwiatkowski, Cordelia Langford, John Sillitoe on behalf of the Wellcome Sanger Institute COVID-19 Surveillance Team |
| EPI_ISL_2962090 | Health Services Laboratories | Wellcome Sanger Institute for the COVID-19 Genomics UK (COG-UK) Consortium | Health Services Laboratories and Alex Alderton, Roberto Amato, Jeffrey Barrett, Sonia Goncalves, Ewan Harrison, David K. Jackson, Ian Johnston, Dominic Kwiatkowski, Cordelia Langford, John Sillitoe on behalf of the Wellcome Sanger Institute COVID-19 Surveillance Team |
| EPI_ISL_2962091, EPI_ISL_2962092, EPI_ISL_2962093, EPI_ISL_2962094 | Lighthouse Lab in Alderley Park | Wellcome Sanger Institute for the COVID-19 Genomics UK (COG-UK) Consortium | Jacquelyn Wynn, Mairead Hyland, The Lighthouse Lab in Alderley Park and Alex Alderton, Roberto Amato, Jeffrey Barrett, Sonia Goncalves, Ewan Harrison, David K. Jackson, Ian Johnston, Dominic Kwiatkowski, Cordelia Langford, John Sillitoe on behalf of the Wellcome Sanger Institute COVID-19 Surveillance Team |
| EPI_ISL_2962095 | Health Services Laboratories | Wellcome Sanger Institute for the COVID-19 Genomics UK (COG-UK) Consortium | Health Services Laboratories and Alex Alderton, Roberto Amato, Jeffrey Barrett, Sonia Goncalves, Ewan Harrison, David K. Jackson, Ian Johnston, Dominic Kwiatkowski, Cordelia Langford, John Sillitoe on behalf of the Wellcome Sanger Institute COVID-19 Surveillance Team |
| EPI_ISL_2962096 | Lighthouse Lab in Alderley Park | Wellcome Sanger Institute for the COVID-19 Genomics UK (COG-UK) Consortium | Jacquelyn Wynn, Mairead Hyland, The Lighthouse Lab in Alderley Park and Alex Alderton, Roberto Amato, Jeffrey Barrett, Sonia Goncalves, Ewan Harrison, David K. Jackson, Ian Johnston, Dominic Kwiatkowski, Cordelia Langford, John Sillitoe on behalf of the Wellcome Sanger Institute COVID-19 Surveillance Team |
| EPI_ISL_2962097 | Health Services Laboratories | Wellcome Sanger Institute for the COVID-19 Genomics UK (COG-UK) Consortium | Health Services Laboratories and Alex Alderton, Roberto Amato, Jeffrey Barrett, Sonia Goncalves, Ewan Harrison, David K. Jackson, Ian Johnston, Dominic Kwiatkowski, Cordelia Langford, John Sillitoe on behalf of the Wellcome Sanger Institute COVID-19 Surveillance Team |
| EPI_ISL_2962098, EPI_ISL_2962099 | Lighthouse Lab in Alderley Park | Wellcome Sanger Institute for the COVID-19 Genomics UK (COG-UK) Consortium | Jacquelyn Wynn, Mairead Hyland, The Lighthouse Lab in Alderley Park and Alex Alderton, Roberto Amato, Jeffrey Barrett, Sonia Goncalves, Ewan Harrison, David K. Jackson, Ian Johnston, Dominic Kwiatkowski, Cordelia Langford, John Sillitoe on behalf of the Wellcome Sanger Institute COVID-19 Surveillance Team |
| EPI_ISL_2962100 | Lighthouse Laboratory Plymouth | Wellcome Sanger Institute for the COVID-19 Genomics UK (COG-UK) Consortium | Lighthouse Laboratory Plymouth and Alex Alderton, Roberto Amato, Jeffrey Barrett, Sonia Goncalves, Ewan Harrison, David K. Jackson, Ian Johnston, Dominic Kwiatkowski, Cordelia Langford, John Sillitoe on behalf of the Wellcome Sanger Institute COVID-19 Surveillance Team |
| EPI_ISL_2962101 | Health Services Laboratories | Wellcome Sanger Institute for the COVID-19 Genomics UK (COG-UK) Consortium | Health Services Laboratories and Alex Alderton, Roberto Amato, Jeffrey Barrett, Sonia Goncalves, Ewan Harrison, David K. Jackson, Ian Johnston, Dominic Kwiatkowski, Cordelia Langford, John Sillitoe on behalf of the Wellcome Sanger Institute COVID-19 Surveillance Team |
| EPI_ISL_2962102, EPI_ISL_2962103, EPI_ISL_2962104, EPI_ISL_2962105, EPI_ISL_2962106, EPI_ISL_2962108 | Lighthouse Lab in Alderley Park | Wellcome Sanger Institute for the COVID-19 Genomics UK (COG-UK) Consortium | Jacquelyn Wynn, Mairead Hyland, The Lighthouse Lab in Alderley Park and Alex Alderton, Roberto Amato, Jeffrey Barrett, Sonia Goncalves, Ewan Harrison, David K. Jackson, Ian Johnston, Dominic Kwiatkowski, Cordelia Langford, John Sillitoe on behalf of the Wellcome Sanger Institute COVID-19 Surveillance Team |
| EPI_ISL_2962109 | Health Services Laboratories | Wellcome Sanger Institute for the COVID-19 Genomics UK (COG-UK) Consortium | Health Services Laboratories and Alex Alderton, Roberto Amato, Jeffrey Barrett, Sonia Goncalves, Ewan Harrison, David K. Jackson, Ian Johnston, Dominic Kwiatkowski, Cordelia Langford, John Sillitoe on behalf of the Wellcome Sanger Institute COVID-19 Surveillance Team |
| EPI_ISL_2962110 | Lighthouse Lab in Alderley Park | Wellcome Sanger Institute for the COVID-19 Genomics UK (COG-UK) Consortium | Jacquelyn Wynn, Mairead Hyland, The Lighthouse Lab in Alderley Park and Alex Alderton, Roberto Amato, Jeffrey Barrett, Sonia Goncalves, Ewan Harrison, David K. Jackson, Ian Johnston, Dominic Kwiatkowski, Cordelia Langford, John Sillitoe on behalf of the Wellcome Sanger Institute COVID-19 Surveillance Team |
| EPI_ISL_2962111 | Health Services Laboratories | Wellcome Sanger Institute for the COVID-19 Genomics UK (COG-UK) Consortium | Health Services Laboratories and Alex Alderton, Roberto Amato, Jeffrey Barrett, Sonia Goncalves, Ewan Harrison, David K. Jackson, Ian Johnston, Dominic Kwiatkowski, Cordelia Langford, John Sillitoe on behalf of the Wellcome Sanger Institute COVID-19 Surveillance Team |
| EPI_ISL_2962112 | Lighthouse Lab in Alderley Park | Wellcome Sanger Institute for the COVID-19 Genomics UK (COG-UK) Consortium | Jacquelyn Wynn, Mairead Hyland, The Lighthouse Lab in Alderley Park and Alex Alderton, Roberto Amato, Jeffrey Barrett, Sonia Goncalves, Ewan Harrison, David K. Jackson, Ian Johnston, Dominic Kwiatkowski, Cordelia Langford, John Sillitoe on behalf of the Wellcome Sanger Institute COVID-19 Surveillance Team |
| EPI_ISL_2962113 | Health Services Laboratories | Wellcome Sanger Institute for the COVID-19 Genomics UK (COG-UK) Consortium | Health Services Laboratories and Alex Alderton, Roberto Amato, Jeffrey Barrett, Sonia Goncalves, Ewan Harrison, David K. Jackson, Ian Johnston, Dominic Kwiatkowski, Cordelia Langford, John Sillitoe on behalf of the Wellcome Sanger Institute COVID-19 Surveillance Team |
| EPI_ISL_2962114, EPI_ISL_2962115, EPI_ISL_2962116, EPI_ISL_2962117, EPI_ISL_2962118 | Lighthouse Lab in Alderley Park | Wellcome Sanger Institute for the COVID-19 Genomics UK (COG-UK) Consortium | Jacquelyn Wynn, Mairead Hyland, The Lighthouse Lab in Alderley Park and Alex Alderton, Roberto Amato, Jeffrey Barrett, Sonia Goncalves, Ewan Harrison, David K. Jackson, Ian Johnston, Dominic Kwiatkowski, Cordelia Langford, John Sillitoe on behalf of the Wellcome Sanger Institute COVID-19 Surveillance Team |
| EPI_ISL_2962119, EPI_ISL_2962120 | Health Services Laboratories | Wellcome Sanger Institute for the COVID-19 Genomics UK (COG-UK) Consortium | Health Services Laboratories and Alex Alderton, Roberto Amato, Jeffrey Barrett, Sonia Goncalves, Ewan Harrison, David K. Jackson, Ian Johnston, Dominic Kwiatkowski, Cordelia Langford, John Sillitoe on behalf of the Wellcome Sanger Institute COVID-19 Surveillance Team |
| EPI_ISL_2962121, EPI_ISL_2962122 | Lighthouse Lab in Alderley Park | Wellcome Sanger Institute for the COVID-19 Genomics UK (COG-UK) Consortium | Jacquelyn Wynn, Mairead Hyland, The Lighthouse Lab in Alderley Park and Alex Alderton, Roberto Amato, Jeffrey Barrett, Sonia Goncalves, Ewan Harrison, David K. Jackson, Ian Johnston, Dominic Kwiatkowski, Cordelia Langford, John Sillitoe on behalf of the Wellcome Sanger Institute COVID-19 Surveillance Team |
| EPI_ISL_2962123, EPI_ISL_2962124 | Lighthouse Laboratory Plymouth | Wellcome Sanger Institute for the COVID-19 Genomics UK (COG-UK) Consortium | Lighthouse Laboratory Plymouth and Alex Alderton, Roberto Amato, Jeffrey Barrett, Sonia Goncalves, Ewan Harrison, David K. Jackson, Ian Johnston, Dominic Kwiatkowski, Cordelia Langford, John Sillitoe on behalf of the Wellcome Sanger Institute COVID-19 Surveillance Team |
| EPI_ISL_2962125, EPI_ISL_2962126, EPI_ISL_2962127 | Lighthouse Lab in Alderley Park | Wellcome Sanger Institute for the COVID-19 Genomics UK (COG-UK) Consortium | Jacquelyn Wynn, Mairead Hyland, The Lighthouse Lab in Alderley Park and Alex Alderton, Roberto Amato, Jeffrey Barrett, Sonia Goncalves, Ewan Harrison, David K. Jackson, Ian Johnston, Dominic Kwiatkowski, Cordelia Langford, John Sillitoe on behalf of the Wellcome Sanger Institute COVID-19 Surveillance Team |
| EPI_ISL_2962128 | Health Services Laboratories | Wellcome Sanger Institute for the COVID-19 Genomics UK (COG-UK) Consortium | Health Services Laboratories and Alex Alderton, Roberto Amato, Jeffrey Barrett, Sonia Goncalves, Ewan Harrison, David K. Jackson, Ian Johnston, Dominic Kwiatkowski, Cordelia Langford, John Sillitoe on behalf of the Wellcome Sanger Institute COVID-19 Surveillance Team |
| EPI_ISL_2962129 | Lighthouse Lab in Alderley Park | Wellcome Sanger Institute for the COVID-19 Genomics UK (COG-UK) Consortium | Jacquelyn Wynn, Mairead Hyland, The Lighthouse Lab in Alderley Park and Alex Alderton, Roberto Amato, Jeffrey Barrett, Sonia Goncalves, Ewan Harrison, David K. Jackson, Ian Johnston, Dominic Kwiatkowski, Cordelia Langford, John Sillitoe on behalf of the Wellcome Sanger Institute COVID-19 Surveillance Team |
| EPI_ISL_2962130 | Lighthouse Laboratory Plymouth | Wellcome Sanger Institute for the COVID-19 Genomics UK (COG-UK) Consortium | Lighthouse Laboratory Plymouth and Alex Alderton, Roberto Amato, Jeffrey Barrett, Sonia Goncalves, Ewan Harrison, David K. Jackson, Ian Johnston, Dominic Kwiatkowski, Cordelia Langford, John Sillitoe on behalf of the Wellcome Sanger Institute COVID-19 Surveillance Team |
| EPI_ISL_2962131 | Health Services Laboratories | Wellcome Sanger Institute for the COVID-19 Genomics UK (COG-UK) Consortium | Health Services Laboratories and Alex Alderton, Roberto Amato, Jeffrey Barrett, Sonia Goncalves, Ewan Harrison, David K. Jackson, Ian Johnston, Dominic Kwiatkowski, Cordelia Langford, John Sillitoe on behalf of the Wellcome Sanger Institute COVID-19 Surveillance Team |
| EPI_ISL_2962132, EPI_ISL_2962133, EPI_ISL_2962134, EPI_ISL_2962135 | Lighthouse Lab in Alderley Park | Wellcome Sanger Institute for the COVID-19 Genomics UK (COG-UK) Consortium | Jacquelyn Wynn, Mairead Hyland, The Lighthouse Lab in Alderley Park and Alex Alderton, Roberto Amato, Jeffrey Barrett, Sonia Goncalves, Ewan Harrison, David K. Jackson, Ian Johnston, Dominic Kwiatkowski, Cordelia Langford, John Sillitoe on behalf of the Wellcome Sanger Institute COVID-19 Surveillance Team |
| EPI_ISL_2962136 | Lighthouse Laboratory Plymouth | Wellcome Sanger Institute for the COVID-19 Genomics UK (COG-UK) Consortium | Lighthouse Laboratory Plymouth and Alex Alderton, Roberto Amato, Jeffrey Barrett, Sonia Goncalves, Ewan Harrison, David K. Jackson, Ian Johnston, Dominic Kwiatkowski, Cordelia Langford, John Sillitoe on behalf of the Wellcome Sanger Institute COVID-19 Surveillance Team |
| EPI_ISL_2962137, EPI_ISL_2962138, EPI_ISL_2962139 | Lighthouse Lab in Alderley Park | Wellcome Sanger Institute for the COVID-19 Genomics UK (COG-UK) Consortium | Jacquelyn Wynn, Mairead Hyland, The Lighthouse Lab in Alderley Park and Alex Alderton, Roberto Amato, Jeffrey Barrett, Sonia Goncalves, Ewan Harrison, David K. Jackson, Ian Johnston, Dominic Kwiatkowski, Cordelia Langford, John Sillitoe on behalf of the Wellcome Sanger Institute COVID-19 Surveillance Team |
| EPI_ISL_2962140, EPI_ISL_2962141, | Lighthouse Laboratory Plymouth | Wellcome Sanger Institute for the COVID-19 Genomics UK (COG-UK) Consortium | Lighthouse Laboratory Plymouth and Alex Alderton, Roberto Amato, Jeffrey Barrett, Sonia Goncalves, Ewan Harrison, David K. Jackson, Ian Johnston, Dominic Kwiatkowski, Cordelia Langford, John Sillitoe on behalf of the Wellcome Sanger Institute COVID-19 Surveillance Team |

| | | | |
|---|---|---|---|
| EPI_ISL_2962142, EPI_ISL_2962143 | | | |
| EPI_ISL_2962144 | Health Services Laboratories | Wellcome Sanger Institute for the COVID-19 Genomics UK (COG-UK) Consortium | Health Services Laboratories and Alex Alderton, Roberto Amato, Jeffrey Barrett, Sonia Goncalves, Ewan Harrison, David K. Jackson, Ian Johnston, Dominic Kwiatkowski, Cordelia Langford, John Sillitoe on behalf of the Wellcome Sanger Institute COVID-19 Surveillance Team |
| EPI_ISL_2962145 | Lighthouse Laboratory Plymouth | Wellcome Sanger Institute for the COVID-19 Genomics UK (COG-UK) Consortium | Lighthouse Laboratory Plymouth and Alex Alderton, Roberto Amato, Jeffrey Barrett, Sonia Goncalves, Ewan Harrison, David K. Jackson, Ian Johnston, Dominic Kwiatkowski, Cordelia Langford, John Sillitoe on behalf of the Wellcome Sanger Institute COVID-19 Surveillance Team |
| EPI_ISL_2962146 | Lighthouse Lab in Milton Keynes | Wellcome Sanger Institute for the COVID-19 Genomics UK (COG-UK) Consortium | The Lighthouse Lab in Milton Keynes and Alex Alderton, Roberto Amato, Jeffrey Barrett, Sonia Goncalves, Ewan Harrison, David K. Jackson, Ian Johnston, Dominic Kwiatkowski, Cordelia Langford, John Sillitoe on behalf of the Wellcome Sanger Institute COVID-19 Surveillance Team |
| EPI_ISL_2962147 | Lighthouse Laboratory Plymouth | Wellcome Sanger Institute for the COVID-19 Genomics UK (COG-UK) Consortium | Lighthouse Laboratory Plymouth and Alex Alderton, Roberto Amato, Jeffrey Barrett, Sonia Goncalves, Ewan Harrison, David K. Jackson, Ian Johnston, Dominic Kwiatkowski, Cordelia Langford, John Sillitoe on behalf of the Wellcome Sanger Institute COVID-19 Surveillance Team |
| EPI_ISL_2962148, EPI_ISL_2962149, EPI_ISL_2962150, EPI_ISL_2962151, EPI_ISL_2962152 | Lighthouse Lab in Milton Keynes | Wellcome Sanger Institute for the COVID-19 Genomics UK (COG-UK) Consortium | The Lighthouse Lab in Milton Keynes and Alex Alderton, Roberto Amato, Jeffrey Barrett, Sonia Goncalves, Ewan Harrison, David K. Jackson, Ian Johnston, Dominic Kwiatkowski, Cordelia Langford, John Sillitoe on behalf of the Wellcome Sanger Institute COVID-19 Surveillance Team |
| EPI_ISL_2962153 | Lighthouse Laboratory Plymouth | Wellcome Sanger Institute for the COVID-19 Genomics UK (COG-UK) Consortium | Lighthouse Laboratory Plymouth and Alex Alderton, Roberto Amato, Jeffrey Barrett, Sonia Goncalves, Ewan Harrison, David K. Jackson, Ian Johnston, Dominic Kwiatkowski, Cordelia Langford, John Sillitoe on behalf of the Wellcome Sanger Institute COVID-19 Surveillance Team |
| EPI_ISL_2962154 | Health Services Laboratories | Wellcome Sanger Institute for the COVID-19 Genomics UK (COG-UK) Consortium | Health Services Laboratories and Alex Alderton, Roberto Amato, Jeffrey Barrett, Sonia Goncalves, Ewan Harrison, David K. Jackson, Ian Johnston, Dominic Kwiatkowski, Cordelia Langford, John Sillitoe on behalf of the Wellcome Sanger Institute COVID-19 Surveillance Team |
| EPI_ISL_2962155, EPI_ISL_2962156, EPI_ISL_2962157, EPI_ISL_2962158 | Lighthouse Lab in Milton Keynes | Wellcome Sanger Institute for the COVID-19 Genomics UK (COG-UK) Consortium | The Lighthouse Lab in Milton Keynes and Alex Alderton, Roberto Amato, Jeffrey Barrett, Sonia Goncalves, Ewan Harrison, David K. Jackson, Ian Johnston, Dominic Kwiatkowski, Cordelia Langford, John Sillitoe on behalf of the Wellcome Sanger Institute COVID-19 Surveillance Team |
| EPI_ISL_2962159 | Lighthouse Laboratory Plymouth | Wellcome Sanger Institute for the COVID-19 Genomics UK (COG-UK) Consortium | Lighthouse Laboratory Plymouth and Alex Alderton, Roberto Amato, Jeffrey Barrett, Sonia Goncalves, Ewan Harrison, David K. Jackson, Ian Johnston, Dominic Kwiatkowski, Cordelia Langford, John Sillitoe on behalf of the Wellcome Sanger Institute COVID-19 Surveillance Team |
| EPI_ISL_2962160, EPI_ISL_2962161, EPI_ISL_2962162, EPI_ISL_2962163 | Lighthouse Lab in Milton Keynes | Wellcome Sanger Institute for the COVID-19 Genomics UK (COG-UK) Consortium | The Lighthouse Lab in Milton Keynes and Alex Alderton, Roberto Amato, Jeffrey Barrett, Sonia Goncalves, Ewan Harrison, David K. Jackson, Ian Johnston, Dominic Kwiatkowski, Cordelia Langford, John Sillitoe on behalf of the Wellcome Sanger Institute COVID-19 Surveillance Team |
| EPI_ISL_2962164 | Lighthouse Laboratory Plymouth | Wellcome Sanger Institute for the COVID-19 Genomics UK (COG-UK) Consortium | Lighthouse Laboratory Plymouth and Alex Alderton, Roberto Amato, Jeffrey Barrett, Sonia Goncalves, Ewan Harrison, David K. Jackson, Ian Johnston, Dominic Kwiatkowski, Cordelia Langford, John Sillitoe on behalf of the Wellcome Sanger Institute COVID-19 Surveillance Team |
| EPI_ISL_2962165 | Lighthouse Lab in Milton Keynes | Wellcome Sanger Institute for the COVID-19 Genomics UK (COG-UK) Consortium | The Lighthouse Lab in Milton Keynes and Alex Alderton, Roberto Amato, Jeffrey Barrett, Sonia Goncalves, Ewan Harrison, David K. Jackson, Ian Johnston, Dominic Kwiatkowski, Cordelia Langford, John Sillitoe on behalf of the Wellcome Sanger Institute COVID-19 Surveillance Team |
| EPI_ISL_2962166, EPI_ISL_2962167 | Health Services Laboratories | Wellcome Sanger Institute for the COVID-19 Genomics UK (COG-UK) Consortium | Health Services Laboratories and Alex Alderton, Roberto Amato, Jeffrey Barrett, Sonia Goncalves, Ewan Harrison, David K. Jackson, Ian Johnston, Dominic Kwiatkowski, Cordelia Langford, John Sillitoe on behalf of the Wellcome Sanger Institute COVID-19 Surveillance Team |
| EPI_ISL_2962168, EPI_ISL_2962169 | Lighthouse Lab in Milton Keynes | Wellcome Sanger Institute for the COVID-19 Genomics UK (COG-UK) Consortium | The Lighthouse Lab in Milton Keynes and Alex Alderton, Roberto Amato, Jeffrey Barrett, Sonia Goncalves, Ewan Harrison, David K. Jackson, Ian Johnston, Dominic Kwiatkowski, Cordelia Langford, John Sillitoe on behalf of the Wellcome Sanger Institute COVID-19 Surveillance Team |
| EPI_ISL_2962170 | Lighthouse Laboratory Plymouth | Wellcome Sanger Institute for the COVID-19 Genomics UK (COG-UK) Consortium | Lighthouse Laboratory Plymouth and Alex Alderton, Roberto Amato, Jeffrey Barrett, Sonia Goncalves, Ewan Harrison, David K. Jackson, Ian Johnston, Dominic Kwiatkowski, Cordelia Langford, John Sillitoe on behalf of the Wellcome Sanger Institute COVID-19 Surveillance Team |
| EPI_ISL_2962171, EPI_ISL_2962172 | Lighthouse Lab in Milton Keynes | Wellcome Sanger Institute for the COVID-19 Genomics UK (COG-UK) Consortium | The Lighthouse Lab in Milton Keynes and Alex Alderton, Roberto Amato, Jeffrey Barrett, Sonia Goncalves, Ewan Harrison, David K. Jackson, Ian Johnston, Dominic Kwiatkowski, Cordelia Langford, John Sillitoe on behalf of the Wellcome Sanger Institute COVID-19 Surveillance Team |
| EPI_ISL_2962173 | Health Services Laboratories | Wellcome Sanger Institute for the COVID-19 Genomics UK (COG-UK) Consortium | Health Services Laboratories and Alex Alderton, Roberto Amato, Jeffrey Barrett, Sonia Goncalves, Ewan Harrison, David K. Jackson, Ian Johnston, Dominic Kwiatkowski, Cordelia Langford, John Sillitoe on behalf of the Wellcome Sanger Institute COVID-19 Surveillance Team |
| EPI_ISL_2962174 | Lighthouse Laboratory Plymouth | Wellcome Sanger Institute for the COVID-19 Genomics UK (COG-UK) Consortium | Lighthouse Laboratory Plymouth and Alex Alderton, Roberto Amato, Jeffrey Barrett, Sonia Goncalves, Ewan Harrison, David K. Jackson, Ian Johnston, Dominic Kwiatkowski, Cordelia Langford, John Sillitoe on behalf of the Wellcome Sanger Institute COVID-19 Surveillance Team |
| EPI_ISL_2962175, EPI_ISL_2962176 | Lighthouse Lab in Milton Keynes | Wellcome Sanger Institute for the COVID-19 Genomics UK (COG-UK) Consortium | The Lighthouse Lab in Milton Keynes and Alex Alderton, Roberto Amato, Jeffrey Barrett, Sonia Goncalves, Ewan Harrison, David K. Jackson, Ian Johnston, Dominic Kwiatkowski, Cordelia Langford, John Sillitoe on behalf of the Wellcome Sanger Institute COVID-19 Surveillance Team |
| EPI_ISL_2962177 | Lighthouse Laboratory Plymouth | Wellcome Sanger Institute for the COVID-19 Genomics UK (COG-UK) Consortium | Lighthouse Laboratory Plymouth and Alex Alderton, Roberto Amato, Jeffrey Barrett, Sonia Goncalves, Ewan Harrison, David K. Jackson, Ian Johnston, Dominic Kwiatkowski, Cordelia Langford, John Sillitoe on behalf of the Wellcome Sanger Institute COVID-19 Surveillance Team |
| EPI_ISL_2962178 | Lighthouse Lab in Milton Keynes | Wellcome Sanger Institute for the COVID-19 Genomics UK (COG-UK) Consortium | The Lighthouse Lab in Milton Keynes and Alex Alderton, Roberto Amato, Jeffrey Barrett, Sonia Goncalves, Ewan Harrison, David K. Jackson, Ian Johnston, Dominic Kwiatkowski, Cordelia Langford, John Sillitoe on behalf of the Wellcome Sanger Institute COVID-19 Surveillance Team |
| EPI_ISL_2962179 | Lighthouse Laboratory Plymouth | Wellcome Sanger Institute for the COVID-19 Genomics UK (COG-UK) Consortium | Lighthouse Laboratory Plymouth and Alex Alderton, Roberto Amato, Jeffrey Barrett, Sonia Goncalves, Ewan Harrison, David K. Jackson, Ian Johnston, Dominic Kwiatkowski, Cordelia Langford, John Sillitoe on behalf of the Wellcome Sanger Institute COVID-19 Surveillance Team |
| EPI_ISL_2962180, EPI_ISL_2962181, EPI_ISL_2962182, EPI_ISL_2962183, EPI_ISL_2962184, EPI_ISL_2962185, EPI_ISL_2962186 | Lighthouse Lab in Milton Keynes | Wellcome Sanger Institute for the COVID-19 Genomics UK (COG-UK) Consortium | The Lighthouse Lab in Milton Keynes and Alex Alderton, Roberto Amato, Jeffrey Barrett, Sonia Goncalves, Ewan Harrison, David K. Jackson, Ian Johnston, Dominic Kwiatkowski, Cordelia Langford, John Sillitoe on behalf of the Wellcome Sanger Institute COVID-19 Surveillance Team |
| EPI_ISL_2962187, EPI_ISL_2962188 | Health Services Laboratories | Wellcome Sanger Institute for the COVID-19 Genomics UK (COG-UK) Consortium | Health Services Laboratories and Alex Alderton, Roberto Amato, Jeffrey Barrett, Sonia Goncalves, Ewan Harrison, David K. Jackson, Ian Johnston, Dominic Kwiatkowski, Cordelia Langford, John Sillitoe on behalf of the Wellcome Sanger Institute COVID-19 Surveillance Team |
| EPI_ISL_2962189 | Lighthouse Lab in Milton Keynes | Wellcome Sanger Institute for the COVID-19 Genomics UK (COG-UK) Consortium | The Lighthouse Lab in Milton Keynes and Alex Alderton, Roberto Amato, Jeffrey Barrett, Sonia Goncalves, Ewan Harrison, David K. Jackson, Ian Johnston, Dominic Kwiatkowski, Cordelia Langford, John Sillitoe on behalf of the Wellcome Sanger Institute COVID-19 Surveillance Team |
| EPI_ISL_2962190 | Health Services Laboratories | Wellcome Sanger Institute for the COVID-19 Genomics UK (COG-UK) Consortium | Health Services Laboratories and Alex Alderton, Roberto Amato, Jeffrey Barrett, Sonia Goncalves, Ewan Harrison, David K. Jackson, Ian Johnston, Dominic Kwiatkowski, Cordelia Langford, John Sillitoe on behalf of the Wellcome Sanger Institute COVID-19 Surveillance Team |
| EPI_ISL_2962191 | Lighthouse Laboratory Plymouth | Wellcome Sanger Institute for the COVID-19 Genomics UK (COG-UK) Consortium | Lighthouse Laboratory Plymouth and Alex Alderton, Roberto Amato, Jeffrey Barrett, Sonia Goncalves, Ewan Harrison, David K. Jackson, Ian Johnston, Dominic Kwiatkowski, Cordelia Langford, John Sillitoe on behalf of the Wellcome Sanger Institute COVID-19 Surveillance Team |
| EPI_ISL_2962192 | Health Services Laboratories | Wellcome Sanger Institute for the COVID-19 Genomics UK (COG-UK) Consortium | Health Services Laboratories and Alex Alderton, Roberto Amato, Jeffrey Barrett, Sonia Goncalves, Ewan Harrison, David K. Jackson, Ian Johnston, Dominic Kwiatkowski, Cordelia Langford, John Sillitoe on behalf of the Wellcome Sanger Institute COVID-19 Surveillance Team |
| EPI_ISL_2962193, EPI_ISL_2962194 | Lighthouse Lab in Milton Keynes | Wellcome Sanger Institute for the COVID-19 Genomics UK (COG-UK) Consortium | The Lighthouse Lab in Milton Keynes and Alex Alderton, Roberto Amato, Jeffrey Barrett, Sonia Goncalves, Ewan Harrison, David K. Jackson, Ian Johnston, Dominic Kwiatkowski, Cordelia Langford, John Sillitoe on behalf of the Wellcome Sanger Institute COVID-19 Surveillance Team |
| EPI_ISL_2962195 | Lighthouse Laboratory Plymouth | Wellcome Sanger Institute for the COVID-19 Genomics UK (COG-UK) Consortium | Lighthouse Laboratory Plymouth and Alex Alderton, Roberto Amato, Jeffrey Barrett, Sonia Goncalves, Ewan Harrison, David K. Jackson, Ian Johnston, Dominic Kwiatkowski, Cordelia Langford, John Sillitoe on behalf of the Wellcome Sanger Institute COVID-19 Surveillance Team |
| EPI_ISL_2962196 | Health Services Laboratories | Wellcome Sanger Institute for the COVID-19 Genomics UK (COG-UK) Consortium | Health Services Laboratories and Alex Alderton, Roberto Amato, Jeffrey Barrett, Sonia Goncalves, Ewan Harrison, David K. Jackson, Ian Johnston, Dominic Kwiatkowski, Cordelia Langford, John Sillitoe on behalf of the Wellcome Sanger Institute COVID-19 Surveillance Team |
| EPI_ISL_2962197 | Lighthouse Lab in Milton Keynes | Wellcome Sanger Institute for the COVID-19 Genomics UK (COG-UK) Consortium | The Lighthouse Lab in Milton Keynes and Alex Alderton, Roberto Amato, Jeffrey Barrett, Sonia Goncalves, Ewan Harrison, David K. Jackson, Ian Johnston, Dominic Kwiatkowski, Cordelia Langford, John Sillitoe on behalf of the Wellcome Sanger Institute COVID-19 Surveillance Team |
| EPI_ISL_2962198 | Lighthouse Laboratory Plymouth | Wellcome Sanger Institute for the COVID-19 Genomics UK (COG-UK) Consortium | Lighthouse Laboratory Plymouth and Alex Alderton, Roberto Amato, Jeffrey Barrett, Sonia Goncalves, Ewan Harrison, David K. Jackson, Ian Johnston, Dominic Kwiatkowski, Cordelia Langford, John Sillitoe on behalf of the Wellcome Sanger Institute COVID-19 Surveillance Team |
| EPI_ISL_2962199 | Lighthouse Lab in Milton Keynes | Wellcome Sanger Institute for the COVID-19 Genomics UK (COG-UK) Consortium | The Lighthouse Lab in Milton Keynes and Alex Alderton, Roberto Amato, Jeffrey Barrett, Sonia Goncalves, Ewan Harrison, David K. Jackson, Ian Johnston, Dominic Kwiatkowski, Cordelia Langford, John Sillitoe on behalf of the Wellcome Sanger Institute COVID-19 Surveillance Team |
| EPI_ISL_2962200 | Health Services Laboratories | Wellcome Sanger Institute for the COVID-19 Genomics UK (COG-UK) Consortium | Health Services Laboratories and Alex Alderton, Roberto Amato, Jeffrey Barrett, Sonia Goncalves, Ewan Harrison, David K. Jackson, Ian Johnston, Dominic Kwiatkowski, Cordelia Langford, John Sillitoe on behalf of the Wellcome Sanger Institute COVID-19 Surveillance Team |
| EPI_ISL_2962201 | Lighthouse Lab in Milton Keynes | Wellcome Sanger Institute for the COVID-19 Genomics UK (COG-UK) Consortium | The Lighthouse Lab in Milton Keynes and Alex Alderton, Roberto Amato, Jeffrey Barrett, Sonia Goncalves, Ewan Harrison, David K. Jackson, Ian Johnston, Dominic Kwiatkowski, Cordelia Langford, John Sillitoe on behalf of the Wellcome Sanger Institute COVID-19 Surveillance Team |
| EPI_ISL_2962202 | Health Services Laboratories | Wellcome Sanger Institute for the COVID-19 Genomics UK (COG-UK) Consortium | Health Services Laboratories and Alex Alderton, Roberto Amato, Jeffrey Barrett, Sonia Goncalves, Ewan Harrison, David K. Jackson, Ian Johnston, Dominic Kwiatkowski, Cordelia Langford, John Sillitoe on behalf of the Wellcome Sanger Institute COVID-19 Surveillance Team |
| EPI_ISL_2962203, EPI_ISL_2962204, EPI_ISL_2962205 | Lighthouse Lab in Milton Keynes | Wellcome Sanger Institute for the COVID-19 Genomics UK (COG-UK) Consortium | The Lighthouse Lab in Milton Keynes and Alex Alderton, Roberto Amato, Jeffrey Barrett, Sonia Goncalves, Ewan Harrison, David K. Jackson, Ian Johnston, Dominic Kwiatkowski, Cordelia Langford, John Sillitoe on behalf of the Wellcome Sanger Institute COVID-19 Surveillance Team |
| EPI_ISL_2962206, | Lighthouse Laboratory Plymouth | Wellcome Sanger Institute for the COVID-19 Genomics UK (COG-UK) | Lighthouse Laboratory Plymouth and Alex Alderton, Roberto Amato, Jeffrey Barrett, Sonia Goncalves, Ewan Harrison, David K. Jackson, Ian Johnston, Dominic Kwiatkowski, Cordelia Langford, John Sillitoe on behalf of the Wellcome Sanger |

| | | | |
|---|---|---|---|
| EPI_ISL_2962207 | | Consortium | Institute COVID-19 Surveillance Team |
| EPI_ISL_2962208 | Lighthouse Lab in Milton Keynes | Wellcome Sanger Institute for the COVID-19 Genomics UK (COG-UK) Consortium | The Lighthouse Lab in Milton Keynes and Alex Alderton, Roberto Amato, Jeffrey Barrett, Sonia Goncalves, Ewan Harrison, David K. Jackson, Ian Johnston, Dominic Kwiatkowski, Cordelia Langford, John Sillitoe on behalf of the Wellcome Sanger Institute COVID-19 Surveillance Team |
| EPI_ISL_2962209 | Lighthouse Laboratory Plymouth | Wellcome Sanger Institute for the COVID-19 Genomics UK (COG-UK) Consortium | Lighthouse Laboratory Plymouth and Alex Alderton, Roberto Amato, Jeffrey Barrett, Sonia Goncalves, Ewan Harrison, David K. Jackson, Ian Johnston, Dominic Kwiatkowski, Cordelia Langford, John Sillitoe on behalf of the Wellcome Sanger Institute COVID-19 Surveillance Team |
| EPI_ISL_2962210 | Lighthouse Lab in Milton Keynes | Wellcome Sanger Institute for the COVID-19 Genomics UK (COG-UK) Consortium | The Lighthouse Lab in Milton Keynes and Alex Alderton, Roberto Amato, Jeffrey Barrett, Sonia Goncalves, Ewan Harrison, David K. Jackson, Ian Johnston, Dominic Kwiatkowski, Cordelia Langford, John Sillitoe on behalf of the Wellcome Sanger Institute COVID-19 Surveillance Team |
| EPI_ISL_2962211, EPI_ISL_2962212 | Lighthouse Laboratory Plymouth | Wellcome Sanger Institute for the COVID-19 Genomics UK (COG-UK) Consortium | Lighthouse Laboratory Plymouth and Alex Alderton, Roberto Amato, Jeffrey Barrett, Sonia Goncalves, Ewan Harrison, David K. Jackson, Ian Johnston, Dominic Kwiatkowski, Cordelia Langford, John Sillitoe on behalf of the Wellcome Sanger Institute COVID-19 Surveillance Team |
| EPI_ISL_2962213 | Health Services Laboratories | Wellcome Sanger Institute for the COVID-19 Genomics UK (COG-UK) Consortium | Health Services Laboratories and Alex Alderton, Roberto Amato, Jeffrey Barrett, Sonia Goncalves, Ewan Harrison, David K. Jackson, Ian Johnston, Dominic Kwiatkowski, Cordelia Langford, John Sillitoe on behalf of the Wellcome Sanger Institute COVID-19 Surveillance Team |
| EPI_ISL_2962214, EPI_ISL_2962215, EPI_ISL_2962216 | Lighthouse Lab in Milton Keynes | Wellcome Sanger Institute for the COVID-19 Genomics UK (COG-UK) Consortium | The Lighthouse Lab in Milton Keynes and Alex Alderton, Roberto Amato, Jeffrey Barrett, Sonia Goncalves, Ewan Harrison, David K. Jackson, Ian Johnston, Dominic Kwiatkowski, Cordelia Langford, John Sillitoe on behalf of the Wellcome Sanger Institute COVID-19 Surveillance Team |
| EPI_ISL_2962217 | Lighthouse Laboratory Plymouth | Wellcome Sanger Institute for the COVID-19 Genomics UK (COG-UK) Consortium | Lighthouse Laboratory Plymouth and Alex Alderton, Roberto Amato, Jeffrey Barrett, Sonia Goncalves, Ewan Harrison, David K. Jackson, Ian Johnston, Dominic Kwiatkowski, Cordelia Langford, John Sillitoe on behalf of the Wellcome Sanger Institute COVID-19 Surveillance Team |
| EPI_ISL_2962218, EPI_ISL_2962219, EPI_ISL_2962220, EPI_ISL_2962221, EPI_ISL_2962222, EPI_ISL_2962223 | Lighthouse Lab in Milton Keynes | Wellcome Sanger Institute for the COVID-19 Genomics UK (COG-UK) Consortium | The Lighthouse Lab in Milton Keynes and Alex Alderton, Roberto Amato, Jeffrey Barrett, Sonia Goncalves, Ewan Harrison, David K. Jackson, Ian Johnston, Dominic Kwiatkowski, Cordelia Langford, John Sillitoe on behalf of the Wellcome Sanger Institute COVID-19 Surveillance Team |
| EPI_ISL_2962224 | Lighthouse Laboratory Plymouth | Wellcome Sanger Institute for the COVID-19 Genomics UK (COG-UK) Consortium | Lighthouse Laboratory Plymouth and Alex Alderton, Roberto Amato, Jeffrey Barrett, Sonia Goncalves, Ewan Harrison, David K. Jackson, Ian Johnston, Dominic Kwiatkowski, Cordelia Langford, John Sillitoe on behalf of the Wellcome Sanger Institute COVID-19 Surveillance Team |
| EPI_ISL_2962225 | Lighthouse Lab in Milton Keynes | Wellcome Sanger Institute for the COVID-19 Genomics UK (COG-UK) Consortium | The Lighthouse Lab in Milton Keynes and Alex Alderton, Roberto Amato, Jeffrey Barrett, Sonia Goncalves, Ewan Harrison, David K. Jackson, Ian Johnston, Dominic Kwiatkowski, Cordelia Langford, John Sillitoe on behalf of the Wellcome Sanger Institute COVID-19 Surveillance Team |
| EPI_ISL_2962226, EPI_ISL_2962227, EPI_ISL_2962228, EPI_ISL_2962229, EPI_ISL_2962230 | Lighthouse Laboratory Plymouth | Wellcome Sanger Institute for the COVID-19 Genomics UK (COG-UK) Consortium | Lighthouse Laboratory Plymouth and Alex Alderton, Roberto Amato, Jeffrey Barrett, Sonia Goncalves, Ewan Harrison, David K. Jackson, Ian Johnston, Dominic Kwiatkowski, Cordelia Langford, John Sillitoe on behalf of the Wellcome Sanger Institute COVID-19 Surveillance Team |
| EPI_ISL_2962231, EPI_ISL_2962232, EPI_ISL_2962233 | Lighthouse Lab in Milton Keynes | Wellcome Sanger Institute for the COVID-19 Genomics UK (COG-UK) Consortium | The Lighthouse Lab in Milton Keynes and Alex Alderton, Roberto Amato, Jeffrey Barrett, Sonia Goncalves, Ewan Harrison, David K. Jackson, Ian Johnston, Dominic Kwiatkowski, Cordelia Langford, John Sillitoe on behalf of the Wellcome Sanger Institute COVID-19 Surveillance Team |
| EPI_ISL_2962234, EPI_ISL_2962235 | Health Services Laboratories | Wellcome Sanger Institute for the COVID-19 Genomics UK (COG-UK) Consortium | Health Services Laboratories and Alex Alderton, Roberto Amato, Jeffrey Barrett, Sonia Goncalves, Ewan Harrison, David K. Jackson, Ian Johnston, Dominic Kwiatkowski, Cordelia Langford, John Sillitoe on behalf of the Wellcome Sanger Institute COVID-19 Surveillance Team |
| EPI_ISL_2962236 | Lighthouse Laboratory Plymouth | Wellcome Sanger Institute for the COVID-19 Genomics UK (COG-UK) Consortium | Lighthouse Laboratory Plymouth and Alex Alderton, Roberto Amato, Jeffrey Barrett, Sonia Goncalves, Ewan Harrison, David K. Jackson, Ian Johnston, Dominic Kwiatkowski, Cordelia Langford, John Sillitoe on behalf of the Wellcome Sanger Institute COVID-19 Surveillance Team |
| EPI_ISL_2962237 | Lighthouse Lab in Milton Keynes | Wellcome Sanger Institute for the COVID-19 Genomics UK (COG-UK) Consortium | The Lighthouse Lab in Milton Keynes and Alex Alderton, Roberto Amato, Jeffrey Barrett, Sonia Goncalves, Ewan Harrison, David K. Jackson, Ian Johnston, Dominic Kwiatkowski, Cordelia Langford, John Sillitoe on behalf of the Wellcome Sanger Institute COVID-19 Surveillance Team |
| EPI_ISL_2962238 | Health Services Laboratories | Wellcome Sanger Institute for the COVID-19 Genomics UK (COG-UK) Consortium | Health Services Laboratories and Alex Alderton, Roberto Amato, Jeffrey Barrett, Sonia Goncalves, Ewan Harrison, David K. Jackson, Ian Johnston, Dominic Kwiatkowski, Cordelia Langford, John Sillitoe on behalf of the Wellcome Sanger Institute COVID-19 Surveillance Team |
| EPI_ISL_2962239, EPI_ISL_2962240 | Lighthouse Lab in Milton Keynes | Wellcome Sanger Institute for the COVID-19 Genomics UK (COG-UK) Consortium | The Lighthouse Lab in Milton Keynes and Alex Alderton, Roberto Amato, Jeffrey Barrett, Sonia Goncalves, Ewan Harrison, David K. Jackson, Ian Johnston, Dominic Kwiatkowski, Cordelia Langford, John Sillitoe on behalf of the Wellcome Sanger Institute COVID-19 Surveillance Team |
| EPI_ISL_2962241 | Lighthouse Laboratory Plymouth | Wellcome Sanger Institute for the COVID-19 Genomics UK (COG-UK) Consortium | Lighthouse Laboratory Plymouth and Alex Alderton, Roberto Amato, Jeffrey Barrett, Sonia Goncalves, Ewan Harrison, David K. Jackson, Ian Johnston, Dominic Kwiatkowski, Cordelia Langford, John Sillitoe on behalf of the Wellcome Sanger Institute COVID-19 Surveillance Team |
| EPI_ISL_2962242, EPI_ISL_2962243 | Lighthouse Lab in Milton Keynes | Wellcome Sanger Institute for the COVID-19 Genomics UK (COG-UK) Consortium | The Lighthouse Lab in Milton Keynes and Alex Alderton, Roberto Amato, Jeffrey Barrett, Sonia Goncalves, Ewan Harrison, David K. Jackson, Ian Johnston, Dominic Kwiatkowski, Cordelia Langford, John Sillitoe on behalf of the Wellcome Sanger Institute COVID-19 Surveillance Team |
| EPI_ISL_2962244, EPI_ISL_2962245 | Lighthouse Laboratory Plymouth | Wellcome Sanger Institute for the COVID-19 Genomics UK (COG-UK) Consortium | Lighthouse Laboratory Plymouth and Alex Alderton, Roberto Amato, Jeffrey Barrett, Sonia Goncalves, Ewan Harrison, David K. Jackson, Ian Johnston, Dominic Kwiatkowski, Cordelia Langford, John Sillitoe on behalf of the Wellcome Sanger Institute COVID-19 Surveillance Team |
| EPI_ISL_2962246 | Lighthouse Lab in Milton Keynes | Wellcome Sanger Institute for the COVID-19 Genomics UK (COG-UK) Consortium | The Lighthouse Lab in Milton Keynes and Alex Alderton, Roberto Amato, Jeffrey Barrett, Sonia Goncalves, Ewan Harrison, David K. Jackson, Ian Johnston, Dominic Kwiatkowski, Cordelia Langford, John Sillitoe on behalf of the Wellcome Sanger Institute COVID-19 Surveillance Team |
| EPI_ISL_2962247 | Lighthouse Laboratory Plymouth | Wellcome Sanger Institute for the COVID-19 Genomics UK (COG-UK) Consortium | Lighthouse Laboratory Plymouth and Alex Alderton, Roberto Amato, Jeffrey Barrett, Sonia Goncalves, Ewan Harrison, David K. Jackson, Ian Johnston, Dominic Kwiatkowski, Cordelia Langford, John Sillitoe on behalf of the Wellcome Sanger Institute COVID-19 Surveillance Team |
| EPI_ISL_2962248 | Health Services Laboratories | Wellcome Sanger Institute for the COVID-19 Genomics UK (COG-UK) Consortium | Health Services Laboratories and Alex Alderton, Roberto Amato, Jeffrey Barrett, Sonia Goncalves, Ewan Harrison, David K. Jackson, Ian Johnston, Dominic Kwiatkowski, Cordelia Langford, John Sillitoe on behalf of the Wellcome Sanger Institute COVID-19 Surveillance Team |
| EPI_ISL_2962249, EPI_ISL_2962250, EPI_ISL_2962251, EPI_ISL_2962252 | Lighthouse Lab in Milton Keynes | Wellcome Sanger Institute for the COVID-19 Genomics UK (COG-UK) Consortium | The Lighthouse Lab in Milton Keynes and Alex Alderton, Roberto Amato, Jeffrey Barrett, Sonia Goncalves, Ewan Harrison, David K. Jackson, Ian Johnston, Dominic Kwiatkowski, Cordelia Langford, John Sillitoe on behalf of the Wellcome Sanger Institute COVID-19 Surveillance Team |
| EPI_ISL_2962253 | Lighthouse Laboratory Plymouth | Wellcome Sanger Institute for the COVID-19 Genomics UK (COG-UK) Consortium | Lighthouse Laboratory Plymouth and Alex Alderton, Roberto Amato, Jeffrey Barrett, Sonia Goncalves, Ewan Harrison, David K. Jackson, Ian Johnston, Dominic Kwiatkowski, Cordelia Langford, John Sillitoe on behalf of the Wellcome Sanger Institute COVID-19 Surveillance Team |
| EPI_ISL_2962254, EPI_ISL_2962255 | Health Services Laboratories | Wellcome Sanger Institute for the COVID-19 Genomics UK (COG-UK) Consortium | Health Services Laboratories and Alex Alderton, Roberto Amato, Jeffrey Barrett, Sonia Goncalves, Ewan Harrison, David K. Jackson, Ian Johnston, Dominic Kwiatkowski, Cordelia Langford, John Sillitoe on behalf of the Wellcome Sanger Institute COVID-19 Surveillance Team |
| EPI_ISL_2962256, EPI_ISL_2962257 | Lighthouse Lab in Milton Keynes | Wellcome Sanger Institute for the COVID-19 Genomics UK (COG-UK) Consortium | The Lighthouse Lab in Milton Keynes and Alex Alderton, Roberto Amato, Jeffrey Barrett, Sonia Goncalves, Ewan Harrison, David K. Jackson, Ian Johnston, Dominic Kwiatkowski, Cordelia Langford, John Sillitoe on behalf of the Wellcome Sanger Institute COVID-19 Surveillance Team |
| EPI_ISL_2962258 | Health Services Laboratories | Wellcome Sanger Institute for the COVID-19 Genomics UK (COG-UK) Consortium | Health Services Laboratories and Alex Alderton, Roberto Amato, Jeffrey Barrett, Sonia Goncalves, Ewan Harrison, David K. Jackson, Ian Johnston, Dominic Kwiatkowski, Cordelia Langford, John Sillitoe on behalf of the Wellcome Sanger Institute COVID-19 Surveillance Team |
| EPI_ISL_2962259 | Lighthouse Lab in Milton Keynes | Wellcome Sanger Institute for the COVID-19 Genomics UK (COG-UK) Consortium | The Lighthouse Lab in Milton Keynes and Alex Alderton, Roberto Amato, Jeffrey Barrett, Sonia Goncalves, Ewan Harrison, David K. Jackson, Ian Johnston, Dominic Kwiatkowski, Cordelia Langford, John Sillitoe on behalf of the Wellcome Sanger Institute COVID-19 Surveillance Team |
| EPI_ISL_2962260 | Health Services Laboratories | Wellcome Sanger Institute for the COVID-19 Genomics UK (COG-UK) Consortium | Health Services Laboratories and Alex Alderton, Roberto Amato, Jeffrey Barrett, Sonia Goncalves, Ewan Harrison, David K. Jackson, Ian Johnston, Dominic Kwiatkowski, Cordelia Langford, John Sillitoe on behalf of the Wellcome Sanger Institute COVID-19 Surveillance Team |
| EPI_ISL_2962261, EPI_ISL_2962262, EPI_ISL_2962263, EPI_ISL_2962264, EPI_ISL_2962265 | Lighthouse Lab in Milton Keynes | Wellcome Sanger Institute for the COVID-19 Genomics UK (COG-UK) Consortium | The Lighthouse Lab in Milton Keynes and Alex Alderton, Roberto Amato, Jeffrey Barrett, Sonia Goncalves, Ewan Harrison, David K. Jackson, Ian Johnston, Dominic Kwiatkowski, Cordelia Langford, John Sillitoe on behalf of the Wellcome Sanger Institute COVID-19 Surveillance Team |
| EPI_ISL_2962266 | Health Services Laboratories | Wellcome Sanger Institute for the COVID-19 Genomics UK (COG-UK) Consortium | Health Services Laboratories and Alex Alderton, Roberto Amato, Jeffrey Barrett, Sonia Goncalves, Ewan Harrison, David K. Jackson, Ian Johnston, Dominic Kwiatkowski, Cordelia Langford, John Sillitoe on behalf of the Wellcome Sanger Institute COVID-19 Surveillance Team |
| EPI_ISL_2962267 | Lighthouse Lab in Milton Keynes | Wellcome Sanger Institute for the COVID-19 Genomics UK (COG-UK) Consortium | The Lighthouse Lab in Milton Keynes and Alex Alderton, Roberto Amato, Jeffrey Barrett, Sonia Goncalves, Ewan Harrison, David K. Jackson, Ian Johnston, Dominic Kwiatkowski, Cordelia Langford, John Sillitoe on behalf of the Wellcome Sanger Institute COVID-19 Surveillance Team |
| EPI_ISL_2962268 | Lighthouse Laboratory Plymouth | Wellcome Sanger Institute for the COVID-19 Genomics UK (COG-UK) Consortium | Lighthouse Laboratory Plymouth and Alex Alderton, Roberto Amato, Jeffrey Barrett, Sonia Goncalves, Ewan Harrison, David K. Jackson, Ian Johnston, Dominic Kwiatkowski, Cordelia Langford, John Sillitoe on behalf of the Wellcome Sanger Institute COVID-19 Surveillance Team |
| EPI_ISL_2962269 | Lighthouse Lab in Milton Keynes | Wellcome Sanger Institute for the COVID-19 Genomics UK (COG-UK) Consortium | The Lighthouse Lab in Milton Keynes and Alex Alderton, Roberto Amato, Jeffrey Barrett, Sonia Goncalves, Ewan Harrison, David K. Jackson, Ian Johnston, Dominic Kwiatkowski, Cordelia Langford, John Sillitoe on behalf of the Wellcome Sanger Institute COVID-19 Surveillance Team |
| EPI_ISL_2962270, EPI_ISL_2962271 | Health Services Laboratories | Wellcome Sanger Institute for the COVID-19 Genomics UK (COG-UK) Consortium | Health Services Laboratories and Alex Alderton, Roberto Amato, Jeffrey Barrett, Sonia Goncalves, Ewan Harrison, David K. Jackson, Ian Johnston, Dominic Kwiatkowski, Cordelia Langford, John Sillitoe on behalf of the Wellcome Sanger Institute COVID-19 Surveillance Team |
| EPI_ISL_2962272 | Lighthouse Laboratory Plymouth | Wellcome Sanger Institute for the COVID-19 Genomics UK (COG-UK) Consortium | Lighthouse Laboratory Plymouth and Alex Alderton, Roberto Amato, Jeffrey Barrett, Sonia Goncalves, Ewan Harrison, David K. Jackson, Ian Johnston, Dominic Kwiatkowski, Cordelia Langford, John Sillitoe on behalf of the Wellcome Sanger Institute COVID-19 Surveillance Team |
| EPI_ISL_2962273 | Lighthouse Lab in Milton Keynes | Wellcome Sanger Institute for the COVID-19 Genomics UK (COG-UK) Consortium | The Lighthouse Lab in Milton Keynes and Alex Alderton, Roberto Amato, Jeffrey Barrett, Sonia Goncalves, Ewan Harrison, David K. Jackson, Ian Johnston, Dominic Kwiatkowski, Cordelia Langford, John Sillitoe on behalf of the Wellcome Sanger Institute COVID-19 Surveillance Team |
| EPI_ISL_2962274 | Health Services Laboratories | Wellcome Sanger Institute for the COVID-19 Genomics UK (COG-UK) | Health Services Laboratories and Alex Alderton, Roberto Amato, Jeffrey Barrett, Sonia Goncalves, Ewan Harrison, David K. Jackson, Ian Johnston, Dominic Kwiatkowski, Cordelia Langford, John Sillitoe on behalf of the Wellcome Sanger Institute |

| | | | |
|---|---|---|---|
| | | Consortium | COVID-19 Surveillance Team |
| EPI_ISL_2962275, EPI_ISL_2962276 | Lighthouse Laboratory Plymouth | Wellcome Sanger Institute for the COVID-19 Genomics UK (COG-UK) Consortium | Lighthouse Laboratory Plymouth and Alex Alderton, Roberto Amato, Jeffrey Barrett, Sonia Goncalves, Ewan Harrison, David K. Jackson, Ian Johnston, Dominic Kwiatkowski, Cordelia Langford, John Sillitoe on behalf of the Wellcome Sanger Institute COVID-19 Surveillance Team |
| EPI_ISL_2962277 | Lighthouse Lab in Milton Keynes | Wellcome Sanger Institute for the COVID-19 Genomics UK (COG-UK) Consortium | The Lighthouse Lab in Milton Keynes and Alex Alderton, Roberto Amato, Jeffrey Barrett, Sonia Goncalves, Ewan Harrison, David K. Jackson, Ian Johnston, Dominic Kwiatkowski, Cordelia Langford, John Sillitoe on behalf of the Wellcome Sanger Institute COVID-19 Surveillance Team |
| EPI_ISL_2962278 | Health Services Laboratories | Wellcome Sanger Institute for the COVID-19 Genomics UK (COG-UK) Consortium | Health Services Laboratories and Alex Alderton, Roberto Amato, Jeffrey Barrett, Sonia Goncalves, Ewan Harrison, David K. Jackson, Ian Johnston, Dominic Kwiatkowski, Cordelia Langford, John Sillitoe on behalf of the Wellcome Sanger Institute COVID-19 Surveillance Team |
| EPI_ISL_2962279 | Lighthouse Lab in Milton Keynes | Wellcome Sanger Institute for the COVID-19 Genomics UK (COG-UK) Consortium | The Lighthouse Lab in Milton Keynes and Alex Alderton, Roberto Amato, Jeffrey Barrett, Sonia Goncalves, Ewan Harrison, David K. Jackson, Ian Johnston, Dominic Kwiatkowski, Cordelia Langford, John Sillitoe on behalf of the Wellcome Sanger Institute COVID-19 Surveillance Team |
| EPI_ISL_2962280 | Health Services Laboratories | Wellcome Sanger Institute for the COVID-19 Genomics UK (COG-UK) Consortium | Health Services Laboratories and Alex Alderton, Roberto Amato, Jeffrey Barrett, Sonia Goncalves, Ewan Harrison, David K. Jackson, Ian Johnston, Dominic Kwiatkowski, Cordelia Langford, John Sillitoe on behalf of the Wellcome Sanger Institute COVID-19 Surveillance Team |
| EPI_ISL_2962281 | Lighthouse Lab in Milton Keynes | Wellcome Sanger Institute for the COVID-19 Genomics UK (COG-UK) Consortium | The Lighthouse Lab in Milton Keynes and Alex Alderton, Roberto Amato, Jeffrey Barrett, Sonia Goncalves, Ewan Harrison, David K. Jackson, Ian Johnston, Dominic Kwiatkowski, Cordelia Langford, John Sillitoe on behalf of the Wellcome Sanger Institute COVID-19 Surveillance Team |
| EPI_ISL_2962282 | Lighthouse Laboratory Plymouth | Wellcome Sanger Institute for the COVID-19 Genomics UK (COG-UK) Consortium | Lighthouse Laboratory Plymouth and Alex Alderton, Roberto Amato, Jeffrey Barrett, Sonia Goncalves, Ewan Harrison, David K. Jackson, Ian Johnston, Dominic Kwiatkowski, Cordelia Langford, John Sillitoe on behalf of the Wellcome Sanger Institute COVID-19 Surveillance Team |
| EPI_ISL_2962283 | Health Services Laboratories | Wellcome Sanger Institute for the COVID-19 Genomics UK (COG-UK) Consortium | Health Services Laboratories and Alex Alderton, Roberto Amato, Jeffrey Barrett, Sonia Goncalves, Ewan Harrison, David K. Jackson, Ian Johnston, Dominic Kwiatkowski, Cordelia Langford, John Sillitoe on behalf of the Wellcome Sanger Institute COVID-19 Surveillance Team |
| EPI_ISL_2962284 | Lighthouse Laboratory Plymouth | Wellcome Sanger Institute for the COVID-19 Genomics UK (COG-UK) Consortium | Lighthouse Laboratory Plymouth and Alex Alderton, Roberto Amato, Jeffrey Barrett, Sonia Goncalves, Ewan Harrison, David K. Jackson, Ian Johnston, Dominic Kwiatkowski, Cordelia Langford, John Sillitoe on behalf of the Wellcome Sanger Institute COVID-19 Surveillance Team |
| EPI_ISL_2962285, EPI_ISL_2962286 | Lighthouse Lab in Milton Keynes | Wellcome Sanger Institute for the COVID-19 Genomics UK (COG-UK) Consortium | The Lighthouse Lab in Milton Keynes and Alex Alderton, Roberto Amato, Jeffrey Barrett, Sonia Goncalves, Ewan Harrison, David K. Jackson, Ian Johnston, Dominic Kwiatkowski, Cordelia Langford, John Sillitoe on behalf of the Wellcome Sanger Institute COVID-19 Surveillance Team |
| EPI_ISL_2962287 | Health Services Laboratories | Wellcome Sanger Institute for the COVID-19 Genomics UK (COG-UK) Consortium | Health Services Laboratories and Alex Alderton, Roberto Amato, Jeffrey Barrett, Sonia Goncalves, Ewan Harrison, David K. Jackson, Ian Johnston, Dominic Kwiatkowski, Cordelia Langford, John Sillitoe on behalf of the Wellcome Sanger Institute COVID-19 Surveillance Team |
| EPI_ISL_2962288 | Lighthouse Laboratory Plymouth | Wellcome Sanger Institute for the COVID-19 Genomics UK (COG-UK) Consortium | Lighthouse Laboratory Plymouth and Alex Alderton, Roberto Amato, Jeffrey Barrett, Sonia Goncalves, Ewan Harrison, David K. Jackson, Ian Johnston, Dominic Kwiatkowski, Cordelia Langford, John Sillitoe on behalf of the Wellcome Sanger Institute COVID-19 Surveillance Team |
| EPI_ISL_2962289 | Health Services Laboratories | Wellcome Sanger Institute for the COVID-19 Genomics UK (COG-UK) Consortium | Health Services Laboratories and Alex Alderton, Roberto Amato, Jeffrey Barrett, Sonia Goncalves, Ewan Harrison, David K. Jackson, Ian Johnston, Dominic Kwiatkowski, Cordelia Langford, John Sillitoe on behalf of the Wellcome Sanger Institute COVID-19 Surveillance Team |
| EPI_ISL_2962290, EPI_ISL_2962291 | Lighthouse Lab in Milton Keynes | Wellcome Sanger Institute for the COVID-19 Genomics UK (COG-UK) Consortium | The Lighthouse Lab in Milton Keynes and Alex Alderton, Roberto Amato, Jeffrey Barrett, Sonia Goncalves, Ewan Harrison, David K. Jackson, Ian Johnston, Dominic Kwiatkowski, Cordelia Langford, John Sillitoe on behalf of the Wellcome Sanger Institute COVID-19 Surveillance Team |
| EPI_ISL_2962292 | Health Services Laboratories | Wellcome Sanger Institute for the COVID-19 Genomics UK (COG-UK) Consortium | Health Services Laboratories and Alex Alderton, Roberto Amato, Jeffrey Barrett, Sonia Goncalves, Ewan Harrison, David K. Jackson, Ian Johnston, Dominic Kwiatkowski, Cordelia Langford, John Sillitoe on behalf of the Wellcome Sanger Institute COVID-19 Surveillance Team |
| EPI_ISL_2962293 | Lighthouse Lab in Milton Keynes | Wellcome Sanger Institute for the COVID-19 Genomics UK (COG-UK) Consortium | The Lighthouse Lab in Milton Keynes and Alex Alderton, Roberto Amato, Jeffrey Barrett, Sonia Goncalves, Ewan Harrison, David K. Jackson, Ian Johnston, Dominic Kwiatkowski, Cordelia Langford, John Sillitoe on behalf of the Wellcome Sanger Institute COVID-19 Surveillance Team |
| EPI_ISL_2962294 | Lighthouse Laboratory Plymouth | Wellcome Sanger Institute for the COVID-19 Genomics UK (COG-UK) Consortium | Lighthouse Laboratory Plymouth and Alex Alderton, Roberto Amato, Jeffrey Barrett, Sonia Goncalves, Ewan Harrison, David K. Jackson, Ian Johnston, Dominic Kwiatkowski, Cordelia Langford, John Sillitoe on behalf of the Wellcome Sanger Institute COVID-19 Surveillance Team |
| EPI_ISL_2962295 | Health Services Laboratories | Wellcome Sanger Institute for the COVID-19 Genomics UK (COG-UK) Consortium | Health Services Laboratories and Alex Alderton, Roberto Amato, Jeffrey Barrett, Sonia Goncalves, Ewan Harrison, David K. Jackson, Ian Johnston, Dominic Kwiatkowski, Cordelia Langford, John Sillitoe on behalf of the Wellcome Sanger Institute COVID-19 Surveillance Team |
| EPI_ISL_2962296, EPI_ISL_2962297, EPI_ISL_2962298, EPI_ISL_2962299 | Lighthouse Lab in Milton Keynes | Wellcome Sanger Institute for the COVID-19 Genomics UK (COG-UK) Consortium | The Lighthouse Lab in Milton Keynes and Alex Alderton, Roberto Amato, Jeffrey Barrett, Sonia Goncalves, Ewan Harrison, David K. Jackson, Ian Johnston, Dominic Kwiatkowski, Cordelia Langford, John Sillitoe on behalf of the Wellcome Sanger Institute COVID-19 Surveillance Team |
| EPI_ISL_2962300 | Lighthouse Laboratory Plymouth | Wellcome Sanger Institute for the COVID-19 Genomics UK (COG-UK) Consortium | Lighthouse Laboratory Plymouth and Alex Alderton, Roberto Amato, Jeffrey Barrett, Sonia Goncalves, Ewan Harrison, David K. Jackson, Ian Johnston, Dominic Kwiatkowski, Cordelia Langford, John Sillitoe on behalf of the Wellcome Sanger Institute COVID-19 Surveillance Team |
| EPI_ISL_2962301 | Lighthouse Lab in Milton Keynes | Wellcome Sanger Institute for the COVID-19 Genomics UK (COG-UK) Consortium | The Lighthouse Lab in Milton Keynes and Alex Alderton, Roberto Amato, Jeffrey Barrett, Sonia Goncalves, Ewan Harrison, David K. Jackson, Ian Johnston, Dominic Kwiatkowski, Cordelia Langford, John Sillitoe on behalf of the Wellcome Sanger Institute COVID-19 Surveillance Team |
| EPI_ISL_2962302 | Lighthouse Laboratory Plymouth | Wellcome Sanger Institute for the COVID-19 Genomics UK (COG-UK) Consortium | Lighthouse Laboratory Plymouth and Alex Alderton, Roberto Amato, Jeffrey Barrett, Sonia Goncalves, Ewan Harrison, David K. Jackson, Ian Johnston, Dominic Kwiatkowski, Cordelia Langford, John Sillitoe on behalf of the Wellcome Sanger Institute COVID-19 Surveillance Team |
| EPI_ISL_2962303 | Lighthouse Lab in Milton Keynes | Wellcome Sanger Institute for the COVID-19 Genomics UK (COG-UK) Consortium | The Lighthouse Lab in Milton Keynes and Alex Alderton, Roberto Amato, Jeffrey Barrett, Sonia Goncalves, Ewan Harrison, David K. Jackson, Ian Johnston, Dominic Kwiatkowski, Cordelia Langford, John Sillitoe on behalf of the Wellcome Sanger Institute COVID-19 Surveillance Team |
| EPI_ISL_2962304 | Lighthouse Laboratory Plymouth | Wellcome Sanger Institute for the COVID-19 Genomics UK (COG-UK) Consortium | Lighthouse Laboratory Plymouth and Alex Alderton, Roberto Amato, Jeffrey Barrett, Sonia Goncalves, Ewan Harrison, David K. Jackson, Ian Johnston, Dominic Kwiatkowski, Cordelia Langford, John Sillitoe on behalf of the Wellcome Sanger Institute COVID-19 Surveillance Team |
| EPI_ISL_2962305 | Health Services Laboratories | Wellcome Sanger Institute for the COVID-19 Genomics UK (COG-UK) Consortium | Health Services Laboratories and Alex Alderton, Roberto Amato, Jeffrey Barrett, Sonia Goncalves, Ewan Harrison, David K. Jackson, Ian Johnston, Dominic Kwiatkowski, Cordelia Langford, John Sillitoe on behalf of the Wellcome Sanger Institute COVID-19 Surveillance Team |
| EPI_ISL_2962306, EPI_ISL_2962307, EPI_ISL_2962308, EPI_ISL_2962309 | Lighthouse Lab in Milton Keynes | Wellcome Sanger Institute for the COVID-19 Genomics UK (COG-UK) Consortium | The Lighthouse Lab in Milton Keynes and Alex Alderton, Roberto Amato, Jeffrey Barrett, Sonia Goncalves, Ewan Harrison, David K. Jackson, Ian Johnston, Dominic Kwiatkowski, Cordelia Langford, John Sillitoe on behalf of the Wellcome Sanger Institute COVID-19 Surveillance Team |
| EPI_ISL_2962310, EPI_ISL_2962311 | Health Services Laboratories | Wellcome Sanger Institute for the COVID-19 Genomics UK (COG-UK) Consortium | Health Services Laboratories and Alex Alderton, Roberto Amato, Jeffrey Barrett, Sonia Goncalves, Ewan Harrison, David K. Jackson, Ian Johnston, Dominic Kwiatkowski, Cordelia Langford, John Sillitoe on behalf of the Wellcome Sanger Institute COVID-19 Surveillance Team |
| EPI_ISL_2962312, EPI_ISL_2962313 | Lighthouse Lab in Milton Keynes | Wellcome Sanger Institute for the COVID-19 Genomics UK (COG-UK) Consortium | The Lighthouse Lab in Milton Keynes and Alex Alderton, Roberto Amato, Jeffrey Barrett, Sonia Goncalves, Ewan Harrison, David K. Jackson, Ian Johnston, Dominic Kwiatkowski, Cordelia Langford, John Sillitoe on behalf of the Wellcome Sanger Institute COVID-19 Surveillance Team |
| EPI_ISL_2962314 | Lighthouse Laboratory Plymouth | Wellcome Sanger Institute for the COVID-19 Genomics UK (COG-UK) Consortium | Lighthouse Laboratory Plymouth and Alex Alderton, Roberto Amato, Jeffrey Barrett, Sonia Goncalves, Ewan Harrison, David K. Jackson, Ian Johnston, Dominic Kwiatkowski, Cordelia Langford, John Sillitoe on behalf of the Wellcome Sanger Institute COVID-19 Surveillance Team |
| EPI_ISL_2962315 | Health Services Laboratories | Wellcome Sanger Institute for the COVID-19 Genomics UK (COG-UK) Consortium | Health Services Laboratories and Alex Alderton, Roberto Amato, Jeffrey Barrett, Sonia Goncalves, Ewan Harrison, David K. Jackson, Ian Johnston, Dominic Kwiatkowski, Cordelia Langford, John Sillitoe on behalf of the Wellcome Sanger Institute COVID-19 Surveillance Team |
| EPI_ISL_2962316 | Lighthouse Laboratory Plymouth | Wellcome Sanger Institute for the COVID-19 Genomics UK (COG-UK) Consortium | Lighthouse Laboratory Plymouth and Alex Alderton, Roberto Amato, Jeffrey Barrett, Sonia Goncalves, Ewan Harrison, David K. Jackson, Ian Johnston, Dominic Kwiatkowski, Cordelia Langford, John Sillitoe on behalf of the Wellcome Sanger Institute COVID-19 Surveillance Team |
| EPI_ISL_2962317, EPI_ISL_2962318, EPI_ISL_2962319 | Lighthouse Lab in Milton Keynes | Wellcome Sanger Institute for the COVID-19 Genomics UK (COG-UK) Consortium | The Lighthouse Lab in Milton Keynes and Alex Alderton, Roberto Amato, Jeffrey Barrett, Sonia Goncalves, Ewan Harrison, David K. Jackson, Ian Johnston, Dominic Kwiatkowski, Cordelia Langford, John Sillitoe on behalf of the Wellcome Sanger Institute COVID-19 Surveillance Team |
| EPI_ISL_2962320, EPI_ISL_2962321 | Health Services Laboratories | Wellcome Sanger Institute for the COVID-19 Genomics UK (COG-UK) Consortium | Health Services Laboratories and Alex Alderton, Roberto Amato, Jeffrey Barrett, Sonia Goncalves, Ewan Harrison, David K. Jackson, Ian Johnston, Dominic Kwiatkowski, Cordelia Langford, John Sillitoe on behalf of the Wellcome Sanger Institute COVID-19 Surveillance Team |
| EPI_ISL_2962322 | Lighthouse Lab in Milton Keynes | Wellcome Sanger Institute for the COVID-19 Genomics UK (COG-UK) Consortium | The Lighthouse Lab in Milton Keynes and Alex Alderton, Roberto Amato, Jeffrey Barrett, Sonia Goncalves, Ewan Harrison, David K. Jackson, Ian Johnston, Dominic Kwiatkowski, Cordelia Langford, John Sillitoe on behalf of the Wellcome Sanger Institute COVID-19 Surveillance Team |
| EPI_ISL_2962323, EPI_ISL_2962324 | Lighthouse Laboratory Plymouth | Wellcome Sanger Institute for the COVID-19 Genomics UK (COG-UK) Consortium | Lighthouse Laboratory Plymouth and Alex Alderton, Roberto Amato, Jeffrey Barrett, Sonia Goncalves, Ewan Harrison, David K. Jackson, Ian Johnston, Dominic Kwiatkowski, Cordelia Langford, John Sillitoe on behalf of the Wellcome Sanger Institute COVID-19 Surveillance Team |
| EPI_ISL_2962325, EPI_ISL_2962326, EPI_ISL_2962327, EPI_ISL_2962328, EPI_ISL_2962329 | Lighthouse Lab in Milton Keynes | Wellcome Sanger Institute for the COVID-19 Genomics UK (COG-UK) Consortium | The Lighthouse Lab in Milton Keynes and Alex Alderton, Roberto Amato, Jeffrey Barrett, Sonia Goncalves, Ewan Harrison, David K. Jackson, Ian Johnston, Dominic Kwiatkowski, Cordelia Langford, John Sillitoe on behalf of the Wellcome Sanger Institute COVID-19 Surveillance Team |
| EPI_ISL_2962330 | Health Services Laboratories | Wellcome Sanger Institute for the COVID-19 Genomics UK (COG-UK) Consortium | Health Services Laboratories and Alex Alderton, Roberto Amato, Jeffrey Barrett, Sonia Goncalves, Ewan Harrison, David K. Jackson, Ian Johnston, Dominic Kwiatkowski, Cordelia Langford, John Sillitoe on behalf of the Wellcome Sanger Institute COVID-19 Surveillance Team |
| EPI_ISL_2962331 | Lighthouse Laboratory Plymouth | Wellcome Sanger Institute for the COVID-19 Genomics UK (COG-UK) Consortium | Lighthouse Laboratory Plymouth and Alex Alderton, Roberto Amato, Jeffrey Barrett, Sonia Goncalves, Ewan Harrison, David K. Jackson, Ian Johnston, Dominic Kwiatkowski, Cordelia Langford, John Sillitoe on behalf of the Wellcome Sanger Institute COVID-19 Surveillance Team |
| EPI_ISL_2962332 | Health Services Laboratories | Wellcome Sanger Institute for the COVID-19 Genomics UK (COG-UK) Consortium | Health Services Laboratories and Alex Alderton, Roberto Amato, Jeffrey Barrett, Sonia Goncalves, Ewan Harrison, David K. Jackson, Ian Johnston, Dominic Kwiatkowski, Cordelia Langford, John Sillitoe on behalf of the Wellcome Sanger Institute COVID-19 Surveillance Team |
| EPI_ISL_2962333 | Lighthouse Lab in Milton Keynes | Wellcome Sanger Institute for the COVID-19 Genomics UK (COG-UK) Consortium | The Lighthouse Lab in Milton Keynes and Alex Alderton, Roberto Amato, Jeffrey Barrett, Sonia Goncalves, Ewan Harrison, David K. Jackson, Ian Johnston, Dominic Kwiatkowski, Cordelia Langford, John Sillitoe on behalf of the Wellcome Sanger Institute COVID-19 Surveillance Team |
| EPI_ISL_2962334 | Lighthouse Laboratory Plymouth | Wellcome Sanger Institute for the COVID-19 Genomics UK (COG-UK) Consortium | Lighthouse Laboratory Plymouth and Alex Alderton, Roberto Amato, Jeffrey Barrett, Sonia Goncalves, Ewan Harrison, David K. Jackson, Ian Johnston, Dominic Kwiatkowski, Cordelia Langford, John Sillitoe on behalf of the Wellcome Sanger Institute COVID-19 Surveillance Team |
| EPI_ISL_2962335 | Health Services Laboratories | Wellcome Sanger Institute for the COVID-19 Genomics UK (COG-UK) | Health Services Laboratories and Alex Alderton, Roberto Amato, Jeffrey Barrett, Sonia Goncalves, Ewan Harrison, David K. Jackson, Ian Johnston, Dominic Kwiatkowski, Cordelia Langford, John Sillitoe on behalf of the Wellcome Sanger Institute |

| | | | |
|---|---|---|---|
| | | Consortium | COVID-19 Surveillance Team |
| EPI_ISL_2962336 | Lighthouse Lab in Milton Keynes | Wellcome Sanger Institute for the COVID-19 Genomics UK (COG-UK) Consortium | The Lighthouse Lab in Milton Keynes and Alex Alderton, Roberto Amato, Jeffrey Barrett, Sonia Goncalves, Ewan Harrison, David K. Jackson, Ian Johnston, Dominic Kwiatkowski, Cordelia Langford, John Sillitoe on behalf of the Wellcome Sanger Institute COVID-19 Surveillance Team |
| EPI_ISL_2962337, EPI_ISL_2962338, EPI_ISL_2962339, EPI_ISL_2962341, EPI_ISL_2962342 | Lighthouse Laboratory Plymouth | Wellcome Sanger Institute for the COVID-19 Genomics UK (COG-UK) Consortium | Lighthouse Laboratory Plymouth and Alex Alderton, Roberto Amato, Jeffrey Barrett, Sonia Goncalves, Ewan Harrison, David K. Jackson, Ian Johnston, Dominic Kwiatkowski, Cordelia Langford, John Sillitoe on behalf of the Wellcome Sanger Institute COVID-19 Surveillance Team |
| EPI_ISL_2962343, EPI_ISL_2962344, EPI_ISL_2962345 | Lighthouse Lab in Milton Keynes | Wellcome Sanger Institute for the COVID-19 Genomics UK (COG-UK) Consortium | The Lighthouse Lab in Milton Keynes and Alex Alderton, Roberto Amato, Jeffrey Barrett, Sonia Goncalves, Ewan Harrison, David K. Jackson, Ian Johnston, Dominic Kwiatkowski, Cordelia Langford, John Sillitoe on behalf of the Wellcome Sanger Institute COVID-19 Surveillance Team |
| EPI_ISL_2962346, EPI_ISL_2962347 | Lighthouse Laboratory Plymouth | Wellcome Sanger Institute for the COVID-19 Genomics UK (COG-UK) Consortium | Lighthouse Laboratory Plymouth and Alex Alderton, Roberto Amato, Jeffrey Barrett, Sonia Goncalves, Ewan Harrison, David K. Jackson, Ian Johnston, Dominic Kwiatkowski, Cordelia Langford, John Sillitoe on behalf of the Wellcome Sanger Institute COVID-19 Surveillance Team |
| EPI_ISL_2962348 | Lighthouse Lab in Milton Keynes | Wellcome Sanger Institute for the COVID-19 Genomics UK (COG-UK) Consortium | The Lighthouse Lab in Milton Keynes and Alex Alderton, Roberto Amato, Jeffrey Barrett, Sonia Goncalves, Ewan Harrison, David K. Jackson, Ian Johnston, Dominic Kwiatkowski, Cordelia Langford, John Sillitoe on behalf of the Wellcome Sanger Institute COVID-19 Surveillance Team |
| EPI_ISL_2962349 | Health Services Laboratories | Wellcome Sanger Institute for the COVID-19 Genomics UK (COG-UK) Consortium | Health Services Laboratories and Alex Alderton, Roberto Amato, Jeffrey Barrett, Sonia Goncalves, Ewan Harrison, David K. Jackson, Ian Johnston, Dominic Kwiatkowski, Cordelia Langford, John Sillitoe on behalf of the Wellcome Sanger Institute COVID-19 Surveillance Team |
| EPI_ISL_2962350, EPI_ISL_2962351 | Lighthouse Lab in Milton Keynes | Wellcome Sanger Institute for the COVID-19 Genomics UK (COG-UK) Consortium | The Lighthouse Lab in Milton Keynes and Alex Alderton, Roberto Amato, Jeffrey Barrett, Sonia Goncalves, Ewan Harrison, David K. Jackson, Ian Johnston, Dominic Kwiatkowski, Cordelia Langford, John Sillitoe on behalf of the Wellcome Sanger Institute COVID-19 Surveillance Team |
| EPI_ISL_2962352 | Lighthouse Laboratory Plymouth | Wellcome Sanger Institute for the COVID-19 Genomics UK (COG-UK) Consortium | Lighthouse Laboratory Plymouth and Alex Alderton, Roberto Amato, Jeffrey Barrett, Sonia Goncalves, Ewan Harrison, David K. Jackson, Ian Johnston, Dominic Kwiatkowski, Cordelia Langford, John Sillitoe on behalf of the Wellcome Sanger Institute COVID-19 Surveillance Team |
| EPI_ISL_2962353 | Lighthouse Lab in Milton Keynes | Wellcome Sanger Institute for the COVID-19 Genomics UK (COG-UK) Consortium | The Lighthouse Lab in Milton Keynes and Alex Alderton, Roberto Amato, Jeffrey Barrett, Sonia Goncalves, Ewan Harrison, David K. Jackson, Ian Johnston, Dominic Kwiatkowski, Cordelia Langford, John Sillitoe on behalf of the Wellcome Sanger Institute COVID-19 Surveillance Team |
| EPI_ISL_2962354 | Lighthouse Laboratory Plymouth | Wellcome Sanger Institute for the COVID-19 Genomics UK (COG-UK) Consortium | Lighthouse Laboratory Plymouth and Alex Alderton, Roberto Amato, Jeffrey Barrett, Sonia Goncalves, Ewan Harrison, David K. Jackson, Ian Johnston, Dominic Kwiatkowski, Cordelia Langford, John Sillitoe on behalf of the Wellcome Sanger Institute COVID-19 Surveillance Team |
| EPI_ISL_2962355 | Lighthouse Lab in Milton Keynes | Wellcome Sanger Institute for the COVID-19 Genomics UK (COG-UK) Consortium | The Lighthouse Lab in Milton Keynes and Alex Alderton, Roberto Amato, Jeffrey Barrett, Sonia Goncalves, Ewan Harrison, David K. Jackson, Ian Johnston, Dominic Kwiatkowski, Cordelia Langford, John Sillitoe on behalf of the Wellcome Sanger Institute COVID-19 Surveillance Team |
| EPI_ISL_2962356 | Health Services Laboratories | Wellcome Sanger Institute for the COVID-19 Genomics UK (COG-UK) Consortium | Health Services Laboratories and Alex Alderton, Roberto Amato, Jeffrey Barrett, Sonia Goncalves, Ewan Harrison, David K. Jackson, Ian Johnston, Dominic Kwiatkowski, Cordelia Langford, John Sillitoe on behalf of the Wellcome Sanger Institute COVID-19 Surveillance Team |
| EPI_ISL_2962357, EPI_ISL_2962358 | Lighthouse Lab in Milton Keynes | Wellcome Sanger Institute for the COVID-19 Genomics UK (COG-UK) Consortium | The Lighthouse Lab in Milton Keynes and Alex Alderton, Roberto Amato, Jeffrey Barrett, Sonia Goncalves, Ewan Harrison, David K. Jackson, Ian Johnston, Dominic Kwiatkowski, Cordelia Langford, John Sillitoe on behalf of the Wellcome Sanger Institute COVID-19 Surveillance Team |
| EPI_ISL_2962359 | Lighthouse Laboratory Plymouth | Wellcome Sanger Institute for the COVID-19 Genomics UK (COG-UK) Consortium | Lighthouse Laboratory Plymouth and Alex Alderton, Roberto Amato, Jeffrey Barrett, Sonia Goncalves, Ewan Harrison, David K. Jackson, Ian Johnston, Dominic Kwiatkowski, Cordelia Langford, John Sillitoe on behalf of the Wellcome Sanger Institute COVID-19 Surveillance Team |
| EPI_ISL_2962360 | Health Services Laboratories | Wellcome Sanger Institute for the COVID-19 Genomics UK (COG-UK) Consortium | Health Services Laboratories and Alex Alderton, Roberto Amato, Jeffrey Barrett, Sonia Goncalves, Ewan Harrison, David K. Jackson, Ian Johnston, Dominic Kwiatkowski, Cordelia Langford, John Sillitoe on behalf of the Wellcome Sanger Institute COVID-19 Surveillance Team |
| EPI_ISL_2962361, EPI_ISL_2962362, EPI_ISL_2962363 | Lighthouse Lab in Milton Keynes | Wellcome Sanger Institute for the COVID-19 Genomics UK (COG-UK) Consortium | The Lighthouse Lab in Milton Keynes and Alex Alderton, Roberto Amato, Jeffrey Barrett, Sonia Goncalves, Ewan Harrison, David K. Jackson, Ian Johnston, Dominic Kwiatkowski, Cordelia Langford, John Sillitoe on behalf of the Wellcome Sanger Institute COVID-19 Surveillance Team |
| EPI_ISL_2962364 | Lighthouse Laboratory Plymouth | Wellcome Sanger Institute for the COVID-19 Genomics UK (COG-UK) Consortium | Lighthouse Laboratory Plymouth and Alex Alderton, Roberto Amato, Jeffrey Barrett, Sonia Goncalves, Ewan Harrison, David K. Jackson, Ian Johnston, Dominic Kwiatkowski, Cordelia Langford, John Sillitoe on behalf of the Wellcome Sanger Institute COVID-19 Surveillance Team |
| EPI_ISL_2962365, EPI_ISL_2962366 | Lighthouse Lab in Milton Keynes | Wellcome Sanger Institute for the COVID-19 Genomics UK (COG-UK) Consortium | The Lighthouse Lab in Milton Keynes and Alex Alderton, Roberto Amato, Jeffrey Barrett, Sonia Goncalves, Ewan Harrison, David K. Jackson, Ian Johnston, Dominic Kwiatkowski, Cordelia Langford, John Sillitoe on behalf of the Wellcome Sanger Institute COVID-19 Surveillance Team |
| EPI_ISL_2962367 | Lighthouse Laboratory Plymouth | Wellcome Sanger Institute for the COVID-19 Genomics UK (COG-UK) Consortium | Lighthouse Laboratory Plymouth and Alex Alderton, Roberto Amato, Jeffrey Barrett, Sonia Goncalves, Ewan Harrison, David K. Jackson, Ian Johnston, Dominic Kwiatkowski, Cordelia Langford, John Sillitoe on behalf of the Wellcome Sanger Institute COVID-19 Surveillance Team |
| EPI_ISL_2962368, EPI_ISL_2962369, EPI_ISL_2962370, EPI_ISL_2962371 | Lighthouse Lab in Milton Keynes | Wellcome Sanger Institute for the COVID-19 Genomics UK (COG-UK) Consortium | The Lighthouse Lab in Milton Keynes and Alex Alderton, Roberto Amato, Jeffrey Barrett, Sonia Goncalves, Ewan Harrison, David K. Jackson, Ian Johnston, Dominic Kwiatkowski, Cordelia Langford, John Sillitoe on behalf of the Wellcome Sanger Institute COVID-19 Surveillance Team |
| EPI_ISL_2962372, EPI_ISL_2962373, EPI_ISL_2962374 | Lighthouse Laboratory Plymouth | Wellcome Sanger Institute for the COVID-19 Genomics UK (COG-UK) Consortium | Lighthouse Laboratory Plymouth and Alex Alderton, Roberto Amato, Jeffrey Barrett, Sonia Goncalves, Ewan Harrison, David K. Jackson, Ian Johnston, Dominic Kwiatkowski, Cordelia Langford, John Sillitoe on behalf of the Wellcome Sanger Institute COVID-19 Surveillance Team |
| EPI_ISL_2962375 | Lighthouse Lab in Milton Keynes | Wellcome Sanger Institute for the COVID-19 Genomics UK (COG-UK) Consortium | The Lighthouse Lab in Milton Keynes and Alex Alderton, Roberto Amato, Jeffrey Barrett, Sonia Goncalves, Ewan Harrison, David K. Jackson, Ian Johnston, Dominic Kwiatkowski, Cordelia Langford, John Sillitoe on behalf of the Wellcome Sanger Institute COVID-19 Surveillance Team |
| EPI_ISL_2962376 | Lighthouse Laboratory Plymouth | Wellcome Sanger Institute for the COVID-19 Genomics UK (COG-UK) Consortium | Lighthouse Laboratory Plymouth and Alex Alderton, Roberto Amato, Jeffrey Barrett, Sonia Goncalves, Ewan Harrison, David K. Jackson, Ian Johnston, Dominic Kwiatkowski, Cordelia Langford, John Sillitoe on behalf of the Wellcome Sanger Institute COVID-19 Surveillance Team |
| EPI_ISL_2962377 | Health Services Laboratories | Wellcome Sanger Institute for the COVID-19 Genomics UK (COG-UK) Consortium | Health Services Laboratories and Alex Alderton, Roberto Amato, Jeffrey Barrett, Sonia Goncalves, Ewan Harrison, David K. Jackson, Ian Johnston, Dominic Kwiatkowski, Cordelia Langford, John Sillitoe on behalf of the Wellcome Sanger Institute COVID-19 Surveillance Team |
| EPI_ISL_2962378, EPI_ISL_2962379, EPI_ISL_2962380, EPI_ISL_2962381, EPI_ISL_2962382 | Lighthouse Lab in Milton Keynes | Wellcome Sanger Institute for the COVID-19 Genomics UK (COG-UK) Consortium | The Lighthouse Lab in Milton Keynes and Alex Alderton, Roberto Amato, Jeffrey Barrett, Sonia Goncalves, Ewan Harrison, David K. Jackson, Ian Johnston, Dominic Kwiatkowski, Cordelia Langford, John Sillitoe on behalf of the Wellcome Sanger Institute COVID-19 Surveillance Team |
| EPI_ISL_2962383 | Health Services Laboratories | Wellcome Sanger Institute for the COVID-19 Genomics UK (COG-UK) Consortium | Health Services Laboratories and Alex Alderton, Roberto Amato, Jeffrey Barrett, Sonia Goncalves, Ewan Harrison, David K. Jackson, Ian Johnston, Dominic Kwiatkowski, Cordelia Langford, John Sillitoe on behalf of the Wellcome Sanger Institute COVID-19 Surveillance Team |
| EPI_ISL_2962384, EPI_ISL_2962385, EPI_ISL_2962386 | Lighthouse Lab in Milton Keynes | Wellcome Sanger Institute for the COVID-19 Genomics UK (COG-UK) Consortium | The Lighthouse Lab in Milton Keynes and Alex Alderton, Roberto Amato, Jeffrey Barrett, Sonia Goncalves, Ewan Harrison, David K. Jackson, Ian Johnston, Dominic Kwiatkowski, Cordelia Langford, John Sillitoe on behalf of the Wellcome Sanger Institute COVID-19 Surveillance Team |
| EPI_ISL_2962387 | Lighthouse Laboratory Plymouth | Wellcome Sanger Institute for the COVID-19 Genomics UK (COG-UK) Consortium | Lighthouse Laboratory Plymouth and Alex Alderton, Roberto Amato, Jeffrey Barrett, Sonia Goncalves, Ewan Harrison, David K. Jackson, Ian Johnston, Dominic Kwiatkowski, Cordelia Langford, John Sillitoe on behalf of the Wellcome Sanger Institute COVID-19 Surveillance Team |
| EPI_ISL_2962388, EPI_ISL_2962389 | Lighthouse Lab in Milton Keynes | Wellcome Sanger Institute for the COVID-19 Genomics UK (COG-UK) Consortium | The Lighthouse Lab in Milton Keynes and Alex Alderton, Roberto Amato, Jeffrey Barrett, Sonia Goncalves, Ewan Harrison, David K. Jackson, Ian Johnston, Dominic Kwiatkowski, Cordelia Langford, John Sillitoe on behalf of the Wellcome Sanger Institute COVID-19 Surveillance Team |
| EPI_ISL_2962390, EPI_ISL_2962391 | Lighthouse Laboratory Plymouth | Wellcome Sanger Institute for the COVID-19 Genomics UK (COG-UK) Consortium | Lighthouse Laboratory Plymouth and Alex Alderton, Roberto Amato, Jeffrey Barrett, Sonia Goncalves, Ewan Harrison, David K. Jackson, Ian Johnston, Dominic Kwiatkowski, Cordelia Langford, John Sillitoe on behalf of the Wellcome Sanger Institute COVID-19 Surveillance Team |
| EPI_ISL_2962392 | Health Services Laboratories | Wellcome Sanger Institute for the COVID-19 Genomics UK (COG-UK) Consortium | Health Services Laboratories and Alex Alderton, Roberto Amato, Jeffrey Barrett, Sonia Goncalves, Ewan Harrison, David K. Jackson, Ian Johnston, Dominic Kwiatkowski, Cordelia Langford, John Sillitoe on behalf of the Wellcome Sanger Institute COVID-19 Surveillance Team |
| EPI_ISL_2962393 | Lighthouse Laboratory Plymouth | Wellcome Sanger Institute for the COVID-19 Genomics UK (COG-UK) Consortium | Lighthouse Laboratory Plymouth and Alex Alderton, Roberto Amato, Jeffrey Barrett, Sonia Goncalves, Ewan Harrison, David K. Jackson, Ian Johnston, Dominic Kwiatkowski, Cordelia Langford, John Sillitoe on behalf of the Wellcome Sanger Institute COVID-19 Surveillance Team |
| EPI_ISL_2962394, EPI_ISL_2962395, EPI_ISL_2962396, EPI_ISL_2962397 | Lighthouse Lab in Milton Keynes | Wellcome Sanger Institute for the COVID-19 Genomics UK (COG-UK) Consortium | The Lighthouse Lab in Milton Keynes and Alex Alderton, Roberto Amato, Jeffrey Barrett, Sonia Goncalves, Ewan Harrison, David K. Jackson, Ian Johnston, Dominic Kwiatkowski, Cordelia Langford, John Sillitoe on behalf of the Wellcome Sanger Institute COVID-19 Surveillance Team |
| EPI_ISL_2962398, EPI_ISL_2962399 | Lighthouse Laboratory Plymouth | Wellcome Sanger Institute for the COVID-19 Genomics UK (COG-UK) Consortium | Lighthouse Laboratory Plymouth and Alex Alderton, Roberto Amato, Jeffrey Barrett, Sonia Goncalves, Ewan Harrison, David K. Jackson, Ian Johnston, Dominic Kwiatkowski, Cordelia Langford, John Sillitoe on behalf of the Wellcome Sanger Institute COVID-19 Surveillance Team |
| EPI_ISL_2962400 | Health Services Laboratories | Wellcome Sanger Institute for the COVID-19 Genomics UK (COG-UK) Consortium | Health Services Laboratories and Alex Alderton, Roberto Amato, Jeffrey Barrett, Sonia Goncalves, Ewan Harrison, David K. Jackson, Ian Johnston, Dominic Kwiatkowski, Cordelia Langford, John Sillitoe on behalf of the Wellcome Sanger Institute COVID-19 Surveillance Team |
| EPI_ISL_2962401 | Lighthouse Lab in Milton Keynes | Wellcome Sanger Institute for the COVID-19 Genomics UK (COG-UK) Consortium | The Lighthouse Lab in Milton Keynes and Alex Alderton, Roberto Amato, Jeffrey Barrett, Sonia Goncalves, Ewan Harrison, David K. Jackson, Ian Johnston, Dominic Kwiatkowski, Cordelia Langford, John Sillitoe on behalf of the Wellcome Sanger Institute COVID-19 Surveillance Team |
| EPI_ISL_2962402 | Lighthouse Laboratory Plymouth | Wellcome Sanger Institute for the COVID-19 Genomics UK (COG-UK) Consortium | Lighthouse Laboratory Plymouth and Alex Alderton, Roberto Amato, Jeffrey Barrett, Sonia Goncalves, Ewan Harrison, David K. Jackson, Ian Johnston, Dominic Kwiatkowski, Cordelia Langford, John Sillitoe on behalf of the Wellcome Sanger Institute COVID-19 Surveillance Team |
| EPI_ISL_2962403 | Lighthouse Lab in Milton Keynes | Wellcome Sanger Institute for the COVID-19 Genomics UK (COG-UK) Consortium | The Lighthouse Lab in Milton Keynes and Alex Alderton, Roberto Amato, Jeffrey Barrett, Sonia Goncalves, Ewan Harrison, David K. Jackson, Ian Johnston, Dominic Kwiatkowski, Cordelia Langford, John Sillitoe on behalf of the Wellcome Sanger Institute COVID-19 Surveillance Team |
| EPI_ISL_2962404 | Health Services Laboratories | Wellcome Sanger Institute for the COVID-19 Genomics UK (COG-UK) Consortium | Health Services Laboratories and Alex Alderton, Roberto Amato, Jeffrey Barrett, Sonia Goncalves, Ewan Harrison, David K. Jackson, Ian Johnston, Dominic Kwiatkowski, Cordelia Langford, John Sillitoe on behalf of the Wellcome Sanger Institute COVID-19 Surveillance Team |

| | | | |
|---|---|---|---|
| EPI_ISL_2962405 | Lighthouse Laboratory Plymouth | Wellcome Sanger Institute for the COVID-19 Genomics UK (COG-UK) Consortium | Lighthouse Laboratory Plymouth and Alex Alderton, Roberto Amato, Jeffrey Barrett, Sonia Goncalves, Ewan Harrison, David K. Jackson, Ian Johnston, Dominic Kwiatkowski, Cordelia Langford, John Sillitoe on behalf of the Wellcome Sanger Institute COVID-19 Surveillance Team |
| EPI_ISL_2962406, EPI_ISL_2962407, EPI_ISL_2962408 | Lighthouse Lab in Milton Keynes | Wellcome Sanger Institute for the COVID-19 Genomics UK (COG-UK) Consortium | The Lighthouse Lab in Milton Keynes and Alex Alderton, Roberto Amato, Jeffrey Barrett, Sonia Goncalves, Ewan Harrison, David K. Jackson, Ian Johnston, Dominic Kwiatkowski, Cordelia Langford, John Sillitoe on behalf of the Wellcome Sanger Institute COVID-19 Surveillance Team |
| EPI_ISL_2962409 | Health Services Laboratories | Wellcome Sanger Institute for the COVID-19 Genomics UK (COG-UK) Consortium | Health Services Laboratories and Alex Alderton, Roberto Amato, Jeffrey Barrett, Sonia Goncalves, Ewan Harrison, David K. Jackson, Ian Johnston, Dominic Kwiatkowski, Cordelia Langford, John Sillitoe on behalf of the Wellcome Sanger Institute COVID-19 Surveillance Team |
| EPI_ISL_2962410 | Lighthouse Lab in Milton Keynes | Wellcome Sanger Institute for the COVID-19 Genomics UK (COG-UK) Consortium | The Lighthouse Lab in Milton Keynes and Alex Alderton, Roberto Amato, Jeffrey Barrett, Sonia Goncalves, Ewan Harrison, David K. Jackson, Ian Johnston, Dominic Kwiatkowski, Cordelia Langford, John Sillitoe on behalf of the Wellcome Sanger Institute COVID-19 Surveillance Team |
| EPI_ISL_2962411, EPI_ISL_2962412 | Health Services Laboratories | Wellcome Sanger Institute for the COVID-19 Genomics UK (COG-UK) Consortium | Health Services Laboratories and Alex Alderton, Roberto Amato, Jeffrey Barrett, Sonia Goncalves, Ewan Harrison, David K. Jackson, Ian Johnston, Dominic Kwiatkowski, Cordelia Langford, John Sillitoe on behalf of the Wellcome Sanger Institute COVID-19 Surveillance Team |
| EPI_ISL_2962413, EPI_ISL_2962414, EPI_ISL_2962415, EPI_ISL_2962416, EPI_ISL_2962417, EPI_ISL_2962418, EPI_ISL_2962419, EPI_ISL_2962420, EPI_ISL_2962421, EPI_ISL_2962422 | Lighthouse Lab in Milton Keynes | Wellcome Sanger Institute for the COVID-19 Genomics UK (COG-UK) Consortium | The Lighthouse Lab in Milton Keynes and Alex Alderton, Roberto Amato, Jeffrey Barrett, Sonia Goncalves, Ewan Harrison, David K. Jackson, Ian Johnston, Dominic Kwiatkowski, Cordelia Langford, John Sillitoe on behalf of the Wellcome Sanger Institute COVID-19 Surveillance Team |
| EPI_ISL_2962423 | Lighthouse Laboratory Plymouth | Wellcome Sanger Institute for the COVID-19 Genomics UK (COG-UK) Consortium | Lighthouse Laboratory Plymouth and Alex Alderton, Roberto Amato, Jeffrey Barrett, Sonia Goncalves, Ewan Harrison, David K. Jackson, Ian Johnston, Dominic Kwiatkowski, Cordelia Langford, John Sillitoe on behalf of the Wellcome Sanger Institute COVID-19 Surveillance Team |
| EPI_ISL_2962424 | Lighthouse Lab in Milton Keynes | Wellcome Sanger Institute for the COVID-19 Genomics UK (COG-UK) Consortium | The Lighthouse Lab in Milton Keynes and Alex Alderton, Roberto Amato, Jeffrey Barrett, Sonia Goncalves, Ewan Harrison, David K. Jackson, Ian Johnston, Dominic Kwiatkowski, Cordelia Langford, John Sillitoe on behalf of the Wellcome Sanger Institute COVID-19 Surveillance Team |
| EPI_ISL_2962425, EPI_ISL_2962426 | Lighthouse Laboratory Plymouth | Wellcome Sanger Institute for the COVID-19 Genomics UK (COG-UK) Consortium | Lighthouse Laboratory Plymouth and Alex Alderton, Roberto Amato, Jeffrey Barrett, Sonia Goncalves, Ewan Harrison, David K. Jackson, Ian Johnston, Dominic Kwiatkowski, Cordelia Langford, John Sillitoe on behalf of the Wellcome Sanger Institute COVID-19 Surveillance Team |
| EPI_ISL_2962427 | Lighthouse Lab in Milton Keynes | Wellcome Sanger Institute for the COVID-19 Genomics UK (COG-UK) Consortium | The Lighthouse Lab in Milton Keynes and Alex Alderton, Roberto Amato, Jeffrey Barrett, Sonia Goncalves, Ewan Harrison, David K. Jackson, Ian Johnston, Dominic Kwiatkowski, Cordelia Langford, John Sillitoe on behalf of the Wellcome Sanger Institute COVID-19 Surveillance Team |
| EPI_ISL_2962428 | Health Services Laboratories | Wellcome Sanger Institute for the COVID-19 Genomics UK (COG-UK) Consortium | Health Services Laboratories and Alex Alderton, Roberto Amato, Jeffrey Barrett, Sonia Goncalves, Ewan Harrison, David K. Jackson, Ian Johnston, Dominic Kwiatkowski, Cordelia Langford, John Sillitoe on behalf of the Wellcome Sanger Institute COVID-19 Surveillance Team |
| EPI_ISL_2962429 | Lighthouse Lab in Milton Keynes | Wellcome Sanger Institute for the COVID-19 Genomics UK (COG-UK) Consortium | The Lighthouse Lab in Milton Keynes and Alex Alderton, Roberto Amato, Jeffrey Barrett, Sonia Goncalves, Ewan Harrison, David K. Jackson, Ian Johnston, Dominic Kwiatkowski, Cordelia Langford, John Sillitoe on behalf of the Wellcome Sanger Institute COVID-19 Surveillance Team |
| EPI_ISL_2962430 | Lighthouse Laboratory Plymouth | Wellcome Sanger Institute for the COVID-19 Genomics UK (COG-UK) Consortium | Lighthouse Laboratory Plymouth and Alex Alderton, Roberto Amato, Jeffrey Barrett, Sonia Goncalves, Ewan Harrison, David K. Jackson, Ian Johnston, Dominic Kwiatkowski, Cordelia Langford, John Sillitoe on behalf of the Wellcome Sanger Institute COVID-19 Surveillance Team |
| EPI_ISL_2962431, EPI_ISL_2962432 | Lighthouse Lab in Milton Keynes | Wellcome Sanger Institute for the COVID-19 Genomics UK (COG-UK) Consortium | The Lighthouse Lab in Milton Keynes and Alex Alderton, Roberto Amato, Jeffrey Barrett, Sonia Goncalves, Ewan Harrison, David K. Jackson, Ian Johnston, Dominic Kwiatkowski, Cordelia Langford, John Sillitoe on behalf of the Wellcome Sanger Institute COVID-19 Surveillance Team |
| EPI_ISL_2962433 | Health Services Laboratories | Wellcome Sanger Institute for the COVID-19 Genomics UK (COG-UK) Consortium | Health Services Laboratories and Alex Alderton, Roberto Amato, Jeffrey Barrett, Sonia Goncalves, Ewan Harrison, David K. Jackson, Ian Johnston, Dominic Kwiatkowski, Cordelia Langford, John Sillitoe on behalf of the Wellcome Sanger Institute COVID-19 Surveillance Team |
| EPI_ISL_2962434, EPI_ISL_2962435 | Lighthouse Lab in Milton Keynes | Wellcome Sanger Institute for the COVID-19 Genomics UK (COG-UK) Consortium | The Lighthouse Lab in Milton Keynes and Alex Alderton, Roberto Amato, Jeffrey Barrett, Sonia Goncalves, Ewan Harrison, David K. Jackson, Ian Johnston, Dominic Kwiatkowski, Cordelia Langford, John Sillitoe on behalf of the Wellcome Sanger Institute COVID-19 Surveillance Team |
| EPI_ISL_2962436 | Health Services Laboratories | Wellcome Sanger Institute for the COVID-19 Genomics UK (COG-UK) Consortium | Health Services Laboratories and Alex Alderton, Roberto Amato, Jeffrey Barrett, Sonia Goncalves, Ewan Harrison, David K. Jackson, Ian Johnston, Dominic Kwiatkowski, Cordelia Langford, John Sillitoe on behalf of the Wellcome Sanger Institute COVID-19 Surveillance Team |
| EPI_ISL_2962437, EPI_ISL_2962438 | Lighthouse Laboratory Plymouth | Wellcome Sanger Institute for the COVID-19 Genomics UK (COG-UK) Consortium | Lighthouse Laboratory Plymouth and Alex Alderton, Roberto Amato, Jeffrey Barrett, Sonia Goncalves, Ewan Harrison, David K. Jackson, Ian Johnston, Dominic Kwiatkowski, Cordelia Langford, John Sillitoe on behalf of the Wellcome Sanger Institute COVID-19 Surveillance Team |
| EPI_ISL_2962439, EPI_ISL_2962440, EPI_ISL_2962441 | Lighthouse Lab in Milton Keynes | Wellcome Sanger Institute for the COVID-19 Genomics UK (COG-UK) Consortium | The Lighthouse Lab in Milton Keynes and Alex Alderton, Roberto Amato, Jeffrey Barrett, Sonia Goncalves, Ewan Harrison, David K. Jackson, Ian Johnston, Dominic Kwiatkowski, Cordelia Langford, John Sillitoe on behalf of the Wellcome Sanger Institute COVID-19 Surveillance Team |
| EPI_ISL_2962443, EPI_ISL_2962444 | Lighthouse Laboratory Plymouth | Wellcome Sanger Institute for the COVID-19 Genomics UK (COG-UK) Consortium | Lighthouse Laboratory Plymouth and Alex Alderton, Roberto Amato, Jeffrey Barrett, Sonia Goncalves, Ewan Harrison, David K. Jackson, Ian Johnston, Dominic Kwiatkowski, Cordelia Langford, John Sillitoe on behalf of the Wellcome Sanger Institute COVID-19 Surveillance Team |
| EPI_ISL_2962445 | Lighthouse Lab in Milton Keynes | Wellcome Sanger Institute for the COVID-19 Genomics UK (COG-UK) Consortium | The Lighthouse Lab in Milton Keynes and Alex Alderton, Roberto Amato, Jeffrey Barrett, Sonia Goncalves, Ewan Harrison, David K. Jackson, Ian Johnston, Dominic Kwiatkowski, Cordelia Langford, John Sillitoe on behalf of the Wellcome Sanger Institute COVID-19 Surveillance Team |
| EPI_ISL_2962446 | Health Services Laboratories | Wellcome Sanger Institute for the COVID-19 Genomics UK (COG-UK) Consortium | Health Services Laboratories and Alex Alderton, Roberto Amato, Jeffrey Barrett, Sonia Goncalves, Ewan Harrison, David K. Jackson, Ian Johnston, Dominic Kwiatkowski, Cordelia Langford, John Sillitoe on behalf of the Wellcome Sanger Institute COVID-19 Surveillance Team |
| EPI_ISL_2962447 | Lighthouse Laboratory Plymouth | Wellcome Sanger Institute for the COVID-19 Genomics UK (COG-UK) Consortium | Lighthouse Laboratory Plymouth and Alex Alderton, Roberto Amato, Jeffrey Barrett, Sonia Goncalves, Ewan Harrison, David K. Jackson, Ian Johnston, Dominic Kwiatkowski, Cordelia Langford, John Sillitoe on behalf of the Wellcome Sanger Institute COVID-19 Surveillance Team |
| EPI_ISL_2962448 | Health Services Laboratories | Wellcome Sanger Institute for the COVID-19 Genomics UK (COG-UK) Consortium | Health Services Laboratories and Alex Alderton, Roberto Amato, Jeffrey Barrett, Sonia Goncalves, Ewan Harrison, David K. Jackson, Ian Johnston, Dominic Kwiatkowski, Cordelia Langford, John Sillitoe on behalf of the Wellcome Sanger Institute COVID-19 Surveillance Team |
| EPI_ISL_2962449 | Lighthouse Lab in Milton Keynes | Wellcome Sanger Institute for the COVID-19 Genomics UK (COG-UK) Consortium | The Lighthouse Lab in Milton Keynes and Alex Alderton, Roberto Amato, Jeffrey Barrett, Sonia Goncalves, Ewan Harrison, David K. Jackson, Ian Johnston, Dominic Kwiatkowski, Cordelia Langford, John Sillitoe on behalf of the Wellcome Sanger Institute COVID-19 Surveillance Team |
| EPI_ISL_2962450 | Lighthouse Laboratory Plymouth | Wellcome Sanger Institute for the COVID-19 Genomics UK (COG-UK) Consortium | Lighthouse Laboratory Plymouth and Alex Alderton, Roberto Amato, Jeffrey Barrett, Sonia Goncalves, Ewan Harrison, David K. Jackson, Ian Johnston, Dominic Kwiatkowski, Cordelia Langford, John Sillitoe on behalf of the Wellcome Sanger Institute COVID-19 Surveillance Team |
| EPI_ISL_2962451, EPI_ISL_2962452, EPI_ISL_2962453, EPI_ISL_2962454 | Lighthouse Lab in Milton Keynes | Wellcome Sanger Institute for the COVID-19 Genomics UK (COG-UK) Consortium | The Lighthouse Lab in Milton Keynes and Alex Alderton, Roberto Amato, Jeffrey Barrett, Sonia Goncalves, Ewan Harrison, David K. Jackson, Ian Johnston, Dominic Kwiatkowski, Cordelia Langford, John Sillitoe on behalf of the Wellcome Sanger Institute COVID-19 Surveillance Team |
| EPI_ISL_2962455 | Health Services Laboratories | Wellcome Sanger Institute for the COVID-19 Genomics UK (COG-UK) Consortium | Health Services Laboratories and Alex Alderton, Roberto Amato, Jeffrey Barrett, Sonia Goncalves, Ewan Harrison, David K. Jackson, Ian Johnston, Dominic Kwiatkowski, Cordelia Langford, John Sillitoe on behalf of the Wellcome Sanger Institute COVID-19 Surveillance Team |
| EPI_ISL_2962456 | Lighthouse Lab in Milton Keynes | Wellcome Sanger Institute for the COVID-19 Genomics UK (COG-UK) Consortium | The Lighthouse Lab in Milton Keynes and Alex Alderton, Roberto Amato, Jeffrey Barrett, Sonia Goncalves, Ewan Harrison, David K. Jackson, Ian Johnston, Dominic Kwiatkowski, Cordelia Langford, John Sillitoe on behalf of the Wellcome Sanger Institute COVID-19 Surveillance Team |
| EPI_ISL_2962457 | Health Services Laboratories | Wellcome Sanger Institute for the COVID-19 Genomics UK (COG-UK) Consortium | Health Services Laboratories and Alex Alderton, Roberto Amato, Jeffrey Barrett, Sonia Goncalves, Ewan Harrison, David K. Jackson, Ian Johnston, Dominic Kwiatkowski, Cordelia Langford, John Sillitoe on behalf of the Wellcome Sanger Institute COVID-19 Surveillance Team |
| EPI_ISL_2962458, EPI_ISL_2962459 | Lighthouse Lab in Milton Keynes | Wellcome Sanger Institute for the COVID-19 Genomics UK (COG-UK) Consortium | The Lighthouse Lab in Milton Keynes and Alex Alderton, Roberto Amato, Jeffrey Barrett, Sonia Goncalves, Ewan Harrison, David K. Jackson, Ian Johnston, Dominic Kwiatkowski, Cordelia Langford, John Sillitoe on behalf of the Wellcome Sanger Institute COVID-19 Surveillance Team |
| EPI_ISL_2962460 | Health Services Laboratories | Wellcome Sanger Institute for the COVID-19 Genomics UK (COG-UK) Consortium | Health Services Laboratories and Alex Alderton, Roberto Amato, Jeffrey Barrett, Sonia Goncalves, Ewan Harrison, David K. Jackson, Ian Johnston, Dominic Kwiatkowski, Cordelia Langford, John Sillitoe on behalf of the Wellcome Sanger Institute COVID-19 Surveillance Team |
| EPI_ISL_2962461, EPI_ISL_2962462, EPI_ISL_2962463 | Lighthouse Lab in Milton Keynes | Wellcome Sanger Institute for the COVID-19 Genomics UK (COG-UK) Consortium | The Lighthouse Lab in Milton Keynes and Alex Alderton, Roberto Amato, Jeffrey Barrett, Sonia Goncalves, Ewan Harrison, David K. Jackson, Ian Johnston, Dominic Kwiatkowski, Cordelia Langford, John Sillitoe on behalf of the Wellcome Sanger Institute COVID-19 Surveillance Team |
| EPI_ISL_2962464 | Health Services Laboratories | Wellcome Sanger Institute for the COVID-19 Genomics UK (COG-UK) Consortium | Health Services Laboratories and Alex Alderton, Roberto Amato, Jeffrey Barrett, Sonia Goncalves, Ewan Harrison, David K. Jackson, Ian Johnston, Dominic Kwiatkowski, Cordelia Langford, John Sillitoe on behalf of the Wellcome Sanger Institute COVID-19 Surveillance Team |
| EPI_ISL_2962465 | Lighthouse Laboratory Plymouth | Wellcome Sanger Institute for the COVID-19 Genomics UK (COG-UK) Consortium | Lighthouse Laboratory Plymouth and Alex Alderton, Roberto Amato, Jeffrey Barrett, Sonia Goncalves, Ewan Harrison, David K. Jackson, Ian Johnston, Dominic Kwiatkowski, Cordelia Langford, John Sillitoe on behalf of the Wellcome Sanger Institute COVID-19 Surveillance Team |
| EPI_ISL_2962466 | Lighthouse Lab in Milton Keynes | Wellcome Sanger Institute for the COVID-19 Genomics UK (COG-UK) Consortium | The Lighthouse Lab in Milton Keynes and Alex Alderton, Roberto Amato, Jeffrey Barrett, Sonia Goncalves, Ewan Harrison, David K. Jackson, Ian Johnston, Dominic Kwiatkowski, Cordelia Langford, John Sillitoe on behalf of the Wellcome Sanger Institute COVID-19 Surveillance Team |
| EPI_ISL_2962467 | Lighthouse Lab in Alderley Park | Wellcome Sanger Institute for the COVID-19 Genomics UK (COG-UK) Consortium | Jacquelyn Wynn, Mairead Hyland, The Lighthouse Lab in Alderley Park and Alex Alderton, Roberto Amato, Jeffrey Barrett, Sonia Goncalves, Ewan Harrison, David K. Jackson, Ian Johnston, Dominic Kwiatkowski, Cordelia Langford, John Sillitoe on behalf of the Wellcome Sanger Institute COVID-19 Surveillance Team |
| EPI_ISL_2962468, EPI_ISL_2962469, EPI_ISL_2962470 | Lighthouse Lab in Milton Keynes | Wellcome Sanger Institute for the COVID-19 Genomics UK (COG-UK) Consortium | The Lighthouse Lab in Milton Keynes and Alex Alderton, Roberto Amato, Jeffrey Barrett, Sonia Goncalves, Ewan Harrison, David K. Jackson, Ian Johnston, Dominic Kwiatkowski, Cordelia Langford, John Sillitoe on behalf of the Wellcome Sanger Institute COVID-19 Surveillance Team |
| EPI_ISL_2962471 | Lighthouse Lab in Alderley Park | Wellcome Sanger Institute for the COVID-19 Genomics UK (COG-UK) Consortium | Jacquelyn Wynn, Mairead Hyland, The Lighthouse Lab in Alderley Park and Alex Alderton, Roberto Amato, Jeffrey Barrett, Sonia Goncalves, Ewan Harrison, David K. Jackson, Ian Johnston, Dominic Kwiatkowski, Cordelia Langford, John Sillitoe on behalf of the Wellcome Sanger Institute COVID-19 Surveillance Team |

| | | | |
|---|---|---|---|
| EPI_ISL_2962472, EPI_ISL_2962473, EPI_ISL_2962474, EPI_ISL_2962475 | Lighthouse Lab in Milton Keynes | Wellcome Sanger Institute for the COVID-19 Genomics UK (COG-UK) Consortium | The Lighthouse Lab in Milton Keynes and Alex Alderton, Roberto Amato, Jeffrey Barrett, Sonia Goncalves, Ewan Harrison, David K. Jackson, Ian Johnston, Dominic Kwiatkowski, Cordelia Langford, John Sillitoe on behalf of the Wellcome Sanger Institute COVID-19 Surveillance Team |
| EPI_ISL_2962476 | Lighthouse Lab in Alderley Park | Wellcome Sanger Institute for the COVID-19 Genomics UK (COG-UK) Consortium | Jacquelyn Wynn, Mairead Hyland, The Lighthouse Lab in Alderley Park and Alex Alderton, Roberto Amato, Jeffrey Barrett, Sonia Goncalves, Ewan Harrison, David K. Jackson, Ian Johnston, Dominic Kwiatkowski, Cordelia Langford, John Sillitoe on behalf of the Wellcome Sanger Institute COVID-19 Surveillance Team |
| EPI_ISL_2962477 | Lighthouse Lab in Milton Keynes | Wellcome Sanger Institute for the COVID-19 Genomics UK (COG-UK) Consortium | The Lighthouse Lab in Milton Keynes and Alex Alderton, Roberto Amato, Jeffrey Barrett, Sonia Goncalves, Ewan Harrison, David K. Jackson, Ian Johnston, Dominic Kwiatkowski, Cordelia Langford, John Sillitoe on behalf of the Wellcome Sanger Institute COVID-19 Surveillance Team |
| EPI_ISL_2962478 | Lighthouse Lab in Alderley Park | Wellcome Sanger Institute for the COVID-19 Genomics UK (COG-UK) Consortium | Jacquelyn Wynn, Mairead Hyland, The Lighthouse Lab in Alderley Park and Alex Alderton, Roberto Amato, Jeffrey Barrett, Sonia Goncalves, Ewan Harrison, David K. Jackson, Ian Johnston, Dominic Kwiatkowski, Cordelia Langford, John Sillitoe on behalf of the Wellcome Sanger Institute COVID-19 Surveillance Team |
| EPI_ISL_2962479 | Lighthouse Lab in Milton Keynes | Wellcome Sanger Institute for the COVID-19 Genomics UK (COG-UK) Consortium | The Lighthouse Lab in Milton Keynes and Alex Alderton, Roberto Amato, Jeffrey Barrett, Sonia Goncalves, Ewan Harrison, David K. Jackson, Ian Johnston, Dominic Kwiatkowski, Cordelia Langford, John Sillitoe on behalf of the Wellcome Sanger Institute COVID-19 Surveillance Team |
| EPI_ISL_2962480 | Lighthouse Lab in Alderley Park | Wellcome Sanger Institute for the COVID-19 Genomics UK (COG-UK) Consortium | Jacquelyn Wynn, Mairead Hyland, The Lighthouse Lab in Alderley Park and Alex Alderton, Roberto Amato, Jeffrey Barrett, Sonia Goncalves, Ewan Harrison, David K. Jackson, Ian Johnston, Dominic Kwiatkowski, Cordelia Langford, John Sillitoe on behalf of the Wellcome Sanger Institute COVID-19 Surveillance Team |
| EPI_ISL_2962481 | Lighthouse Lab in Milton Keynes | Wellcome Sanger Institute for the COVID-19 Genomics UK (COG-UK) Consortium | The Lighthouse Lab in Milton Keynes and Alex Alderton, Roberto Amato, Jeffrey Barrett, Sonia Goncalves, Ewan Harrison, David K. Jackson, Ian Johnston, Dominic Kwiatkowski, Cordelia Langford, John Sillitoe on behalf of the Wellcome Sanger Institute COVID-19 Surveillance Team |
| EPI_ISL_2962482 | Lighthouse Lab in Alderley Park | Wellcome Sanger Institute for the COVID-19 Genomics UK (COG-UK) Consortium | Jacquelyn Wynn, Mairead Hyland, The Lighthouse Lab in Alderley Park and Alex Alderton, Roberto Amato, Jeffrey Barrett, Sonia Goncalves, Ewan Harrison, David K. Jackson, Ian Johnston, Dominic Kwiatkowski, Cordelia Langford, John Sillitoe on behalf of the Wellcome Sanger Institute COVID-19 Surveillance Team |
| EPI_ISL_2962483 | Lighthouse Lab in Milton Keynes | Wellcome Sanger Institute for the COVID-19 Genomics UK (COG-UK) Consortium | The Lighthouse Lab in Milton Keynes and Alex Alderton, Roberto Amato, Jeffrey Barrett, Sonia Goncalves, Ewan Harrison, David K. Jackson, Ian Johnston, Dominic Kwiatkowski, Cordelia Langford, John Sillitoe on behalf of the Wellcome Sanger Institute COVID-19 Surveillance Team |
| EPI_ISL_2962484 | Lighthouse Lab in Alderley Park | Wellcome Sanger Institute for the COVID-19 Genomics UK (COG-UK) Consortium | Jacquelyn Wynn, Mairead Hyland, The Lighthouse Lab in Alderley Park and Alex Alderton, Roberto Amato, Jeffrey Barrett, Sonia Goncalves, Ewan Harrison, David K. Jackson, Ian Johnston, Dominic Kwiatkowski, Cordelia Langford, John Sillitoe on behalf of the Wellcome Sanger Institute COVID-19 Surveillance Team |
| EPI_ISL_2962485, EPI_ISL_2962486, EPI_ISL_2962487 | Lighthouse Lab in Milton Keynes | Wellcome Sanger Institute for the COVID-19 Genomics UK (COG-UK) Consortium | The Lighthouse Lab in Milton Keynes and Alex Alderton, Roberto Amato, Jeffrey Barrett, Sonia Goncalves, Ewan Harrison, David K. Jackson, Ian Johnston, Dominic Kwiatkowski, Cordelia Langford, John Sillitoe on behalf of the Wellcome Sanger Institute COVID-19 Surveillance Team |
| EPI_ISL_2962488, EPI_ISL_2962489, EPI_ISL_2962490 | Lighthouse Lab in Alderley Park | Wellcome Sanger Institute for the COVID-19 Genomics UK (COG-UK) Consortium | Jacquelyn Wynn, Mairead Hyland, The Lighthouse Lab in Alderley Park and Alex Alderton, Roberto Amato, Jeffrey Barrett, Sonia Goncalves, Ewan Harrison, David K. Jackson, Ian Johnston, Dominic Kwiatkowski, Cordelia Langford, John Sillitoe on behalf of the Wellcome Sanger Institute COVID-19 Surveillance Team |
| EPI_ISL_2962491 | Lighthouse Lab in Milton Keynes | Wellcome Sanger Institute for the COVID-19 Genomics UK (COG-UK) Consortium | The Lighthouse Lab in Milton Keynes and Alex Alderton, Roberto Amato, Jeffrey Barrett, Sonia Goncalves, Ewan Harrison, David K. Jackson, Ian Johnston, Dominic Kwiatkowski, Cordelia Langford, John Sillitoe on behalf of the Wellcome Sanger Institute COVID-19 Surveillance Team |
| EPI_ISL_2962492 | Lighthouse Lab in Alderley Park | Wellcome Sanger Institute for the COVID-19 Genomics UK (COG-UK) Consortium | Jacquelyn Wynn, Mairead Hyland, The Lighthouse Lab in Alderley Park and Alex Alderton, Roberto Amato, Jeffrey Barrett, Sonia Goncalves, Ewan Harrison, David K. Jackson, Ian Johnston, Dominic Kwiatkowski, Cordelia Langford, John Sillitoe on behalf of the Wellcome Sanger Institute COVID-19 Surveillance Team |
| EPI_ISL_2962493 | Lighthouse Lab in Milton Keynes | Wellcome Sanger Institute for the COVID-19 Genomics UK (COG-UK) Consortium | The Lighthouse Lab in Milton Keynes and Alex Alderton, Roberto Amato, Jeffrey Barrett, Sonia Goncalves, Ewan Harrison, David K. Jackson, Ian Johnston, Dominic Kwiatkowski, Cordelia Langford, John Sillitoe on behalf of the Wellcome Sanger Institute COVID-19 Surveillance Team |
| EPI_ISL_2962494, EPI_ISL_2962495, EPI_ISL_2962496 | Lighthouse Lab in Alderley Park | Wellcome Sanger Institute for the COVID-19 Genomics UK (COG-UK) Consortium | Jacquelyn Wynn, Mairead Hyland, The Lighthouse Lab in Alderley Park and Alex Alderton, Roberto Amato, Jeffrey Barrett, Sonia Goncalves, Ewan Harrison, David K. Jackson, Ian Johnston, Dominic Kwiatkowski, Cordelia Langford, John Sillitoe on behalf of the Wellcome Sanger Institute COVID-19 Surveillance Team |
| EPI_ISL_2962497, EPI_ISL_2962498 | Lighthouse Lab in Milton Keynes | Wellcome Sanger Institute for the COVID-19 Genomics UK (COG-UK) Consortium | The Lighthouse Lab in Milton Keynes and Alex Alderton, Roberto Amato, Jeffrey Barrett, Sonia Goncalves, Ewan Harrison, David K. Jackson, Ian Johnston, Dominic Kwiatkowski, Cordelia Langford, John Sillitoe on behalf of the Wellcome Sanger Institute COVID-19 Surveillance Team |
| EPI_ISL_2962499 | Lighthouse Lab in Alderley Park | Wellcome Sanger Institute for the COVID-19 Genomics UK (COG-UK) Consortium | Jacquelyn Wynn, Mairead Hyland, The Lighthouse Lab in Alderley Park and Alex Alderton, Roberto Amato, Jeffrey Barrett, Sonia Goncalves, Ewan Harrison, David K. Jackson, Ian Johnston, Dominic Kwiatkowski, Cordelia Langford, John Sillitoe on behalf of the Wellcome Sanger Institute COVID-19 Surveillance Team |
| EPI_ISL_2962500, EPI_ISL_2962501 | Lighthouse Lab in Milton Keynes | Wellcome Sanger Institute for the COVID-19 Genomics UK (COG-UK) Consortium | The Lighthouse Lab in Milton Keynes and Alex Alderton, Roberto Amato, Jeffrey Barrett, Sonia Goncalves, Ewan Harrison, David K. Jackson, Ian Johnston, Dominic Kwiatkowski, Cordelia Langford, John Sillitoe on behalf of the Wellcome Sanger Institute COVID-19 Surveillance Team |
| EPI_ISL_2962502 | Lighthouse Lab in Alderley Park | Wellcome Sanger Institute for the COVID-19 Genomics UK (COG-UK) Consortium | Jacquelyn Wynn, Mairead Hyland, The Lighthouse Lab in Alderley Park and Alex Alderton, Roberto Amato, Jeffrey Barrett, Sonia Goncalves, Ewan Harrison, David K. Jackson, Ian Johnston, Dominic Kwiatkowski, Cordelia Langford, John Sillitoe on behalf of the Wellcome Sanger Institute COVID-19 Surveillance Team |
| EPI_ISL_2962503, EPI_ISL_2962504, EPI_ISL_2962505, EPI_ISL_2962506, EPI_ISL_2962507, EPI_ISL_2962508, EPI_ISL_2962509, EPI_ISL_2962510, EPI_ISL_2962511, EPI_ISL_2962512, EPI_ISL_2962513, EPI_ISL_2962514 | | | |
| see above | Lighthouse Lab in Milton Keynes | Wellcome Sanger Institute for the COVID-19 Genomics UK (COG-UK) Consortium | The Lighthouse Lab in Milton Keynes and Alex Alderton, Roberto Amato, Jeffrey Barrett, Sonia Goncalves, Ewan Harrison, David K. Jackson, Ian Johnston, Dominic Kwiatkowski, Cordelia Langford, John Sillitoe on behalf of the Wellcome Sanger Institute COVID-19 Surveillance Team |
| EPI_ISL_2962515 | Lighthouse Lab in Alderley Park | Wellcome Sanger Institute for the COVID-19 Genomics UK (COG-UK) Consortium | Jacquelyn Wynn, Mairead Hyland, The Lighthouse Lab in Alderley Park and Alex Alderton, Roberto Amato, Jeffrey Barrett, Sonia Goncalves, Ewan Harrison, David K. Jackson, Ian Johnston, Dominic Kwiatkowski, Cordelia Langford, John Sillitoe on behalf of the Wellcome Sanger Institute COVID-19 Surveillance Team |
| EPI_ISL_2962516, EPI_ISL_2962517, EPI_ISL_2962518, EPI_ISL_2962519, EPI_ISL_2962520, EPI_ISL_2962521, EPI_ISL_2962522 | Lighthouse Lab in Milton Keynes | Wellcome Sanger Institute for the COVID-19 Genomics UK (COG-UK) Consortium | The Lighthouse Lab in Milton Keynes and Alex Alderton, Roberto Amato, Jeffrey Barrett, Sonia Goncalves, Ewan Harrison, David K. Jackson, Ian Johnston, Dominic Kwiatkowski, Cordelia Langford, John Sillitoe on behalf of the Wellcome Sanger Institute COVID-19 Surveillance Team |
| EPI_ISL_2962523 | Lighthouse Lab in Alderley Park | Wellcome Sanger Institute for the COVID-19 Genomics UK (COG-UK) Consortium | Jacquelyn Wynn, Mairead Hyland, The Lighthouse Lab in Alderley Park and Alex Alderton, Roberto Amato, Jeffrey Barrett, Sonia Goncalves, Ewan Harrison, David K. Jackson, Ian Johnston, Dominic Kwiatkowski, Cordelia Langford, John Sillitoe on behalf of the Wellcome Sanger Institute COVID-19 Surveillance Team |
| EPI_ISL_2962524 | Lighthouse Lab in Milton Keynes | Wellcome Sanger Institute for the COVID-19 Genomics UK (COG-UK) Consortium | The Lighthouse Lab in Milton Keynes and Alex Alderton, Roberto Amato, Jeffrey Barrett, Sonia Goncalves, Ewan Harrison, David K. Jackson, Ian Johnston, Dominic Kwiatkowski, Cordelia Langford, John Sillitoe on behalf of the Wellcome Sanger Institute COVID-19 Surveillance Team |
| EPI_ISL_2962525 | Lighthouse Lab in Alderley Park | Wellcome Sanger Institute for the COVID-19 Genomics UK (COG-UK) Consortium | Jacquelyn Wynn, Mairead Hyland, The Lighthouse Lab in Alderley Park and Alex Alderton, Roberto Amato, Jeffrey Barrett, Sonia Goncalves, Ewan Harrison, David K. Jackson, Ian Johnston, Dominic Kwiatkowski, Cordelia Langford, John Sillitoe on behalf of the Wellcome Sanger Institute COVID-19 Surveillance Team |
| EPI_ISL_2962526 | Lighthouse Lab in Milton Keynes | Wellcome Sanger Institute for the COVID-19 Genomics UK (COG-UK) Consortium | The Lighthouse Lab in Milton Keynes and Alex Alderton, Roberto Amato, Jeffrey Barrett, Sonia Goncalves, Ewan Harrison, David K. Jackson, Ian Johnston, Dominic Kwiatkowski, Cordelia Langford, John Sillitoe on behalf of the Wellcome Sanger Institute COVID-19 Surveillance Team |
| EPI_ISL_2962527 | Lighthouse Lab in Alderley Park | Wellcome Sanger Institute for the COVID-19 Genomics UK (COG-UK) Consortium | Jacquelyn Wynn, Mairead Hyland, The Lighthouse Lab in Alderley Park and Alex Alderton, Roberto Amato, Jeffrey Barrett, Sonia Goncalves, Ewan Harrison, David K. Jackson, Ian Johnston, Dominic Kwiatkowski, Cordelia Langford, John Sillitoe on behalf of the Wellcome Sanger Institute COVID-19 Surveillance Team |
| EPI_ISL_2962528 | Lighthouse Lab in Milton Keynes | Wellcome Sanger Institute for the COVID-19 Genomics UK (COG-UK) Consortium | The Lighthouse Lab in Milton Keynes and Alex Alderton, Roberto Amato, Jeffrey Barrett, Sonia Goncalves, Ewan Harrison, David K. Jackson, Ian Johnston, Dominic Kwiatkowski, Cordelia Langford, John Sillitoe on behalf of the Wellcome Sanger Institute COVID-19 Surveillance Team |
| EPI_ISL_2962529, EPI_ISL_2962530 | Lighthouse Lab in Alderley Park | Wellcome Sanger Institute for the COVID-19 Genomics UK (COG-UK) Consortium | Jacquelyn Wynn, Mairead Hyland, The Lighthouse Lab in Alderley Park and Alex Alderton, Roberto Amato, Jeffrey Barrett, Sonia Goncalves, Ewan Harrison, David K. Jackson, Ian Johnston, Dominic Kwiatkowski, Cordelia Langford, John Sillitoe on behalf of the Wellcome Sanger Institute COVID-19 Surveillance Team |
| EPI_ISL_2962531, EPI_ISL_2962532, EPI_ISL_2962533, EPI_ISL_2962534, EPI_ISL_2962535 | Lighthouse Lab in Milton Keynes | Wellcome Sanger Institute for the COVID-19 Genomics UK (COG-UK) Consortium | The Lighthouse Lab in Milton Keynes and Alex Alderton, Roberto Amato, Jeffrey Barrett, Sonia Goncalves, Ewan Harrison, David K. Jackson, Ian Johnston, Dominic Kwiatkowski, Cordelia Langford, John Sillitoe on behalf of the Wellcome Sanger Institute COVID-19 Surveillance Team |
| EPI_ISL_2962536 | Lighthouse Lab in Alderley Park | Wellcome Sanger Institute for the COVID-19 Genomics UK (COG-UK) Consortium | Jacquelyn Wynn, Mairead Hyland, The Lighthouse Lab in Alderley Park and Alex Alderton, Roberto Amato, Jeffrey Barrett, Sonia Goncalves, Ewan Harrison, David K. Jackson, Ian Johnston, Dominic Kwiatkowski, Cordelia Langford, John Sillitoe on behalf of the Wellcome Sanger Institute COVID-19 Surveillance Team |
| EPI_ISL_2962537, EPI_ISL_2962538, EPI_ISL_2962539, EPI_ISL_2962540, EPI_ISL_2962541 | Lighthouse Lab in Milton Keynes | Wellcome Sanger Institute for the COVID-19 Genomics UK (COG-UK) Consortium | The Lighthouse Lab in Milton Keynes and Alex Alderton, Roberto Amato, Jeffrey Barrett, Sonia Goncalves, Ewan Harrison, David K. Jackson, Ian Johnston, Dominic Kwiatkowski, Cordelia Langford, John Sillitoe on behalf of the Wellcome Sanger Institute COVID-19 Surveillance Team |
| EPI_ISL_2962542 | Lighthouse Lab in Alderley Park | Wellcome Sanger Institute for the COVID-19 Genomics UK (COG-UK) Consortium | Jacquelyn Wynn, Mairead Hyland, The Lighthouse Lab in Alderley Park and Alex Alderton, Roberto Amato, Jeffrey Barrett, Sonia Goncalves, Ewan Harrison, David K. Jackson, Ian Johnston, Dominic Kwiatkowski, Cordelia Langford, John Sillitoe on behalf of the Wellcome Sanger Institute COVID-19 Surveillance Team |
| EPI_ISL_2962543, EPI_ISL_2962544, EPI_ISL_2962545, EPI_ISL_2962546, EPI_ISL_2962547, EPI_ISL_2962548, EPI_ISL_2962549, EPI_ISL_2962550, EPI_ISL_2962551, EPI_ISL_2962552, EPI_ISL_2962553, EPI_ISL_2962554, EPI_ISL_2962555, EPI_ISL_2962556 | | | |
| see above | Lighthouse Lab in Milton Keynes | Wellcome Sanger Institute for the COVID-19 Genomics UK (COG-UK) Consortium | The Lighthouse Lab in Milton Keynes and Alex Alderton, Roberto Amato, Jeffrey Barrett, Sonia Goncalves, Ewan Harrison, David K. Jackson, Ian Johnston, Dominic Kwiatkowski, Cordelia Langford, John Sillitoe on behalf of the Wellcome Sanger Institute COVID-19 Surveillance Team |
| EPI_ISL_2962557 | Lighthouse Lab in Alderley Park | Wellcome Sanger Institute for the COVID-19 Genomics UK (COG-UK) Consortium | Jacquelyn Wynn, Mairead Hyland, The Lighthouse Lab in Alderley Park and Alex Alderton, Roberto Amato, Jeffrey Barrett, Sonia Goncalves, Ewan Harrison, David K. Jackson, Ian Johnston, Dominic Kwiatkowski, Cordelia Langford, John Sillitoe on behalf of the Wellcome Sanger Institute COVID-19 Surveillance Team |
| EPI_ISL_2962558, EPI_ISL_2962559, EPI_ISL_2962560, | Lighthouse Lab in Milton Keynes | Wellcome Sanger Institute for the COVID-19 Genomics UK (COG-UK) Consortium | The Lighthouse Lab in Milton Keynes and Alex Alderton, Roberto Amato, Jeffrey Barrett, Sonia Goncalves, Ewan Harrison, David K. Jackson, Ian Johnston, Dominic Kwiatkowski, Cordelia Langford, John Sillitoe on behalf of the Wellcome Sanger Institute COVID-19 Surveillance Team |

| | | | |
|---|---|---|---|
| EPI_ISL_2962561, EPI_ISL_2962562, EPI_ISL_2962563, EPI_ISL_2962564, EPI_ISL_2962565, EPI_ISL_2962566, EPI_ISL_2962567 | | | |
| EPI_ISL_2962568 | Lighthouse Lab in Alderley Park | Wellcome Sanger Institute for the COVID-19 Genomics UK (COG-UK) Consortium | Jacquelyn Wynn, Mairead Hyland, The Lighthouse Lab in Alderley Park and Alex Alderton, Roberto Amato, Jeffrey Barrett, Sonia Goncalves, Ewan Harrison, David K. Jackson, Ian Johnston, Dominic Kwiatkowski, Cordelia Langford, John Sillitoe on behalf of the Wellcome Sanger Institute COVID-19 Surveillance Team |
| EPI_ISL_2962569, EPI_ISL_2962570, EPI_ISL_2962571 | Lighthouse Lab in Milton Keynes | Wellcome Sanger Institute for the COVID-19 Genomics UK (COG-UK) Consortium | The Lighthouse Lab in Milton Keynes and Alex Alderton, Roberto Amato, Jeffrey Barrett, Sonia Goncalves, Ewan Harrison, David K. Jackson, Ian Johnston, Dominic Kwiatkowski, Cordelia Langford, John Sillitoe on behalf of the Wellcome Sanger Institute COVID-19 Surveillance Team |
| EPI_ISL_2962572, EPI_ISL_2962573, EPI_ISL_2962574, EPI_ISL_2962575 | Lighthouse Lab in Alderley Park | Wellcome Sanger Institute for the COVID-19 Genomics UK (COG-UK) Consortium | Jacquelyn Wynn, Mairead Hyland, The Lighthouse Lab in Alderley Park and Alex Alderton, Roberto Amato, Jeffrey Barrett, Sonia Goncalves, Ewan Harrison, David K. Jackson, Ian Johnston, Dominic Kwiatkowski, Cordelia Langford, John Sillitoe on behalf of the Wellcome Sanger Institute COVID-19 Surveillance Team |
| EPI_ISL_2962576 | Lighthouse Lab in Milton Keynes | Wellcome Sanger Institute for the COVID-19 Genomics UK (COG-UK) Consortium | The Lighthouse Lab in Milton Keynes and Alex Alderton, Roberto Amato, Jeffrey Barrett, Sonia Goncalves, Ewan Harrison, David K. Jackson, Ian Johnston, Dominic Kwiatkowski, Cordelia Langford, John Sillitoe on behalf of the Wellcome Sanger Institute COVID-19 Surveillance Team |
| EPI_ISL_2962577, EPI_ISL_2962578 | Lighthouse Lab in Alderley Park | Wellcome Sanger Institute for the COVID-19 Genomics UK (COG-UK) Consortium | Jacquelyn Wynn, Mairead Hyland, The Lighthouse Lab in Alderley Park and Alex Alderton, Roberto Amato, Jeffrey Barrett, Sonia Goncalves, Ewan Harrison, David K. Jackson, Ian Johnston, Dominic Kwiatkowski, Cordelia Langford, John Sillitoe on behalf of the Wellcome Sanger Institute COVID-19 Surveillance Team |
| EPI_ISL_2962579, EPI_ISL_2962580, EPI_ISL_2962581, EPI_ISL_2962582, EPI_ISL_2962583 | Lighthouse Lab in Milton Keynes | Wellcome Sanger Institute for the COVID-19 Genomics UK (COG-UK) Consortium | The Lighthouse Lab in Milton Keynes and Alex Alderton, Roberto Amato, Jeffrey Barrett, Sonia Goncalves, Ewan Harrison, David K. Jackson, Ian Johnston, Dominic Kwiatkowski, Cordelia Langford, John Sillitoe on behalf of the Wellcome Sanger Institute COVID-19 Surveillance Team |
| EPI_ISL_2962584, EPI_ISL_2962585 | Lighthouse Lab in Alderley Park | Wellcome Sanger Institute for the COVID-19 Genomics UK (COG-UK) Consortium | Jacquelyn Wynn, Mairead Hyland, The Lighthouse Lab in Alderley Park and Alex Alderton, Roberto Amato, Jeffrey Barrett, Sonia Goncalves, Ewan Harrison, David K. Jackson, Ian Johnston, Dominic Kwiatkowski, Cordelia Langford, John Sillitoe on behalf of the Wellcome Sanger Institute COVID-19 Surveillance Team |
| EPI_ISL_2962586, EPI_ISL_2962587, EPI_ISL_2962588 | Lighthouse Lab in Milton Keynes | Wellcome Sanger Institute for the COVID-19 Genomics UK (COG-UK) Consortium | The Lighthouse Lab in Milton Keynes and Alex Alderton, Roberto Amato, Jeffrey Barrett, Sonia Goncalves, Ewan Harrison, David K. Jackson, Ian Johnston, Dominic Kwiatkowski, Cordelia Langford, John Sillitoe on behalf of the Wellcome Sanger Institute COVID-19 Surveillance Team |
| EPI_ISL_2962589 | Lighthouse Lab in Alderley Park | Wellcome Sanger Institute for the COVID-19 Genomics UK (COG-UK) Consortium | Jacquelyn Wynn, Mairead Hyland, The Lighthouse Lab in Alderley Park and Alex Alderton, Roberto Amato, Jeffrey Barrett, Sonia Goncalves, Ewan Harrison, David K. Jackson, Ian Johnston, Dominic Kwiatkowski, Cordelia Langford, John Sillitoe on behalf of the Wellcome Sanger Institute COVID-19 Surveillance Team |
| EPI_ISL_2962590, EPI_ISL_2962591, EPI_ISL_2962592, EPI_ISL_2962593, EPI_ISL_2962594, EPI_ISL_2962595, EPI_ISL_2962596, EPI_ISL_2962597, EPI_ISL_2962598, EPI_ISL_2962599 | Lighthouse Lab in Milton Keynes | Wellcome Sanger Institute for the COVID-19 Genomics UK (COG-UK) Consortium | The Lighthouse Lab in Milton Keynes and Alex Alderton, Roberto Amato, Jeffrey Barrett, Sonia Goncalves, Ewan Harrison, David K. Jackson, Ian Johnston, Dominic Kwiatkowski, Cordelia Langford, John Sillitoe on behalf of the Wellcome Sanger Institute COVID-19 Surveillance Team |
| EPI_ISL_2962600 | Lighthouse Lab in Alderley Park | Wellcome Sanger Institute for the COVID-19 Genomics UK (COG-UK) Consortium | Jacquelyn Wynn, Mairead Hyland, The Lighthouse Lab in Alderley Park and Alex Alderton, Roberto Amato, Jeffrey Barrett, Sonia Goncalves, Ewan Harrison, David K. Jackson, Ian Johnston, Dominic Kwiatkowski, Cordelia Langford, John Sillitoe on behalf of the Wellcome Sanger Institute COVID-19 Surveillance Team |
| EPI_ISL_2962601, EPI_ISL_2962602, EPI_ISL_2962603, EPI_ISL_2962604, EPI_ISL_2962605 | Lighthouse Lab in Milton Keynes | Wellcome Sanger Institute for the COVID-19 Genomics UK (COG-UK) Consortium | The Lighthouse Lab in Milton Keynes and Alex Alderton, Roberto Amato, Jeffrey Barrett, Sonia Goncalves, Ewan Harrison, David K. Jackson, Ian Johnston, Dominic Kwiatkowski, Cordelia Langford, John Sillitoe on behalf of the Wellcome Sanger Institute COVID-19 Surveillance Team |
| EPI_ISL_2962606, EPI_ISL_2962607 | Lighthouse Lab in Alderley Park | Wellcome Sanger Institute for the COVID-19 Genomics UK (COG-UK) Consortium | Jacquelyn Wynn, Mairead Hyland, The Lighthouse Lab in Alderley Park and Alex Alderton, Roberto Amato, Jeffrey Barrett, Sonia Goncalves, Ewan Harrison, David K. Jackson, Ian Johnston, Dominic Kwiatkowski, Cordelia Langford, John Sillitoe on behalf of the Wellcome Sanger Institute COVID-19 Surveillance Team |
| EPI_ISL_2962608, EPI_ISL_2962609 | Lighthouse Lab in Milton Keynes | Wellcome Sanger Institute for the COVID-19 Genomics UK (COG-UK) Consortium | The Lighthouse Lab in Milton Keynes and Alex Alderton, Roberto Amato, Jeffrey Barrett, Sonia Goncalves, Ewan Harrison, David K. Jackson, Ian Johnston, Dominic Kwiatkowski, Cordelia Langford, John Sillitoe on behalf of the Wellcome Sanger Institute COVID-19 Surveillance Team |
| EPI_ISL_2962610 | Lighthouse Lab in Alderley Park | Wellcome Sanger Institute for the COVID-19 Genomics UK (COG-UK) Consortium | Jacquelyn Wynn, Mairead Hyland, The Lighthouse Lab in Alderley Park and Alex Alderton, Roberto Amato, Jeffrey Barrett, Sonia Goncalves, Ewan Harrison, David K. Jackson, Ian Johnston, Dominic Kwiatkowski, Cordelia Langford, John Sillitoe on behalf of the Wellcome Sanger Institute COVID-19 Surveillance Team |
| EPI_ISL_2962611, EPI_ISL_2962612, EPI_ISL_2962613, EPI_ISL_2962614 | Lighthouse Lab in Milton Keynes | Wellcome Sanger Institute for the COVID-19 Genomics UK (COG-UK) Consortium | The Lighthouse Lab in Milton Keynes and Alex Alderton, Roberto Amato, Jeffrey Barrett, Sonia Goncalves, Ewan Harrison, David K. Jackson, Ian Johnston, Dominic Kwiatkowski, Cordelia Langford, John Sillitoe on behalf of the Wellcome Sanger Institute COVID-19 Surveillance Team |
| EPI_ISL_2962615, EPI_ISL_2962616, EPI_ISL_2962617 | Lighthouse Lab in Alderley Park | Wellcome Sanger Institute for the COVID-19 Genomics UK (COG-UK) Consortium | Jacquelyn Wynn, Mairead Hyland, The Lighthouse Lab in Alderley Park and Alex Alderton, Roberto Amato, Jeffrey Barrett, Sonia Goncalves, Ewan Harrison, David K. Jackson, Ian Johnston, Dominic Kwiatkowski, Cordelia Langford, John Sillitoe on behalf of the Wellcome Sanger Institute COVID-19 Surveillance Team |
| EPI_ISL_2962618, EPI_ISL_2962619, EPI_ISL_2962620, EPI_ISL_2962621, EPI_ISL_2962622 | Lighthouse Lab in Milton Keynes | Wellcome Sanger Institute for the COVID-19 Genomics UK (COG-UK) Consortium | The Lighthouse Lab in Milton Keynes and Alex Alderton, Roberto Amato, Jeffrey Barrett, Sonia Goncalves, Ewan Harrison, David K. Jackson, Ian Johnston, Dominic Kwiatkowski, Cordelia Langford, John Sillitoe on behalf of the Wellcome Sanger Institute COVID-19 Surveillance Team |
| EPI_ISL_2962623 | Lighthouse Lab in Alderley Park | Wellcome Sanger Institute for the COVID-19 Genomics UK (COG-UK) Consortium | Jacquelyn Wynn, Mairead Hyland, The Lighthouse Lab in Alderley Park and Alex Alderton, Roberto Amato, Jeffrey Barrett, Sonia Goncalves, Ewan Harrison, David K. Jackson, Ian Johnston, Dominic Kwiatkowski, Cordelia Langford, John Sillitoe on behalf of the Wellcome Sanger Institute COVID-19 Surveillance Team |
| EPI_ISL_2962624 | Lighthouse Lab in Milton Keynes | Wellcome Sanger Institute for the COVID-19 Genomics UK (COG-UK) Consortium | The Lighthouse Lab in Milton Keynes and Alex Alderton, Roberto Amato, Jeffrey Barrett, Sonia Goncalves, Ewan Harrison, David K. Jackson, Ian Johnston, Dominic Kwiatkowski, Cordelia Langford, John Sillitoe on behalf of the Wellcome Sanger Institute COVID-19 Surveillance Team |
| EPI_ISL_2962625 | Lighthouse Lab in Alderley Park | Wellcome Sanger Institute for the COVID-19 Genomics UK (COG-UK) Consortium | Jacquelyn Wynn, Mairead Hyland, The Lighthouse Lab in Alderley Park and Alex Alderton, Roberto Amato, Jeffrey Barrett, Sonia Goncalves, Ewan Harrison, David K. Jackson, Ian Johnston, Dominic Kwiatkowski, Cordelia Langford, John Sillitoe on behalf of the Wellcome Sanger Institute COVID-19 Surveillance Team |
| EPI_ISL_2962626, EPI_ISL_2962627, EPI_ISL_2962628, EPI_ISL_2962629, EPI_ISL_2962630, EPI_ISL_2962631, EPI_ISL_2962632, EPI_ISL_2962633, EPI_ISL_2962634, EPI_ISL_2962635 | Lighthouse Lab in Milton Keynes | Wellcome Sanger Institute for the COVID-19 Genomics UK (COG-UK) Consortium | The Lighthouse Lab in Milton Keynes and Alex Alderton, Roberto Amato, Jeffrey Barrett, Sonia Goncalves, Ewan Harrison, David K. Jackson, Ian Johnston, Dominic Kwiatkowski, Cordelia Langford, John Sillitoe on behalf of the Wellcome Sanger Institute COVID-19 Surveillance Team |
| EPI_ISL_2962636, EPI_ISL_2962637 | Lighthouse Lab in Alderley Park | Wellcome Sanger Institute for the COVID-19 Genomics UK (COG-UK) Consortium | Jacquelyn Wynn, Mairead Hyland, The Lighthouse Lab in Alderley Park and Alex Alderton, Roberto Amato, Jeffrey Barrett, Sonia Goncalves, Ewan Harrison, David K. Jackson, Ian Johnston, Dominic Kwiatkowski, Cordelia Langford, John Sillitoe on behalf of the Wellcome Sanger Institute COVID-19 Surveillance Team |
| EPI_ISL_2962638 | Lighthouse Lab in Milton Keynes | Wellcome Sanger Institute for the COVID-19 Genomics UK (COG-UK) Consortium | The Lighthouse Lab in Milton Keynes and Alex Alderton, Roberto Amato, Jeffrey Barrett, Sonia Goncalves, Ewan Harrison, David K. Jackson, Ian Johnston, Dominic Kwiatkowski, Cordelia Langford, John Sillitoe on behalf of the Wellcome Sanger Institute COVID-19 Surveillance Team |
| EPI_ISL_2962639 | Lighthouse Lab in Alderley Park | Wellcome Sanger Institute for the COVID-19 Genomics UK (COG-UK) Consortium | Jacquelyn Wynn, Mairead Hyland, The Lighthouse Lab in Alderley Park and Alex Alderton, Roberto Amato, Jeffrey Barrett, Sonia Goncalves, Ewan Harrison, David K. Jackson, Ian Johnston, Dominic Kwiatkowski, Cordelia Langford, John Sillitoe on behalf of the Wellcome Sanger Institute COVID-19 Surveillance Team |
| EPI_ISL_2962640, EPI_ISL_2962641, EPI_ISL_2962642, EPI_ISL_2962643, EPI_ISL_2962644, EPI_ISL_2962645 | Lighthouse Lab in Milton Keynes | Wellcome Sanger Institute for the COVID-19 Genomics UK (COG-UK) Consortium | The Lighthouse Lab in Milton Keynes and Alex Alderton, Roberto Amato, Jeffrey Barrett, Sonia Goncalves, Ewan Harrison, David K. Jackson, Ian Johnston, Dominic Kwiatkowski, Cordelia Langford, John Sillitoe on behalf of the Wellcome Sanger Institute COVID-19 Surveillance Team |
| EPI_ISL_2962646 | Lighthouse Lab in Alderley Park | Wellcome Sanger Institute for the COVID-19 Genomics UK (COG-UK) Consortium | Jacquelyn Wynn, Mairead Hyland, The Lighthouse Lab in Alderley Park and Alex Alderton, Roberto Amato, Jeffrey Barrett, Sonia Goncalves, Ewan Harrison, David K. Jackson, Ian Johnston, Dominic Kwiatkowski, Cordelia Langford, John Sillitoe on behalf of the Wellcome Sanger Institute COVID-19 Surveillance Team |
| EPI_ISL_2962647, | Lighthouse Lab in Milton Keynes | Wellcome Sanger Institute for the COVID-19 Genomics UK (COG-UK) | The Lighthouse Lab in Milton Keynes and Alex Alderton, Roberto Amato, Jeffrey Barrett, Sonia Goncalves, Ewan Harrison, David K. Jackson, Ian Johnston, Dominic Kwiatkowski, Cordelia Langford, John Sillitoe on behalf of the Wellcome Sanger |

| | | | |
|---|---|---|---|
| EPI_ISL_2962648 | | Consortium | Institute COVID-19 Surveillance Team |
| EPI_ISL_2962649, EPI_ISL_2962650 | Lighthouse Lab in Alderley Park | Wellcome Sanger Institute for the COVID-19 Genomics UK (COG-UK) Consortium | Jacquelyn Wynn, Mairead Hyland, The Lighthouse Lab in Alderley Park and Alex Alderton, Roberto Amato, Jeffrey Barrett, Sonia Goncalves, Ewan Harrison, David K. Jackson, Ian Johnston, Dominic Kwiatkowski, Cordelia Langford, John Sillitoe on behalf of the Wellcome Sanger Institute COVID-19 Surveillance Team |
| EPI_ISL_2962651, EPI_ISL_2962652, EPI_ISL_2962653 | Lighthouse Lab in Milton Keynes | Wellcome Sanger Institute for the COVID-19 Genomics UK (COG-UK) Consortium | The Lighthouse Lab in Milton Keynes and Alex Alderton, Roberto Amato, Jeffrey Barrett, Sonia Goncalves, Ewan Harrison, David K. Jackson, Ian Johnston, Dominic Kwiatkowski, Cordelia Langford, John Sillitoe on behalf of the Wellcome Sanger Institute COVID-19 Surveillance Team |
| EPI_ISL_2962654 | Lighthouse Lab in Alderley Park | Wellcome Sanger Institute for the COVID-19 Genomics UK (COG-UK) Consortium | Jacquelyn Wynn, Mairead Hyland, The Lighthouse Lab in Alderley Park and Alex Alderton, Roberto Amato, Jeffrey Barrett, Sonia Goncalves, Ewan Harrison, David K. Jackson, Ian Johnston, Dominic Kwiatkowski, Cordelia Langford, John Sillitoe on behalf of the Wellcome Sanger Institute COVID-19 Surveillance Team |
| EPI_ISL_2962655, EPI_ISL_2962656, EPI_ISL_2962657 | Lighthouse Lab in Milton Keynes | Wellcome Sanger Institute for the COVID-19 Genomics UK (COG-UK) Consortium | The Lighthouse Lab in Milton Keynes and Alex Alderton, Roberto Amato, Jeffrey Barrett, Sonia Goncalves, Ewan Harrison, David K. Jackson, Ian Johnston, Dominic Kwiatkowski, Cordelia Langford, John Sillitoe on behalf of the Wellcome Sanger Institute COVID-19 Surveillance Team |
| EPI_ISL_2962658 | Lighthouse Lab in Alderley Park | Wellcome Sanger Institute for the COVID-19 Genomics UK (COG-UK) Consortium | Jacquelyn Wynn, Mairead Hyland, The Lighthouse Lab in Alderley Park and Alex Alderton, Roberto Amato, Jeffrey Barrett, Sonia Goncalves, Ewan Harrison, David K. Jackson, Ian Johnston, Dominic Kwiatkowski, Cordelia Langford, John Sillitoe on behalf of the Wellcome Sanger Institute COVID-19 Surveillance Team |
| EPI_ISL_2962659 | Lighthouse Lab in Milton Keynes | Wellcome Sanger Institute for the COVID-19 Genomics UK (COG-UK) Consortium | The Lighthouse Lab in Milton Keynes and Alex Alderton, Roberto Amato, Jeffrey Barrett, Sonia Goncalves, Ewan Harrison, David K. Jackson, Ian Johnston, Dominic Kwiatkowski, Cordelia Langford, John Sillitoe on behalf of the Wellcome Sanger Institute COVID-19 Surveillance Team |
| EPI_ISL_2962660 | Lighthouse Lab in Alderley Park | Wellcome Sanger Institute for the COVID-19 Genomics UK (COG-UK) Consortium | Jacquelyn Wynn, Mairead Hyland, The Lighthouse Lab in Alderley Park and Alex Alderton, Roberto Amato, Jeffrey Barrett, Sonia Goncalves, Ewan Harrison, David K. Jackson, Ian Johnston, Dominic Kwiatkowski, Cordelia Langford, John Sillitoe on behalf of the Wellcome Sanger Institute COVID-19 Surveillance Team |
| EPI_ISL_2962661, EPI_ISL_2962662, EPI_ISL_2962663 | Lighthouse Lab in Milton Keynes | Wellcome Sanger Institute for the COVID-19 Genomics UK (COG-UK) Consortium | The Lighthouse Lab in Milton Keynes and Alex Alderton, Roberto Amato, Jeffrey Barrett, Sonia Goncalves, Ewan Harrison, David K. Jackson, Ian Johnston, Dominic Kwiatkowski, Cordelia Langford, John Sillitoe on behalf of the Wellcome Sanger Institute COVID-19 Surveillance Team |
| EPI_ISL_2962664 | Lighthouse Lab in Alderley Park | Wellcome Sanger Institute for the COVID-19 Genomics UK (COG-UK) Consortium | Jacquelyn Wynn, Mairead Hyland, The Lighthouse Lab in Alderley Park and Alex Alderton, Roberto Amato, Jeffrey Barrett, Sonia Goncalves, Ewan Harrison, David K. Jackson, Ian Johnston, Dominic Kwiatkowski, Cordelia Langford, John Sillitoe on behalf of the Wellcome Sanger Institute COVID-19 Surveillance Team |
| EPI_ISL_2962665, EPI_ISL_2962666, EPI_ISL_2962667 | Lighthouse Lab in Milton Keynes | Wellcome Sanger Institute for the COVID-19 Genomics UK (COG-UK) Consortium | The Lighthouse Lab in Milton Keynes and Alex Alderton, Roberto Amato, Jeffrey Barrett, Sonia Goncalves, Ewan Harrison, David K. Jackson, Ian Johnston, Dominic Kwiatkowski, Cordelia Langford, John Sillitoe on behalf of the Wellcome Sanger Institute COVID-19 Surveillance Team |
| EPI_ISL_2962668 | Lighthouse Lab in Alderley Park | Wellcome Sanger Institute for the COVID-19 Genomics UK (COG-UK) Consortium | Jacquelyn Wynn, Mairead Hyland, The Lighthouse Lab in Alderley Park and Alex Alderton, Roberto Amato, Jeffrey Barrett, Sonia Goncalves, Ewan Harrison, David K. Jackson, Ian Johnston, Dominic Kwiatkowski, Cordelia Langford, John Sillitoe on behalf of the Wellcome Sanger Institute COVID-19 Surveillance Team |
| EPI_ISL_2962669, EPI_ISL_2962670, EPI_ISL_2962671, EPI_ISL_2962672, EPI_ISL_2962673 | Lighthouse Lab in Milton Keynes | Wellcome Sanger Institute for the COVID-19 Genomics UK (COG-UK) Consortium | The Lighthouse Lab in Milton Keynes and Alex Alderton, Roberto Amato, Jeffrey Barrett, Sonia Goncalves, Ewan Harrison, David K. Jackson, Ian Johnston, Dominic Kwiatkowski, Cordelia Langford, John Sillitoe on behalf of the Wellcome Sanger Institute COVID-19 Surveillance Team |
| EPI_ISL_2962674 | Lighthouse Lab in Alderley Park | Wellcome Sanger Institute for the COVID-19 Genomics UK (COG-UK) Consortium | Jacquelyn Wynn, Mairead Hyland, The Lighthouse Lab in Alderley Park and Alex Alderton, Roberto Amato, Jeffrey Barrett, Sonia Goncalves, Ewan Harrison, David K. Jackson, Ian Johnston, Dominic Kwiatkowski, Cordelia Langford, John Sillitoe on behalf of the Wellcome Sanger Institute COVID-19 Surveillance Team |
| EPI_ISL_2962675, EPI_ISL_2962676 | Lighthouse Lab in Milton Keynes | Wellcome Sanger Institute for the COVID-19 Genomics UK (COG-UK) Consortium | The Lighthouse Lab in Milton Keynes and Alex Alderton, Roberto Amato, Jeffrey Barrett, Sonia Goncalves, Ewan Harrison, David K. Jackson, Ian Johnston, Dominic Kwiatkowski, Cordelia Langford, John Sillitoe on behalf of the Wellcome Sanger Institute COVID-19 Surveillance Team |
| EPI_ISL_2962677 | Lighthouse Lab in Alderley Park | Wellcome Sanger Institute for the COVID-19 Genomics UK (COG-UK) Consortium | Jacquelyn Wynn, Mairead Hyland, The Lighthouse Lab in Alderley Park and Alex Alderton, Roberto Amato, Jeffrey Barrett, Sonia Goncalves, Ewan Harrison, David K. Jackson, Ian Johnston, Dominic Kwiatkowski, Cordelia Langford, John Sillitoe on behalf of the Wellcome Sanger Institute COVID-19 Surveillance Team |
| EPI_ISL_2962678, EPI_ISL_2962679, EPI_ISL_2962680, EPI_ISL_2962681, EPI_ISL_2962682 | Lighthouse Lab in Milton Keynes | Wellcome Sanger Institute for the COVID-19 Genomics UK (COG-UK) Consortium | The Lighthouse Lab in Milton Keynes and Alex Alderton, Roberto Amato, Jeffrey Barrett, Sonia Goncalves, Ewan Harrison, David K. Jackson, Ian Johnston, Dominic Kwiatkowski, Cordelia Langford, John Sillitoe on behalf of the Wellcome Sanger Institute COVID-19 Surveillance Team |
| EPI_ISL_2962683 | Lighthouse Lab in Alderley Park | Wellcome Sanger Institute for the COVID-19 Genomics UK (COG-UK) Consortium | Jacquelyn Wynn, Mairead Hyland, The Lighthouse Lab in Alderley Park and Alex Alderton, Roberto Amato, Jeffrey Barrett, Sonia Goncalves, Ewan Harrison, David K. Jackson, Ian Johnston, Dominic Kwiatkowski, Cordelia Langford, John Sillitoe on behalf of the Wellcome Sanger Institute COVID-19 Surveillance Team |
| EPI_ISL_2962684, EPI_ISL_2962685, EPI_ISL_2962686, EPI_ISL_2962687, EPI_ISL_2962688 | Lighthouse Lab in Milton Keynes | Wellcome Sanger Institute for the COVID-19 Genomics UK (COG-UK) Consortium | The Lighthouse Lab in Milton Keynes and Alex Alderton, Roberto Amato, Jeffrey Barrett, Sonia Goncalves, Ewan Harrison, David K. Jackson, Ian Johnston, Dominic Kwiatkowski, Cordelia Langford, John Sillitoe on behalf of the Wellcome Sanger Institute COVID-19 Surveillance Team |
| EPI_ISL_2962689 | Lighthouse Lab in Alderley Park | Wellcome Sanger Institute for the COVID-19 Genomics UK (COG-UK) Consortium | Jacquelyn Wynn, Mairead Hyland, The Lighthouse Lab in Alderley Park and Alex Alderton, Roberto Amato, Jeffrey Barrett, Sonia Goncalves, Ewan Harrison, David K. Jackson, Ian Johnston, Dominic Kwiatkowski, Cordelia Langford, John Sillitoe on behalf of the Wellcome Sanger Institute COVID-19 Surveillance Team |
| EPI_ISL_2962690, EPI_ISL_2962691 | Lighthouse Lab in Milton Keynes | Wellcome Sanger Institute for the COVID-19 Genomics UK (COG-UK) Consortium | The Lighthouse Lab in Milton Keynes and Alex Alderton, Roberto Amato, Jeffrey Barrett, Sonia Goncalves, Ewan Harrison, David K. Jackson, Ian Johnston, Dominic Kwiatkowski, Cordelia Langford, John Sillitoe on behalf of the Wellcome Sanger Institute COVID-19 Surveillance Team |
| EPI_ISL_2962692 | Lighthouse Lab in Alderley Park | Wellcome Sanger Institute for the COVID-19 Genomics UK (COG-UK) Consortium | Jacquelyn Wynn, Mairead Hyland, The Lighthouse Lab in Alderley Park and Alex Alderton, Roberto Amato, Jeffrey Barrett, Sonia Goncalves, Ewan Harrison, David K. Jackson, Ian Johnston, Dominic Kwiatkowski, Cordelia Langford, John Sillitoe on behalf of the Wellcome Sanger Institute COVID-19 Surveillance Team |
| EPI_ISL_2962693, EPI_ISL_2962694, EPI_ISL_2962695, EPI_ISL_2962696, EPI_ISL_2962697, EPI_ISL_2962698, EPI_ISL_2962699, EPI_ISL_2962700, EPI_ISL_2962701, EPI_ISL_2962702, EPI_ISL_2962703 | | | |
| see above | Lighthouse Lab in Milton Keynes | Wellcome Sanger Institute for the COVID-19 Genomics UK (COG-UK) Consortium | The Lighthouse Lab in Milton Keynes and Alex Alderton, Roberto Amato, Jeffrey Barrett, Sonia Goncalves, Ewan Harrison, David K. Jackson, Ian Johnston, Dominic Kwiatkowski, Cordelia Langford, John Sillitoe on behalf of the Wellcome Sanger Institute COVID-19 Surveillance Team |
| EPI_ISL_2962704, EPI_ISL_2962705, EPI_ISL_2962706 | Lighthouse Lab in Alderley Park | Wellcome Sanger Institute for the COVID-19 Genomics UK (COG-UK) Consortium | Jacquelyn Wynn, Mairead Hyland, The Lighthouse Lab in Alderley Park and Alex Alderton, Roberto Amato, Jeffrey Barrett, Sonia Goncalves, Ewan Harrison, David K. Jackson, Ian Johnston, Dominic Kwiatkowski, Cordelia Langford, John Sillitoe on behalf of the Wellcome Sanger Institute COVID-19 Surveillance Team |
| EPI_ISL_2962707, EPI_ISL_2962708 | Lighthouse Lab in Milton Keynes | Wellcome Sanger Institute for the COVID-19 Genomics UK (COG-UK) Consortium | The Lighthouse Lab in Milton Keynes and Alex Alderton, Roberto Amato, Jeffrey Barrett, Sonia Goncalves, Ewan Harrison, David K. Jackson, Ian Johnston, Dominic Kwiatkowski, Cordelia Langford, John Sillitoe on behalf of the Wellcome Sanger Institute COVID-19 Surveillance Team |
| EPI_ISL_2962709 | Lighthouse Lab in Alderley Park | Wellcome Sanger Institute for the COVID-19 Genomics UK (COG-UK) Consortium | Jacquelyn Wynn, Mairead Hyland, The Lighthouse Lab in Alderley Park and Alex Alderton, Roberto Amato, Jeffrey Barrett, Sonia Goncalves, Ewan Harrison, David K. Jackson, Ian Johnston, Dominic Kwiatkowski, Cordelia Langford, John Sillitoe on behalf of the Wellcome Sanger Institute COVID-19 Surveillance Team |
| EPI_ISL_2962710, EPI_ISL_2962711, EPI_ISL_2962712, EPI_ISL_2962713 | Lighthouse Lab in Milton Keynes | Wellcome Sanger Institute for the COVID-19 Genomics UK (COG-UK) Consortium | The Lighthouse Lab in Milton Keynes and Alex Alderton, Roberto Amato, Jeffrey Barrett, Sonia Goncalves, Ewan Harrison, David K. Jackson, Ian Johnston, Dominic Kwiatkowski, Cordelia Langford, John Sillitoe on behalf of the Wellcome Sanger Institute COVID-19 Surveillance Team |
| EPI_ISL_2962714, EPI_ISL_2962715, EPI_ISL_2962716, EPI_ISL_2962717, EPI_ISL_2962718 | Lighthouse Lab in Alderley Park | Wellcome Sanger Institute for the COVID-19 Genomics UK (COG-UK) Consortium | Jacquelyn Wynn, Mairead Hyland, The Lighthouse Lab in Alderley Park and Alex Alderton, Roberto Amato, Jeffrey Barrett, Sonia Goncalves, Ewan Harrison, David K. Jackson, Ian Johnston, Dominic Kwiatkowski, Cordelia Langford, John Sillitoe on behalf of the Wellcome Sanger Institute COVID-19 Surveillance Team |
| EPI_ISL_2962719, EPI_ISL_2962720, EPI_ISL_2962721, EPI_ISL_2962722, EPI_ISL_2962723, EPI_ISL_2962724, EPI_ISL_2962725 | Lighthouse Lab in Milton Keynes | Wellcome Sanger Institute for the COVID-19 Genomics UK (COG-UK) Consortium | The Lighthouse Lab in Milton Keynes and Alex Alderton, Roberto Amato, Jeffrey Barrett, Sonia Goncalves, Ewan Harrison, David K. Jackson, Ian Johnston, Dominic Kwiatkowski, Cordelia Langford, John Sillitoe on behalf of the Wellcome Sanger Institute COVID-19 Surveillance Team |
| EPI_ISL_2962726 | Lighthouse Lab in Alderley Park | Wellcome Sanger Institute for the COVID-19 Genomics UK (COG-UK) Consortium | Jacquelyn Wynn, Mairead Hyland, The Lighthouse Lab in Alderley Park and Alex Alderton, Roberto Amato, Jeffrey Barrett, Sonia Goncalves, Ewan Harrison, David K. Jackson, Ian Johnston, Dominic Kwiatkowski, Cordelia Langford, John Sillitoe on behalf of the Wellcome Sanger Institute COVID-19 Surveillance Team |
| EPI_ISL_2962727 | Lighthouse Lab in Milton Keynes | Wellcome Sanger Institute for the COVID-19 Genomics UK (COG-UK) Consortium | The Lighthouse Lab in Milton Keynes and Alex Alderton, Roberto Amato, Jeffrey Barrett, Sonia Goncalves, Ewan Harrison, David K. Jackson, Ian Johnston, Dominic Kwiatkowski, Cordelia Langford, John Sillitoe on behalf of the Wellcome Sanger Institute COVID-19 Surveillance Team |
| EPI_ISL_2962728 | Lighthouse Lab in Alderley Park | Wellcome Sanger Institute for the COVID-19 Genomics UK (COG-UK) Consortium | Jacquelyn Wynn, Mairead Hyland, The Lighthouse Lab in Alderley Park and Alex Alderton, Roberto Amato, Jeffrey Barrett, Sonia Goncalves, Ewan Harrison, David K. Jackson, Ian Johnston, Dominic Kwiatkowski, Cordelia Langford, John Sillitoe on behalf of the Wellcome Sanger Institute COVID-19 Surveillance Team |
| EPI_ISL_2962729, EPI_ISL_2962730, EPI_ISL_2962731, EPI_ISL_2962732, EPI_ISL_2962733 | Lighthouse Lab in Milton Keynes | Wellcome Sanger Institute for the COVID-19 Genomics UK (COG-UK) Consortium | The Lighthouse Lab in Milton Keynes and Alex Alderton, Roberto Amato, Jeffrey Barrett, Sonia Goncalves, Ewan Harrison, David K. Jackson, Ian Johnston, Dominic Kwiatkowski, Cordelia Langford, John Sillitoe on behalf of the Wellcome Sanger Institute COVID-19 Surveillance Team |
| EPI_ISL_2962734, EPI_ISL_2962735, | Lighthouse Lab in Alderley Park | Wellcome Sanger Institute for the COVID-19 Genomics UK (COG-UK) Consortium | Jacquelyn Wynn, Mairead Hyland, The Lighthouse Lab in Alderley Park and Alex Alderton, Roberto Amato, Jeffrey Barrett, Sonia Goncalves, Ewan Harrison, David K. Jackson, Ian Johnston, Dominic Kwiatkowski, Cordelia Langford, John Sillitoe on behalf of the Wellcome Sanger Institute COVID-19 Surveillance Team |

| | | | |
|---|---|---|---|
| EPI_ISL_2962736 | | | |
| EPI_ISL_2962737, EPI_ISL_2962738, EPI_ISL_2962739 | Lighthouse Lab in Milton Keynes | Wellcome Sanger Institute for the COVID-19 Genomics UK (COG-UK) Consortium | The Lighthouse Lab in Milton Keynes and Alex Alderton, Roberto Amato, Jeffrey Barrett, Sonia Goncalves, Ewan Harrison, David K. Jackson, Ian Johnston, Dominic Kwiatkowski, Cordelia Langford, John Sillitoe on behalf of the Wellcome Sanger Institute COVID-19 Surveillance Team |
| EPI_ISL_2962740, EPI_ISL_2962741, EPI_ISL_2962742, EPI_ISL_2962743, EPI_ISL_2962744, EPI_ISL_2962745, EPI_ISL_2962746 | Lighthouse Lab in Alderley Park | Wellcome Sanger Institute for the COVID-19 Genomics UK (COG-UK) Consortium | Jacquelyn Wynn, Mairead Hyland, The Lighthouse Lab in Alderley Park and Alex Alderton, Roberto Amato, Jeffrey Barrett, Sonia Goncalves, Ewan Harrison, David K. Jackson, Ian Johnston, Dominic Kwiatkowski, Cordelia Langford, John Sillitoe on behalf of the Wellcome Sanger Institute COVID-19 Surveillance Team |
| EPI_ISL_2962747 | Lighthouse Lab in Milton Keynes | Wellcome Sanger Institute for the COVID-19 Genomics UK (COG-UK) Consortium | The Lighthouse Lab in Milton Keynes and Alex Alderton, Roberto Amato, Jeffrey Barrett, Sonia Goncalves, Ewan Harrison, David K. Jackson, Ian Johnston, Dominic Kwiatkowski, Cordelia Langford, John Sillitoe on behalf of the Wellcome Sanger Institute COVID-19 Surveillance Team |
| EPI_ISL_2962748 | Lighthouse Lab in Alderley Park | Wellcome Sanger Institute for the COVID-19 Genomics UK (COG-UK) Consortium | Jacquelyn Wynn, Mairead Hyland, The Lighthouse Lab in Alderley Park and Alex Alderton, Roberto Amato, Jeffrey Barrett, Sonia Goncalves, Ewan Harrison, David K. Jackson, Ian Johnston, Dominic Kwiatkowski, Cordelia Langford, John Sillitoe on behalf of the Wellcome Sanger Institute COVID-19 Surveillance Team |
| EPI_ISL_2962749 | Lighthouse Lab in Milton Keynes | Wellcome Sanger Institute for the COVID-19 Genomics UK (COG-UK) Consortium | The Lighthouse Lab in Milton Keynes and Alex Alderton, Roberto Amato, Jeffrey Barrett, Sonia Goncalves, Ewan Harrison, David K. Jackson, Ian Johnston, Dominic Kwiatkowski, Cordelia Langford, John Sillitoe on behalf of the Wellcome Sanger Institute COVID-19 Surveillance Team |
| EPI_ISL_2962750, EPI_ISL_2962751, EPI_ISL_2962752, EPI_ISL_2962753 | Lighthouse Lab in Alderley Park | Wellcome Sanger Institute for the COVID-19 Genomics UK (COG-UK) Consortium | Jacquelyn Wynn, Mairead Hyland, The Lighthouse Lab in Alderley Park and Alex Alderton, Roberto Amato, Jeffrey Barrett, Sonia Goncalves, Ewan Harrison, David K. Jackson, Ian Johnston, Dominic Kwiatkowski, Cordelia Langford, John Sillitoe on behalf of the Wellcome Sanger Institute COVID-19 Surveillance Team |
| EPI_ISL_2962754 | Lighthouse Lab in Milton Keynes | Wellcome Sanger Institute for the COVID-19 Genomics UK (COG-UK) Consortium | The Lighthouse Lab in Milton Keynes and Alex Alderton, Roberto Amato, Jeffrey Barrett, Sonia Goncalves, Ewan Harrison, David K. Jackson, Ian Johnston, Dominic Kwiatkowski, Cordelia Langford, John Sillitoe on behalf of the Wellcome Sanger Institute COVID-19 Surveillance Team |
| EPI_ISL_2962755, EPI_ISL_2962756 | Lighthouse Lab in Alderley Park | Wellcome Sanger Institute for the COVID-19 Genomics UK (COG-UK) Consortium | Jacquelyn Wynn, Mairead Hyland, The Lighthouse Lab in Alderley Park and Alex Alderton, Roberto Amato, Jeffrey Barrett, Sonia Goncalves, Ewan Harrison, David K. Jackson, Ian Johnston, Dominic Kwiatkowski, Cordelia Langford, John Sillitoe on behalf of the Wellcome Sanger Institute COVID-19 Surveillance Team |
| EPI_ISL_2962757 | Lighthouse Lab in Milton Keynes | Wellcome Sanger Institute for the COVID-19 Genomics UK (COG-UK) Consortium | The Lighthouse Lab in Milton Keynes and Alex Alderton, Roberto Amato, Jeffrey Barrett, Sonia Goncalves, Ewan Harrison, David K. Jackson, Ian Johnston, Dominic Kwiatkowski, Cordelia Langford, John Sillitoe on behalf of the Wellcome Sanger Institute COVID-19 Surveillance Team |
| EPI_ISL_2962758, EPI_ISL_2962759, EPI_ISL_2962760, EPI_ISL_2962761, EPI_ISL_2962762, EPI_ISL_2962763 | Lighthouse Lab in Alderley Park | Wellcome Sanger Institute for the COVID-19 Genomics UK (COG-UK) Consortium | Jacquelyn Wynn, Mairead Hyland, The Lighthouse Lab in Alderley Park and Alex Alderton, Roberto Amato, Jeffrey Barrett, Sonia Goncalves, Ewan Harrison, David K. Jackson, Ian Johnston, Dominic Kwiatkowski, Cordelia Langford, John Sillitoe on behalf of the Wellcome Sanger Institute COVID-19 Surveillance Team |
| EPI_ISL_2962764 | Lighthouse Lab in Milton Keynes | Wellcome Sanger Institute for the COVID-19 Genomics UK (COG-UK) Consortium | The Lighthouse Lab in Milton Keynes and Alex Alderton, Roberto Amato, Jeffrey Barrett, Sonia Goncalves, Ewan Harrison, David K. Jackson, Ian Johnston, Dominic Kwiatkowski, Cordelia Langford, John Sillitoe on behalf of the Wellcome Sanger Institute COVID-19 Surveillance Team |
| EPI_ISL_2962765, EPI_ISL_2962766, EPI_ISL_2962767, EPI_ISL_2962768 | Lighthouse Lab in Alderley Park | Wellcome Sanger Institute for the COVID-19 Genomics UK (COG-UK) Consortium | Jacquelyn Wynn, Mairead Hyland, The Lighthouse Lab in Alderley Park and Alex Alderton, Roberto Amato, Jeffrey Barrett, Sonia Goncalves, Ewan Harrison, David K. Jackson, Ian Johnston, Dominic Kwiatkowski, Cordelia Langford, John Sillitoe on behalf of the Wellcome Sanger Institute COVID-19 Surveillance Team |
| EPI_ISL_2962769, EPI_ISL_2962770 | Lighthouse Lab in Milton Keynes | Wellcome Sanger Institute for the COVID-19 Genomics UK (COG-UK) Consortium | The Lighthouse Lab in Milton Keynes and Alex Alderton, Roberto Amato, Jeffrey Barrett, Sonia Goncalves, Ewan Harrison, David K. Jackson, Ian Johnston, Dominic Kwiatkowski, Cordelia Langford, John Sillitoe on behalf of the Wellcome Sanger Institute COVID-19 Surveillance Team |
| EPI_ISL_2962771, EPI_ISL_2962772, EPI_ISL_2962773, EPI_ISL_2962774, EPI_ISL_2962776, EPI_ISL_2962777 | Lighthouse Lab in Alderley Park | Wellcome Sanger Institute for the COVID-19 Genomics UK (COG-UK) Consortium | Jacquelyn Wynn, Mairead Hyland, The Lighthouse Lab in Alderley Park and Alex Alderton, Roberto Amato, Jeffrey Barrett, Sonia Goncalves, Ewan Harrison, David K. Jackson, Ian Johnston, Dominic Kwiatkowski, Cordelia Langford, John Sillitoe on behalf of the Wellcome Sanger Institute COVID-19 Surveillance Team |
| EPI_ISL_2962778, EPI_ISL_2962779 | Lighthouse Lab in Milton Keynes | Wellcome Sanger Institute for the COVID-19 Genomics UK (COG-UK) Consortium | The Lighthouse Lab in Milton Keynes and Alex Alderton, Roberto Amato, Jeffrey Barrett, Sonia Goncalves, Ewan Harrison, David K. Jackson, Ian Johnston, Dominic Kwiatkowski, Cordelia Langford, John Sillitoe on behalf of the Wellcome Sanger Institute COVID-19 Surveillance Team |
| EPI_ISL_2962780 | Lighthouse Lab in Alderley Park | Wellcome Sanger Institute for the COVID-19 Genomics UK (COG-UK) Consortium | Jacquelyn Wynn, Mairead Hyland, The Lighthouse Lab in Alderley Park and Alex Alderton, Roberto Amato, Jeffrey Barrett, Sonia Goncalves, Ewan Harrison, David K. Jackson, Ian Johnston, Dominic Kwiatkowski, Cordelia Langford, John Sillitoe on behalf of the Wellcome Sanger Institute COVID-19 Surveillance Team |
| EPI_ISL_2962781 | Lighthouse Lab in Milton Keynes | Wellcome Sanger Institute for the COVID-19 Genomics UK (COG-UK) Consortium | The Lighthouse Lab in Milton Keynes and Alex Alderton, Roberto Amato, Jeffrey Barrett, Sonia Goncalves, Ewan Harrison, David K. Jackson, Ian Johnston, Dominic Kwiatkowski, Cordelia Langford, John Sillitoe on behalf of the Wellcome Sanger Institute COVID-19 Surveillance Team |
| EPI_ISL_2962782 | Lighthouse Lab in Alderley Park | Wellcome Sanger Institute for the COVID-19 Genomics UK (COG-UK) Consortium | Jacquelyn Wynn, Mairead Hyland, The Lighthouse Lab in Alderley Park and Alex Alderton, Roberto Amato, Jeffrey Barrett, Sonia Goncalves, Ewan Harrison, David K. Jackson, Ian Johnston, Dominic Kwiatkowski, Cordelia Langford, John Sillitoe on behalf of the Wellcome Sanger Institute COVID-19 Surveillance Team |
| EPI_ISL_2962783 | Lighthouse Lab in Milton Keynes | Wellcome Sanger Institute for the COVID-19 Genomics UK (COG-UK) Consortium | The Lighthouse Lab in Milton Keynes and Alex Alderton, Roberto Amato, Jeffrey Barrett, Sonia Goncalves, Ewan Harrison, David K. Jackson, Ian Johnston, Dominic Kwiatkowski, Cordelia Langford, John Sillitoe on behalf of the Wellcome Sanger Institute COVID-19 Surveillance Team |
| EPI_ISL_2962784, EPI_ISL_2962785 | Lighthouse Lab in Alderley Park | Wellcome Sanger Institute for the COVID-19 Genomics UK (COG-UK) Consortium | Jacquelyn Wynn, Mairead Hyland, The Lighthouse Lab in Alderley Park and Alex Alderton, Roberto Amato, Jeffrey Barrett, Sonia Goncalves, Ewan Harrison, David K. Jackson, Ian Johnston, Dominic Kwiatkowski, Cordelia Langford, John Sillitoe on behalf of the Wellcome Sanger Institute COVID-19 Surveillance Team |
| EPI_ISL_2962786 | Lighthouse Lab in Milton Keynes | Wellcome Sanger Institute for the COVID-19 Genomics UK (COG-UK) Consortium | The Lighthouse Lab in Milton Keynes and Alex Alderton, Roberto Amato, Jeffrey Barrett, Sonia Goncalves, Ewan Harrison, David K. Jackson, Ian Johnston, Dominic Kwiatkowski, Cordelia Langford, John Sillitoe on behalf of the Wellcome Sanger Institute COVID-19 Surveillance Team |
| EPI_ISL_2962787, EPI_ISL_2962788, EPI_ISL_2962789, EPI_ISL_2962790 | Lighthouse Lab in Alderley Park | Wellcome Sanger Institute for the COVID-19 Genomics UK (COG-UK) Consortium | Jacquelyn Wynn, Mairead Hyland, The Lighthouse Lab in Alderley Park and Alex Alderton, Roberto Amato, Jeffrey Barrett, Sonia Goncalves, Ewan Harrison, David K. Jackson, Ian Johnston, Dominic Kwiatkowski, Cordelia Langford, John Sillitoe on behalf of the Wellcome Sanger Institute COVID-19 Surveillance Team |
| EPI_ISL_2962791 | Lighthouse Lab in Milton Keynes | Wellcome Sanger Institute for the COVID-19 Genomics UK (COG-UK) Consortium | The Lighthouse Lab in Milton Keynes and Alex Alderton, Roberto Amato, Jeffrey Barrett, Sonia Goncalves, Ewan Harrison, David K. Jackson, Ian Johnston, Dominic Kwiatkowski, Cordelia Langford, John Sillitoe on behalf of the Wellcome Sanger Institute COVID-19 Surveillance Team |
| EPI_ISL_2962792 | Lighthouse Lab in Alderley Park | Wellcome Sanger Institute for the COVID-19 Genomics UK (COG-UK) Consortium | Jacquelyn Wynn, Mairead Hyland, The Lighthouse Lab in Alderley Park and Alex Alderton, Roberto Amato, Jeffrey Barrett, Sonia Goncalves, Ewan Harrison, David K. Jackson, Ian Johnston, Dominic Kwiatkowski, Cordelia Langford, John Sillitoe on behalf of the Wellcome Sanger Institute COVID-19 Surveillance Team |
| EPI_ISL_2962793 | Lighthouse Lab in Milton Keynes | Wellcome Sanger Institute for the COVID-19 Genomics UK (COG-UK) Consortium | The Lighthouse Lab in Milton Keynes and Alex Alderton, Roberto Amato, Jeffrey Barrett, Sonia Goncalves, Ewan Harrison, David K. Jackson, Ian Johnston, Dominic Kwiatkowski, Cordelia Langford, John Sillitoe on behalf of the Wellcome Sanger Institute COVID-19 Surveillance Team |
| EPI_ISL_2962794, EPI_ISL_2962795, EPI_ISL_2962796, EPI_ISL_2962797, EPI_ISL_2962798, EPI_ISL_2962799, EPI_ISL_2962800, EPI_ISL_2962801, EPI_ISL_2962802, EPI_ISL_2962803, EPI_ISL_2962804, EPI_ISL_2962805, EPI_ISL_2962806 | | | |
| see above | Lighthouse Lab in Alderley Park | Wellcome Sanger Institute for the COVID-19 Genomics UK (COG-UK) Consortium | Jacquelyn Wynn, Mairead Hyland, The Lighthouse Lab in Alderley Park and Alex Alderton, Roberto Amato, Jeffrey Barrett, Sonia Goncalves, Ewan Harrison, David K. Jackson, Ian Johnston, Dominic Kwiatkowski, Cordelia Langford, John Sillitoe on behalf of the Wellcome Sanger Institute COVID-19 Surveillance Team |
| EPI_ISL_2962807, EPI_ISL_2962808 | Lighthouse Lab in Milton Keynes | Wellcome Sanger Institute for the COVID-19 Genomics UK (COG-UK) Consortium | The Lighthouse Lab in Milton Keynes and Alex Alderton, Roberto Amato, Jeffrey Barrett, Sonia Goncalves, Ewan Harrison, David K. Jackson, Ian Johnston, Dominic Kwiatkowski, Cordelia Langford, John Sillitoe on behalf of the Wellcome Sanger Institute COVID-19 Surveillance Team |
| EPI_ISL_2962809 | Lighthouse Lab in Alderley Park | Wellcome Sanger Institute for the COVID-19 Genomics UK (COG-UK) Consortium | Jacquelyn Wynn, Mairead Hyland, The Lighthouse Lab in Alderley Park and Alex Alderton, Roberto Amato, Jeffrey Barrett, Sonia Goncalves, Ewan Harrison, David K. Jackson, Ian Johnston, Dominic Kwiatkowski, Cordelia Langford, John Sillitoe on behalf of the Wellcome Sanger Institute COVID-19 Surveillance Team |
| EPI_ISL_2962810, EPI_ISL_2962811 | Lighthouse Lab in Milton Keynes | Wellcome Sanger Institute for the COVID-19 Genomics UK (COG-UK) Consortium | The Lighthouse Lab in Milton Keynes and Alex Alderton, Roberto Amato, Jeffrey Barrett, Sonia Goncalves, Ewan Harrison, David K. Jackson, Ian Johnston, Dominic Kwiatkowski, Cordelia Langford, John Sillitoe on behalf of the Wellcome Sanger Institute COVID-19 Surveillance Team |
| EPI_ISL_2962812, EPI_ISL_2962813 | Lighthouse Lab in Alderley Park | Wellcome Sanger Institute for the COVID-19 Genomics UK (COG-UK) Consortium | Jacquelyn Wynn, Mairead Hyland, The Lighthouse Lab in Alderley Park and Alex Alderton, Roberto Amato, Jeffrey Barrett, Sonia Goncalves, Ewan Harrison, David K. Jackson, Ian Johnston, Dominic Kwiatkowski, Cordelia Langford, John Sillitoe on behalf of the Wellcome Sanger Institute COVID-19 Surveillance Team |
| EPI_ISL_2962814 | Lighthouse Lab in Milton Keynes | Wellcome Sanger Institute for the COVID-19 Genomics UK (COG-UK) Consortium | The Lighthouse Lab in Milton Keynes and Alex Alderton, Roberto Amato, Jeffrey Barrett, Sonia Goncalves, Ewan Harrison, David K. Jackson, Ian Johnston, Dominic Kwiatkowski, Cordelia Langford, John Sillitoe on behalf of the Wellcome Sanger Institute COVID-19 Surveillance Team |
| EPI_ISL_2962815, EPI_ISL_2962816, EPI_ISL_2962817, EPI_ISL_2962818 | Lighthouse Lab in Alderley Park | Wellcome Sanger Institute for the COVID-19 Genomics UK (COG-UK) Consortium | Jacquelyn Wynn, Mairead Hyland, The Lighthouse Lab in Alderley Park and Alex Alderton, Roberto Amato, Jeffrey Barrett, Sonia Goncalves, Ewan Harrison, David K. Jackson, Ian Johnston, Dominic Kwiatkowski, Cordelia Langford, John Sillitoe on behalf of the Wellcome Sanger Institute COVID-19 Surveillance Team |
| EPI_ISL_2962819 | Lighthouse Lab in Milton Keynes | Wellcome Sanger Institute for the COVID-19 Genomics UK (COG-UK) Consortium | The Lighthouse Lab in Milton Keynes and Alex Alderton, Roberto Amato, Jeffrey Barrett, Sonia Goncalves, Ewan Harrison, David K. Jackson, Ian Johnston, Dominic Kwiatkowski, Cordelia Langford, John Sillitoe on behalf of the Wellcome Sanger Institute COVID-19 Surveillance Team |
| EPI_ISL_2962820, EPI_ISL_2962821, EPI_ISL_2962822, EPI_ISL_2962823, EPI_ISL_2962824, | Lighthouse Lab in Alderley Park | Wellcome Sanger Institute for the COVID-19 Genomics UK (COG-UK) Consortium | Jacquelyn Wynn, Mairead Hyland, The Lighthouse Lab in Alderley Park and Alex Alderton, Roberto Amato, Jeffrey Barrett, Sonia Goncalves, Ewan Harrison, David K. Jackson, Ian Johnston, Dominic Kwiatkowski, Cordelia Langford, John Sillitoe on behalf of the Wellcome Sanger Institute COVID-19 Surveillance Team |

| | | | |
|---|---|---|---|
| EPI_ISL_2962825, EPI_ISL_2962826, EPI_ISL_2962827, EPI_ISL_2962828 | | | |
| EPI_ISL_2962829 | Lighthouse Lab in Milton Keynes | Wellcome Sanger Institute for the COVID-19 Genomics UK (COG-UK) Consortium | The Lighthouse Lab in Milton Keynes and Alex Alderton, Roberto Amato, Jeffrey Barrett, Sonia Goncalves, Ewan Harrison, David K. Jackson, Ian Johnston, Dominic Kwiatkowski, Cordelia Langford, John Sillitoe on behalf of the Wellcome Sanger Institute COVID-19 Surveillance Team |
| EPI_ISL_2962830, EPI_ISL_2962831, EPI_ISL_2962832, EPI_ISL_2962833, EPI_ISL_2962834, EPI_ISL_2962835, EPI_ISL_2962836, EPI_ISL_2962837, EPI_ISL_2962838, EPI_ISL_2962839 | Lighthouse Lab in Alderley Park | Wellcome Sanger Institute for the COVID-19 Genomics UK (COG-UK) Consortium | Jacquelyn Wynn, Mairead Hyland, The Lighthouse Lab in Alderley Park and Alex Alderton, Roberto Amato, Jeffrey Barrett, Sonia Goncalves, Ewan Harrison, David K. Jackson, Ian Johnston, Dominic Kwiatkowski, Cordelia Langford, John Sillitoe on behalf of the Wellcome Sanger Institute COVID-19 Surveillance Team |
| EPI_ISL_2962840 | Lighthouse Lab in Milton Keynes | Wellcome Sanger Institute for the COVID-19 Genomics UK (COG-UK) Consortium | The Lighthouse Lab in Milton Keynes and Alex Alderton, Roberto Amato, Jeffrey Barrett, Sonia Goncalves, Ewan Harrison, David K. Jackson, Ian Johnston, Dominic Kwiatkowski, Cordelia Langford, John Sillitoe on behalf of the Wellcome Sanger Institute COVID-19 Surveillance Team |
| EPI_ISL_2962841, EPI_ISL_2962842 | Lighthouse Lab in Alderley Park | Wellcome Sanger Institute for the COVID-19 Genomics UK (COG-UK) Consortium | Jacquelyn Wynn, Mairead Hyland, The Lighthouse Lab in Alderley Park and Alex Alderton, Roberto Amato, Jeffrey Barrett, Sonia Goncalves, Ewan Harrison, David K. Jackson, Ian Johnston, Dominic Kwiatkowski, Cordelia Langford, John Sillitoe on behalf of the Wellcome Sanger Institute COVID-19 Surveillance Team |
| EPI_ISL_2962843, EPI_ISL_2962844 | Lighthouse Lab in Milton Keynes | Wellcome Sanger Institute for the COVID-19 Genomics UK (COG-UK) Consortium | The Lighthouse Lab in Milton Keynes and Alex Alderton, Roberto Amato, Jeffrey Barrett, Sonia Goncalves, Ewan Harrison, David K. Jackson, Ian Johnston, Dominic Kwiatkowski, Cordelia Langford, John Sillitoe on behalf of the Wellcome Sanger Institute COVID-19 Surveillance Team |
| EPI_ISL_2962845, EPI_ISL_2962846, EPI_ISL_2962847, EPI_ISL_2962848, EPI_ISL_2962849, EPI_ISL_2962850, EPI_ISL_2962851, EPI_ISL_2962852, EPI_ISL_2962853, EPI_ISL_2962854, EPI_ISL_2962855, EPI_ISL_2962856, EPI_ISL_2962857, EPI_ISL_2962858, EPI_ISL_2962859, EPI_ISL_2962861 | | | |
| see above | Lighthouse Lab in Alderley Park | Wellcome Sanger Institute for the COVID-19 Genomics UK (COG-UK) Consortium | Jacquelyn Wynn, Mairead Hyland, The Lighthouse Lab in Alderley Park and Alex Alderton, Roberto Amato, Jeffrey Barrett, Sonia Goncalves, Ewan Harrison, David K. Jackson, Ian Johnston, Dominic Kwiatkowski, Cordelia Langford, John Sillitoe on behalf of the Wellcome Sanger Institute COVID-19 Surveillance Team |
| EPI_ISL_2962862 | Lighthouse Lab in Milton Keynes | Wellcome Sanger Institute for the COVID-19 Genomics UK (COG-UK) Consortium | The Lighthouse Lab in Milton Keynes and Alex Alderton, Roberto Amato, Jeffrey Barrett, Sonia Goncalves, Ewan Harrison, David K. Jackson, Ian Johnston, Dominic Kwiatkowski, Cordelia Langford, John Sillitoe on behalf of the Wellcome Sanger Institute COVID-19 Surveillance Team |
| EPI_ISL_2962863, EPI_ISL_2962864 | Lighthouse Lab in Alderley Park | Wellcome Sanger Institute for the COVID-19 Genomics UK (COG-UK) Consortium | Jacquelyn Wynn, Mairead Hyland, The Lighthouse Lab in Alderley Park and Alex Alderton, Roberto Amato, Jeffrey Barrett, Sonia Goncalves, Ewan Harrison, David K. Jackson, Ian Johnston, Dominic Kwiatkowski, Cordelia Langford, John Sillitoe on behalf of the Wellcome Sanger Institute COVID-19 Surveillance Team |
| EPI_ISL_2962865 | Lighthouse Lab in Milton Keynes | Wellcome Sanger Institute for the COVID-19 Genomics UK (COG-UK) Consortium | The Lighthouse Lab in Milton Keynes and Alex Alderton, Roberto Amato, Jeffrey Barrett, Sonia Goncalves, Ewan Harrison, David K. Jackson, Ian Johnston, Dominic Kwiatkowski, Cordelia Langford, John Sillitoe on behalf of the Wellcome Sanger Institute COVID-19 Surveillance Team |
| EPI_ISL_2962866, EPI_ISL_2962867, EPI_ISL_2962868 | Lighthouse Lab in Alderley Park | Wellcome Sanger Institute for the COVID-19 Genomics UK (COG-UK) Consortium | Jacquelyn Wynn, Mairead Hyland, The Lighthouse Lab in Alderley Park and Alex Alderton, Roberto Amato, Jeffrey Barrett, Sonia Goncalves, Ewan Harrison, David K. Jackson, Ian Johnston, Dominic Kwiatkowski, Cordelia Langford, John Sillitoe on behalf of the Wellcome Sanger Institute COVID-19 Surveillance Team |
| EPI_ISL_2962869 | Lighthouse Lab in Milton Keynes | Wellcome Sanger Institute for the COVID-19 Genomics UK (COG-UK) Consortium | The Lighthouse Lab in Milton Keynes and Alex Alderton, Roberto Amato, Jeffrey Barrett, Sonia Goncalves, Ewan Harrison, David K. Jackson, Ian Johnston, Dominic Kwiatkowski, Cordelia Langford, John Sillitoe on behalf of the Wellcome Sanger Institute COVID-19 Surveillance Team |
| EPI_ISL_2962870 | Lighthouse Lab in Alderley Park | Wellcome Sanger Institute for the COVID-19 Genomics UK (COG-UK) Consortium | Jacquelyn Wynn, Mairead Hyland, The Lighthouse Lab in Alderley Park and Alex Alderton, Roberto Amato, Jeffrey Barrett, Sonia Goncalves, Ewan Harrison, David K. Jackson, Ian Johnston, Dominic Kwiatkowski, Cordelia Langford, John Sillitoe on behalf of the Wellcome Sanger Institute COVID-19 Surveillance Team |
| EPI_ISL_2962871, EPI_ISL_2962872, EPI_ISL_2962873 | Lighthouse Lab in Milton Keynes | Wellcome Sanger Institute for the COVID-19 Genomics UK (COG-UK) Consortium | The Lighthouse Lab in Milton Keynes and Alex Alderton, Roberto Amato, Jeffrey Barrett, Sonia Goncalves, Ewan Harrison, David K. Jackson, Ian Johnston, Dominic Kwiatkowski, Cordelia Langford, John Sillitoe on behalf of the Wellcome Sanger Institute COVID-19 Surveillance Team |
| EPI_ISL_2962874 | Lighthouse Lab in Alderley Park | Wellcome Sanger Institute for the COVID-19 Genomics UK (COG-UK) Consortium | Jacquelyn Wynn, Mairead Hyland, The Lighthouse Lab in Alderley Park and Alex Alderton, Roberto Amato, Jeffrey Barrett, Sonia Goncalves, Ewan Harrison, David K. Jackson, Ian Johnston, Dominic Kwiatkowski, Cordelia Langford, John Sillitoe on behalf of the Wellcome Sanger Institute COVID-19 Surveillance Team |
| EPI_ISL_2962875 | Lighthouse Lab in Milton Keynes | Wellcome Sanger Institute for the COVID-19 Genomics UK (COG-UK) Consortium | The Lighthouse Lab in Milton Keynes and Alex Alderton, Roberto Amato, Jeffrey Barrett, Sonia Goncalves, Ewan Harrison, David K. Jackson, Ian Johnston, Dominic Kwiatkowski, Cordelia Langford, John Sillitoe on behalf of the Wellcome Sanger Institute COVID-19 Surveillance Team |
| EPI_ISL_2962876 | Lighthouse Lab in Alderley Park | Wellcome Sanger Institute for the COVID-19 Genomics UK (COG-UK) Consortium | Jacquelyn Wynn, Mairead Hyland, The Lighthouse Lab in Alderley Park and Alex Alderton, Roberto Amato, Jeffrey Barrett, Sonia Goncalves, Ewan Harrison, David K. Jackson, Ian Johnston, Dominic Kwiatkowski, Cordelia Langford, John Sillitoe on behalf of the Wellcome Sanger Institute COVID-19 Surveillance Team |
| EPI_ISL_2962877 | Lighthouse Lab in Milton Keynes | Wellcome Sanger Institute for the COVID-19 Genomics UK (COG-UK) Consortium | The Lighthouse Lab in Milton Keynes and Alex Alderton, Roberto Amato, Jeffrey Barrett, Sonia Goncalves, Ewan Harrison, David K. Jackson, Ian Johnston, Dominic Kwiatkowski, Cordelia Langford, John Sillitoe on behalf of the Wellcome Sanger Institute COVID-19 Surveillance Team |
| EPI_ISL_2962878, EPI_ISL_2962879, EPI_ISL_2962880, EPI_ISL_2962881, EPI_ISL_2962882, EPI_ISL_2962883, EPI_ISL_2962884 | Lighthouse Lab in Alderley Park | Wellcome Sanger Institute for the COVID-19 Genomics UK (COG-UK) Consortium | Jacquelyn Wynn, Mairead Hyland, The Lighthouse Lab in Alderley Park and Alex Alderton, Roberto Amato, Jeffrey Barrett, Sonia Goncalves, Ewan Harrison, David K. Jackson, Ian Johnston, Dominic Kwiatkowski, Cordelia Langford, John Sillitoe on behalf of the Wellcome Sanger Institute COVID-19 Surveillance Team |
| EPI_ISL_2962885 | Lighthouse Lab in Milton Keynes | Wellcome Sanger Institute for the COVID-19 Genomics UK (COG-UK) Consortium | The Lighthouse Lab in Milton Keynes and Alex Alderton, Roberto Amato, Jeffrey Barrett, Sonia Goncalves, Ewan Harrison, David K. Jackson, Ian Johnston, Dominic Kwiatkowski, Cordelia Langford, John Sillitoe on behalf of the Wellcome Sanger Institute COVID-19 Surveillance Team |
| EPI_ISL_2962886 | Lighthouse Lab in Alderley Park | Wellcome Sanger Institute for the COVID-19 Genomics UK (COG-UK) Consortium | Jacquelyn Wynn, Mairead Hyland, The Lighthouse Lab in Alderley Park and Alex Alderton, Roberto Amato, Jeffrey Barrett, Sonia Goncalves, Ewan Harrison, David K. Jackson, Ian Johnston, Dominic Kwiatkowski, Cordelia Langford, John Sillitoe on behalf of the Wellcome Sanger Institute COVID-19 Surveillance Team |
| EPI_ISL_2962887 | Lighthouse Lab in Milton Keynes | Wellcome Sanger Institute for the COVID-19 Genomics UK (COG-UK) Consortium | The Lighthouse Lab in Milton Keynes and Alex Alderton, Roberto Amato, Jeffrey Barrett, Sonia Goncalves, Ewan Harrison, David K. Jackson, Ian Johnston, Dominic Kwiatkowski, Cordelia Langford, John Sillitoe on behalf of the Wellcome Sanger Institute COVID-19 Surveillance Team |
| EPI_ISL_2962888, EPI_ISL_2962889, EPI_ISL_2962890 | Lighthouse Lab in Alderley Park | Wellcome Sanger Institute for the COVID-19 Genomics UK (COG-UK) Consortium | Jacquelyn Wynn, Mairead Hyland, The Lighthouse Lab in Alderley Park and Alex Alderton, Roberto Amato, Jeffrey Barrett, Sonia Goncalves, Ewan Harrison, David K. Jackson, Ian Johnston, Dominic Kwiatkowski, Cordelia Langford, John Sillitoe on behalf of the Wellcome Sanger Institute COVID-19 Surveillance Team |
| EPI_ISL_2962891, EPI_ISL_2962892 | Lighthouse Lab in Milton Keynes | Wellcome Sanger Institute for the COVID-19 Genomics UK (COG-UK) Consortium | The Lighthouse Lab in Milton Keynes and Alex Alderton, Roberto Amato, Jeffrey Barrett, Sonia Goncalves, Ewan Harrison, David K. Jackson, Ian Johnston, Dominic Kwiatkowski, Cordelia Langford, John Sillitoe on behalf of the Wellcome Sanger Institute COVID-19 Surveillance Team |
| EPI_ISL_2962893, EPI_ISL_2962894 | Lighthouse Lab in Alderley Park | Wellcome Sanger Institute for the COVID-19 Genomics UK (COG-UK) Consortium | Jacquelyn Wynn, Mairead Hyland, The Lighthouse Lab in Alderley Park and Alex Alderton, Roberto Amato, Jeffrey Barrett, Sonia Goncalves, Ewan Harrison, David K. Jackson, Ian Johnston, Dominic Kwiatkowski, Cordelia Langford, John Sillitoe on behalf of the Wellcome Sanger Institute COVID-19 Surveillance Team |
| EPI_ISL_2962895 | Lighthouse Lab in Milton Keynes | Wellcome Sanger Institute for the COVID-19 Genomics UK (COG-UK) Consortium | The Lighthouse Lab in Milton Keynes and Alex Alderton, Roberto Amato, Jeffrey Barrett, Sonia Goncalves, Ewan Harrison, David K. Jackson, Ian Johnston, Dominic Kwiatkowski, Cordelia Langford, John Sillitoe on behalf of the Wellcome Sanger Institute COVID-19 Surveillance Team |
| EPI_ISL_2962896 | Lighthouse Lab in Alderley Park | Wellcome Sanger Institute for the COVID-19 Genomics UK (COG-UK) Consortium | Jacquelyn Wynn, Mairead Hyland, The Lighthouse Lab in Alderley Park and Alex Alderton, Roberto Amato, Jeffrey Barrett, Sonia Goncalves, Ewan Harrison, David K. Jackson, Ian Johnston, Dominic Kwiatkowski, Cordelia Langford, John Sillitoe on behalf of the Wellcome Sanger Institute COVID-19 Surveillance Team |
| EPI_ISL_2962897 | Lighthouse Lab in Milton Keynes | Wellcome Sanger Institute for the COVID-19 Genomics UK (COG-UK) Consortium | The Lighthouse Lab in Milton Keynes and Alex Alderton, Roberto Amato, Jeffrey Barrett, Sonia Goncalves, Ewan Harrison, David K. Jackson, Ian Johnston, Dominic Kwiatkowski, Cordelia Langford, John Sillitoe on behalf of the Wellcome Sanger Institute COVID-19 Surveillance Team |
| EPI_ISL_2962898, EPI_ISL_2962899, EPI_ISL_2962900, EPI_ISL_2962901, EPI_ISL_2962902 | Lighthouse Lab in Alderley Park | Wellcome Sanger Institute for the COVID-19 Genomics UK (COG-UK) Consortium | Jacquelyn Wynn, Mairead Hyland, The Lighthouse Lab in Alderley Park and Alex Alderton, Roberto Amato, Jeffrey Barrett, Sonia Goncalves, Ewan Harrison, David K. Jackson, Ian Johnston, Dominic Kwiatkowski, Cordelia Langford, John Sillitoe on behalf of the Wellcome Sanger Institute COVID-19 Surveillance Team |
| EPI_ISL_2962903 | Lighthouse Lab in Milton Keynes | Wellcome Sanger Institute for the COVID-19 Genomics UK (COG-UK) Consortium | The Lighthouse Lab in Milton Keynes and Alex Alderton, Roberto Amato, Jeffrey Barrett, Sonia Goncalves, Ewan Harrison, David K. Jackson, Ian Johnston, Dominic Kwiatkowski, Cordelia Langford, John Sillitoe on behalf of the Wellcome Sanger Institute COVID-19 Surveillance Team |
| EPI_ISL_2962904 | Lighthouse Lab in Alderley Park | Wellcome Sanger Institute for the COVID-19 Genomics UK (COG-UK) Consortium | Jacquelyn Wynn, Mairead Hyland, The Lighthouse Lab in Alderley Park and Alex Alderton, Roberto Amato, Jeffrey Barrett, Sonia Goncalves, Ewan Harrison, David K. Jackson, Ian Johnston, Dominic Kwiatkowski, Cordelia Langford, John Sillitoe on behalf of the Wellcome Sanger Institute COVID-19 Surveillance Team |
| EPI_ISL_2962905 | Lighthouse Lab in Milton Keynes | Wellcome Sanger Institute for the COVID-19 Genomics UK (COG-UK) Consortium | The Lighthouse Lab in Milton Keynes and Alex Alderton, Roberto Amato, Jeffrey Barrett, Sonia Goncalves, Ewan Harrison, David K. Jackson, Ian Johnston, Dominic Kwiatkowski, Cordelia Langford, John Sillitoe on behalf of the Wellcome Sanger Institute COVID-19 Surveillance Team |
| EPI_ISL_2962906, EPI_ISL_2962907 | Lighthouse Lab in Alderley Park | Wellcome Sanger Institute for the COVID-19 Genomics UK (COG-UK) Consortium | Jacquelyn Wynn, Mairead Hyland, The Lighthouse Lab in Alderley Park and Alex Alderton, Roberto Amato, Jeffrey Barrett, Sonia Goncalves, Ewan Harrison, David K. Jackson, Ian Johnston, Dominic Kwiatkowski, Cordelia Langford, John Sillitoe on behalf of the Wellcome Sanger Institute COVID-19 Surveillance Team |
| EPI_ISL_2962908, EPI_ISL_2962909 | Lighthouse Lab in Milton Keynes | Wellcome Sanger Institute for the COVID-19 Genomics UK (COG-UK) Consortium | The Lighthouse Lab in Milton Keynes and Alex Alderton, Roberto Amato, Jeffrey Barrett, Sonia Goncalves, Ewan Harrison, David K. Jackson, Ian Johnston, Dominic Kwiatkowski, Cordelia Langford, John Sillitoe on behalf of the Wellcome Sanger Institute COVID-19 Surveillance Team |
| EPI_ISL_2962910, EPI_ISL_2962911, EPI_ISL_2962912 | Lighthouse Lab in Alderley Park | Wellcome Sanger Institute for the COVID-19 Genomics UK (COG-UK) Consortium | Jacquelyn Wynn, Mairead Hyland, The Lighthouse Lab in Alderley Park and Alex Alderton, Roberto Amato, Jeffrey Barrett, Sonia Goncalves, Ewan Harrison, David K. Jackson, Ian Johnston, Dominic Kwiatkowski, Cordelia Langford, John Sillitoe on behalf of the Wellcome Sanger Institute COVID-19 Surveillance Team |
| EPI_ISL_2962913, | Lighthouse Lab in Milton Keynes | Wellcome Sanger Institute for the COVID-19 Genomics UK (COG-UK) | The Lighthouse Lab in Milton Keynes and Alex Alderton, Roberto Amato, Jeffrey Barrett, Sonia Goncalves, Ewan Harrison, David K. Jackson, Ian Johnston, Dominic Kwiatkowski, Cordelia Langford, John Sillitoe on behalf of the Wellcome Sanger |

| | | | |
|---|---|---|---|
| EPI_ISL_2962914 | | Consortium | Institute COVID-19 Surveillance Team |
| EPI_ISL_2962915 | Lighthouse Lab in Alderley Park | Wellcome Sanger Institute for the COVID-19 Genomics UK (COG-UK) Consortium | Jacquelyn Wynn, Mairead Hyland, The Lighthouse Lab in Alderley Park and Alex Alderton, Roberto Amato, Jeffrey Barrett, Sonia Goncalves, Ewan Harrison, David K. Jackson, Ian Johnston, Dominic Kwiatkowski, Cordelia Langford, John Sillitoe on behalf of the Wellcome Sanger Institute COVID-19 Surveillance Team |
| EPI_ISL_2962916, EPI_ISL_2962917 | Lighthouse Lab in Milton Keynes | Wellcome Sanger Institute for the COVID-19 Genomics UK (COG-UK) Consortium | The Lighthouse Lab in Milton Keynes and Alex Alderton, Roberto Amato, Jeffrey Barrett, Sonia Goncalves, Ewan Harrison, David K. Jackson, Ian Johnston, Dominic Kwiatkowski, Cordelia Langford, John Sillitoe on behalf of the Wellcome Sanger Institute COVID-19 Surveillance Team |
| EPI_ISL_2962918, EPI_ISL_2962919, EPI_ISL_2962920, EPI_ISL_2962921, EPI_ISL_2962922, EPI_ISL_2962923 | Lighthouse Lab in Alderley Park | Wellcome Sanger Institute for the COVID-19 Genomics UK (COG-UK) Consortium | Jacquelyn Wynn, Mairead Hyland, The Lighthouse Lab in Alderley Park and Alex Alderton, Roberto Amato, Jeffrey Barrett, Sonia Goncalves, Ewan Harrison, David K. Jackson, Ian Johnston, Dominic Kwiatkowski, Cordelia Langford, John Sillitoe on behalf of the Wellcome Sanger Institute COVID-19 Surveillance Team |
| EPI_ISL_2962924, EPI_ISL_2962925, EPI_ISL_2962926 | Lighthouse Lab in Milton Keynes | Wellcome Sanger Institute for the COVID-19 Genomics UK (COG-UK) Consortium | The Lighthouse Lab in Milton Keynes and Alex Alderton, Roberto Amato, Jeffrey Barrett, Sonia Goncalves, Ewan Harrison, David K. Jackson, Ian Johnston, Dominic Kwiatkowski, Cordelia Langford, John Sillitoe on behalf of the Wellcome Sanger Institute COVID-19 Surveillance Team |
| EPI_ISL_2962927, EPI_ISL_2962928 | Lighthouse Lab in Alderley Park | Wellcome Sanger Institute for the COVID-19 Genomics UK (COG-UK) Consortium | Jacquelyn Wynn, Mairead Hyland, The Lighthouse Lab in Alderley Park and Alex Alderton, Roberto Amato, Jeffrey Barrett, Sonia Goncalves, Ewan Harrison, David K. Jackson, Ian Johnston, Dominic Kwiatkowski, Cordelia Langford, John Sillitoe on behalf of the Wellcome Sanger Institute COVID-19 Surveillance Team |
| EPI_ISL_2962929 | Lighthouse Lab in Milton Keynes | Wellcome Sanger Institute for the COVID-19 Genomics UK (COG-UK) Consortium | The Lighthouse Lab in Milton Keynes and Alex Alderton, Roberto Amato, Jeffrey Barrett, Sonia Goncalves, Ewan Harrison, David K. Jackson, Ian Johnston, Dominic Kwiatkowski, Cordelia Langford, John Sillitoe on behalf of the Wellcome Sanger Institute COVID-19 Surveillance Team |
| EPI_ISL_2962930, EPI_ISL_2962931, EPI_ISL_2962932, EPI_ISL_2962933, EPI_ISL_2962934, EPI_ISL_2962935 | Lighthouse Lab in Alderley Park | Wellcome Sanger Institute for the COVID-19 Genomics UK (COG-UK) Consortium | Jacquelyn Wynn, Mairead Hyland, The Lighthouse Lab in Alderley Park and Alex Alderton, Roberto Amato, Jeffrey Barrett, Sonia Goncalves, Ewan Harrison, David K. Jackson, Ian Johnston, Dominic Kwiatkowski, Cordelia Langford, John Sillitoe on behalf of the Wellcome Sanger Institute COVID-19 Surveillance Team |
| EPI_ISL_2962936 | Lighthouse Lab in Milton Keynes | Wellcome Sanger Institute for the COVID-19 Genomics UK (COG-UK) Consortium | The Lighthouse Lab in Milton Keynes and Alex Alderton, Roberto Amato, Jeffrey Barrett, Sonia Goncalves, Ewan Harrison, David K. Jackson, Ian Johnston, Dominic Kwiatkowski, Cordelia Langford, John Sillitoe on behalf of the Wellcome Sanger Institute COVID-19 Surveillance Team |
| EPI_ISL_2962937 | Lighthouse Lab in Alderley Park | Wellcome Sanger Institute for the COVID-19 Genomics UK (COG-UK) Consortium | Jacquelyn Wynn, Mairead Hyland, The Lighthouse Lab in Alderley Park and Alex Alderton, Roberto Amato, Jeffrey Barrett, Sonia Goncalves, Ewan Harrison, David K. Jackson, Ian Johnston, Dominic Kwiatkowski, Cordelia Langford, John Sillitoe on behalf of the Wellcome Sanger Institute COVID-19 Surveillance Team |
| EPI_ISL_2962938 | Lighthouse Lab in Milton Keynes | Wellcome Sanger Institute for the COVID-19 Genomics UK (COG-UK) Consortium | The Lighthouse Lab in Milton Keynes and Alex Alderton, Roberto Amato, Jeffrey Barrett, Sonia Goncalves, Ewan Harrison, David K. Jackson, Ian Johnston, Dominic Kwiatkowski, Cordelia Langford, John Sillitoe on behalf of the Wellcome Sanger Institute COVID-19 Surveillance Team |
| EPI_ISL_2962939, EPI_ISL_2962940, EPI_ISL_2962941, EPI_ISL_2962942, EPI_ISL_2962943, EPI_ISL_2962944, EPI_ISL_2962945, EPI_ISL_2962946, EPI_ISL_2962947 | Lighthouse Lab in Alderley Park | Wellcome Sanger Institute for the COVID-19 Genomics UK (COG-UK) Consortium | Jacquelyn Wynn, Mairead Hyland, The Lighthouse Lab in Alderley Park and Alex Alderton, Roberto Amato, Jeffrey Barrett, Sonia Goncalves, Ewan Harrison, David K. Jackson, Ian Johnston, Dominic Kwiatkowski, Cordelia Langford, John Sillitoe on behalf of the Wellcome Sanger Institute COVID-19 Surveillance Team |
| EPI_ISL_2962948 | Lighthouse Lab in Milton Keynes | Wellcome Sanger Institute for the COVID-19 Genomics UK (COG-UK) Consortium | The Lighthouse Lab in Milton Keynes and Alex Alderton, Roberto Amato, Jeffrey Barrett, Sonia Goncalves, Ewan Harrison, David K. Jackson, Ian Johnston, Dominic Kwiatkowski, Cordelia Langford, John Sillitoe on behalf of the Wellcome Sanger Institute COVID-19 Surveillance Team |
| EPI_ISL_2962949 | Lighthouse Lab in Alderley Park | Wellcome Sanger Institute for the COVID-19 Genomics UK (COG-UK) Consortium | Jacquelyn Wynn, Mairead Hyland, The Lighthouse Lab in Alderley Park and Alex Alderton, Roberto Amato, Jeffrey Barrett, Sonia Goncalves, Ewan Harrison, David K. Jackson, Ian Johnston, Dominic Kwiatkowski, Cordelia Langford, John Sillitoe on behalf of the Wellcome Sanger Institute COVID-19 Surveillance Team |
| EPI_ISL_2962950, EPI_ISL_2962951 | Lighthouse Lab in Milton Keynes | Wellcome Sanger Institute for the COVID-19 Genomics UK (COG-UK) Consortium | The Lighthouse Lab in Milton Keynes and Alex Alderton, Roberto Amato, Jeffrey Barrett, Sonia Goncalves, Ewan Harrison, David K. Jackson, Ian Johnston, Dominic Kwiatkowski, Cordelia Langford, John Sillitoe on behalf of the Wellcome Sanger Institute COVID-19 Surveillance Team |
| EPI_ISL_2962952 | Lighthouse Lab in Alderley Park | Wellcome Sanger Institute for the COVID-19 Genomics UK (COG-UK) Consortium | Jacquelyn Wynn, Mairead Hyland, The Lighthouse Lab in Alderley Park and Alex Alderton, Roberto Amato, Jeffrey Barrett, Sonia Goncalves, Ewan Harrison, David K. Jackson, Ian Johnston, Dominic Kwiatkowski, Cordelia Langford, John Sillitoe on behalf of the Wellcome Sanger Institute COVID-19 Surveillance Team |
| EPI_ISL_2962953, EPI_ISL_2962954 | Lighthouse Lab in Milton Keynes | Wellcome Sanger Institute for the COVID-19 Genomics UK (COG-UK) Consortium | The Lighthouse Lab in Milton Keynes and Alex Alderton, Roberto Amato, Jeffrey Barrett, Sonia Goncalves, Ewan Harrison, David K. Jackson, Ian Johnston, Dominic Kwiatkowski, Cordelia Langford, John Sillitoe on behalf of the Wellcome Sanger Institute COVID-19 Surveillance Team |
| EPI_ISL_2962955, EPI_ISL_2962956, EPI_ISL_2962957, EPI_ISL_2962958 | Lighthouse Lab in Alderley Park | Wellcome Sanger Institute for the COVID-19 Genomics UK (COG-UK) Consortium | Jacquelyn Wynn, Mairead Hyland, The Lighthouse Lab in Alderley Park and Alex Alderton, Roberto Amato, Jeffrey Barrett, Sonia Goncalves, Ewan Harrison, David K. Jackson, Ian Johnston, Dominic Kwiatkowski, Cordelia Langford, John Sillitoe on behalf of the Wellcome Sanger Institute COVID-19 Surveillance Team |
| EPI_ISL_2962959 | Lighthouse Lab in Milton Keynes | Wellcome Sanger Institute for the COVID-19 Genomics UK (COG-UK) Consortium | The Lighthouse Lab in Milton Keynes and Alex Alderton, Roberto Amato, Jeffrey Barrett, Sonia Goncalves, Ewan Harrison, David K. Jackson, Ian Johnston, Dominic Kwiatkowski, Cordelia Langford, John Sillitoe on behalf of the Wellcome Sanger Institute COVID-19 Surveillance Team |
| EPI_ISL_2962960, EPI_ISL_2962961, EPI_ISL_2962962, EPI_ISL_2962963, EPI_ISL_2962964, EPI_ISL_2962965, EPI_ISL_2962966 | Lighthouse Lab in Alderley Park | Wellcome Sanger Institute for the COVID-19 Genomics UK (COG-UK) Consortium | Jacquelyn Wynn, Mairead Hyland, The Lighthouse Lab in Alderley Park and Alex Alderton, Roberto Amato, Jeffrey Barrett, Sonia Goncalves, Ewan Harrison, David K. Jackson, Ian Johnston, Dominic Kwiatkowski, Cordelia Langford, John Sillitoe on behalf of the Wellcome Sanger Institute COVID-19 Surveillance Team |
| EPI_ISL_2962967, EPI_ISL_2962968 | Lighthouse Lab in Milton Keynes | Wellcome Sanger Institute for the COVID-19 Genomics UK (COG-UK) Consortium | The Lighthouse Lab in Milton Keynes and Alex Alderton, Roberto Amato, Jeffrey Barrett, Sonia Goncalves, Ewan Harrison, David K. Jackson, Ian Johnston, Dominic Kwiatkowski, Cordelia Langford, John Sillitoe on behalf of the Wellcome Sanger Institute COVID-19 Surveillance Team |
| EPI_ISL_2962969, EPI_ISL_2962970 | Lighthouse Lab in Alderley Park | Wellcome Sanger Institute for the COVID-19 Genomics UK (COG-UK) Consortium | Jacquelyn Wynn, Mairead Hyland, The Lighthouse Lab in Alderley Park and Alex Alderton, Roberto Amato, Jeffrey Barrett, Sonia Goncalves, Ewan Harrison, David K. Jackson, Ian Johnston, Dominic Kwiatkowski, Cordelia Langford, John Sillitoe on behalf of the Wellcome Sanger Institute COVID-19 Surveillance Team |
| EPI_ISL_2962971 | Lighthouse Lab in Milton Keynes | Wellcome Sanger Institute for the COVID-19 Genomics UK (COG-UK) Consortium | The Lighthouse Lab in Milton Keynes and Alex Alderton, Roberto Amato, Jeffrey Barrett, Sonia Goncalves, Ewan Harrison, David K. Jackson, Ian Johnston, Dominic Kwiatkowski, Cordelia Langford, John Sillitoe on behalf of the Wellcome Sanger Institute COVID-19 Surveillance Team |
| EPI_ISL_2962972, EPI_ISL_2962973, EPI_ISL_2962974, EPI_ISL_2962975, EPI_ISL_2962976, EPI_ISL_2962977, EPI_ISL_2962978, EPI_ISL_2962979, EPI_ISL_2962980, EPI_ISL_2962981, EPI_ISL_2962982, EPI_ISL_2962983, EPI_ISL_2962984, EPI_ISL_2962985, EPI_ISL_2962986, EPI_ISL_2962987, EPI_ISL_2962988 | | | |
| see above | Lighthouse Lab in Alderley Park | Wellcome Sanger Institute for the COVID-19 Genomics UK (COG-UK) Consortium | Jacquelyn Wynn, Mairead Hyland, The Lighthouse Lab in Alderley Park and Alex Alderton, Roberto Amato, Jeffrey Barrett, Sonia Goncalves, Ewan Harrison, David K. Jackson, Ian Johnston, Dominic Kwiatkowski, Cordelia Langford, John Sillitoe on behalf of the Wellcome Sanger Institute COVID-19 Surveillance Team |
| EPI_ISL_2962989 | Lighthouse Lab in Milton Keynes | Wellcome Sanger Institute for the COVID-19 Genomics UK (COG-UK) Consortium | The Lighthouse Lab in Milton Keynes and Alex Alderton, Roberto Amato, Jeffrey Barrett, Sonia Goncalves, Ewan Harrison, David K. Jackson, Ian Johnston, Dominic Kwiatkowski, Cordelia Langford, John Sillitoe on behalf of the Wellcome Sanger Institute COVID-19 Surveillance Team |
| EPI_ISL_2962990, EPI_ISL_2962991, EPI_ISL_2962992, EPI_ISL_2962993, EPI_ISL_2962994, EPI_ISL_2962995 | Lighthouse Lab in Alderley Park | Wellcome Sanger Institute for the COVID-19 Genomics UK (COG-UK) Consortium | Jacquelyn Wynn, Mairead Hyland, The Lighthouse Lab in Alderley Park and Alex Alderton, Roberto Amato, Jeffrey Barrett, Sonia Goncalves, Ewan Harrison, David K. Jackson, Ian Johnston, Dominic Kwiatkowski, Cordelia Langford, John Sillitoe on behalf of the Wellcome Sanger Institute COVID-19 Surveillance Team |
| EPI_ISL_2962996 | Lighthouse Lab in Milton Keynes | Wellcome Sanger Institute for the COVID-19 Genomics UK (COG-UK) Consortium | The Lighthouse Lab in Milton Keynes and Alex Alderton, Roberto Amato, Jeffrey Barrett, Sonia Goncalves, Ewan Harrison, David K. Jackson, Ian Johnston, Dominic Kwiatkowski, Cordelia Langford, John Sillitoe on behalf of the Wellcome Sanger Institute COVID-19 Surveillance Team |
| EPI_ISL_2962997 | Lighthouse Lab in Alderley Park | Wellcome Sanger Institute for the COVID-19 Genomics UK (COG-UK) Consortium | Jacquelyn Wynn, Mairead Hyland, The Lighthouse Lab in Alderley Park and Alex Alderton, Roberto Amato, Jeffrey Barrett, Sonia Goncalves, Ewan Harrison, David K. Jackson, Ian Johnston, Dominic Kwiatkowski, Cordelia Langford, John Sillitoe on behalf of the Wellcome Sanger Institute COVID-19 Surveillance Team |
| EPI_ISL_2962998 | Lighthouse Lab in Milton Keynes | Wellcome Sanger Institute for the COVID-19 Genomics UK (COG-UK) Consortium | The Lighthouse Lab in Milton Keynes and Alex Alderton, Roberto Amato, Jeffrey Barrett, Sonia Goncalves, Ewan Harrison, David K. Jackson, Ian Johnston, Dominic Kwiatkowski, Cordelia Langford, John Sillitoe on behalf of the Wellcome Sanger Institute COVID-19 Surveillance Team |
| EPI_ISL_2962999, EPI_ISL_2963000, EPI_ISL_2963001 | Lighthouse Lab in Alderley Park | Wellcome Sanger Institute for the COVID-19 Genomics UK (COG-UK) Consortium | Jacquelyn Wynn, Mairead Hyland, The Lighthouse Lab in Alderley Park and Alex Alderton, Roberto Amato, Jeffrey Barrett, Sonia Goncalves, Ewan Harrison, David K. Jackson, Ian Johnston, Dominic Kwiatkowski, Cordelia Langford, John Sillitoe on behalf of the Wellcome Sanger Institute COVID-19 Surveillance Team |
| EPI_ISL_2963002 | Lighthouse Lab in Milton Keynes | Wellcome Sanger Institute for the COVID-19 Genomics UK (COG-UK) Consortium | The Lighthouse Lab in Milton Keynes and Alex Alderton, Roberto Amato, Jeffrey Barrett, Sonia Goncalves, Ewan Harrison, David K. Jackson, Ian Johnston, Dominic Kwiatkowski, Cordelia Langford, John Sillitoe on behalf of the Wellcome Sanger Institute COVID-19 Surveillance Team |
| EPI_ISL_2963003, EPI_ISL_2963004, EPI_ISL_2963005, EPI_ISL_2963006, EPI_ISL_2963007, EPI_ISL_2963008, | Lighthouse Lab in Alderley Park | Wellcome Sanger Institute for the COVID-19 Genomics UK (COG-UK) Consortium | Jacquelyn Wynn, Mairead Hyland, The Lighthouse Lab in Alderley Park and Alex Alderton, Roberto Amato, Jeffrey Barrett, Sonia Goncalves, Ewan Harrison, David K. Jackson, Ian Johnston, Dominic Kwiatkowski, Cordelia Langford, John Sillitoe on behalf of the Wellcome Sanger Institute COVID-19 Surveillance Team |

| | | | |
|---|---|---|---|
| EPI_ISL_2963009, EPI_ISL_2963010, EPI_ISL_2963011 | | | |
| EPI_ISL_2963012 | Lighthouse Lab in Milton Keynes | Wellcome Sanger Institute for the COVID-19 Genomics UK (COG-UK) Consortium | The Lighthouse Lab in Milton Keynes and Alex Alderton, Roberto Amato, Jeffrey Barrett, Sonia Goncalves, Ewan Harrison, David K. Jackson, Ian Johnston, Dominic Kwiatkowski, Cordelia Langford, John Sillitoe on behalf of the Wellcome Sanger Institute COVID-19 Surveillance Team |
| EPI_ISL_2963013, EPI_ISL_2963014 | Lighthouse Lab in Alderley Park | Wellcome Sanger Institute for the COVID-19 Genomics UK (COG-UK) Consortium | Jacquelyn Wynn, Mairead Hyland, The Lighthouse Lab in Alderley Park and Alex Alderton, Roberto Amato, Jeffrey Barrett, Sonia Goncalves, Ewan Harrison, David K. Jackson, Ian Johnston, Dominic Kwiatkowski, Cordelia Langford, John Sillitoe on behalf of the Wellcome Sanger Institute COVID-19 Surveillance Team |
| EPI_ISL_2963015, EPI_ISL_2963016 | Lighthouse Lab in Milton Keynes | Wellcome Sanger Institute for the COVID-19 Genomics UK (COG-UK) Consortium | The Lighthouse Lab in Milton Keynes and Alex Alderton, Roberto Amato, Jeffrey Barrett, Sonia Goncalves, Ewan Harrison, David K. Jackson, Ian Johnston, Dominic Kwiatkowski, Cordelia Langford, John Sillitoe on behalf of the Wellcome Sanger Institute COVID-19 Surveillance Team |
| EPI_ISL_2963017, EPI_ISL_2963018, EPI_ISL_2963019, EPI_ISL_2963020 | Lighthouse Lab in Alderley Park | Wellcome Sanger Institute for the COVID-19 Genomics UK (COG-UK) Consortium | Jacquelyn Wynn, Mairead Hyland, The Lighthouse Lab in Alderley Park and Alex Alderton, Roberto Amato, Jeffrey Barrett, Sonia Goncalves, Ewan Harrison, David K. Jackson, Ian Johnston, Dominic Kwiatkowski, Cordelia Langford, John Sillitoe on behalf of the Wellcome Sanger Institute COVID-19 Surveillance Team |
| EPI_ISL_2963021 | Lighthouse Lab in Milton Keynes | Wellcome Sanger Institute for the COVID-19 Genomics UK (COG-UK) Consortium | The Lighthouse Lab in Milton Keynes and Alex Alderton, Roberto Amato, Jeffrey Barrett, Sonia Goncalves, Ewan Harrison, David K. Jackson, Ian Johnston, Dominic Kwiatkowski, Cordelia Langford, John Sillitoe on behalf of the Wellcome Sanger Institute COVID-19 Surveillance Team |
| EPI_ISL_2963022 | Lighthouse Lab in Alderley Park | Wellcome Sanger Institute for the COVID-19 Genomics UK (COG-UK) Consortium | Jacquelyn Wynn, Mairead Hyland, The Lighthouse Lab in Alderley Park and Alex Alderton, Roberto Amato, Jeffrey Barrett, Sonia Goncalves, Ewan Harrison, David K. Jackson, Ian Johnston, Dominic Kwiatkowski, Cordelia Langford, John Sillitoe on behalf of the Wellcome Sanger Institute COVID-19 Surveillance Team |
| EPI_ISL_2963023 | Lighthouse Lab in Milton Keynes | Wellcome Sanger Institute for the COVID-19 Genomics UK (COG-UK) Consortium | The Lighthouse Lab in Milton Keynes and Alex Alderton, Roberto Amato, Jeffrey Barrett, Sonia Goncalves, Ewan Harrison, David K. Jackson, Ian Johnston, Dominic Kwiatkowski, Cordelia Langford, John Sillitoe on behalf of the Wellcome Sanger Institute COVID-19 Surveillance Team |
| EPI_ISL_2963024, EPI_ISL_2963025, EPI_ISL_2963026, EPI_ISL_2963027, EPI_ISL_2963028, EPI_ISL_2963029, EPI_ISL_2963030 | Lighthouse Lab in Alderley Park | Wellcome Sanger Institute for the COVID-19 Genomics UK (COG-UK) Consortium | Jacquelyn Wynn, Mairead Hyland, The Lighthouse Lab in Alderley Park and Alex Alderton, Roberto Amato, Jeffrey Barrett, Sonia Goncalves, Ewan Harrison, David K. Jackson, Ian Johnston, Dominic Kwiatkowski, Cordelia Langford, John Sillitoe on behalf of the Wellcome Sanger Institute COVID-19 Surveillance Team |
| EPI_ISL_2963031 | Lighthouse Lab in Milton Keynes | Wellcome Sanger Institute for the COVID-19 Genomics UK (COG-UK) Consortium | The Lighthouse Lab in Milton Keynes and Alex Alderton, Roberto Amato, Jeffrey Barrett, Sonia Goncalves, Ewan Harrison, David K. Jackson, Ian Johnston, Dominic Kwiatkowski, Cordelia Langford, John Sillitoe on behalf of the Wellcome Sanger Institute COVID-19 Surveillance Team |
| EPI_ISL_2963032, EPI_ISL_2963033, EPI_ISL_2963034, EPI_ISL_2963035, EPI_ISL_2963036, EPI_ISL_2963037 | Lighthouse Lab in Alderley Park | Wellcome Sanger Institute for the COVID-19 Genomics UK (COG-UK) Consortium | Jacquelyn Wynn, Mairead Hyland, The Lighthouse Lab in Alderley Park and Alex Alderton, Roberto Amato, Jeffrey Barrett, Sonia Goncalves, Ewan Harrison, David K. Jackson, Ian Johnston, Dominic Kwiatkowski, Cordelia Langford, John Sillitoe on behalf of the Wellcome Sanger Institute COVID-19 Surveillance Team |
| EPI_ISL_2963038 | Lighthouse Lab in Milton Keynes | Wellcome Sanger Institute for the COVID-19 Genomics UK (COG-UK) Consortium | The Lighthouse Lab in Milton Keynes and Alex Alderton, Roberto Amato, Jeffrey Barrett, Sonia Goncalves, Ewan Harrison, David K. Jackson, Ian Johnston, Dominic Kwiatkowski, Cordelia Langford, John Sillitoe on behalf of the Wellcome Sanger Institute COVID-19 Surveillance Team |
| EPI_ISL_2963039 | Lighthouse Lab in Alderley Park | Wellcome Sanger Institute for the COVID-19 Genomics UK (COG-UK) Consortium | Jacquelyn Wynn, Mairead Hyland, The Lighthouse Lab in Alderley Park and Alex Alderton, Roberto Amato, Jeffrey Barrett, Sonia Goncalves, Ewan Harrison, David K. Jackson, Ian Johnston, Dominic Kwiatkowski, Cordelia Langford, John Sillitoe on behalf of the Wellcome Sanger Institute COVID-19 Surveillance Team |
| EPI_ISL_2963040 | Lighthouse Lab in Milton Keynes | Wellcome Sanger Institute for the COVID-19 Genomics UK (COG-UK) Consortium | The Lighthouse Lab in Milton Keynes and Alex Alderton, Roberto Amato, Jeffrey Barrett, Sonia Goncalves, Ewan Harrison, David K. Jackson, Ian Johnston, Dominic Kwiatkowski, Cordelia Langford, John Sillitoe on behalf of the Wellcome Sanger Institute COVID-19 Surveillance Team |
| EPI_ISL_2963041, EPI_ISL_2963042, EPI_ISL_2963043, EPI_ISL_2963044, EPI_ISL_2963045, EPI_ISL_2963046, EPI_ISL_2963047, EPI_ISL_2963048 | Lighthouse Lab in Alderley Park | Wellcome Sanger Institute for the COVID-19 Genomics UK (COG-UK) Consortium | Jacquelyn Wynn, Mairead Hyland, The Lighthouse Lab in Alderley Park and Alex Alderton, Roberto Amato, Jeffrey Barrett, Sonia Goncalves, Ewan Harrison, David K. Jackson, Ian Johnston, Dominic Kwiatkowski, Cordelia Langford, John Sillitoe on behalf of the Wellcome Sanger Institute COVID-19 Surveillance Team |
| EPI_ISL_2963049, EPI_ISL_2963050 | Lighthouse Lab in Milton Keynes | Wellcome Sanger Institute for the COVID-19 Genomics UK (COG-UK) Consortium | The Lighthouse Lab in Milton Keynes and Alex Alderton, Roberto Amato, Jeffrey Barrett, Sonia Goncalves, Ewan Harrison, David K. Jackson, Ian Johnston, Dominic Kwiatkowski, Cordelia Langford, John Sillitoe on behalf of the Wellcome Sanger Institute COVID-19 Surveillance Team |
| EPI_ISL_2963051, EPI_ISL_2963052 | Lighthouse Lab in Alderley Park | Wellcome Sanger Institute for the COVID-19 Genomics UK (COG-UK) Consortium | Jacquelyn Wynn, Mairead Hyland, The Lighthouse Lab in Alderley Park and Alex Alderton, Roberto Amato, Jeffrey Barrett, Sonia Goncalves, Ewan Harrison, David K. Jackson, Ian Johnston, Dominic Kwiatkowski, Cordelia Langford, John Sillitoe on behalf of the Wellcome Sanger Institute COVID-19 Surveillance Team |
| EPI_ISL_2963053 | Lighthouse Lab in Milton Keynes | Wellcome Sanger Institute for the COVID-19 Genomics UK (COG-UK) Consortium | The Lighthouse Lab in Milton Keynes and Alex Alderton, Roberto Amato, Jeffrey Barrett, Sonia Goncalves, Ewan Harrison, David K. Jackson, Ian Johnston, Dominic Kwiatkowski, Cordelia Langford, John Sillitoe on behalf of the Wellcome Sanger Institute COVID-19 Surveillance Team |
| EPI_ISL_2963054, EPI_ISL_2963055, EPI_ISL_2963056, EPI_ISL_2963057, EPI_ISL_2963058, EPI_ISL_2963059 | Lighthouse Lab in Alderley Park | Wellcome Sanger Institute for the COVID-19 Genomics UK (COG-UK) Consortium | Jacquelyn Wynn, Mairead Hyland, The Lighthouse Lab in Alderley Park and Alex Alderton, Roberto Amato, Jeffrey Barrett, Sonia Goncalves, Ewan Harrison, David K. Jackson, Ian Johnston, Dominic Kwiatkowski, Cordelia Langford, John Sillitoe on behalf of the Wellcome Sanger Institute COVID-19 Surveillance Team |
| EPI_ISL_2963060 | Lighthouse Lab in Milton Keynes | Wellcome Sanger Institute for the COVID-19 Genomics UK (COG-UK) Consortium | The Lighthouse Lab in Milton Keynes and Alex Alderton, Roberto Amato, Jeffrey Barrett, Sonia Goncalves, Ewan Harrison, David K. Jackson, Ian Johnston, Dominic Kwiatkowski, Cordelia Langford, John Sillitoe on behalf of the Wellcome Sanger Institute COVID-19 Surveillance Team |
| EPI_ISL_2963061, EPI_ISL_2963062, EPI_ISL_2963063 | Lighthouse Lab in Alderley Park | Wellcome Sanger Institute for the COVID-19 Genomics UK (COG-UK) Consortium | Jacquelyn Wynn, Mairead Hyland, The Lighthouse Lab in Alderley Park and Alex Alderton, Roberto Amato, Jeffrey Barrett, Sonia Goncalves, Ewan Harrison, David K. Jackson, Ian Johnston, Dominic Kwiatkowski, Cordelia Langford, John Sillitoe on behalf of the Wellcome Sanger Institute COVID-19 Surveillance Team |
| EPI_ISL_2963064 | Lighthouse Lab in Milton Keynes | Wellcome Sanger Institute for the COVID-19 Genomics UK (COG-UK) Consortium | The Lighthouse Lab in Milton Keynes and Alex Alderton, Roberto Amato, Jeffrey Barrett, Sonia Goncalves, Ewan Harrison, David K. Jackson, Ian Johnston, Dominic Kwiatkowski, Cordelia Langford, John Sillitoe on behalf of the Wellcome Sanger Institute COVID-19 Surveillance Team |
| EPI_ISL_2963065 | Lighthouse Lab in Alderley Park | Wellcome Sanger Institute for the COVID-19 Genomics UK (COG-UK) Consortium | Jacquelyn Wynn, Mairead Hyland, The Lighthouse Lab in Alderley Park and Alex Alderton, Roberto Amato, Jeffrey Barrett, Sonia Goncalves, Ewan Harrison, David K. Jackson, Ian Johnston, Dominic Kwiatkowski, Cordelia Langford, John Sillitoe on behalf of the Wellcome Sanger Institute COVID-19 Surveillance Team |
| EPI_ISL_2963066 | Lighthouse Lab in Milton Keynes | Wellcome Sanger Institute for the COVID-19 Genomics UK (COG-UK) Consortium | The Lighthouse Lab in Milton Keynes and Alex Alderton, Roberto Amato, Jeffrey Barrett, Sonia Goncalves, Ewan Harrison, David K. Jackson, Ian Johnston, Dominic Kwiatkowski, Cordelia Langford, John Sillitoe on behalf of the Wellcome Sanger Institute COVID-19 Surveillance Team |
| EPI_ISL_2963067, EPI_ISL_2963068, EPI_ISL_2963069, EPI_ISL_2963070, EPI_ISL_2963071, EPI_ISL_2963072 | Lighthouse Lab in Alderley Park | Wellcome Sanger Institute for the COVID-19 Genomics UK (COG-UK) Consortium | Jacquelyn Wynn, Mairead Hyland, The Lighthouse Lab in Alderley Park and Alex Alderton, Roberto Amato, Jeffrey Barrett, Sonia Goncalves, Ewan Harrison, David K. Jackson, Ian Johnston, Dominic Kwiatkowski, Cordelia Langford, John Sillitoe on behalf of the Wellcome Sanger Institute COVID-19 Surveillance Team |
| EPI_ISL_2963073, EPI_ISL_2963074 | Lighthouse Lab in Milton Keynes | Wellcome Sanger Institute for the COVID-19 Genomics UK (COG-UK) Consortium | The Lighthouse Lab in Milton Keynes and Alex Alderton, Roberto Amato, Jeffrey Barrett, Sonia Goncalves, Ewan Harrison, David K. Jackson, Ian Johnston, Dominic Kwiatkowski, Cordelia Langford, John Sillitoe on behalf of the Wellcome Sanger Institute COVID-19 Surveillance Team |
| EPI_ISL_2963075, EPI_ISL_2963076 | Lighthouse Lab in Alderley Park | Wellcome Sanger Institute for the COVID-19 Genomics UK (COG-UK) Consortium | Jacquelyn Wynn, Mairead Hyland, The Lighthouse Lab in Alderley Park and Alex Alderton, Roberto Amato, Jeffrey Barrett, Sonia Goncalves, Ewan Harrison, David K. Jackson, Ian Johnston, Dominic Kwiatkowski, Cordelia Langford, John Sillitoe on behalf of the Wellcome Sanger Institute COVID-19 Surveillance Team |
| EPI_ISL_2963077 | Lighthouse Lab in Milton Keynes | Wellcome Sanger Institute for the COVID-19 Genomics UK (COG-UK) Consortium | The Lighthouse Lab in Milton Keynes and Alex Alderton, Roberto Amato, Jeffrey Barrett, Sonia Goncalves, Ewan Harrison, David K. Jackson, Ian Johnston, Dominic Kwiatkowski, Cordelia Langford, John Sillitoe on behalf of the Wellcome Sanger Institute COVID-19 Surveillance Team |
| EPI_ISL_2963078, EPI_ISL_2963079, EPI_ISL_2963080, EPI_ISL_2963081, EPI_ISL_2963082, EPI_ISL_2963083, EPI_ISL_2963084, EPI_ISL_2963085, EPI_ISL_2963086, EPI_ISL_2963087, EPI_ISL_2963088, EPI_ISL_2963089, EPI_ISL_2963090, EPI_ISL_2963091 | | | |
| see above | Lighthouse Lab in Alderley Park | Wellcome Sanger Institute for the COVID-19 Genomics UK (COG-UK) Consortium | Jacquelyn Wynn, Mairead Hyland, The Lighthouse Lab in Alderley Park and Alex Alderton, Roberto Amato, Jeffrey Barrett, Sonia Goncalves, Ewan Harrison, David K. Jackson, Ian Johnston, Dominic Kwiatkowski, Cordelia Langford, John Sillitoe on behalf of the Wellcome Sanger Institute COVID-19 Surveillance Team |
| EPI_ISL_2963092, EPI_ISL_2963093, EPI_ISL_2963094, EPI_ISL_2963095, EPI_ISL_2963096 | Lighthouse Lab in Milton Keynes | Wellcome Sanger Institute for the COVID-19 Genomics UK (COG-UK) Consortium | The Lighthouse Lab in Milton Keynes and Alex Alderton, Roberto Amato, Jeffrey Barrett, Sonia Goncalves, Ewan Harrison, David K. Jackson, Ian Johnston, Dominic Kwiatkowski, Cordelia Langford, John Sillitoe on behalf of the Wellcome Sanger Institute COVID-19 Surveillance Team |
| EPI_ISL_2963097 | Lighthouse Laboratory Plymouth | Wellcome Sanger Institute for the COVID-19 Genomics UK (COG-UK) Consortium | Lighthouse Laboratory Plymouth and Alex Alderton, Roberto Amato, Jeffrey Barrett, Sonia Goncalves, Ewan Harrison, David K. Jackson, Ian Johnston, Dominic Kwiatkowski, Cordelia Langford, John Sillitoe on behalf of the Wellcome Sanger Institute COVID-19 Surveillance Team |
| EPI_ISL_2963098 | Lighthouse Lab in Milton Keynes | Wellcome Sanger Institute for the COVID-19 Genomics UK (COG-UK) Consortium | The Lighthouse Lab in Milton Keynes and Alex Alderton, Roberto Amato, Jeffrey Barrett, Sonia Goncalves, Ewan Harrison, David K. Jackson, Ian Johnston, Dominic Kwiatkowski, Cordelia Langford, John Sillitoe on behalf of the Wellcome Sanger Institute COVID-19 Surveillance Team |

| | | | |
|---|---|---|---|
| EPI_ISL_2963099 | Health Services Laboratories | Wellcome Sanger Institute for the COVID-19 Genomics UK (COG-UK) Consortium | Health Services Laboratories and Alex Alderton, Roberto Amato, Jeffrey Barrett, Sonia Goncalves, Ewan Harrison, David K. Jackson, Ian Johnston, Dominic Kwiatkowski, Cordelia Langford, John Sillitoe on behalf of the Wellcome Sanger Institute COVID-19 Surveillance Team |
| EPI_ISL_2963100 | Lighthouse Laboratory Plymouth | Wellcome Sanger Institute for the COVID-19 Genomics UK (COG-UK) Consortium | Lighthouse Laboratory Plymouth and Alex Alderton, Roberto Amato, Jeffrey Barrett, Sonia Goncalves, Ewan Harrison, David K. Jackson, Ian Johnston, Dominic Kwiatkowski, Cordelia Langford, John Sillitoe on behalf of the Wellcome Sanger Institute COVID-19 Surveillance Team |
| EPI_ISL_2963101 | Lighthouse Lab in Milton Keynes | Wellcome Sanger Institute for the COVID-19 Genomics UK (COG-UK) Consortium | The Lighthouse Lab in Milton Keynes and Alex Alderton, Roberto Amato, Jeffrey Barrett, Sonia Goncalves, Ewan Harrison, David K. Jackson, Ian Johnston, Dominic Kwiatkowski, Cordelia Langford, John Sillitoe on behalf of the Wellcome Sanger Institute COVID-19 Surveillance Team |
| EPI_ISL_2963102 | Health Services Laboratories | Wellcome Sanger Institute for the COVID-19 Genomics UK (COG-UK) Consortium | Health Services Laboratories and Alex Alderton, Roberto Amato, Jeffrey Barrett, Sonia Goncalves, Ewan Harrison, David K. Jackson, Ian Johnston, Dominic Kwiatkowski, Cordelia Langford, John Sillitoe on behalf of the Wellcome Sanger Institute COVID-19 Surveillance Team |
| EPI_ISL_2963103, EPI_ISL_2963104, EPI_ISL_2963105 | Lighthouse Lab in Milton Keynes | Wellcome Sanger Institute for the COVID-19 Genomics UK (COG-UK) Consortium | The Lighthouse Lab in Milton Keynes and Alex Alderton, Roberto Amato, Jeffrey Barrett, Sonia Goncalves, Ewan Harrison, David K. Jackson, Ian Johnston, Dominic Kwiatkowski, Cordelia Langford, John Sillitoe on behalf of the Wellcome Sanger Institute COVID-19 Surveillance Team |
| EPI_ISL_2963106, EPI_ISL_2963107, EPI_ISL_2963108 | Health Services Laboratories | Wellcome Sanger Institute for the COVID-19 Genomics UK (COG-UK) Consortium | Health Services Laboratories and Alex Alderton, Roberto Amato, Jeffrey Barrett, Sonia Goncalves, Ewan Harrison, David K. Jackson, Ian Johnston, Dominic Kwiatkowski, Cordelia Langford, John Sillitoe on behalf of the Wellcome Sanger Institute COVID-19 Surveillance Team |
| EPI_ISL_2963109 | Lighthouse Laboratory Plymouth | Wellcome Sanger Institute for the COVID-19 Genomics UK (COG-UK) Consortium | Lighthouse Laboratory Plymouth and Alex Alderton, Roberto Amato, Jeffrey Barrett, Sonia Goncalves, Ewan Harrison, David K. Jackson, Ian Johnston, Dominic Kwiatkowski, Cordelia Langford, John Sillitoe on behalf of the Wellcome Sanger Institute COVID-19 Surveillance Team |
| EPI_ISL_2963110 | Health Services Laboratories | Wellcome Sanger Institute for the COVID-19 Genomics UK (COG-UK) Consortium | Health Services Laboratories and Alex Alderton, Roberto Amato, Jeffrey Barrett, Sonia Goncalves, Ewan Harrison, David K. Jackson, Ian Johnston, Dominic Kwiatkowski, Cordelia Langford, John Sillitoe on behalf of the Wellcome Sanger Institute COVID-19 Surveillance Team |
| EPI_ISL_2963111 | Lighthouse Lab in Milton Keynes | Wellcome Sanger Institute for the COVID-19 Genomics UK (COG-UK) Consortium | The Lighthouse Lab in Milton Keynes and Alex Alderton, Roberto Amato, Jeffrey Barrett, Sonia Goncalves, Ewan Harrison, David K. Jackson, Ian Johnston, Dominic Kwiatkowski, Cordelia Langford, John Sillitoe on behalf of the Wellcome Sanger Institute COVID-19 Surveillance Team |
| EPI_ISL_2963112, EPI_ISL_2963113 | Lighthouse Laboratory Plymouth | Wellcome Sanger Institute for the COVID-19 Genomics UK (COG-UK) Consortium | Lighthouse Laboratory Plymouth and Alex Alderton, Roberto Amato, Jeffrey Barrett, Sonia Goncalves, Ewan Harrison, David K. Jackson, Ian Johnston, Dominic Kwiatkowski, Cordelia Langford, John Sillitoe on behalf of the Wellcome Sanger Institute COVID-19 Surveillance Team |
| EPI_ISL_2963114 | Lighthouse Lab in Milton Keynes | Wellcome Sanger Institute for the COVID-19 Genomics UK (COG-UK) Consortium | The Lighthouse Lab in Milton Keynes and Alex Alderton, Roberto Amato, Jeffrey Barrett, Sonia Goncalves, Ewan Harrison, David K. Jackson, Ian Johnston, Dominic Kwiatkowski, Cordelia Langford, John Sillitoe on behalf of the Wellcome Sanger Institute COVID-19 Surveillance Team |
| EPI_ISL_2963115 | Lighthouse Laboratory Plymouth | Wellcome Sanger Institute for the COVID-19 Genomics UK (COG-UK) Consortium | Lighthouse Laboratory Plymouth and Alex Alderton, Roberto Amato, Jeffrey Barrett, Sonia Goncalves, Ewan Harrison, David K. Jackson, Ian Johnston, Dominic Kwiatkowski, Cordelia Langford, John Sillitoe on behalf of the Wellcome Sanger Institute COVID-19 Surveillance Team |
| EPI_ISL_2963116, EPI_ISL_2963117, EPI_ISL_2963118 | Lighthouse Lab in Milton Keynes | Wellcome Sanger Institute for the COVID-19 Genomics UK (COG-UK) Consortium | The Lighthouse Lab in Milton Keynes and Alex Alderton, Roberto Amato, Jeffrey Barrett, Sonia Goncalves, Ewan Harrison, David K. Jackson, Ian Johnston, Dominic Kwiatkowski, Cordelia Langford, John Sillitoe on behalf of the Wellcome Sanger Institute COVID-19 Surveillance Team |
| EPI_ISL_2963119, EPI_ISL_2963120 | Lighthouse Laboratory Plymouth | Wellcome Sanger Institute for the COVID-19 Genomics UK (COG-UK) Consortium | Lighthouse Laboratory Plymouth and Alex Alderton, Roberto Amato, Jeffrey Barrett, Sonia Goncalves, Ewan Harrison, David K. Jackson, Ian Johnston, Dominic Kwiatkowski, Cordelia Langford, John Sillitoe on behalf of the Wellcome Sanger Institute COVID-19 Surveillance Team |
| EPI_ISL_2963121 | Lighthouse Lab in Milton Keynes | Wellcome Sanger Institute for the COVID-19 Genomics UK (COG-UK) Consortium | The Lighthouse Lab in Milton Keynes and Alex Alderton, Roberto Amato, Jeffrey Barrett, Sonia Goncalves, Ewan Harrison, David K. Jackson, Ian Johnston, Dominic Kwiatkowski, Cordelia Langford, John Sillitoe on behalf of the Wellcome Sanger Institute COVID-19 Surveillance Team |
| EPI_ISL_2963122, EPI_ISL_2963123 | Lighthouse Laboratory Plymouth | Wellcome Sanger Institute for the COVID-19 Genomics UK (COG-UK) Consortium | Lighthouse Laboratory Plymouth and Alex Alderton, Roberto Amato, Jeffrey Barrett, Sonia Goncalves, Ewan Harrison, David K. Jackson, Ian Johnston, Dominic Kwiatkowski, Cordelia Langford, John Sillitoe on behalf of the Wellcome Sanger Institute COVID-19 Surveillance Team |
| EPI_ISL_2963124, EPI_ISL_2963125 | Lighthouse Lab in Milton Keynes | Wellcome Sanger Institute for the COVID-19 Genomics UK (COG-UK) Consortium | The Lighthouse Lab in Milton Keynes and Alex Alderton, Roberto Amato, Jeffrey Barrett, Sonia Goncalves, Ewan Harrison, David K. Jackson, Ian Johnston, Dominic Kwiatkowski, Cordelia Langford, John Sillitoe on behalf of the Wellcome Sanger Institute COVID-19 Surveillance Team |
| EPI_ISL_2963126 | Lighthouse Laboratory Plymouth | Wellcome Sanger Institute for the COVID-19 Genomics UK (COG-UK) Consortium | Lighthouse Laboratory Plymouth and Alex Alderton, Roberto Amato, Jeffrey Barrett, Sonia Goncalves, Ewan Harrison, David K. Jackson, Ian Johnston, Dominic Kwiatkowski, Cordelia Langford, John Sillitoe on behalf of the Wellcome Sanger Institute COVID-19 Surveillance Team |
| EPI_ISL_2963127, EPI_ISL_2963128 | Lighthouse Lab in Milton Keynes | Wellcome Sanger Institute for the COVID-19 Genomics UK (COG-UK) Consortium | The Lighthouse Lab in Milton Keynes and Alex Alderton, Roberto Amato, Jeffrey Barrett, Sonia Goncalves, Ewan Harrison, David K. Jackson, Ian Johnston, Dominic Kwiatkowski, Cordelia Langford, John Sillitoe on behalf of the Wellcome Sanger Institute COVID-19 Surveillance Team |
| EPI_ISL_2963129 | Lighthouse Laboratory Plymouth | Wellcome Sanger Institute for the COVID-19 Genomics UK (COG-UK) Consortium | Lighthouse Laboratory Plymouth and Alex Alderton, Roberto Amato, Jeffrey Barrett, Sonia Goncalves, Ewan Harrison, David K. Jackson, Ian Johnston, Dominic Kwiatkowski, Cordelia Langford, John Sillitoe on behalf of the Wellcome Sanger Institute COVID-19 Surveillance Team |
| EPI_ISL_2963130 | Lighthouse Lab in Milton Keynes | Wellcome Sanger Institute for the COVID-19 Genomics UK (COG-UK) Consortium | The Lighthouse Lab in Milton Keynes and Alex Alderton, Roberto Amato, Jeffrey Barrett, Sonia Goncalves, Ewan Harrison, David K. Jackson, Ian Johnston, Dominic Kwiatkowski, Cordelia Langford, John Sillitoe on behalf of the Wellcome Sanger Institute COVID-19 Surveillance Team |
| EPI_ISL_2963131 | Health Services Laboratories | Wellcome Sanger Institute for the COVID-19 Genomics UK (COG-UK) Consortium | Health Services Laboratories and Alex Alderton, Roberto Amato, Jeffrey Barrett, Sonia Goncalves, Ewan Harrison, David K. Jackson, Ian Johnston, Dominic Kwiatkowski, Cordelia Langford, John Sillitoe on behalf of the Wellcome Sanger Institute COVID-19 Surveillance Team |
| EPI_ISL_2963132 | Lighthouse Lab in Milton Keynes | Wellcome Sanger Institute for the COVID-19 Genomics UK (COG-UK) Consortium | The Lighthouse Lab in Milton Keynes and Alex Alderton, Roberto Amato, Jeffrey Barrett, Sonia Goncalves, Ewan Harrison, David K. Jackson, Ian Johnston, Dominic Kwiatkowski, Cordelia Langford, John Sillitoe on behalf of the Wellcome Sanger Institute COVID-19 Surveillance Team |
| EPI_ISL_2963133 | Health Services Laboratories | Wellcome Sanger Institute for the COVID-19 Genomics UK (COG-UK) Consortium | Health Services Laboratories and Alex Alderton, Roberto Amato, Jeffrey Barrett, Sonia Goncalves, Ewan Harrison, David K. Jackson, Ian Johnston, Dominic Kwiatkowski, Cordelia Langford, John Sillitoe on behalf of the Wellcome Sanger Institute COVID-19 Surveillance Team |
| EPI_ISL_2963134 | Lighthouse Laboratory Plymouth | Wellcome Sanger Institute for the COVID-19 Genomics UK (COG-UK) Consortium | Lighthouse Laboratory Plymouth and Alex Alderton, Roberto Amato, Jeffrey Barrett, Sonia Goncalves, Ewan Harrison, David K. Jackson, Ian Johnston, Dominic Kwiatkowski, Cordelia Langford, John Sillitoe on behalf of the Wellcome Sanger Institute COVID-19 Surveillance Team |
| EPI_ISL_2963135 | Health Services Laboratories | Wellcome Sanger Institute for the COVID-19 Genomics UK (COG-UK) Consortium | Health Services Laboratories and Alex Alderton, Roberto Amato, Jeffrey Barrett, Sonia Goncalves, Ewan Harrison, David K. Jackson, Ian Johnston, Dominic Kwiatkowski, Cordelia Langford, John Sillitoe on behalf of the Wellcome Sanger Institute COVID-19 Surveillance Team |
| EPI_ISL_2963136, EPI_ISL_2963137, EPI_ISL_2963138, EPI_ISL_2963139 | Lighthouse Lab in Milton Keynes | Wellcome Sanger Institute for the COVID-19 Genomics UK (COG-UK) Consortium | The Lighthouse Lab in Milton Keynes and Alex Alderton, Roberto Amato, Jeffrey Barrett, Sonia Goncalves, Ewan Harrison, David K. Jackson, Ian Johnston, Dominic Kwiatkowski, Cordelia Langford, John Sillitoe on behalf of the Wellcome Sanger Institute COVID-19 Surveillance Team |
| EPI_ISL_2963140 | Health Services Laboratories | Wellcome Sanger Institute for the COVID-19 Genomics UK (COG-UK) Consortium | Health Services Laboratories and Alex Alderton, Roberto Amato, Jeffrey Barrett, Sonia Goncalves, Ewan Harrison, David K. Jackson, Ian Johnston, Dominic Kwiatkowski, Cordelia Langford, John Sillitoe on behalf of the Wellcome Sanger Institute COVID-19 Surveillance Team |
| EPI_ISL_2963141 | Lighthouse Lab in Milton Keynes | Wellcome Sanger Institute for the COVID-19 Genomics UK (COG-UK) Consortium | The Lighthouse Lab in Milton Keynes and Alex Alderton, Roberto Amato, Jeffrey Barrett, Sonia Goncalves, Ewan Harrison, David K. Jackson, Ian Johnston, Dominic Kwiatkowski, Cordelia Langford, John Sillitoe on behalf of the Wellcome Sanger Institute COVID-19 Surveillance Team |
| EPI_ISL_2963142 | Health Services Laboratories | Wellcome Sanger Institute for the COVID-19 Genomics UK (COG-UK) Consortium | Health Services Laboratories and Alex Alderton, Roberto Amato, Jeffrey Barrett, Sonia Goncalves, Ewan Harrison, David K. Jackson, Ian Johnston, Dominic Kwiatkowski, Cordelia Langford, John Sillitoe on behalf of the Wellcome Sanger Institute COVID-19 Surveillance Team |
| EPI_ISL_2963143, EPI_ISL_2963144 | Lighthouse Laboratory Plymouth | Wellcome Sanger Institute for the COVID-19 Genomics UK (COG-UK) Consortium | Lighthouse Laboratory Plymouth and Alex Alderton, Roberto Amato, Jeffrey Barrett, Sonia Goncalves, Ewan Harrison, David K. Jackson, Ian Johnston, Dominic Kwiatkowski, Cordelia Langford, John Sillitoe on behalf of the Wellcome Sanger Institute COVID-19 Surveillance Team |
| EPI_ISL_2963145 | Lighthouse Lab in Milton Keynes | Wellcome Sanger Institute for the COVID-19 Genomics UK (COG-UK) Consortium | The Lighthouse Lab in Milton Keynes and Alex Alderton, Roberto Amato, Jeffrey Barrett, Sonia Goncalves, Ewan Harrison, David K. Jackson, Ian Johnston, Dominic Kwiatkowski, Cordelia Langford, John Sillitoe on behalf of the Wellcome Sanger Institute COVID-19 Surveillance Team |
| EPI_ISL_2963146, EPI_ISL_2963147 | Lighthouse Laboratory Plymouth | Wellcome Sanger Institute for the COVID-19 Genomics UK (COG-UK) Consortium | Lighthouse Laboratory Plymouth and Alex Alderton, Roberto Amato, Jeffrey Barrett, Sonia Goncalves, Ewan Harrison, David K. Jackson, Ian Johnston, Dominic Kwiatkowski, Cordelia Langford, John Sillitoe on behalf of the Wellcome Sanger Institute COVID-19 Surveillance Team |
| EPI_ISL_2963148 | Health Services Laboratories | Wellcome Sanger Institute for the COVID-19 Genomics UK (COG-UK) Consortium | Health Services Laboratories and Alex Alderton, Roberto Amato, Jeffrey Barrett, Sonia Goncalves, Ewan Harrison, David K. Jackson, Ian Johnston, Dominic Kwiatkowski, Cordelia Langford, John Sillitoe on behalf of the Wellcome Sanger Institute COVID-19 Surveillance Team |
| EPI_ISL_2963149 | Lighthouse Laboratory Plymouth | Wellcome Sanger Institute for the COVID-19 Genomics UK (COG-UK) Consortium | Lighthouse Laboratory Plymouth and Alex Alderton, Roberto Amato, Jeffrey Barrett, Sonia Goncalves, Ewan Harrison, David K. Jackson, Ian Johnston, Dominic Kwiatkowski, Cordelia Langford, John Sillitoe on behalf of the Wellcome Sanger Institute COVID-19 Surveillance Team |
| EPI_ISL_2963150 | Lighthouse Lab in Milton Keynes | Wellcome Sanger Institute for the COVID-19 Genomics UK (COG-UK) Consortium | The Lighthouse Lab in Milton Keynes and Alex Alderton, Roberto Amato, Jeffrey Barrett, Sonia Goncalves, Ewan Harrison, David K. Jackson, Ian Johnston, Dominic Kwiatkowski, Cordelia Langford, John Sillitoe on behalf of the Wellcome Sanger Institute COVID-19 Surveillance Team |
| EPI_ISL_2963151 | Health Services Laboratories | Wellcome Sanger Institute for the COVID-19 Genomics UK (COG-UK) Consortium | Health Services Laboratories and Alex Alderton, Roberto Amato, Jeffrey Barrett, Sonia Goncalves, Ewan Harrison, David K. Jackson, Ian Johnston, Dominic Kwiatkowski, Cordelia Langford, John Sillitoe on behalf of the Wellcome Sanger Institute COVID-19 Surveillance Team |
| EPI_ISL_2963152 | Lighthouse Lab in Milton Keynes | Wellcome Sanger Institute for the COVID-19 Genomics UK (COG-UK) Consortium | The Lighthouse Lab in Milton Keynes and Alex Alderton, Roberto Amato, Jeffrey Barrett, Sonia Goncalves, Ewan Harrison, David K. Jackson, Ian Johnston, Dominic Kwiatkowski, Cordelia Langford, John Sillitoe on behalf of the Wellcome Sanger Institute COVID-19 Surveillance Team |
| EPI_ISL_2963153 | Health Services Laboratories | Wellcome Sanger Institute for the COVID-19 Genomics UK (COG-UK) Consortium | Health Services Laboratories and Alex Alderton, Roberto Amato, Jeffrey Barrett, Sonia Goncalves, Ewan Harrison, David K. Jackson, Ian Johnston, Dominic Kwiatkowski, Cordelia Langford, John Sillitoe on behalf of the Wellcome Sanger Institute COVID-19 Surveillance Team |
| EPI_ISL_2963154, EPI_ISL_2963155 | Lighthouse Laboratory Plymouth | Wellcome Sanger Institute for the COVID-19 Genomics UK (COG-UK) Consortium | Lighthouse Laboratory Plymouth and Alex Alderton, Roberto Amato, Jeffrey Barrett, Sonia Goncalves, Ewan Harrison, David K. Jackson, Ian Johnston, Dominic Kwiatkowski, Cordelia Langford, John Sillitoe on behalf of the Wellcome Sanger Institute COVID-19 Surveillance Team |
| EPI_ISL_2963156 | Health Services Laboratories | Wellcome Sanger Institute for the COVID-19 Genomics UK (COG-UK) Consortium | Health Services Laboratories and Alex Alderton, Roberto Amato, Jeffrey Barrett, Sonia Goncalves, Ewan Harrison, David K. Jackson, Ian Johnston, Dominic Kwiatkowski, Cordelia Langford, John Sillitoe on behalf of the Wellcome Sanger Institute COVID-19 Surveillance Team |
| EPI_ISL_2963157 | Lighthouse Laboratory Plymouth | Wellcome Sanger Institute for the COVID-19 Genomics UK (COG-UK) Consortium | Lighthouse Laboratory Plymouth and Alex Alderton, Roberto Amato, Jeffrey Barrett, Sonia Goncalves, Ewan Harrison, David K. Jackson, Ian Johnston, Dominic Kwiatkowski, Cordelia Langford, John Sillitoe on behalf of the Wellcome Sanger Institute COVID-19 Surveillance Team |
| EPI_ISL_2963158, EPI_ISL_2963159, EPI_ISL_2963160 | Lighthouse Lab in Milton Keynes | Wellcome Sanger Institute for the COVID-19 Genomics UK (COG-UK) Consortium | The Lighthouse Lab in Milton Keynes and Alex Alderton, Roberto Amato, Jeffrey Barrett, Sonia Goncalves, Ewan Harrison, David K. Jackson, Ian Johnston, Dominic Kwiatkowski, Cordelia Langford, John Sillitoe on behalf of the Wellcome Sanger Institute COVID-19 Surveillance Team |

| | | | |
|---|---|---|---|
| EPI_ISL_2963161 | Health Services Laboratories | Wellcome Sanger Institute for the COVID-19 Genomics UK (COG-UK) Consortium | Health Services Laboratories and Alex Alderton, Roberto Amato, Jeffrey Barrett, Sonia Goncalves, Ewan Harrison, David K. Jackson, Ian Johnston, Dominic Kwiatkowski, Cordelia Langford, John Sillitoe on behalf of the Wellcome Sanger Institute COVID-19 Surveillance Team |
| EPI_ISL_2963162 | Lighthouse Lab in Milton Keynes | Wellcome Sanger Institute for the COVID-19 Genomics UK (COG-UK) Consortium | The Lighthouse Lab in Milton Keynes and Alex Alderton, Roberto Amato, Jeffrey Barrett, Sonia Goncalves, Ewan Harrison, David K. Jackson, Ian Johnston, Dominic Kwiatkowski, Cordelia Langford, John Sillitoe on behalf of the Wellcome Sanger Institute COVID-19 Surveillance Team |
| EPI_ISL_2963163, EPI_ISL_2963164 | Lighthouse Laboratory Plymouth | Wellcome Sanger Institute for the COVID-19 Genomics UK (COG-UK) Consortium | Lighthouse Laboratory Plymouth and Alex Alderton, Roberto Amato, Jeffrey Barrett, Sonia Goncalves, Ewan Harrison, David K. Jackson, Ian Johnston, Dominic Kwiatkowski, Cordelia Langford, John Sillitoe on behalf of the Wellcome Sanger Institute COVID-19 Surveillance Team |
| EPI_ISL_2963165 | Lighthouse Lab in Milton Keynes | Wellcome Sanger Institute for the COVID-19 Genomics UK (COG-UK) Consortium | The Lighthouse Lab in Milton Keynes and Alex Alderton, Roberto Amato, Jeffrey Barrett, Sonia Goncalves, Ewan Harrison, David K. Jackson, Ian Johnston, Dominic Kwiatkowski, Cordelia Langford, John Sillitoe on behalf of the Wellcome Sanger Institute COVID-19 Surveillance Team |
| EPI_ISL_2963166 | Lighthouse Laboratory Plymouth | Wellcome Sanger Institute for the COVID-19 Genomics UK (COG-UK) Consortium | Lighthouse Laboratory Plymouth and Alex Alderton, Roberto Amato, Jeffrey Barrett, Sonia Goncalves, Ewan Harrison, David K. Jackson, Ian Johnston, Dominic Kwiatkowski, Cordelia Langford, John Sillitoe on behalf of the Wellcome Sanger Institute COVID-19 Surveillance Team |
| EPI_ISL_2963167 | Health Services Laboratories | Wellcome Sanger Institute for the COVID-19 Genomics UK (COG-UK) Consortium | Health Services Laboratories and Alex Alderton, Roberto Amato, Jeffrey Barrett, Sonia Goncalves, Ewan Harrison, David K. Jackson, Ian Johnston, Dominic Kwiatkowski, Cordelia Langford, John Sillitoe on behalf of the Wellcome Sanger Institute COVID-19 Surveillance Team |
| EPI_ISL_2963168 | Lighthouse Laboratory Plymouth | Wellcome Sanger Institute for the COVID-19 Genomics UK (COG-UK) Consortium | Lighthouse Laboratory Plymouth and Alex Alderton, Roberto Amato, Jeffrey Barrett, Sonia Goncalves, Ewan Harrison, David K. Jackson, Ian Johnston, Dominic Kwiatkowski, Cordelia Langford, John Sillitoe on behalf of the Wellcome Sanger Institute COVID-19 Surveillance Team |
| EPI_ISL_2963169 | Health Services Laboratories | Wellcome Sanger Institute for the COVID-19 Genomics UK (COG-UK) Consortium | Health Services Laboratories and Alex Alderton, Roberto Amato, Jeffrey Barrett, Sonia Goncalves, Ewan Harrison, David K. Jackson, Ian Johnston, Dominic Kwiatkowski, Cordelia Langford, John Sillitoe on behalf of the Wellcome Sanger Institute COVID-19 Surveillance Team |
| EPI_ISL_2963170 | Lighthouse Lab in Milton Keynes | Wellcome Sanger Institute for the COVID-19 Genomics UK (COG-UK) Consortium | The Lighthouse Lab in Milton Keynes and Alex Alderton, Roberto Amato, Jeffrey Barrett, Sonia Goncalves, Ewan Harrison, David K. Jackson, Ian Johnston, Dominic Kwiatkowski, Cordelia Langford, John Sillitoe on behalf of the Wellcome Sanger Institute COVID-19 Surveillance Team |
| EPI_ISL_2963171 | Lighthouse Laboratory Plymouth | Wellcome Sanger Institute for the COVID-19 Genomics UK (COG-UK) Consortium | Lighthouse Laboratory Plymouth and Alex Alderton, Roberto Amato, Jeffrey Barrett, Sonia Goncalves, Ewan Harrison, David K. Jackson, Ian Johnston, Dominic Kwiatkowski, Cordelia Langford, John Sillitoe on behalf of the Wellcome Sanger Institute COVID-19 Surveillance Team |
| EPI_ISL_2963172 | Health Services Laboratories | Wellcome Sanger Institute for the COVID-19 Genomics UK (COG-UK) Consortium | Health Services Laboratories and Alex Alderton, Roberto Amato, Jeffrey Barrett, Sonia Goncalves, Ewan Harrison, David K. Jackson, Ian Johnston, Dominic Kwiatkowski, Cordelia Langford, John Sillitoe on behalf of the Wellcome Sanger Institute COVID-19 Surveillance Team |
| EPI_ISL_2963173, EPI_ISL_2963174 | Lighthouse Lab in Milton Keynes | Wellcome Sanger Institute for the COVID-19 Genomics UK (COG-UK) Consortium | The Lighthouse Lab in Milton Keynes and Alex Alderton, Roberto Amato, Jeffrey Barrett, Sonia Goncalves, Ewan Harrison, David K. Jackson, Ian Johnston, Dominic Kwiatkowski, Cordelia Langford, John Sillitoe on behalf of the Wellcome Sanger Institute COVID-19 Surveillance Team |
| EPI_ISL_2963175 | Health Services Laboratories | Wellcome Sanger Institute for the COVID-19 Genomics UK (COG-UK) Consortium | Health Services Laboratories and Alex Alderton, Roberto Amato, Jeffrey Barrett, Sonia Goncalves, Ewan Harrison, David K. Jackson, Ian Johnston, Dominic Kwiatkowski, Cordelia Langford, John Sillitoe on behalf of the Wellcome Sanger Institute COVID-19 Surveillance Team |
| EPI_ISL_2963176, EPI_ISL_2963177, EPI_ISL_2963178 | Lighthouse Laboratory Plymouth | Wellcome Sanger Institute for the COVID-19 Genomics UK (COG-UK) Consortium | Lighthouse Laboratory Plymouth and Alex Alderton, Roberto Amato, Jeffrey Barrett, Sonia Goncalves, Ewan Harrison, David K. Jackson, Ian Johnston, Dominic Kwiatkowski, Cordelia Langford, John Sillitoe on behalf of the Wellcome Sanger Institute COVID-19 Surveillance Team |
| EPI_ISL_2963179, EPI_ISL_2963180, EPI_ISL_2963181 | Lighthouse Lab in Milton Keynes | Wellcome Sanger Institute for the COVID-19 Genomics UK (COG-UK) Consortium | The Lighthouse Lab in Milton Keynes and Alex Alderton, Roberto Amato, Jeffrey Barrett, Sonia Goncalves, Ewan Harrison, David K. Jackson, Ian Johnston, Dominic Kwiatkowski, Cordelia Langford, John Sillitoe on behalf of the Wellcome Sanger Institute COVID-19 Surveillance Team |
| EPI_ISL_2963182 | Lighthouse Laboratory Plymouth | Wellcome Sanger Institute for the COVID-19 Genomics UK (COG-UK) Consortium | Lighthouse Laboratory Plymouth and Alex Alderton, Roberto Amato, Jeffrey Barrett, Sonia Goncalves, Ewan Harrison, David K. Jackson, Ian Johnston, Dominic Kwiatkowski, Cordelia Langford, John Sillitoe on behalf of the Wellcome Sanger Institute COVID-19 Surveillance Team |
| EPI_ISL_2963183, EPI_ISL_2963184 | Lighthouse Lab in Milton Keynes | Wellcome Sanger Institute for the COVID-19 Genomics UK (COG-UK) Consortium | The Lighthouse Lab in Milton Keynes and Alex Alderton, Roberto Amato, Jeffrey Barrett, Sonia Goncalves, Ewan Harrison, David K. Jackson, Ian Johnston, Dominic Kwiatkowski, Cordelia Langford, John Sillitoe on behalf of the Wellcome Sanger Institute COVID-19 Surveillance Team |
| EPI_ISL_2963185 | Lighthouse Laboratory Plymouth | Wellcome Sanger Institute for the COVID-19 Genomics UK (COG-UK) Consortium | Lighthouse Laboratory Plymouth and Alex Alderton, Roberto Amato, Jeffrey Barrett, Sonia Goncalves, Ewan Harrison, David K. Jackson, Ian Johnston, Dominic Kwiatkowski, Cordelia Langford, John Sillitoe on behalf of the Wellcome Sanger Institute COVID-19 Surveillance Team |
| EPI_ISL_2963186, EPI_ISL_2963187 | Lighthouse Lab in Milton Keynes | Wellcome Sanger Institute for the COVID-19 Genomics UK (COG-UK) Consortium | The Lighthouse Lab in Milton Keynes and Alex Alderton, Roberto Amato, Jeffrey Barrett, Sonia Goncalves, Ewan Harrison, David K. Jackson, Ian Johnston, Dominic Kwiatkowski, Cordelia Langford, John Sillitoe on behalf of the Wellcome Sanger Institute COVID-19 Surveillance Team |
| EPI_ISL_2963188 | Health Services Laboratories | Wellcome Sanger Institute for the COVID-19 Genomics UK (COG-UK) Consortium | Health Services Laboratories and Alex Alderton, Roberto Amato, Jeffrey Barrett, Sonia Goncalves, Ewan Harrison, David K. Jackson, Ian Johnston, Dominic Kwiatkowski, Cordelia Langford, John Sillitoe on behalf of the Wellcome Sanger Institute COVID-19 Surveillance Team |
| EPI_ISL_2963189 | Lighthouse Lab in Milton Keynes | Wellcome Sanger Institute for the COVID-19 Genomics UK (COG-UK) Consortium | The Lighthouse Lab in Milton Keynes and Alex Alderton, Roberto Amato, Jeffrey Barrett, Sonia Goncalves, Ewan Harrison, David K. Jackson, Ian Johnston, Dominic Kwiatkowski, Cordelia Langford, John Sillitoe on behalf of the Wellcome Sanger Institute COVID-19 Surveillance Team |
| EPI_ISL_2963190 | Health Services Laboratories | Wellcome Sanger Institute for the COVID-19 Genomics UK (COG-UK) Consortium | Health Services Laboratories and Alex Alderton, Roberto Amato, Jeffrey Barrett, Sonia Goncalves, Ewan Harrison, David K. Jackson, Ian Johnston, Dominic Kwiatkowski, Cordelia Langford, John Sillitoe on behalf of the Wellcome Sanger Institute COVID-19 Surveillance Team |
| EPI_ISL_2963191 | Lighthouse Lab in Milton Keynes | Wellcome Sanger Institute for the COVID-19 Genomics UK (COG-UK) Consortium | The Lighthouse Lab in Milton Keynes and Alex Alderton, Roberto Amato, Jeffrey Barrett, Sonia Goncalves, Ewan Harrison, David K. Jackson, Ian Johnston, Dominic Kwiatkowski, Cordelia Langford, John Sillitoe on behalf of the Wellcome Sanger Institute COVID-19 Surveillance Team |
| EPI_ISL_2963192 | Lighthouse Laboratory Plymouth | Wellcome Sanger Institute for the COVID-19 Genomics UK (COG-UK) Consortium | Lighthouse Laboratory Plymouth and Alex Alderton, Roberto Amato, Jeffrey Barrett, Sonia Goncalves, Ewan Harrison, David K. Jackson, Ian Johnston, Dominic Kwiatkowski, Cordelia Langford, John Sillitoe on behalf of the Wellcome Sanger Institute COVID-19 Surveillance Team |
| EPI_ISL_2963193, EPI_ISL_2963194 | Lighthouse Lab in Milton Keynes | Wellcome Sanger Institute for the COVID-19 Genomics UK (COG-UK) Consortium | The Lighthouse Lab in Milton Keynes and Alex Alderton, Roberto Amato, Jeffrey Barrett, Sonia Goncalves, Ewan Harrison, David K. Jackson, Ian Johnston, Dominic Kwiatkowski, Cordelia Langford, John Sillitoe on behalf of the Wellcome Sanger Institute COVID-19 Surveillance Team |
| EPI_ISL_2963195 | Health Services Laboratories | Wellcome Sanger Institute for the COVID-19 Genomics UK (COG-UK) Consortium | Health Services Laboratories and Alex Alderton, Roberto Amato, Jeffrey Barrett, Sonia Goncalves, Ewan Harrison, David K. Jackson, Ian Johnston, Dominic Kwiatkowski, Cordelia Langford, John Sillitoe on behalf of the Wellcome Sanger Institute COVID-19 Surveillance Team |
| EPI_ISL_2963196 | Lighthouse Lab in Milton Keynes | Wellcome Sanger Institute for the COVID-19 Genomics UK (COG-UK) Consortium | The Lighthouse Lab in Milton Keynes and Alex Alderton, Roberto Amato, Jeffrey Barrett, Sonia Goncalves, Ewan Harrison, David K. Jackson, Ian Johnston, Dominic Kwiatkowski, Cordelia Langford, John Sillitoe on behalf of the Wellcome Sanger Institute COVID-19 Surveillance Team |
| EPI_ISL_2963197, EPI_ISL_2963198 | Health Services Laboratories | Wellcome Sanger Institute for the COVID-19 Genomics UK (COG-UK) Consortium | Health Services Laboratories and Alex Alderton, Roberto Amato, Jeffrey Barrett, Sonia Goncalves, Ewan Harrison, David K. Jackson, Ian Johnston, Dominic Kwiatkowski, Cordelia Langford, John Sillitoe on behalf of the Wellcome Sanger Institute COVID-19 Surveillance Team |
| EPI_ISL_2963199 | Lighthouse Lab in Milton Keynes | Wellcome Sanger Institute for the COVID-19 Genomics UK (COG-UK) Consortium | The Lighthouse Lab in Milton Keynes and Alex Alderton, Roberto Amato, Jeffrey Barrett, Sonia Goncalves, Ewan Harrison, David K. Jackson, Ian Johnston, Dominic Kwiatkowski, Cordelia Langford, John Sillitoe on behalf of the Wellcome Sanger Institute COVID-19 Surveillance Team |
| EPI_ISL_2963200 | Lighthouse Laboratory Plymouth | Wellcome Sanger Institute for the COVID-19 Genomics UK (COG-UK) Consortium | Lighthouse Laboratory Plymouth and Alex Alderton, Roberto Amato, Jeffrey Barrett, Sonia Goncalves, Ewan Harrison, David K. Jackson, Ian Johnston, Dominic Kwiatkowski, Cordelia Langford, John Sillitoe on behalf of the Wellcome Sanger Institute COVID-19 Surveillance Team |
| EPI_ISL_2963201 | Health Services Laboratories | Wellcome Sanger Institute for the COVID-19 Genomics UK (COG-UK) Consortium | Health Services Laboratories and Alex Alderton, Roberto Amato, Jeffrey Barrett, Sonia Goncalves, Ewan Harrison, David K. Jackson, Ian Johnston, Dominic Kwiatkowski, Cordelia Langford, John Sillitoe on behalf of the Wellcome Sanger Institute COVID-19 Surveillance Team |
| EPI_ISL_2963202 | Lighthouse Laboratory Plymouth | Wellcome Sanger Institute for the COVID-19 Genomics UK (COG-UK) Consortium | Lighthouse Laboratory Plymouth and Alex Alderton, Roberto Amato, Jeffrey Barrett, Sonia Goncalves, Ewan Harrison, David K. Jackson, Ian Johnston, Dominic Kwiatkowski, Cordelia Langford, John Sillitoe on behalf of the Wellcome Sanger Institute COVID-19 Surveillance Team |
| EPI_ISL_2963203, EPI_ISL_2963204, EPI_ISL_2963205 | Lighthouse Lab in Milton Keynes | Wellcome Sanger Institute for the COVID-19 Genomics UK (COG-UK) Consortium | The Lighthouse Lab in Milton Keynes and Alex Alderton, Roberto Amato, Jeffrey Barrett, Sonia Goncalves, Ewan Harrison, David K. Jackson, Ian Johnston, Dominic Kwiatkowski, Cordelia Langford, John Sillitoe on behalf of the Wellcome Sanger Institute COVID-19 Surveillance Team |
| EPI_ISL_2963206 | Health Services Laboratories | Wellcome Sanger Institute for the COVID-19 Genomics UK (COG-UK) Consortium | Health Services Laboratories and Alex Alderton, Roberto Amato, Jeffrey Barrett, Sonia Goncalves, Ewan Harrison, David K. Jackson, Ian Johnston, Dominic Kwiatkowski, Cordelia Langford, John Sillitoe on behalf of the Wellcome Sanger Institute COVID-19 Surveillance Team |
| EPI_ISL_2963207 | Lighthouse Laboratory Plymouth | Wellcome Sanger Institute for the COVID-19 Genomics UK (COG-UK) Consortium | Lighthouse Laboratory Plymouth and Alex Alderton, Roberto Amato, Jeffrey Barrett, Sonia Goncalves, Ewan Harrison, David K. Jackson, Ian Johnston, Dominic Kwiatkowski, Cordelia Langford, John Sillitoe on behalf of the Wellcome Sanger Institute COVID-19 Surveillance Team |
| EPI_ISL_2963208 | Health Services Laboratories | Wellcome Sanger Institute for the COVID-19 Genomics UK (COG-UK) Consortium | Health Services Laboratories and Alex Alderton, Roberto Amato, Jeffrey Barrett, Sonia Goncalves, Ewan Harrison, David K. Jackson, Ian Johnston, Dominic Kwiatkowski, Cordelia Langford, John Sillitoe on behalf of the Wellcome Sanger Institute COVID-19 Surveillance Team |
| EPI_ISL_2963209, EPI_ISL_2963210 | Lighthouse Lab in Milton Keynes | Wellcome Sanger Institute for the COVID-19 Genomics UK (COG-UK) Consortium | The Lighthouse Lab in Milton Keynes and Alex Alderton, Roberto Amato, Jeffrey Barrett, Sonia Goncalves, Ewan Harrison, David K. Jackson, Ian Johnston, Dominic Kwiatkowski, Cordelia Langford, John Sillitoe on behalf of the Wellcome Sanger Institute COVID-19 Surveillance Team |
| EPI_ISL_2963211 | Lighthouse Laboratory Plymouth | Wellcome Sanger Institute for the COVID-19 Genomics UK (COG-UK) Consortium | Lighthouse Laboratory Plymouth and Alex Alderton, Roberto Amato, Jeffrey Barrett, Sonia Goncalves, Ewan Harrison, David K. Jackson, Ian Johnston, Dominic Kwiatkowski, Cordelia Langford, John Sillitoe on behalf of the Wellcome Sanger Institute COVID-19 Surveillance Team |
| EPI_ISL_2963212, EPI_ISL_2963213 | Health Services Laboratories | Wellcome Sanger Institute for the COVID-19 Genomics UK (COG-UK) Consortium | Health Services Laboratories and Alex Alderton, Roberto Amato, Jeffrey Barrett, Sonia Goncalves, Ewan Harrison, David K. Jackson, Ian Johnston, Dominic Kwiatkowski, Cordelia Langford, John Sillitoe on behalf of the Wellcome Sanger Institute COVID-19 Surveillance Team |
| EPI_ISL_2963214 | Lighthouse Laboratory Plymouth | Wellcome Sanger Institute for the COVID-19 Genomics UK (COG-UK) Consortium | Lighthouse Laboratory Plymouth and Alex Alderton, Roberto Amato, Jeffrey Barrett, Sonia Goncalves, Ewan Harrison, David K. Jackson, Ian Johnston, Dominic Kwiatkowski, Cordelia Langford, John Sillitoe on behalf of the Wellcome Sanger Institute COVID-19 Surveillance Team |
| EPI_ISL_2963215 | Lighthouse Lab in Milton Keynes | Wellcome Sanger Institute for the COVID-19 Genomics UK (COG-UK) Consortium | The Lighthouse Lab in Milton Keynes and Alex Alderton, Roberto Amato, Jeffrey Barrett, Sonia Goncalves, Ewan Harrison, David K. Jackson, Ian Johnston, Dominic Kwiatkowski, Cordelia Langford, John Sillitoe on behalf of the Wellcome Sanger Institute COVID-19 Surveillance Team |
| EPI_ISL_2963216 | Health Services Laboratories | Wellcome Sanger Institute for the COVID-19 Genomics UK (COG-UK) Consortium | Health Services Laboratories and Alex Alderton, Roberto Amato, Jeffrey Barrett, Sonia Goncalves, Ewan Harrison, David K. Jackson, Ian Johnston, Dominic Kwiatkowski, Cordelia Langford, John Sillitoe on behalf of the Wellcome Sanger Institute COVID-19 Surveillance Team |
| EPI_ISL_2963217, EPI_ISL_2963218 | Lighthouse Lab in Milton Keynes | Wellcome Sanger Institute for the COVID-19 Genomics UK (COG-UK) Consortium | The Lighthouse Lab in Milton Keynes and Alex Alderton, Roberto Amato, Jeffrey Barrett, Sonia Goncalves, Ewan Harrison, David K. Jackson, Ian Johnston, Dominic Kwiatkowski, Cordelia Langford, John Sillitoe on behalf of the Wellcome Sanger Institute COVID-19 Surveillance Team |
| EPI_ISL_2963219 | Lighthouse Laboratory Plymouth | Wellcome Sanger Institute for the COVID-19 Genomics UK (COG-UK) | Lighthouse Laboratory Plymouth and Alex Alderton, Roberto Amato, Jeffrey Barrett, Sonia Goncalves, Ewan Harrison, David K. Jackson, Ian Johnston, Dominic Kwiatkowski, Cordelia Langford, John Sillitoe on behalf of the Wellcome Sanger |

| | | | |
|---|---|---|---|
| | | Consortium | Institute COVID-19 Surveillance Team |
| EPI_ISL_2963220 | Health Services Laboratories | Wellcome Sanger Institute for the COVID-19 Genomics UK (COG-UK) Consortium | Health Services Laboratories and Alex Alderton, Roberto Amato, Jeffrey Barrett, Sonia Goncalves, Ewan Harrison, David K. Jackson, Ian Johnston, Dominic Kwiatkowski, Cordelia Langford, John Sillitoe on behalf of the Wellcome Sanger Institute COVID-19 Surveillance Team |
| EPI_ISL_2963221 | Lighthouse Laboratory Plymouth | Wellcome Sanger Institute for the COVID-19 Genomics UK (COG-UK) Consortium | Lighthouse Laboratory Plymouth and Alex Alderton, Roberto Amato, Jeffrey Barrett, Sonia Goncalves, Ewan Harrison, David K. Jackson, Ian Johnston, Dominic Kwiatkowski, Cordelia Langford, John Sillitoe on behalf of the Wellcome Sanger Institute COVID-19 Surveillance Team |
| EPI_ISL_2963222 | Health Services Laboratories | Wellcome Sanger Institute for the COVID-19 Genomics UK (COG-UK) Consortium | Health Services Laboratories and Alex Alderton, Roberto Amato, Jeffrey Barrett, Sonia Goncalves, Ewan Harrison, David K. Jackson, Ian Johnston, Dominic Kwiatkowski, Cordelia Langford, John Sillitoe on behalf of the Wellcome Sanger Institute COVID-19 Surveillance Team |
| EPI_ISL_2963223, EPI_ISL_2963224, EPI_ISL_2963226, EPI_ISL_2963227 | Lighthouse Lab in Milton Keynes | Wellcome Sanger Institute for the COVID-19 Genomics UK (COG-UK) Consortium | The Lighthouse Lab in Milton Keynes and Alex Alderton, Roberto Amato, Jeffrey Barrett, Sonia Goncalves, Ewan Harrison, David K. Jackson, Ian Johnston, Dominic Kwiatkowski, Cordelia Langford, John Sillitoe on behalf of the Wellcome Sanger Institute COVID-19 Surveillance Team |
| EPI_ISL_2963228, EPI_ISL_2963229, EPI_ISL_2963230, EPI_ISL_2963231, EPI_ISL_2963232, EPI_ISL_2963233 | Lighthouse Laboratory Plymouth | Wellcome Sanger Institute for the COVID-19 Genomics UK (COG-UK) Consortium | Lighthouse Laboratory Plymouth and Alex Alderton, Roberto Amato, Jeffrey Barrett, Sonia Goncalves, Ewan Harrison, David K. Jackson, Ian Johnston, Dominic Kwiatkowski, Cordelia Langford, John Sillitoe on behalf of the Wellcome Sanger Institute COVID-19 Surveillance Team |
| EPI_ISL_2963234 | Health Services Laboratories | Wellcome Sanger Institute for the COVID-19 Genomics UK (COG-UK) Consortium | Health Services Laboratories and Alex Alderton, Roberto Amato, Jeffrey Barrett, Sonia Goncalves, Ewan Harrison, David K. Jackson, Ian Johnston, Dominic Kwiatkowski, Cordelia Langford, John Sillitoe on behalf of the Wellcome Sanger Institute COVID-19 Surveillance Team |
| EPI_ISL_2963235 | Lighthouse Lab in Milton Keynes | Wellcome Sanger Institute for the COVID-19 Genomics UK (COG-UK) Consortium | The Lighthouse Lab in Milton Keynes and Alex Alderton, Roberto Amato, Jeffrey Barrett, Sonia Goncalves, Ewan Harrison, David K. Jackson, Ian Johnston, Dominic Kwiatkowski, Cordelia Langford, John Sillitoe on behalf of the Wellcome Sanger Institute COVID-19 Surveillance Team |
| EPI_ISL_2963236 | Lighthouse Laboratory Plymouth | Wellcome Sanger Institute for the COVID-19 Genomics UK (COG-UK) Consortium | Lighthouse Laboratory Plymouth and Alex Alderton, Roberto Amato, Jeffrey Barrett, Sonia Goncalves, Ewan Harrison, David K. Jackson, Ian Johnston, Dominic Kwiatkowski, Cordelia Langford, John Sillitoe on behalf of the Wellcome Sanger Institute COVID-19 Surveillance Team |
| EPI_ISL_2963237, EPI_ISL_2963238 | Health Services Laboratories | Wellcome Sanger Institute for the COVID-19 Genomics UK (COG-UK) Consortium | Health Services Laboratories and Alex Alderton, Roberto Amato, Jeffrey Barrett, Sonia Goncalves, Ewan Harrison, David K. Jackson, Ian Johnston, Dominic Kwiatkowski, Cordelia Langford, John Sillitoe on behalf of the Wellcome Sanger Institute COVID-19 Surveillance Team |
| EPI_ISL_2963239, EPI_ISL_2963240, EPI_ISL_2963241, EPI_ISL_2963242 | Lighthouse Lab in Milton Keynes | Wellcome Sanger Institute for the COVID-19 Genomics UK (COG-UK) Consortium | The Lighthouse Lab in Milton Keynes and Alex Alderton, Roberto Amato, Jeffrey Barrett, Sonia Goncalves, Ewan Harrison, David K. Jackson, Ian Johnston, Dominic Kwiatkowski, Cordelia Langford, John Sillitoe on behalf of the Wellcome Sanger Institute COVID-19 Surveillance Team |
| EPI_ISL_2963243 | Health Services Laboratories | Wellcome Sanger Institute for the COVID-19 Genomics UK (COG-UK) Consortium | Health Services Laboratories and Alex Alderton, Roberto Amato, Jeffrey Barrett, Sonia Goncalves, Ewan Harrison, David K. Jackson, Ian Johnston, Dominic Kwiatkowski, Cordelia Langford, John Sillitoe on behalf of the Wellcome Sanger Institute COVID-19 Surveillance Team |
| EPI_ISL_2963244 | Lighthouse Lab in Milton Keynes | Wellcome Sanger Institute for the COVID-19 Genomics UK (COG-UK) Consortium | The Lighthouse Lab in Milton Keynes and Alex Alderton, Roberto Amato, Jeffrey Barrett, Sonia Goncalves, Ewan Harrison, David K. Jackson, Ian Johnston, Dominic Kwiatkowski, Cordelia Langford, John Sillitoe on behalf of the Wellcome Sanger Institute COVID-19 Surveillance Team |
| EPI_ISL_2963245, EPI_ISL_2963246 | Lighthouse Laboratory Plymouth | Wellcome Sanger Institute for the COVID-19 Genomics UK (COG-UK) Consortium | Lighthouse Laboratory Plymouth and Alex Alderton, Roberto Amato, Jeffrey Barrett, Sonia Goncalves, Ewan Harrison, David K. Jackson, Ian Johnston, Dominic Kwiatkowski, Cordelia Langford, John Sillitoe on behalf of the Wellcome Sanger Institute COVID-19 Surveillance Team |
| EPI_ISL_2963247, EPI_ISL_2963248 | Lighthouse Lab in Milton Keynes | Wellcome Sanger Institute for the COVID-19 Genomics UK (COG-UK) Consortium | The Lighthouse Lab in Milton Keynes and Alex Alderton, Roberto Amato, Jeffrey Barrett, Sonia Goncalves, Ewan Harrison, David K. Jackson, Ian Johnston, Dominic Kwiatkowski, Cordelia Langford, John Sillitoe on behalf of the Wellcome Sanger Institute COVID-19 Surveillance Team |
| EPI_ISL_2963249 | Lighthouse Laboratory Plymouth | Wellcome Sanger Institute for the COVID-19 Genomics UK (COG-UK) Consortium | Lighthouse Laboratory Plymouth and Alex Alderton, Roberto Amato, Jeffrey Barrett, Sonia Goncalves, Ewan Harrison, David K. Jackson, Ian Johnston, Dominic Kwiatkowski, Cordelia Langford, John Sillitoe on behalf of the Wellcome Sanger Institute COVID-19 Surveillance Team |
| EPI_ISL_2963250, EPI_ISL_2963251, EPI_ISL_2963252, EPI_ISL_2963253, EPI_ISL_2963254 | Lighthouse Lab in Milton Keynes | Wellcome Sanger Institute for the COVID-19 Genomics UK (COG-UK) Consortium | The Lighthouse Lab in Milton Keynes and Alex Alderton, Roberto Amato, Jeffrey Barrett, Sonia Goncalves, Ewan Harrison, David K. Jackson, Ian Johnston, Dominic Kwiatkowski, Cordelia Langford, John Sillitoe on behalf of the Wellcome Sanger Institute COVID-19 Surveillance Team |
| EPI_ISL_2963255, EPI_ISL_2963256 | Lighthouse Laboratory Plymouth | Wellcome Sanger Institute for the COVID-19 Genomics UK (COG-UK) Consortium | Lighthouse Laboratory Plymouth and Alex Alderton, Roberto Amato, Jeffrey Barrett, Sonia Goncalves, Ewan Harrison, David K. Jackson, Ian Johnston, Dominic Kwiatkowski, Cordelia Langford, John Sillitoe on behalf of the Wellcome Sanger Institute COVID-19 Surveillance Team |
| EPI_ISL_2963257, EPI_ISL_2963258 | Lighthouse Lab in Milton Keynes | Wellcome Sanger Institute for the COVID-19 Genomics UK (COG-UK) Consortium | The Lighthouse Lab in Milton Keynes and Alex Alderton, Roberto Amato, Jeffrey Barrett, Sonia Goncalves, Ewan Harrison, David K. Jackson, Ian Johnston, Dominic Kwiatkowski, Cordelia Langford, John Sillitoe on behalf of the Wellcome Sanger Institute COVID-19 Surveillance Team |
| EPI_ISL_2963259 | Lighthouse Laboratory Plymouth | Wellcome Sanger Institute for the COVID-19 Genomics UK (COG-UK) Consortium | Lighthouse Laboratory Plymouth and Alex Alderton, Roberto Amato, Jeffrey Barrett, Sonia Goncalves, Ewan Harrison, David K. Jackson, Ian Johnston, Dominic Kwiatkowski, Cordelia Langford, John Sillitoe on behalf of the Wellcome Sanger Institute COVID-19 Surveillance Team |
| EPI_ISL_2963260 | Lighthouse Lab in Milton Keynes | Wellcome Sanger Institute for the COVID-19 Genomics UK (COG-UK) Consortium | The Lighthouse Lab in Milton Keynes and Alex Alderton, Roberto Amato, Jeffrey Barrett, Sonia Goncalves, Ewan Harrison, David K. Jackson, Ian Johnston, Dominic Kwiatkowski, Cordelia Langford, John Sillitoe on behalf of the Wellcome Sanger Institute COVID-19 Surveillance Team |
| EPI_ISL_2963261, EPI_ISL_2963262 | Lighthouse Laboratory Plymouth | Wellcome Sanger Institute for the COVID-19 Genomics UK (COG-UK) Consortium | Lighthouse Laboratory Plymouth and Alex Alderton, Roberto Amato, Jeffrey Barrett, Sonia Goncalves, Ewan Harrison, David K. Jackson, Ian Johnston, Dominic Kwiatkowski, Cordelia Langford, John Sillitoe on behalf of the Wellcome Sanger Institute COVID-19 Surveillance Team |
| EPI_ISL_2963263 | Lighthouse Lab in Milton Keynes | Wellcome Sanger Institute for the COVID-19 Genomics UK (COG-UK) Consortium | The Lighthouse Lab in Milton Keynes and Alex Alderton, Roberto Amato, Jeffrey Barrett, Sonia Goncalves, Ewan Harrison, David K. Jackson, Ian Johnston, Dominic Kwiatkowski, Cordelia Langford, John Sillitoe on behalf of the Wellcome Sanger Institute COVID-19 Surveillance Team |
| EPI_ISL_2963264 | Health Services Laboratories | Wellcome Sanger Institute for the COVID-19 Genomics UK (COG-UK) Consortium | Health Services Laboratories and Alex Alderton, Roberto Amato, Jeffrey Barrett, Sonia Goncalves, Ewan Harrison, David K. Jackson, Ian Johnston, Dominic Kwiatkowski, Cordelia Langford, John Sillitoe on behalf of the Wellcome Sanger Institute COVID-19 Surveillance Team |
| EPI_ISL_2963265 | Lighthouse Lab in Milton Keynes | Wellcome Sanger Institute for the COVID-19 Genomics UK (COG-UK) Consortium | The Lighthouse Lab in Milton Keynes and Alex Alderton, Roberto Amato, Jeffrey Barrett, Sonia Goncalves, Ewan Harrison, David K. Jackson, Ian Johnston, Dominic Kwiatkowski, Cordelia Langford, John Sillitoe on behalf of the Wellcome Sanger Institute COVID-19 Surveillance Team |
| EPI_ISL_2963266 | Lighthouse Laboratory Plymouth | Wellcome Sanger Institute for the COVID-19 Genomics UK (COG-UK) Consortium | Lighthouse Laboratory Plymouth and Alex Alderton, Roberto Amato, Jeffrey Barrett, Sonia Goncalves, Ewan Harrison, David K. Jackson, Ian Johnston, Dominic Kwiatkowski, Cordelia Langford, John Sillitoe on behalf of the Wellcome Sanger Institute COVID-19 Surveillance Team |
| EPI_ISL_2963267, EPI_ISL_2963268 | Lighthouse Lab in Milton Keynes | Wellcome Sanger Institute for the COVID-19 Genomics UK (COG-UK) Consortium | The Lighthouse Lab in Milton Keynes and Alex Alderton, Roberto Amato, Jeffrey Barrett, Sonia Goncalves, Ewan Harrison, David K. Jackson, Ian Johnston, Dominic Kwiatkowski, Cordelia Langford, John Sillitoe on behalf of the Wellcome Sanger Institute COVID-19 Surveillance Team |
| EPI_ISL_2963269, EPI_ISL_2963270, EPI_ISL_2963271 | Lighthouse Laboratory Plymouth | Wellcome Sanger Institute for the COVID-19 Genomics UK (COG-UK) Consortium | Lighthouse Laboratory Plymouth and Alex Alderton, Roberto Amato, Jeffrey Barrett, Sonia Goncalves, Ewan Harrison, David K. Jackson, Ian Johnston, Dominic Kwiatkowski, Cordelia Langford, John Sillitoe on behalf of the Wellcome Sanger Institute COVID-19 Surveillance Team |
| EPI_ISL_2963272 | Lighthouse Lab in Milton Keynes | Wellcome Sanger Institute for the COVID-19 Genomics UK (COG-UK) Consortium | The Lighthouse Lab in Milton Keynes and Alex Alderton, Roberto Amato, Jeffrey Barrett, Sonia Goncalves, Ewan Harrison, David K. Jackson, Ian Johnston, Dominic Kwiatkowski, Cordelia Langford, John Sillitoe on behalf of the Wellcome Sanger Institute COVID-19 Surveillance Team |
| EPI_ISL_2963273 | Lighthouse Laboratory Plymouth | Wellcome Sanger Institute for the COVID-19 Genomics UK (COG-UK) Consortium | Lighthouse Laboratory Plymouth and Alex Alderton, Roberto Amato, Jeffrey Barrett, Sonia Goncalves, Ewan Harrison, David K. Jackson, Ian Johnston, Dominic Kwiatkowski, Cordelia Langford, John Sillitoe on behalf of the Wellcome Sanger Institute COVID-19 Surveillance Team |
| EPI_ISL_2963274 | Lighthouse Lab in Milton Keynes | Wellcome Sanger Institute for the COVID-19 Genomics UK (COG-UK) Consortium | The Lighthouse Lab in Milton Keynes and Alex Alderton, Roberto Amato, Jeffrey Barrett, Sonia Goncalves, Ewan Harrison, David K. Jackson, Ian Johnston, Dominic Kwiatkowski, Cordelia Langford, John Sillitoe on behalf of the Wellcome Sanger Institute COVID-19 Surveillance Team |
| EPI_ISL_2963275, EPI_ISL_2963276 | Lighthouse Laboratory Plymouth | Wellcome Sanger Institute for the COVID-19 Genomics UK (COG-UK) Consortium | Lighthouse Laboratory Plymouth and Alex Alderton, Roberto Amato, Jeffrey Barrett, Sonia Goncalves, Ewan Harrison, David K. Jackson, Ian Johnston, Dominic Kwiatkowski, Cordelia Langford, John Sillitoe on behalf of the Wellcome Sanger Institute COVID-19 Surveillance Team |
| EPI_ISL_2963277 | Health Services Laboratories | Wellcome Sanger Institute for the COVID-19 Genomics UK (COG-UK) Consortium | Health Services Laboratories and Alex Alderton, Roberto Amato, Jeffrey Barrett, Sonia Goncalves, Ewan Harrison, David K. Jackson, Ian Johnston, Dominic Kwiatkowski, Cordelia Langford, John Sillitoe on behalf of the Wellcome Sanger Institute COVID-19 Surveillance Team |
| EPI_ISL_2963278 | Lighthouse Lab in Milton Keynes | Wellcome Sanger Institute for the COVID-19 Genomics UK (COG-UK) Consortium | The Lighthouse Lab in Milton Keynes and Alex Alderton, Roberto Amato, Jeffrey Barrett, Sonia Goncalves, Ewan Harrison, David K. Jackson, Ian Johnston, Dominic Kwiatkowski, Cordelia Langford, John Sillitoe on behalf of the Wellcome Sanger Institute COVID-19 Surveillance Team |
| EPI_ISL_2963279 | Health Services Laboratories | Wellcome Sanger Institute for the COVID-19 Genomics UK (COG-UK) Consortium | Health Services Laboratories and Alex Alderton, Roberto Amato, Jeffrey Barrett, Sonia Goncalves, Ewan Harrison, David K. Jackson, Ian Johnston, Dominic Kwiatkowski, Cordelia Langford, John Sillitoe on behalf of the Wellcome Sanger Institute COVID-19 Surveillance Team |
| EPI_ISL_2963280, EPI_ISL_2963281, EPI_ISL_2963282 | Lighthouse Laboratory Plymouth | Wellcome Sanger Institute for the COVID-19 Genomics UK (COG-UK) Consortium | Lighthouse Laboratory Plymouth and Alex Alderton, Roberto Amato, Jeffrey Barrett, Sonia Goncalves, Ewan Harrison, David K. Jackson, Ian Johnston, Dominic Kwiatkowski, Cordelia Langford, John Sillitoe on behalf of the Wellcome Sanger Institute COVID-19 Surveillance Team |
| EPI_ISL_2963283, EPI_ISL_2963284 | Lighthouse Lab in Milton Keynes | Wellcome Sanger Institute for the COVID-19 Genomics UK (COG-UK) Consortium | The Lighthouse Lab in Milton Keynes and Alex Alderton, Roberto Amato, Jeffrey Barrett, Sonia Goncalves, Ewan Harrison, David K. Jackson, Ian Johnston, Dominic Kwiatkowski, Cordelia Langford, John Sillitoe on behalf of the Wellcome Sanger Institute COVID-19 Surveillance Team |
| EPI_ISL_2963285 | Lighthouse Laboratory Plymouth | Wellcome Sanger Institute for the COVID-19 Genomics UK (COG-UK) Consortium | Lighthouse Laboratory Plymouth and Alex Alderton, Roberto Amato, Jeffrey Barrett, Sonia Goncalves, Ewan Harrison, David K. Jackson, Ian Johnston, Dominic Kwiatkowski, Cordelia Langford, John Sillitoe on behalf of the Wellcome Sanger Institute COVID-19 Surveillance Team |
| EPI_ISL_2963286 | Health Services Laboratories | Wellcome Sanger Institute for the COVID-19 Genomics UK (COG-UK) Consortium | Health Services Laboratories and Alex Alderton, Roberto Amato, Jeffrey Barrett, Sonia Goncalves, Ewan Harrison, David K. Jackson, Ian Johnston, Dominic Kwiatkowski, Cordelia Langford, John Sillitoe on behalf of the Wellcome Sanger Institute COVID-19 Surveillance Team |
| EPI_ISL_2963287 | Lighthouse Laboratory Plymouth | Wellcome Sanger Institute for the COVID-19 Genomics UK (COG-UK) Consortium | Lighthouse Laboratory Plymouth and Alex Alderton, Roberto Amato, Jeffrey Barrett, Sonia Goncalves, Ewan Harrison, David K. Jackson, Ian Johnston, Dominic Kwiatkowski, Cordelia Langford, John Sillitoe on behalf of the Wellcome Sanger Institute COVID-19 Surveillance Team |
| EPI_ISL_2963288, | Lighthouse Lab in Milton Keynes | Wellcome Sanger Institute for the COVID-19 Genomics UK (COG-UK) | The Lighthouse Lab in Milton Keynes and Alex Alderton, Roberto Amato, Jeffrey Barrett, Sonia Goncalves, Ewan Harrison, David K. Jackson, Ian Johnston, Dominic Kwiatkowski, Cordelia Langford, John Sillitoe on behalf of the Wellcome Sanger |

| | | | |
|---|---|---|---|
| EPI_ISL_2963289, EPI_ISL_2963290, EPI_ISL_2963291, EPI_ISL_2963292, EPI_ISL_2963293, EPI_ISL_2963294 | | Consortium | Institute COVID-19 Surveillance Team |
| EPI_ISL_2963295 | Health Services Laboratories | Wellcome Sanger Institute for the COVID-19 Genomics UK (COG-UK) Consortium | Health Services Laboratories and Alex Alderton, Roberto Amato, Jeffrey Barrett, Sonia Goncalves, Ewan Harrison, David K. Jackson, Ian Johnston, Dominic Kwiatkowski, Cordelia Langford, John Sillitoe on behalf of the Wellcome Sanger Institute COVID-19 Surveillance Team |
| EPI_ISL_2963296 | Lighthouse Laboratory Plymouth | Wellcome Sanger Institute for the COVID-19 Genomics UK (COG-UK) Consortium | Lighthouse Laboratory Plymouth and Alex Alderton, Roberto Amato, Jeffrey Barrett, Sonia Goncalves, Ewan Harrison, David K. Jackson, Ian Johnston, Dominic Kwiatkowski, Cordelia Langford, John Sillitoe on behalf of the Wellcome Sanger Institute COVID-19 Surveillance Team |
| EPI_ISL_2963297 | Lighthouse Lab in Milton Keynes | Wellcome Sanger Institute for the COVID-19 Genomics UK (COG-UK) Consortium | The Lighthouse Lab in Milton Keynes and Alex Alderton, Roberto Amato, Jeffrey Barrett, Sonia Goncalves, Ewan Harrison, David K. Jackson, Ian Johnston, Dominic Kwiatkowski, Cordelia Langford, John Sillitoe on behalf of the Wellcome Sanger Institute COVID-19 Surveillance Team |
| EPI_ISL_2963298 | Lighthouse Laboratory Plymouth | Wellcome Sanger Institute for the COVID-19 Genomics UK (COG-UK) Consortium | Lighthouse Laboratory Plymouth and Alex Alderton, Roberto Amato, Jeffrey Barrett, Sonia Goncalves, Ewan Harrison, David K. Jackson, Ian Johnston, Dominic Kwiatkowski, Cordelia Langford, John Sillitoe on behalf of the Wellcome Sanger Institute COVID-19 Surveillance Team |
| EPI_ISL_2963299, EPI_ISL_2963300 | Lighthouse Lab in Milton Keynes | Wellcome Sanger Institute for the COVID-19 Genomics UK (COG-UK) Consortium | The Lighthouse Lab in Milton Keynes and Alex Alderton, Roberto Amato, Jeffrey Barrett, Sonia Goncalves, Ewan Harrison, David K. Jackson, Ian Johnston, Dominic Kwiatkowski, Cordelia Langford, John Sillitoe on behalf of the Wellcome Sanger Institute COVID-19 Surveillance Team |
| EPI_ISL_2963301 | Health Services Laboratories | Wellcome Sanger Institute for the COVID-19 Genomics UK (COG-UK) Consortium | Health Services Laboratories and Alex Alderton, Roberto Amato, Jeffrey Barrett, Sonia Goncalves, Ewan Harrison, David K. Jackson, Ian Johnston, Dominic Kwiatkowski, Cordelia Langford, John Sillitoe on behalf of the Wellcome Sanger Institute COVID-19 Surveillance Team |
| EPI_ISL_2963302, EPI_ISL_2963303 | Lighthouse Lab in Milton Keynes | Wellcome Sanger Institute for the COVID-19 Genomics UK (COG-UK) Consortium | The Lighthouse Lab in Milton Keynes and Alex Alderton, Roberto Amato, Jeffrey Barrett, Sonia Goncalves, Ewan Harrison, David K. Jackson, Ian Johnston, Dominic Kwiatkowski, Cordelia Langford, John Sillitoe on behalf of the Wellcome Sanger Institute COVID-19 Surveillance Team |
| EPI_ISL_2963304 | Lighthouse Laboratory Plymouth | Wellcome Sanger Institute for the COVID-19 Genomics UK (COG-UK) Consortium | Lighthouse Laboratory Plymouth and Alex Alderton, Roberto Amato, Jeffrey Barrett, Sonia Goncalves, Ewan Harrison, David K. Jackson, Ian Johnston, Dominic Kwiatkowski, Cordelia Langford, John Sillitoe on behalf of the Wellcome Sanger Institute COVID-19 Surveillance Team |
| EPI_ISL_2963305, EPI_ISL_2963306 | Lighthouse Lab in Milton Keynes | Wellcome Sanger Institute for the COVID-19 Genomics UK (COG-UK) Consortium | The Lighthouse Lab in Milton Keynes and Alex Alderton, Roberto Amato, Jeffrey Barrett, Sonia Goncalves, Ewan Harrison, David K. Jackson, Ian Johnston, Dominic Kwiatkowski, Cordelia Langford, John Sillitoe on behalf of the Wellcome Sanger Institute COVID-19 Surveillance Team |
| EPI_ISL_2963307, EPI_ISL_2963308 | Lighthouse Laboratory Plymouth | Wellcome Sanger Institute for the COVID-19 Genomics UK (COG-UK) Consortium | Lighthouse Laboratory Plymouth and Alex Alderton, Roberto Amato, Jeffrey Barrett, Sonia Goncalves, Ewan Harrison, David K. Jackson, Ian Johnston, Dominic Kwiatkowski, Cordelia Langford, John Sillitoe on behalf of the Wellcome Sanger Institute COVID-19 Surveillance Team |
| EPI_ISL_2963309, EPI_ISL_2963310 | Lighthouse Lab in Milton Keynes | Wellcome Sanger Institute for the COVID-19 Genomics UK (COG-UK) Consortium | The Lighthouse Lab in Milton Keynes and Alex Alderton, Roberto Amato, Jeffrey Barrett, Sonia Goncalves, Ewan Harrison, David K. Jackson, Ian Johnston, Dominic Kwiatkowski, Cordelia Langford, John Sillitoe on behalf of the Wellcome Sanger Institute COVID-19 Surveillance Team |
| EPI_ISL_2963311, EPI_ISL_2963312 | Health Services Laboratories | Wellcome Sanger Institute for the COVID-19 Genomics UK (COG-UK) Consortium | Health Services Laboratories and Alex Alderton, Roberto Amato, Jeffrey Barrett, Sonia Goncalves, Ewan Harrison, David K. Jackson, Ian Johnston, Dominic Kwiatkowski, Cordelia Langford, John Sillitoe on behalf of the Wellcome Sanger Institute COVID-19 Surveillance Team |
| EPI_ISL_2963313 | Lighthouse Lab in Milton Keynes | Wellcome Sanger Institute for the COVID-19 Genomics UK (COG-UK) Consortium | The Lighthouse Lab in Milton Keynes and Alex Alderton, Roberto Amato, Jeffrey Barrett, Sonia Goncalves, Ewan Harrison, David K. Jackson, Ian Johnston, Dominic Kwiatkowski, Cordelia Langford, John Sillitoe on behalf of the Wellcome Sanger Institute COVID-19 Surveillance Team |
| EPI_ISL_2963314 | Lighthouse Laboratory Plymouth | Wellcome Sanger Institute for the COVID-19 Genomics UK (COG-UK) Consortium | Lighthouse Laboratory Plymouth and Alex Alderton, Roberto Amato, Jeffrey Barrett, Sonia Goncalves, Ewan Harrison, David K. Jackson, Ian Johnston, Dominic Kwiatkowski, Cordelia Langford, John Sillitoe on behalf of the Wellcome Sanger Institute COVID-19 Surveillance Team |
| EPI_ISL_2963315 | Lighthouse Lab in Milton Keynes | Wellcome Sanger Institute for the COVID-19 Genomics UK (COG-UK) Consortium | The Lighthouse Lab in Milton Keynes and Alex Alderton, Roberto Amato, Jeffrey Barrett, Sonia Goncalves, Ewan Harrison, David K. Jackson, Ian Johnston, Dominic Kwiatkowski, Cordelia Langford, John Sillitoe on behalf of the Wellcome Sanger Institute COVID-19 Surveillance Team |
| EPI_ISL_2963316 | Lighthouse Laboratory Plymouth | Wellcome Sanger Institute for the COVID-19 Genomics UK (COG-UK) Consortium | Lighthouse Laboratory Plymouth and Alex Alderton, Roberto Amato, Jeffrey Barrett, Sonia Goncalves, Ewan Harrison, David K. Jackson, Ian Johnston, Dominic Kwiatkowski, Cordelia Langford, John Sillitoe on behalf of the Wellcome Sanger Institute COVID-19 Surveillance Team |
| EPI_ISL_2963317, EPI_ISL_2963318 | Lighthouse Lab in Milton Keynes | Wellcome Sanger Institute for the COVID-19 Genomics UK (COG-UK) Consortium | The Lighthouse Lab in Milton Keynes and Alex Alderton, Roberto Amato, Jeffrey Barrett, Sonia Goncalves, Ewan Harrison, David K. Jackson, Ian Johnston, Dominic Kwiatkowski, Cordelia Langford, John Sillitoe on behalf of the Wellcome Sanger Institute COVID-19 Surveillance Team |
| EPI_ISL_2963319 | Health Services Laboratories | Wellcome Sanger Institute for the COVID-19 Genomics UK (COG-UK) Consortium | Health Services Laboratories and Alex Alderton, Roberto Amato, Jeffrey Barrett, Sonia Goncalves, Ewan Harrison, David K. Jackson, Ian Johnston, Dominic Kwiatkowski, Cordelia Langford, John Sillitoe on behalf of the Wellcome Sanger Institute COVID-19 Surveillance Team |
| EPI_ISL_2963320 | Lighthouse Lab in Milton Keynes | Wellcome Sanger Institute for the COVID-19 Genomics UK (COG-UK) Consortium | The Lighthouse Lab in Milton Keynes and Alex Alderton, Roberto Amato, Jeffrey Barrett, Sonia Goncalves, Ewan Harrison, David K. Jackson, Ian Johnston, Dominic Kwiatkowski, Cordelia Langford, John Sillitoe on behalf of the Wellcome Sanger Institute COVID-19 Surveillance Team |
| EPI_ISL_2963321 | Health Services Laboratories | Wellcome Sanger Institute for the COVID-19 Genomics UK (COG-UK) Consortium | Health Services Laboratories and Alex Alderton, Roberto Amato, Jeffrey Barrett, Sonia Goncalves, Ewan Harrison, David K. Jackson, Ian Johnston, Dominic Kwiatkowski, Cordelia Langford, John Sillitoe on behalf of the Wellcome Sanger Institute COVID-19 Surveillance Team |
| EPI_ISL_2963322 | Lighthouse Lab in Milton Keynes | Wellcome Sanger Institute for the COVID-19 Genomics UK (COG-UK) Consortium | The Lighthouse Lab in Milton Keynes and Alex Alderton, Roberto Amato, Jeffrey Barrett, Sonia Goncalves, Ewan Harrison, David K. Jackson, Ian Johnston, Dominic Kwiatkowski, Cordelia Langford, John Sillitoe on behalf of the Wellcome Sanger Institute COVID-19 Surveillance Team |
| EPI_ISL_2963323 | Lighthouse Laboratory Plymouth | Wellcome Sanger Institute for the COVID-19 Genomics UK (COG-UK) Consortium | Lighthouse Laboratory Plymouth and Alex Alderton, Roberto Amato, Jeffrey Barrett, Sonia Goncalves, Ewan Harrison, David K. Jackson, Ian Johnston, Dominic Kwiatkowski, Cordelia Langford, John Sillitoe on behalf of the Wellcome Sanger Institute COVID-19 Surveillance Team |
| EPI_ISL_2963324 | Lighthouse Lab in Milton Keynes | Wellcome Sanger Institute for the COVID-19 Genomics UK (COG-UK) Consortium | The Lighthouse Lab in Milton Keynes and Alex Alderton, Roberto Amato, Jeffrey Barrett, Sonia Goncalves, Ewan Harrison, David K. Jackson, Ian Johnston, Dominic Kwiatkowski, Cordelia Langford, John Sillitoe on behalf of the Wellcome Sanger Institute COVID-19 Surveillance Team |
| EPI_ISL_2963325 | Lighthouse Laboratory Plymouth | Wellcome Sanger Institute for the COVID-19 Genomics UK (COG-UK) Consortium | Lighthouse Laboratory Plymouth and Alex Alderton, Roberto Amato, Jeffrey Barrett, Sonia Goncalves, Ewan Harrison, David K. Jackson, Ian Johnston, Dominic Kwiatkowski, Cordelia Langford, John Sillitoe on behalf of the Wellcome Sanger Institute COVID-19 Surveillance Team |
| EPI_ISL_2963326 | Health Services Laboratories | Wellcome Sanger Institute for the COVID-19 Genomics UK (COG-UK) Consortium | Health Services Laboratories and Alex Alderton, Roberto Amato, Jeffrey Barrett, Sonia Goncalves, Ewan Harrison, David K. Jackson, Ian Johnston, Dominic Kwiatkowski, Cordelia Langford, John Sillitoe on behalf of the Wellcome Sanger Institute COVID-19 Surveillance Team |
| EPI_ISL_2963327 | Lighthouse Laboratory Plymouth | Wellcome Sanger Institute for the COVID-19 Genomics UK (COG-UK) Consortium | Lighthouse Laboratory Plymouth and Alex Alderton, Roberto Amato, Jeffrey Barrett, Sonia Goncalves, Ewan Harrison, David K. Jackson, Ian Johnston, Dominic Kwiatkowski, Cordelia Langford, John Sillitoe on behalf of the Wellcome Sanger Institute COVID-19 Surveillance Team |
| EPI_ISL_2963328 | Health Services Laboratories | Wellcome Sanger Institute for the COVID-19 Genomics UK (COG-UK) Consortium | Health Services Laboratories and Alex Alderton, Roberto Amato, Jeffrey Barrett, Sonia Goncalves, Ewan Harrison, David K. Jackson, Ian Johnston, Dominic Kwiatkowski, Cordelia Langford, John Sillitoe on behalf of the Wellcome Sanger Institute COVID-19 Surveillance Team |
| EPI_ISL_2963329 | Lighthouse Laboratory Plymouth | Wellcome Sanger Institute for the COVID-19 Genomics UK (COG-UK) Consortium | Lighthouse Laboratory Plymouth and Alex Alderton, Roberto Amato, Jeffrey Barrett, Sonia Goncalves, Ewan Harrison, David K. Jackson, Ian Johnston, Dominic Kwiatkowski, Cordelia Langford, John Sillitoe on behalf of the Wellcome Sanger Institute COVID-19 Surveillance Team |
| EPI_ISL_2963330 | Health Services Laboratories | Wellcome Sanger Institute for the COVID-19 Genomics UK (COG-UK) Consortium | Health Services Laboratories and Alex Alderton, Roberto Amato, Jeffrey Barrett, Sonia Goncalves, Ewan Harrison, David K. Jackson, Ian Johnston, Dominic Kwiatkowski, Cordelia Langford, John Sillitoe on behalf of the Wellcome Sanger Institute COVID-19 Surveillance Team |
| EPI_ISL_2963331 | Lighthouse Laboratory Plymouth | Wellcome Sanger Institute for the COVID-19 Genomics UK (COG-UK) Consortium | Lighthouse Laboratory Plymouth and Alex Alderton, Roberto Amato, Jeffrey Barrett, Sonia Goncalves, Ewan Harrison, David K. Jackson, Ian Johnston, Dominic Kwiatkowski, Cordelia Langford, John Sillitoe on behalf of the Wellcome Sanger Institute COVID-19 Surveillance Team |
| EPI_ISL_2963332 | Lighthouse Lab in Milton Keynes | Wellcome Sanger Institute for the COVID-19 Genomics UK (COG-UK) Consortium | The Lighthouse Lab in Milton Keynes and Alex Alderton, Roberto Amato, Jeffrey Barrett, Sonia Goncalves, Ewan Harrison, David K. Jackson, Ian Johnston, Dominic Kwiatkowski, Cordelia Langford, John Sillitoe on behalf of the Wellcome Sanger Institute COVID-19 Surveillance Team |
| EPI_ISL_2963333 | Health Services Laboratories | Wellcome Sanger Institute for the COVID-19 Genomics UK (COG-UK) Consortium | Health Services Laboratories and Alex Alderton, Roberto Amato, Jeffrey Barrett, Sonia Goncalves, Ewan Harrison, David K. Jackson, Ian Johnston, Dominic Kwiatkowski, Cordelia Langford, John Sillitoe on behalf of the Wellcome Sanger Institute COVID-19 Surveillance Team |
| EPI_ISL_2963334, EPI_ISL_2963335, EPI_ISL_2963336 | Lighthouse Lab in Milton Keynes | Wellcome Sanger Institute for the COVID-19 Genomics UK (COG-UK) Consortium | The Lighthouse Lab in Milton Keynes and Alex Alderton, Roberto Amato, Jeffrey Barrett, Sonia Goncalves, Ewan Harrison, David K. Jackson, Ian Johnston, Dominic Kwiatkowski, Cordelia Langford, John Sillitoe on behalf of the Wellcome Sanger Institute COVID-19 Surveillance Team |
| EPI_ISL_2963337 | Lighthouse Laboratory Plymouth | Wellcome Sanger Institute for the COVID-19 Genomics UK (COG-UK) Consortium | Lighthouse Laboratory Plymouth and Alex Alderton, Roberto Amato, Jeffrey Barrett, Sonia Goncalves, Ewan Harrison, David K. Jackson, Ian Johnston, Dominic Kwiatkowski, Cordelia Langford, John Sillitoe on behalf of the Wellcome Sanger Institute COVID-19 Surveillance Team |
| EPI_ISL_2963338, EPI_ISL_2963339 | Lighthouse Lab in Milton Keynes | Wellcome Sanger Institute for the COVID-19 Genomics UK (COG-UK) Consortium | The Lighthouse Lab in Milton Keynes and Alex Alderton, Roberto Amato, Jeffrey Barrett, Sonia Goncalves, Ewan Harrison, David K. Jackson, Ian Johnston, Dominic Kwiatkowski, Cordelia Langford, John Sillitoe on behalf of the Wellcome Sanger Institute COVID-19 Surveillance Team |
| EPI_ISL_2963340 | Lighthouse Laboratory Plymouth | Wellcome Sanger Institute for the COVID-19 Genomics UK (COG-UK) Consortium | Lighthouse Laboratory Plymouth and Alex Alderton, Roberto Amato, Jeffrey Barrett, Sonia Goncalves, Ewan Harrison, David K. Jackson, Ian Johnston, Dominic Kwiatkowski, Cordelia Langford, John Sillitoe on behalf of the Wellcome Sanger Institute COVID-19 Surveillance Team |
| EPI_ISL_2963341, EPI_ISL_2963342, EPI_ISL_2963343, EPI_ISL_2963344 | Lighthouse Lab in Milton Keynes | Wellcome Sanger Institute for the COVID-19 Genomics UK (COG-UK) Consortium | The Lighthouse Lab in Milton Keynes and Alex Alderton, Roberto Amato, Jeffrey Barrett, Sonia Goncalves, Ewan Harrison, David K. Jackson, Ian Johnston, Dominic Kwiatkowski, Cordelia Langford, John Sillitoe on behalf of the Wellcome Sanger Institute COVID-19 Surveillance Team |
| EPI_ISL_2963345 | Health Services Laboratories | Wellcome Sanger Institute for the COVID-19 Genomics UK (COG-UK) Consortium | Health Services Laboratories and Alex Alderton, Roberto Amato, Jeffrey Barrett, Sonia Goncalves, Ewan Harrison, David K. Jackson, Ian Johnston, Dominic Kwiatkowski, Cordelia Langford, John Sillitoe on behalf of the Wellcome Sanger Institute COVID-19 Surveillance Team |
| EPI_ISL_2963346, EPI_ISL_2963347, EPI_ISL_2963348 | Lighthouse Lab in Milton Keynes | Wellcome Sanger Institute for the COVID-19 Genomics UK (COG-UK) Consortium | The Lighthouse Lab in Milton Keynes and Alex Alderton, Roberto Amato, Jeffrey Barrett, Sonia Goncalves, Ewan Harrison, David K. Jackson, Ian Johnston, Dominic Kwiatkowski, Cordelia Langford, John Sillitoe on behalf of the Wellcome Sanger Institute COVID-19 Surveillance Team |
| EPI_ISL_2963349 | Health Services Laboratories | Wellcome Sanger Institute for the COVID-19 Genomics UK (COG-UK) Consortium | Health Services Laboratories and Alex Alderton, Roberto Amato, Jeffrey Barrett, Sonia Goncalves, Ewan Harrison, David K. Jackson, Ian Johnston, Dominic Kwiatkowski, Cordelia Langford, John Sillitoe on behalf of the Wellcome Sanger Institute COVID-19 Surveillance Team |
| EPI_ISL_2963350, EPI_ISL_2963351 | Lighthouse Lab in Milton Keynes | Wellcome Sanger Institute for the COVID-19 Genomics UK (COG-UK) Consortium | The Lighthouse Lab in Milton Keynes and Alex Alderton, Roberto Amato, Jeffrey Barrett, Sonia Goncalves, Ewan Harrison, David K. Jackson, Ian Johnston, Dominic Kwiatkowski, Cordelia Langford, John Sillitoe on behalf of the Wellcome Sanger Institute COVID-19 Surveillance Team |

| | | | |
|---|---|---|---|
| EPI_ISL_2963352 | Lighthouse Laboratory Plymouth | Wellcome Sanger Institute for the COVID-19 Genomics UK (COG-UK) Consortium | Lighthouse Laboratory Plymouth and Alex Alderton, Roberto Amato, Jeffrey Barrett, Sonia Goncalves, Ewan Harrison, David K. Jackson, Ian Johnston, Dominic Kwiatkowski, Cordelia Langford, John Sillitoe on behalf of the Wellcome Sanger Institute COVID-19 Surveillance Team |
| EPI_ISL_2963353, EPI_ISL_2963354, EPI_ISL_2963355, EPI_ISL_2963356, EPI_ISL_2963357 | Lighthouse Lab in Milton Keynes | Wellcome Sanger Institute for the COVID-19 Genomics UK (COG-UK) Consortium | The Lighthouse Lab in Milton Keynes and Alex Alderton, Roberto Amato, Jeffrey Barrett, Sonia Goncalves, Ewan Harrison, David K. Jackson, Ian Johnston, Dominic Kwiatkowski, Cordelia Langford, John Sillitoe on behalf of the Wellcome Sanger Institute COVID-19 Surveillance Team |
| EPI_ISL_2963358 | Health Services Laboratories | Wellcome Sanger Institute for the COVID-19 Genomics UK (COG-UK) Consortium | Health Services Laboratories and Alex Alderton, Roberto Amato, Jeffrey Barrett, Sonia Goncalves, Ewan Harrison, David K. Jackson, Ian Johnston, Dominic Kwiatkowski, Cordelia Langford, John Sillitoe on behalf of the Wellcome Sanger Institute COVID-19 Surveillance Team |
| EPI_ISL_2963359 | Lighthouse Lab in Milton Keynes | Wellcome Sanger Institute for the COVID-19 Genomics UK (COG-UK) Consortium | The Lighthouse Lab in Milton Keynes and Alex Alderton, Roberto Amato, Jeffrey Barrett, Sonia Goncalves, Ewan Harrison, David K. Jackson, Ian Johnston, Dominic Kwiatkowski, Cordelia Langford, John Sillitoe on behalf of the Wellcome Sanger Institute COVID-19 Surveillance Team |
| EPI_ISL_2963360 | Health Services Laboratories | Wellcome Sanger Institute for the COVID-19 Genomics UK (COG-UK) Consortium | Health Services Laboratories and Alex Alderton, Roberto Amato, Jeffrey Barrett, Sonia Goncalves, Ewan Harrison, David K. Jackson, Ian Johnston, Dominic Kwiatkowski, Cordelia Langford, John Sillitoe on behalf of the Wellcome Sanger Institute COVID-19 Surveillance Team |
| EPI_ISL_2963361, EPI_ISL_2963362 | Lighthouse Lab in Milton Keynes | Wellcome Sanger Institute for the COVID-19 Genomics UK (COG-UK) Consortium | The Lighthouse Lab in Milton Keynes and Alex Alderton, Roberto Amato, Jeffrey Barrett, Sonia Goncalves, Ewan Harrison, David K. Jackson, Ian Johnston, Dominic Kwiatkowski, Cordelia Langford, John Sillitoe on behalf of the Wellcome Sanger Institute COVID-19 Surveillance Team |
| EPI_ISL_2963363 | Health Services Laboratories | Wellcome Sanger Institute for the COVID-19 Genomics UK (COG-UK) Consortium | Health Services Laboratories and Alex Alderton, Roberto Amato, Jeffrey Barrett, Sonia Goncalves, Ewan Harrison, David K. Jackson, Ian Johnston, Dominic Kwiatkowski, Cordelia Langford, John Sillitoe on behalf of the Wellcome Sanger Institute COVID-19 Surveillance Team |
| EPI_ISL_2963364 | Lighthouse Lab in Milton Keynes | Wellcome Sanger Institute for the COVID-19 Genomics UK (COG-UK) Consortium | The Lighthouse Lab in Milton Keynes and Alex Alderton, Roberto Amato, Jeffrey Barrett, Sonia Goncalves, Ewan Harrison, David K. Jackson, Ian Johnston, Dominic Kwiatkowski, Cordelia Langford, John Sillitoe on behalf of the Wellcome Sanger Institute COVID-19 Surveillance Team |
| EPI_ISL_2963365 | Lighthouse Laboratory Plymouth | Wellcome Sanger Institute for the COVID-19 Genomics UK (COG-UK) Consortium | Lighthouse Laboratory Plymouth and Alex Alderton, Roberto Amato, Jeffrey Barrett, Sonia Goncalves, Ewan Harrison, David K. Jackson, Ian Johnston, Dominic Kwiatkowski, Cordelia Langford, John Sillitoe on behalf of the Wellcome Sanger Institute COVID-19 Surveillance Team |
| EPI_ISL_2963366 | Health Services Laboratories | Wellcome Sanger Institute for the COVID-19 Genomics UK (COG-UK) Consortium | Health Services Laboratories and Alex Alderton, Roberto Amato, Jeffrey Barrett, Sonia Goncalves, Ewan Harrison, David K. Jackson, Ian Johnston, Dominic Kwiatkowski, Cordelia Langford, John Sillitoe on behalf of the Wellcome Sanger Institute COVID-19 Surveillance Team |
| EPI_ISL_2963367, EPI_ISL_2963368, EPI_ISL_2963369 | Lighthouse Lab in Milton Keynes | Wellcome Sanger Institute for the COVID-19 Genomics UK (COG-UK) Consortium | The Lighthouse Lab in Milton Keynes and Alex Alderton, Roberto Amato, Jeffrey Barrett, Sonia Goncalves, Ewan Harrison, David K. Jackson, Ian Johnston, Dominic Kwiatkowski, Cordelia Langford, John Sillitoe on behalf of the Wellcome Sanger Institute COVID-19 Surveillance Team |
| EPI_ISL_2963370 | Lighthouse Laboratory Plymouth | Wellcome Sanger Institute for the COVID-19 Genomics UK (COG-UK) Consortium | Lighthouse Laboratory Plymouth and Alex Alderton, Roberto Amato, Jeffrey Barrett, Sonia Goncalves, Ewan Harrison, David K. Jackson, Ian Johnston, Dominic Kwiatkowski, Cordelia Langford, John Sillitoe on behalf of the Wellcome Sanger Institute COVID-19 Surveillance Team |
| EPI_ISL_2963371 | Health Services Laboratories | Wellcome Sanger Institute for the COVID-19 Genomics UK (COG-UK) Consortium | Health Services Laboratories and Alex Alderton, Roberto Amato, Jeffrey Barrett, Sonia Goncalves, Ewan Harrison, David K. Jackson, Ian Johnston, Dominic Kwiatkowski, Cordelia Langford, John Sillitoe on behalf of the Wellcome Sanger Institute COVID-19 Surveillance Team |
| EPI_ISL_2963372 | Lighthouse Lab in Milton Keynes | Wellcome Sanger Institute for the COVID-19 Genomics UK (COG-UK) Consortium | The Lighthouse Lab in Milton Keynes and Alex Alderton, Roberto Amato, Jeffrey Barrett, Sonia Goncalves, Ewan Harrison, David K. Jackson, Ian Johnston, Dominic Kwiatkowski, Cordelia Langford, John Sillitoe on behalf of the Wellcome Sanger Institute COVID-19 Surveillance Team |
| EPI_ISL_2963373 | Berkshire and Surrey Pathology Services Lighthouse Laboratory | Wellcome Sanger Institute for the COVID-19 Genomics UK (COG-UK) Consortium | Berkshire and Surrey Pathology Services Lighthouse Laboratory and Alex Alderton, Roberto Amato, Jeffrey Barrett, Sonia Goncalves, Ewan Harrison, David K. Jackson, Ian Johnston, Dominic Kwiatkowski, Cordelia Langford, John Sillitoe on behalf of the Wellcome Sanger Institute COVID-19 Surveillance Team |
| EPI_ISL_2963374, EPI_ISL_2963375, EPI_ISL_2963376, EPI_ISL_2963377 | Lighthouse Lab in Milton Keynes | Wellcome Sanger Institute for the COVID-19 Genomics UK (COG-UK) Consortium | The Lighthouse Lab in Milton Keynes and Alex Alderton, Roberto Amato, Jeffrey Barrett, Sonia Goncalves, Ewan Harrison, David K. Jackson, Ian Johnston, Dominic Kwiatkowski, Cordelia Langford, John Sillitoe on behalf of the Wellcome Sanger Institute COVID-19 Surveillance Team |
| EPI_ISL_2963378, EPI_ISL_2963379 | Lighthouse Lab in Glasgow | Wellcome Sanger Institute for the COVID-19 Genomics UK (COG-UK) Consortium | Harper VanSteenhouse, Yumi Kasai, David Gray, Carol Clugston, Anna Dominiczak and Alex Alderton, Roberto Amato, Jeffrey Barrett, Sonia Goncalves, Ewan Harrison, David K. Jackson, Ian Johnston, Dominic Kwiatkowski, Cordelia Langford, John Sillitoe on behalf of the Wellcome Sanger Institute COVID-19 Surveillance Team |
| EPI_ISL_2963380 | Lighthouse Lab in Milton Keynes | Wellcome Sanger Institute for the COVID-19 Genomics UK (COG-UK) Consortium | The Lighthouse Lab in Milton Keynes and Alex Alderton, Roberto Amato, Jeffrey Barrett, Sonia Goncalves, Ewan Harrison, David K. Jackson, Ian Johnston, Dominic Kwiatkowski, Cordelia Langford, John Sillitoe on behalf of the Wellcome Sanger Institute COVID-19 Surveillance Team |
| EPI_ISL_2963381 | Lighthouse Lab in Glasgow | Wellcome Sanger Institute for the COVID-19 Genomics UK (COG-UK) Consortium | Harper VanSteenhouse, Yumi Kasai, David Gray, Carol Clugston, Anna Dominiczak and Alex Alderton, Roberto Amato, Jeffrey Barrett, Sonia Goncalves, Ewan Harrison, David K. Jackson, Ian Johnston, Dominic Kwiatkowski, Cordelia Langford, John Sillitoe on behalf of the Wellcome Sanger Institute COVID-19 Surveillance Team |
| EPI_ISL_2963382 | Lighthouse Lab in Milton Keynes | Wellcome Sanger Institute for the COVID-19 Genomics UK (COG-UK) Consortium | The Lighthouse Lab in Milton Keynes and Alex Alderton, Roberto Amato, Jeffrey Barrett, Sonia Goncalves, Ewan Harrison, David K. Jackson, Ian Johnston, Dominic Kwiatkowski, Cordelia Langford, John Sillitoe on behalf of the Wellcome Sanger Institute COVID-19 Surveillance Team |
| EPI_ISL_2963383 | Lighthouse Lab in Glasgow | Wellcome Sanger Institute for the COVID-19 Genomics UK (COG-UK) Consortium | Harper VanSteenhouse, Yumi Kasai, David Gray, Carol Clugston, Anna Dominiczak and Alex Alderton, Roberto Amato, Jeffrey Barrett, Sonia Goncalves, Ewan Harrison, David K. Jackson, Ian Johnston, Dominic Kwiatkowski, Cordelia Langford, John Sillitoe on behalf of the Wellcome Sanger Institute COVID-19 Surveillance Team |
| EPI_ISL_2963384, EPI_ISL_2963385, EPI_ISL_2963386, EPI_ISL_2963387, EPI_ISL_2963388, EPI_ISL_2963389, EPI_ISL_2963390, EPI_ISL_2963391, EPI_ISL_2963392, EPI_ISL_2963393, EPI_ISL_2963394, EPI_ISL_2963395, EPI_ISL_2963396, EPI_ISL_2963397 | | | |
| see above | Lighthouse Lab in Milton Keynes | Wellcome Sanger Institute for the COVID-19 Genomics UK (COG-UK) Consortium | The Lighthouse Lab in Milton Keynes and Alex Alderton, Roberto Amato, Jeffrey Barrett, Sonia Goncalves, Ewan Harrison, David K. Jackson, Ian Johnston, Dominic Kwiatkowski, Cordelia Langford, John Sillitoe on behalf of the Wellcome Sanger Institute COVID-19 Surveillance Team |
| EPI_ISL_2963398 | Lighthouse Lab in Alderley Park | Wellcome Sanger Institute for the COVID-19 Genomics UK (COG-UK) Consortium | Jacquelyn Wynn, Mairead Hyland, The Lighthouse Lab in Alderley Park and Alex Alderton, Roberto Amato, Jeffrey Barrett, Sonia Goncalves, Ewan Harrison, David K. Jackson, Ian Johnston, Dominic Kwiatkowski, Cordelia Langford, John Sillitoe on behalf of the Wellcome Sanger Institute COVID-19 Surveillance Team |
| EPI_ISL_2963399, EPI_ISL_2963400 | Lighthouse Lab in Glasgow | Wellcome Sanger Institute for the COVID-19 Genomics UK (COG-UK) Consortium | Harper VanSteenhouse, Yumi Kasai, David Gray, Carol Clugston, Anna Dominiczak and Alex Alderton, Roberto Amato, Jeffrey Barrett, Sonia Goncalves, Ewan Harrison, David K. Jackson, Ian Johnston, Dominic Kwiatkowski, Cordelia Langford, John Sillitoe on behalf of the Wellcome Sanger Institute COVID-19 Surveillance Team |
| EPI_ISL_2963401, EPI_ISL_2963402, EPI_ISL_2963403 | Lighthouse Lab in Milton Keynes | Wellcome Sanger Institute for the COVID-19 Genomics UK (COG-UK) Consortium | The Lighthouse Lab in Milton Keynes and Alex Alderton, Roberto Amato, Jeffrey Barrett, Sonia Goncalves, Ewan Harrison, David K. Jackson, Ian Johnston, Dominic Kwiatkowski, Cordelia Langford, John Sillitoe on behalf of the Wellcome Sanger Institute COVID-19 Surveillance Team |
| EPI_ISL_2963404 | Lighthouse Lab in Alderley Park | Wellcome Sanger Institute for the COVID-19 Genomics UK (COG-UK) Consortium | Jacquelyn Wynn, Mairead Hyland, The Lighthouse Lab in Alderley Park and Alex Alderton, Roberto Amato, Jeffrey Barrett, Sonia Goncalves, Ewan Harrison, David K. Jackson, Ian Johnston, Dominic Kwiatkowski, Cordelia Langford, John Sillitoe on behalf of the Wellcome Sanger Institute COVID-19 Surveillance Team |
| EPI_ISL_2963405 | Lighthouse Lab in Milton Keynes | Wellcome Sanger Institute for the COVID-19 Genomics UK (COG-UK) Consortium | The Lighthouse Lab in Milton Keynes and Alex Alderton, Roberto Amato, Jeffrey Barrett, Sonia Goncalves, Ewan Harrison, David K. Jackson, Ian Johnston, Dominic Kwiatkowski, Cordelia Langford, John Sillitoe on behalf of the Wellcome Sanger Institute COVID-19 Surveillance Team |
| EPI_ISL_2963406 | Lighthouse Lab in Alderley Park | Wellcome Sanger Institute for the COVID-19 Genomics UK (COG-UK) Consortium | Jacquelyn Wynn, Mairead Hyland, The Lighthouse Lab in Alderley Park and Alex Alderton, Roberto Amato, Jeffrey Barrett, Sonia Goncalves, Ewan Harrison, David K. Jackson, Ian Johnston, Dominic Kwiatkowski, Cordelia Langford, John Sillitoe on behalf of the Wellcome Sanger Institute COVID-19 Surveillance Team |
| EPI_ISL_2963407, EPI_ISL_2963408, EPI_ISL_2963409, EPI_ISL_2963410 | Lighthouse Lab in Milton Keynes | Wellcome Sanger Institute for the COVID-19 Genomics UK (COG-UK) Consortium | The Lighthouse Lab in Milton Keynes and Alex Alderton, Roberto Amato, Jeffrey Barrett, Sonia Goncalves, Ewan Harrison, David K. Jackson, Ian Johnston, Dominic Kwiatkowski, Cordelia Langford, John Sillitoe on behalf of the Wellcome Sanger Institute COVID-19 Surveillance Team |
| EPI_ISL_2963411, EPI_ISL_2963412, EPI_ISL_2963413 | Lighthouse Lab in Alderley Park | Wellcome Sanger Institute for the COVID-19 Genomics UK (COG-UK) Consortium | Jacquelyn Wynn, Mairead Hyland, The Lighthouse Lab in Alderley Park and Alex Alderton, Roberto Amato, Jeffrey Barrett, Sonia Goncalves, Ewan Harrison, David K. Jackson, Ian Johnston, Dominic Kwiatkowski, Cordelia Langford, John Sillitoe on behalf of the Wellcome Sanger Institute COVID-19 Surveillance Team |
| EPI_ISL_2963414, EPI_ISL_2963415, EPI_ISL_2963416 | Lighthouse Lab in Milton Keynes | Wellcome Sanger Institute for the COVID-19 Genomics UK (COG-UK) Consortium | The Lighthouse Lab in Milton Keynes and Alex Alderton, Roberto Amato, Jeffrey Barrett, Sonia Goncalves, Ewan Harrison, David K. Jackson, Ian Johnston, Dominic Kwiatkowski, Cordelia Langford, John Sillitoe on behalf of the Wellcome Sanger Institute COVID-19 Surveillance Team |
| EPI_ISL_2963419, EPI_ISL_2963421 | AULSS 2 Marca Trevigiana | Istituto Zooprofilattico Sperimentale delle Venezie | Adelaide Milani, Alessia Schivo, Annalisa Salviato, Elisa Palumbo, Erika Giorgia Quaranta, Luca Tassoni, Ambra Pastori, Edoardo Giussani, Alice Fusaro, Isabella Monne, Calogero Terregino, Antonia Ricci |
| EPI_ISL_2963427, EPI_ISL_2963429, EPI_ISL_2963430, EPI_ISL_2963431 | AOPD | Istituto Zooprofilattico Sperimentale delle Venezie | Adelaide Milani, Alessia Schivo, Annalisa Salviato, Elisa Palumbo, Erika Giorgia Quaranta, Luca Tassoni, Ambra Pastori, Edoardo Giussani, Alice Fusaro, Isabella Monne, Calogero Terregino, Antonia Ricci |
| EPI_ISL_2963444, EPI_ISL_2963445, EPI_ISL_2963448, EPI_ISL_2963451, EPI_ISL_2963452, EPI_ISL_2963453 | AULSS 2 Marca Trevigiana | Istituto Zooprofilattico Sperimentale delle Venezie | Adelaide Milani, Alessia Schivo, Annalisa Salviato, Elisa Palumbo, Erika Giorgia Quaranta, Luca Tassoni, Ambra Pastori, Edoardo Giussani, Alice Fusaro, Isabella Monne, Calogero Terregino, Antonia Ricci |
| EPI_ISL_2963458 | AULSS 6 Euganea | Istituto Zooprofilattico Sperimentale delle Venezie | Adelaide Milani, Alessia Schivo, Annalisa Salviato, Elisa Palumbo, Erika Giorgia Quaranta, Luca Tassoni, Ambra Pastori, Edoardo Giussani, Alice Fusaro, Isabella Monne, Calogero Terregino, Antonia Ricci |
| EPI_ISL_2963477 | AULSS 5 Polesana | Istituto Zooprofilattico Sperimentale delle Venezie | Adelaide Milani, Alessia Schivo, Annalisa Salviato, Elisa Palumbo, Erika Giorgia Quaranta, Luca Tassoni, Ambra Pastori, Edoardo Giussani, Alice Fusaro, Isabella Monne, Calogero Terregino, Antonia Ricci |
| EPI_ISL_2963478 | AULSS 6 Euganea | Istituto Zooprofilattico Sperimentale delle Venezie | Adelaide Milani, Alessia Schivo, Annalisa Salviato, Elisa Palumbo, Erika Giorgia Quaranta, Luca Tassoni, Ambra Pastori, Edoardo Giussani, Alice Fusaro, Isabella Monne, Calogero Terregino, Antonia Ricci |
| EPI_ISL_2963479, EPI_ISL_2963480, EPI_ISL_2963481 | AOPD | Istituto Zooprofilattico Sperimentale delle Venezie | Adelaide Milani, Alessia Schivo, Annalisa Salviato, Elisa Palumbo, Erika Giorgia Quaranta, Luca Tassoni, Ambra Pastori, Edoardo Giussani, Alice Fusaro, Isabella Monne, Calogero Terregino, Antonia Ricci |

| | | | |
|---|---|---|---|
| EPI_ISL_2963521, EPI_ISL_2963522, EPI_ISL_2963523, EPI_ISL_2963524, EPI_ISL_2963525, EPI_ISL_2963526, EPI_ISL_2963527, EPI_ISL_2963528, EPI_ISL_2963529, EPI_ISL_2963530, EPI_ISL_2963531, EPI_ISL_2963532, EPI_ISL_2963533, EPI_ISL_2963534, EPI_ISL_2963535, EPI_ISL_2963536, EPI_ISL_2963537, EPI_ISL_2963538, EPI_ISL_2963539, EPI_ISL_2963540, EPI_ISL_2963541, EPI_ISL_2963542, EPI_ISL_2963543, EPI_ISL_2963544, EPI_ISL_2963545, EPI_ISL_2963546, EPI_ISL_2963547, EPI_ISL_2963548, EPI_ISL_2963549, EPI_ISL_2963551, EPI_ISL_2963552, EPI_ISL_2963553, EPI_ISL_2963554, EPI_ISL_2963555, EPI_ISL_2963556, EPI_ISL_2963557, EPI_ISL_2963558, EPI_ISL_2963559, EPI_ISL_2963560, EPI_ISL_2963561, EPI_ISL_2963562, EPI_ISL_2963563, EPI_ISL_2963564, EPI_ISL_2963565, EPI_ISL_2963566, EPI_ISL_2963567, EPI_ISL_2963568, EPI_ISL_2963569, EPI_ISL_2963570, EPI_ISL_2963571, EPI_ISL_2963572, EPI_ISL_2963573, EPI_ISL_2963574, EPI_ISL_2963575, EPI_ISL_2963576, EPI_ISL_2963577, EPI_ISL_2963578, EPI_ISL_2963579, EPI_ISL_2963580, EPI_ISL_2963581, EPI_ISL_2963582, EPI_ISL_2963583, EPI_ISL_2963584, EPI_ISL_2963585, EPI_ISL_2963586, EPI_ISL_2963587, EPI_ISL_2963588, EPI_ISL_2963589 | | | |
| see above | Department of Bacteria, Parasites and Fungi, Statens Serum Institut, Copenhagen, Denmark | Statens Serum Institut Bioinformatics and Microbial Genomics | Danish Covid-19 Genome Consortium |
| EPI_ISL_2963590 | Molekylær Medicinsk Afdeling, Aarhus University Hospital, Aarhus, Denmark | Statens Serum Institut Bioinformatics and Microbial Genomics | Danish Covid-19 Genome Consortium |
| EPI_ISL_2963591, EPI_ISL_2963592, EPI_ISL_2963593, EPI_ISL_2963594, EPI_ISL_2963595, EPI_ISL_2963596, EPI_ISL_2963597, EPI_ISL_2963598, EPI_ISL_2963599, EPI_ISL_2963600, EPI_ISL_2963601, EPI_ISL_2963602, EPI_ISL_2963603, EPI_ISL_2963604, EPI_ISL_2963605, EPI_ISL_2963606, EPI_ISL_2963607, EPI_ISL_2963608, EPI_ISL_2963609, EPI_ISL_2963610, EPI_ISL_2963611, EPI_ISL_2963612, EPI_ISL_2963613, EPI_ISL_2963614, EPI_ISL_2963615, EPI_ISL_2963616, EPI_ISL_2963617, EPI_ISL_2963618, EPI_ISL_2963619, EPI_ISL_2963620, EPI_ISL_2963621, EPI_ISL_2963622 | | | |
| see above | Department of Bacteria, Parasites and Fungi, Statens Serum Institut, Copenhagen, Denmark | Statens Serum Institut Bioinformatics and Microbial Genomics | Danish Covid-19 Genome Consortium |
| EPI_ISL_2963623 | Molekylær Medicinsk Afdeling, Aarhus University Hospital, Aarhus, Denmark | Statens Serum Institut Bioinformatics and Microbial Genomics | Danish Covid-19 Genome Consortium |
| EPI_ISL_2963625, EPI_ISL_2963626, EPI_ISL_2963627, EPI_ISL_2963628, EPI_ISL_2963629, EPI_ISL_2963630, EPI_ISL_2963631, EPI_ISL_2963632, EPI_ISL_2963633, EPI_ISL_2963635, EPI_ISL_2963636, EPI_ISL_2963637, EPI_ISL_2963638, EPI_ISL_2963639, EPI_ISL_2963640, EPI_ISL_2963641, EPI_ISL_2963642, EPI_ISL_2963643, EPI_ISL_2963644, EPI_ISL_2963645, EPI_ISL_2963646, EPI_ISL_2963647, EPI_ISL_2963648, EPI_ISL_2963649, EPI_ISL_2963650, EPI_ISL_2963651, EPI_ISL_2963652, EPI_ISL_2963653, EPI_ISL_2963654, EPI_ISL_2963656, EPI_ISL_2963657, EPI_ISL_2963658, EPI_ISL_2963659, EPI_ISL_2963660, EPI_ISL_2963662, EPI_ISL_2963663, EPI_ISL_2963664, EPI_ISL_2963665, EPI_ISL_2963666, EPI_ISL_2963667, EPI_ISL_2963668, EPI_ISL_2963669, EPI_ISL_2963670, EPI_ISL_2963671, EPI_ISL_2963672, EPI_ISL_2963673, EPI_ISL_2963674, EPI_ISL_2963675, EPI_ISL_2963676, EPI_ISL_2963677, EPI_ISL_2963678, EPI_ISL_2963679, EPI_ISL_2963680, EPI_ISL_2963682, EPI_ISL_2963683, EPI_ISL_2963684, EPI_ISL_2963685, EPI_ISL_2963686, EPI_ISL_2963687, EPI_ISL_2963688, EPI_ISL_2963689, EPI_ISL_2963690, EPI_ISL_2963691, EPI_ISL_2963692, EPI_ISL_2963693, EPI_ISL_2963694, EPI_ISL_2963695, EPI_ISL_2963696, EPI_ISL_2963697, EPI_ISL_2963699, EPI_ISL_2963700, EPI_ISL_2963701, EPI_ISL_2963702, EPI_ISL_2963703, EPI_ISL_2963704, EPI_ISL_2963705, EPI_ISL_2963706, EPI_ISL_2963707, EPI_ISL_2963708, EPI_ISL_2963709, EPI_ISL_2963710, EPI_ISL_2963711, EPI_ISL_2963712, EPI_ISL_2963713, EPI_ISL_2963714, EPI_ISL_2963715, EPI_ISL_2963716, EPI_ISL_2963717, EPI_ISL_2963718, EPI_ISL_2963719, EPI_ISL_2963720, EPI_ISL_2963721, EPI_ISL_2963722, EPI_ISL_2963723, EPI_ISL_2963724, EPI_ISL_2963725, EPI_ISL_2963726, EPI_ISL_2963727, EPI_ISL_2963728, EPI_ISL_2963729, EPI_ISL_2963730, EPI_ISL_2963731, EPI_ISL_2963732, EPI_ISL_2963733, EPI_ISL_2963734, EPI_ISL_2963735, EPI_ISL_2963736, EPI_ISL_2963737, EPI_ISL_2963738, EPI_ISL_2963739, EPI_ISL_2963740, EPI_ISL_2963741, EPI_ISL_2963742, EPI_ISL_2963744, EPI_ISL_2963745, EPI_ISL_2963746, EPI_ISL_2963747, EPI_ISL_2963748, EPI_ISL_2963749, EPI_ISL_2963750, EPI_ISL_2963751, EPI_ISL_2963752, EPI_ISL_2963753, EPI_ISL_2963754, EPI_ISL_2963755, EPI_ISL_2963756, EPI_ISL_2963757, EPI_ISL_2963758, EPI_ISL_2963760, EPI_ISL_2963761, EPI_ISL_2963762, EPI_ISL_2963763, EPI_ISL_2963764, EPI_ISL_2963765, EPI_ISL_2963766, EPI_ISL_2963767, EPI_ISL_2963768, EPI_ISL_2963769, EPI_ISL_2963770, EPI_ISL_2963771, EPI_ISL_2963772, EPI_ISL_2963773, EPI_ISL_2963774, EPI_ISL_2963775, EPI_ISL_2963776, EPI_ISL_2963777, EPI_ISL_2963778, EPI_ISL_2963779, EPI_ISL_2963780, EPI_ISL_2963781, EPI_ISL_2963782, EPI_ISL_2963783, EPI_ISL_2963785, EPI_ISL_2963786, EPI_ISL_2963787, EPI_ISL_2963788, EPI_ISL_2963789, EPI_ISL_2963790, EPI_ISL_2963791, EPI_ISL_2963792, EPI_ISL_2963793, EPI_ISL_2963794, EPI_ISL_2963795, EPI_ISL_2963796, EPI_ISL_2963797, EPI_ISL_2963798, EPI_ISL_2963799, EPI_ISL_2963800, EPI_ISL_2963801, EPI_ISL_2963803, EPI_ISL_2963804, EPI_ISL_2963805, EPI_ISL_2963806, EPI_ISL_2963807, EPI_ISL_2963808, EPI_ISL_2963809, EPI_ISL_2963810, EPI_ISL_2963811, EPI_ISL_2963812, EPI_ISL_2963813, EPI_ISL_2963814, EPI_ISL_2963815, EPI_ISL_2963816, EPI_ISL_2963817, EPI_ISL_2963818, EPI_ISL_2963819, EPI_ISL_2963820, EPI_ISL_2963821, EPI_ISL_2963822, EPI_ISL_2963823, EPI_ISL_2963824, EPI_ISL_2963825, EPI_ISL_2963826, EPI_ISL_2963827, EPI_ISL_2963828, EPI_ISL_2963829, EPI_ISL_2963830, EPI_ISL_2963831, EPI_ISL_2963832, EPI_ISL_2963833, EPI_ISL_2963834, EPI_ISL_2963835, EPI_ISL_2963836, EPI_ISL_2963837, EPI_ISL_2963838, EPI_ISL_2963840, EPI_ISL_2963841, EPI_ISL_2963842, EPI_ISL_2963843, EPI_ISL_2963844, EPI_ISL_2963845, EPI_ISL_2963846, EPI_ISL_2963847, EPI_ISL_2963848, EPI_ISL_2963850, EPI_ISL_2963851, EPI_ISL_2963852, EPI_ISL_2963853, EPI_ISL_2963854, EPI_ISL_2963855, EPI_ISL_2963856, EPI_ISL_2963857, EPI_ISL_2963858, EPI_ISL_2963859, EPI_ISL_2963860, EPI_ISL_2963861, EPI_ISL_2963862, EPI_ISL_2963863, EPI_ISL_2963864, EPI_ISL_2963865, EPI_ISL_2963867, EPI_ISL_2963869, EPI_ISL_2963870, EPI_ISL_2963871, EPI_ISL_2963872, EPI_ISL_2963873, EPI_ISL_2963874, EPI_ISL_2963875, EPI_ISL_2963876, EPI_ISL_2963877, EPI_ISL_2963878, EPI_ISL_2963879, EPI_ISL_2963880, EPI_ISL_2963881, EPI_ISL_2963882, EPI_ISL_2963883, EPI_ISL_2963884, EPI_ISL_2963885, EPI_ISL_2963886, EPI_ISL_2963887, EPI_ISL_2963888, EPI_ISL_2963889, EPI_ISL_2963890, EPI_ISL_2963891, EPI_ISL_2963892, EPI_ISL_2963894, EPI_ISL_2963895, EPI_ISL_2963896, EPI_ISL_2963897, EPI_ISL_2963898, EPI_ISL_2963899, EPI_ISL_2963900, EPI_ISL_2963901, EPI_ISL_2963902, EPI_ISL_2963903, EPI_ISL_2963904, EPI_ISL_2963905, EPI_ISL_2963906, EPI_ISL_2963907, EPI_ISL_2963908, EPI_ISL_2963909, EPI_ISL_2963910, EPI_ISL_2963911, EPI_ISL_2963912, EPI_ISL_2963913, EPI_ISL_2963914, EPI_ISL_2963915, EPI_ISL_2963916, EPI_ISL_2963917, EPI_ISL_2963918, EPI_ISL_2963919, EPI_ISL_2963921, EPI_ISL_2963922, EPI_ISL_2963923, EPI_ISL_2963925, EPI_ISL_2963926, EPI_ISL_2963927, EPI_ISL_2963928, EPI_ISL_2963929, EPI_ISL_2963930, EPI_ISL_2963931, EPI_ISL_2963932, EPI_ISL_2963933, EPI_ISL_2963934, EPI_ISL_2963935, EPI_ISL_2963936, EPI_ISL_2963937, EPI_ISL_2963938, EPI_ISL_2963939, EPI_ISL_2963940, EPI_ISL_2963941, EPI_ISL_2963942, EPI_ISL_2963943, EPI_ISL_2963944, EPI_ISL_2963945, EPI_ISL_2963946, EPI_ISL_2963947, EPI_ISL_2963948, EPI_ISL_2963949, EPI_ISL_2963950, EPI_ISL_2963952, EPI_ISL_2963953, EPI_ISL_2963955, EPI_ISL_2963956, EPI_ISL_2963957, EPI_ISL_2963958, EPI_ISL_2963960, EPI_ISL_2963962, EPI_ISL_2963963, EPI_ISL_2963964, EPI_ISL_2963965, EPI_ISL_2963966, EPI_ISL_2963967, EPI_ISL_2963968, EPI_ISL_2963969, EPI_ISL_2963970, EPI_ISL_2963971, EPI_ISL_2963973, EPI_ISL_2963974, EPI_ISL_2963975, EPI_ISL_2963976, EPI_ISL_2963977, EPI_ISL_2963978, EPI_ISL_2963979, EPI_ISL_2963980, EPI_ISL_2963981, EPI_ISL_2963982, EPI_ISL_2963983, EPI_ISL_2963984, EPI_ISL_2963985, EPI_ISL_2963986, EPI_ISL_2963987, EPI_ISL_2963988, EPI_ISL_2963989, EPI_ISL_2963990, EPI_ISL_2963991 | | | |
| see above | Department of Bacteria, Parasites and Fungi, Statens Serum Institut, Copenhagen, Denmark | Statens Serum Institut Bioinformatics and Microbial Genomics | Danish Covid-19 Genome Consortium |
| EPI_ISL_2963992 | Molekylær Medicinsk Afdeling, Aarhus University Hospital, Aarhus, Denmark | Statens Serum Institut Bioinformatics and Microbial Genomics | Danish Covid-19 Genome Consortium |
| EPI_ISL_2963993, EPI_ISL_2963994, EPI_ISL_2963995, EPI_ISL_2963997, EPI_ISL_2963998, EPI_ISL_2963999, EPI_ISL_2964000, EPI_ISL_2964001, EPI_ISL_2964002, EPI_ISL_2964003, EPI_ISL_2964005, EPI_ISL_2964006, EPI_ISL_2964007, EPI_ISL_2964008, EPI_ISL_2964009, EPI_ISL_2964010, EPI_ISL_2964011, EPI_ISL_2964012, EPI_ISL_2964013, EPI_ISL_2964014, EPI_ISL_2964015, EPI_ISL_2964016, EPI_ISL_2964017, EPI_ISL_2964018, EPI_ISL_2964019, EPI_ISL_2964020, EPI_ISL_2964021, EPI_ISL_2964022, EPI_ISL_2964023, EPI_ISL_2964024, EPI_ISL_2964025, EPI_ISL_2964026, EPI_ISL_2964027, EPI_ISL_2964028, EPI_ISL_2964029, EPI_ISL_2964030, EPI_ISL_2964031, EPI_ISL_2964032, EPI_ISL_2964033, EPI_ISL_2964034, EPI_ISL_2964035, EPI_ISL_2964036, EPI_ISL_2964037, EPI_ISL_2964038, EPI_ISL_2964039, EPI_ISL_2964040, EPI_ISL_2964041, EPI_ISL_2964042, EPI_ISL_2964043, EPI_ISL_2964045, EPI_ISL_2964046, EPI_ISL_2964047, EPI_ISL_2964048, EPI_ISL_2964049, EPI_ISL_2964050, EPI_ISL_2964051, EPI_ISL_2964052, EPI_ISL_2964053, EPI_ISL_2964054, EPI_ISL_2964056, EPI_ISL_2964057, EPI_ISL_2964058, EPI_ISL_2964059, EPI_ISL_2964060, EPI_ISL_2964061, EPI_ISL_2964062, EPI_ISL_2964063, EPI_ISL_2964064, EPI_ISL_2964065, EPI_ISL_2964066, EPI_ISL_2964067, EPI_ISL_2964068, EPI_ISL_2964069, EPI_ISL_2964070, EPI_ISL_2964071, EPI_ISL_2964072, EPI_ISL_2964073, EPI_ISL_2964074, EPI_ISL_2964075, EPI_ISL_2964076, EPI_ISL_2964077, EPI_ISL_2964078, EPI_ISL_2964079, EPI_ISL_2964080, EPI_ISL_2964081, EPI_ISL_2964082, EPI_ISL_2964083, EPI_ISL_2964084, EPI_ISL_2964085, EPI_ISL_2964086, EPI_ISL_2964087, EPI_ISL_2964088, EPI_ISL_2964089, EPI_ISL_2964090, EPI_ISL_2964092, EPI_ISL_2964093, EPI_ISL_2964094, EPI_ISL_2964095, EPI_ISL_2964096, EPI_ISL_2964097, EPI_ISL_2964098, EPI_ISL_2964099, EPI_ISL_2964100, EPI_ISL_2964102, EPI_ISL_2964103, EPI_ISL_2964105, EPI_ISL_2964106, EPI_ISL_2964107, EPI_ISL_2964109, EPI_ISL_2964110, EPI_ISL_2964111, EPI_ISL_2964112, EPI_ISL_2964113, EPI_ISL_2964114, EPI_ISL_2964115, EPI_ISL_2964116, EPI_ISL_2964117, EPI_ISL_2964118, EPI_ISL_2964119, EPI_ISL_2964120, EPI_ISL_2964121, EPI_ISL_2964122, EPI_ISL_2964123, EPI_ISL_2964125, EPI_ISL_2964126, EPI_ISL_2964127, EPI_ISL_2964128, EPI_ISL_2964129, EPI_ISL_2964130, EPI_ISL_2964131, EPI_ISL_2964132, EPI_ISL_2964133, EPI_ISL_2964134, EPI_ISL_2964135, EPI_ISL_2964136, EPI_ISL_2964137, EPI_ISL_2964138, EPI_ISL_2964139, EPI_ISL_2964140, EPI_ISL_2964141, EPI_ISL_2964142, EPI_ISL_2964143, EPI_ISL_2964144, EPI_ISL_2964145, EPI_ISL_2964146, EPI_ISL_2964147, EPI_ISL_2964148, EPI_ISL_2964149, EPI_ISL_2964150, EPI_ISL_2964151, EPI_ISL_2964152, EPI_ISL_2964154, EPI_ISL_2964155, EPI_ISL_2964156, EPI_ISL_2964157, EPI_ISL_2964158, EPI_ISL_2964159, EPI_ISL_2964160, EPI_ISL_2964161, EPI_ISL_2964162, EPI_ISL_2964163, EPI_ISL_2964164, EPI_ISL_2964165, EPI_ISL_2964166, EPI_ISL_2964167, EPI_ISL_2964168, EPI_ISL_2964169, EPI_ISL_2964170, EPI_ISL_2964171, EPI_ISL_2964172, EPI_ISL_2964174, EPI_ISL_2964175, EPI_ISL_2964176, EPI_ISL_2964177, EPI_ISL_2964178, EPI_ISL_2964179, EPI_ISL_2964180, EPI_ISL_2964181, EPI_ISL_2964182, EPI_ISL_2964183, EPI_ISL_2964184, EPI_ISL_2964185, EPI_ISL_2964186, EPI_ISL_2964187, EPI_ISL_2964188, EPI_ISL_2964189, EPI_ISL_2964190, EPI_ISL_2964191, EPI_ISL_2964192, EPI_ISL_2964193, EPI_ISL_2964194, EPI_ISL_2964195, EPI_ISL_2964196, EPI_ISL_2964197, EPI_ISL_2964198, EPI_ISL_2964199, EPI_ISL_2964200, EPI_ISL_2964201, EPI_ISL_2964204, EPI_ISL_2964205, EPI_ISL_2964206, EPI_ISL_2964207, EPI_ISL_2964208, EPI_ISL_2964209, EPI_ISL_2964210, EPI_ISL_2964211, EPI_ISL_2964212, EPI_ISL_2964213, EPI_ISL_2964214, EPI_ISL_2964215, EPI_ISL_2964216, EPI_ISL_2964217, EPI_ISL_2964218, EPI_ISL_2964219, EPI_ISL_2964220, EPI_ISL_2964221, EPI_ISL_2964222, EPI_ISL_2964225, EPI_ISL_2964227, EPI_ISL_2964228, EPI_ISL_2964229, EPI_ISL_2964230, EPI_ISL_2964231, EPI_ISL_2964232, EPI_ISL_2964233, EPI_ISL_2964234 | | | |
| see above | Department of Bacteria, Parasites and Fungi, Statens Serum Institut, Copenhagen, Denmark | Statens Serum Institut Bioinformatics and Microbial Genomics | Danish Covid-19 Genome Consortium |
| EPI_ISL_2964235 | Molekylær Medicinsk Afdeling, Aarhus University Hospital, Aarhus, Denmark | Statens Serum Institut Bioinformatics and Microbial Genomics | Danish Covid-19 Genome Consortium |
| EPI_ISL_2964236, EPI_ISL_2964237, EPI_ISL_2964239, EPI_ISL_2964240, EPI_ISL_2964241, EPI_ISL_2964242, EPI_ISL_2964243, EPI_ISL_2964244, EPI_ISL_2964245, EPI_ISL_2964246, EPI_ISL_2964247, EPI_ISL_2964248, EPI_ISL_2964249, EPI_ISL_2964250, EPI_ISL_2964251, EPI_ISL_2964252, EPI_ISL_2964253, EPI_ISL_2964254 | | | |
| see above | Department of Bacteria, Parasites and Fungi, Statens Serum Institut, Copenhagen, Denmark | Statens Serum Institut Bioinformatics and Microbial Genomics | Danish Covid-19 Genome Consortium |
| EPI_ISL_2964255, EPI_ISL_2964256 | Molekylær Medicinsk Afdeling, Aarhus University Hospital, Aarhus, Denmark | Statens Serum Institut Bioinformatics and Microbial Genomics | Danish Covid-19 Genome Consortium |
| EPI_ISL_2964257, EPI_ISL_2964258, EPI_ISL_2964259, EPI_ISL_2964260, EPI_ISL_2964261, EPI_ISL_2964262, EPI_ISL_2964263, EPI_ISL_2964264, EPI_ISL_2964265, EPI_ISL_2964266, EPI_ISL_2964267, EPI_ISL_2964268, EPI_ISL_2964269, EPI_ISL_2964270, EPI_ISL_2964271, EPI_ISL_2964272, EPI_ISL_2964274, EPI_ISL_2964275, EPI_ISL_2964276, EPI_ISL_2964277, EPI_ISL_2964278, EPI_ISL_2964279, EPI_ISL_2964280, EPI_ISL_2964282, EPI_ISL_2964283, EPI_ISL_2964284, EPI_ISL_2964285, EPI_ISL_2964286, EPI_ISL_2964287, EPI_ISL_2964288, EPI_ISL_2964289, EPI_ISL_2964290, EPI_ISL_2964291, EPI_ISL_2964292, EPI_ISL_2964293, EPI_ISL_2964294, EPI_ISL_2964295, EPI_ISL_2964296, EPI_ISL_2964297, EPI_ISL_2964298, EPI_ISL_2964299, EPI_ISL_2964300 | | | |
| see above | Department of Bacteria, Parasites and Fungi, Statens Serum Institut, Copenhagen, Denmark | Statens Serum Institut Bioinformatics and Microbial Genomics | Danish Covid-19 Genome Consortium |
| EPI_ISL_2964301 | Molekylær Medicinsk Afdeling, Aarhus University Hospital, Aarhus, Denmark | Statens Serum Institut Bioinformatics and Microbial Genomics | Danish Covid-19 Genome Consortium |
| EPI_ISL_2964302, EPI_ISL_2964303, EPI_ISL_2964304, EPI_ISL_2964305, EPI_ISL_2964306, EPI_ISL_2964307, EPI_ISL_2964308, EPI_ISL_2964309, EPI_ISL_2964310, EPI_ISL_2964311, EPI_ISL_2964312, EPI_ISL_2964313, EPI_ISL_2964314, EPI_ISL_2964315, EPI_ISL_2964316, EPI_ISL_2964317, EPI_ISL_2964318, EPI_ISL_2964319, EPI_ISL_2964320, EPI_ISL_2964321, EPI_ISL_2964322, EPI_ISL_2964323, EPI_ISL_2964324, EPI_ISL_2964325, EPI_ISL_2964326, EPI_ISL_2964327, EPI_ISL_2964328, EPI_ISL_2964330, EPI_ISL_2964331, EPI_ISL_2964332, EPI_ISL_2964333, EPI_ISL_2964334, EPI_ISL_2964335, EPI_ISL_2964336, EPI_ISL_2964337, EPI_ISL_2964338, EPI_ISL_2964339, EPI_ISL_2964340, EPI_ISL_2964341, EPI_ISL_2964342, EPI_ISL_2964343, EPI_ISL_2964344, EPI_ISL_2964346, EPI_ISL_2964347, EPI_ISL_2964348, EPI_ISL_2964349, EPI_ISL_2964350, EPI_ISL_2964351, EPI_ISL_2964352, EPI_ISL_2964353, EPI_ISL_2964354, EPI_ISL_2964355, EPI_ISL_2964356, EPI_ISL_2964357, EPI_ISL_2964358, EPI_ISL_2964359, EPI_ISL_2964360, EPI_ISL_2964361, EPI_ISL_2964362, EPI_ISL_2964363, EPI_ISL_2964364, EPI_ISL_2964365, EPI_ISL_2964366, EPI_ISL_2964367, EPI_ISL_2964368, EPI_ISL_2964369, EPI_ISL_2964370, EPI_ISL_2964371, EPI_ISL_2964372, EPI_ISL_2964373, EPI_ISL_2964374, EPI_ISL_2964375, EPI_ISL_2964376, EPI_ISL_2964377, EPI_ISL_2964378, EPI_ISL_2964379 | | | |
| see above | Department of Bacteria, Parasites and Fungi, Statens Serum Institut, Copenhagen, Denmark | Statens Serum Institut Bioinformatics and Microbial Genomics | Danish Covid-19 Genome Consortium |
| EPI_ISL_2964390, EPI_ISL_2964392, EPI_ISL_2964393, EPI_ISL_2964399, EPI_ISL_2964400, EPI_ISL_2964401, EPI_ISL_2964402 | Krai Clinical Infectious Diseases Hospital | WHO National Influenza Centre Russian Federation | Andrey Komissarov, Artem Fadeev, Kseniya Komissarova, Oula Mansour, Kirill Varchenko, Mikhail Bakaev, Tamila Musaeva, Veronika Eder, Maria Pisareva, Nikita Yolshin, Daria Danilenko, Ksenia Safina, Elena Nabieva, Georgii Bazykin, Dmitry Lioznov |
| EPI_ISL_2964403, EPI_ISL_2964404, EPI_ISL_2964405, EPI_ISL_2964406, EPI_ISL_2964407, EPI_ISL_2964408, EPI_ISL_2964409, EPI_ISL_2964410, EPI_ISL_2964411, EPI_ISL_2964412, EPI_ISL_2964413 | | | |
| see above | Infectious Diseases Hospital No 4 | WHO National Influenza Centre Russian Federation | Andrey Komissarov, Artem Fadeev, Kseniya Komissarova, Oula Mansour, Kirill Varchenko, Mikhail Bakaev, Tamila Musaeva, Veronika Eder, Maria Pisareva, Nikita Yolshin, Daria Danilenko, Ksenia Safina, Elena Nabieva, Georgii Bazykin, Dmitry Lioznov |
| EPI_ISL_2964414, EPI_ISL_2964416 | Krai Clinical Hospital No 2 | WHO National Influenza Centre Russian Federation | Andrey Komissarov, Artem Fadeev, Kseniya Komissarova, Oula Mansour, Kirill Varchenko, Mikhail Bakaev, Tamila Musaeva, Veronika Eder, Maria Pisareva, Nikita Yolshin, Daria Danilenko, Ksenia Safina, Elena Nabieva, Georgii Bazykin, Dmitry Lioznov |
| EPI_ISL_2964417, EPI_ISL_2964418, EPI_ISL_2964420, EPI_ISL_2964421, EPI_ISL_2964422, EPI_ISL_2964423, EPI_ISL_2964424, EPI_ISL_2964425, EPI_ISL_2964426, EPI_ISL_2964427, EPI_ISL_2964428, EPI_ISL_2964429, EPI_ISL_2964430, EPI_ISL_2964431, EPI_ISL_2964432, EPI_ISL_2964433, EPI_ISL_2964434, EPI_ISL_2964435, EPI_ISL_2964436, EPI_ISL_2964437, EPI_ISL_2964438, EPI_ISL_2964439, EPI_ISL_2964440, EPI_ISL_2964441, EPI_ISL_2964442, EPI_ISL_2964443, EPI_ISL_2964444, EPI_ISL_2964445, EPI_ISL_2964446, EPI_ISL_2964448, EPI_ISL_2964449, EPI_ISL_2964451, EPI_ISL_2964452, EPI_ISL_2964453, EPI_ISL_2964454, EPI_ISL_2964455, EPI_ISL_2964456, EPI_ISL_2964457, EPI_ISL_2964458, EPI_ISL_2964459, EPI_ISL_2964460, EPI_ISL_2964461, EPI_ISL_2964462, EPI_ISL_2964463, EPI_ISL_2964464, EPI_ISL_2964465, EPI_ISL_2964466, EPI_ISL_2964467, EPI_ISL_2964468, EPI_ISL_2964469, EPI_ISL_2964470, EPI_ISL_2964471, EPI_ISL_2964472, EPI_ISL_2964473, EPI_ISL_2964474, EPI_ISL_2964475, EPI_ISL_2964476, EPI_ISL_2964477, EPI_ISL_2964478, EPI_ISL_2964479, EPI_ISL_2964480, EPI_ISL_2964481, EPI_ISL_2964482, EPI_ISL_2964484, EPI_ISL_2964485, EPI_ISL_2964486, EPI_ISL_2964488, EPI_ISL_2964489, EPI_ISL_2964490, EPI_ISL_2964491, EPI_ISL_2964492, EPI_ISL_2964494, EPI_ISL_2964496, EPI_ISL_2964497, EPI_ISL_2964498, EPI_ISL_2964499, EPI_ISL_2964500, EPI_ISL_2964501, EPI_ISL_2964502, EPI_ISL_2964503, EPI_ISL_2964504, EPI_ISL_2964505, EPI_ISL_2964506, EPI_ISL_2964507, EPI_ISL_2964508, EPI_ISL_2964509, EPI_ISL_2964510, EPI_ISL_2964511, EPI_ISL_2964514, EPI_ISL_2964515, EPI_ISL_2964516, EPI_ISL_2964517, EPI_ISL_2964518, EPI_ISL_2964519, EPI_ISL_2964520, EPI_ISL_2964521, EPI_ISL_2964522, EPI_ISL_2964523, EPI_ISL_2964524, EPI_ISL_2964525, EPI_ISL_2964526, EPI_ISL_2964527, EPI_ISL_2964529, EPI_ISL_2964530, EPI_ISL_2964531, EPI_ISL_2964532, EPI_ISL_2964533, EPI_ISL_2964534, EPI_ISL_2964535, EPI_ISL_2964536, EPI_ISL_2964537, EPI_ISL_2964538, EPI_ISL_2964539, EPI_ISL_2964540, EPI_ISL_2964541, EPI_ISL_2964542, EPI_ISL_2964543, EPI_ISL_2964544, EPI_ISL_2964545, EPI_ISL_2964546, EPI_ISL_2964547, EPI_ISL_2964548, EPI_ISL_2964549, EPI_ISL_2964550, EPI_ISL_2964551, EPI_ISL_2964552, EPI_ISL_2964553, EPI_ISL_2964554, EPI_ISL_2964555, EPI_ISL_2964556 | | | |
| see above | N.F. Gamaleya Research Center for Epidemiology and Microbiology | WHO National Influenza Centre Russian Federation | Andrey Komissarov, Artem Fadeev, Kseniya Komissarova, Oula Mansour, Kirill Varchenko, Mikhail Bakaev, Tamila Musaeva, Veronika Eder, Maria Pisareva, Nikita Yolshin, Daria Danilenko, Elizaveta Divisenko, Alina Odintsova, Elena Shidlovskaya, Ludmila Vasilchenko, Nadezhda Kuznetsova, Maria Nikiforova, Alexey Shchetinin, Artem Tkachuk, Andrei Pochtovyi, Andrei Siniavin, Denis Kleymenov, Evgeniia Bykonia, Liubov Popova, Elena Mazunina, Inna Dolzhikova, Anna Kovyrshina, Andrei Botikov, Dmitry Shcheblyakov, Vladimir Gushchin, Denis Logunov, Alexander Gintsburg, Ksenia Safina, Elena Nabieva, Georgii Bazykin, Dmitry Lioznov |
| EPI_ISL_2964572, EPI_ISL_2964573, EPI_ISL_2964575, EPI_ISL_2964576, EPI_ISL_2964577, EPI_ISL_2964578, EPI_ISL_2964579, EPI_ISL_2964580, EPI_ISL_2964581, EPI_ISL_2964585, EPI_ISL_2964586 | | | |
| see above | Wojewodzka Stacja Sanitarno-Epidemiologiczna w Olsztynie, Laboratorium Badan Epidemiologiczno-Klinicznych | Wojewodzka Stacja Sanitarno-Epidemiologiczna w Olsztynie, Laboratorium Badan Epidemiologiczno-Klinicznych | Sylwia Krzetowska, Monika Czerminska, Ewa Liszewska, Tomasz Jakubczak, Marta Lukian, Patryk Bielecki, Aleksandra Kobiatko, Emilia Tarabasz, Paulina Rozycka, Barbara Dolinska |
| EPI_ISL_2964642, EPI_ISL_2964643, EPI_ISL_2964644, EPI_ISL_2964645 | Public Health Authority of the Slovak Republic | Laboratory of Genomics and Bioinformatics, Comenius University Science Park | Tatiana Sedláčková, Diana Rusňáková, Miroslav Böhmer, Anna Gičová, Jaroslav Budiš, Tomáš Szemes, Jakub Styk |
| EPI_ISL_2964647 | HELEN JOSEPH LABORATORY | National Institute for Communicable Diseases of the National Health Laboratory Service | Amoako DG, Everatt J, Scheepers C, Mohale T, Ntuli N, Mahlangu B, Mnguni A, Ismail A, Bhiman JN |
| EPI_ISL_2964652 | NILMRC | Genomic Research Laboratory, Bangabandhu Sheikh Mujib Medical | Laila Anjuman Banu, Md.Saydur Rahman, Sharadindu Kanti Sinha, Arifa Akram, Md.Sharfuddin Ahmed |

| | | | |
|---|---|---|---|
| | | University Bangabandhu Sheikh Mujib Medical University (BSMMU) Shahbag, Dhaka_1000 | |
| EPI_ISL_2964677 | RSUD DR. MOEWARDI Surakarta | Genetics Working Group (Pokja Genetik) Faculty of Medicine, Public Health and Nursing Universitas Gadjah Mada (FK-KMK UGM); Disease Investigation Center Wates Ministry of Agriculture Indonesia; Department of Microbiology FK-KMK UGM; Laboratorium Diagnostik Yayasan Tahija World Mosquito Program (WMP) Yogyakarta Center for Tropical Medicine FK-KMK UGM; Integrated Research Center FK-KMK UGM; Department of Computer Science and Electronics FMIPA UGM; RSUP Dr. Sardjito | Gunadi, Hendra Wibawa, Marcellus, Mohamad S. Hakim, Edwin W. Daniwijaya, Ludhang P. Rizki, Endah Supriyati, Eggi Arguni, Titik Nuryastuti, Tri Wibawa, Dwi AA Nugrahaningsih, Afiahayati, Siswanto,Kristy Iskandar,Nungki Anggorowati, Pramesti G Dewi, Dwi Indaryati,Indarto Sulistiyono, Khanza a Vujira, Alvina A Setiawan |
| EPI_ISL_2964681 | Puskesmas Colomadu 1 DKK Karanganyar | Genetics Working Group (Pokja Genetik) Faculty of Medicine, Public Health and Nursing Universitas Gadjah Mada (FK-KMK UGM); Disease Investigation Center Wates Ministry of Agriculture Indonesia; Department of Microbiology FK-KMK UGM; Laboratorium Diagnostik Yayasan Tahija World Mosquito Program (WMP) Yogyakarta Center for Tropical Medicine FK-KMK UGM; Integrated Research Center FK-KMK UGM; Department of Computer Science and Electronics FMIPA UGM; RSUP Dr. Sardjito | Gunadi, Hendra Wibawa, Marcellus, Mohamad S. Hakim, Edwin W. Daniwijaya, Ludhang P. Rizki, Endah Supriyati, Eggi Arguni, Titik Nuryastuti, Tri Wibawa, Dwi AA Nugrahaningsih, Afiahayati, Siswanto,Kristy Iskandar,Nungki Anggorowati, Pramesti G Dewi, Dwi Indaryati,Indarto Sulistiyono, Khanza a Vujira, Alvina A Setiawan |
| EPI_ISL_2964684 | RSUD DR. MOEWARDI | Genetics Working Group (Pokja Genetik) Faculty of Medicine, Public Health and Nursing Universitas Gadjah Mada (FK-KMK UGM); Disease Investigation Center Wates Ministry of Agriculture Indonesia; Department of Microbiology FK-KMK UGM; Laboratorium Diagnostik Yayasan Tahija World Mosquito Program (WMP) Yogyakarta Center for Tropical Medicine FK-KMK UGM; Integrated Research Center FK-KMK UGM; Department of Computer Science and Electronics FMIPA UGM; RSUP Dr. Sardjito | Gunadi, Hendra Wibawa, Marcellus, Mohamad S. Hakim, Edwin W. Daniwijaya, Ludhang P. Rizki, Endah Supriyati, Eggi Arguni, Titik Nuryastuti, Tri Wibawa, Dwi AA Nugrahaningsih, Afiahayati, Siswanto,Kristy Iskandar,Nungki Anggorowati, Pramesti G Dewi, Dwi Indaryati,Indarto Sulistiyono, Cita S Amalia, Alvina A Setiawan |
| EPI_ISL_2964685 | NILMRC | Genomic Research Laboratory, BSMMU Bangabandhu Sheikh Mujib Medical University (BSMMU) i | Laila Anjuman Banu, Md.Saydur Rahman, Sharadindu Kanti Sinha, Arifa Akram, Md.Sharfuddin Ahmed |
| EPI_ISL_2964688 | RSUD DR. MOEWARDI | Genetics Working Group (Pokja Genetik) Faculty of Medicine, Public Health and Nursing Universitas Gadjah Mada (FK-KMK UGM); Disease Investigation Center Wates Ministry of Agriculture Indonesia; Department of Microbiology FK-KMK UGM; Laboratorium Diagnostik Yayasan Tahija World Mosquito Program (WMP) Yogyakarta Center for Tropical Medicine FK-KMK UGM; Integrated Research Center FK-KMK UGM; Department of Computer Science and Electronics FMIPA UGM; RSUP Dr. Sardjito | Gunadi, Hendra Wibawa, Marcellus, Mohamad S. Hakim, Edwin W. Daniwijaya, Ludhang P. Rizki, Endah Supriyati, Eggi Arguni, Titik Nuryastuti, Tri Wibawa, Dwi AA Nugrahaningsih, Afiahayati, Siswanto,Kristy Iskandar,Nungki Anggorowati, Pramesti G Dewi, Dwi Indaryati,Indarto Sulistiyono, Irene Tania, Hana F Hanifin |
| EPI_ISL_2964692 | RSUD DR. MOEWARDI | Genetics Working Group (Pokja Genetik) Faculty of Medicine, Public Health and Nursing Universitas Gadjah Mada (FK-KMK UGM); Disease Investigation Center Wates Ministry of Agriculture Indonesia; Department of Microbiology FK-KMK UGM; Laboratorium Diagnostik Yayasan Tahija World Mosquito Program (WMP) Yogyakarta Center for Tropical Medicine FK-KMK UGM; Integrated Research Center FK-KMK UGM; Department of Computer Science and Electronics FMIPA UGM; RSUP Dr. Sardjito | Gunadi, Hendra Wibawa, Marcellus, Mohamad S. Hakim, Edwin W. Daniwijaya, Ludhang P. Rizki, Endah Supriyati, Eggi Arguni, Titik Nuryastuti, Tri Wibawa, Dwi AA Nugrahaningsih, Afiahayati, Siswanto,Kristy Iskandar,Nungki Anggorowati, Pramesti G Dewi, Dwi Indaryati,Indarto Sulistiyono, Susan Simanjaya, Dyah A Puspitarani |
| EPI_ISL_2964697 | DINKES KAB. JEPARA | Genetics Working Group (Pokja Genetik) Faculty of Medicine, Public Health and Nursing Universitas Gadjah Mada (FK-KMK UGM); Disease Investigation Center Wates Ministry of Agriculture Indonesia; Department of Microbiology FK-KMK UGM; Laboratorium Diagnostik Yayasan Tahija World Mosquito Program (WMP) Yogyakarta Center for Tropical Medicine FK-KMK UGM; Integrated Research Center FK-KMK UGM; Department of Computer Science and Electronics FMIPA UGM; RSUP Dr. Sardjito | Gunadi, Hendra Wibawa, Marcellus, Mohamad S. Hakim, Edwin W. Daniwijaya, Ludhang P. Rizki, Endah Supriyati, Eggi Arguni, Titik Nuryastuti, Tri Wibawa, Dwi AA Nugrahaningsih, Afiahayati, Siswanto,Kristy Iskandar,Nungki Anggorowati, Pramesti G Dewi, Dwi Indaryati,Indarto Sulistiyono, Cita S Amalia, Alvina A Setiawan |
| EPI_ISL_2964700 | DINKES KAB. JEPARA | Genetics Working Group (Pokja Genetik) Faculty of Medicine, Public Health and Nursing Universitas Gadjah Mada (FK-KMK UGM); Disease Investigation Center Wates Ministry of Agriculture Indonesia; Department of Microbiology FK-KMK UGM; Laboratorium Diagnostik Yayasan Tahija World Mosquito Program (WMP) Yogyakarta Center for Tropical Medicine FK-KMK UGM; Integrated Research Center FK-KMK UGM; Department of Computer Science and Electronics FMIPA UGM; RSUP Dr. Sardjito | Gunadi, Hendra Wibawa, Marcellus, Mohamad S. Hakim, Edwin W. Daniwijaya, Ludhang P. Rizki, Endah Supriyati, Eggi Arguni, Titik Nuryastuti, Tri Wibawa, Dwi AA Nugrahaningsih, Afiahayati, Siswanto,Kristy Iskandar,Nungki Anggorowati, Pramesti G Dewi, Dwi Indaryati,Indarto Sulistiyono, Khanza a Vujira, Alvina A Setiawan |
| EPI_ISL_2964704 | DINKES KAB. GROBOGAN | Genetics Working Group (Pokja Genetik) Faculty of Medicine, Public Health and Nursing Universitas Gadjah Mada (FK-KMK UGM); Disease Investigation Center Wates Ministry of Agriculture Indonesia; Department of Microbiology FK-KMK UGM; Laboratorium Diagnostik Yayasan Tahija World Mosquito Program (WMP) Yogyakarta Center for Tropical Medicine FK-KMK UGM; Integrated Research Center FK-KMK UGM; Department of Computer Science and Electronics FMIPA UGM; RSUP Dr. Sardjito | Gunadi, Hendra Wibawa, Marcellus, Mohamad S. Hakim, Edwin W. Daniwijaya, Ludhang P. Rizki, Endah Supriyati, Eggi Arguni, Titik Nuryastuti, Tri Wibawa, Dwi AA Nugrahaningsih, Afiahayati, Siswanto,Kristy Iskandar,Nungki Anggorowati, Pramesti G Dewi, Dwi Indaryati,Indarto Sulistiyono, Susan Simanjaya, Dyah A Puspitarani |
| EPI_ISL_2964706 | DINKES KAB. GROBOGAN | Genetics Working Group (Pokja Genetik) Faculty of Medicine, Public Health and Nursing Universitas Gadjah Mada (FK-KMK UGM); Disease Investigation Center Wates Ministry of Agriculture Indonesia; Department of Microbiology FK-KMK UGM; Laboratorium Diagnostik Yayasan Tahija World Mosquito Program (WMP) Yogyakarta Center for Tropical Medicine FK-KMK UGM; Integrated Research Center FK-KMK UGM; Department of Computer Science and Electronics FMIPA UGM; RSUP Dr. Sardjito | Gunadi, Hendra Wibawa, Marcellus, Mohamad S. Hakim, Edwin W. Daniwijaya, Ludhang P. Rizki, Endah Supriyati, Eggi Arguni, Titik Nuryastuti, Tri Wibawa, Dwi AA Nugrahaningsih, Afiahayati, Siswanto,Kristy Iskandar,Nungki Anggorowati, Pramesti G Dewi, Dwi Indaryati,Indarto Sulistiyono, Irene Tania, Hana F Hanifin |
| EPI_ISL_2964707 | RS Dr. Oen Kandang Sapi Solo | Genetics Working Group (Pokja Genetik) Faculty of Medicine, Public Health and Nursing Universitas Gadjah Mada (FK-KMK UGM); Disease Investigation Center Wates Ministry of Agriculture Indonesia; Department of Microbiology FK-KMK UGM; Laboratorium Diagnostik Yayasan Tahija World Mosquito Program (WMP) Yogyakarta Center for Tropical Medicine FK-KMK UGM; Integrated Research Center FK-KMK UGM; Department of Computer Science and Electronics FMIPA UGM; RSUP Dr. Sardjito | Gunadi, Hendra Wibawa, Marcellus, Mohamad S. Hakim, Edwin W. Daniwijaya, Ludhang P. Rizki, Endah Supriyati, Eggi Arguni, Titik Nuryastuti, Tri Wibawa, Dwi AA Nugrahaningsih, Afiahayati, Siswanto,Kristy Iskandar,Nungki Anggorowati, Pramesti G Dewi, Dwi Indaryati,Indarto Sulistiyono, Susan Simanjaya, Dyah A Puspitarani |
| EPI_ISL_2964753, EPI_ISL_2964762, EPI_ISL_2964765, EPI_ISL_2964766, EPI_ISL_2964772, EPI_ISL_2964782, EPI_ISL_2964786, EPI_ISL_2964788, EPI_ISL_2964842, EPI_ISL_2964843, EPI_ISL_2964844, EPI_ISL_2964845, EPI_ISL_2964857 | | | |
| see above | KU Leuven, Rega Institute, Clinical and Epidemiological Virology | KU Leuven, Rega Institute, Clinical and Epidemiological Virology | Tony Wawina-Bokalanga, Bert Vanmechelen, Joan Marti-Carerras, Piet Maes |
| EPI_ISL_2964872 | RS Dr. Oen Kandang Sapi Solo | Genetics Working Group (Pokja Genetik) Faculty of Medicine, Public Health and Nursing Universitas Gadjah Mada (FK-KMK UGM); Disease Investigation Center Wates Ministry of Agriculture Indonesia; Department of Microbiology FK-KMK UGM; Laboratorium Diagnostik Yayasan Tahija World Mosquito Program (WMP) Yogyakarta Center for Tropical Medicine FK-KMK UGM; Integrated Research Center FK-KMK UGM; Department of Computer Science and Electronics FMIPA UGM; RSUP Dr. Sardjito | Gunadi, Hendra Wibawa, Marcellus, Mohamad S. Hakim, Edwin W. Daniwijaya, Ludhang P. Rizki, Endah Supriyati, Eggi Arguni, Titik Nuryastuti, Tri Wibawa, Dwi AA Nugrahaningsih, Afiahayati, Siswanto,Kristy Iskandar,Nungki Anggorowati, Pramesti G Dewi, Dwi Indaryati,Indarto Sulistiyono, Cita S Amalia, Alvina A Setiawan |
| EPI_ISL_2964917 | RS Dr. OEN KANDANG SAPI SOLO | Genetics Working Group (Pokja Genetik) Faculty of Medicine, Public Health and Nursing Universitas Gadjah Mada (FK-KMK UGM); Disease Investigation Center Wates Ministry of Agriculture Indonesia; Department of Microbiology FK-KMK UGM; Laboratorium Diagnostik Yayasan Tahija World Mosquito Program (WMP) Yogyakarta Center for Tropical Medicine FK-KMK UGM; Integrated Research Center FK-KMK UGM; Department of Computer Science and Electronics FMIPA UGM; RSUP Dr. Sardjito | Gunadi, Hendra Wibawa, Marcellus, Mohamad S. Hakim, Edwin W. Daniwijaya, Ludhang P. Rizki, Endah Supriyati, Eggi Arguni, Titik Nuryastuti, Tri Wibawa, Dwi AA Nugrahaningsih, Afiahayati, Siswanto,Kristy Iskandar,Nungki Anggorowati, Pramesti G Dewi, Dwi Indaryati,Indarto Sulistiyono, Irene Tania, Hana F Hanifin |
| EPI_ISL_2964918, | RS Dr. OEN KANDANG SAPI SOLO | Genetics Working Group (Pokja Genetik) Faculty of Medicine, Public | Gunadi, Hendra Wibawa, Marcellus, Mohamad S. Hakim, Edwin W. Daniwijaya, Ludhang P. Rizki, Endah Supriyati, Eggi Arguni, Titik Nuryastuti, Tri Wibawa, Dwi AA Nugrahaningsih, Afiahayati, Siswanto,Kristy Iskandar,Nungki Anggorowati, |

| | | | |
|---|---|---|---|
| EPI_ISL_2964919 | | Health and Nursing Universitas Gadjah Mada (FK-KMK UGM); Disease Investigation Center Wates Ministry of Agriculture Indonesia; Department of Microbiology FK-KMK UGM; Laboratorium Diagnostik Yayasan Tahija World Mosquito Program (WMP) Yogyakarta Center for Tropical Medicine FK-KMK UGM; Integrated Research Center FK-KMK UGM; Department of Computer Science and Electronics FMIPA UGM; RSUP Dr. Sardjito | Pramesti G Dewi, Dwi Indaryati,Indarto Sulistiyono, Cita S Amalia, Alvina A Setiawan |
| EPI_ISL_2964920 | RS Dr. OEN KANDANG SAPI SOLO | Genetics Working Group (Pokja Genetik) Faculty of Medicine, Public Health and Nursing Universitas Gadjah Mada (FK-KMK UGM); Disease Investigation Center Wates Ministry of Agriculture Indonesia; Department of Microbiology FK-KMK UGM; Laboratorium Diagnostik Yayasan Tahija World Mosquito Program (WMP) Yogyakarta Center for Tropical Medicine FK-KMK UGM; Integrated Research Center FK-KMK UGM; Department of Computer Science and Electronics FMIPA UGM; RSUP Dr. Sardjito | Gunadi, Hendra Wibawa, Marcellus, Mohamad S. Hakim, Edwin W. Daniwijaya, Ludhang P. Rizki, Endah Supriyati, Eggi Arguni, Titik Nuryastuti, Tri Wibawa, Dwi AA Nugrahaningsih, Afiahayati, Siswanto,Kristy Iskandar,Nungki Anggorowati, Pramesti G Dewi, Dwi Indaryati,Indarto Sulistiyono, Susan Simanjaya, Dyah A Puspitarani |
| EPI_ISL_2964921 | RS Dr. OEN KANDANG SAPI SOLO | Genetics Working Group (Pokja Genetik) Faculty of Medicine, Public Health and Nursing Universitas Gadjah Mada (FK-KMK UGM); Disease Investigation Center Wates Ministry of Agriculture Indonesia; Department of Microbiology FK-KMK UGM; Laboratorium Diagnostik Yayasan Tahija World Mosquito Program (WMP) Yogyakarta Center for Tropical Medicine FK-KMK UGM; Integrated Research Center FK-KMK UGM; Department of Computer Science and Electronics FMIPA UGM; RSUP Dr. Sardjito | Gunadi, Hendra Wibawa, Marcellus, Mohamad S. Hakim, Edwin W. Daniwijaya, Ludhang P. Rizki, Endah Supriyati, Eggi Arguni, Titik Nuryastuti, Tri Wibawa, Dwi AA Nugrahaningsih, Afiahayati, Siswanto,Kristy Iskandar,Nungki Anggorowati, Pramesti G Dewi, Dwi Indaryati,Indarto Sulistiyono, Khanza a Vujira, Alvina A Setiawan |
| EPI_ISL_2964922, EPI_ISL_2964923 | Middlemore Hospital | Institute of Environmental Science and Research (ESR) | Rachel Boyle, SallyAnn Harbison, Olivia Stroeven, Xiaoyun Ren, Matt Storey, Nikki Freed, Muhammad Faisal, Jing Wang, Hermes Perez, Anja Werno, Antje van der Linden, Arlo Upton, Chris Mansell, David Hammer, Dragana Drinkovic, Gary McAuliffe, Hana Sofia Andersson, James Ussher, Jill Sherwood, Josh Freeman, Julia Howard, Juliet Elvy, Mary DeAlmeida, Matt Blakiston, Matthew Rogers, Max Bloomfield, Michael Addidle, Michelle Balm, Sally Roberts, Sarah Jefferies, Sharmini Muttaiyah, Susan Morpeth, Susan Taylor, Timothy Blackmore, Vani Sathyendran, Veronica Playle, Virginia Hope, Erasmus Smit, Lauren Jelly, Olin Silander, Joep de Ligt |
| EPI_ISL_2964924, EPI_ISL_2964925, EPI_ISL_2964926, EPI_ISL_2964927, EPI_ISL_2964928, EPI_ISL_2964930 | LabPLUS | Institute of Environmental Science and Research (ESR) | Rachel Boyle, SallyAnn Harbison, Olivia Stroeven, Xiaoyun Ren, Matt Storey, Nikki Freed, Muhammad Faisal, Jing Wang, Hermes Perez, Anja Werno, Antje van der Linden, Arlo Upton, Chris Mansell, David Hammer, Dragana Drinkovic, Gary McAuliffe, Hana Sofia Andersson, James Ussher, Jill Sherwood, Josh Freeman, Julia Howard, Juliet Elvy, Mary DeAlmeida, Matt Blakiston, Matthew Rogers, Max Bloomfield, Michael Addidle, Michelle Balm, Sally Roberts, Sarah Jefferies, Sharmini Muttaiyah, Susan Morpeth, Susan Taylor, Timothy Blackmore, Vani Sathyendran, Veronica Playle, Virginia Hope, Erasmus Smit, Lauren Jelly, Olin Silander, Joep de Ligt |
| EPI_ISL_2964931 | Wellington SCL (WN) | Institute of Environmental Science and Research (ESR) | Xiaoyun Ren, Matt Storey, Nikki Freed, Muhammad Faisal, Jing Wang, Hermes Perez, Anja Werno, Antje van der Linden, Arlo Upton, Chris Mansell, David Hammer, Dragana Drinkovic, Gary McAuliffe, Hana Sofia Andersson, James Ussher, Jill Sherwood, Josh Freeman, Julia Howard, Juliet Elvy, Mary DeAlmeida, Matt Blakiston, Matthew Rogers, Max Bloomfield, Michael Addidle, Michelle Balm, Sally Roberts, Sarah Jefferies, Sharmini Muttaiyah, Susan Morpeth, Susan Taylor, Timothy Blackmore, Vani Sathyendran, Veronica Playle, Virginia Hope, Erasmus Smit, Lauren Jelly, Olin Silander, Joep de Ligt |
| EPI_ISL_2964935 | LabPLUS | Institute of Environmental Science and Research (ESR) | Rachel Boyle, SallyAnn Harbison, Olivia Stroeven, Xiaoyun Ren, Matt Storey, Nikki Freed, Muhammad Faisal, Jing Wang, Hermes Perez, Anja Werno, Antje van der Linden, Arlo Upton, Chris Mansell, David Hammer, Dragana Drinkovic, Gary McAuliffe, Hana Sofia Andersson, James Ussher, Jill Sherwood, Josh Freeman, Julia Howard, Juliet Elvy, Mary DeAlmeida, Matt Blakiston, Matthew Rogers, Max Bloomfield, Michael Addidle, Michelle Balm, Sally Roberts, Sarah Jefferies, Sharmini Muttaiyah, Susan Morpeth, Susan Taylor, Timothy Blackmore, Vani Sathyendran, Veronica Playle, Virginia Hope, Erasmus Smit, Lauren Jelly, Olin Silander, Joep de Ligt |
| EPI_ISL_2964936, EPI_ISL_2964937 | Middlemore Hospital | Institute of Environmental Science and Research (ESR) | Rachel Boyle, SallyAnn Harbison, Olivia Stroeven, Xiaoyun Ren, Matt Storey, Nikki Freed, Muhammad Faisal, Jing Wang, Hermes Perez, Anja Werno, Antje van der Linden, Arlo Upton, Chris Mansell, David Hammer, Dragana Drinkovic, Gary McAuliffe, Hana Sofia Andersson, James Ussher, Jill Sherwood, Josh Freeman, Julia Howard, Juliet Elvy, Mary DeAlmeida, Matt Blakiston, Matthew Rogers, Max Bloomfield, Michael Addidle, Michelle Balm, Sally Roberts, Sarah Jefferies, Sharmini Muttaiyah, Susan Morpeth, Susan Taylor, Timothy Blackmore, Vani Sathyendran, Veronica Playle, Virginia Hope, Erasmus Smit, Lauren Jelly, Olin Silander, Joep de Ligt |
| EPI_ISL_2964938, EPI_ISL_2964939, EPI_ISL_2964940 | Wellington SCL (WN) | Institute of Environmental Science and Research (ESR) | Xiaoyun Ren, Matt Storey, Nikki Freed, Muhammad Faisal, Jing Wang, Hermes Perez, Anja Werno, Antje van der Linden, Arlo Upton, Chris Mansell, David Hammer, Dragana Drinkovic, Gary McAuliffe, Hana Sofia Andersson, James Ussher, Jill Sherwood, Josh Freeman, Julia Howard, Juliet Elvy, Mary DeAlmeida, Matt Blakiston, Matthew Rogers, Max Bloomfield, Michael Addidle, Michelle Balm, Sally Roberts, Sarah Jefferies, Sharmini Muttaiyah, Susan Morpeth, Susan Taylor, Timothy Blackmore, Vani Sathyendran, Veronica Playle, Virginia Hope, Erasmus Smit, Lauren Jelly, Olin Silander, Joep de Ligt |
| EPI_ISL_2964941 | RSUD DR. MOEWARDI | Genetics Working Group (Pokja Genetik) Faculty of Medicine, Public Health and Nursing Universitas Gadjah Mada (FK-KMK UGM); Disease Investigation Center Wates Ministry of Agriculture Indonesia; Department of Microbiology FK-KMK UGM; Laboratorium Diagnostik Yayasan Tahija World Mosquito Program (WMP) Yogyakarta Center for Tropical Medicine FK-KMK UGM; Integrated Research Center FK-KMK UGM; Department of Computer Science and Electronics FMIPA UGM; RSUP Dr. Sardjito | Gunadi, Hendra Wibawa, Marcellus, Mohamad S. Hakim, Edwin W. Daniwijaya, Ludhang P. Rizki, Endah Supriyati, Eggi Arguni, Titik Nuryastuti, Tri Wibawa, Dwi AA Nugrahaningsih, Afiahayati, Siswanto,Kristy Iskandar,Nungki Anggorowati, Pramesti G Dewi, Dwi Indaryati,Indarto Sulistiyono, Cita S Amalia, Alvina A Setiawan |
| EPI_ISL_2964942 | RSUD DR. MOEWARDI | Genetics Working Group (Pokja Genetik) Faculty of Medicine, Public Health and Nursing Universitas Gadjah Mada (FK-KMK UGM); Disease Investigation Center Wates Ministry of Agriculture Indonesia; Department of Microbiology FK-KMK UGM; Laboratorium Diagnostik Yayasan Tahija World Mosquito Program (WMP) Yogyakarta Center for Tropical Medicine FK-KMK UGM; Integrated Research Center FK-KMK UGM; Department of Computer Science and Electronics FMIPA UGM; RSUP Dr. Sardjito | Gunadi, Hendra Wibawa, Marcellus, Mohamad S. Hakim, Edwin W. Daniwijaya, Ludhang P. Rizki, Endah Supriyati, Eggi Arguni, Titik Nuryastuti, Tri Wibawa, Dwi AA Nugrahaningsih, Afiahayati, Siswanto,Kristy Iskandar,Nungki Anggorowati, Pramesti G Dewi, Dwi Indaryati,Indarto Sulistiyono, Irene Tania, Hana F Hanifin |
| EPI_ISL_2964943 | RSUD DR. MOEWARDI | Genetics Working Group (Pokja Genetik) Faculty of Medicine, Public Health and Nursing Universitas Gadjah Mada (FK-KMK UGM); Disease Investigation Center Wates Ministry of Agriculture Indonesia; Department of Microbiology FK-KMK UGM; Laboratorium Diagnostik Yayasan Tahija World Mosquito Program (WMP) Yogyakarta Center for Tropical Medicine FK-KMK UGM; Integrated Research Center FK-KMK UGM; Department of Computer Science and Electronics FMIPA UGM; RSUP Dr. Sardjito | Gunadi, Hendra Wibawa, Marcellus, Mohamad S. Hakim, Edwin W. Daniwijaya, Ludhang P. Rizki, Endah Supriyati, Eggi Arguni, Titik Nuryastuti, Tri Wibawa, Dwi AA Nugrahaningsih, Afiahayati, Siswanto,Kristy Iskandar,Nungki Anggorowati, Pramesti G Dewi, Dwi Indaryati,Indarto Sulistiyono, Susan Simanjaya, Dyah A Puspitarani |
| EPI_ISL_2964944 | RSUD DR. MOEWARDI | Genetics Working Group (Pokja Genetik) Faculty of Medicine, Public Health and Nursing Universitas Gadjah Mada (FK-KMK UGM); Disease Investigation Center Wates Ministry of Agriculture Indonesia; Department of Microbiology FK-KMK UGM; Laboratorium Diagnostik Yayasan Tahija World Mosquito Program (WMP) Yogyakarta Center for Tropical Medicine FK-KMK UGM; Integrated Research Center FK-KMK UGM; Department of Computer Science and Electronics FMIPA UGM; RSUP Dr. Sardjito | Gunadi, Hendra Wibawa, Marcellus, Mohamad S. Hakim, Edwin W. Daniwijaya, Ludhang P. Rizki, Endah Supriyati, Eggi Arguni, Titik Nuryastuti, Tri Wibawa, Dwi AA Nugrahaningsih, Afiahayati, Siswanto,Kristy Iskandar,Nungki Anggorowati, Pramesti G Dewi, Dwi Indaryati,Indarto Sulistiyono, Susan Simanjaya, I Putu Aditio A |
| EPI_ISL_2964955 | Fondation Congolaise pour la recherche medicale (FCRM), Francine Ntoumi | Fondation Congolaise pour la Recherche Médicale | Mfoutou Mapanguy Claujens Chastel; Batchi-Bouyou Armel Landry; Dr. Jean Claude Djontu; Dr. Abel Lissom; Prof. Dr. Thirumalaisamy P. Velavan; Prof. Francine Ntoumi |
| EPI_ISL_2964956 | RSUD DR. MOEWARDI | Genetics Working Group (Pokja Genetik) Faculty of Medicine, Public Health and Nursing Universitas Gadjah Mada (FK-KMK UGM); Disease Investigation Center Wates Ministry of Agriculture Indonesia; Department of Microbiology FK-KMK UGM; Laboratorium Diagnostik Yayasan Tahija World Mosquito Program (WMP) Yogyakarta Center for Tropical Medicine FK-KMK UGM; Integrated Research Center FK-KMK UGM; Department of Computer Science and Electronics FMIPA UGM; RSUP Dr. Sardjito | Gunadi, Hendra Wibawa, Marcellus, Mohamad S. Hakim, Edwin W. Daniwijaya, Ludhang P. Rizki, Endah Supriyati, Eggi Arguni, Titik Nuryastuti, Tri Wibawa, Dwi AA Nugrahaningsih, Afiahayati, Siswanto,Kristy Iskandar,Nungki Anggorowati, Pramesti G Dewi, Dwi Indaryati,Indarto Sulistiyono, Cita S Amalia, Alvina A Setiawan |
| EPI_ISL_2964962, EPI_ISL_2965002, EPI_ISL_2965173 | Pandemic Response Lab - NYC | Pandemic Response Lab, R&D | Isabel Fernandez Escapa, Henry Lee, Michael Hammerling, Melissa Hopkins, Cybill del Castillo, Shinyoung Clair Kang, William Ward, Pradeep Bugga, Sol Rey, Dylan Law, Haiping Hao, Jon Laurent |
| EPI_ISL_2965566, EPI_ISL_2965567 | Zdravotní ústav Ústí nad Labem | Institute of Medical Microbiology and Virology, University Hospital Carl Gustav Carus, TU Dresden | Marlena Stadtmüller, Alexa Laubner, Johanna Beil, Fabian Rost, Sylvia Klemroth, Grit Mehnert, Montserrat Palau de Miquel, Susanne Reinhardt, Leo Büttner, Anett Zabzinski, Sylke Winkler, Alexander Dalpke, Romana Mikešová, Jitka Pohořská, Ivana Stiborová, Eva Patrasová, Lenka Šimůnková |
| EPI_ISL_2965577, EPI_ISL_2965582 | LATE - Laboratório de Técnicas Especiais - Hospital Israelita Albert Einstein | LATE - Laboratório de Técnicas Especiais - Hospital Israelita Albert Einstein | Deyvid Amgarten, Fernanda de Mello Malta, Raquel Riyuzo, Ana Paula Moreira Salles, Pedro Henrique Sebe Rodrigues, Alexandre Hideaki Takara, Anelisie da Silva Santos, Erick Gustavo Dorlass, Marcio Anunciacao Menezes, João Renato Rebello Pinho |
| EPI_ISL_2965590, EPI_ISL_2965591, EPI_ISL_2965592 | Zdravotní ústav Ústí nad Labem | Institute of Medical Microbiology and Virology, University Hospital Carl Gustav Carus, TU Dresden | Marlena Stadtmüller, Alexa Laubner, Johanna Beil, Fabian Rost, Sylvia Klemroth, Grit Mehnert, Montserrat Palau de Miquel, Susanne Reinhardt, Leo Büttner, Anett Zabzinski, Sylke Winkler, Alexander Dalpke, Romana Mikešová, Jitka Pohořská, Ivana Stiborová, Eva Patrasová, Lenka Šimůnková |
| EPI_ISL_2965600, EPI_ISL_2965601, EPI_ISL_2965602, EPI_ISL_2965603, EPI_ISL_2965604 | Fujian Center for Disease Control and Prevention | Fujian Center for Disease Control and Prevention | Lin Qi, Huang Zhimiao, Weng Yuwei |

| | | | |
|---|---|---|---|
| EPI_ISL_2965607, EPI_ISL_2965608, EPI_ISL_2965610, EPI_ISL_2965611, EPI_ISL_2965613, EPI_ISL_2965616, EPI_ISL_2965620, EPI_ISL_2965621, EPI_ISL_2965626, EPI_ISL_2965627, EPI_ISL_2965631, EPI_ISL_2965632, EPI_ISL_2965633, EPI_ISL_2965634, EPI_ISL_2965635 | | | |
| see above | AZDelta | AZ Delta Medical Laboratories in Roeselare, Belgium | Geert Martens, Dieter De Smet, Merijn Vanhee, on behalf of AZ Delta COVID-19 Genomics core (member of Genomic surveillance of SARS-CoV-2 in Belgium network) |
| EPI_ISL_2965645, EPI_ISL_2965646, EPI_ISL_2965647, EPI_ISL_2965648, EPI_ISL_2965649, EPI_ISL_2965650, EPI_ISL_2965651, EPI_ISL_2965652, EPI_ISL_2965653, EPI_ISL_2965654, EPI_ISL_2965656, EPI_ISL_2965657, EPI_ISL_2965659, EPI_ISL_2965660, EPI_ISL_2965661, EPI_ISL_2965662, EPI_ISL_2965663 | | | |
| see above | Institute for Infectious Diseases | Institute for Infectious Diseases | Stefan Neuenschwander, Christian Baumann, Miguel A Terrazos Miani, Cora SÃ¤gesser, Pascal Bittel, Peter Keller, Franziska Suter-Riniker, Stephen L Leib, Alban Ramette |
| EPI_ISL_2965733, EPI_ISL_2965750 | Public Health Authority of the Slovak Republic | Public Health Authority of the Slovak Republic | Barbora Kotvasová, Lucia Ševčíková, Terézia Vrabľová, Anna Gičová, Elena Tichá, Miroslav Böhmer, Tomáš Szemes, Pavol Mišenko |
| EPI_ISL_2965759 | RS Advent Bandar Lampung | National Institute of Health Research and Development | Vivi Setiawaty, Hana Apsari Pawestri, Subangkit, Kartika Dewi Puspa, Arie Ardiansyah Nugraha, Hartanti Dian Ikawati, Krisna Pangesti, Triyani Soekarso, Fajar Nur Sulistiyohadi, Nelly Puspandari |
| EPI_ISL_2965760, EPI_ISL_2965761, EPI_ISL_2965762 | Dinkes Kabupaten Merauke | National Institute of Health Research and Development | Vivi Setiawaty, Hana Apsari Pawestri, Subangkit, Kartika Dewi Puspa, Arie Ardiansyah Nugraha, Hartanti Dian Ikawati, Krisna Pangesti, Triyani Soekarso, Fajar Nur Sulistiyohadi, Nelly Puspandari |
| EPI_ISL_2965763, EPI_ISL_2965764, EPI_ISL_2965765, EPI_ISL_2965766, EPI_ISL_2965767 | Dinkes Kabupaten Merauke | National Institute of Health Research and Development | Arie Ardiansyah Nugraha, Hana Apsari Pawestri, Subangkit, Kartika Dewi Puspa, Hartanti Dian Ikawati, Krisna Pangesti, Triyani Soekarso, Fajar Nur Sulistiyohadi, Nelly Puspandari, Vivi Setiawaty |
| EPI_ISL_2965768, EPI_ISL_2965769 | Dinkes Kabupaten Merauke | National Institute of Health Research and Development | Hana Apsari Pawestri, Subangkit, Arie Ardiansyah Nugraha, Kartika Dewi Puspa, Hartanti Dian Ikawati, Krisna Pangesti, Triyani Soekarso, Fajar Nur Sulistiyohadi, Nelly Puspandari, Vivi Setiawaty |
| EPI_ISL_2965770, EPI_ISL_2965771, EPI_ISL_2965772, EPI_ISL_2965773, EPI_ISL_2965774, EPI_ISL_2965775 | Dinkes Provinsi Kalimantan Utara | National Institute of Health Research and Development | Hana Apsari Pawestri, Subangkit, Arie Ardiansyah Nugraha, Kartika Dewi Puspa, Hartanti Dian Ikawati, Krisna Pangesti, Triyani Soekarso, Fajar Nur Sulistiyohadi, Nelly Puspandari, Vivi Setiawaty |
| EPI_ISL_2965776, EPI_ISL_2965777 | Dinkes Provinsi Kalimantan Utara | National Institute of Health Research and Development | Kartika Dewi Puspa, Arie Ardiansyah Nugraha, Hana Apsari Pawestri, Subangkit, Hartanti Dian Ikawati, Krisna Pangesti, Triyani Soekarso, Fajar Nur Sulistiyohadi, Nelly Puspandari, Vivi Setiawaty |
| EPI_ISL_2965778, EPI_ISL_2965779, EPI_ISL_2965780, EPI_ISL_2965781, EPI_ISL_2965782, EPI_ISL_2965783 | Dinkes Provinsi Kalimantan Timur | National Institute of Health Research and Development | Kartika Dewi Puspa, Arie Ardiansyah Nugraha, Hana Apsari Pawestri, Subangkit, Hartanti Dian Ikawati, Krisna Pangesti, Triyani Soekarso, Fajar Nur Sulistiyohadi, Nelly Puspandari, Vivi Setiawaty |
| EPI_ISL_2965784, EPI_ISL_2965785, EPI_ISL_2965786 | Dinkes Provinsi Kalimantan Timur | National Institute of Health Research and Development | Hartanti Dian Ikawati, Arie Ardiansyah Nugraha, Hana Apsari Pawestri, Subangkit, Kartika Dewi Puspa, Krisna Pangesti, Triyani Soekarso, Fajar Nur Sulistiyohadi, Nelly Puspandari, Vivi Setiawaty |
| EPI_ISL_2965787, EPI_ISL_2965788 | RS Prima Husada Cipta | National Institute of Health Research and Development | Hartanti Dian Ikawati, Arie Ardiansyah Nugraha, Hana Apsari Pawestri, Subangkit, Kartika Dewi Puspa, Krisna Pangesti, Triyani Soekarso, Fajar Nur Sulistiyohadi, Nelly Puspandari, Vivi Setiawaty |
| EPI_ISL_2965789, EPI_ISL_2965790, EPI_ISL_2965791, EPI_ISL_2965792, EPI_ISL_2965793, EPI_ISL_2965794, EPI_ISL_2965795, EPI_ISL_2965796, EPI_ISL_2965797, EPI_ISL_2965798, EPI_ISL_2965799, EPI_ISL_2965800, EPI_ISL_2965801, EPI_ISL_2965802, EPI_ISL_2965803, EPI_ISL_2965804, EPI_ISL_2965805, EPI_ISL_2965807, EPI_ISL_2965808, EPI_ISL_2965809, EPI_ISL_2965810, EPI_ISL_2965811, EPI_ISL_2965812, EPI_ISL_2965813, EPI_ISL_2965814, EPI_ISL_2965815, EPI_ISL_2965816, EPI_ISL_2965817, EPI_ISL_2965818, EPI_ISL_2965819, EPI_ISL_2965820, EPI_ISL_2965821, EPI_ISL_2965822, EPI_ISL_2965823, EPI_ISL_2965824, EPI_ISL_2965825, EPI_ISL_2965826, EPI_ISL_2965827, EPI_ISL_2965828, EPI_ISL_2965829, EPI_ISL_2965830, EPI_ISL_2965831, EPI_ISL_2965832, EPI_ISL_2965833, EPI_ISL_2965834, EPI_ISL_2965835, EPI_ISL_2965836, EPI_ISL_2965837, EPI_ISL_2965838, EPI_ISL_2965839, EPI_ISL_2965840, EPI_ISL_2965841, EPI_ISL_2965842, EPI_ISL_2965843, EPI_ISL_2965844, EPI_ISL_2965845, EPI_ISL_2965846, EPI_ISL_2965847, EPI_ISL_2965848, EPI_ISL_2965849, EPI_ISL_2965850, EPI_ISL_2965851, EPI_ISL_2965852, EPI_ISL_2965853, EPI_ISL_2965854, EPI_ISL_2965855, EPI_ISL_2965856, EPI_ISL_2965857, EPI_ISL_2965858, EPI_ISL_2965859, EPI_ISL_2965860, EPI_ISL_2965861, EPI_ISL_2965862, EPI_ISL_2965863, EPI_ISL_2965864, EPI_ISL_2965865, EPI_ISL_2965866, EPI_ISL_2965867, EPI_ISL_2965868, EPI_ISL_2965869, EPI_ISL_2965870, EPI_ISL_2965871, EPI_ISL_2965872, EPI_ISL_2965873, EPI_ISL_2965874, EPI_ISL_2965875, EPI_ISL_2965876, EPI_ISL_2965877, EPI_ISL_2965878, EPI_ISL_2965879, EPI_ISL_2965880, EPI_ISL_2965881, EPI_ISL_2965882, EPI_ISL_2965883, EPI_ISL_2965884, EPI_ISL_2965885, EPI_ISL_2965886, EPI_ISL_2965887, EPI_ISL_2965888, EPI_ISL_2965889, EPI_ISL_2965890, EPI_ISL_2965891, EPI_ISL_2965892, EPI_ISL_2965893, EPI_ISL_2965894, EPI_ISL_2965895, EPI_ISL_2965896, EPI_ISL_2965897, EPI_ISL_2965898, EPI_ISL_2965899, EPI_ISL_2965900, EPI_ISL_2965901, EPI_ISL_2965902, EPI_ISL_2965903, EPI_ISL_2965904, EPI_ISL_2965905, EPI_ISL_2965906, EPI_ISL_2965907, EPI_ISL_2965908, EPI_ISL_2965909, EPI_ISL_2965910, EPI_ISL_2965911, EPI_ISL_2965912, EPI_ISL_2965913, EPI_ISL_2965914, EPI_ISL_2965915, EPI_ISL_2965916, EPI_ISL_2965917, EPI_ISL_2965918, EPI_ISL_2965919, EPI_ISL_2965920, EPI_ISL_2965921, EPI_ISL_2965922, EPI_ISL_2965923, EPI_ISL_2965924, EPI_ISL_2965925, EPI_ISL_2965926, EPI_ISL_2965927, EPI_ISL_2965929, EPI_ISL_2965930, EPI_ISL_2965931, EPI_ISL_2965932, EPI_ISL_2965933, EPI_ISL_2965934, EPI_ISL_2965935, EPI_ISL_2965936, EPI_ISL_2965937, EPI_ISL_2965938, EPI_ISL_2965939, EPI_ISL_2965940, EPI_ISL_2965941, EPI_ISL_2965942, EPI_ISL_2965943, EPI_ISL_2965944, EPI_ISL_2965945, EPI_ISL_2965946, EPI_ISL_2965947, EPI_ISL_2965948, EPI_ISL_2965949, EPI_ISL_2965950, EPI_ISL_2965951, EPI_ISL_2965952, EPI_ISL_2965953, EPI_ISL_2965954, EPI_ISL_2965955, EPI_ISL_2965956, EPI_ISL_2965957, EPI_ISL_2965958, EPI_ISL_2965959, EPI_ISL_2965960, EPI_ISL_2965961, EPI_ISL_2965962, EPI_ISL_2965963, EPI_ISL_2965964, EPI_ISL_2965965, EPI_ISL_2965966, EPI_ISL_2965967, EPI_ISL_2965968, EPI_ISL_2965969, EPI_ISL_2965970, EPI_ISL_2965971, EPI_ISL_2965972, EPI_ISL_2965973, EPI_ISL_2965974, EPI_ISL_2965975, EPI_ISL_2965976, EPI_ISL_2965977, EPI_ISL_2965978, EPI_ISL_2965979, EPI_ISL_2965980, EPI_ISL_2965981, EPI_ISL_2965982, EPI_ISL_2965983, EPI_ISL_2965985, EPI_ISL_2965986, EPI_ISL_2965987, EPI_ISL_2965988, EPI_ISL_2965989, EPI_ISL_2965990, EPI_ISL_2965991, EPI_ISL_2965992, EPI_ISL_2965993, EPI_ISL_2965994, EPI_ISL_2965995, EPI_ISL_2965996, EPI_ISL_2965997, EPI_ISL_2965998, EPI_ISL_2965999, EPI_ISL_2966000, EPI_ISL_2966001, EPI_ISL_2966002, EPI_ISL_2966003, EPI_ISL_2966004, EPI_ISL_2966005, EPI_ISL_2966006, EPI_ISL_2966008, EPI_ISL_2966009, EPI_ISL_2966010, EPI_ISL_2966011, EPI_ISL_2966012, EPI_ISL_2966013, EPI_ISL_2966014, EPI_ISL_2966015, EPI_ISL_2966016, EPI_ISL_2966017, EPI_ISL_2966018, EPI_ISL_2966019, EPI_ISL_2966020, EPI_ISL_2966021, EPI_ISL_2966022, EPI_ISL_2966023, EPI_ISL_2966024, EPI_ISL_2966025, EPI_ISL_2966026, EPI_ISL_2966027, EPI_ISL_2966028, EPI_ISL_2966029, EPI_ISL_2966030, EPI_ISL_2966031, EPI_ISL_2966032, EPI_ISL_2966033, EPI_ISL_2966034, EPI_ISL_2966035, EPI_ISL_2966036, EPI_ISL_2966037, EPI_ISL_2966038, EPI_ISL_2966039, EPI_ISL_2966040, EPI_ISL_2966041, EPI_ISL_2966042, EPI_ISL_2966043, EPI_ISL_2966044, EPI_ISL_2966045, EPI_ISL_2966046, EPI_ISL_2966047, EPI_ISL_2966048, EPI_ISL_2966049, EPI_ISL_2966050, EPI_ISL_2966051, EPI_ISL_2966052, EPI_ISL_2966053, EPI_ISL_2966055, EPI_ISL_2966056, EPI_ISL_2966057, EPI_ISL_2966058, EPI_ISL_2966059, EPI_ISL_2966060, EPI_ISL_2966061, EPI_ISL_2966062, EPI_ISL_2966063, EPI_ISL_2966064, EPI_ISL_2966066, EPI_ISL_2966067, EPI_ISL_2966068, EPI_ISL_2966069, EPI_ISL_2966070, EPI_ISL_2966071, EPI_ISL_2966072, EPI_ISL_2966073, EPI_ISL_2966074, EPI_ISL_2966075, EPI_ISL_2966076, EPI_ISL_2966077, EPI_ISL_2966078, EPI_ISL_2966079, EPI_ISL_2966080, EPI_ISL_2966081, EPI_ISL_2966082, EPI_ISL_2966083, EPI_ISL_2966084, EPI_ISL_2966085, EPI_ISL_2966087, EPI_ISL_2966088, EPI_ISL_2966089, EPI_ISL_2966090, EPI_ISL_2966091, EPI_ISL_2966092, EPI_ISL_2966093, EPI_ISL_2966094, EPI_ISL_2966095, EPI_ISL_2966096, EPI_ISL_2966097, EPI_ISL_2966098, EPI_ISL_2966099, EPI_ISL_2966100, EPI_ISL_2966101, EPI_ISL_2966102, EPI_ISL_2966103, EPI_ISL_2966104, EPI_ISL_2966105, EPI_ISL_2966106, EPI_ISL_2966107, EPI_ISL_2966108, EPI_ISL_2966109, EPI_ISL_2966110, EPI_ISL_2966111, EPI_ISL_2966112, EPI_ISL_2966113, EPI_ISL_2966114, EPI_ISL_2966115, EPI_ISL_2966116, EPI_ISL_2966117, EPI_ISL_2966118, EPI_ISL_2966119, EPI_ISL_2966120, EPI_ISL_2966121, EPI_ISL_2966122, EPI_ISL_2966123, EPI_ISL_2966124, EPI_ISL_2966125, EPI_ISL_2966126, EPI_ISL_2966127, EPI_ISL_2966128, EPI_ISL_2966129, EPI_ISL_2966130, EPI_ISL_2966131, EPI_ISL_2966132, EPI_ISL_2966133, EPI_ISL_2966134, EPI_ISL_2966135, EPI_ISL_2966136, EPI_ISL_2966137, EPI_ISL_2966138, EPI_ISL_2966139, EPI_ISL_2966140, EPI_ISL_2966141, EPI_ISL_2966142, EPI_ISL_2966143, EPI_ISL_2966144, EPI_ISL_2966145, EPI_ISL_2966146, EPI_ISL_2966147, EPI_ISL_2966148, EPI_ISL_2966149, EPI_ISL_2966150, EPI_ISL_2966151, EPI_ISL_2966152, EPI_ISL_2966153, EPI_ISL_2966154, EPI_ISL_2966155, EPI_ISL_2966156, EPI_ISL_2966157, EPI_ISL_2966158, EPI_ISL_2966159, EPI_ISL_2966160, EPI_ISL_2966161, EPI_ISL_2966162, EPI_ISL_2966163, EPI_ISL_2966164, EPI_ISL_2966165, EPI_ISL_2966166, EPI_ISL_2966167, EPI_ISL_2966168, EPI_ISL_2966169, EPI_ISL_2966170, EPI_ISL_2966171, EPI_ISL_2966172, EPI_ISL_2966173, EPI_ISL_2966174, EPI_ISL_2966175, EPI_ISL_2966177, EPI_ISL_2966178, EPI_ISL_2966179, EPI_ISL_2966180, EPI_ISL_2966181, EPI_ISL_2966182, EPI_ISL_2966183, EPI_ISL_2966184, EPI_ISL_2966185, EPI_ISL_2966186, EPI_ISL_2966187, EPI_ISL_2966188, EPI_ISL_2966189, EPI_ISL_2966190, EPI_ISL_2966191, EPI_ISL_2966192, EPI_ISL_2966193, EPI_ISL_2966194, EPI_ISL_2966195, EPI_ISL_2966196, EPI_ISL_2966197, EPI_ISL_2966198, EPI_ISL_2966199, EPI_ISL_2966200, EPI_ISL_2966201, EPI_ISL_2966202, EPI_ISL_2966203, EPI_ISL_2966204, EPI_ISL_2966205, EPI_ISL_2966206, EPI_ISL_2966207, EPI_ISL_2966208, EPI_ISL_2966209, EPI_ISL_2966210, EPI_ISL_2966211, EPI_ISL_2966212, EPI_ISL_2966213, EPI_ISL_2966214, EPI_ISL_2966215, EPI_ISL_2966216, EPI_ISL_2966217, EPI_ISL_2966218, EPI_ISL_2966219, EPI_ISL_2966220, EPI_ISL_2966221, EPI_ISL_2966222, EPI_ISL_2966223, EPI_ISL_2966224, EPI_ISL_2966225, EPI_ISL_2966226, EPI_ISL_2966227, EPI_ISL_2966228, EPI_ISL_2966229, EPI_ISL_2966230, EPI_ISL_2966231, EPI_ISL_2966232, EPI_ISL_2966233, EPI_ISL_2966234, EPI_ISL_2966235 | | | |
| see above | ICMR-National Institute of Virology - INSACOG | NIV Influenza | Dr. Varsha Potdar and NIC Team |
| EPI_ISL_2966248, EPI_ISL_2966250, EPI_ISL_2966251, EPI_ISL_2966252, EPI_ISL_2966253, EPI_ISL_2966254, EPI_ISL_2966255, EPI_ISL_2966256, EPI_ISL_2966257, EPI_ISL_2966258, EPI_ISL_2966259, EPI_ISL_2966261, EPI_ISL_2966262, EPI_ISL_2966263, EPI_ISL_2966264, EPI_ISL_2966265, EPI_ISL_2966266, EPI_ISL_2966267, EPI_ISL_2966268, EPI_ISL_2966269, EPI_ISL_2966270, EPI_ISL_2966271, EPI_ISL_2966272, EPI_ISL_2966274, EPI_ISL_2966275, EPI_ISL_2966276, EPI_ISL_2966278, EPI_ISL_2966279, EPI_ISL_2966280, EPI_ISL_2966281, EPI_ISL_2966282, EPI_ISL_2966283, EPI_ISL_2966284, EPI_ISL_2966285, EPI_ISL_2966287, EPI_ISL_2966288, EPI_ISL_2966289, EPI_ISL_2966290, EPI_ISL_2966292, EPI_ISL_2966293, EPI_ISL_2966294, EPI_ISL_2966299, EPI_ISL_2966300, EPI_ISL_2966302, EPI_ISL_2966303, EPI_ISL_2966306, EPI_ISL_2966307, EPI_ISL_2966308, EPI_ISL_2966309, EPI_ISL_2966310, EPI_ISL_2966311, EPI_ISL_2966312, EPI_ISL_2966313, EPI_ISL_2966314, EPI_ISL_2966315, EPI_ISL_2966316, EPI_ISL_2966317, EPI_ISL_2966318, EPI_ISL_2966320, EPI_ISL_2966321, EPI_ISL_2966322, EPI_ISL_2966324, EPI_ISL_2966325, EPI_ISL_2966326, EPI_ISL_2966327, EPI_ISL_2966328, EPI_ISL_2966329, EPI_ISL_2966330, EPI_ISL_2966331, EPI_ISL_2966332, EPI_ISL_2966333, EPI_ISL_2966334, EPI_ISL_2966335, EPI_ISL_2966336, EPI_ISL_2966337, EPI_ISL_2966338, EPI_ISL_2966339 | | | |
| see above | Cliniques universitaires Saint-Luc | UCLouvain/IREC/MBLG | Jean Ruelle, Bertrand Bearzatto, Benoit Kabamba Mukadi |
| EPI_ISL_2966416 | LESP Ciudad de Mexico | Instituto de Diagnostico y Referencia Epidemiologicos (INDRE) | Claudia Wong-Arambula, Abril Rodriguez-Maldonado, Vanessa Rivero-Arredondo, Ariadna Medina-Benitez, Joaquin Quiroz-Mercado, Sergio Rangel-Guerrero, Maribel Gonzalez-Villa, Natividad Cruz-Ortiz, Tatiana Nunez-Garcia, Gisela Barrera-Badillo, Lucia Hernandez-Rivas, Irma Lopez-Martinez, Ernesto Ramirez-Gonzalez. |
| EPI_ISL_2966418, EPI_ISL_2966420, EPI_ISL_2966421, EPI_ISL_2966422, EPI_ISL_2966423, EPI_ISL_2966425, EPI_ISL_2966428, EPI_ISL_2966429, EPI_ISL_2966431, EPI_ISL_2966432, EPI_ISL_2966435, EPI_ISL_2966436, EPI_ISL_2966437, EPI_ISL_2966438, EPI_ISL_2966441, EPI_ISL_2966443, EPI_ISL_2966446, EPI_ISL_2966447, EPI_ISL_2966448, EPI_ISL_2966449, EPI_ISL_2966450, EPI_ISL_2966454, EPI_ISL_2966455, EPI_ISL_2966456, EPI_ISL_2966457, EPI_ISL_2966458 | | | |
| see above | Alaska State Virology Laboratory | Alaska State Virology Laboratory | Stephanie DeRonde, Elva House, Jacob Zidek, Lisa Smith, Ph.D., Jack Chen, Ph.D. |
| EPI_ISL_2966459, EPI_ISL_2966460, EPI_ISL_2966461, EPI_ISL_2966462, EPI_ISL_2966463, EPI_ISL_2966464, EPI_ISL_2966466, EPI_ISL_2966467, EPI_ISL_2966469, EPI_ISL_2966470, EPI_ISL_2966471, EPI_ISL_2966472, EPI_ISL_2966474, EPI_ISL_2966475, EPI_ISL_2966476, EPI_ISL_2966477, EPI_ISL_2966478, EPI_ISL_2966479, EPI_ISL_2966480, EPI_ISL_2966481, EPI_ISL_2966482, EPI_ISL_2966483, EPI_ISL_2966484, EPI_ISL_2966485, EPI_ISL_2966486, EPI_ISL_2966487, EPI_ISL_2966488, EPI_ISL_2966489, EPI_ISL_2966490, EPI_ISL_2966491, EPI_ISL_2966492, EPI_ISL_2966493, EPI_ISL_2966494, EPI_ISL_2966495, EPI_ISL_2966496, EPI_ISL_2966497, EPI_ISL_2966498, EPI_ISL_2966499, EPI_ISL_2966500, EPI_ISL_2966501, EPI_ISL_2966503, EPI_ISL_2966504, EPI_ISL_2966505, EPI_ISL_2966508, EPI_ISL_2966510 | | | |
| see above | Nebraska Public Health Laboratory | NPHL COVID-19 Response Team | NPHL COVID-19 Response Team |
| EPI_ISL_2966512 | SECP MHU | The Institute of Molecular Biology and Genetics of NASU | M.Tukalo et al. |
| EPI_ISL_2966514 | Wyoming Public Health Laboratory | Wyoming Public Health Laboratory | Jim Mildenberger, Taylor Fearing, Channing Weber, Ashley Norberg, Chayse Rowley, Marley Goetz, Brian Dominguez, Elliot Thomasson, Sam Britz, Cari Sloma, Robert Petit, and Rob Christensen |
| EPI_ISL_2966537, EPI_ISL_2966538, EPI_ISL_2966539, EPI_ISL_2966540, EPI_ISL_2966541, EPI_ISL_2966542, EPI_ISL_2966543, EPI_ISL_2966544, EPI_ISL_2966545, EPI_ISL_2966546, EPI_ISL_2966547, EPI_ISL_2966548, EPI_ISL_2966549, EPI_ISL_2966550, EPI_ISL_2966551, EPI_ISL_2966552, EPI_ISL_2966553, EPI_ISL_2966554, EPI_ISL_2966555, EPI_ISL_2966556, EPI_ISL_2966559, EPI_ISL_2966560, EPI_ISL_2966561, EPI_ISL_2966562, EPI_ISL_2966564, EPI_ISL_2966565, EPI_ISL_2966566, EPI_ISL_2966567, EPI_ISL_2966568, EPI_ISL_2966569, EPI_ISL_2966570, EPI_ISL_2966571, EPI_ISL_2966572, EPI_ISL_2966573, EPI_ISL_2966574, EPI_ISL_2966575, EPI_ISL_2966576 | | | |
| see above | University of Mississippi Medical Center, Department of Pathology | University of Mississippi Medical Center, Molecular and Genomics Core Facility | Ashley C. Johnson, Wenjie Wu, Ithiel J. Frame, Krishna K. Ayyalasomayajula, Michael R. Garrett, D. Ashley Robinson |
| EPI_ISL_2966577, EPI_ISL_2966578, EPI_ISL_2966579, EPI_ISL_2966580, EPI_ISL_2966581, EPI_ISL_2966582, EPI_ISL_2966584, EPI_ISL_2966585, EPI_ISL_2966586, EPI_ISL_2966588, EPI_ISL_2966589, EPI_ISL_2966590, EPI_ISL_2966592, EPI_ISL_2966593, EPI_ISL_2966594, EPI_ISL_2966595, EPI_ISL_2966596, EPI_ISL_2966597, EPI_ISL_2966598, EPI_ISL_2966599, EPI_ISL_2966601, EPI_ISL_2966602, EPI_ISL_2966603, EPI_ISL_2966604, EPI_ISL_2966605, EPI_ISL_2966606, EPI_ISL_2966607, EPI_ISL_2966608, EPI_ISL_2966610, EPI_ISL_2966611 | | | |
| see above | MS Public Health Laboratory | University of Mississippi Medical Center, Molecular and Genomics Core Facility | Ashley C. Johnson, Wenjie Wu, Ithiel J. Frame, Krishna K. Ayyalasomayajula, Michael R. Garrett, D. Ashley Robinson |
| EPI_ISL_2966613, EPI_ISL_2966614, EPI_ISL_2966617 | Ochsner Health | Bioinfoexperts, LLC | Judy Crabtree, Amy Feehan, Rebecca Rose, David J. Nolan, Tessa LaFleur, Julia-Garcia-Diaz, Lucio Miele, Susanna L. Lamers |
| EPI_ISL_2966620, EPI_ISL_2966621, EPI_ISL_2966622, EPI_ISL_2966623, EPI_ISL_2966624, EPI_ISL_2966625, EPI_ISL_2966626, EPI_ISL_2966627, EPI_ISL_2966628, EPI_ISL_2966629, EPI_ISL_2966630, EPI_ISL_2966631, EPI_ISL_2966632, EPI_ISL_2966633, EPI_ISL_2966634, EPI_ISL_2966635, EPI_ISL_2966636, EPI_ISL_2966638, EPI_ISL_2966639, EPI_ISL_2966640, EPI_ISL_2966644, EPI_ISL_2966645, EPI_ISL_2966646, EPI_ISL_2966647, EPI_ISL_2966649, EPI_ISL_2966651, EPI_ISL_2966652, EPI_ISL_2966653, EPI_ISL_2966655 | | | |
| see above | Viral Respiratory Lab, National Institute for Biomedical Research (INRB) | Pathogen Sequencing Lab, National Institute for Biomedical Research (INRB) | Placide Mbala-Kingebeni, Edith Nkwembe, Eddy Kinganda-Lusamaki, Amuri Aziza, Francisca Muyembe Mawete, Emmanuel Lokilo Lofiko, Jean Claude Makangara, Catherine Pratt, Matthias Pauthner, Josh Quick, Allison Black, James Hadfield, Trevor Bedford, Ian Goodfellow, Andrew Rambaut, Nick Loman, Kristian Andersen, Michael Wiley, Steve Ahuka-Mundeke, Jean-Jacques Muyembe Tamfum |
| EPI_ISL_2966671, EPI_ISL_2966673, EPI_ISL_2966675, EPI_ISL_2966676, EPI_ISL_2966677, EPI_ISL_2966683, EPI_ISL_2966684, EPI_ISL_2966685, EPI_ISL_2966686, EPI_ISL_2966688, EPI_ISL_2966690, EPI_ISL_2966691, EPI_ISL_2966692, EPI_ISL_2966694, EPI_ISL_2966695, EPI_ISL_2966696, EPI_ISL_2966697, EPI_ISL_2966699 | | | |
| see above | Hospital General Universitario Gregorio Marañón | Hospital General Universitario Gregorio Marañón | Sergio Buenestado Serrano, Pedro Sola Campoy, Laura Pérez-Lago, Cristina Rodriguez-Grande, Marta Herranz Martin, Victor Manuel de la Cueva, Julia Suárez, Pilar Catalán, Patricia Muñoz, Darío García de Viedma |
| EPI_ISL_2966701, EPI_ISL_2966702, EPI_ISL_2966703, EPI_ISL_2966704 | Lafene Health Center | Kansas State Veterinary Diagnostic Laboratory | Tyler Doerksen, Andreea Lu, Lance Noll, Elizabeth Porter, Emily Cox, Jianfa Bai, Jamie Henningson, Rachel Palinski |

| | | | |
|---|---|---|---|
| EPI_ISL_2966705 | National Institute of Public Health | State Veterinary Institute Prague | Nagy,A;Cernikov,L;Stara,M;Suri,T;Vecerova,J;Jirincova,H |
| EPI_ISL_2966707, EPI_ISL_2966708, EPI_ISL_2966709, EPI_ISL_2966710, EPI_ISL_2966711, EPI_ISL_2966712, EPI_ISL_2966713, EPI_ISL_2966723, EPI_ISL_2966725, EPI_ISL_2966728, EPI_ISL_2966739, EPI_ISL_2966740, EPI_ISL_2966743, EPI_ISL_2966752, EPI_ISL_2966753, EPI_ISL_2966754, EPI_ISL_2966755, EPI_ISL_2966757, EPI_ISL_2966760 | | | |
| see above | US Air Force School of Aerospace Medicine | US Air Force School of Aerospace Medicine | Anthony Fries, Jennifer Meyer, William Gruner, Amanda Javorina, Carol Garrett, Sarah Purves, Clarise Starr, Elizabeth Macias |
| EPI_ISL_2966761, EPI_ISL_2966763, EPI_ISL_2966764, EPI_ISL_2966765, EPI_ISL_2966766 | US Air Force School of Aerospace Medicine | US Air Force School of Aerospace Medicine | Anthony Fries, Jennifer Meyer, William Gruner, Amanda Javorina, Carol Garrett, Sarah Purves, Clarise Starr, Elizabeth Macias, Fritz Castillo, Jennifer Black, Cole Anderson |
| EPI_ISL_2966776, EPI_ISL_2966777, EPI_ISL_2966778, EPI_ISL_2966779, EPI_ISL_2966780, EPI_ISL_2966781, EPI_ISL_2966782, EPI_ISL_2966783, EPI_ISL_2966784, EPI_ISL_2966785, EPI_ISL_2966786, EPI_ISL_2966787, EPI_ISL_2966788, EPI_ISL_2966789, EPI_ISL_2966790, EPI_ISL_2966791, EPI_ISL_2966792, EPI_ISL_2966793, EPI_ISL_2966794, EPI_ISL_2966795, EPI_ISL_2966796, EPI_ISL_2966797, EPI_ISL_2966798, EPI_ISL_2966799, EPI_ISL_2966800, EPI_ISL_2966801, EPI_ISL_2966802, EPI_ISL_2966803, EPI_ISL_2966804, EPI_ISL_2966805, EPI_ISL_2966806 | | | |
| see above | Viral Respiratory Lab, National Institute for Biomedical Research (INRB) | Pathogen Sequencing Lab, National Institute for Biomedical Research (INRB) | Placide Mbala-Kingebeni, Edith Nkwembe, Eddy Kinganda-Lusamaki, Amuri Aziza, Francisca Muyembe Mawete, Emmanuel Lokilo Lofiko, Jean Claude Makangara, Catherine Pratt, Matthias Pauthner, Josh Quick, Allison Black, James Hadfield, Trevor Bedford, Ian Goodfellow, Andrew Rambaut, Nick Loman, Kristian Andersen, Michael Wiley, Steve Ahuka-Mundeke, Jean-Jacques Muyembe Tamfum |
| EPI_ISL_2966837, EPI_ISL_2966838, EPI_ISL_2966839, EPI_ISL_2966841, EPI_ISL_2966842, EPI_ISL_2966843, EPI_ISL_2966846, EPI_ISL_2966847, EPI_ISL_2966848, EPI_ISL_2966849, EPI_ISL_2966851, EPI_ISL_2966852, EPI_ISL_2966853 | | | |
| see above | Tampa General Hospital Esoteric Lab | Tampa General Hospital Esoteric Research & Development Lab | Jon Faughn, Deanna Becker, Dominic Uy, Jorge Mecias-Frias, Elaine Vendrone, Vicki Healer, Amorce Lima, Suzane Silbert |
| EPI_ISL_2966859, EPI_ISL_2966860, EPI_ISL_2966863, EPI_ISL_2966864 | Nebraska Public Health Laboratory | NPHL COVID-19 Response Team | NPHL COVID-19 Response Team |
| EPI_ISL_2966876, EPI_ISL_2966877, EPI_ISL_2966878, EPI_ISL_2966879, EPI_ISL_2966880, EPI_ISL_2966881, EPI_ISL_2966882, EPI_ISL_2966883, EPI_ISL_2966885, EPI_ISL_2966886, EPI_ISL_2966887, EPI_ISL_2966888, EPI_ISL_2966889, EPI_ISL_2966890, EPI_ISL_2966891, EPI_ISL_2966892, EPI_ISL_2966893, EPI_ISL_2966894, EPI_ISL_2966895, EPI_ISL_2966896, EPI_ISL_2966897, EPI_ISL_2966898, EPI_ISL_2966899 | | | |
| see above | Kansas Health and Environmental Lab | Kansas Health and Environmental Lab | Mike Grose, Katherine Wiggins, Jonathan Barnell, Ben Olsen, and Phil Adam |
| EPI_ISL_2966919, EPI_ISL_2966923, EPI_ISL_2966924, EPI_ISL_2966926, EPI_ISL_2966928, EPI_ISL_2966931, EPI_ISL_2966938, EPI_ISL_2966940, EPI_ISL_2966945, EPI_ISL_2966984 | Institute of Microbiology and Immunology, Faculty of Medicine, University of Ljubljana | Institute of Microbiology and Immunology, Faculty of Medicine, University of Ljubljana | Alen Suljič, Samo Zakotnik, Tomaž Mark Zorec, Matic Brvar, Doroteja Vlaj, Andraž Celar, Dominika Šturm, Patricija Pozvek, Špela Pleh, Miša Korva, Mario Poljak, Tatjana Avšič – Županc |
| EPI_ISL_2966993, EPI_ISL_2966994, EPI_ISL_2966995, EPI_ISL_2966996, EPI_ISL_2966997, EPI_ISL_2967001, EPI_ISL_2967004, EPI_ISL_2967006, EPI_ISL_2967007, EPI_ISL_2967008, EPI_ISL_2967009, EPI_ISL_2967010, EPI_ISL_2967011, EPI_ISL_2967012, EPI_ISL_2967013, EPI_ISL_2967015, EPI_ISL_2967016, EPI_ISL_2967017, EPI_ISL_2967018, EPI_ISL_2967019, EPI_ISL_2967020, EPI_ISL_2967021, EPI_ISL_2967022, EPI_ISL_2967023, EPI_ISL_2967024, EPI_ISL_2967025, EPI_ISL_2967026, EPI_ISL_2967027, EPI_ISL_2967028, EPI_ISL_2967029, EPI_ISL_2967030 | | | |
| see above | IN State Department of Health Laboratory Services | IN State Department of Health Laboratory Services | Cassandra Campion, Jamie Yeadon, Brian Pope, Lixia Liu, Kyle Brownlee, Melissa Hindenlang, Mark Glazier |
| EPI_ISL_2967038, EPI_ISL_2967042, EPI_ISL_2967058, EPI_ISL_2967059, EPI_ISL_2967060, EPI_ISL_2967062, EPI_ISL_2967064, EPI_ISL_2967066, EPI_ISL_2967067, EPI_ISL_2967071, EPI_ISL_2967076, EPI_ISL_2967078, EPI_ISL_2967079, EPI_ISL_2967081, EPI_ISL_2967082, EPI_ISL_2967083, EPI_ISL_2967092, EPI_ISL_2967096, EPI_ISL_2967108, EPI_ISL_2967110, EPI_ISL_2967113, EPI_ISL_2967114, EPI_ISL_2967120, EPI_ISL_2967122, EPI_ISL_2967125, EPI_ISL_2967132, EPI_ISL_2967136, EPI_ISL_2967139, EPI_ISL_2967141, EPI_ISL_2967145, EPI_ISL_2967148, EPI_ISL_2967150, EPI_ISL_2967153, EPI_ISL_2967154, EPI_ISL_2967157, EPI_ISL_2967165, EPI_ISL_2967166, EPI_ISL_2967167, EPI_ISL_2967168, EPI_ISL_2967169, EPI_ISL_2967170, EPI_ISL_2967171, EPI_ISL_2967172, EPI_ISL_2967173, EPI_ISL_2967174, EPI_ISL_2967175, EPI_ISL_2967176, EPI_ISL_2967177, EPI_ISL_2967185, EPI_ISL_2967187, EPI_ISL_2967189, EPI_ISL_2967190, EPI_ISL_2967191, EPI_ISL_2967192, EPI_ISL_2967193, EPI_ISL_2967194, EPI_ISL_2967195, EPI_ISL_2967196, EPI_ISL_2967197, EPI_ISL_2967198, EPI_ISL_2967199, EPI_ISL_2967200, EPI_ISL_2967201 | | | |
| see above | National Center of Infectious and Parasitic Diseases | National Center of Infectious and Parasitic Diseases | Alexiev et al |
| EPI_ISL_2967203, EPI_ISL_2967205, EPI_ISL_2967206, EPI_ISL_2967208, EPI_ISL_2967213, EPI_ISL_2967215, EPI_ISL_2967216 | Sri Ganga Ram Hospital, New Delhi, India | NCDC Delhi, Biotechnology Division | Kalaiarasan Ponnusamy, Meena Datta, Priyanka Singh, Uma Sharma, Manoj K Singh, Radhakrishnan V. S, Robin Marwal, Mahesh S Dhar, Hemlata Lall, Hema Gogia, Preeti Madan, Sandhya Kabra, Sujeet K Singh, Partha Rakshit |
| EPI_ISL_2967218, EPI_ISL_2967219, EPI_ISL_2967224, EPI_ISL_2967225 | Max von Pettenkofer Institute, Virology, National Reference Center for Retroviruses, LMU Munich | Laboratory for Functional Genome Analysis; Dept. Genomics; Gene Center of the LMU Munich | Max Muenchhoff; Stefan Krebs; Alexander Graf; Oliver Keppler; Helmut Blum |
| EPI_ISL_2967226, EPI_ISL_2967227, EPI_ISL_2967228, EPI_ISL_2967229, EPI_ISL_2967230, EPI_ISL_2967231, EPI_ISL_2967232, EPI_ISL_2967233, EPI_ISL_2967234, EPI_ISL_2967235, EPI_ISL_2967236, EPI_ISL_2967237, EPI_ISL_2967240, EPI_ISL_2967242, EPI_ISL_2967243, EPI_ISL_2967244, EPI_ISL_2967245, EPI_ISL_2967246, EPI_ISL_2967247, EPI_ISL_2967248 | | | |
| see above | Kansas Health and Environmental Lab | Kansas Health and Environmental Lab | Katherine Wiggins, Mike Grose, Jonathan Barnell, Ben Olsen, and Phil Adam |
| EPI_ISL_2967250, EPI_ISL_2967251, EPI_ISL_2967259, EPI_ISL_2967279, EPI_ISL_2967298, EPI_ISL_2967307, EPI_ISL_2967308, EPI_ISL_2967312, EPI_ISL_2967323, EPI_ISL_2967325, EPI_ISL_2967326, EPI_ISL_2967327, EPI_ISL_2967328, EPI_ISL_2967329, EPI_ISL_2967331, EPI_ISL_2967338, EPI_ISL_2967346, EPI_ISL_2967349, EPI_ISL_2967350, EPI_ISL_2967351, EPI_ISL_2967352, EPI_ISL_2967353, EPI_ISL_2967355, EPI_ISL_2967356, EPI_ISL_2967357, EPI_ISL_2967359, EPI_ISL_2967361, EPI_ISL_2967369, EPI_ISL_2967371, EPI_ISL_2967374, EPI_ISL_2967376, EPI_ISL_2967377, EPI_ISL_2967378, EPI_ISL_2967379, EPI_ISL_2967380, EPI_ISL_2967381, EPI_ISL_2967382, EPI_ISL_2967383, EPI_ISL_2967385, EPI_ISL_2967387, EPI_ISL_2967389, EPI_ISL_2967391, EPI_ISL_2967392, EPI_ISL_2967393, EPI_ISL_2967394, EPI_ISL_2967395, EPI_ISL_2967396, EPI_ISL_2967397, EPI_ISL_2967398, EPI_ISL_2967399, EPI_ISL_2967400, EPI_ISL_2967401, EPI_ISL_2967402, EPI_ISL_2967404, EPI_ISL_2967412, EPI_ISL_2967413, EPI_ISL_2967423, EPI_ISL_2967424, EPI_ISL_2967426, EPI_ISL_2967434, EPI_ISL_2967437, EPI_ISL_2967447 | | | |
| see above | Division of Emerging Infectious Diseases, Bureau of Infectious Diseases Diagnosis Control, Korea Disease Control and Prevention Agency | Division of Emerging Infectious Diseases, Bureau of Infectious Diseases Diagnosis Control, Korea Disease Control and Prevention Agency | Jeong-Ah Kim, Ae Kyung Park, Il-Hwan Kim, Heui Man Kim, Jeong-Min Kim, Chae Young Lee, Eun-Jin Kim |
| EPI_ISL_2967582, EPI_ISL_2967583, EPI_ISL_2967584, EPI_ISL_2967585, EPI_ISL_2967586, EPI_ISL_2967587, EPI_ISL_2967588, EPI_ISL_2967589, EPI_ISL_2967590, EPI_ISL_2967592, EPI_ISL_2967593, EPI_ISL_2967594, EPI_ISL_2967595, EPI_ISL_2967596, EPI_ISL_2967598, EPI_ISL_2967602, EPI_ISL_2967603, EPI_ISL_2967604, EPI_ISL_2967605, EPI_ISL_2967606, EPI_ISL_2967607, EPI_ISL_2967609, EPI_ISL_2967610, EPI_ISL_2967611, EPI_ISL_2967612, EPI_ISL_2967613, EPI_ISL_2967615, EPI_ISL_2967617, EPI_ISL_2967618, EPI_ISL_2967619, EPI_ISL_2967620, EPI_ISL_2967621, EPI_ISL_2967622, EPI_ISL_2967623, EPI_ISL_2967624, EPI_ISL_2967626, EPI_ISL_2967627, EPI_ISL_2967628, EPI_ISL_2967629, EPI_ISL_2967630, EPI_ISL_2967631, EPI_ISL_2967632, EPI_ISL_2967633, EPI_ISL_2967634, EPI_ISL_2967635, EPI_ISL_2967636, EPI_ISL_2967638, EPI_ISL_2967639, EPI_ISL_2967640, EPI_ISL_2967641, EPI_ISL_2967643, EPI_ISL_2967644, EPI_ISL_2967645, EPI_ISL_2967646, EPI_ISL_2967647, EPI_ISL_2967648, EPI_ISL_2967650, EPI_ISL_2967651, EPI_ISL_2967652, EPI_ISL_2967653, EPI_ISL_2967654, EPI_ISL_2967655, EPI_ISL_2967656, EPI_ISL_2967657, EPI_ISL_2967659, EPI_ISL_2967660, EPI_ISL_2967661, EPI_ISL_2967662, EPI_ISL_2967664, EPI_ISL_2967665, EPI_ISL_2967666, EPI_ISL_2967667, EPI_ISL_2967668, EPI_ISL_2967670, EPI_ISL_2967673, EPI_ISL_2967674, EPI_ISL_2967675, EPI_ISL_2967676, EPI_ISL_2967677, EPI_ISL_2967679, EPI_ISL_2967680, EPI_ISL_2967681, EPI_ISL_2967682, EPI_ISL_2967683, EPI_ISL_2967684, EPI_ISL_2967685, EPI_ISL_2967686, EPI_ISL_2967687, EPI_ISL_2967688, EPI_ISL_2967689, EPI_ISL_2967690, EPI_ISL_2967693, EPI_ISL_2967694, EPI_ISL_2967696, EPI_ISL_2967697, EPI_ISL_2967698, EPI_ISL_2967699, EPI_ISL_2967701, EPI_ISL_2967702, EPI_ISL_2967703, EPI_ISL_2967704, EPI_ISL_2967705, EPI_ISL_2967706, EPI_ISL_2967709, EPI_ISL_2967710, EPI_ISL_2967711, EPI_ISL_2967713, EPI_ISL_2967714, EPI_ISL_2967716, EPI_ISL_2967717, EPI_ISL_2967718, EPI_ISL_2967720, EPI_ISL_2967721, EPI_ISL_2967723, EPI_ISL_2967724, EPI_ISL_2967726, EPI_ISL_2967727, EPI_ISL_2967728, EPI_ISL_2967730, EPI_ISL_2967731, EPI_ISL_2967734, EPI_ISL_2967735, EPI_ISL_2967736, EPI_ISL_2967737, EPI_ISL_2967739, EPI_ISL_2967740, EPI_ISL_2967741, EPI_ISL_2967742, EPI_ISL_2967746, EPI_ISL_2967747, EPI_ISL_2967749, EPI_ISL_2967751, EPI_ISL_2967752, EPI_ISL_2967753, EPI_ISL_2967756, EPI_ISL_2967757, EPI_ISL_2967758, EPI_ISL_2967759, EPI_ISL_2967760, EPI_ISL_2967761, EPI_ISL_2967762, EPI_ISL_2967765, EPI_ISL_2967767, EPI_ISL_2967769, EPI_ISL_2967770, EPI_ISL_2967772 | | | |
| see above | UW Virology Lab | UW Virology Lab | Pavitra Roychoudhury, Hong Xie, Lasata Shrestha, Tien V. Nguyen, Shah Mohamed Bakhash, Michelle Lin, Noah R. Baker, Ricardo Perez, Sean Ellis, Nathan Breit, Robert J. Livingston, Meei-Li Huang, Keith R Jerome, Patrick Mathias, Alexander Greninger |
| EPI_ISL_2967781, EPI_ISL_2967782, EPI_ISL_2967783, EPI_ISL_2967784, EPI_ISL_2967785, EPI_ISL_2967786, EPI_ISL_2967787, EPI_ISL_2967788 | Salzkammergutklinikum Vöcklabruck, Institut für Pathologie | Salzkammergutklinikum Vöcklabruck, Institut für Pathologie | Regina Stitz, Penka Lechner, Senka Rohregger, René Silye |
| EPI_ISL_2967804, EPI_ISL_2967819, EPI_ISL_2967833, EPI_ISL_2967834, EPI_ISL_2967835, EPI_ISL_2967836, EPI_ISL_2967842, EPI_ISL_2967855, EPI_ISL_2967870, EPI_ISL_2967871, EPI_ISL_2967872, EPI_ISL_2967873, EPI_ISL_2967874, EPI_ISL_2967875, EPI_ISL_2967876, EPI_ISL_2967877, EPI_ISL_2967886, EPI_ISL_2967887, EPI_ISL_2967894, EPI_ISL_2967895, EPI_ISL_2967896 | | | |
| see above | Microbiology Department. Complexo Hospitalario Universitario de Vigo | Microbiology Department. Complexo Hospitalario Universitario de Vigo | Alfaya N, Alonso I, Alvarez M, Cabrera JJ, Carballo R, Cores O, Cortizo S, del-Campo V, Martinez L, Mediero G, Perez S, Davina C, Potel C, Regueiro B, Rey S, Vasallo FJ |
| EPI_ISL_2967897, EPI_ISL_2967898, EPI_ISL_2967899, EPI_ISL_2967900, EPI_ISL_2967901, EPI_ISL_2967902, EPI_ISL_2967903, EPI_ISL_2967904, EPI_ISL_2967905, EPI_ISL_2967906, EPI_ISL_2967907, EPI_ISL_2967908, EPI_ISL_2967909, EPI_ISL_2967910, EPI_ISL_2967911, EPI_ISL_2967912, EPI_ISL_2967913, EPI_ISL_2967914, EPI_ISL_2967915, EPI_ISL_2967916, EPI_ISL_2967917, EPI_ISL_2967918, EPI_ISL_2967919, EPI_ISL_2967920, EPI_ISL_2967921, EPI_ISL_2967922, EPI_ISL_2967923, EPI_ISL_2967924, EPI_ISL_2967925 | | | |
| see above | Nebraska Public Health Laboratory | NPHL COVID-19 Response Team | NPHL COVID-19 Response Team |
| EPI_ISL_2967930 | Santa Clara Valley Medical Center | Santa Clara County public Health Laboratory | Santa Clara County Public Health Department |
| EPI_ISL_2967931 | Santa Clara County Public Health Laboratory | Santa Clara County public Health Laboratory | Santa Clara County Public Health Department |
| EPI_ISL_2967932, EPI_ISL_2967933, EPI_ISL_2967934, EPI_ISL_2967938, EPI_ISL_2967941, EPI_ISL_2967942, EPI_ISL_2967946, EPI_ISL_2967947, EPI_ISL_2967948, EPI_ISL_2967949, EPI_ISL_2967950, EPI_ISL_2967951, EPI_ISL_2967952, EPI_ISL_2967953, EPI_ISL_2967955, EPI_ISL_2967956, EPI_ISL_2967958, EPI_ISL_2967960, EPI_ISL_2967963 | | | |
| see above | Santa Clara Valley Medical Center | Santa Clara County public Health Laboratory | Santa Clara County Public Health Department |
| EPI_ISL_2967964 | Kaiser Santa Clara | Santa Clara County public Health Laboratory | Santa Clara County Public Health Department |
| EPI_ISL_2967966 | Good Samaritan Hospital | Santa Clara County public Health Laboratory | Santa Clara County Public Health Department |
| EPI_ISL_2967967, EPI_ISL_2967968 | El Camino Hospital | Santa Clara County public Health Laboratory | Santa Clara County Public Health Department |
| EPI_ISL_2967969, EPI_ISL_2967970, EPI_ISL_2967971 | Kaiser San Jose | Santa Clara County public Health Laboratory | Santa Clara County Public Health Department |
| EPI_ISL_2967973 | PAMF Palo Alto | Santa Clara County public Health Laboratory | Santa Clara County Public Health Department |
| EPI_ISL_2967991, | The Caribbean Public Health Agency | Carrington Lab, Department of Preclinical Sciences, Faculty of | Nikita S. D. Sahadeo, Arianne Brown-Jordan, Sarah Hill, Vernie Ramkissoon, Anushka Ramjag, Kenneth George, Avery Hinds, Risha Singh, SueMin Nathaniel, Nuno Faria, Oliver Pybus, Gabriel Escobar, Christine V. F. Carrington |

| | | | |
|---|---|---|---|
| EPI_ISL_2967992 | | Medical Sciences, The University of the West Indies, St Augustine Campus | |
| EPI_ISL_2967997, EPI_ISL_2968004, EPI_ISL_2968009, EPI_ISL_2968010, EPI_ISL_2968015, EPI_ISL_2968016, EPI_ISL_2968025, EPI_ISL_2968031 | WACCBIP, University of Ghana, Accra, Ghana | WACCBIP, University of Ghana, Volta Road, Legon-Accra, Ghana | Collins M. Morang'a, Joyce M. Ngoi, Dominic S. Y. Amuzu, Vincent Appiah, Evelyn B. Quansah, Philip M. Soglo, Violette V. M'cormack, Samirah Saiid, Bright K. Yemi, Israel Osei-Wusu, Deborah N. A. Mettle, Oliver Commey, Samuel Armoo, Nicaise T. Ndam, Frederick Tei-Maya, Joe K. Mutungi, William K. Ampofo, Yaw Bediako, Peter K Quashie, Lucas N. Amenga-Etego and Gordon A. Awandare |
| EPI_ISL_2968070, EPI_ISL_2968071, EPI_ISL_2968072, EPI_ISL_2968073, EPI_ISL_2968074, EPI_ISL_2968075, EPI_ISL_2968076, EPI_ISL_2968077, EPI_ISL_2968078, EPI_ISL_2968079, EPI_ISL_2968080, EPI_ISL_2968081, EPI_ISL_2968082, EPI_ISL_2968083, EPI_ISL_2968084, EPI_ISL_2968085, EPI_ISL_2968086, EPI_ISL_2968087, EPI_ISL_2968088, EPI_ISL_2968089, EPI_ISL_2968090, EPI_ISL_2968091, EPI_ISL_2968092, EPI_ISL_2968093 | | | |
| see above | NILMRC | Genomic Research Laboratory, Bangabandhu Sheikh Mujib Medical University | Laila Anjuman Banu, Md.Saydur Rahman, Sharadindu Kanti Sinha, Arifa Akram, Md.Sharfuddin Ahmed |
| EPI_ISL_2968094, EPI_ISL_2968095, EPI_ISL_2968096, EPI_ISL_2968097, EPI_ISL_2968098, EPI_ISL_2968099, EPI_ISL_2968100, EPI_ISL_2968101, EPI_ISL_2968102, EPI_ISL_2968103, EPI_ISL_2968104, EPI_ISL_2968105, EPI_ISL_2968106, EPI_ISL_2968107, EPI_ISL_2968108, EPI_ISL_2968109, EPI_ISL_2968110, EPI_ISL_2968111, EPI_ISL_2968112, EPI_ISL_2968113, EPI_ISL_2968114, EPI_ISL_2968115, EPI_ISL_2968116, EPI_ISL_2968118, EPI_ISL_2968119, EPI_ISL_2968120, EPI_ISL_2968121, EPI_ISL_2968122, EPI_ISL_2968123, EPI_ISL_2968124, EPI_ISL_2968125 | | | |
| see above | ASST MONZA | ASST MONZA | Sergio Maria Ivano Malandrin |
| EPI_ISL_2968126, EPI_ISL_2968127, EPI_ISL_2968129, EPI_ISL_2968130, EPI_ISL_2968131, EPI_ISL_2968132, EPI_ISL_2968133, EPI_ISL_2968134, EPI_ISL_2968135, EPI_ISL_2968136, EPI_ISL_2968137, EPI_ISL_2968139, EPI_ISL_2968141, EPI_ISL_2968142, EPI_ISL_2968143, EPI_ISL_2968144, EPI_ISL_2968145, EPI_ISL_2968148, EPI_ISL_2968150, EPI_ISL_2968152, EPI_ISL_2968153, EPI_ISL_2968154, EPI_ISL_2968156, EPI_ISL_2968157, EPI_ISL_2968158, EPI_ISL_2968159, EPI_ISL_2968160, EPI_ISL_2968161, EPI_ISL_2968162, EPI_ISL_2968163, EPI_ISL_2968164, EPI_ISL_2968165, EPI_ISL_2968166, EPI_ISL_2968167, EPI_ISL_2968168, EPI_ISL_2968169, EPI_ISL_2968170, EPI_ISL_2968171, EPI_ISL_2968172, EPI_ISL_2968173, EPI_ISL_2968174, EPI_ISL_2968175, EPI_ISL_2968176, EPI_ISL_2968177, EPI_ISL_2968178, EPI_ISL_2968179, EPI_ISL_2968180, EPI_ISL_2968181, EPI_ISL_2968182, EPI_ISL_2968183, EPI_ISL_2968184, EPI_ISL_2968185, EPI_ISL_2968186, EPI_ISL_2968187, EPI_ISL_2968188, EPI_ISL_2968189, EPI_ISL_2968190, EPI_ISL_2968191, EPI_ISL_2968192, EPI_ISL_2968193, EPI_ISL_2968194, EPI_ISL_2968195, EPI_ISL_2968198, EPI_ISL_2968199, EPI_ISL_2968200, EPI_ISL_2968201, EPI_ISL_2968202, EPI_ISL_2968203, EPI_ISL_2968204, EPI_ISL_2968205, EPI_ISL_2968206, EPI_ISL_2968207, EPI_ISL_2968208, EPI_ISL_2968209, EPI_ISL_2968210, EPI_ISL_2968211, EPI_ISL_2968214, EPI_ISL_2968220, EPI_ISL_2968223, EPI_ISL_2968224, EPI_ISL_2968225, EPI_ISL_2968226, EPI_ISL_2968227, EPI_ISL_2968228, EPI_ISL_2968229, EPI_ISL_2968230, EPI_ISL_2968231, EPI_ISL_2968232, EPI_ISL_2968233, EPI_ISL_2968234, EPI_ISL_2968235, EPI_ISL_2968236, EPI_ISL_2968237, EPI_ISL_2968238, EPI_ISL_2968239, EPI_ISL_2968240, EPI_ISL_2968241, EPI_ISL_2968242, EPI_ISL_2968243, EPI_ISL_2968244, EPI_ISL_2968245, EPI_ISL_2968246, EPI_ISL_2968247, EPI_ISL_2968248, EPI_ISL_2968249, EPI_ISL_2968250, EPI_ISL_2968251, EPI_ISL_2968252, EPI_ISL_2968253, EPI_ISL_2968254, EPI_ISL_2968255, EPI_ISL_2968256, EPI_ISL_2968257, EPI_ISL_2968258, EPI_ISL_2968259, EPI_ISL_2968260, EPI_ISL_2968261, EPI_ISL_2968262, EPI_ISL_2968263, EPI_ISL_2968264, EPI_ISL_2968265, EPI_ISL_2968266, EPI_ISL_2968268, EPI_ISL_2968269, EPI_ISL_2968270, EPI_ISL_2968271, EPI_ISL_2968272, EPI_ISL_2968273, EPI_ISL_2968274, EPI_ISL_2968275, EPI_ISL_2968276, EPI_ISL_2968277, EPI_ISL_2968278, EPI_ISL_2968279, EPI_ISL_2968280, EPI_ISL_2968281, EPI_ISL_2968282, EPI_ISL_2968283, EPI_ISL_2968285 | | | |
| see above | National Virus Reference Laboratory | National Virus Reference Laboratory | Zoe Yandle, Charlene Bennett, Gabriel Gonzalez, Michael Carr, Jonathan Dean, Cillian F De Gascun |
| EPI_ISL_2968537, EPI_ISL_2968538, EPI_ISL_2968539, EPI_ISL_2968540, EPI_ISL_2968541, EPI_ISL_2968542, EPI_ISL_2968544, EPI_ISL_2968545, EPI_ISL_2968546, EPI_ISL_2968547, EPI_ISL_2968548, EPI_ISL_2968550, EPI_ISL_2968552, EPI_ISL_2968553, EPI_ISL_2968554, EPI_ISL_2968555, EPI_ISL_2968556, EPI_ISL_2968557, EPI_ISL_2968558, EPI_ISL_2968559, EPI_ISL_2968561, EPI_ISL_2968562, EPI_ISL_2968563, EPI_ISL_2968564, EPI_ISL_2968565, EPI_ISL_2968566, EPI_ISL_2968567, EPI_ISL_2968568, EPI_ISL_2968569, EPI_ISL_2968570, EPI_ISL_2968571, EPI_ISL_2968572, EPI_ISL_2968573, EPI_ISL_2968575, EPI_ISL_2968576, EPI_ISL_2968577, EPI_ISL_2968578, EPI_ISL_2968579, EPI_ISL_2968580, EPI_ISL_2968581, EPI_ISL_2968582, EPI_ISL_2968583, EPI_ISL_2968585, EPI_ISL_2968586, EPI_ISL_2968587, EPI_ISL_2968588, EPI_ISL_2968589, EPI_ISL_2968590, EPI_ISL_2968591, EPI_ISL_2968592, EPI_ISL_2968593, EPI_ISL_2968594, EPI_ISL_2968595 | | | |
| see above | Viral Respiratory Lab, National Institute for Biomedical Research (INRB) | Pathogen Sequencing Lab, National Institute for Biomedical Research (INRB) | Placide Mbala-Kingebeni, Edith Nkwembe, Eddy Kinganda-Lusamaki, Amuri Aziza, Francisca Muyembe Mawete, Emmanuel Lokilo Lofiko, Jean Claude Makangara, Catherine Pratt, Matthias Pauthner, Josh Quick, Allison Black, James Hadfield, Trevor Bedford, Ian Goodfellow, Andrew Rambaut, Nick Loman, Kristian Andersen, Michael Wiley, Steve Ahuka-Mundeke, Jean-Jacques Muyembe Tamfum |
| EPI_ISL_2968596, EPI_ISL_2968597, EPI_ISL_2968599, EPI_ISL_2968600, EPI_ISL_2968601, EPI_ISL_2968605, EPI_ISL_2968606, EPI_ISL_2968607, EPI_ISL_2968608, EPI_ISL_2968609, EPI_ISL_2968610, EPI_ISL_2968611, EPI_ISL_2968612, EPI_ISL_2968614, EPI_ISL_2968615, EPI_ISL_2968616, EPI_ISL_2968618, EPI_ISL_2968622, EPI_ISL_2968623, EPI_ISL_2968624, EPI_ISL_2968626, EPI_ISL_2968627, EPI_ISL_2968628, EPI_ISL_2968629, EPI_ISL_2968630, EPI_ISL_2968638 | | | |
| see above | Houston Health Dept. | Houston Health Dept. | Ryker Penn, Pamela Brown, Adolpho Lara, Yanlai Lai |
| EPI_ISL_2968639, EPI_ISL_2968640, EPI_ISL_2968641, EPI_ISL_2968642, EPI_ISL_2968643, EPI_ISL_2968645, EPI_ISL_2968646, EPI_ISL_2968648, EPI_ISL_2968649, EPI_ISL_2968650, EPI_ISL_2968653, EPI_ISL_2968655, EPI_ISL_2968656, EPI_ISL_2968658, EPI_ISL_2968659, EPI_ISL_2968661 | | | |
| see above | Arizona State University | Arizona State University | Peter T. Skidmore, LaRinda A. Holland, Matthew F. Smith, Rabia Maqsood, Nicholas J. Mellor, Joy M. Blain, Valerie Harris, Joshua LaBaer, Vel Murugan, Efrem S. Lim |
| EPI_ISL_2968662, EPI_ISL_2968666, EPI_ISL_2968667, EPI_ISL_2968668, EPI_ISL_2968669, EPI_ISL_2968670, EPI_ISL_2968671, EPI_ISL_2968672, EPI_ISL_2968673, EPI_ISL_2968674, EPI_ISL_2968675, EPI_ISL_2968676, EPI_ISL_2968677, EPI_ISL_2968678, EPI_ISL_2968679, EPI_ISL_2968680, EPI_ISL_2968681, EPI_ISL_2968682, EPI_ISL_2968685, EPI_ISL_2968686, EPI_ISL_2968687, EPI_ISL_2968688 | | | |
| see above | Houston Health Dept. | Houston Health Dept. | Ryker Penn, Pamela Brown, Adolpho Lara, Yanlai Lai |
| EPI_ISL_2968692, EPI_ISL_2968698, EPI_ISL_2968699, EPI_ISL_2968701, EPI_ISL_2968704, EPI_ISL_2968705, EPI_ISL_2968707, EPI_ISL_2968708, EPI_ISL_2968709, EPI_ISL_2968712, EPI_ISL_2968713, EPI_ISL_2968715, EPI_ISL_2968718, EPI_ISL_2968719, EPI_ISL_2968720, EPI_ISL_2968721, EPI_ISL_2968722, EPI_ISL_2968723, EPI_ISL_2968724, EPI_ISL_2968725, EPI_ISL_2968726, EPI_ISL_2968727, EPI_ISL_2968728, EPI_ISL_2968729, EPI_ISL_2968730, EPI_ISL_2968732, EPI_ISL_2968733, EPI_ISL_2968734, EPI_ISL_2968735, EPI_ISL_2968736, EPI_ISL_2968737, EPI_ISL_2968738, EPI_ISL_2968740, EPI_ISL_2968741, EPI_ISL_2968742, EPI_ISL_2968743, EPI_ISL_2968744, EPI_ISL_2968745, EPI_ISL_2968746, EPI_ISL_2968747, EPI_ISL_2968748, EPI_ISL_2968749, EPI_ISL_2968750, EPI_ISL_2968752, EPI_ISL_2968753, EPI_ISL_2968754, EPI_ISL_2968755, EPI_ISL_2968756, EPI_ISL_2968757, EPI_ISL_2968758, EPI_ISL_2968759, EPI_ISL_2968760, EPI_ISL_2968762, EPI_ISL_2968763, EPI_ISL_2968764, EPI_ISL_2968765, EPI_ISL_2968766, EPI_ISL_2969520, EPI_ISL_2969525, EPI_ISL_2969526, EPI_ISL_2969529, EPI_ISL_2969530, EPI_ISL_2969531 | | | |
| see above | Viral Respiratory Lab, National Institute for Biomedical Research (INRB) | Pathogen Sequencing Lab, National Institute for Biomedical Research (INRB) | Placide Mbala-Kingebeni, Edith Nkwembe, Eddy Kinganda-Lusamaki, Amuri Aziza, Francisca Muyembe Mawete, Emmanuel Lokilo Lofiko, Jean Claude Makangara, Catherine Pratt, Matthias Pauthner, Josh Quick, Allison Black, James Hadfield, Trevor Bedford, Ian Goodfellow, Andrew Rambaut, Nick Loman, Kristian Andersen, Michael Wiley, Steve Ahuka-Mundeke, Jean-Jacques Muyembe Tamfum |
| EPI_ISL_2969533, EPI_ISL_2969534, EPI_ISL_2969536, EPI_ISL_2969538, EPI_ISL_2969539, EPI_ISL_2969540, EPI_ISL_2969541, EPI_ISL_2969542, EPI_ISL_2969543, EPI_ISL_2969544, EPI_ISL_2969546, EPI_ISL_2969548, EPI_ISL_2969549, EPI_ISL_2969554, EPI_ISL_2969555, EPI_ISL_2969558, EPI_ISL_2969560, EPI_ISL_2969561, EPI_ISL_2969564, EPI_ISL_2969565, EPI_ISL_2969566, EPI_ISL_2969567, EPI_ISL_2969568, EPI_ISL_2969572, EPI_ISL_2969573, EPI_ISL_2969575, EPI_ISL_2969576, EPI_ISL_2969577, EPI_ISL_2969579 | | | |
| see above | Clinical Microbiology, Infection Prevention and Control | Section for Molecular Diagnostics | Björn Hallström, Jonas Björkman |
| EPI_ISL_2969582, EPI_ISL_2969583, EPI_ISL_2969584, EPI_ISL_2969586, EPI_ISL_2969592, EPI_ISL_2969593, EPI_ISL_2969596, EPI_ISL_2969597, EPI_ISL_2969602, EPI_ISL_2969603, EPI_ISL_2969605, EPI_ISL_2969606, EPI_ISL_2969607, EPI_ISL_2969612 | | | |
| see above | Alaska State Virology Laboratory | Alaska State Virology Laboratory | Stephanie DeRonde, Elva House, Jacob Zidek, Lisa Smith, Ph.D., Jack Chen, Ph.D. |
| EPI_ISL_2969616, EPI_ISL_2969617, EPI_ISL_2969618, EPI_ISL_2969620, EPI_ISL_2969621, EPI_ISL_2969623, EPI_ISL_2969625, EPI_ISL_2969626 | Gundersen Clinical Microbiology Laboratory | Kabara Cancer Research Institute | Craig S. Richmond, Paraic A. Kenny |
| EPI_ISL_2969657 | A. Krumbholz, Labor Dr. Krause und Kollegen MVZ GmbH, Kiel | Charité Universitätsmedizin Berlin, Institute of Virology | Victor M Corman, Barbara Mühlemann, Julia Schneider, Jörn Beheim-Schwarzbach, Tobias Bleicker, Julia Tesch, Talitha Veith, Andi Krumbholz, Terry Jones, Christian Drosten |
| EPI_ISL_2969658, EPI_ISL_2969659, EPI_ISL_2969858, EPI_ISL_2969860 | National Platform bis UMONS/Jolimont | National Platform bis UMONS/Jolimont | François Dufrasne, Guillaume Bayon-Vicente, Florian Juszczak, Eric Tarantino, Gautier Detry, Ruddy Wattiez |
| EPI_ISL_2969894 | Centro de Investigacion Biomedica del Noreste (CIBIN) | Unidad de Genomica Avanzada | Consorcio Mexicano de Vigilancia Genomica (CoViGen-Mex). Authors (in alphabetical order): Julio Elias Alvarado-Yaah, Carlos F. Arias, Santiago avila-Rios, Eduardo Becerril-Vargas, Victor Hugo Borja-Aburto, Celia Boukadida, Cristobal Chaidez-Quiroz, Juan Bautista Chale-Dzul, Ricardo Ciria Merce, Andreu Comas-Garcia, Celida Duque Molina, Julissa Enciso-Ibarra, Jose Antonio Enciso-Moreno, Gloria Elena Espinosa-Ayala, Fernando Fontove-Herrera, Daniel Fregoso-Rueda, Victor Eduardo Garcia-Arias, Alejandra Garcia-Gasca, Bruno Gomez-Gil, Jean Pierre Gonzalez, Irvin Gonzalez-Lopez, Concepcion Grajales-Muñiz, Ricardo Grande, Rosa Maria Gutierrez Rios, Jesus Hernandez, Alejandra Hernandez-Teran, Alfredo Herrera-Estrella, Carla Ivon Herrera-Najera, Pavel Isa, Veronica Mata-Haro, Daniel Lira Morales, Susana Lopez, Antonio Loza Roman, Brenda Irasema Maldonado-Meza, Bernardo Martinez-Miguel, Jose Arturo Martinez-Orozco, Celida Martinez-Rodriguez, Margarita Matias-Florentino, Fidencio Mejia-Nepomuceno, Maria Guadalupe de Jesus Mireles-Rivera, Gloria Maria Molina-Salinas, Hector Montoya-Fuentes, , Mario Mujica-Sanchez, Jose Esteban Muñoz-Medina, Jose de Jesus Nuñez-Contreras, Alicia Ocaña-Mondragon, Luis Alberto Ochoa-Carrera, Hector Esteban Paz-Juarez, Marissa Perez-Garcia, Francisco Pulido, Helen Haydee Fernanda Ramirez-Plascencia, Jorge Salas-Hernandez, Angel Gustavo Salas-Lais, Alejandro Sanchez-Flores, Clara Esperanza Santacruz-Tinoco, Maria Guadalupe Santiago-Mauricio, Selene Zarate, Nelly Selem-Mojica, Blanca Taboada, Gloria Vazquez, Joel Armando Vazquez-Perez. |
| EPI_ISL_2969963, EPI_ISL_2969964 | Laboratorio Central de Epidemiologia (LCE) | Unidad de Genomica Avanzada | Consorcio Mexicano de Vigilancia Genomica (CoViGen-Mex). Authors (in alphabetical order): Julio Elias Alvarado-Yaah, Carlos F. Arias, Santiago avila-Rios, Eduardo Becerril-Vargas, Victor Hugo Borja-Aburto, Celia Boukadida, Cristobal Chaidez-Quiroz, Juan Bautista Chale-Dzul, Ricardo Ciria Merce, Andreu Comas-Garcia, Celida Duque Molina, Julissa Enciso-Ibarra, Jose Antonio Enciso-Moreno, Gloria Elena Espinosa-Ayala, Fernando Fontove-Herrera, Daniel Fregoso-Rueda, Victor Eduardo Garcia-Arias, Alejandra Garcia-Gasca, Bruno Gomez-Gil, Jean Pierre Gonzalez, Irvin Gonzalez-Lopez, Concepcion Grajales-Muñiz, Ricardo Grande, Rosa Maria Gutierrez Rios, Jesus Hernandez, Alejandra Hernandez-Teran, Alfredo Herrera-Estrella, Carla Ivon Herrera-Najera, Pavel Isa, Veronica Mata-Haro, Daniel Lira Morales, Susana Lopez, Antonio Loza Roman, Brenda Irasema Maldonado-Meza, Bernardo Martinez-Miguel, Jose Arturo Martinez-Orozco, Celida Martinez-Rodriguez, Margarita Matias-Florentino, Fidencio Mejia-Nepomuceno, Maria Guadalupe de Jesus Mireles-Rivera, Gloria Maria Molina-Salinas, Hector Montoya-Fuentes, , Mario Mujica-Sanchez, Jose Esteban Muñoz-Medina, Jose de Jesus Nuñez-Contreras, Alicia Ocaña-Mondragon, Luis Alberto Ochoa-Carrera, Hector Esteban Paz-Juarez, Marissa Perez-Garcia, Francisco Pulido, Helen Haydee Fernanda Ramirez-Plascencia, Jorge Salas-Hernandez, Angel Gustavo Salas-Lais, Alejandro Sanchez-Flores, Clara Esperanza Santacruz-Tinoco, Maria Guadalupe Santiago-Mauricio, Selene Zarate, Nelly Selem-Mojica, Blanca Taboada, Gloria Vazquez, Joel Armando Vazquez-Perez. |
| EPI_ISL_2969975 | National Platform bis UMONS/Jolimont | National Platform bis UMONS/Jolimont | François Dufrasne, Guillaume Bayon-Vicente, Florian Juszczak, Eric Tarantino, Gautier Detry, Ruddy Wattiez |
| EPI_ISL_2970008 | Centro de Investigacion Biomedica del Noreste (CIBIN) | Unidad de Genomica Avanzada | Consorcio Mexicano de Vigilancia Genomica (CoViGen-Mex). Authors (in alphabetical order): Julio Elias Alvarado-Yaah, Carlos F. Arias, Santiago avila-Rios, Eduardo Becerril-Vargas, Victor Hugo Borja-Aburto, Celia Boukadida, Cristobal Chaidez-Quiroz, Juan Bautista Chale-Dzul, Ricardo Ciria Merce, Andreu Comas-Garcia, Celida Duque Molina, Julissa Enciso-Ibarra, Jose Antonio Enciso-Moreno, Gloria Elena Espinosa-Ayala, Fernando Fontove-Herrera, Daniel Fregoso-Rueda, Victor Eduardo Garcia-Arias, Alejandra Garcia-Gasca, Bruno Gomez-Gil, Jean Pierre Gonzalez, Irvin Gonzalez-Lopez, Concepcion Grajales-Muñiz, Ricardo Grande, Rosa Maria Gutierrez Rios, Jesus Hernandez, Alejandra Hernandez-Teran, Alfredo Herrera-Estrella, Carla Ivon Herrera-Najera, Pavel Isa, Veronica Mata-Haro, Daniel Lira Morales, Susana Lopez, Antonio Loza Roman, Brenda Irasema Maldonado-Meza, Bernardo Martinez-Miguel, Jose Arturo Martinez-Orozco, Celida Martinez-Rodriguez, Margarita Matias-Florentino, Fidencio Mejia-Nepomuceno, Maria Guadalupe de Jesus Mireles-Rivera, Gloria Maria Molina-Salinas, Hector Montoya-Fuentes, , Mario Mujica-Sanchez, Jose Esteban Muñoz-Medina, Jose de Jesus Nuñez-Contreras, Alicia Ocaña-Mondragon, Luis Alberto Ochoa-Carrera, Hector Esteban Paz-Juarez, Marissa Perez-Garcia, Francisco Pulido, Helen Haydee Fernanda Ramirez-Plascencia, Jorge Salas-Hernandez, Angel Gustavo Salas-Lais, Alejandro Sanchez-Flores, Clara Esperanza Santacruz-Tinoco, Maria Guadalupe Santiago-Mauricio, Selene Zarate, Nelly Selem-Mojica, Blanca Taboada, Gloria Vazquez, Joel Armando Vazquez-Perez. |
| EPI_ISL_2970030, EPI_ISL_2970082 | Unidad de Investigacion Medica de Yucatan (UIMY) | Unidad de Genomica Avanzada | Consorcio Mexicano de Vigilancia Genomica (CoViGen-Mex). Authors (in alphabetical order): Julio Elias Alvarado-Yaah, Carlos F. Arias, Santiago avila-Rios, Eduardo Becerril-Vargas, Victor Hugo Borja-Aburto, Celia Boukadida, Cristobal Chaidez-Quiroz, Juan Bautista Chale-Dzul, Ricardo Ciria Merce, Andreu Comas-Garcia, Celida Duque Molina, Julissa Enciso-Ibarra, Jose Antonio Enciso-Moreno, Gloria Elena Espinosa-Ayala, Fernando Fontove-Herrera, Daniel Fregoso-Rueda, Victor Eduardo Garcia-Arias, Alejandra Garcia-Gasca, Bruno Gomez-Gil, Jean Pierre Gonzalez, Irvin Gonzalez-Lopez, Concepcion Grajales-Muñiz, Ricardo Grande, Rosa Maria Gutierrez Rios, Jesus Hernandez, Alejandra Hernandez-Teran, Alfredo Herrera-Estrella, Carla Ivon Herrera-Najera, Pavel Isa, Veronica Mata-Haro, Daniel Lira Morales, Susana Lopez, Antonio Loza Roman, Brenda Irasema Maldonado-Meza, Bernardo Martinez-Miguel, Jose Arturo Martinez-Orozco, Celida Martinez-Rodriguez, Margarita Matias-Florentino, Fidencio Mejia-Nepomuceno, Maria Guadalupe de Jesus Mireles-Rivera, Gloria Maria Molina-Salinas, Hector Montoya-Fuentes, , Mario Mujica-Sanchez, Jose Esteban Muñoz-Medina, Jose de Jesus Nuñez-Contreras, Alicia Ocaña-Mondragon, Luis Alberto Ochoa-Carrera, Hector Esteban Paz-Juarez, Marissa Perez-Garcia, Francisco Pulido, Helen Haydee Fernanda Ramirez-Plascencia, Jorge Salas-Hernandez, Angel Gustavo Salas-Lais, Alejandro Sanchez-Flores, Clara Esperanza Santacruz-Tinoco, Maria Guadalupe Santiago-Mauricio, Selene Zarate, Nelly Selem-Mojica, Blanca Taboada, Gloria Vazquez, Joel Armando Vazquez-Perez. |
| EPI_ISL_2970108, EPI_ISL_2970109, EPI_ISL_2970110, EPI_ISL_2970111 | Ospedale Regionale Umberto Parini | U.O. Genomica - Istituto Zooprofilattico Sperimentale del Piemonte, Liguria e Valle d'Aosta | Simone Peletto |
| EPI_ISL_2970370, EPI_ISL_2970371, EPI_ISL_2970376, | Molecular Diagnostics Pathology Department Mater Dei Hospital Malta | Molecular Diagnostics Pathology Department Mater Dei Hospital Malta | G Zahra, R Borg, C Cilia, L Grech, M Briffa |

| | | | |
|---|---|---|---|
| EPI_ISL_2970379 | | | |
| EPI_ISL_2970383, EPI_ISL_2970392, EPI_ISL_2970400 | Instituto Nacional de Medicina Genomica | Instituto Nacional de Medicina Genomica | Hidalgo-Miranda A, Cedro-Tanda A, Cisneros-Villanueva M, Blancas S, Orjuela-Rodríguez M, Mendoza-Vargas A, Reyes-Grajeda JP, Cisneros- Villanueva M, Gonzalez-Barrera D, Rangel-DeLeon D, Munguia-Garza P, Ramirez-Vega O, Escobar-Arrazola, M, Herrera-Montalvo LA. |
| EPI_ISL_2970561, EPI_ISL_2970562, EPI_ISL_2970563, EPI_ISL_2970827, EPI_ISL_2970828, EPI_ISL_2970829, EPI_ISL_2970962, EPI_ISL_2970963, EPI_ISL_2970965 | Molecular Diagnostics Pathology Department Mater Dei Hospital Malta | Molecular Diagnostics Pathology Department Mater Dei Hospital Malta | G Zahra, R Borg, C Cilia, L Grech, M Briffa |
| EPI_ISL_2970972, EPI_ISL_2970974, EPI_ISL_2970976 | Lighthouse Lab in Alderley Park | Wellcome Sanger Institute for the COVID-19 Genomics UK (COG-UK) Consortium | Jacquelyn Wynn, Mairead Hyland, The Lighthouse Lab in Alderley Park and Alex Alderton, Roberto Amato, Jeffrey Barrett, Sonia Goncalves, Ewan Harrison, David K. Jackson, Ian Johnston, Dominic Kwiatkowski, Cordelia Langford, John Sillitoe on behalf of the Wellcome Sanger Institute COVID-19 Surveillance Team |
| EPI_ISL_2970977 | Lighthouse Laboratory Plymouth | Wellcome Sanger Institute for the COVID-19 Genomics UK (COG-UK) Consortium | Lighthouse Laboratory Plymouth and Alex Alderton, Roberto Amato, Jeffrey Barrett, Sonia Goncalves, Ewan Harrison, David K. Jackson, Ian Johnston, Dominic Kwiatkowski, Cordelia Langford, John Sillitoe on behalf of the Wellcome Sanger Institute COVID-19 Surveillance Team |
| EPI_ISL_2970980 | Lighthouse Lab in Alderley Park | Wellcome Sanger Institute for the COVID-19 Genomics UK (COG-UK) Consortium | Jacquelyn Wynn, Mairead Hyland, The Lighthouse Lab in Alderley Park and Alex Alderton, Roberto Amato, Jeffrey Barrett, Sonia Goncalves, Ewan Harrison, David K. Jackson, Ian Johnston, Dominic Kwiatkowski, Cordelia Langford, John Sillitoe on behalf of the Wellcome Sanger Institute COVID-19 Surveillance Team |
| EPI_ISL_2970981 | Lighthouse Laboratory Plymouth | Wellcome Sanger Institute for the COVID-19 Genomics UK (COG-UK) Consortium | Lighthouse Laboratory Plymouth and Alex Alderton, Roberto Amato, Jeffrey Barrett, Sonia Goncalves, Ewan Harrison, David K. Jackson, Ian Johnston, Dominic Kwiatkowski, Cordelia Langford, John Sillitoe on behalf of the Wellcome Sanger Institute COVID-19 Surveillance Team |
| EPI_ISL_2970983 | Lighthouse Lab in Alderley Park | Wellcome Sanger Institute for the COVID-19 Genomics UK (COG-UK) Consortium | Jacquelyn Wynn, Mairead Hyland, The Lighthouse Lab in Alderley Park and Alex Alderton, Roberto Amato, Jeffrey Barrett, Sonia Goncalves, Ewan Harrison, David K. Jackson, Ian Johnston, Dominic Kwiatkowski, Cordelia Langford, John Sillitoe on behalf of the Wellcome Sanger Institute COVID-19 Surveillance Team |
| EPI_ISL_2970986 | Lighthouse Laboratory Plymouth | Wellcome Sanger Institute for the COVID-19 Genomics UK (COG-UK) Consortium | Lighthouse Laboratory Plymouth and Alex Alderton, Roberto Amato, Jeffrey Barrett, Sonia Goncalves, Ewan Harrison, David K. Jackson, Ian Johnston, Dominic Kwiatkowski, Cordelia Langford, John Sillitoe on behalf of the Wellcome Sanger Institute COVID-19 Surveillance Team |
| EPI_ISL_2970988 | Lighthouse Lab in Milton Keynes | Wellcome Sanger Institute for the COVID-19 Genomics UK (COG-UK) Consortium | The Lighthouse Lab in Milton Keynes and Alex Alderton, Roberto Amato, Jeffrey Barrett, Sonia Goncalves, Ewan Harrison, David K. Jackson, Ian Johnston, Dominic Kwiatkowski, Cordelia Langford, John Sillitoe on behalf of the Wellcome Sanger Institute COVID-19 Surveillance Team |
| EPI_ISL_2970989, EPI_ISL_2970990, EPI_ISL_2970996, EPI_ISL_2970999, EPI_ISL_2971001 | Lighthouse Lab in Alderley Park | Wellcome Sanger Institute for the COVID-19 Genomics UK (COG-UK) Consortium | Jacquelyn Wynn, Mairead Hyland, The Lighthouse Lab in Alderley Park and Alex Alderton, Roberto Amato, Jeffrey Barrett, Sonia Goncalves, Ewan Harrison, David K. Jackson, Ian Johnston, Dominic Kwiatkowski, Cordelia Langford, John Sillitoe on behalf of the Wellcome Sanger Institute COVID-19 Surveillance Team |
| EPI_ISL_2971003 | Lighthouse Lab in Milton Keynes | Wellcome Sanger Institute for the COVID-19 Genomics UK (COG-UK) Consortium | The Lighthouse Lab in Milton Keynes and Alex Alderton, Roberto Amato, Jeffrey Barrett, Sonia Goncalves, Ewan Harrison, David K. Jackson, Ian Johnston, Dominic Kwiatkowski, Cordelia Langford, John Sillitoe on behalf of the Wellcome Sanger Institute COVID-19 Surveillance Team |
| EPI_ISL_2971011 | Lighthouse Lab in Alderley Park | Wellcome Sanger Institute for the COVID-19 Genomics UK (COG-UK) Consortium | Jacquelyn Wynn, Mairead Hyland, The Lighthouse Lab in Alderley Park and Alex Alderton, Roberto Amato, Jeffrey Barrett, Sonia Goncalves, Ewan Harrison, David K. Jackson, Ian Johnston, Dominic Kwiatkowski, Cordelia Langford, John Sillitoe on behalf of the Wellcome Sanger Institute COVID-19 Surveillance Team |
| EPI_ISL_2971012, EPI_ISL_2971014 | Lighthouse Lab in Milton Keynes | Wellcome Sanger Institute for the COVID-19 Genomics UK (COG-UK) Consortium | The Lighthouse Lab in Milton Keynes and Alex Alderton, Roberto Amato, Jeffrey Barrett, Sonia Goncalves, Ewan Harrison, David K. Jackson, Ian Johnston, Dominic Kwiatkowski, Cordelia Langford, John Sillitoe on behalf of the Wellcome Sanger Institute COVID-19 Surveillance Team |
| EPI_ISL_2971018, EPI_ISL_2971022, EPI_ISL_2971029 | Lighthouse Lab in Alderley Park | Wellcome Sanger Institute for the COVID-19 Genomics UK (COG-UK) Consortium | Jacquelyn Wynn, Mairead Hyland, The Lighthouse Lab in Alderley Park and Alex Alderton, Roberto Amato, Jeffrey Barrett, Sonia Goncalves, Ewan Harrison, David K. Jackson, Ian Johnston, Dominic Kwiatkowski, Cordelia Langford, John Sillitoe on behalf of the Wellcome Sanger Institute COVID-19 Surveillance Team |
| EPI_ISL_2971039 | Lighthouse Lab in Milton Keynes | Wellcome Sanger Institute for the COVID-19 Genomics UK (COG-UK) Consortium | The Lighthouse Lab in Milton Keynes and Alex Alderton, Roberto Amato, Jeffrey Barrett, Sonia Goncalves, Ewan Harrison, David K. Jackson, Ian Johnston, Dominic Kwiatkowski, Cordelia Langford, John Sillitoe on behalf of the Wellcome Sanger Institute COVID-19 Surveillance Team |
| EPI_ISL_2971042 | Lighthouse Laboratory Plymouth | Wellcome Sanger Institute for the COVID-19 Genomics UK (COG-UK) Consortium | Lighthouse Laboratory Plymouth and Alex Alderton, Roberto Amato, Jeffrey Barrett, Sonia Goncalves, Ewan Harrison, David K. Jackson, Ian Johnston, Dominic Kwiatkowski, Cordelia Langford, John Sillitoe on behalf of the Wellcome Sanger Institute COVID-19 Surveillance Team |
| EPI_ISL_2971050, EPI_ISL_2971051 | Lighthouse Lab in Milton Keynes | Wellcome Sanger Institute for the COVID-19 Genomics UK (COG-UK) Consortium | The Lighthouse Lab in Milton Keynes and Alex Alderton, Roberto Amato, Jeffrey Barrett, Sonia Goncalves, Ewan Harrison, David K. Jackson, Ian Johnston, Dominic Kwiatkowski, Cordelia Langford, John Sillitoe on behalf of the Wellcome Sanger Institute COVID-19 Surveillance Team |
| EPI_ISL_2971057 | Lighthouse Lab in Glasgow | Wellcome Sanger Institute for the COVID-19 Genomics UK (COG-UK) Consortium | Harper VanSteenhouse, Yumi Kasai, David Gray, Carol Clugston, Anna Dominiczak and Alex Alderton, Roberto Amato, Jeffrey Barrett, Sonia Goncalves, Ewan Harrison, David K. Jackson, Ian Johnston, Dominic Kwiatkowski, Cordelia Langford, John Sillitoe on behalf of the Wellcome Sanger Institute COVID-19 Surveillance Team |
| EPI_ISL_2971061 | Lighthouse Lab in Alderley Park | Wellcome Sanger Institute for the COVID-19 Genomics UK (COG-UK) Consortium | Jacquelyn Wynn, Mairead Hyland, The Lighthouse Lab in Alderley Park and Alex Alderton, Roberto Amato, Jeffrey Barrett, Sonia Goncalves, Ewan Harrison, David K. Jackson, Ian Johnston, Dominic Kwiatkowski, Cordelia Langford, John Sillitoe on behalf of the Wellcome Sanger Institute COVID-19 Surveillance Team |
| EPI_ISL_2971063, EPI_ISL_2971065 | Lighthouse Lab in Milton Keynes | Wellcome Sanger Institute for the COVID-19 Genomics UK (COG-UK) Consortium | The Lighthouse Lab in Milton Keynes and Alex Alderton, Roberto Amato, Jeffrey Barrett, Sonia Goncalves, Ewan Harrison, David K. Jackson, Ian Johnston, Dominic Kwiatkowski, Cordelia Langford, John Sillitoe on behalf of the Wellcome Sanger Institute COVID-19 Surveillance Team |
| EPI_ISL_2971068 | Lighthouse Laboratory Plymouth | Wellcome Sanger Institute for the COVID-19 Genomics UK (COG-UK) Consortium | Lighthouse Laboratory Plymouth and Alex Alderton, Roberto Amato, Jeffrey Barrett, Sonia Goncalves, Ewan Harrison, David K. Jackson, Ian Johnston, Dominic Kwiatkowski, Cordelia Langford, John Sillitoe on behalf of the Wellcome Sanger Institute COVID-19 Surveillance Team |
| EPI_ISL_2971077, EPI_ISL_2971085 | Lighthouse Lab in Alderley Park | Wellcome Sanger Institute for the COVID-19 Genomics UK (COG-UK) Consortium | Jacquelyn Wynn, Mairead Hyland, The Lighthouse Lab in Alderley Park and Alex Alderton, Roberto Amato, Jeffrey Barrett, Sonia Goncalves, Ewan Harrison, David K. Jackson, Ian Johnston, Dominic Kwiatkowski, Cordelia Langford, John Sillitoe on behalf of the Wellcome Sanger Institute COVID-19 Surveillance Team |
| EPI_ISL_2971086 | Lighthouse Laboratory Plymouth | Wellcome Sanger Institute for the COVID-19 Genomics UK (COG-UK) Consortium | Lighthouse Laboratory Plymouth and Alex Alderton, Roberto Amato, Jeffrey Barrett, Sonia Goncalves, Ewan Harrison, David K. Jackson, Ian Johnston, Dominic Kwiatkowski, Cordelia Langford, John Sillitoe on behalf of the Wellcome Sanger Institute COVID-19 Surveillance Team |
| EPI_ISL_2971089 | Lighthouse Lab in Milton Keynes | Wellcome Sanger Institute for the COVID-19 Genomics UK (COG-UK) Consortium | The Lighthouse Lab in Milton Keynes and Alex Alderton, Roberto Amato, Jeffrey Barrett, Sonia Goncalves, Ewan Harrison, David K. Jackson, Ian Johnston, Dominic Kwiatkowski, Cordelia Langford, John Sillitoe on behalf of the Wellcome Sanger Institute COVID-19 Surveillance Team |
| EPI_ISL_2971093 | Lighthouse Lab in Alderley Park | Wellcome Sanger Institute for the COVID-19 Genomics UK (COG-UK) Consortium | Jacquelyn Wynn, Mairead Hyland, The Lighthouse Lab in Alderley Park and Alex Alderton, Roberto Amato, Jeffrey Barrett, Sonia Goncalves, Ewan Harrison, David K. Jackson, Ian Johnston, Dominic Kwiatkowski, Cordelia Langford, John Sillitoe on behalf of the Wellcome Sanger Institute COVID-19 Surveillance Team |
| EPI_ISL_2971099 | Lighthouse Lab in Milton Keynes | Wellcome Sanger Institute for the COVID-19 Genomics UK (COG-UK) Consortium | The Lighthouse Lab in Milton Keynes and Alex Alderton, Roberto Amato, Jeffrey Barrett, Sonia Goncalves, Ewan Harrison, David K. Jackson, Ian Johnston, Dominic Kwiatkowski, Cordelia Langford, John Sillitoe on behalf of the Wellcome Sanger Institute COVID-19 Surveillance Team |
| EPI_ISL_2971118 | Lighthouse Laboratory Plymouth | Wellcome Sanger Institute for the COVID-19 Genomics UK (COG-UK) Consortium | Lighthouse Laboratory Plymouth and Alex Alderton, Roberto Amato, Jeffrey Barrett, Sonia Goncalves, Ewan Harrison, David K. Jackson, Ian Johnston, Dominic Kwiatkowski, Cordelia Langford, John Sillitoe on behalf of the Wellcome Sanger Institute COVID-19 Surveillance Team |
| EPI_ISL_2971121, EPI_ISL_2971123, EPI_ISL_2971141, EPI_ISL_2971145 | Lighthouse Lab in Milton Keynes | Wellcome Sanger Institute for the COVID-19 Genomics UK (COG-UK) Consortium | The Lighthouse Lab in Milton Keynes and Alex Alderton, Roberto Amato, Jeffrey Barrett, Sonia Goncalves, Ewan Harrison, David K. Jackson, Ian Johnston, Dominic Kwiatkowski, Cordelia Langford, John Sillitoe on behalf of the Wellcome Sanger Institute COVID-19 Surveillance Team |
| EPI_ISL_2971157 | Lighthouse Lab in Alderley Park | Wellcome Sanger Institute for the COVID-19 Genomics UK (COG-UK) Consortium | Jacquelyn Wynn, Mairead Hyland, The Lighthouse Lab in Alderley Park and Alex Alderton, Roberto Amato, Jeffrey Barrett, Sonia Goncalves, Ewan Harrison, David K. Jackson, Ian Johnston, Dominic Kwiatkowski, Cordelia Langford, John Sillitoe on behalf of the Wellcome Sanger Institute COVID-19 Surveillance Team |
| EPI_ISL_2971159, EPI_ISL_2971164 | Lighthouse Lab in Milton Keynes | Wellcome Sanger Institute for the COVID-19 Genomics UK (COG-UK) Consortium | The Lighthouse Lab in Milton Keynes and Alex Alderton, Roberto Amato, Jeffrey Barrett, Sonia Goncalves, Ewan Harrison, David K. Jackson, Ian Johnston, Dominic Kwiatkowski, Cordelia Langford, John Sillitoe on behalf of the Wellcome Sanger Institute COVID-19 Surveillance Team |
| EPI_ISL_2971165, EPI_ISL_2971167, EPI_ISL_2971172 | Lighthouse Laboratory Plymouth | Wellcome Sanger Institute for the COVID-19 Genomics UK (COG-UK) Consortium | Lighthouse Laboratory Plymouth and Alex Alderton, Roberto Amato, Jeffrey Barrett, Sonia Goncalves, Ewan Harrison, David K. Jackson, Ian Johnston, Dominic Kwiatkowski, Cordelia Langford, John Sillitoe on behalf of the Wellcome Sanger Institute COVID-19 Surveillance Team |
| EPI_ISL_2971191 | Lighthouse Lab in Milton Keynes | Wellcome Sanger Institute for the COVID-19 Genomics UK (COG-UK) Consortium | The Lighthouse Lab in Milton Keynes and Alex Alderton, Roberto Amato, Jeffrey Barrett, Sonia Goncalves, Ewan Harrison, David K. Jackson, Ian Johnston, Dominic Kwiatkowski, Cordelia Langford, John Sillitoe on behalf of the Wellcome Sanger Institute COVID-19 Surveillance Team |
| EPI_ISL_2971207 | Lighthouse Lab in Alderley Park | Wellcome Sanger Institute for the COVID-19 Genomics UK (COG-UK) Consortium | Jacquelyn Wynn, Mairead Hyland, The Lighthouse Lab in Alderley Park and Alex Alderton, Roberto Amato, Jeffrey Barrett, Sonia Goncalves, Ewan Harrison, David K. Jackson, Ian Johnston, Dominic Kwiatkowski, Cordelia Langford, John Sillitoe on behalf of the Wellcome Sanger Institute COVID-19 Surveillance Team |
| EPI_ISL_2971209 | Lighthouse Laboratory Plymouth | Wellcome Sanger Institute for the COVID-19 Genomics UK (COG-UK) Consortium | Lighthouse Laboratory Plymouth and Alex Alderton, Roberto Amato, Jeffrey Barrett, Sonia Goncalves, Ewan Harrison, David K. Jackson, Ian Johnston, Dominic Kwiatkowski, Cordelia Langford, John Sillitoe on behalf of the Wellcome Sanger Institute COVID-19 Surveillance Team |
| EPI_ISL_2971210, EPI_ISL_2971211 | Berkshire and Surrey Pathology Services Lighthouse Laboratory | Wellcome Sanger Institute for the COVID-19 Genomics UK (COG-UK) Consortium | Berkshire and Surrey Pathology Services Lighthouse Laboratory and Alex Alderton, Roberto Amato, Jeffrey Barrett, Sonia Goncalves, Ewan Harrison, David K. Jackson, Ian Johnston, Dominic Kwiatkowski, Cordelia Langford, John Sillitoe on behalf of the Wellcome Sanger Institute COVID-19 Surveillance Team |
| EPI_ISL_2971212, EPI_ISL_2971213, EPI_ISL_2971214, EPI_ISL_2971215, EPI_ISL_2971216 | Lighthouse Lab in Alderley Park | Wellcome Sanger Institute for the COVID-19 Genomics UK (COG-UK) Consortium | Jacquelyn Wynn, Mairead Hyland, The Lighthouse Lab in Alderley Park and Alex Alderton, Roberto Amato, Jeffrey Barrett, Sonia Goncalves, Ewan Harrison, David K. Jackson, Ian Johnston, Dominic Kwiatkowski, Cordelia Langford, John Sillitoe on behalf of the Wellcome Sanger Institute COVID-19 Surveillance Team |
| EPI_ISL_2971217 | Lighthouse Laboratory Plymouth | Wellcome Sanger Institute for the COVID-19 Genomics UK (COG-UK) Consortium | Lighthouse Laboratory Plymouth and Alex Alderton, Roberto Amato, Jeffrey Barrett, Sonia Goncalves, Ewan Harrison, David K. Jackson, Ian Johnston, Dominic Kwiatkowski, Cordelia Langford, John Sillitoe on behalf of the Wellcome Sanger Institute COVID-19 Surveillance Team |

| | | | |
|---|---|---|---|
| EPI_ISL_2971218, EPI_ISL_2971219, EPI_ISL_2971220, EPI_ISL_2971221, EPI_ISL_2971222, EPI_ISL_2971223, EPI_ISL_2971224, EPI_ISL_2971225 | Lighthouse Lab in Alderley Park | Wellcome Sanger Institute for the COVID-19 Genomics UK (COG-UK) Consortium | Jacquelyn Wynn, Mairead Hyland, The Lighthouse Lab in Alderley Park and Alex Alderton, Roberto Amato, Jeffrey Barrett, Sonia Goncalves, Ewan Harrison, David K. Jackson, Ian Johnston, Dominic Kwiatkowski, Cordelia Langford, John Sillitoe on behalf of the Wellcome Sanger Institute COVID-19 Surveillance Team |
| EPI_ISL_2971226 | Berkshire and Surrey Pathology Services Lighthouse Laboratory | Wellcome Sanger Institute for the COVID-19 Genomics UK (COG-UK) Consortium | Berkshire and Surrey Pathology Services Lighthouse Laboratory and Alex Alderton, Roberto Amato, Jeffrey Barrett, Sonia Goncalves, Ewan Harrison, David K. Jackson, Ian Johnston, Dominic Kwiatkowski, Cordelia Langford, John Sillitoe on behalf of the Wellcome Sanger Institute COVID-19 Surveillance Team |
| EPI_ISL_2971227 | Lighthouse Lab in Alderley Park | Wellcome Sanger Institute for the COVID-19 Genomics UK (COG-UK) Consortium | Jacquelyn Wynn, Mairead Hyland, The Lighthouse Lab in Alderley Park and Alex Alderton, Roberto Amato, Jeffrey Barrett, Sonia Goncalves, Ewan Harrison, David K. Jackson, Ian Johnston, Dominic Kwiatkowski, Cordelia Langford, John Sillitoe on behalf of the Wellcome Sanger Institute COVID-19 Surveillance Team |
| EPI_ISL_2971228 | Berkshire and Surrey Pathology Services Lighthouse Laboratory | Wellcome Sanger Institute for the COVID-19 Genomics UK (COG-UK) Consortium | Berkshire and Surrey Pathology Services Lighthouse Laboratory and Alex Alderton, Roberto Amato, Jeffrey Barrett, Sonia Goncalves, Ewan Harrison, David K. Jackson, Ian Johnston, Dominic Kwiatkowski, Cordelia Langford, John Sillitoe on behalf of the Wellcome Sanger Institute COVID-19 Surveillance Team |
| EPI_ISL_2971229 | Lighthouse Lab in Alderley Park | Wellcome Sanger Institute for the COVID-19 Genomics UK (COG-UK) Consortium | Jacquelyn Wynn, Mairead Hyland, The Lighthouse Lab in Alderley Park and Alex Alderton, Roberto Amato, Jeffrey Barrett, Sonia Goncalves, Ewan Harrison, David K. Jackson, Ian Johnston, Dominic Kwiatkowski, Cordelia Langford, John Sillitoe on behalf of the Wellcome Sanger Institute COVID-19 Surveillance Team |
| EPI_ISL_2971230 | Lighthouse Laboratory Plymouth | Wellcome Sanger Institute for the COVID-19 Genomics UK (COG-UK) Consortium | Lighthouse Laboratory Plymouth and Alex Alderton, Roberto Amato, Jeffrey Barrett, Sonia Goncalves, Ewan Harrison, David K. Jackson, Ian Johnston, Dominic Kwiatkowski, Cordelia Langford, John Sillitoe on behalf of the Wellcome Sanger Institute COVID-19 Surveillance Team |
| EPI_ISL_2971231, EPI_ISL_2971232 | Lighthouse Lab in Alderley Park | Wellcome Sanger Institute for the COVID-19 Genomics UK (COG-UK) Consortium | Jacquelyn Wynn, Mairead Hyland, The Lighthouse Lab in Alderley Park and Alex Alderton, Roberto Amato, Jeffrey Barrett, Sonia Goncalves, Ewan Harrison, David K. Jackson, Ian Johnston, Dominic Kwiatkowski, Cordelia Langford, John Sillitoe on behalf of the Wellcome Sanger Institute COVID-19 Surveillance Team |
| EPI_ISL_2971233, EPI_ISL_2971234 | Lighthouse Laboratory Plymouth | Wellcome Sanger Institute for the COVID-19 Genomics UK (COG-UK) Consortium | Lighthouse Laboratory Plymouth and Alex Alderton, Roberto Amato, Jeffrey Barrett, Sonia Goncalves, Ewan Harrison, David K. Jackson, Ian Johnston, Dominic Kwiatkowski, Cordelia Langford, John Sillitoe on behalf of the Wellcome Sanger Institute COVID-19 Surveillance Team |
| EPI_ISL_2971235 | Lighthouse Lab in Alderley Park | Wellcome Sanger Institute for the COVID-19 Genomics UK (COG-UK) Consortium | Jacquelyn Wynn, Mairead Hyland, The Lighthouse Lab in Alderley Park and Alex Alderton, Roberto Amato, Jeffrey Barrett, Sonia Goncalves, Ewan Harrison, David K. Jackson, Ian Johnston, Dominic Kwiatkowski, Cordelia Langford, John Sillitoe on behalf of the Wellcome Sanger Institute COVID-19 Surveillance Team |
| EPI_ISL_2971236 | Berkshire and Surrey Pathology Services Lighthouse Laboratory | Wellcome Sanger Institute for the COVID-19 Genomics UK (COG-UK) Consortium | Berkshire and Surrey Pathology Services Lighthouse Laboratory and Alex Alderton, Roberto Amato, Jeffrey Barrett, Sonia Goncalves, Ewan Harrison, David K. Jackson, Ian Johnston, Dominic Kwiatkowski, Cordelia Langford, John Sillitoe on behalf of the Wellcome Sanger Institute COVID-19 Surveillance Team |
| EPI_ISL_2971237 | Lighthouse Lab in Alderley Park | Wellcome Sanger Institute for the COVID-19 Genomics UK (COG-UK) Consortium | Jacquelyn Wynn, Mairead Hyland, The Lighthouse Lab in Alderley Park and Alex Alderton, Roberto Amato, Jeffrey Barrett, Sonia Goncalves, Ewan Harrison, David K. Jackson, Ian Johnston, Dominic Kwiatkowski, Cordelia Langford, John Sillitoe on behalf of the Wellcome Sanger Institute COVID-19 Surveillance Team |
| EPI_ISL_2971238, EPI_ISL_2971239, EPI_ISL_2971240, EPI_ISL_2971241 | Berkshire and Surrey Pathology Services Lighthouse Laboratory | Wellcome Sanger Institute for the COVID-19 Genomics UK (COG-UK) Consortium | Berkshire and Surrey Pathology Services Lighthouse Laboratory and Alex Alderton, Roberto Amato, Jeffrey Barrett, Sonia Goncalves, Ewan Harrison, David K. Jackson, Ian Johnston, Dominic Kwiatkowski, Cordelia Langford, John Sillitoe on behalf of the Wellcome Sanger Institute COVID-19 Surveillance Team |
| EPI_ISL_2971242 | Lighthouse Lab in Alderley Park | Wellcome Sanger Institute for the COVID-19 Genomics UK (COG-UK) Consortium | Jacquelyn Wynn, Mairead Hyland, The Lighthouse Lab in Alderley Park and Alex Alderton, Roberto Amato, Jeffrey Barrett, Sonia Goncalves, Ewan Harrison, David K. Jackson, Ian Johnston, Dominic Kwiatkowski, Cordelia Langford, John Sillitoe on behalf of the Wellcome Sanger Institute COVID-19 Surveillance Team |
| EPI_ISL_2971243 | Berkshire and Surrey Pathology Services Lighthouse Laboratory | Wellcome Sanger Institute for the COVID-19 Genomics UK (COG-UK) Consortium | Berkshire and Surrey Pathology Services Lighthouse Laboratory and Alex Alderton, Roberto Amato, Jeffrey Barrett, Sonia Goncalves, Ewan Harrison, David K. Jackson, Ian Johnston, Dominic Kwiatkowski, Cordelia Langford, John Sillitoe on behalf of the Wellcome Sanger Institute COVID-19 Surveillance Team |
| EPI_ISL_2971244 | Lighthouse Laboratory Plymouth | Wellcome Sanger Institute for the COVID-19 Genomics UK (COG-UK) Consortium | Lighthouse Laboratory Plymouth and Alex Alderton, Roberto Amato, Jeffrey Barrett, Sonia Goncalves, Ewan Harrison, David K. Jackson, Ian Johnston, Dominic Kwiatkowski, Cordelia Langford, John Sillitoe on behalf of the Wellcome Sanger Institute COVID-19 Surveillance Team |
| EPI_ISL_2971245, EPI_ISL_2971246 | Lighthouse Lab in Alderley Park | Wellcome Sanger Institute for the COVID-19 Genomics UK (COG-UK) Consortium | Jacquelyn Wynn, Mairead Hyland, The Lighthouse Lab in Alderley Park and Alex Alderton, Roberto Amato, Jeffrey Barrett, Sonia Goncalves, Ewan Harrison, David K. Jackson, Ian Johnston, Dominic Kwiatkowski, Cordelia Langford, John Sillitoe on behalf of the Wellcome Sanger Institute COVID-19 Surveillance Team |
| EPI_ISL_2971247, EPI_ISL_2971248 | Berkshire and Surrey Pathology Services Lighthouse Laboratory | Wellcome Sanger Institute for the COVID-19 Genomics UK (COG-UK) Consortium | Berkshire and Surrey Pathology Services Lighthouse Laboratory and Alex Alderton, Roberto Amato, Jeffrey Barrett, Sonia Goncalves, Ewan Harrison, David K. Jackson, Ian Johnston, Dominic Kwiatkowski, Cordelia Langford, John Sillitoe on behalf of the Wellcome Sanger Institute COVID-19 Surveillance Team |
| EPI_ISL_2971249 | Lighthouse Lab in Alderley Park | Wellcome Sanger Institute for the COVID-19 Genomics UK (COG-UK) Consortium | Jacquelyn Wynn, Mairead Hyland, The Lighthouse Lab in Alderley Park and Alex Alderton, Roberto Amato, Jeffrey Barrett, Sonia Goncalves, Ewan Harrison, David K. Jackson, Ian Johnston, Dominic Kwiatkowski, Cordelia Langford, John Sillitoe on behalf of the Wellcome Sanger Institute COVID-19 Surveillance Team |
| EPI_ISL_2971250 | Lighthouse Laboratory Plymouth | Wellcome Sanger Institute for the COVID-19 Genomics UK (COG-UK) Consortium | Lighthouse Laboratory Plymouth and Alex Alderton, Roberto Amato, Jeffrey Barrett, Sonia Goncalves, Ewan Harrison, David K. Jackson, Ian Johnston, Dominic Kwiatkowski, Cordelia Langford, John Sillitoe on behalf of the Wellcome Sanger Institute COVID-19 Surveillance Team |
| EPI_ISL_2971251 | Lighthouse Lab in Alderley Park | Wellcome Sanger Institute for the COVID-19 Genomics UK (COG-UK) Consortium | Jacquelyn Wynn, Mairead Hyland, The Lighthouse Lab in Alderley Park and Alex Alderton, Roberto Amato, Jeffrey Barrett, Sonia Goncalves, Ewan Harrison, David K. Jackson, Ian Johnston, Dominic Kwiatkowski, Cordelia Langford, John Sillitoe on behalf of the Wellcome Sanger Institute COVID-19 Surveillance Team |
| EPI_ISL_2971252 | Lighthouse Laboratory Plymouth | Wellcome Sanger Institute for the COVID-19 Genomics UK (COG-UK) Consortium | Lighthouse Laboratory Plymouth and Alex Alderton, Roberto Amato, Jeffrey Barrett, Sonia Goncalves, Ewan Harrison, David K. Jackson, Ian Johnston, Dominic Kwiatkowski, Cordelia Langford, John Sillitoe on behalf of the Wellcome Sanger Institute COVID-19 Surveillance Team |
| EPI_ISL_2971253, EPI_ISL_2971254 | Lighthouse Lab in Alderley Park | Wellcome Sanger Institute for the COVID-19 Genomics UK (COG-UK) Consortium | Jacquelyn Wynn, Mairead Hyland, The Lighthouse Lab in Alderley Park and Alex Alderton, Roberto Amato, Jeffrey Barrett, Sonia Goncalves, Ewan Harrison, David K. Jackson, Ian Johnston, Dominic Kwiatkowski, Cordelia Langford, John Sillitoe on behalf of the Wellcome Sanger Institute COVID-19 Surveillance Team |
| EPI_ISL_2971255, EPI_ISL_2971256 | Lighthouse Laboratory Plymouth | Wellcome Sanger Institute for the COVID-19 Genomics UK (COG-UK) Consortium | Lighthouse Laboratory Plymouth and Alex Alderton, Roberto Amato, Jeffrey Barrett, Sonia Goncalves, Ewan Harrison, David K. Jackson, Ian Johnston, Dominic Kwiatkowski, Cordelia Langford, John Sillitoe on behalf of the Wellcome Sanger Institute COVID-19 Surveillance Team |
| EPI_ISL_2971257, EPI_ISL_2971258, EPI_ISL_2971259, EPI_ISL_2971260 | Berkshire and Surrey Pathology Services Lighthouse Laboratory | Wellcome Sanger Institute for the COVID-19 Genomics UK (COG-UK) Consortium | Berkshire and Surrey Pathology Services Lighthouse Laboratory and Alex Alderton, Roberto Amato, Jeffrey Barrett, Sonia Goncalves, Ewan Harrison, David K. Jackson, Ian Johnston, Dominic Kwiatkowski, Cordelia Langford, John Sillitoe on behalf of the Wellcome Sanger Institute COVID-19 Surveillance Team |
| EPI_ISL_2971261, EPI_ISL_2971262, EPI_ISL_2971263 | Lighthouse Lab in Alderley Park | Wellcome Sanger Institute for the COVID-19 Genomics UK (COG-UK) Consortium | Jacquelyn Wynn, Mairead Hyland, The Lighthouse Lab in Alderley Park and Alex Alderton, Roberto Amato, Jeffrey Barrett, Sonia Goncalves, Ewan Harrison, David K. Jackson, Ian Johnston, Dominic Kwiatkowski, Cordelia Langford, John Sillitoe on behalf of the Wellcome Sanger Institute COVID-19 Surveillance Team |
| EPI_ISL_2971264 | Lighthouse Laboratory Plymouth | Wellcome Sanger Institute for the COVID-19 Genomics UK (COG-UK) Consortium | Lighthouse Laboratory Plymouth and Alex Alderton, Roberto Amato, Jeffrey Barrett, Sonia Goncalves, Ewan Harrison, David K. Jackson, Ian Johnston, Dominic Kwiatkowski, Cordelia Langford, John Sillitoe on behalf of the Wellcome Sanger Institute COVID-19 Surveillance Team |
| EPI_ISL_2971265, EPI_ISL_2971266 | Lighthouse Lab in Alderley Park | Wellcome Sanger Institute for the COVID-19 Genomics UK (COG-UK) Consortium | Jacquelyn Wynn, Mairead Hyland, The Lighthouse Lab in Alderley Park and Alex Alderton, Roberto Amato, Jeffrey Barrett, Sonia Goncalves, Ewan Harrison, David K. Jackson, Ian Johnston, Dominic Kwiatkowski, Cordelia Langford, John Sillitoe on behalf of the Wellcome Sanger Institute COVID-19 Surveillance Team |
| EPI_ISL_2971267 | Berkshire and Surrey Pathology Services Lighthouse Laboratory | Wellcome Sanger Institute for the COVID-19 Genomics UK (COG-UK) Consortium | Berkshire and Surrey Pathology Services Lighthouse Laboratory and Alex Alderton, Roberto Amato, Jeffrey Barrett, Sonia Goncalves, Ewan Harrison, David K. Jackson, Ian Johnston, Dominic Kwiatkowski, Cordelia Langford, John Sillitoe on behalf of the Wellcome Sanger Institute COVID-19 Surveillance Team |
| EPI_ISL_2971268, EPI_ISL_2971269 | Lighthouse Lab in Alderley Park | Wellcome Sanger Institute for the COVID-19 Genomics UK (COG-UK) Consortium | Jacquelyn Wynn, Mairead Hyland, The Lighthouse Lab in Alderley Park and Alex Alderton, Roberto Amato, Jeffrey Barrett, Sonia Goncalves, Ewan Harrison, David K. Jackson, Ian Johnston, Dominic Kwiatkowski, Cordelia Langford, John Sillitoe on behalf of the Wellcome Sanger Institute COVID-19 Surveillance Team |
| EPI_ISL_2971270, EPI_ISL_2971271 | Berkshire and Surrey Pathology Services Lighthouse Laboratory | Wellcome Sanger Institute for the COVID-19 Genomics UK (COG-UK) Consortium | Berkshire and Surrey Pathology Services Lighthouse Laboratory and Alex Alderton, Roberto Amato, Jeffrey Barrett, Sonia Goncalves, Ewan Harrison, David K. Jackson, Ian Johnston, Dominic Kwiatkowski, Cordelia Langford, John Sillitoe on behalf of the Wellcome Sanger Institute COVID-19 Surveillance Team |
| EPI_ISL_2971272 | Lighthouse Lab in Alderley Park | Wellcome Sanger Institute for the COVID-19 Genomics UK (COG-UK) Consortium | Jacquelyn Wynn, Mairead Hyland, The Lighthouse Lab in Alderley Park and Alex Alderton, Roberto Amato, Jeffrey Barrett, Sonia Goncalves, Ewan Harrison, David K. Jackson, Ian Johnston, Dominic Kwiatkowski, Cordelia Langford, John Sillitoe on behalf of the Wellcome Sanger Institute COVID-19 Surveillance Team |
| EPI_ISL_2971273 | Lighthouse Laboratory Plymouth | Wellcome Sanger Institute for the COVID-19 Genomics UK (COG-UK) Consortium | Lighthouse Laboratory Plymouth and Alex Alderton, Roberto Amato, Jeffrey Barrett, Sonia Goncalves, Ewan Harrison, David K. Jackson, Ian Johnston, Dominic Kwiatkowski, Cordelia Langford, John Sillitoe on behalf of the Wellcome Sanger Institute COVID-19 Surveillance Team |
| EPI_ISL_2971274 | Berkshire and Surrey Pathology Services Lighthouse Laboratory | Wellcome Sanger Institute for the COVID-19 Genomics UK (COG-UK) Consortium | Berkshire and Surrey Pathology Services Lighthouse Laboratory and Alex Alderton, Roberto Amato, Jeffrey Barrett, Sonia Goncalves, Ewan Harrison, David K. Jackson, Ian Johnston, Dominic Kwiatkowski, Cordelia Langford, John Sillitoe on behalf of the Wellcome Sanger Institute COVID-19 Surveillance Team |
| EPI_ISL_2971275 | Lighthouse Laboratory Plymouth | Wellcome Sanger Institute for the COVID-19 Genomics UK (COG-UK) Consortium | Lighthouse Laboratory Plymouth and Alex Alderton, Roberto Amato, Jeffrey Barrett, Sonia Goncalves, Ewan Harrison, David K. Jackson, Ian Johnston, Dominic Kwiatkowski, Cordelia Langford, John Sillitoe on behalf of the Wellcome Sanger Institute COVID-19 Surveillance Team |
| EPI_ISL_2971276 | Lighthouse Lab in Alderley Park | Wellcome Sanger Institute for the COVID-19 Genomics UK (COG-UK) Consortium | Jacquelyn Wynn, Mairead Hyland, The Lighthouse Lab in Alderley Park and Alex Alderton, Roberto Amato, Jeffrey Barrett, Sonia Goncalves, Ewan Harrison, David K. Jackson, Ian Johnston, Dominic Kwiatkowski, Cordelia Langford, John Sillitoe on behalf of the Wellcome Sanger Institute COVID-19 Surveillance Team |
| EPI_ISL_2971277 | Berkshire and Surrey Pathology Services Lighthouse Laboratory | Wellcome Sanger Institute for the COVID-19 Genomics UK (COG-UK) Consortium | Berkshire and Surrey Pathology Services Lighthouse Laboratory and Alex Alderton, Roberto Amato, Jeffrey Barrett, Sonia Goncalves, Ewan Harrison, David K. Jackson, Ian Johnston, Dominic Kwiatkowski, Cordelia Langford, John Sillitoe on behalf of the Wellcome Sanger Institute COVID-19 Surveillance Team |
| EPI_ISL_2971278 | Lighthouse Lab in Alderley Park | Wellcome Sanger Institute for the COVID-19 Genomics UK (COG-UK) Consortium | Jacquelyn Wynn, Mairead Hyland, The Lighthouse Lab in Alderley Park and Alex Alderton, Roberto Amato, Jeffrey Barrett, Sonia Goncalves, Ewan Harrison, David K. Jackson, Ian Johnston, Dominic Kwiatkowski, Cordelia Langford, John Sillitoe on behalf of the Wellcome Sanger Institute COVID-19 Surveillance Team |
| EPI_ISL_2971279 | Berkshire and Surrey Pathology Services Lighthouse Laboratory | Wellcome Sanger Institute for the COVID-19 Genomics UK (COG-UK) Consortium | Berkshire and Surrey Pathology Services Lighthouse Laboratory and Alex Alderton, Roberto Amato, Jeffrey Barrett, Sonia Goncalves, Ewan Harrison, David K. Jackson, Ian Johnston, Dominic Kwiatkowski, Cordelia Langford, John Sillitoe on behalf of the Wellcome Sanger Institute COVID-19 Surveillance Team |
| EPI_ISL_2971280, EPI_ISL_2971281, EPI_ISL_2971282, EPI_ISL_2971283 | Lighthouse Lab in Alderley Park | Wellcome Sanger Institute for the COVID-19 Genomics UK (COG-UK) Consortium | Jacquelyn Wynn, Mairead Hyland, The Lighthouse Lab in Alderley Park and Alex Alderton, Roberto Amato, Jeffrey Barrett, Sonia Goncalves, Ewan Harrison, David K. Jackson, Ian Johnston, Dominic Kwiatkowski, Cordelia Langford, John Sillitoe on behalf of the Wellcome Sanger Institute COVID-19 Surveillance Team |
| EPI_ISL_2971284 | Lighthouse Laboratory Plymouth | Wellcome Sanger Institute for the COVID-19 Genomics UK (COG-UK) | Lighthouse Laboratory Plymouth and Alex Alderton, Roberto Amato, Jeffrey Barrett, Sonia Goncalves, Ewan Harrison, David K. Jackson, Ian Johnston, Dominic Kwiatkowski, Cordelia Langford, John Sillitoe on behalf of the Wellcome Sanger |

| | | | |
|---|---|---|---|
| | | Consortium | Institute COVID-19 Surveillance Team |
| EPI_ISL_2971285, EPI_ISL_2971286 | Lighthouse Lab in Alderley Park | Wellcome Sanger Institute for the COVID-19 Genomics UK (COG-UK) Consortium | Jacquelyn Wynn, Mairead Hyland, The Lighthouse Lab in Alderley Park and Alex Alderton, Roberto Amato, Jeffrey Barrett, Sonia Goncalves, Ewan Harrison, David K. Jackson, Ian Johnston, Dominic Kwiatkowski, Cordelia Langford, John Sillitoe on behalf of the Wellcome Sanger Institute COVID-19 Surveillance Team |
| EPI_ISL_2971287, EPI_ISL_2971288 | Berkshire and Surrey Pathology Services Lighthouse Laboratory | Wellcome Sanger Institute for the COVID-19 Genomics UK (COG-UK) Consortium | Berkshire and Surrey Pathology Services Lighthouse Laboratory and Alex Alderton, Roberto Amato, Jeffrey Barrett, Sonia Goncalves, Ewan Harrison, David K. Jackson, Ian Johnston, Dominic Kwiatkowski, Cordelia Langford, John Sillitoe on behalf of the Wellcome Sanger Institute COVID-19 Surveillance Team |
| EPI_ISL_2971289, EPI_ISL_2971290, EPI_ISL_2971291, EPI_ISL_2971292, EPI_ISL_2971293 | Lighthouse Lab in Alderley Park | Wellcome Sanger Institute for the COVID-19 Genomics UK (COG-UK) Consortium | Jacquelyn Wynn, Mairead Hyland, The Lighthouse Lab in Alderley Park and Alex Alderton, Roberto Amato, Jeffrey Barrett, Sonia Goncalves, Ewan Harrison, David K. Jackson, Ian Johnston, Dominic Kwiatkowski, Cordelia Langford, John Sillitoe on behalf of the Wellcome Sanger Institute COVID-19 Surveillance Team |
| EPI_ISL_2971294, EPI_ISL_2971295, EPI_ISL_2971296, EPI_ISL_2971297, EPI_ISL_2971298 | Lighthouse Laboratory Plymouth | Wellcome Sanger Institute for the COVID-19 Genomics UK (COG-UK) Consortium | Lighthouse Laboratory Plymouth and Alex Alderton, Roberto Amato, Jeffrey Barrett, Sonia Goncalves, Ewan Harrison, David K. Jackson, Ian Johnston, Dominic Kwiatkowski, Cordelia Langford, John Sillitoe on behalf of the Wellcome Sanger Institute COVID-19 Surveillance Team |
| EPI_ISL_2971299 | Lighthouse Lab in Alderley Park | Wellcome Sanger Institute for the COVID-19 Genomics UK (COG-UK) Consortium | Jacquelyn Wynn, Mairead Hyland, The Lighthouse Lab in Alderley Park and Alex Alderton, Roberto Amato, Jeffrey Barrett, Sonia Goncalves, Ewan Harrison, David K. Jackson, Ian Johnston, Dominic Kwiatkowski, Cordelia Langford, John Sillitoe on behalf of the Wellcome Sanger Institute COVID-19 Surveillance Team |
| EPI_ISL_2971300 | Berkshire and Surrey Pathology Services Lighthouse Laboratory | Wellcome Sanger Institute for the COVID-19 Genomics UK (COG-UK) Consortium | Berkshire and Surrey Pathology Services Lighthouse Laboratory and Alex Alderton, Roberto Amato, Jeffrey Barrett, Sonia Goncalves, Ewan Harrison, David K. Jackson, Ian Johnston, Dominic Kwiatkowski, Cordelia Langford, John Sillitoe on behalf of the Wellcome Sanger Institute COVID-19 Surveillance Team |
| EPI_ISL_2971301 | Lighthouse Laboratory Plymouth | Wellcome Sanger Institute for the COVID-19 Genomics UK (COG-UK) Consortium | Lighthouse Laboratory Plymouth and Alex Alderton, Roberto Amato, Jeffrey Barrett, Sonia Goncalves, Ewan Harrison, David K. Jackson, Ian Johnston, Dominic Kwiatkowski, Cordelia Langford, John Sillitoe on behalf of the Wellcome Sanger Institute COVID-19 Surveillance Team |
| EPI_ISL_2971302 | Berkshire and Surrey Pathology Services Lighthouse Laboratory | Wellcome Sanger Institute for the COVID-19 Genomics UK (COG-UK) Consortium | Berkshire and Surrey Pathology Services Lighthouse Laboratory and Alex Alderton, Roberto Amato, Jeffrey Barrett, Sonia Goncalves, Ewan Harrison, David K. Jackson, Ian Johnston, Dominic Kwiatkowski, Cordelia Langford, John Sillitoe on behalf of the Wellcome Sanger Institute COVID-19 Surveillance Team |
| EPI_ISL_2971303 | Lighthouse Lab in Alderley Park | Wellcome Sanger Institute for the COVID-19 Genomics UK (COG-UK) Consortium | Jacquelyn Wynn, Mairead Hyland, The Lighthouse Lab in Alderley Park and Alex Alderton, Roberto Amato, Jeffrey Barrett, Sonia Goncalves, Ewan Harrison, David K. Jackson, Ian Johnston, Dominic Kwiatkowski, Cordelia Langford, John Sillitoe on behalf of the Wellcome Sanger Institute COVID-19 Surveillance Team |
| EPI_ISL_2971304 | Lighthouse Laboratory Plymouth | Wellcome Sanger Institute for the COVID-19 Genomics UK (COG-UK) Consortium | Lighthouse Laboratory Plymouth and Alex Alderton, Roberto Amato, Jeffrey Barrett, Sonia Goncalves, Ewan Harrison, David K. Jackson, Ian Johnston, Dominic Kwiatkowski, Cordelia Langford, John Sillitoe on behalf of the Wellcome Sanger Institute COVID-19 Surveillance Team |
| EPI_ISL_2971305 | Lighthouse Lab in Alderley Park | Wellcome Sanger Institute for the COVID-19 Genomics UK (COG-UK) Consortium | Jacquelyn Wynn, Mairead Hyland, The Lighthouse Lab in Alderley Park and Alex Alderton, Roberto Amato, Jeffrey Barrett, Sonia Goncalves, Ewan Harrison, David K. Jackson, Ian Johnston, Dominic Kwiatkowski, Cordelia Langford, John Sillitoe on behalf of the Wellcome Sanger Institute COVID-19 Surveillance Team |
| EPI_ISL_2971306, EPI_ISL_2971307, EPI_ISL_2971308 | Lighthouse Laboratory Plymouth | Wellcome Sanger Institute for the COVID-19 Genomics UK (COG-UK) Consortium | Lighthouse Laboratory Plymouth and Alex Alderton, Roberto Amato, Jeffrey Barrett, Sonia Goncalves, Ewan Harrison, David K. Jackson, Ian Johnston, Dominic Kwiatkowski, Cordelia Langford, John Sillitoe on behalf of the Wellcome Sanger Institute COVID-19 Surveillance Team |
| EPI_ISL_2971309 | Lighthouse Lab in Alderley Park | Wellcome Sanger Institute for the COVID-19 Genomics UK (COG-UK) Consortium | Jacquelyn Wynn, Mairead Hyland, The Lighthouse Lab in Alderley Park and Alex Alderton, Roberto Amato, Jeffrey Barrett, Sonia Goncalves, Ewan Harrison, David K. Jackson, Ian Johnston, Dominic Kwiatkowski, Cordelia Langford, John Sillitoe on behalf of the Wellcome Sanger Institute COVID-19 Surveillance Team |
| EPI_ISL_2971310 | Berkshire and Surrey Pathology Services Lighthouse Laboratory | Wellcome Sanger Institute for the COVID-19 Genomics UK (COG-UK) Consortium | Berkshire and Surrey Pathology Services Lighthouse Laboratory and Alex Alderton, Roberto Amato, Jeffrey Barrett, Sonia Goncalves, Ewan Harrison, David K. Jackson, Ian Johnston, Dominic Kwiatkowski, Cordelia Langford, John Sillitoe on behalf of the Wellcome Sanger Institute COVID-19 Surveillance Team |
| EPI_ISL_2971311, EPI_ISL_2971312 | Lighthouse Lab in Alderley Park | Wellcome Sanger Institute for the COVID-19 Genomics UK (COG-UK) Consortium | Jacquelyn Wynn, Mairead Hyland, The Lighthouse Lab in Alderley Park and Alex Alderton, Roberto Amato, Jeffrey Barrett, Sonia Goncalves, Ewan Harrison, David K. Jackson, Ian Johnston, Dominic Kwiatkowski, Cordelia Langford, John Sillitoe on behalf of the Wellcome Sanger Institute COVID-19 Surveillance Team |
| EPI_ISL_2971313 | Berkshire and Surrey Pathology Services Lighthouse Laboratory | Wellcome Sanger Institute for the COVID-19 Genomics UK (COG-UK) Consortium | Berkshire and Surrey Pathology Services Lighthouse Laboratory and Alex Alderton, Roberto Amato, Jeffrey Barrett, Sonia Goncalves, Ewan Harrison, David K. Jackson, Ian Johnston, Dominic Kwiatkowski, Cordelia Langford, John Sillitoe on behalf of the Wellcome Sanger Institute COVID-19 Surveillance Team |
| EPI_ISL_2971314 | Lighthouse Lab in Alderley Park | Wellcome Sanger Institute for the COVID-19 Genomics UK (COG-UK) Consortium | Jacquelyn Wynn, Mairead Hyland, The Lighthouse Lab in Alderley Park and Alex Alderton, Roberto Amato, Jeffrey Barrett, Sonia Goncalves, Ewan Harrison, David K. Jackson, Ian Johnston, Dominic Kwiatkowski, Cordelia Langford, John Sillitoe on behalf of the Wellcome Sanger Institute COVID-19 Surveillance Team |
| EPI_ISL_2971315, EPI_ISL_2971316 | Berkshire and Surrey Pathology Services Lighthouse Laboratory | Wellcome Sanger Institute for the COVID-19 Genomics UK (COG-UK) Consortium | Berkshire and Surrey Pathology Services Lighthouse Laboratory and Alex Alderton, Roberto Amato, Jeffrey Barrett, Sonia Goncalves, Ewan Harrison, David K. Jackson, Ian Johnston, Dominic Kwiatkowski, Cordelia Langford, John Sillitoe on behalf of the Wellcome Sanger Institute COVID-19 Surveillance Team |
| EPI_ISL_2971317 | Lighthouse Laboratory Plymouth | Wellcome Sanger Institute for the COVID-19 Genomics UK (COG-UK) Consortium | Lighthouse Laboratory Plymouth and Alex Alderton, Roberto Amato, Jeffrey Barrett, Sonia Goncalves, Ewan Harrison, David K. Jackson, Ian Johnston, Dominic Kwiatkowski, Cordelia Langford, John Sillitoe on behalf of the Wellcome Sanger Institute COVID-19 Surveillance Team |
| EPI_ISL_2971318 | Berkshire and Surrey Pathology Services Lighthouse Laboratory | Wellcome Sanger Institute for the COVID-19 Genomics UK (COG-UK) Consortium | Berkshire and Surrey Pathology Services Lighthouse Laboratory and Alex Alderton, Roberto Amato, Jeffrey Barrett, Sonia Goncalves, Ewan Harrison, David K. Jackson, Ian Johnston, Dominic Kwiatkowski, Cordelia Langford, John Sillitoe on behalf of the Wellcome Sanger Institute COVID-19 Surveillance Team |
| EPI_ISL_2971319 | Lighthouse Laboratory Plymouth | Wellcome Sanger Institute for the COVID-19 Genomics UK (COG-UK) Consortium | Lighthouse Laboratory Plymouth and Alex Alderton, Roberto Amato, Jeffrey Barrett, Sonia Goncalves, Ewan Harrison, David K. Jackson, Ian Johnston, Dominic Kwiatkowski, Cordelia Langford, John Sillitoe on behalf of the Wellcome Sanger Institute COVID-19 Surveillance Team |
| EPI_ISL_2971320 | Lighthouse Lab in Alderley Park | Wellcome Sanger Institute for the COVID-19 Genomics UK (COG-UK) Consortium | Jacquelyn Wynn, Mairead Hyland, The Lighthouse Lab in Alderley Park and Alex Alderton, Roberto Amato, Jeffrey Barrett, Sonia Goncalves, Ewan Harrison, David K. Jackson, Ian Johnston, Dominic Kwiatkowski, Cordelia Langford, John Sillitoe on behalf of the Wellcome Sanger Institute COVID-19 Surveillance Team |
| EPI_ISL_2971321, EPI_ISL_2971322 | Lighthouse Laboratory Plymouth | Wellcome Sanger Institute for the COVID-19 Genomics UK (COG-UK) Consortium | Lighthouse Laboratory Plymouth and Alex Alderton, Roberto Amato, Jeffrey Barrett, Sonia Goncalves, Ewan Harrison, David K. Jackson, Ian Johnston, Dominic Kwiatkowski, Cordelia Langford, John Sillitoe on behalf of the Wellcome Sanger Institute COVID-19 Surveillance Team |
| EPI_ISL_2971323 | Berkshire and Surrey Pathology Services Lighthouse Laboratory | Wellcome Sanger Institute for the COVID-19 Genomics UK (COG-UK) Consortium | Berkshire and Surrey Pathology Services Lighthouse Laboratory and Alex Alderton, Roberto Amato, Jeffrey Barrett, Sonia Goncalves, Ewan Harrison, David K. Jackson, Ian Johnston, Dominic Kwiatkowski, Cordelia Langford, John Sillitoe on behalf of the Wellcome Sanger Institute COVID-19 Surveillance Team |
| EPI_ISL_2971324, EPI_ISL_2971325 | Lighthouse Laboratory Plymouth | Wellcome Sanger Institute for the COVID-19 Genomics UK (COG-UK) Consortium | Lighthouse Laboratory Plymouth and Alex Alderton, Roberto Amato, Jeffrey Barrett, Sonia Goncalves, Ewan Harrison, David K. Jackson, Ian Johnston, Dominic Kwiatkowski, Cordelia Langford, John Sillitoe on behalf of the Wellcome Sanger Institute COVID-19 Surveillance Team |
| EPI_ISL_2971326 | Lighthouse Lab in Alderley Park | Wellcome Sanger Institute for the COVID-19 Genomics UK (COG-UK) Consortium | Jacquelyn Wynn, Mairead Hyland, The Lighthouse Lab in Alderley Park and Alex Alderton, Roberto Amato, Jeffrey Barrett, Sonia Goncalves, Ewan Harrison, David K. Jackson, Ian Johnston, Dominic Kwiatkowski, Cordelia Langford, John Sillitoe on behalf of the Wellcome Sanger Institute COVID-19 Surveillance Team |
| EPI_ISL_2971327 | Lighthouse Laboratory Plymouth | Wellcome Sanger Institute for the COVID-19 Genomics UK (COG-UK) Consortium | Lighthouse Laboratory Plymouth and Alex Alderton, Roberto Amato, Jeffrey Barrett, Sonia Goncalves, Ewan Harrison, David K. Jackson, Ian Johnston, Dominic Kwiatkowski, Cordelia Langford, John Sillitoe on behalf of the Wellcome Sanger Institute COVID-19 Surveillance Team |
| EPI_ISL_2971328 | Berkshire and Surrey Pathology Services Lighthouse Laboratory | Wellcome Sanger Institute for the COVID-19 Genomics UK (COG-UK) Consortium | Berkshire and Surrey Pathology Services Lighthouse Laboratory and Alex Alderton, Roberto Amato, Jeffrey Barrett, Sonia Goncalves, Ewan Harrison, David K. Jackson, Ian Johnston, Dominic Kwiatkowski, Cordelia Langford, John Sillitoe on behalf of the Wellcome Sanger Institute COVID-19 Surveillance Team |
| EPI_ISL_2971329 | Lighthouse Lab in Alderley Park | Wellcome Sanger Institute for the COVID-19 Genomics UK (COG-UK) Consortium | Jacquelyn Wynn, Mairead Hyland, The Lighthouse Lab in Alderley Park and Alex Alderton, Roberto Amato, Jeffrey Barrett, Sonia Goncalves, Ewan Harrison, David K. Jackson, Ian Johnston, Dominic Kwiatkowski, Cordelia Langford, John Sillitoe on behalf of the Wellcome Sanger Institute COVID-19 Surveillance Team |
| EPI_ISL_2971330, EPI_ISL_2971331 | Lighthouse Laboratory Plymouth | Wellcome Sanger Institute for the COVID-19 Genomics UK (COG-UK) Consortium | Lighthouse Laboratory Plymouth and Alex Alderton, Roberto Amato, Jeffrey Barrett, Sonia Goncalves, Ewan Harrison, David K. Jackson, Ian Johnston, Dominic Kwiatkowski, Cordelia Langford, John Sillitoe on behalf of the Wellcome Sanger Institute COVID-19 Surveillance Team |
| EPI_ISL_2971332 | Berkshire and Surrey Pathology Services Lighthouse Laboratory | Wellcome Sanger Institute for the COVID-19 Genomics UK (COG-UK) Consortium | Berkshire and Surrey Pathology Services Lighthouse Laboratory and Alex Alderton, Roberto Amato, Jeffrey Barrett, Sonia Goncalves, Ewan Harrison, David K. Jackson, Ian Johnston, Dominic Kwiatkowski, Cordelia Langford, John Sillitoe on behalf of the Wellcome Sanger Institute COVID-19 Surveillance Team |
| EPI_ISL_2971333 | Lighthouse Laboratory Plymouth | Wellcome Sanger Institute for the COVID-19 Genomics UK (COG-UK) Consortium | Lighthouse Laboratory Plymouth and Alex Alderton, Roberto Amato, Jeffrey Barrett, Sonia Goncalves, Ewan Harrison, David K. Jackson, Ian Johnston, Dominic Kwiatkowski, Cordelia Langford, John Sillitoe on behalf of the Wellcome Sanger Institute COVID-19 Surveillance Team |
| EPI_ISL_2971334, EPI_ISL_2971335 | Lighthouse Lab in Alderley Park | Wellcome Sanger Institute for the COVID-19 Genomics UK (COG-UK) Consortium | Jacquelyn Wynn, Mairead Hyland, The Lighthouse Lab in Alderley Park and Alex Alderton, Roberto Amato, Jeffrey Barrett, Sonia Goncalves, Ewan Harrison, David K. Jackson, Ian Johnston, Dominic Kwiatkowski, Cordelia Langford, John Sillitoe on behalf of the Wellcome Sanger Institute COVID-19 Surveillance Team |
| EPI_ISL_2971336 | Berkshire and Surrey Pathology Services Lighthouse Laboratory | Wellcome Sanger Institute for the COVID-19 Genomics UK (COG-UK) Consortium | Berkshire and Surrey Pathology Services Lighthouse Laboratory and Alex Alderton, Roberto Amato, Jeffrey Barrett, Sonia Goncalves, Ewan Harrison, David K. Jackson, Ian Johnston, Dominic Kwiatkowski, Cordelia Langford, John Sillitoe on behalf of the Wellcome Sanger Institute COVID-19 Surveillance Team |
| EPI_ISL_2971337 | Lighthouse Lab in Alderley Park | Wellcome Sanger Institute for the COVID-19 Genomics UK (COG-UK) Consortium | Jacquelyn Wynn, Mairead Hyland, The Lighthouse Lab in Alderley Park and Alex Alderton, Roberto Amato, Jeffrey Barrett, Sonia Goncalves, Ewan Harrison, David K. Jackson, Ian Johnston, Dominic Kwiatkowski, Cordelia Langford, John Sillitoe on behalf of the Wellcome Sanger Institute COVID-19 Surveillance Team |
| EPI_ISL_2971338 | Berkshire and Surrey Pathology Services Lighthouse Laboratory | Wellcome Sanger Institute for the COVID-19 Genomics UK (COG-UK) Consortium | Berkshire and Surrey Pathology Services Lighthouse Laboratory and Alex Alderton, Roberto Amato, Jeffrey Barrett, Sonia Goncalves, Ewan Harrison, David K. Jackson, Ian Johnston, Dominic Kwiatkowski, Cordelia Langford, John Sillitoe on behalf of the Wellcome Sanger Institute COVID-19 Surveillance Team |
| EPI_ISL_2971339, EPI_ISL_2971340, EPI_ISL_2971341 | Lighthouse Lab in Alderley Park | Wellcome Sanger Institute for the COVID-19 Genomics UK (COG-UK) Consortium | Jacquelyn Wynn, Mairead Hyland, The Lighthouse Lab in Alderley Park and Alex Alderton, Roberto Amato, Jeffrey Barrett, Sonia Goncalves, Ewan Harrison, David K. Jackson, Ian Johnston, Dominic Kwiatkowski, Cordelia Langford, John Sillitoe on behalf of the Wellcome Sanger Institute COVID-19 Surveillance Team |
| EPI_ISL_2971342 | Lighthouse Laboratory Plymouth | Wellcome Sanger Institute for the COVID-19 Genomics UK (COG-UK) Consortium | Lighthouse Laboratory Plymouth and Alex Alderton, Roberto Amato, Jeffrey Barrett, Sonia Goncalves, Ewan Harrison, David K. Jackson, Ian Johnston, Dominic Kwiatkowski, Cordelia Langford, John Sillitoe on behalf of the Wellcome Sanger Institute COVID-19 Surveillance Team |
| EPI_ISL_2971343, EPI_ISL_2971344 | Lighthouse Lab in Alderley Park | Wellcome Sanger Institute for the COVID-19 Genomics UK (COG-UK) Consortium | Jacquelyn Wynn, Mairead Hyland, The Lighthouse Lab in Alderley Park and Alex Alderton, Roberto Amato, Jeffrey Barrett, Sonia Goncalves, Ewan Harrison, David K. Jackson, Ian Johnston, Dominic Kwiatkowski, Cordelia Langford, John Sillitoe on behalf of the Wellcome Sanger Institute COVID-19 Surveillance Team |
| EPI_ISL_2971345 | Berkshire and Surrey Pathology Services Lighthouse Laboratory | Wellcome Sanger Institute for the COVID-19 Genomics UK (COG-UK) Consortium | Berkshire and Surrey Pathology Services Lighthouse Laboratory and Alex Alderton, Roberto Amato, Jeffrey Barrett, Sonia Goncalves, Ewan Harrison, David K. Jackson, Ian Johnston, Dominic Kwiatkowski, Cordelia Langford, John Sillitoe on behalf of the Wellcome Sanger Institute COVID-19 Surveillance Team |
| EPI_ISL_2971346, EPI_ISL_2971347 | Lighthouse Lab in Alderley Park | Wellcome Sanger Institute for the COVID-19 Genomics UK (COG-UK) Consortium | Jacquelyn Wynn, Mairead Hyland, The Lighthouse Lab in Alderley Park and Alex Alderton, Roberto Amato, Jeffrey Barrett, Sonia Goncalves, Ewan Harrison, David K. Jackson, Ian Johnston, Dominic Kwiatkowski, Cordelia Langford, John Sillitoe on behalf of the Wellcome Sanger Institute COVID-19 Surveillance Team |

| | | | |
|---|---|---|---|
| EPI_ISL_2971348 | Lighthouse Laboratory Plymouth | Wellcome Sanger Institute for the COVID-19 Genomics UK (COG-UK) Consortium | Lighthouse Laboratory Plymouth and Alex Alderton, Roberto Amato, Jeffrey Barrett, Sonia Goncalves, Ewan Harrison, David K. Jackson, Ian Johnston, Dominic Kwiatkowski, Cordelia Langford, John Sillitoe on behalf of the Wellcome Sanger Institute COVID-19 Surveillance Team |
| EPI_ISL_2971349 | Lighthouse Lab in Alderley Park | Wellcome Sanger Institute for the COVID-19 Genomics UK (COG-UK) Consortium | Jacquelyn Wynn, Mairead Hyland, The Lighthouse Lab in Alderley Park and Alex Alderton, Roberto Amato, Jeffrey Barrett, Sonia Goncalves, Ewan Harrison, David K. Jackson, Ian Johnston, Dominic Kwiatkowski, Cordelia Langford, John Sillitoe on behalf of the Wellcome Sanger Institute COVID-19 Surveillance Team |
| EPI_ISL_2971350, EPI_ISL_2971351, EPI_ISL_2971352, EPI_ISL_2971353 | Lighthouse Laboratory Plymouth | Wellcome Sanger Institute for the COVID-19 Genomics UK (COG-UK) Consortium | Lighthouse Laboratory Plymouth and Alex Alderton, Roberto Amato, Jeffrey Barrett, Sonia Goncalves, Ewan Harrison, David K. Jackson, Ian Johnston, Dominic Kwiatkowski, Cordelia Langford, John Sillitoe on behalf of the Wellcome Sanger Institute COVID-19 Surveillance Team |
| EPI_ISL_2971354, EPI_ISL_2971355, EPI_ISL_2971356, EPI_ISL_2971357 | Lighthouse Lab in Alderley Park | Wellcome Sanger Institute for the COVID-19 Genomics UK (COG-UK) Consortium | Jacquelyn Wynn, Mairead Hyland, The Lighthouse Lab in Alderley Park and Alex Alderton, Roberto Amato, Jeffrey Barrett, Sonia Goncalves, Ewan Harrison, David K. Jackson, Ian Johnston, Dominic Kwiatkowski, Cordelia Langford, John Sillitoe on behalf of the Wellcome Sanger Institute COVID-19 Surveillance Team |
| EPI_ISL_2971358 | Berkshire and Surrey Pathology Services Lighthouse Laboratory | Wellcome Sanger Institute for the COVID-19 Genomics UK (COG-UK) Consortium | Berkshire and Surrey Pathology Services Lighthouse Laboratory and Alex Alderton, Roberto Amato, Jeffrey Barrett, Sonia Goncalves, Ewan Harrison, David K. Jackson, Ian Johnston, Dominic Kwiatkowski, Cordelia Langford, John Sillitoe on behalf of the Wellcome Sanger Institute COVID-19 Surveillance Team |
| EPI_ISL_2971359, EPI_ISL_2971360 | Lighthouse Laboratory Plymouth | Wellcome Sanger Institute for the COVID-19 Genomics UK (COG-UK) Consortium | Lighthouse Laboratory Plymouth and Alex Alderton, Roberto Amato, Jeffrey Barrett, Sonia Goncalves, Ewan Harrison, David K. Jackson, Ian Johnston, Dominic Kwiatkowski, Cordelia Langford, John Sillitoe on behalf of the Wellcome Sanger Institute COVID-19 Surveillance Team |
| EPI_ISL_2971361, EPI_ISL_2971362 | Berkshire and Surrey Pathology Services Lighthouse Laboratory | Wellcome Sanger Institute for the COVID-19 Genomics UK (COG-UK) Consortium | Berkshire and Surrey Pathology Services Lighthouse Laboratory and Alex Alderton, Roberto Amato, Jeffrey Barrett, Sonia Goncalves, Ewan Harrison, David K. Jackson, Ian Johnston, Dominic Kwiatkowski, Cordelia Langford, John Sillitoe on behalf of the Wellcome Sanger Institute COVID-19 Surveillance Team |
| EPI_ISL_2971363 | Lighthouse Lab in Alderley Park | Wellcome Sanger Institute for the COVID-19 Genomics UK (COG-UK) Consortium | Jacquelyn Wynn, Mairead Hyland, The Lighthouse Lab in Alderley Park and Alex Alderton, Roberto Amato, Jeffrey Barrett, Sonia Goncalves, Ewan Harrison, David K. Jackson, Ian Johnston, Dominic Kwiatkowski, Cordelia Langford, John Sillitoe on behalf of the Wellcome Sanger Institute COVID-19 Surveillance Team |
| EPI_ISL_2971364 | Berkshire and Surrey Pathology Services Lighthouse Laboratory | Wellcome Sanger Institute for the COVID-19 Genomics UK (COG-UK) Consortium | Berkshire and Surrey Pathology Services Lighthouse Laboratory and Alex Alderton, Roberto Amato, Jeffrey Barrett, Sonia Goncalves, Ewan Harrison, David K. Jackson, Ian Johnston, Dominic Kwiatkowski, Cordelia Langford, John Sillitoe on behalf of the Wellcome Sanger Institute COVID-19 Surveillance Team |
| EPI_ISL_2971365 | Lighthouse Lab in Alderley Park | Wellcome Sanger Institute for the COVID-19 Genomics UK (COG-UK) Consortium | Jacquelyn Wynn, Mairead Hyland, The Lighthouse Lab in Alderley Park and Alex Alderton, Roberto Amato, Jeffrey Barrett, Sonia Goncalves, Ewan Harrison, David K. Jackson, Ian Johnston, Dominic Kwiatkowski, Cordelia Langford, John Sillitoe on behalf of the Wellcome Sanger Institute COVID-19 Surveillance Team |
| EPI_ISL_2971366 | Lighthouse Laboratory Plymouth | Wellcome Sanger Institute for the COVID-19 Genomics UK (COG-UK) Consortium | Lighthouse Laboratory Plymouth and Alex Alderton, Roberto Amato, Jeffrey Barrett, Sonia Goncalves, Ewan Harrison, David K. Jackson, Ian Johnston, Dominic Kwiatkowski, Cordelia Langford, John Sillitoe on behalf of the Wellcome Sanger Institute COVID-19 Surveillance Team |
| EPI_ISL_2971367 | Lighthouse Lab in Alderley Park | Wellcome Sanger Institute for the COVID-19 Genomics UK (COG-UK) Consortium | Jacquelyn Wynn, Mairead Hyland, The Lighthouse Lab in Alderley Park and Alex Alderton, Roberto Amato, Jeffrey Barrett, Sonia Goncalves, Ewan Harrison, David K. Jackson, Ian Johnston, Dominic Kwiatkowski, Cordelia Langford, John Sillitoe on behalf of the Wellcome Sanger Institute COVID-19 Surveillance Team |
| EPI_ISL_2971368 | Berkshire and Surrey Pathology Services Lighthouse Laboratory | Wellcome Sanger Institute for the COVID-19 Genomics UK (COG-UK) Consortium | Berkshire and Surrey Pathology Services Lighthouse Laboratory and Alex Alderton, Roberto Amato, Jeffrey Barrett, Sonia Goncalves, Ewan Harrison, David K. Jackson, Ian Johnston, Dominic Kwiatkowski, Cordelia Langford, John Sillitoe on behalf of the Wellcome Sanger Institute COVID-19 Surveillance Team |
| EPI_ISL_2971369 | Department of Bacteria, Parasites and Fungi, Statens Serum Institut, Copenhagen, Denmark | Statens Serum Institut Bioinformatics and Microbial Genomics | Danish Covid-19 Genome Consortium |
| EPI_ISL_2971370, EPI_ISL_2971371 | Lighthouse Lab in Alderley Park | Wellcome Sanger Institute for the COVID-19 Genomics UK (COG-UK) Consortium | Jacquelyn Wynn, Mairead Hyland, The Lighthouse Lab in Alderley Park and Alex Alderton, Roberto Amato, Jeffrey Barrett, Sonia Goncalves, Ewan Harrison, David K. Jackson, Ian Johnston, Dominic Kwiatkowski, Cordelia Langford, John Sillitoe on behalf of the Wellcome Sanger Institute COVID-19 Surveillance Team |
| EPI_ISL_2971372 | Department of Bacteria, Parasites and Fungi, Statens Serum Institut, Copenhagen, Denmark | Statens Serum Institut Bioinformatics and Microbial Genomics | Danish Covid-19 Genome Consortium |
| EPI_ISL_2971373, EPI_ISL_2971374 | Lighthouse Lab in Alderley Park | Wellcome Sanger Institute for the COVID-19 Genomics UK (COG-UK) Consortium | Jacquelyn Wynn, Mairead Hyland, The Lighthouse Lab in Alderley Park and Alex Alderton, Roberto Amato, Jeffrey Barrett, Sonia Goncalves, Ewan Harrison, David K. Jackson, Ian Johnston, Dominic Kwiatkowski, Cordelia Langford, John Sillitoe on behalf of the Wellcome Sanger Institute COVID-19 Surveillance Team |
| EPI_ISL_2971375 | Department of Bacteria, Parasites and Fungi, Statens Serum Institut, Copenhagen, Denmark | Statens Serum Institut Bioinformatics and Microbial Genomics | Danish Covid-19 Genome Consortium |
| EPI_ISL_2971376 | Lighthouse Laboratory Plymouth | Wellcome Sanger Institute for the COVID-19 Genomics UK (COG-UK) Consortium | Lighthouse Laboratory Plymouth and Alex Alderton, Roberto Amato, Jeffrey Barrett, Sonia Goncalves, Ewan Harrison, David K. Jackson, Ian Johnston, Dominic Kwiatkowski, Cordelia Langford, John Sillitoe on behalf of the Wellcome Sanger Institute COVID-19 Surveillance Team |
| EPI_ISL_2971377, EPI_ISL_2971378, EPI_ISL_2971380, EPI_ISL_2971381 | Lighthouse Lab in Alderley Park | Wellcome Sanger Institute for the COVID-19 Genomics UK (COG-UK) Consortium | Jacquelyn Wynn, Mairead Hyland, The Lighthouse Lab in Alderley Park and Alex Alderton, Roberto Amato, Jeffrey Barrett, Sonia Goncalves, Ewan Harrison, David K. Jackson, Ian Johnston, Dominic Kwiatkowski, Cordelia Langford, John Sillitoe on behalf of the Wellcome Sanger Institute COVID-19 Surveillance Team |
| EPI_ISL_2971382 | Department of Bacteria, Parasites and Fungi, Statens Serum Institut, Copenhagen, Denmark | Statens Serum Institut Bioinformatics and Microbial Genomics | Danish Covid-19 Genome Consortium |
| EPI_ISL_2971383 | Berkshire and Surrey Pathology Services Lighthouse Laboratory | Wellcome Sanger Institute for the COVID-19 Genomics UK (COG-UK) Consortium | Berkshire and Surrey Pathology Services Lighthouse Laboratory and Alex Alderton, Roberto Amato, Jeffrey Barrett, Sonia Goncalves, Ewan Harrison, David K. Jackson, Ian Johnston, Dominic Kwiatkowski, Cordelia Langford, John Sillitoe on behalf of the Wellcome Sanger Institute COVID-19 Surveillance Team |
| EPI_ISL_2971384, EPI_ISL_2971385 | Lighthouse Lab in Alderley Park | Wellcome Sanger Institute for the COVID-19 Genomics UK (COG-UK) Consortium | Jacquelyn Wynn, Mairead Hyland, The Lighthouse Lab in Alderley Park and Alex Alderton, Roberto Amato, Jeffrey Barrett, Sonia Goncalves, Ewan Harrison, David K. Jackson, Ian Johnston, Dominic Kwiatkowski, Cordelia Langford, John Sillitoe on behalf of the Wellcome Sanger Institute COVID-19 Surveillance Team |
| EPI_ISL_2971386 | Department of Bacteria, Parasites and Fungi, Statens Serum Institut, Copenhagen, Denmark | Statens Serum Institut Bioinformatics and Microbial Genomics | Danish Covid-19 Genome Consortium |
| EPI_ISL_2971387, EPI_ISL_2971388 | Berkshire and Surrey Pathology Services Lighthouse Laboratory | Wellcome Sanger Institute for the COVID-19 Genomics UK (COG-UK) Consortium | Berkshire and Surrey Pathology Services Lighthouse Laboratory and Alex Alderton, Roberto Amato, Jeffrey Barrett, Sonia Goncalves, Ewan Harrison, David K. Jackson, Ian Johnston, Dominic Kwiatkowski, Cordelia Langford, John Sillitoe on behalf of the Wellcome Sanger Institute COVID-19 Surveillance Team |
| EPI_ISL_2971389 | Department of Bacteria, Parasites and Fungi, Statens Serum Institut, Copenhagen, Denmark | Statens Serum Institut Bioinformatics and Microbial Genomics | Danish Covid-19 Genome Consortium |
| EPI_ISL_2971390 | Berkshire and Surrey Pathology Services Lighthouse Laboratory | Wellcome Sanger Institute for the COVID-19 Genomics UK (COG-UK) Consortium | Berkshire and Surrey Pathology Services Lighthouse Laboratory and Alex Alderton, Roberto Amato, Jeffrey Barrett, Sonia Goncalves, Ewan Harrison, David K. Jackson, Ian Johnston, Dominic Kwiatkowski, Cordelia Langford, John Sillitoe on behalf of the Wellcome Sanger Institute COVID-19 Surveillance Team |
| EPI_ISL_2971391 | Molecular Diagnostics Pathology Department Mater Dei Hospital Malta | Molecular Diagnostics Pathology Department Mater Dei Hospital Malta | G Zahra, R Borg, C Cilia, L Grech, M Briffa |
| EPI_ISL_2971392 | Lighthouse Laboratory Plymouth | Wellcome Sanger Institute for the COVID-19 Genomics UK (COG-UK) Consortium | Lighthouse Laboratory Plymouth and Alex Alderton, Roberto Amato, Jeffrey Barrett, Sonia Goncalves, Ewan Harrison, David K. Jackson, Ian Johnston, Dominic Kwiatkowski, Cordelia Langford, John Sillitoe on behalf of the Wellcome Sanger Institute COVID-19 Surveillance Team |
| EPI_ISL_2971393 | Lighthouse Lab in Alderley Park | Wellcome Sanger Institute for the COVID-19 Genomics UK (COG-UK) Consortium | Jacquelyn Wynn, Mairead Hyland, The Lighthouse Lab in Alderley Park and Alex Alderton, Roberto Amato, Jeffrey Barrett, Sonia Goncalves, Ewan Harrison, David K. Jackson, Ian Johnston, Dominic Kwiatkowski, Cordelia Langford, John Sillitoe on behalf of the Wellcome Sanger Institute COVID-19 Surveillance Team |
| EPI_ISL_2971394 | Department of Bacteria, Parasites and Fungi, Statens Serum Institut, Copenhagen, Denmark | Statens Serum Institut Bioinformatics and Microbial Genomics | Danish Covid-19 Genome Consortium |
| EPI_ISL_2971395, EPI_ISL_2971396 | Lighthouse Lab in Alderley Park | Wellcome Sanger Institute for the COVID-19 Genomics UK (COG-UK) Consortium | Jacquelyn Wynn, Mairead Hyland, The Lighthouse Lab in Alderley Park and Alex Alderton, Roberto Amato, Jeffrey Barrett, Sonia Goncalves, Ewan Harrison, David K. Jackson, Ian Johnston, Dominic Kwiatkowski, Cordelia Langford, John Sillitoe on behalf of the Wellcome Sanger Institute COVID-19 Surveillance Team |
| EPI_ISL_2971397 | Department of Bacteria, Parasites and Fungi, Statens Serum Institut, Copenhagen, Denmark | Statens Serum Institut Bioinformatics and Microbial Genomics | Danish Covid-19 Genome Consortium |
| EPI_ISL_2971398, EPI_ISL_2971399 | Lighthouse Lab in Alderley Park | Wellcome Sanger Institute for the COVID-19 Genomics UK (COG-UK) Consortium | Jacquelyn Wynn, Mairead Hyland, The Lighthouse Lab in Alderley Park and Alex Alderton, Roberto Amato, Jeffrey Barrett, Sonia Goncalves, Ewan Harrison, David K. Jackson, Ian Johnston, Dominic Kwiatkowski, Cordelia Langford, John Sillitoe on behalf of the Wellcome Sanger Institute COVID-19 Surveillance Team |
| EPI_ISL_2971400 | Department of Bacteria, Parasites and Fungi, Statens Serum Institut, Copenhagen, Denmark | Statens Serum Institut Bioinformatics and Microbial Genomics | Danish Covid-19 Genome Consortium |
| EPI_ISL_2971401, EPI_ISL_2971402 | Lighthouse Lab in Alderley Park | Wellcome Sanger Institute for the COVID-19 Genomics UK (COG-UK) Consortium | Jacquelyn Wynn, Mairead Hyland, The Lighthouse Lab in Alderley Park and Alex Alderton, Roberto Amato, Jeffrey Barrett, Sonia Goncalves, Ewan Harrison, David K. Jackson, Ian Johnston, Dominic Kwiatkowski, Cordelia Langford, John Sillitoe on behalf of the Wellcome Sanger Institute COVID-19 Surveillance Team |
| EPI_ISL_2971403 | Department of Bacteria, Parasites and Fungi, Statens Serum Institut, Copenhagen, Denmark | Statens Serum Institut Bioinformatics and Microbial Genomics | Danish Covid-19 Genome Consortium |
| EPI_ISL_2971404, EPI_ISL_2971405 | Lighthouse Lab in Alderley Park | Wellcome Sanger Institute for the COVID-19 Genomics UK (COG-UK) Consortium | Jacquelyn Wynn, Mairead Hyland, The Lighthouse Lab in Alderley Park and Alex Alderton, Roberto Amato, Jeffrey Barrett, Sonia Goncalves, Ewan Harrison, David K. Jackson, Ian Johnston, Dominic Kwiatkowski, Cordelia Langford, John Sillitoe on behalf of the Wellcome Sanger Institute COVID-19 Surveillance Team |
| EPI_ISL_2971406 | Department of Bacteria, Parasites and Fungi, Statens Serum Institut, Copenhagen, Denmark | Statens Serum Institut Bioinformatics and Microbial Genomics | Danish Covid-19 Genome Consortium |
| EPI_ISL_2971407 | Lighthouse Lab in Alderley Park | Wellcome Sanger Institute for the COVID-19 Genomics UK (COG-UK) Consortium | Jacquelyn Wynn, Mairead Hyland, The Lighthouse Lab in Alderley Park and Alex Alderton, Roberto Amato, Jeffrey Barrett, Sonia Goncalves, Ewan Harrison, David K. Jackson, Ian Johnston, Dominic Kwiatkowski, Cordelia Langford, John Sillitoe on behalf of the Wellcome Sanger Institute COVID-19 Surveillance Team |
| EPI_ISL_2971408 | Berkshire and Surrey Pathology Services Lighthouse Laboratory | Wellcome Sanger Institute for the COVID-19 Genomics UK (COG-UK) Consortium | Berkshire and Surrey Pathology Services Lighthouse Laboratory and Alex Alderton, Roberto Amato, Jeffrey Barrett, Sonia Goncalves, Ewan Harrison, David K. Jackson, Ian Johnston, Dominic Kwiatkowski, Cordelia Langford, John Sillitoe on behalf of the Wellcome Sanger Institute COVID-19 Surveillance Team |
| EPI_ISL_2971409 | Lighthouse Lab in Alderley Park | Wellcome Sanger Institute for the COVID-19 Genomics UK (COG-UK) Consortium | Jacquelyn Wynn, Mairead Hyland, The Lighthouse Lab in Alderley Park and Alex Alderton, Roberto Amato, Jeffrey Barrett, Sonia Goncalves, Ewan Harrison, David K. Jackson, Ian Johnston, Dominic Kwiatkowski, Cordelia Langford, John Sillitoe on behalf of the Wellcome Sanger Institute COVID-19 Surveillance Team |
| EPI_ISL_2971411 | Berkshire and Surrey Pathology Services Lighthouse Laboratory | Wellcome Sanger Institute for the COVID-19 Genomics UK (COG-UK) | Berkshire and Surrey Pathology Services Lighthouse Laboratory and Alex Alderton, Roberto Amato, Jeffrey Barrett, Sonia Goncalves, Ewan Harrison, David K. Jackson, Ian Johnston, Dominic Kwiatkowski, Cordelia Langford, John Sillitoe on behalf |

| | | | |
|---|---|---|---|
| | | Consortium | of the Wellcome Sanger Institute COVID-19 Surveillance Team |
| EPI_ISL_2971412 | Lighthouse Laboratory Plymouth | Wellcome Sanger Institute for the COVID-19 Genomics UK (COG-UK) Consortium | Lighthouse Laboratory Plymouth and Alex Alderton, Roberto Amato, Jeffrey Barrett, Sonia Goncalves, Ewan Harrison, David K. Jackson, Ian Johnston, Dominic Kwiatkowski, Cordelia Langford, John Sillitoe on behalf of the Wellcome Sanger Institute COVID-19 Surveillance Team |
| EPI_ISL_2971413 | Department of Bacteria, Parasites and Fungi, Statens Serum Institut, Copenhagen, Denmark | Statens Serum Institut Bioinformatics and Microbial Genomics | Danish Covid-19 Genome Consortium |
| EPI_ISL_2971414, EPI_ISL_2971415 | Lighthouse Laboratory Plymouth | Wellcome Sanger Institute for the COVID-19 Genomics UK (COG-UK) Consortium | Lighthouse Laboratory Plymouth and Alex Alderton, Roberto Amato, Jeffrey Barrett, Sonia Goncalves, Ewan Harrison, David K. Jackson, Ian Johnston, Dominic Kwiatkowski, Cordelia Langford, John Sillitoe on behalf of the Wellcome Sanger Institute COVID-19 Surveillance Team |
| EPI_ISL_2971416 | Department of Bacteria, Parasites and Fungi, Statens Serum Institut, Copenhagen, Denmark | Statens Serum Institut Bioinformatics and Microbial Genomics | Danish Covid-19 Genome Consortium |
| EPI_ISL_2971417, EPI_ISL_2971418 | Lighthouse Lab in Alderley Park | Wellcome Sanger Institute for the COVID-19 Genomics UK (COG-UK) Consortium | Jacquelyn Wynn, Mairead Hyland, The Lighthouse Lab in Alderley Park and Alex Alderton, Roberto Amato, Jeffrey Barrett, Sonia Goncalves, Ewan Harrison, David K. Jackson, Ian Johnston, Dominic Kwiatkowski, Cordelia Langford, John Sillitoe on behalf of the Wellcome Sanger Institute COVID-19 Surveillance Team |
| EPI_ISL_2971419 | Lighthouse Laboratory Plymouth | Wellcome Sanger Institute for the COVID-19 Genomics UK (COG-UK) Consortium | Lighthouse Laboratory Plymouth and Alex Alderton, Roberto Amato, Jeffrey Barrett, Sonia Goncalves, Ewan Harrison, David K. Jackson, Ian Johnston, Dominic Kwiatkowski, Cordelia Langford, John Sillitoe on behalf of the Wellcome Sanger Institute COVID-19 Surveillance Team |
| EPI_ISL_2971420 | Department of Bacteria, Parasites and Fungi, Statens Serum Institut, Copenhagen, Denmark | Statens Serum Institut Bioinformatics and Microbial Genomics | Danish Covid-19 Genome Consortium |
| EPI_ISL_2971421, EPI_ISL_2971422 | Lighthouse Lab in Alderley Park | Wellcome Sanger Institute for the COVID-19 Genomics UK (COG-UK) Consortium | Jacquelyn Wynn, Mairead Hyland, The Lighthouse Lab in Alderley Park and Alex Alderton, Roberto Amato, Jeffrey Barrett, Sonia Goncalves, Ewan Harrison, David K. Jackson, Ian Johnston, Dominic Kwiatkowski, Cordelia Langford, John Sillitoe on behalf of the Wellcome Sanger Institute COVID-19 Surveillance Team |
| EPI_ISL_2971423 | Department of Bacteria, Parasites and Fungi, Statens Serum Institut, Copenhagen, Denmark | Statens Serum Institut Bioinformatics and Microbial Genomics | Danish Covid-19 Genome Consortium |
| EPI_ISL_2971424 | Lighthouse Laboratory Plymouth | Wellcome Sanger Institute for the COVID-19 Genomics UK (COG-UK) Consortium | Lighthouse Laboratory Plymouth and Alex Alderton, Roberto Amato, Jeffrey Barrett, Sonia Goncalves, Ewan Harrison, David K. Jackson, Ian Johnston, Dominic Kwiatkowski, Cordelia Langford, John Sillitoe on behalf of the Wellcome Sanger Institute COVID-19 Surveillance Team |
| EPI_ISL_2971425 | Berkshire and Surrey Pathology Services Lighthouse Laboratory | Wellcome Sanger Institute for the COVID-19 Genomics UK (COG-UK) Consortium | Berkshire and Surrey Pathology Services Lighthouse Laboratory and Alex Alderton, Roberto Amato, Jeffrey Barrett, Sonia Goncalves, Ewan Harrison, David K. Jackson, Ian Johnston, Dominic Kwiatkowski, Cordelia Langford, John Sillitoe on behalf of the Wellcome Sanger Institute COVID-19 Surveillance Team |
| EPI_ISL_2971426 | Department of Bacteria, Parasites and Fungi, Statens Serum Institut, Copenhagen, Denmark | Statens Serum Institut Bioinformatics and Microbial Genomics | Danish Covid-19 Genome Consortium |
| EPI_ISL_2971427 | Lighthouse Laboratory Plymouth | Wellcome Sanger Institute for the COVID-19 Genomics UK (COG-UK) Consortium | Lighthouse Laboratory Plymouth and Alex Alderton, Roberto Amato, Jeffrey Barrett, Sonia Goncalves, Ewan Harrison, David K. Jackson, Ian Johnston, Dominic Kwiatkowski, Cordelia Langford, John Sillitoe on behalf of the Wellcome Sanger Institute COVID-19 Surveillance Team |
| EPI_ISL_2971428 | Lighthouse Lab in Alderley Park | Wellcome Sanger Institute for the COVID-19 Genomics UK (COG-UK) Consortium | Jacquelyn Wynn, Mairead Hyland, The Lighthouse Lab in Alderley Park and Alex Alderton, Roberto Amato, Jeffrey Barrett, Sonia Goncalves, Ewan Harrison, David K. Jackson, Ian Johnston, Dominic Kwiatkowski, Cordelia Langford, John Sillitoe on behalf of the Wellcome Sanger Institute COVID-19 Surveillance Team |
| EPI_ISL_2971429 | Lighthouse Laboratory Plymouth | Wellcome Sanger Institute for the COVID-19 Genomics UK (COG-UK) Consortium | Lighthouse Laboratory Plymouth and Alex Alderton, Roberto Amato, Jeffrey Barrett, Sonia Goncalves, Ewan Harrison, David K. Jackson, Ian Johnston, Dominic Kwiatkowski, Cordelia Langford, John Sillitoe on behalf of the Wellcome Sanger Institute COVID-19 Surveillance Team |
| EPI_ISL_2971430 | Department of Bacteria, Parasites and Fungi, Statens Serum Institut, Copenhagen, Denmark | Statens Serum Institut Bioinformatics and Microbial Genomics | Danish Covid-19 Genome Consortium |
| EPI_ISL_2971431 | Berkshire and Surrey Pathology Services Lighthouse Laboratory | Wellcome Sanger Institute for the COVID-19 Genomics UK (COG-UK) Consortium | Berkshire and Surrey Pathology Services Lighthouse Laboratory and Alex Alderton, Roberto Amato, Jeffrey Barrett, Sonia Goncalves, Ewan Harrison, David K. Jackson, Ian Johnston, Dominic Kwiatkowski, Cordelia Langford, John Sillitoe on behalf of the Wellcome Sanger Institute COVID-19 Surveillance Team |
| EPI_ISL_2971432 | Lighthouse Lab in Alderley Park | Wellcome Sanger Institute for the COVID-19 Genomics UK (COG-UK) Consortium | Jacquelyn Wynn, Mairead Hyland, The Lighthouse Lab in Alderley Park and Alex Alderton, Roberto Amato, Jeffrey Barrett, Sonia Goncalves, Ewan Harrison, David K. Jackson, Ian Johnston, Dominic Kwiatkowski, Cordelia Langford, John Sillitoe on behalf of the Wellcome Sanger Institute COVID-19 Surveillance Team |
| EPI_ISL_2971433 | Department of Bacteria, Parasites and Fungi, Statens Serum Institut, Copenhagen, Denmark | Statens Serum Institut Bioinformatics and Microbial Genomics | Danish Covid-19 Genome Consortium |
| EPI_ISL_2971434 | Lighthouse Lab in Alderley Park | Wellcome Sanger Institute for the COVID-19 Genomics UK (COG-UK) Consortium | Jacquelyn Wynn, Mairead Hyland, The Lighthouse Lab in Alderley Park and Alex Alderton, Roberto Amato, Jeffrey Barrett, Sonia Goncalves, Ewan Harrison, David K. Jackson, Ian Johnston, Dominic Kwiatkowski, Cordelia Langford, John Sillitoe on behalf of the Wellcome Sanger Institute COVID-19 Surveillance Team |
| EPI_ISL_2971435 | Lighthouse Laboratory Plymouth | Wellcome Sanger Institute for the COVID-19 Genomics UK (COG-UK) Consortium | Lighthouse Laboratory Plymouth and Alex Alderton, Roberto Amato, Jeffrey Barrett, Sonia Goncalves, Ewan Harrison, David K. Jackson, Ian Johnston, Dominic Kwiatkowski, Cordelia Langford, John Sillitoe on behalf of the Wellcome Sanger Institute COVID-19 Surveillance Team |
| EPI_ISL_2971436 | Lighthouse Lab in Alderley Park | Wellcome Sanger Institute for the COVID-19 Genomics UK (COG-UK) Consortium | Jacquelyn Wynn, Mairead Hyland, The Lighthouse Lab in Alderley Park and Alex Alderton, Roberto Amato, Jeffrey Barrett, Sonia Goncalves, Ewan Harrison, David K. Jackson, Ian Johnston, Dominic Kwiatkowski, Cordelia Langford, John Sillitoe on behalf of the Wellcome Sanger Institute COVID-19 Surveillance Team |
| EPI_ISL_2971437 | Department of Bacteria, Parasites and Fungi, Statens Serum Institut, Copenhagen, Denmark | Statens Serum Institut Bioinformatics and Microbial Genomics | Danish Covid-19 Genome Consortium |
| EPI_ISL_2971438 | Lighthouse Laboratory Plymouth | Wellcome Sanger Institute for the COVID-19 Genomics UK (COG-UK) Consortium | Lighthouse Laboratory Plymouth and Alex Alderton, Roberto Amato, Jeffrey Barrett, Sonia Goncalves, Ewan Harrison, David K. Jackson, Ian Johnston, Dominic Kwiatkowski, Cordelia Langford, John Sillitoe on behalf of the Wellcome Sanger Institute COVID-19 Surveillance Team |
| EPI_ISL_2971439 | Department of Bacteria, Parasites and Fungi, Statens Serum Institut, Copenhagen, Denmark | Statens Serum Institut Bioinformatics and Microbial Genomics | Danish Covid-19 Genome Consortium |
| EPI_ISL_2971440 | Berkshire and Surrey Pathology Services Lighthouse Laboratory | Wellcome Sanger Institute for the COVID-19 Genomics UK (COG-UK) Consortium | Berkshire and Surrey Pathology Services Lighthouse Laboratory and Alex Alderton, Roberto Amato, Jeffrey Barrett, Sonia Goncalves, Ewan Harrison, David K. Jackson, Ian Johnston, Dominic Kwiatkowski, Cordelia Langford, John Sillitoe on behalf of the Wellcome Sanger Institute COVID-19 Surveillance Team |
| EPI_ISL_2971441, EPI_ISL_2971442 | Lighthouse Laboratory Plymouth | Wellcome Sanger Institute for the COVID-19 Genomics UK (COG-UK) Consortium | Lighthouse Laboratory Plymouth and Alex Alderton, Roberto Amato, Jeffrey Barrett, Sonia Goncalves, Ewan Harrison, David K. Jackson, Ian Johnston, Dominic Kwiatkowski, Cordelia Langford, John Sillitoe on behalf of the Wellcome Sanger Institute COVID-19 Surveillance Team |
| EPI_ISL_2971443 | Department of Bacteria, Parasites and Fungi, Statens Serum Institut, Copenhagen, Denmark | Statens Serum Institut Bioinformatics and Microbial Genomics | Danish Covid-19 Genome Consortium |
| EPI_ISL_2971444, EPI_ISL_2971445 | Lighthouse Lab in Alderley Park | Wellcome Sanger Institute for the COVID-19 Genomics UK (COG-UK) Consortium | Jacquelyn Wynn, Mairead Hyland, The Lighthouse Lab in Alderley Park and Alex Alderton, Roberto Amato, Jeffrey Barrett, Sonia Goncalves, Ewan Harrison, David K. Jackson, Ian Johnston, Dominic Kwiatkowski, Cordelia Langford, John Sillitoe on behalf of the Wellcome Sanger Institute COVID-19 Surveillance Team |
| EPI_ISL_2971446 | Department of Bacteria, Parasites and Fungi, Statens Serum Institut, Copenhagen, Denmark | Statens Serum Institut Bioinformatics and Microbial Genomics | Danish Covid-19 Genome Consortium |
| EPI_ISL_2971447 | Berkshire and Surrey Pathology Services Lighthouse Laboratory | Wellcome Sanger Institute for the COVID-19 Genomics UK (COG-UK) Consortium | Berkshire and Surrey Pathology Services Lighthouse Laboratory and Alex Alderton, Roberto Amato, Jeffrey Barrett, Sonia Goncalves, Ewan Harrison, David K. Jackson, Ian Johnston, Dominic Kwiatkowski, Cordelia Langford, John Sillitoe on behalf of the Wellcome Sanger Institute COVID-19 Surveillance Team |
| EPI_ISL_2971448 | Lighthouse Lab in Alderley Park | Wellcome Sanger Institute for the COVID-19 Genomics UK (COG-UK) Consortium | Jacquelyn Wynn, Mairead Hyland, The Lighthouse Lab in Alderley Park and Alex Alderton, Roberto Amato, Jeffrey Barrett, Sonia Goncalves, Ewan Harrison, David K. Jackson, Ian Johnston, Dominic Kwiatkowski, Cordelia Langford, John Sillitoe on behalf of the Wellcome Sanger Institute COVID-19 Surveillance Team |
| EPI_ISL_2971449 | Department of Bacteria, Parasites and Fungi, Statens Serum Institut, Copenhagen, Denmark | Statens Serum Institut Bioinformatics and Microbial Genomics | Danish Covid-19 Genome Consortium |
| EPI_ISL_2971450 | Lighthouse Lab in Alderley Park | Wellcome Sanger Institute for the COVID-19 Genomics UK (COG-UK) Consortium | Jacquelyn Wynn, Mairead Hyland, The Lighthouse Lab in Alderley Park and Alex Alderton, Roberto Amato, Jeffrey Barrett, Sonia Goncalves, Ewan Harrison, David K. Jackson, Ian Johnston, Dominic Kwiatkowski, Cordelia Langford, John Sillitoe on behalf of the Wellcome Sanger Institute COVID-19 Surveillance Team |
| EPI_ISL_2971451 | Lighthouse Laboratory Plymouth | Wellcome Sanger Institute for the COVID-19 Genomics UK (COG-UK) Consortium | Lighthouse Laboratory Plymouth and Alex Alderton, Roberto Amato, Jeffrey Barrett, Sonia Goncalves, Ewan Harrison, David K. Jackson, Ian Johnston, Dominic Kwiatkowski, Cordelia Langford, John Sillitoe on behalf of the Wellcome Sanger Institute COVID-19 Surveillance Team |
| EPI_ISL_2971452 | Department of Bacteria, Parasites and Fungi, Statens Serum Institut, Copenhagen, Denmark | Statens Serum Institut Bioinformatics and Microbial Genomics | Danish Covid-19 Genome Consortium |
| EPI_ISL_2971453, EPI_ISL_2971454 | Berkshire and Surrey Pathology Services Lighthouse Laboratory | Wellcome Sanger Institute for the COVID-19 Genomics UK (COG-UK) Consortium | Berkshire and Surrey Pathology Services Lighthouse Laboratory and Alex Alderton, Roberto Amato, Jeffrey Barrett, Sonia Goncalves, Ewan Harrison, David K. Jackson, Ian Johnston, Dominic Kwiatkowski, Cordelia Langford, John Sillitoe on behalf of the Wellcome Sanger Institute COVID-19 Surveillance Team |
| EPI_ISL_2971455 | Department of Bacteria, Parasites and Fungi, Statens Serum Institut, Copenhagen, Denmark | Statens Serum Institut Bioinformatics and Microbial Genomics | Danish Covid-19 Genome Consortium |
| EPI_ISL_2971456, EPI_ISL_2971457 | Lighthouse Laboratory Plymouth | Wellcome Sanger Institute for the COVID-19 Genomics UK (COG-UK) Consortium | Lighthouse Laboratory Plymouth and Alex Alderton, Roberto Amato, Jeffrey Barrett, Sonia Goncalves, Ewan Harrison, David K. Jackson, Ian Johnston, Dominic Kwiatkowski, Cordelia Langford, John Sillitoe on behalf of the Wellcome Sanger Institute COVID-19 Surveillance Team |
| EPI_ISL_2971458 | Lighthouse Lab in Alderley Park | Wellcome Sanger Institute for the COVID-19 Genomics UK (COG-UK) Consortium | Jacquelyn Wynn, Mairead Hyland, The Lighthouse Lab in Alderley Park and Alex Alderton, Roberto Amato, Jeffrey Barrett, Sonia Goncalves, Ewan Harrison, David K. Jackson, Ian Johnston, Dominic Kwiatkowski, Cordelia Langford, John Sillitoe on behalf of the Wellcome Sanger Institute COVID-19 Surveillance Team |
| EPI_ISL_2971459 | Department of Bacteria, Parasites and Fungi, Statens Serum Institut, Copenhagen, Denmark | Statens Serum Institut Bioinformatics and Microbial Genomics | Danish Covid-19 Genome Consortium |
| EPI_ISL_2971460 | Berkshire and Surrey Pathology Services Lighthouse Laboratory | Wellcome Sanger Institute for the COVID-19 Genomics UK (COG-UK) Consortium | Berkshire and Surrey Pathology Services Lighthouse Laboratory and Alex Alderton, Roberto Amato, Jeffrey Barrett, Sonia Goncalves, Ewan Harrison, David K. Jackson, Ian Johnston, Dominic Kwiatkowski, Cordelia Langford, John Sillitoe on behalf of the Wellcome Sanger Institute COVID-19 Surveillance Team |
| EPI_ISL_2971461 | Lighthouse Lab in Alderley Park | Wellcome Sanger Institute for the COVID-19 Genomics UK (COG-UK) Consortium | Jacquelyn Wynn, Mairead Hyland, The Lighthouse Lab in Alderley Park and Alex Alderton, Roberto Amato, Jeffrey Barrett, Sonia Goncalves, Ewan Harrison, David K. Jackson, Ian Johnston, Dominic Kwiatkowski, Cordelia Langford, John Sillitoe on behalf of the Wellcome Sanger Institute COVID-19 Surveillance Team |
| EPI_ISL_2971462 | Department of Bacteria, Parasites and Fungi, Statens Serum Institut, Copenhagen, Denmark | Statens Serum Institut Bioinformatics and Microbial Genomics | Danish Covid-19 Genome Consortium |
| EPI_ISL_2971463 | Lighthouse Laboratory Plymouth | Wellcome Sanger Institute for the COVID-19 Genomics UK (COG-UK) | Lighthouse Laboratory Plymouth and Alex Alderton, Roberto Amato, Jeffrey Barrett, Sonia Goncalves, Ewan Harrison, David K. Jackson, Ian Johnston, Dominic Kwiatkowski, Cordelia Langford, John Sillitoe on behalf of the Wellcome Sanger |

| | | | |
|---|---|---|---|
| | | Consortium | Institute COVID-19 Surveillance Team |
| EPI_ISL_2971464 | Lighthouse Lab in Alderley Park | Wellcome Sanger Institute for the COVID-19 Genomics UK (COG-UK) Consortium | Jacquelyn Wynn, Mairead Hyland, The Lighthouse Lab in Alderley Park and Alex Alderton, Roberto Amato, Jeffrey Barrett, Sonia Goncalves, Ewan Harrison, David K. Jackson, Ian Johnston, Dominic Kwiatkowski, Cordelia Langford, John Sillitoe on behalf of the Wellcome Sanger Institute COVID-19 Surveillance Team |
| EPI_ISL_2971465 | Department of Bacteria, Parasites and Fungi, Statens Serum Institut, Copenhagen, Denmark | Statens Serum Institut Bioinformatics and Microbial Genomics | Danish Covid-19 Genome Consortium |
| EPI_ISL_2971466, EPI_ISL_2971467, EPI_ISL_2971468 | Lighthouse Lab in Alderley Park | Wellcome Sanger Institute for the COVID-19 Genomics UK (COG-UK) Consortium | Jacquelyn Wynn, Mairead Hyland, The Lighthouse Lab in Alderley Park and Alex Alderton, Roberto Amato, Jeffrey Barrett, Sonia Goncalves, Ewan Harrison, David K. Jackson, Ian Johnston, Dominic Kwiatkowski, Cordelia Langford, John Sillitoe on behalf of the Wellcome Sanger Institute COVID-19 Surveillance Team |
| EPI_ISL_2971469 | Department of Bacteria, Parasites and Fungi, Statens Serum Institut, Copenhagen, Denmark | Statens Serum Institut Bioinformatics and Microbial Genomics | Danish Covid-19 Genome Consortium |
| EPI_ISL_2971470 | Lighthouse Lab in Alderley Park | Wellcome Sanger Institute for the COVID-19 Genomics UK (COG-UK) Consortium | Jacquelyn Wynn, Mairead Hyland, The Lighthouse Lab in Alderley Park and Alex Alderton, Roberto Amato, Jeffrey Barrett, Sonia Goncalves, Ewan Harrison, David K. Jackson, Ian Johnston, Dominic Kwiatkowski, Cordelia Langford, John Sillitoe on behalf of the Wellcome Sanger Institute COVID-19 Surveillance Team |
| EPI_ISL_2971471 | Lighthouse Laboratory Plymouth | Wellcome Sanger Institute for the COVID-19 Genomics UK (COG-UK) Consortium | Lighthouse Laboratory Plymouth and Alex Alderton, Roberto Amato, Jeffrey Barrett, Sonia Goncalves, Ewan Harrison, David K. Jackson, Ian Johnston, Dominic Kwiatkowski, Cordelia Langford, John Sillitoe on behalf of the Wellcome Sanger Institute COVID-19 Surveillance Team |
| EPI_ISL_2971472 | Department of Bacteria, Parasites and Fungi, Statens Serum Institut, Copenhagen, Denmark | Statens Serum Institut Bioinformatics and Microbial Genomics | Danish Covid-19 Genome Consortium |
| EPI_ISL_2971473 | Lighthouse Lab in Alderley Park | Wellcome Sanger Institute for the COVID-19 Genomics UK (COG-UK) Consortium | Jacquelyn Wynn, Mairead Hyland, The Lighthouse Lab in Alderley Park and Alex Alderton, Roberto Amato, Jeffrey Barrett, Sonia Goncalves, Ewan Harrison, David K. Jackson, Ian Johnston, Dominic Kwiatkowski, Cordelia Langford, John Sillitoe on behalf of the Wellcome Sanger Institute COVID-19 Surveillance Team |
| EPI_ISL_2971474 | Lighthouse Laboratory Plymouth | Wellcome Sanger Institute for the COVID-19 Genomics UK (COG-UK) Consortium | Lighthouse Laboratory Plymouth and Alex Alderton, Roberto Amato, Jeffrey Barrett, Sonia Goncalves, Ewan Harrison, David K. Jackson, Ian Johnston, Dominic Kwiatkowski, Cordelia Langford, John Sillitoe on behalf of the Wellcome Sanger Institute COVID-19 Surveillance Team |
| EPI_ISL_2971475 | Department of Bacteria, Parasites and Fungi, Statens Serum Institut, Copenhagen, Denmark | Statens Serum Institut Bioinformatics and Microbial Genomics | Danish Covid-19 Genome Consortium |
| EPI_ISL_2971476, EPI_ISL_2971477 | Lighthouse Lab in Alderley Park | Wellcome Sanger Institute for the COVID-19 Genomics UK (COG-UK) Consortium | Jacquelyn Wynn, Mairead Hyland, The Lighthouse Lab in Alderley Park and Alex Alderton, Roberto Amato, Jeffrey Barrett, Sonia Goncalves, Ewan Harrison, David K. Jackson, Ian Johnston, Dominic Kwiatkowski, Cordelia Langford, John Sillitoe on behalf of the Wellcome Sanger Institute COVID-19 Surveillance Team |
| EPI_ISL_2971479 | Berkshire and Surrey Pathology Services Lighthouse Laboratory | Wellcome Sanger Institute for the COVID-19 Genomics UK (COG-UK) Consortium | Berkshire and Surrey Pathology Services Lighthouse Laboratory and Alex Alderton, Roberto Amato, Jeffrey Barrett, Sonia Goncalves, Ewan Harrison, David K. Jackson, Ian Johnston, Dominic Kwiatkowski, Cordelia Langford, John Sillitoe on behalf of the Wellcome Sanger Institute COVID-19 Surveillance Team |
| EPI_ISL_2971480 | Lighthouse Lab in Alderley Park | Wellcome Sanger Institute for the COVID-19 Genomics UK (COG-UK) Consortium | Jacquelyn Wynn, Mairead Hyland, The Lighthouse Lab in Alderley Park and Alex Alderton, Roberto Amato, Jeffrey Barrett, Sonia Goncalves, Ewan Harrison, David K. Jackson, Ian Johnston, Dominic Kwiatkowski, Cordelia Langford, John Sillitoe on behalf of the Wellcome Sanger Institute COVID-19 Surveillance Team |
| EPI_ISL_2971481 | Department of Bacteria, Parasites and Fungi, Statens Serum Institut, Copenhagen, Denmark | Statens Serum Institut Bioinformatics and Microbial Genomics | Danish Covid-19 Genome Consortium |
| EPI_ISL_2971482 | Lighthouse Laboratory Plymouth | Wellcome Sanger Institute for the COVID-19 Genomics UK (COG-UK) Consortium | Lighthouse Laboratory Plymouth and Alex Alderton, Roberto Amato, Jeffrey Barrett, Sonia Goncalves, Ewan Harrison, David K. Jackson, Ian Johnston, Dominic Kwiatkowski, Cordelia Langford, John Sillitoe on behalf of the Wellcome Sanger Institute COVID-19 Surveillance Team |
| EPI_ISL_2971483 | Berkshire and Surrey Pathology Services Lighthouse Laboratory | Wellcome Sanger Institute for the COVID-19 Genomics UK (COG-UK) Consortium | Berkshire and Surrey Pathology Services Lighthouse Laboratory and Alex Alderton, Roberto Amato, Jeffrey Barrett, Sonia Goncalves, Ewan Harrison, David K. Jackson, Ian Johnston, Dominic Kwiatkowski, Cordelia Langford, John Sillitoe on behalf of the Wellcome Sanger Institute COVID-19 Surveillance Team |
| EPI_ISL_2971484 | Lighthouse Lab in Alderley Park | Wellcome Sanger Institute for the COVID-19 Genomics UK (COG-UK) Consortium | Jacquelyn Wynn, Mairead Hyland, The Lighthouse Lab in Alderley Park and Alex Alderton, Roberto Amato, Jeffrey Barrett, Sonia Goncalves, Ewan Harrison, David K. Jackson, Ian Johnston, Dominic Kwiatkowski, Cordelia Langford, John Sillitoe on behalf of the Wellcome Sanger Institute COVID-19 Surveillance Team |
| EPI_ISL_2971485 | Department of Bacteria, Parasites and Fungi, Statens Serum Institut, Copenhagen, Denmark | Statens Serum Institut Bioinformatics and Microbial Genomics | Danish Covid-19 Genome Consortium |
| EPI_ISL_2971486, EPI_ISL_2971487 | Lighthouse Laboratory Plymouth | Wellcome Sanger Institute for the COVID-19 Genomics UK (COG-UK) Consortium | Lighthouse Laboratory Plymouth and Alex Alderton, Roberto Amato, Jeffrey Barrett, Sonia Goncalves, Ewan Harrison, David K. Jackson, Ian Johnston, Dominic Kwiatkowski, Cordelia Langford, John Sillitoe on behalf of the Wellcome Sanger Institute COVID-19 Surveillance Team |
| EPI_ISL_2971488 | Department of Bacteria, Parasites and Fungi, Statens Serum Institut, Copenhagen, Denmark | Statens Serum Institut Bioinformatics and Microbial Genomics | Danish Covid-19 Genome Consortium |
| EPI_ISL_2971489, EPI_ISL_2971490 | Lighthouse Lab in Alderley Park | Wellcome Sanger Institute for the COVID-19 Genomics UK (COG-UK) Consortium | Jacquelyn Wynn, Mairead Hyland, The Lighthouse Lab in Alderley Park and Alex Alderton, Roberto Amato, Jeffrey Barrett, Sonia Goncalves, Ewan Harrison, David K. Jackson, Ian Johnston, Dominic Kwiatkowski, Cordelia Langford, John Sillitoe on behalf of the Wellcome Sanger Institute COVID-19 Surveillance Team |
| EPI_ISL_2971491 | Department of Bacteria, Parasites and Fungi, Statens Serum Institut, Copenhagen, Denmark | Statens Serum Institut Bioinformatics and Microbial Genomics | Danish Covid-19 Genome Consortium |
| EPI_ISL_2971492 | Berkshire and Surrey Pathology Services Lighthouse Laboratory | Wellcome Sanger Institute for the COVID-19 Genomics UK (COG-UK) Consortium | Berkshire and Surrey Pathology Services Lighthouse Laboratory and Alex Alderton, Roberto Amato, Jeffrey Barrett, Sonia Goncalves, Ewan Harrison, David K. Jackson, Ian Johnston, Dominic Kwiatkowski, Cordelia Langford, John Sillitoe on behalf of the Wellcome Sanger Institute COVID-19 Surveillance Team |
| EPI_ISL_2971493 | Lighthouse Laboratory Plymouth | Wellcome Sanger Institute for the COVID-19 Genomics UK (COG-UK) Consortium | Lighthouse Laboratory Plymouth and Alex Alderton, Roberto Amato, Jeffrey Barrett, Sonia Goncalves, Ewan Harrison, David K. Jackson, Ian Johnston, Dominic Kwiatkowski, Cordelia Langford, John Sillitoe on behalf of the Wellcome Sanger Institute COVID-19 Surveillance Team |
| EPI_ISL_2971494 | Department of Bacteria, Parasites and Fungi, Statens Serum Institut, Copenhagen, Denmark | Statens Serum Institut Bioinformatics and Microbial Genomics | Danish Covid-19 Genome Consortium |
| EPI_ISL_2971495 | Lighthouse Lab in Alderley Park | Wellcome Sanger Institute for the COVID-19 Genomics UK (COG-UK) Consortium | Jacquelyn Wynn, Mairead Hyland, The Lighthouse Lab in Alderley Park and Alex Alderton, Roberto Amato, Jeffrey Barrett, Sonia Goncalves, Ewan Harrison, David K. Jackson, Ian Johnston, Dominic Kwiatkowski, Cordelia Langford, John Sillitoe on behalf of the Wellcome Sanger Institute COVID-19 Surveillance Team |
| EPI_ISL_2971496 | Lighthouse Laboratory Plymouth | Wellcome Sanger Institute for the COVID-19 Genomics UK (COG-UK) Consortium | Lighthouse Laboratory Plymouth and Alex Alderton, Roberto Amato, Jeffrey Barrett, Sonia Goncalves, Ewan Harrison, David K. Jackson, Ian Johnston, Dominic Kwiatkowski, Cordelia Langford, John Sillitoe on behalf of the Wellcome Sanger Institute COVID-19 Surveillance Team |
| EPI_ISL_2971497 | Berkshire and Surrey Pathology Services Lighthouse Laboratory | Wellcome Sanger Institute for the COVID-19 Genomics UK (COG-UK) Consortium | Berkshire and Surrey Pathology Services Lighthouse Laboratory and Alex Alderton, Roberto Amato, Jeffrey Barrett, Sonia Goncalves, Ewan Harrison, David K. Jackson, Ian Johnston, Dominic Kwiatkowski, Cordelia Langford, John Sillitoe on behalf of the Wellcome Sanger Institute COVID-19 Surveillance Team |
| EPI_ISL_2971498 | Department of Bacteria, Parasites and Fungi, Statens Serum Institut, Copenhagen, Denmark | Statens Serum Institut Bioinformatics and Microbial Genomics | Danish Covid-19 Genome Consortium |
| EPI_ISL_2971499 | Lighthouse Laboratory Plymouth | Wellcome Sanger Institute for the COVID-19 Genomics UK (COG-UK) Consortium | Lighthouse Laboratory Plymouth and Alex Alderton, Roberto Amato, Jeffrey Barrett, Sonia Goncalves, Ewan Harrison, David K. Jackson, Ian Johnston, Dominic Kwiatkowski, Cordelia Langford, John Sillitoe on behalf of the Wellcome Sanger Institute COVID-19 Surveillance Team |
| EPI_ISL_2971500 | Lighthouse Lab in Alderley Park | Wellcome Sanger Institute for the COVID-19 Genomics UK (COG-UK) Consortium | Jacquelyn Wynn, Mairead Hyland, The Lighthouse Lab in Alderley Park and Alex Alderton, Roberto Amato, Jeffrey Barrett, Sonia Goncalves, Ewan Harrison, David K. Jackson, Ian Johnston, Dominic Kwiatkowski, Cordelia Langford, John Sillitoe on behalf of the Wellcome Sanger Institute COVID-19 Surveillance Team |
| EPI_ISL_2971501 | Department of Bacteria, Parasites and Fungi, Statens Serum Institut, Copenhagen, Denmark | Statens Serum Institut Bioinformatics and Microbial Genomics | Danish Covid-19 Genome Consortium |
| EPI_ISL_2971502, EPI_ISL_2971503 | Lighthouse Lab in Alderley Park | Wellcome Sanger Institute for the COVID-19 Genomics UK (COG-UK) Consortium | Jacquelyn Wynn, Mairead Hyland, The Lighthouse Lab in Alderley Park and Alex Alderton, Roberto Amato, Jeffrey Barrett, Sonia Goncalves, Ewan Harrison, David K. Jackson, Ian Johnston, Dominic Kwiatkowski, Cordelia Langford, John Sillitoe on behalf of the Wellcome Sanger Institute COVID-19 Surveillance Team |
| EPI_ISL_2971504 | Department of Bacteria, Parasites and Fungi, Statens Serum Institut, Copenhagen, Denmark | Statens Serum Institut Bioinformatics and Microbial Genomics | Danish Covid-19 Genome Consortium |
| EPI_ISL_2971505 | Lighthouse Laboratory Plymouth | Wellcome Sanger Institute for the COVID-19 Genomics UK (COG-UK) Consortium | Lighthouse Laboratory Plymouth and Alex Alderton, Roberto Amato, Jeffrey Barrett, Sonia Goncalves, Ewan Harrison, David K. Jackson, Ian Johnston, Dominic Kwiatkowski, Cordelia Langford, John Sillitoe on behalf of the Wellcome Sanger Institute COVID-19 Surveillance Team |
| EPI_ISL_2971506 | Lighthouse Lab in Alderley Park | Wellcome Sanger Institute for the COVID-19 Genomics UK (COG-UK) Consortium | Jacquelyn Wynn, Mairead Hyland, The Lighthouse Lab in Alderley Park and Alex Alderton, Roberto Amato, Jeffrey Barrett, Sonia Goncalves, Ewan Harrison, David K. Jackson, Ian Johnston, Dominic Kwiatkowski, Cordelia Langford, John Sillitoe on behalf of the Wellcome Sanger Institute COVID-19 Surveillance Team |
| EPI_ISL_2971507 | Lighthouse Laboratory Plymouth | Wellcome Sanger Institute for the COVID-19 Genomics UK (COG-UK) Consortium | Lighthouse Laboratory Plymouth and Alex Alderton, Roberto Amato, Jeffrey Barrett, Sonia Goncalves, Ewan Harrison, David K. Jackson, Ian Johnston, Dominic Kwiatkowski, Cordelia Langford, John Sillitoe on behalf of the Wellcome Sanger Institute COVID-19 Surveillance Team |
| EPI_ISL_2971508 | Lighthouse Lab in Alderley Park | Wellcome Sanger Institute for the COVID-19 Genomics UK (COG-UK) Consortium | Jacquelyn Wynn, Mairead Hyland, The Lighthouse Lab in Alderley Park and Alex Alderton, Roberto Amato, Jeffrey Barrett, Sonia Goncalves, Ewan Harrison, David K. Jackson, Ian Johnston, Dominic Kwiatkowski, Cordelia Langford, John Sillitoe on behalf of the Wellcome Sanger Institute COVID-19 Surveillance Team |
| EPI_ISL_2971509, EPI_ISL_2971511, EPI_ISL_2971512, EPI_ISL_2971513, EPI_ISL_2971514, EPI_ISL_2971515, EPI_ISL_2971516, EPI_ISL_2971517, EPI_ISL_2971519, EPI_ISL_2971520, EPI_ISL_2971521, EPI_ISL_2971522, EPI_ISL_2971524, EPI_ISL_2971525, EPI_ISL_2971526, EPI_ISL_2971527, EPI_ISL_2971528, EPI_ISL_2971529, EPI_ISL_2971530, EPI_ISL_2971531, EPI_ISL_2971532, EPI_ISL_2971533, EPI_ISL_2971534, EPI_ISL_2971535, EPI_ISL_2971536, EPI_ISL_2971537, EPI_ISL_2971538, EPI_ISL_2971539, EPI_ISL_2971540, EPI_ISL_2971541, EPI_ISL_2971542, EPI_ISL_2971543, EPI_ISL_2971544, EPI_ISL_2971545, EPI_ISL_2971546, EPI_ISL_2971547, EPI_ISL_2971548, EPI_ISL_2971549, EPI_ISL_2971550, EPI_ISL_2971551, EPI_ISL_2971552, EPI_ISL_2971553, EPI_ISL_2971554, EPI_ISL_2971555, EPI_ISL_2971556, EPI_ISL_2971557, EPI_ISL_2971558, EPI_ISL_2971559, EPI_ISL_2971560, EPI_ISL_2971561 | | | |
| see above | Department of Bacteria, Parasites and Fungi, Statens Serum Institut, Copenhagen, Denmark | Statens Serum Institut Bioinformatics and Microbial Genomics | Danish Covid-19 Genome Consortium |
| EPI_ISL_2971562, EPI_ISL_2971563 | Lighthouse Lab in Alderley Park | Wellcome Sanger Institute for the COVID-19 Genomics UK (COG-UK) Consortium | Jacquelyn Wynn, Mairead Hyland, The Lighthouse Lab in Alderley Park and Alex Alderton, Roberto Amato, Jeffrey Barrett, Sonia Goncalves, Ewan Harrison, David K. Jackson, Ian Johnston, Dominic Kwiatkowski, Cordelia Langford, John Sillitoe on behalf of the Wellcome Sanger Institute COVID-19 Surveillance Team |
| EPI_ISL_2971564 | Department of Bacteria, Parasites and Fungi, Statens Serum Institut, Copenhagen, Denmark | Statens Serum Institut Bioinformatics and Microbial Genomics | Danish Covid-19 Genome Consortium |
| EPI_ISL_2971565 | Berkshire and Surrey Pathology Services Lighthouse Laboratory | Wellcome Sanger Institute for the COVID-19 Genomics UK (COG-UK) Consortium | Berkshire and Surrey Pathology Services Lighthouse Laboratory and Alex Alderton, Roberto Amato, Jeffrey Barrett, Sonia Goncalves, Ewan Harrison, David K. Jackson, Ian Johnston, Dominic Kwiatkowski, Cordelia Langford, John Sillitoe on behalf of the Wellcome Sanger Institute COVID-19 Surveillance Team |
| EPI_ISL_2971566 | Department of Bacteria, Parasites and Fungi, Statens Serum Institut, Copenhagen, Denmark | Statens Serum Institut Bioinformatics and Microbial Genomics | Danish Covid-19 Genome Consortium |

| | | | |
|---|---|---|---|
| EPI_ISL_2971567 | Lighthouse Laboratory Plymouth | Wellcome Sanger Institute for the COVID-19 Genomics UK (COG-UK) Consortium | Lighthouse Laboratory Plymouth and Alex Alderton, Roberto Amato, Jeffrey Barrett, Sonia Goncalves, Ewan Harrison, David K. Jackson, Ian Johnston, Dominic Kwiatkowski, Cordelia Langford, John Sillitoe on behalf of the Wellcome Sanger Institute COVID-19 Surveillance Team |
| EPI_ISL_2971568 | Berkshire and Surrey Pathology Services Lighthouse Laboratory | Wellcome Sanger Institute for the COVID-19 Genomics UK (COG-UK) Consortium | Berkshire and Surrey Pathology Services Lighthouse Laboratory and Alex Alderton, Roberto Amato, Jeffrey Barrett, Sonia Goncalves, Ewan Harrison, David K. Jackson, Ian Johnston, Dominic Kwiatkowski, Cordelia Langford, John Sillitoe on behalf of the Wellcome Sanger Institute COVID-19 Surveillance Team |
| EPI_ISL_2971569 | Lighthouse Lab in Alderley Park | Wellcome Sanger Institute for the COVID-19 Genomics UK (COG-UK) Consortium | Jacquelyn Wynn, Mairead Hyland, The Lighthouse Lab in Alderley Park and Alex Alderton, Roberto Amato, Jeffrey Barrett, Sonia Goncalves, Ewan Harrison, David K. Jackson, Ian Johnston, Dominic Kwiatkowski, Cordelia Langford, John Sillitoe on behalf of the Wellcome Sanger Institute COVID-19 Surveillance Team |
| EPI_ISL_2971570 | Department of Bacteria, Parasites and Fungi, Statens Serum Institut, Copenhagen, Denmark | Statens Serum Institut Bioinformatics and Microbial Genomics | Danish Covid-19 Genome Consortium |
| EPI_ISL_2971571 | Berkshire and Surrey Pathology Services Lighthouse Laboratory | Wellcome Sanger Institute for the COVID-19 Genomics UK (COG-UK) Consortium | Berkshire and Surrey Pathology Services Lighthouse Laboratory and Alex Alderton, Roberto Amato, Jeffrey Barrett, Sonia Goncalves, Ewan Harrison, David K. Jackson, Ian Johnston, Dominic Kwiatkowski, Cordelia Langford, John Sillitoe on behalf of the Wellcome Sanger Institute COVID-19 Surveillance Team |
| EPI_ISL_2971572 | Lighthouse Laboratory Plymouth | Wellcome Sanger Institute for the COVID-19 Genomics UK (COG-UK) Consortium | Lighthouse Laboratory Plymouth and Alex Alderton, Roberto Amato, Jeffrey Barrett, Sonia Goncalves, Ewan Harrison, David K. Jackson, Ian Johnston, Dominic Kwiatkowski, Cordelia Langford, John Sillitoe on behalf of the Wellcome Sanger Institute COVID-19 Surveillance Team |
| EPI_ISL_2971573 | Department of Bacteria, Parasites and Fungi, Statens Serum Institut, Copenhagen, Denmark | Statens Serum Institut Bioinformatics and Microbial Genomics | Danish Covid-19 Genome Consortium |
| EPI_ISL_2971574 | Lighthouse Laboratory Plymouth | Wellcome Sanger Institute for the COVID-19 Genomics UK (COG-UK) Consortium | Lighthouse Laboratory Plymouth and Alex Alderton, Roberto Amato, Jeffrey Barrett, Sonia Goncalves, Ewan Harrison, David K. Jackson, Ian Johnston, Dominic Kwiatkowski, Cordelia Langford, John Sillitoe on behalf of the Wellcome Sanger Institute COVID-19 Surveillance Team |
| EPI_ISL_2971575 | Lighthouse Lab in Alderley Park | Wellcome Sanger Institute for the COVID-19 Genomics UK (COG-UK) Consortium | Jacquelyn Wynn, Mairead Hyland, The Lighthouse Lab in Alderley Park and Alex Alderton, Roberto Amato, Jeffrey Barrett, Sonia Goncalves, Ewan Harrison, David K. Jackson, Ian Johnston, Dominic Kwiatkowski, Cordelia Langford, John Sillitoe on behalf of the Wellcome Sanger Institute COVID-19 Surveillance Team |
| EPI_ISL_2971576 | Department of Bacteria, Parasites and Fungi, Statens Serum Institut, Copenhagen, Denmark | Statens Serum Institut Bioinformatics and Microbial Genomics | Danish Covid-19 Genome Consortium |
| EPI_ISL_2971577, EPI_ISL_2971578 | Berkshire and Surrey Pathology Services Lighthouse Laboratory | Wellcome Sanger Institute for the COVID-19 Genomics UK (COG-UK) Consortium | Berkshire and Surrey Pathology Services Lighthouse Laboratory and Alex Alderton, Roberto Amato, Jeffrey Barrett, Sonia Goncalves, Ewan Harrison, David K. Jackson, Ian Johnston, Dominic Kwiatkowski, Cordelia Langford, John Sillitoe on behalf of the Wellcome Sanger Institute COVID-19 Surveillance Team |
| EPI_ISL_2971579 | Department of Bacteria, Parasites and Fungi, Statens Serum Institut, Copenhagen, Denmark | Statens Serum Institut Bioinformatics and Microbial Genomics | Danish Covid-19 Genome Consortium |
| EPI_ISL_2971580 | Lighthouse Laboratory Plymouth | Wellcome Sanger Institute for the COVID-19 Genomics UK (COG-UK) Consortium | Lighthouse Laboratory Plymouth and Alex Alderton, Roberto Amato, Jeffrey Barrett, Sonia Goncalves, Ewan Harrison, David K. Jackson, Ian Johnston, Dominic Kwiatkowski, Cordelia Langford, John Sillitoe on behalf of the Wellcome Sanger Institute COVID-19 Surveillance Team |
| EPI_ISL_2971581 | Department of Bacteria, Parasites and Fungi, Statens Serum Institut, Copenhagen, Denmark | Statens Serum Institut Bioinformatics and Microbial Genomics | Danish Covid-19 Genome Consortium |
| EPI_ISL_2971582, EPI_ISL_2971583, EPI_ISL_2971584 | Lighthouse Lab in Alderley Park | Wellcome Sanger Institute for the COVID-19 Genomics UK (COG-UK) Consortium | Jacquelyn Wynn, Mairead Hyland, The Lighthouse Lab in Alderley Park and Alex Alderton, Roberto Amato, Jeffrey Barrett, Sonia Goncalves, Ewan Harrison, David K. Jackson, Ian Johnston, Dominic Kwiatkowski, Cordelia Langford, John Sillitoe on behalf of the Wellcome Sanger Institute COVID-19 Surveillance Team |
| EPI_ISL_2971585 | Department of Bacteria, Parasites and Fungi, Statens Serum Institut, Copenhagen, Denmark | Statens Serum Institut Bioinformatics and Microbial Genomics | Danish Covid-19 Genome Consortium |
| EPI_ISL_2971586, EPI_ISL_2971587 | Lighthouse Lab in Alderley Park | Wellcome Sanger Institute for the COVID-19 Genomics UK (COG-UK) Consortium | Jacquelyn Wynn, Mairead Hyland, The Lighthouse Lab in Alderley Park and Alex Alderton, Roberto Amato, Jeffrey Barrett, Sonia Goncalves, Ewan Harrison, David K. Jackson, Ian Johnston, Dominic Kwiatkowski, Cordelia Langford, John Sillitoe on behalf of the Wellcome Sanger Institute COVID-19 Surveillance Team |
| EPI_ISL_2971588 | Department of Bacteria, Parasites and Fungi, Statens Serum Institut, Copenhagen, Denmark | Statens Serum Institut Bioinformatics and Microbial Genomics | Danish Covid-19 Genome Consortium |
| EPI_ISL_2971589 | Lighthouse Lab in Alderley Park | Wellcome Sanger Institute for the COVID-19 Genomics UK (COG-UK) Consortium | Jacquelyn Wynn, Mairead Hyland, The Lighthouse Lab in Alderley Park and Alex Alderton, Roberto Amato, Jeffrey Barrett, Sonia Goncalves, Ewan Harrison, David K. Jackson, Ian Johnston, Dominic Kwiatkowski, Cordelia Langford, John Sillitoe on behalf of the Wellcome Sanger Institute COVID-19 Surveillance Team |
| EPI_ISL_2971590 | Lighthouse Laboratory Plymouth | Wellcome Sanger Institute for the COVID-19 Genomics UK (COG-UK) Consortium | Lighthouse Laboratory Plymouth and Alex Alderton, Roberto Amato, Jeffrey Barrett, Sonia Goncalves, Ewan Harrison, David K. Jackson, Ian Johnston, Dominic Kwiatkowski, Cordelia Langford, John Sillitoe on behalf of the Wellcome Sanger Institute COVID-19 Surveillance Team |
| EPI_ISL_2971591 | Department of Bacteria, Parasites and Fungi, Statens Serum Institut, Copenhagen, Denmark | Statens Serum Institut Bioinformatics and Microbial Genomics | Danish Covid-19 Genome Consortium |
| EPI_ISL_2971592 | Lighthouse Lab in Alderley Park | Wellcome Sanger Institute for the COVID-19 Genomics UK (COG-UK) Consortium | Jacquelyn Wynn, Mairead Hyland, The Lighthouse Lab in Alderley Park and Alex Alderton, Roberto Amato, Jeffrey Barrett, Sonia Goncalves, Ewan Harrison, David K. Jackson, Ian Johnston, Dominic Kwiatkowski, Cordelia Langford, John Sillitoe on behalf of the Wellcome Sanger Institute COVID-19 Surveillance Team |
| EPI_ISL_2971593, EPI_ISL_2971594 | Lighthouse Laboratory Plymouth | Wellcome Sanger Institute for the COVID-19 Genomics UK (COG-UK) Consortium | Lighthouse Laboratory Plymouth and Alex Alderton, Roberto Amato, Jeffrey Barrett, Sonia Goncalves, Ewan Harrison, David K. Jackson, Ian Johnston, Dominic Kwiatkowski, Cordelia Langford, John Sillitoe on behalf of the Wellcome Sanger Institute COVID-19 Surveillance Team |
| EPI_ISL_2971595 | Department of Bacteria, Parasites and Fungi, Statens Serum Institut, Copenhagen, Denmark | Statens Serum Institut Bioinformatics and Microbial Genomics | Danish Covid-19 Genome Consortium |
| EPI_ISL_2971596, EPI_ISL_2971597 | Berkshire and Surrey Pathology Services Lighthouse Laboratory | Wellcome Sanger Institute for the COVID-19 Genomics UK (COG-UK) Consortium | Berkshire and Surrey Pathology Services Lighthouse Laboratory and Alex Alderton, Roberto Amato, Jeffrey Barrett, Sonia Goncalves, Ewan Harrison, David K. Jackson, Ian Johnston, Dominic Kwiatkowski, Cordelia Langford, John Sillitoe on behalf of the Wellcome Sanger Institute COVID-19 Surveillance Team |
| EPI_ISL_2971598 | Department of Bacteria, Parasites and Fungi, Statens Serum Institut, Copenhagen, Denmark | Statens Serum Institut Bioinformatics and Microbial Genomics | Danish Covid-19 Genome Consortium |
| EPI_ISL_2971599, EPI_ISL_2971600 | Lighthouse Lab in Alderley Park | Wellcome Sanger Institute for the COVID-19 Genomics UK (COG-UK) Consortium | Jacquelyn Wynn, Mairead Hyland, The Lighthouse Lab in Alderley Park and Alex Alderton, Roberto Amato, Jeffrey Barrett, Sonia Goncalves, Ewan Harrison, David K. Jackson, Ian Johnston, Dominic Kwiatkowski, Cordelia Langford, John Sillitoe on behalf of the Wellcome Sanger Institute COVID-19 Surveillance Team |
| EPI_ISL_2971601 | Department of Bacteria, Parasites and Fungi, Statens Serum Institut, Copenhagen, Denmark | Statens Serum Institut Bioinformatics and Microbial Genomics | Danish Covid-19 Genome Consortium |
| EPI_ISL_2971602 | Berkshire and Surrey Pathology Services Lighthouse Laboratory | Wellcome Sanger Institute for the COVID-19 Genomics UK (COG-UK) Consortium | Berkshire and Surrey Pathology Services Lighthouse Laboratory and Alex Alderton, Roberto Amato, Jeffrey Barrett, Sonia Goncalves, Ewan Harrison, David K. Jackson, Ian Johnston, Dominic Kwiatkowski, Cordelia Langford, John Sillitoe on behalf of the Wellcome Sanger Institute COVID-19 Surveillance Team |
| EPI_ISL_2971603 | Lighthouse Laboratory Plymouth | Wellcome Sanger Institute for the COVID-19 Genomics UK (COG-UK) Consortium | Lighthouse Laboratory Plymouth and Alex Alderton, Roberto Amato, Jeffrey Barrett, Sonia Goncalves, Ewan Harrison, David K. Jackson, Ian Johnston, Dominic Kwiatkowski, Cordelia Langford, John Sillitoe on behalf of the Wellcome Sanger Institute COVID-19 Surveillance Team |
| EPI_ISL_2971604 | Lighthouse Lab in Alderley Park | Wellcome Sanger Institute for the COVID-19 Genomics UK (COG-UK) Consortium | Jacquelyn Wynn, Mairead Hyland, The Lighthouse Lab in Alderley Park and Alex Alderton, Roberto Amato, Jeffrey Barrett, Sonia Goncalves, Ewan Harrison, David K. Jackson, Ian Johnston, Dominic Kwiatkowski, Cordelia Langford, John Sillitoe on behalf of the Wellcome Sanger Institute COVID-19 Surveillance Team |
| EPI_ISL_2971605 | Department of Bacteria, Parasites and Fungi, Statens Serum Institut, Copenhagen, Denmark | Statens Serum Institut Bioinformatics and Microbial Genomics | Danish Covid-19 Genome Consortium |
| EPI_ISL_2971606 | Lighthouse Lab in Alderley Park | Wellcome Sanger Institute for the COVID-19 Genomics UK (COG-UK) Consortium | Jacquelyn Wynn, Mairead Hyland, The Lighthouse Lab in Alderley Park and Alex Alderton, Roberto Amato, Jeffrey Barrett, Sonia Goncalves, Ewan Harrison, David K. Jackson, Ian Johnston, Dominic Kwiatkowski, Cordelia Langford, John Sillitoe on behalf of the Wellcome Sanger Institute COVID-19 Surveillance Team |
| EPI_ISL_2971607 | Lighthouse Lab in Glasgow | Wellcome Sanger Institute for the COVID-19 Genomics UK (COG-UK) Consortium | Harper VanSteenhouse, Yumi Kasai, David Gray, Carol Clugston, Anna Dominiczak and Alex Alderton, Roberto Amato, Jeffrey Barrett, Sonia Goncalves, Ewan Harrison, David K. Jackson, Ian Johnston, Dominic Kwiatkowski, Cordelia Langford, John Sillitoe on behalf of the Wellcome Sanger Institute COVID-19 Surveillance Team |
| EPI_ISL_2971608 | Department of Bacteria, Parasites and Fungi, Statens Serum Institut, Copenhagen, Denmark | Statens Serum Institut Bioinformatics and Microbial Genomics | Danish Covid-19 Genome Consortium |
| EPI_ISL_2971609, EPI_ISL_2971610 | Lighthouse Laboratory Plymouth | Wellcome Sanger Institute for the COVID-19 Genomics UK (COG-UK) Consortium | Lighthouse Laboratory Plymouth and Alex Alderton, Roberto Amato, Jeffrey Barrett, Sonia Goncalves, Ewan Harrison, David K. Jackson, Ian Johnston, Dominic Kwiatkowski, Cordelia Langford, John Sillitoe on behalf of the Wellcome Sanger Institute COVID-19 Surveillance Team |
| EPI_ISL_2971611 | Randox Laboratories | Wellcome Sanger Institute for the COVID-19 Genomics UK (COG-UK) Consortium | Randox Laboratories and Alex Alderton, Roberto Amato, Jeffrey Barrett, Sonia Goncalves, Ewan Harrison, David K. Jackson, Ian Johnston, Dominic Kwiatkowski, Cordelia Langford, John Sillitoe on behalf of the Wellcome Sanger Institute COVID-19 Surveillance Team |
| EPI_ISL_2971612 | Department of Bacteria, Parasites and Fungi, Statens Serum Institut, Copenhagen, Denmark | Statens Serum Institut Bioinformatics and Microbial Genomics | Danish Covid-19 Genome Consortium |
| EPI_ISL_2971613, EPI_ISL_2971614, EPI_ISL_2971616, EPI_ISL_2971617, EPI_ISL_2971618 | Lighthouse Laboratory Plymouth | Wellcome Sanger Institute for the COVID-19 Genomics UK (COG-UK) Consortium | Lighthouse Laboratory Plymouth and Alex Alderton, Roberto Amato, Jeffrey Barrett, Sonia Goncalves, Ewan Harrison, David K. Jackson, Ian Johnston, Dominic Kwiatkowski, Cordelia Langford, John Sillitoe on behalf of the Wellcome Sanger Institute COVID-19 Surveillance Team |
| EPI_ISL_2971619 | Department of Bacteria, Parasites and Fungi, Statens Serum Institut, Copenhagen, Denmark | Statens Serum Institut Bioinformatics and Microbial Genomics | Danish Covid-19 Genome Consortium |
| EPI_ISL_2971620 | Lighthouse Lab in Glasgow | Wellcome Sanger Institute for the COVID-19 Genomics UK (COG-UK) Consortium | Harper VanSteenhouse, Yumi Kasai, David Gray, Carol Clugston, Anna Dominiczak and Alex Alderton, Roberto Amato, Jeffrey Barrett, Sonia Goncalves, Ewan Harrison, David K. Jackson, Ian Johnston, Dominic Kwiatkowski, Cordelia Langford, John Sillitoe on behalf of the Wellcome Sanger Institute COVID-19 Surveillance Team |
| EPI_ISL_2971621 | Lighthouse Laboratory Plymouth | Wellcome Sanger Institute for the COVID-19 Genomics UK (COG-UK) Consortium | Lighthouse Laboratory Plymouth and Alex Alderton, Roberto Amato, Jeffrey Barrett, Sonia Goncalves, Ewan Harrison, David K. Jackson, Ian Johnston, Dominic Kwiatkowski, Cordelia Langford, John Sillitoe on behalf of the Wellcome Sanger Institute COVID-19 Surveillance Team |
| EPI_ISL_2971622 | Randox Laboratories | Wellcome Sanger Institute for the COVID-19 Genomics UK (COG-UK) Consortium | Randox Laboratories and Alex Alderton, Roberto Amato, Jeffrey Barrett, Sonia Goncalves, Ewan Harrison, David K. Jackson, Ian Johnston, Dominic Kwiatkowski, Cordelia Langford, John Sillitoe on behalf of the Wellcome Sanger Institute COVID-19 Surveillance Team |

| | | | |
|---|---|---|---|
| EPI_ISL_2971623 | Department of Bacteria, Parasites and Fungi, Statens Serum Institut, Copenhagen, Denmark | Statens Serum Institut Bioinformatics and Microbial Genomics | Danish Covid-19 Genome Consortium |
| EPI_ISL_2971624 | Lighthouse Laboratory Plymouth | Wellcome Sanger Institute for the COVID-19 Genomics UK (COG-UK) Consortium | Lighthouse Laboratory Plymouth and Alex Alderton, Roberto Amato, Jeffrey Barrett, Sonia Goncalves, Ewan Harrison, David K. Jackson, Ian Johnston, Dominic Kwiatkowski, Cordelia Langford, John Sillitoe on behalf of the Wellcome Sanger Institute COVID-19 Surveillance Team |
| EPI_ISL_2971625 | Lighthouse Lab in Glasgow | Wellcome Sanger Institute for the COVID-19 Genomics UK (COG-UK) Consortium | Harper VanSteenhouse, Yumi Kasai, David Gray, Carol Clugston, Anna Dominiczak and Alex Alderton, Roberto Amato, Jeffrey Barrett, Sonia Goncalves, Ewan Harrison, David K. Jackson, Ian Johnston, Dominic Kwiatkowski, Cordelia Langford, John Sillitoe on behalf of the Wellcome Sanger Institute COVID-19 Surveillance Team |
| EPI_ISL_2971626 | Department of Bacteria, Parasites and Fungi, Statens Serum Institut, Copenhagen, Denmark | Statens Serum Institut Bioinformatics and Microbial Genomics | Danish Covid-19 Genome Consortium |
| EPI_ISL_2971627 | Lighthouse Lab in Glasgow | Wellcome Sanger Institute for the COVID-19 Genomics UK (COG-UK) Consortium | Harper VanSteenhouse, Yumi Kasai, David Gray, Carol Clugston, Anna Dominiczak and Alex Alderton, Roberto Amato, Jeffrey Barrett, Sonia Goncalves, Ewan Harrison, David K. Jackson, Ian Johnston, Dominic Kwiatkowski, Cordelia Langford, John Sillitoe on behalf of the Wellcome Sanger Institute COVID-19 Surveillance Team |
| EPI_ISL_2971628 | Lighthouse Laboratory Plymouth | Wellcome Sanger Institute for the COVID-19 Genomics UK (COG-UK) Consortium | Lighthouse Laboratory Plymouth and Alex Alderton, Roberto Amato, Jeffrey Barrett, Sonia Goncalves, Ewan Harrison, David K. Jackson, Ian Johnston, Dominic Kwiatkowski, Cordelia Langford, John Sillitoe on behalf of the Wellcome Sanger Institute COVID-19 Surveillance Team |
| EPI_ISL_2971629 | Department of Bacteria, Parasites and Fungi, Statens Serum Institut, Copenhagen, Denmark | Statens Serum Institut Bioinformatics and Microbial Genomics | Danish Covid-19 Genome Consortium |
| EPI_ISL_2971630 | Randox Laboratories | Wellcome Sanger Institute for the COVID-19 Genomics UK (COG-UK) Consortium | Randox Laboratories and Alex Alderton, Roberto Amato, Jeffrey Barrett, Sonia Goncalves, Ewan Harrison, David K. Jackson, Ian Johnston, Dominic Kwiatkowski, Cordelia Langford, John Sillitoe on behalf of the Wellcome Sanger Institute COVID-19 Surveillance Team |
| EPI_ISL_2971631 | Lighthouse Laboratory Plymouth | Wellcome Sanger Institute for the COVID-19 Genomics UK (COG-UK) Consortium | Lighthouse Laboratory Plymouth and Alex Alderton, Roberto Amato, Jeffrey Barrett, Sonia Goncalves, Ewan Harrison, David K. Jackson, Ian Johnston, Dominic Kwiatkowski, Cordelia Langford, John Sillitoe on behalf of the Wellcome Sanger Institute COVID-19 Surveillance Team |
| EPI_ISL_2971632 | Department of Bacteria, Parasites and Fungi, Statens Serum Institut, Copenhagen, Denmark | Statens Serum Institut Bioinformatics and Microbial Genomics | Danish Covid-19 Genome Consortium |
| EPI_ISL_2971633 | Lighthouse Laboratory Plymouth | Wellcome Sanger Institute for the COVID-19 Genomics UK (COG-UK) Consortium | Lighthouse Laboratory Plymouth and Alex Alderton, Roberto Amato, Jeffrey Barrett, Sonia Goncalves, Ewan Harrison, David K. Jackson, Ian Johnston, Dominic Kwiatkowski, Cordelia Langford, John Sillitoe on behalf of the Wellcome Sanger Institute COVID-19 Surveillance Team |
| EPI_ISL_2971634 | Lighthouse Lab in Glasgow | Wellcome Sanger Institute for the COVID-19 Genomics UK (COG-UK) Consortium | Harper VanSteenhouse, Yumi Kasai, David Gray, Carol Clugston, Anna Dominiczak and Alex Alderton, Roberto Amato, Jeffrey Barrett, Sonia Goncalves, Ewan Harrison, David K. Jackson, Ian Johnston, Dominic Kwiatkowski, Cordelia Langford, John Sillitoe on behalf of the Wellcome Sanger Institute COVID-19 Surveillance Team |
| EPI_ISL_2971635 | Department of Bacteria, Parasites and Fungi, Statens Serum Institut, Copenhagen, Denmark | Statens Serum Institut Bioinformatics and Microbial Genomics | Danish Covid-19 Genome Consortium |
| EPI_ISL_2971636 | Lighthouse Laboratory Plymouth | Wellcome Sanger Institute for the COVID-19 Genomics UK (COG-UK) Consortium | Lighthouse Laboratory Plymouth and Alex Alderton, Roberto Amato, Jeffrey Barrett, Sonia Goncalves, Ewan Harrison, David K. Jackson, Ian Johnston, Dominic Kwiatkowski, Cordelia Langford, John Sillitoe on behalf of the Wellcome Sanger Institute COVID-19 Surveillance Team |
| EPI_ISL_2971637 | Lighthouse Lab in Glasgow | Wellcome Sanger Institute for the COVID-19 Genomics UK (COG-UK) Consortium | Harper VanSteenhouse, Yumi Kasai, David Gray, Carol Clugston, Anna Dominiczak and Alex Alderton, Roberto Amato, Jeffrey Barrett, Sonia Goncalves, Ewan Harrison, David K. Jackson, Ian Johnston, Dominic Kwiatkowski, Cordelia Langford, John Sillitoe on behalf of the Wellcome Sanger Institute COVID-19 Surveillance Team |
| EPI_ISL_2971638 | Department of Bacteria, Parasites and Fungi, Statens Serum Institut, Copenhagen, Denmark | Statens Serum Institut Bioinformatics and Microbial Genomics | Danish Covid-19 Genome Consortium |
| EPI_ISL_2971639, EPI_ISL_2971640 | Lighthouse Laboratory Plymouth | Wellcome Sanger Institute for the COVID-19 Genomics UK (COG-UK) Consortium | Lighthouse Laboratory Plymouth and Alex Alderton, Roberto Amato, Jeffrey Barrett, Sonia Goncalves, Ewan Harrison, David K. Jackson, Ian Johnston, Dominic Kwiatkowski, Cordelia Langford, John Sillitoe on behalf of the Wellcome Sanger Institute COVID-19 Surveillance Team |
| EPI_ISL_2971641 | Lighthouse Lab in Glasgow | Wellcome Sanger Institute for the COVID-19 Genomics UK (COG-UK) Consortium | Harper VanSteenhouse, Yumi Kasai, David Gray, Carol Clugston, Anna Dominiczak and Alex Alderton, Roberto Amato, Jeffrey Barrett, Sonia Goncalves, Ewan Harrison, David K. Jackson, Ian Johnston, Dominic Kwiatkowski, Cordelia Langford, John Sillitoe on behalf of the Wellcome Sanger Institute COVID-19 Surveillance Team |
| EPI_ISL_2971642 | Department of Bacteria, Parasites and Fungi, Statens Serum Institut, Copenhagen, Denmark | Statens Serum Institut Bioinformatics and Microbial Genomics | Danish Covid-19 Genome Consortium |
| EPI_ISL_2971643 | Lighthouse Laboratory Plymouth | Wellcome Sanger Institute for the COVID-19 Genomics UK (COG-UK) Consortium | Lighthouse Laboratory Plymouth and Alex Alderton, Roberto Amato, Jeffrey Barrett, Sonia Goncalves, Ewan Harrison, David K. Jackson, Ian Johnston, Dominic Kwiatkowski, Cordelia Langford, John Sillitoe on behalf of the Wellcome Sanger Institute COVID-19 Surveillance Team |
| EPI_ISL_2971644 | Randox Laboratories | Wellcome Sanger Institute for the COVID-19 Genomics UK (COG-UK) Consortium | Randox Laboratories and Alex Alderton, Roberto Amato, Jeffrey Barrett, Sonia Goncalves, Ewan Harrison, David K. Jackson, Ian Johnston, Dominic Kwiatkowski, Cordelia Langford, John Sillitoe on behalf of the Wellcome Sanger Institute COVID-19 Surveillance Team |
| EPI_ISL_2971645 | Department of Bacteria, Parasites and Fungi, Statens Serum Institut, Copenhagen, Denmark | Statens Serum Institut Bioinformatics and Microbial Genomics | Danish Covid-19 Genome Consortium |
| EPI_ISL_2971646, EPI_ISL_2971647, EPI_ISL_2971648, EPI_ISL_2971650, EPI_ISL_2971651 | Lighthouse Laboratory Plymouth | Wellcome Sanger Institute for the COVID-19 Genomics UK (COG-UK) Consortium | Lighthouse Laboratory Plymouth and Alex Alderton, Roberto Amato, Jeffrey Barrett, Sonia Goncalves, Ewan Harrison, David K. Jackson, Ian Johnston, Dominic Kwiatkowski, Cordelia Langford, John Sillitoe on behalf of the Wellcome Sanger Institute COVID-19 Surveillance Team |
| EPI_ISL_2971652 | Department of Bacteria, Parasites and Fungi, Statens Serum Institut, Copenhagen, Denmark | Statens Serum Institut Bioinformatics and Microbial Genomics | Danish Covid-19 Genome Consortium |
| EPI_ISL_2971653, EPI_ISL_2971654 | Lighthouse Laboratory Plymouth | Wellcome Sanger Institute for the COVID-19 Genomics UK (COG-UK) Consortium | Lighthouse Laboratory Plymouth and Alex Alderton, Roberto Amato, Jeffrey Barrett, Sonia Goncalves, Ewan Harrison, David K. Jackson, Ian Johnston, Dominic Kwiatkowski, Cordelia Langford, John Sillitoe on behalf of the Wellcome Sanger Institute COVID-19 Surveillance Team |
| EPI_ISL_2971655 | Department of Bacteria, Parasites and Fungi, Statens Serum Institut, Copenhagen, Denmark | Statens Serum Institut Bioinformatics and Microbial Genomics | Danish Covid-19 Genome Consortium |
| EPI_ISL_2971656 | Lighthouse Laboratory Plymouth | Wellcome Sanger Institute for the COVID-19 Genomics UK (COG-UK) Consortium | Lighthouse Laboratory Plymouth and Alex Alderton, Roberto Amato, Jeffrey Barrett, Sonia Goncalves, Ewan Harrison, David K. Jackson, Ian Johnston, Dominic Kwiatkowski, Cordelia Langford, John Sillitoe on behalf of the Wellcome Sanger Institute COVID-19 Surveillance Team |
| EPI_ISL_2971657 | Department of Bacteria, Parasites and Fungi, Statens Serum Institut, Copenhagen, Denmark | Statens Serum Institut Bioinformatics and Microbial Genomics | Danish Covid-19 Genome Consortium |
| EPI_ISL_2971658, EPI_ISL_2971659, EPI_ISL_2971660 | Lighthouse Laboratory Plymouth | Wellcome Sanger Institute for the COVID-19 Genomics UK (COG-UK) Consortium | Lighthouse Laboratory Plymouth and Alex Alderton, Roberto Amato, Jeffrey Barrett, Sonia Goncalves, Ewan Harrison, David K. Jackson, Ian Johnston, Dominic Kwiatkowski, Cordelia Langford, John Sillitoe on behalf of the Wellcome Sanger Institute COVID-19 Surveillance Team |
| EPI_ISL_2971661 | Department of Bacteria, Parasites and Fungi, Statens Serum Institut, Copenhagen, Denmark | Statens Serum Institut Bioinformatics and Microbial Genomics | Danish Covid-19 Genome Consortium |
| EPI_ISL_2971662 | Lighthouse Laboratory Plymouth | Wellcome Sanger Institute for the COVID-19 Genomics UK (COG-UK) Consortium | Lighthouse Laboratory Plymouth and Alex Alderton, Roberto Amato, Jeffrey Barrett, Sonia Goncalves, Ewan Harrison, David K. Jackson, Ian Johnston, Dominic Kwiatkowski, Cordelia Langford, John Sillitoe on behalf of the Wellcome Sanger Institute COVID-19 Surveillance Team |
| EPI_ISL_2971663 | Lighthouse Lab in Glasgow | Wellcome Sanger Institute for the COVID-19 Genomics UK (COG-UK) Consortium | Harper VanSteenhouse, Yumi Kasai, David Gray, Carol Clugston, Anna Dominiczak and Alex Alderton, Roberto Amato, Jeffrey Barrett, Sonia Goncalves, Ewan Harrison, David K. Jackson, Ian Johnston, Dominic Kwiatkowski, Cordelia Langford, John Sillitoe on behalf of the Wellcome Sanger Institute COVID-19 Surveillance Team |
| EPI_ISL_2971664 | Department of Bacteria, Parasites and Fungi, Statens Serum Institut, Copenhagen, Denmark | Statens Serum Institut Bioinformatics and Microbial Genomics | Danish Covid-19 Genome Consortium |
| EPI_ISL_2971665, EPI_ISL_2971666 | Lighthouse Laboratory Plymouth | Wellcome Sanger Institute for the COVID-19 Genomics UK (COG-UK) Consortium | Lighthouse Laboratory Plymouth and Alex Alderton, Roberto Amato, Jeffrey Barrett, Sonia Goncalves, Ewan Harrison, David K. Jackson, Ian Johnston, Dominic Kwiatkowski, Cordelia Langford, John Sillitoe on behalf of the Wellcome Sanger Institute COVID-19 Surveillance Team |
| EPI_ISL_2971667 | Department of Bacteria, Parasites and Fungi, Statens Serum Institut, Copenhagen, Denmark | Statens Serum Institut Bioinformatics and Microbial Genomics | Danish Covid-19 Genome Consortium |
| EPI_ISL_2971668 | Randox Laboratories | Wellcome Sanger Institute for the COVID-19 Genomics UK (COG-UK) Consortium | Randox Laboratories and Alex Alderton, Roberto Amato, Jeffrey Barrett, Sonia Goncalves, Ewan Harrison, David K. Jackson, Ian Johnston, Dominic Kwiatkowski, Cordelia Langford, John Sillitoe on behalf of the Wellcome Sanger Institute COVID-19 Surveillance Team |
| EPI_ISL_2971669 | Lighthouse Laboratory Plymouth | Wellcome Sanger Institute for the COVID-19 Genomics UK (COG-UK) Consortium | Lighthouse Laboratory Plymouth and Alex Alderton, Roberto Amato, Jeffrey Barrett, Sonia Goncalves, Ewan Harrison, David K. Jackson, Ian Johnston, Dominic Kwiatkowski, Cordelia Langford, John Sillitoe on behalf of the Wellcome Sanger Institute COVID-19 Surveillance Team |
| EPI_ISL_2971670 | Department of Bacteria, Parasites and Fungi, Statens Serum Institut, Copenhagen, Denmark | Statens Serum Institut Bioinformatics and Microbial Genomics | Danish Covid-19 Genome Consortium |
| EPI_ISL_2971671, EPI_ISL_2971672, EPI_ISL_2971673 | Lighthouse Laboratory Plymouth | Wellcome Sanger Institute for the COVID-19 Genomics UK (COG-UK) Consortium | Lighthouse Laboratory Plymouth and Alex Alderton, Roberto Amato, Jeffrey Barrett, Sonia Goncalves, Ewan Harrison, David K. Jackson, Ian Johnston, Dominic Kwiatkowski, Cordelia Langford, John Sillitoe on behalf of the Wellcome Sanger Institute COVID-19 Surveillance Team |
| EPI_ISL_2971674 | Department of Bacteria, Parasites and Fungi, Statens Serum Institut, Copenhagen, Denmark | Statens Serum Institut Bioinformatics and Microbial Genomics | Danish Covid-19 Genome Consortium |
| EPI_ISL_2971675 | Lighthouse Laboratory Plymouth | Wellcome Sanger Institute for the COVID-19 Genomics UK (COG-UK) Consortium | Lighthouse Laboratory Plymouth and Alex Alderton, Roberto Amato, Jeffrey Barrett, Sonia Goncalves, Ewan Harrison, David K. Jackson, Ian Johnston, Dominic Kwiatkowski, Cordelia Langford, John Sillitoe on behalf of the Wellcome Sanger Institute COVID-19 Surveillance Team |
| EPI_ISL_2971676 | Department of Bacteria, Parasites and Fungi, Statens Serum Institut, Copenhagen, Denmark | Statens Serum Institut Bioinformatics and Microbial Genomics | Danish Covid-19 Genome Consortium |
| EPI_ISL_2971677 | Lighthouse Lab in Glasgow | Wellcome Sanger Institute for the COVID-19 Genomics UK (COG-UK) Consortium | Harper VanSteenhouse, Yumi Kasai, David Gray, Carol Clugston, Anna Dominiczak and Alex Alderton, Roberto Amato, Jeffrey Barrett, Sonia Goncalves, Ewan Harrison, David K. Jackson, Ian Johnston, Dominic Kwiatkowski, Cordelia Langford, John Sillitoe on behalf of the Wellcome Sanger Institute COVID-19 Surveillance Team |

| | | | |
|---|---|---|---|
| EPI_ISL_2971678 | Lighthouse Laboratory Plymouth | Wellcome Sanger Institute for the COVID-19 Genomics UK (COG-UK) Consortium | Lighthouse Laboratory Plymouth and Alex Alderton, Roberto Amato, Jeffrey Barrett, Sonia Goncalves, Ewan Harrison, David K. Jackson, Ian Johnston, Dominic Kwiatkowski, Cordelia Langford, John Sillitoe on behalf of the Wellcome Sanger Institute COVID-19 Surveillance Team |
| EPI_ISL_2971679 | Department of Bacteria, Parasites and Fungi, Statens Serum Institut, Copenhagen, Denmark | Statens Serum Institut Bioinformatics and Microbial Genomics | Danish Covid-19 Genome Consortium |
| EPI_ISL_2971680, EPI_ISL_2971681 | Lighthouse Laboratory Plymouth | Wellcome Sanger Institute for the COVID-19 Genomics UK (COG-UK) Consortium | Lighthouse Laboratory Plymouth and Alex Alderton, Roberto Amato, Jeffrey Barrett, Sonia Goncalves, Ewan Harrison, David K. Jackson, Ian Johnston, Dominic Kwiatkowski, Cordelia Langford, John Sillitoe on behalf of the Wellcome Sanger Institute COVID-19 Surveillance Team |
| EPI_ISL_2971682 | Department of Bacteria, Parasites and Fungi, Statens Serum Institut, Copenhagen, Denmark | Statens Serum Institut Bioinformatics and Microbial Genomics | Danish Covid-19 Genome Consortium |
| EPI_ISL_2971683, EPI_ISL_2971684 | Lighthouse Laboratory Plymouth | Wellcome Sanger Institute for the COVID-19 Genomics UK (COG-UK) Consortium | Lighthouse Laboratory Plymouth and Alex Alderton, Roberto Amato, Jeffrey Barrett, Sonia Goncalves, Ewan Harrison, David K. Jackson, Ian Johnston, Dominic Kwiatkowski, Cordelia Langford, John Sillitoe on behalf of the Wellcome Sanger Institute COVID-19 Surveillance Team |
| EPI_ISL_2971685 | Department of Bacteria, Parasites and Fungi, Statens Serum Institut, Copenhagen, Denmark | Statens Serum Institut Bioinformatics and Microbial Genomics | Danish Covid-19 Genome Consortium |
| EPI_ISL_2971686, EPI_ISL_2971687 | Randox Laboratories | Wellcome Sanger Institute for the COVID-19 Genomics UK (COG-UK) Consortium | Randox Laboratories and Alex Alderton, Roberto Amato, Jeffrey Barrett, Sonia Goncalves, Ewan Harrison, David K. Jackson, Ian Johnston, Dominic Kwiatkowski, Cordelia Langford, John Sillitoe on behalf of the Wellcome Sanger Institute COVID-19 Surveillance Team |
| EPI_ISL_2971688 | Lighthouse Laboratory Plymouth | Wellcome Sanger Institute for the COVID-19 Genomics UK (COG-UK) Consortium | Lighthouse Laboratory Plymouth and Alex Alderton, Roberto Amato, Jeffrey Barrett, Sonia Goncalves, Ewan Harrison, David K. Jackson, Ian Johnston, Dominic Kwiatkowski, Cordelia Langford, John Sillitoe on behalf of the Wellcome Sanger Institute COVID-19 Surveillance Team |
| EPI_ISL_2971689 | Department of Bacteria, Parasites and Fungi, Statens Serum Institut, Copenhagen, Denmark | Statens Serum Institut Bioinformatics and Microbial Genomics | Danish Covid-19 Genome Consortium |
| EPI_ISL_2971690 | Lighthouse Lab in Glasgow | Wellcome Sanger Institute for the COVID-19 Genomics UK (COG-UK) Consortium | Harper VanSteenhouse, Yumi Kasai, David Gray, Carol Clugston, Anna Dominiczak and Alex Alderton, Roberto Amato, Jeffrey Barrett, Sonia Goncalves, Ewan Harrison, David K. Jackson, Ian Johnston, Dominic Kwiatkowski, Cordelia Langford, John Sillitoe on behalf of the Wellcome Sanger Institute COVID-19 Surveillance Team |
| EPI_ISL_2971691 | Lighthouse Laboratory Plymouth | Wellcome Sanger Institute for the COVID-19 Genomics UK (COG-UK) Consortium | Lighthouse Laboratory Plymouth and Alex Alderton, Roberto Amato, Jeffrey Barrett, Sonia Goncalves, Ewan Harrison, David K. Jackson, Ian Johnston, Dominic Kwiatkowski, Cordelia Langford, John Sillitoe on behalf of the Wellcome Sanger Institute COVID-19 Surveillance Team |
| EPI_ISL_2971692 | Department of Bacteria, Parasites and Fungi, Statens Serum Institut, Copenhagen, Denmark | Statens Serum Institut Bioinformatics and Microbial Genomics | Danish Covid-19 Genome Consortium |
| EPI_ISL_2971693, EPI_ISL_2971694 | Lighthouse Laboratory Plymouth | Wellcome Sanger Institute for the COVID-19 Genomics UK (COG-UK) Consortium | Lighthouse Laboratory Plymouth and Alex Alderton, Roberto Amato, Jeffrey Barrett, Sonia Goncalves, Ewan Harrison, David K. Jackson, Ian Johnston, Dominic Kwiatkowski, Cordelia Langford, John Sillitoe on behalf of the Wellcome Sanger Institute COVID-19 Surveillance Team |
| EPI_ISL_2971695 | Department of Bacteria, Parasites and Fungi, Statens Serum Institut, Copenhagen, Denmark | Statens Serum Institut Bioinformatics and Microbial Genomics | Danish Covid-19 Genome Consortium |
| EPI_ISL_2971696, EPI_ISL_2971697 | Lighthouse Laboratory Plymouth | Wellcome Sanger Institute for the COVID-19 Genomics UK (COG-UK) Consortium | Lighthouse Laboratory Plymouth and Alex Alderton, Roberto Amato, Jeffrey Barrett, Sonia Goncalves, Ewan Harrison, David K. Jackson, Ian Johnston, Dominic Kwiatkowski, Cordelia Langford, John Sillitoe on behalf of the Wellcome Sanger Institute COVID-19 Surveillance Team |
| EPI_ISL_2971698 | Department of Bacteria, Parasites and Fungi, Statens Serum Institut, Copenhagen, Denmark | Statens Serum Institut Bioinformatics and Microbial Genomics | Danish Covid-19 Genome Consortium |
| EPI_ISL_2971699, EPI_ISL_2971700 | Lighthouse Laboratory Plymouth | Wellcome Sanger Institute for the COVID-19 Genomics UK (COG-UK) Consortium | Lighthouse Laboratory Plymouth and Alex Alderton, Roberto Amato, Jeffrey Barrett, Sonia Goncalves, Ewan Harrison, David K. Jackson, Ian Johnston, Dominic Kwiatkowski, Cordelia Langford, John Sillitoe on behalf of the Wellcome Sanger Institute COVID-19 Surveillance Team |
| EPI_ISL_2971701 | Department of Bacteria, Parasites and Fungi, Statens Serum Institut, Copenhagen, Denmark | Statens Serum Institut Bioinformatics and Microbial Genomics | Danish Covid-19 Genome Consortium |
| EPI_ISL_2971702, EPI_ISL_2971703 | Lighthouse Laboratory Plymouth | Wellcome Sanger Institute for the COVID-19 Genomics UK (COG-UK) Consortium | Lighthouse Laboratory Plymouth and Alex Alderton, Roberto Amato, Jeffrey Barrett, Sonia Goncalves, Ewan Harrison, David K. Jackson, Ian Johnston, Dominic Kwiatkowski, Cordelia Langford, John Sillitoe on behalf of the Wellcome Sanger Institute COVID-19 Surveillance Team |
| EPI_ISL_2971704 | Department of Bacteria, Parasites and Fungi, Statens Serum Institut, Copenhagen, Denmark | Statens Serum Institut Bioinformatics and Microbial Genomics | Danish Covid-19 Genome Consortium |
| EPI_ISL_2971705 | Randox Laboratories | Wellcome Sanger Institute for the COVID-19 Genomics UK (COG-UK) Consortium | Randox Laboratories and Alex Alderton, Roberto Amato, Jeffrey Barrett, Sonia Goncalves, Ewan Harrison, David K. Jackson, Ian Johnston, Dominic Kwiatkowski, Cordelia Langford, John Sillitoe on behalf of the Wellcome Sanger Institute COVID-19 Surveillance Team |
| EPI_ISL_2971706 | Lighthouse Laboratory Plymouth | Wellcome Sanger Institute for the COVID-19 Genomics UK (COG-UK) Consortium | Lighthouse Laboratory Plymouth and Alex Alderton, Roberto Amato, Jeffrey Barrett, Sonia Goncalves, Ewan Harrison, David K. Jackson, Ian Johnston, Dominic Kwiatkowski, Cordelia Langford, John Sillitoe on behalf of the Wellcome Sanger Institute COVID-19 Surveillance Team |
| EPI_ISL_2971707 | Department of Bacteria, Parasites and Fungi, Statens Serum Institut, Copenhagen, Denmark | Statens Serum Institut Bioinformatics and Microbial Genomics | Danish Covid-19 Genome Consortium |
| EPI_ISL_2971708 | Lighthouse Laboratory Plymouth | Wellcome Sanger Institute for the COVID-19 Genomics UK (COG-UK) Consortium | Lighthouse Laboratory Plymouth and Alex Alderton, Roberto Amato, Jeffrey Barrett, Sonia Goncalves, Ewan Harrison, David K. Jackson, Ian Johnston, Dominic Kwiatkowski, Cordelia Langford, John Sillitoe on behalf of the Wellcome Sanger Institute COVID-19 Surveillance Team |
| EPI_ISL_2971709 | Lighthouse Lab in Glasgow | Wellcome Sanger Institute for the COVID-19 Genomics UK (COG-UK) Consortium | Harper VanSteenhouse, Yumi Kasai, David Gray, Carol Clugston, Anna Dominiczak and Alex Alderton, Roberto Amato, Jeffrey Barrett, Sonia Goncalves, Ewan Harrison, David K. Jackson, Ian Johnston, Dominic Kwiatkowski, Cordelia Langford, John Sillitoe on behalf of the Wellcome Sanger Institute COVID-19 Surveillance Team |
| EPI_ISL_2971710 | Lighthouse Laboratory Plymouth | Wellcome Sanger Institute for the COVID-19 Genomics UK (COG-UK) Consortium | Lighthouse Laboratory Plymouth and Alex Alderton, Roberto Amato, Jeffrey Barrett, Sonia Goncalves, Ewan Harrison, David K. Jackson, Ian Johnston, Dominic Kwiatkowski, Cordelia Langford, John Sillitoe on behalf of the Wellcome Sanger Institute COVID-19 Surveillance Team |
| EPI_ISL_2971711 | Department of Bacteria, Parasites and Fungi, Statens Serum Institut, Copenhagen, Denmark | Statens Serum Institut Bioinformatics and Microbial Genomics | Danish Covid-19 Genome Consortium |
| EPI_ISL_2971712, EPI_ISL_2971713 | Lighthouse Laboratory Plymouth | Wellcome Sanger Institute for the COVID-19 Genomics UK (COG-UK) Consortium | Lighthouse Laboratory Plymouth and Alex Alderton, Roberto Amato, Jeffrey Barrett, Sonia Goncalves, Ewan Harrison, David K. Jackson, Ian Johnston, Dominic Kwiatkowski, Cordelia Langford, John Sillitoe on behalf of the Wellcome Sanger Institute COVID-19 Surveillance Team |
| EPI_ISL_2971714 | Department of Bacteria, Parasites and Fungi, Statens Serum Institut, Copenhagen, Denmark | Statens Serum Institut Bioinformatics and Microbial Genomics | Danish Covid-19 Genome Consortium |
| EPI_ISL_2971715 | Randox Laboratories | Wellcome Sanger Institute for the COVID-19 Genomics UK (COG-UK) Consortium | Randox Laboratories and Alex Alderton, Roberto Amato, Jeffrey Barrett, Sonia Goncalves, Ewan Harrison, David K. Jackson, Ian Johnston, Dominic Kwiatkowski, Cordelia Langford, John Sillitoe on behalf of the Wellcome Sanger Institute COVID-19 Surveillance Team |
| EPI_ISL_2971716 | Lighthouse Laboratory Plymouth | Wellcome Sanger Institute for the COVID-19 Genomics UK (COG-UK) Consortium | Lighthouse Laboratory Plymouth and Alex Alderton, Roberto Amato, Jeffrey Barrett, Sonia Goncalves, Ewan Harrison, David K. Jackson, Ian Johnston, Dominic Kwiatkowski, Cordelia Langford, John Sillitoe on behalf of the Wellcome Sanger Institute COVID-19 Surveillance Team |
| EPI_ISL_2971717 | Department of Bacteria, Parasites and Fungi, Statens Serum Institut, Copenhagen, Denmark | Statens Serum Institut Bioinformatics and Microbial Genomics | Danish Covid-19 Genome Consortium |
| EPI_ISL_2971718, EPI_ISL_2971719 | Lighthouse Laboratory Plymouth | Wellcome Sanger Institute for the COVID-19 Genomics UK (COG-UK) Consortium | Lighthouse Laboratory Plymouth and Alex Alderton, Roberto Amato, Jeffrey Barrett, Sonia Goncalves, Ewan Harrison, David K. Jackson, Ian Johnston, Dominic Kwiatkowski, Cordelia Langford, John Sillitoe on behalf of the Wellcome Sanger Institute COVID-19 Surveillance Team |
| EPI_ISL_2971720 | Department of Bacteria, Parasites and Fungi, Statens Serum Institut, Copenhagen, Denmark | Statens Serum Institut Bioinformatics and Microbial Genomics | Danish Covid-19 Genome Consortium |
| EPI_ISL_2971721 | Lighthouse Lab in Glasgow | Wellcome Sanger Institute for the COVID-19 Genomics UK (COG-UK) Consortium | Harper VanSteenhouse, Yumi Kasai, David Gray, Carol Clugston, Anna Dominiczak and Alex Alderton, Roberto Amato, Jeffrey Barrett, Sonia Goncalves, Ewan Harrison, David K. Jackson, Ian Johnston, Dominic Kwiatkowski, Cordelia Langford, John Sillitoe on behalf of the Wellcome Sanger Institute COVID-19 Surveillance Team |
| EPI_ISL_2971722, EPI_ISL_2971723 | Lighthouse Laboratory Plymouth | Wellcome Sanger Institute for the COVID-19 Genomics UK (COG-UK) Consortium | Lighthouse Laboratory Plymouth and Alex Alderton, Roberto Amato, Jeffrey Barrett, Sonia Goncalves, Ewan Harrison, David K. Jackson, Ian Johnston, Dominic Kwiatkowski, Cordelia Langford, John Sillitoe on behalf of the Wellcome Sanger Institute COVID-19 Surveillance Team |
| EPI_ISL_2971724 | Department of Bacteria, Parasites and Fungi, Statens Serum Institut, Copenhagen, Denmark | Statens Serum Institut Bioinformatics and Microbial Genomics | Danish Covid-19 Genome Consortium |
| EPI_ISL_2971725 | Lighthouse Laboratory Plymouth | Wellcome Sanger Institute for the COVID-19 Genomics UK (COG-UK) Consortium | Lighthouse Laboratory Plymouth and Alex Alderton, Roberto Amato, Jeffrey Barrett, Sonia Goncalves, Ewan Harrison, David K. Jackson, Ian Johnston, Dominic Kwiatkowski, Cordelia Langford, John Sillitoe on behalf of the Wellcome Sanger Institute COVID-19 Surveillance Team |
| EPI_ISL_2971726 | Department of Bacteria, Parasites and Fungi, Statens Serum Institut, Copenhagen, Denmark | Statens Serum Institut Bioinformatics and Microbial Genomics | Danish Covid-19 Genome Consortium |
| EPI_ISL_2971727 | Randox Laboratories | Wellcome Sanger Institute for the COVID-19 Genomics UK (COG-UK) Consortium | Randox Laboratories and Alex Alderton, Roberto Amato, Jeffrey Barrett, Sonia Goncalves, Ewan Harrison, David K. Jackson, Ian Johnston, Dominic Kwiatkowski, Cordelia Langford, John Sillitoe on behalf of the Wellcome Sanger Institute COVID-19 Surveillance Team |
| EPI_ISL_2971728, EPI_ISL_2971729 | Lighthouse Laboratory Plymouth | Wellcome Sanger Institute for the COVID-19 Genomics UK (COG-UK) Consortium | Lighthouse Laboratory Plymouth and Alex Alderton, Roberto Amato, Jeffrey Barrett, Sonia Goncalves, Ewan Harrison, David K. Jackson, Ian Johnston, Dominic Kwiatkowski, Cordelia Langford, John Sillitoe on behalf of the Wellcome Sanger Institute COVID-19 Surveillance Team |
| EPI_ISL_2971730 | Department of Bacteria, Parasites and Fungi, Statens Serum Institut, Copenhagen, Denmark | Statens Serum Institut Bioinformatics and Microbial Genomics | Danish Covid-19 Genome Consortium |
| EPI_ISL_2971731, EPI_ISL_2971732 | Lighthouse Laboratory Plymouth | Wellcome Sanger Institute for the COVID-19 Genomics UK (COG-UK) Consortium | Lighthouse Laboratory Plymouth and Alex Alderton, Roberto Amato, Jeffrey Barrett, Sonia Goncalves, Ewan Harrison, David K. Jackson, Ian Johnston, Dominic Kwiatkowski, Cordelia Langford, John Sillitoe on behalf of the Wellcome Sanger Institute COVID-19 Surveillance Team |
| EPI_ISL_2971733 | Department of Bacteria, Parasites and Fungi, Statens Serum Institut, Copenhagen, Denmark | Statens Serum Institut Bioinformatics and Microbial Genomics | Danish Covid-19 Genome Consortium |
| EPI_ISL_2971734 | Lighthouse Lab in Glasgow | Wellcome Sanger Institute for the COVID-19 Genomics UK (COG-UK) Consortium | Harper VanSteenhouse, Yumi Kasai, David Gray, Carol Clugston, Anna Dominiczak and Alex Alderton, Roberto Amato, Jeffrey Barrett, Sonia Goncalves, Ewan Harrison, David K. Jackson, Ian Johnston, Dominic Kwiatkowski, Cordelia Langford, John Sillitoe on behalf of the Wellcome Sanger Institute COVID-19 Surveillance Team |

| | | | |
|---|---|---|---|
| EPI_ISL_2971735, EPI_ISL_2971736 | Lighthouse Laboratory Plymouth | Wellcome Sanger Institute for the COVID-19 Genomics UK (COG-UK) Consortium | Lighthouse Laboratory Plymouth and Alex Alderton, Roberto Amato, Jeffrey Barrett, Sonia Goncalves, Ewan Harrison, David K. Jackson, Ian Johnston, Dominic Kwiatkowski, Cordelia Langford, John Sillitoe on behalf of the Wellcome Sanger Institute COVID-19 Surveillance Team |
| EPI_ISL_2971737 | Department of Bacteria, Parasites and Fungi, Statens Serum Institut, Copenhagen, Denmark | Statens Serum Institut Bioinformatics and Microbial Genomics | Danish Covid-19 Genome Consortium |
| EPI_ISL_2971738, EPI_ISL_2971739 | Lighthouse Laboratory Plymouth | Wellcome Sanger Institute for the COVID-19 Genomics UK (COG-UK) Consortium | Lighthouse Laboratory Plymouth and Alex Alderton, Roberto Amato, Jeffrey Barrett, Sonia Goncalves, Ewan Harrison, David K. Jackson, Ian Johnston, Dominic Kwiatkowski, Cordelia Langford, John Sillitoe on behalf of the Wellcome Sanger Institute COVID-19 Surveillance Team |
| EPI_ISL_2971740 | Department of Bacteria, Parasites and Fungi, Statens Serum Institut, Copenhagen, Denmark | Statens Serum Institut Bioinformatics and Microbial Genomics | Danish Covid-19 Genome Consortium |
| EPI_ISL_2971741, EPI_ISL_2971742 | Lighthouse Laboratory Plymouth | Wellcome Sanger Institute for the COVID-19 Genomics UK (COG-UK) Consortium | Lighthouse Laboratory Plymouth and Alex Alderton, Roberto Amato, Jeffrey Barrett, Sonia Goncalves, Ewan Harrison, David K. Jackson, Ian Johnston, Dominic Kwiatkowski, Cordelia Langford, John Sillitoe on behalf of the Wellcome Sanger Institute COVID-19 Surveillance Team |
| EPI_ISL_2971743 | Department of Bacteria, Parasites and Fungi, Statens Serum Institut, Copenhagen, Denmark | Statens Serum Institut Bioinformatics and Microbial Genomics | Danish Covid-19 Genome Consortium |
| EPI_ISL_2971744, EPI_ISL_2971745 | Lighthouse Laboratory Plymouth | Wellcome Sanger Institute for the COVID-19 Genomics UK (COG-UK) Consortium | Lighthouse Laboratory Plymouth and Alex Alderton, Roberto Amato, Jeffrey Barrett, Sonia Goncalves, Ewan Harrison, David K. Jackson, Ian Johnston, Dominic Kwiatkowski, Cordelia Langford, John Sillitoe on behalf of the Wellcome Sanger Institute COVID-19 Surveillance Team |
| EPI_ISL_2971746 | Department of Bacteria, Parasites and Fungi, Statens Serum Institut, Copenhagen, Denmark | Statens Serum Institut Bioinformatics and Microbial Genomics | Danish Covid-19 Genome Consortium |
| EPI_ISL_2971747, EPI_ISL_2971748, EPI_ISL_2971749 | Lighthouse Laboratory Plymouth | Wellcome Sanger Institute for the COVID-19 Genomics UK (COG-UK) Consortium | Lighthouse Laboratory Plymouth and Alex Alderton, Roberto Amato, Jeffrey Barrett, Sonia Goncalves, Ewan Harrison, David K. Jackson, Ian Johnston, Dominic Kwiatkowski, Cordelia Langford, John Sillitoe on behalf of the Wellcome Sanger Institute COVID-19 Surveillance Team |
| EPI_ISL_2971750 | Department of Bacteria, Parasites and Fungi, Statens Serum Institut, Copenhagen, Denmark | Statens Serum Institut Bioinformatics and Microbial Genomics | Danish Covid-19 Genome Consortium |
| EPI_ISL_2971751, EPI_ISL_2971752 | Lighthouse Laboratory Plymouth | Wellcome Sanger Institute for the COVID-19 Genomics UK (COG-UK) Consortium | Lighthouse Laboratory Plymouth and Alex Alderton, Roberto Amato, Jeffrey Barrett, Sonia Goncalves, Ewan Harrison, David K. Jackson, Ian Johnston, Dominic Kwiatkowski, Cordelia Langford, John Sillitoe on behalf of the Wellcome Sanger Institute COVID-19 Surveillance Team |
| EPI_ISL_2971753 | Department of Bacteria, Parasites and Fungi, Statens Serum Institut, Copenhagen, Denmark | Statens Serum Institut Bioinformatics and Microbial Genomics | Danish Covid-19 Genome Consortium |
| EPI_ISL_2971754 | Randox Laboratories | Wellcome Sanger Institute for the COVID-19 Genomics UK (COG-UK) Consortium | Randox Laboratories and Alex Alderton, Roberto Amato, Jeffrey Barrett, Sonia Goncalves, Ewan Harrison, David K. Jackson, Ian Johnston, Dominic Kwiatkowski, Cordelia Langford, John Sillitoe on behalf of the Wellcome Sanger Institute COVID-19 Surveillance Team |
| EPI_ISL_2971755, EPI_ISL_2971757 | Lighthouse Laboratory Plymouth | Wellcome Sanger Institute for the COVID-19 Genomics UK (COG-UK) Consortium | Lighthouse Laboratory Plymouth and Alex Alderton, Roberto Amato, Jeffrey Barrett, Sonia Goncalves, Ewan Harrison, David K. Jackson, Ian Johnston, Dominic Kwiatkowski, Cordelia Langford, John Sillitoe on behalf of the Wellcome Sanger Institute COVID-19 Surveillance Team |
| EPI_ISL_2971758 | Lighthouse Lab in Glasgow | Wellcome Sanger Institute for the COVID-19 Genomics UK (COG-UK) Consortium | Harper VanSteenhouse, Yumi Kasai, David Gray, Carol Clugston, Anna Dominiczak and Alex Alderton, Roberto Amato, Jeffrey Barrett, Sonia Goncalves, Ewan Harrison, David K. Jackson, Ian Johnston, Dominic Kwiatkowski, Cordelia Langford, John Sillitoe on behalf of the Wellcome Sanger Institute COVID-19 Surveillance Team |
| EPI_ISL_2971759 | Department of Bacteria, Parasites and Fungi, Statens Serum Institut, Copenhagen, Denmark | Statens Serum Institut Bioinformatics and Microbial Genomics | Danish Covid-19 Genome Consortium |
| EPI_ISL_2971760 | Lighthouse Lab in Glasgow | Wellcome Sanger Institute for the COVID-19 Genomics UK (COG-UK) Consortium | Harper VanSteenhouse, Yumi Kasai, David Gray, Carol Clugston, Anna Dominiczak and Alex Alderton, Roberto Amato, Jeffrey Barrett, Sonia Goncalves, Ewan Harrison, David K. Jackson, Ian Johnston, Dominic Kwiatkowski, Cordelia Langford, John Sillitoe on behalf of the Wellcome Sanger Institute COVID-19 Surveillance Team |
| EPI_ISL_2971761 | Lighthouse Laboratory Plymouth | Wellcome Sanger Institute for the COVID-19 Genomics UK (COG-UK) Consortium | Lighthouse Laboratory Plymouth and Alex Alderton, Roberto Amato, Jeffrey Barrett, Sonia Goncalves, Ewan Harrison, David K. Jackson, Ian Johnston, Dominic Kwiatkowski, Cordelia Langford, John Sillitoe on behalf of the Wellcome Sanger Institute COVID-19 Surveillance Team |
| EPI_ISL_2971762 | Department of Bacteria, Parasites and Fungi, Statens Serum Institut, Copenhagen, Denmark | Statens Serum Institut Bioinformatics and Microbial Genomics | Danish Covid-19 Genome Consortium |
| EPI_ISL_2971763 | Lighthouse Lab in Glasgow | Wellcome Sanger Institute for the COVID-19 Genomics UK (COG-UK) Consortium | Harper VanSteenhouse, Yumi Kasai, David Gray, Carol Clugston, Anna Dominiczak and Alex Alderton, Roberto Amato, Jeffrey Barrett, Sonia Goncalves, Ewan Harrison, David K. Jackson, Ian Johnston, Dominic Kwiatkowski, Cordelia Langford, John Sillitoe on behalf of the Wellcome Sanger Institute COVID-19 Surveillance Team |
| EPI_ISL_2971764 | Lighthouse Laboratory Plymouth | Wellcome Sanger Institute for the COVID-19 Genomics UK (COG-UK) Consortium | Lighthouse Laboratory Plymouth and Alex Alderton, Roberto Amato, Jeffrey Barrett, Sonia Goncalves, Ewan Harrison, David K. Jackson, Ian Johnston, Dominic Kwiatkowski, Cordelia Langford, John Sillitoe on behalf of the Wellcome Sanger Institute COVID-19 Surveillance Team |
| EPI_ISL_2971765 | Randox Laboratories | Wellcome Sanger Institute for the COVID-19 Genomics UK (COG-UK) Consortium | Randox Laboratories and Alex Alderton, Roberto Amato, Jeffrey Barrett, Sonia Goncalves, Ewan Harrison, David K. Jackson, Ian Johnston, Dominic Kwiatkowski, Cordelia Langford, John Sillitoe on behalf of the Wellcome Sanger Institute COVID-19 Surveillance Team |
| EPI_ISL_2971766 | Department of Bacteria, Parasites and Fungi, Statens Serum Institut, Copenhagen, Denmark | Statens Serum Institut Bioinformatics and Microbial Genomics | Danish Covid-19 Genome Consortium |
| EPI_ISL_2971767, EPI_ISL_2971768 | Lighthouse Laboratory Plymouth | Wellcome Sanger Institute for the COVID-19 Genomics UK (COG-UK) Consortium | Lighthouse Laboratory Plymouth and Alex Alderton, Roberto Amato, Jeffrey Barrett, Sonia Goncalves, Ewan Harrison, David K. Jackson, Ian Johnston, Dominic Kwiatkowski, Cordelia Langford, John Sillitoe on behalf of the Wellcome Sanger Institute COVID-19 Surveillance Team |
| EPI_ISL_2971769 | Department of Bacteria, Parasites and Fungi, Statens Serum Institut, Copenhagen, Denmark | Statens Serum Institut Bioinformatics and Microbial Genomics | Danish Covid-19 Genome Consortium |
| EPI_ISL_2971770, EPI_ISL_2971771 | Lighthouse Laboratory Plymouth | Wellcome Sanger Institute for the COVID-19 Genomics UK (COG-UK) Consortium | Lighthouse Laboratory Plymouth and Alex Alderton, Roberto Amato, Jeffrey Barrett, Sonia Goncalves, Ewan Harrison, David K. Jackson, Ian Johnston, Dominic Kwiatkowski, Cordelia Langford, John Sillitoe on behalf of the Wellcome Sanger Institute COVID-19 Surveillance Team |
| EPI_ISL_2971772 | Department of Bacteria, Parasites and Fungi, Statens Serum Institut, Copenhagen, Denmark | Statens Serum Institut Bioinformatics and Microbial Genomics | Danish Covid-19 Genome Consortium |
| EPI_ISL_2971773, EPI_ISL_2971774 | Lighthouse Laboratory Plymouth | Wellcome Sanger Institute for the COVID-19 Genomics UK (COG-UK) Consortium | Lighthouse Laboratory Plymouth and Alex Alderton, Roberto Amato, Jeffrey Barrett, Sonia Goncalves, Ewan Harrison, David K. Jackson, Ian Johnston, Dominic Kwiatkowski, Cordelia Langford, John Sillitoe on behalf of the Wellcome Sanger Institute COVID-19 Surveillance Team |
| EPI_ISL_2971775 | Department of Bacteria, Parasites and Fungi, Statens Serum Institut, Copenhagen, Denmark | Statens Serum Institut Bioinformatics and Microbial Genomics | Danish Covid-19 Genome Consortium |
| EPI_ISL_2971776 | Lighthouse Laboratory Plymouth | Wellcome Sanger Institute for the COVID-19 Genomics UK (COG-UK) Consortium | Lighthouse Laboratory Plymouth and Alex Alderton, Roberto Amato, Jeffrey Barrett, Sonia Goncalves, Ewan Harrison, David K. Jackson, Ian Johnston, Dominic Kwiatkowski, Cordelia Langford, John Sillitoe on behalf of the Wellcome Sanger Institute COVID-19 Surveillance Team |
| EPI_ISL_2971777 | Lighthouse Lab in Glasgow | Wellcome Sanger Institute for the COVID-19 Genomics UK (COG-UK) Consortium | Harper VanSteenhouse, Yumi Kasai, David Gray, Carol Clugston, Anna Dominiczak and Alex Alderton, Roberto Amato, Jeffrey Barrett, Sonia Goncalves, Ewan Harrison, David K. Jackson, Ian Johnston, Dominic Kwiatkowski, Cordelia Langford, John Sillitoe on behalf of the Wellcome Sanger Institute COVID-19 Surveillance Team |
| EPI_ISL_2971778 | Department of Bacteria, Parasites and Fungi, Statens Serum Institut, Copenhagen, Denmark | Statens Serum Institut Bioinformatics and Microbial Genomics | Danish Covid-19 Genome Consortium |
| EPI_ISL_2971779, EPI_ISL_2971780 | Lighthouse Laboratory Plymouth | Wellcome Sanger Institute for the COVID-19 Genomics UK (COG-UK) Consortium | Lighthouse Laboratory Plymouth and Alex Alderton, Roberto Amato, Jeffrey Barrett, Sonia Goncalves, Ewan Harrison, David K. Jackson, Ian Johnston, Dominic Kwiatkowski, Cordelia Langford, John Sillitoe on behalf of the Wellcome Sanger Institute COVID-19 Surveillance Team |
| EPI_ISL_2971781 | Department of Bacteria, Parasites and Fungi, Statens Serum Institut, Copenhagen, Denmark | Statens Serum Institut Bioinformatics and Microbial Genomics | Danish Covid-19 Genome Consortium |
| EPI_ISL_2971782 | Randox Laboratories | Wellcome Sanger Institute for the COVID-19 Genomics UK (COG-UK) Consortium | Randox Laboratories and Alex Alderton, Roberto Amato, Jeffrey Barrett, Sonia Goncalves, Ewan Harrison, David K. Jackson, Ian Johnston, Dominic Kwiatkowski, Cordelia Langford, John Sillitoe on behalf of the Wellcome Sanger Institute COVID-19 Surveillance Team |
| EPI_ISL_2971783 | Lighthouse Lab in Glasgow | Wellcome Sanger Institute for the COVID-19 Genomics UK (COG-UK) Consortium | Harper VanSteenhouse, Yumi Kasai, David Gray, Carol Clugston, Anna Dominiczak and Alex Alderton, Roberto Amato, Jeffrey Barrett, Sonia Goncalves, Ewan Harrison, David K. Jackson, Ian Johnston, Dominic Kwiatkowski, Cordelia Langford, John Sillitoe on behalf of the Wellcome Sanger Institute COVID-19 Surveillance Team |
| EPI_ISL_2971784 | Department of Bacteria, Parasites and Fungi, Statens Serum Institut, Copenhagen, Denmark | Statens Serum Institut Bioinformatics and Microbial Genomics | Danish Covid-19 Genome Consortium |
| EPI_ISL_2971785, EPI_ISL_2971786 | Lighthouse Laboratory Plymouth | Wellcome Sanger Institute for the COVID-19 Genomics UK (COG-UK) Consortium | Lighthouse Laboratory Plymouth and Alex Alderton, Roberto Amato, Jeffrey Barrett, Sonia Goncalves, Ewan Harrison, David K. Jackson, Ian Johnston, Dominic Kwiatkowski, Cordelia Langford, John Sillitoe on behalf of the Wellcome Sanger Institute COVID-19 Surveillance Team |
| EPI_ISL_2971787 | Lighthouse Lab in Glasgow | Wellcome Sanger Institute for the COVID-19 Genomics UK (COG-UK) Consortium | Harper VanSteenhouse, Yumi Kasai, David Gray, Carol Clugston, Anna Dominiczak and Alex Alderton, Roberto Amato, Jeffrey Barrett, Sonia Goncalves, Ewan Harrison, David K. Jackson, Ian Johnston, Dominic Kwiatkowski, Cordelia Langford, John Sillitoe on behalf of the Wellcome Sanger Institute COVID-19 Surveillance Team |
| EPI_ISL_2971788 | Department of Bacteria, Parasites and Fungi, Statens Serum Institut, Copenhagen, Denmark | Statens Serum Institut Bioinformatics and Microbial Genomics | Danish Covid-19 Genome Consortium |
| EPI_ISL_2971789, EPI_ISL_2971790 | Lighthouse Laboratory Plymouth | Wellcome Sanger Institute for the COVID-19 Genomics UK (COG-UK) Consortium | Lighthouse Laboratory Plymouth and Alex Alderton, Roberto Amato, Jeffrey Barrett, Sonia Goncalves, Ewan Harrison, David K. Jackson, Ian Johnston, Dominic Kwiatkowski, Cordelia Langford, John Sillitoe on behalf of the Wellcome Sanger Institute COVID-19 Surveillance Team |
| EPI_ISL_2971791 | Department of Bacteria, Parasites and Fungi, Statens Serum Institut, Copenhagen, Denmark | Statens Serum Institut Bioinformatics and Microbial Genomics | Danish Covid-19 Genome Consortium |
| EPI_ISL_2971792, EPI_ISL_2971793 | Lighthouse Laboratory Plymouth | Wellcome Sanger Institute for the COVID-19 Genomics UK (COG-UK) Consortium | Lighthouse Laboratory Plymouth and Alex Alderton, Roberto Amato, Jeffrey Barrett, Sonia Goncalves, Ewan Harrison, David K. Jackson, Ian Johnston, Dominic Kwiatkowski, Cordelia Langford, John Sillitoe on behalf of the Wellcome Sanger Institute COVID-19 Surveillance Team |
| EPI_ISL_2971794 | Department of Bacteria, Parasites and Fungi, Statens Serum Institut, Copenhagen, Denmark | Statens Serum Institut Bioinformatics and Microbial Genomics | Danish Covid-19 Genome Consortium |
| EPI_ISL_2971795, EPI_ISL_2971796 | Lighthouse Laboratory Plymouth | Wellcome Sanger Institute for the COVID-19 Genomics UK (COG-UK) Consortium | Lighthouse Laboratory Plymouth and Alex Alderton, Roberto Amato, Jeffrey Barrett, Sonia Goncalves, Ewan Harrison, David K. Jackson, Ian Johnston, Dominic Kwiatkowski, Cordelia Langford, John Sillitoe on behalf of the Wellcome Sanger Institute COVID-19 Surveillance Team |

| | | | |
|---|---|---|---|
| EPI_ISL_2971797 | Department of Bacteria, Parasites and Fungi, Statens Serum Institut, Copenhagen, Denmark | Statens Serum Institut Bioinformatics and Microbial Genomics | Danish Covid-19 Genome Consortium |
| EPI_ISL_2971798, EPI_ISL_2971799, EPI_ISL_2971801, EPI_ISL_2971802 | Lighthouse Laboratory Plymouth | Wellcome Sanger Institute for the COVID-19 Genomics UK (COG-UK) Consortium | Lighthouse Laboratory Plymouth and Alex Alderton, Roberto Amato, Jeffrey Barrett, Sonia Goncalves, Ewan Harrison, David K. Jackson, Ian Johnston, Dominic Kwiatkowski, Cordelia Langford, John Sillitoe on behalf of the Wellcome Sanger Institute COVID-19 Surveillance Team |
| EPI_ISL_2971803 | Lighthouse Lab in Glasgow | Wellcome Sanger Institute for the COVID-19 Genomics UK (COG-UK) Consortium | Harper VanSteenhouse, Yumi Kasai, David Gray, Carol Clugston, Anna Dominiczak and Alex Alderton, Roberto Amato, Jeffrey Barrett, Sonia Goncalves, Ewan Harrison, David K. Jackson, Ian Johnston, Dominic Kwiatkowski, Cordelia Langford, John Sillitoe on behalf of the Wellcome Sanger Institute COVID-19 Surveillance Team |
| EPI_ISL_2971804 | Department of Bacteria, Parasites and Fungi, Statens Serum Institut, Copenhagen, Denmark | Statens Serum Institut Bioinformatics and Microbial Genomics | Danish Covid-19 Genome Consortium |
| EPI_ISL_2971805 | Randox Laboratories | Wellcome Sanger Institute for the COVID-19 Genomics UK (COG-UK) Consortium | Randox Laboratories and Alex Alderton, Roberto Amato, Jeffrey Barrett, Sonia Goncalves, Ewan Harrison, David K. Jackson, Ian Johnston, Dominic Kwiatkowski, Cordelia Langford, John Sillitoe on behalf of the Wellcome Sanger Institute COVID-19 Surveillance Team |
| EPI_ISL_2971806 | Lighthouse Laboratory Plymouth | Wellcome Sanger Institute for the COVID-19 Genomics UK (COG-UK) Consortium | Lighthouse Laboratory Plymouth and Alex Alderton, Roberto Amato, Jeffrey Barrett, Sonia Goncalves, Ewan Harrison, David K. Jackson, Ian Johnston, Dominic Kwiatkowski, Cordelia Langford, John Sillitoe on behalf of the Wellcome Sanger Institute COVID-19 Surveillance Team |
| EPI_ISL_2971807 | Department of Bacteria, Parasites and Fungi, Statens Serum Institut, Copenhagen, Denmark | Statens Serum Institut Bioinformatics and Microbial Genomics | Danish Covid-19 Genome Consortium |
| EPI_ISL_2971808 | Randox Laboratories | Wellcome Sanger Institute for the COVID-19 Genomics UK (COG-UK) Consortium | Randox Laboratories and Alex Alderton, Roberto Amato, Jeffrey Barrett, Sonia Goncalves, Ewan Harrison, David K. Jackson, Ian Johnston, Dominic Kwiatkowski, Cordelia Langford, John Sillitoe on behalf of the Wellcome Sanger Institute COVID-19 Surveillance Team |
| EPI_ISL_2971809, EPI_ISL_2971810 | Lighthouse Laboratory Plymouth | Wellcome Sanger Institute for the COVID-19 Genomics UK (COG-UK) Consortium | Lighthouse Laboratory Plymouth and Alex Alderton, Roberto Amato, Jeffrey Barrett, Sonia Goncalves, Ewan Harrison, David K. Jackson, Ian Johnston, Dominic Kwiatkowski, Cordelia Langford, John Sillitoe on behalf of the Wellcome Sanger Institute COVID-19 Surveillance Team |
| EPI_ISL_2971811 | Department of Bacteria, Parasites and Fungi, Statens Serum Institut, Copenhagen, Denmark | Statens Serum Institut Bioinformatics and Microbial Genomics | Danish Covid-19 Genome Consortium |
| EPI_ISL_2971812, EPI_ISL_2971813 | Lighthouse Laboratory Plymouth | Wellcome Sanger Institute for the COVID-19 Genomics UK (COG-UK) Consortium | Lighthouse Laboratory Plymouth and Alex Alderton, Roberto Amato, Jeffrey Barrett, Sonia Goncalves, Ewan Harrison, David K. Jackson, Ian Johnston, Dominic Kwiatkowski, Cordelia Langford, John Sillitoe on behalf of the Wellcome Sanger Institute COVID-19 Surveillance Team |
| EPI_ISL_2971814 | Department of Bacteria, Parasites and Fungi, Statens Serum Institut, Copenhagen, Denmark | Statens Serum Institut Bioinformatics and Microbial Genomics | Danish Covid-19 Genome Consortium |
| EPI_ISL_2971815, EPI_ISL_2971816 | Lighthouse Laboratory Plymouth | Wellcome Sanger Institute for the COVID-19 Genomics UK (COG-UK) Consortium | Lighthouse Laboratory Plymouth and Alex Alderton, Roberto Amato, Jeffrey Barrett, Sonia Goncalves, Ewan Harrison, David K. Jackson, Ian Johnston, Dominic Kwiatkowski, Cordelia Langford, John Sillitoe on behalf of the Wellcome Sanger Institute COVID-19 Surveillance Team |
| EPI_ISL_2971817 | Department of Bacteria, Parasites and Fungi, Statens Serum Institut, Copenhagen, Denmark | Statens Serum Institut Bioinformatics and Microbial Genomics | Danish Covid-19 Genome Consortium |
| EPI_ISL_2971818, EPI_ISL_2971819 | Lighthouse Laboratory Plymouth | Wellcome Sanger Institute for the COVID-19 Genomics UK (COG-UK) Consortium | Lighthouse Laboratory Plymouth and Alex Alderton, Roberto Amato, Jeffrey Barrett, Sonia Goncalves, Ewan Harrison, David K. Jackson, Ian Johnston, Dominic Kwiatkowski, Cordelia Langford, John Sillitoe on behalf of the Wellcome Sanger Institute COVID-19 Surveillance Team |
| EPI_ISL_2971820 | Department of Bacteria, Parasites and Fungi, Statens Serum Institut, Copenhagen, Denmark | Statens Serum Institut Bioinformatics and Microbial Genomics | Danish Covid-19 Genome Consortium |
| EPI_ISL_2971821, EPI_ISL_2971822 | Lighthouse Laboratory Plymouth | Wellcome Sanger Institute for the COVID-19 Genomics UK (COG-UK) Consortium | Lighthouse Laboratory Plymouth and Alex Alderton, Roberto Amato, Jeffrey Barrett, Sonia Goncalves, Ewan Harrison, David K. Jackson, Ian Johnston, Dominic Kwiatkowski, Cordelia Langford, John Sillitoe on behalf of the Wellcome Sanger Institute COVID-19 Surveillance Team |
| EPI_ISL_2971823 | Department of Bacteria, Parasites and Fungi, Statens Serum Institut, Copenhagen, Denmark | Statens Serum Institut Bioinformatics and Microbial Genomics | Danish Covid-19 Genome Consortium |
| EPI_ISL_2971824, EPI_ISL_2971825, EPI_ISL_2971827 | Lighthouse Laboratory Plymouth | Wellcome Sanger Institute for the COVID-19 Genomics UK (COG-UK) Consortium | Lighthouse Laboratory Plymouth and Alex Alderton, Roberto Amato, Jeffrey Barrett, Sonia Goncalves, Ewan Harrison, David K. Jackson, Ian Johnston, Dominic Kwiatkowski, Cordelia Langford, John Sillitoe on behalf of the Wellcome Sanger Institute COVID-19 Surveillance Team |
| EPI_ISL_2971828 | Randox Laboratories | Wellcome Sanger Institute for the COVID-19 Genomics UK (COG-UK) Consortium | Randox Laboratories and Alex Alderton, Roberto Amato, Jeffrey Barrett, Sonia Goncalves, Ewan Harrison, David K. Jackson, Ian Johnston, Dominic Kwiatkowski, Cordelia Langford, John Sillitoe on behalf of the Wellcome Sanger Institute COVID-19 Surveillance Team |
| EPI_ISL_2971829 | Department of Bacteria, Parasites and Fungi, Statens Serum Institut, Copenhagen, Denmark | Statens Serum Institut Bioinformatics and Microbial Genomics | Danish Covid-19 Genome Consortium |
| EPI_ISL_2971830 | Lighthouse Lab in Glasgow | Wellcome Sanger Institute for the COVID-19 Genomics UK (COG-UK) Consortium | Harper VanSteenhouse, Yumi Kasai, David Gray, Carol Clugston, Anna Dominiczak and Alex Alderton, Roberto Amato, Jeffrey Barrett, Sonia Goncalves, Ewan Harrison, David K. Jackson, Ian Johnston, Dominic Kwiatkowski, Cordelia Langford, John Sillitoe on behalf of the Wellcome Sanger Institute COVID-19 Surveillance Team |
| EPI_ISL_2971831 | Lighthouse Laboratory Plymouth | Wellcome Sanger Institute for the COVID-19 Genomics UK (COG-UK) Consortium | Lighthouse Laboratory Plymouth and Alex Alderton, Roberto Amato, Jeffrey Barrett, Sonia Goncalves, Ewan Harrison, David K. Jackson, Ian Johnston, Dominic Kwiatkowski, Cordelia Langford, John Sillitoe on behalf of the Wellcome Sanger Institute COVID-19 Surveillance Team |
| EPI_ISL_2971832 | Department of Bacteria, Parasites and Fungi, Statens Serum Institut, Copenhagen, Denmark | Statens Serum Institut Bioinformatics and Microbial Genomics | Danish Covid-19 Genome Consortium |
| EPI_ISL_2971833 | Lighthouse Lab in Glasgow | Wellcome Sanger Institute for the COVID-19 Genomics UK (COG-UK) Consortium | Harper VanSteenhouse, Yumi Kasai, David Gray, Carol Clugston, Anna Dominiczak and Alex Alderton, Roberto Amato, Jeffrey Barrett, Sonia Goncalves, Ewan Harrison, David K. Jackson, Ian Johnston, Dominic Kwiatkowski, Cordelia Langford, John Sillitoe on behalf of the Wellcome Sanger Institute COVID-19 Surveillance Team |
| EPI_ISL_2971834, EPI_ISL_2971835 | Lighthouse Laboratory Plymouth | Wellcome Sanger Institute for the COVID-19 Genomics UK (COG-UK) Consortium | Lighthouse Laboratory Plymouth and Alex Alderton, Roberto Amato, Jeffrey Barrett, Sonia Goncalves, Ewan Harrison, David K. Jackson, Ian Johnston, Dominic Kwiatkowski, Cordelia Langford, John Sillitoe on behalf of the Wellcome Sanger Institute COVID-19 Surveillance Team |
| EPI_ISL_2971836 | Department of Bacteria, Parasites and Fungi, Statens Serum Institut, Copenhagen, Denmark | Statens Serum Institut Bioinformatics and Microbial Genomics | Danish Covid-19 Genome Consortium |
| EPI_ISL_2971837, EPI_ISL_2971838 | Lighthouse Laboratory Plymouth | Wellcome Sanger Institute for the COVID-19 Genomics UK (COG-UK) Consortium | Lighthouse Laboratory Plymouth and Alex Alderton, Roberto Amato, Jeffrey Barrett, Sonia Goncalves, Ewan Harrison, David K. Jackson, Ian Johnston, Dominic Kwiatkowski, Cordelia Langford, John Sillitoe on behalf of the Wellcome Sanger Institute COVID-19 Surveillance Team |
| EPI_ISL_2971839 | Department of Bacteria, Parasites and Fungi, Statens Serum Institut, Copenhagen, Denmark | Statens Serum Institut Bioinformatics and Microbial Genomics | Danish Covid-19 Genome Consortium |
| EPI_ISL_2971840, EPI_ISL_2971841, EPI_ISL_2971843, EPI_ISL_2971844 | Lighthouse Laboratory Plymouth | Wellcome Sanger Institute for the COVID-19 Genomics UK (COG-UK) Consortium | Lighthouse Laboratory Plymouth and Alex Alderton, Roberto Amato, Jeffrey Barrett, Sonia Goncalves, Ewan Harrison, David K. Jackson, Ian Johnston, Dominic Kwiatkowski, Cordelia Langford, John Sillitoe on behalf of the Wellcome Sanger Institute COVID-19 Surveillance Team |
| EPI_ISL_2971845 | Department of Bacteria, Parasites and Fungi, Statens Serum Institut, Copenhagen, Denmark | Statens Serum Institut Bioinformatics and Microbial Genomics | Danish Covid-19 Genome Consortium |
| EPI_ISL_2971846, EPI_ISL_2971847 | Lighthouse Laboratory Plymouth | Wellcome Sanger Institute for the COVID-19 Genomics UK (COG-UK) Consortium | Lighthouse Laboratory Plymouth and Alex Alderton, Roberto Amato, Jeffrey Barrett, Sonia Goncalves, Ewan Harrison, David K. Jackson, Ian Johnston, Dominic Kwiatkowski, Cordelia Langford, John Sillitoe on behalf of the Wellcome Sanger Institute COVID-19 Surveillance Team |
| EPI_ISL_2971848 | Randox Laboratories | Wellcome Sanger Institute for the COVID-19 Genomics UK (COG-UK) Consortium | Randox Laboratories and Alex Alderton, Roberto Amato, Jeffrey Barrett, Sonia Goncalves, Ewan Harrison, David K. Jackson, Ian Johnston, Dominic Kwiatkowski, Cordelia Langford, John Sillitoe on behalf of the Wellcome Sanger Institute COVID-19 Surveillance Team |
| EPI_ISL_2971849 | Department of Bacteria, Parasites and Fungi, Statens Serum Institut, Copenhagen, Denmark | Statens Serum Institut Bioinformatics and Microbial Genomics | Danish Covid-19 Genome Consortium |
| EPI_ISL_2971850 | Lighthouse Lab in Glasgow | Wellcome Sanger Institute for the COVID-19 Genomics UK (COG-UK) Consortium | Harper VanSteenhouse, Yumi Kasai, David Gray, Carol Clugston, Anna Dominiczak and Alex Alderton, Roberto Amato, Jeffrey Barrett, Sonia Goncalves, Ewan Harrison, David K. Jackson, Ian Johnston, Dominic Kwiatkowski, Cordelia Langford, John Sillitoe on behalf of the Wellcome Sanger Institute COVID-19 Surveillance Team |
| EPI_ISL_2971851, EPI_ISL_2971852 | Lighthouse Laboratory Plymouth | Wellcome Sanger Institute for the COVID-19 Genomics UK (COG-UK) Consortium | Lighthouse Laboratory Plymouth and Alex Alderton, Roberto Amato, Jeffrey Barrett, Sonia Goncalves, Ewan Harrison, David K. Jackson, Ian Johnston, Dominic Kwiatkowski, Cordelia Langford, John Sillitoe on behalf of the Wellcome Sanger Institute COVID-19 Surveillance Team |
| EPI_ISL_2971853 | Department of Bacteria, Parasites and Fungi, Statens Serum Institut, Copenhagen, Denmark | Statens Serum Institut Bioinformatics and Microbial Genomics | Danish Covid-19 Genome Consortium |
| EPI_ISL_2971854 | Randox Laboratories | Wellcome Sanger Institute for the COVID-19 Genomics UK (COG-UK) Consortium | Randox Laboratories and Alex Alderton, Roberto Amato, Jeffrey Barrett, Sonia Goncalves, Ewan Harrison, David K. Jackson, Ian Johnston, Dominic Kwiatkowski, Cordelia Langford, John Sillitoe on behalf of the Wellcome Sanger Institute COVID-19 Surveillance Team |
| EPI_ISL_2971855 | Lighthouse Laboratory Plymouth | Wellcome Sanger Institute for the COVID-19 Genomics UK (COG-UK) Consortium | Lighthouse Laboratory Plymouth and Alex Alderton, Roberto Amato, Jeffrey Barrett, Sonia Goncalves, Ewan Harrison, David K. Jackson, Ian Johnston, Dominic Kwiatkowski, Cordelia Langford, John Sillitoe on behalf of the Wellcome Sanger Institute COVID-19 Surveillance Team |
| EPI_ISL_2971856 | Department of Bacteria, Parasites and Fungi, Statens Serum Institut, Copenhagen, Denmark | Statens Serum Institut Bioinformatics and Microbial Genomics | Danish Covid-19 Genome Consortium |
| EPI_ISL_2971857, EPI_ISL_2971858, EPI_ISL_2971860, EPI_ISL_2971861, EPI_ISL_2971862 | Lighthouse Laboratory Plymouth | Wellcome Sanger Institute for the COVID-19 Genomics UK (COG-UK) Consortium | Lighthouse Laboratory Plymouth and Alex Alderton, Roberto Amato, Jeffrey Barrett, Sonia Goncalves, Ewan Harrison, David K. Jackson, Ian Johnston, Dominic Kwiatkowski, Cordelia Langford, John Sillitoe on behalf of the Wellcome Sanger Institute COVID-19 Surveillance Team |
| EPI_ISL_2971863 | Department of Bacteria, Parasites and Fungi, Statens Serum | Statens Serum Institut Bioinformatics and Microbial Genomics | Danish Covid-19 Genome Consortium |

| | | | |
|---|---|---|---|
| | Institut, Copenhagen, Denmark | | |
| EPI_ISL_2971864 | Randox Laboratories | Wellcome Sanger Institute for the COVID-19 Genomics UK (COG-UK) Consortium | Randox Laboratories and Alex Alderton, Roberto Amato, Jeffrey Barrett, Sonia Goncalves, Ewan Harrison, David K. Jackson, Ian Johnston, Dominic Kwiatkowski, Cordelia Langford, John Sillitoe on behalf of the Wellcome Sanger Institute COVID-19 Surveillance Team |
| EPI_ISL_2971865 | Lighthouse Laboratory Plymouth | Wellcome Sanger Institute for the COVID-19 Genomics UK (COG-UK) Consortium | Lighthouse Laboratory Plymouth and Alex Alderton, Roberto Amato, Jeffrey Barrett, Sonia Goncalves, Ewan Harrison, David K. Jackson, Ian Johnston, Dominic Kwiatkowski, Cordelia Langford, John Sillitoe on behalf of the Wellcome Sanger Institute COVID-19 Surveillance Team |
| EPI_ISL_2971866 | Department of Bacteria, Parasites and Fungi, Statens Serum Institut, Copenhagen, Denmark | Statens Serum Institut Bioinformatics and Microbial Genomics | Danish Covid-19 Genome Consortium |
| EPI_ISL_2971867 | Lighthouse Lab in Glasgow | Wellcome Sanger Institute for the COVID-19 Genomics UK (COG-UK) Consortium | Harper VanSteenhouse, Yumi Kasai, David Gray, Carol Clugston, Anna Dominiczak and Alex Alderton, Roberto Amato, Jeffrey Barrett, Sonia Goncalves, Ewan Harrison, David K. Jackson, Ian Johnston, Dominic Kwiatkowski, Cordelia Langford, John Sillitoe on behalf of the Wellcome Sanger Institute COVID-19 Surveillance Team |
| EPI_ISL_2971868 | Randox Laboratories | Wellcome Sanger Institute for the COVID-19 Genomics UK (COG-UK) Consortium | Randox Laboratories and Alex Alderton, Roberto Amato, Jeffrey Barrett, Sonia Goncalves, Ewan Harrison, David K. Jackson, Ian Johnston, Dominic Kwiatkowski, Cordelia Langford, John Sillitoe on behalf of the Wellcome Sanger Institute COVID-19 Surveillance Team |
| EPI_ISL_2971869 | Department of Bacteria, Parasites and Fungi, Statens Serum Institut, Copenhagen, Denmark | Statens Serum Institut Bioinformatics and Microbial Genomics | Danish Covid-19 Genome Consortium |
| EPI_ISL_2971870 | Lighthouse Laboratory Plymouth | Wellcome Sanger Institute for the COVID-19 Genomics UK (COG-UK) Consortium | Lighthouse Laboratory Plymouth and Alex Alderton, Roberto Amato, Jeffrey Barrett, Sonia Goncalves, Ewan Harrison, David K. Jackson, Ian Johnston, Dominic Kwiatkowski, Cordelia Langford, John Sillitoe on behalf of the Wellcome Sanger Institute COVID-19 Surveillance Team |
| EPI_ISL_2971871 | Lighthouse Lab in Glasgow | Wellcome Sanger Institute for the COVID-19 Genomics UK (COG-UK) Consortium | Harper VanSteenhouse, Yumi Kasai, David Gray, Carol Clugston, Anna Dominiczak and Alex Alderton, Roberto Amato, Jeffrey Barrett, Sonia Goncalves, Ewan Harrison, David K. Jackson, Ian Johnston, Dominic Kwiatkowski, Cordelia Langford, John Sillitoe on behalf of the Wellcome Sanger Institute COVID-19 Surveillance Team |
| EPI_ISL_2971872 | Department of Bacteria, Parasites and Fungi, Statens Serum Institut, Copenhagen, Denmark | Statens Serum Institut Bioinformatics and Microbial Genomics | Danish Covid-19 Genome Consortium |
| EPI_ISL_2971873, EPI_ISL_2971874 | Lighthouse Laboratory Plymouth | Wellcome Sanger Institute for the COVID-19 Genomics UK (COG-UK) Consortium | Lighthouse Laboratory Plymouth and Alex Alderton, Roberto Amato, Jeffrey Barrett, Sonia Goncalves, Ewan Harrison, David K. Jackson, Ian Johnston, Dominic Kwiatkowski, Cordelia Langford, John Sillitoe on behalf of the Wellcome Sanger Institute COVID-19 Surveillance Team |
| EPI_ISL_2971875 | Department of Bacteria, Parasites and Fungi, Statens Serum Institut, Copenhagen, Denmark | Statens Serum Institut Bioinformatics and Microbial Genomics | Danish Covid-19 Genome Consortium |
| EPI_ISL_2971876, EPI_ISL_2971877 | Lighthouse Laboratory Plymouth | Wellcome Sanger Institute for the COVID-19 Genomics UK (COG-UK) Consortium | Lighthouse Laboratory Plymouth and Alex Alderton, Roberto Amato, Jeffrey Barrett, Sonia Goncalves, Ewan Harrison, David K. Jackson, Ian Johnston, Dominic Kwiatkowski, Cordelia Langford, John Sillitoe on behalf of the Wellcome Sanger Institute COVID-19 Surveillance Team |
| EPI_ISL_2971878 | Department of Bacteria, Parasites and Fungi, Statens Serum Institut, Copenhagen, Denmark | Statens Serum Institut Bioinformatics and Microbial Genomics | Danish Covid-19 Genome Consortium |
| EPI_ISL_2971879, EPI_ISL_2971880 | Lighthouse Laboratory Plymouth | Wellcome Sanger Institute for the COVID-19 Genomics UK (COG-UK) Consortium | Lighthouse Laboratory Plymouth and Alex Alderton, Roberto Amato, Jeffrey Barrett, Sonia Goncalves, Ewan Harrison, David K. Jackson, Ian Johnston, Dominic Kwiatkowski, Cordelia Langford, John Sillitoe on behalf of the Wellcome Sanger Institute COVID-19 Surveillance Team |
| EPI_ISL_2971881 | Department of Bacteria, Parasites and Fungi, Statens Serum Institut, Copenhagen, Denmark | Statens Serum Institut Bioinformatics and Microbial Genomics | Danish Covid-19 Genome Consortium |
| EPI_ISL_2971882, EPI_ISL_2971883 | Lighthouse Laboratory Plymouth | Wellcome Sanger Institute for the COVID-19 Genomics UK (COG-UK) Consortium | Lighthouse Laboratory Plymouth and Alex Alderton, Roberto Amato, Jeffrey Barrett, Sonia Goncalves, Ewan Harrison, David K. Jackson, Ian Johnston, Dominic Kwiatkowski, Cordelia Langford, John Sillitoe on behalf of the Wellcome Sanger Institute COVID-19 Surveillance Team |
| EPI_ISL_2971884 | Department of Bacteria, Parasites and Fungi, Statens Serum Institut, Copenhagen, Denmark | Statens Serum Institut Bioinformatics and Microbial Genomics | Danish Covid-19 Genome Consortium |
| EPI_ISL_2971885 | Randox Laboratories | Wellcome Sanger Institute for the COVID-19 Genomics UK (COG-UK) Consortium | Randox Laboratories and Alex Alderton, Roberto Amato, Jeffrey Barrett, Sonia Goncalves, Ewan Harrison, David K. Jackson, Ian Johnston, Dominic Kwiatkowski, Cordelia Langford, John Sillitoe on behalf of the Wellcome Sanger Institute COVID-19 Surveillance Team |
| EPI_ISL_2971886 | Lighthouse Laboratory Plymouth | Wellcome Sanger Institute for the COVID-19 Genomics UK (COG-UK) Consortium | Lighthouse Laboratory Plymouth and Alex Alderton, Roberto Amato, Jeffrey Barrett, Sonia Goncalves, Ewan Harrison, David K. Jackson, Ian Johnston, Dominic Kwiatkowski, Cordelia Langford, John Sillitoe on behalf of the Wellcome Sanger Institute COVID-19 Surveillance Team |
| EPI_ISL_2971887 | Department of Bacteria, Parasites and Fungi, Statens Serum Institut, Copenhagen, Denmark | Statens Serum Institut Bioinformatics and Microbial Genomics | Danish Covid-19 Genome Consortium |
| EPI_ISL_2971888, EPI_ISL_2971889 | Lighthouse Laboratory Plymouth | Wellcome Sanger Institute for the COVID-19 Genomics UK (COG-UK) Consortium | Lighthouse Laboratory Plymouth and Alex Alderton, Roberto Amato, Jeffrey Barrett, Sonia Goncalves, Ewan Harrison, David K. Jackson, Ian Johnston, Dominic Kwiatkowski, Cordelia Langford, John Sillitoe on behalf of the Wellcome Sanger Institute COVID-19 Surveillance Team |
| EPI_ISL_2971890 | Department of Bacteria, Parasites and Fungi, Statens Serum Institut, Copenhagen, Denmark | Statens Serum Institut Bioinformatics and Microbial Genomics | Danish Covid-19 Genome Consortium |
| EPI_ISL_2971891, EPI_ISL_2971892 | Lighthouse Laboratory Plymouth | Wellcome Sanger Institute for the COVID-19 Genomics UK (COG-UK) Consortium | Lighthouse Laboratory Plymouth and Alex Alderton, Roberto Amato, Jeffrey Barrett, Sonia Goncalves, Ewan Harrison, David K. Jackson, Ian Johnston, Dominic Kwiatkowski, Cordelia Langford, John Sillitoe on behalf of the Wellcome Sanger Institute COVID-19 Surveillance Team |
| EPI_ISL_2971893 | Department of Bacteria, Parasites and Fungi, Statens Serum Institut, Copenhagen, Denmark | Statens Serum Institut Bioinformatics and Microbial Genomics | Danish Covid-19 Genome Consortium |
| EPI_ISL_2971894, EPI_ISL_2971895 | Lighthouse Laboratory Plymouth | Wellcome Sanger Institute for the COVID-19 Genomics UK (COG-UK) Consortium | Lighthouse Laboratory Plymouth and Alex Alderton, Roberto Amato, Jeffrey Barrett, Sonia Goncalves, Ewan Harrison, David K. Jackson, Ian Johnston, Dominic Kwiatkowski, Cordelia Langford, John Sillitoe on behalf of the Wellcome Sanger Institute COVID-19 Surveillance Team |
| EPI_ISL_2971896 | Lighthouse Lab in Glasgow | Wellcome Sanger Institute for the COVID-19 Genomics UK (COG-UK) Consortium | Harper VanSteenhouse, Yumi Kasai, David Gray, Carol Clugston, Anna Dominiczak and Alex Alderton, Roberto Amato, Jeffrey Barrett, Sonia Goncalves, Ewan Harrison, David K. Jackson, Ian Johnston, Dominic Kwiatkowski, Cordelia Langford, John Sillitoe on behalf of the Wellcome Sanger Institute COVID-19 Surveillance Team |
| EPI_ISL_2971897 | Randox Laboratories | Wellcome Sanger Institute for the COVID-19 Genomics UK (COG-UK) Consortium | Randox Laboratories and Alex Alderton, Roberto Amato, Jeffrey Barrett, Sonia Goncalves, Ewan Harrison, David K. Jackson, Ian Johnston, Dominic Kwiatkowski, Cordelia Langford, John Sillitoe on behalf of the Wellcome Sanger Institute COVID-19 Surveillance Team |
| EPI_ISL_2971898 | Department of Bacteria, Parasites and Fungi, Statens Serum Institut, Copenhagen, Denmark | Statens Serum Institut Bioinformatics and Microbial Genomics | Danish Covid-19 Genome Consortium |
| EPI_ISL_2971900, EPI_ISL_2971901 | Lighthouse Laboratory Plymouth | Wellcome Sanger Institute for the COVID-19 Genomics UK (COG-UK) Consortium | Lighthouse Laboratory Plymouth and Alex Alderton, Roberto Amato, Jeffrey Barrett, Sonia Goncalves, Ewan Harrison, David K. Jackson, Ian Johnston, Dominic Kwiatkowski, Cordelia Langford, John Sillitoe on behalf of the Wellcome Sanger Institute COVID-19 Surveillance Team |
| EPI_ISL_2971902 | Department of Bacteria, Parasites and Fungi, Statens Serum Institut, Copenhagen, Denmark | Statens Serum Institut Bioinformatics and Microbial Genomics | Danish Covid-19 Genome Consortium |
| EPI_ISL_2971903, EPI_ISL_2971904 | Lighthouse Laboratory Plymouth | Wellcome Sanger Institute for the COVID-19 Genomics UK (COG-UK) Consortium | Lighthouse Laboratory Plymouth and Alex Alderton, Roberto Amato, Jeffrey Barrett, Sonia Goncalves, Ewan Harrison, David K. Jackson, Ian Johnston, Dominic Kwiatkowski, Cordelia Langford, John Sillitoe on behalf of the Wellcome Sanger Institute COVID-19 Surveillance Team |
| EPI_ISL_2971905 | Department of Bacteria, Parasites and Fungi, Statens Serum Institut, Copenhagen, Denmark | Statens Serum Institut Bioinformatics and Microbial Genomics | Danish Covid-19 Genome Consortium |
| EPI_ISL_2971906, EPI_ISL_2971907 | Lighthouse Laboratory Plymouth | Wellcome Sanger Institute for the COVID-19 Genomics UK (COG-UK) Consortium | Lighthouse Laboratory Plymouth and Alex Alderton, Roberto Amato, Jeffrey Barrett, Sonia Goncalves, Ewan Harrison, David K. Jackson, Ian Johnston, Dominic Kwiatkowski, Cordelia Langford, John Sillitoe on behalf of the Wellcome Sanger Institute COVID-19 Surveillance Team |
| EPI_ISL_2971908 | Department of Bacteria, Parasites and Fungi, Statens Serum Institut, Copenhagen, Denmark | Statens Serum Institut Bioinformatics and Microbial Genomics | Danish Covid-19 Genome Consortium |
| EPI_ISL_2971909, EPI_ISL_2971910 | Lighthouse Laboratory Plymouth | Wellcome Sanger Institute for the COVID-19 Genomics UK (COG-UK) Consortium | Lighthouse Laboratory Plymouth and Alex Alderton, Roberto Amato, Jeffrey Barrett, Sonia Goncalves, Ewan Harrison, David K. Jackson, Ian Johnston, Dominic Kwiatkowski, Cordelia Langford, John Sillitoe on behalf of the Wellcome Sanger Institute COVID-19 Surveillance Team |
| EPI_ISL_2971911 | Department of Bacteria, Parasites and Fungi, Statens Serum Institut, Copenhagen, Denmark | Statens Serum Institut Bioinformatics and Microbial Genomics | Danish Covid-19 Genome Consortium |
| EPI_ISL_2971912, EPI_ISL_2971913, EPI_ISL_2971914 | Lighthouse Laboratory Plymouth | Wellcome Sanger Institute for the COVID-19 Genomics UK (COG-UK) Consortium | Lighthouse Laboratory Plymouth and Alex Alderton, Roberto Amato, Jeffrey Barrett, Sonia Goncalves, Ewan Harrison, David K. Jackson, Ian Johnston, Dominic Kwiatkowski, Cordelia Langford, John Sillitoe on behalf of the Wellcome Sanger Institute COVID-19 Surveillance Team |
| EPI_ISL_2971915 | Department of Bacteria, Parasites and Fungi, Statens Serum Institut, Copenhagen, Denmark | Statens Serum Institut Bioinformatics and Microbial Genomics | Danish Covid-19 Genome Consortium |
| EPI_ISL_2971916, EPI_ISL_2971917 | Lighthouse Laboratory Plymouth | Wellcome Sanger Institute for the COVID-19 Genomics UK (COG-UK) Consortium | Lighthouse Laboratory Plymouth and Alex Alderton, Roberto Amato, Jeffrey Barrett, Sonia Goncalves, Ewan Harrison, David K. Jackson, Ian Johnston, Dominic Kwiatkowski, Cordelia Langford, John Sillitoe on behalf of the Wellcome Sanger Institute COVID-19 Surveillance Team |
| EPI_ISL_2971918 | Department of Bacteria, Parasites and Fungi, Statens Serum Institut, Copenhagen, Denmark | Statens Serum Institut Bioinformatics and Microbial Genomics | Danish Covid-19 Genome Consortium |
| EPI_ISL_2971919, EPI_ISL_2971920 | Lighthouse Laboratory Plymouth | Wellcome Sanger Institute for the COVID-19 Genomics UK (COG-UK) Consortium | Lighthouse Laboratory Plymouth and Alex Alderton, Roberto Amato, Jeffrey Barrett, Sonia Goncalves, Ewan Harrison, David K. Jackson, Ian Johnston, Dominic Kwiatkowski, Cordelia Langford, John Sillitoe on behalf of the Wellcome Sanger Institute COVID-19 Surveillance Team |
| EPI_ISL_2971921 | Department of Bacteria, Parasites and Fungi, Statens Serum Institut, Copenhagen, Denmark | Statens Serum Institut Bioinformatics and Microbial Genomics | Danish Covid-19 Genome Consortium |
| EPI_ISL_2971922 | Lighthouse Laboratory Plymouth | Wellcome Sanger Institute for the COVID-19 Genomics UK (COG-UK) Consortium | Lighthouse Laboratory Plymouth and Alex Alderton, Roberto Amato, Jeffrey Barrett, Sonia Goncalves, Ewan Harrison, David K. Jackson, Ian Johnston, Dominic Kwiatkowski, Cordelia Langford, John Sillitoe on behalf of the Wellcome Sanger Institute COVID-19 Surveillance Team |
| EPI_ISL_2971923 | Randox Laboratories | Wellcome Sanger Institute for the COVID-19 Genomics UK (COG-UK) Consortium | Randox Laboratories and Alex Alderton, Roberto Amato, Jeffrey Barrett, Sonia Goncalves, Ewan Harrison, David K. Jackson, Ian Johnston, Dominic Kwiatkowski, Cordelia Langford, John Sillitoe on behalf of the Wellcome Sanger Institute COVID-19 Surveillance Team |

| | | | |
|---|---|---|---|
| EPI_ISL_2971924 | Department of Bacteria, Parasites and Fungi, Statens Serum Institut, Copenhagen, Denmark | Statens Serum Institut Bioinformatics and Microbial Genomics | Danish Covid-19 Genome Consortium |
| EPI_ISL_2971925 | Lighthouse Lab in Glasgow | Wellcome Sanger Institute for the COVID-19 Genomics UK (COG-UK) Consortium | Harper VanSteenhouse, Yumi Kasai, David Gray, Carol Clugston, Anna Dominiczak and Alex Alderton, Roberto Amato, Jeffrey Barrett, Sonia Goncalves, Ewan Harrison, David K. Jackson, Ian Johnston, Dominic Kwiatkowski, Cordelia Langford, John Sillitoe on behalf of the Wellcome Sanger Institute COVID-19 Surveillance Team |
| EPI_ISL_2971926, EPI_ISL_2971927 | Lighthouse Laboratory Plymouth | Wellcome Sanger Institute for the COVID-19 Genomics UK (COG-UK) Consortium | Lighthouse Laboratory Plymouth and Alex Alderton, Roberto Amato, Jeffrey Barrett, Sonia Goncalves, Ewan Harrison, David K. Jackson, Ian Johnston, Dominic Kwiatkowski, Cordelia Langford, John Sillitoe on behalf of the Wellcome Sanger Institute COVID-19 Surveillance Team |
| EPI_ISL_2971928 | Department of Bacteria, Parasites and Fungi, Statens Serum Institut, Copenhagen, Denmark | Statens Serum Institut Bioinformatics and Microbial Genomics | Danish Covid-19 Genome Consortium |
| EPI_ISL_2971929 | Randox Laboratories | Wellcome Sanger Institute for the COVID-19 Genomics UK (COG-UK) Consortium | Randox Laboratories and Alex Alderton, Roberto Amato, Jeffrey Barrett, Sonia Goncalves, Ewan Harrison, David K. Jackson, Ian Johnston, Dominic Kwiatkowski, Cordelia Langford, John Sillitoe on behalf of the Wellcome Sanger Institute COVID-19 Surveillance Team |
| EPI_ISL_2971930 | Lighthouse Laboratory Plymouth | Wellcome Sanger Institute for the COVID-19 Genomics UK (COG-UK) Consortium | Lighthouse Laboratory Plymouth and Alex Alderton, Roberto Amato, Jeffrey Barrett, Sonia Goncalves, Ewan Harrison, David K. Jackson, Ian Johnston, Dominic Kwiatkowski, Cordelia Langford, John Sillitoe on behalf of the Wellcome Sanger Institute COVID-19 Surveillance Team |
| EPI_ISL_2971931 | Department of Bacteria, Parasites and Fungi, Statens Serum Institut, Copenhagen, Denmark | Statens Serum Institut Bioinformatics and Microbial Genomics | Danish Covid-19 Genome Consortium |
| EPI_ISL_2971932, EPI_ISL_2971933 | Lighthouse Laboratory Plymouth | Wellcome Sanger Institute for the COVID-19 Genomics UK (COG-UK) Consortium | Lighthouse Laboratory Plymouth and Alex Alderton, Roberto Amato, Jeffrey Barrett, Sonia Goncalves, Ewan Harrison, David K. Jackson, Ian Johnston, Dominic Kwiatkowski, Cordelia Langford, John Sillitoe on behalf of the Wellcome Sanger Institute COVID-19 Surveillance Team |
| EPI_ISL_2971934 | Department of Bacteria, Parasites and Fungi, Statens Serum Institut, Copenhagen, Denmark | Statens Serum Institut Bioinformatics and Microbial Genomics | Danish Covid-19 Genome Consortium |
| EPI_ISL_2971935, EPI_ISL_2971936, EPI_ISL_2971937 | Lighthouse Laboratory Plymouth | Wellcome Sanger Institute for the COVID-19 Genomics UK (COG-UK) Consortium | Lighthouse Laboratory Plymouth and Alex Alderton, Roberto Amato, Jeffrey Barrett, Sonia Goncalves, Ewan Harrison, David K. Jackson, Ian Johnston, Dominic Kwiatkowski, Cordelia Langford, John Sillitoe on behalf of the Wellcome Sanger Institute COVID-19 Surveillance Team |
| EPI_ISL_2971938 | Department of Bacteria, Parasites and Fungi, Statens Serum Institut, Copenhagen, Denmark | Statens Serum Institut Bioinformatics and Microbial Genomics | Danish Covid-19 Genome Consortium |
| EPI_ISL_2971939, EPI_ISL_2971940 | Lighthouse Laboratory Plymouth | Wellcome Sanger Institute for the COVID-19 Genomics UK (COG-UK) Consortium | Lighthouse Laboratory Plymouth and Alex Alderton, Roberto Amato, Jeffrey Barrett, Sonia Goncalves, Ewan Harrison, David K. Jackson, Ian Johnston, Dominic Kwiatkowski, Cordelia Langford, John Sillitoe on behalf of the Wellcome Sanger Institute COVID-19 Surveillance Team |
| EPI_ISL_2971941 | Department of Bacteria, Parasites and Fungi, Statens Serum Institut, Copenhagen, Denmark | Statens Serum Institut Bioinformatics and Microbial Genomics | Danish Covid-19 Genome Consortium |
| EPI_ISL_2971942, EPI_ISL_2971943, EPI_ISL_2971944 | Lighthouse Laboratory Plymouth | Wellcome Sanger Institute for the COVID-19 Genomics UK (COG-UK) Consortium | Lighthouse Laboratory Plymouth and Alex Alderton, Roberto Amato, Jeffrey Barrett, Sonia Goncalves, Ewan Harrison, David K. Jackson, Ian Johnston, Dominic Kwiatkowski, Cordelia Langford, John Sillitoe on behalf of the Wellcome Sanger Institute COVID-19 Surveillance Team |
| EPI_ISL_2971945 | Department of Bacteria, Parasites and Fungi, Statens Serum Institut, Copenhagen, Denmark | Statens Serum Institut Bioinformatics and Microbial Genomics | Danish Covid-19 Genome Consortium |
| EPI_ISL_2971946, EPI_ISL_2971947 | Lighthouse Laboratory Plymouth | Wellcome Sanger Institute for the COVID-19 Genomics UK (COG-UK) Consortium | Lighthouse Laboratory Plymouth and Alex Alderton, Roberto Amato, Jeffrey Barrett, Sonia Goncalves, Ewan Harrison, David K. Jackson, Ian Johnston, Dominic Kwiatkowski, Cordelia Langford, John Sillitoe on behalf of the Wellcome Sanger Institute COVID-19 Surveillance Team |
| EPI_ISL_2971948 | Department of Bacteria, Parasites and Fungi, Statens Serum Institut, Copenhagen, Denmark | Statens Serum Institut Bioinformatics and Microbial Genomics | Danish Covid-19 Genome Consortium |
| EPI_ISL_2971949, EPI_ISL_2971950, EPI_ISL_2971951 | Lighthouse Laboratory Plymouth | Wellcome Sanger Institute for the COVID-19 Genomics UK (COG-UK) Consortium | Lighthouse Laboratory Plymouth and Alex Alderton, Roberto Amato, Jeffrey Barrett, Sonia Goncalves, Ewan Harrison, David K. Jackson, Ian Johnston, Dominic Kwiatkowski, Cordelia Langford, John Sillitoe on behalf of the Wellcome Sanger Institute COVID-19 Surveillance Team |
| EPI_ISL_2971952 | Department of Bacteria, Parasites and Fungi, Statens Serum Institut, Copenhagen, Denmark | Statens Serum Institut Bioinformatics and Microbial Genomics | Danish Covid-19 Genome Consortium |
| EPI_ISL_2971953 | Lighthouse Laboratory Plymouth | Wellcome Sanger Institute for the COVID-19 Genomics UK (COG-UK) Consortium | Lighthouse Laboratory Plymouth and Alex Alderton, Roberto Amato, Jeffrey Barrett, Sonia Goncalves, Ewan Harrison, David K. Jackson, Ian Johnston, Dominic Kwiatkowski, Cordelia Langford, John Sillitoe on behalf of the Wellcome Sanger Institute COVID-19 Surveillance Team |
| EPI_ISL_2971954 | Lighthouse Lab in Glasgow | Wellcome Sanger Institute for the COVID-19 Genomics UK (COG-UK) Consortium | Harper VanSteenhouse, Yumi Kasai, David Gray, Carol Clugston, Anna Dominiczak and Alex Alderton, Roberto Amato, Jeffrey Barrett, Sonia Goncalves, Ewan Harrison, David K. Jackson, Ian Johnston, Dominic Kwiatkowski, Cordelia Langford, John Sillitoe on behalf of the Wellcome Sanger Institute COVID-19 Surveillance Team |
| EPI_ISL_2971955 | Department of Bacteria, Parasites and Fungi, Statens Serum Institut, Copenhagen, Denmark | Statens Serum Institut Bioinformatics and Microbial Genomics | Danish Covid-19 Genome Consortium |
| EPI_ISL_2971956, EPI_ISL_2971957 | Lighthouse Laboratory Plymouth | Wellcome Sanger Institute for the COVID-19 Genomics UK (COG-UK) Consortium | Lighthouse Laboratory Plymouth and Alex Alderton, Roberto Amato, Jeffrey Barrett, Sonia Goncalves, Ewan Harrison, David K. Jackson, Ian Johnston, Dominic Kwiatkowski, Cordelia Langford, John Sillitoe on behalf of the Wellcome Sanger Institute COVID-19 Surveillance Team |
| EPI_ISL_2971958 | Department of Bacteria, Parasites and Fungi, Statens Serum Institut, Copenhagen, Denmark | Statens Serum Institut Bioinformatics and Microbial Genomics | Danish Covid-19 Genome Consortium |
| EPI_ISL_2971959 | Lighthouse Lab in Glasgow | Wellcome Sanger Institute for the COVID-19 Genomics UK (COG-UK) Consortium | Harper VanSteenhouse, Yumi Kasai, David Gray, Carol Clugston, Anna Dominiczak and Alex Alderton, Roberto Amato, Jeffrey Barrett, Sonia Goncalves, Ewan Harrison, David K. Jackson, Ian Johnston, Dominic Kwiatkowski, Cordelia Langford, John Sillitoe on behalf of the Wellcome Sanger Institute COVID-19 Surveillance Team |
| EPI_ISL_2971960 | Lighthouse Laboratory Plymouth | Wellcome Sanger Institute for the COVID-19 Genomics UK (COG-UK) Consortium | Lighthouse Laboratory Plymouth and Alex Alderton, Roberto Amato, Jeffrey Barrett, Sonia Goncalves, Ewan Harrison, David K. Jackson, Ian Johnston, Dominic Kwiatkowski, Cordelia Langford, John Sillitoe on behalf of the Wellcome Sanger Institute COVID-19 Surveillance Team |
| EPI_ISL_2971961 | Department of Bacteria, Parasites and Fungi, Statens Serum Institut, Copenhagen, Denmark | Statens Serum Institut Bioinformatics and Microbial Genomics | Danish Covid-19 Genome Consortium |
| EPI_ISL_2971962 | Lighthouse Laboratory Plymouth | Wellcome Sanger Institute for the COVID-19 Genomics UK (COG-UK) Consortium | Lighthouse Laboratory Plymouth and Alex Alderton, Roberto Amato, Jeffrey Barrett, Sonia Goncalves, Ewan Harrison, David K. Jackson, Ian Johnston, Dominic Kwiatkowski, Cordelia Langford, John Sillitoe on behalf of the Wellcome Sanger Institute COVID-19 Surveillance Team |
| EPI_ISL_2971963 | Lighthouse Lab in Glasgow | Wellcome Sanger Institute for the COVID-19 Genomics UK (COG-UK) Consortium | Harper VanSteenhouse, Yumi Kasai, David Gray, Carol Clugston, Anna Dominiczak and Alex Alderton, Roberto Amato, Jeffrey Barrett, Sonia Goncalves, Ewan Harrison, David K. Jackson, Ian Johnston, Dominic Kwiatkowski, Cordelia Langford, John Sillitoe on behalf of the Wellcome Sanger Institute COVID-19 Surveillance Team |
| EPI_ISL_2971964 | Department of Bacteria, Parasites and Fungi, Statens Serum Institut, Copenhagen, Denmark | Statens Serum Institut Bioinformatics and Microbial Genomics | Danish Covid-19 Genome Consortium |
| EPI_ISL_2971965, EPI_ISL_2971966, EPI_ISL_2971967 | Lighthouse Laboratory Plymouth | Wellcome Sanger Institute for the COVID-19 Genomics UK (COG-UK) Consortium | Lighthouse Laboratory Plymouth and Alex Alderton, Roberto Amato, Jeffrey Barrett, Sonia Goncalves, Ewan Harrison, David K. Jackson, Ian Johnston, Dominic Kwiatkowski, Cordelia Langford, John Sillitoe on behalf of the Wellcome Sanger Institute COVID-19 Surveillance Team |
| EPI_ISL_2971968 | Department of Bacteria, Parasites and Fungi, Statens Serum Institut, Copenhagen, Denmark | Statens Serum Institut Bioinformatics and Microbial Genomics | Danish Covid-19 Genome Consortium |
| EPI_ISL_2971969 | Lighthouse Laboratory Plymouth | Wellcome Sanger Institute for the COVID-19 Genomics UK (COG-UK) Consortium | Lighthouse Laboratory Plymouth and Alex Alderton, Roberto Amato, Jeffrey Barrett, Sonia Goncalves, Ewan Harrison, David K. Jackson, Ian Johnston, Dominic Kwiatkowski, Cordelia Langford, John Sillitoe on behalf of the Wellcome Sanger Institute COVID-19 Surveillance Team |
| EPI_ISL_2971970 | Randox Laboratories | Wellcome Sanger Institute for the COVID-19 Genomics UK (COG-UK) Consortium | Randox Laboratories and Alex Alderton, Roberto Amato, Jeffrey Barrett, Sonia Goncalves, Ewan Harrison, David K. Jackson, Ian Johnston, Dominic Kwiatkowski, Cordelia Langford, John Sillitoe on behalf of the Wellcome Sanger Institute COVID-19 Surveillance Team |
| EPI_ISL_2971971 | Department of Bacteria, Parasites and Fungi, Statens Serum Institut, Copenhagen, Denmark | Statens Serum Institut Bioinformatics and Microbial Genomics | Danish Covid-19 Genome Consortium |
| EPI_ISL_2971972, EPI_ISL_2971973, EPI_ISL_2971974 | Lighthouse Laboratory Plymouth | Wellcome Sanger Institute for the COVID-19 Genomics UK (COG-UK) Consortium | Lighthouse Laboratory Plymouth and Alex Alderton, Roberto Amato, Jeffrey Barrett, Sonia Goncalves, Ewan Harrison, David K. Jackson, Ian Johnston, Dominic Kwiatkowski, Cordelia Langford, John Sillitoe on behalf of the Wellcome Sanger Institute COVID-19 Surveillance Team |
| EPI_ISL_2971975 | Department of Bacteria, Parasites and Fungi, Statens Serum Institut, Copenhagen, Denmark | Statens Serum Institut Bioinformatics and Microbial Genomics | Danish Covid-19 Genome Consortium |
| EPI_ISL_2971976 | Lighthouse Laboratory Plymouth | Wellcome Sanger Institute for the COVID-19 Genomics UK (COG-UK) Consortium | Lighthouse Laboratory Plymouth and Alex Alderton, Roberto Amato, Jeffrey Barrett, Sonia Goncalves, Ewan Harrison, David K. Jackson, Ian Johnston, Dominic Kwiatkowski, Cordelia Langford, John Sillitoe on behalf of the Wellcome Sanger Institute COVID-19 Surveillance Team |
| EPI_ISL_2971977 | Lighthouse Lab in Glasgow | Wellcome Sanger Institute for the COVID-19 Genomics UK (COG-UK) Consortium | Harper VanSteenhouse, Yumi Kasai, David Gray, Carol Clugston, Anna Dominiczak and Alex Alderton, Roberto Amato, Jeffrey Barrett, Sonia Goncalves, Ewan Harrison, David K. Jackson, Ian Johnston, Dominic Kwiatkowski, Cordelia Langford, John Sillitoe on behalf of the Wellcome Sanger Institute COVID-19 Surveillance Team |
| EPI_ISL_2971978 | Department of Bacteria, Parasites and Fungi, Statens Serum Institut, Copenhagen, Denmark | Statens Serum Institut Bioinformatics and Microbial Genomics | Danish Covid-19 Genome Consortium |
| EPI_ISL_2971979 | Lighthouse Laboratory Plymouth | Wellcome Sanger Institute for the COVID-19 Genomics UK (COG-UK) Consortium | Lighthouse Laboratory Plymouth and Alex Alderton, Roberto Amato, Jeffrey Barrett, Sonia Goncalves, Ewan Harrison, David K. Jackson, Ian Johnston, Dominic Kwiatkowski, Cordelia Langford, John Sillitoe on behalf of the Wellcome Sanger Institute COVID-19 Surveillance Team |
| EPI_ISL_2971980 | Randox Laboratories | Wellcome Sanger Institute for the COVID-19 Genomics UK (COG-UK) Consortium | Randox Laboratories and Alex Alderton, Roberto Amato, Jeffrey Barrett, Sonia Goncalves, Ewan Harrison, David K. Jackson, Ian Johnston, Dominic Kwiatkowski, Cordelia Langford, John Sillitoe on behalf of the Wellcome Sanger Institute COVID-19 Surveillance Team |
| EPI_ISL_2971982, EPI_ISL_2971983, | Lighthouse Laboratory Plymouth | Wellcome Sanger Institute for the COVID-19 Genomics UK (COG-UK) Consortium | Lighthouse Laboratory Plymouth and Alex Alderton, Roberto Amato, Jeffrey Barrett, Sonia Goncalves, Ewan Harrison, David K. Jackson, Ian Johnston, Dominic Kwiatkowski, Cordelia Langford, John Sillitoe on behalf of the Wellcome Sanger Institute COVID-19 Surveillance Team |

| | | | |
|---|---|---|---|
| EPI_ISL_2971985, EPI_ISL_2971986 | | | |
| EPI_ISL_2971987 | Randox Laboratories | Wellcome Sanger Institute for the COVID-19 Genomics UK (COG-UK) Consortium | Randox Laboratories and Alex Alderton, Roberto Amato, Jeffrey Barrett, Sonia Goncalves, Ewan Harrison, David K. Jackson, Ian Johnston, Dominic Kwiatkowski, Cordelia Langford, John Sillitoe on behalf of the Wellcome Sanger Institute COVID-19 Surveillance Team |
| EPI_ISL_2971988 | Department of Bacteria, Parasites and Fungi, Statens Serum Institut, Copenhagen, Denmark | Statens Serum Institut Bioinformatics and Microbial Genomics | Danish Covid-19 Genome Consortium |
| EPI_ISL_2971989 | Lighthouse Laboratory Plymouth | Wellcome Sanger Institute for the COVID-19 Genomics UK (COG-UK) Consortium | Lighthouse Laboratory Plymouth and Alex Alderton, Roberto Amato, Jeffrey Barrett, Sonia Goncalves, Ewan Harrison, David K. Jackson, Ian Johnston, Dominic Kwiatkowski, Cordelia Langford, John Sillitoe on behalf of the Wellcome Sanger Institute COVID-19 Surveillance Team |
| EPI_ISL_2971990 | Randox Laboratories | Wellcome Sanger Institute for the COVID-19 Genomics UK (COG-UK) Consortium | Randox Laboratories and Alex Alderton, Roberto Amato, Jeffrey Barrett, Sonia Goncalves, Ewan Harrison, David K. Jackson, Ian Johnston, Dominic Kwiatkowski, Cordelia Langford, John Sillitoe on behalf of the Wellcome Sanger Institute COVID-19 Surveillance Team |
| EPI_ISL_2971991 | Department of Bacteria, Parasites and Fungi, Statens Serum Institut, Copenhagen, Denmark | Statens Serum Institut Bioinformatics and Microbial Genomics | Danish Covid-19 Genome Consortium |
| EPI_ISL_2971992 | Lighthouse Laboratory Plymouth | Wellcome Sanger Institute for the COVID-19 Genomics UK (COG-UK) Consortium | Lighthouse Laboratory Plymouth and Alex Alderton, Roberto Amato, Jeffrey Barrett, Sonia Goncalves, Ewan Harrison, David K. Jackson, Ian Johnston, Dominic Kwiatkowski, Cordelia Langford, John Sillitoe on behalf of the Wellcome Sanger Institute COVID-19 Surveillance Team |
| EPI_ISL_2971993 | Randox Laboratories | Wellcome Sanger Institute for the COVID-19 Genomics UK (COG-UK) Consortium | Randox Laboratories and Alex Alderton, Roberto Amato, Jeffrey Barrett, Sonia Goncalves, Ewan Harrison, David K. Jackson, Ian Johnston, Dominic Kwiatkowski, Cordelia Langford, John Sillitoe on behalf of the Wellcome Sanger Institute COVID-19 Surveillance Team |
| EPI_ISL_2971994 | Department of Bacteria, Parasites and Fungi, Statens Serum Institut, Copenhagen, Denmark | Statens Serum Institut Bioinformatics and Microbial Genomics | Danish Covid-19 Genome Consortium |
| EPI_ISL_2971995 | Lighthouse Laboratory Plymouth | Wellcome Sanger Institute for the COVID-19 Genomics UK (COG-UK) Consortium | Lighthouse Laboratory Plymouth and Alex Alderton, Roberto Amato, Jeffrey Barrett, Sonia Goncalves, Ewan Harrison, David K. Jackson, Ian Johnston, Dominic Kwiatkowski, Cordelia Langford, John Sillitoe on behalf of the Wellcome Sanger Institute COVID-19 Surveillance Team |
| EPI_ISL_2971996 | Lighthouse Lab in Glasgow | Wellcome Sanger Institute for the COVID-19 Genomics UK (COG-UK) Consortium | Harper VanSteenhouse, Yumi Kasai, David Gray, Carol Clugston, Anna Dominiczak and Alex Alderton, Roberto Amato, Jeffrey Barrett, Sonia Goncalves, Ewan Harrison, David K. Jackson, Ian Johnston, Dominic Kwiatkowski, Cordelia Langford, John Sillitoe on behalf of the Wellcome Sanger Institute COVID-19 Surveillance Team |
| EPI_ISL_2971998, EPI_ISL_2971999 | Lighthouse Laboratory Plymouth | Wellcome Sanger Institute for the COVID-19 Genomics UK (COG-UK) Consortium | Lighthouse Laboratory Plymouth and Alex Alderton, Roberto Amato, Jeffrey Barrett, Sonia Goncalves, Ewan Harrison, David K. Jackson, Ian Johnston, Dominic Kwiatkowski, Cordelia Langford, John Sillitoe on behalf of the Wellcome Sanger Institute COVID-19 Surveillance Team |
| EPI_ISL_2972000 | Department of Bacteria, Parasites and Fungi, Statens Serum Institut, Copenhagen, Denmark | Statens Serum Institut Bioinformatics and Microbial Genomics | Danish Covid-19 Genome Consortium |
| EPI_ISL_2972001 | Lighthouse Laboratory Plymouth | Wellcome Sanger Institute for the COVID-19 Genomics UK (COG-UK) Consortium | Lighthouse Laboratory Plymouth and Alex Alderton, Roberto Amato, Jeffrey Barrett, Sonia Goncalves, Ewan Harrison, David K. Jackson, Ian Johnston, Dominic Kwiatkowski, Cordelia Langford, John Sillitoe on behalf of the Wellcome Sanger Institute COVID-19 Surveillance Team |
| EPI_ISL_2972002 | Randox Laboratories | Wellcome Sanger Institute for the COVID-19 Genomics UK (COG-UK) Consortium | Randox Laboratories and Alex Alderton, Roberto Amato, Jeffrey Barrett, Sonia Goncalves, Ewan Harrison, David K. Jackson, Ian Johnston, Dominic Kwiatkowski, Cordelia Langford, John Sillitoe on behalf of the Wellcome Sanger Institute COVID-19 Surveillance Team |
| EPI_ISL_2972003 | Department of Bacteria, Parasites and Fungi, Statens Serum Institut, Copenhagen, Denmark | Statens Serum Institut Bioinformatics and Microbial Genomics | Danish Covid-19 Genome Consortium |
| EPI_ISL_2972004, EPI_ISL_2972005, EPI_ISL_2972006 | Lighthouse Laboratory Plymouth | Wellcome Sanger Institute for the COVID-19 Genomics UK (COG-UK) Consortium | Lighthouse Laboratory Plymouth and Alex Alderton, Roberto Amato, Jeffrey Barrett, Sonia Goncalves, Ewan Harrison, David K. Jackson, Ian Johnston, Dominic Kwiatkowski, Cordelia Langford, John Sillitoe on behalf of the Wellcome Sanger Institute COVID-19 Surveillance Team |
| EPI_ISL_2972007 | Department of Bacteria, Parasites and Fungi, Statens Serum Institut, Copenhagen, Denmark | Statens Serum Institut Bioinformatics and Microbial Genomics | Danish Covid-19 Genome Consortium |
| EPI_ISL_2972008, EPI_ISL_2972009 | Lighthouse Laboratory Plymouth | Wellcome Sanger Institute for the COVID-19 Genomics UK (COG-UK) Consortium | Lighthouse Laboratory Plymouth and Alex Alderton, Roberto Amato, Jeffrey Barrett, Sonia Goncalves, Ewan Harrison, David K. Jackson, Ian Johnston, Dominic Kwiatkowski, Cordelia Langford, John Sillitoe on behalf of the Wellcome Sanger Institute COVID-19 Surveillance Team |
| EPI_ISL_2972010 | Department of Bacteria, Parasites and Fungi, Statens Serum Institut, Copenhagen, Denmark | Statens Serum Institut Bioinformatics and Microbial Genomics | Danish Covid-19 Genome Consortium |
| EPI_ISL_2972011, EPI_ISL_2972012 | Lighthouse Laboratory Plymouth | Wellcome Sanger Institute for the COVID-19 Genomics UK (COG-UK) Consortium | Lighthouse Laboratory Plymouth and Alex Alderton, Roberto Amato, Jeffrey Barrett, Sonia Goncalves, Ewan Harrison, David K. Jackson, Ian Johnston, Dominic Kwiatkowski, Cordelia Langford, John Sillitoe on behalf of the Wellcome Sanger Institute COVID-19 Surveillance Team |
| EPI_ISL_2972013 | Department of Bacteria, Parasites and Fungi, Statens Serum Institut, Copenhagen, Denmark | Statens Serum Institut Bioinformatics and Microbial Genomics | Danish Covid-19 Genome Consortium |
| EPI_ISL_2972014 | Lighthouse Lab in Glasgow | Wellcome Sanger Institute for the COVID-19 Genomics UK (COG-UK) Consortium | Harper VanSteenhouse, Yumi Kasai, David Gray, Carol Clugston, Anna Dominiczak and Alex Alderton, Roberto Amato, Jeffrey Barrett, Sonia Goncalves, Ewan Harrison, David K. Jackson, Ian Johnston, Dominic Kwiatkowski, Cordelia Langford, John Sillitoe on behalf of the Wellcome Sanger Institute COVID-19 Surveillance Team |
| EPI_ISL_2972015, EPI_ISL_2972016 | Lighthouse Laboratory Plymouth | Wellcome Sanger Institute for the COVID-19 Genomics UK (COG-UK) Consortium | Lighthouse Laboratory Plymouth and Alex Alderton, Roberto Amato, Jeffrey Barrett, Sonia Goncalves, Ewan Harrison, David K. Jackson, Ian Johnston, Dominic Kwiatkowski, Cordelia Langford, John Sillitoe on behalf of the Wellcome Sanger Institute COVID-19 Surveillance Team |
| EPI_ISL_2972017 | Department of Bacteria, Parasites and Fungi, Statens Serum Institut, Copenhagen, Denmark | Statens Serum Institut Bioinformatics and Microbial Genomics | Danish Covid-19 Genome Consortium |
| EPI_ISL_2972018, EPI_ISL_2972019, EPI_ISL_2972020 | Lighthouse Laboratory Plymouth | Wellcome Sanger Institute for the COVID-19 Genomics UK (COG-UK) Consortium | Lighthouse Laboratory Plymouth and Alex Alderton, Roberto Amato, Jeffrey Barrett, Sonia Goncalves, Ewan Harrison, David K. Jackson, Ian Johnston, Dominic Kwiatkowski, Cordelia Langford, John Sillitoe on behalf of the Wellcome Sanger Institute COVID-19 Surveillance Team |
| EPI_ISL_2972021 | Department of Bacteria, Parasites and Fungi, Statens Serum Institut, Copenhagen, Denmark | Statens Serum Institut Bioinformatics and Microbial Genomics | Danish Covid-19 Genome Consortium |
| EPI_ISL_2972022, EPI_ISL_2972023 | Lighthouse Laboratory Plymouth | Wellcome Sanger Institute for the COVID-19 Genomics UK (COG-UK) Consortium | Lighthouse Laboratory Plymouth and Alex Alderton, Roberto Amato, Jeffrey Barrett, Sonia Goncalves, Ewan Harrison, David K. Jackson, Ian Johnston, Dominic Kwiatkowski, Cordelia Langford, John Sillitoe on behalf of the Wellcome Sanger Institute COVID-19 Surveillance Team |
| EPI_ISL_2972024 | Department of Bacteria, Parasites and Fungi, Statens Serum Institut, Copenhagen, Denmark | Statens Serum Institut Bioinformatics and Microbial Genomics | Danish Covid-19 Genome Consortium |
| EPI_ISL_2972025, EPI_ISL_2972026 | Lighthouse Laboratory Plymouth | Wellcome Sanger Institute for the COVID-19 Genomics UK (COG-UK) Consortium | Lighthouse Laboratory Plymouth and Alex Alderton, Roberto Amato, Jeffrey Barrett, Sonia Goncalves, Ewan Harrison, David K. Jackson, Ian Johnston, Dominic Kwiatkowski, Cordelia Langford, John Sillitoe on behalf of the Wellcome Sanger Institute COVID-19 Surveillance Team |
| EPI_ISL_2972027 | Lighthouse Lab in Glasgow | Wellcome Sanger Institute for the COVID-19 Genomics UK (COG-UK) Consortium | Harper VanSteenhouse, Yumi Kasai, David Gray, Carol Clugston, Anna Dominiczak and Alex Alderton, Roberto Amato, Jeffrey Barrett, Sonia Goncalves, Ewan Harrison, David K. Jackson, Ian Johnston, Dominic Kwiatkowski, Cordelia Langford, John Sillitoe on behalf of the Wellcome Sanger Institute COVID-19 Surveillance Team |
| EPI_ISL_2972028 | Department of Bacteria, Parasites and Fungi, Statens Serum Institut, Copenhagen, Denmark | Statens Serum Institut Bioinformatics and Microbial Genomics | Danish Covid-19 Genome Consortium |
| EPI_ISL_2972029, EPI_ISL_2972030 | Lighthouse Laboratory Plymouth | Wellcome Sanger Institute for the COVID-19 Genomics UK (COG-UK) Consortium | Lighthouse Laboratory Plymouth and Alex Alderton, Roberto Amato, Jeffrey Barrett, Sonia Goncalves, Ewan Harrison, David K. Jackson, Ian Johnston, Dominic Kwiatkowski, Cordelia Langford, John Sillitoe on behalf of the Wellcome Sanger Institute COVID-19 Surveillance Team |
| EPI_ISL_2972031 | Department of Bacteria, Parasites and Fungi, Statens Serum Institut, Copenhagen, Denmark | Statens Serum Institut Bioinformatics and Microbial Genomics | Danish Covid-19 Genome Consortium |
| EPI_ISL_2972032, EPI_ISL_2972033, EPI_ISL_2972034, EPI_ISL_2972036, EPI_ISL_2972037 | Lighthouse Laboratory Plymouth | Wellcome Sanger Institute for the COVID-19 Genomics UK (COG-UK) Consortium | Lighthouse Laboratory Plymouth and Alex Alderton, Roberto Amato, Jeffrey Barrett, Sonia Goncalves, Ewan Harrison, David K. Jackson, Ian Johnston, Dominic Kwiatkowski, Cordelia Langford, John Sillitoe on behalf of the Wellcome Sanger Institute COVID-19 Surveillance Team |
| EPI_ISL_2972038 | Department of Bacteria, Parasites and Fungi, Statens Serum Institut, Copenhagen, Denmark | Statens Serum Institut Bioinformatics and Microbial Genomics | Danish Covid-19 Genome Consortium |
| EPI_ISL_2972039 | Lighthouse Lab in Glasgow | Wellcome Sanger Institute for the COVID-19 Genomics UK (COG-UK) Consortium | Harper VanSteenhouse, Yumi Kasai, David Gray, Carol Clugston, Anna Dominiczak and Alex Alderton, Roberto Amato, Jeffrey Barrett, Sonia Goncalves, Ewan Harrison, David K. Jackson, Ian Johnston, Dominic Kwiatkowski, Cordelia Langford, John Sillitoe on behalf of the Wellcome Sanger Institute COVID-19 Surveillance Team |
| EPI_ISL_2972040 | Lighthouse Laboratory Plymouth | Wellcome Sanger Institute for the COVID-19 Genomics UK (COG-UK) Consortium | Lighthouse Laboratory Plymouth and Alex Alderton, Roberto Amato, Jeffrey Barrett, Sonia Goncalves, Ewan Harrison, David K. Jackson, Ian Johnston, Dominic Kwiatkowski, Cordelia Langford, John Sillitoe on behalf of the Wellcome Sanger Institute COVID-19 Surveillance Team |
| EPI_ISL_2972041 | Department of Bacteria, Parasites and Fungi, Statens Serum Institut, Copenhagen, Denmark | Statens Serum Institut Bioinformatics and Microbial Genomics | Danish Covid-19 Genome Consortium |
| EPI_ISL_2972042 | Lighthouse Laboratory Plymouth | Wellcome Sanger Institute for the COVID-19 Genomics UK (COG-UK) Consortium | Lighthouse Laboratory Plymouth and Alex Alderton, Roberto Amato, Jeffrey Barrett, Sonia Goncalves, Ewan Harrison, David K. Jackson, Ian Johnston, Dominic Kwiatkowski, Cordelia Langford, John Sillitoe on behalf of the Wellcome Sanger Institute COVID-19 Surveillance Team |
| EPI_ISL_2972043 | Randox Laboratories | Wellcome Sanger Institute for the COVID-19 Genomics UK (COG-UK) Consortium | Randox Laboratories and Alex Alderton, Roberto Amato, Jeffrey Barrett, Sonia Goncalves, Ewan Harrison, David K. Jackson, Ian Johnston, Dominic Kwiatkowski, Cordelia Langford, John Sillitoe on behalf of the Wellcome Sanger Institute COVID-19 Surveillance Team |
| EPI_ISL_2972044 | Department of Bacteria, Parasites and Fungi, Statens Serum Institut, Copenhagen, Denmark | Statens Serum Institut Bioinformatics and Microbial Genomics | Danish Covid-19 Genome Consortium |
| EPI_ISL_2972045, EPI_ISL_2972046 | Lighthouse Laboratory Plymouth | Wellcome Sanger Institute for the COVID-19 Genomics UK (COG-UK) Consortium | Lighthouse Laboratory Plymouth and Alex Alderton, Roberto Amato, Jeffrey Barrett, Sonia Goncalves, Ewan Harrison, David K. Jackson, Ian Johnston, Dominic Kwiatkowski, Cordelia Langford, John Sillitoe on behalf of the Wellcome Sanger Institute COVID-19 Surveillance Team |

| | | | |
|---|---|---|---|
| EPI_ISL_2972047 | Department of Bacteria, Parasites and Fungi, Statens Serum Institut, Copenhagen, Denmark | Statens Serum Institut Bioinformatics and Microbial Genomics | Danish Covid-19 Genome Consortium |
| EPI_ISL_2972048 | Randox Laboratories | Wellcome Sanger Institute for the COVID-19 Genomics UK (COG-UK) Consortium | Randox Laboratories and Alex Alderton, Roberto Amato, Jeffrey Barrett, Sonia Goncalves, Ewan Harrison, David K. Jackson, Ian Johnston, Dominic Kwiatkowski, Cordelia Langford, John Sillitoe on behalf of the Wellcome Sanger Institute COVID-19 Surveillance Team |
| EPI_ISL_2972049 | Lighthouse Laboratory Plymouth | Wellcome Sanger Institute for the COVID-19 Genomics UK (COG-UK) Consortium | Lighthouse Laboratory Plymouth and Alex Alderton, Roberto Amato, Jeffrey Barrett, Sonia Goncalves, Ewan Harrison, David K. Jackson, Ian Johnston, Dominic Kwiatkowski, Cordelia Langford, John Sillitoe on behalf of the Wellcome Sanger Institute COVID-19 Surveillance Team |
| EPI_ISL_2972050 | Randox Laboratories | Wellcome Sanger Institute for the COVID-19 Genomics UK (COG-UK) Consortium | Randox Laboratories and Alex Alderton, Roberto Amato, Jeffrey Barrett, Sonia Goncalves, Ewan Harrison, David K. Jackson, Ian Johnston, Dominic Kwiatkowski, Cordelia Langford, John Sillitoe on behalf of the Wellcome Sanger Institute COVID-19 Surveillance Team |
| EPI_ISL_2972051 | Department of Bacteria, Parasites and Fungi, Statens Serum Institut, Copenhagen, Denmark | Statens Serum Institut Bioinformatics and Microbial Genomics | Danish Covid-19 Genome Consortium |
| EPI_ISL_2972052, EPI_ISL_2972053 | Lighthouse Laboratory Plymouth | Wellcome Sanger Institute for the COVID-19 Genomics UK (COG-UK) Consortium | Lighthouse Laboratory Plymouth and Alex Alderton, Roberto Amato, Jeffrey Barrett, Sonia Goncalves, Ewan Harrison, David K. Jackson, Ian Johnston, Dominic Kwiatkowski, Cordelia Langford, John Sillitoe on behalf of the Wellcome Sanger Institute COVID-19 Surveillance Team |
| EPI_ISL_2972054 | Department of Bacteria, Parasites and Fungi, Statens Serum Institut, Copenhagen, Denmark | Statens Serum Institut Bioinformatics and Microbial Genomics | Danish Covid-19 Genome Consortium |
| EPI_ISL_2972055 | Lighthouse Laboratory Plymouth | Wellcome Sanger Institute for the COVID-19 Genomics UK (COG-UK) Consortium | Lighthouse Laboratory Plymouth and Alex Alderton, Roberto Amato, Jeffrey Barrett, Sonia Goncalves, Ewan Harrison, David K. Jackson, Ian Johnston, Dominic Kwiatkowski, Cordelia Langford, John Sillitoe on behalf of the Wellcome Sanger Institute COVID-19 Surveillance Team |
| EPI_ISL_2972056 | Department of Bacteria, Parasites and Fungi, Statens Serum Institut, Copenhagen, Denmark | Statens Serum Institut Bioinformatics and Microbial Genomics | Danish Covid-19 Genome Consortium |
| EPI_ISL_2972057 | Lighthouse Lab in Glasgow | Wellcome Sanger Institute for the COVID-19 Genomics UK (COG-UK) Consortium | Harper VanSteenhouse, Yumi Kasai, David Gray, Carol Clugston, Anna Dominiczak and Alex Alderton, Roberto Amato, Jeffrey Barrett, Sonia Goncalves, Ewan Harrison, David K. Jackson, Ian Johnston, Dominic Kwiatkowski, Cordelia Langford, John Sillitoe on behalf of the Wellcome Sanger Institute COVID-19 Surveillance Team |
| EPI_ISL_2972058, EPI_ISL_2972059 | Lighthouse Laboratory Plymouth | Wellcome Sanger Institute for the COVID-19 Genomics UK (COG-UK) Consortium | Lighthouse Laboratory Plymouth and Alex Alderton, Roberto Amato, Jeffrey Barrett, Sonia Goncalves, Ewan Harrison, David K. Jackson, Ian Johnston, Dominic Kwiatkowski, Cordelia Langford, John Sillitoe on behalf of the Wellcome Sanger Institute COVID-19 Surveillance Team |
| EPI_ISL_2972060 | Department of Bacteria, Parasites and Fungi, Statens Serum Institut, Copenhagen, Denmark | Statens Serum Institut Bioinformatics and Microbial Genomics | Danish Covid-19 Genome Consortium |
| EPI_ISL_2972061 | Lighthouse Laboratory Plymouth | Wellcome Sanger Institute for the COVID-19 Genomics UK (COG-UK) Consortium | Lighthouse Laboratory Plymouth and Alex Alderton, Roberto Amato, Jeffrey Barrett, Sonia Goncalves, Ewan Harrison, David K. Jackson, Ian Johnston, Dominic Kwiatkowski, Cordelia Langford, John Sillitoe on behalf of the Wellcome Sanger Institute COVID-19 Surveillance Team |
| EPI_ISL_2972062 | Randox Laboratories | Wellcome Sanger Institute for the COVID-19 Genomics UK (COG-UK) Consortium | Randox Laboratories and Alex Alderton, Roberto Amato, Jeffrey Barrett, Sonia Goncalves, Ewan Harrison, David K. Jackson, Ian Johnston, Dominic Kwiatkowski, Cordelia Langford, John Sillitoe on behalf of the Wellcome Sanger Institute COVID-19 Surveillance Team |
| EPI_ISL_2972063 | Department of Bacteria, Parasites and Fungi, Statens Serum Institut, Copenhagen, Denmark | Statens Serum Institut Bioinformatics and Microbial Genomics | Danish Covid-19 Genome Consortium |
| EPI_ISL_2972064, EPI_ISL_2972065 | Lighthouse Laboratory Plymouth | Wellcome Sanger Institute for the COVID-19 Genomics UK (COG-UK) Consortium | Lighthouse Laboratory Plymouth and Alex Alderton, Roberto Amato, Jeffrey Barrett, Sonia Goncalves, Ewan Harrison, David K. Jackson, Ian Johnston, Dominic Kwiatkowski, Cordelia Langford, John Sillitoe on behalf of the Wellcome Sanger Institute COVID-19 Surveillance Team |
| EPI_ISL_2972066 | Department of Bacteria, Parasites and Fungi, Statens Serum Institut, Copenhagen, Denmark | Statens Serum Institut Bioinformatics and Microbial Genomics | Danish Covid-19 Genome Consortium |
| EPI_ISL_2972067, EPI_ISL_2972068 | Lighthouse Laboratory Plymouth | Wellcome Sanger Institute for the COVID-19 Genomics UK (COG-UK) Consortium | Lighthouse Laboratory Plymouth and Alex Alderton, Roberto Amato, Jeffrey Barrett, Sonia Goncalves, Ewan Harrison, David K. Jackson, Ian Johnston, Dominic Kwiatkowski, Cordelia Langford, John Sillitoe on behalf of the Wellcome Sanger Institute COVID-19 Surveillance Team |
| EPI_ISL_2972069 | Department of Bacteria, Parasites and Fungi, Statens Serum Institut, Copenhagen, Denmark | Statens Serum Institut Bioinformatics and Microbial Genomics | Danish Covid-19 Genome Consortium |
| EPI_ISL_2972070 | Lighthouse Laboratory Plymouth | Wellcome Sanger Institute for the COVID-19 Genomics UK (COG-UK) Consortium | Lighthouse Laboratory Plymouth and Alex Alderton, Roberto Amato, Jeffrey Barrett, Sonia Goncalves, Ewan Harrison, David K. Jackson, Ian Johnston, Dominic Kwiatkowski, Cordelia Langford, John Sillitoe on behalf of the Wellcome Sanger Institute COVID-19 Surveillance Team |
| EPI_ISL_2972071 | Randox Laboratories | Wellcome Sanger Institute for the COVID-19 Genomics UK (COG-UK) Consortium | Randox Laboratories and Alex Alderton, Roberto Amato, Jeffrey Barrett, Sonia Goncalves, Ewan Harrison, David K. Jackson, Ian Johnston, Dominic Kwiatkowski, Cordelia Langford, John Sillitoe on behalf of the Wellcome Sanger Institute COVID-19 Surveillance Team |
| EPI_ISL_2972072 | Lighthouse Laboratory Plymouth | Wellcome Sanger Institute for the COVID-19 Genomics UK (COG-UK) Consortium | Lighthouse Laboratory Plymouth and Alex Alderton, Roberto Amato, Jeffrey Barrett, Sonia Goncalves, Ewan Harrison, David K. Jackson, Ian Johnston, Dominic Kwiatkowski, Cordelia Langford, John Sillitoe on behalf of the Wellcome Sanger Institute COVID-19 Surveillance Team |
| EPI_ISL_2972073 | Department of Bacteria, Parasites and Fungi, Statens Serum Institut, Copenhagen, Denmark | Statens Serum Institut Bioinformatics and Microbial Genomics | Danish Covid-19 Genome Consortium |
| EPI_ISL_2972074, EPI_ISL_2972075 | Lighthouse Laboratory Plymouth | Wellcome Sanger Institute for the COVID-19 Genomics UK (COG-UK) Consortium | Lighthouse Laboratory Plymouth and Alex Alderton, Roberto Amato, Jeffrey Barrett, Sonia Goncalves, Ewan Harrison, David K. Jackson, Ian Johnston, Dominic Kwiatkowski, Cordelia Langford, John Sillitoe on behalf of the Wellcome Sanger Institute COVID-19 Surveillance Team |
| EPI_ISL_2972076 | Department of Bacteria, Parasites and Fungi, Statens Serum Institut, Copenhagen, Denmark | Statens Serum Institut Bioinformatics and Microbial Genomics | Danish Covid-19 Genome Consortium |
| EPI_ISL_2972077, EPI_ISL_2972078 | Lighthouse Laboratory Plymouth | Wellcome Sanger Institute for the COVID-19 Genomics UK (COG-UK) Consortium | Lighthouse Laboratory Plymouth and Alex Alderton, Roberto Amato, Jeffrey Barrett, Sonia Goncalves, Ewan Harrison, David K. Jackson, Ian Johnston, Dominic Kwiatkowski, Cordelia Langford, John Sillitoe on behalf of the Wellcome Sanger Institute COVID-19 Surveillance Team |
| EPI_ISL_2972079 | Department of Bacteria, Parasites and Fungi, Statens Serum Institut, Copenhagen, Denmark | Statens Serum Institut Bioinformatics and Microbial Genomics | Danish Covid-19 Genome Consortium |
| EPI_ISL_2972080, EPI_ISL_2972081 | Lighthouse Laboratory Plymouth | Wellcome Sanger Institute for the COVID-19 Genomics UK (COG-UK) Consortium | Lighthouse Laboratory Plymouth and Alex Alderton, Roberto Amato, Jeffrey Barrett, Sonia Goncalves, Ewan Harrison, David K. Jackson, Ian Johnston, Dominic Kwiatkowski, Cordelia Langford, John Sillitoe on behalf of the Wellcome Sanger Institute COVID-19 Surveillance Team |
| EPI_ISL_2972082 | Lighthouse Lab in Glasgow | Wellcome Sanger Institute for the COVID-19 Genomics UK (COG-UK) Consortium | Harper VanSteenhouse, Yumi Kasai, David Gray, Carol Clugston, Anna Dominiczak and Alex Alderton, Roberto Amato, Jeffrey Barrett, Sonia Goncalves, Ewan Harrison, David K. Jackson, Ian Johnston, Dominic Kwiatkowski, Cordelia Langford, John Sillitoe on behalf of the Wellcome Sanger Institute COVID-19 Surveillance Team |
| EPI_ISL_2972083 | Lighthouse Laboratory Plymouth | Wellcome Sanger Institute for the COVID-19 Genomics UK (COG-UK) Consortium | Lighthouse Laboratory Plymouth and Alex Alderton, Roberto Amato, Jeffrey Barrett, Sonia Goncalves, Ewan Harrison, David K. Jackson, Ian Johnston, Dominic Kwiatkowski, Cordelia Langford, John Sillitoe on behalf of the Wellcome Sanger Institute COVID-19 Surveillance Team |
| EPI_ISL_2972084 | Lighthouse Lab in Alderley Park | Wellcome Sanger Institute for the COVID-19 Genomics UK (COG-UK) Consortium | Jacquelyn Wynn, Mairead Hyland, The Lighthouse Lab in Alderley Park and Alex Alderton, Roberto Amato, Jeffrey Barrett, Sonia Goncalves, Ewan Harrison, David K. Jackson, Ian Johnston, Dominic Kwiatkowski, Cordelia Langford, John Sillitoe on behalf of the Wellcome Sanger Institute COVID-19 Surveillance Team |
| EPI_ISL_2972085 | Department of Bacteria, Parasites and Fungi, Statens Serum Institut, Copenhagen, Denmark | Statens Serum Institut Bioinformatics and Microbial Genomics | Danish Covid-19 Genome Consortium |
| EPI_ISL_2972086, EPI_ISL_2972087 | Lighthouse Lab in Alderley Park | Wellcome Sanger Institute for the COVID-19 Genomics UK (COG-UK) Consortium | Jacquelyn Wynn, Mairead Hyland, The Lighthouse Lab in Alderley Park and Alex Alderton, Roberto Amato, Jeffrey Barrett, Sonia Goncalves, Ewan Harrison, David K. Jackson, Ian Johnston, Dominic Kwiatkowski, Cordelia Langford, John Sillitoe on behalf of the Wellcome Sanger Institute COVID-19 Surveillance Team |
| EPI_ISL_2972088, EPI_ISL_2972090 | Lighthouse Lab in Milton Keynes | Wellcome Sanger Institute for the COVID-19 Genomics UK (COG-UK) Consortium | The Lighthouse Lab in Milton Keynes and Alex Alderton, Roberto Amato, Jeffrey Barrett, Sonia Goncalves, Ewan Harrison, David K. Jackson, Ian Johnston, Dominic Kwiatkowski, Cordelia Langford, John Sillitoe on behalf of the Wellcome Sanger Institute COVID-19 Surveillance Team |
| EPI_ISL_2972091 | Lighthouse Lab in Alderley Park | Wellcome Sanger Institute for the COVID-19 Genomics UK (COG-UK) Consortium | Jacquelyn Wynn, Mairead Hyland, The Lighthouse Lab in Alderley Park and Alex Alderton, Roberto Amato, Jeffrey Barrett, Sonia Goncalves, Ewan Harrison, David K. Jackson, Ian Johnston, Dominic Kwiatkowski, Cordelia Langford, John Sillitoe on behalf of the Wellcome Sanger Institute COVID-19 Surveillance Team |
| EPI_ISL_2972092 | Department of Bacteria, Parasites and Fungi, Statens Serum Institut, Copenhagen, Denmark | Statens Serum Institut Bioinformatics and Microbial Genomics | Danish Covid-19 Genome Consortium |
| EPI_ISL_2972093 | Lighthouse Lab in Alderley Park | Wellcome Sanger Institute for the COVID-19 Genomics UK (COG-UK) Consortium | Jacquelyn Wynn, Mairead Hyland, The Lighthouse Lab in Alderley Park and Alex Alderton, Roberto Amato, Jeffrey Barrett, Sonia Goncalves, Ewan Harrison, David K. Jackson, Ian Johnston, Dominic Kwiatkowski, Cordelia Langford, John Sillitoe on behalf of the Wellcome Sanger Institute COVID-19 Surveillance Team |
| EPI_ISL_2972094 | Lighthouse Lab in Milton Keynes | Wellcome Sanger Institute for the COVID-19 Genomics UK (COG-UK) Consortium | The Lighthouse Lab in Milton Keynes and Alex Alderton, Roberto Amato, Jeffrey Barrett, Sonia Goncalves, Ewan Harrison, David K. Jackson, Ian Johnston, Dominic Kwiatkowski, Cordelia Langford, John Sillitoe on behalf of the Wellcome Sanger Institute COVID-19 Surveillance Team |
| EPI_ISL_2972095 | Department of Bacteria, Parasites and Fungi, Statens Serum Institut, Copenhagen, Denmark | Statens Serum Institut Bioinformatics and Microbial Genomics | Danish Covid-19 Genome Consortium |
| EPI_ISL_2972096, EPI_ISL_2972097, EPI_ISL_2972098 | Lighthouse Lab in Alderley Park | Wellcome Sanger Institute for the COVID-19 Genomics UK (COG-UK) Consortium | Jacquelyn Wynn, Mairead Hyland, The Lighthouse Lab in Alderley Park and Alex Alderton, Roberto Amato, Jeffrey Barrett, Sonia Goncalves, Ewan Harrison, David K. Jackson, Ian Johnston, Dominic Kwiatkowski, Cordelia Langford, John Sillitoe on behalf of the Wellcome Sanger Institute COVID-19 Surveillance Team |
| EPI_ISL_2972099 | Department of Bacteria, Parasites and Fungi, Statens Serum Institut, Copenhagen, Denmark | Statens Serum Institut Bioinformatics and Microbial Genomics | Danish Covid-19 Genome Consortium |
| EPI_ISL_2972100 | Lighthouse Lab in Alderley Park | Wellcome Sanger Institute for the COVID-19 Genomics UK (COG-UK) Consortium | Jacquelyn Wynn, Mairead Hyland, The Lighthouse Lab in Alderley Park and Alex Alderton, Roberto Amato, Jeffrey Barrett, Sonia Goncalves, Ewan Harrison, David K. Jackson, Ian Johnston, Dominic Kwiatkowski, Cordelia Langford, John Sillitoe on behalf of the Wellcome Sanger Institute COVID-19 Surveillance Team |
| EPI_ISL_2972101 | Lighthouse Lab in Milton Keynes | Wellcome Sanger Institute for the COVID-19 Genomics UK (COG-UK) Consortium | The Lighthouse Lab in Milton Keynes and Alex Alderton, Roberto Amato, Jeffrey Barrett, Sonia Goncalves, Ewan Harrison, David K. Jackson, Ian Johnston, Dominic Kwiatkowski, Cordelia Langford, John Sillitoe on behalf of the Wellcome Sanger Institute COVID-19 Surveillance Team |
| EPI_ISL_2972102 | Department of Bacteria, Parasites and Fungi, Statens Serum Institut, Copenhagen, Denmark | Statens Serum Institut Bioinformatics and Microbial Genomics | Danish Covid-19 Genome Consortium |
| EPI_ISL_2972103, EPI_ISL_2972104, | Lighthouse Lab in Milton Keynes | Wellcome Sanger Institute for the COVID-19 Genomics UK (COG-UK) Consortium | The Lighthouse Lab in Milton Keynes and Alex Alderton, Roberto Amato, Jeffrey Barrett, Sonia Goncalves, Ewan Harrison, David K. Jackson, Ian Johnston, Dominic Kwiatkowski, Cordelia Langford, John Sillitoe on behalf of the Wellcome Sanger Institute COVID-19 Surveillance Team |

| | | | |
|---|---|---|---|
| EPI_ISL_2972105 | | | |
| EPI_ISL_2972106 | Department of Bacteria, Parasites and Fungi, Statens Serum Institut, Copenhagen, Denmark | Statens Serum Institut Bioinformatics and Microbial Genomics | Danish Covid-19 Genome Consortium |
| EPI_ISL_2972107 | Lighthouse Lab in Alderley Park | Wellcome Sanger Institute for the COVID-19 Genomics UK (COG-UK) Consortium | Jacquelyn Wynn, Mairead Hyland, The Lighthouse Lab in Alderley Park and Alex Alderton, Roberto Amato, Jeffrey Barrett, Sonia Goncalves, Ewan Harrison, David K. Jackson, Ian Johnston, Dominic Kwiatkowski, Cordelia Langford, John Sillitoe on behalf of the Wellcome Sanger Institute COVID-19 Surveillance Team |
| EPI_ISL_2972108, EPI_ISL_2972109 | Lighthouse Lab in Milton Keynes | Wellcome Sanger Institute for the COVID-19 Genomics UK (COG-UK) Consortium | The Lighthouse Lab in Milton Keynes and Alex Alderton, Roberto Amato, Jeffrey Barrett, Sonia Goncalves, Ewan Harrison, David K. Jackson, Ian Johnston, Dominic Kwiatkowski, Cordelia Langford, John Sillitoe on behalf of the Wellcome Sanger Institute COVID-19 Surveillance Team |
| EPI_ISL_2972110 | Department of Bacteria, Parasites and Fungi, Statens Serum Institut, Copenhagen, Denmark | Statens Serum Institut Bioinformatics and Microbial Genomics | Danish Covid-19 Genome Consortium |
| EPI_ISL_2972111 | Lighthouse Lab in Alderley Park | Wellcome Sanger Institute for the COVID-19 Genomics UK (COG-UK) Consortium | Jacquelyn Wynn, Mairead Hyland, The Lighthouse Lab in Alderley Park and Alex Alderton, Roberto Amato, Jeffrey Barrett, Sonia Goncalves, Ewan Harrison, David K. Jackson, Ian Johnston, Dominic Kwiatkowski, Cordelia Langford, John Sillitoe on behalf of the Wellcome Sanger Institute COVID-19 Surveillance Team |
| EPI_ISL_2972112 | Lighthouse Lab in Milton Keynes | Wellcome Sanger Institute for the COVID-19 Genomics UK (COG-UK) Consortium | The Lighthouse Lab in Milton Keynes and Alex Alderton, Roberto Amato, Jeffrey Barrett, Sonia Goncalves, Ewan Harrison, David K. Jackson, Ian Johnston, Dominic Kwiatkowski, Cordelia Langford, John Sillitoe on behalf of the Wellcome Sanger Institute COVID-19 Surveillance Team |
| EPI_ISL_2972113 | Department of Bacteria, Parasites and Fungi, Statens Serum Institut, Copenhagen, Denmark | Statens Serum Institut Bioinformatics and Microbial Genomics | Danish Covid-19 Genome Consortium |
| EPI_ISL_2972114, EPI_ISL_2972115 | Lighthouse Lab in Alderley Park | Wellcome Sanger Institute for the COVID-19 Genomics UK (COG-UK) Consortium | Jacquelyn Wynn, Mairead Hyland, The Lighthouse Lab in Alderley Park and Alex Alderton, Roberto Amato, Jeffrey Barrett, Sonia Goncalves, Ewan Harrison, David K. Jackson, Ian Johnston, Dominic Kwiatkowski, Cordelia Langford, John Sillitoe on behalf of the Wellcome Sanger Institute COVID-19 Surveillance Team |
| EPI_ISL_2972116 | Department of Bacteria, Parasites and Fungi, Statens Serum Institut, Copenhagen, Denmark | Statens Serum Institut Bioinformatics and Microbial Genomics | Danish Covid-19 Genome Consortium |
| EPI_ISL_2972117, EPI_ISL_2972118 | Lighthouse Lab in Alderley Park | Wellcome Sanger Institute for the COVID-19 Genomics UK (COG-UK) Consortium | Jacquelyn Wynn, Mairead Hyland, The Lighthouse Lab in Alderley Park and Alex Alderton, Roberto Amato, Jeffrey Barrett, Sonia Goncalves, Ewan Harrison, David K. Jackson, Ian Johnston, Dominic Kwiatkowski, Cordelia Langford, John Sillitoe on behalf of the Wellcome Sanger Institute COVID-19 Surveillance Team |
| EPI_ISL_2972119 | Lighthouse Lab in Milton Keynes | Wellcome Sanger Institute for the COVID-19 Genomics UK (COG-UK) Consortium | The Lighthouse Lab in Milton Keynes and Alex Alderton, Roberto Amato, Jeffrey Barrett, Sonia Goncalves, Ewan Harrison, David K. Jackson, Ian Johnston, Dominic Kwiatkowski, Cordelia Langford, John Sillitoe on behalf of the Wellcome Sanger Institute COVID-19 Surveillance Team |
| EPI_ISL_2972120 | Department of Bacteria, Parasites and Fungi, Statens Serum Institut, Copenhagen, Denmark | Statens Serum Institut Bioinformatics and Microbial Genomics | Danish Covid-19 Genome Consortium |
| EPI_ISL_2972121, EPI_ISL_2972122, EPI_ISL_2972124 | Lighthouse Lab in Alderley Park | Wellcome Sanger Institute for the COVID-19 Genomics UK (COG-UK) Consortium | Jacquelyn Wynn, Mairead Hyland, The Lighthouse Lab in Alderley Park and Alex Alderton, Roberto Amato, Jeffrey Barrett, Sonia Goncalves, Ewan Harrison, David K. Jackson, Ian Johnston, Dominic Kwiatkowski, Cordelia Langford, John Sillitoe on behalf of the Wellcome Sanger Institute COVID-19 Surveillance Team |
| EPI_ISL_2972125 | Lighthouse Lab in Milton Keynes | Wellcome Sanger Institute for the COVID-19 Genomics UK (COG-UK) Consortium | The Lighthouse Lab in Milton Keynes and Alex Alderton, Roberto Amato, Jeffrey Barrett, Sonia Goncalves, Ewan Harrison, David K. Jackson, Ian Johnston, Dominic Kwiatkowski, Cordelia Langford, John Sillitoe on behalf of the Wellcome Sanger Institute COVID-19 Surveillance Team |
| EPI_ISL_2972126 | Department of Bacteria, Parasites and Fungi, Statens Serum Institut, Copenhagen, Denmark | Statens Serum Institut Bioinformatics and Microbial Genomics | Danish Covid-19 Genome Consortium |
| EPI_ISL_2972127 | Lighthouse Lab in Alderley Park | Wellcome Sanger Institute for the COVID-19 Genomics UK (COG-UK) Consortium | Jacquelyn Wynn, Mairead Hyland, The Lighthouse Lab in Alderley Park and Alex Alderton, Roberto Amato, Jeffrey Barrett, Sonia Goncalves, Ewan Harrison, David K. Jackson, Ian Johnston, Dominic Kwiatkowski, Cordelia Langford, John Sillitoe on behalf of the Wellcome Sanger Institute COVID-19 Surveillance Team |
| EPI_ISL_2972128 | Lighthouse Lab in Milton Keynes | Wellcome Sanger Institute for the COVID-19 Genomics UK (COG-UK) Consortium | The Lighthouse Lab in Milton Keynes and Alex Alderton, Roberto Amato, Jeffrey Barrett, Sonia Goncalves, Ewan Harrison, David K. Jackson, Ian Johnston, Dominic Kwiatkowski, Cordelia Langford, John Sillitoe on behalf of the Wellcome Sanger Institute COVID-19 Surveillance Team |
| EPI_ISL_2972129 | Lighthouse Lab in Alderley Park | Wellcome Sanger Institute for the COVID-19 Genomics UK (COG-UK) Consortium | Jacquelyn Wynn, Mairead Hyland, The Lighthouse Lab in Alderley Park and Alex Alderton, Roberto Amato, Jeffrey Barrett, Sonia Goncalves, Ewan Harrison, David K. Jackson, Ian Johnston, Dominic Kwiatkowski, Cordelia Langford, John Sillitoe on behalf of the Wellcome Sanger Institute COVID-19 Surveillance Team |
| EPI_ISL_2972130 | Department of Bacteria, Parasites and Fungi, Statens Serum Institut, Copenhagen, Denmark | Statens Serum Institut Bioinformatics and Microbial Genomics | Danish Covid-19 Genome Consortium |
| EPI_ISL_2972131 | Lighthouse Lab in Milton Keynes | Wellcome Sanger Institute for the COVID-19 Genomics UK (COG-UK) Consortium | The Lighthouse Lab in Milton Keynes and Alex Alderton, Roberto Amato, Jeffrey Barrett, Sonia Goncalves, Ewan Harrison, David K. Jackson, Ian Johnston, Dominic Kwiatkowski, Cordelia Langford, John Sillitoe on behalf of the Wellcome Sanger Institute COVID-19 Surveillance Team |
| EPI_ISL_2972132 | Lighthouse Lab in Alderley Park | Wellcome Sanger Institute for the COVID-19 Genomics UK (COG-UK) Consortium | Jacquelyn Wynn, Mairead Hyland, The Lighthouse Lab in Alderley Park and Alex Alderton, Roberto Amato, Jeffrey Barrett, Sonia Goncalves, Ewan Harrison, David K. Jackson, Ian Johnston, Dominic Kwiatkowski, Cordelia Langford, John Sillitoe on behalf of the Wellcome Sanger Institute COVID-19 Surveillance Team |
| EPI_ISL_2972133 | Department of Bacteria, Parasites and Fungi, Statens Serum Institut, Copenhagen, Denmark | Statens Serum Institut Bioinformatics and Microbial Genomics | Danish Covid-19 Genome Consortium |
| EPI_ISL_2972134, EPI_ISL_2972135, EPI_ISL_2972137, EPI_ISL_2972138 | Lighthouse Lab in Alderley Park | Wellcome Sanger Institute for the COVID-19 Genomics UK (COG-UK) Consortium | Jacquelyn Wynn, Mairead Hyland, The Lighthouse Lab in Alderley Park and Alex Alderton, Roberto Amato, Jeffrey Barrett, Sonia Goncalves, Ewan Harrison, David K. Jackson, Ian Johnston, Dominic Kwiatkowski, Cordelia Langford, John Sillitoe on behalf of the Wellcome Sanger Institute COVID-19 Surveillance Team |
| EPI_ISL_2972139 | Department of Bacteria, Parasites and Fungi, Statens Serum Institut, Copenhagen, Denmark | Statens Serum Institut Bioinformatics and Microbial Genomics | Danish Covid-19 Genome Consortium |
| EPI_ISL_2972140, EPI_ISL_2972141 | Lighthouse Lab in Milton Keynes | Wellcome Sanger Institute for the COVID-19 Genomics UK (COG-UK) Consortium | The Lighthouse Lab in Milton Keynes and Alex Alderton, Roberto Amato, Jeffrey Barrett, Sonia Goncalves, Ewan Harrison, David K. Jackson, Ian Johnston, Dominic Kwiatkowski, Cordelia Langford, John Sillitoe on behalf of the Wellcome Sanger Institute COVID-19 Surveillance Team |
| EPI_ISL_2972142 | Department of Bacteria, Parasites and Fungi, Statens Serum Institut, Copenhagen, Denmark | Statens Serum Institut Bioinformatics and Microbial Genomics | Danish Covid-19 Genome Consortium |
| EPI_ISL_2972143, EPI_ISL_2972144, EPI_ISL_2972145 | Lighthouse Lab in Alderley Park | Wellcome Sanger Institute for the COVID-19 Genomics UK (COG-UK) Consortium | Jacquelyn Wynn, Mairead Hyland, The Lighthouse Lab in Alderley Park and Alex Alderton, Roberto Amato, Jeffrey Barrett, Sonia Goncalves, Ewan Harrison, David K. Jackson, Ian Johnston, Dominic Kwiatkowski, Cordelia Langford, John Sillitoe on behalf of the Wellcome Sanger Institute COVID-19 Surveillance Team |
| EPI_ISL_2972146 | Department of Bacteria, Parasites and Fungi, Statens Serum Institut, Copenhagen, Denmark | Statens Serum Institut Bioinformatics and Microbial Genomics | Danish Covid-19 Genome Consortium |
| EPI_ISL_2972147, EPI_ISL_2972148 | Lighthouse Lab in Alderley Park | Wellcome Sanger Institute for the COVID-19 Genomics UK (COG-UK) Consortium | Jacquelyn Wynn, Mairead Hyland, The Lighthouse Lab in Alderley Park and Alex Alderton, Roberto Amato, Jeffrey Barrett, Sonia Goncalves, Ewan Harrison, David K. Jackson, Ian Johnston, Dominic Kwiatkowski, Cordelia Langford, John Sillitoe on behalf of the Wellcome Sanger Institute COVID-19 Surveillance Team |
| EPI_ISL_2972150, EPI_ISL_2972151 | Lighthouse Lab in Milton Keynes | Wellcome Sanger Institute for the COVID-19 Genomics UK (COG-UK) Consortium | The Lighthouse Lab in Milton Keynes and Alex Alderton, Roberto Amato, Jeffrey Barrett, Sonia Goncalves, Ewan Harrison, David K. Jackson, Ian Johnston, Dominic Kwiatkowski, Cordelia Langford, John Sillitoe on behalf of the Wellcome Sanger Institute COVID-19 Surveillance Team |
| EPI_ISL_2972152 | Department of Bacteria, Parasites and Fungi, Statens Serum Institut, Copenhagen, Denmark | Statens Serum Institut Bioinformatics and Microbial Genomics | Danish Covid-19 Genome Consortium |
| EPI_ISL_2972153 | Lighthouse Lab in Alderley Park | Wellcome Sanger Institute for the COVID-19 Genomics UK (COG-UK) Consortium | Jacquelyn Wynn, Mairead Hyland, The Lighthouse Lab in Alderley Park and Alex Alderton, Roberto Amato, Jeffrey Barrett, Sonia Goncalves, Ewan Harrison, David K. Jackson, Ian Johnston, Dominic Kwiatkowski, Cordelia Langford, John Sillitoe on behalf of the Wellcome Sanger Institute COVID-19 Surveillance Team |
| EPI_ISL_2972154 | Lighthouse Lab in Milton Keynes | Wellcome Sanger Institute for the COVID-19 Genomics UK (COG-UK) Consortium | The Lighthouse Lab in Milton Keynes and Alex Alderton, Roberto Amato, Jeffrey Barrett, Sonia Goncalves, Ewan Harrison, David K. Jackson, Ian Johnston, Dominic Kwiatkowski, Cordelia Langford, John Sillitoe on behalf of the Wellcome Sanger Institute COVID-19 Surveillance Team |
| EPI_ISL_2972155 | Department of Bacteria, Parasites and Fungi, Statens Serum Institut, Copenhagen, Denmark | Statens Serum Institut Bioinformatics and Microbial Genomics | Danish Covid-19 Genome Consortium |
| EPI_ISL_2972156, EPI_ISL_2972157 | Lighthouse Lab in Alderley Park | Wellcome Sanger Institute for the COVID-19 Genomics UK (COG-UK) Consortium | Jacquelyn Wynn, Mairead Hyland, The Lighthouse Lab in Alderley Park and Alex Alderton, Roberto Amato, Jeffrey Barrett, Sonia Goncalves, Ewan Harrison, David K. Jackson, Ian Johnston, Dominic Kwiatkowski, Cordelia Langford, John Sillitoe on behalf of the Wellcome Sanger Institute COVID-19 Surveillance Team |
| EPI_ISL_2972158 | Department of Bacteria, Parasites and Fungi, Statens Serum Institut, Copenhagen, Denmark | Statens Serum Institut Bioinformatics and Microbial Genomics | Danish Covid-19 Genome Consortium |
| EPI_ISL_2972159, EPI_ISL_2972160 | Lighthouse Lab in Milton Keynes | Wellcome Sanger Institute for the COVID-19 Genomics UK (COG-UK) Consortium | The Lighthouse Lab in Milton Keynes and Alex Alderton, Roberto Amato, Jeffrey Barrett, Sonia Goncalves, Ewan Harrison, David K. Jackson, Ian Johnston, Dominic Kwiatkowski, Cordelia Langford, John Sillitoe on behalf of the Wellcome Sanger Institute COVID-19 Surveillance Team |
| EPI_ISL_2972161 | Lighthouse Lab in Alderley Park | Wellcome Sanger Institute for the COVID-19 Genomics UK (COG-UK) Consortium | Jacquelyn Wynn, Mairead Hyland, The Lighthouse Lab in Alderley Park and Alex Alderton, Roberto Amato, Jeffrey Barrett, Sonia Goncalves, Ewan Harrison, David K. Jackson, Ian Johnston, Dominic Kwiatkowski, Cordelia Langford, John Sillitoe on behalf of the Wellcome Sanger Institute COVID-19 Surveillance Team |
| EPI_ISL_2972162 | Department of Bacteria, Parasites and Fungi, Statens Serum Institut, Copenhagen, Denmark | Statens Serum Institut Bioinformatics and Microbial Genomics | Danish Covid-19 Genome Consortium |
| EPI_ISL_2972163, EPI_ISL_2972164 | Lighthouse Lab in Alderley Park | Wellcome Sanger Institute for the COVID-19 Genomics UK (COG-UK) Consortium | Jacquelyn Wynn, Mairead Hyland, The Lighthouse Lab in Alderley Park and Alex Alderton, Roberto Amato, Jeffrey Barrett, Sonia Goncalves, Ewan Harrison, David K. Jackson, Ian Johnston, Dominic Kwiatkowski, Cordelia Langford, John Sillitoe on behalf of the Wellcome Sanger Institute COVID-19 Surveillance Team |
| EPI_ISL_2972165 | Department of Bacteria, Parasites and Fungi, Statens Serum Institut, Copenhagen, Denmark | Statens Serum Institut Bioinformatics and Microbial Genomics | Danish Covid-19 Genome Consortium |
| EPI_ISL_2972166 | Lighthouse Lab in Alderley Park | Wellcome Sanger Institute for the COVID-19 Genomics UK (COG-UK) Consortium | Jacquelyn Wynn, Mairead Hyland, The Lighthouse Lab in Alderley Park and Alex Alderton, Roberto Amato, Jeffrey Barrett, Sonia Goncalves, Ewan Harrison, David K. Jackson, Ian Johnston, Dominic Kwiatkowski, Cordelia Langford, John Sillitoe on behalf of the Wellcome Sanger Institute COVID-19 Surveillance Team |
| EPI_ISL_2972167 | Lighthouse Lab in Milton Keynes | Wellcome Sanger Institute for the COVID-19 Genomics UK (COG-UK) Consortium | The Lighthouse Lab in Milton Keynes and Alex Alderton, Roberto Amato, Jeffrey Barrett, Sonia Goncalves, Ewan Harrison, David K. Jackson, Ian Johnston, Dominic Kwiatkowski, Cordelia Langford, John Sillitoe on behalf of the Wellcome Sanger Institute COVID-19 Surveillance Team |

| | | | |
|---|---|---|---|
| EPI_ISL_2972168 | Department of Bacteria, Parasites and Fungi, Statens Serum Institut, Copenhagen, Denmark | Statens Serum Institut Bioinformatics and Microbial Genomics | Danish Covid-19 Genome Consortium |
| EPI_ISL_2972169, EPI_ISL_2972170 | Lighthouse Lab in Alderley Park | Wellcome Sanger Institute for the COVID-19 Genomics UK (COG-UK) Consortium | Jacquelyn Wynn, Mairead Hyland, The Lighthouse Lab in Alderley Park and Alex Alderton, Roberto Amato, Jeffrey Barrett, Sonia Goncalves, Ewan Harrison, David K. Jackson, Ian Johnston, Dominic Kwiatkowski, Cordelia Langford, John Sillitoe on behalf of the Wellcome Sanger Institute COVID-19 Surveillance Team |
| EPI_ISL_2972171 | Department of Bacteria, Parasites and Fungi, Statens Serum Institut, Copenhagen, Denmark | Statens Serum Institut Bioinformatics and Microbial Genomics | Danish Covid-19 Genome Consortium |
| EPI_ISL_2972172 | Lighthouse Lab in Alderley Park | Wellcome Sanger Institute for the COVID-19 Genomics UK (COG-UK) Consortium | Jacquelyn Wynn, Mairead Hyland, The Lighthouse Lab in Alderley Park and Alex Alderton, Roberto Amato, Jeffrey Barrett, Sonia Goncalves, Ewan Harrison, David K. Jackson, Ian Johnston, Dominic Kwiatkowski, Cordelia Langford, John Sillitoe on behalf of the Wellcome Sanger Institute COVID-19 Surveillance Team |
| EPI_ISL_2972173 | Lighthouse Lab in Milton Keynes | Wellcome Sanger Institute for the COVID-19 Genomics UK (COG-UK) Consortium | The Lighthouse Lab in Milton Keynes and Alex Alderton, Roberto Amato, Jeffrey Barrett, Sonia Goncalves, Ewan Harrison, David K. Jackson, Ian Johnston, Dominic Kwiatkowski, Cordelia Langford, John Sillitoe on behalf of the Wellcome Sanger Institute COVID-19 Surveillance Team |
| EPI_ISL_2972174 | Lighthouse Lab in Alderley Park | Wellcome Sanger Institute for the COVID-19 Genomics UK (COG-UK) Consortium | Jacquelyn Wynn, Mairead Hyland, The Lighthouse Lab in Alderley Park and Alex Alderton, Roberto Amato, Jeffrey Barrett, Sonia Goncalves, Ewan Harrison, David K. Jackson, Ian Johnston, Dominic Kwiatkowski, Cordelia Langford, John Sillitoe on behalf of the Wellcome Sanger Institute COVID-19 Surveillance Team |
| EPI_ISL_2972175 | Department of Bacteria, Parasites and Fungi, Statens Serum Institut, Copenhagen, Denmark | Statens Serum Institut Bioinformatics and Microbial Genomics | Danish Covid-19 Genome Consortium |
| EPI_ISL_2972176, EPI_ISL_2972177, EPI_ISL_2972178 | Lighthouse Lab in Alderley Park | Wellcome Sanger Institute for the COVID-19 Genomics UK (COG-UK) Consortium | Jacquelyn Wynn, Mairead Hyland, The Lighthouse Lab in Alderley Park and Alex Alderton, Roberto Amato, Jeffrey Barrett, Sonia Goncalves, Ewan Harrison, David K. Jackson, Ian Johnston, Dominic Kwiatkowski, Cordelia Langford, John Sillitoe on behalf of the Wellcome Sanger Institute COVID-19 Surveillance Team |
| EPI_ISL_2972179 | Department of Bacteria, Parasites and Fungi, Statens Serum Institut, Copenhagen, Denmark | Statens Serum Institut Bioinformatics and Microbial Genomics | Danish Covid-19 Genome Consortium |
| EPI_ISL_2972180 | Lighthouse Lab in Milton Keynes | Wellcome Sanger Institute for the COVID-19 Genomics UK (COG-UK) Consortium | The Lighthouse Lab in Milton Keynes and Alex Alderton, Roberto Amato, Jeffrey Barrett, Sonia Goncalves, Ewan Harrison, David K. Jackson, Ian Johnston, Dominic Kwiatkowski, Cordelia Langford, John Sillitoe on behalf of the Wellcome Sanger Institute COVID-19 Surveillance Team |
| EPI_ISL_2972181 | Lighthouse Lab in Alderley Park | Wellcome Sanger Institute for the COVID-19 Genomics UK (COG-UK) Consortium | Jacquelyn Wynn, Mairead Hyland, The Lighthouse Lab in Alderley Park and Alex Alderton, Roberto Amato, Jeffrey Barrett, Sonia Goncalves, Ewan Harrison, David K. Jackson, Ian Johnston, Dominic Kwiatkowski, Cordelia Langford, John Sillitoe on behalf of the Wellcome Sanger Institute COVID-19 Surveillance Team |
| EPI_ISL_2972182 | Department of Bacteria, Parasites and Fungi, Statens Serum Institut, Copenhagen, Denmark | Statens Serum Institut Bioinformatics and Microbial Genomics | Danish Covid-19 Genome Consortium |
| EPI_ISL_2972183, EPI_ISL_2972184 | Lighthouse Lab in Alderley Park | Wellcome Sanger Institute for the COVID-19 Genomics UK (COG-UK) Consortium | Jacquelyn Wynn, Mairead Hyland, The Lighthouse Lab in Alderley Park and Alex Alderton, Roberto Amato, Jeffrey Barrett, Sonia Goncalves, Ewan Harrison, David K. Jackson, Ian Johnston, Dominic Kwiatkowski, Cordelia Langford, John Sillitoe on behalf of the Wellcome Sanger Institute COVID-19 Surveillance Team |
| EPI_ISL_2972186 | Lighthouse Lab in Milton Keynes | Wellcome Sanger Institute for the COVID-19 Genomics UK (COG-UK) Consortium | The Lighthouse Lab in Milton Keynes and Alex Alderton, Roberto Amato, Jeffrey Barrett, Sonia Goncalves, Ewan Harrison, David K. Jackson, Ian Johnston, Dominic Kwiatkowski, Cordelia Langford, John Sillitoe on behalf of the Wellcome Sanger Institute COVID-19 Surveillance Team |
| EPI_ISL_2972187 | Lighthouse Lab in Alderley Park | Wellcome Sanger Institute for the COVID-19 Genomics UK (COG-UK) Consortium | Jacquelyn Wynn, Mairead Hyland, The Lighthouse Lab in Alderley Park and Alex Alderton, Roberto Amato, Jeffrey Barrett, Sonia Goncalves, Ewan Harrison, David K. Jackson, Ian Johnston, Dominic Kwiatkowski, Cordelia Langford, John Sillitoe on behalf of the Wellcome Sanger Institute COVID-19 Surveillance Team |
| EPI_ISL_2972188 | Department of Bacteria, Parasites and Fungi, Statens Serum Institut, Copenhagen, Denmark | Statens Serum Institut Bioinformatics and Microbial Genomics | Danish Covid-19 Genome Consortium |
| EPI_ISL_2972189 | Lighthouse Lab in Alderley Park | Wellcome Sanger Institute for the COVID-19 Genomics UK (COG-UK) Consortium | Jacquelyn Wynn, Mairead Hyland, The Lighthouse Lab in Alderley Park and Alex Alderton, Roberto Amato, Jeffrey Barrett, Sonia Goncalves, Ewan Harrison, David K. Jackson, Ian Johnston, Dominic Kwiatkowski, Cordelia Langford, John Sillitoe on behalf of the Wellcome Sanger Institute COVID-19 Surveillance Team |
| EPI_ISL_2972190 | Lighthouse Lab in Milton Keynes | Wellcome Sanger Institute for the COVID-19 Genomics UK (COG-UK) Consortium | The Lighthouse Lab in Milton Keynes and Alex Alderton, Roberto Amato, Jeffrey Barrett, Sonia Goncalves, Ewan Harrison, David K. Jackson, Ian Johnston, Dominic Kwiatkowski, Cordelia Langford, John Sillitoe on behalf of the Wellcome Sanger Institute COVID-19 Surveillance Team |
| EPI_ISL_2972191 | Lighthouse Lab in Alderley Park | Wellcome Sanger Institute for the COVID-19 Genomics UK (COG-UK) Consortium | Jacquelyn Wynn, Mairead Hyland, The Lighthouse Lab in Alderley Park and Alex Alderton, Roberto Amato, Jeffrey Barrett, Sonia Goncalves, Ewan Harrison, David K. Jackson, Ian Johnston, Dominic Kwiatkowski, Cordelia Langford, John Sillitoe on behalf of the Wellcome Sanger Institute COVID-19 Surveillance Team |
| EPI_ISL_2972192 | Department of Bacteria, Parasites and Fungi, Statens Serum Institut, Copenhagen, Denmark | Statens Serum Institut Bioinformatics and Microbial Genomics | Danish Covid-19 Genome Consortium |
| EPI_ISL_2972193 | Lighthouse Lab in Milton Keynes | Wellcome Sanger Institute for the COVID-19 Genomics UK (COG-UK) Consortium | The Lighthouse Lab in Milton Keynes and Alex Alderton, Roberto Amato, Jeffrey Barrett, Sonia Goncalves, Ewan Harrison, David K. Jackson, Ian Johnston, Dominic Kwiatkowski, Cordelia Langford, John Sillitoe on behalf of the Wellcome Sanger Institute COVID-19 Surveillance Team |
| EPI_ISL_2972194 | Lighthouse Lab in Alderley Park | Wellcome Sanger Institute for the COVID-19 Genomics UK (COG-UK) Consortium | Jacquelyn Wynn, Mairead Hyland, The Lighthouse Lab in Alderley Park and Alex Alderton, Roberto Amato, Jeffrey Barrett, Sonia Goncalves, Ewan Harrison, David K. Jackson, Ian Johnston, Dominic Kwiatkowski, Cordelia Langford, John Sillitoe on behalf of the Wellcome Sanger Institute COVID-19 Surveillance Team |
| EPI_ISL_2972195 | Department of Bacteria, Parasites and Fungi, Statens Serum Institut, Copenhagen, Denmark | Statens Serum Institut Bioinformatics and Microbial Genomics | Danish Covid-19 Genome Consortium |
| EPI_ISL_2972196 | Lighthouse Lab in Alderley Park | Wellcome Sanger Institute for the COVID-19 Genomics UK (COG-UK) Consortium | Jacquelyn Wynn, Mairead Hyland, The Lighthouse Lab in Alderley Park and Alex Alderton, Roberto Amato, Jeffrey Barrett, Sonia Goncalves, Ewan Harrison, David K. Jackson, Ian Johnston, Dominic Kwiatkowski, Cordelia Langford, John Sillitoe on behalf of the Wellcome Sanger Institute COVID-19 Surveillance Team |
| EPI_ISL_2972197 | Lighthouse Lab in Milton Keynes | Wellcome Sanger Institute for the COVID-19 Genomics UK (COG-UK) Consortium | The Lighthouse Lab in Milton Keynes and Alex Alderton, Roberto Amato, Jeffrey Barrett, Sonia Goncalves, Ewan Harrison, David K. Jackson, Ian Johnston, Dominic Kwiatkowski, Cordelia Langford, John Sillitoe on behalf of the Wellcome Sanger Institute COVID-19 Surveillance Team |
| EPI_ISL_2972198 | Department of Bacteria, Parasites and Fungi, Statens Serum Institut, Copenhagen, Denmark | Statens Serum Institut Bioinformatics and Microbial Genomics | Danish Covid-19 Genome Consortium |
| EPI_ISL_2972199 | Lighthouse Lab in Milton Keynes | Wellcome Sanger Institute for the COVID-19 Genomics UK (COG-UK) Consortium | The Lighthouse Lab in Milton Keynes and Alex Alderton, Roberto Amato, Jeffrey Barrett, Sonia Goncalves, Ewan Harrison, David K. Jackson, Ian Johnston, Dominic Kwiatkowski, Cordelia Langford, John Sillitoe on behalf of the Wellcome Sanger Institute COVID-19 Surveillance Team |
| EPI_ISL_2972200 | Lighthouse Lab in Alderley Park | Wellcome Sanger Institute for the COVID-19 Genomics UK (COG-UK) Consortium | Jacquelyn Wynn, Mairead Hyland, The Lighthouse Lab in Alderley Park and Alex Alderton, Roberto Amato, Jeffrey Barrett, Sonia Goncalves, Ewan Harrison, David K. Jackson, Ian Johnston, Dominic Kwiatkowski, Cordelia Langford, John Sillitoe on behalf of the Wellcome Sanger Institute COVID-19 Surveillance Team |
| EPI_ISL_2972201 | Department of Bacteria, Parasites and Fungi, Statens Serum Institut, Copenhagen, Denmark | Statens Serum Institut Bioinformatics and Microbial Genomics | Danish Covid-19 Genome Consortium |
| EPI_ISL_2972202, EPI_ISL_2972203, EPI_ISL_2972204 | Lighthouse Lab in Alderley Park | Wellcome Sanger Institute for the COVID-19 Genomics UK (COG-UK) Consortium | Jacquelyn Wynn, Mairead Hyland, The Lighthouse Lab in Alderley Park and Alex Alderton, Roberto Amato, Jeffrey Barrett, Sonia Goncalves, Ewan Harrison, David K. Jackson, Ian Johnston, Dominic Kwiatkowski, Cordelia Langford, John Sillitoe on behalf of the Wellcome Sanger Institute COVID-19 Surveillance Team |
| EPI_ISL_2972205 | Department of Bacteria, Parasites and Fungi, Statens Serum Institut, Copenhagen, Denmark | Statens Serum Institut Bioinformatics and Microbial Genomics | Danish Covid-19 Genome Consortium |
| EPI_ISL_2972206 | Lighthouse Lab in Alderley Park | Wellcome Sanger Institute for the COVID-19 Genomics UK (COG-UK) Consortium | Jacquelyn Wynn, Mairead Hyland, The Lighthouse Lab in Alderley Park and Alex Alderton, Roberto Amato, Jeffrey Barrett, Sonia Goncalves, Ewan Harrison, David K. Jackson, Ian Johnston, Dominic Kwiatkowski, Cordelia Langford, John Sillitoe on behalf of the Wellcome Sanger Institute COVID-19 Surveillance Team |
| EPI_ISL_2972207 | Lighthouse Lab in Milton Keynes | Wellcome Sanger Institute for the COVID-19 Genomics UK (COG-UK) Consortium | The Lighthouse Lab in Milton Keynes and Alex Alderton, Roberto Amato, Jeffrey Barrett, Sonia Goncalves, Ewan Harrison, David K. Jackson, Ian Johnston, Dominic Kwiatkowski, Cordelia Langford, John Sillitoe on behalf of the Wellcome Sanger Institute COVID-19 Surveillance Team |
| EPI_ISL_2972208 | Lighthouse Lab in Alderley Park | Wellcome Sanger Institute for the COVID-19 Genomics UK (COG-UK) Consortium | Jacquelyn Wynn, Mairead Hyland, The Lighthouse Lab in Alderley Park and Alex Alderton, Roberto Amato, Jeffrey Barrett, Sonia Goncalves, Ewan Harrison, David K. Jackson, Ian Johnston, Dominic Kwiatkowski, Cordelia Langford, John Sillitoe on behalf of the Wellcome Sanger Institute COVID-19 Surveillance Team |
| EPI_ISL_2972209 | Department of Bacteria, Parasites and Fungi, Statens Serum Institut, Copenhagen, Denmark | Statens Serum Institut Bioinformatics and Microbial Genomics | Danish Covid-19 Genome Consortium |
| EPI_ISL_2972210, EPI_ISL_2972211 | Lighthouse Lab in Alderley Park | Wellcome Sanger Institute for the COVID-19 Genomics UK (COG-UK) Consortium | Jacquelyn Wynn, Mairead Hyland, The Lighthouse Lab in Alderley Park and Alex Alderton, Roberto Amato, Jeffrey Barrett, Sonia Goncalves, Ewan Harrison, David K. Jackson, Ian Johnston, Dominic Kwiatkowski, Cordelia Langford, John Sillitoe on behalf of the Wellcome Sanger Institute COVID-19 Surveillance Team |
| EPI_ISL_2972212 | Department of Bacteria, Parasites and Fungi, Statens Serum Institut, Copenhagen, Denmark | Statens Serum Institut Bioinformatics and Microbial Genomics | Danish Covid-19 Genome Consortium |
| EPI_ISL_2972213 | Lighthouse Lab in Alderley Park | Wellcome Sanger Institute for the COVID-19 Genomics UK (COG-UK) Consortium | Jacquelyn Wynn, Mairead Hyland, The Lighthouse Lab in Alderley Park and Alex Alderton, Roberto Amato, Jeffrey Barrett, Sonia Goncalves, Ewan Harrison, David K. Jackson, Ian Johnston, Dominic Kwiatkowski, Cordelia Langford, John Sillitoe on behalf of the Wellcome Sanger Institute COVID-19 Surveillance Team |
| EPI_ISL_2972214, EPI_ISL_2972215 | Lighthouse Lab in Milton Keynes | Wellcome Sanger Institute for the COVID-19 Genomics UK (COG-UK) Consortium | The Lighthouse Lab in Milton Keynes and Alex Alderton, Roberto Amato, Jeffrey Barrett, Sonia Goncalves, Ewan Harrison, David K. Jackson, Ian Johnston, Dominic Kwiatkowski, Cordelia Langford, John Sillitoe on behalf of the Wellcome Sanger Institute COVID-19 Surveillance Team |
| EPI_ISL_2972216 | Department of Bacteria, Parasites and Fungi, Statens Serum Institut, Copenhagen, Denmark | Statens Serum Institut Bioinformatics and Microbial Genomics | Danish Covid-19 Genome Consortium |
| EPI_ISL_2972217, EPI_ISL_2972218 | Lighthouse Lab in Alderley Park | Wellcome Sanger Institute for the COVID-19 Genomics UK (COG-UK) Consortium | Jacquelyn Wynn, Mairead Hyland, The Lighthouse Lab in Alderley Park and Alex Alderton, Roberto Amato, Jeffrey Barrett, Sonia Goncalves, Ewan Harrison, David K. Jackson, Ian Johnston, Dominic Kwiatkowski, Cordelia Langford, John Sillitoe on behalf of the Wellcome Sanger Institute COVID-19 Surveillance Team |
| EPI_ISL_2972220 | Lighthouse Lab in Milton Keynes | Wellcome Sanger Institute for the COVID-19 Genomics UK (COG-UK) Consortium | The Lighthouse Lab in Milton Keynes and Alex Alderton, Roberto Amato, Jeffrey Barrett, Sonia Goncalves, Ewan Harrison, David K. Jackson, Ian Johnston, Dominic Kwiatkowski, Cordelia Langford, John Sillitoe on behalf of the Wellcome Sanger Institute COVID-19 Surveillance Team |
| EPI_ISL_2972221, EPI_ISL_2972223, EPI_ISL_2972224 | Lighthouse Lab in Alderley Park | Wellcome Sanger Institute for the COVID-19 Genomics UK (COG-UK) Consortium | Jacquelyn Wynn, Mairead Hyland, The Lighthouse Lab in Alderley Park and Alex Alderton, Roberto Amato, Jeffrey Barrett, Sonia Goncalves, Ewan Harrison, David K. Jackson, Ian Johnston, Dominic Kwiatkowski, Cordelia Langford, John Sillitoe on behalf of the Wellcome Sanger Institute COVID-19 Surveillance Team |
| EPI_ISL_2972225 | Department of Bacteria, Parasites and Fungi, Statens Serum Institut, Copenhagen, Denmark | Statens Serum Institut Bioinformatics and Microbial Genomics | Danish Covid-19 Genome Consortium |

| | | | |
|---|---|---|---|
| EPI_ISL_2972226, EPI_ISL_2972227, EPI_ISL_2972228 | Lighthouse Lab in Alderley Park | Wellcome Sanger Institute for the COVID-19 Genomics UK (COG-UK) Consortium | Jacquelyn Wynn, Mairead Hyland, The Lighthouse Lab in Alderley Park and Alex Alderton, Roberto Amato, Jeffrey Barrett, Sonia Goncalves, Ewan Harrison, David K. Jackson, Ian Johnston, Dominic Kwiatkowski, Cordelia Langford, John Sillitoe on behalf of the Wellcome Sanger Institute COVID-19 Surveillance Team |
| EPI_ISL_2972229 | Department of Bacteria, Parasites and Fungi, Statens Serum Institut, Copenhagen, Denmark | Statens Serum Institut Bioinformatics and Microbial Genomics | Danish Covid-19 Genome Consortium |
| EPI_ISL_2972230 | Lighthouse Lab in Alderley Park | Wellcome Sanger Institute for the COVID-19 Genomics UK (COG-UK) Consortium | Jacquelyn Wynn, Mairead Hyland, The Lighthouse Lab in Alderley Park and Alex Alderton, Roberto Amato, Jeffrey Barrett, Sonia Goncalves, Ewan Harrison, David K. Jackson, Ian Johnston, Dominic Kwiatkowski, Cordelia Langford, John Sillitoe on behalf of the Wellcome Sanger Institute COVID-19 Surveillance Team |
| EPI_ISL_2972231 | Lighthouse Lab in Milton Keynes | Wellcome Sanger Institute for the COVID-19 Genomics UK (COG-UK) Consortium | The Lighthouse Lab in Milton Keynes and Alex Alderton, Roberto Amato, Jeffrey Barrett, Sonia Goncalves, Ewan Harrison, David K. Jackson, Ian Johnston, Dominic Kwiatkowski, Cordelia Langford, John Sillitoe on behalf of the Wellcome Sanger Institute COVID-19 Surveillance Team |
| EPI_ISL_2972232 | Department of Bacteria, Parasites and Fungi, Statens Serum Institut, Copenhagen, Denmark | Statens Serum Institut Bioinformatics and Microbial Genomics | Danish Covid-19 Genome Consortium |
| EPI_ISL_2972233 | Lighthouse Lab in Milton Keynes | Wellcome Sanger Institute for the COVID-19 Genomics UK (COG-UK) Consortium | The Lighthouse Lab in Milton Keynes and Alex Alderton, Roberto Amato, Jeffrey Barrett, Sonia Goncalves, Ewan Harrison, David K. Jackson, Ian Johnston, Dominic Kwiatkowski, Cordelia Langford, John Sillitoe on behalf of the Wellcome Sanger Institute COVID-19 Surveillance Team |
| EPI_ISL_2972234, EPI_ISL_2972235 | Lighthouse Lab in Alderley Park | Wellcome Sanger Institute for the COVID-19 Genomics UK (COG-UK) Consortium | Jacquelyn Wynn, Mairead Hyland, The Lighthouse Lab in Alderley Park and Alex Alderton, Roberto Amato, Jeffrey Barrett, Sonia Goncalves, Ewan Harrison, David K. Jackson, Ian Johnston, Dominic Kwiatkowski, Cordelia Langford, John Sillitoe on behalf of the Wellcome Sanger Institute COVID-19 Surveillance Team |
| EPI_ISL_2972236 | Department of Bacteria, Parasites and Fungi, Statens Serum Institut, Copenhagen, Denmark | Statens Serum Institut Bioinformatics and Microbial Genomics | Danish Covid-19 Genome Consortium |
| EPI_ISL_2972237, EPI_ISL_2972238 | Lighthouse Lab in Alderley Park | Wellcome Sanger Institute for the COVID-19 Genomics UK (COG-UK) Consortium | Jacquelyn Wynn, Mairead Hyland, The Lighthouse Lab in Alderley Park and Alex Alderton, Roberto Amato, Jeffrey Barrett, Sonia Goncalves, Ewan Harrison, David K. Jackson, Ian Johnston, Dominic Kwiatkowski, Cordelia Langford, John Sillitoe on behalf of the Wellcome Sanger Institute COVID-19 Surveillance Team |
| EPI_ISL_2972239 | Department of Clinical Microbiology, Odense University Hospital, Odense, Denmark | Statens Serum Institut Bioinformatics and Microbial Genomics | Danish Covid-19 Genome Consortium |
| EPI_ISL_2972240 | Lighthouse Lab in Alderley Park | Wellcome Sanger Institute for the COVID-19 Genomics UK (COG-UK) Consortium | Jacquelyn Wynn, Mairead Hyland, The Lighthouse Lab in Alderley Park and Alex Alderton, Roberto Amato, Jeffrey Barrett, Sonia Goncalves, Ewan Harrison, David K. Jackson, Ian Johnston, Dominic Kwiatkowski, Cordelia Langford, John Sillitoe on behalf of the Wellcome Sanger Institute COVID-19 Surveillance Team |
| EPI_ISL_2972241 | Lighthouse Lab in Milton Keynes | Wellcome Sanger Institute for the COVID-19 Genomics UK (COG-UK) Consortium | The Lighthouse Lab in Milton Keynes and Alex Alderton, Roberto Amato, Jeffrey Barrett, Sonia Goncalves, Ewan Harrison, David K. Jackson, Ian Johnston, Dominic Kwiatkowski, Cordelia Langford, John Sillitoe on behalf of the Wellcome Sanger Institute COVID-19 Surveillance Team |
| EPI_ISL_2972242 | Department of Bacteria, Parasites and Fungi, Statens Serum Institut, Copenhagen, Denmark | Statens Serum Institut Bioinformatics and Microbial Genomics | Danish Covid-19 Genome Consortium |
| EPI_ISL_2972243 | Lighthouse Lab in Milton Keynes | Wellcome Sanger Institute for the COVID-19 Genomics UK (COG-UK) Consortium | The Lighthouse Lab in Milton Keynes and Alex Alderton, Roberto Amato, Jeffrey Barrett, Sonia Goncalves, Ewan Harrison, David K. Jackson, Ian Johnston, Dominic Kwiatkowski, Cordelia Langford, John Sillitoe on behalf of the Wellcome Sanger Institute COVID-19 Surveillance Team |
| EPI_ISL_2972244, EPI_ISL_2972245 | Lighthouse Lab in Alderley Park | Wellcome Sanger Institute for the COVID-19 Genomics UK (COG-UK) Consortium | Jacquelyn Wynn, Mairead Hyland, The Lighthouse Lab in Alderley Park and Alex Alderton, Roberto Amato, Jeffrey Barrett, Sonia Goncalves, Ewan Harrison, David K. Jackson, Ian Johnston, Dominic Kwiatkowski, Cordelia Langford, John Sillitoe on behalf of the Wellcome Sanger Institute COVID-19 Surveillance Team |
| EPI_ISL_2972246 | Department of Bacteria, Parasites and Fungi, Statens Serum Institut, Copenhagen, Denmark | Statens Serum Institut Bioinformatics and Microbial Genomics | Danish Covid-19 Genome Consortium |
| EPI_ISL_2972247, EPI_ISL_2972248 | Lighthouse Lab in Alderley Park | Wellcome Sanger Institute for the COVID-19 Genomics UK (COG-UK) Consortium | Jacquelyn Wynn, Mairead Hyland, The Lighthouse Lab in Alderley Park and Alex Alderton, Roberto Amato, Jeffrey Barrett, Sonia Goncalves, Ewan Harrison, David K. Jackson, Ian Johnston, Dominic Kwiatkowski, Cordelia Langford, John Sillitoe on behalf of the Wellcome Sanger Institute COVID-19 Surveillance Team |
| EPI_ISL_2972249 | Department of Bacteria, Parasites and Fungi, Statens Serum Institut, Copenhagen, Denmark | Statens Serum Institut Bioinformatics and Microbial Genomics | Danish Covid-19 Genome Consortium |
| EPI_ISL_2972250 | Lighthouse Lab in Alderley Park | Wellcome Sanger Institute for the COVID-19 Genomics UK (COG-UK) Consortium | Jacquelyn Wynn, Mairead Hyland, The Lighthouse Lab in Alderley Park and Alex Alderton, Roberto Amato, Jeffrey Barrett, Sonia Goncalves, Ewan Harrison, David K. Jackson, Ian Johnston, Dominic Kwiatkowski, Cordelia Langford, John Sillitoe on behalf of the Wellcome Sanger Institute COVID-19 Surveillance Team |
| EPI_ISL_2972251 | Lighthouse Lab in Milton Keynes | Wellcome Sanger Institute for the COVID-19 Genomics UK (COG-UK) Consortium | The Lighthouse Lab in Milton Keynes and Alex Alderton, Roberto Amato, Jeffrey Barrett, Sonia Goncalves, Ewan Harrison, David K. Jackson, Ian Johnston, Dominic Kwiatkowski, Cordelia Langford, John Sillitoe on behalf of the Wellcome Sanger Institute COVID-19 Surveillance Team |
| EPI_ISL_2972252 | Department of Bacteria, Parasites and Fungi, Statens Serum Institut, Copenhagen, Denmark | Statens Serum Institut Bioinformatics and Microbial Genomics | Danish Covid-19 Genome Consortium |
| EPI_ISL_2972253 | Lighthouse Lab in Alderley Park | Wellcome Sanger Institute for the COVID-19 Genomics UK (COG-UK) Consortium | Jacquelyn Wynn, Mairead Hyland, The Lighthouse Lab in Alderley Park and Alex Alderton, Roberto Amato, Jeffrey Barrett, Sonia Goncalves, Ewan Harrison, David K. Jackson, Ian Johnston, Dominic Kwiatkowski, Cordelia Langford, John Sillitoe on behalf of the Wellcome Sanger Institute COVID-19 Surveillance Team |
| EPI_ISL_2972254 | Lighthouse Lab in Milton Keynes | Wellcome Sanger Institute for the COVID-19 Genomics UK (COG-UK) Consortium | The Lighthouse Lab in Milton Keynes and Alex Alderton, Roberto Amato, Jeffrey Barrett, Sonia Goncalves, Ewan Harrison, David K. Jackson, Ian Johnston, Dominic Kwiatkowski, Cordelia Langford, John Sillitoe on behalf of the Wellcome Sanger Institute COVID-19 Surveillance Team |
| EPI_ISL_2972255 | Department of Bacteria, Parasites and Fungi, Statens Serum Institut, Copenhagen, Denmark | Statens Serum Institut Bioinformatics and Microbial Genomics | Danish Covid-19 Genome Consortium |
| EPI_ISL_2972256, EPI_ISL_2972257, EPI_ISL_2972258 | Lighthouse Lab in Alderley Park | Wellcome Sanger Institute for the COVID-19 Genomics UK (COG-UK) Consortium | Jacquelyn Wynn, Mairead Hyland, The Lighthouse Lab in Alderley Park and Alex Alderton, Roberto Amato, Jeffrey Barrett, Sonia Goncalves, Ewan Harrison, David K. Jackson, Ian Johnston, Dominic Kwiatkowski, Cordelia Langford, John Sillitoe on behalf of the Wellcome Sanger Institute COVID-19 Surveillance Team |
| EPI_ISL_2972259 | Department of Bacteria, Parasites and Fungi, Statens Serum Institut, Copenhagen, Denmark | Statens Serum Institut Bioinformatics and Microbial Genomics | Danish Covid-19 Genome Consortium |
| EPI_ISL_2972260, EPI_ISL_2972261 | Lighthouse Lab in Milton Keynes | Wellcome Sanger Institute for the COVID-19 Genomics UK (COG-UK) Consortium | The Lighthouse Lab in Milton Keynes and Alex Alderton, Roberto Amato, Jeffrey Barrett, Sonia Goncalves, Ewan Harrison, David K. Jackson, Ian Johnston, Dominic Kwiatkowski, Cordelia Langford, John Sillitoe on behalf of the Wellcome Sanger Institute COVID-19 Surveillance Team |
| EPI_ISL_2972262 | Department of Bacteria, Parasites and Fungi, Statens Serum Institut, Copenhagen, Denmark | Statens Serum Institut Bioinformatics and Microbial Genomics | Danish Covid-19 Genome Consortium |
| EPI_ISL_2972263, EPI_ISL_2972264 | Lighthouse Lab in Alderley Park | Wellcome Sanger Institute for the COVID-19 Genomics UK (COG-UK) Consortium | Jacquelyn Wynn, Mairead Hyland, The Lighthouse Lab in Alderley Park and Alex Alderton, Roberto Amato, Jeffrey Barrett, Sonia Goncalves, Ewan Harrison, David K. Jackson, Ian Johnston, Dominic Kwiatkowski, Cordelia Langford, John Sillitoe on behalf of the Wellcome Sanger Institute COVID-19 Surveillance Team |
| EPI_ISL_2972265 | Lighthouse Lab in Milton Keynes | Wellcome Sanger Institute for the COVID-19 Genomics UK (COG-UK) Consortium | The Lighthouse Lab in Milton Keynes and Alex Alderton, Roberto Amato, Jeffrey Barrett, Sonia Goncalves, Ewan Harrison, David K. Jackson, Ian Johnston, Dominic Kwiatkowski, Cordelia Langford, John Sillitoe on behalf of the Wellcome Sanger Institute COVID-19 Surveillance Team |
| EPI_ISL_2972266 | Department of Bacteria, Parasites and Fungi, Statens Serum Institut, Copenhagen, Denmark | Statens Serum Institut Bioinformatics and Microbial Genomics | Danish Covid-19 Genome Consortium |
| EPI_ISL_2972267 | Lighthouse Lab in Alderley Park | Wellcome Sanger Institute for the COVID-19 Genomics UK (COG-UK) Consortium | Jacquelyn Wynn, Mairead Hyland, The Lighthouse Lab in Alderley Park and Alex Alderton, Roberto Amato, Jeffrey Barrett, Sonia Goncalves, Ewan Harrison, David K. Jackson, Ian Johnston, Dominic Kwiatkowski, Cordelia Langford, John Sillitoe on behalf of the Wellcome Sanger Institute COVID-19 Surveillance Team |
| EPI_ISL_2972268 | Lighthouse Lab in Milton Keynes | Wellcome Sanger Institute for the COVID-19 Genomics UK (COG-UK) Consortium | The Lighthouse Lab in Milton Keynes and Alex Alderton, Roberto Amato, Jeffrey Barrett, Sonia Goncalves, Ewan Harrison, David K. Jackson, Ian Johnston, Dominic Kwiatkowski, Cordelia Langford, John Sillitoe on behalf of the Wellcome Sanger Institute COVID-19 Surveillance Team |
| EPI_ISL_2972269 | Department of Bacteria, Parasites and Fungi, Statens Serum Institut, Copenhagen, Denmark | Statens Serum Institut Bioinformatics and Microbial Genomics | Danish Covid-19 Genome Consortium |
| EPI_ISL_2972270 | Lighthouse Lab in Alderley Park | Wellcome Sanger Institute for the COVID-19 Genomics UK (COG-UK) Consortium | Jacquelyn Wynn, Mairead Hyland, The Lighthouse Lab in Alderley Park and Alex Alderton, Roberto Amato, Jeffrey Barrett, Sonia Goncalves, Ewan Harrison, David K. Jackson, Ian Johnston, Dominic Kwiatkowski, Cordelia Langford, John Sillitoe on behalf of the Wellcome Sanger Institute COVID-19 Surveillance Team |
| EPI_ISL_2972271 | Lighthouse Lab in Milton Keynes | Wellcome Sanger Institute for the COVID-19 Genomics UK (COG-UK) Consortium | The Lighthouse Lab in Milton Keynes and Alex Alderton, Roberto Amato, Jeffrey Barrett, Sonia Goncalves, Ewan Harrison, David K. Jackson, Ian Johnston, Dominic Kwiatkowski, Cordelia Langford, John Sillitoe on behalf of the Wellcome Sanger Institute COVID-19 Surveillance Team |
| EPI_ISL_2972272 | Department of Bacteria, Parasites and Fungi, Statens Serum Institut, Copenhagen, Denmark | Statens Serum Institut Bioinformatics and Microbial Genomics | Danish Covid-19 Genome Consortium |
| EPI_ISL_2972273 | Lighthouse Lab in Alderley Park | Wellcome Sanger Institute for the COVID-19 Genomics UK (COG-UK) Consortium | Jacquelyn Wynn, Mairead Hyland, The Lighthouse Lab in Alderley Park and Alex Alderton, Roberto Amato, Jeffrey Barrett, Sonia Goncalves, Ewan Harrison, David K. Jackson, Ian Johnston, Dominic Kwiatkowski, Cordelia Langford, John Sillitoe on behalf of the Wellcome Sanger Institute COVID-19 Surveillance Team |
| EPI_ISL_2972274 | Lighthouse Lab in Milton Keynes | Wellcome Sanger Institute for the COVID-19 Genomics UK (COG-UK) Consortium | The Lighthouse Lab in Milton Keynes and Alex Alderton, Roberto Amato, Jeffrey Barrett, Sonia Goncalves, Ewan Harrison, David K. Jackson, Ian Johnston, Dominic Kwiatkowski, Cordelia Langford, John Sillitoe on behalf of the Wellcome Sanger Institute COVID-19 Surveillance Team |
| EPI_ISL_2972275 | Department of Bacteria, Parasites and Fungi, Statens Serum Institut, Copenhagen, Denmark | Statens Serum Institut Bioinformatics and Microbial Genomics | Danish Covid-19 Genome Consortium |
| EPI_ISL_2972276 | Lighthouse Lab in Alderley Park | Wellcome Sanger Institute for the COVID-19 Genomics UK (COG-UK) Consortium | Jacquelyn Wynn, Mairead Hyland, The Lighthouse Lab in Alderley Park and Alex Alderton, Roberto Amato, Jeffrey Barrett, Sonia Goncalves, Ewan Harrison, David K. Jackson, Ian Johnston, Dominic Kwiatkowski, Cordelia Langford, John Sillitoe on behalf of the Wellcome Sanger Institute COVID-19 Surveillance Team |
| EPI_ISL_2972277, EPI_ISL_2972278 | Lighthouse Lab in Milton Keynes | Wellcome Sanger Institute for the COVID-19 Genomics UK (COG-UK) Consortium | The Lighthouse Lab in Milton Keynes and Alex Alderton, Roberto Amato, Jeffrey Barrett, Sonia Goncalves, Ewan Harrison, David K. Jackson, Ian Johnston, Dominic Kwiatkowski, Cordelia Langford, John Sillitoe on behalf of the Wellcome Sanger Institute COVID-19 Surveillance Team |
| EPI_ISL_2972279 | Department of Bacteria, Parasites and Fungi, Statens Serum Institut, Copenhagen, Denmark | Statens Serum Institut Bioinformatics and Microbial Genomics | Danish Covid-19 Genome Consortium |
| EPI_ISL_2972280 | Lighthouse Lab in Alderley Park | Wellcome Sanger Institute for the COVID-19 Genomics UK (COG-UK) Consortium | Jacquelyn Wynn, Mairead Hyland, The Lighthouse Lab in Alderley Park and Alex Alderton, Roberto Amato, Jeffrey Barrett, Sonia Goncalves, Ewan Harrison, David K. Jackson, Ian Johnston, Dominic Kwiatkowski, Cordelia Langford, John Sillitoe on behalf of the Wellcome Sanger Institute COVID-19 Surveillance Team |

| | | | |
|---|---|---|---|
| EPI_ISL_2972281 | Lighthouse Lab in Milton Keynes | Wellcome Sanger Institute for the COVID-19 Genomics UK (COG-UK) Consortium | The Lighthouse Lab in Milton Keynes and Alex Alderton, Roberto Amato, Jeffrey Barrett, Sonia Goncalves, Ewan Harrison, David K. Jackson, Ian Johnston, Dominic Kwiatkowski, Cordelia Langford, John Sillitoe on behalf of the Wellcome Sanger Institute COVID-19 Surveillance Team |
| EPI_ISL_2972282, EPI_ISL_2972283, EPI_ISL_2972284, EPI_ISL_2972285, EPI_ISL_2972286, EPI_ISL_2972287, EPI_ISL_2972288, EPI_ISL_2972289, EPI_ISL_2972290, EPI_ISL_2972291, EPI_ISL_2972292, EPI_ISL_2972293, EPI_ISL_2972294, EPI_ISL_2972295, EPI_ISL_2972296, EPI_ISL_2972297, EPI_ISL_2972298, EPI_ISL_2972300, EPI_ISL_2972301, EPI_ISL_2972302, EPI_ISL_2972303, EPI_ISL_2972304, EPI_ISL_2972305, EPI_ISL_2972306, EPI_ISL_2972307, EPI_ISL_2972308, EPI_ISL_2972309, EPI_ISL_2972310, EPI_ISL_2972311, EPI_ISL_2972312, EPI_ISL_2972313, EPI_ISL_2972315, EPI_ISL_2972316, EPI_ISL_2972317, EPI_ISL_2972318, EPI_ISL_2972319, EPI_ISL_2972320, EPI_ISL_2972321, EPI_ISL_2972322, EPI_ISL_2972323, EPI_ISL_2972324, EPI_ISL_2972325, EPI_ISL_2972326, EPI_ISL_2972327, EPI_ISL_2972328, EPI_ISL_2972329, EPI_ISL_2972330, EPI_ISL_2972331, EPI_ISL_2972332, EPI_ISL_2972333, EPI_ISL_2972334, EPI_ISL_2972336 | | | |
| see above | Department of Bacteria, Parasites and Fungi, Statens Serum Institut, Copenhagen, Denmark | Statens Serum Institut Bioinformatics and Microbial Genomics | Danish Covid-19 Genome Consortium |
| EPI_ISL_2972337, EPI_ISL_2972338 | Lighthouse Lab in Alderley Park | Wellcome Sanger Institute for the COVID-19 Genomics UK (COG-UK) Consortium | Jacquelyn Wynn, Mairead Hyland, The Lighthouse Lab in Alderley Park and Alex Alderton, Roberto Amato, Jeffrey Barrett, Sonia Goncalves, Ewan Harrison, David K. Jackson, Ian Johnston, Dominic Kwiatkowski, Cordelia Langford, John Sillitoe on behalf of the Wellcome Sanger Institute COVID-19 Surveillance Team |
| EPI_ISL_2972339 | Lighthouse Lab in Milton Keynes | Wellcome Sanger Institute for the COVID-19 Genomics UK (COG-UK) Consortium | The Lighthouse Lab in Milton Keynes and Alex Alderton, Roberto Amato, Jeffrey Barrett, Sonia Goncalves, Ewan Harrison, David K. Jackson, Ian Johnston, Dominic Kwiatkowski, Cordelia Langford, John Sillitoe on behalf of the Wellcome Sanger Institute COVID-19 Surveillance Team |
| EPI_ISL_2972340 | Department of Bacteria, Parasites and Fungi, Statens Serum Institut, Copenhagen, Denmark | Statens Serum Institut Bioinformatics and Microbial Genomics | Danish Covid-19 Genome Consortium |
| EPI_ISL_2972341, EPI_ISL_2972342 | Lighthouse Lab in Alderley Park | Wellcome Sanger Institute for the COVID-19 Genomics UK (COG-UK) Consortium | Jacquelyn Wynn, Mairead Hyland, The Lighthouse Lab in Alderley Park and Alex Alderton, Roberto Amato, Jeffrey Barrett, Sonia Goncalves, Ewan Harrison, David K. Jackson, Ian Johnston, Dominic Kwiatkowski, Cordelia Langford, John Sillitoe on behalf of the Wellcome Sanger Institute COVID-19 Surveillance Team |
| EPI_ISL_2972343 | Department of Bacteria, Parasites and Fungi, Statens Serum Institut, Copenhagen, Denmark | Statens Serum Institut Bioinformatics and Microbial Genomics | Danish Covid-19 Genome Consortium |
| EPI_ISL_2972344 | Lighthouse Lab in Milton Keynes | Wellcome Sanger Institute for the COVID-19 Genomics UK (COG-UK) Consortium | The Lighthouse Lab in Milton Keynes and Alex Alderton, Roberto Amato, Jeffrey Barrett, Sonia Goncalves, Ewan Harrison, David K. Jackson, Ian Johnston, Dominic Kwiatkowski, Cordelia Langford, John Sillitoe on behalf of the Wellcome Sanger Institute COVID-19 Surveillance Team |
| EPI_ISL_2972345 | Department of Bacteria, Parasites and Fungi, Statens Serum Institut, Copenhagen, Denmark | Statens Serum Institut Bioinformatics and Microbial Genomics | Danish Covid-19 Genome Consortium |
| EPI_ISL_2972346, EPI_ISL_2972347, EPI_ISL_2972348 | Lighthouse Lab in Alderley Park | Wellcome Sanger Institute for the COVID-19 Genomics UK (COG-UK) Consortium | Jacquelyn Wynn, Mairead Hyland, The Lighthouse Lab in Alderley Park and Alex Alderton, Roberto Amato, Jeffrey Barrett, Sonia Goncalves, Ewan Harrison, David K. Jackson, Ian Johnston, Dominic Kwiatkowski, Cordelia Langford, John Sillitoe on behalf of the Wellcome Sanger Institute COVID-19 Surveillance Team |
| EPI_ISL_2972349 | Department of Bacteria, Parasites and Fungi, Statens Serum Institut, Copenhagen, Denmark | Statens Serum Institut Bioinformatics and Microbial Genomics | Danish Covid-19 Genome Consortium |
| EPI_ISL_2972350, EPI_ISL_2972351 | Lighthouse Lab in Alderley Park | Wellcome Sanger Institute for the COVID-19 Genomics UK (COG-UK) Consortium | Jacquelyn Wynn, Mairead Hyland, The Lighthouse Lab in Alderley Park and Alex Alderton, Roberto Amato, Jeffrey Barrett, Sonia Goncalves, Ewan Harrison, David K. Jackson, Ian Johnston, Dominic Kwiatkowski, Cordelia Langford, John Sillitoe on behalf of the Wellcome Sanger Institute COVID-19 Surveillance Team |
| EPI_ISL_2972352 | Department of Bacteria, Parasites and Fungi, Statens Serum Institut, Copenhagen, Denmark | Statens Serum Institut Bioinformatics and Microbial Genomics | Danish Covid-19 Genome Consortium |
| EPI_ISL_2972353, EPI_ISL_2972355 | Lighthouse Lab in Milton Keynes | Wellcome Sanger Institute for the COVID-19 Genomics UK (COG-UK) Consortium | The Lighthouse Lab in Milton Keynes and Alex Alderton, Roberto Amato, Jeffrey Barrett, Sonia Goncalves, Ewan Harrison, David K. Jackson, Ian Johnston, Dominic Kwiatkowski, Cordelia Langford, John Sillitoe on behalf of the Wellcome Sanger Institute COVID-19 Surveillance Team |
| EPI_ISL_2972356, EPI_ISL_2972357 | Lighthouse Lab in Alderley Park | Wellcome Sanger Institute for the COVID-19 Genomics UK (COG-UK) Consortium | Jacquelyn Wynn, Mairead Hyland, The Lighthouse Lab in Alderley Park and Alex Alderton, Roberto Amato, Jeffrey Barrett, Sonia Goncalves, Ewan Harrison, David K. Jackson, Ian Johnston, Dominic Kwiatkowski, Cordelia Langford, John Sillitoe on behalf of the Wellcome Sanger Institute COVID-19 Surveillance Team |
| EPI_ISL_2972358 | Department of Bacteria, Parasites and Fungi, Statens Serum Institut, Copenhagen, Denmark | Statens Serum Institut Bioinformatics and Microbial Genomics | Danish Covid-19 Genome Consortium |
| EPI_ISL_2972359, EPI_ISL_2972360 | Lighthouse Lab in Milton Keynes | Wellcome Sanger Institute for the COVID-19 Genomics UK (COG-UK) Consortium | The Lighthouse Lab in Milton Keynes and Alex Alderton, Roberto Amato, Jeffrey Barrett, Sonia Goncalves, Ewan Harrison, David K. Jackson, Ian Johnston, Dominic Kwiatkowski, Cordelia Langford, John Sillitoe on behalf of the Wellcome Sanger Institute COVID-19 Surveillance Team |
| EPI_ISL_2972361 | Department of Bacteria, Parasites and Fungi, Statens Serum Institut, Copenhagen, Denmark | Statens Serum Institut Bioinformatics and Microbial Genomics | Danish Covid-19 Genome Consortium |
| EPI_ISL_2972362 | Lighthouse Lab in Milton Keynes | Wellcome Sanger Institute for the COVID-19 Genomics UK (COG-UK) Consortium | The Lighthouse Lab in Milton Keynes and Alex Alderton, Roberto Amato, Jeffrey Barrett, Sonia Goncalves, Ewan Harrison, David K. Jackson, Ian Johnston, Dominic Kwiatkowski, Cordelia Langford, John Sillitoe on behalf of the Wellcome Sanger Institute COVID-19 Surveillance Team |
| EPI_ISL_2972363 | Lighthouse Lab in Alderley Park | Wellcome Sanger Institute for the COVID-19 Genomics UK (COG-UK) Consortium | Jacquelyn Wynn, Mairead Hyland, The Lighthouse Lab in Alderley Park and Alex Alderton, Roberto Amato, Jeffrey Barrett, Sonia Goncalves, Ewan Harrison, David K. Jackson, Ian Johnston, Dominic Kwiatkowski, Cordelia Langford, John Sillitoe on behalf of the Wellcome Sanger Institute COVID-19 Surveillance Team |
| EPI_ISL_2972364 | Department of Bacteria, Parasites and Fungi, Statens Serum Institut, Copenhagen, Denmark | Statens Serum Institut Bioinformatics and Microbial Genomics | Danish Covid-19 Genome Consortium |
| EPI_ISL_2972365, EPI_ISL_2972366, EPI_ISL_2972367 | Lighthouse Lab in Alderley Park | Wellcome Sanger Institute for the COVID-19 Genomics UK (COG-UK) Consortium | Jacquelyn Wynn, Mairead Hyland, The Lighthouse Lab in Alderley Park and Alex Alderton, Roberto Amato, Jeffrey Barrett, Sonia Goncalves, Ewan Harrison, David K. Jackson, Ian Johnston, Dominic Kwiatkowski, Cordelia Langford, John Sillitoe on behalf of the Wellcome Sanger Institute COVID-19 Surveillance Team |
| EPI_ISL_2972368 | Department of Bacteria, Parasites and Fungi, Statens Serum Institut, Copenhagen, Denmark | Statens Serum Institut Bioinformatics and Microbial Genomics | Danish Covid-19 Genome Consortium |
| EPI_ISL_2972369 | Lighthouse Lab in Milton Keynes | Wellcome Sanger Institute for the COVID-19 Genomics UK (COG-UK) Consortium | The Lighthouse Lab in Milton Keynes and Alex Alderton, Roberto Amato, Jeffrey Barrett, Sonia Goncalves, Ewan Harrison, David K. Jackson, Ian Johnston, Dominic Kwiatkowski, Cordelia Langford, John Sillitoe on behalf of the Wellcome Sanger Institute COVID-19 Surveillance Team |
| EPI_ISL_2972370 | Department of Bacteria, Parasites and Fungi, Statens Serum Institut, Copenhagen, Denmark | Statens Serum Institut Bioinformatics and Microbial Genomics | Danish Covid-19 Genome Consortium |
| EPI_ISL_2972371 | Lighthouse Lab in Alderley Park | Wellcome Sanger Institute for the COVID-19 Genomics UK (COG-UK) Consortium | Jacquelyn Wynn, Mairead Hyland, The Lighthouse Lab in Alderley Park and Alex Alderton, Roberto Amato, Jeffrey Barrett, Sonia Goncalves, Ewan Harrison, David K. Jackson, Ian Johnston, Dominic Kwiatkowski, Cordelia Langford, John Sillitoe on behalf of the Wellcome Sanger Institute COVID-19 Surveillance Team |
| EPI_ISL_2972372 | Lighthouse Lab in Milton Keynes | Wellcome Sanger Institute for the COVID-19 Genomics UK (COG-UK) Consortium | The Lighthouse Lab in Milton Keynes and Alex Alderton, Roberto Amato, Jeffrey Barrett, Sonia Goncalves, Ewan Harrison, David K. Jackson, Ian Johnston, Dominic Kwiatkowski, Cordelia Langford, John Sillitoe on behalf of the Wellcome Sanger Institute COVID-19 Surveillance Team |
| EPI_ISL_2972373 | Department of Bacteria, Parasites and Fungi, Statens Serum Institut, Copenhagen, Denmark | Statens Serum Institut Bioinformatics and Microbial Genomics | Danish Covid-19 Genome Consortium |
| EPI_ISL_2972374 | Lighthouse Lab in Alderley Park | Wellcome Sanger Institute for the COVID-19 Genomics UK (COG-UK) Consortium | Jacquelyn Wynn, Mairead Hyland, The Lighthouse Lab in Alderley Park and Alex Alderton, Roberto Amato, Jeffrey Barrett, Sonia Goncalves, Ewan Harrison, David K. Jackson, Ian Johnston, Dominic Kwiatkowski, Cordelia Langford, John Sillitoe on behalf of the Wellcome Sanger Institute COVID-19 Surveillance Team |
| EPI_ISL_2972375 | Lighthouse Lab in Milton Keynes | Wellcome Sanger Institute for the COVID-19 Genomics UK (COG-UK) Consortium | The Lighthouse Lab in Milton Keynes and Alex Alderton, Roberto Amato, Jeffrey Barrett, Sonia Goncalves, Ewan Harrison, David K. Jackson, Ian Johnston, Dominic Kwiatkowski, Cordelia Langford, John Sillitoe on behalf of the Wellcome Sanger Institute COVID-19 Surveillance Team |
| EPI_ISL_2972376 | Department of Bacteria, Parasites and Fungi, Statens Serum Institut, Copenhagen, Denmark | Statens Serum Institut Bioinformatics and Microbial Genomics | Danish Covid-19 Genome Consortium |
| EPI_ISL_2972377, EPI_ISL_2972378 | Lighthouse Lab in Alderley Park | Wellcome Sanger Institute for the COVID-19 Genomics UK (COG-UK) Consortium | Jacquelyn Wynn, Mairead Hyland, The Lighthouse Lab in Alderley Park and Alex Alderton, Roberto Amato, Jeffrey Barrett, Sonia Goncalves, Ewan Harrison, David K. Jackson, Ian Johnston, Dominic Kwiatkowski, Cordelia Langford, John Sillitoe on behalf of the Wellcome Sanger Institute COVID-19 Surveillance Team |
| EPI_ISL_2972379 | Department of Bacteria, Parasites and Fungi, Statens Serum Institut, Copenhagen, Denmark | Statens Serum Institut Bioinformatics and Microbial Genomics | Danish Covid-19 Genome Consortium |
| EPI_ISL_2972380, EPI_ISL_2972381, EPI_ISL_2972382 | Lighthouse Lab in Milton Keynes | Wellcome Sanger Institute for the COVID-19 Genomics UK (COG-UK) Consortium | The Lighthouse Lab in Milton Keynes and Alex Alderton, Roberto Amato, Jeffrey Barrett, Sonia Goncalves, Ewan Harrison, David K. Jackson, Ian Johnston, Dominic Kwiatkowski, Cordelia Langford, John Sillitoe on behalf of the Wellcome Sanger Institute COVID-19 Surveillance Team |
| EPI_ISL_2972383 | Department of Bacteria, Parasites and Fungi, Statens Serum Institut, Copenhagen, Denmark | Statens Serum Institut Bioinformatics and Microbial Genomics | Danish Covid-19 Genome Consortium |
| EPI_ISL_2972384, EPI_ISL_2972385, EPI_ISL_2972386 | Lighthouse Lab in Alderley Park | Wellcome Sanger Institute for the COVID-19 Genomics UK (COG-UK) Consortium | Jacquelyn Wynn, Mairead Hyland, The Lighthouse Lab in Alderley Park and Alex Alderton, Roberto Amato, Jeffrey Barrett, Sonia Goncalves, Ewan Harrison, David K. Jackson, Ian Johnston, Dominic Kwiatkowski, Cordelia Langford, John Sillitoe on behalf of the Wellcome Sanger Institute COVID-19 Surveillance Team |
| EPI_ISL_2972387 | Department of Bacteria, Parasites and Fungi, Statens Serum Institut, Copenhagen, Denmark | Statens Serum Institut Bioinformatics and Microbial Genomics | Danish Covid-19 Genome Consortium |
| EPI_ISL_2972388, EPI_ISL_2972389 | Lighthouse Lab in Alderley Park | Wellcome Sanger Institute for the COVID-19 Genomics UK (COG-UK) Consortium | Jacquelyn Wynn, Mairead Hyland, The Lighthouse Lab in Alderley Park and Alex Alderton, Roberto Amato, Jeffrey Barrett, Sonia Goncalves, Ewan Harrison, David K. Jackson, Ian Johnston, Dominic Kwiatkowski, Cordelia Langford, John Sillitoe on behalf of the Wellcome Sanger Institute COVID-19 Surveillance Team |
| EPI_ISL_2972390 | Department of Bacteria, Parasites and Fungi, Statens Serum Institut, Copenhagen, Denmark | Statens Serum Institut Bioinformatics and Microbial Genomics | Danish Covid-19 Genome Consortium |
| EPI_ISL_2972391, EPI_ISL_2972392 | Lighthouse Lab in Alderley Park | Wellcome Sanger Institute for the COVID-19 Genomics UK (COG-UK) Consortium | Jacquelyn Wynn, Mairead Hyland, The Lighthouse Lab in Alderley Park and Alex Alderton, Roberto Amato, Jeffrey Barrett, Sonia Goncalves, Ewan Harrison, David K. Jackson, Ian Johnston, Dominic Kwiatkowski, Cordelia Langford, John Sillitoe on behalf of the Wellcome Sanger Institute COVID-19 Surveillance Team |
| EPI_ISL_2972393 | Department of Bacteria, Parasites and Fungi, Statens Serum Institut, Copenhagen, Denmark | Statens Serum Institut Bioinformatics and Microbial Genomics | Danish Covid-19 Genome Consortium |
| EPI_ISL_2972394, EPI_ISL_2972395 | Lighthouse Lab in Alderley Park | Wellcome Sanger Institute for the COVID-19 Genomics UK (COG-UK) Consortium | Jacquelyn Wynn, Mairead Hyland, The Lighthouse Lab in Alderley Park and Alex Alderton, Roberto Amato, Jeffrey Barrett, Sonia Goncalves, Ewan Harrison, David K. Jackson, Ian Johnston, Dominic Kwiatkowski, Cordelia Langford, John Sillitoe on behalf of the Wellcome Sanger Institute COVID-19 Surveillance Team |

| | | | |
|---|---|---|---|
| EPI_ISL_2972396 | Department of Bacteria, Parasites and Fungi, Statens Serum Institut, Copenhagen, Denmark | Statens Serum Institut Bioinformatics and Microbial Genomics | Danish Covid-19 Genome Consortium |
| EPI_ISL_2972397 | Lighthouse Lab in Alderley Park | Wellcome Sanger Institute for the COVID-19 Genomics UK (COG-UK) Consortium | Jacquelyn Wynn, Mairead Hyland, The Lighthouse Lab in Alderley Park and Alex Alderton, Roberto Amato, Jeffrey Barrett, Sonia Goncalves, Ewan Harrison, David K. Jackson, Ian Johnston, Dominic Kwiatkowski, Cordelia Langford, John Sillitoe on behalf of the Wellcome Sanger Institute COVID-19 Surveillance Team |
| EPI_ISL_2972398 | Lighthouse Lab in Milton Keynes | Wellcome Sanger Institute for the COVID-19 Genomics UK (COG-UK) Consortium | The Lighthouse Lab in Milton Keynes and Alex Alderton, Roberto Amato, Jeffrey Barrett, Sonia Goncalves, Ewan Harrison, David K. Jackson, Ian Johnston, Dominic Kwiatkowski, Cordelia Langford, John Sillitoe on behalf of the Wellcome Sanger Institute COVID-19 Surveillance Team |
| EPI_ISL_2972399 | Lighthouse Lab in Alderley Park | Wellcome Sanger Institute for the COVID-19 Genomics UK (COG-UK) Consortium | Jacquelyn Wynn, Mairead Hyland, The Lighthouse Lab in Alderley Park and Alex Alderton, Roberto Amato, Jeffrey Barrett, Sonia Goncalves, Ewan Harrison, David K. Jackson, Ian Johnston, Dominic Kwiatkowski, Cordelia Langford, John Sillitoe on behalf of the Wellcome Sanger Institute COVID-19 Surveillance Team |
| EPI_ISL_2972400 | Department of Bacteria, Parasites and Fungi, Statens Serum Institut, Copenhagen, Denmark | Statens Serum Institut Bioinformatics and Microbial Genomics | Danish Covid-19 Genome Consortium |
| EPI_ISL_2972401, EPI_ISL_2972402 | Lighthouse Lab in Alderley Park | Wellcome Sanger Institute for the COVID-19 Genomics UK (COG-UK) Consortium | Jacquelyn Wynn, Mairead Hyland, The Lighthouse Lab in Alderley Park and Alex Alderton, Roberto Amato, Jeffrey Barrett, Sonia Goncalves, Ewan Harrison, David K. Jackson, Ian Johnston, Dominic Kwiatkowski, Cordelia Langford, John Sillitoe on behalf of the Wellcome Sanger Institute COVID-19 Surveillance Team |
| EPI_ISL_2972403 | Department of Bacteria, Parasites and Fungi, Statens Serum Institut, Copenhagen, Denmark | Statens Serum Institut Bioinformatics and Microbial Genomics | Danish Covid-19 Genome Consortium |
| EPI_ISL_2972404, EPI_ISL_2972405 | Lighthouse Lab in Alderley Park | Wellcome Sanger Institute for the COVID-19 Genomics UK (COG-UK) Consortium | Jacquelyn Wynn, Mairead Hyland, The Lighthouse Lab in Alderley Park and Alex Alderton, Roberto Amato, Jeffrey Barrett, Sonia Goncalves, Ewan Harrison, David K. Jackson, Ian Johnston, Dominic Kwiatkowski, Cordelia Langford, John Sillitoe on behalf of the Wellcome Sanger Institute COVID-19 Surveillance Team |
| EPI_ISL_2972406 | Department of Bacteria, Parasites and Fungi, Statens Serum Institut, Copenhagen, Denmark | Statens Serum Institut Bioinformatics and Microbial Genomics | Danish Covid-19 Genome Consortium |
| EPI_ISL_2972407 | Lighthouse Lab in Alderley Park | Wellcome Sanger Institute for the COVID-19 Genomics UK (COG-UK) Consortium | Jacquelyn Wynn, Mairead Hyland, The Lighthouse Lab in Alderley Park and Alex Alderton, Roberto Amato, Jeffrey Barrett, Sonia Goncalves, Ewan Harrison, David K. Jackson, Ian Johnston, Dominic Kwiatkowski, Cordelia Langford, John Sillitoe on behalf of the Wellcome Sanger Institute COVID-19 Surveillance Team |
| EPI_ISL_2972408 | Lighthouse Lab in Milton Keynes | Wellcome Sanger Institute for the COVID-19 Genomics UK (COG-UK) Consortium | The Lighthouse Lab in Milton Keynes and Alex Alderton, Roberto Amato, Jeffrey Barrett, Sonia Goncalves, Ewan Harrison, David K. Jackson, Ian Johnston, Dominic Kwiatkowski, Cordelia Langford, John Sillitoe on behalf of the Wellcome Sanger Institute COVID-19 Surveillance Team |
| EPI_ISL_2972409 | Department of Bacteria, Parasites and Fungi, Statens Serum Institut, Copenhagen, Denmark | Statens Serum Institut Bioinformatics and Microbial Genomics | Danish Covid-19 Genome Consortium |
| EPI_ISL_2972410 | Lighthouse Lab in Alderley Park | Wellcome Sanger Institute for the COVID-19 Genomics UK (COG-UK) Consortium | Jacquelyn Wynn, Mairead Hyland, The Lighthouse Lab in Alderley Park and Alex Alderton, Roberto Amato, Jeffrey Barrett, Sonia Goncalves, Ewan Harrison, David K. Jackson, Ian Johnston, Dominic Kwiatkowski, Cordelia Langford, John Sillitoe on behalf of the Wellcome Sanger Institute COVID-19 Surveillance Team |
| EPI_ISL_2972411 | Lighthouse Lab in Milton Keynes | Wellcome Sanger Institute for the COVID-19 Genomics UK (COG-UK) Consortium | The Lighthouse Lab in Milton Keynes and Alex Alderton, Roberto Amato, Jeffrey Barrett, Sonia Goncalves, Ewan Harrison, David K. Jackson, Ian Johnston, Dominic Kwiatkowski, Cordelia Langford, John Sillitoe on behalf of the Wellcome Sanger Institute COVID-19 Surveillance Team |
| EPI_ISL_2972412 | Department of Bacteria, Parasites and Fungi, Statens Serum Institut, Copenhagen, Denmark | Statens Serum Institut Bioinformatics and Microbial Genomics | Danish Covid-19 Genome Consortium |
| EPI_ISL_2972413, EPI_ISL_2972414 | Lighthouse Lab in Alderley Park | Wellcome Sanger Institute for the COVID-19 Genomics UK (COG-UK) Consortium | Jacquelyn Wynn, Mairead Hyland, The Lighthouse Lab in Alderley Park and Alex Alderton, Roberto Amato, Jeffrey Barrett, Sonia Goncalves, Ewan Harrison, David K. Jackson, Ian Johnston, Dominic Kwiatkowski, Cordelia Langford, John Sillitoe on behalf of the Wellcome Sanger Institute COVID-19 Surveillance Team |
| EPI_ISL_2972415 | Department of Bacteria, Parasites and Fungi, Statens Serum Institut, Copenhagen, Denmark | Statens Serum Institut Bioinformatics and Microbial Genomics | Danish Covid-19 Genome Consortium |
| EPI_ISL_2972416 | Lighthouse Lab in Alderley Park | Wellcome Sanger Institute for the COVID-19 Genomics UK (COG-UK) Consortium | Jacquelyn Wynn, Mairead Hyland, The Lighthouse Lab in Alderley Park and Alex Alderton, Roberto Amato, Jeffrey Barrett, Sonia Goncalves, Ewan Harrison, David K. Jackson, Ian Johnston, Dominic Kwiatkowski, Cordelia Langford, John Sillitoe on behalf of the Wellcome Sanger Institute COVID-19 Surveillance Team |
| EPI_ISL_2972417 | Lighthouse Lab in Milton Keynes | Wellcome Sanger Institute for the COVID-19 Genomics UK (COG-UK) Consortium | The Lighthouse Lab in Milton Keynes and Alex Alderton, Roberto Amato, Jeffrey Barrett, Sonia Goncalves, Ewan Harrison, David K. Jackson, Ian Johnston, Dominic Kwiatkowski, Cordelia Langford, John Sillitoe on behalf of the Wellcome Sanger Institute COVID-19 Surveillance Team |
| EPI_ISL_2972418 | Department of Bacteria, Parasites and Fungi, Statens Serum Institut, Copenhagen, Denmark | Statens Serum Institut Bioinformatics and Microbial Genomics | Danish Covid-19 Genome Consortium |
| EPI_ISL_2972419 | Lighthouse Lab in Milton Keynes | Wellcome Sanger Institute for the COVID-19 Genomics UK (COG-UK) Consortium | The Lighthouse Lab in Milton Keynes and Alex Alderton, Roberto Amato, Jeffrey Barrett, Sonia Goncalves, Ewan Harrison, David K. Jackson, Ian Johnston, Dominic Kwiatkowski, Cordelia Langford, John Sillitoe on behalf of the Wellcome Sanger Institute COVID-19 Surveillance Team |
| EPI_ISL_2972420 | Lighthouse Lab in Alderley Park | Wellcome Sanger Institute for the COVID-19 Genomics UK (COG-UK) Consortium | Jacquelyn Wynn, Mairead Hyland, The Lighthouse Lab in Alderley Park and Alex Alderton, Roberto Amato, Jeffrey Barrett, Sonia Goncalves, Ewan Harrison, David K. Jackson, Ian Johnston, Dominic Kwiatkowski, Cordelia Langford, John Sillitoe on behalf of the Wellcome Sanger Institute COVID-19 Surveillance Team |
| EPI_ISL_2972421 | Department of Bacteria, Parasites and Fungi, Statens Serum Institut, Copenhagen, Denmark | Statens Serum Institut Bioinformatics and Microbial Genomics | Danish Covid-19 Genome Consortium |
| EPI_ISL_2972422 | Lighthouse Lab in Milton Keynes | Wellcome Sanger Institute for the COVID-19 Genomics UK (COG-UK) Consortium | The Lighthouse Lab in Milton Keynes and Alex Alderton, Roberto Amato, Jeffrey Barrett, Sonia Goncalves, Ewan Harrison, David K. Jackson, Ian Johnston, Dominic Kwiatkowski, Cordelia Langford, John Sillitoe on behalf of the Wellcome Sanger Institute COVID-19 Surveillance Team |
| EPI_ISL_2972423, EPI_ISL_2972424 | Lighthouse Lab in Alderley Park | Wellcome Sanger Institute for the COVID-19 Genomics UK (COG-UK) Consortium | Jacquelyn Wynn, Mairead Hyland, The Lighthouse Lab in Alderley Park and Alex Alderton, Roberto Amato, Jeffrey Barrett, Sonia Goncalves, Ewan Harrison, David K. Jackson, Ian Johnston, Dominic Kwiatkowski, Cordelia Langford, John Sillitoe on behalf of the Wellcome Sanger Institute COVID-19 Surveillance Team |
| EPI_ISL_2972425 | Department of Bacteria, Parasites and Fungi, Statens Serum Institut, Copenhagen, Denmark | Statens Serum Institut Bioinformatics and Microbial Genomics | Danish Covid-19 Genome Consortium |
| EPI_ISL_2972426, EPI_ISL_2972427 | Lighthouse Lab in Alderley Park | Wellcome Sanger Institute for the COVID-19 Genomics UK (COG-UK) Consortium | Jacquelyn Wynn, Mairead Hyland, The Lighthouse Lab in Alderley Park and Alex Alderton, Roberto Amato, Jeffrey Barrett, Sonia Goncalves, Ewan Harrison, David K. Jackson, Ian Johnston, Dominic Kwiatkowski, Cordelia Langford, John Sillitoe on behalf of the Wellcome Sanger Institute COVID-19 Surveillance Team |
| EPI_ISL_2972428 | Department of Bacteria, Parasites and Fungi, Statens Serum Institut, Copenhagen, Denmark | Statens Serum Institut Bioinformatics and Microbial Genomics | Danish Covid-19 Genome Consortium |
| EPI_ISL_2972429 | Lighthouse Lab in Milton Keynes | Wellcome Sanger Institute for the COVID-19 Genomics UK (COG-UK) Consortium | The Lighthouse Lab in Milton Keynes and Alex Alderton, Roberto Amato, Jeffrey Barrett, Sonia Goncalves, Ewan Harrison, David K. Jackson, Ian Johnston, Dominic Kwiatkowski, Cordelia Langford, John Sillitoe on behalf of the Wellcome Sanger Institute COVID-19 Surveillance Team |
| EPI_ISL_2972430 | Lighthouse Lab in Alderley Park | Wellcome Sanger Institute for the COVID-19 Genomics UK (COG-UK) Consortium | Jacquelyn Wynn, Mairead Hyland, The Lighthouse Lab in Alderley Park and Alex Alderton, Roberto Amato, Jeffrey Barrett, Sonia Goncalves, Ewan Harrison, David K. Jackson, Ian Johnston, Dominic Kwiatkowski, Cordelia Langford, John Sillitoe on behalf of the Wellcome Sanger Institute COVID-19 Surveillance Team |
| EPI_ISL_2972431 | Lighthouse Lab in Milton Keynes | Wellcome Sanger Institute for the COVID-19 Genomics UK (COG-UK) Consortium | The Lighthouse Lab in Milton Keynes and Alex Alderton, Roberto Amato, Jeffrey Barrett, Sonia Goncalves, Ewan Harrison, David K. Jackson, Ian Johnston, Dominic Kwiatkowski, Cordelia Langford, John Sillitoe on behalf of the Wellcome Sanger Institute COVID-19 Surveillance Team |
| EPI_ISL_2972432 | Department of Bacteria, Parasites and Fungi, Statens Serum Institut, Copenhagen, Denmark | Statens Serum Institut Bioinformatics and Microbial Genomics | Danish Covid-19 Genome Consortium |
| EPI_ISL_2972433, EPI_ISL_2972434 | Lighthouse Lab in Alderley Park | Wellcome Sanger Institute for the COVID-19 Genomics UK (COG-UK) Consortium | Jacquelyn Wynn, Mairead Hyland, The Lighthouse Lab in Alderley Park and Alex Alderton, Roberto Amato, Jeffrey Barrett, Sonia Goncalves, Ewan Harrison, David K. Jackson, Ian Johnston, Dominic Kwiatkowski, Cordelia Langford, John Sillitoe on behalf of the Wellcome Sanger Institute COVID-19 Surveillance Team |
| EPI_ISL_2972435 | Department of Bacteria, Parasites and Fungi, Statens Serum Institut, Copenhagen, Denmark | Statens Serum Institut Bioinformatics and Microbial Genomics | Danish Covid-19 Genome Consortium |
| EPI_ISL_2972436, EPI_ISL_2972437 | Lighthouse Lab in Alderley Park | Wellcome Sanger Institute for the COVID-19 Genomics UK (COG-UK) Consortium | Jacquelyn Wynn, Mairead Hyland, The Lighthouse Lab in Alderley Park and Alex Alderton, Roberto Amato, Jeffrey Barrett, Sonia Goncalves, Ewan Harrison, David K. Jackson, Ian Johnston, Dominic Kwiatkowski, Cordelia Langford, John Sillitoe on behalf of the Wellcome Sanger Institute COVID-19 Surveillance Team |
| EPI_ISL_2972438 | Department of Bacteria, Parasites and Fungi, Statens Serum Institut, Copenhagen, Denmark | Statens Serum Institut Bioinformatics and Microbial Genomics | Danish Covid-19 Genome Consortium |
| EPI_ISL_2972439 | Lighthouse Lab in Alderley Park | Wellcome Sanger Institute for the COVID-19 Genomics UK (COG-UK) Consortium | Jacquelyn Wynn, Mairead Hyland, The Lighthouse Lab in Alderley Park and Alex Alderton, Roberto Amato, Jeffrey Barrett, Sonia Goncalves, Ewan Harrison, David K. Jackson, Ian Johnston, Dominic Kwiatkowski, Cordelia Langford, John Sillitoe on behalf of the Wellcome Sanger Institute COVID-19 Surveillance Team |
| EPI_ISL_2972440 | Lighthouse Lab in Milton Keynes | Wellcome Sanger Institute for the COVID-19 Genomics UK (COG-UK) Consortium | The Lighthouse Lab in Milton Keynes and Alex Alderton, Roberto Amato, Jeffrey Barrett, Sonia Goncalves, Ewan Harrison, David K. Jackson, Ian Johnston, Dominic Kwiatkowski, Cordelia Langford, John Sillitoe on behalf of the Wellcome Sanger Institute COVID-19 Surveillance Team |
| EPI_ISL_2972441 | Lighthouse Lab in Alderley Park | Wellcome Sanger Institute for the COVID-19 Genomics UK (COG-UK) Consortium | Jacquelyn Wynn, Mairead Hyland, The Lighthouse Lab in Alderley Park and Alex Alderton, Roberto Amato, Jeffrey Barrett, Sonia Goncalves, Ewan Harrison, David K. Jackson, Ian Johnston, Dominic Kwiatkowski, Cordelia Langford, John Sillitoe on behalf of the Wellcome Sanger Institute COVID-19 Surveillance Team |
| EPI_ISL_2972442 | Department of Bacteria, Parasites and Fungi, Statens Serum Institut, Copenhagen, Denmark | Statens Serum Institut Bioinformatics and Microbial Genomics | Danish Covid-19 Genome Consortium |
| EPI_ISL_2972443, EPI_ISL_2972444 | Lighthouse Lab in Alderley Park | Wellcome Sanger Institute for the COVID-19 Genomics UK (COG-UK) Consortium | Jacquelyn Wynn, Mairead Hyland, The Lighthouse Lab in Alderley Park and Alex Alderton, Roberto Amato, Jeffrey Barrett, Sonia Goncalves, Ewan Harrison, David K. Jackson, Ian Johnston, Dominic Kwiatkowski, Cordelia Langford, John Sillitoe on behalf of the Wellcome Sanger Institute COVID-19 Surveillance Team |
| EPI_ISL_2972445 | Department of Bacteria, Parasites and Fungi, Statens Serum Institut, Copenhagen, Denmark | Statens Serum Institut Bioinformatics and Microbial Genomics | Danish Covid-19 Genome Consortium |
| EPI_ISL_2972446, EPI_ISL_2972447, EPI_ISL_2972448 | Lighthouse Lab in Alderley Park | Wellcome Sanger Institute for the COVID-19 Genomics UK (COG-UK) Consortium | Jacquelyn Wynn, Mairead Hyland, The Lighthouse Lab in Alderley Park and Alex Alderton, Roberto Amato, Jeffrey Barrett, Sonia Goncalves, Ewan Harrison, David K. Jackson, Ian Johnston, Dominic Kwiatkowski, Cordelia Langford, John Sillitoe on behalf of the Wellcome Sanger Institute COVID-19 Surveillance Team |
| EPI_ISL_2972449 | Department of Bacteria, Parasites and Fungi, Statens Serum Institut, Copenhagen, Denmark | Statens Serum Institut Bioinformatics and Microbial Genomics | Danish Covid-19 Genome Consortium |
| EPI_ISL_2972450, EPI_ISL_2972451 | Lighthouse Lab in Alderley Park | Wellcome Sanger Institute for the COVID-19 Genomics UK (COG-UK) Consortium | Jacquelyn Wynn, Mairead Hyland, The Lighthouse Lab in Alderley Park and Alex Alderton, Roberto Amato, Jeffrey Barrett, Sonia Goncalves, Ewan Harrison, David K. Jackson, Ian Johnston, Dominic Kwiatkowski, Cordelia Langford, John Sillitoe on behalf of the Wellcome Sanger Institute COVID-19 Surveillance Team |

| | | | |
|---|---|---|---|
| EPI_ISL_2972452 | Department of Bacteria, Parasites and Fungi, Statens Serum Institut, Copenhagen, Denmark | Statens Serum Institut Bioinformatics and Microbial Genomics | Danish Covid-19 Genome Consortium |
| EPI_ISL_2972453, EPI_ISL_2972454 | Lighthouse Lab in Alderley Park | Wellcome Sanger Institute for the COVID-19 Genomics UK (COG-UK) Consortium | Jacquelyn Wynn, Mairead Hyland, The Lighthouse Lab in Alderley Park and Alex Alderton, Roberto Amato, Jeffrey Barrett, Sonia Goncalves, Ewan Harrison, David K. Jackson, Ian Johnston, Dominic Kwiatkowski, Cordelia Langford, John Sillitoe on behalf of the Wellcome Sanger Institute COVID-19 Surveillance Team |
| EPI_ISL_2972455 | Lighthouse Lab in Milton Keynes | Wellcome Sanger Institute for the COVID-19 Genomics UK (COG-UK) Consortium | The Lighthouse Lab in Milton Keynes and Alex Alderton, Roberto Amato, Jeffrey Barrett, Sonia Goncalves, Ewan Harrison, David K. Jackson, Ian Johnston, Dominic Kwiatkowski, Cordelia Langford, John Sillitoe on behalf of the Wellcome Sanger Institute COVID-19 Surveillance Team |
| EPI_ISL_2972456 | Department of Bacteria, Parasites and Fungi, Statens Serum Institut, Copenhagen, Denmark | Statens Serum Institut Bioinformatics and Microbial Genomics | Danish Covid-19 Genome Consortium |
| EPI_ISL_2972457 | Lighthouse Lab in Alderley Park | Wellcome Sanger Institute for the COVID-19 Genomics UK (COG-UK) Consortium | Jacquelyn Wynn, Mairead Hyland, The Lighthouse Lab in Alderley Park and Alex Alderton, Roberto Amato, Jeffrey Barrett, Sonia Goncalves, Ewan Harrison, David K. Jackson, Ian Johnston, Dominic Kwiatkowski, Cordelia Langford, John Sillitoe on behalf of the Wellcome Sanger Institute COVID-19 Surveillance Team |
| EPI_ISL_2972458 | Lighthouse Lab in Milton Keynes | Wellcome Sanger Institute for the COVID-19 Genomics UK (COG-UK) Consortium | The Lighthouse Lab in Milton Keynes and Alex Alderton, Roberto Amato, Jeffrey Barrett, Sonia Goncalves, Ewan Harrison, David K. Jackson, Ian Johnston, Dominic Kwiatkowski, Cordelia Langford, John Sillitoe on behalf of the Wellcome Sanger Institute COVID-19 Surveillance Team |
| EPI_ISL_2972459 | Department of Bacteria, Parasites and Fungi, Statens Serum Institut, Copenhagen, Denmark | Statens Serum Institut Bioinformatics and Microbial Genomics | Danish Covid-19 Genome Consortium |
| EPI_ISL_2972460 | Lighthouse Lab in Milton Keynes | Wellcome Sanger Institute for the COVID-19 Genomics UK (COG-UK) Consortium | The Lighthouse Lab in Milton Keynes and Alex Alderton, Roberto Amato, Jeffrey Barrett, Sonia Goncalves, Ewan Harrison, David K. Jackson, Ian Johnston, Dominic Kwiatkowski, Cordelia Langford, John Sillitoe on behalf of the Wellcome Sanger Institute COVID-19 Surveillance Team |
| EPI_ISL_2972461 | Lighthouse Lab in Alderley Park | Wellcome Sanger Institute for the COVID-19 Genomics UK (COG-UK) Consortium | Jacquelyn Wynn, Mairead Hyland, The Lighthouse Lab in Alderley Park and Alex Alderton, Roberto Amato, Jeffrey Barrett, Sonia Goncalves, Ewan Harrison, David K. Jackson, Ian Johnston, Dominic Kwiatkowski, Cordelia Langford, John Sillitoe on behalf of the Wellcome Sanger Institute COVID-19 Surveillance Team |
| EPI_ISL_2972462 | Department of Bacteria, Parasites and Fungi, Statens Serum Institut, Copenhagen, Denmark | Statens Serum Institut Bioinformatics and Microbial Genomics | Danish Covid-19 Genome Consortium |
| EPI_ISL_2972463, EPI_ISL_2972464, EPI_ISL_2972465 | Lighthouse Lab in Alderley Park | Wellcome Sanger Institute for the COVID-19 Genomics UK (COG-UK) Consortium | Jacquelyn Wynn, Mairead Hyland, The Lighthouse Lab in Alderley Park and Alex Alderton, Roberto Amato, Jeffrey Barrett, Sonia Goncalves, Ewan Harrison, David K. Jackson, Ian Johnston, Dominic Kwiatkowski, Cordelia Langford, John Sillitoe on behalf of the Wellcome Sanger Institute COVID-19 Surveillance Team |
| EPI_ISL_2972466 | Department of Bacteria, Parasites and Fungi, Statens Serum Institut, Copenhagen, Denmark | Statens Serum Institut Bioinformatics and Microbial Genomics | Danish Covid-19 Genome Consortium |
| EPI_ISL_2972467 | Lighthouse Lab in Alderley Park | Wellcome Sanger Institute for the COVID-19 Genomics UK (COG-UK) Consortium | Jacquelyn Wynn, Mairead Hyland, The Lighthouse Lab in Alderley Park and Alex Alderton, Roberto Amato, Jeffrey Barrett, Sonia Goncalves, Ewan Harrison, David K. Jackson, Ian Johnston, Dominic Kwiatkowski, Cordelia Langford, John Sillitoe on behalf of the Wellcome Sanger Institute COVID-19 Surveillance Team |
| EPI_ISL_2972468 | Lighthouse Lab in Milton Keynes | Wellcome Sanger Institute for the COVID-19 Genomics UK (COG-UK) Consortium | The Lighthouse Lab in Milton Keynes and Alex Alderton, Roberto Amato, Jeffrey Barrett, Sonia Goncalves, Ewan Harrison, David K. Jackson, Ian Johnston, Dominic Kwiatkowski, Cordelia Langford, John Sillitoe on behalf of the Wellcome Sanger Institute COVID-19 Surveillance Team |
| EPI_ISL_2972469 | Department of Bacteria, Parasites and Fungi, Statens Serum Institut, Copenhagen, Denmark | Statens Serum Institut Bioinformatics and Microbial Genomics | Danish Covid-19 Genome Consortium |
| EPI_ISL_2972470 | Lighthouse Lab in Alderley Park | Wellcome Sanger Institute for the COVID-19 Genomics UK (COG-UK) Consortium | Jacquelyn Wynn, Mairead Hyland, The Lighthouse Lab in Alderley Park and Alex Alderton, Roberto Amato, Jeffrey Barrett, Sonia Goncalves, Ewan Harrison, David K. Jackson, Ian Johnston, Dominic Kwiatkowski, Cordelia Langford, John Sillitoe on behalf of the Wellcome Sanger Institute COVID-19 Surveillance Team |
| EPI_ISL_2972471 | Lighthouse Lab in Milton Keynes | Wellcome Sanger Institute for the COVID-19 Genomics UK (COG-UK) Consortium | The Lighthouse Lab in Milton Keynes and Alex Alderton, Roberto Amato, Jeffrey Barrett, Sonia Goncalves, Ewan Harrison, David K. Jackson, Ian Johnston, Dominic Kwiatkowski, Cordelia Langford, John Sillitoe on behalf of the Wellcome Sanger Institute COVID-19 Surveillance Team |
| EPI_ISL_2972472 | Department of Bacteria, Parasites and Fungi, Statens Serum Institut, Copenhagen, Denmark | Statens Serum Institut Bioinformatics and Microbial Genomics | Danish Covid-19 Genome Consortium |
| EPI_ISL_2972473 | Lighthouse Lab in Alderley Park | Wellcome Sanger Institute for the COVID-19 Genomics UK (COG-UK) Consortium | Jacquelyn Wynn, Mairead Hyland, The Lighthouse Lab in Alderley Park and Alex Alderton, Roberto Amato, Jeffrey Barrett, Sonia Goncalves, Ewan Harrison, David K. Jackson, Ian Johnston, Dominic Kwiatkowski, Cordelia Langford, John Sillitoe on behalf of the Wellcome Sanger Institute COVID-19 Surveillance Team |
| EPI_ISL_2972474 | Lighthouse Lab in Milton Keynes | Wellcome Sanger Institute for the COVID-19 Genomics UK (COG-UK) Consortium | The Lighthouse Lab in Milton Keynes and Alex Alderton, Roberto Amato, Jeffrey Barrett, Sonia Goncalves, Ewan Harrison, David K. Jackson, Ian Johnston, Dominic Kwiatkowski, Cordelia Langford, John Sillitoe on behalf of the Wellcome Sanger Institute COVID-19 Surveillance Team |
| EPI_ISL_2972475 | Department of Bacteria, Parasites and Fungi, Statens Serum Institut, Copenhagen, Denmark | Statens Serum Institut Bioinformatics and Microbial Genomics | Danish Covid-19 Genome Consortium |
| EPI_ISL_2972476 | Lighthouse Lab in Milton Keynes | Wellcome Sanger Institute for the COVID-19 Genomics UK (COG-UK) Consortium | The Lighthouse Lab in Milton Keynes and Alex Alderton, Roberto Amato, Jeffrey Barrett, Sonia Goncalves, Ewan Harrison, David K. Jackson, Ian Johnston, Dominic Kwiatkowski, Cordelia Langford, John Sillitoe on behalf of the Wellcome Sanger Institute COVID-19 Surveillance Team |
| EPI_ISL_2972477, EPI_ISL_2972479 | Lighthouse Lab in Alderley Park | Wellcome Sanger Institute for the COVID-19 Genomics UK (COG-UK) Consortium | Jacquelyn Wynn, Mairead Hyland, The Lighthouse Lab in Alderley Park and Alex Alderton, Roberto Amato, Jeffrey Barrett, Sonia Goncalves, Ewan Harrison, David K. Jackson, Ian Johnston, Dominic Kwiatkowski, Cordelia Langford, John Sillitoe on behalf of the Wellcome Sanger Institute COVID-19 Surveillance Team |
| EPI_ISL_2972480 | Lighthouse Lab in Milton Keynes | Wellcome Sanger Institute for the COVID-19 Genomics UK (COG-UK) Consortium | The Lighthouse Lab in Milton Keynes and Alex Alderton, Roberto Amato, Jeffrey Barrett, Sonia Goncalves, Ewan Harrison, David K. Jackson, Ian Johnston, Dominic Kwiatkowski, Cordelia Langford, John Sillitoe on behalf of the Wellcome Sanger Institute COVID-19 Surveillance Team |
| EPI_ISL_2972481 | Lighthouse Lab in Alderley Park | Wellcome Sanger Institute for the COVID-19 Genomics UK (COG-UK) Consortium | Jacquelyn Wynn, Mairead Hyland, The Lighthouse Lab in Alderley Park and Alex Alderton, Roberto Amato, Jeffrey Barrett, Sonia Goncalves, Ewan Harrison, David K. Jackson, Ian Johnston, Dominic Kwiatkowski, Cordelia Langford, John Sillitoe on behalf of the Wellcome Sanger Institute COVID-19 Surveillance Team |
| EPI_ISL_2972482 | Department of Bacteria, Parasites and Fungi, Statens Serum Institut, Copenhagen, Denmark | Statens Serum Institut Bioinformatics and Microbial Genomics | Danish Covid-19 Genome Consortium |
| EPI_ISL_2972483, EPI_ISL_2972484 | Lighthouse Laboratory Plymouth | Wellcome Sanger Institute for the COVID-19 Genomics UK (COG-UK) Consortium | Lighthouse Laboratory Plymouth and Alex Alderton, Roberto Amato, Jeffrey Barrett, Sonia Goncalves, Ewan Harrison, David K. Jackson, Ian Johnston, Dominic Kwiatkowski, Cordelia Langford, John Sillitoe on behalf of the Wellcome Sanger Institute COVID-19 Surveillance Team |
| EPI_ISL_2972485 | Department of Bacteria, Parasites and Fungi, Statens Serum Institut, Copenhagen, Denmark | Statens Serum Institut Bioinformatics and Microbial Genomics | Danish Covid-19 Genome Consortium |
| EPI_ISL_2972486, EPI_ISL_2972487 | Lighthouse Lab in Milton Keynes | Wellcome Sanger Institute for the COVID-19 Genomics UK (COG-UK) Consortium | The Lighthouse Lab in Milton Keynes and Alex Alderton, Roberto Amato, Jeffrey Barrett, Sonia Goncalves, Ewan Harrison, David K. Jackson, Ian Johnston, Dominic Kwiatkowski, Cordelia Langford, John Sillitoe on behalf of the Wellcome Sanger Institute COVID-19 Surveillance Team |
| EPI_ISL_2972488 | Department of Bacteria, Parasites and Fungi, Statens Serum Institut, Copenhagen, Denmark | Statens Serum Institut Bioinformatics and Microbial Genomics | Danish Covid-19 Genome Consortium |
| EPI_ISL_2972489 | Lighthouse Lab in Alderley Park | Wellcome Sanger Institute for the COVID-19 Genomics UK (COG-UK) Consortium | Jacquelyn Wynn, Mairead Hyland, The Lighthouse Lab in Alderley Park and Alex Alderton, Roberto Amato, Jeffrey Barrett, Sonia Goncalves, Ewan Harrison, David K. Jackson, Ian Johnston, Dominic Kwiatkowski, Cordelia Langford, John Sillitoe on behalf of the Wellcome Sanger Institute COVID-19 Surveillance Team |
| EPI_ISL_2972490 | Lighthouse Laboratory Plymouth | Wellcome Sanger Institute for the COVID-19 Genomics UK (COG-UK) Consortium | Lighthouse Laboratory Plymouth and Alex Alderton, Roberto Amato, Jeffrey Barrett, Sonia Goncalves, Ewan Harrison, David K. Jackson, Ian Johnston, Dominic Kwiatkowski, Cordelia Langford, John Sillitoe on behalf of the Wellcome Sanger Institute COVID-19 Surveillance Team |
| EPI_ISL_2972491 | Lighthouse Lab in Alderley Park | Wellcome Sanger Institute for the COVID-19 Genomics UK (COG-UK) Consortium | Jacquelyn Wynn, Mairead Hyland, The Lighthouse Lab in Alderley Park and Alex Alderton, Roberto Amato, Jeffrey Barrett, Sonia Goncalves, Ewan Harrison, David K. Jackson, Ian Johnston, Dominic Kwiatkowski, Cordelia Langford, John Sillitoe on behalf of the Wellcome Sanger Institute COVID-19 Surveillance Team |
| EPI_ISL_2972492 | Department of Bacteria, Parasites and Fungi, Statens Serum Institut, Copenhagen, Denmark | Statens Serum Institut Bioinformatics and Microbial Genomics | Danish Covid-19 Genome Consortium |
| EPI_ISL_2972493 | Lighthouse Laboratory Plymouth | Wellcome Sanger Institute for the COVID-19 Genomics UK (COG-UK) Consortium | Lighthouse Laboratory Plymouth and Alex Alderton, Roberto Amato, Jeffrey Barrett, Sonia Goncalves, Ewan Harrison, David K. Jackson, Ian Johnston, Dominic Kwiatkowski, Cordelia Langford, John Sillitoe on behalf of the Wellcome Sanger Institute COVID-19 Surveillance Team |
| EPI_ISL_2972494 | Lighthouse Lab in Alderley Park | Wellcome Sanger Institute for the COVID-19 Genomics UK (COG-UK) Consortium | Jacquelyn Wynn, Mairead Hyland, The Lighthouse Lab in Alderley Park and Alex Alderton, Roberto Amato, Jeffrey Barrett, Sonia Goncalves, Ewan Harrison, David K. Jackson, Ian Johnston, Dominic Kwiatkowski, Cordelia Langford, John Sillitoe on behalf of the Wellcome Sanger Institute COVID-19 Surveillance Team |
| EPI_ISL_2972495 | Department of Bacteria, Parasites and Fungi, Statens Serum Institut, Copenhagen, Denmark | Statens Serum Institut Bioinformatics and Microbial Genomics | Danish Covid-19 Genome Consortium |
| EPI_ISL_2972496 | Lighthouse Lab in Milton Keynes | Wellcome Sanger Institute for the COVID-19 Genomics UK (COG-UK) Consortium | The Lighthouse Lab in Milton Keynes and Alex Alderton, Roberto Amato, Jeffrey Barrett, Sonia Goncalves, Ewan Harrison, David K. Jackson, Ian Johnston, Dominic Kwiatkowski, Cordelia Langford, John Sillitoe on behalf of the Wellcome Sanger Institute COVID-19 Surveillance Team |
| EPI_ISL_2972497 | Lighthouse Laboratory Plymouth | Wellcome Sanger Institute for the COVID-19 Genomics UK (COG-UK) Consortium | Lighthouse Laboratory Plymouth and Alex Alderton, Roberto Amato, Jeffrey Barrett, Sonia Goncalves, Ewan Harrison, David K. Jackson, Ian Johnston, Dominic Kwiatkowski, Cordelia Langford, John Sillitoe on behalf of the Wellcome Sanger Institute COVID-19 Surveillance Team |
| EPI_ISL_2972498 | Department of Bacteria, Parasites and Fungi, Statens Serum Institut, Copenhagen, Denmark | Statens Serum Institut Bioinformatics and Microbial Genomics | Danish Covid-19 Genome Consortium |
| EPI_ISL_2972499 | Lighthouse Laboratory Plymouth | Wellcome Sanger Institute for the COVID-19 Genomics UK (COG-UK) Consortium | Lighthouse Laboratory Plymouth and Alex Alderton, Roberto Amato, Jeffrey Barrett, Sonia Goncalves, Ewan Harrison, David K. Jackson, Ian Johnston, Dominic Kwiatkowski, Cordelia Langford, John Sillitoe on behalf of the Wellcome Sanger Institute COVID-19 Surveillance Team |
| EPI_ISL_2972500 | Lighthouse Lab in Alderley Park | Wellcome Sanger Institute for the COVID-19 Genomics UK (COG-UK) Consortium | Jacquelyn Wynn, Mairead Hyland, The Lighthouse Lab in Alderley Park and Alex Alderton, Roberto Amato, Jeffrey Barrett, Sonia Goncalves, Ewan Harrison, David K. Jackson, Ian Johnston, Dominic Kwiatkowski, Cordelia Langford, John Sillitoe on behalf of the Wellcome Sanger Institute COVID-19 Surveillance Team |
| EPI_ISL_2972501 | Department of Bacteria, Parasites and Fungi, Statens Serum Institut, Copenhagen, Denmark | Statens Serum Institut Bioinformatics and Microbial Genomics | Danish Covid-19 Genome Consortium |
| EPI_ISL_2972502 | Lighthouse Laboratory Plymouth | Wellcome Sanger Institute for the COVID-19 Genomics UK (COG-UK) Consortium | Lighthouse Laboratory Plymouth and Alex Alderton, Roberto Amato, Jeffrey Barrett, Sonia Goncalves, Ewan Harrison, David K. Jackson, Ian Johnston, Dominic Kwiatkowski, Cordelia Langford, John Sillitoe on behalf of the Wellcome Sanger Institute COVID-19 Surveillance Team |
| EPI_ISL_2972503, | Lighthouse Lab in Alderley Park | Wellcome Sanger Institute for the COVID-19 Genomics UK (COG-UK) | Jacquelyn Wynn, Mairead Hyland, The Lighthouse Lab in Alderley Park and Alex Alderton, Roberto Amato, Jeffrey Barrett, Sonia Goncalves, Ewan Harrison, David K. Jackson, Ian Johnston, Dominic Kwiatkowski, Cordelia Langford, John Sillitoe on |

| | | | |
|---|---|---|---|
| EPI_ISL_2972505 | | Consortium | behalf of the Wellcome Sanger Institute COVID-19 Surveillance Team |
| EPI_ISL_2972506 | Lighthouse Lab in Milton Keynes | Wellcome Sanger Institute for the COVID-19 Genomics UK (COG-UK) Consortium | The Lighthouse Lab in Milton Keynes and Alex Alderton, Roberto Amato, Jeffrey Barrett, Sonia Goncalves, Ewan Harrison, David K. Jackson, Ian Johnston, Dominic Kwiatkowski, Cordelia Langford, John Sillitoe on behalf of the Wellcome Sanger Institute COVID-19 Surveillance Team |
| EPI_ISL_2972507 | Lighthouse Lab in Alderley Park | Wellcome Sanger Institute for the COVID-19 Genomics UK (COG-UK) Consortium | Jacquelyn Wynn, Mairead Hyland, The Lighthouse Lab in Alderley Park and Alex Alderton, Roberto Amato, Jeffrey Barrett, Sonia Goncalves, Ewan Harrison, David K. Jackson, Ian Johnston, Dominic Kwiatkowski, Cordelia Langford, John Sillitoe on behalf of the Wellcome Sanger Institute COVID-19 Surveillance Team |
| EPI_ISL_2972509 | Lighthouse Lab in Milton Keynes | Wellcome Sanger Institute for the COVID-19 Genomics UK (COG-UK) Consortium | The Lighthouse Lab in Milton Keynes and Alex Alderton, Roberto Amato, Jeffrey Barrett, Sonia Goncalves, Ewan Harrison, David K. Jackson, Ian Johnston, Dominic Kwiatkowski, Cordelia Langford, John Sillitoe on behalf of the Wellcome Sanger Institute COVID-19 Surveillance Team |
| EPI_ISL_2972510 | Lighthouse Lab in Alderley Park | Wellcome Sanger Institute for the COVID-19 Genomics UK (COG-UK) Consortium | Jacquelyn Wynn, Mairead Hyland, The Lighthouse Lab in Alderley Park and Alex Alderton, Roberto Amato, Jeffrey Barrett, Sonia Goncalves, Ewan Harrison, David K. Jackson, Ian Johnston, Dominic Kwiatkowski, Cordelia Langford, John Sillitoe on behalf of the Wellcome Sanger Institute COVID-19 Surveillance Team |
| EPI_ISL_2972511 | Department of Bacteria, Parasites and Fungi, Statens Serum Institut, Copenhagen, Denmark | Statens Serum Institut Bioinformatics and Microbial Genomics | Danish Covid-19 Genome Consortium |
| EPI_ISL_2972512 | Lighthouse Lab in Alderley Park | Wellcome Sanger Institute for the COVID-19 Genomics UK (COG-UK) Consortium | Jacquelyn Wynn, Mairead Hyland, The Lighthouse Lab in Alderley Park and Alex Alderton, Roberto Amato, Jeffrey Barrett, Sonia Goncalves, Ewan Harrison, David K. Jackson, Ian Johnston, Dominic Kwiatkowski, Cordelia Langford, John Sillitoe on behalf of the Wellcome Sanger Institute COVID-19 Surveillance Team |
| EPI_ISL_2972513 | Lighthouse Lab in Milton Keynes | Wellcome Sanger Institute for the COVID-19 Genomics UK (COG-UK) Consortium | The Lighthouse Lab in Milton Keynes and Alex Alderton, Roberto Amato, Jeffrey Barrett, Sonia Goncalves, Ewan Harrison, David K. Jackson, Ian Johnston, Dominic Kwiatkowski, Cordelia Langford, John Sillitoe on behalf of the Wellcome Sanger Institute COVID-19 Surveillance Team |
| EPI_ISL_2972514 | Department of Bacteria, Parasites and Fungi, Statens Serum Institut, Copenhagen, Denmark | Statens Serum Institut Bioinformatics and Microbial Genomics | Danish Covid-19 Genome Consortium |
| EPI_ISL_2972515, EPI_ISL_2972516 | Lighthouse Lab in Alderley Park | Wellcome Sanger Institute for the COVID-19 Genomics UK (COG-UK) Consortium | Jacquelyn Wynn, Mairead Hyland, The Lighthouse Lab in Alderley Park and Alex Alderton, Roberto Amato, Jeffrey Barrett, Sonia Goncalves, Ewan Harrison, David K. Jackson, Ian Johnston, Dominic Kwiatkowski, Cordelia Langford, John Sillitoe on behalf of the Wellcome Sanger Institute COVID-19 Surveillance Team |
| EPI_ISL_2972517 | Lighthouse Laboratory Plymouth | Wellcome Sanger Institute for the COVID-19 Genomics UK (COG-UK) Consortium | Lighthouse Laboratory Plymouth and Alex Alderton, Roberto Amato, Jeffrey Barrett, Sonia Goncalves, Ewan Harrison, David K. Jackson, Ian Johnston, Dominic Kwiatkowski, Cordelia Langford, John Sillitoe on behalf of the Wellcome Sanger Institute COVID-19 Surveillance Team |
| EPI_ISL_2972518 | Department of Bacteria, Parasites and Fungi, Statens Serum Institut, Copenhagen, Denmark | Statens Serum Institut Bioinformatics and Microbial Genomics | Danish Covid-19 Genome Consortium |
| EPI_ISL_2972519 | Lighthouse Lab in Milton Keynes | Wellcome Sanger Institute for the COVID-19 Genomics UK (COG-UK) Consortium | The Lighthouse Lab in Milton Keynes and Alex Alderton, Roberto Amato, Jeffrey Barrett, Sonia Goncalves, Ewan Harrison, David K. Jackson, Ian Johnston, Dominic Kwiatkowski, Cordelia Langford, John Sillitoe on behalf of the Wellcome Sanger Institute COVID-19 Surveillance Team |
| EPI_ISL_2972520 | Lighthouse Lab in Alderley Park | Wellcome Sanger Institute for the COVID-19 Genomics UK (COG-UK) Consortium | Jacquelyn Wynn, Mairead Hyland, The Lighthouse Lab in Alderley Park and Alex Alderton, Roberto Amato, Jeffrey Barrett, Sonia Goncalves, Ewan Harrison, David K. Jackson, Ian Johnston, Dominic Kwiatkowski, Cordelia Langford, John Sillitoe on behalf of the Wellcome Sanger Institute COVID-19 Surveillance Team |
| EPI_ISL_2972521 | Department of Bacteria, Parasites and Fungi, Statens Serum Institut, Copenhagen, Denmark | Statens Serum Institut Bioinformatics and Microbial Genomics | Danish Covid-19 Genome Consortium |
| EPI_ISL_2972522, EPI_ISL_2972523 | Lighthouse Lab in Milton Keynes | Wellcome Sanger Institute for the COVID-19 Genomics UK (COG-UK) Consortium | The Lighthouse Lab in Milton Keynes and Alex Alderton, Roberto Amato, Jeffrey Barrett, Sonia Goncalves, Ewan Harrison, David K. Jackson, Ian Johnston, Dominic Kwiatkowski, Cordelia Langford, John Sillitoe on behalf of the Wellcome Sanger Institute COVID-19 Surveillance Team |
| EPI_ISL_2972524 | Lighthouse Lab in Alderley Park | Wellcome Sanger Institute for the COVID-19 Genomics UK (COG-UK) Consortium | Jacquelyn Wynn, Mairead Hyland, The Lighthouse Lab in Alderley Park and Alex Alderton, Roberto Amato, Jeffrey Barrett, Sonia Goncalves, Ewan Harrison, David K. Jackson, Ian Johnston, Dominic Kwiatkowski, Cordelia Langford, John Sillitoe on behalf of the Wellcome Sanger Institute COVID-19 Surveillance Team |
| EPI_ISL_2972525 | Department of Bacteria, Parasites and Fungi, Statens Serum Institut, Copenhagen, Denmark | Statens Serum Institut Bioinformatics and Microbial Genomics | Danish Covid-19 Genome Consortium |
| EPI_ISL_2972526, EPI_ISL_2972527, EPI_ISL_2972529 | Lighthouse Lab in Alderley Park | Wellcome Sanger Institute for the COVID-19 Genomics UK (COG-UK) Consortium | Jacquelyn Wynn, Mairead Hyland, The Lighthouse Lab in Alderley Park and Alex Alderton, Roberto Amato, Jeffrey Barrett, Sonia Goncalves, Ewan Harrison, David K. Jackson, Ian Johnston, Dominic Kwiatkowski, Cordelia Langford, John Sillitoe on behalf of the Wellcome Sanger Institute COVID-19 Surveillance Team |
| EPI_ISL_2972530 | Lighthouse Laboratory Plymouth | Wellcome Sanger Institute for the COVID-19 Genomics UK (COG-UK) Consortium | Lighthouse Laboratory Plymouth and Alex Alderton, Roberto Amato, Jeffrey Barrett, Sonia Goncalves, Ewan Harrison, David K. Jackson, Ian Johnston, Dominic Kwiatkowski, Cordelia Langford, John Sillitoe on behalf of the Wellcome Sanger Institute COVID-19 Surveillance Team |
| EPI_ISL_2972531 | Lighthouse Lab in Alderley Park | Wellcome Sanger Institute for the COVID-19 Genomics UK (COG-UK) Consortium | Jacquelyn Wynn, Mairead Hyland, The Lighthouse Lab in Alderley Park and Alex Alderton, Roberto Amato, Jeffrey Barrett, Sonia Goncalves, Ewan Harrison, David K. Jackson, Ian Johnston, Dominic Kwiatkowski, Cordelia Langford, John Sillitoe on behalf of the Wellcome Sanger Institute COVID-19 Surveillance Team |
| EPI_ISL_2972532 | Department of Bacteria, Parasites and Fungi, Statens Serum Institut, Copenhagen, Denmark | Statens Serum Institut Bioinformatics and Microbial Genomics | Danish Covid-19 Genome Consortium |
| EPI_ISL_2972533 | Lighthouse Lab in Milton Keynes | Wellcome Sanger Institute for the COVID-19 Genomics UK (COG-UK) Consortium | The Lighthouse Lab in Milton Keynes and Alex Alderton, Roberto Amato, Jeffrey Barrett, Sonia Goncalves, Ewan Harrison, David K. Jackson, Ian Johnston, Dominic Kwiatkowski, Cordelia Langford, John Sillitoe on behalf of the Wellcome Sanger Institute COVID-19 Surveillance Team |
| EPI_ISL_2972534 | Lighthouse Lab in Alderley Park | Wellcome Sanger Institute for the COVID-19 Genomics UK (COG-UK) Consortium | Jacquelyn Wynn, Mairead Hyland, The Lighthouse Lab in Alderley Park and Alex Alderton, Roberto Amato, Jeffrey Barrett, Sonia Goncalves, Ewan Harrison, David K. Jackson, Ian Johnston, Dominic Kwiatkowski, Cordelia Langford, John Sillitoe on behalf of the Wellcome Sanger Institute COVID-19 Surveillance Team |
| EPI_ISL_2972535 | Department of Bacteria, Parasites and Fungi, Statens Serum Institut, Copenhagen, Denmark | Statens Serum Institut Bioinformatics and Microbial Genomics | Danish Covid-19 Genome Consortium |
| EPI_ISL_2972536, EPI_ISL_2972537 | Lighthouse Lab in Alderley Park | Wellcome Sanger Institute for the COVID-19 Genomics UK (COG-UK) Consortium | Jacquelyn Wynn, Mairead Hyland, The Lighthouse Lab in Alderley Park and Alex Alderton, Roberto Amato, Jeffrey Barrett, Sonia Goncalves, Ewan Harrison, David K. Jackson, Ian Johnston, Dominic Kwiatkowski, Cordelia Langford, John Sillitoe on behalf of the Wellcome Sanger Institute COVID-19 Surveillance Team |
| EPI_ISL_2972538 | Department of Bacteria, Parasites and Fungi, Statens Serum Institut, Copenhagen, Denmark | Statens Serum Institut Bioinformatics and Microbial Genomics | Danish Covid-19 Genome Consortium |
| EPI_ISL_2972539 | Lighthouse Laboratory Plymouth | Wellcome Sanger Institute for the COVID-19 Genomics UK (COG-UK) Consortium | Lighthouse Laboratory Plymouth and Alex Alderton, Roberto Amato, Jeffrey Barrett, Sonia Goncalves, Ewan Harrison, David K. Jackson, Ian Johnston, Dominic Kwiatkowski, Cordelia Langford, John Sillitoe on behalf of the Wellcome Sanger Institute COVID-19 Surveillance Team |
| EPI_ISL_2972540 | Lighthouse Lab in Milton Keynes | Wellcome Sanger Institute for the COVID-19 Genomics UK (COG-UK) Consortium | The Lighthouse Lab in Milton Keynes and Alex Alderton, Roberto Amato, Jeffrey Barrett, Sonia Goncalves, Ewan Harrison, David K. Jackson, Ian Johnston, Dominic Kwiatkowski, Cordelia Langford, John Sillitoe on behalf of the Wellcome Sanger Institute COVID-19 Surveillance Team |
| EPI_ISL_2972542, EPI_ISL_2972543, EPI_ISL_2972544 | Lighthouse Lab in Alderley Park | Wellcome Sanger Institute for the COVID-19 Genomics UK (COG-UK) Consortium | Jacquelyn Wynn, Mairead Hyland, The Lighthouse Lab in Alderley Park and Alex Alderton, Roberto Amato, Jeffrey Barrett, Sonia Goncalves, Ewan Harrison, David K. Jackson, Ian Johnston, Dominic Kwiatkowski, Cordelia Langford, John Sillitoe on behalf of the Wellcome Sanger Institute COVID-19 Surveillance Team |
| EPI_ISL_2972545 | Department of Bacteria, Parasites and Fungi, Statens Serum Institut, Copenhagen, Denmark | Statens Serum Institut Bioinformatics and Microbial Genomics | Danish Covid-19 Genome Consortium |
| EPI_ISL_2972546 | Lighthouse Lab in Alderley Park | Wellcome Sanger Institute for the COVID-19 Genomics UK (COG-UK) Consortium | Jacquelyn Wynn, Mairead Hyland, The Lighthouse Lab in Alderley Park and Alex Alderton, Roberto Amato, Jeffrey Barrett, Sonia Goncalves, Ewan Harrison, David K. Jackson, Ian Johnston, Dominic Kwiatkowski, Cordelia Langford, John Sillitoe on behalf of the Wellcome Sanger Institute COVID-19 Surveillance Team |
| EPI_ISL_2972547 | Lighthouse Lab in Milton Keynes | Wellcome Sanger Institute for the COVID-19 Genomics UK (COG-UK) Consortium | The Lighthouse Lab in Milton Keynes and Alex Alderton, Roberto Amato, Jeffrey Barrett, Sonia Goncalves, Ewan Harrison, David K. Jackson, Ian Johnston, Dominic Kwiatkowski, Cordelia Langford, John Sillitoe on behalf of the Wellcome Sanger Institute COVID-19 Surveillance Team |
| EPI_ISL_2972548 | Department of Bacteria, Parasites and Fungi, Statens Serum Institut, Copenhagen, Denmark | Statens Serum Institut Bioinformatics and Microbial Genomics | Danish Covid-19 Genome Consortium |
| EPI_ISL_2972549 | Lighthouse Laboratory Plymouth | Wellcome Sanger Institute for the COVID-19 Genomics UK (COG-UK) Consortium | Lighthouse Laboratory Plymouth and Alex Alderton, Roberto Amato, Jeffrey Barrett, Sonia Goncalves, Ewan Harrison, David K. Jackson, Ian Johnston, Dominic Kwiatkowski, Cordelia Langford, John Sillitoe on behalf of the Wellcome Sanger Institute COVID-19 Surveillance Team |
| EPI_ISL_2972550 | Lighthouse Lab in Alderley Park | Wellcome Sanger Institute for the COVID-19 Genomics UK (COG-UK) Consortium | Jacquelyn Wynn, Mairead Hyland, The Lighthouse Lab in Alderley Park and Alex Alderton, Roberto Amato, Jeffrey Barrett, Sonia Goncalves, Ewan Harrison, David K. Jackson, Ian Johnston, Dominic Kwiatkowski, Cordelia Langford, John Sillitoe on behalf of the Wellcome Sanger Institute COVID-19 Surveillance Team |
| EPI_ISL_2972551 | Lighthouse Laboratory Plymouth | Wellcome Sanger Institute for the COVID-19 Genomics UK (COG-UK) Consortium | Lighthouse Laboratory Plymouth and Alex Alderton, Roberto Amato, Jeffrey Barrett, Sonia Goncalves, Ewan Harrison, David K. Jackson, Ian Johnston, Dominic Kwiatkowski, Cordelia Langford, John Sillitoe on behalf of the Wellcome Sanger Institute COVID-19 Surveillance Team |
| EPI_ISL_2972552 | Department of Bacteria, Parasites and Fungi, Statens Serum Institut, Copenhagen, Denmark | Statens Serum Institut Bioinformatics and Microbial Genomics | Danish Covid-19 Genome Consortium |
| EPI_ISL_2972553, EPI_ISL_2972554 | Lighthouse Lab in Alderley Park | Wellcome Sanger Institute for the COVID-19 Genomics UK (COG-UK) Consortium | Jacquelyn Wynn, Mairead Hyland, The Lighthouse Lab in Alderley Park and Alex Alderton, Roberto Amato, Jeffrey Barrett, Sonia Goncalves, Ewan Harrison, David K. Jackson, Ian Johnston, Dominic Kwiatkowski, Cordelia Langford, John Sillitoe on behalf of the Wellcome Sanger Institute COVID-19 Surveillance Team |
| EPI_ISL_2972556 | Lighthouse Laboratory Plymouth | Wellcome Sanger Institute for the COVID-19 Genomics UK (COG-UK) Consortium | Lighthouse Laboratory Plymouth and Alex Alderton, Roberto Amato, Jeffrey Barrett, Sonia Goncalves, Ewan Harrison, David K. Jackson, Ian Johnston, Dominic Kwiatkowski, Cordelia Langford, John Sillitoe on behalf of the Wellcome Sanger Institute COVID-19 Surveillance Team |
| EPI_ISL_2972557 | Lighthouse Lab in Alderley Park | Wellcome Sanger Institute for the COVID-19 Genomics UK (COG-UK) Consortium | Jacquelyn Wynn, Mairead Hyland, The Lighthouse Lab in Alderley Park and Alex Alderton, Roberto Amato, Jeffrey Barrett, Sonia Goncalves, Ewan Harrison, David K. Jackson, Ian Johnston, Dominic Kwiatkowski, Cordelia Langford, John Sillitoe on behalf of the Wellcome Sanger Institute COVID-19 Surveillance Team |
| EPI_ISL_2972558 | Department of Bacteria, Parasites and Fungi, Statens Serum Institut, Copenhagen, Denmark | Statens Serum Institut Bioinformatics and Microbial Genomics | Danish Covid-19 Genome Consortium |
| EPI_ISL_2972559 | Lighthouse Lab in Alderley Park | Wellcome Sanger Institute for the COVID-19 Genomics UK (COG-UK) Consortium | Jacquelyn Wynn, Mairead Hyland, The Lighthouse Lab in Alderley Park and Alex Alderton, Roberto Amato, Jeffrey Barrett, Sonia Goncalves, Ewan Harrison, David K. Jackson, Ian Johnston, Dominic Kwiatkowski, Cordelia Langford, John Sillitoe on behalf of the Wellcome Sanger Institute COVID-19 Surveillance Team |
| EPI_ISL_2972560 | Lighthouse Lab in Milton Keynes | Wellcome Sanger Institute for the COVID-19 Genomics UK (COG-UK) Consortium | The Lighthouse Lab in Milton Keynes and Alex Alderton, Roberto Amato, Jeffrey Barrett, Sonia Goncalves, Ewan Harrison, David K. Jackson, Ian Johnston, Dominic Kwiatkowski, Cordelia Langford, John Sillitoe on behalf of the Wellcome Sanger Institute COVID-19 Surveillance Team |
| EPI_ISL_2972561 | Lighthouse Lab in Alderley Park | Wellcome Sanger Institute for the COVID-19 Genomics UK (COG-UK) | Jacquelyn Wynn, Mairead Hyland, The Lighthouse Lab in Alderley Park and Alex Alderton, Roberto Amato, Jeffrey Barrett, Sonia Goncalves, Ewan Harrison, David K. Jackson, Ian Johnston, Dominic Kwiatkowski, Cordelia Langford, John Sillitoe on |

| | | | |
|---|---|---|---|
| | | Consortium | behalf of the Wellcome Sanger Institute COVID-19 Surveillance Team |
| EPI_ISL_2972562 | Department of Bacteria, Parasites and Fungi, Statens Serum Institut, Copenhagen, Denmark | Statens Serum Institut Bioinformatics and Microbial Genomics | Danish Covid-19 Genome Consortium |
| EPI_ISL_2972563, EPI_ISL_2972564 | Lighthouse Laboratory Plymouth | Wellcome Sanger Institute for the COVID-19 Genomics UK (COG-UK) Consortium | Lighthouse Laboratory Plymouth and Alex Alderton, Roberto Amato, Jeffrey Barrett, Sonia Goncalves, Ewan Harrison, David K. Jackson, Ian Johnston, Dominic Kwiatkowski, Cordelia Langford, John Sillitoe on behalf of the Wellcome Sanger Institute COVID-19 Surveillance Team |
| EPI_ISL_2972565 | Department of Bacteria, Parasites and Fungi, Statens Serum Institut, Copenhagen, Denmark | Statens Serum Institut Bioinformatics and Microbial Genomics | Danish Covid-19 Genome Consortium |
| EPI_ISL_2972566, EPI_ISL_2972567 | Lighthouse Lab in Milton Keynes | Wellcome Sanger Institute for the COVID-19 Genomics UK (COG-UK) Consortium | The Lighthouse Lab in Milton Keynes and Alex Alderton, Roberto Amato, Jeffrey Barrett, Sonia Goncalves, Ewan Harrison, David K. Jackson, Ian Johnston, Dominic Kwiatkowski, Cordelia Langford, John Sillitoe on behalf of the Wellcome Sanger Institute COVID-19 Surveillance Team |
| EPI_ISL_2972568 | Department of Bacteria, Parasites and Fungi, Statens Serum Institut, Copenhagen, Denmark | Statens Serum Institut Bioinformatics and Microbial Genomics | Danish Covid-19 Genome Consortium |
| EPI_ISL_2972569 | Lighthouse Lab in Alderley Park | Wellcome Sanger Institute for the COVID-19 Genomics UK (COG-UK) Consortium | Jacquelyn Wynn, Mairead Hyland, The Lighthouse Lab in Alderley Park and Alex Alderton, Roberto Amato, Jeffrey Barrett, Sonia Goncalves, Ewan Harrison, David K. Jackson, Ian Johnston, Dominic Kwiatkowski, Cordelia Langford, John Sillitoe on behalf of the Wellcome Sanger Institute COVID-19 Surveillance Team |
| EPI_ISL_2972570 | Lighthouse Lab in Milton Keynes | Wellcome Sanger Institute for the COVID-19 Genomics UK (COG-UK) Consortium | The Lighthouse Lab in Milton Keynes and Alex Alderton, Roberto Amato, Jeffrey Barrett, Sonia Goncalves, Ewan Harrison, David K. Jackson, Ian Johnston, Dominic Kwiatkowski, Cordelia Langford, John Sillitoe on behalf of the Wellcome Sanger Institute COVID-19 Surveillance Team |
| EPI_ISL_2972571 | Lighthouse Laboratory Plymouth | Wellcome Sanger Institute for the COVID-19 Genomics UK (COG-UK) Consortium | Lighthouse Laboratory Plymouth and Alex Alderton, Roberto Amato, Jeffrey Barrett, Sonia Goncalves, Ewan Harrison, David K. Jackson, Ian Johnston, Dominic Kwiatkowski, Cordelia Langford, John Sillitoe on behalf of the Wellcome Sanger Institute COVID-19 Surveillance Team |
| EPI_ISL_2972572 | Department of Bacteria, Parasites and Fungi, Statens Serum Institut, Copenhagen, Denmark | Statens Serum Institut Bioinformatics and Microbial Genomics | Danish Covid-19 Genome Consortium |
| EPI_ISL_2972573 | Lighthouse Lab in Milton Keynes | Wellcome Sanger Institute for the COVID-19 Genomics UK (COG-UK) Consortium | The Lighthouse Lab in Milton Keynes and Alex Alderton, Roberto Amato, Jeffrey Barrett, Sonia Goncalves, Ewan Harrison, David K. Jackson, Ian Johnston, Dominic Kwiatkowski, Cordelia Langford, John Sillitoe on behalf of the Wellcome Sanger Institute COVID-19 Surveillance Team |
| EPI_ISL_2972574 | Lighthouse Lab in Alderley Park | Wellcome Sanger Institute for the COVID-19 Genomics UK (COG-UK) Consortium | Jacquelyn Wynn, Mairead Hyland, The Lighthouse Lab in Alderley Park and Alex Alderton, Roberto Amato, Jeffrey Barrett, Sonia Goncalves, Ewan Harrison, David K. Jackson, Ian Johnston, Dominic Kwiatkowski, Cordelia Langford, John Sillitoe on behalf of the Wellcome Sanger Institute COVID-19 Surveillance Team |
| EPI_ISL_2972575 | Department of Bacteria, Parasites and Fungi, Statens Serum Institut, Copenhagen, Denmark | Statens Serum Institut Bioinformatics and Microbial Genomics | Danish Covid-19 Genome Consortium |
| EPI_ISL_2972576 | Lighthouse Lab in Alderley Park | Wellcome Sanger Institute for the COVID-19 Genomics UK (COG-UK) Consortium | Jacquelyn Wynn, Mairead Hyland, The Lighthouse Lab in Alderley Park and Alex Alderton, Roberto Amato, Jeffrey Barrett, Sonia Goncalves, Ewan Harrison, David K. Jackson, Ian Johnston, Dominic Kwiatkowski, Cordelia Langford, John Sillitoe on behalf of the Wellcome Sanger Institute COVID-19 Surveillance Team |
| EPI_ISL_2972577 | Lighthouse Laboratory Plymouth | Wellcome Sanger Institute for the COVID-19 Genomics UK (COG-UK) Consortium | Lighthouse Laboratory Plymouth and Alex Alderton, Roberto Amato, Jeffrey Barrett, Sonia Goncalves, Ewan Harrison, David K. Jackson, Ian Johnston, Dominic Kwiatkowski, Cordelia Langford, John Sillitoe on behalf of the Wellcome Sanger Institute COVID-19 Surveillance Team |
| EPI_ISL_2972578 | Lighthouse Lab in Alderley Park | Wellcome Sanger Institute for the COVID-19 Genomics UK (COG-UK) Consortium | Jacquelyn Wynn, Mairead Hyland, The Lighthouse Lab in Alderley Park and Alex Alderton, Roberto Amato, Jeffrey Barrett, Sonia Goncalves, Ewan Harrison, David K. Jackson, Ian Johnston, Dominic Kwiatkowski, Cordelia Langford, John Sillitoe on behalf of the Wellcome Sanger Institute COVID-19 Surveillance Team |
| EPI_ISL_2972579 | Department of Bacteria, Parasites and Fungi, Statens Serum Institut, Copenhagen, Denmark | Statens Serum Institut Bioinformatics and Microbial Genomics | Danish Covid-19 Genome Consortium |
| EPI_ISL_2972580 | Lighthouse Lab in Alderley Park | Wellcome Sanger Institute for the COVID-19 Genomics UK (COG-UK) Consortium | Jacquelyn Wynn, Mairead Hyland, The Lighthouse Lab in Alderley Park and Alex Alderton, Roberto Amato, Jeffrey Barrett, Sonia Goncalves, Ewan Harrison, David K. Jackson, Ian Johnston, Dominic Kwiatkowski, Cordelia Langford, John Sillitoe on behalf of the Wellcome Sanger Institute COVID-19 Surveillance Team |
| EPI_ISL_2972581 | Lighthouse Laboratory Plymouth | Wellcome Sanger Institute for the COVID-19 Genomics UK (COG-UK) Consortium | Lighthouse Laboratory Plymouth and Alex Alderton, Roberto Amato, Jeffrey Barrett, Sonia Goncalves, Ewan Harrison, David K. Jackson, Ian Johnston, Dominic Kwiatkowski, Cordelia Langford, John Sillitoe on behalf of the Wellcome Sanger Institute COVID-19 Surveillance Team |
| EPI_ISL_2972582 | Department of Bacteria, Parasites and Fungi, Statens Serum Institut, Copenhagen, Denmark | Statens Serum Institut Bioinformatics and Microbial Genomics | Danish Covid-19 Genome Consortium |
| EPI_ISL_2972583, EPI_ISL_2972584 | Lighthouse Lab in Milton Keynes | Wellcome Sanger Institute for the COVID-19 Genomics UK (COG-UK) Consortium | The Lighthouse Lab in Milton Keynes and Alex Alderton, Roberto Amato, Jeffrey Barrett, Sonia Goncalves, Ewan Harrison, David K. Jackson, Ian Johnston, Dominic Kwiatkowski, Cordelia Langford, John Sillitoe on behalf of the Wellcome Sanger Institute COVID-19 Surveillance Team |
| EPI_ISL_2972585 | Lighthouse Laboratory Plymouth | Wellcome Sanger Institute for the COVID-19 Genomics UK (COG-UK) Consortium | Lighthouse Laboratory Plymouth and Alex Alderton, Roberto Amato, Jeffrey Barrett, Sonia Goncalves, Ewan Harrison, David K. Jackson, Ian Johnston, Dominic Kwiatkowski, Cordelia Langford, John Sillitoe on behalf of the Wellcome Sanger Institute COVID-19 Surveillance Team |
| EPI_ISL_2972586 | Department of Bacteria, Parasites and Fungi, Statens Serum Institut, Copenhagen, Denmark | Statens Serum Institut Bioinformatics and Microbial Genomics | Danish Covid-19 Genome Consortium |
| EPI_ISL_2972587, EPI_ISL_2972588 | Lighthouse Lab in Alderley Park | Wellcome Sanger Institute for the COVID-19 Genomics UK (COG-UK) Consortium | Jacquelyn Wynn, Mairead Hyland, The Lighthouse Lab in Alderley Park and Alex Alderton, Roberto Amato, Jeffrey Barrett, Sonia Goncalves, Ewan Harrison, David K. Jackson, Ian Johnston, Dominic Kwiatkowski, Cordelia Langford, John Sillitoe on behalf of the Wellcome Sanger Institute COVID-19 Surveillance Team |
| EPI_ISL_2972589 | Department of Bacteria, Parasites and Fungi, Statens Serum Institut, Copenhagen, Denmark | Statens Serum Institut Bioinformatics and Microbial Genomics | Danish Covid-19 Genome Consortium |
| EPI_ISL_2972590 | Lighthouse Laboratory Plymouth | Wellcome Sanger Institute for the COVID-19 Genomics UK (COG-UK) Consortium | Lighthouse Laboratory Plymouth and Alex Alderton, Roberto Amato, Jeffrey Barrett, Sonia Goncalves, Ewan Harrison, David K. Jackson, Ian Johnston, Dominic Kwiatkowski, Cordelia Langford, John Sillitoe on behalf of the Wellcome Sanger Institute COVID-19 Surveillance Team |
| EPI_ISL_2972591 | Lighthouse Lab in Milton Keynes | Wellcome Sanger Institute for the COVID-19 Genomics UK (COG-UK) Consortium | The Lighthouse Lab in Milton Keynes and Alex Alderton, Roberto Amato, Jeffrey Barrett, Sonia Goncalves, Ewan Harrison, David K. Jackson, Ian Johnston, Dominic Kwiatkowski, Cordelia Langford, John Sillitoe on behalf of the Wellcome Sanger Institute COVID-19 Surveillance Team |
| EPI_ISL_2972592 | Department of Bacteria, Parasites and Fungi, Statens Serum Institut, Copenhagen, Denmark | Statens Serum Institut Bioinformatics and Microbial Genomics | Danish Covid-19 Genome Consortium |
| EPI_ISL_2972593 | Lighthouse Lab in Milton Keynes | Wellcome Sanger Institute for the COVID-19 Genomics UK (COG-UK) Consortium | The Lighthouse Lab in Milton Keynes and Alex Alderton, Roberto Amato, Jeffrey Barrett, Sonia Goncalves, Ewan Harrison, David K. Jackson, Ian Johnston, Dominic Kwiatkowski, Cordelia Langford, John Sillitoe on behalf of the Wellcome Sanger Institute COVID-19 Surveillance Team |
| EPI_ISL_2972594 | Lighthouse Laboratory Plymouth | Wellcome Sanger Institute for the COVID-19 Genomics UK (COG-UK) Consortium | Lighthouse Laboratory Plymouth and Alex Alderton, Roberto Amato, Jeffrey Barrett, Sonia Goncalves, Ewan Harrison, David K. Jackson, Ian Johnston, Dominic Kwiatkowski, Cordelia Langford, John Sillitoe on behalf of the Wellcome Sanger Institute COVID-19 Surveillance Team |
| EPI_ISL_2972595 | Department of Bacteria, Parasites and Fungi, Statens Serum Institut, Copenhagen, Denmark | Statens Serum Institut Bioinformatics and Microbial Genomics | Danish Covid-19 Genome Consortium |
| EPI_ISL_2972596, EPI_ISL_2972597 | Lighthouse Lab in Alderley Park | Wellcome Sanger Institute for the COVID-19 Genomics UK (COG-UK) Consortium | Jacquelyn Wynn, Mairead Hyland, The Lighthouse Lab in Alderley Park and Alex Alderton, Roberto Amato, Jeffrey Barrett, Sonia Goncalves, Ewan Harrison, David K. Jackson, Ian Johnston, Dominic Kwiatkowski, Cordelia Langford, John Sillitoe on behalf of the Wellcome Sanger Institute COVID-19 Surveillance Team |
| EPI_ISL_2972598 | Department of Bacteria, Parasites and Fungi, Statens Serum Institut, Copenhagen, Denmark | Statens Serum Institut Bioinformatics and Microbial Genomics | Danish Covid-19 Genome Consortium |
| EPI_ISL_2972599, EPI_ISL_2972600 | Lighthouse Lab in Alderley Park | Wellcome Sanger Institute for the COVID-19 Genomics UK (COG-UK) Consortium | Jacquelyn Wynn, Mairead Hyland, The Lighthouse Lab in Alderley Park and Alex Alderton, Roberto Amato, Jeffrey Barrett, Sonia Goncalves, Ewan Harrison, David K. Jackson, Ian Johnston, Dominic Kwiatkowski, Cordelia Langford, John Sillitoe on behalf of the Wellcome Sanger Institute COVID-19 Surveillance Team |
| EPI_ISL_2972601 | Lighthouse Lab in Milton Keynes | Wellcome Sanger Institute for the COVID-19 Genomics UK (COG-UK) Consortium | The Lighthouse Lab in Milton Keynes and Alex Alderton, Roberto Amato, Jeffrey Barrett, Sonia Goncalves, Ewan Harrison, David K. Jackson, Ian Johnston, Dominic Kwiatkowski, Cordelia Langford, John Sillitoe on behalf of the Wellcome Sanger Institute COVID-19 Surveillance Team |
| EPI_ISL_2972603, EPI_ISL_2972604 | Lighthouse Lab in Alderley Park | Wellcome Sanger Institute for the COVID-19 Genomics UK (COG-UK) Consortium | Jacquelyn Wynn, Mairead Hyland, The Lighthouse Lab in Alderley Park and Alex Alderton, Roberto Amato, Jeffrey Barrett, Sonia Goncalves, Ewan Harrison, David K. Jackson, Ian Johnston, Dominic Kwiatkowski, Cordelia Langford, John Sillitoe on behalf of the Wellcome Sanger Institute COVID-19 Surveillance Team |
| EPI_ISL_2972605 | Department of Bacteria, Parasites and Fungi, Statens Serum Institut, Copenhagen, Denmark | Statens Serum Institut Bioinformatics and Microbial Genomics | Danish Covid-19 Genome Consortium |
| EPI_ISL_2972606 | Lighthouse Lab in Alderley Park | Wellcome Sanger Institute for the COVID-19 Genomics UK (COG-UK) Consortium | Jacquelyn Wynn, Mairead Hyland, The Lighthouse Lab in Alderley Park and Alex Alderton, Roberto Amato, Jeffrey Barrett, Sonia Goncalves, Ewan Harrison, David K. Jackson, Ian Johnston, Dominic Kwiatkowski, Cordelia Langford, John Sillitoe on behalf of the Wellcome Sanger Institute COVID-19 Surveillance Team |
| EPI_ISL_2972607 | Lighthouse Laboratory Plymouth | Wellcome Sanger Institute for the COVID-19 Genomics UK (COG-UK) Consortium | Lighthouse Laboratory Plymouth and Alex Alderton, Roberto Amato, Jeffrey Barrett, Sonia Goncalves, Ewan Harrison, David K. Jackson, Ian Johnston, Dominic Kwiatkowski, Cordelia Langford, John Sillitoe on behalf of the Wellcome Sanger Institute COVID-19 Surveillance Team |
| EPI_ISL_2972608 | Department of Bacteria, Parasites and Fungi, Statens Serum Institut, Copenhagen, Denmark | Statens Serum Institut Bioinformatics and Microbial Genomics | Danish Covid-19 Genome Consortium |
| EPI_ISL_2972609, EPI_ISL_2972610 | Lighthouse Lab in Alderley Park | Wellcome Sanger Institute for the COVID-19 Genomics UK (COG-UK) Consortium | Jacquelyn Wynn, Mairead Hyland, The Lighthouse Lab in Alderley Park and Alex Alderton, Roberto Amato, Jeffrey Barrett, Sonia Goncalves, Ewan Harrison, David K. Jackson, Ian Johnston, Dominic Kwiatkowski, Cordelia Langford, John Sillitoe on behalf of the Wellcome Sanger Institute COVID-19 Surveillance Team |
| EPI_ISL_2972611 | Department of Bacteria, Parasites and Fungi, Statens Serum Institut, Copenhagen, Denmark | Statens Serum Institut Bioinformatics and Microbial Genomics | Danish Covid-19 Genome Consortium |
| EPI_ISL_2972612 | Lighthouse Laboratory Plymouth | Wellcome Sanger Institute for the COVID-19 Genomics UK (COG-UK) Consortium | Lighthouse Laboratory Plymouth and Alex Alderton, Roberto Amato, Jeffrey Barrett, Sonia Goncalves, Ewan Harrison, David K. Jackson, Ian Johnston, Dominic Kwiatkowski, Cordelia Langford, John Sillitoe on behalf of the Wellcome Sanger Institute COVID-19 Surveillance Team |
| EPI_ISL_2972613 | Lighthouse Lab in Alderley Park | Wellcome Sanger Institute for the COVID-19 Genomics UK (COG-UK) Consortium | Jacquelyn Wynn, Mairead Hyland, The Lighthouse Lab in Alderley Park and Alex Alderton, Roberto Amato, Jeffrey Barrett, Sonia Goncalves, Ewan Harrison, David K. Jackson, Ian Johnston, Dominic Kwiatkowski, Cordelia Langford, John Sillitoe on behalf of the Wellcome Sanger Institute COVID-19 Surveillance Team |
| EPI_ISL_2972614 | Lighthouse Laboratory Plymouth | Wellcome Sanger Institute for the COVID-19 Genomics UK (COG-UK) Consortium | Lighthouse Laboratory Plymouth and Alex Alderton, Roberto Amato, Jeffrey Barrett, Sonia Goncalves, Ewan Harrison, David K. Jackson, Ian Johnston, Dominic Kwiatkowski, Cordelia Langford, John Sillitoe on behalf of the Wellcome Sanger Institute COVID-19 Surveillance Team |
| EPI_ISL_2972615 | Department of Bacteria, Parasites and Fungi, Statens Serum | Statens Serum Institut Bioinformatics and Microbial Genomics | Danish Covid-19 Genome Consortium |

| | | | |
|---|---|---|---|
| | Institut, Copenhagen, Denmark | | |
| EPI_ISL_2972616 | Lighthouse Laboratory Plymouth | Wellcome Sanger Institute for the COVID-19 Genomics UK (COG-UK) Consortium | Lighthouse Laboratory Plymouth and Alex Alderton, Roberto Amato, Jeffrey Barrett, Sonia Goncalves, Ewan Harrison, David K. Jackson, Ian Johnston, Dominic Kwiatkowski, Cordelia Langford, John Sillitoe on behalf of the Wellcome Sanger Institute COVID-19 Surveillance Team |
| EPI_ISL_2972617 | Lighthouse Lab in Milton Keynes | Wellcome Sanger Institute for the COVID-19 Genomics UK (COG-UK) Consortium | The Lighthouse Lab in Milton Keynes and Alex Alderton, Roberto Amato, Jeffrey Barrett, Sonia Goncalves, Ewan Harrison, David K. Jackson, Ian Johnston, Dominic Kwiatkowski, Cordelia Langford, John Sillitoe on behalf of the Wellcome Sanger Institute COVID-19 Surveillance Team |
| EPI_ISL_2972618 | Department of Bacteria, Parasites and Fungi, Statens Serum Institut, Copenhagen, Denmark | Statens Serum Institut Bioinformatics and Microbial Genomics | Danish Covid-19 Genome Consortium |
| EPI_ISL_2972619 | Lighthouse Laboratory Plymouth | Wellcome Sanger Institute for the COVID-19 Genomics UK (COG-UK) Consortium | Lighthouse Laboratory Plymouth and Alex Alderton, Roberto Amato, Jeffrey Barrett, Sonia Goncalves, Ewan Harrison, David K. Jackson, Ian Johnston, Dominic Kwiatkowski, Cordelia Langford, John Sillitoe on behalf of the Wellcome Sanger Institute COVID-19 Surveillance Team |
| EPI_ISL_2972620 | Lighthouse Lab in Alderley Park | Wellcome Sanger Institute for the COVID-19 Genomics UK (COG-UK) Consortium | Jacquelyn Wynn, Mairead Hyland, The Lighthouse Lab in Alderley Park and Alex Alderton, Roberto Amato, Jeffrey Barrett, Sonia Goncalves, Ewan Harrison, David K. Jackson, Ian Johnston, Dominic Kwiatkowski, Cordelia Langford, John Sillitoe on behalf of the Wellcome Sanger Institute COVID-19 Surveillance Team |
| EPI_ISL_2972621 | Department of Bacteria, Parasites and Fungi, Statens Serum Institut, Copenhagen, Denmark | Statens Serum Institut Bioinformatics and Microbial Genomics | Danish Covid-19 Genome Consortium |
| EPI_ISL_2972622 | Lighthouse Lab in Milton Keynes | Wellcome Sanger Institute for the COVID-19 Genomics UK (COG-UK) Consortium | The Lighthouse Lab in Milton Keynes and Alex Alderton, Roberto Amato, Jeffrey Barrett, Sonia Goncalves, Ewan Harrison, David K. Jackson, Ian Johnston, Dominic Kwiatkowski, Cordelia Langford, John Sillitoe on behalf of the Wellcome Sanger Institute COVID-19 Surveillance Team |
| EPI_ISL_2972623, EPI_ISL_2972624 | Lighthouse Laboratory Plymouth | Wellcome Sanger Institute for the COVID-19 Genomics UK (COG-UK) Consortium | Lighthouse Laboratory Plymouth and Alex Alderton, Roberto Amato, Jeffrey Barrett, Sonia Goncalves, Ewan Harrison, David K. Jackson, Ian Johnston, Dominic Kwiatkowski, Cordelia Langford, John Sillitoe on behalf of the Wellcome Sanger Institute COVID-19 Surveillance Team |
| EPI_ISL_2972626 | Lighthouse Lab in Alderley Park | Wellcome Sanger Institute for the COVID-19 Genomics UK (COG-UK) Consortium | Jacquelyn Wynn, Mairead Hyland, The Lighthouse Lab in Alderley Park and Alex Alderton, Roberto Amato, Jeffrey Barrett, Sonia Goncalves, Ewan Harrison, David K. Jackson, Ian Johnston, Dominic Kwiatkowski, Cordelia Langford, John Sillitoe on behalf of the Wellcome Sanger Institute COVID-19 Surveillance Team |
| EPI_ISL_2972627 | Lighthouse Lab in Milton Keynes | Wellcome Sanger Institute for the COVID-19 Genomics UK (COG-UK) Consortium | The Lighthouse Lab in Milton Keynes and Alex Alderton, Roberto Amato, Jeffrey Barrett, Sonia Goncalves, Ewan Harrison, David K. Jackson, Ian Johnston, Dominic Kwiatkowski, Cordelia Langford, John Sillitoe on behalf of the Wellcome Sanger Institute COVID-19 Surveillance Team |
| EPI_ISL_2972628 | Lighthouse Lab in Alderley Park | Wellcome Sanger Institute for the COVID-19 Genomics UK (COG-UK) Consortium | Jacquelyn Wynn, Mairead Hyland, The Lighthouse Lab in Alderley Park and Alex Alderton, Roberto Amato, Jeffrey Barrett, Sonia Goncalves, Ewan Harrison, David K. Jackson, Ian Johnston, Dominic Kwiatkowski, Cordelia Langford, John Sillitoe on behalf of the Wellcome Sanger Institute COVID-19 Surveillance Team |
| EPI_ISL_2972629 | Department of Bacteria, Parasites and Fungi, Statens Serum Institut, Copenhagen, Denmark | Statens Serum Institut Bioinformatics and Microbial Genomics | Danish Covid-19 Genome Consortium |
| EPI_ISL_2972630 | Lighthouse Lab in Alderley Park | Wellcome Sanger Institute for the COVID-19 Genomics UK (COG-UK) Consortium | Jacquelyn Wynn, Mairead Hyland, The Lighthouse Lab in Alderley Park and Alex Alderton, Roberto Amato, Jeffrey Barrett, Sonia Goncalves, Ewan Harrison, David K. Jackson, Ian Johnston, Dominic Kwiatkowski, Cordelia Langford, John Sillitoe on behalf of the Wellcome Sanger Institute COVID-19 Surveillance Team |
| EPI_ISL_2972631, EPI_ISL_2972632 | Lighthouse Lab in Milton Keynes | Wellcome Sanger Institute for the COVID-19 Genomics UK (COG-UK) Consortium | The Lighthouse Lab in Milton Keynes and Alex Alderton, Roberto Amato, Jeffrey Barrett, Sonia Goncalves, Ewan Harrison, David K. Jackson, Ian Johnston, Dominic Kwiatkowski, Cordelia Langford, John Sillitoe on behalf of the Wellcome Sanger Institute COVID-19 Surveillance Team |
| EPI_ISL_2972633 | Department of Bacteria, Parasites and Fungi, Statens Serum Institut, Copenhagen, Denmark | Statens Serum Institut Bioinformatics and Microbial Genomics | Danish Covid-19 Genome Consortium |
| EPI_ISL_2972634 | Lighthouse Lab in Milton Keynes | Wellcome Sanger Institute for the COVID-19 Genomics UK (COG-UK) Consortium | The Lighthouse Lab in Milton Keynes and Alex Alderton, Roberto Amato, Jeffrey Barrett, Sonia Goncalves, Ewan Harrison, David K. Jackson, Ian Johnston, Dominic Kwiatkowski, Cordelia Langford, John Sillitoe on behalf of the Wellcome Sanger Institute COVID-19 Surveillance Team |
| EPI_ISL_2972635 | Lighthouse Lab in Alderley Park | Wellcome Sanger Institute for the COVID-19 Genomics UK (COG-UK) Consortium | Jacquelyn Wynn, Mairead Hyland, The Lighthouse Lab in Alderley Park and Alex Alderton, Roberto Amato, Jeffrey Barrett, Sonia Goncalves, Ewan Harrison, David K. Jackson, Ian Johnston, Dominic Kwiatkowski, Cordelia Langford, John Sillitoe on behalf of the Wellcome Sanger Institute COVID-19 Surveillance Team |
| EPI_ISL_2972636 | Department of Bacteria, Parasites and Fungi, Statens Serum Institut, Copenhagen, Denmark | Statens Serum Institut Bioinformatics and Microbial Genomics | Danish Covid-19 Genome Consortium |
| EPI_ISL_2972637 | Lighthouse Laboratory Plymouth | Wellcome Sanger Institute for the COVID-19 Genomics UK (COG-UK) Consortium | Lighthouse Laboratory Plymouth and Alex Alderton, Roberto Amato, Jeffrey Barrett, Sonia Goncalves, Ewan Harrison, David K. Jackson, Ian Johnston, Dominic Kwiatkowski, Cordelia Langford, John Sillitoe on behalf of the Wellcome Sanger Institute COVID-19 Surveillance Team |
| EPI_ISL_2972638 | Lighthouse Lab in Alderley Park | Wellcome Sanger Institute for the COVID-19 Genomics UK (COG-UK) Consortium | Jacquelyn Wynn, Mairead Hyland, The Lighthouse Lab in Alderley Park and Alex Alderton, Roberto Amato, Jeffrey Barrett, Sonia Goncalves, Ewan Harrison, David K. Jackson, Ian Johnston, Dominic Kwiatkowski, Cordelia Langford, John Sillitoe on behalf of the Wellcome Sanger Institute COVID-19 Surveillance Team |
| EPI_ISL_2972639 | Lighthouse Lab in Milton Keynes | Wellcome Sanger Institute for the COVID-19 Genomics UK (COG-UK) Consortium | The Lighthouse Lab in Milton Keynes and Alex Alderton, Roberto Amato, Jeffrey Barrett, Sonia Goncalves, Ewan Harrison, David K. Jackson, Ian Johnston, Dominic Kwiatkowski, Cordelia Langford, John Sillitoe on behalf of the Wellcome Sanger Institute COVID-19 Surveillance Team |
| EPI_ISL_2972640 | Department of Bacteria, Parasites and Fungi, Statens Serum Institut, Copenhagen, Denmark | Statens Serum Institut Bioinformatics and Microbial Genomics | Danish Covid-19 Genome Consortium |
| EPI_ISL_2972641, EPI_ISL_2972642 | Lighthouse Lab in Alderley Park | Wellcome Sanger Institute for the COVID-19 Genomics UK (COG-UK) Consortium | Jacquelyn Wynn, Mairead Hyland, The Lighthouse Lab in Alderley Park and Alex Alderton, Roberto Amato, Jeffrey Barrett, Sonia Goncalves, Ewan Harrison, David K. Jackson, Ian Johnston, Dominic Kwiatkowski, Cordelia Langford, John Sillitoe on behalf of the Wellcome Sanger Institute COVID-19 Surveillance Team |
| EPI_ISL_2972643 | Department of Bacteria, Parasites and Fungi, Statens Serum Institut, Copenhagen, Denmark | Statens Serum Institut Bioinformatics and Microbial Genomics | Danish Covid-19 Genome Consortium |
| EPI_ISL_2972644 | Lighthouse Lab in Alderley Park | Wellcome Sanger Institute for the COVID-19 Genomics UK (COG-UK) Consortium | Jacquelyn Wynn, Mairead Hyland, The Lighthouse Lab in Alderley Park and Alex Alderton, Roberto Amato, Jeffrey Barrett, Sonia Goncalves, Ewan Harrison, David K. Jackson, Ian Johnston, Dominic Kwiatkowski, Cordelia Langford, John Sillitoe on behalf of the Wellcome Sanger Institute COVID-19 Surveillance Team |
| EPI_ISL_2972645 | Lighthouse Laboratory Plymouth | Wellcome Sanger Institute for the COVID-19 Genomics UK (COG-UK) Consortium | Lighthouse Laboratory Plymouth and Alex Alderton, Roberto Amato, Jeffrey Barrett, Sonia Goncalves, Ewan Harrison, David K. Jackson, Ian Johnston, Dominic Kwiatkowski, Cordelia Langford, John Sillitoe on behalf of the Wellcome Sanger Institute COVID-19 Surveillance Team |
| EPI_ISL_2972646 | Department of Bacteria, Parasites and Fungi, Statens Serum Institut, Copenhagen, Denmark | Statens Serum Institut Bioinformatics and Microbial Genomics | Danish Covid-19 Genome Consortium |
| EPI_ISL_2972647, EPI_ISL_2972648, EPI_ISL_2972649 | Lighthouse Lab in Alderley Park | Wellcome Sanger Institute for the COVID-19 Genomics UK (COG-UK) Consortium | Jacquelyn Wynn, Mairead Hyland, The Lighthouse Lab in Alderley Park and Alex Alderton, Roberto Amato, Jeffrey Barrett, Sonia Goncalves, Ewan Harrison, David K. Jackson, Ian Johnston, Dominic Kwiatkowski, Cordelia Langford, John Sillitoe on behalf of the Wellcome Sanger Institute COVID-19 Surveillance Team |
| EPI_ISL_2972650 | Department of Bacteria, Parasites and Fungi, Statens Serum Institut, Copenhagen, Denmark | Statens Serum Institut Bioinformatics and Microbial Genomics | Danish Covid-19 Genome Consortium |
| EPI_ISL_2972651 | Lighthouse Lab in Alderley Park | Wellcome Sanger Institute for the COVID-19 Genomics UK (COG-UK) Consortium | Jacquelyn Wynn, Mairead Hyland, The Lighthouse Lab in Alderley Park and Alex Alderton, Roberto Amato, Jeffrey Barrett, Sonia Goncalves, Ewan Harrison, David K. Jackson, Ian Johnston, Dominic Kwiatkowski, Cordelia Langford, John Sillitoe on behalf of the Wellcome Sanger Institute COVID-19 Surveillance Team |
| EPI_ISL_2972652 | Department of Bacteria, Parasites and Fungi, Statens Serum Institut, Copenhagen, Denmark | Statens Serum Institut Bioinformatics and Microbial Genomics | Danish Covid-19 Genome Consortium |
| EPI_ISL_2972653 | Lighthouse Laboratory Plymouth | Wellcome Sanger Institute for the COVID-19 Genomics UK (COG-UK) Consortium | Lighthouse Laboratory Plymouth and Alex Alderton, Roberto Amato, Jeffrey Barrett, Sonia Goncalves, Ewan Harrison, David K. Jackson, Ian Johnston, Dominic Kwiatkowski, Cordelia Langford, John Sillitoe on behalf of the Wellcome Sanger Institute COVID-19 Surveillance Team |
| EPI_ISL_2972654, EPI_ISL_2972655 | Lighthouse Lab in Alderley Park | Wellcome Sanger Institute for the COVID-19 Genomics UK (COG-UK) Consortium | Jacquelyn Wynn, Mairead Hyland, The Lighthouse Lab in Alderley Park and Alex Alderton, Roberto Amato, Jeffrey Barrett, Sonia Goncalves, Ewan Harrison, David K. Jackson, Ian Johnston, Dominic Kwiatkowski, Cordelia Langford, John Sillitoe on behalf of the Wellcome Sanger Institute COVID-19 Surveillance Team |
| EPI_ISL_2972656 | Department of Bacteria, Parasites and Fungi, Statens Serum Institut, Copenhagen, Denmark | Statens Serum Institut Bioinformatics and Microbial Genomics | Danish Covid-19 Genome Consortium |
| EPI_ISL_2972657, EPI_ISL_2972658 | Lighthouse Lab in Alderley Park | Wellcome Sanger Institute for the COVID-19 Genomics UK (COG-UK) Consortium | Jacquelyn Wynn, Mairead Hyland, The Lighthouse Lab in Alderley Park and Alex Alderton, Roberto Amato, Jeffrey Barrett, Sonia Goncalves, Ewan Harrison, David K. Jackson, Ian Johnston, Dominic Kwiatkowski, Cordelia Langford, John Sillitoe on behalf of the Wellcome Sanger Institute COVID-19 Surveillance Team |
| EPI_ISL_2972659 | Department of Bacteria, Parasites and Fungi, Statens Serum Institut, Copenhagen, Denmark | Statens Serum Institut Bioinformatics and Microbial Genomics | Danish Covid-19 Genome Consortium |
| EPI_ISL_2972660 | Lighthouse Lab in Alderley Park | Wellcome Sanger Institute for the COVID-19 Genomics UK (COG-UK) Consortium | Jacquelyn Wynn, Mairead Hyland, The Lighthouse Lab in Alderley Park and Alex Alderton, Roberto Amato, Jeffrey Barrett, Sonia Goncalves, Ewan Harrison, David K. Jackson, Ian Johnston, Dominic Kwiatkowski, Cordelia Langford, John Sillitoe on behalf of the Wellcome Sanger Institute COVID-19 Surveillance Team |
| EPI_ISL_2972661 | Lighthouse Lab in Milton Keynes | Wellcome Sanger Institute for the COVID-19 Genomics UK (COG-UK) Consortium | The Lighthouse Lab in Milton Keynes and Alex Alderton, Roberto Amato, Jeffrey Barrett, Sonia Goncalves, Ewan Harrison, David K. Jackson, Ian Johnston, Dominic Kwiatkowski, Cordelia Langford, John Sillitoe on behalf of the Wellcome Sanger Institute COVID-19 Surveillance Team |
| EPI_ISL_2972662 | Department of Bacteria, Parasites and Fungi, Statens Serum Institut, Copenhagen, Denmark | Statens Serum Institut Bioinformatics and Microbial Genomics | Danish Covid-19 Genome Consortium |
| EPI_ISL_2972663 | Lighthouse Lab in Alderley Park | Wellcome Sanger Institute for the COVID-19 Genomics UK (COG-UK) Consortium | Jacquelyn Wynn, Mairead Hyland, The Lighthouse Lab in Alderley Park and Alex Alderton, Roberto Amato, Jeffrey Barrett, Sonia Goncalves, Ewan Harrison, David K. Jackson, Ian Johnston, Dominic Kwiatkowski, Cordelia Langford, John Sillitoe on behalf of the Wellcome Sanger Institute COVID-19 Surveillance Team |
| EPI_ISL_2972664 | Lighthouse Laboratory Plymouth | Wellcome Sanger Institute for the COVID-19 Genomics UK (COG-UK) Consortium | Lighthouse Laboratory Plymouth and Alex Alderton, Roberto Amato, Jeffrey Barrett, Sonia Goncalves, Ewan Harrison, David K. Jackson, Ian Johnston, Dominic Kwiatkowski, Cordelia Langford, John Sillitoe on behalf of the Wellcome Sanger Institute COVID-19 Surveillance Team |
| EPI_ISL_2972665 | Lighthouse Lab in Alderley Park | Wellcome Sanger Institute for the COVID-19 Genomics UK (COG-UK) Consortium | Jacquelyn Wynn, Mairead Hyland, The Lighthouse Lab in Alderley Park and Alex Alderton, Roberto Amato, Jeffrey Barrett, Sonia Goncalves, Ewan Harrison, David K. Jackson, Ian Johnston, Dominic Kwiatkowski, Cordelia Langford, John Sillitoe on behalf of the Wellcome Sanger Institute COVID-19 Surveillance Team |
| EPI_ISL_2972666 | Department of Bacteria, Parasites and Fungi, Statens Serum Institut, Copenhagen, Denmark | Statens Serum Institut Bioinformatics and Microbial Genomics | Danish Covid-19 Genome Consortium |
| EPI_ISL_2972667, EPI_ISL_2972668 | Lighthouse Lab in Alderley Park | Wellcome Sanger Institute for the COVID-19 Genomics UK (COG-UK) Consortium | Jacquelyn Wynn, Mairead Hyland, The Lighthouse Lab in Alderley Park and Alex Alderton, Roberto Amato, Jeffrey Barrett, Sonia Goncalves, Ewan Harrison, David K. Jackson, Ian Johnston, Dominic Kwiatkowski, Cordelia Langford, John Sillitoe on behalf of the Wellcome Sanger Institute COVID-19 Surveillance Team |

| | | | |
|---|---|---|---|
| EPI_ISL_2972669 | Department of Bacteria, Parasites and Fungi, Statens Serum Institut, Copenhagen, Denmark | Statens Serum Institut Bioinformatics and Microbial Genomics | Danish Covid-19 Genome Consortium |
| EPI_ISL_2972670 | Lighthouse Laboratory Plymouth | Wellcome Sanger Institute for the COVID-19 Genomics UK (COG-UK) Consortium | Lighthouse Laboratory Plymouth and Alex Alderton, Roberto Amato, Jeffrey Barrett, Sonia Goncalves, Ewan Harrison, David K. Jackson, Ian Johnston, Dominic Kwiatkowski, Cordelia Langford, John Sillitoe on behalf of the Wellcome Sanger Institute COVID-19 Surveillance Team |
| EPI_ISL_2972671 | Molecular Diagnostics Pathology Department Mater Dei Hospital Malta | Molecular Diagnostics Pathology Department Mater Dei Hospital Malta | G Zahra, R Borg, C Cilia, L Grech, M Briffa |
| EPI_ISL_2972672, EPI_ISL_2972673 | Lighthouse Lab in Alderley Park | Wellcome Sanger Institute for the COVID-19 Genomics UK (COG-UK) Consortium | Jacquelyn Wynn, Mairead Hyland, The Lighthouse Lab in Alderley Park and Alex Alderton, Roberto Amato, Jeffrey Barrett, Sonia Goncalves, Ewan Harrison, David K. Jackson, Ian Johnston, Dominic Kwiatkowski, Cordelia Langford, John Sillitoe on behalf of the Wellcome Sanger Institute COVID-19 Surveillance Team |
| EPI_ISL_2972674 | Department of Bacteria, Parasites and Fungi, Statens Serum Institut, Copenhagen, Denmark | Statens Serum Institut Bioinformatics and Microbial Genomics | Danish Covid-19 Genome Consortium |
| EPI_ISL_2972675 | Lighthouse Laboratory Plymouth | Wellcome Sanger Institute for the COVID-19 Genomics UK (COG-UK) Consortium | Lighthouse Laboratory Plymouth and Alex Alderton, Roberto Amato, Jeffrey Barrett, Sonia Goncalves, Ewan Harrison, David K. Jackson, Ian Johnston, Dominic Kwiatkowski, Cordelia Langford, John Sillitoe on behalf of the Wellcome Sanger Institute COVID-19 Surveillance Team |
| EPI_ISL_2972676 | Lighthouse Lab in Alderley Park | Wellcome Sanger Institute for the COVID-19 Genomics UK (COG-UK) Consortium | Jacquelyn Wynn, Mairead Hyland, The Lighthouse Lab in Alderley Park and Alex Alderton, Roberto Amato, Jeffrey Barrett, Sonia Goncalves, Ewan Harrison, David K. Jackson, Ian Johnston, Dominic Kwiatkowski, Cordelia Langford, John Sillitoe on behalf of the Wellcome Sanger Institute COVID-19 Surveillance Team |
| EPI_ISL_2972677 | Department of Bacteria, Parasites and Fungi, Statens Serum Institut, Copenhagen, Denmark | Statens Serum Institut Bioinformatics and Microbial Genomics | Danish Covid-19 Genome Consortium |
| EPI_ISL_2972678 | Lighthouse Laboratory Plymouth | Wellcome Sanger Institute for the COVID-19 Genomics UK (COG-UK) Consortium | Lighthouse Laboratory Plymouth and Alex Alderton, Roberto Amato, Jeffrey Barrett, Sonia Goncalves, Ewan Harrison, David K. Jackson, Ian Johnston, Dominic Kwiatkowski, Cordelia Langford, John Sillitoe on behalf of the Wellcome Sanger Institute COVID-19 Surveillance Team |
| EPI_ISL_2972679, EPI_ISL_2972680 | Lighthouse Lab in Alderley Park | Wellcome Sanger Institute for the COVID-19 Genomics UK (COG-UK) Consortium | Jacquelyn Wynn, Mairead Hyland, The Lighthouse Lab in Alderley Park and Alex Alderton, Roberto Amato, Jeffrey Barrett, Sonia Goncalves, Ewan Harrison, David K. Jackson, Ian Johnston, Dominic Kwiatkowski, Cordelia Langford, John Sillitoe on behalf of the Wellcome Sanger Institute COVID-19 Surveillance Team |
| EPI_ISL_2972681 | Department of Bacteria, Parasites and Fungi, Statens Serum Institut, Copenhagen, Denmark | Statens Serum Institut Bioinformatics and Microbial Genomics | Danish Covid-19 Genome Consortium |
| EPI_ISL_2972682, EPI_ISL_2972683 | Lighthouse Laboratory Plymouth | Wellcome Sanger Institute for the COVID-19 Genomics UK (COG-UK) Consortium | Lighthouse Laboratory Plymouth and Alex Alderton, Roberto Amato, Jeffrey Barrett, Sonia Goncalves, Ewan Harrison, David K. Jackson, Ian Johnston, Dominic Kwiatkowski, Cordelia Langford, John Sillitoe on behalf of the Wellcome Sanger Institute COVID-19 Surveillance Team |
| EPI_ISL_2972684 | Department of Bacteria, Parasites and Fungi, Statens Serum Institut, Copenhagen, Denmark | Statens Serum Institut Bioinformatics and Microbial Genomics | Danish Covid-19 Genome Consortium |
| EPI_ISL_2972685 | Lighthouse Lab in Alderley Park | Wellcome Sanger Institute for the COVID-19 Genomics UK (COG-UK) Consortium | Jacquelyn Wynn, Mairead Hyland, The Lighthouse Lab in Alderley Park and Alex Alderton, Roberto Amato, Jeffrey Barrett, Sonia Goncalves, Ewan Harrison, David K. Jackson, Ian Johnston, Dominic Kwiatkowski, Cordelia Langford, John Sillitoe on behalf of the Wellcome Sanger Institute COVID-19 Surveillance Team |
| EPI_ISL_2972686 | Lighthouse Laboratory Plymouth | Wellcome Sanger Institute for the COVID-19 Genomics UK (COG-UK) Consortium | Lighthouse Laboratory Plymouth and Alex Alderton, Roberto Amato, Jeffrey Barrett, Sonia Goncalves, Ewan Harrison, David K. Jackson, Ian Johnston, Dominic Kwiatkowski, Cordelia Langford, John Sillitoe on behalf of the Wellcome Sanger Institute COVID-19 Surveillance Team |
| EPI_ISL_2972687 | Department of Bacteria, Parasites and Fungi, Statens Serum Institut, Copenhagen, Denmark | Statens Serum Institut Bioinformatics and Microbial Genomics | Danish Covid-19 Genome Consortium |
| EPI_ISL_2972688 | Lighthouse Lab in Alderley Park | Wellcome Sanger Institute for the COVID-19 Genomics UK (COG-UK) Consortium | Jacquelyn Wynn, Mairead Hyland, The Lighthouse Lab in Alderley Park and Alex Alderton, Roberto Amato, Jeffrey Barrett, Sonia Goncalves, Ewan Harrison, David K. Jackson, Ian Johnston, Dominic Kwiatkowski, Cordelia Langford, John Sillitoe on behalf of the Wellcome Sanger Institute COVID-19 Surveillance Team |
| EPI_ISL_2972689 | Lighthouse Laboratory Plymouth | Wellcome Sanger Institute for the COVID-19 Genomics UK (COG-UK) Consortium | Lighthouse Laboratory Plymouth and Alex Alderton, Roberto Amato, Jeffrey Barrett, Sonia Goncalves, Ewan Harrison, David K. Jackson, Ian Johnston, Dominic Kwiatkowski, Cordelia Langford, John Sillitoe on behalf of the Wellcome Sanger Institute COVID-19 Surveillance Team |
| EPI_ISL_2972690 | Lighthouse Lab in Milton Keynes | Wellcome Sanger Institute for the COVID-19 Genomics UK (COG-UK) Consortium | The Lighthouse Lab in Milton Keynes and Alex Alderton, Roberto Amato, Jeffrey Barrett, Sonia Goncalves, Ewan Harrison, David K. Jackson, Ian Johnston, Dominic Kwiatkowski, Cordelia Langford, John Sillitoe on behalf of the Wellcome Sanger Institute COVID-19 Surveillance Team |
| EPI_ISL_2972691 | Department of Bacteria, Parasites and Fungi, Statens Serum Institut, Copenhagen, Denmark | Statens Serum Institut Bioinformatics and Microbial Genomics | Danish Covid-19 Genome Consortium |
| EPI_ISL_2972692, EPI_ISL_2972693, EPI_ISL_2972694 | Lighthouse Lab in Alderley Park | Wellcome Sanger Institute for the COVID-19 Genomics UK (COG-UK) Consortium | Jacquelyn Wynn, Mairead Hyland, The Lighthouse Lab in Alderley Park and Alex Alderton, Roberto Amato, Jeffrey Barrett, Sonia Goncalves, Ewan Harrison, David K. Jackson, Ian Johnston, Dominic Kwiatkowski, Cordelia Langford, John Sillitoe on behalf of the Wellcome Sanger Institute COVID-19 Surveillance Team |
| EPI_ISL_2972695 | Department of Bacteria, Parasites and Fungi, Statens Serum Institut, Copenhagen, Denmark | Statens Serum Institut Bioinformatics and Microbial Genomics | Danish Covid-19 Genome Consortium |
| EPI_ISL_2972696 | Lighthouse Lab in Milton Keynes | Wellcome Sanger Institute for the COVID-19 Genomics UK (COG-UK) Consortium | The Lighthouse Lab in Milton Keynes and Alex Alderton, Roberto Amato, Jeffrey Barrett, Sonia Goncalves, Ewan Harrison, David K. Jackson, Ian Johnston, Dominic Kwiatkowski, Cordelia Langford, John Sillitoe on behalf of the Wellcome Sanger Institute COVID-19 Surveillance Team |
| EPI_ISL_2972697 | Department of Bacteria, Parasites and Fungi, Statens Serum Institut, Copenhagen, Denmark | Statens Serum Institut Bioinformatics and Microbial Genomics | Danish Covid-19 Genome Consortium |
| EPI_ISL_2972698, EPI_ISL_2972699 | Lighthouse Lab in Alderley Park | Wellcome Sanger Institute for the COVID-19 Genomics UK (COG-UK) Consortium | Jacquelyn Wynn, Mairead Hyland, The Lighthouse Lab in Alderley Park and Alex Alderton, Roberto Amato, Jeffrey Barrett, Sonia Goncalves, Ewan Harrison, David K. Jackson, Ian Johnston, Dominic Kwiatkowski, Cordelia Langford, John Sillitoe on behalf of the Wellcome Sanger Institute COVID-19 Surveillance Team |
| EPI_ISL_2972700 | Department of Bacteria, Parasites and Fungi, Statens Serum Institut, Copenhagen, Denmark | Statens Serum Institut Bioinformatics and Microbial Genomics | Danish Covid-19 Genome Consortium |
| EPI_ISL_2972701 | Lighthouse Laboratory Plymouth | Wellcome Sanger Institute for the COVID-19 Genomics UK (COG-UK) Consortium | Lighthouse Laboratory Plymouth and Alex Alderton, Roberto Amato, Jeffrey Barrett, Sonia Goncalves, Ewan Harrison, David K. Jackson, Ian Johnston, Dominic Kwiatkowski, Cordelia Langford, John Sillitoe on behalf of the Wellcome Sanger Institute COVID-19 Surveillance Team |
| EPI_ISL_2972702 | Lighthouse Lab in Alderley Park | Wellcome Sanger Institute for the COVID-19 Genomics UK (COG-UK) Consortium | Jacquelyn Wynn, Mairead Hyland, The Lighthouse Lab in Alderley Park and Alex Alderton, Roberto Amato, Jeffrey Barrett, Sonia Goncalves, Ewan Harrison, David K. Jackson, Ian Johnston, Dominic Kwiatkowski, Cordelia Langford, John Sillitoe on behalf of the Wellcome Sanger Institute COVID-19 Surveillance Team |
| EPI_ISL_2972703 | Lighthouse Lab in Milton Keynes | Wellcome Sanger Institute for the COVID-19 Genomics UK (COG-UK) Consortium | The Lighthouse Lab in Milton Keynes and Alex Alderton, Roberto Amato, Jeffrey Barrett, Sonia Goncalves, Ewan Harrison, David K. Jackson, Ian Johnston, Dominic Kwiatkowski, Cordelia Langford, John Sillitoe on behalf of the Wellcome Sanger Institute COVID-19 Surveillance Team |
| EPI_ISL_2972704 | Department of Bacteria, Parasites and Fungi, Statens Serum Institut, Copenhagen, Denmark | Statens Serum Institut Bioinformatics and Microbial Genomics | Danish Covid-19 Genome Consortium |
| EPI_ISL_2972705 | Lighthouse Lab in Milton Keynes | Wellcome Sanger Institute for the COVID-19 Genomics UK (COG-UK) Consortium | The Lighthouse Lab in Milton Keynes and Alex Alderton, Roberto Amato, Jeffrey Barrett, Sonia Goncalves, Ewan Harrison, David K. Jackson, Ian Johnston, Dominic Kwiatkowski, Cordelia Langford, John Sillitoe on behalf of the Wellcome Sanger Institute COVID-19 Surveillance Team |
| EPI_ISL_2972706 | Lighthouse Lab in Alderley Park | Wellcome Sanger Institute for the COVID-19 Genomics UK (COG-UK) Consortium | Jacquelyn Wynn, Mairead Hyland, The Lighthouse Lab in Alderley Park and Alex Alderton, Roberto Amato, Jeffrey Barrett, Sonia Goncalves, Ewan Harrison, David K. Jackson, Ian Johnston, Dominic Kwiatkowski, Cordelia Langford, John Sillitoe on behalf of the Wellcome Sanger Institute COVID-19 Surveillance Team |
| EPI_ISL_2972707 | Department of Bacteria, Parasites and Fungi, Statens Serum Institut, Copenhagen, Denmark | Statens Serum Institut Bioinformatics and Microbial Genomics | Danish Covid-19 Genome Consortium |
| EPI_ISL_2972708, EPI_ISL_2972709 | Lighthouse Lab in Milton Keynes | Wellcome Sanger Institute for the COVID-19 Genomics UK (COG-UK) Consortium | The Lighthouse Lab in Milton Keynes and Alex Alderton, Roberto Amato, Jeffrey Barrett, Sonia Goncalves, Ewan Harrison, David K. Jackson, Ian Johnston, Dominic Kwiatkowski, Cordelia Langford, John Sillitoe on behalf of the Wellcome Sanger Institute COVID-19 Surveillance Team |
| EPI_ISL_2972710 | Department of Bacteria, Parasites and Fungi, Statens Serum Institut, Copenhagen, Denmark | Statens Serum Institut Bioinformatics and Microbial Genomics | Danish Covid-19 Genome Consortium |
| EPI_ISL_2972711 | Lighthouse Laboratory Plymouth | Wellcome Sanger Institute for the COVID-19 Genomics UK (COG-UK) Consortium | Lighthouse Laboratory Plymouth and Alex Alderton, Roberto Amato, Jeffrey Barrett, Sonia Goncalves, Ewan Harrison, David K. Jackson, Ian Johnston, Dominic Kwiatkowski, Cordelia Langford, John Sillitoe on behalf of the Wellcome Sanger Institute COVID-19 Surveillance Team |
| EPI_ISL_2972712 | Lighthouse Lab in Milton Keynes | Wellcome Sanger Institute for the COVID-19 Genomics UK (COG-UK) Consortium | The Lighthouse Lab in Milton Keynes and Alex Alderton, Roberto Amato, Jeffrey Barrett, Sonia Goncalves, Ewan Harrison, David K. Jackson, Ian Johnston, Dominic Kwiatkowski, Cordelia Langford, John Sillitoe on behalf of the Wellcome Sanger Institute COVID-19 Surveillance Team |
| EPI_ISL_2972714 | Lighthouse Laboratory Plymouth | Wellcome Sanger Institute for the COVID-19 Genomics UK (COG-UK) Consortium | Lighthouse Laboratory Plymouth and Alex Alderton, Roberto Amato, Jeffrey Barrett, Sonia Goncalves, Ewan Harrison, David K. Jackson, Ian Johnston, Dominic Kwiatkowski, Cordelia Langford, John Sillitoe on behalf of the Wellcome Sanger Institute COVID-19 Surveillance Team |
| EPI_ISL_2972715 | Lighthouse Lab in Alderley Park | Wellcome Sanger Institute for the COVID-19 Genomics UK (COG-UK) Consortium | Jacquelyn Wynn, Mairead Hyland, The Lighthouse Lab in Alderley Park and Alex Alderton, Roberto Amato, Jeffrey Barrett, Sonia Goncalves, Ewan Harrison, David K. Jackson, Ian Johnston, Dominic Kwiatkowski, Cordelia Langford, John Sillitoe on behalf of the Wellcome Sanger Institute COVID-19 Surveillance Team |
| EPI_ISL_2972716 | Department of Bacteria, Parasites and Fungi, Statens Serum Institut, Copenhagen, Denmark | Statens Serum Institut Bioinformatics and Microbial Genomics | Danish Covid-19 Genome Consortium |
| EPI_ISL_2972717 | Lighthouse Laboratory Plymouth | Wellcome Sanger Institute for the COVID-19 Genomics UK (COG-UK) Consortium | Lighthouse Laboratory Plymouth and Alex Alderton, Roberto Amato, Jeffrey Barrett, Sonia Goncalves, Ewan Harrison, David K. Jackson, Ian Johnston, Dominic Kwiatkowski, Cordelia Langford, John Sillitoe on behalf of the Wellcome Sanger Institute COVID-19 Surveillance Team |
| EPI_ISL_2972718, EPI_ISL_2972720 | Lighthouse Lab in Alderley Park | Wellcome Sanger Institute for the COVID-19 Genomics UK (COG-UK) Consortium | Jacquelyn Wynn, Mairead Hyland, The Lighthouse Lab in Alderley Park and Alex Alderton, Roberto Amato, Jeffrey Barrett, Sonia Goncalves, Ewan Harrison, David K. Jackson, Ian Johnston, Dominic Kwiatkowski, Cordelia Langford, John Sillitoe on behalf of the Wellcome Sanger Institute COVID-19 Surveillance Team |
| EPI_ISL_2972721 | Lighthouse Lab in Milton Keynes | Wellcome Sanger Institute for the COVID-19 Genomics UK (COG-UK) Consortium | The Lighthouse Lab in Milton Keynes and Alex Alderton, Roberto Amato, Jeffrey Barrett, Sonia Goncalves, Ewan Harrison, David K. Jackson, Ian Johnston, Dominic Kwiatkowski, Cordelia Langford, John Sillitoe on behalf of the Wellcome Sanger Institute COVID-19 Surveillance Team |
| EPI_ISL_2972722 | Department of Bacteria, Parasites and Fungi, Statens Serum Institut, Copenhagen, Denmark | Statens Serum Institut Bioinformatics and Microbial Genomics | Danish Covid-19 Genome Consortium |
| EPI_ISL_2972723 | Lighthouse Lab in Milton Keynes | Wellcome Sanger Institute for the COVID-19 Genomics UK (COG-UK) | The Lighthouse Lab in Milton Keynes and Alex Alderton, Roberto Amato, Jeffrey Barrett, Sonia Goncalves, Ewan Harrison, David K. Jackson, Ian Johnston, Dominic Kwiatkowski, Cordelia Langford, John Sillitoe on behalf of the Wellcome Sanger |

| | | | |
|---|---|---|---|
| | | Consortium | Institute COVID-19 Surveillance Team |
| EPI_ISL_2972724, EPI_ISL_2972725 | Lighthouse Lab in Alderley Park | Wellcome Sanger Institute for the COVID-19 Genomics UK (COG-UK) Consortium | Jacquelyn Wynn, Mairead Hyland, The Lighthouse Lab in Alderley Park and Alex Alderton, Roberto Amato, Jeffrey Barrett, Sonia Goncalves, Ewan Harrison, David K. Jackson, Ian Johnston, Dominic Kwiatkowski, Cordelia Langford, John Sillitoe on behalf of the Wellcome Sanger Institute COVID-19 Surveillance Team |
| EPI_ISL_2972726 | Department of Bacteria, Parasites and Fungi, Statens Serum Institut, Copenhagen, Denmark | Statens Serum Institut Bioinformatics and Microbial Genomics | Danish Covid-19 Genome Consortium |
| EPI_ISL_2972727, EPI_ISL_2972728 | Lighthouse Lab in Alderley Park | Wellcome Sanger Institute for the COVID-19 Genomics UK (COG-UK) Consortium | Jacquelyn Wynn, Mairead Hyland, The Lighthouse Lab in Alderley Park and Alex Alderton, Roberto Amato, Jeffrey Barrett, Sonia Goncalves, Ewan Harrison, David K. Jackson, Ian Johnston, Dominic Kwiatkowski, Cordelia Langford, John Sillitoe on behalf of the Wellcome Sanger Institute COVID-19 Surveillance Team |
| EPI_ISL_2972729 | Department of Bacteria, Parasites and Fungi, Statens Serum Institut, Copenhagen, Denmark | Statens Serum Institut Bioinformatics and Microbial Genomics | Danish Covid-19 Genome Consortium |
| EPI_ISL_2972730 | Lighthouse Laboratory Plymouth | Wellcome Sanger Institute for the COVID-19 Genomics UK (COG-UK) Consortium | Lighthouse Laboratory Plymouth and Alex Alderton, Roberto Amato, Jeffrey Barrett, Sonia Goncalves, Ewan Harrison, David K. Jackson, Ian Johnston, Dominic Kwiatkowski, Cordelia Langford, John Sillitoe on behalf of the Wellcome Sanger Institute COVID-19 Surveillance Team |
| EPI_ISL_2972731 | Lighthouse Lab in Milton Keynes | Wellcome Sanger Institute for the COVID-19 Genomics UK (COG-UK) Consortium | The Lighthouse Lab in Milton Keynes and Alex Alderton, Roberto Amato, Jeffrey Barrett, Sonia Goncalves, Ewan Harrison, David K. Jackson, Ian Johnston, Dominic Kwiatkowski, Cordelia Langford, John Sillitoe on behalf of the Wellcome Sanger Institute COVID-19 Surveillance Team |
| EPI_ISL_2972732 | Department of Bacteria, Parasites and Fungi, Statens Serum Institut, Copenhagen, Denmark | Statens Serum Institut Bioinformatics and Microbial Genomics | Danish Covid-19 Genome Consortium |
| EPI_ISL_2972733 | Lighthouse Lab in Alderley Park | Wellcome Sanger Institute for the COVID-19 Genomics UK (COG-UK) Consortium | Jacquelyn Wynn, Mairead Hyland, The Lighthouse Lab in Alderley Park and Alex Alderton, Roberto Amato, Jeffrey Barrett, Sonia Goncalves, Ewan Harrison, David K. Jackson, Ian Johnston, Dominic Kwiatkowski, Cordelia Langford, John Sillitoe on behalf of the Wellcome Sanger Institute COVID-19 Surveillance Team |
| EPI_ISL_2972734 | Department of Bacteria, Parasites and Fungi, Statens Serum Institut, Copenhagen, Denmark | Statens Serum Institut Bioinformatics and Microbial Genomics | Danish Covid-19 Genome Consortium |
| EPI_ISL_2972735 | Lighthouse Lab in Milton Keynes | Wellcome Sanger Institute for the COVID-19 Genomics UK (COG-UK) Consortium | The Lighthouse Lab in Milton Keynes and Alex Alderton, Roberto Amato, Jeffrey Barrett, Sonia Goncalves, Ewan Harrison, David K. Jackson, Ian Johnston, Dominic Kwiatkowski, Cordelia Langford, John Sillitoe on behalf of the Wellcome Sanger Institute COVID-19 Surveillance Team |
| EPI_ISL_2972736 | Lighthouse Lab in Alderley Park | Wellcome Sanger Institute for the COVID-19 Genomics UK (COG-UK) Consortium | Jacquelyn Wynn, Mairead Hyland, The Lighthouse Lab in Alderley Park and Alex Alderton, Roberto Amato, Jeffrey Barrett, Sonia Goncalves, Ewan Harrison, David K. Jackson, Ian Johnston, Dominic Kwiatkowski, Cordelia Langford, John Sillitoe on behalf of the Wellcome Sanger Institute COVID-19 Surveillance Team |
| EPI_ISL_2972737 | Lighthouse Lab in Milton Keynes | Wellcome Sanger Institute for the COVID-19 Genomics UK (COG-UK) Consortium | The Lighthouse Lab in Milton Keynes and Alex Alderton, Roberto Amato, Jeffrey Barrett, Sonia Goncalves, Ewan Harrison, David K. Jackson, Ian Johnston, Dominic Kwiatkowski, Cordelia Langford, John Sillitoe on behalf of the Wellcome Sanger Institute COVID-19 Surveillance Team |
| EPI_ISL_2972738 | Department of Bacteria, Parasites and Fungi, Statens Serum Institut, Copenhagen, Denmark | Statens Serum Institut Bioinformatics and Microbial Genomics | Danish Covid-19 Genome Consortium |
| EPI_ISL_2972739 | Lighthouse Lab in Alderley Park | Wellcome Sanger Institute for the COVID-19 Genomics UK (COG-UK) Consortium | Jacquelyn Wynn, Mairead Hyland, The Lighthouse Lab in Alderley Park and Alex Alderton, Roberto Amato, Jeffrey Barrett, Sonia Goncalves, Ewan Harrison, David K. Jackson, Ian Johnston, Dominic Kwiatkowski, Cordelia Langford, John Sillitoe on behalf of the Wellcome Sanger Institute COVID-19 Surveillance Team |
| EPI_ISL_2972740 | Lighthouse Laboratory Plymouth | Wellcome Sanger Institute for the COVID-19 Genomics UK (COG-UK) Consortium | Lighthouse Laboratory Plymouth and Alex Alderton, Roberto Amato, Jeffrey Barrett, Sonia Goncalves, Ewan Harrison, David K. Jackson, Ian Johnston, Dominic Kwiatkowski, Cordelia Langford, John Sillitoe on behalf of the Wellcome Sanger Institute COVID-19 Surveillance Team |
| EPI_ISL_2972741 | Lighthouse Lab in Alderley Park | Wellcome Sanger Institute for the COVID-19 Genomics UK (COG-UK) Consortium | Jacquelyn Wynn, Mairead Hyland, The Lighthouse Lab in Alderley Park and Alex Alderton, Roberto Amato, Jeffrey Barrett, Sonia Goncalves, Ewan Harrison, David K. Jackson, Ian Johnston, Dominic Kwiatkowski, Cordelia Langford, John Sillitoe on behalf of the Wellcome Sanger Institute COVID-19 Surveillance Team |
| EPI_ISL_2972742 | Department of Bacteria, Parasites and Fungi, Statens Serum Institut, Copenhagen, Denmark | Statens Serum Institut Bioinformatics and Microbial Genomics | Danish Covid-19 Genome Consortium |
| EPI_ISL_2972743 | Lighthouse Lab in Milton Keynes | Wellcome Sanger Institute for the COVID-19 Genomics UK (COG-UK) Consortium | The Lighthouse Lab in Milton Keynes and Alex Alderton, Roberto Amato, Jeffrey Barrett, Sonia Goncalves, Ewan Harrison, David K. Jackson, Ian Johnston, Dominic Kwiatkowski, Cordelia Langford, John Sillitoe on behalf of the Wellcome Sanger Institute COVID-19 Surveillance Team |
| EPI_ISL_2972744 | Lighthouse Laboratory Plymouth | Wellcome Sanger Institute for the COVID-19 Genomics UK (COG-UK) Consortium | Lighthouse Laboratory Plymouth and Alex Alderton, Roberto Amato, Jeffrey Barrett, Sonia Goncalves, Ewan Harrison, David K. Jackson, Ian Johnston, Dominic Kwiatkowski, Cordelia Langford, John Sillitoe on behalf of the Wellcome Sanger Institute COVID-19 Surveillance Team |
| EPI_ISL_2972745 | Department of Bacteria, Parasites and Fungi, Statens Serum Institut, Copenhagen, Denmark | Statens Serum Institut Bioinformatics and Microbial Genomics | Danish Covid-19 Genome Consortium |
| EPI_ISL_2972746 | Lighthouse Laboratory Plymouth | Wellcome Sanger Institute for the COVID-19 Genomics UK (COG-UK) Consortium | Lighthouse Laboratory Plymouth and Alex Alderton, Roberto Amato, Jeffrey Barrett, Sonia Goncalves, Ewan Harrison, David K. Jackson, Ian Johnston, Dominic Kwiatkowski, Cordelia Langford, John Sillitoe on behalf of the Wellcome Sanger Institute COVID-19 Surveillance Team |
| EPI_ISL_2972747 | Lighthouse Lab in Alderley Park | Wellcome Sanger Institute for the COVID-19 Genomics UK (COG-UK) Consortium | Jacquelyn Wynn, Mairead Hyland, The Lighthouse Lab in Alderley Park and Alex Alderton, Roberto Amato, Jeffrey Barrett, Sonia Goncalves, Ewan Harrison, David K. Jackson, Ian Johnston, Dominic Kwiatkowski, Cordelia Langford, John Sillitoe on behalf of the Wellcome Sanger Institute COVID-19 Surveillance Team |
| EPI_ISL_2972748 | Department of Bacteria, Parasites and Fungi, Statens Serum Institut, Copenhagen, Denmark | Statens Serum Institut Bioinformatics and Microbial Genomics | Danish Covid-19 Genome Consortium |
| EPI_ISL_2972749 | Lighthouse Lab in Milton Keynes | Wellcome Sanger Institute for the COVID-19 Genomics UK (COG-UK) Consortium | The Lighthouse Lab in Milton Keynes and Alex Alderton, Roberto Amato, Jeffrey Barrett, Sonia Goncalves, Ewan Harrison, David K. Jackson, Ian Johnston, Dominic Kwiatkowski, Cordelia Langford, John Sillitoe on behalf of the Wellcome Sanger Institute COVID-19 Surveillance Team |
| EPI_ISL_2972750 | Lighthouse Lab in Alderley Park | Wellcome Sanger Institute for the COVID-19 Genomics UK (COG-UK) Consortium | Jacquelyn Wynn, Mairead Hyland, The Lighthouse Lab in Alderley Park and Alex Alderton, Roberto Amato, Jeffrey Barrett, Sonia Goncalves, Ewan Harrison, David K. Jackson, Ian Johnston, Dominic Kwiatkowski, Cordelia Langford, John Sillitoe on behalf of the Wellcome Sanger Institute COVID-19 Surveillance Team |
| EPI_ISL_2972751 | Department of Bacteria, Parasites and Fungi, Statens Serum Institut, Copenhagen, Denmark | Statens Serum Institut Bioinformatics and Microbial Genomics | Danish Covid-19 Genome Consortium |
| EPI_ISL_2972752 | Lighthouse Laboratory Plymouth | Wellcome Sanger Institute for the COVID-19 Genomics UK (COG-UK) Consortium | Lighthouse Laboratory Plymouth and Alex Alderton, Roberto Amato, Jeffrey Barrett, Sonia Goncalves, Ewan Harrison, David K. Jackson, Ian Johnston, Dominic Kwiatkowski, Cordelia Langford, John Sillitoe on behalf of the Wellcome Sanger Institute COVID-19 Surveillance Team |
| EPI_ISL_2972753 | Lighthouse Lab in Alderley Park | Wellcome Sanger Institute for the COVID-19 Genomics UK (COG-UK) Consortium | Jacquelyn Wynn, Mairead Hyland, The Lighthouse Lab in Alderley Park and Alex Alderton, Roberto Amato, Jeffrey Barrett, Sonia Goncalves, Ewan Harrison, David K. Jackson, Ian Johnston, Dominic Kwiatkowski, Cordelia Langford, John Sillitoe on behalf of the Wellcome Sanger Institute COVID-19 Surveillance Team |
| EPI_ISL_2972754 | Department of Bacteria, Parasites and Fungi, Statens Serum Institut, Copenhagen, Denmark | Statens Serum Institut Bioinformatics and Microbial Genomics | Danish Covid-19 Genome Consortium |
| EPI_ISL_2972755 | Lighthouse Lab in Alderley Park | Wellcome Sanger Institute for the COVID-19 Genomics UK (COG-UK) Consortium | Jacquelyn Wynn, Mairead Hyland, The Lighthouse Lab in Alderley Park and Alex Alderton, Roberto Amato, Jeffrey Barrett, Sonia Goncalves, Ewan Harrison, David K. Jackson, Ian Johnston, Dominic Kwiatkowski, Cordelia Langford, John Sillitoe on behalf of the Wellcome Sanger Institute COVID-19 Surveillance Team |
| EPI_ISL_2972756 | Lighthouse Laboratory Plymouth | Wellcome Sanger Institute for the COVID-19 Genomics UK (COG-UK) Consortium | Lighthouse Laboratory Plymouth and Alex Alderton, Roberto Amato, Jeffrey Barrett, Sonia Goncalves, Ewan Harrison, David K. Jackson, Ian Johnston, Dominic Kwiatkowski, Cordelia Langford, John Sillitoe on behalf of the Wellcome Sanger Institute COVID-19 Surveillance Team |
| EPI_ISL_2972758, EPI_ISL_2972759 | Lighthouse Lab in Alderley Park | Wellcome Sanger Institute for the COVID-19 Genomics UK (COG-UK) Consortium | Jacquelyn Wynn, Mairead Hyland, The Lighthouse Lab in Alderley Park and Alex Alderton, Roberto Amato, Jeffrey Barrett, Sonia Goncalves, Ewan Harrison, David K. Jackson, Ian Johnston, Dominic Kwiatkowski, Cordelia Langford, John Sillitoe on behalf of the Wellcome Sanger Institute COVID-19 Surveillance Team |
| EPI_ISL_2972760 | Department of Bacteria, Parasites and Fungi, Statens Serum Institut, Copenhagen, Denmark | Statens Serum Institut Bioinformatics and Microbial Genomics | Danish Covid-19 Genome Consortium |
| EPI_ISL_2972761 | Lighthouse Laboratory Plymouth | Wellcome Sanger Institute for the COVID-19 Genomics UK (COG-UK) Consortium | Lighthouse Laboratory Plymouth and Alex Alderton, Roberto Amato, Jeffrey Barrett, Sonia Goncalves, Ewan Harrison, David K. Jackson, Ian Johnston, Dominic Kwiatkowski, Cordelia Langford, John Sillitoe on behalf of the Wellcome Sanger Institute COVID-19 Surveillance Team |
| EPI_ISL_2972762 | Lighthouse Lab in Alderley Park | Wellcome Sanger Institute for the COVID-19 Genomics UK (COG-UK) Consortium | Jacquelyn Wynn, Mairead Hyland, The Lighthouse Lab in Alderley Park and Alex Alderton, Roberto Amato, Jeffrey Barrett, Sonia Goncalves, Ewan Harrison, David K. Jackson, Ian Johnston, Dominic Kwiatkowski, Cordelia Langford, John Sillitoe on behalf of the Wellcome Sanger Institute COVID-19 Surveillance Team |
| EPI_ISL_2972763 | Lighthouse Lab in Milton Keynes | Wellcome Sanger Institute for the COVID-19 Genomics UK (COG-UK) Consortium | The Lighthouse Lab in Milton Keynes and Alex Alderton, Roberto Amato, Jeffrey Barrett, Sonia Goncalves, Ewan Harrison, David K. Jackson, Ian Johnston, Dominic Kwiatkowski, Cordelia Langford, John Sillitoe on behalf of the Wellcome Sanger Institute COVID-19 Surveillance Team |
| EPI_ISL_2972764 | Department of Bacteria, Parasites and Fungi, Statens Serum Institut, Copenhagen, Denmark | Statens Serum Institut Bioinformatics and Microbial Genomics | Danish Covid-19 Genome Consortium |
| EPI_ISL_2972765 | Lighthouse Lab in Alderley Park | Wellcome Sanger Institute for the COVID-19 Genomics UK (COG-UK) Consortium | Jacquelyn Wynn, Mairead Hyland, The Lighthouse Lab in Alderley Park and Alex Alderton, Roberto Amato, Jeffrey Barrett, Sonia Goncalves, Ewan Harrison, David K. Jackson, Ian Johnston, Dominic Kwiatkowski, Cordelia Langford, John Sillitoe on behalf of the Wellcome Sanger Institute COVID-19 Surveillance Team |
| EPI_ISL_2972766 | Lighthouse Lab in Milton Keynes | Wellcome Sanger Institute for the COVID-19 Genomics UK (COG-UK) Consortium | The Lighthouse Lab in Milton Keynes and Alex Alderton, Roberto Amato, Jeffrey Barrett, Sonia Goncalves, Ewan Harrison, David K. Jackson, Ian Johnston, Dominic Kwiatkowski, Cordelia Langford, John Sillitoe on behalf of the Wellcome Sanger Institute COVID-19 Surveillance Team |
| EPI_ISL_2972767 | Department of Bacteria, Parasites and Fungi, Statens Serum Institut, Copenhagen, Denmark | Statens Serum Institut Bioinformatics and Microbial Genomics | Danish Covid-19 Genome Consortium |
| EPI_ISL_2972768 | Lighthouse Lab in Alderley Park | Wellcome Sanger Institute for the COVID-19 Genomics UK (COG-UK) Consortium | Jacquelyn Wynn, Mairead Hyland, The Lighthouse Lab in Alderley Park and Alex Alderton, Roberto Amato, Jeffrey Barrett, Sonia Goncalves, Ewan Harrison, David K. Jackson, Ian Johnston, Dominic Kwiatkowski, Cordelia Langford, John Sillitoe on behalf of the Wellcome Sanger Institute COVID-19 Surveillance Team |
| EPI_ISL_2972769 | Lighthouse Laboratory Plymouth | Wellcome Sanger Institute for the COVID-19 Genomics UK (COG-UK) Consortium | Lighthouse Laboratory Plymouth and Alex Alderton, Roberto Amato, Jeffrey Barrett, Sonia Goncalves, Ewan Harrison, David K. Jackson, Ian Johnston, Dominic Kwiatkowski, Cordelia Langford, John Sillitoe on behalf of the Wellcome Sanger Institute COVID-19 Surveillance Team |
| EPI_ISL_2972770, EPI_ISL_2972771 | Lighthouse Lab in Milton Keynes | Wellcome Sanger Institute for the COVID-19 Genomics UK (COG-UK) Consortium | The Lighthouse Lab in Milton Keynes and Alex Alderton, Roberto Amato, Jeffrey Barrett, Sonia Goncalves, Ewan Harrison, David K. Jackson, Ian Johnston, Dominic Kwiatkowski, Cordelia Langford, John Sillitoe on behalf of the Wellcome Sanger Institute COVID-19 Surveillance Team |
| EPI_ISL_2972772, EPI_ISL_2972773 | Lighthouse Lab in Alderley Park | Wellcome Sanger Institute for the COVID-19 Genomics UK (COG-UK) Consortium | Jacquelyn Wynn, Mairead Hyland, The Lighthouse Lab in Alderley Park and Alex Alderton, Roberto Amato, Jeffrey Barrett, Sonia Goncalves, Ewan Harrison, David K. Jackson, Ian Johnston, Dominic Kwiatkowski, Cordelia Langford, John Sillitoe on behalf of the Wellcome Sanger Institute COVID-19 Surveillance Team |
| EPI_ISL_2972774 | Lighthouse Lab in Milton Keynes | Wellcome Sanger Institute for the COVID-19 Genomics UK (COG-UK) | The Lighthouse Lab in Milton Keynes and Alex Alderton, Roberto Amato, Jeffrey Barrett, Sonia Goncalves, Ewan Harrison, David K. Jackson, Ian Johnston, Dominic Kwiatkowski, Cordelia Langford, John Sillitoe on behalf of the Wellcome Sanger |

| | | | |
|---|---|---|---|
| | | Consortium | Institute COVID-19 Surveillance Team |
| EPI_ISL_2972775, EPI_ISL_2972776 | Lighthouse Lab in Alderley Park | Wellcome Sanger Institute for the COVID-19 Genomics UK (COG-UK) Consortium | Jacquelyn Wynn, Mairead Hyland, The Lighthouse Lab in Alderley Park and Alex Alderton, Roberto Amato, Jeffrey Barrett, Sonia Goncalves, Ewan Harrison, David K. Jackson, Ian Johnston, Dominic Kwiatkowski, Cordelia Langford, John Sillitoe on behalf of the Wellcome Sanger Institute COVID-19 Surveillance Team |
| EPI_ISL_2972777 | Department of Bacteria, Parasites and Fungi, Statens Serum Institut, Copenhagen, Denmark | Statens Serum Institut Bioinformatics and Microbial Genomics | Danish Covid-19 Genome Consortium |
| EPI_ISL_2972778, EPI_ISL_2972779 | Lighthouse Lab in Alderley Park | Wellcome Sanger Institute for the COVID-19 Genomics UK (COG-UK) Consortium | Jacquelyn Wynn, Mairead Hyland, The Lighthouse Lab in Alderley Park and Alex Alderton, Roberto Amato, Jeffrey Barrett, Sonia Goncalves, Ewan Harrison, David K. Jackson, Ian Johnston, Dominic Kwiatkowski, Cordelia Langford, John Sillitoe on behalf of the Wellcome Sanger Institute COVID-19 Surveillance Team |
| EPI_ISL_2972780 | Department of Bacteria, Parasites and Fungi, Statens Serum Institut, Copenhagen, Denmark | Statens Serum Institut Bioinformatics and Microbial Genomics | Danish Covid-19 Genome Consortium |
| EPI_ISL_2972781, EPI_ISL_2972782 | Lighthouse Lab in Alderley Park | Wellcome Sanger Institute for the COVID-19 Genomics UK (COG-UK) Consortium | Jacquelyn Wynn, Mairead Hyland, The Lighthouse Lab in Alderley Park and Alex Alderton, Roberto Amato, Jeffrey Barrett, Sonia Goncalves, Ewan Harrison, David K. Jackson, Ian Johnston, Dominic Kwiatkowski, Cordelia Langford, John Sillitoe on behalf of the Wellcome Sanger Institute COVID-19 Surveillance Team |
| EPI_ISL_2972783 | Lighthouse Laboratory Plymouth | Wellcome Sanger Institute for the COVID-19 Genomics UK (COG-UK) Consortium | Lighthouse Laboratory Plymouth and Alex Alderton, Roberto Amato, Jeffrey Barrett, Sonia Goncalves, Ewan Harrison, David K. Jackson, Ian Johnston, Dominic Kwiatkowski, Cordelia Langford, John Sillitoe on behalf of the Wellcome Sanger Institute COVID-19 Surveillance Team |
| EPI_ISL_2972784 | Department of Bacteria, Parasites and Fungi, Statens Serum Institut, Copenhagen, Denmark | Statens Serum Institut Bioinformatics and Microbial Genomics | Danish Covid-19 Genome Consortium |
| EPI_ISL_2972785, EPI_ISL_2972786, EPI_ISL_2972788 | Lighthouse Lab in Alderley Park | Wellcome Sanger Institute for the COVID-19 Genomics UK (COG-UK) Consortium | Jacquelyn Wynn, Mairead Hyland, The Lighthouse Lab in Alderley Park and Alex Alderton, Roberto Amato, Jeffrey Barrett, Sonia Goncalves, Ewan Harrison, David K. Jackson, Ian Johnston, Dominic Kwiatkowski, Cordelia Langford, John Sillitoe on behalf of the Wellcome Sanger Institute COVID-19 Surveillance Team |
| EPI_ISL_2972789, EPI_ISL_2972790 | Lighthouse Lab in Milton Keynes | Wellcome Sanger Institute for the COVID-19 Genomics UK (COG-UK) Consortium | The Lighthouse Lab in Milton Keynes and Alex Alderton, Roberto Amato, Jeffrey Barrett, Sonia Goncalves, Ewan Harrison, David K. Jackson, Ian Johnston, Dominic Kwiatkowski, Cordelia Langford, John Sillitoe on behalf of the Wellcome Sanger Institute COVID-19 Surveillance Team |
| EPI_ISL_2972791 | Department of Bacteria, Parasites and Fungi, Statens Serum Institut, Copenhagen, Denmark | Statens Serum Institut Bioinformatics and Microbial Genomics | Danish Covid-19 Genome Consortium |
| EPI_ISL_2972792, EPI_ISL_2972793, EPI_ISL_2972794 | Lighthouse Laboratory Plymouth | Wellcome Sanger Institute for the COVID-19 Genomics UK (COG-UK) Consortium | Lighthouse Laboratory Plymouth and Alex Alderton, Roberto Amato, Jeffrey Barrett, Sonia Goncalves, Ewan Harrison, David K. Jackson, Ian Johnston, Dominic Kwiatkowski, Cordelia Langford, John Sillitoe on behalf of the Wellcome Sanger Institute COVID-19 Surveillance Team |
| EPI_ISL_2972795 | Department of Bacteria, Parasites and Fungi, Statens Serum Institut, Copenhagen, Denmark | Statens Serum Institut Bioinformatics and Microbial Genomics | Danish Covid-19 Genome Consortium |
| EPI_ISL_2972796 | Lighthouse Lab in Alderley Park | Wellcome Sanger Institute for the COVID-19 Genomics UK (COG-UK) Consortium | Jacquelyn Wynn, Mairead Hyland, The Lighthouse Lab in Alderley Park and Alex Alderton, Roberto Amato, Jeffrey Barrett, Sonia Goncalves, Ewan Harrison, David K. Jackson, Ian Johnston, Dominic Kwiatkowski, Cordelia Langford, John Sillitoe on behalf of the Wellcome Sanger Institute COVID-19 Surveillance Team |
| EPI_ISL_2972797 | Lighthouse Lab in Milton Keynes | Wellcome Sanger Institute for the COVID-19 Genomics UK (COG-UK) Consortium | The Lighthouse Lab in Milton Keynes and Alex Alderton, Roberto Amato, Jeffrey Barrett, Sonia Goncalves, Ewan Harrison, David K. Jackson, Ian Johnston, Dominic Kwiatkowski, Cordelia Langford, John Sillitoe on behalf of the Wellcome Sanger Institute COVID-19 Surveillance Team |
| EPI_ISL_2972798 | Department of Bacteria, Parasites and Fungi, Statens Serum Institut, Copenhagen, Denmark | Statens Serum Institut Bioinformatics and Microbial Genomics | Danish Covid-19 Genome Consortium |
| EPI_ISL_2972799, EPI_ISL_2972800 | Lighthouse Lab in Milton Keynes | Wellcome Sanger Institute for the COVID-19 Genomics UK (COG-UK) Consortium | The Lighthouse Lab in Milton Keynes and Alex Alderton, Roberto Amato, Jeffrey Barrett, Sonia Goncalves, Ewan Harrison, David K. Jackson, Ian Johnston, Dominic Kwiatkowski, Cordelia Langford, John Sillitoe on behalf of the Wellcome Sanger Institute COVID-19 Surveillance Team |
| EPI_ISL_2972801 | Department of Bacteria, Parasites and Fungi, Statens Serum Institut, Copenhagen, Denmark | Statens Serum Institut Bioinformatics and Microbial Genomics | Danish Covid-19 Genome Consortium |
| EPI_ISL_2972802 | Lighthouse Lab in Milton Keynes | Wellcome Sanger Institute for the COVID-19 Genomics UK (COG-UK) Consortium | The Lighthouse Lab in Milton Keynes and Alex Alderton, Roberto Amato, Jeffrey Barrett, Sonia Goncalves, Ewan Harrison, David K. Jackson, Ian Johnston, Dominic Kwiatkowski, Cordelia Langford, John Sillitoe on behalf of the Wellcome Sanger Institute COVID-19 Surveillance Team |
| EPI_ISL_2972803 | Lighthouse Lab in Alderley Park | Wellcome Sanger Institute for the COVID-19 Genomics UK (COG-UK) Consortium | Jacquelyn Wynn, Mairead Hyland, The Lighthouse Lab in Alderley Park and Alex Alderton, Roberto Amato, Jeffrey Barrett, Sonia Goncalves, Ewan Harrison, David K. Jackson, Ian Johnston, Dominic Kwiatkowski, Cordelia Langford, John Sillitoe on behalf of the Wellcome Sanger Institute COVID-19 Surveillance Team |
| EPI_ISL_2972804 | Lighthouse Laboratory Plymouth | Wellcome Sanger Institute for the COVID-19 Genomics UK (COG-UK) Consortium | Lighthouse Laboratory Plymouth and Alex Alderton, Roberto Amato, Jeffrey Barrett, Sonia Goncalves, Ewan Harrison, David K. Jackson, Ian Johnston, Dominic Kwiatkowski, Cordelia Langford, John Sillitoe on behalf of the Wellcome Sanger Institute COVID-19 Surveillance Team |
| EPI_ISL_2972805 | Department of Bacteria, Parasites and Fungi, Statens Serum Institut, Copenhagen, Denmark | Statens Serum Institut Bioinformatics and Microbial Genomics | Danish Covid-19 Genome Consortium |
| EPI_ISL_2972806 | Lighthouse Lab in Alderley Park | Wellcome Sanger Institute for the COVID-19 Genomics UK (COG-UK) Consortium | Jacquelyn Wynn, Mairead Hyland, The Lighthouse Lab in Alderley Park and Alex Alderton, Roberto Amato, Jeffrey Barrett, Sonia Goncalves, Ewan Harrison, David K. Jackson, Ian Johnston, Dominic Kwiatkowski, Cordelia Langford, John Sillitoe on behalf of the Wellcome Sanger Institute COVID-19 Surveillance Team |
| EPI_ISL_2972807 | Lighthouse Laboratory Plymouth | Wellcome Sanger Institute for the COVID-19 Genomics UK (COG-UK) Consortium | Lighthouse Laboratory Plymouth and Alex Alderton, Roberto Amato, Jeffrey Barrett, Sonia Goncalves, Ewan Harrison, David K. Jackson, Ian Johnston, Dominic Kwiatkowski, Cordelia Langford, John Sillitoe on behalf of the Wellcome Sanger Institute COVID-19 Surveillance Team |
| EPI_ISL_2972808 | Lighthouse Lab in Alderley Park | Wellcome Sanger Institute for the COVID-19 Genomics UK (COG-UK) Consortium | Jacquelyn Wynn, Mairead Hyland, The Lighthouse Lab in Alderley Park and Alex Alderton, Roberto Amato, Jeffrey Barrett, Sonia Goncalves, Ewan Harrison, David K. Jackson, Ian Johnston, Dominic Kwiatkowski, Cordelia Langford, John Sillitoe on behalf of the Wellcome Sanger Institute COVID-19 Surveillance Team |
| EPI_ISL_2972809 | Department of Bacteria, Parasites and Fungi, Statens Serum Institut, Copenhagen, Denmark | Statens Serum Institut Bioinformatics and Microbial Genomics | Danish Covid-19 Genome Consortium |
| EPI_ISL_2972810 | Lighthouse Lab in Alderley Park | Wellcome Sanger Institute for the COVID-19 Genomics UK (COG-UK) Consortium | Jacquelyn Wynn, Mairead Hyland, The Lighthouse Lab in Alderley Park and Alex Alderton, Roberto Amato, Jeffrey Barrett, Sonia Goncalves, Ewan Harrison, David K. Jackson, Ian Johnston, Dominic Kwiatkowski, Cordelia Langford, John Sillitoe on behalf of the Wellcome Sanger Institute COVID-19 Surveillance Team |
| EPI_ISL_2972811 | Lighthouse Laboratory Plymouth | Wellcome Sanger Institute for the COVID-19 Genomics UK (COG-UK) Consortium | Lighthouse Laboratory Plymouth and Alex Alderton, Roberto Amato, Jeffrey Barrett, Sonia Goncalves, Ewan Harrison, David K. Jackson, Ian Johnston, Dominic Kwiatkowski, Cordelia Langford, John Sillitoe on behalf of the Wellcome Sanger Institute COVID-19 Surveillance Team |
| EPI_ISL_2972812 | Department of Bacteria, Parasites and Fungi, Statens Serum Institut, Copenhagen, Denmark | Statens Serum Institut Bioinformatics and Microbial Genomics | Danish Covid-19 Genome Consortium |
| EPI_ISL_2972813, EPI_ISL_2972814 | Lighthouse Lab in Alderley Park | Wellcome Sanger Institute for the COVID-19 Genomics UK (COG-UK) Consortium | Jacquelyn Wynn, Mairead Hyland, The Lighthouse Lab in Alderley Park and Alex Alderton, Roberto Amato, Jeffrey Barrett, Sonia Goncalves, Ewan Harrison, David K. Jackson, Ian Johnston, Dominic Kwiatkowski, Cordelia Langford, John Sillitoe on behalf of the Wellcome Sanger Institute COVID-19 Surveillance Team |
| EPI_ISL_2972815 | Lighthouse Lab in Milton Keynes | Wellcome Sanger Institute for the COVID-19 Genomics UK (COG-UK) Consortium | The Lighthouse Lab in Milton Keynes and Alex Alderton, Roberto Amato, Jeffrey Barrett, Sonia Goncalves, Ewan Harrison, David K. Jackson, Ian Johnston, Dominic Kwiatkowski, Cordelia Langford, John Sillitoe on behalf of the Wellcome Sanger Institute COVID-19 Surveillance Team |
| EPI_ISL_2972816 | Department of Bacteria, Parasites and Fungi, Statens Serum Institut, Copenhagen, Denmark | Statens Serum Institut Bioinformatics and Microbial Genomics | Danish Covid-19 Genome Consortium |
| EPI_ISL_2972817 | Lighthouse Lab in Milton Keynes | Wellcome Sanger Institute for the COVID-19 Genomics UK (COG-UK) Consortium | The Lighthouse Lab in Milton Keynes and Alex Alderton, Roberto Amato, Jeffrey Barrett, Sonia Goncalves, Ewan Harrison, David K. Jackson, Ian Johnston, Dominic Kwiatkowski, Cordelia Langford, John Sillitoe on behalf of the Wellcome Sanger Institute COVID-19 Surveillance Team |
| EPI_ISL_2972818 | Lighthouse Lab in Alderley Park | Wellcome Sanger Institute for the COVID-19 Genomics UK (COG-UK) Consortium | Jacquelyn Wynn, Mairead Hyland, The Lighthouse Lab in Alderley Park and Alex Alderton, Roberto Amato, Jeffrey Barrett, Sonia Goncalves, Ewan Harrison, David K. Jackson, Ian Johnston, Dominic Kwiatkowski, Cordelia Langford, John Sillitoe on behalf of the Wellcome Sanger Institute COVID-19 Surveillance Team |
| EPI_ISL_2972819 | Department of Bacteria, Parasites and Fungi, Statens Serum Institut, Copenhagen, Denmark | Statens Serum Institut Bioinformatics and Microbial Genomics | Danish Covid-19 Genome Consortium |
| EPI_ISL_2972820 | Lighthouse Lab in Alderley Park | Wellcome Sanger Institute for the COVID-19 Genomics UK (COG-UK) Consortium | Jacquelyn Wynn, Mairead Hyland, The Lighthouse Lab in Alderley Park and Alex Alderton, Roberto Amato, Jeffrey Barrett, Sonia Goncalves, Ewan Harrison, David K. Jackson, Ian Johnston, Dominic Kwiatkowski, Cordelia Langford, John Sillitoe on behalf of the Wellcome Sanger Institute COVID-19 Surveillance Team |
| EPI_ISL_2972821 | Lighthouse Laboratory Plymouth | Wellcome Sanger Institute for the COVID-19 Genomics UK (COG-UK) Consortium | Lighthouse Laboratory Plymouth and Alex Alderton, Roberto Amato, Jeffrey Barrett, Sonia Goncalves, Ewan Harrison, David K. Jackson, Ian Johnston, Dominic Kwiatkowski, Cordelia Langford, John Sillitoe on behalf of the Wellcome Sanger Institute COVID-19 Surveillance Team |
| EPI_ISL_2972822 | Lighthouse Lab in Alderley Park | Wellcome Sanger Institute for the COVID-19 Genomics UK (COG-UK) Consortium | Jacquelyn Wynn, Mairead Hyland, The Lighthouse Lab in Alderley Park and Alex Alderton, Roberto Amato, Jeffrey Barrett, Sonia Goncalves, Ewan Harrison, David K. Jackson, Ian Johnston, Dominic Kwiatkowski, Cordelia Langford, John Sillitoe on behalf of the Wellcome Sanger Institute COVID-19 Surveillance Team |
| EPI_ISL_2972823 | Department of Bacteria, Parasites and Fungi, Statens Serum Institut, Copenhagen, Denmark | Statens Serum Institut Bioinformatics and Microbial Genomics | Danish Covid-19 Genome Consortium |
| EPI_ISL_2972824 | Lighthouse Lab in Milton Keynes | Wellcome Sanger Institute for the COVID-19 Genomics UK (COG-UK) Consortium | The Lighthouse Lab in Milton Keynes and Alex Alderton, Roberto Amato, Jeffrey Barrett, Sonia Goncalves, Ewan Harrison, David K. Jackson, Ian Johnston, Dominic Kwiatkowski, Cordelia Langford, John Sillitoe on behalf of the Wellcome Sanger Institute COVID-19 Surveillance Team |
| EPI_ISL_2972825 | Lighthouse Lab in Alderley Park | Wellcome Sanger Institute for the COVID-19 Genomics UK (COG-UK) Consortium | Jacquelyn Wynn, Mairead Hyland, The Lighthouse Lab in Alderley Park and Alex Alderton, Roberto Amato, Jeffrey Barrett, Sonia Goncalves, Ewan Harrison, David K. Jackson, Ian Johnston, Dominic Kwiatkowski, Cordelia Langford, John Sillitoe on behalf of the Wellcome Sanger Institute COVID-19 Surveillance Team |
| EPI_ISL_2972826 | Department of Bacteria, Parasites and Fungi, Statens Serum Institut, Copenhagen, Denmark | Statens Serum Institut Bioinformatics and Microbial Genomics | Danish Covid-19 Genome Consortium |
| EPI_ISL_2972827, EPI_ISL_2972828 | Lighthouse Laboratory Plymouth | Wellcome Sanger Institute for the COVID-19 Genomics UK (COG-UK) Consortium | Lighthouse Laboratory Plymouth and Alex Alderton, Roberto Amato, Jeffrey Barrett, Sonia Goncalves, Ewan Harrison, David K. Jackson, Ian Johnston, Dominic Kwiatkowski, Cordelia Langford, John Sillitoe on behalf of the Wellcome Sanger Institute COVID-19 Surveillance Team |
| EPI_ISL_2972829 | Department of Bacteria, Parasites and Fungi, Statens Serum Institut, Copenhagen, Denmark | Statens Serum Institut Bioinformatics and Microbial Genomics | Danish Covid-19 Genome Consortium |
| EPI_ISL_2972830 | Lighthouse Laboratory Plymouth | Wellcome Sanger Institute for the COVID-19 Genomics UK (COG-UK) Consortium | Lighthouse Laboratory Plymouth and Alex Alderton, Roberto Amato, Jeffrey Barrett, Sonia Goncalves, Ewan Harrison, David K. Jackson, Ian Johnston, Dominic Kwiatkowski, Cordelia Langford, John Sillitoe on behalf of the Wellcome Sanger Institute COVID-19 Surveillance Team |

| | | | |
|---|---|---|---|
| EPI_ISL_2972831, EPI_ISL_2972832 | Lighthouse Lab in Alderley Park | Wellcome Sanger Institute for the COVID-19 Genomics UK (COG-UK) Consortium | Jacquelyn Wynn, Mairead Hyland, The Lighthouse Lab in Alderley Park and Alex Alderton, Roberto Amato, Jeffrey Barrett, Sonia Goncalves, Ewan Harrison, David K. Jackson, Ian Johnston, Dominic Kwiatkowski, Cordelia Langford, John Sillitoe on behalf of the Wellcome Sanger Institute COVID-19 Surveillance Team |
| EPI_ISL_2972833 | Department of Bacteria, Parasites and Fungi, Statens Serum Institut, Copenhagen, Denmark | Statens Serum Institut Bioinformatics and Microbial Genomics | Danish Covid-19 Genome Consortium |
| EPI_ISL_2972834 | Lighthouse Laboratory Plymouth | Wellcome Sanger Institute for the COVID-19 Genomics UK (COG-UK) Consortium | Lighthouse Laboratory Plymouth and Alex Alderton, Roberto Amato, Jeffrey Barrett, Sonia Goncalves, Ewan Harrison, David K. Jackson, Ian Johnston, Dominic Kwiatkowski, Cordelia Langford, John Sillitoe on behalf of the Wellcome Sanger Institute COVID-19 Surveillance Team |
| EPI_ISL_2972835 | Lighthouse Lab in Milton Keynes | Wellcome Sanger Institute for the COVID-19 Genomics UK (COG-UK) Consortium | The Lighthouse Lab in Milton Keynes and Alex Alderton, Roberto Amato, Jeffrey Barrett, Sonia Goncalves, Ewan Harrison, David K. Jackson, Ian Johnston, Dominic Kwiatkowski, Cordelia Langford, John Sillitoe on behalf of the Wellcome Sanger Institute COVID-19 Surveillance Team |
| EPI_ISL_2972836 | Department of Bacteria, Parasites and Fungi, Statens Serum Institut, Copenhagen, Denmark | Statens Serum Institut Bioinformatics and Microbial Genomics | Danish Covid-19 Genome Consortium |
| EPI_ISL_2972837 | Lighthouse Lab in Milton Keynes | Wellcome Sanger Institute for the COVID-19 Genomics UK (COG-UK) Consortium | The Lighthouse Lab in Milton Keynes and Alex Alderton, Roberto Amato, Jeffrey Barrett, Sonia Goncalves, Ewan Harrison, David K. Jackson, Ian Johnston, Dominic Kwiatkowski, Cordelia Langford, John Sillitoe on behalf of the Wellcome Sanger Institute COVID-19 Surveillance Team |
| EPI_ISL_2972838, EPI_ISL_2972839 | Lighthouse Lab in Alderley Park | Wellcome Sanger Institute for the COVID-19 Genomics UK (COG-UK) Consortium | Jacquelyn Wynn, Mairead Hyland, The Lighthouse Lab in Alderley Park and Alex Alderton, Roberto Amato, Jeffrey Barrett, Sonia Goncalves, Ewan Harrison, David K. Jackson, Ian Johnston, Dominic Kwiatkowski, Cordelia Langford, John Sillitoe on behalf of the Wellcome Sanger Institute COVID-19 Surveillance Team |
| EPI_ISL_2972840 | Department of Bacteria, Parasites and Fungi, Statens Serum Institut, Copenhagen, Denmark | Statens Serum Institut Bioinformatics and Microbial Genomics | Danish Covid-19 Genome Consortium |
| EPI_ISL_2972841, EPI_ISL_2972842, EPI_ISL_2972844, EPI_ISL_2972845 | Lighthouse Lab in Alderley Park | Wellcome Sanger Institute for the COVID-19 Genomics UK (COG-UK) Consortium | Jacquelyn Wynn, Mairead Hyland, The Lighthouse Lab in Alderley Park and Alex Alderton, Roberto Amato, Jeffrey Barrett, Sonia Goncalves, Ewan Harrison, David K. Jackson, Ian Johnston, Dominic Kwiatkowski, Cordelia Langford, John Sillitoe on behalf of the Wellcome Sanger Institute COVID-19 Surveillance Team |
| EPI_ISL_2972846 | Department of Bacteria, Parasites and Fungi, Statens Serum Institut, Copenhagen, Denmark | Statens Serum Institut Bioinformatics and Microbial Genomics | Danish Covid-19 Genome Consortium |
| EPI_ISL_2972847, EPI_ISL_2972848 | Lighthouse Lab in Alderley Park | Wellcome Sanger Institute for the COVID-19 Genomics UK (COG-UK) Consortium | Jacquelyn Wynn, Mairead Hyland, The Lighthouse Lab in Alderley Park and Alex Alderton, Roberto Amato, Jeffrey Barrett, Sonia Goncalves, Ewan Harrison, David K. Jackson, Ian Johnston, Dominic Kwiatkowski, Cordelia Langford, John Sillitoe on behalf of the Wellcome Sanger Institute COVID-19 Surveillance Team |
| EPI_ISL_2972849 | Lighthouse Lab in Milton Keynes | Wellcome Sanger Institute for the COVID-19 Genomics UK (COG-UK) Consortium | The Lighthouse Lab in Milton Keynes and Alex Alderton, Roberto Amato, Jeffrey Barrett, Sonia Goncalves, Ewan Harrison, David K. Jackson, Ian Johnston, Dominic Kwiatkowski, Cordelia Langford, John Sillitoe on behalf of the Wellcome Sanger Institute COVID-19 Surveillance Team |
| EPI_ISL_2972850 | Department of Bacteria, Parasites and Fungi, Statens Serum Institut, Copenhagen, Denmark | Statens Serum Institut Bioinformatics and Microbial Genomics | Danish Covid-19 Genome Consortium |
| EPI_ISL_2972851 | Lighthouse Lab in Alderley Park | Wellcome Sanger Institute for the COVID-19 Genomics UK (COG-UK) Consortium | Jacquelyn Wynn, Mairead Hyland, The Lighthouse Lab in Alderley Park and Alex Alderton, Roberto Amato, Jeffrey Barrett, Sonia Goncalves, Ewan Harrison, David K. Jackson, Ian Johnston, Dominic Kwiatkowski, Cordelia Langford, John Sillitoe on behalf of the Wellcome Sanger Institute COVID-19 Surveillance Team |
| EPI_ISL_2972852 | Lighthouse Laboratory Plymouth | Wellcome Sanger Institute for the COVID-19 Genomics UK (COG-UK) Consortium | Lighthouse Laboratory Plymouth and Alex Alderton, Roberto Amato, Jeffrey Barrett, Sonia Goncalves, Ewan Harrison, David K. Jackson, Ian Johnston, Dominic Kwiatkowski, Cordelia Langford, John Sillitoe on behalf of the Wellcome Sanger Institute COVID-19 Surveillance Team |
| EPI_ISL_2972853 | Department of Bacteria, Parasites and Fungi, Statens Serum Institut, Copenhagen, Denmark | Statens Serum Institut Bioinformatics and Microbial Genomics | Danish Covid-19 Genome Consortium |
| EPI_ISL_2972854, EPI_ISL_2972855 | Lighthouse Laboratory Plymouth | Wellcome Sanger Institute for the COVID-19 Genomics UK (COG-UK) Consortium | Lighthouse Laboratory Plymouth and Alex Alderton, Roberto Amato, Jeffrey Barrett, Sonia Goncalves, Ewan Harrison, David K. Jackson, Ian Johnston, Dominic Kwiatkowski, Cordelia Langford, John Sillitoe on behalf of the Wellcome Sanger Institute COVID-19 Surveillance Team |
| EPI_ISL_2972856, EPI_ISL_2972858, EPI_ISL_2972859 | Lighthouse Lab in Alderley Park | Wellcome Sanger Institute for the COVID-19 Genomics UK (COG-UK) Consortium | Jacquelyn Wynn, Mairead Hyland, The Lighthouse Lab in Alderley Park and Alex Alderton, Roberto Amato, Jeffrey Barrett, Sonia Goncalves, Ewan Harrison, David K. Jackson, Ian Johnston, Dominic Kwiatkowski, Cordelia Langford, John Sillitoe on behalf of the Wellcome Sanger Institute COVID-19 Surveillance Team |
| EPI_ISL_2972860 | Department of Bacteria, Parasites and Fungi, Statens Serum Institut, Copenhagen, Denmark | Statens Serum Institut Bioinformatics and Microbial Genomics | Danish Covid-19 Genome Consortium |
| EPI_ISL_2972861 | Lighthouse Laboratory Plymouth | Wellcome Sanger Institute for the COVID-19 Genomics UK (COG-UK) Consortium | Lighthouse Laboratory Plymouth and Alex Alderton, Roberto Amato, Jeffrey Barrett, Sonia Goncalves, Ewan Harrison, David K. Jackson, Ian Johnston, Dominic Kwiatkowski, Cordelia Langford, John Sillitoe on behalf of the Wellcome Sanger Institute COVID-19 Surveillance Team |
| EPI_ISL_2972862 | Lighthouse Lab in Alderley Park | Wellcome Sanger Institute for the COVID-19 Genomics UK (COG-UK) Consortium | Jacquelyn Wynn, Mairead Hyland, The Lighthouse Lab in Alderley Park and Alex Alderton, Roberto Amato, Jeffrey Barrett, Sonia Goncalves, Ewan Harrison, David K. Jackson, Ian Johnston, Dominic Kwiatkowski, Cordelia Langford, John Sillitoe on behalf of the Wellcome Sanger Institute COVID-19 Surveillance Team |
| EPI_ISL_2972863 | Department of Bacteria, Parasites and Fungi, Statens Serum Institut, Copenhagen, Denmark | Statens Serum Institut Bioinformatics and Microbial Genomics | Danish Covid-19 Genome Consortium |
| EPI_ISL_2972864 | Lighthouse Lab in Alderley Park | Wellcome Sanger Institute for the COVID-19 Genomics UK (COG-UK) Consortium | Jacquelyn Wynn, Mairead Hyland, The Lighthouse Lab in Alderley Park and Alex Alderton, Roberto Amato, Jeffrey Barrett, Sonia Goncalves, Ewan Harrison, David K. Jackson, Ian Johnston, Dominic Kwiatkowski, Cordelia Langford, John Sillitoe on behalf of the Wellcome Sanger Institute COVID-19 Surveillance Team |
| EPI_ISL_2972865 | Lighthouse Lab in Milton Keynes | Wellcome Sanger Institute for the COVID-19 Genomics UK (COG-UK) Consortium | The Lighthouse Lab in Milton Keynes and Alex Alderton, Roberto Amato, Jeffrey Barrett, Sonia Goncalves, Ewan Harrison, David K. Jackson, Ian Johnston, Dominic Kwiatkowski, Cordelia Langford, John Sillitoe on behalf of the Wellcome Sanger Institute COVID-19 Surveillance Team |
| EPI_ISL_2972866 | Department of Bacteria, Parasites and Fungi, Statens Serum Institut, Copenhagen, Denmark | Statens Serum Institut Bioinformatics and Microbial Genomics | Danish Covid-19 Genome Consortium |
| EPI_ISL_2972867, EPI_ISL_2972868 | Lighthouse Laboratory Plymouth | Wellcome Sanger Institute for the COVID-19 Genomics UK (COG-UK) Consortium | Lighthouse Laboratory Plymouth and Alex Alderton, Roberto Amato, Jeffrey Barrett, Sonia Goncalves, Ewan Harrison, David K. Jackson, Ian Johnston, Dominic Kwiatkowski, Cordelia Langford, John Sillitoe on behalf of the Wellcome Sanger Institute COVID-19 Surveillance Team |
| EPI_ISL_2972870 | Lighthouse Lab in Alderley Park | Wellcome Sanger Institute for the COVID-19 Genomics UK (COG-UK) Consortium | Jacquelyn Wynn, Mairead Hyland, The Lighthouse Lab in Alderley Park and Alex Alderton, Roberto Amato, Jeffrey Barrett, Sonia Goncalves, Ewan Harrison, David K. Jackson, Ian Johnston, Dominic Kwiatkowski, Cordelia Langford, John Sillitoe on behalf of the Wellcome Sanger Institute COVID-19 Surveillance Team |
| EPI_ISL_2972871 | Lighthouse Lab in Milton Keynes | Wellcome Sanger Institute for the COVID-19 Genomics UK (COG-UK) Consortium | The Lighthouse Lab in Milton Keynes and Alex Alderton, Roberto Amato, Jeffrey Barrett, Sonia Goncalves, Ewan Harrison, David K. Jackson, Ian Johnston, Dominic Kwiatkowski, Cordelia Langford, John Sillitoe on behalf of the Wellcome Sanger Institute COVID-19 Surveillance Team |
| EPI_ISL_2972872 | Lighthouse Lab in Alderley Park | Wellcome Sanger Institute for the COVID-19 Genomics UK (COG-UK) Consortium | Jacquelyn Wynn, Mairead Hyland, The Lighthouse Lab in Alderley Park and Alex Alderton, Roberto Amato, Jeffrey Barrett, Sonia Goncalves, Ewan Harrison, David K. Jackson, Ian Johnston, Dominic Kwiatkowski, Cordelia Langford, John Sillitoe on behalf of the Wellcome Sanger Institute COVID-19 Surveillance Team |
| EPI_ISL_2972873 | Department of Bacteria, Parasites and Fungi, Statens Serum Institut, Copenhagen, Denmark | Statens Serum Institut Bioinformatics and Microbial Genomics | Danish Covid-19 Genome Consortium |
| EPI_ISL_2972874 | Lighthouse Lab in Alderley Park | Wellcome Sanger Institute for the COVID-19 Genomics UK (COG-UK) Consortium | Jacquelyn Wynn, Mairead Hyland, The Lighthouse Lab in Alderley Park and Alex Alderton, Roberto Amato, Jeffrey Barrett, Sonia Goncalves, Ewan Harrison, David K. Jackson, Ian Johnston, Dominic Kwiatkowski, Cordelia Langford, John Sillitoe on behalf of the Wellcome Sanger Institute COVID-19 Surveillance Team |
| EPI_ISL_2972875 | Lighthouse Laboratory Plymouth | Wellcome Sanger Institute for the COVID-19 Genomics UK (COG-UK) Consortium | Lighthouse Laboratory Plymouth and Alex Alderton, Roberto Amato, Jeffrey Barrett, Sonia Goncalves, Ewan Harrison, David K. Jackson, Ian Johnston, Dominic Kwiatkowski, Cordelia Langford, John Sillitoe on behalf of the Wellcome Sanger Institute COVID-19 Surveillance Team |
| EPI_ISL_2972877, EPI_ISL_2972878 | Lighthouse Lab in Alderley Park | Wellcome Sanger Institute for the COVID-19 Genomics UK (COG-UK) Consortium | Jacquelyn Wynn, Mairead Hyland, The Lighthouse Lab in Alderley Park and Alex Alderton, Roberto Amato, Jeffrey Barrett, Sonia Goncalves, Ewan Harrison, David K. Jackson, Ian Johnston, Dominic Kwiatkowski, Cordelia Langford, John Sillitoe on behalf of the Wellcome Sanger Institute COVID-19 Surveillance Team |
| EPI_ISL_2972879 | Department of Bacteria, Parasites and Fungi, Statens Serum Institut, Copenhagen, Denmark | Statens Serum Institut Bioinformatics and Microbial Genomics | Danish Covid-19 Genome Consortium |
| EPI_ISL_2972880, EPI_ISL_2972881 | Lighthouse Lab in Milton Keynes | Wellcome Sanger Institute for the COVID-19 Genomics UK (COG-UK) Consortium | The Lighthouse Lab in Milton Keynes and Alex Alderton, Roberto Amato, Jeffrey Barrett, Sonia Goncalves, Ewan Harrison, David K. Jackson, Ian Johnston, Dominic Kwiatkowski, Cordelia Langford, John Sillitoe on behalf of the Wellcome Sanger Institute COVID-19 Surveillance Team |
| EPI_ISL_2972882 | Lighthouse Lab in Alderley Park | Wellcome Sanger Institute for the COVID-19 Genomics UK (COG-UK) Consortium | Jacquelyn Wynn, Mairead Hyland, The Lighthouse Lab in Alderley Park and Alex Alderton, Roberto Amato, Jeffrey Barrett, Sonia Goncalves, Ewan Harrison, David K. Jackson, Ian Johnston, Dominic Kwiatkowski, Cordelia Langford, John Sillitoe on behalf of the Wellcome Sanger Institute COVID-19 Surveillance Team |
| EPI_ISL_2972883 | Department of Bacteria, Parasites and Fungi, Statens Serum Institut, Copenhagen, Denmark | Statens Serum Institut Bioinformatics and Microbial Genomics | Danish Covid-19 Genome Consortium |
| EPI_ISL_2972884 | Lighthouse Laboratory Plymouth | Wellcome Sanger Institute for the COVID-19 Genomics UK (COG-UK) Consortium | Lighthouse Laboratory Plymouth and Alex Alderton, Roberto Amato, Jeffrey Barrett, Sonia Goncalves, Ewan Harrison, David K. Jackson, Ian Johnston, Dominic Kwiatkowski, Cordelia Langford, John Sillitoe on behalf of the Wellcome Sanger Institute COVID-19 Surveillance Team |
| EPI_ISL_2972885, EPI_ISL_2972886 | Lighthouse Lab in Alderley Park | Wellcome Sanger Institute for the COVID-19 Genomics UK (COG-UK) Consortium | Jacquelyn Wynn, Mairead Hyland, The Lighthouse Lab in Alderley Park and Alex Alderton, Roberto Amato, Jeffrey Barrett, Sonia Goncalves, Ewan Harrison, David K. Jackson, Ian Johnston, Dominic Kwiatkowski, Cordelia Langford, John Sillitoe on behalf of the Wellcome Sanger Institute COVID-19 Surveillance Team |
| EPI_ISL_2972887 | Department of Bacteria, Parasites and Fungi, Statens Serum Institut, Copenhagen, Denmark | Statens Serum Institut Bioinformatics and Microbial Genomics | Danish Covid-19 Genome Consortium |
| EPI_ISL_2972888 | Lighthouse Lab in Alderley Park | Wellcome Sanger Institute for the COVID-19 Genomics UK (COG-UK) Consortium | Jacquelyn Wynn, Mairead Hyland, The Lighthouse Lab in Alderley Park and Alex Alderton, Roberto Amato, Jeffrey Barrett, Sonia Goncalves, Ewan Harrison, David K. Jackson, Ian Johnston, Dominic Kwiatkowski, Cordelia Langford, John Sillitoe on behalf of the Wellcome Sanger Institute COVID-19 Surveillance Team |
| EPI_ISL_2972889 | Lighthouse Laboratory Plymouth | Wellcome Sanger Institute for the COVID-19 Genomics UK (COG-UK) Consortium | Lighthouse Laboratory Plymouth and Alex Alderton, Roberto Amato, Jeffrey Barrett, Sonia Goncalves, Ewan Harrison, David K. Jackson, Ian Johnston, Dominic Kwiatkowski, Cordelia Langford, John Sillitoe on behalf of the Wellcome Sanger Institute COVID-19 Surveillance Team |
| EPI_ISL_2972891 | Department of Bacteria, Parasites and Fungi, Statens Serum Institut, Copenhagen, Denmark | Statens Serum Institut Bioinformatics and Microbial Genomics | Danish Covid-19 Genome Consortium |
| EPI_ISL_2972892, EPI_ISL_2972893 | Lighthouse Lab in Alderley Park | Wellcome Sanger Institute for the COVID-19 Genomics UK (COG-UK) Consortium | Jacquelyn Wynn, Mairead Hyland, The Lighthouse Lab in Alderley Park and Alex Alderton, Roberto Amato, Jeffrey Barrett, Sonia Goncalves, Ewan Harrison, David K. Jackson, Ian Johnston, Dominic Kwiatkowski, Cordelia Langford, John Sillitoe on behalf of the Wellcome Sanger Institute COVID-19 Surveillance Team |

| | | | |
|---|---|---|---|
| EPI_ISL_2972894 | Department of Bacteria, Parasites and Fungi, Statens Serum Institut, Copenhagen, Denmark | Statens Serum Institut Bioinformatics and Microbial Genomics | Danish Covid-19 Genome Consortium |
| EPI_ISL_2972895 | Lighthouse Lab in Milton Keynes | Wellcome Sanger Institute for the COVID-19 Genomics UK (COG-UK) Consortium | The Lighthouse Lab in Milton Keynes and Alex Alderton, Roberto Amato, Jeffrey Barrett, Sonia Goncalves, Ewan Harrison, David K. Jackson, Ian Johnston, Dominic Kwiatkowski, Cordelia Langford, John Sillitoe on behalf of the Wellcome Sanger Institute COVID-19 Surveillance Team |
| EPI_ISL_2972896 | Lighthouse Lab in Alderley Park | Wellcome Sanger Institute for the COVID-19 Genomics UK (COG-UK) Consortium | Jacquelyn Wynn, Mairead Hyland, The Lighthouse Lab in Alderley Park and Alex Alderton, Roberto Amato, Jeffrey Barrett, Sonia Goncalves, Ewan Harrison, David K. Jackson, Ian Johnston, Dominic Kwiatkowski, Cordelia Langford, John Sillitoe on behalf of the Wellcome Sanger Institute COVID-19 Surveillance Team |
| EPI_ISL_2972897 | Department of Bacteria, Parasites and Fungi, Statens Serum Institut, Copenhagen, Denmark | Statens Serum Institut Bioinformatics and Microbial Genomics | Danish Covid-19 Genome Consortium |
| EPI_ISL_2972898 | Lighthouse Lab in Alderley Park | Wellcome Sanger Institute for the COVID-19 Genomics UK (COG-UK) Consortium | Jacquelyn Wynn, Mairead Hyland, The Lighthouse Lab in Alderley Park and Alex Alderton, Roberto Amato, Jeffrey Barrett, Sonia Goncalves, Ewan Harrison, David K. Jackson, Ian Johnston, Dominic Kwiatkowski, Cordelia Langford, John Sillitoe on behalf of the Wellcome Sanger Institute COVID-19 Surveillance Team |
| EPI_ISL_2972899 | Lighthouse Laboratory Plymouth | Wellcome Sanger Institute for the COVID-19 Genomics UK (COG-UK) Consortium | Lighthouse Laboratory Plymouth and Alex Alderton, Roberto Amato, Jeffrey Barrett, Sonia Goncalves, Ewan Harrison, David K. Jackson, Ian Johnston, Dominic Kwiatkowski, Cordelia Langford, John Sillitoe on behalf of the Wellcome Sanger Institute COVID-19 Surveillance Team |
| EPI_ISL_2972900 | Lighthouse Lab in Alderley Park | Wellcome Sanger Institute for the COVID-19 Genomics UK (COG-UK) Consortium | Jacquelyn Wynn, Mairead Hyland, The Lighthouse Lab in Alderley Park and Alex Alderton, Roberto Amato, Jeffrey Barrett, Sonia Goncalves, Ewan Harrison, David K. Jackson, Ian Johnston, Dominic Kwiatkowski, Cordelia Langford, John Sillitoe on behalf of the Wellcome Sanger Institute COVID-19 Surveillance Team |
| EPI_ISL_2972901 | Department of Bacteria, Parasites and Fungi, Statens Serum Institut, Copenhagen, Denmark | Statens Serum Institut Bioinformatics and Microbial Genomics | Danish Covid-19 Genome Consortium |
| EPI_ISL_2972902 | Lighthouse Laboratory Plymouth | Wellcome Sanger Institute for the COVID-19 Genomics UK (COG-UK) Consortium | Lighthouse Laboratory Plymouth and Alex Alderton, Roberto Amato, Jeffrey Barrett, Sonia Goncalves, Ewan Harrison, David K. Jackson, Ian Johnston, Dominic Kwiatkowski, Cordelia Langford, John Sillitoe on behalf of the Wellcome Sanger Institute COVID-19 Surveillance Team |
| EPI_ISL_2972903 | Lighthouse Lab in Alderley Park | Wellcome Sanger Institute for the COVID-19 Genomics UK (COG-UK) Consortium | Jacquelyn Wynn, Mairead Hyland, The Lighthouse Lab in Alderley Park and Alex Alderton, Roberto Amato, Jeffrey Barrett, Sonia Goncalves, Ewan Harrison, David K. Jackson, Ian Johnston, Dominic Kwiatkowski, Cordelia Langford, John Sillitoe on behalf of the Wellcome Sanger Institute COVID-19 Surveillance Team |
| EPI_ISL_2972904 | Lighthouse Lab in Milton Keynes | Wellcome Sanger Institute for the COVID-19 Genomics UK (COG-UK) Consortium | The Lighthouse Lab in Milton Keynes and Alex Alderton, Roberto Amato, Jeffrey Barrett, Sonia Goncalves, Ewan Harrison, David K. Jackson, Ian Johnston, Dominic Kwiatkowski, Cordelia Langford, John Sillitoe on behalf of the Wellcome Sanger Institute COVID-19 Surveillance Team |
| EPI_ISL_2972905 | Department of Bacteria, Parasites and Fungi, Statens Serum Institut, Copenhagen, Denmark | Statens Serum Institut Bioinformatics and Microbial Genomics | Danish Covid-19 Genome Consortium |
| EPI_ISL_2972906, EPI_ISL_2972907 | Lighthouse Laboratory Plymouth | Wellcome Sanger Institute for the COVID-19 Genomics UK (COG-UK) Consortium | Lighthouse Laboratory Plymouth and Alex Alderton, Roberto Amato, Jeffrey Barrett, Sonia Goncalves, Ewan Harrison, David K. Jackson, Ian Johnston, Dominic Kwiatkowski, Cordelia Langford, John Sillitoe on behalf of the Wellcome Sanger Institute COVID-19 Surveillance Team |
| EPI_ISL_2972908 | Department of Bacteria, Parasites and Fungi, Statens Serum Institut, Copenhagen, Denmark | Statens Serum Institut Bioinformatics and Microbial Genomics | Danish Covid-19 Genome Consortium |
| EPI_ISL_2972909 | Lighthouse Lab in Alderley Park | Wellcome Sanger Institute for the COVID-19 Genomics UK (COG-UK) Consortium | Jacquelyn Wynn, Mairead Hyland, The Lighthouse Lab in Alderley Park and Alex Alderton, Roberto Amato, Jeffrey Barrett, Sonia Goncalves, Ewan Harrison, David K. Jackson, Ian Johnston, Dominic Kwiatkowski, Cordelia Langford, John Sillitoe on behalf of the Wellcome Sanger Institute COVID-19 Surveillance Team |
| EPI_ISL_2972910 | Lighthouse Lab in Milton Keynes | Wellcome Sanger Institute for the COVID-19 Genomics UK (COG-UK) Consortium | The Lighthouse Lab in Milton Keynes and Alex Alderton, Roberto Amato, Jeffrey Barrett, Sonia Goncalves, Ewan Harrison, David K. Jackson, Ian Johnston, Dominic Kwiatkowski, Cordelia Langford, John Sillitoe on behalf of the Wellcome Sanger Institute COVID-19 Surveillance Team |
| EPI_ISL_2972911 | Department of Bacteria, Parasites and Fungi, Statens Serum Institut, Copenhagen, Denmark | Statens Serum Institut Bioinformatics and Microbial Genomics | Danish Covid-19 Genome Consortium |
| EPI_ISL_2972912, EPI_ISL_2972913 | Lighthouse Lab in Milton Keynes | Wellcome Sanger Institute for the COVID-19 Genomics UK (COG-UK) Consortium | The Lighthouse Lab in Milton Keynes and Alex Alderton, Roberto Amato, Jeffrey Barrett, Sonia Goncalves, Ewan Harrison, David K. Jackson, Ian Johnston, Dominic Kwiatkowski, Cordelia Langford, John Sillitoe on behalf of the Wellcome Sanger Institute COVID-19 Surveillance Team |
| EPI_ISL_2972914 | Lighthouse Lab in Alderley Park | Wellcome Sanger Institute for the COVID-19 Genomics UK (COG-UK) Consortium | Jacquelyn Wynn, Mairead Hyland, The Lighthouse Lab in Alderley Park and Alex Alderton, Roberto Amato, Jeffrey Barrett, Sonia Goncalves, Ewan Harrison, David K. Jackson, Ian Johnston, Dominic Kwiatkowski, Cordelia Langford, John Sillitoe on behalf of the Wellcome Sanger Institute COVID-19 Surveillance Team |
| EPI_ISL_2972915 | Department of Bacteria, Parasites and Fungi, Statens Serum Institut, Copenhagen, Denmark | Statens Serum Institut Bioinformatics and Microbial Genomics | Danish Covid-19 Genome Consortium |
| EPI_ISL_2972916 | Lighthouse Lab in Alderley Park | Wellcome Sanger Institute for the COVID-19 Genomics UK (COG-UK) Consortium | Jacquelyn Wynn, Mairead Hyland, The Lighthouse Lab in Alderley Park and Alex Alderton, Roberto Amato, Jeffrey Barrett, Sonia Goncalves, Ewan Harrison, David K. Jackson, Ian Johnston, Dominic Kwiatkowski, Cordelia Langford, John Sillitoe on behalf of the Wellcome Sanger Institute COVID-19 Surveillance Team |
| EPI_ISL_2972917 | Lighthouse Laboratory Plymouth | Wellcome Sanger Institute for the COVID-19 Genomics UK (COG-UK) Consortium | Lighthouse Laboratory Plymouth and Alex Alderton, Roberto Amato, Jeffrey Barrett, Sonia Goncalves, Ewan Harrison, David K. Jackson, Ian Johnston, Dominic Kwiatkowski, Cordelia Langford, John Sillitoe on behalf of the Wellcome Sanger Institute COVID-19 Surveillance Team |
| EPI_ISL_2972918 | Department of Bacteria, Parasites and Fungi, Statens Serum Institut, Copenhagen, Denmark | Statens Serum Institut Bioinformatics and Microbial Genomics | Danish Covid-19 Genome Consortium |
| EPI_ISL_2972919 | Lighthouse Lab in Milton Keynes | Wellcome Sanger Institute for the COVID-19 Genomics UK (COG-UK) Consortium | The Lighthouse Lab in Milton Keynes and Alex Alderton, Roberto Amato, Jeffrey Barrett, Sonia Goncalves, Ewan Harrison, David K. Jackson, Ian Johnston, Dominic Kwiatkowski, Cordelia Langford, John Sillitoe on behalf of the Wellcome Sanger Institute COVID-19 Surveillance Team |
| EPI_ISL_2972920, EPI_ISL_2972921 | Lighthouse Laboratory Plymouth | Wellcome Sanger Institute for the COVID-19 Genomics UK (COG-UK) Consortium | Lighthouse Laboratory Plymouth and Alex Alderton, Roberto Amato, Jeffrey Barrett, Sonia Goncalves, Ewan Harrison, David K. Jackson, Ian Johnston, Dominic Kwiatkowski, Cordelia Langford, John Sillitoe on behalf of the Wellcome Sanger Institute COVID-19 Surveillance Team |
| EPI_ISL_2972922 | Department of Bacteria, Parasites and Fungi, Statens Serum Institut, Copenhagen, Denmark | Statens Serum Institut Bioinformatics and Microbial Genomics | Danish Covid-19 Genome Consortium |
| EPI_ISL_2972923 | Lighthouse Laboratory Plymouth | Wellcome Sanger Institute for the COVID-19 Genomics UK (COG-UK) Consortium | Lighthouse Laboratory Plymouth and Alex Alderton, Roberto Amato, Jeffrey Barrett, Sonia Goncalves, Ewan Harrison, David K. Jackson, Ian Johnston, Dominic Kwiatkowski, Cordelia Langford, John Sillitoe on behalf of the Wellcome Sanger Institute COVID-19 Surveillance Team |
| EPI_ISL_2972924, EPI_ISL_2972925 | Lighthouse Lab in Alderley Park | Wellcome Sanger Institute for the COVID-19 Genomics UK (COG-UK) Consortium | Jacquelyn Wynn, Mairead Hyland, The Lighthouse Lab in Alderley Park and Alex Alderton, Roberto Amato, Jeffrey Barrett, Sonia Goncalves, Ewan Harrison, David K. Jackson, Ian Johnston, Dominic Kwiatkowski, Cordelia Langford, John Sillitoe on behalf of the Wellcome Sanger Institute COVID-19 Surveillance Team |
| EPI_ISL_2972926 | Department of Bacteria, Parasites and Fungi, Statens Serum Institut, Copenhagen, Denmark | Statens Serum Institut Bioinformatics and Microbial Genomics | Danish Covid-19 Genome Consortium |
| EPI_ISL_2972927 | Lighthouse Lab in Alderley Park | Wellcome Sanger Institute for the COVID-19 Genomics UK (COG-UK) Consortium | Jacquelyn Wynn, Mairead Hyland, The Lighthouse Lab in Alderley Park and Alex Alderton, Roberto Amato, Jeffrey Barrett, Sonia Goncalves, Ewan Harrison, David K. Jackson, Ian Johnston, Dominic Kwiatkowski, Cordelia Langford, John Sillitoe on behalf of the Wellcome Sanger Institute COVID-19 Surveillance Team |
| EPI_ISL_2972928 | Lighthouse Lab in Milton Keynes | Wellcome Sanger Institute for the COVID-19 Genomics UK (COG-UK) Consortium | The Lighthouse Lab in Milton Keynes and Alex Alderton, Roberto Amato, Jeffrey Barrett, Sonia Goncalves, Ewan Harrison, David K. Jackson, Ian Johnston, Dominic Kwiatkowski, Cordelia Langford, John Sillitoe on behalf of the Wellcome Sanger Institute COVID-19 Surveillance Team |
| EPI_ISL_2972929 | Department of Bacteria, Parasites and Fungi, Statens Serum Institut, Copenhagen, Denmark | Statens Serum Institut Bioinformatics and Microbial Genomics | Danish Covid-19 Genome Consortium |
| EPI_ISL_2972930 | Lighthouse Lab in Alderley Park | Wellcome Sanger Institute for the COVID-19 Genomics UK (COG-UK) Consortium | Jacquelyn Wynn, Mairead Hyland, The Lighthouse Lab in Alderley Park and Alex Alderton, Roberto Amato, Jeffrey Barrett, Sonia Goncalves, Ewan Harrison, David K. Jackson, Ian Johnston, Dominic Kwiatkowski, Cordelia Langford, John Sillitoe on behalf of the Wellcome Sanger Institute COVID-19 Surveillance Team |
| EPI_ISL_2972931 | Lighthouse Laboratory Plymouth | Wellcome Sanger Institute for the COVID-19 Genomics UK (COG-UK) Consortium | Lighthouse Laboratory Plymouth and Alex Alderton, Roberto Amato, Jeffrey Barrett, Sonia Goncalves, Ewan Harrison, David K. Jackson, Ian Johnston, Dominic Kwiatkowski, Cordelia Langford, John Sillitoe on behalf of the Wellcome Sanger Institute COVID-19 Surveillance Team |
| EPI_ISL_2972932 | Lighthouse Lab in Alderley Park | Wellcome Sanger Institute for the COVID-19 Genomics UK (COG-UK) Consortium | Jacquelyn Wynn, Mairead Hyland, The Lighthouse Lab in Alderley Park and Alex Alderton, Roberto Amato, Jeffrey Barrett, Sonia Goncalves, Ewan Harrison, David K. Jackson, Ian Johnston, Dominic Kwiatkowski, Cordelia Langford, John Sillitoe on behalf of the Wellcome Sanger Institute COVID-19 Surveillance Team |
| EPI_ISL_2972933 | Department of Bacteria, Parasites and Fungi, Statens Serum Institut, Copenhagen, Denmark | Statens Serum Institut Bioinformatics and Microbial Genomics | Danish Covid-19 Genome Consortium |
| EPI_ISL_2972934, EPI_ISL_2972935 | Lighthouse Lab in Alderley Park | Wellcome Sanger Institute for the COVID-19 Genomics UK (COG-UK) Consortium | Jacquelyn Wynn, Mairead Hyland, The Lighthouse Lab in Alderley Park and Alex Alderton, Roberto Amato, Jeffrey Barrett, Sonia Goncalves, Ewan Harrison, David K. Jackson, Ian Johnston, Dominic Kwiatkowski, Cordelia Langford, John Sillitoe on behalf of the Wellcome Sanger Institute COVID-19 Surveillance Team |
| EPI_ISL_2972936 | Lighthouse Laboratory Plymouth | Wellcome Sanger Institute for the COVID-19 Genomics UK (COG-UK) Consortium | Lighthouse Laboratory Plymouth and Alex Alderton, Roberto Amato, Jeffrey Barrett, Sonia Goncalves, Ewan Harrison, David K. Jackson, Ian Johnston, Dominic Kwiatkowski, Cordelia Langford, John Sillitoe on behalf of the Wellcome Sanger Institute COVID-19 Surveillance Team |
| EPI_ISL_2972937 | Department of Bacteria, Parasites and Fungi, Statens Serum Institut, Copenhagen, Denmark | Statens Serum Institut Bioinformatics and Microbial Genomics | Danish Covid-19 Genome Consortium |
| EPI_ISL_2972938 | Lighthouse Lab in Alderley Park | Wellcome Sanger Institute for the COVID-19 Genomics UK (COG-UK) Consortium | Jacquelyn Wynn, Mairead Hyland, The Lighthouse Lab in Alderley Park and Alex Alderton, Roberto Amato, Jeffrey Barrett, Sonia Goncalves, Ewan Harrison, David K. Jackson, Ian Johnston, Dominic Kwiatkowski, Cordelia Langford, John Sillitoe on behalf of the Wellcome Sanger Institute COVID-19 Surveillance Team |
| EPI_ISL_2972939 | Lighthouse Lab in Milton Keynes | Wellcome Sanger Institute for the COVID-19 Genomics UK (COG-UK) Consortium | The Lighthouse Lab in Milton Keynes and Alex Alderton, Roberto Amato, Jeffrey Barrett, Sonia Goncalves, Ewan Harrison, David K. Jackson, Ian Johnston, Dominic Kwiatkowski, Cordelia Langford, John Sillitoe on behalf of the Wellcome Sanger Institute COVID-19 Surveillance Team |
| EPI_ISL_2972940 | Department of Bacteria, Parasites and Fungi, Statens Serum Institut, Copenhagen, Denmark | Statens Serum Institut Bioinformatics and Microbial Genomics | Danish Covid-19 Genome Consortium |
| EPI_ISL_2972941 | Lighthouse Lab in Milton Keynes | Wellcome Sanger Institute for the COVID-19 Genomics UK (COG-UK) Consortium | The Lighthouse Lab in Milton Keynes and Alex Alderton, Roberto Amato, Jeffrey Barrett, Sonia Goncalves, Ewan Harrison, David K. Jackson, Ian Johnston, Dominic Kwiatkowski, Cordelia Langford, John Sillitoe on behalf of the Wellcome Sanger Institute COVID-19 Surveillance Team |
| EPI_ISL_2972942 | Lighthouse Lab in Alderley Park | Wellcome Sanger Institute for the COVID-19 Genomics UK (COG-UK) Consortium | Jacquelyn Wynn, Mairead Hyland, The Lighthouse Lab in Alderley Park and Alex Alderton, Roberto Amato, Jeffrey Barrett, Sonia Goncalves, Ewan Harrison, David K. Jackson, Ian Johnston, Dominic Kwiatkowski, Cordelia Langford, John Sillitoe on behalf of the Wellcome Sanger Institute COVID-19 Surveillance Team |
| EPI_ISL_2972943 | Department of Bacteria, Parasites and Fungi, Statens Serum Institut, Copenhagen, Denmark | Statens Serum Institut Bioinformatics and Microbial Genomics | Danish Covid-19 Genome Consortium |

| | | | |
|---|---|---|---|
| EPI_ISL_2972944 | Berkshire and Surrey Pathology Services Lighthouse Laboratory | Wellcome Sanger Institute for the COVID-19 Genomics UK (COG-UK) Consortium | Berkshire and Surrey Pathology Services Lighthouse Laboratory and Alex Alderton, Roberto Amato, Jeffrey Barrett, Sonia Goncalves, Ewan Harrison, David K. Jackson, Ian Johnston, Dominic Kwiatkowski, Cordelia Langford, John Sillitoe on behalf of the Wellcome Sanger Institute COVID-19 Surveillance Team |
| EPI_ISL_2972945 | Lighthouse Laboratory Plymouth | Wellcome Sanger Institute for the COVID-19 Genomics UK (COG-UK) Consortium | Lighthouse Laboratory Plymouth and Alex Alderton, Roberto Amato, Jeffrey Barrett, Sonia Goncalves, Ewan Harrison, David K. Jackson, Ian Johnston, Dominic Kwiatkowski, Cordelia Langford, John Sillitoe on behalf of the Wellcome Sanger Institute COVID-19 Surveillance Team |
| EPI_ISL_2972946 | Lighthouse Lab in Milton Keynes | Wellcome Sanger Institute for the COVID-19 Genomics UK (COG-UK) Consortium | The Lighthouse Lab in Milton Keynes and Alex Alderton, Roberto Amato, Jeffrey Barrett, Sonia Goncalves, Ewan Harrison, David K. Jackson, Ian Johnston, Dominic Kwiatkowski, Cordelia Langford, John Sillitoe on behalf of the Wellcome Sanger Institute COVID-19 Surveillance Team |
| EPI_ISL_2972947 | Department of Bacteria, Parasites and Fungi, Statens Serum Institut, Copenhagen, Denmark | Statens Serum Institut Bioinformatics and Microbial Genomics | Danish Covid-19 Genome Consortium |
| EPI_ISL_2972948 | Berkshire and Surrey Pathology Services Lighthouse Laboratory | Wellcome Sanger Institute for the COVID-19 Genomics UK (COG-UK) Consortium | Berkshire and Surrey Pathology Services Lighthouse Laboratory and Alex Alderton, Roberto Amato, Jeffrey Barrett, Sonia Goncalves, Ewan Harrison, David K. Jackson, Ian Johnston, Dominic Kwiatkowski, Cordelia Langford, John Sillitoe on behalf of the Wellcome Sanger Institute COVID-19 Surveillance Team |
| EPI_ISL_2972949 | Lighthouse Laboratory Plymouth | Wellcome Sanger Institute for the COVID-19 Genomics UK (COG-UK) Consortium | Lighthouse Laboratory Plymouth and Alex Alderton, Roberto Amato, Jeffrey Barrett, Sonia Goncalves, Ewan Harrison, David K. Jackson, Ian Johnston, Dominic Kwiatkowski, Cordelia Langford, John Sillitoe on behalf of the Wellcome Sanger Institute COVID-19 Surveillance Team |
| EPI_ISL_2972950 | Department of Bacteria, Parasites and Fungi, Statens Serum Institut, Copenhagen, Denmark | Statens Serum Institut Bioinformatics and Microbial Genomics | Danish Covid-19 Genome Consortium |
| EPI_ISL_2972951 | Lighthouse Lab in Alderley Park | Wellcome Sanger Institute for the COVID-19 Genomics UK (COG-UK) Consortium | Jacquelyn Wynn, Mairead Hyland, The Lighthouse Lab in Alderley Park and Alex Alderton, Roberto Amato, Jeffrey Barrett, Sonia Goncalves, Ewan Harrison, David K. Jackson, Ian Johnston, Dominic Kwiatkowski, Cordelia Langford, John Sillitoe on behalf of the Wellcome Sanger Institute COVID-19 Surveillance Team |
| EPI_ISL_2972952 | Lighthouse Lab in Milton Keynes | Wellcome Sanger Institute for the COVID-19 Genomics UK (COG-UK) Consortium | The Lighthouse Lab in Milton Keynes and Alex Alderton, Roberto Amato, Jeffrey Barrett, Sonia Goncalves, Ewan Harrison, David K. Jackson, Ian Johnston, Dominic Kwiatkowski, Cordelia Langford, John Sillitoe on behalf of the Wellcome Sanger Institute COVID-19 Surveillance Team |
| EPI_ISL_2972953 | Department of Bacteria, Parasites and Fungi, Statens Serum Institut, Copenhagen, Denmark | Statens Serum Institut Bioinformatics and Microbial Genomics | Danish Covid-19 Genome Consortium |
| EPI_ISL_2972954, EPI_ISL_2972955 | Lighthouse Lab in Alderley Park | Wellcome Sanger Institute for the COVID-19 Genomics UK (COG-UK) Consortium | Jacquelyn Wynn, Mairead Hyland, The Lighthouse Lab in Alderley Park and Alex Alderton, Roberto Amato, Jeffrey Barrett, Sonia Goncalves, Ewan Harrison, David K. Jackson, Ian Johnston, Dominic Kwiatkowski, Cordelia Langford, John Sillitoe on behalf of the Wellcome Sanger Institute COVID-19 Surveillance Team |
| EPI_ISL_2972956 | Lighthouse Lab in Milton Keynes | Wellcome Sanger Institute for the COVID-19 Genomics UK (COG-UK) Consortium | The Lighthouse Lab in Milton Keynes and Alex Alderton, Roberto Amato, Jeffrey Barrett, Sonia Goncalves, Ewan Harrison, David K. Jackson, Ian Johnston, Dominic Kwiatkowski, Cordelia Langford, John Sillitoe on behalf of the Wellcome Sanger Institute COVID-19 Surveillance Team |
| EPI_ISL_2972957 | Department of Bacteria, Parasites and Fungi, Statens Serum Institut, Copenhagen, Denmark | Statens Serum Institut Bioinformatics and Microbial Genomics | Danish Covid-19 Genome Consortium |
| EPI_ISL_2972958 | Lighthouse Lab in Alderley Park | Wellcome Sanger Institute for the COVID-19 Genomics UK (COG-UK) Consortium | Jacquelyn Wynn, Mairead Hyland, The Lighthouse Lab in Alderley Park and Alex Alderton, Roberto Amato, Jeffrey Barrett, Sonia Goncalves, Ewan Harrison, David K. Jackson, Ian Johnston, Dominic Kwiatkowski, Cordelia Langford, John Sillitoe on behalf of the Wellcome Sanger Institute COVID-19 Surveillance Team |
| EPI_ISL_2972959 | Lighthouse Lab in Milton Keynes | Wellcome Sanger Institute for the COVID-19 Genomics UK (COG-UK) Consortium | The Lighthouse Lab in Milton Keynes and Alex Alderton, Roberto Amato, Jeffrey Barrett, Sonia Goncalves, Ewan Harrison, David K. Jackson, Ian Johnston, Dominic Kwiatkowski, Cordelia Langford, John Sillitoe on behalf of the Wellcome Sanger Institute COVID-19 Surveillance Team |
| EPI_ISL_2972960 | Department of Bacteria, Parasites and Fungi, Statens Serum Institut, Copenhagen, Denmark | Statens Serum Institut Bioinformatics and Microbial Genomics | Danish Covid-19 Genome Consortium |
| EPI_ISL_2972962 | Lighthouse Lab in Alderley Park | Wellcome Sanger Institute for the COVID-19 Genomics UK (COG-UK) Consortium | Jacquelyn Wynn, Mairead Hyland, The Lighthouse Lab in Alderley Park and Alex Alderton, Roberto Amato, Jeffrey Barrett, Sonia Goncalves, Ewan Harrison, David K. Jackson, Ian Johnston, Dominic Kwiatkowski, Cordelia Langford, John Sillitoe on behalf of the Wellcome Sanger Institute COVID-19 Surveillance Team |
| EPI_ISL_2972963 | Department of Bacteria, Parasites and Fungi, Statens Serum Institut, Copenhagen, Denmark | Statens Serum Institut Bioinformatics and Microbial Genomics | Danish Covid-19 Genome Consortium |
| EPI_ISL_2972964 | Lighthouse Laboratory Plymouth | Wellcome Sanger Institute for the COVID-19 Genomics UK (COG-UK) Consortium | Lighthouse Laboratory Plymouth and Alex Alderton, Roberto Amato, Jeffrey Barrett, Sonia Goncalves, Ewan Harrison, David K. Jackson, Ian Johnston, Dominic Kwiatkowski, Cordelia Langford, John Sillitoe on behalf of the Wellcome Sanger Institute COVID-19 Surveillance Team |
| EPI_ISL_2972965 | Lighthouse Lab in Milton Keynes | Wellcome Sanger Institute for the COVID-19 Genomics UK (COG-UK) Consortium | The Lighthouse Lab in Milton Keynes and Alex Alderton, Roberto Amato, Jeffrey Barrett, Sonia Goncalves, Ewan Harrison, David K. Jackson, Ian Johnston, Dominic Kwiatkowski, Cordelia Langford, John Sillitoe on behalf of the Wellcome Sanger Institute COVID-19 Surveillance Team |
| EPI_ISL_2972966 | Department of Bacteria, Parasites and Fungi, Statens Serum Institut, Copenhagen, Denmark | Statens Serum Institut Bioinformatics and Microbial Genomics | Danish Covid-19 Genome Consortium |
| EPI_ISL_2972967 | Lighthouse Lab in Milton Keynes | Wellcome Sanger Institute for the COVID-19 Genomics UK (COG-UK) Consortium | The Lighthouse Lab in Milton Keynes and Alex Alderton, Roberto Amato, Jeffrey Barrett, Sonia Goncalves, Ewan Harrison, David K. Jackson, Ian Johnston, Dominic Kwiatkowski, Cordelia Langford, John Sillitoe on behalf of the Wellcome Sanger Institute COVID-19 Surveillance Team |
| EPI_ISL_2972968 | Lighthouse Lab in Alderley Park | Wellcome Sanger Institute for the COVID-19 Genomics UK (COG-UK) Consortium | Jacquelyn Wynn, Mairead Hyland, The Lighthouse Lab in Alderley Park and Alex Alderton, Roberto Amato, Jeffrey Barrett, Sonia Goncalves, Ewan Harrison, David K. Jackson, Ian Johnston, Dominic Kwiatkowski, Cordelia Langford, John Sillitoe on behalf of the Wellcome Sanger Institute COVID-19 Surveillance Team |
| EPI_ISL_2972969 | Lighthouse Laboratory Plymouth | Wellcome Sanger Institute for the COVID-19 Genomics UK (COG-UK) Consortium | Lighthouse Laboratory Plymouth and Alex Alderton, Roberto Amato, Jeffrey Barrett, Sonia Goncalves, Ewan Harrison, David K. Jackson, Ian Johnston, Dominic Kwiatkowski, Cordelia Langford, John Sillitoe on behalf of the Wellcome Sanger Institute COVID-19 Surveillance Team |
| EPI_ISL_2972970 | Department of Bacteria, Parasites and Fungi, Statens Serum Institut, Copenhagen, Denmark | Statens Serum Institut Bioinformatics and Microbial Genomics | Danish Covid-19 Genome Consortium |
| EPI_ISL_2972971, EPI_ISL_2972972 | Lighthouse Lab in Alderley Park | Wellcome Sanger Institute for the COVID-19 Genomics UK (COG-UK) Consortium | Jacquelyn Wynn, Mairead Hyland, The Lighthouse Lab in Alderley Park and Alex Alderton, Roberto Amato, Jeffrey Barrett, Sonia Goncalves, Ewan Harrison, David K. Jackson, Ian Johnston, Dominic Kwiatkowski, Cordelia Langford, John Sillitoe on behalf of the Wellcome Sanger Institute COVID-19 Surveillance Team |
| EPI_ISL_2972973 | Department of Bacteria, Parasites and Fungi, Statens Serum Institut, Copenhagen, Denmark | Statens Serum Institut Bioinformatics and Microbial Genomics | Danish Covid-19 Genome Consortium |
| EPI_ISL_2972974 | Lighthouse Laboratory Plymouth | Wellcome Sanger Institute for the COVID-19 Genomics UK (COG-UK) Consortium | Lighthouse Laboratory Plymouth and Alex Alderton, Roberto Amato, Jeffrey Barrett, Sonia Goncalves, Ewan Harrison, David K. Jackson, Ian Johnston, Dominic Kwiatkowski, Cordelia Langford, John Sillitoe on behalf of the Wellcome Sanger Institute COVID-19 Surveillance Team |
| EPI_ISL_2972975 | Lighthouse Lab in Milton Keynes | Wellcome Sanger Institute for the COVID-19 Genomics UK (COG-UK) Consortium | The Lighthouse Lab in Milton Keynes and Alex Alderton, Roberto Amato, Jeffrey Barrett, Sonia Goncalves, Ewan Harrison, David K. Jackson, Ian Johnston, Dominic Kwiatkowski, Cordelia Langford, John Sillitoe on behalf of the Wellcome Sanger Institute COVID-19 Surveillance Team |
| EPI_ISL_2972976 | Department of Bacteria, Parasites and Fungi, Statens Serum Institut, Copenhagen, Denmark | Statens Serum Institut Bioinformatics and Microbial Genomics | Danish Covid-19 Genome Consortium |
| EPI_ISL_2972977 | Lighthouse Laboratory Plymouth | Wellcome Sanger Institute for the COVID-19 Genomics UK (COG-UK) Consortium | Lighthouse Laboratory Plymouth and Alex Alderton, Roberto Amato, Jeffrey Barrett, Sonia Goncalves, Ewan Harrison, David K. Jackson, Ian Johnston, Dominic Kwiatkowski, Cordelia Langford, John Sillitoe on behalf of the Wellcome Sanger Institute COVID-19 Surveillance Team |
| EPI_ISL_2972978 | Lighthouse Lab in Alderley Park | Wellcome Sanger Institute for the COVID-19 Genomics UK (COG-UK) Consortium | Jacquelyn Wynn, Mairead Hyland, The Lighthouse Lab in Alderley Park and Alex Alderton, Roberto Amato, Jeffrey Barrett, Sonia Goncalves, Ewan Harrison, David K. Jackson, Ian Johnston, Dominic Kwiatkowski, Cordelia Langford, John Sillitoe on behalf of the Wellcome Sanger Institute COVID-19 Surveillance Team |
| EPI_ISL_2972979 | Department of Bacteria, Parasites and Fungi, Statens Serum Institut, Copenhagen, Denmark | Statens Serum Institut Bioinformatics and Microbial Genomics | Danish Covid-19 Genome Consortium |
| EPI_ISL_2972980 | Lighthouse Laboratory Plymouth | Wellcome Sanger Institute for the COVID-19 Genomics UK (COG-UK) Consortium | Lighthouse Laboratory Plymouth and Alex Alderton, Roberto Amato, Jeffrey Barrett, Sonia Goncalves, Ewan Harrison, David K. Jackson, Ian Johnston, Dominic Kwiatkowski, Cordelia Langford, John Sillitoe on behalf of the Wellcome Sanger Institute COVID-19 Surveillance Team |
| EPI_ISL_2972981, EPI_ISL_2972983, EPI_ISL_2972984 | Lighthouse Lab in Alderley Park | Wellcome Sanger Institute for the COVID-19 Genomics UK (COG-UK) Consortium | Jacquelyn Wynn, Mairead Hyland, The Lighthouse Lab in Alderley Park and Alex Alderton, Roberto Amato, Jeffrey Barrett, Sonia Goncalves, Ewan Harrison, David K. Jackson, Ian Johnston, Dominic Kwiatkowski, Cordelia Langford, John Sillitoe on behalf of the Wellcome Sanger Institute COVID-19 Surveillance Team |
| EPI_ISL_2972985 | Department of Bacteria, Parasites and Fungi, Statens Serum Institut, Copenhagen, Denmark | Statens Serum Institut Bioinformatics and Microbial Genomics | Danish Covid-19 Genome Consortium |
| EPI_ISL_2972986 | Lighthouse Lab in Milton Keynes | Wellcome Sanger Institute for the COVID-19 Genomics UK (COG-UK) Consortium | The Lighthouse Lab in Milton Keynes and Alex Alderton, Roberto Amato, Jeffrey Barrett, Sonia Goncalves, Ewan Harrison, David K. Jackson, Ian Johnston, Dominic Kwiatkowski, Cordelia Langford, John Sillitoe on behalf of the Wellcome Sanger Institute COVID-19 Surveillance Team |
| EPI_ISL_2972987 | Lighthouse Lab in Alderley Park | Wellcome Sanger Institute for the COVID-19 Genomics UK (COG-UK) Consortium | Jacquelyn Wynn, Mairead Hyland, The Lighthouse Lab in Alderley Park and Alex Alderton, Roberto Amato, Jeffrey Barrett, Sonia Goncalves, Ewan Harrison, David K. Jackson, Ian Johnston, Dominic Kwiatkowski, Cordelia Langford, John Sillitoe on behalf of the Wellcome Sanger Institute COVID-19 Surveillance Team |
| EPI_ISL_2972988 | Lighthouse Lab in Milton Keynes | Wellcome Sanger Institute for the COVID-19 Genomics UK (COG-UK) Consortium | The Lighthouse Lab in Milton Keynes and Alex Alderton, Roberto Amato, Jeffrey Barrett, Sonia Goncalves, Ewan Harrison, David K. Jackson, Ian Johnston, Dominic Kwiatkowski, Cordelia Langford, John Sillitoe on behalf of the Wellcome Sanger Institute COVID-19 Surveillance Team |
| EPI_ISL_2972989 | Department of Bacteria, Parasites and Fungi, Statens Serum Institut, Copenhagen, Denmark | Statens Serum Institut Bioinformatics and Microbial Genomics | Danish Covid-19 Genome Consortium |
| EPI_ISL_2972990 | Lighthouse Laboratory Plymouth | Wellcome Sanger Institute for the COVID-19 Genomics UK (COG-UK) Consortium | Lighthouse Laboratory Plymouth and Alex Alderton, Roberto Amato, Jeffrey Barrett, Sonia Goncalves, Ewan Harrison, David K. Jackson, Ian Johnston, Dominic Kwiatkowski, Cordelia Langford, John Sillitoe on behalf of the Wellcome Sanger Institute COVID-19 Surveillance Team |
| EPI_ISL_2972991 | Lighthouse Lab in Alderley Park | Wellcome Sanger Institute for the COVID-19 Genomics UK (COG-UK) Consortium | Jacquelyn Wynn, Mairead Hyland, The Lighthouse Lab in Alderley Park and Alex Alderton, Roberto Amato, Jeffrey Barrett, Sonia Goncalves, Ewan Harrison, David K. Jackson, Ian Johnston, Dominic Kwiatkowski, Cordelia Langford, John Sillitoe on behalf of the Wellcome Sanger Institute COVID-19 Surveillance Team |
| EPI_ISL_2972992 | Department of Bacteria, Parasites and Fungi, Statens Serum Institut, Copenhagen, Denmark | Statens Serum Institut Bioinformatics and Microbial Genomics | Danish Covid-19 Genome Consortium |
| EPI_ISL_2972993 | Lighthouse Laboratory Plymouth | Wellcome Sanger Institute for the COVID-19 Genomics UK (COG-UK) Consortium | Lighthouse Laboratory Plymouth and Alex Alderton, Roberto Amato, Jeffrey Barrett, Sonia Goncalves, Ewan Harrison, David K. Jackson, Ian Johnston, Dominic Kwiatkowski, Cordelia Langford, John Sillitoe on behalf of the Wellcome Sanger Institute COVID-19 Surveillance Team |
| EPI_ISL_2972994 | Lighthouse Lab in Alderley Park | Wellcome Sanger Institute for the COVID-19 Genomics UK (COG-UK) | Jacquelyn Wynn, Mairead Hyland, The Lighthouse Lab in Alderley Park and Alex Alderton, Roberto Amato, Jeffrey Barrett, Sonia Goncalves, Ewan Harrison, David K. Jackson, Ian Johnston, Dominic Kwiatkowski, Cordelia Langford, John Sillitoe on |

| | | | |
|---|---|---|---|
| | | Consortium | behalf of the Wellcome Sanger Institute COVID-19 Surveillance Team |
| EPI_ISL_2972995 | Department of Bacteria, Parasites and Fungi, Statens Serum Institut, Copenhagen, Denmark | Statens Serum Institut Bioinformatics and Microbial Genomics | Danish Covid-19 Genome Consortium |
| EPI_ISL_2972996 | Lighthouse Lab in Alderley Park | Wellcome Sanger Institute for the COVID-19 Genomics UK (COG-UK) Consortium | Jacquelyn Wynn, Mairead Hyland, The Lighthouse Lab in Alderley Park and Alex Alderton, Roberto Amato, Jeffrey Barrett, Sonia Goncalves, Ewan Harrison, David K. Jackson, Ian Johnston, Dominic Kwiatkowski, Cordelia Langford, John Sillitoe on behalf of the Wellcome Sanger Institute COVID-19 Surveillance Team |
| EPI_ISL_2972997 | Department of Bacteria, Parasites and Fungi, Statens Serum Institut, Copenhagen, Denmark | Statens Serum Institut Bioinformatics and Microbial Genomics | Danish Covid-19 Genome Consortium |
| EPI_ISL_2972998, EPI_ISL_2972999, EPI_ISL_2973000 | Lighthouse Laboratory Plymouth | Wellcome Sanger Institute for the COVID-19 Genomics UK (COG-UK) Consortium | Lighthouse Laboratory Plymouth and Alex Alderton, Roberto Amato, Jeffrey Barrett, Sonia Goncalves, Ewan Harrison, David K. Jackson, Ian Johnston, Dominic Kwiatkowski, Cordelia Langford, John Sillitoe on behalf of the Wellcome Sanger Institute COVID-19 Surveillance Team |
| EPI_ISL_2973001 | Department of Bacteria, Parasites and Fungi, Statens Serum Institut, Copenhagen, Denmark | Statens Serum Institut Bioinformatics and Microbial Genomics | Danish Covid-19 Genome Consortium |
| EPI_ISL_2973002, EPI_ISL_2973003 | Lighthouse Lab in Alderley Park | Wellcome Sanger Institute for the COVID-19 Genomics UK (COG-UK) Consortium | Jacquelyn Wynn, Mairead Hyland, The Lighthouse Lab in Alderley Park and Alex Alderton, Roberto Amato, Jeffrey Barrett, Sonia Goncalves, Ewan Harrison, David K. Jackson, Ian Johnston, Dominic Kwiatkowski, Cordelia Langford, John Sillitoe on behalf of the Wellcome Sanger Institute COVID-19 Surveillance Team |
| EPI_ISL_2973004 | Department of Bacteria, Parasites and Fungi, Statens Serum Institut, Copenhagen, Denmark | Statens Serum Institut Bioinformatics and Microbial Genomics | Danish Covid-19 Genome Consortium |
| EPI_ISL_2973005 | Lighthouse Laboratory Plymouth | Wellcome Sanger Institute for the COVID-19 Genomics UK (COG-UK) Consortium | Lighthouse Laboratory Plymouth and Alex Alderton, Roberto Amato, Jeffrey Barrett, Sonia Goncalves, Ewan Harrison, David K. Jackson, Ian Johnston, Dominic Kwiatkowski, Cordelia Langford, John Sillitoe on behalf of the Wellcome Sanger Institute COVID-19 Surveillance Team |
| EPI_ISL_2973006 | Berkshire and Surrey Pathology Services Lighthouse Laboratory | Wellcome Sanger Institute for the COVID-19 Genomics UK (COG-UK) Consortium | Berkshire and Surrey Pathology Services Lighthouse Laboratory and Alex Alderton, Roberto Amato, Jeffrey Barrett, Sonia Goncalves, Ewan Harrison, David K. Jackson, Ian Johnston, Dominic Kwiatkowski, Cordelia Langford, John Sillitoe on behalf of the Wellcome Sanger Institute COVID-19 Surveillance Team |
| EPI_ISL_2973007 | Department of Bacteria, Parasites and Fungi, Statens Serum Institut, Copenhagen, Denmark | Statens Serum Institut Bioinformatics and Microbial Genomics | Danish Covid-19 Genome Consortium |
| EPI_ISL_2973008, EPI_ISL_2973009 | Lighthouse Laboratory Plymouth | Wellcome Sanger Institute for the COVID-19 Genomics UK (COG-UK) Consortium | Lighthouse Laboratory Plymouth and Alex Alderton, Roberto Amato, Jeffrey Barrett, Sonia Goncalves, Ewan Harrison, David K. Jackson, Ian Johnston, Dominic Kwiatkowski, Cordelia Langford, John Sillitoe on behalf of the Wellcome Sanger Institute COVID-19 Surveillance Team |
| EPI_ISL_2973010 | Department of Bacteria, Parasites and Fungi, Statens Serum Institut, Copenhagen, Denmark | Statens Serum Institut Bioinformatics and Microbial Genomics | Danish Covid-19 Genome Consortium |
| EPI_ISL_2973011 | Berkshire and Surrey Pathology Services Lighthouse Laboratory | Wellcome Sanger Institute for the COVID-19 Genomics UK (COG-UK) Consortium | Berkshire and Surrey Pathology Services Lighthouse Laboratory and Alex Alderton, Roberto Amato, Jeffrey Barrett, Sonia Goncalves, Ewan Harrison, David K. Jackson, Ian Johnston, Dominic Kwiatkowski, Cordelia Langford, John Sillitoe on behalf of the Wellcome Sanger Institute COVID-19 Surveillance Team |
| EPI_ISL_2973012 | Lighthouse Lab in Alderley Park | Wellcome Sanger Institute for the COVID-19 Genomics UK (COG-UK) Consortium | Jacquelyn Wynn, Mairead Hyland, The Lighthouse Lab in Alderley Park and Alex Alderton, Roberto Amato, Jeffrey Barrett, Sonia Goncalves, Ewan Harrison, David K. Jackson, Ian Johnston, Dominic Kwiatkowski, Cordelia Langford, John Sillitoe on behalf of the Wellcome Sanger Institute COVID-19 Surveillance Team |
| EPI_ISL_2973014, EPI_ISL_2973015 | Lighthouse Laboratory Plymouth | Wellcome Sanger Institute for the COVID-19 Genomics UK (COG-UK) Consortium | Lighthouse Laboratory Plymouth and Alex Alderton, Roberto Amato, Jeffrey Barrett, Sonia Goncalves, Ewan Harrison, David K. Jackson, Ian Johnston, Dominic Kwiatkowski, Cordelia Langford, John Sillitoe on behalf of the Wellcome Sanger Institute COVID-19 Surveillance Team |
| EPI_ISL_2973016 | Lighthouse Lab in Alderley Park | Wellcome Sanger Institute for the COVID-19 Genomics UK (COG-UK) Consortium | Jacquelyn Wynn, Mairead Hyland, The Lighthouse Lab in Alderley Park and Alex Alderton, Roberto Amato, Jeffrey Barrett, Sonia Goncalves, Ewan Harrison, David K. Jackson, Ian Johnston, Dominic Kwiatkowski, Cordelia Langford, John Sillitoe on behalf of the Wellcome Sanger Institute COVID-19 Surveillance Team |
| EPI_ISL_2973017 | Department of Bacteria, Parasites and Fungi, Statens Serum Institut, Copenhagen, Denmark | Statens Serum Institut Bioinformatics and Microbial Genomics | Danish Covid-19 Genome Consortium |
| EPI_ISL_2973018 | Lighthouse Laboratory Plymouth | Wellcome Sanger Institute for the COVID-19 Genomics UK (COG-UK) Consortium | Lighthouse Laboratory Plymouth and Alex Alderton, Roberto Amato, Jeffrey Barrett, Sonia Goncalves, Ewan Harrison, David K. Jackson, Ian Johnston, Dominic Kwiatkowski, Cordelia Langford, John Sillitoe on behalf of the Wellcome Sanger Institute COVID-19 Surveillance Team |
| EPI_ISL_2973019 | Lighthouse Lab in Alderley Park | Wellcome Sanger Institute for the COVID-19 Genomics UK (COG-UK) Consortium | Jacquelyn Wynn, Mairead Hyland, The Lighthouse Lab in Alderley Park and Alex Alderton, Roberto Amato, Jeffrey Barrett, Sonia Goncalves, Ewan Harrison, David K. Jackson, Ian Johnston, Dominic Kwiatkowski, Cordelia Langford, John Sillitoe on behalf of the Wellcome Sanger Institute COVID-19 Surveillance Team |
| EPI_ISL_2973020 | Department of Bacteria, Parasites and Fungi, Statens Serum Institut, Copenhagen, Denmark | Statens Serum Institut Bioinformatics and Microbial Genomics | Danish Covid-19 Genome Consortium |
| EPI_ISL_2973021 | Lighthouse Laboratory Plymouth | Wellcome Sanger Institute for the COVID-19 Genomics UK (COG-UK) Consortium | Lighthouse Laboratory Plymouth and Alex Alderton, Roberto Amato, Jeffrey Barrett, Sonia Goncalves, Ewan Harrison, David K. Jackson, Ian Johnston, Dominic Kwiatkowski, Cordelia Langford, John Sillitoe on behalf of the Wellcome Sanger Institute COVID-19 Surveillance Team |
| EPI_ISL_2973022 | Lighthouse Lab in Milton Keynes | Wellcome Sanger Institute for the COVID-19 Genomics UK (COG-UK) Consortium | The Lighthouse Lab in Milton Keynes and Alex Alderton, Roberto Amato, Jeffrey Barrett, Sonia Goncalves, Ewan Harrison, David K. Jackson, Ian Johnston, Dominic Kwiatkowski, Cordelia Langford, John Sillitoe on behalf of the Wellcome Sanger Institute COVID-19 Surveillance Team |
| EPI_ISL_2973023 | Department of Bacteria, Parasites and Fungi, Statens Serum Institut, Copenhagen, Denmark | Statens Serum Institut Bioinformatics and Microbial Genomics | Danish Covid-19 Genome Consortium |
| EPI_ISL_2973024 | Lighthouse Lab in Milton Keynes | Wellcome Sanger Institute for the COVID-19 Genomics UK (COG-UK) Consortium | The Lighthouse Lab in Milton Keynes and Alex Alderton, Roberto Amato, Jeffrey Barrett, Sonia Goncalves, Ewan Harrison, David K. Jackson, Ian Johnston, Dominic Kwiatkowski, Cordelia Langford, John Sillitoe on behalf of the Wellcome Sanger Institute COVID-19 Surveillance Team |
| EPI_ISL_2973025, EPI_ISL_2973026 | Lighthouse Laboratory Plymouth | Wellcome Sanger Institute for the COVID-19 Genomics UK (COG-UK) Consortium | Lighthouse Laboratory Plymouth and Alex Alderton, Roberto Amato, Jeffrey Barrett, Sonia Goncalves, Ewan Harrison, David K. Jackson, Ian Johnston, Dominic Kwiatkowski, Cordelia Langford, John Sillitoe on behalf of the Wellcome Sanger Institute COVID-19 Surveillance Team |
| EPI_ISL_2973027 | Department of Bacteria, Parasites and Fungi, Statens Serum Institut, Copenhagen, Denmark | Statens Serum Institut Bioinformatics and Microbial Genomics | Danish Covid-19 Genome Consortium |
| EPI_ISL_2973028, EPI_ISL_2973029 | Lighthouse Lab in Alderley Park | Wellcome Sanger Institute for the COVID-19 Genomics UK (COG-UK) Consortium | Jacquelyn Wynn, Mairead Hyland, The Lighthouse Lab in Alderley Park and Alex Alderton, Roberto Amato, Jeffrey Barrett, Sonia Goncalves, Ewan Harrison, David K. Jackson, Ian Johnston, Dominic Kwiatkowski, Cordelia Langford, John Sillitoe on behalf of the Wellcome Sanger Institute COVID-19 Surveillance Team |
| EPI_ISL_2973030 | Department of Bacteria, Parasites and Fungi, Statens Serum Institut, Copenhagen, Denmark | Statens Serum Institut Bioinformatics and Microbial Genomics | Danish Covid-19 Genome Consortium |
| EPI_ISL_2973031, EPI_ISL_2973032 | Lighthouse Lab in Alderley Park | Wellcome Sanger Institute for the COVID-19 Genomics UK (COG-UK) Consortium | Jacquelyn Wynn, Mairead Hyland, The Lighthouse Lab in Alderley Park and Alex Alderton, Roberto Amato, Jeffrey Barrett, Sonia Goncalves, Ewan Harrison, David K. Jackson, Ian Johnston, Dominic Kwiatkowski, Cordelia Langford, John Sillitoe on behalf of the Wellcome Sanger Institute COVID-19 Surveillance Team |
| EPI_ISL_2973034 | Berkshire and Surrey Pathology Services Lighthouse Laboratory | Wellcome Sanger Institute for the COVID-19 Genomics UK (COG-UK) Consortium | Berkshire and Surrey Pathology Services Lighthouse Laboratory and Alex Alderton, Roberto Amato, Jeffrey Barrett, Sonia Goncalves, Ewan Harrison, David K. Jackson, Ian Johnston, Dominic Kwiatkowski, Cordelia Langford, John Sillitoe on behalf of the Wellcome Sanger Institute COVID-19 Surveillance Team |
| EPI_ISL_2973035 | Lighthouse Lab in Milton Keynes | Wellcome Sanger Institute for the COVID-19 Genomics UK (COG-UK) Consortium | The Lighthouse Lab in Milton Keynes and Alex Alderton, Roberto Amato, Jeffrey Barrett, Sonia Goncalves, Ewan Harrison, David K. Jackson, Ian Johnston, Dominic Kwiatkowski, Cordelia Langford, John Sillitoe on behalf of the Wellcome Sanger Institute COVID-19 Surveillance Team |
| EPI_ISL_2973036 | Department of Bacteria, Parasites and Fungi, Statens Serum Institut, Copenhagen, Denmark | Statens Serum Institut Bioinformatics and Microbial Genomics | Danish Covid-19 Genome Consortium |
| EPI_ISL_2973037, EPI_ISL_2973038 | Lighthouse Lab in Alderley Park | Wellcome Sanger Institute for the COVID-19 Genomics UK (COG-UK) Consortium | Jacquelyn Wynn, Mairead Hyland, The Lighthouse Lab in Alderley Park and Alex Alderton, Roberto Amato, Jeffrey Barrett, Sonia Goncalves, Ewan Harrison, David K. Jackson, Ian Johnston, Dominic Kwiatkowski, Cordelia Langford, John Sillitoe on behalf of the Wellcome Sanger Institute COVID-19 Surveillance Team |
| EPI_ISL_2973039 | Department of Bacteria, Parasites and Fungi, Statens Serum Institut, Copenhagen, Denmark | Statens Serum Institut Bioinformatics and Microbial Genomics | Danish Covid-19 Genome Consortium |
| EPI_ISL_2973040, EPI_ISL_2973041 | Lighthouse Lab in Milton Keynes | Wellcome Sanger Institute for the COVID-19 Genomics UK (COG-UK) Consortium | The Lighthouse Lab in Milton Keynes and Alex Alderton, Roberto Amato, Jeffrey Barrett, Sonia Goncalves, Ewan Harrison, David K. Jackson, Ian Johnston, Dominic Kwiatkowski, Cordelia Langford, John Sillitoe on behalf of the Wellcome Sanger Institute COVID-19 Surveillance Team |
| EPI_ISL_2973042 | Department of Bacteria, Parasites and Fungi, Statens Serum Institut, Copenhagen, Denmark | Statens Serum Institut Bioinformatics and Microbial Genomics | Danish Covid-19 Genome Consortium |
| EPI_ISL_2973043, EPI_ISL_2973044 | Lighthouse Lab in Alderley Park | Wellcome Sanger Institute for the COVID-19 Genomics UK (COG-UK) Consortium | Jacquelyn Wynn, Mairead Hyland, The Lighthouse Lab in Alderley Park and Alex Alderton, Roberto Amato, Jeffrey Barrett, Sonia Goncalves, Ewan Harrison, David K. Jackson, Ian Johnston, Dominic Kwiatkowski, Cordelia Langford, John Sillitoe on behalf of the Wellcome Sanger Institute COVID-19 Surveillance Team |
| EPI_ISL_2973045 | Department of Bacteria, Parasites and Fungi, Statens Serum Institut, Copenhagen, Denmark | Statens Serum Institut Bioinformatics and Microbial Genomics | Danish Covid-19 Genome Consortium |
| EPI_ISL_2973046 | Lighthouse Laboratory Plymouth | Wellcome Sanger Institute for the COVID-19 Genomics UK (COG-UK) Consortium | Lighthouse Laboratory Plymouth and Alex Alderton, Roberto Amato, Jeffrey Barrett, Sonia Goncalves, Ewan Harrison, David K. Jackson, Ian Johnston, Dominic Kwiatkowski, Cordelia Langford, John Sillitoe on behalf of the Wellcome Sanger Institute COVID-19 Surveillance Team |
| EPI_ISL_2973047 | Lighthouse Lab in Alderley Park | Wellcome Sanger Institute for the COVID-19 Genomics UK (COG-UK) Consortium | Jacquelyn Wynn, Mairead Hyland, The Lighthouse Lab in Alderley Park and Alex Alderton, Roberto Amato, Jeffrey Barrett, Sonia Goncalves, Ewan Harrison, David K. Jackson, Ian Johnston, Dominic Kwiatkowski, Cordelia Langford, John Sillitoe on behalf of the Wellcome Sanger Institute COVID-19 Surveillance Team |
| EPI_ISL_2973048 | Department of Bacteria, Parasites and Fungi, Statens Serum Institut, Copenhagen, Denmark | Statens Serum Institut Bioinformatics and Microbial Genomics | Danish Covid-19 Genome Consortium |
| EPI_ISL_2973049, EPI_ISL_2973050 | Lighthouse Laboratory Plymouth | Wellcome Sanger Institute for the COVID-19 Genomics UK (COG-UK) Consortium | Lighthouse Laboratory Plymouth and Alex Alderton, Roberto Amato, Jeffrey Barrett, Sonia Goncalves, Ewan Harrison, David K. Jackson, Ian Johnston, Dominic Kwiatkowski, Cordelia Langford, John Sillitoe on behalf of the Wellcome Sanger Institute COVID-19 Surveillance Team |
| EPI_ISL_2973051, EPI_ISL_2973052, EPI_ISL_2973053, EPI_ISL_2973054, EPI_ISL_2973055, EPI_ISL_2973056, EPI_ISL_2973057, EPI_ISL_2973058, EPI_ISL_2973059, EPI_ISL_2973060, EPI_ISL_2973061, EPI_ISL_2973062, EPI_ISL_2973063, EPI_ISL_2973064, EPI_ISL_2973065, EPI_ISL_2973066, EPI_ISL_2973067, EPI_ISL_2973068, EPI_ISL_2973069, EPI_ISL_2973070, EPI_ISL_2973071, EPI_ISL_2973073, EPI_ISL_2973074, EPI_ISL_2973075, EPI_ISL_2973076, EPI_ISL_2973077, EPI_ISL_2973078, EPI_ISL_2973079, EPI_ISL_2973080, EPI_ISL_2973081, EPI_ISL_2973082, EPI_ISL_2973083, EPI_ISL_2973084, EPI_ISL_2973085, EPI_ISL_2973086, EPI_ISL_2973087, EPI_ISL_2973088, EPI_ISL_2973089, EPI_ISL_2973090, EPI_ISL_2973091, EPI_ISL_2973092, EPI_ISL_2973093, EPI_ISL_2973094, EPI_ISL_2973095, EPI_ISL_2973097, EPI_ISL_2973098, EPI_ISL_2973099, EPI_ISL_2973100, EPI_ISL_2973102 | | | |
| see above | Department of Bacteria, Parasites and Fungi, Statens Serum Institut, Copenhagen, Denmark | Statens Serum Institut Bioinformatics and Microbial Genomics | Danish Covid-19 Genome Consortium |

| | | | |
|---|---|---|---|
| EPI_ISL_2973103 | Lighthouse Lab in Milton Keynes | Wellcome Sanger Institute for the COVID-19 Genomics UK (COG-UK) Consortium | The Lighthouse Lab in Milton Keynes and Alex Alderton, Roberto Amato, Jeffrey Barrett, Sonia Goncalves, Ewan Harrison, David K. Jackson, Ian Johnston, Dominic Kwiatkowski, Cordelia Langford, John Sillitoe on behalf of the Wellcome Sanger Institute COVID-19 Surveillance Team |
| EPI_ISL_2973104 | Lighthouse Lab in Alderley Park | Wellcome Sanger Institute for the COVID-19 Genomics UK (COG-UK) Consortium | Jacquelyn Wynn, Mairead Hyland, The Lighthouse Lab in Alderley Park and Alex Alderton, Roberto Amato, Jeffrey Barrett, Sonia Goncalves, Ewan Harrison, David K. Jackson, Ian Johnston, Dominic Kwiatkowski, Cordelia Langford, John Sillitoe on behalf of the Wellcome Sanger Institute COVID-19 Surveillance Team |
| EPI_ISL_2973105 | Department of Bacteria, Parasites and Fungi, Statens Serum Institut, Copenhagen, Denmark | Statens Serum Institut Bioinformatics and Microbial Genomics | Danish Covid-19 Genome Consortium |
| EPI_ISL_2973106 | Lighthouse Lab in Milton Keynes | Wellcome Sanger Institute for the COVID-19 Genomics UK (COG-UK) Consortium | The Lighthouse Lab in Milton Keynes and Alex Alderton, Roberto Amato, Jeffrey Barrett, Sonia Goncalves, Ewan Harrison, David K. Jackson, Ian Johnston, Dominic Kwiatkowski, Cordelia Langford, John Sillitoe on behalf of the Wellcome Sanger Institute COVID-19 Surveillance Team |
| EPI_ISL_2973107 | Lighthouse Laboratory Plymouth | Wellcome Sanger Institute for the COVID-19 Genomics UK (COG-UK) Consortium | Lighthouse Laboratory Plymouth and Alex Alderton, Roberto Amato, Jeffrey Barrett, Sonia Goncalves, Ewan Harrison, David K. Jackson, Ian Johnston, Dominic Kwiatkowski, Cordelia Langford, John Sillitoe on behalf of the Wellcome Sanger Institute COVID-19 Surveillance Team |
| EPI_ISL_2973108 | Department of Bacteria, Parasites and Fungi, Statens Serum Institut, Copenhagen, Denmark | Statens Serum Institut Bioinformatics and Microbial Genomics | Danish Covid-19 Genome Consortium |
| EPI_ISL_2973109 | Berkshire and Surrey Pathology Services Lighthouse Laboratory | Wellcome Sanger Institute for the COVID-19 Genomics UK (COG-UK) Consortium | Berkshire and Surrey Pathology Services Lighthouse Laboratory and Alex Alderton, Roberto Amato, Jeffrey Barrett, Sonia Goncalves, Ewan Harrison, David K. Jackson, Ian Johnston, Dominic Kwiatkowski, Cordelia Langford, John Sillitoe on behalf of the Wellcome Sanger Institute COVID-19 Surveillance Team |
| EPI_ISL_2973110 | Lighthouse Lab in Milton Keynes | Wellcome Sanger Institute for the COVID-19 Genomics UK (COG-UK) Consortium | The Lighthouse Lab in Milton Keynes and Alex Alderton, Roberto Amato, Jeffrey Barrett, Sonia Goncalves, Ewan Harrison, David K. Jackson, Ian Johnston, Dominic Kwiatkowski, Cordelia Langford, John Sillitoe on behalf of the Wellcome Sanger Institute COVID-19 Surveillance Team |
| EPI_ISL_2973111 | Department of Bacteria, Parasites and Fungi, Statens Serum Institut, Copenhagen, Denmark | Statens Serum Institut Bioinformatics and Microbial Genomics | Danish Covid-19 Genome Consortium |
| EPI_ISL_2973112 | Berkshire and Surrey Pathology Services Lighthouse Laboratory | Wellcome Sanger Institute for the COVID-19 Genomics UK (COG-UK) Consortium | Berkshire and Surrey Pathology Services Lighthouse Laboratory and Alex Alderton, Roberto Amato, Jeffrey Barrett, Sonia Goncalves, Ewan Harrison, David K. Jackson, Ian Johnston, Dominic Kwiatkowski, Cordelia Langford, John Sillitoe on behalf of the Wellcome Sanger Institute COVID-19 Surveillance Team |
| EPI_ISL_2973113 | Lighthouse Laboratory Plymouth | Wellcome Sanger Institute for the COVID-19 Genomics UK (COG-UK) Consortium | Lighthouse Laboratory Plymouth and Alex Alderton, Roberto Amato, Jeffrey Barrett, Sonia Goncalves, Ewan Harrison, David K. Jackson, Ian Johnston, Dominic Kwiatkowski, Cordelia Langford, John Sillitoe on behalf of the Wellcome Sanger Institute COVID-19 Surveillance Team |
| EPI_ISL_2973114 | Department of Bacteria, Parasites and Fungi, Statens Serum Institut, Copenhagen, Denmark | Statens Serum Institut Bioinformatics and Microbial Genomics | Danish Covid-19 Genome Consortium |
| EPI_ISL_2973115 | Lighthouse Lab in Alderley Park | Wellcome Sanger Institute for the COVID-19 Genomics UK (COG-UK) Consortium | Jacquelyn Wynn, Mairead Hyland, The Lighthouse Lab in Alderley Park and Alex Alderton, Roberto Amato, Jeffrey Barrett, Sonia Goncalves, Ewan Harrison, David K. Jackson, Ian Johnston, Dominic Kwiatkowski, Cordelia Langford, John Sillitoe on behalf of the Wellcome Sanger Institute COVID-19 Surveillance Team |
| EPI_ISL_2973116 | Lighthouse Laboratory Plymouth | Wellcome Sanger Institute for the COVID-19 Genomics UK (COG-UK) Consortium | Lighthouse Laboratory Plymouth and Alex Alderton, Roberto Amato, Jeffrey Barrett, Sonia Goncalves, Ewan Harrison, David K. Jackson, Ian Johnston, Dominic Kwiatkowski, Cordelia Langford, John Sillitoe on behalf of the Wellcome Sanger Institute COVID-19 Surveillance Team |
| EPI_ISL_2973117 | Department of Bacteria, Parasites and Fungi, Statens Serum Institut, Copenhagen, Denmark | Statens Serum Institut Bioinformatics and Microbial Genomics | Danish Covid-19 Genome Consortium |
| EPI_ISL_2973118 | Lighthouse Laboratory Plymouth | Wellcome Sanger Institute for the COVID-19 Genomics UK (COG-UK) Consortium | Lighthouse Laboratory Plymouth and Alex Alderton, Roberto Amato, Jeffrey Barrett, Sonia Goncalves, Ewan Harrison, David K. Jackson, Ian Johnston, Dominic Kwiatkowski, Cordelia Langford, John Sillitoe on behalf of the Wellcome Sanger Institute COVID-19 Surveillance Team |
| EPI_ISL_2973119 | Department of Bacteria, Parasites and Fungi, Statens Serum Institut, Copenhagen, Denmark | Statens Serum Institut Bioinformatics and Microbial Genomics | Danish Covid-19 Genome Consortium |
| EPI_ISL_2973120 | Berkshire and Surrey Pathology Services Lighthouse Laboratory | Wellcome Sanger Institute for the COVID-19 Genomics UK (COG-UK) Consortium | Berkshire and Surrey Pathology Services Lighthouse Laboratory and Alex Alderton, Roberto Amato, Jeffrey Barrett, Sonia Goncalves, Ewan Harrison, David K. Jackson, Ian Johnston, Dominic Kwiatkowski, Cordelia Langford, John Sillitoe on behalf of the Wellcome Sanger Institute COVID-19 Surveillance Team |
| EPI_ISL_2973121, EPI_ISL_2973122 | Lighthouse Laboratory Plymouth | Wellcome Sanger Institute for the COVID-19 Genomics UK (COG-UK) Consortium | Lighthouse Laboratory Plymouth and Alex Alderton, Roberto Amato, Jeffrey Barrett, Sonia Goncalves, Ewan Harrison, David K. Jackson, Ian Johnston, Dominic Kwiatkowski, Cordelia Langford, John Sillitoe on behalf of the Wellcome Sanger Institute COVID-19 Surveillance Team |
| EPI_ISL_2973123 | Department of Bacteria, Parasites and Fungi, Statens Serum Institut, Copenhagen, Denmark | Statens Serum Institut Bioinformatics and Microbial Genomics | Danish Covid-19 Genome Consortium |
| EPI_ISL_2973124, EPI_ISL_2973125 | Lighthouse Lab in Milton Keynes | Wellcome Sanger Institute for the COVID-19 Genomics UK (COG-UK) Consortium | The Lighthouse Lab in Milton Keynes and Alex Alderton, Roberto Amato, Jeffrey Barrett, Sonia Goncalves, Ewan Harrison, David K. Jackson, Ian Johnston, Dominic Kwiatkowski, Cordelia Langford, John Sillitoe on behalf of the Wellcome Sanger Institute COVID-19 Surveillance Team |
| EPI_ISL_2973126 | Department of Bacteria, Parasites and Fungi, Statens Serum Institut, Copenhagen, Denmark | Statens Serum Institut Bioinformatics and Microbial Genomics | Danish Covid-19 Genome Consortium |
| EPI_ISL_2973127 | Lighthouse Laboratory Plymouth | Wellcome Sanger Institute for the COVID-19 Genomics UK (COG-UK) Consortium | Lighthouse Laboratory Plymouth and Alex Alderton, Roberto Amato, Jeffrey Barrett, Sonia Goncalves, Ewan Harrison, David K. Jackson, Ian Johnston, Dominic Kwiatkowski, Cordelia Langford, John Sillitoe on behalf of the Wellcome Sanger Institute COVID-19 Surveillance Team |
| EPI_ISL_2973128 | Lighthouse Lab in Milton Keynes | Wellcome Sanger Institute for the COVID-19 Genomics UK (COG-UK) Consortium | The Lighthouse Lab in Milton Keynes and Alex Alderton, Roberto Amato, Jeffrey Barrett, Sonia Goncalves, Ewan Harrison, David K. Jackson, Ian Johnston, Dominic Kwiatkowski, Cordelia Langford, John Sillitoe on behalf of the Wellcome Sanger Institute COVID-19 Surveillance Team |
| EPI_ISL_2973130 | Lighthouse Laboratory Plymouth | Wellcome Sanger Institute for the COVID-19 Genomics UK (COG-UK) Consortium | Lighthouse Laboratory Plymouth and Alex Alderton, Roberto Amato, Jeffrey Barrett, Sonia Goncalves, Ewan Harrison, David K. Jackson, Ian Johnston, Dominic Kwiatkowski, Cordelia Langford, John Sillitoe on behalf of the Wellcome Sanger Institute COVID-19 Surveillance Team |
| EPI_ISL_2973131 | Lighthouse Lab in Alderley Park | Wellcome Sanger Institute for the COVID-19 Genomics UK (COG-UK) Consortium | Jacquelyn Wynn, Mairead Hyland, The Lighthouse Lab in Alderley Park and Alex Alderton, Roberto Amato, Jeffrey Barrett, Sonia Goncalves, Ewan Harrison, David K. Jackson, Ian Johnston, Dominic Kwiatkowski, Cordelia Langford, John Sillitoe on behalf of the Wellcome Sanger Institute COVID-19 Surveillance Team |
| EPI_ISL_2973132 | Lighthouse Lab in Milton Keynes | Wellcome Sanger Institute for the COVID-19 Genomics UK (COG-UK) Consortium | The Lighthouse Lab in Milton Keynes and Alex Alderton, Roberto Amato, Jeffrey Barrett, Sonia Goncalves, Ewan Harrison, David K. Jackson, Ian Johnston, Dominic Kwiatkowski, Cordelia Langford, John Sillitoe on behalf of the Wellcome Sanger Institute COVID-19 Surveillance Team |
| EPI_ISL_2973133 | Department of Bacteria, Parasites and Fungi, Statens Serum Institut, Copenhagen, Denmark | Statens Serum Institut Bioinformatics and Microbial Genomics | Danish Covid-19 Genome Consortium |
| EPI_ISL_2973134 | Lighthouse Laboratory Plymouth | Wellcome Sanger Institute for the COVID-19 Genomics UK (COG-UK) Consortium | Lighthouse Laboratory Plymouth and Alex Alderton, Roberto Amato, Jeffrey Barrett, Sonia Goncalves, Ewan Harrison, David K. Jackson, Ian Johnston, Dominic Kwiatkowski, Cordelia Langford, John Sillitoe on behalf of the Wellcome Sanger Institute COVID-19 Surveillance Team |
| EPI_ISL_2973135 | Berkshire and Surrey Pathology Services Lighthouse Laboratory | Wellcome Sanger Institute for the COVID-19 Genomics UK (COG-UK) Consortium | Berkshire and Surrey Pathology Services Lighthouse Laboratory and Alex Alderton, Roberto Amato, Jeffrey Barrett, Sonia Goncalves, Ewan Harrison, David K. Jackson, Ian Johnston, Dominic Kwiatkowski, Cordelia Langford, John Sillitoe on behalf of the Wellcome Sanger Institute COVID-19 Surveillance Team |
| EPI_ISL_2973136 | Department of Bacteria, Parasites and Fungi, Statens Serum Institut, Copenhagen, Denmark | Statens Serum Institut Bioinformatics and Microbial Genomics | Danish Covid-19 Genome Consortium |
| EPI_ISL_2973137, EPI_ISL_2973138 | Lighthouse Laboratory Plymouth | Wellcome Sanger Institute for the COVID-19 Genomics UK (COG-UK) Consortium | Lighthouse Laboratory Plymouth and Alex Alderton, Roberto Amato, Jeffrey Barrett, Sonia Goncalves, Ewan Harrison, David K. Jackson, Ian Johnston, Dominic Kwiatkowski, Cordelia Langford, John Sillitoe on behalf of the Wellcome Sanger Institute COVID-19 Surveillance Team |
| EPI_ISL_2973139 | Department of Bacteria, Parasites and Fungi, Statens Serum Institut, Copenhagen, Denmark | Statens Serum Institut Bioinformatics and Microbial Genomics | Danish Covid-19 Genome Consortium |
| EPI_ISL_2973140 | Lighthouse Laboratory Plymouth | Wellcome Sanger Institute for the COVID-19 Genomics UK (COG-UK) Consortium | Lighthouse Laboratory Plymouth and Alex Alderton, Roberto Amato, Jeffrey Barrett, Sonia Goncalves, Ewan Harrison, David K. Jackson, Ian Johnston, Dominic Kwiatkowski, Cordelia Langford, John Sillitoe on behalf of the Wellcome Sanger Institute COVID-19 Surveillance Team |
| EPI_ISL_2973141 | Berkshire and Surrey Pathology Services Lighthouse Laboratory | Wellcome Sanger Institute for the COVID-19 Genomics UK (COG-UK) Consortium | Berkshire and Surrey Pathology Services Lighthouse Laboratory and Alex Alderton, Roberto Amato, Jeffrey Barrett, Sonia Goncalves, Ewan Harrison, David K. Jackson, Ian Johnston, Dominic Kwiatkowski, Cordelia Langford, John Sillitoe on behalf of the Wellcome Sanger Institute COVID-19 Surveillance Team |
| EPI_ISL_2973142 | Department of Bacteria, Parasites and Fungi, Statens Serum Institut, Copenhagen, Denmark | Statens Serum Institut Bioinformatics and Microbial Genomics | Danish Covid-19 Genome Consortium |
| EPI_ISL_2973143 | Lighthouse Laboratory Plymouth | Wellcome Sanger Institute for the COVID-19 Genomics UK (COG-UK) Consortium | Lighthouse Laboratory Plymouth and Alex Alderton, Roberto Amato, Jeffrey Barrett, Sonia Goncalves, Ewan Harrison, David K. Jackson, Ian Johnston, Dominic Kwiatkowski, Cordelia Langford, John Sillitoe on behalf of the Wellcome Sanger Institute COVID-19 Surveillance Team |
| EPI_ISL_2973144 | Lighthouse Lab in Milton Keynes | Wellcome Sanger Institute for the COVID-19 Genomics UK (COG-UK) Consortium | The Lighthouse Lab in Milton Keynes and Alex Alderton, Roberto Amato, Jeffrey Barrett, Sonia Goncalves, Ewan Harrison, David K. Jackson, Ian Johnston, Dominic Kwiatkowski, Cordelia Langford, John Sillitoe on behalf of the Wellcome Sanger Institute COVID-19 Surveillance Team |
| EPI_ISL_2973146, EPI_ISL_2973147 | Lighthouse Lab in Alderley Park | Wellcome Sanger Institute for the COVID-19 Genomics UK (COG-UK) Consortium | Jacquelyn Wynn, Mairead Hyland, The Lighthouse Lab in Alderley Park and Alex Alderton, Roberto Amato, Jeffrey Barrett, Sonia Goncalves, Ewan Harrison, David K. Jackson, Ian Johnston, Dominic Kwiatkowski, Cordelia Langford, John Sillitoe on behalf of the Wellcome Sanger Institute COVID-19 Surveillance Team |
| EPI_ISL_2973148 | Department of Bacteria, Parasites and Fungi, Statens Serum Institut, Copenhagen, Denmark | Statens Serum Institut Bioinformatics and Microbial Genomics | Danish Covid-19 Genome Consortium |
| EPI_ISL_2973149 | Lighthouse Lab in Alderley Park | Wellcome Sanger Institute for the COVID-19 Genomics UK (COG-UK) Consortium | Jacquelyn Wynn, Mairead Hyland, The Lighthouse Lab in Alderley Park and Alex Alderton, Roberto Amato, Jeffrey Barrett, Sonia Goncalves, Ewan Harrison, David K. Jackson, Ian Johnston, Dominic Kwiatkowski, Cordelia Langford, John Sillitoe on behalf of the Wellcome Sanger Institute COVID-19 Surveillance Team |
| EPI_ISL_2973150 | Lighthouse Lab in Milton Keynes | Wellcome Sanger Institute for the COVID-19 Genomics UK (COG-UK) Consortium | The Lighthouse Lab in Milton Keynes and Alex Alderton, Roberto Amato, Jeffrey Barrett, Sonia Goncalves, Ewan Harrison, David K. Jackson, Ian Johnston, Dominic Kwiatkowski, Cordelia Langford, John Sillitoe on behalf of the Wellcome Sanger Institute COVID-19 Surveillance Team |
| EPI_ISL_2973151 | Lighthouse Lab in Alderley Park | Wellcome Sanger Institute for the COVID-19 Genomics UK (COG-UK) Consortium | Jacquelyn Wynn, Mairead Hyland, The Lighthouse Lab in Alderley Park and Alex Alderton, Roberto Amato, Jeffrey Barrett, Sonia Goncalves, Ewan Harrison, David K. Jackson, Ian Johnston, Dominic Kwiatkowski, Cordelia Langford, John Sillitoe on behalf of the Wellcome Sanger Institute COVID-19 Surveillance Team |
| EPI_ISL_2973152 | Department of Bacteria, Parasites and Fungi, Statens Serum Institut, Copenhagen, Denmark | Statens Serum Institut Bioinformatics and Microbial Genomics | Danish Covid-19 Genome Consortium |
| EPI_ISL_2973153 | Lighthouse Lab in Alderley Park | Wellcome Sanger Institute for the COVID-19 Genomics UK (COG-UK) Consortium | Jacquelyn Wynn, Mairead Hyland, The Lighthouse Lab in Alderley Park and Alex Alderton, Roberto Amato, Jeffrey Barrett, Sonia Goncalves, Ewan Harrison, David K. Jackson, Ian Johnston, Dominic Kwiatkowski, Cordelia Langford, John Sillitoe on behalf of the Wellcome Sanger Institute COVID-19 Surveillance Team |

| | | | |
|---|---|---|---|
| EPI_ISL_2973154 | Department of Bacteria, Parasites and Fungi, Statens Serum Institut, Copenhagen, Denmark | Statens Serum Institut Bioinformatics and Microbial Genomics | Danish Covid-19 Genome Consortium |
| EPI_ISL_2973156 | Berkshire and Surrey Pathology Services Lighthouse Laboratory | Wellcome Sanger Institute for the COVID-19 Genomics UK (COG-UK) Consortium | Berkshire and Surrey Pathology Services Lighthouse Laboratory and Alex Alderton, Roberto Amato, Jeffrey Barrett, Sonia Goncalves, Ewan Harrison, David K. Jackson, Ian Johnston, Dominic Kwiatkowski, Cordelia Langford, John Sillitoe on behalf of the Wellcome Sanger Institute COVID-19 Surveillance Team |
| EPI_ISL_2973157 | Department of Bacteria, Parasites and Fungi, Statens Serum Institut, Copenhagen, Denmark | Statens Serum Institut Bioinformatics and Microbial Genomics | Danish Covid-19 Genome Consortium |
| EPI_ISL_2973158 | Lighthouse Lab in Alderley Park | Wellcome Sanger Institute for the COVID-19 Genomics UK (COG-UK) Consortium | Jacquelyn Wynn, Mairead Hyland, The Lighthouse Lab in Alderley Park and Alex Alderton, Roberto Amato, Jeffrey Barrett, Sonia Goncalves, Ewan Harrison, David K. Jackson, Ian Johnston, Dominic Kwiatkowski, Cordelia Langford, John Sillitoe on behalf of the Wellcome Sanger Institute COVID-19 Surveillance Team |
| EPI_ISL_2973159, EPI_ISL_2973160 | Lighthouse Laboratory Plymouth | Wellcome Sanger Institute for the COVID-19 Genomics UK (COG-UK) Consortium | Lighthouse Laboratory Plymouth and Alex Alderton, Roberto Amato, Jeffrey Barrett, Sonia Goncalves, Ewan Harrison, David K. Jackson, Ian Johnston, Dominic Kwiatkowski, Cordelia Langford, John Sillitoe on behalf of the Wellcome Sanger Institute COVID-19 Surveillance Team |
| EPI_ISL_2973161 | Department of Bacteria, Parasites and Fungi, Statens Serum Institut, Copenhagen, Denmark | Statens Serum Institut Bioinformatics and Microbial Genomics | Danish Covid-19 Genome Consortium |
| EPI_ISL_2973162 | Lighthouse Lab in Milton Keynes | Wellcome Sanger Institute for the COVID-19 Genomics UK (COG-UK) Consortium | The Lighthouse Lab in Milton Keynes and Alex Alderton, Roberto Amato, Jeffrey Barrett, Sonia Goncalves, Ewan Harrison, David K. Jackson, Ian Johnston, Dominic Kwiatkowski, Cordelia Langford, John Sillitoe on behalf of the Wellcome Sanger Institute COVID-19 Surveillance Team |
| EPI_ISL_2973163 | Lighthouse Lab in Alderley Park | Wellcome Sanger Institute for the COVID-19 Genomics UK (COG-UK) Consortium | Jacquelyn Wynn, Mairead Hyland, The Lighthouse Lab in Alderley Park and Alex Alderton, Roberto Amato, Jeffrey Barrett, Sonia Goncalves, Ewan Harrison, David K. Jackson, Ian Johnston, Dominic Kwiatkowski, Cordelia Langford, John Sillitoe on behalf of the Wellcome Sanger Institute COVID-19 Surveillance Team |
| EPI_ISL_2973164 | Department of Bacteria, Parasites and Fungi, Statens Serum Institut, Copenhagen, Denmark | Statens Serum Institut Bioinformatics and Microbial Genomics | Danish Covid-19 Genome Consortium |
| EPI_ISL_2973165 | Lighthouse Laboratory Plymouth | Wellcome Sanger Institute for the COVID-19 Genomics UK (COG-UK) Consortium | Lighthouse Laboratory Plymouth and Alex Alderton, Roberto Amato, Jeffrey Barrett, Sonia Goncalves, Ewan Harrison, David K. Jackson, Ian Johnston, Dominic Kwiatkowski, Cordelia Langford, John Sillitoe on behalf of the Wellcome Sanger Institute COVID-19 Surveillance Team |
| EPI_ISL_2973166 | Lighthouse Lab in Alderley Park | Wellcome Sanger Institute for the COVID-19 Genomics UK (COG-UK) Consortium | Jacquelyn Wynn, Mairead Hyland, The Lighthouse Lab in Alderley Park and Alex Alderton, Roberto Amato, Jeffrey Barrett, Sonia Goncalves, Ewan Harrison, David K. Jackson, Ian Johnston, Dominic Kwiatkowski, Cordelia Langford, John Sillitoe on behalf of the Wellcome Sanger Institute COVID-19 Surveillance Team |
| EPI_ISL_2973167 | Department of Bacteria, Parasites and Fungi, Statens Serum Institut, Copenhagen, Denmark | Statens Serum Institut Bioinformatics and Microbial Genomics | Danish Covid-19 Genome Consortium |
| EPI_ISL_2973168 | Lighthouse Lab in Milton Keynes | Wellcome Sanger Institute for the COVID-19 Genomics UK (COG-UK) Consortium | The Lighthouse Lab in Milton Keynes and Alex Alderton, Roberto Amato, Jeffrey Barrett, Sonia Goncalves, Ewan Harrison, David K. Jackson, Ian Johnston, Dominic Kwiatkowski, Cordelia Langford, John Sillitoe on behalf of the Wellcome Sanger Institute COVID-19 Surveillance Team |
| EPI_ISL_2973169 | Lighthouse Lab in Alderley Park | Wellcome Sanger Institute for the COVID-19 Genomics UK (COG-UK) Consortium | Jacquelyn Wynn, Mairead Hyland, The Lighthouse Lab in Alderley Park and Alex Alderton, Roberto Amato, Jeffrey Barrett, Sonia Goncalves, Ewan Harrison, David K. Jackson, Ian Johnston, Dominic Kwiatkowski, Cordelia Langford, John Sillitoe on behalf of the Wellcome Sanger Institute COVID-19 Surveillance Team |
| EPI_ISL_2973170 | Department of Bacteria, Parasites and Fungi, Statens Serum Institut, Copenhagen, Denmark | Statens Serum Institut Bioinformatics and Microbial Genomics | Danish Covid-19 Genome Consortium |
| EPI_ISL_2973171 | Lighthouse Lab in Alderley Park | Wellcome Sanger Institute for the COVID-19 Genomics UK (COG-UK) Consortium | Jacquelyn Wynn, Mairead Hyland, The Lighthouse Lab in Alderley Park and Alex Alderton, Roberto Amato, Jeffrey Barrett, Sonia Goncalves, Ewan Harrison, David K. Jackson, Ian Johnston, Dominic Kwiatkowski, Cordelia Langford, John Sillitoe on behalf of the Wellcome Sanger Institute COVID-19 Surveillance Team |
| EPI_ISL_2973172 | Lighthouse Laboratory Plymouth | Wellcome Sanger Institute for the COVID-19 Genomics UK (COG-UK) Consortium | Lighthouse Laboratory Plymouth and Alex Alderton, Roberto Amato, Jeffrey Barrett, Sonia Goncalves, Ewan Harrison, David K. Jackson, Ian Johnston, Dominic Kwiatkowski, Cordelia Langford, John Sillitoe on behalf of the Wellcome Sanger Institute COVID-19 Surveillance Team |
| EPI_ISL_2973173 | Lighthouse Lab in Milton Keynes | Wellcome Sanger Institute for the COVID-19 Genomics UK (COG-UK) Consortium | The Lighthouse Lab in Milton Keynes and Alex Alderton, Roberto Amato, Jeffrey Barrett, Sonia Goncalves, Ewan Harrison, David K. Jackson, Ian Johnston, Dominic Kwiatkowski, Cordelia Langford, John Sillitoe on behalf of the Wellcome Sanger Institute COVID-19 Surveillance Team |
| EPI_ISL_2973174 | Lighthouse Lab in Alderley Park | Wellcome Sanger Institute for the COVID-19 Genomics UK (COG-UK) Consortium | Jacquelyn Wynn, Mairead Hyland, The Lighthouse Lab in Alderley Park and Alex Alderton, Roberto Amato, Jeffrey Barrett, Sonia Goncalves, Ewan Harrison, David K. Jackson, Ian Johnston, Dominic Kwiatkowski, Cordelia Langford, John Sillitoe on behalf of the Wellcome Sanger Institute COVID-19 Surveillance Team |
| EPI_ISL_2973175 | Lighthouse Laboratory Plymouth | Wellcome Sanger Institute for the COVID-19 Genomics UK (COG-UK) Consortium | Lighthouse Laboratory Plymouth and Alex Alderton, Roberto Amato, Jeffrey Barrett, Sonia Goncalves, Ewan Harrison, David K. Jackson, Ian Johnston, Dominic Kwiatkowski, Cordelia Langford, John Sillitoe on behalf of the Wellcome Sanger Institute COVID-19 Surveillance Team |
| EPI_ISL_2973176 | Lighthouse Lab in Alderley Park | Wellcome Sanger Institute for the COVID-19 Genomics UK (COG-UK) Consortium | Jacquelyn Wynn, Mairead Hyland, The Lighthouse Lab in Alderley Park and Alex Alderton, Roberto Amato, Jeffrey Barrett, Sonia Goncalves, Ewan Harrison, David K. Jackson, Ian Johnston, Dominic Kwiatkowski, Cordelia Langford, John Sillitoe on behalf of the Wellcome Sanger Institute COVID-19 Surveillance Team |
| EPI_ISL_2973177, EPI_ISL_2973178 | Lighthouse Laboratory Plymouth | Wellcome Sanger Institute for the COVID-19 Genomics UK (COG-UK) Consortium | Lighthouse Laboratory Plymouth and Alex Alderton, Roberto Amato, Jeffrey Barrett, Sonia Goncalves, Ewan Harrison, David K. Jackson, Ian Johnston, Dominic Kwiatkowski, Cordelia Langford, John Sillitoe on behalf of the Wellcome Sanger Institute COVID-19 Surveillance Team |
| EPI_ISL_2973179 | Lighthouse Lab in Milton Keynes | Wellcome Sanger Institute for the COVID-19 Genomics UK (COG-UK) Consortium | The Lighthouse Lab in Milton Keynes and Alex Alderton, Roberto Amato, Jeffrey Barrett, Sonia Goncalves, Ewan Harrison, David K. Jackson, Ian Johnston, Dominic Kwiatkowski, Cordelia Langford, John Sillitoe on behalf of the Wellcome Sanger Institute COVID-19 Surveillance Team |
| EPI_ISL_2973180 | Lighthouse Laboratory Plymouth | Wellcome Sanger Institute for the COVID-19 Genomics UK (COG-UK) Consortium | Lighthouse Laboratory Plymouth and Alex Alderton, Roberto Amato, Jeffrey Barrett, Sonia Goncalves, Ewan Harrison, David K. Jackson, Ian Johnston, Dominic Kwiatkowski, Cordelia Langford, John Sillitoe on behalf of the Wellcome Sanger Institute COVID-19 Surveillance Team |
| EPI_ISL_2973181 | Lighthouse Lab in Milton Keynes | Wellcome Sanger Institute for the COVID-19 Genomics UK (COG-UK) Consortium | The Lighthouse Lab in Milton Keynes and Alex Alderton, Roberto Amato, Jeffrey Barrett, Sonia Goncalves, Ewan Harrison, David K. Jackson, Ian Johnston, Dominic Kwiatkowski, Cordelia Langford, John Sillitoe on behalf of the Wellcome Sanger Institute COVID-19 Surveillance Team |
| EPI_ISL_2973182 | Lighthouse Lab in Alderley Park | Wellcome Sanger Institute for the COVID-19 Genomics UK (COG-UK) Consortium | Jacquelyn Wynn, Mairead Hyland, The Lighthouse Lab in Alderley Park and Alex Alderton, Roberto Amato, Jeffrey Barrett, Sonia Goncalves, Ewan Harrison, David K. Jackson, Ian Johnston, Dominic Kwiatkowski, Cordelia Langford, John Sillitoe on behalf of the Wellcome Sanger Institute COVID-19 Surveillance Team |
| EPI_ISL_2973183 | Lighthouse Laboratory Plymouth | Wellcome Sanger Institute for the COVID-19 Genomics UK (COG-UK) Consortium | Lighthouse Laboratory Plymouth and Alex Alderton, Roberto Amato, Jeffrey Barrett, Sonia Goncalves, Ewan Harrison, David K. Jackson, Ian Johnston, Dominic Kwiatkowski, Cordelia Langford, John Sillitoe on behalf of the Wellcome Sanger Institute COVID-19 Surveillance Team |
| EPI_ISL_2973184 | Lighthouse Lab in Alderley Park | Wellcome Sanger Institute for the COVID-19 Genomics UK (COG-UK) Consortium | Jacquelyn Wynn, Mairead Hyland, The Lighthouse Lab in Alderley Park and Alex Alderton, Roberto Amato, Jeffrey Barrett, Sonia Goncalves, Ewan Harrison, David K. Jackson, Ian Johnston, Dominic Kwiatkowski, Cordelia Langford, John Sillitoe on behalf of the Wellcome Sanger Institute COVID-19 Surveillance Team |
| EPI_ISL_2973185, EPI_ISL_2973186, EPI_ISL_2973187 | Lighthouse Laboratory Plymouth | Wellcome Sanger Institute for the COVID-19 Genomics UK (COG-UK) Consortium | Lighthouse Laboratory Plymouth and Alex Alderton, Roberto Amato, Jeffrey Barrett, Sonia Goncalves, Ewan Harrison, David K. Jackson, Ian Johnston, Dominic Kwiatkowski, Cordelia Langford, John Sillitoe on behalf of the Wellcome Sanger Institute COVID-19 Surveillance Team |
| EPI_ISL_2973188 | Lighthouse Lab in Alderley Park | Wellcome Sanger Institute for the COVID-19 Genomics UK (COG-UK) Consortium | Jacquelyn Wynn, Mairead Hyland, The Lighthouse Lab in Alderley Park and Alex Alderton, Roberto Amato, Jeffrey Barrett, Sonia Goncalves, Ewan Harrison, David K. Jackson, Ian Johnston, Dominic Kwiatkowski, Cordelia Langford, John Sillitoe on behalf of the Wellcome Sanger Institute COVID-19 Surveillance Team |
| EPI_ISL_2973189, EPI_ISL_2973190 | Lighthouse Lab in Milton Keynes | Wellcome Sanger Institute for the COVID-19 Genomics UK (COG-UK) Consortium | The Lighthouse Lab in Milton Keynes and Alex Alderton, Roberto Amato, Jeffrey Barrett, Sonia Goncalves, Ewan Harrison, David K. Jackson, Ian Johnston, Dominic Kwiatkowski, Cordelia Langford, John Sillitoe on behalf of the Wellcome Sanger Institute COVID-19 Surveillance Team |
| EPI_ISL_2973191 | Berkshire and Surrey Pathology Services Lighthouse Laboratory | Wellcome Sanger Institute for the COVID-19 Genomics UK (COG-UK) Consortium | Berkshire and Surrey Pathology Services Lighthouse Laboratory and Alex Alderton, Roberto Amato, Jeffrey Barrett, Sonia Goncalves, Ewan Harrison, David K. Jackson, Ian Johnston, Dominic Kwiatkowski, Cordelia Langford, John Sillitoe on behalf of the Wellcome Sanger Institute COVID-19 Surveillance Team |
| EPI_ISL_2973192 | Lighthouse Lab in Alderley Park | Wellcome Sanger Institute for the COVID-19 Genomics UK (COG-UK) Consortium | Jacquelyn Wynn, Mairead Hyland, The Lighthouse Lab in Alderley Park and Alex Alderton, Roberto Amato, Jeffrey Barrett, Sonia Goncalves, Ewan Harrison, David K. Jackson, Ian Johnston, Dominic Kwiatkowski, Cordelia Langford, John Sillitoe on behalf of the Wellcome Sanger Institute COVID-19 Surveillance Team |
| EPI_ISL_2973193 | Lighthouse Laboratory Plymouth | Wellcome Sanger Institute for the COVID-19 Genomics UK (COG-UK) Consortium | Lighthouse Laboratory Plymouth and Alex Alderton, Roberto Amato, Jeffrey Barrett, Sonia Goncalves, Ewan Harrison, David K. Jackson, Ian Johnston, Dominic Kwiatkowski, Cordelia Langford, John Sillitoe on behalf of the Wellcome Sanger Institute COVID-19 Surveillance Team |
| EPI_ISL_2973194 | Lighthouse Lab in Alderley Park | Wellcome Sanger Institute for the COVID-19 Genomics UK (COG-UK) Consortium | Jacquelyn Wynn, Mairead Hyland, The Lighthouse Lab in Alderley Park and Alex Alderton, Roberto Amato, Jeffrey Barrett, Sonia Goncalves, Ewan Harrison, David K. Jackson, Ian Johnston, Dominic Kwiatkowski, Cordelia Langford, John Sillitoe on behalf of the Wellcome Sanger Institute COVID-19 Surveillance Team |
| EPI_ISL_2973195, EPI_ISL_2973196, EPI_ISL_2973197, EPI_ISL_2973198 | Lighthouse Laboratory Plymouth | Wellcome Sanger Institute for the COVID-19 Genomics UK (COG-UK) Consortium | Lighthouse Laboratory Plymouth and Alex Alderton, Roberto Amato, Jeffrey Barrett, Sonia Goncalves, Ewan Harrison, David K. Jackson, Ian Johnston, Dominic Kwiatkowski, Cordelia Langford, John Sillitoe on behalf of the Wellcome Sanger Institute COVID-19 Surveillance Team |
| EPI_ISL_2973199 | Lighthouse Lab in Alderley Park | Wellcome Sanger Institute for the COVID-19 Genomics UK (COG-UK) Consortium | Jacquelyn Wynn, Mairead Hyland, The Lighthouse Lab in Alderley Park and Alex Alderton, Roberto Amato, Jeffrey Barrett, Sonia Goncalves, Ewan Harrison, David K. Jackson, Ian Johnston, Dominic Kwiatkowski, Cordelia Langford, John Sillitoe on behalf of the Wellcome Sanger Institute COVID-19 Surveillance Team |
| EPI_ISL_2973200 | Lighthouse Laboratory Plymouth | Wellcome Sanger Institute for the COVID-19 Genomics UK (COG-UK) Consortium | Lighthouse Laboratory Plymouth and Alex Alderton, Roberto Amato, Jeffrey Barrett, Sonia Goncalves, Ewan Harrison, David K. Jackson, Ian Johnston, Dominic Kwiatkowski, Cordelia Langford, John Sillitoe on behalf of the Wellcome Sanger Institute COVID-19 Surveillance Team |
| EPI_ISL_2973201, EPI_ISL_2973202 | Lighthouse Lab in Milton Keynes | Wellcome Sanger Institute for the COVID-19 Genomics UK (COG-UK) Consortium | The Lighthouse Lab in Milton Keynes and Alex Alderton, Roberto Amato, Jeffrey Barrett, Sonia Goncalves, Ewan Harrison, David K. Jackson, Ian Johnston, Dominic Kwiatkowski, Cordelia Langford, John Sillitoe on behalf of the Wellcome Sanger Institute COVID-19 Surveillance Team |
| EPI_ISL_2973203 | Berkshire and Surrey Pathology Services Lighthouse Laboratory | Wellcome Sanger Institute for the COVID-19 Genomics UK (COG-UK) Consortium | Berkshire and Surrey Pathology Services Lighthouse Laboratory and Alex Alderton, Roberto Amato, Jeffrey Barrett, Sonia Goncalves, Ewan Harrison, David K. Jackson, Ian Johnston, Dominic Kwiatkowski, Cordelia Langford, John Sillitoe on behalf of the Wellcome Sanger Institute COVID-19 Surveillance Team |
| EPI_ISL_2973204 | Lighthouse Lab in Alderley Park | Wellcome Sanger Institute for the COVID-19 Genomics UK (COG-UK) Consortium | Jacquelyn Wynn, Mairead Hyland, The Lighthouse Lab in Alderley Park and Alex Alderton, Roberto Amato, Jeffrey Barrett, Sonia Goncalves, Ewan Harrison, David K. Jackson, Ian Johnston, Dominic Kwiatkowski, Cordelia Langford, John Sillitoe on behalf of the Wellcome Sanger Institute COVID-19 Surveillance Team |
| EPI_ISL_2973205 | Berkshire and Surrey Pathology Services Lighthouse Laboratory | Wellcome Sanger Institute for the COVID-19 Genomics UK (COG-UK) Consortium | Berkshire and Surrey Pathology Services Lighthouse Laboratory and Alex Alderton, Roberto Amato, Jeffrey Barrett, Sonia Goncalves, Ewan Harrison, David K. Jackson, Ian Johnston, Dominic Kwiatkowski, Cordelia Langford, John Sillitoe on behalf of the Wellcome Sanger Institute COVID-19 Surveillance Team |
| EPI_ISL_2973206, EPI_ISL_2973207, EPI_ISL_2973208, EPI_ISL_2973209 | Lighthouse Lab in Milton Keynes | Wellcome Sanger Institute for the COVID-19 Genomics UK (COG-UK) Consortium | The Lighthouse Lab in Milton Keynes and Alex Alderton, Roberto Amato, Jeffrey Barrett, Sonia Goncalves, Ewan Harrison, David K. Jackson, Ian Johnston, Dominic Kwiatkowski, Cordelia Langford, John Sillitoe on behalf of the Wellcome Sanger Institute COVID-19 Surveillance Team |

| | | | |
|---|---|---|---|
| EPI_ISL_2973210 | Lighthouse Lab in Alderley Park | Wellcome Sanger Institute for the COVID-19 Genomics UK (COG-UK) Consortium | Jacquelyn Wynn, Mairead Hyland, The Lighthouse Lab in Alderley Park and Alex Alderton, Roberto Amato, Jeffrey Barrett, Sonia Goncalves, Ewan Harrison, David K. Jackson, Ian Johnston, Dominic Kwiatkowski, Cordelia Langford, John Sillitoe on behalf of the Wellcome Sanger Institute COVID-19 Surveillance Team |
| EPI_ISL_2973211 | Lighthouse Laboratory Plymouth | Wellcome Sanger Institute for the COVID-19 Genomics UK (COG-UK) Consortium | Lighthouse Laboratory Plymouth and Alex Alderton, Roberto Amato, Jeffrey Barrett, Sonia Goncalves, Ewan Harrison, David K. Jackson, Ian Johnston, Dominic Kwiatkowski, Cordelia Langford, John Sillitoe on behalf of the Wellcome Sanger Institute COVID-19 Surveillance Team |
| EPI_ISL_2973212 | Berkshire and Surrey Pathology Services Lighthouse Laboratory | Wellcome Sanger Institute for the COVID-19 Genomics UK (COG-UK) Consortium | Berkshire and Surrey Pathology Services Lighthouse Laboratory and Alex Alderton, Roberto Amato, Jeffrey Barrett, Sonia Goncalves, Ewan Harrison, David K. Jackson, Ian Johnston, Dominic Kwiatkowski, Cordelia Langford, John Sillitoe on behalf of the Wellcome Sanger Institute COVID-19 Surveillance Team |
| EPI_ISL_2973213, EPI_ISL_2973214 | Lighthouse Laboratory Plymouth | Wellcome Sanger Institute for the COVID-19 Genomics UK (COG-UK) Consortium | Lighthouse Laboratory Plymouth and Alex Alderton, Roberto Amato, Jeffrey Barrett, Sonia Goncalves, Ewan Harrison, David K. Jackson, Ian Johnston, Dominic Kwiatkowski, Cordelia Langford, John Sillitoe on behalf of the Wellcome Sanger Institute COVID-19 Surveillance Team |
| EPI_ISL_2973215 | Berkshire and Surrey Pathology Services Lighthouse Laboratory | Wellcome Sanger Institute for the COVID-19 Genomics UK (COG-UK) Consortium | Berkshire and Surrey Pathology Services Lighthouse Laboratory and Alex Alderton, Roberto Amato, Jeffrey Barrett, Sonia Goncalves, Ewan Harrison, David K. Jackson, Ian Johnston, Dominic Kwiatkowski, Cordelia Langford, John Sillitoe on behalf of the Wellcome Sanger Institute COVID-19 Surveillance Team |
| EPI_ISL_2973216 | Lighthouse Lab in Milton Keynes | Wellcome Sanger Institute for the COVID-19 Genomics UK (COG-UK) Consortium | The Lighthouse Lab in Milton Keynes and Alex Alderton, Roberto Amato, Jeffrey Barrett, Sonia Goncalves, Ewan Harrison, David K. Jackson, Ian Johnston, Dominic Kwiatkowski, Cordelia Langford, John Sillitoe on behalf of the Wellcome Sanger Institute COVID-19 Surveillance Team |
| EPI_ISL_2973217, EPI_ISL_2973218, EPI_ISL_2973219, EPI_ISL_2973220, EPI_ISL_2973221 | Lighthouse Laboratory Plymouth | Wellcome Sanger Institute for the COVID-19 Genomics UK (COG-UK) Consortium | Lighthouse Laboratory Plymouth and Alex Alderton, Roberto Amato, Jeffrey Barrett, Sonia Goncalves, Ewan Harrison, David K. Jackson, Ian Johnston, Dominic Kwiatkowski, Cordelia Langford, John Sillitoe on behalf of the Wellcome Sanger Institute COVID-19 Surveillance Team |
| EPI_ISL_2973222, EPI_ISL_2973223 | Lighthouse Lab in Alderley Park | Wellcome Sanger Institute for the COVID-19 Genomics UK (COG-UK) Consortium | Jacquelyn Wynn, Mairead Hyland, The Lighthouse Lab in Alderley Park and Alex Alderton, Roberto Amato, Jeffrey Barrett, Sonia Goncalves, Ewan Harrison, David K. Jackson, Ian Johnston, Dominic Kwiatkowski, Cordelia Langford, John Sillitoe on behalf of the Wellcome Sanger Institute COVID-19 Surveillance Team |
| EPI_ISL_2973224, EPI_ISL_2973225 | Lighthouse Laboratory Plymouth | Wellcome Sanger Institute for the COVID-19 Genomics UK (COG-UK) Consortium | Lighthouse Laboratory Plymouth and Alex Alderton, Roberto Amato, Jeffrey Barrett, Sonia Goncalves, Ewan Harrison, David K. Jackson, Ian Johnston, Dominic Kwiatkowski, Cordelia Langford, John Sillitoe on behalf of the Wellcome Sanger Institute COVID-19 Surveillance Team |
| EPI_ISL_2973226 | Berkshire and Surrey Pathology Services Lighthouse Laboratory | Wellcome Sanger Institute for the COVID-19 Genomics UK (COG-UK) Consortium | Berkshire and Surrey Pathology Services Lighthouse Laboratory and Alex Alderton, Roberto Amato, Jeffrey Barrett, Sonia Goncalves, Ewan Harrison, David K. Jackson, Ian Johnston, Dominic Kwiatkowski, Cordelia Langford, John Sillitoe on behalf of the Wellcome Sanger Institute COVID-19 Surveillance Team |
| EPI_ISL_2973227 | Lighthouse Laboratory Plymouth | Wellcome Sanger Institute for the COVID-19 Genomics UK (COG-UK) Consortium | Lighthouse Laboratory Plymouth and Alex Alderton, Roberto Amato, Jeffrey Barrett, Sonia Goncalves, Ewan Harrison, David K. Jackson, Ian Johnston, Dominic Kwiatkowski, Cordelia Langford, John Sillitoe on behalf of the Wellcome Sanger Institute COVID-19 Surveillance Team |
| EPI_ISL_2973228, EPI_ISL_2973229 | Lighthouse Lab in Alderley Park | Wellcome Sanger Institute for the COVID-19 Genomics UK (COG-UK) Consortium | Jacquelyn Wynn, Mairead Hyland, The Lighthouse Lab in Alderley Park and Alex Alderton, Roberto Amato, Jeffrey Barrett, Sonia Goncalves, Ewan Harrison, David K. Jackson, Ian Johnston, Dominic Kwiatkowski, Cordelia Langford, John Sillitoe on behalf of the Wellcome Sanger Institute COVID-19 Surveillance Team |
| EPI_ISL_2973230 | Lighthouse Laboratory Plymouth | Wellcome Sanger Institute for the COVID-19 Genomics UK (COG-UK) Consortium | Lighthouse Laboratory Plymouth and Alex Alderton, Roberto Amato, Jeffrey Barrett, Sonia Goncalves, Ewan Harrison, David K. Jackson, Ian Johnston, Dominic Kwiatkowski, Cordelia Langford, John Sillitoe on behalf of the Wellcome Sanger Institute COVID-19 Surveillance Team |
| EPI_ISL_2973231 | Lighthouse Lab in Alderley Park | Wellcome Sanger Institute for the COVID-19 Genomics UK (COG-UK) Consortium | Jacquelyn Wynn, Mairead Hyland, The Lighthouse Lab in Alderley Park and Alex Alderton, Roberto Amato, Jeffrey Barrett, Sonia Goncalves, Ewan Harrison, David K. Jackson, Ian Johnston, Dominic Kwiatkowski, Cordelia Langford, John Sillitoe on behalf of the Wellcome Sanger Institute COVID-19 Surveillance Team |
| EPI_ISL_2973232 | Lighthouse Lab in Milton Keynes | Wellcome Sanger Institute for the COVID-19 Genomics UK (COG-UK) Consortium | The Lighthouse Lab in Milton Keynes and Alex Alderton, Roberto Amato, Jeffrey Barrett, Sonia Goncalves, Ewan Harrison, David K. Jackson, Ian Johnston, Dominic Kwiatkowski, Cordelia Langford, John Sillitoe on behalf of the Wellcome Sanger Institute COVID-19 Surveillance Team |
| EPI_ISL_2973233, EPI_ISL_2973234 | Lighthouse Laboratory Plymouth | Wellcome Sanger Institute for the COVID-19 Genomics UK (COG-UK) Consortium | Lighthouse Laboratory Plymouth and Alex Alderton, Roberto Amato, Jeffrey Barrett, Sonia Goncalves, Ewan Harrison, David K. Jackson, Ian Johnston, Dominic Kwiatkowski, Cordelia Langford, John Sillitoe on behalf of the Wellcome Sanger Institute COVID-19 Surveillance Team |
| EPI_ISL_2973235 | Lighthouse Lab in Alderley Park | Wellcome Sanger Institute for the COVID-19 Genomics UK (COG-UK) Consortium | Jacquelyn Wynn, Mairead Hyland, The Lighthouse Lab in Alderley Park and Alex Alderton, Roberto Amato, Jeffrey Barrett, Sonia Goncalves, Ewan Harrison, David K. Jackson, Ian Johnston, Dominic Kwiatkowski, Cordelia Langford, John Sillitoe on behalf of the Wellcome Sanger Institute COVID-19 Surveillance Team |
| EPI_ISL_2973236, EPI_ISL_2973237 | Berkshire and Surrey Pathology Services Lighthouse Laboratory | Wellcome Sanger Institute for the COVID-19 Genomics UK (COG-UK) Consortium | Berkshire and Surrey Pathology Services Lighthouse Laboratory and Alex Alderton, Roberto Amato, Jeffrey Barrett, Sonia Goncalves, Ewan Harrison, David K. Jackson, Ian Johnston, Dominic Kwiatkowski, Cordelia Langford, John Sillitoe on behalf of the Wellcome Sanger Institute COVID-19 Surveillance Team |
| EPI_ISL_2973238 | Lighthouse Laboratory Plymouth | Wellcome Sanger Institute for the COVID-19 Genomics UK (COG-UK) Consortium | Lighthouse Laboratory Plymouth and Alex Alderton, Roberto Amato, Jeffrey Barrett, Sonia Goncalves, Ewan Harrison, David K. Jackson, Ian Johnston, Dominic Kwiatkowski, Cordelia Langford, John Sillitoe on behalf of the Wellcome Sanger Institute COVID-19 Surveillance Team |
| EPI_ISL_2973239 | Lighthouse Lab in Alderley Park | Wellcome Sanger Institute for the COVID-19 Genomics UK (COG-UK) Consortium | Jacquelyn Wynn, Mairead Hyland, The Lighthouse Lab in Alderley Park and Alex Alderton, Roberto Amato, Jeffrey Barrett, Sonia Goncalves, Ewan Harrison, David K. Jackson, Ian Johnston, Dominic Kwiatkowski, Cordelia Langford, John Sillitoe on behalf of the Wellcome Sanger Institute COVID-19 Surveillance Team |
| EPI_ISL_2973240 | Lighthouse Laboratory Plymouth | Wellcome Sanger Institute for the COVID-19 Genomics UK (COG-UK) Consortium | Lighthouse Laboratory Plymouth and Alex Alderton, Roberto Amato, Jeffrey Barrett, Sonia Goncalves, Ewan Harrison, David K. Jackson, Ian Johnston, Dominic Kwiatkowski, Cordelia Langford, John Sillitoe on behalf of the Wellcome Sanger Institute COVID-19 Surveillance Team |
| EPI_ISL_2973241, EPI_ISL_2973242, EPI_ISL_2973243 | Lighthouse Lab in Alderley Park | Wellcome Sanger Institute for the COVID-19 Genomics UK (COG-UK) Consortium | Jacquelyn Wynn, Mairead Hyland, The Lighthouse Lab in Alderley Park and Alex Alderton, Roberto Amato, Jeffrey Barrett, Sonia Goncalves, Ewan Harrison, David K. Jackson, Ian Johnston, Dominic Kwiatkowski, Cordelia Langford, John Sillitoe on behalf of the Wellcome Sanger Institute COVID-19 Surveillance Team |
| EPI_ISL_2973244 | Lighthouse Lab in Milton Keynes | Wellcome Sanger Institute for the COVID-19 Genomics UK (COG-UK) Consortium | The Lighthouse Lab in Milton Keynes and Alex Alderton, Roberto Amato, Jeffrey Barrett, Sonia Goncalves, Ewan Harrison, David K. Jackson, Ian Johnston, Dominic Kwiatkowski, Cordelia Langford, John Sillitoe on behalf of the Wellcome Sanger Institute COVID-19 Surveillance Team |
| EPI_ISL_2973245 | Lighthouse Lab in Alderley Park | Wellcome Sanger Institute for the COVID-19 Genomics UK (COG-UK) Consortium | Jacquelyn Wynn, Mairead Hyland, The Lighthouse Lab in Alderley Park and Alex Alderton, Roberto Amato, Jeffrey Barrett, Sonia Goncalves, Ewan Harrison, David K. Jackson, Ian Johnston, Dominic Kwiatkowski, Cordelia Langford, John Sillitoe on behalf of the Wellcome Sanger Institute COVID-19 Surveillance Team |
| EPI_ISL_2973246, EPI_ISL_2973247, EPI_ISL_2973248 | Lighthouse Laboratory Plymouth | Wellcome Sanger Institute for the COVID-19 Genomics UK (COG-UK) Consortium | Lighthouse Laboratory Plymouth and Alex Alderton, Roberto Amato, Jeffrey Barrett, Sonia Goncalves, Ewan Harrison, David K. Jackson, Ian Johnston, Dominic Kwiatkowski, Cordelia Langford, John Sillitoe on behalf of the Wellcome Sanger Institute COVID-19 Surveillance Team |
| EPI_ISL_2973249, EPI_ISL_2973250 | Lighthouse Lab in Alderley Park | Wellcome Sanger Institute for the COVID-19 Genomics UK (COG-UK) Consortium | Jacquelyn Wynn, Mairead Hyland, The Lighthouse Lab in Alderley Park and Alex Alderton, Roberto Amato, Jeffrey Barrett, Sonia Goncalves, Ewan Harrison, David K. Jackson, Ian Johnston, Dominic Kwiatkowski, Cordelia Langford, John Sillitoe on behalf of the Wellcome Sanger Institute COVID-19 Surveillance Team |
| EPI_ISL_2973251 | Lighthouse Lab in Milton Keynes | Wellcome Sanger Institute for the COVID-19 Genomics UK (COG-UK) Consortium | The Lighthouse Lab in Milton Keynes and Alex Alderton, Roberto Amato, Jeffrey Barrett, Sonia Goncalves, Ewan Harrison, David K. Jackson, Ian Johnston, Dominic Kwiatkowski, Cordelia Langford, John Sillitoe on behalf of the Wellcome Sanger Institute COVID-19 Surveillance Team |
| EPI_ISL_2973252 | Lighthouse Lab in Alderley Park | Wellcome Sanger Institute for the COVID-19 Genomics UK (COG-UK) Consortium | Jacquelyn Wynn, Mairead Hyland, The Lighthouse Lab in Alderley Park and Alex Alderton, Roberto Amato, Jeffrey Barrett, Sonia Goncalves, Ewan Harrison, David K. Jackson, Ian Johnston, Dominic Kwiatkowski, Cordelia Langford, John Sillitoe on behalf of the Wellcome Sanger Institute COVID-19 Surveillance Team |
| EPI_ISL_2973253 | Berkshire and Surrey Pathology Services Lighthouse Laboratory | Wellcome Sanger Institute for the COVID-19 Genomics UK (COG-UK) Consortium | Berkshire and Surrey Pathology Services Lighthouse Laboratory and Alex Alderton, Roberto Amato, Jeffrey Barrett, Sonia Goncalves, Ewan Harrison, David K. Jackson, Ian Johnston, Dominic Kwiatkowski, Cordelia Langford, John Sillitoe on behalf of the Wellcome Sanger Institute COVID-19 Surveillance Team |
| EPI_ISL_2973254 | Lighthouse Laboratory Plymouth | Wellcome Sanger Institute for the COVID-19 Genomics UK (COG-UK) Consortium | Lighthouse Laboratory Plymouth and Alex Alderton, Roberto Amato, Jeffrey Barrett, Sonia Goncalves, Ewan Harrison, David K. Jackson, Ian Johnston, Dominic Kwiatkowski, Cordelia Langford, John Sillitoe on behalf of the Wellcome Sanger Institute COVID-19 Surveillance Team |
| EPI_ISL_2973255 | Lighthouse Lab in Milton Keynes | Wellcome Sanger Institute for the COVID-19 Genomics UK (COG-UK) Consortium | The Lighthouse Lab in Milton Keynes and Alex Alderton, Roberto Amato, Jeffrey Barrett, Sonia Goncalves, Ewan Harrison, David K. Jackson, Ian Johnston, Dominic Kwiatkowski, Cordelia Langford, John Sillitoe on behalf of the Wellcome Sanger Institute COVID-19 Surveillance Team |
| EPI_ISL_2973256 | Lighthouse Laboratory Plymouth | Wellcome Sanger Institute for the COVID-19 Genomics UK (COG-UK) Consortium | Lighthouse Laboratory Plymouth and Alex Alderton, Roberto Amato, Jeffrey Barrett, Sonia Goncalves, Ewan Harrison, David K. Jackson, Ian Johnston, Dominic Kwiatkowski, Cordelia Langford, John Sillitoe on behalf of the Wellcome Sanger Institute COVID-19 Surveillance Team |
| EPI_ISL_2973257 | Lighthouse Lab in Milton Keynes | Wellcome Sanger Institute for the COVID-19 Genomics UK (COG-UK) Consortium | The Lighthouse Lab in Milton Keynes and Alex Alderton, Roberto Amato, Jeffrey Barrett, Sonia Goncalves, Ewan Harrison, David K. Jackson, Ian Johnston, Dominic Kwiatkowski, Cordelia Langford, John Sillitoe on behalf of the Wellcome Sanger Institute COVID-19 Surveillance Team |
| EPI_ISL_2973258 | Lighthouse Lab in Alderley Park | Wellcome Sanger Institute for the COVID-19 Genomics UK (COG-UK) Consortium | Jacquelyn Wynn, Mairead Hyland, The Lighthouse Lab in Alderley Park and Alex Alderton, Roberto Amato, Jeffrey Barrett, Sonia Goncalves, Ewan Harrison, David K. Jackson, Ian Johnston, Dominic Kwiatkowski, Cordelia Langford, John Sillitoe on behalf of the Wellcome Sanger Institute COVID-19 Surveillance Team |
| EPI_ISL_2973259, EPI_ISL_2973260 | Lighthouse Lab in Milton Keynes | Wellcome Sanger Institute for the COVID-19 Genomics UK (COG-UK) Consortium | The Lighthouse Lab in Milton Keynes and Alex Alderton, Roberto Amato, Jeffrey Barrett, Sonia Goncalves, Ewan Harrison, David K. Jackson, Ian Johnston, Dominic Kwiatkowski, Cordelia Langford, John Sillitoe on behalf of the Wellcome Sanger Institute COVID-19 Surveillance Team |
| EPI_ISL_2973261, EPI_ISL_2973262, EPI_ISL_2973263 | Lighthouse Lab in Alderley Park | Wellcome Sanger Institute for the COVID-19 Genomics UK (COG-UK) Consortium | Jacquelyn Wynn, Mairead Hyland, The Lighthouse Lab in Alderley Park and Alex Alderton, Roberto Amato, Jeffrey Barrett, Sonia Goncalves, Ewan Harrison, David K. Jackson, Ian Johnston, Dominic Kwiatkowski, Cordelia Langford, John Sillitoe on behalf of the Wellcome Sanger Institute COVID-19 Surveillance Team |
| EPI_ISL_2973264 | Lighthouse Lab in Milton Keynes | Wellcome Sanger Institute for the COVID-19 Genomics UK (COG-UK) Consortium | The Lighthouse Lab in Milton Keynes and Alex Alderton, Roberto Amato, Jeffrey Barrett, Sonia Goncalves, Ewan Harrison, David K. Jackson, Ian Johnston, Dominic Kwiatkowski, Cordelia Langford, John Sillitoe on behalf of the Wellcome Sanger Institute COVID-19 Surveillance Team |
| EPI_ISL_2973265 | Lighthouse Laboratory Plymouth | Wellcome Sanger Institute for the COVID-19 Genomics UK (COG-UK) Consortium | Lighthouse Laboratory Plymouth and Alex Alderton, Roberto Amato, Jeffrey Barrett, Sonia Goncalves, Ewan Harrison, David K. Jackson, Ian Johnston, Dominic Kwiatkowski, Cordelia Langford, John Sillitoe on behalf of the Wellcome Sanger Institute COVID-19 Surveillance Team |
| EPI_ISL_2973266 | Lighthouse Lab in Milton Keynes | Wellcome Sanger Institute for the COVID-19 Genomics UK (COG-UK) Consortium | The Lighthouse Lab in Milton Keynes and Alex Alderton, Roberto Amato, Jeffrey Barrett, Sonia Goncalves, Ewan Harrison, David K. Jackson, Ian Johnston, Dominic Kwiatkowski, Cordelia Langford, John Sillitoe on behalf of the Wellcome Sanger Institute COVID-19 Surveillance Team |
| EPI_ISL_2973267 | Lighthouse Lab in Alderley Park | Wellcome Sanger Institute for the COVID-19 Genomics UK (COG-UK) Consortium | Jacquelyn Wynn, Mairead Hyland, The Lighthouse Lab in Alderley Park and Alex Alderton, Roberto Amato, Jeffrey Barrett, Sonia Goncalves, Ewan Harrison, David K. Jackson, Ian Johnston, Dominic Kwiatkowski, Cordelia Langford, John Sillitoe on behalf of the Wellcome Sanger Institute COVID-19 Surveillance Team |
| EPI_ISL_2973268 | Lighthouse Lab in Milton Keynes | Wellcome Sanger Institute for the COVID-19 Genomics UK (COG-UK) Consortium | The Lighthouse Lab in Milton Keynes and Alex Alderton, Roberto Amato, Jeffrey Barrett, Sonia Goncalves, Ewan Harrison, David K. Jackson, Ian Johnston, Dominic Kwiatkowski, Cordelia Langford, John Sillitoe on behalf of the Wellcome Sanger Institute COVID-19 Surveillance Team |
| EPI_ISL_2973269 | Lighthouse Lab in Alderley Park | Wellcome Sanger Institute for the COVID-19 Genomics UK (COG-UK) Consortium | Jacquelyn Wynn, Mairead Hyland, The Lighthouse Lab in Alderley Park and Alex Alderton, Roberto Amato, Jeffrey Barrett, Sonia Goncalves, Ewan Harrison, David K. Jackson, Ian Johnston, Dominic Kwiatkowski, Cordelia Langford, John Sillitoe on behalf of the Wellcome Sanger Institute COVID-19 Surveillance Team |

| | | | |
|---|---|---|---|
| EPI_ISL_2973270 | Berkshire and Surrey Pathology Services Lighthouse Laboratory | Wellcome Sanger Institute for the COVID-19 Genomics UK (COG-UK) Consortium | Berkshire and Surrey Pathology Services Lighthouse Laboratory and Alex Alderton, Roberto Amato, Jeffrey Barrett, Sonia Goncalves, Ewan Harrison, David K. Jackson, Ian Johnston, Dominic Kwiatkowski, Cordelia Langford, John Sillitoe on behalf of the Wellcome Sanger Institute COVID-19 Surveillance Team |
| EPI_ISL_2973271 | Lighthouse Lab in Alderley Park | Wellcome Sanger Institute for the COVID-19 Genomics UK (COG-UK) Consortium | Jacquelyn Wynn, Mairead Hyland, The Lighthouse Lab in Alderley Park and Alex Alderton, Roberto Amato, Jeffrey Barrett, Sonia Goncalves, Ewan Harrison, David K. Jackson, Ian Johnston, Dominic Kwiatkowski, Cordelia Langford, John Sillitoe on behalf of the Wellcome Sanger Institute COVID-19 Surveillance Team |
| EPI_ISL_2973272 | Berkshire and Surrey Pathology Services Lighthouse Laboratory | Wellcome Sanger Institute for the COVID-19 Genomics UK (COG-UK) Consortium | Berkshire and Surrey Pathology Services Lighthouse Laboratory and Alex Alderton, Roberto Amato, Jeffrey Barrett, Sonia Goncalves, Ewan Harrison, David K. Jackson, Ian Johnston, Dominic Kwiatkowski, Cordelia Langford, John Sillitoe on behalf of the Wellcome Sanger Institute COVID-19 Surveillance Team |
| EPI_ISL_2973273 | Lighthouse Lab in Alderley Park | Wellcome Sanger Institute for the COVID-19 Genomics UK (COG-UK) Consortium | Jacquelyn Wynn, Mairead Hyland, The Lighthouse Lab in Alderley Park and Alex Alderton, Roberto Amato, Jeffrey Barrett, Sonia Goncalves, Ewan Harrison, David K. Jackson, Ian Johnston, Dominic Kwiatkowski, Cordelia Langford, John Sillitoe on behalf of the Wellcome Sanger Institute COVID-19 Surveillance Team |
| EPI_ISL_2973274 | Lighthouse Laboratory Plymouth | Wellcome Sanger Institute for the COVID-19 Genomics UK (COG-UK) Consortium | Lighthouse Laboratory Plymouth and Alex Alderton, Roberto Amato, Jeffrey Barrett, Sonia Goncalves, Ewan Harrison, David K. Jackson, Ian Johnston, Dominic Kwiatkowski, Cordelia Langford, John Sillitoe on behalf of the Wellcome Sanger Institute COVID-19 Surveillance Team |
| EPI_ISL_2973275, EPI_ISL_2973276, EPI_ISL_2973277 | Lighthouse Lab in Milton Keynes | Wellcome Sanger Institute for the COVID-19 Genomics UK (COG-UK) Consortium | The Lighthouse Lab in Milton Keynes and Alex Alderton, Roberto Amato, Jeffrey Barrett, Sonia Goncalves, Ewan Harrison, David K. Jackson, Ian Johnston, Dominic Kwiatkowski, Cordelia Langford, John Sillitoe on behalf of the Wellcome Sanger Institute COVID-19 Surveillance Team |
| EPI_ISL_2973278 | Lighthouse Laboratory Plymouth | Wellcome Sanger Institute for the COVID-19 Genomics UK (COG-UK) Consortium | Lighthouse Laboratory Plymouth and Alex Alderton, Roberto Amato, Jeffrey Barrett, Sonia Goncalves, Ewan Harrison, David K. Jackson, Ian Johnston, Dominic Kwiatkowski, Cordelia Langford, John Sillitoe on behalf of the Wellcome Sanger Institute COVID-19 Surveillance Team |
| EPI_ISL_2973279 | Lighthouse Lab in Alderley Park | Wellcome Sanger Institute for the COVID-19 Genomics UK (COG-UK) Consortium | Jacquelyn Wynn, Mairead Hyland, The Lighthouse Lab in Alderley Park and Alex Alderton, Roberto Amato, Jeffrey Barrett, Sonia Goncalves, Ewan Harrison, David K. Jackson, Ian Johnston, Dominic Kwiatkowski, Cordelia Langford, John Sillitoe on behalf of the Wellcome Sanger Institute COVID-19 Surveillance Team |
| EPI_ISL_2973280 | Lighthouse Lab in Milton Keynes | Wellcome Sanger Institute for the COVID-19 Genomics UK (COG-UK) Consortium | The Lighthouse Lab in Milton Keynes and Alex Alderton, Roberto Amato, Jeffrey Barrett, Sonia Goncalves, Ewan Harrison, David K. Jackson, Ian Johnston, Dominic Kwiatkowski, Cordelia Langford, John Sillitoe on behalf of the Wellcome Sanger Institute COVID-19 Surveillance Team |
| EPI_ISL_2973281 | Lighthouse Laboratory Plymouth | Wellcome Sanger Institute for the COVID-19 Genomics UK (COG-UK) Consortium | Lighthouse Laboratory Plymouth and Alex Alderton, Roberto Amato, Jeffrey Barrett, Sonia Goncalves, Ewan Harrison, David K. Jackson, Ian Johnston, Dominic Kwiatkowski, Cordelia Langford, John Sillitoe on behalf of the Wellcome Sanger Institute COVID-19 Surveillance Team |
| EPI_ISL_2973282 | Lighthouse Lab in Milton Keynes | Wellcome Sanger Institute for the COVID-19 Genomics UK (COG-UK) Consortium | The Lighthouse Lab in Milton Keynes and Alex Alderton, Roberto Amato, Jeffrey Barrett, Sonia Goncalves, Ewan Harrison, David K. Jackson, Ian Johnston, Dominic Kwiatkowski, Cordelia Langford, John Sillitoe on behalf of the Wellcome Sanger Institute COVID-19 Surveillance Team |
| EPI_ISL_2973283, EPI_ISL_2973284 | Berkshire and Surrey Pathology Services Lighthouse Laboratory | Wellcome Sanger Institute for the COVID-19 Genomics UK (COG-UK) Consortium | Berkshire and Surrey Pathology Services Lighthouse Laboratory and Alex Alderton, Roberto Amato, Jeffrey Barrett, Sonia Goncalves, Ewan Harrison, David K. Jackson, Ian Johnston, Dominic Kwiatkowski, Cordelia Langford, John Sillitoe on behalf of the Wellcome Sanger Institute COVID-19 Surveillance Team |
| EPI_ISL_2973285, EPI_ISL_2973286, EPI_ISL_2973287, EPI_ISL_2973289 | Lighthouse Laboratory Plymouth | Wellcome Sanger Institute for the COVID-19 Genomics UK (COG-UK) Consortium | Lighthouse Laboratory Plymouth and Alex Alderton, Roberto Amato, Jeffrey Barrett, Sonia Goncalves, Ewan Harrison, David K. Jackson, Ian Johnston, Dominic Kwiatkowski, Cordelia Langford, John Sillitoe on behalf of the Wellcome Sanger Institute COVID-19 Surveillance Team |
| EPI_ISL_2973290 | Lighthouse Lab in Milton Keynes | Wellcome Sanger Institute for the COVID-19 Genomics UK (COG-UK) Consortium | The Lighthouse Lab in Milton Keynes and Alex Alderton, Roberto Amato, Jeffrey Barrett, Sonia Goncalves, Ewan Harrison, David K. Jackson, Ian Johnston, Dominic Kwiatkowski, Cordelia Langford, John Sillitoe on behalf of the Wellcome Sanger Institute COVID-19 Surveillance Team |
| EPI_ISL_2973291 | Berkshire and Surrey Pathology Services Lighthouse Laboratory | Wellcome Sanger Institute for the COVID-19 Genomics UK (COG-UK) Consortium | Berkshire and Surrey Pathology Services Lighthouse Laboratory and Alex Alderton, Roberto Amato, Jeffrey Barrett, Sonia Goncalves, Ewan Harrison, David K. Jackson, Ian Johnston, Dominic Kwiatkowski, Cordelia Langford, John Sillitoe on behalf of the Wellcome Sanger Institute COVID-19 Surveillance Team |
| EPI_ISL_2973292 | Lighthouse Laboratory Plymouth | Wellcome Sanger Institute for the COVID-19 Genomics UK (COG-UK) Consortium | Lighthouse Laboratory Plymouth and Alex Alderton, Roberto Amato, Jeffrey Barrett, Sonia Goncalves, Ewan Harrison, David K. Jackson, Ian Johnston, Dominic Kwiatkowski, Cordelia Langford, John Sillitoe on behalf of the Wellcome Sanger Institute COVID-19 Surveillance Team |
| EPI_ISL_2973293 | Berkshire and Surrey Pathology Services Lighthouse Laboratory | Wellcome Sanger Institute for the COVID-19 Genomics UK (COG-UK) Consortium | Berkshire and Surrey Pathology Services Lighthouse Laboratory and Alex Alderton, Roberto Amato, Jeffrey Barrett, Sonia Goncalves, Ewan Harrison, David K. Jackson, Ian Johnston, Dominic Kwiatkowski, Cordelia Langford, John Sillitoe on behalf of the Wellcome Sanger Institute COVID-19 Surveillance Team |
| EPI_ISL_2973294 | Lighthouse Lab in Milton Keynes | Wellcome Sanger Institute for the COVID-19 Genomics UK (COG-UK) Consortium | The Lighthouse Lab in Milton Keynes and Alex Alderton, Roberto Amato, Jeffrey Barrett, Sonia Goncalves, Ewan Harrison, David K. Jackson, Ian Johnston, Dominic Kwiatkowski, Cordelia Langford, John Sillitoe on behalf of the Wellcome Sanger Institute COVID-19 Surveillance Team |
| EPI_ISL_2973295, EPI_ISL_2973296 | Lighthouse Lab in Alderley Park | Wellcome Sanger Institute for the COVID-19 Genomics UK (COG-UK) Consortium | Jacquelyn Wynn, Mairead Hyland, The Lighthouse Lab in Alderley Park and Alex Alderton, Roberto Amato, Jeffrey Barrett, Sonia Goncalves, Ewan Harrison, David K. Jackson, Ian Johnston, Dominic Kwiatkowski, Cordelia Langford, John Sillitoe on behalf of the Wellcome Sanger Institute COVID-19 Surveillance Team |
| EPI_ISL_2973297 | Lighthouse Lab in Milton Keynes | Wellcome Sanger Institute for the COVID-19 Genomics UK (COG-UK) Consortium | The Lighthouse Lab in Milton Keynes and Alex Alderton, Roberto Amato, Jeffrey Barrett, Sonia Goncalves, Ewan Harrison, David K. Jackson, Ian Johnston, Dominic Kwiatkowski, Cordelia Langford, John Sillitoe on behalf of the Wellcome Sanger Institute COVID-19 Surveillance Team |
| EPI_ISL_2973298, EPI_ISL_2973299 | Lighthouse Laboratory Plymouth | Wellcome Sanger Institute for the COVID-19 Genomics UK (COG-UK) Consortium | Lighthouse Laboratory Plymouth and Alex Alderton, Roberto Amato, Jeffrey Barrett, Sonia Goncalves, Ewan Harrison, David K. Jackson, Ian Johnston, Dominic Kwiatkowski, Cordelia Langford, John Sillitoe on behalf of the Wellcome Sanger Institute COVID-19 Surveillance Team |
| EPI_ISL_2973300 | Molecular Diagnostics Pathology Department Mater Dei Hospital Malta | Molecular Diagnostics Pathology Department Mater Dei Hospital Malta | G Zahra, R Borg, C Cilia, L Grech, M Briffa |
| EPI_ISL_2973301 | Lighthouse Lab in Alderley Park | Wellcome Sanger Institute for the COVID-19 Genomics UK (COG-UK) Consortium | Jacquelyn Wynn, Mairead Hyland, The Lighthouse Lab in Alderley Park and Alex Alderton, Roberto Amato, Jeffrey Barrett, Sonia Goncalves, Ewan Harrison, David K. Jackson, Ian Johnston, Dominic Kwiatkowski, Cordelia Langford, John Sillitoe on behalf of the Wellcome Sanger Institute COVID-19 Surveillance Team |
| EPI_ISL_2973302 | Lighthouse Lab in Milton Keynes | Wellcome Sanger Institute for the COVID-19 Genomics UK (COG-UK) Consortium | The Lighthouse Lab in Milton Keynes and Alex Alderton, Roberto Amato, Jeffrey Barrett, Sonia Goncalves, Ewan Harrison, David K. Jackson, Ian Johnston, Dominic Kwiatkowski, Cordelia Langford, John Sillitoe on behalf of the Wellcome Sanger Institute COVID-19 Surveillance Team |
| EPI_ISL_2973303 | Lighthouse Laboratory Plymouth | Wellcome Sanger Institute for the COVID-19 Genomics UK (COG-UK) Consortium | Lighthouse Laboratory Plymouth and Alex Alderton, Roberto Amato, Jeffrey Barrett, Sonia Goncalves, Ewan Harrison, David K. Jackson, Ian Johnston, Dominic Kwiatkowski, Cordelia Langford, John Sillitoe on behalf of the Wellcome Sanger Institute COVID-19 Surveillance Team |
| EPI_ISL_2973304 | Lighthouse Lab in Alderley Park | Wellcome Sanger Institute for the COVID-19 Genomics UK (COG-UK) Consortium | Jacquelyn Wynn, Mairead Hyland, The Lighthouse Lab in Alderley Park and Alex Alderton, Roberto Amato, Jeffrey Barrett, Sonia Goncalves, Ewan Harrison, David K. Jackson, Ian Johnston, Dominic Kwiatkowski, Cordelia Langford, John Sillitoe on behalf of the Wellcome Sanger Institute COVID-19 Surveillance Team |
| EPI_ISL_2973305 | Lighthouse Lab in Milton Keynes | Wellcome Sanger Institute for the COVID-19 Genomics UK (COG-UK) Consortium | The Lighthouse Lab in Milton Keynes and Alex Alderton, Roberto Amato, Jeffrey Barrett, Sonia Goncalves, Ewan Harrison, David K. Jackson, Ian Johnston, Dominic Kwiatkowski, Cordelia Langford, John Sillitoe on behalf of the Wellcome Sanger Institute COVID-19 Surveillance Team |
| EPI_ISL_2973306, EPI_ISL_2973307 | Lighthouse Lab in Alderley Park | Wellcome Sanger Institute for the COVID-19 Genomics UK (COG-UK) Consortium | Jacquelyn Wynn, Mairead Hyland, The Lighthouse Lab in Alderley Park and Alex Alderton, Roberto Amato, Jeffrey Barrett, Sonia Goncalves, Ewan Harrison, David K. Jackson, Ian Johnston, Dominic Kwiatkowski, Cordelia Langford, John Sillitoe on behalf of the Wellcome Sanger Institute COVID-19 Surveillance Team |
| EPI_ISL_2973308, EPI_ISL_2973309, EPI_ISL_2973310, EPI_ISL_2973311, EPI_ISL_2973312, EPI_ISL_2973313, EPI_ISL_2973314, EPI_ISL_2973315, EPI_ISL_2973316, EPI_ISL_2973317, EPI_ISL_2973318, EPI_ISL_2973319, EPI_ISL_2973320, EPI_ISL_2973321, EPI_ISL_2973322, EPI_ISL_2973323, EPI_ISL_2973324, EPI_ISL_2973325, EPI_ISL_2973326, EPI_ISL_2973327, EPI_ISL_2973328, EPI_ISL_2973329, EPI_ISL_2973330, EPI_ISL_2973331, EPI_ISL_2973332, EPI_ISL_2973333, EPI_ISL_2973334, EPI_ISL_2973335, EPI_ISL_2973336, EPI_ISL_2973337, EPI_ISL_2973338, EPI_ISL_2973339, EPI_ISL_2973340, EPI_ISL_2973341, EPI_ISL_2973342, EPI_ISL_2973343, EPI_ISL_2973344, EPI_ISL_2973345, EPI_ISL_2973346, EPI_ISL_2973347, EPI_ISL_2973348, EPI_ISL_2973349, EPI_ISL_2973350, EPI_ISL_2973351, EPI_ISL_2973352, EPI_ISL_2973353, EPI_ISL_2973354, EPI_ISL_2973355, EPI_ISL_2973356, EPI_ISL_2973357, EPI_ISL_2973358, EPI_ISL_2973359, EPI_ISL_2973360, EPI_ISL_2973361, EPI_ISL_2973362, EPI_ISL_2973363, EPI_ISL_2973364, EPI_ISL_2973365, EPI_ISL_2973366, EPI_ISL_2973367, EPI_ISL_2973368, EPI_ISL_2973369, EPI_ISL_2973370, EPI_ISL_2973371, EPI_ISL_2973372, EPI_ISL_2973373, EPI_ISL_2973374, EPI_ISL_2973375, EPI_ISL_2973376, EPI_ISL_2973377, EPI_ISL_2973378, EPI_ISL_2973379, EPI_ISL_2973380, EPI_ISL_2973381, EPI_ISL_2973382, EPI_ISL_2973383, EPI_ISL_2973384, EPI_ISL_2973385, EPI_ISL_2973386, EPI_ISL_2973387, EPI_ISL_2973388, EPI_ISL_2973389, EPI_ISL_2973390, EPI_ISL_2973391, EPI_ISL_2973392, EPI_ISL_2973393, EPI_ISL_2973394, EPI_ISL_2973395, EPI_ISL_2973396, EPI_ISL_2973397, EPI_ISL_2973398, EPI_ISL_2973399, EPI_ISL_2973400, EPI_ISL_2973401, EPI_ISL_2973402, EPI_ISL_2973403, EPI_ISL_2973404, EPI_ISL_2973405, EPI_ISL_2973406, EPI_ISL_2973407, EPI_ISL_2973408, EPI_ISL_2973409, EPI_ISL_2973410, EPI_ISL_2973412, EPI_ISL_2973413, EPI_ISL_2973414, EPI_ISL_2973415, EPI_ISL_2973416, EPI_ISL_2973417, EPI_ISL_2973418, EPI_ISL_2973419, EPI_ISL_2973420, EPI_ISL_2973421, EPI_ISL_2973422, EPI_ISL_2973423, EPI_ISL_2973424, EPI_ISL_2973425, EPI_ISL_2973426, EPI_ISL_2973427, EPI_ISL_2973428, EPI_ISL_2973429, EPI_ISL_2973430, EPI_ISL_2973431, EPI_ISL_2973432, EPI_ISL_2973433, EPI_ISL_2973434, EPI_ISL_2973435, EPI_ISL_2973436, EPI_ISL_2973437, EPI_ISL_2973438, EPI_ISL_2973439, EPI_ISL_2973440, EPI_ISL_2973441, EPI_ISL_2973442, EPI_ISL_2973443, EPI_ISL_2973444, EPI_ISL_2973445, EPI_ISL_2973446, EPI_ISL_2973447, EPI_ISL_2973448, EPI_ISL_2973449, EPI_ISL_2973450, EPI_ISL_2973451, EPI_ISL_2973452, EPI_ISL_2973453, EPI_ISL_2973454, EPI_ISL_2973455, EPI_ISL_2973456, EPI_ISL_2973457, EPI_ISL_2973458, EPI_ISL_2973459, EPI_ISL_2973460, EPI_ISL_2973461, EPI_ISL_2973462, EPI_ISL_2973463, EPI_ISL_2973464, EPI_ISL_2973465, EPI_ISL_2973466, EPI_ISL_2973467, EPI_ISL_2973468, EPI_ISL_2973469, EPI_ISL_2973470, EPI_ISL_2973471, EPI_ISL_2973472, EPI_ISL_2973473, EPI_ISL_2973474, EPI_ISL_2973475, EPI_ISL_2973476, EPI_ISL_2973477, EPI_ISL_2973478, EPI_ISL_2973479, EPI_ISL_2973480, EPI_ISL_2973481, EPI_ISL_2973482, EPI_ISL_2973483, EPI_ISL_2973484, EPI_ISL_2973485, EPI_ISL_2973486, EPI_ISL_2973487, EPI_ISL_2973488, EPI_ISL_2973489, EPI_ISL_2973490, EPI_ISL_2973491, EPI_ISL_2973492, EPI_ISL_2973493, EPI_ISL_2973494, EPI_ISL_2973495, EPI_ISL_2973496, EPI_ISL_2973497, EPI_ISL_2973498, EPI_ISL_2973499, EPI_ISL_2973500, EPI_ISL_2973501 | | | |
| see above | Lighthouse Lab in Glasgow | Wellcome Sanger Institute for the COVID-19 Genomics UK (COG-UK) Consortium | Harper VanSteenhouse, Yumi Kasai, David Gray, Carol Clugston, Anna Dominiczak and Alex Alderton, Roberto Amato, Jeffrey Barrett, Sonia Goncalves, Ewan Harrison, David K. Jackson, Ian Johnston, Dominic Kwiatkowski, Cordelia Langford, John Sillitoe on behalf of the Wellcome Sanger Institute COVID-19 Surveillance Team |
| EPI_ISL_2973502 | Lighthouse Lab in Alderley Park | Wellcome Sanger Institute for the COVID-19 Genomics UK (COG-UK) Consortium | Jacquelyn Wynn, Mairead Hyland, The Lighthouse Lab in Alderley Park and Alex Alderton, Roberto Amato, Jeffrey Barrett, Sonia Goncalves, Ewan Harrison, David K. Jackson, Ian Johnston, Dominic Kwiatkowski, Cordelia Langford, John Sillitoe on behalf of the Wellcome Sanger Institute COVID-19 Surveillance Team |
| EPI_ISL_2973503, EPI_ISL_2973504, EPI_ISL_2973505 | Lighthouse Lab in Glasgow | Wellcome Sanger Institute for the COVID-19 Genomics UK (COG-UK) Consortium | Harper VanSteenhouse, Yumi Kasai, David Gray, Carol Clugston, Anna Dominiczak and Alex Alderton, Roberto Amato, Jeffrey Barrett, Sonia Goncalves, Ewan Harrison, David K. Jackson, Ian Johnston, Dominic Kwiatkowski, Cordelia Langford, John Sillitoe on behalf of the Wellcome Sanger Institute COVID-19 Surveillance Team |
| EPI_ISL_2973506 | Lighthouse Lab in Alderley Park | Wellcome Sanger Institute for the COVID-19 Genomics UK (COG-UK) Consortium | Jacquelyn Wynn, Mairead Hyland, The Lighthouse Lab in Alderley Park and Alex Alderton, Roberto Amato, Jeffrey Barrett, Sonia Goncalves, Ewan Harrison, David K. Jackson, Ian Johnston, Dominic Kwiatkowski, Cordelia Langford, John Sillitoe on behalf of the Wellcome Sanger Institute COVID-19 Surveillance Team |
| EPI_ISL_2973507, EPI_ISL_2973508 | Lighthouse Lab in Glasgow | Wellcome Sanger Institute for the COVID-19 Genomics UK (COG-UK) Consortium | Harper VanSteenhouse, Yumi Kasai, David Gray, Carol Clugston, Anna Dominiczak and Alex Alderton, Roberto Amato, Jeffrey Barrett, Sonia Goncalves, Ewan Harrison, David K. Jackson, Ian Johnston, Dominic Kwiatkowski, Cordelia Langford, John Sillitoe on behalf of the Wellcome Sanger Institute COVID-19 Surveillance Team |
| EPI_ISL_2973509 | Lighthouse Lab in Alderley Park | Wellcome Sanger Institute for the COVID-19 Genomics UK (COG-UK) Consortium | Jacquelyn Wynn, Mairead Hyland, The Lighthouse Lab in Alderley Park and Alex Alderton, Roberto Amato, Jeffrey Barrett, Sonia Goncalves, Ewan Harrison, David K. Jackson, Ian Johnston, Dominic Kwiatkowski, Cordelia Langford, John Sillitoe on behalf of the Wellcome Sanger Institute COVID-19 Surveillance Team |
| EPI_ISL_2973510 | Lighthouse Lab in Glasgow | Wellcome Sanger Institute for the COVID-19 Genomics UK (COG-UK) Consortium | Harper VanSteenhouse, Yumi Kasai, David Gray, Carol Clugston, Anna Dominiczak and Alex Alderton, Roberto Amato, Jeffrey Barrett, Sonia Goncalves, Ewan Harrison, David K. Jackson, Ian Johnston, Dominic Kwiatkowski, Cordelia Langford, John Sillitoe on behalf of the Wellcome Sanger Institute COVID-19 Surveillance Team |
| EPI_ISL_2973511, EPI_ISL_2973512, EPI_ISL_2973513 | Lighthouse Lab in Alderley Park | Wellcome Sanger Institute for the COVID-19 Genomics UK (COG-UK) Consortium | Jacquelyn Wynn, Mairead Hyland, The Lighthouse Lab in Alderley Park and Alex Alderton, Roberto Amato, Jeffrey Barrett, Sonia Goncalves, Ewan Harrison, David K. Jackson, Ian Johnston, Dominic Kwiatkowski, Cordelia Langford, John Sillitoe on behalf of the Wellcome Sanger Institute COVID-19 Surveillance Team |
| EPI_ISL_2973514, EPI_ISL_2973515, EPI_ISL_2973516, EPI_ISL_2973517, EPI_ISL_2973518 | Lighthouse Lab in Glasgow | Wellcome Sanger Institute for the COVID-19 Genomics UK (COG-UK) Consortium | Harper VanSteenhouse, Yumi Kasai, David Gray, Carol Clugston, Anna Dominiczak and Alex Alderton, Roberto Amato, Jeffrey Barrett, Sonia Goncalves, Ewan Harrison, David K. Jackson, Ian Johnston, Dominic Kwiatkowski, Cordelia Langford, John Sillitoe on behalf of the Wellcome Sanger Institute COVID-19 Surveillance Team |
| EPI_ISL_2973519 | Lighthouse Lab in Alderley Park | Wellcome Sanger Institute for the COVID-19 Genomics UK (COG-UK) Consortium | Jacquelyn Wynn, Mairead Hyland, The Lighthouse Lab in Alderley Park and Alex Alderton, Roberto Amato, Jeffrey Barrett, Sonia Goncalves, Ewan Harrison, David K. Jackson, Ian Johnston, Dominic Kwiatkowski, Cordelia Langford, John Sillitoe on behalf of the Wellcome Sanger Institute COVID-19 Surveillance Team |

| | | | |
|---|---|---|---|
| EPI_ISL_2973520, EPI_ISL_2973521 | Lighthouse Lab in Glasgow | Wellcome Sanger Institute for the COVID-19 Genomics UK (COG-UK) Consortium | Harper VanSteenhouse, Yumi Kasai, David Gray, Carol Clugston, Anna Dominiczak and Alex Alderton, Roberto Amato, Jeffrey Barrett, Sonia Goncalves, Ewan Harrison, David K. Jackson, Ian Johnston, Dominic Kwiatkowski, Cordelia Langford, John Sillitoe on behalf of the Wellcome Sanger Institute COVID-19 Surveillance Team |
| EPI_ISL_2973522, EPI_ISL_2973523 | Lighthouse Lab in Alderley Park | Wellcome Sanger Institute for the COVID-19 Genomics UK (COG-UK) Consortium | Jacquelyn Wynn, Mairead Hyland, The Lighthouse Lab in Alderley Park and Alex Alderton, Roberto Amato, Jeffrey Barrett, Sonia Goncalves, Ewan Harrison, David K. Jackson, Ian Johnston, Dominic Kwiatkowski, Cordelia Langford, John Sillitoe on behalf of the Wellcome Sanger Institute COVID-19 Surveillance Team |
| EPI_ISL_2973524 | Lighthouse Lab in Glasgow | Wellcome Sanger Institute for the COVID-19 Genomics UK (COG-UK) Consortium | Harper VanSteenhouse, Yumi Kasai, David Gray, Carol Clugston, Anna Dominiczak and Alex Alderton, Roberto Amato, Jeffrey Barrett, Sonia Goncalves, Ewan Harrison, David K. Jackson, Ian Johnston, Dominic Kwiatkowski, Cordelia Langford, John Sillitoe on behalf of the Wellcome Sanger Institute COVID-19 Surveillance Team |
| EPI_ISL_2973525 | Lighthouse Lab in Alderley Park | Wellcome Sanger Institute for the COVID-19 Genomics UK (COG-UK) Consortium | Jacquelyn Wynn, Mairead Hyland, The Lighthouse Lab in Alderley Park and Alex Alderton, Roberto Amato, Jeffrey Barrett, Sonia Goncalves, Ewan Harrison, David K. Jackson, Ian Johnston, Dominic Kwiatkowski, Cordelia Langford, John Sillitoe on behalf of the Wellcome Sanger Institute COVID-19 Surveillance Team |
| EPI_ISL_2973526, EPI_ISL_2973527, EPI_ISL_2973528, EPI_ISL_2973529, EPI_ISL_2973530, EPI_ISL_2973531 | Lighthouse Lab in Glasgow | Wellcome Sanger Institute for the COVID-19 Genomics UK (COG-UK) Consortium | Harper VanSteenhouse, Yumi Kasai, David Gray, Carol Clugston, Anna Dominiczak and Alex Alderton, Roberto Amato, Jeffrey Barrett, Sonia Goncalves, Ewan Harrison, David K. Jackson, Ian Johnston, Dominic Kwiatkowski, Cordelia Langford, John Sillitoe on behalf of the Wellcome Sanger Institute COVID-19 Surveillance Team |
| EPI_ISL_2973532, EPI_ISL_2973533, EPI_ISL_2973534, EPI_ISL_2973535, EPI_ISL_2973536 | Lighthouse Lab in Alderley Park | Wellcome Sanger Institute for the COVID-19 Genomics UK (COG-UK) Consortium | Jacquelyn Wynn, Mairead Hyland, The Lighthouse Lab in Alderley Park and Alex Alderton, Roberto Amato, Jeffrey Barrett, Sonia Goncalves, Ewan Harrison, David K. Jackson, Ian Johnston, Dominic Kwiatkowski, Cordelia Langford, John Sillitoe on behalf of the Wellcome Sanger Institute COVID-19 Surveillance Team |
| EPI_ISL_2973537, EPI_ISL_2973538, EPI_ISL_2973539, EPI_ISL_2973540 | Lighthouse Lab in Glasgow | Wellcome Sanger Institute for the COVID-19 Genomics UK (COG-UK) Consortium | Harper VanSteenhouse, Yumi Kasai, David Gray, Carol Clugston, Anna Dominiczak and Alex Alderton, Roberto Amato, Jeffrey Barrett, Sonia Goncalves, Ewan Harrison, David K. Jackson, Ian Johnston, Dominic Kwiatkowski, Cordelia Langford, John Sillitoe on behalf of the Wellcome Sanger Institute COVID-19 Surveillance Team |
| EPI_ISL_2973541 | Lighthouse Lab in Alderley Park | Wellcome Sanger Institute for the COVID-19 Genomics UK (COG-UK) Consortium | Jacquelyn Wynn, Mairead Hyland, The Lighthouse Lab in Alderley Park and Alex Alderton, Roberto Amato, Jeffrey Barrett, Sonia Goncalves, Ewan Harrison, David K. Jackson, Ian Johnston, Dominic Kwiatkowski, Cordelia Langford, John Sillitoe on behalf of the Wellcome Sanger Institute COVID-19 Surveillance Team |
| EPI_ISL_2973542, EPI_ISL_2973543 | Lighthouse Lab in Glasgow | Wellcome Sanger Institute for the COVID-19 Genomics UK (COG-UK) Consortium | Harper VanSteenhouse, Yumi Kasai, David Gray, Carol Clugston, Anna Dominiczak and Alex Alderton, Roberto Amato, Jeffrey Barrett, Sonia Goncalves, Ewan Harrison, David K. Jackson, Ian Johnston, Dominic Kwiatkowski, Cordelia Langford, John Sillitoe on behalf of the Wellcome Sanger Institute COVID-19 Surveillance Team |
| EPI_ISL_2973545, EPI_ISL_2973546 | Lighthouse Lab in Alderley Park | Wellcome Sanger Institute for the COVID-19 Genomics UK (COG-UK) Consortium | Jacquelyn Wynn, Mairead Hyland, The Lighthouse Lab in Alderley Park and Alex Alderton, Roberto Amato, Jeffrey Barrett, Sonia Goncalves, Ewan Harrison, David K. Jackson, Ian Johnston, Dominic Kwiatkowski, Cordelia Langford, John Sillitoe on behalf of the Wellcome Sanger Institute COVID-19 Surveillance Team |
| EPI_ISL_2973547, EPI_ISL_2973548, EPI_ISL_2973549, EPI_ISL_2973550 | Lighthouse Lab in Glasgow | Wellcome Sanger Institute for the COVID-19 Genomics UK (COG-UK) Consortium | Harper VanSteenhouse, Yumi Kasai, David Gray, Carol Clugston, Anna Dominiczak and Alex Alderton, Roberto Amato, Jeffrey Barrett, Sonia Goncalves, Ewan Harrison, David K. Jackson, Ian Johnston, Dominic Kwiatkowski, Cordelia Langford, John Sillitoe on behalf of the Wellcome Sanger Institute COVID-19 Surveillance Team |
| EPI_ISL_2973551, EPI_ISL_2973552, EPI_ISL_2973553, EPI_ISL_2973554, EPI_ISL_2973555 | Lighthouse Lab in Alderley Park | Wellcome Sanger Institute for the COVID-19 Genomics UK (COG-UK) Consortium | Jacquelyn Wynn, Mairead Hyland, The Lighthouse Lab in Alderley Park and Alex Alderton, Roberto Amato, Jeffrey Barrett, Sonia Goncalves, Ewan Harrison, David K. Jackson, Ian Johnston, Dominic Kwiatkowski, Cordelia Langford, John Sillitoe on behalf of the Wellcome Sanger Institute COVID-19 Surveillance Team |
| EPI_ISL_2973556, EPI_ISL_2973557 | Lighthouse Lab in Glasgow | Wellcome Sanger Institute for the COVID-19 Genomics UK (COG-UK) Consortium | Harper VanSteenhouse, Yumi Kasai, David Gray, Carol Clugston, Anna Dominiczak and Alex Alderton, Roberto Amato, Jeffrey Barrett, Sonia Goncalves, Ewan Harrison, David K. Jackson, Ian Johnston, Dominic Kwiatkowski, Cordelia Langford, John Sillitoe on behalf of the Wellcome Sanger Institute COVID-19 Surveillance Team |
| EPI_ISL_2973558 | Lighthouse Lab in Alderley Park | Wellcome Sanger Institute for the COVID-19 Genomics UK (COG-UK) Consortium | Jacquelyn Wynn, Mairead Hyland, The Lighthouse Lab in Alderley Park and Alex Alderton, Roberto Amato, Jeffrey Barrett, Sonia Goncalves, Ewan Harrison, David K. Jackson, Ian Johnston, Dominic Kwiatkowski, Cordelia Langford, John Sillitoe on behalf of the Wellcome Sanger Institute COVID-19 Surveillance Team |
| EPI_ISL_2973559 | Lighthouse Lab in Glasgow | Wellcome Sanger Institute for the COVID-19 Genomics UK (COG-UK) Consortium | Harper VanSteenhouse, Yumi Kasai, David Gray, Carol Clugston, Anna Dominiczak and Alex Alderton, Roberto Amato, Jeffrey Barrett, Sonia Goncalves, Ewan Harrison, David K. Jackson, Ian Johnston, Dominic Kwiatkowski, Cordelia Langford, John Sillitoe on behalf of the Wellcome Sanger Institute COVID-19 Surveillance Team |
| EPI_ISL_2973560 | Lighthouse Lab in Alderley Park | Wellcome Sanger Institute for the COVID-19 Genomics UK (COG-UK) Consortium | Jacquelyn Wynn, Mairead Hyland, The Lighthouse Lab in Alderley Park and Alex Alderton, Roberto Amato, Jeffrey Barrett, Sonia Goncalves, Ewan Harrison, David K. Jackson, Ian Johnston, Dominic Kwiatkowski, Cordelia Langford, John Sillitoe on behalf of the Wellcome Sanger Institute COVID-19 Surveillance Team |
| EPI_ISL_2973561, EPI_ISL_2973562, EPI_ISL_2973563 | Lighthouse Lab in Glasgow | Wellcome Sanger Institute for the COVID-19 Genomics UK (COG-UK) Consortium | Harper VanSteenhouse, Yumi Kasai, David Gray, Carol Clugston, Anna Dominiczak and Alex Alderton, Roberto Amato, Jeffrey Barrett, Sonia Goncalves, Ewan Harrison, David K. Jackson, Ian Johnston, Dominic Kwiatkowski, Cordelia Langford, John Sillitoe on behalf of the Wellcome Sanger Institute COVID-19 Surveillance Team |
| EPI_ISL_2973564, EPI_ISL_2973565 | Lighthouse Lab in Alderley Park | Wellcome Sanger Institute for the COVID-19 Genomics UK (COG-UK) Consortium | Jacquelyn Wynn, Mairead Hyland, The Lighthouse Lab in Alderley Park and Alex Alderton, Roberto Amato, Jeffrey Barrett, Sonia Goncalves, Ewan Harrison, David K. Jackson, Ian Johnston, Dominic Kwiatkowski, Cordelia Langford, John Sillitoe on behalf of the Wellcome Sanger Institute COVID-19 Surveillance Team |
| EPI_ISL_2973566, EPI_ISL_2973567 | Lighthouse Lab in Glasgow | Wellcome Sanger Institute for the COVID-19 Genomics UK (COG-UK) Consortium | Harper VanSteenhouse, Yumi Kasai, David Gray, Carol Clugston, Anna Dominiczak and Alex Alderton, Roberto Amato, Jeffrey Barrett, Sonia Goncalves, Ewan Harrison, David K. Jackson, Ian Johnston, Dominic Kwiatkowski, Cordelia Langford, John Sillitoe on behalf of the Wellcome Sanger Institute COVID-19 Surveillance Team |
| EPI_ISL_2973568 | Lighthouse Lab in Alderley Park | Wellcome Sanger Institute for the COVID-19 Genomics UK (COG-UK) Consortium | Jacquelyn Wynn, Mairead Hyland, The Lighthouse Lab in Alderley Park and Alex Alderton, Roberto Amato, Jeffrey Barrett, Sonia Goncalves, Ewan Harrison, David K. Jackson, Ian Johnston, Dominic Kwiatkowski, Cordelia Langford, John Sillitoe on behalf of the Wellcome Sanger Institute COVID-19 Surveillance Team |
| EPI_ISL_2973569 | Lighthouse Lab in Glasgow | Wellcome Sanger Institute for the COVID-19 Genomics UK (COG-UK) Consortium | Harper VanSteenhouse, Yumi Kasai, David Gray, Carol Clugston, Anna Dominiczak and Alex Alderton, Roberto Amato, Jeffrey Barrett, Sonia Goncalves, Ewan Harrison, David K. Jackson, Ian Johnston, Dominic Kwiatkowski, Cordelia Langford, John Sillitoe on behalf of the Wellcome Sanger Institute COVID-19 Surveillance Team |
| EPI_ISL_2973570, EPI_ISL_2973571 | Lighthouse Lab in Alderley Park | Wellcome Sanger Institute for the COVID-19 Genomics UK (COG-UK) Consortium | Jacquelyn Wynn, Mairead Hyland, The Lighthouse Lab in Alderley Park and Alex Alderton, Roberto Amato, Jeffrey Barrett, Sonia Goncalves, Ewan Harrison, David K. Jackson, Ian Johnston, Dominic Kwiatkowski, Cordelia Langford, John Sillitoe on behalf of the Wellcome Sanger Institute COVID-19 Surveillance Team |
| EPI_ISL_2973572 | Lighthouse Lab in Glasgow | Wellcome Sanger Institute for the COVID-19 Genomics UK (COG-UK) Consortium | Harper VanSteenhouse, Yumi Kasai, David Gray, Carol Clugston, Anna Dominiczak and Alex Alderton, Roberto Amato, Jeffrey Barrett, Sonia Goncalves, Ewan Harrison, David K. Jackson, Ian Johnston, Dominic Kwiatkowski, Cordelia Langford, John Sillitoe on behalf of the Wellcome Sanger Institute COVID-19 Surveillance Team |
| EPI_ISL_2973573, EPI_ISL_2973574, EPI_ISL_2973575, EPI_ISL_2973576 | Lighthouse Lab in Alderley Park | Wellcome Sanger Institute for the COVID-19 Genomics UK (COG-UK) Consortium | Jacquelyn Wynn, Mairead Hyland, The Lighthouse Lab in Alderley Park and Alex Alderton, Roberto Amato, Jeffrey Barrett, Sonia Goncalves, Ewan Harrison, David K. Jackson, Ian Johnston, Dominic Kwiatkowski, Cordelia Langford, John Sillitoe on behalf of the Wellcome Sanger Institute COVID-19 Surveillance Team |
| EPI_ISL_2973577 | Lighthouse Lab in Glasgow | Wellcome Sanger Institute for the COVID-19 Genomics UK (COG-UK) Consortium | Harper VanSteenhouse, Yumi Kasai, David Gray, Carol Clugston, Anna Dominiczak and Alex Alderton, Roberto Amato, Jeffrey Barrett, Sonia Goncalves, Ewan Harrison, David K. Jackson, Ian Johnston, Dominic Kwiatkowski, Cordelia Langford, John Sillitoe on behalf of the Wellcome Sanger Institute COVID-19 Surveillance Team |
| EPI_ISL_2973578 | Lighthouse Lab in Alderley Park | Wellcome Sanger Institute for the COVID-19 Genomics UK (COG-UK) Consortium | Jacquelyn Wynn, Mairead Hyland, The Lighthouse Lab in Alderley Park and Alex Alderton, Roberto Amato, Jeffrey Barrett, Sonia Goncalves, Ewan Harrison, David K. Jackson, Ian Johnston, Dominic Kwiatkowski, Cordelia Langford, John Sillitoe on behalf of the Wellcome Sanger Institute COVID-19 Surveillance Team |
| EPI_ISL_2973579, EPI_ISL_2973580, EPI_ISL_2973581, EPI_ISL_2973582, EPI_ISL_2973583, EPI_ISL_2973584 | Lighthouse Lab in Glasgow | Wellcome Sanger Institute for the COVID-19 Genomics UK (COG-UK) Consortium | Harper VanSteenhouse, Yumi Kasai, David Gray, Carol Clugston, Anna Dominiczak and Alex Alderton, Roberto Amato, Jeffrey Barrett, Sonia Goncalves, Ewan Harrison, David K. Jackson, Ian Johnston, Dominic Kwiatkowski, Cordelia Langford, John Sillitoe on behalf of the Wellcome Sanger Institute COVID-19 Surveillance Team |
| EPI_ISL_2973585 | Lighthouse Lab in Alderley Park | Wellcome Sanger Institute for the COVID-19 Genomics UK (COG-UK) Consortium | Jacquelyn Wynn, Mairead Hyland, The Lighthouse Lab in Alderley Park and Alex Alderton, Roberto Amato, Jeffrey Barrett, Sonia Goncalves, Ewan Harrison, David K. Jackson, Ian Johnston, Dominic Kwiatkowski, Cordelia Langford, John Sillitoe on behalf of the Wellcome Sanger Institute COVID-19 Surveillance Team |
| EPI_ISL_2973586, EPI_ISL_2973587 | Lighthouse Lab in Glasgow | Wellcome Sanger Institute for the COVID-19 Genomics UK (COG-UK) Consortium | Harper VanSteenhouse, Yumi Kasai, David Gray, Carol Clugston, Anna Dominiczak and Alex Alderton, Roberto Amato, Jeffrey Barrett, Sonia Goncalves, Ewan Harrison, David K. Jackson, Ian Johnston, Dominic Kwiatkowski, Cordelia Langford, John Sillitoe on behalf of the Wellcome Sanger Institute COVID-19 Surveillance Team |
| EPI_ISL_2973588 | Lighthouse Lab in Alderley Park | Wellcome Sanger Institute for the COVID-19 Genomics UK (COG-UK) Consortium | Jacquelyn Wynn, Mairead Hyland, The Lighthouse Lab in Alderley Park and Alex Alderton, Roberto Amato, Jeffrey Barrett, Sonia Goncalves, Ewan Harrison, David K. Jackson, Ian Johnston, Dominic Kwiatkowski, Cordelia Langford, John Sillitoe on behalf of the Wellcome Sanger Institute COVID-19 Surveillance Team |
| EPI_ISL_2973589 | Lighthouse Lab in Glasgow | Wellcome Sanger Institute for the COVID-19 Genomics UK (COG-UK) Consortium | Harper VanSteenhouse, Yumi Kasai, David Gray, Carol Clugston, Anna Dominiczak and Alex Alderton, Roberto Amato, Jeffrey Barrett, Sonia Goncalves, Ewan Harrison, David K. Jackson, Ian Johnston, Dominic Kwiatkowski, Cordelia Langford, John Sillitoe on behalf of the Wellcome Sanger Institute COVID-19 Surveillance Team |
| EPI_ISL_2973590, EPI_ISL_2973591 | Lighthouse Lab in Alderley Park | Wellcome Sanger Institute for the COVID-19 Genomics UK (COG-UK) Consortium | Jacquelyn Wynn, Mairead Hyland, The Lighthouse Lab in Alderley Park and Alex Alderton, Roberto Amato, Jeffrey Barrett, Sonia Goncalves, Ewan Harrison, David K. Jackson, Ian Johnston, Dominic Kwiatkowski, Cordelia Langford, John Sillitoe on behalf of the Wellcome Sanger Institute COVID-19 Surveillance Team |
| EPI_ISL_2973592, EPI_ISL_2973593, EPI_ISL_2973594, EPI_ISL_2973595 | Lighthouse Lab in Glasgow | Wellcome Sanger Institute for the COVID-19 Genomics UK (COG-UK) Consortium | Harper VanSteenhouse, Yumi Kasai, David Gray, Carol Clugston, Anna Dominiczak and Alex Alderton, Roberto Amato, Jeffrey Barrett, Sonia Goncalves, Ewan Harrison, David K. Jackson, Ian Johnston, Dominic Kwiatkowski, Cordelia Langford, John Sillitoe on behalf of the Wellcome Sanger Institute COVID-19 Surveillance Team |
| EPI_ISL_2973596, EPI_ISL_2973597, EPI_ISL_2973598 | Lighthouse Lab in Alderley Park | Wellcome Sanger Institute for the COVID-19 Genomics UK (COG-UK) Consortium | Jacquelyn Wynn, Mairead Hyland, The Lighthouse Lab in Alderley Park and Alex Alderton, Roberto Amato, Jeffrey Barrett, Sonia Goncalves, Ewan Harrison, David K. Jackson, Ian Johnston, Dominic Kwiatkowski, Cordelia Langford, John Sillitoe on behalf of the Wellcome Sanger Institute COVID-19 Surveillance Team |
| EPI_ISL_2973599 | Lighthouse Lab in Glasgow | Wellcome Sanger Institute for the COVID-19 Genomics UK (COG-UK) Consortium | Harper VanSteenhouse, Yumi Kasai, David Gray, Carol Clugston, Anna Dominiczak and Alex Alderton, Roberto Amato, Jeffrey Barrett, Sonia Goncalves, Ewan Harrison, David K. Jackson, Ian Johnston, Dominic Kwiatkowski, Cordelia Langford, John Sillitoe on behalf of the Wellcome Sanger Institute COVID-19 Surveillance Team |

| | | | |
|---|---|---|---|
| EPI_ISL_2973600 | Lighthouse Lab in Alderley Park | Wellcome Sanger Institute for the COVID-19 Genomics UK (COG-UK) Consortium | Jacquelyn Wynn, Mairead Hyland, The Lighthouse Lab in Alderley Park and Alex Alderton, Roberto Amato, Jeffrey Barrett, Sonia Goncalves, Ewan Harrison, David K. Jackson, Ian Johnston, Dominic Kwiatkowski, Cordelia Langford, John Sillitoe on behalf of the Wellcome Sanger Institute COVID-19 Surveillance Team |
| EPI_ISL_2973601, EPI_ISL_2973602, EPI_ISL_2973603, EPI_ISL_2973604, EPI_ISL_2973605 | Lighthouse Lab in Glasgow | Wellcome Sanger Institute for the COVID-19 Genomics UK (COG-UK) Consortium | Harper VanSteenhouse, Yumi Kasai, David Gray, Carol Clugston, Anna Dominiczak and Alex Alderton, Roberto Amato, Jeffrey Barrett, Sonia Goncalves, Ewan Harrison, David K. Jackson, Ian Johnston, Dominic Kwiatkowski, Cordelia Langford, John Sillitoe on behalf of the Wellcome Sanger Institute COVID-19 Surveillance Team |
| EPI_ISL_2973606, EPI_ISL_2973607 | Lighthouse Lab in Alderley Park | Wellcome Sanger Institute for the COVID-19 Genomics UK (COG-UK) Consortium | Jacquelyn Wynn, Mairead Hyland, The Lighthouse Lab in Alderley Park and Alex Alderton, Roberto Amato, Jeffrey Barrett, Sonia Goncalves, Ewan Harrison, David K. Jackson, Ian Johnston, Dominic Kwiatkowski, Cordelia Langford, John Sillitoe on behalf of the Wellcome Sanger Institute COVID-19 Surveillance Team |
| EPI_ISL_2973608, EPI_ISL_2973609, EPI_ISL_2973610, EPI_ISL_2973611, EPI_ISL_2973612 | Lighthouse Lab in Glasgow | Wellcome Sanger Institute for the COVID-19 Genomics UK (COG-UK) Consortium | Harper VanSteenhouse, Yumi Kasai, David Gray, Carol Clugston, Anna Dominiczak and Alex Alderton, Roberto Amato, Jeffrey Barrett, Sonia Goncalves, Ewan Harrison, David K. Jackson, Ian Johnston, Dominic Kwiatkowski, Cordelia Langford, John Sillitoe on behalf of the Wellcome Sanger Institute COVID-19 Surveillance Team |
| EPI_ISL_2973613, EPI_ISL_2973614 | Lighthouse Lab in Alderley Park | Wellcome Sanger Institute for the COVID-19 Genomics UK (COG-UK) Consortium | Jacquelyn Wynn, Mairead Hyland, The Lighthouse Lab in Alderley Park and Alex Alderton, Roberto Amato, Jeffrey Barrett, Sonia Goncalves, Ewan Harrison, David K. Jackson, Ian Johnston, Dominic Kwiatkowski, Cordelia Langford, John Sillitoe on behalf of the Wellcome Sanger Institute COVID-19 Surveillance Team |
| EPI_ISL_2973615, EPI_ISL_2973616 | Lighthouse Lab in Glasgow | Wellcome Sanger Institute for the COVID-19 Genomics UK (COG-UK) Consortium | Harper VanSteenhouse, Yumi Kasai, David Gray, Carol Clugston, Anna Dominiczak and Alex Alderton, Roberto Amato, Jeffrey Barrett, Sonia Goncalves, Ewan Harrison, David K. Jackson, Ian Johnston, Dominic Kwiatkowski, Cordelia Langford, John Sillitoe on behalf of the Wellcome Sanger Institute COVID-19 Surveillance Team |
| EPI_ISL_2973617, EPI_ISL_2973618 | Lighthouse Lab in Alderley Park | Wellcome Sanger Institute for the COVID-19 Genomics UK (COG-UK) Consortium | Jacquelyn Wynn, Mairead Hyland, The Lighthouse Lab in Alderley Park and Alex Alderton, Roberto Amato, Jeffrey Barrett, Sonia Goncalves, Ewan Harrison, David K. Jackson, Ian Johnston, Dominic Kwiatkowski, Cordelia Langford, John Sillitoe on behalf of the Wellcome Sanger Institute COVID-19 Surveillance Team |
| EPI_ISL_2973619, EPI_ISL_2973620 | Lighthouse Lab in Glasgow | Wellcome Sanger Institute for the COVID-19 Genomics UK (COG-UK) Consortium | Harper VanSteenhouse, Yumi Kasai, David Gray, Carol Clugston, Anna Dominiczak and Alex Alderton, Roberto Amato, Jeffrey Barrett, Sonia Goncalves, Ewan Harrison, David K. Jackson, Ian Johnston, Dominic Kwiatkowski, Cordelia Langford, John Sillitoe on behalf of the Wellcome Sanger Institute COVID-19 Surveillance Team |
| EPI_ISL_2973621 | Lighthouse Lab in Alderley Park | Wellcome Sanger Institute for the COVID-19 Genomics UK (COG-UK) Consortium | Jacquelyn Wynn, Mairead Hyland, The Lighthouse Lab in Alderley Park and Alex Alderton, Roberto Amato, Jeffrey Barrett, Sonia Goncalves, Ewan Harrison, David K. Jackson, Ian Johnston, Dominic Kwiatkowski, Cordelia Langford, John Sillitoe on behalf of the Wellcome Sanger Institute COVID-19 Surveillance Team |
| EPI_ISL_2973622, EPI_ISL_2973623, EPI_ISL_2973624 | Lighthouse Lab in Glasgow | Wellcome Sanger Institute for the COVID-19 Genomics UK (COG-UK) Consortium | Harper VanSteenhouse, Yumi Kasai, David Gray, Carol Clugston, Anna Dominiczak and Alex Alderton, Roberto Amato, Jeffrey Barrett, Sonia Goncalves, Ewan Harrison, David K. Jackson, Ian Johnston, Dominic Kwiatkowski, Cordelia Langford, John Sillitoe on behalf of the Wellcome Sanger Institute COVID-19 Surveillance Team |
| EPI_ISL_2973625 | Lighthouse Lab in Alderley Park | Wellcome Sanger Institute for the COVID-19 Genomics UK (COG-UK) Consortium | Jacquelyn Wynn, Mairead Hyland, The Lighthouse Lab in Alderley Park and Alex Alderton, Roberto Amato, Jeffrey Barrett, Sonia Goncalves, Ewan Harrison, David K. Jackson, Ian Johnston, Dominic Kwiatkowski, Cordelia Langford, John Sillitoe on behalf of the Wellcome Sanger Institute COVID-19 Surveillance Team |
| EPI_ISL_2973626, EPI_ISL_2973627, EPI_ISL_2973628, EPI_ISL_2973629, EPI_ISL_2973630, EPI_ISL_2973631 | Lighthouse Lab in Glasgow | Wellcome Sanger Institute for the COVID-19 Genomics UK (COG-UK) Consortium | Harper VanSteenhouse, Yumi Kasai, David Gray, Carol Clugston, Anna Dominiczak and Alex Alderton, Roberto Amato, Jeffrey Barrett, Sonia Goncalves, Ewan Harrison, David K. Jackson, Ian Johnston, Dominic Kwiatkowski, Cordelia Langford, John Sillitoe on behalf of the Wellcome Sanger Institute COVID-19 Surveillance Team |
| EPI_ISL_2973632, EPI_ISL_2973633, EPI_ISL_2973634 | Lighthouse Lab in Alderley Park | Wellcome Sanger Institute for the COVID-19 Genomics UK (COG-UK) Consortium | Jacquelyn Wynn, Mairead Hyland, The Lighthouse Lab in Alderley Park and Alex Alderton, Roberto Amato, Jeffrey Barrett, Sonia Goncalves, Ewan Harrison, David K. Jackson, Ian Johnston, Dominic Kwiatkowski, Cordelia Langford, John Sillitoe on behalf of the Wellcome Sanger Institute COVID-19 Surveillance Team |
| EPI_ISL_2973635 | Lighthouse Lab in Glasgow | Wellcome Sanger Institute for the COVID-19 Genomics UK (COG-UK) Consortium | Harper VanSteenhouse, Yumi Kasai, David Gray, Carol Clugston, Anna Dominiczak and Alex Alderton, Roberto Amato, Jeffrey Barrett, Sonia Goncalves, Ewan Harrison, David K. Jackson, Ian Johnston, Dominic Kwiatkowski, Cordelia Langford, John Sillitoe on behalf of the Wellcome Sanger Institute COVID-19 Surveillance Team |
| EPI_ISL_2973636 | Lighthouse Lab in Alderley Park | Wellcome Sanger Institute for the COVID-19 Genomics UK (COG-UK) Consortium | Jacquelyn Wynn, Mairead Hyland, The Lighthouse Lab in Alderley Park and Alex Alderton, Roberto Amato, Jeffrey Barrett, Sonia Goncalves, Ewan Harrison, David K. Jackson, Ian Johnston, Dominic Kwiatkowski, Cordelia Langford, John Sillitoe on behalf of the Wellcome Sanger Institute COVID-19 Surveillance Team |
| EPI_ISL_2973637, EPI_ISL_2973638 | Lighthouse Lab in Glasgow | Wellcome Sanger Institute for the COVID-19 Genomics UK (COG-UK) Consortium | Harper VanSteenhouse, Yumi Kasai, David Gray, Carol Clugston, Anna Dominiczak and Alex Alderton, Roberto Amato, Jeffrey Barrett, Sonia Goncalves, Ewan Harrison, David K. Jackson, Ian Johnston, Dominic Kwiatkowski, Cordelia Langford, John Sillitoe on behalf of the Wellcome Sanger Institute COVID-19 Surveillance Team |
| EPI_ISL_2973639, EPI_ISL_2973640 | Lighthouse Lab in Alderley Park | Wellcome Sanger Institute for the COVID-19 Genomics UK (COG-UK) Consortium | Jacquelyn Wynn, Mairead Hyland, The Lighthouse Lab in Alderley Park and Alex Alderton, Roberto Amato, Jeffrey Barrett, Sonia Goncalves, Ewan Harrison, David K. Jackson, Ian Johnston, Dominic Kwiatkowski, Cordelia Langford, John Sillitoe on behalf of the Wellcome Sanger Institute COVID-19 Surveillance Team |
| EPI_ISL_2973641, EPI_ISL_2973642, EPI_ISL_2973643 | Lighthouse Lab in Glasgow | Wellcome Sanger Institute for the COVID-19 Genomics UK (COG-UK) Consortium | Harper VanSteenhouse, Yumi Kasai, David Gray, Carol Clugston, Anna Dominiczak and Alex Alderton, Roberto Amato, Jeffrey Barrett, Sonia Goncalves, Ewan Harrison, David K. Jackson, Ian Johnston, Dominic Kwiatkowski, Cordelia Langford, John Sillitoe on behalf of the Wellcome Sanger Institute COVID-19 Surveillance Team |
| EPI_ISL_2973644, EPI_ISL_2973645, EPI_ISL_2973646, EPI_ISL_2973647 | Lighthouse Lab in Alderley Park | Wellcome Sanger Institute for the COVID-19 Genomics UK (COG-UK) Consortium | Jacquelyn Wynn, Mairead Hyland, The Lighthouse Lab in Alderley Park and Alex Alderton, Roberto Amato, Jeffrey Barrett, Sonia Goncalves, Ewan Harrison, David K. Jackson, Ian Johnston, Dominic Kwiatkowski, Cordelia Langford, John Sillitoe on behalf of the Wellcome Sanger Institute COVID-19 Surveillance Team |
| EPI_ISL_2973648 | Lighthouse Lab in Glasgow | Wellcome Sanger Institute for the COVID-19 Genomics UK (COG-UK) Consortium | Harper VanSteenhouse, Yumi Kasai, David Gray, Carol Clugston, Anna Dominiczak and Alex Alderton, Roberto Amato, Jeffrey Barrett, Sonia Goncalves, Ewan Harrison, David K. Jackson, Ian Johnston, Dominic Kwiatkowski, Cordelia Langford, John Sillitoe on behalf of the Wellcome Sanger Institute COVID-19 Surveillance Team |
| EPI_ISL_2973649, EPI_ISL_2973650 | Lighthouse Lab in Alderley Park | Wellcome Sanger Institute for the COVID-19 Genomics UK (COG-UK) Consortium | Jacquelyn Wynn, Mairead Hyland, The Lighthouse Lab in Alderley Park and Alex Alderton, Roberto Amato, Jeffrey Barrett, Sonia Goncalves, Ewan Harrison, David K. Jackson, Ian Johnston, Dominic Kwiatkowski, Cordelia Langford, John Sillitoe on behalf of the Wellcome Sanger Institute COVID-19 Surveillance Team |
| EPI_ISL_2973651, EPI_ISL_2973652 | Lighthouse Lab in Glasgow | Wellcome Sanger Institute for the COVID-19 Genomics UK (COG-UK) Consortium | Harper VanSteenhouse, Yumi Kasai, David Gray, Carol Clugston, Anna Dominiczak and Alex Alderton, Roberto Amato, Jeffrey Barrett, Sonia Goncalves, Ewan Harrison, David K. Jackson, Ian Johnston, Dominic Kwiatkowski, Cordelia Langford, John Sillitoe on behalf of the Wellcome Sanger Institute COVID-19 Surveillance Team |
| EPI_ISL_2973653 | Lighthouse Lab in Alderley Park | Wellcome Sanger Institute for the COVID-19 Genomics UK (COG-UK) Consortium | Jacquelyn Wynn, Mairead Hyland, The Lighthouse Lab in Alderley Park and Alex Alderton, Roberto Amato, Jeffrey Barrett, Sonia Goncalves, Ewan Harrison, David K. Jackson, Ian Johnston, Dominic Kwiatkowski, Cordelia Langford, John Sillitoe on behalf of the Wellcome Sanger Institute COVID-19 Surveillance Team |
| EPI_ISL_2973654, EPI_ISL_2973655, EPI_ISL_2973656, EPI_ISL_2973657 | Lighthouse Lab in Glasgow | Wellcome Sanger Institute for the COVID-19 Genomics UK (COG-UK) Consortium | Harper VanSteenhouse, Yumi Kasai, David Gray, Carol Clugston, Anna Dominiczak and Alex Alderton, Roberto Amato, Jeffrey Barrett, Sonia Goncalves, Ewan Harrison, David K. Jackson, Ian Johnston, Dominic Kwiatkowski, Cordelia Langford, John Sillitoe on behalf of the Wellcome Sanger Institute COVID-19 Surveillance Team |
| EPI_ISL_2973658 | Lighthouse Lab in Alderley Park | Wellcome Sanger Institute for the COVID-19 Genomics UK (COG-UK) Consortium | Jacquelyn Wynn, Mairead Hyland, The Lighthouse Lab in Alderley Park and Alex Alderton, Roberto Amato, Jeffrey Barrett, Sonia Goncalves, Ewan Harrison, David K. Jackson, Ian Johnston, Dominic Kwiatkowski, Cordelia Langford, John Sillitoe on behalf of the Wellcome Sanger Institute COVID-19 Surveillance Team |
| EPI_ISL_2973659, EPI_ISL_2973660 | Lighthouse Lab in Glasgow | Wellcome Sanger Institute for the COVID-19 Genomics UK (COG-UK) Consortium | Harper VanSteenhouse, Yumi Kasai, David Gray, Carol Clugston, Anna Dominiczak and Alex Alderton, Roberto Amato, Jeffrey Barrett, Sonia Goncalves, Ewan Harrison, David K. Jackson, Ian Johnston, Dominic Kwiatkowski, Cordelia Langford, John Sillitoe on behalf of the Wellcome Sanger Institute COVID-19 Surveillance Team |
| EPI_ISL_2973661 | Lighthouse Lab in Alderley Park | Wellcome Sanger Institute for the COVID-19 Genomics UK (COG-UK) Consortium | Jacquelyn Wynn, Mairead Hyland, The Lighthouse Lab in Alderley Park and Alex Alderton, Roberto Amato, Jeffrey Barrett, Sonia Goncalves, Ewan Harrison, David K. Jackson, Ian Johnston, Dominic Kwiatkowski, Cordelia Langford, John Sillitoe on behalf of the Wellcome Sanger Institute COVID-19 Surveillance Team |
| EPI_ISL_2973662, EPI_ISL_2973663, EPI_ISL_2973664 | Lighthouse Lab in Glasgow | Wellcome Sanger Institute for the COVID-19 Genomics UK (COG-UK) Consortium | Harper VanSteenhouse, Yumi Kasai, David Gray, Carol Clugston, Anna Dominiczak and Alex Alderton, Roberto Amato, Jeffrey Barrett, Sonia Goncalves, Ewan Harrison, David K. Jackson, Ian Johnston, Dominic Kwiatkowski, Cordelia Langford, John Sillitoe on behalf of the Wellcome Sanger Institute COVID-19 Surveillance Team |
| EPI_ISL_2973665, EPI_ISL_2973666, EPI_ISL_2973667, EPI_ISL_2973668, EPI_ISL_2973669, EPI_ISL_2973670, EPI_ISL_2973671 | Lighthouse Lab in Alderley Park | Wellcome Sanger Institute for the COVID-19 Genomics UK (COG-UK) Consortium | Jacquelyn Wynn, Mairead Hyland, The Lighthouse Lab in Alderley Park and Alex Alderton, Roberto Amato, Jeffrey Barrett, Sonia Goncalves, Ewan Harrison, David K. Jackson, Ian Johnston, Dominic Kwiatkowski, Cordelia Langford, John Sillitoe on behalf of the Wellcome Sanger Institute COVID-19 Surveillance Team |
| EPI_ISL_2973672 | Lighthouse Lab in Glasgow | Wellcome Sanger Institute for the COVID-19 Genomics UK (COG-UK) Consortium | Harper VanSteenhouse, Yumi Kasai, David Gray, Carol Clugston, Anna Dominiczak and Alex Alderton, Roberto Amato, Jeffrey Barrett, Sonia Goncalves, Ewan Harrison, David K. Jackson, Ian Johnston, Dominic Kwiatkowski, Cordelia Langford, John Sillitoe on behalf of the Wellcome Sanger Institute COVID-19 Surveillance Team |
| EPI_ISL_2973673, EPI_ISL_2973674 | Lighthouse Lab in Alderley Park | Wellcome Sanger Institute for the COVID-19 Genomics UK (COG-UK) Consortium | Jacquelyn Wynn, Mairead Hyland, The Lighthouse Lab in Alderley Park and Alex Alderton, Roberto Amato, Jeffrey Barrett, Sonia Goncalves, Ewan Harrison, David K. Jackson, Ian Johnston, Dominic Kwiatkowski, Cordelia Langford, John Sillitoe on behalf of the Wellcome Sanger Institute COVID-19 Surveillance Team |
| EPI_ISL_2973675, EPI_ISL_2973676 | Lighthouse Lab in Glasgow | Wellcome Sanger Institute for the COVID-19 Genomics UK (COG-UK) Consortium | Harper VanSteenhouse, Yumi Kasai, David Gray, Carol Clugston, Anna Dominiczak and Alex Alderton, Roberto Amato, Jeffrey Barrett, Sonia Goncalves, Ewan Harrison, David K. Jackson, Ian Johnston, Dominic Kwiatkowski, Cordelia Langford, John Sillitoe on behalf of the Wellcome Sanger Institute COVID-19 Surveillance Team |
| EPI_ISL_2973677 | Lighthouse Lab in Alderley Park | Wellcome Sanger Institute for the COVID-19 Genomics UK (COG-UK) Consortium | Jacquelyn Wynn, Mairead Hyland, The Lighthouse Lab in Alderley Park and Alex Alderton, Roberto Amato, Jeffrey Barrett, Sonia Goncalves, Ewan Harrison, David K. Jackson, Ian Johnston, Dominic Kwiatkowski, Cordelia Langford, John Sillitoe on behalf of the Wellcome Sanger Institute COVID-19 Surveillance Team |
| EPI_ISL_2973678 | Lighthouse Lab in Glasgow | Wellcome Sanger Institute for the COVID-19 Genomics UK (COG-UK) Consortium | Harper VanSteenhouse, Yumi Kasai, David Gray, Carol Clugston, Anna Dominiczak and Alex Alderton, Roberto Amato, Jeffrey Barrett, Sonia Goncalves, Ewan Harrison, David K. Jackson, Ian Johnston, Dominic Kwiatkowski, Cordelia Langford, John Sillitoe on behalf of the Wellcome Sanger Institute COVID-19 Surveillance Team |
| EPI_ISL_2973679, EPI_ISL_2973680 | Lighthouse Lab in Alderley Park | Wellcome Sanger Institute for the COVID-19 Genomics UK (COG-UK) Consortium | Jacquelyn Wynn, Mairead Hyland, The Lighthouse Lab in Alderley Park and Alex Alderton, Roberto Amato, Jeffrey Barrett, Sonia Goncalves, Ewan Harrison, David K. Jackson, Ian Johnston, Dominic Kwiatkowski, Cordelia Langford, John Sillitoe on behalf of the Wellcome Sanger Institute COVID-19 Surveillance Team |
| EPI_ISL_2973681, EPI_ISL_2973682, | Lighthouse Lab in Glasgow | Wellcome Sanger Institute for the COVID-19 Genomics UK (COG-UK) Consortium | Harper VanSteenhouse, Yumi Kasai, David Gray, Carol Clugston, Anna Dominiczak and Alex Alderton, Roberto Amato, Jeffrey Barrett, Sonia Goncalves, Ewan Harrison, David K. Jackson, Ian Johnston, Dominic Kwiatkowski, Cordelia Langford, John Sillitoe on behalf of the Wellcome Sanger Institute COVID-19 Surveillance Team |

| | | | |
|---|---|---|---|
| EPI_ISL_2973683 | | | |
| EPI_ISL_2973684 | Lighthouse Lab in Alderley Park | Wellcome Sanger Institute for the COVID-19 Genomics UK (COG-UK) Consortium | Jacquelyn Wynn, Mairead Hyland, The Lighthouse Lab in Alderley Park and Alex Alderton, Roberto Amato, Jeffrey Barrett, Sonia Goncalves, Ewan Harrison, David K. Jackson, Ian Johnston, Dominic Kwiatkowski, Cordelia Langford, John Sillitoe on behalf of the Wellcome Sanger Institute COVID-19 Surveillance Team |
| EPI_ISL_2973685, EPI_ISL_2973686, EPI_ISL_2973687, EPI_ISL_2973688 | Lighthouse Lab in Glasgow | Wellcome Sanger Institute for the COVID-19 Genomics UK (COG-UK) Consortium | Harper VanSteenhouse, Yumi Kasai, David Gray, Carol Clugston, Anna Dominiczak and Alex Alderton, Roberto Amato, Jeffrey Barrett, Sonia Goncalves, Ewan Harrison, David K. Jackson, Ian Johnston, Dominic Kwiatkowski, Cordelia Langford, John Sillitoe on behalf of the Wellcome Sanger Institute COVID-19 Surveillance Team |
| EPI_ISL_2973689 | Lighthouse Lab in Alderley Park | Wellcome Sanger Institute for the COVID-19 Genomics UK (COG-UK) Consortium | Jacquelyn Wynn, Mairead Hyland, The Lighthouse Lab in Alderley Park and Alex Alderton, Roberto Amato, Jeffrey Barrett, Sonia Goncalves, Ewan Harrison, David K. Jackson, Ian Johnston, Dominic Kwiatkowski, Cordelia Langford, John Sillitoe on behalf of the Wellcome Sanger Institute COVID-19 Surveillance Team |
| EPI_ISL_2973690, EPI_ISL_2973691 | Lighthouse Lab in Glasgow | Wellcome Sanger Institute for the COVID-19 Genomics UK (COG-UK) Consortium | Harper VanSteenhouse, Yumi Kasai, David Gray, Carol Clugston, Anna Dominiczak and Alex Alderton, Roberto Amato, Jeffrey Barrett, Sonia Goncalves, Ewan Harrison, David K. Jackson, Ian Johnston, Dominic Kwiatkowski, Cordelia Langford, John Sillitoe on behalf of the Wellcome Sanger Institute COVID-19 Surveillance Team |
| EPI_ISL_2973692, EPI_ISL_2973693 | Lighthouse Lab in Alderley Park | Wellcome Sanger Institute for the COVID-19 Genomics UK (COG-UK) Consortium | Jacquelyn Wynn, Mairead Hyland, The Lighthouse Lab in Alderley Park and Alex Alderton, Roberto Amato, Jeffrey Barrett, Sonia Goncalves, Ewan Harrison, David K. Jackson, Ian Johnston, Dominic Kwiatkowski, Cordelia Langford, John Sillitoe on behalf of the Wellcome Sanger Institute COVID-19 Surveillance Team |
| EPI_ISL_2973694 | Lighthouse Lab in Glasgow | Wellcome Sanger Institute for the COVID-19 Genomics UK (COG-UK) Consortium | Harper VanSteenhouse, Yumi Kasai, David Gray, Carol Clugston, Anna Dominiczak and Alex Alderton, Roberto Amato, Jeffrey Barrett, Sonia Goncalves, Ewan Harrison, David K. Jackson, Ian Johnston, Dominic Kwiatkowski, Cordelia Langford, John Sillitoe on behalf of the Wellcome Sanger Institute COVID-19 Surveillance Team |
| EPI_ISL_2973695, EPI_ISL_2973696, EPI_ISL_2973697, EPI_ISL_2973698, EPI_ISL_2973699 | Lighthouse Lab in Alderley Park | Wellcome Sanger Institute for the COVID-19 Genomics UK (COG-UK) Consortium | Jacquelyn Wynn, Mairead Hyland, The Lighthouse Lab in Alderley Park and Alex Alderton, Roberto Amato, Jeffrey Barrett, Sonia Goncalves, Ewan Harrison, David K. Jackson, Ian Johnston, Dominic Kwiatkowski, Cordelia Langford, John Sillitoe on behalf of the Wellcome Sanger Institute COVID-19 Surveillance Team |
| EPI_ISL_2973700, EPI_ISL_2973701, EPI_ISL_2973702, EPI_ISL_2973703 | Lighthouse Lab in Glasgow | Wellcome Sanger Institute for the COVID-19 Genomics UK (COG-UK) Consortium | Harper VanSteenhouse, Yumi Kasai, David Gray, Carol Clugston, Anna Dominiczak and Alex Alderton, Roberto Amato, Jeffrey Barrett, Sonia Goncalves, Ewan Harrison, David K. Jackson, Ian Johnston, Dominic Kwiatkowski, Cordelia Langford, John Sillitoe on behalf of the Wellcome Sanger Institute COVID-19 Surveillance Team |
| EPI_ISL_2973704, EPI_ISL_2973705, EPI_ISL_2973706, EPI_ISL_2973707 | Lighthouse Lab in Alderley Park | Wellcome Sanger Institute for the COVID-19 Genomics UK (COG-UK) Consortium | Jacquelyn Wynn, Mairead Hyland, The Lighthouse Lab in Alderley Park and Alex Alderton, Roberto Amato, Jeffrey Barrett, Sonia Goncalves, Ewan Harrison, David K. Jackson, Ian Johnston, Dominic Kwiatkowski, Cordelia Langford, John Sillitoe on behalf of the Wellcome Sanger Institute COVID-19 Surveillance Team |
| EPI_ISL_2973708 | Lighthouse Lab in Glasgow | Wellcome Sanger Institute for the COVID-19 Genomics UK (COG-UK) Consortium | Harper VanSteenhouse, Yumi Kasai, David Gray, Carol Clugston, Anna Dominiczak and Alex Alderton, Roberto Amato, Jeffrey Barrett, Sonia Goncalves, Ewan Harrison, David K. Jackson, Ian Johnston, Dominic Kwiatkowski, Cordelia Langford, John Sillitoe on behalf of the Wellcome Sanger Institute COVID-19 Surveillance Team |
| EPI_ISL_2973709, EPI_ISL_2973710 | Lighthouse Lab in Alderley Park | Wellcome Sanger Institute for the COVID-19 Genomics UK (COG-UK) Consortium | Jacquelyn Wynn, Mairead Hyland, The Lighthouse Lab in Alderley Park and Alex Alderton, Roberto Amato, Jeffrey Barrett, Sonia Goncalves, Ewan Harrison, David K. Jackson, Ian Johnston, Dominic Kwiatkowski, Cordelia Langford, John Sillitoe on behalf of the Wellcome Sanger Institute COVID-19 Surveillance Team |
| EPI_ISL_2973711 | Lighthouse Lab in Glasgow | Wellcome Sanger Institute for the COVID-19 Genomics UK (COG-UK) Consortium | Harper VanSteenhouse, Yumi Kasai, David Gray, Carol Clugston, Anna Dominiczak and Alex Alderton, Roberto Amato, Jeffrey Barrett, Sonia Goncalves, Ewan Harrison, David K. Jackson, Ian Johnston, Dominic Kwiatkowski, Cordelia Langford, John Sillitoe on behalf of the Wellcome Sanger Institute COVID-19 Surveillance Team |
| EPI_ISL_2973712 | Lighthouse Lab in Alderley Park | Wellcome Sanger Institute for the COVID-19 Genomics UK (COG-UK) Consortium | Jacquelyn Wynn, Mairead Hyland, The Lighthouse Lab in Alderley Park and Alex Alderton, Roberto Amato, Jeffrey Barrett, Sonia Goncalves, Ewan Harrison, David K. Jackson, Ian Johnston, Dominic Kwiatkowski, Cordelia Langford, John Sillitoe on behalf of the Wellcome Sanger Institute COVID-19 Surveillance Team |
| EPI_ISL_2973713 | Lighthouse Lab in Glasgow | Wellcome Sanger Institute for the COVID-19 Genomics UK (COG-UK) Consortium | Harper VanSteenhouse, Yumi Kasai, David Gray, Carol Clugston, Anna Dominiczak and Alex Alderton, Roberto Amato, Jeffrey Barrett, Sonia Goncalves, Ewan Harrison, David K. Jackson, Ian Johnston, Dominic Kwiatkowski, Cordelia Langford, John Sillitoe on behalf of the Wellcome Sanger Institute COVID-19 Surveillance Team |
| EPI_ISL_2973714, EPI_ISL_2973715, EPI_ISL_2973716 | Lighthouse Lab in Alderley Park | Wellcome Sanger Institute for the COVID-19 Genomics UK (COG-UK) Consortium | Jacquelyn Wynn, Mairead Hyland, The Lighthouse Lab in Alderley Park and Alex Alderton, Roberto Amato, Jeffrey Barrett, Sonia Goncalves, Ewan Harrison, David K. Jackson, Ian Johnston, Dominic Kwiatkowski, Cordelia Langford, John Sillitoe on behalf of the Wellcome Sanger Institute COVID-19 Surveillance Team |
| EPI_ISL_2973717 | Lighthouse Lab in Glasgow | Wellcome Sanger Institute for the COVID-19 Genomics UK (COG-UK) Consortium | Harper VanSteenhouse, Yumi Kasai, David Gray, Carol Clugston, Anna Dominiczak and Alex Alderton, Roberto Amato, Jeffrey Barrett, Sonia Goncalves, Ewan Harrison, David K. Jackson, Ian Johnston, Dominic Kwiatkowski, Cordelia Langford, John Sillitoe on behalf of the Wellcome Sanger Institute COVID-19 Surveillance Team |
| EPI_ISL_2973718, EPI_ISL_2973719 | Lighthouse Lab in Alderley Park | Wellcome Sanger Institute for the COVID-19 Genomics UK (COG-UK) Consortium | Jacquelyn Wynn, Mairead Hyland, The Lighthouse Lab in Alderley Park and Alex Alderton, Roberto Amato, Jeffrey Barrett, Sonia Goncalves, Ewan Harrison, David K. Jackson, Ian Johnston, Dominic Kwiatkowski, Cordelia Langford, John Sillitoe on behalf of the Wellcome Sanger Institute COVID-19 Surveillance Team |
| EPI_ISL_2973720 | Lighthouse Lab in Glasgow | Wellcome Sanger Institute for the COVID-19 Genomics UK (COG-UK) Consortium | Harper VanSteenhouse, Yumi Kasai, David Gray, Carol Clugston, Anna Dominiczak and Alex Alderton, Roberto Amato, Jeffrey Barrett, Sonia Goncalves, Ewan Harrison, David K. Jackson, Ian Johnston, Dominic Kwiatkowski, Cordelia Langford, John Sillitoe on behalf of the Wellcome Sanger Institute COVID-19 Surveillance Team |
| EPI_ISL_2973721 | Lighthouse Lab in Alderley Park | Wellcome Sanger Institute for the COVID-19 Genomics UK (COG-UK) Consortium | Jacquelyn Wynn, Mairead Hyland, The Lighthouse Lab in Alderley Park and Alex Alderton, Roberto Amato, Jeffrey Barrett, Sonia Goncalves, Ewan Harrison, David K. Jackson, Ian Johnston, Dominic Kwiatkowski, Cordelia Langford, John Sillitoe on behalf of the Wellcome Sanger Institute COVID-19 Surveillance Team |
| EPI_ISL_2973722, EPI_ISL_2973723, EPI_ISL_2973724 | Lighthouse Lab in Glasgow | Wellcome Sanger Institute for the COVID-19 Genomics UK (COG-UK) Consortium | Harper VanSteenhouse, Yumi Kasai, David Gray, Carol Clugston, Anna Dominiczak and Alex Alderton, Roberto Amato, Jeffrey Barrett, Sonia Goncalves, Ewan Harrison, David K. Jackson, Ian Johnston, Dominic Kwiatkowski, Cordelia Langford, John Sillitoe on behalf of the Wellcome Sanger Institute COVID-19 Surveillance Team |
| EPI_ISL_2973725, EPI_ISL_2973726, EPI_ISL_2973727, EPI_ISL_2973728, EPI_ISL_2973729 | Lighthouse Lab in Alderley Park | Wellcome Sanger Institute for the COVID-19 Genomics UK (COG-UK) Consortium | Jacquelyn Wynn, Mairead Hyland, The Lighthouse Lab in Alderley Park and Alex Alderton, Roberto Amato, Jeffrey Barrett, Sonia Goncalves, Ewan Harrison, David K. Jackson, Ian Johnston, Dominic Kwiatkowski, Cordelia Langford, John Sillitoe on behalf of the Wellcome Sanger Institute COVID-19 Surveillance Team |
| EPI_ISL_2973730 | Lighthouse Lab in Glasgow | Wellcome Sanger Institute for the COVID-19 Genomics UK (COG-UK) Consortium | Harper VanSteenhouse, Yumi Kasai, David Gray, Carol Clugston, Anna Dominiczak and Alex Alderton, Roberto Amato, Jeffrey Barrett, Sonia Goncalves, Ewan Harrison, David K. Jackson, Ian Johnston, Dominic Kwiatkowski, Cordelia Langford, John Sillitoe on behalf of the Wellcome Sanger Institute COVID-19 Surveillance Team |
| EPI_ISL_2973731, EPI_ISL_2973732 | Lighthouse Lab in Alderley Park | Wellcome Sanger Institute for the COVID-19 Genomics UK (COG-UK) Consortium | Jacquelyn Wynn, Mairead Hyland, The Lighthouse Lab in Alderley Park and Alex Alderton, Roberto Amato, Jeffrey Barrett, Sonia Goncalves, Ewan Harrison, David K. Jackson, Ian Johnston, Dominic Kwiatkowski, Cordelia Langford, John Sillitoe on behalf of the Wellcome Sanger Institute COVID-19 Surveillance Team |
| EPI_ISL_2973733 | Lighthouse Lab in Glasgow | Wellcome Sanger Institute for the COVID-19 Genomics UK (COG-UK) Consortium | Harper VanSteenhouse, Yumi Kasai, David Gray, Carol Clugston, Anna Dominiczak and Alex Alderton, Roberto Amato, Jeffrey Barrett, Sonia Goncalves, Ewan Harrison, David K. Jackson, Ian Johnston, Dominic Kwiatkowski, Cordelia Langford, John Sillitoe on behalf of the Wellcome Sanger Institute COVID-19 Surveillance Team |
| EPI_ISL_2973734 | Lighthouse Lab in Alderley Park | Wellcome Sanger Institute for the COVID-19 Genomics UK (COG-UK) Consortium | Jacquelyn Wynn, Mairead Hyland, The Lighthouse Lab in Alderley Park and Alex Alderton, Roberto Amato, Jeffrey Barrett, Sonia Goncalves, Ewan Harrison, David K. Jackson, Ian Johnston, Dominic Kwiatkowski, Cordelia Langford, John Sillitoe on behalf of the Wellcome Sanger Institute COVID-19 Surveillance Team |
| EPI_ISL_2973735, EPI_ISL_2973736, EPI_ISL_2973737, EPI_ISL_2973738, EPI_ISL_2973739 | Lighthouse Lab in Glasgow | Wellcome Sanger Institute for the COVID-19 Genomics UK (COG-UK) Consortium | Harper VanSteenhouse, Yumi Kasai, David Gray, Carol Clugston, Anna Dominiczak and Alex Alderton, Roberto Amato, Jeffrey Barrett, Sonia Goncalves, Ewan Harrison, David K. Jackson, Ian Johnston, Dominic Kwiatkowski, Cordelia Langford, John Sillitoe on behalf of the Wellcome Sanger Institute COVID-19 Surveillance Team |
| EPI_ISL_2973740 | Lighthouse Lab in Alderley Park | Wellcome Sanger Institute for the COVID-19 Genomics UK (COG-UK) Consortium | Jacquelyn Wynn, Mairead Hyland, The Lighthouse Lab in Alderley Park and Alex Alderton, Roberto Amato, Jeffrey Barrett, Sonia Goncalves, Ewan Harrison, David K. Jackson, Ian Johnston, Dominic Kwiatkowski, Cordelia Langford, John Sillitoe on behalf of the Wellcome Sanger Institute COVID-19 Surveillance Team |
| EPI_ISL_2973741, EPI_ISL_2973742 | Lighthouse Lab in Glasgow | Wellcome Sanger Institute for the COVID-19 Genomics UK (COG-UK) Consortium | Harper VanSteenhouse, Yumi Kasai, David Gray, Carol Clugston, Anna Dominiczak and Alex Alderton, Roberto Amato, Jeffrey Barrett, Sonia Goncalves, Ewan Harrison, David K. Jackson, Ian Johnston, Dominic Kwiatkowski, Cordelia Langford, John Sillitoe on behalf of the Wellcome Sanger Institute COVID-19 Surveillance Team |
| EPI_ISL_2973743, EPI_ISL_2973744 | Lighthouse Lab in Alderley Park | Wellcome Sanger Institute for the COVID-19 Genomics UK (COG-UK) Consortium | Jacquelyn Wynn, Mairead Hyland, The Lighthouse Lab in Alderley Park and Alex Alderton, Roberto Amato, Jeffrey Barrett, Sonia Goncalves, Ewan Harrison, David K. Jackson, Ian Johnston, Dominic Kwiatkowski, Cordelia Langford, John Sillitoe on behalf of the Wellcome Sanger Institute COVID-19 Surveillance Team |
| EPI_ISL_2973745 | Lighthouse Lab in Glasgow | Wellcome Sanger Institute for the COVID-19 Genomics UK (COG-UK) Consortium | Harper VanSteenhouse, Yumi Kasai, David Gray, Carol Clugston, Anna Dominiczak and Alex Alderton, Roberto Amato, Jeffrey Barrett, Sonia Goncalves, Ewan Harrison, David K. Jackson, Ian Johnston, Dominic Kwiatkowski, Cordelia Langford, John Sillitoe on behalf of the Wellcome Sanger Institute COVID-19 Surveillance Team |
| EPI_ISL_2973746 | Lighthouse Lab in Alderley Park | Wellcome Sanger Institute for the COVID-19 Genomics UK (COG-UK) Consortium | Jacquelyn Wynn, Mairead Hyland, The Lighthouse Lab in Alderley Park and Alex Alderton, Roberto Amato, Jeffrey Barrett, Sonia Goncalves, Ewan Harrison, David K. Jackson, Ian Johnston, Dominic Kwiatkowski, Cordelia Langford, John Sillitoe on behalf of the Wellcome Sanger Institute COVID-19 Surveillance Team |
| EPI_ISL_2973747 | Lighthouse Lab in Glasgow | Wellcome Sanger Institute for the COVID-19 Genomics UK (COG-UK) Consortium | Harper VanSteenhouse, Yumi Kasai, David Gray, Carol Clugston, Anna Dominiczak and Alex Alderton, Roberto Amato, Jeffrey Barrett, Sonia Goncalves, Ewan Harrison, David K. Jackson, Ian Johnston, Dominic Kwiatkowski, Cordelia Langford, John Sillitoe on behalf of the Wellcome Sanger Institute COVID-19 Surveillance Team |
| EPI_ISL_2973748, EPI_ISL_2973749 | Lighthouse Lab in Alderley Park | Wellcome Sanger Institute for the COVID-19 Genomics UK (COG-UK) Consortium | Jacquelyn Wynn, Mairead Hyland, The Lighthouse Lab in Alderley Park and Alex Alderton, Roberto Amato, Jeffrey Barrett, Sonia Goncalves, Ewan Harrison, David K. Jackson, Ian Johnston, Dominic Kwiatkowski, Cordelia Langford, John Sillitoe on behalf of the Wellcome Sanger Institute COVID-19 Surveillance Team |
| EPI_ISL_2973750 | Lighthouse Lab in Glasgow | Wellcome Sanger Institute for the COVID-19 Genomics UK (COG-UK) Consortium | Harper VanSteenhouse, Yumi Kasai, David Gray, Carol Clugston, Anna Dominiczak and Alex Alderton, Roberto Amato, Jeffrey Barrett, Sonia Goncalves, Ewan Harrison, David K. Jackson, Ian Johnston, Dominic Kwiatkowski, Cordelia Langford, John Sillitoe on behalf of the Wellcome Sanger Institute COVID-19 Surveillance Team |
| EPI_ISL_2973751, EPI_ISL_2973752, EPI_ISL_2973753, EPI_ISL_2973754 | Lighthouse Lab in Alderley Park | Wellcome Sanger Institute for the COVID-19 Genomics UK (COG-UK) Consortium | Jacquelyn Wynn, Mairead Hyland, The Lighthouse Lab in Alderley Park and Alex Alderton, Roberto Amato, Jeffrey Barrett, Sonia Goncalves, Ewan Harrison, David K. Jackson, Ian Johnston, Dominic Kwiatkowski, Cordelia Langford, John Sillitoe on behalf of the Wellcome Sanger Institute COVID-19 Surveillance Team |
| EPI_ISL_2973755 | Lighthouse Lab in Glasgow | Wellcome Sanger Institute for the COVID-19 Genomics UK (COG-UK) Consortium | Harper VanSteenhouse, Yumi Kasai, David Gray, Carol Clugston, Anna Dominiczak and Alex Alderton, Roberto Amato, Jeffrey Barrett, Sonia Goncalves, Ewan Harrison, David K. Jackson, Ian Johnston, Dominic Kwiatkowski, Cordelia Langford, John Sillitoe on behalf of the Wellcome Sanger Institute COVID-19 Surveillance Team |
| EPI_ISL_2973756, | Lighthouse Lab in Alderley Park | Wellcome Sanger Institute for the COVID-19 Genomics UK (COG-UK) | Jacquelyn Wynn, Mairead Hyland, The Lighthouse Lab in Alderley Park and Alex Alderton, Roberto Amato, Jeffrey Barrett, Sonia Goncalves, Ewan Harrison, David K. Jackson, Ian Johnston, Dominic Kwiatkowski, Cordelia Langford, John Sillitoe on |

| | | | |
|---|---|---|---|
| EPI_ISL_2973757, EPI_ISL_2973758 | | Consortium | behalf of the Wellcome Sanger Institute COVID-19 Surveillance Team |
| EPI_ISL_2973759, EPI_ISL_2973760 | Lighthouse Lab in Glasgow | Wellcome Sanger Institute for the COVID-19 Genomics UK (COG-UK) Consortium | Harper VanSteenhouse, Yumi Kasai, David Gray, Carol Clugston, Anna Dominiczak and Alex Alderton, Roberto Amato, Jeffrey Barrett, Sonia Goncalves, Ewan Harrison, David K. Jackson, Ian Johnston, Dominic Kwiatkowski, Cordelia Langford, John Sillitoe on behalf of the Wellcome Sanger Institute COVID-19 Surveillance Team |
| EPI_ISL_2973761, EPI_ISL_2973762, EPI_ISL_2973763 | Lighthouse Lab in Alderley Park | Wellcome Sanger Institute for the COVID-19 Genomics UK (COG-UK) Consortium | Jacquelyn Wynn, Mairead Hyland, The Lighthouse Lab in Alderley Park and Alex Alderton, Roberto Amato, Jeffrey Barrett, Sonia Goncalves, Ewan Harrison, David K. Jackson, Ian Johnston, Dominic Kwiatkowski, Cordelia Langford, John Sillitoe on behalf of the Wellcome Sanger Institute COVID-19 Surveillance Team |
| EPI_ISL_2973764, EPI_ISL_2973765 | Lighthouse Lab in Glasgow | Wellcome Sanger Institute for the COVID-19 Genomics UK (COG-UK) Consortium | Harper VanSteenhouse, Yumi Kasai, David Gray, Carol Clugston, Anna Dominiczak and Alex Alderton, Roberto Amato, Jeffrey Barrett, Sonia Goncalves, Ewan Harrison, David K. Jackson, Ian Johnston, Dominic Kwiatkowski, Cordelia Langford, John Sillitoe on behalf of the Wellcome Sanger Institute COVID-19 Surveillance Team |
| EPI_ISL_2973766 | Lighthouse Lab in Alderley Park | Wellcome Sanger Institute for the COVID-19 Genomics UK (COG-UK) Consortium | Jacquelyn Wynn, Mairead Hyland, The Lighthouse Lab in Alderley Park and Alex Alderton, Roberto Amato, Jeffrey Barrett, Sonia Goncalves, Ewan Harrison, David K. Jackson, Ian Johnston, Dominic Kwiatkowski, Cordelia Langford, John Sillitoe on behalf of the Wellcome Sanger Institute COVID-19 Surveillance Team |
| EPI_ISL_2973767, EPI_ISL_2973768, EPI_ISL_2973769, EPI_ISL_2973770 | Lighthouse Lab in Glasgow | Wellcome Sanger Institute for the COVID-19 Genomics UK (COG-UK) Consortium | Harper VanSteenhouse, Yumi Kasai, David Gray, Carol Clugston, Anna Dominiczak and Alex Alderton, Roberto Amato, Jeffrey Barrett, Sonia Goncalves, Ewan Harrison, David K. Jackson, Ian Johnston, Dominic Kwiatkowski, Cordelia Langford, John Sillitoe on behalf of the Wellcome Sanger Institute COVID-19 Surveillance Team |
| EPI_ISL_2973771, EPI_ISL_2973772 | Lighthouse Lab in Alderley Park | Wellcome Sanger Institute for the COVID-19 Genomics UK (COG-UK) Consortium | Jacquelyn Wynn, Mairead Hyland, The Lighthouse Lab in Alderley Park and Alex Alderton, Roberto Amato, Jeffrey Barrett, Sonia Goncalves, Ewan Harrison, David K. Jackson, Ian Johnston, Dominic Kwiatkowski, Cordelia Langford, John Sillitoe on behalf of the Wellcome Sanger Institute COVID-19 Surveillance Team |
| EPI_ISL_2973773, EPI_ISL_2973774 | Lighthouse Lab in Glasgow | Wellcome Sanger Institute for the COVID-19 Genomics UK (COG-UK) Consortium | Harper VanSteenhouse, Yumi Kasai, David Gray, Carol Clugston, Anna Dominiczak and Alex Alderton, Roberto Amato, Jeffrey Barrett, Sonia Goncalves, Ewan Harrison, David K. Jackson, Ian Johnston, Dominic Kwiatkowski, Cordelia Langford, John Sillitoe on behalf of the Wellcome Sanger Institute COVID-19 Surveillance Team |
| EPI_ISL_2973775 | Lighthouse Lab in Alderley Park | Wellcome Sanger Institute for the COVID-19 Genomics UK (COG-UK) Consortium | Jacquelyn Wynn, Mairead Hyland, The Lighthouse Lab in Alderley Park and Alex Alderton, Roberto Amato, Jeffrey Barrett, Sonia Goncalves, Ewan Harrison, David K. Jackson, Ian Johnston, Dominic Kwiatkowski, Cordelia Langford, John Sillitoe on behalf of the Wellcome Sanger Institute COVID-19 Surveillance Team |
| EPI_ISL_2973776 | Lighthouse Lab in Glasgow | Wellcome Sanger Institute for the COVID-19 Genomics UK (COG-UK) Consortium | Harper VanSteenhouse, Yumi Kasai, David Gray, Carol Clugston, Anna Dominiczak and Alex Alderton, Roberto Amato, Jeffrey Barrett, Sonia Goncalves, Ewan Harrison, David K. Jackson, Ian Johnston, Dominic Kwiatkowski, Cordelia Langford, John Sillitoe on behalf of the Wellcome Sanger Institute COVID-19 Surveillance Team |
| EPI_ISL_2973777, EPI_ISL_2973778, EPI_ISL_2973779, EPI_ISL_2973780 | Lighthouse Lab in Alderley Park | Wellcome Sanger Institute for the COVID-19 Genomics UK (COG-UK) Consortium | Jacquelyn Wynn, Mairead Hyland, The Lighthouse Lab in Alderley Park and Alex Alderton, Roberto Amato, Jeffrey Barrett, Sonia Goncalves, Ewan Harrison, David K. Jackson, Ian Johnston, Dominic Kwiatkowski, Cordelia Langford, John Sillitoe on behalf of the Wellcome Sanger Institute COVID-19 Surveillance Team |
| EPI_ISL_2973781, EPI_ISL_2973782, EPI_ISL_2973783, EPI_ISL_2973784 | Lighthouse Lab in Glasgow | Wellcome Sanger Institute for the COVID-19 Genomics UK (COG-UK) Consortium | Harper VanSteenhouse, Yumi Kasai, David Gray, Carol Clugston, Anna Dominiczak and Alex Alderton, Roberto Amato, Jeffrey Barrett, Sonia Goncalves, Ewan Harrison, David K. Jackson, Ian Johnston, Dominic Kwiatkowski, Cordelia Langford, John Sillitoe on behalf of the Wellcome Sanger Institute COVID-19 Surveillance Team |
| EPI_ISL_2973785, EPI_ISL_2973786, EPI_ISL_2973787, EPI_ISL_2973788, EPI_ISL_2973789, EPI_ISL_2973790, EPI_ISL_2973791, EPI_ISL_2973792, EPI_ISL_2973793 | Lighthouse Lab in Alderley Park | Wellcome Sanger Institute for the COVID-19 Genomics UK (COG-UK) Consortium | Jacquelyn Wynn, Mairead Hyland, The Lighthouse Lab in Alderley Park and Alex Alderton, Roberto Amato, Jeffrey Barrett, Sonia Goncalves, Ewan Harrison, David K. Jackson, Ian Johnston, Dominic Kwiatkowski, Cordelia Langford, John Sillitoe on behalf of the Wellcome Sanger Institute COVID-19 Surveillance Team |
| EPI_ISL_2973794 | Lighthouse Lab in Glasgow | Wellcome Sanger Institute for the COVID-19 Genomics UK (COG-UK) Consortium | Harper VanSteenhouse, Yumi Kasai, David Gray, Carol Clugston, Anna Dominiczak and Alex Alderton, Roberto Amato, Jeffrey Barrett, Sonia Goncalves, Ewan Harrison, David K. Jackson, Ian Johnston, Dominic Kwiatkowski, Cordelia Langford, John Sillitoe on behalf of the Wellcome Sanger Institute COVID-19 Surveillance Team |
| EPI_ISL_2973795, EPI_ISL_2973796, EPI_ISL_2973797, EPI_ISL_2973798, EPI_ISL_2973799 | Lighthouse Lab in Alderley Park | Wellcome Sanger Institute for the COVID-19 Genomics UK (COG-UK) Consortium | Jacquelyn Wynn, Mairead Hyland, The Lighthouse Lab in Alderley Park and Alex Alderton, Roberto Amato, Jeffrey Barrett, Sonia Goncalves, Ewan Harrison, David K. Jackson, Ian Johnston, Dominic Kwiatkowski, Cordelia Langford, John Sillitoe on behalf of the Wellcome Sanger Institute COVID-19 Surveillance Team |
| EPI_ISL_2973800, EPI_ISL_2973801, EPI_ISL_2973802 | Lighthouse Lab in Glasgow | Wellcome Sanger Institute for the COVID-19 Genomics UK (COG-UK) Consortium | Harper VanSteenhouse, Yumi Kasai, David Gray, Carol Clugston, Anna Dominiczak and Alex Alderton, Roberto Amato, Jeffrey Barrett, Sonia Goncalves, Ewan Harrison, David K. Jackson, Ian Johnston, Dominic Kwiatkowski, Cordelia Langford, John Sillitoe on behalf of the Wellcome Sanger Institute COVID-19 Surveillance Team |
| EPI_ISL_2973803 | Lighthouse Lab in Alderley Park | Wellcome Sanger Institute for the COVID-19 Genomics UK (COG-UK) Consortium | Jacquelyn Wynn, Mairead Hyland, The Lighthouse Lab in Alderley Park and Alex Alderton, Roberto Amato, Jeffrey Barrett, Sonia Goncalves, Ewan Harrison, David K. Jackson, Ian Johnston, Dominic Kwiatkowski, Cordelia Langford, John Sillitoe on behalf of the Wellcome Sanger Institute COVID-19 Surveillance Team |
| EPI_ISL_2973804 | Lighthouse Lab in Glasgow | Wellcome Sanger Institute for the COVID-19 Genomics UK (COG-UK) Consortium | Harper VanSteenhouse, Yumi Kasai, David Gray, Carol Clugston, Anna Dominiczak and Alex Alderton, Roberto Amato, Jeffrey Barrett, Sonia Goncalves, Ewan Harrison, David K. Jackson, Ian Johnston, Dominic Kwiatkowski, Cordelia Langford, John Sillitoe on behalf of the Wellcome Sanger Institute COVID-19 Surveillance Team |
| EPI_ISL_2973805 | Lighthouse Lab in Alderley Park | Wellcome Sanger Institute for the COVID-19 Genomics UK (COG-UK) Consortium | Jacquelyn Wynn, Mairead Hyland, The Lighthouse Lab in Alderley Park and Alex Alderton, Roberto Amato, Jeffrey Barrett, Sonia Goncalves, Ewan Harrison, David K. Jackson, Ian Johnston, Dominic Kwiatkowski, Cordelia Langford, John Sillitoe on behalf of the Wellcome Sanger Institute COVID-19 Surveillance Team |
| EPI_ISL_2973806 | Lighthouse Lab in Glasgow | Wellcome Sanger Institute for the COVID-19 Genomics UK (COG-UK) Consortium | Harper VanSteenhouse, Yumi Kasai, David Gray, Carol Clugston, Anna Dominiczak and Alex Alderton, Roberto Amato, Jeffrey Barrett, Sonia Goncalves, Ewan Harrison, David K. Jackson, Ian Johnston, Dominic Kwiatkowski, Cordelia Langford, John Sillitoe on behalf of the Wellcome Sanger Institute COVID-19 Surveillance Team |
| EPI_ISL_2973807, EPI_ISL_2973808 | Lighthouse Lab in Alderley Park | Wellcome Sanger Institute for the COVID-19 Genomics UK (COG-UK) Consortium | Jacquelyn Wynn, Mairead Hyland, The Lighthouse Lab in Alderley Park and Alex Alderton, Roberto Amato, Jeffrey Barrett, Sonia Goncalves, Ewan Harrison, David K. Jackson, Ian Johnston, Dominic Kwiatkowski, Cordelia Langford, John Sillitoe on behalf of the Wellcome Sanger Institute COVID-19 Surveillance Team |
| EPI_ISL_2973809 | Lighthouse Lab in Glasgow | Wellcome Sanger Institute for the COVID-19 Genomics UK (COG-UK) Consortium | Harper VanSteenhouse, Yumi Kasai, David Gray, Carol Clugston, Anna Dominiczak and Alex Alderton, Roberto Amato, Jeffrey Barrett, Sonia Goncalves, Ewan Harrison, David K. Jackson, Ian Johnston, Dominic Kwiatkowski, Cordelia Langford, John Sillitoe on behalf of the Wellcome Sanger Institute COVID-19 Surveillance Team |
| EPI_ISL_2973810, EPI_ISL_2973811, EPI_ISL_2973812 | Lighthouse Lab in Alderley Park | Wellcome Sanger Institute for the COVID-19 Genomics UK (COG-UK) Consortium | Jacquelyn Wynn, Mairead Hyland, The Lighthouse Lab in Alderley Park and Alex Alderton, Roberto Amato, Jeffrey Barrett, Sonia Goncalves, Ewan Harrison, David K. Jackson, Ian Johnston, Dominic Kwiatkowski, Cordelia Langford, John Sillitoe on behalf of the Wellcome Sanger Institute COVID-19 Surveillance Team |
| EPI_ISL_2973813, EPI_ISL_2973814, EPI_ISL_2973815 | Lighthouse Lab in Glasgow | Wellcome Sanger Institute for the COVID-19 Genomics UK (COG-UK) Consortium | Harper VanSteenhouse, Yumi Kasai, David Gray, Carol Clugston, Anna Dominiczak and Alex Alderton, Roberto Amato, Jeffrey Barrett, Sonia Goncalves, Ewan Harrison, David K. Jackson, Ian Johnston, Dominic Kwiatkowski, Cordelia Langford, John Sillitoe on behalf of the Wellcome Sanger Institute COVID-19 Surveillance Team |
| EPI_ISL_2973816, EPI_ISL_2973817 | Lighthouse Lab in Alderley Park | Wellcome Sanger Institute for the COVID-19 Genomics UK (COG-UK) Consortium | Jacquelyn Wynn, Mairead Hyland, The Lighthouse Lab in Alderley Park and Alex Alderton, Roberto Amato, Jeffrey Barrett, Sonia Goncalves, Ewan Harrison, David K. Jackson, Ian Johnston, Dominic Kwiatkowski, Cordelia Langford, John Sillitoe on behalf of the Wellcome Sanger Institute COVID-19 Surveillance Team |
| EPI_ISL_2973818 | Lighthouse Lab in Glasgow | Wellcome Sanger Institute for the COVID-19 Genomics UK (COG-UK) Consortium | Harper VanSteenhouse, Yumi Kasai, David Gray, Carol Clugston, Anna Dominiczak and Alex Alderton, Roberto Amato, Jeffrey Barrett, Sonia Goncalves, Ewan Harrison, David K. Jackson, Ian Johnston, Dominic Kwiatkowski, Cordelia Langford, John Sillitoe on behalf of the Wellcome Sanger Institute COVID-19 Surveillance Team |
| EPI_ISL_2973819 | Lighthouse Lab in Alderley Park | Wellcome Sanger Institute for the COVID-19 Genomics UK (COG-UK) Consortium | Jacquelyn Wynn, Mairead Hyland, The Lighthouse Lab in Alderley Park and Alex Alderton, Roberto Amato, Jeffrey Barrett, Sonia Goncalves, Ewan Harrison, David K. Jackson, Ian Johnston, Dominic Kwiatkowski, Cordelia Langford, John Sillitoe on behalf of the Wellcome Sanger Institute COVID-19 Surveillance Team |
| EPI_ISL_2973820, EPI_ISL_2973821, EPI_ISL_2973822 | Lighthouse Lab in Glasgow | Wellcome Sanger Institute for the COVID-19 Genomics UK (COG-UK) Consortium | Harper VanSteenhouse, Yumi Kasai, David Gray, Carol Clugston, Anna Dominiczak and Alex Alderton, Roberto Amato, Jeffrey Barrett, Sonia Goncalves, Ewan Harrison, David K. Jackson, Ian Johnston, Dominic Kwiatkowski, Cordelia Langford, John Sillitoe on behalf of the Wellcome Sanger Institute COVID-19 Surveillance Team |
| EPI_ISL_2973823, EPI_ISL_2973824, EPI_ISL_2973825, EPI_ISL_2973826, EPI_ISL_2973827, EPI_ISL_2973828 | Lighthouse Lab in Alderley Park | Wellcome Sanger Institute for the COVID-19 Genomics UK (COG-UK) Consortium | Jacquelyn Wynn, Mairead Hyland, The Lighthouse Lab in Alderley Park and Alex Alderton, Roberto Amato, Jeffrey Barrett, Sonia Goncalves, Ewan Harrison, David K. Jackson, Ian Johnston, Dominic Kwiatkowski, Cordelia Langford, John Sillitoe on behalf of the Wellcome Sanger Institute COVID-19 Surveillance Team |
| EPI_ISL_2973829 | Lighthouse Lab in Glasgow | Wellcome Sanger Institute for the COVID-19 Genomics UK (COG-UK) Consortium | Harper VanSteenhouse, Yumi Kasai, David Gray, Carol Clugston, Anna Dominiczak and Alex Alderton, Roberto Amato, Jeffrey Barrett, Sonia Goncalves, Ewan Harrison, David K. Jackson, Ian Johnston, Dominic Kwiatkowski, Cordelia Langford, John Sillitoe on behalf of the Wellcome Sanger Institute COVID-19 Surveillance Team |
| EPI_ISL_2973830 | Lighthouse Lab in Alderley Park | Wellcome Sanger Institute for the COVID-19 Genomics UK (COG-UK) Consortium | Jacquelyn Wynn, Mairead Hyland, The Lighthouse Lab in Alderley Park and Alex Alderton, Roberto Amato, Jeffrey Barrett, Sonia Goncalves, Ewan Harrison, David K. Jackson, Ian Johnston, Dominic Kwiatkowski, Cordelia Langford, John Sillitoe on behalf of the Wellcome Sanger Institute COVID-19 Surveillance Team |
| EPI_ISL_2973831 | Lighthouse Lab in Glasgow | Wellcome Sanger Institute for the COVID-19 Genomics UK (COG-UK) Consortium | Harper VanSteenhouse, Yumi Kasai, David Gray, Carol Clugston, Anna Dominiczak and Alex Alderton, Roberto Amato, Jeffrey Barrett, Sonia Goncalves, Ewan Harrison, David K. Jackson, Ian Johnston, Dominic Kwiatkowski, Cordelia Langford, John Sillitoe on behalf of the Wellcome Sanger Institute COVID-19 Surveillance Team |
| EPI_ISL_2973832 | Lighthouse Lab in Alderley Park | Wellcome Sanger Institute for the COVID-19 Genomics UK (COG-UK) Consortium | Jacquelyn Wynn, Mairead Hyland, The Lighthouse Lab in Alderley Park and Alex Alderton, Roberto Amato, Jeffrey Barrett, Sonia Goncalves, Ewan Harrison, David K. Jackson, Ian Johnston, Dominic Kwiatkowski, Cordelia Langford, John Sillitoe on behalf of the Wellcome Sanger Institute COVID-19 Surveillance Team |
| EPI_ISL_2973833 | Lighthouse Lab in Glasgow | Wellcome Sanger Institute for the COVID-19 Genomics UK (COG-UK) Consortium | Harper VanSteenhouse, Yumi Kasai, David Gray, Carol Clugston, Anna Dominiczak and Alex Alderton, Roberto Amato, Jeffrey Barrett, Sonia Goncalves, Ewan Harrison, David K. Jackson, Ian Johnston, Dominic Kwiatkowski, Cordelia Langford, John Sillitoe on behalf of the Wellcome Sanger Institute COVID-19 Surveillance Team |
| EPI_ISL_2973834, EPI_ISL_2973835 | Lighthouse Lab in Alderley Park | Wellcome Sanger Institute for the COVID-19 Genomics UK (COG-UK) Consortium | Jacquelyn Wynn, Mairead Hyland, The Lighthouse Lab in Alderley Park and Alex Alderton, Roberto Amato, Jeffrey Barrett, Sonia Goncalves, Ewan Harrison, David K. Jackson, Ian Johnston, Dominic Kwiatkowski, Cordelia Langford, John Sillitoe on behalf of the Wellcome Sanger Institute COVID-19 Surveillance Team |

| | | | |
|---|---|---|---|
| EPI_ISL_2973836, EPI_ISL_2973837 | Lighthouse Lab in Glasgow | Wellcome Sanger Institute for the COVID-19 Genomics UK (COG-UK) Consortium | Harper VanSteenhouse, Yumi Kasai, David Gray, Carol Clugston, Anna Dominiczak and Alex Alderton, Roberto Amato, Jeffrey Barrett, Sonia Goncalves, Ewan Harrison, David K. Jackson, Ian Johnston, Dominic Kwiatkowski, Cordelia Langford, John Sillitoe on behalf of the Wellcome Sanger Institute COVID-19 Surveillance Team |
| EPI_ISL_2973838, EPI_ISL_2973839, EPI_ISL_2973840, EPI_ISL_2973841, EPI_ISL_2973842, EPI_ISL_2973843 | Lighthouse Lab in Alderley Park | Wellcome Sanger Institute for the COVID-19 Genomics UK (COG-UK) Consortium | Jacquelyn Wynn, Mairead Hyland, The Lighthouse Lab in Alderley Park and Alex Alderton, Roberto Amato, Jeffrey Barrett, Sonia Goncalves, Ewan Harrison, David K. Jackson, Ian Johnston, Dominic Kwiatkowski, Cordelia Langford, John Sillitoe on behalf of the Wellcome Sanger Institute COVID-19 Surveillance Team |
| EPI_ISL_2973844, EPI_ISL_2973845, EPI_ISL_2973846, EPI_ISL_2973847, EPI_ISL_2973848, EPI_ISL_2973849 | Lighthouse Lab in Glasgow | Wellcome Sanger Institute for the COVID-19 Genomics UK (COG-UK) Consortium | Harper VanSteenhouse, Yumi Kasai, David Gray, Carol Clugston, Anna Dominiczak and Alex Alderton, Roberto Amato, Jeffrey Barrett, Sonia Goncalves, Ewan Harrison, David K. Jackson, Ian Johnston, Dominic Kwiatkowski, Cordelia Langford, John Sillitoe on behalf of the Wellcome Sanger Institute COVID-19 Surveillance Team |
| EPI_ISL_2973850, EPI_ISL_2973851, EPI_ISL_2973852 | Lighthouse Lab in Alderley Park | Wellcome Sanger Institute for the COVID-19 Genomics UK (COG-UK) Consortium | Jacquelyn Wynn, Mairead Hyland, The Lighthouse Lab in Alderley Park and Alex Alderton, Roberto Amato, Jeffrey Barrett, Sonia Goncalves, Ewan Harrison, David K. Jackson, Ian Johnston, Dominic Kwiatkowski, Cordelia Langford, John Sillitoe on behalf of the Wellcome Sanger Institute COVID-19 Surveillance Team |
| EPI_ISL_2973853, EPI_ISL_2973854, EPI_ISL_2973855 | Lighthouse Lab in Glasgow | Wellcome Sanger Institute for the COVID-19 Genomics UK (COG-UK) Consortium | Harper VanSteenhouse, Yumi Kasai, David Gray, Carol Clugston, Anna Dominiczak and Alex Alderton, Roberto Amato, Jeffrey Barrett, Sonia Goncalves, Ewan Harrison, David K. Jackson, Ian Johnston, Dominic Kwiatkowski, Cordelia Langford, John Sillitoe on behalf of the Wellcome Sanger Institute COVID-19 Surveillance Team |
| EPI_ISL_2973856, EPI_ISL_2973857 | Lighthouse Lab in Alderley Park | Wellcome Sanger Institute for the COVID-19 Genomics UK (COG-UK) Consortium | Jacquelyn Wynn, Mairead Hyland, The Lighthouse Lab in Alderley Park and Alex Alderton, Roberto Amato, Jeffrey Barrett, Sonia Goncalves, Ewan Harrison, David K. Jackson, Ian Johnston, Dominic Kwiatkowski, Cordelia Langford, John Sillitoe on behalf of the Wellcome Sanger Institute COVID-19 Surveillance Team |
| EPI_ISL_2973858 | Lighthouse Lab in Glasgow | Wellcome Sanger Institute for the COVID-19 Genomics UK (COG-UK) Consortium | Harper VanSteenhouse, Yumi Kasai, David Gray, Carol Clugston, Anna Dominiczak and Alex Alderton, Roberto Amato, Jeffrey Barrett, Sonia Goncalves, Ewan Harrison, David K. Jackson, Ian Johnston, Dominic Kwiatkowski, Cordelia Langford, John Sillitoe on behalf of the Wellcome Sanger Institute COVID-19 Surveillance Team |
| EPI_ISL_2973859 | Lighthouse Lab in Alderley Park | Wellcome Sanger Institute for the COVID-19 Genomics UK (COG-UK) Consortium | Jacquelyn Wynn, Mairead Hyland, The Lighthouse Lab in Alderley Park and Alex Alderton, Roberto Amato, Jeffrey Barrett, Sonia Goncalves, Ewan Harrison, David K. Jackson, Ian Johnston, Dominic Kwiatkowski, Cordelia Langford, John Sillitoe on behalf of the Wellcome Sanger Institute COVID-19 Surveillance Team |
| EPI_ISL_2973860 | Lighthouse Lab in Glasgow | Wellcome Sanger Institute for the COVID-19 Genomics UK (COG-UK) Consortium | Harper VanSteenhouse, Yumi Kasai, David Gray, Carol Clugston, Anna Dominiczak and Alex Alderton, Roberto Amato, Jeffrey Barrett, Sonia Goncalves, Ewan Harrison, David K. Jackson, Ian Johnston, Dominic Kwiatkowski, Cordelia Langford, John Sillitoe on behalf of the Wellcome Sanger Institute COVID-19 Surveillance Team |
| EPI_ISL_2973861 | Lighthouse Lab in Alderley Park | Wellcome Sanger Institute for the COVID-19 Genomics UK (COG-UK) Consortium | Jacquelyn Wynn, Mairead Hyland, The Lighthouse Lab in Alderley Park and Alex Alderton, Roberto Amato, Jeffrey Barrett, Sonia Goncalves, Ewan Harrison, David K. Jackson, Ian Johnston, Dominic Kwiatkowski, Cordelia Langford, John Sillitoe on behalf of the Wellcome Sanger Institute COVID-19 Surveillance Team |
| EPI_ISL_2973862 | Lighthouse Lab in Glasgow | Wellcome Sanger Institute for the COVID-19 Genomics UK (COG-UK) Consortium | Harper VanSteenhouse, Yumi Kasai, David Gray, Carol Clugston, Anna Dominiczak and Alex Alderton, Roberto Amato, Jeffrey Barrett, Sonia Goncalves, Ewan Harrison, David K. Jackson, Ian Johnston, Dominic Kwiatkowski, Cordelia Langford, John Sillitoe on behalf of the Wellcome Sanger Institute COVID-19 Surveillance Team |
| EPI_ISL_2973863 | Lighthouse Lab in Alderley Park | Wellcome Sanger Institute for the COVID-19 Genomics UK (COG-UK) Consortium | Jacquelyn Wynn, Mairead Hyland, The Lighthouse Lab in Alderley Park and Alex Alderton, Roberto Amato, Jeffrey Barrett, Sonia Goncalves, Ewan Harrison, David K. Jackson, Ian Johnston, Dominic Kwiatkowski, Cordelia Langford, John Sillitoe on behalf of the Wellcome Sanger Institute COVID-19 Surveillance Team |
| EPI_ISL_2973864, EPI_ISL_2973865 | Lighthouse Lab in Glasgow | Wellcome Sanger Institute for the COVID-19 Genomics UK (COG-UK) Consortium | Harper VanSteenhouse, Yumi Kasai, David Gray, Carol Clugston, Anna Dominiczak and Alex Alderton, Roberto Amato, Jeffrey Barrett, Sonia Goncalves, Ewan Harrison, David K. Jackson, Ian Johnston, Dominic Kwiatkowski, Cordelia Langford, John Sillitoe on behalf of the Wellcome Sanger Institute COVID-19 Surveillance Team |
| EPI_ISL_2973866 | Lighthouse Lab in Alderley Park | Wellcome Sanger Institute for the COVID-19 Genomics UK (COG-UK) Consortium | Jacquelyn Wynn, Mairead Hyland, The Lighthouse Lab in Alderley Park and Alex Alderton, Roberto Amato, Jeffrey Barrett, Sonia Goncalves, Ewan Harrison, David K. Jackson, Ian Johnston, Dominic Kwiatkowski, Cordelia Langford, John Sillitoe on behalf of the Wellcome Sanger Institute COVID-19 Surveillance Team |
| EPI_ISL_2973867, EPI_ISL_2973868 | Lighthouse Lab in Glasgow | Wellcome Sanger Institute for the COVID-19 Genomics UK (COG-UK) Consortium | Harper VanSteenhouse, Yumi Kasai, David Gray, Carol Clugston, Anna Dominiczak and Alex Alderton, Roberto Amato, Jeffrey Barrett, Sonia Goncalves, Ewan Harrison, David K. Jackson, Ian Johnston, Dominic Kwiatkowski, Cordelia Langford, John Sillitoe on behalf of the Wellcome Sanger Institute COVID-19 Surveillance Team |
| EPI_ISL_2973869, EPI_ISL_2973870, EPI_ISL_2973871, EPI_ISL_2973872, EPI_ISL_2973873, EPI_ISL_2973874, EPI_ISL_2973875 | Lighthouse Lab in Alderley Park | Wellcome Sanger Institute for the COVID-19 Genomics UK (COG-UK) Consortium | Jacquelyn Wynn, Mairead Hyland, The Lighthouse Lab in Alderley Park and Alex Alderton, Roberto Amato, Jeffrey Barrett, Sonia Goncalves, Ewan Harrison, David K. Jackson, Ian Johnston, Dominic Kwiatkowski, Cordelia Langford, John Sillitoe on behalf of the Wellcome Sanger Institute COVID-19 Surveillance Team |
| EPI_ISL_2973876 | Lighthouse Lab in Glasgow | Wellcome Sanger Institute for the COVID-19 Genomics UK (COG-UK) Consortium | Harper VanSteenhouse, Yumi Kasai, David Gray, Carol Clugston, Anna Dominiczak and Alex Alderton, Roberto Amato, Jeffrey Barrett, Sonia Goncalves, Ewan Harrison, David K. Jackson, Ian Johnston, Dominic Kwiatkowski, Cordelia Langford, John Sillitoe on behalf of the Wellcome Sanger Institute COVID-19 Surveillance Team |
| EPI_ISL_2973877, EPI_ISL_2973878, EPI_ISL_2973879, EPI_ISL_2973880, EPI_ISL_2973881, EPI_ISL_2973882, EPI_ISL_2973883 | Lighthouse Lab in Alderley Park | Wellcome Sanger Institute for the COVID-19 Genomics UK (COG-UK) Consortium | Jacquelyn Wynn, Mairead Hyland, The Lighthouse Lab in Alderley Park and Alex Alderton, Roberto Amato, Jeffrey Barrett, Sonia Goncalves, Ewan Harrison, David K. Jackson, Ian Johnston, Dominic Kwiatkowski, Cordelia Langford, John Sillitoe on behalf of the Wellcome Sanger Institute COVID-19 Surveillance Team |
| EPI_ISL_2973884 | Lighthouse Lab in Glasgow | Wellcome Sanger Institute for the COVID-19 Genomics UK (COG-UK) Consortium | Harper VanSteenhouse, Yumi Kasai, David Gray, Carol Clugston, Anna Dominiczak and Alex Alderton, Roberto Amato, Jeffrey Barrett, Sonia Goncalves, Ewan Harrison, David K. Jackson, Ian Johnston, Dominic Kwiatkowski, Cordelia Langford, John Sillitoe on behalf of the Wellcome Sanger Institute COVID-19 Surveillance Team |
| EPI_ISL_2973885 | Lighthouse Lab in Alderley Park | Wellcome Sanger Institute for the COVID-19 Genomics UK (COG-UK) Consortium | Jacquelyn Wynn, Mairead Hyland, The Lighthouse Lab in Alderley Park and Alex Alderton, Roberto Amato, Jeffrey Barrett, Sonia Goncalves, Ewan Harrison, David K. Jackson, Ian Johnston, Dominic Kwiatkowski, Cordelia Langford, John Sillitoe on behalf of the Wellcome Sanger Institute COVID-19 Surveillance Team |
| EPI_ISL_2973886, EPI_ISL_2973887, EPI_ISL_2973888 | Lighthouse Lab in Glasgow | Wellcome Sanger Institute for the COVID-19 Genomics UK (COG-UK) Consortium | Harper VanSteenhouse, Yumi Kasai, David Gray, Carol Clugston, Anna Dominiczak and Alex Alderton, Roberto Amato, Jeffrey Barrett, Sonia Goncalves, Ewan Harrison, David K. Jackson, Ian Johnston, Dominic Kwiatkowski, Cordelia Langford, John Sillitoe on behalf of the Wellcome Sanger Institute COVID-19 Surveillance Team |
| EPI_ISL_2973889 | Lighthouse Lab in Alderley Park | Wellcome Sanger Institute for the COVID-19 Genomics UK (COG-UK) Consortium | Jacquelyn Wynn, Mairead Hyland, The Lighthouse Lab in Alderley Park and Alex Alderton, Roberto Amato, Jeffrey Barrett, Sonia Goncalves, Ewan Harrison, David K. Jackson, Ian Johnston, Dominic Kwiatkowski, Cordelia Langford, John Sillitoe on behalf of the Wellcome Sanger Institute COVID-19 Surveillance Team |
| EPI_ISL_2973890, EPI_ISL_2973891 | Lighthouse Lab in Glasgow | Wellcome Sanger Institute for the COVID-19 Genomics UK (COG-UK) Consortium | Harper VanSteenhouse, Yumi Kasai, David Gray, Carol Clugston, Anna Dominiczak and Alex Alderton, Roberto Amato, Jeffrey Barrett, Sonia Goncalves, Ewan Harrison, David K. Jackson, Ian Johnston, Dominic Kwiatkowski, Cordelia Langford, John Sillitoe on behalf of the Wellcome Sanger Institute COVID-19 Surveillance Team |
| EPI_ISL_2973892 | Lighthouse Lab in Alderley Park | Wellcome Sanger Institute for the COVID-19 Genomics UK (COG-UK) Consortium | Jacquelyn Wynn, Mairead Hyland, The Lighthouse Lab in Alderley Park and Alex Alderton, Roberto Amato, Jeffrey Barrett, Sonia Goncalves, Ewan Harrison, David K. Jackson, Ian Johnston, Dominic Kwiatkowski, Cordelia Langford, John Sillitoe on behalf of the Wellcome Sanger Institute COVID-19 Surveillance Team |
| EPI_ISL_2973893, EPI_ISL_2973894, EPI_ISL_2973895 | Lighthouse Lab in Glasgow | Wellcome Sanger Institute for the COVID-19 Genomics UK (COG-UK) Consortium | Harper VanSteenhouse, Yumi Kasai, David Gray, Carol Clugston, Anna Dominiczak and Alex Alderton, Roberto Amato, Jeffrey Barrett, Sonia Goncalves, Ewan Harrison, David K. Jackson, Ian Johnston, Dominic Kwiatkowski, Cordelia Langford, John Sillitoe on behalf of the Wellcome Sanger Institute COVID-19 Surveillance Team |
| EPI_ISL_2973896, EPI_ISL_2973897, EPI_ISL_2973898 | Lighthouse Lab in Alderley Park | Wellcome Sanger Institute for the COVID-19 Genomics UK (COG-UK) Consortium | Jacquelyn Wynn, Mairead Hyland, The Lighthouse Lab in Alderley Park and Alex Alderton, Roberto Amato, Jeffrey Barrett, Sonia Goncalves, Ewan Harrison, David K. Jackson, Ian Johnston, Dominic Kwiatkowski, Cordelia Langford, John Sillitoe on behalf of the Wellcome Sanger Institute COVID-19 Surveillance Team |
| EPI_ISL_2973899 | Lighthouse Lab in Glasgow | Wellcome Sanger Institute for the COVID-19 Genomics UK (COG-UK) Consortium | Harper VanSteenhouse, Yumi Kasai, David Gray, Carol Clugston, Anna Dominiczak and Alex Alderton, Roberto Amato, Jeffrey Barrett, Sonia Goncalves, Ewan Harrison, David K. Jackson, Ian Johnston, Dominic Kwiatkowski, Cordelia Langford, John Sillitoe on behalf of the Wellcome Sanger Institute COVID-19 Surveillance Team |
| EPI_ISL_2973900, EPI_ISL_2973901 | Lighthouse Lab in Alderley Park | Wellcome Sanger Institute for the COVID-19 Genomics UK (COG-UK) Consortium | Jacquelyn Wynn, Mairead Hyland, The Lighthouse Lab in Alderley Park and Alex Alderton, Roberto Amato, Jeffrey Barrett, Sonia Goncalves, Ewan Harrison, David K. Jackson, Ian Johnston, Dominic Kwiatkowski, Cordelia Langford, John Sillitoe on behalf of the Wellcome Sanger Institute COVID-19 Surveillance Team |
| EPI_ISL_2973902, EPI_ISL_2973903, EPI_ISL_2973904, EPI_ISL_2973905, EPI_ISL_2973906 | Lighthouse Lab in Glasgow | Wellcome Sanger Institute for the COVID-19 Genomics UK (COG-UK) Consortium | Harper VanSteenhouse, Yumi Kasai, David Gray, Carol Clugston, Anna Dominiczak and Alex Alderton, Roberto Amato, Jeffrey Barrett, Sonia Goncalves, Ewan Harrison, David K. Jackson, Ian Johnston, Dominic Kwiatkowski, Cordelia Langford, John Sillitoe on behalf of the Wellcome Sanger Institute COVID-19 Surveillance Team |
| EPI_ISL_2973907, EPI_ISL_2973908, EPI_ISL_2973909, EPI_ISL_2973910, EPI_ISL_2973911, EPI_ISL_2973912, EPI_ISL_2973913 | Lighthouse Lab in Alderley Park | Wellcome Sanger Institute for the COVID-19 Genomics UK (COG-UK) Consortium | Jacquelyn Wynn, Mairead Hyland, The Lighthouse Lab in Alderley Park and Alex Alderton, Roberto Amato, Jeffrey Barrett, Sonia Goncalves, Ewan Harrison, David K. Jackson, Ian Johnston, Dominic Kwiatkowski, Cordelia Langford, John Sillitoe on behalf of the Wellcome Sanger Institute COVID-19 Surveillance Team |
| EPI_ISL_2973914 | Lighthouse Lab in Glasgow | Wellcome Sanger Institute for the COVID-19 Genomics UK (COG-UK) Consortium | Harper VanSteenhouse, Yumi Kasai, David Gray, Carol Clugston, Anna Dominiczak and Alex Alderton, Roberto Amato, Jeffrey Barrett, Sonia Goncalves, Ewan Harrison, David K. Jackson, Ian Johnston, Dominic Kwiatkowski, Cordelia Langford, John Sillitoe on behalf of the Wellcome Sanger Institute COVID-19 Surveillance Team |
| EPI_ISL_2973915, EPI_ISL_2973916 | Lighthouse Lab in Alderley Park | Wellcome Sanger Institute for the COVID-19 Genomics UK (COG-UK) Consortium | Jacquelyn Wynn, Mairead Hyland, The Lighthouse Lab in Alderley Park and Alex Alderton, Roberto Amato, Jeffrey Barrett, Sonia Goncalves, Ewan Harrison, David K. Jackson, Ian Johnston, Dominic Kwiatkowski, Cordelia Langford, John Sillitoe on behalf of the Wellcome Sanger Institute COVID-19 Surveillance Team |
| EPI_ISL_2973917, | Lighthouse Lab in Glasgow | Wellcome Sanger Institute for the COVID-19 Genomics UK (COG-UK) | Harper VanSteenhouse, Yumi Kasai, David Gray, Carol Clugston, Anna Dominiczak and Alex Alderton, Roberto Amato, Jeffrey Barrett, Sonia Goncalves, Ewan Harrison, David K. Jackson, Ian Johnston, Dominic Kwiatkowski, Cordelia Langford, |

| | | | |
|---|---|---|---|
| EPI_ISL_2973918 | | Consortium | John Sillitoe on behalf of the Wellcome Sanger Institute COVID-19 Surveillance Team |
| EPI_ISL_2973919 | Lighthouse Lab in Alderley Park | Wellcome Sanger Institute for the COVID-19 Genomics UK (COG-UK) Consortium | Jacquelyn Wynn, Mairead Hyland, The Lighthouse Lab in Alderley Park and Alex Alderton, Roberto Amato, Jeffrey Barrett, Sonia Goncalves, Ewan Harrison, David K. Jackson, Ian Johnston, Dominic Kwiatkowski, Cordelia Langford, John Sillitoe on behalf of the Wellcome Sanger Institute COVID-19 Surveillance Team |
| EPI_ISL_2973920, EPI_ISL_2973921, EPI_ISL_2973922, EPI_ISL_2973923, EPI_ISL_2973924, EPI_ISL_2973925 | Lighthouse Lab in Glasgow | Wellcome Sanger Institute for the COVID-19 Genomics UK (COG-UK) Consortium | Harper VanSteenhouse, Yumi Kasai, David Gray, Carol Clugston, Anna Dominiczak and Alex Alderton, Roberto Amato, Jeffrey Barrett, Sonia Goncalves, Ewan Harrison, David K. Jackson, Ian Johnston, Dominic Kwiatkowski, Cordelia Langford, John Sillitoe on behalf of the Wellcome Sanger Institute COVID-19 Surveillance Team |
| EPI_ISL_2973926, EPI_ISL_2973927 | Lighthouse Lab in Alderley Park | Wellcome Sanger Institute for the COVID-19 Genomics UK (COG-UK) Consortium | Jacquelyn Wynn, Mairead Hyland, The Lighthouse Lab in Alderley Park and Alex Alderton, Roberto Amato, Jeffrey Barrett, Sonia Goncalves, Ewan Harrison, David K. Jackson, Ian Johnston, Dominic Kwiatkowski, Cordelia Langford, John Sillitoe on behalf of the Wellcome Sanger Institute COVID-19 Surveillance Team |
| EPI_ISL_2973928, EPI_ISL_2973929 | Lighthouse Lab in Glasgow | Wellcome Sanger Institute for the COVID-19 Genomics UK (COG-UK) Consortium | Harper VanSteenhouse, Yumi Kasai, David Gray, Carol Clugston, Anna Dominiczak and Alex Alderton, Roberto Amato, Jeffrey Barrett, Sonia Goncalves, Ewan Harrison, David K. Jackson, Ian Johnston, Dominic Kwiatkowski, Cordelia Langford, John Sillitoe on behalf of the Wellcome Sanger Institute COVID-19 Surveillance Team |
| EPI_ISL_2973930, EPI_ISL_2973931, EPI_ISL_2973932, EPI_ISL_2973933, EPI_ISL_2973934 | Lighthouse Lab in Alderley Park | Wellcome Sanger Institute for the COVID-19 Genomics UK (COG-UK) Consortium | Jacquelyn Wynn, Mairead Hyland, The Lighthouse Lab in Alderley Park and Alex Alderton, Roberto Amato, Jeffrey Barrett, Sonia Goncalves, Ewan Harrison, David K. Jackson, Ian Johnston, Dominic Kwiatkowski, Cordelia Langford, John Sillitoe on behalf of the Wellcome Sanger Institute COVID-19 Surveillance Team |
| EPI_ISL_2973935 | Lighthouse Lab in Glasgow | Wellcome Sanger Institute for the COVID-19 Genomics UK (COG-UK) Consortium | Harper VanSteenhouse, Yumi Kasai, David Gray, Carol Clugston, Anna Dominiczak and Alex Alderton, Roberto Amato, Jeffrey Barrett, Sonia Goncalves, Ewan Harrison, David K. Jackson, Ian Johnston, Dominic Kwiatkowski, Cordelia Langford, John Sillitoe on behalf of the Wellcome Sanger Institute COVID-19 Surveillance Team |
| EPI_ISL_2973936, EPI_ISL_2973937, EPI_ISL_2973938 | Lighthouse Lab in Alderley Park | Wellcome Sanger Institute for the COVID-19 Genomics UK (COG-UK) Consortium | Jacquelyn Wynn, Mairead Hyland, The Lighthouse Lab in Alderley Park and Alex Alderton, Roberto Amato, Jeffrey Barrett, Sonia Goncalves, Ewan Harrison, David K. Jackson, Ian Johnston, Dominic Kwiatkowski, Cordelia Langford, John Sillitoe on behalf of the Wellcome Sanger Institute COVID-19 Surveillance Team |
| EPI_ISL_2973939 | Lighthouse Lab in Glasgow | Wellcome Sanger Institute for the COVID-19 Genomics UK (COG-UK) Consortium | Harper VanSteenhouse, Yumi Kasai, David Gray, Carol Clugston, Anna Dominiczak and Alex Alderton, Roberto Amato, Jeffrey Barrett, Sonia Goncalves, Ewan Harrison, David K. Jackson, Ian Johnston, Dominic Kwiatkowski, Cordelia Langford, John Sillitoe on behalf of the Wellcome Sanger Institute COVID-19 Surveillance Team |
| EPI_ISL_2973940 | Molecular Diagnostics Pathology Department Mater Dei Hospital Malta | Molecular Diagnostics Pathology Department Mater Dei Hospital Malta | G Zahra, R Borg, C Cilia, L Grech, M Briffa |
| EPI_ISL_2973941, EPI_ISL_2973942 | Lighthouse Lab in Glasgow | Wellcome Sanger Institute for the COVID-19 Genomics UK (COG-UK) Consortium | Harper VanSteenhouse, Yumi Kasai, David Gray, Carol Clugston, Anna Dominiczak and Alex Alderton, Roberto Amato, Jeffrey Barrett, Sonia Goncalves, Ewan Harrison, David K. Jackson, Ian Johnston, Dominic Kwiatkowski, Cordelia Langford, John Sillitoe on behalf of the Wellcome Sanger Institute COVID-19 Surveillance Team |
| EPI_ISL_2973943, EPI_ISL_2973944 | Lighthouse Lab in Alderley Park | Wellcome Sanger Institute for the COVID-19 Genomics UK (COG-UK) Consortium | Jacquelyn Wynn, Mairead Hyland, The Lighthouse Lab in Alderley Park and Alex Alderton, Roberto Amato, Jeffrey Barrett, Sonia Goncalves, Ewan Harrison, David K. Jackson, Ian Johnston, Dominic Kwiatkowski, Cordelia Langford, John Sillitoe on behalf of the Wellcome Sanger Institute COVID-19 Surveillance Team |
| EPI_ISL_2973945, EPI_ISL_2973946 | Lighthouse Lab in Glasgow | Wellcome Sanger Institute for the COVID-19 Genomics UK (COG-UK) Consortium | Harper VanSteenhouse, Yumi Kasai, David Gray, Carol Clugston, Anna Dominiczak and Alex Alderton, Roberto Amato, Jeffrey Barrett, Sonia Goncalves, Ewan Harrison, David K. Jackson, Ian Johnston, Dominic Kwiatkowski, Cordelia Langford, John Sillitoe on behalf of the Wellcome Sanger Institute COVID-19 Surveillance Team |
| EPI_ISL_2973947 | Lighthouse Lab in Alderley Park | Wellcome Sanger Institute for the COVID-19 Genomics UK (COG-UK) Consortium | Jacquelyn Wynn, Mairead Hyland, The Lighthouse Lab in Alderley Park and Alex Alderton, Roberto Amato, Jeffrey Barrett, Sonia Goncalves, Ewan Harrison, David K. Jackson, Ian Johnston, Dominic Kwiatkowski, Cordelia Langford, John Sillitoe on behalf of the Wellcome Sanger Institute COVID-19 Surveillance Team |
| EPI_ISL_2973948 | Lighthouse Lab in Glasgow | Wellcome Sanger Institute for the COVID-19 Genomics UK (COG-UK) Consortium | Harper VanSteenhouse, Yumi Kasai, David Gray, Carol Clugston, Anna Dominiczak and Alex Alderton, Roberto Amato, Jeffrey Barrett, Sonia Goncalves, Ewan Harrison, David K. Jackson, Ian Johnston, Dominic Kwiatkowski, Cordelia Langford, John Sillitoe on behalf of the Wellcome Sanger Institute COVID-19 Surveillance Team |
| EPI_ISL_2973949, EPI_ISL_2973950 | Lighthouse Lab in Alderley Park | Wellcome Sanger Institute for the COVID-19 Genomics UK (COG-UK) Consortium | Jacquelyn Wynn, Mairead Hyland, The Lighthouse Lab in Alderley Park and Alex Alderton, Roberto Amato, Jeffrey Barrett, Sonia Goncalves, Ewan Harrison, David K. Jackson, Ian Johnston, Dominic Kwiatkowski, Cordelia Langford, John Sillitoe on behalf of the Wellcome Sanger Institute COVID-19 Surveillance Team |
| EPI_ISL_2973951, EPI_ISL_2973952, EPI_ISL_2973953, EPI_ISL_2973954 | Lighthouse Lab in Glasgow | Wellcome Sanger Institute for the COVID-19 Genomics UK (COG-UK) Consortium | Harper VanSteenhouse, Yumi Kasai, David Gray, Carol Clugston, Anna Dominiczak and Alex Alderton, Roberto Amato, Jeffrey Barrett, Sonia Goncalves, Ewan Harrison, David K. Jackson, Ian Johnston, Dominic Kwiatkowski, Cordelia Langford, John Sillitoe on behalf of the Wellcome Sanger Institute COVID-19 Surveillance Team |
| EPI_ISL_2973955 | Lighthouse Lab in Alderley Park | Wellcome Sanger Institute for the COVID-19 Genomics UK (COG-UK) Consortium | Jacquelyn Wynn, Mairead Hyland, The Lighthouse Lab in Alderley Park and Alex Alderton, Roberto Amato, Jeffrey Barrett, Sonia Goncalves, Ewan Harrison, David K. Jackson, Ian Johnston, Dominic Kwiatkowski, Cordelia Langford, John Sillitoe on behalf of the Wellcome Sanger Institute COVID-19 Surveillance Team |
| EPI_ISL_2973956, EPI_ISL_2973957 | Lighthouse Lab in Glasgow | Wellcome Sanger Institute for the COVID-19 Genomics UK (COG-UK) Consortium | Harper VanSteenhouse, Yumi Kasai, David Gray, Carol Clugston, Anna Dominiczak and Alex Alderton, Roberto Amato, Jeffrey Barrett, Sonia Goncalves, Ewan Harrison, David K. Jackson, Ian Johnston, Dominic Kwiatkowski, Cordelia Langford, John Sillitoe on behalf of the Wellcome Sanger Institute COVID-19 Surveillance Team |
| EPI_ISL_2973958 | Lighthouse Lab in Alderley Park | Wellcome Sanger Institute for the COVID-19 Genomics UK (COG-UK) Consortium | Jacquelyn Wynn, Mairead Hyland, The Lighthouse Lab in Alderley Park and Alex Alderton, Roberto Amato, Jeffrey Barrett, Sonia Goncalves, Ewan Harrison, David K. Jackson, Ian Johnston, Dominic Kwiatkowski, Cordelia Langford, John Sillitoe on behalf of the Wellcome Sanger Institute COVID-19 Surveillance Team |
| EPI_ISL_2973959 | Lighthouse Lab in Glasgow | Wellcome Sanger Institute for the COVID-19 Genomics UK (COG-UK) Consortium | Harper VanSteenhouse, Yumi Kasai, David Gray, Carol Clugston, Anna Dominiczak and Alex Alderton, Roberto Amato, Jeffrey Barrett, Sonia Goncalves, Ewan Harrison, David K. Jackson, Ian Johnston, Dominic Kwiatkowski, Cordelia Langford, John Sillitoe on behalf of the Wellcome Sanger Institute COVID-19 Surveillance Team |
| EPI_ISL_2973960, EPI_ISL_2973961 | Lighthouse Lab in Alderley Park | Wellcome Sanger Institute for the COVID-19 Genomics UK (COG-UK) Consortium | Jacquelyn Wynn, Mairead Hyland, The Lighthouse Lab in Alderley Park and Alex Alderton, Roberto Amato, Jeffrey Barrett, Sonia Goncalves, Ewan Harrison, David K. Jackson, Ian Johnston, Dominic Kwiatkowski, Cordelia Langford, John Sillitoe on behalf of the Wellcome Sanger Institute COVID-19 Surveillance Team |
| EPI_ISL_2973962 | Lighthouse Lab in Glasgow | Wellcome Sanger Institute for the COVID-19 Genomics UK (COG-UK) Consortium | Harper VanSteenhouse, Yumi Kasai, David Gray, Carol Clugston, Anna Dominiczak and Alex Alderton, Roberto Amato, Jeffrey Barrett, Sonia Goncalves, Ewan Harrison, David K. Jackson, Ian Johnston, Dominic Kwiatkowski, Cordelia Langford, John Sillitoe on behalf of the Wellcome Sanger Institute COVID-19 Surveillance Team |
| EPI_ISL_2973963 | Lighthouse Lab in Alderley Park | Wellcome Sanger Institute for the COVID-19 Genomics UK (COG-UK) Consortium | Jacquelyn Wynn, Mairead Hyland, The Lighthouse Lab in Alderley Park and Alex Alderton, Roberto Amato, Jeffrey Barrett, Sonia Goncalves, Ewan Harrison, David K. Jackson, Ian Johnston, Dominic Kwiatkowski, Cordelia Langford, John Sillitoe on behalf of the Wellcome Sanger Institute COVID-19 Surveillance Team |
| EPI_ISL_2973964 | Lighthouse Lab in Glasgow | Wellcome Sanger Institute for the COVID-19 Genomics UK (COG-UK) Consortium | Harper VanSteenhouse, Yumi Kasai, David Gray, Carol Clugston, Anna Dominiczak and Alex Alderton, Roberto Amato, Jeffrey Barrett, Sonia Goncalves, Ewan Harrison, David K. Jackson, Ian Johnston, Dominic Kwiatkowski, Cordelia Langford, John Sillitoe on behalf of the Wellcome Sanger Institute COVID-19 Surveillance Team |
| EPI_ISL_2973965, EPI_ISL_2973966 | Lighthouse Lab in Alderley Park | Wellcome Sanger Institute for the COVID-19 Genomics UK (COG-UK) Consortium | Jacquelyn Wynn, Mairead Hyland, The Lighthouse Lab in Alderley Park and Alex Alderton, Roberto Amato, Jeffrey Barrett, Sonia Goncalves, Ewan Harrison, David K. Jackson, Ian Johnston, Dominic Kwiatkowski, Cordelia Langford, John Sillitoe on behalf of the Wellcome Sanger Institute COVID-19 Surveillance Team |
| EPI_ISL_2973967, EPI_ISL_2973968 | Lighthouse Lab in Glasgow | Wellcome Sanger Institute for the COVID-19 Genomics UK (COG-UK) Consortium | Harper VanSteenhouse, Yumi Kasai, David Gray, Carol Clugston, Anna Dominiczak and Alex Alderton, Roberto Amato, Jeffrey Barrett, Sonia Goncalves, Ewan Harrison, David K. Jackson, Ian Johnston, Dominic Kwiatkowski, Cordelia Langford, John Sillitoe on behalf of the Wellcome Sanger Institute COVID-19 Surveillance Team |
| EPI_ISL_2973969 | Lighthouse Lab in Alderley Park | Wellcome Sanger Institute for the COVID-19 Genomics UK (COG-UK) Consortium | Jacquelyn Wynn, Mairead Hyland, The Lighthouse Lab in Alderley Park and Alex Alderton, Roberto Amato, Jeffrey Barrett, Sonia Goncalves, Ewan Harrison, David K. Jackson, Ian Johnston, Dominic Kwiatkowski, Cordelia Langford, John Sillitoe on behalf of the Wellcome Sanger Institute COVID-19 Surveillance Team |
| EPI_ISL_2973970 | Lighthouse Lab in Glasgow | Wellcome Sanger Institute for the COVID-19 Genomics UK (COG-UK) Consortium | Harper VanSteenhouse, Yumi Kasai, David Gray, Carol Clugston, Anna Dominiczak and Alex Alderton, Roberto Amato, Jeffrey Barrett, Sonia Goncalves, Ewan Harrison, David K. Jackson, Ian Johnston, Dominic Kwiatkowski, Cordelia Langford, John Sillitoe on behalf of the Wellcome Sanger Institute COVID-19 Surveillance Team |
| EPI_ISL_2973971, EPI_ISL_2973972 | Lighthouse Lab in Alderley Park | Wellcome Sanger Institute for the COVID-19 Genomics UK (COG-UK) Consortium | Jacquelyn Wynn, Mairead Hyland, The Lighthouse Lab in Alderley Park and Alex Alderton, Roberto Amato, Jeffrey Barrett, Sonia Goncalves, Ewan Harrison, David K. Jackson, Ian Johnston, Dominic Kwiatkowski, Cordelia Langford, John Sillitoe on behalf of the Wellcome Sanger Institute COVID-19 Surveillance Team |
| EPI_ISL_2973973 | Lighthouse Lab in Glasgow | Wellcome Sanger Institute for the COVID-19 Genomics UK (COG-UK) Consortium | Harper VanSteenhouse, Yumi Kasai, David Gray, Carol Clugston, Anna Dominiczak and Alex Alderton, Roberto Amato, Jeffrey Barrett, Sonia Goncalves, Ewan Harrison, David K. Jackson, Ian Johnston, Dominic Kwiatkowski, Cordelia Langford, John Sillitoe on behalf of the Wellcome Sanger Institute COVID-19 Surveillance Team |
| EPI_ISL_2973974, EPI_ISL_2973975 | Lighthouse Lab in Alderley Park | Wellcome Sanger Institute for the COVID-19 Genomics UK (COG-UK) Consortium | Jacquelyn Wynn, Mairead Hyland, The Lighthouse Lab in Alderley Park and Alex Alderton, Roberto Amato, Jeffrey Barrett, Sonia Goncalves, Ewan Harrison, David K. Jackson, Ian Johnston, Dominic Kwiatkowski, Cordelia Langford, John Sillitoe on behalf of the Wellcome Sanger Institute COVID-19 Surveillance Team |
| EPI_ISL_2973976 | Lighthouse Lab in Glasgow | Wellcome Sanger Institute for the COVID-19 Genomics UK (COG-UK) Consortium | Harper VanSteenhouse, Yumi Kasai, David Gray, Carol Clugston, Anna Dominiczak and Alex Alderton, Roberto Amato, Jeffrey Barrett, Sonia Goncalves, Ewan Harrison, David K. Jackson, Ian Johnston, Dominic Kwiatkowski, Cordelia Langford, John Sillitoe on behalf of the Wellcome Sanger Institute COVID-19 Surveillance Team |
| EPI_ISL_2973977 | Lighthouse Lab in Alderley Park | Wellcome Sanger Institute for the COVID-19 Genomics UK (COG-UK) Consortium | Jacquelyn Wynn, Mairead Hyland, The Lighthouse Lab in Alderley Park and Alex Alderton, Roberto Amato, Jeffrey Barrett, Sonia Goncalves, Ewan Harrison, David K. Jackson, Ian Johnston, Dominic Kwiatkowski, Cordelia Langford, John Sillitoe on behalf of the Wellcome Sanger Institute COVID-19 Surveillance Team |
| EPI_ISL_2973978, EPI_ISL_2973979, EPI_ISL_2973980, EPI_ISL_2973981 | Lighthouse Lab in Glasgow | Wellcome Sanger Institute for the COVID-19 Genomics UK (COG-UK) Consortium | Harper VanSteenhouse, Yumi Kasai, David Gray, Carol Clugston, Anna Dominiczak and Alex Alderton, Roberto Amato, Jeffrey Barrett, Sonia Goncalves, Ewan Harrison, David K. Jackson, Ian Johnston, Dominic Kwiatkowski, Cordelia Langford, John Sillitoe on behalf of the Wellcome Sanger Institute COVID-19 Surveillance Team |
| EPI_ISL_2973982 | Lighthouse Lab in Alderley Park | Wellcome Sanger Institute for the COVID-19 Genomics UK (COG-UK) Consortium | Jacquelyn Wynn, Mairead Hyland, The Lighthouse Lab in Alderley Park and Alex Alderton, Roberto Amato, Jeffrey Barrett, Sonia Goncalves, Ewan Harrison, David K. Jackson, Ian Johnston, Dominic Kwiatkowski, Cordelia Langford, John Sillitoe on behalf of the Wellcome Sanger Institute COVID-19 Surveillance Team |
| EPI_ISL_2973983, EPI_ISL_2973984 | Lighthouse Lab in Glasgow | Wellcome Sanger Institute for the COVID-19 Genomics UK (COG-UK) Consortium | Harper VanSteenhouse, Yumi Kasai, David Gray, Carol Clugston, Anna Dominiczak and Alex Alderton, Roberto Amato, Jeffrey Barrett, Sonia Goncalves, Ewan Harrison, David K. Jackson, Ian Johnston, Dominic Kwiatkowski, Cordelia Langford, John Sillitoe on behalf of the Wellcome Sanger Institute COVID-19 Surveillance Team |
| EPI_ISL_2973985, EPI_ISL_2973986, EPI_ISL_2973987 | Lighthouse Lab in Alderley Park | Wellcome Sanger Institute for the COVID-19 Genomics UK (COG-UK) Consortium | Jacquelyn Wynn, Mairead Hyland, The Lighthouse Lab in Alderley Park and Alex Alderton, Roberto Amato, Jeffrey Barrett, Sonia Goncalves, Ewan Harrison, David K. Jackson, Ian Johnston, Dominic Kwiatkowski, Cordelia Langford, John Sillitoe on behalf of the Wellcome Sanger Institute COVID-19 Surveillance Team |
| EPI_ISL_2973988, EPI_ISL_2973989 | Lighthouse Lab in Glasgow | Wellcome Sanger Institute for the COVID-19 Genomics UK (COG-UK) Consortium | Harper VanSteenhouse, Yumi Kasai, David Gray, Carol Clugston, Anna Dominiczak and Alex Alderton, Roberto Amato, Jeffrey Barrett, Sonia Goncalves, Ewan Harrison, David K. Jackson, Ian Johnston, Dominic Kwiatkowski, Cordelia Langford, John Sillitoe on behalf of the Wellcome Sanger Institute COVID-19 Surveillance Team |
| EPI_ISL_2973990 | Lighthouse Lab in Alderley Park | Wellcome Sanger Institute for the COVID-19 Genomics UK (COG-UK) Consortium | Jacquelyn Wynn, Mairead Hyland, The Lighthouse Lab in Alderley Park and Alex Alderton, Roberto Amato, Jeffrey Barrett, Sonia Goncalves, Ewan Harrison, David K. Jackson, Ian Johnston, Dominic Kwiatkowski, Cordelia Langford, John Sillitoe on behalf of the Wellcome Sanger Institute COVID-19 Surveillance Team |
| EPI_ISL_2973991 | Lighthouse Lab in Glasgow | Wellcome Sanger Institute for the COVID-19 Genomics UK (COG-UK) | Harper VanSteenhouse, Yumi Kasai, David Gray, Carol Clugston, Anna Dominiczak and Alex Alderton, Roberto Amato, Jeffrey Barrett, Sonia Goncalves, Ewan Harrison, David K. Jackson, Ian Johnston, Dominic Kwiatkowski, Cordelia Langford, |

| | | | |
|---|---|---|---|
| | | Consortium | John Sillitoe on behalf of the Wellcome Sanger Institute COVID-19 Surveillance Team |
| EPI_ISL_2973992, EPI_ISL_2973993, EPI_ISL_2973994, EPI_ISL_2973995, EPI_ISL_2973996, EPI_ISL_2973997, EPI_ISL_2973998, EPI_ISL_2973999, EPI_ISL_2974000, EPI_ISL_2974001, EPI_ISL_2974002 | | | |
| see above | Lighthouse Lab in Alderley Park | Wellcome Sanger Institute for the COVID-19 Genomics UK (COG-UK) Consortium | Jacquelyn Wynn, Mairead Hyland, The Lighthouse Lab in Alderley Park and Alex Alderton, Roberto Amato, Jeffrey Barrett, Sonia Goncalves, Ewan Harrison, David K. Jackson, Ian Johnston, Dominic Kwiatkowski, Cordelia Langford, John Sillitoe on behalf of the Wellcome Sanger Institute COVID-19 Surveillance Team |
| EPI_ISL_2974003, EPI_ISL_2974004 | Lighthouse Lab in Glasgow | Wellcome Sanger Institute for the COVID-19 Genomics UK (COG-UK) Consortium | Harper VanSteenhouse, Yumi Kasai, David Gray, Carol Clugston, Anna Dominiczak and Alex Alderton, Roberto Amato, Jeffrey Barrett, Sonia Goncalves, Ewan Harrison, David K. Jackson, Ian Johnston, Dominic Kwiatkowski, Cordelia Langford, John Sillitoe on behalf of the Wellcome Sanger Institute COVID-19 Surveillance Team |
| EPI_ISL_2974005 | Lighthouse Lab in Alderley Park | Wellcome Sanger Institute for the COVID-19 Genomics UK (COG-UK) Consortium | Jacquelyn Wynn, Mairead Hyland, The Lighthouse Lab in Alderley Park and Alex Alderton, Roberto Amato, Jeffrey Barrett, Sonia Goncalves, Ewan Harrison, David K. Jackson, Ian Johnston, Dominic Kwiatkowski, Cordelia Langford, John Sillitoe on behalf of the Wellcome Sanger Institute COVID-19 Surveillance Team |
| EPI_ISL_2974006, EPI_ISL_2974007, EPI_ISL_2974008 | Lighthouse Lab in Glasgow | Wellcome Sanger Institute for the COVID-19 Genomics UK (COG-UK) Consortium | Harper VanSteenhouse, Yumi Kasai, David Gray, Carol Clugston, Anna Dominiczak and Alex Alderton, Roberto Amato, Jeffrey Barrett, Sonia Goncalves, Ewan Harrison, David K. Jackson, Ian Johnston, Dominic Kwiatkowski, Cordelia Langford, John Sillitoe on behalf of the Wellcome Sanger Institute COVID-19 Surveillance Team |
| EPI_ISL_2974009 | Randox Laboratories | Wellcome Sanger Institute for the COVID-19 Genomics UK (COG-UK) Consortium | Randox Laboratories and Alex Alderton, Roberto Amato, Jeffrey Barrett, Sonia Goncalves, Ewan Harrison, David K. Jackson, Ian Johnston, Dominic Kwiatkowski, Cordelia Langford, John Sillitoe on behalf of the Wellcome Sanger Institute COVID-19 Surveillance Team |
| EPI_ISL_2974010 | Lighthouse Lab in Glasgow | Wellcome Sanger Institute for the COVID-19 Genomics UK (COG-UK) Consortium | Harper VanSteenhouse, Yumi Kasai, David Gray, Carol Clugston, Anna Dominiczak and Alex Alderton, Roberto Amato, Jeffrey Barrett, Sonia Goncalves, Ewan Harrison, David K. Jackson, Ian Johnston, Dominic Kwiatkowski, Cordelia Langford, John Sillitoe on behalf of the Wellcome Sanger Institute COVID-19 Surveillance Team |
| EPI_ISL_2974011, EPI_ISL_2974012 | Randox Laboratories | Wellcome Sanger Institute for the COVID-19 Genomics UK (COG-UK) Consortium | Randox Laboratories and Alex Alderton, Roberto Amato, Jeffrey Barrett, Sonia Goncalves, Ewan Harrison, David K. Jackson, Ian Johnston, Dominic Kwiatkowski, Cordelia Langford, John Sillitoe on behalf of the Wellcome Sanger Institute COVID-19 Surveillance Team |
| EPI_ISL_2974013 | Lighthouse Lab in Glasgow | Wellcome Sanger Institute for the COVID-19 Genomics UK (COG-UK) Consortium | Harper VanSteenhouse, Yumi Kasai, David Gray, Carol Clugston, Anna Dominiczak and Alex Alderton, Roberto Amato, Jeffrey Barrett, Sonia Goncalves, Ewan Harrison, David K. Jackson, Ian Johnston, Dominic Kwiatkowski, Cordelia Langford, John Sillitoe on behalf of the Wellcome Sanger Institute COVID-19 Surveillance Team |
| EPI_ISL_2974014, EPI_ISL_2974015 | Randox Laboratories | Wellcome Sanger Institute for the COVID-19 Genomics UK (COG-UK) Consortium | Randox Laboratories and Alex Alderton, Roberto Amato, Jeffrey Barrett, Sonia Goncalves, Ewan Harrison, David K. Jackson, Ian Johnston, Dominic Kwiatkowski, Cordelia Langford, John Sillitoe on behalf of the Wellcome Sanger Institute COVID-19 Surveillance Team |
| EPI_ISL_2974016, EPI_ISL_2974017, EPI_ISL_2974018 | Lighthouse Lab in Glasgow | Wellcome Sanger Institute for the COVID-19 Genomics UK (COG-UK) Consortium | Harper VanSteenhouse, Yumi Kasai, David Gray, Carol Clugston, Anna Dominiczak and Alex Alderton, Roberto Amato, Jeffrey Barrett, Sonia Goncalves, Ewan Harrison, David K. Jackson, Ian Johnston, Dominic Kwiatkowski, Cordelia Langford, John Sillitoe on behalf of the Wellcome Sanger Institute COVID-19 Surveillance Team |
| EPI_ISL_2974019 | Randox Laboratories | Wellcome Sanger Institute for the COVID-19 Genomics UK (COG-UK) Consortium | Randox Laboratories and Alex Alderton, Roberto Amato, Jeffrey Barrett, Sonia Goncalves, Ewan Harrison, David K. Jackson, Ian Johnston, Dominic Kwiatkowski, Cordelia Langford, John Sillitoe on behalf of the Wellcome Sanger Institute COVID-19 Surveillance Team |
| EPI_ISL_2974020, EPI_ISL_2974021, EPI_ISL_2974022, EPI_ISL_2974023 | Lighthouse Lab in Glasgow | Wellcome Sanger Institute for the COVID-19 Genomics UK (COG-UK) Consortium | Harper VanSteenhouse, Yumi Kasai, David Gray, Carol Clugston, Anna Dominiczak and Alex Alderton, Roberto Amato, Jeffrey Barrett, Sonia Goncalves, Ewan Harrison, David K. Jackson, Ian Johnston, Dominic Kwiatkowski, Cordelia Langford, John Sillitoe on behalf of the Wellcome Sanger Institute COVID-19 Surveillance Team |
| EPI_ISL_2974024, EPI_ISL_2974025, EPI_ISL_2974026, EPI_ISL_2974027 | Randox Laboratories | Wellcome Sanger Institute for the COVID-19 Genomics UK (COG-UK) Consortium | Randox Laboratories and Alex Alderton, Roberto Amato, Jeffrey Barrett, Sonia Goncalves, Ewan Harrison, David K. Jackson, Ian Johnston, Dominic Kwiatkowski, Cordelia Langford, John Sillitoe on behalf of the Wellcome Sanger Institute COVID-19 Surveillance Team |
| EPI_ISL_2974028 | Lighthouse Lab in Glasgow | Wellcome Sanger Institute for the COVID-19 Genomics UK (COG-UK) Consortium | Harper VanSteenhouse, Yumi Kasai, David Gray, Carol Clugston, Anna Dominiczak and Alex Alderton, Roberto Amato, Jeffrey Barrett, Sonia Goncalves, Ewan Harrison, David K. Jackson, Ian Johnston, Dominic Kwiatkowski, Cordelia Langford, John Sillitoe on behalf of the Wellcome Sanger Institute COVID-19 Surveillance Team |
| EPI_ISL_2974029 | Randox Laboratories | Wellcome Sanger Institute for the COVID-19 Genomics UK (COG-UK) Consortium | Randox Laboratories and Alex Alderton, Roberto Amato, Jeffrey Barrett, Sonia Goncalves, Ewan Harrison, David K. Jackson, Ian Johnston, Dominic Kwiatkowski, Cordelia Langford, John Sillitoe on behalf of the Wellcome Sanger Institute COVID-19 Surveillance Team |
| EPI_ISL_2974030, EPI_ISL_2974031, EPI_ISL_2974032 | Lighthouse Lab in Glasgow | Wellcome Sanger Institute for the COVID-19 Genomics UK (COG-UK) Consortium | Harper VanSteenhouse, Yumi Kasai, David Gray, Carol Clugston, Anna Dominiczak and Alex Alderton, Roberto Amato, Jeffrey Barrett, Sonia Goncalves, Ewan Harrison, David K. Jackson, Ian Johnston, Dominic Kwiatkowski, Cordelia Langford, John Sillitoe on behalf of the Wellcome Sanger Institute COVID-19 Surveillance Team |
| EPI_ISL_2974033, EPI_ISL_2974034, EPI_ISL_2974035 | Randox Laboratories | Wellcome Sanger Institute for the COVID-19 Genomics UK (COG-UK) Consortium | Randox Laboratories and Alex Alderton, Roberto Amato, Jeffrey Barrett, Sonia Goncalves, Ewan Harrison, David K. Jackson, Ian Johnston, Dominic Kwiatkowski, Cordelia Langford, John Sillitoe on behalf of the Wellcome Sanger Institute COVID-19 Surveillance Team |
| EPI_ISL_2974036, EPI_ISL_2974037 | Lighthouse Lab in Glasgow | Wellcome Sanger Institute for the COVID-19 Genomics UK (COG-UK) Consortium | Harper VanSteenhouse, Yumi Kasai, David Gray, Carol Clugston, Anna Dominiczak and Alex Alderton, Roberto Amato, Jeffrey Barrett, Sonia Goncalves, Ewan Harrison, David K. Jackson, Ian Johnston, Dominic Kwiatkowski, Cordelia Langford, John Sillitoe on behalf of the Wellcome Sanger Institute COVID-19 Surveillance Team |
| EPI_ISL_2974038 | Randox Laboratories | Wellcome Sanger Institute for the COVID-19 Genomics UK (COG-UK) Consortium | Randox Laboratories and Alex Alderton, Roberto Amato, Jeffrey Barrett, Sonia Goncalves, Ewan Harrison, David K. Jackson, Ian Johnston, Dominic Kwiatkowski, Cordelia Langford, John Sillitoe on behalf of the Wellcome Sanger Institute COVID-19 Surveillance Team |
| EPI_ISL_2974039 | Lighthouse Lab in Glasgow | Wellcome Sanger Institute for the COVID-19 Genomics UK (COG-UK) Consortium | Harper VanSteenhouse, Yumi Kasai, David Gray, Carol Clugston, Anna Dominiczak and Alex Alderton, Roberto Amato, Jeffrey Barrett, Sonia Goncalves, Ewan Harrison, David K. Jackson, Ian Johnston, Dominic Kwiatkowski, Cordelia Langford, John Sillitoe on behalf of the Wellcome Sanger Institute COVID-19 Surveillance Team |
| EPI_ISL_2974040, EPI_ISL_2974041, EPI_ISL_2974042 | Randox Laboratories | Wellcome Sanger Institute for the COVID-19 Genomics UK (COG-UK) Consortium | Randox Laboratories and Alex Alderton, Roberto Amato, Jeffrey Barrett, Sonia Goncalves, Ewan Harrison, David K. Jackson, Ian Johnston, Dominic Kwiatkowski, Cordelia Langford, John Sillitoe on behalf of the Wellcome Sanger Institute COVID-19 Surveillance Team |
| EPI_ISL_2974043, EPI_ISL_2974044 | Lighthouse Lab in Glasgow | Wellcome Sanger Institute for the COVID-19 Genomics UK (COG-UK) Consortium | Harper VanSteenhouse, Yumi Kasai, David Gray, Carol Clugston, Anna Dominiczak and Alex Alderton, Roberto Amato, Jeffrey Barrett, Sonia Goncalves, Ewan Harrison, David K. Jackson, Ian Johnston, Dominic Kwiatkowski, Cordelia Langford, John Sillitoe on behalf of the Wellcome Sanger Institute COVID-19 Surveillance Team |
| EPI_ISL_2974045, EPI_ISL_2974046 | Randox Laboratories | Wellcome Sanger Institute for the COVID-19 Genomics UK (COG-UK) Consortium | Randox Laboratories and Alex Alderton, Roberto Amato, Jeffrey Barrett, Sonia Goncalves, Ewan Harrison, David K. Jackson, Ian Johnston, Dominic Kwiatkowski, Cordelia Langford, John Sillitoe on behalf of the Wellcome Sanger Institute COVID-19 Surveillance Team |
| EPI_ISL_2974047, EPI_ISL_2974048, EPI_ISL_2974049 | Lighthouse Lab in Glasgow | Wellcome Sanger Institute for the COVID-19 Genomics UK (COG-UK) Consortium | Harper VanSteenhouse, Yumi Kasai, David Gray, Carol Clugston, Anna Dominiczak and Alex Alderton, Roberto Amato, Jeffrey Barrett, Sonia Goncalves, Ewan Harrison, David K. Jackson, Ian Johnston, Dominic Kwiatkowski, Cordelia Langford, John Sillitoe on behalf of the Wellcome Sanger Institute COVID-19 Surveillance Team |
| EPI_ISL_2974050, EPI_ISL_2974051 | Randox Laboratories | Wellcome Sanger Institute for the COVID-19 Genomics UK (COG-UK) Consortium | Randox Laboratories and Alex Alderton, Roberto Amato, Jeffrey Barrett, Sonia Goncalves, Ewan Harrison, David K. Jackson, Ian Johnston, Dominic Kwiatkowski, Cordelia Langford, John Sillitoe on behalf of the Wellcome Sanger Institute COVID-19 Surveillance Team |
| EPI_ISL_2974052, EPI_ISL_2974053 | Lighthouse Lab in Glasgow | Wellcome Sanger Institute for the COVID-19 Genomics UK (COG-UK) Consortium | Harper VanSteenhouse, Yumi Kasai, David Gray, Carol Clugston, Anna Dominiczak and Alex Alderton, Roberto Amato, Jeffrey Barrett, Sonia Goncalves, Ewan Harrison, David K. Jackson, Ian Johnston, Dominic Kwiatkowski, Cordelia Langford, John Sillitoe on behalf of the Wellcome Sanger Institute COVID-19 Surveillance Team |
| EPI_ISL_2974054, EPI_ISL_2974055, EPI_ISL_2974056, EPI_ISL_2974057 | Randox Laboratories | Wellcome Sanger Institute for the COVID-19 Genomics UK (COG-UK) Consortium | Randox Laboratories and Alex Alderton, Roberto Amato, Jeffrey Barrett, Sonia Goncalves, Ewan Harrison, David K. Jackson, Ian Johnston, Dominic Kwiatkowski, Cordelia Langford, John Sillitoe on behalf of the Wellcome Sanger Institute COVID-19 Surveillance Team |
| EPI_ISL_2974058, EPI_ISL_2974059, EPI_ISL_2974060 | Lighthouse Lab in Glasgow | Wellcome Sanger Institute for the COVID-19 Genomics UK (COG-UK) Consortium | Harper VanSteenhouse, Yumi Kasai, David Gray, Carol Clugston, Anna Dominiczak and Alex Alderton, Roberto Amato, Jeffrey Barrett, Sonia Goncalves, Ewan Harrison, David K. Jackson, Ian Johnston, Dominic Kwiatkowski, Cordelia Langford, John Sillitoe on behalf of the Wellcome Sanger Institute COVID-19 Surveillance Team |
| EPI_ISL_2974061, EPI_ISL_2974062, EPI_ISL_2974063 | Randox Laboratories | Wellcome Sanger Institute for the COVID-19 Genomics UK (COG-UK) Consortium | Randox Laboratories and Alex Alderton, Roberto Amato, Jeffrey Barrett, Sonia Goncalves, Ewan Harrison, David K. Jackson, Ian Johnston, Dominic Kwiatkowski, Cordelia Langford, John Sillitoe on behalf of the Wellcome Sanger Institute COVID-19 Surveillance Team |
| EPI_ISL_2974064 | Lighthouse Lab in Glasgow | Wellcome Sanger Institute for the COVID-19 Genomics UK (COG-UK) Consortium | Harper VanSteenhouse, Yumi Kasai, David Gray, Carol Clugston, Anna Dominiczak and Alex Alderton, Roberto Amato, Jeffrey Barrett, Sonia Goncalves, Ewan Harrison, David K. Jackson, Ian Johnston, Dominic Kwiatkowski, Cordelia Langford, John Sillitoe on behalf of the Wellcome Sanger Institute COVID-19 Surveillance Team |
| EPI_ISL_2974065 | Randox Laboratories | Wellcome Sanger Institute for the COVID-19 Genomics UK (COG-UK) Consortium | Randox Laboratories and Alex Alderton, Roberto Amato, Jeffrey Barrett, Sonia Goncalves, Ewan Harrison, David K. Jackson, Ian Johnston, Dominic Kwiatkowski, Cordelia Langford, John Sillitoe on behalf of the Wellcome Sanger Institute COVID-19 Surveillance Team |
| EPI_ISL_2974066 | Lighthouse Lab in Glasgow | Wellcome Sanger Institute for the COVID-19 Genomics UK (COG-UK) Consortium | Harper VanSteenhouse, Yumi Kasai, David Gray, Carol Clugston, Anna Dominiczak and Alex Alderton, Roberto Amato, Jeffrey Barrett, Sonia Goncalves, Ewan Harrison, David K. Jackson, Ian Johnston, Dominic Kwiatkowski, Cordelia Langford, John Sillitoe on behalf of the Wellcome Sanger Institute COVID-19 Surveillance Team |
| EPI_ISL_2974067, EPI_ISL_2974068, EPI_ISL_2974069 | Randox Laboratories | Wellcome Sanger Institute for the COVID-19 Genomics UK (COG-UK) Consortium | Randox Laboratories and Alex Alderton, Roberto Amato, Jeffrey Barrett, Sonia Goncalves, Ewan Harrison, David K. Jackson, Ian Johnston, Dominic Kwiatkowski, Cordelia Langford, John Sillitoe on behalf of the Wellcome Sanger Institute COVID-19 Surveillance Team |
| EPI_ISL_2974070, EPI_ISL_2974071 | Lighthouse Lab in Glasgow | Wellcome Sanger Institute for the COVID-19 Genomics UK (COG-UK) Consortium | Harper VanSteenhouse, Yumi Kasai, David Gray, Carol Clugston, Anna Dominiczak and Alex Alderton, Roberto Amato, Jeffrey Barrett, Sonia Goncalves, Ewan Harrison, David K. Jackson, Ian Johnston, Dominic Kwiatkowski, Cordelia Langford, John Sillitoe on behalf of the Wellcome Sanger Institute COVID-19 Surveillance Team |
| EPI_ISL_2974072 | Randox Laboratories | Wellcome Sanger Institute for the COVID-19 Genomics UK (COG-UK) Consortium | Randox Laboratories and Alex Alderton, Roberto Amato, Jeffrey Barrett, Sonia Goncalves, Ewan Harrison, David K. Jackson, Ian Johnston, Dominic Kwiatkowski, Cordelia Langford, John Sillitoe on behalf of the Wellcome Sanger Institute COVID-19 Surveillance Team |
| EPI_ISL_2974073 | Lighthouse Lab in Glasgow | Wellcome Sanger Institute for the COVID-19 Genomics UK (COG-UK) Consortium | Harper VanSteenhouse, Yumi Kasai, David Gray, Carol Clugston, Anna Dominiczak and Alex Alderton, Roberto Amato, Jeffrey Barrett, Sonia Goncalves, Ewan Harrison, David K. Jackson, Ian Johnston, Dominic Kwiatkowski, Cordelia Langford, John Sillitoe on behalf of the Wellcome Sanger Institute COVID-19 Surveillance Team |
| EPI_ISL_2974074 | Randox Laboratories | Wellcome Sanger Institute for the COVID-19 Genomics UK (COG-UK) Consortium | Randox Laboratories and Alex Alderton, Roberto Amato, Jeffrey Barrett, Sonia Goncalves, Ewan Harrison, David K. Jackson, Ian Johnston, Dominic Kwiatkowski, Cordelia Langford, John Sillitoe on behalf of the Wellcome Sanger Institute COVID-19 Surveillance Team |
| EPI_ISL_2974075, EPI_ISL_2974076 | Lighthouse Lab in Glasgow | Wellcome Sanger Institute for the COVID-19 Genomics UK (COG-UK) Consortium | Harper VanSteenhouse, Yumi Kasai, David Gray, Carol Clugston, Anna Dominiczak and Alex Alderton, Roberto Amato, Jeffrey Barrett, Sonia Goncalves, Ewan Harrison, David K. Jackson, Ian Johnston, Dominic Kwiatkowski, Cordelia Langford, John Sillitoe on behalf of the Wellcome Sanger Institute COVID-19 Surveillance Team |

| | | | |
|---|---|---|---|
| EPI_ISL_2974077, EPI_ISL_2974078 | Randox Laboratories | Wellcome Sanger Institute for the COVID-19 Genomics UK (COG-UK) Consortium | Randox Laboratories and Alex Alderton, Roberto Amato, Jeffrey Barrett, Sonia Goncalves, Ewan Harrison, David K. Jackson, Ian Johnston, Dominic Kwiatkowski, Cordelia Langford, John Sillitoe on behalf of the Wellcome Sanger Institute COVID-19 Surveillance Team |
| EPI_ISL_2974079, EPI_ISL_2974080 | Lighthouse Lab in Milton Keynes | Wellcome Sanger Institute for the COVID-19 Genomics UK (COG-UK) Consortium | The Lighthouse Lab in Milton Keynes and Alex Alderton, Roberto Amato, Jeffrey Barrett, Sonia Goncalves, Ewan Harrison, David K. Jackson, Ian Johnston, Dominic Kwiatkowski, Cordelia Langford, John Sillitoe on behalf of the Wellcome Sanger Institute COVID-19 Surveillance Team |
| EPI_ISL_2974081 | Lighthouse Lab in Alderley Park | Wellcome Sanger Institute for the COVID-19 Genomics UK (COG-UK) Consortium | Jacquelyn Wynn, Mairead Hyland, The Lighthouse Lab in Alderley Park and Alex Alderton, Roberto Amato, Jeffrey Barrett, Sonia Goncalves, Ewan Harrison, David K. Jackson, Ian Johnston, Dominic Kwiatkowski, Cordelia Langford, John Sillitoe on behalf of the Wellcome Sanger Institute COVID-19 Surveillance Team |
| EPI_ISL_2974082, EPI_ISL_2974083 | Lighthouse Lab in Milton Keynes | Wellcome Sanger Institute for the COVID-19 Genomics UK (COG-UK) Consortium | The Lighthouse Lab in Milton Keynes and Alex Alderton, Roberto Amato, Jeffrey Barrett, Sonia Goncalves, Ewan Harrison, David K. Jackson, Ian Johnston, Dominic Kwiatkowski, Cordelia Langford, John Sillitoe on behalf of the Wellcome Sanger Institute COVID-19 Surveillance Team |
| EPI_ISL_2974084 | Lighthouse Laboratory Plymouth | Wellcome Sanger Institute for the COVID-19 Genomics UK (COG-UK) Consortium | Lighthouse Laboratory Plymouth and Alex Alderton, Roberto Amato, Jeffrey Barrett, Sonia Goncalves, Ewan Harrison, David K. Jackson, Ian Johnston, Dominic Kwiatkowski, Cordelia Langford, John Sillitoe on behalf of the Wellcome Sanger Institute COVID-19 Surveillance Team |
| EPI_ISL_2974085 | Lighthouse Lab in Milton Keynes | Wellcome Sanger Institute for the COVID-19 Genomics UK (COG-UK) Consortium | The Lighthouse Lab in Milton Keynes and Alex Alderton, Roberto Amato, Jeffrey Barrett, Sonia Goncalves, Ewan Harrison, David K. Jackson, Ian Johnston, Dominic Kwiatkowski, Cordelia Langford, John Sillitoe on behalf of the Wellcome Sanger Institute COVID-19 Surveillance Team |
| EPI_ISL_2974086 | Lighthouse Lab in Alderley Park | Wellcome Sanger Institute for the COVID-19 Genomics UK (COG-UK) Consortium | Jacquelyn Wynn, Mairead Hyland, The Lighthouse Lab in Alderley Park and Alex Alderton, Roberto Amato, Jeffrey Barrett, Sonia Goncalves, Ewan Harrison, David K. Jackson, Ian Johnston, Dominic Kwiatkowski, Cordelia Langford, John Sillitoe on behalf of the Wellcome Sanger Institute COVID-19 Surveillance Team |
| EPI_ISL_2974087, EPI_ISL_2974088 | Lighthouse Laboratory Plymouth | Wellcome Sanger Institute for the COVID-19 Genomics UK (COG-UK) Consortium | Lighthouse Laboratory Plymouth and Alex Alderton, Roberto Amato, Jeffrey Barrett, Sonia Goncalves, Ewan Harrison, David K. Jackson, Ian Johnston, Dominic Kwiatkowski, Cordelia Langford, John Sillitoe on behalf of the Wellcome Sanger Institute COVID-19 Surveillance Team |
| EPI_ISL_2974089 | Lighthouse Lab in Milton Keynes | Wellcome Sanger Institute for the COVID-19 Genomics UK (COG-UK) Consortium | The Lighthouse Lab in Milton Keynes and Alex Alderton, Roberto Amato, Jeffrey Barrett, Sonia Goncalves, Ewan Harrison, David K. Jackson, Ian Johnston, Dominic Kwiatkowski, Cordelia Langford, John Sillitoe on behalf of the Wellcome Sanger Institute COVID-19 Surveillance Team |
| EPI_ISL_2974090 | Lighthouse Laboratory Plymouth | Wellcome Sanger Institute for the COVID-19 Genomics UK (COG-UK) Consortium | Lighthouse Laboratory Plymouth and Alex Alderton, Roberto Amato, Jeffrey Barrett, Sonia Goncalves, Ewan Harrison, David K. Jackson, Ian Johnston, Dominic Kwiatkowski, Cordelia Langford, John Sillitoe on behalf of the Wellcome Sanger Institute COVID-19 Surveillance Team |
| EPI_ISL_2974091 | Lighthouse Lab in Milton Keynes | Wellcome Sanger Institute for the COVID-19 Genomics UK (COG-UK) Consortium | The Lighthouse Lab in Milton Keynes and Alex Alderton, Roberto Amato, Jeffrey Barrett, Sonia Goncalves, Ewan Harrison, David K. Jackson, Ian Johnston, Dominic Kwiatkowski, Cordelia Langford, John Sillitoe on behalf of the Wellcome Sanger Institute COVID-19 Surveillance Team |
| EPI_ISL_2974092, EPI_ISL_2974093 | Lighthouse Laboratory Plymouth | Wellcome Sanger Institute for the COVID-19 Genomics UK (COG-UK) Consortium | Lighthouse Laboratory Plymouth and Alex Alderton, Roberto Amato, Jeffrey Barrett, Sonia Goncalves, Ewan Harrison, David K. Jackson, Ian Johnston, Dominic Kwiatkowski, Cordelia Langford, John Sillitoe on behalf of the Wellcome Sanger Institute COVID-19 Surveillance Team |
| EPI_ISL_2974094 | Lighthouse Lab in Milton Keynes | Wellcome Sanger Institute for the COVID-19 Genomics UK (COG-UK) Consortium | The Lighthouse Lab in Milton Keynes and Alex Alderton, Roberto Amato, Jeffrey Barrett, Sonia Goncalves, Ewan Harrison, David K. Jackson, Ian Johnston, Dominic Kwiatkowski, Cordelia Langford, John Sillitoe on behalf of the Wellcome Sanger Institute COVID-19 Surveillance Team |
| EPI_ISL_2974095 | Lighthouse Lab in Alderley Park | Wellcome Sanger Institute for the COVID-19 Genomics UK (COG-UK) Consortium | Jacquelyn Wynn, Mairead Hyland, The Lighthouse Lab in Alderley Park and Alex Alderton, Roberto Amato, Jeffrey Barrett, Sonia Goncalves, Ewan Harrison, David K. Jackson, Ian Johnston, Dominic Kwiatkowski, Cordelia Langford, John Sillitoe on behalf of the Wellcome Sanger Institute COVID-19 Surveillance Team |
| EPI_ISL_2974096 | Lighthouse Laboratory Plymouth | Wellcome Sanger Institute for the COVID-19 Genomics UK (COG-UK) Consortium | Lighthouse Laboratory Plymouth and Alex Alderton, Roberto Amato, Jeffrey Barrett, Sonia Goncalves, Ewan Harrison, David K. Jackson, Ian Johnston, Dominic Kwiatkowski, Cordelia Langford, John Sillitoe on behalf of the Wellcome Sanger Institute COVID-19 Surveillance Team |
| EPI_ISL_2974097 | Lighthouse Lab in Milton Keynes | Wellcome Sanger Institute for the COVID-19 Genomics UK (COG-UK) Consortium | The Lighthouse Lab in Milton Keynes and Alex Alderton, Roberto Amato, Jeffrey Barrett, Sonia Goncalves, Ewan Harrison, David K. Jackson, Ian Johnston, Dominic Kwiatkowski, Cordelia Langford, John Sillitoe on behalf of the Wellcome Sanger Institute COVID-19 Surveillance Team |
| EPI_ISL_2974098, EPI_ISL_2974099, EPI_ISL_2974100 | Lighthouse Laboratory Plymouth | Wellcome Sanger Institute for the COVID-19 Genomics UK (COG-UK) Consortium | Lighthouse Laboratory Plymouth and Alex Alderton, Roberto Amato, Jeffrey Barrett, Sonia Goncalves, Ewan Harrison, David K. Jackson, Ian Johnston, Dominic Kwiatkowski, Cordelia Langford, John Sillitoe on behalf of the Wellcome Sanger Institute COVID-19 Surveillance Team |
| EPI_ISL_2974101 | Lighthouse Lab in Milton Keynes | Wellcome Sanger Institute for the COVID-19 Genomics UK (COG-UK) Consortium | The Lighthouse Lab in Milton Keynes and Alex Alderton, Roberto Amato, Jeffrey Barrett, Sonia Goncalves, Ewan Harrison, David K. Jackson, Ian Johnston, Dominic Kwiatkowski, Cordelia Langford, John Sillitoe on behalf of the Wellcome Sanger Institute COVID-19 Surveillance Team |
| EPI_ISL_2974102 | Lighthouse Laboratory Plymouth | Wellcome Sanger Institute for the COVID-19 Genomics UK (COG-UK) Consortium | Lighthouse Laboratory Plymouth and Alex Alderton, Roberto Amato, Jeffrey Barrett, Sonia Goncalves, Ewan Harrison, David K. Jackson, Ian Johnston, Dominic Kwiatkowski, Cordelia Langford, John Sillitoe on behalf of the Wellcome Sanger Institute COVID-19 Surveillance Team |
| EPI_ISL_2974103 | Lighthouse Lab in Alderley Park | Wellcome Sanger Institute for the COVID-19 Genomics UK (COG-UK) Consortium | Jacquelyn Wynn, Mairead Hyland, The Lighthouse Lab in Alderley Park and Alex Alderton, Roberto Amato, Jeffrey Barrett, Sonia Goncalves, Ewan Harrison, David K. Jackson, Ian Johnston, Dominic Kwiatkowski, Cordelia Langford, John Sillitoe on behalf of the Wellcome Sanger Institute COVID-19 Surveillance Team |
| EPI_ISL_2974104 | Lighthouse Laboratory Plymouth | Wellcome Sanger Institute for the COVID-19 Genomics UK (COG-UK) Consortium | Lighthouse Laboratory Plymouth and Alex Alderton, Roberto Amato, Jeffrey Barrett, Sonia Goncalves, Ewan Harrison, David K. Jackson, Ian Johnston, Dominic Kwiatkowski, Cordelia Langford, John Sillitoe on behalf of the Wellcome Sanger Institute COVID-19 Surveillance Team |
| EPI_ISL_2974105 | Lighthouse Lab in Milton Keynes | Wellcome Sanger Institute for the COVID-19 Genomics UK (COG-UK) Consortium | The Lighthouse Lab in Milton Keynes and Alex Alderton, Roberto Amato, Jeffrey Barrett, Sonia Goncalves, Ewan Harrison, David K. Jackson, Ian Johnston, Dominic Kwiatkowski, Cordelia Langford, John Sillitoe on behalf of the Wellcome Sanger Institute COVID-19 Surveillance Team |
| EPI_ISL_2974106, EPI_ISL_2974107 | Lighthouse Lab in Alderley Park | Wellcome Sanger Institute for the COVID-19 Genomics UK (COG-UK) Consortium | Jacquelyn Wynn, Mairead Hyland, The Lighthouse Lab in Alderley Park and Alex Alderton, Roberto Amato, Jeffrey Barrett, Sonia Goncalves, Ewan Harrison, David K. Jackson, Ian Johnston, Dominic Kwiatkowski, Cordelia Langford, John Sillitoe on behalf of the Wellcome Sanger Institute COVID-19 Surveillance Team |
| EPI_ISL_2974108 | Lighthouse Lab in Milton Keynes | Wellcome Sanger Institute for the COVID-19 Genomics UK (COG-UK) Consortium | The Lighthouse Lab in Milton Keynes and Alex Alderton, Roberto Amato, Jeffrey Barrett, Sonia Goncalves, Ewan Harrison, David K. Jackson, Ian Johnston, Dominic Kwiatkowski, Cordelia Langford, John Sillitoe on behalf of the Wellcome Sanger Institute COVID-19 Surveillance Team |
| EPI_ISL_2974109 | Lighthouse Laboratory Plymouth | Wellcome Sanger Institute for the COVID-19 Genomics UK (COG-UK) Consortium | Lighthouse Laboratory Plymouth and Alex Alderton, Roberto Amato, Jeffrey Barrett, Sonia Goncalves, Ewan Harrison, David K. Jackson, Ian Johnston, Dominic Kwiatkowski, Cordelia Langford, John Sillitoe on behalf of the Wellcome Sanger Institute COVID-19 Surveillance Team |
| EPI_ISL_2974110 | Lighthouse Lab in Alderley Park | Wellcome Sanger Institute for the COVID-19 Genomics UK (COG-UK) Consortium | Jacquelyn Wynn, Mairead Hyland, The Lighthouse Lab in Alderley Park and Alex Alderton, Roberto Amato, Jeffrey Barrett, Sonia Goncalves, Ewan Harrison, David K. Jackson, Ian Johnston, Dominic Kwiatkowski, Cordelia Langford, John Sillitoe on behalf of the Wellcome Sanger Institute COVID-19 Surveillance Team |
| EPI_ISL_2974111, EPI_ISL_2974112 | Lighthouse Lab in Milton Keynes | Wellcome Sanger Institute for the COVID-19 Genomics UK (COG-UK) Consortium | The Lighthouse Lab in Milton Keynes and Alex Alderton, Roberto Amato, Jeffrey Barrett, Sonia Goncalves, Ewan Harrison, David K. Jackson, Ian Johnston, Dominic Kwiatkowski, Cordelia Langford, John Sillitoe on behalf of the Wellcome Sanger Institute COVID-19 Surveillance Team |
| EPI_ISL_2974113, EPI_ISL_2974114, EPI_ISL_2974115 | Lighthouse Laboratory Plymouth | Wellcome Sanger Institute for the COVID-19 Genomics UK (COG-UK) Consortium | Lighthouse Laboratory Plymouth and Alex Alderton, Roberto Amato, Jeffrey Barrett, Sonia Goncalves, Ewan Harrison, David K. Jackson, Ian Johnston, Dominic Kwiatkowski, Cordelia Langford, John Sillitoe on behalf of the Wellcome Sanger Institute COVID-19 Surveillance Team |
| EPI_ISL_2974116, EPI_ISL_2974117 | Lighthouse Lab in Alderley Park | Wellcome Sanger Institute for the COVID-19 Genomics UK (COG-UK) Consortium | Jacquelyn Wynn, Mairead Hyland, The Lighthouse Lab in Alderley Park and Alex Alderton, Roberto Amato, Jeffrey Barrett, Sonia Goncalves, Ewan Harrison, David K. Jackson, Ian Johnston, Dominic Kwiatkowski, Cordelia Langford, John Sillitoe on behalf of the Wellcome Sanger Institute COVID-19 Surveillance Team |
| EPI_ISL_2974118 | Lighthouse Laboratory Plymouth | Wellcome Sanger Institute for the COVID-19 Genomics UK (COG-UK) Consortium | Lighthouse Laboratory Plymouth and Alex Alderton, Roberto Amato, Jeffrey Barrett, Sonia Goncalves, Ewan Harrison, David K. Jackson, Ian Johnston, Dominic Kwiatkowski, Cordelia Langford, John Sillitoe on behalf of the Wellcome Sanger Institute COVID-19 Surveillance Team |
| EPI_ISL_2974119 | Lighthouse Lab in Alderley Park | Wellcome Sanger Institute for the COVID-19 Genomics UK (COG-UK) Consortium | Jacquelyn Wynn, Mairead Hyland, The Lighthouse Lab in Alderley Park and Alex Alderton, Roberto Amato, Jeffrey Barrett, Sonia Goncalves, Ewan Harrison, David K. Jackson, Ian Johnston, Dominic Kwiatkowski, Cordelia Langford, John Sillitoe on behalf of the Wellcome Sanger Institute COVID-19 Surveillance Team |
| EPI_ISL_2974120 | Lighthouse Laboratory Plymouth | Wellcome Sanger Institute for the COVID-19 Genomics UK (COG-UK) Consortium | Lighthouse Laboratory Plymouth and Alex Alderton, Roberto Amato, Jeffrey Barrett, Sonia Goncalves, Ewan Harrison, David K. Jackson, Ian Johnston, Dominic Kwiatkowski, Cordelia Langford, John Sillitoe on behalf of the Wellcome Sanger Institute COVID-19 Surveillance Team |
| EPI_ISL_2974121, EPI_ISL_2974122 | Lighthouse Lab in Milton Keynes | Wellcome Sanger Institute for the COVID-19 Genomics UK (COG-UK) Consortium | The Lighthouse Lab in Milton Keynes and Alex Alderton, Roberto Amato, Jeffrey Barrett, Sonia Goncalves, Ewan Harrison, David K. Jackson, Ian Johnston, Dominic Kwiatkowski, Cordelia Langford, John Sillitoe on behalf of the Wellcome Sanger Institute COVID-19 Surveillance Team |
| EPI_ISL_2974123 | Lighthouse Lab in Alderley Park | Wellcome Sanger Institute for the COVID-19 Genomics UK (COG-UK) Consortium | Jacquelyn Wynn, Mairead Hyland, The Lighthouse Lab in Alderley Park and Alex Alderton, Roberto Amato, Jeffrey Barrett, Sonia Goncalves, Ewan Harrison, David K. Jackson, Ian Johnston, Dominic Kwiatkowski, Cordelia Langford, John Sillitoe on behalf of the Wellcome Sanger Institute COVID-19 Surveillance Team |
| EPI_ISL_2974124 | Lighthouse Laboratory Plymouth | Wellcome Sanger Institute for the COVID-19 Genomics UK (COG-UK) Consortium | Lighthouse Laboratory Plymouth and Alex Alderton, Roberto Amato, Jeffrey Barrett, Sonia Goncalves, Ewan Harrison, David K. Jackson, Ian Johnston, Dominic Kwiatkowski, Cordelia Langford, John Sillitoe on behalf of the Wellcome Sanger Institute COVID-19 Surveillance Team |
| EPI_ISL_2974125 | Lighthouse Lab in Alderley Park | Wellcome Sanger Institute for the COVID-19 Genomics UK (COG-UK) Consortium | Jacquelyn Wynn, Mairead Hyland, The Lighthouse Lab in Alderley Park and Alex Alderton, Roberto Amato, Jeffrey Barrett, Sonia Goncalves, Ewan Harrison, David K. Jackson, Ian Johnston, Dominic Kwiatkowski, Cordelia Langford, John Sillitoe on behalf of the Wellcome Sanger Institute COVID-19 Surveillance Team |
| EPI_ISL_2974126 | Lighthouse Lab in Milton Keynes | Wellcome Sanger Institute for the COVID-19 Genomics UK (COG-UK) Consortium | The Lighthouse Lab in Milton Keynes and Alex Alderton, Roberto Amato, Jeffrey Barrett, Sonia Goncalves, Ewan Harrison, David K. Jackson, Ian Johnston, Dominic Kwiatkowski, Cordelia Langford, John Sillitoe on behalf of the Wellcome Sanger Institute COVID-19 Surveillance Team |
| EPI_ISL_2974127, EPI_ISL_2974128, EPI_ISL_2974129, EPI_ISL_2974130 | Lighthouse Lab in Alderley Park | Wellcome Sanger Institute for the COVID-19 Genomics UK (COG-UK) Consortium | Jacquelyn Wynn, Mairead Hyland, The Lighthouse Lab in Alderley Park and Alex Alderton, Roberto Amato, Jeffrey Barrett, Sonia Goncalves, Ewan Harrison, David K. Jackson, Ian Johnston, Dominic Kwiatkowski, Cordelia Langford, John Sillitoe on behalf of the Wellcome Sanger Institute COVID-19 Surveillance Team |
| EPI_ISL_2974131 | Lighthouse Lab in Milton Keynes | Wellcome Sanger Institute for the COVID-19 Genomics UK (COG-UK) Consortium | The Lighthouse Lab in Milton Keynes and Alex Alderton, Roberto Amato, Jeffrey Barrett, Sonia Goncalves, Ewan Harrison, David K. Jackson, Ian Johnston, Dominic Kwiatkowski, Cordelia Langford, John Sillitoe on behalf of the Wellcome Sanger Institute COVID-19 Surveillance Team |
| EPI_ISL_2974132, EPI_ISL_2974133 | Lighthouse Lab in Alderley Park | Wellcome Sanger Institute for the COVID-19 Genomics UK (COG-UK) Consortium | Jacquelyn Wynn, Mairead Hyland, The Lighthouse Lab in Alderley Park and Alex Alderton, Roberto Amato, Jeffrey Barrett, Sonia Goncalves, Ewan Harrison, David K. Jackson, Ian Johnston, Dominic Kwiatkowski, Cordelia Langford, John Sillitoe on behalf of the Wellcome Sanger Institute COVID-19 Surveillance Team |
| EPI_ISL_2974134 | Lighthouse Lab in Milton Keynes | Wellcome Sanger Institute for the COVID-19 Genomics UK (COG-UK) Consortium | The Lighthouse Lab in Milton Keynes and Alex Alderton, Roberto Amato, Jeffrey Barrett, Sonia Goncalves, Ewan Harrison, David K. Jackson, Ian Johnston, Dominic Kwiatkowski, Cordelia Langford, John Sillitoe on behalf of the Wellcome Sanger Institute COVID-19 Surveillance Team |
| EPI_ISL_2974135 | Lighthouse Laboratory Plymouth | Wellcome Sanger Institute for the COVID-19 Genomics UK (COG-UK) Consortium | Lighthouse Laboratory Plymouth and Alex Alderton, Roberto Amato, Jeffrey Barrett, Sonia Goncalves, Ewan Harrison, David K. Jackson, Ian Johnston, Dominic Kwiatkowski, Cordelia Langford, John Sillitoe on behalf of the Wellcome Sanger Institute COVID-19 Surveillance Team |
| EPI_ISL_2974136, EPI_ISL_2974137, EPI_ISL_2974138, | Lighthouse Lab in Milton Keynes | Wellcome Sanger Institute for the COVID-19 Genomics UK (COG-UK) Consortium | The Lighthouse Lab in Milton Keynes and Alex Alderton, Roberto Amato, Jeffrey Barrett, Sonia Goncalves, Ewan Harrison, David K. Jackson, Ian Johnston, Dominic Kwiatkowski, Cordelia Langford, John Sillitoe on behalf of the Wellcome Sanger Institute COVID-19 Surveillance Team |

| | | | |
|---|---|---|---|
| EPI_ISL_2974139 | | | |
| EPI_ISL_2974140 | Lighthouse Laboratory Plymouth | Wellcome Sanger Institute for the COVID-19 Genomics UK (COG-UK) Consortium | Lighthouse Laboratory Plymouth and Alex Alderton, Roberto Amato, Jeffrey Barrett, Sonia Goncalves, Ewan Harrison, David K. Jackson, Ian Johnston, Dominic Kwiatkowski, Cordelia Langford, John Sillitoe on behalf of the Wellcome Sanger Institute COVID-19 Surveillance Team |
| EPI_ISL_2974141 | Lighthouse Lab in Milton Keynes | Wellcome Sanger Institute for the COVID-19 Genomics UK (COG-UK) Consortium | The Lighthouse Lab in Milton Keynes and Alex Alderton, Roberto Amato, Jeffrey Barrett, Sonia Goncalves, Ewan Harrison, David K. Jackson, Ian Johnston, Dominic Kwiatkowski, Cordelia Langford, John Sillitoe on behalf of the Wellcome Sanger Institute COVID-19 Surveillance Team |
| EPI_ISL_2974142, EPI_ISL_2974143 | Lighthouse Lab in Alderley Park | Wellcome Sanger Institute for the COVID-19 Genomics UK (COG-UK) Consortium | Jacquelyn Wynn, Mairead Hyland, The Lighthouse Lab in Alderley Park and Alex Alderton, Roberto Amato, Jeffrey Barrett, Sonia Goncalves, Ewan Harrison, David K. Jackson, Ian Johnston, Dominic Kwiatkowski, Cordelia Langford, John Sillitoe on behalf of the Wellcome Sanger Institute COVID-19 Surveillance Team |
| EPI_ISL_2974144, EPI_ISL_2974145, EPI_ISL_2974146 | Lighthouse Lab in Milton Keynes | Wellcome Sanger Institute for the COVID-19 Genomics UK (COG-UK) Consortium | The Lighthouse Lab in Milton Keynes and Alex Alderton, Roberto Amato, Jeffrey Barrett, Sonia Goncalves, Ewan Harrison, David K. Jackson, Ian Johnston, Dominic Kwiatkowski, Cordelia Langford, John Sillitoe on behalf of the Wellcome Sanger Institute COVID-19 Surveillance Team |
| EPI_ISL_2974147, EPI_ISL_2974148 | Lighthouse Lab in Alderley Park | Wellcome Sanger Institute for the COVID-19 Genomics UK (COG-UK) Consortium | Jacquelyn Wynn, Mairead Hyland, The Lighthouse Lab in Alderley Park and Alex Alderton, Roberto Amato, Jeffrey Barrett, Sonia Goncalves, Ewan Harrison, David K. Jackson, Ian Johnston, Dominic Kwiatkowski, Cordelia Langford, John Sillitoe on behalf of the Wellcome Sanger Institute COVID-19 Surveillance Team |
| EPI_ISL_2974149, EPI_ISL_2974150 | Lighthouse Lab in Milton Keynes | Wellcome Sanger Institute for the COVID-19 Genomics UK (COG-UK) Consortium | The Lighthouse Lab in Milton Keynes and Alex Alderton, Roberto Amato, Jeffrey Barrett, Sonia Goncalves, Ewan Harrison, David K. Jackson, Ian Johnston, Dominic Kwiatkowski, Cordelia Langford, John Sillitoe on behalf of the Wellcome Sanger Institute COVID-19 Surveillance Team |
| EPI_ISL_2974151 | Lighthouse Laboratory Plymouth | Wellcome Sanger Institute for the COVID-19 Genomics UK (COG-UK) Consortium | Lighthouse Laboratory Plymouth and Alex Alderton, Roberto Amato, Jeffrey Barrett, Sonia Goncalves, Ewan Harrison, David K. Jackson, Ian Johnston, Dominic Kwiatkowski, Cordelia Langford, John Sillitoe on behalf of the Wellcome Sanger Institute COVID-19 Surveillance Team |
| EPI_ISL_2974152, EPI_ISL_2974153 | Lighthouse Lab in Alderley Park | Wellcome Sanger Institute for the COVID-19 Genomics UK (COG-UK) Consortium | Jacquelyn Wynn, Mairead Hyland, The Lighthouse Lab in Alderley Park and Alex Alderton, Roberto Amato, Jeffrey Barrett, Sonia Goncalves, Ewan Harrison, David K. Jackson, Ian Johnston, Dominic Kwiatkowski, Cordelia Langford, John Sillitoe on behalf of the Wellcome Sanger Institute COVID-19 Surveillance Team |
| EPI_ISL_2974154 | Lighthouse Lab in Milton Keynes | Wellcome Sanger Institute for the COVID-19 Genomics UK (COG-UK) Consortium | The Lighthouse Lab in Milton Keynes and Alex Alderton, Roberto Amato, Jeffrey Barrett, Sonia Goncalves, Ewan Harrison, David K. Jackson, Ian Johnston, Dominic Kwiatkowski, Cordelia Langford, John Sillitoe on behalf of the Wellcome Sanger Institute COVID-19 Surveillance Team |
| EPI_ISL_2974155, EPI_ISL_2974156 | Lighthouse Lab in Alderley Park | Wellcome Sanger Institute for the COVID-19 Genomics UK (COG-UK) Consortium | Jacquelyn Wynn, Mairead Hyland, The Lighthouse Lab in Alderley Park and Alex Alderton, Roberto Amato, Jeffrey Barrett, Sonia Goncalves, Ewan Harrison, David K. Jackson, Ian Johnston, Dominic Kwiatkowski, Cordelia Langford, John Sillitoe on behalf of the Wellcome Sanger Institute COVID-19 Surveillance Team |
| EPI_ISL_2974157 | Lighthouse Laboratory Plymouth | Wellcome Sanger Institute for the COVID-19 Genomics UK (COG-UK) Consortium | Lighthouse Laboratory Plymouth and Alex Alderton, Roberto Amato, Jeffrey Barrett, Sonia Goncalves, Ewan Harrison, David K. Jackson, Ian Johnston, Dominic Kwiatkowski, Cordelia Langford, John Sillitoe on behalf of the Wellcome Sanger Institute COVID-19 Surveillance Team |
| EPI_ISL_2974158 | Lighthouse Lab in Alderley Park | Wellcome Sanger Institute for the COVID-19 Genomics UK (COG-UK) Consortium | Jacquelyn Wynn, Mairead Hyland, The Lighthouse Lab in Alderley Park and Alex Alderton, Roberto Amato, Jeffrey Barrett, Sonia Goncalves, Ewan Harrison, David K. Jackson, Ian Johnston, Dominic Kwiatkowski, Cordelia Langford, John Sillitoe on behalf of the Wellcome Sanger Institute COVID-19 Surveillance Team |
| EPI_ISL_2974159, EPI_ISL_2974160 | Lighthouse Laboratory Plymouth | Wellcome Sanger Institute for the COVID-19 Genomics UK (COG-UK) Consortium | Lighthouse Laboratory Plymouth and Alex Alderton, Roberto Amato, Jeffrey Barrett, Sonia Goncalves, Ewan Harrison, David K. Jackson, Ian Johnston, Dominic Kwiatkowski, Cordelia Langford, John Sillitoe on behalf of the Wellcome Sanger Institute COVID-19 Surveillance Team |
| EPI_ISL_2974161 | Lighthouse Lab in Alderley Park | Wellcome Sanger Institute for the COVID-19 Genomics UK (COG-UK) Consortium | Jacquelyn Wynn, Mairead Hyland, The Lighthouse Lab in Alderley Park and Alex Alderton, Roberto Amato, Jeffrey Barrett, Sonia Goncalves, Ewan Harrison, David K. Jackson, Ian Johnston, Dominic Kwiatkowski, Cordelia Langford, John Sillitoe on behalf of the Wellcome Sanger Institute COVID-19 Surveillance Team |
| EPI_ISL_2974162, EPI_ISL_2974163 | Lighthouse Lab in Milton Keynes | Wellcome Sanger Institute for the COVID-19 Genomics UK (COG-UK) Consortium | The Lighthouse Lab in Milton Keynes and Alex Alderton, Roberto Amato, Jeffrey Barrett, Sonia Goncalves, Ewan Harrison, David K. Jackson, Ian Johnston, Dominic Kwiatkowski, Cordelia Langford, John Sillitoe on behalf of the Wellcome Sanger Institute COVID-19 Surveillance Team |
| EPI_ISL_2974164 | Lighthouse Laboratory Plymouth | Wellcome Sanger Institute for the COVID-19 Genomics UK (COG-UK) Consortium | Lighthouse Laboratory Plymouth and Alex Alderton, Roberto Amato, Jeffrey Barrett, Sonia Goncalves, Ewan Harrison, David K. Jackson, Ian Johnston, Dominic Kwiatkowski, Cordelia Langford, John Sillitoe on behalf of the Wellcome Sanger Institute COVID-19 Surveillance Team |
| EPI_ISL_2974165, EPI_ISL_2974166 | Lighthouse Lab in Milton Keynes | Wellcome Sanger Institute for the COVID-19 Genomics UK (COG-UK) Consortium | The Lighthouse Lab in Milton Keynes and Alex Alderton, Roberto Amato, Jeffrey Barrett, Sonia Goncalves, Ewan Harrison, David K. Jackson, Ian Johnston, Dominic Kwiatkowski, Cordelia Langford, John Sillitoe on behalf of the Wellcome Sanger Institute COVID-19 Surveillance Team |
| EPI_ISL_2974167 | Lighthouse Laboratory Plymouth | Wellcome Sanger Institute for the COVID-19 Genomics UK (COG-UK) Consortium | Lighthouse Laboratory Plymouth and Alex Alderton, Roberto Amato, Jeffrey Barrett, Sonia Goncalves, Ewan Harrison, David K. Jackson, Ian Johnston, Dominic Kwiatkowski, Cordelia Langford, John Sillitoe on behalf of the Wellcome Sanger Institute COVID-19 Surveillance Team |
| EPI_ISL_2974168, EPI_ISL_2974169 | Lighthouse Lab in Milton Keynes | Wellcome Sanger Institute for the COVID-19 Genomics UK (COG-UK) Consortium | The Lighthouse Lab in Milton Keynes and Alex Alderton, Roberto Amato, Jeffrey Barrett, Sonia Goncalves, Ewan Harrison, David K. Jackson, Ian Johnston, Dominic Kwiatkowski, Cordelia Langford, John Sillitoe on behalf of the Wellcome Sanger Institute COVID-19 Surveillance Team |
| EPI_ISL_2974170 | Lighthouse Laboratory Plymouth | Wellcome Sanger Institute for the COVID-19 Genomics UK (COG-UK) Consortium | Lighthouse Laboratory Plymouth and Alex Alderton, Roberto Amato, Jeffrey Barrett, Sonia Goncalves, Ewan Harrison, David K. Jackson, Ian Johnston, Dominic Kwiatkowski, Cordelia Langford, John Sillitoe on behalf of the Wellcome Sanger Institute COVID-19 Surveillance Team |
| EPI_ISL_2974171, EPI_ISL_2974172, EPI_ISL_2974173, EPI_ISL_2974174, EPI_ISL_2974175, EPI_ISL_2974176 | Lighthouse Lab in Milton Keynes | Wellcome Sanger Institute for the COVID-19 Genomics UK (COG-UK) Consortium | The Lighthouse Lab in Milton Keynes and Alex Alderton, Roberto Amato, Jeffrey Barrett, Sonia Goncalves, Ewan Harrison, David K. Jackson, Ian Johnston, Dominic Kwiatkowski, Cordelia Langford, John Sillitoe on behalf of the Wellcome Sanger Institute COVID-19 Surveillance Team |
| EPI_ISL_2974177, EPI_ISL_2974178 | Lighthouse Laboratory Plymouth | Wellcome Sanger Institute for the COVID-19 Genomics UK (COG-UK) Consortium | Lighthouse Laboratory Plymouth and Alex Alderton, Roberto Amato, Jeffrey Barrett, Sonia Goncalves, Ewan Harrison, David K. Jackson, Ian Johnston, Dominic Kwiatkowski, Cordelia Langford, John Sillitoe on behalf of the Wellcome Sanger Institute COVID-19 Surveillance Team |
| EPI_ISL_2974179 | Lighthouse Lab in Milton Keynes | Wellcome Sanger Institute for the COVID-19 Genomics UK (COG-UK) Consortium | The Lighthouse Lab in Milton Keynes and Alex Alderton, Roberto Amato, Jeffrey Barrett, Sonia Goncalves, Ewan Harrison, David K. Jackson, Ian Johnston, Dominic Kwiatkowski, Cordelia Langford, John Sillitoe on behalf of the Wellcome Sanger Institute COVID-19 Surveillance Team |
| EPI_ISL_2974180 | Lighthouse Laboratory Plymouth | Wellcome Sanger Institute for the COVID-19 Genomics UK (COG-UK) Consortium | Lighthouse Laboratory Plymouth and Alex Alderton, Roberto Amato, Jeffrey Barrett, Sonia Goncalves, Ewan Harrison, David K. Jackson, Ian Johnston, Dominic Kwiatkowski, Cordelia Langford, John Sillitoe on behalf of the Wellcome Sanger Institute COVID-19 Surveillance Team |
| EPI_ISL_2974181 | Lighthouse Lab in Alderley Park | Wellcome Sanger Institute for the COVID-19 Genomics UK (COG-UK) Consortium | Jacquelyn Wynn, Mairead Hyland, The Lighthouse Lab in Alderley Park and Alex Alderton, Roberto Amato, Jeffrey Barrett, Sonia Goncalves, Ewan Harrison, David K. Jackson, Ian Johnston, Dominic Kwiatkowski, Cordelia Langford, John Sillitoe on behalf of the Wellcome Sanger Institute COVID-19 Surveillance Team |
| EPI_ISL_2974182 | Lighthouse Laboratory Plymouth | Wellcome Sanger Institute for the COVID-19 Genomics UK (COG-UK) Consortium | Lighthouse Laboratory Plymouth and Alex Alderton, Roberto Amato, Jeffrey Barrett, Sonia Goncalves, Ewan Harrison, David K. Jackson, Ian Johnston, Dominic Kwiatkowski, Cordelia Langford, John Sillitoe on behalf of the Wellcome Sanger Institute COVID-19 Surveillance Team |
| EPI_ISL_2974183 | Lighthouse Lab in Milton Keynes | Wellcome Sanger Institute for the COVID-19 Genomics UK (COG-UK) Consortium | The Lighthouse Lab in Milton Keynes and Alex Alderton, Roberto Amato, Jeffrey Barrett, Sonia Goncalves, Ewan Harrison, David K. Jackson, Ian Johnston, Dominic Kwiatkowski, Cordelia Langford, John Sillitoe on behalf of the Wellcome Sanger Institute COVID-19 Surveillance Team |
| EPI_ISL_2974184 | Lighthouse Lab in Alderley Park | Wellcome Sanger Institute for the COVID-19 Genomics UK (COG-UK) Consortium | Jacquelyn Wynn, Mairead Hyland, The Lighthouse Lab in Alderley Park and Alex Alderton, Roberto Amato, Jeffrey Barrett, Sonia Goncalves, Ewan Harrison, David K. Jackson, Ian Johnston, Dominic Kwiatkowski, Cordelia Langford, John Sillitoe on behalf of the Wellcome Sanger Institute COVID-19 Surveillance Team |
| EPI_ISL_2974185 | Lighthouse Lab in Milton Keynes | Wellcome Sanger Institute for the COVID-19 Genomics UK (COG-UK) Consortium | The Lighthouse Lab in Milton Keynes and Alex Alderton, Roberto Amato, Jeffrey Barrett, Sonia Goncalves, Ewan Harrison, David K. Jackson, Ian Johnston, Dominic Kwiatkowski, Cordelia Langford, John Sillitoe on behalf of the Wellcome Sanger Institute COVID-19 Surveillance Team |
| EPI_ISL_2974186, EPI_ISL_2974187, EPI_ISL_2974188 | Lighthouse Lab in Alderley Park | Wellcome Sanger Institute for the COVID-19 Genomics UK (COG-UK) Consortium | Jacquelyn Wynn, Mairead Hyland, The Lighthouse Lab in Alderley Park and Alex Alderton, Roberto Amato, Jeffrey Barrett, Sonia Goncalves, Ewan Harrison, David K. Jackson, Ian Johnston, Dominic Kwiatkowski, Cordelia Langford, John Sillitoe on behalf of the Wellcome Sanger Institute COVID-19 Surveillance Team |
| EPI_ISL_2974189 | Lighthouse Lab in Milton Keynes | Wellcome Sanger Institute for the COVID-19 Genomics UK (COG-UK) Consortium | The Lighthouse Lab in Milton Keynes and Alex Alderton, Roberto Amato, Jeffrey Barrett, Sonia Goncalves, Ewan Harrison, David K. Jackson, Ian Johnston, Dominic Kwiatkowski, Cordelia Langford, John Sillitoe on behalf of the Wellcome Sanger Institute COVID-19 Surveillance Team |
| EPI_ISL_2974190 | Lighthouse Lab in Alderley Park | Wellcome Sanger Institute for the COVID-19 Genomics UK (COG-UK) Consortium | Jacquelyn Wynn, Mairead Hyland, The Lighthouse Lab in Alderley Park and Alex Alderton, Roberto Amato, Jeffrey Barrett, Sonia Goncalves, Ewan Harrison, David K. Jackson, Ian Johnston, Dominic Kwiatkowski, Cordelia Langford, John Sillitoe on behalf of the Wellcome Sanger Institute COVID-19 Surveillance Team |
| EPI_ISL_2974191 | Lighthouse Laboratory Plymouth | Wellcome Sanger Institute for the COVID-19 Genomics UK (COG-UK) Consortium | Lighthouse Laboratory Plymouth and Alex Alderton, Roberto Amato, Jeffrey Barrett, Sonia Goncalves, Ewan Harrison, David K. Jackson, Ian Johnston, Dominic Kwiatkowski, Cordelia Langford, John Sillitoe on behalf of the Wellcome Sanger Institute COVID-19 Surveillance Team |
| EPI_ISL_2974192 | Lighthouse Lab in Alderley Park | Wellcome Sanger Institute for the COVID-19 Genomics UK (COG-UK) Consortium | Jacquelyn Wynn, Mairead Hyland, The Lighthouse Lab in Alderley Park and Alex Alderton, Roberto Amato, Jeffrey Barrett, Sonia Goncalves, Ewan Harrison, David K. Jackson, Ian Johnston, Dominic Kwiatkowski, Cordelia Langford, John Sillitoe on behalf of the Wellcome Sanger Institute COVID-19 Surveillance Team |
| EPI_ISL_2974193, EPI_ISL_2974194, EPI_ISL_2974195, EPI_ISL_2974196 | Lighthouse Laboratory Plymouth | Wellcome Sanger Institute for the COVID-19 Genomics UK (COG-UK) Consortium | Lighthouse Laboratory Plymouth and Alex Alderton, Roberto Amato, Jeffrey Barrett, Sonia Goncalves, Ewan Harrison, David K. Jackson, Ian Johnston, Dominic Kwiatkowski, Cordelia Langford, John Sillitoe on behalf of the Wellcome Sanger Institute COVID-19 Surveillance Team |
| EPI_ISL_2974197 | Lighthouse Lab in Milton Keynes | Wellcome Sanger Institute for the COVID-19 Genomics UK (COG-UK) Consortium | The Lighthouse Lab in Milton Keynes and Alex Alderton, Roberto Amato, Jeffrey Barrett, Sonia Goncalves, Ewan Harrison, David K. Jackson, Ian Johnston, Dominic Kwiatkowski, Cordelia Langford, John Sillitoe on behalf of the Wellcome Sanger Institute COVID-19 Surveillance Team |
| EPI_ISL_2974198 | Lighthouse Lab in Alderley Park | Wellcome Sanger Institute for the COVID-19 Genomics UK (COG-UK) Consortium | Jacquelyn Wynn, Mairead Hyland, The Lighthouse Lab in Alderley Park and Alex Alderton, Roberto Amato, Jeffrey Barrett, Sonia Goncalves, Ewan Harrison, David K. Jackson, Ian Johnston, Dominic Kwiatkowski, Cordelia Langford, John Sillitoe on behalf of the Wellcome Sanger Institute COVID-19 Surveillance Team |
| EPI_ISL_2974199, EPI_ISL_2974200 | Lighthouse Lab in Milton Keynes | Wellcome Sanger Institute for the COVID-19 Genomics UK (COG-UK) Consortium | The Lighthouse Lab in Milton Keynes and Alex Alderton, Roberto Amato, Jeffrey Barrett, Sonia Goncalves, Ewan Harrison, David K. Jackson, Ian Johnston, Dominic Kwiatkowski, Cordelia Langford, John Sillitoe on behalf of the Wellcome Sanger Institute COVID-19 Surveillance Team |
| EPI_ISL_2974201 | Lighthouse Laboratory Plymouth | Wellcome Sanger Institute for the COVID-19 Genomics UK (COG-UK) Consortium | Lighthouse Laboratory Plymouth and Alex Alderton, Roberto Amato, Jeffrey Barrett, Sonia Goncalves, Ewan Harrison, David K. Jackson, Ian Johnston, Dominic Kwiatkowski, Cordelia Langford, John Sillitoe on behalf of the Wellcome Sanger Institute COVID-19 Surveillance Team |
| EPI_ISL_2974202, EPI_ISL_2974203 | Lighthouse Lab in Milton Keynes | Wellcome Sanger Institute for the COVID-19 Genomics UK (COG-UK) Consortium | The Lighthouse Lab in Milton Keynes and Alex Alderton, Roberto Amato, Jeffrey Barrett, Sonia Goncalves, Ewan Harrison, David K. Jackson, Ian Johnston, Dominic Kwiatkowski, Cordelia Langford, John Sillitoe on behalf of the Wellcome Sanger Institute COVID-19 Surveillance Team |
| EPI_ISL_2974204, EPI_ISL_2974205 | Lighthouse Lab in Alderley Park | Wellcome Sanger Institute for the COVID-19 Genomics UK (COG-UK) Consortium | Jacquelyn Wynn, Mairead Hyland, The Lighthouse Lab in Alderley Park and Alex Alderton, Roberto Amato, Jeffrey Barrett, Sonia Goncalves, Ewan Harrison, David K. Jackson, Ian Johnston, Dominic Kwiatkowski, Cordelia Langford, John Sillitoe on behalf of the Wellcome Sanger Institute COVID-19 Surveillance Team |

| | | | |
|---|---|---|---|
| EPI_ISL_2974206, EPI_ISL_2974207, EPI_ISL_2974208, EPI_ISL_2974209 | Lighthouse Lab in Milton Keynes | Wellcome Sanger Institute for the COVID-19 Genomics UK (COG-UK) Consortium | The Lighthouse Lab in Milton Keynes and Alex Alderton, Roberto Amato, Jeffrey Barrett, Sonia Goncalves, Ewan Harrison, David K. Jackson, Ian Johnston, Dominic Kwiatkowski, Cordelia Langford, John Sillitoe on behalf of the Wellcome Sanger Institute COVID-19 Surveillance Team |
| EPI_ISL_2974210 | Lighthouse Lab in Alderley Park | Wellcome Sanger Institute for the COVID-19 Genomics UK (COG-UK) Consortium | Jacquelyn Wynn, Mairead Hyland, The Lighthouse Lab in Alderley Park and Alex Alderton, Roberto Amato, Jeffrey Barrett, Sonia Goncalves, Ewan Harrison, David K. Jackson, Ian Johnston, Dominic Kwiatkowski, Cordelia Langford, John Sillitoe on behalf of the Wellcome Sanger Institute COVID-19 Surveillance Team |
| EPI_ISL_2974211, EPI_ISL_2974212 | Lighthouse Lab in Milton Keynes | Wellcome Sanger Institute for the COVID-19 Genomics UK (COG-UK) Consortium | The Lighthouse Lab in Milton Keynes and Alex Alderton, Roberto Amato, Jeffrey Barrett, Sonia Goncalves, Ewan Harrison, David K. Jackson, Ian Johnston, Dominic Kwiatkowski, Cordelia Langford, John Sillitoe on behalf of the Wellcome Sanger Institute COVID-19 Surveillance Team |
| EPI_ISL_2974213 | Lighthouse Lab in Alderley Park | Wellcome Sanger Institute for the COVID-19 Genomics UK (COG-UK) Consortium | Jacquelyn Wynn, Mairead Hyland, The Lighthouse Lab in Alderley Park and Alex Alderton, Roberto Amato, Jeffrey Barrett, Sonia Goncalves, Ewan Harrison, David K. Jackson, Ian Johnston, Dominic Kwiatkowski, Cordelia Langford, John Sillitoe on behalf of the Wellcome Sanger Institute COVID-19 Surveillance Team |
| EPI_ISL_2974214, EPI_ISL_2974215, EPI_ISL_2974216, EPI_ISL_2974217 | Lighthouse Laboratory Plymouth | Wellcome Sanger Institute for the COVID-19 Genomics UK (COG-UK) Consortium | Lighthouse Laboratory Plymouth and Alex Alderton, Roberto Amato, Jeffrey Barrett, Sonia Goncalves, Ewan Harrison, David K. Jackson, Ian Johnston, Dominic Kwiatkowski, Cordelia Langford, John Sillitoe on behalf of the Wellcome Sanger Institute COVID-19 Surveillance Team |
| EPI_ISL_2974218 | Lighthouse Lab in Alderley Park | Wellcome Sanger Institute for the COVID-19 Genomics UK (COG-UK) Consortium | Jacquelyn Wynn, Mairead Hyland, The Lighthouse Lab in Alderley Park and Alex Alderton, Roberto Amato, Jeffrey Barrett, Sonia Goncalves, Ewan Harrison, David K. Jackson, Ian Johnston, Dominic Kwiatkowski, Cordelia Langford, John Sillitoe on behalf of the Wellcome Sanger Institute COVID-19 Surveillance Team |
| EPI_ISL_2974219 | Lighthouse Lab in Milton Keynes | Wellcome Sanger Institute for the COVID-19 Genomics UK (COG-UK) Consortium | The Lighthouse Lab in Milton Keynes and Alex Alderton, Roberto Amato, Jeffrey Barrett, Sonia Goncalves, Ewan Harrison, David K. Jackson, Ian Johnston, Dominic Kwiatkowski, Cordelia Langford, John Sillitoe on behalf of the Wellcome Sanger Institute COVID-19 Surveillance Team |
| EPI_ISL_2974220 | Lighthouse Lab in Alderley Park | Wellcome Sanger Institute for the COVID-19 Genomics UK (COG-UK) Consortium | Jacquelyn Wynn, Mairead Hyland, The Lighthouse Lab in Alderley Park and Alex Alderton, Roberto Amato, Jeffrey Barrett, Sonia Goncalves, Ewan Harrison, David K. Jackson, Ian Johnston, Dominic Kwiatkowski, Cordelia Langford, John Sillitoe on behalf of the Wellcome Sanger Institute COVID-19 Surveillance Team |
| EPI_ISL_2974221, EPI_ISL_2974222, EPI_ISL_2974223 | Lighthouse Lab in Milton Keynes | Wellcome Sanger Institute for the COVID-19 Genomics UK (COG-UK) Consortium | The Lighthouse Lab in Milton Keynes and Alex Alderton, Roberto Amato, Jeffrey Barrett, Sonia Goncalves, Ewan Harrison, David K. Jackson, Ian Johnston, Dominic Kwiatkowski, Cordelia Langford, John Sillitoe on behalf of the Wellcome Sanger Institute COVID-19 Surveillance Team |
| EPI_ISL_2974224, EPI_ISL_2974225 | Lighthouse Laboratory Plymouth | Wellcome Sanger Institute for the COVID-19 Genomics UK (COG-UK) Consortium | Lighthouse Laboratory Plymouth and Alex Alderton, Roberto Amato, Jeffrey Barrett, Sonia Goncalves, Ewan Harrison, David K. Jackson, Ian Johnston, Dominic Kwiatkowski, Cordelia Langford, John Sillitoe on behalf of the Wellcome Sanger Institute COVID-19 Surveillance Team |
| EPI_ISL_2974226, EPI_ISL_2974227, EPI_ISL_2974228, EPI_ISL_2974229, EPI_ISL_2974230, EPI_ISL_2974231, EPI_ISL_2974232 | Lighthouse Lab in Milton Keynes | Wellcome Sanger Institute for the COVID-19 Genomics UK (COG-UK) Consortium | The Lighthouse Lab in Milton Keynes and Alex Alderton, Roberto Amato, Jeffrey Barrett, Sonia Goncalves, Ewan Harrison, David K. Jackson, Ian Johnston, Dominic Kwiatkowski, Cordelia Langford, John Sillitoe on behalf of the Wellcome Sanger Institute COVID-19 Surveillance Team |
| EPI_ISL_2974233 | Lighthouse Laboratory Plymouth | Wellcome Sanger Institute for the COVID-19 Genomics UK (COG-UK) Consortium | Lighthouse Laboratory Plymouth and Alex Alderton, Roberto Amato, Jeffrey Barrett, Sonia Goncalves, Ewan Harrison, David K. Jackson, Ian Johnston, Dominic Kwiatkowski, Cordelia Langford, John Sillitoe on behalf of the Wellcome Sanger Institute COVID-19 Surveillance Team |
| EPI_ISL_2974234 | Lighthouse Lab in Milton Keynes | Wellcome Sanger Institute for the COVID-19 Genomics UK (COG-UK) Consortium | The Lighthouse Lab in Milton Keynes and Alex Alderton, Roberto Amato, Jeffrey Barrett, Sonia Goncalves, Ewan Harrison, David K. Jackson, Ian Johnston, Dominic Kwiatkowski, Cordelia Langford, John Sillitoe on behalf of the Wellcome Sanger Institute COVID-19 Surveillance Team |
| EPI_ISL_2974235 | Lighthouse Lab in Alderley Park | Wellcome Sanger Institute for the COVID-19 Genomics UK (COG-UK) Consortium | Jacquelyn Wynn, Mairead Hyland, The Lighthouse Lab in Alderley Park and Alex Alderton, Roberto Amato, Jeffrey Barrett, Sonia Goncalves, Ewan Harrison, David K. Jackson, Ian Johnston, Dominic Kwiatkowski, Cordelia Langford, John Sillitoe on behalf of the Wellcome Sanger Institute COVID-19 Surveillance Team |
| EPI_ISL_2974236 | Lighthouse Lab in Milton Keynes | Wellcome Sanger Institute for the COVID-19 Genomics UK (COG-UK) Consortium | The Lighthouse Lab in Milton Keynes and Alex Alderton, Roberto Amato, Jeffrey Barrett, Sonia Goncalves, Ewan Harrison, David K. Jackson, Ian Johnston, Dominic Kwiatkowski, Cordelia Langford, John Sillitoe on behalf of the Wellcome Sanger Institute COVID-19 Surveillance Team |
| EPI_ISL_2974237 | Lighthouse Laboratory Plymouth | Wellcome Sanger Institute for the COVID-19 Genomics UK (COG-UK) Consortium | Lighthouse Laboratory Plymouth and Alex Alderton, Roberto Amato, Jeffrey Barrett, Sonia Goncalves, Ewan Harrison, David K. Jackson, Ian Johnston, Dominic Kwiatkowski, Cordelia Langford, John Sillitoe on behalf of the Wellcome Sanger Institute COVID-19 Surveillance Team |
| EPI_ISL_2974238 | Lighthouse Lab in Alderley Park | Wellcome Sanger Institute for the COVID-19 Genomics UK (COG-UK) Consortium | Jacquelyn Wynn, Mairead Hyland, The Lighthouse Lab in Alderley Park and Alex Alderton, Roberto Amato, Jeffrey Barrett, Sonia Goncalves, Ewan Harrison, David K. Jackson, Ian Johnston, Dominic Kwiatkowski, Cordelia Langford, John Sillitoe on behalf of the Wellcome Sanger Institute COVID-19 Surveillance Team |
| EPI_ISL_2974239 | Lighthouse Laboratory Plymouth | Wellcome Sanger Institute for the COVID-19 Genomics UK (COG-UK) Consortium | Lighthouse Laboratory Plymouth and Alex Alderton, Roberto Amato, Jeffrey Barrett, Sonia Goncalves, Ewan Harrison, David K. Jackson, Ian Johnston, Dominic Kwiatkowski, Cordelia Langford, John Sillitoe on behalf of the Wellcome Sanger Institute COVID-19 Surveillance Team |
| EPI_ISL_2974240 | Lighthouse Lab in Milton Keynes | Wellcome Sanger Institute for the COVID-19 Genomics UK (COG-UK) Consortium | The Lighthouse Lab in Milton Keynes and Alex Alderton, Roberto Amato, Jeffrey Barrett, Sonia Goncalves, Ewan Harrison, David K. Jackson, Ian Johnston, Dominic Kwiatkowski, Cordelia Langford, John Sillitoe on behalf of the Wellcome Sanger Institute COVID-19 Surveillance Team |
| EPI_ISL_2974241 | Lighthouse Laboratory Plymouth | Wellcome Sanger Institute for the COVID-19 Genomics UK (COG-UK) Consortium | Lighthouse Laboratory Plymouth and Alex Alderton, Roberto Amato, Jeffrey Barrett, Sonia Goncalves, Ewan Harrison, David K. Jackson, Ian Johnston, Dominic Kwiatkowski, Cordelia Langford, John Sillitoe on behalf of the Wellcome Sanger Institute COVID-19 Surveillance Team |
| EPI_ISL_2974242 | Lighthouse Lab in Alderley Park | Wellcome Sanger Institute for the COVID-19 Genomics UK (COG-UK) Consortium | Jacquelyn Wynn, Mairead Hyland, The Lighthouse Lab in Alderley Park and Alex Alderton, Roberto Amato, Jeffrey Barrett, Sonia Goncalves, Ewan Harrison, David K. Jackson, Ian Johnston, Dominic Kwiatkowski, Cordelia Langford, John Sillitoe on behalf of the Wellcome Sanger Institute COVID-19 Surveillance Team |
| EPI_ISL_2974243 | Lighthouse Lab in Milton Keynes | Wellcome Sanger Institute for the COVID-19 Genomics UK (COG-UK) Consortium | The Lighthouse Lab in Milton Keynes and Alex Alderton, Roberto Amato, Jeffrey Barrett, Sonia Goncalves, Ewan Harrison, David K. Jackson, Ian Johnston, Dominic Kwiatkowski, Cordelia Langford, John Sillitoe on behalf of the Wellcome Sanger Institute COVID-19 Surveillance Team |
| EPI_ISL_2974244 | Lighthouse Laboratory Plymouth | Wellcome Sanger Institute for the COVID-19 Genomics UK (COG-UK) Consortium | Lighthouse Laboratory Plymouth and Alex Alderton, Roberto Amato, Jeffrey Barrett, Sonia Goncalves, Ewan Harrison, David K. Jackson, Ian Johnston, Dominic Kwiatkowski, Cordelia Langford, John Sillitoe on behalf of the Wellcome Sanger Institute COVID-19 Surveillance Team |
| EPI_ISL_2974245 | Lighthouse Lab in Alderley Park | Wellcome Sanger Institute for the COVID-19 Genomics UK (COG-UK) Consortium | Jacquelyn Wynn, Mairead Hyland, The Lighthouse Lab in Alderley Park and Alex Alderton, Roberto Amato, Jeffrey Barrett, Sonia Goncalves, Ewan Harrison, David K. Jackson, Ian Johnston, Dominic Kwiatkowski, Cordelia Langford, John Sillitoe on behalf of the Wellcome Sanger Institute COVID-19 Surveillance Team |
| EPI_ISL_2974246 | Lighthouse Laboratory Plymouth | Wellcome Sanger Institute for the COVID-19 Genomics UK (COG-UK) Consortium | Lighthouse Laboratory Plymouth and Alex Alderton, Roberto Amato, Jeffrey Barrett, Sonia Goncalves, Ewan Harrison, David K. Jackson, Ian Johnston, Dominic Kwiatkowski, Cordelia Langford, John Sillitoe on behalf of the Wellcome Sanger Institute COVID-19 Surveillance Team |
| EPI_ISL_2974247 | Lighthouse Lab in Alderley Park | Wellcome Sanger Institute for the COVID-19 Genomics UK (COG-UK) Consortium | Jacquelyn Wynn, Mairead Hyland, The Lighthouse Lab in Alderley Park and Alex Alderton, Roberto Amato, Jeffrey Barrett, Sonia Goncalves, Ewan Harrison, David K. Jackson, Ian Johnston, Dominic Kwiatkowski, Cordelia Langford, John Sillitoe on behalf of the Wellcome Sanger Institute COVID-19 Surveillance Team |
| EPI_ISL_2974248 | Lighthouse Lab in Milton Keynes | Wellcome Sanger Institute for the COVID-19 Genomics UK (COG-UK) Consortium | The Lighthouse Lab in Milton Keynes and Alex Alderton, Roberto Amato, Jeffrey Barrett, Sonia Goncalves, Ewan Harrison, David K. Jackson, Ian Johnston, Dominic Kwiatkowski, Cordelia Langford, John Sillitoe on behalf of the Wellcome Sanger Institute COVID-19 Surveillance Team |
| EPI_ISL_2974249 | Lighthouse Laboratory Plymouth | Wellcome Sanger Institute for the COVID-19 Genomics UK (COG-UK) Consortium | Lighthouse Laboratory Plymouth and Alex Alderton, Roberto Amato, Jeffrey Barrett, Sonia Goncalves, Ewan Harrison, David K. Jackson, Ian Johnston, Dominic Kwiatkowski, Cordelia Langford, John Sillitoe on behalf of the Wellcome Sanger Institute COVID-19 Surveillance Team |
| EPI_ISL_2974250, EPI_ISL_2974251 | Lighthouse Lab in Milton Keynes | Wellcome Sanger Institute for the COVID-19 Genomics UK (COG-UK) Consortium | The Lighthouse Lab in Milton Keynes and Alex Alderton, Roberto Amato, Jeffrey Barrett, Sonia Goncalves, Ewan Harrison, David K. Jackson, Ian Johnston, Dominic Kwiatkowski, Cordelia Langford, John Sillitoe on behalf of the Wellcome Sanger Institute COVID-19 Surveillance Team |
| EPI_ISL_2974252 | Lighthouse Laboratory Plymouth | Wellcome Sanger Institute for the COVID-19 Genomics UK (COG-UK) Consortium | Lighthouse Laboratory Plymouth and Alex Alderton, Roberto Amato, Jeffrey Barrett, Sonia Goncalves, Ewan Harrison, David K. Jackson, Ian Johnston, Dominic Kwiatkowski, Cordelia Langford, John Sillitoe on behalf of the Wellcome Sanger Institute COVID-19 Surveillance Team |
| EPI_ISL_2974253 | Lighthouse Lab in Milton Keynes | Wellcome Sanger Institute for the COVID-19 Genomics UK (COG-UK) Consortium | The Lighthouse Lab in Milton Keynes and Alex Alderton, Roberto Amato, Jeffrey Barrett, Sonia Goncalves, Ewan Harrison, David K. Jackson, Ian Johnston, Dominic Kwiatkowski, Cordelia Langford, John Sillitoe on behalf of the Wellcome Sanger Institute COVID-19 Surveillance Team |
| EPI_ISL_2974254 | Lighthouse Laboratory Plymouth | Wellcome Sanger Institute for the COVID-19 Genomics UK (COG-UK) Consortium | Lighthouse Laboratory Plymouth and Alex Alderton, Roberto Amato, Jeffrey Barrett, Sonia Goncalves, Ewan Harrison, David K. Jackson, Ian Johnston, Dominic Kwiatkowski, Cordelia Langford, John Sillitoe on behalf of the Wellcome Sanger Institute COVID-19 Surveillance Team |
| EPI_ISL_2974255 | Lighthouse Lab in Milton Keynes | Wellcome Sanger Institute for the COVID-19 Genomics UK (COG-UK) Consortium | The Lighthouse Lab in Milton Keynes and Alex Alderton, Roberto Amato, Jeffrey Barrett, Sonia Goncalves, Ewan Harrison, David K. Jackson, Ian Johnston, Dominic Kwiatkowski, Cordelia Langford, John Sillitoe on behalf of the Wellcome Sanger Institute COVID-19 Surveillance Team |
| EPI_ISL_2974256 | Lighthouse Lab in Alderley Park | Wellcome Sanger Institute for the COVID-19 Genomics UK (COG-UK) Consortium | Jacquelyn Wynn, Mairead Hyland, The Lighthouse Lab in Alderley Park and Alex Alderton, Roberto Amato, Jeffrey Barrett, Sonia Goncalves, Ewan Harrison, David K. Jackson, Ian Johnston, Dominic Kwiatkowski, Cordelia Langford, John Sillitoe on behalf of the Wellcome Sanger Institute COVID-19 Surveillance Team |
| EPI_ISL_2974257 | Lighthouse Laboratory Plymouth | Wellcome Sanger Institute for the COVID-19 Genomics UK (COG-UK) Consortium | Lighthouse Laboratory Plymouth and Alex Alderton, Roberto Amato, Jeffrey Barrett, Sonia Goncalves, Ewan Harrison, David K. Jackson, Ian Johnston, Dominic Kwiatkowski, Cordelia Langford, John Sillitoe on behalf of the Wellcome Sanger Institute COVID-19 Surveillance Team |
| EPI_ISL_2974258 | Lighthouse Lab in Milton Keynes | Wellcome Sanger Institute for the COVID-19 Genomics UK (COG-UK) Consortium | The Lighthouse Lab in Milton Keynes and Alex Alderton, Roberto Amato, Jeffrey Barrett, Sonia Goncalves, Ewan Harrison, David K. Jackson, Ian Johnston, Dominic Kwiatkowski, Cordelia Langford, John Sillitoe on behalf of the Wellcome Sanger Institute COVID-19 Surveillance Team |
| EPI_ISL_2974259 | Lighthouse Lab in Alderley Park | Wellcome Sanger Institute for the COVID-19 Genomics UK (COG-UK) Consortium | Jacquelyn Wynn, Mairead Hyland, The Lighthouse Lab in Alderley Park and Alex Alderton, Roberto Amato, Jeffrey Barrett, Sonia Goncalves, Ewan Harrison, David K. Jackson, Ian Johnston, Dominic Kwiatkowski, Cordelia Langford, John Sillitoe on behalf of the Wellcome Sanger Institute COVID-19 Surveillance Team |
| EPI_ISL_2974260 | Lighthouse Lab in Milton Keynes | Wellcome Sanger Institute for the COVID-19 Genomics UK (COG-UK) Consortium | The Lighthouse Lab in Milton Keynes and Alex Alderton, Roberto Amato, Jeffrey Barrett, Sonia Goncalves, Ewan Harrison, David K. Jackson, Ian Johnston, Dominic Kwiatkowski, Cordelia Langford, John Sillitoe on behalf of the Wellcome Sanger Institute COVID-19 Surveillance Team |
| EPI_ISL_2974261 | Lighthouse Lab in Alderley Park | Wellcome Sanger Institute for the COVID-19 Genomics UK (COG-UK) Consortium | Jacquelyn Wynn, Mairead Hyland, The Lighthouse Lab in Alderley Park and Alex Alderton, Roberto Amato, Jeffrey Barrett, Sonia Goncalves, Ewan Harrison, David K. Jackson, Ian Johnston, Dominic Kwiatkowski, Cordelia Langford, John Sillitoe on behalf of the Wellcome Sanger Institute COVID-19 Surveillance Team |
| EPI_ISL_2974262 | Lighthouse Laboratory Plymouth | Wellcome Sanger Institute for the COVID-19 Genomics UK (COG-UK) Consortium | Lighthouse Laboratory Plymouth and Alex Alderton, Roberto Amato, Jeffrey Barrett, Sonia Goncalves, Ewan Harrison, David K. Jackson, Ian Johnston, Dominic Kwiatkowski, Cordelia Langford, John Sillitoe on behalf of the Wellcome Sanger Institute COVID-19 Surveillance Team |
| EPI_ISL_2974263, EPI_ISL_2974264 | Lighthouse Lab in Alderley Park | Wellcome Sanger Institute for the COVID-19 Genomics UK (COG-UK) Consortium | Jacquelyn Wynn, Mairead Hyland, The Lighthouse Lab in Alderley Park and Alex Alderton, Roberto Amato, Jeffrey Barrett, Sonia Goncalves, Ewan Harrison, David K. Jackson, Ian Johnston, Dominic Kwiatkowski, Cordelia Langford, John Sillitoe on behalf of the Wellcome Sanger Institute COVID-19 Surveillance Team |

| | | | |
|---|---|---|---|
| EPI_ISL_2974265 | Lighthouse Lab in Milton Keynes | Wellcome Sanger Institute for the COVID-19 Genomics UK (COG-UK) Consortium | The Lighthouse Lab in Milton Keynes and Alex Alderton, Roberto Amato, Jeffrey Barrett, Sonia Goncalves, Ewan Harrison, David K. Jackson, Ian Johnston, Dominic Kwiatkowski, Cordelia Langford, John Sillitoe on behalf of the Wellcome Sanger Institute COVID-19 Surveillance Team |
| EPI_ISL_2974266 | Lighthouse Laboratory Plymouth | Wellcome Sanger Institute for the COVID-19 Genomics UK (COG-UK) Consortium | Lighthouse Laboratory Plymouth and Alex Alderton, Roberto Amato, Jeffrey Barrett, Sonia Goncalves, Ewan Harrison, David K. Jackson, Ian Johnston, Dominic Kwiatkowski, Cordelia Langford, John Sillitoe on behalf of the Wellcome Sanger Institute COVID-19 Surveillance Team |
| EPI_ISL_2974267, EPI_ISL_2974268 | Lighthouse Lab in Milton Keynes | Wellcome Sanger Institute for the COVID-19 Genomics UK (COG-UK) Consortium | The Lighthouse Lab in Milton Keynes and Alex Alderton, Roberto Amato, Jeffrey Barrett, Sonia Goncalves, Ewan Harrison, David K. Jackson, Ian Johnston, Dominic Kwiatkowski, Cordelia Langford, John Sillitoe on behalf of the Wellcome Sanger Institute COVID-19 Surveillance Team |
| EPI_ISL_2974269 | Lighthouse Lab in Alderley Park | Wellcome Sanger Institute for the COVID-19 Genomics UK (COG-UK) Consortium | Jacquelyn Wynn, Mairead Hyland, The Lighthouse Lab in Alderley Park and Alex Alderton, Roberto Amato, Jeffrey Barrett, Sonia Goncalves, Ewan Harrison, David K. Jackson, Ian Johnston, Dominic Kwiatkowski, Cordelia Langford, John Sillitoe on behalf of the Wellcome Sanger Institute COVID-19 Surveillance Team |
| EPI_ISL_2974270 | Lighthouse Laboratory Plymouth | Wellcome Sanger Institute for the COVID-19 Genomics UK (COG-UK) Consortium | Lighthouse Laboratory Plymouth and Alex Alderton, Roberto Amato, Jeffrey Barrett, Sonia Goncalves, Ewan Harrison, David K. Jackson, Ian Johnston, Dominic Kwiatkowski, Cordelia Langford, John Sillitoe on behalf of the Wellcome Sanger Institute COVID-19 Surveillance Team |
| EPI_ISL_2974271 | Lighthouse Lab in Milton Keynes | Wellcome Sanger Institute for the COVID-19 Genomics UK (COG-UK) Consortium | The Lighthouse Lab in Milton Keynes and Alex Alderton, Roberto Amato, Jeffrey Barrett, Sonia Goncalves, Ewan Harrison, David K. Jackson, Ian Johnston, Dominic Kwiatkowski, Cordelia Langford, John Sillitoe on behalf of the Wellcome Sanger Institute COVID-19 Surveillance Team |
| EPI_ISL_2974272 | Lighthouse Laboratory Plymouth | Wellcome Sanger Institute for the COVID-19 Genomics UK (COG-UK) Consortium | Lighthouse Laboratory Plymouth and Alex Alderton, Roberto Amato, Jeffrey Barrett, Sonia Goncalves, Ewan Harrison, David K. Jackson, Ian Johnston, Dominic Kwiatkowski, Cordelia Langford, John Sillitoe on behalf of the Wellcome Sanger Institute COVID-19 Surveillance Team |
| EPI_ISL_2974273 | Lighthouse Lab in Milton Keynes | Wellcome Sanger Institute for the COVID-19 Genomics UK (COG-UK) Consortium | The Lighthouse Lab in Milton Keynes and Alex Alderton, Roberto Amato, Jeffrey Barrett, Sonia Goncalves, Ewan Harrison, David K. Jackson, Ian Johnston, Dominic Kwiatkowski, Cordelia Langford, John Sillitoe on behalf of the Wellcome Sanger Institute COVID-19 Surveillance Team |
| EPI_ISL_2974274 | Lighthouse Lab in Alderley Park | Wellcome Sanger Institute for the COVID-19 Genomics UK (COG-UK) Consortium | Jacquelyn Wynn, Mairead Hyland, The Lighthouse Lab in Alderley Park and Alex Alderton, Roberto Amato, Jeffrey Barrett, Sonia Goncalves, Ewan Harrison, David K. Jackson, Ian Johnston, Dominic Kwiatkowski, Cordelia Langford, John Sillitoe on behalf of the Wellcome Sanger Institute COVID-19 Surveillance Team |
| EPI_ISL_2974275 | Lighthouse Laboratory Plymouth | Wellcome Sanger Institute for the COVID-19 Genomics UK (COG-UK) Consortium | Lighthouse Laboratory Plymouth and Alex Alderton, Roberto Amato, Jeffrey Barrett, Sonia Goncalves, Ewan Harrison, David K. Jackson, Ian Johnston, Dominic Kwiatkowski, Cordelia Langford, John Sillitoe on behalf of the Wellcome Sanger Institute COVID-19 Surveillance Team |
| EPI_ISL_2974276, EPI_ISL_2974277 | Lighthouse Lab in Milton Keynes | Wellcome Sanger Institute for the COVID-19 Genomics UK (COG-UK) Consortium | The Lighthouse Lab in Milton Keynes and Alex Alderton, Roberto Amato, Jeffrey Barrett, Sonia Goncalves, Ewan Harrison, David K. Jackson, Ian Johnston, Dominic Kwiatkowski, Cordelia Langford, John Sillitoe on behalf of the Wellcome Sanger Institute COVID-19 Surveillance Team |
| EPI_ISL_2974278 | Lighthouse Laboratory Plymouth | Wellcome Sanger Institute for the COVID-19 Genomics UK (COG-UK) Consortium | Lighthouse Laboratory Plymouth and Alex Alderton, Roberto Amato, Jeffrey Barrett, Sonia Goncalves, Ewan Harrison, David K. Jackson, Ian Johnston, Dominic Kwiatkowski, Cordelia Langford, John Sillitoe on behalf of the Wellcome Sanger Institute COVID-19 Surveillance Team |
| EPI_ISL_2974279 | Lighthouse Lab in Milton Keynes | Wellcome Sanger Institute for the COVID-19 Genomics UK (COG-UK) Consortium | The Lighthouse Lab in Milton Keynes and Alex Alderton, Roberto Amato, Jeffrey Barrett, Sonia Goncalves, Ewan Harrison, David K. Jackson, Ian Johnston, Dominic Kwiatkowski, Cordelia Langford, John Sillitoe on behalf of the Wellcome Sanger Institute COVID-19 Surveillance Team |
| EPI_ISL_2974280 | Lighthouse Laboratory Plymouth | Wellcome Sanger Institute for the COVID-19 Genomics UK (COG-UK) Consortium | Lighthouse Laboratory Plymouth and Alex Alderton, Roberto Amato, Jeffrey Barrett, Sonia Goncalves, Ewan Harrison, David K. Jackson, Ian Johnston, Dominic Kwiatkowski, Cordelia Langford, John Sillitoe on behalf of the Wellcome Sanger Institute COVID-19 Surveillance Team |
| EPI_ISL_2974281, EPI_ISL_2974282, EPI_ISL_2974283 | Lighthouse Lab in Milton Keynes | Wellcome Sanger Institute for the COVID-19 Genomics UK (COG-UK) Consortium | The Lighthouse Lab in Milton Keynes and Alex Alderton, Roberto Amato, Jeffrey Barrett, Sonia Goncalves, Ewan Harrison, David K. Jackson, Ian Johnston, Dominic Kwiatkowski, Cordelia Langford, John Sillitoe on behalf of the Wellcome Sanger Institute COVID-19 Surveillance Team |
| EPI_ISL_2974284 | Lighthouse Lab in Alderley Park | Wellcome Sanger Institute for the COVID-19 Genomics UK (COG-UK) Consortium | Jacquelyn Wynn, Mairead Hyland, The Lighthouse Lab in Alderley Park and Alex Alderton, Roberto Amato, Jeffrey Barrett, Sonia Goncalves, Ewan Harrison, David K. Jackson, Ian Johnston, Dominic Kwiatkowski, Cordelia Langford, John Sillitoe on behalf of the Wellcome Sanger Institute COVID-19 Surveillance Team |
| EPI_ISL_2974285 | Lighthouse Lab in Milton Keynes | Wellcome Sanger Institute for the COVID-19 Genomics UK (COG-UK) Consortium | The Lighthouse Lab in Milton Keynes and Alex Alderton, Roberto Amato, Jeffrey Barrett, Sonia Goncalves, Ewan Harrison, David K. Jackson, Ian Johnston, Dominic Kwiatkowski, Cordelia Langford, John Sillitoe on behalf of the Wellcome Sanger Institute COVID-19 Surveillance Team |
| EPI_ISL_2974286 | Lighthouse Lab in Alderley Park | Wellcome Sanger Institute for the COVID-19 Genomics UK (COG-UK) Consortium | Jacquelyn Wynn, Mairead Hyland, The Lighthouse Lab in Alderley Park and Alex Alderton, Roberto Amato, Jeffrey Barrett, Sonia Goncalves, Ewan Harrison, David K. Jackson, Ian Johnston, Dominic Kwiatkowski, Cordelia Langford, John Sillitoe on behalf of the Wellcome Sanger Institute COVID-19 Surveillance Team |
| EPI_ISL_2974287, EPI_ISL_2974288 | Lighthouse Lab in Milton Keynes | Wellcome Sanger Institute for the COVID-19 Genomics UK (COG-UK) Consortium | The Lighthouse Lab in Milton Keynes and Alex Alderton, Roberto Amato, Jeffrey Barrett, Sonia Goncalves, Ewan Harrison, David K. Jackson, Ian Johnston, Dominic Kwiatkowski, Cordelia Langford, John Sillitoe on behalf of the Wellcome Sanger Institute COVID-19 Surveillance Team |
| EPI_ISL_2974289 | Lighthouse Laboratory Plymouth | Wellcome Sanger Institute for the COVID-19 Genomics UK (COG-UK) Consortium | Lighthouse Laboratory Plymouth and Alex Alderton, Roberto Amato, Jeffrey Barrett, Sonia Goncalves, Ewan Harrison, David K. Jackson, Ian Johnston, Dominic Kwiatkowski, Cordelia Langford, John Sillitoe on behalf of the Wellcome Sanger Institute COVID-19 Surveillance Team |
| EPI_ISL_2974290 | Lighthouse Lab in Milton Keynes | Wellcome Sanger Institute for the COVID-19 Genomics UK (COG-UK) Consortium | The Lighthouse Lab in Milton Keynes and Alex Alderton, Roberto Amato, Jeffrey Barrett, Sonia Goncalves, Ewan Harrison, David K. Jackson, Ian Johnston, Dominic Kwiatkowski, Cordelia Langford, John Sillitoe on behalf of the Wellcome Sanger Institute COVID-19 Surveillance Team |
| EPI_ISL_2974291 | Lighthouse Laboratory Plymouth | Wellcome Sanger Institute for the COVID-19 Genomics UK (COG-UK) Consortium | Lighthouse Laboratory Plymouth and Alex Alderton, Roberto Amato, Jeffrey Barrett, Sonia Goncalves, Ewan Harrison, David K. Jackson, Ian Johnston, Dominic Kwiatkowski, Cordelia Langford, John Sillitoe on behalf of the Wellcome Sanger Institute COVID-19 Surveillance Team |
| EPI_ISL_2974292 | Lighthouse Lab in Alderley Park | Wellcome Sanger Institute for the COVID-19 Genomics UK (COG-UK) Consortium | Jacquelyn Wynn, Mairead Hyland, The Lighthouse Lab in Alderley Park and Alex Alderton, Roberto Amato, Jeffrey Barrett, Sonia Goncalves, Ewan Harrison, David K. Jackson, Ian Johnston, Dominic Kwiatkowski, Cordelia Langford, John Sillitoe on behalf of the Wellcome Sanger Institute COVID-19 Surveillance Team |
| EPI_ISL_2974293 | Lighthouse Lab in Milton Keynes | Wellcome Sanger Institute for the COVID-19 Genomics UK (COG-UK) Consortium | The Lighthouse Lab in Milton Keynes and Alex Alderton, Roberto Amato, Jeffrey Barrett, Sonia Goncalves, Ewan Harrison, David K. Jackson, Ian Johnston, Dominic Kwiatkowski, Cordelia Langford, John Sillitoe on behalf of the Wellcome Sanger Institute COVID-19 Surveillance Team |
| EPI_ISL_2974294, EPI_ISL_2974295 | Lighthouse Laboratory Plymouth | Wellcome Sanger Institute for the COVID-19 Genomics UK (COG-UK) Consortium | Lighthouse Laboratory Plymouth and Alex Alderton, Roberto Amato, Jeffrey Barrett, Sonia Goncalves, Ewan Harrison, David K. Jackson, Ian Johnston, Dominic Kwiatkowski, Cordelia Langford, John Sillitoe on behalf of the Wellcome Sanger Institute COVID-19 Surveillance Team |
| EPI_ISL_2974296 | Lighthouse Lab in Milton Keynes | Wellcome Sanger Institute for the COVID-19 Genomics UK (COG-UK) Consortium | The Lighthouse Lab in Milton Keynes and Alex Alderton, Roberto Amato, Jeffrey Barrett, Sonia Goncalves, Ewan Harrison, David K. Jackson, Ian Johnston, Dominic Kwiatkowski, Cordelia Langford, John Sillitoe on behalf of the Wellcome Sanger Institute COVID-19 Surveillance Team |
| EPI_ISL_2974297, EPI_ISL_2974298 | Lighthouse Lab in Alderley Park | Wellcome Sanger Institute for the COVID-19 Genomics UK (COG-UK) Consortium | Jacquelyn Wynn, Mairead Hyland, The Lighthouse Lab in Alderley Park and Alex Alderton, Roberto Amato, Jeffrey Barrett, Sonia Goncalves, Ewan Harrison, David K. Jackson, Ian Johnston, Dominic Kwiatkowski, Cordelia Langford, John Sillitoe on behalf of the Wellcome Sanger Institute COVID-19 Surveillance Team |
| EPI_ISL_2974299, EPI_ISL_2974300, EPI_ISL_2974301 | Lighthouse Lab in Milton Keynes | Wellcome Sanger Institute for the COVID-19 Genomics UK (COG-UK) Consortium | The Lighthouse Lab in Milton Keynes and Alex Alderton, Roberto Amato, Jeffrey Barrett, Sonia Goncalves, Ewan Harrison, David K. Jackson, Ian Johnston, Dominic Kwiatkowski, Cordelia Langford, John Sillitoe on behalf of the Wellcome Sanger Institute COVID-19 Surveillance Team |
| EPI_ISL_2974302 | Lighthouse Lab in Alderley Park | Wellcome Sanger Institute for the COVID-19 Genomics UK (COG-UK) Consortium | Jacquelyn Wynn, Mairead Hyland, The Lighthouse Lab in Alderley Park and Alex Alderton, Roberto Amato, Jeffrey Barrett, Sonia Goncalves, Ewan Harrison, David K. Jackson, Ian Johnston, Dominic Kwiatkowski, Cordelia Langford, John Sillitoe on behalf of the Wellcome Sanger Institute COVID-19 Surveillance Team |
| EPI_ISL_2974303 | Lighthouse Lab in Milton Keynes | Wellcome Sanger Institute for the COVID-19 Genomics UK (COG-UK) Consortium | The Lighthouse Lab in Milton Keynes and Alex Alderton, Roberto Amato, Jeffrey Barrett, Sonia Goncalves, Ewan Harrison, David K. Jackson, Ian Johnston, Dominic Kwiatkowski, Cordelia Langford, John Sillitoe on behalf of the Wellcome Sanger Institute COVID-19 Surveillance Team |
| EPI_ISL_2974304 | Lighthouse Laboratory Plymouth | Wellcome Sanger Institute for the COVID-19 Genomics UK (COG-UK) Consortium | Lighthouse Laboratory Plymouth and Alex Alderton, Roberto Amato, Jeffrey Barrett, Sonia Goncalves, Ewan Harrison, David K. Jackson, Ian Johnston, Dominic Kwiatkowski, Cordelia Langford, John Sillitoe on behalf of the Wellcome Sanger Institute COVID-19 Surveillance Team |
| EPI_ISL_2974305 | Lighthouse Lab in Milton Keynes | Wellcome Sanger Institute for the COVID-19 Genomics UK (COG-UK) Consortium | The Lighthouse Lab in Milton Keynes and Alex Alderton, Roberto Amato, Jeffrey Barrett, Sonia Goncalves, Ewan Harrison, David K. Jackson, Ian Johnston, Dominic Kwiatkowski, Cordelia Langford, John Sillitoe on behalf of the Wellcome Sanger Institute COVID-19 Surveillance Team |
| EPI_ISL_2974306 | Lighthouse Lab in Alderley Park | Wellcome Sanger Institute for the COVID-19 Genomics UK (COG-UK) Consortium | Jacquelyn Wynn, Mairead Hyland, The Lighthouse Lab in Alderley Park and Alex Alderton, Roberto Amato, Jeffrey Barrett, Sonia Goncalves, Ewan Harrison, David K. Jackson, Ian Johnston, Dominic Kwiatkowski, Cordelia Langford, John Sillitoe on behalf of the Wellcome Sanger Institute COVID-19 Surveillance Team |
| EPI_ISL_2974307 | Lighthouse Lab in Milton Keynes | Wellcome Sanger Institute for the COVID-19 Genomics UK (COG-UK) Consortium | The Lighthouse Lab in Milton Keynes and Alex Alderton, Roberto Amato, Jeffrey Barrett, Sonia Goncalves, Ewan Harrison, David K. Jackson, Ian Johnston, Dominic Kwiatkowski, Cordelia Langford, John Sillitoe on behalf of the Wellcome Sanger Institute COVID-19 Surveillance Team |
| EPI_ISL_2974308 | Lighthouse Laboratory Plymouth | Wellcome Sanger Institute for the COVID-19 Genomics UK (COG-UK) Consortium | Lighthouse Laboratory Plymouth and Alex Alderton, Roberto Amato, Jeffrey Barrett, Sonia Goncalves, Ewan Harrison, David K. Jackson, Ian Johnston, Dominic Kwiatkowski, Cordelia Langford, John Sillitoe on behalf of the Wellcome Sanger Institute COVID-19 Surveillance Team |
| EPI_ISL_2974309 | Lighthouse Lab in Milton Keynes | Wellcome Sanger Institute for the COVID-19 Genomics UK (COG-UK) Consortium | The Lighthouse Lab in Milton Keynes and Alex Alderton, Roberto Amato, Jeffrey Barrett, Sonia Goncalves, Ewan Harrison, David K. Jackson, Ian Johnston, Dominic Kwiatkowski, Cordelia Langford, John Sillitoe on behalf of the Wellcome Sanger Institute COVID-19 Surveillance Team |
| EPI_ISL_2974310, EPI_ISL_2974311, EPI_ISL_2974312, EPI_ISL_2974313 | Lighthouse Lab in Alderley Park | Wellcome Sanger Institute for the COVID-19 Genomics UK (COG-UK) Consortium | Jacquelyn Wynn, Mairead Hyland, The Lighthouse Lab in Alderley Park and Alex Alderton, Roberto Amato, Jeffrey Barrett, Sonia Goncalves, Ewan Harrison, David K. Jackson, Ian Johnston, Dominic Kwiatkowski, Cordelia Langford, John Sillitoe on behalf of the Wellcome Sanger Institute COVID-19 Surveillance Team |
| EPI_ISL_2974314 | Lighthouse Laboratory Plymouth | Wellcome Sanger Institute for the COVID-19 Genomics UK (COG-UK) Consortium | Lighthouse Laboratory Plymouth and Alex Alderton, Roberto Amato, Jeffrey Barrett, Sonia Goncalves, Ewan Harrison, David K. Jackson, Ian Johnston, Dominic Kwiatkowski, Cordelia Langford, John Sillitoe on behalf of the Wellcome Sanger Institute COVID-19 Surveillance Team |
| EPI_ISL_2974315, EPI_ISL_2974316 | Lighthouse Lab in Milton Keynes | Wellcome Sanger Institute for the COVID-19 Genomics UK (COG-UK) Consortium | The Lighthouse Lab in Milton Keynes and Alex Alderton, Roberto Amato, Jeffrey Barrett, Sonia Goncalves, Ewan Harrison, David K. Jackson, Ian Johnston, Dominic Kwiatkowski, Cordelia Langford, John Sillitoe on behalf of the Wellcome Sanger Institute COVID-19 Surveillance Team |
| EPI_ISL_2974317 | Lighthouse Lab in Alderley Park | Wellcome Sanger Institute for the COVID-19 Genomics UK (COG-UK) Consortium | Jacquelyn Wynn, Mairead Hyland, The Lighthouse Lab in Alderley Park and Alex Alderton, Roberto Amato, Jeffrey Barrett, Sonia Goncalves, Ewan Harrison, David K. Jackson, Ian Johnston, Dominic Kwiatkowski, Cordelia Langford, John Sillitoe on behalf of the Wellcome Sanger Institute COVID-19 Surveillance Team |
| EPI_ISL_2974318, EPI_ISL_2974319, EPI_ISL_2974320 | Lighthouse Lab in Milton Keynes | Wellcome Sanger Institute for the COVID-19 Genomics UK (COG-UK) Consortium | The Lighthouse Lab in Milton Keynes and Alex Alderton, Roberto Amato, Jeffrey Barrett, Sonia Goncalves, Ewan Harrison, David K. Jackson, Ian Johnston, Dominic Kwiatkowski, Cordelia Langford, John Sillitoe on behalf of the Wellcome Sanger Institute COVID-19 Surveillance Team |
| EPI_ISL_2974321 | Lighthouse Laboratory Plymouth | Wellcome Sanger Institute for the COVID-19 Genomics UK (COG-UK) Consortium | Lighthouse Laboratory Plymouth and Alex Alderton, Roberto Amato, Jeffrey Barrett, Sonia Goncalves, Ewan Harrison, David K. Jackson, Ian Johnston, Dominic Kwiatkowski, Cordelia Langford, John Sillitoe on behalf of the Wellcome Sanger Institute COVID-19 Surveillance Team |
| EPI_ISL_2974322 | Lighthouse Lab in Milton Keynes | Wellcome Sanger Institute for the COVID-19 Genomics UK (COG-UK) Consortium | The Lighthouse Lab in Milton Keynes and Alex Alderton, Roberto Amato, Jeffrey Barrett, Sonia Goncalves, Ewan Harrison, David K. Jackson, Ian Johnston, Dominic Kwiatkowski, Cordelia Langford, John Sillitoe on behalf of the Wellcome Sanger Institute COVID-19 Surveillance Team |

| | | | |
|---|---|---|---|
| EPI_ISL_2974323 | Lighthouse Laboratory Plymouth | Wellcome Sanger Institute for the COVID-19 Genomics UK (COG-UK) Consortium | Lighthouse Laboratory Plymouth and Alex Alderton, Roberto Amato, Jeffrey Barrett, Sonia Goncalves, Ewan Harrison, David K. Jackson, Ian Johnston, Dominic Kwiatkowski, Cordelia Langford, John Sillitoe on behalf of the Wellcome Sanger Institute COVID-19 Surveillance Team |
| EPI_ISL_2974324 | Lighthouse Lab in Milton Keynes | Wellcome Sanger Institute for the COVID-19 Genomics UK (COG-UK) Consortium | The Lighthouse Lab in Milton Keynes and Alex Alderton, Roberto Amato, Jeffrey Barrett, Sonia Goncalves, Ewan Harrison, David K. Jackson, Ian Johnston, Dominic Kwiatkowski, Cordelia Langford, John Sillitoe on behalf of the Wellcome Sanger Institute COVID-19 Surveillance Team |
| EPI_ISL_2974325 | Lighthouse Lab in Alderley Park | Wellcome Sanger Institute for the COVID-19 Genomics UK (COG-UK) Consortium | Jacquelyn Wynn, Mairead Hyland, The Lighthouse Lab in Alderley Park and Alex Alderton, Roberto Amato, Jeffrey Barrett, Sonia Goncalves, Ewan Harrison, David K. Jackson, Ian Johnston, Dominic Kwiatkowski, Cordelia Langford, John Sillitoe on behalf of the Wellcome Sanger Institute COVID-19 Surveillance Team |
| EPI_ISL_2974326 | Lighthouse Lab in Milton Keynes | Wellcome Sanger Institute for the COVID-19 Genomics UK (COG-UK) Consortium | The Lighthouse Lab in Milton Keynes and Alex Alderton, Roberto Amato, Jeffrey Barrett, Sonia Goncalves, Ewan Harrison, David K. Jackson, Ian Johnston, Dominic Kwiatkowski, Cordelia Langford, John Sillitoe on behalf of the Wellcome Sanger Institute COVID-19 Surveillance Team |
| EPI_ISL_2974327 | Lighthouse Lab in Alderley Park | Wellcome Sanger Institute for the COVID-19 Genomics UK (COG-UK) Consortium | Jacquelyn Wynn, Mairead Hyland, The Lighthouse Lab in Alderley Park and Alex Alderton, Roberto Amato, Jeffrey Barrett, Sonia Goncalves, Ewan Harrison, David K. Jackson, Ian Johnston, Dominic Kwiatkowski, Cordelia Langford, John Sillitoe on behalf of the Wellcome Sanger Institute COVID-19 Surveillance Team |
| EPI_ISL_2974328 | Lighthouse Laboratory Plymouth | Wellcome Sanger Institute for the COVID-19 Genomics UK (COG-UK) Consortium | Lighthouse Laboratory Plymouth and Alex Alderton, Roberto Amato, Jeffrey Barrett, Sonia Goncalves, Ewan Harrison, David K. Jackson, Ian Johnston, Dominic Kwiatkowski, Cordelia Langford, John Sillitoe on behalf of the Wellcome Sanger Institute COVID-19 Surveillance Team |
| EPI_ISL_2974329 | Lighthouse Lab in Alderley Park | Wellcome Sanger Institute for the COVID-19 Genomics UK (COG-UK) Consortium | Jacquelyn Wynn, Mairead Hyland, The Lighthouse Lab in Alderley Park and Alex Alderton, Roberto Amato, Jeffrey Barrett, Sonia Goncalves, Ewan Harrison, David K. Jackson, Ian Johnston, Dominic Kwiatkowski, Cordelia Langford, John Sillitoe on behalf of the Wellcome Sanger Institute COVID-19 Surveillance Team |
| EPI_ISL_2974330 | Lighthouse Lab in Milton Keynes | Wellcome Sanger Institute for the COVID-19 Genomics UK (COG-UK) Consortium | The Lighthouse Lab in Milton Keynes and Alex Alderton, Roberto Amato, Jeffrey Barrett, Sonia Goncalves, Ewan Harrison, David K. Jackson, Ian Johnston, Dominic Kwiatkowski, Cordelia Langford, John Sillitoe on behalf of the Wellcome Sanger Institute COVID-19 Surveillance Team |
| EPI_ISL_2974331 | Lighthouse Lab in Alderley Park | Wellcome Sanger Institute for the COVID-19 Genomics UK (COG-UK) Consortium | Jacquelyn Wynn, Mairead Hyland, The Lighthouse Lab in Alderley Park and Alex Alderton, Roberto Amato, Jeffrey Barrett, Sonia Goncalves, Ewan Harrison, David K. Jackson, Ian Johnston, Dominic Kwiatkowski, Cordelia Langford, John Sillitoe on behalf of the Wellcome Sanger Institute COVID-19 Surveillance Team |
| EPI_ISL_2974332 | Lighthouse Lab in Milton Keynes | Wellcome Sanger Institute for the COVID-19 Genomics UK (COG-UK) Consortium | The Lighthouse Lab in Milton Keynes and Alex Alderton, Roberto Amato, Jeffrey Barrett, Sonia Goncalves, Ewan Harrison, David K. Jackson, Ian Johnston, Dominic Kwiatkowski, Cordelia Langford, John Sillitoe on behalf of the Wellcome Sanger Institute COVID-19 Surveillance Team |
| EPI_ISL_2974333 | Lighthouse Laboratory Plymouth | Wellcome Sanger Institute for the COVID-19 Genomics UK (COG-UK) Consortium | Lighthouse Laboratory Plymouth and Alex Alderton, Roberto Amato, Jeffrey Barrett, Sonia Goncalves, Ewan Harrison, David K. Jackson, Ian Johnston, Dominic Kwiatkowski, Cordelia Langford, John Sillitoe on behalf of the Wellcome Sanger Institute COVID-19 Surveillance Team |
| EPI_ISL_2974334 | Lighthouse Lab in Alderley Park | Wellcome Sanger Institute for the COVID-19 Genomics UK (COG-UK) Consortium | Jacquelyn Wynn, Mairead Hyland, The Lighthouse Lab in Alderley Park and Alex Alderton, Roberto Amato, Jeffrey Barrett, Sonia Goncalves, Ewan Harrison, David K. Jackson, Ian Johnston, Dominic Kwiatkowski, Cordelia Langford, John Sillitoe on behalf of the Wellcome Sanger Institute COVID-19 Surveillance Team |
| EPI_ISL_2974335, EPI_ISL_2974336 | Lighthouse Lab in Milton Keynes | Wellcome Sanger Institute for the COVID-19 Genomics UK (COG-UK) Consortium | The Lighthouse Lab in Milton Keynes and Alex Alderton, Roberto Amato, Jeffrey Barrett, Sonia Goncalves, Ewan Harrison, David K. Jackson, Ian Johnston, Dominic Kwiatkowski, Cordelia Langford, John Sillitoe on behalf of the Wellcome Sanger Institute COVID-19 Surveillance Team |
| EPI_ISL_2974337 | Lighthouse Lab in Alderley Park | Wellcome Sanger Institute for the COVID-19 Genomics UK (COG-UK) Consortium | Jacquelyn Wynn, Mairead Hyland, The Lighthouse Lab in Alderley Park and Alex Alderton, Roberto Amato, Jeffrey Barrett, Sonia Goncalves, Ewan Harrison, David K. Jackson, Ian Johnston, Dominic Kwiatkowski, Cordelia Langford, John Sillitoe on behalf of the Wellcome Sanger Institute COVID-19 Surveillance Team |
| EPI_ISL_2974338, EPI_ISL_2974339 | Lighthouse Lab in Milton Keynes | Wellcome Sanger Institute for the COVID-19 Genomics UK (COG-UK) Consortium | The Lighthouse Lab in Milton Keynes and Alex Alderton, Roberto Amato, Jeffrey Barrett, Sonia Goncalves, Ewan Harrison, David K. Jackson, Ian Johnston, Dominic Kwiatkowski, Cordelia Langford, John Sillitoe on behalf of the Wellcome Sanger Institute COVID-19 Surveillance Team |
| EPI_ISL_2974340, EPI_ISL_2974341, EPI_ISL_2974342 | Lighthouse Lab in Alderley Park | Wellcome Sanger Institute for the COVID-19 Genomics UK (COG-UK) Consortium | Jacquelyn Wynn, Mairead Hyland, The Lighthouse Lab in Alderley Park and Alex Alderton, Roberto Amato, Jeffrey Barrett, Sonia Goncalves, Ewan Harrison, David K. Jackson, Ian Johnston, Dominic Kwiatkowski, Cordelia Langford, John Sillitoe on behalf of the Wellcome Sanger Institute COVID-19 Surveillance Team |
| EPI_ISL_2974343, EPI_ISL_2974344, EPI_ISL_2974345 | Lighthouse Lab in Milton Keynes | Wellcome Sanger Institute for the COVID-19 Genomics UK (COG-UK) Consortium | The Lighthouse Lab in Milton Keynes and Alex Alderton, Roberto Amato, Jeffrey Barrett, Sonia Goncalves, Ewan Harrison, David K. Jackson, Ian Johnston, Dominic Kwiatkowski, Cordelia Langford, John Sillitoe on behalf of the Wellcome Sanger Institute COVID-19 Surveillance Team |
| EPI_ISL_2974346 | Lighthouse Laboratory Plymouth | Wellcome Sanger Institute for the COVID-19 Genomics UK (COG-UK) Consortium | Lighthouse Laboratory Plymouth and Alex Alderton, Roberto Amato, Jeffrey Barrett, Sonia Goncalves, Ewan Harrison, David K. Jackson, Ian Johnston, Dominic Kwiatkowski, Cordelia Langford, John Sillitoe on behalf of the Wellcome Sanger Institute COVID-19 Surveillance Team |
| EPI_ISL_2974347 | Lighthouse Lab in Milton Keynes | Wellcome Sanger Institute for the COVID-19 Genomics UK (COG-UK) Consortium | The Lighthouse Lab in Milton Keynes and Alex Alderton, Roberto Amato, Jeffrey Barrett, Sonia Goncalves, Ewan Harrison, David K. Jackson, Ian Johnston, Dominic Kwiatkowski, Cordelia Langford, John Sillitoe on behalf of the Wellcome Sanger Institute COVID-19 Surveillance Team |
| EPI_ISL_2974348, EPI_ISL_2974349 | Lighthouse Laboratory Plymouth | Wellcome Sanger Institute for the COVID-19 Genomics UK (COG-UK) Consortium | Lighthouse Laboratory Plymouth and Alex Alderton, Roberto Amato, Jeffrey Barrett, Sonia Goncalves, Ewan Harrison, David K. Jackson, Ian Johnston, Dominic Kwiatkowski, Cordelia Langford, John Sillitoe on behalf of the Wellcome Sanger Institute COVID-19 Surveillance Team |
| EPI_ISL_2974350, EPI_ISL_2974351 | Lighthouse Lab in Milton Keynes | Wellcome Sanger Institute for the COVID-19 Genomics UK (COG-UK) Consortium | The Lighthouse Lab in Milton Keynes and Alex Alderton, Roberto Amato, Jeffrey Barrett, Sonia Goncalves, Ewan Harrison, David K. Jackson, Ian Johnston, Dominic Kwiatkowski, Cordelia Langford, John Sillitoe on behalf of the Wellcome Sanger Institute COVID-19 Surveillance Team |
| EPI_ISL_2974352, EPI_ISL_2974353, EPI_ISL_2974354, EPI_ISL_2974355 | Lighthouse Laboratory Plymouth | Wellcome Sanger Institute for the COVID-19 Genomics UK (COG-UK) Consortium | Lighthouse Laboratory Plymouth and Alex Alderton, Roberto Amato, Jeffrey Barrett, Sonia Goncalves, Ewan Harrison, David K. Jackson, Ian Johnston, Dominic Kwiatkowski, Cordelia Langford, John Sillitoe on behalf of the Wellcome Sanger Institute COVID-19 Surveillance Team |
| EPI_ISL_2974356, EPI_ISL_2974357 | Lighthouse Lab in Alderley Park | Wellcome Sanger Institute for the COVID-19 Genomics UK (COG-UK) Consortium | Jacquelyn Wynn, Mairead Hyland, The Lighthouse Lab in Alderley Park and Alex Alderton, Roberto Amato, Jeffrey Barrett, Sonia Goncalves, Ewan Harrison, David K. Jackson, Ian Johnston, Dominic Kwiatkowski, Cordelia Langford, John Sillitoe on behalf of the Wellcome Sanger Institute COVID-19 Surveillance Team |
| EPI_ISL_2974358, EPI_ISL_2974359 | Lighthouse Lab in Milton Keynes | Wellcome Sanger Institute for the COVID-19 Genomics UK (COG-UK) Consortium | The Lighthouse Lab in Milton Keynes and Alex Alderton, Roberto Amato, Jeffrey Barrett, Sonia Goncalves, Ewan Harrison, David K. Jackson, Ian Johnston, Dominic Kwiatkowski, Cordelia Langford, John Sillitoe on behalf of the Wellcome Sanger Institute COVID-19 Surveillance Team |
| EPI_ISL_2974360 | Lighthouse Laboratory Plymouth | Wellcome Sanger Institute for the COVID-19 Genomics UK (COG-UK) Consortium | Lighthouse Laboratory Plymouth and Alex Alderton, Roberto Amato, Jeffrey Barrett, Sonia Goncalves, Ewan Harrison, David K. Jackson, Ian Johnston, Dominic Kwiatkowski, Cordelia Langford, John Sillitoe on behalf of the Wellcome Sanger Institute COVID-19 Surveillance Team |
| EPI_ISL_2974361 | Lighthouse Lab in Milton Keynes | Wellcome Sanger Institute for the COVID-19 Genomics UK (COG-UK) Consortium | The Lighthouse Lab in Milton Keynes and Alex Alderton, Roberto Amato, Jeffrey Barrett, Sonia Goncalves, Ewan Harrison, David K. Jackson, Ian Johnston, Dominic Kwiatkowski, Cordelia Langford, John Sillitoe on behalf of the Wellcome Sanger Institute COVID-19 Surveillance Team |
| EPI_ISL_2974362 | Lighthouse Lab in Alderley Park | Wellcome Sanger Institute for the COVID-19 Genomics UK (COG-UK) Consortium | Jacquelyn Wynn, Mairead Hyland, The Lighthouse Lab in Alderley Park and Alex Alderton, Roberto Amato, Jeffrey Barrett, Sonia Goncalves, Ewan Harrison, David K. Jackson, Ian Johnston, Dominic Kwiatkowski, Cordelia Langford, John Sillitoe on behalf of the Wellcome Sanger Institute COVID-19 Surveillance Team |
| EPI_ISL_2974363 | Lighthouse Laboratory Plymouth | Wellcome Sanger Institute for the COVID-19 Genomics UK (COG-UK) Consortium | Lighthouse Laboratory Plymouth and Alex Alderton, Roberto Amato, Jeffrey Barrett, Sonia Goncalves, Ewan Harrison, David K. Jackson, Ian Johnston, Dominic Kwiatkowski, Cordelia Langford, John Sillitoe on behalf of the Wellcome Sanger Institute COVID-19 Surveillance Team |
| EPI_ISL_2974364 | Lighthouse Lab in Alderley Park | Wellcome Sanger Institute for the COVID-19 Genomics UK (COG-UK) Consortium | Jacquelyn Wynn, Mairead Hyland, The Lighthouse Lab in Alderley Park and Alex Alderton, Roberto Amato, Jeffrey Barrett, Sonia Goncalves, Ewan Harrison, David K. Jackson, Ian Johnston, Dominic Kwiatkowski, Cordelia Langford, John Sillitoe on behalf of the Wellcome Sanger Institute COVID-19 Surveillance Team |
| EPI_ISL_2974365 | Lighthouse Lab in Milton Keynes | Wellcome Sanger Institute for the COVID-19 Genomics UK (COG-UK) Consortium | The Lighthouse Lab in Milton Keynes and Alex Alderton, Roberto Amato, Jeffrey Barrett, Sonia Goncalves, Ewan Harrison, David K. Jackson, Ian Johnston, Dominic Kwiatkowski, Cordelia Langford, John Sillitoe on behalf of the Wellcome Sanger Institute COVID-19 Surveillance Team |
| EPI_ISL_2974366, EPI_ISL_2974367 | Lighthouse Lab in Alderley Park | Wellcome Sanger Institute for the COVID-19 Genomics UK (COG-UK) Consortium | Jacquelyn Wynn, Mairead Hyland, The Lighthouse Lab in Alderley Park and Alex Alderton, Roberto Amato, Jeffrey Barrett, Sonia Goncalves, Ewan Harrison, David K. Jackson, Ian Johnston, Dominic Kwiatkowski, Cordelia Langford, John Sillitoe on behalf of the Wellcome Sanger Institute COVID-19 Surveillance Team |
| EPI_ISL_2974368 | Lighthouse Laboratory Plymouth | Wellcome Sanger Institute for the COVID-19 Genomics UK (COG-UK) Consortium | Lighthouse Laboratory Plymouth and Alex Alderton, Roberto Amato, Jeffrey Barrett, Sonia Goncalves, Ewan Harrison, David K. Jackson, Ian Johnston, Dominic Kwiatkowski, Cordelia Langford, John Sillitoe on behalf of the Wellcome Sanger Institute COVID-19 Surveillance Team |
| EPI_ISL_2974369 | Lighthouse Lab in Alderley Park | Wellcome Sanger Institute for the COVID-19 Genomics UK (COG-UK) Consortium | Jacquelyn Wynn, Mairead Hyland, The Lighthouse Lab in Alderley Park and Alex Alderton, Roberto Amato, Jeffrey Barrett, Sonia Goncalves, Ewan Harrison, David K. Jackson, Ian Johnston, Dominic Kwiatkowski, Cordelia Langford, John Sillitoe on behalf of the Wellcome Sanger Institute COVID-19 Surveillance Team |
| EPI_ISL_2974370 | Lighthouse Lab in Milton Keynes | Wellcome Sanger Institute for the COVID-19 Genomics UK (COG-UK) Consortium | The Lighthouse Lab in Milton Keynes and Alex Alderton, Roberto Amato, Jeffrey Barrett, Sonia Goncalves, Ewan Harrison, David K. Jackson, Ian Johnston, Dominic Kwiatkowski, Cordelia Langford, John Sillitoe on behalf of the Wellcome Sanger Institute COVID-19 Surveillance Team |
| EPI_ISL_2974371 | Fimlab Laboratories | Fimlab Laboratories | Minna Paloniemi, Leena Huhti, Sara Lehtinen, Bruno Luukinen, Tapio Seiskari, Mauri Keinänen |
| EPI_ISL_2974372 | Lighthouse Laboratory Plymouth | Wellcome Sanger Institute for the COVID-19 Genomics UK (COG-UK) Consortium | Lighthouse Laboratory Plymouth and Alex Alderton, Roberto Amato, Jeffrey Barrett, Sonia Goncalves, Ewan Harrison, David K. Jackson, Ian Johnston, Dominic Kwiatkowski, Cordelia Langford, John Sillitoe on behalf of the Wellcome Sanger Institute COVID-19 Surveillance Team |
| EPI_ISL_2974373 | Lighthouse Lab in Milton Keynes | Wellcome Sanger Institute for the COVID-19 Genomics UK (COG-UK) Consortium | The Lighthouse Lab in Milton Keynes and Alex Alderton, Roberto Amato, Jeffrey Barrett, Sonia Goncalves, Ewan Harrison, David K. Jackson, Ian Johnston, Dominic Kwiatkowski, Cordelia Langford, John Sillitoe on behalf of the Wellcome Sanger Institute COVID-19 Surveillance Team |
| EPI_ISL_2974374, EPI_ISL_2974375, EPI_ISL_2974376 | Lighthouse Lab in Alderley Park | Wellcome Sanger Institute for the COVID-19 Genomics UK (COG-UK) Consortium | Jacquelyn Wynn, Mairead Hyland, The Lighthouse Lab in Alderley Park and Alex Alderton, Roberto Amato, Jeffrey Barrett, Sonia Goncalves, Ewan Harrison, David K. Jackson, Ian Johnston, Dominic Kwiatkowski, Cordelia Langford, John Sillitoe on behalf of the Wellcome Sanger Institute COVID-19 Surveillance Team |
| EPI_ISL_2974377 | Lighthouse Lab in Milton Keynes | Wellcome Sanger Institute for the COVID-19 Genomics UK (COG-UK) Consortium | The Lighthouse Lab in Milton Keynes and Alex Alderton, Roberto Amato, Jeffrey Barrett, Sonia Goncalves, Ewan Harrison, David K. Jackson, Ian Johnston, Dominic Kwiatkowski, Cordelia Langford, John Sillitoe on behalf of the Wellcome Sanger Institute COVID-19 Surveillance Team |
| EPI_ISL_2974378 | Lighthouse Lab in Alderley Park | Wellcome Sanger Institute for the COVID-19 Genomics UK (COG-UK) Consortium | Jacquelyn Wynn, Mairead Hyland, The Lighthouse Lab in Alderley Park and Alex Alderton, Roberto Amato, Jeffrey Barrett, Sonia Goncalves, Ewan Harrison, David K. Jackson, Ian Johnston, Dominic Kwiatkowski, Cordelia Langford, John Sillitoe on behalf of the Wellcome Sanger Institute COVID-19 Surveillance Team |
| EPI_ISL_2974379, EPI_ISL_2974380 | Lighthouse Lab in Milton Keynes | Wellcome Sanger Institute for the COVID-19 Genomics UK (COG-UK) Consortium | The Lighthouse Lab in Milton Keynes and Alex Alderton, Roberto Amato, Jeffrey Barrett, Sonia Goncalves, Ewan Harrison, David K. Jackson, Ian Johnston, Dominic Kwiatkowski, Cordelia Langford, John Sillitoe on behalf of the Wellcome Sanger Institute COVID-19 Surveillance Team |
| EPI_ISL_2974381, EPI_ISL_2974382, EPI_ISL_2974383, EPI_ISL_2974384, EPI_ISL_2974385, EPI_ISL_2974386, EPI_ISL_2974387, EPI_ISL_2974388, EPI_ISL_2974389, EPI_ISL_2974390, EPI_ISL_2974391, EPI_ISL_2974392, EPI_ISL_2974393, EPI_ISL_2974394 | | | |
| see above | Lighthouse Lab in Glasgow | Wellcome Sanger Institute for the COVID-19 Genomics UK (COG-UK) Consortium | Harper VanSteenhouse, Yumi Kasai, David Gray, Carol Clugston, Anna Dominiczak and Alex Alderton, Roberto Amato, Jeffrey Barrett, Sonia Goncalves, Ewan Harrison, David K. Jackson, Ian Johnston, Dominic Kwiatkowski, Cordelia Langford, John Sillitoe on behalf of the Wellcome Sanger Institute COVID-19 Surveillance Team |
| EPI_ISL_2974395 | Lighthouse Lab in Alderley Park | Wellcome Sanger Institute for the COVID-19 Genomics UK (COG-UK) Consortium | Jacquelyn Wynn, Mairead Hyland, The Lighthouse Lab in Alderley Park and Alex Alderton, Roberto Amato, Jeffrey Barrett, Sonia Goncalves, Ewan Harrison, David K. Jackson, Ian Johnston, Dominic Kwiatkowski, Cordelia Langford, John Sillitoe on behalf of the Wellcome Sanger Institute COVID-19 Surveillance Team |

| | | | |
|---|---|---|---|
| EPI_ISL_2974396, EPI_ISL_2974397, EPI_ISL_2974398, EPI_ISL_2974399, EPI_ISL_2974400, EPI_ISL_2974401, EPI_ISL_2974402, EPI_ISL_2974403, EPI_ISL_2974404, EPI_ISL_2974405, EPI_ISL_2974406, EPI_ISL_2974407, EPI_ISL_2974408, EPI_ISL_2974409, EPI_ISL_2974410 | | | |
| see above | Lighthouse Lab in Glasgow | Wellcome Sanger Institute for the COVID-19 Genomics UK (COG-UK) Consortium | Harper VanSteenhouse, Yumi Kasai, David Gray, Carol Clugston, Anna Dominiczak and Alex Alderton, Roberto Amato, Jeffrey Barrett, Sonia Goncalves, Ewan Harrison, David K. Jackson, Ian Johnston, Dominic Kwiatkowski, Cordelia Langford, John Sillitoe on behalf of the Wellcome Sanger Institute COVID-19 Surveillance Team |
| EPI_ISL_2974411 | Lighthouse Lab in Alderley Park | Wellcome Sanger Institute for the COVID-19 Genomics UK (COG-UK) Consortium | Jacquelyn Wynn, Mairead Hyland, The Lighthouse Lab in Alderley Park and Alex Alderton, Roberto Amato, Jeffrey Barrett, Sonia Goncalves, Ewan Harrison, David K. Jackson, Ian Johnston, Dominic Kwiatkowski, Cordelia Langford, John Sillitoe on behalf of the Wellcome Sanger Institute COVID-19 Surveillance Team |
| EPI_ISL_2974412, EPI_ISL_2974413, EPI_ISL_2974414, EPI_ISL_2974415, EPI_ISL_2974416, EPI_ISL_2974417, EPI_ISL_2974418, EPI_ISL_2974419, EPI_ISL_2974420, EPI_ISL_2974421, EPI_ISL_2974422, EPI_ISL_2974423, EPI_ISL_2974424, EPI_ISL_2974425, EPI_ISL_2974426, EPI_ISL_2974427, EPI_ISL_2974428, EPI_ISL_2974429, EPI_ISL_2974430, EPI_ISL_2974431 | | | |
| see above | Lighthouse Lab in Glasgow | Wellcome Sanger Institute for the COVID-19 Genomics UK (COG-UK) Consortium | Harper VanSteenhouse, Yumi Kasai, David Gray, Carol Clugston, Anna Dominiczak and Alex Alderton, Roberto Amato, Jeffrey Barrett, Sonia Goncalves, Ewan Harrison, David K. Jackson, Ian Johnston, Dominic Kwiatkowski, Cordelia Langford, John Sillitoe on behalf of the Wellcome Sanger Institute COVID-19 Surveillance Team |
| EPI_ISL_2974432 | Lighthouse Lab in Alderley Park | Wellcome Sanger Institute for the COVID-19 Genomics UK (COG-UK) Consortium | Jacquelyn Wynn, Mairead Hyland, The Lighthouse Lab in Alderley Park and Alex Alderton, Roberto Amato, Jeffrey Barrett, Sonia Goncalves, Ewan Harrison, David K. Jackson, Ian Johnston, Dominic Kwiatkowski, Cordelia Langford, John Sillitoe on behalf of the Wellcome Sanger Institute COVID-19 Surveillance Team |
| EPI_ISL_2974433, EPI_ISL_2974434, EPI_ISL_2974435 | Lighthouse Lab in Glasgow | Wellcome Sanger Institute for the COVID-19 Genomics UK (COG-UK) Consortium | Harper VanSteenhouse, Yumi Kasai, David Gray, Carol Clugston, Anna Dominiczak and Alex Alderton, Roberto Amato, Jeffrey Barrett, Sonia Goncalves, Ewan Harrison, David K. Jackson, Ian Johnston, Dominic Kwiatkowski, Cordelia Langford, John Sillitoe on behalf of the Wellcome Sanger Institute COVID-19 Surveillance Team |
| EPI_ISL_2974436, EPI_ISL_2974437, EPI_ISL_2974438, EPI_ISL_2974439, EPI_ISL_2974440, EPI_ISL_2974441, EPI_ISL_2974442, EPI_ISL_2974443, EPI_ISL_2974444, EPI_ISL_2974445, EPI_ISL_2974446, EPI_ISL_2974447 | | | |
| see above | Lighthouse Lab in Alderley Park | Wellcome Sanger Institute for the COVID-19 Genomics UK (COG-UK) Consortium | Jacquelyn Wynn, Mairead Hyland, The Lighthouse Lab in Alderley Park and Alex Alderton, Roberto Amato, Jeffrey Barrett, Sonia Goncalves, Ewan Harrison, David K. Jackson, Ian Johnston, Dominic Kwiatkowski, Cordelia Langford, John Sillitoe on behalf of the Wellcome Sanger Institute COVID-19 Surveillance Team |
| EPI_ISL_2974448 | Berkshire and Surrey Pathology Services Lighthouse Laboratory | Wellcome Sanger Institute for the COVID-19 Genomics UK (COG-UK) Consortium | Berkshire and Surrey Pathology Services Lighthouse Laboratory and Alex Alderton, Roberto Amato, Jeffrey Barrett, Sonia Goncalves, Ewan Harrison, David K. Jackson, Ian Johnston, Dominic Kwiatkowski, Cordelia Langford, John Sillitoe on behalf of the Wellcome Sanger Institute COVID-19 Surveillance Team |
| EPI_ISL_2974449, EPI_ISL_2974450, EPI_ISL_2974451, EPI_ISL_2974452, EPI_ISL_2974453, EPI_ISL_2974454 | Lighthouse Lab in Glasgow | Wellcome Sanger Institute for the COVID-19 Genomics UK (COG-UK) Consortium | Harper VanSteenhouse, Yumi Kasai, David Gray, Carol Clugston, Anna Dominiczak and Alex Alderton, Roberto Amato, Jeffrey Barrett, Sonia Goncalves, Ewan Harrison, David K. Jackson, Ian Johnston, Dominic Kwiatkowski, Cordelia Langford, John Sillitoe on behalf of the Wellcome Sanger Institute COVID-19 Surveillance Team |
| EPI_ISL_2974455 | Lighthouse Lab in Alderley Park | Wellcome Sanger Institute for the COVID-19 Genomics UK (COG-UK) Consortium | Jacquelyn Wynn, Mairead Hyland, The Lighthouse Lab in Alderley Park and Alex Alderton, Roberto Amato, Jeffrey Barrett, Sonia Goncalves, Ewan Harrison, David K. Jackson, Ian Johnston, Dominic Kwiatkowski, Cordelia Langford, John Sillitoe on behalf of the Wellcome Sanger Institute COVID-19 Surveillance Team |
| EPI_ISL_2974456, EPI_ISL_2974457, EPI_ISL_2974458, EPI_ISL_2974459, EPI_ISL_2974460, EPI_ISL_2974461, EPI_ISL_2974462, EPI_ISL_2974463, EPI_ISL_2974464, EPI_ISL_2974465, EPI_ISL_2974466 | | | |
| see above | Lighthouse Lab in Glasgow | Wellcome Sanger Institute for the COVID-19 Genomics UK (COG-UK) Consortium | Harper VanSteenhouse, Yumi Kasai, David Gray, Carol Clugston, Anna Dominiczak and Alex Alderton, Roberto Amato, Jeffrey Barrett, Sonia Goncalves, Ewan Harrison, David K. Jackson, Ian Johnston, Dominic Kwiatkowski, Cordelia Langford, John Sillitoe on behalf of the Wellcome Sanger Institute COVID-19 Surveillance Team |
| EPI_ISL_2974467, EPI_ISL_2974468 | Lighthouse Lab in Alderley Park | Wellcome Sanger Institute for the COVID-19 Genomics UK (COG-UK) Consortium | Jacquelyn Wynn, Mairead Hyland, The Lighthouse Lab in Alderley Park and Alex Alderton, Roberto Amato, Jeffrey Barrett, Sonia Goncalves, Ewan Harrison, David K. Jackson, Ian Johnston, Dominic Kwiatkowski, Cordelia Langford, John Sillitoe on behalf of the Wellcome Sanger Institute COVID-19 Surveillance Team |
| EPI_ISL_2974469, EPI_ISL_2974470, EPI_ISL_2974471, EPI_ISL_2974472, EPI_ISL_2974473, EPI_ISL_2974474 | Lighthouse Lab in Glasgow | Wellcome Sanger Institute for the COVID-19 Genomics UK (COG-UK) Consortium | Harper VanSteenhouse, Yumi Kasai, David Gray, Carol Clugston, Anna Dominiczak and Alex Alderton, Roberto Amato, Jeffrey Barrett, Sonia Goncalves, Ewan Harrison, David K. Jackson, Ian Johnston, Dominic Kwiatkowski, Cordelia Langford, John Sillitoe on behalf of the Wellcome Sanger Institute COVID-19 Surveillance Team |
| EPI_ISL_2974475 | Berkshire and Surrey Pathology Services Lighthouse Laboratory | Wellcome Sanger Institute for the COVID-19 Genomics UK (COG-UK) Consortium | Berkshire and Surrey Pathology Services Lighthouse Laboratory and Alex Alderton, Roberto Amato, Jeffrey Barrett, Sonia Goncalves, Ewan Harrison, David K. Jackson, Ian Johnston, Dominic Kwiatkowski, Cordelia Langford, John Sillitoe on behalf of the Wellcome Sanger Institute COVID-19 Surveillance Team |
| EPI_ISL_2974476, EPI_ISL_2974477, EPI_ISL_2974478, EPI_ISL_2974479, EPI_ISL_2974480, EPI_ISL_2974481, EPI_ISL_2974482 | Lighthouse Lab in Glasgow | Wellcome Sanger Institute for the COVID-19 Genomics UK (COG-UK) Consortium | Harper VanSteenhouse, Yumi Kasai, David Gray, Carol Clugston, Anna Dominiczak and Alex Alderton, Roberto Amato, Jeffrey Barrett, Sonia Goncalves, Ewan Harrison, David K. Jackson, Ian Johnston, Dominic Kwiatkowski, Cordelia Langford, John Sillitoe on behalf of the Wellcome Sanger Institute COVID-19 Surveillance Team |
| EPI_ISL_2974483 | Lighthouse Lab in Alderley Park | Wellcome Sanger Institute for the COVID-19 Genomics UK (COG-UK) Consortium | Jacquelyn Wynn, Mairead Hyland, The Lighthouse Lab in Alderley Park and Alex Alderton, Roberto Amato, Jeffrey Barrett, Sonia Goncalves, Ewan Harrison, David K. Jackson, Ian Johnston, Dominic Kwiatkowski, Cordelia Langford, John Sillitoe on behalf of the Wellcome Sanger Institute COVID-19 Surveillance Team |
| EPI_ISL_2974484 | Lighthouse Lab in Glasgow | Wellcome Sanger Institute for the COVID-19 Genomics UK (COG-UK) Consortium | Harper VanSteenhouse, Yumi Kasai, David Gray, Carol Clugston, Anna Dominiczak and Alex Alderton, Roberto Amato, Jeffrey Barrett, Sonia Goncalves, Ewan Harrison, David K. Jackson, Ian Johnston, Dominic Kwiatkowski, Cordelia Langford, John Sillitoe on behalf of the Wellcome Sanger Institute COVID-19 Surveillance Team |
| EPI_ISL_2974485 | Lighthouse Lab in Alderley Park | Wellcome Sanger Institute for the COVID-19 Genomics UK (COG-UK) Consortium | Jacquelyn Wynn, Mairead Hyland, The Lighthouse Lab in Alderley Park and Alex Alderton, Roberto Amato, Jeffrey Barrett, Sonia Goncalves, Ewan Harrison, David K. Jackson, Ian Johnston, Dominic Kwiatkowski, Cordelia Langford, John Sillitoe on behalf of the Wellcome Sanger Institute COVID-19 Surveillance Team |
| EPI_ISL_2974486 | Lighthouse Lab in Glasgow | Wellcome Sanger Institute for the COVID-19 Genomics UK (COG-UK) Consortium | Harper VanSteenhouse, Yumi Kasai, David Gray, Carol Clugston, Anna Dominiczak and Alex Alderton, Roberto Amato, Jeffrey Barrett, Sonia Goncalves, Ewan Harrison, David K. Jackson, Ian Johnston, Dominic Kwiatkowski, Cordelia Langford, John Sillitoe on behalf of the Wellcome Sanger Institute COVID-19 Surveillance Team |
| EPI_ISL_2974488 | Berkshire and Surrey Pathology Services Lighthouse Laboratory | Wellcome Sanger Institute for the COVID-19 Genomics UK (COG-UK) Consortium | Berkshire and Surrey Pathology Services Lighthouse Laboratory and Alex Alderton, Roberto Amato, Jeffrey Barrett, Sonia Goncalves, Ewan Harrison, David K. Jackson, Ian Johnston, Dominic Kwiatkowski, Cordelia Langford, John Sillitoe on behalf of the Wellcome Sanger Institute COVID-19 Surveillance Team |
| EPI_ISL_2974489 | Lighthouse Lab in Glasgow | Wellcome Sanger Institute for the COVID-19 Genomics UK (COG-UK) Consortium | Harper VanSteenhouse, Yumi Kasai, David Gray, Carol Clugston, Anna Dominiczak and Alex Alderton, Roberto Amato, Jeffrey Barrett, Sonia Goncalves, Ewan Harrison, David K. Jackson, Ian Johnston, Dominic Kwiatkowski, Cordelia Langford, John Sillitoe on behalf of the Wellcome Sanger Institute COVID-19 Surveillance Team |
| EPI_ISL_2974490, EPI_ISL_2974491 | Berkshire and Surrey Pathology Services Lighthouse Laboratory | Wellcome Sanger Institute for the COVID-19 Genomics UK (COG-UK) Consortium | Berkshire and Surrey Pathology Services Lighthouse Laboratory and Alex Alderton, Roberto Amato, Jeffrey Barrett, Sonia Goncalves, Ewan Harrison, David K. Jackson, Ian Johnston, Dominic Kwiatkowski, Cordelia Langford, John Sillitoe on behalf of the Wellcome Sanger Institute COVID-19 Surveillance Team |
| EPI_ISL_2974492, EPI_ISL_2974493, EPI_ISL_2974494, EPI_ISL_2974495, EPI_ISL_2974496, EPI_ISL_2974497, EPI_ISL_2974498, EPI_ISL_2974499, EPI_ISL_2974500, EPI_ISL_2974501, EPI_ISL_2974502, EPI_ISL_2974503, EPI_ISL_2974504, EPI_ISL_2974505, EPI_ISL_2974506, EPI_ISL_2974507, EPI_ISL_2974508, EPI_ISL_2974509 | | | |
| see above | Lighthouse Lab in Glasgow | Wellcome Sanger Institute for the COVID-19 Genomics UK (COG-UK) Consortium | Harper VanSteenhouse, Yumi Kasai, David Gray, Carol Clugston, Anna Dominiczak and Alex Alderton, Roberto Amato, Jeffrey Barrett, Sonia Goncalves, Ewan Harrison, David K. Jackson, Ian Johnston, Dominic Kwiatkowski, Cordelia Langford, John Sillitoe on behalf of the Wellcome Sanger Institute COVID-19 Surveillance Team |
| EPI_ISL_2974510 | Lighthouse Lab in Alderley Park | Wellcome Sanger Institute for the COVID-19 Genomics UK (COG-UK) Consortium | Jacquelyn Wynn, Mairead Hyland, The Lighthouse Lab in Alderley Park and Alex Alderton, Roberto Amato, Jeffrey Barrett, Sonia Goncalves, Ewan Harrison, David K. Jackson, Ian Johnston, Dominic Kwiatkowski, Cordelia Langford, John Sillitoe on behalf of the Wellcome Sanger Institute COVID-19 Surveillance Team |
| EPI_ISL_2974511, EPI_ISL_2974512, EPI_ISL_2974513, EPI_ISL_2974514 | Lighthouse Lab in Glasgow | Wellcome Sanger Institute for the COVID-19 Genomics UK (COG-UK) Consortium | Harper VanSteenhouse, Yumi Kasai, David Gray, Carol Clugston, Anna Dominiczak and Alex Alderton, Roberto Amato, Jeffrey Barrett, Sonia Goncalves, Ewan Harrison, David K. Jackson, Ian Johnston, Dominic Kwiatkowski, Cordelia Langford, John Sillitoe on behalf of the Wellcome Sanger Institute COVID-19 Surveillance Team |
| EPI_ISL_2974515, EPI_ISL_2974516 | Berkshire and Surrey Pathology Services Lighthouse Laboratory | Wellcome Sanger Institute for the COVID-19 Genomics UK (COG-UK) Consortium | Berkshire and Surrey Pathology Services Lighthouse Laboratory and Alex Alderton, Roberto Amato, Jeffrey Barrett, Sonia Goncalves, Ewan Harrison, David K. Jackson, Ian Johnston, Dominic Kwiatkowski, Cordelia Langford, John Sillitoe on behalf of the Wellcome Sanger Institute COVID-19 Surveillance Team |
| EPI_ISL_2974517 | Lighthouse Lab in Alderley Park | Wellcome Sanger Institute for the COVID-19 Genomics UK (COG-UK) Consortium | Jacquelyn Wynn, Mairead Hyland, The Lighthouse Lab in Alderley Park and Alex Alderton, Roberto Amato, Jeffrey Barrett, Sonia Goncalves, Ewan Harrison, David K. Jackson, Ian Johnston, Dominic Kwiatkowski, Cordelia Langford, John Sillitoe on behalf of the Wellcome Sanger Institute COVID-19 Surveillance Team |
| EPI_ISL_2974518, EPI_ISL_2974519, EPI_ISL_2974520 | Berkshire and Surrey Pathology Services Lighthouse Laboratory | Wellcome Sanger Institute for the COVID-19 Genomics UK (COG-UK) Consortium | Berkshire and Surrey Pathology Services Lighthouse Laboratory and Alex Alderton, Roberto Amato, Jeffrey Barrett, Sonia Goncalves, Ewan Harrison, David K. Jackson, Ian Johnston, Dominic Kwiatkowski, Cordelia Langford, John Sillitoe on behalf of the Wellcome Sanger Institute COVID-19 Surveillance Team |
| EPI_ISL_2974521, EPI_ISL_2974522 | Lighthouse Lab in Alderley Park | Wellcome Sanger Institute for the COVID-19 Genomics UK (COG-UK) Consortium | Jacquelyn Wynn, Mairead Hyland, The Lighthouse Lab in Alderley Park and Alex Alderton, Roberto Amato, Jeffrey Barrett, Sonia Goncalves, Ewan Harrison, David K. Jackson, Ian Johnston, Dominic Kwiatkowski, Cordelia Langford, John Sillitoe on behalf of the Wellcome Sanger Institute COVID-19 Surveillance Team |
| EPI_ISL_2974523 | Lighthouse Laboratory Plymouth | Wellcome Sanger Institute for the COVID-19 Genomics UK (COG-UK) Consortium | Lighthouse Laboratory Plymouth and Alex Alderton, Roberto Amato, Jeffrey Barrett, Sonia Goncalves, Ewan Harrison, David K. Jackson, Ian Johnston, Dominic Kwiatkowski, Cordelia Langford, John Sillitoe on behalf of the Wellcome Sanger Institute COVID-19 Surveillance Team |
| EPI_ISL_2974524, EPI_ISL_2974525, EPI_ISL_2974526, EPI_ISL_2974527, EPI_ISL_2974528, EPI_ISL_2974529, EPI_ISL_2974530, EPI_ISL_2974531, EPI_ISL_2974532, EPI_ISL_2974533, EPI_ISL_2974534 | | | |
| see above | Lighthouse Lab in Alderley Park | Wellcome Sanger Institute for the COVID-19 Genomics UK (COG-UK) Consortium | Jacquelyn Wynn, Mairead Hyland, The Lighthouse Lab in Alderley Park and Alex Alderton, Roberto Amato, Jeffrey Barrett, Sonia Goncalves, Ewan Harrison, David K. Jackson, Ian Johnston, Dominic Kwiatkowski, Cordelia Langford, John Sillitoe on behalf of the Wellcome Sanger Institute COVID-19 Surveillance Team |
| EPI_ISL_2974535 | Lighthouse Laboratory Plymouth | Wellcome Sanger Institute for the COVID-19 Genomics UK (COG-UK) Consortium | Lighthouse Laboratory Plymouth and Alex Alderton, Roberto Amato, Jeffrey Barrett, Sonia Goncalves, Ewan Harrison, David K. Jackson, Ian Johnston, Dominic Kwiatkowski, Cordelia Langford, John Sillitoe on behalf of the Wellcome Sanger Institute COVID-19 Surveillance Team |
| EPI_ISL_2974536, EPI_ISL_2974537 | Lighthouse Lab in Alderley Park | Wellcome Sanger Institute for the COVID-19 Genomics UK (COG-UK) Consortium | Jacquelyn Wynn, Mairead Hyland, The Lighthouse Lab in Alderley Park and Alex Alderton, Roberto Amato, Jeffrey Barrett, Sonia Goncalves, Ewan Harrison, David K. Jackson, Ian Johnston, Dominic Kwiatkowski, Cordelia Langford, John Sillitoe on behalf of the Wellcome Sanger Institute COVID-19 Surveillance Team |
| EPI_ISL_2974538 | Lighthouse Laboratory Plymouth | Wellcome Sanger Institute for the COVID-19 Genomics UK (COG-UK) Consortium | Lighthouse Laboratory Plymouth and Alex Alderton, Roberto Amato, Jeffrey Barrett, Sonia Goncalves, Ewan Harrison, David K. Jackson, Ian Johnston, Dominic Kwiatkowski, Cordelia Langford, John Sillitoe on behalf of the Wellcome Sanger Institute COVID-19 Surveillance Team |
| EPI_ISL_2974539 | Health Services Laboratories | Wellcome Sanger Institute for the COVID-19 Genomics UK (COG-UK) Consortium | Health Services Laboratories and Alex Alderton, Roberto Amato, Jeffrey Barrett, Sonia Goncalves, Ewan Harrison, David K. Jackson, Ian Johnston, Dominic Kwiatkowski, Cordelia Langford, John Sillitoe on behalf of the Wellcome Sanger Institute COVID-19 Surveillance Team |
| EPI_ISL_2974540 | Lighthouse Lab in Alderley Park | Wellcome Sanger Institute for the COVID-19 Genomics UK (COG-UK) Consortium | Jacquelyn Wynn, Mairead Hyland, The Lighthouse Lab in Alderley Park and Alex Alderton, Roberto Amato, Jeffrey Barrett, Sonia Goncalves, Ewan Harrison, David K. Jackson, Ian Johnston, Dominic Kwiatkowski, Cordelia Langford, John Sillitoe on |

| | | | |
|---|---|---|---|
| | | Consortium | behalf of the Wellcome Sanger Institute COVID-19 Surveillance Team |
| EPI_ISL_2974541, EPI_ISL_2974542 | Health Services Laboratories | Wellcome Sanger Institute for the COVID-19 Genomics UK (COG-UK) Consortium | Health Services Laboratories and Alex Alderton, Roberto Amato, Jeffrey Barrett, Sonia Goncalves, Ewan Harrison, David K. Jackson, Ian Johnston, Dominic Kwiatkowski, Cordelia Langford, John Sillitoe on behalf of the Wellcome Sanger Institute COVID-19 Surveillance Team |
| EPI_ISL_2974543, EPI_ISL_2974544 | Lighthouse Lab in Alderley Park | Wellcome Sanger Institute for the COVID-19 Genomics UK (COG-UK) Consortium | Jacquelyn Wynn, Mairead Hyland, The Lighthouse Lab in Alderley Park and Alex Alderton, Roberto Amato, Jeffrey Barrett, Sonia Goncalves, Ewan Harrison, David K. Jackson, Ian Johnston, Dominic Kwiatkowski, Cordelia Langford, John Sillitoe on behalf of the Wellcome Sanger Institute COVID-19 Surveillance Team |
| EPI_ISL_2974545 | Health Services Laboratories | Wellcome Sanger Institute for the COVID-19 Genomics UK (COG-UK) Consortium | Health Services Laboratories and Alex Alderton, Roberto Amato, Jeffrey Barrett, Sonia Goncalves, Ewan Harrison, David K. Jackson, Ian Johnston, Dominic Kwiatkowski, Cordelia Langford, John Sillitoe on behalf of the Wellcome Sanger Institute COVID-19 Surveillance Team |
| EPI_ISL_2974546 | Lighthouse Lab in Alderley Park | Wellcome Sanger Institute for the COVID-19 Genomics UK (COG-UK) Consortium | Jacquelyn Wynn, Mairead Hyland, The Lighthouse Lab in Alderley Park and Alex Alderton, Roberto Amato, Jeffrey Barrett, Sonia Goncalves, Ewan Harrison, David K. Jackson, Ian Johnston, Dominic Kwiatkowski, Cordelia Langford, John Sillitoe on behalf of the Wellcome Sanger Institute COVID-19 Surveillance Team |
| EPI_ISL_2974547 | Health Services Laboratories | Wellcome Sanger Institute for the COVID-19 Genomics UK (COG-UK) Consortium | Health Services Laboratories and Alex Alderton, Roberto Amato, Jeffrey Barrett, Sonia Goncalves, Ewan Harrison, David K. Jackson, Ian Johnston, Dominic Kwiatkowski, Cordelia Langford, John Sillitoe on behalf of the Wellcome Sanger Institute COVID-19 Surveillance Team |
| EPI_ISL_2974548 | Lighthouse Lab in Alderley Park | Wellcome Sanger Institute for the COVID-19 Genomics UK (COG-UK) Consortium | Jacquelyn Wynn, Mairead Hyland, The Lighthouse Lab in Alderley Park and Alex Alderton, Roberto Amato, Jeffrey Barrett, Sonia Goncalves, Ewan Harrison, David K. Jackson, Ian Johnston, Dominic Kwiatkowski, Cordelia Langford, John Sillitoe on behalf of the Wellcome Sanger Institute COVID-19 Surveillance Team |
| EPI_ISL_2974549, EPI_ISL_2974550, EPI_ISL_2974551, EPI_ISL_2974552, EPI_ISL_2974553, EPI_ISL_2974554, EPI_ISL_2974555, EPI_ISL_2974556 | Health Services Laboratories | Wellcome Sanger Institute for the COVID-19 Genomics UK (COG-UK) Consortium | Health Services Laboratories and Alex Alderton, Roberto Amato, Jeffrey Barrett, Sonia Goncalves, Ewan Harrison, David K. Jackson, Ian Johnston, Dominic Kwiatkowski, Cordelia Langford, John Sillitoe on behalf of the Wellcome Sanger Institute COVID-19 Surveillance Team |
| EPI_ISL_2974557 | Lighthouse Lab in Alderley Park | Wellcome Sanger Institute for the COVID-19 Genomics UK (COG-UK) Consortium | Jacquelyn Wynn, Mairead Hyland, The Lighthouse Lab in Alderley Park and Alex Alderton, Roberto Amato, Jeffrey Barrett, Sonia Goncalves, Ewan Harrison, David K. Jackson, Ian Johnston, Dominic Kwiatkowski, Cordelia Langford, John Sillitoe on behalf of the Wellcome Sanger Institute COVID-19 Surveillance Team |
| EPI_ISL_2974558 | Health Services Laboratories | Wellcome Sanger Institute for the COVID-19 Genomics UK (COG-UK) Consortium | Health Services Laboratories and Alex Alderton, Roberto Amato, Jeffrey Barrett, Sonia Goncalves, Ewan Harrison, David K. Jackson, Ian Johnston, Dominic Kwiatkowski, Cordelia Langford, John Sillitoe on behalf of the Wellcome Sanger Institute COVID-19 Surveillance Team |
| EPI_ISL_2974559 | Lighthouse Lab in Milton Keynes | Wellcome Sanger Institute for the COVID-19 Genomics UK (COG-UK) Consortium | The Lighthouse Lab in Milton Keynes and Alex Alderton, Roberto Amato, Jeffrey Barrett, Sonia Goncalves, Ewan Harrison, David K. Jackson, Ian Johnston, Dominic Kwiatkowski, Cordelia Langford, John Sillitoe on behalf of the Wellcome Sanger Institute COVID-19 Surveillance Team |
| EPI_ISL_2974560, EPI_ISL_2974561 | Health Services Laboratories | Wellcome Sanger Institute for the COVID-19 Genomics UK (COG-UK) Consortium | Health Services Laboratories and Alex Alderton, Roberto Amato, Jeffrey Barrett, Sonia Goncalves, Ewan Harrison, David K. Jackson, Ian Johnston, Dominic Kwiatkowski, Cordelia Langford, John Sillitoe on behalf of the Wellcome Sanger Institute COVID-19 Surveillance Team |
| EPI_ISL_2974562 | Lighthouse Laboratory Plymouth | Wellcome Sanger Institute for the COVID-19 Genomics UK (COG-UK) Consortium | Lighthouse Laboratory Plymouth and Alex Alderton, Roberto Amato, Jeffrey Barrett, Sonia Goncalves, Ewan Harrison, David K. Jackson, Ian Johnston, Dominic Kwiatkowski, Cordelia Langford, John Sillitoe on behalf of the Wellcome Sanger Institute COVID-19 Surveillance Team |
| EPI_ISL_2974563, EPI_ISL_2974564, EPI_ISL_2974565, EPI_ISL_2974566, EPI_ISL_2974567, EPI_ISL_2974568, EPI_ISL_2974569, EPI_ISL_2974570, EPI_ISL_2974571, EPI_ISL_2974572, EPI_ISL_2974573, EPI_ISL_2974574, EPI_ISL_2974575, EPI_ISL_2974576, EPI_ISL_2974577, EPI_ISL_2974578, EPI_ISL_2974579, EPI_ISL_2974580, EPI_ISL_2974581, EPI_ISL_2974582, EPI_ISL_2974583 | | | |
| see above | Health Services Laboratories | Wellcome Sanger Institute for the COVID-19 Genomics UK (COG-UK) Consortium | Health Services Laboratories and Alex Alderton, Roberto Amato, Jeffrey Barrett, Sonia Goncalves, Ewan Harrison, David K. Jackson, Ian Johnston, Dominic Kwiatkowski, Cordelia Langford, John Sillitoe on behalf of the Wellcome Sanger Institute COVID-19 Surveillance Team |
| EPI_ISL_2974584 | Lighthouse Lab in Milton Keynes | Wellcome Sanger Institute for the COVID-19 Genomics UK (COG-UK) Consortium | The Lighthouse Lab in Milton Keynes and Alex Alderton, Roberto Amato, Jeffrey Barrett, Sonia Goncalves, Ewan Harrison, David K. Jackson, Ian Johnston, Dominic Kwiatkowski, Cordelia Langford, John Sillitoe on behalf of the Wellcome Sanger Institute COVID-19 Surveillance Team |
| EPI_ISL_2974585, EPI_ISL_2974586, EPI_ISL_2974587, EPI_ISL_2974588, EPI_ISL_2974589, EPI_ISL_2974590 | Health Services Laboratories | Wellcome Sanger Institute for the COVID-19 Genomics UK (COG-UK) Consortium | Health Services Laboratories and Alex Alderton, Roberto Amato, Jeffrey Barrett, Sonia Goncalves, Ewan Harrison, David K. Jackson, Ian Johnston, Dominic Kwiatkowski, Cordelia Langford, John Sillitoe on behalf of the Wellcome Sanger Institute COVID-19 Surveillance Team |
| EPI_ISL_2974591 | Lighthouse Lab in Milton Keynes | Wellcome Sanger Institute for the COVID-19 Genomics UK (COG-UK) Consortium | The Lighthouse Lab in Milton Keynes and Alex Alderton, Roberto Amato, Jeffrey Barrett, Sonia Goncalves, Ewan Harrison, David K. Jackson, Ian Johnston, Dominic Kwiatkowski, Cordelia Langford, John Sillitoe on behalf of the Wellcome Sanger Institute COVID-19 Surveillance Team |
| EPI_ISL_2974592 | Health Services Laboratories | Wellcome Sanger Institute for the COVID-19 Genomics UK (COG-UK) Consortium | Health Services Laboratories and Alex Alderton, Roberto Amato, Jeffrey Barrett, Sonia Goncalves, Ewan Harrison, David K. Jackson, Ian Johnston, Dominic Kwiatkowski, Cordelia Langford, John Sillitoe on behalf of the Wellcome Sanger Institute COVID-19 Surveillance Team |
| EPI_ISL_2974593, EPI_ISL_2974594, EPI_ISL_2974595, EPI_ISL_2974596, EPI_ISL_2974597, EPI_ISL_2974598, EPI_ISL_2974599, EPI_ISL_2974600 | Lighthouse Lab in Alderley Park | Wellcome Sanger Institute for the COVID-19 Genomics UK (COG-UK) Consortium | Jacquelyn Wynn, Mairead Hyland, The Lighthouse Lab in Alderley Park and Alex Alderton, Roberto Amato, Jeffrey Barrett, Sonia Goncalves, Ewan Harrison, David K. Jackson, Ian Johnston, Dominic Kwiatkowski, Cordelia Langford, John Sillitoe on behalf of the Wellcome Sanger Institute COVID-19 Surveillance Team |
| EPI_ISL_2974601, EPI_ISL_2974602, EPI_ISL_2974603, EPI_ISL_2974604, EPI_ISL_2974605 | Health Services Laboratories | Wellcome Sanger Institute for the COVID-19 Genomics UK (COG-UK) Consortium | Health Services Laboratories and Alex Alderton, Roberto Amato, Jeffrey Barrett, Sonia Goncalves, Ewan Harrison, David K. Jackson, Ian Johnston, Dominic Kwiatkowski, Cordelia Langford, John Sillitoe on behalf of the Wellcome Sanger Institute COVID-19 Surveillance Team |
| EPI_ISL_2974606 | Lighthouse Lab in Milton Keynes | Wellcome Sanger Institute for the COVID-19 Genomics UK (COG-UK) Consortium | The Lighthouse Lab in Milton Keynes and Alex Alderton, Roberto Amato, Jeffrey Barrett, Sonia Goncalves, Ewan Harrison, David K. Jackson, Ian Johnston, Dominic Kwiatkowski, Cordelia Langford, John Sillitoe on behalf of the Wellcome Sanger Institute COVID-19 Surveillance Team |
| EPI_ISL_2974607, EPI_ISL_2974608, EPI_ISL_2974609, EPI_ISL_2974610, EPI_ISL_2974611 | Health Services Laboratories | Wellcome Sanger Institute for the COVID-19 Genomics UK (COG-UK) Consortium | Health Services Laboratories and Alex Alderton, Roberto Amato, Jeffrey Barrett, Sonia Goncalves, Ewan Harrison, David K. Jackson, Ian Johnston, Dominic Kwiatkowski, Cordelia Langford, John Sillitoe on behalf of the Wellcome Sanger Institute COVID-19 Surveillance Team |
| EPI_ISL_2974612 | Lighthouse Lab in Milton Keynes | Wellcome Sanger Institute for the COVID-19 Genomics UK (COG-UK) Consortium | The Lighthouse Lab in Milton Keynes and Alex Alderton, Roberto Amato, Jeffrey Barrett, Sonia Goncalves, Ewan Harrison, David K. Jackson, Ian Johnston, Dominic Kwiatkowski, Cordelia Langford, John Sillitoe on behalf of the Wellcome Sanger Institute COVID-19 Surveillance Team |
| EPI_ISL_2974613 | Health Services Laboratories | Wellcome Sanger Institute for the COVID-19 Genomics UK (COG-UK) Consortium | Health Services Laboratories and Alex Alderton, Roberto Amato, Jeffrey Barrett, Sonia Goncalves, Ewan Harrison, David K. Jackson, Ian Johnston, Dominic Kwiatkowski, Cordelia Langford, John Sillitoe on behalf of the Wellcome Sanger Institute COVID-19 Surveillance Team |
| EPI_ISL_2974614 | Lighthouse Lab in Milton Keynes | Wellcome Sanger Institute for the COVID-19 Genomics UK (COG-UK) Consortium | The Lighthouse Lab in Milton Keynes and Alex Alderton, Roberto Amato, Jeffrey Barrett, Sonia Goncalves, Ewan Harrison, David K. Jackson, Ian Johnston, Dominic Kwiatkowski, Cordelia Langford, John Sillitoe on behalf of the Wellcome Sanger Institute COVID-19 Surveillance Team |
| EPI_ISL_2974615, EPI_ISL_2974616, EPI_ISL_2974617, EPI_ISL_2974618, EPI_ISL_2974619, EPI_ISL_2974620, EPI_ISL_2974621, EPI_ISL_2974622, EPI_ISL_2974623, EPI_ISL_2974624, EPI_ISL_2974625, EPI_ISL_2974626 | | | |
| see above | Health Services Laboratories | Wellcome Sanger Institute for the COVID-19 Genomics UK (COG-UK) Consortium | Health Services Laboratories and Alex Alderton, Roberto Amato, Jeffrey Barrett, Sonia Goncalves, Ewan Harrison, David K. Jackson, Ian Johnston, Dominic Kwiatkowski, Cordelia Langford, John Sillitoe on behalf of the Wellcome Sanger Institute COVID-19 Surveillance Team |
| EPI_ISL_2974627 | Lighthouse Lab in Milton Keynes | Wellcome Sanger Institute for the COVID-19 Genomics UK (COG-UK) Consortium | The Lighthouse Lab in Milton Keynes and Alex Alderton, Roberto Amato, Jeffrey Barrett, Sonia Goncalves, Ewan Harrison, David K. Jackson, Ian Johnston, Dominic Kwiatkowski, Cordelia Langford, John Sillitoe on behalf of the Wellcome Sanger Institute COVID-19 Surveillance Team |
| EPI_ISL_2974628, EPI_ISL_2974629 | Health Services Laboratories | Wellcome Sanger Institute for the COVID-19 Genomics UK (COG-UK) Consortium | Health Services Laboratories and Alex Alderton, Roberto Amato, Jeffrey Barrett, Sonia Goncalves, Ewan Harrison, David K. Jackson, Ian Johnston, Dominic Kwiatkowski, Cordelia Langford, John Sillitoe on behalf of the Wellcome Sanger Institute COVID-19 Surveillance Team |
| EPI_ISL_2974630 | Lighthouse Lab in Milton Keynes | Wellcome Sanger Institute for the COVID-19 Genomics UK (COG-UK) Consortium | The Lighthouse Lab in Milton Keynes and Alex Alderton, Roberto Amato, Jeffrey Barrett, Sonia Goncalves, Ewan Harrison, David K. Jackson, Ian Johnston, Dominic Kwiatkowski, Cordelia Langford, John Sillitoe on behalf of the Wellcome Sanger Institute COVID-19 Surveillance Team |
| EPI_ISL_2974631, EPI_ISL_2974632, EPI_ISL_2974633, EPI_ISL_2974634, EPI_ISL_2974635, EPI_ISL_2974636 | Health Services Laboratories | Wellcome Sanger Institute for the COVID-19 Genomics UK (COG-UK) Consortium | Health Services Laboratories and Alex Alderton, Roberto Amato, Jeffrey Barrett, Sonia Goncalves, Ewan Harrison, David K. Jackson, Ian Johnston, Dominic Kwiatkowski, Cordelia Langford, John Sillitoe on behalf of the Wellcome Sanger Institute COVID-19 Surveillance Team |
| EPI_ISL_2974637, EPI_ISL_2974638, EPI_ISL_2974639, EPI_ISL_2974640, EPI_ISL_2974641, EPI_ISL_2974642, EPI_ISL_2974643, EPI_ISL_2974644, EPI_ISL_2974645, EPI_ISL_2974646, EPI_ISL_2974647, EPI_ISL_2974648, EPI_ISL_2974649, EPI_ISL_2974650, EPI_ISL_2974651, EPI_ISL_2974652, EPI_ISL_2974653, EPI_ISL_2974654, EPI_ISL_2974655, EPI_ISL_2974656, EPI_ISL_2974657, EPI_ISL_2974658, EPI_ISL_2974659, EPI_ISL_2974660, EPI_ISL_2974661, EPI_ISL_2974662, EPI_ISL_2974663, EPI_ISL_2974664, EPI_ISL_2974665, EPI_ISL_2974666, EPI_ISL_2974667, EPI_ISL_2974668, EPI_ISL_2974669, EPI_ISL_2974670, EPI_ISL_2974671, EPI_ISL_2974672, EPI_ISL_2974673, EPI_ISL_2974674, EPI_ISL_2974675, EPI_ISL_2974676, EPI_ISL_2974677, EPI_ISL_2974678, EPI_ISL_2974679, EPI_ISL_2974680, EPI_ISL_2974681, EPI_ISL_2974682, EPI_ISL_2974683, EPI_ISL_2974684, EPI_ISL_2974685, EPI_ISL_2974686, EPI_ISL_2974687, EPI_ISL_2974688, EPI_ISL_2974689, EPI_ISL_2974690, EPI_ISL_2974691, EPI_ISL_2974692, EPI_ISL_2974693, EPI_ISL_2974694, EPI_ISL_2974695, EPI_ISL_2974696, EPI_ISL_2974697, EPI_ISL_2974698, EPI_ISL_2974699, EPI_ISL_2974700, EPI_ISL_2974701, EPI_ISL_2974702, EPI_ISL_2974703, EPI_ISL_2974704, EPI_ISL_2974705, EPI_ISL_2974706, EPI_ISL_2974707, EPI_ISL_2974708, EPI_ISL_2974709, EPI_ISL_2974710, EPI_ISL_2974711, EPI_ISL_2974712, EPI_ISL_2974713, EPI_ISL_2974714, EPI_ISL_2974715, EPI_ISL_2974716, EPI_ISL_2974717, EPI_ISL_2974718, EPI_ISL_2974719, EPI_ISL_2974720, EPI_ISL_2974721, EPI_ISL_2974722, EPI_ISL_2974723, EPI_ISL_2974724, EPI_ISL_2974725, EPI_ISL_2974726, EPI_ISL_2974727, EPI_ISL_2974728, EPI_ISL_2974729, EPI_ISL_2974730, EPI_ISL_2974731, EPI_ISL_2974732, EPI_ISL_2974733, EPI_ISL_2974734, EPI_ISL_2974735, EPI_ISL_2974736, EPI_ISL_2974737, EPI_ISL_2974738, EPI_ISL_2974739, EPI_ISL_2974740, EPI_ISL_2974741, EPI_ISL_2974742, EPI_ISL_2974743, EPI_ISL_2974744, EPI_ISL_2974745, EPI_ISL_2974746, EPI_ISL_2974747, EPI_ISL_2974748, EPI_ISL_2974749, EPI_ISL_2974750, EPI_ISL_2974751, EPI_ISL_2974752, EPI_ISL_2974753, EPI_ISL_2974754, EPI_ISL_2974755, EPI_ISL_2974756, EPI_ISL_2974757, EPI_ISL_2974758, EPI_ISL_2974759, EPI_ISL_2974760, EPI_ISL_2974761, EPI_ISL_2974762, EPI_ISL_2974763, EPI_ISL_2974764, EPI_ISL_2974765, EPI_ISL_2974766, EPI_ISL_2974767, EPI_ISL_2974768, EPI_ISL_2974769, EPI_ISL_2974770, EPI_ISL_2974771, EPI_ISL_2974772, EPI_ISL_2974773, EPI_ISL_2974774, EPI_ISL_2974775, EPI_ISL_2974776, EPI_ISL_2974777, EPI_ISL_2974778, EPI_ISL_2974779, EPI_ISL_2974780, EPI_ISL_2974781, EPI_ISL_2974782, EPI_ISL_2974783, EPI_ISL_2974784, EPI_ISL_2974785, EPI_ISL_2974786, EPI_ISL_2974787, EPI_ISL_2974788, EPI_ISL_2974789, EPI_ISL_2974790, EPI_ISL_2974791, EPI_ISL_2974792, EPI_ISL_2974793, EPI_ISL_2974794, EPI_ISL_2974795, EPI_ISL_2974796, EPI_ISL_2974797, EPI_ISL_2974798, EPI_ISL_2974799, EPI_ISL_2974800, EPI_ISL_2974801, EPI_ISL_2974802, EPI_ISL_2974803, EPI_ISL_2974804, EPI_ISL_2974805, EPI_ISL_2974807, EPI_ISL_2974808, EPI_ISL_2974809, EPI_ISL_2974810, EPI_ISL_2974811, EPI_ISL_2974812, EPI_ISL_2974813, EPI_ISL_2974814, EPI_ISL_2974815, EPI_ISL_2974816, EPI_ISL_2974817, EPI_ISL_2974818, EPI_ISL_2974819, EPI_ISL_2974820, EPI_ISL_2974821, EPI_ISL_2974822, EPI_ISL_2974823, EPI_ISL_2974824, EPI_ISL_2974825, EPI_ISL_2974826, EPI_ISL_2974827, EPI_ISL_2974828, EPI_ISL_2974829, EPI_ISL_2974830, EPI_ISL_2974831, EPI_ISL_2974832, EPI_ISL_2974833, EPI_ISL_2974834, EPI_ISL_2974835, | | | |

| | | | |
|---|---|---|---|
| EPI_ISL_2974836, EPI_ISL_2974837, EPI_ISL_2974838, EPI_ISL_2974839 | | | |
| see above | Lighthouse Lab in Glasgow | Wellcome Sanger Institute for the COVID-19 Genomics UK (COG-UK) Consortium | Harper VanSteenhouse, Yumi Kasai, David Gray, Carol Clugston, Anna Dominiczak and Alex Alderton, Roberto Amato, Jeffrey Barrett, Sonia Goncalves, Ewan Harrison, David K. Jackson, Ian Johnston, Dominic Kwiatkowski, Cordelia Langford, John Sillitoe on behalf of the Wellcome Sanger Institute COVID-19 Surveillance Team |
| EPI_ISL_2974840 | Lighthouse Lab in Milton Keynes | Wellcome Sanger Institute for the COVID-19 Genomics UK (COG-UK) Consortium | The Lighthouse Lab in Milton Keynes and Alex Alderton, Roberto Amato, Jeffrey Barrett, Sonia Goncalves, Ewan Harrison, David K. Jackson, Ian Johnston, Dominic Kwiatkowski, Cordelia Langford, John Sillitoe on behalf of the Wellcome Sanger Institute COVID-19 Surveillance Team |
| EPI_ISL_2974841, EPI_ISL_2974842, EPI_ISL_2974843, EPI_ISL_2974844, EPI_ISL_2974845, EPI_ISL_2974846, EPI_ISL_2974847, EPI_ISL_2974848, EPI_ISL_2974849, EPI_ISL_2974850, EPI_ISL_2974851, EPI_ISL_2974852, EPI_ISL_2974853, EPI_ISL_2974855, EPI_ISL_2974856, EPI_ISL_2974857, EPI_ISL_2974858, EPI_ISL_2974859, EPI_ISL_2974860, EPI_ISL_2974861, EPI_ISL_2974862, EPI_ISL_2974863, EPI_ISL_2974864, EPI_ISL_2974865, EPI_ISL_2974866, EPI_ISL_2974867, EPI_ISL_2974868, EPI_ISL_2974869, EPI_ISL_2974870, EPI_ISL_2974871, EPI_ISL_2974872, EPI_ISL_2974873, EPI_ISL_2974874, EPI_ISL_2974875, EPI_ISL_2974876, EPI_ISL_2974877, EPI_ISL_2974878, EPI_ISL_2974879, EPI_ISL_2974880, EPI_ISL_2974881, EPI_ISL_2974882, EPI_ISL_2974883, EPI_ISL_2974884, EPI_ISL_2974885, EPI_ISL_2974886, EPI_ISL_2974887, EPI_ISL_2974888, EPI_ISL_2974889, EPI_ISL_2974890, EPI_ISL_2974891, EPI_ISL_2974892, EPI_ISL_2974893, EPI_ISL_2974894, EPI_ISL_2974895, EPI_ISL_2974896, EPI_ISL_2974897, EPI_ISL_2974898, EPI_ISL_2974899, EPI_ISL_2974900, EPI_ISL_2974901, EPI_ISL_2974902, EPI_ISL_2974903, EPI_ISL_2974904, EPI_ISL_2974905, EPI_ISL_2974906, EPI_ISL_2974907, EPI_ISL_2974908, EPI_ISL_2974909, EPI_ISL_2974910, EPI_ISL_2974911, EPI_ISL_2974912, EPI_ISL_2974913, EPI_ISL_2974914, EPI_ISL_2974915, EPI_ISL_2974916, EPI_ISL_2974917, EPI_ISL_2974918 | | | |
| see above | Lighthouse Lab in Glasgow | Wellcome Sanger Institute for the COVID-19 Genomics UK (COG-UK) Consortium | Harper VanSteenhouse, Yumi Kasai, David Gray, Carol Clugston, Anna Dominiczak and Alex Alderton, Roberto Amato, Jeffrey Barrett, Sonia Goncalves, Ewan Harrison, David K. Jackson, Ian Johnston, Dominic Kwiatkowski, Cordelia Langford, John Sillitoe on behalf of the Wellcome Sanger Institute COVID-19 Surveillance Team |
| EPI_ISL_2974919, EPI_ISL_2974920, EPI_ISL_2974921, EPI_ISL_2974922, EPI_ISL_2974923, EPI_ISL_2974924, EPI_ISL_2974925, EPI_ISL_2974926, EPI_ISL_2974927, EPI_ISL_2974928, EPI_ISL_2974929, EPI_ISL_2974930, EPI_ISL_2974931, EPI_ISL_2974932, EPI_ISL_2974933, EPI_ISL_2974934, EPI_ISL_2974935, EPI_ISL_2974936, EPI_ISL_2974937, EPI_ISL_2974938, EPI_ISL_2974939, EPI_ISL_2974940, EPI_ISL_2974941, EPI_ISL_2974942, EPI_ISL_2974943, EPI_ISL_2974945, EPI_ISL_2974946, EPI_ISL_2974948, EPI_ISL_2974950, EPI_ISL_2974951, EPI_ISL_2974952, EPI_ISL_2974953, EPI_ISL_2974954, EPI_ISL_2974955, EPI_ISL_2974956, EPI_ISL_2974957, EPI_ISL_2974958, EPI_ISL_2974959, EPI_ISL_2974960, EPI_ISL_2974961, EPI_ISL_2974962, EPI_ISL_2974963, EPI_ISL_2974964, EPI_ISL_2974965, EPI_ISL_2974966, EPI_ISL_2974967, EPI_ISL_2974969, EPI_ISL_2974970, EPI_ISL_2974971, EPI_ISL_2974972, EPI_ISL_2974973, EPI_ISL_2974975, EPI_ISL_2974976, EPI_ISL_2974977, EPI_ISL_2974978, EPI_ISL_2974979, EPI_ISL_2974981, EPI_ISL_2974983, EPI_ISL_2974984, EPI_ISL_2974985, EPI_ISL_2974986, EPI_ISL_2974987, EPI_ISL_2974988, EPI_ISL_2974990, EPI_ISL_2974991, EPI_ISL_2974992, EPI_ISL_2974993, EPI_ISL_2974994, EPI_ISL_2974995, EPI_ISL_2974996, EPI_ISL_2974997, EPI_ISL_2974998, EPI_ISL_2974999, EPI_ISL_2975000, EPI_ISL_2975001, EPI_ISL_2975002, EPI_ISL_2975003, EPI_ISL_2975004, EPI_ISL_2975005, EPI_ISL_2975006, EPI_ISL_2975007, EPI_ISL_2975008, EPI_ISL_2975009 | | | |
| see above | Randox Laboratories | Wellcome Sanger Institute for the COVID-19 Genomics UK (COG-UK) Consortium | Randox Laboratories and Alex Alderton, Roberto Amato, Jeffrey Barrett, Sonia Goncalves, Ewan Harrison, David K. Jackson, Ian Johnston, Dominic Kwiatkowski, Cordelia Langford, John Sillitoe on behalf of the Wellcome Sanger Institute COVID-19 Surveillance Team |
| EPI_ISL_2975010, EPI_ISL_2975011, EPI_ISL_2975012, EPI_ISL_2975013, EPI_ISL_2975014, EPI_ISL_2975015, EPI_ISL_2975016, EPI_ISL_2975017, EPI_ISL_2975018, EPI_ISL_2975019, EPI_ISL_2975020, EPI_ISL_2975021, EPI_ISL_2975022, EPI_ISL_2975023, EPI_ISL_2975024, EPI_ISL_2975025, EPI_ISL_2975026, EPI_ISL_2975027, EPI_ISL_2975028, EPI_ISL_2975029, EPI_ISL_2975030, EPI_ISL_2975031, EPI_ISL_2975032, EPI_ISL_2975033, EPI_ISL_2975034, EPI_ISL_2975035, EPI_ISL_2975036, EPI_ISL_2975037, EPI_ISL_2975038, EPI_ISL_2975039, EPI_ISL_2975040, EPI_ISL_2975041, EPI_ISL_2975042, EPI_ISL_2975043, EPI_ISL_2975044, EPI_ISL_2975045, EPI_ISL_2975046 | | | |
| see above | Lighthouse Lab in Glasgow | Wellcome Sanger Institute for the COVID-19 Genomics UK (COG-UK) Consortium | Harper VanSteenhouse, Yumi Kasai, David Gray, Carol Clugston, Anna Dominiczak and Alex Alderton, Roberto Amato, Jeffrey Barrett, Sonia Goncalves, Ewan Harrison, David K. Jackson, Ian Johnston, Dominic Kwiatkowski, Cordelia Langford, John Sillitoe on behalf of the Wellcome Sanger Institute COVID-19 Surveillance Team |
| EPI_ISL_2975047 | Lighthouse Lab in Alderley Park | Wellcome Sanger Institute for the COVID-19 Genomics UK (COG-UK) Consortium | Jacquelyn Wynn, Mairead Hyland, The Lighthouse Lab in Alderley Park and Alex Alderton, Roberto Amato, Jeffrey Barrett, Sonia Goncalves, Ewan Harrison, David K. Jackson, Ian Johnston, Dominic Kwiatkowski, Cordelia Langford, John Sillitoe on behalf of the Wellcome Sanger Institute COVID-19 Surveillance Team |
| EPI_ISL_2975048, EPI_ISL_2975049, EPI_ISL_2975050, EPI_ISL_2975051, EPI_ISL_2975052, EPI_ISL_2975053, EPI_ISL_2975054, EPI_ISL_2975055, EPI_ISL_2975056, EPI_ISL_2975057, EPI_ISL_2975058, EPI_ISL_2975059, EPI_ISL_2975060, EPI_ISL_2975061, EPI_ISL_2975062, EPI_ISL_2975063 | | | |
| see above | Lighthouse Lab in Glasgow | Wellcome Sanger Institute for the COVID-19 Genomics UK (COG-UK) Consortium | Harper VanSteenhouse, Yumi Kasai, David Gray, Carol Clugston, Anna Dominiczak and Alex Alderton, Roberto Amato, Jeffrey Barrett, Sonia Goncalves, Ewan Harrison, David K. Jackson, Ian Johnston, Dominic Kwiatkowski, Cordelia Langford, John Sillitoe on behalf of the Wellcome Sanger Institute COVID-19 Surveillance Team |
| EPI_ISL_2975064 | Berkshire and Surrey Pathology Services Lighthouse Laboratory | Wellcome Sanger Institute for the COVID-19 Genomics UK (COG-UK) Consortium | Berkshire and Surrey Pathology Services Lighthouse Laboratory and Alex Alderton, Roberto Amato, Jeffrey Barrett, Sonia Goncalves, Ewan Harrison, David K. Jackson, Ian Johnston, Dominic Kwiatkowski, Cordelia Langford, John Sillitoe on behalf of the Wellcome Sanger Institute COVID-19 Surveillance Team |
| EPI_ISL_2975066, EPI_ISL_2975067, EPI_ISL_2975068, EPI_ISL_2975069, EPI_ISL_2975070, EPI_ISL_2975071, EPI_ISL_2975072, EPI_ISL_2975073, EPI_ISL_2975074, EPI_ISL_2975075, EPI_ISL_2975077, EPI_ISL_2975079, EPI_ISL_2975080, EPI_ISL_2975081, EPI_ISL_2975082, EPI_ISL_2975084, EPI_ISL_2975085, EPI_ISL_2975089, EPI_ISL_2975090, EPI_ISL_2975091, EPI_ISL_2975092, EPI_ISL_2975094, EPI_ISL_2975095, EPI_ISL_2975096, EPI_ISL_2975097, EPI_ISL_2975098, EPI_ISL_2975099, EPI_ISL_2975100, EPI_ISL_2975101, EPI_ISL_2975102, EPI_ISL_2975103, EPI_ISL_2975104, EPI_ISL_2975105, EPI_ISL_2975107, EPI_ISL_2975108, EPI_ISL_2975109, EPI_ISL_2975110, EPI_ISL_2975111, EPI_ISL_2975113, EPI_ISL_2975114, EPI_ISL_2975116, EPI_ISL_2975117, EPI_ISL_2975118, EPI_ISL_2975119, EPI_ISL_2975121, EPI_ISL_2975122, EPI_ISL_2975123, EPI_ISL_2975125, EPI_ISL_2975126 | | | |
| see above | Randox Laboratories | Wellcome Sanger Institute for the COVID-19 Genomics UK (COG-UK) Consortium | Randox Laboratories and Alex Alderton, Roberto Amato, Jeffrey Barrett, Sonia Goncalves, Ewan Harrison, David K. Jackson, Ian Johnston, Dominic Kwiatkowski, Cordelia Langford, John Sillitoe on behalf of the Wellcome Sanger Institute COVID-19 Surveillance Team |
| EPI_ISL_2975127 | Lighthouse Lab in Alderley Park | Wellcome Sanger Institute for the COVID-19 Genomics UK (COG-UK) Consortium | Jacquelyn Wynn, Mairead Hyland, The Lighthouse Lab in Alderley Park and Alex Alderton, Roberto Amato, Jeffrey Barrett, Sonia Goncalves, Ewan Harrison, David K. Jackson, Ian Johnston, Dominic Kwiatkowski, Cordelia Langford, John Sillitoe on behalf of the Wellcome Sanger Institute COVID-19 Surveillance Team |
| EPI_ISL_2975128, EPI_ISL_2975131, EPI_ISL_2975132, EPI_ISL_2975133, EPI_ISL_2975134 | Fimlab Laboratories | Fimlab Laboratories | Minna Paloniemi, Leena Huhti, Sara Lehtinen, Bruno Luukinen, Tapio Seiskari, Mauri Keinänen |
| EPI_ISL_2975141, EPI_ISL_2975143, EPI_ISL_2975144, EPI_ISL_2975145, EPI_ISL_2975146, EPI_ISL_2975148, EPI_ISL_2975150 | AULSS 2 Marca Trevigiana | Istituto Zooprofilattico Sperimentale delle Venezie | Adelaide Milani, Alessia Schivo, Annalisa Salviato, Elisa Palumbo, Erika Giorgia Quaranta, Luca Tassoni, Ambra Pastori, Edoardo Giussani, Alice Fusaro, Isabella Monne, Calogero Terregino, Antonia Ricci |
| EPI_ISL_2975153 | AULSS 8 Berica | Istituto Zooprofilattico Sperimentale delle Venezie | Adelaide Milani, Alessia Schivo, Annalisa Salviato, Elisa Palumbo, Erika Giorgia Quaranta, Luca Tassoni, Ambra Pastori, Edoardo Giussani, Alice Fusaro, Isabella Monne, Calogero Terregino, Antonia Ricci |
| EPI_ISL_2975159 | AULSS 2 Marca Trevigiana | Istituto Zooprofilattico Sperimentale delle Venezie | Adelaide Milani, Alessia Schivo, Annalisa Salviato, Elisa Palumbo, Erika Giorgia Quaranta, Luca Tassoni, Ambra Pastori, Edoardo Giussani, Alice Fusaro, Isabella Monne, Calogero Terregino, Antonia Ricci |
| EPI_ISL_2975160 | AULSS 6 Euganea | Istituto Zooprofilattico Sperimentale delle Venezie | Adelaide Milani, Alessia Schivo, Annalisa Salviato, Elisa Palumbo, Erika Giorgia Quaranta, Luca Tassoni, Ambra Pastori, Edoardo Giussani, Alice Fusaro, Isabella Monne, Calogero Terregino, Antonia Ricci |
| EPI_ISL_2975162, EPI_ISL_2975163, EPI_ISL_2975165, EPI_ISL_2975168 | AULSS 2 Marca Trevigiana | Istituto Zooprofilattico Sperimentale delle Venezie | Adelaide Milani, Alessia Schivo, Annalisa Salviato, Elisa Palumbo, Erika Giorgia Quaranta, Luca Tassoni, Ambra Pastori, Edoardo Giussani, Alice Fusaro, Isabella Monne, Calogero Terregino, Antonia Ricci |
| EPI_ISL_2975189 | AULSS 1 Dolomiti | Istituto Zooprofilattico Sperimentale delle Venezie | Adelaide Milani, Alessia Schivo, Annalisa Salviato, Elisa Palumbo, Erika Giorgia Quaranta, Luca Tassoni, Ambra Pastori, Edoardo Giussani, Alice Fusaro, Isabella Monne, Calogero Terregino, Antonia Ricci |
| EPI_ISL_2975192, EPI_ISL_2975194, EPI_ISL_2975195, EPI_ISL_2975197, EPI_ISL_2975199, EPI_ISL_2975202, EPI_ISL_2975204, EPI_ISL_2975206 | Laboratorio di Riferimento Regionale della Sicilia Occidentale per l'Emergenza COVID-19 | Laboratorio di Riferimento Regionale della Sicilia Occidentale per l’Emergenza COVID-19 | Fabio Tramuto, Carmelo Massimo Maida, Daniela Di Naro, Giulia Randazzo, Walter Mazzucco, Giorgio Graziano, Vincenzo Restivo, Claudio Costantino, Francesco Vitale |
| EPI_ISL_2975207, EPI_ISL_2975209 | AULSS 8 Berica | Istituto Zooprofilattico Sperimentale delle Venezie | Adelaide Milani, Alessia Schivo, Annalisa Salviato, Elisa Palumbo, Erika Giorgia Quaranta, Luca Tassoni, Ambra Pastori, Edoardo Giussani, Alice Fusaro, Isabella Monne, Calogero Terregino, Antonia Ricci |
| EPI_ISL_2975210 | Laboratorio di Riferimento Regionale della Sicilia Occidentale per l'Emergenza COVID-19 | Laboratorio di Riferimento Regionale della Sicilia Occidentale per l’Emergenza COVID-19 | Fabio Tramuto, Carmelo Massimo Maida, Daniela Di Naro, Giulia Randazzo, Walter Mazzucco, Giorgio Graziano, Vincenzo Restivo, Claudio Costantino, Francesco Vitale |
| EPI_ISL_2975211 | AULSS 8 Berica | Istituto Zooprofilattico Sperimentale delle Venezie | Adelaide Milani, Alessia Schivo, Annalisa Salviato, Elisa Palumbo, Erika Giorgia Quaranta, Luca Tassoni, Ambra Pastori, Edoardo Giussani, Alice Fusaro, Isabella Monne, Calogero Terregino, Antonia Ricci |
| EPI_ISL_2975212 | Laboratorio di Riferimento Regionale della Sicilia Occidentale per l'Emergenza COVID-19 | Laboratorio di Riferimento Regionale della Sicilia Occidentale per l’Emergenza COVID-19 | Fabio Tramuto, Carmelo Massimo Maida, Daniela Di Naro, Giulia Randazzo, Walter Mazzucco, Giorgio Graziano, Vincenzo Restivo, Claudio Costantino, Francesco Vitale |
| EPI_ISL_2975213 | AULSS 8 Berica | Istituto Zooprofilattico Sperimentale delle Venezie | Adelaide Milani, Alessia Schivo, Annalisa Salviato, Elisa Palumbo, Erika Giorgia Quaranta, Luca Tassoni, Ambra Pastori, Edoardo Giussani, Alice Fusaro, Isabella Monne, Calogero Terregino, Antonia Ricci |
| EPI_ISL_2975215, EPI_ISL_2975219, EPI_ISL_2975221 | AULSS 6 Euganea | Istituto Zooprofilattico Sperimentale delle Venezie | Adelaide Milani, Alessia Schivo, Annalisa Salviato, Elisa Palumbo, Erika Giorgia Quaranta, Luca Tassoni, Ambra Pastori, Edoardo Giussani, Alice Fusaro, Isabella Monne, Calogero Terregino, Antonia Ricci |
| EPI_ISL_2975223, EPI_ISL_2975231, EPI_ISL_2975233, EPI_ISL_2975240, EPI_ISL_2975242 | Laboratorio di Riferimento Regionale della Sicilia Occidentale per l'Emergenza COVID-19 | Laboratorio di Riferimento Regionale della Sicilia Occidentale per l’Emergenza COVID-19 | Fabio Tramuto, Carmelo Massimo Maida, Daniela Di Naro, Giulia Randazzo, Walter Mazzucco, Giorgio Graziano, Vincenzo Restivo, Claudio Costantino, Francesco Vitale |
| EPI_ISL_2975247, EPI_ISL_2975249, EPI_ISL_2975250 | AULSS 5 Polesana | Istituto Zooprofilattico Sperimentale delle Venezie | Adelaide Milani, Alessia Schivo, Annalisa Salviato, Elisa Palumbo, Erika Giorgia Quaranta, Luca Tassoni, Ambra Pastori, Edoardo Giussani, Alice Fusaro, Isabella Monne, Calogero Terregino, Antonia Ricci |
| EPI_ISL_2975257, EPI_ISL_2975258, EPI_ISL_2975259, EPI_ISL_2975260 | AULSS 9 Scaligera | Istituto Zooprofilattico Sperimentale delle Venezie | Adelaide Milani, Alessia Schivo, Annalisa Salviato, Elisa Palumbo, Erika Giorgia Quaranta, Luca Tassoni, Ambra Pastori, Edoardo Giussani, Alice Fusaro, Isabella Monne, Calogero Terregino, Antonia Ricci |
| EPI_ISL_2975262, EPI_ISL_2975263 | AULSS 1 Dolomiti | Istituto Zooprofilattico Sperimentale delle Venezie | Adelaide Milani, Alessia Schivo, Annalisa Salviato, Elisa Palumbo, Erika Giorgia Quaranta, Luca Tassoni, Ambra Pastori, Edoardo Giussani, Alice Fusaro, Isabella Monne, Calogero Terregino, Antonia Ricci |
| EPI_ISL_2975265 | AULSS 7 Pedemontana | Istituto Zooprofilattico Sperimentale delle Venezie | Adelaide Milani, Alessia Schivo, Annalisa Salviato, Elisa Palumbo, Erika Giorgia Quaranta, Luca Tassoni, Ambra Pastori, Edoardo Giussani, Alice Fusaro, Isabella Monne, Calogero Terregino, Antonia Ricci |
| EPI_ISL_2975270 | Fimlab Laboratories | Fimlab Laboratories | Minna Paloniemi, Leena Huhti, Sara Lehtinen, Bruno Luukinen, Tapio Seiskari, Mauri Keinänen |
| EPI_ISL_2975272 | Usansolo-Galdakao University Hospital | Cruces University Hospital | Ana Gual-de-Torrella, Izaskun Alejo-Cancho, Mikel Gallego, Ana Belén de la Hoz |
| EPI_ISL_2975273 | Fimlab Laboratories | Fimlab Laboratories | Minna Paloniemi, Leena Huhti, Sara Lehtinen, Bruno Luukinen, Tapio Seiskari, Mauri Keinänen |
| EPI_ISL_2975274 | Usansolo-Galdakao University Hospital | Cruces University Hospital | Ana Gual-de-Torrella, Izaskun Alejo-Cancho, Mikel Gallego, Ana Belén de la Hoz |

| | | | |
|---|---|---|---|
| EPI_ISL_2975292, EPI_ISL_2975293, EPI_ISL_2975294, EPI_ISL_2975295, EPI_ISL_2975296 | AULSS 9 Scaligera | Istituto Zooprofilattico Sperimentale delle Venezie | Adelaide Milani, Alessia Schivo, Annalisa Salviato, Elisa Palumbo, Erika Giorgia Quaranta, Luca Tassoni, Ambra Pastori, Edoardo Giussani, Alice Fusaro, Isabella Monne, Calogero Terregino, Antonia Ricci |
| EPI_ISL_2975297, EPI_ISL_2975298, EPI_ISL_2975299, EPI_ISL_2975300 | AULSS 6 Euganea | Istituto Zooprofilattico Sperimentale delle Venezie | Adelaide Milani, Alessia Schivo, Annalisa Salviato, Elisa Palumbo, Erika Giorgia Quaranta, Luca Tassoni, Ambra Pastori, Edoardo Giussani, Alice Fusaro, Isabella Monne, Calogero Terregino, Antonia Ricci |
| EPI_ISL_2975301, EPI_ISL_2975302 | AULSS 9 Scaligera | Istituto Zooprofilattico Sperimentale delle Venezie | Adelaide Milani, Alessia Schivo, Annalisa Salviato, Elisa Palumbo, Erika Giorgia Quaranta, Luca Tassoni, Ambra Pastori, Edoardo Giussani, Alice Fusaro, Isabella Monne, Calogero Terregino, Antonia Ricci |
| EPI_ISL_2975303 | AULSS 6 Euganea | Istituto Zooprofilattico Sperimentale delle Venezie | Adelaide Milani, Alessia Schivo, Annalisa Salviato, Elisa Palumbo, Erika Giorgia Quaranta, Luca Tassoni, Ambra Pastori, Edoardo Giussani, Alice Fusaro, Isabella Monne, Calogero Terregino, Antonia Ricci |
| EPI_ISL_2975304, EPI_ISL_2975305, EPI_ISL_2975306, EPI_ISL_2975307, EPI_ISL_2975308 | AULSS 9 Scaligera | Istituto Zooprofilattico Sperimentale delle Venezie | Adelaide Milani, Alessia Schivo, Annalisa Salviato, Elisa Palumbo, Erika Giorgia Quaranta, Luca Tassoni, Ambra Pastori, Edoardo Giussani, Alice Fusaro, Isabella Monne, Calogero Terregino, Antonia Ricci |
| EPI_ISL_2975309 | AULSS 6 Euganea | Istituto Zooprofilattico Sperimentale delle Venezie | Adelaide Milani, Alessia Schivo, Annalisa Salviato, Elisa Palumbo, Erika Giorgia Quaranta, Luca Tassoni, Ambra Pastori, Edoardo Giussani, Alice Fusaro, Isabella Monne, Calogero Terregino, Antonia Ricci |
| EPI_ISL_2975310 | AULSS 9 Scaligera | Istituto Zooprofilattico Sperimentale delle Venezie | Adelaide Milani, Alessia Schivo, Annalisa Salviato, Elisa Palumbo, Erika Giorgia Quaranta, Luca Tassoni, Ambra Pastori, Edoardo Giussani, Alice Fusaro, Isabella Monne, Calogero Terregino, Antonia Ricci |
| EPI_ISL_2975311 | AULSS 8 Berica | Istituto Zooprofilattico Sperimentale delle Venezie | Adelaide Milani, Alessia Schivo, Annalisa Salviato, Elisa Palumbo, Erika Giorgia Quaranta, Luca Tassoni, Ambra Pastori, Edoardo Giussani, Alice Fusaro, Isabella Monne, Calogero Terregino, Antonia Ricci |
| EPI_ISL_2975312, EPI_ISL_2975313, EPI_ISL_2975314, EPI_ISL_2975315, EPI_ISL_2975316, EPI_ISL_2975317, EPI_ISL_2975318, EPI_ISL_2975319, EPI_ISL_2975320, EPI_ISL_2975321 | AULSS 9 Scaligera | Istituto Zooprofilattico Sperimentale delle Venezie | Adelaide Milani, Alessia Schivo, Annalisa Salviato, Elisa Palumbo, Erika Giorgia Quaranta, Luca Tassoni, Ambra Pastori, Edoardo Giussani, Alice Fusaro, Isabella Monne, Calogero Terregino, Antonia Ricci |
| EPI_ISL_2975322 | AULSS 8 Berica | Istituto Zooprofilattico Sperimentale delle Venezie | Adelaide Milani, Alessia Schivo, Annalisa Salviato, Elisa Palumbo, Erika Giorgia Quaranta, Luca Tassoni, Ambra Pastori, Edoardo Giussani, Alice Fusaro, Isabella Monne, Calogero Terregino, Antonia Ricci |
| EPI_ISL_2975323, EPI_ISL_2975324 | AULSS 9 Scaligera | Istituto Zooprofilattico Sperimentale delle Venezie | Adelaide Milani, Alessia Schivo, Annalisa Salviato, Elisa Palumbo, Erika Giorgia Quaranta, Luca Tassoni, Ambra Pastori, Edoardo Giussani, Alice Fusaro, Isabella Monne, Calogero Terregino, Antonia Ricci |
| EPI_ISL_2975325 | AULSS 8 Berica | Istituto Zooprofilattico Sperimentale delle Venezie | Adelaide Milani, Alessia Schivo, Annalisa Salviato, Elisa Palumbo, Erika Giorgia Quaranta, Luca Tassoni, Ambra Pastori, Edoardo Giussani, Alice Fusaro, Isabella Monne, Calogero Terregino, Antonia Ricci |
| EPI_ISL_2975326, EPI_ISL_2975327, EPI_ISL_2975328, EPI_ISL_2975329, EPI_ISL_2975331, EPI_ISL_2975333, EPI_ISL_2975334, EPI_ISL_2975335, EPI_ISL_2975336, EPI_ISL_2975337, EPI_ISL_2975338, EPI_ISL_2975339 | | | |
| see above | AULSS 9 Scaligera | Istituto Zooprofilattico Sperimentale delle Venezie | Adelaide Milani, Alessia Schivo, Annalisa Salviato, Elisa Palumbo, Erika Giorgia Quaranta, Luca Tassoni, Ambra Pastori, Edoardo Giussani, Alice Fusaro, Isabella Monne, Calogero Terregino, Antonia Ricci |
| EPI_ISL_2975340, EPI_ISL_2975341 | AULSS 6 Euganea | Istituto Zooprofilattico Sperimentale delle Venezie | Adelaide Milani, Alessia Schivo, Annalisa Salviato, Elisa Palumbo, Erika Giorgia Quaranta, Luca Tassoni, Ambra Pastori, Edoardo Giussani, Alice Fusaro, Isabella Monne, Calogero Terregino, Antonia Ricci |
| EPI_ISL_2975342, EPI_ISL_2975343 | AULSS 8 Berica | Istituto Zooprofilattico Sperimentale delle Venezie | Adelaide Milani, Alessia Schivo, Annalisa Salviato, Elisa Palumbo, Erika Giorgia Quaranta, Luca Tassoni, Ambra Pastori, Edoardo Giussani, Alice Fusaro, Isabella Monne, Calogero Terregino, Antonia Ricci |
| EPI_ISL_2975344, EPI_ISL_2975345, EPI_ISL_2975346, EPI_ISL_2975347, EPI_ISL_2975348, EPI_ISL_2975349 | AULSS 6 Euganea | Istituto Zooprofilattico Sperimentale delle Venezie | Adelaide Milani, Alessia Schivo, Annalisa Salviato, Elisa Palumbo, Erika Giorgia Quaranta, Luca Tassoni, Ambra Pastori, Edoardo Giussani, Alice Fusaro, Isabella Monne, Calogero Terregino, Antonia Ricci |
| EPI_ISL_2976148, EPI_ISL_2976151, EPI_ISL_2976152, EPI_ISL_2976153, EPI_ISL_2976154, EPI_ISL_2976155, EPI_ISL_2976156, EPI_ISL_2976157, EPI_ISL_2976158, EPI_ISL_2976159, EPI_ISL_2976160, EPI_ISL_2976161, EPI_ISL_2976162, EPI_ISL_2976164, EPI_ISL_2976165, EPI_ISL_2976166, EPI_ISL_2976167, EPI_ISL_2976168, EPI_ISL_2976169, EPI_ISL_2976170, EPI_ISL_2976171, EPI_ISL_2976172, EPI_ISL_2976173, EPI_ISL_2976174, EPI_ISL_2976175, EPI_ISL_2976176, EPI_ISL_2976177, EPI_ISL_2976178, EPI_ISL_2976179, EPI_ISL_2976180, EPI_ISL_2976181, EPI_ISL_2976182, EPI_ISL_2976183, EPI_ISL_2976186, EPI_ISL_2976187, EPI_ISL_2976188 | | | |
| see above | Southern Nevada Public Health Laboratory | Southern Nevada Public Health Laboratory | Michael Picker |
| EPI_ISL_2976945, EPI_ISL_2977058, EPI_ISL_2977128, EPI_ISL_2977221, EPI_ISL_2977222, EPI_ISL_2977227, EPI_ISL_2977258, EPI_ISL_2977351, EPI_ISL_2977366, EPI_ISL_2977370, EPI_ISL_2977376, EPI_ISL_2977381, EPI_ISL_2977388, EPI_ISL_2977421, EPI_ISL_2977438, EPI_ISL_2977439, EPI_ISL_2977443, EPI_ISL_2977469, EPI_ISL_2978159, EPI_ISL_2978163, EPI_ISL_2978165 | | | |
| see above | CDPH VBL | California Department of Public Health | CDPH-COVIDNet,UCSD EXCITE lab |
| EPI_ISL_2978177, EPI_ISL_2978178, EPI_ISL_2978179, EPI_ISL_2978180, EPI_ISL_2978181 | LSUHS Emerging Viral Threat Laboratory | LSUHS Emerging Viral Threat Laboratory | Gregory L. Ware, Alexander Mijalis, Jeremy P. Kamil, Jennifer L. Carroll, Maarten Van Diest, Rona S. Scott, Andrew D. Yurochko, Christopher G. Kevil, John A. Vanchiere |
| EPI_ISL_2978238 | EHNV | Laboratory of genomics and metagenomics | Trestan Pillonel, Damien Jacot, SÃ©bastien Aeby, Gilbert Greub, Claire Bertelli |
| EPI_ISL_2978239, EPI_ISL_2978241, EPI_ISL_2978244, EPI_ISL_2978245, EPI_ISL_2978247 | CHUV | Laboratory of genomics and metagenomics | Trestan Pillonel, Damien Jacot, SÃ©bastien Aeby, Gilbert Greub, Claire Bertelli |
| EPI_ISL_2978250, EPI_ISL_2978253 | VIDYMED LAUSANNE | Laboratory of genomics and metagenomics | Trestan Pillonel, Damien Jacot, Sébastien Aeby, Gilbert Greub, Claire Bertelli |
| EPI_ISL_2978254 | CHUV | Laboratory of genomics and metagenomics | Trestan Pillonel, Damien Jacot, Sébastien Aeby, Gilbert Greub, Claire Bertelli |
| EPI_ISL_2978256 | VIDYMED EPALINGES | Laboratory of genomics and metagenomics | Trestan Pillonel, Damien Jacot, Sébastien Aeby, Gilbert Greub, Claire Bertelli |
| EPI_ISL_2978257 | CHUV | Laboratory of genomics and metagenomics | Trestan Pillonel, Damien Jacot, Sébastien Aeby, Gilbert Greub, Claire Bertelli |
| EPI_ISL_2978259, EPI_ISL_2978261, EPI_ISL_2978262 | EHNV | Laboratory of genomics and metagenomics | Trestan Pillonel, Damien Jacot, Sébastien Aeby, Gilbert Greub, Claire Bertelli |
| EPI_ISL_2978264 | UNILABS - COPPET | Laboratory of genomics and metagenomics | Trestan Pillonel, Damien Jacot, Sébastien Aeby, Gilbert Greub, Claire Bertelli |
| EPI_ISL_2978265, EPI_ISL_2978267, EPI_ISL_2978268 | CHUV | Laboratory of genomics and metagenomics | Trestan Pillonel, Damien Jacot, Sébastien Aeby, Gilbert Greub, Claire Bertelli |
| EPI_ISL_2978269 | EHC MORGES | Laboratory of genomics and metagenomics | Trestan Pillonel, Damien Jacot, Sébastien Aeby, Gilbert Greub, Claire Bertelli |
| EPI_ISL_2978270, EPI_ISL_2978273, EPI_ISL_2978274, EPI_ISL_2978275, EPI_ISL_2978276, EPI_ISL_2978277, EPI_ISL_2978278 | CHUV | Laboratory of genomics and metagenomics | Trestan Pillonel, Damien Jacot, Sébastien Aeby, Gilbert Greub, Claire Bertelli |
| EPI_ISL_2978279 | VIDYMED LAUSANNE | Laboratory of genomics and metagenomics | Trestan Pillonel, Damien Jacot, Sébastien Aeby, Gilbert Greub, Claire Bertelli |
| EPI_ISL_2978280, EPI_ISL_2978283, EPI_ISL_2978284, EPI_ISL_2978285 | CHUV | Laboratory of genomics and metagenomics | Trestan Pillonel, Damien Jacot, Sébastien Aeby, Gilbert Greub, Claire Bertelli |
| EPI_ISL_2978286, EPI_ISL_2978287, EPI_ISL_2978288, EPI_ISL_2978290, EPI_ISL_2978292 | EHNV | Laboratory of genomics and metagenomics | Trestan Pillonel, Damien Jacot, Sébastien Aeby, Gilbert Greub, Claire Bertelli |
| EPI_ISL_2978293, EPI_ISL_2978294 | CHUV | Laboratory of genomics and metagenomics | Trestan Pillonel, Damien Jacot, Sébastien Aeby, Gilbert Greub, Claire Bertelli |

| | | | |
|---|---|---|---|
| EPI_ISL_2978295 | EHNV | Laboratory of genomics and metagenomics | Trestan Pillonel, Damien Jacot, Sébastien Aeby, Gilbert Greub, Claire Bertelli |
| EPI_ISL_2978298 | VIDYMED LAUSANNE | Laboratory of genomics and metagenomics | Trestan Pillonel, Damien Jacot, Sébastien Aeby, Gilbert Greub, Claire Bertelli |
| EPI_ISL_2978300, EPI_ISL_2978301 | CHUV | Laboratory of genomics and metagenomics | Trestan Pillonel, Damien Jacot, Sébastien Aeby, Gilbert Greub, Claire Bertelli |
| EPI_ISL_2978303 | VIDYMED LAUSANNE | Laboratory of genomics and metagenomics | Trestan Pillonel, Damien Jacot, Sébastien Aeby, Gilbert Greub, Claire Bertelli |
| EPI_ISL_2978304 | CHUV | Laboratory of genomics and metagenomics | Trestan Pillonel, Damien Jacot, Sébastien Aeby, Gilbert Greub, Claire Bertelli |
| EPI_ISL_2978305, EPI_ISL_2978306 | EHNV | Laboratory of genomics and metagenomics | Trestan Pillonel, Damien Jacot, Sébastien Aeby, Gilbert Greub, Claire Bertelli |
| EPI_ISL_2978307, EPI_ISL_2978308 | Spital Männedorf AG | Institute of Medical Virology | Verena Kufner, Gabriela Ziltener, Maryam Zaheri, Stefan Schmutz, Annette Audigé, Maria Grünberg, Kevin Steiner, Jon Huder, Cyril Shah, Riccarda Capaul, Guido Bloemberg, Jürg Böni, Michael Huber, Alexandra Trkola |
| EPI_ISL_2978309 | Spital Limmattal | Institute of Medical Virology | Verena Kufner, Gabriela Ziltener, Maryam Zaheri, Stefan Schmutz, Annette Audigé, Maria Grünberg, Kevin Steiner, Jon Huder, Cyril Shah, Riccarda Capaul, Guido Bloemberg, Jürg Böni, Michael Huber, Alexandra Trkola |
| EPI_ISL_2978310, EPI_ISL_2978311, EPI_ISL_2978312, EPI_ISL_2978313 | Stadtspital Triemli | Institute of Medical Virology | Verena Kufner, Gabriela Ziltener, Maryam Zaheri, Stefan Schmutz, Annette Audigé, Maria Grünberg, Kevin Steiner, Jon Huder, Cyril Shah, Riccarda Capaul, Guido Bloemberg, Jürg Böni, Michael Huber, Alexandra Trkola |
| EPI_ISL_2978314, EPI_ISL_2978315 | UniversitätsSpital Zürich | Institute of Medical Virology | Verena Kufner, Gabriela Ziltener, Maryam Zaheri, Stefan Schmutz, Annette Audigé, Maria Grünberg, Kevin Steiner, Jon Huder, Cyril Shah, Riccarda Capaul, Guido Bloemberg, Jürg Böni, Michael Huber, Alexandra Trkola |
| EPI_ISL_2978316 | Spital Limmattal | Institute of Medical Virology | Verena Kufner, Gabriela Ziltener, Maryam Zaheri, Stefan Schmutz, Annette Audigé, Maria Grünberg, Kevin Steiner, Jon Huder, Cyril Shah, Riccarda Capaul, Guido Bloemberg, Jürg Böni, Michael Huber, Alexandra Trkola |
| EPI_ISL_2978317, EPI_ISL_2978318 | UniversitätsSpital Zürich | Institute of Medical Virology | Verena Kufner, Gabriela Ziltener, Maryam Zaheri, Stefan Schmutz, Annette Audigé, Maria Grünberg, Kevin Steiner, Jon Huder, Cyril Shah, Riccarda Capaul, Guido Bloemberg, Jürg Böni, Michael Huber, Alexandra Trkola |
| EPI_ISL_2978319, EPI_ISL_2978320 | Spital Limmattal | Institute of Medical Virology | Verena Kufner, Gabriela Ziltener, Maryam Zaheri, Stefan Schmutz, Annette Audigé, Maria Grünberg, Kevin Steiner, Jon Huder, Cyril Shah, Riccarda Capaul, Guido Bloemberg, Jürg Böni, Michael Huber, Alexandra Trkola |
| EPI_ISL_2978321 | Stadtspital Triemli | Institute of Medical Virology | Verena Kufner, Gabriela Ziltener, Maryam Zaheri, Stefan Schmutz, Annette Audigé, Maria Grünberg, Kevin Steiner, Jon Huder, Cyril Shah, Riccarda Capaul, Guido Bloemberg, Jürg Böni, Michael Huber, Alexandra Trkola |
| EPI_ISL_2978322, EPI_ISL_2978323, EPI_ISL_2978324, EPI_ISL_2978326, EPI_ISL_2978327, EPI_ISL_2978328 | UniversitätsSpital Zürich | Institute of Medical Virology | Verena Kufner, Gabriela Ziltener, Maryam Zaheri, Stefan Schmutz, Annette Audigé, Maria Grünberg, Kevin Steiner, Jon Huder, Cyril Shah, Riccarda Capaul, Guido Bloemberg, Jürg Böni, Michael Huber, Alexandra Trkola |
| EPI_ISL_2978330 | UniversitätsSpital Zürich 009 | Institute of Medical Virology | Verena Kufner, Gabriela Ziltener, Maryam Zaheri, Stefan Schmutz, Annette Audigé, Maria Grünberg, Kevin Steiner, Jon Huder, Cyril Shah, Riccarda Capaul, Guido Bloemberg, Jürg Böni, Michael Huber, Alexandra Trkola |
| EPI_ISL_2978331, EPI_ISL_2978332, EPI_ISL_2978333, EPI_ISL_2978334 | Stadtspital Triemli | Institute of Medical Virology | Verena Kufner, Gabriela Ziltener, Maryam Zaheri, Stefan Schmutz, Annette Audigé, Maria Grünberg, Kevin Steiner, Jon Huder, Cyril Shah, Riccarda Capaul, Guido Bloemberg, Jürg Böni, Michael Huber, Alexandra Trkola |
| EPI_ISL_2978335, EPI_ISL_2978336 | Medica | Institute of Medical Virology | Verena Kufner, Gabriela Ziltener, Maryam Zaheri, Stefan Schmutz, Annette Audigé, Maria Grünberg, Kevin Steiner, Jon Huder, Cyril Shah, Riccarda Capaul, Guido Bloemberg, Jürg Böni, Michael Huber, Alexandra Trkola |
| EPI_ISL_2978337 | Kantonsspital Aarau | Institute of Medical Virology | Verena Kufner, Gabriela Ziltener, Maryam Zaheri, Stefan Schmutz, Annette Audigé, Maria Grünberg, Kevin Steiner, Jon Huder, Cyril Shah, Riccarda Capaul, Guido Bloemberg, Jürg Böni, Michael Huber, Alexandra Trkola |
| EPI_ISL_2978338 | Unilabs | Institute of Medical Virology | Verena Kufner, Gabriela Ziltener, Maryam Zaheri, Stefan Schmutz, Annette Audigé, Maria Grünberg, Kevin Steiner, Jon Huder, Cyril Shah, Riccarda Capaul, Guido Bloemberg, Jürg Böni, Michael Huber, Alexandra Trkola |
| EPI_ISL_2978339 | Universitätsspital Zürich 046 | Institute of Medical Virology | Verena Kufner, Gabriela Ziltener, Maryam Zaheri, Stefan Schmutz, Annette Audigé, Maria Grünberg, Kevin Steiner, Jon Huder, Cyril Shah, Riccarda Capaul, Guido Bloemberg, Jürg Böni, Michael Huber, Alexandra Trkola |
| EPI_ISL_2978340, EPI_ISL_2978341 | UniversitätsSpital Zürich 009 | Institute of Medical Virology | Verena Kufner, Gabriela Ziltener, Maryam Zaheri, Stefan Schmutz, Annette Audigé, Maria Grünberg, Kevin Steiner, Jon Huder, Cyril Shah, Riccarda Capaul, Guido Bloemberg, Jürg Böni, Michael Huber, Alexandra Trkola |
| EPI_ISL_2978342, EPI_ISL_2978343, EPI_ISL_2978344 | Spital Limmattal | Institute of Medical Virology | Verena Kufner, Gabriela Ziltener, Maryam Zaheri, Stefan Schmutz, Annette Audigé, Maria Grünberg, Kevin Steiner, Jon Huder, Cyril Shah, Riccarda Capaul, Guido Bloemberg, Jürg Böni, Michael Huber, Alexandra Trkola |
| EPI_ISL_2978345, EPI_ISL_2978346 | Stadtspital Triemli | Institute of Medical Virology | Verena Kufner, Gabriela Ziltener, Maryam Zaheri, Stefan Schmutz, Annette Audigé, Maria Grünberg, Kevin Steiner, Jon Huder, Cyril Shah, Riccarda Capaul, Guido Bloemberg, Jürg Böni, Michael Huber, Alexandra Trkola |
| EPI_ISL_2978347 | NJDOH, Public Health and Environmental Laboratories | NJ_PHEL | Lindsey Bodnar, Shiv K. Verma, Ryan Pachucki, Chelsea San Filippo, Jacquelyn Deverell, Dana Woell, Allison Roder, Byeong Jeong |
| EPI_ISL_2978348, EPI_ISL_2978349, EPI_ISL_2978350, EPI_ISL_2978351 | Victorian Infectious Diseases Reference Laboratory (VIDRL) | VIDRL and MDU-PHL | Caly L., Seemann T., Sait, M.L., Druce J., Sherry, N.L. |
| EPI_ISL_2978362, EPI_ISL_2978364, EPI_ISL_2978365, EPI_ISL_2978366, EPI_ISL_2978367, EPI_ISL_2978368, EPI_ISL_2978369, EPI_ISL_2978370, EPI_ISL_2978372, EPI_ISL_2978373, EPI_ISL_2978374, EPI_ISL_2978375, EPI_ISL_2978377, EPI_ISL_2978378, EPI_ISL_2978379, EPI_ISL_2978380, EPI_ISL_2978381, EPI_ISL_2978382, EPI_ISL_2978383, EPI_ISL_2978384, EPI_ISL_2978385, EPI_ISL_2978386, EPI_ISL_2978387, EPI_ISL_2978388, EPI_ISL_2978390, EPI_ISL_2978392, EPI_ISL_2978393, EPI_ISL_2978394, EPI_ISL_2978395, EPI_ISL_2978397, EPI_ISL_2978398, EPI_ISL_2978399, EPI_ISL_2978400, EPI_ISL_2978401, EPI_ISL_2978402, EPI_ISL_2978403 | | | |
| see above | Salud Digna | Instituto Nacional de Medicina Genomica | Cedro-Tanda A, Hidalgo-Miranda A, Mendoza-Vargas A, Reyes-Grajeda JP, Abraham Campos-Romero, Moreno-Camacho José Luis, Rodriguez-Gallegos Jorge, Luna-Ruiz Marco, Gonzalez-Barrera D, Rangel-DeLeon D, Munguia-Garza P, Ramirez-Vega O, Escobar-Arrazola, M, Herrera-Montalvo LA. |
| EPI_ISL_2978405, EPI_ISL_2978406, EPI_ISL_2978407, EPI_ISL_2978408, EPI_ISL_2978409, EPI_ISL_2978410, EPI_ISL_2978411, EPI_ISL_2978412, EPI_ISL_2978413, EPI_ISL_2978414, EPI_ISL_2978415, EPI_ISL_2978416, EPI_ISL_2978417, EPI_ISL_2978418, EPI_ISL_2978419, EPI_ISL_2978420, EPI_ISL_2978421, EPI_ISL_2978422, EPI_ISL_2978424, EPI_ISL_2978425, EPI_ISL_2978426, EPI_ISL_2978427, EPI_ISL_2978428, EPI_ISL_2978429, EPI_ISL_2978430, EPI_ISL_2978431, EPI_ISL_2978432, EPI_ISL_2978433, EPI_ISL_2978434, EPI_ISL_2978435, EPI_ISL_2978436, EPI_ISL_2978437, EPI_ISL_2978438, EPI_ISL_2978439, EPI_ISL_2978440 | | | |
| see above | Instituto Nacional de Medicina Genomica | Instituto Nacional de Medicina Genomica | Cedro-Tanda A, Hidalgo-Miranda A, Mendoza-Vargas A, Reyes-Grajeda JP, Cisneros- Villanueva M, Gonzalez-Barrera D, Rangel-DeLeon D, Munguia-Garza P, Ramirez-Vega O, Escobar-Arrazola, M, Herrera-Montalvo LA. |
| EPI_ISL_2978443, EPI_ISL_2978446, EPI_ISL_2978447, EPI_ISL_2978448, EPI_ISL_2978450, EPI_ISL_2978451, EPI_ISL_2978452, EPI_ISL_2978453, EPI_ISL_2978454, EPI_ISL_2978455, EPI_ISL_2978457, EPI_ISL_2978459, EPI_ISL_2978460, EPI_ISL_2978461, EPI_ISL_2978463, EPI_ISL_2978464, EPI_ISL_2978465, EPI_ISL_2978466, EPI_ISL_2978467, EPI_ISL_2978468, EPI_ISL_2978470, EPI_ISL_2978471, EPI_ISL_2978472, EPI_ISL_2978474, EPI_ISL_2978475, EPI_ISL_2978476, EPI_ISL_2978477, EPI_ISL_2978478, EPI_ISL_2978479, EPI_ISL_2978480, EPI_ISL_2978481, EPI_ISL_2978482, EPI_ISL_2978483, EPI_ISL_2978486, EPI_ISL_2978487, EPI_ISL_2978488, EPI_ISL_2978490, EPI_ISL_2978492, EPI_ISL_2978493, EPI_ISL_2978494, EPI_ISL_2978495, EPI_ISL_2978496, EPI_ISL_2978497, EPI_ISL_2978498, EPI_ISL_2978500, EPI_ISL_2978501, EPI_ISL_2978502, EPI_ISL_2978504, EPI_ISL_2978505, EPI_ISL_2978506, EPI_ISL_2978507, EPI_ISL_2978508, EPI_ISL_2978509, EPI_ISL_2978510, EPI_ISL_2978511, EPI_ISL_2978512, EPI_ISL_2978513, EPI_ISL_2978514, EPI_ISL_2978515, EPI_ISL_2978516, EPI_ISL_2978518, EPI_ISL_2978519, EPI_ISL_2978520, EPI_ISL_2978521, EPI_ISL_2978522, EPI_ISL_2978523, EPI_ISL_2978524, EPI_ISL_2978525 | | | |
| see above | Salud Digna | Instituto Nacional de Medicina Genomica | Cedro-Tanda A, Hidalgo-Miranda A, Mendoza-Vargas A, Reyes-Grajeda JP, Abraham Campos-Romero, Moreno-Camacho José Luis, Rodriguez-Gallegos Jorge, Luna-Ruiz Marco, Gonzalez-Barrera D, Rangel-DeLeon D, Munguia-Garza P, Ramirez-Vega O, Escobar-Arrazola, M, Herrera-Montalvo LA. |
| EPI_ISL_2978526, EPI_ISL_2978527, EPI_ISL_2978528, EPI_ISL_2978529, EPI_ISL_2978530, EPI_ISL_2978531, EPI_ISL_2978532, EPI_ISL_2978533, EPI_ISL_2978534, EPI_ISL_2978535, EPI_ISL_2978536, EPI_ISL_2978537, EPI_ISL_2978538, EPI_ISL_2978539 | | | |
| see above | Instituto Nacional de Medicina Genomica | Instituto Nacional de Medicina Genomica | Cedro-Tanda A, Hidalgo-Miranda A, Mendoza-Vargas A, Reyes-Grajeda JP, Cisneros- Villanueva M, Gonzalez-Barrera D, Rangel-DeLeon D, Munguia-Garza P, Ramirez-Vega O, Escobar-Arrazola, M, Herrera-Montalvo LA. |
| EPI_ISL_2978540, EPI_ISL_2978541, EPI_ISL_2978542, EPI_ISL_2978543, EPI_ISL_2978544, EPI_ISL_2978545, EPI_ISL_2978546, EPI_ISL_2978547, EPI_ISL_2978548, EPI_ISL_2978549, EPI_ISL_2978550, EPI_ISL_2978551, EPI_ISL_2978553, EPI_ISL_2978554, EPI_ISL_2978555, EPI_ISL_2978556, EPI_ISL_2978557, EPI_ISL_2978558, EPI_ISL_2978559, EPI_ISL_2978560, EPI_ISL_2978561, EPI_ISL_2978562, EPI_ISL_2978564, EPI_ISL_2978565, EPI_ISL_2978566, EPI_ISL_2978567, EPI_ISL_2978568, EPI_ISL_2978570, EPI_ISL_2978571, EPI_ISL_2978572, EPI_ISL_2978573, EPI_ISL_2978575, EPI_ISL_2978578, EPI_ISL_2978579, EPI_ISL_2978580, EPI_ISL_2978581, EPI_ISL_2978582, EPI_ISL_2978583, EPI_ISL_2978584, EPI_ISL_2978585, EPI_ISL_2978586, EPI_ISL_2978587, EPI_ISL_2978588, EPI_ISL_2978589, EPI_ISL_2978590, EPI_ISL_2978591, EPI_ISL_2978592, EPI_ISL_2978593, EPI_ISL_2978602, EPI_ISL_2978607, EPI_ISL_2978613, EPI_ISL_2978625, EPI_ISL_2978631, EPI_ISL_2978639, EPI_ISL_2978648, EPI_ISL_2978649, EPI_ISL_2978650, EPI_ISL_2978655, EPI_ISL_2978669, EPI_ISL_2978672, EPI_ISL_2978677 | | | |
| see above | Salud Digna | Instituto Nacional de Medicina Genomica | Cedro-Tanda A, Hidalgo-Miranda A, Mendoza-Vargas A, Reyes-Grajeda JP, Abraham Campos-Romero, Moreno-Camacho José Luis, Rodriguez-Gallegos Jorge, Luna-Ruiz Marco, Gonzalez-Barrera D, Rangel-DeLeon D, Munguia-Garza P, Ramirez-Vega O, Escobar-Arrazola, M, Herrera-Montalvo LA. |
| EPI_ISL_2978680 | National Public Health Laboratory, National Centre for Infectious Diseases | National Public Health Laboratory, National Centre for Infectious Diseases | Tze Minn Mak, Zhenyang Zhou, Royce Ang, Lin Cui, Raymond Tzer Pin Lin |
| EPI_ISL_2978685 | A.S.L. BIELLA | Fondazione del Piemonte per l'Oncologia IRCCS | Antonino Sottile, Silvia Brossa, Paola Marino, Giorgio Giardina |
| EPI_ISL_2978687 | A.S.L. NOVARA-BORGOMANERO | Fondazione del Piemonte per l'Oncologia IRCCS | Antonino Sottile, Silvia Brossa, Paola Marino, Giorgio Giardina |
| EPI_ISL_2978690 | SA Pathology | SA Pathology | Caitlin Selway, Lex Leong, Chuan Kok Lim, Mark Turra, Ivan Bastian, Geoff Higgins |
| EPI_ISL_2978691 | A.S.L. CN1 | Fondazione del Piemonte per l'Oncologia IRCCS | Antonino Sottile, Silvia Brossa, Paola Marino, Giorgio Giardina |
| EPI_ISL_2978692 | SA Pathology | SA Pathology | Caitlin Selway, Lex Leong, Chuan Kok Lim, Mark Turra, Ivan Bastian, Geoff Higgins |
| EPI_ISL_2978693 | A.S.L. CN1 | Fondazione del Piemonte per l'Oncologia IRCCS | Antonino Sottile, Silvia Brossa, Paola Marino, Giorgio Giardina |
| EPI_ISL_2978694 | SA Pathology | SA Pathology | Caitlin Selway, Lex Leong, Chuan Kok Lim, Mark Turra, Ivan Bastian, Geoff Higgins |
| EPI_ISL_2978695 | A.S.L. TO4 | Fondazione del Piemonte per l'Oncologia IRCCS | Antonino Sottile, Silvia Brossa, Paola Marino, Giorgio Giardina |
| EPI_ISL_2978696 | SA Pathology | SA Pathology | Caitlin Selway, Lex Leong, Chuan Kok Lim, Mark Turra, Ivan Bastian, Geoff Higgins |
| EPI_ISL_2978697 | A.S.L. CN1 | Fondazione del Piemonte per l'Oncologia IRCCS | Antonino Sottile, Silvia Brossa, Paola Marino, Giorgio Giardina |
| EPI_ISL_2978698 | SA Pathology | SA Pathology | Caitlin Selway, Lex Leong, Chuan Kok Lim, Mark Turra, Ivan Bastian, Geoff Higgins |
| EPI_ISL_2978699 | A.S.L. CN1 | Fondazione del Piemonte per l'Oncologia IRCCS | Antonino Sottile, Silvia Brossa, Paola Marino, Giorgio Giardina |
| EPI_ISL_2978700 | SA Pathology | SA Pathology | Caitlin Selway, Lex Leong, Chuan Kok Lim, Mark Turra, Ivan Bastian, Geoff Higgins |
| EPI_ISL_2978701, | A.S.L. TO4 | Fondazione del Piemonte per l'Oncologia IRCCS | Antonino Sottile, Silvia Brossa, Paola Marino, Giorgio Giardina |

| | | | |
|---|---|---|---|
| EPI_ISL_2978702 | | | |
| EPI_ISL_2978703, EPI_ISL_2978704 | A.S.L.VCO | Fondazione del Piemonte per l'Oncologia IRCCS | Antonino Sottile, Silvia Brossa, Paola Marino, Giorgio Giardina |
| EPI_ISL_2978705, EPI_ISL_2978706 | A.S.L. TO4 | Fondazione del Piemonte per l'Oncologia IRCCS | Antonino Sottile, Silvia Brossa, Paola Marino, Giorgio Giardina |
| EPI_ISL_2978707, EPI_ISL_2978709, EPI_ISL_2978711, EPI_ISL_2978712 | A.S.L.VCO | Fondazione del Piemonte per l'Oncologia IRCCS | Antonino Sottile, Silvia Brossa, Paola Marino, Giorgio Giardina |
| EPI_ISL_2978713 | A.S.L. NOVARA-BORGOMANERO | Fondazione del Piemonte per l'Oncologia IRCCS | Antonino Sottile, Silvia Brossa, Paola Marino, Giorgio Giardina |
| EPI_ISL_2978714 | A.S.L.VCO | Fondazione del Piemonte per l'Oncologia IRCCS | Antonino Sottile, Silvia Brossa, Paola Marino, Giorgio Giardina |
| EPI_ISL_2978715, EPI_ISL_2978716, EPI_ISL_2978717, EPI_ISL_2978719, EPI_ISL_2978721, EPI_ISL_2978722, EPI_ISL_2978725, EPI_ISL_2978727 | A.S.L. NOVARA-BORGOMANERO | Fondazione del Piemonte per l'Oncologia IRCCS | Antonino Sottile, Silvia Brossa, Paola Marino, Giorgio Giardina |
| EPI_ISL_2978728 | U.O. Microbiologia Laboratorio Unico Centro Servizi AUSL della Romagna | U.O. Microbiologia Laboratorio Unico Centro Servizi AUSL della Romagna | Giorgio Dirani, Silvia Zannoli, Francesca Taddei, Giulia Gatti, Ilaria Poggianti, Vittorio Sambri |
| EPI_ISL_2980020, EPI_ISL_2980021, EPI_ISL_2980022, EPI_ISL_2980023, EPI_ISL_2980025, EPI_ISL_2980026, EPI_ISL_2980027, EPI_ISL_2980028, EPI_ISL_2980034, EPI_ISL_2980035, EPI_ISL_2980036, EPI_ISL_2980048, EPI_ISL_2980053, EPI_ISL_2980054, EPI_ISL_2980055, EPI_ISL_2980058, EPI_ISL_2980060 | | | |
| see above | MD PHL | MD PHL | Maryland Department of Health Laboratories Administration |
| EPI_ISL_2980169, EPI_ISL_2980170, EPI_ISL_2980172, EPI_ISL_2980173, EPI_ISL_2980175, EPI_ISL_2980176, EPI_ISL_2980177, EPI_ISL_2980178, EPI_ISL_2980179, EPI_ISL_2980181, EPI_ISL_2980182, EPI_ISL_2980185, EPI_ISL_2980186, EPI_ISL_2980189, EPI_ISL_2980190, EPI_ISL_2980191, EPI_ISL_2980192, EPI_ISL_2980193, EPI_ISL_2980194, EPI_ISL_2980195, EPI_ISL_2980196, EPI_ISL_2980197, EPI_ISL_2980199, EPI_ISL_2980200, EPI_ISL_2980201, EPI_ISL_2980202, EPI_ISL_2980203, EPI_ISL_2980204, EPI_ISL_2980205, EPI_ISL_2980206, EPI_ISL_2980207, EPI_ISL_2980208, EPI_ISL_2980211, EPI_ISL_2980212, EPI_ISL_2980213, EPI_ISL_2980214, EPI_ISL_2980216, EPI_ISL_2980217, EPI_ISL_2980218, EPI_ISL_2980219, EPI_ISL_2980220, EPI_ISL_2980221, EPI_ISL_2980223, EPI_ISL_2980225, EPI_ISL_2980226, EPI_ISL_2980227, EPI_ISL_2980228, EPI_ISL_2980230, EPI_ISL_2980231, EPI_ISL_2980232, EPI_ISL_2980233, EPI_ISL_2980234, EPI_ISL_2980235, EPI_ISL_2980236, EPI_ISL_2980238, EPI_ISL_2980239, EPI_ISL_2980240, EPI_ISL_2980241, EPI_ISL_2980242, EPI_ISL_2980243, EPI_ISL_2980244, EPI_ISL_2980245, EPI_ISL_2980246, EPI_ISL_2980247, EPI_ISL_2980248, EPI_ISL_2980249, EPI_ISL_2980250, EPI_ISL_2980252, EPI_ISL_2980253, EPI_ISL_2980255, EPI_ISL_2980256, EPI_ISL_2980258, EPI_ISL_2980259, EPI_ISL_2980260, EPI_ISL_2980261, EPI_ISL_2980262, EPI_ISL_2980263, EPI_ISL_2980264, EPI_ISL_2980265, EPI_ISL_2980267, EPI_ISL_2980268, EPI_ISL_2980270, EPI_ISL_2980271, EPI_ISL_2980272, EPI_ISL_2980275, EPI_ISL_2980278, EPI_ISL_2980279, EPI_ISL_2980280, EPI_ISL_2980281, EPI_ISL_2980282, EPI_ISL_2980283, EPI_ISL_2980284, EPI_ISL_2980285, EPI_ISL_2980286, EPI_ISL_2980289, EPI_ISL_2980290, EPI_ISL_2980291, EPI_ISL_2980292, EPI_ISL_2980293, EPI_ISL_2980304, EPI_ISL_2980306, EPI_ISL_2980307, EPI_ISL_2980308, EPI_ISL_2980309, EPI_ISL_2980310, EPI_ISL_2980311, EPI_ISL_2980313, EPI_ISL_2980315, EPI_ISL_2980318, EPI_ISL_2980319, EPI_ISL_2980323, EPI_ISL_2980327, EPI_ISL_2980330, EPI_ISL_2980333, EPI_ISL_2980334, EPI_ISL_2980336, EPI_ISL_2980339, EPI_ISL_2980340, EPI_ISL_2980342, EPI_ISL_2980343, EPI_ISL_2980345, EPI_ISL_2980350, EPI_ISL_2980352, EPI_ISL_2980354, EPI_ISL_2980355, EPI_ISL_2980356, EPI_ISL_2980357, EPI_ISL_2980358, EPI_ISL_2980359, EPI_ISL_2980360, EPI_ISL_2980361, EPI_ISL_2980362, EPI_ISL_2980363, EPI_ISL_2980364, EPI_ISL_2980365, EPI_ISL_2980366, EPI_ISL_2980367, EPI_ISL_2980368, EPI_ISL_2980369, EPI_ISL_2980370, EPI_ISL_2980371, EPI_ISL_2980372, EPI_ISL_2980373, EPI_ISL_2980374, EPI_ISL_2980375, EPI_ISL_2980376, EPI_ISL_2980377, EPI_ISL_2980378, EPI_ISL_2980379, EPI_ISL_2980381, EPI_ISL_2980383, EPI_ISL_2980384, EPI_ISL_2980385, EPI_ISL_2980386, EPI_ISL_2980387, EPI_ISL_2980388, EPI_ISL_2980389, EPI_ISL_2980390, EPI_ISL_2980391, EPI_ISL_2980392, EPI_ISL_2980393, EPI_ISL_2980394, EPI_ISL_2980395, EPI_ISL_2980396, EPI_ISL_2980397, EPI_ISL_2980398, EPI_ISL_2980399, EPI_ISL_2980400, EPI_ISL_2980488, EPI_ISL_2980489, EPI_ISL_2980490, EPI_ISL_2980491, EPI_ISL_2980492, EPI_ISL_2980493, EPI_ISL_2980494, EPI_ISL_2980495, EPI_ISL_2980496, EPI_ISL_2980497, EPI_ISL_2980498, EPI_ISL_2980499, EPI_ISL_2980500, EPI_ISL_2980501, EPI_ISL_2980502, EPI_ISL_2980503, EPI_ISL_2980504, EPI_ISL_2980505, EPI_ISL_2980506, EPI_ISL_2980507, EPI_ISL_2980508, EPI_ISL_2980509, EPI_ISL_2980510, EPI_ISL_2980511, EPI_ISL_2980512, EPI_ISL_2980513, EPI_ISL_2980514, EPI_ISL_2980515, EPI_ISL_2980516, EPI_ISL_2980517, EPI_ISL_2980518, EPI_ISL_2980519, EPI_ISL_2980520, EPI_ISL_2980521, EPI_ISL_2980522, EPI_ISL_2980523, EPI_ISL_2980524, EPI_ISL_2980525, EPI_ISL_2980526, EPI_ISL_2980527, EPI_ISL_2980528, EPI_ISL_2980529, EPI_ISL_2980530, EPI_ISL_2980531, EPI_ISL_2980532, EPI_ISL_2980533, EPI_ISL_2980534, EPI_ISL_2980535, EPI_ISL_2980536, EPI_ISL_2980537, EPI_ISL_2980538, EPI_ISL_2980539, EPI_ISL_2980540, EPI_ISL_2980541, EPI_ISL_2980542, EPI_ISL_2980543, EPI_ISL_2980544, EPI_ISL_2980545, EPI_ISL_2980546, EPI_ISL_2980547, EPI_ISL_2980548, EPI_ISL_2980549, EPI_ISL_2980550, EPI_ISL_2980551, EPI_ISL_2980552, EPI_ISL_2980553, EPI_ISL_2980554, EPI_ISL_2980555, EPI_ISL_2980556, EPI_ISL_2980557, EPI_ISL_2980558, EPI_ISL_2980559, EPI_ISL_2980560, EPI_ISL_2980561, EPI_ISL_2980562, EPI_ISL_2980563, EPI_ISL_2980564, EPI_ISL_2980565, EPI_ISL_2980566, EPI_ISL_2980567, EPI_ISL_2980568, EPI_ISL_2980569, EPI_ISL_2980570, EPI_ISL_2980571, EPI_ISL_2980572, EPI_ISL_2980573, EPI_ISL_2980574, EPI_ISL_2980575, EPI_ISL_2980577, EPI_ISL_2980578, EPI_ISL_2980579, EPI_ISL_2980580, EPI_ISL_2980581, EPI_ISL_2980582, EPI_ISL_2980583, EPI_ISL_2980584, EPI_ISL_2980585, EPI_ISL_2980586, EPI_ISL_2980588, EPI_ISL_2980589, EPI_ISL_2980590, EPI_ISL_2980591, EPI_ISL_2980592, EPI_ISL_2980593, EPI_ISL_2980594, EPI_ISL_2980595, EPI_ISL_2980596, EPI_ISL_2980597, EPI_ISL_2980598, EPI_ISL_2980599, EPI_ISL_2980600, EPI_ISL_2980601, EPI_ISL_2980602, EPI_ISL_2980603, EPI_ISL_2980604, EPI_ISL_2980613, EPI_ISL_2980614, EPI_ISL_2980615, EPI_ISL_2980616, EPI_ISL_2980617, EPI_ISL_2980618, EPI_ISL_2980619, EPI_ISL_2980621, EPI_ISL_2980622, EPI_ISL_2980623, EPI_ISL_2980624, EPI_ISL_2980625, EPI_ISL_2980626, EPI_ISL_2980627, EPI_ISL_2980628, EPI_ISL_2980629, EPI_ISL_2980630, EPI_ISL_2980631, EPI_ISL_2980632, EPI_ISL_2980633, EPI_ISL_2980634, EPI_ISL_2980635, EPI_ISL_2980636, EPI_ISL_2980637, EPI_ISL_2980638, EPI_ISL_2980639, EPI_ISL_2980640, EPI_ISL_2980641, EPI_ISL_2980642, EPI_ISL_2980643, EPI_ISL_2980644, EPI_ISL_2980645, EPI_ISL_2980646, EPI_ISL_2980647, EPI_ISL_2980648, EPI_ISL_2980649, EPI_ISL_2980651, EPI_ISL_2980653, EPI_ISL_2980654, EPI_ISL_2980655, EPI_ISL_2980656, EPI_ISL_2980657, EPI_ISL_2980658, EPI_ISL_2980659, EPI_ISL_2980660, EPI_ISL_2980661, EPI_ISL_2980662, EPI_ISL_2980663, EPI_ISL_2980664, EPI_ISL_2980665, EPI_ISL_2980666, EPI_ISL_2980667, EPI_ISL_2980669, EPI_ISL_2980670, EPI_ISL_2980671, EPI_ISL_2980672, EPI_ISL_2980673, EPI_ISL_2980674, EPI_ISL_2980675, EPI_ISL_2980676, EPI_ISL_2980677, EPI_ISL_2980678, EPI_ISL_2980679, EPI_ISL_2980680, EPI_ISL_2980681, EPI_ISL_2980682, EPI_ISL_2980683, EPI_ISL_2980684, EPI_ISL_2980685, EPI_ISL_2980686, EPI_ISL_2980687, EPI_ISL_2980688, EPI_ISL_2980689, EPI_ISL_2980690, EPI_ISL_2980692, EPI_ISL_2980693, EPI_ISL_2980694, EPI_ISL_2980695, EPI_ISL_2980696, EPI_ISL_2980697, EPI_ISL_2980698, EPI_ISL_2980699, EPI_ISL_2980700, EPI_ISL_2980701, EPI_ISL_2980702, EPI_ISL_2980703, EPI_ISL_2980704, EPI_ISL_2980705, EPI_ISL_2980706, EPI_ISL_2980707, EPI_ISL_2980708, EPI_ISL_2980709, EPI_ISL_2980710, EPI_ISL_2980711, EPI_ISL_2980712, EPI_ISL_2980713, EPI_ISL_2980714, EPI_ISL_2980715, EPI_ISL_2980716, EPI_ISL_2980717, EPI_ISL_2980718, EPI_ISL_2980719, EPI_ISL_2980720, EPI_ISL_2980721, EPI_ISL_2980722, EPI_ISL_2980724, EPI_ISL_2980725, EPI_ISL_2980726, EPI_ISL_2980727, EPI_ISL_2980728, EPI_ISL_2980729, EPI_ISL_2980730, EPI_ISL_2980731, EPI_ISL_2980732, EPI_ISL_2980733, EPI_ISL_2980734, EPI_ISL_2980735, EPI_ISL_2980736, EPI_ISL_2980737, EPI_ISL_2980738, EPI_ISL_2980739, EPI_ISL_2980740, EPI_ISL_2980741, EPI_ISL_2980742, EPI_ISL_2980743, EPI_ISL_2980744, EPI_ISL_2980745, EPI_ISL_2980746, EPI_ISL_2980747, EPI_ISL_2980748, EPI_ISL_2980749, EPI_ISL_2980750, EPI_ISL_2980751, EPI_ISL_2980752, EPI_ISL_2980753, EPI_ISL_2980754, EPI_ISL_2980755, EPI_ISL_2980756, EPI_ISL_2980758, EPI_ISL_2980759, EPI_ISL_2980760, EPI_ISL_2980761, EPI_ISL_2980762, EPI_ISL_2980763, EPI_ISL_2980764, EPI_ISL_2980765, EPI_ISL_2980766, EPI_ISL_2980767, EPI_ISL_2980768, EPI_ISL_2980769, EPI_ISL_2980770, EPI_ISL_2980771, EPI_ISL_2980772, EPI_ISL_2980773, EPI_ISL_2980774, EPI_ISL_2980775, EPI_ISL_2980776, EPI_ISL_2980777, EPI_ISL_2980778, EPI_ISL_2980779, EPI_ISL_2980780, EPI_ISL_2980781, EPI_ISL_2980782, EPI_ISL_2980783, EPI_ISL_2980784, EPI_ISL_2980785, EPI_ISL_2980786, EPI_ISL_2980787, EPI_ISL_2980788, EPI_ISL_2980789, EPI_ISL_2980790, EPI_ISL_2980791, EPI_ISL_2980792, EPI_ISL_2980793, EPI_ISL_2980794, EPI_ISL_2980795, EPI_ISL_2980796, EPI_ISL_2980798, EPI_ISL_2980799, EPI_ISL_2980800, EPI_ISL_2980801, EPI_ISL_2980816, EPI_ISL_2980822, EPI_ISL_2980823, EPI_ISL_2980824, EPI_ISL_2980825, EPI_ISL_2980826, EPI_ISL_2980827, EPI_ISL_2980835, EPI_ISL_2980836, EPI_ISL_2980837, EPI_ISL_2980838, EPI_ISL_2980839, EPI_ISL_2980840, EPI_ISL_2980841, EPI_ISL_2980842, EPI_ISL_2980843, EPI_ISL_2980844, EPI_ISL_2980845, EPI_ISL_2980846, EPI_ISL_2980847, EPI_ISL_2980848, EPI_ISL_2980849, EPI_ISL_2980850, EPI_ISL_2980851, EPI_ISL_2980852, EPI_ISL_2980880, EPI_ISL_2980881, EPI_ISL_2980882, EPI_ISL_2980900, EPI_ISL_2980907, EPI_ISL_2980911, EPI_ISL_2980917, EPI_ISL_2980918, EPI_ISL_2980920, EPI_ISL_2980922, EPI_ISL_2980930, EPI_ISL_2980932, EPI_ISL_2980935, EPI_ISL_2980939, EPI_ISL_2980955, EPI_ISL_2980956, EPI_ISL_2980957, EPI_ISL_2980958, EPI_ISL_2980959, EPI_ISL_2980960, EPI_ISL_2980961, EPI_ISL_2980962, EPI_ISL_2980963, EPI_ISL_2980964, EPI_ISL_2980965, EPI_ISL_2980966, EPI_ISL_2980987, EPI_ISL_2980988, EPI_ISL_2980989, EPI_ISL_2980990, EPI_ISL_2980991, EPI_ISL_2980992, EPI_ISL_2980993, EPI_ISL_2980994, EPI_ISL_2980995, EPI_ISL_2980996, EPI_ISL_2980997, EPI_ISL_2980998, EPI_ISL_2980999, EPI_ISL_2981000, EPI_ISL_2981001, EPI_ISL_2981002, EPI_ISL_2981003, EPI_ISL_2981004, EPI_ISL_2981005, EPI_ISL_2981006, EPI_ISL_2981033, EPI_ISL_2981034, EPI_ISL_2981038, EPI_ISL_2981039, EPI_ISL_2981041, EPI_ISL_2981042, EPI_ISL_2981043, EPI_ISL_2981044, EPI_ISL_2981047, EPI_ISL_2981048, EPI_ISL_2981049, EPI_ISL_2981050, EPI_ISL_2981051 | | | |
| see above | Swedish national genomic surveillance program of SARS-CoV-2 | The Public Health Agency of Sweden | Maximilian Riess, Maria Lind Karlberg, Alma Brolund, Swedish national genomic surveillance program of SARS-CoV-2 |
| EPI_ISL_2981055, EPI_ISL_2981056, EPI_ISL_2981066, EPI_ISL_2981073, EPI_ISL_2981074, EPI_ISL_2981077, EPI_ISL_2981080, EPI_ISL_2981082, EPI_ISL_2981086, EPI_ISL_2981089, EPI_ISL_2981094, EPI_ISL_2981095, EPI_ISL_2981096, EPI_ISL_2981097, EPI_ISL_2981102, EPI_ISL_2981103, EPI_ISL_2981104, EPI_ISL_2981108, EPI_ISL_2981111, EPI_ISL_2981112, EPI_ISL_2981113, EPI_ISL_2981114, EPI_ISL_2981119, EPI_ISL_2981120, EPI_ISL_2981121, EPI_ISL_2981122, EPI_ISL_2981124, EPI_ISL_2981126, EPI_ISL_2981127, EPI_ISL_2981133, EPI_ISL_2981134, EPI_ISL_2981137, EPI_ISL_2981139, EPI_ISL_2981140, EPI_ISL_2981142, EPI_ISL_2981143, EPI_ISL_2981144, EPI_ISL_2981145, EPI_ISL_2981146, EPI_ISL_2981147, EPI_ISL_2981148, EPI_ISL_2981149, EPI_ISL_2981152, EPI_ISL_2981153, EPI_ISL_2981154, EPI_ISL_2981155, EPI_ISL_2981156, EPI_ISL_2981157, EPI_ISL_2981158, EPI_ISL_2981159, EPI_ISL_2981162, EPI_ISL_2981163, EPI_ISL_2981164, EPI_ISL_2981165, EPI_ISL_2981166, EPI_ISL_2981167, EPI_ISL_2981170, EPI_ISL_2981171, EPI_ISL_2981172, EPI_ISL_2981178, EPI_ISL_2981187, EPI_ISL_2981190, EPI_ISL_2981192, EPI_ISL_2981195, EPI_ISL_2981196, EPI_ISL_2981197, EPI_ISL_2981199, EPI_ISL_2981201, EPI_ISL_2981205, EPI_ISL_2981207, EPI_ISL_2981208, EPI_ISL_2981209, EPI_ISL_2981210, EPI_ISL_2981211, EPI_ISL_2981214, EPI_ISL_2981215, EPI_ISL_2981216, EPI_ISL_2981250, EPI_ISL_2981252, EPI_ISL_2981264, EPI_ISL_2981265, EPI_ISL_2981266, EPI_ISL_2981270, EPI_ISL_2981286, EPI_ISL_2981290, EPI_ISL_2981293, EPI_ISL_2981294, EPI_ISL_2981298, EPI_ISL_2981299, EPI_ISL_2981300, EPI_ISL_2981301, EPI_ISL_2981303, EPI_ISL_2981304, EPI_ISL_2981307, EPI_ISL_2981312, EPI_ISL_2981314, EPI_ISL_2981316, EPI_ISL_2981319, EPI_ISL_2981320, EPI_ISL_2981321, EPI_ISL_2981322, EPI_ISL_2981323, EPI_ISL_2981324, EPI_ISL_2981326, EPI_ISL_2981327, EPI_ISL_2981328, EPI_ISL_2981329, EPI_ISL_2981332, EPI_ISL_2981333, EPI_ISL_2981335, EPI_ISL_2981336, EPI_ISL_2981337, EPI_ISL_2981338, EPI_ISL_2981346, EPI_ISL_2981348, EPI_ISL_2981353, EPI_ISL_2981355, EPI_ISL_2981363, EPI_ISL_2981364, EPI_ISL_2981365, EPI_ISL_2981366, EPI_ISL_2981367, EPI_ISL_2981369, EPI_ISL_2981370, EPI_ISL_2981371, EPI_ISL_2981373, EPI_ISL_2981374, EPI_ISL_2981376, EPI_ISL_2981377, EPI_ISL_2981379, EPI_ISL_2981381, EPI_ISL_2981383, EPI_ISL_2981384, EPI_ISL_2981385, EPI_ISL_2981387, EPI_ISL_2981389, EPI_ISL_2981390, EPI_ISL_2981391, EPI_ISL_2981393, EPI_ISL_2981394, EPI_ISL_2981395, EPI_ISL_2981398, EPI_ISL_2981399, EPI_ISL_2981401, EPI_ISL_2981402, EPI_ISL_2981403, EPI_ISL_2981404, EPI_ISL_2981405, EPI_ISL_2981406, EPI_ISL_2981407, EPI_ISL_2981410, EPI_ISL_2981411, EPI_ISL_2981412, EPI_ISL_2981413, EPI_ISL_2981415, EPI_ISL_2981416, EPI_ISL_2981418, EPI_ISL_2981419, EPI_ISL_2981421, EPI_ISL_2981422, EPI_ISL_2981424, EPI_ISL_2981428, EPI_ISL_2981429, EPI_ISL_2981431, EPI_ISL_2981432, EPI_ISL_2981435, EPI_ISL_2981436, EPI_ISL_2981437, EPI_ISL_2981440, EPI_ISL_2981441, EPI_ISL_2981442, EPI_ISL_2981450, EPI_ISL_2981453, EPI_ISL_2981454, EPI_ISL_2981459, EPI_ISL_2981460, EPI_ISL_2981461, EPI_ISL_2981462, EPI_ISL_2981463, EPI_ISL_2981464, EPI_ISL_2981468, EPI_ISL_2981469, EPI_ISL_2981470, EPI_ISL_2981471, EPI_ISL_2981472, EPI_ISL_2981474, EPI_ISL_2981475, EPI_ISL_2981476, EPI_ISL_2981477, EPI_ISL_2981480, EPI_ISL_2981482, EPI_ISL_2981483, EPI_ISL_2981484, EPI_ISL_2981486, EPI_ISL_2981487, EPI_ISL_2981488, EPI_ISL_2981497, EPI_ISL_2981499, EPI_ISL_2981500, EPI_ISL_2981501, EPI_ISL_2981502, EPI_ISL_2981503, EPI_ISL_2981504, EPI_ISL_2981505, EPI_ISL_2981509, EPI_ISL_2981510, EPI_ISL_2981512, EPI_ISL_2981513, EPI_ISL_2981523, EPI_ISL_2981526, EPI_ISL_2981534, EPI_ISL_2981543, EPI_ISL_2981552, EPI_ISL_2981556, EPI_ISL_2981569, EPI_ISL_2981573, EPI_ISL_2981578, EPI_ISL_2981580, EPI_ISL_2981581, EPI_ISL_2981587, EPI_ISL_2981592, EPI_ISL_2981594, EPI_ISL_2981599, EPI_ISL_2981600, EPI_ISL_2981602, EPI_ISL_2981603, EPI_ISL_2981605, EPI_ISL_2981615, EPI_ISL_2981617, EPI_ISL_2981620, EPI_ISL_2981622, EPI_ISL_2981623, EPI_ISL_2981630, EPI_ISL_2981633, EPI_ISL_2981634, EPI_ISL_2981635, EPI_ISL_2981636, EPI_ISL_2981637, EPI_ISL_2981638, EPI_ISL_2981641, EPI_ISL_2981642, EPI_ISL_2981643, EPI_ISL_2981644, EPI_ISL_2981647, EPI_ISL_2981650, EPI_ISL_2981651, EPI_ISL_2981652, EPI_ISL_2981654, EPI_ISL_2981655, EPI_ISL_2981656, EPI_ISL_2981657, EPI_ISL_2981659, EPI_ISL_2981662, EPI_ISL_2981669, EPI_ISL_2981674, EPI_ISL_2981679, EPI_ISL_2981682, EPI_ISL_2981684, EPI_ISL_2981689, EPI_ISL_2981690, EPI_ISL_2981695, EPI_ISL_2981697, EPI_ISL_2981698, EPI_ISL_2981700, EPI_ISL_2981703, EPI_ISL_2981704, EPI_ISL_2981705, EPI_ISL_2981707, EPI_ISL_2981708, EPI_ISL_2981716, EPI_ISL_2981718, EPI_ISL_2981722, EPI_ISL_2981724, EPI_ISL_2981725, EPI_ISL_2981729, EPI_ISL_2981732, EPI_ISL_2981735, EPI_ISL_2981737, EPI_ISL_2981739, EPI_ISL_2981741, EPI_ISL_2981742, EPI_ISL_2981743, EPI_ISL_2981744, EPI_ISL_2981747, EPI_ISL_2981748, EPI_ISL_2981750, EPI_ISL_2981752, EPI_ISL_2981754, EPI_ISL_2981759, EPI_ISL_2981762, EPI_ISL_2981763, EPI_ISL_2981764, EPI_ISL_2981766, EPI_ISL_2981769, EPI_ISL_2981772, EPI_ISL_2981773, EPI_ISL_2981774, EPI_ISL_2981776, EPI_ISL_2981779, EPI_ISL_2981780, EPI_ISL_2981781, EPI_ISL_2981782, EPI_ISL_2981784, EPI_ISL_2981786, EPI_ISL_2981788, EPI_ISL_2981789, EPI_ISL_2981790, EPI_ISL_2981791, EPI_ISL_2981793, EPI_ISL_2981795, EPI_ISL_2981796, EPI_ISL_2981797, EPI_ISL_2981798, EPI_ISL_2981800, EPI_ISL_2981802, EPI_ISL_2981804, EPI_ISL_2981805, EPI_ISL_2981806, EPI_ISL_2981808, EPI_ISL_2981810, EPI_ISL_2981811, EPI_ISL_2981812, EPI_ISL_2981813, EPI_ISL_2981814, EPI_ISL_2981815, EPI_ISL_2981816, EPI_ISL_2981817, EPI_ISL_2981820, EPI_ISL_2981821, EPI_ISL_2981822, EPI_ISL_2981823, EPI_ISL_2981824, EPI_ISL_2981825, EPI_ISL_2981828, EPI_ISL_2981832, EPI_ISL_2981835, EPI_ISL_2981837, EPI_ISL_2981839, EPI_ISL_2981843, EPI_ISL_2981844, EPI_ISL_2981845, EPI_ISL_2981847, EPI_ISL_2981848, EPI_ISL_2981861, EPI_ISL_2981864, EPI_ISL_2981867, EPI_ISL_2981872, EPI_ISL_2981880, EPI_ISL_2981892 | | | |
| see above | Dutch COVID-19 response team | National Institute for Public Health and the Environment (RIVM) | Adam Meijer, Harry Vennema, Dirk Eggink, Jeroen Cremer, Sharon van den Brink, Bas van der Veer, AnneMarie van den Brandt, Lisa Wijsman, Kim Freriks, Ryanne Jaarsma, Euníce Then, Lynn Aarts, Sanne Bos, Melissa van Tuil, Florian Zwagemaker, Dennis Schmitz, Annelies Kroneman, Karim Hajji, Chantal Reusken, on behalf of the national COVID-19 response team |
| EPI_ISL_2981906, EPI_ISL_2981907, EPI_ISL_2981908, EPI_ISL_2981909, EPI_ISL_2981910, EPI_ISL_2981911, EPI_ISL_2981912, EPI_ISL_2981913, EPI_ISL_2981914, EPI_ISL_2981915, EPI_ISL_2981916, EPI_ISL_2981917, EPI_ISL_2981918 | | | |
| see above | Arizona State Public Health Laboratory | Arizona State Public Health Laboratory | Trung Huynh, Jessica Escobar, Katherine Fullerton, Nobuko Fukushima, Matthew Contursi, Stacy White, Linda Getsinger, Victor Waddell |
| EPI_ISL_2981920, EPI_ISL_2981921, EPI_ISL_2981922, EPI_ISL_2981923, EPI_ISL_2981924, EPI_ISL_2981925, EPI_ISL_2981926 | University of Wisconsin-Madison AIDS Vaccine Research Laboratories | University of Wisconsin-Madison AIDS Vaccine Research Laboratories | Gage Moreno, Katarina Braun, et al. AIDS Vaccine Research Laboratories |
| EPI_ISL_2981928, EPI_ISL_2981945, EPI_ISL_2981949, EPI_ISL_2981950, EPI_ISL_2981951, EPI_ISL_2981952, EPI_ISL_2981953, EPI_ISL_2981954, EPI_ISL_2981955, EPI_ISL_2981956, EPI_ISL_2981957, EPI_ISL_2981958, EPI_ISL_2981959, EPI_ISL_2981960, EPI_ISL_2981961, EPI_ISL_2981962, EPI_ISL_2981963, EPI_ISL_2981964, EPI_ISL_2981965, EPI_ISL_2981966, EPI_ISL_2981967, EPI_ISL_2981968, EPI_ISL_2981969, EPI_ISL_2981970, EPI_ISL_2981971, EPI_ISL_2981972, EPI_ISL_2981973, EPI_ISL_2981974, EPI_ISL_2981975, EPI_ISL_2981976, EPI_ISL_2981977, EPI_ISL_2981978, EPI_ISL_2981979, EPI_ISL_2981980, EPI_ISL_2981981, EPI_ISL_2981982, EPI_ISL_2981983, EPI_ISL_2981985, EPI_ISL_2981986, EPI_ISL_2981987, EPI_ISL_2981988, EPI_ISL_2981989, EPI_ISL_2981990, EPI_ISL_2981991, EPI_ISL_2981993, EPI_ISL_2981994, EPI_ISL_2981995, EPI_ISL_2981996, EPI_ISL_2981997, EPI_ISL_2981998, EPI_ISL_2981999, EPI_ISL_2982000 | | | |
| see above | University Hospitals of Geneva, Laboratory of Virology | HUG, Laboratory of Virology and the Health2030 Genome Center | Samuel Cordey, Ana Rita Goncalves, Laurent Kaiser, Lorenzo Cerutti, Henri Pegeot, Melyssa Elies, Deborah Penet, Keith Harshman, Ioannis Xenarios, Emmanouil Dermitzakis |
| EPI_ISL_2982011, EPI_ISL_2982013, EPI_ISL_2982014, EPI_ISL_2982015, EPI_ISL_2982016, EPI_ISL_2982017 | University of Wisconsin-Madison AIDS Vaccine Research Laboratories | University of Wisconsin-Madison AIDS Vaccine Research Laboratories | Gage Moreno, Katarina Braun, et al. AIDS Vaccine Research Laboratories |
| EPI_ISL_2982020, EPI_ISL_2982021, EPI_ISL_2982022, EPI_ISL_2982023 | UCSC Genomics Institute | UCSC Genomics Institute | Mark Akeson, Ikenna Anigbogu, Eric Beraut, Isabel Bjork, Molly Cassatt-Johnstone, Terren Chang, Russell Corbett-Detig, Namrita Dhillon, Maximilian Haeussler, David Haussler, Angie Hinrichs, Miten Jain, Joshua Kapp, A. Marm Kilpatrick, Jakob McBroome, Hugh Olsen, Jeremy Sanford, Beth Shapiro, Michael Stone, Bryan Thornlow, Yatish Turakhia, Ciara Wanket |
| EPI_ISL_2982121, EPI_ISL_2982122, EPI_ISL_2982123, EPI_ISL_2982124, EPI_ISL_2982125, EPI_ISL_2982126, EPI_ISL_2982127, EPI_ISL_2982128, EPI_ISL_2982129, EPI_ISL_2982130, EPI_ISL_2982131, EPI_ISL_2982132, EPI_ISL_2982133, EPI_ISL_2982135 | | | |
| see above | Microbiology Department. Complexo Hospitalario Universitario de Vigo | Microbiology Department. Complexo Hospitalario Universitario de Vigo | Alfaya N, Alonso I, Alvarez M, Cabrera JJ, Carballo R, Cores O, Cortizo S, del-Campo V, Martinez L, Mediero G, Perez S, Davina C, Potel C, Regueiro B, Rey S, Vasallo FJ |
| EPI_ISL_2982136, EPI_ISL_2982137, EPI_ISL_2982138, EPI_ISL_2982139, EPI_ISL_2982140, EPI_ISL_2982141, EPI_ISL_2982142, EPI_ISL_2982143, EPI_ISL_2982144, EPI_ISL_2982145, EPI_ISL_2982146, EPI_ISL_2982147, EPI_ISL_2982148, EPI_ISL_2982150, EPI_ISL_2982151, EPI_ISL_2982152, EPI_ISL_2982153, EPI_ISL_2982154 | | | |
| see above | OSPEDALE CIVILE TERAMO - CENTRO TRASFUSIONALE | Istituto Zooprofilattico Sperimentale dell'Abruzzo e Molise "G. Caporale" | Lorusso A, Marcacci M, Di Domenico M, Ancora M, Curini V, Di Lollo Valeria, Mangone I, Rinaldi A, Delli Compagni E, Scialabba S, Caporale M, Di Pasquale A, Cammà C, Puglia I, Calistri P, Savini G |
| EPI_ISL_2982155 | OSPEDALE SAN SALVATORE - U.O.C. Malattie infettive | Istituto Zooprofilattico Sperimentale dell'Abruzzo e Molise "G. Caporale" | Lorusso A, Marcacci M, Di Domenico M, Ancora M, Curini V, Di Lollo Valeria, Mangone I, Rinaldi A, Delli Compagni E, Scialabba S, Caporale M, Di Pasquale A, Cammà C, Puglia I, Calistri P, Savini G |
| EPI_ISL_2982156 | OSPEDALE CIVILE TERAMO - CENTRO TRASFUSIONALE | Istituto Zooprofilattico Sperimentale dell'Abruzzo e Molise "G. | Lorusso A, Marcacci M, Di Domenico M, Ancora M, Curini V, Di Lollo Valeria, Mangone I, Rinaldi A, Delli Compagni E, Scialabba S, Caporale M, Di Pasquale A, Cammà C, Puglia I, Calistri P, Savini G |

| | | | |
|---|---|---|---|
| | | Caporale" | |
| EPI_ISL_2982157, EPI_ISL_2982158, EPI_ISL_2982159 | OSPEDALE SAN SALVATORE - MEDICINA DI LABORATORIO | Istituto Zooprofilattico Sperimentale dell'Abruzzo e Molise "G. Caporale" | Lorusso A, Marcacci M, Di Domenico M, Ancora M, Curini V, Di Lollo Valeria, Mangone I, Rinaldi A, Delli Compagni E, Scialabba S, Caporale M, Di Pasquale A, Cammà C, Puglia I, Calistri P, Savini G |
| EPI_ISL_2982170, EPI_ISL_2982171, EPI_ISL_2982172, EPI_ISL_2982174, EPI_ISL_2982175, EPI_ISL_2982176 | PO AVEZZANO - U.O.C. Laboratorio Analisi | Istituto Zooprofilattico Sperimentale dell'Abruzzo e Molise "G. Caporale" | Lorusso A, Marcacci M, Di Domenico M, Ancora M, Curini V, Di Lollo Valeria, Mangone I, Rinaldi A, Delli Compagni E, Scialabba S, Caporale M, Di Pasquale A, Cammà C, Puglia I, Calistri P, Savini G |
| EPI_ISL_2982177, EPI_ISL_2982178, EPI_ISL_2982179, EPI_ISL_2982180, EPI_ISL_2982181, EPI_ISL_2982182, EPI_ISL_2982183, EPI_ISL_2982184, EPI_ISL_2982185, EPI_ISL_2982186, EPI_ISL_2982188, EPI_ISL_2982189, EPI_ISL_2982190, EPI_ISL_2982194, EPI_ISL_2982196, EPI_ISL_2982197, EPI_ISL_2982198, EPI_ISL_2982199, EPI_ISL_2982200, EPI_ISL_2982201, EPI_ISL_2982203, EPI_ISL_2982204, EPI_ISL_2982205, EPI_ISL_2982207, EPI_ISL_2982208, EPI_ISL_2982209 | | | |
| see above | OSPEDALE CIVILE TERAMO - CENTRO TRASFUSIONALE | Istituto Zooprofilattico Sperimentale dell'Abruzzo e Molise "G. Caporale" | Lorusso A, Marcacci M, Di Domenico M, Ancora M, Curini V, Di Lollo Valeria, Mangone I, Rinaldi A, Delli Compagni E, Scialabba S, Caporale M, Di Pasquale A, Cammà C, Puglia I, Calistri P, Savini G |
| EPI_ISL_2982210, EPI_ISL_2982211 | OSPEDALE SAN SALVATORE - MEDICINA DI LABORATORIO | Istituto Zooprofilattico Sperimentale dell'Abruzzo e Molise "G. Caporale" | Lorusso A, Marcacci M, Di Domenico M, Ancora M, Curini V, Di Lollo Valeria, Mangone I, Rinaldi A, Delli Compagni E, Scialabba S, Caporale M, Di Pasquale A, Cammà C, Puglia I, Calistri P, Savini G |
| EPI_ISL_2982215 | PO AVEZZANO - U.O.C. Laboratorio Analisi | Istituto Zooprofilattico Sperimentale dell'Abruzzo e Molise "G. Caporale" | Lorusso A, Marcacci M, Di Domenico M, Ancora M, Curini V, Di Lollo Valeria, Mangone I, Rinaldi A, Delli Compagni E, Scialabba S, Caporale M, Di Pasquale A, Cammà C, Puglia I, Calistri P, Savini G |
| EPI_ISL_2982216, EPI_ISL_2982217, EPI_ISL_2982218, EPI_ISL_2982219, EPI_ISL_2982220, EPI_ISL_2982221, EPI_ISL_2982222, EPI_ISL_2982223, EPI_ISL_2982225 | OSPEDALE CIVILE TERAMO - CENTRO TRASFUSIONALE | Istituto Zooprofilattico Sperimentale dell'Abruzzo e Molise "G. Caporale" | Lorusso A, Marcacci M, Di Domenico M, Ancora M, Curini V, Di Lollo Valeria, Mangone I, Rinaldi A, Delli Compagni E, Scialabba S, Caporale M, Di Pasquale A, Cammà C, Puglia I, Calistri P, Savini G |
| EPI_ISL_2982227, EPI_ISL_2982228, EPI_ISL_2982229, EPI_ISL_2982230, EPI_ISL_2982231, EPI_ISL_2982232, EPI_ISL_2982233, EPI_ISL_2982234, EPI_ISL_2982235, EPI_ISL_2982236, EPI_ISL_2982237, EPI_ISL_2982238, EPI_ISL_2982239, EPI_ISL_2982240, EPI_ISL_2982241, EPI_ISL_2982242, EPI_ISL_2982243, EPI_ISL_2982244, EPI_ISL_2982245, EPI_ISL_2982246, EPI_ISL_2982247, EPI_ISL_2982248, EPI_ISL_2982249, EPI_ISL_2982250, EPI_ISL_2982251, EPI_ISL_2982252, EPI_ISL_2982253, EPI_ISL_2982254, EPI_ISL_2982255, EPI_ISL_2982256 | | | |
| see above | National Public Health Laboratory, National Centre for Infectious Diseases | National Public Health Laboratory, National Centre for Infectious Diseases | Tze Minn Mak, Katherine Ching, Zhenyang Zhou, Royce Ang, Lin Cui, Raymond Tzer Pin Lin |
| EPI_ISL_2982260, EPI_ISL_2982262 | Heilig hart Lier | Imelda Ziekenhuis | Johan Frans, Hanne Valgaeren, Dagmar Obbels |
| EPI_ISL_2982263 | AZ Jan Portaels | Imelda Ziekenhuis | Johan Frans, Hanne Valgaeren, Dagmar Obbels |
| EPI_ISL_2982264, EPI_ISL_2982266 | Imelda Ziekenhuis | Imelda Ziekenhuis | Johan Frans, Hanne Valgaeren, Dagmar Obbels |
| EPI_ISL_2982268 | Salzkammergutklinikum Vöcklabruck, Institut für Pathologie | Salzkammergutklinikum Vöcklabruck, Institut für Pathologie | Regina Stitz, Penka Lechner, Senka Rohregger, René Silye |
| EPI_ISL_2982269, EPI_ISL_2982271, EPI_ISL_2982272, EPI_ISL_2982274, EPI_ISL_2982276, EPI_ISL_2982277, EPI_ISL_2982278, EPI_ISL_2982279, EPI_ISL_2982280, EPI_ISL_2982281, EPI_ISL_2982282, EPI_ISL_2982283, EPI_ISL_2982284, EPI_ISL_2982285, EPI_ISL_2982286, EPI_ISL_2982287, EPI_ISL_2982288, EPI_ISL_2982289, EPI_ISL_2982290, EPI_ISL_2982291, EPI_ISL_2982292, EPI_ISL_2982293, EPI_ISL_2982294, EPI_ISL_2982295, EPI_ISL_2982296, EPI_ISL_2982297, EPI_ISL_2982298, EPI_ISL_2982299, EPI_ISL_2982300, EPI_ISL_2982301, EPI_ISL_2982302 | | | |
| see above | Dept. of Microbiology and Infection Control, Akershus University Hospital HF | Dept. of Microbiology and Infection Control, Akershus University Hospital HF | Hege Vangstein Aamot, Alexander Hesselberg Løvestad |
| EPI_ISL_2982303 | PathWest Laboratory Medicine WA | PathWest Laboratory Medicine WA Microbial Surveillance Unit | PathWest Laboratory Medicine WA Microbial Surveillance Unit |
| EPI_ISL_2982306, EPI_ISL_2982307, EPI_ISL_2982308, EPI_ISL_2982310, EPI_ISL_2982311, EPI_ISL_2982312, EPI_ISL_2982313, EPI_ISL_2982314, EPI_ISL_2982315, EPI_ISL_2982316, EPI_ISL_2982318, EPI_ISL_2982319, EPI_ISL_2982320, EPI_ISL_2982321, EPI_ISL_2982322, EPI_ISL_2982323, EPI_ISL_2982324, EPI_ISL_2982325, EPI_ISL_2982326, EPI_ISL_2982327, EPI_ISL_2982328, EPI_ISL_2982329, EPI_ISL_2982330, EPI_ISL_2982332, EPI_ISL_2982334, EPI_ISL_2982335, EPI_ISL_2982336, EPI_ISL_2982337, EPI_ISL_2982338, EPI_ISL_2982339, EPI_ISL_2982340, EPI_ISL_2982341, EPI_ISL_2982342, EPI_ISL_2982343, EPI_ISL_2982344, EPI_ISL_2982345, EPI_ISL_2982346, EPI_ISL_2982347, EPI_ISL_2982348, EPI_ISL_2982349, EPI_ISL_2982350, EPI_ISL_2982351, EPI_ISL_2982352, EPI_ISL_2982353, EPI_ISL_2982354, EPI_ISL_2982355, EPI_ISL_2982356, EPI_ISL_2982357, EPI_ISL_2982358, EPI_ISL_2982359, EPI_ISL_2982360, EPI_ISL_2982361, EPI_ISL_2982362, EPI_ISL_2982363, EPI_ISL_2982364, EPI_ISL_2982365, EPI_ISL_2982366, EPI_ISL_2982367, EPI_ISL_2982368, EPI_ISL_2982369, EPI_ISL_2982370, EPI_ISL_2982371, EPI_ISL_2982372, EPI_ISL_2982373, EPI_ISL_2982374, EPI_ISL_2982375, EPI_ISL_2982376, EPI_ISL_2982377, EPI_ISL_2982378, EPI_ISL_2982379, EPI_ISL_2982380, EPI_ISL_2982381, EPI_ISL_2982382, EPI_ISL_2982383, EPI_ISL_2982384, EPI_ISL_2982385, EPI_ISL_2982386, EPI_ISL_2982388, EPI_ISL_2982389, EPI_ISL_2982390, EPI_ISL_2982391, EPI_ISL_2982392, EPI_ISL_2982393, EPI_ISL_2982394, EPI_ISL_2982395, EPI_ISL_2982396, EPI_ISL_2982397, EPI_ISL_2982398, EPI_ISL_2982399, EPI_ISL_2982400, EPI_ISL_2982401, EPI_ISL_2982402, EPI_ISL_2982403, EPI_ISL_2982404, EPI_ISL_2982405, EPI_ISL_2982406, EPI_ISL_2982407, EPI_ISL_2982408, EPI_ISL_2982409, EPI_ISL_2982410, EPI_ISL_2982411, EPI_ISL_2982412, EPI_ISL_2982413, EPI_ISL_2982414, EPI_ISL_2982415, EPI_ISL_2982416, EPI_ISL_2982417, EPI_ISL_2982419 | | | |
| see above | Hospital Universitari Vall d'Hebron - Vall d'Hebron Institut de Recerca | Hospital Universitari Vall d'Hebron - Vall d'Hebron Institut de Recerca | Cristina Andrés, Maria Piñana, Alejandra González-Sánchez, Damir Garcia-Cehic, Ariadna Rando, Juliana Esperalba, Maria Gema Codina, Carla Castillo, Karen García, Rodrigo Vásquez, Maria Carmen Martin, Tomàs Pumarola, Josep Quer, Andrés Antón |
| EPI_ISL_2982423, EPI_ISL_2982424, EPI_ISL_2982425, EPI_ISL_2982426, EPI_ISL_2982427, EPI_ISL_2982428, EPI_ISL_2982429, EPI_ISL_2982430, EPI_ISL_2982431, EPI_ISL_2982432, EPI_ISL_2982433, EPI_ISL_2982434, EPI_ISL_2982435, EPI_ISL_2982436, EPI_ISL_2982437, EPI_ISL_2982438, EPI_ISL_2982439, EPI_ISL_2982440, EPI_ISL_2982441, EPI_ISL_2982442, EPI_ISL_2982443, EPI_ISL_2982444, EPI_ISL_2982445, EPI_ISL_2982446 | | | |
| see above | Hospital Universitari Arnau de Vilanova | Hospital Universitari Vall d'Hebron - Vall d'Hebron Institut de Recerca | Cristina Andrés, Maria Piñana, Alejandra González-Sánchez, Damir Garcia-Cehic, Ariadna Rando, Juliana Esperalba, Maria Gema Codina, Carla Castillo, Karen García, Rodrigo Vásquez, Maria Carmen Martin, Tomàs Pumarola, Josep Quer, Andrés Antón |
| EPI_ISL_2982447, EPI_ISL_2982448, EPI_ISL_2982449 | Department of Microbiology, University Hospital in Pilsen | Department of Microbiology, University Hospital in Pilsen | Jaroslav Hrabak |
| EPI_ISL_2982453, EPI_ISL_2982455, EPI_ISL_2982457, EPI_ISL_2982459, EPI_ISL_2982460, EPI_ISL_2982461, EPI_ISL_2982462 | IN State Department of Health Laboratory Services | IN State Department of Health Laboratory Services | Cassandra Campion, Jamie Yeadon, Brian Pope, Lixia Liu, Kyle Brownlee, Melissa Hindenlang, Mark Glazier |
| EPI_ISL_2982468, EPI_ISL_2982469, EPI_ISL_2982470 | ADPH | ADPH | Adrianna Walker |
| EPI_ISL_2982629, EPI_ISL_2982630, EPI_ISL_2982631, EPI_ISL_2982632 | UW Virology Lab | UW Virology Lab | Pavitra Roychoudhury, Hong Xie, Lasata Shrestha, Shah Mohamed Bakhash, Tien V. Nguyen, Noah R. Baker, Sean Ellis, Meei-Li Huang, Keith R Jerome, Alexander Greninger |
| EPI_ISL_2982633 | UCSC Genomics Institute | UCSC Genomics Institute | Mark Akeson, Ikenna Anigbogu, Eric Beraut, Isabel Bjork, Molly Cassatt-Johnstone, Terren Chang, Russell Corbett-Detig, Namrita Dhillon, Maximilian Haeussler, David Haussler, Angie Hinrichs, Miten Jain, Joshua Kapp, A. Marm Kilpatrick, Jakob McBroome, Hugh Olsen, Jeremy Sanford, Beth Shapiro, Michael Stone, Bryan Thornlow, Yatish Turakhia, Ciara Wanket |
| EPI_ISL_2982634, EPI_ISL_2982635, EPI_ISL_2982637, EPI_ISL_2982638, EPI_ISL_2982642, EPI_ISL_2982643, EPI_ISL_2982644, EPI_ISL_2982645, EPI_ISL_2982648, EPI_ISL_2982649, EPI_ISL_2982650, EPI_ISL_2982651, EPI_ISL_2982652, EPI_ISL_2982655, EPI_ISL_2982658, EPI_ISL_2982661, EPI_ISL_2982662, EPI_ISL_2982664, EPI_ISL_2982666, EPI_ISL_2982667, EPI_ISL_2982669, EPI_ISL_2982674, EPI_ISL_2982675, EPI_ISL_2982683, EPI_ISL_2982684, EPI_ISL_2982685, EPI_ISL_2982687, EPI_ISL_2982689, EPI_ISL_2982690, EPI_ISL_2982691 | | | |
| see above | Virginia Division of Consolidated Laboratory Services | Virginia Division of Consolidated Laboratory Services | Virginia Division of Consolidated Laboratory Services |
| EPI_ISL_2982693, EPI_ISL_2982694, EPI_ISL_2982695, EPI_ISL_2982697, EPI_ISL_2982698, EPI_ISL_2982699, EPI_ISL_2982702, EPI_ISL_2982703, EPI_ISL_2982704, EPI_ISL_2982705, EPI_ISL_2982706, EPI_ISL_2982710, EPI_ISL_2982713, EPI_ISL_2982714, EPI_ISL_2982715 | | | |
| see above | Montana Public Health Laboratory | Montana Public Health Laboratory | Joy Ritter, Michelle Mozer, Carrie Biskupiak, Deborah Gibson, Michael Dills |
| EPI_ISL_2982735 | Laboratory of Respiratory Viruses and Measles, Oswaldo Cruz Institute, FIOCRUZ | Laboratory of Respiratory Viruses and Measles, Oswaldo Cruz Institute, FIOCRUZ | Paola Resende, Luciana Appolinario, Fernando Motta, Anna Carolina Paixao, Ana Carolina Mendonca, Ighor Leonardo Arantes Gomes, Agatha Cristinne Prudencio, Alice Sampaio Rocha, Taina Venas, Elisa Cavalcante Pereira, Renata Serrano Lopes, Marilda Siqueira on behalf of the Fiocruz COVID-19 Genomic Surveillance Network |
| EPI_ISL_2982743 | Laboratorio Central de Saude Publica do Estado do Rio de Janeiro (LACEN-RJ) | Laboratory of Respiratory Viruses and Measles, Oswaldo Cruz Institute, FIOCRUZ | Paola Resende, Luciana Appolinario, Fernando Motta, Anna Carolina Paixao, Ighor Leonardo Arantes Gomes, Agatha Cristinne Prudencio, Ana Carolina Mendonca, Alice Sampaio Rocha, Taina Venas, Elisa Cavalcante Pereira, Renata Serrano Lopes, Andrea Cony Cavalcanti, Marilda Siqueira on behalf of the Fiocruz COVID-19 Genomic Surveillance Network |
| EPI_ISL_2982914, EPI_ISL_2982921, EPI_ISL_2982925 | National Laboratory for Health, Environment and Food, OMM, Maribor | NLZOH (National Laboratory for Health, Environment and Food) / CISLD (Clinical Institute of Special Laboratory Diagnostics), University Children's Hospital, University Medical Center Ljubljana | Sandra Janezic, Aleksander Mahnic, Maja Rupnik, Tjaša Žohar Čretnik, Alenka Štorman, Nika Gobec, Aleksander Kocuvan, Kaja Tominc, Maša Jarčič, Mojca Cimerman, Nika Volmajer, Andrej Golle / Jernej Kovač, Barbara Jenko Bizjan, Tine Tesovnik, Robert Šket, Katarina Kozmos, Ana Grom, Maruša Debeljak, Marko Pokorn, Tadej Battelino |
| EPI_ISL_2982940, EPI_ISL_2982941, EPI_ISL_2982944, EPI_ISL_2982956, EPI_ISL_2982959, EPI_ISL_2982960 | National Laboratory for Health, Environment and Food, OMM, Kranj | NLZOH (National Laboratory for Health, Environment and Food) / CISLD (Clinical Institute of Special Laboratory Diagnostics), University Children's Hospital, University Medical Center Ljubljana | Sandra Janezic, Aleksander Mahnic, Maja Rupnik, Tjaša Žohar Čretnik, Alenka Štorman, Nika Gobec, Aleksander Kocuvan, Kaja Tominc, Maša Jarčič, Mateja Ravnik, Monika Korošec, Marjana Petrevčič / Jernej Kovač, Barbara Jenko Bizjan, Tine Tesovnik, Robert Šket, Katarina Kozmos, Ana Grom, Maruša Debeljak, Marko Pokorn, Tadej Battelino |
| EPI_ISL_2982971, EPI_ISL_2982972, EPI_ISL_2982976, EPI_ISL_2982992 | National Laboratory for Health, Environment and Food, OMM, Celje | NLZOH (National Laboratory for Health, Environment and Food) / CISLD (Clinical Institute of Special Laboratory Diagnostics), University Children's Hospital, University Medical Center Ljubljana | Sandra Janezic, Aleksander Mahnic, Maja Rupnik, Tjaša Žohar Čretnik, Alenka Štorman, Nika Gobec, Aleksander Kocuvan, Kaja Tominc, Maša Jarčič, Mateja Borinc, Daša Kavka / Jernej Kovač, Barbara Jenko Bizjan, Tine Tesovnik, Robert Šket, Katarina Kozmos, Ana Grom, Maruša Debeljak, Marko Pokorn, Tadej Battelino |
| EPI_ISL_2983000, | National Laboratory for Health, Environment and Food, OMM, | NLZOH (National Laboratory for Health, Environment and Food) / | Sandra Janezic, Aleksander Mahnic, Maja Rupnik, Tjaša Žohar Čretnik, Alenka Štorman, Nika Gobec, Aleksander Kocuvan, Kaja Tominc, Maša Jarčič, Gašper Strugar, Tina Cvetković, Mitja Rak / Jernej Kovač, Barbara Jenko Bizjan, Tine Tesovnik, |

| | | | |
|---|---|---|---|
| EPI_ISL_2983002, EPI_ISL_2983014, EPI_ISL_2983015, EPI_ISL_2983016, EPI_ISL_2983017, EPI_ISL_2983018, EPI_ISL_2983022, EPI_ISL_2983026, EPI_ISL_2983028 | Koper | CISLD (Clinical Institute of Special Laboratory Diagnostics), University Children's Hospital, University Medical Center Ljubljana | Robert Šket, Katarina Kozmos, Ana Grom, Maruša Debeljak, Marko Pokorn, Tadej Battelino |
| EPI_ISL_2983060, EPI_ISL_2983062 | National Laboratory for Health, Environment and Food, OMM, Novo mesto | NLZOH (National Laboratory for Health, Environment and Food) / CISLD (Clinical Institute of Special Laboratory Diagnostics), University Children's Hospital, University Medical Center Ljubljana | Sandra Janezic, Aleksander Mahnic, Maja Rupnik, Tjaša Žohar Čretnik, Alenka Štorman, Nika Gobec, Aleksander Kocuvan, Kaja Tominc, Maša Jarčič, David Cvetko, Tatjana Harlander, Matjaž Retelj / Jernej Kovač, Barbara Jenko Bizjan, Tine Tesovnik, Robert Šket, Katarina Kozmos, Ana Grom, Maruša Debeljak, Marko Pokorn, Tadej Battelino |
| EPI_ISL_2983219, EPI_ISL_2983220 | IN State Department of Health Laboratory Services | IN State Department of Health Laboratory Services | Cassandra Campion, Jamie Yeadon, Brian Pope, Lixia Liu, Kyle Brownlee, Melissa Hindenlang, Mark Glazier |
| EPI_ISL_2983221, EPI_ISL_2983222, EPI_ISL_2983223, EPI_ISL_2983224 | DNA Labs | National Reference Laboratory, Nigeria Centre for Disease Control | Dr Ndodo Nnaemeka, Olusola Anuoluwapo Akanbi, Chimaobi Chukwu, Dr Omoare Adesuyi, Anthony Ahumibe, Catherine Okoi, Nwando Mba, Dr Chikwe Ihekweazu |
| EPI_ISL_2983373 | Laboratorio Central de Saude Publica do Estado do Parana (LACEN/PR) | Laboratory of Respiratory Viruses and Measles, Oswaldo Cruz Institute, FIOCRUZ | Paola Resende, Luciana Appolinario, Fernando Motta, Anna Carolina Paixao, Ana Carolina Mendonca, Alice Sampaio Rocha, Taina Venas, Elisa Cavalcante Pereira, Renata Serrano Lopes, Irina Riediger, Marilda Siqueira on behalf of the Fiocruz COVID-19 Genomic Surveillance Network |
| EPI_ISL_2983377, EPI_ISL_2983378 | IRCCS San Gallicano Dermatological Institute | IRCCS Regina Elena National Cancer Institute | Fulvia Pimpinelli, Martina Pontone, Elena Papa, Francesco Marchesi |
| EPI_ISL_2983452 | Laboratorio Central de Saude Publica do Estado do Parana (LACEN/PR) | Laboratory of Respiratory Viruses and Measles, Oswaldo Cruz Institute, FIOCRUZ | Paola Resende, Luciana Appolinario, Fernando Motta, Anna Carolina Paixao, Ana Carolina Mendonca, Alice Sampaio Rocha, Taina Venas, Elisa Cavalcante Pereira, Renata Serrano Lopes, Irina Riediger, Marilda Siqueira on behalf of the Fiocruz COVID-19 Genomic Surveillance Network |
| EPI_ISL_2983482, EPI_ISL_2983483, EPI_ISL_2983484, EPI_ISL_2983485, EPI_ISL_2983486, EPI_ISL_2983488, EPI_ISL_2983491, EPI_ISL_2983492 | TXDSHS | TXDSHS | Rashmi Tuladhar, Bonnie Oh, Jenny Zhang, Maliha Rahman, Mayela Pedrueza, Anita Pokharel, Karen Bobier, Lorraine Rodriguez, Myong Koag, Chun Wang, Rachel Lee, Grace Kubin |
| EPI_ISL_2983494, EPI_ISL_2983495, EPI_ISL_2983496, EPI_ISL_2983497, EPI_ISL_2983498, EPI_ISL_2983499, EPI_ISL_2983500, EPI_ISL_2983501, EPI_ISL_2983502, EPI_ISL_2983503, EPI_ISL_2983504, EPI_ISL_2983505, EPI_ISL_2983506, EPI_ISL_2983507, EPI_ISL_2983508, EPI_ISL_2983509, EPI_ISL_2983510, EPI_ISL_2983511 | | | |
| see above | Kansas Health and Environmental Lab | Kansas Health and Environmental Lab | Mike Grose, Katherine Wiggins, Jonathan Barnell, Ben Olsen, and Phil Adam |
| EPI_ISL_2983512, EPI_ISL_2983513, EPI_ISL_2983514, EPI_ISL_2983515, EPI_ISL_2983516 | Wichita State University - Molecular Diagnostics Lab | Kansas Health and Environmental Lab | Mike Grose, Katherine Wiggins, Jonathan Barnell, Ben Olsen, and Phil Adam |
| EPI_ISL_2983517, EPI_ISL_2983518, EPI_ISL_2983519, EPI_ISL_2983520, EPI_ISL_2983521, EPI_ISL_2983522 | Kansas Health and Environmental Lab | Kansas Health and Environmental Lab | Mike Grose, Katherine Wiggins, Jonathan Barnell, Ben Olsen, and Phil Adam |
| EPI_ISL_2983557, EPI_ISL_2983558, EPI_ISL_2983559, EPI_ISL_2983560, EPI_ISL_2983561, EPI_ISL_2983562, EPI_ISL_2983563, EPI_ISL_2983564, EPI_ISL_2983565, EPI_ISL_2983566, EPI_ISL_2983567, EPI_ISL_2983568, EPI_ISL_2983569, EPI_ISL_2983570, EPI_ISL_2983571, EPI_ISL_2983572, EPI_ISL_2983573, EPI_ISL_2983574, EPI_ISL_2983575, EPI_ISL_2983576, EPI_ISL_2983577, EPI_ISL_2983578, EPI_ISL_2983579, EPI_ISL_2983580, EPI_ISL_2983581 | | | |
| see above | Pathology and Laboratory Medicine Institute, Cleveland Clinic, Ohio, USA | Pathology and Laboratory Medicine Institute, Cleveland Clinic, Ohio, USA | Concetta Peck, Jennifer Starbuck, Joy Nakitandwe, Kristen McDonnell, David Plunkett, Zheng Jin Tu, Jay Brock, Yu-Wei Cheng, Gary Procop, Daniel Rhoads, Daniel H. Farkas, David Bosler |
| EPI_ISL_2983582, EPI_ISL_2983583, EPI_ISL_2983585, EPI_ISL_2983586, EPI_ISL_2983587, EPI_ISL_2983588, EPI_ISL_2983589, EPI_ISL_2983590, EPI_ISL_2983591, EPI_ISL_2983592, EPI_ISL_2983593, EPI_ISL_2983595, EPI_ISL_2983596, EPI_ISL_2983597, EPI_ISL_2983598, EPI_ISL_2983599, EPI_ISL_2983600, EPI_ISL_2983601, EPI_ISL_2983602, EPI_ISL_2983603, EPI_ISL_2983604, EPI_ISL_2983605, EPI_ISL_2983606, EPI_ISL_2983607, EPI_ISL_2983608, EPI_ISL_2983609, EPI_ISL_2983610, EPI_ISL_2983611, EPI_ISL_2983612, EPI_ISL_2983613 | | | |
| see above | Wichita State University - Molecular Diagnostics Lab | Kansas Health and Environmental Lab | Mike Grose, Katherine Wiggins, Jonathan Barnell, Ben Olsen, and Phil Adam |
| EPI_ISL_2983615, EPI_ISL_2983616, EPI_ISL_2983617, EPI_ISL_2983618, EPI_ISL_2983619, EPI_ISL_2983620, EPI_ISL_2983621, EPI_ISL_2983622, EPI_ISL_2983623, EPI_ISL_2983624, EPI_ISL_2983625, EPI_ISL_2983626, EPI_ISL_2983627, EPI_ISL_2983628, EPI_ISL_2983633, EPI_ISL_2983635, EPI_ISL_2983636, EPI_ISL_2983637, EPI_ISL_2983638, EPI_ISL_2983640, EPI_ISL_2983641, EPI_ISL_2983643, EPI_ISL_2983644, EPI_ISL_2983645, EPI_ISL_2983646, EPI_ISL_2983647, EPI_ISL_2983648, EPI_ISL_2983649, EPI_ISL_2983650, EPI_ISL_2983651, EPI_ISL_2983652, EPI_ISL_2983654, EPI_ISL_2983655, EPI_ISL_2983656, EPI_ISL_2983657, EPI_ISL_2983658, EPI_ISL_2983660, EPI_ISL_2983661, EPI_ISL_2983663, EPI_ISL_2983664, EPI_ISL_2983665, EPI_ISL_2983666, EPI_ISL_2983667, EPI_ISL_2983668, EPI_ISL_2983669, EPI_ISL_2983670, EPI_ISL_2983671, EPI_ISL_2983672, EPI_ISL_2983673, EPI_ISL_2983674, EPI_ISL_2983675, EPI_ISL_2983676, EPI_ISL_2983677, EPI_ISL_2983678, EPI_ISL_2983679, EPI_ISL_2983681, EPI_ISL_2983683, EPI_ISL_2983684, EPI_ISL_2983686, EPI_ISL_2983687, EPI_ISL_2983688, EPI_ISL_2983689, EPI_ISL_2983690, EPI_ISL_2983691, EPI_ISL_2983692, EPI_ISL_2983693, EPI_ISL_2983696, EPI_ISL_2983697, EPI_ISL_2983698, EPI_ISL_2983699, EPI_ISL_2983701, EPI_ISL_2983702, EPI_ISL_2983703, EPI_ISL_2983704 | | | |
| see above | Illinois Department of Public Health - Springfield Lab | Illinois Department of Public Health - Springfield Lab | Bryan Sim, Gordon McCall |
| EPI_ISL_2983729, EPI_ISL_2983730, EPI_ISL_2983731, EPI_ISL_2983734, EPI_ISL_2983735, EPI_ISL_2983736, EPI_ISL_2983738, EPI_ISL_2983740, EPI_ISL_2983741, EPI_ISL_2983743, EPI_ISL_2983744, EPI_ISL_2983746, EPI_ISL_2983748, EPI_ISL_2983749, EPI_ISL_2983750, EPI_ISL_2983751, EPI_ISL_2983752, EPI_ISL_2983753, EPI_ISL_2983754, EPI_ISL_2983755, EPI_ISL_2983756, EPI_ISL_2983757, EPI_ISL_2983758, EPI_ISL_2983760, EPI_ISL_2983761, EPI_ISL_2983762, EPI_ISL_2983763, EPI_ISL_2983764, EPI_ISL_2983765, EPI_ISL_2983766, EPI_ISL_2983767, EPI_ISL_2983769, EPI_ISL_2983770, EPI_ISL_2983771, EPI_ISL_2983772, EPI_ISL_2983773 | | | |
| see above | UMC Groningen, Clinical Virology, Department of Medical Microbiology and Infection Prevention | UMC Groningen, Clinical Virology, Department of Medical Microbiology and Infection Prevention | Hubert Niesters, Alexander Friedrich, Erley Lizarazo-Forero, Monika Fliss, Lilli Gard, Sigrid Rosema, Coretta Van Leer-Buter, Xuewei Zhou, Marjolein Knoester |
| EPI_ISL_2983774, EPI_ISL_2983775, EPI_ISL_2983776, EPI_ISL_2983777, EPI_ISL_2983778, EPI_ISL_2983779, EPI_ISL_2983780, EPI_ISL_2983781, EPI_ISL_2983782, EPI_ISL_2983783, EPI_ISL_2983785, EPI_ISL_2983786, EPI_ISL_2983787, EPI_ISL_2983788, EPI_ISL_2983789, EPI_ISL_2983790, EPI_ISL_2983791, EPI_ISL_2983792, EPI_ISL_2983793, EPI_ISL_2983794, EPI_ISL_2983795, EPI_ISL_2983796, EPI_ISL_2983797, EPI_ISL_2983798, EPI_ISL_2983799, EPI_ISL_2983800, EPI_ISL_2983801, EPI_ISL_2983802, EPI_ISL_2983804, EPI_ISL_2983805, EPI_ISL_2983806, EPI_ISL_2983807, EPI_ISL_2983808, EPI_ISL_2983809 | | | |
| see above | IN State Department of Health Laboratory Services | IN State Department of Health Laboratory Services | Cassandra Campion, Jamie Yeadon, Brian Pope, Lixia Liu, Kyle Brownlee, Melissa Hindenlang, Mark Glazier |
| EPI_ISL_2983812, EPI_ISL_2983814, EPI_ISL_2983815 | SELAS MEDILYS | Department of Virology, Henri Mondor University Hospital, Assistance Publique Hôpitaux de Paris, Université Paris-Est Créteil, INSERM U955 | Christophe Rodriguez, Slim Fourati, Vanessa Demontant, Guillaume Gricourt, Melissa N'Debi, Alexandre Soulier, Elisabeth Trawinski, Jean-Michel Pawlotsky |
| EPI_ISL_2983817 | GH de l'Est Francilien | Department of Virology, Henri Mondor University Hospital, Assistance Publique Hôpitaux de Paris, Université Paris-Est Créteil, INSERM U955 | Christophe Rodriguez, Slim Fourati, Vanessa Demontant, Guillaume Gricourt, Melissa N'Debi, Alexandre Soulier, Elisabeth Trawinski, Jean-Michel Pawlotsky |
| EPI_ISL_2983818 | BIO67-BIOSPHERE | Department of Virology, Henri Mondor University Hospital, Assistance Publique Hôpitaux de Paris, Université Paris-Est Créteil, INSERM U955 | Christophe Rodriguez, Slim Fourati, Vanessa Demontant, Guillaume Gricourt, Melissa N'Debi, Alexandre Soulier, Elisabeth Trawinski, Jean-Michel Pawlotsky |
| EPI_ISL_2983820 | Hôpital Saint Antoine | Department of Virology, Henri Mondor University Hospital, Assistance Publique Hôpitaux de Paris, Université Paris-Est Créteil, INSERM U955 | Christophe Rodriguez, Slim Fourati, Vanessa Demontant, Guillaume Gricourt, Melissa N'Debi, Alexandre Soulier, Elisabeth Trawinski, Jean-Michel Pawlotsky |
| EPI_ISL_2983821 | Laboratoire multi-site du Diaconat | Department of Virology, Henri Mondor University Hospital, Assistance Publique Hôpitaux de Paris, Université Paris-Est Créteil, INSERM U955 | Christophe Rodriguez, Slim Fourati, Vanessa Demontant, Guillaume Gricourt, Melissa N'Debi, Alexandre Soulier, Elisabeth Trawinski, Jean-Michel Pawlotsky |
| EPI_ISL_2983822, EPI_ISL_2983823 | BIO67-BIOSPHERE | Department of Virology, Henri Mondor University Hospital, Assistance Publique Hôpitaux de Paris, Université Paris-Est Créteil, INSERM U955 | Christophe Rodriguez, Slim Fourati, Vanessa Demontant, Guillaume Gricourt, Melissa N'Debi, Alexandre Soulier, Elisabeth Trawinski, Jean-Michel Pawlotsky |
| EPI_ISL_2983824 | Hôpital Avicenne | Department of Virology, Henri Mondor University Hospital, Assistance Publique Hôpitaux de Paris, Université Paris-Est Créteil, INSERM U955 | Christophe Rodriguez, Slim Fourati, Vanessa Demontant, Guillaume Gricourt, Melissa N'Debi, Alexandre Soulier, Elisabeth Trawinski, Jean-Michel Pawlotsky |
| EPI_ISL_2983825 | SELAS MEDILYS | Department of Virology, Henri Mondor University Hospital, Assistance Publique Hôpitaux de Paris, Université Paris-Est Créteil, INSERM U955 | Christophe Rodriguez, Slim Fourati, Vanessa Demontant, Guillaume Gricourt, Melissa N'Debi, Alexandre Soulier, Elisabeth Trawinski, Jean-Michel Pawlotsky |
| EPI_ISL_2983826 | GH de l'Est Francilien | Department of Virology, Henri Mondor University Hospital, Assistance Publique Hôpitaux de Paris, Université Paris-Est Créteil, INSERM U955 | Christophe Rodriguez, Slim Fourati, Vanessa Demontant, Guillaume Gricourt, Melissa N'Debi, Alexandre Soulier, Elisabeth Trawinski, Jean-Michel Pawlotsky |
| EPI_ISL_2983827 | BIO67-BIOSPHERE | Department of Virology, Henri Mondor University Hospital, Assistance Publique Hôpitaux de Paris, Université Paris-Est Créteil, INSERM U955 | Christophe Rodriguez, Slim Fourati, Vanessa Demontant, Guillaume Gricourt, Melissa N'Debi, Alexandre Soulier, Elisabeth Trawinski, Jean-Michel Pawlotsky |
| EPI_ISL_2983828 | SELAS MEDILYS | Department of Virology, Henri Mondor University Hospital, Assistance Publique Hôpitaux de Paris, Université Paris-Est Créteil, | Christophe Rodriguez, Slim Fourati, Vanessa Demontant, Guillaume Gricourt, Melissa N'Debi, Alexandre Soulier, Elisabeth Trawinski, Jean-Michel Pawlotsky |

| | | | |
|---|---|---|---|
| | | INSERM U955 | |
| EPI_ISL_2983834 | BROUSSAIS | Department of Virology, Henri Mondor University Hospital, Assistance Publique Hôpitaux de Paris, Université Paris-Est Créteil, INSERM U955 | Christophe Rodriguez, Slim Fourati, Vanessa Demontant, Guillaume Gricourt, Melissa N'Debi, Alexandre Soulier, Elisabeth Trawinski, Jean-Michel Pawlotsky |
| EPI_ISL_2983835, EPI_ISL_2983837, EPI_ISL_2983838 | BIOMNIS EUROFINS IVRY | Department of Virology, Henri Mondor University Hospital, Assistance Publique Hôpitaux de Paris, Université Paris-Est Créteil, INSERM U955 | Christophe Rodriguez, Slim Fourati, Vanessa Demontant, Guillaume Gricourt, Melissa N'Debi, Alexandre Soulier, Elisabeth Trawinski, Jean-Michel Pawlotsky |
| EPI_ISL_2983839 | Hôpital Avicenne | Department of Virology, Henri Mondor University Hospital, Assistance Publique Hôpitaux de Paris, Université Paris-Est Créteil, INSERM U955 | Christophe Rodriguez, Slim Fourati, Vanessa Demontant, Guillaume Gricourt, Melissa N'Debi, Alexandre Soulier, Elisabeth Trawinski, Jean-Michel Pawlotsky |
| EPI_ISL_2983843 | Laboratoire LENYS | Department of Virology, Henri Mondor University Hospital, Assistance Publique Hôpitaux de Paris, Université Paris-Est Créteil, INSERM U955 | Christophe Rodriguez, Slim Fourati, Vanessa Demontant, Guillaume Gricourt, Melissa N'Debi, Alexandre Soulier, Elisabeth Trawinski, Jean-Michel Pawlotsky |
| EPI_ISL_2983845 | SELAS MEDILYS | Department of Virology, Henri Mondor University Hospital, Assistance Publique Hôpitaux de Paris, Université Paris-Est Créteil, INSERM U955 | Christophe Rodriguez, Slim Fourati, Vanessa Demontant, Guillaume Gricourt, Melissa N'Debi, Alexandre Soulier, Elisabeth Trawinski, Jean-Michel Pawlotsky |
| EPI_ISL_2983848 | CH. ANDRE GREGOIRE | Department of Virology, Henri Mondor University Hospital, Assistance Publique Hôpitaux de Paris, Université Paris-Est Créteil, INSERM U955 | Christophe Rodriguez, Slim Fourati, Vanessa Demontant, Guillaume Gricourt, Melissa N'Debi, Alexandre Soulier, Elisabeth Trawinski, Jean-Michel Pawlotsky |
| EPI_ISL_2983850 | SELAS MEDILYS | Department of Virology, Henri Mondor University Hospital, Assistance Publique Hôpitaux de Paris, Université Paris-Est Créteil, INSERM U955 | Christophe Rodriguez, Slim Fourati, Vanessa Demontant, Guillaume Gricourt, Melissa N'Debi, Alexandre Soulier, Elisabeth Trawinski, Jean-Michel Pawlotsky |
| EPI_ISL_2983851 | BIO67-BIOSPHERE | Department of Virology, Henri Mondor University Hospital, Assistance Publique Hôpitaux de Paris, Université Paris-Est Créteil, INSERM U955 | Christophe Rodriguez, Slim Fourati, Vanessa Demontant, Guillaume Gricourt, Melissa N'Debi, Alexandre Soulier, Elisabeth Trawinski, Jean-Michel Pawlotsky |
| EPI_ISL_2983854 | SELAS MEDILYS | Department of Virology, Henri Mondor University Hospital, Assistance Publique Hôpitaux de Paris, Université Paris-Est Créteil, INSERM U955 | Christophe Rodriguez, Slim Fourati, Vanessa Demontant, Guillaume Gricourt, Melissa N'Debi, Alexandre Soulier, Elisabeth Trawinski, Jean-Michel Pawlotsky |
| EPI_ISL_2983857, EPI_ISL_2983858 | BIO67-BIOSPHERE | Department of Virology, Henri Mondor University Hospital, Assistance Publique Hôpitaux de Paris, Université Paris-Est Créteil, INSERM U955 | Christophe Rodriguez, Slim Fourati, Vanessa Demontant, Guillaume Gricourt, Melissa N'Debi, Alexandre Soulier, Elisabeth Trawinski, Jean-Michel Pawlotsky |
| EPI_ISL_2983862 | SELAS MEDILYS | Department of Virology, Henri Mondor University Hospital, Assistance Publique Hôpitaux de Paris, Université Paris-Est Créteil, INSERM U955 | Christophe Rodriguez, Slim Fourati, Vanessa Demontant, Guillaume Gricourt, Melissa N'Debi, Alexandre Soulier, Elisabeth Trawinski, Jean-Michel Pawlotsky |
| EPI_ISL_2983863 | BIO67-BIOSPHERE | Department of Virology, Henri Mondor University Hospital, Assistance Publique Hôpitaux de Paris, Université Paris-Est Créteil, INSERM U955 | Christophe Rodriguez, Slim Fourati, Vanessa Demontant, Guillaume Gricourt, Melissa N'Debi, Alexandre Soulier, Elisabeth Trawinski, Jean-Michel Pawlotsky |
| EPI_ISL_2983864, EPI_ISL_2983865 | SELAS MEDILYS | Department of Virology, Henri Mondor University Hospital, Assistance Publique Hôpitaux de Paris, Université Paris-Est Créteil, INSERM U955 | Christophe Rodriguez, Slim Fourati, Vanessa Demontant, Guillaume Gricourt, Melissa N'Debi, Alexandre Soulier, Elisabeth Trawinski, Jean-Michel Pawlotsky |
| EPI_ISL_2983866 | GH A.CHENEVIER-H.MONDOR | Department of Virology, Henri Mondor University Hospital, Assistance Publique Hôpitaux de Paris, Université Paris-Est Créteil, INSERM U955 | Christophe Rodriguez, Slim Fourati, Vanessa Demontant, Guillaume Gricourt, Melissa N'Debi, Alexandre Soulier, Elisabeth Trawinski, Jean-Michel Pawlotsky |
| EPI_ISL_2983868, EPI_ISL_2983869 | BIOMNIS EUROFINS IVRY | Department of Virology, Henri Mondor University Hospital, Assistance Publique Hôpitaux de Paris, Université Paris-Est Créteil, INSERM U955 | Christophe Rodriguez, Slim Fourati, Vanessa Demontant, Guillaume Gricourt, Melissa N'Debi, Alexandre Soulier, Elisabeth Trawinski, Jean-Michel Pawlotsky |
| EPI_ISL_2983870 | SELAS MEDILYS | Department of Virology, Henri Mondor University Hospital, Assistance Publique Hôpitaux de Paris, Université Paris-Est Créteil, INSERM U955 | Christophe Rodriguez, Slim Fourati, Vanessa Demontant, Guillaume Gricourt, Melissa N'Debi, Alexandre Soulier, Elisabeth Trawinski, Jean-Michel Pawlotsky |
| EPI_ISL_2983873, EPI_ISL_2983874 | BROUSSAIS | Department of Virology, Henri Mondor University Hospital, Assistance Publique Hôpitaux de Paris, Université Paris-Est Créteil, INSERM U955 | Christophe Rodriguez, Slim Fourati, Vanessa Demontant, Guillaume Gricourt, Melissa N'Debi, Alexandre Soulier, Elisabeth Trawinski, Jean-Michel Pawlotsky |
| EPI_ISL_2983878 | BIO ARD'AISNE | Department of Virology, Henri Mondor University Hospital, Assistance Publique Hôpitaux de Paris, Université Paris-Est Créteil, INSERM U955 | Christophe Rodriguez, Slim Fourati, Vanessa Demontant, Guillaume Gricourt, Melissa N'Debi, Alexandre Soulier, Elisabeth Trawinski, Jean-Michel Pawlotsky |
| EPI_ISL_2983879 | SELAS MEDILYS | Department of Virology, Henri Mondor University Hospital, Assistance Publique Hôpitaux de Paris, Université Paris-Est Créteil, INSERM U955 | Christophe Rodriguez, Slim Fourati, Vanessa Demontant, Guillaume Gricourt, Melissa N'Debi, Alexandre Soulier, Elisabeth Trawinski, Jean-Michel Pawlotsky |
| EPI_ISL_2983881 | BIO ARD'AISNE | Department of Virology, Henri Mondor University Hospital, Assistance Publique Hôpitaux de Paris, Université Paris-Est Créteil, INSERM U955 | Christophe Rodriguez, Slim Fourati, Vanessa Demontant, Guillaume Gricourt, Melissa N'Debi, Alexandre Soulier, Elisabeth Trawinski, Jean-Michel Pawlotsky |
| EPI_ISL_2983883 | Laboratoire ATOUTBIO JDC | Department of Virology, Henri Mondor University Hospital, Assistance Publique Hôpitaux de Paris, Université Paris-Est Créteil, INSERM U955 | Christophe Rodriguez, Slim Fourati, Vanessa Demontant, Guillaume Gricourt, Melissa N'Debi, Alexandre Soulier, Elisabeth Trawinski, Jean-Michel Pawlotsky |
| EPI_ISL_2983886 | GH A.CHENEVIER-H.MONDOR | Department of Virology, Henri Mondor University Hospital, Assistance Publique Hôpitaux de Paris, Université Paris-Est Créteil, INSERM U955 | Christophe Rodriguez, Slim Fourati, Vanessa Demontant, Guillaume Gricourt, Melissa N'Debi, Alexandre Soulier, Elisabeth Trawinski, Jean-Michel Pawlotsky |
| EPI_ISL_2983887 | SELAS MEDILYS | Department of Virology, Henri Mondor University Hospital, Assistance Publique Hôpitaux de Paris, Université Paris-Est Créteil, INSERM U955 | Christophe Rodriguez, Slim Fourati, Vanessa Demontant, Guillaume Gricourt, Melissa N'Debi, Alexandre Soulier, Elisabeth Trawinski, Jean-Michel Pawlotsky |
| EPI_ISL_2983888 | GH de l'Est Francilien | Department of Virology, Henri Mondor University Hospital, Assistance Publique Hôpitaux de Paris, Université Paris-Est Créteil, INSERM U955 | Christophe Rodriguez, Slim Fourati, Vanessa Demontant, Guillaume Gricourt, Melissa N'Debi, Alexandre Soulier, Elisabeth Trawinski, Jean-Michel Pawlotsky |
| EPI_ISL_2983890 | Laboratoire multi-site du Diaconat | Department of Virology, Henri Mondor University Hospital, Assistance Publique Hôpitaux de Paris, Université Paris-Est Créteil, INSERM U955 | Christophe Rodriguez, Slim Fourati, Vanessa Demontant, Guillaume Gricourt, Melissa N'Debi, Alexandre Soulier, Elisabeth Trawinski, Jean-Michel Pawlotsky |
| EPI_ISL_2983891, EPI_ISL_2983892 | BIO67-BIOSPHERE | Department of Virology, Henri Mondor University Hospital, Assistance Publique Hôpitaux de Paris, Université Paris-Est Créteil, INSERM U955 | Christophe Rodriguez, Slim Fourati, Vanessa Demontant, Guillaume Gricourt, Melissa N'Debi, Alexandre Soulier, Elisabeth Trawinski, Jean-Michel Pawlotsky |
| EPI_ISL_2983893 | Cerballiance Paris IDF est Saint-Denis | Department of Virology, Henri Mondor University Hospital, Assistance Publique Hôpitaux de Paris, Université Paris-Est Créteil, INSERM U955 | Christophe Rodriguez, Slim Fourati, Vanessa Demontant, Guillaume Gricourt, Melissa N'Debi, Alexandre Soulier, Elisabeth Trawinski, Jean-Michel Pawlotsky |
| EPI_ISL_2983895 | BIO67-BIOSPHERE | Department of Virology, Henri Mondor University Hospital, Assistance Publique Hôpitaux de Paris, Université Paris-Est Créteil, INSERM U955 | Christophe Rodriguez, Slim Fourati, Vanessa Demontant, Guillaume Gricourt, Melissa N'Debi, Alexandre Soulier, Elisabeth Trawinski, Jean-Michel Pawlotsky |
| EPI_ISL_2983896 | CHRU de NANCY | Department of Virology, Henri Mondor University Hospital, Assistance Publique Hôpitaux de Paris, Université Paris-Est Créteil, INSERM U955 | Christophe Rodriguez, Slim Fourati, Vanessa Demontant, Guillaume Gricourt, Melissa N'Debi, Alexandre Soulier, Elisabeth Trawinski, Jean-Michel Pawlotsky |
| EPI_ISL_2983897 | SELAS MEDILYS | Department of Virology, Henri Mondor University Hospital, Assistance Publique Hôpitaux de Paris, Université Paris-Est Créteil, INSERM U955 | Christophe Rodriguez, Slim Fourati, Vanessa Demontant, Guillaume Gricourt, Melissa N'Debi, Alexandre Soulier, Elisabeth Trawinski, Jean-Michel Pawlotsky |
| EPI_ISL_2983898 | BIO ARD'AISNE | Department of Virology, Henri Mondor University Hospital, Assistance Publique Hôpitaux de Paris, Université Paris-Est Créteil, INSERM U955 | Christophe Rodriguez, Slim Fourati, Vanessa Demontant, Guillaume Gricourt, Melissa N'Debi, Alexandre Soulier, Elisabeth Trawinski, Jean-Michel Pawlotsky |
| EPI_ISL_2983899, EPI_ISL_2983900 | SELAS MEDILYS | Department of Virology, Henri Mondor University Hospital, Assistance Publique Hôpitaux de Paris, Université Paris-Est Créteil, | Christophe Rodriguez, Slim Fourati, Vanessa Demontant, Guillaume Gricourt, Melissa N'Debi, Alexandre Soulier, Elisabeth Trawinski, Jean-Michel Pawlotsky |

| | | | |
|---|---|---|---|
| | | INSERM U955 | |
| EPI_ISL_2983901 | BIO67-BIOSPHERE | Department of Virology, Henri Mondor University Hospital, Assistance Publique Hôpitaux de Paris, Université Paris-Est Créteil, INSERM U955 | Christophe Rodriguez, Slim Fourati, Vanessa Demontant, Guillaume Gricourt, Melissa N'Debi, Alexandre Soulier, Elisabeth Trawinski, Jean-Michel Pawlotsky |
| EPI_ISL_2983903 | Laboratoire Bonhoure | Department of Virology, Henri Mondor University Hospital, Assistance Publique Hôpitaux de Paris, Université Paris-Est Créteil, INSERM U955 | Christophe Rodriguez, Slim Fourati, Vanessa Demontant, Guillaume Gricourt, Melissa N'Debi, Alexandre Soulier, Elisabeth Trawinski, Jean-Michel Pawlotsky |
| EPI_ISL_2983905 | BROUSSAIS | Department of Virology, Henri Mondor University Hospital, Assistance Publique Hôpitaux de Paris, Université Paris-Est Créteil, INSERM U955 | Christophe Rodriguez, Slim Fourati, Vanessa Demontant, Guillaume Gricourt, Melissa N'Debi, Alexandre Soulier, Elisabeth Trawinski, Jean-Michel Pawlotsky |
| EPI_ISL_2983906 | BIO67-BIOSPHERE | Department of Virology, Henri Mondor University Hospital, Assistance Publique Hôpitaux de Paris, Université Paris-Est Créteil, INSERM U955 | Christophe Rodriguez, Slim Fourati, Vanessa Demontant, Guillaume Gricourt, Melissa N'Debi, Alexandre Soulier, Elisabeth Trawinski, Jean-Michel Pawlotsky |
| EPI_ISL_2983910 | SELAS MEDILYS | Department of Virology, Henri Mondor University Hospital, Assistance Publique Hôpitaux de Paris, Université Paris-Est Créteil, INSERM U955 | Christophe Rodriguez, Slim Fourati, Vanessa Demontant, Guillaume Gricourt, Melissa N'Debi, Alexandre Soulier, Elisabeth Trawinski, Jean-Michel Pawlotsky |
| EPI_ISL_2983911 | Cerballiance Paris IDF est Saint-Denis | Department of Virology, Henri Mondor University Hospital, Assistance Publique Hôpitaux de Paris, Université Paris-Est Créteil, INSERM U955 | Christophe Rodriguez, Slim Fourati, Vanessa Demontant, Guillaume Gricourt, Melissa N'Debi, Alexandre Soulier, Elisabeth Trawinski, Jean-Michel Pawlotsky |
| EPI_ISL_2983912 | BIO67-BIOSPHERE | Department of Virology, Henri Mondor University Hospital, Assistance Publique Hôpitaux de Paris, Université Paris-Est Créteil, INSERM U955 | Christophe Rodriguez, Slim Fourati, Vanessa Demontant, Guillaume Gricourt, Melissa N'Debi, Alexandre Soulier, Elisabeth Trawinski, Jean-Michel Pawlotsky |
| EPI_ISL_2983913 | BROUSSAIS | Department of Virology, Henri Mondor University Hospital, Assistance Publique Hôpitaux de Paris, Université Paris-Est Créteil, INSERM U955 | Christophe Rodriguez, Slim Fourati, Vanessa Demontant, Guillaume Gricourt, Melissa N'Debi, Alexandre Soulier, Elisabeth Trawinski, Jean-Michel Pawlotsky |
| EPI_ISL_2983914 | SELAS MEDILYS | Department of Virology, Henri Mondor University Hospital, Assistance Publique Hôpitaux de Paris, Université Paris-Est Créteil, INSERM U955 | Christophe Rodriguez, Slim Fourati, Vanessa Demontant, Guillaume Gricourt, Melissa N'Debi, Alexandre Soulier, Elisabeth Trawinski, Jean-Michel Pawlotsky |
| EPI_ISL_2983915 | BIOMNIS EUROFINS IVRY | Department of Virology, Henri Mondor University Hospital, Assistance Publique Hôpitaux de Paris, Université Paris-Est Créteil, INSERM U955 | Christophe Rodriguez, Slim Fourati, Vanessa Demontant, Guillaume Gricourt, Melissa N'Debi, Alexandre Soulier, Elisabeth Trawinski, Jean-Michel Pawlotsky |
| EPI_ISL_2983916 | BROUSSAIS | Department of Virology, Henri Mondor University Hospital, Assistance Publique Hôpitaux de Paris, Université Paris-Est Créteil, INSERM U955 | Christophe Rodriguez, Slim Fourati, Vanessa Demontant, Guillaume Gricourt, Melissa N'Debi, Alexandre Soulier, Elisabeth Trawinski, Jean-Michel Pawlotsky |
| EPI_ISL_2983918 | SELAS MEDILYS | Department of Virology, Henri Mondor University Hospital, Assistance Publique Hôpitaux de Paris, Université Paris-Est Créteil, INSERM U955 | Christophe Rodriguez, Slim Fourati, Vanessa Demontant, Guillaume Gricourt, Melissa N'Debi, Alexandre Soulier, Elisabeth Trawinski, Jean-Michel Pawlotsky |
| EPI_ISL_2983920 | BIO67-BIOSPHERE | Department of Virology, Henri Mondor University Hospital, Assistance Publique Hôpitaux de Paris, Université Paris-Est Créteil, INSERM U955 | Christophe Rodriguez, Slim Fourati, Vanessa Demontant, Guillaume Gricourt, Melissa N'Debi, Alexandre Soulier, Elisabeth Trawinski, Jean-Michel Pawlotsky |
| EPI_ISL_2983922 | BIOMNIS EUROFINS IVRY | Department of Virology, Henri Mondor University Hospital, Assistance Publique Hôpitaux de Paris, Université Paris-Est Créteil, INSERM U955 | Christophe Rodriguez, Slim Fourati, Vanessa Demontant, Guillaume Gricourt, Melissa N'Debi, Alexandre Soulier, Elisabeth Trawinski, Jean-Michel Pawlotsky |
| EPI_ISL_2983923, EPI_ISL_2983924 | SELAS MEDILYS | Department of Virology, Henri Mondor University Hospital, Assistance Publique Hôpitaux de Paris, Université Paris-Est Créteil, INSERM U955 | Christophe Rodriguez, Slim Fourati, Vanessa Demontant, Guillaume Gricourt, Melissa N'Debi, Alexandre Soulier, Elisabeth Trawinski, Jean-Michel Pawlotsky |
| EPI_ISL_2983925 | BIOMNIS EUROFINS IVRY | Department of Virology, Henri Mondor University Hospital, Assistance Publique Hôpitaux de Paris, Université Paris-Est Créteil, INSERM U955 | Christophe Rodriguez, Slim Fourati, Vanessa Demontant, Guillaume Gricourt, Melissa N'Debi, Alexandre Soulier, Elisabeth Trawinski, Jean-Michel Pawlotsky |
| EPI_ISL_2983929 | BIO67-BIOSPHERE | Department of Virology, Henri Mondor University Hospital, Assistance Publique Hôpitaux de Paris, Université Paris-Est Créteil, INSERM U955 | Christophe Rodriguez, Slim Fourati, Vanessa Demontant, Guillaume Gricourt, Melissa N'Debi, Alexandre Soulier, Elisabeth Trawinski, Jean-Michel Pawlotsky |
| EPI_ISL_2983931 | Hôpital Avicenne | Department of Virology, Henri Mondor University Hospital, Assistance Publique Hôpitaux de Paris, Université Paris-Est Créteil, INSERM U955 | Christophe Rodriguez, Slim Fourati, Vanessa Demontant, Guillaume Gricourt, Melissa N'Debi, Alexandre Soulier, Elisabeth Trawinski, Jean-Michel Pawlotsky |
| EPI_ISL_2983934, EPI_ISL_2983935 | SELAS MEDILYS | Department of Virology, Henri Mondor University Hospital, Assistance Publique Hôpitaux de Paris, Université Paris-Est Créteil, INSERM U955 | Christophe Rodriguez, Slim Fourati, Vanessa Demontant, Guillaume Gricourt, Melissa N'Debi, Alexandre Soulier, Elisabeth Trawinski, Jean-Michel Pawlotsky |
| EPI_ISL_2983936, EPI_ISL_2983939, EPI_ISL_2983940 | BIO67-BIOSPHERE | Department of Virology, Henri Mondor University Hospital, Assistance Publique Hôpitaux de Paris, Université Paris-Est Créteil, INSERM U955 | Christophe Rodriguez, Slim Fourati, Vanessa Demontant, Guillaume Gricourt, Melissa N'Debi, Alexandre Soulier, Elisabeth Trawinski, Jean-Michel Pawlotsky |
| EPI_ISL_2983942 | SELAS MEDILYS | Department of Virology, Henri Mondor University Hospital, Assistance Publique Hôpitaux de Paris, Université Paris-Est Créteil, INSERM U955 | Christophe Rodriguez, Slim Fourati, Vanessa Demontant, Guillaume Gricourt, Melissa N'Debi, Alexandre Soulier, Elisabeth Trawinski, Jean-Michel Pawlotsky |
| EPI_ISL_2983945 | BIO67-BIOSPHERE | Department of Virology, Henri Mondor University Hospital, Assistance Publique Hôpitaux de Paris, Université Paris-Est Créteil, INSERM U955 | Christophe Rodriguez, Slim Fourati, Vanessa Demontant, Guillaume Gricourt, Melissa N'Debi, Alexandre Soulier, Elisabeth Trawinski, Jean-Michel Pawlotsky |
| EPI_ISL_2983950 | SELAS MEDILYS | Department of Virology, Henri Mondor University Hospital, Assistance Publique Hôpitaux de Paris, Université Paris-Est Créteil, INSERM U955 | Christophe Rodriguez, Slim Fourati, Vanessa Demontant, Guillaume Gricourt, Melissa N'Debi, Alexandre Soulier, Elisabeth Trawinski, Jean-Michel Pawlotsky |
| EPI_ISL_2983951, EPI_ISL_2983952 | BIOMNIS EUROFINS IVRY | Department of Virology, Henri Mondor University Hospital, Assistance Publique Hôpitaux de Paris, Université Paris-Est Créteil, INSERM U955 | Christophe Rodriguez, Slim Fourati, Vanessa Demontant, Guillaume Gricourt, Melissa N'Debi, Alexandre Soulier, Elisabeth Trawinski, Jean-Michel Pawlotsky |
| EPI_ISL_2983953 | Plateau technique Biomer Metz Verriers | Department of Virology, Henri Mondor University Hospital, Assistance Publique Hôpitaux de Paris, Université Paris-Est Créteil, INSERM U955 | Christophe Rodriguez, Slim Fourati, Vanessa Demontant, Guillaume Gricourt, Melissa N'Debi, Alexandre Soulier, Elisabeth Trawinski, Jean-Michel Pawlotsky |
| EPI_ISL_2983955 | BIOMNIS EUROFINS IVRY | Department of Virology, Henri Mondor University Hospital, Assistance Publique Hôpitaux de Paris, Université Paris-Est Créteil, INSERM U955 | Christophe Rodriguez, Slim Fourati, Vanessa Demontant, Guillaume Gricourt, Melissa N'Debi, Alexandre Soulier, Elisabeth Trawinski, Jean-Michel Pawlotsky |
| EPI_ISL_2983957 | Plateau technique Biomer Metz Verriers | Department of Virology, Henri Mondor University Hospital, Assistance Publique Hôpitaux de Paris, Université Paris-Est Créteil, INSERM U955 | Christophe Rodriguez, Slim Fourati, Vanessa Demontant, Guillaume Gricourt, Melissa N'Debi, Alexandre Soulier, Elisabeth Trawinski, Jean-Michel Pawlotsky |
| EPI_ISL_2983959 | Biogroup Bio Lam-LCD Saint-Denis | Department of Virology, Henri Mondor University Hospital, Assistance Publique Hôpitaux de Paris, Université Paris-Est Créteil, INSERM U955 | Christophe Rodriguez, Slim Fourati, Vanessa Demontant, Guillaume Gricourt, Melissa N'Debi, Alexandre Soulier, Elisabeth Trawinski, Jean-Michel Pawlotsky |
| EPI_ISL_2983960 | Laboratoire LENYS | Department of Virology, Henri Mondor University Hospital, Assistance Publique Hôpitaux de Paris, Université Paris-Est Créteil, INSERM U955 | Christophe Rodriguez, Slim Fourati, Vanessa Demontant, Guillaume Gricourt, Melissa N'Debi, Alexandre Soulier, Elisabeth Trawinski, Jean-Michel Pawlotsky |
| EPI_ISL_2983962 | Cerballiance Paris IDF est Saint-Denis | Department of Virology, Henri Mondor University Hospital, Assistance Publique Hôpitaux de Paris, Université Paris-Est Créteil, INSERM U955 | Christophe Rodriguez, Slim Fourati, Vanessa Demontant, Guillaume Gricourt, Melissa N'Debi, Alexandre Soulier, Elisabeth Trawinski, Jean-Michel Pawlotsky |
| EPI_ISL_2983963 | Biogroup Bio Lam-LCD Saint-Denis | Department of Virology, Henri Mondor University Hospital, Assistance Publique Hôpitaux de Paris, Université Paris-Est Créteil, INSERM U955 | Christophe Rodriguez, Slim Fourati, Vanessa Demontant, Guillaume Gricourt, Melissa N'Debi, Alexandre Soulier, Elisabeth Trawinski, Jean-Michel Pawlotsky |
| EPI_ISL_2983964 | LBM CARRON SELAS | Department of Virology, Henri Mondor University Hospital, Assistance Publique Hôpitaux de Paris, Université Paris-Est Créteil, | Christophe Rodriguez, Slim Fourati, Vanessa Demontant, Guillaume Gricourt, Melissa N'Debi, Alexandre Soulier, Elisabeth Trawinski, Jean-Michel Pawlotsky |

| | | | |
|---|---|---|---|
| | | INSERM U955 | |
| EPI_ISL_2983965 | BIOMNIS EUROFINS IVRY | Department of Virology, Henri Mondor University Hospital, Assistance Publique Hôpitaux de Paris, Université Paris-Est Créteil, INSERM U955 | Christophe Rodriguez, Slim Fourati, Vanessa Demontant, Guillaume Gricourt, Melissa N'Debi, Alexandre Soulier, Elisabeth Trawinski, Jean-Michel Pawlotsky |
| EPI_ISL_2983966 | Plateau technique Biomer Metz Verriers | Department of Virology, Henri Mondor University Hospital, Assistance Publique Hôpitaux de Paris, Université Paris-Est Créteil, INSERM U955 | Christophe Rodriguez, Slim Fourati, Vanessa Demontant, Guillaume Gricourt, Melissa N'Debi, Alexandre Soulier, Elisabeth Trawinski, Jean-Michel Pawlotsky |
| EPI_ISL_2983970 | Biogroup Bio Lam-LCD Saint-Denis | Department of Virology, Henri Mondor University Hospital, Assistance Publique Hôpitaux de Paris, Université Paris-Est Créteil, INSERM U955 | Christophe Rodriguez, Slim Fourati, Vanessa Demontant, Guillaume Gricourt, Melissa N'Debi, Alexandre Soulier, Elisabeth Trawinski, Jean-Michel Pawlotsky |
| EPI_ISL_2983971 | Laboratoire LENYS | Department of Virology, Henri Mondor University Hospital, Assistance Publique Hôpitaux de Paris, Université Paris-Est Créteil, INSERM U955 | Christophe Rodriguez, Slim Fourati, Vanessa Demontant, Guillaume Gricourt, Melissa N'Debi, Alexandre Soulier, Elisabeth Trawinski, Jean-Michel Pawlotsky |
| EPI_ISL_2983972 | Cerballiance Paris IDF est Saint-Denis | Department of Virology, Henri Mondor University Hospital, Assistance Publique Hôpitaux de Paris, Université Paris-Est Créteil, INSERM U955 | Christophe Rodriguez, Slim Fourati, Vanessa Demontant, Guillaume Gricourt, Melissa N'Debi, Alexandre Soulier, Elisabeth Trawinski, Jean-Michel Pawlotsky |
| EPI_ISL_2983973 | SELAFA SANTE LABO | Department of Virology, Henri Mondor University Hospital, Assistance Publique Hôpitaux de Paris, Université Paris-Est Créteil, INSERM U955 | Christophe Rodriguez, Slim Fourati, Vanessa Demontant, Guillaume Gricourt, Melissa N'Debi, Alexandre Soulier, Elisabeth Trawinski, Jean-Michel Pawlotsky |
| EPI_ISL_2983974 | Biogroup Bio Lam-LCD Saint-Denis | Department of Virology, Henri Mondor University Hospital, Assistance Publique Hôpitaux de Paris, Université Paris-Est Créteil, INSERM U955 | Christophe Rodriguez, Slim Fourati, Vanessa Demontant, Guillaume Gricourt, Melissa N'Debi, Alexandre Soulier, Elisabeth Trawinski, Jean-Michel Pawlotsky |
| EPI_ISL_2983977 | Cerballiance Paris IDF est Saint-Denis | Department of Virology, Henri Mondor University Hospital, Assistance Publique Hôpitaux de Paris, Université Paris-Est Créteil, INSERM U955 | Christophe Rodriguez, Slim Fourati, Vanessa Demontant, Guillaume Gricourt, Melissa N'Debi, Alexandre Soulier, Elisabeth Trawinski, Jean-Michel Pawlotsky |
| EPI_ISL_2983981 | Plateau technique Biomer Metz Verriers | Department of Virology, Henri Mondor University Hospital, Assistance Publique Hôpitaux de Paris, Université Paris-Est Créteil, INSERM U955 | Christophe Rodriguez, Slim Fourati, Vanessa Demontant, Guillaume Gricourt, Melissa N'Debi, Alexandre Soulier, Elisabeth Trawinski, Jean-Michel Pawlotsky |
| EPI_ISL_2983982, EPI_ISL_2983983, EPI_ISL_2983984 | Biogroup Bio Lam-LCD Saint-Denis | Department of Virology, Henri Mondor University Hospital, Assistance Publique Hôpitaux de Paris, Université Paris-Est Créteil, INSERM U955 | Christophe Rodriguez, Slim Fourati, Vanessa Demontant, Guillaume Gricourt, Melissa N'Debi, Alexandre Soulier, Elisabeth Trawinski, Jean-Michel Pawlotsky |
| EPI_ISL_2983985 | BIOMNIS EUROFINS IVRY | Department of Virology, Henri Mondor University Hospital, Assistance Publique Hôpitaux de Paris, Université Paris-Est Créteil, INSERM U955 | Christophe Rodriguez, Slim Fourati, Vanessa Demontant, Guillaume Gricourt, Melissa N'Debi, Alexandre Soulier, Elisabeth Trawinski, Jean-Michel Pawlotsky |
| EPI_ISL_2983986 | Cerballiance Paris IDF est Saint-Denis | Department of Virology, Henri Mondor University Hospital, Assistance Publique Hôpitaux de Paris, Université Paris-Est Créteil, INSERM U955 | Christophe Rodriguez, Slim Fourati, Vanessa Demontant, Guillaume Gricourt, Melissa N'Debi, Alexandre Soulier, Elisabeth Trawinski, Jean-Michel Pawlotsky |
| EPI_ISL_2983987, EPI_ISL_2983988 | BIOMNIS EUROFINS IVRY | Department of Virology, Henri Mondor University Hospital, Assistance Publique Hôpitaux de Paris, Université Paris-Est Créteil, INSERM U955 | Christophe Rodriguez, Slim Fourati, Vanessa Demontant, Guillaume Gricourt, Melissa N'Debi, Alexandre Soulier, Elisabeth Trawinski, Jean-Michel Pawlotsky |
| EPI_ISL_2983989 | Plateau technique Biomer Metz Verriers | Department of Virology, Henri Mondor University Hospital, Assistance Publique Hôpitaux de Paris, Université Paris-Est Créteil, INSERM U955 | Christophe Rodriguez, Slim Fourati, Vanessa Demontant, Guillaume Gricourt, Melissa N'Debi, Alexandre Soulier, Elisabeth Trawinski, Jean-Michel Pawlotsky |
| EPI_ISL_2983991 | Biogroup Bio Lam-LCD Saint-Denis | Department of Virology, Henri Mondor University Hospital, Assistance Publique Hôpitaux de Paris, Université Paris-Est Créteil, INSERM U955 | Christophe Rodriguez, Slim Fourati, Vanessa Demontant, Guillaume Gricourt, Melissa N'Debi, Alexandre Soulier, Elisabeth Trawinski, Jean-Michel Pawlotsky |
| EPI_ISL_2983992, EPI_ISL_2983994 | Plateau technique Biomer Metz Verriers | Department of Virology, Henri Mondor University Hospital, Assistance Publique Hôpitaux de Paris, Université Paris-Est Créteil, INSERM U955 | Christophe Rodriguez, Slim Fourati, Vanessa Demontant, Guillaume Gricourt, Melissa N'Debi, Alexandre Soulier, Elisabeth Trawinski, Jean-Michel Pawlotsky |
| EPI_ISL_2983996 | Biogroup Bio Lam-LCD Saint-Denis | Department of Virology, Henri Mondor University Hospital, Assistance Publique Hôpitaux de Paris, Université Paris-Est Créteil, INSERM U955 | Christophe Rodriguez, Slim Fourati, Vanessa Demontant, Guillaume Gricourt, Melissa N'Debi, Alexandre Soulier, Elisabeth Trawinski, Jean-Michel Pawlotsky |
| EPI_ISL_2983999, EPI_ISL_2984000 | BIOMNIS EUROFINS IVRY | Department of Virology, Henri Mondor University Hospital, Assistance Publique Hôpitaux de Paris, Université Paris-Est Créteil, INSERM U955 | Christophe Rodriguez, Slim Fourati, Vanessa Demontant, Guillaume Gricourt, Melissa N'Debi, Alexandre Soulier, Elisabeth Trawinski, Jean-Michel Pawlotsky |
| EPI_ISL_2984001, EPI_ISL_2984002 | Plateau technique Biomer Metz Verriers | Department of Virology, Henri Mondor University Hospital, Assistance Publique Hôpitaux de Paris, Université Paris-Est Créteil, INSERM U955 | Christophe Rodriguez, Slim Fourati, Vanessa Demontant, Guillaume Gricourt, Melissa N'Debi, Alexandre Soulier, Elisabeth Trawinski, Jean-Michel Pawlotsky |
| EPI_ISL_2984004 | Laboratoire LENYS | Department of Virology, Henri Mondor University Hospital, Assistance Publique Hôpitaux de Paris, Université Paris-Est Créteil, INSERM U955 | Christophe Rodriguez, Slim Fourati, Vanessa Demontant, Guillaume Gricourt, Melissa N'Debi, Alexandre Soulier, Elisabeth Trawinski, Jean-Michel Pawlotsky |
| EPI_ISL_2984006 | Biogroup Bio Lam-LCD Saint-Denis | Department of Virology, Henri Mondor University Hospital, Assistance Publique Hôpitaux de Paris, Université Paris-Est Créteil, INSERM U955 | Christophe Rodriguez, Slim Fourati, Vanessa Demontant, Guillaume Gricourt, Melissa N'Debi, Alexandre Soulier, Elisabeth Trawinski, Jean-Michel Pawlotsky |
| EPI_ISL_2984007 | SELAS MEDILYS | Department of Virology, Henri Mondor University Hospital, Assistance Publique Hôpitaux de Paris, Université Paris-Est Créteil, INSERM U955 | Christophe Rodriguez, Slim Fourati, Vanessa Demontant, Guillaume Gricourt, Melissa N'Debi, Alexandre Soulier, Elisabeth Trawinski, Jean-Michel Pawlotsky |
| EPI_ISL_2984008, EPI_ISL_2984009 | Biogroup Bio Lam-LCD Saint-Denis | Department of Virology, Henri Mondor University Hospital, Assistance Publique Hôpitaux de Paris, Université Paris-Est Créteil, INSERM U955 | Christophe Rodriguez, Slim Fourati, Vanessa Demontant, Guillaume Gricourt, Melissa N'Debi, Alexandre Soulier, Elisabeth Trawinski, Jean-Michel Pawlotsky |
| EPI_ISL_2984010 | Plateau technique Biomer Metz Verriers | Department of Virology, Henri Mondor University Hospital, Assistance Publique Hôpitaux de Paris, Université Paris-Est Créteil, INSERM U955 | Christophe Rodriguez, Slim Fourati, Vanessa Demontant, Guillaume Gricourt, Melissa N'Debi, Alexandre Soulier, Elisabeth Trawinski, Jean-Michel Pawlotsky |
| EPI_ISL_2984011 | Biogroup Bio Lam-LCD Saint-Denis | Department of Virology, Henri Mondor University Hospital, Assistance Publique Hôpitaux de Paris, Université Paris-Est Créteil, INSERM U955 | Christophe Rodriguez, Slim Fourati, Vanessa Demontant, Guillaume Gricourt, Melissa N'Debi, Alexandre Soulier, Elisabeth Trawinski, Jean-Michel Pawlotsky |
| EPI_ISL_2984012 | BIOMNIS EUROFINS IVRY | Department of Virology, Henri Mondor University Hospital, Assistance Publique Hôpitaux de Paris, Université Paris-Est Créteil, INSERM U955 | Christophe Rodriguez, Slim Fourati, Vanessa Demontant, Guillaume Gricourt, Melissa N'Debi, Alexandre Soulier, Elisabeth Trawinski, Jean-Michel Pawlotsky |
| EPI_ISL_2984013 | LBM CARRON SELAS | Department of Virology, Henri Mondor University Hospital, Assistance Publique Hôpitaux de Paris, Université Paris-Est Créteil, INSERM U955 | Christophe Rodriguez, Slim Fourati, Vanessa Demontant, Guillaume Gricourt, Melissa N'Debi, Alexandre Soulier, Elisabeth Trawinski, Jean-Michel Pawlotsky |
| EPI_ISL_2984014 | Laboratoire Ana-L | Department of Virology, Henri Mondor University Hospital, Assistance Publique Hôpitaux de Paris, Université Paris-Est Créteil, INSERM U955 | Christophe Rodriguez, Slim Fourati, Vanessa Demontant, Guillaume Gricourt, Melissa N'Debi, Alexandre Soulier, Elisabeth Trawinski, Jean-Michel Pawlotsky |
| EPI_ISL_2984015, EPI_ISL_2984017 | Biogroup Bio Lam-LCD Saint-Denis | Department of Virology, Henri Mondor University Hospital, Assistance Publique Hôpitaux de Paris, Université Paris-Est Créteil, INSERM U955 | Christophe Rodriguez, Slim Fourati, Vanessa Demontant, Guillaume Gricourt, Melissa N'Debi, Alexandre Soulier, Elisabeth Trawinski, Jean-Michel Pawlotsky |
| EPI_ISL_2984019 | Cerballiance Paris IDF est Saint-Denis | Department of Virology, Henri Mondor University Hospital, Assistance Publique Hôpitaux de Paris, Université Paris-Est Créteil, INSERM U955 | Christophe Rodriguez, Slim Fourati, Vanessa Demontant, Guillaume Gricourt, Melissa N'Debi, Alexandre Soulier, Elisabeth Trawinski, Jean-Michel Pawlotsky |
| EPI_ISL_2984021 | Biogroup Bio Lam-LCD Saint-Denis | Department of Virology, Henri Mondor University Hospital, Assistance Publique Hôpitaux de Paris, Université Paris-Est Créteil, INSERM U955 | Christophe Rodriguez, Slim Fourati, Vanessa Demontant, Guillaume Gricourt, Melissa N'Debi, Alexandre Soulier, Elisabeth Trawinski, Jean-Michel Pawlotsky |
| EPI_ISL_2984022 | BIOMNIS EUROFINS IVRY | Department of Virology, Henri Mondor University Hospital, Assistance Publique Hôpitaux de Paris, Université Paris-Est Créteil, | Christophe Rodriguez, Slim Fourati, Vanessa Demontant, Guillaume Gricourt, Melissa N'Debi, Alexandre Soulier, Elisabeth Trawinski, Jean-Michel Pawlotsky |

| | | | |
|---|---|---|---|
| | | INSERM U955 | |
| EPI_ISL_2984023 | Biogroup Bio Lam-LCD Saint-Denis | Department of Virology, Henri Mondor University Hospital, Assistance Publique Hôpitaux de Paris, Université Paris-Est Créteil, INSERM U955 | Christophe Rodriguez, Slim Fourati, Vanessa Demontant, Guillaume Gricourt, Melissa N'Debi, Alexandre Soulier, Elisabeth Trawinski, Jean-Michel Pawlotsky |
| EPI_ISL_2984024 | Laboratoire LENYS | Department of Virology, Henri Mondor University Hospital, Assistance Publique Hôpitaux de Paris, Université Paris-Est Créteil, INSERM U955 | Christophe Rodriguez, Slim Fourati, Vanessa Demontant, Guillaume Gricourt, Melissa N'Debi, Alexandre Soulier, Elisabeth Trawinski, Jean-Michel Pawlotsky |
| EPI_ISL_2984026, EPI_ISL_2984027 | Biogroup Bio Lam-LCD Saint-Denis | Department of Virology, Henri Mondor University Hospital, Assistance Publique Hôpitaux de Paris, Université Paris-Est Créteil, INSERM U955 | Christophe Rodriguez, Slim Fourati, Vanessa Demontant, Guillaume Gricourt, Melissa N'Debi, Alexandre Soulier, Elisabeth Trawinski, Jean-Michel Pawlotsky |
| EPI_ISL_2984029 | Cerballiance Paris IDF est Saint-Denis | Department of Virology, Henri Mondor University Hospital, Assistance Publique Hôpitaux de Paris, Université Paris-Est Créteil, INSERM U955 | Christophe Rodriguez, Slim Fourati, Vanessa Demontant, Guillaume Gricourt, Melissa N'Debi, Alexandre Soulier, Elisabeth Trawinski, Jean-Michel Pawlotsky |
| EPI_ISL_2984032 | Biogroup Bio Lam-LCD Saint-Denis | Department of Virology, Henri Mondor University Hospital, Assistance Publique Hôpitaux de Paris, Université Paris-Est Créteil, INSERM U955 | Christophe Rodriguez, Slim Fourati, Vanessa Demontant, Guillaume Gricourt, Melissa N'Debi, Alexandre Soulier, Elisabeth Trawinski, Jean-Michel Pawlotsky |
| EPI_ISL_2984033 | Laboratoire LENYS | Department of Virology, Henri Mondor University Hospital, Assistance Publique Hôpitaux de Paris, Université Paris-Est Créteil, INSERM U955 | Christophe Rodriguez, Slim Fourati, Vanessa Demontant, Guillaume Gricourt, Melissa N'Debi, Alexandre Soulier, Elisabeth Trawinski, Jean-Michel Pawlotsky |
| EPI_ISL_2984034 | Laboratoire Ana-L | Department of Virology, Henri Mondor University Hospital, Assistance Publique Hôpitaux de Paris, Université Paris-Est Créteil, INSERM U955 | Christophe Rodriguez, Slim Fourati, Vanessa Demontant, Guillaume Gricourt, Melissa N'Debi, Alexandre Soulier, Elisabeth Trawinski, Jean-Michel Pawlotsky |
| EPI_ISL_2984036 | LBM CARRON SELAS | Department of Virology, Henri Mondor University Hospital, Assistance Publique Hôpitaux de Paris, Université Paris-Est Créteil, INSERM U955 | Christophe Rodriguez, Slim Fourati, Vanessa Demontant, Guillaume Gricourt, Melissa N'Debi, Alexandre Soulier, Elisabeth Trawinski, Jean-Michel Pawlotsky |
| EPI_ISL_2984037 | Biogroup Bio Lam-LCD Saint-Denis | Department of Virology, Henri Mondor University Hospital, Assistance Publique Hôpitaux de Paris, Université Paris-Est Créteil, INSERM U955 | Christophe Rodriguez, Slim Fourati, Vanessa Demontant, Guillaume Gricourt, Melissa N'Debi, Alexandre Soulier, Elisabeth Trawinski, Jean-Michel Pawlotsky |
| EPI_ISL_2984039 | BIOMNIS EUROFINS IVRY | Department of Virology, Henri Mondor University Hospital, Assistance Publique Hôpitaux de Paris, Université Paris-Est Créteil, INSERM U955 | Christophe Rodriguez, Slim Fourati, Vanessa Demontant, Guillaume Gricourt, Melissa N'Debi, Alexandre Soulier, Elisabeth Trawinski, Jean-Michel Pawlotsky |
| EPI_ISL_2984041 | Laboratoire Ana-L | Department of Virology, Henri Mondor University Hospital, Assistance Publique Hôpitaux de Paris, Université Paris-Est Créteil, INSERM U955 | Christophe Rodriguez, Slim Fourati, Vanessa Demontant, Guillaume Gricourt, Melissa N'Debi, Alexandre Soulier, Elisabeth Trawinski, Jean-Michel Pawlotsky |
| EPI_ISL_2984042 | BIOMNIS EUROFINS IVRY | Department of Virology, Henri Mondor University Hospital, Assistance Publique Hôpitaux de Paris, Université Paris-Est Créteil, INSERM U955 | Christophe Rodriguez, Slim Fourati, Vanessa Demontant, Guillaume Gricourt, Melissa N'Debi, Alexandre Soulier, Elisabeth Trawinski, Jean-Michel Pawlotsky |
| EPI_ISL_2984043 | Biogroup Bio Lam-LCD Saint-Denis | Department of Virology, Henri Mondor University Hospital, Assistance Publique Hôpitaux de Paris, Université Paris-Est Créteil, INSERM U955 | Christophe Rodriguez, Slim Fourati, Vanessa Demontant, Guillaume Gricourt, Melissa N'Debi, Alexandre Soulier, Elisabeth Trawinski, Jean-Michel Pawlotsky |
| EPI_ISL_2984045 | Laboratoire LENYS | Department of Virology, Henri Mondor University Hospital, Assistance Publique Hôpitaux de Paris, Université Paris-Est Créteil, INSERM U955 | Christophe Rodriguez, Slim Fourati, Vanessa Demontant, Guillaume Gricourt, Melissa N'Debi, Alexandre Soulier, Elisabeth Trawinski, Jean-Michel Pawlotsky |
| EPI_ISL_2984046 | Laboratoire Ana-L | Department of Virology, Henri Mondor University Hospital, Assistance Publique Hôpitaux de Paris, Université Paris-Est Créteil, INSERM U955 | Christophe Rodriguez, Slim Fourati, Vanessa Demontant, Guillaume Gricourt, Melissa N'Debi, Alexandre Soulier, Elisabeth Trawinski, Jean-Michel Pawlotsky |
| EPI_ISL_2984047 | BIOMNIS EUROFINS IVRY | Department of Virology, Henri Mondor University Hospital, Assistance Publique Hôpitaux de Paris, Université Paris-Est Créteil, INSERM U955 | Christophe Rodriguez, Slim Fourati, Vanessa Demontant, Guillaume Gricourt, Melissa N'Debi, Alexandre Soulier, Elisabeth Trawinski, Jean-Michel Pawlotsky |
| EPI_ISL_2984048 | Biogroup Bio Lam-LCD Saint-Denis | Department of Virology, Henri Mondor University Hospital, Assistance Publique Hôpitaux de Paris, Université Paris-Est Créteil, INSERM U955 | Christophe Rodriguez, Slim Fourati, Vanessa Demontant, Guillaume Gricourt, Melissa N'Debi, Alexandre Soulier, Elisabeth Trawinski, Jean-Michel Pawlotsky |
| EPI_ISL_2984049, EPI_ISL_2984050 | Cerballiance Paris IDF est Saint-Denis | Department of Virology, Henri Mondor University Hospital, Assistance Publique Hôpitaux de Paris, Université Paris-Est Créteil, INSERM U955 | Christophe Rodriguez, Slim Fourati, Vanessa Demontant, Guillaume Gricourt, Melissa N'Debi, Alexandre Soulier, Elisabeth Trawinski, Jean-Michel Pawlotsky |
| EPI_ISL_2984052 | Laboratoire LENYS | Department of Virology, Henri Mondor University Hospital, Assistance Publique Hôpitaux de Paris, Université Paris-Est Créteil, INSERM U955 | Christophe Rodriguez, Slim Fourati, Vanessa Demontant, Guillaume Gricourt, Melissa N'Debi, Alexandre Soulier, Elisabeth Trawinski, Jean-Michel Pawlotsky |
| EPI_ISL_2984053, EPI_ISL_2984056 | BIOMNIS EUROFINS IVRY | Department of Virology, Henri Mondor University Hospital, Assistance Publique Hôpitaux de Paris, Université Paris-Est Créteil, INSERM U955 | Christophe Rodriguez, Slim Fourati, Vanessa Demontant, Guillaume Gricourt, Melissa N'Debi, Alexandre Soulier, Elisabeth Trawinski, Jean-Michel Pawlotsky |
| EPI_ISL_2984057, EPI_ISL_2984058 | Biogroup Bio Lam-LCD Saint-Denis | Department of Virology, Henri Mondor University Hospital, Assistance Publique Hôpitaux de Paris, Université Paris-Est Créteil, INSERM U955 | Christophe Rodriguez, Slim Fourati, Vanessa Demontant, Guillaume Gricourt, Melissa N'Debi, Alexandre Soulier, Elisabeth Trawinski, Jean-Michel Pawlotsky |
| EPI_ISL_2984059 | SELAS MEDILYS | Department of Virology, Henri Mondor University Hospital, Assistance Publique Hôpitaux de Paris, Université Paris-Est Créteil, INSERM U955 | Christophe Rodriguez, Slim Fourati, Vanessa Demontant, Guillaume Gricourt, Melissa N'Debi, Alexandre Soulier, Elisabeth Trawinski, Jean-Michel Pawlotsky |
| EPI_ISL_2984060 | Plateau technique Biomer Metz Verriers | Department of Virology, Henri Mondor University Hospital, Assistance Publique Hôpitaux de Paris, Université Paris-Est Créteil, INSERM U955 | Christophe Rodriguez, Slim Fourati, Vanessa Demontant, Guillaume Gricourt, Melissa N'Debi, Alexandre Soulier, Elisabeth Trawinski, Jean-Michel Pawlotsky |
| EPI_ISL_2984061 | BIOLAB UNILABS | Department of Virology, Henri Mondor University Hospital, Assistance Publique Hôpitaux de Paris, Université Paris-Est Créteil, INSERM U955 | Christophe Rodriguez, Slim Fourati, Vanessa Demontant, Guillaume Gricourt, Melissa N'Debi, Alexandre Soulier, Elisabeth Trawinski, Jean-Michel Pawlotsky |
| EPI_ISL_2984062 | Cerballiance Paris IDF est Saint-Denis | Department of Virology, Henri Mondor University Hospital, Assistance Publique Hôpitaux de Paris, Université Paris-Est Créteil, INSERM U955 | Christophe Rodriguez, Slim Fourati, Vanessa Demontant, Guillaume Gricourt, Melissa N'Debi, Alexandre Soulier, Elisabeth Trawinski, Jean-Michel Pawlotsky |
| EPI_ISL_2984064 | BIOMNIS EUROFINS IVRY | Department of Virology, Henri Mondor University Hospital, Assistance Publique Hôpitaux de Paris, Université Paris-Est Créteil, INSERM U955 | Christophe Rodriguez, Slim Fourati, Vanessa Demontant, Guillaume Gricourt, Melissa N'Debi, Alexandre Soulier, Elisabeth Trawinski, Jean-Michel Pawlotsky |
| EPI_ISL_2984065 | Biogroup Bio Lam-LCD Saint-Denis | Department of Virology, Henri Mondor University Hospital, Assistance Publique Hôpitaux de Paris, Université Paris-Est Créteil, INSERM U955 | Christophe Rodriguez, Slim Fourati, Vanessa Demontant, Guillaume Gricourt, Melissa N'Debi, Alexandre Soulier, Elisabeth Trawinski, Jean-Michel Pawlotsky |
| EPI_ISL_2984066 | Plateau technique Biomer Metz Verriers | Department of Virology, Henri Mondor University Hospital, Assistance Publique Hôpitaux de Paris, Université Paris-Est Créteil, INSERM U955 | Christophe Rodriguez, Slim Fourati, Vanessa Demontant, Guillaume Gricourt, Melissa N'Debi, Alexandre Soulier, Elisabeth Trawinski, Jean-Michel Pawlotsky |
| EPI_ISL_2984068 | Cerballiance Paris IDF est Saint-Denis | Department of Virology, Henri Mondor University Hospital, Assistance Publique Hôpitaux de Paris, Université Paris-Est Créteil, INSERM U955 | Christophe Rodriguez, Slim Fourati, Vanessa Demontant, Guillaume Gricourt, Melissa N'Debi, Alexandre Soulier, Elisabeth Trawinski, Jean-Michel Pawlotsky |
| EPI_ISL_2984070 | Biogroup Bio Lam-LCD Saint-Denis | Department of Virology, Henri Mondor University Hospital, Assistance Publique Hôpitaux de Paris, Université Paris-Est Créteil, INSERM U955 | Christophe Rodriguez, Slim Fourati, Vanessa Demontant, Guillaume Gricourt, Melissa N'Debi, Alexandre Soulier, Elisabeth Trawinski, Jean-Michel Pawlotsky |
| EPI_ISL_2984072 | BIOMNIS EUROFINS IVRY | Department of Virology, Henri Mondor University Hospital, Assistance Publique Hôpitaux de Paris, Université Paris-Est Créteil, INSERM U955 | Christophe Rodriguez, Slim Fourati, Vanessa Demontant, Guillaume Gricourt, Melissa N'Debi, Alexandre Soulier, Elisabeth Trawinski, Jean-Michel Pawlotsky |
| EPI_ISL_2984073, EPI_ISL_2984076, | Biogroup Bio Lam-LCD Saint-Denis | Department of Virology, Henri Mondor University Hospital, Assistance Publique Hôpitaux de Paris, Université Paris-Est Créteil, | Christophe Rodriguez, Slim Fourati, Vanessa Demontant, Guillaume Gricourt, Melissa N'Debi, Alexandre Soulier, Elisabeth Trawinski, Jean-Michel Pawlotsky |

| | | | |
|---|---|---|---|
| EPI_ISL_2984078 | | INSERM U955 | |
| EPI_ISL_2984079 | Laboratoire Ana-L | Department of Virology, Henri Mondor University Hospital, Assistance Publique Hôpitaux de Paris, Université Paris-Est Créteil, INSERM U955 | Christophe Rodriguez, Slim Fourati, Vanessa Demontant, Guillaume Gricourt, Melissa N'Debi, Alexandre Soulier, Elisabeth Trawinski, Jean-Michel Pawlotsky |
| EPI_ISL_2984080, EPI_ISL_2984081 | Biogroup Bio Lam-LCD Saint-Denis | Department of Virology, Henri Mondor University Hospital, Assistance Publique Hôpitaux de Paris, Université Paris-Est Créteil, INSERM U955 | Christophe Rodriguez, Slim Fourati, Vanessa Demontant, Guillaume Gricourt, Melissa N'Debi, Alexandre Soulier, Elisabeth Trawinski, Jean-Michel Pawlotsky |
| EPI_ISL_2984082, EPI_ISL_2984084 | Cerballiance Paris IDF est Saint-Denis | Department of Virology, Henri Mondor University Hospital, Assistance Publique Hôpitaux de Paris, Université Paris-Est Créteil, INSERM U955 | Christophe Rodriguez, Slim Fourati, Vanessa Demontant, Guillaume Gricourt, Melissa N'Debi, Alexandre Soulier, Elisabeth Trawinski, Jean-Michel Pawlotsky |
| EPI_ISL_2984085 | Laboratoire Ana-L | Department of Virology, Henri Mondor University Hospital, Assistance Publique Hôpitaux de Paris, Université Paris-Est Créteil, INSERM U955 | Christophe Rodriguez, Slim Fourati, Vanessa Demontant, Guillaume Gricourt, Melissa N'Debi, Alexandre Soulier, Elisabeth Trawinski, Jean-Michel Pawlotsky |
| EPI_ISL_2984086 | Biogroup Bio Lam-LCD Saint-Denis | Department of Virology, Henri Mondor University Hospital, Assistance Publique Hôpitaux de Paris, Université Paris-Est Créteil, INSERM U955 | Christophe Rodriguez, Slim Fourati, Vanessa Demontant, Guillaume Gricourt, Melissa N'Debi, Alexandre Soulier, Elisabeth Trawinski, Jean-Michel Pawlotsky |
| EPI_ISL_2984087 | Cerballiance Paris IDF est Saint-Denis | Department of Virology, Henri Mondor University Hospital, Assistance Publique Hôpitaux de Paris, Université Paris-Est Créteil, INSERM U955 | Christophe Rodriguez, Slim Fourati, Vanessa Demontant, Guillaume Gricourt, Melissa N'Debi, Alexandre Soulier, Elisabeth Trawinski, Jean-Michel Pawlotsky |
| EPI_ISL_2984088 | LBM CARRON SELAS | Department of Virology, Henri Mondor University Hospital, Assistance Publique Hôpitaux de Paris, Université Paris-Est Créteil, INSERM U955 | Christophe Rodriguez, Slim Fourati, Vanessa Demontant, Guillaume Gricourt, Melissa N'Debi, Alexandre Soulier, Elisabeth Trawinski, Jean-Michel Pawlotsky |
| EPI_ISL_2984089 | CH AUXERRE | Department of Virology, Henri Mondor University Hospital, Assistance Publique Hôpitaux de Paris, Université Paris-Est Créteil, INSERM U955 | Christophe Rodriguez, Slim Fourati, Vanessa Demontant, Guillaume Gricourt, Melissa N'Debi, Alexandre Soulier, Elisabeth Trawinski, Jean-Michel Pawlotsky |
| EPI_ISL_2984090 | ASST MONZA | ASST MONZA | Sergio Maria Ivano Malandrin |
| EPI_ISL_2984091 | Laboratoire Ana-L | Department of Virology, Henri Mondor University Hospital, Assistance Publique Hôpitaux de Paris, Université Paris-Est Créteil, INSERM U955 | Christophe Rodriguez, Slim Fourati, Vanessa Demontant, Guillaume Gricourt, Melissa N'Debi, Alexandre Soulier, Elisabeth Trawinski, Jean-Michel Pawlotsky |
| EPI_ISL_2984092 | BIOMNIS EUROFINS IVRY | Department of Virology, Henri Mondor University Hospital, Assistance Publique Hôpitaux de Paris, Université Paris-Est Créteil, INSERM U955 | Christophe Rodriguez, Slim Fourati, Vanessa Demontant, Guillaume Gricourt, Melissa N'Debi, Alexandre Soulier, Elisabeth Trawinski, Jean-Michel Pawlotsky |
| EPI_ISL_2984093 | ASST MONZA | ASST MONZA | Sergio Maria Ivano Malandrin |
| EPI_ISL_2984094 | BIOMNIS EUROFINS IVRY | Department of Virology, Henri Mondor University Hospital, Assistance Publique Hôpitaux de Paris, Université Paris-Est Créteil, INSERM U955 | Christophe Rodriguez, Slim Fourati, Vanessa Demontant, Guillaume Gricourt, Melissa N'Debi, Alexandre Soulier, Elisabeth Trawinski, Jean-Michel Pawlotsky |
| EPI_ISL_2984095 | ASST MONZA | ASST MONZA | Sergio Maria Ivano Malandrin |
| EPI_ISL_2984096 | Biogroup Bio Lam-LCD Saint-Denis | Department of Virology, Henri Mondor University Hospital, Assistance Publique Hôpitaux de Paris, Université Paris-Est Créteil, INSERM U955 | Christophe Rodriguez, Slim Fourati, Vanessa Demontant, Guillaume Gricourt, Melissa N'Debi, Alexandre Soulier, Elisabeth Trawinski, Jean-Michel Pawlotsky |
| EPI_ISL_2984097 | ASST MONZA | ASST MONZA | Sergio Maria Ivano Malandrin |
| EPI_ISL_2984098 | Laboratoire LENYS | Department of Virology, Henri Mondor University Hospital, Assistance Publique Hôpitaux de Paris, Université Paris-Est Créteil, INSERM U955 | Christophe Rodriguez, Slim Fourati, Vanessa Demontant, Guillaume Gricourt, Melissa N'Debi, Alexandre Soulier, Elisabeth Trawinski, Jean-Michel Pawlotsky |
| EPI_ISL_2984099 | ASST MONZA | ASST MONZA | Sergio Maria Ivano Malandrin |
| EPI_ISL_2984100 | BIOMNIS EUROFINS IVRY | Department of Virology, Henri Mondor University Hospital, Assistance Publique Hôpitaux de Paris, Université Paris-Est Créteil, INSERM U955 | Christophe Rodriguez, Slim Fourati, Vanessa Demontant, Guillaume Gricourt, Melissa N'Debi, Alexandre Soulier, Elisabeth Trawinski, Jean-Michel Pawlotsky |
| EPI_ISL_2984101 | ASST MONZA | ASST MONZA | Sergio Maria Ivano Malandrin |
| EPI_ISL_2984102 | Laboratoire Ana-L | Department of Virology, Henri Mondor University Hospital, Assistance Publique Hôpitaux de Paris, Université Paris-Est Créteil, INSERM U955 | Christophe Rodriguez, Slim Fourati, Vanessa Demontant, Guillaume Gricourt, Melissa N'Debi, Alexandre Soulier, Elisabeth Trawinski, Jean-Michel Pawlotsky |
| EPI_ISL_2984104, EPI_ISL_2984105 | Cerballiance Paris IDF est Saint-Denis | Department of Virology, Henri Mondor University Hospital, Assistance Publique Hôpitaux de Paris, Université Paris-Est Créteil, INSERM U955 | Christophe Rodriguez, Slim Fourati, Vanessa Demontant, Guillaume Gricourt, Melissa N'Debi, Alexandre Soulier, Elisabeth Trawinski, Jean-Michel Pawlotsky |
| EPI_ISL_2984106 | ASST MONZA | ASST MONZA | Sergio Maria Ivano Malandrin |
| EPI_ISL_2984107 | Biogroup Bio Lam-LCD Saint-Denis | Department of Virology, Henri Mondor University Hospital, Assistance Publique Hôpitaux de Paris, Université Paris-Est Créteil, INSERM U955 | Christophe Rodriguez, Slim Fourati, Vanessa Demontant, Guillaume Gricourt, Melissa N'Debi, Alexandre Soulier, Elisabeth Trawinski, Jean-Michel Pawlotsky |
| EPI_ISL_2984108, EPI_ISL_2984110 | ASST MONZA | ASST MONZA | Sergio Maria Ivano Malandrin |
| EPI_ISL_2984111 | Laboratoire Ana-L | Department of Virology, Henri Mondor University Hospital, Assistance Publique Hôpitaux de Paris, Université Paris-Est Créteil, INSERM U955 | Christophe Rodriguez, Slim Fourati, Vanessa Demontant, Guillaume Gricourt, Melissa N'Debi, Alexandre Soulier, Elisabeth Trawinski, Jean-Michel Pawlotsky |
| EPI_ISL_2984112 | ASST MONZA | ASST MONZA | Sergio Maria Ivano Malandrin |
| EPI_ISL_2984113 | Biogroup Bio Lam-LCD Saint-Denis | Department of Virology, Henri Mondor University Hospital, Assistance Publique Hôpitaux de Paris, Université Paris-Est Créteil, INSERM U955 | Christophe Rodriguez, Slim Fourati, Vanessa Demontant, Guillaume Gricourt, Melissa N'Debi, Alexandre Soulier, Elisabeth Trawinski, Jean-Michel Pawlotsky |
| EPI_ISL_2984114 | ASST MONZA | ASST MONZA | Sergio Maria Ivano Malandrin |
| EPI_ISL_2984115 | Biogroup Bio Lam-LCD Saint-Denis | Department of Virology, Henri Mondor University Hospital, Assistance Publique Hôpitaux de Paris, Université Paris-Est Créteil, INSERM U955 | Christophe Rodriguez, Slim Fourati, Vanessa Demontant, Guillaume Gricourt, Melissa N'Debi, Alexandre Soulier, Elisabeth Trawinski, Jean-Michel Pawlotsky |
| EPI_ISL_2984116, EPI_ISL_2984118 | ASST MONZA | ASST MONZA | Sergio Maria Ivano Malandrin |
| EPI_ISL_2984119 | BIOMNIS EUROFINS IVRY | Department of Virology, Henri Mondor University Hospital, Assistance Publique Hôpitaux de Paris, Université Paris-Est Créteil, INSERM U955 | Christophe Rodriguez, Slim Fourati, Vanessa Demontant, Guillaume Gricourt, Melissa N'Debi, Alexandre Soulier, Elisabeth Trawinski, Jean-Michel Pawlotsky |
| EPI_ISL_2984120 | ASST MONZA | ASST MONZA | Sergio Maria Ivano Malandrin |
| EPI_ISL_2984121 | SELAS MEDILYS | Department of Virology, Henri Mondor University Hospital, Assistance Publique Hôpitaux de Paris, Université Paris-Est Créteil, INSERM U955 | Christophe Rodriguez, Slim Fourati, Vanessa Demontant, Guillaume Gricourt, Melissa N'Debi, Alexandre Soulier, Elisabeth Trawinski, Jean-Michel Pawlotsky |
| EPI_ISL_2984122 | ASST MONZA | ASST MONZA | Sergio Maria Ivano Malandrin |
| EPI_ISL_2984123 | Laboratoire LENYS | Department of Virology, Henri Mondor University Hospital, Assistance Publique Hôpitaux de Paris, Université Paris-Est Créteil, INSERM U955 | Christophe Rodriguez, Slim Fourati, Vanessa Demontant, Guillaume Gricourt, Melissa N'Debi, Alexandre Soulier, Elisabeth Trawinski, Jean-Michel Pawlotsky |
| EPI_ISL_2984124 | ASST MONZA | ASST MONZA | Sergio Maria Ivano Malandrin |
| EPI_ISL_2984125 | LBM CARRON SELAS | Department of Virology, Henri Mondor University Hospital, Assistance Publique Hôpitaux de Paris, Université Paris-Est Créteil, INSERM U955 | Christophe Rodriguez, Slim Fourati, Vanessa Demontant, Guillaume Gricourt, Melissa N'Debi, Alexandre Soulier, Elisabeth Trawinski, Jean-Michel Pawlotsky |
| EPI_ISL_2984126, EPI_ISL_2984128 | ASST MONZA | ASST MONZA | Sergio Maria Ivano Malandrin |
| EPI_ISL_2984129 | Laboratoire LENYS | Department of Virology, Henri Mondor University Hospital, | Christophe Rodriguez, Slim Fourati, Vanessa Demontant, Guillaume Gricourt, Melissa N'Debi, Alexandre Soulier, Elisabeth Trawinski, Jean-Michel Pawlotsky |

| | | | |
|---|---|---|---|
| | | Assistance Publique Hôpitaux de Paris, Université Paris-Est Créteil, INSERM U955 | |
| EPI_ISL_2984130 | ASST MONZA | ASST MONZA | Sergio Maria Ivano Malandrin |
| EPI_ISL_2984131 | Laboratoire LENYS | Department of Virology, Henri Mondor University Hospital, Assistance Publique Hôpitaux de Paris, Université Paris-Est Créteil, INSERM U955 | Christophe Rodriguez, Slim Fourati, Vanessa Demontant, Guillaume Gricourt, Melissa N'Debi, Alexandre Soulier, Elisabeth Trawinski, Jean-Michel Pawlotsky |
| EPI_ISL_2984132 | ASST MONZA | ASST MONZA | Sergio Maria Ivano Malandrin |
| EPI_ISL_2984133 | BIOMNIS EUROFINS IVRY | Department of Virology, Henri Mondor University Hospital, Assistance Publique Hôpitaux de Paris, Université Paris-Est Créteil, INSERM U955 | Christophe Rodriguez, Slim Fourati, Vanessa Demontant, Guillaume Gricourt, Melissa N'Debi, Alexandre Soulier, Elisabeth Trawinski, Jean-Michel Pawlotsky |
| EPI_ISL_2984134 | ASST MONZA | ASST MONZA | Sergio Maria Ivano Malandrin |
| EPI_ISL_2984135 | Plateau technique Biomer Metz Verriers | Department of Virology, Henri Mondor University Hospital, Assistance Publique Hôpitaux de Paris, Université Paris-Est Créteil, INSERM U955 | Christophe Rodriguez, Slim Fourati, Vanessa Demontant, Guillaume Gricourt, Melissa N'Debi, Alexandre Soulier, Elisabeth Trawinski, Jean-Michel Pawlotsky |
| EPI_ISL_2984136 | Cerballiance Paris IDF est Saint-Denis | Department of Virology, Henri Mondor University Hospital, Assistance Publique Hôpitaux de Paris, Université Paris-Est Créteil, INSERM U955 | Christophe Rodriguez, Slim Fourati, Vanessa Demontant, Guillaume Gricourt, Melissa N'Debi, Alexandre Soulier, Elisabeth Trawinski, Jean-Michel Pawlotsky |
| EPI_ISL_2984137 | ASST MONZA | ASST MONZA | Sergio Maria Ivano Malandrin |
| EPI_ISL_2984138 | Cerballiance Paris IDF est Saint-Denis | Department of Virology, Henri Mondor University Hospital, Assistance Publique Hôpitaux de Paris, Université Paris-Est Créteil, INSERM U955 | Christophe Rodriguez, Slim Fourati, Vanessa Demontant, Guillaume Gricourt, Melissa N'Debi, Alexandre Soulier, Elisabeth Trawinski, Jean-Michel Pawlotsky |
| EPI_ISL_2984139 | ASST MONZA | ASST MONZA | Sergio Maria Ivano Malandrin |
| EPI_ISL_2984140 | Laboratoire LENYS | Department of Virology, Henri Mondor University Hospital, Assistance Publique Hôpitaux de Paris, Université Paris-Est Créteil, INSERM U955 | Christophe Rodriguez, Slim Fourati, Vanessa Demontant, Guillaume Gricourt, Melissa N'Debi, Alexandre Soulier, Elisabeth Trawinski, Jean-Michel Pawlotsky |
| EPI_ISL_2984141 | ASST MONZA | ASST MONZA | Sergio Maria Ivano Malandrin |
| EPI_ISL_2984142 | BIOMNIS EUROFINS IVRY | Department of Virology, Henri Mondor University Hospital, Assistance Publique Hôpitaux de Paris, Université Paris-Est Créteil, INSERM U955 | Christophe Rodriguez, Slim Fourati, Vanessa Demontant, Guillaume Gricourt, Melissa N'Debi, Alexandre Soulier, Elisabeth Trawinski, Jean-Michel Pawlotsky |
| EPI_ISL_2984143 | ASST MONZA | ASST MONZA | Sergio Maria Ivano Malandrin |
| EPI_ISL_2984144 | Plateau technique Biomer Metz Verriers | Department of Virology, Henri Mondor University Hospital, Assistance Publique Hôpitaux de Paris, Université Paris-Est Créteil, INSERM U955 | Christophe Rodriguez, Slim Fourati, Vanessa Demontant, Guillaume Gricourt, Melissa N'Debi, Alexandre Soulier, Elisabeth Trawinski, Jean-Michel Pawlotsky |
| EPI_ISL_2984145 | ASST MONZA | ASST MONZA | Sergio Maria Ivano Malandrin |
| EPI_ISL_2984146 | Plateau technique Biomer Metz Verriers | Department of Virology, Henri Mondor University Hospital, Assistance Publique Hôpitaux de Paris, Université Paris-Est Créteil, INSERM U955 | Christophe Rodriguez, Slim Fourati, Vanessa Demontant, Guillaume Gricourt, Melissa N'Debi, Alexandre Soulier, Elisabeth Trawinski, Jean-Michel Pawlotsky |
| EPI_ISL_2984147 | BIOMNIS EUROFINS IVRY | Department of Virology, Henri Mondor University Hospital, Assistance Publique Hôpitaux de Paris, Université Paris-Est Créteil, INSERM U955 | Christophe Rodriguez, Slim Fourati, Vanessa Demontant, Guillaume Gricourt, Melissa N'Debi, Alexandre Soulier, Elisabeth Trawinski, Jean-Michel Pawlotsky |
| EPI_ISL_2984148 | ASST MONZA | ASST MONZA | Sergio Maria Ivano Malandrin |
| EPI_ISL_2984149 | Biogroup Bio Lam-LCD Saint-Denis | Department of Virology, Henri Mondor University Hospital, Assistance Publique Hôpitaux de Paris, Université Paris-Est Créteil, INSERM U955 | Christophe Rodriguez, Slim Fourati, Vanessa Demontant, Guillaume Gricourt, Melissa N'Debi, Alexandre Soulier, Elisabeth Trawinski, Jean-Michel Pawlotsky |
| EPI_ISL_2984150 | Cerballiance Paris IDF est Saint-Denis | Department of Virology, Henri Mondor University Hospital, Assistance Publique Hôpitaux de Paris, Université Paris-Est Créteil, INSERM U955 | Christophe Rodriguez, Slim Fourati, Vanessa Demontant, Guillaume Gricourt, Melissa N'Debi, Alexandre Soulier, Elisabeth Trawinski, Jean-Michel Pawlotsky |
| EPI_ISL_2984152, EPI_ISL_2984153, EPI_ISL_2984154 | Biogroup Bio Lam-LCD Saint-Denis | Department of Virology, Henri Mondor University Hospital, Assistance Publique Hôpitaux de Paris, Université Paris-Est Créteil, INSERM U955 | Christophe Rodriguez, Slim Fourati, Vanessa Demontant, Guillaume Gricourt, Melissa N'Debi, Alexandre Soulier, Elisabeth Trawinski, Jean-Michel Pawlotsky |
| EPI_ISL_2984155 | CH. ROBERT BISSON | Department of Virology, Henri Mondor University Hospital, Assistance Publique Hôpitaux de Paris, Université Paris-Est Créteil, INSERM U955 | Christophe Rodriguez, Slim Fourati, Vanessa Demontant, Guillaume Gricourt, Melissa N'Debi, Alexandre Soulier, Elisabeth Trawinski, Jean-Michel Pawlotsky |
| EPI_ISL_2984156, EPI_ISL_2984158 | Biogroup Bio Lam-LCD Saint-Denis | Department of Virology, Henri Mondor University Hospital, Assistance Publique Hôpitaux de Paris, Université Paris-Est Créteil, INSERM U955 | Christophe Rodriguez, Slim Fourati, Vanessa Demontant, Guillaume Gricourt, Melissa N'Debi, Alexandre Soulier, Elisabeth Trawinski, Jean-Michel Pawlotsky |
| EPI_ISL_2984159 | BIOMNIS EUROFINS IVRY | Department of Virology, Henri Mondor University Hospital, Assistance Publique Hôpitaux de Paris, Université Paris-Est Créteil, INSERM U955 | Christophe Rodriguez, Slim Fourati, Vanessa Demontant, Guillaume Gricourt, Melissa N'Debi, Alexandre Soulier, Elisabeth Trawinski, Jean-Michel Pawlotsky |
| EPI_ISL_2984160 | Plateau technique Biomer Metz Verriers | Department of Virology, Henri Mondor University Hospital, Assistance Publique Hôpitaux de Paris, Université Paris-Est Créteil, INSERM U955 | Christophe Rodriguez, Slim Fourati, Vanessa Demontant, Guillaume Gricourt, Melissa N'Debi, Alexandre Soulier, Elisabeth Trawinski, Jean-Michel Pawlotsky |
| EPI_ISL_2984161 | Biogroup Bio Lam-LCD Saint-Denis | Department of Virology, Henri Mondor University Hospital, Assistance Publique Hôpitaux de Paris, Université Paris-Est Créteil, INSERM U955 | Christophe Rodriguez, Slim Fourati, Vanessa Demontant, Guillaume Gricourt, Melissa N'Debi, Alexandre Soulier, Elisabeth Trawinski, Jean-Michel Pawlotsky |
| EPI_ISL_2984163 | CH. ROBERT BISSON | Department of Virology, Henri Mondor University Hospital, Assistance Publique Hôpitaux de Paris, Université Paris-Est Créteil, INSERM U955 | Christophe Rodriguez, Slim Fourati, Vanessa Demontant, Guillaume Gricourt, Melissa N'Debi, Alexandre Soulier, Elisabeth Trawinski, Jean-Michel Pawlotsky |
| EPI_ISL_2984164, EPI_ISL_2984166 | BIOMNIS EUROFINS IVRY | Department of Virology, Henri Mondor University Hospital, Assistance Publique Hôpitaux de Paris, Université Paris-Est Créteil, INSERM U955 | Christophe Rodriguez, Slim Fourati, Vanessa Demontant, Guillaume Gricourt, Melissa N'Debi, Alexandre Soulier, Elisabeth Trawinski, Jean-Michel Pawlotsky |
| EPI_ISL_2984167 | Cerballiance Paris IDF est Saint-Denis | Department of Virology, Henri Mondor University Hospital, Assistance Publique Hôpitaux de Paris, Université Paris-Est Créteil, INSERM U955 | Christophe Rodriguez, Slim Fourati, Vanessa Demontant, Guillaume Gricourt, Melissa N'Debi, Alexandre Soulier, Elisabeth Trawinski, Jean-Michel Pawlotsky |
| EPI_ISL_2984169 | Laboratoire Ana-L | Department of Virology, Henri Mondor University Hospital, Assistance Publique Hôpitaux de Paris, Université Paris-Est Créteil, INSERM U955 | Christophe Rodriguez, Slim Fourati, Vanessa Demontant, Guillaume Gricourt, Melissa N'Debi, Alexandre Soulier, Elisabeth Trawinski, Jean-Michel Pawlotsky |
| EPI_ISL_2984171 | Cerballiance Paris IDF est Saint-Denis | Department of Virology, Henri Mondor University Hospital, Assistance Publique Hôpitaux de Paris, Université Paris-Est Créteil, INSERM U955 | Christophe Rodriguez, Slim Fourati, Vanessa Demontant, Guillaume Gricourt, Melissa N'Debi, Alexandre Soulier, Elisabeth Trawinski, Jean-Michel Pawlotsky |
| EPI_ISL_2984172 | BIOMNIS EUROFINS IVRY | Department of Virology, Henri Mondor University Hospital, Assistance Publique Hôpitaux de Paris, Université Paris-Est Créteil, INSERM U955 | Christophe Rodriguez, Slim Fourati, Vanessa Demontant, Guillaume Gricourt, Melissa N'Debi, Alexandre Soulier, Elisabeth Trawinski, Jean-Michel Pawlotsky |
| EPI_ISL_2984173 | Biogroup Bio Lam-LCD Saint-Denis | Department of Virology, Henri Mondor University Hospital, Assistance Publique Hôpitaux de Paris, Université Paris-Est Créteil, INSERM U955 | Christophe Rodriguez, Slim Fourati, Vanessa Demontant, Guillaume Gricourt, Melissa N'Debi, Alexandre Soulier, Elisabeth Trawinski, Jean-Michel Pawlotsky |
| EPI_ISL_2984174 | Laboratoire LENYS | Department of Virology, Henri Mondor University Hospital, Assistance Publique Hôpitaux de Paris, Université Paris-Est Créteil, INSERM U955 | Christophe Rodriguez, Slim Fourati, Vanessa Demontant, Guillaume Gricourt, Melissa N'Debi, Alexandre Soulier, Elisabeth Trawinski, Jean-Michel Pawlotsky |
| EPI_ISL_2984175, EPI_ISL_2984177, EPI_ISL_2984178 | Biogroup Bio Lam-LCD Saint-Denis | Department of Virology, Henri Mondor University Hospital, Assistance Publique Hôpitaux de Paris, Université Paris-Est Créteil, INSERM U955 | Christophe Rodriguez, Slim Fourati, Vanessa Demontant, Guillaume Gricourt, Melissa N'Debi, Alexandre Soulier, Elisabeth Trawinski, Jean-Michel Pawlotsky |
| EPI_ISL_2984179 | Laboratoire CBM 25 TERRE ROUGE | Department of Virology, Henri Mondor University Hospital, | Christophe Rodriguez, Slim Fourati, Vanessa Demontant, Guillaume Gricourt, Melissa N'Debi, Alexandre Soulier, Elisabeth Trawinski, Jean-Michel Pawlotsky |

| | | | |
|---|---|---|---|
| | | Assistance Publique Hôpitaux de Paris, Université Paris-Est Créteil, INSERM U955 | |
| EPI_ISL_2984180 | BIOMNIS EUROFINS IVRY | Department of Virology, Henri Mondor University Hospital, Assistance Publique Hôpitaux de Paris, Université Paris-Est Créteil, INSERM U955 | Christophe Rodriguez, Slim Fourati, Vanessa Demontant, Guillaume Gricourt, Melissa N'Debi, Alexandre Soulier, Elisabeth Trawinski, Jean-Michel Pawlotsky |
| EPI_ISL_2984181 | Biogroup Bio Lam-LCD Saint-Denis | Department of Virology, Henri Mondor University Hospital, Assistance Publique Hôpitaux de Paris, Université Paris-Est Créteil, INSERM U955 | Christophe Rodriguez, Slim Fourati, Vanessa Demontant, Guillaume Gricourt, Melissa N'Debi, Alexandre Soulier, Elisabeth Trawinski, Jean-Michel Pawlotsky |
| EPI_ISL_2984183 | LBM BIOEMERAUDE | Department of Virology, Henri Mondor University Hospital, Assistance Publique Hôpitaux de Paris, Université Paris-Est Créteil, INSERM U955 | Christophe Rodriguez, Slim Fourati, Vanessa Demontant, Guillaume Gricourt, Melissa N'Debi, Alexandre Soulier, Elisabeth Trawinski, Jean-Michel Pawlotsky |
| EPI_ISL_2984184 | Biogroup Bio Lam-LCD Saint-Denis | Department of Virology, Henri Mondor University Hospital, Assistance Publique Hôpitaux de Paris, Université Paris-Est Créteil, INSERM U955 | Christophe Rodriguez, Slim Fourati, Vanessa Demontant, Guillaume Gricourt, Melissa N'Debi, Alexandre Soulier, Elisabeth Trawinski, Jean-Michel Pawlotsky |
| EPI_ISL_2984185 | LBM CARRON SELAS | Department of Virology, Henri Mondor University Hospital, Assistance Publique Hôpitaux de Paris, Université Paris-Est Créteil, INSERM U955 | Christophe Rodriguez, Slim Fourati, Vanessa Demontant, Guillaume Gricourt, Melissa N'Debi, Alexandre Soulier, Elisabeth Trawinski, Jean-Michel Pawlotsky |
| EPI_ISL_2984188 | Laboratoire LENYS | Department of Virology, Henri Mondor University Hospital, Assistance Publique Hôpitaux de Paris, Université Paris-Est Créteil, INSERM U955 | Christophe Rodriguez, Slim Fourati, Vanessa Demontant, Guillaume Gricourt, Melissa N'Debi, Alexandre Soulier, Elisabeth Trawinski, Jean-Michel Pawlotsky |
| EPI_ISL_2984189, EPI_ISL_2984190 | Biogroup Bio Lam-LCD Saint-Denis | Department of Virology, Henri Mondor University Hospital, Assistance Publique Hôpitaux de Paris, Université Paris-Est Créteil, INSERM U955 | Christophe Rodriguez, Slim Fourati, Vanessa Demontant, Guillaume Gricourt, Melissa N'Debi, Alexandre Soulier, Elisabeth Trawinski, Jean-Michel Pawlotsky |
| EPI_ISL_2984191, EPI_ISL_2984192 | Plateau technique Biomer Metz Verriers | Department of Virology, Henri Mondor University Hospital, Assistance Publique Hôpitaux de Paris, Université Paris-Est Créteil, INSERM U955 | Christophe Rodriguez, Slim Fourati, Vanessa Demontant, Guillaume Gricourt, Melissa N'Debi, Alexandre Soulier, Elisabeth Trawinski, Jean-Michel Pawlotsky |
| EPI_ISL_2984193 | Biogroup Bio Lam-LCD Saint-Denis | Department of Virology, Henri Mondor University Hospital, Assistance Publique Hôpitaux de Paris, Université Paris-Est Créteil, INSERM U955 | Christophe Rodriguez, Slim Fourati, Vanessa Demontant, Guillaume Gricourt, Melissa N'Debi, Alexandre Soulier, Elisabeth Trawinski, Jean-Michel Pawlotsky |
| EPI_ISL_2984194 | Laboratoire LENYS | Department of Virology, Henri Mondor University Hospital, Assistance Publique Hôpitaux de Paris, Université Paris-Est Créteil, INSERM U955 | Christophe Rodriguez, Slim Fourati, Vanessa Demontant, Guillaume Gricourt, Melissa N'Debi, Alexandre Soulier, Elisabeth Trawinski, Jean-Michel Pawlotsky |
| EPI_ISL_2984195 | Biogroup Bio Lam-LCD Saint-Denis | Department of Virology, Henri Mondor University Hospital, Assistance Publique Hôpitaux de Paris, Université Paris-Est Créteil, INSERM U955 | Christophe Rodriguez, Slim Fourati, Vanessa Demontant, Guillaume Gricourt, Melissa N'Debi, Alexandre Soulier, Elisabeth Trawinski, Jean-Michel Pawlotsky |
| EPI_ISL_2984197 | Laboratoire LENYS | Department of Virology, Henri Mondor University Hospital, Assistance Publique Hôpitaux de Paris, Université Paris-Est Créteil, INSERM U955 | Christophe Rodriguez, Slim Fourati, Vanessa Demontant, Guillaume Gricourt, Melissa N'Debi, Alexandre Soulier, Elisabeth Trawinski, Jean-Michel Pawlotsky |
| EPI_ISL_2984198, EPI_ISL_2984199 | Biogroup Bio Lam-LCD Saint-Denis | Department of Virology, Henri Mondor University Hospital, Assistance Publique Hôpitaux de Paris, Université Paris-Est Créteil, INSERM U955 | Christophe Rodriguez, Slim Fourati, Vanessa Demontant, Guillaume Gricourt, Melissa N'Debi, Alexandre Soulier, Elisabeth Trawinski, Jean-Michel Pawlotsky |
| EPI_ISL_2984200 | SELAS MEDILYS | Department of Virology, Henri Mondor University Hospital, Assistance Publique Hôpitaux de Paris, Université Paris-Est Créteil, INSERM U955 | Christophe Rodriguez, Slim Fourati, Vanessa Demontant, Guillaume Gricourt, Melissa N'Debi, Alexandre Soulier, Elisabeth Trawinski, Jean-Michel Pawlotsky |
| EPI_ISL_2984201 | BIOMNIS EUROFINS IVRY | Department of Virology, Henri Mondor University Hospital, Assistance Publique Hôpitaux de Paris, Université Paris-Est Créteil, INSERM U955 | Christophe Rodriguez, Slim Fourati, Vanessa Demontant, Guillaume Gricourt, Melissa N'Debi, Alexandre Soulier, Elisabeth Trawinski, Jean-Michel Pawlotsky |
| EPI_ISL_2984202 | Biogroup Bio Lam-LCD Saint-Denis | Department of Virology, Henri Mondor University Hospital, Assistance Publique Hôpitaux de Paris, Université Paris-Est Créteil, INSERM U955 | Christophe Rodriguez, Slim Fourati, Vanessa Demontant, Guillaume Gricourt, Melissa N'Debi, Alexandre Soulier, Elisabeth Trawinski, Jean-Michel Pawlotsky |
| EPI_ISL_2984204 | Plateau technique Biomer Metz Verriers | Department of Virology, Henri Mondor University Hospital, Assistance Publique Hôpitaux de Paris, Université Paris-Est Créteil, INSERM U955 | Christophe Rodriguez, Slim Fourati, Vanessa Demontant, Guillaume Gricourt, Melissa N'Debi, Alexandre Soulier, Elisabeth Trawinski, Jean-Michel Pawlotsky |
| EPI_ISL_2984205, EPI_ISL_2984206 | Biogroup Bio Lam-LCD Saint-Denis | Department of Virology, Henri Mondor University Hospital, Assistance Publique Hôpitaux de Paris, Université Paris-Est Créteil, INSERM U955 | Christophe Rodriguez, Slim Fourati, Vanessa Demontant, Guillaume Gricourt, Melissa N'Debi, Alexandre Soulier, Elisabeth Trawinski, Jean-Michel Pawlotsky |
| EPI_ISL_2984207 | SELAFA SANTE LABO | Department of Virology, Henri Mondor University Hospital, Assistance Publique Hôpitaux de Paris, Université Paris-Est Créteil, INSERM U955 | Christophe Rodriguez, Slim Fourati, Vanessa Demontant, Guillaume Gricourt, Melissa N'Debi, Alexandre Soulier, Elisabeth Trawinski, Jean-Michel Pawlotsky |
| EPI_ISL_2984208 | BIOMNIS EUROFINS IVRY | Department of Virology, Henri Mondor University Hospital, Assistance Publique Hôpitaux de Paris, Université Paris-Est Créteil, INSERM U955 | Christophe Rodriguez, Slim Fourati, Vanessa Demontant, Guillaume Gricourt, Melissa N'Debi, Alexandre Soulier, Elisabeth Trawinski, Jean-Michel Pawlotsky |
| EPI_ISL_2984209 | Laboratoire Ana-L | Department of Virology, Henri Mondor University Hospital, Assistance Publique Hôpitaux de Paris, Université Paris-Est Créteil, INSERM U955 | Christophe Rodriguez, Slim Fourati, Vanessa Demontant, Guillaume Gricourt, Melissa N'Debi, Alexandre Soulier, Elisabeth Trawinski, Jean-Michel Pawlotsky |
| EPI_ISL_2984210 | SELAS MEDILYS | Department of Virology, Henri Mondor University Hospital, Assistance Publique Hôpitaux de Paris, Université Paris-Est Créteil, INSERM U955 | Christophe Rodriguez, Slim Fourati, Vanessa Demontant, Guillaume Gricourt, Melissa N'Debi, Alexandre Soulier, Elisabeth Trawinski, Jean-Michel Pawlotsky |
| EPI_ISL_2984211 | Biogroup Bio Lam-LCD Saint-Denis | Department of Virology, Henri Mondor University Hospital, Assistance Publique Hôpitaux de Paris, Université Paris-Est Créteil, INSERM U955 | Christophe Rodriguez, Slim Fourati, Vanessa Demontant, Guillaume Gricourt, Melissa N'Debi, Alexandre Soulier, Elisabeth Trawinski, Jean-Michel Pawlotsky |
| EPI_ISL_2984212 | BIOMNIS EUROFINS IVRY | Department of Virology, Henri Mondor University Hospital, Assistance Publique Hôpitaux de Paris, Université Paris-Est Créteil, INSERM U955 | Christophe Rodriguez, Slim Fourati, Vanessa Demontant, Guillaume Gricourt, Melissa N'Debi, Alexandre Soulier, Elisabeth Trawinski, Jean-Michel Pawlotsky |
| EPI_ISL_2984215 | Biogroup Bio Lam-LCD Saint-Denis | Department of Virology, Henri Mondor University Hospital, Assistance Publique Hôpitaux de Paris, Université Paris-Est Créteil, INSERM U955 | Christophe Rodriguez, Slim Fourati, Vanessa Demontant, Guillaume Gricourt, Melissa N'Debi, Alexandre Soulier, Elisabeth Trawinski, Jean-Michel Pawlotsky |
| EPI_ISL_2984218 | BIOMNIS EUROFINS IVRY | Department of Virology, Henri Mondor University Hospital, Assistance Publique Hôpitaux de Paris, Université Paris-Est Créteil, INSERM U955 | Christophe Rodriguez, Slim Fourati, Vanessa Demontant, Guillaume Gricourt, Melissa N'Debi, Alexandre Soulier, Elisabeth Trawinski, Jean-Michel Pawlotsky |
| EPI_ISL_2984219, EPI_ISL_2984221 | Biogroup Bio Lam-LCD Saint-Denis | Department of Virology, Henri Mondor University Hospital, Assistance Publique Hôpitaux de Paris, Université Paris-Est Créteil, INSERM U955 | Christophe Rodriguez, Slim Fourati, Vanessa Demontant, Guillaume Gricourt, Melissa N'Debi, Alexandre Soulier, Elisabeth Trawinski, Jean-Michel Pawlotsky |
| EPI_ISL_2984223 | BIOMNIS EUROFINS IVRY | Department of Virology, Henri Mondor University Hospital, Assistance Publique Hôpitaux de Paris, Université Paris-Est Créteil, INSERM U955 | Christophe Rodriguez, Slim Fourati, Vanessa Demontant, Guillaume Gricourt, Melissa N'Debi, Alexandre Soulier, Elisabeth Trawinski, Jean-Michel Pawlotsky |
| EPI_ISL_2984224, EPI_ISL_2984226, EPI_ISL_2984227, EPI_ISL_2984228, EPI_ISL_2984229, EPI_ISL_2984230, EPI_ISL_2984231, EPI_ISL_2984232, EPI_ISL_2984233, EPI_ISL_2984234, EPI_ISL_2984235, EPI_ISL_2984236, EPI_ISL_2984237, EPI_ISL_2984238, EPI_ISL_2984239, EPI_ISL_2984240, EPI_ISL_2984242, EPI_ISL_2984243, EPI_ISL_2984244, EPI_ISL_2984245, EPI_ISL_2984246 | | | |
| see above | Jahra Hospital | Kuwait Cancer Control Center | Shakir Bahzad, Mona Alateeqi |
| EPI_ISL_2984247, EPI_ISL_2984252, EPI_ISL_2984253, EPI_ISL_2984255, EPI_ISL_2984256, EPI_ISL_2984257, EPI_ISL_2984258, EPI_ISL_2984259, EPI_ISL_2984260, EPI_ISL_2984261, EPI_ISL_2984262, EPI_ISL_2984263, EPI_ISL_2984264, EPI_ISL_2984265, EPI_ISL_2984266, EPI_ISL_2984267, EPI_ISL_2984268, EPI_ISL_2984269, EPI_ISL_2984270, EPI_ISL_2984271, EPI_ISL_2984272, EPI_ISL_2984273, EPI_ISL_2984274, EPI_ISL_2984275, EPI_ISL_2984276, EPI_ISL_2984277, EPI_ISL_2984278, EPI_ISL_2984279, EPI_ISL_2984280, EPI_ISL_2984281, EPI_ISL_2984282, EPI_ISL_2984283, EPI_ISL_2984284, EPI_ISL_2984285, EPI_ISL_2984286, EPI_ISL_2984287, EPI_ISL_2984288, EPI_ISL_2984289, EPI_ISL_2984290, EPI_ISL_2984291, EPI_ISL_2984292, EPI_ISL_2984293, EPI_ISL_2984294, EPI_ISL_2984295, EPI_ISL_2984296, EPI_ISL_2984297, EPI_ISL_2984298, EPI_ISL_2984299, EPI_ISL_2984300, EPI_ISL_2984301, EPI_ISL_2984302, EPI_ISL_2984303, EPI_ISL_2984304, EPI_ISL_2984305, EPI_ISL_2984306, EPI_ISL_2984307, EPI_ISL_2984308, EPI_ISL_2984309, EPI_ISL_2984310, EPI_ISL_2984311, EPI_ISL_2984312, EPI_ISL_2984313, EPI_ISL_2984314, EPI_ISL_2984315, EPI_ISL_2984316, EPI_ISL_2984317, EPI_ISL_2984318, EPI_ISL_2984319, EPI_ISL_2984320, EPI_ISL_2984321, EPI_ISL_2984322, EPI_ISL_2984323, EPI_ISL_2984324, EPI_ISL_2984325, EPI_ISL_2984326, EPI_ISL_2984327, EPI_ISL_2984328, EPI_ISL_2984329, EPI_ISL_2984330, EPI_ISL_2984331, EPI_ISL_2984332, EPI_ISL_2984333, EPI_ISL_2984334, EPI_ISL_2984335, EPI_ISL_2984336, EPI_ISL_2984337, EPI_ISL_2984338, EPI_ISL_2984339, EPI_ISL_2984340, EPI_ISL_2984341, EPI_ISL_2984342, EPI_ISL_2984344, EPI_ISL_2984345, EPI_ISL_2984346, EPI_ISL_2984348, EPI_ISL_2984350, EPI_ISL_2984364, EPI_ISL_2984366, EPI_ISL_2984367, EPI_ISL_2984368, EPI_ISL_2984377, EPI_ISL_2984378, EPI_ISL_2984379, EPI_ISL_2984380, EPI_ISL_2984382, EPI_ISL_2984383, EPI_ISL_2984384, EPI_ISL_2984385, EPI_ISL_2984386, EPI_ISL_2984387, EPI_ISL_2984388, EPI_ISL_2984389, EPI_ISL_2984390, EPI_ISL_2984391, EPI_ISL_2984392, EPI_ISL_2984393, EPI_ISL_2984394, EPI_ISL_2984395, EPI_ISL_2984396, EPI_ISL_2984397, EPI_ISL_2984398, EPI_ISL_2984399, EPI_ISL_2984400, EPI_ISL_2984401, EPI_ISL_2984402, EPI_ISL_2984403, EPI_ISL_2984404, EPI_ISL_2984405, EPI_ISL_2984406, EPI_ISL_2984407, EPI_ISL_2984408, EPI_ISL_2984409, EPI_ISL_2984411, EPI_ISL_2984412, EPI_ISL_2984413, EPI_ISL_2984415, EPI_ISL_2984416, EPI_ISL_2984417, EPI_ISL_2984418, EPI_ISL_2984419, EPI_ISL_2984420, EPI_ISL_2984422, EPI_ISL_2984423, EPI_ISL_2984424, EPI_ISL_2984425, EPI_ISL_2984426, EPI_ISL_2984427, EPI_ISL_2984428 | | | |
| see above | National Virus Reference Laboratory | National Virus Reference Laboratory | Zoe Yandle, Charlene Bennett, Gabriel Gonzalez, Michael Carr, Jonathan Dean, Cillian F De Gascun |

| | | | |
|---|---|---|---|
| EPI_ISL_2984429, EPI_ISL_2984430, EPI_ISL_2984431, EPI_ISL_2984438, EPI_ISL_2984439, EPI_ISL_2984440, EPI_ISL_2984441, EPI_ISL_2984442, EPI_ISL_2984443, EPI_ISL_2984444, EPI_ISL_2984445, EPI_ISL_2984449, EPI_ISL_2984450, EPI_ISL_2984451, EPI_ISL_2984452, EPI_ISL_2984453, EPI_ISL_2984455, EPI_ISL_2984456, EPI_ISL_2984463, EPI_ISL_2984464, EPI_ISL_2984465, EPI_ISL_2984482, EPI_ISL_2984485, EPI_ISL_2984489, EPI_ISL_2984491, EPI_ISL_2984492, EPI_ISL_2984493, EPI_ISL_2984494, EPI_ISL_2984497, EPI_ISL_2984498, EPI_ISL_2984499, EPI_ISL_2984500, EPI_ISL_2984501, EPI_ISL_2984502, EPI_ISL_2984503, EPI_ISL_2984504, EPI_ISL_2984505, EPI_ISL_2984507, EPI_ISL_2984508, EPI_ISL_2984509, EPI_ISL_2984510, EPI_ISL_2984511 | | | |
| see above | National Virus Reference Laboratory | National Virus Reference Laboratory | Guerrino Macori, Gabriel Gonzalez, Michael Carr, Zoe Yandle, Charlene Bennett, Jonathan Dean, Seamus Fanning, Cillian F De Gascun |
| EPI_ISL_2984513, EPI_ISL_2984514, EPI_ISL_2984515, EPI_ISL_2984516, EPI_ISL_2984517, EPI_ISL_2984518, EPI_ISL_2984520, EPI_ISL_2984526, EPI_ISL_2984527, EPI_ISL_2984530, EPI_ISL_2984534, EPI_ISL_2984537, EPI_ISL_2984546, EPI_ISL_2984547, EPI_ISL_2984552, EPI_ISL_2984557, EPI_ISL_2984560, EPI_ISL_2984561, EPI_ISL_2984562, EPI_ISL_2984563, EPI_ISL_2984578, EPI_ISL_2984584, EPI_ISL_2984585, EPI_ISL_2984591, EPI_ISL_2984592, EPI_ISL_2984594, EPI_ISL_2984595, EPI_ISL_2984605, EPI_ISL_2984606, EPI_ISL_2984609, EPI_ISL_2984610, EPI_ISL_2984612, EPI_ISL_2984613, EPI_ISL_2984614, EPI_ISL_2984615, EPI_ISL_2984617, EPI_ISL_2984618, EPI_ISL_2984619, EPI_ISL_2984620, EPI_ISL_2984621, EPI_ISL_2984628, EPI_ISL_2984629, EPI_ISL_2984632, EPI_ISL_2984634, EPI_ISL_2984636, EPI_ISL_2984637, EPI_ISL_2984640, EPI_ISL_2984641, EPI_ISL_2984642, EPI_ISL_2984645, EPI_ISL_2984646, EPI_ISL_2984648, EPI_ISL_2984650, EPI_ISL_2984652, EPI_ISL_2984653, EPI_ISL_2984655, EPI_ISL_2984656, EPI_ISL_2984658, EPI_ISL_2984659, EPI_ISL_2984660, EPI_ISL_2984662, EPI_ISL_2984667, EPI_ISL_2984669, EPI_ISL_2984670, EPI_ISL_2984671, EPI_ISL_2984673, EPI_ISL_2984675, EPI_ISL_2984676, EPI_ISL_2984678, EPI_ISL_2984679, EPI_ISL_2984680, EPI_ISL_2984681, EPI_ISL_2984682, EPI_ISL_2984683, EPI_ISL_2984684, EPI_ISL_2984685, EPI_ISL_2984687, EPI_ISL_2984692, EPI_ISL_2984693, EPI_ISL_2984695, EPI_ISL_2984705, EPI_ISL_2984706, EPI_ISL_2984707, EPI_ISL_2984708, EPI_ISL_2984710, EPI_ISL_2984712, EPI_ISL_2984714, EPI_ISL_2984715, EPI_ISL_2984716, EPI_ISL_2984719, EPI_ISL_2984720, EPI_ISL_2984721, EPI_ISL_2984722, EPI_ISL_2984724, EPI_ISL_2984755, EPI_ISL_2984765, EPI_ISL_2984785 | | | |
| see above | Ministry of Health Turkey | Ministry of Health Turkey | Fatma Bayrakdar, Yasemin Cosgun, Suleyman Yalcin, Gulay Korukluoglu |
| EPI_ISL_2984804, EPI_ISL_2984805, EPI_ISL_2984807, EPI_ISL_2984808, EPI_ISL_2984809 | KIMBERLEY LABORATORY | National Institute for Communicable Diseases of the National Health Laboratory Service | Amoako DG, Everatt J, Scheepers C, Mohale T, Ntuli N, Mahlangu B, Mnguni A, Ismail A, Bhiman JN |
| EPI_ISL_2984810, EPI_ISL_2984811 | MANKWENG LABORATORY | National Institute for Communicable Diseases of the National Health Laboratory Service | Amoako DG, Everatt J, Scheepers C, Mohale T, Ntuli N, Mahlangu B, Mnguni A, Ismail A, Bhiman JN |
| EPI_ISL_2984812, EPI_ISL_2984813, EPI_ISL_2984814, EPI_ISL_2984815, EPI_ISL_2984816, EPI_ISL_2984817, EPI_ISL_2984818, EPI_ISL_2984819, EPI_ISL_2984820, EPI_ISL_2984821, EPI_ISL_2984822, EPI_ISL_2984825, EPI_ISL_2984826, EPI_ISL_2984827, EPI_ISL_2984828, EPI_ISL_2984830, EPI_ISL_2984831, EPI_ISL_2984832, EPI_ISL_2984833, EPI_ISL_2984834, EPI_ISL_2984835 | | | |
| see above | HELEN JOSEPH LABORATORY | National Institute for Communicable Diseases of the National Health Laboratory Service | Amoako DG, Everatt J, Scheepers C, Mohale T, Ntuli N, Mahlangu B, Mnguni A, Ismail A, Bhiman JN |
| EPI_ISL_2984836, EPI_ISL_2984837, EPI_ISL_2984838, EPI_ISL_2984839, EPI_ISL_2984840, EPI_ISL_2984841, EPI_ISL_2984842, EPI_ISL_2984843, EPI_ISL_2984844, EPI_ISL_2984845, EPI_ISL_2984846, EPI_ISL_2984847, EPI_ISL_2984848, EPI_ISL_2984849, EPI_ISL_2984850, EPI_ISL_2984851, EPI_ISL_2984852, EPI_ISL_2984853, EPI_ISL_2984854, EPI_ISL_2984855, EPI_ISL_2984856, EPI_ISL_2984857, EPI_ISL_2984859, EPI_ISL_2984860, EPI_ISL_2984861, EPI_ISL_2984862 | | | |
| see above | MANKWENG LABORATORY | National Institute for Communicable Diseases of the National Health Laboratory Service | Amoako DG, Everatt J, Scheepers C, Mohale T, Ntuli N, Mahlangu B, Mnguni A, Ismail A, Bhiman JN |
| EPI_ISL_2984863, EPI_ISL_2984864, EPI_ISL_2984865 | PORT ELIZABETH LABOARTORY | National Institute for Communicable Diseases of the National Health Laboratory Service | Amoako DG, Everatt J, Scheepers C, Mohale T, Ntuli N, Mahlangu B, Mnguni A, Ismail A, Bhiman JN |
| EPI_ISL_2984866, EPI_ISL_2984867, EPI_ISL_2984868, EPI_ISL_2984869, EPI_ISL_2984870, EPI_ISL_2984872, EPI_ISL_2984873 | Lancet Laboratory | National Institute for Communicable Diseases of the National Health Laboratory Service | Amoako DG, Everatt J, Scheepers C, Mohale T, Ntuli N, Mahlangu B, Mnguni A, Ismail A, Bhiman JN |
| EPI_ISL_2984874, EPI_ISL_2984875, EPI_ISL_2984877, EPI_ISL_2984878, EPI_ISL_2984879, EPI_ISL_2984881, EPI_ISL_2984882, EPI_ISL_2984883, EPI_ISL_2984885, EPI_ISL_2984886, EPI_ISL_2984887, EPI_ISL_2984888, EPI_ISL_2984889, EPI_ISL_2984892 | | | |
| see above | TSHEPONG LABORATORY | National Institute for Communicable Diseases of the National Health Laboratory Service | Amoako DG, Everatt J, Scheepers C, Mohale T, Ntuli N, Mahlangu B, Mnguni A, Ismail A, Bhiman JN |
| EPI_ISL_2984893, EPI_ISL_2984894, EPI_ISL_2984895, EPI_ISL_2984896, EPI_ISL_2984898, EPI_ISL_2984899, EPI_ISL_2984900, EPI_ISL_2984901, EPI_ISL_2984903, EPI_ISL_2984904, EPI_ISL_2984905, EPI_ISL_2984906, EPI_ISL_2984907, EPI_ISL_2984908, EPI_ISL_2984909, EPI_ISL_2984910 | | | |
| see above | HELEN JOSEPH LABORATORY | National Institute for Communicable Diseases of the National Health Laboratory Service | Amoako DG, Everatt J, Scheepers C, Mohale T, Ntuli N, Mahlangu B, Mnguni A, Ismail A, Bhiman JN |
| EPI_ISL_2984913, EPI_ISL_2984915, EPI_ISL_2984916, EPI_ISL_2984917, EPI_ISL_2984918, EPI_ISL_2984919, EPI_ISL_2984921 | TSHEPONG LABORATORY | National Institute for Communicable Diseases of the National Health Laboratory Service | Amoako DG, Everatt J, Scheepers C, Mohale T, Ntuli N, Mahlangu B, Mnguni A, Ismail A, Bhiman JN |
| EPI_ISL_2984922 | POTCHEFSTROOM | National Institute for Communicable Diseases of the National Health Laboratory Service | Amoako DG, Everatt J, Scheepers C, Mohale T, Ntuli N, Mahlangu B, Mnguni A, Ismail A, Bhiman JN |
| EPI_ISL_2984924, EPI_ISL_2984925, EPI_ISL_2984929, EPI_ISL_2984930, EPI_ISL_2984932, EPI_ISL_2984933, EPI_ISL_2984934, EPI_ISL_2984935, EPI_ISL_2984938 | TSHEPONG LABORATORY | National Institute for Communicable Diseases of the National Health Laboratory Service | Amoako DG, Everatt J, Scheepers C, Mohale T, Ntuli N, Mahlangu B, Mnguni A, Ismail A, Bhiman JN |
| EPI_ISL_2984962 | AOU Polclinico "G Rodolico-S.Marco" PO Rodolico CATANIA | AOU Polclinico "G Rodolico-S.Marco" PO Rodolico and BIOMETEC UNICT CATANIA | STEFANI AND SCALIA |
| EPI_ISL_2984971, EPI_ISL_2984975, EPI_ISL_2984978, EPI_ISL_2984982, EPI_ISL_2984985, EPI_ISL_2984989, EPI_ISL_2984990, EPI_ISL_2984991, EPI_ISL_2984992, EPI_ISL_2984996, EPI_ISL_2985000, EPI_ISL_2985003, EPI_ISL_2985004, EPI_ISL_2985008, EPI_ISL_2985009, EPI_ISL_2985010, EPI_ISL_2985011, EPI_ISL_2985015 | | | |
| see above | Allergy, Immunology and Cell Biology Unit (AICBU) | Allergy, Immunology and Cell Biology Unit (AICBU) | Chandima Jeewandara, Deshni Jayathilaka, Dumni Gunasinghe, Tibutius Thanesh, Dinuka Ariyarathne, Ayesha Wijesinghe, Diyanath Ranasinghe, Gathsaurie Neelika Malavige |
| EPI_ISL_2985016 | AOU Polclinico "G Rodolico-S.Marco" PO Rodolico CATANIA | AOU Polclinico "G Rodolico-S.Marco" PO Rodolico and BIOMETEC UNICT CATANIA | STEFANI AND SCALIA |
| EPI_ISL_2985017 | IRCCS San Gallicano Dermatological Institute | IRCCS Regina Elena National Cancer Institute | Sabrina Strano, Sara Donzelli, Matteo Pallocca, Grazia Prignano, Fabrizio Ensoli, Fulvia Pimpinelli, Maurizio Fanciulli, Giovanni Blandino, Aldo Morrone, Gennaro Ciliberto |
| EPI_ISL_2985019 | IRCCS San Gallicano Dermatological Institute | IRCCS Regina Elena National Cancer Institute | Ludovica Ciuffreda, Sara Donzelli, Alice Massacci, Arianna Mastrofrancesco, Fabrizio Ensoli, Fulvia Pimpinelli, Maurizio Fanciulli, Giovanni Blandino, Aldo Morrone, Gennaro Ciliberto |
| EPI_ISL_2985020 | IRCCS San Gallicano Dermatological Institute | IRCCS Regina Elena National Cancer Institute | Frauke Goeman, Giulia Orlandi, Matteo Pallocca, Giovanna D'agosto, Fabrizio Ensoli, Fulvia Pimpinelli, Maurizio Fanciulli, Giovanni Blandino, Aldo Morrone, Gennaro Ciliberto |
| EPI_ISL_2985021 | IRCCS San Gallicano Dermatological Institute | IRCCS Regina Elena National Cancer Institute | Sabrina Strano, Sara Donzelli, Eleonora Sperandio, Carla Mottini, Fabrizio Ensoli, Fulvia Pimpinelli, Maurizio Fanciulli, Giovanni Blandino, Aldo Morrone, Gennaro Ciliberto |
| EPI_ISL_2985022 | IRCCS San Gallicano Dermatological Institute | IRCCS Regina Elena National Cancer Institute | Francesca De Nicola, Giulia Orlandi, Alice Massacci, Martina Pontone, Fabrizio Ensoli, Fulvia Pimpinelli, Maurizio Fanciulli, Giovanni Blandino, Aldo Morrone, Gennaro Ciliberto |
| EPI_ISL_2985024 | IRCCS San Gallicano Dermatological Institute | IRCCS Regina Elena National Cancer Institute | Frauke Goeman, Giulia Orlandi, Eleonora Sperandio, Ilaria Cavallo, Fabrizio Ensoli, Fulvia Pimpinelli, Maurizio Fanciulli, Giovanni Blandino, Aldo Morrone, Gennaro Ciliberto |
| EPI_ISL_2985025 | IRCCS San Gallicano Dermatological Institute | IRCCS Regina Elena National Cancer Institute | Sabrina Strano, Sara Donzelli, Alice Massacci, Francesca Sivori, Fabrizio Ensoli, Fulvia Pimpinelli, Maurizio Fanciulli, Giovanni Blandino, Aldo Morrone, Gennaro Ciliberto |
| EPI_ISL_2985026 | IRCCS San Gallicano Dermatological Institute | IRCCS Regina Elena National Cancer Institute | Francesca De Nicola, Giulia Orlandi, Matteo Pallocca, Andrea Cazzani, Fabrizio Ensoli, Fulvia Pimpinelli, Maurizio Fanciulli, Giovanni Blandino, Aldo Morrone, Gennaro Ciliberto |
| EPI_ISL_2985027 | IRCCS San Gallicano Dermatological Institute | IRCCS Regina Elena National Cancer Institute | Ludovica Ciuffreda, Sara Donzelli, Eleonora Sperandio, Ilaria Celesti, Fabrizio Ensoli, Fulvia Pimpinelli, Maurizio Fanciulli, Giovanni Blandino, Aldo Morrone, Gennaro Ciliberto |
| EPI_ISL_2985028 | IRCCS San Gallicano Dermatological Institute | IRCCS Regina Elena National Cancer Institute | Frauke Goeman, Giulia Orlandi, Alice Massacci, Martina Diano, Fabrizio Ensoli, Fulvia Pimpinelli, Maurizio Fanciulli, Giovanni Blandino, Aldo Morrone, Gennaro Ciliberto |
| EPI_ISL_2985029 | IRCCS San Gallicano Dermatological Institute | IRCCS Regina Elena National Cancer Institute | Sabrina Strano, Sara Donzelli, Matteo Pallocca, Antonio Federico, Fabrizio Ensoli, Fulvia Pimpinelli, Maurizio Fanciulli, Giovanni Blandino, Aldo Morrone, Gennaro Ciliberto |
| EPI_ISL_2985030 | IRCCS San Gallicano Dermatological Institute | IRCCS Regina Elena National Cancer Institute | Francesca De Nicola, Giulia Orlandi, Eleonora Sperandio, Fulvia Fraticelli, Fabrizio Ensoli, Fulvia Pimpinelli, Maurizio Fanciulli, Giovanni Blandino, Aldo Morrone, Gennaro Ciliberto |
| EPI_ISL_2985031 | IRCCS San Gallicano Dermatological Institute | IRCCS Regina Elena National Cancer Institute | Ludovica Ciuffreda, Sara Donzelli, Alice Massacci, Lorenzo Furzi, Fabrizio Ensoli, Fulvia Pimpinelli, Maurizio Fanciulli, Giovanni Blandino, Aldo Morrone, Gennaro Ciliberto |
| EPI_ISL_2985032 | IRCCS San Gallicano Dermatological Institute | IRCCS Regina Elena National Cancer Institute | Frauke Goeman, Giulia Orlandi, Matteo Pallocca, Alessia Lauretti, Fabrizio Ensoli, Fulvia Pimpinelli, Maurizio Fanciulli, Giovanni Blandino, Aldo Morrone, Gennaro Ciliberto |
| EPI_ISL_2985033 | IRCCS San Gallicano Dermatological Institute | IRCCS Regina Elena National Cancer Institute | Sabrina Strano, Sara Donzelli, Eleonora Sperandio, Francesca Maione, Fabrizio Ensoli, Fulvia Pimpinelli, Maurizio Fanciulli, Giovanni Blandino, Aldo Morrone, Gennaro Ciliberto |
| EPI_ISL_2985034 | IRCCS San Gallicano Dermatological Institute | IRCCS Regina Elena National Cancer Institute | Francesca De Nicola, Giulia Orlandi, Alice Massacci, Francisco Obregon, Fabrizio Ensoli, Fulvia Pimpinelli, Maurizio Fanciulli, Giovanni Blandino, Aldo Morrone, Gennaro Ciliberto |
| EPI_ISL_2985035 | IRCCS San Gallicano Dermatological Institute | IRCCS Regina Elena National Cancer Institute | Ludovica Ciuffreda, Sara Donzelli, Matteo Pallocca, Silvia Paluzzi, Fabrizio Ensoli, Fulvia Pimpinelli, Maurizio Fanciulli, Giovanni Blandino, Aldo Morrone, Gennaro Ciliberto |
| EPI_ISL_2985036 | IRCCS San Gallicano Dermatological Institute | IRCCS Regina Elena National Cancer Institute | Frauke Goeman, Giulia Orlandi, Eleonora Sperandio, Sara Petrolo, Fabrizio Ensoli, Fulvia Pimpinelli, Maurizio Fanciulli, Giovanni Blandino, Aldo Morrone, Gennaro Ciliberto |
| EPI_ISL_2985037 | IRCCS San Gallicano Dermatological Institute | IRCCS Regina Elena National Cancer Institute | Sabrina Strano, Sara Donzelli, Alice Massacci, Valentina Ricca, Fabrizio Ensoli, Fulvia Pimpinelli, Maurizio Fanciulli, Giovanni Blandino, Aldo Morrone, Gennaro Ciliberto |
| EPI_ISL_2985038 | IRCCS San Gallicano Dermatological Institute | IRCCS Regina Elena National Cancer Institute | Francesca De Nicola, Giulia Orlandi, Matteo Pallocca, Serena Salvo, Fabrizio Ensoli, Fulvia Pimpinelli, Maurizio Fanciulli, Giovanni Blandino, Aldo Morrone, Gennaro Ciliberto |
| EPI_ISL_2985039 | IRCCS San Gallicano Dermatological Institute | IRCCS Regina Elena National Cancer Institute | Ludovica Ciuffreda, Sara Donzelli, Eleonora Sperandio, Grazia Prignano, Fabrizio Ensoli, Fulvia Pimpinelli, Maurizio Fanciulli, Giovanni Blandino, Aldo Morrone, Gennaro Ciliberto |
| EPI_ISL_2985041 | IRCCS San Gallicano Dermatological Institute | IRCCS Regina Elena National Cancer Institute | Sabrina Strano, Sara Donzelli, Matteo Pallocca, Arianna Mastrofrancesco, Fabrizio Ensoli, Fulvia Pimpinelli, Maurizio Fanciulli, Giovanni Blandino, Aldo Morrone, Gennaro Ciliberto |
| EPI_ISL_2985042 | IRCCS San Gallicano Dermatological Institute | IRCCS Regina Elena National Cancer Institute | Francesca De Nicola, Giulia Orlandi, Eleonora Sperandio, Giovanna D'agosto, Fabrizio Ensoli, Fulvia Pimpinelli, Maurizio Fanciulli, Giovanni Blandino, Aldo Morrone, Gennaro Ciliberto |
| EPI_ISL_2985043 | IRCCS San Gallicano Dermatological Institute | IRCCS Regina Elena National Cancer Institute | Ludovica Ciuffreda, Sara Donzelli, Alice Massacci, Carla Mottini, Fabrizio Ensoli, Fulvia Pimpinelli, Maurizio Fanciulli, Giovanni Blandino, Aldo Morrone, Gennaro Ciliberto |
| EPI_ISL_2985044 | IRCCS San Gallicano Dermatological Institute | IRCCS Regina Elena National Cancer Institute | Frauke Goeman, Giulia Orlandi, Matteo Pallocca, Martina Pontone, Fabrizio Ensoli, Fulvia Pimpinelli, Maurizio Fanciulli, Giovanni Blandino, Aldo Morrone, Gennaro Ciliberto |
| EPI_ISL_2985045 | IRCCS San Gallicano Dermatological Institute | IRCCS Regina Elena National Cancer Institute | Sabrina Strano, Sara Donzelli, Eleonora Sperandio, di Domenico Enea Gino, Fabrizio Ensoli, Fulvia Pimpinelli, Maurizio Fanciulli, Giovanni Blandino, Aldo Morrone, Gennaro Ciliberto |

| | | | |
|---|---|---|---|
| EPI_ISL_2985046 | IRCCS San Gallicano Dermatological Institute | IRCCS Regina Elena National Cancer Institute | Francesca De Nicola, Giulia Orlandi, Alice Massacci, Ilaria Cavallo, Fabrizio Ensoli, Fulvia Pimpinelli, Maurizio Fanciulli, Giovanni Blandino, Aldo Morrone, Gennaro Ciliberto |
| EPI_ISL_2985047 | IRCCS San Gallicano Dermatological Institute | IRCCS Regina Elena National Cancer Institute | Ludovica Ciuffreda, Sara Donzelli, Matteo Pallocca, Francesca Sivori, Fabrizio Ensoli, Fulvia Pimpinelli, Maurizio Fanciulli, Giovanni Blandino, Aldo Morrone, Gennaro Ciliberto |
| EPI_ISL_2985048 | IRCCS San Gallicano Dermatological Institute | IRCCS Regina Elena National Cancer Institute | Frauke Goeman, Giulia Orlandi, Eleonora Sperandio, Andrea Cazzani, Fabrizio Ensoli, Fulvia Pimpinelli, Maurizio Fanciulli, Giovanni Blandino, Aldo Morrone, Gennaro Ciliberto |
| EPI_ISL_2985049 | IRCCS San Gallicano Dermatological Institute | IRCCS Regina Elena National Cancer Institute | Sabrina Strano, Sara Donzelli, Alice Massacci, Ilaria Celesti, Fabrizio Ensoli, Fulvia Pimpinelli, Maurizio Fanciulli, Giovanni Blandino, Aldo Morrone, Gennaro Ciliberto |
| EPI_ISL_2985050 | IRCCS San Gallicano Dermatological Institute | IRCCS Regina Elena National Cancer Institute | Francesca De Nicola, Giulia Orlandi, Matteo Pallocca, Martina Diano, Fabrizio Ensoli, Fulvia Pimpinelli, Maurizio Fanciulli, Giovanni Blandino, Aldo Morrone, Gennaro Ciliberto |
| EPI_ISL_2985051 | IRCCS San Gallicano Dermatological Institute | IRCCS Regina Elena National Cancer Institute | Ludovica Ciuffreda, Sara Donzelli, Eleonora Sperandio, Antonio Federico, Fabrizio Ensoli, Fulvia Pimpinelli, Maurizio Fanciulli, Giovanni Blandino, Aldo Morrone, Gennaro Ciliberto |
| EPI_ISL_2985052 | IRCCS San Gallicano Dermatological Institute | IRCCS Regina Elena National Cancer Institute | Frauke Goeman, Giulia Orlandi, Alice Massacci, Fulvia Fraticelli, Fabrizio Ensoli, Fulvia Pimpinelli, Maurizio Fanciulli, Giovanni Blandino, Aldo Morrone, Gennaro Ciliberto |
| EPI_ISL_2985053 | IRCCS San Gallicano Dermatological Institute | IRCCS Regina Elena National Cancer Institute | Sabrina Strano, Sara Donzelli, Matteo Pallocca, Lorenzo Furzi, Fabrizio Ensoli, Fulvia Pimpinelli, Maurizio Fanciulli, Giovanni Blandino, Aldo Morrone, Gennaro Ciliberto |
| EPI_ISL_2985054 | IRCCS San Gallicano Dermatological Institute | IRCCS Regina Elena National Cancer Institute | Francesca De Nicola, Giulia Orlandi, Eleonora Sperandio, Alessia Lauretti, Fabrizio Ensoli, Fulvia Pimpinelli, Maurizio Fanciulli, Giovanni Blandino, Aldo Morrone, Gennaro Ciliberto |
| EPI_ISL_2985055 | IRCCS San Gallicano Dermatological Institute | IRCCS Regina Elena National Cancer Institute | Ludovica Ciuffreda, Sara Donzelli, Alice Massacci, Francesca Maione, Fabrizio Ensoli, Fulvia Pimpinelli, Maurizio Fanciulli, Giovanni Blandino, Aldo Morrone, Gennaro Ciliberto |
| EPI_ISL_2985056 | IRCCS San Gallicano Dermatological Institute | IRCCS Regina Elena National Cancer Institute | Frauke Goeman, Giulia Orlandi, Matteo Pallocca, Francisco Obregon, Fabrizio Ensoli, Fulvia Pimpinelli, Maurizio Fanciulli, Giovanni Blandino, Aldo Morrone, Gennaro Ciliberto |
| EPI_ISL_2985057 | IRCCS San Gallicano Dermatological Institute | IRCCS Regina Elena National Cancer Institute | Sabrina Strano, Sara Donzelli, Eleonora Sperandio, Silvia Paluzzi, Fabrizio Ensoli, Fulvia Pimpinelli, Maurizio Fanciulli, Giovanni Blandino, Aldo Morrone, Gennaro Ciliberto |
| EPI_ISL_2985058 | IRCCS San Gallicano Dermatological Institute | IRCCS Regina Elena National Cancer Institute | Francesca De Nicola, Giulia Orlandi, Alice Massacci, Sara Petrolo, Fabrizio Ensoli, Fulvia Pimpinelli, Maurizio Fanciulli, Giovanni Blandino, Aldo Morrone, Gennaro Ciliberto |
| EPI_ISL_2985059 | IRCCS San Gallicano Dermatological Institute | IRCCS Regina Elena National Cancer Institute | Ludovica Ciuffreda, Sara Donzelli, Matteo Pallocca, Valentina Ricca, Fabrizio Ensoli, Fulvia Pimpinelli, Maurizio Fanciulli, Giovanni Blandino, Aldo Morrone, Gennaro Ciliberto |
| EPI_ISL_2985060 | IRCCS San Gallicano Dermatological Institute | IRCCS Regina Elena National Cancer Institute | Frauke Goeman, Giulia Orlandi, Eleonora Sperandio, Serena Salvo, Fabrizio Ensoli, Fulvia Pimpinelli, Maurizio Fanciulli, Giovanni Blandino, Aldo Morrone, Gennaro Ciliberto |
| EPI_ISL_2985061 | IRCCS San Gallicano Dermatological Institute | IRCCS Regina Elena National Cancer Institute | Sabrina Strano, Sara Donzelli, Alice Massacci, Grazia Prignano, Fabrizio Ensoli, Fulvia Pimpinelli, Maurizio Fanciulli, Giovanni Blandino, Aldo Morrone, Gennaro Ciliberto |
| EPI_ISL_2985062 | IRCCS San Gallicano Dermatological Institute | IRCCS Regina Elena National Cancer Institute | Francesca De Nicola, Giulia Orlandi, Matteo Pallocca, Elisabetta Trento, Fabrizio Ensoli, Fulvia Pimpinelli, Maurizio Fanciulli, Giovanni Blandino, Aldo Morrone, Gennaro Ciliberto |
| EPI_ISL_2985063 | IRCCS San Gallicano Dermatological Institute | IRCCS Regina Elena National Cancer Institute | Ludovica Ciuffreda, Sara Donzelli, Eleonora Sperandio, Arianna Mastrofrancesco, Fabrizio Ensoli, Fulvia Pimpinelli, Maurizio Fanciulli, Giovanni Blandino, Aldo Morrone, Gennaro Ciliberto |
| EPI_ISL_2985064 | IRCCS San Gallicano Dermatological Institute | IRCCS Regina Elena National Cancer Institute | Frauke Goeman, Giulia Orlandi, Alice Massacci, Giovanna D’agosto, Fabrizio Ensoli, Fulvia Pimpinelli, Maurizio Fanciulli, Giovanni Blandino, Aldo Morrone, Gennaro Ciliberto |
| EPI_ISL_2985065 | IRCCS San Gallicano Dermatological Institute | IRCCS Regina Elena National Cancer Institute | Francesca De Nicola, Giulia Orlandi, Eleonora Sperandio, Martina Pontone, Fabrizio Ensoli, Fulvia Pimpinelli, Maurizio Fanciulli, Giovanni Blandino, Aldo Morrone, Gennaro Ciliberto |
| EPI_ISL_2985066 | IRCCS San Gallicano Dermatological Institute | IRCCS Regina Elena National Cancer Institute | Ludovica Ciuffreda, Sara Donzelli, Alice Massacci, di Domenico Enea Gino, Fabrizio Ensoli, Fulvia Pimpinelli, Maurizio Fanciulli, Giovanni Blandino, Aldo Morrone, Gennaro Ciliberto |
| EPI_ISL_2985067 | IRCCS San Gallicano Dermatological Institute | IRCCS Regina Elena National Cancer Institute | Frauke Goeman, Giulia Orlandi, Matteo Pallocca, Ilaria Cavallo, Fabrizio Ensoli, Fulvia Pimpinelli, Maurizio Fanciulli, Giovanni Blandino, Aldo Morrone, Gennaro Ciliberto |
| EPI_ISL_2985068 | IRCCS San Gallicano Dermatological Institute | IRCCS Regina Elena National Cancer Institute | Sabrina Strano, Sara Donzelli, Eleonora Sperandio, Francesca Sivori, Fabrizio Ensoli, Fulvia Pimpinelli, Maurizio Fanciulli, Giovanni Blandino, Aldo Morrone, Gennaro Ciliberto |
| EPI_ISL_2985070 | IRCCS San Gallicano Dermatological Institute | IRCCS Regina Elena National Cancer Institute | Ludovica Ciuffreda, Sara Donzelli, Matteo Pallocca, Ilaria Celesti, Fabrizio Ensoli, Fulvia Pimpinelli, Maurizio Fanciulli, Giovanni Blandino, Aldo Morrone, Gennaro Ciliberto |
| EPI_ISL_2985071 | IRCCS San Gallicano Dermatological Institute | IRCCS Regina Elena National Cancer Institute | Frauke Goeman, Giulia Orlandi, Eleonora Sperandio, Martina Diano, Fabrizio Ensoli, Fulvia Pimpinelli, Maurizio Fanciulli, Giovanni Blandino, Aldo Morrone, Gennaro Ciliberto |
| EPI_ISL_2985072 | IRCCS San Gallicano Dermatological Institute | IRCCS Regina Elena National Cancer Institute | Sabrina Strano, Sara Donzelli, Alice Massacci, Antonio Federico, Fabrizio Ensoli, Fulvia Pimpinelli, Maurizio Fanciulli, Giovanni Blandino, Aldo Morrone, Gennaro Ciliberto |
| EPI_ISL_2985075 | IRCCS San Gallicano Dermatological Institute | IRCCS Regina Elena National Cancer Institute | Frauke Goeman, Giulia Orlandi, Alice Massacci, Alessia Lauretti, Fabrizio Ensoli, Fulvia Pimpinelli, Maurizio Fanciulli, Giovanni Blandino, Aldo Morrone, Gennaro Ciliberto |
| EPI_ISL_2985076 | IRCCS San Gallicano Dermatological Institute | IRCCS Regina Elena National Cancer Institute | Sabrina Strano, Sara Donzelli, Matteo Pallocca, Francesca Maione, Fabrizio Ensoli, Fulvia Pimpinelli, Maurizio Fanciulli, Giovanni Blandino, Aldo Morrone, Gennaro Ciliberto |
| EPI_ISL_2985078 | IRCCS San Gallicano Dermatological Institute | IRCCS Regina Elena National Cancer Institute | Ludovica Ciuffreda, Sara Donzelli, Alice Massacci, Silvia Paluzzi, Fabrizio Ensoli, Fulvia Pimpinelli, Maurizio Fanciulli, Giovanni Blandino, Aldo Morrone, Gennaro Ciliberto |
| EPI_ISL_2985080 | IRCCS San Gallicano Dermatological Institute | IRCCS Regina Elena National Cancer Institute | Sabrina Strano, Sara Donzelli, Eleonora Sperandio, Valentina Ricca, Fabrizio Ensoli, Fulvia Pimpinelli, Maurizio Fanciulli, Giovanni Blandino, Aldo Morrone, Gennaro Ciliberto |
| EPI_ISL_2985082 | IRCCS San Gallicano Dermatological Institute | IRCCS Regina Elena National Cancer Institute | Ludovica Ciuffreda, Sara Donzelli, Matteo Pallocca, Grazia Prignano, Fabrizio Ensoli, Fulvia Pimpinelli, Maurizio Fanciulli, Giovanni Blandino, Aldo Morrone, Gennaro Ciliberto |
| EPI_ISL_2985084 | IRCCS San Gallicano Dermatological Institute | IRCCS Regina Elena National Cancer Institute | Sabrina Strano, Sara Donzelli, Alice Massacci, Arianna Mastrofrancesco, Fabrizio Ensoli, Fulvia Pimpinelli, Maurizio Fanciulli, Giovanni Blandino, Aldo Morrone, Gennaro Ciliberto |
| EPI_ISL_2985085 | IRCCS San Gallicano Dermatological Institute | IRCCS Regina Elena National Cancer Institute | Francesca De Nicola, Giulia Orlandi, Matteo Pallocca, Giovanna D’agosto, Fabrizio Ensoli, Fulvia Pimpinelli, Maurizio Fanciulli, Giovanni Blandino, Aldo Morrone, Gennaro Ciliberto |
| EPI_ISL_2985086 | IRCCS San Gallicano Dermatological Institute | IRCCS Regina Elena National Cancer Institute | Ludovica Ciuffreda, Sara Donzelli, Eleonora Sperandio, Carla Mottini, Fabrizio Ensoli, Fulvia Pimpinelli, Maurizio Fanciulli, Giovanni Blandino, Aldo Morrone, Gennaro Ciliberto |
| EPI_ISL_2985089 | IRCCS San Gallicano Dermatological Institute | IRCCS Regina Elena National Cancer Institute | Francesca De Nicola, Giulia Orlandi, Eleonora Sperandio, Ilaria Cavallo, Fabrizio Ensoli, Fulvia Pimpinelli, Maurizio Fanciulli, Giovanni Blandino, Aldo Morrone, Gennaro Ciliberto |
| EPI_ISL_2985090 | IRCCS San Gallicano Dermatological Institute | IRCCS Regina Elena National Cancer Institute | Ludovica Ciuffreda, Sara Donzelli, Alice Massacci, Francesca Sivori, Fabrizio Ensoli, Fulvia Pimpinelli, Maurizio Fanciulli, Giovanni Blandino, Aldo Morrone, Gennaro Ciliberto |
| EPI_ISL_2985092 | IRCCS San Gallicano Dermatological Institute | IRCCS Regina Elena National Cancer Institute | Francesca De Nicola, Giulia Orlandi, Alice Massacci, Martina Diano, Fabrizio Ensoli, Fulvia Pimpinelli, Maurizio Fanciulli, Giovanni Blandino, Aldo Morrone, Gennaro Ciliberto |
| EPI_ISL_2985093 | IRCCS San Gallicano Dermatological Institute | IRCCS Regina Elena National Cancer Institute | Ludovica Ciuffreda, Sara Donzelli, Matteo Pallocca, Antonio Federico, Fabrizio Ensoli, Fulvia Pimpinelli, Maurizio Fanciulli, Giovanni Blandino, Aldo Morrone, Gennaro Ciliberto |
| EPI_ISL_2985094 | IRCCS San Gallicano Dermatological Institute | IRCCS Regina Elena National Cancer Institute | Frauke Goeman, Giulia Orlandi, Eleonora Sperandio, Fulvia Fraticelli, Fabrizio Ensoli, Fulvia Pimpinelli, Maurizio Fanciulli, Giovanni Blandino, Aldo Morrone, Gennaro Ciliberto |
| EPI_ISL_2985095 | IRCCS San Gallicano Dermatological Institute | IRCCS Regina Elena National Cancer Institute | Sabrina Strano, Sara Donzelli, Alice Massacci, Lorenzo Furzi, Fabrizio Ensoli, Fulvia Pimpinelli, Maurizio Fanciulli, Giovanni Blandino, Aldo Morrone, Gennaro Ciliberto |
| EPI_ISL_2985096 | IRCCS San Gallicano Dermatological Institute | IRCCS Regina Elena National Cancer Institute | Francesca De Nicola, Giulia Orlandi, Matteo Pallocca, Alessia Lauretti, Fabrizio Ensoli, Fulvia Pimpinelli, Maurizio Fanciulli, Giovanni Blandino, Aldo Morrone, Gennaro Ciliberto |
| EPI_ISL_2985097 | IRCCS San Gallicano Dermatological Institute | IRCCS Regina Elena National Cancer Institute | Ludovica Ciuffreda, Sara Donzelli, Eleonora Sperandio, Francesca Maione, Fabrizio Ensoli, Fulvia Pimpinelli, Maurizio Fanciulli, Giovanni Blandino, Aldo Morrone, Gennaro Ciliberto |
| EPI_ISL_2985098 | IRCCS San Gallicano Dermatological Institute | IRCCS Regina Elena National Cancer Institute | Sabrina Strano, Sara Donzelli, Matteo Pallocca, Silvia Paluzzi, Fabrizio Ensoli, Fulvia Pimpinelli, Maurizio Fanciulli, Giovanni Blandino, Aldo Morrone, Gennaro Ciliberto |
| EPI_ISL_2985099 | IRCCS San Gallicano Dermatological Institute | IRCCS Regina Elena National Cancer Institute | Francesca De Nicola, Giulia Orlandi, Eleonora Sperandio, Sara Petrolo, Fabrizio Ensoli, Fulvia Pimpinelli, Maurizio Fanciulli, Giovanni Blandino, Aldo Morrone, Gennaro Ciliberto |
| EPI_ISL_2985101 | IRCCS San Gallicano Dermatological Institute | IRCCS Regina Elena National Cancer Institute | Frauke Goeman, Giulia Orlandi, Matteo Pallocca, Serena Salvo, Fabrizio Ensoli, Fulvia Pimpinelli, Maurizio Fanciulli, Giovanni Blandino, Aldo Morrone, Gennaro Ciliberto |
| EPI_ISL_2985102 | IRCCS San Gallicano Dermatological Institute | IRCCS Regina Elena National Cancer Institute | Sabrina Strano, Sara Donzelli, Eleonora Sperandio, Grazia Prignano, Fabrizio Ensoli, Fulvia Pimpinelli, Maurizio Fanciulli, Giovanni Blandino, Aldo Morrone, Gennaro Ciliberto |
| EPI_ISL_2985103 | IRCCS San Gallicano Dermatological Institute | IRCCS Regina Elena National Cancer Institute | Francesca De Nicola, Giulia Orlandi, Alice Massacci, Elisabetta Trento, Fabrizio Ensoli, Fulvia Pimpinelli, Maurizio Fanciulli, Giovanni Blandino, Aldo Morrone, Gennaro Ciliberto |
| EPI_ISL_2985104 | IRCCS San Gallicano Dermatological Institute | IRCCS Regina Elena National Cancer Institute | Ludovica Ciuffreda, Sara Donzelli, Matteo Pallocca, Arianna Mastrofrancesco, Fabrizio Ensoli, Fulvia Pimpinelli, Maurizio Fanciulli, Giovanni Blandino, Aldo Morrone, Gennaro Ciliberto |
| EPI_ISL_2985105 | IRCCS San Gallicano Dermatological Institute | IRCCS Regina Elena National Cancer Institute | Frauke Goeman, Giulia Orlandi, Eleonora Sperandio, Giovanna D’agosto, Fabrizio Ensoli, Fulvia Pimpinelli, Maurizio Fanciulli, Giovanni Blandino, Aldo Morrone, Gennaro Ciliberto |
| EPI_ISL_2985106 | IRCCS San Gallicano Dermatological Institute | IRCCS Regina Elena National Cancer Institute | Sabrina Strano, Sara Donzelli, Alice Massacci, Carla Mottini, Fabrizio Ensoli, Fulvia Pimpinelli, Maurizio Fanciulli, Giovanni Blandino, Aldo Morrone, Gennaro Ciliberto |
| EPI_ISL_2985107 | IRCCS San Gallicano Dermatological Institute | IRCCS Regina Elena National Cancer Institute | Francesca De Nicola, Giulia Orlandi, Matteo Pallocca, Martina Pontone, Fabrizio Ensoli, Fulvia Pimpinelli, Maurizio Fanciulli, Giovanni Blandino, Aldo Morrone, Gennaro Ciliberto |
| EPI_ISL_2985108 | IRCCS San Gallicano Dermatological Institute | IRCCS Regina Elena National Cancer Institute | Ludovica Ciuffreda, Sara Donzelli, Eleonora Sperandio, di Domenico Enea Gino, Fabrizio Ensoli, Fulvia Pimpinelli, Maurizio Fanciulli, Giovanni Blandino, Aldo Morrone, Gennaro Ciliberto |
| EPI_ISL_2985109 | IRCCS San Gallicano Dermatological Institute | IRCCS Regina Elena National Cancer Institute | Frauke Goeman, Giulia Orlandi, Alice Massacci, Ilaria Cavallo, Fabrizio Ensoli, Fulvia Pimpinelli, Maurizio Fanciulli, Giovanni Blandino, Aldo Morrone, Gennaro Ciliberto |
| EPI_ISL_2985110 | IRCCS San Gallicano Dermatological Institute | IRCCS Regina Elena National Cancer Institute | Sabrina Strano, Sara Donzelli, Matteo Pallocca, Francesca Sivori, Fabrizio Ensoli, Fulvia Pimpinelli, Maurizio Fanciulli, Giovanni Blandino, Aldo Morrone, Gennaro Ciliberto |
| EPI_ISL_2985112 | IRCCS San Gallicano Dermatological Institute | IRCCS Regina Elena National Cancer Institute | Ludovica Ciuffreda, Sara Donzelli, Alice Massacci, Ilaria Celesti, Fabrizio Ensoli, Fulvia Pimpinelli, Maurizio Fanciulli, Giovanni Blandino, Aldo Morrone, Gennaro Ciliberto |
| EPI_ISL_2985113 | IRCCS San Gallicano Dermatological Institute | IRCCS Regina Elena National Cancer Institute | Frauke Goeman, Giulia Orlandi, Matteo Pallocca, Martina Diano, Fabrizio Ensoli, Fulvia Pimpinelli, Maurizio Fanciulli, Giovanni Blandino, Aldo Morrone, Gennaro Ciliberto |
| EPI_ISL_2985114 | IRCCS San Gallicano Dermatological Institute | IRCCS Regina Elena National Cancer Institute | Sabrina Strano, Sara Donzelli, Eleonora Sperandio, Antonio Federico, Fabrizio Ensoli, Fulvia Pimpinelli, Maurizio Fanciulli, Giovanni Blandino, Aldo Morrone, Gennaro Ciliberto |
| EPI_ISL_2985118 | IRCCS San Gallicano Dermatological Institute | IRCCS Regina Elena National Cancer Institute | Sabrina Strano, Sara Donzelli, Alice Massacci, Francesca Maione, Fabrizio Ensoli, Fulvia Pimpinelli, Maurizio Fanciulli, Giovanni Blandino, Aldo Morrone, Gennaro Ciliberto |
| EPI_ISL_2985119 | IRCCS San Gallicano Dermatological Institute | IRCCS Regina Elena National Cancer Institute | Francesca De Nicola, Giulia Orlandi, Matteo Pallocca, Francisco Obregon, Fabrizio Ensoli, Fulvia Pimpinelli, Maurizio Fanciulli, Giovanni Blandino, Aldo Morrone, Gennaro Ciliberto |
| EPI_ISL_2985120 | IRCCS San Gallicano Dermatological Institute | IRCCS Regina Elena National Cancer Institute | Ludovica Ciuffreda, Sara Donzelli, Eleonora Sperandio, Silvia Paluzzi, Fabrizio Ensoli, Fulvia Pimpinelli, Maurizio Fanciulli, Giovanni Blandino, Aldo Morrone, Gennaro Ciliberto |
| EPI_ISL_2985122 | IRCCS San Gallicano Dermatological Institute | IRCCS Regina Elena National Cancer Institute | Sabrina Strano, Sara Donzelli, Matteo Pallocca, Valentina Ricca, Fabrizio Ensoli, Fulvia Pimpinelli, Maurizio Fanciulli, Giovanni Blandino, Aldo Morrone, Gennaro Ciliberto |
| EPI_ISL_2985123 | IRCCS San Gallicano Dermatological Institute | IRCCS Regina Elena National Cancer Institute | Francesca De Nicola, Giulia Orlandi, Eleonora Sperandio, Serena Salvo, Fabrizio Ensoli, Fulvia Pimpinelli, Maurizio Fanciulli, Giovanni Blandino, Aldo Morrone, Gennaro Ciliberto |
| EPI_ISL_2985124 | IRCCS San Gallicano Dermatological Institute | IRCCS Regina Elena National Cancer Institute | Ludovica Ciuffreda, Sara Donzelli, Alice Massacci, Grazia Prignano, Fabrizio Ensoli, Fulvia Pimpinelli, Maurizio Fanciulli, Giovanni Blandino, Aldo Morrone, Gennaro Ciliberto |
| EPI_ISL_2985125 | IRCCS San Gallicano Dermatological Institute | IRCCS Regina Elena National Cancer Institute | Frauke Goeman, Giulia Orlandi, Matteo Pallocca, Elisabetta Trento, Fabrizio Ensoli, Fulvia Pimpinelli, Maurizio Fanciulli, Giovanni Blandino, Aldo Morrone, Gennaro Ciliberto |
| EPI_ISL_2985126 | IRCCS San Gallicano Dermatological Institute | IRCCS Regina Elena National Cancer Institute | Sabrina Strano, Sara Donzelli, Eleonora Sperandio, Arianna Mastrofrancesco, Fabrizio Ensoli, Fulvia Pimpinelli, Maurizio Fanciulli, Giovanni Blandino, Aldo Morrone, Gennaro Ciliberto |
| EPI_ISL_2985129 | IRCCS San Gallicano Dermatological Institute | IRCCS Regina Elena National Cancer Institute | Frauke Goeman, Giulia Orlandi, Eleonora Sperandio, Martina Pontone, Fabrizio Ensoli, Fulvia Pimpinelli, Maurizio Fanciulli, Giovanni Blandino, Aldo Morrone, Gennaro Ciliberto |
| EPI_ISL_2985130 | IRCCS San Gallicano Dermatological Institute | IRCCS Regina Elena National Cancer Institute | Sabrina Strano, Sara Donzelli, Alice Massacci, di Domenico Enea Gino, Fabrizio Ensoli, Fulvia Pimpinelli, Maurizio Fanciulli, Giovanni Blandino, Aldo Morrone, Gennaro Ciliberto |
| EPI_ISL_2985131 | IRCCS San Gallicano Dermatological Institute | IRCCS Regina Elena National Cancer Institute | Francesca De Nicola, Giulia Orlandi, Matteo Pallocca, Ilaria Cavallo, Fabrizio Ensoli, Fulvia Pimpinelli, Maurizio Fanciulli, Giovanni Blandino, Aldo Morrone, Gennaro Ciliberto |
| EPI_ISL_2985132 | IRCCS San Gallicano Dermatological Institute | IRCCS Regina Elena National Cancer Institute | Ludovica Ciuffreda, Sara Donzelli, Eleonora Sperandio, Francesca Sivori, Fabrizio Ensoli, Fulvia Pimpinelli, Maurizio Fanciulli, Giovanni Blandino, Aldo Morrone, Gennaro Ciliberto |
| EPI_ISL_2985133 | IRCCS San Gallicano Dermatological Institute | IRCCS Regina Elena National Cancer Institute | Frauke Goeman, Giulia Orlandi, Alice Massacci, Andrea Cazzani, Fabrizio Ensoli, Fulvia Pimpinelli, Maurizio Fanciulli, Giovanni Blandino, Aldo Morrone, Gennaro Ciliberto |
| EPI_ISL_2985134 | IRCCS San Gallicano Dermatological Institute | IRCCS Regina Elena National Cancer Institute | Sabrina Strano, Sara Donzelli, Matteo Pallocca, Ilaria Celesti, Fabrizio Ensoli, Fulvia Pimpinelli, Maurizio Fanciulli, Giovanni Blandino, Aldo Morrone, Gennaro Ciliberto |
| EPI_ISL_2985135 | IRCCS San Gallicano Dermatological Institute | IRCCS Regina Elena National Cancer Institute | Francesca De Nicola, Giulia Orlandi, Eleonora Sperandio, Martina Diano, Fabrizio Ensoli, Fulvia Pimpinelli, Maurizio Fanciulli, Giovanni Blandino, Aldo Morrone, Gennaro Ciliberto |
| EPI_ISL_2985136 | IRCCS San Gallicano Dermatological Institute | IRCCS Regina Elena National Cancer Institute | Ludovica Ciuffreda, Sara Donzelli, Alice Massacci, Antonio Federico, Fabrizio Ensoli, Fulvia Pimpinelli, Maurizio Fanciulli, Giovanni Blandino, Aldo Morrone, Gennaro Ciliberto |
| EPI_ISL_2985137 | IRCCS San Gallicano Dermatological Institute | IRCCS Regina Elena National Cancer Institute | Frauke Goeman, Giulia Orlandi, Matteo Pallocca, Fulvia Fraticelli, Fabrizio Ensoli, Fulvia Pimpinelli, Maurizio Fanciulli, Giovanni Blandino, Aldo Morrone, Gennaro Ciliberto |
| EPI_ISL_2985138 | IRCCS San Gallicano Dermatological Institute | IRCCS Regina Elena National Cancer Institute | Sabrina Strano, Sara Donzelli, Eleonora Sperandio, Lorenzo Furzi, Fabrizio Ensoli, Fulvia Pimpinelli, Maurizio Fanciulli, Giovanni Blandino, Aldo Morrone, Gennaro Ciliberto |
| EPI_ISL_2985139 | IRCCS San Gallicano Dermatological Institute | IRCCS Regina Elena National Cancer Institute | Francesca De Nicola, Giulia Orlandi, Alice Massacci, Alessia Lauretti, Fabrizio Ensoli, Fulvia Pimpinelli, Maurizio Fanciulli, Giovanni Blandino, Aldo Morrone, Gennaro Ciliberto |

| | | | |
|---|---|---|---|
| EPI_ISL_2985140 | IRCCS San Gallicano Dermatological Institute | IRCCS Regina Elena National Cancer Institute | Ludovica Ciuffreda, Sara Donzelli, Matteo Pallocca, Francesca Maione, Fabrizio Ensoli, Fulvia Pimpinelli, Maurizio Fanciulli, Giovanni Blandino, Aldo Morrone, Gennaro Ciliberto |
| EPI_ISL_2985141 | IRCCS San Gallicano Dermatological Institute | IRCCS Regina Elena National Cancer Institute | Frauke Goeman, Giulia Orlandi, Eleonora Sperandio, Francisco Obregon, Fabrizio Ensoli, Fulvia Pimpinelli, Maurizio Fanciulli, Giovanni Blandino, Aldo Morrone, Gennaro Ciliberto |
| EPI_ISL_2985142 | IRCCS San Gallicano Dermatological Institute | IRCCS Regina Elena National Cancer Institute | Sabrina Strano, Sara Donzelli, Alice Massacci, Silvia Paluzzi, Fabrizio Ensoli, Fulvia Pimpinelli, Maurizio Fanciulli, Giovanni Blandino, Aldo Morrone, Gennaro Ciliberto |
| EPI_ISL_2985143 | IRCCS San Gallicano Dermatological Institute | IRCCS Regina Elena National Cancer Institute | Francesca De Nicola, Giulia Orlandi, Matteo Pallocca, Sara Petrolo, Fabrizio Ensoli, Fulvia Pimpinelli, Maurizio Fanciulli, Giovanni Blandino, Aldo Morrone, Gennaro Ciliberto |
| EPI_ISL_2985144 | IRCCS San Gallicano Dermatological Institute | IRCCS Regina Elena National Cancer Institute | Ludovica Ciuffreda, Sara Donzelli, Eleonora Sperandio, Valentina Ricca, Fabrizio Ensoli, Fulvia Pimpinelli, Maurizio Fanciulli, Giovanni Blandino, Aldo Morrone, Gennaro Ciliberto |
| EPI_ISL_2985146 | IRCCS San Gallicano Dermatological Institute | IRCCS Regina Elena National Cancer Institute | Sabrina Strano, Sara Donzelli, Matteo Pallocca, Grazia Prignano, Fabrizio Ensoli, Fulvia Pimpinelli, Maurizio Fanciulli, Giovanni Blandino, Aldo Morrone, Gennaro Ciliberto |
| EPI_ISL_2985148 | IRCCS San Gallicano Dermatological Institute | IRCCS Regina Elena National Cancer Institute | Ludovica Ciuffreda, Sara Donzelli, Alice Massacci, Arianna Mastrofrancesco, Fabrizio Ensoli, Fulvia Pimpinelli, Maurizio Fanciulli, Giovanni Blandino, Aldo Morrone, Gennaro Ciliberto |
| EPI_ISL_2985149 | IRCCS San Gallicano Dermatological Institute | IRCCS Regina Elena National Cancer Institute | Frauke Goeman, Giulia Orlandi, Matteo Pallocca, Giovanna D'agosto, Fabrizio Ensoli, Fulvia Pimpinelli, Maurizio Fanciulli, Giovanni Blandino, Aldo Morrone, Gennaro Ciliberto |
| EPI_ISL_2985150 | IRCCS San Gallicano Dermatological Institute | IRCCS Regina Elena National Cancer Institute | Sabrina Strano, Sara Donzelli, Eleonora Sperandio, Carla Mottini, Fabrizio Ensoli, Fulvia Pimpinelli, Maurizio Fanciulli, Giovanni Blandino, Aldo Morrone, Gennaro Ciliberto |
| EPI_ISL_2985152 | IRCCS San Gallicano Dermatological Institute | IRCCS Regina Elena National Cancer Institute | Ludovica Ciuffreda, Sara Donzelli, Matteo Pallocca, di Domenico Enea Gino, Fabrizio Ensoli, Fulvia Pimpinelli, Maurizio Fanciulli, Giovanni Blandino, Aldo Morrone, Gennaro Ciliberto |
| EPI_ISL_2985153 | IRCCS San Gallicano Dermatological Institute | IRCCS Regina Elena National Cancer Institute | Frauke Goeman, Giulia Orlandi, Eleonora Sperandio, Ilaria Cavallo, Fabrizio Ensoli, Fulvia Pimpinelli, Maurizio Fanciulli, Giovanni Blandino, Aldo Morrone, Gennaro Ciliberto |
| EPI_ISL_2985154 | IRCCS San Gallicano Dermatological Institute | IRCCS Regina Elena National Cancer Institute | Sabrina Strano, Sara Donzelli, Alice Massacci, Francesca Sivori, Fabrizio Ensoli, Fulvia Pimpinelli, Maurizio Fanciulli, Giovanni Blandino, Aldo Morrone, Gennaro Ciliberto |
| EPI_ISL_2985155 | IRCCS San Gallicano Dermatological Institute | IRCCS Regina Elena National Cancer Institute | Francesca De Nicola, Giulia Orlandi, Matteo Pallocca, Andrea Cazzani, Fabrizio Ensoli, Fulvia Pimpinelli, Maurizio Fanciulli, Giovanni Blandino, Aldo Morrone, Gennaro Ciliberto |
| EPI_ISL_2985156 | IRCCS San Gallicano Dermatological Institute | IRCCS Regina Elena National Cancer Institute | Ludovica Ciuffreda, Sara Donzelli, Eleonora Sperandio, Ilaria Celesti, Fabrizio Ensoli, Fulvia Pimpinelli, Maurizio Fanciulli, Giovanni Blandino, Aldo Morrone, Gennaro Ciliberto |
| EPI_ISL_2985157 | IRCCS San Gallicano Dermatological Institute | IRCCS Regina Elena National Cancer Institute | Frauke Goeman, Giulia Orlandi, Alice Massacci, Martina Diano, Fabrizio Ensoli, Fulvia Pimpinelli, Maurizio Fanciulli, Giovanni Blandino, Aldo Morrone, Gennaro Ciliberto |
| EPI_ISL_2985158 | IRCCS San Gallicano Dermatological Institute | IRCCS Regina Elena National Cancer Institute | Sabrina Strano, Sara Donzelli, Matteo Pallocca, Antonio Federico, Fabrizio Ensoli, Fulvia Pimpinelli, Maurizio Fanciulli, Giovanni Blandino, Aldo Morrone, Gennaro Ciliberto |
| EPI_ISL_2985159 | IRCCS San Gallicano Dermatological Institute | IRCCS Regina Elena National Cancer Institute | Francesca De Nicola, Giulia Orlandi, Eleonora Sperandio, Fulvia Fraticelli, Fabrizio Ensoli, Fulvia Pimpinelli, Maurizio Fanciulli, Giovanni Blandino, Aldo Morrone, Gennaro Ciliberto |
| EPI_ISL_2985162 | IRCCS San Gallicano Dermatological Institute | IRCCS Regina Elena National Cancer Institute | Sabrina Strano, Sara Donzelli, Eleonora Sperandio, Francesca Maione, Fabrizio Ensoli, Fulvia Pimpinelli, Maurizio Fanciulli, Giovanni Blandino, Aldo Morrone, Gennaro Ciliberto |
| EPI_ISL_2985163 | IRCCS San Gallicano Dermatological Institute | IRCCS Regina Elena National Cancer Institute | Francesca De Nicola, Giulia Orlandi, Alice Massacci, Francisco Obregon, Fabrizio Ensoli, Fulvia Pimpinelli, Maurizio Fanciulli, Giovanni Blandino, Aldo Morrone, Gennaro Ciliberto |
| EPI_ISL_2985164 | IRCCS San Gallicano Dermatological Institute | IRCCS Regina Elena National Cancer Institute | Ludovica Ciuffreda, Sara Donzelli, Matteo Pallocca, Silvia Paluzzi, Fabrizio Ensoli, Fulvia Pimpinelli, Maurizio Fanciulli, Giovanni Blandino, Aldo Morrone, Gennaro Ciliberto |
| EPI_ISL_2985165 | IRCCS San Gallicano Dermatological Institute | IRCCS Regina Elena National Cancer Institute | Frauke Goeman, Giulia Orlandi, Eleonora Sperandio, Sara Petrolo, Fabrizio Ensoli, Fulvia Pimpinelli, Maurizio Fanciulli, Giovanni Blandino, Aldo Morrone, Gennaro Ciliberto |
| EPI_ISL_2985166 | IRCCS San Gallicano Dermatological Institute | IRCCS Regina Elena National Cancer Institute | Sabrina Strano, Sara Donzelli, Alice Massacci, Valentina Ricca, Fabrizio Ensoli, Fulvia Pimpinelli, Maurizio Fanciulli, Giovanni Blandino, Aldo Morrone, Gennaro Ciliberto |
| EPI_ISL_2985168 | IRCCS San Gallicano Dermatological Institute | IRCCS Regina Elena National Cancer Institute | Ludovica Ciuffreda, Sara Donzelli, Eleonora Sperandio, Grazia Prignano, Fabrizio Ensoli, Fulvia Pimpinelli, Maurizio Fanciulli, Giovanni Blandino, Aldo Morrone, Gennaro Ciliberto |
| EPI_ISL_2985169 | IRCCS San Gallicano Dermatological Institute | IRCCS Regina Elena National Cancer Institute | Frauke Goeman, Giulia Orlandi, Alice Massacci, Elisabetta Trento, Fabrizio Ensoli, Fulvia Pimpinelli, Maurizio Fanciulli, Giovanni Blandino, Aldo Morrone, Gennaro Ciliberto |
| EPI_ISL_2985170 | IRCCS San Gallicano Dermatological Institute | IRCCS Regina Elena National Cancer Institute | Sabrina Strano, Sara Donzelli, Matteo Pallocca, Arianna Mastrofrancesco, Fabrizio Ensoli, Fulvia Pimpinelli, Maurizio Fanciulli, Giovanni Blandino, Aldo Morrone, Gennaro Ciliberto |
| EPI_ISL_2985171 | IRCCS San Gallicano Dermatological Institute | IRCCS Regina Elena National Cancer Institute | Francesca De Nicola, Giulia Orlandi, Eleonora Sperandio, Giovanna D'agosto, Fabrizio Ensoli, Fulvia Pimpinelli, Maurizio Fanciulli, Giovanni Blandino, Aldo Morrone, Gennaro Ciliberto |
| EPI_ISL_2985172 | IRCCS San Gallicano Dermatological Institute | IRCCS Regina Elena National Cancer Institute | Ludovica Ciuffreda, Sara Donzelli, Alice Massacci, Carla Mottini, Fabrizio Ensoli, Fulvia Pimpinelli, Maurizio Fanciulli, Giovanni Blandino, Aldo Morrone, Gennaro Ciliberto |
| EPI_ISL_2985173 | IRCCS San Gallicano Dermatological Institute | IRCCS Regina Elena National Cancer Institute | Frauke Goeman, Giulia Orlandi, Matteo Pallocca, Martina Pontone, Fabrizio Ensoli, Fulvia Pimpinelli, Maurizio Fanciulli, Giovanni Blandino, Aldo Morrone, Gennaro Ciliberto |
| EPI_ISL_2985174 | IRCCS San Gallicano Dermatological Institute | IRCCS Regina Elena National Cancer Institute | Sabrina Strano, Sara Donzelli, Eleonora Sperandio, di Domenico Enea Gino, Fabrizio Ensoli, Fulvia Pimpinelli, Maurizio Fanciulli, Giovanni Blandino, Aldo Morrone, Gennaro Ciliberto |
| EPI_ISL_2985175 | IRCCS San Gallicano Dermatological Institute | IRCCS Regina Elena National Cancer Institute | Francesca De Nicola, Giulia Orlandi, Alice Massacci, Ilaria Cavallo, Fabrizio Ensoli, Fulvia Pimpinelli, Maurizio Fanciulli, Giovanni Blandino, Aldo Morrone, Gennaro Ciliberto |
| EPI_ISL_2985176 | IRCCS San Gallicano Dermatological Institute | IRCCS Regina Elena National Cancer Institute | Ludovica Ciuffreda, Sara Donzelli, Matteo Pallocca, Francesca Sivori, Fabrizio Ensoli, Fulvia Pimpinelli, Maurizio Fanciulli, Giovanni Blandino, Aldo Morrone, Gennaro Ciliberto |
| EPI_ISL_2985178 | IRCCS San Gallicano Dermatological Institute | IRCCS Regina Elena National Cancer Institute | Francesca De Nicola, Giulia Orlandi, Matteo Pallocca, Martina Diano, Fabrizio Ensoli, Fulvia Pimpinelli, Maurizio Fanciulli, Giovanni Blandino, Aldo Morrone, Gennaro Ciliberto |
| EPI_ISL_2985180 | IRCCS San Gallicano Dermatological Institute | IRCCS Regina Elena National Cancer Institute | Frauke Goeman, Giulia Orlandi, Alice Massacci, Fulvia Fraticelli, Fabrizio Ensoli, Fulvia Pimpinelli, Maurizio Fanciulli, Giovanni Blandino, Aldo Morrone, Gennaro Ciliberto |
| EPI_ISL_2985181 | IRCCS San Gallicano Dermatological Institute | IRCCS Regina Elena National Cancer Institute | Sabrina Strano, Sara Donzelli, Matteo Pallocca, Lorenzo Furzi, Fabrizio Ensoli, Fulvia Pimpinelli, Maurizio Fanciulli, Giovanni Blandino, Aldo Morrone, Gennaro Ciliberto |
| EPI_ISL_2985182 | IRCCS San Gallicano Dermatological Institute | IRCCS Regina Elena National Cancer Institute | Francesca De Nicola, Giulia Orlandi, Eleonora Sperandio, Alessia Lauretti, Fabrizio Ensoli, Fulvia Pimpinelli, Maurizio Fanciulli, Giovanni Blandino, Aldo Morrone, Gennaro Ciliberto |
| EPI_ISL_2985183 | IRCCS San Gallicano Dermatological Institute | IRCCS Regina Elena National Cancer Institute | Ludovica Ciuffreda, Sara Donzelli, Alice Massacci, Francesca Maione, Fabrizio Ensoli, Fulvia Pimpinelli, Maurizio Fanciulli, Giovanni Blandino, Aldo Morrone, Gennaro Ciliberto |
| EPI_ISL_2985185 | IRCCS San Gallicano Dermatological Institute | IRCCS Regina Elena National Cancer Institute | Sabrina Strano, Sara Donzelli, Eleonora Sperandio, Silvia Paluzzi, Fabrizio Ensoli, Fulvia Pimpinelli, Maurizio Fanciulli, Giovanni Blandino, Aldo Morrone, Gennaro Ciliberto |
| EPI_ISL_2985186 | IRCCS San Gallicano Dermatological Institute | IRCCS Regina Elena National Cancer Institute | Francesca De Nicola, Giulia Orlandi, Alice Massacci, Sara Petrolo, Fabrizio Ensoli, Fulvia Pimpinelli, Maurizio Fanciulli, Giovanni Blandino, Aldo Morrone, Gennaro Ciliberto |
| EPI_ISL_2985187 | IRCCS San Gallicano Dermatological Institute | IRCCS Regina Elena National Cancer Institute | Ludovica Ciuffreda, Sara Donzelli, Matteo Pallocca, Valentina Ricca, Fabrizio Ensoli, Fulvia Pimpinelli, Maurizio Fanciulli, Giovanni Blandino, Aldo Morrone, Gennaro Ciliberto |
| EPI_ISL_2985188 | IRCCS San Gallicano Dermatological Institute | IRCCS Regina Elena National Cancer Institute | Frauke Goeman, Giulia Orlandi, Eleonora Sperandio, Serena Salvo, Fabrizio Ensoli, Fulvia Pimpinelli, Maurizio Fanciulli, Giovanni Blandino, Aldo Morrone, Gennaro Ciliberto |
| EPI_ISL_2985189 | IRCCS San Gallicano Dermatological Institute | IRCCS Regina Elena National Cancer Institute | Sabrina Strano, Sara Donzelli, Alice Massacci, Grazia Prignano, Fabrizio Ensoli, Fulvia Pimpinelli, Maurizio Fanciulli, Giovanni Blandino, Aldo Morrone, Gennaro Ciliberto |
| EPI_ISL_2985190 | IRCCS San Gallicano Dermatological Institute | IRCCS Regina Elena National Cancer Institute | Francesca De Nicola, Giulia Orlandi, Matteo Pallocca, Elisabetta Trento, Fabrizio Ensoli, Fulvia Pimpinelli, Maurizio Fanciulli, Giovanni Blandino, Aldo Morrone, Gennaro Ciliberto |
| EPI_ISL_2985191 | IRCCS San Gallicano Dermatological Institute | IRCCS Regina Elena National Cancer Institute | Ludovica Ciuffreda, Sara Donzelli, Eleonora Sperandio, Arianna Mastrofrancesco, Fabrizio Ensoli, Fulvia Pimpinelli, Maurizio Fanciulli, Giovanni Blandino, Aldo Morrone, Gennaro Ciliberto |
| EPI_ISL_2985192 | IRCCS San Gallicano Dermatological Institute | IRCCS Regina Elena National Cancer Institute | Frauke Goeman, Giulia Orlandi, Alice Massacci, Giovanna D'agosto, Fabrizio Ensoli, Fulvia Pimpinelli, Maurizio Fanciulli, Giovanni Blandino, Aldo Morrone, Gennaro Ciliberto |
| EPI_ISL_2985193 | IRCCS San Gallicano Dermatological Institute | IRCCS Regina Elena National Cancer Institute | Sabrina Strano, Sara Donzelli, Matteo Pallocca, Carla Mottini, Fabrizio Ensoli, Fulvia Pimpinelli, Maurizio Fanciulli, Giovanni Blandino, Aldo Morrone, Gennaro Ciliberto |
| EPI_ISL_2985195 | IRCCS San Gallicano Dermatological Institute | IRCCS Regina Elena National Cancer Institute | Ludovica Ciuffreda, Sara Donzelli, Alice Massacci, di Domenico Enea Gino, Fabrizio Ensoli, Fulvia Pimpinelli, Maurizio Fanciulli, Giovanni Blandino, Aldo Morrone, Gennaro Ciliberto |
| EPI_ISL_2985196 | IRCCS San Gallicano Dermatological Institute | IRCCS Regina Elena National Cancer Institute | Frauke Goeman, Giulia Orlandi, Matteo Pallocca, Ilaria Cavallo, Fabrizio Ensoli, Fulvia Pimpinelli, Maurizio Fanciulli, Giovanni Blandino, Aldo Morrone, Gennaro Ciliberto |
| EPI_ISL_2985197 | IRCCS San Gallicano Dermatological Institute | IRCCS Regina Elena National Cancer Institute | Sabrina Strano, Sara Donzelli, Eleonora Sperandio, Francesca Sivori, Fabrizio Ensoli, Fulvia Pimpinelli, Maurizio Fanciulli, Giovanni Blandino, Aldo Morrone, Gennaro Ciliberto |
| EPI_ISL_2985198 | IRCCS San Gallicano Dermatological Institute | IRCCS Regina Elena National Cancer Institute | Francesca De Nicola, Giulia Orlandi, Alice Massacci, Andrea Cazzani, Fabrizio Ensoli, Fulvia Pimpinelli, Maurizio Fanciulli, Giovanni Blandino, Aldo Morrone, Gennaro Ciliberto |
| EPI_ISL_2985199 | IRCCS San Gallicano Dermatological Institute | IRCCS Regina Elena National Cancer Institute | Ludovica Ciuffreda, Sara Donzelli, Matteo Pallocca, Ilaria Celesti, Fabrizio Ensoli, Fulvia Pimpinelli, Maurizio Fanciulli, Giovanni Blandino, Aldo Morrone, Gennaro Ciliberto |
| EPI_ISL_2985200 | IRCCS San Gallicano Dermatological Institute | IRCCS Regina Elena National Cancer Institute | Frauke Goeman, Giulia Orlandi, Eleonora Sperandio, Martina Diano, Fabrizio Ensoli, Fulvia Pimpinelli, Maurizio Fanciulli, Giovanni Blandino, Aldo Morrone, Gennaro Ciliberto |
| EPI_ISL_2985201 | IRCCS San Gallicano Dermatological Institute | IRCCS Regina Elena National Cancer Institute | Sabrina Strano, Sara Donzelli, Alice Massacci, Antonio Federico, Fabrizio Ensoli, Fulvia Pimpinelli, Maurizio Fanciulli, Giovanni Blandino, Aldo Morrone, Gennaro Ciliberto |
| EPI_ISL_2985202 | IRCCS San Gallicano Dermatological Institute | IRCCS Regina Elena National Cancer Institute | Francesca De Nicola, Giulia Orlandi, Matteo Pallocca, Fulvia Fraticelli, Fabrizio Ensoli, Fulvia Pimpinelli, Maurizio Fanciulli, Giovanni Blandino, Aldo Morrone, Gennaro Ciliberto |
| EPI_ISL_2985203 | IRCCS San Gallicano Dermatological Institute | IRCCS Regina Elena National Cancer Institute | Ludovica Ciuffreda, Sara Donzelli, Eleonora Sperandio, Lorenzo Furzi, Fabrizio Ensoli, Fulvia Pimpinelli, Maurizio Fanciulli, Giovanni Blandino, Aldo Morrone, Gennaro Ciliberto |
| EPI_ISL_2985205 | IRCCS San Gallicano Dermatological Institute | IRCCS Regina Elena National Cancer Institute | Sabrina Strano, Sara Donzelli, Matteo Pallocca, Francesca Maione, Fabrizio Ensoli, Fulvia Pimpinelli, Maurizio Fanciulli, Giovanni Blandino, Aldo Morrone, Gennaro Ciliberto |
| EPI_ISL_2985206 | IRCCS San Gallicano Dermatological Institute | IRCCS Regina Elena National Cancer Institute | Francesca De Nicola, Giulia Orlandi, Eleonora Sperandio, Francisco Obregon, Fabrizio Ensoli, Fulvia Pimpinelli, Maurizio Fanciulli, Giovanni Blandino, Aldo Morrone, Gennaro Ciliberto |
| EPI_ISL_2985207 | IRCCS San Gallicano Dermatological Institute | IRCCS Regina Elena National Cancer Institute | Ludovica Ciuffreda, Sara Donzelli, Alice Massacci, Silvia Paluzzi, Fabrizio Ensoli, Fulvia Pimpinelli, Maurizio Fanciulli, Giovanni Blandino, Aldo Morrone, Gennaro Ciliberto |
| EPI_ISL_2985208 | IRCCS San Gallicano Dermatological Institute | IRCCS Regina Elena National Cancer Institute | Frauke Goeman, Giulia Orlandi, Matteo Pallocca, Sara Petrolo, Fabrizio Ensoli, Fulvia Pimpinelli, Maurizio Fanciulli, Giovanni Blandino, Aldo Morrone, Gennaro Ciliberto |
| EPI_ISL_2985209 | IRCCS San Gallicano Dermatological Institute | IRCCS Regina Elena National Cancer Institute | Sabrina Strano, Sara Donzelli, Eleonora Sperandio, Valentina Ricca, Fabrizio Ensoli, Fulvia Pimpinelli, Maurizio Fanciulli, Giovanni Blandino, Aldo Morrone, Gennaro Ciliberto |
| EPI_ISL_2985210 | IRCCS San Gallicano Dermatological Institute | IRCCS Regina Elena National Cancer Institute | Francesca De Nicola, Giulia Orlandi, Alice Massacci, Serena Salvo, Fabrizio Ensoli, Fulvia Pimpinelli, Maurizio Fanciulli, Giovanni Blandino, Aldo Morrone, Gennaro Ciliberto |
| EPI_ISL_2985211 | IRCCS San Gallicano Dermatological Institute | IRCCS Regina Elena National Cancer Institute | Ludovica Ciuffreda, Sara Donzelli, Matteo Pallocca, Grazia Prignano, Fabrizio Ensoli, Fulvia Pimpinelli, Maurizio Fanciulli, Giovanni Blandino, Aldo Morrone, Gennaro Ciliberto |
| EPI_ISL_2985212 | IRCCS San Gallicano Dermatological Institute | IRCCS Regina Elena National Cancer Institute | Frauke Goeman, Giulia Orlandi, Eleonora Sperandio, Elisabetta Trento, Fabrizio Ensoli, Fulvia Pimpinelli, Maurizio Fanciulli, Giovanni Blandino, Aldo Morrone, Gennaro Ciliberto |
| EPI_ISL_2985213 | IRCCS San Gallicano Dermatological Institute | IRCCS Regina Elena National Cancer Institute | Sabrina Strano, Sara Donzelli, Alice Massacci, Arianna Mastrofrancesco, Fabrizio Ensoli, Fulvia Pimpinelli, Maurizio Fanciulli, Giovanni Blandino, Aldo Morrone, Gennaro Ciliberto |
| EPI_ISL_2985214 | IRCCS San Gallicano Dermatological Institute | IRCCS Regina Elena National Cancer Institute | Francesca De Nicola, Giulia Orlandi, Matteo Pallocca, Giovanna D'agosto, Fabrizio Ensoli, Fulvia Pimpinelli, Maurizio Fanciulli, Giovanni Blandino, Aldo Morrone, Gennaro Ciliberto |
| EPI_ISL_2985215 | IRCCS San Gallicano Dermatological Institute | IRCCS Regina Elena National Cancer Institute | Ludovica Ciuffreda, Sara Donzelli, Eleonora Sperandio, Carla Mottini, Fabrizio Ensoli, Fulvia Pimpinelli, Maurizio Fanciulli, Giovanni Blandino, Aldo Morrone, Gennaro Ciliberto |
| EPI_ISL_2985216 | IRCCS San Gallicano Dermatological Institute | IRCCS Regina Elena National Cancer Institute | Frauke Goeman, Giulia Orlandi, Alice Massacci, Martina Pontone, Fabrizio Ensoli, Fulvia Pimpinelli, Maurizio Fanciulli, Giovanni Blandino, Aldo Morrone, Gennaro Ciliberto |
| EPI_ISL_2985217 | IRCCS San Gallicano Dermatological Institute | IRCCS Regina Elena National Cancer Institute | Sabrina Strano, Sara Donzelli, Matteo Pallocca, di Domenico Enea Gino, Fabrizio Ensoli, Fulvia Pimpinelli, Maurizio Fanciulli, Giovanni Blandino, Aldo Morrone, Gennaro Ciliberto |
| EPI_ISL_2985218 | IRCCS San Gallicano Dermatological Institute | IRCCS Regina Elena National Cancer Institute | Francesca De Nicola, Giulia Orlandi, Eleonora Sperandio, Ilaria Cavallo, Fabrizio Ensoli, Fulvia Pimpinelli, Maurizio Fanciulli, Giovanni Blandino, Aldo Morrone, Gennaro Ciliberto |
| EPI_ISL_2985219 | IRCCS San Gallicano Dermatological Institute | IRCCS Regina Elena National Cancer Institute | Ludovica Ciuffreda, Sara Donzelli, Alice Massacci, Francesca Sivori, Fabrizio Ensoli, Fulvia Pimpinelli, Maurizio Fanciulli, Giovanni Blandino, Aldo Morrone, Gennaro Ciliberto |
| EPI_ISL_2985221 | IRCCS San Gallicano Dermatological Institute | IRCCS Regina Elena National Cancer Institute | Sabrina Strano, Sara Donzelli, Eleonora Sperandio, Ilaria Celesti, Fabrizio Ensoli, Fulvia Pimpinelli, Maurizio Fanciulli, Giovanni Blandino, Aldo Morrone, Gennaro Ciliberto |
| EPI_ISL_2985224 | IRCCS San Gallicano Dermatological Institute | IRCCS Regina Elena National Cancer Institute | Frauke Goeman, Giulia Orlandi, Eleonora Sperandio, Fulvia Fraticelli, Fabrizio Ensoli, Fulvia Pimpinelli, Maurizio Fanciulli, Giovanni Blandino, Aldo Morrone, Gennaro Ciliberto |
| EPI_ISL_2985225 | IRCCS San Gallicano Dermatological Institute | IRCCS Regina Elena National Cancer Institute | Sabrina Strano, Sara Donzelli, Alice Massacci, Lorenzo Furzi, Fabrizio Ensoli, Fulvia Pimpinelli, Maurizio Fanciulli, Giovanni Blandino, Aldo Morrone, Gennaro Ciliberto |
| EPI_ISL_2985226 | IRCCS San Gallicano Dermatological Institute | IRCCS Regina Elena National Cancer Institute | Francesca De Nicola, Giulia Orlandi, Matteo Pallocca, Alessia Lauretti, Fabrizio Ensoli, Fulvia Pimpinelli, Maurizio Fanciulli, Giovanni Blandino, Aldo Morrone, Gennaro Ciliberto |
| EPI_ISL_2985227 | IRCCS San Gallicano Dermatological Institute | IRCCS Regina Elena National Cancer Institute | Ludovica Ciuffreda, Sara Donzelli, Eleonora Sperandio, Francesca Maione, Fabrizio Ensoli, Fulvia Pimpinelli, Maurizio Fanciulli, Giovanni Blandino, Aldo Morrone, Gennaro Ciliberto |
| EPI_ISL_2985228 | IRCCS San Gallicano Dermatological Institute | IRCCS Regina Elena National Cancer Institute | Frauke Goeman, Giulia Orlandi, Alice Massacci, Francisco Obregon, Fabrizio Ensoli, Fulvia Pimpinelli, Maurizio Fanciulli, Giovanni Blandino, Aldo Morrone, Gennaro Ciliberto |
| EPI_ISL_2985229 | IRCCS San Gallicano Dermatological Institute | IRCCS Regina Elena National Cancer Institute | Sabrina Strano, Sara Donzelli, Matteo Pallocca, Silvia Paluzzi, Fabrizio Ensoli, Fulvia Pimpinelli, Maurizio Fanciulli, Giovanni Blandino, Aldo Morrone, Gennaro Ciliberto |
| EPI_ISL_2985230 | IRCCS San Gallicano Dermatological Institute | IRCCS Regina Elena National Cancer Institute | Francesca De Nicola, Giulia Orlandi, Eleonora Sperandio, Sara Petrolo, Fabrizio Ensoli, Fulvia Pimpinelli, Maurizio Fanciulli, Giovanni Blandino, Aldo Morrone, Gennaro Ciliberto |

| | | | |
|---|---|---|---|
| EPI_ISL_2985231 | IRCCS San Gallicano Dermatological Institute | IRCCS Regina Elena National Cancer Institute | Ludovica Ciuffreda, Sara Donzelli, Alice Massacci, Valentina Ricca, Fabrizio Ensoli, Fulvia Pimpinelli, Maurizio Fanciulli, Giovanni Blandino, Aldo Morrone, Gennaro Ciliberto |
| EPI_ISL_2985232 | IRCCS San Gallicano Dermatological Institute | IRCCS Regina Elena National Cancer Institute | Frauke Goeman, Giulia Orlandi, Matteo Pallocca, Serena Salvo, Fabrizio Ensoli, Fulvia Pimpinelli, Maurizio Fanciulli, Giovanni Blandino, Aldo Morrone, Gennaro Ciliberto |
| EPI_ISL_2985233 | IRCCS San Gallicano Dermatological Institute | IRCCS Regina Elena National Cancer Institute | Sabrina Strano, Sara Donzelli, Eleonora Sperandio, Grazia Prignano, Fabrizio Ensoli, Fulvia Pimpinelli, Maurizio Fanciulli, Giovanni Blandino, Aldo Morrone, Gennaro Ciliberto |
| EPI_ISL_2985234 | IRCCS San Gallicano Dermatological Institute | IRCCS Regina Elena National Cancer Institute | Francesca De Nicola, Giulia Orlandi, Alice Massacci, Elisabetta Trento, Fabrizio Ensoli, Fulvia Pimpinelli, Maurizio Fanciulli, Giovanni Blandino, Aldo Morrone, Gennaro Ciliberto |
| EPI_ISL_2985235 | IRCCS San Gallicano Dermatological Institute | IRCCS Regina Elena National Cancer Institute | Ludovica Ciuffreda, Sara Donzelli, Matteo Pallocca, Arianna Mastrofrancesco, Fabrizio Ensoli, Fulvia Pimpinelli, Maurizio Fanciulli, Giovanni Blandino, Aldo Morrone, Gennaro Ciliberto |
| EPI_ISL_2985236 | IRCCS San Gallicano Dermatological Institute | IRCCS Regina Elena National Cancer Institute | Sabrina Strano, Sara Donzelli, Alice Massacci, Carla Mottini, Fabrizio Ensoli, Fulvia Pimpinelli, Maurizio Fanciulli, Giovanni Blandino, Aldo Morrone, Gennaro Ciliberto |
| EPI_ISL_2985237 | IRCCS San Gallicano Dermatological Institute | IRCCS Regina Elena National Cancer Institute | Francesca De Nicola, Giulia Orlandi, Matteo Pallocca, Martina Pontone, Fabrizio Ensoli, Fulvia Pimpinelli, Maurizio Fanciulli, Giovanni Blandino, Aldo Morrone, Gennaro Ciliberto |
| EPI_ISL_2985238 | IRCCS San Gallicano Dermatological Institute | IRCCS Regina Elena National Cancer Institute | Ludovica Ciuffreda, Sara Donzelli, Eleonora Sperandio, di Domenico Enea Gino, Fabrizio Ensoli, Fulvia Pimpinelli, Maurizio Fanciulli, Giovanni Blandino, Aldo Morrone, Gennaro Ciliberto |
| EPI_ISL_2985239 | IRCCS San Gallicano Dermatological Institute | IRCCS Regina Elena National Cancer Institute | Frauke Goeman, Giulia Orlandi, Alice Massacci, Ilaria Cavallo, Fabrizio Ensoli, Fulvia Pimpinelli, Maurizio Fanciulli, Giovanni Blandino, Aldo Morrone, Gennaro Ciliberto |
| EPI_ISL_2985240 | IRCCS San Gallicano Dermatological Institute | IRCCS Regina Elena National Cancer Institute | Sabrina Strano, Sara Donzelli, Matteo Pallocca, Francesca Sivori, Fabrizio Ensoli, Fulvia Pimpinelli, Maurizio Fanciulli, Giovanni Blandino, Aldo Morrone, Gennaro Ciliberto |
| EPI_ISL_2985241 | IRCCS San Gallicano Dermatological Institute | IRCCS Regina Elena National Cancer Institute | Francesca De Nicola, Giulia Orlandi, Eleonora Sperandio, Andrea Cazzani, Fabrizio Ensoli, Fulvia Pimpinelli, Maurizio Fanciulli, Giovanni Blandino, Aldo Morrone, Gennaro Ciliberto |
| EPI_ISL_2985243 | IRCCS San Gallicano Dermatological Institute | IRCCS Regina Elena National Cancer Institute | Frauke Goeman, Giulia Orlandi, Matteo Pallocca, Martina Diano, Fabrizio Ensoli, Fulvia Pimpinelli, Maurizio Fanciulli, Giovanni Blandino, Aldo Morrone, Gennaro Ciliberto |
| EPI_ISL_2985244 | IRCCS San Gallicano Dermatological Institute | IRCCS Regina Elena National Cancer Institute | Sabrina Strano, Sara Donzelli, Eleonora Sperandio, Antonio Federico, Fabrizio Ensoli, Fulvia Pimpinelli, Maurizio Fanciulli, Giovanni Blandino, Aldo Morrone, Gennaro Ciliberto |
| EPI_ISL_2985245 | IRCCS San Gallicano Dermatological Institute | IRCCS Regina Elena National Cancer Institute | Francesca De Nicola, Giulia Orlandi, Alice Massacci, Fulvia Fraticelli, Fabrizio Ensoli, Fulvia Pimpinelli, Maurizio Fanciulli, Giovanni Blandino, Aldo Morrone, Gennaro Ciliberto |
| EPI_ISL_2985246 | IRCCS San Gallicano Dermatological Institute | IRCCS Regina Elena National Cancer Institute | Ludovica Ciuffreda, Sara Donzelli, Matteo Pallocca, Lorenzo Furzi, Fabrizio Ensoli, Fulvia Pimpinelli, Maurizio Fanciulli, Giovanni Blandino, Aldo Morrone, Gennaro Ciliberto |
| EPI_ISL_2985247 | IRCCS San Gallicano Dermatological Institute | IRCCS Regina Elena National Cancer Institute | Frauke Goeman, Giulia Orlandi, Eleonora Sperandio, Alessia Lauretti, Fabrizio Ensoli, Fulvia Pimpinelli, Maurizio Fanciulli, Giovanni Blandino, Aldo Morrone, Gennaro Ciliberto |
| EPI_ISL_2985248 | IRCCS San Gallicano Dermatological Institute | IRCCS Regina Elena National Cancer Institute | Sabrina Strano, Sara Donzelli, Alice Massacci, Francesca Maione, Fabrizio Ensoli, Fulvia Pimpinelli, Maurizio Fanciulli, Giovanni Blandino, Aldo Morrone, Gennaro Ciliberto |
| EPI_ISL_2985249 | IRCCS San Gallicano Dermatological Institute | IRCCS Regina Elena National Cancer Institute | Francesca De Nicola, Giulia Orlandi, Matteo Pallocca, Francisco Obregon, Fabrizio Ensoli, Fulvia Pimpinelli, Maurizio Fanciulli, Giovanni Blandino, Aldo Morrone, Gennaro Ciliberto |
| EPI_ISL_2985250 | IRCCS San Gallicano Dermatological Institute | IRCCS Regina Elena National Cancer Institute | Ludovica Ciuffreda, Sara Donzelli, Eleonora Sperandio, Silvia Paluzzi, Fabrizio Ensoli, Fulvia Pimpinelli, Maurizio Fanciulli, Giovanni Blandino, Aldo Morrone, Gennaro Ciliberto |
| EPI_ISL_2985251 | IRCCS San Gallicano Dermatological Institute | IRCCS Regina Elena National Cancer Institute | Frauke Goeman, Giulia Orlandi, Alice Massacci, Sara Petrolo, Fabrizio Ensoli, Fulvia Pimpinelli, Maurizio Fanciulli, Giovanni Blandino, Aldo Morrone, Gennaro Ciliberto |
| EPI_ISL_2985252 | IRCCS San Gallicano Dermatological Institute | IRCCS Regina Elena National Cancer Institute | Sabrina Strano, Sara Donzelli, Matteo Pallocca, Valentina Ricca, Fabrizio Ensoli, Fulvia Pimpinelli, Maurizio Fanciulli, Giovanni Blandino, Aldo Morrone, Gennaro Ciliberto |
| EPI_ISL_2985253 | IRCCS San Gallicano Dermatological Institute | IRCCS Regina Elena National Cancer Institute | Francesca De Nicola, Giulia Orlandi, Eleonora Sperandio, Serena Salvo, Fabrizio Ensoli, Fulvia Pimpinelli, Maurizio Fanciulli, Giovanni Blandino, Aldo Morrone, Gennaro Ciliberto |
| EPI_ISL_2985254 | IRCCS San Gallicano Dermatological Institute | IRCCS Regina Elena National Cancer Institute | Frauke Goeman, Giulia Orlandi, Matteo Pallocca, Elisabetta Trento, Fabrizio Ensoli, Fulvia Pimpinelli, Maurizio Fanciulli, Giovanni Blandino, Aldo Morrone, Gennaro Ciliberto |
| EPI_ISL_2985255 | IRCCS San Gallicano Dermatological Institute | IRCCS Regina Elena National Cancer Institute | Sabrina Strano, Sara Donzelli, Eleonora Sperandio, Arianna Mastrofrancesco, Fabrizio Ensoli, Fulvia Pimpinelli, Maurizio Fanciulli, Giovanni Blandino, Aldo Morrone, Gennaro Ciliberto |
| EPI_ISL_2985257 | IRCCS San Gallicano Dermatological Institute | IRCCS Regina Elena National Cancer Institute | Ludovica Ciuffreda, Sara Donzelli, Matteo Pallocca, Carla Mottini, Fabrizio Ensoli, Fulvia Pimpinelli, Maurizio Fanciulli, Giovanni Blandino, Aldo Morrone, Gennaro Ciliberto |
| EPI_ISL_2985258 | IRCCS San Gallicano Dermatological Institute | IRCCS Regina Elena National Cancer Institute | Frauke Goeman, Giulia Orlandi, Eleonora Sperandio, Martina Pontone, Fabrizio Ensoli, Fulvia Pimpinelli, Maurizio Fanciulli, Giovanni Blandino, Aldo Morrone, Gennaro Ciliberto |
| EPI_ISL_2985259 | IRCCS San Gallicano Dermatological Institute | IRCCS Regina Elena National Cancer Institute | Sabrina Strano, Sara Donzelli, Alice Massacci, di Domenico Enea Gino, Fabrizio Ensoli, Fulvia Pimpinelli, Maurizio Fanciulli, Giovanni Blandino, Aldo Morrone, Gennaro Ciliberto |
| EPI_ISL_2985260 | IRCCS San Gallicano Dermatological Institute | IRCCS Regina Elena National Cancer Institute | Francesca De Nicola, Giulia Orlandi, Matteo Pallocca, Ilaria Cavallo, Fabrizio Ensoli, Fulvia Pimpinelli, Maurizio Fanciulli, Giovanni Blandino, Aldo Morrone, Gennaro Ciliberto |
| EPI_ISL_2985261 | IRCCS San Gallicano Dermatological Institute | IRCCS Regina Elena National Cancer Institute | Ludovica Ciuffreda, Sara Donzelli, Eleonora Sperandio, Francesca Sivori, Fabrizio Ensoli, Fulvia Pimpinelli, Maurizio Fanciulli, Giovanni Blandino, Aldo Morrone, Gennaro Ciliberto |
| EPI_ISL_2985262 | IRCCS San Gallicano Dermatological Institute | IRCCS Regina Elena National Cancer Institute | Frauke Goeman, Giulia Orlandi, Alice Massacci, Andrea Cazzani, Fabrizio Ensoli, Fulvia Pimpinelli, Maurizio Fanciulli, Giovanni Blandino, Aldo Morrone, Gennaro Ciliberto |
| EPI_ISL_2985263 | IRCCS San Gallicano Dermatological Institute | IRCCS Regina Elena National Cancer Institute | Sabrina Strano, Sara Donzelli, Matteo Pallocca, Ilaria Celesti, Fabrizio Ensoli, Fulvia Pimpinelli, Maurizio Fanciulli, Giovanni Blandino, Aldo Morrone, Gennaro Ciliberto |
| EPI_ISL_2985264 | IRCCS San Gallicano Dermatological Institute | IRCCS Regina Elena National Cancer Institute | Francesca De Nicola, Giulia Orlandi, Eleonora Sperandio, Martina Diano, Fabrizio Ensoli, Fulvia Pimpinelli, Maurizio Fanciulli, Giovanni Blandino, Aldo Morrone, Gennaro Ciliberto |
| EPI_ISL_2985265 | IRCCS San Gallicano Dermatological Institute | IRCCS Regina Elena National Cancer Institute | Ludovica Ciuffreda, Sara Donzelli, Alice Massacci, Antonio Federico, Fabrizio Ensoli, Fulvia Pimpinelli, Maurizio Fanciulli, Giovanni Blandino, Aldo Morrone, Gennaro Ciliberto |
| EPI_ISL_2985266 | IRCCS San Gallicano Dermatological Institute | IRCCS Regina Elena National Cancer Institute | Frauke Goeman, Giulia Orlandi, Matteo Pallocca, Fulvia Fraticelli, Fabrizio Ensoli, Fulvia Pimpinelli, Maurizio Fanciulli, Giovanni Blandino, Aldo Morrone, Gennaro Ciliberto |
| EPI_ISL_2985267 | IRCCS San Gallicano Dermatological Institute | IRCCS Regina Elena National Cancer Institute | Sabrina Strano, Sara Donzelli, Eleonora Sperandio, Lorenzo Furzi, Fabrizio Ensoli, Fulvia Pimpinelli, Maurizio Fanciulli, Giovanni Blandino, Aldo Morrone, Gennaro Ciliberto |
| EPI_ISL_2985268 | IRCCS San Gallicano Dermatological Institute | IRCCS Regina Elena National Cancer Institute | Francesca De Nicola, Giulia Orlandi, Alice Massacci, Alessia Lauretti, Fabrizio Ensoli, Fulvia Pimpinelli, Maurizio Fanciulli, Giovanni Blandino, Aldo Morrone, Gennaro Ciliberto |
| EPI_ISL_2985269 | IRCCS San Gallicano Dermatological Institute | IRCCS Regina Elena National Cancer Institute | Ludovica Ciuffreda, Sara Donzelli, Matteo Pallocca, Francesca Maione, Fabrizio Ensoli, Fulvia Pimpinelli, Maurizio Fanciulli, Giovanni Blandino, Aldo Morrone, Gennaro Ciliberto |
| EPI_ISL_2985270 | IRCCS San Gallicano Dermatological Institute | IRCCS Regina Elena National Cancer Institute | Frauke Goeman, Giulia Orlandi, Eleonora Sperandio, Francisco Obregon, Fabrizio Ensoli, Fulvia Pimpinelli, Maurizio Fanciulli, Giovanni Blandino, Aldo Morrone, Gennaro Ciliberto |
| EPI_ISL_2985271 | IRCCS San Gallicano Dermatological Institute | IRCCS Regina Elena National Cancer Institute | Sabrina Strano, Sara Donzelli, Alice Massacci, Silvia Paluzzi, Fabrizio Ensoli, Fulvia Pimpinelli, Maurizio Fanciulli, Giovanni Blandino, Aldo Morrone, Gennaro Ciliberto |
| EPI_ISL_2985272 | IRCCS San Gallicano Dermatological Institute | IRCCS Regina Elena National Cancer Institute | Francesca De Nicola, Giulia Orlandi, Matteo Pallocca, Sara Petrolo, Fabrizio Ensoli, Fulvia Pimpinelli, Maurizio Fanciulli, Giovanni Blandino, Aldo Morrone, Gennaro Ciliberto |
| EPI_ISL_2985274 | IRCCS San Gallicano Dermatological Institute | IRCCS Regina Elena National Cancer Institute | Frauke Goeman, Giulia Orlandi, Alice Massacci, Serena Salvo, Fabrizio Ensoli, Fulvia Pimpinelli, Maurizio Fanciulli, Giovanni Blandino, Aldo Morrone, Gennaro Ciliberto |
| EPI_ISL_2985275 | IRCCS San Gallicano Dermatological Institute | IRCCS Regina Elena National Cancer Institute | Sabrina Strano, Sara Donzelli, Matteo Pallocca, Grazia Prignano, Fabrizio Ensoli, Fulvia Pimpinelli, Maurizio Fanciulli, Giovanni Blandino, Aldo Morrone, Gennaro Ciliberto |
| EPI_ISL_2985276 | IRCCS San Gallicano Dermatological Institute | IRCCS Regina Elena National Cancer Institute | Francesca De Nicola, Giulia Orlandi, Eleonora Sperandio, Elisabetta Trento, Fabrizio Ensoli, Fulvia Pimpinelli, Maurizio Fanciulli, Giovanni Blandino, Aldo Morrone, Gennaro Ciliberto |
| EPI_ISL_2985277 | IRCCS San Gallicano Dermatological Institute | IRCCS Regina Elena National Cancer Institute | Ludovica Ciuffreda, Sara Donzelli, Alice Massacci, Arianna Mastrofrancesco, Fabrizio Ensoli, Fulvia Pimpinelli, Maurizio Fanciulli, Giovanni Blandino, Aldo Morrone, Gennaro Ciliberto |
| EPI_ISL_2985279 | IRCCS San Gallicano Dermatological Institute | IRCCS Regina Elena National Cancer Institute | Sabrina Strano, Sara Donzelli, Eleonora Sperandio, Carla Mottini, Fabrizio Ensoli, Fulvia Pimpinelli, Maurizio Fanciulli, Giovanni Blandino, Aldo Morrone, Gennaro Ciliberto |
| EPI_ISL_2985280 | IRCCS San Gallicano Dermatological Institute | IRCCS Regina Elena National Cancer Institute | Francesca De Nicola, Giulia Orlandi, Alice Massacci, Martina Pontone, Fabrizio Ensoli, Fulvia Pimpinelli, Maurizio Fanciulli, Giovanni Blandino, Aldo Morrone, Gennaro Ciliberto |
| EPI_ISL_2985281 | IRCCS San Gallicano Dermatological Institute | IRCCS Regina Elena National Cancer Institute | Ludovica Ciuffreda, Sara Donzelli, Matteo Pallocca, di Domenico Enea Gino, Fabrizio Ensoli, Fulvia Pimpinelli, Maurizio Fanciulli, Giovanni Blandino, Aldo Morrone, Gennaro Ciliberto |
| EPI_ISL_2985282 | IRCCS San Gallicano Dermatological Institute | IRCCS Regina Elena National Cancer Institute | Frauke Goeman, Giulia Orlandi, Eleonora Sperandio, Ilaria Cavallo, Fabrizio Ensoli, Fulvia Pimpinelli, Maurizio Fanciulli, Giovanni Blandino, Aldo Morrone, Gennaro Ciliberto |
| EPI_ISL_2985283 | IRCCS San Gallicano Dermatological Institute | IRCCS Regina Elena National Cancer Institute | Sabrina Strano, Sara Donzelli, Alice Massacci, Francesca Sivori, Fabrizio Ensoli, Fulvia Pimpinelli, Maurizio Fanciulli, Giovanni Blandino, Aldo Morrone, Gennaro Ciliberto |
| EPI_ISL_2985284 | IRCCS San Gallicano Dermatological Institute | IRCCS Regina Elena National Cancer Institute | Francesca De Nicola, Giulia Orlandi, Matteo Pallocca, Andrea Cazzani, Fabrizio Ensoli, Fulvia Pimpinelli, Maurizio Fanciulli, Giovanni Blandino, Aldo Morrone, Gennaro Ciliberto |
| EPI_ISL_2985285 | IRCCS San Gallicano Dermatological Institute | IRCCS Regina Elena National Cancer Institute | Ludovica Ciuffreda, Sara Donzelli, Eleonora Sperandio, Ilaria Celesti, Fabrizio Ensoli, Fulvia Pimpinelli, Maurizio Fanciulli, Giovanni Blandino, Aldo Morrone, Gennaro Ciliberto |
| EPI_ISL_2985286 | IRCCS San Gallicano Dermatological Institute | IRCCS Regina Elena National Cancer Institute | Frauke Goeman, Giulia Orlandi, Alice Massacci, Martina Diano, Fabrizio Ensoli, Fulvia Pimpinelli, Maurizio Fanciulli, Giovanni Blandino, Aldo Morrone, Gennaro Ciliberto |
| EPI_ISL_2985287 | IRCCS San Gallicano Dermatological Institute | IRCCS Regina Elena National Cancer Institute | Sabrina Strano, Sara Donzelli, Matteo Pallocca, Antonio Federico, Fabrizio Ensoli, Fulvia Pimpinelli, Maurizio Fanciulli, Giovanni Blandino, Aldo Morrone, Gennaro Ciliberto |
| EPI_ISL_2985288 | IRCCS San Gallicano Dermatological Institute | IRCCS Regina Elena National Cancer Institute | Francesca De Nicola, Giulia Orlandi, Eleonora Sperandio, Fulvia Fraticelli, Fabrizio Ensoli, Fulvia Pimpinelli, Maurizio Fanciulli, Giovanni Blandino, Aldo Morrone, Gennaro Ciliberto |
| EPI_ISL_2985290 | IRCCS San Gallicano Dermatological Institute | IRCCS Regina Elena National Cancer Institute | Frauke Goeman, Giulia Orlandi, Matteo Pallocca, Alessia Lauretti, Fabrizio Ensoli, Fulvia Pimpinelli, Maurizio Fanciulli, Giovanni Blandino, Aldo Morrone, Gennaro Ciliberto |
| EPI_ISL_2985292 | IRCCS San Gallicano Dermatological Institute | IRCCS Regina Elena National Cancer Institute | Francesca De Nicola, Giulia Orlandi, Alice Massacci, Francisco Obregon, Fabrizio Ensoli, Fulvia Pimpinelli, Maurizio Fanciulli, Giovanni Blandino, Aldo Morrone, Gennaro Ciliberto |
| EPI_ISL_2985293 | IRCCS San Gallicano Dermatological Institute | IRCCS Regina Elena National Cancer Institute | Ludovica Ciuffreda, Sara Donzelli, Matteo Pallocca, Silvia Paluzzi, Fabrizio Ensoli, Fulvia Pimpinelli, Maurizio Fanciulli, Giovanni Blandino, Aldo Morrone, Gennaro Ciliberto |
| EPI_ISL_2985294 | IRCCS San Gallicano Dermatological Institute | IRCCS Regina Elena National Cancer Institute | Frauke Goeman, Giulia Orlandi, Eleonora Sperandio, Sara Petrolo, Fabrizio Ensoli, Fulvia Pimpinelli, Maurizio Fanciulli, Giovanni Blandino, Aldo Morrone, Gennaro Ciliberto |
| EPI_ISL_2985297 | IRCCS San Gallicano Dermatological Institute | IRCCS Regina Elena National Cancer Institute | Ludovica Ciuffreda, Sara Donzelli, Eleonora Sperandio, Grazia Prignano, Fabrizio Ensoli, Fulvia Pimpinelli, Maurizio Fanciulli, Giovanni Blandino, Aldo Morrone, Gennaro Ciliberto |
| EPI_ISL_2985299 | IRCCS San Gallicano Dermatological Institute | IRCCS Regina Elena National Cancer Institute | Sabrina Strano, Sara Donzelli, Matteo Pallocca, Arianna Mastrofrancesco, Fabrizio Ensoli, Fulvia Pimpinelli, Maurizio Fanciulli, Giovanni Blandino, Aldo Morrone, Gennaro Ciliberto |
| EPI_ISL_2985300 | IRCCS San Gallicano Dermatological Institute | IRCCS Regina Elena National Cancer Institute | Francesca De Nicola, Giulia Orlandi, Eleonora Sperandio, Giovanna D'agosto, Fabrizio Ensoli, Fulvia Pimpinelli, Maurizio Fanciulli, Giovanni Blandino, Aldo Morrone, Gennaro Ciliberto |
| EPI_ISL_2985301 | IRCCS San Gallicano Dermatological Institute | IRCCS Regina Elena National Cancer Institute | Ludovica Ciuffreda, Sara Donzelli, Alice Massacci, Carla Mottini, Fabrizio Ensoli, Fulvia Pimpinelli, Maurizio Fanciulli, Giovanni Blandino, Aldo Morrone, Gennaro Ciliberto |
| EPI_ISL_2985303 | IRCCS San Gallicano Dermatological Institute | IRCCS Regina Elena National Cancer Institute | Sabrina Strano, Sara Donzelli, Eleonora Sperandio, di Domenico Enea Gino, Fabrizio Ensoli, Fulvia Pimpinelli, Maurizio Fanciulli, Giovanni Blandino, Aldo Morrone, Gennaro Ciliberto |
| EPI_ISL_2985305 | IRCCS San Gallicano Dermatological Institute | IRCCS Regina Elena National Cancer Institute | Ludovica Ciuffreda, Sara Donzelli, Matteo Pallocca, Francesca Sivori, Fabrizio Ensoli, Fulvia Pimpinelli, Maurizio Fanciulli, Giovanni Blandino, Aldo Morrone, Gennaro Ciliberto |
| EPI_ISL_2985306 | IRCCS San Gallicano Dermatological Institute | IRCCS Regina Elena National Cancer Institute | Frauke Goeman, Giulia Orlandi, Eleonora Sperandio, Andrea Cazzani, Fabrizio Ensoli, Fulvia Pimpinelli, Maurizio Fanciulli, Giovanni Blandino, Aldo Morrone, Gennaro Ciliberto |
| EPI_ISL_2985307 | IRCCS San Gallicano Dermatological Institute | IRCCS Regina Elena National Cancer Institute | Sabrina Strano, Sara Donzelli, Alice Massacci, Ilaria Celesti, Fabrizio Ensoli, Fulvia Pimpinelli, Maurizio Fanciulli, Giovanni Blandino, Aldo Morrone, Gennaro Ciliberto |
| EPI_ISL_2985308 | IRCCS San Gallicano Dermatological Institute | IRCCS Regina Elena National Cancer Institute | Francesca De Nicola, Giulia Orlandi, Matteo Pallocca, Martina Diano, Fabrizio Ensoli, Fulvia Pimpinelli, Maurizio Fanciulli, Giovanni Blandino, Aldo Morrone, Gennaro Ciliberto |
| EPI_ISL_2985309 | IRCCS San Gallicano Dermatological Institute | IRCCS Regina Elena National Cancer Institute | Ludovica Ciuffreda, Sara Donzelli, Eleonora Sperandio, Antonio Federico, Fabrizio Ensoli, Fulvia Pimpinelli, Maurizio Fanciulli, Giovanni Blandino, Aldo Morrone, Gennaro Ciliberto |
| EPI_ISL_2985310 | IRCCS San Gallicano Dermatological Institute | IRCCS Regina Elena National Cancer Institute | Frauke Goeman, Giulia Orlandi, Alice Massacci, Fulvia Fraticelli, Fabrizio Ensoli, Fulvia Pimpinelli, Maurizio Fanciulli, Giovanni Blandino, Aldo Morrone, Gennaro Ciliberto |
| EPI_ISL_2985311 | IRCCS San Gallicano Dermatological Institute | IRCCS Regina Elena National Cancer Institute | Sabrina Strano, Sara Donzelli, Matteo Pallocca, Lorenzo Furzi, Fabrizio Ensoli, Fulvia Pimpinelli, Maurizio Fanciulli, Giovanni Blandino, Aldo Morrone, Gennaro Ciliberto |
| EPI_ISL_2985312 | IRCCS San Gallicano Dermatological Institute | IRCCS Regina Elena National Cancer Institute | Francesca De Nicola, Giulia Orlandi, Eleonora Sperandio, Alessia Lauretti, Fabrizio Ensoli, Fulvia Pimpinelli, Maurizio Fanciulli, Giovanni Blandino, Aldo Morrone, Gennaro Ciliberto |
| EPI_ISL_2985313 | IRCCS San Gallicano Dermatological Institute | IRCCS Regina Elena National Cancer Institute | Ludovica Ciuffreda, Sara Donzelli, Alice Massacci, Francesca Maione, Fabrizio Ensoli, Fulvia Pimpinelli, Maurizio Fanciulli, Giovanni Blandino, Aldo Morrone, Gennaro Ciliberto |
| EPI_ISL_2985314 | IRCCS San Gallicano Dermatological Institute | IRCCS Regina Elena National Cancer Institute | Frauke Goeman, Giulia Orlandi, Matteo Pallocca, Francisco Obregon, Fabrizio Ensoli, Fulvia Pimpinelli, Maurizio Fanciulli, Giovanni Blandino, Aldo Morrone, Gennaro Ciliberto |
| EPI_ISL_2985315 | IRCCS San Gallicano Dermatological Institute | IRCCS Regina Elena National Cancer Institute | Sabrina Strano, Sara Donzelli, Eleonora Sperandio, Silvia Paluzzi, Fabrizio Ensoli, Fulvia Pimpinelli, Maurizio Fanciulli, Giovanni Blandino, Aldo Morrone, Gennaro Ciliberto |
| EPI_ISL_2985316 | IRCCS San Gallicano Dermatological Institute | IRCCS Regina Elena National Cancer Institute | Francesca De Nicola, Giulia Orlandi, Alice Massacci, Sara Petrolo, Fabrizio Ensoli, Fulvia Pimpinelli, Maurizio Fanciulli, Giovanni Blandino, Aldo Morrone, Gennaro Ciliberto |
| EPI_ISL_2985317 | IRCCS San Gallicano Dermatological Institute | IRCCS Regina Elena National Cancer Institute | Ludovica Ciuffreda, Sara Donzelli, Matteo Pallocca, Valentina Ricca, Fabrizio Ensoli, Fulvia Pimpinelli, Maurizio Fanciulli, Giovanni Blandino, Aldo Morrone, Gennaro Ciliberto |

| | | | |
|---|---|---|---|
| EPI_ISL_2985318 | IRCCS San Gallicano Dermatological Institute | IRCCS Regina Elena National Cancer Institute | Frauke Goeman, Giulia Orlandi, Eleonora Sperandio, Serena Salvo, Fabrizio Ensoli, Fulvia Pimpinelli, Maurizio Fanciulli, Giovanni Blandino, Aldo Morrone, Gennaro Ciliberto |
| EPI_ISL_2985319 | IRCCS San Gallicano Dermatological Institute | IRCCS Regina Elena National Cancer Institute | Sabrina Strano, Sara Donzelli, Alice Massacci, Grazia Prignano, Fabrizio Ensoli, Fulvia Pimpinelli, Maurizio Fanciulli, Giovanni Blandino, Aldo Morrone, Gennaro Ciliberto |
| EPI_ISL_2985323 | IRCCS San Gallicano Dermatological Institute | IRCCS Regina Elena National Cancer Institute | Sabrina Strano, Sara Donzelli, Matteo Pallocca, Carla Mottini, Fabrizio Ensoli, Fulvia Pimpinelli, Maurizio Fanciulli, Giovanni Blandino, Aldo Morrone, Gennaro Ciliberto |
| EPI_ISL_2985324 | IRCCS San Gallicano Dermatological Institute | IRCCS Regina Elena National Cancer Institute | Francesca De Nicola, Giulia Orlandi, Eleonora Sperandio, Martina Pontone, Fabrizio Ensoli, Fulvia Pimpinelli, Maurizio Fanciulli, Giovanni Blandino, Aldo Morrone, Gennaro Ciliberto |
| EPI_ISL_2985325 | IRCCS San Gallicano Dermatological Institute | IRCCS Regina Elena National Cancer Institute | Frauke Goeman, Giulia Orlandi, Matteo Pallocca, Ilaria Cavallo, Fabrizio Ensoli, Fulvia Pimpinelli, Maurizio Fanciulli, Giovanni Blandino, Aldo Morrone, Gennaro Ciliberto |
| EPI_ISL_2985326 | IRCCS San Gallicano Dermatological Institute | IRCCS Regina Elena National Cancer Institute | Sabrina Strano, Sara Donzelli, Eleonora Sperandio, Francesca Sivori, Fabrizio Ensoli, Fulvia Pimpinelli, Maurizio Fanciulli, Giovanni Blandino, Aldo Morrone, Gennaro Ciliberto |
| EPI_ISL_2985327 | IRCCS San Gallicano Dermatological Institute | IRCCS Regina Elena National Cancer Institute | Francesca De Nicola, Giulia Orlandi, Alice Massacci, Andrea Cazzani, Fabrizio Ensoli, Fulvia Pimpinelli, Maurizio Fanciulli, Giovanni Blandino, Aldo Morrone, Gennaro Ciliberto |
| EPI_ISL_2985328 | IRCCS San Gallicano Dermatological Institute | IRCCS Regina Elena National Cancer Institute | Ludovica Ciuffreda, Sara Donzelli, Matteo Pallocca, Ilaria Celesti, Fabrizio Ensoli, Fulvia Pimpinelli, Maurizio Fanciulli, Giovanni Blandino, Aldo Morrone, Gennaro Ciliberto |
| EPI_ISL_2985329 | IRCCS San Gallicano Dermatological Institute | IRCCS Regina Elena National Cancer Institute | Frauke Goeman, Giulia Orlandi, Eleonora Sperandio, Martina Diano, Fabrizio Ensoli, Fulvia Pimpinelli, Maurizio Fanciulli, Giovanni Blandino, Aldo Morrone, Gennaro Ciliberto |
| EPI_ISL_2985330 | IRCCS San Gallicano Dermatological Institute | IRCCS Regina Elena National Cancer Institute | Sabrina Strano, Sara Donzelli, Alice Massacci, Antonio Federico, Fabrizio Ensoli, Fulvia Pimpinelli, Maurizio Fanciulli, Giovanni Blandino, Aldo Morrone, Gennaro Ciliberto |
| EPI_ISL_2985331 | IRCCS San Gallicano Dermatological Institute | IRCCS Regina Elena National Cancer Institute | Francesca De Nicola, Giulia Orlandi, Matteo Pallocca, Fulvia Fraticelli, Fabrizio Ensoli, Fulvia Pimpinelli, Maurizio Fanciulli, Giovanni Blandino, Aldo Morrone, Gennaro Ciliberto |
| EPI_ISL_2985332 | IRCCS San Gallicano Dermatological Institute | IRCCS Regina Elena National Cancer Institute | Ludovica Ciuffreda, Sara Donzelli, Eleonora Sperandio, Lorenzo Furzi, Fabrizio Ensoli, Fulvia Pimpinelli, Maurizio Fanciulli, Giovanni Blandino, Aldo Morrone, Gennaro Ciliberto |
| EPI_ISL_2985333 | IRCCS San Gallicano Dermatological Institute | IRCCS Regina Elena National Cancer Institute | Frauke Goeman, Giulia Orlandi, Alice Massacci, Alessia Lauretti, Fabrizio Ensoli, Fulvia Pimpinelli, Maurizio Fanciulli, Giovanni Blandino, Aldo Morrone, Gennaro Ciliberto |
| EPI_ISL_2985334 | IRCCS San Gallicano Dermatological Institute | IRCCS Regina Elena National Cancer Institute | Sabrina Strano, Sara Donzelli, Matteo Pallocca, Francesca Maione, Fabrizio Ensoli, Fulvia Pimpinelli, Maurizio Fanciulli, Giovanni Blandino, Aldo Morrone, Gennaro Ciliberto |
| EPI_ISL_2985335 | IRCCS San Gallicano Dermatological Institute | IRCCS Regina Elena National Cancer Institute | Francesca De Nicola, Giulia Orlandi, Eleonora Sperandio, Francisco Obregon, Fabrizio Ensoli, Fulvia Pimpinelli, Maurizio Fanciulli, Giovanni Blandino, Aldo Morrone, Gennaro Ciliberto |
| EPI_ISL_2985336 | IRCCS San Gallicano Dermatological Institute | IRCCS Regina Elena National Cancer Institute | Ludovica Ciuffreda, Sara Donzelli, Alice Massacci, Silvia Paluzzi, Fabrizio Ensoli, Fulvia Pimpinelli, Maurizio Fanciulli, Giovanni Blandino, Aldo Morrone, Gennaro Ciliberto |
| EPI_ISL_2985338 | IRCCS San Gallicano Dermatological Institute | IRCCS Regina Elena National Cancer Institute | Sabrina Strano, Sara Donzelli, Eleonora Sperandio, Valentina Ricca, Fabrizio Ensoli, Fulvia Pimpinelli, Maurizio Fanciulli, Giovanni Blandino, Aldo Morrone, Gennaro Ciliberto |
| EPI_ISL_2985339 | IRCCS San Gallicano Dermatological Institute | IRCCS Regina Elena National Cancer Institute | Francesca De Nicola, Giulia Orlandi, Alice Massacci, Serena Salvo, Fabrizio Ensoli, Fulvia Pimpinelli, Maurizio Fanciulli, Giovanni Blandino, Aldo Morrone, Gennaro Ciliberto |
| EPI_ISL_2985340 | IRCCS San Gallicano Dermatological Institute | IRCCS Regina Elena National Cancer Institute | Ludovica Ciuffreda, Sara Donzelli, Matteo Pallocca, Grazia Prignano, Fabrizio Ensoli, Fulvia Pimpinelli, Maurizio Fanciulli, Giovanni Blandino, Aldo Morrone, Gennaro Ciliberto |
| EPI_ISL_2985341 | IRCCS San Gallicano Dermatological Institute | IRCCS Regina Elena National Cancer Institute | Frauke Goeman, Giulia Orlandi, Eleonora Sperandio, Elisabetta Trento, Fabrizio Ensoli, Fulvia Pimpinelli, Maurizio Fanciulli, Giovanni Blandino, Aldo Morrone, Gennaro Ciliberto |
| EPI_ISL_2985342 | IRCCS San Gallicano Dermatological Institute | IRCCS Regina Elena National Cancer Institute | Sabrina Strano, Sara Donzelli, Alice Massacci, Arianna Mastrofrancesco, Fabrizio Ensoli, Fulvia Pimpinelli, Maurizio Fanciulli, Giovanni Blandino, Aldo Morrone, Gennaro Ciliberto |
| EPI_ISL_2985345 | IRCCS San Gallicano Dermatological Institute | IRCCS Regina Elena National Cancer Institute | Frauke Goeman, Giulia Orlandi, Alice Massacci, Martina Pontone, Fabrizio Ensoli, Fulvia Pimpinelli, Maurizio Fanciulli, Giovanni Blandino, Aldo Morrone, Gennaro Ciliberto |
| EPI_ISL_2985347 | IRCCS San Gallicano Dermatological Institute | IRCCS Regina Elena National Cancer Institute | Ludovica Ciuffreda, Sara Donzelli, Alice Massacci, Francesca Sivori, Fabrizio Ensoli, Fulvia Pimpinelli, Maurizio Fanciulli, Giovanni Blandino, Aldo Morrone, Gennaro Ciliberto |
| EPI_ISL_2985348 | Department of Virology, Istituto Zooprofilattico Sperimentale del Lazio e della Toscana (IZSLT) | Department of General Diagnostics; Department of Virology; Istituto Zooprofilattico Sperimentale del Lazio e della Toscana (IZSLT) | Patricia Alba, Giuseppe Manna, Elena L. Diaconu, Fabiola Feltrin, Raffaela Conti, Teresa Scicluna, Virginia Carfora, Antonella Cersini, Alessia Franco, Antonio Battisti. |
| EPI_ISL_2985351, EPI_ISL_2985353, EPI_ISL_2985354, EPI_ISL_2985355, EPI_ISL_2985356, EPI_ISL_2985357, EPI_ISL_2985358, EPI_ISL_2985359, EPI_ISL_2985360, EPI_ISL_2985362, EPI_ISL_2985363, EPI_ISL_2985364, EPI_ISL_2985365, EPI_ISL_2985366, EPI_ISL_2985367, EPI_ISL_2985368, EPI_ISL_2985369, EPI_ISL_2985370, EPI_ISL_2985371, EPI_ISL_2985372, EPI_ISL_2985373, EPI_ISL_2985374, EPI_ISL_2985375, EPI_ISL_2985376, EPI_ISL_2985377, EPI_ISL_2985378 | | | |
| see above | Ospedale San Filippo Neri | Department of General Diagnostics; Department of Virology; Istituto Zooprofilattico Sperimentale del Lazio e della Toscana (IZSLT) | Patricia Alba, Giuseppe Manna, Elena L. Diaconu, Fabiola Feltrin, Raffaela Conti, Teresa Scicluna, Virginia Carfora, Antonella Cersini, Alessia Franco, Antonio Battisti. |
| EPI_ISL_2985379 | Policlinico Universitario Campus Bio-Medico | Department of General Diagnostics; Department of Virology; Istituto Zooprofilattico Sperimentale del Lazio e della Toscana (IZSLT) | Patricia Alba, Giuseppe Manna, Elena L. Diaconu, Fabiola Feltrin, Raffaela Conti, Teresa Scicluna, Virginia Carfora, Antonella Cersini, Alessia Franco, Antonio Battisti. |
| EPI_ISL_2985380, EPI_ISL_2985381, EPI_ISL_2985382, EPI_ISL_2985383, EPI_ISL_2985386, EPI_ISL_2985387, EPI_ISL_2985388, EPI_ISL_2985389, EPI_ISL_2985390, EPI_ISL_2985391, EPI_ISL_2985392, EPI_ISL_2985393, EPI_ISL_2985394, EPI_ISL_2985395 | | | |
| see above | Ospedale Sandro Pertini ASL Roma2 | Department of General Diagnostics; Department of Virology; Istituto Zooprofilattico Sperimentale del Lazio e della Toscana (IZSLT) | Patricia Alba, Giuseppe Manna, Elena L. Diaconu, Fabiola Feltrin, Raffaela Conti, Teresa Scicluna, Virginia Carfora, Antonella Cersini, Alessia Franco, Antonio Battisti. |
| EPI_ISL_2985399, EPI_ISL_2985400, EPI_ISL_2985401, EPI_ISL_2985402, EPI_ISL_2985403, EPI_ISL_2985404, EPI_ISL_2985405, EPI_ISL_2985406, EPI_ISL_2985408, EPI_ISL_2985409, EPI_ISL_2985410, EPI_ISL_2985411, EPI_ISL_2985413, EPI_ISL_2985414, EPI_ISL_2985415, EPI_ISL_2985416, EPI_ISL_2985418, EPI_ISL_2985419, EPI_ISL_2985420, EPI_ISL_2985421, EPI_ISL_2985423, EPI_ISL_2985424, EPI_ISL_2985425, EPI_ISL_2985426, EPI_ISL_2985428, EPI_ISL_2985429, EPI_ISL_2985430, EPI_ISL_2985431, EPI_ISL_2985432, EPI_ISL_2985433, EPI_ISL_2985434, EPI_ISL_2985435, EPI_ISL_2985436, EPI_ISL_2985437, EPI_ISL_2985438, EPI_ISL_2985439, EPI_ISL_2985440, EPI_ISL_2985441, EPI_ISL_2985442, EPI_ISL_2985443, EPI_ISL_2985444, EPI_ISL_2985445, EPI_ISL_2985446, EPI_ISL_2985447, EPI_ISL_2985448, EPI_ISL_2985449, EPI_ISL_2985450, EPI_ISL_2985451, EPI_ISL_2985452, EPI_ISL_2985453, EPI_ISL_2985454, EPI_ISL_2985455, EPI_ISL_2985456, EPI_ISL_2985457, EPI_ISL_2985458, EPI_ISL_2985459, EPI_ISL_2985460, EPI_ISL_2985461, EPI_ISL_2985462, EPI_ISL_2985463, EPI_ISL_2985464, EPI_ISL_2985465, EPI_ISL_2985466, EPI_ISL_2985467, EPI_ISL_2985469, EPI_ISL_2985470, EPI_ISL_2985471, EPI_ISL_2985472, EPI_ISL_2985474, EPI_ISL_2985475, EPI_ISL_2985476, EPI_ISL_2985477, EPI_ISL_2985478, EPI_ISL_2985479, EPI_ISL_2985480, EPI_ISL_2985481, EPI_ISL_2985482, EPI_ISL_2985483, EPI_ISL_2985484, EPI_ISL_2985485, EPI_ISL_2985486, EPI_ISL_2985487, EPI_ISL_2985488, EPI_ISL_2985489, EPI_ISL_2985490, EPI_ISL_2985491, EPI_ISL_2985492, EPI_ISL_2985493, EPI_ISL_2985494, EPI_ISL_2985495, EPI_ISL_2985497, EPI_ISL_2985498, EPI_ISL_2985499, EPI_ISL_2985500, EPI_ISL_2985501, EPI_ISL_2985502, EPI_ISL_2985503, EPI_ISL_2985504, EPI_ISL_2985505, EPI_ISL_2985506, EPI_ISL_2985507, EPI_ISL_2985508, EPI_ISL_2985509, EPI_ISL_2985510, EPI_ISL_2985511, EPI_ISL_2985512, EPI_ISL_2985513, EPI_ISL_2985514, EPI_ISL_2985515, EPI_ISL_2985517, EPI_ISL_2985518, EPI_ISL_2985519, EPI_ISL_2985520, EPI_ISL_2985521, EPI_ISL_2985522, EPI_ISL_2985523, EPI_ISL_2985524, EPI_ISL_2985525, EPI_ISL_2985526, EPI_ISL_2985527, EPI_ISL_2985528, EPI_ISL_2985529, EPI_ISL_2985530, EPI_ISL_2985531, EPI_ISL_2985533, EPI_ISL_2985535, EPI_ISL_2985536, EPI_ISL_2985538, EPI_ISL_2985545, EPI_ISL_2985546, EPI_ISL_2985547, EPI_ISL_2985551, EPI_ISL_2985556, EPI_ISL_2985557, EPI_ISL_2985558, EPI_ISL_2985566, EPI_ISL_2985567, EPI_ISL_2985568, EPI_ISL_2985569, EPI_ISL_2985577, EPI_ISL_2985578, EPI_ISL_2985579, EPI_ISL_2985580, EPI_ISL_2985587, EPI_ISL_2985588, EPI_ISL_2985589, EPI_ISL_2985592, EPI_ISL_2985596, EPI_ISL_2985598, EPI_ISL_2985599, EPI_ISL_2985600 | | | |
| see above | Austrian Agency for Health and Food Safety (AGES) | Bergthaler laboratory, CeMM Research Center for Molecular Medicine of the Austrian Academy of Sciences | Lukas Endler, Anna Schedl, Fabian Amman, Petr Triska, Matthew Thornton, Thomas Penz, Benedikt Agerer, Maelle Le Moing, Michael Schuster, Bekir Erguner, Jan Laine, Martin Senekowitsch, Christoph Bock, Andreas Bergthaler |
| EPI_ISL_2985605 | Universität Innsbruck, Institut für Mikrobiologie | Bergthaler laboratory, CeMM Research Center for Molecular Medicine of the Austrian Academy of Sciences | Lukas Endler, Anna Schedl, Fabian Amman, Petr Triska, Matthew Thornton, Thomas Penz, Benedikt Agerer, Maelle Le Moing, Michael Schuster, Bekir Erguner, Jan Laine, Martin Senekowitsch, Christoph Bock, Andreas Bergthaler |
| EPI_ISL_2985606 | Johannes Kepler Universtität Linz | Bergthaler laboratory, CeMM Research Center for Molecular Medicine of the Austrian Academy of Sciences | Lukas Endler, Anna Schedl, Fabian Amman, Petr Triska, Matthew Thornton, Thomas Penz, Benedikt Agerer, Maelle Le Moing, Michael Schuster, Bekir Erguner, Jan Laine, Martin Senekowitsch, Christoph Bock, Andreas Bergthaler |
| EPI_ISL_2985628, EPI_ISL_2985629, EPI_ISL_2985639, EPI_ISL_2985645, EPI_ISL_2985651, EPI_ISL_2985652, EPI_ISL_2985653, EPI_ISL_2985654, EPI_ISL_2985659, EPI_ISL_2985667, EPI_ISL_2985670, EPI_ISL_2985671, EPI_ISL_2985673, EPI_ISL_2985676, EPI_ISL_2985679, EPI_ISL_2985680, EPI_ISL_2985681, EPI_ISL_2985685, EPI_ISL_2985687, EPI_ISL_2985688, EPI_ISL_2985698, EPI_ISL_2985701, EPI_ISL_2985702, EPI_ISL_2985703, EPI_ISL_2985704, EPI_ISL_2985705, EPI_ISL_2985707, EPI_ISL_2985708, EPI_ISL_2985712, EPI_ISL_2985713, EPI_ISL_2985715, EPI_ISL_2985719, EPI_ISL_2985722, EPI_ISL_2985725, EPI_ISL_2985726, EPI_ISL_2985729, EPI_ISL_2985730, EPI_ISL_2985732, EPI_ISL_2985735, EPI_ISL_2985742, EPI_ISL_2985744, EPI_ISL_2985746, EPI_ISL_2985747, EPI_ISL_2985748, EPI_ISL_2985749, EPI_ISL_2985750, EPI_ISL_2985752, EPI_ISL_2985753, EPI_ISL_2985754, EPI_ISL_2985755, EPI_ISL_2985762, EPI_ISL_2985763, EPI_ISL_2985767, EPI_ISL_2985774, EPI_ISL_2985775, EPI_ISL_2985778, EPI_ISL_2985780, EPI_ISL_2985781, EPI_ISL_2985783, EPI_ISL_2985784, EPI_ISL_2985785, EPI_ISL_2985787, EPI_ISL_2985788, EPI_ISL_2985789, EPI_ISL_2985790, EPI_ISL_2985791, EPI_ISL_2985792, EPI_ISL_2985794 | | | |
| see above | Gravity Diagnostics, LLC | Gravity Diagnostics, LLC | Gravity Diagnostics |
| EPI_ISL_2985873, EPI_ISL_2985874, EPI_ISL_2985875, EPI_ISL_2985876, EPI_ISL_2985877, EPI_ISL_2985878, EPI_ISL_2985880, EPI_ISL_2985883, EPI_ISL_2985884, EPI_ISL_2985887, EPI_ISL_2985888, EPI_ISL_2985890, EPI_ISL_2985891, EPI_ISL_2985892, EPI_ISL_2985893, EPI_ISL_2985894, EPI_ISL_2985895, EPI_ISL_2985897, EPI_ISL_2985898, EPI_ISL_2985899 | | | |
| see above | ZOL | Jessa | Severine Berden et al. on behalf of the Jessa_cmdLab |
| EPI_ISL_2985901, EPI_ISL_2985902, EPI_ISL_2985903, EPI_ISL_2985904, EPI_ISL_2985905, EPI_ISL_2985906, EPI_ISL_2985907, EPI_ISL_2985908, EPI_ISL_2985909, EPI_ISL_2985910, EPI_ISL_2985911, EPI_ISL_2985912, EPI_ISL_2985913, EPI_ISL_2985914, EPI_ISL_2985915, EPI_ISL_2985916, EPI_ISL_2985917, EPI_ISL_2985918, EPI_ISL_2985919, EPI_ISL_2985920, EPI_ISL_2985921, EPI_ISL_2985922 | | | |
| see above | Jessa | Jessa | Severine Berden et al. on behalf of the Jessa_cmdLab |
| EPI_ISL_2985961, EPI_ISL_2985986, EPI_ISL_2986056, EPI_ISL_2986061, EPI_ISL_2986064, EPI_ISL_2986072, EPI_ISL_2986073, EPI_ISL_2986074, EPI_ISL_2986082, EPI_ISL_2986086, EPI_ISL_2986091, EPI_ISL_2986237, EPI_ISL_2986238, EPI_ISL_2986251, EPI_ISL_2986287, EPI_ISL_2986360, EPI_ISL_2986416, EPI_ISL_2986424, EPI_ISL_2986425, EPI_ISL_2986427, EPI_ISL_2986430, EPI_ISL_2986431, EPI_ISL_2986435, EPI_ISL_2986438, EPI_ISL_2986477, EPI_ISL_2986483, EPI_ISL_2986503, EPI_ISL_2986555, EPI_ISL_2986582, EPI_ISL_2986584, EPI_ISL_2986585, EPI_ISL_2986588, EPI_ISL_2986589, EPI_ISL_2986590, EPI_ISL_2986591, EPI_ISL_2986592, EPI_ISL_2986593, EPI_ISL_2986595, EPI_ISL_2986596, EPI_ISL_2986597, EPI_ISL_2986598, EPI_ISL_2986599, EPI_ISL_2986600, EPI_ISL_2986601, EPI_ISL_2986602, EPI_ISL_2986659, EPI_ISL_2986660, EPI_ISL_2986664, EPI_ISL_2986665, EPI_ISL_2986667, EPI_ISL_2986668, EPI_ISL_2986670, EPI_ISL_2986694, EPI_ISL_2986695, EPI_ISL_2986761, EPI_ISL_2986777, EPI_ISL_2986938 | | | |
| see above | SARS-CoV-2 Sequencing Castilla y Leon-Spain Consortium | SARS-CoV-2 Sequencing Castilla y Leon-Spain Consortium | Antonio Orduña-Domingo, Marta Hernandez, David Abad, Marta Dominguez-Gil, Silvia Rojo, Gabriel March Rossello, Sonsoles Garcinuño Pérez, Carmen Aldea-Mansilla, Mª Fe Brezmes-Valdivieso, Gregoria Megías Lobón, María Antonia García Castro, Carmen Gimeno Crespo, Noelia Arenal Andrés, Carlos Fuster Foz, M. Isabel Fernandez-Natal, Jose María Eiros Bouza |
| EPI_ISL_2986954, EPI_ISL_2986974, EPI_ISL_2986980, EPI_ISL_2986981 | SARS-CoV-2 Sequencing Castilla y Leon-Spain Consortium | SARS-CoV-2 Sequencing Castilla y Leon-Spain Consortium | Antonio Orduña-Domingo, Marta Hernandez, David Abad, Marta Dominguez-Gil, Silvia Rojo, Laura Sánchez de Prada, Sonsoles Garcinuño Pérez, Carmen Aldea-Mansilla, Mª Fe Brezmes-Valdivieso, Gregoria Megías Lobón, María Antonia García Castro, Carmen Gimeno Crespo, Noelia Arenal Andrés, Carlos Fuster Foz, M. Isabel Fernandez-Natal, Jose María Eiros Bouza |
| EPI_ISL_2986985, EPI_ISL_2986986 | Ayass Bioscience LLC | Ayass Bioscience LLC | Mohamad Ammar Ayass, Natalya Griko, Kevin Zhu, Nazanin Taheri, Lina Abi Mosleh |
| EPI_ISL_2986987 | Institute of Public Health Kraljevo | Veterinary Specialized Institute Kraljevo | Vidanovic,D., Tesovic,B., Knezevic,A., Jovanovic,T., Jankovic,M., Sekler,M., Djukic V.,. Banovic Djeri,B., Petrovic,T., Volkening,J., Afonso,C. |
| EPI_ISL_2986988 | "CO Dept. of Public Health and Environment, Lab Services Division" | Centers for Disease Control and Prevention Division of Viral Diseases, Pathogen Discovery | Mili Sheth, Sarah Nobles, Jasmine Padilla, Mark Burroughs, Shoshona Le, Katie Dillon, Peter Cook, Clinton R. Paden, Dhwani Batra, Krista Queen, Kristen Knipe, Dakota Howard, Yvette Unoarumhi, Darlene Wagner, Matthew Schmerer, Ben L. Rambo-Martin, Kristine Lacek, Sam Shepard, Alison Laufer Halpin, Dave Wentworth, Vivien Dugan, Suxiang Tong, Justin Lee |
| EPI_ISL_2986989 | WY Public Health Laboratory | Centers for Disease Control and Prevention Division of Viral Diseases, Pathogen Discovery | Mili Sheth, Sarah Nobles, Jasmine Padilla, Mark Burroughs, Shoshona Le, Katie Dillon, Peter Cook, Clinton R. Paden, Dhwani Batra, Krista Queen, Kristen Knipe, Dakota Howard, Yvette Unoarumhi, Darlene Wagner, Matthew Schmerer, Ben L. Rambo-Martin, Kristine Lacek, Sam Shepard, Alison Laufer Halpin, Dave Wentworth, Vivien Dugan, Suxiang Tong, Justin Lee |
| EPI_ISL_2986990, EPI_ISL_2986991 | "CDPH, Viral and Rickettsial Disease Laboratory" | Centers for Disease Control and Prevention Division of Viral Diseases, Pathogen Discovery | Mili Sheth, Sarah Nobles, Jasmine Padilla, Mark Burroughs, Shoshona Le, Katie Dillon, Peter Cook, Clinton R. Paden, Dhwani Batra, Krista Queen, Kristen Knipe, Dakota Howard, Yvette Unoarumhi, Darlene Wagner, Matthew Schmerer, Ben L. Rambo-Martin, Kristine Lacek, Sam Shepard, Alison Laufer Halpin, Dave Wentworth, Vivien Dugan, Suxiang Tong, Justin Lee |
| EPI_ISL_2986993, EPI_ISL_2986995, EPI_ISL_2986996, EPI_ISL_2986997, EPI_ISL_2987000 | NV-Southern Nevada Public Health Laboratory | Centers for Disease Control and Prevention Division of Viral Diseases, Pathogen Discovery | Mili Sheth, Sarah Nobles, Jasmine Padilla, Mark Burroughs, Shoshona Le, Katie Dillon, Peter Cook, Clinton R. Paden, Dhwani Batra, Krista Queen, Kristen Knipe, Dakota Howard, Yvette Unoarumhi, Darlene Wagner, Matthew Schmerer, Ben L. Rambo-Martin, Kristine Lacek, Sam Shepard, Alison Laufer Halpin, Dave Wentworth, Vivien Dugan, Suxiang Tong, Justin Lee |
| EPI_ISL_2987001, EPI_ISL_2987002, EPI_ISL_2987003, EPI_ISL_2987006, EPI_ISL_2987008, EPI_ISL_2987009 | NE Public Health Laboratory | Centers for Disease Control and Prevention Division of Viral Diseases, Pathogen Discovery | Mili Sheth, Sarah Nobles, Jasmine Padilla, Mark Burroughs, Shoshona Le, Katie Dillon, Peter Cook, Clinton R. Paden, Dhwani Batra, Krista Queen, Kristen Knipe, Dakota Howard, Yvette Unoarumhi, Darlene Wagner, Matthew Schmerer, Ben L. Rambo-Martin, Kristine Lacek, Sam Shepard, Alison Laufer Halpin, Dave Wentworth, Vivien Dugan, Suxiang Tong, Justin Lee |
| EPI_ISL_2987011 | OR State PHL-Virology/Immunology Section | Centers for Disease Control and Prevention Division of Viral Diseases, Pathogen Discovery | Mili Sheth, Sarah Nobles, Jasmine Padilla, Mark Burroughs, Shoshona Le, Katie Dillon, Peter Cook, Clinton R. Paden, Dhwani Batra, Krista Queen, Kristen Knipe, Dakota Howard, Yvette Unoarumhi, Darlene Wagner, Matthew Schmerer, Ben L. Rambo-Martin, Kristine Lacek, Sam Shepard, Alison Laufer Halpin, Dave Wentworth, Vivien Dugan, Suxiang Tong, Justin Lee |
| EPI_ISL_2987021, | "CO Dept. of Public Health and Environment, Lab Services | Centers for Disease Control and Prevention Division of Viral | Mili Sheth, Sarah Nobles, Jasmine Padilla, Mark Burroughs, Shoshona Le, Katie Dillon, Peter Cook, Clinton R. Paden, Dhwani Batra, Krista Queen, Kristen Knipe, Dakota Howard, Yvette Unoarumhi, Darlene Wagner, Matthew Schmerer, Ben L. |

| | | | |
|---|---|---|---|
| EPI_ISL_2987022, EPI_ISL_2987023, EPI_ISL_2987026, EPI_ISL_2987034 | Division" | Diseases, Pathogen Discovery | Rambo-Martin, Kristine Lacek, Sam Shepard, Alison Laufer Halpin, Dave Wentworth, Vivien Dugan, Suxiang Tong, Justin Lee |
| EPI_ISL_2987036 | SD Public Health Laboratory | Centers for Disease Control and Prevention Division of Viral Diseases, Pathogen Discovery | Mili Sheth, Sarah Nobles, Jasmine Padilla, Mark Burroughs, Shoshona Le, Katie Dillon, Peter Cook, Clinton R. Paden, Dhwani Batra, Krista Queen, Kristen Knipe, Dakota Howard, Yvette Unoarumhi, Darlene Wagner, Matthew Schmerer, Ben L. Rambo-Martin, Kristine Lacek, Sam Shepard, Alison Laufer Halpin, Dave Wentworth, Vivien Dugan, Suxiang Tong, Justin Lee |
| EPI_ISL_2987042, EPI_ISL_2987055 | ID Bureau of Laboratories | Centers for Disease Control and Prevention Division of Viral Diseases, Pathogen Discovery | Mili Sheth, Sarah Nobles, Jasmine Padilla, Mark Burroughs, Shoshona Le, Katie Dillon, Peter Cook, Clinton R. Paden, Dhwani Batra, Krista Queen, Kristen Knipe, Dakota Howard, Yvette Unoarumhi, Darlene Wagner, Matthew Schmerer, Ben L. Rambo-Martin, Kristine Lacek, Sam Shepard, Alison Laufer Halpin, Dave Wentworth, Vivien Dugan, Suxiang Tong, Justin Lee |
| EPI_ISL_2987071 | ME Health and Environmental Testing Laboratory | Centers for Disease Control and Prevention Division of Viral Diseases, Pathogen Discovery | Mili Sheth, Sarah Nobles, Jasmine Padilla, Mark Burroughs, Shoshona Le, Katie Dillon, Peter Cook, Clinton R. Paden, Dhwani Batra, Krista Queen, Kristen Knipe, Dakota Howard, Yvette Unoarumhi, Darlene Wagner, Matthew Schmerer, Ben L. Rambo-Martin, Kristine Lacek, Sam Shepard, Alison Laufer Halpin, Dave Wentworth, Vivien Dugan, Suxiang Tong, Justin Lee |
| EPI_ISL_2987076, EPI_ISL_2987079, EPI_ISL_2987081, EPI_ISL_2987082 | OH Department of Health Laboratory | Centers for Disease Control and Prevention Division of Viral Diseases, Pathogen Discovery | Mili Sheth, Sarah Nobles, Jasmine Padilla, Mark Burroughs, Shoshona Le, Katie Dillon, Peter Cook, Clinton R. Paden, Dhwani Batra, Krista Queen, Kristen Knipe, Dakota Howard, Yvette Unoarumhi, Darlene Wagner, Matthew Schmerer, Ben L. Rambo-Martin, Kristine Lacek, Sam Shepard, Alison Laufer Halpin, Dave Wentworth, Vivien Dugan, Suxiang Tong, Justin Lee |
| EPI_ISL_2987083, EPI_ISL_2987084, EPI_ISL_2987085, EPI_ISL_2987086, EPI_ISL_2987087, EPI_ISL_2987088, EPI_ISL_2987089 | WY Public Health Laboratory | Centers for Disease Control and Prevention Division of Viral Diseases, Pathogen Discovery | Mili Sheth, Sarah Nobles, Jasmine Padilla, Mark Burroughs, Shoshona Le, Katie Dillon, Peter Cook, Clinton R. Paden, Dhwani Batra, Krista Queen, Kristen Knipe, Dakota Howard, Yvette Unoarumhi, Darlene Wagner, Matthew Schmerer, Ben L. Rambo-Martin, Kristine Lacek, Sam Shepard, Alison Laufer Halpin, Dave Wentworth, Vivien Dugan, Suxiang Tong, Justin Lee |
| EPI_ISL_2987097, EPI_ISL_2987102 | RI State Health Laboratories | Centers for Disease Control and Prevention Division of Viral Diseases, Pathogen Discovery | Mili Sheth, Sarah Nobles, Jasmine Padilla, Mark Burroughs, Shoshona Le, Katie Dillon, Peter Cook, Clinton R. Paden, Dhwani Batra, Krista Queen, Kristen Knipe, Dakota Howard, Yvette Unoarumhi, Darlene Wagner, Matthew Schmerer, Ben L. Rambo-Martin, Kristine Lacek, Sam Shepard, Alison Laufer Halpin, Dave Wentworth, Vivien Dugan, Suxiang Tong, Justin Lee |
| EPI_ISL_2987106, EPI_ISL_2987108, EPI_ISL_2987109, EPI_ISL_2987110 | "CDPH, Viral and Rickettsial Disease Laboratory" | Centers for Disease Control and Prevention Division of Viral Diseases, Pathogen Discovery | Mili Sheth, Sarah Nobles, Jasmine Padilla, Mark Burroughs, Shoshona Le, Katie Dillon, Peter Cook, Clinton R. Paden, Dhwani Batra, Krista Queen, Kristen Knipe, Dakota Howard, Yvette Unoarumhi, Darlene Wagner, Matthew Schmerer, Ben L. Rambo-Martin, Kristine Lacek, Sam Shepard, Alison Laufer Halpin, Dave Wentworth, Vivien Dugan, Suxiang Tong, Justin Lee |
| EPI_ISL_2987117, EPI_ISL_2987118, EPI_ISL_2987119, EPI_ISL_2987120, EPI_ISL_2987121, EPI_ISL_2987122, EPI_ISL_2987123, EPI_ISL_2987124, EPI_ISL_2987125, EPI_ISL_2987126, EPI_ISL_2987127, EPI_ISL_2987129, EPI_ISL_2987130, EPI_ISL_2987131, EPI_ISL_2987132, EPI_ISL_2987133, EPI_ISL_2987134, EPI_ISL_2987135 | | | |
| see above | Southern Nevada Public Health Laboratory | Southern Nevada Public Health Laboratory | Michael Picker |
| EPI_ISL_2987136 | Pathology and Laboratory Medicine Institute, Cleveland Clinic, Ohio, USA | Pathology and Laboratory Medicine Institute, Cleveland Clinic, Ohio, USA | Concetta Peck, Jennifer Starbuck, Joy Nakitandwe, Kristen McDonnell, David Plunkett, Zheng Jin Tu, Jay Brock, Yu-Wei Cheng, Gary Procop, Daniel Rhoads, Daniel H. Farkas, David Bosler |
| EPI_ISL_2987140, EPI_ISL_2987141, EPI_ISL_2987142 | Stanford Health Care | Stanford University School of Medicine, Clinical Virology Laboratory | Mamdouh Sibai, Daniel Solis, Selamawit Bihon, Bernadette Troung, Becky Jiang, Nathan Hammond, James Zehnder, Malaya K. Sahoo, and Benjamin A. Pinsky |
| EPI_ISL_2987143, EPI_ISL_2987144, EPI_ISL_2987145, EPI_ISL_2987146, EPI_ISL_2987147, EPI_ISL_2987148, EPI_ISL_2987149, EPI_ISL_2987150, EPI_ISL_2987167, EPI_ISL_2987168, EPI_ISL_2987169, EPI_ISL_2987170, EPI_ISL_2987171, EPI_ISL_2987172, EPI_ISL_2987173, EPI_ISL_2987174 | | | |
| see above | Alaska State Virology Laboratory | Alaska State Virology Laboratory | Stephanie DeRonde, Elva House, Jacob Zidek, Lisa Smith, Ph.D., Jack Chen, Ph.D. |
| EPI_ISL_2987211, EPI_ISL_2987214, EPI_ISL_2987218 | DNA Labs | National Reference Laboratory, Nigeria Centre for Disease Control | Dr Ndodo Nnaemeka, Olusola Anuoluwapo Akanbi, Chimaobi Chukwu, Dr Omoare Adesuyi, Anthony Ahumibe, Catherine Okoi, Nwando Mba, Dr Chikwe Ihekweazu |
| EPI_ISL_2987219, EPI_ISL_2987220, EPI_ISL_2987221, EPI_ISL_2987222, EPI_ISL_2987223, EPI_ISL_2987224, EPI_ISL_2987225, EPI_ISL_2987226, EPI_ISL_2987227, EPI_ISL_2987228, EPI_ISL_2987229, EPI_ISL_2987230, EPI_ISL_2987231, EPI_ISL_2987232 | | | |
| see above | Center of Advanced Studies and Technology, Molecular Genetics Laboratory | Center of Advanced Studies and Technology, Molecular Genetics Laboratory | Anaclerio Federico, De Fabritiis Simone |
| EPI_ISL_2987233 | TSHEPONG LABORATORY | National Institute for Communicable Diseases of the National Health Laboratory Service | Amoako DG, Everatt J, Scheepers C, Mohale T, Ntuli N, Mahlangu B, Mnguni A, Ismail A, Bhiman JN |
| EPI_ISL_2987234 | HELEN JOSEPH LABORATORY | National Institute for Communicable Diseases of the National Health Laboratory Service | Amoako DG, Everatt J, Scheepers C, Mohale T, Ntuli N, Mahlangu B, Mnguni A, Ismail A, Bhiman JN |
| EPI_ISL_2987235 | ERMELO LABORATORY | National Institute for Communicable Diseases of the National Health Laboratory Service | Amoako DG, Everatt J, Scheepers C, Mohale T, Ntuli N, Mahlangu B, Mnguni A, Ismail A, Bhiman JN |
| EPI_ISL_2987239, EPI_ISL_2987240, EPI_ISL_2987241, EPI_ISL_2987242, EPI_ISL_2987244 | TSHEPONG LABORATORY | National Institute for Communicable Diseases of the National Health Laboratory Service | Amoako DG, Everatt J, Scheepers C, Mohale T, Ntuli N, Mahlangu B, Mnguni A, Ismail A, Bhiman JN |
| EPI_ISL_2987246 | ERMELO LABORATORY | National Institute for Communicable Diseases of the National Health Laboratory Service | Amoako DG, Everatt J, Scheepers C, Mohale T, Ntuli N, Mahlangu B, Mnguni A, Ismail A, Bhiman JN |
| EPI_ISL_2987249 | TSHEPONG LABORATORY | National Institute for Communicable Diseases of the National Health Laboratory Service | Amoako DG, Everatt J, Scheepers C, Mohale T, Ntuli N, Mahlangu B, Mnguni A, Ismail A, Bhiman JN |
| EPI_ISL_2987258, EPI_ISL_2987265 | ERMELO LABORATORY | National Institute for Communicable Diseases of the National Health Laboratory Service | Amoako DG, Everatt J, Scheepers C, Mohale T, Ntuli N, Mahlangu B, Mnguni A, Ismail A, Bhiman JN |
| EPI_ISL_2987274, EPI_ISL_2987275, EPI_ISL_2987279 | TSHEPONG LABORATORY | National Institute for Communicable Diseases of the National Health Laboratory Service | Amoako DG, Everatt J, Scheepers C, Mohale T, Ntuli N, Mahlangu B, Mnguni A, Ismail A, Bhiman JN |
| EPI_ISL_2987294 | ERMELO LABORATORY | National Institute for Communicable Diseases of the National Health Laboratory Service | Amoako DG, Everatt J, Scheepers C, Mohale T, Ntuli N, Mahlangu B, Mnguni A, Ismail A, Bhiman JN |
| EPI_ISL_2987309, EPI_ISL_2987314, EPI_ISL_2987315, EPI_ISL_2987316 | TSHEPONG LABORATORY | National Institute for Communicable Diseases of the National Health Laboratory Service | Amoako DG, Everatt J, Scheepers C, Mohale T, Ntuli N, Mahlangu B, Mnguni A, Ismail A, Bhiman JN |
| EPI_ISL_2987317, EPI_ISL_2987318 | ERMELO LABORATORY | National Institute for Communicable Diseases of the National Health Laboratory Service | Amoako DG, Everatt J, Scheepers C, Mohale T, Ntuli N, Mahlangu B, Mnguni A, Ismail A, Bhiman JN |
| EPI_ISL_2987319, EPI_ISL_2987320, EPI_ISL_2987322, EPI_ISL_2987323, EPI_ISL_2987324, EPI_ISL_2987325, EPI_ISL_2987326, EPI_ISL_2987327, EPI_ISL_2987328, EPI_ISL_2987329, EPI_ISL_2987330, EPI_ISL_2987331, EPI_ISL_2987332, EPI_ISL_2987333, EPI_ISL_2987334 | | | |
| see above | TSHEPONG LABORATORY | National Institute for Communicable Diseases of the National Health Laboratory Service | Amoako DG, Everatt J, Scheepers C, Mohale T, Ntuli N, Mahlangu B, Mnguni A, Ismail A, Bhiman JN |
| EPI_ISL_2987335 | ERMELO LABORATORY | National Institute for Communicable Diseases of the National Health Laboratory Service | Amoako DG, Everatt J, Scheepers C, Mohale T, Ntuli N, Mahlangu B, Mnguni A, Ismail A, Bhiman JN |
| EPI_ISL_2987336, EPI_ISL_2987337, EPI_ISL_2987338, EPI_ISL_2987339, EPI_ISL_2987340, EPI_ISL_2987341, EPI_ISL_2987342, EPI_ISL_2987343, EPI_ISL_2987344 | TSHEPONG LABORATORY | National Institute for Communicable Diseases of the National Health Laboratory Service | Amoako DG, Everatt J, Scheepers C, Mohale T, Ntuli N, Mahlangu B, Mnguni A, Ismail A, Bhiman JN |
| EPI_ISL_2987398, EPI_ISL_2987399, EPI_ISL_2987400, EPI_ISL_2987401, EPI_ISL_2987402, EPI_ISL_2987403, EPI_ISL_2987404, EPI_ISL_2987405, EPI_ISL_2987406, EPI_ISL_2987407, EPI_ISL_2987408 | | | |
| see above | Colorado Department of Public Health and Environment | Colorado Department of Public Health and Environment | Laura Bankers, Molly C. Hetherington-Rauth, Diana Ir, Alexandria Rossheim, Michael Martin, Mandy Waters, Shannon R. Matzinger, Sarah Elizabeth Totten, Emily A. Travanty |
| EPI_ISL_2987420, EPI_ISL_2987421, EPI_ISL_2987422, EPI_ISL_2987423, EPI_ISL_2987424, EPI_ISL_2987425, EPI_ISL_2987426, EPI_ISL_2987427, EPI_ISL_2987428, EPI_ISL_2987430, EPI_ISL_2987431, EPI_ISL_2987433, EPI_ISL_2987435, EPI_ISL_2987436, EPI_ISL_2987437, EPI_ISL_2987438, EPI_ISL_2987439 | | | |
| see above | Plateforme de testing Namuroise | Plateforme de testing Namuroise | Lesly Nyinkeu Kemamen; Demars Aurore ; Maschietto Céline ; Nobis Chloé ; Otto Gaetan ; Denis Olivier ; Degosserie Jonathan ; Mullier François |
| EPI_ISL_2987440, EPI_ISL_2987441, EPI_ISL_2987442 | Chittagong General Hospital/ Disease Biology & Molecular Epidemiology Research Lab, University of Chittagong | Virology Laboratory, International Centre for Diarrhoeal Disease Research, Bangladesh (ICDDR,B) | Adnan Mannan, H.M. Hamidullah Mehedi, Md. Abdur Rob, Sanjoy Biswas, Nahid Sultana, Md. Minhazul Hoque, Rajdeep Biswas, Fahim Reza, Mojnu Miah, Rashedul Hasan, Mustafizur Rahman, Mohammad Enayet Hossain |
| EPI_ISL_2987443, EPI_ISL_2987444, EPI_ISL_2987445, | Chattogram Maa-O-Shishu Hospital / Disease Biology & Molecular Epidemiology Research Lab, University of Chittagong | Virology Laboratory, International Centre for Diarrhoeal Disease Research, Bangladesh (ICDDR,B) | Adnan Mannan, H.M. Hamidullah Mehedi, Md. Abdur Rob, Sanjoy Biswas, Nahid Sultana, Md. Minhazul Hoque, Rajdeep Biswas, Fahim Reza, Mojnu Miah, Rashedul Hasan, Mustafizur Rahman, Mohammad Enayet Hossain |

| | | | |
|---|---|---|---|
| EPI_ISL_2987446, EPI_ISL_2987447, EPI_ISL_2987448, EPI_ISL_2987449 | | | |
| EPI_ISL_2987450 | Chittagong General Hospital/ Disease Biology & Molecular Epidemiology Research Lab, University of Chittagong | Virology Laboratory, International Centre for Diarrhoeal Disease Research, Bangladesh (ICDDR,B) | Adnan Mannan, H.M. Hamidullah Mehedi, Md. Abdur Rob, Sanjoy Biswas, Nahid Sultana, Md. Minhazul Hoque, Rajdeep Biswas, Fahim Reza, Mojnu Miah, Rashedul Hasan, Mustafizur Rahman, Mohammad Enayet Hossain |
| EPI_ISL_2987451 | Chattogram Maa-O-Shishu Hospital / Disease Biology & Molecular Epidemiology Research Lab, University of Chittagong | Virology Laboratory, International Centre for Diarrhoeal Disease Research, Bangladesh (ICDDR,B) | Adnan Mannan, H.M. Hamidullah Mehedi, Md. Abdur Rob, Sanjoy Biswas, Nahid Sultana, Md. Minhazul Hoque, Rajdeep Biswas, Fahim Reza, Mojnu Miah, Rashedul Hasan, Mustafizur Rahman, Mohammad Enayet Hossain |
| EPI_ISL_2987453, EPI_ISL_2987454, EPI_ISL_2987456, EPI_ISL_2987457 | Chittagong General Hospital/ Disease Biology & Molecular Epidemiology Research Lab, University of Chittagong | Virology Laboratory, International Centre for Diarrhoeal Disease Research, Bangladesh (ICDDR,B) | Adnan Mannan, H.M. Hamidullah Mehedi, Md. Abdur Rob, Sanjoy Biswas, Nahid Sultana, Md. Minhazul Hoque, Rajdeep Biswas, Fahim Reza, Mojnu Miah, Rashedul Hasan, Mustafizur Rahman, Mohammad Enayet Hossain |
| EPI_ISL_2987458, EPI_ISL_2987459, EPI_ISL_2987460, EPI_ISL_2987461, EPI_ISL_2987462, EPI_ISL_2987463 | Chattogram Maa-O-Shishu Hospital / Disease Biology & Molecular Epidemiology Research Lab, University of Chittagong | Virology Laboratory, International Centre for Diarrhoeal Disease Research, Bangladesh (ICDDR,B) | Adnan Mannan, H.M. Hamidullah Mehedi, Md. Abdur Rob, Sanjoy Biswas, Nahid Sultana, Md. Minhazul Hoque, Rajdeep Biswas, Fahim Reza, Mojnu Miah, Rashedul Hasan, Mustafizur Rahman, Mohammad Enayet Hossain |
| EPI_ISL_2987464, EPI_ISL_2987465, EPI_ISL_2987466, EPI_ISL_2987467, EPI_ISL_2987468, EPI_ISL_2987469, EPI_ISL_2987470, EPI_ISL_2987471, EPI_ISL_2987472, EPI_ISL_2987474, EPI_ISL_2987475, EPI_ISL_2987476, EPI_ISL_2987477, EPI_ISL_2987478, EPI_ISL_2987479, EPI_ISL_2987480, EPI_ISL_2987482, EPI_ISL_2987483, EPI_ISL_2987484, EPI_ISL_2987485, EPI_ISL_2987486, EPI_ISL_2987487, EPI_ISL_2987488, EPI_ISL_2987489, EPI_ISL_2987490, EPI_ISL_2987491, EPI_ISL_2987492, EPI_ISL_2987493, EPI_ISL_2987494, EPI_ISL_2987495, EPI_ISL_2987496, EPI_ISL_2987497, EPI_ISL_2987498, EPI_ISL_2987499, EPI_ISL_2987501, EPI_ISL_2987502, EPI_ISL_2987503, EPI_ISL_2987504, EPI_ISL_2987505, EPI_ISL_2987506, EPI_ISL_2987507, EPI_ISL_2987508, EPI_ISL_2987509, EPI_ISL_2987510, EPI_ISL_2987511, EPI_ISL_2987512, EPI_ISL_2987513, EPI_ISL_2987514, EPI_ISL_2987515, EPI_ISL_2987516, EPI_ISL_2987517, EPI_ISL_2987518, EPI_ISL_2987519, EPI_ISL_2987520, EPI_ISL_2987521, EPI_ISL_2987522, EPI_ISL_2987523, EPI_ISL_2987524, EPI_ISL_2987525, EPI_ISL_2987526, EPI_ISL_2987527, EPI_ISL_2987528, EPI_ISL_2987529, EPI_ISL_2987530, EPI_ISL_2987531, EPI_ISL_2987532, EPI_ISL_2987533, EPI_ISL_2987534, EPI_ISL_2987535, EPI_ISL_2987536, EPI_ISL_2987537, EPI_ISL_2987538, EPI_ISL_2987539, EPI_ISL_2987540, EPI_ISL_2987541 | | | |
| see above | Virology Laboratory, Department of Microbiology, King George's Medical University | Virology Laboratory, Department of Microbiology, King George's Medical University | Shantanu Prakash, Hricha Mishra, Suruchi Shukla, Amita Jain |
| EPI_ISL_2987585, EPI_ISL_2987588, EPI_ISL_2987589 | RSUP Sanglah | National Institute of Health Research and Development | Hana Apsari Pawestri, Subangkit, Arie Ardiansyah Nugraha, Kartika Dewi Puspa, Hartanti Dian Ikawati, Krisna Pangesti, Triyani Soekarso, Fajar Nur Sulistiyohadi, Nelly Puspandari, Vivi Setiawaty |
| EPI_ISL_2987591 | RSUD Kabupaten Buleleng | National Institute of Health Research and Development | Hana Apsari Pawestri, Subangkit, Arie Ardiansyah Nugraha, Kartika Dewi Puspa, Hartanti Dian Ikawati, Krisna Pangesti, Triyani Soekarso, Fajar Nur Sulistiyohadi, Nelly Puspandari, Vivi Setiawaty |
| EPI_ISL_2987597 | Labkesda DKI Jakarta | National Institute of Health Research and Development | Arie Ardiansyah Nugraha, Hana Apsari Pawestri, Subangkit, Kartika Dewi Puspa, Hartanti Dian Ikawati, Krisna Pangesti, Triyani Soekarso, Fajar Nur Sulistiyohadi, Nelly Puspandari, Vivi Setiawaty |
| EPI_ISL_2987599 | LAB RSUD Koja Jakut | National Institute of Health Research and Development | Hana Apsari Pawestri, Subangkit, Arie Ardiansyah Nugraha, Kartika Dewi Puspa, Hartanti Dian Ikawati, Krisna Pangesti, Triyani Soekarso, Fajar Nur Sulistiyohadi, Nelly Puspandari, Vivi Setiawaty |
| EPI_ISL_2987600 | RS Mitra Keluarga | National Institute of Health Research and Development | Hana Apsari Pawestri, Subangkit, Arie Ardiansyah Nugraha, Kartika Dewi Puspa, Hartanti Dian Ikawati, Krisna Pangesti, Triyani Soekarso, Fajar Nur Sulistiyohadi, Nelly Puspandari, Vivi Setiawaty |
| EPI_ISL_2987601 | Labkesda DKI Jakarta | National Institute of Health Research and Development | Hana Apsari Pawestri, Subangkit, Arie Ardiansyah Nugraha, Kartika Dewi Puspa, Hartanti Dian Ikawati, Krisna Pangesti, Triyani Soekarso, Fajar Nur Sulistiyohadi, Nelly Puspandari, Vivi Setiawaty |
| EPI_ISL_2987602 | RS Mitra Keluarga | National Institute of Health Research and Development | Hana Apsari Pawestri, Subangkit, Arie Ardiansyah Nugraha, Kartika Dewi Puspa, Hartanti Dian Ikawati, Krisna Pangesti, Triyani Soekarso, Fajar Nur Sulistiyohadi, Nelly Puspandari, Vivi Setiawaty |
| EPI_ISL_2987603 | RSDC Wisma Atlet Kemayoran | National Institute of Health Research and Development | Hana Apsari Pawestri, Subangkit, Arie Ardiansyah Nugraha, Kartika Dewi Puspa, Hartanti Dian Ikawati, Krisna Pangesti, Triyani Soekarso, Fajar Nur Sulistiyohadi, Nelly Puspandari, Vivi Setiawaty |
| EPI_ISL_2987605, EPI_ISL_2987606 | Labkesda DKI Jakarta | National Institute of Health Research and Development | Hana Apsari Pawestri, Subangkit, Arie Ardiansyah Nugraha, Kartika Dewi Puspa, Hartanti Dian Ikawati, Krisna Pangesti, Triyani Soekarso, Fajar Nur Sulistiyohadi, Nelly Puspandari, Vivi Setiawaty |
| EPI_ISL_2987607, EPI_ISL_2987609 | RSDC Wisma Atlet Kemayoran | National Institute of Health Research and Development | Hana Apsari Pawestri, Subangkit, Arie Ardiansyah Nugraha, Kartika Dewi Puspa, Hartanti Dian Ikawati, Krisna Pangesti, Triyani Soekarso, Fajar Nur Sulistiyohadi, Nelly Puspandari, Vivi Setiawaty |
| EPI_ISL_2987611 | RSU Hermina Kemayoran | National Institute of Health Research and Development | Kartika Dewi Puspa, Arie Ardiansyah Nugraha, Hana Apsari Pawestri, Subangkit, Hartanti Dian Ikawati, Krisna Pangesti, Triyani Soekarso, Fajar Nur Sulistiyohadi, Nelly Puspandari, Vivi Setiawaty |
| EPI_ISL_2987612 | PUSKESMAS SINDANG BARANG | National Institute of Health Research and Development | Hana Apsari Pawestri, Subangkit, Arie Ardiansyah Nugraha, Kartika Dewi Puspa, Hartanti Dian Ikawati, Krisna Pangesti, Triyani Soekarso, Fajar Nur Sulistiyohadi, Nelly Puspandari, Vivi Setiawaty |
| EPI_ISL_2987613 | PUSKESMAS BOJONGSARI | National Institute of Health Research and Development | Hana Apsari Pawestri, Subangkit, Arie Ardiansyah Nugraha, Kartika Dewi Puspa, Hartanti Dian Ikawati, Krisna Pangesti, Triyani Soekarso, Fajar Nur Sulistiyohadi, Nelly Puspandari, Vivi Setiawaty |
| EPI_ISL_2987614 | PUSKESMAS TANA SEREAL | National Institute of Health Research and Development | Hana Apsari Pawestri, Subangkit, Arie Ardiansyah Nugraha, Kartika Dewi Puspa, Hartanti Dian Ikawati, Krisna Pangesti, Triyani Soekarso, Fajar Nur Sulistiyohadi, Nelly Puspandari, Vivi Setiawaty |
| EPI_ISL_2987615 | RS Mata Cicendo Bandung | National Institute of Health Research and Development | Hana Apsari Pawestri, Subangkit, Arie Ardiansyah Nugraha, Kartika Dewi Puspa, Hartanti Dian Ikawati, Krisna Pangesti, Triyani Soekarso, Fajar Nur Sulistiyohadi, Nelly Puspandari, Vivi Setiawaty |
| EPI_ISL_2987617 | PUSKESMAS BOJONGSARI | National Institute of Health Research and Development | Kartika Dewi Puspa, Arie Ardiansyah Nugraha, Hana Apsari Pawestri, Subangkit, Hartanti Dian Ikawati, Krisna Pangesti, Triyani Soekarso, Fajar Nur Sulistiyohadi, Nelly Puspandari, Vivi Setiawaty |
| EPI_ISL_2987620 | PUSKESMAS SINDANG BARANG | National Institute of Health Research and Development | Arie Ardiansyah Nugraha, Hana Apsari Pawestri, Subangkit, Kartika Dewi Puspa, Hartanti Dian Ikawati, Krisna Pangesti, Triyani Soekarso, Fajar Nur Sulistiyohadi, Nelly Puspandari, Vivi Setiawaty |
| EPI_ISL_2987621 | PUSKESMAS BOJONGSARI | National Institute of Health Research and Development | Arie Ardiansyah Nugraha, Hana Apsari Pawestri, Subangkit, Kartika Dewi Puspa, Hartanti Dian Ikawati, Krisna Pangesti, Triyani Soekarso, Fajar Nur Sulistiyohadi, Nelly Puspandari, Vivi Setiawaty |
| EPI_ISL_2987622, EPI_ISL_2987623 | PUSKESMAS BOJONGSARI | National Institute of Health Research and Development | Hana Apsari Pawestri, Subangkit, Arie Ardiansyah Nugraha, Kartika Dewi Puspa, Hartanti Dian Ikawati, Krisna Pangesti, Triyani Soekarso, Fajar Nur Sulistiyohadi, Nelly Puspandari, Vivi Setiawaty |
| EPI_ISL_2987624 | PUSKESMAS BOJONGSARI | National Institute of Health Research and Development | Kartika Dewi Puspa, Arie Ardiansyah Nugraha, Hana Apsari Pawestri, Subangkit, Hartanti Dian Ikawati, Krisna Pangesti, Triyani Soekarso, Fajar Nur Sulistiyohadi, Nelly Puspandari, Vivi Setiawaty |
| EPI_ISL_2987625 | RSUD dr Slamet Garut | National Institute of Health Research and Development | Hana Apsari Pawestri, Subangkit, Arie Ardiansyah Nugraha, Kartika Dewi Puspa, Hartanti Dian Ikawati, Krisna Pangesti, Triyani Soekarso, Fajar Nur Sulistiyohadi, Nelly Puspandari, Vivi Setiawaty |
| EPI_ISL_2987626, EPI_ISL_2987627, EPI_ISL_2987628 | RSUD dr Slamet Garut | National Institute of Health Research and Development | Kartika Dewi Puspa, Arie Ardiansyah Nugraha, Hana Apsari Pawestri, Subangkit, Hartanti Dian Ikawati, Krisna Pangesti, Triyani Soekarso, Fajar Nur Sulistiyohadi, Nelly Puspandari, Vivi Setiawaty |
| EPI_ISL_2987631 | RSUD KRMT Wongsonegoro | National Institute of Health Research and Development | Kartika Dewi Puspa, Arie Ardiansyah Nugraha, Hana Apsari Pawestri, Subangkit, Hartanti Dian Ikawati, Krisna Pangesti, Triyani Soekarso, Fajar Nur Sulistiyohadi, Nelly Puspandari, Vivi Setiawaty |
| EPI_ISL_2987634 | RS Premier Surabaya | National Institute of Health Research and Development | Hana Apsari Pawestri, Subangkit, Arie Ardiansyah Nugraha, Kartika Dewi Puspa, Hartanti Dian Ikawati, Krisna Pangesti, Triyani Soekarso, Fajar Nur Sulistiyohadi, Nelly Puspandari, Vivi Setiawaty |
| EPI_ISL_2987654, EPI_ISL_2987655, EPI_ISL_2987656 | Lab Genetik Sumbawa | National Institute of Health Research and Development | Kartika Dewi Puspa, Arie Ardiansyah Nugraha, Hana Apsari Pawestri, Subangkit, Hartanti Dian Ikawati, Krisna Pangesti, Triyani Soekarso, Fajar Nur Sulistiyohadi, Nelly Puspandari, Arief Budi Witarto, Rian Adha Ardinata, Lalu Hamzi Fikri, Nurhandini Eka Dewi, Laily Indrayanti, Vivi Setiawaty |
| EPI_ISL_2987658, EPI_ISL_2987660 | RSUD Dr. W.Z. Johanes kupang | National Institute of Health Research and Development | Hana Apsari Pawestri, Subangkit, Arie Ardiansyah Nugraha, Kartika Dewi Puspa, Hartanti Dian Ikawati, Krisna Pangesti, Triyani Soekarso, Fajar Nur Sulistiyohadi, Nelly Puspandari, Vivi Setiawaty |
| EPI_ISL_2987675, EPI_ISL_2987676, EPI_ISL_2987677, EPI_ISL_2987678, EPI_ISL_2987679, EPI_ISL_2987680, EPI_ISL_2987681, EPI_ISL_2987682, EPI_ISL_2987683, EPI_ISL_2987684, EPI_ISL_2987685 | | | |
| see above | Laboratorio nacional-LANIIA-CIAD | Laboratorio nacional-LANIIA-CIAD | Consorcio Mexicano de Vigilancia Genómica (CoViGen-Mex). Authors (in alphabetical order): Julio Elias Alvarado-Yaah, Carlos F. Arias, Santiago Ávila-Ríos, Eduardo Becerril-Vargas, Víctor Hugo Borja-Aburto, Celia Boukadida, Cristóbal Cháidez-Quiróz, Juan Bautista Chale-Dzul, Ricardo Ciria Merce, Andreu Comas-García, Célida Duque Molina, Julissa Enciso-Ibarra, José Antonio Enciso-Moreno, Gloria Elena Espinosa-Ayala, Fernando Fontove-Herrera, Daniel Fregoso-Rueda, Victor Eduardo García-Arias, Alejandra García-Gasca, Bruno Gómez-Gil, Jean Pierre González, Irvin González-López, Concepción Grajales-Muñiz, Ricardo Grande, Rosa María Gutiérrez Rios, Jesús Hernández, Alejandra Hernández-Terán, Alfredo Herrera-Estrella, Carla Ivón Herrera-Najera, Pavel Isa, Verónica Mata-Haro, Daniel Lira Morales, Susana Lopez, Antonio Loza Román, Brenda Irasema Maldonado-Meza, Bernardo Martínez-Miguel, José Arturo Martínez-Orozco, Célida Martínez-Rodríguez, Margarita Matías-Florentino, Fidencio Mejía-Nepomuceno, María Guadalupe de Jesús Mireles-Rivera, Gloria María Molina-Salinas, Hector Montoya-Fuentes, , Mario Mújica-Sánchez, José Esteban Muñoz-Medina, José de Jesús Nuñez-Contreras, Alicia Ocaña-Mondragón, Luis Alberto Ochoa-Carrera, Hector Esteban Paz-Juárez, Marissa Perez-Garcia, Francisco Pulido, Helen Haydee Fernanda Ramírez-Plascencia, Jorge Salas-Hernández, Angel Gustavo Salas-Lais, Alejandro Sánchez-Flores, Clara Esperanza Santacruz-Tinoco, María Guadalupe Santiago-Mauricio, Selene Zárate, Nelly Sélem-Mojica, Blanca Taboada, Gloria Vazquez, Joel Armando Vázquez-Pérez. |
| EPI_ISL_2987690 | Laboratorios Delia Barraza | Laboratorio nacional-LANIIA-CIAD | Consorcio Mexicano de Vigilancia Genómica (CoViGen-Mex). Authors (in alphabetical order): Julio Elias Alvarado-Yaah, Carlos F. Arias, Santiago Ávila-Ríos, Eduardo Becerril-Vargas, Víctor Hugo Borja-Aburto, Celia Boukadida, Cristóbal Cháidez-Quiróz, Juan Bautista Chale-Dzul, Ricardo Ciria Merce, Andreu Comas-García, Célida Duque Molina, Julissa Enciso-Ibarra, José Antonio Enciso-Moreno, Gloria Elena Espinosa-Ayala, Fernando Fontove-Herrera, Daniel Fregoso-Rueda, Victor Eduardo García-Arias, Alejandra García-Gasca, Bruno Gómez-Gil, Jean Pierre González, Irvin González-López, Concepción Grajales-Muñiz, Ricardo Grande, Rosa María Gutiérrez Rios, Jesús Hernández, Alejandra Hernández-Terán, Alfredo Herrera-Estrella, Carla Ivón Herrera-Najera, Pavel Isa, Verónica Mata-Haro, Daniel Lira Morales, Susana Lopez, Antonio Loza Román, Brenda Irasema Maldonado-Meza, Bernardo Martínez-Miguel, José Arturo Martínez-Orozco, Célida Martínez-Rodríguez, Margarita Matías-Florentino, Fidencio Mejía-Nepomuceno, María Guadalupe de Jesús Mireles-Rivera, Gloria María Molina-Salinas, Hector Montoya-Fuentes, , Mario Mújica-Sánchez, José Esteban Muñoz-Medina, José de Jesús Nuñez-Contreras, Alicia Ocaña-Mondragón, Luis Alberto Ochoa-Carrera, Hector Esteban Paz-Juárez, Marissa Perez-Garcia, Francisco Pulido, Helen Haydee Fernanda Ramírez-Plascencia, Jorge Salas-Hernández, Angel Gustavo Salas-Lais, Alejandro Sánchez-Flores, Clara Esperanza Santacruz-Tinoco, María Guadalupe Santiago-Mauricio, Selene Zárate, Nelly Sélem-Mojica, Blanca Taboada, Gloria Vazquez, Joel Armando Vázquez-Pérez. Laboratorios Delia Barraza: María Delia Barraza Sámano, Claudia Soto Félix, Melissa García Angulo. |
| EPI_ISL_2987695 | Hospital Fátima S.A. de C.V. | Laboratorio nacional-LANIIA-CIAD | Consorcio Mexicano de Vigilancia Genómica (CoViGen-Mex). Authors (in alphabetical order): Julio Elias Alvarado-Yaah, Carlos F. Arias, Santiago Ávila-Ríos, Eduardo Becerril-Vargas, Víctor Hugo Borja-Aburto, Celia Boukadida, Cristóbal Cháidez-Quiróz, Juan Bautista Chale-Dzul, Ricardo Ciria Merce, Andreu Comas-García, Célida Duque Molina, Julissa Enciso-Ibarra, José Antonio Enciso-Moreno, Gloria Elena Espinosa-Ayala, Fernando Fontove-Herrera, Daniel Fregoso-Rueda, Victor Eduardo García-Arias, Alejandra García-Gasca, Bruno Gómez-Gil, Jean Pierre González, Irvin González-López, Concepción Grajales-Muñiz, Ricardo Grande, Rosa María Gutiérrez Rios, Jesús Hernández, Alejandra Hernández-Terán, Alfredo Herrera-Estrella, Carla Ivón Herrera-Najera, Pavel Isa, Verónica Mata-Haro, Daniel Lira Morales, Susana Lopez, Antonio Loza Román, Brenda Irasema Maldonado-Meza, Bernardo Martínez-Miguel, José Arturo Martínez-Orozco, Célida Martínez-Rodríguez, Margarita Matías-Florentino, Fidencio Mejía-Nepomuceno, María Guadalupe de Jesús Mireles-Rivera, Gloria María Molina-Salinas, Hector Montoya-Fuentes, , Mario Mújica-Sánchez, José Esteban Muñoz-Medina, José de Jesús Nuñez-Contreras, Alicia Ocaña-Mondragón, Luis Alberto Ochoa-Carrera, Hector Esteban Paz-Juárez, Marissa Perez-Garcia, Francisco Pulido, Helen Haydee Fernanda Ramírez-Plascencia, Jorge Salas-Hernández, Angel Gustavo Salas-Lais, Alejandro Sánchez-Flores, Clara Esperanza Santacruz-Tinoco, María Guadalupe Santiago-Mauricio, Selene Zárate, Nelly Sélem-Mojica, Blanca Taboada, Gloria Vazquez, Joel Armando Vázquez-Pérez. |
| EPI_ISL_2987699, EPI_ISL_2987704, EPI_ISL_2987705, EPI_ISL_2987706, EPI_ISL_2987708, EPI_ISL_2987709, EPI_ISL_2987710 | Hospital Margarita Maza de Juárez | Microbial Genomics Laboratory | Consorcio Mexicano de Vigilancia Genómica (CoViGen-Mex). Authors (in alphabetical order): Julio Elias Alvarado-Yaah, Carlos F. Arias, Santiago Ávila-Ríos, Eduardo Becerril-Vargas, Víctor Hugo Borja-Aburto, Celia Boukadida, Cristóbal Cháidez-Quiróz, Juan Bautista Chale-Dzul, Ricardo Ciria Merce, Andreu Comas-García, Célida Duque Molina, Julissa Enciso-Ibarra, José Antonio Enciso-Moreno, Gloria Elena Espinosa-Ayala, Fernando Fontove-Herrera, Daniel Fregoso-Rueda, Victor Eduardo García-Arias, Alejandra García-Gasca, Bruno Gómez-Gil, Jean Pierre González, Irvin González-López, Concepción Grajales-Muñiz, Ricardo Grande, Rosa María Gutiérrez Rios, Jesús Hernández, Alejandra Hernández-Terán, Alfredo Herrera-Estrella, Carla Ivón Herrera-Najera, Pavel Isa, Verónica Mata-Haro, Daniel Lira Morales, Susana Lopez, Antonio Loza Román, Brenda Irasema Maldonado-Meza, Bernardo Martínez-Miguel, José Arturo Martínez-Orozco, Célida Martínez-Rodríguez, Margarita Matías-Florentino, Fidencio Mejía-Nepomuceno, María Guadalupe de Jesús Mireles-Rivera, Gloria María Molina-Salinas, Hector Montoya-Fuentes, , Mario Mújica-Sánchez, José Esteban Muñoz-Medina, José de Jesús Nuñez-Contreras, Alicia Ocaña-Mondragón, Luis Alberto Ochoa-Carrera, Hector Esteban Paz-Juárez, Marissa Perez-Garcia, Francisco Pulido, Helen Haydee Fernanda Ramírez-Plascencia, Jorge Salas-Hernández, Angel Gustavo Salas-Lais, Alejandro Sánchez-Flores, Clara Esperanza Santacruz-Tinoco, María Guadalupe Santiago-Mauricio, Selene Zárate, Nelly Sélem-Mojica, Blanca Taboada, Gloria Vazquez, Joel Armando Vázquez-Pérez. |
| EPI_ISL_2987722 | Hospital Sharp | Microbial Genomics Laboratory | Consorcio Mexicano de Vigilancia Genómica (CoViGen-Mex). Authors (in alphabetical order): Julio Elias Alvarado-Yaah, Carlos F. Arias, Santiago Ávila-Ríos, Eduardo Becerril-Vargas, Víctor Hugo Borja-Aburto, Celia Boukadida, Cristóbal Cháidez-Quiróz, Juan Bautista Chale-Dzul, Ricardo Ciria Merce, Andreu Comas-García, Célida Duque Molina, Julissa Enciso-Ibarra, José Antonio Enciso-Moreno, Gloria Elena Espinosa-Ayala, Fernando Fontove-Herrera, Daniel Fregoso-Rueda, Victor Eduardo García-Arias, Alejandra García-Gasca, Bruno Gómez-Gil, Jean Pierre González, Irvin González-López, Concepción Grajales-Muñiz, Ricardo Grande, Rosa María Gutiérrez Rios, Jesús Hernández, Alejandra Hernández-Terán, Alfredo |

| | | | |
|---|---|---|---|
| | | | Herrera-Estrella, Carla Ivón Herrera-Najera, Pavel Isa, Verónica Mata-Haro, Daniel Lira Morales, Susana Lopez, Antonio Loza Román, Brenda Irasema Maldonado-Meza, Bernardo Martínez-Miguel, José Arturo Martínez-Orozco, Célida Martínez-Rodríguez, Margarita Matías-Florentino, Fidencio Mejía-Nepomuceno, María Guadalupe de Jesús Mireles-Rivera, Gloria María Molina-Salinas, Hector Montoya-Fuentes, , Mario Mújica-Sánchez, José Esteban Muñoz-Medina, José de Jesús Nuñez-Contreras, Alicia Ocaña-Mondragón, Luis Alberto Ochoa-Carrera, Hector Esteban Paz-Juárez, Marissa Perez-Garcia, Francisco Pulido, Helen Haydee Fernanda Ramírez-Plascencia, Jorge Salas-Hernández, Angel Gustavo Salas-Lais, Alejandro Sánchez-Flores, Clara Esperanza Santacruz-Tinoco, María Guadalupe Santiago-Mauricio, Selene Zárate, Nelly Sélem-Mojica, Blanca Taboada, Gloria Vazquez, Joel Armando Vázquez-Pérez. |
| EPI_ISL_2987723, EPI_ISL_2987727, EPI_ISL_2987729, EPI_ISL_2987732, EPI_ISL_2987737, EPI_ISL_2987748, EPI_ISL_2987758, EPI_ISL_2987764, EPI_ISL_2987765, EPI_ISL_2987768, EPI_ISL_2987771, EPI_ISL_2987772, EPI_ISL_2987776, EPI_ISL_2987781, EPI_ISL_2987782, EPI_ISL_2987783, EPI_ISL_2987784, EPI_ISL_2987785, EPI_ISL_2987786, EPI_ISL_2987787, EPI_ISL_2987801, EPI_ISL_2987802, EPI_ISL_2987803, EPI_ISL_2987804, EPI_ISL_2987805, EPI_ISL_2987807, EPI_ISL_2987808, EPI_ISL_2987809, EPI_ISL_2987821, EPI_ISL_2987822, EPI_ISL_2987823, EPI_ISL_2987824, EPI_ISL_2987825, EPI_ISL_2987826, EPI_ISL_2987827, EPI_ISL_2987828, EPI_ISL_2987829, EPI_ISL_2987830, EPI_ISL_2987831, EPI_ISL_2987833, EPI_ISL_2987834, EPI_ISL_2987835, EPI_ISL_2987836, EPI_ISL_2987837, EPI_ISL_2987838, EPI_ISL_2987840, EPI_ISL_2987841, EPI_ISL_2987842, EPI_ISL_2987843, EPI_ISL_2987844, EPI_ISL_2987845, EPI_ISL_2987846, EPI_ISL_2987847, EPI_ISL_2987848, EPI_ISL_2987850, EPI_ISL_2987851, EPI_ISL_2987852, EPI_ISL_2987853, EPI_ISL_2987854, EPI_ISL_2987855, EPI_ISL_2987856, EPI_ISL_2987857, EPI_ISL_2987858, EPI_ISL_2987859, EPI_ISL_2987860, EPI_ISL_2987862, EPI_ISL_2987863, EPI_ISL_2987864, EPI_ISL_2987865, EPI_ISL_2987866, EPI_ISL_2987867, EPI_ISL_2987869, EPI_ISL_2987870, EPI_ISL_2987871, EPI_ISL_2987872, EPI_ISL_2987873, EPI_ISL_2987874, EPI_ISL_2987875, EPI_ISL_2987876, EPI_ISL_2987877, EPI_ISL_2987878, EPI_ISL_2987879, EPI_ISL_2987880, EPI_ISL_2987881, EPI_ISL_2987882, EPI_ISL_2987883, EPI_ISL_2987884, EPI_ISL_2987885, EPI_ISL_2987886, EPI_ISL_2987888, EPI_ISL_2987889, EPI_ISL_2987891, EPI_ISL_2987893, EPI_ISL_2987894, EPI_ISL_2987896, EPI_ISL_2987897, EPI_ISL_2987898, EPI_ISL_2987899, EPI_ISL_2987900, EPI_ISL_2987901, EPI_ISL_2987902, EPI_ISL_2987903, EPI_ISL_2987904, EPI_ISL_2987905, EPI_ISL_2987906, EPI_ISL_2987907, EPI_ISL_2987909, EPI_ISL_2987910, EPI_ISL_2987911, EPI_ISL_2987912, EPI_ISL_2987913, EPI_ISL_2987914, EPI_ISL_2987915, EPI_ISL_2987916, EPI_ISL_2987917, EPI_ISL_2987918, EPI_ISL_2987919, EPI_ISL_2987921, EPI_ISL_2987922, EPI_ISL_2987924, EPI_ISL_2987925, EPI_ISL_2987926, EPI_ISL_2987927, EPI_ISL_2987928, EPI_ISL_2987929, EPI_ISL_2987930, EPI_ISL_2987931, EPI_ISL_2987932, EPI_ISL_2987933, EPI_ISL_2987951, EPI_ISL_2988005, EPI_ISL_2988012, EPI_ISL_2988014, EPI_ISL_2988016, EPI_ISL_2988018, EPI_ISL_2988036, EPI_ISL_2988040, EPI_ISL_2988046, EPI_ISL_2988047, EPI_ISL_2988048, EPI_ISL_2988049, EPI_ISL_2988050, EPI_ISL_2988051, EPI_ISL_2988052, EPI_ISL_2988053, EPI_ISL_2988054, EPI_ISL_2988055, EPI_ISL_2988056, EPI_ISL_2988057, EPI_ISL_2988058, EPI_ISL_2988059, EPI_ISL_2988060, EPI_ISL_2988061, EPI_ISL_2988062, EPI_ISL_2988063, EPI_ISL_2988064, EPI_ISL_2988065, EPI_ISL_2988066, EPI_ISL_2988067, EPI_ISL_2988068, EPI_ISL_2988069, EPI_ISL_2988070, EPI_ISL_2988071, EPI_ISL_2988072, EPI_ISL_2988073, EPI_ISL_2988074, EPI_ISL_2988075, EPI_ISL_2988076, EPI_ISL_2988077, EPI_ISL_2988078, EPI_ISL_2988079, EPI_ISL_2988080, EPI_ISL_2988081, EPI_ISL_2988082, EPI_ISL_2988083, EPI_ISL_2988084, EPI_ISL_2988085, EPI_ISL_2988086, EPI_ISL_2988087, EPI_ISL_2988088, EPI_ISL_2988090, EPI_ISL_2988091, EPI_ISL_2988092, EPI_ISL_2988093, EPI_ISL_2988094, EPI_ISL_2988095, EPI_ISL_2988096, EPI_ISL_2988097, EPI_ISL_2988098, EPI_ISL_2988099, EPI_ISL_2988100, EPI_ISL_2988101, EPI_ISL_2988102, EPI_ISL_2988103, EPI_ISL_2988104, EPI_ISL_2988105, EPI_ISL_2988106, EPI_ISL_2988107, EPI_ISL_2988108, EPI_ISL_2988109, EPI_ISL_2988110, EPI_ISL_2988111, EPI_ISL_2988112, EPI_ISL_2988113, EPI_ISL_2988114, EPI_ISL_2988115, EPI_ISL_2988116, EPI_ISL_2988117, EPI_ISL_2988118, EPI_ISL_2988119, EPI_ISL_2988120, EPI_ISL_2988121, EPI_ISL_2988122, EPI_ISL_2988123, EPI_ISL_2988124, EPI_ISL_2988125, EPI_ISL_2988126, EPI_ISL_2988127, EPI_ISL_2988128, EPI_ISL_2988129, EPI_ISL_2988130, EPI_ISL_2988131, EPI_ISL_2988132, EPI_ISL_2988133, EPI_ISL_2988134, EPI_ISL_2988135, EPI_ISL_2988136, EPI_ISL_2988137, EPI_ISL_2988138, EPI_ISL_2988139, EPI_ISL_2988140, EPI_ISL_2988141, EPI_ISL_2988142, EPI_ISL_2988143, EPI_ISL_2988144, EPI_ISL_2988145, EPI_ISL_2988146, EPI_ISL_2988147, EPI_ISL_2988148, EPI_ISL_2988149, EPI_ISL_2988150, EPI_ISL_2988151, EPI_ISL_2988152, EPI_ISL_2988154, EPI_ISL_2988155, EPI_ISL_2988156, EPI_ISL_2988157, EPI_ISL_2988158, EPI_ISL_2988159, EPI_ISL_2988160, EPI_ISL_2988162, EPI_ISL_2988164, EPI_ISL_2988165, EPI_ISL_2988166, EPI_ISL_2988167, EPI_ISL_2988168, EPI_ISL_2988170, EPI_ISL_2988171, EPI_ISL_2988172, EPI_ISL_2988173, EPI_ISL_2988174, EPI_ISL_2988175, EPI_ISL_2988176, EPI_ISL_2988177, EPI_ISL_2988178, EPI_ISL_2988179, EPI_ISL_2988180, EPI_ISL_2988181, EPI_ISL_2988182, EPI_ISL_2988183, EPI_ISL_2988184, EPI_ISL_2988185, EPI_ISL_2988186, EPI_ISL_2988187 | | | |
| see above | Ministry of Health Turkey | Ministry of Health Turkey | Fatma Bayrakdar, Yasemin Cosgun, Suleyman Yalcin, Gulay Korukluoglu |
| EPI_ISL_2988188, EPI_ISL_2988189, EPI_ISL_2988190, EPI_ISL_2988191, EPI_ISL_2988192, EPI_ISL_2988193, EPI_ISL_2988194, EPI_ISL_2988195 | Advance Virology Lab, National Botanical Research Institute | Virology Laboratory, Department of Microbiology, King George's Medical University | Samir Sawant, Shantanu Prakash, Suruchi Shukla, Amita Jain |
| EPI_ISL_2988196, EPI_ISL_2988197, EPI_ISL_2988198, EPI_ISL_2988199 | Department of Microbiology, Dr. Ram Manohar Lohia Institute of Medical Sciences | Virology Laboratory, Department of Microbiology, King George's Medical University | Jyotsana Agarwal, Shantanu Prakash, Suruchi Shukla, Amita Jain |
| EPI_ISL_2988200, EPI_ISL_2988201, EPI_ISL_2988202, EPI_ISL_2988203, EPI_ISL_2988204, EPI_ISL_2988205, EPI_ISL_2988206, EPI_ISL_2988207, EPI_ISL_2988208, EPI_ISL_2988209, EPI_ISL_2988210, EPI_ISL_2988211, EPI_ISL_2988213, EPI_ISL_2988215, EPI_ISL_2988218, EPI_ISL_2988219, EPI_ISL_2988224, EPI_ISL_2988234, EPI_ISL_2988235, EPI_ISL_2988236, EPI_ISL_2988237, EPI_ISL_2988238, EPI_ISL_2988239, EPI_ISL_2988240, EPI_ISL_2988241, EPI_ISL_2988242, EPI_ISL_2988251, EPI_ISL_2988253, EPI_ISL_2988254, EPI_ISL_2988255, EPI_ISL_2988256, EPI_ISL_2988257, EPI_ISL_2988258, EPI_ISL_2988259, EPI_ISL_2988260, EPI_ISL_2988263, EPI_ISL_2988265, EPI_ISL_2988267, EPI_ISL_2988268, EPI_ISL_2988269, EPI_ISL_2988270, EPI_ISL_2988271, EPI_ISL_2988273, EPI_ISL_2988274, EPI_ISL_2988275, EPI_ISL_2988276, EPI_ISL_2988278, EPI_ISL_2988280, EPI_ISL_2988281, EPI_ISL_2988285, EPI_ISL_2988286, EPI_ISL_2988288, EPI_ISL_2988289, EPI_ISL_2988290, EPI_ISL_2988291, EPI_ISL_2988292, EPI_ISL_2988293, EPI_ISL_2988294, EPI_ISL_2988295, EPI_ISL_2988296, EPI_ISL_2988297, EPI_ISL_2988298, EPI_ISL_2988299, EPI_ISL_2988300, EPI_ISL_2988301, EPI_ISL_2988302, EPI_ISL_2988303, EPI_ISL_2988304, EPI_ISL_2988305, EPI_ISL_2988306, EPI_ISL_2988307, EPI_ISL_2988308, EPI_ISL_2988309, EPI_ISL_2988311, EPI_ISL_2988312, EPI_ISL_2988313, EPI_ISL_2988314, EPI_ISL_2988316, EPI_ISL_2988318, EPI_ISL_2988319, EPI_ISL_2988320, EPI_ISL_2988321, EPI_ISL_2988322, EPI_ISL_2988323, EPI_ISL_2988324, EPI_ISL_2988325, EPI_ISL_2988326, EPI_ISL_2988328, EPI_ISL_2988329, EPI_ISL_2988330, EPI_ISL_2988331, EPI_ISL_2988332, EPI_ISL_2988333, EPI_ISL_2988334, EPI_ISL_2988335, EPI_ISL_2988336, EPI_ISL_2988337, EPI_ISL_2988338, EPI_ISL_2988339, EPI_ISL_2988340, EPI_ISL_2988341, EPI_ISL_2988342, EPI_ISL_2988343, EPI_ISL_2988344, EPI_ISL_2988345, EPI_ISL_2988346, EPI_ISL_2988347, EPI_ISL_2988348, EPI_ISL_2988349, EPI_ISL_2988350, EPI_ISL_2988351, EPI_ISL_2988352, EPI_ISL_2988353, EPI_ISL_2988354, EPI_ISL_2988355, EPI_ISL_2988356, EPI_ISL_2988357, EPI_ISL_2988358, EPI_ISL_2988359, EPI_ISL_2988360, EPI_ISL_2988361, EPI_ISL_2988362, EPI_ISL_2988363, EPI_ISL_2988365 | | | |
| see above | Platform BIS UZA/UAntwerpen | Labo Klinische Biologie, UZA | Marie Le Mercier, Jasmine Coppens, Kathleen Holemans, Ines Verbesselt, Basil Britto Xavier, Christine Lammens, Veerle Matheeussen, Herman Goossens |
| EPI_ISL_2988366, EPI_ISL_2988367, EPI_ISL_2988368, EPI_ISL_2988369, EPI_ISL_2988370, EPI_ISL_2988373, EPI_ISL_2988374, EPI_ISL_2988376, EPI_ISL_2988377, EPI_ISL_2988378, EPI_ISL_2988379, EPI_ISL_2988380, EPI_ISL_2988381, EPI_ISL_2988382, EPI_ISL_2988383, EPI_ISL_2988384, EPI_ISL_2988385 | | | |
| see above | Algemeen Medisch Labo | Labo Klinische Biologie, UZA | Marie Le Mercier, Jasmine Coppens, Kathleen Holemans, Ines Verbesselt, Basil Britto Xavier, Christine Lammens, Veerle Matheeussen, Herman Goossens |
| EPI_ISL_2988406 | MANKWENG LABORATORY | National Institute for Communicable Diseases of the National Health Laboratory Service | Amoako DG, Everatt J, Scheepers C, Mohale T, Ntuli N, Mahlangu B, Mnguni A, Ismail A, Bhiman JN |
| EPI_ISL_2988417 | TSHEPONG LABORATORY | National Institute for Communicable Diseases of the National Health Laboratory Service | Amoako DG, Everatt J, Scheepers C, Mohale T, Ntuli N, Mahlangu B, Mnguni A, Ismail A, Bhiman JN |
| EPI_ISL_2988419, EPI_ISL_2988420, EPI_ISL_2988423, EPI_ISL_2988425, EPI_ISL_2988426, EPI_ISL_2988427, EPI_ISL_2988428, EPI_ISL_2988430, EPI_ISL_2988431, EPI_ISL_2988432, EPI_ISL_2988433, EPI_ISL_2988435, EPI_ISL_2988437, EPI_ISL_2988438, EPI_ISL_2988442, EPI_ISL_2988443, EPI_ISL_2988445, EPI_ISL_2988446 | | | |
| see above | Montana Public Health Laboratory | Montana Public Health Laboratory | Joy Ritter, Michelle Mozer, Carrie Biskupiak, Deborah Gibson, Michael Dills |
| EPI_ISL_2988617, EPI_ISL_2988618, EPI_ISL_2988619, EPI_ISL_2988620, EPI_ISL_2988621, EPI_ISL_2988622, EPI_ISL_2988623, EPI_ISL_2988624, EPI_ISL_2988625, EPI_ISL_2988626, EPI_ISL_2988627 | | | |
| see above | Tampa General Hospital Esoteric Lab | Tampa General Hospital Esoteric Research & Development Lab | Amorce Lima, Jon Faughn, Deanna Becker, Elaine Vendrone, Suzane Silbert |
| EPI_ISL_2988629, EPI_ISL_2988630, EPI_ISL_2988631, EPI_ISL_2988632, EPI_ISL_2988633, EPI_ISL_2988634 | Wyoming Public Health Laboratory | Wyoming Public Health Laboratory | Jim Mildenberger, Taylor Fearing, Channing Weber, Ashley Norberg, Chayse Rowley, Marley Goetz, Brian Dominguez, Elliot Thomasson, Sam Britz, Cari Sloma, Robert Petit, and Rob Christensen |
| EPI_ISL_2988649, EPI_ISL_2988650, EPI_ISL_2988651, EPI_ISL_2988652, EPI_ISL_2988653, EPI_ISL_2988654, EPI_ISL_2988655, EPI_ISL_2988656, EPI_ISL_2988657 | Molecular Diagnostics Pathology Department Mater Dei Hospital Malta | Molecular Diagnostics Pathology Department Mater Dei Hospital Malta | G Zahra, R Borg, C Cilia, L Grech, M Briffa |
| EPI_ISL_2988658 | Molecular Diagnostics Pathology Department Mater Dei Hospital Malta | Molecular Diagnostics Pathology Department Mater Dei Hospital Malta | G Zahra, R Borg, C Cilia,L Grech, M Briffa |
| EPI_ISL_2988659, EPI_ISL_2988660, EPI_ISL_2988661, EPI_ISL_2988663, EPI_ISL_2988664, EPI_ISL_2988665, EPI_ISL_2988666, EPI_ISL_2988667, EPI_ISL_2988668, EPI_ISL_2988669, EPI_ISL_2988670, EPI_ISL_2988672, EPI_ISL_2988673, EPI_ISL_2988674, EPI_ISL_2988675, EPI_ISL_2988676, EPI_ISL_2988677, EPI_ISL_2988678, EPI_ISL_2988679, EPI_ISL_2988681, EPI_ISL_2988682, EPI_ISL_2988683, EPI_ISL_2988684, EPI_ISL_2988685, EPI_ISL_2988686, EPI_ISL_2988688, EPI_ISL_2988689, EPI_ISL_2988690, EPI_ISL_2988691, EPI_ISL_2988692, EPI_ISL_2988693, EPI_ISL_2988694, EPI_ISL_2988695, EPI_ISL_2988696, EPI_ISL_2988697, EPI_ISL_2988698 | | | |
| see above | Clinical Microbiology, Infection Prevention and Control | Section for Molecular Diagnostics | Björn Hallström, Jonas Björkman |
| EPI_ISL_2988701 | 1 Wojskowy Szpital Kliniczny z Poliklinką SP ZOZ | 1. Academic Center for Pathomorphological and Genetic-Molecular Diagnostics ltd, Bialystok, Poland 2. National Institute of Public Health - National Institute of Hygiene, Warsaw, Poland | Radosław Charkiewicz, Jacek Nikliński, Przemysław Biecek. Joanna Reszeć, Piotr Majewski, Anetta Sulewska, Piotr Karabowicz, Joanna Kiśluk, Janusz Dzięcioł, Konrad Raczkowski, Katarzyna Zacharczuk, Magdalena Nowakowska, Małgorzata Sadkowska-Todys, Tomasz Wołkowicz |
| EPI_ISL_2988707, EPI_ISL_2988710, EPI_ISL_2988711, EPI_ISL_2988714, EPI_ISL_2988715, EPI_ISL_2988721 | Wojewódzki Szpital Specjalistyczny Laboratorium Mikrobiologiczne | 1. Academic Center for Pathomorphological and Genetic-Molecular Diagnostics ltd, Bialystok, Poland 2. National Institute of Public Health - National Institute of Hygiene, Warsaw, Poland | Radosław Charkiewicz, Jacek Nikliński, Przemysław Biecek. Joanna Reszeć, Piotr Majewski, Anetta Sulewska, Piotr Karabowicz, Joanna Kiśluk, Janusz Dzięcioł, Konrad Raczkowski, Katarzyna Zacharczuk, Magdalena Nowakowska, Małgorzata Sadkowska-Todys, Tomasz Wołkowicz |
| EPI_ISL_2988725 | SPZOZ K-KOŻLE | 1. Academic Center for Pathomorphological and Genetic-Molecular Diagnostics ltd, Bialystok, Poland 2. National Institute of Public Health - National Institute of Hygiene, Warsaw, Poland | Radosław Charkiewicz, Jacek Nikliński, Przemysław Biecek. Joanna Reszeć, Piotr Majewski, Anetta Sulewska, Piotr Karabowicz, Joanna Kiśluk, Janusz Dzięcioł, Konrad Raczkowski, Katarzyna Zacharczuk, Magdalena Nowakowska, Małgorzata Sadkowska-Todys, Tomasz Wołkowicz |
| EPI_ISL_2988728, EPI_ISL_2988729 | Labolatorium Biologi Molekularnej USK | 1. Academic Center for Pathomorphological and Genetic-Molecular Diagnostics ltd, Bialystok, Poland 2. National Institute of Public Health - National Institute of Hygiene, Warsaw, Poland | Radosław Charkiewicz, Jacek Nikliński, Przemysław Biecek. Joanna Reszeć, Piotr Majewski, Anetta Sulewska, Piotr Karabowicz, Joanna Kiśluk, Janusz Dzięcioł, Konrad Raczkowski, Katarzyna Zacharczuk, Magdalena Nowakowska, Małgorzata Sadkowska-Todys, Tomasz Wołkowicz |
| EPI_ISL_2988730, EPI_ISL_2988731 | LAB-MED. | 1. Academic Center for Pathomorphological and Genetic-Molecular Diagnostics ltd, Bialystok, Poland 2. National Institute of Public Health - National Institute of Hygiene, Warsaw, Poland | Radosław Charkiewicz, Jacek Nikliński, Przemysław Biecek. Joanna Reszeć, Piotr Majewski, Anetta Sulewska, Piotr Karabowicz, Joanna Kiśluk, Janusz Dzięcioł, Konrad Raczkowski, Katarzyna Zacharczuk, Magdalena Nowakowska, Małgorzata Sadkowska-Todys, Tomasz Wołkowicz |
| EPI_ISL_2988840, EPI_ISL_2988841, EPI_ISL_2988842, EPI_ISL_2988843, EPI_ISL_2988844, EPI_ISL_2988847, EPI_ISL_2988848, EPI_ISL_2988849, EPI_ISL_2988850, EPI_ISL_2988851, EPI_ISL_2988852, EPI_ISL_2988853, EPI_ISL_2988854, EPI_ISL_2988856, EPI_ISL_2988857, EPI_ISL_2988858 | | | |
| see above | Sharp HealthCare Laboratory | Andersen lab at Scripps Research | SEARCH Alliance San Diego with Aaron Harding, Jacquelyn Berumen, Cathy Woerle, Liam McGinnis, Art Mendoza, Omid Bakhtar |
| EPI_ISL_2988881, EPI_ISL_2988882, EPI_ISL_2988883, EPI_ISL_2988884, EPI_ISL_2988886, EPI_ISL_2988887, EPI_ISL_2988888, EPI_ISL_2988890, EPI_ISL_2988891, EPI_ISL_2988892, EPI_ISL_2988894, EPI_ISL_2988895, EPI_ISL_2988896, EPI_ISL_2988897, EPI_ISL_2988898, EPI_ISL_2988900, EPI_ISL_2988901, EPI_ISL_2988902, EPI_ISL_2988903, EPI_ISL_2988904, EPI_ISL_2988905, EPI_ISL_2988906, EPI_ISL_2988907, EPI_ISL_2988908, EPI_ISL_2988909, EPI_ISL_2988910, EPI_ISL_2988911, EPI_ISL_2988912, EPI_ISL_2988914, EPI_ISL_2988915, EPI_ISL_2988917, EPI_ISL_2988918, EPI_ISL_2988920, EPI_ISL_2988921, EPI_ISL_2988922, EPI_ISL_2988924, EPI_ISL_2988925, EPI_ISL_2988927, EPI_ISL_2988928, EPI_ISL_2988929, EPI_ISL_2988931, EPI_ISL_2988932, EPI_ISL_2988933, EPI_ISL_2988934, EPI_ISL_2988935, EPI_ISL_2988936, EPI_ISL_2988937, EPI_ISL_2988938, EPI_ISL_2988939, EPI_ISL_2988940, EPI_ISL_2988941, EPI_ISL_2988942, EPI_ISL_2988943, EPI_ISL_2988944, EPI_ISL_2988945, EPI_ISL_2988946, EPI_ISL_2988947, EPI_ISL_2988949, EPI_ISL_2988950, EPI_ISL_2988951, EPI_ISL_2988952, EPI_ISL_2988953, EPI_ISL_2988954, EPI_ISL_2988955, EPI_ISL_2988956, EPI_ISL_2988957, EPI_ISL_2988958, EPI_ISL_2988959, EPI_ISL_2988960, EPI_ISL_2988961, EPI_ISL_2988962, EPI_ISL_2988963 | | | |
| see above | UMC Groningen, Clinical Virology, Department of Medical Microbiology and Infection Prevention | UMC Groningen, Clinical Virology, Department of Medical Microbiology and Infection Prevention | Hubert Niesters, Alexander Friedrich, Erley Lizarazo-Forero, Monika Fliss, Lilli Gard, Sigrid Rosema, Coretta Van Leer-Buter, Xuewei Zhou, Marjolein Knoester |
| EPI_ISL_2988964 | CHLN | Instituto Nacional de Saude (INSA) | Borges et al |
| EPI_ISL_2988965 | SYNLAB | Instituto Nacional de Saude (INSA) | Borges et al |
| EPI_ISL_2988966 | CHLC | Instituto Nacional de Saude (INSA) | Borges et al |

| | | | |
|---|---|---|---|
| EPI_ISL_2988967 | CHEDV | Instituto Nacional de Saude (INSA) | Borges et al |
| EPI_ISL_2988968 | SYNLAB | Instituto Nacional de Saude (INSA) | Borges et al |
| EPI_ISL_2988969 | H Divino Espirito Santo - Ponta Delgada | Instituto Nacional de Saude (INSA) | Borges et al |
| EPI_ISL_2988970 | Avelab | Instituto Nacional de Saude (INSA) | Borges et al |
| EPI_ISL_2988971 | CH Tondela Viseu | Instituto Nacional de Saude (INSA) | Borges et al |
| EPI_ISL_2988972 | H Braga | Instituto Nacional de Saude (INSA) | Borges et al |
| EPI_ISL_2988973 | UNILABS | Instituto Nacional de Saude (INSA) | Borges et al |
| EPI_ISL_2988974 | INSA | Instituto Nacional de Saude (INSA) | Borges et al |
| EPI_ISL_2988975 | SYNLAB | Instituto Nacional de Saude (INSA) | Borges et al |
| EPI_ISL_2988976 | H Divino Espirito Santo - Ponta Delgada | Instituto Nacional de Saude (INSA) | Borges et al |
| EPI_ISL_2988977 | CHU Sao Joao, Porto | Instituto Nacional de Saude (INSA) | Borges et al |
| EPI_ISL_2988978 | CH Setubal | Instituto Nacional de Saude (INSA) | Borges et al |
| EPI_ISL_2988979 | CHLO | Instituto Nacional de Saude (INSA) | Borges et al |
| EPI_ISL_2988980 | CHTMAD | Instituto Nacional de Saude (INSA) | Borges et al |
| EPI_ISL_2988982 | H Divino Espirito Santo - Ponta Delgada | Instituto Nacional de Saude (INSA) | Borges et al |
| EPI_ISL_2988983, EPI_ISL_2988984 | UNILABS | Instituto Nacional de Saude (INSA) | Borges et al |
| EPI_ISL_2988985 | CH Setubal | Instituto Nacional de Saude (INSA) | Borges et al |
| EPI_ISL_2988986 | H Braga | Instituto Nacional de Saude (INSA) | Borges et al |
| EPI_ISL_2988987 | H Divino Espirito Santo - Ponta Delgada | Instituto Nacional de Saude (INSA) | Borges et al |
| EPI_ISL_2988988, EPI_ISL_2988989 | CHLO | Instituto Nacional de Saude (INSA) | Borges et al |
| EPI_ISL_2988990 | UNILABS | Instituto Nacional de Saude (INSA) | Borges et al |
| EPI_ISL_2988991 | CHLO | Instituto Nacional de Saude (INSA) | Borges et al |
| EPI_ISL_2988992 | H Braga | Instituto Nacional de Saude (INSA) | Borges et al |
| EPI_ISL_2988993 | H Garcia de Orta | Instituto Nacional de Saude (INSA) | Borges et al |
| EPI_ISL_2988994 | Avelab | Instituto Nacional de Saude (INSA) | Borges et al |
| EPI_ISL_2988996 | CHLO | Instituto Nacional de Saude (INSA) | Borges et al |
| EPI_ISL_2988997, EPI_ISL_2988998 | UNILABS | Instituto Nacional de Saude (INSA) | Borges et al |
| EPI_ISL_2988999 | Avelab | Instituto Nacional de Saude (INSA) | Borges et al |
| EPI_ISL_2989000 | H Evora | Instituto Nacional de Saude (INSA) | Borges et al |
| EPI_ISL_2989001 | H Divino Espirito Santo - Ponta Delgada | Instituto Nacional de Saude (INSA) | Borges et al |
| EPI_ISL_2989002 | CH Tondela Viseu | Instituto Nacional de Saude (INSA) | Borges et al |
| EPI_ISL_2989003 | CHLC | Instituto Nacional de Saude (INSA) | Borges et al |
| EPI_ISL_2989004 | H Vila Franca Xira | Instituto Nacional de Saude (INSA) | Borges et al |
| EPI_ISL_2989005 | H Divino Espirito Santo - Ponta Delgada | Instituto Nacional de Saude (INSA) | Borges et al |
| EPI_ISL_2989006 | ULS Norte Alentejano | Instituto Nacional de Saude (INSA) | Borges et al |
| EPI_ISL_2989007 | CHTMAD | Instituto Nacional de Saude (INSA) | Borges et al |
| EPI_ISL_2989008 | ULS Litoral Alentejano | Instituto Nacional de Saude (INSA) | Borges et al |
| EPI_ISL_2989009 | Avelab | Instituto Nacional de Saude (INSA) | Borges et al |
| EPI_ISL_2989010 | H Divino Espirito Santo - Ponta Delgada | Instituto Nacional de Saude (INSA) | Borges et al |
| EPI_ISL_2989011 | H Evora | Instituto Nacional de Saude (INSA) | Borges et al |
| EPI_ISL_2989012 | CHLO | Instituto Nacional de Saude (INSA) | Borges et al |
| EPI_ISL_2989013 | H Braga | Instituto Nacional de Saude (INSA) | Borges et al |
| EPI_ISL_2989015 | ULS Norte Alentejano | Instituto Nacional de Saude (INSA) | Borges et al |
| EPI_ISL_2989016 | H Divino Espirito Santo - Ponta Delgada | Instituto Nacional de Saude (INSA) | Borges et al |
| EPI_ISL_2989017 | CH Porto - H Sto Antonio | Instituto Nacional de Saude (INSA) | Borges et al |
| EPI_ISL_2989018, EPI_ISL_2989019 | INSA | Instituto Nacional de Saude (INSA) | Borges et al |
| EPI_ISL_2989020 | H Garcia de Orta | Instituto Nacional de Saude (INSA) | Borges et al |
| EPI_ISL_2989022 | Avelab | Instituto Nacional de Saude (INSA) | Borges et al |
| EPI_ISL_2989023 | H Divino Espirito Santo - Ponta Delgada | Instituto Nacional de Saude (INSA) | Borges et al |
| EPI_ISL_2989025 | H Vila Franca Xira | Instituto Nacional de Saude (INSA) | Borges et al |
| EPI_ISL_2989026, EPI_ISL_2989027 | UNILABS | Instituto Nacional de Saude (INSA) | Borges et al |
| EPI_ISL_2989028 | Avelab | Instituto Nacional de Saude (INSA) | Borges et al |
| EPI_ISL_2989030 | H Evora | Instituto Nacional de Saude (INSA) | Borges et al |
| EPI_ISL_2989031 | H Braga | Instituto Nacional de Saude (INSA) | Borges et al |
| EPI_ISL_2989032 | CHU Sao Joao, Porto | Instituto Nacional de Saude (INSA) | Borges et al |
| EPI_ISL_2989033 | CH Setubal | Instituto Nacional de Saude (INSA) | Borges et al |
| EPI_ISL_2989034, EPI_ISL_2989035 | H Divino Espirito Santo - Ponta Delgada | Instituto Nacional de Saude (INSA) | Borges et al |
| EPI_ISL_2989036 | CHLO | Instituto Nacional de Saude (INSA) | Borges et al |
| EPI_ISL_2989037 | UNILABS | Instituto Nacional de Saude (INSA) | Borges et al |
| EPI_ISL_2989038 | ULS Litoral Alentejano | Instituto Nacional de Saude (INSA) | Borges et al |
| EPI_ISL_2989039 | CHU Sao Joao, Porto | Instituto Nacional de Saude (INSA) | Borges et al |
| EPI_ISL_2989053, EPI_ISL_2989054 | CH Porto - H Sto Antonio | Instituto Nacional de Saude (INSA) | Borges et al |
| EPI_ISL_2989055, EPI_ISL_2989056, EPI_ISL_2989057, EPI_ISL_2989058 | ULS Castelo Branco | Instituto Nacional de Saude (INSA) | Borges et al |
| EPI_ISL_2989059, EPI_ISL_2989060 | UNILABS | Instituto Nacional de Saude (INSA) | Borges et al |
| EPI_ISL_2989075 | SYNLAB | Instituto Nacional de Saude (INSA) | Borges et al |
| EPI_ISL_2989076 | CHEDV | Instituto Nacional de Saude (INSA) | Borges et al |
| EPI_ISL_2989077 | ULS Castelo Branco | Instituto Nacional de Saude (INSA) | Borges et al |
| EPI_ISL_2989078 | ULS Norte Alentejano | Instituto Nacional de Saude (INSA) | Borges et al |
| EPI_ISL_2989079 | H Vila Franca Xira | Instituto Nacional de Saude (INSA) | Borges et al |

| | | | |
|---|---|---|---|
| EPI_ISL_2989080 | SYNLAB | Instituto Nacional de Saude (INSA) | Borges et al |
| EPI_ISL_2989081 | UNILABS | Instituto Nacional de Saude (INSA) | Borges et al |
| EPI_ISL_2989082 | SYNLAB | Instituto Nacional de Saude (INSA) | Borges et al |
| EPI_ISL_2989083, EPI_ISL_2989084 | UNILABS | Instituto Nacional de Saude (INSA) | Borges et al |
| EPI_ISL_2989085 | SYNLAB | Instituto Nacional de Saude (INSA) | Borges et al |
| EPI_ISL_2989086 | UNILABS | Instituto Nacional de Saude (INSA) | Borges et al |
| EPI_ISL_2989087, EPI_ISL_2989088 | CHLC | Instituto Nacional de Saude (INSA) | Borges et al |
| EPI_ISL_2989089 | LAC UCoimbra | Instituto Nacional de Saude (INSA) | Borges et al |
| EPI_ISL_2989090 | INSA | Instituto Nacional de Saude (INSA) | Borges et al |
| EPI_ISL_2989091 | UNILABS | Instituto Nacional de Saude (INSA) | Borges et al |
| EPI_ISL_2989093 | ABC Algarve | Instituto Nacional de Saude (INSA) | Borges et al |
| EPI_ISL_2989094, EPI_ISL_2989095, EPI_ISL_2989096 | H Fernando Fonseca | Instituto Nacional de Saude (INSA) | Borges et al |
| EPI_ISL_2989097 | INSA | Instituto Nacional de Saude (INSA) | Borges et al |
| EPI_ISL_2989098, EPI_ISL_2989099 | SYNLAB | Instituto Nacional de Saude (INSA) | Borges et al |
| EPI_ISL_2989100 | CHMT | Instituto Nacional de Saude (INSA) | Borges et al |
| EPI_ISL_2989101, EPI_ISL_2989102, EPI_ISL_2989103 | CHTMAD | Instituto Nacional de Saude (INSA) | Borges et al |
| EPI_ISL_2989104, EPI_ISL_2989105 | H Divino Espirito Santo - Ponta Delgada | Instituto Nacional de Saude (INSA) | Borges et al |
| EPI_ISL_2989106 | SESARAM | Instituto Nacional de Saude (INSA) | Borges et al |
| EPI_ISL_2989107 | H Fernando Fonseca | Instituto Nacional de Saude (INSA) | Borges et al |
| EPI_ISL_2989109 | H Santarem | Instituto Nacional de Saude (INSA) | Borges et al |
| EPI_ISL_2989110 | SYNLAB | Instituto Nacional de Saude (INSA) | Borges et al |
| EPI_ISL_2989111, EPI_ISL_2989112 | CHLN | Instituto Nacional de Saude (INSA) | Borges et al |
| EPI_ISL_2989114, EPI_ISL_2989115 | SYNLAB | Instituto Nacional de Saude (INSA) | Borges et al |
| EPI_ISL_2989116 | H Fernando Fonseca | Instituto Nacional de Saude (INSA) | Borges et al |
| EPI_ISL_2989117 | CHLC | Instituto Nacional de Saude (INSA) | Borges et al |
| EPI_ISL_2989118, EPI_ISL_2989119, EPI_ISL_2989120, EPI_ISL_2989121, EPI_ISL_2989122, EPI_ISL_2989123, EPI_ISL_2989124, EPI_ISL_2989125, EPI_ISL_2989126, EPI_ISL_2989127, EPI_ISL_2989128, EPI_ISL_2989129 | | | |
| see above | ABC Algarve | Instituto Nacional de Saude (INSA) | Borges et al |
| EPI_ISL_2989130, EPI_ISL_2989131, EPI_ISL_2989132, EPI_ISL_2989133, EPI_ISL_2989134, EPI_ISL_2989135 | CHEDV | Instituto Nacional de Saude (INSA) | Borges et al |
| EPI_ISL_2989136, EPI_ISL_2989137, EPI_ISL_2989138, EPI_ISL_2989139, EPI_ISL_2989140, EPI_ISL_2989141, EPI_ISL_2989142, EPI_ISL_2989143, EPI_ISL_2989144, EPI_ISL_2989145 | CHLC | Instituto Nacional de Saude (INSA) | Borges et al |
| EPI_ISL_2989146, EPI_ISL_2989147, EPI_ISL_2989148 | CH Setubal | Instituto Nacional de Saude (INSA) | Borges et al |
| EPI_ISL_2989149, EPI_ISL_2989150, EPI_ISL_2989151 | CH Tondela Viseu | Instituto Nacional de Saude (INSA) | Borges et al |
| EPI_ISL_2989152, EPI_ISL_2989153, EPI_ISL_2989154, EPI_ISL_2989155, EPI_ISL_2989156, EPI_ISL_2989157, EPI_ISL_2989158, EPI_ISL_2989159 | CH Porto - H Sto Antonio | Instituto Nacional de Saude (INSA) | Borges et al |
| EPI_ISL_2989160, EPI_ISL_2989161, EPI_ISL_2989162, EPI_ISL_2989163, EPI_ISL_2989164, EPI_ISL_2989165, EPI_ISL_2989166, EPI_ISL_2989167, EPI_ISL_2989168, EPI_ISL_2989169 | H Braga | Instituto Nacional de Saude (INSA) | Borges et al |
| EPI_ISL_2989170, EPI_ISL_2989171, EPI_ISL_2989172, EPI_ISL_2989173, EPI_ISL_2989174, EPI_ISL_2989175, EPI_ISL_2989176, EPI_ISL_2989177, EPI_ISL_2989178 | H Santarem | Instituto Nacional de Saude (INSA) | Borges et al |
| EPI_ISL_2989179, EPI_ISL_2989180, EPI_ISL_2989181, EPI_ISL_2989182, EPI_ISL_2989183, EPI_ISL_2989184, EPI_ISL_2989185, EPI_ISL_2989186, EPI_ISL_2989187, EPI_ISL_2989188, EPI_ISL_2989189, EPI_ISL_2989190 | | | |
| see above | H Evora | Instituto Nacional de Saude (INSA) | Borges et al |
| EPI_ISL_2989191, EPI_ISL_2989192, EPI_ISL_2989193, EPI_ISL_2989194, EPI_ISL_2989195, EPI_ISL_2989196, | H Fernando Fonseca | Instituto Nacional de Saude (INSA) | Borges et al |

| | | | |
|---|---|---|---|
| EPI_ISL_2989197, EPI_ISL_2989198, EPI_ISL_2989199, EPI_ISL_2989200 | | | |
| EPI_ISL_2989201, EPI_ISL_2989202, EPI_ISL_2989203, EPI_ISL_2989204, EPI_ISL_2989205, EPI_ISL_2989206, EPI_ISL_2989207, EPI_ISL_2989208 | H Vila Franca Xira | Instituto Nacional de Saude (INSA) | Borges et al |
| EPI_ISL_2989209, EPI_ISL_2989210, EPI_ISL_2989211, EPI_ISL_2989212, EPI_ISL_2989213 | INSA | Instituto Nacional de Saude (INSA) | Borges et al |
| EPI_ISL_2989214, EPI_ISL_2989215, EPI_ISL_2989216, EPI_ISL_2989217, EPI_ISL_2989218, EPI_ISL_2989219, EPI_ISL_2989220, EPI_ISL_2989221, EPI_ISL_2989222, EPI_ISL_2989223, EPI_ISL_2989224, EPI_ISL_2989225, EPI_ISL_2989226, EPI_ISL_2989227, EPI_ISL_2989228, EPI_ISL_2989229, EPI_ISL_2989230, EPI_ISL_2989231, EPI_ISL_2989232, EPI_ISL_2989233, EPI_ISL_2989234, EPI_ISL_2989235, EPI_ISL_2989236, EPI_ISL_2989237, EPI_ISL_2989238, EPI_ISL_2989239, EPI_ISL_2989240, EPI_ISL_2989241, EPI_ISL_2989242, EPI_ISL_2989243, EPI_ISL_2989244, EPI_ISL_2989245, EPI_ISL_2989246, EPI_ISL_2989247, EPI_ISL_2989248, EPI_ISL_2989249, EPI_ISL_2989250, EPI_ISL_2989251, EPI_ISL_2989252, EPI_ISL_2989253, EPI_ISL_2989254, EPI_ISL_2989255 | | | |
| see above | SYNLAB | Instituto Nacional de Saude (INSA) | Borges et al |
| EPI_ISL_2989256, EPI_ISL_2989257, EPI_ISL_2989258, EPI_ISL_2989259, EPI_ISL_2989260, EPI_ISL_2989261, EPI_ISL_2989262, EPI_ISL_2989263, EPI_ISL_2989264, EPI_ISL_2989265, EPI_ISL_2989266, EPI_ISL_2989267, EPI_ISL_2989268, EPI_ISL_2989269, EPI_ISL_2989270, EPI_ISL_2989271 | | | |
| see above | LAC UCoimbra | Instituto Nacional de Saude (INSA) | Borges et al |
| EPI_ISL_2989272, EPI_ISL_2989273, EPI_ISL_2989274, EPI_ISL_2989275, EPI_ISL_2989276, EPI_ISL_2989277, EPI_ISL_2989278 | ULS Castelo Branco | Instituto Nacional de Saude (INSA) | Borges et al |
| EPI_ISL_2989279, EPI_ISL_2989280, EPI_ISL_2989281, EPI_ISL_2989282, EPI_ISL_2989283, EPI_ISL_2989284, EPI_ISL_2989285, EPI_ISL_2989286, EPI_ISL_2989287 | ULS Litoral Alentejano | Instituto Nacional de Saude (INSA) | Borges et al |
| EPI_ISL_2989288, EPI_ISL_2989289, EPI_ISL_2989290, EPI_ISL_2989291, EPI_ISL_2989292, EPI_ISL_2989293, EPI_ISL_2989294, EPI_ISL_2989295, EPI_ISL_2989296, EPI_ISL_2989297, EPI_ISL_2989298, EPI_ISL_2989299, EPI_ISL_2989300, EPI_ISL_2989301, EPI_ISL_2989302, EPI_ISL_2989303 | | | |
| see above | ULS Norte Alentejano | Instituto Nacional de Saude (INSA) | Borges et al |
| EPI_ISL_2989304, EPI_ISL_2989305, EPI_ISL_2989306, EPI_ISL_2989307, EPI_ISL_2989308, EPI_ISL_2989309, EPI_ISL_2989310, EPI_ISL_2989311, EPI_ISL_2989312, EPI_ISL_2989313, EPI_ISL_2989314, EPI_ISL_2989315, EPI_ISL_2989316, EPI_ISL_2989317, EPI_ISL_2989318, EPI_ISL_2989319, EPI_ISL_2989320, EPI_ISL_2989321, EPI_ISL_2989322, EPI_ISL_2989323, EPI_ISL_2989324, EPI_ISL_2989325, EPI_ISL_2989326 | | | |
| see above | ARS Algarve - Laboratorio Laura Ayres | Instituto Nacional de Saude (INSA) | Borges et al |
| EPI_ISL_2989327, EPI_ISL_2989328, EPI_ISL_2989329, EPI_ISL_2989330, EPI_ISL_2989331, EPI_ISL_2989332, EPI_ISL_2989333, EPI_ISL_2989334 | Avelab | Instituto Nacional de Saude (INSA) | Borges et al |
| EPI_ISL_2989335, EPI_ISL_2989336, EPI_ISL_2989337, EPI_ISL_2989338, EPI_ISL_2989339, EPI_ISL_2989340, EPI_ISL_2989341, EPI_ISL_2989342, EPI_ISL_2989343, EPI_ISL_2989344, EPI_ISL_2989345 | | | |
| see above | CHLN | Instituto Nacional de Saude (INSA) | Borges et al |
| EPI_ISL_2989346, EPI_ISL_2989347, EPI_ISL_2989348, EPI_ISL_2989349, EPI_ISL_2989350, EPI_ISL_2989351, EPI_ISL_2989352, EPI_ISL_2989353 | CHLO | Instituto Nacional de Saude (INSA) | Borges et al |
| EPI_ISL_2989354, EPI_ISL_2989355 | CHMT | Instituto Nacional de Saude (INSA) | Borges et al |
| EPI_ISL_2989356, EPI_ISL_2989357, EPI_ISL_2989358, EPI_ISL_2989359, EPI_ISL_2989360, EPI_ISL_2989361, EPI_ISL_2989362, EPI_ISL_2989363, EPI_ISL_2989364, EPI_ISL_2989365, EPI_ISL_2989366, EPI_ISL_2989367, EPI_ISL_2989368 | | | |
| see above | CHTMAD | Instituto Nacional de Saude (INSA) | Borges et al |
| EPI_ISL_2989369 | CH Tondela Viseu | Instituto Nacional de Saude (INSA) | Borges et al |
| EPI_ISL_2989370, EPI_ISL_2989371, EPI_ISL_2989372, EPI_ISL_2989373, EPI_ISL_2989374, EPI_ISL_2989375, EPI_ISL_2989376, EPI_ISL_2989377, EPI_ISL_2989378 | CHU Sao Joao, Porto | Instituto Nacional de Saude (INSA) | Borges et al |
| EPI_ISL_2989379, EPI_ISL_2989380, EPI_ISL_2989381, EPI_ISL_2989382, EPI_ISL_2989383, EPI_ISL_2989384, EPI_ISL_2989385, EPI_ISL_2989386, EPI_ISL_2989387, EPI_ISL_2989388, EPI_ISL_2989389, EPI_ISL_2989390, EPI_ISL_2989391, EPI_ISL_2989392 | | | |
| see above | H Divino Espirito Santo - Ponta Delgada | Instituto Nacional de Saude (INSA) | Borges et al |
| EPI_ISL_2989393, EPI_ISL_2989394, EPI_ISL_2989395, EPI_ISL_2989396, EPI_ISL_2989397, EPI_ISL_2989398, EPI_ISL_2989399, EPI_ISL_2989400, EPI_ISL_2989401, EPI_ISL_2989402, EPI_ISL_2989403, EPI_ISL_2989404, EPI_ISL_2989405, EPI_ISL_2989406, EPI_ISL_2989407, EPI_ISL_2989408, EPI_ISL_2989409, EPI_ISL_2989410, EPI_ISL_2989411, EPI_ISL_2989412, EPI_ISL_2989413, EPI_ISL_2989414 | | | |
| see above | H Garcia de Orta | Instituto Nacional de Saude (INSA) | Borges et al |
| EPI_ISL_2989415, EPI_ISL_2989416, EPI_ISL_2989417, EPI_ISL_2989418, EPI_ISL_2989419, EPI_ISL_2989420, EPI_ISL_2989421, EPI_ISL_2989422, EPI_ISL_2989423, EPI_ISL_2989424, EPI_ISL_2989425, EPI_ISL_2989426, EPI_ISL_2989427, EPI_ISL_2989428, EPI_ISL_2989429, EPI_ISL_2989430, EPI_ISL_2989431, EPI_ISL_2989432, EPI_ISL_2989433, EPI_ISL_2989434, EPI_ISL_2989435 | | | |
| see above | SEEBMO | Instituto Nacional de Saude (INSA) | Borges et al |
| EPI_ISL_2989436, EPI_ISL_2989437, EPI_ISL_2989438, EPI_ISL_2989439, EPI_ISL_2989440, EPI_ISL_2989441, EPI_ISL_2989442, EPI_ISL_2989443, EPI_ISL_2989444, EPI_ISL_2989445, EPI_ISL_2989446, EPI_ISL_2989447, EPI_ISL_2989448, EPI_ISL_2989449, EPI_ISL_2989450, EPI_ISL_2989451, EPI_ISL_2989452, EPI_ISL_2989453, EPI_ISL_2989454, EPI_ISL_2989455, EPI_ISL_2989456, EPI_ISL_2989457, EPI_ISL_2989458, EPI_ISL_2989459, EPI_ISL_2989460, EPI_ISL_2989461, EPI_ISL_2989462, EPI_ISL_2989463, EPI_ISL_2989464, EPI_ISL_2989465 | | | |
| see above | SESARAM | Instituto Nacional de Saude (INSA) | Borges et al |
| EPI_ISL_2989466, EPI_ISL_2989467, EPI_ISL_2989468, EPI_ISL_2989469, EPI_ISL_2989470, EPI_ISL_2989471, EPI_ISL_2989472, EPI_ISL_2989473, EPI_ISL_2989474, EPI_ISL_2989475, EPI_ISL_2989476, EPI_ISL_2989477, EPI_ISL_2989478, EPI_ISL_2989479, EPI_ISL_2989480, EPI_ISL_2989481, EPI_ISL_2989482, EPI_ISL_2989483, EPI_ISL_2989484, EPI_ISL_2989485, EPI_ISL_2989486, EPI_ISL_2989487, EPI_ISL_2989488, EPI_ISL_2989489, EPI_ISL_2989490, EPI_ISL_2989491, EPI_ISL_2989492, EPI_ISL_2989493, EPI_ISL_2989494, EPI_ISL_2989495, EPI_ISL_2989496, EPI_ISL_2989497, EPI_ISL_2989498, EPI_ISL_2989499, EPI_ISL_2989500 | | | |
| see above | UNILABS | Instituto Nacional de Saude (INSA) | Borges et al |
| EPI_ISL_2989501 | CHMT | Instituto Nacional de Saude (INSA) | Borges et al |
| EPI_ISL_2989504, EPI_ISL_2989505 | H Vila Franca Xira | Instituto Nacional de Saude (INSA) | Borges et al |
| EPI_ISL_2989506 | CHLN | Instituto Nacional de Saude (INSA) | Borges et al |
| EPI_ISL_2991494 | Utah Public Health Laboratory, Utah Public Health Laboratory Infectious Disease submission group | Utah Public Health Laboratory, Utah Public Health Laboratory Infectious Disease submission group | Linares-Perdomo,O., Oakeson,K., Gallagher,T., Young,E. |
| EPI_ISL_2991508 | Infectious Disease Program, Broad Institute of Harvard and MIT | Infectious Disease Program, Broad Institute of Harvard and MIT | Lemieux,J.E., Siddle,K.J., Shaw,B., Adams,G., Pierce,V., Turbett,S., Anahtar,M., Branda,J., Slater,D., Harris,J., Lin,A.E., Gladden-Young,A., Lagerborg,K., Rudy,M., DeRuff,K., Carter,A., Normandin,E., Bauer,M., Reilly,S., Tomkins-Tinch,C., Loreth,C., |

| | | | |
|---|---|---|---|
| | | | Chaluvadi,S., Neumann,A., Cusick,C., Chapman,S.B., Gnirke,A., Flowers,K., Cerrato,F., Birren,B.W., Gallagher,G., Smole,S., Park,D.J., MacInnis,B.L., Ryan,E., LaRocque,R., Rosenberg,E., Sabeti,P.C. |
| EPI_ISL_2991528, EPI_ISL_2991532, EPI_ISL_2991542, EPI_ISL_2991545, EPI_ISL_2991548, EPI_ISL_2991553, EPI_ISL_2991572, EPI_ISL_2991578, EPI_ISL_2991581, EPI_ISL_2991589, EPI_ISL_2991603, EPI_ISL_2991604, EPI_ISL_2991666, EPI_ISL_2991674, EPI_ISL_2991677, EPI_ISL_2991691, EPI_ISL_2991693, EPI_ISL_2991695, EPI_ISL_2991702, EPI_ISL_2991716, EPI_ISL_2991726, EPI_ISL_2991731, EPI_ISL_2991740, EPI_ISL_2991748, EPI_ISL_2991754, EPI_ISL_2991771, EPI_ISL_2991783, EPI_ISL_2991784, EPI_ISL_2991807, EPI_ISL_2991811, EPI_ISL_2991837, EPI_ISL_2991843, EPI_ISL_2991847, EPI_ISL_2991855, EPI_ISL_2991872, EPI_ISL_2991877, EPI_ISL_2991883, EPI_ISL_2991884, EPI_ISL_2991890, EPI_ISL_2991899, EPI_ISL_2991900, EPI_ISL_2991902, EPI_ISL_2991903, EPI_ISL_2991904, EPI_ISL_2991909, EPI_ISL_2991916, EPI_ISL_2991922, EPI_ISL_2991932, EPI_ISL_2991937, EPI_ISL_2991938, EPI_ISL_2991950, EPI_ISL_2991963, EPI_ISL_2991966 | | | |
| see above | Broad Institute Clinical Research Sequencing Platform | Infectious Disease Program, Broad Institute of Harvard and MIT | Lemieux,J.E., Siddle,K.J., Shaw,B., Adams,G., Pierce,V., Turbett,S., Anahtar,M., Branda,J., Slater,D., Harris,J., Lin,A.E., Gladden-Young,A., Lagerborg,K., Rudy,M., DeRuff,K., Carter,A., Normandin,E., Bauer,M., Reilly,S., Tomkins-Tinch,C., Loreth,C., Chaluvadi,S., Neumann,A., Cusick,C., Chapman,S.B., Gnirke,A., Flowers,K., Cerrato,F., Birren,B.W., Gallagher,G., Smole,S., Park,D.J., MacInnis,B.L., Ryan,E., LaRocque,R., Rosenberg,E., Sabeti,P.C. |
| EPI_ISL_2992019 | UNR School of Medicine, Nevada State Public Health Laboratory | UNR School of Medicine, Nevada State Public Health Laboratory | Pandori,M., Gorzalski,A.J. |
| EPI_ISL_2992072, EPI_ISL_2992078 | Infectious Disease Program, Broad Institute of Harvard and MIT | Infectious Disease Program, Broad Institute of Harvard and MIT | Lemieux,J.E., Siddle,K.J., Shaw,B., Adams,G., Pierce,V., Turbett,S., Anahtar,M., Branda,J., Slater,D., Harris,J., Lin,A.E., Gladden-Young,A., Lagerborg,K., Rudy,M., DeRuff,K., Carter,A., Normandin,E., Bauer,M., Reilly,S., Tomkins-Tinch,C., Loreth,C., Chaluvadi,S., Neumann,A., Cusick,C., Chapman,S.B., Gnirke,A., Flowers,K., Cerrato,F., Birren,B.W., Gallagher,G., Smole,S., Park,D.J., MacInnis,B.L., Ryan,E., LaRocque,R., Rosenberg,E., Sabeti,P.C. |
| EPI_ISL_2992080, EPI_ISL_2992081, EPI_ISL_2992082, EPI_ISL_2992083, EPI_ISL_2992089, EPI_ISL_2992090, EPI_ISL_2992091, EPI_ISL_2992092, EPI_ISL_2992093, EPI_ISL_2992094, EPI_ISL_2992095, EPI_ISL_2992096, EPI_ISL_2992097, EPI_ISL_2992098, EPI_ISL_2992099, EPI_ISL_2992100, EPI_ISL_2992101, EPI_ISL_2992102, EPI_ISL_2992103, EPI_ISL_2992104, EPI_ISL_2992105, EPI_ISL_2992106, EPI_ISL_2992107, EPI_ISL_2992108, EPI_ISL_2992110 | | | |
| see above | Communicable Disease Laboratory, Public Health Directorate | Communicable Disease Laboratory, Public Health Directorate | Alwasti,H., AlHujairi,Z., AlAbbas,Z., Marhoon,A., Almoamen,G. |
| EPI_ISL_2992112, EPI_ISL_2992113, EPI_ISL_2992114, EPI_ISL_2992115, EPI_ISL_2992116, EPI_ISL_2992117, EPI_ISL_2992118, EPI_ISL_2992119, EPI_ISL_2992120, EPI_ISL_2992121, EPI_ISL_2992122, EPI_ISL_2992123, EPI_ISL_2992124, EPI_ISL_2992125, EPI_ISL_2992126, EPI_ISL_2992127, EPI_ISL_2992128, EPI_ISL_2992129, EPI_ISL_2992130, EPI_ISL_2992131, EPI_ISL_2992133, EPI_ISL_2992134, EPI_ISL_2992135, EPI_ISL_2992136, EPI_ISL_2992137, EPI_ISL_2992138, EPI_ISL_2992139, EPI_ISL_2992140, EPI_ISL_2992141, EPI_ISL_2992142, EPI_ISL_2992143, EPI_ISL_2992144, EPI_ISL_2992145, EPI_ISL_2992146, EPI_ISL_2992147, EPI_ISL_2992148, EPI_ISL_2992149, EPI_ISL_2992151, EPI_ISL_2992152, EPI_ISL_2992153, EPI_ISL_2992154, EPI_ISL_2992155, EPI_ISL_2992156, EPI_ISL_2992157, EPI_ISL_2992158, EPI_ISL_2992159, EPI_ISL_2992160, EPI_ISL_2992161, EPI_ISL_2992162, EPI_ISL_2992163, EPI_ISL_2992164, EPI_ISL_2992165, EPI_ISL_2992166, EPI_ISL_2992167, EPI_ISL_2992168, EPI_ISL_2992169, EPI_ISL_2992170, EPI_ISL_2992171, EPI_ISL_2992172, EPI_ISL_2992173, EPI_ISL_2992174, EPI_ISL_2992175, EPI_ISL_2992176, EPI_ISL_2992177, EPI_ISL_2992178, EPI_ISL_2992179, EPI_ISL_2992180, EPI_ISL_2992181, EPI_ISL_2992182, EPI_ISL_2992183, EPI_ISL_2992184, EPI_ISL_2992185, EPI_ISL_2992186, EPI_ISL_2992187, EPI_ISL_2992188, EPI_ISL_2992189, EPI_ISL_2992190, EPI_ISL_2992191, EPI_ISL_2992192, EPI_ISL_2992193, EPI_ISL_2992194, EPI_ISL_2992195, EPI_ISL_2992196, EPI_ISL_2992197, EPI_ISL_2992198, EPI_ISL_2992199, EPI_ISL_2992200, EPI_ISL_2992201, EPI_ISL_2992202, EPI_ISL_2992203, EPI_ISL_2992204, EPI_ISL_2992205, EPI_ISL_2992206, EPI_ISL_2992207, EPI_ISL_2992208, EPI_ISL_2992209, EPI_ISL_2992210, EPI_ISL_2992211, EPI_ISL_2992212, EPI_ISL_2992214, EPI_ISL_2992215, EPI_ISL_2992216, EPI_ISL_2992217, EPI_ISL_2992218, EPI_ISL_2992219, EPI_ISL_2992220, EPI_ISL_2992221, EPI_ISL_2992222, EPI_ISL_2992223, EPI_ISL_2992224 | | | |
| see above | Department of Bacteria, Parasites and Fungi, Statens Serum Institut, Copenhagen, Denmark | Statens Serum Institut Bioinformatics and Microbial Genomics | Danish Covid-19 Genome Consortium |
| EPI_ISL_2992225 | Molekylær Medicinsk Afdeling, Aarhus University Hospital, Aarhus, Denmark | Statens Serum Institut Bioinformatics and Microbial Genomics | Danish Covid-19 Genome Consortium |
| EPI_ISL_2992226, EPI_ISL_2992227, EPI_ISL_2992228, EPI_ISL_2992229, EPI_ISL_2992230, EPI_ISL_2992231, EPI_ISL_2992235, EPI_ISL_2992236, EPI_ISL_2992237, EPI_ISL_2992238, EPI_ISL_2992239, EPI_ISL_2992240, EPI_ISL_2992241, EPI_ISL_2992242, EPI_ISL_2992243, EPI_ISL_2992244, EPI_ISL_2992245, EPI_ISL_2992246, EPI_ISL_2992247, EPI_ISL_2992248, EPI_ISL_2992249, EPI_ISL_2992250, EPI_ISL_2992251, EPI_ISL_2992252, EPI_ISL_2992253, EPI_ISL_2992254 | | | |
| see above | Department of Bacteria, Parasites and Fungi, Statens Serum Institut, Copenhagen, Denmark | Statens Serum Institut Bioinformatics and Microbial Genomics | Danish Covid-19 Genome Consortium |
| EPI_ISL_2992255 | Molekylær Medicinsk Afdeling, Aarhus University Hospital, Aarhus, Denmark | Statens Serum Institut Bioinformatics and Microbial Genomics | Danish Covid-19 Genome Consortium |
| EPI_ISL_2992256 | Department of Bacteria, Parasites and Fungi, Statens Serum Institut, Copenhagen, Denmark | Statens Serum Institut Bioinformatics and Microbial Genomics | Danish Covid-19 Genome Consortium |
| EPI_ISL_2992257 | Department of Clinical Microbiology, Odense University Hospital, Odense, Denmark | Statens Serum Institut Bioinformatics and Microbial Genomics | Danish Covid-19 Genome Consortium |
| EPI_ISL_2992258, EPI_ISL_2992259, EPI_ISL_2992260, EPI_ISL_2992261, EPI_ISL_2992262, EPI_ISL_2992263, EPI_ISL_2992264, EPI_ISL_2992265, EPI_ISL_2992266, EPI_ISL_2992267, EPI_ISL_2992268, EPI_ISL_2992269, EPI_ISL_2992270, EPI_ISL_2992271, EPI_ISL_2992272, EPI_ISL_2992273, EPI_ISL_2992274, EPI_ISL_2992275, EPI_ISL_2992276, EPI_ISL_2992277, EPI_ISL_2992278, EPI_ISL_2992279, EPI_ISL_2992280, EPI_ISL_2992281, EPI_ISL_2992282, EPI_ISL_2992283, EPI_ISL_2992284, EPI_ISL_2992285, EPI_ISL_2992286, EPI_ISL_2992287, EPI_ISL_2992288, EPI_ISL_2992289, EPI_ISL_2992290, EPI_ISL_2992291, EPI_ISL_2992292, EPI_ISL_2992293, EPI_ISL_2992294, EPI_ISL_2992295, EPI_ISL_2992296, EPI_ISL_2992297, EPI_ISL_2992298 | | | |
| see above | Department of Bacteria, Parasites and Fungi, Statens Serum Institut, Copenhagen, Denmark | Statens Serum Institut Bioinformatics and Microbial Genomics | Danish Covid-19 Genome Consortium |
| EPI_ISL_2992299 | Molekylær Medicinsk Afdeling, Aarhus University Hospital, Aarhus, Denmark | Statens Serum Institut Bioinformatics and Microbial Genomics | Danish Covid-19 Genome Consortium |
| EPI_ISL_2992300, EPI_ISL_2992301, EPI_ISL_2992302, EPI_ISL_2992303, EPI_ISL_2992304, EPI_ISL_2992305, EPI_ISL_2992306, EPI_ISL_2992307, EPI_ISL_2992308, EPI_ISL_2992309, EPI_ISL_2992310, EPI_ISL_2992311, EPI_ISL_2992313, EPI_ISL_2992314, EPI_ISL_2992315, EPI_ISL_2992316, EPI_ISL_2992317, EPI_ISL_2992318 | | | |
| see above | Department of Bacteria, Parasites and Fungi, Statens Serum Institut, Copenhagen, Denmark | Statens Serum Institut Bioinformatics and Microbial Genomics | Danish Covid-19 Genome Consortium |
| EPI_ISL_2992319 | Molekylær Medicinsk Afdeling, Aarhus University Hospital, Aarhus, Denmark | Statens Serum Institut Bioinformatics and Microbial Genomics | Danish Covid-19 Genome Consortium |
| EPI_ISL_2992320 | Department of Bacteria, Parasites and Fungi, Statens Serum Institut, Copenhagen, Denmark | Statens Serum Institut Bioinformatics and Microbial Genomics | Danish Covid-19 Genome Consortium |
| EPI_ISL_2992321 | Molekylær Medicinsk Afdeling, Aarhus University Hospital, Aarhus, Denmark | Statens Serum Institut Bioinformatics and Microbial Genomics | Danish Covid-19 Genome Consortium |
| EPI_ISL_2992322, EPI_ISL_2992323, EPI_ISL_2992324, EPI_ISL_2992326, EPI_ISL_2992327, EPI_ISL_2992328, EPI_ISL_2992329, EPI_ISL_2992330, EPI_ISL_2992331, EPI_ISL_2992332, EPI_ISL_2992333, EPI_ISL_2992334, EPI_ISL_2992335, EPI_ISL_2992336, EPI_ISL_2992337, EPI_ISL_2992338, EPI_ISL_2992339, EPI_ISL_2992340, EPI_ISL_2992341, EPI_ISL_2992342, EPI_ISL_2992343, EPI_ISL_2992344, EPI_ISL_2992345, EPI_ISL_2992346, EPI_ISL_2992347, EPI_ISL_2992348, EPI_ISL_2992349, EPI_ISL_2992350, EPI_ISL_2992351, EPI_ISL_2992352, EPI_ISL_2992353, EPI_ISL_2992354, EPI_ISL_2992355, EPI_ISL_2992356, EPI_ISL_2992357, EPI_ISL_2992358, EPI_ISL_2992359, EPI_ISL_2992360, EPI_ISL_2992361, EPI_ISL_2992362, EPI_ISL_2992363, EPI_ISL_2992364, EPI_ISL_2992366, EPI_ISL_2992367, EPI_ISL_2992368, EPI_ISL_2992369, EPI_ISL_2992370, EPI_ISL_2992371, EPI_ISL_2992372, EPI_ISL_2992373, EPI_ISL_2992374, EPI_ISL_2992375, EPI_ISL_2992376, EPI_ISL_2992377, EPI_ISL_2992378, EPI_ISL_2992379, EPI_ISL_2992380, EPI_ISL_2992381, EPI_ISL_2992382, EPI_ISL_2992383, EPI_ISL_2992384, EPI_ISL_2992385, EPI_ISL_2992386, EPI_ISL_2992387, EPI_ISL_2992388, EPI_ISL_2992389, EPI_ISL_2992390, EPI_ISL_2992391, EPI_ISL_2992392, EPI_ISL_2992393, EPI_ISL_2992394, EPI_ISL_2992395, EPI_ISL_2992396, EPI_ISL_2992397, EPI_ISL_2992398, EPI_ISL_2992399, EPI_ISL_2992400, EPI_ISL_2992401, EPI_ISL_2992402, EPI_ISL_2992403, EPI_ISL_2992404, EPI_ISL_2992405, EPI_ISL_2992406, EPI_ISL_2992407, EPI_ISL_2992408, EPI_ISL_2992409, EPI_ISL_2992410, EPI_ISL_2992411, EPI_ISL_2992412, EPI_ISL_2992413, EPI_ISL_2992414, EPI_ISL_2992415, EPI_ISL_2992416, EPI_ISL_2992418, EPI_ISL_2992419, EPI_ISL_2992420, EPI_ISL_2992421, EPI_ISL_2992422, EPI_ISL_2992423, EPI_ISL_2992424, EPI_ISL_2992425, EPI_ISL_2992426, EPI_ISL_2992427, EPI_ISL_2992428, EPI_ISL_2992429, EPI_ISL_2992430, EPI_ISL_2992431, EPI_ISL_2992432, EPI_ISL_2992433, EPI_ISL_2992434, EPI_ISL_2992435, EPI_ISL_2992436, EPI_ISL_2992437, EPI_ISL_2992438, EPI_ISL_2992439, EPI_ISL_2992440, EPI_ISL_2992441, EPI_ISL_2992442, EPI_ISL_2992443, EPI_ISL_2992445, EPI_ISL_2992446, EPI_ISL_2992447, EPI_ISL_2992448, EPI_ISL_2992449, EPI_ISL_2992450, EPI_ISL_2992451, EPI_ISL_2992452, EPI_ISL_2992453, EPI_ISL_2992454, EPI_ISL_2992455, EPI_ISL_2992456, EPI_ISL_2992457, EPI_ISL_2992458, EPI_ISL_2992459, EPI_ISL_2992460, EPI_ISL_2992461, EPI_ISL_2992462, EPI_ISL_2992463, EPI_ISL_2992464, EPI_ISL_2992465, EPI_ISL_2992466, EPI_ISL_2992467, EPI_ISL_2992468, EPI_ISL_2992469, EPI_ISL_2992470, EPI_ISL_2992471, EPI_ISL_2992472, EPI_ISL_2992473, EPI_ISL_2992474, EPI_ISL_2992475, EPI_ISL_2992476, EPI_ISL_2992477, EPI_ISL_2992478, EPI_ISL_2992479, EPI_ISL_2992480, EPI_ISL_2992481, EPI_ISL_2992482, EPI_ISL_2992483, EPI_ISL_2992484, EPI_ISL_2992485, EPI_ISL_2992486, EPI_ISL_2992487, EPI_ISL_2992488, EPI_ISL_2992489, EPI_ISL_2992490, EPI_ISL_2992491, EPI_ISL_2992492, EPI_ISL_2992493, EPI_ISL_2992494, EPI_ISL_2992495, EPI_ISL_2992496 | | | |
| see above | Department of Bacteria, Parasites and Fungi, Statens Serum Institut, Copenhagen, Denmark | Statens Serum Institut Bioinformatics and Microbial Genomics | Danish Covid-19 Genome Consortium |
| EPI_ISL_2992497 | Department of Clinical Microbiology, Odense University Hospital, Odense, Denmark | Statens Serum Institut Bioinformatics and Microbial Genomics | Danish Covid-19 Genome Consortium |
| EPI_ISL_2992499, EPI_ISL_2992500, EPI_ISL_2992501, EPI_ISL_2992502, EPI_ISL_2992503, EPI_ISL_2992504, EPI_ISL_2992505, EPI_ISL_2992506, EPI_ISL_2992507, EPI_ISL_2992508, EPI_ISL_2992509, EPI_ISL_2992510, EPI_ISL_2992511, EPI_ISL_2992512, EPI_ISL_2992513, EPI_ISL_2992514, EPI_ISL_2992515, EPI_ISL_2992516, EPI_ISL_2992517, EPI_ISL_2992518, EPI_ISL_2992519, EPI_ISL_2992520, EPI_ISL_2992521, EPI_ISL_2992522, EPI_ISL_2992524, EPI_ISL_2992525, EPI_ISL_2992526, EPI_ISL_2992527, EPI_ISL_2992528, EPI_ISL_2992529, EPI_ISL_2992530, EPI_ISL_2992531, EPI_ISL_2992532, EPI_ISL_2992533, EPI_ISL_2992534, EPI_ISL_2992535, EPI_ISL_2992536, EPI_ISL_2992537, EPI_ISL_2992538, EPI_ISL_2992539, EPI_ISL_2992540, EPI_ISL_2992541, EPI_ISL_2992542, EPI_ISL_2992543, EPI_ISL_2992544, EPI_ISL_2992545, EPI_ISL_2992546, EPI_ISL_2992547, EPI_ISL_2992548, EPI_ISL_2992549, EPI_ISL_2992550, EPI_ISL_2992551, EPI_ISL_2992552, EPI_ISL_2992553, EPI_ISL_2992554, EPI_ISL_2992556, EPI_ISL_2992557, EPI_ISL_2992558, EPI_ISL_2992559, EPI_ISL_2992560, EPI_ISL_2992561, EPI_ISL_2992562, EPI_ISL_2992563, EPI_ISL_2992564, EPI_ISL_2992565, EPI_ISL_2992566, EPI_ISL_2992567, EPI_ISL_2992568, EPI_ISL_2992569, EPI_ISL_2992570, EPI_ISL_2992571, EPI_ISL_2992572, EPI_ISL_2992573, EPI_ISL_2992574, EPI_ISL_2992575, EPI_ISL_2992576, EPI_ISL_2992577, EPI_ISL_2992578, EPI_ISL_2992579, EPI_ISL_2992580, EPI_ISL_2992581, EPI_ISL_2992582, EPI_ISL_2992583, EPI_ISL_2992584, EPI_ISL_2992585, EPI_ISL_2992586, EPI_ISL_2992587, EPI_ISL_2992588, EPI_ISL_2992589, EPI_ISL_2992590, EPI_ISL_2992591, EPI_ISL_2992592, EPI_ISL_2992593, EPI_ISL_2992594, EPI_ISL_2992595, EPI_ISL_2992596, EPI_ISL_2992597, EPI_ISL_2992598, EPI_ISL_2992599, EPI_ISL_2992600 | | | |
| see above | Department of Bacteria, Parasites and Fungi, Statens Serum Institut, Copenhagen, Denmark | Statens Serum Institut Bioinformatics and Microbial Genomics | Danish Covid-19 Genome Consortium |
| EPI_ISL_2992601 | Department of Clinical Microbiology and Center for Genomic Medicine, Rigshospitalet, Copenhagen, Denmark | Statens Serum Institut Bioinformatics and Microbial Genomics | Danish Covid-19 Genome Consortium |
| EPI_ISL_2992602, EPI_ISL_2992603, EPI_ISL_2992604, EPI_ISL_2992605, EPI_ISL_2992606, EPI_ISL_2992607, EPI_ISL_2992608, EPI_ISL_2992609, EPI_ISL_2992610, EPI_ISL_2992611, EPI_ISL_2992612, EPI_ISL_2992613, EPI_ISL_2992614, EPI_ISL_2992615, EPI_ISL_2992616, EPI_ISL_2992617, EPI_ISL_2992618, EPI_ISL_2992619, EPI_ISL_2992620, EPI_ISL_2992621, EPI_ISL_2992623, EPI_ISL_2992624, EPI_ISL_2992625, EPI_ISL_2992626, EPI_ISL_2992627, EPI_ISL_2992628, EPI_ISL_2992629, EPI_ISL_2992630, EPI_ISL_2992631, EPI_ISL_2992632, EPI_ISL_2992633, EPI_ISL_2992634, EPI_ISL_2992635, EPI_ISL_2992636, EPI_ISL_2992637, EPI_ISL_2992638, EPI_ISL_2992639, EPI_ISL_2992640, EPI_ISL_2992641, EPI_ISL_2992642, EPI_ISL_2992643, EPI_ISL_2992644, EPI_ISL_2992645, EPI_ISL_2992646, EPI_ISL_2992647, EPI_ISL_2992648, EPI_ISL_2992649, EPI_ISL_2992650, EPI_ISL_2992651, EPI_ISL_2992652, EPI_ISL_2992653 | | | |
| see above | Department of Bacteria, Parasites and Fungi, Statens Serum Institut, Copenhagen, Denmark | Statens Serum Institut Bioinformatics and Microbial Genomics | Danish Covid-19 Genome Consortium |
| EPI_ISL_2992654 | Department of Clinical Microbiology, Odense University Hospital, Odense, Denmark | Statens Serum Institut Bioinformatics and Microbial Genomics | Danish Covid-19 Genome Consortium |
| EPI_ISL_2992655, EPI_ISL_2992656, EPI_ISL_2992657, EPI_ISL_2992658 | Department of Bacteria, Parasites and Fungi, Statens Serum Institut, Copenhagen, Denmark | Statens Serum Institut Bioinformatics and Microbial Genomics | Danish Covid-19 Genome Consortium |
| EPI_ISL_2992659 | Department of Clinical Microbiology, Odense University Hospital, Odense, Denmark | Statens Serum Institut Bioinformatics and Microbial Genomics | Danish Covid-19 Genome Consortium |
| EPI_ISL_2992660, EPI_ISL_2992661, EPI_ISL_2992662, EPI_ISL_2992663, EPI_ISL_2992664, EPI_ISL_2992665, EPI_ISL_2992666, EPI_ISL_2992667, EPI_ISL_2992668 | Department of Bacteria, Parasites and Fungi, Statens Serum Institut, Copenhagen, Denmark | Statens Serum Institut Bioinformatics and Microbial Genomics | Danish Covid-19 Genome Consortium |
| EPI_ISL_2992669 | Department of Clinical Microbiology and Center for Genomic Medicine, Rigshospitalet, Copenhagen, Denmark | Statens Serum Institut Bioinformatics and Microbial Genomics | Danish Covid-19 Genome Consortium |
| EPI_ISL_2992670, EPI_ISL_2992671, EPI_ISL_2992672, EPI_ISL_2992673, EPI_ISL_2992674, EPI_ISL_2992675, EPI_ISL_2992676, EPI_ISL_2992677, EPI_ISL_2992678, EPI_ISL_2992679, EPI_ISL_2992680, EPI_ISL_2992682, EPI_ISL_2992683, EPI_ISL_2992684, EPI_ISL_2992685, EPI_ISL_2992686, EPI_ISL_2992687, EPI_ISL_2992688, EPI_ISL_2992689, EPI_ISL_2992690 | | | |
| see above | Department of Bacteria, Parasites and Fungi, Statens Serum Institut, Copenhagen, Denmark | Statens Serum Institut Bioinformatics and Microbial Genomics | Danish Covid-19 Genome Consortium |

| | | | |
|---|---|---|---|
| EPI_ISL_2992691 | Molekylær Medicinsk Afdeling, Aarhus University Hospital, Aarhus, Denmark | Statens Serum Institut Bioinformatics and Microbial Genomics | Danish Covid-19 Genome Consortium |
| EPI_ISL_2992692, EPI_ISL_2992693, EPI_ISL_2992694, EPI_ISL_2992695, EPI_ISL_2992696, EPI_ISL_2992697, EPI_ISL_2992698, EPI_ISL_2992699, EPI_ISL_2992700, EPI_ISL_2992701 | Department of Bacteria, Parasites and Fungi, Statens Serum Institut, Copenhagen, Denmark | Statens Serum Institut Bioinformatics and Microbial Genomics | Danish Covid-19 Genome Consortium |
| EPI_ISL_2992702 | Molekylær Medicinsk Afdeling, Aarhus University Hospital, Aarhus, Denmark | Statens Serum Institut Bioinformatics and Microbial Genomics | Danish Covid-19 Genome Consortium |

EPI_ISL_2992703, EPI_ISL_2992704, EPI_ISL_2992705, EPI_ISL_2992706, EPI_ISL_2992708, EPI_ISL_2992709, EPI_ISL_2992710, EPI_ISL_2992711, EPI_ISL_2992712, EPI_ISL_2992713, EPI_ISL_2992714, EPI_ISL_2992715, EPI_ISL_2992716, EPI_ISL_2992717, EPI_ISL_2992718, EPI_ISL_2992719, EPI_ISL_2992720, EPI_ISL_2992721, EPI_ISL_2992722, EPI_ISL_2992723, EPI_ISL_2992724, EPI_ISL_2992725, EPI_ISL_2992726, EPI_ISL_2992727, EPI_ISL_2992728, EPI_ISL_2992729, EPI_ISL_2992730, EPI_ISL_2992731, EPI_ISL_2992732, EPI_ISL_2992733, EPI_ISL_2992734, EPI_ISL_2992735, EPI_ISL_2992736, EPI_ISL_2992737, EPI_ISL_2992738, EPI_ISL_2992741

| | | | |
|---|---|---|---|
| see above | Department of Bacteria, Parasites and Fungi, Statens Serum Institut, Copenhagen, Denmark | Statens Serum Institut Bioinformatics and Microbial Genomics | Danish Covid-19 Genome Consortium |
| EPI_ISL_2992742 | Department of Clinical Microbiology, Odense University Hospital, Odense, Denmark | Statens Serum Institut Bioinformatics and Microbial Genomics | Danish Covid-19 Genome Consortium |

EPI_ISL_2992743, EPI_ISL_2992744, EPI_ISL_2992745, EPI_ISL_2992746, EPI_ISL_2992747, EPI_ISL_2992748, EPI_ISL_2992749, EPI_ISL_2992750, EPI_ISL_2992751, EPI_ISL_2992752, EPI_ISL_2992753, EPI_ISL_2992754, EPI_ISL_2992755, EPI_ISL_2992756, EPI_ISL_2992757, EPI_ISL_2992758, EPI_ISL_2992759, EPI_ISL_2992760, EPI_ISL_2992761, EPI_ISL_2992762, EPI_ISL_2992763, EPI_ISL_2992764, EPI_ISL_2992765, EPI_ISL_2992766, EPI_ISL_2992767, EPI_ISL_2992768, EPI_ISL_2992769, EPI_ISL_2992770, EPI_ISL_2992771, EPI_ISL_2992772, EPI_ISL_2992773, EPI_ISL_2992774, EPI_ISL_2992775, EPI_ISL_2992777, EPI_ISL_2992778

| | | | |
|---|---|---|---|
| see above | Department of Bacteria, Parasites and Fungi, Statens Serum Institut, Copenhagen, Denmark | Statens Serum Institut Bioinformatics and Microbial Genomics | Danish Covid-19 Genome Consortium |
| EPI_ISL_2992779 | Molekylær Medicinsk Afdeling, Aarhus University Hospital, Aarhus, Denmark | Statens Serum Institut Bioinformatics and Microbial Genomics | Danish Covid-19 Genome Consortium |

EPI_ISL_2992780, EPI_ISL_2992781, EPI_ISL_2992782, EPI_ISL_2992783, EPI_ISL_2992784, EPI_ISL_2992785, EPI_ISL_2992786, EPI_ISL_2992787, EPI_ISL_2992788, EPI_ISL_2992789, EPI_ISL_2992790, EPI_ISL_2992791, EPI_ISL_2992792, EPI_ISL_2992793, EPI_ISL_2992794, EPI_ISL_2992795, EPI_ISL_2992796, EPI_ISL_2992797, EPI_ISL_2992798, EPI_ISL_2992799, EPI_ISL_2992800, EPI_ISL_2992801, EPI_ISL_2992802, EPI_ISL_2992803, EPI_ISL_2992804, EPI_ISL_2992805, EPI_ISL_2992806, EPI_ISL_2992807, EPI_ISL_2992808, EPI_ISL_2992809, EPI_ISL_2992810, EPI_ISL_2992811, EPI_ISL_2992812, EPI_ISL_2992813, EPI_ISL_2992814, EPI_ISL_2992815, EPI_ISL_2992816, EPI_ISL_2992817, EPI_ISL_2992818, EPI_ISL_2992819, EPI_ISL_2992820, EPI_ISL_2992821, EPI_ISL_2992822, EPI_ISL_2992823, EPI_ISL_2992824

| | | | |
|---|---|---|---|
| see above | Department of Bacteria, Parasites and Fungi, Statens Serum Institut, Copenhagen, Denmark | Statens Serum Institut Bioinformatics and Microbial Genomics | Danish Covid-19 Genome Consortium |
| EPI_ISL_2992825 | Molekylær Medicinsk Afdeling, Aarhus University Hospital, Aarhus, Denmark | Statens Serum Institut Bioinformatics and Microbial Genomics | Danish Covid-19 Genome Consortium |

EPI_ISL_2992826, EPI_ISL_2992827, EPI_ISL_2992828, EPI_ISL_2992829, EPI_ISL_2992830, EPI_ISL_2992831, EPI_ISL_2992832, EPI_ISL_2992833, EPI_ISL_2992834, EPI_ISL_2992835, EPI_ISL_2992836, EPI_ISL_2992837, EPI_ISL_2992838, EPI_ISL_2992840, EPI_ISL_2992841, EPI_ISL_2992842, EPI_ISL_2992843, EPI_ISL_2992844, EPI_ISL_2992845, EPI_ISL_2992846, EPI_ISL_2992847, EPI_ISL_2992848, EPI_ISL_2992849, EPI_ISL_2992850, EPI_ISL_2992851, EPI_ISL_2992852, EPI_ISL_2992853, EPI_ISL_2992854, EPI_ISL_2992855, EPI_ISL_2992856, EPI_ISL_2992857, EPI_ISL_2992858, EPI_ISL_2992859, EPI_ISL_2992860, EPI_ISL_2992861, EPI_ISL_2992862, EPI_ISL_2992863, EPI_ISL_2992864, EPI_ISL_2992865, EPI_ISL_2992866, EPI_ISL_2992867, EPI_ISL_2992868, EPI_ISL_2992869, EPI_ISL_2992870, EPI_ISL_2992871, EPI_ISL_2992872, EPI_ISL_2992873, EPI_ISL_2992874, EPI_ISL_2992875, EPI_ISL_2992876, EPI_ISL_2992877, EPI_ISL_2992878, EPI_ISL_2992879, EPI_ISL_2992880, EPI_ISL_2992881, EPI_ISL_2992882, EPI_ISL_2992883, EPI_ISL_2992884, EPI_ISL_2992885, EPI_ISL_2992886, EPI_ISL_2992887, EPI_ISL_2992888, EPI_ISL_2992889, EPI_ISL_2992890, EPI_ISL_2992891, EPI_ISL_2992892, EPI_ISL_2992893, EPI_ISL_2992894, EPI_ISL_2992895, EPI_ISL_2992896, EPI_ISL_2992897, EPI_ISL_2992898, EPI_ISL_2992899, EPI_ISL_2992900, EPI_ISL_2992901, EPI_ISL_2992902, EPI_ISL_2992903, EPI_ISL_2992904, EPI_ISL_2992905, EPI_ISL_2992906, EPI_ISL_2992907, EPI_ISL_2992908, EPI_ISL_2992909, EPI_ISL_2992910, EPI_ISL_2992911, EPI_ISL_2992912, EPI_ISL_2992913, EPI_ISL_2992914, EPI_ISL_2992915, EPI_ISL_2992916, EPI_ISL_2992917, EPI_ISL_2992919, EPI_ISL_2992920, EPI_ISL_2992921, EPI_ISL_2992922, EPI_ISL_2992923, EPI_ISL_2992924, EPI_ISL_2992925, EPI_ISL_2992926, EPI_ISL_2992927, EPI_ISL_2992928, EPI_ISL_2992929, EPI_ISL_2992930, EPI_ISL_2992931, EPI_ISL_2992932, EPI_ISL_2992933, EPI_ISL_2992934, EPI_ISL_2992935, EPI_ISL_2992936, EPI_ISL_2992937, EPI_ISL_2992938, EPI_ISL_2992939, EPI_ISL_2992940, EPI_ISL_2992941, EPI_ISL_2992942, EPI_ISL_2992943, EPI_ISL_2992944, EPI_ISL_2992946, EPI_ISL_2992947, EPI_ISL_2992948, EPI_ISL_2992949, EPI_ISL_2992950, EPI_ISL_2992951, EPI_ISL_2992952, EPI_ISL_2992953, EPI_ISL_2992954, EPI_ISL_2992955, EPI_ISL_2992956, EPI_ISL_2992957, EPI_ISL_2992958, EPI_ISL_2992959, EPI_ISL_2992960, EPI_ISL_2992961, EPI_ISL_2992962, EPI_ISL_2992963, EPI_ISL_2992964, EPI_ISL_2992965, EPI_ISL_2992966, EPI_ISL_2992967, EPI_ISL_2992968, EPI_ISL_2992969, EPI_ISL_2992970, EPI_ISL_2992971, EPI_ISL_2992972, EPI_ISL_2992973, EPI_ISL_2992974

| | | | |
|---|---|---|---|
| see above | Department of Bacteria, Parasites and Fungi, Statens Serum Institut, Copenhagen, Denmark | Statens Serum Institut Bioinformatics and Microbial Genomics | Danish Covid-19 Genome Consortium |
| EPI_ISL_2992975 | Department of Clinical Microbiology, Odense University Hospital, Odense, Denmark | Statens Serum Institut Bioinformatics and Microbial Genomics | Danish Covid-19 Genome Consortium |

EPI_ISL_2992976, EPI_ISL_2992977, EPI_ISL_2992978, EPI_ISL_2992979, EPI_ISL_2992980, EPI_ISL_2992981, EPI_ISL_2992982, EPI_ISL_2992983, EPI_ISL_2992984, EPI_ISL_2992985, EPI_ISL_2992986, EPI_ISL_2992987, EPI_ISL_2992988, EPI_ISL_2992989, EPI_ISL_2992990, EPI_ISL_2992991, EPI_ISL_2992992, EPI_ISL_2992993, EPI_ISL_2992994, EPI_ISL_2992995

| | | | |
|---|---|---|---|
| see above | Department of Bacteria, Parasites and Fungi, Statens Serum Institut, Copenhagen, Denmark | Statens Serum Institut Bioinformatics and Microbial Genomics | Danish Covid-19 Genome Consortium |
| EPI_ISL_2992996 | Department of Clinical Microbiology, Odense University Hospital, Odense, Denmark | Statens Serum Institut Bioinformatics and Microbial Genomics | Danish Covid-19 Genome Consortium |

EPI_ISL_2992997, EPI_ISL_2992998, EPI_ISL_2992999, EPI_ISL_2993000, EPI_ISL_2993001, EPI_ISL_2993002, EPI_ISL_2993003, EPI_ISL_2993004, EPI_ISL_2993005, EPI_ISL_2993006, EPI_ISL_2993007, EPI_ISL_2993008, EPI_ISL_2993009, EPI_ISL_2993010, EPI_ISL_2993011, EPI_ISL_2993012, EPI_ISL_2993013

| | | | |
|---|---|---|---|
| see above | Department of Bacteria, Parasites and Fungi, Statens Serum Institut, Copenhagen, Denmark | Statens Serum Institut Bioinformatics and Microbial Genomics | Danish Covid-19 Genome Consortium |
| EPI_ISL_2993014 | Molekylær Medicinsk Afdeling, Aarhus University Hospital, Aarhus, Denmark | Statens Serum Institut Bioinformatics and Microbial Genomics | Danish Covid-19 Genome Consortium |

EPI_ISL_2993015, EPI_ISL_2993016, EPI_ISL_2993018, EPI_ISL_2993019, EPI_ISL_2993020, EPI_ISL_2993021, EPI_ISL_2993022, EPI_ISL_2993023, EPI_ISL_2993024, EPI_ISL_2993025, EPI_ISL_2993026, EPI_ISL_2993028, EPI_ISL_2993029, EPI_ISL_2993030, EPI_ISL_2993031, EPI_ISL_2993032, EPI_ISL_2993033, EPI_ISL_2993034, EPI_ISL_2993035, EPI_ISL_2993036, EPI_ISL_2993037, EPI_ISL_2993038, EPI_ISL_2993039, EPI_ISL_2993040, EPI_ISL_2993041, EPI_ISL_2993042, EPI_ISL_2993043, EPI_ISL_2993044, EPI_ISL_2993045, EPI_ISL_2993046, EPI_ISL_2993047, EPI_ISL_2993048, EPI_ISL_2993049, EPI_ISL_2993050, EPI_ISL_2993051, EPI_ISL_2993052, EPI_ISL_2993053, EPI_ISL_2993054, EPI_ISL_2993055, EPI_ISL_2993056, EPI_ISL_2993057, EPI_ISL_2993058, EPI_ISL_2993059, EPI_ISL_2993060, EPI_ISL_2993061, EPI_ISL_2993062, EPI_ISL_2993063, EPI_ISL_2993064, EPI_ISL_2993066, EPI_ISL_2993067, EPI_ISL_2993068, EPI_ISL_2993069, EPI_ISL_2993070, EPI_ISL_2993071, EPI_ISL_2993072, EPI_ISL_2993073, EPI_ISL_2993074, EPI_ISL_2993075, EPI_ISL_2993076, EPI_ISL_2993077, EPI_ISL_2993078, EPI_ISL_2993079, EPI_ISL_2993080, EPI_ISL_2993081, EPI_ISL_2993082, EPI_ISL_2993083, EPI_ISL_2993084, EPI_ISL_2993085, EPI_ISL_2993086, EPI_ISL_2993088, EPI_ISL_2993089, EPI_ISL_2993090, EPI_ISL_2993091, EPI_ISL_2993092, EPI_ISL_2993093, EPI_ISL_2993094, EPI_ISL_2993095, EPI_ISL_2993096, EPI_ISL_2993097, EPI_ISL_2993098, EPI_ISL_2993099, EPI_ISL_2993100, EPI_ISL_2993101, EPI_ISL_2993102, EPI_ISL_2993103, EPI_ISL_2993104, EPI_ISL_2993105, EPI_ISL_2993106, EPI_ISL_2993107

| | | | |
|---|---|---|---|
| see above | Department of Bacteria, Parasites and Fungi, Statens Serum Institut, Copenhagen, Denmark | Statens Serum Institut Bioinformatics and Microbial Genomics | Danish Covid-19 Genome Consortium |
| EPI_ISL_2993108 | Molekylær Medicinsk Afdeling, Aarhus University Hospital, Aarhus, Denmark | Statens Serum Institut Bioinformatics and Microbial Genomics | Danish Covid-19 Genome Consortium |
| EPI_ISL_2993109 | Department of Clinical Microbiology, Odense University Hospital, Odense, Denmark | Statens Serum Institut Bioinformatics and Microbial Genomics | Danish Covid-19 Genome Consortium |

EPI_ISL_2993110, EPI_ISL_2993111, EPI_ISL_2993112, EPI_ISL_2993113, EPI_ISL_2993114, EPI_ISL_2993115, EPI_ISL_2993116, EPI_ISL_2993117, EPI_ISL_2993118, EPI_ISL_2993119, EPI_ISL_2993120, EPI_ISL_2993122, EPI_ISL_2993123, EPI_ISL_2993124, EPI_ISL_2993125, EPI_ISL_2993126, EPI_ISL_2993127, EPI_ISL_2993128, EPI_ISL_2993130, EPI_ISL_2993131, EPI_ISL_2993132, EPI_ISL_2993133, EPI_ISL_2993134, EPI_ISL_2993135, EPI_ISL_2993136, EPI_ISL_2993137, EPI_ISL_2993138, EPI_ISL_2993139, EPI_ISL_2993140, EPI_ISL_2993141, EPI_ISL_2993142, EPI_ISL_2993143, EPI_ISL_2993144, EPI_ISL_2993145, EPI_ISL_2993146, EPI_ISL_2993147

| | | | |
|---|---|---|---|
| see above | Department of Bacteria, Parasites and Fungi, Statens Serum Institut, Copenhagen, Denmark | Statens Serum Institut Bioinformatics and Microbial Genomics | Danish Covid-19 Genome Consortium |
| EPI_ISL_2993148 | Department of Clinical Microbiology, Odense University Hospital, Odense, Denmark | Statens Serum Institut Bioinformatics and Microbial Genomics | Danish Covid-19 Genome Consortium |
| EPI_ISL_2993149, EPI_ISL_2993150, EPI_ISL_2993151, EPI_ISL_2993152, EPI_ISL_2993153 | Department of Bacteria, Parasites and Fungi, Statens Serum Institut, Copenhagen, Denmark | Statens Serum Institut Bioinformatics and Microbial Genomics | Danish Covid-19 Genome Consortium |
| EPI_ISL_2993154 | Department of Clinical Microbiology, Odense University Hospital, Odense, Denmark | Statens Serum Institut Bioinformatics and Microbial Genomics | Danish Covid-19 Genome Consortium |

EPI_ISL_2993156, EPI_ISL_2993157, EPI_ISL_2993158, EPI_ISL_2993159, EPI_ISL_2993160, EPI_ISL_2993161, EPI_ISL_2993162, EPI_ISL_2993163, EPI_ISL_2993164, EPI_ISL_2993165, EPI_ISL_2993166, EPI_ISL_2993167, EPI_ISL_2993168, EPI_ISL_2993169, EPI_ISL_2993170, EPI_ISL_2993171, EPI_ISL_2993172, EPI_ISL_2993173, EPI_ISL_2993174, EPI_ISL_2993175, EPI_ISL_2993176, EPI_ISL_2993177, EPI_ISL_2993178, EPI_ISL_2993179, EPI_ISL_2993180, EPI_ISL_2993181, EPI_ISL_2993182, EPI_ISL_2993183, EPI_ISL_2993184, EPI_ISL_2993185, EPI_ISL_2993186, EPI_ISL_2993187

| | | | |
|---|---|---|---|
| see above | Department of Bacteria, Parasites and Fungi, Statens Serum Institut, Copenhagen, Denmark | Statens Serum Institut Bioinformatics and Microbial Genomics | Danish Covid-19 Genome Consortium |
| EPI_ISL_2993188 | Molekylær Medicinsk Afdeling, Aarhus University Hospital, Aarhus, Denmark | Statens Serum Institut Bioinformatics and Microbial Genomics | Danish Covid-19 Genome Consortium |

EPI_ISL_2993190, EPI_ISL_2993191, EPI_ISL_2993192, EPI_ISL_2993193, EPI_ISL_2993194, EPI_ISL_2993195, EPI_ISL_2993196, EPI_ISL_2993198, EPI_ISL_2993199, EPI_ISL_2993200, EPI_ISL_2993201, EPI_ISL_2993202, EPI_ISL_2993203, EPI_ISL_2993204, EPI_ISL_2993205, EPI_ISL_2993206, EPI_ISL_2993207, EPI_ISL_2993208, EPI_ISL_2993209, EPI_ISL_2993210, EPI_ISL_2993211, EPI_ISL_2993212, EPI_ISL_2993213, EPI_ISL_2993214, EPI_ISL_2993215, EPI_ISL_2993216, EPI_ISL_2993217

| | | | |
|---|---|---|---|
| see above | Department of Bacteria, Parasites and Fungi, Statens Serum Institut, Copenhagen, Denmark | Statens Serum Institut Bioinformatics and Microbial Genomics | Danish Covid-19 Genome Consortium |
| EPI_ISL_2993218 | Molekylær Medicinsk Afdeling, Aarhus University Hospital, Aarhus, Denmark | Statens Serum Institut Bioinformatics and Microbial Genomics | Danish Covid-19 Genome Consortium |
| EPI_ISL_2993219, EPI_ISL_2993220, EPI_ISL_2993221, EPI_ISL_2993222, | Department of Bacteria, Parasites and Fungi, Statens Serum Institut, Copenhagen, Denmark | Statens Serum Institut Bioinformatics and Microbial Genomics | Danish Covid-19 Genome Consortium |

| | | | |
|---|---|---|---|
| EPI_ISL_2993224, EPI_ISL_2993225 | | | |
| EPI_ISL_2993226 | Department of Clinical Microbiology, Odense University Hospital, Odense, Denmark | Statens Serum Institut Bioinformatics and Microbial Genomics | Danish Covid-19 Genome Consortium |
| EPI_ISL_2993228, EPI_ISL_2993229, EPI_ISL_2993230, EPI_ISL_2993231 | Department of Bacteria, Parasites and Fungi, Statens Serum Institut, Copenhagen, Denmark | Statens Serum Institut Bioinformatics and Microbial Genomics | Danish Covid-19 Genome Consortium |
| EPI_ISL_2993232 | Department of Clinical Microbiology and Center for Genomic Medicine, Rigshospitalet, Copenhagen, Denmark | Statens Serum Institut Bioinformatics and Microbial Genomics | Danish Covid-19 Genome Consortium |
| EPI_ISL_2993233, EPI_ISL_2993234, EPI_ISL_2993235, EPI_ISL_2993236, EPI_ISL_2993237, EPI_ISL_2993238, EPI_ISL_2993239, EPI_ISL_2993240, EPI_ISL_2993241, EPI_ISL_2993242, EPI_ISL_2993243, EPI_ISL_2993244, EPI_ISL_2993245, EPI_ISL_2993246, EPI_ISL_2993247, EPI_ISL_2993248, EPI_ISL_2993249, EPI_ISL_2993250, EPI_ISL_2993251, EPI_ISL_2993252, EPI_ISL_2993253, EPI_ISL_2993254, EPI_ISL_2993255, EPI_ISL_2993256, EPI_ISL_2993257, EPI_ISL_2993258, EPI_ISL_2993259, EPI_ISL_2993260, EPI_ISL_2993261, EPI_ISL_2993262, EPI_ISL_2993263, EPI_ISL_2993264, EPI_ISL_2993265, EPI_ISL_2993266, EPI_ISL_2993267, EPI_ISL_2993268, EPI_ISL_2993269, EPI_ISL_2993270, EPI_ISL_2993271, EPI_ISL_2993272, EPI_ISL_2993273, EPI_ISL_2993274, EPI_ISL_2993275, EPI_ISL_2993276, EPI_ISL_2993277, EPI_ISL_2993278, EPI_ISL_2993279, EPI_ISL_2993281, EPI_ISL_2993282, EPI_ISL_2993283, EPI_ISL_2993285, EPI_ISL_2993286, EPI_ISL_2993287, EPI_ISL_2993288, EPI_ISL_2993289, EPI_ISL_2993290 | | | |
| see above | Department of Bacteria, Parasites and Fungi, Statens Serum Institut, Copenhagen, Denmark | Statens Serum Institut Bioinformatics and Microbial Genomics | Danish Covid-19 Genome Consortium |
| EPI_ISL_2993291 | Department of Clinical Microbiology, Odense University Hospital, Odense, Denmark | Statens Serum Institut Bioinformatics and Microbial Genomics | Danish Covid-19 Genome Consortium |
| EPI_ISL_2993292, EPI_ISL_2993293, EPI_ISL_2993294, EPI_ISL_2993295, EPI_ISL_2993296, EPI_ISL_2993297, EPI_ISL_2993298, EPI_ISL_2993299, EPI_ISL_2993300, EPI_ISL_2993301, EPI_ISL_2993302, EPI_ISL_2993303, EPI_ISL_2993304, EPI_ISL_2993305, EPI_ISL_2993306, EPI_ISL_2993307, EPI_ISL_2993308, EPI_ISL_2993309, EPI_ISL_2993310, EPI_ISL_2993311, EPI_ISL_2993312, EPI_ISL_2993313, EPI_ISL_2993314, EPI_ISL_2993315, EPI_ISL_2993316, EPI_ISL_2993317, EPI_ISL_2993319, EPI_ISL_2993321, EPI_ISL_2993322, EPI_ISL_2993323, EPI_ISL_2993324, EPI_ISL_2993325, EPI_ISL_2993326, EPI_ISL_2993327, EPI_ISL_2993328, EPI_ISL_2993329, EPI_ISL_2993330, EPI_ISL_2993331, EPI_ISL_2993332, EPI_ISL_2993333, EPI_ISL_2993334, EPI_ISL_2993335, EPI_ISL_2993336, EPI_ISL_2993337, EPI_ISL_2993338 | | | |
| see above | Department of Bacteria, Parasites and Fungi, Statens Serum Institut, Copenhagen, Denmark | Statens Serum Institut Bioinformatics and Microbial Genomics | Danish Covid-19 Genome Consortium |
| EPI_ISL_2993340, EPI_ISL_2993341, EPI_ISL_2993432 | Clinical Microbiology Laboratory, Faculty of Medicine, Universitas Indonesia | Clinical Microbiology Laboratory, Faculty of Medicine, Universitas Indonesia | Ibrahim F, Natasha A, Yasmon A, Fitrhiyah, Saharman YR, Karuniawati A, Sudarmono P. |
| EPI_ISL_2993496 | Pathology, UAB | Pathology, UAB | Moates,D., Hendrickson,R.C., Hernandez,J., Lefkowitz,E.J., Leal,S.M. |
| EPI_ISL_2993636, EPI_ISL_2993637, EPI_ISL_2993638, EPI_ISL_2993639, EPI_ISL_2993640, EPI_ISL_2993641, EPI_ISL_2993642, EPI_ISL_2993643, EPI_ISL_2993644, EPI_ISL_2993645, EPI_ISL_2993646, EPI_ISL_2993647, EPI_ISL_2993648, EPI_ISL_2993649, EPI_ISL_2993652, EPI_ISL_2993653, EPI_ISL_2993655, EPI_ISL_2993656, EPI_ISL_2993657 | | | |
| see above | Medical Laboratories Duesseldorf | Center of Medical Microbiology, Virology, and Hospital Hygiene, University of Duesseldorf | Maximilian Damagnez;Alexander Dilthey;Angelika Helmer;Torsten Houwaart;Lisanna Hülse;Christian Lange;Malte Kohns Vasconcelos;Nadine Lübke;Jessica Nicolai;Klaus Pfeffer;Daniel Strelow;Jörg Timm;Andreas Walker;Tobias Wienemann |
| EPI_ISL_2993690, EPI_ISL_2993692, EPI_ISL_2993693, EPI_ISL_2993695, EPI_ISL_2993698, EPI_ISL_2993699, EPI_ISL_2993701, EPI_ISL_2993705, EPI_ISL_2993707, EPI_ISL_2993710, EPI_ISL_2993711, EPI_ISL_2993713, EPI_ISL_2993718, EPI_ISL_2993719 | | | |
| see above | Laboratorio CQRC | CQRC_QUALITY CONTROL CHEMICAL BIOLOGICAL RISK_AOOR Villa Sofia Cervello Palermo | Di Gaudio F., Contino F., Brunacci G. |
| EPI_ISL_2993728, EPI_ISL_2993797 | Pathology, UAB | Pathology, UAB | Moates,D., Hendrickson,R.C., Hernandez,J., Lefkowitz,E.J., Leal,S.M. |
| EPI_ISL_2993798 | Lighthouse Lab in Glasgow | Wellcome Sanger Institute for the COVID-19 Genomics UK (COG-UK) Consortium | Harper VanSteenhouse, Yumi Kasai, David Gray, Carol Clugston, Anna Dominiczak and Alex Alderton, Roberto Amato, Jeffrey Barrett, Sonia Goncalves, Ewan Harrison, David K. Jackson, Ian Johnston, Dominic Kwiatkowski, Cordelia Langford, John Sillitoe on behalf of the Wellcome Sanger Institute COVID-19 Surveillance Team |
| EPI_ISL_2993799 | Lighthouse Lab in Alderley Park | Wellcome Sanger Institute for the COVID-19 Genomics UK (COG-UK) Consortium | Jacquelyn Wynn, Mairead Hyland, The Lighthouse Lab in Alderley Park and Alex Alderton, Roberto Amato, Jeffrey Barrett, Sonia Goncalves, Ewan Harrison, David K. Jackson, Ian Johnston, Dominic Kwiatkowski, Cordelia Langford, John Sillitoe on behalf of the Wellcome Sanger Institute COVID-19 Surveillance Team |
| EPI_ISL_2993800, EPI_ISL_2993801 | Lighthouse Laboratory Plymouth | Wellcome Sanger Institute for the COVID-19 Genomics UK (COG-UK) Consortium | Lighthouse Laboratory Plymouth and Alex Alderton, Roberto Amato, Jeffrey Barrett, Sonia Goncalves, Ewan Harrison, David K. Jackson, Ian Johnston, Dominic Kwiatkowski, Cordelia Langford, John Sillitoe on behalf of the Wellcome Sanger Institute COVID-19 Surveillance Team |
| EPI_ISL_2993802 | Lighthouse Lab in Alderley Park | Wellcome Sanger Institute for the COVID-19 Genomics UK (COG-UK) Consortium | Jacquelyn Wynn, Mairead Hyland, The Lighthouse Lab in Alderley Park and Alex Alderton, Roberto Amato, Jeffrey Barrett, Sonia Goncalves, Ewan Harrison, David K. Jackson, Ian Johnston, Dominic Kwiatkowski, Cordelia Langford, John Sillitoe on behalf of the Wellcome Sanger Institute COVID-19 Surveillance Team |
| EPI_ISL_2993803 | Health Services Laboratories | Wellcome Sanger Institute for the COVID-19 Genomics UK (COG-UK) Consortium | Health Services Laboratories and Alex Alderton, Roberto Amato, Jeffrey Barrett, Sonia Goncalves, Ewan Harrison, David K. Jackson, Ian Johnston, Dominic Kwiatkowski, Cordelia Langford, John Sillitoe on behalf of the Wellcome Sanger Institute COVID-19 Surveillance Team |
| EPI_ISL_2993804, EPI_ISL_2993805 | Lighthouse Laboratory Plymouth | Wellcome Sanger Institute for the COVID-19 Genomics UK (COG-UK) Consortium | Lighthouse Laboratory Plymouth and Alex Alderton, Roberto Amato, Jeffrey Barrett, Sonia Goncalves, Ewan Harrison, David K. Jackson, Ian Johnston, Dominic Kwiatkowski, Cordelia Langford, John Sillitoe on behalf of the Wellcome Sanger Institute COVID-19 Surveillance Team |
| EPI_ISL_2993806 | Lighthouse Lab in Glasgow | Wellcome Sanger Institute for the COVID-19 Genomics UK (COG-UK) Consortium | Harper VanSteenhouse, Yumi Kasai, David Gray, Carol Clugston, Anna Dominiczak and Alex Alderton, Roberto Amato, Jeffrey Barrett, Sonia Goncalves, Ewan Harrison, David K. Jackson, Ian Johnston, Dominic Kwiatkowski, Cordelia Langford, John Sillitoe on behalf of the Wellcome Sanger Institute COVID-19 Surveillance Team |
| EPI_ISL_2993807 | Lighthouse Lab in Alderley Park | Wellcome Sanger Institute for the COVID-19 Genomics UK (COG-UK) Consortium | Jacquelyn Wynn, Mairead Hyland, The Lighthouse Lab in Alderley Park and Alex Alderton, Roberto Amato, Jeffrey Barrett, Sonia Goncalves, Ewan Harrison, David K. Jackson, Ian Johnston, Dominic Kwiatkowski, Cordelia Langford, John Sillitoe on behalf of the Wellcome Sanger Institute COVID-19 Surveillance Team |
| EPI_ISL_2993808 | Lighthouse Laboratory Plymouth | Wellcome Sanger Institute for the COVID-19 Genomics UK (COG-UK) Consortium | Lighthouse Laboratory Plymouth and Alex Alderton, Roberto Amato, Jeffrey Barrett, Sonia Goncalves, Ewan Harrison, David K. Jackson, Ian Johnston, Dominic Kwiatkowski, Cordelia Langford, John Sillitoe on behalf of the Wellcome Sanger Institute COVID-19 Surveillance Team |
| EPI_ISL_2993809 | Lighthouse Lab in Alderley Park | Wellcome Sanger Institute for the COVID-19 Genomics UK (COG-UK) Consortium | Jacquelyn Wynn, Mairead Hyland, The Lighthouse Lab in Alderley Park and Alex Alderton, Roberto Amato, Jeffrey Barrett, Sonia Goncalves, Ewan Harrison, David K. Jackson, Ian Johnston, Dominic Kwiatkowski, Cordelia Langford, John Sillitoe on behalf of the Wellcome Sanger Institute COVID-19 Surveillance Team |
| EPI_ISL_2993810 | Lighthouse Laboratory Plymouth | Wellcome Sanger Institute for the COVID-19 Genomics UK (COG-UK) Consortium | Lighthouse Laboratory Plymouth and Alex Alderton, Roberto Amato, Jeffrey Barrett, Sonia Goncalves, Ewan Harrison, David K. Jackson, Ian Johnston, Dominic Kwiatkowski, Cordelia Langford, John Sillitoe on behalf of the Wellcome Sanger Institute COVID-19 Surveillance Team |
| EPI_ISL_2993811 | Health Services Laboratories | Wellcome Sanger Institute for the COVID-19 Genomics UK (COG-UK) Consortium | Health Services Laboratories and Alex Alderton, Roberto Amato, Jeffrey Barrett, Sonia Goncalves, Ewan Harrison, David K. Jackson, Ian Johnston, Dominic Kwiatkowski, Cordelia Langford, John Sillitoe on behalf of the Wellcome Sanger Institute COVID-19 Surveillance Team |
| EPI_ISL_2993812 | Lighthouse Lab in Alderley Park | Wellcome Sanger Institute for the COVID-19 Genomics UK (COG-UK) Consortium | Jacquelyn Wynn, Mairead Hyland, The Lighthouse Lab in Alderley Park and Alex Alderton, Roberto Amato, Jeffrey Barrett, Sonia Goncalves, Ewan Harrison, David K. Jackson, Ian Johnston, Dominic Kwiatkowski, Cordelia Langford, John Sillitoe on behalf of the Wellcome Sanger Institute COVID-19 Surveillance Team |
| EPI_ISL_2993813 | Lighthouse Laboratory Plymouth | Wellcome Sanger Institute for the COVID-19 Genomics UK (COG-UK) Consortium | Lighthouse Laboratory Plymouth and Alex Alderton, Roberto Amato, Jeffrey Barrett, Sonia Goncalves, Ewan Harrison, David K. Jackson, Ian Johnston, Dominic Kwiatkowski, Cordelia Langford, John Sillitoe on behalf of the Wellcome Sanger Institute COVID-19 Surveillance Team |
| EPI_ISL_2993814 | Health Services Laboratories | Wellcome Sanger Institute for the COVID-19 Genomics UK (COG-UK) Consortium | Health Services Laboratories and Alex Alderton, Roberto Amato, Jeffrey Barrett, Sonia Goncalves, Ewan Harrison, David K. Jackson, Ian Johnston, Dominic Kwiatkowski, Cordelia Langford, John Sillitoe on behalf of the Wellcome Sanger Institute COVID-19 Surveillance Team |
| EPI_ISL_2993815, EPI_ISL_2993816, EPI_ISL_2993817 | Lighthouse Laboratory Plymouth | Wellcome Sanger Institute for the COVID-19 Genomics UK (COG-UK) Consortium | Lighthouse Laboratory Plymouth and Alex Alderton, Roberto Amato, Jeffrey Barrett, Sonia Goncalves, Ewan Harrison, David K. Jackson, Ian Johnston, Dominic Kwiatkowski, Cordelia Langford, John Sillitoe on behalf of the Wellcome Sanger Institute COVID-19 Surveillance Team |
| EPI_ISL_2993818 | Lighthouse Lab in Alderley Park | Wellcome Sanger Institute for the COVID-19 Genomics UK (COG-UK) Consortium | Jacquelyn Wynn, Mairead Hyland, The Lighthouse Lab in Alderley Park and Alex Alderton, Roberto Amato, Jeffrey Barrett, Sonia Goncalves, Ewan Harrison, David K. Jackson, Ian Johnston, Dominic Kwiatkowski, Cordelia Langford, John Sillitoe on behalf of the Wellcome Sanger Institute COVID-19 Surveillance Team |
| EPI_ISL_2993819 | Lighthouse Laboratory Plymouth | Wellcome Sanger Institute for the COVID-19 Genomics UK (COG-UK) Consortium | Lighthouse Laboratory Plymouth and Alex Alderton, Roberto Amato, Jeffrey Barrett, Sonia Goncalves, Ewan Harrison, David K. Jackson, Ian Johnston, Dominic Kwiatkowski, Cordelia Langford, John Sillitoe on behalf of the Wellcome Sanger Institute COVID-19 Surveillance Team |
| EPI_ISL_2993820 | Health Services Laboratories | Wellcome Sanger Institute for the COVID-19 Genomics UK (COG-UK) Consortium | Health Services Laboratories and Alex Alderton, Roberto Amato, Jeffrey Barrett, Sonia Goncalves, Ewan Harrison, David K. Jackson, Ian Johnston, Dominic Kwiatkowski, Cordelia Langford, John Sillitoe on behalf of the Wellcome Sanger Institute COVID-19 Surveillance Team |
| EPI_ISL_2993821 | Lighthouse Laboratory Plymouth | Wellcome Sanger Institute for the COVID-19 Genomics UK (COG-UK) Consortium | Lighthouse Laboratory Plymouth and Alex Alderton, Roberto Amato, Jeffrey Barrett, Sonia Goncalves, Ewan Harrison, David K. Jackson, Ian Johnston, Dominic Kwiatkowski, Cordelia Langford, John Sillitoe on behalf of the Wellcome Sanger Institute COVID-19 Surveillance Team |
| EPI_ISL_2993822 | Lighthouse Lab in Glasgow | Wellcome Sanger Institute for the COVID-19 Genomics UK (COG-UK) Consortium | Harper VanSteenhouse, Yumi Kasai, David Gray, Carol Clugston, Anna Dominiczak and Alex Alderton, Roberto Amato, Jeffrey Barrett, Sonia Goncalves, Ewan Harrison, David K. Jackson, Ian Johnston, Dominic Kwiatkowski, Cordelia Langford, John Sillitoe on behalf of the Wellcome Sanger Institute COVID-19 Surveillance Team |
| EPI_ISL_2993823, EPI_ISL_2993824 | Lighthouse Laboratory Plymouth | Wellcome Sanger Institute for the COVID-19 Genomics UK (COG-UK) Consortium | Lighthouse Laboratory Plymouth and Alex Alderton, Roberto Amato, Jeffrey Barrett, Sonia Goncalves, Ewan Harrison, David K. Jackson, Ian Johnston, Dominic Kwiatkowski, Cordelia Langford, John Sillitoe on behalf of the Wellcome Sanger Institute COVID-19 Surveillance Team |
| EPI_ISL_2993825 | Health Services Laboratories | Wellcome Sanger Institute for the COVID-19 Genomics UK (COG-UK) Consortium | Health Services Laboratories and Alex Alderton, Roberto Amato, Jeffrey Barrett, Sonia Goncalves, Ewan Harrison, David K. Jackson, Ian Johnston, Dominic Kwiatkowski, Cordelia Langford, John Sillitoe on behalf of the Wellcome Sanger Institute COVID-19 Surveillance Team |
| EPI_ISL_2993826 | Lighthouse Laboratory Plymouth | Wellcome Sanger Institute for the COVID-19 Genomics UK (COG-UK) Consortium | Lighthouse Laboratory Plymouth and Alex Alderton, Roberto Amato, Jeffrey Barrett, Sonia Goncalves, Ewan Harrison, David K. Jackson, Ian Johnston, Dominic Kwiatkowski, Cordelia Langford, John Sillitoe on behalf of the Wellcome Sanger Institute COVID-19 Surveillance Team |
| EPI_ISL_2993827 | Lighthouse Lab in Alderley Park | Wellcome Sanger Institute for the COVID-19 Genomics UK (COG-UK) Consortium | Jacquelyn Wynn, Mairead Hyland, The Lighthouse Lab in Alderley Park and Alex Alderton, Roberto Amato, Jeffrey Barrett, Sonia Goncalves, Ewan Harrison, David K. Jackson, Ian Johnston, Dominic Kwiatkowski, Cordelia Langford, John Sillitoe on behalf of the Wellcome Sanger Institute COVID-19 Surveillance Team |
| EPI_ISL_2993828 | Lighthouse Lab in Glasgow | Wellcome Sanger Institute for the COVID-19 Genomics UK (COG-UK) Consortium | Harper VanSteenhouse, Yumi Kasai, David Gray, Carol Clugston, Anna Dominiczak and Alex Alderton, Roberto Amato, Jeffrey Barrett, Sonia Goncalves, Ewan Harrison, David K. Jackson, Ian Johnston, Dominic Kwiatkowski, Cordelia Langford, John Sillitoe on behalf of the Wellcome Sanger Institute COVID-19 Surveillance Team |
| EPI_ISL_2993829 | Lighthouse Lab in Alderley Park | Wellcome Sanger Institute for the COVID-19 Genomics UK (COG-UK) Consortium | Jacquelyn Wynn, Mairead Hyland, The Lighthouse Lab in Alderley Park and Alex Alderton, Roberto Amato, Jeffrey Barrett, Sonia Goncalves, Ewan Harrison, David K. Jackson, Ian Johnston, Dominic Kwiatkowski, Cordelia Langford, John Sillitoe on behalf of the Wellcome Sanger Institute COVID-19 Surveillance Team |
| EPI_ISL_2993830 | Lighthouse Lab in Glasgow | Wellcome Sanger Institute for the COVID-19 Genomics UK (COG-UK) Consortium | Harper VanSteenhouse, Yumi Kasai, David Gray, Carol Clugston, Anna Dominiczak and Alex Alderton, Roberto Amato, Jeffrey Barrett, Sonia Goncalves, Ewan Harrison, David K. Jackson, Ian Johnston, Dominic Kwiatkowski, Cordelia Langford, John Sillitoe on behalf of the Wellcome Sanger Institute COVID-19 Surveillance Team |

| | | | |
|---|---|---|---|
| EPI_ISL_2993831 | Lighthouse Lab in Alderley Park | Wellcome Sanger Institute for the COVID-19 Genomics UK (COG-UK) Consortium | Jacquelyn Wynn, Mairead Hyland, The Lighthouse Lab in Alderley Park and Alex Alderton, Roberto Amato, Jeffrey Barrett, Sonia Goncalves, Ewan Harrison, David K. Jackson, Ian Johnston, Dominic Kwiatkowski, Cordelia Langford, John Sillitoe on behalf of the Wellcome Sanger Institute COVID-19 Surveillance Team |
| EPI_ISL_2993832 | Lighthouse Laboratory Plymouth | Wellcome Sanger Institute for the COVID-19 Genomics UK (COG-UK) Consortium | Lighthouse Laboratory Plymouth and Alex Alderton, Roberto Amato, Jeffrey Barrett, Sonia Goncalves, Ewan Harrison, David K. Jackson, Ian Johnston, Dominic Kwiatkowski, Cordelia Langford, John Sillitoe on behalf of the Wellcome Sanger Institute COVID-19 Surveillance Team |
| EPI_ISL_2993833 | Lighthouse Lab in Alderley Park | Wellcome Sanger Institute for the COVID-19 Genomics UK (COG-UK) Consortium | Jacquelyn Wynn, Mairead Hyland, The Lighthouse Lab in Alderley Park and Alex Alderton, Roberto Amato, Jeffrey Barrett, Sonia Goncalves, Ewan Harrison, David K. Jackson, Ian Johnston, Dominic Kwiatkowski, Cordelia Langford, John Sillitoe on behalf of the Wellcome Sanger Institute COVID-19 Surveillance Team |
| EPI_ISL_2993834, EPI_ISL_2993835, EPI_ISL_2993836, EPI_ISL_2993837 | Lighthouse Laboratory Plymouth | Wellcome Sanger Institute for the COVID-19 Genomics UK (COG-UK) Consortium | Lighthouse Laboratory Plymouth and Alex Alderton, Roberto Amato, Jeffrey Barrett, Sonia Goncalves, Ewan Harrison, David K. Jackson, Ian Johnston, Dominic Kwiatkowski, Cordelia Langford, John Sillitoe on behalf of the Wellcome Sanger Institute COVID-19 Surveillance Team |
| EPI_ISL_2993838 | Lighthouse Lab in Alderley Park | Wellcome Sanger Institute for the COVID-19 Genomics UK (COG-UK) Consortium | Jacquelyn Wynn, Mairead Hyland, The Lighthouse Lab in Alderley Park and Alex Alderton, Roberto Amato, Jeffrey Barrett, Sonia Goncalves, Ewan Harrison, David K. Jackson, Ian Johnston, Dominic Kwiatkowski, Cordelia Langford, John Sillitoe on behalf of the Wellcome Sanger Institute COVID-19 Surveillance Team |
| EPI_ISL_2993839 | Health Services Laboratories | Wellcome Sanger Institute for the COVID-19 Genomics UK (COG-UK) Consortium | Health Services Laboratories and Alex Alderton, Roberto Amato, Jeffrey Barrett, Sonia Goncalves, Ewan Harrison, David K. Jackson, Ian Johnston, Dominic Kwiatkowski, Cordelia Langford, John Sillitoe on behalf of the Wellcome Sanger Institute COVID-19 Surveillance Team |
| EPI_ISL_2993840 | Lighthouse Lab in Alderley Park | Wellcome Sanger Institute for the COVID-19 Genomics UK (COG-UK) Consortium | Jacquelyn Wynn, Mairead Hyland, The Lighthouse Lab in Alderley Park and Alex Alderton, Roberto Amato, Jeffrey Barrett, Sonia Goncalves, Ewan Harrison, David K. Jackson, Ian Johnston, Dominic Kwiatkowski, Cordelia Langford, John Sillitoe on behalf of the Wellcome Sanger Institute COVID-19 Surveillance Team |
| EPI_ISL_2993841, EPI_ISL_2993842, EPI_ISL_2993843 | Lighthouse Laboratory Plymouth | Wellcome Sanger Institute for the COVID-19 Genomics UK (COG-UK) Consortium | Lighthouse Laboratory Plymouth and Alex Alderton, Roberto Amato, Jeffrey Barrett, Sonia Goncalves, Ewan Harrison, David K. Jackson, Ian Johnston, Dominic Kwiatkowski, Cordelia Langford, John Sillitoe on behalf of the Wellcome Sanger Institute COVID-19 Surveillance Team |
| EPI_ISL_2993844 | Lighthouse Lab in Alderley Park | Wellcome Sanger Institute for the COVID-19 Genomics UK (COG-UK) Consortium | Jacquelyn Wynn, Mairead Hyland, The Lighthouse Lab in Alderley Park and Alex Alderton, Roberto Amato, Jeffrey Barrett, Sonia Goncalves, Ewan Harrison, David K. Jackson, Ian Johnston, Dominic Kwiatkowski, Cordelia Langford, John Sillitoe on behalf of the Wellcome Sanger Institute COVID-19 Surveillance Team |
| EPI_ISL_2993845 | Health Services Laboratories | Wellcome Sanger Institute for the COVID-19 Genomics UK (COG-UK) Consortium | Health Services Laboratories and Alex Alderton, Roberto Amato, Jeffrey Barrett, Sonia Goncalves, Ewan Harrison, David K. Jackson, Ian Johnston, Dominic Kwiatkowski, Cordelia Langford, John Sillitoe on behalf of the Wellcome Sanger Institute COVID-19 Surveillance Team |
| EPI_ISL_2993846, EPI_ISL_2993847, EPI_ISL_2993848 | Lighthouse Lab in Alderley Park | Wellcome Sanger Institute for the COVID-19 Genomics UK (COG-UK) Consortium | Jacquelyn Wynn, Mairead Hyland, The Lighthouse Lab in Alderley Park and Alex Alderton, Roberto Amato, Jeffrey Barrett, Sonia Goncalves, Ewan Harrison, David K. Jackson, Ian Johnston, Dominic Kwiatkowski, Cordelia Langford, John Sillitoe on behalf of the Wellcome Sanger Institute COVID-19 Surveillance Team |
| EPI_ISL_2993849 | Lighthouse Lab in Glasgow | Wellcome Sanger Institute for the COVID-19 Genomics UK (COG-UK) Consortium | Harper VanSteenhouse, Yumi Kasai, David Gray, Carol Clugston, Anna Dominiczak and Alex Alderton, Roberto Amato, Jeffrey Barrett, Sonia Goncalves, Ewan Harrison, David K. Jackson, Ian Johnston, Dominic Kwiatkowski, Cordelia Langford, John Sillitoe on behalf of the Wellcome Sanger Institute COVID-19 Surveillance Team |
| EPI_ISL_2993850 | Lighthouse Laboratory Plymouth | Wellcome Sanger Institute for the COVID-19 Genomics UK (COG-UK) Consortium | Lighthouse Laboratory Plymouth and Alex Alderton, Roberto Amato, Jeffrey Barrett, Sonia Goncalves, Ewan Harrison, David K. Jackson, Ian Johnston, Dominic Kwiatkowski, Cordelia Langford, John Sillitoe on behalf of the Wellcome Sanger Institute COVID-19 Surveillance Team |
| EPI_ISL_2993851, EPI_ISL_2993852 | Lighthouse Lab in Alderley Park | Wellcome Sanger Institute for the COVID-19 Genomics UK (COG-UK) Consortium | Jacquelyn Wynn, Mairead Hyland, The Lighthouse Lab in Alderley Park and Alex Alderton, Roberto Amato, Jeffrey Barrett, Sonia Goncalves, Ewan Harrison, David K. Jackson, Ian Johnston, Dominic Kwiatkowski, Cordelia Langford, John Sillitoe on behalf of the Wellcome Sanger Institute COVID-19 Surveillance Team |
| EPI_ISL_2993853, EPI_ISL_2993854 | Lighthouse Lab in Glasgow | Wellcome Sanger Institute for the COVID-19 Genomics UK (COG-UK) Consortium | Harper VanSteenhouse, Yumi Kasai, David Gray, Carol Clugston, Anna Dominiczak and Alex Alderton, Roberto Amato, Jeffrey Barrett, Sonia Goncalves, Ewan Harrison, David K. Jackson, Ian Johnston, Dominic Kwiatkowski, Cordelia Langford, John Sillitoe on behalf of the Wellcome Sanger Institute COVID-19 Surveillance Team |
| EPI_ISL_2993855 | Lighthouse Laboratory Plymouth | Wellcome Sanger Institute for the COVID-19 Genomics UK (COG-UK) Consortium | Lighthouse Laboratory Plymouth and Alex Alderton, Roberto Amato, Jeffrey Barrett, Sonia Goncalves, Ewan Harrison, David K. Jackson, Ian Johnston, Dominic Kwiatkowski, Cordelia Langford, John Sillitoe on behalf of the Wellcome Sanger Institute COVID-19 Surveillance Team |
| EPI_ISL_2993856, EPI_ISL_2993857 | Lighthouse Lab in Alderley Park | Wellcome Sanger Institute for the COVID-19 Genomics UK (COG-UK) Consortium | Jacquelyn Wynn, Mairead Hyland, The Lighthouse Lab in Alderley Park and Alex Alderton, Roberto Amato, Jeffrey Barrett, Sonia Goncalves, Ewan Harrison, David K. Jackson, Ian Johnston, Dominic Kwiatkowski, Cordelia Langford, John Sillitoe on behalf of the Wellcome Sanger Institute COVID-19 Surveillance Team |
| EPI_ISL_2993858 | Health Services Laboratories | Wellcome Sanger Institute for the COVID-19 Genomics UK (COG-UK) Consortium | Health Services Laboratories and Alex Alderton, Roberto Amato, Jeffrey Barrett, Sonia Goncalves, Ewan Harrison, David K. Jackson, Ian Johnston, Dominic Kwiatkowski, Cordelia Langford, John Sillitoe on behalf of the Wellcome Sanger Institute COVID-19 Surveillance Team |
| EPI_ISL_2993859 | Lighthouse Lab in Glasgow | Wellcome Sanger Institute for the COVID-19 Genomics UK (COG-UK) Consortium | Harper VanSteenhouse, Yumi Kasai, David Gray, Carol Clugston, Anna Dominiczak and Alex Alderton, Roberto Amato, Jeffrey Barrett, Sonia Goncalves, Ewan Harrison, David K. Jackson, Ian Johnston, Dominic Kwiatkowski, Cordelia Langford, John Sillitoe on behalf of the Wellcome Sanger Institute COVID-19 Surveillance Team |
| EPI_ISL_2993860 | Lighthouse Laboratory Plymouth | Wellcome Sanger Institute for the COVID-19 Genomics UK (COG-UK) Consortium | Lighthouse Laboratory Plymouth and Alex Alderton, Roberto Amato, Jeffrey Barrett, Sonia Goncalves, Ewan Harrison, David K. Jackson, Ian Johnston, Dominic Kwiatkowski, Cordelia Langford, John Sillitoe on behalf of the Wellcome Sanger Institute COVID-19 Surveillance Team |
| EPI_ISL_2993861, EPI_ISL_2993862 | Lighthouse Lab in Alderley Park | Wellcome Sanger Institute for the COVID-19 Genomics UK (COG-UK) Consortium | Jacquelyn Wynn, Mairead Hyland, The Lighthouse Lab in Alderley Park and Alex Alderton, Roberto Amato, Jeffrey Barrett, Sonia Goncalves, Ewan Harrison, David K. Jackson, Ian Johnston, Dominic Kwiatkowski, Cordelia Langford, John Sillitoe on behalf of the Wellcome Sanger Institute COVID-19 Surveillance Team |
| EPI_ISL_2993863, EPI_ISL_2993864 | Health Services Laboratories | Wellcome Sanger Institute for the COVID-19 Genomics UK (COG-UK) Consortium | Health Services Laboratories and Alex Alderton, Roberto Amato, Jeffrey Barrett, Sonia Goncalves, Ewan Harrison, David K. Jackson, Ian Johnston, Dominic Kwiatkowski, Cordelia Langford, John Sillitoe on behalf of the Wellcome Sanger Institute COVID-19 Surveillance Team |
| EPI_ISL_2993865 | Lighthouse Laboratory Plymouth | Wellcome Sanger Institute for the COVID-19 Genomics UK (COG-UK) Consortium | Lighthouse Laboratory Plymouth and Alex Alderton, Roberto Amato, Jeffrey Barrett, Sonia Goncalves, Ewan Harrison, David K. Jackson, Ian Johnston, Dominic Kwiatkowski, Cordelia Langford, John Sillitoe on behalf of the Wellcome Sanger Institute COVID-19 Surveillance Team |
| EPI_ISL_2993866 | Lighthouse Lab in Alderley Park | Wellcome Sanger Institute for the COVID-19 Genomics UK (COG-UK) Consortium | Jacquelyn Wynn, Mairead Hyland, The Lighthouse Lab in Alderley Park and Alex Alderton, Roberto Amato, Jeffrey Barrett, Sonia Goncalves, Ewan Harrison, David K. Jackson, Ian Johnston, Dominic Kwiatkowski, Cordelia Langford, John Sillitoe on behalf of the Wellcome Sanger Institute COVID-19 Surveillance Team |
| EPI_ISL_2993867 | Lighthouse Laboratory Plymouth | Wellcome Sanger Institute for the COVID-19 Genomics UK (COG-UK) Consortium | Lighthouse Laboratory Plymouth and Alex Alderton, Roberto Amato, Jeffrey Barrett, Sonia Goncalves, Ewan Harrison, David K. Jackson, Ian Johnston, Dominic Kwiatkowski, Cordelia Langford, John Sillitoe on behalf of the Wellcome Sanger Institute COVID-19 Surveillance Team |
| EPI_ISL_2993868 | Health Services Laboratories | Wellcome Sanger Institute for the COVID-19 Genomics UK (COG-UK) Consortium | Health Services Laboratories and Alex Alderton, Roberto Amato, Jeffrey Barrett, Sonia Goncalves, Ewan Harrison, David K. Jackson, Ian Johnston, Dominic Kwiatkowski, Cordelia Langford, John Sillitoe on behalf of the Wellcome Sanger Institute COVID-19 Surveillance Team |
| EPI_ISL_2993869 | Lighthouse Lab in Glasgow | Wellcome Sanger Institute for the COVID-19 Genomics UK (COG-UK) Consortium | Harper VanSteenhouse, Yumi Kasai, David Gray, Carol Clugston, Anna Dominiczak and Alex Alderton, Roberto Amato, Jeffrey Barrett, Sonia Goncalves, Ewan Harrison, David K. Jackson, Ian Johnston, Dominic Kwiatkowski, Cordelia Langford, John Sillitoe on behalf of the Wellcome Sanger Institute COVID-19 Surveillance Team |
| EPI_ISL_2993870 | Health Services Laboratories | Wellcome Sanger Institute for the COVID-19 Genomics UK (COG-UK) Consortium | Health Services Laboratories and Alex Alderton, Roberto Amato, Jeffrey Barrett, Sonia Goncalves, Ewan Harrison, David K. Jackson, Ian Johnston, Dominic Kwiatkowski, Cordelia Langford, John Sillitoe on behalf of the Wellcome Sanger Institute COVID-19 Surveillance Team |
| EPI_ISL_2993871 | Lighthouse Lab in Glasgow | Wellcome Sanger Institute for the COVID-19 Genomics UK (COG-UK) Consortium | Harper VanSteenhouse, Yumi Kasai, David Gray, Carol Clugston, Anna Dominiczak and Alex Alderton, Roberto Amato, Jeffrey Barrett, Sonia Goncalves, Ewan Harrison, David K. Jackson, Ian Johnston, Dominic Kwiatkowski, Cordelia Langford, John Sillitoe on behalf of the Wellcome Sanger Institute COVID-19 Surveillance Team |
| EPI_ISL_2993872 | Health Services Laboratories | Wellcome Sanger Institute for the COVID-19 Genomics UK (COG-UK) Consortium | Health Services Laboratories and Alex Alderton, Roberto Amato, Jeffrey Barrett, Sonia Goncalves, Ewan Harrison, David K. Jackson, Ian Johnston, Dominic Kwiatkowski, Cordelia Langford, John Sillitoe on behalf of the Wellcome Sanger Institute COVID-19 Surveillance Team |
| EPI_ISL_2993873 | Lighthouse Lab in Alderley Park | Wellcome Sanger Institute for the COVID-19 Genomics UK (COG-UK) Consortium | Jacquelyn Wynn, Mairead Hyland, The Lighthouse Lab in Alderley Park and Alex Alderton, Roberto Amato, Jeffrey Barrett, Sonia Goncalves, Ewan Harrison, David K. Jackson, Ian Johnston, Dominic Kwiatkowski, Cordelia Langford, John Sillitoe on behalf of the Wellcome Sanger Institute COVID-19 Surveillance Team |
| EPI_ISL_2993874 | Lighthouse Laboratory Plymouth | Wellcome Sanger Institute for the COVID-19 Genomics UK (COG-UK) Consortium | Lighthouse Laboratory Plymouth and Alex Alderton, Roberto Amato, Jeffrey Barrett, Sonia Goncalves, Ewan Harrison, David K. Jackson, Ian Johnston, Dominic Kwiatkowski, Cordelia Langford, John Sillitoe on behalf of the Wellcome Sanger Institute COVID-19 Surveillance Team |
| EPI_ISL_2993875, EPI_ISL_2993876 | Health Services Laboratories | Wellcome Sanger Institute for the COVID-19 Genomics UK (COG-UK) Consortium | Health Services Laboratories and Alex Alderton, Roberto Amato, Jeffrey Barrett, Sonia Goncalves, Ewan Harrison, David K. Jackson, Ian Johnston, Dominic Kwiatkowski, Cordelia Langford, John Sillitoe on behalf of the Wellcome Sanger Institute COVID-19 Surveillance Team |
| EPI_ISL_2993877 | Lighthouse Lab in Glasgow | Wellcome Sanger Institute for the COVID-19 Genomics UK (COG-UK) Consortium | Harper VanSteenhouse, Yumi Kasai, David Gray, Carol Clugston, Anna Dominiczak and Alex Alderton, Roberto Amato, Jeffrey Barrett, Sonia Goncalves, Ewan Harrison, David K. Jackson, Ian Johnston, Dominic Kwiatkowski, Cordelia Langford, John Sillitoe on behalf of the Wellcome Sanger Institute COVID-19 Surveillance Team |
| EPI_ISL_2993878, EPI_ISL_2993879, EPI_ISL_2993880, EPI_ISL_2993881 | Health Services Laboratories | Wellcome Sanger Institute for the COVID-19 Genomics UK (COG-UK) Consortium | Health Services Laboratories and Alex Alderton, Roberto Amato, Jeffrey Barrett, Sonia Goncalves, Ewan Harrison, David K. Jackson, Ian Johnston, Dominic Kwiatkowski, Cordelia Langford, John Sillitoe on behalf of the Wellcome Sanger Institute COVID-19 Surveillance Team |
| EPI_ISL_2993882 | Lighthouse Lab in Alderley Park | Wellcome Sanger Institute for the COVID-19 Genomics UK (COG-UK) Consortium | Jacquelyn Wynn, Mairead Hyland, The Lighthouse Lab in Alderley Park and Alex Alderton, Roberto Amato, Jeffrey Barrett, Sonia Goncalves, Ewan Harrison, David K. Jackson, Ian Johnston, Dominic Kwiatkowski, Cordelia Langford, John Sillitoe on behalf of the Wellcome Sanger Institute COVID-19 Surveillance Team |
| EPI_ISL_2993883 | Lighthouse Laboratory Plymouth | Wellcome Sanger Institute for the COVID-19 Genomics UK (COG-UK) Consortium | Lighthouse Laboratory Plymouth and Alex Alderton, Roberto Amato, Jeffrey Barrett, Sonia Goncalves, Ewan Harrison, David K. Jackson, Ian Johnston, Dominic Kwiatkowski, Cordelia Langford, John Sillitoe on behalf of the Wellcome Sanger Institute COVID-19 Surveillance Team |
| EPI_ISL_2993884, EPI_ISL_2993885 | Lighthouse Lab in Alderley Park | Wellcome Sanger Institute for the COVID-19 Genomics UK (COG-UK) Consortium | Jacquelyn Wynn, Mairead Hyland, The Lighthouse Lab in Alderley Park and Alex Alderton, Roberto Amato, Jeffrey Barrett, Sonia Goncalves, Ewan Harrison, David K. Jackson, Ian Johnston, Dominic Kwiatkowski, Cordelia Langford, John Sillitoe on behalf of the Wellcome Sanger Institute COVID-19 Surveillance Team |
| EPI_ISL_2993886 | Lighthouse Lab in Glasgow | Wellcome Sanger Institute for the COVID-19 Genomics UK (COG-UK) Consortium | Harper VanSteenhouse, Yumi Kasai, David Gray, Carol Clugston, Anna Dominiczak and Alex Alderton, Roberto Amato, Jeffrey Barrett, Sonia Goncalves, Ewan Harrison, David K. Jackson, Ian Johnston, Dominic Kwiatkowski, Cordelia Langford, John Sillitoe on behalf of the Wellcome Sanger Institute COVID-19 Surveillance Team |
| EPI_ISL_2993887 | Lighthouse Laboratory Plymouth | Wellcome Sanger Institute for the COVID-19 Genomics UK (COG-UK) Consortium | Lighthouse Laboratory Plymouth and Alex Alderton, Roberto Amato, Jeffrey Barrett, Sonia Goncalves, Ewan Harrison, David K. Jackson, Ian Johnston, Dominic Kwiatkowski, Cordelia Langford, John Sillitoe on behalf of the Wellcome Sanger Institute COVID-19 Surveillance Team |
| EPI_ISL_2993888 | Health Services Laboratories | Wellcome Sanger Institute for the COVID-19 Genomics UK (COG-UK) Consortium | Health Services Laboratories and Alex Alderton, Roberto Amato, Jeffrey Barrett, Sonia Goncalves, Ewan Harrison, David K. Jackson, Ian Johnston, Dominic Kwiatkowski, Cordelia Langford, John Sillitoe on behalf of the Wellcome Sanger Institute COVID-19 Surveillance Team |
| EPI_ISL_2993889 | Lighthouse Lab in Glasgow | Wellcome Sanger Institute for the COVID-19 Genomics UK (COG-UK) Consortium | Harper VanSteenhouse, Yumi Kasai, David Gray, Carol Clugston, Anna Dominiczak and Alex Alderton, Roberto Amato, Jeffrey Barrett, Sonia Goncalves, Ewan Harrison, David K. Jackson, Ian Johnston, Dominic Kwiatkowski, Cordelia Langford, John Sillitoe on behalf of the Wellcome Sanger Institute COVID-19 Surveillance Team |
| EPI_ISL_2993890, EPI_ISL_2993891 | Lighthouse Laboratory Plymouth | Wellcome Sanger Institute for the COVID-19 Genomics UK (COG-UK) Consortium | Lighthouse Laboratory Plymouth and Alex Alderton, Roberto Amato, Jeffrey Barrett, Sonia Goncalves, Ewan Harrison, David K. Jackson, Ian Johnston, Dominic Kwiatkowski, Cordelia Langford, John Sillitoe on behalf of the Wellcome Sanger Institute COVID-19 Surveillance Team |

| | | | |
|---|---|---|---|
| EPI_ISL_2993892 | Health Services Laboratories | Wellcome Sanger Institute for the COVID-19 Genomics UK (COG-UK) Consortium | Health Services Laboratories and Alex Alderton, Roberto Amato, Jeffrey Barrett, Sonia Goncalves, Ewan Harrison, David K. Jackson, Ian Johnston, Dominic Kwiatkowski, Cordelia Langford, John Sillitoe on behalf of the Wellcome Sanger Institute COVID-19 Surveillance Team |
| EPI_ISL_2993893 | Lighthouse Lab in Alderley Park | Wellcome Sanger Institute for the COVID-19 Genomics UK (COG-UK) Consortium | Jacquelyn Wynn, Mairead Hyland, The Lighthouse Lab in Alderley Park and Alex Alderton, Roberto Amato, Jeffrey Barrett, Sonia Goncalves, Ewan Harrison, David K. Jackson, Ian Johnston, Dominic Kwiatkowski, Cordelia Langford, John Sillitoe on behalf of the Wellcome Sanger Institute COVID-19 Surveillance Team |
| EPI_ISL_2993894 | Health Services Laboratories | Wellcome Sanger Institute for the COVID-19 Genomics UK (COG-UK) Consortium | Health Services Laboratories and Alex Alderton, Roberto Amato, Jeffrey Barrett, Sonia Goncalves, Ewan Harrison, David K. Jackson, Ian Johnston, Dominic Kwiatkowski, Cordelia Langford, John Sillitoe on behalf of the Wellcome Sanger Institute COVID-19 Surveillance Team |
| EPI_ISL_2993895 | Lighthouse Lab in Glasgow | Wellcome Sanger Institute for the COVID-19 Genomics UK (COG-UK) Consortium | Harper VanSteenhouse, Yumi Kasai, David Gray, Carol Clugston, Anna Dominiczak and Alex Alderton, Roberto Amato, Jeffrey Barrett, Sonia Goncalves, Ewan Harrison, David K. Jackson, Ian Johnston, Dominic Kwiatkowski, Cordelia Langford, John Sillitoe on behalf of the Wellcome Sanger Institute COVID-19 Surveillance Team |
| EPI_ISL_2993896 | Health Services Laboratories | Wellcome Sanger Institute for the COVID-19 Genomics UK (COG-UK) Consortium | Health Services Laboratories and Alex Alderton, Roberto Amato, Jeffrey Barrett, Sonia Goncalves, Ewan Harrison, David K. Jackson, Ian Johnston, Dominic Kwiatkowski, Cordelia Langford, John Sillitoe on behalf of the Wellcome Sanger Institute COVID-19 Surveillance Team |
| EPI_ISL_2993897 | Lighthouse Lab in Alderley Park | Wellcome Sanger Institute for the COVID-19 Genomics UK (COG-UK) Consortium | Jacquelyn Wynn, Mairead Hyland, The Lighthouse Lab in Alderley Park and Alex Alderton, Roberto Amato, Jeffrey Barrett, Sonia Goncalves, Ewan Harrison, David K. Jackson, Ian Johnston, Dominic Kwiatkowski, Cordelia Langford, John Sillitoe on behalf of the Wellcome Sanger Institute COVID-19 Surveillance Team |
| EPI_ISL_2993898 | Lighthouse Laboratory Plymouth | Wellcome Sanger Institute for the COVID-19 Genomics UK (COG-UK) Consortium | Lighthouse Laboratory Plymouth and Alex Alderton, Roberto Amato, Jeffrey Barrett, Sonia Goncalves, Ewan Harrison, David K. Jackson, Ian Johnston, Dominic Kwiatkowski, Cordelia Langford, John Sillitoe on behalf of the Wellcome Sanger Institute COVID-19 Surveillance Team |
| EPI_ISL_2993899 | Lighthouse Lab in Alderley Park | Wellcome Sanger Institute for the COVID-19 Genomics UK (COG-UK) Consortium | Jacquelyn Wynn, Mairead Hyland, The Lighthouse Lab in Alderley Park and Alex Alderton, Roberto Amato, Jeffrey Barrett, Sonia Goncalves, Ewan Harrison, David K. Jackson, Ian Johnston, Dominic Kwiatkowski, Cordelia Langford, John Sillitoe on behalf of the Wellcome Sanger Institute COVID-19 Surveillance Team |
| EPI_ISL_2993900 | Lighthouse Laboratory Plymouth | Wellcome Sanger Institute for the COVID-19 Genomics UK (COG-UK) Consortium | Lighthouse Laboratory Plymouth and Alex Alderton, Roberto Amato, Jeffrey Barrett, Sonia Goncalves, Ewan Harrison, David K. Jackson, Ian Johnston, Dominic Kwiatkowski, Cordelia Langford, John Sillitoe on behalf of the Wellcome Sanger Institute COVID-19 Surveillance Team |
| EPI_ISL_2993901 | Lighthouse Lab in Glasgow | Wellcome Sanger Institute for the COVID-19 Genomics UK (COG-UK) Consortium | Harper VanSteenhouse, Yumi Kasai, David Gray, Carol Clugston, Anna Dominiczak and Alex Alderton, Roberto Amato, Jeffrey Barrett, Sonia Goncalves, Ewan Harrison, David K. Jackson, Ian Johnston, Dominic Kwiatkowski, Cordelia Langford, John Sillitoe on behalf of the Wellcome Sanger Institute COVID-19 Surveillance Team |
| EPI_ISL_2993902 | Health Services Laboratories | Wellcome Sanger Institute for the COVID-19 Genomics UK (COG-UK) Consortium | Health Services Laboratories and Alex Alderton, Roberto Amato, Jeffrey Barrett, Sonia Goncalves, Ewan Harrison, David K. Jackson, Ian Johnston, Dominic Kwiatkowski, Cordelia Langford, John Sillitoe on behalf of the Wellcome Sanger Institute COVID-19 Surveillance Team |
| EPI_ISL_2993903, EPI_ISL_2993904 | Lighthouse Laboratory Plymouth | Wellcome Sanger Institute for the COVID-19 Genomics UK (COG-UK) Consortium | Lighthouse Laboratory Plymouth and Alex Alderton, Roberto Amato, Jeffrey Barrett, Sonia Goncalves, Ewan Harrison, David K. Jackson, Ian Johnston, Dominic Kwiatkowski, Cordelia Langford, John Sillitoe on behalf of the Wellcome Sanger Institute COVID-19 Surveillance Team |
| EPI_ISL_2993905, EPI_ISL_2993906 | Lighthouse Lab in Alderley Park | Wellcome Sanger Institute for the COVID-19 Genomics UK (COG-UK) Consortium | Jacquelyn Wynn, Mairead Hyland, The Lighthouse Lab in Alderley Park and Alex Alderton, Roberto Amato, Jeffrey Barrett, Sonia Goncalves, Ewan Harrison, David K. Jackson, Ian Johnston, Dominic Kwiatkowski, Cordelia Langford, John Sillitoe on behalf of the Wellcome Sanger Institute COVID-19 Surveillance Team |
| EPI_ISL_2993907 | Lighthouse Laboratory Plymouth | Wellcome Sanger Institute for the COVID-19 Genomics UK (COG-UK) Consortium | Lighthouse Laboratory Plymouth and Alex Alderton, Roberto Amato, Jeffrey Barrett, Sonia Goncalves, Ewan Harrison, David K. Jackson, Ian Johnston, Dominic Kwiatkowski, Cordelia Langford, John Sillitoe on behalf of the Wellcome Sanger Institute COVID-19 Surveillance Team |
| EPI_ISL_2993908 | Lighthouse Lab in Alderley Park | Wellcome Sanger Institute for the COVID-19 Genomics UK (COG-UK) Consortium | Jacquelyn Wynn, Mairead Hyland, The Lighthouse Lab in Alderley Park and Alex Alderton, Roberto Amato, Jeffrey Barrett, Sonia Goncalves, Ewan Harrison, David K. Jackson, Ian Johnston, Dominic Kwiatkowski, Cordelia Langford, John Sillitoe on behalf of the Wellcome Sanger Institute COVID-19 Surveillance Team |
| EPI_ISL_2993910 | Lighthouse Lab in Glasgow | Wellcome Sanger Institute for the COVID-19 Genomics UK (COG-UK) Consortium | Harper VanSteenhouse, Yumi Kasai, David Gray, Carol Clugston, Anna Dominiczak and Alex Alderton, Roberto Amato, Jeffrey Barrett, Sonia Goncalves, Ewan Harrison, David K. Jackson, Ian Johnston, Dominic Kwiatkowski, Cordelia Langford, John Sillitoe on behalf of the Wellcome Sanger Institute COVID-19 Surveillance Team |
| EPI_ISL_2993911 | Lighthouse Lab in Alderley Park | Wellcome Sanger Institute for the COVID-19 Genomics UK (COG-UK) Consortium | Jacquelyn Wynn, Mairead Hyland, The Lighthouse Lab in Alderley Park and Alex Alderton, Roberto Amato, Jeffrey Barrett, Sonia Goncalves, Ewan Harrison, David K. Jackson, Ian Johnston, Dominic Kwiatkowski, Cordelia Langford, John Sillitoe on behalf of the Wellcome Sanger Institute COVID-19 Surveillance Team |
| EPI_ISL_2993912 | Health Services Laboratories | Wellcome Sanger Institute for the COVID-19 Genomics UK (COG-UK) Consortium | Health Services Laboratories and Alex Alderton, Roberto Amato, Jeffrey Barrett, Sonia Goncalves, Ewan Harrison, David K. Jackson, Ian Johnston, Dominic Kwiatkowski, Cordelia Langford, John Sillitoe on behalf of the Wellcome Sanger Institute COVID-19 Surveillance Team |
| EPI_ISL_2993913 | Lighthouse Laboratory Plymouth | Wellcome Sanger Institute for the COVID-19 Genomics UK (COG-UK) Consortium | Lighthouse Laboratory Plymouth and Alex Alderton, Roberto Amato, Jeffrey Barrett, Sonia Goncalves, Ewan Harrison, David K. Jackson, Ian Johnston, Dominic Kwiatkowski, Cordelia Langford, John Sillitoe on behalf of the Wellcome Sanger Institute COVID-19 Surveillance Team |
| EPI_ISL_2993914 | Health Services Laboratories | Wellcome Sanger Institute for the COVID-19 Genomics UK (COG-UK) Consortium | Health Services Laboratories and Alex Alderton, Roberto Amato, Jeffrey Barrett, Sonia Goncalves, Ewan Harrison, David K. Jackson, Ian Johnston, Dominic Kwiatkowski, Cordelia Langford, John Sillitoe on behalf of the Wellcome Sanger Institute COVID-19 Surveillance Team |
| EPI_ISL_2993915 | Lighthouse Lab in Glasgow | Wellcome Sanger Institute for the COVID-19 Genomics UK (COG-UK) Consortium | Harper VanSteenhouse, Yumi Kasai, David Gray, Carol Clugston, Anna Dominiczak and Alex Alderton, Roberto Amato, Jeffrey Barrett, Sonia Goncalves, Ewan Harrison, David K. Jackson, Ian Johnston, Dominic Kwiatkowski, Cordelia Langford, John Sillitoe on behalf of the Wellcome Sanger Institute COVID-19 Surveillance Team |
| EPI_ISL_2993916, EPI_ISL_2993917, EPI_ISL_2993918, EPI_ISL_2993919 | Lighthouse Laboratory Plymouth | Wellcome Sanger Institute for the COVID-19 Genomics UK (COG-UK) Consortium | Lighthouse Laboratory Plymouth and Alex Alderton, Roberto Amato, Jeffrey Barrett, Sonia Goncalves, Ewan Harrison, David K. Jackson, Ian Johnston, Dominic Kwiatkowski, Cordelia Langford, John Sillitoe on behalf of the Wellcome Sanger Institute COVID-19 Surveillance Team |
| EPI_ISL_2993920 | Lighthouse Lab in Alderley Park | Wellcome Sanger Institute for the COVID-19 Genomics UK (COG-UK) Consortium | Jacquelyn Wynn, Mairead Hyland, The Lighthouse Lab in Alderley Park and Alex Alderton, Roberto Amato, Jeffrey Barrett, Sonia Goncalves, Ewan Harrison, David K. Jackson, Ian Johnston, Dominic Kwiatkowski, Cordelia Langford, John Sillitoe on behalf of the Wellcome Sanger Institute COVID-19 Surveillance Team |
| EPI_ISL_2993921, EPI_ISL_2993922 | Health Services Laboratories | Wellcome Sanger Institute for the COVID-19 Genomics UK (COG-UK) Consortium | Health Services Laboratories and Alex Alderton, Roberto Amato, Jeffrey Barrett, Sonia Goncalves, Ewan Harrison, David K. Jackson, Ian Johnston, Dominic Kwiatkowski, Cordelia Langford, John Sillitoe on behalf of the Wellcome Sanger Institute COVID-19 Surveillance Team |
| EPI_ISL_2993923 | Lighthouse Lab in Alderley Park | Wellcome Sanger Institute for the COVID-19 Genomics UK (COG-UK) Consortium | Jacquelyn Wynn, Mairead Hyland, The Lighthouse Lab in Alderley Park and Alex Alderton, Roberto Amato, Jeffrey Barrett, Sonia Goncalves, Ewan Harrison, David K. Jackson, Ian Johnston, Dominic Kwiatkowski, Cordelia Langford, John Sillitoe on behalf of the Wellcome Sanger Institute COVID-19 Surveillance Team |
| EPI_ISL_2993924 | Lighthouse Laboratory Plymouth | Wellcome Sanger Institute for the COVID-19 Genomics UK (COG-UK) Consortium | Lighthouse Laboratory Plymouth and Alex Alderton, Roberto Amato, Jeffrey Barrett, Sonia Goncalves, Ewan Harrison, David K. Jackson, Ian Johnston, Dominic Kwiatkowski, Cordelia Langford, John Sillitoe on behalf of the Wellcome Sanger Institute COVID-19 Surveillance Team |
| EPI_ISL_2993925 | Health Services Laboratories | Wellcome Sanger Institute for the COVID-19 Genomics UK (COG-UK) Consortium | Health Services Laboratories and Alex Alderton, Roberto Amato, Jeffrey Barrett, Sonia Goncalves, Ewan Harrison, David K. Jackson, Ian Johnston, Dominic Kwiatkowski, Cordelia Langford, John Sillitoe on behalf of the Wellcome Sanger Institute COVID-19 Surveillance Team |
| EPI_ISL_2993926 | Lighthouse Lab in Glasgow | Wellcome Sanger Institute for the COVID-19 Genomics UK (COG-UK) Consortium | Harper VanSteenhouse, Yumi Kasai, David Gray, Carol Clugston, Anna Dominiczak and Alex Alderton, Roberto Amato, Jeffrey Barrett, Sonia Goncalves, Ewan Harrison, David K. Jackson, Ian Johnston, Dominic Kwiatkowski, Cordelia Langford, John Sillitoe on behalf of the Wellcome Sanger Institute COVID-19 Surveillance Team |
| EPI_ISL_2993928, EPI_ISL_2993929, EPI_ISL_2993930 | Lighthouse Laboratory Plymouth | Wellcome Sanger Institute for the COVID-19 Genomics UK (COG-UK) Consortium | Lighthouse Laboratory Plymouth and Alex Alderton, Roberto Amato, Jeffrey Barrett, Sonia Goncalves, Ewan Harrison, David K. Jackson, Ian Johnston, Dominic Kwiatkowski, Cordelia Langford, John Sillitoe on behalf of the Wellcome Sanger Institute COVID-19 Surveillance Team |
| EPI_ISL_2993931 | Lighthouse Lab in Alderley Park | Wellcome Sanger Institute for the COVID-19 Genomics UK (COG-UK) Consortium | Jacquelyn Wynn, Mairead Hyland, The Lighthouse Lab in Alderley Park and Alex Alderton, Roberto Amato, Jeffrey Barrett, Sonia Goncalves, Ewan Harrison, David K. Jackson, Ian Johnston, Dominic Kwiatkowski, Cordelia Langford, John Sillitoe on behalf of the Wellcome Sanger Institute COVID-19 Surveillance Team |
| EPI_ISL_2993932 | Health Services Laboratories | Wellcome Sanger Institute for the COVID-19 Genomics UK (COG-UK) Consortium | Health Services Laboratories and Alex Alderton, Roberto Amato, Jeffrey Barrett, Sonia Goncalves, Ewan Harrison, David K. Jackson, Ian Johnston, Dominic Kwiatkowski, Cordelia Langford, John Sillitoe on behalf of the Wellcome Sanger Institute COVID-19 Surveillance Team |
| EPI_ISL_2993933, EPI_ISL_2993934 | Lighthouse Lab in Alderley Park | Wellcome Sanger Institute for the COVID-19 Genomics UK (COG-UK) Consortium | Jacquelyn Wynn, Mairead Hyland, The Lighthouse Lab in Alderley Park and Alex Alderton, Roberto Amato, Jeffrey Barrett, Sonia Goncalves, Ewan Harrison, David K. Jackson, Ian Johnston, Dominic Kwiatkowski, Cordelia Langford, John Sillitoe on behalf of the Wellcome Sanger Institute COVID-19 Surveillance Team |
| EPI_ISL_2993935, EPI_ISL_2993936, EPI_ISL_2993937 | Health Services Laboratories | Wellcome Sanger Institute for the COVID-19 Genomics UK (COG-UK) Consortium | Health Services Laboratories and Alex Alderton, Roberto Amato, Jeffrey Barrett, Sonia Goncalves, Ewan Harrison, David K. Jackson, Ian Johnston, Dominic Kwiatkowski, Cordelia Langford, John Sillitoe on behalf of the Wellcome Sanger Institute COVID-19 Surveillance Team |
| EPI_ISL_2993938, EPI_ISL_2993939 | Lighthouse Lab in Alderley Park | Wellcome Sanger Institute for the COVID-19 Genomics UK (COG-UK) Consortium | Jacquelyn Wynn, Mairead Hyland, The Lighthouse Lab in Alderley Park and Alex Alderton, Roberto Amato, Jeffrey Barrett, Sonia Goncalves, Ewan Harrison, David K. Jackson, Ian Johnston, Dominic Kwiatkowski, Cordelia Langford, John Sillitoe on behalf of the Wellcome Sanger Institute COVID-19 Surveillance Team |
| EPI_ISL_2993940, EPI_ISL_2993941 | Lighthouse Laboratory Plymouth | Wellcome Sanger Institute for the COVID-19 Genomics UK (COG-UK) Consortium | Lighthouse Laboratory Plymouth and Alex Alderton, Roberto Amato, Jeffrey Barrett, Sonia Goncalves, Ewan Harrison, David K. Jackson, Ian Johnston, Dominic Kwiatkowski, Cordelia Langford, John Sillitoe on behalf of the Wellcome Sanger Institute COVID-19 Surveillance Team |
| EPI_ISL_2993942 | Lighthouse Lab in Alderley Park | Wellcome Sanger Institute for the COVID-19 Genomics UK (COG-UK) Consortium | Jacquelyn Wynn, Mairead Hyland, The Lighthouse Lab in Alderley Park and Alex Alderton, Roberto Amato, Jeffrey Barrett, Sonia Goncalves, Ewan Harrison, David K. Jackson, Ian Johnston, Dominic Kwiatkowski, Cordelia Langford, John Sillitoe on behalf of the Wellcome Sanger Institute COVID-19 Surveillance Team |
| EPI_ISL_2993943 | Lighthouse Laboratory Plymouth | Wellcome Sanger Institute for the COVID-19 Genomics UK (COG-UK) Consortium | Lighthouse Laboratory Plymouth and Alex Alderton, Roberto Amato, Jeffrey Barrett, Sonia Goncalves, Ewan Harrison, David K. Jackson, Ian Johnston, Dominic Kwiatkowski, Cordelia Langford, John Sillitoe on behalf of the Wellcome Sanger Institute COVID-19 Surveillance Team |
| EPI_ISL_2993944 | Lighthouse Lab in Alderley Park | Wellcome Sanger Institute for the COVID-19 Genomics UK (COG-UK) Consortium | Jacquelyn Wynn, Mairead Hyland, The Lighthouse Lab in Alderley Park and Alex Alderton, Roberto Amato, Jeffrey Barrett, Sonia Goncalves, Ewan Harrison, David K. Jackson, Ian Johnston, Dominic Kwiatkowski, Cordelia Langford, John Sillitoe on behalf of the Wellcome Sanger Institute COVID-19 Surveillance Team |
| EPI_ISL_2993945 | Health Services Laboratories | Wellcome Sanger Institute for the COVID-19 Genomics UK (COG-UK) Consortium | Health Services Laboratories and Alex Alderton, Roberto Amato, Jeffrey Barrett, Sonia Goncalves, Ewan Harrison, David K. Jackson, Ian Johnston, Dominic Kwiatkowski, Cordelia Langford, John Sillitoe on behalf of the Wellcome Sanger Institute COVID-19 Surveillance Team |
| EPI_ISL_2993946 | Lighthouse Lab in Glasgow | Wellcome Sanger Institute for the COVID-19 Genomics UK (COG-UK) Consortium | Harper VanSteenhouse, Yumi Kasai, David Gray, Carol Clugston, Anna Dominiczak and Alex Alderton, Roberto Amato, Jeffrey Barrett, Sonia Goncalves, Ewan Harrison, David K. Jackson, Ian Johnston, Dominic Kwiatkowski, Cordelia Langford, John Sillitoe on behalf of the Wellcome Sanger Institute COVID-19 Surveillance Team |
| EPI_ISL_2993947 | Lighthouse Laboratory Plymouth | Wellcome Sanger Institute for the COVID-19 Genomics UK (COG-UK) Consortium | Lighthouse Laboratory Plymouth and Alex Alderton, Roberto Amato, Jeffrey Barrett, Sonia Goncalves, Ewan Harrison, David K. Jackson, Ian Johnston, Dominic Kwiatkowski, Cordelia Langford, John Sillitoe on behalf of the Wellcome Sanger Institute COVID-19 Surveillance Team |
| EPI_ISL_2993948 | Lighthouse Lab in Alderley Park | Wellcome Sanger Institute for the COVID-19 Genomics UK (COG-UK) Consortium | Jacquelyn Wynn, Mairead Hyland, The Lighthouse Lab in Alderley Park and Alex Alderton, Roberto Amato, Jeffrey Barrett, Sonia Goncalves, Ewan Harrison, David K. Jackson, Ian Johnston, Dominic Kwiatkowski, Cordelia Langford, John Sillitoe on behalf of the Wellcome Sanger Institute COVID-19 Surveillance Team |
| EPI_ISL_2993949 | Lighthouse Lab in Glasgow | Wellcome Sanger Institute for the COVID-19 Genomics UK (COG-UK) Consortium | Harper VanSteenhouse, Yumi Kasai, David Gray, Carol Clugston, Anna Dominiczak and Alex Alderton, Roberto Amato, Jeffrey Barrett, Sonia Goncalves, Ewan Harrison, David K. Jackson, Ian Johnston, Dominic Kwiatkowski, Cordelia Langford, John Sillitoe on behalf of the Wellcome Sanger Institute COVID-19 Surveillance Team |
| EPI_ISL_2993950 | Lighthouse Lab in Alderley Park | Wellcome Sanger Institute for the COVID-19 Genomics UK (COG-UK) | Jacquelyn Wynn, Mairead Hyland, The Lighthouse Lab in Alderley Park and Alex Alderton, Roberto Amato, Jeffrey Barrett, Sonia Goncalves, Ewan Harrison, David K. Jackson, Ian Johnston, Dominic Kwiatkowski, Cordelia Langford, John Sillitoe on |

| | | | |
|---|---|---|---|
| | | Consortium | behalf of the Wellcome Sanger Institute COVID-19 Surveillance Team |
| EPI_ISL_2993951, EPI_ISL_2993952 | Lighthouse Laboratory Plymouth | Wellcome Sanger Institute for the COVID-19 Genomics UK (COG-UK) Consortium | Lighthouse Laboratory Plymouth and Alex Alderton, Roberto Amato, Jeffrey Barrett, Sonia Goncalves, Ewan Harrison, David K. Jackson, Ian Johnston, Dominic Kwiatkowski, Cordelia Langford, John Sillitoe on behalf of the Wellcome Sanger Institute COVID-19 Surveillance Team |
| EPI_ISL_2993953 | Lighthouse Lab in Alderley Park | Wellcome Sanger Institute for the COVID-19 Genomics UK (COG-UK) Consortium | Jacquelyn Wynn, Mairead Hyland, The Lighthouse Lab in Alderley Park and Alex Alderton, Roberto Amato, Jeffrey Barrett, Sonia Goncalves, Ewan Harrison, David K. Jackson, Ian Johnston, Dominic Kwiatkowski, Cordelia Langford, John Sillitoe on behalf of the Wellcome Sanger Institute COVID-19 Surveillance Team |
| EPI_ISL_2993954 | Lighthouse Laboratory Plymouth | Wellcome Sanger Institute for the COVID-19 Genomics UK (COG-UK) Consortium | Lighthouse Laboratory Plymouth and Alex Alderton, Roberto Amato, Jeffrey Barrett, Sonia Goncalves, Ewan Harrison, David K. Jackson, Ian Johnston, Dominic Kwiatkowski, Cordelia Langford, John Sillitoe on behalf of the Wellcome Sanger Institute COVID-19 Surveillance Team |
| EPI_ISL_2993955 | Health Services Laboratories | Wellcome Sanger Institute for the COVID-19 Genomics UK (COG-UK) Consortium | Health Services Laboratories and Alex Alderton, Roberto Amato, Jeffrey Barrett, Sonia Goncalves, Ewan Harrison, David K. Jackson, Ian Johnston, Dominic Kwiatkowski, Cordelia Langford, John Sillitoe on behalf of the Wellcome Sanger Institute COVID-19 Surveillance Team |
| EPI_ISL_2993956, EPI_ISL_2993957 | Lighthouse Laboratory Plymouth | Wellcome Sanger Institute for the COVID-19 Genomics UK (COG-UK) Consortium | Lighthouse Laboratory Plymouth and Alex Alderton, Roberto Amato, Jeffrey Barrett, Sonia Goncalves, Ewan Harrison, David K. Jackson, Ian Johnston, Dominic Kwiatkowski, Cordelia Langford, John Sillitoe on behalf of the Wellcome Sanger Institute COVID-19 Surveillance Team |
| EPI_ISL_2993958 | Health Services Laboratories | Wellcome Sanger Institute for the COVID-19 Genomics UK (COG-UK) Consortium | Health Services Laboratories and Alex Alderton, Roberto Amato, Jeffrey Barrett, Sonia Goncalves, Ewan Harrison, David K. Jackson, Ian Johnston, Dominic Kwiatkowski, Cordelia Langford, John Sillitoe on behalf of the Wellcome Sanger Institute COVID-19 Surveillance Team |
| EPI_ISL_2993959 | Lighthouse Lab in Glasgow | Wellcome Sanger Institute for the COVID-19 Genomics UK (COG-UK) Consortium | Harper VanSteenhouse, Yumi Kasai, David Gray, Carol Clugston, Anna Dominiczak and Alex Alderton, Roberto Amato, Jeffrey Barrett, Sonia Goncalves, Ewan Harrison, David K. Jackson, Ian Johnston, Dominic Kwiatkowski, Cordelia Langford, John Sillitoe on behalf of the Wellcome Sanger Institute COVID-19 Surveillance Team |
| EPI_ISL_2993960, EPI_ISL_2993961 | Health Services Laboratories | Wellcome Sanger Institute for the COVID-19 Genomics UK (COG-UK) Consortium | Health Services Laboratories and Alex Alderton, Roberto Amato, Jeffrey Barrett, Sonia Goncalves, Ewan Harrison, David K. Jackson, Ian Johnston, Dominic Kwiatkowski, Cordelia Langford, John Sillitoe on behalf of the Wellcome Sanger Institute COVID-19 Surveillance Team |
| EPI_ISL_2993962 | Lighthouse Laboratory Plymouth | Wellcome Sanger Institute for the COVID-19 Genomics UK (COG-UK) Consortium | Lighthouse Laboratory Plymouth and Alex Alderton, Roberto Amato, Jeffrey Barrett, Sonia Goncalves, Ewan Harrison, David K. Jackson, Ian Johnston, Dominic Kwiatkowski, Cordelia Langford, John Sillitoe on behalf of the Wellcome Sanger Institute COVID-19 Surveillance Team |
| EPI_ISL_2993963 | Health Services Laboratories | Wellcome Sanger Institute for the COVID-19 Genomics UK (COG-UK) Consortium | Health Services Laboratories and Alex Alderton, Roberto Amato, Jeffrey Barrett, Sonia Goncalves, Ewan Harrison, David K. Jackson, Ian Johnston, Dominic Kwiatkowski, Cordelia Langford, John Sillitoe on behalf of the Wellcome Sanger Institute COVID-19 Surveillance Team |
| EPI_ISL_2993964, EPI_ISL_2993965 | Lighthouse Laboratory Plymouth | Wellcome Sanger Institute for the COVID-19 Genomics UK (COG-UK) Consortium | Lighthouse Laboratory Plymouth and Alex Alderton, Roberto Amato, Jeffrey Barrett, Sonia Goncalves, Ewan Harrison, David K. Jackson, Ian Johnston, Dominic Kwiatkowski, Cordelia Langford, John Sillitoe on behalf of the Wellcome Sanger Institute COVID-19 Surveillance Team |
| EPI_ISL_2993966 | Lighthouse Lab in Alderley Park | Wellcome Sanger Institute for the COVID-19 Genomics UK (COG-UK) Consortium | Jacquelyn Wynn, Mairead Hyland, The Lighthouse Lab in Alderley Park and Alex Alderton, Roberto Amato, Jeffrey Barrett, Sonia Goncalves, Ewan Harrison, David K. Jackson, Ian Johnston, Dominic Kwiatkowski, Cordelia Langford, John Sillitoe on behalf of the Wellcome Sanger Institute COVID-19 Surveillance Team |
| EPI_ISL_2993967 | Lighthouse Lab in Glasgow | Wellcome Sanger Institute for the COVID-19 Genomics UK (COG-UK) Consortium | Harper VanSteenhouse, Yumi Kasai, David Gray, Carol Clugston, Anna Dominiczak and Alex Alderton, Roberto Amato, Jeffrey Barrett, Sonia Goncalves, Ewan Harrison, David K. Jackson, Ian Johnston, Dominic Kwiatkowski, Cordelia Langford, John Sillitoe on behalf of the Wellcome Sanger Institute COVID-19 Surveillance Team |
| EPI_ISL_2993968 | Lighthouse Laboratory Plymouth | Wellcome Sanger Institute for the COVID-19 Genomics UK (COG-UK) Consortium | Lighthouse Laboratory Plymouth and Alex Alderton, Roberto Amato, Jeffrey Barrett, Sonia Goncalves, Ewan Harrison, David K. Jackson, Ian Johnston, Dominic Kwiatkowski, Cordelia Langford, John Sillitoe on behalf of the Wellcome Sanger Institute COVID-19 Surveillance Team |
| EPI_ISL_2993969 | Lighthouse Lab in Glasgow | Wellcome Sanger Institute for the COVID-19 Genomics UK (COG-UK) Consortium | Harper VanSteenhouse, Yumi Kasai, David Gray, Carol Clugston, Anna Dominiczak and Alex Alderton, Roberto Amato, Jeffrey Barrett, Sonia Goncalves, Ewan Harrison, David K. Jackson, Ian Johnston, Dominic Kwiatkowski, Cordelia Langford, John Sillitoe on behalf of the Wellcome Sanger Institute COVID-19 Surveillance Team |
| EPI_ISL_2993970 | Health Services Laboratories | Wellcome Sanger Institute for the COVID-19 Genomics UK (COG-UK) Consortium | Health Services Laboratories and Alex Alderton, Roberto Amato, Jeffrey Barrett, Sonia Goncalves, Ewan Harrison, David K. Jackson, Ian Johnston, Dominic Kwiatkowski, Cordelia Langford, John Sillitoe on behalf of the Wellcome Sanger Institute COVID-19 Surveillance Team |
| EPI_ISL_2993971 | Lighthouse Lab in Glasgow | Wellcome Sanger Institute for the COVID-19 Genomics UK (COG-UK) Consortium | Harper VanSteenhouse, Yumi Kasai, David Gray, Carol Clugston, Anna Dominiczak and Alex Alderton, Roberto Amato, Jeffrey Barrett, Sonia Goncalves, Ewan Harrison, David K. Jackson, Ian Johnston, Dominic Kwiatkowski, Cordelia Langford, John Sillitoe on behalf of the Wellcome Sanger Institute COVID-19 Surveillance Team |
| EPI_ISL_2993972 | Health Services Laboratories | Wellcome Sanger Institute for the COVID-19 Genomics UK (COG-UK) Consortium | Health Services Laboratories and Alex Alderton, Roberto Amato, Jeffrey Barrett, Sonia Goncalves, Ewan Harrison, David K. Jackson, Ian Johnston, Dominic Kwiatkowski, Cordelia Langford, John Sillitoe on behalf of the Wellcome Sanger Institute COVID-19 Surveillance Team |
| EPI_ISL_2993973 | Lighthouse Lab in Glasgow | Wellcome Sanger Institute for the COVID-19 Genomics UK (COG-UK) Consortium | Harper VanSteenhouse, Yumi Kasai, David Gray, Carol Clugston, Anna Dominiczak and Alex Alderton, Roberto Amato, Jeffrey Barrett, Sonia Goncalves, Ewan Harrison, David K. Jackson, Ian Johnston, Dominic Kwiatkowski, Cordelia Langford, John Sillitoe on behalf of the Wellcome Sanger Institute COVID-19 Surveillance Team |
| EPI_ISL_2993974, EPI_ISL_2993975, EPI_ISL_2993976 | Lighthouse Lab in Alderley Park | Wellcome Sanger Institute for the COVID-19 Genomics UK (COG-UK) Consortium | Jacquelyn Wynn, Mairead Hyland, The Lighthouse Lab in Alderley Park and Alex Alderton, Roberto Amato, Jeffrey Barrett, Sonia Goncalves, Ewan Harrison, David K. Jackson, Ian Johnston, Dominic Kwiatkowski, Cordelia Langford, John Sillitoe on behalf of the Wellcome Sanger Institute COVID-19 Surveillance Team |
| EPI_ISL_2993977, EPI_ISL_2993978 | Health Services Laboratories | Wellcome Sanger Institute for the COVID-19 Genomics UK (COG-UK) Consortium | Health Services Laboratories and Alex Alderton, Roberto Amato, Jeffrey Barrett, Sonia Goncalves, Ewan Harrison, David K. Jackson, Ian Johnston, Dominic Kwiatkowski, Cordelia Langford, John Sillitoe on behalf of the Wellcome Sanger Institute COVID-19 Surveillance Team |
| EPI_ISL_2993979 | Lighthouse Laboratory Plymouth | Wellcome Sanger Institute for the COVID-19 Genomics UK (COG-UK) Consortium | Lighthouse Laboratory Plymouth and Alex Alderton, Roberto Amato, Jeffrey Barrett, Sonia Goncalves, Ewan Harrison, David K. Jackson, Ian Johnston, Dominic Kwiatkowski, Cordelia Langford, John Sillitoe on behalf of the Wellcome Sanger Institute COVID-19 Surveillance Team |
| EPI_ISL_2993980 | Health Services Laboratories | Wellcome Sanger Institute for the COVID-19 Genomics UK (COG-UK) Consortium | Health Services Laboratories and Alex Alderton, Roberto Amato, Jeffrey Barrett, Sonia Goncalves, Ewan Harrison, David K. Jackson, Ian Johnston, Dominic Kwiatkowski, Cordelia Langford, John Sillitoe on behalf of the Wellcome Sanger Institute COVID-19 Surveillance Team |
| EPI_ISL_2993981 | Lighthouse Lab in Alderley Park | Wellcome Sanger Institute for the COVID-19 Genomics UK (COG-UK) Consortium | Jacquelyn Wynn, Mairead Hyland, The Lighthouse Lab in Alderley Park and Alex Alderton, Roberto Amato, Jeffrey Barrett, Sonia Goncalves, Ewan Harrison, David K. Jackson, Ian Johnston, Dominic Kwiatkowski, Cordelia Langford, John Sillitoe on behalf of the Wellcome Sanger Institute COVID-19 Surveillance Team |
| EPI_ISL_2993982 | Health Services Laboratories | Wellcome Sanger Institute for the COVID-19 Genomics UK (COG-UK) Consortium | Health Services Laboratories and Alex Alderton, Roberto Amato, Jeffrey Barrett, Sonia Goncalves, Ewan Harrison, David K. Jackson, Ian Johnston, Dominic Kwiatkowski, Cordelia Langford, John Sillitoe on behalf of the Wellcome Sanger Institute COVID-19 Surveillance Team |
| EPI_ISL_2993983 | Lighthouse Laboratory Plymouth | Wellcome Sanger Institute for the COVID-19 Genomics UK (COG-UK) Consortium | Lighthouse Laboratory Plymouth and Alex Alderton, Roberto Amato, Jeffrey Barrett, Sonia Goncalves, Ewan Harrison, David K. Jackson, Ian Johnston, Dominic Kwiatkowski, Cordelia Langford, John Sillitoe on behalf of the Wellcome Sanger Institute COVID-19 Surveillance Team |
| EPI_ISL_2993984 | Lighthouse Lab in Glasgow | Wellcome Sanger Institute for the COVID-19 Genomics UK (COG-UK) Consortium | Harper VanSteenhouse, Yumi Kasai, David Gray, Carol Clugston, Anna Dominiczak and Alex Alderton, Roberto Amato, Jeffrey Barrett, Sonia Goncalves, Ewan Harrison, David K. Jackson, Ian Johnston, Dominic Kwiatkowski, Cordelia Langford, John Sillitoe on behalf of the Wellcome Sanger Institute COVID-19 Surveillance Team |
| EPI_ISL_2993985, EPI_ISL_2993986 | Health Services Laboratories | Wellcome Sanger Institute for the COVID-19 Genomics UK (COG-UK) Consortium | Health Services Laboratories and Alex Alderton, Roberto Amato, Jeffrey Barrett, Sonia Goncalves, Ewan Harrison, David K. Jackson, Ian Johnston, Dominic Kwiatkowski, Cordelia Langford, John Sillitoe on behalf of the Wellcome Sanger Institute COVID-19 Surveillance Team |
| EPI_ISL_2993987 | Lighthouse Lab in Glasgow | Wellcome Sanger Institute for the COVID-19 Genomics UK (COG-UK) Consortium | Harper VanSteenhouse, Yumi Kasai, David Gray, Carol Clugston, Anna Dominiczak and Alex Alderton, Roberto Amato, Jeffrey Barrett, Sonia Goncalves, Ewan Harrison, David K. Jackson, Ian Johnston, Dominic Kwiatkowski, Cordelia Langford, John Sillitoe on behalf of the Wellcome Sanger Institute COVID-19 Surveillance Team |
| EPI_ISL_2993988, EPI_ISL_2993989, EPI_ISL_2993990 | Lighthouse Lab in Alderley Park | Wellcome Sanger Institute for the COVID-19 Genomics UK (COG-UK) Consortium | Jacquelyn Wynn, Mairead Hyland, The Lighthouse Lab in Alderley Park and Alex Alderton, Roberto Amato, Jeffrey Barrett, Sonia Goncalves, Ewan Harrison, David K. Jackson, Ian Johnston, Dominic Kwiatkowski, Cordelia Langford, John Sillitoe on behalf of the Wellcome Sanger Institute COVID-19 Surveillance Team |
| EPI_ISL_2993991 | Lighthouse Laboratory Plymouth | Wellcome Sanger Institute for the COVID-19 Genomics UK (COG-UK) Consortium | Lighthouse Laboratory Plymouth and Alex Alderton, Roberto Amato, Jeffrey Barrett, Sonia Goncalves, Ewan Harrison, David K. Jackson, Ian Johnston, Dominic Kwiatkowski, Cordelia Langford, John Sillitoe on behalf of the Wellcome Sanger Institute COVID-19 Surveillance Team |
| EPI_ISL_2993992, EPI_ISL_2993993 | Lighthouse Lab in Alderley Park | Wellcome Sanger Institute for the COVID-19 Genomics UK (COG-UK) Consortium | Jacquelyn Wynn, Mairead Hyland, The Lighthouse Lab in Alderley Park and Alex Alderton, Roberto Amato, Jeffrey Barrett, Sonia Goncalves, Ewan Harrison, David K. Jackson, Ian Johnston, Dominic Kwiatkowski, Cordelia Langford, John Sillitoe on behalf of the Wellcome Sanger Institute COVID-19 Surveillance Team |
| EPI_ISL_2993994 | Lighthouse Lab in Glasgow | Wellcome Sanger Institute for the COVID-19 Genomics UK (COG-UK) Consortium | Harper VanSteenhouse, Yumi Kasai, David Gray, Carol Clugston, Anna Dominiczak and Alex Alderton, Roberto Amato, Jeffrey Barrett, Sonia Goncalves, Ewan Harrison, David K. Jackson, Ian Johnston, Dominic Kwiatkowski, Cordelia Langford, John Sillitoe on behalf of the Wellcome Sanger Institute COVID-19 Surveillance Team |
| EPI_ISL_2993995, EPI_ISL_2993996 | Lighthouse Lab in Alderley Park | Wellcome Sanger Institute for the COVID-19 Genomics UK (COG-UK) Consortium | Jacquelyn Wynn, Mairead Hyland, The Lighthouse Lab in Alderley Park and Alex Alderton, Roberto Amato, Jeffrey Barrett, Sonia Goncalves, Ewan Harrison, David K. Jackson, Ian Johnston, Dominic Kwiatkowski, Cordelia Langford, John Sillitoe on behalf of the Wellcome Sanger Institute COVID-19 Surveillance Team |
| EPI_ISL_2993997, EPI_ISL_2993998 | Lighthouse Laboratory Plymouth | Wellcome Sanger Institute for the COVID-19 Genomics UK (COG-UK) Consortium | Lighthouse Laboratory Plymouth and Alex Alderton, Roberto Amato, Jeffrey Barrett, Sonia Goncalves, Ewan Harrison, David K. Jackson, Ian Johnston, Dominic Kwiatkowski, Cordelia Langford, John Sillitoe on behalf of the Wellcome Sanger Institute COVID-19 Surveillance Team |
| EPI_ISL_2993999 | Health Services Laboratories | Wellcome Sanger Institute for the COVID-19 Genomics UK (COG-UK) Consortium | Health Services Laboratories and Alex Alderton, Roberto Amato, Jeffrey Barrett, Sonia Goncalves, Ewan Harrison, David K. Jackson, Ian Johnston, Dominic Kwiatkowski, Cordelia Langford, John Sillitoe on behalf of the Wellcome Sanger Institute COVID-19 Surveillance Team |
| EPI_ISL_2994000 | Lighthouse Lab in Alderley Park | Wellcome Sanger Institute for the COVID-19 Genomics UK (COG-UK) Consortium | Jacquelyn Wynn, Mairead Hyland, The Lighthouse Lab in Alderley Park and Alex Alderton, Roberto Amato, Jeffrey Barrett, Sonia Goncalves, Ewan Harrison, David K. Jackson, Ian Johnston, Dominic Kwiatkowski, Cordelia Langford, John Sillitoe on behalf of the Wellcome Sanger Institute COVID-19 Surveillance Team |
| EPI_ISL_2994001, EPI_ISL_2994002, EPI_ISL_2994003, EPI_ISL_2994004 | Lighthouse Laboratory Plymouth | Wellcome Sanger Institute for the COVID-19 Genomics UK (COG-UK) Consortium | Lighthouse Laboratory Plymouth and Alex Alderton, Roberto Amato, Jeffrey Barrett, Sonia Goncalves, Ewan Harrison, David K. Jackson, Ian Johnston, Dominic Kwiatkowski, Cordelia Langford, John Sillitoe on behalf of the Wellcome Sanger Institute COVID-19 Surveillance Team |
| EPI_ISL_2994005, EPI_ISL_2994006 | Lighthouse Lab in Alderley Park | Wellcome Sanger Institute for the COVID-19 Genomics UK (COG-UK) Consortium | Jacquelyn Wynn, Mairead Hyland, The Lighthouse Lab in Alderley Park and Alex Alderton, Roberto Amato, Jeffrey Barrett, Sonia Goncalves, Ewan Harrison, David K. Jackson, Ian Johnston, Dominic Kwiatkowski, Cordelia Langford, John Sillitoe on behalf of the Wellcome Sanger Institute COVID-19 Surveillance Team |
| EPI_ISL_2994007 | Lighthouse Laboratory Plymouth | Wellcome Sanger Institute for the COVID-19 Genomics UK (COG-UK) Consortium | Lighthouse Laboratory Plymouth and Alex Alderton, Roberto Amato, Jeffrey Barrett, Sonia Goncalves, Ewan Harrison, David K. Jackson, Ian Johnston, Dominic Kwiatkowski, Cordelia Langford, John Sillitoe on behalf of the Wellcome Sanger Institute COVID-19 Surveillance Team |
| EPI_ISL_2994008, EPI_ISL_2994009 | Lighthouse Lab in Alderley Park | Wellcome Sanger Institute for the COVID-19 Genomics UK (COG-UK) Consortium | Jacquelyn Wynn, Mairead Hyland, The Lighthouse Lab in Alderley Park and Alex Alderton, Roberto Amato, Jeffrey Barrett, Sonia Goncalves, Ewan Harrison, David K. Jackson, Ian Johnston, Dominic Kwiatkowski, Cordelia Langford, John Sillitoe on behalf of the Wellcome Sanger Institute COVID-19 Surveillance Team |
| EPI_ISL_2994010 | Health Services Laboratories | Wellcome Sanger Institute for the COVID-19 Genomics UK (COG-UK) Consortium | Health Services Laboratories and Alex Alderton, Roberto Amato, Jeffrey Barrett, Sonia Goncalves, Ewan Harrison, David K. Jackson, Ian Johnston, Dominic Kwiatkowski, Cordelia Langford, John Sillitoe on behalf of the Wellcome Sanger Institute COVID-19 Surveillance Team |
| EPI_ISL_2994011 | Lighthouse Laboratory Plymouth | Wellcome Sanger Institute for the COVID-19 Genomics UK (COG-UK) Consortium | Lighthouse Laboratory Plymouth and Alex Alderton, Roberto Amato, Jeffrey Barrett, Sonia Goncalves, Ewan Harrison, David K. Jackson, Ian Johnston, Dominic Kwiatkowski, Cordelia Langford, John Sillitoe on behalf of the Wellcome Sanger Institute COVID-19 Surveillance Team |

| | | | |
|---|---|---|---|
| EPI_ISL_2994012 | Lighthouse Lab in Alderley Park | Wellcome Sanger Institute for the COVID-19 Genomics UK (COG-UK) Consortium | Jacquelyn Wynn, Mairead Hyland, The Lighthouse Lab in Alderley Park and Alex Alderton, Roberto Amato, Jeffrey Barrett, Sonia Goncalves, Ewan Harrison, David K. Jackson, Ian Johnston, Dominic Kwiatkowski, Cordelia Langford, John Sillitoe on behalf of the Wellcome Sanger Institute COVID-19 Surveillance Team |
| EPI_ISL_2994013 | Lighthouse Lab in Glasgow | Wellcome Sanger Institute for the COVID-19 Genomics UK (COG-UK) Consortium | Harper VanSteenhouse, Yumi Kasai, David Gray, Carol Clugston, Anna Dominiczak and Alex Alderton, Roberto Amato, Jeffrey Barrett, Sonia Goncalves, Ewan Harrison, David K. Jackson, Ian Johnston, Dominic Kwiatkowski, Cordelia Langford, John Sillitoe on behalf of the Wellcome Sanger Institute COVID-19 Surveillance Team |
| EPI_ISL_2994014 | Health Services Laboratories | Wellcome Sanger Institute for the COVID-19 Genomics UK (COG-UK) Consortium | Health Services Laboratories and Alex Alderton, Roberto Amato, Jeffrey Barrett, Sonia Goncalves, Ewan Harrison, David K. Jackson, Ian Johnston, Dominic Kwiatkowski, Cordelia Langford, John Sillitoe on behalf of the Wellcome Sanger Institute COVID-19 Surveillance Team |
| EPI_ISL_2994015 | Lighthouse Lab in Alderley Park | Wellcome Sanger Institute for the COVID-19 Genomics UK (COG-UK) Consortium | Jacquelyn Wynn, Mairead Hyland, The Lighthouse Lab in Alderley Park and Alex Alderton, Roberto Amato, Jeffrey Barrett, Sonia Goncalves, Ewan Harrison, David K. Jackson, Ian Johnston, Dominic Kwiatkowski, Cordelia Langford, John Sillitoe on behalf of the Wellcome Sanger Institute COVID-19 Surveillance Team |
| EPI_ISL_2994016 | Lighthouse Laboratory Plymouth | Wellcome Sanger Institute for the COVID-19 Genomics UK (COG-UK) Consortium | Lighthouse Laboratory Plymouth and Alex Alderton, Roberto Amato, Jeffrey Barrett, Sonia Goncalves, Ewan Harrison, David K. Jackson, Ian Johnston, Dominic Kwiatkowski, Cordelia Langford, John Sillitoe on behalf of the Wellcome Sanger Institute COVID-19 Surveillance Team |
| EPI_ISL_2994017 | Lighthouse Lab in Alderley Park | Wellcome Sanger Institute for the COVID-19 Genomics UK (COG-UK) Consortium | Jacquelyn Wynn, Mairead Hyland, The Lighthouse Lab in Alderley Park and Alex Alderton, Roberto Amato, Jeffrey Barrett, Sonia Goncalves, Ewan Harrison, David K. Jackson, Ian Johnston, Dominic Kwiatkowski, Cordelia Langford, John Sillitoe on behalf of the Wellcome Sanger Institute COVID-19 Surveillance Team |
| EPI_ISL_2994018 | Health Services Laboratories | Wellcome Sanger Institute for the COVID-19 Genomics UK (COG-UK) Consortium | Health Services Laboratories and Alex Alderton, Roberto Amato, Jeffrey Barrett, Sonia Goncalves, Ewan Harrison, David K. Jackson, Ian Johnston, Dominic Kwiatkowski, Cordelia Langford, John Sillitoe on behalf of the Wellcome Sanger Institute COVID-19 Surveillance Team |
| EPI_ISL_2994019 | Lighthouse Lab in Glasgow | Wellcome Sanger Institute for the COVID-19 Genomics UK (COG-UK) Consortium | Harper VanSteenhouse, Yumi Kasai, David Gray, Carol Clugston, Anna Dominiczak and Alex Alderton, Roberto Amato, Jeffrey Barrett, Sonia Goncalves, Ewan Harrison, David K. Jackson, Ian Johnston, Dominic Kwiatkowski, Cordelia Langford, John Sillitoe on behalf of the Wellcome Sanger Institute COVID-19 Surveillance Team |
| EPI_ISL_2994020 | Health Services Laboratories | Wellcome Sanger Institute for the COVID-19 Genomics UK (COG-UK) Consortium | Health Services Laboratories and Alex Alderton, Roberto Amato, Jeffrey Barrett, Sonia Goncalves, Ewan Harrison, David K. Jackson, Ian Johnston, Dominic Kwiatkowski, Cordelia Langford, John Sillitoe on behalf of the Wellcome Sanger Institute COVID-19 Surveillance Team |
| EPI_ISL_2994021 | Lighthouse Lab in Alderley Park | Wellcome Sanger Institute for the COVID-19 Genomics UK (COG-UK) Consortium | Jacquelyn Wynn, Mairead Hyland, The Lighthouse Lab in Alderley Park and Alex Alderton, Roberto Amato, Jeffrey Barrett, Sonia Goncalves, Ewan Harrison, David K. Jackson, Ian Johnston, Dominic Kwiatkowski, Cordelia Langford, John Sillitoe on behalf of the Wellcome Sanger Institute COVID-19 Surveillance Team |
| EPI_ISL_2994022, EPI_ISL_2994023, EPI_ISL_2994024 | Health Services Laboratories | Wellcome Sanger Institute for the COVID-19 Genomics UK (COG-UK) Consortium | Health Services Laboratories and Alex Alderton, Roberto Amato, Jeffrey Barrett, Sonia Goncalves, Ewan Harrison, David K. Jackson, Ian Johnston, Dominic Kwiatkowski, Cordelia Langford, John Sillitoe on behalf of the Wellcome Sanger Institute COVID-19 Surveillance Team |
| EPI_ISL_2994025 | Lighthouse Laboratory Plymouth | Wellcome Sanger Institute for the COVID-19 Genomics UK (COG-UK) Consortium | Lighthouse Laboratory Plymouth and Alex Alderton, Roberto Amato, Jeffrey Barrett, Sonia Goncalves, Ewan Harrison, David K. Jackson, Ian Johnston, Dominic Kwiatkowski, Cordelia Langford, John Sillitoe on behalf of the Wellcome Sanger Institute COVID-19 Surveillance Team |
| EPI_ISL_2994026, EPI_ISL_2994027 | Lighthouse Lab in Alderley Park | Wellcome Sanger Institute for the COVID-19 Genomics UK (COG-UK) Consortium | Jacquelyn Wynn, Mairead Hyland, The Lighthouse Lab in Alderley Park and Alex Alderton, Roberto Amato, Jeffrey Barrett, Sonia Goncalves, Ewan Harrison, David K. Jackson, Ian Johnston, Dominic Kwiatkowski, Cordelia Langford, John Sillitoe on behalf of the Wellcome Sanger Institute COVID-19 Surveillance Team |
| EPI_ISL_2994028 | Health Services Laboratories | Wellcome Sanger Institute for the COVID-19 Genomics UK (COG-UK) Consortium | Health Services Laboratories and Alex Alderton, Roberto Amato, Jeffrey Barrett, Sonia Goncalves, Ewan Harrison, David K. Jackson, Ian Johnston, Dominic Kwiatkowski, Cordelia Langford, John Sillitoe on behalf of the Wellcome Sanger Institute COVID-19 Surveillance Team |
| EPI_ISL_2994029 | Lighthouse Laboratory Plymouth | Wellcome Sanger Institute for the COVID-19 Genomics UK (COG-UK) Consortium | Lighthouse Laboratory Plymouth and Alex Alderton, Roberto Amato, Jeffrey Barrett, Sonia Goncalves, Ewan Harrison, David K. Jackson, Ian Johnston, Dominic Kwiatkowski, Cordelia Langford, John Sillitoe on behalf of the Wellcome Sanger Institute COVID-19 Surveillance Team |
| EPI_ISL_2994030 | Health Services Laboratories | Wellcome Sanger Institute for the COVID-19 Genomics UK (COG-UK) Consortium | Health Services Laboratories and Alex Alderton, Roberto Amato, Jeffrey Barrett, Sonia Goncalves, Ewan Harrison, David K. Jackson, Ian Johnston, Dominic Kwiatkowski, Cordelia Langford, John Sillitoe on behalf of the Wellcome Sanger Institute COVID-19 Surveillance Team |
| EPI_ISL_2994031 | Lighthouse Laboratory Plymouth | Wellcome Sanger Institute for the COVID-19 Genomics UK (COG-UK) Consortium | Lighthouse Laboratory Plymouth and Alex Alderton, Roberto Amato, Jeffrey Barrett, Sonia Goncalves, Ewan Harrison, David K. Jackson, Ian Johnston, Dominic Kwiatkowski, Cordelia Langford, John Sillitoe on behalf of the Wellcome Sanger Institute COVID-19 Surveillance Team |
| EPI_ISL_2994032 | Health Services Laboratories | Wellcome Sanger Institute for the COVID-19 Genomics UK (COG-UK) Consortium | Health Services Laboratories and Alex Alderton, Roberto Amato, Jeffrey Barrett, Sonia Goncalves, Ewan Harrison, David K. Jackson, Ian Johnston, Dominic Kwiatkowski, Cordelia Langford, John Sillitoe on behalf of the Wellcome Sanger Institute COVID-19 Surveillance Team |
| EPI_ISL_2994033 | Lighthouse Lab in Glasgow | Wellcome Sanger Institute for the COVID-19 Genomics UK (COG-UK) Consortium | Harper VanSteenhouse, Yumi Kasai, David Gray, Carol Clugston, Anna Dominiczak and Alex Alderton, Roberto Amato, Jeffrey Barrett, Sonia Goncalves, Ewan Harrison, David K. Jackson, Ian Johnston, Dominic Kwiatkowski, Cordelia Langford, John Sillitoe on behalf of the Wellcome Sanger Institute COVID-19 Surveillance Team |
| EPI_ISL_2994034 | Lighthouse Laboratory Plymouth | Wellcome Sanger Institute for the COVID-19 Genomics UK (COG-UK) Consortium | Lighthouse Laboratory Plymouth and Alex Alderton, Roberto Amato, Jeffrey Barrett, Sonia Goncalves, Ewan Harrison, David K. Jackson, Ian Johnston, Dominic Kwiatkowski, Cordelia Langford, John Sillitoe on behalf of the Wellcome Sanger Institute COVID-19 Surveillance Team |
| EPI_ISL_2994035 | Lighthouse Lab in Glasgow | Wellcome Sanger Institute for the COVID-19 Genomics UK (COG-UK) Consortium | Harper VanSteenhouse, Yumi Kasai, David Gray, Carol Clugston, Anna Dominiczak and Alex Alderton, Roberto Amato, Jeffrey Barrett, Sonia Goncalves, Ewan Harrison, David K. Jackson, Ian Johnston, Dominic Kwiatkowski, Cordelia Langford, John Sillitoe on behalf of the Wellcome Sanger Institute COVID-19 Surveillance Team |
| EPI_ISL_2994036, EPI_ISL_2994037 | Health Services Laboratories | Wellcome Sanger Institute for the COVID-19 Genomics UK (COG-UK) Consortium | Health Services Laboratories and Alex Alderton, Roberto Amato, Jeffrey Barrett, Sonia Goncalves, Ewan Harrison, David K. Jackson, Ian Johnston, Dominic Kwiatkowski, Cordelia Langford, John Sillitoe on behalf of the Wellcome Sanger Institute COVID-19 Surveillance Team |
| EPI_ISL_2994038 | Lighthouse Laboratory Plymouth | Wellcome Sanger Institute for the COVID-19 Genomics UK (COG-UK) Consortium | Lighthouse Laboratory Plymouth and Alex Alderton, Roberto Amato, Jeffrey Barrett, Sonia Goncalves, Ewan Harrison, David K. Jackson, Ian Johnston, Dominic Kwiatkowski, Cordelia Langford, John Sillitoe on behalf of the Wellcome Sanger Institute COVID-19 Surveillance Team |
| EPI_ISL_2994039, EPI_ISL_2994040, EPI_ISL_2994041, EPI_ISL_2994042, EPI_ISL_2994043 | Lighthouse Lab in Alderley Park | Wellcome Sanger Institute for the COVID-19 Genomics UK (COG-UK) Consortium | Jacquelyn Wynn, Mairead Hyland, The Lighthouse Lab in Alderley Park and Alex Alderton, Roberto Amato, Jeffrey Barrett, Sonia Goncalves, Ewan Harrison, David K. Jackson, Ian Johnston, Dominic Kwiatkowski, Cordelia Langford, John Sillitoe on behalf of the Wellcome Sanger Institute COVID-19 Surveillance Team |
| EPI_ISL_2994044, EPI_ISL_2994045 | Lighthouse Laboratory Plymouth | Wellcome Sanger Institute for the COVID-19 Genomics UK (COG-UK) Consortium | Lighthouse Laboratory Plymouth and Alex Alderton, Roberto Amato, Jeffrey Barrett, Sonia Goncalves, Ewan Harrison, David K. Jackson, Ian Johnston, Dominic Kwiatkowski, Cordelia Langford, John Sillitoe on behalf of the Wellcome Sanger Institute COVID-19 Surveillance Team |
| EPI_ISL_2994046 | Lighthouse Lab in Glasgow | Wellcome Sanger Institute for the COVID-19 Genomics UK (COG-UK) Consortium | Harper VanSteenhouse, Yumi Kasai, David Gray, Carol Clugston, Anna Dominiczak and Alex Alderton, Roberto Amato, Jeffrey Barrett, Sonia Goncalves, Ewan Harrison, David K. Jackson, Ian Johnston, Dominic Kwiatkowski, Cordelia Langford, John Sillitoe on behalf of the Wellcome Sanger Institute COVID-19 Surveillance Team |
| EPI_ISL_2994047, EPI_ISL_2994048 | Lighthouse Lab in Alderley Park | Wellcome Sanger Institute for the COVID-19 Genomics UK (COG-UK) Consortium | Jacquelyn Wynn, Mairead Hyland, The Lighthouse Lab in Alderley Park and Alex Alderton, Roberto Amato, Jeffrey Barrett, Sonia Goncalves, Ewan Harrison, David K. Jackson, Ian Johnston, Dominic Kwiatkowski, Cordelia Langford, John Sillitoe on behalf of the Wellcome Sanger Institute COVID-19 Surveillance Team |
| EPI_ISL_2994049, EPI_ISL_2994050 | Health Services Laboratories | Wellcome Sanger Institute for the COVID-19 Genomics UK (COG-UK) Consortium | Health Services Laboratories and Alex Alderton, Roberto Amato, Jeffrey Barrett, Sonia Goncalves, Ewan Harrison, David K. Jackson, Ian Johnston, Dominic Kwiatkowski, Cordelia Langford, John Sillitoe on behalf of the Wellcome Sanger Institute COVID-19 Surveillance Team |
| EPI_ISL_2994051, EPI_ISL_2994052 | Lighthouse Laboratory Plymouth | Wellcome Sanger Institute for the COVID-19 Genomics UK (COG-UK) Consortium | Lighthouse Laboratory Plymouth and Alex Alderton, Roberto Amato, Jeffrey Barrett, Sonia Goncalves, Ewan Harrison, David K. Jackson, Ian Johnston, Dominic Kwiatkowski, Cordelia Langford, John Sillitoe on behalf of the Wellcome Sanger Institute COVID-19 Surveillance Team |
| EPI_ISL_2994053, EPI_ISL_2994054 | Lighthouse Lab in Alderley Park | Wellcome Sanger Institute for the COVID-19 Genomics UK (COG-UK) Consortium | Jacquelyn Wynn, Mairead Hyland, The Lighthouse Lab in Alderley Park and Alex Alderton, Roberto Amato, Jeffrey Barrett, Sonia Goncalves, Ewan Harrison, David K. Jackson, Ian Johnston, Dominic Kwiatkowski, Cordelia Langford, John Sillitoe on behalf of the Wellcome Sanger Institute COVID-19 Surveillance Team |
| EPI_ISL_2994055, EPI_ISL_2994056 | Lighthouse Lab in Milton Keynes | Wellcome Sanger Institute for the COVID-19 Genomics UK (COG-UK) Consortium | The Lighthouse Lab in Milton Keynes and Alex Alderton, Roberto Amato, Jeffrey Barrett, Sonia Goncalves, Ewan Harrison, David K. Jackson, Ian Johnston, Dominic Kwiatkowski, Cordelia Langford, John Sillitoe on behalf of the Wellcome Sanger Institute COVID-19 Surveillance Team |
| EPI_ISL_2994057 | Berkshire and Surrey Pathology Services Lighthouse Laboratory | Wellcome Sanger Institute for the COVID-19 Genomics UK (COG-UK) Consortium | Berkshire and Surrey Pathology Services Lighthouse Laboratory and Alex Alderton, Roberto Amato, Jeffrey Barrett, Sonia Goncalves, Ewan Harrison, David K. Jackson, Ian Johnston, Dominic Kwiatkowski, Cordelia Langford, John Sillitoe on behalf of the Wellcome Sanger Institute COVID-19 Surveillance Team |
| EPI_ISL_2994058 | Lighthouse Laboratory Plymouth | Wellcome Sanger Institute for the COVID-19 Genomics UK (COG-UK) Consortium | Lighthouse Laboratory Plymouth and Alex Alderton, Roberto Amato, Jeffrey Barrett, Sonia Goncalves, Ewan Harrison, David K. Jackson, Ian Johnston, Dominic Kwiatkowski, Cordelia Langford, John Sillitoe on behalf of the Wellcome Sanger Institute COVID-19 Surveillance Team |
| EPI_ISL_2994059, EPI_ISL_2994060 | Lighthouse Lab in Alderley Park | Wellcome Sanger Institute for the COVID-19 Genomics UK (COG-UK) Consortium | Jacquelyn Wynn, Mairead Hyland, The Lighthouse Lab in Alderley Park and Alex Alderton, Roberto Amato, Jeffrey Barrett, Sonia Goncalves, Ewan Harrison, David K. Jackson, Ian Johnston, Dominic Kwiatkowski, Cordelia Langford, John Sillitoe on behalf of the Wellcome Sanger Institute COVID-19 Surveillance Team |
| EPI_ISL_2994061 | Lighthouse Laboratory Plymouth | Wellcome Sanger Institute for the COVID-19 Genomics UK (COG-UK) Consortium | Lighthouse Laboratory Plymouth and Alex Alderton, Roberto Amato, Jeffrey Barrett, Sonia Goncalves, Ewan Harrison, David K. Jackson, Ian Johnston, Dominic Kwiatkowski, Cordelia Langford, John Sillitoe on behalf of the Wellcome Sanger Institute COVID-19 Surveillance Team |
| EPI_ISL_2994062, EPI_ISL_2994063 | Lighthouse Lab in Alderley Park | Wellcome Sanger Institute for the COVID-19 Genomics UK (COG-UK) Consortium | Jacquelyn Wynn, Mairead Hyland, The Lighthouse Lab in Alderley Park and Alex Alderton, Roberto Amato, Jeffrey Barrett, Sonia Goncalves, Ewan Harrison, David K. Jackson, Ian Johnston, Dominic Kwiatkowski, Cordelia Langford, John Sillitoe on behalf of the Wellcome Sanger Institute COVID-19 Surveillance Team |
| EPI_ISL_2994064 | Lighthouse Laboratory Plymouth | Wellcome Sanger Institute for the COVID-19 Genomics UK (COG-UK) Consortium | Lighthouse Laboratory Plymouth and Alex Alderton, Roberto Amato, Jeffrey Barrett, Sonia Goncalves, Ewan Harrison, David K. Jackson, Ian Johnston, Dominic Kwiatkowski, Cordelia Langford, John Sillitoe on behalf of the Wellcome Sanger Institute COVID-19 Surveillance Team |
| EPI_ISL_2994065 | Lighthouse Lab in Alderley Park | Wellcome Sanger Institute for the COVID-19 Genomics UK (COG-UK) Consortium | Jacquelyn Wynn, Mairead Hyland, The Lighthouse Lab in Alderley Park and Alex Alderton, Roberto Amato, Jeffrey Barrett, Sonia Goncalves, Ewan Harrison, David K. Jackson, Ian Johnston, Dominic Kwiatkowski, Cordelia Langford, John Sillitoe on behalf of the Wellcome Sanger Institute COVID-19 Surveillance Team |
| EPI_ISL_2994066 | Berkshire and Surrey Pathology Services Lighthouse Laboratory | Wellcome Sanger Institute for the COVID-19 Genomics UK (COG-UK) Consortium | Berkshire and Surrey Pathology Services Lighthouse Laboratory and Alex Alderton, Roberto Amato, Jeffrey Barrett, Sonia Goncalves, Ewan Harrison, David K. Jackson, Ian Johnston, Dominic Kwiatkowski, Cordelia Langford, John Sillitoe on behalf of the Wellcome Sanger Institute COVID-19 Surveillance Team |
| EPI_ISL_2994067, EPI_ISL_2994068 | Lighthouse Laboratory Plymouth | Wellcome Sanger Institute for the COVID-19 Genomics UK (COG-UK) Consortium | Lighthouse Laboratory Plymouth and Alex Alderton, Roberto Amato, Jeffrey Barrett, Sonia Goncalves, Ewan Harrison, David K. Jackson, Ian Johnston, Dominic Kwiatkowski, Cordelia Langford, John Sillitoe on behalf of the Wellcome Sanger Institute COVID-19 Surveillance Team |
| EPI_ISL_2994069 | Lighthouse Lab in Milton Keynes | Wellcome Sanger Institute for the COVID-19 Genomics UK (COG-UK) Consortium | The Lighthouse Lab in Milton Keynes and Alex Alderton, Roberto Amato, Jeffrey Barrett, Sonia Goncalves, Ewan Harrison, David K. Jackson, Ian Johnston, Dominic Kwiatkowski, Cordelia Langford, John Sillitoe on behalf of the Wellcome Sanger Institute COVID-19 Surveillance Team |
| EPI_ISL_2994070 | Lighthouse Laboratory Plymouth | Wellcome Sanger Institute for the COVID-19 Genomics UK (COG-UK) Consortium | Lighthouse Laboratory Plymouth and Alex Alderton, Roberto Amato, Jeffrey Barrett, Sonia Goncalves, Ewan Harrison, David K. Jackson, Ian Johnston, Dominic Kwiatkowski, Cordelia Langford, John Sillitoe on behalf of the Wellcome Sanger Institute COVID-19 Surveillance Team |
| EPI_ISL_2994071 | Lighthouse Lab in Alderley Park | Wellcome Sanger Institute for the COVID-19 Genomics UK (COG-UK) Consortium | Jacquelyn Wynn, Mairead Hyland, The Lighthouse Lab in Alderley Park and Alex Alderton, Roberto Amato, Jeffrey Barrett, Sonia Goncalves, Ewan Harrison, David K. Jackson, Ian Johnston, Dominic Kwiatkowski, Cordelia Langford, John Sillitoe on behalf of the Wellcome Sanger Institute COVID-19 Surveillance Team |
| EPI_ISL_2994072 | Berkshire and Surrey Pathology Services Lighthouse Laboratory | Wellcome Sanger Institute for the COVID-19 Genomics UK (COG-UK) | Berkshire and Surrey Pathology Services Lighthouse Laboratory and Alex Alderton, Roberto Amato, Jeffrey Barrett, Sonia Goncalves, Ewan Harrison, David K. Jackson, Ian Johnston, Dominic Kwiatkowski, Cordelia Langford, John Sillitoe on behalf |

| | | | |
|---|---|---|---|
| | | Consortium | of the Wellcome Sanger Institute COVID-19 Surveillance Team |
| EPI_ISL_2994073 | Lighthouse Lab in Milton Keynes | Wellcome Sanger Institute for the COVID-19 Genomics UK (COG-UK) Consortium | The Lighthouse Lab in Milton Keynes and Alex Alderton, Roberto Amato, Jeffrey Barrett, Sonia Goncalves, Ewan Harrison, David K. Jackson, Ian Johnston, Dominic Kwiatkowski, Cordelia Langford, John Sillitoe on behalf of the Wellcome Sanger Institute COVID-19 Surveillance Team |
| EPI_ISL_2994074, EPI_ISL_2994075 | Berkshire and Surrey Pathology Services Lighthouse Laboratory | Wellcome Sanger Institute for the COVID-19 Genomics UK (COG-UK) Consortium | Berkshire and Surrey Pathology Services Lighthouse Laboratory and Alex Alderton, Roberto Amato, Jeffrey Barrett, Sonia Goncalves, Ewan Harrison, David K. Jackson, Ian Johnston, Dominic Kwiatkowski, Cordelia Langford, John Sillitoe on behalf of the Wellcome Sanger Institute COVID-19 Surveillance Team |
| EPI_ISL_2994076 | Lighthouse Lab in Milton Keynes | Wellcome Sanger Institute for the COVID-19 Genomics UK (COG-UK) Consortium | The Lighthouse Lab in Milton Keynes and Alex Alderton, Roberto Amato, Jeffrey Barrett, Sonia Goncalves, Ewan Harrison, David K. Jackson, Ian Johnston, Dominic Kwiatkowski, Cordelia Langford, John Sillitoe on behalf of the Wellcome Sanger Institute COVID-19 Surveillance Team |
| EPI_ISL_2994077 | Lighthouse Lab in Alderley Park | Wellcome Sanger Institute for the COVID-19 Genomics UK (COG-UK) Consortium | Jacquelyn Wynn, Mairead Hyland, The Lighthouse Lab in Alderley Park and Alex Alderton, Roberto Amato, Jeffrey Barrett, Sonia Goncalves, Ewan Harrison, David K. Jackson, Ian Johnston, Dominic Kwiatkowski, Cordelia Langford, John Sillitoe on behalf of the Wellcome Sanger Institute COVID-19 Surveillance Team |
| EPI_ISL_2994078 | Lighthouse Lab in Milton Keynes | Wellcome Sanger Institute for the COVID-19 Genomics UK (COG-UK) Consortium | The Lighthouse Lab in Milton Keynes and Alex Alderton, Roberto Amato, Jeffrey Barrett, Sonia Goncalves, Ewan Harrison, David K. Jackson, Ian Johnston, Dominic Kwiatkowski, Cordelia Langford, John Sillitoe on behalf of the Wellcome Sanger Institute COVID-19 Surveillance Team |
| EPI_ISL_2994079 | Lighthouse Lab in Alderley Park | Wellcome Sanger Institute for the COVID-19 Genomics UK (COG-UK) Consortium | Jacquelyn Wynn, Mairead Hyland, The Lighthouse Lab in Alderley Park and Alex Alderton, Roberto Amato, Jeffrey Barrett, Sonia Goncalves, Ewan Harrison, David K. Jackson, Ian Johnston, Dominic Kwiatkowski, Cordelia Langford, John Sillitoe on behalf of the Wellcome Sanger Institute COVID-19 Surveillance Team |
| EPI_ISL_2994080 | Berkshire and Surrey Pathology Services Lighthouse Laboratory | Wellcome Sanger Institute for the COVID-19 Genomics UK (COG-UK) Consortium | Berkshire and Surrey Pathology Services Lighthouse Laboratory and Alex Alderton, Roberto Amato, Jeffrey Barrett, Sonia Goncalves, Ewan Harrison, David K. Jackson, Ian Johnston, Dominic Kwiatkowski, Cordelia Langford, John Sillitoe on behalf of the Wellcome Sanger Institute COVID-19 Surveillance Team |
| EPI_ISL_2994081 | Lighthouse Lab in Alderley Park | Wellcome Sanger Institute for the COVID-19 Genomics UK (COG-UK) Consortium | Jacquelyn Wynn, Mairead Hyland, The Lighthouse Lab in Alderley Park and Alex Alderton, Roberto Amato, Jeffrey Barrett, Sonia Goncalves, Ewan Harrison, David K. Jackson, Ian Johnston, Dominic Kwiatkowski, Cordelia Langford, John Sillitoe on behalf of the Wellcome Sanger Institute COVID-19 Surveillance Team |
| EPI_ISL_2994082, EPI_ISL_2994083, EPI_ISL_2994084 | Lighthouse Lab in Milton Keynes | Wellcome Sanger Institute for the COVID-19 Genomics UK (COG-UK) Consortium | The Lighthouse Lab in Milton Keynes and Alex Alderton, Roberto Amato, Jeffrey Barrett, Sonia Goncalves, Ewan Harrison, David K. Jackson, Ian Johnston, Dominic Kwiatkowski, Cordelia Langford, John Sillitoe on behalf of the Wellcome Sanger Institute COVID-19 Surveillance Team |
| EPI_ISL_2994085 | Lighthouse Lab in Alderley Park | Wellcome Sanger Institute for the COVID-19 Genomics UK (COG-UK) Consortium | Jacquelyn Wynn, Mairead Hyland, The Lighthouse Lab in Alderley Park and Alex Alderton, Roberto Amato, Jeffrey Barrett, Sonia Goncalves, Ewan Harrison, David K. Jackson, Ian Johnston, Dominic Kwiatkowski, Cordelia Langford, John Sillitoe on behalf of the Wellcome Sanger Institute COVID-19 Surveillance Team |
| EPI_ISL_2994086 | Lighthouse Lab in Milton Keynes | Wellcome Sanger Institute for the COVID-19 Genomics UK (COG-UK) Consortium | The Lighthouse Lab in Milton Keynes and Alex Alderton, Roberto Amato, Jeffrey Barrett, Sonia Goncalves, Ewan Harrison, David K. Jackson, Ian Johnston, Dominic Kwiatkowski, Cordelia Langford, John Sillitoe on behalf of the Wellcome Sanger Institute COVID-19 Surveillance Team |
| EPI_ISL_2994087, EPI_ISL_2994088 | Berkshire and Surrey Pathology Services Lighthouse Laboratory | Wellcome Sanger Institute for the COVID-19 Genomics UK (COG-UK) Consortium | Berkshire and Surrey Pathology Services Lighthouse Laboratory and Alex Alderton, Roberto Amato, Jeffrey Barrett, Sonia Goncalves, Ewan Harrison, David K. Jackson, Ian Johnston, Dominic Kwiatkowski, Cordelia Langford, John Sillitoe on behalf of the Wellcome Sanger Institute COVID-19 Surveillance Team |
| EPI_ISL_2994089 | Lighthouse Lab in Milton Keynes | Wellcome Sanger Institute for the COVID-19 Genomics UK (COG-UK) Consortium | The Lighthouse Lab in Milton Keynes and Alex Alderton, Roberto Amato, Jeffrey Barrett, Sonia Goncalves, Ewan Harrison, David K. Jackson, Ian Johnston, Dominic Kwiatkowski, Cordelia Langford, John Sillitoe on behalf of the Wellcome Sanger Institute COVID-19 Surveillance Team |
| EPI_ISL_2994090 | Berkshire and Surrey Pathology Services Lighthouse Laboratory | Wellcome Sanger Institute for the COVID-19 Genomics UK (COG-UK) Consortium | Berkshire and Surrey Pathology Services Lighthouse Laboratory and Alex Alderton, Roberto Amato, Jeffrey Barrett, Sonia Goncalves, Ewan Harrison, David K. Jackson, Ian Johnston, Dominic Kwiatkowski, Cordelia Langford, John Sillitoe on behalf of the Wellcome Sanger Institute COVID-19 Surveillance Team |
| EPI_ISL_2994091 | Lighthouse Laboratory Plymouth | Wellcome Sanger Institute for the COVID-19 Genomics UK (COG-UK) Consortium | Lighthouse Laboratory Plymouth and Alex Alderton, Roberto Amato, Jeffrey Barrett, Sonia Goncalves, Ewan Harrison, David K. Jackson, Ian Johnston, Dominic Kwiatkowski, Cordelia Langford, John Sillitoe on behalf of the Wellcome Sanger Institute COVID-19 Surveillance Team |
| EPI_ISL_2994092 | Berkshire and Surrey Pathology Services Lighthouse Laboratory | Wellcome Sanger Institute for the COVID-19 Genomics UK (COG-UK) Consortium | Berkshire and Surrey Pathology Services Lighthouse Laboratory and Alex Alderton, Roberto Amato, Jeffrey Barrett, Sonia Goncalves, Ewan Harrison, David K. Jackson, Ian Johnston, Dominic Kwiatkowski, Cordelia Langford, John Sillitoe on behalf of the Wellcome Sanger Institute COVID-19 Surveillance Team |
| EPI_ISL_2994093 | Lighthouse Lab in Alderley Park | Wellcome Sanger Institute for the COVID-19 Genomics UK (COG-UK) Consortium | Jacquelyn Wynn, Mairead Hyland, The Lighthouse Lab in Alderley Park and Alex Alderton, Roberto Amato, Jeffrey Barrett, Sonia Goncalves, Ewan Harrison, David K. Jackson, Ian Johnston, Dominic Kwiatkowski, Cordelia Langford, John Sillitoe on behalf of the Wellcome Sanger Institute COVID-19 Surveillance Team |
| EPI_ISL_2994094 | Lighthouse Lab in Milton Keynes | Wellcome Sanger Institute for the COVID-19 Genomics UK (COG-UK) Consortium | The Lighthouse Lab in Milton Keynes and Alex Alderton, Roberto Amato, Jeffrey Barrett, Sonia Goncalves, Ewan Harrison, David K. Jackson, Ian Johnston, Dominic Kwiatkowski, Cordelia Langford, John Sillitoe on behalf of the Wellcome Sanger Institute COVID-19 Surveillance Team |
| EPI_ISL_2994095 | Lighthouse Lab in Alderley Park | Wellcome Sanger Institute for the COVID-19 Genomics UK (COG-UK) Consortium | Jacquelyn Wynn, Mairead Hyland, The Lighthouse Lab in Alderley Park and Alex Alderton, Roberto Amato, Jeffrey Barrett, Sonia Goncalves, Ewan Harrison, David K. Jackson, Ian Johnston, Dominic Kwiatkowski, Cordelia Langford, John Sillitoe on behalf of the Wellcome Sanger Institute COVID-19 Surveillance Team |
| EPI_ISL_2994096 | Lighthouse Laboratory Plymouth | Wellcome Sanger Institute for the COVID-19 Genomics UK (COG-UK) Consortium | Lighthouse Laboratory Plymouth and Alex Alderton, Roberto Amato, Jeffrey Barrett, Sonia Goncalves, Ewan Harrison, David K. Jackson, Ian Johnston, Dominic Kwiatkowski, Cordelia Langford, John Sillitoe on behalf of the Wellcome Sanger Institute COVID-19 Surveillance Team |
| EPI_ISL_2994097, EPI_ISL_2994098 | Lighthouse Lab in Milton Keynes | Wellcome Sanger Institute for the COVID-19 Genomics UK (COG-UK) Consortium | The Lighthouse Lab in Milton Keynes and Alex Alderton, Roberto Amato, Jeffrey Barrett, Sonia Goncalves, Ewan Harrison, David K. Jackson, Ian Johnston, Dominic Kwiatkowski, Cordelia Langford, John Sillitoe on behalf of the Wellcome Sanger Institute COVID-19 Surveillance Team |
| EPI_ISL_2994099 | Berkshire and Surrey Pathology Services Lighthouse Laboratory | Wellcome Sanger Institute for the COVID-19 Genomics UK (COG-UK) Consortium | Berkshire and Surrey Pathology Services Lighthouse Laboratory and Alex Alderton, Roberto Amato, Jeffrey Barrett, Sonia Goncalves, Ewan Harrison, David K. Jackson, Ian Johnston, Dominic Kwiatkowski, Cordelia Langford, John Sillitoe on behalf of the Wellcome Sanger Institute COVID-19 Surveillance Team |
| EPI_ISL_2994100 | Lighthouse Lab in Alderley Park | Wellcome Sanger Institute for the COVID-19 Genomics UK (COG-UK) Consortium | Jacquelyn Wynn, Mairead Hyland, The Lighthouse Lab in Alderley Park and Alex Alderton, Roberto Amato, Jeffrey Barrett, Sonia Goncalves, Ewan Harrison, David K. Jackson, Ian Johnston, Dominic Kwiatkowski, Cordelia Langford, John Sillitoe on behalf of the Wellcome Sanger Institute COVID-19 Surveillance Team |
| EPI_ISL_2994101, EPI_ISL_2994102 | Berkshire and Surrey Pathology Services Lighthouse Laboratory | Wellcome Sanger Institute for the COVID-19 Genomics UK (COG-UK) Consortium | Berkshire and Surrey Pathology Services Lighthouse Laboratory and Alex Alderton, Roberto Amato, Jeffrey Barrett, Sonia Goncalves, Ewan Harrison, David K. Jackson, Ian Johnston, Dominic Kwiatkowski, Cordelia Langford, John Sillitoe on behalf of the Wellcome Sanger Institute COVID-19 Surveillance Team |
| EPI_ISL_2994103 | Lighthouse Lab in Milton Keynes | Wellcome Sanger Institute for the COVID-19 Genomics UK (COG-UK) Consortium | The Lighthouse Lab in Milton Keynes and Alex Alderton, Roberto Amato, Jeffrey Barrett, Sonia Goncalves, Ewan Harrison, David K. Jackson, Ian Johnston, Dominic Kwiatkowski, Cordelia Langford, John Sillitoe on behalf of the Wellcome Sanger Institute COVID-19 Surveillance Team |
| EPI_ISL_2994104 | Lighthouse Laboratory Plymouth | Wellcome Sanger Institute for the COVID-19 Genomics UK (COG-UK) Consortium | Lighthouse Laboratory Plymouth and Alex Alderton, Roberto Amato, Jeffrey Barrett, Sonia Goncalves, Ewan Harrison, David K. Jackson, Ian Johnston, Dominic Kwiatkowski, Cordelia Langford, John Sillitoe on behalf of the Wellcome Sanger Institute COVID-19 Surveillance Team |
| EPI_ISL_2994105, EPI_ISL_2994106 | Lighthouse Lab in Alderley Park | Wellcome Sanger Institute for the COVID-19 Genomics UK (COG-UK) Consortium | Jacquelyn Wynn, Mairead Hyland, The Lighthouse Lab in Alderley Park and Alex Alderton, Roberto Amato, Jeffrey Barrett, Sonia Goncalves, Ewan Harrison, David K. Jackson, Ian Johnston, Dominic Kwiatkowski, Cordelia Langford, John Sillitoe on behalf of the Wellcome Sanger Institute COVID-19 Surveillance Team |
| EPI_ISL_2994107 | Lighthouse Laboratory Plymouth | Wellcome Sanger Institute for the COVID-19 Genomics UK (COG-UK) Consortium | Lighthouse Laboratory Plymouth and Alex Alderton, Roberto Amato, Jeffrey Barrett, Sonia Goncalves, Ewan Harrison, David K. Jackson, Ian Johnston, Dominic Kwiatkowski, Cordelia Langford, John Sillitoe on behalf of the Wellcome Sanger Institute COVID-19 Surveillance Team |
| EPI_ISL_2994108, EPI_ISL_2994109 | Lighthouse Lab in Milton Keynes | Wellcome Sanger Institute for the COVID-19 Genomics UK (COG-UK) Consortium | The Lighthouse Lab in Milton Keynes and Alex Alderton, Roberto Amato, Jeffrey Barrett, Sonia Goncalves, Ewan Harrison, David K. Jackson, Ian Johnston, Dominic Kwiatkowski, Cordelia Langford, John Sillitoe on behalf of the Wellcome Sanger Institute COVID-19 Surveillance Team |
| EPI_ISL_2994110 | Lighthouse Lab in Alderley Park | Wellcome Sanger Institute for the COVID-19 Genomics UK (COG-UK) Consortium | Jacquelyn Wynn, Mairead Hyland, The Lighthouse Lab in Alderley Park and Alex Alderton, Roberto Amato, Jeffrey Barrett, Sonia Goncalves, Ewan Harrison, David K. Jackson, Ian Johnston, Dominic Kwiatkowski, Cordelia Langford, John Sillitoe on behalf of the Wellcome Sanger Institute COVID-19 Surveillance Team |
| EPI_ISL_2994111 | Lighthouse Laboratory Plymouth | Wellcome Sanger Institute for the COVID-19 Genomics UK (COG-UK) Consortium | Lighthouse Laboratory Plymouth and Alex Alderton, Roberto Amato, Jeffrey Barrett, Sonia Goncalves, Ewan Harrison, David K. Jackson, Ian Johnston, Dominic Kwiatkowski, Cordelia Langford, John Sillitoe on behalf of the Wellcome Sanger Institute COVID-19 Surveillance Team |
| EPI_ISL_2994112 | Lighthouse Lab in Alderley Park | Wellcome Sanger Institute for the COVID-19 Genomics UK (COG-UK) Consortium | Jacquelyn Wynn, Mairead Hyland, The Lighthouse Lab in Alderley Park and Alex Alderton, Roberto Amato, Jeffrey Barrett, Sonia Goncalves, Ewan Harrison, David K. Jackson, Ian Johnston, Dominic Kwiatkowski, Cordelia Langford, John Sillitoe on behalf of the Wellcome Sanger Institute COVID-19 Surveillance Team |
| EPI_ISL_2994113 | Lighthouse Lab in Milton Keynes | Wellcome Sanger Institute for the COVID-19 Genomics UK (COG-UK) Consortium | The Lighthouse Lab in Milton Keynes and Alex Alderton, Roberto Amato, Jeffrey Barrett, Sonia Goncalves, Ewan Harrison, David K. Jackson, Ian Johnston, Dominic Kwiatkowski, Cordelia Langford, John Sillitoe on behalf of the Wellcome Sanger Institute COVID-19 Surveillance Team |
| EPI_ISL_2994114, EPI_ISL_2994115 | Lighthouse Laboratory Plymouth | Wellcome Sanger Institute for the COVID-19 Genomics UK (COG-UK) Consortium | Lighthouse Laboratory Plymouth and Alex Alderton, Roberto Amato, Jeffrey Barrett, Sonia Goncalves, Ewan Harrison, David K. Jackson, Ian Johnston, Dominic Kwiatkowski, Cordelia Langford, John Sillitoe on behalf of the Wellcome Sanger Institute COVID-19 Surveillance Team |
| EPI_ISL_2994116 | Lighthouse Lab in Alderley Park | Wellcome Sanger Institute for the COVID-19 Genomics UK (COG-UK) Consortium | Jacquelyn Wynn, Mairead Hyland, The Lighthouse Lab in Alderley Park and Alex Alderton, Roberto Amato, Jeffrey Barrett, Sonia Goncalves, Ewan Harrison, David K. Jackson, Ian Johnston, Dominic Kwiatkowski, Cordelia Langford, John Sillitoe on behalf of the Wellcome Sanger Institute COVID-19 Surveillance Team |
| EPI_ISL_2994117 | Berkshire and Surrey Pathology Services Lighthouse Laboratory | Wellcome Sanger Institute for the COVID-19 Genomics UK (COG-UK) Consortium | Berkshire and Surrey Pathology Services Lighthouse Laboratory and Alex Alderton, Roberto Amato, Jeffrey Barrett, Sonia Goncalves, Ewan Harrison, David K. Jackson, Ian Johnston, Dominic Kwiatkowski, Cordelia Langford, John Sillitoe on behalf of the Wellcome Sanger Institute COVID-19 Surveillance Team |
| EPI_ISL_2994118 | Lighthouse Lab in Milton Keynes | Wellcome Sanger Institute for the COVID-19 Genomics UK (COG-UK) Consortium | The Lighthouse Lab in Milton Keynes and Alex Alderton, Roberto Amato, Jeffrey Barrett, Sonia Goncalves, Ewan Harrison, David K. Jackson, Ian Johnston, Dominic Kwiatkowski, Cordelia Langford, John Sillitoe on behalf of the Wellcome Sanger Institute COVID-19 Surveillance Team |
| EPI_ISL_2994119 | Lighthouse Laboratory Plymouth | Wellcome Sanger Institute for the COVID-19 Genomics UK (COG-UK) Consortium | Lighthouse Laboratory Plymouth and Alex Alderton, Roberto Amato, Jeffrey Barrett, Sonia Goncalves, Ewan Harrison, David K. Jackson, Ian Johnston, Dominic Kwiatkowski, Cordelia Langford, John Sillitoe on behalf of the Wellcome Sanger Institute COVID-19 Surveillance Team |
| EPI_ISL_2994120 | Lighthouse Lab in Milton Keynes | Wellcome Sanger Institute for the COVID-19 Genomics UK (COG-UK) Consortium | The Lighthouse Lab in Milton Keynes and Alex Alderton, Roberto Amato, Jeffrey Barrett, Sonia Goncalves, Ewan Harrison, David K. Jackson, Ian Johnston, Dominic Kwiatkowski, Cordelia Langford, John Sillitoe on behalf of the Wellcome Sanger Institute COVID-19 Surveillance Team |
| EPI_ISL_2994121, EPI_ISL_2994122, EPI_ISL_2994123, EPI_ISL_2994124 | Lighthouse Lab in Alderley Park | Wellcome Sanger Institute for the COVID-19 Genomics UK (COG-UK) Consortium | Jacquelyn Wynn, Mairead Hyland, The Lighthouse Lab in Alderley Park and Alex Alderton, Roberto Amato, Jeffrey Barrett, Sonia Goncalves, Ewan Harrison, David K. Jackson, Ian Johnston, Dominic Kwiatkowski, Cordelia Langford, John Sillitoe on behalf of the Wellcome Sanger Institute COVID-19 Surveillance Team |
| EPI_ISL_2994125, EPI_ISL_2994126 | Lighthouse Lab in Milton Keynes | Wellcome Sanger Institute for the COVID-19 Genomics UK (COG-UK) Consortium | The Lighthouse Lab in Milton Keynes and Alex Alderton, Roberto Amato, Jeffrey Barrett, Sonia Goncalves, Ewan Harrison, David K. Jackson, Ian Johnston, Dominic Kwiatkowski, Cordelia Langford, John Sillitoe on behalf of the Wellcome Sanger Institute COVID-19 Surveillance Team |
| EPI_ISL_2994127 | Lighthouse Lab in Alderley Park | Wellcome Sanger Institute for the COVID-19 Genomics UK (COG-UK) Consortium | Jacquelyn Wynn, Mairead Hyland, The Lighthouse Lab in Alderley Park and Alex Alderton, Roberto Amato, Jeffrey Barrett, Sonia Goncalves, Ewan Harrison, David K. Jackson, Ian Johnston, Dominic Kwiatkowski, Cordelia Langford, John Sillitoe on behalf of the Wellcome Sanger Institute COVID-19 Surveillance Team |
| EPI_ISL_2994128 | Berkshire and Surrey Pathology Services Lighthouse Laboratory | Wellcome Sanger Institute for the COVID-19 Genomics UK (COG-UK) Consortium | Berkshire and Surrey Pathology Services Lighthouse Laboratory and Alex Alderton, Roberto Amato, Jeffrey Barrett, Sonia Goncalves, Ewan Harrison, David K. Jackson, Ian Johnston, Dominic Kwiatkowski, Cordelia Langford, John Sillitoe on behalf of the Wellcome Sanger Institute COVID-19 Surveillance Team |
| EPI_ISL_2994129, | Lighthouse Lab in Milton Keynes | Wellcome Sanger Institute for the COVID-19 Genomics UK (COG-UK) | The Lighthouse Lab in Milton Keynes and Alex Alderton, Roberto Amato, Jeffrey Barrett, Sonia Goncalves, Ewan Harrison, David K. Jackson, Ian Johnston, Dominic Kwiatkowski, Cordelia Langford, John Sillitoe on behalf of the Wellcome Sanger |

| | | | |
|---|---|---|---|
| EPI_ISL_2994130 | | Consortium | Institute COVID-19 Surveillance Team |
| EPI_ISL_2994131, EPI_ISL_2994132 | Lighthouse Lab in Alderley Park | Wellcome Sanger Institute for the COVID-19 Genomics UK (COG-UK) Consortium | Jacquelyn Wynn, Mairead Hyland, The Lighthouse Lab in Alderley Park and Alex Alderton, Roberto Amato, Jeffrey Barrett, Sonia Goncalves, Ewan Harrison, David K. Jackson, Ian Johnston, Dominic Kwiatkowski, Cordelia Langford, John Sillitoe on behalf of the Wellcome Sanger Institute COVID-19 Surveillance Team |
| EPI_ISL_2994133 | Lighthouse Lab in Milton Keynes | Wellcome Sanger Institute for the COVID-19 Genomics UK (COG-UK) Consortium | The Lighthouse Lab in Milton Keynes and Alex Alderton, Roberto Amato, Jeffrey Barrett, Sonia Goncalves, Ewan Harrison, David K. Jackson, Ian Johnston, Dominic Kwiatkowski, Cordelia Langford, John Sillitoe on behalf of the Wellcome Sanger Institute COVID-19 Surveillance Team |
| EPI_ISL_2994134 | Lighthouse Laboratory Plymouth | Wellcome Sanger Institute for the COVID-19 Genomics UK (COG-UK) Consortium | Lighthouse Laboratory Plymouth and Alex Alderton, Roberto Amato, Jeffrey Barrett, Sonia Goncalves, Ewan Harrison, David K. Jackson, Ian Johnston, Dominic Kwiatkowski, Cordelia Langford, John Sillitoe on behalf of the Wellcome Sanger Institute COVID-19 Surveillance Team |
| EPI_ISL_2994135, EPI_ISL_2994136, EPI_ISL_2994137, EPI_ISL_2994138, EPI_ISL_2994139 | Lighthouse Lab in Milton Keynes | Wellcome Sanger Institute for the COVID-19 Genomics UK (COG-UK) Consortium | The Lighthouse Lab in Milton Keynes and Alex Alderton, Roberto Amato, Jeffrey Barrett, Sonia Goncalves, Ewan Harrison, David K. Jackson, Ian Johnston, Dominic Kwiatkowski, Cordelia Langford, John Sillitoe on behalf of the Wellcome Sanger Institute COVID-19 Surveillance Team |
| EPI_ISL_2994140 | Lighthouse Lab in Alderley Park | Wellcome Sanger Institute for the COVID-19 Genomics UK (COG-UK) Consortium | Jacquelyn Wynn, Mairead Hyland, The Lighthouse Lab in Alderley Park and Alex Alderton, Roberto Amato, Jeffrey Barrett, Sonia Goncalves, Ewan Harrison, David K. Jackson, Ian Johnston, Dominic Kwiatkowski, Cordelia Langford, John Sillitoe on behalf of the Wellcome Sanger Institute COVID-19 Surveillance Team |
| EPI_ISL_2994141 | Berkshire and Surrey Pathology Services Lighthouse Laboratory | Wellcome Sanger Institute for the COVID-19 Genomics UK (COG-UK) Consortium | Berkshire and Surrey Pathology Services Lighthouse Laboratory and Alex Alderton, Roberto Amato, Jeffrey Barrett, Sonia Goncalves, Ewan Harrison, David K. Jackson, Ian Johnston, Dominic Kwiatkowski, Cordelia Langford, John Sillitoe on behalf of the Wellcome Sanger Institute COVID-19 Surveillance Team |
| EPI_ISL_2994142, EPI_ISL_2994143 | Lighthouse Lab in Milton Keynes | Wellcome Sanger Institute for the COVID-19 Genomics UK (COG-UK) Consortium | The Lighthouse Lab in Milton Keynes and Alex Alderton, Roberto Amato, Jeffrey Barrett, Sonia Goncalves, Ewan Harrison, David K. Jackson, Ian Johnston, Dominic Kwiatkowski, Cordelia Langford, John Sillitoe on behalf of the Wellcome Sanger Institute COVID-19 Surveillance Team |
| EPI_ISL_2994144, EPI_ISL_2994145 | Berkshire and Surrey Pathology Services Lighthouse Laboratory | Wellcome Sanger Institute for the COVID-19 Genomics UK (COG-UK) Consortium | Berkshire and Surrey Pathology Services Lighthouse Laboratory and Alex Alderton, Roberto Amato, Jeffrey Barrett, Sonia Goncalves, Ewan Harrison, David K. Jackson, Ian Johnston, Dominic Kwiatkowski, Cordelia Langford, John Sillitoe on behalf of the Wellcome Sanger Institute COVID-19 Surveillance Team |
| EPI_ISL_2994146 | Lighthouse Lab in Milton Keynes | Wellcome Sanger Institute for the COVID-19 Genomics UK (COG-UK) Consortium | The Lighthouse Lab in Milton Keynes and Alex Alderton, Roberto Amato, Jeffrey Barrett, Sonia Goncalves, Ewan Harrison, David K. Jackson, Ian Johnston, Dominic Kwiatkowski, Cordelia Langford, John Sillitoe on behalf of the Wellcome Sanger Institute COVID-19 Surveillance Team |
| EPI_ISL_2994147 | Berkshire and Surrey Pathology Services Lighthouse Laboratory | Wellcome Sanger Institute for the COVID-19 Genomics UK (COG-UK) Consortium | Berkshire and Surrey Pathology Services Lighthouse Laboratory and Alex Alderton, Roberto Amato, Jeffrey Barrett, Sonia Goncalves, Ewan Harrison, David K. Jackson, Ian Johnston, Dominic Kwiatkowski, Cordelia Langford, John Sillitoe on behalf of the Wellcome Sanger Institute COVID-19 Surveillance Team |
| EPI_ISL_2994148 | Lighthouse Laboratory Plymouth | Wellcome Sanger Institute for the COVID-19 Genomics UK (COG-UK) Consortium | Lighthouse Laboratory Plymouth and Alex Alderton, Roberto Amato, Jeffrey Barrett, Sonia Goncalves, Ewan Harrison, David K. Jackson, Ian Johnston, Dominic Kwiatkowski, Cordelia Langford, John Sillitoe on behalf of the Wellcome Sanger Institute COVID-19 Surveillance Team |
| EPI_ISL_2994149 | Lighthouse Lab in Milton Keynes | Wellcome Sanger Institute for the COVID-19 Genomics UK (COG-UK) Consortium | The Lighthouse Lab in Milton Keynes and Alex Alderton, Roberto Amato, Jeffrey Barrett, Sonia Goncalves, Ewan Harrison, David K. Jackson, Ian Johnston, Dominic Kwiatkowski, Cordelia Langford, John Sillitoe on behalf of the Wellcome Sanger Institute COVID-19 Surveillance Team |
| EPI_ISL_2994150 | Lighthouse Lab in Alderley Park | Wellcome Sanger Institute for the COVID-19 Genomics UK (COG-UK) Consortium | Jacquelyn Wynn, Mairead Hyland, The Lighthouse Lab in Alderley Park and Alex Alderton, Roberto Amato, Jeffrey Barrett, Sonia Goncalves, Ewan Harrison, David K. Jackson, Ian Johnston, Dominic Kwiatkowski, Cordelia Langford, John Sillitoe on behalf of the Wellcome Sanger Institute COVID-19 Surveillance Team |
| EPI_ISL_2994151 | Lighthouse Lab in Milton Keynes | Wellcome Sanger Institute for the COVID-19 Genomics UK (COG-UK) Consortium | The Lighthouse Lab in Milton Keynes and Alex Alderton, Roberto Amato, Jeffrey Barrett, Sonia Goncalves, Ewan Harrison, David K. Jackson, Ian Johnston, Dominic Kwiatkowski, Cordelia Langford, John Sillitoe on behalf of the Wellcome Sanger Institute COVID-19 Surveillance Team |
| EPI_ISL_2994152 | Lighthouse Lab in Alderley Park | Wellcome Sanger Institute for the COVID-19 Genomics UK (COG-UK) Consortium | Jacquelyn Wynn, Mairead Hyland, The Lighthouse Lab in Alderley Park and Alex Alderton, Roberto Amato, Jeffrey Barrett, Sonia Goncalves, Ewan Harrison, David K. Jackson, Ian Johnston, Dominic Kwiatkowski, Cordelia Langford, John Sillitoe on behalf of the Wellcome Sanger Institute COVID-19 Surveillance Team |
| EPI_ISL_2994153 | Lighthouse Laboratory Plymouth | Wellcome Sanger Institute for the COVID-19 Genomics UK (COG-UK) Consortium | Lighthouse Laboratory Plymouth and Alex Alderton, Roberto Amato, Jeffrey Barrett, Sonia Goncalves, Ewan Harrison, David K. Jackson, Ian Johnston, Dominic Kwiatkowski, Cordelia Langford, John Sillitoe on behalf of the Wellcome Sanger Institute COVID-19 Surveillance Team |
| EPI_ISL_2994154 | Lighthouse Lab in Alderley Park | Wellcome Sanger Institute for the COVID-19 Genomics UK (COG-UK) Consortium | Jacquelyn Wynn, Mairead Hyland, The Lighthouse Lab in Alderley Park and Alex Alderton, Roberto Amato, Jeffrey Barrett, Sonia Goncalves, Ewan Harrison, David K. Jackson, Ian Johnston, Dominic Kwiatkowski, Cordelia Langford, John Sillitoe on behalf of the Wellcome Sanger Institute COVID-19 Surveillance Team |
| EPI_ISL_2994155 | Lighthouse Lab in Milton Keynes | Wellcome Sanger Institute for the COVID-19 Genomics UK (COG-UK) Consortium | The Lighthouse Lab in Milton Keynes and Alex Alderton, Roberto Amato, Jeffrey Barrett, Sonia Goncalves, Ewan Harrison, David K. Jackson, Ian Johnston, Dominic Kwiatkowski, Cordelia Langford, John Sillitoe on behalf of the Wellcome Sanger Institute COVID-19 Surveillance Team |
| EPI_ISL_2994156 | Lighthouse Laboratory Plymouth | Wellcome Sanger Institute for the COVID-19 Genomics UK (COG-UK) Consortium | Lighthouse Laboratory Plymouth and Alex Alderton, Roberto Amato, Jeffrey Barrett, Sonia Goncalves, Ewan Harrison, David K. Jackson, Ian Johnston, Dominic Kwiatkowski, Cordelia Langford, John Sillitoe on behalf of the Wellcome Sanger Institute COVID-19 Surveillance Team |
| EPI_ISL_2994157 | Berkshire and Surrey Pathology Services Lighthouse Laboratory | Wellcome Sanger Institute for the COVID-19 Genomics UK (COG-UK) Consortium | Berkshire and Surrey Pathology Services Lighthouse Laboratory and Alex Alderton, Roberto Amato, Jeffrey Barrett, Sonia Goncalves, Ewan Harrison, David K. Jackson, Ian Johnston, Dominic Kwiatkowski, Cordelia Langford, John Sillitoe on behalf of the Wellcome Sanger Institute COVID-19 Surveillance Team |
| EPI_ISL_2994158, EPI_ISL_2994159 | Lighthouse Lab in Alderley Park | Wellcome Sanger Institute for the COVID-19 Genomics UK (COG-UK) Consortium | Jacquelyn Wynn, Mairead Hyland, The Lighthouse Lab in Alderley Park and Alex Alderton, Roberto Amato, Jeffrey Barrett, Sonia Goncalves, Ewan Harrison, David K. Jackson, Ian Johnston, Dominic Kwiatkowski, Cordelia Langford, John Sillitoe on behalf of the Wellcome Sanger Institute COVID-19 Surveillance Team |
| EPI_ISL_2994160 | Berkshire and Surrey Pathology Services Lighthouse Laboratory | Wellcome Sanger Institute for the COVID-19 Genomics UK (COG-UK) Consortium | Berkshire and Surrey Pathology Services Lighthouse Laboratory and Alex Alderton, Roberto Amato, Jeffrey Barrett, Sonia Goncalves, Ewan Harrison, David K. Jackson, Ian Johnston, Dominic Kwiatkowski, Cordelia Langford, John Sillitoe on behalf of the Wellcome Sanger Institute COVID-19 Surveillance Team |
| EPI_ISL_2994161 | Lighthouse Laboratory Plymouth | Wellcome Sanger Institute for the COVID-19 Genomics UK (COG-UK) Consortium | Lighthouse Laboratory Plymouth and Alex Alderton, Roberto Amato, Jeffrey Barrett, Sonia Goncalves, Ewan Harrison, David K. Jackson, Ian Johnston, Dominic Kwiatkowski, Cordelia Langford, John Sillitoe on behalf of the Wellcome Sanger Institute COVID-19 Surveillance Team |
| EPI_ISL_2994162 | Berkshire and Surrey Pathology Services Lighthouse Laboratory | Wellcome Sanger Institute for the COVID-19 Genomics UK (COG-UK) Consortium | Berkshire and Surrey Pathology Services Lighthouse Laboratory and Alex Alderton, Roberto Amato, Jeffrey Barrett, Sonia Goncalves, Ewan Harrison, David K. Jackson, Ian Johnston, Dominic Kwiatkowski, Cordelia Langford, John Sillitoe on behalf of the Wellcome Sanger Institute COVID-19 Surveillance Team |
| EPI_ISL_2994163 | Lighthouse Laboratory Plymouth | Wellcome Sanger Institute for the COVID-19 Genomics UK (COG-UK) Consortium | Lighthouse Laboratory Plymouth and Alex Alderton, Roberto Amato, Jeffrey Barrett, Sonia Goncalves, Ewan Harrison, David K. Jackson, Ian Johnston, Dominic Kwiatkowski, Cordelia Langford, John Sillitoe on behalf of the Wellcome Sanger Institute COVID-19 Surveillance Team |
| EPI_ISL_2994164 | Lighthouse Lab in Milton Keynes | Wellcome Sanger Institute for the COVID-19 Genomics UK (COG-UK) Consortium | The Lighthouse Lab in Milton Keynes and Alex Alderton, Roberto Amato, Jeffrey Barrett, Sonia Goncalves, Ewan Harrison, David K. Jackson, Ian Johnston, Dominic Kwiatkowski, Cordelia Langford, John Sillitoe on behalf of the Wellcome Sanger Institute COVID-19 Surveillance Team |
| EPI_ISL_2994165, EPI_ISL_2994166 | Lighthouse Laboratory Plymouth | Wellcome Sanger Institute for the COVID-19 Genomics UK (COG-UK) Consortium | Lighthouse Laboratory Plymouth and Alex Alderton, Roberto Amato, Jeffrey Barrett, Sonia Goncalves, Ewan Harrison, David K. Jackson, Ian Johnston, Dominic Kwiatkowski, Cordelia Langford, John Sillitoe on behalf of the Wellcome Sanger Institute COVID-19 Surveillance Team |
| EPI_ISL_2994167, EPI_ISL_2994168 | Berkshire and Surrey Pathology Services Lighthouse Laboratory | Wellcome Sanger Institute for the COVID-19 Genomics UK (COG-UK) Consortium | Berkshire and Surrey Pathology Services Lighthouse Laboratory and Alex Alderton, Roberto Amato, Jeffrey Barrett, Sonia Goncalves, Ewan Harrison, David K. Jackson, Ian Johnston, Dominic Kwiatkowski, Cordelia Langford, John Sillitoe on behalf of the Wellcome Sanger Institute COVID-19 Surveillance Team |
| EPI_ISL_2994169 | Lighthouse Lab in Alderley Park | Wellcome Sanger Institute for the COVID-19 Genomics UK (COG-UK) Consortium | Jacquelyn Wynn, Mairead Hyland, The Lighthouse Lab in Alderley Park and Alex Alderton, Roberto Amato, Jeffrey Barrett, Sonia Goncalves, Ewan Harrison, David K. Jackson, Ian Johnston, Dominic Kwiatkowski, Cordelia Langford, John Sillitoe on behalf of the Wellcome Sanger Institute COVID-19 Surveillance Team |
| EPI_ISL_2994170 | Lighthouse Laboratory Plymouth | Wellcome Sanger Institute for the COVID-19 Genomics UK (COG-UK) Consortium | Lighthouse Laboratory Plymouth and Alex Alderton, Roberto Amato, Jeffrey Barrett, Sonia Goncalves, Ewan Harrison, David K. Jackson, Ian Johnston, Dominic Kwiatkowski, Cordelia Langford, John Sillitoe on behalf of the Wellcome Sanger Institute COVID-19 Surveillance Team |
| EPI_ISL_2994171, EPI_ISL_2994172 | Lighthouse Lab in Alderley Park | Wellcome Sanger Institute for the COVID-19 Genomics UK (COG-UK) Consortium | Jacquelyn Wynn, Mairead Hyland, The Lighthouse Lab in Alderley Park and Alex Alderton, Roberto Amato, Jeffrey Barrett, Sonia Goncalves, Ewan Harrison, David K. Jackson, Ian Johnston, Dominic Kwiatkowski, Cordelia Langford, John Sillitoe on behalf of the Wellcome Sanger Institute COVID-19 Surveillance Team |
| EPI_ISL_2994173 | Berkshire and Surrey Pathology Services Lighthouse Laboratory | Wellcome Sanger Institute for the COVID-19 Genomics UK (COG-UK) Consortium | Berkshire and Surrey Pathology Services Lighthouse Laboratory and Alex Alderton, Roberto Amato, Jeffrey Barrett, Sonia Goncalves, Ewan Harrison, David K. Jackson, Ian Johnston, Dominic Kwiatkowski, Cordelia Langford, John Sillitoe on behalf of the Wellcome Sanger Institute COVID-19 Surveillance Team |
| EPI_ISL_2994174 | Lighthouse Lab in Alderley Park | Wellcome Sanger Institute for the COVID-19 Genomics UK (COG-UK) Consortium | Jacquelyn Wynn, Mairead Hyland, The Lighthouse Lab in Alderley Park and Alex Alderton, Roberto Amato, Jeffrey Barrett, Sonia Goncalves, Ewan Harrison, David K. Jackson, Ian Johnston, Dominic Kwiatkowski, Cordelia Langford, John Sillitoe on behalf of the Wellcome Sanger Institute COVID-19 Surveillance Team |
| EPI_ISL_2994175, EPI_ISL_2994176 | Berkshire and Surrey Pathology Services Lighthouse Laboratory | Wellcome Sanger Institute for the COVID-19 Genomics UK (COG-UK) Consortium | Berkshire and Surrey Pathology Services Lighthouse Laboratory and Alex Alderton, Roberto Amato, Jeffrey Barrett, Sonia Goncalves, Ewan Harrison, David K. Jackson, Ian Johnston, Dominic Kwiatkowski, Cordelia Langford, John Sillitoe on behalf of the Wellcome Sanger Institute COVID-19 Surveillance Team |
| EPI_ISL_2994177, EPI_ISL_2994178 | Lighthouse Laboratory Plymouth | Wellcome Sanger Institute for the COVID-19 Genomics UK (COG-UK) Consortium | Lighthouse Laboratory Plymouth and Alex Alderton, Roberto Amato, Jeffrey Barrett, Sonia Goncalves, Ewan Harrison, David K. Jackson, Ian Johnston, Dominic Kwiatkowski, Cordelia Langford, John Sillitoe on behalf of the Wellcome Sanger Institute COVID-19 Surveillance Team |
| EPI_ISL_2994179, EPI_ISL_2994180, EPI_ISL_2994181 | Lighthouse Lab in Milton Keynes | Wellcome Sanger Institute for the COVID-19 Genomics UK (COG-UK) Consortium | The Lighthouse Lab in Milton Keynes and Alex Alderton, Roberto Amato, Jeffrey Barrett, Sonia Goncalves, Ewan Harrison, David K. Jackson, Ian Johnston, Dominic Kwiatkowski, Cordelia Langford, John Sillitoe on behalf of the Wellcome Sanger Institute COVID-19 Surveillance Team |
| EPI_ISL_2994182, EPI_ISL_2994183 | Lighthouse Lab in Alderley Park | Wellcome Sanger Institute for the COVID-19 Genomics UK (COG-UK) Consortium | Jacquelyn Wynn, Mairead Hyland, The Lighthouse Lab in Alderley Park and Alex Alderton, Roberto Amato, Jeffrey Barrett, Sonia Goncalves, Ewan Harrison, David K. Jackson, Ian Johnston, Dominic Kwiatkowski, Cordelia Langford, John Sillitoe on behalf of the Wellcome Sanger Institute COVID-19 Surveillance Team |
| EPI_ISL_2994184 | Lighthouse Laboratory Plymouth | Wellcome Sanger Institute for the COVID-19 Genomics UK (COG-UK) Consortium | Lighthouse Laboratory Plymouth and Alex Alderton, Roberto Amato, Jeffrey Barrett, Sonia Goncalves, Ewan Harrison, David K. Jackson, Ian Johnston, Dominic Kwiatkowski, Cordelia Langford, John Sillitoe on behalf of the Wellcome Sanger Institute COVID-19 Surveillance Team |
| EPI_ISL_2994185, EPI_ISL_2994186 | Berkshire and Surrey Pathology Services Lighthouse Laboratory | Wellcome Sanger Institute for the COVID-19 Genomics UK (COG-UK) Consortium | Berkshire and Surrey Pathology Services Lighthouse Laboratory and Alex Alderton, Roberto Amato, Jeffrey Barrett, Sonia Goncalves, Ewan Harrison, David K. Jackson, Ian Johnston, Dominic Kwiatkowski, Cordelia Langford, John Sillitoe on behalf of the Wellcome Sanger Institute COVID-19 Surveillance Team |
| EPI_ISL_2994187 | Lighthouse Lab in Alderley Park | Wellcome Sanger Institute for the COVID-19 Genomics UK (COG-UK) Consortium | Jacquelyn Wynn, Mairead Hyland, The Lighthouse Lab in Alderley Park and Alex Alderton, Roberto Amato, Jeffrey Barrett, Sonia Goncalves, Ewan Harrison, David K. Jackson, Ian Johnston, Dominic Kwiatkowski, Cordelia Langford, John Sillitoe on behalf of the Wellcome Sanger Institute COVID-19 Surveillance Team |
| EPI_ISL_2994188 | Berkshire and Surrey Pathology Services Lighthouse Laboratory | Wellcome Sanger Institute for the COVID-19 Genomics UK (COG-UK) Consortium | Berkshire and Surrey Pathology Services Lighthouse Laboratory and Alex Alderton, Roberto Amato, Jeffrey Barrett, Sonia Goncalves, Ewan Harrison, David K. Jackson, Ian Johnston, Dominic Kwiatkowski, Cordelia Langford, John Sillitoe on behalf of the Wellcome Sanger Institute COVID-19 Surveillance Team |
| EPI_ISL_2994189 | Lighthouse Lab in Milton Keynes | Wellcome Sanger Institute for the COVID-19 Genomics UK (COG-UK) Consortium | The Lighthouse Lab in Milton Keynes and Alex Alderton, Roberto Amato, Jeffrey Barrett, Sonia Goncalves, Ewan Harrison, David K. Jackson, Ian Johnston, Dominic Kwiatkowski, Cordelia Langford, John Sillitoe on behalf of the Wellcome Sanger Institute COVID-19 Surveillance Team |
| EPI_ISL_2994190, EPI_ISL_2994191 | Lighthouse Laboratory Plymouth | Wellcome Sanger Institute for the COVID-19 Genomics UK (COG-UK) Consortium | Lighthouse Laboratory Plymouth and Alex Alderton, Roberto Amato, Jeffrey Barrett, Sonia Goncalves, Ewan Harrison, David K. Jackson, Ian Johnston, Dominic Kwiatkowski, Cordelia Langford, John Sillitoe on behalf of the Wellcome Sanger Institute COVID-19 Surveillance Team |

| | | | |
|---|---|---|---|
| EPI_ISL_2994192 | Lighthouse Lab in Milton Keynes | Wellcome Sanger Institute for the COVID-19 Genomics UK (COG-UK) Consortium | The Lighthouse Lab in Milton Keynes and Alex Alderton, Roberto Amato, Jeffrey Barrett, Sonia Goncalves, Ewan Harrison, David K. Jackson, Ian Johnston, Dominic Kwiatkowski, Cordelia Langford, John Sillitoe on behalf of the Wellcome Sanger Institute COVID-19 Surveillance Team |
| EPI_ISL_2994193 | Lighthouse Lab in Alderley Park | Wellcome Sanger Institute for the COVID-19 Genomics UK (COG-UK) Consortium | Jacquelyn Wynn, Mairead Hyland, The Lighthouse Lab in Alderley Park and Alex Alderton, Roberto Amato, Jeffrey Barrett, Sonia Goncalves, Ewan Harrison, David K. Jackson, Ian Johnston, Dominic Kwiatkowski, Cordelia Langford, John Sillitoe on behalf of the Wellcome Sanger Institute COVID-19 Surveillance Team |
| EPI_ISL_2994194 | Lighthouse Lab in Milton Keynes | Wellcome Sanger Institute for the COVID-19 Genomics UK (COG-UK) Consortium | The Lighthouse Lab in Milton Keynes and Alex Alderton, Roberto Amato, Jeffrey Barrett, Sonia Goncalves, Ewan Harrison, David K. Jackson, Ian Johnston, Dominic Kwiatkowski, Cordelia Langford, John Sillitoe on behalf of the Wellcome Sanger Institute COVID-19 Surveillance Team |
| EPI_ISL_2994195 | Berkshire and Surrey Pathology Services Lighthouse Laboratory | Wellcome Sanger Institute for the COVID-19 Genomics UK (COG-UK) Consortium | Berkshire and Surrey Pathology Services Lighthouse Laboratory and Alex Alderton, Roberto Amato, Jeffrey Barrett, Sonia Goncalves, Ewan Harrison, David K. Jackson, Ian Johnston, Dominic Kwiatkowski, Cordelia Langford, John Sillitoe on behalf of the Wellcome Sanger Institute COVID-19 Surveillance Team |
| EPI_ISL_2994196 | Lighthouse Lab in Alderley Park | Wellcome Sanger Institute for the COVID-19 Genomics UK (COG-UK) Consortium | Jacquelyn Wynn, Mairead Hyland, The Lighthouse Lab in Alderley Park and Alex Alderton, Roberto Amato, Jeffrey Barrett, Sonia Goncalves, Ewan Harrison, David K. Jackson, Ian Johnston, Dominic Kwiatkowski, Cordelia Langford, John Sillitoe on behalf of the Wellcome Sanger Institute COVID-19 Surveillance Team |
| EPI_ISL_2994197 | Berkshire and Surrey Pathology Services Lighthouse Laboratory | Wellcome Sanger Institute for the COVID-19 Genomics UK (COG-UK) Consortium | Berkshire and Surrey Pathology Services Lighthouse Laboratory and Alex Alderton, Roberto Amato, Jeffrey Barrett, Sonia Goncalves, Ewan Harrison, David K. Jackson, Ian Johnston, Dominic Kwiatkowski, Cordelia Langford, John Sillitoe on behalf of the Wellcome Sanger Institute COVID-19 Surveillance Team |
| EPI_ISL_2994198 | Lighthouse Laboratory Plymouth | Wellcome Sanger Institute for the COVID-19 Genomics UK (COG-UK) Consortium | Lighthouse Laboratory Plymouth and Alex Alderton, Roberto Amato, Jeffrey Barrett, Sonia Goncalves, Ewan Harrison, David K. Jackson, Ian Johnston, Dominic Kwiatkowski, Cordelia Langford, John Sillitoe on behalf of the Wellcome Sanger Institute COVID-19 Surveillance Team |
| EPI_ISL_2994199, EPI_ISL_2994200 | Lighthouse Lab in Alderley Park | Wellcome Sanger Institute for the COVID-19 Genomics UK (COG-UK) Consortium | Jacquelyn Wynn, Mairead Hyland, The Lighthouse Lab in Alderley Park and Alex Alderton, Roberto Amato, Jeffrey Barrett, Sonia Goncalves, Ewan Harrison, David K. Jackson, Ian Johnston, Dominic Kwiatkowski, Cordelia Langford, John Sillitoe on behalf of the Wellcome Sanger Institute COVID-19 Surveillance Team |
| EPI_ISL_2994201 | Lighthouse Laboratory Plymouth | Wellcome Sanger Institute for the COVID-19 Genomics UK (COG-UK) Consortium | Lighthouse Laboratory Plymouth and Alex Alderton, Roberto Amato, Jeffrey Barrett, Sonia Goncalves, Ewan Harrison, David K. Jackson, Ian Johnston, Dominic Kwiatkowski, Cordelia Langford, John Sillitoe on behalf of the Wellcome Sanger Institute COVID-19 Surveillance Team |
| EPI_ISL_2994202, EPI_ISL_2994203 | Berkshire and Surrey Pathology Services Lighthouse Laboratory | Wellcome Sanger Institute for the COVID-19 Genomics UK (COG-UK) Consortium | Berkshire and Surrey Pathology Services Lighthouse Laboratory and Alex Alderton, Roberto Amato, Jeffrey Barrett, Sonia Goncalves, Ewan Harrison, David K. Jackson, Ian Johnston, Dominic Kwiatkowski, Cordelia Langford, John Sillitoe on behalf of the Wellcome Sanger Institute COVID-19 Surveillance Team |
| EPI_ISL_2994204, EPI_ISL_2994205, EPI_ISL_2994206 | Lighthouse Lab in Milton Keynes | Wellcome Sanger Institute for the COVID-19 Genomics UK (COG-UK) Consortium | The Lighthouse Lab in Milton Keynes and Alex Alderton, Roberto Amato, Jeffrey Barrett, Sonia Goncalves, Ewan Harrison, David K. Jackson, Ian Johnston, Dominic Kwiatkowski, Cordelia Langford, John Sillitoe on behalf of the Wellcome Sanger Institute COVID-19 Surveillance Team |
| EPI_ISL_2994207, EPI_ISL_2994208, EPI_ISL_2994209, EPI_ISL_2994210 | Lighthouse Lab in Alderley Park | Wellcome Sanger Institute for the COVID-19 Genomics UK (COG-UK) Consortium | Jacquelyn Wynn, Mairead Hyland, The Lighthouse Lab in Alderley Park and Alex Alderton, Roberto Amato, Jeffrey Barrett, Sonia Goncalves, Ewan Harrison, David K. Jackson, Ian Johnston, Dominic Kwiatkowski, Cordelia Langford, John Sillitoe on behalf of the Wellcome Sanger Institute COVID-19 Surveillance Team |
| EPI_ISL_2994211 | Lighthouse Lab in Milton Keynes | Wellcome Sanger Institute for the COVID-19 Genomics UK (COG-UK) Consortium | The Lighthouse Lab in Milton Keynes and Alex Alderton, Roberto Amato, Jeffrey Barrett, Sonia Goncalves, Ewan Harrison, David K. Jackson, Ian Johnston, Dominic Kwiatkowski, Cordelia Langford, John Sillitoe on behalf of the Wellcome Sanger Institute COVID-19 Surveillance Team |
| EPI_ISL_2994212 | Lighthouse Laboratory Plymouth | Wellcome Sanger Institute for the COVID-19 Genomics UK (COG-UK) Consortium | Lighthouse Laboratory Plymouth and Alex Alderton, Roberto Amato, Jeffrey Barrett, Sonia Goncalves, Ewan Harrison, David K. Jackson, Ian Johnston, Dominic Kwiatkowski, Cordelia Langford, John Sillitoe on behalf of the Wellcome Sanger Institute COVID-19 Surveillance Team |
| EPI_ISL_2994213 | Lighthouse Lab in Alderley Park | Wellcome Sanger Institute for the COVID-19 Genomics UK (COG-UK) Consortium | Jacquelyn Wynn, Mairead Hyland, The Lighthouse Lab in Alderley Park and Alex Alderton, Roberto Amato, Jeffrey Barrett, Sonia Goncalves, Ewan Harrison, David K. Jackson, Ian Johnston, Dominic Kwiatkowski, Cordelia Langford, John Sillitoe on behalf of the Wellcome Sanger Institute COVID-19 Surveillance Team |
| EPI_ISL_2994214 | Lighthouse Laboratory Plymouth | Wellcome Sanger Institute for the COVID-19 Genomics UK (COG-UK) Consortium | Lighthouse Laboratory Plymouth and Alex Alderton, Roberto Amato, Jeffrey Barrett, Sonia Goncalves, Ewan Harrison, David K. Jackson, Ian Johnston, Dominic Kwiatkowski, Cordelia Langford, John Sillitoe on behalf of the Wellcome Sanger Institute COVID-19 Surveillance Team |
| EPI_ISL_2994215 | Lighthouse Lab in Milton Keynes | Wellcome Sanger Institute for the COVID-19 Genomics UK (COG-UK) Consortium | The Lighthouse Lab in Milton Keynes and Alex Alderton, Roberto Amato, Jeffrey Barrett, Sonia Goncalves, Ewan Harrison, David K. Jackson, Ian Johnston, Dominic Kwiatkowski, Cordelia Langford, John Sillitoe on behalf of the Wellcome Sanger Institute COVID-19 Surveillance Team |
| EPI_ISL_2994216 | Lighthouse Laboratory Plymouth | Wellcome Sanger Institute for the COVID-19 Genomics UK (COG-UK) Consortium | Lighthouse Laboratory Plymouth and Alex Alderton, Roberto Amato, Jeffrey Barrett, Sonia Goncalves, Ewan Harrison, David K. Jackson, Ian Johnston, Dominic Kwiatkowski, Cordelia Langford, John Sillitoe on behalf of the Wellcome Sanger Institute COVID-19 Surveillance Team |
| EPI_ISL_2994217 | Berkshire and Surrey Pathology Services Lighthouse Laboratory | Wellcome Sanger Institute for the COVID-19 Genomics UK (COG-UK) Consortium | Berkshire and Surrey Pathology Services Lighthouse Laboratory and Alex Alderton, Roberto Amato, Jeffrey Barrett, Sonia Goncalves, Ewan Harrison, David K. Jackson, Ian Johnston, Dominic Kwiatkowski, Cordelia Langford, John Sillitoe on behalf of the Wellcome Sanger Institute COVID-19 Surveillance Team |
| EPI_ISL_2994218, EPI_ISL_2994219 | Lighthouse Laboratory Plymouth | Wellcome Sanger Institute for the COVID-19 Genomics UK (COG-UK) Consortium | Lighthouse Laboratory Plymouth and Alex Alderton, Roberto Amato, Jeffrey Barrett, Sonia Goncalves, Ewan Harrison, David K. Jackson, Ian Johnston, Dominic Kwiatkowski, Cordelia Langford, John Sillitoe on behalf of the Wellcome Sanger Institute COVID-19 Surveillance Team |
| EPI_ISL_2994220 | Lighthouse Lab in Milton Keynes | Wellcome Sanger Institute for the COVID-19 Genomics UK (COG-UK) Consortium | The Lighthouse Lab in Milton Keynes and Alex Alderton, Roberto Amato, Jeffrey Barrett, Sonia Goncalves, Ewan Harrison, David K. Jackson, Ian Johnston, Dominic Kwiatkowski, Cordelia Langford, John Sillitoe on behalf of the Wellcome Sanger Institute COVID-19 Surveillance Team |
| EPI_ISL_2994221, EPI_ISL_2994222 | Berkshire and Surrey Pathology Services Lighthouse Laboratory | Wellcome Sanger Institute for the COVID-19 Genomics UK (COG-UK) Consortium | Berkshire and Surrey Pathology Services Lighthouse Laboratory and Alex Alderton, Roberto Amato, Jeffrey Barrett, Sonia Goncalves, Ewan Harrison, David K. Jackson, Ian Johnston, Dominic Kwiatkowski, Cordelia Langford, John Sillitoe on behalf of the Wellcome Sanger Institute COVID-19 Surveillance Team |
| EPI_ISL_2994223 | Lighthouse Laboratory Plymouth | Wellcome Sanger Institute for the COVID-19 Genomics UK (COG-UK) Consortium | Lighthouse Laboratory Plymouth and Alex Alderton, Roberto Amato, Jeffrey Barrett, Sonia Goncalves, Ewan Harrison, David K. Jackson, Ian Johnston, Dominic Kwiatkowski, Cordelia Langford, John Sillitoe on behalf of the Wellcome Sanger Institute COVID-19 Surveillance Team |
| EPI_ISL_2994224, EPI_ISL_2994225, EPI_ISL_2994226 | Lighthouse Lab in Milton Keynes | Wellcome Sanger Institute for the COVID-19 Genomics UK (COG-UK) Consortium | The Lighthouse Lab in Milton Keynes and Alex Alderton, Roberto Amato, Jeffrey Barrett, Sonia Goncalves, Ewan Harrison, David K. Jackson, Ian Johnston, Dominic Kwiatkowski, Cordelia Langford, John Sillitoe on behalf of the Wellcome Sanger Institute COVID-19 Surveillance Team |
| EPI_ISL_2994227 | Berkshire and Surrey Pathology Services Lighthouse Laboratory | Wellcome Sanger Institute for the COVID-19 Genomics UK (COG-UK) Consortium | Berkshire and Surrey Pathology Services Lighthouse Laboratory and Alex Alderton, Roberto Amato, Jeffrey Barrett, Sonia Goncalves, Ewan Harrison, David K. Jackson, Ian Johnston, Dominic Kwiatkowski, Cordelia Langford, John Sillitoe on behalf of the Wellcome Sanger Institute COVID-19 Surveillance Team |
| EPI_ISL_2994228, EPI_ISL_2994229 | Lighthouse Lab in Alderley Park | Wellcome Sanger Institute for the COVID-19 Genomics UK (COG-UK) Consortium | Jacquelyn Wynn, Mairead Hyland, The Lighthouse Lab in Alderley Park and Alex Alderton, Roberto Amato, Jeffrey Barrett, Sonia Goncalves, Ewan Harrison, David K. Jackson, Ian Johnston, Dominic Kwiatkowski, Cordelia Langford, John Sillitoe on behalf of the Wellcome Sanger Institute COVID-19 Surveillance Team |
| EPI_ISL_2994230 | Lighthouse Lab in Milton Keynes | Wellcome Sanger Institute for the COVID-19 Genomics UK (COG-UK) Consortium | The Lighthouse Lab in Milton Keynes and Alex Alderton, Roberto Amato, Jeffrey Barrett, Sonia Goncalves, Ewan Harrison, David K. Jackson, Ian Johnston, Dominic Kwiatkowski, Cordelia Langford, John Sillitoe on behalf of the Wellcome Sanger Institute COVID-19 Surveillance Team |
| EPI_ISL_2994231 | Lighthouse Lab in Alderley Park | Wellcome Sanger Institute for the COVID-19 Genomics UK (COG-UK) Consortium | Jacquelyn Wynn, Mairead Hyland, The Lighthouse Lab in Alderley Park and Alex Alderton, Roberto Amato, Jeffrey Barrett, Sonia Goncalves, Ewan Harrison, David K. Jackson, Ian Johnston, Dominic Kwiatkowski, Cordelia Langford, John Sillitoe on behalf of the Wellcome Sanger Institute COVID-19 Surveillance Team |
| EPI_ISL_2994232 | Lighthouse Lab in Milton Keynes | Wellcome Sanger Institute for the COVID-19 Genomics UK (COG-UK) Consortium | The Lighthouse Lab in Milton Keynes and Alex Alderton, Roberto Amato, Jeffrey Barrett, Sonia Goncalves, Ewan Harrison, David K. Jackson, Ian Johnston, Dominic Kwiatkowski, Cordelia Langford, John Sillitoe on behalf of the Wellcome Sanger Institute COVID-19 Surveillance Team |
| EPI_ISL_2994233 | Lighthouse Lab in Alderley Park | Wellcome Sanger Institute for the COVID-19 Genomics UK (COG-UK) Consortium | Jacquelyn Wynn, Mairead Hyland, The Lighthouse Lab in Alderley Park and Alex Alderton, Roberto Amato, Jeffrey Barrett, Sonia Goncalves, Ewan Harrison, David K. Jackson, Ian Johnston, Dominic Kwiatkowski, Cordelia Langford, John Sillitoe on behalf of the Wellcome Sanger Institute COVID-19 Surveillance Team |
| EPI_ISL_2994234 | Lighthouse Lab in Milton Keynes | Wellcome Sanger Institute for the COVID-19 Genomics UK (COG-UK) Consortium | The Lighthouse Lab in Milton Keynes and Alex Alderton, Roberto Amato, Jeffrey Barrett, Sonia Goncalves, Ewan Harrison, David K. Jackson, Ian Johnston, Dominic Kwiatkowski, Cordelia Langford, John Sillitoe on behalf of the Wellcome Sanger Institute COVID-19 Surveillance Team |
| EPI_ISL_2994235 | Lighthouse Lab in Alderley Park | Wellcome Sanger Institute for the COVID-19 Genomics UK (COG-UK) Consortium | Jacquelyn Wynn, Mairead Hyland, The Lighthouse Lab in Alderley Park and Alex Alderton, Roberto Amato, Jeffrey Barrett, Sonia Goncalves, Ewan Harrison, David K. Jackson, Ian Johnston, Dominic Kwiatkowski, Cordelia Langford, John Sillitoe on behalf of the Wellcome Sanger Institute COVID-19 Surveillance Team |
| EPI_ISL_2994236, EPI_ISL_2994237 | Lighthouse Lab in Milton Keynes | Wellcome Sanger Institute for the COVID-19 Genomics UK (COG-UK) Consortium | The Lighthouse Lab in Milton Keynes and Alex Alderton, Roberto Amato, Jeffrey Barrett, Sonia Goncalves, Ewan Harrison, David K. Jackson, Ian Johnston, Dominic Kwiatkowski, Cordelia Langford, John Sillitoe on behalf of the Wellcome Sanger Institute COVID-19 Surveillance Team |
| EPI_ISL_2994238 | Lighthouse Lab in Alderley Park | Wellcome Sanger Institute for the COVID-19 Genomics UK (COG-UK) Consortium | Jacquelyn Wynn, Mairead Hyland, The Lighthouse Lab in Alderley Park and Alex Alderton, Roberto Amato, Jeffrey Barrett, Sonia Goncalves, Ewan Harrison, David K. Jackson, Ian Johnston, Dominic Kwiatkowski, Cordelia Langford, John Sillitoe on behalf of the Wellcome Sanger Institute COVID-19 Surveillance Team |
| EPI_ISL_2994239, EPI_ISL_2994240 | Lighthouse Laboratory Plymouth | Wellcome Sanger Institute for the COVID-19 Genomics UK (COG-UK) Consortium | Lighthouse Laboratory Plymouth and Alex Alderton, Roberto Amato, Jeffrey Barrett, Sonia Goncalves, Ewan Harrison, David K. Jackson, Ian Johnston, Dominic Kwiatkowski, Cordelia Langford, John Sillitoe on behalf of the Wellcome Sanger Institute COVID-19 Surveillance Team |
| EPI_ISL_2994241 | Lighthouse Lab in Milton Keynes | Wellcome Sanger Institute for the COVID-19 Genomics UK (COG-UK) Consortium | The Lighthouse Lab in Milton Keynes and Alex Alderton, Roberto Amato, Jeffrey Barrett, Sonia Goncalves, Ewan Harrison, David K. Jackson, Ian Johnston, Dominic Kwiatkowski, Cordelia Langford, John Sillitoe on behalf of the Wellcome Sanger Institute COVID-19 Surveillance Team |
| EPI_ISL_2994242 | Lighthouse Lab in Alderley Park | Wellcome Sanger Institute for the COVID-19 Genomics UK (COG-UK) Consortium | Jacquelyn Wynn, Mairead Hyland, The Lighthouse Lab in Alderley Park and Alex Alderton, Roberto Amato, Jeffrey Barrett, Sonia Goncalves, Ewan Harrison, David K. Jackson, Ian Johnston, Dominic Kwiatkowski, Cordelia Langford, John Sillitoe on behalf of the Wellcome Sanger Institute COVID-19 Surveillance Team |
| EPI_ISL_2994243, EPI_ISL_2994244 | Berkshire and Surrey Pathology Services Lighthouse Laboratory | Wellcome Sanger Institute for the COVID-19 Genomics UK (COG-UK) Consortium | Berkshire and Surrey Pathology Services Lighthouse Laboratory and Alex Alderton, Roberto Amato, Jeffrey Barrett, Sonia Goncalves, Ewan Harrison, David K. Jackson, Ian Johnston, Dominic Kwiatkowski, Cordelia Langford, John Sillitoe on behalf of the Wellcome Sanger Institute COVID-19 Surveillance Team |
| EPI_ISL_2994245 | Lighthouse Lab in Alderley Park | Wellcome Sanger Institute for the COVID-19 Genomics UK (COG-UK) Consortium | Jacquelyn Wynn, Mairead Hyland, The Lighthouse Lab in Alderley Park and Alex Alderton, Roberto Amato, Jeffrey Barrett, Sonia Goncalves, Ewan Harrison, David K. Jackson, Ian Johnston, Dominic Kwiatkowski, Cordelia Langford, John Sillitoe on behalf of the Wellcome Sanger Institute COVID-19 Surveillance Team |
| EPI_ISL_2994246, EPI_ISL_2994247 | Lighthouse Laboratory Plymouth | Wellcome Sanger Institute for the COVID-19 Genomics UK (COG-UK) Consortium | Lighthouse Laboratory Plymouth and Alex Alderton, Roberto Amato, Jeffrey Barrett, Sonia Goncalves, Ewan Harrison, David K. Jackson, Ian Johnston, Dominic Kwiatkowski, Cordelia Langford, John Sillitoe on behalf of the Wellcome Sanger Institute COVID-19 Surveillance Team |
| EPI_ISL_2994248 | Lighthouse Lab in Alderley Park | Wellcome Sanger Institute for the COVID-19 Genomics UK (COG-UK) Consortium | Jacquelyn Wynn, Mairead Hyland, The Lighthouse Lab in Alderley Park and Alex Alderton, Roberto Amato, Jeffrey Barrett, Sonia Goncalves, Ewan Harrison, David K. Jackson, Ian Johnston, Dominic Kwiatkowski, Cordelia Langford, John Sillitoe on behalf of the Wellcome Sanger Institute COVID-19 Surveillance Team |
| EPI_ISL_2994249, EPI_ISL_2994250, EPI_ISL_2994251 | Lighthouse Laboratory Plymouth | Wellcome Sanger Institute for the COVID-19 Genomics UK (COG-UK) Consortium | Lighthouse Laboratory Plymouth and Alex Alderton, Roberto Amato, Jeffrey Barrett, Sonia Goncalves, Ewan Harrison, David K. Jackson, Ian Johnston, Dominic Kwiatkowski, Cordelia Langford, John Sillitoe on behalf of the Wellcome Sanger Institute COVID-19 Surveillance Team |
| EPI_ISL_2994252 | Lighthouse Lab in Alderley Park | Wellcome Sanger Institute for the COVID-19 Genomics UK (COG-UK) Consortium | Jacquelyn Wynn, Mairead Hyland, The Lighthouse Lab in Alderley Park and Alex Alderton, Roberto Amato, Jeffrey Barrett, Sonia Goncalves, Ewan Harrison, David K. Jackson, Ian Johnston, Dominic Kwiatkowski, Cordelia Langford, John Sillitoe on behalf of the Wellcome Sanger Institute COVID-19 Surveillance Team |

| | | | |
|---|---|---|---|
| EPI_ISL_2994253 | Lighthouse Laboratory Plymouth | Wellcome Sanger Institute for the COVID-19 Genomics UK (COG-UK) Consortium | Lighthouse Laboratory Plymouth and Alex Alderton, Roberto Amato, Jeffrey Barrett, Sonia Goncalves, Ewan Harrison, David K. Jackson, Ian Johnston, Dominic Kwiatkowski, Cordelia Langford, John Sillitoe on behalf of the Wellcome Sanger Institute COVID-19 Surveillance Team |
| EPI_ISL_2994254 | Berkshire and Surrey Pathology Services Lighthouse Laboratory | Wellcome Sanger Institute for the COVID-19 Genomics UK (COG-UK) Consortium | Berkshire and Surrey Pathology Services Lighthouse Laboratory and Alex Alderton, Roberto Amato, Jeffrey Barrett, Sonia Goncalves, Ewan Harrison, David K. Jackson, Ian Johnston, Dominic Kwiatkowski, Cordelia Langford, John Sillitoe on behalf of the Wellcome Sanger Institute COVID-19 Surveillance Team |
| EPI_ISL_2994255 | Lighthouse Lab in Milton Keynes | Wellcome Sanger Institute for the COVID-19 Genomics UK (COG-UK) Consortium | The Lighthouse Lab in Milton Keynes and Alex Alderton, Roberto Amato, Jeffrey Barrett, Sonia Goncalves, Ewan Harrison, David K. Jackson, Ian Johnston, Dominic Kwiatkowski, Cordelia Langford, John Sillitoe on behalf of the Wellcome Sanger Institute COVID-19 Surveillance Team |
| EPI_ISL_2994256 | Lighthouse Laboratory Plymouth | Wellcome Sanger Institute for the COVID-19 Genomics UK (COG-UK) Consortium | Lighthouse Laboratory Plymouth and Alex Alderton, Roberto Amato, Jeffrey Barrett, Sonia Goncalves, Ewan Harrison, David K. Jackson, Ian Johnston, Dominic Kwiatkowski, Cordelia Langford, John Sillitoe on behalf of the Wellcome Sanger Institute COVID-19 Surveillance Team |
| EPI_ISL_2994257 | Lighthouse Lab in Alderley Park | Wellcome Sanger Institute for the COVID-19 Genomics UK (COG-UK) Consortium | Jacquelyn Wynn, Mairead Hyland, The Lighthouse Lab in Alderley Park and Alex Alderton, Roberto Amato, Jeffrey Barrett, Sonia Goncalves, Ewan Harrison, David K. Jackson, Ian Johnston, Dominic Kwiatkowski, Cordelia Langford, John Sillitoe on behalf of the Wellcome Sanger Institute COVID-19 Surveillance Team |
| EPI_ISL_2994258 | Berkshire and Surrey Pathology Services Lighthouse Laboratory | Wellcome Sanger Institute for the COVID-19 Genomics UK (COG-UK) Consortium | Berkshire and Surrey Pathology Services Lighthouse Laboratory and Alex Alderton, Roberto Amato, Jeffrey Barrett, Sonia Goncalves, Ewan Harrison, David K. Jackson, Ian Johnston, Dominic Kwiatkowski, Cordelia Langford, John Sillitoe on behalf of the Wellcome Sanger Institute COVID-19 Surveillance Team |
| EPI_ISL_2994259, EPI_ISL_2994260 | Lighthouse Lab in Milton Keynes | Wellcome Sanger Institute for the COVID-19 Genomics UK (COG-UK) Consortium | The Lighthouse Lab in Milton Keynes and Alex Alderton, Roberto Amato, Jeffrey Barrett, Sonia Goncalves, Ewan Harrison, David K. Jackson, Ian Johnston, Dominic Kwiatkowski, Cordelia Langford, John Sillitoe on behalf of the Wellcome Sanger Institute COVID-19 Surveillance Team |
| EPI_ISL_2994261, EPI_ISL_2994262 | Lighthouse Laboratory Plymouth | Wellcome Sanger Institute for the COVID-19 Genomics UK (COG-UK) Consortium | Lighthouse Laboratory Plymouth and Alex Alderton, Roberto Amato, Jeffrey Barrett, Sonia Goncalves, Ewan Harrison, David K. Jackson, Ian Johnston, Dominic Kwiatkowski, Cordelia Langford, John Sillitoe on behalf of the Wellcome Sanger Institute COVID-19 Surveillance Team |
| EPI_ISL_2994263 | Lighthouse Lab in Alderley Park | Wellcome Sanger Institute for the COVID-19 Genomics UK (COG-UK) Consortium | Jacquelyn Wynn, Mairead Hyland, The Lighthouse Lab in Alderley Park and Alex Alderton, Roberto Amato, Jeffrey Barrett, Sonia Goncalves, Ewan Harrison, David K. Jackson, Ian Johnston, Dominic Kwiatkowski, Cordelia Langford, John Sillitoe on behalf of the Wellcome Sanger Institute COVID-19 Surveillance Team |
| EPI_ISL_2994264 | Lighthouse Laboratory Plymouth | Wellcome Sanger Institute for the COVID-19 Genomics UK (COG-UK) Consortium | Lighthouse Laboratory Plymouth and Alex Alderton, Roberto Amato, Jeffrey Barrett, Sonia Goncalves, Ewan Harrison, David K. Jackson, Ian Johnston, Dominic Kwiatkowski, Cordelia Langford, John Sillitoe on behalf of the Wellcome Sanger Institute COVID-19 Surveillance Team |
| EPI_ISL_2994265 | Lighthouse Lab in Alderley Park | Wellcome Sanger Institute for the COVID-19 Genomics UK (COG-UK) Consortium | Jacquelyn Wynn, Mairead Hyland, The Lighthouse Lab in Alderley Park and Alex Alderton, Roberto Amato, Jeffrey Barrett, Sonia Goncalves, Ewan Harrison, David K. Jackson, Ian Johnston, Dominic Kwiatkowski, Cordelia Langford, John Sillitoe on behalf of the Wellcome Sanger Institute COVID-19 Surveillance Team |
| EPI_ISL_2994266 | Lighthouse Lab in Milton Keynes | Wellcome Sanger Institute for the COVID-19 Genomics UK (COG-UK) Consortium | The Lighthouse Lab in Milton Keynes and Alex Alderton, Roberto Amato, Jeffrey Barrett, Sonia Goncalves, Ewan Harrison, David K. Jackson, Ian Johnston, Dominic Kwiatkowski, Cordelia Langford, John Sillitoe on behalf of the Wellcome Sanger Institute COVID-19 Surveillance Team |
| EPI_ISL_2994267, EPI_ISL_2994268, EPI_ISL_2994269, EPI_ISL_2994270 | Berkshire and Surrey Pathology Services Lighthouse Laboratory | Wellcome Sanger Institute for the COVID-19 Genomics UK (COG-UK) Consortium | Berkshire and Surrey Pathology Services Lighthouse Laboratory and Alex Alderton, Roberto Amato, Jeffrey Barrett, Sonia Goncalves, Ewan Harrison, David K. Jackson, Ian Johnston, Dominic Kwiatkowski, Cordelia Langford, John Sillitoe on behalf of the Wellcome Sanger Institute COVID-19 Surveillance Team |
| EPI_ISL_2994271, EPI_ISL_2994272 | Lighthouse Laboratory Plymouth | Wellcome Sanger Institute for the COVID-19 Genomics UK (COG-UK) Consortium | Lighthouse Laboratory Plymouth and Alex Alderton, Roberto Amato, Jeffrey Barrett, Sonia Goncalves, Ewan Harrison, David K. Jackson, Ian Johnston, Dominic Kwiatkowski, Cordelia Langford, John Sillitoe on behalf of the Wellcome Sanger Institute COVID-19 Surveillance Team |
| EPI_ISL_2994273, EPI_ISL_2994274 | Lighthouse Lab in Alderley Park | Wellcome Sanger Institute for the COVID-19 Genomics UK (COG-UK) Consortium | Jacquelyn Wynn, Mairead Hyland, The Lighthouse Lab in Alderley Park and Alex Alderton, Roberto Amato, Jeffrey Barrett, Sonia Goncalves, Ewan Harrison, David K. Jackson, Ian Johnston, Dominic Kwiatkowski, Cordelia Langford, John Sillitoe on behalf of the Wellcome Sanger Institute COVID-19 Surveillance Team |
| EPI_ISL_2994275 | Berkshire and Surrey Pathology Services Lighthouse Laboratory | Wellcome Sanger Institute for the COVID-19 Genomics UK (COG-UK) Consortium | Berkshire and Surrey Pathology Services Lighthouse Laboratory and Alex Alderton, Roberto Amato, Jeffrey Barrett, Sonia Goncalves, Ewan Harrison, David K. Jackson, Ian Johnston, Dominic Kwiatkowski, Cordelia Langford, John Sillitoe on behalf of the Wellcome Sanger Institute COVID-19 Surveillance Team |
| EPI_ISL_2994276 | Lighthouse Lab in Alderley Park | Wellcome Sanger Institute for the COVID-19 Genomics UK (COG-UK) Consortium | Jacquelyn Wynn, Mairead Hyland, The Lighthouse Lab in Alderley Park and Alex Alderton, Roberto Amato, Jeffrey Barrett, Sonia Goncalves, Ewan Harrison, David K. Jackson, Ian Johnston, Dominic Kwiatkowski, Cordelia Langford, John Sillitoe on behalf of the Wellcome Sanger Institute COVID-19 Surveillance Team |
| EPI_ISL_2994277 | Berkshire and Surrey Pathology Services Lighthouse Laboratory | Wellcome Sanger Institute for the COVID-19 Genomics UK (COG-UK) Consortium | Berkshire and Surrey Pathology Services Lighthouse Laboratory and Alex Alderton, Roberto Amato, Jeffrey Barrett, Sonia Goncalves, Ewan Harrison, David K. Jackson, Ian Johnston, Dominic Kwiatkowski, Cordelia Langford, John Sillitoe on behalf of the Wellcome Sanger Institute COVID-19 Surveillance Team |
| EPI_ISL_2994278 | Lighthouse Lab in Milton Keynes | Wellcome Sanger Institute for the COVID-19 Genomics UK (COG-UK) Consortium | The Lighthouse Lab in Milton Keynes and Alex Alderton, Roberto Amato, Jeffrey Barrett, Sonia Goncalves, Ewan Harrison, David K. Jackson, Ian Johnston, Dominic Kwiatkowski, Cordelia Langford, John Sillitoe on behalf of the Wellcome Sanger Institute COVID-19 Surveillance Team |
| EPI_ISL_2994279 | Berkshire and Surrey Pathology Services Lighthouse Laboratory | Wellcome Sanger Institute for the COVID-19 Genomics UK (COG-UK) Consortium | Berkshire and Surrey Pathology Services Lighthouse Laboratory and Alex Alderton, Roberto Amato, Jeffrey Barrett, Sonia Goncalves, Ewan Harrison, David K. Jackson, Ian Johnston, Dominic Kwiatkowski, Cordelia Langford, John Sillitoe on behalf of the Wellcome Sanger Institute COVID-19 Surveillance Team |
| EPI_ISL_2994280, EPI_ISL_2994281 | Lighthouse Lab in Alderley Park | Wellcome Sanger Institute for the COVID-19 Genomics UK (COG-UK) Consortium | Jacquelyn Wynn, Mairead Hyland, The Lighthouse Lab in Alderley Park and Alex Alderton, Roberto Amato, Jeffrey Barrett, Sonia Goncalves, Ewan Harrison, David K. Jackson, Ian Johnston, Dominic Kwiatkowski, Cordelia Langford, John Sillitoe on behalf of the Wellcome Sanger Institute COVID-19 Surveillance Team |
| EPI_ISL_2994282 | Lighthouse Laboratory Plymouth | Wellcome Sanger Institute for the COVID-19 Genomics UK (COG-UK) Consortium | Lighthouse Laboratory Plymouth and Alex Alderton, Roberto Amato, Jeffrey Barrett, Sonia Goncalves, Ewan Harrison, David K. Jackson, Ian Johnston, Dominic Kwiatkowski, Cordelia Langford, John Sillitoe on behalf of the Wellcome Sanger Institute COVID-19 Surveillance Team |
| EPI_ISL_2994283, EPI_ISL_2994284 | Lighthouse Lab in Milton Keynes | Wellcome Sanger Institute for the COVID-19 Genomics UK (COG-UK) Consortium | The Lighthouse Lab in Milton Keynes and Alex Alderton, Roberto Amato, Jeffrey Barrett, Sonia Goncalves, Ewan Harrison, David K. Jackson, Ian Johnston, Dominic Kwiatkowski, Cordelia Langford, John Sillitoe on behalf of the Wellcome Sanger Institute COVID-19 Surveillance Team |
| EPI_ISL_2994285 | Lighthouse Lab in Alderley Park | Wellcome Sanger Institute for the COVID-19 Genomics UK (COG-UK) Consortium | Jacquelyn Wynn, Mairead Hyland, The Lighthouse Lab in Alderley Park and Alex Alderton, Roberto Amato, Jeffrey Barrett, Sonia Goncalves, Ewan Harrison, David K. Jackson, Ian Johnston, Dominic Kwiatkowski, Cordelia Langford, John Sillitoe on behalf of the Wellcome Sanger Institute COVID-19 Surveillance Team |
| EPI_ISL_2994286 | Berkshire and Surrey Pathology Services Lighthouse Laboratory | Wellcome Sanger Institute for the COVID-19 Genomics UK (COG-UK) Consortium | Berkshire and Surrey Pathology Services Lighthouse Laboratory and Alex Alderton, Roberto Amato, Jeffrey Barrett, Sonia Goncalves, Ewan Harrison, David K. Jackson, Ian Johnston, Dominic Kwiatkowski, Cordelia Langford, John Sillitoe on behalf of the Wellcome Sanger Institute COVID-19 Surveillance Team |
| EPI_ISL_2994287, EPI_ISL_2994288 | Lighthouse Laboratory Plymouth | Wellcome Sanger Institute for the COVID-19 Genomics UK (COG-UK) Consortium | Lighthouse Laboratory Plymouth and Alex Alderton, Roberto Amato, Jeffrey Barrett, Sonia Goncalves, Ewan Harrison, David K. Jackson, Ian Johnston, Dominic Kwiatkowski, Cordelia Langford, John Sillitoe on behalf of the Wellcome Sanger Institute COVID-19 Surveillance Team |
| EPI_ISL_2994289 | Berkshire and Surrey Pathology Services Lighthouse Laboratory | Wellcome Sanger Institute for the COVID-19 Genomics UK (COG-UK) Consortium | Berkshire and Surrey Pathology Services Lighthouse Laboratory and Alex Alderton, Roberto Amato, Jeffrey Barrett, Sonia Goncalves, Ewan Harrison, David K. Jackson, Ian Johnston, Dominic Kwiatkowski, Cordelia Langford, John Sillitoe on behalf of the Wellcome Sanger Institute COVID-19 Surveillance Team |
| EPI_ISL_2994290 | Lighthouse Lab in Alderley Park | Wellcome Sanger Institute for the COVID-19 Genomics UK (COG-UK) Consortium | Jacquelyn Wynn, Mairead Hyland, The Lighthouse Lab in Alderley Park and Alex Alderton, Roberto Amato, Jeffrey Barrett, Sonia Goncalves, Ewan Harrison, David K. Jackson, Ian Johnston, Dominic Kwiatkowski, Cordelia Langford, John Sillitoe on behalf of the Wellcome Sanger Institute COVID-19 Surveillance Team |
| EPI_ISL_2994291 | Lighthouse Lab in Milton Keynes | Wellcome Sanger Institute for the COVID-19 Genomics UK (COG-UK) Consortium | The Lighthouse Lab in Milton Keynes and Alex Alderton, Roberto Amato, Jeffrey Barrett, Sonia Goncalves, Ewan Harrison, David K. Jackson, Ian Johnston, Dominic Kwiatkowski, Cordelia Langford, John Sillitoe on behalf of the Wellcome Sanger Institute COVID-19 Surveillance Team |
| EPI_ISL_2994292 | Berkshire and Surrey Pathology Services Lighthouse Laboratory | Wellcome Sanger Institute for the COVID-19 Genomics UK (COG-UK) Consortium | Berkshire and Surrey Pathology Services Lighthouse Laboratory and Alex Alderton, Roberto Amato, Jeffrey Barrett, Sonia Goncalves, Ewan Harrison, David K. Jackson, Ian Johnston, Dominic Kwiatkowski, Cordelia Langford, John Sillitoe on behalf of the Wellcome Sanger Institute COVID-19 Surveillance Team |
| EPI_ISL_2994293, EPI_ISL_2994294 | Lighthouse Laboratory Plymouth | Wellcome Sanger Institute for the COVID-19 Genomics UK (COG-UK) Consortium | Lighthouse Laboratory Plymouth and Alex Alderton, Roberto Amato, Jeffrey Barrett, Sonia Goncalves, Ewan Harrison, David K. Jackson, Ian Johnston, Dominic Kwiatkowski, Cordelia Langford, John Sillitoe on behalf of the Wellcome Sanger Institute COVID-19 Surveillance Team |
| EPI_ISL_2994295 | Berkshire and Surrey Pathology Services Lighthouse Laboratory | Wellcome Sanger Institute for the COVID-19 Genomics UK (COG-UK) Consortium | Berkshire and Surrey Pathology Services Lighthouse Laboratory and Alex Alderton, Roberto Amato, Jeffrey Barrett, Sonia Goncalves, Ewan Harrison, David K. Jackson, Ian Johnston, Dominic Kwiatkowski, Cordelia Langford, John Sillitoe on behalf of the Wellcome Sanger Institute COVID-19 Surveillance Team |
| EPI_ISL_2994296 | Lighthouse Lab in Milton Keynes | Wellcome Sanger Institute for the COVID-19 Genomics UK (COG-UK) Consortium | The Lighthouse Lab in Milton Keynes and Alex Alderton, Roberto Amato, Jeffrey Barrett, Sonia Goncalves, Ewan Harrison, David K. Jackson, Ian Johnston, Dominic Kwiatkowski, Cordelia Langford, John Sillitoe on behalf of the Wellcome Sanger Institute COVID-19 Surveillance Team |
| EPI_ISL_2994297 | Lighthouse Laboratory Plymouth | Wellcome Sanger Institute for the COVID-19 Genomics UK (COG-UK) Consortium | Lighthouse Laboratory Plymouth and Alex Alderton, Roberto Amato, Jeffrey Barrett, Sonia Goncalves, Ewan Harrison, David K. Jackson, Ian Johnston, Dominic Kwiatkowski, Cordelia Langford, John Sillitoe on behalf of the Wellcome Sanger Institute COVID-19 Surveillance Team |
| EPI_ISL_2994298 | Lighthouse Lab in Milton Keynes | Wellcome Sanger Institute for the COVID-19 Genomics UK (COG-UK) Consortium | The Lighthouse Lab in Milton Keynes and Alex Alderton, Roberto Amato, Jeffrey Barrett, Sonia Goncalves, Ewan Harrison, David K. Jackson, Ian Johnston, Dominic Kwiatkowski, Cordelia Langford, John Sillitoe on behalf of the Wellcome Sanger Institute COVID-19 Surveillance Team |
| EPI_ISL_2994299 | Lighthouse Laboratory Plymouth | Wellcome Sanger Institute for the COVID-19 Genomics UK (COG-UK) Consortium | Lighthouse Laboratory Plymouth and Alex Alderton, Roberto Amato, Jeffrey Barrett, Sonia Goncalves, Ewan Harrison, David K. Jackson, Ian Johnston, Dominic Kwiatkowski, Cordelia Langford, John Sillitoe on behalf of the Wellcome Sanger Institute COVID-19 Surveillance Team |
| EPI_ISL_2994300 | Lighthouse Lab in Alderley Park | Wellcome Sanger Institute for the COVID-19 Genomics UK (COG-UK) Consortium | Jacquelyn Wynn, Mairead Hyland, The Lighthouse Lab in Alderley Park and Alex Alderton, Roberto Amato, Jeffrey Barrett, Sonia Goncalves, Ewan Harrison, David K. Jackson, Ian Johnston, Dominic Kwiatkowski, Cordelia Langford, John Sillitoe on behalf of the Wellcome Sanger Institute COVID-19 Surveillance Team |
| EPI_ISL_2994301 | Lighthouse Laboratory Plymouth | Wellcome Sanger Institute for the COVID-19 Genomics UK (COG-UK) Consortium | Lighthouse Laboratory Plymouth and Alex Alderton, Roberto Amato, Jeffrey Barrett, Sonia Goncalves, Ewan Harrison, David K. Jackson, Ian Johnston, Dominic Kwiatkowski, Cordelia Langford, John Sillitoe on behalf of the Wellcome Sanger Institute COVID-19 Surveillance Team |
| EPI_ISL_2994302 | Lighthouse Lab in Alderley Park | Wellcome Sanger Institute for the COVID-19 Genomics UK (COG-UK) Consortium | Jacquelyn Wynn, Mairead Hyland, The Lighthouse Lab in Alderley Park and Alex Alderton, Roberto Amato, Jeffrey Barrett, Sonia Goncalves, Ewan Harrison, David K. Jackson, Ian Johnston, Dominic Kwiatkowski, Cordelia Langford, John Sillitoe on behalf of the Wellcome Sanger Institute COVID-19 Surveillance Team |
| EPI_ISL_2994303 | Berkshire and Surrey Pathology Services Lighthouse Laboratory | Wellcome Sanger Institute for the COVID-19 Genomics UK (COG-UK) Consortium | Berkshire and Surrey Pathology Services Lighthouse Laboratory and Alex Alderton, Roberto Amato, Jeffrey Barrett, Sonia Goncalves, Ewan Harrison, David K. Jackson, Ian Johnston, Dominic Kwiatkowski, Cordelia Langford, John Sillitoe on behalf of the Wellcome Sanger Institute COVID-19 Surveillance Team |
| EPI_ISL_2994304 | Lighthouse Laboratory Plymouth | Wellcome Sanger Institute for the COVID-19 Genomics UK (COG-UK) Consortium | Lighthouse Laboratory Plymouth and Alex Alderton, Roberto Amato, Jeffrey Barrett, Sonia Goncalves, Ewan Harrison, David K. Jackson, Ian Johnston, Dominic Kwiatkowski, Cordelia Langford, John Sillitoe on behalf of the Wellcome Sanger Institute COVID-19 Surveillance Team |
| EPI_ISL_2994305, EPI_ISL_2994306 | Lighthouse Lab in Milton Keynes | Wellcome Sanger Institute for the COVID-19 Genomics UK (COG-UK) Consortium | The Lighthouse Lab in Milton Keynes and Alex Alderton, Roberto Amato, Jeffrey Barrett, Sonia Goncalves, Ewan Harrison, David K. Jackson, Ian Johnston, Dominic Kwiatkowski, Cordelia Langford, John Sillitoe on behalf of the Wellcome Sanger Institute COVID-19 Surveillance Team |
| EPI_ISL_2994307 | Lighthouse Lab in Alderley Park | Wellcome Sanger Institute for the COVID-19 Genomics UK (COG-UK) Consortium | Jacquelyn Wynn, Mairead Hyland, The Lighthouse Lab in Alderley Park and Alex Alderton, Roberto Amato, Jeffrey Barrett, Sonia Goncalves, Ewan Harrison, David K. Jackson, Ian Johnston, Dominic Kwiatkowski, Cordelia Langford, John Sillitoe on behalf of the Wellcome Sanger Institute COVID-19 Surveillance Team |
| EPI_ISL_2994308 | Berkshire and Surrey Pathology Services Lighthouse Laboratory | Wellcome Sanger Institute for the COVID-19 Genomics UK (COG-UK) Consortium | Berkshire and Surrey Pathology Services Lighthouse Laboratory and Alex Alderton, Roberto Amato, Jeffrey Barrett, Sonia Goncalves, Ewan Harrison, David K. Jackson, Ian Johnston, Dominic Kwiatkowski, Cordelia Langford, John Sillitoe on behalf of the Wellcome Sanger Institute COVID-19 Surveillance Team |

| | | | |
|---|---|---|---|
| EPI_ISL_2994309 | Lighthouse Laboratory Plymouth | Wellcome Sanger Institute for the COVID-19 Genomics UK (COG-UK) Consortium | Lighthouse Laboratory Plymouth and Alex Alderton, Roberto Amato, Jeffrey Barrett, Sonia Goncalves, Ewan Harrison, David K. Jackson, Ian Johnston, Dominic Kwiatkowski, Cordelia Langford, John Sillitoe on behalf of the Wellcome Sanger Institute COVID-19 Surveillance Team |
| EPI_ISL_2994310, EPI_ISL_2994311 | Lighthouse Lab in Milton Keynes | Wellcome Sanger Institute for the COVID-19 Genomics UK (COG-UK) Consortium | The Lighthouse Lab in Milton Keynes and Alex Alderton, Roberto Amato, Jeffrey Barrett, Sonia Goncalves, Ewan Harrison, David K. Jackson, Ian Johnston, Dominic Kwiatkowski, Cordelia Langford, John Sillitoe on behalf of the Wellcome Sanger Institute COVID-19 Surveillance Team |
| EPI_ISL_2994312 | Lighthouse Laboratory Plymouth | Wellcome Sanger Institute for the COVID-19 Genomics UK (COG-UK) Consortium | Lighthouse Laboratory Plymouth and Alex Alderton, Roberto Amato, Jeffrey Barrett, Sonia Goncalves, Ewan Harrison, David K. Jackson, Ian Johnston, Dominic Kwiatkowski, Cordelia Langford, John Sillitoe on behalf of the Wellcome Sanger Institute COVID-19 Surveillance Team |
| EPI_ISL_2994313 | Lighthouse Lab in Alderley Park | Wellcome Sanger Institute for the COVID-19 Genomics UK (COG-UK) Consortium | Jacquelyn Wynn, Mairead Hyland, The Lighthouse Lab in Alderley Park and Alex Alderton, Roberto Amato, Jeffrey Barrett, Sonia Goncalves, Ewan Harrison, David K. Jackson, Ian Johnston, Dominic Kwiatkowski, Cordelia Langford, John Sillitoe on behalf of the Wellcome Sanger Institute COVID-19 Surveillance Team |
| EPI_ISL_2994314 | Lighthouse Laboratory Plymouth | Wellcome Sanger Institute for the COVID-19 Genomics UK (COG-UK) Consortium | Lighthouse Laboratory Plymouth and Alex Alderton, Roberto Amato, Jeffrey Barrett, Sonia Goncalves, Ewan Harrison, David K. Jackson, Ian Johnston, Dominic Kwiatkowski, Cordelia Langford, John Sillitoe on behalf of the Wellcome Sanger Institute COVID-19 Surveillance Team |
| EPI_ISL_2994315 | Berkshire and Surrey Pathology Services Lighthouse Laboratory | Wellcome Sanger Institute for the COVID-19 Genomics UK (COG-UK) Consortium | Berkshire and Surrey Pathology Services Lighthouse Laboratory and Alex Alderton, Roberto Amato, Jeffrey Barrett, Sonia Goncalves, Ewan Harrison, David K. Jackson, Ian Johnston, Dominic Kwiatkowski, Cordelia Langford, John Sillitoe on behalf of the Wellcome Sanger Institute COVID-19 Surveillance Team |
| EPI_ISL_2994316 | Lighthouse Lab in Milton Keynes | Wellcome Sanger Institute for the COVID-19 Genomics UK (COG-UK) Consortium | The Lighthouse Lab in Milton Keynes and Alex Alderton, Roberto Amato, Jeffrey Barrett, Sonia Goncalves, Ewan Harrison, David K. Jackson, Ian Johnston, Dominic Kwiatkowski, Cordelia Langford, John Sillitoe on behalf of the Wellcome Sanger Institute COVID-19 Surveillance Team |
| EPI_ISL_2994317 | Lighthouse Laboratory Plymouth | Wellcome Sanger Institute for the COVID-19 Genomics UK (COG-UK) Consortium | Lighthouse Laboratory Plymouth and Alex Alderton, Roberto Amato, Jeffrey Barrett, Sonia Goncalves, Ewan Harrison, David K. Jackson, Ian Johnston, Dominic Kwiatkowski, Cordelia Langford, John Sillitoe on behalf of the Wellcome Sanger Institute COVID-19 Surveillance Team |
| EPI_ISL_2994318 | Lighthouse Lab in Alderley Park | Wellcome Sanger Institute for the COVID-19 Genomics UK (COG-UK) Consortium | Jacquelyn Wynn, Mairead Hyland, The Lighthouse Lab in Alderley Park and Alex Alderton, Roberto Amato, Jeffrey Barrett, Sonia Goncalves, Ewan Harrison, David K. Jackson, Ian Johnston, Dominic Kwiatkowski, Cordelia Langford, John Sillitoe on behalf of the Wellcome Sanger Institute COVID-19 Surveillance Team |
| EPI_ISL_2994319 | Lighthouse Lab in Milton Keynes | Wellcome Sanger Institute for the COVID-19 Genomics UK (COG-UK) Consortium | The Lighthouse Lab in Milton Keynes and Alex Alderton, Roberto Amato, Jeffrey Barrett, Sonia Goncalves, Ewan Harrison, David K. Jackson, Ian Johnston, Dominic Kwiatkowski, Cordelia Langford, John Sillitoe on behalf of the Wellcome Sanger Institute COVID-19 Surveillance Team |
| EPI_ISL_2994320 | Lighthouse Laboratory Plymouth | Wellcome Sanger Institute for the COVID-19 Genomics UK (COG-UK) Consortium | Lighthouse Laboratory Plymouth and Alex Alderton, Roberto Amato, Jeffrey Barrett, Sonia Goncalves, Ewan Harrison, David K. Jackson, Ian Johnston, Dominic Kwiatkowski, Cordelia Langford, John Sillitoe on behalf of the Wellcome Sanger Institute COVID-19 Surveillance Team |
| EPI_ISL_2994321, EPI_ISL_2994322 | Lighthouse Lab in Alderley Park | Wellcome Sanger Institute for the COVID-19 Genomics UK (COG-UK) Consortium | Jacquelyn Wynn, Mairead Hyland, The Lighthouse Lab in Alderley Park and Alex Alderton, Roberto Amato, Jeffrey Barrett, Sonia Goncalves, Ewan Harrison, David K. Jackson, Ian Johnston, Dominic Kwiatkowski, Cordelia Langford, John Sillitoe on behalf of the Wellcome Sanger Institute COVID-19 Surveillance Team |
| EPI_ISL_2994323 | Lighthouse Lab in Milton Keynes | Wellcome Sanger Institute for the COVID-19 Genomics UK (COG-UK) Consortium | The Lighthouse Lab in Milton Keynes and Alex Alderton, Roberto Amato, Jeffrey Barrett, Sonia Goncalves, Ewan Harrison, David K. Jackson, Ian Johnston, Dominic Kwiatkowski, Cordelia Langford, John Sillitoe on behalf of the Wellcome Sanger Institute COVID-19 Surveillance Team |
| EPI_ISL_2994324 | Berkshire and Surrey Pathology Services Lighthouse Laboratory | Wellcome Sanger Institute for the COVID-19 Genomics UK (COG-UK) Consortium | Berkshire and Surrey Pathology Services Lighthouse Laboratory and Alex Alderton, Roberto Amato, Jeffrey Barrett, Sonia Goncalves, Ewan Harrison, David K. Jackson, Ian Johnston, Dominic Kwiatkowski, Cordelia Langford, John Sillitoe on behalf of the Wellcome Sanger Institute COVID-19 Surveillance Team |
| EPI_ISL_2994325 | Lighthouse Lab in Milton Keynes | Wellcome Sanger Institute for the COVID-19 Genomics UK (COG-UK) Consortium | The Lighthouse Lab in Milton Keynes and Alex Alderton, Roberto Amato, Jeffrey Barrett, Sonia Goncalves, Ewan Harrison, David K. Jackson, Ian Johnston, Dominic Kwiatkowski, Cordelia Langford, John Sillitoe on behalf of the Wellcome Sanger Institute COVID-19 Surveillance Team |
| EPI_ISL_2994326 | Berkshire and Surrey Pathology Services Lighthouse Laboratory | Wellcome Sanger Institute for the COVID-19 Genomics UK (COG-UK) Consortium | Berkshire and Surrey Pathology Services Lighthouse Laboratory and Alex Alderton, Roberto Amato, Jeffrey Barrett, Sonia Goncalves, Ewan Harrison, David K. Jackson, Ian Johnston, Dominic Kwiatkowski, Cordelia Langford, John Sillitoe on behalf of the Wellcome Sanger Institute COVID-19 Surveillance Team |
| EPI_ISL_2994327 | Lighthouse Lab in Milton Keynes | Wellcome Sanger Institute for the COVID-19 Genomics UK (COG-UK) Consortium | The Lighthouse Lab in Milton Keynes and Alex Alderton, Roberto Amato, Jeffrey Barrett, Sonia Goncalves, Ewan Harrison, David K. Jackson, Ian Johnston, Dominic Kwiatkowski, Cordelia Langford, John Sillitoe on behalf of the Wellcome Sanger Institute COVID-19 Surveillance Team |
| EPI_ISL_2994328 | Berkshire and Surrey Pathology Services Lighthouse Laboratory | Wellcome Sanger Institute for the COVID-19 Genomics UK (COG-UK) Consortium | Berkshire and Surrey Pathology Services Lighthouse Laboratory and Alex Alderton, Roberto Amato, Jeffrey Barrett, Sonia Goncalves, Ewan Harrison, David K. Jackson, Ian Johnston, Dominic Kwiatkowski, Cordelia Langford, John Sillitoe on behalf of the Wellcome Sanger Institute COVID-19 Surveillance Team |
| EPI_ISL_2994329 | Lighthouse Laboratory Plymouth | Wellcome Sanger Institute for the COVID-19 Genomics UK (COG-UK) Consortium | Lighthouse Laboratory Plymouth and Alex Alderton, Roberto Amato, Jeffrey Barrett, Sonia Goncalves, Ewan Harrison, David K. Jackson, Ian Johnston, Dominic Kwiatkowski, Cordelia Langford, John Sillitoe on behalf of the Wellcome Sanger Institute COVID-19 Surveillance Team |
| EPI_ISL_2994330 | Lighthouse Lab in Milton Keynes | Wellcome Sanger Institute for the COVID-19 Genomics UK (COG-UK) Consortium | The Lighthouse Lab in Milton Keynes and Alex Alderton, Roberto Amato, Jeffrey Barrett, Sonia Goncalves, Ewan Harrison, David K. Jackson, Ian Johnston, Dominic Kwiatkowski, Cordelia Langford, John Sillitoe on behalf of the Wellcome Sanger Institute COVID-19 Surveillance Team |
| EPI_ISL_2994331 | Lighthouse Laboratory Plymouth | Wellcome Sanger Institute for the COVID-19 Genomics UK (COG-UK) Consortium | Lighthouse Laboratory Plymouth and Alex Alderton, Roberto Amato, Jeffrey Barrett, Sonia Goncalves, Ewan Harrison, David K. Jackson, Ian Johnston, Dominic Kwiatkowski, Cordelia Langford, John Sillitoe on behalf of the Wellcome Sanger Institute COVID-19 Surveillance Team |
| EPI_ISL_2994332 | Lighthouse Lab in Alderley Park | Wellcome Sanger Institute for the COVID-19 Genomics UK (COG-UK) Consortium | Jacquelyn Wynn, Mairead Hyland, The Lighthouse Lab in Alderley Park and Alex Alderton, Roberto Amato, Jeffrey Barrett, Sonia Goncalves, Ewan Harrison, David K. Jackson, Ian Johnston, Dominic Kwiatkowski, Cordelia Langford, John Sillitoe on behalf of the Wellcome Sanger Institute COVID-19 Surveillance Team |
| EPI_ISL_2994333, EPI_ISL_2994334 | Lighthouse Laboratory Plymouth | Wellcome Sanger Institute for the COVID-19 Genomics UK (COG-UK) Consortium | Lighthouse Laboratory Plymouth and Alex Alderton, Roberto Amato, Jeffrey Barrett, Sonia Goncalves, Ewan Harrison, David K. Jackson, Ian Johnston, Dominic Kwiatkowski, Cordelia Langford, John Sillitoe on behalf of the Wellcome Sanger Institute COVID-19 Surveillance Team |
| EPI_ISL_2994335 | Berkshire and Surrey Pathology Services Lighthouse Laboratory | Wellcome Sanger Institute for the COVID-19 Genomics UK (COG-UK) Consortium | Berkshire and Surrey Pathology Services Lighthouse Laboratory and Alex Alderton, Roberto Amato, Jeffrey Barrett, Sonia Goncalves, Ewan Harrison, David K. Jackson, Ian Johnston, Dominic Kwiatkowski, Cordelia Langford, John Sillitoe on behalf of the Wellcome Sanger Institute COVID-19 Surveillance Team |
| EPI_ISL_2994336 | Lighthouse Lab in Alderley Park | Wellcome Sanger Institute for the COVID-19 Genomics UK (COG-UK) Consortium | Jacquelyn Wynn, Mairead Hyland, The Lighthouse Lab in Alderley Park and Alex Alderton, Roberto Amato, Jeffrey Barrett, Sonia Goncalves, Ewan Harrison, David K. Jackson, Ian Johnston, Dominic Kwiatkowski, Cordelia Langford, John Sillitoe on behalf of the Wellcome Sanger Institute COVID-19 Surveillance Team |
| EPI_ISL_2994337, EPI_ISL_2994338, EPI_ISL_2994339, EPI_ISL_2994340, EPI_ISL_2994341 | Lighthouse Laboratory Plymouth | Wellcome Sanger Institute for the COVID-19 Genomics UK (COG-UK) Consortium | Lighthouse Laboratory Plymouth and Alex Alderton, Roberto Amato, Jeffrey Barrett, Sonia Goncalves, Ewan Harrison, David K. Jackson, Ian Johnston, Dominic Kwiatkowski, Cordelia Langford, John Sillitoe on behalf of the Wellcome Sanger Institute COVID-19 Surveillance Team |
| EPI_ISL_2994342, EPI_ISL_2994343 | Lighthouse Lab in Alderley Park | Wellcome Sanger Institute for the COVID-19 Genomics UK (COG-UK) Consortium | Jacquelyn Wynn, Mairead Hyland, The Lighthouse Lab in Alderley Park and Alex Alderton, Roberto Amato, Jeffrey Barrett, Sonia Goncalves, Ewan Harrison, David K. Jackson, Ian Johnston, Dominic Kwiatkowski, Cordelia Langford, John Sillitoe on behalf of the Wellcome Sanger Institute COVID-19 Surveillance Team |
| EPI_ISL_2994344 | Berkshire and Surrey Pathology Services Lighthouse Laboratory | Wellcome Sanger Institute for the COVID-19 Genomics UK (COG-UK) Consortium | Berkshire and Surrey Pathology Services Lighthouse Laboratory and Alex Alderton, Roberto Amato, Jeffrey Barrett, Sonia Goncalves, Ewan Harrison, David K. Jackson, Ian Johnston, Dominic Kwiatkowski, Cordelia Langford, John Sillitoe on behalf of the Wellcome Sanger Institute COVID-19 Surveillance Team |
| EPI_ISL_2994345 | Lighthouse Lab in Alderley Park | Wellcome Sanger Institute for the COVID-19 Genomics UK (COG-UK) Consortium | Jacquelyn Wynn, Mairead Hyland, The Lighthouse Lab in Alderley Park and Alex Alderton, Roberto Amato, Jeffrey Barrett, Sonia Goncalves, Ewan Harrison, David K. Jackson, Ian Johnston, Dominic Kwiatkowski, Cordelia Langford, John Sillitoe on behalf of the Wellcome Sanger Institute COVID-19 Surveillance Team |
| EPI_ISL_2994346 | Lighthouse Laboratory Plymouth | Wellcome Sanger Institute for the COVID-19 Genomics UK (COG-UK) Consortium | Lighthouse Laboratory Plymouth and Alex Alderton, Roberto Amato, Jeffrey Barrett, Sonia Goncalves, Ewan Harrison, David K. Jackson, Ian Johnston, Dominic Kwiatkowski, Cordelia Langford, John Sillitoe on behalf of the Wellcome Sanger Institute COVID-19 Surveillance Team |
| EPI_ISL_2994347 | Lighthouse Lab in Milton Keynes | Wellcome Sanger Institute for the COVID-19 Genomics UK (COG-UK) Consortium | The Lighthouse Lab in Milton Keynes and Alex Alderton, Roberto Amato, Jeffrey Barrett, Sonia Goncalves, Ewan Harrison, David K. Jackson, Ian Johnston, Dominic Kwiatkowski, Cordelia Langford, John Sillitoe on behalf of the Wellcome Sanger Institute COVID-19 Surveillance Team |
| EPI_ISL_2994348 | Lighthouse Laboratory Plymouth | Wellcome Sanger Institute for the COVID-19 Genomics UK (COG-UK) Consortium | Lighthouse Laboratory Plymouth and Alex Alderton, Roberto Amato, Jeffrey Barrett, Sonia Goncalves, Ewan Harrison, David K. Jackson, Ian Johnston, Dominic Kwiatkowski, Cordelia Langford, John Sillitoe on behalf of the Wellcome Sanger Institute COVID-19 Surveillance Team |
| EPI_ISL_2994349, EPI_ISL_2994350 | Berkshire and Surrey Pathology Services Lighthouse Laboratory | Wellcome Sanger Institute for the COVID-19 Genomics UK (COG-UK) Consortium | Berkshire and Surrey Pathology Services Lighthouse Laboratory and Alex Alderton, Roberto Amato, Jeffrey Barrett, Sonia Goncalves, Ewan Harrison, David K. Jackson, Ian Johnston, Dominic Kwiatkowski, Cordelia Langford, John Sillitoe on behalf of the Wellcome Sanger Institute COVID-19 Surveillance Team |
| EPI_ISL_2994351, EPI_ISL_2994352, EPI_ISL_2994353 | Lighthouse Lab in Alderley Park | Wellcome Sanger Institute for the COVID-19 Genomics UK (COG-UK) Consortium | Jacquelyn Wynn, Mairead Hyland, The Lighthouse Lab in Alderley Park and Alex Alderton, Roberto Amato, Jeffrey Barrett, Sonia Goncalves, Ewan Harrison, David K. Jackson, Ian Johnston, Dominic Kwiatkowski, Cordelia Langford, John Sillitoe on behalf of the Wellcome Sanger Institute COVID-19 Surveillance Team |
| EPI_ISL_2994354 | Lighthouse Laboratory Plymouth | Wellcome Sanger Institute for the COVID-19 Genomics UK (COG-UK) Consortium | Lighthouse Laboratory Plymouth and Alex Alderton, Roberto Amato, Jeffrey Barrett, Sonia Goncalves, Ewan Harrison, David K. Jackson, Ian Johnston, Dominic Kwiatkowski, Cordelia Langford, John Sillitoe on behalf of the Wellcome Sanger Institute COVID-19 Surveillance Team |
| EPI_ISL_2994355 | Berkshire and Surrey Pathology Services Lighthouse Laboratory | Wellcome Sanger Institute for the COVID-19 Genomics UK (COG-UK) Consortium | Berkshire and Surrey Pathology Services Lighthouse Laboratory and Alex Alderton, Roberto Amato, Jeffrey Barrett, Sonia Goncalves, Ewan Harrison, David K. Jackson, Ian Johnston, Dominic Kwiatkowski, Cordelia Langford, John Sillitoe on behalf of the Wellcome Sanger Institute COVID-19 Surveillance Team |
| EPI_ISL_2994356 | Lighthouse Laboratory Plymouth | Wellcome Sanger Institute for the COVID-19 Genomics UK (COG-UK) Consortium | Lighthouse Laboratory Plymouth and Alex Alderton, Roberto Amato, Jeffrey Barrett, Sonia Goncalves, Ewan Harrison, David K. Jackson, Ian Johnston, Dominic Kwiatkowski, Cordelia Langford, John Sillitoe on behalf of the Wellcome Sanger Institute COVID-19 Surveillance Team |
| EPI_ISL_2994357, EPI_ISL_2994358, EPI_ISL_2994359 | Lighthouse Lab in Alderley Park | Wellcome Sanger Institute for the COVID-19 Genomics UK (COG-UK) Consortium | Jacquelyn Wynn, Mairead Hyland, The Lighthouse Lab in Alderley Park and Alex Alderton, Roberto Amato, Jeffrey Barrett, Sonia Goncalves, Ewan Harrison, David K. Jackson, Ian Johnston, Dominic Kwiatkowski, Cordelia Langford, John Sillitoe on behalf of the Wellcome Sanger Institute COVID-19 Surveillance Team |
| EPI_ISL_2994360 | Lighthouse Laboratory Plymouth | Wellcome Sanger Institute for the COVID-19 Genomics UK (COG-UK) Consortium | Lighthouse Laboratory Plymouth and Alex Alderton, Roberto Amato, Jeffrey Barrett, Sonia Goncalves, Ewan Harrison, David K. Jackson, Ian Johnston, Dominic Kwiatkowski, Cordelia Langford, John Sillitoe on behalf of the Wellcome Sanger Institute COVID-19 Surveillance Team |
| EPI_ISL_2994361 | Berkshire and Surrey Pathology Services Lighthouse Laboratory | Wellcome Sanger Institute for the COVID-19 Genomics UK (COG-UK) Consortium | Berkshire and Surrey Pathology Services Lighthouse Laboratory and Alex Alderton, Roberto Amato, Jeffrey Barrett, Sonia Goncalves, Ewan Harrison, David K. Jackson, Ian Johnston, Dominic Kwiatkowski, Cordelia Langford, John Sillitoe on behalf of the Wellcome Sanger Institute COVID-19 Surveillance Team |
| EPI_ISL_2994362 | Lighthouse Lab in Milton Keynes | Wellcome Sanger Institute for the COVID-19 Genomics UK (COG-UK) Consortium | The Lighthouse Lab in Milton Keynes and Alex Alderton, Roberto Amato, Jeffrey Barrett, Sonia Goncalves, Ewan Harrison, David K. Jackson, Ian Johnston, Dominic Kwiatkowski, Cordelia Langford, John Sillitoe on behalf of the Wellcome Sanger Institute COVID-19 Surveillance Team |
| EPI_ISL_2994363 | Lighthouse Lab in Alderley Park | Wellcome Sanger Institute for the COVID-19 Genomics UK (COG-UK) Consortium | Jacquelyn Wynn, Mairead Hyland, The Lighthouse Lab in Alderley Park and Alex Alderton, Roberto Amato, Jeffrey Barrett, Sonia Goncalves, Ewan Harrison, David K. Jackson, Ian Johnston, Dominic Kwiatkowski, Cordelia Langford, John Sillitoe on behalf of the Wellcome Sanger Institute COVID-19 Surveillance Team |
| EPI_ISL_2994364, EPI_ISL_2994365 | Lighthouse Laboratory Plymouth | Wellcome Sanger Institute for the COVID-19 Genomics UK (COG-UK) Consortium | Lighthouse Laboratory Plymouth and Alex Alderton, Roberto Amato, Jeffrey Barrett, Sonia Goncalves, Ewan Harrison, David K. Jackson, Ian Johnston, Dominic Kwiatkowski, Cordelia Langford, John Sillitoe on behalf of the Wellcome Sanger Institute COVID-19 Surveillance Team |

| | | | |
|---|---|---|---|
| EPI_ISL_2994366 | Lighthouse Lab in Milton Keynes | Wellcome Sanger Institute for the COVID-19 Genomics UK (COG-UK) Consortium | The Lighthouse Lab in Milton Keynes and Alex Alderton, Roberto Amato, Jeffrey Barrett, Sonia Goncalves, Ewan Harrison, David K. Jackson, Ian Johnston, Dominic Kwiatkowski, Cordelia Langford, John Sillitoe on behalf of the Wellcome Sanger Institute COVID-19 Surveillance Team |
| EPI_ISL_2994367 | Berkshire and Surrey Pathology Services Lighthouse Laboratory | Wellcome Sanger Institute for the COVID-19 Genomics UK (COG-UK) Consortium | Berkshire and Surrey Pathology Services Lighthouse Laboratory and Alex Alderton, Roberto Amato, Jeffrey Barrett, Sonia Goncalves, Ewan Harrison, David K. Jackson, Ian Johnston, Dominic Kwiatkowski, Cordelia Langford, John Sillitoe on behalf of the Wellcome Sanger Institute COVID-19 Surveillance Team |
| EPI_ISL_2994368, EPI_ISL_2994369 | Lighthouse Laboratory Plymouth | Wellcome Sanger Institute for the COVID-19 Genomics UK (COG-UK) Consortium | Lighthouse Laboratory Plymouth and Alex Alderton, Roberto Amato, Jeffrey Barrett, Sonia Goncalves, Ewan Harrison, David K. Jackson, Ian Johnston, Dominic Kwiatkowski, Cordelia Langford, John Sillitoe on behalf of the Wellcome Sanger Institute COVID-19 Surveillance Team |
| EPI_ISL_2994370 | Lighthouse Lab in Alderley Park | Wellcome Sanger Institute for the COVID-19 Genomics UK (COG-UK) Consortium | Jacquelyn Wynn, Mairead Hyland, The Lighthouse Lab in Alderley Park and Alex Alderton, Roberto Amato, Jeffrey Barrett, Sonia Goncalves, Ewan Harrison, David K. Jackson, Ian Johnston, Dominic Kwiatkowski, Cordelia Langford, John Sillitoe on behalf of the Wellcome Sanger Institute COVID-19 Surveillance Team |
| EPI_ISL_2994371, EPI_ISL_2994372 | Lighthouse Lab in Milton Keynes | Wellcome Sanger Institute for the COVID-19 Genomics UK (COG-UK) Consortium | The Lighthouse Lab in Milton Keynes and Alex Alderton, Roberto Amato, Jeffrey Barrett, Sonia Goncalves, Ewan Harrison, David K. Jackson, Ian Johnston, Dominic Kwiatkowski, Cordelia Langford, John Sillitoe on behalf of the Wellcome Sanger Institute COVID-19 Surveillance Team |
| EPI_ISL_2994373, EPI_ISL_2994374 | Lighthouse Laboratory Plymouth | Wellcome Sanger Institute for the COVID-19 Genomics UK (COG-UK) Consortium | Lighthouse Laboratory Plymouth and Alex Alderton, Roberto Amato, Jeffrey Barrett, Sonia Goncalves, Ewan Harrison, David K. Jackson, Ian Johnston, Dominic Kwiatkowski, Cordelia Langford, John Sillitoe on behalf of the Wellcome Sanger Institute COVID-19 Surveillance Team |
| EPI_ISL_2994375 | Berkshire and Surrey Pathology Services Lighthouse Laboratory | Wellcome Sanger Institute for the COVID-19 Genomics UK (COG-UK) Consortium | Berkshire and Surrey Pathology Services Lighthouse Laboratory and Alex Alderton, Roberto Amato, Jeffrey Barrett, Sonia Goncalves, Ewan Harrison, David K. Jackson, Ian Johnston, Dominic Kwiatkowski, Cordelia Langford, John Sillitoe on behalf of the Wellcome Sanger Institute COVID-19 Surveillance Team |
| EPI_ISL_2994376 | Lighthouse Lab in Milton Keynes | Wellcome Sanger Institute for the COVID-19 Genomics UK (COG-UK) Consortium | The Lighthouse Lab in Milton Keynes and Alex Alderton, Roberto Amato, Jeffrey Barrett, Sonia Goncalves, Ewan Harrison, David K. Jackson, Ian Johnston, Dominic Kwiatkowski, Cordelia Langford, John Sillitoe on behalf of the Wellcome Sanger Institute COVID-19 Surveillance Team |
| EPI_ISL_2994377, EPI_ISL_2994378 | Lighthouse Lab in Glasgow | Wellcome Sanger Institute for the COVID-19 Genomics UK (COG-UK) Consortium | Harper VanSteenhouse, Yumi Kasai, David Gray, Carol Clugston, Anna Dominiczak and Alex Alderton, Roberto Amato, Jeffrey Barrett, Sonia Goncalves, Ewan Harrison, David K. Jackson, Ian Johnston, Dominic Kwiatkowski, Cordelia Langford, John Sillitoe on behalf of the Wellcome Sanger Institute COVID-19 Surveillance Team |
| EPI_ISL_2994379, EPI_ISL_2994380, EPI_ISL_2994381 | Lighthouse Lab in Milton Keynes | Wellcome Sanger Institute for the COVID-19 Genomics UK (COG-UK) Consortium | The Lighthouse Lab in Milton Keynes and Alex Alderton, Roberto Amato, Jeffrey Barrett, Sonia Goncalves, Ewan Harrison, David K. Jackson, Ian Johnston, Dominic Kwiatkowski, Cordelia Langford, John Sillitoe on behalf of the Wellcome Sanger Institute COVID-19 Surveillance Team |
| EPI_ISL_2994382 | Berkshire and Surrey Pathology Services Lighthouse Laboratory | Wellcome Sanger Institute for the COVID-19 Genomics UK (COG-UK) Consortium | Berkshire and Surrey Pathology Services Lighthouse Laboratory and Alex Alderton, Roberto Amato, Jeffrey Barrett, Sonia Goncalves, Ewan Harrison, David K. Jackson, Ian Johnston, Dominic Kwiatkowski, Cordelia Langford, John Sillitoe on behalf of the Wellcome Sanger Institute COVID-19 Surveillance Team |
| EPI_ISL_2994383, EPI_ISL_2994384 | Lighthouse Lab in Alderley Park | Wellcome Sanger Institute for the COVID-19 Genomics UK (COG-UK) Consortium | Jacquelyn Wynn, Mairead Hyland, The Lighthouse Lab in Alderley Park and Alex Alderton, Roberto Amato, Jeffrey Barrett, Sonia Goncalves, Ewan Harrison, David K. Jackson, Ian Johnston, Dominic Kwiatkowski, Cordelia Langford, John Sillitoe on behalf of the Wellcome Sanger Institute COVID-19 Surveillance Team |
| EPI_ISL_2994385, EPI_ISL_2994386 | Berkshire and Surrey Pathology Services Lighthouse Laboratory | Wellcome Sanger Institute for the COVID-19 Genomics UK (COG-UK) Consortium | Berkshire and Surrey Pathology Services Lighthouse Laboratory and Alex Alderton, Roberto Amato, Jeffrey Barrett, Sonia Goncalves, Ewan Harrison, David K. Jackson, Ian Johnston, Dominic Kwiatkowski, Cordelia Langford, John Sillitoe on behalf of the Wellcome Sanger Institute COVID-19 Surveillance Team |
| EPI_ISL_2994387, EPI_ISL_2994388 | Lighthouse Lab in Glasgow | Wellcome Sanger Institute for the COVID-19 Genomics UK (COG-UK) Consortium | Harper VanSteenhouse, Yumi Kasai, David Gray, Carol Clugston, Anna Dominiczak and Alex Alderton, Roberto Amato, Jeffrey Barrett, Sonia Goncalves, Ewan Harrison, David K. Jackson, Ian Johnston, Dominic Kwiatkowski, Cordelia Langford, John Sillitoe on behalf of the Wellcome Sanger Institute COVID-19 Surveillance Team |
| EPI_ISL_2994389 | Berkshire and Surrey Pathology Services Lighthouse Laboratory | Wellcome Sanger Institute for the COVID-19 Genomics UK (COG-UK) Consortium | Berkshire and Surrey Pathology Services Lighthouse Laboratory and Alex Alderton, Roberto Amato, Jeffrey Barrett, Sonia Goncalves, Ewan Harrison, David K. Jackson, Ian Johnston, Dominic Kwiatkowski, Cordelia Langford, John Sillitoe on behalf of the Wellcome Sanger Institute COVID-19 Surveillance Team |
| EPI_ISL_2994390 | Lighthouse Lab in Milton Keynes | Wellcome Sanger Institute for the COVID-19 Genomics UK (COG-UK) Consortium | The Lighthouse Lab in Milton Keynes and Alex Alderton, Roberto Amato, Jeffrey Barrett, Sonia Goncalves, Ewan Harrison, David K. Jackson, Ian Johnston, Dominic Kwiatkowski, Cordelia Langford, John Sillitoe on behalf of the Wellcome Sanger Institute COVID-19 Surveillance Team |
| EPI_ISL_2994391 | Lighthouse Lab in Alderley Park | Wellcome Sanger Institute for the COVID-19 Genomics UK (COG-UK) Consortium | Jacquelyn Wynn, Mairead Hyland, The Lighthouse Lab in Alderley Park and Alex Alderton, Roberto Amato, Jeffrey Barrett, Sonia Goncalves, Ewan Harrison, David K. Jackson, Ian Johnston, Dominic Kwiatkowski, Cordelia Langford, John Sillitoe on behalf of the Wellcome Sanger Institute COVID-19 Surveillance Team |
| EPI_ISL_2994392 | Lighthouse Lab in Glasgow | Wellcome Sanger Institute for the COVID-19 Genomics UK (COG-UK) Consortium | Harper VanSteenhouse, Yumi Kasai, David Gray, Carol Clugston, Anna Dominiczak and Alex Alderton, Roberto Amato, Jeffrey Barrett, Sonia Goncalves, Ewan Harrison, David K. Jackson, Ian Johnston, Dominic Kwiatkowski, Cordelia Langford, John Sillitoe on behalf of the Wellcome Sanger Institute COVID-19 Surveillance Team |
| EPI_ISL_2994393 | Lighthouse Lab in Milton Keynes | Wellcome Sanger Institute for the COVID-19 Genomics UK (COG-UK) Consortium | The Lighthouse Lab in Milton Keynes and Alex Alderton, Roberto Amato, Jeffrey Barrett, Sonia Goncalves, Ewan Harrison, David K. Jackson, Ian Johnston, Dominic Kwiatkowski, Cordelia Langford, John Sillitoe on behalf of the Wellcome Sanger Institute COVID-19 Surveillance Team |
| EPI_ISL_2994394 | Berkshire and Surrey Pathology Services Lighthouse Laboratory | Wellcome Sanger Institute for the COVID-19 Genomics UK (COG-UK) Consortium | Berkshire and Surrey Pathology Services Lighthouse Laboratory and Alex Alderton, Roberto Amato, Jeffrey Barrett, Sonia Goncalves, Ewan Harrison, David K. Jackson, Ian Johnston, Dominic Kwiatkowski, Cordelia Langford, John Sillitoe on behalf of the Wellcome Sanger Institute COVID-19 Surveillance Team |
| EPI_ISL_2994395 | Lighthouse Lab in Milton Keynes | Wellcome Sanger Institute for the COVID-19 Genomics UK (COG-UK) Consortium | The Lighthouse Lab in Milton Keynes and Alex Alderton, Roberto Amato, Jeffrey Barrett, Sonia Goncalves, Ewan Harrison, David K. Jackson, Ian Johnston, Dominic Kwiatkowski, Cordelia Langford, John Sillitoe on behalf of the Wellcome Sanger Institute COVID-19 Surveillance Team |
| EPI_ISL_2994396 | Berkshire and Surrey Pathology Services Lighthouse Laboratory | Wellcome Sanger Institute for the COVID-19 Genomics UK (COG-UK) Consortium | Berkshire and Surrey Pathology Services Lighthouse Laboratory and Alex Alderton, Roberto Amato, Jeffrey Barrett, Sonia Goncalves, Ewan Harrison, David K. Jackson, Ian Johnston, Dominic Kwiatkowski, Cordelia Langford, John Sillitoe on behalf of the Wellcome Sanger Institute COVID-19 Surveillance Team |
| EPI_ISL_2994397 | Lighthouse Lab in Milton Keynes | Wellcome Sanger Institute for the COVID-19 Genomics UK (COG-UK) Consortium | The Lighthouse Lab in Milton Keynes and Alex Alderton, Roberto Amato, Jeffrey Barrett, Sonia Goncalves, Ewan Harrison, David K. Jackson, Ian Johnston, Dominic Kwiatkowski, Cordelia Langford, John Sillitoe on behalf of the Wellcome Sanger Institute COVID-19 Surveillance Team |
| EPI_ISL_2994398 | Berkshire and Surrey Pathology Services Lighthouse Laboratory | Wellcome Sanger Institute for the COVID-19 Genomics UK (COG-UK) Consortium | Berkshire and Surrey Pathology Services Lighthouse Laboratory and Alex Alderton, Roberto Amato, Jeffrey Barrett, Sonia Goncalves, Ewan Harrison, David K. Jackson, Ian Johnston, Dominic Kwiatkowski, Cordelia Langford, John Sillitoe on behalf of the Wellcome Sanger Institute COVID-19 Surveillance Team |
| EPI_ISL_2994399 | Lighthouse Lab in Glasgow | Wellcome Sanger Institute for the COVID-19 Genomics UK (COG-UK) Consortium | Harper VanSteenhouse, Yumi Kasai, David Gray, Carol Clugston, Anna Dominiczak and Alex Alderton, Roberto Amato, Jeffrey Barrett, Sonia Goncalves, Ewan Harrison, David K. Jackson, Ian Johnston, Dominic Kwiatkowski, Cordelia Langford, John Sillitoe on behalf of the Wellcome Sanger Institute COVID-19 Surveillance Team |
| EPI_ISL_2994400 | Lighthouse Lab in Alderley Park | Wellcome Sanger Institute for the COVID-19 Genomics UK (COG-UK) Consortium | Jacquelyn Wynn, Mairead Hyland, The Lighthouse Lab in Alderley Park and Alex Alderton, Roberto Amato, Jeffrey Barrett, Sonia Goncalves, Ewan Harrison, David K. Jackson, Ian Johnston, Dominic Kwiatkowski, Cordelia Langford, John Sillitoe on behalf of the Wellcome Sanger Institute COVID-19 Surveillance Team |
| EPI_ISL_2994401, EPI_ISL_2994402 | Berkshire and Surrey Pathology Services Lighthouse Laboratory | Wellcome Sanger Institute for the COVID-19 Genomics UK (COG-UK) Consortium | Berkshire and Surrey Pathology Services Lighthouse Laboratory and Alex Alderton, Roberto Amato, Jeffrey Barrett, Sonia Goncalves, Ewan Harrison, David K. Jackson, Ian Johnston, Dominic Kwiatkowski, Cordelia Langford, John Sillitoe on behalf of the Wellcome Sanger Institute COVID-19 Surveillance Team |
| EPI_ISL_2994403 | Lighthouse Lab in Milton Keynes | Wellcome Sanger Institute for the COVID-19 Genomics UK (COG-UK) Consortium | The Lighthouse Lab in Milton Keynes and Alex Alderton, Roberto Amato, Jeffrey Barrett, Sonia Goncalves, Ewan Harrison, David K. Jackson, Ian Johnston, Dominic Kwiatkowski, Cordelia Langford, John Sillitoe on behalf of the Wellcome Sanger Institute COVID-19 Surveillance Team |
| EPI_ISL_2994404 | Berkshire and Surrey Pathology Services Lighthouse Laboratory | Wellcome Sanger Institute for the COVID-19 Genomics UK (COG-UK) Consortium | Berkshire and Surrey Pathology Services Lighthouse Laboratory and Alex Alderton, Roberto Amato, Jeffrey Barrett, Sonia Goncalves, Ewan Harrison, David K. Jackson, Ian Johnston, Dominic Kwiatkowski, Cordelia Langford, John Sillitoe on behalf of the Wellcome Sanger Institute COVID-19 Surveillance Team |
| EPI_ISL_2994405 | Lighthouse Lab in Alderley Park | Wellcome Sanger Institute for the COVID-19 Genomics UK (COG-UK) Consortium | Jacquelyn Wynn, Mairead Hyland, The Lighthouse Lab in Alderley Park and Alex Alderton, Roberto Amato, Jeffrey Barrett, Sonia Goncalves, Ewan Harrison, David K. Jackson, Ian Johnston, Dominic Kwiatkowski, Cordelia Langford, John Sillitoe on behalf of the Wellcome Sanger Institute COVID-19 Surveillance Team |
| EPI_ISL_2994406 | Lighthouse Lab in Milton Keynes | Wellcome Sanger Institute for the COVID-19 Genomics UK (COG-UK) Consortium | The Lighthouse Lab in Milton Keynes and Alex Alderton, Roberto Amato, Jeffrey Barrett, Sonia Goncalves, Ewan Harrison, David K. Jackson, Ian Johnston, Dominic Kwiatkowski, Cordelia Langford, John Sillitoe on behalf of the Wellcome Sanger Institute COVID-19 Surveillance Team |
| EPI_ISL_2994407 | Berkshire and Surrey Pathology Services Lighthouse Laboratory | Wellcome Sanger Institute for the COVID-19 Genomics UK (COG-UK) Consortium | Berkshire and Surrey Pathology Services Lighthouse Laboratory and Alex Alderton, Roberto Amato, Jeffrey Barrett, Sonia Goncalves, Ewan Harrison, David K. Jackson, Ian Johnston, Dominic Kwiatkowski, Cordelia Langford, John Sillitoe on behalf of the Wellcome Sanger Institute COVID-19 Surveillance Team |
| EPI_ISL_2994408 | Lighthouse Lab in Milton Keynes | Wellcome Sanger Institute for the COVID-19 Genomics UK (COG-UK) Consortium | The Lighthouse Lab in Milton Keynes and Alex Alderton, Roberto Amato, Jeffrey Barrett, Sonia Goncalves, Ewan Harrison, David K. Jackson, Ian Johnston, Dominic Kwiatkowski, Cordelia Langford, John Sillitoe on behalf of the Wellcome Sanger Institute COVID-19 Surveillance Team |
| EPI_ISL_2994409 | Berkshire and Surrey Pathology Services Lighthouse Laboratory | Wellcome Sanger Institute for the COVID-19 Genomics UK (COG-UK) Consortium | Berkshire and Surrey Pathology Services Lighthouse Laboratory and Alex Alderton, Roberto Amato, Jeffrey Barrett, Sonia Goncalves, Ewan Harrison, David K. Jackson, Ian Johnston, Dominic Kwiatkowski, Cordelia Langford, John Sillitoe on behalf of the Wellcome Sanger Institute COVID-19 Surveillance Team |
| EPI_ISL_2994410 | Lighthouse Lab in Alderley Park | Wellcome Sanger Institute for the COVID-19 Genomics UK (COG-UK) Consortium | Jacquelyn Wynn, Mairead Hyland, The Lighthouse Lab in Alderley Park and Alex Alderton, Roberto Amato, Jeffrey Barrett, Sonia Goncalves, Ewan Harrison, David K. Jackson, Ian Johnston, Dominic Kwiatkowski, Cordelia Langford, John Sillitoe on behalf of the Wellcome Sanger Institute COVID-19 Surveillance Team |
| EPI_ISL_2994411, EPI_ISL_2994412, EPI_ISL_2994413 | Lighthouse Lab in Milton Keynes | Wellcome Sanger Institute for the COVID-19 Genomics UK (COG-UK) Consortium | The Lighthouse Lab in Milton Keynes and Alex Alderton, Roberto Amato, Jeffrey Barrett, Sonia Goncalves, Ewan Harrison, David K. Jackson, Ian Johnston, Dominic Kwiatkowski, Cordelia Langford, John Sillitoe on behalf of the Wellcome Sanger Institute COVID-19 Surveillance Team |
| EPI_ISL_2994414 | Lighthouse Lab in Alderley Park | Wellcome Sanger Institute for the COVID-19 Genomics UK (COG-UK) Consortium | Jacquelyn Wynn, Mairead Hyland, The Lighthouse Lab in Alderley Park and Alex Alderton, Roberto Amato, Jeffrey Barrett, Sonia Goncalves, Ewan Harrison, David K. Jackson, Ian Johnston, Dominic Kwiatkowski, Cordelia Langford, John Sillitoe on behalf of the Wellcome Sanger Institute COVID-19 Surveillance Team |
| EPI_ISL_2994415 | Berkshire and Surrey Pathology Services Lighthouse Laboratory | Wellcome Sanger Institute for the COVID-19 Genomics UK (COG-UK) Consortium | Berkshire and Surrey Pathology Services Lighthouse Laboratory and Alex Alderton, Roberto Amato, Jeffrey Barrett, Sonia Goncalves, Ewan Harrison, David K. Jackson, Ian Johnston, Dominic Kwiatkowski, Cordelia Langford, John Sillitoe on behalf of the Wellcome Sanger Institute COVID-19 Surveillance Team |
| EPI_ISL_2994416 | Lighthouse Lab in Alderley Park | Wellcome Sanger Institute for the COVID-19 Genomics UK (COG-UK) Consortium | Jacquelyn Wynn, Mairead Hyland, The Lighthouse Lab in Alderley Park and Alex Alderton, Roberto Amato, Jeffrey Barrett, Sonia Goncalves, Ewan Harrison, David K. Jackson, Ian Johnston, Dominic Kwiatkowski, Cordelia Langford, John Sillitoe on behalf of the Wellcome Sanger Institute COVID-19 Surveillance Team |
| EPI_ISL_2994417 | Berkshire and Surrey Pathology Services Lighthouse Laboratory | Wellcome Sanger Institute for the COVID-19 Genomics UK (COG-UK) Consortium | Berkshire and Surrey Pathology Services Lighthouse Laboratory and Alex Alderton, Roberto Amato, Jeffrey Barrett, Sonia Goncalves, Ewan Harrison, David K. Jackson, Ian Johnston, Dominic Kwiatkowski, Cordelia Langford, John Sillitoe on behalf of the Wellcome Sanger Institute COVID-19 Surveillance Team |
| EPI_ISL_2994418 | Lighthouse Lab in Alderley Park | Wellcome Sanger Institute for the COVID-19 Genomics UK (COG-UK) Consortium | Jacquelyn Wynn, Mairead Hyland, The Lighthouse Lab in Alderley Park and Alex Alderton, Roberto Amato, Jeffrey Barrett, Sonia Goncalves, Ewan Harrison, David K. Jackson, Ian Johnston, Dominic Kwiatkowski, Cordelia Langford, John Sillitoe on behalf of the Wellcome Sanger Institute COVID-19 Surveillance Team |
| EPI_ISL_2994419 | Berkshire and Surrey Pathology Services Lighthouse Laboratory | Wellcome Sanger Institute for the COVID-19 Genomics UK (COG-UK) Consortium | Berkshire and Surrey Pathology Services Lighthouse Laboratory and Alex Alderton, Roberto Amato, Jeffrey Barrett, Sonia Goncalves, Ewan Harrison, David K. Jackson, Ian Johnston, Dominic Kwiatkowski, Cordelia Langford, John Sillitoe on behalf of the Wellcome Sanger Institute COVID-19 Surveillance Team |
| EPI_ISL_2994420 | Lighthouse Lab in Alderley Park | Wellcome Sanger Institute for the COVID-19 Genomics UK (COG-UK) Consortium | Jacquelyn Wynn, Mairead Hyland, The Lighthouse Lab in Alderley Park and Alex Alderton, Roberto Amato, Jeffrey Barrett, Sonia Goncalves, Ewan Harrison, David K. Jackson, Ian Johnston, Dominic Kwiatkowski, Cordelia Langford, John Sillitoe on behalf of the Wellcome Sanger Institute COVID-19 Surveillance Team |
| EPI_ISL_2994421, EPI_ISL_2994422 | Lighthouse Lab in Milton Keynes | Wellcome Sanger Institute for the COVID-19 Genomics UK (COG-UK) Consortium | The Lighthouse Lab in Milton Keynes and Alex Alderton, Roberto Amato, Jeffrey Barrett, Sonia Goncalves, Ewan Harrison, David K. Jackson, Ian Johnston, Dominic Kwiatkowski, Cordelia Langford, John Sillitoe on behalf of the Wellcome Sanger Institute COVID-19 Surveillance Team |

| | | | |
|---|---|---|---|
| EPI_ISL_2994423, EPI_ISL_2994424, EPI_ISL_2994425, EPI_ISL_2994426, EPI_ISL_2994427 | Lighthouse Lab in Glasgow | Wellcome Sanger Institute for the COVID-19 Genomics UK (COG-UK) Consortium | Harper VanSteenhouse, Yumi Kasai, David Gray, Carol Clugston, Anna Dominiczak and Alex Alderton, Roberto Amato, Jeffrey Barrett, Sonia Goncalves, Ewan Harrison, David K. Jackson, Ian Johnston, Dominic Kwiatkowski, Cordelia Langford, John Sillitoe on behalf of the Wellcome Sanger Institute COVID-19 Surveillance Team |
| EPI_ISL_2994428 | Lighthouse Lab in Alderley Park | Wellcome Sanger Institute for the COVID-19 Genomics UK (COG-UK) Consortium | Jacquelyn Wynn, Mairead Hyland, The Lighthouse Lab in Alderley Park and Alex Alderton, Roberto Amato, Jeffrey Barrett, Sonia Goncalves, Ewan Harrison, David K. Jackson, Ian Johnston, Dominic Kwiatkowski, Cordelia Langford, John Sillitoe on behalf of the Wellcome Sanger Institute COVID-19 Surveillance Team |
| EPI_ISL_2994429 | Lighthouse Lab in Milton Keynes | Wellcome Sanger Institute for the COVID-19 Genomics UK (COG-UK) Consortium | The Lighthouse Lab in Milton Keynes and Alex Alderton, Roberto Amato, Jeffrey Barrett, Sonia Goncalves, Ewan Harrison, David K. Jackson, Ian Johnston, Dominic Kwiatkowski, Cordelia Langford, John Sillitoe on behalf of the Wellcome Sanger Institute COVID-19 Surveillance Team |
| EPI_ISL_2994430 | Lighthouse Lab in Glasgow | Wellcome Sanger Institute for the COVID-19 Genomics UK (COG-UK) Consortium | Harper VanSteenhouse, Yumi Kasai, David Gray, Carol Clugston, Anna Dominiczak and Alex Alderton, Roberto Amato, Jeffrey Barrett, Sonia Goncalves, Ewan Harrison, David K. Jackson, Ian Johnston, Dominic Kwiatkowski, Cordelia Langford, John Sillitoe on behalf of the Wellcome Sanger Institute COVID-19 Surveillance Team |
| EPI_ISL_2994431 | Lighthouse Lab in Alderley Park | Wellcome Sanger Institute for the COVID-19 Genomics UK (COG-UK) Consortium | Jacquelyn Wynn, Mairead Hyland, The Lighthouse Lab in Alderley Park and Alex Alderton, Roberto Amato, Jeffrey Barrett, Sonia Goncalves, Ewan Harrison, David K. Jackson, Ian Johnston, Dominic Kwiatkowski, Cordelia Langford, John Sillitoe on behalf of the Wellcome Sanger Institute COVID-19 Surveillance Team |
| EPI_ISL_2994432, EPI_ISL_2994433 | Berkshire and Surrey Pathology Services Lighthouse Laboratory | Wellcome Sanger Institute for the COVID-19 Genomics UK (COG-UK) Consortium | Berkshire and Surrey Pathology Services Lighthouse Laboratory and Alex Alderton, Roberto Amato, Jeffrey Barrett, Sonia Goncalves, Ewan Harrison, David K. Jackson, Ian Johnston, Dominic Kwiatkowski, Cordelia Langford, John Sillitoe on behalf of the Wellcome Sanger Institute COVID-19 Surveillance Team |
| EPI_ISL_2994434, EPI_ISL_2994435 | Lighthouse Lab in Glasgow | Wellcome Sanger Institute for the COVID-19 Genomics UK (COG-UK) Consortium | Harper VanSteenhouse, Yumi Kasai, David Gray, Carol Clugston, Anna Dominiczak and Alex Alderton, Roberto Amato, Jeffrey Barrett, Sonia Goncalves, Ewan Harrison, David K. Jackson, Ian Johnston, Dominic Kwiatkowski, Cordelia Langford, John Sillitoe on behalf of the Wellcome Sanger Institute COVID-19 Surveillance Team |
| EPI_ISL_2994436 | Berkshire and Surrey Pathology Services Lighthouse Laboratory | Wellcome Sanger Institute for the COVID-19 Genomics UK (COG-UK) Consortium | Berkshire and Surrey Pathology Services Lighthouse Laboratory and Alex Alderton, Roberto Amato, Jeffrey Barrett, Sonia Goncalves, Ewan Harrison, David K. Jackson, Ian Johnston, Dominic Kwiatkowski, Cordelia Langford, John Sillitoe on behalf of the Wellcome Sanger Institute COVID-19 Surveillance Team |
| EPI_ISL_2994437 | Lighthouse Lab in Alderley Park | Wellcome Sanger Institute for the COVID-19 Genomics UK (COG-UK) Consortium | Jacquelyn Wynn, Mairead Hyland, The Lighthouse Lab in Alderley Park and Alex Alderton, Roberto Amato, Jeffrey Barrett, Sonia Goncalves, Ewan Harrison, David K. Jackson, Ian Johnston, Dominic Kwiatkowski, Cordelia Langford, John Sillitoe on behalf of the Wellcome Sanger Institute COVID-19 Surveillance Team |
| EPI_ISL_2994438 | Berkshire and Surrey Pathology Services Lighthouse Laboratory | Wellcome Sanger Institute for the COVID-19 Genomics UK (COG-UK) Consortium | Berkshire and Surrey Pathology Services Lighthouse Laboratory and Alex Alderton, Roberto Amato, Jeffrey Barrett, Sonia Goncalves, Ewan Harrison, David K. Jackson, Ian Johnston, Dominic Kwiatkowski, Cordelia Langford, John Sillitoe on behalf of the Wellcome Sanger Institute COVID-19 Surveillance Team |
| EPI_ISL_2994439 | Lighthouse Lab in Glasgow | Wellcome Sanger Institute for the COVID-19 Genomics UK (COG-UK) Consortium | Harper VanSteenhouse, Yumi Kasai, David Gray, Carol Clugston, Anna Dominiczak and Alex Alderton, Roberto Amato, Jeffrey Barrett, Sonia Goncalves, Ewan Harrison, David K. Jackson, Ian Johnston, Dominic Kwiatkowski, Cordelia Langford, John Sillitoe on behalf of the Wellcome Sanger Institute COVID-19 Surveillance Team |
| EPI_ISL_2994440 | Berkshire and Surrey Pathology Services Lighthouse Laboratory | Wellcome Sanger Institute for the COVID-19 Genomics UK (COG-UK) Consortium | Berkshire and Surrey Pathology Services Lighthouse Laboratory and Alex Alderton, Roberto Amato, Jeffrey Barrett, Sonia Goncalves, Ewan Harrison, David K. Jackson, Ian Johnston, Dominic Kwiatkowski, Cordelia Langford, John Sillitoe on behalf of the Wellcome Sanger Institute COVID-19 Surveillance Team |
| EPI_ISL_2994441 | Lighthouse Lab in Glasgow | Wellcome Sanger Institute for the COVID-19 Genomics UK (COG-UK) Consortium | Harper VanSteenhouse, Yumi Kasai, David Gray, Carol Clugston, Anna Dominiczak and Alex Alderton, Roberto Amato, Jeffrey Barrett, Sonia Goncalves, Ewan Harrison, David K. Jackson, Ian Johnston, Dominic Kwiatkowski, Cordelia Langford, John Sillitoe on behalf of the Wellcome Sanger Institute COVID-19 Surveillance Team |
| EPI_ISL_2994442 | Lighthouse Lab in Milton Keynes | Wellcome Sanger Institute for the COVID-19 Genomics UK (COG-UK) Consortium | The Lighthouse Lab in Milton Keynes and Alex Alderton, Roberto Amato, Jeffrey Barrett, Sonia Goncalves, Ewan Harrison, David K. Jackson, Ian Johnston, Dominic Kwiatkowski, Cordelia Langford, John Sillitoe on behalf of the Wellcome Sanger Institute COVID-19 Surveillance Team |
| EPI_ISL_2994443 | Lighthouse Lab in Alderley Park | Wellcome Sanger Institute for the COVID-19 Genomics UK (COG-UK) Consortium | Jacquelyn Wynn, Mairead Hyland, The Lighthouse Lab in Alderley Park and Alex Alderton, Roberto Amato, Jeffrey Barrett, Sonia Goncalves, Ewan Harrison, David K. Jackson, Ian Johnston, Dominic Kwiatkowski, Cordelia Langford, John Sillitoe on behalf of the Wellcome Sanger Institute COVID-19 Surveillance Team |
| EPI_ISL_2994444 | Berkshire and Surrey Pathology Services Lighthouse Laboratory | Wellcome Sanger Institute for the COVID-19 Genomics UK (COG-UK) Consortium | Berkshire and Surrey Pathology Services Lighthouse Laboratory and Alex Alderton, Roberto Amato, Jeffrey Barrett, Sonia Goncalves, Ewan Harrison, David K. Jackson, Ian Johnston, Dominic Kwiatkowski, Cordelia Langford, John Sillitoe on behalf of the Wellcome Sanger Institute COVID-19 Surveillance Team |
| EPI_ISL_2994445 | Lighthouse Lab in Alderley Park | Wellcome Sanger Institute for the COVID-19 Genomics UK (COG-UK) Consortium | Jacquelyn Wynn, Mairead Hyland, The Lighthouse Lab in Alderley Park and Alex Alderton, Roberto Amato, Jeffrey Barrett, Sonia Goncalves, Ewan Harrison, David K. Jackson, Ian Johnston, Dominic Kwiatkowski, Cordelia Langford, John Sillitoe on behalf of the Wellcome Sanger Institute COVID-19 Surveillance Team |
| EPI_ISL_2994446 | Berkshire and Surrey Pathology Services Lighthouse Laboratory | Wellcome Sanger Institute for the COVID-19 Genomics UK (COG-UK) Consortium | Berkshire and Surrey Pathology Services Lighthouse Laboratory and Alex Alderton, Roberto Amato, Jeffrey Barrett, Sonia Goncalves, Ewan Harrison, David K. Jackson, Ian Johnston, Dominic Kwiatkowski, Cordelia Langford, John Sillitoe on behalf of the Wellcome Sanger Institute COVID-19 Surveillance Team |
| EPI_ISL_2994447 | Lighthouse Lab in Milton Keynes | Wellcome Sanger Institute for the COVID-19 Genomics UK (COG-UK) Consortium | The Lighthouse Lab in Milton Keynes and Alex Alderton, Roberto Amato, Jeffrey Barrett, Sonia Goncalves, Ewan Harrison, David K. Jackson, Ian Johnston, Dominic Kwiatkowski, Cordelia Langford, John Sillitoe on behalf of the Wellcome Sanger Institute COVID-19 Surveillance Team |
| EPI_ISL_2994448 | Berkshire and Surrey Pathology Services Lighthouse Laboratory | Wellcome Sanger Institute for the COVID-19 Genomics UK (COG-UK) Consortium | Berkshire and Surrey Pathology Services Lighthouse Laboratory and Alex Alderton, Roberto Amato, Jeffrey Barrett, Sonia Goncalves, Ewan Harrison, David K. Jackson, Ian Johnston, Dominic Kwiatkowski, Cordelia Langford, John Sillitoe on behalf of the Wellcome Sanger Institute COVID-19 Surveillance Team |
| EPI_ISL_2994449, EPI_ISL_2994450 | Lighthouse Lab in Milton Keynes | Wellcome Sanger Institute for the COVID-19 Genomics UK (COG-UK) Consortium | The Lighthouse Lab in Milton Keynes and Alex Alderton, Roberto Amato, Jeffrey Barrett, Sonia Goncalves, Ewan Harrison, David K. Jackson, Ian Johnston, Dominic Kwiatkowski, Cordelia Langford, John Sillitoe on behalf of the Wellcome Sanger Institute COVID-19 Surveillance Team |
| EPI_ISL_2994451 | Berkshire and Surrey Pathology Services Lighthouse Laboratory | Wellcome Sanger Institute for the COVID-19 Genomics UK (COG-UK) Consortium | Berkshire and Surrey Pathology Services Lighthouse Laboratory and Alex Alderton, Roberto Amato, Jeffrey Barrett, Sonia Goncalves, Ewan Harrison, David K. Jackson, Ian Johnston, Dominic Kwiatkowski, Cordelia Langford, John Sillitoe on behalf of the Wellcome Sanger Institute COVID-19 Surveillance Team |
| EPI_ISL_2994452, EPI_ISL_2994453 | Lighthouse Lab in Glasgow | Wellcome Sanger Institute for the COVID-19 Genomics UK (COG-UK) Consortium | Harper VanSteenhouse, Yumi Kasai, David Gray, Carol Clugston, Anna Dominiczak and Alex Alderton, Roberto Amato, Jeffrey Barrett, Sonia Goncalves, Ewan Harrison, David K. Jackson, Ian Johnston, Dominic Kwiatkowski, Cordelia Langford, John Sillitoe on behalf of the Wellcome Sanger Institute COVID-19 Surveillance Team |
| EPI_ISL_2994454 | Berkshire and Surrey Pathology Services Lighthouse Laboratory | Wellcome Sanger Institute for the COVID-19 Genomics UK (COG-UK) Consortium | Berkshire and Surrey Pathology Services Lighthouse Laboratory and Alex Alderton, Roberto Amato, Jeffrey Barrett, Sonia Goncalves, Ewan Harrison, David K. Jackson, Ian Johnston, Dominic Kwiatkowski, Cordelia Langford, John Sillitoe on behalf of the Wellcome Sanger Institute COVID-19 Surveillance Team |
| EPI_ISL_2994455 | Lighthouse Lab in Glasgow | Wellcome Sanger Institute for the COVID-19 Genomics UK (COG-UK) Consortium | Harper VanSteenhouse, Yumi Kasai, David Gray, Carol Clugston, Anna Dominiczak and Alex Alderton, Roberto Amato, Jeffrey Barrett, Sonia Goncalves, Ewan Harrison, David K. Jackson, Ian Johnston, Dominic Kwiatkowski, Cordelia Langford, John Sillitoe on behalf of the Wellcome Sanger Institute COVID-19 Surveillance Team |
| EPI_ISL_2994456 | Lighthouse Lab in Alderley Park | Wellcome Sanger Institute for the COVID-19 Genomics UK (COG-UK) Consortium | Jacquelyn Wynn, Mairead Hyland, The Lighthouse Lab in Alderley Park and Alex Alderton, Roberto Amato, Jeffrey Barrett, Sonia Goncalves, Ewan Harrison, David K. Jackson, Ian Johnston, Dominic Kwiatkowski, Cordelia Langford, John Sillitoe on behalf of the Wellcome Sanger Institute COVID-19 Surveillance Team |
| EPI_ISL_2994457 | Berkshire and Surrey Pathology Services Lighthouse Laboratory | Wellcome Sanger Institute for the COVID-19 Genomics UK (COG-UK) Consortium | Berkshire and Surrey Pathology Services Lighthouse Laboratory and Alex Alderton, Roberto Amato, Jeffrey Barrett, Sonia Goncalves, Ewan Harrison, David K. Jackson, Ian Johnston, Dominic Kwiatkowski, Cordelia Langford, John Sillitoe on behalf of the Wellcome Sanger Institute COVID-19 Surveillance Team |
| EPI_ISL_2994458, EPI_ISL_2994459, EPI_ISL_2994460 | Lighthouse Lab in Alderley Park | Wellcome Sanger Institute for the COVID-19 Genomics UK (COG-UK) Consortium | Jacquelyn Wynn, Mairead Hyland, The Lighthouse Lab in Alderley Park and Alex Alderton, Roberto Amato, Jeffrey Barrett, Sonia Goncalves, Ewan Harrison, David K. Jackson, Ian Johnston, Dominic Kwiatkowski, Cordelia Langford, John Sillitoe on behalf of the Wellcome Sanger Institute COVID-19 Surveillance Team |
| EPI_ISL_2994461 | Lighthouse Lab in Milton Keynes | Wellcome Sanger Institute for the COVID-19 Genomics UK (COG-UK) Consortium | The Lighthouse Lab in Milton Keynes and Alex Alderton, Roberto Amato, Jeffrey Barrett, Sonia Goncalves, Ewan Harrison, David K. Jackson, Ian Johnston, Dominic Kwiatkowski, Cordelia Langford, John Sillitoe on behalf of the Wellcome Sanger Institute COVID-19 Surveillance Team |
| EPI_ISL_2994462 | Lighthouse Lab in Glasgow | Wellcome Sanger Institute for the COVID-19 Genomics UK (COG-UK) Consortium | Harper VanSteenhouse, Yumi Kasai, David Gray, Carol Clugston, Anna Dominiczak and Alex Alderton, Roberto Amato, Jeffrey Barrett, Sonia Goncalves, Ewan Harrison, David K. Jackson, Ian Johnston, Dominic Kwiatkowski, Cordelia Langford, John Sillitoe on behalf of the Wellcome Sanger Institute COVID-19 Surveillance Team |
| EPI_ISL_2994463 | Berkshire and Surrey Pathology Services Lighthouse Laboratory | Wellcome Sanger Institute for the COVID-19 Genomics UK (COG-UK) Consortium | Berkshire and Surrey Pathology Services Lighthouse Laboratory and Alex Alderton, Roberto Amato, Jeffrey Barrett, Sonia Goncalves, Ewan Harrison, David K. Jackson, Ian Johnston, Dominic Kwiatkowski, Cordelia Langford, John Sillitoe on behalf of the Wellcome Sanger Institute COVID-19 Surveillance Team |
| EPI_ISL_2994464 | Lighthouse Lab in Alderley Park | Wellcome Sanger Institute for the COVID-19 Genomics UK (COG-UK) Consortium | Jacquelyn Wynn, Mairead Hyland, The Lighthouse Lab in Alderley Park and Alex Alderton, Roberto Amato, Jeffrey Barrett, Sonia Goncalves, Ewan Harrison, David K. Jackson, Ian Johnston, Dominic Kwiatkowski, Cordelia Langford, John Sillitoe on behalf of the Wellcome Sanger Institute COVID-19 Surveillance Team |
| EPI_ISL_2994465 | Lighthouse Lab in Milton Keynes | Wellcome Sanger Institute for the COVID-19 Genomics UK (COG-UK) Consortium | The Lighthouse Lab in Milton Keynes and Alex Alderton, Roberto Amato, Jeffrey Barrett, Sonia Goncalves, Ewan Harrison, David K. Jackson, Ian Johnston, Dominic Kwiatkowski, Cordelia Langford, John Sillitoe on behalf of the Wellcome Sanger Institute COVID-19 Surveillance Team |
| EPI_ISL_2994466 | Lighthouse Lab in Glasgow | Wellcome Sanger Institute for the COVID-19 Genomics UK (COG-UK) Consortium | Harper VanSteenhouse, Yumi Kasai, David Gray, Carol Clugston, Anna Dominiczak and Alex Alderton, Roberto Amato, Jeffrey Barrett, Sonia Goncalves, Ewan Harrison, David K. Jackson, Ian Johnston, Dominic Kwiatkowski, Cordelia Langford, John Sillitoe on behalf of the Wellcome Sanger Institute COVID-19 Surveillance Team |
| EPI_ISL_2994467 | Berkshire and Surrey Pathology Services Lighthouse Laboratory | Wellcome Sanger Institute for the COVID-19 Genomics UK (COG-UK) Consortium | Berkshire and Surrey Pathology Services Lighthouse Laboratory and Alex Alderton, Roberto Amato, Jeffrey Barrett, Sonia Goncalves, Ewan Harrison, David K. Jackson, Ian Johnston, Dominic Kwiatkowski, Cordelia Langford, John Sillitoe on behalf of the Wellcome Sanger Institute COVID-19 Surveillance Team |
| EPI_ISL_2994468 | Lighthouse Lab in Milton Keynes | Wellcome Sanger Institute for the COVID-19 Genomics UK (COG-UK) Consortium | The Lighthouse Lab in Milton Keynes and Alex Alderton, Roberto Amato, Jeffrey Barrett, Sonia Goncalves, Ewan Harrison, David K. Jackson, Ian Johnston, Dominic Kwiatkowski, Cordelia Langford, John Sillitoe on behalf of the Wellcome Sanger Institute COVID-19 Surveillance Team |
| EPI_ISL_2994469 | Berkshire and Surrey Pathology Services Lighthouse Laboratory | Wellcome Sanger Institute for the COVID-19 Genomics UK (COG-UK) Consortium | Berkshire and Surrey Pathology Services Lighthouse Laboratory and Alex Alderton, Roberto Amato, Jeffrey Barrett, Sonia Goncalves, Ewan Harrison, David K. Jackson, Ian Johnston, Dominic Kwiatkowski, Cordelia Langford, John Sillitoe on behalf of the Wellcome Sanger Institute COVID-19 Surveillance Team |
| EPI_ISL_2994470 | Lighthouse Lab in Glasgow | Wellcome Sanger Institute for the COVID-19 Genomics UK (COG-UK) Consortium | Harper VanSteenhouse, Yumi Kasai, David Gray, Carol Clugston, Anna Dominiczak and Alex Alderton, Roberto Amato, Jeffrey Barrett, Sonia Goncalves, Ewan Harrison, David K. Jackson, Ian Johnston, Dominic Kwiatkowski, Cordelia Langford, John Sillitoe on behalf of the Wellcome Sanger Institute COVID-19 Surveillance Team |
| EPI_ISL_2994471 | Lighthouse Lab in Milton Keynes | Wellcome Sanger Institute for the COVID-19 Genomics UK (COG-UK) Consortium | The Lighthouse Lab in Milton Keynes and Alex Alderton, Roberto Amato, Jeffrey Barrett, Sonia Goncalves, Ewan Harrison, David K. Jackson, Ian Johnston, Dominic Kwiatkowski, Cordelia Langford, John Sillitoe on behalf of the Wellcome Sanger Institute COVID-19 Surveillance Team |
| EPI_ISL_2994472 | Lighthouse Lab in Glasgow | Wellcome Sanger Institute for the COVID-19 Genomics UK (COG-UK) Consortium | Harper VanSteenhouse, Yumi Kasai, David Gray, Carol Clugston, Anna Dominiczak and Alex Alderton, Roberto Amato, Jeffrey Barrett, Sonia Goncalves, Ewan Harrison, David K. Jackson, Ian Johnston, Dominic Kwiatkowski, Cordelia Langford, John Sillitoe on behalf of the Wellcome Sanger Institute COVID-19 Surveillance Team |
| EPI_ISL_2994473 | Lighthouse Lab in Milton Keynes | Wellcome Sanger Institute for the COVID-19 Genomics UK (COG-UK) Consortium | The Lighthouse Lab in Milton Keynes and Alex Alderton, Roberto Amato, Jeffrey Barrett, Sonia Goncalves, Ewan Harrison, David K. Jackson, Ian Johnston, Dominic Kwiatkowski, Cordelia Langford, John Sillitoe on behalf of the Wellcome Sanger Institute COVID-19 Surveillance Team |
| EPI_ISL_2994474 | Lighthouse Lab in Alderley Park | Wellcome Sanger Institute for the COVID-19 Genomics UK (COG-UK) Consortium | Jacquelyn Wynn, Mairead Hyland, The Lighthouse Lab in Alderley Park and Alex Alderton, Roberto Amato, Jeffrey Barrett, Sonia Goncalves, Ewan Harrison, David K. Jackson, Ian Johnston, Dominic Kwiatkowski, Cordelia Langford, John Sillitoe on behalf of the Wellcome Sanger Institute COVID-19 Surveillance Team |
| EPI_ISL_2994475, EPI_ISL_2994476 | Lighthouse Lab in Glasgow | Wellcome Sanger Institute for the COVID-19 Genomics UK (COG-UK) Consortium | Harper VanSteenhouse, Yumi Kasai, David Gray, Carol Clugston, Anna Dominiczak and Alex Alderton, Roberto Amato, Jeffrey Barrett, Sonia Goncalves, Ewan Harrison, David K. Jackson, Ian Johnston, Dominic Kwiatkowski, Cordelia Langford, John Sillitoe on behalf of the Wellcome Sanger Institute COVID-19 Surveillance Team |
| EPI_ISL_2994477 | Berkshire and Surrey Pathology Services Lighthouse Laboratory | Wellcome Sanger Institute for the COVID-19 Genomics UK (COG-UK) | Berkshire and Surrey Pathology Services Lighthouse Laboratory and Alex Alderton, Roberto Amato, Jeffrey Barrett, Sonia Goncalves, Ewan Harrison, David K. Jackson, Ian Johnston, Dominic Kwiatkowski, Cordelia Langford, John Sillitoe on behalf |

| | | | |
|---|---|---|---|
| | | Consortium | of the Wellcome Sanger Institute COVID-19 Surveillance Team |
| EPI_ISL_2994478, EPI_ISL_2994479 | Lighthouse Lab in Milton Keynes | Wellcome Sanger Institute for the COVID-19 Genomics UK (COG-UK) Consortium | The Lighthouse Lab in Milton Keynes and Alex Alderton, Roberto Amato, Jeffrey Barrett, Sonia Goncalves, Ewan Harrison, David K. Jackson, Ian Johnston, Dominic Kwiatkowski, Cordelia Langford, John Sillitoe on behalf of the Wellcome Sanger Institute COVID-19 Surveillance Team |
| EPI_ISL_2994480, EPI_ISL_2994481 | Berkshire and Surrey Pathology Services Lighthouse Laboratory | Wellcome Sanger Institute for the COVID-19 Genomics UK (COG-UK) Consortium | Berkshire and Surrey Pathology Services Lighthouse Laboratory and Alex Alderton, Roberto Amato, Jeffrey Barrett, Sonia Goncalves, Ewan Harrison, David K. Jackson, Ian Johnston, Dominic Kwiatkowski, Cordelia Langford, John Sillitoe on behalf of the Wellcome Sanger Institute COVID-19 Surveillance Team |
| EPI_ISL_2994482, EPI_ISL_2994483, EPI_ISL_2994484 | Lighthouse Lab in Alderley Park | Wellcome Sanger Institute for the COVID-19 Genomics UK (COG-UK) Consortium | Jacquelyn Wynn, Mairead Hyland, The Lighthouse Lab in Alderley Park and Alex Alderton, Roberto Amato, Jeffrey Barrett, Sonia Goncalves, Ewan Harrison, David K. Jackson, Ian Johnston, Dominic Kwiatkowski, Cordelia Langford, John Sillitoe on behalf of the Wellcome Sanger Institute COVID-19 Surveillance Team |
| EPI_ISL_2994485 | Lighthouse Lab in Milton Keynes | Wellcome Sanger Institute for the COVID-19 Genomics UK (COG-UK) Consortium | The Lighthouse Lab in Milton Keynes and Alex Alderton, Roberto Amato, Jeffrey Barrett, Sonia Goncalves, Ewan Harrison, David K. Jackson, Ian Johnston, Dominic Kwiatkowski, Cordelia Langford, John Sillitoe on behalf of the Wellcome Sanger Institute COVID-19 Surveillance Team |
| EPI_ISL_2994486, EPI_ISL_2994487 | Berkshire and Surrey Pathology Services Lighthouse Laboratory | Wellcome Sanger Institute for the COVID-19 Genomics UK (COG-UK) Consortium | Berkshire and Surrey Pathology Services Lighthouse Laboratory and Alex Alderton, Roberto Amato, Jeffrey Barrett, Sonia Goncalves, Ewan Harrison, David K. Jackson, Ian Johnston, Dominic Kwiatkowski, Cordelia Langford, John Sillitoe on behalf of the Wellcome Sanger Institute COVID-19 Surveillance Team |
| EPI_ISL_2994488 | Lighthouse Lab in Milton Keynes | Wellcome Sanger Institute for the COVID-19 Genomics UK (COG-UK) Consortium | The Lighthouse Lab in Milton Keynes and Alex Alderton, Roberto Amato, Jeffrey Barrett, Sonia Goncalves, Ewan Harrison, David K. Jackson, Ian Johnston, Dominic Kwiatkowski, Cordelia Langford, John Sillitoe on behalf of the Wellcome Sanger Institute COVID-19 Surveillance Team |
| EPI_ISL_2994489 | Lighthouse Lab in Alderley Park | Wellcome Sanger Institute for the COVID-19 Genomics UK (COG-UK) Consortium | Jacquelyn Wynn, Mairead Hyland, The Lighthouse Lab in Alderley Park and Alex Alderton, Roberto Amato, Jeffrey Barrett, Sonia Goncalves, Ewan Harrison, David K. Jackson, Ian Johnston, Dominic Kwiatkowski, Cordelia Langford, John Sillitoe on behalf of the Wellcome Sanger Institute COVID-19 Surveillance Team |
| EPI_ISL_2994490, EPI_ISL_2994491 | Lighthouse Lab in Milton Keynes | Wellcome Sanger Institute for the COVID-19 Genomics UK (COG-UK) Consortium | The Lighthouse Lab in Milton Keynes and Alex Alderton, Roberto Amato, Jeffrey Barrett, Sonia Goncalves, Ewan Harrison, David K. Jackson, Ian Johnston, Dominic Kwiatkowski, Cordelia Langford, John Sillitoe on behalf of the Wellcome Sanger Institute COVID-19 Surveillance Team |
| EPI_ISL_2994492 | Lighthouse Lab in Alderley Park | Wellcome Sanger Institute for the COVID-19 Genomics UK (COG-UK) Consortium | Jacquelyn Wynn, Mairead Hyland, The Lighthouse Lab in Alderley Park and Alex Alderton, Roberto Amato, Jeffrey Barrett, Sonia Goncalves, Ewan Harrison, David K. Jackson, Ian Johnston, Dominic Kwiatkowski, Cordelia Langford, John Sillitoe on behalf of the Wellcome Sanger Institute COVID-19 Surveillance Team |
| EPI_ISL_2994493 | Berkshire and Surrey Pathology Services Lighthouse Laboratory | Wellcome Sanger Institute for the COVID-19 Genomics UK (COG-UK) Consortium | Berkshire and Surrey Pathology Services Lighthouse Laboratory and Alex Alderton, Roberto Amato, Jeffrey Barrett, Sonia Goncalves, Ewan Harrison, David K. Jackson, Ian Johnston, Dominic Kwiatkowski, Cordelia Langford, John Sillitoe on behalf of the Wellcome Sanger Institute COVID-19 Surveillance Team |
| EPI_ISL_2994494 | Lighthouse Lab in Alderley Park | Wellcome Sanger Institute for the COVID-19 Genomics UK (COG-UK) Consortium | Jacquelyn Wynn, Mairead Hyland, The Lighthouse Lab in Alderley Park and Alex Alderton, Roberto Amato, Jeffrey Barrett, Sonia Goncalves, Ewan Harrison, David K. Jackson, Ian Johnston, Dominic Kwiatkowski, Cordelia Langford, John Sillitoe on behalf of the Wellcome Sanger Institute COVID-19 Surveillance Team |
| EPI_ISL_2994495 | Lighthouse Lab in Milton Keynes | Wellcome Sanger Institute for the COVID-19 Genomics UK (COG-UK) Consortium | The Lighthouse Lab in Milton Keynes and Alex Alderton, Roberto Amato, Jeffrey Barrett, Sonia Goncalves, Ewan Harrison, David K. Jackson, Ian Johnston, Dominic Kwiatkowski, Cordelia Langford, John Sillitoe on behalf of the Wellcome Sanger Institute COVID-19 Surveillance Team |
| EPI_ISL_2994496 | Berkshire and Surrey Pathology Services Lighthouse Laboratory | Wellcome Sanger Institute for the COVID-19 Genomics UK (COG-UK) Consortium | Berkshire and Surrey Pathology Services Lighthouse Laboratory and Alex Alderton, Roberto Amato, Jeffrey Barrett, Sonia Goncalves, Ewan Harrison, David K. Jackson, Ian Johnston, Dominic Kwiatkowski, Cordelia Langford, John Sillitoe on behalf of the Wellcome Sanger Institute COVID-19 Surveillance Team |
| EPI_ISL_2994497 | Lighthouse Lab in Alderley Park | Wellcome Sanger Institute for the COVID-19 Genomics UK (COG-UK) Consortium | Jacquelyn Wynn, Mairead Hyland, The Lighthouse Lab in Alderley Park and Alex Alderton, Roberto Amato, Jeffrey Barrett, Sonia Goncalves, Ewan Harrison, David K. Jackson, Ian Johnston, Dominic Kwiatkowski, Cordelia Langford, John Sillitoe on behalf of the Wellcome Sanger Institute COVID-19 Surveillance Team |
| EPI_ISL_2994498, EPI_ISL_2994499 | Lighthouse Lab in Milton Keynes | Wellcome Sanger Institute for the COVID-19 Genomics UK (COG-UK) Consortium | The Lighthouse Lab in Milton Keynes and Alex Alderton, Roberto Amato, Jeffrey Barrett, Sonia Goncalves, Ewan Harrison, David K. Jackson, Ian Johnston, Dominic Kwiatkowski, Cordelia Langford, John Sillitoe on behalf of the Wellcome Sanger Institute COVID-19 Surveillance Team |
| EPI_ISL_2994500 | Lighthouse Lab in Glasgow | Wellcome Sanger Institute for the COVID-19 Genomics UK (COG-UK) Consortium | Harper VanSteenhouse, Yumi Kasai, David Gray, Carol Clugston, Anna Dominiczak and Alex Alderton, Roberto Amato, Jeffrey Barrett, Sonia Goncalves, Ewan Harrison, David K. Jackson, Ian Johnston, Dominic Kwiatkowski, Cordelia Langford, John Sillitoe on behalf of the Wellcome Sanger Institute COVID-19 Surveillance Team |
| EPI_ISL_2994501 | Berkshire and Surrey Pathology Services Lighthouse Laboratory | Wellcome Sanger Institute for the COVID-19 Genomics UK (COG-UK) Consortium | Berkshire and Surrey Pathology Services Lighthouse Laboratory and Alex Alderton, Roberto Amato, Jeffrey Barrett, Sonia Goncalves, Ewan Harrison, David K. Jackson, Ian Johnston, Dominic Kwiatkowski, Cordelia Langford, John Sillitoe on behalf of the Wellcome Sanger Institute COVID-19 Surveillance Team |
| EPI_ISL_2994502 | Lighthouse Lab in Glasgow | Wellcome Sanger Institute for the COVID-19 Genomics UK (COG-UK) Consortium | Harper VanSteenhouse, Yumi Kasai, David Gray, Carol Clugston, Anna Dominiczak and Alex Alderton, Roberto Amato, Jeffrey Barrett, Sonia Goncalves, Ewan Harrison, David K. Jackson, Ian Johnston, Dominic Kwiatkowski, Cordelia Langford, John Sillitoe on behalf of the Wellcome Sanger Institute COVID-19 Surveillance Team |
| EPI_ISL_2994503, EPI_ISL_2994504, EPI_ISL_2994505, EPI_ISL_2994506 | Lighthouse Lab in Alderley Park | Wellcome Sanger Institute for the COVID-19 Genomics UK (COG-UK) Consortium | Jacquelyn Wynn, Mairead Hyland, The Lighthouse Lab in Alderley Park and Alex Alderton, Roberto Amato, Jeffrey Barrett, Sonia Goncalves, Ewan Harrison, David K. Jackson, Ian Johnston, Dominic Kwiatkowski, Cordelia Langford, John Sillitoe on behalf of the Wellcome Sanger Institute COVID-19 Surveillance Team |
| EPI_ISL_2994507, EPI_ISL_2994508 | Berkshire and Surrey Pathology Services Lighthouse Laboratory | Wellcome Sanger Institute for the COVID-19 Genomics UK (COG-UK) Consortium | Berkshire and Surrey Pathology Services Lighthouse Laboratory and Alex Alderton, Roberto Amato, Jeffrey Barrett, Sonia Goncalves, Ewan Harrison, David K. Jackson, Ian Johnston, Dominic Kwiatkowski, Cordelia Langford, John Sillitoe on behalf of the Wellcome Sanger Institute COVID-19 Surveillance Team |
| EPI_ISL_2994509, EPI_ISL_2994510, EPI_ISL_2994511, EPI_ISL_2994512 | Lighthouse Lab in Glasgow | Wellcome Sanger Institute for the COVID-19 Genomics UK (COG-UK) Consortium | Harper VanSteenhouse, Yumi Kasai, David Gray, Carol Clugston, Anna Dominiczak and Alex Alderton, Roberto Amato, Jeffrey Barrett, Sonia Goncalves, Ewan Harrison, David K. Jackson, Ian Johnston, Dominic Kwiatkowski, Cordelia Langford, John Sillitoe on behalf of the Wellcome Sanger Institute COVID-19 Surveillance Team |
| EPI_ISL_2994513, EPI_ISL_2994514 | Berkshire and Surrey Pathology Services Lighthouse Laboratory | Wellcome Sanger Institute for the COVID-19 Genomics UK (COG-UK) Consortium | Berkshire and Surrey Pathology Services Lighthouse Laboratory and Alex Alderton, Roberto Amato, Jeffrey Barrett, Sonia Goncalves, Ewan Harrison, David K. Jackson, Ian Johnston, Dominic Kwiatkowski, Cordelia Langford, John Sillitoe on behalf of the Wellcome Sanger Institute COVID-19 Surveillance Team |
| EPI_ISL_2994515, EPI_ISL_2994516 | Lighthouse Lab in Alderley Park | Wellcome Sanger Institute for the COVID-19 Genomics UK (COG-UK) Consortium | Jacquelyn Wynn, Mairead Hyland, The Lighthouse Lab in Alderley Park and Alex Alderton, Roberto Amato, Jeffrey Barrett, Sonia Goncalves, Ewan Harrison, David K. Jackson, Ian Johnston, Dominic Kwiatkowski, Cordelia Langford, John Sillitoe on behalf of the Wellcome Sanger Institute COVID-19 Surveillance Team |
| EPI_ISL_2994517 | Lighthouse Lab in Milton Keynes | Wellcome Sanger Institute for the COVID-19 Genomics UK (COG-UK) Consortium | The Lighthouse Lab in Milton Keynes and Alex Alderton, Roberto Amato, Jeffrey Barrett, Sonia Goncalves, Ewan Harrison, David K. Jackson, Ian Johnston, Dominic Kwiatkowski, Cordelia Langford, John Sillitoe on behalf of the Wellcome Sanger Institute COVID-19 Surveillance Team |
| EPI_ISL_2994518 | Lighthouse Lab in Glasgow | Wellcome Sanger Institute for the COVID-19 Genomics UK (COG-UK) Consortium | Harper VanSteenhouse, Yumi Kasai, David Gray, Carol Clugston, Anna Dominiczak and Alex Alderton, Roberto Amato, Jeffrey Barrett, Sonia Goncalves, Ewan Harrison, David K. Jackson, Ian Johnston, Dominic Kwiatkowski, Cordelia Langford, John Sillitoe on behalf of the Wellcome Sanger Institute COVID-19 Surveillance Team |
| EPI_ISL_2994519 | Lighthouse Lab in Alderley Park | Wellcome Sanger Institute for the COVID-19 Genomics UK (COG-UK) Consortium | Jacquelyn Wynn, Mairead Hyland, The Lighthouse Lab in Alderley Park and Alex Alderton, Roberto Amato, Jeffrey Barrett, Sonia Goncalves, Ewan Harrison, David K. Jackson, Ian Johnston, Dominic Kwiatkowski, Cordelia Langford, John Sillitoe on behalf of the Wellcome Sanger Institute COVID-19 Surveillance Team |
| EPI_ISL_2994520 | Lighthouse Lab in Milton Keynes | Wellcome Sanger Institute for the COVID-19 Genomics UK (COG-UK) Consortium | The Lighthouse Lab in Milton Keynes and Alex Alderton, Roberto Amato, Jeffrey Barrett, Sonia Goncalves, Ewan Harrison, David K. Jackson, Ian Johnston, Dominic Kwiatkowski, Cordelia Langford, John Sillitoe on behalf of the Wellcome Sanger Institute COVID-19 Surveillance Team |
| EPI_ISL_2994521 | Berkshire and Surrey Pathology Services Lighthouse Laboratory | Wellcome Sanger Institute for the COVID-19 Genomics UK (COG-UK) Consortium | Berkshire and Surrey Pathology Services Lighthouse Laboratory and Alex Alderton, Roberto Amato, Jeffrey Barrett, Sonia Goncalves, Ewan Harrison, David K. Jackson, Ian Johnston, Dominic Kwiatkowski, Cordelia Langford, John Sillitoe on behalf of the Wellcome Sanger Institute COVID-19 Surveillance Team |
| EPI_ISL_2994522 | Lighthouse Lab in Milton Keynes | Wellcome Sanger Institute for the COVID-19 Genomics UK (COG-UK) Consortium | The Lighthouse Lab in Milton Keynes and Alex Alderton, Roberto Amato, Jeffrey Barrett, Sonia Goncalves, Ewan Harrison, David K. Jackson, Ian Johnston, Dominic Kwiatkowski, Cordelia Langford, John Sillitoe on behalf of the Wellcome Sanger Institute COVID-19 Surveillance Team |
| EPI_ISL_2994523 | Lighthouse Lab in Glasgow | Wellcome Sanger Institute for the COVID-19 Genomics UK (COG-UK) Consortium | Harper VanSteenhouse, Yumi Kasai, David Gray, Carol Clugston, Anna Dominiczak and Alex Alderton, Roberto Amato, Jeffrey Barrett, Sonia Goncalves, Ewan Harrison, David K. Jackson, Ian Johnston, Dominic Kwiatkowski, Cordelia Langford, John Sillitoe on behalf of the Wellcome Sanger Institute COVID-19 Surveillance Team |
| EPI_ISL_2994524, EPI_ISL_2994525, EPI_ISL_2994526 | Lighthouse Lab in Alderley Park | Wellcome Sanger Institute for the COVID-19 Genomics UK (COG-UK) Consortium | Jacquelyn Wynn, Mairead Hyland, The Lighthouse Lab in Alderley Park and Alex Alderton, Roberto Amato, Jeffrey Barrett, Sonia Goncalves, Ewan Harrison, David K. Jackson, Ian Johnston, Dominic Kwiatkowski, Cordelia Langford, John Sillitoe on behalf of the Wellcome Sanger Institute COVID-19 Surveillance Team |
| EPI_ISL_2994527 | Lighthouse Lab in Glasgow | Wellcome Sanger Institute for the COVID-19 Genomics UK (COG-UK) Consortium | Harper VanSteenhouse, Yumi Kasai, David Gray, Carol Clugston, Anna Dominiczak and Alex Alderton, Roberto Amato, Jeffrey Barrett, Sonia Goncalves, Ewan Harrison, David K. Jackson, Ian Johnston, Dominic Kwiatkowski, Cordelia Langford, John Sillitoe on behalf of the Wellcome Sanger Institute COVID-19 Surveillance Team |
| EPI_ISL_2994528 | Lighthouse Lab in Milton Keynes | Wellcome Sanger Institute for the COVID-19 Genomics UK (COG-UK) Consortium | The Lighthouse Lab in Milton Keynes and Alex Alderton, Roberto Amato, Jeffrey Barrett, Sonia Goncalves, Ewan Harrison, David K. Jackson, Ian Johnston, Dominic Kwiatkowski, Cordelia Langford, John Sillitoe on behalf of the Wellcome Sanger Institute COVID-19 Surveillance Team |
| EPI_ISL_2994529 | Lighthouse Lab in Alderley Park | Wellcome Sanger Institute for the COVID-19 Genomics UK (COG-UK) Consortium | Jacquelyn Wynn, Mairead Hyland, The Lighthouse Lab in Alderley Park and Alex Alderton, Roberto Amato, Jeffrey Barrett, Sonia Goncalves, Ewan Harrison, David K. Jackson, Ian Johnston, Dominic Kwiatkowski, Cordelia Langford, John Sillitoe on behalf of the Wellcome Sanger Institute COVID-19 Surveillance Team |
| EPI_ISL_2994530 | Lighthouse Lab in Glasgow | Wellcome Sanger Institute for the COVID-19 Genomics UK (COG-UK) Consortium | Harper VanSteenhouse, Yumi Kasai, David Gray, Carol Clugston, Anna Dominiczak and Alex Alderton, Roberto Amato, Jeffrey Barrett, Sonia Goncalves, Ewan Harrison, David K. Jackson, Ian Johnston, Dominic Kwiatkowski, Cordelia Langford, John Sillitoe on behalf of the Wellcome Sanger Institute COVID-19 Surveillance Team |
| EPI_ISL_2994531 | Berkshire and Surrey Pathology Services Lighthouse Laboratory | Wellcome Sanger Institute for the COVID-19 Genomics UK (COG-UK) Consortium | Berkshire and Surrey Pathology Services Lighthouse Laboratory and Alex Alderton, Roberto Amato, Jeffrey Barrett, Sonia Goncalves, Ewan Harrison, David K. Jackson, Ian Johnston, Dominic Kwiatkowski, Cordelia Langford, John Sillitoe on behalf of the Wellcome Sanger Institute COVID-19 Surveillance Team |
| EPI_ISL_2994532, EPI_ISL_2994533 | Lighthouse Lab in Glasgow | Wellcome Sanger Institute for the COVID-19 Genomics UK (COG-UK) Consortium | Harper VanSteenhouse, Yumi Kasai, David Gray, Carol Clugston, Anna Dominiczak and Alex Alderton, Roberto Amato, Jeffrey Barrett, Sonia Goncalves, Ewan Harrison, David K. Jackson, Ian Johnston, Dominic Kwiatkowski, Cordelia Langford, John Sillitoe on behalf of the Wellcome Sanger Institute COVID-19 Surveillance Team |
| EPI_ISL_2994534 | Lighthouse Lab in Milton Keynes | Wellcome Sanger Institute for the COVID-19 Genomics UK (COG-UK) Consortium | The Lighthouse Lab in Milton Keynes and Alex Alderton, Roberto Amato, Jeffrey Barrett, Sonia Goncalves, Ewan Harrison, David K. Jackson, Ian Johnston, Dominic Kwiatkowski, Cordelia Langford, John Sillitoe on behalf of the Wellcome Sanger Institute COVID-19 Surveillance Team |
| EPI_ISL_2994535 | Lighthouse Lab in Glasgow | Wellcome Sanger Institute for the COVID-19 Genomics UK (COG-UK) Consortium | Harper VanSteenhouse, Yumi Kasai, David Gray, Carol Clugston, Anna Dominiczak and Alex Alderton, Roberto Amato, Jeffrey Barrett, Sonia Goncalves, Ewan Harrison, David K. Jackson, Ian Johnston, Dominic Kwiatkowski, Cordelia Langford, John Sillitoe on behalf of the Wellcome Sanger Institute COVID-19 Surveillance Team |
| EPI_ISL_2994536 | Berkshire and Surrey Pathology Services Lighthouse Laboratory | Wellcome Sanger Institute for the COVID-19 Genomics UK (COG-UK) Consortium | Berkshire and Surrey Pathology Services Lighthouse Laboratory and Alex Alderton, Roberto Amato, Jeffrey Barrett, Sonia Goncalves, Ewan Harrison, David K. Jackson, Ian Johnston, Dominic Kwiatkowski, Cordelia Langford, John Sillitoe on behalf of the Wellcome Sanger Institute COVID-19 Surveillance Team |
| EPI_ISL_2994537 | Lighthouse Lab in Glasgow | Wellcome Sanger Institute for the COVID-19 Genomics UK (COG-UK) Consortium | Harper VanSteenhouse, Yumi Kasai, David Gray, Carol Clugston, Anna Dominiczak and Alex Alderton, Roberto Amato, Jeffrey Barrett, Sonia Goncalves, Ewan Harrison, David K. Jackson, Ian Johnston, Dominic Kwiatkowski, Cordelia Langford, John Sillitoe on behalf of the Wellcome Sanger Institute COVID-19 Surveillance Team |
| EPI_ISL_2994538 | Lighthouse Lab in Alderley Park | Wellcome Sanger Institute for the COVID-19 Genomics UK (COG-UK) Consortium | Jacquelyn Wynn, Mairead Hyland, The Lighthouse Lab in Alderley Park and Alex Alderton, Roberto Amato, Jeffrey Barrett, Sonia Goncalves, Ewan Harrison, David K. Jackson, Ian Johnston, Dominic Kwiatkowski, Cordelia Langford, John Sillitoe on behalf of the Wellcome Sanger Institute COVID-19 Surveillance Team |

| | | | |
|---|---|---|---|
| EPI_ISL_2994539 | Berkshire and Surrey Pathology Services Lighthouse Laboratory | Wellcome Sanger Institute for the COVID-19 Genomics UK (COG-UK) Consortium | Berkshire and Surrey Pathology Services Lighthouse Laboratory and Alex Alderton, Roberto Amato, Jeffrey Barrett, Sonia Goncalves, Ewan Harrison, David K. Jackson, Ian Johnston, Dominic Kwiatkowski, Cordelia Langford, John Sillitoe on behalf of the Wellcome Sanger Institute COVID-19 Surveillance Team |
| EPI_ISL_2994540 | Lighthouse Lab in Alderley Park | Wellcome Sanger Institute for the COVID-19 Genomics UK (COG-UK) Consortium | Jacquelyn Wynn, Mairead Hyland, The Lighthouse Lab in Alderley Park and Alex Alderton, Roberto Amato, Jeffrey Barrett, Sonia Goncalves, Ewan Harrison, David K. Jackson, Ian Johnston, Dominic Kwiatkowski, Cordelia Langford, John Sillitoe on behalf of the Wellcome Sanger Institute COVID-19 Surveillance Team |
| EPI_ISL_2994541, EPI_ISL_2994542 | Berkshire and Surrey Pathology Services Lighthouse Laboratory | Wellcome Sanger Institute for the COVID-19 Genomics UK (COG-UK) Consortium | Berkshire and Surrey Pathology Services Lighthouse Laboratory and Alex Alderton, Roberto Amato, Jeffrey Barrett, Sonia Goncalves, Ewan Harrison, David K. Jackson, Ian Johnston, Dominic Kwiatkowski, Cordelia Langford, John Sillitoe on behalf of the Wellcome Sanger Institute COVID-19 Surveillance Team |
| EPI_ISL_2994543 | Lighthouse Lab in Glasgow | Wellcome Sanger Institute for the COVID-19 Genomics UK (COG-UK) Consortium | Harper VanSteenhouse, Yumi Kasai, David Gray, Carol Clugston, Anna Dominiczak and Alex Alderton, Roberto Amato, Jeffrey Barrett, Sonia Goncalves, Ewan Harrison, David K. Jackson, Ian Johnston, Dominic Kwiatkowski, Cordelia Langford, John Sillitoe on behalf of the Wellcome Sanger Institute COVID-19 Surveillance Team |
| EPI_ISL_2994544, EPI_ISL_2994545 | Lighthouse Lab in Alderley Park | Wellcome Sanger Institute for the COVID-19 Genomics UK (COG-UK) Consortium | Jacquelyn Wynn, Mairead Hyland, The Lighthouse Lab in Alderley Park and Alex Alderton, Roberto Amato, Jeffrey Barrett, Sonia Goncalves, Ewan Harrison, David K. Jackson, Ian Johnston, Dominic Kwiatkowski, Cordelia Langford, John Sillitoe on behalf of the Wellcome Sanger Institute COVID-19 Surveillance Team |
| EPI_ISL_2994546 | Lighthouse Lab in Milton Keynes | Wellcome Sanger Institute for the COVID-19 Genomics UK (COG-UK) Consortium | The Lighthouse Lab in Milton Keynes and Alex Alderton, Roberto Amato, Jeffrey Barrett, Sonia Goncalves, Ewan Harrison, David K. Jackson, Ian Johnston, Dominic Kwiatkowski, Cordelia Langford, John Sillitoe on behalf of the Wellcome Sanger Institute COVID-19 Surveillance Team |
| EPI_ISL_2994547 | Lighthouse Lab in Glasgow | Wellcome Sanger Institute for the COVID-19 Genomics UK (COG-UK) Consortium | Harper VanSteenhouse, Yumi Kasai, David Gray, Carol Clugston, Anna Dominiczak and Alex Alderton, Roberto Amato, Jeffrey Barrett, Sonia Goncalves, Ewan Harrison, David K. Jackson, Ian Johnston, Dominic Kwiatkowski, Cordelia Langford, John Sillitoe on behalf of the Wellcome Sanger Institute COVID-19 Surveillance Team |
| EPI_ISL_2994548, EPI_ISL_2994549 | Lighthouse Lab in Milton Keynes | Wellcome Sanger Institute for the COVID-19 Genomics UK (COG-UK) Consortium | The Lighthouse Lab in Milton Keynes and Alex Alderton, Roberto Amato, Jeffrey Barrett, Sonia Goncalves, Ewan Harrison, David K. Jackson, Ian Johnston, Dominic Kwiatkowski, Cordelia Langford, John Sillitoe on behalf of the Wellcome Sanger Institute COVID-19 Surveillance Team |
| EPI_ISL_2994550 | Lighthouse Lab in Glasgow | Wellcome Sanger Institute for the COVID-19 Genomics UK (COG-UK) Consortium | Harper VanSteenhouse, Yumi Kasai, David Gray, Carol Clugston, Anna Dominiczak and Alex Alderton, Roberto Amato, Jeffrey Barrett, Sonia Goncalves, Ewan Harrison, David K. Jackson, Ian Johnston, Dominic Kwiatkowski, Cordelia Langford, John Sillitoe on behalf of the Wellcome Sanger Institute COVID-19 Surveillance Team |
| EPI_ISL_2994551 | Lighthouse Lab in Milton Keynes | Wellcome Sanger Institute for the COVID-19 Genomics UK (COG-UK) Consortium | The Lighthouse Lab in Milton Keynes and Alex Alderton, Roberto Amato, Jeffrey Barrett, Sonia Goncalves, Ewan Harrison, David K. Jackson, Ian Johnston, Dominic Kwiatkowski, Cordelia Langford, John Sillitoe on behalf of the Wellcome Sanger Institute COVID-19 Surveillance Team |
| EPI_ISL_2994552, EPI_ISL_2994553 | Lighthouse Lab in Alderley Park | Wellcome Sanger Institute for the COVID-19 Genomics UK (COG-UK) Consortium | Jacquelyn Wynn, Mairead Hyland, The Lighthouse Lab in Alderley Park and Alex Alderton, Roberto Amato, Jeffrey Barrett, Sonia Goncalves, Ewan Harrison, David K. Jackson, Ian Johnston, Dominic Kwiatkowski, Cordelia Langford, John Sillitoe on behalf of the Wellcome Sanger Institute COVID-19 Surveillance Team |
| EPI_ISL_2994554 | Berkshire and Surrey Pathology Services Lighthouse Laboratory | Wellcome Sanger Institute for the COVID-19 Genomics UK (COG-UK) Consortium | Berkshire and Surrey Pathology Services Lighthouse Laboratory and Alex Alderton, Roberto Amato, Jeffrey Barrett, Sonia Goncalves, Ewan Harrison, David K. Jackson, Ian Johnston, Dominic Kwiatkowski, Cordelia Langford, John Sillitoe on behalf of the Wellcome Sanger Institute COVID-19 Surveillance Team |
| EPI_ISL_2994555 | Lighthouse Lab in Alderley Park | Wellcome Sanger Institute for the COVID-19 Genomics UK (COG-UK) Consortium | Jacquelyn Wynn, Mairead Hyland, The Lighthouse Lab in Alderley Park and Alex Alderton, Roberto Amato, Jeffrey Barrett, Sonia Goncalves, Ewan Harrison, David K. Jackson, Ian Johnston, Dominic Kwiatkowski, Cordelia Langford, John Sillitoe on behalf of the Wellcome Sanger Institute COVID-19 Surveillance Team |
| EPI_ISL_2994556 | Berkshire and Surrey Pathology Services Lighthouse Laboratory | Wellcome Sanger Institute for the COVID-19 Genomics UK (COG-UK) Consortium | Berkshire and Surrey Pathology Services Lighthouse Laboratory and Alex Alderton, Roberto Amato, Jeffrey Barrett, Sonia Goncalves, Ewan Harrison, David K. Jackson, Ian Johnston, Dominic Kwiatkowski, Cordelia Langford, John Sillitoe on behalf of the Wellcome Sanger Institute COVID-19 Surveillance Team |
| EPI_ISL_2994557 | Lighthouse Lab in Glasgow | Wellcome Sanger Institute for the COVID-19 Genomics UK (COG-UK) Consortium | Harper VanSteenhouse, Yumi Kasai, David Gray, Carol Clugston, Anna Dominiczak and Alex Alderton, Roberto Amato, Jeffrey Barrett, Sonia Goncalves, Ewan Harrison, David K. Jackson, Ian Johnston, Dominic Kwiatkowski, Cordelia Langford, John Sillitoe on behalf of the Wellcome Sanger Institute COVID-19 Surveillance Team |
| EPI_ISL_2994558, EPI_ISL_2994559, EPI_ISL_2994560 | Berkshire and Surrey Pathology Services Lighthouse Laboratory | Wellcome Sanger Institute for the COVID-19 Genomics UK (COG-UK) Consortium | Berkshire and Surrey Pathology Services Lighthouse Laboratory and Alex Alderton, Roberto Amato, Jeffrey Barrett, Sonia Goncalves, Ewan Harrison, David K. Jackson, Ian Johnston, Dominic Kwiatkowski, Cordelia Langford, John Sillitoe on behalf of the Wellcome Sanger Institute COVID-19 Surveillance Team |
| EPI_ISL_2994561, EPI_ISL_2994562 | Lighthouse Lab in Glasgow | Wellcome Sanger Institute for the COVID-19 Genomics UK (COG-UK) Consortium | Harper VanSteenhouse, Yumi Kasai, David Gray, Carol Clugston, Anna Dominiczak and Alex Alderton, Roberto Amato, Jeffrey Barrett, Sonia Goncalves, Ewan Harrison, David K. Jackson, Ian Johnston, Dominic Kwiatkowski, Cordelia Langford, John Sillitoe on behalf of the Wellcome Sanger Institute COVID-19 Surveillance Team |
| EPI_ISL_2994563, EPI_ISL_2994564 | Berkshire and Surrey Pathology Services Lighthouse Laboratory | Wellcome Sanger Institute for the COVID-19 Genomics UK (COG-UK) Consortium | Berkshire and Surrey Pathology Services Lighthouse Laboratory and Alex Alderton, Roberto Amato, Jeffrey Barrett, Sonia Goncalves, Ewan Harrison, David K. Jackson, Ian Johnston, Dominic Kwiatkowski, Cordelia Langford, John Sillitoe on behalf of the Wellcome Sanger Institute COVID-19 Surveillance Team |
| EPI_ISL_2994565 | Lighthouse Lab in Milton Keynes | Wellcome Sanger Institute for the COVID-19 Genomics UK (COG-UK) Consortium | The Lighthouse Lab in Milton Keynes and Alex Alderton, Roberto Amato, Jeffrey Barrett, Sonia Goncalves, Ewan Harrison, David K. Jackson, Ian Johnston, Dominic Kwiatkowski, Cordelia Langford, John Sillitoe on behalf of the Wellcome Sanger Institute COVID-19 Surveillance Team |
| EPI_ISL_2994566 | Lighthouse Lab in Alderley Park | Wellcome Sanger Institute for the COVID-19 Genomics UK (COG-UK) Consortium | Jacquelyn Wynn, Mairead Hyland, The Lighthouse Lab in Alderley Park and Alex Alderton, Roberto Amato, Jeffrey Barrett, Sonia Goncalves, Ewan Harrison, David K. Jackson, Ian Johnston, Dominic Kwiatkowski, Cordelia Langford, John Sillitoe on behalf of the Wellcome Sanger Institute COVID-19 Surveillance Team |
| EPI_ISL_2994567 | Berkshire and Surrey Pathology Services Lighthouse Laboratory | Wellcome Sanger Institute for the COVID-19 Genomics UK (COG-UK) Consortium | Berkshire and Surrey Pathology Services Lighthouse Laboratory and Alex Alderton, Roberto Amato, Jeffrey Barrett, Sonia Goncalves, Ewan Harrison, David K. Jackson, Ian Johnston, Dominic Kwiatkowski, Cordelia Langford, John Sillitoe on behalf of the Wellcome Sanger Institute COVID-19 Surveillance Team |
| EPI_ISL_2994568, EPI_ISL_2994569 | Lighthouse Lab in Milton Keynes | Wellcome Sanger Institute for the COVID-19 Genomics UK (COG-UK) Consortium | The Lighthouse Lab in Milton Keynes and Alex Alderton, Roberto Amato, Jeffrey Barrett, Sonia Goncalves, Ewan Harrison, David K. Jackson, Ian Johnston, Dominic Kwiatkowski, Cordelia Langford, John Sillitoe on behalf of the Wellcome Sanger Institute COVID-19 Surveillance Team |
| EPI_ISL_2994570 | Lighthouse Lab in Alderley Park | Wellcome Sanger Institute for the COVID-19 Genomics UK (COG-UK) Consortium | Jacquelyn Wynn, Mairead Hyland, The Lighthouse Lab in Alderley Park and Alex Alderton, Roberto Amato, Jeffrey Barrett, Sonia Goncalves, Ewan Harrison, David K. Jackson, Ian Johnston, Dominic Kwiatkowski, Cordelia Langford, John Sillitoe on behalf of the Wellcome Sanger Institute COVID-19 Surveillance Team |
| EPI_ISL_2994571 | Lighthouse Lab in Milton Keynes | Wellcome Sanger Institute for the COVID-19 Genomics UK (COG-UK) Consortium | The Lighthouse Lab in Milton Keynes and Alex Alderton, Roberto Amato, Jeffrey Barrett, Sonia Goncalves, Ewan Harrison, David K. Jackson, Ian Johnston, Dominic Kwiatkowski, Cordelia Langford, John Sillitoe on behalf of the Wellcome Sanger Institute COVID-19 Surveillance Team |
| EPI_ISL_2994572 | Lighthouse Lab in Glasgow | Wellcome Sanger Institute for the COVID-19 Genomics UK (COG-UK) Consortium | Harper VanSteenhouse, Yumi Kasai, David Gray, Carol Clugston, Anna Dominiczak and Alex Alderton, Roberto Amato, Jeffrey Barrett, Sonia Goncalves, Ewan Harrison, David K. Jackson, Ian Johnston, Dominic Kwiatkowski, Cordelia Langford, John Sillitoe on behalf of the Wellcome Sanger Institute COVID-19 Surveillance Team |
| EPI_ISL_2994573 | Lighthouse Lab in Milton Keynes | Wellcome Sanger Institute for the COVID-19 Genomics UK (COG-UK) Consortium | The Lighthouse Lab in Milton Keynes and Alex Alderton, Roberto Amato, Jeffrey Barrett, Sonia Goncalves, Ewan Harrison, David K. Jackson, Ian Johnston, Dominic Kwiatkowski, Cordelia Langford, John Sillitoe on behalf of the Wellcome Sanger Institute COVID-19 Surveillance Team |
| EPI_ISL_2994574 | Lighthouse Lab in Glasgow | Wellcome Sanger Institute for the COVID-19 Genomics UK (COG-UK) Consortium | Harper VanSteenhouse, Yumi Kasai, David Gray, Carol Clugston, Anna Dominiczak and Alex Alderton, Roberto Amato, Jeffrey Barrett, Sonia Goncalves, Ewan Harrison, David K. Jackson, Ian Johnston, Dominic Kwiatkowski, Cordelia Langford, John Sillitoe on behalf of the Wellcome Sanger Institute COVID-19 Surveillance Team |
| EPI_ISL_2994575, EPI_ISL_2994576 | Lighthouse Lab in Milton Keynes | Wellcome Sanger Institute for the COVID-19 Genomics UK (COG-UK) Consortium | The Lighthouse Lab in Milton Keynes and Alex Alderton, Roberto Amato, Jeffrey Barrett, Sonia Goncalves, Ewan Harrison, David K. Jackson, Ian Johnston, Dominic Kwiatkowski, Cordelia Langford, John Sillitoe on behalf of the Wellcome Sanger Institute COVID-19 Surveillance Team |
| EPI_ISL_2994577 | Berkshire and Surrey Pathology Services Lighthouse Laboratory | Wellcome Sanger Institute for the COVID-19 Genomics UK (COG-UK) Consortium | Berkshire and Surrey Pathology Services Lighthouse Laboratory and Alex Alderton, Roberto Amato, Jeffrey Barrett, Sonia Goncalves, Ewan Harrison, David K. Jackson, Ian Johnston, Dominic Kwiatkowski, Cordelia Langford, John Sillitoe on behalf of the Wellcome Sanger Institute COVID-19 Surveillance Team |
| EPI_ISL_2994578 | Lighthouse Lab in Milton Keynes | Wellcome Sanger Institute for the COVID-19 Genomics UK (COG-UK) Consortium | The Lighthouse Lab in Milton Keynes and Alex Alderton, Roberto Amato, Jeffrey Barrett, Sonia Goncalves, Ewan Harrison, David K. Jackson, Ian Johnston, Dominic Kwiatkowski, Cordelia Langford, John Sillitoe on behalf of the Wellcome Sanger Institute COVID-19 Surveillance Team |
| EPI_ISL_2994579, EPI_ISL_2994580 | Berkshire and Surrey Pathology Services Lighthouse Laboratory | Wellcome Sanger Institute for the COVID-19 Genomics UK (COG-UK) Consortium | Berkshire and Surrey Pathology Services Lighthouse Laboratory and Alex Alderton, Roberto Amato, Jeffrey Barrett, Sonia Goncalves, Ewan Harrison, David K. Jackson, Ian Johnston, Dominic Kwiatkowski, Cordelia Langford, John Sillitoe on behalf of the Wellcome Sanger Institute COVID-19 Surveillance Team |
| EPI_ISL_2994581 | Lighthouse Lab in Alderley Park | Wellcome Sanger Institute for the COVID-19 Genomics UK (COG-UK) Consortium | Jacquelyn Wynn, Mairead Hyland, The Lighthouse Lab in Alderley Park and Alex Alderton, Roberto Amato, Jeffrey Barrett, Sonia Goncalves, Ewan Harrison, David K. Jackson, Ian Johnston, Dominic Kwiatkowski, Cordelia Langford, John Sillitoe on behalf of the Wellcome Sanger Institute COVID-19 Surveillance Team |
| EPI_ISL_2994582 | Berkshire and Surrey Pathology Services Lighthouse Laboratory | Wellcome Sanger Institute for the COVID-19 Genomics UK (COG-UK) Consortium | Berkshire and Surrey Pathology Services Lighthouse Laboratory and Alex Alderton, Roberto Amato, Jeffrey Barrett, Sonia Goncalves, Ewan Harrison, David K. Jackson, Ian Johnston, Dominic Kwiatkowski, Cordelia Langford, John Sillitoe on behalf of the Wellcome Sanger Institute COVID-19 Surveillance Team |
| EPI_ISL_2994583, EPI_ISL_2994584 | Lighthouse Lab in Alderley Park | Wellcome Sanger Institute for the COVID-19 Genomics UK (COG-UK) Consortium | Jacquelyn Wynn, Mairead Hyland, The Lighthouse Lab in Alderley Park and Alex Alderton, Roberto Amato, Jeffrey Barrett, Sonia Goncalves, Ewan Harrison, David K. Jackson, Ian Johnston, Dominic Kwiatkowski, Cordelia Langford, John Sillitoe on behalf of the Wellcome Sanger Institute COVID-19 Surveillance Team |
| EPI_ISL_2994585, EPI_ISL_2994586 | Lighthouse Lab in Glasgow | Wellcome Sanger Institute for the COVID-19 Genomics UK (COG-UK) Consortium | Harper VanSteenhouse, Yumi Kasai, David Gray, Carol Clugston, Anna Dominiczak and Alex Alderton, Roberto Amato, Jeffrey Barrett, Sonia Goncalves, Ewan Harrison, David K. Jackson, Ian Johnston, Dominic Kwiatkowski, Cordelia Langford, John Sillitoe on behalf of the Wellcome Sanger Institute COVID-19 Surveillance Team |
| EPI_ISL_2994587, EPI_ISL_2994588, EPI_ISL_2994589, EPI_ISL_2994590, EPI_ISL_2994591 | Lighthouse Lab in Alderley Park | Wellcome Sanger Institute for the COVID-19 Genomics UK (COG-UK) Consortium | Jacquelyn Wynn, Mairead Hyland, The Lighthouse Lab in Alderley Park and Alex Alderton, Roberto Amato, Jeffrey Barrett, Sonia Goncalves, Ewan Harrison, David K. Jackson, Ian Johnston, Dominic Kwiatkowski, Cordelia Langford, John Sillitoe on behalf of the Wellcome Sanger Institute COVID-19 Surveillance Team |
| EPI_ISL_2994592, EPI_ISL_2994593 | Lighthouse Lab in Glasgow | Wellcome Sanger Institute for the COVID-19 Genomics UK (COG-UK) Consortium | Harper VanSteenhouse, Yumi Kasai, David Gray, Carol Clugston, Anna Dominiczak and Alex Alderton, Roberto Amato, Jeffrey Barrett, Sonia Goncalves, Ewan Harrison, David K. Jackson, Ian Johnston, Dominic Kwiatkowski, Cordelia Langford, John Sillitoe on behalf of the Wellcome Sanger Institute COVID-19 Surveillance Team |
| EPI_ISL_2994594 | Berkshire and Surrey Pathology Services Lighthouse Laboratory | Wellcome Sanger Institute for the COVID-19 Genomics UK (COG-UK) Consortium | Berkshire and Surrey Pathology Services Lighthouse Laboratory and Alex Alderton, Roberto Amato, Jeffrey Barrett, Sonia Goncalves, Ewan Harrison, David K. Jackson, Ian Johnston, Dominic Kwiatkowski, Cordelia Langford, John Sillitoe on behalf of the Wellcome Sanger Institute COVID-19 Surveillance Team |
| EPI_ISL_2994595, EPI_ISL_2994596 | Lighthouse Lab in Alderley Park | Wellcome Sanger Institute for the COVID-19 Genomics UK (COG-UK) Consortium | Jacquelyn Wynn, Mairead Hyland, The Lighthouse Lab in Alderley Park and Alex Alderton, Roberto Amato, Jeffrey Barrett, Sonia Goncalves, Ewan Harrison, David K. Jackson, Ian Johnston, Dominic Kwiatkowski, Cordelia Langford, John Sillitoe on behalf of the Wellcome Sanger Institute COVID-19 Surveillance Team |
| EPI_ISL_2994597 | Lighthouse Lab in Glasgow | Wellcome Sanger Institute for the COVID-19 Genomics UK (COG-UK) Consortium | Harper VanSteenhouse, Yumi Kasai, David Gray, Carol Clugston, Anna Dominiczak and Alex Alderton, Roberto Amato, Jeffrey Barrett, Sonia Goncalves, Ewan Harrison, David K. Jackson, Ian Johnston, Dominic Kwiatkowski, Cordelia Langford, John Sillitoe on behalf of the Wellcome Sanger Institute COVID-19 Surveillance Team |
| EPI_ISL_2994598 | Lighthouse Lab in Milton Keynes | Wellcome Sanger Institute for the COVID-19 Genomics UK (COG-UK) Consortium | The Lighthouse Lab in Milton Keynes and Alex Alderton, Roberto Amato, Jeffrey Barrett, Sonia Goncalves, Ewan Harrison, David K. Jackson, Ian Johnston, Dominic Kwiatkowski, Cordelia Langford, John Sillitoe on behalf of the Wellcome Sanger Institute COVID-19 Surveillance Team |
| EPI_ISL_2994599, EPI_ISL_2994600, EPI_ISL_2994601, EPI_ISL_2994602 | Lighthouse Lab in Glasgow | Wellcome Sanger Institute for the COVID-19 Genomics UK (COG-UK) Consortium | Harper VanSteenhouse, Yumi Kasai, David Gray, Carol Clugston, Anna Dominiczak and Alex Alderton, Roberto Amato, Jeffrey Barrett, Sonia Goncalves, Ewan Harrison, David K. Jackson, Ian Johnston, Dominic Kwiatkowski, Cordelia Langford, John Sillitoe on behalf of the Wellcome Sanger Institute COVID-19 Surveillance Team |
| EPI_ISL_2994603 | Lighthouse Lab in Milton Keynes | Wellcome Sanger Institute for the COVID-19 Genomics UK (COG-UK) | The Lighthouse Lab in Milton Keynes and Alex Alderton, Roberto Amato, Jeffrey Barrett, Sonia Goncalves, Ewan Harrison, David K. Jackson, Ian Johnston, Dominic Kwiatkowski, Cordelia Langford, John Sillitoe on behalf of the Wellcome Sanger |

| | | | |
|---|---|---|---|
| | | Consortium | Institute COVID-19 Surveillance Team |
| EPI_ISL_2994604, EPI_ISL_2994605 | Berkshire and Surrey Pathology Services Lighthouse Laboratory | Wellcome Sanger Institute for the COVID-19 Genomics UK (COG-UK) Consortium | Berkshire and Surrey Pathology Services Lighthouse Laboratory and Alex Alderton, Roberto Amato, Jeffrey Barrett, Sonia Goncalves, Ewan Harrison, David K. Jackson, Ian Johnston, Dominic Kwiatkowski, Cordelia Langford, John Sillitoe on behalf of the Wellcome Sanger Institute COVID-19 Surveillance Team |
| EPI_ISL_2994606 | Lighthouse Lab in Glasgow | Wellcome Sanger Institute for the COVID-19 Genomics UK (COG-UK) Consortium | Harper VanSteenhouse, Yumi Kasai, David Gray, Carol Clugston, Anna Dominiczak and Alex Alderton, Roberto Amato, Jeffrey Barrett, Sonia Goncalves, Ewan Harrison, David K. Jackson, Ian Johnston, Dominic Kwiatkowski, Cordelia Langford, John Sillitoe on behalf of the Wellcome Sanger Institute COVID-19 Surveillance Team |
| EPI_ISL_2994607, EPI_ISL_2994608, EPI_ISL_2994609 | Lighthouse Lab in Milton Keynes | Wellcome Sanger Institute for the COVID-19 Genomics UK (COG-UK) Consortium | The Lighthouse Lab in Milton Keynes and Alex Alderton, Roberto Amato, Jeffrey Barrett, Sonia Goncalves, Ewan Harrison, David K. Jackson, Ian Johnston, Dominic Kwiatkowski, Cordelia Langford, John Sillitoe on behalf of the Wellcome Sanger Institute COVID-19 Surveillance Team |
| EPI_ISL_2994610 | Berkshire and Surrey Pathology Services Lighthouse Laboratory | Wellcome Sanger Institute for the COVID-19 Genomics UK (COG-UK) Consortium | Berkshire and Surrey Pathology Services Lighthouse Laboratory and Alex Alderton, Roberto Amato, Jeffrey Barrett, Sonia Goncalves, Ewan Harrison, David K. Jackson, Ian Johnston, Dominic Kwiatkowski, Cordelia Langford, John Sillitoe on behalf of the Wellcome Sanger Institute COVID-19 Surveillance Team |
| EPI_ISL_2994611, EPI_ISL_2994612, EPI_ISL_2994613, EPI_ISL_2994614, EPI_ISL_2994615, EPI_ISL_2994616 | Lighthouse Lab in Milton Keynes | Wellcome Sanger Institute for the COVID-19 Genomics UK (COG-UK) Consortium | The Lighthouse Lab in Milton Keynes and Alex Alderton, Roberto Amato, Jeffrey Barrett, Sonia Goncalves, Ewan Harrison, David K. Jackson, Ian Johnston, Dominic Kwiatkowski, Cordelia Langford, John Sillitoe on behalf of the Wellcome Sanger Institute COVID-19 Surveillance Team |
| EPI_ISL_2994617, EPI_ISL_2994618, EPI_ISL_2994619 | Lighthouse Lab in Alderley Park | Wellcome Sanger Institute for the COVID-19 Genomics UK (COG-UK) Consortium | Jacquelyn Wynn, Mairead Hyland, The Lighthouse Lab in Alderley Park and Alex Alderton, Roberto Amato, Jeffrey Barrett, Sonia Goncalves, Ewan Harrison, David K. Jackson, Ian Johnston, Dominic Kwiatkowski, Cordelia Langford, John Sillitoe on behalf of the Wellcome Sanger Institute COVID-19 Surveillance Team |
| EPI_ISL_2994620, EPI_ISL_2994621 | Lighthouse Lab in Glasgow | Wellcome Sanger Institute for the COVID-19 Genomics UK (COG-UK) Consortium | Harper VanSteenhouse, Yumi Kasai, David Gray, Carol Clugston, Anna Dominiczak and Alex Alderton, Roberto Amato, Jeffrey Barrett, Sonia Goncalves, Ewan Harrison, David K. Jackson, Ian Johnston, Dominic Kwiatkowski, Cordelia Langford, John Sillitoe on behalf of the Wellcome Sanger Institute COVID-19 Surveillance Team |
| EPI_ISL_2994622 | Lighthouse Lab in Milton Keynes | Wellcome Sanger Institute for the COVID-19 Genomics UK (COG-UK) Consortium | The Lighthouse Lab in Milton Keynes and Alex Alderton, Roberto Amato, Jeffrey Barrett, Sonia Goncalves, Ewan Harrison, David K. Jackson, Ian Johnston, Dominic Kwiatkowski, Cordelia Langford, John Sillitoe on behalf of the Wellcome Sanger Institute COVID-19 Surveillance Team |
| EPI_ISL_2994623 | Lighthouse Lab in Glasgow | Wellcome Sanger Institute for the COVID-19 Genomics UK (COG-UK) Consortium | Harper VanSteenhouse, Yumi Kasai, David Gray, Carol Clugston, Anna Dominiczak and Alex Alderton, Roberto Amato, Jeffrey Barrett, Sonia Goncalves, Ewan Harrison, David K. Jackson, Ian Johnston, Dominic Kwiatkowski, Cordelia Langford, John Sillitoe on behalf of the Wellcome Sanger Institute COVID-19 Surveillance Team |
| EPI_ISL_2994624 | Lighthouse Lab in Alderley Park | Wellcome Sanger Institute for the COVID-19 Genomics UK (COG-UK) Consortium | Jacquelyn Wynn, Mairead Hyland, The Lighthouse Lab in Alderley Park and Alex Alderton, Roberto Amato, Jeffrey Barrett, Sonia Goncalves, Ewan Harrison, David K. Jackson, Ian Johnston, Dominic Kwiatkowski, Cordelia Langford, John Sillitoe on behalf of the Wellcome Sanger Institute COVID-19 Surveillance Team |
| EPI_ISL_2994625, EPI_ISL_2994626 | Lighthouse Lab in Glasgow | Wellcome Sanger Institute for the COVID-19 Genomics UK (COG-UK) Consortium | Harper VanSteenhouse, Yumi Kasai, David Gray, Carol Clugston, Anna Dominiczak and Alex Alderton, Roberto Amato, Jeffrey Barrett, Sonia Goncalves, Ewan Harrison, David K. Jackson, Ian Johnston, Dominic Kwiatkowski, Cordelia Langford, John Sillitoe on behalf of the Wellcome Sanger Institute COVID-19 Surveillance Team |
| EPI_ISL_2994627 | Lighthouse Lab in Alderley Park | Wellcome Sanger Institute for the COVID-19 Genomics UK (COG-UK) Consortium | Jacquelyn Wynn, Mairead Hyland, The Lighthouse Lab in Alderley Park and Alex Alderton, Roberto Amato, Jeffrey Barrett, Sonia Goncalves, Ewan Harrison, David K. Jackson, Ian Johnston, Dominic Kwiatkowski, Cordelia Langford, John Sillitoe on behalf of the Wellcome Sanger Institute COVID-19 Surveillance Team |
| EPI_ISL_2994628 | Lighthouse Lab in Milton Keynes | Wellcome Sanger Institute for the COVID-19 Genomics UK (COG-UK) Consortium | The Lighthouse Lab in Milton Keynes and Alex Alderton, Roberto Amato, Jeffrey Barrett, Sonia Goncalves, Ewan Harrison, David K. Jackson, Ian Johnston, Dominic Kwiatkowski, Cordelia Langford, John Sillitoe on behalf of the Wellcome Sanger Institute COVID-19 Surveillance Team |
| EPI_ISL_2994629 | Berkshire and Surrey Pathology Services Lighthouse Laboratory | Wellcome Sanger Institute for the COVID-19 Genomics UK (COG-UK) Consortium | Berkshire and Surrey Pathology Services Lighthouse Laboratory and Alex Alderton, Roberto Amato, Jeffrey Barrett, Sonia Goncalves, Ewan Harrison, David K. Jackson, Ian Johnston, Dominic Kwiatkowski, Cordelia Langford, John Sillitoe on behalf of the Wellcome Sanger Institute COVID-19 Surveillance Team |
| EPI_ISL_2994630, EPI_ISL_2994631 | Lighthouse Lab in Alderley Park | Wellcome Sanger Institute for the COVID-19 Genomics UK (COG-UK) Consortium | Jacquelyn Wynn, Mairead Hyland, The Lighthouse Lab in Alderley Park and Alex Alderton, Roberto Amato, Jeffrey Barrett, Sonia Goncalves, Ewan Harrison, David K. Jackson, Ian Johnston, Dominic Kwiatkowski, Cordelia Langford, John Sillitoe on behalf of the Wellcome Sanger Institute COVID-19 Surveillance Team |
| EPI_ISL_2994632 | Lighthouse Lab in Milton Keynes | Wellcome Sanger Institute for the COVID-19 Genomics UK (COG-UK) Consortium | The Lighthouse Lab in Milton Keynes and Alex Alderton, Roberto Amato, Jeffrey Barrett, Sonia Goncalves, Ewan Harrison, David K. Jackson, Ian Johnston, Dominic Kwiatkowski, Cordelia Langford, John Sillitoe on behalf of the Wellcome Sanger Institute COVID-19 Surveillance Team |
| EPI_ISL_2994633, EPI_ISL_2994634 | Lighthouse Lab in Glasgow | Wellcome Sanger Institute for the COVID-19 Genomics UK (COG-UK) Consortium | Harper VanSteenhouse, Yumi Kasai, David Gray, Carol Clugston, Anna Dominiczak and Alex Alderton, Roberto Amato, Jeffrey Barrett, Sonia Goncalves, Ewan Harrison, David K. Jackson, Ian Johnston, Dominic Kwiatkowski, Cordelia Langford, John Sillitoe on behalf of the Wellcome Sanger Institute COVID-19 Surveillance Team |
| EPI_ISL_2994635 | Berkshire and Surrey Pathology Services Lighthouse Laboratory | Wellcome Sanger Institute for the COVID-19 Genomics UK (COG-UK) Consortium | Berkshire and Surrey Pathology Services Lighthouse Laboratory and Alex Alderton, Roberto Amato, Jeffrey Barrett, Sonia Goncalves, Ewan Harrison, David K. Jackson, Ian Johnston, Dominic Kwiatkowski, Cordelia Langford, John Sillitoe on behalf of the Wellcome Sanger Institute COVID-19 Surveillance Team |
| EPI_ISL_2994636 | Lighthouse Lab in Alderley Park | Wellcome Sanger Institute for the COVID-19 Genomics UK (COG-UK) Consortium | Jacquelyn Wynn, Mairead Hyland, The Lighthouse Lab in Alderley Park and Alex Alderton, Roberto Amato, Jeffrey Barrett, Sonia Goncalves, Ewan Harrison, David K. Jackson, Ian Johnston, Dominic Kwiatkowski, Cordelia Langford, John Sillitoe on behalf of the Wellcome Sanger Institute COVID-19 Surveillance Team |
| EPI_ISL_2994637, EPI_ISL_2994638, EPI_ISL_2994639 | Lighthouse Lab in Glasgow | Wellcome Sanger Institute for the COVID-19 Genomics UK (COG-UK) Consortium | Harper VanSteenhouse, Yumi Kasai, David Gray, Carol Clugston, Anna Dominiczak and Alex Alderton, Roberto Amato, Jeffrey Barrett, Sonia Goncalves, Ewan Harrison, David K. Jackson, Ian Johnston, Dominic Kwiatkowski, Cordelia Langford, John Sillitoe on behalf of the Wellcome Sanger Institute COVID-19 Surveillance Team |
| EPI_ISL_2994640 | Berkshire and Surrey Pathology Services Lighthouse Laboratory | Wellcome Sanger Institute for the COVID-19 Genomics UK (COG-UK) Consortium | Berkshire and Surrey Pathology Services Lighthouse Laboratory and Alex Alderton, Roberto Amato, Jeffrey Barrett, Sonia Goncalves, Ewan Harrison, David K. Jackson, Ian Johnston, Dominic Kwiatkowski, Cordelia Langford, John Sillitoe on behalf of the Wellcome Sanger Institute COVID-19 Surveillance Team |
| EPI_ISL_2994641 | Lighthouse Lab in Alderley Park | Wellcome Sanger Institute for the COVID-19 Genomics UK (COG-UK) Consortium | Jacquelyn Wynn, Mairead Hyland, The Lighthouse Lab in Alderley Park and Alex Alderton, Roberto Amato, Jeffrey Barrett, Sonia Goncalves, Ewan Harrison, David K. Jackson, Ian Johnston, Dominic Kwiatkowski, Cordelia Langford, John Sillitoe on behalf of the Wellcome Sanger Institute COVID-19 Surveillance Team |
| EPI_ISL_2994642, EPI_ISL_2994643, EPI_ISL_2994644 | Lighthouse Lab in Milton Keynes | Wellcome Sanger Institute for the COVID-19 Genomics UK (COG-UK) Consortium | The Lighthouse Lab in Milton Keynes and Alex Alderton, Roberto Amato, Jeffrey Barrett, Sonia Goncalves, Ewan Harrison, David K. Jackson, Ian Johnston, Dominic Kwiatkowski, Cordelia Langford, John Sillitoe on behalf of the Wellcome Sanger Institute COVID-19 Surveillance Team |
| EPI_ISL_2994645, EPI_ISL_2994646 | Lighthouse Lab in Glasgow | Wellcome Sanger Institute for the COVID-19 Genomics UK (COG-UK) Consortium | Harper VanSteenhouse, Yumi Kasai, David Gray, Carol Clugston, Anna Dominiczak and Alex Alderton, Roberto Amato, Jeffrey Barrett, Sonia Goncalves, Ewan Harrison, David K. Jackson, Ian Johnston, Dominic Kwiatkowski, Cordelia Langford, John Sillitoe on behalf of the Wellcome Sanger Institute COVID-19 Surveillance Team |
| EPI_ISL_2994647 | Berkshire and Surrey Pathology Services Lighthouse Laboratory | Wellcome Sanger Institute for the COVID-19 Genomics UK (COG-UK) Consortium | Berkshire and Surrey Pathology Services Lighthouse Laboratory and Alex Alderton, Roberto Amato, Jeffrey Barrett, Sonia Goncalves, Ewan Harrison, David K. Jackson, Ian Johnston, Dominic Kwiatkowski, Cordelia Langford, John Sillitoe on behalf of the Wellcome Sanger Institute COVID-19 Surveillance Team |
| EPI_ISL_2994648, EPI_ISL_2994649 | Lighthouse Lab in Milton Keynes | Wellcome Sanger Institute for the COVID-19 Genomics UK (COG-UK) Consortium | The Lighthouse Lab in Milton Keynes and Alex Alderton, Roberto Amato, Jeffrey Barrett, Sonia Goncalves, Ewan Harrison, David K. Jackson, Ian Johnston, Dominic Kwiatkowski, Cordelia Langford, John Sillitoe on behalf of the Wellcome Sanger Institute COVID-19 Surveillance Team |
| EPI_ISL_2994650 | Lighthouse Lab in Glasgow | Wellcome Sanger Institute for the COVID-19 Genomics UK (COG-UK) Consortium | Harper VanSteenhouse, Yumi Kasai, David Gray, Carol Clugston, Anna Dominiczak and Alex Alderton, Roberto Amato, Jeffrey Barrett, Sonia Goncalves, Ewan Harrison, David K. Jackson, Ian Johnston, Dominic Kwiatkowski, Cordelia Langford, John Sillitoe on behalf of the Wellcome Sanger Institute COVID-19 Surveillance Team |
| EPI_ISL_2994651 | Lighthouse Lab in Milton Keynes | Wellcome Sanger Institute for the COVID-19 Genomics UK (COG-UK) Consortium | The Lighthouse Lab in Milton Keynes and Alex Alderton, Roberto Amato, Jeffrey Barrett, Sonia Goncalves, Ewan Harrison, David K. Jackson, Ian Johnston, Dominic Kwiatkowski, Cordelia Langford, John Sillitoe on behalf of the Wellcome Sanger Institute COVID-19 Surveillance Team |
| EPI_ISL_2994652 | Lighthouse Lab in Alderley Park | Wellcome Sanger Institute for the COVID-19 Genomics UK (COG-UK) Consortium | Jacquelyn Wynn, Mairead Hyland, The Lighthouse Lab in Alderley Park and Alex Alderton, Roberto Amato, Jeffrey Barrett, Sonia Goncalves, Ewan Harrison, David K. Jackson, Ian Johnston, Dominic Kwiatkowski, Cordelia Langford, John Sillitoe on behalf of the Wellcome Sanger Institute COVID-19 Surveillance Team |
| EPI_ISL_2994653 | Berkshire and Surrey Pathology Services Lighthouse Laboratory | Wellcome Sanger Institute for the COVID-19 Genomics UK (COG-UK) Consortium | Berkshire and Surrey Pathology Services Lighthouse Laboratory and Alex Alderton, Roberto Amato, Jeffrey Barrett, Sonia Goncalves, Ewan Harrison, David K. Jackson, Ian Johnston, Dominic Kwiatkowski, Cordelia Langford, John Sillitoe on behalf of the Wellcome Sanger Institute COVID-19 Surveillance Team |
| EPI_ISL_2994654 | Lighthouse Lab in Glasgow | Wellcome Sanger Institute for the COVID-19 Genomics UK (COG-UK) Consortium | Harper VanSteenhouse, Yumi Kasai, David Gray, Carol Clugston, Anna Dominiczak and Alex Alderton, Roberto Amato, Jeffrey Barrett, Sonia Goncalves, Ewan Harrison, David K. Jackson, Ian Johnston, Dominic Kwiatkowski, Cordelia Langford, John Sillitoe on behalf of the Wellcome Sanger Institute COVID-19 Surveillance Team |
| EPI_ISL_2994655 | Berkshire and Surrey Pathology Services Lighthouse Laboratory | Wellcome Sanger Institute for the COVID-19 Genomics UK (COG-UK) Consortium | Berkshire and Surrey Pathology Services Lighthouse Laboratory and Alex Alderton, Roberto Amato, Jeffrey Barrett, Sonia Goncalves, Ewan Harrison, David K. Jackson, Ian Johnston, Dominic Kwiatkowski, Cordelia Langford, John Sillitoe on behalf of the Wellcome Sanger Institute COVID-19 Surveillance Team |
| EPI_ISL_2994656 | Lighthouse Lab in Alderley Park | Wellcome Sanger Institute for the COVID-19 Genomics UK (COG-UK) Consortium | Jacquelyn Wynn, Mairead Hyland, The Lighthouse Lab in Alderley Park and Alex Alderton, Roberto Amato, Jeffrey Barrett, Sonia Goncalves, Ewan Harrison, David K. Jackson, Ian Johnston, Dominic Kwiatkowski, Cordelia Langford, John Sillitoe on behalf of the Wellcome Sanger Institute COVID-19 Surveillance Team |
| EPI_ISL_2994657, EPI_ISL_2994658 | Berkshire and Surrey Pathology Services Lighthouse Laboratory | Wellcome Sanger Institute for the COVID-19 Genomics UK (COG-UK) Consortium | Berkshire and Surrey Pathology Services Lighthouse Laboratory and Alex Alderton, Roberto Amato, Jeffrey Barrett, Sonia Goncalves, Ewan Harrison, David K. Jackson, Ian Johnston, Dominic Kwiatkowski, Cordelia Langford, John Sillitoe on behalf of the Wellcome Sanger Institute COVID-19 Surveillance Team |
| EPI_ISL_2994659 | Lighthouse Lab in Milton Keynes | Wellcome Sanger Institute for the COVID-19 Genomics UK (COG-UK) Consortium | The Lighthouse Lab in Milton Keynes and Alex Alderton, Roberto Amato, Jeffrey Barrett, Sonia Goncalves, Ewan Harrison, David K. Jackson, Ian Johnston, Dominic Kwiatkowski, Cordelia Langford, John Sillitoe on behalf of the Wellcome Sanger Institute COVID-19 Surveillance Team |
| EPI_ISL_2994660 | Lighthouse Lab in Glasgow | Wellcome Sanger Institute for the COVID-19 Genomics UK (COG-UK) Consortium | Harper VanSteenhouse, Yumi Kasai, David Gray, Carol Clugston, Anna Dominiczak and Alex Alderton, Roberto Amato, Jeffrey Barrett, Sonia Goncalves, Ewan Harrison, David K. Jackson, Ian Johnston, Dominic Kwiatkowski, Cordelia Langford, John Sillitoe on behalf of the Wellcome Sanger Institute COVID-19 Surveillance Team |
| EPI_ISL_2994661 | Berkshire and Surrey Pathology Services Lighthouse Laboratory | Wellcome Sanger Institute for the COVID-19 Genomics UK (COG-UK) Consortium | Berkshire and Surrey Pathology Services Lighthouse Laboratory and Alex Alderton, Roberto Amato, Jeffrey Barrett, Sonia Goncalves, Ewan Harrison, David K. Jackson, Ian Johnston, Dominic Kwiatkowski, Cordelia Langford, John Sillitoe on behalf of the Wellcome Sanger Institute COVID-19 Surveillance Team |
| EPI_ISL_2994662, EPI_ISL_2994663, EPI_ISL_2994664, EPI_ISL_2994665 | Lighthouse Lab in Alderley Park | Wellcome Sanger Institute for the COVID-19 Genomics UK (COG-UK) Consortium | Jacquelyn Wynn, Mairead Hyland, The Lighthouse Lab in Alderley Park and Alex Alderton, Roberto Amato, Jeffrey Barrett, Sonia Goncalves, Ewan Harrison, David K. Jackson, Ian Johnston, Dominic Kwiatkowski, Cordelia Langford, John Sillitoe on behalf of the Wellcome Sanger Institute COVID-19 Surveillance Team |
| EPI_ISL_2994666 | Lighthouse Laboratory Plymouth | Wellcome Sanger Institute for the COVID-19 Genomics UK (COG-UK) Consortium | Lighthouse Laboratory Plymouth and Alex Alderton, Roberto Amato, Jeffrey Barrett, Sonia Goncalves, Ewan Harrison, David K. Jackson, Ian Johnston, Dominic Kwiatkowski, Cordelia Langford, John Sillitoe on behalf of the Wellcome Sanger Institute COVID-19 Surveillance Team |
| EPI_ISL_2994667 | Lighthouse Lab in Milton Keynes | Wellcome Sanger Institute for the COVID-19 Genomics UK (COG-UK) Consortium | The Lighthouse Lab in Milton Keynes and Alex Alderton, Roberto Amato, Jeffrey Barrett, Sonia Goncalves, Ewan Harrison, David K. Jackson, Ian Johnston, Dominic Kwiatkowski, Cordelia Langford, John Sillitoe on behalf of the Wellcome Sanger Institute COVID-19 Surveillance Team |
| EPI_ISL_2994668, | Lighthouse Laboratory Plymouth | Wellcome Sanger Institute for the COVID-19 Genomics UK (COG-UK) | Lighthouse Laboratory Plymouth and Alex Alderton, Roberto Amato, Jeffrey Barrett, Sonia Goncalves, Ewan Harrison, David K. Jackson, Ian Johnston, Dominic Kwiatkowski, Cordelia Langford, John Sillitoe on behalf of the Wellcome Sanger |

| | | | |
|---|---|---|---|
| EPI_ISL_2994669, EPI_ISL_2994670, EPI_ISL_2994671, EPI_ISL_2994672, EPI_ISL_2994673 | | Consortium | Institute COVID-19 Surveillance Team |
| EPI_ISL_2994674 | Lighthouse Lab in Milton Keynes | Wellcome Sanger Institute for the COVID-19 Genomics UK (COG-UK) Consortium | The Lighthouse Lab in Milton Keynes and Alex Alderton, Roberto Amato, Jeffrey Barrett, Sonia Goncalves, Ewan Harrison, David K. Jackson, Ian Johnston, Dominic Kwiatkowski, Cordelia Langford, John Sillitoe on behalf of the Wellcome Sanger Institute COVID-19 Surveillance Team |
| EPI_ISL_2994675 | Lighthouse Lab in Glasgow | Wellcome Sanger Institute for the COVID-19 Genomics UK (COG-UK) Consortium | Harper VanSteenhouse, Yumi Kasai, David Gray, Carol Clugston, Anna Dominiczak and Alex Alderton, Roberto Amato, Jeffrey Barrett, Sonia Goncalves, Ewan Harrison, David K. Jackson, Ian Johnston, Dominic Kwiatkowski, Cordelia Langford, John Sillitoe on behalf of the Wellcome Sanger Institute COVID-19 Surveillance Team |
| EPI_ISL_2994676, EPI_ISL_2994677, EPI_ISL_2994678, EPI_ISL_2994679, EPI_ISL_2994680 | Lighthouse Laboratory Plymouth | Wellcome Sanger Institute for the COVID-19 Genomics UK (COG-UK) Consortium | Lighthouse Laboratory Plymouth and Alex Alderton, Roberto Amato, Jeffrey Barrett, Sonia Goncalves, Ewan Harrison, David K. Jackson, Ian Johnston, Dominic Kwiatkowski, Cordelia Langford, John Sillitoe on behalf of the Wellcome Sanger Institute COVID-19 Surveillance Team |
| EPI_ISL_2994681, EPI_ISL_2994682 | Lighthouse Lab in Glasgow | Wellcome Sanger Institute for the COVID-19 Genomics UK (COG-UK) Consortium | Harper VanSteenhouse, Yumi Kasai, David Gray, Carol Clugston, Anna Dominiczak and Alex Alderton, Roberto Amato, Jeffrey Barrett, Sonia Goncalves, Ewan Harrison, David K. Jackson, Ian Johnston, Dominic Kwiatkowski, Cordelia Langford, John Sillitoe on behalf of the Wellcome Sanger Institute COVID-19 Surveillance Team |
| EPI_ISL_2994683 | Lighthouse Laboratory Plymouth | Wellcome Sanger Institute for the COVID-19 Genomics UK (COG-UK) Consortium | Lighthouse Laboratory Plymouth and Alex Alderton, Roberto Amato, Jeffrey Barrett, Sonia Goncalves, Ewan Harrison, David K. Jackson, Ian Johnston, Dominic Kwiatkowski, Cordelia Langford, John Sillitoe on behalf of the Wellcome Sanger Institute COVID-19 Surveillance Team |
| EPI_ISL_2994684 | Lighthouse Lab in Glasgow | Wellcome Sanger Institute for the COVID-19 Genomics UK (COG-UK) Consortium | Harper VanSteenhouse, Yumi Kasai, David Gray, Carol Clugston, Anna Dominiczak and Alex Alderton, Roberto Amato, Jeffrey Barrett, Sonia Goncalves, Ewan Harrison, David K. Jackson, Ian Johnston, Dominic Kwiatkowski, Cordelia Langford, John Sillitoe on behalf of the Wellcome Sanger Institute COVID-19 Surveillance Team |
| EPI_ISL_2994685 | Lighthouse Lab in Milton Keynes | Wellcome Sanger Institute for the COVID-19 Genomics UK (COG-UK) Consortium | The Lighthouse Lab in Milton Keynes and Alex Alderton, Roberto Amato, Jeffrey Barrett, Sonia Goncalves, Ewan Harrison, David K. Jackson, Ian Johnston, Dominic Kwiatkowski, Cordelia Langford, John Sillitoe on behalf of the Wellcome Sanger Institute COVID-19 Surveillance Team |
| EPI_ISL_2994686 | Lighthouse Laboratory Plymouth | Wellcome Sanger Institute for the COVID-19 Genomics UK (COG-UK) Consortium | Lighthouse Laboratory Plymouth and Alex Alderton, Roberto Amato, Jeffrey Barrett, Sonia Goncalves, Ewan Harrison, David K. Jackson, Ian Johnston, Dominic Kwiatkowski, Cordelia Langford, John Sillitoe on behalf of the Wellcome Sanger Institute COVID-19 Surveillance Team |
| EPI_ISL_2994687, EPI_ISL_2994688 | Lighthouse Lab in Milton Keynes | Wellcome Sanger Institute for the COVID-19 Genomics UK (COG-UK) Consortium | The Lighthouse Lab in Milton Keynes and Alex Alderton, Roberto Amato, Jeffrey Barrett, Sonia Goncalves, Ewan Harrison, David K. Jackson, Ian Johnston, Dominic Kwiatkowski, Cordelia Langford, John Sillitoe on behalf of the Wellcome Sanger Institute COVID-19 Surveillance Team |
| EPI_ISL_2994689 | Lighthouse Laboratory Plymouth | Wellcome Sanger Institute for the COVID-19 Genomics UK (COG-UK) Consortium | Lighthouse Laboratory Plymouth and Alex Alderton, Roberto Amato, Jeffrey Barrett, Sonia Goncalves, Ewan Harrison, David K. Jackson, Ian Johnston, Dominic Kwiatkowski, Cordelia Langford, John Sillitoe on behalf of the Wellcome Sanger Institute COVID-19 Surveillance Team |
| EPI_ISL_2994690 | Lighthouse Lab in Milton Keynes | Wellcome Sanger Institute for the COVID-19 Genomics UK (COG-UK) Consortium | The Lighthouse Lab in Milton Keynes and Alex Alderton, Roberto Amato, Jeffrey Barrett, Sonia Goncalves, Ewan Harrison, David K. Jackson, Ian Johnston, Dominic Kwiatkowski, Cordelia Langford, John Sillitoe on behalf of the Wellcome Sanger Institute COVID-19 Surveillance Team |
| EPI_ISL_2994691 | Lighthouse Lab in Glasgow | Wellcome Sanger Institute for the COVID-19 Genomics UK (COG-UK) Consortium | Harper VanSteenhouse, Yumi Kasai, David Gray, Carol Clugston, Anna Dominiczak and Alex Alderton, Roberto Amato, Jeffrey Barrett, Sonia Goncalves, Ewan Harrison, David K. Jackson, Ian Johnston, Dominic Kwiatkowski, Cordelia Langford, John Sillitoe on behalf of the Wellcome Sanger Institute COVID-19 Surveillance Team |
| EPI_ISL_2994692, EPI_ISL_2994693 | Lighthouse Laboratory Plymouth | Wellcome Sanger Institute for the COVID-19 Genomics UK (COG-UK) Consortium | Lighthouse Laboratory Plymouth and Alex Alderton, Roberto Amato, Jeffrey Barrett, Sonia Goncalves, Ewan Harrison, David K. Jackson, Ian Johnston, Dominic Kwiatkowski, Cordelia Langford, John Sillitoe on behalf of the Wellcome Sanger Institute COVID-19 Surveillance Team |
| EPI_ISL_2994694 | Lighthouse Lab in Milton Keynes | Wellcome Sanger Institute for the COVID-19 Genomics UK (COG-UK) Consortium | The Lighthouse Lab in Milton Keynes and Alex Alderton, Roberto Amato, Jeffrey Barrett, Sonia Goncalves, Ewan Harrison, David K. Jackson, Ian Johnston, Dominic Kwiatkowski, Cordelia Langford, John Sillitoe on behalf of the Wellcome Sanger Institute COVID-19 Surveillance Team |
| EPI_ISL_2994695, EPI_ISL_2994696 | Lighthouse Lab in Glasgow | Wellcome Sanger Institute for the COVID-19 Genomics UK (COG-UK) Consortium | Harper VanSteenhouse, Yumi Kasai, David Gray, Carol Clugston, Anna Dominiczak and Alex Alderton, Roberto Amato, Jeffrey Barrett, Sonia Goncalves, Ewan Harrison, David K. Jackson, Ian Johnston, Dominic Kwiatkowski, Cordelia Langford, John Sillitoe on behalf of the Wellcome Sanger Institute COVID-19 Surveillance Team |
| EPI_ISL_2994697 | Lighthouse Laboratory Plymouth | Wellcome Sanger Institute for the COVID-19 Genomics UK (COG-UK) Consortium | Lighthouse Laboratory Plymouth and Alex Alderton, Roberto Amato, Jeffrey Barrett, Sonia Goncalves, Ewan Harrison, David K. Jackson, Ian Johnston, Dominic Kwiatkowski, Cordelia Langford, John Sillitoe on behalf of the Wellcome Sanger Institute COVID-19 Surveillance Team |
| EPI_ISL_2994698, EPI_ISL_2994699, EPI_ISL_2994700, EPI_ISL_2994701 | Lighthouse Lab in Glasgow | Wellcome Sanger Institute for the COVID-19 Genomics UK (COG-UK) Consortium | Harper VanSteenhouse, Yumi Kasai, David Gray, Carol Clugston, Anna Dominiczak and Alex Alderton, Roberto Amato, Jeffrey Barrett, Sonia Goncalves, Ewan Harrison, David K. Jackson, Ian Johnston, Dominic Kwiatkowski, Cordelia Langford, John Sillitoe on behalf of the Wellcome Sanger Institute COVID-19 Surveillance Team |
| EPI_ISL_2994702 | Lighthouse Laboratory Plymouth | Wellcome Sanger Institute for the COVID-19 Genomics UK (COG-UK) Consortium | Lighthouse Laboratory Plymouth and Alex Alderton, Roberto Amato, Jeffrey Barrett, Sonia Goncalves, Ewan Harrison, David K. Jackson, Ian Johnston, Dominic Kwiatkowski, Cordelia Langford, John Sillitoe on behalf of the Wellcome Sanger Institute COVID-19 Surveillance Team |
| EPI_ISL_2994703, EPI_ISL_2994704, EPI_ISL_2994705, EPI_ISL_2994706, EPI_ISL_2994707 | Lighthouse Lab in Milton Keynes | Wellcome Sanger Institute for the COVID-19 Genomics UK (COG-UK) Consortium | The Lighthouse Lab in Milton Keynes and Alex Alderton, Roberto Amato, Jeffrey Barrett, Sonia Goncalves, Ewan Harrison, David K. Jackson, Ian Johnston, Dominic Kwiatkowski, Cordelia Langford, John Sillitoe on behalf of the Wellcome Sanger Institute COVID-19 Surveillance Team |
| EPI_ISL_2994708, EPI_ISL_2994709, EPI_ISL_2994710, EPI_ISL_2994711, EPI_ISL_2994712 | Lighthouse Laboratory Plymouth | Wellcome Sanger Institute for the COVID-19 Genomics UK (COG-UK) Consortium | Lighthouse Laboratory Plymouth and Alex Alderton, Roberto Amato, Jeffrey Barrett, Sonia Goncalves, Ewan Harrison, David K. Jackson, Ian Johnston, Dominic Kwiatkowski, Cordelia Langford, John Sillitoe on behalf of the Wellcome Sanger Institute COVID-19 Surveillance Team |
| EPI_ISL_2994713 | Lighthouse Lab in Milton Keynes | Wellcome Sanger Institute for the COVID-19 Genomics UK (COG-UK) Consortium | The Lighthouse Lab in Milton Keynes and Alex Alderton, Roberto Amato, Jeffrey Barrett, Sonia Goncalves, Ewan Harrison, David K. Jackson, Ian Johnston, Dominic Kwiatkowski, Cordelia Langford, John Sillitoe on behalf of the Wellcome Sanger Institute COVID-19 Surveillance Team |
| EPI_ISL_2994714 | Lighthouse Lab in Glasgow | Wellcome Sanger Institute for the COVID-19 Genomics UK (COG-UK) Consortium | Harper VanSteenhouse, Yumi Kasai, David Gray, Carol Clugston, Anna Dominiczak and Alex Alderton, Roberto Amato, Jeffrey Barrett, Sonia Goncalves, Ewan Harrison, David K. Jackson, Ian Johnston, Dominic Kwiatkowski, Cordelia Langford, John Sillitoe on behalf of the Wellcome Sanger Institute COVID-19 Surveillance Team |
| EPI_ISL_2994715, EPI_ISL_2994716, EPI_ISL_2994717, EPI_ISL_2994718 | Lighthouse Laboratory Plymouth | Wellcome Sanger Institute for the COVID-19 Genomics UK (COG-UK) Consortium | Lighthouse Laboratory Plymouth and Alex Alderton, Roberto Amato, Jeffrey Barrett, Sonia Goncalves, Ewan Harrison, David K. Jackson, Ian Johnston, Dominic Kwiatkowski, Cordelia Langford, John Sillitoe on behalf of the Wellcome Sanger Institute COVID-19 Surveillance Team |
| EPI_ISL_2994719 | Lighthouse Lab in Glasgow | Wellcome Sanger Institute for the COVID-19 Genomics UK (COG-UK) Consortium | Harper VanSteenhouse, Yumi Kasai, David Gray, Carol Clugston, Anna Dominiczak and Alex Alderton, Roberto Amato, Jeffrey Barrett, Sonia Goncalves, Ewan Harrison, David K. Jackson, Ian Johnston, Dominic Kwiatkowski, Cordelia Langford, John Sillitoe on behalf of the Wellcome Sanger Institute COVID-19 Surveillance Team |
| EPI_ISL_2994720 | Lighthouse Laboratory Plymouth | Wellcome Sanger Institute for the COVID-19 Genomics UK (COG-UK) Consortium | Lighthouse Laboratory Plymouth and Alex Alderton, Roberto Amato, Jeffrey Barrett, Sonia Goncalves, Ewan Harrison, David K. Jackson, Ian Johnston, Dominic Kwiatkowski, Cordelia Langford, John Sillitoe on behalf of the Wellcome Sanger Institute COVID-19 Surveillance Team |
| EPI_ISL_2994721 | Lighthouse Lab in Glasgow | Wellcome Sanger Institute for the COVID-19 Genomics UK (COG-UK) Consortium | Harper VanSteenhouse, Yumi Kasai, David Gray, Carol Clugston, Anna Dominiczak and Alex Alderton, Roberto Amato, Jeffrey Barrett, Sonia Goncalves, Ewan Harrison, David K. Jackson, Ian Johnston, Dominic Kwiatkowski, Cordelia Langford, John Sillitoe on behalf of the Wellcome Sanger Institute COVID-19 Surveillance Team |
| EPI_ISL_2994722 | Lighthouse Laboratory Plymouth | Wellcome Sanger Institute for the COVID-19 Genomics UK (COG-UK) Consortium | Lighthouse Laboratory Plymouth and Alex Alderton, Roberto Amato, Jeffrey Barrett, Sonia Goncalves, Ewan Harrison, David K. Jackson, Ian Johnston, Dominic Kwiatkowski, Cordelia Langford, John Sillitoe on behalf of the Wellcome Sanger Institute COVID-19 Surveillance Team |
| EPI_ISL_2994723 | Lighthouse Lab in Glasgow | Wellcome Sanger Institute for the COVID-19 Genomics UK (COG-UK) Consortium | Harper VanSteenhouse, Yumi Kasai, David Gray, Carol Clugston, Anna Dominiczak and Alex Alderton, Roberto Amato, Jeffrey Barrett, Sonia Goncalves, Ewan Harrison, David K. Jackson, Ian Johnston, Dominic Kwiatkowski, Cordelia Langford, John Sillitoe on behalf of the Wellcome Sanger Institute COVID-19 Surveillance Team |
| EPI_ISL_2994724 | Lighthouse Laboratory Plymouth | Wellcome Sanger Institute for the COVID-19 Genomics UK (COG-UK) Consortium | Lighthouse Laboratory Plymouth and Alex Alderton, Roberto Amato, Jeffrey Barrett, Sonia Goncalves, Ewan Harrison, David K. Jackson, Ian Johnston, Dominic Kwiatkowski, Cordelia Langford, John Sillitoe on behalf of the Wellcome Sanger Institute COVID-19 Surveillance Team |
| EPI_ISL_2994725 | Lighthouse Lab in Milton Keynes | Wellcome Sanger Institute for the COVID-19 Genomics UK (COG-UK) Consortium | The Lighthouse Lab in Milton Keynes and Alex Alderton, Roberto Amato, Jeffrey Barrett, Sonia Goncalves, Ewan Harrison, David K. Jackson, Ian Johnston, Dominic Kwiatkowski, Cordelia Langford, John Sillitoe on behalf of the Wellcome Sanger Institute COVID-19 Surveillance Team |
| EPI_ISL_2994726 | Lighthouse Laboratory Plymouth | Wellcome Sanger Institute for the COVID-19 Genomics UK (COG-UK) Consortium | Lighthouse Laboratory Plymouth and Alex Alderton, Roberto Amato, Jeffrey Barrett, Sonia Goncalves, Ewan Harrison, David K. Jackson, Ian Johnston, Dominic Kwiatkowski, Cordelia Langford, John Sillitoe on behalf of the Wellcome Sanger Institute COVID-19 Surveillance Team |
| EPI_ISL_2994727 | Lighthouse Lab in Glasgow | Wellcome Sanger Institute for the COVID-19 Genomics UK (COG-UK) Consortium | Harper VanSteenhouse, Yumi Kasai, David Gray, Carol Clugston, Anna Dominiczak and Alex Alderton, Roberto Amato, Jeffrey Barrett, Sonia Goncalves, Ewan Harrison, David K. Jackson, Ian Johnston, Dominic Kwiatkowski, Cordelia Langford, John Sillitoe on behalf of the Wellcome Sanger Institute COVID-19 Surveillance Team |
| EPI_ISL_2994728 | Lighthouse Lab in Milton Keynes | Wellcome Sanger Institute for the COVID-19 Genomics UK (COG-UK) Consortium | The Lighthouse Lab in Milton Keynes and Alex Alderton, Roberto Amato, Jeffrey Barrett, Sonia Goncalves, Ewan Harrison, David K. Jackson, Ian Johnston, Dominic Kwiatkowski, Cordelia Langford, John Sillitoe on behalf of the Wellcome Sanger Institute COVID-19 Surveillance Team |
| EPI_ISL_2994729 | Lighthouse Lab in Glasgow | Wellcome Sanger Institute for the COVID-19 Genomics UK (COG-UK) Consortium | Harper VanSteenhouse, Yumi Kasai, David Gray, Carol Clugston, Anna Dominiczak and Alex Alderton, Roberto Amato, Jeffrey Barrett, Sonia Goncalves, Ewan Harrison, David K. Jackson, Ian Johnston, Dominic Kwiatkowski, Cordelia Langford, John Sillitoe on behalf of the Wellcome Sanger Institute COVID-19 Surveillance Team |
| EPI_ISL_2994730 | Lighthouse Laboratory Plymouth | Wellcome Sanger Institute for the COVID-19 Genomics UK (COG-UK) Consortium | Lighthouse Laboratory Plymouth and Alex Alderton, Roberto Amato, Jeffrey Barrett, Sonia Goncalves, Ewan Harrison, David K. Jackson, Ian Johnston, Dominic Kwiatkowski, Cordelia Langford, John Sillitoe on behalf of the Wellcome Sanger Institute COVID-19 Surveillance Team |
| EPI_ISL_2994731 | Lighthouse Lab in Milton Keynes | Wellcome Sanger Institute for the COVID-19 Genomics UK (COG-UK) Consortium | The Lighthouse Lab in Milton Keynes and Alex Alderton, Roberto Amato, Jeffrey Barrett, Sonia Goncalves, Ewan Harrison, David K. Jackson, Ian Johnston, Dominic Kwiatkowski, Cordelia Langford, John Sillitoe on behalf of the Wellcome Sanger Institute COVID-19 Surveillance Team |
| EPI_ISL_2994732, EPI_ISL_2994733, EPI_ISL_2994734 | Lighthouse Laboratory Plymouth | Wellcome Sanger Institute for the COVID-19 Genomics UK (COG-UK) Consortium | Lighthouse Laboratory Plymouth and Alex Alderton, Roberto Amato, Jeffrey Barrett, Sonia Goncalves, Ewan Harrison, David K. Jackson, Ian Johnston, Dominic Kwiatkowski, Cordelia Langford, John Sillitoe on behalf of the Wellcome Sanger Institute COVID-19 Surveillance Team |

| | | | |
|---|---|---|---|
| EPI_ISL_2994735, EPI_ISL_2994736 | Lighthouse Lab in Glasgow | Wellcome Sanger Institute for the COVID-19 Genomics UK (COG-UK) Consortium | Harper VanSteenhouse, Yumi Kasai, David Gray, Carol Clugston, Anna Dominiczak and Alex Alderton, Roberto Amato, Jeffrey Barrett, Sonia Goncalves, Ewan Harrison, David K. Jackson, Ian Johnston, Dominic Kwiatkowski, Cordelia Langford, John Sillitoe on behalf of the Wellcome Sanger Institute COVID-19 Surveillance Team |
| EPI_ISL_2994737 | Lighthouse Lab in Milton Keynes | Wellcome Sanger Institute for the COVID-19 Genomics UK (COG-UK) Consortium | The Lighthouse Lab in Milton Keynes and Alex Alderton, Roberto Amato, Jeffrey Barrett, Sonia Goncalves, Ewan Harrison, David K. Jackson, Ian Johnston, Dominic Kwiatkowski, Cordelia Langford, John Sillitoe on behalf of the Wellcome Sanger Institute COVID-19 Surveillance Team |
| EPI_ISL_2994738 | Lighthouse Laboratory Plymouth | Wellcome Sanger Institute for the COVID-19 Genomics UK (COG-UK) Consortium | Lighthouse Laboratory Plymouth and Alex Alderton, Roberto Amato, Jeffrey Barrett, Sonia Goncalves, Ewan Harrison, David K. Jackson, Ian Johnston, Dominic Kwiatkowski, Cordelia Langford, John Sillitoe on behalf of the Wellcome Sanger Institute COVID-19 Surveillance Team |
| EPI_ISL_2994739 | Lighthouse Lab in Glasgow | Wellcome Sanger Institute for the COVID-19 Genomics UK (COG-UK) Consortium | Harper VanSteenhouse, Yumi Kasai, David Gray, Carol Clugston, Anna Dominiczak and Alex Alderton, Roberto Amato, Jeffrey Barrett, Sonia Goncalves, Ewan Harrison, David K. Jackson, Ian Johnston, Dominic Kwiatkowski, Cordelia Langford, John Sillitoe on behalf of the Wellcome Sanger Institute COVID-19 Surveillance Team |
| EPI_ISL_2994740 | Lighthouse Laboratory Plymouth | Wellcome Sanger Institute for the COVID-19 Genomics UK (COG-UK) Consortium | Lighthouse Laboratory Plymouth and Alex Alderton, Roberto Amato, Jeffrey Barrett, Sonia Goncalves, Ewan Harrison, David K. Jackson, Ian Johnston, Dominic Kwiatkowski, Cordelia Langford, John Sillitoe on behalf of the Wellcome Sanger Institute COVID-19 Surveillance Team |
| EPI_ISL_2994741 | Lighthouse Lab in Milton Keynes | Wellcome Sanger Institute for the COVID-19 Genomics UK (COG-UK) Consortium | The Lighthouse Lab in Milton Keynes and Alex Alderton, Roberto Amato, Jeffrey Barrett, Sonia Goncalves, Ewan Harrison, David K. Jackson, Ian Johnston, Dominic Kwiatkowski, Cordelia Langford, John Sillitoe on behalf of the Wellcome Sanger Institute COVID-19 Surveillance Team |
| EPI_ISL_2994742 | Lighthouse Laboratory Plymouth | Wellcome Sanger Institute for the COVID-19 Genomics UK (COG-UK) Consortium | Lighthouse Laboratory Plymouth and Alex Alderton, Roberto Amato, Jeffrey Barrett, Sonia Goncalves, Ewan Harrison, David K. Jackson, Ian Johnston, Dominic Kwiatkowski, Cordelia Langford, John Sillitoe on behalf of the Wellcome Sanger Institute COVID-19 Surveillance Team |
| EPI_ISL_2994743 | Lighthouse Lab in Milton Keynes | Wellcome Sanger Institute for the COVID-19 Genomics UK (COG-UK) Consortium | The Lighthouse Lab in Milton Keynes and Alex Alderton, Roberto Amato, Jeffrey Barrett, Sonia Goncalves, Ewan Harrison, David K. Jackson, Ian Johnston, Dominic Kwiatkowski, Cordelia Langford, John Sillitoe on behalf of the Wellcome Sanger Institute COVID-19 Surveillance Team |
| EPI_ISL_2994744, EPI_ISL_2994745, EPI_ISL_2994746 | Lighthouse Laboratory Plymouth | Wellcome Sanger Institute for the COVID-19 Genomics UK (COG-UK) Consortium | Lighthouse Laboratory Plymouth and Alex Alderton, Roberto Amato, Jeffrey Barrett, Sonia Goncalves, Ewan Harrison, David K. Jackson, Ian Johnston, Dominic Kwiatkowski, Cordelia Langford, John Sillitoe on behalf of the Wellcome Sanger Institute COVID-19 Surveillance Team |
| EPI_ISL_2994747, EPI_ISL_2994748 | Lighthouse Lab in Glasgow | Wellcome Sanger Institute for the COVID-19 Genomics UK (COG-UK) Consortium | Harper VanSteenhouse, Yumi Kasai, David Gray, Carol Clugston, Anna Dominiczak and Alex Alderton, Roberto Amato, Jeffrey Barrett, Sonia Goncalves, Ewan Harrison, David K. Jackson, Ian Johnston, Dominic Kwiatkowski, Cordelia Langford, John Sillitoe on behalf of the Wellcome Sanger Institute COVID-19 Surveillance Team |
| EPI_ISL_2994749 | Lighthouse Laboratory Plymouth | Wellcome Sanger Institute for the COVID-19 Genomics UK (COG-UK) Consortium | Lighthouse Laboratory Plymouth and Alex Alderton, Roberto Amato, Jeffrey Barrett, Sonia Goncalves, Ewan Harrison, David K. Jackson, Ian Johnston, Dominic Kwiatkowski, Cordelia Langford, John Sillitoe on behalf of the Wellcome Sanger Institute COVID-19 Surveillance Team |
| EPI_ISL_2994750, EPI_ISL_2994751 | Lighthouse Lab in Milton Keynes | Wellcome Sanger Institute for the COVID-19 Genomics UK (COG-UK) Consortium | The Lighthouse Lab in Milton Keynes and Alex Alderton, Roberto Amato, Jeffrey Barrett, Sonia Goncalves, Ewan Harrison, David K. Jackson, Ian Johnston, Dominic Kwiatkowski, Cordelia Langford, John Sillitoe on behalf of the Wellcome Sanger Institute COVID-19 Surveillance Team |
| EPI_ISL_2994752, EPI_ISL_2994753, EPI_ISL_2994754 | Lighthouse Laboratory Plymouth | Wellcome Sanger Institute for the COVID-19 Genomics UK (COG-UK) Consortium | Lighthouse Laboratory Plymouth and Alex Alderton, Roberto Amato, Jeffrey Barrett, Sonia Goncalves, Ewan Harrison, David K. Jackson, Ian Johnston, Dominic Kwiatkowski, Cordelia Langford, John Sillitoe on behalf of the Wellcome Sanger Institute COVID-19 Surveillance Team |
| EPI_ISL_2994755 | Lighthouse Lab in Glasgow | Wellcome Sanger Institute for the COVID-19 Genomics UK (COG-UK) Consortium | Harper VanSteenhouse, Yumi Kasai, David Gray, Carol Clugston, Anna Dominiczak and Alex Alderton, Roberto Amato, Jeffrey Barrett, Sonia Goncalves, Ewan Harrison, David K. Jackson, Ian Johnston, Dominic Kwiatkowski, Cordelia Langford, John Sillitoe on behalf of the Wellcome Sanger Institute COVID-19 Surveillance Team |
| EPI_ISL_2994756 | Lighthouse Laboratory Plymouth | Wellcome Sanger Institute for the COVID-19 Genomics UK (COG-UK) Consortium | Lighthouse Laboratory Plymouth and Alex Alderton, Roberto Amato, Jeffrey Barrett, Sonia Goncalves, Ewan Harrison, David K. Jackson, Ian Johnston, Dominic Kwiatkowski, Cordelia Langford, John Sillitoe on behalf of the Wellcome Sanger Institute COVID-19 Surveillance Team |
| EPI_ISL_2994757 | Lighthouse Lab in Glasgow | Wellcome Sanger Institute for the COVID-19 Genomics UK (COG-UK) Consortium | Harper VanSteenhouse, Yumi Kasai, David Gray, Carol Clugston, Anna Dominiczak and Alex Alderton, Roberto Amato, Jeffrey Barrett, Sonia Goncalves, Ewan Harrison, David K. Jackson, Ian Johnston, Dominic Kwiatkowski, Cordelia Langford, John Sillitoe on behalf of the Wellcome Sanger Institute COVID-19 Surveillance Team |
| EPI_ISL_2994758 | Lighthouse Laboratory Plymouth | Wellcome Sanger Institute for the COVID-19 Genomics UK (COG-UK) Consortium | Lighthouse Laboratory Plymouth and Alex Alderton, Roberto Amato, Jeffrey Barrett, Sonia Goncalves, Ewan Harrison, David K. Jackson, Ian Johnston, Dominic Kwiatkowski, Cordelia Langford, John Sillitoe on behalf of the Wellcome Sanger Institute COVID-19 Surveillance Team |
| EPI_ISL_2994759, EPI_ISL_2994760 | Lighthouse Lab in Milton Keynes | Wellcome Sanger Institute for the COVID-19 Genomics UK (COG-UK) Consortium | The Lighthouse Lab in Milton Keynes and Alex Alderton, Roberto Amato, Jeffrey Barrett, Sonia Goncalves, Ewan Harrison, David K. Jackson, Ian Johnston, Dominic Kwiatkowski, Cordelia Langford, John Sillitoe on behalf of the Wellcome Sanger Institute COVID-19 Surveillance Team |
| EPI_ISL_2994761 | Lighthouse Lab in Glasgow | Wellcome Sanger Institute for the COVID-19 Genomics UK (COG-UK) Consortium | Harper VanSteenhouse, Yumi Kasai, David Gray, Carol Clugston, Anna Dominiczak and Alex Alderton, Roberto Amato, Jeffrey Barrett, Sonia Goncalves, Ewan Harrison, David K. Jackson, Ian Johnston, Dominic Kwiatkowski, Cordelia Langford, John Sillitoe on behalf of the Wellcome Sanger Institute COVID-19 Surveillance Team |
| EPI_ISL_2994762, EPI_ISL_2994763, EPI_ISL_2994764, EPI_ISL_2994765, EPI_ISL_2994766 | Lighthouse Laboratory Plymouth | Wellcome Sanger Institute for the COVID-19 Genomics UK (COG-UK) Consortium | Lighthouse Laboratory Plymouth and Alex Alderton, Roberto Amato, Jeffrey Barrett, Sonia Goncalves, Ewan Harrison, David K. Jackson, Ian Johnston, Dominic Kwiatkowski, Cordelia Langford, John Sillitoe on behalf of the Wellcome Sanger Institute COVID-19 Surveillance Team |
| EPI_ISL_2994767 | Lighthouse Lab in Glasgow | Wellcome Sanger Institute for the COVID-19 Genomics UK (COG-UK) Consortium | Harper VanSteenhouse, Yumi Kasai, David Gray, Carol Clugston, Anna Dominiczak and Alex Alderton, Roberto Amato, Jeffrey Barrett, Sonia Goncalves, Ewan Harrison, David K. Jackson, Ian Johnston, Dominic Kwiatkowski, Cordelia Langford, John Sillitoe on behalf of the Wellcome Sanger Institute COVID-19 Surveillance Team |
| EPI_ISL_2994768, EPI_ISL_2994769 | Lighthouse Laboratory Plymouth | Wellcome Sanger Institute for the COVID-19 Genomics UK (COG-UK) Consortium | Lighthouse Laboratory Plymouth and Alex Alderton, Roberto Amato, Jeffrey Barrett, Sonia Goncalves, Ewan Harrison, David K. Jackson, Ian Johnston, Dominic Kwiatkowski, Cordelia Langford, John Sillitoe on behalf of the Wellcome Sanger Institute COVID-19 Surveillance Team |
| EPI_ISL_2994770 | Lighthouse Lab in Glasgow | Wellcome Sanger Institute for the COVID-19 Genomics UK (COG-UK) Consortium | Harper VanSteenhouse, Yumi Kasai, David Gray, Carol Clugston, Anna Dominiczak and Alex Alderton, Roberto Amato, Jeffrey Barrett, Sonia Goncalves, Ewan Harrison, David K. Jackson, Ian Johnston, Dominic Kwiatkowski, Cordelia Langford, John Sillitoe on behalf of the Wellcome Sanger Institute COVID-19 Surveillance Team |
| EPI_ISL_2994771 | Lighthouse Laboratory Plymouth | Wellcome Sanger Institute for the COVID-19 Genomics UK (COG-UK) Consortium | Lighthouse Laboratory Plymouth and Alex Alderton, Roberto Amato, Jeffrey Barrett, Sonia Goncalves, Ewan Harrison, David K. Jackson, Ian Johnston, Dominic Kwiatkowski, Cordelia Langford, John Sillitoe on behalf of the Wellcome Sanger Institute COVID-19 Surveillance Team |
| EPI_ISL_2994772 | Lighthouse Lab in Glasgow | Wellcome Sanger Institute for the COVID-19 Genomics UK (COG-UK) Consortium | Harper VanSteenhouse, Yumi Kasai, David Gray, Carol Clugston, Anna Dominiczak and Alex Alderton, Roberto Amato, Jeffrey Barrett, Sonia Goncalves, Ewan Harrison, David K. Jackson, Ian Johnston, Dominic Kwiatkowski, Cordelia Langford, John Sillitoe on behalf of the Wellcome Sanger Institute COVID-19 Surveillance Team |
| EPI_ISL_2994773, EPI_ISL_2994774 | Lighthouse Laboratory Plymouth | Wellcome Sanger Institute for the COVID-19 Genomics UK (COG-UK) Consortium | Lighthouse Laboratory Plymouth and Alex Alderton, Roberto Amato, Jeffrey Barrett, Sonia Goncalves, Ewan Harrison, David K. Jackson, Ian Johnston, Dominic Kwiatkowski, Cordelia Langford, John Sillitoe on behalf of the Wellcome Sanger Institute COVID-19 Surveillance Team |
| EPI_ISL_2994775 | Lighthouse Lab in Glasgow | Wellcome Sanger Institute for the COVID-19 Genomics UK (COG-UK) Consortium | Harper VanSteenhouse, Yumi Kasai, David Gray, Carol Clugston, Anna Dominiczak and Alex Alderton, Roberto Amato, Jeffrey Barrett, Sonia Goncalves, Ewan Harrison, David K. Jackson, Ian Johnston, Dominic Kwiatkowski, Cordelia Langford, John Sillitoe on behalf of the Wellcome Sanger Institute COVID-19 Surveillance Team |
| EPI_ISL_2994776, EPI_ISL_2994777 | Lighthouse Laboratory Plymouth | Wellcome Sanger Institute for the COVID-19 Genomics UK (COG-UK) Consortium | Lighthouse Laboratory Plymouth and Alex Alderton, Roberto Amato, Jeffrey Barrett, Sonia Goncalves, Ewan Harrison, David K. Jackson, Ian Johnston, Dominic Kwiatkowski, Cordelia Langford, John Sillitoe on behalf of the Wellcome Sanger Institute COVID-19 Surveillance Team |
| EPI_ISL_2994778, EPI_ISL_2994779 | Lighthouse Lab in Glasgow | Wellcome Sanger Institute for the COVID-19 Genomics UK (COG-UK) Consortium | Harper VanSteenhouse, Yumi Kasai, David Gray, Carol Clugston, Anna Dominiczak and Alex Alderton, Roberto Amato, Jeffrey Barrett, Sonia Goncalves, Ewan Harrison, David K. Jackson, Ian Johnston, Dominic Kwiatkowski, Cordelia Langford, John Sillitoe on behalf of the Wellcome Sanger Institute COVID-19 Surveillance Team |
| EPI_ISL_2994780, EPI_ISL_2994781, EPI_ISL_2994782 | Lighthouse Laboratory Plymouth | Wellcome Sanger Institute for the COVID-19 Genomics UK (COG-UK) Consortium | Lighthouse Laboratory Plymouth and Alex Alderton, Roberto Amato, Jeffrey Barrett, Sonia Goncalves, Ewan Harrison, David K. Jackson, Ian Johnston, Dominic Kwiatkowski, Cordelia Langford, John Sillitoe on behalf of the Wellcome Sanger Institute COVID-19 Surveillance Team |
| EPI_ISL_2994783 | Lighthouse Lab in Milton Keynes | Wellcome Sanger Institute for the COVID-19 Genomics UK (COG-UK) Consortium | The Lighthouse Lab in Milton Keynes and Alex Alderton, Roberto Amato, Jeffrey Barrett, Sonia Goncalves, Ewan Harrison, David K. Jackson, Ian Johnston, Dominic Kwiatkowski, Cordelia Langford, John Sillitoe on behalf of the Wellcome Sanger Institute COVID-19 Surveillance Team |
| EPI_ISL_2994784, EPI_ISL_2994785, EPI_ISL_2994786 | Lighthouse Laboratory Plymouth | Wellcome Sanger Institute for the COVID-19 Genomics UK (COG-UK) Consortium | Lighthouse Laboratory Plymouth and Alex Alderton, Roberto Amato, Jeffrey Barrett, Sonia Goncalves, Ewan Harrison, David K. Jackson, Ian Johnston, Dominic Kwiatkowski, Cordelia Langford, John Sillitoe on behalf of the Wellcome Sanger Institute COVID-19 Surveillance Team |
| EPI_ISL_2994787, EPI_ISL_2994788 | Lighthouse Lab in Glasgow | Wellcome Sanger Institute for the COVID-19 Genomics UK (COG-UK) Consortium | Harper VanSteenhouse, Yumi Kasai, David Gray, Carol Clugston, Anna Dominiczak and Alex Alderton, Roberto Amato, Jeffrey Barrett, Sonia Goncalves, Ewan Harrison, David K. Jackson, Ian Johnston, Dominic Kwiatkowski, Cordelia Langford, John Sillitoe on behalf of the Wellcome Sanger Institute COVID-19 Surveillance Team |
| EPI_ISL_2994789, EPI_ISL_2994790, EPI_ISL_2994791, EPI_ISL_2994792, EPI_ISL_2994793 | Lighthouse Laboratory Plymouth | Wellcome Sanger Institute for the COVID-19 Genomics UK (COG-UK) Consortium | Lighthouse Laboratory Plymouth and Alex Alderton, Roberto Amato, Jeffrey Barrett, Sonia Goncalves, Ewan Harrison, David K. Jackson, Ian Johnston, Dominic Kwiatkowski, Cordelia Langford, John Sillitoe on behalf of the Wellcome Sanger Institute COVID-19 Surveillance Team |
| EPI_ISL_2994794 | Lighthouse Lab in Glasgow | Wellcome Sanger Institute for the COVID-19 Genomics UK (COG-UK) Consortium | Harper VanSteenhouse, Yumi Kasai, David Gray, Carol Clugston, Anna Dominiczak and Alex Alderton, Roberto Amato, Jeffrey Barrett, Sonia Goncalves, Ewan Harrison, David K. Jackson, Ian Johnston, Dominic Kwiatkowski, Cordelia Langford, John Sillitoe on behalf of the Wellcome Sanger Institute COVID-19 Surveillance Team |
| EPI_ISL_2994795 | Lighthouse Laboratory Plymouth | Wellcome Sanger Institute for the COVID-19 Genomics UK (COG-UK) Consortium | Lighthouse Laboratory Plymouth and Alex Alderton, Roberto Amato, Jeffrey Barrett, Sonia Goncalves, Ewan Harrison, David K. Jackson, Ian Johnston, Dominic Kwiatkowski, Cordelia Langford, John Sillitoe on behalf of the Wellcome Sanger Institute COVID-19 Surveillance Team |
| EPI_ISL_2994796 | Lighthouse Lab in Milton Keynes | Wellcome Sanger Institute for the COVID-19 Genomics UK (COG-UK) Consortium | The Lighthouse Lab in Milton Keynes and Alex Alderton, Roberto Amato, Jeffrey Barrett, Sonia Goncalves, Ewan Harrison, David K. Jackson, Ian Johnston, Dominic Kwiatkowski, Cordelia Langford, John Sillitoe on behalf of the Wellcome Sanger Institute COVID-19 Surveillance Team |
| EPI_ISL_2994797 | Lighthouse Lab in Glasgow | Wellcome Sanger Institute for the COVID-19 Genomics UK (COG-UK) Consortium | Harper VanSteenhouse, Yumi Kasai, David Gray, Carol Clugston, Anna Dominiczak and Alex Alderton, Roberto Amato, Jeffrey Barrett, Sonia Goncalves, Ewan Harrison, David K. Jackson, Ian Johnston, Dominic Kwiatkowski, Cordelia Langford, John Sillitoe on behalf of the Wellcome Sanger Institute COVID-19 Surveillance Team |
| EPI_ISL_2994798, EPI_ISL_2994799, EPI_ISL_2994800, EPI_ISL_2994801 | Lighthouse Lab in Milton Keynes | Wellcome Sanger Institute for the COVID-19 Genomics UK (COG-UK) Consortium | The Lighthouse Lab in Milton Keynes and Alex Alderton, Roberto Amato, Jeffrey Barrett, Sonia Goncalves, Ewan Harrison, David K. Jackson, Ian Johnston, Dominic Kwiatkowski, Cordelia Langford, John Sillitoe on behalf of the Wellcome Sanger Institute COVID-19 Surveillance Team |
| EPI_ISL_2994802, EPI_ISL_2994803 | Lighthouse Laboratory Plymouth | Wellcome Sanger Institute for the COVID-19 Genomics UK (COG-UK) Consortium | Lighthouse Laboratory Plymouth and Alex Alderton, Roberto Amato, Jeffrey Barrett, Sonia Goncalves, Ewan Harrison, David K. Jackson, Ian Johnston, Dominic Kwiatkowski, Cordelia Langford, John Sillitoe on behalf of the Wellcome Sanger Institute COVID-19 Surveillance Team |

| | | | |
|---|---|---|---|
| EPI_ISL_2994804 | Lighthouse Lab in Milton Keynes | Wellcome Sanger Institute for the COVID-19 Genomics UK (COG-UK) Consortium | The Lighthouse Lab in Milton Keynes and Alex Alderton, Roberto Amato, Jeffrey Barrett, Sonia Goncalves, Ewan Harrison, David K. Jackson, Ian Johnston, Dominic Kwiatkowski, Cordelia Langford, John Sillitoe on behalf of the Wellcome Sanger Institute COVID-19 Surveillance Team |
| EPI_ISL_2994805 | Lighthouse Laboratory Plymouth | Wellcome Sanger Institute for the COVID-19 Genomics UK (COG-UK) Consortium | Lighthouse Laboratory Plymouth and Alex Alderton, Roberto Amato, Jeffrey Barrett, Sonia Goncalves, Ewan Harrison, David K. Jackson, Ian Johnston, Dominic Kwiatkowski, Cordelia Langford, John Sillitoe on behalf of the Wellcome Sanger Institute COVID-19 Surveillance Team |
| EPI_ISL_2994806, EPI_ISL_2994807 | Lighthouse Lab in Glasgow | Wellcome Sanger Institute for the COVID-19 Genomics UK (COG-UK) Consortium | Harper VanSteenhouse, Yumi Kasai, David Gray, Carol Clugston, Anna Dominiczak and Alex Alderton, Roberto Amato, Jeffrey Barrett, Sonia Goncalves, Ewan Harrison, David K. Jackson, Ian Johnston, Dominic Kwiatkowski, Cordelia Langford, John Sillitoe on behalf of the Wellcome Sanger Institute COVID-19 Surveillance Team |
| EPI_ISL_2994808, EPI_ISL_2994809 | Lighthouse Lab in Milton Keynes | Wellcome Sanger Institute for the COVID-19 Genomics UK (COG-UK) Consortium | The Lighthouse Lab in Milton Keynes and Alex Alderton, Roberto Amato, Jeffrey Barrett, Sonia Goncalves, Ewan Harrison, David K. Jackson, Ian Johnston, Dominic Kwiatkowski, Cordelia Langford, John Sillitoe on behalf of the Wellcome Sanger Institute COVID-19 Surveillance Team |
| EPI_ISL_2994810 | Lighthouse Laboratory Plymouth | Wellcome Sanger Institute for the COVID-19 Genomics UK (COG-UK) Consortium | Lighthouse Laboratory Plymouth and Alex Alderton, Roberto Amato, Jeffrey Barrett, Sonia Goncalves, Ewan Harrison, David K. Jackson, Ian Johnston, Dominic Kwiatkowski, Cordelia Langford, John Sillitoe on behalf of the Wellcome Sanger Institute COVID-19 Surveillance Team |
| EPI_ISL_2994811 | Lighthouse Lab in Milton Keynes | Wellcome Sanger Institute for the COVID-19 Genomics UK (COG-UK) Consortium | The Lighthouse Lab in Milton Keynes and Alex Alderton, Roberto Amato, Jeffrey Barrett, Sonia Goncalves, Ewan Harrison, David K. Jackson, Ian Johnston, Dominic Kwiatkowski, Cordelia Langford, John Sillitoe on behalf of the Wellcome Sanger Institute COVID-19 Surveillance Team |
| EPI_ISL_2994812, EPI_ISL_2994813, EPI_ISL_2994814 | Lighthouse Laboratory Plymouth | Wellcome Sanger Institute for the COVID-19 Genomics UK (COG-UK) Consortium | Lighthouse Laboratory Plymouth and Alex Alderton, Roberto Amato, Jeffrey Barrett, Sonia Goncalves, Ewan Harrison, David K. Jackson, Ian Johnston, Dominic Kwiatkowski, Cordelia Langford, John Sillitoe on behalf of the Wellcome Sanger Institute COVID-19 Surveillance Team |
| EPI_ISL_2994815 | Lighthouse Lab in Milton Keynes | Wellcome Sanger Institute for the COVID-19 Genomics UK (COG-UK) Consortium | The Lighthouse Lab in Milton Keynes and Alex Alderton, Roberto Amato, Jeffrey Barrett, Sonia Goncalves, Ewan Harrison, David K. Jackson, Ian Johnston, Dominic Kwiatkowski, Cordelia Langford, John Sillitoe on behalf of the Wellcome Sanger Institute COVID-19 Surveillance Team |
| EPI_ISL_2994816, EPI_ISL_2994817 | Lighthouse Laboratory Plymouth | Wellcome Sanger Institute for the COVID-19 Genomics UK (COG-UK) Consortium | Lighthouse Laboratory Plymouth and Alex Alderton, Roberto Amato, Jeffrey Barrett, Sonia Goncalves, Ewan Harrison, David K. Jackson, Ian Johnston, Dominic Kwiatkowski, Cordelia Langford, John Sillitoe on behalf of the Wellcome Sanger Institute COVID-19 Surveillance Team |
| EPI_ISL_2994818 | Lighthouse Lab in Glasgow | Wellcome Sanger Institute for the COVID-19 Genomics UK (COG-UK) Consortium | Harper VanSteenhouse, Yumi Kasai, David Gray, Carol Clugston, Anna Dominiczak and Alex Alderton, Roberto Amato, Jeffrey Barrett, Sonia Goncalves, Ewan Harrison, David K. Jackson, Ian Johnston, Dominic Kwiatkowski, Cordelia Langford, John Sillitoe on behalf of the Wellcome Sanger Institute COVID-19 Surveillance Team |
| EPI_ISL_2994819 | Lighthouse Lab in Milton Keynes | Wellcome Sanger Institute for the COVID-19 Genomics UK (COG-UK) Consortium | The Lighthouse Lab in Milton Keynes and Alex Alderton, Roberto Amato, Jeffrey Barrett, Sonia Goncalves, Ewan Harrison, David K. Jackson, Ian Johnston, Dominic Kwiatkowski, Cordelia Langford, John Sillitoe on behalf of the Wellcome Sanger Institute COVID-19 Surveillance Team |
| EPI_ISL_2994820 | Lighthouse Lab in Glasgow | Wellcome Sanger Institute for the COVID-19 Genomics UK (COG-UK) Consortium | Harper VanSteenhouse, Yumi Kasai, David Gray, Carol Clugston, Anna Dominiczak and Alex Alderton, Roberto Amato, Jeffrey Barrett, Sonia Goncalves, Ewan Harrison, David K. Jackson, Ian Johnston, Dominic Kwiatkowski, Cordelia Langford, John Sillitoe on behalf of the Wellcome Sanger Institute COVID-19 Surveillance Team |
| EPI_ISL_2994821, EPI_ISL_2994822 | Lighthouse Laboratory Plymouth | Wellcome Sanger Institute for the COVID-19 Genomics UK (COG-UK) Consortium | Lighthouse Laboratory Plymouth and Alex Alderton, Roberto Amato, Jeffrey Barrett, Sonia Goncalves, Ewan Harrison, David K. Jackson, Ian Johnston, Dominic Kwiatkowski, Cordelia Langford, John Sillitoe on behalf of the Wellcome Sanger Institute COVID-19 Surveillance Team |
| EPI_ISL_2994823, EPI_ISL_2994824, EPI_ISL_2994825, EPI_ISL_2994826 | Lighthouse Lab in Glasgow | Wellcome Sanger Institute for the COVID-19 Genomics UK (COG-UK) Consortium | Harper VanSteenhouse, Yumi Kasai, David Gray, Carol Clugston, Anna Dominiczak and Alex Alderton, Roberto Amato, Jeffrey Barrett, Sonia Goncalves, Ewan Harrison, David K. Jackson, Ian Johnston, Dominic Kwiatkowski, Cordelia Langford, John Sillitoe on behalf of the Wellcome Sanger Institute COVID-19 Surveillance Team |
| EPI_ISL_2994827 | Lighthouse Lab in Milton Keynes | Wellcome Sanger Institute for the COVID-19 Genomics UK (COG-UK) Consortium | The Lighthouse Lab in Milton Keynes and Alex Alderton, Roberto Amato, Jeffrey Barrett, Sonia Goncalves, Ewan Harrison, David K. Jackson, Ian Johnston, Dominic Kwiatkowski, Cordelia Langford, John Sillitoe on behalf of the Wellcome Sanger Institute COVID-19 Surveillance Team |
| EPI_ISL_2994828 | Lighthouse Laboratory Plymouth | Wellcome Sanger Institute for the COVID-19 Genomics UK (COG-UK) Consortium | Lighthouse Laboratory Plymouth and Alex Alderton, Roberto Amato, Jeffrey Barrett, Sonia Goncalves, Ewan Harrison, David K. Jackson, Ian Johnston, Dominic Kwiatkowski, Cordelia Langford, John Sillitoe on behalf of the Wellcome Sanger Institute COVID-19 Surveillance Team |
| EPI_ISL_2994829 | Lighthouse Lab in Milton Keynes | Wellcome Sanger Institute for the COVID-19 Genomics UK (COG-UK) Consortium | The Lighthouse Lab in Milton Keynes and Alex Alderton, Roberto Amato, Jeffrey Barrett, Sonia Goncalves, Ewan Harrison, David K. Jackson, Ian Johnston, Dominic Kwiatkowski, Cordelia Langford, John Sillitoe on behalf of the Wellcome Sanger Institute COVID-19 Surveillance Team |
| EPI_ISL_2994830, EPI_ISL_2994831 | Lighthouse Laboratory Plymouth | Wellcome Sanger Institute for the COVID-19 Genomics UK (COG-UK) Consortium | Lighthouse Laboratory Plymouth and Alex Alderton, Roberto Amato, Jeffrey Barrett, Sonia Goncalves, Ewan Harrison, David K. Jackson, Ian Johnston, Dominic Kwiatkowski, Cordelia Langford, John Sillitoe on behalf of the Wellcome Sanger Institute COVID-19 Surveillance Team |
| EPI_ISL_2994832, EPI_ISL_2994833 | Lighthouse Lab in Milton Keynes | Wellcome Sanger Institute for the COVID-19 Genomics UK (COG-UK) Consortium | The Lighthouse Lab in Milton Keynes and Alex Alderton, Roberto Amato, Jeffrey Barrett, Sonia Goncalves, Ewan Harrison, David K. Jackson, Ian Johnston, Dominic Kwiatkowski, Cordelia Langford, John Sillitoe on behalf of the Wellcome Sanger Institute COVID-19 Surveillance Team |
| EPI_ISL_2994834, EPI_ISL_2994835, EPI_ISL_2994836 | Lighthouse Laboratory Plymouth | Wellcome Sanger Institute for the COVID-19 Genomics UK (COG-UK) Consortium | Lighthouse Laboratory Plymouth and Alex Alderton, Roberto Amato, Jeffrey Barrett, Sonia Goncalves, Ewan Harrison, David K. Jackson, Ian Johnston, Dominic Kwiatkowski, Cordelia Langford, John Sillitoe on behalf of the Wellcome Sanger Institute COVID-19 Surveillance Team |
| EPI_ISL_2994837 | Lighthouse Lab in Glasgow | Wellcome Sanger Institute for the COVID-19 Genomics UK (COG-UK) Consortium | Harper VanSteenhouse, Yumi Kasai, David Gray, Carol Clugston, Anna Dominiczak and Alex Alderton, Roberto Amato, Jeffrey Barrett, Sonia Goncalves, Ewan Harrison, David K. Jackson, Ian Johnston, Dominic Kwiatkowski, Cordelia Langford, John Sillitoe on behalf of the Wellcome Sanger Institute COVID-19 Surveillance Team |
| EPI_ISL_2994838, EPI_ISL_2994839 | Lighthouse Lab in Milton Keynes | Wellcome Sanger Institute for the COVID-19 Genomics UK (COG-UK) Consortium | The Lighthouse Lab in Milton Keynes and Alex Alderton, Roberto Amato, Jeffrey Barrett, Sonia Goncalves, Ewan Harrison, David K. Jackson, Ian Johnston, Dominic Kwiatkowski, Cordelia Langford, John Sillitoe on behalf of the Wellcome Sanger Institute COVID-19 Surveillance Team |
| EPI_ISL_2994840, EPI_ISL_2994841, EPI_ISL_2994842, EPI_ISL_2994843 | Lighthouse Laboratory Plymouth | Wellcome Sanger Institute for the COVID-19 Genomics UK (COG-UK) Consortium | Lighthouse Laboratory Plymouth and Alex Alderton, Roberto Amato, Jeffrey Barrett, Sonia Goncalves, Ewan Harrison, David K. Jackson, Ian Johnston, Dominic Kwiatkowski, Cordelia Langford, John Sillitoe on behalf of the Wellcome Sanger Institute COVID-19 Surveillance Team |
| EPI_ISL_2994844 | Lighthouse Lab in Glasgow | Wellcome Sanger Institute for the COVID-19 Genomics UK (COG-UK) Consortium | Harper VanSteenhouse, Yumi Kasai, David Gray, Carol Clugston, Anna Dominiczak and Alex Alderton, Roberto Amato, Jeffrey Barrett, Sonia Goncalves, Ewan Harrison, David K. Jackson, Ian Johnston, Dominic Kwiatkowski, Cordelia Langford, John Sillitoe on behalf of the Wellcome Sanger Institute COVID-19 Surveillance Team |
| EPI_ISL_2994845 | Lighthouse Lab in Milton Keynes | Wellcome Sanger Institute for the COVID-19 Genomics UK (COG-UK) Consortium | The Lighthouse Lab in Milton Keynes and Alex Alderton, Roberto Amato, Jeffrey Barrett, Sonia Goncalves, Ewan Harrison, David K. Jackson, Ian Johnston, Dominic Kwiatkowski, Cordelia Langford, John Sillitoe on behalf of the Wellcome Sanger Institute COVID-19 Surveillance Team |
| EPI_ISL_2994846 | Lighthouse Lab in Glasgow | Wellcome Sanger Institute for the COVID-19 Genomics UK (COG-UK) Consortium | Harper VanSteenhouse, Yumi Kasai, David Gray, Carol Clugston, Anna Dominiczak and Alex Alderton, Roberto Amato, Jeffrey Barrett, Sonia Goncalves, Ewan Harrison, David K. Jackson, Ian Johnston, Dominic Kwiatkowski, Cordelia Langford, John Sillitoe on behalf of the Wellcome Sanger Institute COVID-19 Surveillance Team |
| EPI_ISL_2994847 | Lighthouse Laboratory Plymouth | Wellcome Sanger Institute for the COVID-19 Genomics UK (COG-UK) Consortium | Lighthouse Laboratory Plymouth and Alex Alderton, Roberto Amato, Jeffrey Barrett, Sonia Goncalves, Ewan Harrison, David K. Jackson, Ian Johnston, Dominic Kwiatkowski, Cordelia Langford, John Sillitoe on behalf of the Wellcome Sanger Institute COVID-19 Surveillance Team |
| EPI_ISL_2994848, EPI_ISL_2994849 | Lighthouse Lab in Milton Keynes | Wellcome Sanger Institute for the COVID-19 Genomics UK (COG-UK) Consortium | The Lighthouse Lab in Milton Keynes and Alex Alderton, Roberto Amato, Jeffrey Barrett, Sonia Goncalves, Ewan Harrison, David K. Jackson, Ian Johnston, Dominic Kwiatkowski, Cordelia Langford, John Sillitoe on behalf of the Wellcome Sanger Institute COVID-19 Surveillance Team |
| EPI_ISL_2994850 | Lighthouse Lab in Glasgow | Wellcome Sanger Institute for the COVID-19 Genomics UK (COG-UK) Consortium | Harper VanSteenhouse, Yumi Kasai, David Gray, Carol Clugston, Anna Dominiczak and Alex Alderton, Roberto Amato, Jeffrey Barrett, Sonia Goncalves, Ewan Harrison, David K. Jackson, Ian Johnston, Dominic Kwiatkowski, Cordelia Langford, John Sillitoe on behalf of the Wellcome Sanger Institute COVID-19 Surveillance Team |
| EPI_ISL_2994851 | Lighthouse Lab in Milton Keynes | Wellcome Sanger Institute for the COVID-19 Genomics UK (COG-UK) Consortium | The Lighthouse Lab in Milton Keynes and Alex Alderton, Roberto Amato, Jeffrey Barrett, Sonia Goncalves, Ewan Harrison, David K. Jackson, Ian Johnston, Dominic Kwiatkowski, Cordelia Langford, John Sillitoe on behalf of the Wellcome Sanger Institute COVID-19 Surveillance Team |
| EPI_ISL_2994852 | Lighthouse Lab in Glasgow | Wellcome Sanger Institute for the COVID-19 Genomics UK (COG-UK) Consortium | Harper VanSteenhouse, Yumi Kasai, David Gray, Carol Clugston, Anna Dominiczak and Alex Alderton, Roberto Amato, Jeffrey Barrett, Sonia Goncalves, Ewan Harrison, David K. Jackson, Ian Johnston, Dominic Kwiatkowski, Cordelia Langford, John Sillitoe on behalf of the Wellcome Sanger Institute COVID-19 Surveillance Team |
| EPI_ISL_2994853, EPI_ISL_2994854 | Lighthouse Laboratory Plymouth | Wellcome Sanger Institute for the COVID-19 Genomics UK (COG-UK) Consortium | Lighthouse Laboratory Plymouth and Alex Alderton, Roberto Amato, Jeffrey Barrett, Sonia Goncalves, Ewan Harrison, David K. Jackson, Ian Johnston, Dominic Kwiatkowski, Cordelia Langford, John Sillitoe on behalf of the Wellcome Sanger Institute COVID-19 Surveillance Team |
| EPI_ISL_2994855 | Lighthouse Lab in Milton Keynes | Wellcome Sanger Institute for the COVID-19 Genomics UK (COG-UK) Consortium | The Lighthouse Lab in Milton Keynes and Alex Alderton, Roberto Amato, Jeffrey Barrett, Sonia Goncalves, Ewan Harrison, David K. Jackson, Ian Johnston, Dominic Kwiatkowski, Cordelia Langford, John Sillitoe on behalf of the Wellcome Sanger Institute COVID-19 Surveillance Team |
| EPI_ISL_2994856 | Lighthouse Lab in Glasgow | Wellcome Sanger Institute for the COVID-19 Genomics UK (COG-UK) Consortium | Harper VanSteenhouse, Yumi Kasai, David Gray, Carol Clugston, Anna Dominiczak and Alex Alderton, Roberto Amato, Jeffrey Barrett, Sonia Goncalves, Ewan Harrison, David K. Jackson, Ian Johnston, Dominic Kwiatkowski, Cordelia Langford, John Sillitoe on behalf of the Wellcome Sanger Institute COVID-19 Surveillance Team |
| EPI_ISL_2994857 | Lighthouse Laboratory Plymouth | Wellcome Sanger Institute for the COVID-19 Genomics UK (COG-UK) Consortium | Lighthouse Laboratory Plymouth and Alex Alderton, Roberto Amato, Jeffrey Barrett, Sonia Goncalves, Ewan Harrison, David K. Jackson, Ian Johnston, Dominic Kwiatkowski, Cordelia Langford, John Sillitoe on behalf of the Wellcome Sanger Institute COVID-19 Surveillance Team |
| EPI_ISL_2994858, EPI_ISL_2994859 | Lighthouse Lab in Glasgow | Wellcome Sanger Institute for the COVID-19 Genomics UK (COG-UK) Consortium | Harper VanSteenhouse, Yumi Kasai, David Gray, Carol Clugston, Anna Dominiczak and Alex Alderton, Roberto Amato, Jeffrey Barrett, Sonia Goncalves, Ewan Harrison, David K. Jackson, Ian Johnston, Dominic Kwiatkowski, Cordelia Langford, John Sillitoe on behalf of the Wellcome Sanger Institute COVID-19 Surveillance Team |
| EPI_ISL_2994860, EPI_ISL_2994861 | Lighthouse Laboratory Plymouth | Wellcome Sanger Institute for the COVID-19 Genomics UK (COG-UK) Consortium | Lighthouse Laboratory Plymouth and Alex Alderton, Roberto Amato, Jeffrey Barrett, Sonia Goncalves, Ewan Harrison, David K. Jackson, Ian Johnston, Dominic Kwiatkowski, Cordelia Langford, John Sillitoe on behalf of the Wellcome Sanger Institute COVID-19 Surveillance Team |
| EPI_ISL_2994862, EPI_ISL_2994863 | Lighthouse Lab in Milton Keynes | Wellcome Sanger Institute for the COVID-19 Genomics UK (COG-UK) Consortium | The Lighthouse Lab in Milton Keynes and Alex Alderton, Roberto Amato, Jeffrey Barrett, Sonia Goncalves, Ewan Harrison, David K. Jackson, Ian Johnston, Dominic Kwiatkowski, Cordelia Langford, John Sillitoe on behalf of the Wellcome Sanger Institute COVID-19 Surveillance Team |
| EPI_ISL_2994864 | Lighthouse Laboratory Plymouth | Wellcome Sanger Institute for the COVID-19 Genomics UK (COG-UK) Consortium | Lighthouse Laboratory Plymouth and Alex Alderton, Roberto Amato, Jeffrey Barrett, Sonia Goncalves, Ewan Harrison, David K. Jackson, Ian Johnston, Dominic Kwiatkowski, Cordelia Langford, John Sillitoe on behalf of the Wellcome Sanger Institute COVID-19 Surveillance Team |
| EPI_ISL_2994865, EPI_ISL_2994866 | Lighthouse Lab in Milton Keynes | Wellcome Sanger Institute for the COVID-19 Genomics UK (COG-UK) Consortium | The Lighthouse Lab in Milton Keynes and Alex Alderton, Roberto Amato, Jeffrey Barrett, Sonia Goncalves, Ewan Harrison, David K. Jackson, Ian Johnston, Dominic Kwiatkowski, Cordelia Langford, John Sillitoe on behalf of the Wellcome Sanger Institute COVID-19 Surveillance Team |
| EPI_ISL_2994867 | Lighthouse Laboratory Plymouth | Wellcome Sanger Institute for the COVID-19 Genomics UK (COG-UK) Consortium | Lighthouse Laboratory Plymouth and Alex Alderton, Roberto Amato, Jeffrey Barrett, Sonia Goncalves, Ewan Harrison, David K. Jackson, Ian Johnston, Dominic Kwiatkowski, Cordelia Langford, John Sillitoe on behalf of the Wellcome Sanger Institute COVID-19 Surveillance Team |
| EPI_ISL_2994868, EPI_ISL_2994869 | Lighthouse Lab in Glasgow | Wellcome Sanger Institute for the COVID-19 Genomics UK (COG-UK) Consortium | Harper VanSteenhouse, Yumi Kasai, David Gray, Carol Clugston, Anna Dominiczak and Alex Alderton, Roberto Amato, Jeffrey Barrett, Sonia Goncalves, Ewan Harrison, David K. Jackson, Ian Johnston, Dominic Kwiatkowski, Cordelia Langford, John Sillitoe on behalf of the Wellcome Sanger Institute COVID-19 Surveillance Team |
| EPI_ISL_2994870 | Lighthouse Lab in Milton Keynes | Wellcome Sanger Institute for the COVID-19 Genomics UK (COG-UK) Consortium | The Lighthouse Lab in Milton Keynes and Alex Alderton, Roberto Amato, Jeffrey Barrett, Sonia Goncalves, Ewan Harrison, David K. Jackson, Ian Johnston, Dominic Kwiatkowski, Cordelia Langford, John Sillitoe on behalf of the Wellcome Sanger Institute COVID-19 Surveillance Team |

| | | | |
|---|---|---|---|
| EPI_ISL_2994871 | Lighthouse Lab in Glasgow | Wellcome Sanger Institute for the COVID-19 Genomics UK (COG-UK) Consortium | Harper VanSteenhouse, Yumi Kasai, David Gray, Carol Clugston, Anna Dominiczak and Alex Alderton, Roberto Amato, Jeffrey Barrett, Sonia Goncalves, Ewan Harrison, David K. Jackson, Ian Johnston, Dominic Kwiatkowski, Cordelia Langford, John Sillitoe on behalf of the Wellcome Sanger Institute COVID-19 Surveillance Team |
| EPI_ISL_2994872 | Lighthouse Laboratory Plymouth | Wellcome Sanger Institute for the COVID-19 Genomics UK (COG-UK) Consortium | Lighthouse Laboratory Plymouth and Alex Alderton, Roberto Amato, Jeffrey Barrett, Sonia Goncalves, Ewan Harrison, David K. Jackson, Ian Johnston, Dominic Kwiatkowski, Cordelia Langford, John Sillitoe on behalf of the Wellcome Sanger Institute COVID-19 Surveillance Team |
| EPI_ISL_2994873, EPI_ISL_2994874 | Lighthouse Lab in Glasgow | Wellcome Sanger Institute for the COVID-19 Genomics UK (COG-UK) Consortium | Harper VanSteenhouse, Yumi Kasai, David Gray, Carol Clugston, Anna Dominiczak and Alex Alderton, Roberto Amato, Jeffrey Barrett, Sonia Goncalves, Ewan Harrison, David K. Jackson, Ian Johnston, Dominic Kwiatkowski, Cordelia Langford, John Sillitoe on behalf of the Wellcome Sanger Institute COVID-19 Surveillance Team |
| EPI_ISL_2994875, EPI_ISL_2994876, EPI_ISL_2994877 | Lighthouse Laboratory Plymouth | Wellcome Sanger Institute for the COVID-19 Genomics UK (COG-UK) Consortium | Lighthouse Laboratory Plymouth and Alex Alderton, Roberto Amato, Jeffrey Barrett, Sonia Goncalves, Ewan Harrison, David K. Jackson, Ian Johnston, Dominic Kwiatkowski, Cordelia Langford, John Sillitoe on behalf of the Wellcome Sanger Institute COVID-19 Surveillance Team |
| EPI_ISL_2994878 | Lighthouse Lab in Milton Keynes | Wellcome Sanger Institute for the COVID-19 Genomics UK (COG-UK) Consortium | The Lighthouse Lab in Milton Keynes and Alex Alderton, Roberto Amato, Jeffrey Barrett, Sonia Goncalves, Ewan Harrison, David K. Jackson, Ian Johnston, Dominic Kwiatkowski, Cordelia Langford, John Sillitoe on behalf of the Wellcome Sanger Institute COVID-19 Surveillance Team |
| EPI_ISL_2994879 | Lighthouse Lab in Glasgow | Wellcome Sanger Institute for the COVID-19 Genomics UK (COG-UK) Consortium | Harper VanSteenhouse, Yumi Kasai, David Gray, Carol Clugston, Anna Dominiczak and Alex Alderton, Roberto Amato, Jeffrey Barrett, Sonia Goncalves, Ewan Harrison, David K. Jackson, Ian Johnston, Dominic Kwiatkowski, Cordelia Langford, John Sillitoe on behalf of the Wellcome Sanger Institute COVID-19 Surveillance Team |
| EPI_ISL_2994880 | Lighthouse Lab in Milton Keynes | Wellcome Sanger Institute for the COVID-19 Genomics UK (COG-UK) Consortium | The Lighthouse Lab in Milton Keynes and Alex Alderton, Roberto Amato, Jeffrey Barrett, Sonia Goncalves, Ewan Harrison, David K. Jackson, Ian Johnston, Dominic Kwiatkowski, Cordelia Langford, John Sillitoe on behalf of the Wellcome Sanger Institute COVID-19 Surveillance Team |
| EPI_ISL_2994881 | Lighthouse Laboratory Plymouth | Wellcome Sanger Institute for the COVID-19 Genomics UK (COG-UK) Consortium | Lighthouse Laboratory Plymouth and Alex Alderton, Roberto Amato, Jeffrey Barrett, Sonia Goncalves, Ewan Harrison, David K. Jackson, Ian Johnston, Dominic Kwiatkowski, Cordelia Langford, John Sillitoe on behalf of the Wellcome Sanger Institute COVID-19 Surveillance Team |
| EPI_ISL_2994882 | Lighthouse Lab in Glasgow | Wellcome Sanger Institute for the COVID-19 Genomics UK (COG-UK) Consortium | Harper VanSteenhouse, Yumi Kasai, David Gray, Carol Clugston, Anna Dominiczak and Alex Alderton, Roberto Amato, Jeffrey Barrett, Sonia Goncalves, Ewan Harrison, David K. Jackson, Ian Johnston, Dominic Kwiatkowski, Cordelia Langford, John Sillitoe on behalf of the Wellcome Sanger Institute COVID-19 Surveillance Team |
| EPI_ISL_2994883 | Lighthouse Laboratory Plymouth | Wellcome Sanger Institute for the COVID-19 Genomics UK (COG-UK) Consortium | Lighthouse Laboratory Plymouth and Alex Alderton, Roberto Amato, Jeffrey Barrett, Sonia Goncalves, Ewan Harrison, David K. Jackson, Ian Johnston, Dominic Kwiatkowski, Cordelia Langford, John Sillitoe on behalf of the Wellcome Sanger Institute COVID-19 Surveillance Team |
| EPI_ISL_2994884 | Lighthouse Lab in Milton Keynes | Wellcome Sanger Institute for the COVID-19 Genomics UK (COG-UK) Consortium | The Lighthouse Lab in Milton Keynes and Alex Alderton, Roberto Amato, Jeffrey Barrett, Sonia Goncalves, Ewan Harrison, David K. Jackson, Ian Johnston, Dominic Kwiatkowski, Cordelia Langford, John Sillitoe on behalf of the Wellcome Sanger Institute COVID-19 Surveillance Team |
| EPI_ISL_2994885 | Lighthouse Laboratory Plymouth | Wellcome Sanger Institute for the COVID-19 Genomics UK (COG-UK) Consortium | Lighthouse Laboratory Plymouth and Alex Alderton, Roberto Amato, Jeffrey Barrett, Sonia Goncalves, Ewan Harrison, David K. Jackson, Ian Johnston, Dominic Kwiatkowski, Cordelia Langford, John Sillitoe on behalf of the Wellcome Sanger Institute COVID-19 Surveillance Team |
| EPI_ISL_2994886 | Lighthouse Lab in Milton Keynes | Wellcome Sanger Institute for the COVID-19 Genomics UK (COG-UK) Consortium | The Lighthouse Lab in Milton Keynes and Alex Alderton, Roberto Amato, Jeffrey Barrett, Sonia Goncalves, Ewan Harrison, David K. Jackson, Ian Johnston, Dominic Kwiatkowski, Cordelia Langford, John Sillitoe on behalf of the Wellcome Sanger Institute COVID-19 Surveillance Team |
| EPI_ISL_2994887 | Lighthouse Laboratory Plymouth | Wellcome Sanger Institute for the COVID-19 Genomics UK (COG-UK) Consortium | Lighthouse Laboratory Plymouth and Alex Alderton, Roberto Amato, Jeffrey Barrett, Sonia Goncalves, Ewan Harrison, David K. Jackson, Ian Johnston, Dominic Kwiatkowski, Cordelia Langford, John Sillitoe on behalf of the Wellcome Sanger Institute COVID-19 Surveillance Team |
| EPI_ISL_2994888 | Lighthouse Lab in Glasgow | Wellcome Sanger Institute for the COVID-19 Genomics UK (COG-UK) Consortium | Harper VanSteenhouse, Yumi Kasai, David Gray, Carol Clugston, Anna Dominiczak and Alex Alderton, Roberto Amato, Jeffrey Barrett, Sonia Goncalves, Ewan Harrison, David K. Jackson, Ian Johnston, Dominic Kwiatkowski, Cordelia Langford, John Sillitoe on behalf of the Wellcome Sanger Institute COVID-19 Surveillance Team |
| EPI_ISL_2994889, EPI_ISL_2994890, EPI_ISL_2994891, EPI_ISL_2994892 | Lighthouse Laboratory Plymouth | Wellcome Sanger Institute for the COVID-19 Genomics UK (COG-UK) Consortium | Lighthouse Laboratory Plymouth and Alex Alderton, Roberto Amato, Jeffrey Barrett, Sonia Goncalves, Ewan Harrison, David K. Jackson, Ian Johnston, Dominic Kwiatkowski, Cordelia Langford, John Sillitoe on behalf of the Wellcome Sanger Institute COVID-19 Surveillance Team |
| EPI_ISL_2994893, EPI_ISL_2994894 | Lighthouse Lab in Glasgow | Wellcome Sanger Institute for the COVID-19 Genomics UK (COG-UK) Consortium | Harper VanSteenhouse, Yumi Kasai, David Gray, Carol Clugston, Anna Dominiczak and Alex Alderton, Roberto Amato, Jeffrey Barrett, Sonia Goncalves, Ewan Harrison, David K. Jackson, Ian Johnston, Dominic Kwiatkowski, Cordelia Langford, John Sillitoe on behalf of the Wellcome Sanger Institute COVID-19 Surveillance Team |
| EPI_ISL_2994895 | Lighthouse Lab in Milton Keynes | Wellcome Sanger Institute for the COVID-19 Genomics UK (COG-UK) Consortium | The Lighthouse Lab in Milton Keynes and Alex Alderton, Roberto Amato, Jeffrey Barrett, Sonia Goncalves, Ewan Harrison, David K. Jackson, Ian Johnston, Dominic Kwiatkowski, Cordelia Langford, John Sillitoe on behalf of the Wellcome Sanger Institute COVID-19 Surveillance Team |
| EPI_ISL_2994896 | Lighthouse Lab in Glasgow | Wellcome Sanger Institute for the COVID-19 Genomics UK (COG-UK) Consortium | Harper VanSteenhouse, Yumi Kasai, David Gray, Carol Clugston, Anna Dominiczak and Alex Alderton, Roberto Amato, Jeffrey Barrett, Sonia Goncalves, Ewan Harrison, David K. Jackson, Ian Johnston, Dominic Kwiatkowski, Cordelia Langford, John Sillitoe on behalf of the Wellcome Sanger Institute COVID-19 Surveillance Team |
| EPI_ISL_2994897 | Lighthouse Lab in Milton Keynes | Wellcome Sanger Institute for the COVID-19 Genomics UK (COG-UK) Consortium | The Lighthouse Lab in Milton Keynes and Alex Alderton, Roberto Amato, Jeffrey Barrett, Sonia Goncalves, Ewan Harrison, David K. Jackson, Ian Johnston, Dominic Kwiatkowski, Cordelia Langford, John Sillitoe on behalf of the Wellcome Sanger Institute COVID-19 Surveillance Team |
| EPI_ISL_2994898, EPI_ISL_2994899 | Lighthouse Lab in Glasgow | Wellcome Sanger Institute for the COVID-19 Genomics UK (COG-UK) Consortium | Harper VanSteenhouse, Yumi Kasai, David Gray, Carol Clugston, Anna Dominiczak and Alex Alderton, Roberto Amato, Jeffrey Barrett, Sonia Goncalves, Ewan Harrison, David K. Jackson, Ian Johnston, Dominic Kwiatkowski, Cordelia Langford, John Sillitoe on behalf of the Wellcome Sanger Institute COVID-19 Surveillance Team |
| EPI_ISL_2994900 | Lighthouse Laboratory Plymouth | Wellcome Sanger Institute for the COVID-19 Genomics UK (COG-UK) Consortium | Lighthouse Laboratory Plymouth and Alex Alderton, Roberto Amato, Jeffrey Barrett, Sonia Goncalves, Ewan Harrison, David K. Jackson, Ian Johnston, Dominic Kwiatkowski, Cordelia Langford, John Sillitoe on behalf of the Wellcome Sanger Institute COVID-19 Surveillance Team |
| EPI_ISL_2994901 | Lighthouse Lab in Milton Keynes | Wellcome Sanger Institute for the COVID-19 Genomics UK (COG-UK) Consortium | The Lighthouse Lab in Milton Keynes and Alex Alderton, Roberto Amato, Jeffrey Barrett, Sonia Goncalves, Ewan Harrison, David K. Jackson, Ian Johnston, Dominic Kwiatkowski, Cordelia Langford, John Sillitoe on behalf of the Wellcome Sanger Institute COVID-19 Surveillance Team |
| EPI_ISL_2994902 | Lighthouse Laboratory Plymouth | Wellcome Sanger Institute for the COVID-19 Genomics UK (COG-UK) Consortium | Lighthouse Laboratory Plymouth and Alex Alderton, Roberto Amato, Jeffrey Barrett, Sonia Goncalves, Ewan Harrison, David K. Jackson, Ian Johnston, Dominic Kwiatkowski, Cordelia Langford, John Sillitoe on behalf of the Wellcome Sanger Institute COVID-19 Surveillance Team |
| EPI_ISL_2994903 | Lighthouse Lab in Milton Keynes | Wellcome Sanger Institute for the COVID-19 Genomics UK (COG-UK) Consortium | The Lighthouse Lab in Milton Keynes and Alex Alderton, Roberto Amato, Jeffrey Barrett, Sonia Goncalves, Ewan Harrison, David K. Jackson, Ian Johnston, Dominic Kwiatkowski, Cordelia Langford, John Sillitoe on behalf of the Wellcome Sanger Institute COVID-19 Surveillance Team |
| EPI_ISL_2994904, EPI_ISL_2994905 | Lighthouse Laboratory Plymouth | Wellcome Sanger Institute for the COVID-19 Genomics UK (COG-UK) Consortium | Lighthouse Laboratory Plymouth and Alex Alderton, Roberto Amato, Jeffrey Barrett, Sonia Goncalves, Ewan Harrison, David K. Jackson, Ian Johnston, Dominic Kwiatkowski, Cordelia Langford, John Sillitoe on behalf of the Wellcome Sanger Institute COVID-19 Surveillance Team |
| EPI_ISL_2994906 | Lighthouse Lab in Milton Keynes | Wellcome Sanger Institute for the COVID-19 Genomics UK (COG-UK) Consortium | The Lighthouse Lab in Milton Keynes and Alex Alderton, Roberto Amato, Jeffrey Barrett, Sonia Goncalves, Ewan Harrison, David K. Jackson, Ian Johnston, Dominic Kwiatkowski, Cordelia Langford, John Sillitoe on behalf of the Wellcome Sanger Institute COVID-19 Surveillance Team |
| EPI_ISL_2994907, EPI_ISL_2994908 | Lighthouse Laboratory Plymouth | Wellcome Sanger Institute for the COVID-19 Genomics UK (COG-UK) Consortium | Lighthouse Laboratory Plymouth and Alex Alderton, Roberto Amato, Jeffrey Barrett, Sonia Goncalves, Ewan Harrison, David K. Jackson, Ian Johnston, Dominic Kwiatkowski, Cordelia Langford, John Sillitoe on behalf of the Wellcome Sanger Institute COVID-19 Surveillance Team |
| EPI_ISL_2994909 | Lighthouse Lab in Milton Keynes | Wellcome Sanger Institute for the COVID-19 Genomics UK (COG-UK) Consortium | The Lighthouse Lab in Milton Keynes and Alex Alderton, Roberto Amato, Jeffrey Barrett, Sonia Goncalves, Ewan Harrison, David K. Jackson, Ian Johnston, Dominic Kwiatkowski, Cordelia Langford, John Sillitoe on behalf of the Wellcome Sanger Institute COVID-19 Surveillance Team |
| EPI_ISL_2994910, EPI_ISL_2994911, EPI_ISL_2994912 | Lighthouse Laboratory Plymouth | Wellcome Sanger Institute for the COVID-19 Genomics UK (COG-UK) Consortium | Lighthouse Laboratory Plymouth and Alex Alderton, Roberto Amato, Jeffrey Barrett, Sonia Goncalves, Ewan Harrison, David K. Jackson, Ian Johnston, Dominic Kwiatkowski, Cordelia Langford, John Sillitoe on behalf of the Wellcome Sanger Institute COVID-19 Surveillance Team |
| EPI_ISL_2994913 | Lighthouse Lab in Milton Keynes | Wellcome Sanger Institute for the COVID-19 Genomics UK (COG-UK) Consortium | The Lighthouse Lab in Milton Keynes and Alex Alderton, Roberto Amato, Jeffrey Barrett, Sonia Goncalves, Ewan Harrison, David K. Jackson, Ian Johnston, Dominic Kwiatkowski, Cordelia Langford, John Sillitoe on behalf of the Wellcome Sanger Institute COVID-19 Surveillance Team |
| EPI_ISL_2994914 | Lighthouse Lab in Glasgow | Wellcome Sanger Institute for the COVID-19 Genomics UK (COG-UK) Consortium | Harper VanSteenhouse, Yumi Kasai, David Gray, Carol Clugston, Anna Dominiczak and Alex Alderton, Roberto Amato, Jeffrey Barrett, Sonia Goncalves, Ewan Harrison, David K. Jackson, Ian Johnston, Dominic Kwiatkowski, Cordelia Langford, John Sillitoe on behalf of the Wellcome Sanger Institute COVID-19 Surveillance Team |
| EPI_ISL_2994915 | Lighthouse Laboratory Plymouth | Wellcome Sanger Institute for the COVID-19 Genomics UK (COG-UK) Consortium | Lighthouse Laboratory Plymouth and Alex Alderton, Roberto Amato, Jeffrey Barrett, Sonia Goncalves, Ewan Harrison, David K. Jackson, Ian Johnston, Dominic Kwiatkowski, Cordelia Langford, John Sillitoe on behalf of the Wellcome Sanger Institute COVID-19 Surveillance Team |
| EPI_ISL_2994916 | Lighthouse Lab in Glasgow | Wellcome Sanger Institute for the COVID-19 Genomics UK (COG-UK) Consortium | Harper VanSteenhouse, Yumi Kasai, David Gray, Carol Clugston, Anna Dominiczak and Alex Alderton, Roberto Amato, Jeffrey Barrett, Sonia Goncalves, Ewan Harrison, David K. Jackson, Ian Johnston, Dominic Kwiatkowski, Cordelia Langford, John Sillitoe on behalf of the Wellcome Sanger Institute COVID-19 Surveillance Team |
| EPI_ISL_2994917 | Lighthouse Laboratory Plymouth | Wellcome Sanger Institute for the COVID-19 Genomics UK (COG-UK) Consortium | Lighthouse Laboratory Plymouth and Alex Alderton, Roberto Amato, Jeffrey Barrett, Sonia Goncalves, Ewan Harrison, David K. Jackson, Ian Johnston, Dominic Kwiatkowski, Cordelia Langford, John Sillitoe on behalf of the Wellcome Sanger Institute COVID-19 Surveillance Team |
| EPI_ISL_2994918 | Lighthouse Lab in Glasgow | Wellcome Sanger Institute for the COVID-19 Genomics UK (COG-UK) Consortium | Harper VanSteenhouse, Yumi Kasai, David Gray, Carol Clugston, Anna Dominiczak and Alex Alderton, Roberto Amato, Jeffrey Barrett, Sonia Goncalves, Ewan Harrison, David K. Jackson, Ian Johnston, Dominic Kwiatkowski, Cordelia Langford, John Sillitoe on behalf of the Wellcome Sanger Institute COVID-19 Surveillance Team |
| EPI_ISL_2994919 | Lighthouse Laboratory Plymouth | Wellcome Sanger Institute for the COVID-19 Genomics UK (COG-UK) Consortium | Lighthouse Laboratory Plymouth and Alex Alderton, Roberto Amato, Jeffrey Barrett, Sonia Goncalves, Ewan Harrison, David K. Jackson, Ian Johnston, Dominic Kwiatkowski, Cordelia Langford, John Sillitoe on behalf of the Wellcome Sanger Institute COVID-19 Surveillance Team |
| EPI_ISL_2994920 | Lighthouse Lab in Glasgow | Wellcome Sanger Institute for the COVID-19 Genomics UK (COG-UK) Consortium | Harper VanSteenhouse, Yumi Kasai, David Gray, Carol Clugston, Anna Dominiczak and Alex Alderton, Roberto Amato, Jeffrey Barrett, Sonia Goncalves, Ewan Harrison, David K. Jackson, Ian Johnston, Dominic Kwiatkowski, Cordelia Langford, John Sillitoe on behalf of the Wellcome Sanger Institute COVID-19 Surveillance Team |
| EPI_ISL_2994921, EPI_ISL_2994922, EPI_ISL_2994923 | Lighthouse Laboratory Plymouth | Wellcome Sanger Institute for the COVID-19 Genomics UK (COG-UK) Consortium | Lighthouse Laboratory Plymouth and Alex Alderton, Roberto Amato, Jeffrey Barrett, Sonia Goncalves, Ewan Harrison, David K. Jackson, Ian Johnston, Dominic Kwiatkowski, Cordelia Langford, John Sillitoe on behalf of the Wellcome Sanger Institute COVID-19 Surveillance Team |
| EPI_ISL_2994924, EPI_ISL_2994925 | Lighthouse Lab in Milton Keynes | Wellcome Sanger Institute for the COVID-19 Genomics UK (COG-UK) Consortium | The Lighthouse Lab in Milton Keynes and Alex Alderton, Roberto Amato, Jeffrey Barrett, Sonia Goncalves, Ewan Harrison, David K. Jackson, Ian Johnston, Dominic Kwiatkowski, Cordelia Langford, John Sillitoe on behalf of the Wellcome Sanger Institute COVID-19 Surveillance Team |
| EPI_ISL_2994926 | Lighthouse Laboratory Plymouth | Wellcome Sanger Institute for the COVID-19 Genomics UK (COG-UK) Consortium | Lighthouse Laboratory Plymouth and Alex Alderton, Roberto Amato, Jeffrey Barrett, Sonia Goncalves, Ewan Harrison, David K. Jackson, Ian Johnston, Dominic Kwiatkowski, Cordelia Langford, John Sillitoe on behalf of the Wellcome Sanger Institute COVID-19 Surveillance Team |
| EPI_ISL_2994927, EPI_ISL_2994928 | Lighthouse Lab in Glasgow | Wellcome Sanger Institute for the COVID-19 Genomics UK (COG-UK) Consortium | Harper VanSteenhouse, Yumi Kasai, David Gray, Carol Clugston, Anna Dominiczak and Alex Alderton, Roberto Amato, Jeffrey Barrett, Sonia Goncalves, Ewan Harrison, David K. Jackson, Ian Johnston, Dominic Kwiatkowski, Cordelia Langford, John Sillitoe on behalf of the Wellcome Sanger Institute COVID-19 Surveillance Team |
| EPI_ISL_2994929, EPI_ISL_2994930 | Lighthouse Lab in Milton Keynes | Wellcome Sanger Institute for the COVID-19 Genomics UK (COG-UK) Consortium | The Lighthouse Lab in Milton Keynes and Alex Alderton, Roberto Amato, Jeffrey Barrett, Sonia Goncalves, Ewan Harrison, David K. Jackson, Ian Johnston, Dominic Kwiatkowski, Cordelia Langford, John Sillitoe on behalf of the Wellcome Sanger Institute COVID-19 Surveillance Team |

| | | | |
|---|---|---|---|
| EPI_ISL_2994931, EPI_ISL_2994932 | Lighthouse Laboratory Plymouth | Wellcome Sanger Institute for the COVID-19 Genomics UK (COG-UK) Consortium | Lighthouse Laboratory Plymouth and Alex Alderton, Roberto Amato, Jeffrey Barrett, Sonia Goncalves, Ewan Harrison, David K. Jackson, Ian Johnston, Dominic Kwiatkowski, Cordelia Langford, John Sillitoe on behalf of the Wellcome Sanger Institute COVID-19 Surveillance Team |
| EPI_ISL_2994933 | Lighthouse Lab in Glasgow | Wellcome Sanger Institute for the COVID-19 Genomics UK (COG-UK) Consortium | Harper VanSteenhouse, Yumi Kasai, David Gray, Carol Clugston, Anna Dominiczak and Alex Alderton, Roberto Amato, Jeffrey Barrett, Sonia Goncalves, Ewan Harrison, David K. Jackson, Ian Johnston, Dominic Kwiatkowski, Cordelia Langford, John Sillitoe on behalf of the Wellcome Sanger Institute COVID-19 Surveillance Team |
| EPI_ISL_2994934, EPI_ISL_2994935 | Lighthouse Lab in Milton Keynes | Wellcome Sanger Institute for the COVID-19 Genomics UK (COG-UK) Consortium | The Lighthouse Lab in Milton Keynes and Alex Alderton, Roberto Amato, Jeffrey Barrett, Sonia Goncalves, Ewan Harrison, David K. Jackson, Ian Johnston, Dominic Kwiatkowski, Cordelia Langford, John Sillitoe on behalf of the Wellcome Sanger Institute COVID-19 Surveillance Team |
| EPI_ISL_2994936, EPI_ISL_2994937 | Lighthouse Laboratory Plymouth | Wellcome Sanger Institute for the COVID-19 Genomics UK (COG-UK) Consortium | Lighthouse Laboratory Plymouth and Alex Alderton, Roberto Amato, Jeffrey Barrett, Sonia Goncalves, Ewan Harrison, David K. Jackson, Ian Johnston, Dominic Kwiatkowski, Cordelia Langford, John Sillitoe on behalf of the Wellcome Sanger Institute COVID-19 Surveillance Team |
| EPI_ISL_2994938 | Lighthouse Lab in Glasgow | Wellcome Sanger Institute for the COVID-19 Genomics UK (COG-UK) Consortium | Harper VanSteenhouse, Yumi Kasai, David Gray, Carol Clugston, Anna Dominiczak and Alex Alderton, Roberto Amato, Jeffrey Barrett, Sonia Goncalves, Ewan Harrison, David K. Jackson, Ian Johnston, Dominic Kwiatkowski, Cordelia Langford, John Sillitoe on behalf of the Wellcome Sanger Institute COVID-19 Surveillance Team |
| EPI_ISL_2994939 | Lighthouse Laboratory Plymouth | Wellcome Sanger Institute for the COVID-19 Genomics UK (COG-UK) Consortium | Lighthouse Laboratory Plymouth and Alex Alderton, Roberto Amato, Jeffrey Barrett, Sonia Goncalves, Ewan Harrison, David K. Jackson, Ian Johnston, Dominic Kwiatkowski, Cordelia Langford, John Sillitoe on behalf of the Wellcome Sanger Institute COVID-19 Surveillance Team |
| EPI_ISL_2994940 | Lighthouse Lab in Glasgow | Wellcome Sanger Institute for the COVID-19 Genomics UK (COG-UK) Consortium | Harper VanSteenhouse, Yumi Kasai, David Gray, Carol Clugston, Anna Dominiczak and Alex Alderton, Roberto Amato, Jeffrey Barrett, Sonia Goncalves, Ewan Harrison, David K. Jackson, Ian Johnston, Dominic Kwiatkowski, Cordelia Langford, John Sillitoe on behalf of the Wellcome Sanger Institute COVID-19 Surveillance Team |
| EPI_ISL_2994941 | Lighthouse Laboratory Plymouth | Wellcome Sanger Institute for the COVID-19 Genomics UK (COG-UK) Consortium | Lighthouse Laboratory Plymouth and Alex Alderton, Roberto Amato, Jeffrey Barrett, Sonia Goncalves, Ewan Harrison, David K. Jackson, Ian Johnston, Dominic Kwiatkowski, Cordelia Langford, John Sillitoe on behalf of the Wellcome Sanger Institute COVID-19 Surveillance Team |
| EPI_ISL_2994942 | Lighthouse Lab in Glasgow | Wellcome Sanger Institute for the COVID-19 Genomics UK (COG-UK) Consortium | Harper VanSteenhouse, Yumi Kasai, David Gray, Carol Clugston, Anna Dominiczak and Alex Alderton, Roberto Amato, Jeffrey Barrett, Sonia Goncalves, Ewan Harrison, David K. Jackson, Ian Johnston, Dominic Kwiatkowski, Cordelia Langford, John Sillitoe on behalf of the Wellcome Sanger Institute COVID-19 Surveillance Team |
| EPI_ISL_2994943 | Lighthouse Lab in Milton Keynes | Wellcome Sanger Institute for the COVID-19 Genomics UK (COG-UK) Consortium | The Lighthouse Lab in Milton Keynes and Alex Alderton, Roberto Amato, Jeffrey Barrett, Sonia Goncalves, Ewan Harrison, David K. Jackson, Ian Johnston, Dominic Kwiatkowski, Cordelia Langford, John Sillitoe on behalf of the Wellcome Sanger Institute COVID-19 Surveillance Team |
| EPI_ISL_2994944 | Lighthouse Laboratory Plymouth | Wellcome Sanger Institute for the COVID-19 Genomics UK (COG-UK) Consortium | Lighthouse Laboratory Plymouth and Alex Alderton, Roberto Amato, Jeffrey Barrett, Sonia Goncalves, Ewan Harrison, David K. Jackson, Ian Johnston, Dominic Kwiatkowski, Cordelia Langford, John Sillitoe on behalf of the Wellcome Sanger Institute COVID-19 Surveillance Team |
| EPI_ISL_2994945, EPI_ISL_2994946, EPI_ISL_2994947 | Lighthouse Lab in Milton Keynes | Wellcome Sanger Institute for the COVID-19 Genomics UK (COG-UK) Consortium | The Lighthouse Lab in Milton Keynes and Alex Alderton, Roberto Amato, Jeffrey Barrett, Sonia Goncalves, Ewan Harrison, David K. Jackson, Ian Johnston, Dominic Kwiatkowski, Cordelia Langford, John Sillitoe on behalf of the Wellcome Sanger Institute COVID-19 Surveillance Team |
| EPI_ISL_2994948 | Lighthouse Lab in Glasgow | Wellcome Sanger Institute for the COVID-19 Genomics UK (COG-UK) Consortium | Harper VanSteenhouse, Yumi Kasai, David Gray, Carol Clugston, Anna Dominiczak and Alex Alderton, Roberto Amato, Jeffrey Barrett, Sonia Goncalves, Ewan Harrison, David K. Jackson, Ian Johnston, Dominic Kwiatkowski, Cordelia Langford, John Sillitoe on behalf of the Wellcome Sanger Institute COVID-19 Surveillance Team |
| EPI_ISL_2994949 | Lighthouse Laboratory Plymouth | Wellcome Sanger Institute for the COVID-19 Genomics UK (COG-UK) Consortium | Lighthouse Laboratory Plymouth and Alex Alderton, Roberto Amato, Jeffrey Barrett, Sonia Goncalves, Ewan Harrison, David K. Jackson, Ian Johnston, Dominic Kwiatkowski, Cordelia Langford, John Sillitoe on behalf of the Wellcome Sanger Institute COVID-19 Surveillance Team |
| EPI_ISL_2994950 | Lighthouse Lab in Milton Keynes | Wellcome Sanger Institute for the COVID-19 Genomics UK (COG-UK) Consortium | The Lighthouse Lab in Milton Keynes and Alex Alderton, Roberto Amato, Jeffrey Barrett, Sonia Goncalves, Ewan Harrison, David K. Jackson, Ian Johnston, Dominic Kwiatkowski, Cordelia Langford, John Sillitoe on behalf of the Wellcome Sanger Institute COVID-19 Surveillance Team |
| EPI_ISL_2994951 | Lighthouse Laboratory Plymouth | Wellcome Sanger Institute for the COVID-19 Genomics UK (COG-UK) Consortium | Lighthouse Laboratory Plymouth and Alex Alderton, Roberto Amato, Jeffrey Barrett, Sonia Goncalves, Ewan Harrison, David K. Jackson, Ian Johnston, Dominic Kwiatkowski, Cordelia Langford, John Sillitoe on behalf of the Wellcome Sanger Institute COVID-19 Surveillance Team |
| EPI_ISL_2994952 | Lighthouse Lab in Glasgow | Wellcome Sanger Institute for the COVID-19 Genomics UK (COG-UK) Consortium | Harper VanSteenhouse, Yumi Kasai, David Gray, Carol Clugston, Anna Dominiczak and Alex Alderton, Roberto Amato, Jeffrey Barrett, Sonia Goncalves, Ewan Harrison, David K. Jackson, Ian Johnston, Dominic Kwiatkowski, Cordelia Langford, John Sillitoe on behalf of the Wellcome Sanger Institute COVID-19 Surveillance Team |
| EPI_ISL_2994953 | Lighthouse Laboratory Plymouth | Wellcome Sanger Institute for the COVID-19 Genomics UK (COG-UK) Consortium | Lighthouse Laboratory Plymouth and Alex Alderton, Roberto Amato, Jeffrey Barrett, Sonia Goncalves, Ewan Harrison, David K. Jackson, Ian Johnston, Dominic Kwiatkowski, Cordelia Langford, John Sillitoe on behalf of the Wellcome Sanger Institute COVID-19 Surveillance Team |
| EPI_ISL_2994954 | Lighthouse Lab in Milton Keynes | Wellcome Sanger Institute for the COVID-19 Genomics UK (COG-UK) Consortium | The Lighthouse Lab in Milton Keynes and Alex Alderton, Roberto Amato, Jeffrey Barrett, Sonia Goncalves, Ewan Harrison, David K. Jackson, Ian Johnston, Dominic Kwiatkowski, Cordelia Langford, John Sillitoe on behalf of the Wellcome Sanger Institute COVID-19 Surveillance Team |
| EPI_ISL_2994955, EPI_ISL_2994956 | Lighthouse Laboratory Plymouth | Wellcome Sanger Institute for the COVID-19 Genomics UK (COG-UK) Consortium | Lighthouse Laboratory Plymouth and Alex Alderton, Roberto Amato, Jeffrey Barrett, Sonia Goncalves, Ewan Harrison, David K. Jackson, Ian Johnston, Dominic Kwiatkowski, Cordelia Langford, John Sillitoe on behalf of the Wellcome Sanger Institute COVID-19 Surveillance Team |
| EPI_ISL_2994957 | Lighthouse Lab in Milton Keynes | Wellcome Sanger Institute for the COVID-19 Genomics UK (COG-UK) Consortium | The Lighthouse Lab in Milton Keynes and Alex Alderton, Roberto Amato, Jeffrey Barrett, Sonia Goncalves, Ewan Harrison, David K. Jackson, Ian Johnston, Dominic Kwiatkowski, Cordelia Langford, John Sillitoe on behalf of the Wellcome Sanger Institute COVID-19 Surveillance Team |
| EPI_ISL_2994958, EPI_ISL_2994959, EPI_ISL_2994960 | Lighthouse Laboratory Plymouth | Wellcome Sanger Institute for the COVID-19 Genomics UK (COG-UK) Consortium | Lighthouse Laboratory Plymouth and Alex Alderton, Roberto Amato, Jeffrey Barrett, Sonia Goncalves, Ewan Harrison, David K. Jackson, Ian Johnston, Dominic Kwiatkowski, Cordelia Langford, John Sillitoe on behalf of the Wellcome Sanger Institute COVID-19 Surveillance Team |
| EPI_ISL_2994961 | Lighthouse Lab in Glasgow | Wellcome Sanger Institute for the COVID-19 Genomics UK (COG-UK) Consortium | Harper VanSteenhouse, Yumi Kasai, David Gray, Carol Clugston, Anna Dominiczak and Alex Alderton, Roberto Amato, Jeffrey Barrett, Sonia Goncalves, Ewan Harrison, David K. Jackson, Ian Johnston, Dominic Kwiatkowski, Cordelia Langford, John Sillitoe on behalf of the Wellcome Sanger Institute COVID-19 Surveillance Team |
| EPI_ISL_2994962 | Lighthouse Laboratory Plymouth | Wellcome Sanger Institute for the COVID-19 Genomics UK (COG-UK) Consortium | Lighthouse Laboratory Plymouth and Alex Alderton, Roberto Amato, Jeffrey Barrett, Sonia Goncalves, Ewan Harrison, David K. Jackson, Ian Johnston, Dominic Kwiatkowski, Cordelia Langford, John Sillitoe on behalf of the Wellcome Sanger Institute COVID-19 Surveillance Team |
| EPI_ISL_2994963 | Lighthouse Lab in Glasgow | Wellcome Sanger Institute for the COVID-19 Genomics UK (COG-UK) Consortium | Harper VanSteenhouse, Yumi Kasai, David Gray, Carol Clugston, Anna Dominiczak and Alex Alderton, Roberto Amato, Jeffrey Barrett, Sonia Goncalves, Ewan Harrison, David K. Jackson, Ian Johnston, Dominic Kwiatkowski, Cordelia Langford, John Sillitoe on behalf of the Wellcome Sanger Institute COVID-19 Surveillance Team |
| EPI_ISL_2994964 | Lighthouse Lab in Milton Keynes | Wellcome Sanger Institute for the COVID-19 Genomics UK (COG-UK) Consortium | The Lighthouse Lab in Milton Keynes and Alex Alderton, Roberto Amato, Jeffrey Barrett, Sonia Goncalves, Ewan Harrison, David K. Jackson, Ian Johnston, Dominic Kwiatkowski, Cordelia Langford, John Sillitoe on behalf of the Wellcome Sanger Institute COVID-19 Surveillance Team |
| EPI_ISL_2994965, EPI_ISL_2994966 | Lighthouse Lab in Glasgow | Wellcome Sanger Institute for the COVID-19 Genomics UK (COG-UK) Consortium | Harper VanSteenhouse, Yumi Kasai, David Gray, Carol Clugston, Anna Dominiczak and Alex Alderton, Roberto Amato, Jeffrey Barrett, Sonia Goncalves, Ewan Harrison, David K. Jackson, Ian Johnston, Dominic Kwiatkowski, Cordelia Langford, John Sillitoe on behalf of the Wellcome Sanger Institute COVID-19 Surveillance Team |
| EPI_ISL_2994967, EPI_ISL_2994968, EPI_ISL_2994969, EPI_ISL_2994970 | Lighthouse Laboratory Plymouth | Wellcome Sanger Institute for the COVID-19 Genomics UK (COG-UK) Consortium | Lighthouse Laboratory Plymouth and Alex Alderton, Roberto Amato, Jeffrey Barrett, Sonia Goncalves, Ewan Harrison, David K. Jackson, Ian Johnston, Dominic Kwiatkowski, Cordelia Langford, John Sillitoe on behalf of the Wellcome Sanger Institute COVID-19 Surveillance Team |
| EPI_ISL_2994971 | Lighthouse Lab in Glasgow | Wellcome Sanger Institute for the COVID-19 Genomics UK (COG-UK) Consortium | Harper VanSteenhouse, Yumi Kasai, David Gray, Carol Clugston, Anna Dominiczak and Alex Alderton, Roberto Amato, Jeffrey Barrett, Sonia Goncalves, Ewan Harrison, David K. Jackson, Ian Johnston, Dominic Kwiatkowski, Cordelia Langford, John Sillitoe on behalf of the Wellcome Sanger Institute COVID-19 Surveillance Team |
| EPI_ISL_2994972 | Lighthouse Lab in Milton Keynes | Wellcome Sanger Institute for the COVID-19 Genomics UK (COG-UK) Consortium | The Lighthouse Lab in Milton Keynes and Alex Alderton, Roberto Amato, Jeffrey Barrett, Sonia Goncalves, Ewan Harrison, David K. Jackson, Ian Johnston, Dominic Kwiatkowski, Cordelia Langford, John Sillitoe on behalf of the Wellcome Sanger Institute COVID-19 Surveillance Team |
| EPI_ISL_2994973, EPI_ISL_2994974 | Lighthouse Laboratory Plymouth | Wellcome Sanger Institute for the COVID-19 Genomics UK (COG-UK) Consortium | Lighthouse Laboratory Plymouth and Alex Alderton, Roberto Amato, Jeffrey Barrett, Sonia Goncalves, Ewan Harrison, David K. Jackson, Ian Johnston, Dominic Kwiatkowski, Cordelia Langford, John Sillitoe on behalf of the Wellcome Sanger Institute COVID-19 Surveillance Team |
| EPI_ISL_2994975 | Lighthouse Lab in Milton Keynes | Wellcome Sanger Institute for the COVID-19 Genomics UK (COG-UK) Consortium | The Lighthouse Lab in Milton Keynes and Alex Alderton, Roberto Amato, Jeffrey Barrett, Sonia Goncalves, Ewan Harrison, David K. Jackson, Ian Johnston, Dominic Kwiatkowski, Cordelia Langford, John Sillitoe on behalf of the Wellcome Sanger Institute COVID-19 Surveillance Team |
| EPI_ISL_2994976, EPI_ISL_2994977, EPI_ISL_2994978 | Lighthouse Laboratory Plymouth | Wellcome Sanger Institute for the COVID-19 Genomics UK (COG-UK) Consortium | Lighthouse Laboratory Plymouth and Alex Alderton, Roberto Amato, Jeffrey Barrett, Sonia Goncalves, Ewan Harrison, David K. Jackson, Ian Johnston, Dominic Kwiatkowski, Cordelia Langford, John Sillitoe on behalf of the Wellcome Sanger Institute COVID-19 Surveillance Team |
| EPI_ISL_2994979 | Lighthouse Lab in Milton Keynes | Wellcome Sanger Institute for the COVID-19 Genomics UK (COG-UK) Consortium | The Lighthouse Lab in Milton Keynes and Alex Alderton, Roberto Amato, Jeffrey Barrett, Sonia Goncalves, Ewan Harrison, David K. Jackson, Ian Johnston, Dominic Kwiatkowski, Cordelia Langford, John Sillitoe on behalf of the Wellcome Sanger Institute COVID-19 Surveillance Team |
| EPI_ISL_2994980, EPI_ISL_2994981, EPI_ISL_2994982 | Lighthouse Laboratory Plymouth | Wellcome Sanger Institute for the COVID-19 Genomics UK (COG-UK) Consortium | Lighthouse Laboratory Plymouth and Alex Alderton, Roberto Amato, Jeffrey Barrett, Sonia Goncalves, Ewan Harrison, David K. Jackson, Ian Johnston, Dominic Kwiatkowski, Cordelia Langford, John Sillitoe on behalf of the Wellcome Sanger Institute COVID-19 Surveillance Team |
| EPI_ISL_2994983 | Lighthouse Lab in Milton Keynes | Wellcome Sanger Institute for the COVID-19 Genomics UK (COG-UK) Consortium | The Lighthouse Lab in Milton Keynes and Alex Alderton, Roberto Amato, Jeffrey Barrett, Sonia Goncalves, Ewan Harrison, David K. Jackson, Ian Johnston, Dominic Kwiatkowski, Cordelia Langford, John Sillitoe on behalf of the Wellcome Sanger Institute COVID-19 Surveillance Team |
| EPI_ISL_2994984, EPI_ISL_2994985 | Lighthouse Laboratory Plymouth | Wellcome Sanger Institute for the COVID-19 Genomics UK (COG-UK) Consortium | Lighthouse Laboratory Plymouth and Alex Alderton, Roberto Amato, Jeffrey Barrett, Sonia Goncalves, Ewan Harrison, David K. Jackson, Ian Johnston, Dominic Kwiatkowski, Cordelia Langford, John Sillitoe on behalf of the Wellcome Sanger Institute COVID-19 Surveillance Team |
| EPI_ISL_2994986, EPI_ISL_2994987, EPI_ISL_2994988 | Lighthouse Lab in Milton Keynes | Wellcome Sanger Institute for the COVID-19 Genomics UK (COG-UK) Consortium | The Lighthouse Lab in Milton Keynes and Alex Alderton, Roberto Amato, Jeffrey Barrett, Sonia Goncalves, Ewan Harrison, David K. Jackson, Ian Johnston, Dominic Kwiatkowski, Cordelia Langford, John Sillitoe on behalf of the Wellcome Sanger Institute COVID-19 Surveillance Team |
| EPI_ISL_2994989, EPI_ISL_2994990 | Lighthouse Lab in Glasgow | Wellcome Sanger Institute for the COVID-19 Genomics UK (COG-UK) Consortium | Harper VanSteenhouse, Yumi Kasai, David Gray, Carol Clugston, Anna Dominiczak and Alex Alderton, Roberto Amato, Jeffrey Barrett, Sonia Goncalves, Ewan Harrison, David K. Jackson, Ian Johnston, Dominic Kwiatkowski, Cordelia Langford, John Sillitoe on behalf of the Wellcome Sanger Institute COVID-19 Surveillance Team |
| EPI_ISL_2994991 | Lighthouse Laboratory Plymouth | Wellcome Sanger Institute for the COVID-19 Genomics UK (COG-UK) Consortium | Lighthouse Laboratory Plymouth and Alex Alderton, Roberto Amato, Jeffrey Barrett, Sonia Goncalves, Ewan Harrison, David K. Jackson, Ian Johnston, Dominic Kwiatkowski, Cordelia Langford, John Sillitoe on behalf of the Wellcome Sanger Institute COVID-19 Surveillance Team |
| EPI_ISL_2994992, EPI_ISL_2994993 | Lighthouse Lab in Glasgow | Wellcome Sanger Institute for the COVID-19 Genomics UK (COG-UK) Consortium | Harper VanSteenhouse, Yumi Kasai, David Gray, Carol Clugston, Anna Dominiczak and Alex Alderton, Roberto Amato, Jeffrey Barrett, Sonia Goncalves, Ewan Harrison, David K. Jackson, Ian Johnston, Dominic Kwiatkowski, Cordelia Langford, John Sillitoe on behalf of the Wellcome Sanger Institute COVID-19 Surveillance Team |
| EPI_ISL_2994994 | Lighthouse Laboratory Plymouth | Wellcome Sanger Institute for the COVID-19 Genomics UK (COG-UK) Consortium | Lighthouse Laboratory Plymouth and Alex Alderton, Roberto Amato, Jeffrey Barrett, Sonia Goncalves, Ewan Harrison, David K. Jackson, Ian Johnston, Dominic Kwiatkowski, Cordelia Langford, John Sillitoe on behalf of the Wellcome Sanger Institute COVID-19 Surveillance Team |
| EPI_ISL_2994995 | Lighthouse Lab in Milton Keynes | Wellcome Sanger Institute for the COVID-19 Genomics UK (COG-UK) | The Lighthouse Lab in Milton Keynes and Alex Alderton, Roberto Amato, Jeffrey Barrett, Sonia Goncalves, Ewan Harrison, David K. Jackson, Ian Johnston, Dominic Kwiatkowski, Cordelia Langford, John Sillitoe on behalf of the Wellcome Sanger |

| | | | |
|---|---|---|---|
| | | Consortium | Institute COVID-19 Surveillance Team |
| EPI_ISL_2994996, EPI_ISL_2994997 | Lighthouse Laboratory Plymouth | Wellcome Sanger Institute for the COVID-19 Genomics UK (COG-UK) Consortium | Lighthouse Laboratory Plymouth and Alex Alderton, Roberto Amato, Jeffrey Barrett, Sonia Goncalves, Ewan Harrison, David K. Jackson, Ian Johnston, Dominic Kwiatkowski, Cordelia Langford, John Sillitoe on behalf of the Wellcome Sanger Institute COVID-19 Surveillance Team |
| EPI_ISL_2994998 | Lighthouse Lab in Glasgow | Wellcome Sanger Institute for the COVID-19 Genomics UK (COG-UK) Consortium | Harper VanSteenhouse, Yumi Kasai, David Gray, Carol Clugston, Anna Dominiczak and Alex Alderton, Roberto Amato, Jeffrey Barrett, Sonia Goncalves, Ewan Harrison, David K. Jackson, Ian Johnston, Dominic Kwiatkowski, Cordelia Langford, John Sillitoe on behalf of the Wellcome Sanger Institute COVID-19 Surveillance Team |
| EPI_ISL_2994999 | Lighthouse Laboratory Plymouth | Wellcome Sanger Institute for the COVID-19 Genomics UK (COG-UK) Consortium | Lighthouse Laboratory Plymouth and Alex Alderton, Roberto Amato, Jeffrey Barrett, Sonia Goncalves, Ewan Harrison, David K. Jackson, Ian Johnston, Dominic Kwiatkowski, Cordelia Langford, John Sillitoe on behalf of the Wellcome Sanger Institute COVID-19 Surveillance Team |
| EPI_ISL_2995000, EPI_ISL_2995001, EPI_ISL_2995002 | Lighthouse Lab in Milton Keynes | Wellcome Sanger Institute for the COVID-19 Genomics UK (COG-UK) Consortium | The Lighthouse Lab in Milton Keynes and Alex Alderton, Roberto Amato, Jeffrey Barrett, Sonia Goncalves, Ewan Harrison, David K. Jackson, Ian Johnston, Dominic Kwiatkowski, Cordelia Langford, John Sillitoe on behalf of the Wellcome Sanger Institute COVID-19 Surveillance Team |
| EPI_ISL_2995003 | Lighthouse Lab in Glasgow | Wellcome Sanger Institute for the COVID-19 Genomics UK (COG-UK) Consortium | Harper VanSteenhouse, Yumi Kasai, David Gray, Carol Clugston, Anna Dominiczak and Alex Alderton, Roberto Amato, Jeffrey Barrett, Sonia Goncalves, Ewan Harrison, David K. Jackson, Ian Johnston, Dominic Kwiatkowski, Cordelia Langford, John Sillitoe on behalf of the Wellcome Sanger Institute COVID-19 Surveillance Team |
| EPI_ISL_2995004, EPI_ISL_2995005 | Lighthouse Laboratory Plymouth | Wellcome Sanger Institute for the COVID-19 Genomics UK (COG-UK) Consortium | Lighthouse Laboratory Plymouth and Alex Alderton, Roberto Amato, Jeffrey Barrett, Sonia Goncalves, Ewan Harrison, David K. Jackson, Ian Johnston, Dominic Kwiatkowski, Cordelia Langford, John Sillitoe on behalf of the Wellcome Sanger Institute COVID-19 Surveillance Team |
| EPI_ISL_2995006, EPI_ISL_2995007 | Lighthouse Lab in Milton Keynes | Wellcome Sanger Institute for the COVID-19 Genomics UK (COG-UK) Consortium | The Lighthouse Lab in Milton Keynes and Alex Alderton, Roberto Amato, Jeffrey Barrett, Sonia Goncalves, Ewan Harrison, David K. Jackson, Ian Johnston, Dominic Kwiatkowski, Cordelia Langford, John Sillitoe on behalf of the Wellcome Sanger Institute COVID-19 Surveillance Team |
| EPI_ISL_2995008 | Lighthouse Lab in Glasgow | Wellcome Sanger Institute for the COVID-19 Genomics UK (COG-UK) Consortium | Harper VanSteenhouse, Yumi Kasai, David Gray, Carol Clugston, Anna Dominiczak and Alex Alderton, Roberto Amato, Jeffrey Barrett, Sonia Goncalves, Ewan Harrison, David K. Jackson, Ian Johnston, Dominic Kwiatkowski, Cordelia Langford, John Sillitoe on behalf of the Wellcome Sanger Institute COVID-19 Surveillance Team |
| EPI_ISL_2995009, EPI_ISL_2995010 | Lighthouse Laboratory Plymouth | Wellcome Sanger Institute for the COVID-19 Genomics UK (COG-UK) Consortium | Lighthouse Laboratory Plymouth and Alex Alderton, Roberto Amato, Jeffrey Barrett, Sonia Goncalves, Ewan Harrison, David K. Jackson, Ian Johnston, Dominic Kwiatkowski, Cordelia Langford, John Sillitoe on behalf of the Wellcome Sanger Institute COVID-19 Surveillance Team |
| EPI_ISL_2995011 | Lighthouse Lab in Glasgow | Wellcome Sanger Institute for the COVID-19 Genomics UK (COG-UK) Consortium | Harper VanSteenhouse, Yumi Kasai, David Gray, Carol Clugston, Anna Dominiczak and Alex Alderton, Roberto Amato, Jeffrey Barrett, Sonia Goncalves, Ewan Harrison, David K. Jackson, Ian Johnston, Dominic Kwiatkowski, Cordelia Langford, John Sillitoe on behalf of the Wellcome Sanger Institute COVID-19 Surveillance Team |
| EPI_ISL_2995012, EPI_ISL_2995013 | Lighthouse Laboratory Plymouth | Wellcome Sanger Institute for the COVID-19 Genomics UK (COG-UK) Consortium | Lighthouse Laboratory Plymouth and Alex Alderton, Roberto Amato, Jeffrey Barrett, Sonia Goncalves, Ewan Harrison, David K. Jackson, Ian Johnston, Dominic Kwiatkowski, Cordelia Langford, John Sillitoe on behalf of the Wellcome Sanger Institute COVID-19 Surveillance Team |
| EPI_ISL_2995014 | Lighthouse Lab in Milton Keynes | Wellcome Sanger Institute for the COVID-19 Genomics UK (COG-UK) Consortium | The Lighthouse Lab in Milton Keynes and Alex Alderton, Roberto Amato, Jeffrey Barrett, Sonia Goncalves, Ewan Harrison, David K. Jackson, Ian Johnston, Dominic Kwiatkowski, Cordelia Langford, John Sillitoe on behalf of the Wellcome Sanger Institute COVID-19 Surveillance Team |
| EPI_ISL_2995015 | Lighthouse Laboratory Plymouth | Wellcome Sanger Institute for the COVID-19 Genomics UK (COG-UK) Consortium | Lighthouse Laboratory Plymouth and Alex Alderton, Roberto Amato, Jeffrey Barrett, Sonia Goncalves, Ewan Harrison, David K. Jackson, Ian Johnston, Dominic Kwiatkowski, Cordelia Langford, John Sillitoe on behalf of the Wellcome Sanger Institute COVID-19 Surveillance Team |
| EPI_ISL_2995016 | Lighthouse Lab in Glasgow | Wellcome Sanger Institute for the COVID-19 Genomics UK (COG-UK) Consortium | Harper VanSteenhouse, Yumi Kasai, David Gray, Carol Clugston, Anna Dominiczak and Alex Alderton, Roberto Amato, Jeffrey Barrett, Sonia Goncalves, Ewan Harrison, David K. Jackson, Ian Johnston, Dominic Kwiatkowski, Cordelia Langford, John Sillitoe on behalf of the Wellcome Sanger Institute COVID-19 Surveillance Team |
| EPI_ISL_2995017 | Lighthouse Laboratory Plymouth | Wellcome Sanger Institute for the COVID-19 Genomics UK (COG-UK) Consortium | Lighthouse Laboratory Plymouth and Alex Alderton, Roberto Amato, Jeffrey Barrett, Sonia Goncalves, Ewan Harrison, David K. Jackson, Ian Johnston, Dominic Kwiatkowski, Cordelia Langford, John Sillitoe on behalf of the Wellcome Sanger Institute COVID-19 Surveillance Team |
| EPI_ISL_2995018 | Lighthouse Lab in Glasgow | Wellcome Sanger Institute for the COVID-19 Genomics UK (COG-UK) Consortium | Harper VanSteenhouse, Yumi Kasai, David Gray, Carol Clugston, Anna Dominiczak and Alex Alderton, Roberto Amato, Jeffrey Barrett, Sonia Goncalves, Ewan Harrison, David K. Jackson, Ian Johnston, Dominic Kwiatkowski, Cordelia Langford, John Sillitoe on behalf of the Wellcome Sanger Institute COVID-19 Surveillance Team |
| EPI_ISL_2995019, EPI_ISL_2995020 | Lighthouse Laboratory Plymouth | Wellcome Sanger Institute for the COVID-19 Genomics UK (COG-UK) Consortium | Lighthouse Laboratory Plymouth and Alex Alderton, Roberto Amato, Jeffrey Barrett, Sonia Goncalves, Ewan Harrison, David K. Jackson, Ian Johnston, Dominic Kwiatkowski, Cordelia Langford, John Sillitoe on behalf of the Wellcome Sanger Institute COVID-19 Surveillance Team |
| EPI_ISL_2995021 | Lighthouse Lab in Milton Keynes | Wellcome Sanger Institute for the COVID-19 Genomics UK (COG-UK) Consortium | The Lighthouse Lab in Milton Keynes and Alex Alderton, Roberto Amato, Jeffrey Barrett, Sonia Goncalves, Ewan Harrison, David K. Jackson, Ian Johnston, Dominic Kwiatkowski, Cordelia Langford, John Sillitoe on behalf of the Wellcome Sanger Institute COVID-19 Surveillance Team |
| EPI_ISL_2995022 | Lighthouse Lab in Glasgow | Wellcome Sanger Institute for the COVID-19 Genomics UK (COG-UK) Consortium | Harper VanSteenhouse, Yumi Kasai, David Gray, Carol Clugston, Anna Dominiczak and Alex Alderton, Roberto Amato, Jeffrey Barrett, Sonia Goncalves, Ewan Harrison, David K. Jackson, Ian Johnston, Dominic Kwiatkowski, Cordelia Langford, John Sillitoe on behalf of the Wellcome Sanger Institute COVID-19 Surveillance Team |
| EPI_ISL_2995023 | Lighthouse Laboratory Plymouth | Wellcome Sanger Institute for the COVID-19 Genomics UK (COG-UK) Consortium | Lighthouse Laboratory Plymouth and Alex Alderton, Roberto Amato, Jeffrey Barrett, Sonia Goncalves, Ewan Harrison, David K. Jackson, Ian Johnston, Dominic Kwiatkowski, Cordelia Langford, John Sillitoe on behalf of the Wellcome Sanger Institute COVID-19 Surveillance Team |
| EPI_ISL_2995024, EPI_ISL_2995025 | Lighthouse Lab in Glasgow | Wellcome Sanger Institute for the COVID-19 Genomics UK (COG-UK) Consortium | Harper VanSteenhouse, Yumi Kasai, David Gray, Carol Clugston, Anna Dominiczak and Alex Alderton, Roberto Amato, Jeffrey Barrett, Sonia Goncalves, Ewan Harrison, David K. Jackson, Ian Johnston, Dominic Kwiatkowski, Cordelia Langford, John Sillitoe on behalf of the Wellcome Sanger Institute COVID-19 Surveillance Team |
| EPI_ISL_2995026 | Lighthouse Laboratory Plymouth | Wellcome Sanger Institute for the COVID-19 Genomics UK (COG-UK) Consortium | Lighthouse Laboratory Plymouth and Alex Alderton, Roberto Amato, Jeffrey Barrett, Sonia Goncalves, Ewan Harrison, David K. Jackson, Ian Johnston, Dominic Kwiatkowski, Cordelia Langford, John Sillitoe on behalf of the Wellcome Sanger Institute COVID-19 Surveillance Team |
| EPI_ISL_2995027 | Lighthouse Lab in Milton Keynes | Wellcome Sanger Institute for the COVID-19 Genomics UK (COG-UK) Consortium | The Lighthouse Lab in Milton Keynes and Alex Alderton, Roberto Amato, Jeffrey Barrett, Sonia Goncalves, Ewan Harrison, David K. Jackson, Ian Johnston, Dominic Kwiatkowski, Cordelia Langford, John Sillitoe on behalf of the Wellcome Sanger Institute COVID-19 Surveillance Team |
| EPI_ISL_2995028 | Berkshire and Surrey Pathology Services Lighthouse Laboratory | Wellcome Sanger Institute for the COVID-19 Genomics UK (COG-UK) Consortium | Berkshire and Surrey Pathology Services Lighthouse Laboratory and Alex Alderton, Roberto Amato, Jeffrey Barrett, Sonia Goncalves, Ewan Harrison, David K. Jackson, Ian Johnston, Dominic Kwiatkowski, Cordelia Langford, John Sillitoe on behalf of the Wellcome Sanger Institute COVID-19 Surveillance Team |
| EPI_ISL_2995029 | Lighthouse Lab in Glasgow | Wellcome Sanger Institute for the COVID-19 Genomics UK (COG-UK) Consortium | Harper VanSteenhouse, Yumi Kasai, David Gray, Carol Clugston, Anna Dominiczak and Alex Alderton, Roberto Amato, Jeffrey Barrett, Sonia Goncalves, Ewan Harrison, David K. Jackson, Ian Johnston, Dominic Kwiatkowski, Cordelia Langford, John Sillitoe on behalf of the Wellcome Sanger Institute COVID-19 Surveillance Team |
| EPI_ISL_2995030 | Berkshire and Surrey Pathology Services Lighthouse Laboratory | Wellcome Sanger Institute for the COVID-19 Genomics UK (COG-UK) Consortium | Berkshire and Surrey Pathology Services Lighthouse Laboratory and Alex Alderton, Roberto Amato, Jeffrey Barrett, Sonia Goncalves, Ewan Harrison, David K. Jackson, Ian Johnston, Dominic Kwiatkowski, Cordelia Langford, John Sillitoe on behalf of the Wellcome Sanger Institute COVID-19 Surveillance Team |
| EPI_ISL_2995031 | Lighthouse Lab in Alderley Park | Wellcome Sanger Institute for the COVID-19 Genomics UK (COG-UK) Consortium | Jacquelyn Wynn, Mairead Hyland, The Lighthouse Lab in Alderley Park and Alex Alderton, Roberto Amato, Jeffrey Barrett, Sonia Goncalves, Ewan Harrison, David K. Jackson, Ian Johnston, Dominic Kwiatkowski, Cordelia Langford, John Sillitoe on behalf of the Wellcome Sanger Institute COVID-19 Surveillance Team |
| EPI_ISL_2995032 | Berkshire and Surrey Pathology Services Lighthouse Laboratory | Wellcome Sanger Institute for the COVID-19 Genomics UK (COG-UK) Consortium | Berkshire and Surrey Pathology Services Lighthouse Laboratory and Alex Alderton, Roberto Amato, Jeffrey Barrett, Sonia Goncalves, Ewan Harrison, David K. Jackson, Ian Johnston, Dominic Kwiatkowski, Cordelia Langford, John Sillitoe on behalf of the Wellcome Sanger Institute COVID-19 Surveillance Team |
| EPI_ISL_2995033 | Lighthouse Lab in Alderley Park | Wellcome Sanger Institute for the COVID-19 Genomics UK (COG-UK) Consortium | Jacquelyn Wynn, Mairead Hyland, The Lighthouse Lab in Alderley Park and Alex Alderton, Roberto Amato, Jeffrey Barrett, Sonia Goncalves, Ewan Harrison, David K. Jackson, Ian Johnston, Dominic Kwiatkowski, Cordelia Langford, John Sillitoe on behalf of the Wellcome Sanger Institute COVID-19 Surveillance Team |
| EPI_ISL_2995034 | Berkshire and Surrey Pathology Services Lighthouse Laboratory | Wellcome Sanger Institute for the COVID-19 Genomics UK (COG-UK) Consortium | Berkshire and Surrey Pathology Services Lighthouse Laboratory and Alex Alderton, Roberto Amato, Jeffrey Barrett, Sonia Goncalves, Ewan Harrison, David K. Jackson, Ian Johnston, Dominic Kwiatkowski, Cordelia Langford, John Sillitoe on behalf of the Wellcome Sanger Institute COVID-19 Surveillance Team |
| EPI_ISL_2995035 | Lighthouse Lab in Glasgow | Wellcome Sanger Institute for the COVID-19 Genomics UK (COG-UK) Consortium | Harper VanSteenhouse, Yumi Kasai, David Gray, Carol Clugston, Anna Dominiczak and Alex Alderton, Roberto Amato, Jeffrey Barrett, Sonia Goncalves, Ewan Harrison, David K. Jackson, Ian Johnston, Dominic Kwiatkowski, Cordelia Langford, John Sillitoe on behalf of the Wellcome Sanger Institute COVID-19 Surveillance Team |
| EPI_ISL_2995036 | Lighthouse Lab in Milton Keynes | Wellcome Sanger Institute for the COVID-19 Genomics UK (COG-UK) Consortium | The Lighthouse Lab in Milton Keynes and Alex Alderton, Roberto Amato, Jeffrey Barrett, Sonia Goncalves, Ewan Harrison, David K. Jackson, Ian Johnston, Dominic Kwiatkowski, Cordelia Langford, John Sillitoe on behalf of the Wellcome Sanger Institute COVID-19 Surveillance Team |
| EPI_ISL_2995037, EPI_ISL_2995038 | Berkshire and Surrey Pathology Services Lighthouse Laboratory | Wellcome Sanger Institute for the COVID-19 Genomics UK (COG-UK) Consortium | Berkshire and Surrey Pathology Services Lighthouse Laboratory and Alex Alderton, Roberto Amato, Jeffrey Barrett, Sonia Goncalves, Ewan Harrison, David K. Jackson, Ian Johnston, Dominic Kwiatkowski, Cordelia Langford, John Sillitoe on behalf of the Wellcome Sanger Institute COVID-19 Surveillance Team |
| EPI_ISL_2995039 | Lighthouse Lab in Alderley Park | Wellcome Sanger Institute for the COVID-19 Genomics UK (COG-UK) Consortium | Jacquelyn Wynn, Mairead Hyland, The Lighthouse Lab in Alderley Park and Alex Alderton, Roberto Amato, Jeffrey Barrett, Sonia Goncalves, Ewan Harrison, David K. Jackson, Ian Johnston, Dominic Kwiatkowski, Cordelia Langford, John Sillitoe on behalf of the Wellcome Sanger Institute COVID-19 Surveillance Team |
| EPI_ISL_2995040 | Lighthouse Lab in Glasgow | Wellcome Sanger Institute for the COVID-19 Genomics UK (COG-UK) Consortium | Harper VanSteenhouse, Yumi Kasai, David Gray, Carol Clugston, Anna Dominiczak and Alex Alderton, Roberto Amato, Jeffrey Barrett, Sonia Goncalves, Ewan Harrison, David K. Jackson, Ian Johnston, Dominic Kwiatkowski, Cordelia Langford, John Sillitoe on behalf of the Wellcome Sanger Institute COVID-19 Surveillance Team |
| EPI_ISL_2995041, EPI_ISL_2995042 | Lighthouse Lab in Alderley Park | Wellcome Sanger Institute for the COVID-19 Genomics UK (COG-UK) Consortium | Jacquelyn Wynn, Mairead Hyland, The Lighthouse Lab in Alderley Park and Alex Alderton, Roberto Amato, Jeffrey Barrett, Sonia Goncalves, Ewan Harrison, David K. Jackson, Ian Johnston, Dominic Kwiatkowski, Cordelia Langford, John Sillitoe on behalf of the Wellcome Sanger Institute COVID-19 Surveillance Team |
| EPI_ISL_2995043 | Lighthouse Lab in Glasgow | Wellcome Sanger Institute for the COVID-19 Genomics UK (COG-UK) Consortium | Harper VanSteenhouse, Yumi Kasai, David Gray, Carol Clugston, Anna Dominiczak and Alex Alderton, Roberto Amato, Jeffrey Barrett, Sonia Goncalves, Ewan Harrison, David K. Jackson, Ian Johnston, Dominic Kwiatkowski, Cordelia Langford, John Sillitoe on behalf of the Wellcome Sanger Institute COVID-19 Surveillance Team |
| EPI_ISL_2995044 | Berkshire and Surrey Pathology Services Lighthouse Laboratory | Wellcome Sanger Institute for the COVID-19 Genomics UK (COG-UK) Consortium | Berkshire and Surrey Pathology Services Lighthouse Laboratory and Alex Alderton, Roberto Amato, Jeffrey Barrett, Sonia Goncalves, Ewan Harrison, David K. Jackson, Ian Johnston, Dominic Kwiatkowski, Cordelia Langford, John Sillitoe on behalf of the Wellcome Sanger Institute COVID-19 Surveillance Team |
| EPI_ISL_2995045 | Lighthouse Lab in Alderley Park | Wellcome Sanger Institute for the COVID-19 Genomics UK (COG-UK) Consortium | Jacquelyn Wynn, Mairead Hyland, The Lighthouse Lab in Alderley Park and Alex Alderton, Roberto Amato, Jeffrey Barrett, Sonia Goncalves, Ewan Harrison, David K. Jackson, Ian Johnston, Dominic Kwiatkowski, Cordelia Langford, John Sillitoe on behalf of the Wellcome Sanger Institute COVID-19 Surveillance Team |
| EPI_ISL_2995046 | Lighthouse Lab in Milton Keynes | Wellcome Sanger Institute for the COVID-19 Genomics UK (COG-UK) Consortium | The Lighthouse Lab in Milton Keynes and Alex Alderton, Roberto Amato, Jeffrey Barrett, Sonia Goncalves, Ewan Harrison, David K. Jackson, Ian Johnston, Dominic Kwiatkowski, Cordelia Langford, John Sillitoe on behalf of the Wellcome Sanger Institute COVID-19 Surveillance Team |
| EPI_ISL_2995047 | Berkshire and Surrey Pathology Services Lighthouse Laboratory | Wellcome Sanger Institute for the COVID-19 Genomics UK (COG-UK) Consortium | Berkshire and Surrey Pathology Services Lighthouse Laboratory and Alex Alderton, Roberto Amato, Jeffrey Barrett, Sonia Goncalves, Ewan Harrison, David K. Jackson, Ian Johnston, Dominic Kwiatkowski, Cordelia Langford, John Sillitoe on behalf of the Wellcome Sanger Institute COVID-19 Surveillance Team |
| EPI_ISL_2995048 | Lighthouse Lab in Alderley Park | Wellcome Sanger Institute for the COVID-19 Genomics UK (COG-UK) Consortium | Jacquelyn Wynn, Mairead Hyland, The Lighthouse Lab in Alderley Park and Alex Alderton, Roberto Amato, Jeffrey Barrett, Sonia Goncalves, Ewan Harrison, David K. Jackson, Ian Johnston, Dominic Kwiatkowski, Cordelia Langford, John Sillitoe on behalf of the Wellcome Sanger Institute COVID-19 Surveillance Team |
| EPI_ISL_2995049 | Lighthouse Lab in Milton Keynes | Wellcome Sanger Institute for the COVID-19 Genomics UK (COG-UK) Consortium | The Lighthouse Lab in Milton Keynes and Alex Alderton, Roberto Amato, Jeffrey Barrett, Sonia Goncalves, Ewan Harrison, David K. Jackson, Ian Johnston, Dominic Kwiatkowski, Cordelia Langford, John Sillitoe on behalf of the Wellcome Sanger Institute COVID-19 Surveillance Team |
| EPI_ISL_2995050 | Lighthouse Lab in Glasgow | Wellcome Sanger Institute for the COVID-19 Genomics UK (COG-UK) Consortium | Harper VanSteenhouse, Yumi Kasai, David Gray, Carol Clugston, Anna Dominiczak and Alex Alderton, Roberto Amato, Jeffrey Barrett, Sonia Goncalves, Ewan Harrison, David K. Jackson, Ian Johnston, Dominic Kwiatkowski, Cordelia Langford, John Sillitoe on behalf of the Wellcome Sanger Institute COVID-19 Surveillance Team |

| | | | |
|---|---|---|---|
| EPI_ISL_2995051 | Lighthouse Lab in Milton Keynes | Wellcome Sanger Institute for the COVID-19 Genomics UK (COG-UK) Consortium | The Lighthouse Lab in Milton Keynes and Alex Alderton, Roberto Amato, Jeffrey Barrett, Sonia Goncalves, Ewan Harrison, David K. Jackson, Ian Johnston, Dominic Kwiatkowski, Cordelia Langford, John Sillitoe on behalf of the Wellcome Sanger Institute COVID-19 Surveillance Team |
| EPI_ISL_2995052 | Lighthouse Lab in Glasgow | Wellcome Sanger Institute for the COVID-19 Genomics UK (COG-UK) Consortium | Harper VanSteenhouse, Yumi Kasai, David Gray, Carol Clugston, Anna Dominiczak and Alex Alderton, Roberto Amato, Jeffrey Barrett, Sonia Goncalves, Ewan Harrison, David K. Jackson, Ian Johnston, Dominic Kwiatkowski, Cordelia Langford, John Sillitoe on behalf of the Wellcome Sanger Institute COVID-19 Surveillance Team |
| EPI_ISL_2995053 | Berkshire and Surrey Pathology Services Lighthouse Laboratory | Wellcome Sanger Institute for the COVID-19 Genomics UK (COG-UK) Consortium | Berkshire and Surrey Pathology Services Lighthouse Laboratory and Alex Alderton, Roberto Amato, Jeffrey Barrett, Sonia Goncalves, Ewan Harrison, David K. Jackson, Ian Johnston, Dominic Kwiatkowski, Cordelia Langford, John Sillitoe on behalf of the Wellcome Sanger Institute COVID-19 Surveillance Team |
| EPI_ISL_2995054 | Lighthouse Lab in Glasgow | Wellcome Sanger Institute for the COVID-19 Genomics UK (COG-UK) Consortium | Harper VanSteenhouse, Yumi Kasai, David Gray, Carol Clugston, Anna Dominiczak and Alex Alderton, Roberto Amato, Jeffrey Barrett, Sonia Goncalves, Ewan Harrison, David K. Jackson, Ian Johnston, Dominic Kwiatkowski, Cordelia Langford, John Sillitoe on behalf of the Wellcome Sanger Institute COVID-19 Surveillance Team |
| EPI_ISL_2995055 | Lighthouse Lab in Alderley Park | Wellcome Sanger Institute for the COVID-19 Genomics UK (COG-UK) Consortium | Jacquelyn Wynn, Mairead Hyland, The Lighthouse Lab in Alderley Park and Alex Alderton, Roberto Amato, Jeffrey Barrett, Sonia Goncalves, Ewan Harrison, David K. Jackson, Ian Johnston, Dominic Kwiatkowski, Cordelia Langford, John Sillitoe on behalf of the Wellcome Sanger Institute COVID-19 Surveillance Team |
| EPI_ISL_2995056 | Lighthouse Lab in Glasgow | Wellcome Sanger Institute for the COVID-19 Genomics UK (COG-UK) Consortium | Harper VanSteenhouse, Yumi Kasai, David Gray, Carol Clugston, Anna Dominiczak and Alex Alderton, Roberto Amato, Jeffrey Barrett, Sonia Goncalves, Ewan Harrison, David K. Jackson, Ian Johnston, Dominic Kwiatkowski, Cordelia Langford, John Sillitoe on behalf of the Wellcome Sanger Institute COVID-19 Surveillance Team |
| EPI_ISL_2995057 | Berkshire and Surrey Pathology Services Lighthouse Laboratory | Wellcome Sanger Institute for the COVID-19 Genomics UK (COG-UK) Consortium | Berkshire and Surrey Pathology Services Lighthouse Laboratory and Alex Alderton, Roberto Amato, Jeffrey Barrett, Sonia Goncalves, Ewan Harrison, David K. Jackson, Ian Johnston, Dominic Kwiatkowski, Cordelia Langford, John Sillitoe on behalf of the Wellcome Sanger Institute COVID-19 Surveillance Team |
| EPI_ISL_2995058 | Lighthouse Lab in Alderley Park | Wellcome Sanger Institute for the COVID-19 Genomics UK (COG-UK) Consortium | Jacquelyn Wynn, Mairead Hyland, The Lighthouse Lab in Alderley Park and Alex Alderton, Roberto Amato, Jeffrey Barrett, Sonia Goncalves, Ewan Harrison, David K. Jackson, Ian Johnston, Dominic Kwiatkowski, Cordelia Langford, John Sillitoe on behalf of the Wellcome Sanger Institute COVID-19 Surveillance Team |
| EPI_ISL_2995059 | Lighthouse Lab in Milton Keynes | Wellcome Sanger Institute for the COVID-19 Genomics UK (COG-UK) Consortium | The Lighthouse Lab in Milton Keynes and Alex Alderton, Roberto Amato, Jeffrey Barrett, Sonia Goncalves, Ewan Harrison, David K. Jackson, Ian Johnston, Dominic Kwiatkowski, Cordelia Langford, John Sillitoe on behalf of the Wellcome Sanger Institute COVID-19 Surveillance Team |
| EPI_ISL_2995060 | Berkshire and Surrey Pathology Services Lighthouse Laboratory | Wellcome Sanger Institute for the COVID-19 Genomics UK (COG-UK) Consortium | Berkshire and Surrey Pathology Services Lighthouse Laboratory and Alex Alderton, Roberto Amato, Jeffrey Barrett, Sonia Goncalves, Ewan Harrison, David K. Jackson, Ian Johnston, Dominic Kwiatkowski, Cordelia Langford, John Sillitoe on behalf of the Wellcome Sanger Institute COVID-19 Surveillance Team |
| EPI_ISL_2995061, EPI_ISL_2995062 | Lighthouse Lab in Milton Keynes | Wellcome Sanger Institute for the COVID-19 Genomics UK (COG-UK) Consortium | The Lighthouse Lab in Milton Keynes and Alex Alderton, Roberto Amato, Jeffrey Barrett, Sonia Goncalves, Ewan Harrison, David K. Jackson, Ian Johnston, Dominic Kwiatkowski, Cordelia Langford, John Sillitoe on behalf of the Wellcome Sanger Institute COVID-19 Surveillance Team |
| EPI_ISL_2995063, EPI_ISL_2995064 | Lighthouse Lab in Alderley Park | Wellcome Sanger Institute for the COVID-19 Genomics UK (COG-UK) Consortium | Jacquelyn Wynn, Mairead Hyland, The Lighthouse Lab in Alderley Park and Alex Alderton, Roberto Amato, Jeffrey Barrett, Sonia Goncalves, Ewan Harrison, David K. Jackson, Ian Johnston, Dominic Kwiatkowski, Cordelia Langford, John Sillitoe on behalf of the Wellcome Sanger Institute COVID-19 Surveillance Team |
| EPI_ISL_2995065 | Lighthouse Lab in Glasgow | Wellcome Sanger Institute for the COVID-19 Genomics UK (COG-UK) Consortium | Harper VanSteenhouse, Yumi Kasai, David Gray, Carol Clugston, Anna Dominiczak and Alex Alderton, Roberto Amato, Jeffrey Barrett, Sonia Goncalves, Ewan Harrison, David K. Jackson, Ian Johnston, Dominic Kwiatkowski, Cordelia Langford, John Sillitoe on behalf of the Wellcome Sanger Institute COVID-19 Surveillance Team |
| EPI_ISL_2995066 | Berkshire and Surrey Pathology Services Lighthouse Laboratory | Wellcome Sanger Institute for the COVID-19 Genomics UK (COG-UK) Consortium | Berkshire and Surrey Pathology Services Lighthouse Laboratory and Alex Alderton, Roberto Amato, Jeffrey Barrett, Sonia Goncalves, Ewan Harrison, David K. Jackson, Ian Johnston, Dominic Kwiatkowski, Cordelia Langford, John Sillitoe on behalf of the Wellcome Sanger Institute COVID-19 Surveillance Team |
| EPI_ISL_2995067, EPI_ISL_2995068 | Lighthouse Lab in Alderley Park | Wellcome Sanger Institute for the COVID-19 Genomics UK (COG-UK) Consortium | Jacquelyn Wynn, Mairead Hyland, The Lighthouse Lab in Alderley Park and Alex Alderton, Roberto Amato, Jeffrey Barrett, Sonia Goncalves, Ewan Harrison, David K. Jackson, Ian Johnston, Dominic Kwiatkowski, Cordelia Langford, John Sillitoe on behalf of the Wellcome Sanger Institute COVID-19 Surveillance Team |
| EPI_ISL_2995069, EPI_ISL_2995070 | Lighthouse Lab in Glasgow | Wellcome Sanger Institute for the COVID-19 Genomics UK (COG-UK) Consortium | Harper VanSteenhouse, Yumi Kasai, David Gray, Carol Clugston, Anna Dominiczak and Alex Alderton, Roberto Amato, Jeffrey Barrett, Sonia Goncalves, Ewan Harrison, David K. Jackson, Ian Johnston, Dominic Kwiatkowski, Cordelia Langford, John Sillitoe on behalf of the Wellcome Sanger Institute COVID-19 Surveillance Team |
| EPI_ISL_2995071 | Lighthouse Lab in Alderley Park | Wellcome Sanger Institute for the COVID-19 Genomics UK (COG-UK) Consortium | Jacquelyn Wynn, Mairead Hyland, The Lighthouse Lab in Alderley Park and Alex Alderton, Roberto Amato, Jeffrey Barrett, Sonia Goncalves, Ewan Harrison, David K. Jackson, Ian Johnston, Dominic Kwiatkowski, Cordelia Langford, John Sillitoe on behalf of the Wellcome Sanger Institute COVID-19 Surveillance Team |
| EPI_ISL_2995072 | Lighthouse Lab in Glasgow | Wellcome Sanger Institute for the COVID-19 Genomics UK (COG-UK) Consortium | Harper VanSteenhouse, Yumi Kasai, David Gray, Carol Clugston, Anna Dominiczak and Alex Alderton, Roberto Amato, Jeffrey Barrett, Sonia Goncalves, Ewan Harrison, David K. Jackson, Ian Johnston, Dominic Kwiatkowski, Cordelia Langford, John Sillitoe on behalf of the Wellcome Sanger Institute COVID-19 Surveillance Team |
| EPI_ISL_2995073 | Lighthouse Lab in Alderley Park | Wellcome Sanger Institute for the COVID-19 Genomics UK (COG-UK) Consortium | Jacquelyn Wynn, Mairead Hyland, The Lighthouse Lab in Alderley Park and Alex Alderton, Roberto Amato, Jeffrey Barrett, Sonia Goncalves, Ewan Harrison, David K. Jackson, Ian Johnston, Dominic Kwiatkowski, Cordelia Langford, John Sillitoe on behalf of the Wellcome Sanger Institute COVID-19 Surveillance Team |
| EPI_ISL_2995074 | Berkshire and Surrey Pathology Services Lighthouse Laboratory | Wellcome Sanger Institute for the COVID-19 Genomics UK (COG-UK) Consortium | Berkshire and Surrey Pathology Services Lighthouse Laboratory and Alex Alderton, Roberto Amato, Jeffrey Barrett, Sonia Goncalves, Ewan Harrison, David K. Jackson, Ian Johnston, Dominic Kwiatkowski, Cordelia Langford, John Sillitoe on behalf of the Wellcome Sanger Institute COVID-19 Surveillance Team |
| EPI_ISL_2995075 | Lighthouse Lab in Glasgow | Wellcome Sanger Institute for the COVID-19 Genomics UK (COG-UK) Consortium | Harper VanSteenhouse, Yumi Kasai, David Gray, Carol Clugston, Anna Dominiczak and Alex Alderton, Roberto Amato, Jeffrey Barrett, Sonia Goncalves, Ewan Harrison, David K. Jackson, Ian Johnston, Dominic Kwiatkowski, Cordelia Langford, John Sillitoe on behalf of the Wellcome Sanger Institute COVID-19 Surveillance Team |
| EPI_ISL_2995076, EPI_ISL_2995077 | Berkshire and Surrey Pathology Services Lighthouse Laboratory | Wellcome Sanger Institute for the COVID-19 Genomics UK (COG-UK) Consortium | Berkshire and Surrey Pathology Services Lighthouse Laboratory and Alex Alderton, Roberto Amato, Jeffrey Barrett, Sonia Goncalves, Ewan Harrison, David K. Jackson, Ian Johnston, Dominic Kwiatkowski, Cordelia Langford, John Sillitoe on behalf of the Wellcome Sanger Institute COVID-19 Surveillance Team |
| EPI_ISL_2995078 | Lighthouse Lab in Glasgow | Wellcome Sanger Institute for the COVID-19 Genomics UK (COG-UK) Consortium | Harper VanSteenhouse, Yumi Kasai, David Gray, Carol Clugston, Anna Dominiczak and Alex Alderton, Roberto Amato, Jeffrey Barrett, Sonia Goncalves, Ewan Harrison, David K. Jackson, Ian Johnston, Dominic Kwiatkowski, Cordelia Langford, John Sillitoe on behalf of the Wellcome Sanger Institute COVID-19 Surveillance Team |
| EPI_ISL_2995079 | Lighthouse Lab in Milton Keynes | Wellcome Sanger Institute for the COVID-19 Genomics UK (COG-UK) Consortium | The Lighthouse Lab in Milton Keynes and Alex Alderton, Roberto Amato, Jeffrey Barrett, Sonia Goncalves, Ewan Harrison, David K. Jackson, Ian Johnston, Dominic Kwiatkowski, Cordelia Langford, John Sillitoe on behalf of the Wellcome Sanger Institute COVID-19 Surveillance Team |
| EPI_ISL_2995080 | Lighthouse Lab in Glasgow | Wellcome Sanger Institute for the COVID-19 Genomics UK (COG-UK) Consortium | Harper VanSteenhouse, Yumi Kasai, David Gray, Carol Clugston, Anna Dominiczak and Alex Alderton, Roberto Amato, Jeffrey Barrett, Sonia Goncalves, Ewan Harrison, David K. Jackson, Ian Johnston, Dominic Kwiatkowski, Cordelia Langford, John Sillitoe on behalf of the Wellcome Sanger Institute COVID-19 Surveillance Team |
| EPI_ISL_2995081 | Lighthouse Lab in Alderley Park | Wellcome Sanger Institute for the COVID-19 Genomics UK (COG-UK) Consortium | Jacquelyn Wynn, Mairead Hyland, The Lighthouse Lab in Alderley Park and Alex Alderton, Roberto Amato, Jeffrey Barrett, Sonia Goncalves, Ewan Harrison, David K. Jackson, Ian Johnston, Dominic Kwiatkowski, Cordelia Langford, John Sillitoe on behalf of the Wellcome Sanger Institute COVID-19 Surveillance Team |
| EPI_ISL_2995082 | Lighthouse Lab in Glasgow | Wellcome Sanger Institute for the COVID-19 Genomics UK (COG-UK) Consortium | Harper VanSteenhouse, Yumi Kasai, David Gray, Carol Clugston, Anna Dominiczak and Alex Alderton, Roberto Amato, Jeffrey Barrett, Sonia Goncalves, Ewan Harrison, David K. Jackson, Ian Johnston, Dominic Kwiatkowski, Cordelia Langford, John Sillitoe on behalf of the Wellcome Sanger Institute COVID-19 Surveillance Team |
| EPI_ISL_2995083, EPI_ISL_2995084 | Lighthouse Lab in Milton Keynes | Wellcome Sanger Institute for the COVID-19 Genomics UK (COG-UK) Consortium | The Lighthouse Lab in Milton Keynes and Alex Alderton, Roberto Amato, Jeffrey Barrett, Sonia Goncalves, Ewan Harrison, David K. Jackson, Ian Johnston, Dominic Kwiatkowski, Cordelia Langford, John Sillitoe on behalf of the Wellcome Sanger Institute COVID-19 Surveillance Team |
| EPI_ISL_2995085 | Lighthouse Lab in Alderley Park | Wellcome Sanger Institute for the COVID-19 Genomics UK (COG-UK) Consortium | Jacquelyn Wynn, Mairead Hyland, The Lighthouse Lab in Alderley Park and Alex Alderton, Roberto Amato, Jeffrey Barrett, Sonia Goncalves, Ewan Harrison, David K. Jackson, Ian Johnston, Dominic Kwiatkowski, Cordelia Langford, John Sillitoe on behalf of the Wellcome Sanger Institute COVID-19 Surveillance Team |
| EPI_ISL_2995086, EPI_ISL_2995087 | Lighthouse Lab in Glasgow | Wellcome Sanger Institute for the COVID-19 Genomics UK (COG-UK) Consortium | Harper VanSteenhouse, Yumi Kasai, David Gray, Carol Clugston, Anna Dominiczak and Alex Alderton, Roberto Amato, Jeffrey Barrett, Sonia Goncalves, Ewan Harrison, David K. Jackson, Ian Johnston, Dominic Kwiatkowski, Cordelia Langford, John Sillitoe on behalf of the Wellcome Sanger Institute COVID-19 Surveillance Team |
| EPI_ISL_2995088 | Berkshire and Surrey Pathology Services Lighthouse Laboratory | Wellcome Sanger Institute for the COVID-19 Genomics UK (COG-UK) Consortium | Berkshire and Surrey Pathology Services Lighthouse Laboratory and Alex Alderton, Roberto Amato, Jeffrey Barrett, Sonia Goncalves, Ewan Harrison, David K. Jackson, Ian Johnston, Dominic Kwiatkowski, Cordelia Langford, John Sillitoe on behalf of the Wellcome Sanger Institute COVID-19 Surveillance Team |
| EPI_ISL_2995089 | Lighthouse Lab in Alderley Park | Wellcome Sanger Institute for the COVID-19 Genomics UK (COG-UK) Consortium | Jacquelyn Wynn, Mairead Hyland, The Lighthouse Lab in Alderley Park and Alex Alderton, Roberto Amato, Jeffrey Barrett, Sonia Goncalves, Ewan Harrison, David K. Jackson, Ian Johnston, Dominic Kwiatkowski, Cordelia Langford, John Sillitoe on behalf of the Wellcome Sanger Institute COVID-19 Surveillance Team |
| EPI_ISL_2995090, EPI_ISL_2995091 | Lighthouse Lab in Milton Keynes | Wellcome Sanger Institute for the COVID-19 Genomics UK (COG-UK) Consortium | The Lighthouse Lab in Milton Keynes and Alex Alderton, Roberto Amato, Jeffrey Barrett, Sonia Goncalves, Ewan Harrison, David K. Jackson, Ian Johnston, Dominic Kwiatkowski, Cordelia Langford, John Sillitoe on behalf of the Wellcome Sanger Institute COVID-19 Surveillance Team |
| EPI_ISL_2995092 | Berkshire and Surrey Pathology Services Lighthouse Laboratory | Wellcome Sanger Institute for the COVID-19 Genomics UK (COG-UK) Consortium | Berkshire and Surrey Pathology Services Lighthouse Laboratory and Alex Alderton, Roberto Amato, Jeffrey Barrett, Sonia Goncalves, Ewan Harrison, David K. Jackson, Ian Johnston, Dominic Kwiatkowski, Cordelia Langford, John Sillitoe on behalf of the Wellcome Sanger Institute COVID-19 Surveillance Team |
| EPI_ISL_2995093 | Lighthouse Lab in Glasgow | Wellcome Sanger Institute for the COVID-19 Genomics UK (COG-UK) Consortium | Harper VanSteenhouse, Yumi Kasai, David Gray, Carol Clugston, Anna Dominiczak and Alex Alderton, Roberto Amato, Jeffrey Barrett, Sonia Goncalves, Ewan Harrison, David K. Jackson, Ian Johnston, Dominic Kwiatkowski, Cordelia Langford, John Sillitoe on behalf of the Wellcome Sanger Institute COVID-19 Surveillance Team |
| EPI_ISL_2995094, EPI_ISL_2995095, EPI_ISL_2995096 | Berkshire and Surrey Pathology Services Lighthouse Laboratory | Wellcome Sanger Institute for the COVID-19 Genomics UK (COG-UK) Consortium | Berkshire and Surrey Pathology Services Lighthouse Laboratory and Alex Alderton, Roberto Amato, Jeffrey Barrett, Sonia Goncalves, Ewan Harrison, David K. Jackson, Ian Johnston, Dominic Kwiatkowski, Cordelia Langford, John Sillitoe on behalf of the Wellcome Sanger Institute COVID-19 Surveillance Team |
| EPI_ISL_2995097 | Lighthouse Lab in Glasgow | Wellcome Sanger Institute for the COVID-19 Genomics UK (COG-UK) Consortium | Harper VanSteenhouse, Yumi Kasai, David Gray, Carol Clugston, Anna Dominiczak and Alex Alderton, Roberto Amato, Jeffrey Barrett, Sonia Goncalves, Ewan Harrison, David K. Jackson, Ian Johnston, Dominic Kwiatkowski, Cordelia Langford, John Sillitoe on behalf of the Wellcome Sanger Institute COVID-19 Surveillance Team |
| EPI_ISL_2995098 | Berkshire and Surrey Pathology Services Lighthouse Laboratory | Wellcome Sanger Institute for the COVID-19 Genomics UK (COG-UK) Consortium | Berkshire and Surrey Pathology Services Lighthouse Laboratory and Alex Alderton, Roberto Amato, Jeffrey Barrett, Sonia Goncalves, Ewan Harrison, David K. Jackson, Ian Johnston, Dominic Kwiatkowski, Cordelia Langford, John Sillitoe on behalf of the Wellcome Sanger Institute COVID-19 Surveillance Team |
| EPI_ISL_2995099 | Lighthouse Lab in Milton Keynes | Wellcome Sanger Institute for the COVID-19 Genomics UK (COG-UK) Consortium | The Lighthouse Lab in Milton Keynes and Alex Alderton, Roberto Amato, Jeffrey Barrett, Sonia Goncalves, Ewan Harrison, David K. Jackson, Ian Johnston, Dominic Kwiatkowski, Cordelia Langford, John Sillitoe on behalf of the Wellcome Sanger Institute COVID-19 Surveillance Team |
| EPI_ISL_2995100, EPI_ISL_2995101 | Lighthouse Lab in Glasgow | Wellcome Sanger Institute for the COVID-19 Genomics UK (COG-UK) Consortium | Harper VanSteenhouse, Yumi Kasai, David Gray, Carol Clugston, Anna Dominiczak and Alex Alderton, Roberto Amato, Jeffrey Barrett, Sonia Goncalves, Ewan Harrison, David K. Jackson, Ian Johnston, Dominic Kwiatkowski, Cordelia Langford, John Sillitoe on behalf of the Wellcome Sanger Institute COVID-19 Surveillance Team |
| EPI_ISL_2995102 | Lighthouse Lab in Milton Keynes | Wellcome Sanger Institute for the COVID-19 Genomics UK (COG-UK) Consortium | The Lighthouse Lab in Milton Keynes and Alex Alderton, Roberto Amato, Jeffrey Barrett, Sonia Goncalves, Ewan Harrison, David K. Jackson, Ian Johnston, Dominic Kwiatkowski, Cordelia Langford, John Sillitoe on behalf of the Wellcome Sanger Institute COVID-19 Surveillance Team |
| EPI_ISL_2995103 | Lighthouse Lab in Glasgow | Wellcome Sanger Institute for the COVID-19 Genomics UK (COG-UK) Consortium | Harper VanSteenhouse, Yumi Kasai, David Gray, Carol Clugston, Anna Dominiczak and Alex Alderton, Roberto Amato, Jeffrey Barrett, Sonia Goncalves, Ewan Harrison, David K. Jackson, Ian Johnston, Dominic Kwiatkowski, Cordelia Langford, John Sillitoe on behalf of the Wellcome Sanger Institute COVID-19 Surveillance Team |
| EPI_ISL_2995104 | Lighthouse Lab in Alderley Park | Wellcome Sanger Institute for the COVID-19 Genomics UK (COG-UK) Consortium | Jacquelyn Wynn, Mairead Hyland, The Lighthouse Lab in Alderley Park and Alex Alderton, Roberto Amato, Jeffrey Barrett, Sonia Goncalves, Ewan Harrison, David K. Jackson, Ian Johnston, Dominic Kwiatkowski, Cordelia Langford, John Sillitoe on behalf of the Wellcome Sanger Institute COVID-19 Surveillance Team |
| EPI_ISL_2995105 | Lighthouse Lab in Milton Keynes | Wellcome Sanger Institute for the COVID-19 Genomics UK (COG-UK) Consortium | The Lighthouse Lab in Milton Keynes and Alex Alderton, Roberto Amato, Jeffrey Barrett, Sonia Goncalves, Ewan Harrison, David K. Jackson, Ian Johnston, Dominic Kwiatkowski, Cordelia Langford, John Sillitoe on behalf of the Wellcome Sanger Institute COVID-19 Surveillance Team |
| EPI_ISL_2995106, EPI_ISL_2995107 | Lighthouse Lab in Glasgow | Wellcome Sanger Institute for the COVID-19 Genomics UK (COG-UK) Consortium | Harper VanSteenhouse, Yumi Kasai, David Gray, Carol Clugston, Anna Dominiczak and Alex Alderton, Roberto Amato, Jeffrey Barrett, Sonia Goncalves, Ewan Harrison, David K. Jackson, Ian Johnston, Dominic Kwiatkowski, Cordelia Langford, John Sillitoe on behalf of the Wellcome Sanger Institute COVID-19 Surveillance Team |

| | | | |
|---|---|---|---|
| EPI_ISL_2995108 | Berkshire and Surrey Pathology Services Lighthouse Laboratory | Wellcome Sanger Institute for the COVID-19 Genomics UK (COG-UK) Consortium | Berkshire and Surrey Pathology Services Lighthouse Laboratory and Alex Alderton, Roberto Amato, Jeffrey Barrett, Sonia Goncalves, Ewan Harrison, David K. Jackson, Ian Johnston, Dominic Kwiatkowski, Cordelia Langford, John Sillitoe on behalf of the Wellcome Sanger Institute COVID-19 Surveillance Team |
| EPI_ISL_2995109 | Lighthouse Lab in Alderley Park | Wellcome Sanger Institute for the COVID-19 Genomics UK (COG-UK) Consortium | Jacquelyn Wynn, Mairead Hyland, The Lighthouse Lab in Alderley Park and Alex Alderton, Roberto Amato, Jeffrey Barrett, Sonia Goncalves, Ewan Harrison, David K. Jackson, Ian Johnston, Dominic Kwiatkowski, Cordelia Langford, John Sillitoe on behalf of the Wellcome Sanger Institute COVID-19 Surveillance Team |
| EPI_ISL_2995110, EPI_ISL_2995111 | Lighthouse Lab in Milton Keynes | Wellcome Sanger Institute for the COVID-19 Genomics UK (COG-UK) Consortium | The Lighthouse Lab in Milton Keynes and Alex Alderton, Roberto Amato, Jeffrey Barrett, Sonia Goncalves, Ewan Harrison, David K. Jackson, Ian Johnston, Dominic Kwiatkowski, Cordelia Langford, John Sillitoe on behalf of the Wellcome Sanger Institute COVID-19 Surveillance Team |
| EPI_ISL_2995112, EPI_ISL_2995113, EPI_ISL_2995114 | Lighthouse Lab in Glasgow | Wellcome Sanger Institute for the COVID-19 Genomics UK (COG-UK) Consortium | Harper VanSteenhouse, Yumi Kasai, David Gray, Carol Clugston, Anna Dominiczak and Alex Alderton, Roberto Amato, Jeffrey Barrett, Sonia Goncalves, Ewan Harrison, David K. Jackson, Ian Johnston, Dominic Kwiatkowski, Cordelia Langford, John Sillitoe on behalf of the Wellcome Sanger Institute COVID-19 Surveillance Team |
| EPI_ISL_2995115 | Lighthouse Lab in Milton Keynes | Wellcome Sanger Institute for the COVID-19 Genomics UK (COG-UK) Consortium | The Lighthouse Lab in Milton Keynes and Alex Alderton, Roberto Amato, Jeffrey Barrett, Sonia Goncalves, Ewan Harrison, David K. Jackson, Ian Johnston, Dominic Kwiatkowski, Cordelia Langford, John Sillitoe on behalf of the Wellcome Sanger Institute COVID-19 Surveillance Team |
| EPI_ISL_2995116 | Lighthouse Lab in Alderley Park | Wellcome Sanger Institute for the COVID-19 Genomics UK (COG-UK) Consortium | Jacquelyn Wynn, Mairead Hyland, The Lighthouse Lab in Alderley Park and Alex Alderton, Roberto Amato, Jeffrey Barrett, Sonia Goncalves, Ewan Harrison, David K. Jackson, Ian Johnston, Dominic Kwiatkowski, Cordelia Langford, John Sillitoe on behalf of the Wellcome Sanger Institute COVID-19 Surveillance Team |
| EPI_ISL_2995117 | Lighthouse Lab in Glasgow | Wellcome Sanger Institute for the COVID-19 Genomics UK (COG-UK) Consortium | Harper VanSteenhouse, Yumi Kasai, David Gray, Carol Clugston, Anna Dominiczak and Alex Alderton, Roberto Amato, Jeffrey Barrett, Sonia Goncalves, Ewan Harrison, David K. Jackson, Ian Johnston, Dominic Kwiatkowski, Cordelia Langford, John Sillitoe on behalf of the Wellcome Sanger Institute COVID-19 Surveillance Team |
| EPI_ISL_2995118 | Berkshire and Surrey Pathology Services Lighthouse Laboratory | Wellcome Sanger Institute for the COVID-19 Genomics UK (COG-UK) Consortium | Berkshire and Surrey Pathology Services Lighthouse Laboratory and Alex Alderton, Roberto Amato, Jeffrey Barrett, Sonia Goncalves, Ewan Harrison, David K. Jackson, Ian Johnston, Dominic Kwiatkowski, Cordelia Langford, John Sillitoe on behalf of the Wellcome Sanger Institute COVID-19 Surveillance Team |
| EPI_ISL_2995119 | Lighthouse Lab in Milton Keynes | Wellcome Sanger Institute for the COVID-19 Genomics UK (COG-UK) Consortium | The Lighthouse Lab in Milton Keynes and Alex Alderton, Roberto Amato, Jeffrey Barrett, Sonia Goncalves, Ewan Harrison, David K. Jackson, Ian Johnston, Dominic Kwiatkowski, Cordelia Langford, John Sillitoe on behalf of the Wellcome Sanger Institute COVID-19 Surveillance Team |
| EPI_ISL_2995120 | Berkshire and Surrey Pathology Services Lighthouse Laboratory | Wellcome Sanger Institute for the COVID-19 Genomics UK (COG-UK) Consortium | Berkshire and Surrey Pathology Services Lighthouse Laboratory and Alex Alderton, Roberto Amato, Jeffrey Barrett, Sonia Goncalves, Ewan Harrison, David K. Jackson, Ian Johnston, Dominic Kwiatkowski, Cordelia Langford, John Sillitoe on behalf of the Wellcome Sanger Institute COVID-19 Surveillance Team |
| EPI_ISL_2995121, EPI_ISL_2995122, EPI_ISL_2995123, EPI_ISL_2995124, EPI_ISL_2995125 | Lighthouse Lab in Glasgow | Wellcome Sanger Institute for the COVID-19 Genomics UK (COG-UK) Consortium | Harper VanSteenhouse, Yumi Kasai, David Gray, Carol Clugston, Anna Dominiczak and Alex Alderton, Roberto Amato, Jeffrey Barrett, Sonia Goncalves, Ewan Harrison, David K. Jackson, Ian Johnston, Dominic Kwiatkowski, Cordelia Langford, John Sillitoe on behalf of the Wellcome Sanger Institute COVID-19 Surveillance Team |
| EPI_ISL_2995126 | Lighthouse Lab in Milton Keynes | Wellcome Sanger Institute for the COVID-19 Genomics UK (COG-UK) Consortium | The Lighthouse Lab in Milton Keynes and Alex Alderton, Roberto Amato, Jeffrey Barrett, Sonia Goncalves, Ewan Harrison, David K. Jackson, Ian Johnston, Dominic Kwiatkowski, Cordelia Langford, John Sillitoe on behalf of the Wellcome Sanger Institute COVID-19 Surveillance Team |
| EPI_ISL_2995127 | Berkshire and Surrey Pathology Services Lighthouse Laboratory | Wellcome Sanger Institute for the COVID-19 Genomics UK (COG-UK) Consortium | Berkshire and Surrey Pathology Services Lighthouse Laboratory and Alex Alderton, Roberto Amato, Jeffrey Barrett, Sonia Goncalves, Ewan Harrison, David K. Jackson, Ian Johnston, Dominic Kwiatkowski, Cordelia Langford, John Sillitoe on behalf of the Wellcome Sanger Institute COVID-19 Surveillance Team |
| EPI_ISL_2995128 | Lighthouse Lab in Glasgow | Wellcome Sanger Institute for the COVID-19 Genomics UK (COG-UK) Consortium | Harper VanSteenhouse, Yumi Kasai, David Gray, Carol Clugston, Anna Dominiczak and Alex Alderton, Roberto Amato, Jeffrey Barrett, Sonia Goncalves, Ewan Harrison, David K. Jackson, Ian Johnston, Dominic Kwiatkowski, Cordelia Langford, John Sillitoe on behalf of the Wellcome Sanger Institute COVID-19 Surveillance Team |
| EPI_ISL_2995129, EPI_ISL_2995130 | Lighthouse Lab in Alderley Park | Wellcome Sanger Institute for the COVID-19 Genomics UK (COG-UK) Consortium | Jacquelyn Wynn, Mairead Hyland, The Lighthouse Lab in Alderley Park and Alex Alderton, Roberto Amato, Jeffrey Barrett, Sonia Goncalves, Ewan Harrison, David K. Jackson, Ian Johnston, Dominic Kwiatkowski, Cordelia Langford, John Sillitoe on behalf of the Wellcome Sanger Institute COVID-19 Surveillance Team |
| EPI_ISL_2995131, EPI_ISL_2995132 | Lighthouse Lab in Glasgow | Wellcome Sanger Institute for the COVID-19 Genomics UK (COG-UK) Consortium | Harper VanSteenhouse, Yumi Kasai, David Gray, Carol Clugston, Anna Dominiczak and Alex Alderton, Roberto Amato, Jeffrey Barrett, Sonia Goncalves, Ewan Harrison, David K. Jackson, Ian Johnston, Dominic Kwiatkowski, Cordelia Langford, John Sillitoe on behalf of the Wellcome Sanger Institute COVID-19 Surveillance Team |
| EPI_ISL_2995133 | Lighthouse Lab in Milton Keynes | Wellcome Sanger Institute for the COVID-19 Genomics UK (COG-UK) Consortium | The Lighthouse Lab in Milton Keynes and Alex Alderton, Roberto Amato, Jeffrey Barrett, Sonia Goncalves, Ewan Harrison, David K. Jackson, Ian Johnston, Dominic Kwiatkowski, Cordelia Langford, John Sillitoe on behalf of the Wellcome Sanger Institute COVID-19 Surveillance Team |
| EPI_ISL_2995134 | Lighthouse Lab in Glasgow | Wellcome Sanger Institute for the COVID-19 Genomics UK (COG-UK) Consortium | Harper VanSteenhouse, Yumi Kasai, David Gray, Carol Clugston, Anna Dominiczak and Alex Alderton, Roberto Amato, Jeffrey Barrett, Sonia Goncalves, Ewan Harrison, David K. Jackson, Ian Johnston, Dominic Kwiatkowski, Cordelia Langford, John Sillitoe on behalf of the Wellcome Sanger Institute COVID-19 Surveillance Team |
| EPI_ISL_2995135 | Lighthouse Lab in Alderley Park | Wellcome Sanger Institute for the COVID-19 Genomics UK (COG-UK) Consortium | Jacquelyn Wynn, Mairead Hyland, The Lighthouse Lab in Alderley Park and Alex Alderton, Roberto Amato, Jeffrey Barrett, Sonia Goncalves, Ewan Harrison, David K. Jackson, Ian Johnston, Dominic Kwiatkowski, Cordelia Langford, John Sillitoe on behalf of the Wellcome Sanger Institute COVID-19 Surveillance Team |
| EPI_ISL_2995136 | Lighthouse Lab in Milton Keynes | Wellcome Sanger Institute for the COVID-19 Genomics UK (COG-UK) Consortium | The Lighthouse Lab in Milton Keynes and Alex Alderton, Roberto Amato, Jeffrey Barrett, Sonia Goncalves, Ewan Harrison, David K. Jackson, Ian Johnston, Dominic Kwiatkowski, Cordelia Langford, John Sillitoe on behalf of the Wellcome Sanger Institute COVID-19 Surveillance Team |
| EPI_ISL_2995137 | Berkshire and Surrey Pathology Services Lighthouse Laboratory | Wellcome Sanger Institute for the COVID-19 Genomics UK (COG-UK) Consortium | Berkshire and Surrey Pathology Services Lighthouse Laboratory and Alex Alderton, Roberto Amato, Jeffrey Barrett, Sonia Goncalves, Ewan Harrison, David K. Jackson, Ian Johnston, Dominic Kwiatkowski, Cordelia Langford, John Sillitoe on behalf of the Wellcome Sanger Institute COVID-19 Surveillance Team |
| EPI_ISL_2995138 | Lighthouse Lab in Glasgow | Wellcome Sanger Institute for the COVID-19 Genomics UK (COG-UK) Consortium | Harper VanSteenhouse, Yumi Kasai, David Gray, Carol Clugston, Anna Dominiczak and Alex Alderton, Roberto Amato, Jeffrey Barrett, Sonia Goncalves, Ewan Harrison, David K. Jackson, Ian Johnston, Dominic Kwiatkowski, Cordelia Langford, John Sillitoe on behalf of the Wellcome Sanger Institute COVID-19 Surveillance Team |
| EPI_ISL_2995139 | Lighthouse Lab in Alderley Park | Wellcome Sanger Institute for the COVID-19 Genomics UK (COG-UK) Consortium | Jacquelyn Wynn, Mairead Hyland, The Lighthouse Lab in Alderley Park and Alex Alderton, Roberto Amato, Jeffrey Barrett, Sonia Goncalves, Ewan Harrison, David K. Jackson, Ian Johnston, Dominic Kwiatkowski, Cordelia Langford, John Sillitoe on behalf of the Wellcome Sanger Institute COVID-19 Surveillance Team |
| EPI_ISL_2995140 | Lighthouse Lab in Milton Keynes | Wellcome Sanger Institute for the COVID-19 Genomics UK (COG-UK) Consortium | The Lighthouse Lab in Milton Keynes and Alex Alderton, Roberto Amato, Jeffrey Barrett, Sonia Goncalves, Ewan Harrison, David K. Jackson, Ian Johnston, Dominic Kwiatkowski, Cordelia Langford, John Sillitoe on behalf of the Wellcome Sanger Institute COVID-19 Surveillance Team |
| EPI_ISL_2995141 | Lighthouse Lab in Alderley Park | Wellcome Sanger Institute for the COVID-19 Genomics UK (COG-UK) Consortium | Jacquelyn Wynn, Mairead Hyland, The Lighthouse Lab in Alderley Park and Alex Alderton, Roberto Amato, Jeffrey Barrett, Sonia Goncalves, Ewan Harrison, David K. Jackson, Ian Johnston, Dominic Kwiatkowski, Cordelia Langford, John Sillitoe on behalf of the Wellcome Sanger Institute COVID-19 Surveillance Team |
| EPI_ISL_2995142 | Berkshire and Surrey Pathology Services Lighthouse Laboratory | Wellcome Sanger Institute for the COVID-19 Genomics UK (COG-UK) Consortium | Berkshire and Surrey Pathology Services Lighthouse Laboratory and Alex Alderton, Roberto Amato, Jeffrey Barrett, Sonia Goncalves, Ewan Harrison, David K. Jackson, Ian Johnston, Dominic Kwiatkowski, Cordelia Langford, John Sillitoe on behalf of the Wellcome Sanger Institute COVID-19 Surveillance Team |
| EPI_ISL_2995143, EPI_ISL_2995144 | Lighthouse Lab in Glasgow | Wellcome Sanger Institute for the COVID-19 Genomics UK (COG-UK) Consortium | Harper VanSteenhouse, Yumi Kasai, David Gray, Carol Clugston, Anna Dominiczak and Alex Alderton, Roberto Amato, Jeffrey Barrett, Sonia Goncalves, Ewan Harrison, David K. Jackson, Ian Johnston, Dominic Kwiatkowski, Cordelia Langford, John Sillitoe on behalf of the Wellcome Sanger Institute COVID-19 Surveillance Team |
| EPI_ISL_2995145 | Berkshire and Surrey Pathology Services Lighthouse Laboratory | Wellcome Sanger Institute for the COVID-19 Genomics UK (COG-UK) Consortium | Berkshire and Surrey Pathology Services Lighthouse Laboratory and Alex Alderton, Roberto Amato, Jeffrey Barrett, Sonia Goncalves, Ewan Harrison, David K. Jackson, Ian Johnston, Dominic Kwiatkowski, Cordelia Langford, John Sillitoe on behalf of the Wellcome Sanger Institute COVID-19 Surveillance Team |
| EPI_ISL_2995146 | Lighthouse Lab in Milton Keynes | Wellcome Sanger Institute for the COVID-19 Genomics UK (COG-UK) Consortium | The Lighthouse Lab in Milton Keynes and Alex Alderton, Roberto Amato, Jeffrey Barrett, Sonia Goncalves, Ewan Harrison, David K. Jackson, Ian Johnston, Dominic Kwiatkowski, Cordelia Langford, John Sillitoe on behalf of the Wellcome Sanger Institute COVID-19 Surveillance Team |
| EPI_ISL_2995147 | Berkshire and Surrey Pathology Services Lighthouse Laboratory | Wellcome Sanger Institute for the COVID-19 Genomics UK (COG-UK) Consortium | Berkshire and Surrey Pathology Services Lighthouse Laboratory and Alex Alderton, Roberto Amato, Jeffrey Barrett, Sonia Goncalves, Ewan Harrison, David K. Jackson, Ian Johnston, Dominic Kwiatkowski, Cordelia Langford, John Sillitoe on behalf of the Wellcome Sanger Institute COVID-19 Surveillance Team |
| EPI_ISL_2995148 | Lighthouse Lab in Glasgow | Wellcome Sanger Institute for the COVID-19 Genomics UK (COG-UK) Consortium | Harper VanSteenhouse, Yumi Kasai, David Gray, Carol Clugston, Anna Dominiczak and Alex Alderton, Roberto Amato, Jeffrey Barrett, Sonia Goncalves, Ewan Harrison, David K. Jackson, Ian Johnston, Dominic Kwiatkowski, Cordelia Langford, John Sillitoe on behalf of the Wellcome Sanger Institute COVID-19 Surveillance Team |
| EPI_ISL_2995149 | Lighthouse Lab in Milton Keynes | Wellcome Sanger Institute for the COVID-19 Genomics UK (COG-UK) Consortium | The Lighthouse Lab in Milton Keynes and Alex Alderton, Roberto Amato, Jeffrey Barrett, Sonia Goncalves, Ewan Harrison, David K. Jackson, Ian Johnston, Dominic Kwiatkowski, Cordelia Langford, John Sillitoe on behalf of the Wellcome Sanger Institute COVID-19 Surveillance Team |
| EPI_ISL_2995150 | Lighthouse Lab in Alderley Park | Wellcome Sanger Institute for the COVID-19 Genomics UK (COG-UK) Consortium | Jacquelyn Wynn, Mairead Hyland, The Lighthouse Lab in Alderley Park and Alex Alderton, Roberto Amato, Jeffrey Barrett, Sonia Goncalves, Ewan Harrison, David K. Jackson, Ian Johnston, Dominic Kwiatkowski, Cordelia Langford, John Sillitoe on behalf of the Wellcome Sanger Institute COVID-19 Surveillance Team |
| EPI_ISL_2995151 | Lighthouse Lab in Glasgow | Wellcome Sanger Institute for the COVID-19 Genomics UK (COG-UK) Consortium | Harper VanSteenhouse, Yumi Kasai, David Gray, Carol Clugston, Anna Dominiczak and Alex Alderton, Roberto Amato, Jeffrey Barrett, Sonia Goncalves, Ewan Harrison, David K. Jackson, Ian Johnston, Dominic Kwiatkowski, Cordelia Langford, John Sillitoe on behalf of the Wellcome Sanger Institute COVID-19 Surveillance Team |
| EPI_ISL_2995152 | Lighthouse Lab in Alderley Park | Wellcome Sanger Institute for the COVID-19 Genomics UK (COG-UK) Consortium | Jacquelyn Wynn, Mairead Hyland, The Lighthouse Lab in Alderley Park and Alex Alderton, Roberto Amato, Jeffrey Barrett, Sonia Goncalves, Ewan Harrison, David K. Jackson, Ian Johnston, Dominic Kwiatkowski, Cordelia Langford, John Sillitoe on behalf of the Wellcome Sanger Institute COVID-19 Surveillance Team |
| EPI_ISL_2995153, EPI_ISL_2995154 | Berkshire and Surrey Pathology Services Lighthouse Laboratory | Wellcome Sanger Institute for the COVID-19 Genomics UK (COG-UK) Consortium | Berkshire and Surrey Pathology Services Lighthouse Laboratory and Alex Alderton, Roberto Amato, Jeffrey Barrett, Sonia Goncalves, Ewan Harrison, David K. Jackson, Ian Johnston, Dominic Kwiatkowski, Cordelia Langford, John Sillitoe on behalf of the Wellcome Sanger Institute COVID-19 Surveillance Team |
| EPI_ISL_2995155 | Lighthouse Lab in Milton Keynes | Wellcome Sanger Institute for the COVID-19 Genomics UK (COG-UK) Consortium | The Lighthouse Lab in Milton Keynes and Alex Alderton, Roberto Amato, Jeffrey Barrett, Sonia Goncalves, Ewan Harrison, David K. Jackson, Ian Johnston, Dominic Kwiatkowski, Cordelia Langford, John Sillitoe on behalf of the Wellcome Sanger Institute COVID-19 Surveillance Team |
| EPI_ISL_2995156, EPI_ISL_2995157 | Lighthouse Lab in Glasgow | Wellcome Sanger Institute for the COVID-19 Genomics UK (COG-UK) Consortium | Harper VanSteenhouse, Yumi Kasai, David Gray, Carol Clugston, Anna Dominiczak and Alex Alderton, Roberto Amato, Jeffrey Barrett, Sonia Goncalves, Ewan Harrison, David K. Jackson, Ian Johnston, Dominic Kwiatkowski, Cordelia Langford, John Sillitoe on behalf of the Wellcome Sanger Institute COVID-19 Surveillance Team |
| EPI_ISL_2995158 | Lighthouse Lab in Alderley Park | Wellcome Sanger Institute for the COVID-19 Genomics UK (COG-UK) Consortium | Jacquelyn Wynn, Mairead Hyland, The Lighthouse Lab in Alderley Park and Alex Alderton, Roberto Amato, Jeffrey Barrett, Sonia Goncalves, Ewan Harrison, David K. Jackson, Ian Johnston, Dominic Kwiatkowski, Cordelia Langford, John Sillitoe on behalf of the Wellcome Sanger Institute COVID-19 Surveillance Team |
| EPI_ISL_2995159, EPI_ISL_2995160 | Lighthouse Lab in Milton Keynes | Wellcome Sanger Institute for the COVID-19 Genomics UK (COG-UK) Consortium | The Lighthouse Lab in Milton Keynes and Alex Alderton, Roberto Amato, Jeffrey Barrett, Sonia Goncalves, Ewan Harrison, David K. Jackson, Ian Johnston, Dominic Kwiatkowski, Cordelia Langford, John Sillitoe on behalf of the Wellcome Sanger Institute COVID-19 Surveillance Team |
| EPI_ISL_2995161 | Lighthouse Lab in Alderley Park | Wellcome Sanger Institute for the COVID-19 Genomics UK (COG-UK) Consortium | Jacquelyn Wynn, Mairead Hyland, The Lighthouse Lab in Alderley Park and Alex Alderton, Roberto Amato, Jeffrey Barrett, Sonia Goncalves, Ewan Harrison, David K. Jackson, Ian Johnston, Dominic Kwiatkowski, Cordelia Langford, John Sillitoe on behalf of the Wellcome Sanger Institute COVID-19 Surveillance Team |
| EPI_ISL_2995162, EPI_ISL_2995163, EPI_ISL_2995164, EPI_ISL_2995165 | Berkshire and Surrey Pathology Services Lighthouse Laboratory | Wellcome Sanger Institute for the COVID-19 Genomics UK (COG-UK) Consortium | Berkshire and Surrey Pathology Services Lighthouse Laboratory and Alex Alderton, Roberto Amato, Jeffrey Barrett, Sonia Goncalves, Ewan Harrison, David K. Jackson, Ian Johnston, Dominic Kwiatkowski, Cordelia Langford, John Sillitoe on behalf of the Wellcome Sanger Institute COVID-19 Surveillance Team |
| EPI_ISL_2995166 | Lighthouse Lab in Glasgow | Wellcome Sanger Institute for the COVID-19 Genomics UK (COG-UK) | Harper VanSteenhouse, Yumi Kasai, David Gray, Carol Clugston, Anna Dominiczak and Alex Alderton, Roberto Amato, Jeffrey Barrett, Sonia Goncalves, Ewan Harrison, David K. Jackson, Ian Johnston, Dominic Kwiatkowski, Cordelia Langford, |

| | | | |
|---|---|---|---|
| | | Consortium | John Sillitoe on behalf of the Wellcome Sanger Institute COVID-19 Surveillance Team |
| EPI_ISL_2995167 | Lighthouse Lab in Milton Keynes | Wellcome Sanger Institute for the COVID-19 Genomics UK (COG-UK) Consortium | The Lighthouse Lab in Milton Keynes and Alex Alderton, Roberto Amato, Jeffrey Barrett, Sonia Goncalves, Ewan Harrison, David K. Jackson, Ian Johnston, Dominic Kwiatkowski, Cordelia Langford, John Sillitoe on behalf of the Wellcome Sanger Institute COVID-19 Surveillance Team |
| EPI_ISL_2995168 | Berkshire and Surrey Pathology Services Lighthouse Laboratory | Wellcome Sanger Institute for the COVID-19 Genomics UK (COG-UK) Consortium | Berkshire and Surrey Pathology Services Lighthouse Laboratory and Alex Alderton, Roberto Amato, Jeffrey Barrett, Sonia Goncalves, Ewan Harrison, David K. Jackson, Ian Johnston, Dominic Kwiatkowski, Cordelia Langford, John Sillitoe on behalf of the Wellcome Sanger Institute COVID-19 Surveillance Team |
| EPI_ISL_2995169 | Lighthouse Lab in Glasgow | Wellcome Sanger Institute for the COVID-19 Genomics UK (COG-UK) Consortium | Harper VanSteenhouse, Yumi Kasai, David Gray, Carol Clugston, Anna Dominiczak and Alex Alderton, Roberto Amato, Jeffrey Barrett, Sonia Goncalves, Ewan Harrison, David K. Jackson, Ian Johnston, Dominic Kwiatkowski, Cordelia Langford, John Sillitoe on behalf of the Wellcome Sanger Institute COVID-19 Surveillance Team |
| EPI_ISL_2995170 | Lighthouse Lab in Alderley Park | Wellcome Sanger Institute for the COVID-19 Genomics UK (COG-UK) Consortium | Jacquelyn Wynn, Mairead Hyland, The Lighthouse Lab in Alderley Park and Alex Alderton, Roberto Amato, Jeffrey Barrett, Sonia Goncalves, Ewan Harrison, David K. Jackson, Ian Johnston, Dominic Kwiatkowski, Cordelia Langford, John Sillitoe on behalf of the Wellcome Sanger Institute COVID-19 Surveillance Team |
| EPI_ISL_2995171 | Berkshire and Surrey Pathology Services Lighthouse Laboratory | Wellcome Sanger Institute for the COVID-19 Genomics UK (COG-UK) Consortium | Berkshire and Surrey Pathology Services Lighthouse Laboratory and Alex Alderton, Roberto Amato, Jeffrey Barrett, Sonia Goncalves, Ewan Harrison, David K. Jackson, Ian Johnston, Dominic Kwiatkowski, Cordelia Langford, John Sillitoe on behalf of the Wellcome Sanger Institute COVID-19 Surveillance Team |
| EPI_ISL_2995172 | Lighthouse Lab in Alderley Park | Wellcome Sanger Institute for the COVID-19 Genomics UK (COG-UK) Consortium | Jacquelyn Wynn, Mairead Hyland, The Lighthouse Lab in Alderley Park and Alex Alderton, Roberto Amato, Jeffrey Barrett, Sonia Goncalves, Ewan Harrison, David K. Jackson, Ian Johnston, Dominic Kwiatkowski, Cordelia Langford, John Sillitoe on behalf of the Wellcome Sanger Institute COVID-19 Surveillance Team |
| EPI_ISL_2995173, EPI_ISL_2995174 | Berkshire and Surrey Pathology Services Lighthouse Laboratory | Wellcome Sanger Institute for the COVID-19 Genomics UK (COG-UK) Consortium | Berkshire and Surrey Pathology Services Lighthouse Laboratory and Alex Alderton, Roberto Amato, Jeffrey Barrett, Sonia Goncalves, Ewan Harrison, David K. Jackson, Ian Johnston, Dominic Kwiatkowski, Cordelia Langford, John Sillitoe on behalf of the Wellcome Sanger Institute COVID-19 Surveillance Team |
| EPI_ISL_2995175 | Lighthouse Lab in Milton Keynes | Wellcome Sanger Institute for the COVID-19 Genomics UK (COG-UK) Consortium | The Lighthouse Lab in Milton Keynes and Alex Alderton, Roberto Amato, Jeffrey Barrett, Sonia Goncalves, Ewan Harrison, David K. Jackson, Ian Johnston, Dominic Kwiatkowski, Cordelia Langford, John Sillitoe on behalf of the Wellcome Sanger Institute COVID-19 Surveillance Team |
| EPI_ISL_2995176 | Lighthouse Lab in Glasgow | Wellcome Sanger Institute for the COVID-19 Genomics UK (COG-UK) Consortium | Harper VanSteenhouse, Yumi Kasai, David Gray, Carol Clugston, Anna Dominiczak and Alex Alderton, Roberto Amato, Jeffrey Barrett, Sonia Goncalves, Ewan Harrison, David K. Jackson, Ian Johnston, Dominic Kwiatkowski, Cordelia Langford, John Sillitoe on behalf of the Wellcome Sanger Institute COVID-19 Surveillance Team |
| EPI_ISL_2995177, EPI_ISL_2995178 | Berkshire and Surrey Pathology Services Lighthouse Laboratory | Wellcome Sanger Institute for the COVID-19 Genomics UK (COG-UK) Consortium | Berkshire and Surrey Pathology Services Lighthouse Laboratory and Alex Alderton, Roberto Amato, Jeffrey Barrett, Sonia Goncalves, Ewan Harrison, David K. Jackson, Ian Johnston, Dominic Kwiatkowski, Cordelia Langford, John Sillitoe on behalf of the Wellcome Sanger Institute COVID-19 Surveillance Team |
| EPI_ISL_2995179, EPI_ISL_2995180 | Lighthouse Lab in Alderley Park | Wellcome Sanger Institute for the COVID-19 Genomics UK (COG-UK) Consortium | Jacquelyn Wynn, Mairead Hyland, The Lighthouse Lab in Alderley Park and Alex Alderton, Roberto Amato, Jeffrey Barrett, Sonia Goncalves, Ewan Harrison, David K. Jackson, Ian Johnston, Dominic Kwiatkowski, Cordelia Langford, John Sillitoe on behalf of the Wellcome Sanger Institute COVID-19 Surveillance Team |
| EPI_ISL_2995181 | Lighthouse Lab in Glasgow | Wellcome Sanger Institute for the COVID-19 Genomics UK (COG-UK) Consortium | Harper VanSteenhouse, Yumi Kasai, David Gray, Carol Clugston, Anna Dominiczak and Alex Alderton, Roberto Amato, Jeffrey Barrett, Sonia Goncalves, Ewan Harrison, David K. Jackson, Ian Johnston, Dominic Kwiatkowski, Cordelia Langford, John Sillitoe on behalf of the Wellcome Sanger Institute COVID-19 Surveillance Team |
| EPI_ISL_2995182 | Lighthouse Lab in Milton Keynes | Wellcome Sanger Institute for the COVID-19 Genomics UK (COG-UK) Consortium | The Lighthouse Lab in Milton Keynes and Alex Alderton, Roberto Amato, Jeffrey Barrett, Sonia Goncalves, Ewan Harrison, David K. Jackson, Ian Johnston, Dominic Kwiatkowski, Cordelia Langford, John Sillitoe on behalf of the Wellcome Sanger Institute COVID-19 Surveillance Team |
| EPI_ISL_2995183 | Berkshire and Surrey Pathology Services Lighthouse Laboratory | Wellcome Sanger Institute for the COVID-19 Genomics UK (COG-UK) Consortium | Berkshire and Surrey Pathology Services Lighthouse Laboratory and Alex Alderton, Roberto Amato, Jeffrey Barrett, Sonia Goncalves, Ewan Harrison, David K. Jackson, Ian Johnston, Dominic Kwiatkowski, Cordelia Langford, John Sillitoe on behalf of the Wellcome Sanger Institute COVID-19 Surveillance Team |
| EPI_ISL_2995184 | Lighthouse Lab in Glasgow | Wellcome Sanger Institute for the COVID-19 Genomics UK (COG-UK) Consortium | Harper VanSteenhouse, Yumi Kasai, David Gray, Carol Clugston, Anna Dominiczak and Alex Alderton, Roberto Amato, Jeffrey Barrett, Sonia Goncalves, Ewan Harrison, David K. Jackson, Ian Johnston, Dominic Kwiatkowski, Cordelia Langford, John Sillitoe on behalf of the Wellcome Sanger Institute COVID-19 Surveillance Team |
| EPI_ISL_2995185 | Lighthouse Lab in Milton Keynes | Wellcome Sanger Institute for the COVID-19 Genomics UK (COG-UK) Consortium | The Lighthouse Lab in Milton Keynes and Alex Alderton, Roberto Amato, Jeffrey Barrett, Sonia Goncalves, Ewan Harrison, David K. Jackson, Ian Johnston, Dominic Kwiatkowski, Cordelia Langford, John Sillitoe on behalf of the Wellcome Sanger Institute COVID-19 Surveillance Team |
| EPI_ISL_2995186, EPI_ISL_2995187 | Lighthouse Lab in Alderley Park | Wellcome Sanger Institute for the COVID-19 Genomics UK (COG-UK) Consortium | Jacquelyn Wynn, Mairead Hyland, The Lighthouse Lab in Alderley Park and Alex Alderton, Roberto Amato, Jeffrey Barrett, Sonia Goncalves, Ewan Harrison, David K. Jackson, Ian Johnston, Dominic Kwiatkowski, Cordelia Langford, John Sillitoe on behalf of the Wellcome Sanger Institute COVID-19 Surveillance Team |
| EPI_ISL_2995188 | Lighthouse Lab in Glasgow | Wellcome Sanger Institute for the COVID-19 Genomics UK (COG-UK) Consortium | Harper VanSteenhouse, Yumi Kasai, David Gray, Carol Clugston, Anna Dominiczak and Alex Alderton, Roberto Amato, Jeffrey Barrett, Sonia Goncalves, Ewan Harrison, David K. Jackson, Ian Johnston, Dominic Kwiatkowski, Cordelia Langford, John Sillitoe on behalf of the Wellcome Sanger Institute COVID-19 Surveillance Team |
| EPI_ISL_2995189 | Lighthouse Lab in Alderley Park | Wellcome Sanger Institute for the COVID-19 Genomics UK (COG-UK) Consortium | Jacquelyn Wynn, Mairead Hyland, The Lighthouse Lab in Alderley Park and Alex Alderton, Roberto Amato, Jeffrey Barrett, Sonia Goncalves, Ewan Harrison, David K. Jackson, Ian Johnston, Dominic Kwiatkowski, Cordelia Langford, John Sillitoe on behalf of the Wellcome Sanger Institute COVID-19 Surveillance Team |
| EPI_ISL_2995190 | Lighthouse Lab in Milton Keynes | Wellcome Sanger Institute for the COVID-19 Genomics UK (COG-UK) Consortium | The Lighthouse Lab in Milton Keynes and Alex Alderton, Roberto Amato, Jeffrey Barrett, Sonia Goncalves, Ewan Harrison, David K. Jackson, Ian Johnston, Dominic Kwiatkowski, Cordelia Langford, John Sillitoe on behalf of the Wellcome Sanger Institute COVID-19 Surveillance Team |
| EPI_ISL_2995191 | Lighthouse Lab in Alderley Park | Wellcome Sanger Institute for the COVID-19 Genomics UK (COG-UK) Consortium | Jacquelyn Wynn, Mairead Hyland, The Lighthouse Lab in Alderley Park and Alex Alderton, Roberto Amato, Jeffrey Barrett, Sonia Goncalves, Ewan Harrison, David K. Jackson, Ian Johnston, Dominic Kwiatkowski, Cordelia Langford, John Sillitoe on behalf of the Wellcome Sanger Institute COVID-19 Surveillance Team |
| EPI_ISL_2995192 | Berkshire and Surrey Pathology Services Lighthouse Laboratory | Wellcome Sanger Institute for the COVID-19 Genomics UK (COG-UK) Consortium | Berkshire and Surrey Pathology Services Lighthouse Laboratory and Alex Alderton, Roberto Amato, Jeffrey Barrett, Sonia Goncalves, Ewan Harrison, David K. Jackson, Ian Johnston, Dominic Kwiatkowski, Cordelia Langford, John Sillitoe on behalf of the Wellcome Sanger Institute COVID-19 Surveillance Team |
| EPI_ISL_2995193 | Lighthouse Lab in Glasgow | Wellcome Sanger Institute for the COVID-19 Genomics UK (COG-UK) Consortium | Harper VanSteenhouse, Yumi Kasai, David Gray, Carol Clugston, Anna Dominiczak and Alex Alderton, Roberto Amato, Jeffrey Barrett, Sonia Goncalves, Ewan Harrison, David K. Jackson, Ian Johnston, Dominic Kwiatkowski, Cordelia Langford, John Sillitoe on behalf of the Wellcome Sanger Institute COVID-19 Surveillance Team |
| EPI_ISL_2995194, EPI_ISL_2995195, EPI_ISL_2995196, EPI_ISL_2995197 | Lighthouse Lab in Alderley Park | Wellcome Sanger Institute for the COVID-19 Genomics UK (COG-UK) Consortium | Jacquelyn Wynn, Mairead Hyland, The Lighthouse Lab in Alderley Park and Alex Alderton, Roberto Amato, Jeffrey Barrett, Sonia Goncalves, Ewan Harrison, David K. Jackson, Ian Johnston, Dominic Kwiatkowski, Cordelia Langford, John Sillitoe on behalf of the Wellcome Sanger Institute COVID-19 Surveillance Team |
| EPI_ISL_2995198 | Lighthouse Lab in Milton Keynes | Wellcome Sanger Institute for the COVID-19 Genomics UK (COG-UK) Consortium | The Lighthouse Lab in Milton Keynes and Alex Alderton, Roberto Amato, Jeffrey Barrett, Sonia Goncalves, Ewan Harrison, David K. Jackson, Ian Johnston, Dominic Kwiatkowski, Cordelia Langford, John Sillitoe on behalf of the Wellcome Sanger Institute COVID-19 Surveillance Team |
| EPI_ISL_2995199 | Berkshire and Surrey Pathology Services Lighthouse Laboratory | Wellcome Sanger Institute for the COVID-19 Genomics UK (COG-UK) Consortium | Berkshire and Surrey Pathology Services Lighthouse Laboratory and Alex Alderton, Roberto Amato, Jeffrey Barrett, Sonia Goncalves, Ewan Harrison, David K. Jackson, Ian Johnston, Dominic Kwiatkowski, Cordelia Langford, John Sillitoe on behalf of the Wellcome Sanger Institute COVID-19 Surveillance Team |
| EPI_ISL_2995200, EPI_ISL_2995201 | Lighthouse Lab in Glasgow | Wellcome Sanger Institute for the COVID-19 Genomics UK (COG-UK) Consortium | Harper VanSteenhouse, Yumi Kasai, David Gray, Carol Clugston, Anna Dominiczak and Alex Alderton, Roberto Amato, Jeffrey Barrett, Sonia Goncalves, Ewan Harrison, David K. Jackson, Ian Johnston, Dominic Kwiatkowski, Cordelia Langford, John Sillitoe on behalf of the Wellcome Sanger Institute COVID-19 Surveillance Team |
| EPI_ISL_2995202 | Lighthouse Lab in Milton Keynes | Wellcome Sanger Institute for the COVID-19 Genomics UK (COG-UK) Consortium | The Lighthouse Lab in Milton Keynes and Alex Alderton, Roberto Amato, Jeffrey Barrett, Sonia Goncalves, Ewan Harrison, David K. Jackson, Ian Johnston, Dominic Kwiatkowski, Cordelia Langford, John Sillitoe on behalf of the Wellcome Sanger Institute COVID-19 Surveillance Team |
| EPI_ISL_2995203 | Lighthouse Lab in Alderley Park | Wellcome Sanger Institute for the COVID-19 Genomics UK (COG-UK) Consortium | Jacquelyn Wynn, Mairead Hyland, The Lighthouse Lab in Alderley Park and Alex Alderton, Roberto Amato, Jeffrey Barrett, Sonia Goncalves, Ewan Harrison, David K. Jackson, Ian Johnston, Dominic Kwiatkowski, Cordelia Langford, John Sillitoe on behalf of the Wellcome Sanger Institute COVID-19 Surveillance Team |
| EPI_ISL_2995204 | Lighthouse Lab in Milton Keynes | Wellcome Sanger Institute for the COVID-19 Genomics UK (COG-UK) Consortium | The Lighthouse Lab in Milton Keynes and Alex Alderton, Roberto Amato, Jeffrey Barrett, Sonia Goncalves, Ewan Harrison, David K. Jackson, Ian Johnston, Dominic Kwiatkowski, Cordelia Langford, John Sillitoe on behalf of the Wellcome Sanger Institute COVID-19 Surveillance Team |
| EPI_ISL_2995205, EPI_ISL_2995206 | Berkshire and Surrey Pathology Services Lighthouse Laboratory | Wellcome Sanger Institute for the COVID-19 Genomics UK (COG-UK) Consortium | Berkshire and Surrey Pathology Services Lighthouse Laboratory and Alex Alderton, Roberto Amato, Jeffrey Barrett, Sonia Goncalves, Ewan Harrison, David K. Jackson, Ian Johnston, Dominic Kwiatkowski, Cordelia Langford, John Sillitoe on behalf of the Wellcome Sanger Institute COVID-19 Surveillance Team |
| EPI_ISL_2995207 | Lighthouse Lab in Alderley Park | Wellcome Sanger Institute for the COVID-19 Genomics UK (COG-UK) Consortium | Jacquelyn Wynn, Mairead Hyland, The Lighthouse Lab in Alderley Park and Alex Alderton, Roberto Amato, Jeffrey Barrett, Sonia Goncalves, Ewan Harrison, David K. Jackson, Ian Johnston, Dominic Kwiatkowski, Cordelia Langford, John Sillitoe on behalf of the Wellcome Sanger Institute COVID-19 Surveillance Team |
| EPI_ISL_2995208 | Lighthouse Lab in Milton Keynes | Wellcome Sanger Institute for the COVID-19 Genomics UK (COG-UK) Consortium | The Lighthouse Lab in Milton Keynes and Alex Alderton, Roberto Amato, Jeffrey Barrett, Sonia Goncalves, Ewan Harrison, David K. Jackson, Ian Johnston, Dominic Kwiatkowski, Cordelia Langford, John Sillitoe on behalf of the Wellcome Sanger Institute COVID-19 Surveillance Team |
| EPI_ISL_2995209 | Lighthouse Lab in Glasgow | Wellcome Sanger Institute for the COVID-19 Genomics UK (COG-UK) Consortium | Harper VanSteenhouse, Yumi Kasai, David Gray, Carol Clugston, Anna Dominiczak and Alex Alderton, Roberto Amato, Jeffrey Barrett, Sonia Goncalves, Ewan Harrison, David K. Jackson, Ian Johnston, Dominic Kwiatkowski, Cordelia Langford, John Sillitoe on behalf of the Wellcome Sanger Institute COVID-19 Surveillance Team |
| EPI_ISL_2995210, EPI_ISL_2995211 | Berkshire and Surrey Pathology Services Lighthouse Laboratory | Wellcome Sanger Institute for the COVID-19 Genomics UK (COG-UK) Consortium | Berkshire and Surrey Pathology Services Lighthouse Laboratory and Alex Alderton, Roberto Amato, Jeffrey Barrett, Sonia Goncalves, Ewan Harrison, David K. Jackson, Ian Johnston, Dominic Kwiatkowski, Cordelia Langford, John Sillitoe on behalf of the Wellcome Sanger Institute COVID-19 Surveillance Team |
| EPI_ISL_2995212 | Lighthouse Lab in Milton Keynes | Wellcome Sanger Institute for the COVID-19 Genomics UK (COG-UK) Consortium | The Lighthouse Lab in Milton Keynes and Alex Alderton, Roberto Amato, Jeffrey Barrett, Sonia Goncalves, Ewan Harrison, David K. Jackson, Ian Johnston, Dominic Kwiatkowski, Cordelia Langford, John Sillitoe on behalf of the Wellcome Sanger Institute COVID-19 Surveillance Team |
| EPI_ISL_2995213, EPI_ISL_2995214 | Lighthouse Lab in Alderley Park | Wellcome Sanger Institute for the COVID-19 Genomics UK (COG-UK) Consortium | Jacquelyn Wynn, Mairead Hyland, The Lighthouse Lab in Alderley Park and Alex Alderton, Roberto Amato, Jeffrey Barrett, Sonia Goncalves, Ewan Harrison, David K. Jackson, Ian Johnston, Dominic Kwiatkowski, Cordelia Langford, John Sillitoe on behalf of the Wellcome Sanger Institute COVID-19 Surveillance Team |
| EPI_ISL_2995215 | Berkshire and Surrey Pathology Services Lighthouse Laboratory | Wellcome Sanger Institute for the COVID-19 Genomics UK (COG-UK) Consortium | Berkshire and Surrey Pathology Services Lighthouse Laboratory and Alex Alderton, Roberto Amato, Jeffrey Barrett, Sonia Goncalves, Ewan Harrison, David K. Jackson, Ian Johnston, Dominic Kwiatkowski, Cordelia Langford, John Sillitoe on behalf of the Wellcome Sanger Institute COVID-19 Surveillance Team |
| EPI_ISL_2995216 | Lighthouse Lab in Milton Keynes | Wellcome Sanger Institute for the COVID-19 Genomics UK (COG-UK) Consortium | The Lighthouse Lab in Milton Keynes and Alex Alderton, Roberto Amato, Jeffrey Barrett, Sonia Goncalves, Ewan Harrison, David K. Jackson, Ian Johnston, Dominic Kwiatkowski, Cordelia Langford, John Sillitoe on behalf of the Wellcome Sanger Institute COVID-19 Surveillance Team |
| EPI_ISL_2995217 | Berkshire and Surrey Pathology Services Lighthouse Laboratory | Wellcome Sanger Institute for the COVID-19 Genomics UK (COG-UK) Consortium | Berkshire and Surrey Pathology Services Lighthouse Laboratory and Alex Alderton, Roberto Amato, Jeffrey Barrett, Sonia Goncalves, Ewan Harrison, David K. Jackson, Ian Johnston, Dominic Kwiatkowski, Cordelia Langford, John Sillitoe on behalf of the Wellcome Sanger Institute COVID-19 Surveillance Team |
| EPI_ISL_2995218 | Lighthouse Lab in Milton Keynes | Wellcome Sanger Institute for the COVID-19 Genomics UK (COG-UK) Consortium | The Lighthouse Lab in Milton Keynes and Alex Alderton, Roberto Amato, Jeffrey Barrett, Sonia Goncalves, Ewan Harrison, David K. Jackson, Ian Johnston, Dominic Kwiatkowski, Cordelia Langford, John Sillitoe on behalf of the Wellcome Sanger Institute COVID-19 Surveillance Team |
| EPI_ISL_2995219, EPI_ISL_2995220 | Lighthouse Lab in Alderley Park | Wellcome Sanger Institute for the COVID-19 Genomics UK (COG-UK) Consortium | Jacquelyn Wynn, Mairead Hyland, The Lighthouse Lab in Alderley Park and Alex Alderton, Roberto Amato, Jeffrey Barrett, Sonia Goncalves, Ewan Harrison, David K. Jackson, Ian Johnston, Dominic Kwiatkowski, Cordelia Langford, John Sillitoe on behalf of the Wellcome Sanger Institute COVID-19 Surveillance Team |
| EPI_ISL_2995221 | Lighthouse Lab in Milton Keynes | Wellcome Sanger Institute for the COVID-19 Genomics UK (COG-UK) Consortium | The Lighthouse Lab in Milton Keynes and Alex Alderton, Roberto Amato, Jeffrey Barrett, Sonia Goncalves, Ewan Harrison, David K. Jackson, Ian Johnston, Dominic Kwiatkowski, Cordelia Langford, John Sillitoe on behalf of the Wellcome Sanger Institute COVID-19 Surveillance Team |

| | | | |
|---|---|---|---|
| EPI_ISL_2995222 | Lighthouse Lab in Glasgow | Wellcome Sanger Institute for the COVID-19 Genomics UK (COG-UK) Consortium | Harper VanSteenhouse, Yumi Kasai, David Gray, Carol Clugston, Anna Dominiczak and Alex Alderton, Roberto Amato, Jeffrey Barrett, Sonia Goncalves, Ewan Harrison, David K. Jackson, Ian Johnston, Dominic Kwiatkowski, Cordelia Langford, John Sillitoe on behalf of the Wellcome Sanger Institute COVID-19 Surveillance Team |
| EPI_ISL_2995223, EPI_ISL_2995224, EPI_ISL_2995225, EPI_ISL_2995226, EPI_ISL_2995227 | Lighthouse Lab in Alderley Park | Wellcome Sanger Institute for the COVID-19 Genomics UK (COG-UK) Consortium | Jacquelyn Wynn, Mairead Hyland, The Lighthouse Lab in Alderley Park and Alex Alderton, Roberto Amato, Jeffrey Barrett, Sonia Goncalves, Ewan Harrison, David K. Jackson, Ian Johnston, Dominic Kwiatkowski, Cordelia Langford, John Sillitoe on behalf of the Wellcome Sanger Institute COVID-19 Surveillance Team |
| EPI_ISL_2995228, EPI_ISL_2995229 | Lighthouse Lab in Glasgow | Wellcome Sanger Institute for the COVID-19 Genomics UK (COG-UK) Consortium | Harper VanSteenhouse, Yumi Kasai, David Gray, Carol Clugston, Anna Dominiczak and Alex Alderton, Roberto Amato, Jeffrey Barrett, Sonia Goncalves, Ewan Harrison, David K. Jackson, Ian Johnston, Dominic Kwiatkowski, Cordelia Langford, John Sillitoe on behalf of the Wellcome Sanger Institute COVID-19 Surveillance Team |
| EPI_ISL_2995230 | Lighthouse Lab in Alderley Park | Wellcome Sanger Institute for the COVID-19 Genomics UK (COG-UK) Consortium | Jacquelyn Wynn, Mairead Hyland, The Lighthouse Lab in Alderley Park and Alex Alderton, Roberto Amato, Jeffrey Barrett, Sonia Goncalves, Ewan Harrison, David K. Jackson, Ian Johnston, Dominic Kwiatkowski, Cordelia Langford, John Sillitoe on behalf of the Wellcome Sanger Institute COVID-19 Surveillance Team |
| EPI_ISL_2995231 | Lighthouse Lab in Milton Keynes | Wellcome Sanger Institute for the COVID-19 Genomics UK (COG-UK) Consortium | The Lighthouse Lab in Milton Keynes and Alex Alderton, Roberto Amato, Jeffrey Barrett, Sonia Goncalves, Ewan Harrison, David K. Jackson, Ian Johnston, Dominic Kwiatkowski, Cordelia Langford, John Sillitoe on behalf of the Wellcome Sanger Institute COVID-19 Surveillance Team |
| EPI_ISL_2995232 | Lighthouse Lab in Alderley Park | Wellcome Sanger Institute for the COVID-19 Genomics UK (COG-UK) Consortium | Jacquelyn Wynn, Mairead Hyland, The Lighthouse Lab in Alderley Park and Alex Alderton, Roberto Amato, Jeffrey Barrett, Sonia Goncalves, Ewan Harrison, David K. Jackson, Ian Johnston, Dominic Kwiatkowski, Cordelia Langford, John Sillitoe on behalf of the Wellcome Sanger Institute COVID-19 Surveillance Team |
| EPI_ISL_2995233 | Lighthouse Lab in Glasgow | Wellcome Sanger Institute for the COVID-19 Genomics UK (COG-UK) Consortium | Harper VanSteenhouse, Yumi Kasai, David Gray, Carol Clugston, Anna Dominiczak and Alex Alderton, Roberto Amato, Jeffrey Barrett, Sonia Goncalves, Ewan Harrison, David K. Jackson, Ian Johnston, Dominic Kwiatkowski, Cordelia Langford, John Sillitoe on behalf of the Wellcome Sanger Institute COVID-19 Surveillance Team |
| EPI_ISL_2995234 | Berkshire and Surrey Pathology Services Lighthouse Laboratory | Wellcome Sanger Institute for the COVID-19 Genomics UK (COG-UK) Consortium | Berkshire and Surrey Pathology Services Lighthouse Laboratory and Alex Alderton, Roberto Amato, Jeffrey Barrett, Sonia Goncalves, Ewan Harrison, David K. Jackson, Ian Johnston, Dominic Kwiatkowski, Cordelia Langford, John Sillitoe on behalf of the Wellcome Sanger Institute COVID-19 Surveillance Team |
| EPI_ISL_2995235, EPI_ISL_2995236 | Lighthouse Lab in Glasgow | Wellcome Sanger Institute for the COVID-19 Genomics UK (COG-UK) Consortium | Harper VanSteenhouse, Yumi Kasai, David Gray, Carol Clugston, Anna Dominiczak and Alex Alderton, Roberto Amato, Jeffrey Barrett, Sonia Goncalves, Ewan Harrison, David K. Jackson, Ian Johnston, Dominic Kwiatkowski, Cordelia Langford, John Sillitoe on behalf of the Wellcome Sanger Institute COVID-19 Surveillance Team |
| EPI_ISL_2995237 | Lighthouse Lab in Milton Keynes | Wellcome Sanger Institute for the COVID-19 Genomics UK (COG-UK) Consortium | The Lighthouse Lab in Milton Keynes and Alex Alderton, Roberto Amato, Jeffrey Barrett, Sonia Goncalves, Ewan Harrison, David K. Jackson, Ian Johnston, Dominic Kwiatkowski, Cordelia Langford, John Sillitoe on behalf of the Wellcome Sanger Institute COVID-19 Surveillance Team |
| EPI_ISL_2995238 | Berkshire and Surrey Pathology Services Lighthouse Laboratory | Wellcome Sanger Institute for the COVID-19 Genomics UK (COG-UK) Consortium | Berkshire and Surrey Pathology Services Lighthouse Laboratory and Alex Alderton, Roberto Amato, Jeffrey Barrett, Sonia Goncalves, Ewan Harrison, David K. Jackson, Ian Johnston, Dominic Kwiatkowski, Cordelia Langford, John Sillitoe on behalf of the Wellcome Sanger Institute COVID-19 Surveillance Team |
| EPI_ISL_2995239, EPI_ISL_2995240, EPI_ISL_2995241 | Lighthouse Lab in Milton Keynes | Wellcome Sanger Institute for the COVID-19 Genomics UK (COG-UK) Consortium | The Lighthouse Lab in Milton Keynes and Alex Alderton, Roberto Amato, Jeffrey Barrett, Sonia Goncalves, Ewan Harrison, David K. Jackson, Ian Johnston, Dominic Kwiatkowski, Cordelia Langford, John Sillitoe on behalf of the Wellcome Sanger Institute COVID-19 Surveillance Team |
| EPI_ISL_2995242, EPI_ISL_2995243 | Lighthouse Lab in Alderley Park | Wellcome Sanger Institute for the COVID-19 Genomics UK (COG-UK) Consortium | Jacquelyn Wynn, Mairead Hyland, The Lighthouse Lab in Alderley Park and Alex Alderton, Roberto Amato, Jeffrey Barrett, Sonia Goncalves, Ewan Harrison, David K. Jackson, Ian Johnston, Dominic Kwiatkowski, Cordelia Langford, John Sillitoe on behalf of the Wellcome Sanger Institute COVID-19 Surveillance Team |
| EPI_ISL_2995244, EPI_ISL_2995245 | Lighthouse Lab in Milton Keynes | Wellcome Sanger Institute for the COVID-19 Genomics UK (COG-UK) Consortium | The Lighthouse Lab in Milton Keynes and Alex Alderton, Roberto Amato, Jeffrey Barrett, Sonia Goncalves, Ewan Harrison, David K. Jackson, Ian Johnston, Dominic Kwiatkowski, Cordelia Langford, John Sillitoe on behalf of the Wellcome Sanger Institute COVID-19 Surveillance Team |
| EPI_ISL_2995246, EPI_ISL_2995247 | Berkshire and Surrey Pathology Services Lighthouse Laboratory | Wellcome Sanger Institute for the COVID-19 Genomics UK (COG-UK) Consortium | Berkshire and Surrey Pathology Services Lighthouse Laboratory and Alex Alderton, Roberto Amato, Jeffrey Barrett, Sonia Goncalves, Ewan Harrison, David K. Jackson, Ian Johnston, Dominic Kwiatkowski, Cordelia Langford, John Sillitoe on behalf of the Wellcome Sanger Institute COVID-19 Surveillance Team |
| EPI_ISL_2995248, EPI_ISL_2995249 | Lighthouse Lab in Glasgow | Wellcome Sanger Institute for the COVID-19 Genomics UK (COG-UK) Consortium | Harper VanSteenhouse, Yumi Kasai, David Gray, Carol Clugston, Anna Dominiczak and Alex Alderton, Roberto Amato, Jeffrey Barrett, Sonia Goncalves, Ewan Harrison, David K. Jackson, Ian Johnston, Dominic Kwiatkowski, Cordelia Langford, John Sillitoe on behalf of the Wellcome Sanger Institute COVID-19 Surveillance Team |
| EPI_ISL_2995250, EPI_ISL_2995251 | Lighthouse Lab in Alderley Park | Wellcome Sanger Institute for the COVID-19 Genomics UK (COG-UK) Consortium | Jacquelyn Wynn, Mairead Hyland, The Lighthouse Lab in Alderley Park and Alex Alderton, Roberto Amato, Jeffrey Barrett, Sonia Goncalves, Ewan Harrison, David K. Jackson, Ian Johnston, Dominic Kwiatkowski, Cordelia Langford, John Sillitoe on behalf of the Wellcome Sanger Institute COVID-19 Surveillance Team |
| EPI_ISL_2995252 | Lighthouse Lab in Milton Keynes | Wellcome Sanger Institute for the COVID-19 Genomics UK (COG-UK) Consortium | The Lighthouse Lab in Milton Keynes and Alex Alderton, Roberto Amato, Jeffrey Barrett, Sonia Goncalves, Ewan Harrison, David K. Jackson, Ian Johnston, Dominic Kwiatkowski, Cordelia Langford, John Sillitoe on behalf of the Wellcome Sanger Institute COVID-19 Surveillance Team |
| EPI_ISL_2995253 | Lighthouse Lab in Glasgow | Wellcome Sanger Institute for the COVID-19 Genomics UK (COG-UK) Consortium | Harper VanSteenhouse, Yumi Kasai, David Gray, Carol Clugston, Anna Dominiczak and Alex Alderton, Roberto Amato, Jeffrey Barrett, Sonia Goncalves, Ewan Harrison, David K. Jackson, Ian Johnston, Dominic Kwiatkowski, Cordelia Langford, John Sillitoe on behalf of the Wellcome Sanger Institute COVID-19 Surveillance Team |
| EPI_ISL_2995254, EPI_ISL_2995255, EPI_ISL_2995256, EPI_ISL_2995257 | Lighthouse Lab in Milton Keynes | Wellcome Sanger Institute for the COVID-19 Genomics UK (COG-UK) Consortium | The Lighthouse Lab in Milton Keynes and Alex Alderton, Roberto Amato, Jeffrey Barrett, Sonia Goncalves, Ewan Harrison, David K. Jackson, Ian Johnston, Dominic Kwiatkowski, Cordelia Langford, John Sillitoe on behalf of the Wellcome Sanger Institute COVID-19 Surveillance Team |
| EPI_ISL_2995258 | Berkshire and Surrey Pathology Services Lighthouse Laboratory | Wellcome Sanger Institute for the COVID-19 Genomics UK (COG-UK) Consortium | Berkshire and Surrey Pathology Services Lighthouse Laboratory and Alex Alderton, Roberto Amato, Jeffrey Barrett, Sonia Goncalves, Ewan Harrison, David K. Jackson, Ian Johnston, Dominic Kwiatkowski, Cordelia Langford, John Sillitoe on behalf of the Wellcome Sanger Institute COVID-19 Surveillance Team |
| EPI_ISL_2995259 | Lighthouse Lab in Alderley Park | Wellcome Sanger Institute for the COVID-19 Genomics UK (COG-UK) Consortium | Jacquelyn Wynn, Mairead Hyland, The Lighthouse Lab in Alderley Park and Alex Alderton, Roberto Amato, Jeffrey Barrett, Sonia Goncalves, Ewan Harrison, David K. Jackson, Ian Johnston, Dominic Kwiatkowski, Cordelia Langford, John Sillitoe on behalf of the Wellcome Sanger Institute COVID-19 Surveillance Team |
| EPI_ISL_2995260 | Lighthouse Lab in Glasgow | Wellcome Sanger Institute for the COVID-19 Genomics UK (COG-UK) Consortium | Harper VanSteenhouse, Yumi Kasai, David Gray, Carol Clugston, Anna Dominiczak and Alex Alderton, Roberto Amato, Jeffrey Barrett, Sonia Goncalves, Ewan Harrison, David K. Jackson, Ian Johnston, Dominic Kwiatkowski, Cordelia Langford, John Sillitoe on behalf of the Wellcome Sanger Institute COVID-19 Surveillance Team |
| EPI_ISL_2995261 | Lighthouse Lab in Alderley Park | Wellcome Sanger Institute for the COVID-19 Genomics UK (COG-UK) Consortium | Jacquelyn Wynn, Mairead Hyland, The Lighthouse Lab in Alderley Park and Alex Alderton, Roberto Amato, Jeffrey Barrett, Sonia Goncalves, Ewan Harrison, David K. Jackson, Ian Johnston, Dominic Kwiatkowski, Cordelia Langford, John Sillitoe on behalf of the Wellcome Sanger Institute COVID-19 Surveillance Team |
| EPI_ISL_2995262, EPI_ISL_2995263 | Lighthouse Lab in Glasgow | Wellcome Sanger Institute for the COVID-19 Genomics UK (COG-UK) Consortium | Harper VanSteenhouse, Yumi Kasai, David Gray, Carol Clugston, Anna Dominiczak and Alex Alderton, Roberto Amato, Jeffrey Barrett, Sonia Goncalves, Ewan Harrison, David K. Jackson, Ian Johnston, Dominic Kwiatkowski, Cordelia Langford, John Sillitoe on behalf of the Wellcome Sanger Institute COVID-19 Surveillance Team |
| EPI_ISL_2995264 | Lighthouse Lab in Milton Keynes | Wellcome Sanger Institute for the COVID-19 Genomics UK (COG-UK) Consortium | The Lighthouse Lab in Milton Keynes and Alex Alderton, Roberto Amato, Jeffrey Barrett, Sonia Goncalves, Ewan Harrison, David K. Jackson, Ian Johnston, Dominic Kwiatkowski, Cordelia Langford, John Sillitoe on behalf of the Wellcome Sanger Institute COVID-19 Surveillance Team |
| EPI_ISL_2995265 | Berkshire and Surrey Pathology Services Lighthouse Laboratory | Wellcome Sanger Institute for the COVID-19 Genomics UK (COG-UK) Consortium | Berkshire and Surrey Pathology Services Lighthouse Laboratory and Alex Alderton, Roberto Amato, Jeffrey Barrett, Sonia Goncalves, Ewan Harrison, David K. Jackson, Ian Johnston, Dominic Kwiatkowski, Cordelia Langford, John Sillitoe on behalf of the Wellcome Sanger Institute COVID-19 Surveillance Team |
| EPI_ISL_2995266, EPI_ISL_2995267, EPI_ISL_2995268 | Lighthouse Lab in Milton Keynes | Wellcome Sanger Institute for the COVID-19 Genomics UK (COG-UK) Consortium | The Lighthouse Lab in Milton Keynes and Alex Alderton, Roberto Amato, Jeffrey Barrett, Sonia Goncalves, Ewan Harrison, David K. Jackson, Ian Johnston, Dominic Kwiatkowski, Cordelia Langford, John Sillitoe on behalf of the Wellcome Sanger Institute COVID-19 Surveillance Team |
| EPI_ISL_2995269, EPI_ISL_2995270, EPI_ISL_2995271 | Lighthouse Lab in Alderley Park | Wellcome Sanger Institute for the COVID-19 Genomics UK (COG-UK) Consortium | Jacquelyn Wynn, Mairead Hyland, The Lighthouse Lab in Alderley Park and Alex Alderton, Roberto Amato, Jeffrey Barrett, Sonia Goncalves, Ewan Harrison, David K. Jackson, Ian Johnston, Dominic Kwiatkowski, Cordelia Langford, John Sillitoe on behalf of the Wellcome Sanger Institute COVID-19 Surveillance Team |
| EPI_ISL_2995272 | Lighthouse Lab in Milton Keynes | Wellcome Sanger Institute for the COVID-19 Genomics UK (COG-UK) Consortium | The Lighthouse Lab in Milton Keynes and Alex Alderton, Roberto Amato, Jeffrey Barrett, Sonia Goncalves, Ewan Harrison, David K. Jackson, Ian Johnston, Dominic Kwiatkowski, Cordelia Langford, John Sillitoe on behalf of the Wellcome Sanger Institute COVID-19 Surveillance Team |
| EPI_ISL_2995273 | Berkshire and Surrey Pathology Services Lighthouse Laboratory | Wellcome Sanger Institute for the COVID-19 Genomics UK (COG-UK) Consortium | Berkshire and Surrey Pathology Services Lighthouse Laboratory and Alex Alderton, Roberto Amato, Jeffrey Barrett, Sonia Goncalves, Ewan Harrison, David K. Jackson, Ian Johnston, Dominic Kwiatkowski, Cordelia Langford, John Sillitoe on behalf of the Wellcome Sanger Institute COVID-19 Surveillance Team |
| EPI_ISL_2995274 | Lighthouse Lab in Milton Keynes | Wellcome Sanger Institute for the COVID-19 Genomics UK (COG-UK) Consortium | The Lighthouse Lab in Milton Keynes and Alex Alderton, Roberto Amato, Jeffrey Barrett, Sonia Goncalves, Ewan Harrison, David K. Jackson, Ian Johnston, Dominic Kwiatkowski, Cordelia Langford, John Sillitoe on behalf of the Wellcome Sanger Institute COVID-19 Surveillance Team |
| EPI_ISL_2995275 | Berkshire and Surrey Pathology Services Lighthouse Laboratory | Wellcome Sanger Institute for the COVID-19 Genomics UK (COG-UK) Consortium | Berkshire and Surrey Pathology Services Lighthouse Laboratory and Alex Alderton, Roberto Amato, Jeffrey Barrett, Sonia Goncalves, Ewan Harrison, David K. Jackson, Ian Johnston, Dominic Kwiatkowski, Cordelia Langford, John Sillitoe on behalf of the Wellcome Sanger Institute COVID-19 Surveillance Team |
| EPI_ISL_2995276 | Lighthouse Lab in Milton Keynes | Wellcome Sanger Institute for the COVID-19 Genomics UK (COG-UK) Consortium | The Lighthouse Lab in Milton Keynes and Alex Alderton, Roberto Amato, Jeffrey Barrett, Sonia Goncalves, Ewan Harrison, David K. Jackson, Ian Johnston, Dominic Kwiatkowski, Cordelia Langford, John Sillitoe on behalf of the Wellcome Sanger Institute COVID-19 Surveillance Team |
| EPI_ISL_2995277 | Lighthouse Lab in Glasgow | Wellcome Sanger Institute for the COVID-19 Genomics UK (COG-UK) Consortium | Harper VanSteenhouse, Yumi Kasai, David Gray, Carol Clugston, Anna Dominiczak and Alex Alderton, Roberto Amato, Jeffrey Barrett, Sonia Goncalves, Ewan Harrison, David K. Jackson, Ian Johnston, Dominic Kwiatkowski, Cordelia Langford, John Sillitoe on behalf of the Wellcome Sanger Institute COVID-19 Surveillance Team |
| EPI_ISL_2995278 | Lighthouse Lab in Alderley Park | Wellcome Sanger Institute for the COVID-19 Genomics UK (COG-UK) Consortium | Jacquelyn Wynn, Mairead Hyland, The Lighthouse Lab in Alderley Park and Alex Alderton, Roberto Amato, Jeffrey Barrett, Sonia Goncalves, Ewan Harrison, David K. Jackson, Ian Johnston, Dominic Kwiatkowski, Cordelia Langford, John Sillitoe on behalf of the Wellcome Sanger Institute COVID-19 Surveillance Team |
| EPI_ISL_2995279, EPI_ISL_2995280 | Lighthouse Lab in Milton Keynes | Wellcome Sanger Institute for the COVID-19 Genomics UK (COG-UK) Consortium | The Lighthouse Lab in Milton Keynes and Alex Alderton, Roberto Amato, Jeffrey Barrett, Sonia Goncalves, Ewan Harrison, David K. Jackson, Ian Johnston, Dominic Kwiatkowski, Cordelia Langford, John Sillitoe on behalf of the Wellcome Sanger Institute COVID-19 Surveillance Team |
| EPI_ISL_2995281 | Berkshire and Surrey Pathology Services Lighthouse Laboratory | Wellcome Sanger Institute for the COVID-19 Genomics UK (COG-UK) Consortium | Berkshire and Surrey Pathology Services Lighthouse Laboratory and Alex Alderton, Roberto Amato, Jeffrey Barrett, Sonia Goncalves, Ewan Harrison, David K. Jackson, Ian Johnston, Dominic Kwiatkowski, Cordelia Langford, John Sillitoe on behalf of the Wellcome Sanger Institute COVID-19 Surveillance Team |
| EPI_ISL_2995282 | Lighthouse Lab in Alderley Park | Wellcome Sanger Institute for the COVID-19 Genomics UK (COG-UK) Consortium | Jacquelyn Wynn, Mairead Hyland, The Lighthouse Lab in Alderley Park and Alex Alderton, Roberto Amato, Jeffrey Barrett, Sonia Goncalves, Ewan Harrison, David K. Jackson, Ian Johnston, Dominic Kwiatkowski, Cordelia Langford, John Sillitoe on behalf of the Wellcome Sanger Institute COVID-19 Surveillance Team |
| EPI_ISL_2995283 | Berkshire and Surrey Pathology Services Lighthouse Laboratory | Wellcome Sanger Institute for the COVID-19 Genomics UK (COG-UK) Consortium | Berkshire and Surrey Pathology Services Lighthouse Laboratory and Alex Alderton, Roberto Amato, Jeffrey Barrett, Sonia Goncalves, Ewan Harrison, David K. Jackson, Ian Johnston, Dominic Kwiatkowski, Cordelia Langford, John Sillitoe on behalf of the Wellcome Sanger Institute COVID-19 Surveillance Team |
| EPI_ISL_2995284 | Lighthouse Lab in Alderley Park | Wellcome Sanger Institute for the COVID-19 Genomics UK (COG-UK) Consortium | Jacquelyn Wynn, Mairead Hyland, The Lighthouse Lab in Alderley Park and Alex Alderton, Roberto Amato, Jeffrey Barrett, Sonia Goncalves, Ewan Harrison, David K. Jackson, Ian Johnston, Dominic Kwiatkowski, Cordelia Langford, John Sillitoe on behalf of the Wellcome Sanger Institute COVID-19 Surveillance Team |
| EPI_ISL_2995285 | Berkshire and Surrey Pathology Services Lighthouse Laboratory | Wellcome Sanger Institute for the COVID-19 Genomics UK (COG-UK) | Berkshire and Surrey Pathology Services Lighthouse Laboratory and Alex Alderton, Roberto Amato, Jeffrey Barrett, Sonia Goncalves, Ewan Harrison, David K. Jackson, Ian Johnston, Dominic Kwiatkowski, Cordelia Langford, John Sillitoe on behalf |

| | | | |
|---|---|---|---|
| | | Consortium | of the Wellcome Sanger Institute COVID-19 Surveillance Team |
| EPI_ISL_2995286, EPI_ISL_2995287 | Lighthouse Lab in Milton Keynes | Wellcome Sanger Institute for the COVID-19 Genomics UK (COG-UK) Consortium | The Lighthouse Lab in Milton Keynes and Alex Alderton, Roberto Amato, Jeffrey Barrett, Sonia Goncalves, Ewan Harrison, David K. Jackson, Ian Johnston, Dominic Kwiatkowski, Cordelia Langford, John Sillitoe on behalf of the Wellcome Sanger Institute COVID-19 Surveillance Team |
| EPI_ISL_2995288 | Lighthouse Lab in Glasgow | Wellcome Sanger Institute for the COVID-19 Genomics UK (COG-UK) Consortium | Harper VanSteenhouse, Yumi Kasai, David Gray, Carol Clugston, Anna Dominiczak and Alex Alderton, Roberto Amato, Jeffrey Barrett, Sonia Goncalves, Ewan Harrison, David K. Jackson, Ian Johnston, Dominic Kwiatkowski, Cordelia Langford, John Sillitoe on behalf of the Wellcome Sanger Institute COVID-19 Surveillance Team |
| EPI_ISL_2995289, EPI_ISL_2995290 | Lighthouse Lab in Alderley Park | Wellcome Sanger Institute for the COVID-19 Genomics UK (COG-UK) Consortium | Jacquelyn Wynn, Mairead Hyland, The Lighthouse Lab in Alderley Park and Alex Alderton, Roberto Amato, Jeffrey Barrett, Sonia Goncalves, Ewan Harrison, David K. Jackson, Ian Johnston, Dominic Kwiatkowski, Cordelia Langford, John Sillitoe on behalf of the Wellcome Sanger Institute COVID-19 Surveillance Team |
| EPI_ISL_2995291 | Lighthouse Lab in Milton Keynes | Wellcome Sanger Institute for the COVID-19 Genomics UK (COG-UK) Consortium | The Lighthouse Lab in Milton Keynes and Alex Alderton, Roberto Amato, Jeffrey Barrett, Sonia Goncalves, Ewan Harrison, David K. Jackson, Ian Johnston, Dominic Kwiatkowski, Cordelia Langford, John Sillitoe on behalf of the Wellcome Sanger Institute COVID-19 Surveillance Team |
| EPI_ISL_2995292, EPI_ISL_2995293 | Lighthouse Lab in Alderley Park | Wellcome Sanger Institute for the COVID-19 Genomics UK (COG-UK) Consortium | Jacquelyn Wynn, Mairead Hyland, The Lighthouse Lab in Alderley Park and Alex Alderton, Roberto Amato, Jeffrey Barrett, Sonia Goncalves, Ewan Harrison, David K. Jackson, Ian Johnston, Dominic Kwiatkowski, Cordelia Langford, John Sillitoe on behalf of the Wellcome Sanger Institute COVID-19 Surveillance Team |
| EPI_ISL_2995294 | Berkshire and Surrey Pathology Services Lighthouse Laboratory | Wellcome Sanger Institute for the COVID-19 Genomics UK (COG-UK) Consortium | Berkshire and Surrey Pathology Services Lighthouse Laboratory and Alex Alderton, Roberto Amato, Jeffrey Barrett, Sonia Goncalves, Ewan Harrison, David K. Jackson, Ian Johnston, Dominic Kwiatkowski, Cordelia Langford, John Sillitoe on behalf of the Wellcome Sanger Institute COVID-19 Surveillance Team |
| EPI_ISL_2995295 | Lighthouse Lab in Milton Keynes | Wellcome Sanger Institute for the COVID-19 Genomics UK (COG-UK) Consortium | The Lighthouse Lab in Milton Keynes and Alex Alderton, Roberto Amato, Jeffrey Barrett, Sonia Goncalves, Ewan Harrison, David K. Jackson, Ian Johnston, Dominic Kwiatkowski, Cordelia Langford, John Sillitoe on behalf of the Wellcome Sanger Institute COVID-19 Surveillance Team |
| EPI_ISL_2995296 | Berkshire and Surrey Pathology Services Lighthouse Laboratory | Wellcome Sanger Institute for the COVID-19 Genomics UK (COG-UK) Consortium | Berkshire and Surrey Pathology Services Lighthouse Laboratory and Alex Alderton, Roberto Amato, Jeffrey Barrett, Sonia Goncalves, Ewan Harrison, David K. Jackson, Ian Johnston, Dominic Kwiatkowski, Cordelia Langford, John Sillitoe on behalf of the Wellcome Sanger Institute COVID-19 Surveillance Team |
| EPI_ISL_2995297 | Lighthouse Lab in Milton Keynes | Wellcome Sanger Institute for the COVID-19 Genomics UK (COG-UK) Consortium | The Lighthouse Lab in Milton Keynes and Alex Alderton, Roberto Amato, Jeffrey Barrett, Sonia Goncalves, Ewan Harrison, David K. Jackson, Ian Johnston, Dominic Kwiatkowski, Cordelia Langford, John Sillitoe on behalf of the Wellcome Sanger Institute COVID-19 Surveillance Team |
| EPI_ISL_2995298 | Lighthouse Lab in Glasgow | Wellcome Sanger Institute for the COVID-19 Genomics UK (COG-UK) Consortium | Harper VanSteenhouse, Yumi Kasai, David Gray, Carol Clugston, Anna Dominiczak and Alex Alderton, Roberto Amato, Jeffrey Barrett, Sonia Goncalves, Ewan Harrison, David K. Jackson, Ian Johnston, Dominic Kwiatkowski, Cordelia Langford, John Sillitoe on behalf of the Wellcome Sanger Institute COVID-19 Surveillance Team |
| EPI_ISL_2995299 | Lighthouse Lab in Alderley Park | Wellcome Sanger Institute for the COVID-19 Genomics UK (COG-UK) Consortium | Jacquelyn Wynn, Mairead Hyland, The Lighthouse Lab in Alderley Park and Alex Alderton, Roberto Amato, Jeffrey Barrett, Sonia Goncalves, Ewan Harrison, David K. Jackson, Ian Johnston, Dominic Kwiatkowski, Cordelia Langford, John Sillitoe on behalf of the Wellcome Sanger Institute COVID-19 Surveillance Team |
| EPI_ISL_2995300 | Lighthouse Lab in Glasgow | Wellcome Sanger Institute for the COVID-19 Genomics UK (COG-UK) Consortium | Harper VanSteenhouse, Yumi Kasai, David Gray, Carol Clugston, Anna Dominiczak and Alex Alderton, Roberto Amato, Jeffrey Barrett, Sonia Goncalves, Ewan Harrison, David K. Jackson, Ian Johnston, Dominic Kwiatkowski, Cordelia Langford, John Sillitoe on behalf of the Wellcome Sanger Institute COVID-19 Surveillance Team |
| EPI_ISL_2995301 | Berkshire and Surrey Pathology Services Lighthouse Laboratory | Wellcome Sanger Institute for the COVID-19 Genomics UK (COG-UK) Consortium | Berkshire and Surrey Pathology Services Lighthouse Laboratory and Alex Alderton, Roberto Amato, Jeffrey Barrett, Sonia Goncalves, Ewan Harrison, David K. Jackson, Ian Johnston, Dominic Kwiatkowski, Cordelia Langford, John Sillitoe on behalf of the Wellcome Sanger Institute COVID-19 Surveillance Team |
| EPI_ISL_2995302, EPI_ISL_2995303 | Lighthouse Lab in Milton Keynes | Wellcome Sanger Institute for the COVID-19 Genomics UK (COG-UK) Consortium | The Lighthouse Lab in Milton Keynes and Alex Alderton, Roberto Amato, Jeffrey Barrett, Sonia Goncalves, Ewan Harrison, David K. Jackson, Ian Johnston, Dominic Kwiatkowski, Cordelia Langford, John Sillitoe on behalf of the Wellcome Sanger Institute COVID-19 Surveillance Team |
| EPI_ISL_2995304 | Lighthouse Lab in Alderley Park | Wellcome Sanger Institute for the COVID-19 Genomics UK (COG-UK) Consortium | Jacquelyn Wynn, Mairead Hyland, The Lighthouse Lab in Alderley Park and Alex Alderton, Roberto Amato, Jeffrey Barrett, Sonia Goncalves, Ewan Harrison, David K. Jackson, Ian Johnston, Dominic Kwiatkowski, Cordelia Langford, John Sillitoe on behalf of the Wellcome Sanger Institute COVID-19 Surveillance Team |
| EPI_ISL_2995305 | Berkshire and Surrey Pathology Services Lighthouse Laboratory | Wellcome Sanger Institute for the COVID-19 Genomics UK (COG-UK) Consortium | Berkshire and Surrey Pathology Services Lighthouse Laboratory and Alex Alderton, Roberto Amato, Jeffrey Barrett, Sonia Goncalves, Ewan Harrison, David K. Jackson, Ian Johnston, Dominic Kwiatkowski, Cordelia Langford, John Sillitoe on behalf of the Wellcome Sanger Institute COVID-19 Surveillance Team |
| EPI_ISL_2995306 | Lighthouse Lab in Milton Keynes | Wellcome Sanger Institute for the COVID-19 Genomics UK (COG-UK) Consortium | The Lighthouse Lab in Milton Keynes and Alex Alderton, Roberto Amato, Jeffrey Barrett, Sonia Goncalves, Ewan Harrison, David K. Jackson, Ian Johnston, Dominic Kwiatkowski, Cordelia Langford, John Sillitoe on behalf of the Wellcome Sanger Institute COVID-19 Surveillance Team |
| EPI_ISL_2995307 | Lighthouse Lab in Glasgow | Wellcome Sanger Institute for the COVID-19 Genomics UK (COG-UK) Consortium | Harper VanSteenhouse, Yumi Kasai, David Gray, Carol Clugston, Anna Dominiczak and Alex Alderton, Roberto Amato, Jeffrey Barrett, Sonia Goncalves, Ewan Harrison, David K. Jackson, Ian Johnston, Dominic Kwiatkowski, Cordelia Langford, John Sillitoe on behalf of the Wellcome Sanger Institute COVID-19 Surveillance Team |
| EPI_ISL_2995308 | Lighthouse Lab in Milton Keynes | Wellcome Sanger Institute for the COVID-19 Genomics UK (COG-UK) Consortium | The Lighthouse Lab in Milton Keynes and Alex Alderton, Roberto Amato, Jeffrey Barrett, Sonia Goncalves, Ewan Harrison, David K. Jackson, Ian Johnston, Dominic Kwiatkowski, Cordelia Langford, John Sillitoe on behalf of the Wellcome Sanger Institute COVID-19 Surveillance Team |
| EPI_ISL_2995309 | Berkshire and Surrey Pathology Services Lighthouse Laboratory | Wellcome Sanger Institute for the COVID-19 Genomics UK (COG-UK) Consortium | Berkshire and Surrey Pathology Services Lighthouse Laboratory and Alex Alderton, Roberto Amato, Jeffrey Barrett, Sonia Goncalves, Ewan Harrison, David K. Jackson, Ian Johnston, Dominic Kwiatkowski, Cordelia Langford, John Sillitoe on behalf of the Wellcome Sanger Institute COVID-19 Surveillance Team |
| EPI_ISL_2995310 | Lighthouse Lab in Alderley Park | Wellcome Sanger Institute for the COVID-19 Genomics UK (COG-UK) Consortium | Jacquelyn Wynn, Mairead Hyland, The Lighthouse Lab in Alderley Park and Alex Alderton, Roberto Amato, Jeffrey Barrett, Sonia Goncalves, Ewan Harrison, David K. Jackson, Ian Johnston, Dominic Kwiatkowski, Cordelia Langford, John Sillitoe on behalf of the Wellcome Sanger Institute COVID-19 Surveillance Team |
| EPI_ISL_2995311 | Berkshire and Surrey Pathology Services Lighthouse Laboratory | Wellcome Sanger Institute for the COVID-19 Genomics UK (COG-UK) Consortium | Berkshire and Surrey Pathology Services Lighthouse Laboratory and Alex Alderton, Roberto Amato, Jeffrey Barrett, Sonia Goncalves, Ewan Harrison, David K. Jackson, Ian Johnston, Dominic Kwiatkowski, Cordelia Langford, John Sillitoe on behalf of the Wellcome Sanger Institute COVID-19 Surveillance Team |
| EPI_ISL_2995312, EPI_ISL_2995313, EPI_ISL_2995314 | Lighthouse Lab in Glasgow | Wellcome Sanger Institute for the COVID-19 Genomics UK (COG-UK) Consortium | Harper VanSteenhouse, Yumi Kasai, David Gray, Carol Clugston, Anna Dominiczak and Alex Alderton, Roberto Amato, Jeffrey Barrett, Sonia Goncalves, Ewan Harrison, David K. Jackson, Ian Johnston, Dominic Kwiatkowski, Cordelia Langford, John Sillitoe on behalf of the Wellcome Sanger Institute COVID-19 Surveillance Team |
| EPI_ISL_2995315 | Lighthouse Lab in Alderley Park | Wellcome Sanger Institute for the COVID-19 Genomics UK (COG-UK) Consortium | Jacquelyn Wynn, Mairead Hyland, The Lighthouse Lab in Alderley Park and Alex Alderton, Roberto Amato, Jeffrey Barrett, Sonia Goncalves, Ewan Harrison, David K. Jackson, Ian Johnston, Dominic Kwiatkowski, Cordelia Langford, John Sillitoe on behalf of the Wellcome Sanger Institute COVID-19 Surveillance Team |
| EPI_ISL_2995316 | Lighthouse Lab in Glasgow | Wellcome Sanger Institute for the COVID-19 Genomics UK (COG-UK) Consortium | Harper VanSteenhouse, Yumi Kasai, David Gray, Carol Clugston, Anna Dominiczak and Alex Alderton, Roberto Amato, Jeffrey Barrett, Sonia Goncalves, Ewan Harrison, David K. Jackson, Ian Johnston, Dominic Kwiatkowski, Cordelia Langford, John Sillitoe on behalf of the Wellcome Sanger Institute COVID-19 Surveillance Team |
| EPI_ISL_2995317 | Lighthouse Lab in Milton Keynes | Wellcome Sanger Institute for the COVID-19 Genomics UK (COG-UK) Consortium | The Lighthouse Lab in Milton Keynes and Alex Alderton, Roberto Amato, Jeffrey Barrett, Sonia Goncalves, Ewan Harrison, David K. Jackson, Ian Johnston, Dominic Kwiatkowski, Cordelia Langford, John Sillitoe on behalf of the Wellcome Sanger Institute COVID-19 Surveillance Team |
| EPI_ISL_2995318, EPI_ISL_2995319 | Lighthouse Lab in Alderley Park | Wellcome Sanger Institute for the COVID-19 Genomics UK (COG-UK) Consortium | Jacquelyn Wynn, Mairead Hyland, The Lighthouse Lab in Alderley Park and Alex Alderton, Roberto Amato, Jeffrey Barrett, Sonia Goncalves, Ewan Harrison, David K. Jackson, Ian Johnston, Dominic Kwiatkowski, Cordelia Langford, John Sillitoe on behalf of the Wellcome Sanger Institute COVID-19 Surveillance Team |
| EPI_ISL_2995320 | Lighthouse Lab in Glasgow | Wellcome Sanger Institute for the COVID-19 Genomics UK (COG-UK) Consortium | Harper VanSteenhouse, Yumi Kasai, David Gray, Carol Clugston, Anna Dominiczak and Alex Alderton, Roberto Amato, Jeffrey Barrett, Sonia Goncalves, Ewan Harrison, David K. Jackson, Ian Johnston, Dominic Kwiatkowski, Cordelia Langford, John Sillitoe on behalf of the Wellcome Sanger Institute COVID-19 Surveillance Team |
| EPI_ISL_2995321, EPI_ISL_2995322, EPI_ISL_2995323 | Lighthouse Lab in Alderley Park | Wellcome Sanger Institute for the COVID-19 Genomics UK (COG-UK) Consortium | Jacquelyn Wynn, Mairead Hyland, The Lighthouse Lab in Alderley Park and Alex Alderton, Roberto Amato, Jeffrey Barrett, Sonia Goncalves, Ewan Harrison, David K. Jackson, Ian Johnston, Dominic Kwiatkowski, Cordelia Langford, John Sillitoe on behalf of the Wellcome Sanger Institute COVID-19 Surveillance Team |
| EPI_ISL_2995324 | Lighthouse Lab in Glasgow | Wellcome Sanger Institute for the COVID-19 Genomics UK (COG-UK) Consortium | Harper VanSteenhouse, Yumi Kasai, David Gray, Carol Clugston, Anna Dominiczak and Alex Alderton, Roberto Amato, Jeffrey Barrett, Sonia Goncalves, Ewan Harrison, David K. Jackson, Ian Johnston, Dominic Kwiatkowski, Cordelia Langford, John Sillitoe on behalf of the Wellcome Sanger Institute COVID-19 Surveillance Team |
| EPI_ISL_2995325 | Lighthouse Lab in Milton Keynes | Wellcome Sanger Institute for the COVID-19 Genomics UK (COG-UK) Consortium | The Lighthouse Lab in Milton Keynes and Alex Alderton, Roberto Amato, Jeffrey Barrett, Sonia Goncalves, Ewan Harrison, David K. Jackson, Ian Johnston, Dominic Kwiatkowski, Cordelia Langford, John Sillitoe on behalf of the Wellcome Sanger Institute COVID-19 Surveillance Team |
| EPI_ISL_2995326 | Lighthouse Lab in Alderley Park | Wellcome Sanger Institute for the COVID-19 Genomics UK (COG-UK) Consortium | Jacquelyn Wynn, Mairead Hyland, The Lighthouse Lab in Alderley Park and Alex Alderton, Roberto Amato, Jeffrey Barrett, Sonia Goncalves, Ewan Harrison, David K. Jackson, Ian Johnston, Dominic Kwiatkowski, Cordelia Langford, John Sillitoe on behalf of the Wellcome Sanger Institute COVID-19 Surveillance Team |
| EPI_ISL_2995327 | Lighthouse Lab in Glasgow | Wellcome Sanger Institute for the COVID-19 Genomics UK (COG-UK) Consortium | Harper VanSteenhouse, Yumi Kasai, David Gray, Carol Clugston, Anna Dominiczak and Alex Alderton, Roberto Amato, Jeffrey Barrett, Sonia Goncalves, Ewan Harrison, David K. Jackson, Ian Johnston, Dominic Kwiatkowski, Cordelia Langford, John Sillitoe on behalf of the Wellcome Sanger Institute COVID-19 Surveillance Team |
| EPI_ISL_2995328 | Lighthouse Lab in Alderley Park | Wellcome Sanger Institute for the COVID-19 Genomics UK (COG-UK) Consortium | Jacquelyn Wynn, Mairead Hyland, The Lighthouse Lab in Alderley Park and Alex Alderton, Roberto Amato, Jeffrey Barrett, Sonia Goncalves, Ewan Harrison, David K. Jackson, Ian Johnston, Dominic Kwiatkowski, Cordelia Langford, John Sillitoe on behalf of the Wellcome Sanger Institute COVID-19 Surveillance Team |
| EPI_ISL_2995329 | Berkshire and Surrey Pathology Services Lighthouse Laboratory | Wellcome Sanger Institute for the COVID-19 Genomics UK (COG-UK) Consortium | Berkshire and Surrey Pathology Services Lighthouse Laboratory and Alex Alderton, Roberto Amato, Jeffrey Barrett, Sonia Goncalves, Ewan Harrison, David K. Jackson, Ian Johnston, Dominic Kwiatkowski, Cordelia Langford, John Sillitoe on behalf of the Wellcome Sanger Institute COVID-19 Surveillance Team |
| EPI_ISL_2995330 | Lighthouse Lab in Milton Keynes | Wellcome Sanger Institute for the COVID-19 Genomics UK (COG-UK) Consortium | The Lighthouse Lab in Milton Keynes and Alex Alderton, Roberto Amato, Jeffrey Barrett, Sonia Goncalves, Ewan Harrison, David K. Jackson, Ian Johnston, Dominic Kwiatkowski, Cordelia Langford, John Sillitoe on behalf of the Wellcome Sanger Institute COVID-19 Surveillance Team |
| EPI_ISL_2995331 | Berkshire and Surrey Pathology Services Lighthouse Laboratory | Wellcome Sanger Institute for the COVID-19 Genomics UK (COG-UK) Consortium | Berkshire and Surrey Pathology Services Lighthouse Laboratory and Alex Alderton, Roberto Amato, Jeffrey Barrett, Sonia Goncalves, Ewan Harrison, David K. Jackson, Ian Johnston, Dominic Kwiatkowski, Cordelia Langford, John Sillitoe on behalf of the Wellcome Sanger Institute COVID-19 Surveillance Team |
| EPI_ISL_2995332 | Lighthouse Lab in Milton Keynes | Wellcome Sanger Institute for the COVID-19 Genomics UK (COG-UK) Consortium | The Lighthouse Lab in Milton Keynes and Alex Alderton, Roberto Amato, Jeffrey Barrett, Sonia Goncalves, Ewan Harrison, David K. Jackson, Ian Johnston, Dominic Kwiatkowski, Cordelia Langford, John Sillitoe on behalf of the Wellcome Sanger Institute COVID-19 Surveillance Team |
| EPI_ISL_2995333, EPI_ISL_2995334, EPI_ISL_2995335 | Lighthouse Lab in Alderley Park | Wellcome Sanger Institute for the COVID-19 Genomics UK (COG-UK) Consortium | Jacquelyn Wynn, Mairead Hyland, The Lighthouse Lab in Alderley Park and Alex Alderton, Roberto Amato, Jeffrey Barrett, Sonia Goncalves, Ewan Harrison, David K. Jackson, Ian Johnston, Dominic Kwiatkowski, Cordelia Langford, John Sillitoe on behalf of the Wellcome Sanger Institute COVID-19 Surveillance Team |
| EPI_ISL_2995336 | Lighthouse Lab in Milton Keynes | Wellcome Sanger Institute for the COVID-19 Genomics UK (COG-UK) Consortium | The Lighthouse Lab in Milton Keynes and Alex Alderton, Roberto Amato, Jeffrey Barrett, Sonia Goncalves, Ewan Harrison, David K. Jackson, Ian Johnston, Dominic Kwiatkowski, Cordelia Langford, John Sillitoe on behalf of the Wellcome Sanger Institute COVID-19 Surveillance Team |
| EPI_ISL_2995337 | Berkshire and Surrey Pathology Services Lighthouse Laboratory | Wellcome Sanger Institute for the COVID-19 Genomics UK (COG-UK) Consortium | Berkshire and Surrey Pathology Services Lighthouse Laboratory and Alex Alderton, Roberto Amato, Jeffrey Barrett, Sonia Goncalves, Ewan Harrison, David K. Jackson, Ian Johnston, Dominic Kwiatkowski, Cordelia Langford, John Sillitoe on behalf of the Wellcome Sanger Institute COVID-19 Surveillance Team |
| EPI_ISL_2995338 | Lighthouse Lab in Milton Keynes | Wellcome Sanger Institute for the COVID-19 Genomics UK (COG-UK) Consortium | The Lighthouse Lab in Milton Keynes and Alex Alderton, Roberto Amato, Jeffrey Barrett, Sonia Goncalves, Ewan Harrison, David K. Jackson, Ian Johnston, Dominic Kwiatkowski, Cordelia Langford, John Sillitoe on behalf of the Wellcome Sanger Institute COVID-19 Surveillance Team |
| EPI_ISL_2995339 | Berkshire and Surrey Pathology Services Lighthouse Laboratory | Wellcome Sanger Institute for the COVID-19 Genomics UK (COG-UK) Consortium | Berkshire and Surrey Pathology Services Lighthouse Laboratory and Alex Alderton, Roberto Amato, Jeffrey Barrett, Sonia Goncalves, Ewan Harrison, David K. Jackson, Ian Johnston, Dominic Kwiatkowski, Cordelia Langford, John Sillitoe on behalf of the Wellcome Sanger Institute COVID-19 Surveillance Team |

| | | | |
|---|---|---|---|
| EPI_ISL_2995340 | Lighthouse Lab in Alderley Park | Wellcome Sanger Institute for the COVID-19 Genomics UK (COG-UK) Consortium | Jacquelyn Wynn, Mairead Hyland, The Lighthouse Lab in Alderley Park and Alex Alderton, Roberto Amato, Jeffrey Barrett, Sonia Goncalves, Ewan Harrison, David K. Jackson, Ian Johnston, Dominic Kwiatkowski, Cordelia Langford, John Sillitoe on behalf of the Wellcome Sanger Institute COVID-19 Surveillance Team |
| EPI_ISL_2995341, EPI_ISL_2995342 | Lighthouse Lab in Glasgow | Wellcome Sanger Institute for the COVID-19 Genomics UK (COG-UK) Consortium | Harper VanSteenhouse, Yumi Kasai, David Gray, Carol Clugston, Anna Dominiczak and Alex Alderton, Roberto Amato, Jeffrey Barrett, Sonia Goncalves, Ewan Harrison, David K. Jackson, Ian Johnston, Dominic Kwiatkowski, Cordelia Langford, John Sillitoe on behalf of the Wellcome Sanger Institute COVID-19 Surveillance Team |
| EPI_ISL_2995343 | Lighthouse Lab in Milton Keynes | Wellcome Sanger Institute for the COVID-19 Genomics UK (COG-UK) Consortium | The Lighthouse Lab in Milton Keynes and Alex Alderton, Roberto Amato, Jeffrey Barrett, Sonia Goncalves, Ewan Harrison, David K. Jackson, Ian Johnston, Dominic Kwiatkowski, Cordelia Langford, John Sillitoe on behalf of the Wellcome Sanger Institute COVID-19 Surveillance Team |
| EPI_ISL_2995344, EPI_ISL_2995345 | Lighthouse Lab in Glasgow | Wellcome Sanger Institute for the COVID-19 Genomics UK (COG-UK) Consortium | Harper VanSteenhouse, Yumi Kasai, David Gray, Carol Clugston, Anna Dominiczak and Alex Alderton, Roberto Amato, Jeffrey Barrett, Sonia Goncalves, Ewan Harrison, David K. Jackson, Ian Johnston, Dominic Kwiatkowski, Cordelia Langford, John Sillitoe on behalf of the Wellcome Sanger Institute COVID-19 Surveillance Team |
| EPI_ISL_2995346, EPI_ISL_2995347 | Lighthouse Lab in Milton Keynes | Wellcome Sanger Institute for the COVID-19 Genomics UK (COG-UK) Consortium | The Lighthouse Lab in Milton Keynes and Alex Alderton, Roberto Amato, Jeffrey Barrett, Sonia Goncalves, Ewan Harrison, David K. Jackson, Ian Johnston, Dominic Kwiatkowski, Cordelia Langford, John Sillitoe on behalf of the Wellcome Sanger Institute COVID-19 Surveillance Team |
| EPI_ISL_2995348 | Lighthouse Lab in Glasgow | Wellcome Sanger Institute for the COVID-19 Genomics UK (COG-UK) Consortium | Harper VanSteenhouse, Yumi Kasai, David Gray, Carol Clugston, Anna Dominiczak and Alex Alderton, Roberto Amato, Jeffrey Barrett, Sonia Goncalves, Ewan Harrison, David K. Jackson, Ian Johnston, Dominic Kwiatkowski, Cordelia Langford, John Sillitoe on behalf of the Wellcome Sanger Institute COVID-19 Surveillance Team |
| EPI_ISL_2995349, EPI_ISL_2995350 | Lighthouse Laboratory Plymouth | Wellcome Sanger Institute for the COVID-19 Genomics UK (COG-UK) Consortium | Lighthouse Laboratory Plymouth and Alex Alderton, Roberto Amato, Jeffrey Barrett, Sonia Goncalves, Ewan Harrison, David K. Jackson, Ian Johnston, Dominic Kwiatkowski, Cordelia Langford, John Sillitoe on behalf of the Wellcome Sanger Institute COVID-19 Surveillance Team |
| EPI_ISL_2995351, EPI_ISL_2995352 | Lighthouse Lab in Glasgow | Wellcome Sanger Institute for the COVID-19 Genomics UK (COG-UK) Consortium | Harper VanSteenhouse, Yumi Kasai, David Gray, Carol Clugston, Anna Dominiczak and Alex Alderton, Roberto Amato, Jeffrey Barrett, Sonia Goncalves, Ewan Harrison, David K. Jackson, Ian Johnston, Dominic Kwiatkowski, Cordelia Langford, John Sillitoe on behalf of the Wellcome Sanger Institute COVID-19 Surveillance Team |
| EPI_ISL_2995353 | Lighthouse Laboratory Plymouth | Wellcome Sanger Institute for the COVID-19 Genomics UK (COG-UK) Consortium | Lighthouse Laboratory Plymouth and Alex Alderton, Roberto Amato, Jeffrey Barrett, Sonia Goncalves, Ewan Harrison, David K. Jackson, Ian Johnston, Dominic Kwiatkowski, Cordelia Langford, John Sillitoe on behalf of the Wellcome Sanger Institute COVID-19 Surveillance Team |
| EPI_ISL_2995354, EPI_ISL_2995355, EPI_ISL_2995356, EPI_ISL_2995357 | Lighthouse Lab in Glasgow | Wellcome Sanger Institute for the COVID-19 Genomics UK (COG-UK) Consortium | Harper VanSteenhouse, Yumi Kasai, David Gray, Carol Clugston, Anna Dominiczak and Alex Alderton, Roberto Amato, Jeffrey Barrett, Sonia Goncalves, Ewan Harrison, David K. Jackson, Ian Johnston, Dominic Kwiatkowski, Cordelia Langford, John Sillitoe on behalf of the Wellcome Sanger Institute COVID-19 Surveillance Team |
| EPI_ISL_2995358 | Lighthouse Laboratory Plymouth | Wellcome Sanger Institute for the COVID-19 Genomics UK (COG-UK) Consortium | Lighthouse Laboratory Plymouth and Alex Alderton, Roberto Amato, Jeffrey Barrett, Sonia Goncalves, Ewan Harrison, David K. Jackson, Ian Johnston, Dominic Kwiatkowski, Cordelia Langford, John Sillitoe on behalf of the Wellcome Sanger Institute COVID-19 Surveillance Team |
| EPI_ISL_2995359, EPI_ISL_2995360, EPI_ISL_2995361 | Lighthouse Lab in Glasgow | Wellcome Sanger Institute for the COVID-19 Genomics UK (COG-UK) Consortium | Harper VanSteenhouse, Yumi Kasai, David Gray, Carol Clugston, Anna Dominiczak and Alex Alderton, Roberto Amato, Jeffrey Barrett, Sonia Goncalves, Ewan Harrison, David K. Jackson, Ian Johnston, Dominic Kwiatkowski, Cordelia Langford, John Sillitoe on behalf of the Wellcome Sanger Institute COVID-19 Surveillance Team |
| EPI_ISL_2995362 | Lighthouse Laboratory Plymouth | Wellcome Sanger Institute for the COVID-19 Genomics UK (COG-UK) Consortium | Lighthouse Laboratory Plymouth and Alex Alderton, Roberto Amato, Jeffrey Barrett, Sonia Goncalves, Ewan Harrison, David K. Jackson, Ian Johnston, Dominic Kwiatkowski, Cordelia Langford, John Sillitoe on behalf of the Wellcome Sanger Institute COVID-19 Surveillance Team |
| EPI_ISL_2995363 | Lighthouse Lab in Glasgow | Wellcome Sanger Institute for the COVID-19 Genomics UK (COG-UK) Consortium | Harper VanSteenhouse, Yumi Kasai, David Gray, Carol Clugston, Anna Dominiczak and Alex Alderton, Roberto Amato, Jeffrey Barrett, Sonia Goncalves, Ewan Harrison, David K. Jackson, Ian Johnston, Dominic Kwiatkowski, Cordelia Langford, John Sillitoe on behalf of the Wellcome Sanger Institute COVID-19 Surveillance Team |
| EPI_ISL_2995364, EPI_ISL_2995365, EPI_ISL_2995366 | Lighthouse Laboratory Plymouth | Wellcome Sanger Institute for the COVID-19 Genomics UK (COG-UK) Consortium | Lighthouse Laboratory Plymouth and Alex Alderton, Roberto Amato, Jeffrey Barrett, Sonia Goncalves, Ewan Harrison, David K. Jackson, Ian Johnston, Dominic Kwiatkowski, Cordelia Langford, John Sillitoe on behalf of the Wellcome Sanger Institute COVID-19 Surveillance Team |
| EPI_ISL_2995367, EPI_ISL_2995368, EPI_ISL_2995369, EPI_ISL_2995370, EPI_ISL_2995371, EPI_ISL_2995372, EPI_ISL_2995373 | Lighthouse Lab in Glasgow | Wellcome Sanger Institute for the COVID-19 Genomics UK (COG-UK) Consortium | Harper VanSteenhouse, Yumi Kasai, David Gray, Carol Clugston, Anna Dominiczak and Alex Alderton, Roberto Amato, Jeffrey Barrett, Sonia Goncalves, Ewan Harrison, David K. Jackson, Ian Johnston, Dominic Kwiatkowski, Cordelia Langford, John Sillitoe on behalf of the Wellcome Sanger Institute COVID-19 Surveillance Team |
| EPI_ISL_2995374 | Lighthouse Laboratory Plymouth | Wellcome Sanger Institute for the COVID-19 Genomics UK (COG-UK) Consortium | Lighthouse Laboratory Plymouth and Alex Alderton, Roberto Amato, Jeffrey Barrett, Sonia Goncalves, Ewan Harrison, David K. Jackson, Ian Johnston, Dominic Kwiatkowski, Cordelia Langford, John Sillitoe on behalf of the Wellcome Sanger Institute COVID-19 Surveillance Team |
| EPI_ISL_2995375 | Lighthouse Lab in Glasgow | Wellcome Sanger Institute for the COVID-19 Genomics UK (COG-UK) Consortium | Harper VanSteenhouse, Yumi Kasai, David Gray, Carol Clugston, Anna Dominiczak and Alex Alderton, Roberto Amato, Jeffrey Barrett, Sonia Goncalves, Ewan Harrison, David K. Jackson, Ian Johnston, Dominic Kwiatkowski, Cordelia Langford, John Sillitoe on behalf of the Wellcome Sanger Institute COVID-19 Surveillance Team |
| EPI_ISL_2995376 | Lighthouse Laboratory Plymouth | Wellcome Sanger Institute for the COVID-19 Genomics UK (COG-UK) Consortium | Lighthouse Laboratory Plymouth and Alex Alderton, Roberto Amato, Jeffrey Barrett, Sonia Goncalves, Ewan Harrison, David K. Jackson, Ian Johnston, Dominic Kwiatkowski, Cordelia Langford, John Sillitoe on behalf of the Wellcome Sanger Institute COVID-19 Surveillance Team |
| EPI_ISL_2995377 | Lighthouse Lab in Glasgow | Wellcome Sanger Institute for the COVID-19 Genomics UK (COG-UK) Consortium | Harper VanSteenhouse, Yumi Kasai, David Gray, Carol Clugston, Anna Dominiczak and Alex Alderton, Roberto Amato, Jeffrey Barrett, Sonia Goncalves, Ewan Harrison, David K. Jackson, Ian Johnston, Dominic Kwiatkowski, Cordelia Langford, John Sillitoe on behalf of the Wellcome Sanger Institute COVID-19 Surveillance Team |
| EPI_ISL_2995378 | Lighthouse Laboratory Plymouth | Wellcome Sanger Institute for the COVID-19 Genomics UK (COG-UK) Consortium | Lighthouse Laboratory Plymouth and Alex Alderton, Roberto Amato, Jeffrey Barrett, Sonia Goncalves, Ewan Harrison, David K. Jackson, Ian Johnston, Dominic Kwiatkowski, Cordelia Langford, John Sillitoe on behalf of the Wellcome Sanger Institute COVID-19 Surveillance Team |
| EPI_ISL_2995379 | Lighthouse Lab in Glasgow | Wellcome Sanger Institute for the COVID-19 Genomics UK (COG-UK) Consortium | Harper VanSteenhouse, Yumi Kasai, David Gray, Carol Clugston, Anna Dominiczak and Alex Alderton, Roberto Amato, Jeffrey Barrett, Sonia Goncalves, Ewan Harrison, David K. Jackson, Ian Johnston, Dominic Kwiatkowski, Cordelia Langford, John Sillitoe on behalf of the Wellcome Sanger Institute COVID-19 Surveillance Team |
| EPI_ISL_2995380 | Lighthouse Laboratory Plymouth | Wellcome Sanger Institute for the COVID-19 Genomics UK (COG-UK) Consortium | Lighthouse Laboratory Plymouth and Alex Alderton, Roberto Amato, Jeffrey Barrett, Sonia Goncalves, Ewan Harrison, David K. Jackson, Ian Johnston, Dominic Kwiatkowski, Cordelia Langford, John Sillitoe on behalf of the Wellcome Sanger Institute COVID-19 Surveillance Team |
| EPI_ISL_2995381, EPI_ISL_2995382, EPI_ISL_2995383, EPI_ISL_2995384, EPI_ISL_2995385 | Lighthouse Lab in Glasgow | Wellcome Sanger Institute for the COVID-19 Genomics UK (COG-UK) Consortium | Harper VanSteenhouse, Yumi Kasai, David Gray, Carol Clugston, Anna Dominiczak and Alex Alderton, Roberto Amato, Jeffrey Barrett, Sonia Goncalves, Ewan Harrison, David K. Jackson, Ian Johnston, Dominic Kwiatkowski, Cordelia Langford, John Sillitoe on behalf of the Wellcome Sanger Institute COVID-19 Surveillance Team |
| EPI_ISL_2995386 | Lighthouse Laboratory Plymouth | Wellcome Sanger Institute for the COVID-19 Genomics UK (COG-UK) Consortium | Lighthouse Laboratory Plymouth and Alex Alderton, Roberto Amato, Jeffrey Barrett, Sonia Goncalves, Ewan Harrison, David K. Jackson, Ian Johnston, Dominic Kwiatkowski, Cordelia Langford, John Sillitoe on behalf of the Wellcome Sanger Institute COVID-19 Surveillance Team |
| EPI_ISL_2995387, EPI_ISL_2995388 | Lighthouse Lab in Glasgow | Wellcome Sanger Institute for the COVID-19 Genomics UK (COG-UK) Consortium | Harper VanSteenhouse, Yumi Kasai, David Gray, Carol Clugston, Anna Dominiczak and Alex Alderton, Roberto Amato, Jeffrey Barrett, Sonia Goncalves, Ewan Harrison, David K. Jackson, Ian Johnston, Dominic Kwiatkowski, Cordelia Langford, John Sillitoe on behalf of the Wellcome Sanger Institute COVID-19 Surveillance Team |
| EPI_ISL_2995389 | Lighthouse Laboratory Plymouth | Wellcome Sanger Institute for the COVID-19 Genomics UK (COG-UK) Consortium | Lighthouse Laboratory Plymouth and Alex Alderton, Roberto Amato, Jeffrey Barrett, Sonia Goncalves, Ewan Harrison, David K. Jackson, Ian Johnston, Dominic Kwiatkowski, Cordelia Langford, John Sillitoe on behalf of the Wellcome Sanger Institute COVID-19 Surveillance Team |
| EPI_ISL_2995390, EPI_ISL_2995391, EPI_ISL_2995392, EPI_ISL_2995393, EPI_ISL_2995394, EPI_ISL_2995395, EPI_ISL_2995396, EPI_ISL_2995397 | Lighthouse Lab in Glasgow | Wellcome Sanger Institute for the COVID-19 Genomics UK (COG-UK) Consortium | Harper VanSteenhouse, Yumi Kasai, David Gray, Carol Clugston, Anna Dominiczak and Alex Alderton, Roberto Amato, Jeffrey Barrett, Sonia Goncalves, Ewan Harrison, David K. Jackson, Ian Johnston, Dominic Kwiatkowski, Cordelia Langford, John Sillitoe on behalf of the Wellcome Sanger Institute COVID-19 Surveillance Team |
| EPI_ISL_2995398 | Lighthouse Laboratory Plymouth | Wellcome Sanger Institute for the COVID-19 Genomics UK (COG-UK) Consortium | Lighthouse Laboratory Plymouth and Alex Alderton, Roberto Amato, Jeffrey Barrett, Sonia Goncalves, Ewan Harrison, David K. Jackson, Ian Johnston, Dominic Kwiatkowski, Cordelia Langford, John Sillitoe on behalf of the Wellcome Sanger Institute COVID-19 Surveillance Team |
| EPI_ISL_2995399, EPI_ISL_2995400, EPI_ISL_2995401, EPI_ISL_2995402, EPI_ISL_2995403 | Lighthouse Lab in Glasgow | Wellcome Sanger Institute for the COVID-19 Genomics UK (COG-UK) Consortium | Harper VanSteenhouse, Yumi Kasai, David Gray, Carol Clugston, Anna Dominiczak and Alex Alderton, Roberto Amato, Jeffrey Barrett, Sonia Goncalves, Ewan Harrison, David K. Jackson, Ian Johnston, Dominic Kwiatkowski, Cordelia Langford, John Sillitoe on behalf of the Wellcome Sanger Institute COVID-19 Surveillance Team |
| EPI_ISL_2995404 | Lighthouse Laboratory Plymouth | Wellcome Sanger Institute for the COVID-19 Genomics UK (COG-UK) Consortium | Lighthouse Laboratory Plymouth and Alex Alderton, Roberto Amato, Jeffrey Barrett, Sonia Goncalves, Ewan Harrison, David K. Jackson, Ian Johnston, Dominic Kwiatkowski, Cordelia Langford, John Sillitoe on behalf of the Wellcome Sanger Institute COVID-19 Surveillance Team |
| EPI_ISL_2995405 | Lighthouse Lab in Glasgow | Wellcome Sanger Institute for the COVID-19 Genomics UK (COG-UK) Consortium | Harper VanSteenhouse, Yumi Kasai, David Gray, Carol Clugston, Anna Dominiczak and Alex Alderton, Roberto Amato, Jeffrey Barrett, Sonia Goncalves, Ewan Harrison, David K. Jackson, Ian Johnston, Dominic Kwiatkowski, Cordelia Langford, John Sillitoe on behalf of the Wellcome Sanger Institute COVID-19 Surveillance Team |
| EPI_ISL_2995406, EPI_ISL_2995407 | Lighthouse Laboratory Plymouth | Wellcome Sanger Institute for the COVID-19 Genomics UK (COG-UK) Consortium | Lighthouse Laboratory Plymouth and Alex Alderton, Roberto Amato, Jeffrey Barrett, Sonia Goncalves, Ewan Harrison, David K. Jackson, Ian Johnston, Dominic Kwiatkowski, Cordelia Langford, John Sillitoe on behalf of the Wellcome Sanger Institute COVID-19 Surveillance Team |
| EPI_ISL_2995408, EPI_ISL_2995409, EPI_ISL_2995410 | Lighthouse Lab in Glasgow | Wellcome Sanger Institute for the COVID-19 Genomics UK (COG-UK) Consortium | Harper VanSteenhouse, Yumi Kasai, David Gray, Carol Clugston, Anna Dominiczak and Alex Alderton, Roberto Amato, Jeffrey Barrett, Sonia Goncalves, Ewan Harrison, David K. Jackson, Ian Johnston, Dominic Kwiatkowski, Cordelia Langford, John Sillitoe on behalf of the Wellcome Sanger Institute COVID-19 Surveillance Team |
| EPI_ISL_2995411 | Lighthouse Laboratory Plymouth | Wellcome Sanger Institute for the COVID-19 Genomics UK (COG-UK) Consortium | Lighthouse Laboratory Plymouth and Alex Alderton, Roberto Amato, Jeffrey Barrett, Sonia Goncalves, Ewan Harrison, David K. Jackson, Ian Johnston, Dominic Kwiatkowski, Cordelia Langford, John Sillitoe on behalf of the Wellcome Sanger Institute COVID-19 Surveillance Team |
| EPI_ISL_2995412, EPI_ISL_2995413 | Lighthouse Lab in Glasgow | Wellcome Sanger Institute for the COVID-19 Genomics UK (COG-UK) Consortium | Harper VanSteenhouse, Yumi Kasai, David Gray, Carol Clugston, Anna Dominiczak and Alex Alderton, Roberto Amato, Jeffrey Barrett, Sonia Goncalves, Ewan Harrison, David K. Jackson, Ian Johnston, Dominic Kwiatkowski, Cordelia Langford, John Sillitoe on behalf of the Wellcome Sanger Institute COVID-19 Surveillance Team |

| | | | |
|---|---|---|---|
| EPI_ISL_2995414 | Lighthouse Laboratory Plymouth | Wellcome Sanger Institute for the COVID-19 Genomics UK (COG-UK) Consortium | Lighthouse Laboratory Plymouth and Alex Alderton, Roberto Amato, Jeffrey Barrett, Sonia Goncalves, Ewan Harrison, David K. Jackson, Ian Johnston, Dominic Kwiatkowski, Cordelia Langford, John Sillitoe on behalf of the Wellcome Sanger Institute COVID-19 Surveillance Team |
| EPI_ISL_2995415 | Lighthouse Lab in Glasgow | Wellcome Sanger Institute for the COVID-19 Genomics UK (COG-UK) Consortium | Harper VanSteenhouse, Yumi Kasai, David Gray, Carol Clugston, Anna Dominiczak and Alex Alderton, Roberto Amato, Jeffrey Barrett, Sonia Goncalves, Ewan Harrison, David K. Jackson, Ian Johnston, Dominic Kwiatkowski, Cordelia Langford, John Sillitoe on behalf of the Wellcome Sanger Institute COVID-19 Surveillance Team |
| EPI_ISL_2995416 | Lighthouse Laboratory Plymouth | Wellcome Sanger Institute for the COVID-19 Genomics UK (COG-UK) Consortium | Lighthouse Laboratory Plymouth and Alex Alderton, Roberto Amato, Jeffrey Barrett, Sonia Goncalves, Ewan Harrison, David K. Jackson, Ian Johnston, Dominic Kwiatkowski, Cordelia Langford, John Sillitoe on behalf of the Wellcome Sanger Institute COVID-19 Surveillance Team |
| EPI_ISL_2995417 | Lighthouse Lab in Glasgow | Wellcome Sanger Institute for the COVID-19 Genomics UK (COG-UK) Consortium | Harper VanSteenhouse, Yumi Kasai, David Gray, Carol Clugston, Anna Dominiczak and Alex Alderton, Roberto Amato, Jeffrey Barrett, Sonia Goncalves, Ewan Harrison, David K. Jackson, Ian Johnston, Dominic Kwiatkowski, Cordelia Langford, John Sillitoe on behalf of the Wellcome Sanger Institute COVID-19 Surveillance Team |
| EPI_ISL_2995418, EPI_ISL_2995419, EPI_ISL_2995420 | Lighthouse Laboratory Plymouth | Wellcome Sanger Institute for the COVID-19 Genomics UK (COG-UK) Consortium | Lighthouse Laboratory Plymouth and Alex Alderton, Roberto Amato, Jeffrey Barrett, Sonia Goncalves, Ewan Harrison, David K. Jackson, Ian Johnston, Dominic Kwiatkowski, Cordelia Langford, John Sillitoe on behalf of the Wellcome Sanger Institute COVID-19 Surveillance Team |
| EPI_ISL_2995421 | Lighthouse Lab in Glasgow | Wellcome Sanger Institute for the COVID-19 Genomics UK (COG-UK) Consortium | Harper VanSteenhouse, Yumi Kasai, David Gray, Carol Clugston, Anna Dominiczak and Alex Alderton, Roberto Amato, Jeffrey Barrett, Sonia Goncalves, Ewan Harrison, David K. Jackson, Ian Johnston, Dominic Kwiatkowski, Cordelia Langford, John Sillitoe on behalf of the Wellcome Sanger Institute COVID-19 Surveillance Team |
| EPI_ISL_2995422, EPI_ISL_2995423 | Lighthouse Laboratory Plymouth | Wellcome Sanger Institute for the COVID-19 Genomics UK (COG-UK) Consortium | Lighthouse Laboratory Plymouth and Alex Alderton, Roberto Amato, Jeffrey Barrett, Sonia Goncalves, Ewan Harrison, David K. Jackson, Ian Johnston, Dominic Kwiatkowski, Cordelia Langford, John Sillitoe on behalf of the Wellcome Sanger Institute COVID-19 Surveillance Team |
| EPI_ISL_2995424, EPI_ISL_2995425, EPI_ISL_2995426 | Lighthouse Lab in Glasgow | Wellcome Sanger Institute for the COVID-19 Genomics UK (COG-UK) Consortium | Harper VanSteenhouse, Yumi Kasai, David Gray, Carol Clugston, Anna Dominiczak and Alex Alderton, Roberto Amato, Jeffrey Barrett, Sonia Goncalves, Ewan Harrison, David K. Jackson, Ian Johnston, Dominic Kwiatkowski, Cordelia Langford, John Sillitoe on behalf of the Wellcome Sanger Institute COVID-19 Surveillance Team |
| EPI_ISL_2995427, EPI_ISL_2995428 | Lighthouse Laboratory Plymouth | Wellcome Sanger Institute for the COVID-19 Genomics UK (COG-UK) Consortium | Lighthouse Laboratory Plymouth and Alex Alderton, Roberto Amato, Jeffrey Barrett, Sonia Goncalves, Ewan Harrison, David K. Jackson, Ian Johnston, Dominic Kwiatkowski, Cordelia Langford, John Sillitoe on behalf of the Wellcome Sanger Institute COVID-19 Surveillance Team |
| EPI_ISL_2995429, EPI_ISL_2995430, EPI_ISL_2995431, EPI_ISL_2995432, EPI_ISL_2995433, EPI_ISL_2995434, EPI_ISL_2995435 | Lighthouse Lab in Glasgow | Wellcome Sanger Institute for the COVID-19 Genomics UK (COG-UK) Consortium | Harper VanSteenhouse, Yumi Kasai, David Gray, Carol Clugston, Anna Dominiczak and Alex Alderton, Roberto Amato, Jeffrey Barrett, Sonia Goncalves, Ewan Harrison, David K. Jackson, Ian Johnston, Dominic Kwiatkowski, Cordelia Langford, John Sillitoe on behalf of the Wellcome Sanger Institute COVID-19 Surveillance Team |
| EPI_ISL_2995436 | Lighthouse Laboratory Plymouth | Wellcome Sanger Institute for the COVID-19 Genomics UK (COG-UK) Consortium | Lighthouse Laboratory Plymouth and Alex Alderton, Roberto Amato, Jeffrey Barrett, Sonia Goncalves, Ewan Harrison, David K. Jackson, Ian Johnston, Dominic Kwiatkowski, Cordelia Langford, John Sillitoe on behalf of the Wellcome Sanger Institute COVID-19 Surveillance Team |
| EPI_ISL_2995437, EPI_ISL_2995438, EPI_ISL_2995439, EPI_ISL_2995440 | Lighthouse Lab in Glasgow | Wellcome Sanger Institute for the COVID-19 Genomics UK (COG-UK) Consortium | Harper VanSteenhouse, Yumi Kasai, David Gray, Carol Clugston, Anna Dominiczak and Alex Alderton, Roberto Amato, Jeffrey Barrett, Sonia Goncalves, Ewan Harrison, David K. Jackson, Ian Johnston, Dominic Kwiatkowski, Cordelia Langford, John Sillitoe on behalf of the Wellcome Sanger Institute COVID-19 Surveillance Team |
| EPI_ISL_2995441 | Lighthouse Laboratory Plymouth | Wellcome Sanger Institute for the COVID-19 Genomics UK (COG-UK) Consortium | Lighthouse Laboratory Plymouth and Alex Alderton, Roberto Amato, Jeffrey Barrett, Sonia Goncalves, Ewan Harrison, David K. Jackson, Ian Johnston, Dominic Kwiatkowski, Cordelia Langford, John Sillitoe on behalf of the Wellcome Sanger Institute COVID-19 Surveillance Team |
| EPI_ISL_2995442, EPI_ISL_2995443, EPI_ISL_2995444, EPI_ISL_2995445, EPI_ISL_2995446, EPI_ISL_2995447, EPI_ISL_2995448, EPI_ISL_2995449, EPI_ISL_2995450, EPI_ISL_2995451, EPI_ISL_2995452, EPI_ISL_2995453, EPI_ISL_2995454 | | | |
| see above | Lighthouse Lab in Glasgow | Wellcome Sanger Institute for the COVID-19 Genomics UK (COG-UK) Consortium | Harper VanSteenhouse, Yumi Kasai, David Gray, Carol Clugston, Anna Dominiczak and Alex Alderton, Roberto Amato, Jeffrey Barrett, Sonia Goncalves, Ewan Harrison, David K. Jackson, Ian Johnston, Dominic Kwiatkowski, Cordelia Langford, John Sillitoe on behalf of the Wellcome Sanger Institute COVID-19 Surveillance Team |
| EPI_ISL_2995455, EPI_ISL_2995456 | Lighthouse Laboratory Plymouth | Wellcome Sanger Institute for the COVID-19 Genomics UK (COG-UK) Consortium | Lighthouse Laboratory Plymouth and Alex Alderton, Roberto Amato, Jeffrey Barrett, Sonia Goncalves, Ewan Harrison, David K. Jackson, Ian Johnston, Dominic Kwiatkowski, Cordelia Langford, John Sillitoe on behalf of the Wellcome Sanger Institute COVID-19 Surveillance Team |
| EPI_ISL_2995457, EPI_ISL_2995458, EPI_ISL_2995459 | Lighthouse Lab in Glasgow | Wellcome Sanger Institute for the COVID-19 Genomics UK (COG-UK) Consortium | Harper VanSteenhouse, Yumi Kasai, David Gray, Carol Clugston, Anna Dominiczak and Alex Alderton, Roberto Amato, Jeffrey Barrett, Sonia Goncalves, Ewan Harrison, David K. Jackson, Ian Johnston, Dominic Kwiatkowski, Cordelia Langford, John Sillitoe on behalf of the Wellcome Sanger Institute COVID-19 Surveillance Team |
| EPI_ISL_2995460, EPI_ISL_2995461 | Lighthouse Laboratory Plymouth | Wellcome Sanger Institute for the COVID-19 Genomics UK (COG-UK) Consortium | Lighthouse Laboratory Plymouth and Alex Alderton, Roberto Amato, Jeffrey Barrett, Sonia Goncalves, Ewan Harrison, David K. Jackson, Ian Johnston, Dominic Kwiatkowski, Cordelia Langford, John Sillitoe on behalf of the Wellcome Sanger Institute COVID-19 Surveillance Team |
| EPI_ISL_2995462, EPI_ISL_2995463, EPI_ISL_2995464, EPI_ISL_2995465, EPI_ISL_2995466, EPI_ISL_2995467, EPI_ISL_2995468 | Lighthouse Lab in Glasgow | Wellcome Sanger Institute for the COVID-19 Genomics UK (COG-UK) Consortium | Harper VanSteenhouse, Yumi Kasai, David Gray, Carol Clugston, Anna Dominiczak and Alex Alderton, Roberto Amato, Jeffrey Barrett, Sonia Goncalves, Ewan Harrison, David K. Jackson, Ian Johnston, Dominic Kwiatkowski, Cordelia Langford, John Sillitoe on behalf of the Wellcome Sanger Institute COVID-19 Surveillance Team |
| EPI_ISL_2995469 | Lighthouse Laboratory Plymouth | Wellcome Sanger Institute for the COVID-19 Genomics UK (COG-UK) Consortium | Lighthouse Laboratory Plymouth and Alex Alderton, Roberto Amato, Jeffrey Barrett, Sonia Goncalves, Ewan Harrison, David K. Jackson, Ian Johnston, Dominic Kwiatkowski, Cordelia Langford, John Sillitoe on behalf of the Wellcome Sanger Institute COVID-19 Surveillance Team |
| EPI_ISL_2995470 | Lighthouse Lab in Glasgow | Wellcome Sanger Institute for the COVID-19 Genomics UK (COG-UK) Consortium | Harper VanSteenhouse, Yumi Kasai, David Gray, Carol Clugston, Anna Dominiczak and Alex Alderton, Roberto Amato, Jeffrey Barrett, Sonia Goncalves, Ewan Harrison, David K. Jackson, Ian Johnston, Dominic Kwiatkowski, Cordelia Langford, John Sillitoe on behalf of the Wellcome Sanger Institute COVID-19 Surveillance Team |
| EPI_ISL_2995471 | Lighthouse Laboratory Plymouth | Wellcome Sanger Institute for the COVID-19 Genomics UK (COG-UK) Consortium | Lighthouse Laboratory Plymouth and Alex Alderton, Roberto Amato, Jeffrey Barrett, Sonia Goncalves, Ewan Harrison, David K. Jackson, Ian Johnston, Dominic Kwiatkowski, Cordelia Langford, John Sillitoe on behalf of the Wellcome Sanger Institute COVID-19 Surveillance Team |
| EPI_ISL_2995472, EPI_ISL_2995473, EPI_ISL_2995474, EPI_ISL_2995475, EPI_ISL_2995476, EPI_ISL_2995477, EPI_ISL_2995478 | Lighthouse Lab in Glasgow | Wellcome Sanger Institute for the COVID-19 Genomics UK (COG-UK) Consortium | Harper VanSteenhouse, Yumi Kasai, David Gray, Carol Clugston, Anna Dominiczak and Alex Alderton, Roberto Amato, Jeffrey Barrett, Sonia Goncalves, Ewan Harrison, David K. Jackson, Ian Johnston, Dominic Kwiatkowski, Cordelia Langford, John Sillitoe on behalf of the Wellcome Sanger Institute COVID-19 Surveillance Team |
| EPI_ISL_2995479 | Lighthouse Laboratory Plymouth | Wellcome Sanger Institute for the COVID-19 Genomics UK (COG-UK) Consortium | Lighthouse Laboratory Plymouth and Alex Alderton, Roberto Amato, Jeffrey Barrett, Sonia Goncalves, Ewan Harrison, David K. Jackson, Ian Johnston, Dominic Kwiatkowski, Cordelia Langford, John Sillitoe on behalf of the Wellcome Sanger Institute COVID-19 Surveillance Team |
| EPI_ISL_2995480, EPI_ISL_2995481 | Lighthouse Lab in Glasgow | Wellcome Sanger Institute for the COVID-19 Genomics UK (COG-UK) Consortium | Harper VanSteenhouse, Yumi Kasai, David Gray, Carol Clugston, Anna Dominiczak and Alex Alderton, Roberto Amato, Jeffrey Barrett, Sonia Goncalves, Ewan Harrison, David K. Jackson, Ian Johnston, Dominic Kwiatkowski, Cordelia Langford, John Sillitoe on behalf of the Wellcome Sanger Institute COVID-19 Surveillance Team |
| EPI_ISL_2995482 | Lighthouse Laboratory Plymouth | Wellcome Sanger Institute for the COVID-19 Genomics UK (COG-UK) Consortium | Lighthouse Laboratory Plymouth and Alex Alderton, Roberto Amato, Jeffrey Barrett, Sonia Goncalves, Ewan Harrison, David K. Jackson, Ian Johnston, Dominic Kwiatkowski, Cordelia Langford, John Sillitoe on behalf of the Wellcome Sanger Institute COVID-19 Surveillance Team |
| EPI_ISL_2995483, EPI_ISL_2995484, EPI_ISL_2995485, EPI_ISL_2995486 | Lighthouse Lab in Glasgow | Wellcome Sanger Institute for the COVID-19 Genomics UK (COG-UK) Consortium | Harper VanSteenhouse, Yumi Kasai, David Gray, Carol Clugston, Anna Dominiczak and Alex Alderton, Roberto Amato, Jeffrey Barrett, Sonia Goncalves, Ewan Harrison, David K. Jackson, Ian Johnston, Dominic Kwiatkowski, Cordelia Langford, John Sillitoe on behalf of the Wellcome Sanger Institute COVID-19 Surveillance Team |
| EPI_ISL_2995487, EPI_ISL_2995488, EPI_ISL_2995489 | Lighthouse Laboratory Plymouth | Wellcome Sanger Institute for the COVID-19 Genomics UK (COG-UK) Consortium | Lighthouse Laboratory Plymouth and Alex Alderton, Roberto Amato, Jeffrey Barrett, Sonia Goncalves, Ewan Harrison, David K. Jackson, Ian Johnston, Dominic Kwiatkowski, Cordelia Langford, John Sillitoe on behalf of the Wellcome Sanger Institute COVID-19 Surveillance Team |
| EPI_ISL_2995490, EPI_ISL_2995491 | Lighthouse Lab in Glasgow | Wellcome Sanger Institute for the COVID-19 Genomics UK (COG-UK) Consortium | Harper VanSteenhouse, Yumi Kasai, David Gray, Carol Clugston, Anna Dominiczak and Alex Alderton, Roberto Amato, Jeffrey Barrett, Sonia Goncalves, Ewan Harrison, David K. Jackson, Ian Johnston, Dominic Kwiatkowski, Cordelia Langford, John Sillitoe on behalf of the Wellcome Sanger Institute COVID-19 Surveillance Team |
| EPI_ISL_2995492 | Lighthouse Laboratory Plymouth | Wellcome Sanger Institute for the COVID-19 Genomics UK (COG-UK) Consortium | Lighthouse Laboratory Plymouth and Alex Alderton, Roberto Amato, Jeffrey Barrett, Sonia Goncalves, Ewan Harrison, David K. Jackson, Ian Johnston, Dominic Kwiatkowski, Cordelia Langford, John Sillitoe on behalf of the Wellcome Sanger Institute COVID-19 Surveillance Team |
| EPI_ISL_2995493, EPI_ISL_2995494 | Lighthouse Lab in Glasgow | Wellcome Sanger Institute for the COVID-19 Genomics UK (COG-UK) Consortium | Harper VanSteenhouse, Yumi Kasai, David Gray, Carol Clugston, Anna Dominiczak and Alex Alderton, Roberto Amato, Jeffrey Barrett, Sonia Goncalves, Ewan Harrison, David K. Jackson, Ian Johnston, Dominic Kwiatkowski, Cordelia Langford, John Sillitoe on behalf of the Wellcome Sanger Institute COVID-19 Surveillance Team |
| EPI_ISL_2995495 | Lighthouse Laboratory Plymouth | Wellcome Sanger Institute for the COVID-19 Genomics UK (COG-UK) Consortium | Lighthouse Laboratory Plymouth and Alex Alderton, Roberto Amato, Jeffrey Barrett, Sonia Goncalves, Ewan Harrison, David K. Jackson, Ian Johnston, Dominic Kwiatkowski, Cordelia Langford, John Sillitoe on behalf of the Wellcome Sanger Institute COVID-19 Surveillance Team |
| EPI_ISL_2995496 | Lighthouse Lab in Glasgow | Wellcome Sanger Institute for the COVID-19 Genomics UK (COG-UK) Consortium | Harper VanSteenhouse, Yumi Kasai, David Gray, Carol Clugston, Anna Dominiczak and Alex Alderton, Roberto Amato, Jeffrey Barrett, Sonia Goncalves, Ewan Harrison, David K. Jackson, Ian Johnston, Dominic Kwiatkowski, Cordelia Langford, John Sillitoe on behalf of the Wellcome Sanger Institute COVID-19 Surveillance Team |
| EPI_ISL_2995497 | Lighthouse Laboratory Plymouth | Wellcome Sanger Institute for the COVID-19 Genomics UK (COG-UK) Consortium | Lighthouse Laboratory Plymouth and Alex Alderton, Roberto Amato, Jeffrey Barrett, Sonia Goncalves, Ewan Harrison, David K. Jackson, Ian Johnston, Dominic Kwiatkowski, Cordelia Langford, John Sillitoe on behalf of the Wellcome Sanger Institute COVID-19 Surveillance Team |
| EPI_ISL_2995498, EPI_ISL_2995499, EPI_ISL_2995500, EPI_ISL_2995501, | Lighthouse Lab in Glasgow | Wellcome Sanger Institute for the COVID-19 Genomics UK (COG-UK) Consortium | Harper VanSteenhouse, Yumi Kasai, David Gray, Carol Clugston, Anna Dominiczak and Alex Alderton, Roberto Amato, Jeffrey Barrett, Sonia Goncalves, Ewan Harrison, David K. Jackson, Ian Johnston, Dominic Kwiatkowski, Cordelia Langford, John Sillitoe on behalf of the Wellcome Sanger Institute COVID-19 Surveillance Team |

| | | | |
|---|---|---|---|
| EPI_ISL_2995502, EPI_ISL_2995503, EPI_ISL_2995504 | | | |
| EPI_ISL_2995505, EPI_ISL_2995506 | Lighthouse Laboratory Plymouth | Wellcome Sanger Institute for the COVID-19 Genomics UK (COG-UK) Consortium | Lighthouse Laboratory Plymouth and Alex Alderton, Roberto Amato, Jeffrey Barrett, Sonia Goncalves, Ewan Harrison, David K. Jackson, Ian Johnston, Dominic Kwiatkowski, Cordelia Langford, John Sillitoe on behalf of the Wellcome Sanger Institute COVID-19 Surveillance Team |
| EPI_ISL_2995507, EPI_ISL_2995508, EPI_ISL_2995509, EPI_ISL_2995510 | Lighthouse Lab in Glasgow | Wellcome Sanger Institute for the COVID-19 Genomics UK (COG-UK) Consortium | Harper VanSteenhouse, Yumi Kasai, David Gray, Carol Clugston, Anna Dominiczak and Alex Alderton, Roberto Amato, Jeffrey Barrett, Sonia Goncalves, Ewan Harrison, David K. Jackson, Ian Johnston, Dominic Kwiatkowski, Cordelia Langford, John Sillitoe on behalf of the Wellcome Sanger Institute COVID-19 Surveillance Team |
| EPI_ISL_2995511 | Lighthouse Laboratory Plymouth | Wellcome Sanger Institute for the COVID-19 Genomics UK (COG-UK) Consortium | Lighthouse Laboratory Plymouth and Alex Alderton, Roberto Amato, Jeffrey Barrett, Sonia Goncalves, Ewan Harrison, David K. Jackson, Ian Johnston, Dominic Kwiatkowski, Cordelia Langford, John Sillitoe on behalf of the Wellcome Sanger Institute COVID-19 Surveillance Team |
| EPI_ISL_2995512, EPI_ISL_2995513, EPI_ISL_2995514, EPI_ISL_2995515, EPI_ISL_2995516, EPI_ISL_2995517, EPI_ISL_2995518 | Lighthouse Lab in Glasgow | Wellcome Sanger Institute for the COVID-19 Genomics UK (COG-UK) Consortium | Harper VanSteenhouse, Yumi Kasai, David Gray, Carol Clugston, Anna Dominiczak and Alex Alderton, Roberto Amato, Jeffrey Barrett, Sonia Goncalves, Ewan Harrison, David K. Jackson, Ian Johnston, Dominic Kwiatkowski, Cordelia Langford, John Sillitoe on behalf of the Wellcome Sanger Institute COVID-19 Surveillance Team |
| EPI_ISL_2995519, EPI_ISL_2995520 | Lighthouse Laboratory Plymouth | Wellcome Sanger Institute for the COVID-19 Genomics UK (COG-UK) Consortium | Lighthouse Laboratory Plymouth and Alex Alderton, Roberto Amato, Jeffrey Barrett, Sonia Goncalves, Ewan Harrison, David K. Jackson, Ian Johnston, Dominic Kwiatkowski, Cordelia Langford, John Sillitoe on behalf of the Wellcome Sanger Institute COVID-19 Surveillance Team |
| EPI_ISL_2995521 | Lighthouse Lab in Glasgow | Wellcome Sanger Institute for the COVID-19 Genomics UK (COG-UK) Consortium | Harper VanSteenhouse, Yumi Kasai, David Gray, Carol Clugston, Anna Dominiczak and Alex Alderton, Roberto Amato, Jeffrey Barrett, Sonia Goncalves, Ewan Harrison, David K. Jackson, Ian Johnston, Dominic Kwiatkowski, Cordelia Langford, John Sillitoe on behalf of the Wellcome Sanger Institute COVID-19 Surveillance Team |
| EPI_ISL_2995522 | Lighthouse Laboratory Plymouth | Wellcome Sanger Institute for the COVID-19 Genomics UK (COG-UK) Consortium | Lighthouse Laboratory Plymouth and Alex Alderton, Roberto Amato, Jeffrey Barrett, Sonia Goncalves, Ewan Harrison, David K. Jackson, Ian Johnston, Dominic Kwiatkowski, Cordelia Langford, John Sillitoe on behalf of the Wellcome Sanger Institute COVID-19 Surveillance Team |
| EPI_ISL_2995523, EPI_ISL_2995524, EPI_ISL_2995525 | Lighthouse Lab in Glasgow | Wellcome Sanger Institute for the COVID-19 Genomics UK (COG-UK) Consortium | Harper VanSteenhouse, Yumi Kasai, David Gray, Carol Clugston, Anna Dominiczak and Alex Alderton, Roberto Amato, Jeffrey Barrett, Sonia Goncalves, Ewan Harrison, David K. Jackson, Ian Johnston, Dominic Kwiatkowski, Cordelia Langford, John Sillitoe on behalf of the Wellcome Sanger Institute COVID-19 Surveillance Team |
| EPI_ISL_2995526 | Lighthouse Laboratory Plymouth | Wellcome Sanger Institute for the COVID-19 Genomics UK (COG-UK) Consortium | Lighthouse Laboratory Plymouth and Alex Alderton, Roberto Amato, Jeffrey Barrett, Sonia Goncalves, Ewan Harrison, David K. Jackson, Ian Johnston, Dominic Kwiatkowski, Cordelia Langford, John Sillitoe on behalf of the Wellcome Sanger Institute COVID-19 Surveillance Team |
| EPI_ISL_2995527, EPI_ISL_2995528, EPI_ISL_2995529, EPI_ISL_2995530 | Lighthouse Lab in Glasgow | Wellcome Sanger Institute for the COVID-19 Genomics UK (COG-UK) Consortium | Harper VanSteenhouse, Yumi Kasai, David Gray, Carol Clugston, Anna Dominiczak and Alex Alderton, Roberto Amato, Jeffrey Barrett, Sonia Goncalves, Ewan Harrison, David K. Jackson, Ian Johnston, Dominic Kwiatkowski, Cordelia Langford, John Sillitoe on behalf of the Wellcome Sanger Institute COVID-19 Surveillance Team |
| EPI_ISL_2995531 | Lighthouse Laboratory Plymouth | Wellcome Sanger Institute for the COVID-19 Genomics UK (COG-UK) Consortium | Lighthouse Laboratory Plymouth and Alex Alderton, Roberto Amato, Jeffrey Barrett, Sonia Goncalves, Ewan Harrison, David K. Jackson, Ian Johnston, Dominic Kwiatkowski, Cordelia Langford, John Sillitoe on behalf of the Wellcome Sanger Institute COVID-19 Surveillance Team |
| EPI_ISL_2995532, EPI_ISL_2995533 | Lighthouse Lab in Glasgow | Wellcome Sanger Institute for the COVID-19 Genomics UK (COG-UK) Consortium | Harper VanSteenhouse, Yumi Kasai, David Gray, Carol Clugston, Anna Dominiczak and Alex Alderton, Roberto Amato, Jeffrey Barrett, Sonia Goncalves, Ewan Harrison, David K. Jackson, Ian Johnston, Dominic Kwiatkowski, Cordelia Langford, John Sillitoe on behalf of the Wellcome Sanger Institute COVID-19 Surveillance Team |
| EPI_ISL_2995534 | Lighthouse Laboratory Plymouth | Wellcome Sanger Institute for the COVID-19 Genomics UK (COG-UK) Consortium | Lighthouse Laboratory Plymouth and Alex Alderton, Roberto Amato, Jeffrey Barrett, Sonia Goncalves, Ewan Harrison, David K. Jackson, Ian Johnston, Dominic Kwiatkowski, Cordelia Langford, John Sillitoe on behalf of the Wellcome Sanger Institute COVID-19 Surveillance Team |
| EPI_ISL_2995535, EPI_ISL_2995536 | Lighthouse Lab in Glasgow | Wellcome Sanger Institute for the COVID-19 Genomics UK (COG-UK) Consortium | Harper VanSteenhouse, Yumi Kasai, David Gray, Carol Clugston, Anna Dominiczak and Alex Alderton, Roberto Amato, Jeffrey Barrett, Sonia Goncalves, Ewan Harrison, David K. Jackson, Ian Johnston, Dominic Kwiatkowski, Cordelia Langford, John Sillitoe on behalf of the Wellcome Sanger Institute COVID-19 Surveillance Team |
| EPI_ISL_2995537 | Lighthouse Laboratory Plymouth | Wellcome Sanger Institute for the COVID-19 Genomics UK (COG-UK) Consortium | Lighthouse Laboratory Plymouth and Alex Alderton, Roberto Amato, Jeffrey Barrett, Sonia Goncalves, Ewan Harrison, David K. Jackson, Ian Johnston, Dominic Kwiatkowski, Cordelia Langford, John Sillitoe on behalf of the Wellcome Sanger Institute COVID-19 Surveillance Team |
| EPI_ISL_2995538 | Lighthouse Lab in Glasgow | Wellcome Sanger Institute for the COVID-19 Genomics UK (COG-UK) Consortium | Harper VanSteenhouse, Yumi Kasai, David Gray, Carol Clugston, Anna Dominiczak and Alex Alderton, Roberto Amato, Jeffrey Barrett, Sonia Goncalves, Ewan Harrison, David K. Jackson, Ian Johnston, Dominic Kwiatkowski, Cordelia Langford, John Sillitoe on behalf of the Wellcome Sanger Institute COVID-19 Surveillance Team |
| EPI_ISL_2995539 | Lighthouse Laboratory Plymouth | Wellcome Sanger Institute for the COVID-19 Genomics UK (COG-UK) Consortium | Lighthouse Laboratory Plymouth and Alex Alderton, Roberto Amato, Jeffrey Barrett, Sonia Goncalves, Ewan Harrison, David K. Jackson, Ian Johnston, Dominic Kwiatkowski, Cordelia Langford, John Sillitoe on behalf of the Wellcome Sanger Institute COVID-19 Surveillance Team |
| EPI_ISL_2995540, EPI_ISL_2995541, EPI_ISL_2995542, EPI_ISL_2995543, EPI_ISL_2995544, EPI_ISL_2995545, EPI_ISL_2995546, EPI_ISL_2995547, EPI_ISL_2995548, EPI_ISL_2995549, EPI_ISL_2995550 | | | |
| see above | Lighthouse Lab in Glasgow | Wellcome Sanger Institute for the COVID-19 Genomics UK (COG-UK) Consortium | Harper VanSteenhouse, Yumi Kasai, David Gray, Carol Clugston, Anna Dominiczak and Alex Alderton, Roberto Amato, Jeffrey Barrett, Sonia Goncalves, Ewan Harrison, David K. Jackson, Ian Johnston, Dominic Kwiatkowski, Cordelia Langford, John Sillitoe on behalf of the Wellcome Sanger Institute COVID-19 Surveillance Team |
| EPI_ISL_2995551 | Lighthouse Laboratory Plymouth | Wellcome Sanger Institute for the COVID-19 Genomics UK (COG-UK) Consortium | Lighthouse Laboratory Plymouth and Alex Alderton, Roberto Amato, Jeffrey Barrett, Sonia Goncalves, Ewan Harrison, David K. Jackson, Ian Johnston, Dominic Kwiatkowski, Cordelia Langford, John Sillitoe on behalf of the Wellcome Sanger Institute COVID-19 Surveillance Team |
| EPI_ISL_2995552, EPI_ISL_2995553 | Lighthouse Lab in Glasgow | Wellcome Sanger Institute for the COVID-19 Genomics UK (COG-UK) Consortium | Harper VanSteenhouse, Yumi Kasai, David Gray, Carol Clugston, Anna Dominiczak and Alex Alderton, Roberto Amato, Jeffrey Barrett, Sonia Goncalves, Ewan Harrison, David K. Jackson, Ian Johnston, Dominic Kwiatkowski, Cordelia Langford, John Sillitoe on behalf of the Wellcome Sanger Institute COVID-19 Surveillance Team |
| EPI_ISL_2995554 | Lighthouse Laboratory Plymouth | Wellcome Sanger Institute for the COVID-19 Genomics UK (COG-UK) Consortium | Lighthouse Laboratory Plymouth and Alex Alderton, Roberto Amato, Jeffrey Barrett, Sonia Goncalves, Ewan Harrison, David K. Jackson, Ian Johnston, Dominic Kwiatkowski, Cordelia Langford, John Sillitoe on behalf of the Wellcome Sanger Institute COVID-19 Surveillance Team |
| EPI_ISL_2995555, EPI_ISL_2995556 | Lighthouse Lab in Glasgow | Wellcome Sanger Institute for the COVID-19 Genomics UK (COG-UK) Consortium | Harper VanSteenhouse, Yumi Kasai, David Gray, Carol Clugston, Anna Dominiczak and Alex Alderton, Roberto Amato, Jeffrey Barrett, Sonia Goncalves, Ewan Harrison, David K. Jackson, Ian Johnston, Dominic Kwiatkowski, Cordelia Langford, John Sillitoe on behalf of the Wellcome Sanger Institute COVID-19 Surveillance Team |
| EPI_ISL_2995557 | Lighthouse Laboratory Plymouth | Wellcome Sanger Institute for the COVID-19 Genomics UK (COG-UK) Consortium | Lighthouse Laboratory Plymouth and Alex Alderton, Roberto Amato, Jeffrey Barrett, Sonia Goncalves, Ewan Harrison, David K. Jackson, Ian Johnston, Dominic Kwiatkowski, Cordelia Langford, John Sillitoe on behalf of the Wellcome Sanger Institute COVID-19 Surveillance Team |
| EPI_ISL_2995558, EPI_ISL_2995559 | Lighthouse Lab in Glasgow | Wellcome Sanger Institute for the COVID-19 Genomics UK (COG-UK) Consortium | Harper VanSteenhouse, Yumi Kasai, David Gray, Carol Clugston, Anna Dominiczak and Alex Alderton, Roberto Amato, Jeffrey Barrett, Sonia Goncalves, Ewan Harrison, David K. Jackson, Ian Johnston, Dominic Kwiatkowski, Cordelia Langford, John Sillitoe on behalf of the Wellcome Sanger Institute COVID-19 Surveillance Team |
| EPI_ISL_2995560 | Lighthouse Laboratory Plymouth | Wellcome Sanger Institute for the COVID-19 Genomics UK (COG-UK) Consortium | Lighthouse Laboratory Plymouth and Alex Alderton, Roberto Amato, Jeffrey Barrett, Sonia Goncalves, Ewan Harrison, David K. Jackson, Ian Johnston, Dominic Kwiatkowski, Cordelia Langford, John Sillitoe on behalf of the Wellcome Sanger Institute COVID-19 Surveillance Team |
| EPI_ISL_2995561, EPI_ISL_2995562, EPI_ISL_2995563, EPI_ISL_2995564, EPI_ISL_2995565 | Lighthouse Lab in Glasgow | Wellcome Sanger Institute for the COVID-19 Genomics UK (COG-UK) Consortium | Harper VanSteenhouse, Yumi Kasai, David Gray, Carol Clugston, Anna Dominiczak and Alex Alderton, Roberto Amato, Jeffrey Barrett, Sonia Goncalves, Ewan Harrison, David K. Jackson, Ian Johnston, Dominic Kwiatkowski, Cordelia Langford, John Sillitoe on behalf of the Wellcome Sanger Institute COVID-19 Surveillance Team |
| EPI_ISL_2995566, EPI_ISL_2995567 | Lighthouse Laboratory Plymouth | Wellcome Sanger Institute for the COVID-19 Genomics UK (COG-UK) Consortium | Lighthouse Laboratory Plymouth and Alex Alderton, Roberto Amato, Jeffrey Barrett, Sonia Goncalves, Ewan Harrison, David K. Jackson, Ian Johnston, Dominic Kwiatkowski, Cordelia Langford, John Sillitoe on behalf of the Wellcome Sanger Institute COVID-19 Surveillance Team |
| EPI_ISL_2995568, EPI_ISL_2995569, EPI_ISL_2995570, EPI_ISL_2995571, EPI_ISL_2995572, EPI_ISL_2995573, EPI_ISL_2995574, EPI_ISL_2995575, EPI_ISL_2995576, EPI_ISL_2995577, EPI_ISL_2995578 | | | |
| see above | Lighthouse Lab in Glasgow | Wellcome Sanger Institute for the COVID-19 Genomics UK (COG-UK) Consortium | Harper VanSteenhouse, Yumi Kasai, David Gray, Carol Clugston, Anna Dominiczak and Alex Alderton, Roberto Amato, Jeffrey Barrett, Sonia Goncalves, Ewan Harrison, David K. Jackson, Ian Johnston, Dominic Kwiatkowski, Cordelia Langford, John Sillitoe on behalf of the Wellcome Sanger Institute COVID-19 Surveillance Team |
| EPI_ISL_2995579 | Lighthouse Laboratory Plymouth | Wellcome Sanger Institute for the COVID-19 Genomics UK (COG-UK) Consortium | Lighthouse Laboratory Plymouth and Alex Alderton, Roberto Amato, Jeffrey Barrett, Sonia Goncalves, Ewan Harrison, David K. Jackson, Ian Johnston, Dominic Kwiatkowski, Cordelia Langford, John Sillitoe on behalf of the Wellcome Sanger Institute COVID-19 Surveillance Team |
| EPI_ISL_2995580 | Lighthouse Lab in Glasgow | Wellcome Sanger Institute for the COVID-19 Genomics UK (COG-UK) Consortium | Harper VanSteenhouse, Yumi Kasai, David Gray, Carol Clugston, Anna Dominiczak and Alex Alderton, Roberto Amato, Jeffrey Barrett, Sonia Goncalves, Ewan Harrison, David K. Jackson, Ian Johnston, Dominic Kwiatkowski, Cordelia Langford, John Sillitoe on behalf of the Wellcome Sanger Institute COVID-19 Surveillance Team |
| EPI_ISL_2995581 | Lighthouse Laboratory Plymouth | Wellcome Sanger Institute for the COVID-19 Genomics UK (COG-UK) Consortium | Lighthouse Laboratory Plymouth and Alex Alderton, Roberto Amato, Jeffrey Barrett, Sonia Goncalves, Ewan Harrison, David K. Jackson, Ian Johnston, Dominic Kwiatkowski, Cordelia Langford, John Sillitoe on behalf of the Wellcome Sanger Institute COVID-19 Surveillance Team |
| EPI_ISL_2995582, EPI_ISL_2995583, EPI_ISL_2995584, EPI_ISL_2995585, EPI_ISL_2995586 | Lighthouse Lab in Glasgow | Wellcome Sanger Institute for the COVID-19 Genomics UK (COG-UK) Consortium | Harper VanSteenhouse, Yumi Kasai, David Gray, Carol Clugston, Anna Dominiczak and Alex Alderton, Roberto Amato, Jeffrey Barrett, Sonia Goncalves, Ewan Harrison, David K. Jackson, Ian Johnston, Dominic Kwiatkowski, Cordelia Langford, John Sillitoe on behalf of the Wellcome Sanger Institute COVID-19 Surveillance Team |
| EPI_ISL_2995587 | Lighthouse Laboratory Plymouth | Wellcome Sanger Institute for the COVID-19 Genomics UK (COG-UK) Consortium | Lighthouse Laboratory Plymouth and Alex Alderton, Roberto Amato, Jeffrey Barrett, Sonia Goncalves, Ewan Harrison, David K. Jackson, Ian Johnston, Dominic Kwiatkowski, Cordelia Langford, John Sillitoe on behalf of the Wellcome Sanger Institute COVID-19 Surveillance Team |
| EPI_ISL_2995588 | Lighthouse Lab in Glasgow | Wellcome Sanger Institute for the COVID-19 Genomics UK (COG-UK) Consortium | Harper VanSteenhouse, Yumi Kasai, David Gray, Carol Clugston, Anna Dominiczak and Alex Alderton, Roberto Amato, Jeffrey Barrett, Sonia Goncalves, Ewan Harrison, David K. Jackson, Ian Johnston, Dominic Kwiatkowski, Cordelia Langford, John Sillitoe on behalf of the Wellcome Sanger Institute COVID-19 Surveillance Team |
| EPI_ISL_2995589, EPI_ISL_2995590 | Lighthouse Laboratory Plymouth | Wellcome Sanger Institute for the COVID-19 Genomics UK (COG-UK) Consortium | Lighthouse Laboratory Plymouth and Alex Alderton, Roberto Amato, Jeffrey Barrett, Sonia Goncalves, Ewan Harrison, David K. Jackson, Ian Johnston, Dominic Kwiatkowski, Cordelia Langford, John Sillitoe on behalf of the Wellcome Sanger Institute COVID-19 Surveillance Team |
| EPI_ISL_2995591, EPI_ISL_2995592, | Lighthouse Lab in Glasgow | Wellcome Sanger Institute for the COVID-19 Genomics UK (COG-UK) Consortium | Harper VanSteenhouse, Yumi Kasai, David Gray, Carol Clugston, Anna Dominiczak and Alex Alderton, Roberto Amato, Jeffrey Barrett, Sonia Goncalves, Ewan Harrison, David K. Jackson, Ian Johnston, Dominic Kwiatkowski, Cordelia Langford, John Sillitoe on behalf of the Wellcome Sanger Institute COVID-19 Surveillance Team |

| | | | |
|---|---|---|---|
| EPI_ISL_2995593, EPI_ISL_2995594, EPI_ISL_2995595 | | | |
| EPI_ISL_2995596 | Lighthouse Laboratory Plymouth | Wellcome Sanger Institute for the COVID-19 Genomics UK (COG-UK) Consortium | Lighthouse Laboratory Plymouth and Alex Alderton, Roberto Amato, Jeffrey Barrett, Sonia Goncalves, Ewan Harrison, David K. Jackson, Ian Johnston, Dominic Kwiatkowski, Cordelia Langford, John Sillitoe on behalf of the Wellcome Sanger Institute COVID-19 Surveillance Team |
| EPI_ISL_2995597, EPI_ISL_2995598, EPI_ISL_2995599, EPI_ISL_2995600 | Lighthouse Lab in Glasgow | Wellcome Sanger Institute for the COVID-19 Genomics UK (COG-UK) Consortium | Harper VanSteenhouse, Yumi Kasai, David Gray, Carol Clugston, Anna Dominiczak and Alex Alderton, Roberto Amato, Jeffrey Barrett, Sonia Goncalves, Ewan Harrison, David K. Jackson, Ian Johnston, Dominic Kwiatkowski, Cordelia Langford, John Sillitoe on behalf of the Wellcome Sanger Institute COVID-19 Surveillance Team |
| EPI_ISL_2995601 | Lighthouse Laboratory Plymouth | Wellcome Sanger Institute for the COVID-19 Genomics UK (COG-UK) Consortium | Lighthouse Laboratory Plymouth and Alex Alderton, Roberto Amato, Jeffrey Barrett, Sonia Goncalves, Ewan Harrison, David K. Jackson, Ian Johnston, Dominic Kwiatkowski, Cordelia Langford, John Sillitoe on behalf of the Wellcome Sanger Institute COVID-19 Surveillance Team |
| EPI_ISL_2995602, EPI_ISL_2995603 | Lighthouse Lab in Glasgow | Wellcome Sanger Institute for the COVID-19 Genomics UK (COG-UK) Consortium | Harper VanSteenhouse, Yumi Kasai, David Gray, Carol Clugston, Anna Dominiczak and Alex Alderton, Roberto Amato, Jeffrey Barrett, Sonia Goncalves, Ewan Harrison, David K. Jackson, Ian Johnston, Dominic Kwiatkowski, Cordelia Langford, John Sillitoe on behalf of the Wellcome Sanger Institute COVID-19 Surveillance Team |
| EPI_ISL_2995604 | Lighthouse Laboratory Plymouth | Wellcome Sanger Institute for the COVID-19 Genomics UK (COG-UK) Consortium | Lighthouse Laboratory Plymouth and Alex Alderton, Roberto Amato, Jeffrey Barrett, Sonia Goncalves, Ewan Harrison, David K. Jackson, Ian Johnston, Dominic Kwiatkowski, Cordelia Langford, John Sillitoe on behalf of the Wellcome Sanger Institute COVID-19 Surveillance Team |
| EPI_ISL_2995605, EPI_ISL_2995606, EPI_ISL_2995607 | Lighthouse Lab in Glasgow | Wellcome Sanger Institute for the COVID-19 Genomics UK (COG-UK) Consortium | Harper VanSteenhouse, Yumi Kasai, David Gray, Carol Clugston, Anna Dominiczak and Alex Alderton, Roberto Amato, Jeffrey Barrett, Sonia Goncalves, Ewan Harrison, David K. Jackson, Ian Johnston, Dominic Kwiatkowski, Cordelia Langford, John Sillitoe on behalf of the Wellcome Sanger Institute COVID-19 Surveillance Team |
| EPI_ISL_2995608, EPI_ISL_2995609, EPI_ISL_2995610 | Lighthouse Laboratory Plymouth | Wellcome Sanger Institute for the COVID-19 Genomics UK (COG-UK) Consortium | Lighthouse Laboratory Plymouth and Alex Alderton, Roberto Amato, Jeffrey Barrett, Sonia Goncalves, Ewan Harrison, David K. Jackson, Ian Johnston, Dominic Kwiatkowski, Cordelia Langford, John Sillitoe on behalf of the Wellcome Sanger Institute COVID-19 Surveillance Team |
| EPI_ISL_2995611, EPI_ISL_2995612, EPI_ISL_2995613, EPI_ISL_2995614, EPI_ISL_2995615, EPI_ISL_2995616, EPI_ISL_2995617, EPI_ISL_2995618, EPI_ISL_2995619, EPI_ISL_2995620, EPI_ISL_2995621, EPI_ISL_2995622, EPI_ISL_2995623, EPI_ISL_2995624, EPI_ISL_2995625 | | | |
| see above | Lighthouse Lab in Glasgow | Wellcome Sanger Institute for the COVID-19 Genomics UK (COG-UK) Consortium | Harper VanSteenhouse, Yumi Kasai, David Gray, Carol Clugston, Anna Dominiczak and Alex Alderton, Roberto Amato, Jeffrey Barrett, Sonia Goncalves, Ewan Harrison, David K. Jackson, Ian Johnston, Dominic Kwiatkowski, Cordelia Langford, John Sillitoe on behalf of the Wellcome Sanger Institute COVID-19 Surveillance Team |
| EPI_ISL_2995626, EPI_ISL_2995627 | Lighthouse Laboratory Plymouth | Wellcome Sanger Institute for the COVID-19 Genomics UK (COG-UK) Consortium | Lighthouse Laboratory Plymouth and Alex Alderton, Roberto Amato, Jeffrey Barrett, Sonia Goncalves, Ewan Harrison, David K. Jackson, Ian Johnston, Dominic Kwiatkowski, Cordelia Langford, John Sillitoe on behalf of the Wellcome Sanger Institute COVID-19 Surveillance Team |
| EPI_ISL_2995628, EPI_ISL_2995629, EPI_ISL_2995630, EPI_ISL_2995631 | Lighthouse Lab in Glasgow | Wellcome Sanger Institute for the COVID-19 Genomics UK (COG-UK) Consortium | Harper VanSteenhouse, Yumi Kasai, David Gray, Carol Clugston, Anna Dominiczak and Alex Alderton, Roberto Amato, Jeffrey Barrett, Sonia Goncalves, Ewan Harrison, David K. Jackson, Ian Johnston, Dominic Kwiatkowski, Cordelia Langford, John Sillitoe on behalf of the Wellcome Sanger Institute COVID-19 Surveillance Team |
| EPI_ISL_2995632 | Lighthouse Laboratory Plymouth | Wellcome Sanger Institute for the COVID-19 Genomics UK (COG-UK) Consortium | Lighthouse Laboratory Plymouth and Alex Alderton, Roberto Amato, Jeffrey Barrett, Sonia Goncalves, Ewan Harrison, David K. Jackson, Ian Johnston, Dominic Kwiatkowski, Cordelia Langford, John Sillitoe on behalf of the Wellcome Sanger Institute COVID-19 Surveillance Team |
| EPI_ISL_2995633 | Lighthouse Lab in Glasgow | Wellcome Sanger Institute for the COVID-19 Genomics UK (COG-UK) Consortium | Harper VanSteenhouse, Yumi Kasai, David Gray, Carol Clugston, Anna Dominiczak and Alex Alderton, Roberto Amato, Jeffrey Barrett, Sonia Goncalves, Ewan Harrison, David K. Jackson, Ian Johnston, Dominic Kwiatkowski, Cordelia Langford, John Sillitoe on behalf of the Wellcome Sanger Institute COVID-19 Surveillance Team |
| EPI_ISL_2995634 | Lighthouse Laboratory Plymouth | Wellcome Sanger Institute for the COVID-19 Genomics UK (COG-UK) Consortium | Lighthouse Laboratory Plymouth and Alex Alderton, Roberto Amato, Jeffrey Barrett, Sonia Goncalves, Ewan Harrison, David K. Jackson, Ian Johnston, Dominic Kwiatkowski, Cordelia Langford, John Sillitoe on behalf of the Wellcome Sanger Institute COVID-19 Surveillance Team |
| EPI_ISL_2995635 | Lighthouse Lab in Glasgow | Wellcome Sanger Institute for the COVID-19 Genomics UK (COG-UK) Consortium | Harper VanSteenhouse, Yumi Kasai, David Gray, Carol Clugston, Anna Dominiczak and Alex Alderton, Roberto Amato, Jeffrey Barrett, Sonia Goncalves, Ewan Harrison, David K. Jackson, Ian Johnston, Dominic Kwiatkowski, Cordelia Langford, John Sillitoe on behalf of the Wellcome Sanger Institute COVID-19 Surveillance Team |
| EPI_ISL_2995636 | Lighthouse Laboratory Plymouth | Wellcome Sanger Institute for the COVID-19 Genomics UK (COG-UK) Consortium | Lighthouse Laboratory Plymouth and Alex Alderton, Roberto Amato, Jeffrey Barrett, Sonia Goncalves, Ewan Harrison, David K. Jackson, Ian Johnston, Dominic Kwiatkowski, Cordelia Langford, John Sillitoe on behalf of the Wellcome Sanger Institute COVID-19 Surveillance Team |
| EPI_ISL_2995637, EPI_ISL_2995638, EPI_ISL_2995639, EPI_ISL_2995640, EPI_ISL_2995641, EPI_ISL_2995642, EPI_ISL_2995643 | Lighthouse Lab in Glasgow | Wellcome Sanger Institute for the COVID-19 Genomics UK (COG-UK) Consortium | Harper VanSteenhouse, Yumi Kasai, David Gray, Carol Clugston, Anna Dominiczak and Alex Alderton, Roberto Amato, Jeffrey Barrett, Sonia Goncalves, Ewan Harrison, David K. Jackson, Ian Johnston, Dominic Kwiatkowski, Cordelia Langford, John Sillitoe on behalf of the Wellcome Sanger Institute COVID-19 Surveillance Team |
| EPI_ISL_2995644, EPI_ISL_2995645, EPI_ISL_2995646 | Lighthouse Laboratory Plymouth | Wellcome Sanger Institute for the COVID-19 Genomics UK (COG-UK) Consortium | Lighthouse Laboratory Plymouth and Alex Alderton, Roberto Amato, Jeffrey Barrett, Sonia Goncalves, Ewan Harrison, David K. Jackson, Ian Johnston, Dominic Kwiatkowski, Cordelia Langford, John Sillitoe on behalf of the Wellcome Sanger Institute COVID-19 Surveillance Team |
| EPI_ISL_2995647, EPI_ISL_2995648, EPI_ISL_2995649, EPI_ISL_2995650, EPI_ISL_2995651, EPI_ISL_2995652, EPI_ISL_2995653 | Lighthouse Lab in Glasgow | Wellcome Sanger Institute for the COVID-19 Genomics UK (COG-UK) Consortium | Harper VanSteenhouse, Yumi Kasai, David Gray, Carol Clugston, Anna Dominiczak and Alex Alderton, Roberto Amato, Jeffrey Barrett, Sonia Goncalves, Ewan Harrison, David K. Jackson, Ian Johnston, Dominic Kwiatkowski, Cordelia Langford, John Sillitoe on behalf of the Wellcome Sanger Institute COVID-19 Surveillance Team |
| EPI_ISL_2995654, EPI_ISL_2995655 | Lighthouse Laboratory Plymouth | Wellcome Sanger Institute for the COVID-19 Genomics UK (COG-UK) Consortium | Lighthouse Laboratory Plymouth and Alex Alderton, Roberto Amato, Jeffrey Barrett, Sonia Goncalves, Ewan Harrison, David K. Jackson, Ian Johnston, Dominic Kwiatkowski, Cordelia Langford, John Sillitoe on behalf of the Wellcome Sanger Institute COVID-19 Surveillance Team |
| EPI_ISL_2995656, EPI_ISL_2995657, EPI_ISL_2995658, EPI_ISL_2995659, EPI_ISL_2995660, EPI_ISL_2995661, EPI_ISL_2995662 | Lighthouse Lab in Glasgow | Wellcome Sanger Institute for the COVID-19 Genomics UK (COG-UK) Consortium | Harper VanSteenhouse, Yumi Kasai, David Gray, Carol Clugston, Anna Dominiczak and Alex Alderton, Roberto Amato, Jeffrey Barrett, Sonia Goncalves, Ewan Harrison, David K. Jackson, Ian Johnston, Dominic Kwiatkowski, Cordelia Langford, John Sillitoe on behalf of the Wellcome Sanger Institute COVID-19 Surveillance Team |
| EPI_ISL_2995663, EPI_ISL_2995664 | Lighthouse Laboratory Plymouth | Wellcome Sanger Institute for the COVID-19 Genomics UK (COG-UK) Consortium | Lighthouse Laboratory Plymouth and Alex Alderton, Roberto Amato, Jeffrey Barrett, Sonia Goncalves, Ewan Harrison, David K. Jackson, Ian Johnston, Dominic Kwiatkowski, Cordelia Langford, John Sillitoe on behalf of the Wellcome Sanger Institute COVID-19 Surveillance Team |
| EPI_ISL_2995665, EPI_ISL_2995666, EPI_ISL_2995667 | Lighthouse Lab in Glasgow | Wellcome Sanger Institute for the COVID-19 Genomics UK (COG-UK) Consortium | Harper VanSteenhouse, Yumi Kasai, David Gray, Carol Clugston, Anna Dominiczak and Alex Alderton, Roberto Amato, Jeffrey Barrett, Sonia Goncalves, Ewan Harrison, David K. Jackson, Ian Johnston, Dominic Kwiatkowski, Cordelia Langford, John Sillitoe on behalf of the Wellcome Sanger Institute COVID-19 Surveillance Team |
| EPI_ISL_2995668 | Lighthouse Laboratory Plymouth | Wellcome Sanger Institute for the COVID-19 Genomics UK (COG-UK) Consortium | Lighthouse Laboratory Plymouth and Alex Alderton, Roberto Amato, Jeffrey Barrett, Sonia Goncalves, Ewan Harrison, David K. Jackson, Ian Johnston, Dominic Kwiatkowski, Cordelia Langford, John Sillitoe on behalf of the Wellcome Sanger Institute COVID-19 Surveillance Team |
| EPI_ISL_2995669, EPI_ISL_2995670, EPI_ISL_2995671, EPI_ISL_2995672, EPI_ISL_2995673, EPI_ISL_2995674, EPI_ISL_2995675, EPI_ISL_2995676, EPI_ISL_2995677 | Lighthouse Lab in Glasgow | Wellcome Sanger Institute for the COVID-19 Genomics UK (COG-UK) Consortium | Harper VanSteenhouse, Yumi Kasai, David Gray, Carol Clugston, Anna Dominiczak and Alex Alderton, Roberto Amato, Jeffrey Barrett, Sonia Goncalves, Ewan Harrison, David K. Jackson, Ian Johnston, Dominic Kwiatkowski, Cordelia Langford, John Sillitoe on behalf of the Wellcome Sanger Institute COVID-19 Surveillance Team |
| EPI_ISL_2995678 | Lighthouse Laboratory Plymouth | Wellcome Sanger Institute for the COVID-19 Genomics UK (COG-UK) Consortium | Lighthouse Laboratory Plymouth and Alex Alderton, Roberto Amato, Jeffrey Barrett, Sonia Goncalves, Ewan Harrison, David K. Jackson, Ian Johnston, Dominic Kwiatkowski, Cordelia Langford, John Sillitoe on behalf of the Wellcome Sanger Institute COVID-19 Surveillance Team |
| EPI_ISL_2995679, EPI_ISL_2995680, EPI_ISL_2995681 | Lighthouse Lab in Glasgow | Wellcome Sanger Institute for the COVID-19 Genomics UK (COG-UK) Consortium | Harper VanSteenhouse, Yumi Kasai, David Gray, Carol Clugston, Anna Dominiczak and Alex Alderton, Roberto Amato, Jeffrey Barrett, Sonia Goncalves, Ewan Harrison, David K. Jackson, Ian Johnston, Dominic Kwiatkowski, Cordelia Langford, John Sillitoe on behalf of the Wellcome Sanger Institute COVID-19 Surveillance Team |
| EPI_ISL_2995682 | Berkshire and Surrey Pathology Services Lighthouse Laboratory | Wellcome Sanger Institute for the COVID-19 Genomics UK (COG-UK) Consortium | Berkshire and Surrey Pathology Services Lighthouse Laboratory and Alex Alderton, Roberto Amato, Jeffrey Barrett, Sonia Goncalves, Ewan Harrison, David K. Jackson, Ian Johnston, Dominic Kwiatkowski, Cordelia Langford, John Sillitoe on behalf of the Wellcome Sanger Institute COVID-19 Surveillance Team |
| EPI_ISL_2995683 | Lighthouse Lab in Milton Keynes | Wellcome Sanger Institute for the COVID-19 Genomics UK (COG-UK) Consortium | The Lighthouse Lab in Milton Keynes and Alex Alderton, Roberto Amato, Jeffrey Barrett, Sonia Goncalves, Ewan Harrison, David K. Jackson, Ian Johnston, Dominic Kwiatkowski, Cordelia Langford, John Sillitoe on behalf of the Wellcome Sanger Institute COVID-19 Surveillance Team |
| EPI_ISL_2995684 | Berkshire and Surrey Pathology Services Lighthouse Laboratory | Wellcome Sanger Institute for the COVID-19 Genomics UK (COG-UK) Consortium | Berkshire and Surrey Pathology Services Lighthouse Laboratory and Alex Alderton, Roberto Amato, Jeffrey Barrett, Sonia Goncalves, Ewan Harrison, David K. Jackson, Ian Johnston, Dominic Kwiatkowski, Cordelia Langford, John Sillitoe on behalf of the Wellcome Sanger Institute COVID-19 Surveillance Team |
| EPI_ISL_2995685, EPI_ISL_2995686, EPI_ISL_2995687 | Lighthouse Lab in Glasgow | Wellcome Sanger Institute for the COVID-19 Genomics UK (COG-UK) Consortium | Harper VanSteenhouse, Yumi Kasai, David Gray, Carol Clugston, Anna Dominiczak and Alex Alderton, Roberto Amato, Jeffrey Barrett, Sonia Goncalves, Ewan Harrison, David K. Jackson, Ian Johnston, Dominic Kwiatkowski, Cordelia Langford, John Sillitoe on behalf of the Wellcome Sanger Institute COVID-19 Surveillance Team |

| | | | |
|---|---|---|---|
| EPI_ISL_2995688 | Lighthouse Lab in Milton Keynes | Wellcome Sanger Institute for the COVID-19 Genomics UK (COG-UK) Consortium | The Lighthouse Lab in Milton Keynes and Alex Alderton, Roberto Amato, Jeffrey Barrett, Sonia Goncalves, Ewan Harrison, David K. Jackson, Ian Johnston, Dominic Kwiatkowski, Cordelia Langford, John Sillitoe on behalf of the Wellcome Sanger Institute COVID-19 Surveillance Team |
| EPI_ISL_2995689 | Lighthouse Lab in Glasgow | Wellcome Sanger Institute for the COVID-19 Genomics UK (COG-UK) Consortium | Harper VanSteenhouse, Yumi Kasai, David Gray, Carol Clugston, Anna Dominiczak and Alex Alderton, Roberto Amato, Jeffrey Barrett, Sonia Goncalves, Ewan Harrison, David K. Jackson, Ian Johnston, Dominic Kwiatkowski, Cordelia Langford, John Sillitoe on behalf of the Wellcome Sanger Institute COVID-19 Surveillance Team |
| EPI_ISL_2995690, EPI_ISL_2995691, EPI_ISL_2995692 | Berkshire and Surrey Pathology Services Lighthouse Laboratory | Wellcome Sanger Institute for the COVID-19 Genomics UK (COG-UK) Consortium | Berkshire and Surrey Pathology Services Lighthouse Laboratory and Alex Alderton, Roberto Amato, Jeffrey Barrett, Sonia Goncalves, Ewan Harrison, David K. Jackson, Ian Johnston, Dominic Kwiatkowski, Cordelia Langford, John Sillitoe on behalf of the Wellcome Sanger Institute COVID-19 Surveillance Team |
| EPI_ISL_2995693 | Lighthouse Lab in Milton Keynes | Wellcome Sanger Institute for the COVID-19 Genomics UK (COG-UK) Consortium | The Lighthouse Lab in Milton Keynes and Alex Alderton, Roberto Amato, Jeffrey Barrett, Sonia Goncalves, Ewan Harrison, David K. Jackson, Ian Johnston, Dominic Kwiatkowski, Cordelia Langford, John Sillitoe on behalf of the Wellcome Sanger Institute COVID-19 Surveillance Team |
| EPI_ISL_2995694 | Berkshire and Surrey Pathology Services Lighthouse Laboratory | Wellcome Sanger Institute for the COVID-19 Genomics UK (COG-UK) Consortium | Berkshire and Surrey Pathology Services Lighthouse Laboratory and Alex Alderton, Roberto Amato, Jeffrey Barrett, Sonia Goncalves, Ewan Harrison, David K. Jackson, Ian Johnston, Dominic Kwiatkowski, Cordelia Langford, John Sillitoe on behalf of the Wellcome Sanger Institute COVID-19 Surveillance Team |
| EPI_ISL_2995695, EPI_ISL_2995696 | Lighthouse Lab in Milton Keynes | Wellcome Sanger Institute for the COVID-19 Genomics UK (COG-UK) Consortium | The Lighthouse Lab in Milton Keynes and Alex Alderton, Roberto Amato, Jeffrey Barrett, Sonia Goncalves, Ewan Harrison, David K. Jackson, Ian Johnston, Dominic Kwiatkowski, Cordelia Langford, John Sillitoe on behalf of the Wellcome Sanger Institute COVID-19 Surveillance Team |
| EPI_ISL_2995697 | Lighthouse Lab in Glasgow | Wellcome Sanger Institute for the COVID-19 Genomics UK (COG-UK) Consortium | Harper VanSteenhouse, Yumi Kasai, David Gray, Carol Clugston, Anna Dominiczak and Alex Alderton, Roberto Amato, Jeffrey Barrett, Sonia Goncalves, Ewan Harrison, David K. Jackson, Ian Johnston, Dominic Kwiatkowski, Cordelia Langford, John Sillitoe on behalf of the Wellcome Sanger Institute COVID-19 Surveillance Team |
| EPI_ISL_2995698 | Lighthouse Lab in Milton Keynes | Wellcome Sanger Institute for the COVID-19 Genomics UK (COG-UK) Consortium | The Lighthouse Lab in Milton Keynes and Alex Alderton, Roberto Amato, Jeffrey Barrett, Sonia Goncalves, Ewan Harrison, David K. Jackson, Ian Johnston, Dominic Kwiatkowski, Cordelia Langford, John Sillitoe on behalf of the Wellcome Sanger Institute COVID-19 Surveillance Team |
| EPI_ISL_2995699, EPI_ISL_2995700 | Berkshire and Surrey Pathology Services Lighthouse Laboratory | Wellcome Sanger Institute for the COVID-19 Genomics UK (COG-UK) Consortium | Berkshire and Surrey Pathology Services Lighthouse Laboratory and Alex Alderton, Roberto Amato, Jeffrey Barrett, Sonia Goncalves, Ewan Harrison, David K. Jackson, Ian Johnston, Dominic Kwiatkowski, Cordelia Langford, John Sillitoe on behalf of the Wellcome Sanger Institute COVID-19 Surveillance Team |
| EPI_ISL_2995701 | Lighthouse Lab in Milton Keynes | Wellcome Sanger Institute for the COVID-19 Genomics UK (COG-UK) Consortium | The Lighthouse Lab in Milton Keynes and Alex Alderton, Roberto Amato, Jeffrey Barrett, Sonia Goncalves, Ewan Harrison, David K. Jackson, Ian Johnston, Dominic Kwiatkowski, Cordelia Langford, John Sillitoe on behalf of the Wellcome Sanger Institute COVID-19 Surveillance Team |
| EPI_ISL_2995702, EPI_ISL_2995703 | Lighthouse Lab in Glasgow | Wellcome Sanger Institute for the COVID-19 Genomics UK (COG-UK) Consortium | Harper VanSteenhouse, Yumi Kasai, David Gray, Carol Clugston, Anna Dominiczak and Alex Alderton, Roberto Amato, Jeffrey Barrett, Sonia Goncalves, Ewan Harrison, David K. Jackson, Ian Johnston, Dominic Kwiatkowski, Cordelia Langford, John Sillitoe on behalf of the Wellcome Sanger Institute COVID-19 Surveillance Team |
| EPI_ISL_2995704 | Berkshire and Surrey Pathology Services Lighthouse Laboratory | Wellcome Sanger Institute for the COVID-19 Genomics UK (COG-UK) Consortium | Berkshire and Surrey Pathology Services Lighthouse Laboratory and Alex Alderton, Roberto Amato, Jeffrey Barrett, Sonia Goncalves, Ewan Harrison, David K. Jackson, Ian Johnston, Dominic Kwiatkowski, Cordelia Langford, John Sillitoe on behalf of the Wellcome Sanger Institute COVID-19 Surveillance Team |
| EPI_ISL_2995705, EPI_ISL_2995706 | Lighthouse Lab in Milton Keynes | Wellcome Sanger Institute for the COVID-19 Genomics UK (COG-UK) Consortium | The Lighthouse Lab in Milton Keynes and Alex Alderton, Roberto Amato, Jeffrey Barrett, Sonia Goncalves, Ewan Harrison, David K. Jackson, Ian Johnston, Dominic Kwiatkowski, Cordelia Langford, John Sillitoe on behalf of the Wellcome Sanger Institute COVID-19 Surveillance Team |
| EPI_ISL_2995707, EPI_ISL_2995708 | Berkshire and Surrey Pathology Services Lighthouse Laboratory | Wellcome Sanger Institute for the COVID-19 Genomics UK (COG-UK) Consortium | Berkshire and Surrey Pathology Services Lighthouse Laboratory and Alex Alderton, Roberto Amato, Jeffrey Barrett, Sonia Goncalves, Ewan Harrison, David K. Jackson, Ian Johnston, Dominic Kwiatkowski, Cordelia Langford, John Sillitoe on behalf of the Wellcome Sanger Institute COVID-19 Surveillance Team |
| EPI_ISL_2995709, EPI_ISL_2995710 | Lighthouse Lab in Milton Keynes | Wellcome Sanger Institute for the COVID-19 Genomics UK (COG-UK) Consortium | The Lighthouse Lab in Milton Keynes and Alex Alderton, Roberto Amato, Jeffrey Barrett, Sonia Goncalves, Ewan Harrison, David K. Jackson, Ian Johnston, Dominic Kwiatkowski, Cordelia Langford, John Sillitoe on behalf of the Wellcome Sanger Institute COVID-19 Surveillance Team |
| EPI_ISL_2995711 | Berkshire and Surrey Pathology Services Lighthouse Laboratory | Wellcome Sanger Institute for the COVID-19 Genomics UK (COG-UK) Consortium | Berkshire and Surrey Pathology Services Lighthouse Laboratory and Alex Alderton, Roberto Amato, Jeffrey Barrett, Sonia Goncalves, Ewan Harrison, David K. Jackson, Ian Johnston, Dominic Kwiatkowski, Cordelia Langford, John Sillitoe on behalf of the Wellcome Sanger Institute COVID-19 Surveillance Team |
| EPI_ISL_2995712, EPI_ISL_2995713 | Lighthouse Lab in Milton Keynes | Wellcome Sanger Institute for the COVID-19 Genomics UK (COG-UK) Consortium | The Lighthouse Lab in Milton Keynes and Alex Alderton, Roberto Amato, Jeffrey Barrett, Sonia Goncalves, Ewan Harrison, David K. Jackson, Ian Johnston, Dominic Kwiatkowski, Cordelia Langford, John Sillitoe on behalf of the Wellcome Sanger Institute COVID-19 Surveillance Team |
| EPI_ISL_2995714 | Berkshire and Surrey Pathology Services Lighthouse Laboratory | Wellcome Sanger Institute for the COVID-19 Genomics UK (COG-UK) Consortium | Berkshire and Surrey Pathology Services Lighthouse Laboratory and Alex Alderton, Roberto Amato, Jeffrey Barrett, Sonia Goncalves, Ewan Harrison, David K. Jackson, Ian Johnston, Dominic Kwiatkowski, Cordelia Langford, John Sillitoe on behalf of the Wellcome Sanger Institute COVID-19 Surveillance Team |
| EPI_ISL_2995715 | Lighthouse Lab in Milton Keynes | Wellcome Sanger Institute for the COVID-19 Genomics UK (COG-UK) Consortium | The Lighthouse Lab in Milton Keynes and Alex Alderton, Roberto Amato, Jeffrey Barrett, Sonia Goncalves, Ewan Harrison, David K. Jackson, Ian Johnston, Dominic Kwiatkowski, Cordelia Langford, John Sillitoe on behalf of the Wellcome Sanger Institute COVID-19 Surveillance Team |
| EPI_ISL_2995716, EPI_ISL_2995717, EPI_ISL_2995718 | Berkshire and Surrey Pathology Services Lighthouse Laboratory | Wellcome Sanger Institute for the COVID-19 Genomics UK (COG-UK) Consortium | Berkshire and Surrey Pathology Services Lighthouse Laboratory and Alex Alderton, Roberto Amato, Jeffrey Barrett, Sonia Goncalves, Ewan Harrison, David K. Jackson, Ian Johnston, Dominic Kwiatkowski, Cordelia Langford, John Sillitoe on behalf of the Wellcome Sanger Institute COVID-19 Surveillance Team |
| EPI_ISL_2995719, EPI_ISL_2995720, EPI_ISL_2995721 | Lighthouse Lab in Milton Keynes | Wellcome Sanger Institute for the COVID-19 Genomics UK (COG-UK) Consortium | The Lighthouse Lab in Milton Keynes and Alex Alderton, Roberto Amato, Jeffrey Barrett, Sonia Goncalves, Ewan Harrison, David K. Jackson, Ian Johnston, Dominic Kwiatkowski, Cordelia Langford, John Sillitoe on behalf of the Wellcome Sanger Institute COVID-19 Surveillance Team |
| EPI_ISL_2995722 | Lighthouse Lab in Glasgow | Wellcome Sanger Institute for the COVID-19 Genomics UK (COG-UK) Consortium | Harper VanSteenhouse, Yumi Kasai, David Gray, Carol Clugston, Anna Dominiczak and Alex Alderton, Roberto Amato, Jeffrey Barrett, Sonia Goncalves, Ewan Harrison, David K. Jackson, Ian Johnston, Dominic Kwiatkowski, Cordelia Langford, John Sillitoe on behalf of the Wellcome Sanger Institute COVID-19 Surveillance Team |
| EPI_ISL_2995723, EPI_ISL_2995724 | Berkshire and Surrey Pathology Services Lighthouse Laboratory | Wellcome Sanger Institute for the COVID-19 Genomics UK (COG-UK) Consortium | Berkshire and Surrey Pathology Services Lighthouse Laboratory and Alex Alderton, Roberto Amato, Jeffrey Barrett, Sonia Goncalves, Ewan Harrison, David K. Jackson, Ian Johnston, Dominic Kwiatkowski, Cordelia Langford, John Sillitoe on behalf of the Wellcome Sanger Institute COVID-19 Surveillance Team |
| EPI_ISL_2995725 | Lighthouse Lab in Milton Keynes | Wellcome Sanger Institute for the COVID-19 Genomics UK (COG-UK) Consortium | The Lighthouse Lab in Milton Keynes and Alex Alderton, Roberto Amato, Jeffrey Barrett, Sonia Goncalves, Ewan Harrison, David K. Jackson, Ian Johnston, Dominic Kwiatkowski, Cordelia Langford, John Sillitoe on behalf of the Wellcome Sanger Institute COVID-19 Surveillance Team |
| EPI_ISL_2995726, EPI_ISL_2995727 | Berkshire and Surrey Pathology Services Lighthouse Laboratory | Wellcome Sanger Institute for the COVID-19 Genomics UK (COG-UK) Consortium | Berkshire and Surrey Pathology Services Lighthouse Laboratory and Alex Alderton, Roberto Amato, Jeffrey Barrett, Sonia Goncalves, Ewan Harrison, David K. Jackson, Ian Johnston, Dominic Kwiatkowski, Cordelia Langford, John Sillitoe on behalf of the Wellcome Sanger Institute COVID-19 Surveillance Team |
| EPI_ISL_2995728 | Lighthouse Lab in Glasgow | Wellcome Sanger Institute for the COVID-19 Genomics UK (COG-UK) Consortium | Harper VanSteenhouse, Yumi Kasai, David Gray, Carol Clugston, Anna Dominiczak and Alex Alderton, Roberto Amato, Jeffrey Barrett, Sonia Goncalves, Ewan Harrison, David K. Jackson, Ian Johnston, Dominic Kwiatkowski, Cordelia Langford, John Sillitoe on behalf of the Wellcome Sanger Institute COVID-19 Surveillance Team |
| EPI_ISL_2995729 | Berkshire and Surrey Pathology Services Lighthouse Laboratory | Wellcome Sanger Institute for the COVID-19 Genomics UK (COG-UK) Consortium | Berkshire and Surrey Pathology Services Lighthouse Laboratory and Alex Alderton, Roberto Amato, Jeffrey Barrett, Sonia Goncalves, Ewan Harrison, David K. Jackson, Ian Johnston, Dominic Kwiatkowski, Cordelia Langford, John Sillitoe on behalf of the Wellcome Sanger Institute COVID-19 Surveillance Team |
| EPI_ISL_2995730 | Lighthouse Lab in Glasgow | Wellcome Sanger Institute for the COVID-19 Genomics UK (COG-UK) Consortium | Harper VanSteenhouse, Yumi Kasai, David Gray, Carol Clugston, Anna Dominiczak and Alex Alderton, Roberto Amato, Jeffrey Barrett, Sonia Goncalves, Ewan Harrison, David K. Jackson, Ian Johnston, Dominic Kwiatkowski, Cordelia Langford, John Sillitoe on behalf of the Wellcome Sanger Institute COVID-19 Surveillance Team |
| EPI_ISL_2995731 | Lighthouse Lab in Milton Keynes | Wellcome Sanger Institute for the COVID-19 Genomics UK (COG-UK) Consortium | The Lighthouse Lab in Milton Keynes and Alex Alderton, Roberto Amato, Jeffrey Barrett, Sonia Goncalves, Ewan Harrison, David K. Jackson, Ian Johnston, Dominic Kwiatkowski, Cordelia Langford, John Sillitoe on behalf of the Wellcome Sanger Institute COVID-19 Surveillance Team |
| EPI_ISL_2995732, EPI_ISL_2995733 | Berkshire and Surrey Pathology Services Lighthouse Laboratory | Wellcome Sanger Institute for the COVID-19 Genomics UK (COG-UK) Consortium | Berkshire and Surrey Pathology Services Lighthouse Laboratory and Alex Alderton, Roberto Amato, Jeffrey Barrett, Sonia Goncalves, Ewan Harrison, David K. Jackson, Ian Johnston, Dominic Kwiatkowski, Cordelia Langford, John Sillitoe on behalf of the Wellcome Sanger Institute COVID-19 Surveillance Team |
| EPI_ISL_2995734 | Lighthouse Lab in Milton Keynes | Wellcome Sanger Institute for the COVID-19 Genomics UK (COG-UK) Consortium | The Lighthouse Lab in Milton Keynes and Alex Alderton, Roberto Amato, Jeffrey Barrett, Sonia Goncalves, Ewan Harrison, David K. Jackson, Ian Johnston, Dominic Kwiatkowski, Cordelia Langford, John Sillitoe on behalf of the Wellcome Sanger Institute COVID-19 Surveillance Team |
| EPI_ISL_2995735, EPI_ISL_2995736 | Lighthouse Lab in Glasgow | Wellcome Sanger Institute for the COVID-19 Genomics UK (COG-UK) Consortium | Harper VanSteenhouse, Yumi Kasai, David Gray, Carol Clugston, Anna Dominiczak and Alex Alderton, Roberto Amato, Jeffrey Barrett, Sonia Goncalves, Ewan Harrison, David K. Jackson, Ian Johnston, Dominic Kwiatkowski, Cordelia Langford, John Sillitoe on behalf of the Wellcome Sanger Institute COVID-19 Surveillance Team |
| EPI_ISL_2995737, EPI_ISL_2995738 | Berkshire and Surrey Pathology Services Lighthouse Laboratory | Wellcome Sanger Institute for the COVID-19 Genomics UK (COG-UK) Consortium | Berkshire and Surrey Pathology Services Lighthouse Laboratory and Alex Alderton, Roberto Amato, Jeffrey Barrett, Sonia Goncalves, Ewan Harrison, David K. Jackson, Ian Johnston, Dominic Kwiatkowski, Cordelia Langford, John Sillitoe on behalf of the Wellcome Sanger Institute COVID-19 Surveillance Team |
| EPI_ISL_2995739 | Lighthouse Lab in Milton Keynes | Wellcome Sanger Institute for the COVID-19 Genomics UK (COG-UK) Consortium | The Lighthouse Lab in Milton Keynes and Alex Alderton, Roberto Amato, Jeffrey Barrett, Sonia Goncalves, Ewan Harrison, David K. Jackson, Ian Johnston, Dominic Kwiatkowski, Cordelia Langford, John Sillitoe on behalf of the Wellcome Sanger Institute COVID-19 Surveillance Team |
| EPI_ISL_2995740 | Berkshire and Surrey Pathology Services Lighthouse Laboratory | Wellcome Sanger Institute for the COVID-19 Genomics UK (COG-UK) Consortium | Berkshire and Surrey Pathology Services Lighthouse Laboratory and Alex Alderton, Roberto Amato, Jeffrey Barrett, Sonia Goncalves, Ewan Harrison, David K. Jackson, Ian Johnston, Dominic Kwiatkowski, Cordelia Langford, John Sillitoe on behalf of the Wellcome Sanger Institute COVID-19 Surveillance Team |
| EPI_ISL_2995741 | Lighthouse Lab in Milton Keynes | Wellcome Sanger Institute for the COVID-19 Genomics UK (COG-UK) Consortium | The Lighthouse Lab in Milton Keynes and Alex Alderton, Roberto Amato, Jeffrey Barrett, Sonia Goncalves, Ewan Harrison, David K. Jackson, Ian Johnston, Dominic Kwiatkowski, Cordelia Langford, John Sillitoe on behalf of the Wellcome Sanger Institute COVID-19 Surveillance Team |
| EPI_ISL_2995742, EPI_ISL_2995743, EPI_ISL_2995744 | Berkshire and Surrey Pathology Services Lighthouse Laboratory | Wellcome Sanger Institute for the COVID-19 Genomics UK (COG-UK) Consortium | Berkshire and Surrey Pathology Services Lighthouse Laboratory and Alex Alderton, Roberto Amato, Jeffrey Barrett, Sonia Goncalves, Ewan Harrison, David K. Jackson, Ian Johnston, Dominic Kwiatkowski, Cordelia Langford, John Sillitoe on behalf of the Wellcome Sanger Institute COVID-19 Surveillance Team |
| EPI_ISL_2995745, EPI_ISL_2995746 | Lighthouse Lab in Glasgow | Wellcome Sanger Institute for the COVID-19 Genomics UK (COG-UK) Consortium | Harper VanSteenhouse, Yumi Kasai, David Gray, Carol Clugston, Anna Dominiczak and Alex Alderton, Roberto Amato, Jeffrey Barrett, Sonia Goncalves, Ewan Harrison, David K. Jackson, Ian Johnston, Dominic Kwiatkowski, Cordelia Langford, John Sillitoe on behalf of the Wellcome Sanger Institute COVID-19 Surveillance Team |
| EPI_ISL_2995747, EPI_ISL_2995748 | Berkshire and Surrey Pathology Services Lighthouse Laboratory | Wellcome Sanger Institute for the COVID-19 Genomics UK (COG-UK) Consortium | Berkshire and Surrey Pathology Services Lighthouse Laboratory and Alex Alderton, Roberto Amato, Jeffrey Barrett, Sonia Goncalves, Ewan Harrison, David K. Jackson, Ian Johnston, Dominic Kwiatkowski, Cordelia Langford, John Sillitoe on behalf of the Wellcome Sanger Institute COVID-19 Surveillance Team |
| EPI_ISL_2995749 | Lighthouse Lab in Milton Keynes | Wellcome Sanger Institute for the COVID-19 Genomics UK (COG-UK) Consortium | The Lighthouse Lab in Milton Keynes and Alex Alderton, Roberto Amato, Jeffrey Barrett, Sonia Goncalves, Ewan Harrison, David K. Jackson, Ian Johnston, Dominic Kwiatkowski, Cordelia Langford, John Sillitoe on behalf of the Wellcome Sanger Institute COVID-19 Surveillance Team |
| EPI_ISL_2995750, EPI_ISL_2995751 | Berkshire and Surrey Pathology Services Lighthouse Laboratory | Wellcome Sanger Institute for the COVID-19 Genomics UK (COG-UK) Consortium | Berkshire and Surrey Pathology Services Lighthouse Laboratory and Alex Alderton, Roberto Amato, Jeffrey Barrett, Sonia Goncalves, Ewan Harrison, David K. Jackson, Ian Johnston, Dominic Kwiatkowski, Cordelia Langford, John Sillitoe on behalf of the Wellcome Sanger Institute COVID-19 Surveillance Team |
| EPI_ISL_2995752 | Lighthouse Lab in Milton Keynes | Wellcome Sanger Institute for the COVID-19 Genomics UK (COG-UK) Consortium | The Lighthouse Lab in Milton Keynes and Alex Alderton, Roberto Amato, Jeffrey Barrett, Sonia Goncalves, Ewan Harrison, David K. Jackson, Ian Johnston, Dominic Kwiatkowski, Cordelia Langford, John Sillitoe on behalf of the Wellcome Sanger Institute COVID-19 Surveillance Team |
| EPI_ISL_2995753 | Berkshire and Surrey Pathology Services Lighthouse Laboratory | Wellcome Sanger Institute for the COVID-19 Genomics UK (COG-UK) Consortium | Berkshire and Surrey Pathology Services Lighthouse Laboratory and Alex Alderton, Roberto Amato, Jeffrey Barrett, Sonia Goncalves, Ewan Harrison, David K. Jackson, Ian Johnston, Dominic Kwiatkowski, Cordelia Langford, John Sillitoe on behalf of the Wellcome Sanger Institute COVID-19 Surveillance Team |
| EPI_ISL_2995754 | Lighthouse Lab in Milton Keynes | Wellcome Sanger Institute for the COVID-19 Genomics UK (COG-UK) | The Lighthouse Lab in Milton Keynes and Alex Alderton, Roberto Amato, Jeffrey Barrett, Sonia Goncalves, Ewan Harrison, David K. Jackson, Ian Johnston, Dominic Kwiatkowski, Cordelia Langford, John Sillitoe on behalf of the Wellcome Sanger |

| | | | |
|---|---|---|---|
| | | Consortium | Institute COVID-19 Surveillance Team |
| EPI_ISL_2995755 | Lighthouse Lab in Glasgow | Wellcome Sanger Institute for the COVID-19 Genomics UK (COG-UK) Consortium | Harper VanSteenhouse, Yumi Kasai, David Gray, Carol Clugston, Anna Dominiczak and Alex Alderton, Roberto Amato, Jeffrey Barrett, Sonia Goncalves, Ewan Harrison, David K. Jackson, Ian Johnston, Dominic Kwiatkowski, Cordelia Langford, John Sillitoe on behalf of the Wellcome Sanger Institute COVID-19 Surveillance Team |
| EPI_ISL_2995756 | Lighthouse Lab in Milton Keynes | Wellcome Sanger Institute for the COVID-19 Genomics UK (COG-UK) Consortium | The Lighthouse Lab in Milton Keynes and Alex Alderton, Roberto Amato, Jeffrey Barrett, Sonia Goncalves, Ewan Harrison, David K. Jackson, Ian Johnston, Dominic Kwiatkowski, Cordelia Langford, John Sillitoe on behalf of the Wellcome Sanger Institute COVID-19 Surveillance Team |
| EPI_ISL_2995757 | Lighthouse Lab in Glasgow | Wellcome Sanger Institute for the COVID-19 Genomics UK (COG-UK) Consortium | Harper VanSteenhouse, Yumi Kasai, David Gray, Carol Clugston, Anna Dominiczak and Alex Alderton, Roberto Amato, Jeffrey Barrett, Sonia Goncalves, Ewan Harrison, David K. Jackson, Ian Johnston, Dominic Kwiatkowski, Cordelia Langford, John Sillitoe on behalf of the Wellcome Sanger Institute COVID-19 Surveillance Team |
| EPI_ISL_2995758, EPI_ISL_2995759 | Berkshire and Surrey Pathology Services Lighthouse Laboratory | Wellcome Sanger Institute for the COVID-19 Genomics UK (COG-UK) Consortium | Berkshire and Surrey Pathology Services Lighthouse Laboratory and Alex Alderton, Roberto Amato, Jeffrey Barrett, Sonia Goncalves, Ewan Harrison, David K. Jackson, Ian Johnston, Dominic Kwiatkowski, Cordelia Langford, John Sillitoe on behalf of the Wellcome Sanger Institute COVID-19 Surveillance Team |
| EPI_ISL_2995760 | Lighthouse Lab in Glasgow | Wellcome Sanger Institute for the COVID-19 Genomics UK (COG-UK) Consortium | Harper VanSteenhouse, Yumi Kasai, David Gray, Carol Clugston, Anna Dominiczak and Alex Alderton, Roberto Amato, Jeffrey Barrett, Sonia Goncalves, Ewan Harrison, David K. Jackson, Ian Johnston, Dominic Kwiatkowski, Cordelia Langford, John Sillitoe on behalf of the Wellcome Sanger Institute COVID-19 Surveillance Team |
| EPI_ISL_2995761, EPI_ISL_2995762, EPI_ISL_2995763 | Berkshire and Surrey Pathology Services Lighthouse Laboratory | Wellcome Sanger Institute for the COVID-19 Genomics UK (COG-UK) Consortium | Berkshire and Surrey Pathology Services Lighthouse Laboratory and Alex Alderton, Roberto Amato, Jeffrey Barrett, Sonia Goncalves, Ewan Harrison, David K. Jackson, Ian Johnston, Dominic Kwiatkowski, Cordelia Langford, John Sillitoe on behalf of the Wellcome Sanger Institute COVID-19 Surveillance Team |
| EPI_ISL_2995764 | Lighthouse Lab in Milton Keynes | Wellcome Sanger Institute for the COVID-19 Genomics UK (COG-UK) Consortium | The Lighthouse Lab in Milton Keynes and Alex Alderton, Roberto Amato, Jeffrey Barrett, Sonia Goncalves, Ewan Harrison, David K. Jackson, Ian Johnston, Dominic Kwiatkowski, Cordelia Langford, John Sillitoe on behalf of the Wellcome Sanger Institute COVID-19 Surveillance Team |
| EPI_ISL_2995765, EPI_ISL_2995766 | Berkshire and Surrey Pathology Services Lighthouse Laboratory | Wellcome Sanger Institute for the COVID-19 Genomics UK (COG-UK) Consortium | Berkshire and Surrey Pathology Services Lighthouse Laboratory and Alex Alderton, Roberto Amato, Jeffrey Barrett, Sonia Goncalves, Ewan Harrison, David K. Jackson, Ian Johnston, Dominic Kwiatkowski, Cordelia Langford, John Sillitoe on behalf of the Wellcome Sanger Institute COVID-19 Surveillance Team |
| EPI_ISL_2995767 | Lighthouse Lab in Glasgow | Wellcome Sanger Institute for the COVID-19 Genomics UK (COG-UK) Consortium | Harper VanSteenhouse, Yumi Kasai, David Gray, Carol Clugston, Anna Dominiczak and Alex Alderton, Roberto Amato, Jeffrey Barrett, Sonia Goncalves, Ewan Harrison, David K. Jackson, Ian Johnston, Dominic Kwiatkowski, Cordelia Langford, John Sillitoe on behalf of the Wellcome Sanger Institute COVID-19 Surveillance Team |
| EPI_ISL_2995768 | Lighthouse Lab in Milton Keynes | Wellcome Sanger Institute for the COVID-19 Genomics UK (COG-UK) Consortium | The Lighthouse Lab in Milton Keynes and Alex Alderton, Roberto Amato, Jeffrey Barrett, Sonia Goncalves, Ewan Harrison, David K. Jackson, Ian Johnston, Dominic Kwiatkowski, Cordelia Langford, John Sillitoe on behalf of the Wellcome Sanger Institute COVID-19 Surveillance Team |
| EPI_ISL_2995769, EPI_ISL_2995770 | Lighthouse Lab in Glasgow | Wellcome Sanger Institute for the COVID-19 Genomics UK (COG-UK) Consortium | Harper VanSteenhouse, Yumi Kasai, David Gray, Carol Clugston, Anna Dominiczak and Alex Alderton, Roberto Amato, Jeffrey Barrett, Sonia Goncalves, Ewan Harrison, David K. Jackson, Ian Johnston, Dominic Kwiatkowski, Cordelia Langford, John Sillitoe on behalf of the Wellcome Sanger Institute COVID-19 Surveillance Team |
| EPI_ISL_2995771 | Lighthouse Lab in Milton Keynes | Wellcome Sanger Institute for the COVID-19 Genomics UK (COG-UK) Consortium | The Lighthouse Lab in Milton Keynes and Alex Alderton, Roberto Amato, Jeffrey Barrett, Sonia Goncalves, Ewan Harrison, David K. Jackson, Ian Johnston, Dominic Kwiatkowski, Cordelia Langford, John Sillitoe on behalf of the Wellcome Sanger Institute COVID-19 Surveillance Team |
| EPI_ISL_2995772, EPI_ISL_2995773 | Berkshire and Surrey Pathology Services Lighthouse Laboratory | Wellcome Sanger Institute for the COVID-19 Genomics UK (COG-UK) Consortium | Berkshire and Surrey Pathology Services Lighthouse Laboratory and Alex Alderton, Roberto Amato, Jeffrey Barrett, Sonia Goncalves, Ewan Harrison, David K. Jackson, Ian Johnston, Dominic Kwiatkowski, Cordelia Langford, John Sillitoe on behalf of the Wellcome Sanger Institute COVID-19 Surveillance Team |
| EPI_ISL_2995774 | Lighthouse Lab in Glasgow | Wellcome Sanger Institute for the COVID-19 Genomics UK (COG-UK) Consortium | Harper VanSteenhouse, Yumi Kasai, David Gray, Carol Clugston, Anna Dominiczak and Alex Alderton, Roberto Amato, Jeffrey Barrett, Sonia Goncalves, Ewan Harrison, David K. Jackson, Ian Johnston, Dominic Kwiatkowski, Cordelia Langford, John Sillitoe on behalf of the Wellcome Sanger Institute COVID-19 Surveillance Team |
| EPI_ISL_2995775 | Lighthouse Lab in Milton Keynes | Wellcome Sanger Institute for the COVID-19 Genomics UK (COG-UK) Consortium | The Lighthouse Lab in Milton Keynes and Alex Alderton, Roberto Amato, Jeffrey Barrett, Sonia Goncalves, Ewan Harrison, David K. Jackson, Ian Johnston, Dominic Kwiatkowski, Cordelia Langford, John Sillitoe on behalf of the Wellcome Sanger Institute COVID-19 Surveillance Team |
| EPI_ISL_2995776 | Berkshire and Surrey Pathology Services Lighthouse Laboratory | Wellcome Sanger Institute for the COVID-19 Genomics UK (COG-UK) Consortium | Berkshire and Surrey Pathology Services Lighthouse Laboratory and Alex Alderton, Roberto Amato, Jeffrey Barrett, Sonia Goncalves, Ewan Harrison, David K. Jackson, Ian Johnston, Dominic Kwiatkowski, Cordelia Langford, John Sillitoe on behalf of the Wellcome Sanger Institute COVID-19 Surveillance Team |
| EPI_ISL_2995777, EPI_ISL_2995778 | Lighthouse Lab in Glasgow | Wellcome Sanger Institute for the COVID-19 Genomics UK (COG-UK) Consortium | Harper VanSteenhouse, Yumi Kasai, David Gray, Carol Clugston, Anna Dominiczak and Alex Alderton, Roberto Amato, Jeffrey Barrett, Sonia Goncalves, Ewan Harrison, David K. Jackson, Ian Johnston, Dominic Kwiatkowski, Cordelia Langford, John Sillitoe on behalf of the Wellcome Sanger Institute COVID-19 Surveillance Team |
| EPI_ISL_2995779 | Lighthouse Lab in Milton Keynes | Wellcome Sanger Institute for the COVID-19 Genomics UK (COG-UK) Consortium | The Lighthouse Lab in Milton Keynes and Alex Alderton, Roberto Amato, Jeffrey Barrett, Sonia Goncalves, Ewan Harrison, David K. Jackson, Ian Johnston, Dominic Kwiatkowski, Cordelia Langford, John Sillitoe on behalf of the Wellcome Sanger Institute COVID-19 Surveillance Team |
| EPI_ISL_2995780 | Lighthouse Lab in Glasgow | Wellcome Sanger Institute for the COVID-19 Genomics UK (COG-UK) Consortium | Harper VanSteenhouse, Yumi Kasai, David Gray, Carol Clugston, Anna Dominiczak and Alex Alderton, Roberto Amato, Jeffrey Barrett, Sonia Goncalves, Ewan Harrison, David K. Jackson, Ian Johnston, Dominic Kwiatkowski, Cordelia Langford, John Sillitoe on behalf of the Wellcome Sanger Institute COVID-19 Surveillance Team |
| EPI_ISL_2995781 | Lighthouse Lab in Milton Keynes | Wellcome Sanger Institute for the COVID-19 Genomics UK (COG-UK) Consortium | The Lighthouse Lab in Milton Keynes and Alex Alderton, Roberto Amato, Jeffrey Barrett, Sonia Goncalves, Ewan Harrison, David K. Jackson, Ian Johnston, Dominic Kwiatkowski, Cordelia Langford, John Sillitoe on behalf of the Wellcome Sanger Institute COVID-19 Surveillance Team |
| EPI_ISL_2995782 | Lighthouse Lab in Glasgow | Wellcome Sanger Institute for the COVID-19 Genomics UK (COG-UK) Consortium | Harper VanSteenhouse, Yumi Kasai, David Gray, Carol Clugston, Anna Dominiczak and Alex Alderton, Roberto Amato, Jeffrey Barrett, Sonia Goncalves, Ewan Harrison, David K. Jackson, Ian Johnston, Dominic Kwiatkowski, Cordelia Langford, John Sillitoe on behalf of the Wellcome Sanger Institute COVID-19 Surveillance Team |
| EPI_ISL_2995783 | Lighthouse Lab in Milton Keynes | Wellcome Sanger Institute for the COVID-19 Genomics UK (COG-UK) Consortium | The Lighthouse Lab in Milton Keynes and Alex Alderton, Roberto Amato, Jeffrey Barrett, Sonia Goncalves, Ewan Harrison, David K. Jackson, Ian Johnston, Dominic Kwiatkowski, Cordelia Langford, John Sillitoe on behalf of the Wellcome Sanger Institute COVID-19 Surveillance Team |
| EPI_ISL_2995784 | Berkshire and Surrey Pathology Services Lighthouse Laboratory | Wellcome Sanger Institute for the COVID-19 Genomics UK (COG-UK) Consortium | Berkshire and Surrey Pathology Services Lighthouse Laboratory and Alex Alderton, Roberto Amato, Jeffrey Barrett, Sonia Goncalves, Ewan Harrison, David K. Jackson, Ian Johnston, Dominic Kwiatkowski, Cordelia Langford, John Sillitoe on behalf of the Wellcome Sanger Institute COVID-19 Surveillance Team |
| EPI_ISL_2995785, EPI_ISL_2995786, EPI_ISL_2995787 | Lighthouse Lab in Glasgow | Wellcome Sanger Institute for the COVID-19 Genomics UK (COG-UK) Consortium | Harper VanSteenhouse, Yumi Kasai, David Gray, Carol Clugston, Anna Dominiczak and Alex Alderton, Roberto Amato, Jeffrey Barrett, Sonia Goncalves, Ewan Harrison, David K. Jackson, Ian Johnston, Dominic Kwiatkowski, Cordelia Langford, John Sillitoe on behalf of the Wellcome Sanger Institute COVID-19 Surveillance Team |
| EPI_ISL_2995788 | Berkshire and Surrey Pathology Services Lighthouse Laboratory | Wellcome Sanger Institute for the COVID-19 Genomics UK (COG-UK) Consortium | Berkshire and Surrey Pathology Services Lighthouse Laboratory and Alex Alderton, Roberto Amato, Jeffrey Barrett, Sonia Goncalves, Ewan Harrison, David K. Jackson, Ian Johnston, Dominic Kwiatkowski, Cordelia Langford, John Sillitoe on behalf of the Wellcome Sanger Institute COVID-19 Surveillance Team |
| EPI_ISL_2995789 | Lighthouse Lab in Glasgow | Wellcome Sanger Institute for the COVID-19 Genomics UK (COG-UK) Consortium | Harper VanSteenhouse, Yumi Kasai, David Gray, Carol Clugston, Anna Dominiczak and Alex Alderton, Roberto Amato, Jeffrey Barrett, Sonia Goncalves, Ewan Harrison, David K. Jackson, Ian Johnston, Dominic Kwiatkowski, Cordelia Langford, John Sillitoe on behalf of the Wellcome Sanger Institute COVID-19 Surveillance Team |
| EPI_ISL_2995790 | Lighthouse Lab in Milton Keynes | Wellcome Sanger Institute for the COVID-19 Genomics UK (COG-UK) Consortium | The Lighthouse Lab in Milton Keynes and Alex Alderton, Roberto Amato, Jeffrey Barrett, Sonia Goncalves, Ewan Harrison, David K. Jackson, Ian Johnston, Dominic Kwiatkowski, Cordelia Langford, John Sillitoe on behalf of the Wellcome Sanger Institute COVID-19 Surveillance Team |
| EPI_ISL_2995791 | Berkshire and Surrey Pathology Services Lighthouse Laboratory | Wellcome Sanger Institute for the COVID-19 Genomics UK (COG-UK) Consortium | Berkshire and Surrey Pathology Services Lighthouse Laboratory and Alex Alderton, Roberto Amato, Jeffrey Barrett, Sonia Goncalves, Ewan Harrison, David K. Jackson, Ian Johnston, Dominic Kwiatkowski, Cordelia Langford, John Sillitoe on behalf of the Wellcome Sanger Institute COVID-19 Surveillance Team |
| EPI_ISL_2995792 | Lighthouse Lab in Glasgow | Wellcome Sanger Institute for the COVID-19 Genomics UK (COG-UK) Consortium | Harper VanSteenhouse, Yumi Kasai, David Gray, Carol Clugston, Anna Dominiczak and Alex Alderton, Roberto Amato, Jeffrey Barrett, Sonia Goncalves, Ewan Harrison, David K. Jackson, Ian Johnston, Dominic Kwiatkowski, Cordelia Langford, John Sillitoe on behalf of the Wellcome Sanger Institute COVID-19 Surveillance Team |
| EPI_ISL_2995793 | Berkshire and Surrey Pathology Services Lighthouse Laboratory | Wellcome Sanger Institute for the COVID-19 Genomics UK (COG-UK) Consortium | Berkshire and Surrey Pathology Services Lighthouse Laboratory and Alex Alderton, Roberto Amato, Jeffrey Barrett, Sonia Goncalves, Ewan Harrison, David K. Jackson, Ian Johnston, Dominic Kwiatkowski, Cordelia Langford, John Sillitoe on behalf of the Wellcome Sanger Institute COVID-19 Surveillance Team |
| EPI_ISL_2995794 | Lighthouse Lab in Glasgow | Wellcome Sanger Institute for the COVID-19 Genomics UK (COG-UK) Consortium | Harper VanSteenhouse, Yumi Kasai, David Gray, Carol Clugston, Anna Dominiczak and Alex Alderton, Roberto Amato, Jeffrey Barrett, Sonia Goncalves, Ewan Harrison, David K. Jackson, Ian Johnston, Dominic Kwiatkowski, Cordelia Langford, John Sillitoe on behalf of the Wellcome Sanger Institute COVID-19 Surveillance Team |
| EPI_ISL_2995795 | Berkshire and Surrey Pathology Services Lighthouse Laboratory | Wellcome Sanger Institute for the COVID-19 Genomics UK (COG-UK) Consortium | Berkshire and Surrey Pathology Services Lighthouse Laboratory and Alex Alderton, Roberto Amato, Jeffrey Barrett, Sonia Goncalves, Ewan Harrison, David K. Jackson, Ian Johnston, Dominic Kwiatkowski, Cordelia Langford, John Sillitoe on behalf of the Wellcome Sanger Institute COVID-19 Surveillance Team |
| EPI_ISL_2995796 | Lighthouse Lab in Milton Keynes | Wellcome Sanger Institute for the COVID-19 Genomics UK (COG-UK) Consortium | The Lighthouse Lab in Milton Keynes and Alex Alderton, Roberto Amato, Jeffrey Barrett, Sonia Goncalves, Ewan Harrison, David K. Jackson, Ian Johnston, Dominic Kwiatkowski, Cordelia Langford, John Sillitoe on behalf of the Wellcome Sanger Institute COVID-19 Surveillance Team |
| EPI_ISL_2995797, EPI_ISL_2995798, EPI_ISL_2995799, EPI_ISL_2995800, EPI_ISL_2995801 | Berkshire and Surrey Pathology Services Lighthouse Laboratory | Wellcome Sanger Institute for the COVID-19 Genomics UK (COG-UK) Consortium | Berkshire and Surrey Pathology Services Lighthouse Laboratory and Alex Alderton, Roberto Amato, Jeffrey Barrett, Sonia Goncalves, Ewan Harrison, David K. Jackson, Ian Johnston, Dominic Kwiatkowski, Cordelia Langford, John Sillitoe on behalf of the Wellcome Sanger Institute COVID-19 Surveillance Team |
| EPI_ISL_2995802, EPI_ISL_2995803 | Lighthouse Lab in Milton Keynes | Wellcome Sanger Institute for the COVID-19 Genomics UK (COG-UK) Consortium | The Lighthouse Lab in Milton Keynes and Alex Alderton, Roberto Amato, Jeffrey Barrett, Sonia Goncalves, Ewan Harrison, David K. Jackson, Ian Johnston, Dominic Kwiatkowski, Cordelia Langford, John Sillitoe on behalf of the Wellcome Sanger Institute COVID-19 Surveillance Team |
| EPI_ISL_2995804, EPI_ISL_2995805 | Berkshire and Surrey Pathology Services Lighthouse Laboratory | Wellcome Sanger Institute for the COVID-19 Genomics UK (COG-UK) Consortium | Berkshire and Surrey Pathology Services Lighthouse Laboratory and Alex Alderton, Roberto Amato, Jeffrey Barrett, Sonia Goncalves, Ewan Harrison, David K. Jackson, Ian Johnston, Dominic Kwiatkowski, Cordelia Langford, John Sillitoe on behalf of the Wellcome Sanger Institute COVID-19 Surveillance Team |
| EPI_ISL_2995806 | Lighthouse Lab in Glasgow | Wellcome Sanger Institute for the COVID-19 Genomics UK (COG-UK) Consortium | Harper VanSteenhouse, Yumi Kasai, David Gray, Carol Clugston, Anna Dominiczak and Alex Alderton, Roberto Amato, Jeffrey Barrett, Sonia Goncalves, Ewan Harrison, David K. Jackson, Ian Johnston, Dominic Kwiatkowski, Cordelia Langford, John Sillitoe on behalf of the Wellcome Sanger Institute COVID-19 Surveillance Team |
| EPI_ISL_2995807, EPI_ISL_2995808, EPI_ISL_2995809, EPI_ISL_2995810, EPI_ISL_2995811, EPI_ISL_2995812 | Berkshire and Surrey Pathology Services Lighthouse Laboratory | Wellcome Sanger Institute for the COVID-19 Genomics UK (COG-UK) Consortium | Berkshire and Surrey Pathology Services Lighthouse Laboratory and Alex Alderton, Roberto Amato, Jeffrey Barrett, Sonia Goncalves, Ewan Harrison, David K. Jackson, Ian Johnston, Dominic Kwiatkowski, Cordelia Langford, John Sillitoe on behalf of the Wellcome Sanger Institute COVID-19 Surveillance Team |
| EPI_ISL_2995813, EPI_ISL_2995814, EPI_ISL_2995815 | Lighthouse Lab in Glasgow | Wellcome Sanger Institute for the COVID-19 Genomics UK (COG-UK) Consortium | Harper VanSteenhouse, Yumi Kasai, David Gray, Carol Clugston, Anna Dominiczak and Alex Alderton, Roberto Amato, Jeffrey Barrett, Sonia Goncalves, Ewan Harrison, David K. Jackson, Ian Johnston, Dominic Kwiatkowski, Cordelia Langford, John Sillitoe on behalf of the Wellcome Sanger Institute COVID-19 Surveillance Team |
| EPI_ISL_2995816, EPI_ISL_2995817 | Berkshire and Surrey Pathology Services Lighthouse Laboratory | Wellcome Sanger Institute for the COVID-19 Genomics UK (COG-UK) Consortium | Berkshire and Surrey Pathology Services Lighthouse Laboratory and Alex Alderton, Roberto Amato, Jeffrey Barrett, Sonia Goncalves, Ewan Harrison, David K. Jackson, Ian Johnston, Dominic Kwiatkowski, Cordelia Langford, John Sillitoe on behalf of the Wellcome Sanger Institute COVID-19 Surveillance Team |

| | | | |
|---|---|---|---|
| EPI_ISL_2995818 | Lighthouse Lab in Glasgow | Wellcome Sanger Institute for the COVID-19 Genomics UK (COG-UK) Consortium | Harper VanSteenhouse, Yumi Kasai, David Gray, Carol Clugston, Anna Dominiczak and Alex Alderton, Roberto Amato, Jeffrey Barrett, Sonia Goncalves, Ewan Harrison, David K. Jackson, Ian Johnston, Dominic Kwiatkowski, Cordelia Langford, John Sillitoe on behalf of the Wellcome Sanger Institute COVID-19 Surveillance Team |
| EPI_ISL_2995819 | Lighthouse Lab in Milton Keynes | Wellcome Sanger Institute for the COVID-19 Genomics UK (COG-UK) Consortium | The Lighthouse Lab in Milton Keynes and Alex Alderton, Roberto Amato, Jeffrey Barrett, Sonia Goncalves, Ewan Harrison, David K. Jackson, Ian Johnston, Dominic Kwiatkowski, Cordelia Langford, John Sillitoe on behalf of the Wellcome Sanger Institute COVID-19 Surveillance Team |
| EPI_ISL_2995820 | Berkshire and Surrey Pathology Services Lighthouse Laboratory | Wellcome Sanger Institute for the COVID-19 Genomics UK (COG-UK) Consortium | Berkshire and Surrey Pathology Services Lighthouse Laboratory and Alex Alderton, Roberto Amato, Jeffrey Barrett, Sonia Goncalves, Ewan Harrison, David K. Jackson, Ian Johnston, Dominic Kwiatkowski, Cordelia Langford, John Sillitoe on behalf of the Wellcome Sanger Institute COVID-19 Surveillance Team |
| EPI_ISL_2995821 | Lighthouse Lab in Glasgow | Wellcome Sanger Institute for the COVID-19 Genomics UK (COG-UK) Consortium | Harper VanSteenhouse, Yumi Kasai, David Gray, Carol Clugston, Anna Dominiczak and Alex Alderton, Roberto Amato, Jeffrey Barrett, Sonia Goncalves, Ewan Harrison, David K. Jackson, Ian Johnston, Dominic Kwiatkowski, Cordelia Langford, John Sillitoe on behalf of the Wellcome Sanger Institute COVID-19 Surveillance Team |
| EPI_ISL_2995822 | Berkshire and Surrey Pathology Services Lighthouse Laboratory | Wellcome Sanger Institute for the COVID-19 Genomics UK (COG-UK) Consortium | Berkshire and Surrey Pathology Services Lighthouse Laboratory and Alex Alderton, Roberto Amato, Jeffrey Barrett, Sonia Goncalves, Ewan Harrison, David K. Jackson, Ian Johnston, Dominic Kwiatkowski, Cordelia Langford, John Sillitoe on behalf of the Wellcome Sanger Institute COVID-19 Surveillance Team |
| EPI_ISL_2995823 | Lighthouse Lab in Milton Keynes | Wellcome Sanger Institute for the COVID-19 Genomics UK (COG-UK) Consortium | The Lighthouse Lab in Milton Keynes and Alex Alderton, Roberto Amato, Jeffrey Barrett, Sonia Goncalves, Ewan Harrison, David K. Jackson, Ian Johnston, Dominic Kwiatkowski, Cordelia Langford, John Sillitoe on behalf of the Wellcome Sanger Institute COVID-19 Surveillance Team |
| EPI_ISL_2995824 | Berkshire and Surrey Pathology Services Lighthouse Laboratory | Wellcome Sanger Institute for the COVID-19 Genomics UK (COG-UK) Consortium | Berkshire and Surrey Pathology Services Lighthouse Laboratory and Alex Alderton, Roberto Amato, Jeffrey Barrett, Sonia Goncalves, Ewan Harrison, David K. Jackson, Ian Johnston, Dominic Kwiatkowski, Cordelia Langford, John Sillitoe on behalf of the Wellcome Sanger Institute COVID-19 Surveillance Team |
| EPI_ISL_2995825 | Lighthouse Lab in Milton Keynes | Wellcome Sanger Institute for the COVID-19 Genomics UK (COG-UK) Consortium | The Lighthouse Lab in Milton Keynes and Alex Alderton, Roberto Amato, Jeffrey Barrett, Sonia Goncalves, Ewan Harrison, David K. Jackson, Ian Johnston, Dominic Kwiatkowski, Cordelia Langford, John Sillitoe on behalf of the Wellcome Sanger Institute COVID-19 Surveillance Team |
| EPI_ISL_2995826 | Lighthouse Lab in Glasgow | Wellcome Sanger Institute for the COVID-19 Genomics UK (COG-UK) Consortium | Harper VanSteenhouse, Yumi Kasai, David Gray, Carol Clugston, Anna Dominiczak and Alex Alderton, Roberto Amato, Jeffrey Barrett, Sonia Goncalves, Ewan Harrison, David K. Jackson, Ian Johnston, Dominic Kwiatkowski, Cordelia Langford, John Sillitoe on behalf of the Wellcome Sanger Institute COVID-19 Surveillance Team |
| EPI_ISL_2995827 | Berkshire and Surrey Pathology Services Lighthouse Laboratory | Wellcome Sanger Institute for the COVID-19 Genomics UK (COG-UK) Consortium | Berkshire and Surrey Pathology Services Lighthouse Laboratory and Alex Alderton, Roberto Amato, Jeffrey Barrett, Sonia Goncalves, Ewan Harrison, David K. Jackson, Ian Johnston, Dominic Kwiatkowski, Cordelia Langford, John Sillitoe on behalf of the Wellcome Sanger Institute COVID-19 Surveillance Team |
| EPI_ISL_2995828 | Lighthouse Lab in Glasgow | Wellcome Sanger Institute for the COVID-19 Genomics UK (COG-UK) Consortium | Harper VanSteenhouse, Yumi Kasai, David Gray, Carol Clugston, Anna Dominiczak and Alex Alderton, Roberto Amato, Jeffrey Barrett, Sonia Goncalves, Ewan Harrison, David K. Jackson, Ian Johnston, Dominic Kwiatkowski, Cordelia Langford, John Sillitoe on behalf of the Wellcome Sanger Institute COVID-19 Surveillance Team |
| EPI_ISL_2995829 | Lighthouse Lab in Milton Keynes | Wellcome Sanger Institute for the COVID-19 Genomics UK (COG-UK) Consortium | The Lighthouse Lab in Milton Keynes and Alex Alderton, Roberto Amato, Jeffrey Barrett, Sonia Goncalves, Ewan Harrison, David K. Jackson, Ian Johnston, Dominic Kwiatkowski, Cordelia Langford, John Sillitoe on behalf of the Wellcome Sanger Institute COVID-19 Surveillance Team |
| EPI_ISL_2995830 | Berkshire and Surrey Pathology Services Lighthouse Laboratory | Wellcome Sanger Institute for the COVID-19 Genomics UK (COG-UK) Consortium | Berkshire and Surrey Pathology Services Lighthouse Laboratory and Alex Alderton, Roberto Amato, Jeffrey Barrett, Sonia Goncalves, Ewan Harrison, David K. Jackson, Ian Johnston, Dominic Kwiatkowski, Cordelia Langford, John Sillitoe on behalf of the Wellcome Sanger Institute COVID-19 Surveillance Team |
| EPI_ISL_2995831 | Lighthouse Lab in Milton Keynes | Wellcome Sanger Institute for the COVID-19 Genomics UK (COG-UK) Consortium | The Lighthouse Lab in Milton Keynes and Alex Alderton, Roberto Amato, Jeffrey Barrett, Sonia Goncalves, Ewan Harrison, David K. Jackson, Ian Johnston, Dominic Kwiatkowski, Cordelia Langford, John Sillitoe on behalf of the Wellcome Sanger Institute COVID-19 Surveillance Team |
| EPI_ISL_2995832 | Berkshire and Surrey Pathology Services Lighthouse Laboratory | Wellcome Sanger Institute for the COVID-19 Genomics UK (COG-UK) Consortium | Berkshire and Surrey Pathology Services Lighthouse Laboratory and Alex Alderton, Roberto Amato, Jeffrey Barrett, Sonia Goncalves, Ewan Harrison, David K. Jackson, Ian Johnston, Dominic Kwiatkowski, Cordelia Langford, John Sillitoe on behalf of the Wellcome Sanger Institute COVID-19 Surveillance Team |
| EPI_ISL_2995833, EPI_ISL_2995834 | Lighthouse Lab in Milton Keynes | Wellcome Sanger Institute for the COVID-19 Genomics UK (COG-UK) Consortium | The Lighthouse Lab in Milton Keynes and Alex Alderton, Roberto Amato, Jeffrey Barrett, Sonia Goncalves, Ewan Harrison, David K. Jackson, Ian Johnston, Dominic Kwiatkowski, Cordelia Langford, John Sillitoe on behalf of the Wellcome Sanger Institute COVID-19 Surveillance Team |
| EPI_ISL_2995835 | Berkshire and Surrey Pathology Services Lighthouse Laboratory | Wellcome Sanger Institute for the COVID-19 Genomics UK (COG-UK) Consortium | Berkshire and Surrey Pathology Services Lighthouse Laboratory and Alex Alderton, Roberto Amato, Jeffrey Barrett, Sonia Goncalves, Ewan Harrison, David K. Jackson, Ian Johnston, Dominic Kwiatkowski, Cordelia Langford, John Sillitoe on behalf of the Wellcome Sanger Institute COVID-19 Surveillance Team |
| EPI_ISL_2995836 | Lighthouse Lab in Glasgow | Wellcome Sanger Institute for the COVID-19 Genomics UK (COG-UK) Consortium | Harper VanSteenhouse, Yumi Kasai, David Gray, Carol Clugston, Anna Dominiczak and Alex Alderton, Roberto Amato, Jeffrey Barrett, Sonia Goncalves, Ewan Harrison, David K. Jackson, Ian Johnston, Dominic Kwiatkowski, Cordelia Langford, John Sillitoe on behalf of the Wellcome Sanger Institute COVID-19 Surveillance Team |
| EPI_ISL_2995837 | Berkshire and Surrey Pathology Services Lighthouse Laboratory | Wellcome Sanger Institute for the COVID-19 Genomics UK (COG-UK) Consortium | Berkshire and Surrey Pathology Services Lighthouse Laboratory and Alex Alderton, Roberto Amato, Jeffrey Barrett, Sonia Goncalves, Ewan Harrison, David K. Jackson, Ian Johnston, Dominic Kwiatkowski, Cordelia Langford, John Sillitoe on behalf of the Wellcome Sanger Institute COVID-19 Surveillance Team |
| EPI_ISL_2995838, EPI_ISL_2995839 | Lighthouse Lab in Glasgow | Wellcome Sanger Institute for the COVID-19 Genomics UK (COG-UK) Consortium | Harper VanSteenhouse, Yumi Kasai, David Gray, Carol Clugston, Anna Dominiczak and Alex Alderton, Roberto Amato, Jeffrey Barrett, Sonia Goncalves, Ewan Harrison, David K. Jackson, Ian Johnston, Dominic Kwiatkowski, Cordelia Langford, John Sillitoe on behalf of the Wellcome Sanger Institute COVID-19 Surveillance Team |
| EPI_ISL_2995840, EPI_ISL_2995841, EPI_ISL_2995842 | Berkshire and Surrey Pathology Services Lighthouse Laboratory | Wellcome Sanger Institute for the COVID-19 Genomics UK (COG-UK) Consortium | Berkshire and Surrey Pathology Services Lighthouse Laboratory and Alex Alderton, Roberto Amato, Jeffrey Barrett, Sonia Goncalves, Ewan Harrison, David K. Jackson, Ian Johnston, Dominic Kwiatkowski, Cordelia Langford, John Sillitoe on behalf of the Wellcome Sanger Institute COVID-19 Surveillance Team |
| EPI_ISL_2995843 | Lighthouse Lab in Glasgow | Wellcome Sanger Institute for the COVID-19 Genomics UK (COG-UK) Consortium | Harper VanSteenhouse, Yumi Kasai, David Gray, Carol Clugston, Anna Dominiczak and Alex Alderton, Roberto Amato, Jeffrey Barrett, Sonia Goncalves, Ewan Harrison, David K. Jackson, Ian Johnston, Dominic Kwiatkowski, Cordelia Langford, John Sillitoe on behalf of the Wellcome Sanger Institute COVID-19 Surveillance Team |
| EPI_ISL_2995844 | Lighthouse Lab in Milton Keynes | Wellcome Sanger Institute for the COVID-19 Genomics UK (COG-UK) Consortium | The Lighthouse Lab in Milton Keynes and Alex Alderton, Roberto Amato, Jeffrey Barrett, Sonia Goncalves, Ewan Harrison, David K. Jackson, Ian Johnston, Dominic Kwiatkowski, Cordelia Langford, John Sillitoe on behalf of the Wellcome Sanger Institute COVID-19 Surveillance Team |
| EPI_ISL_2995845 | Lighthouse Lab in Glasgow | Wellcome Sanger Institute for the COVID-19 Genomics UK (COG-UK) Consortium | Harper VanSteenhouse, Yumi Kasai, David Gray, Carol Clugston, Anna Dominiczak and Alex Alderton, Roberto Amato, Jeffrey Barrett, Sonia Goncalves, Ewan Harrison, David K. Jackson, Ian Johnston, Dominic Kwiatkowski, Cordelia Langford, John Sillitoe on behalf of the Wellcome Sanger Institute COVID-19 Surveillance Team |
| EPI_ISL_2995846 | Berkshire and Surrey Pathology Services Lighthouse Laboratory | Wellcome Sanger Institute for the COVID-19 Genomics UK (COG-UK) Consortium | Berkshire and Surrey Pathology Services Lighthouse Laboratory and Alex Alderton, Roberto Amato, Jeffrey Barrett, Sonia Goncalves, Ewan Harrison, David K. Jackson, Ian Johnston, Dominic Kwiatkowski, Cordelia Langford, John Sillitoe on behalf of the Wellcome Sanger Institute COVID-19 Surveillance Team |
| EPI_ISL_2995847 | Lighthouse Lab in Milton Keynes | Wellcome Sanger Institute for the COVID-19 Genomics UK (COG-UK) Consortium | The Lighthouse Lab in Milton Keynes and Alex Alderton, Roberto Amato, Jeffrey Barrett, Sonia Goncalves, Ewan Harrison, David K. Jackson, Ian Johnston, Dominic Kwiatkowski, Cordelia Langford, John Sillitoe on behalf of the Wellcome Sanger Institute COVID-19 Surveillance Team |
| EPI_ISL_2995848 | Berkshire and Surrey Pathology Services Lighthouse Laboratory | Wellcome Sanger Institute for the COVID-19 Genomics UK (COG-UK) Consortium | Berkshire and Surrey Pathology Services Lighthouse Laboratory and Alex Alderton, Roberto Amato, Jeffrey Barrett, Sonia Goncalves, Ewan Harrison, David K. Jackson, Ian Johnston, Dominic Kwiatkowski, Cordelia Langford, John Sillitoe on behalf of the Wellcome Sanger Institute COVID-19 Surveillance Team |
| EPI_ISL_2995849 | Lighthouse Lab in Milton Keynes | Wellcome Sanger Institute for the COVID-19 Genomics UK (COG-UK) Consortium | The Lighthouse Lab in Milton Keynes and Alex Alderton, Roberto Amato, Jeffrey Barrett, Sonia Goncalves, Ewan Harrison, David K. Jackson, Ian Johnston, Dominic Kwiatkowski, Cordelia Langford, John Sillitoe on behalf of the Wellcome Sanger Institute COVID-19 Surveillance Team |
| EPI_ISL_2995850 | Lighthouse Lab in Glasgow | Wellcome Sanger Institute for the COVID-19 Genomics UK (COG-UK) Consortium | Harper VanSteenhouse, Yumi Kasai, David Gray, Carol Clugston, Anna Dominiczak and Alex Alderton, Roberto Amato, Jeffrey Barrett, Sonia Goncalves, Ewan Harrison, David K. Jackson, Ian Johnston, Dominic Kwiatkowski, Cordelia Langford, John Sillitoe on behalf of the Wellcome Sanger Institute COVID-19 Surveillance Team |
| EPI_ISL_2995851 | Berkshire and Surrey Pathology Services Lighthouse Laboratory | Wellcome Sanger Institute for the COVID-19 Genomics UK (COG-UK) Consortium | Berkshire and Surrey Pathology Services Lighthouse Laboratory and Alex Alderton, Roberto Amato, Jeffrey Barrett, Sonia Goncalves, Ewan Harrison, David K. Jackson, Ian Johnston, Dominic Kwiatkowski, Cordelia Langford, John Sillitoe on behalf of the Wellcome Sanger Institute COVID-19 Surveillance Team |
| EPI_ISL_2995852 | Lighthouse Lab in Glasgow | Wellcome Sanger Institute for the COVID-19 Genomics UK (COG-UK) Consortium | Harper VanSteenhouse, Yumi Kasai, David Gray, Carol Clugston, Anna Dominiczak and Alex Alderton, Roberto Amato, Jeffrey Barrett, Sonia Goncalves, Ewan Harrison, David K. Jackson, Ian Johnston, Dominic Kwiatkowski, Cordelia Langford, John Sillitoe on behalf of the Wellcome Sanger Institute COVID-19 Surveillance Team |
| EPI_ISL_2995853 | Berkshire and Surrey Pathology Services Lighthouse Laboratory | Wellcome Sanger Institute for the COVID-19 Genomics UK (COG-UK) Consortium | Berkshire and Surrey Pathology Services Lighthouse Laboratory and Alex Alderton, Roberto Amato, Jeffrey Barrett, Sonia Goncalves, Ewan Harrison, David K. Jackson, Ian Johnston, Dominic Kwiatkowski, Cordelia Langford, John Sillitoe on behalf of the Wellcome Sanger Institute COVID-19 Surveillance Team |
| EPI_ISL_2995854, EPI_ISL_2995855, EPI_ISL_2995856 | Lighthouse Lab in Glasgow | Wellcome Sanger Institute for the COVID-19 Genomics UK (COG-UK) Consortium | Harper VanSteenhouse, Yumi Kasai, David Gray, Carol Clugston, Anna Dominiczak and Alex Alderton, Roberto Amato, Jeffrey Barrett, Sonia Goncalves, Ewan Harrison, David K. Jackson, Ian Johnston, Dominic Kwiatkowski, Cordelia Langford, John Sillitoe on behalf of the Wellcome Sanger Institute COVID-19 Surveillance Team |
| EPI_ISL_2995857, EPI_ISL_2995858 | Berkshire and Surrey Pathology Services Lighthouse Laboratory | Wellcome Sanger Institute for the COVID-19 Genomics UK (COG-UK) Consortium | Berkshire and Surrey Pathology Services Lighthouse Laboratory and Alex Alderton, Roberto Amato, Jeffrey Barrett, Sonia Goncalves, Ewan Harrison, David K. Jackson, Ian Johnston, Dominic Kwiatkowski, Cordelia Langford, John Sillitoe on behalf of the Wellcome Sanger Institute COVID-19 Surveillance Team |
| EPI_ISL_2995859 | Lighthouse Lab in Glasgow | Wellcome Sanger Institute for the COVID-19 Genomics UK (COG-UK) Consortium | Harper VanSteenhouse, Yumi Kasai, David Gray, Carol Clugston, Anna Dominiczak and Alex Alderton, Roberto Amato, Jeffrey Barrett, Sonia Goncalves, Ewan Harrison, David K. Jackson, Ian Johnston, Dominic Kwiatkowski, Cordelia Langford, John Sillitoe on behalf of the Wellcome Sanger Institute COVID-19 Surveillance Team |
| EPI_ISL_2995860 | Berkshire and Surrey Pathology Services Lighthouse Laboratory | Wellcome Sanger Institute for the COVID-19 Genomics UK (COG-UK) Consortium | Berkshire and Surrey Pathology Services Lighthouse Laboratory and Alex Alderton, Roberto Amato, Jeffrey Barrett, Sonia Goncalves, Ewan Harrison, David K. Jackson, Ian Johnston, Dominic Kwiatkowski, Cordelia Langford, John Sillitoe on behalf of the Wellcome Sanger Institute COVID-19 Surveillance Team |
| EPI_ISL_2995861 | Lighthouse Lab in Glasgow | Wellcome Sanger Institute for the COVID-19 Genomics UK (COG-UK) Consortium | Harper VanSteenhouse, Yumi Kasai, David Gray, Carol Clugston, Anna Dominiczak and Alex Alderton, Roberto Amato, Jeffrey Barrett, Sonia Goncalves, Ewan Harrison, David K. Jackson, Ian Johnston, Dominic Kwiatkowski, Cordelia Langford, John Sillitoe on behalf of the Wellcome Sanger Institute COVID-19 Surveillance Team |
| EPI_ISL_2995862, EPI_ISL_2995863, EPI_ISL_2995864, EPI_ISL_2995865, EPI_ISL_2995866 | Berkshire and Surrey Pathology Services Lighthouse Laboratory | Wellcome Sanger Institute for the COVID-19 Genomics UK (COG-UK) Consortium | Berkshire and Surrey Pathology Services Lighthouse Laboratory and Alex Alderton, Roberto Amato, Jeffrey Barrett, Sonia Goncalves, Ewan Harrison, David K. Jackson, Ian Johnston, Dominic Kwiatkowski, Cordelia Langford, John Sillitoe on behalf of the Wellcome Sanger Institute COVID-19 Surveillance Team |
| EPI_ISL_2995867 | Lighthouse Lab in Milton Keynes | Wellcome Sanger Institute for the COVID-19 Genomics UK (COG-UK) Consortium | The Lighthouse Lab in Milton Keynes and Alex Alderton, Roberto Amato, Jeffrey Barrett, Sonia Goncalves, Ewan Harrison, David K. Jackson, Ian Johnston, Dominic Kwiatkowski, Cordelia Langford, John Sillitoe on behalf of the Wellcome Sanger Institute COVID-19 Surveillance Team |
| EPI_ISL_2995868 | Berkshire and Surrey Pathology Services Lighthouse Laboratory | Wellcome Sanger Institute for the COVID-19 Genomics UK (COG-UK) Consortium | Berkshire and Surrey Pathology Services Lighthouse Laboratory and Alex Alderton, Roberto Amato, Jeffrey Barrett, Sonia Goncalves, Ewan Harrison, David K. Jackson, Ian Johnston, Dominic Kwiatkowski, Cordelia Langford, John Sillitoe on behalf of the Wellcome Sanger Institute COVID-19 Surveillance Team |
| EPI_ISL_2995869 | Lighthouse Lab in Glasgow | Wellcome Sanger Institute for the COVID-19 Genomics UK (COG-UK) Consortium | Harper VanSteenhouse, Yumi Kasai, David Gray, Carol Clugston, Anna Dominiczak and Alex Alderton, Roberto Amato, Jeffrey Barrett, Sonia Goncalves, Ewan Harrison, David K. Jackson, Ian Johnston, Dominic Kwiatkowski, Cordelia Langford, John Sillitoe on behalf of the Wellcome Sanger Institute COVID-19 Surveillance Team |
| EPI_ISL_2995870 | Berkshire and Surrey Pathology Services Lighthouse Laboratory | Wellcome Sanger Institute for the COVID-19 Genomics UK (COG-UK) Consortium | Berkshire and Surrey Pathology Services Lighthouse Laboratory and Alex Alderton, Roberto Amato, Jeffrey Barrett, Sonia Goncalves, Ewan Harrison, David K. Jackson, Ian Johnston, Dominic Kwiatkowski, Cordelia Langford, John Sillitoe on behalf of the Wellcome Sanger Institute COVID-19 Surveillance Team |
| EPI_ISL_2995871 | Lighthouse Lab in Milton Keynes | Wellcome Sanger Institute for the COVID-19 Genomics UK (COG-UK) Consortium | The Lighthouse Lab in Milton Keynes and Alex Alderton, Roberto Amato, Jeffrey Barrett, Sonia Goncalves, Ewan Harrison, David K. Jackson, Ian Johnston, Dominic Kwiatkowski, Cordelia Langford, John Sillitoe on behalf of the Wellcome Sanger Institute COVID-19 Surveillance Team |

| | | | |
|---|---|---|---|
| EPI_ISL_2995872, EPI_ISL_2995873 | Berkshire and Surrey Pathology Services Lighthouse Laboratory | Wellcome Sanger Institute for the COVID-19 Genomics UK (COG-UK) Consortium | Berkshire and Surrey Pathology Services Lighthouse Laboratory and Alex Alderton, Roberto Amato, Jeffrey Barrett, Sonia Goncalves, Ewan Harrison, David K. Jackson, Ian Johnston, Dominic Kwiatkowski, Cordelia Langford, John Sillitoe on behalf of the Wellcome Sanger Institute COVID-19 Surveillance Team |
| EPI_ISL_2995874 | Lighthouse Lab in Milton Keynes | Wellcome Sanger Institute for the COVID-19 Genomics UK (COG-UK) Consortium | The Lighthouse Lab in Milton Keynes and Alex Alderton, Roberto Amato, Jeffrey Barrett, Sonia Goncalves, Ewan Harrison, David K. Jackson, Ian Johnston, Dominic Kwiatkowski, Cordelia Langford, John Sillitoe on behalf of the Wellcome Sanger Institute COVID-19 Surveillance Team |
| EPI_ISL_2995875, EPI_ISL_2995876 | Lighthouse Lab in Glasgow | Wellcome Sanger Institute for the COVID-19 Genomics UK (COG-UK) Consortium | Harper VanSteenhouse, Yumi Kasai, David Gray, Carol Clugston, Anna Dominiczak and Alex Alderton, Roberto Amato, Jeffrey Barrett, Sonia Goncalves, Ewan Harrison, David K. Jackson, Ian Johnston, Dominic Kwiatkowski, Cordelia Langford, John Sillitoe on behalf of the Wellcome Sanger Institute COVID-19 Surveillance Team |
| EPI_ISL_2995877 | Berkshire and Surrey Pathology Services Lighthouse Laboratory | Wellcome Sanger Institute for the COVID-19 Genomics UK (COG-UK) Consortium | Berkshire and Surrey Pathology Services Lighthouse Laboratory and Alex Alderton, Roberto Amato, Jeffrey Barrett, Sonia Goncalves, Ewan Harrison, David K. Jackson, Ian Johnston, Dominic Kwiatkowski, Cordelia Langford, John Sillitoe on behalf of the Wellcome Sanger Institute COVID-19 Surveillance Team |
| EPI_ISL_2995878 | Lighthouse Lab in Glasgow | Wellcome Sanger Institute for the COVID-19 Genomics UK (COG-UK) Consortium | Harper VanSteenhouse, Yumi Kasai, David Gray, Carol Clugston, Anna Dominiczak and Alex Alderton, Roberto Amato, Jeffrey Barrett, Sonia Goncalves, Ewan Harrison, David K. Jackson, Ian Johnston, Dominic Kwiatkowski, Cordelia Langford, John Sillitoe on behalf of the Wellcome Sanger Institute COVID-19 Surveillance Team |
| EPI_ISL_2995879 | Lighthouse Lab in Milton Keynes | Wellcome Sanger Institute for the COVID-19 Genomics UK (COG-UK) Consortium | The Lighthouse Lab in Milton Keynes and Alex Alderton, Roberto Amato, Jeffrey Barrett, Sonia Goncalves, Ewan Harrison, David K. Jackson, Ian Johnston, Dominic Kwiatkowski, Cordelia Langford, John Sillitoe on behalf of the Wellcome Sanger Institute COVID-19 Surveillance Team |
| EPI_ISL_2995880, EPI_ISL_2995881 | Berkshire and Surrey Pathology Services Lighthouse Laboratory | Wellcome Sanger Institute for the COVID-19 Genomics UK (COG-UK) Consortium | Berkshire and Surrey Pathology Services Lighthouse Laboratory and Alex Alderton, Roberto Amato, Jeffrey Barrett, Sonia Goncalves, Ewan Harrison, David K. Jackson, Ian Johnston, Dominic Kwiatkowski, Cordelia Langford, John Sillitoe on behalf of the Wellcome Sanger Institute COVID-19 Surveillance Team |
| EPI_ISL_2995882 | Lighthouse Lab in Milton Keynes | Wellcome Sanger Institute for the COVID-19 Genomics UK (COG-UK) Consortium | The Lighthouse Lab in Milton Keynes and Alex Alderton, Roberto Amato, Jeffrey Barrett, Sonia Goncalves, Ewan Harrison, David K. Jackson, Ian Johnston, Dominic Kwiatkowski, Cordelia Langford, John Sillitoe on behalf of the Wellcome Sanger Institute COVID-19 Surveillance Team |
| EPI_ISL_2995883 | Lighthouse Lab in Glasgow | Wellcome Sanger Institute for the COVID-19 Genomics UK (COG-UK) Consortium | Harper VanSteenhouse, Yumi Kasai, David Gray, Carol Clugston, Anna Dominiczak and Alex Alderton, Roberto Amato, Jeffrey Barrett, Sonia Goncalves, Ewan Harrison, David K. Jackson, Ian Johnston, Dominic Kwiatkowski, Cordelia Langford, John Sillitoe on behalf of the Wellcome Sanger Institute COVID-19 Surveillance Team |
| EPI_ISL_2995884 | Berkshire and Surrey Pathology Services Lighthouse Laboratory | Wellcome Sanger Institute for the COVID-19 Genomics UK (COG-UK) Consortium | Berkshire and Surrey Pathology Services Lighthouse Laboratory and Alex Alderton, Roberto Amato, Jeffrey Barrett, Sonia Goncalves, Ewan Harrison, David K. Jackson, Ian Johnston, Dominic Kwiatkowski, Cordelia Langford, John Sillitoe on behalf of the Wellcome Sanger Institute COVID-19 Surveillance Team |
| EPI_ISL_2995885 | Lighthouse Lab in Milton Keynes | Wellcome Sanger Institute for the COVID-19 Genomics UK (COG-UK) Consortium | The Lighthouse Lab in Milton Keynes and Alex Alderton, Roberto Amato, Jeffrey Barrett, Sonia Goncalves, Ewan Harrison, David K. Jackson, Ian Johnston, Dominic Kwiatkowski, Cordelia Langford, John Sillitoe on behalf of the Wellcome Sanger Institute COVID-19 Surveillance Team |
| EPI_ISL_2995886, EPI_ISL_2995887, EPI_ISL_2995888, EPI_ISL_2995889 | Berkshire and Surrey Pathology Services Lighthouse Laboratory | Wellcome Sanger Institute for the COVID-19 Genomics UK (COG-UK) Consortium | Berkshire and Surrey Pathology Services Lighthouse Laboratory and Alex Alderton, Roberto Amato, Jeffrey Barrett, Sonia Goncalves, Ewan Harrison, David K. Jackson, Ian Johnston, Dominic Kwiatkowski, Cordelia Langford, John Sillitoe on behalf of the Wellcome Sanger Institute COVID-19 Surveillance Team |
| EPI_ISL_2995890 | Lighthouse Lab in Glasgow | Wellcome Sanger Institute for the COVID-19 Genomics UK (COG-UK) Consortium | Harper VanSteenhouse, Yumi Kasai, David Gray, Carol Clugston, Anna Dominiczak and Alex Alderton, Roberto Amato, Jeffrey Barrett, Sonia Goncalves, Ewan Harrison, David K. Jackson, Ian Johnston, Dominic Kwiatkowski, Cordelia Langford, John Sillitoe on behalf of the Wellcome Sanger Institute COVID-19 Surveillance Team |
| EPI_ISL_2995891 | Berkshire and Surrey Pathology Services Lighthouse Laboratory | Wellcome Sanger Institute for the COVID-19 Genomics UK (COG-UK) Consortium | Berkshire and Surrey Pathology Services Lighthouse Laboratory and Alex Alderton, Roberto Amato, Jeffrey Barrett, Sonia Goncalves, Ewan Harrison, David K. Jackson, Ian Johnston, Dominic Kwiatkowski, Cordelia Langford, John Sillitoe on behalf of the Wellcome Sanger Institute COVID-19 Surveillance Team |
| EPI_ISL_2995892, EPI_ISL_2995893, EPI_ISL_2995894 | Lighthouse Lab in Milton Keynes | Wellcome Sanger Institute for the COVID-19 Genomics UK (COG-UK) Consortium | The Lighthouse Lab in Milton Keynes and Alex Alderton, Roberto Amato, Jeffrey Barrett, Sonia Goncalves, Ewan Harrison, David K. Jackson, Ian Johnston, Dominic Kwiatkowski, Cordelia Langford, John Sillitoe on behalf of the Wellcome Sanger Institute COVID-19 Surveillance Team |
| EPI_ISL_2995895, EPI_ISL_2995896, EPI_ISL_2995897 | Berkshire and Surrey Pathology Services Lighthouse Laboratory | Wellcome Sanger Institute for the COVID-19 Genomics UK (COG-UK) Consortium | Berkshire and Surrey Pathology Services Lighthouse Laboratory and Alex Alderton, Roberto Amato, Jeffrey Barrett, Sonia Goncalves, Ewan Harrison, David K. Jackson, Ian Johnston, Dominic Kwiatkowski, Cordelia Langford, John Sillitoe on behalf of the Wellcome Sanger Institute COVID-19 Surveillance Team |
| EPI_ISL_2995898 | Lighthouse Lab in Milton Keynes | Wellcome Sanger Institute for the COVID-19 Genomics UK (COG-UK) Consortium | The Lighthouse Lab in Milton Keynes and Alex Alderton, Roberto Amato, Jeffrey Barrett, Sonia Goncalves, Ewan Harrison, David K. Jackson, Ian Johnston, Dominic Kwiatkowski, Cordelia Langford, John Sillitoe on behalf of the Wellcome Sanger Institute COVID-19 Surveillance Team |
| EPI_ISL_2995899 | Berkshire and Surrey Pathology Services Lighthouse Laboratory | Wellcome Sanger Institute for the COVID-19 Genomics UK (COG-UK) Consortium | Berkshire and Surrey Pathology Services Lighthouse Laboratory and Alex Alderton, Roberto Amato, Jeffrey Barrett, Sonia Goncalves, Ewan Harrison, David K. Jackson, Ian Johnston, Dominic Kwiatkowski, Cordelia Langford, John Sillitoe on behalf of the Wellcome Sanger Institute COVID-19 Surveillance Team |
| EPI_ISL_2995900 | Lighthouse Lab in Glasgow | Wellcome Sanger Institute for the COVID-19 Genomics UK (COG-UK) Consortium | Harper VanSteenhouse, Yumi Kasai, David Gray, Carol Clugston, Anna Dominiczak and Alex Alderton, Roberto Amato, Jeffrey Barrett, Sonia Goncalves, Ewan Harrison, David K. Jackson, Ian Johnston, Dominic Kwiatkowski, Cordelia Langford, John Sillitoe on behalf of the Wellcome Sanger Institute COVID-19 Surveillance Team |
| EPI_ISL_2995901, EPI_ISL_2995902 | Berkshire and Surrey Pathology Services Lighthouse Laboratory | Wellcome Sanger Institute for the COVID-19 Genomics UK (COG-UK) Consortium | Berkshire and Surrey Pathology Services Lighthouse Laboratory and Alex Alderton, Roberto Amato, Jeffrey Barrett, Sonia Goncalves, Ewan Harrison, David K. Jackson, Ian Johnston, Dominic Kwiatkowski, Cordelia Langford, John Sillitoe on behalf of the Wellcome Sanger Institute COVID-19 Surveillance Team |
| EPI_ISL_2995903, EPI_ISL_2995904 | Lighthouse Lab in Glasgow | Wellcome Sanger Institute for the COVID-19 Genomics UK (COG-UK) Consortium | Harper VanSteenhouse, Yumi Kasai, David Gray, Carol Clugston, Anna Dominiczak and Alex Alderton, Roberto Amato, Jeffrey Barrett, Sonia Goncalves, Ewan Harrison, David K. Jackson, Ian Johnston, Dominic Kwiatkowski, Cordelia Langford, John Sillitoe on behalf of the Wellcome Sanger Institute COVID-19 Surveillance Team |
| EPI_ISL_2995905 | Lighthouse Lab in Milton Keynes | Wellcome Sanger Institute for the COVID-19 Genomics UK (COG-UK) Consortium | The Lighthouse Lab in Milton Keynes and Alex Alderton, Roberto Amato, Jeffrey Barrett, Sonia Goncalves, Ewan Harrison, David K. Jackson, Ian Johnston, Dominic Kwiatkowski, Cordelia Langford, John Sillitoe on behalf of the Wellcome Sanger Institute COVID-19 Surveillance Team |
| EPI_ISL_2995906 | Lighthouse Lab in Glasgow | Wellcome Sanger Institute for the COVID-19 Genomics UK (COG-UK) Consortium | Harper VanSteenhouse, Yumi Kasai, David Gray, Carol Clugston, Anna Dominiczak and Alex Alderton, Roberto Amato, Jeffrey Barrett, Sonia Goncalves, Ewan Harrison, David K. Jackson, Ian Johnston, Dominic Kwiatkowski, Cordelia Langford, John Sillitoe on behalf of the Wellcome Sanger Institute COVID-19 Surveillance Team |
| EPI_ISL_2995907 | Berkshire and Surrey Pathology Services Lighthouse Laboratory | Wellcome Sanger Institute for the COVID-19 Genomics UK (COG-UK) Consortium | Berkshire and Surrey Pathology Services Lighthouse Laboratory and Alex Alderton, Roberto Amato, Jeffrey Barrett, Sonia Goncalves, Ewan Harrison, David K. Jackson, Ian Johnston, Dominic Kwiatkowski, Cordelia Langford, John Sillitoe on behalf of the Wellcome Sanger Institute COVID-19 Surveillance Team |
| EPI_ISL_2995908 | Lighthouse Lab in Milton Keynes | Wellcome Sanger Institute for the COVID-19 Genomics UK (COG-UK) Consortium | The Lighthouse Lab in Milton Keynes and Alex Alderton, Roberto Amato, Jeffrey Barrett, Sonia Goncalves, Ewan Harrison, David K. Jackson, Ian Johnston, Dominic Kwiatkowski, Cordelia Langford, John Sillitoe on behalf of the Wellcome Sanger Institute COVID-19 Surveillance Team |
| EPI_ISL_2995909, EPI_ISL_2995910 | Berkshire and Surrey Pathology Services Lighthouse Laboratory | Wellcome Sanger Institute for the COVID-19 Genomics UK (COG-UK) Consortium | Berkshire and Surrey Pathology Services Lighthouse Laboratory and Alex Alderton, Roberto Amato, Jeffrey Barrett, Sonia Goncalves, Ewan Harrison, David K. Jackson, Ian Johnston, Dominic Kwiatkowski, Cordelia Langford, John Sillitoe on behalf of the Wellcome Sanger Institute COVID-19 Surveillance Team |
| EPI_ISL_2995911 | Lighthouse Lab in Glasgow | Wellcome Sanger Institute for the COVID-19 Genomics UK (COG-UK) Consortium | Harper VanSteenhouse, Yumi Kasai, David Gray, Carol Clugston, Anna Dominiczak and Alex Alderton, Roberto Amato, Jeffrey Barrett, Sonia Goncalves, Ewan Harrison, David K. Jackson, Ian Johnston, Dominic Kwiatkowski, Cordelia Langford, John Sillitoe on behalf of the Wellcome Sanger Institute COVID-19 Surveillance Team |
| EPI_ISL_2995912 | Lighthouse Lab in Milton Keynes | Wellcome Sanger Institute for the COVID-19 Genomics UK (COG-UK) Consortium | The Lighthouse Lab in Milton Keynes and Alex Alderton, Roberto Amato, Jeffrey Barrett, Sonia Goncalves, Ewan Harrison, David K. Jackson, Ian Johnston, Dominic Kwiatkowski, Cordelia Langford, John Sillitoe on behalf of the Wellcome Sanger Institute COVID-19 Surveillance Team |
| EPI_ISL_2995913, EPI_ISL_2995914 | Berkshire and Surrey Pathology Services Lighthouse Laboratory | Wellcome Sanger Institute for the COVID-19 Genomics UK (COG-UK) Consortium | Berkshire and Surrey Pathology Services Lighthouse Laboratory and Alex Alderton, Roberto Amato, Jeffrey Barrett, Sonia Goncalves, Ewan Harrison, David K. Jackson, Ian Johnston, Dominic Kwiatkowski, Cordelia Langford, John Sillitoe on behalf of the Wellcome Sanger Institute COVID-19 Surveillance Team |
| EPI_ISL_2995915 | Lighthouse Lab in Milton Keynes | Wellcome Sanger Institute for the COVID-19 Genomics UK (COG-UK) Consortium | The Lighthouse Lab in Milton Keynes and Alex Alderton, Roberto Amato, Jeffrey Barrett, Sonia Goncalves, Ewan Harrison, David K. Jackson, Ian Johnston, Dominic Kwiatkowski, Cordelia Langford, John Sillitoe on behalf of the Wellcome Sanger Institute COVID-19 Surveillance Team |
| EPI_ISL_2995916, EPI_ISL_2995917 | Berkshire and Surrey Pathology Services Lighthouse Laboratory | Wellcome Sanger Institute for the COVID-19 Genomics UK (COG-UK) Consortium | Berkshire and Surrey Pathology Services Lighthouse Laboratory and Alex Alderton, Roberto Amato, Jeffrey Barrett, Sonia Goncalves, Ewan Harrison, David K. Jackson, Ian Johnston, Dominic Kwiatkowski, Cordelia Langford, John Sillitoe on behalf of the Wellcome Sanger Institute COVID-19 Surveillance Team |
| EPI_ISL_2995918 | Lighthouse Lab in Glasgow | Wellcome Sanger Institute for the COVID-19 Genomics UK (COG-UK) Consortium | Harper VanSteenhouse, Yumi Kasai, David Gray, Carol Clugston, Anna Dominiczak and Alex Alderton, Roberto Amato, Jeffrey Barrett, Sonia Goncalves, Ewan Harrison, David K. Jackson, Ian Johnston, Dominic Kwiatkowski, Cordelia Langford, John Sillitoe on behalf of the Wellcome Sanger Institute COVID-19 Surveillance Team |
| EPI_ISL_2995919, EPI_ISL_2995920 | Lighthouse Lab in Milton Keynes | Wellcome Sanger Institute for the COVID-19 Genomics UK (COG-UK) Consortium | The Lighthouse Lab in Milton Keynes and Alex Alderton, Roberto Amato, Jeffrey Barrett, Sonia Goncalves, Ewan Harrison, David K. Jackson, Ian Johnston, Dominic Kwiatkowski, Cordelia Langford, John Sillitoe on behalf of the Wellcome Sanger Institute COVID-19 Surveillance Team |
| EPI_ISL_2995921, EPI_ISL_2995922 | Lighthouse Lab in Glasgow | Wellcome Sanger Institute for the COVID-19 Genomics UK (COG-UK) Consortium | Harper VanSteenhouse, Yumi Kasai, David Gray, Carol Clugston, Anna Dominiczak and Alex Alderton, Roberto Amato, Jeffrey Barrett, Sonia Goncalves, Ewan Harrison, David K. Jackson, Ian Johnston, Dominic Kwiatkowski, Cordelia Langford, John Sillitoe on behalf of the Wellcome Sanger Institute COVID-19 Surveillance Team |
| EPI_ISL_2995923, EPI_ISL_2995924 | Berkshire and Surrey Pathology Services Lighthouse Laboratory | Wellcome Sanger Institute for the COVID-19 Genomics UK (COG-UK) Consortium | Berkshire and Surrey Pathology Services Lighthouse Laboratory and Alex Alderton, Roberto Amato, Jeffrey Barrett, Sonia Goncalves, Ewan Harrison, David K. Jackson, Ian Johnston, Dominic Kwiatkowski, Cordelia Langford, John Sillitoe on behalf of the Wellcome Sanger Institute COVID-19 Surveillance Team |
| EPI_ISL_2995925, EPI_ISL_2995926 | Lighthouse Lab in Glasgow | Wellcome Sanger Institute for the COVID-19 Genomics UK (COG-UK) Consortium | Harper VanSteenhouse, Yumi Kasai, David Gray, Carol Clugston, Anna Dominiczak and Alex Alderton, Roberto Amato, Jeffrey Barrett, Sonia Goncalves, Ewan Harrison, David K. Jackson, Ian Johnston, Dominic Kwiatkowski, Cordelia Langford, John Sillitoe on behalf of the Wellcome Sanger Institute COVID-19 Surveillance Team |
| EPI_ISL_2995927 | Lighthouse Lab in Milton Keynes | Wellcome Sanger Institute for the COVID-19 Genomics UK (COG-UK) Consortium | The Lighthouse Lab in Milton Keynes and Alex Alderton, Roberto Amato, Jeffrey Barrett, Sonia Goncalves, Ewan Harrison, David K. Jackson, Ian Johnston, Dominic Kwiatkowski, Cordelia Langford, John Sillitoe on behalf of the Wellcome Sanger Institute COVID-19 Surveillance Team |
| EPI_ISL_2995928, EPI_ISL_2995929 | Berkshire and Surrey Pathology Services Lighthouse Laboratory | Wellcome Sanger Institute for the COVID-19 Genomics UK (COG-UK) Consortium | Berkshire and Surrey Pathology Services Lighthouse Laboratory and Alex Alderton, Roberto Amato, Jeffrey Barrett, Sonia Goncalves, Ewan Harrison, David K. Jackson, Ian Johnston, Dominic Kwiatkowski, Cordelia Langford, John Sillitoe on behalf of the Wellcome Sanger Institute COVID-19 Surveillance Team |
| EPI_ISL_2995930 | Lighthouse Lab in Glasgow | Wellcome Sanger Institute for the COVID-19 Genomics UK (COG-UK) Consortium | Harper VanSteenhouse, Yumi Kasai, David Gray, Carol Clugston, Anna Dominiczak and Alex Alderton, Roberto Amato, Jeffrey Barrett, Sonia Goncalves, Ewan Harrison, David K. Jackson, Ian Johnston, Dominic Kwiatkowski, Cordelia Langford, John Sillitoe on behalf of the Wellcome Sanger Institute COVID-19 Surveillance Team |
| EPI_ISL_2995931 | Lighthouse Lab in Milton Keynes | Wellcome Sanger Institute for the COVID-19 Genomics UK (COG-UK) Consortium | The Lighthouse Lab in Milton Keynes and Alex Alderton, Roberto Amato, Jeffrey Barrett, Sonia Goncalves, Ewan Harrison, David K. Jackson, Ian Johnston, Dominic Kwiatkowski, Cordelia Langford, John Sillitoe on behalf of the Wellcome Sanger Institute COVID-19 Surveillance Team |
| EPI_ISL_2995932 | Berkshire and Surrey Pathology Services Lighthouse Laboratory | Wellcome Sanger Institute for the COVID-19 Genomics UK (COG-UK) Consortium | Berkshire and Surrey Pathology Services Lighthouse Laboratory and Alex Alderton, Roberto Amato, Jeffrey Barrett, Sonia Goncalves, Ewan Harrison, David K. Jackson, Ian Johnston, Dominic Kwiatkowski, Cordelia Langford, John Sillitoe on behalf of the Wellcome Sanger Institute COVID-19 Surveillance Team |
| EPI_ISL_2995933 | Lighthouse Lab in Glasgow | Wellcome Sanger Institute for the COVID-19 Genomics UK (COG-UK) Consortium | Harper VanSteenhouse, Yumi Kasai, David Gray, Carol Clugston, Anna Dominiczak and Alex Alderton, Roberto Amato, Jeffrey Barrett, Sonia Goncalves, Ewan Harrison, David K. Jackson, Ian Johnston, Dominic Kwiatkowski, Cordelia Langford, John Sillitoe on behalf of the Wellcome Sanger Institute COVID-19 Surveillance Team |
| EPI_ISL_2995934 | Berkshire and Surrey Pathology Services Lighthouse Laboratory | Wellcome Sanger Institute for the COVID-19 Genomics UK (COG-UK) Consortium | Berkshire and Surrey Pathology Services Lighthouse Laboratory and Alex Alderton, Roberto Amato, Jeffrey Barrett, Sonia Goncalves, Ewan Harrison, David K. Jackson, Ian Johnston, Dominic Kwiatkowski, Cordelia Langford, John Sillitoe on behalf of the Wellcome Sanger Institute COVID-19 Surveillance Team |
| EPI_ISL_2995935 | Lighthouse Lab in Glasgow | Wellcome Sanger Institute for the COVID-19 Genomics UK (COG-UK) | Harper VanSteenhouse, Yumi Kasai, David Gray, Carol Clugston, Anna Dominiczak and Alex Alderton, Roberto Amato, Jeffrey Barrett, Sonia Goncalves, Ewan Harrison, David K. Jackson, Ian Johnston, Dominic Kwiatkowski, Cordelia Langford, |

| | | | |
|---|---|---|---|
| | | Consortium | John Sillitoe on behalf of the Wellcome Sanger Institute COVID-19 Surveillance Team |
| EPI_ISL_2995936, EPI_ISL_2995937 | Berkshire and Surrey Pathology Services Lighthouse Laboratory | Wellcome Sanger Institute for the COVID-19 Genomics UK (COG-UK) Consortium | Berkshire and Surrey Pathology Services Lighthouse Laboratory and Alex Alderton, Roberto Amato, Jeffrey Barrett, Sonia Goncalves, Ewan Harrison, David K. Jackson, Ian Johnston, Dominic Kwiatkowski, Cordelia Langford, John Sillitoe on behalf of the Wellcome Sanger Institute COVID-19 Surveillance Team |
| EPI_ISL_2995938, EPI_ISL_2995939, EPI_ISL_2995940, EPI_ISL_2995941 | Lighthouse Lab in Milton Keynes | Wellcome Sanger Institute for the COVID-19 Genomics UK (COG-UK) Consortium | The Lighthouse Lab in Milton Keynes and Alex Alderton, Roberto Amato, Jeffrey Barrett, Sonia Goncalves, Ewan Harrison, David K. Jackson, Ian Johnston, Dominic Kwiatkowski, Cordelia Langford, John Sillitoe on behalf of the Wellcome Sanger Institute COVID-19 Surveillance Team |
| EPI_ISL_2995942 | Berkshire and Surrey Pathology Services Lighthouse Laboratory | Wellcome Sanger Institute for the COVID-19 Genomics UK (COG-UK) Consortium | Berkshire and Surrey Pathology Services Lighthouse Laboratory and Alex Alderton, Roberto Amato, Jeffrey Barrett, Sonia Goncalves, Ewan Harrison, David K. Jackson, Ian Johnston, Dominic Kwiatkowski, Cordelia Langford, John Sillitoe on behalf of the Wellcome Sanger Institute COVID-19 Surveillance Team |
| EPI_ISL_2995943 | Lighthouse Lab in Milton Keynes | Wellcome Sanger Institute for the COVID-19 Genomics UK (COG-UK) Consortium | The Lighthouse Lab in Milton Keynes and Alex Alderton, Roberto Amato, Jeffrey Barrett, Sonia Goncalves, Ewan Harrison, David K. Jackson, Ian Johnston, Dominic Kwiatkowski, Cordelia Langford, John Sillitoe on behalf of the Wellcome Sanger Institute COVID-19 Surveillance Team |
| EPI_ISL_2995944 | Berkshire and Surrey Pathology Services Lighthouse Laboratory | Wellcome Sanger Institute for the COVID-19 Genomics UK (COG-UK) Consortium | Berkshire and Surrey Pathology Services Lighthouse Laboratory and Alex Alderton, Roberto Amato, Jeffrey Barrett, Sonia Goncalves, Ewan Harrison, David K. Jackson, Ian Johnston, Dominic Kwiatkowski, Cordelia Langford, John Sillitoe on behalf of the Wellcome Sanger Institute COVID-19 Surveillance Team |
| EPI_ISL_2995945 | Lighthouse Lab in Glasgow | Wellcome Sanger Institute for the COVID-19 Genomics UK (COG-UK) Consortium | Harper VanSteenhouse, Yumi Kasai, David Gray, Carol Clugston, Anna Dominiczak and Alex Alderton, Roberto Amato, Jeffrey Barrett, Sonia Goncalves, Ewan Harrison, David K. Jackson, Ian Johnston, Dominic Kwiatkowski, Cordelia Langford, John Sillitoe on behalf of the Wellcome Sanger Institute COVID-19 Surveillance Team |
| EPI_ISL_2995946 | Lighthouse Lab in Milton Keynes | Wellcome Sanger Institute for the COVID-19 Genomics UK (COG-UK) Consortium | The Lighthouse Lab in Milton Keynes and Alex Alderton, Roberto Amato, Jeffrey Barrett, Sonia Goncalves, Ewan Harrison, David K. Jackson, Ian Johnston, Dominic Kwiatkowski, Cordelia Langford, John Sillitoe on behalf of the Wellcome Sanger Institute COVID-19 Surveillance Team |
| EPI_ISL_2995947 | Lighthouse Lab in Glasgow | Wellcome Sanger Institute for the COVID-19 Genomics UK (COG-UK) Consortium | Harper VanSteenhouse, Yumi Kasai, David Gray, Carol Clugston, Anna Dominiczak and Alex Alderton, Roberto Amato, Jeffrey Barrett, Sonia Goncalves, Ewan Harrison, David K. Jackson, Ian Johnston, Dominic Kwiatkowski, Cordelia Langford, John Sillitoe on behalf of the Wellcome Sanger Institute COVID-19 Surveillance Team |
| EPI_ISL_2995948, EPI_ISL_2995949, EPI_ISL_2995950, EPI_ISL_2995951 | Berkshire and Surrey Pathology Services Lighthouse Laboratory | Wellcome Sanger Institute for the COVID-19 Genomics UK (COG-UK) Consortium | Berkshire and Surrey Pathology Services Lighthouse Laboratory and Alex Alderton, Roberto Amato, Jeffrey Barrett, Sonia Goncalves, Ewan Harrison, David K. Jackson, Ian Johnston, Dominic Kwiatkowski, Cordelia Langford, John Sillitoe on behalf of the Wellcome Sanger Institute COVID-19 Surveillance Team |
| EPI_ISL_2995952 | Lighthouse Lab in Glasgow | Wellcome Sanger Institute for the COVID-19 Genomics UK (COG-UK) Consortium | Harper VanSteenhouse, Yumi Kasai, David Gray, Carol Clugston, Anna Dominiczak and Alex Alderton, Roberto Amato, Jeffrey Barrett, Sonia Goncalves, Ewan Harrison, David K. Jackson, Ian Johnston, Dominic Kwiatkowski, Cordelia Langford, John Sillitoe on behalf of the Wellcome Sanger Institute COVID-19 Surveillance Team |
| EPI_ISL_2995953 | Lighthouse Lab in Milton Keynes | Wellcome Sanger Institute for the COVID-19 Genomics UK (COG-UK) Consortium | The Lighthouse Lab in Milton Keynes and Alex Alderton, Roberto Amato, Jeffrey Barrett, Sonia Goncalves, Ewan Harrison, David K. Jackson, Ian Johnston, Dominic Kwiatkowski, Cordelia Langford, John Sillitoe on behalf of the Wellcome Sanger Institute COVID-19 Surveillance Team |
| EPI_ISL_2995954 | Berkshire and Surrey Pathology Services Lighthouse Laboratory | Wellcome Sanger Institute for the COVID-19 Genomics UK (COG-UK) Consortium | Berkshire and Surrey Pathology Services Lighthouse Laboratory and Alex Alderton, Roberto Amato, Jeffrey Barrett, Sonia Goncalves, Ewan Harrison, David K. Jackson, Ian Johnston, Dominic Kwiatkowski, Cordelia Langford, John Sillitoe on behalf of the Wellcome Sanger Institute COVID-19 Surveillance Team |
| EPI_ISL_2995955, EPI_ISL_2995956, EPI_ISL_2995957 | Lighthouse Lab in Milton Keynes | Wellcome Sanger Institute for the COVID-19 Genomics UK (COG-UK) Consortium | The Lighthouse Lab in Milton Keynes and Alex Alderton, Roberto Amato, Jeffrey Barrett, Sonia Goncalves, Ewan Harrison, David K. Jackson, Ian Johnston, Dominic Kwiatkowski, Cordelia Langford, John Sillitoe on behalf of the Wellcome Sanger Institute COVID-19 Surveillance Team |
| EPI_ISL_2995958, EPI_ISL_2995959 | Lighthouse Lab in Glasgow | Wellcome Sanger Institute for the COVID-19 Genomics UK (COG-UK) Consortium | Harper VanSteenhouse, Yumi Kasai, David Gray, Carol Clugston, Anna Dominiczak and Alex Alderton, Roberto Amato, Jeffrey Barrett, Sonia Goncalves, Ewan Harrison, David K. Jackson, Ian Johnston, Dominic Kwiatkowski, Cordelia Langford, John Sillitoe on behalf of the Wellcome Sanger Institute COVID-19 Surveillance Team |
| EPI_ISL_2995960, EPI_ISL_2995961, EPI_ISL_2995962, EPI_ISL_2995963 | Berkshire and Surrey Pathology Services Lighthouse Laboratory | Wellcome Sanger Institute for the COVID-19 Genomics UK (COG-UK) Consortium | Berkshire and Surrey Pathology Services Lighthouse Laboratory and Alex Alderton, Roberto Amato, Jeffrey Barrett, Sonia Goncalves, Ewan Harrison, David K. Jackson, Ian Johnston, Dominic Kwiatkowski, Cordelia Langford, John Sillitoe on behalf of the Wellcome Sanger Institute COVID-19 Surveillance Team |
| EPI_ISL_2995964 | Lighthouse Lab in Milton Keynes | Wellcome Sanger Institute for the COVID-19 Genomics UK (COG-UK) Consortium | The Lighthouse Lab in Milton Keynes and Alex Alderton, Roberto Amato, Jeffrey Barrett, Sonia Goncalves, Ewan Harrison, David K. Jackson, Ian Johnston, Dominic Kwiatkowski, Cordelia Langford, John Sillitoe on behalf of the Wellcome Sanger Institute COVID-19 Surveillance Team |
| EPI_ISL_2995965 | Lighthouse Lab in Glasgow | Wellcome Sanger Institute for the COVID-19 Genomics UK (COG-UK) Consortium | Harper VanSteenhouse, Yumi Kasai, David Gray, Carol Clugston, Anna Dominiczak and Alex Alderton, Roberto Amato, Jeffrey Barrett, Sonia Goncalves, Ewan Harrison, David K. Jackson, Ian Johnston, Dominic Kwiatkowski, Cordelia Langford, John Sillitoe on behalf of the Wellcome Sanger Institute COVID-19 Surveillance Team |
| EPI_ISL_2995966, EPI_ISL_2995967 | Berkshire and Surrey Pathology Services Lighthouse Laboratory | Wellcome Sanger Institute for the COVID-19 Genomics UK (COG-UK) Consortium | Berkshire and Surrey Pathology Services Lighthouse Laboratory and Alex Alderton, Roberto Amato, Jeffrey Barrett, Sonia Goncalves, Ewan Harrison, David K. Jackson, Ian Johnston, Dominic Kwiatkowski, Cordelia Langford, John Sillitoe on behalf of the Wellcome Sanger Institute COVID-19 Surveillance Team |
| EPI_ISL_2995968, EPI_ISL_2995969 | Lighthouse Lab in Milton Keynes | Wellcome Sanger Institute for the COVID-19 Genomics UK (COG-UK) Consortium | The Lighthouse Lab in Milton Keynes and Alex Alderton, Roberto Amato, Jeffrey Barrett, Sonia Goncalves, Ewan Harrison, David K. Jackson, Ian Johnston, Dominic Kwiatkowski, Cordelia Langford, John Sillitoe on behalf of the Wellcome Sanger Institute COVID-19 Surveillance Team |
| EPI_ISL_2995970, EPI_ISL_2995971, EPI_ISL_2995972 | Berkshire and Surrey Pathology Services Lighthouse Laboratory | Wellcome Sanger Institute for the COVID-19 Genomics UK (COG-UK) Consortium | Berkshire and Surrey Pathology Services Lighthouse Laboratory and Alex Alderton, Roberto Amato, Jeffrey Barrett, Sonia Goncalves, Ewan Harrison, David K. Jackson, Ian Johnston, Dominic Kwiatkowski, Cordelia Langford, John Sillitoe on behalf of the Wellcome Sanger Institute COVID-19 Surveillance Team |
| EPI_ISL_2995973 | Lighthouse Lab in Glasgow | Wellcome Sanger Institute for the COVID-19 Genomics UK (COG-UK) Consortium | Harper VanSteenhouse, Yumi Kasai, David Gray, Carol Clugston, Anna Dominiczak and Alex Alderton, Roberto Amato, Jeffrey Barrett, Sonia Goncalves, Ewan Harrison, David K. Jackson, Ian Johnston, Dominic Kwiatkowski, Cordelia Langford, John Sillitoe on behalf of the Wellcome Sanger Institute COVID-19 Surveillance Team |
| EPI_ISL_2995974 | Lighthouse Lab in Milton Keynes | Wellcome Sanger Institute for the COVID-19 Genomics UK (COG-UK) Consortium | The Lighthouse Lab in Milton Keynes and Alex Alderton, Roberto Amato, Jeffrey Barrett, Sonia Goncalves, Ewan Harrison, David K. Jackson, Ian Johnston, Dominic Kwiatkowski, Cordelia Langford, John Sillitoe on behalf of the Wellcome Sanger Institute COVID-19 Surveillance Team |
| EPI_ISL_2995975, EPI_ISL_2995976 | Lighthouse Lab in Glasgow | Wellcome Sanger Institute for the COVID-19 Genomics UK (COG-UK) Consortium | Harper VanSteenhouse, Yumi Kasai, David Gray, Carol Clugston, Anna Dominiczak and Alex Alderton, Roberto Amato, Jeffrey Barrett, Sonia Goncalves, Ewan Harrison, David K. Jackson, Ian Johnston, Dominic Kwiatkowski, Cordelia Langford, John Sillitoe on behalf of the Wellcome Sanger Institute COVID-19 Surveillance Team |
| EPI_ISL_2995977 | Berkshire and Surrey Pathology Services Lighthouse Laboratory | Wellcome Sanger Institute for the COVID-19 Genomics UK (COG-UK) Consortium | Berkshire and Surrey Pathology Services Lighthouse Laboratory and Alex Alderton, Roberto Amato, Jeffrey Barrett, Sonia Goncalves, Ewan Harrison, David K. Jackson, Ian Johnston, Dominic Kwiatkowski, Cordelia Langford, John Sillitoe on behalf of the Wellcome Sanger Institute COVID-19 Surveillance Team |
| EPI_ISL_2995978 | Lighthouse Lab in Glasgow | Wellcome Sanger Institute for the COVID-19 Genomics UK (COG-UK) Consortium | Harper VanSteenhouse, Yumi Kasai, David Gray, Carol Clugston, Anna Dominiczak and Alex Alderton, Roberto Amato, Jeffrey Barrett, Sonia Goncalves, Ewan Harrison, David K. Jackson, Ian Johnston, Dominic Kwiatkowski, Cordelia Langford, John Sillitoe on behalf of the Wellcome Sanger Institute COVID-19 Surveillance Team |
| EPI_ISL_2995979 | Lighthouse Lab in Milton Keynes | Wellcome Sanger Institute for the COVID-19 Genomics UK (COG-UK) Consortium | The Lighthouse Lab in Milton Keynes and Alex Alderton, Roberto Amato, Jeffrey Barrett, Sonia Goncalves, Ewan Harrison, David K. Jackson, Ian Johnston, Dominic Kwiatkowski, Cordelia Langford, John Sillitoe on behalf of the Wellcome Sanger Institute COVID-19 Surveillance Team |
| EPI_ISL_2995980, EPI_ISL_2995981, EPI_ISL_2995982 | Berkshire and Surrey Pathology Services Lighthouse Laboratory | Wellcome Sanger Institute for the COVID-19 Genomics UK (COG-UK) Consortium | Berkshire and Surrey Pathology Services Lighthouse Laboratory and Alex Alderton, Roberto Amato, Jeffrey Barrett, Sonia Goncalves, Ewan Harrison, David K. Jackson, Ian Johnston, Dominic Kwiatkowski, Cordelia Langford, John Sillitoe on behalf of the Wellcome Sanger Institute COVID-19 Surveillance Team |
| EPI_ISL_2995983, EPI_ISL_2995984 | Lighthouse Lab in Glasgow | Wellcome Sanger Institute for the COVID-19 Genomics UK (COG-UK) Consortium | Harper VanSteenhouse, Yumi Kasai, David Gray, Carol Clugston, Anna Dominiczak and Alex Alderton, Roberto Amato, Jeffrey Barrett, Sonia Goncalves, Ewan Harrison, David K. Jackson, Ian Johnston, Dominic Kwiatkowski, Cordelia Langford, John Sillitoe on behalf of the Wellcome Sanger Institute COVID-19 Surveillance Team |
| EPI_ISL_2995985, EPI_ISL_2995986, EPI_ISL_2995987 | Lighthouse Lab in Milton Keynes | Wellcome Sanger Institute for the COVID-19 Genomics UK (COG-UK) Consortium | The Lighthouse Lab in Milton Keynes and Alex Alderton, Roberto Amato, Jeffrey Barrett, Sonia Goncalves, Ewan Harrison, David K. Jackson, Ian Johnston, Dominic Kwiatkowski, Cordelia Langford, John Sillitoe on behalf of the Wellcome Sanger Institute COVID-19 Surveillance Team |
| EPI_ISL_2995988, EPI_ISL_2995989, EPI_ISL_2995990 | Lighthouse Lab in Glasgow | Wellcome Sanger Institute for the COVID-19 Genomics UK (COG-UK) Consortium | Harper VanSteenhouse, Yumi Kasai, David Gray, Carol Clugston, Anna Dominiczak and Alex Alderton, Roberto Amato, Jeffrey Barrett, Sonia Goncalves, Ewan Harrison, David K. Jackson, Ian Johnston, Dominic Kwiatkowski, Cordelia Langford, John Sillitoe on behalf of the Wellcome Sanger Institute COVID-19 Surveillance Team |
| EPI_ISL_2995991 | Lighthouse Lab in Milton Keynes | Wellcome Sanger Institute for the COVID-19 Genomics UK (COG-UK) Consortium | The Lighthouse Lab in Milton Keynes and Alex Alderton, Roberto Amato, Jeffrey Barrett, Sonia Goncalves, Ewan Harrison, David K. Jackson, Ian Johnston, Dominic Kwiatkowski, Cordelia Langford, John Sillitoe on behalf of the Wellcome Sanger Institute COVID-19 Surveillance Team |
| EPI_ISL_2995992 | Berkshire and Surrey Pathology Services Lighthouse Laboratory | Wellcome Sanger Institute for the COVID-19 Genomics UK (COG-UK) Consortium | Berkshire and Surrey Pathology Services Lighthouse Laboratory and Alex Alderton, Roberto Amato, Jeffrey Barrett, Sonia Goncalves, Ewan Harrison, David K. Jackson, Ian Johnston, Dominic Kwiatkowski, Cordelia Langford, John Sillitoe on behalf of the Wellcome Sanger Institute COVID-19 Surveillance Team |
| EPI_ISL_2995993 | Lighthouse Lab in Milton Keynes | Wellcome Sanger Institute for the COVID-19 Genomics UK (COG-UK) Consortium | The Lighthouse Lab in Milton Keynes and Alex Alderton, Roberto Amato, Jeffrey Barrett, Sonia Goncalves, Ewan Harrison, David K. Jackson, Ian Johnston, Dominic Kwiatkowski, Cordelia Langford, John Sillitoe on behalf of the Wellcome Sanger Institute COVID-19 Surveillance Team |
| EPI_ISL_2995994 | Lighthouse Lab in Glasgow | Wellcome Sanger Institute for the COVID-19 Genomics UK (COG-UK) Consortium | Harper VanSteenhouse, Yumi Kasai, David Gray, Carol Clugston, Anna Dominiczak and Alex Alderton, Roberto Amato, Jeffrey Barrett, Sonia Goncalves, Ewan Harrison, David K. Jackson, Ian Johnston, Dominic Kwiatkowski, Cordelia Langford, John Sillitoe on behalf of the Wellcome Sanger Institute COVID-19 Surveillance Team |
| EPI_ISL_2995995 | Lighthouse Lab in Milton Keynes | Wellcome Sanger Institute for the COVID-19 Genomics UK (COG-UK) Consortium | The Lighthouse Lab in Milton Keynes and Alex Alderton, Roberto Amato, Jeffrey Barrett, Sonia Goncalves, Ewan Harrison, David K. Jackson, Ian Johnston, Dominic Kwiatkowski, Cordelia Langford, John Sillitoe on behalf of the Wellcome Sanger Institute COVID-19 Surveillance Team |
| EPI_ISL_2995996, EPI_ISL_2995997 | Lighthouse Lab in Glasgow | Wellcome Sanger Institute for the COVID-19 Genomics UK (COG-UK) Consortium | Harper VanSteenhouse, Yumi Kasai, David Gray, Carol Clugston, Anna Dominiczak and Alex Alderton, Roberto Amato, Jeffrey Barrett, Sonia Goncalves, Ewan Harrison, David K. Jackson, Ian Johnston, Dominic Kwiatkowski, Cordelia Langford, John Sillitoe on behalf of the Wellcome Sanger Institute COVID-19 Surveillance Team |
| EPI_ISL_2995998 | Lighthouse Laboratory Plymouth | Wellcome Sanger Institute for the COVID-19 Genomics UK (COG-UK) Consortium | Lighthouse Laboratory Plymouth and Alex Alderton, Roberto Amato, Jeffrey Barrett, Sonia Goncalves, Ewan Harrison, David K. Jackson, Ian Johnston, Dominic Kwiatkowski, Cordelia Langford, John Sillitoe on behalf of the Wellcome Sanger Institute COVID-19 Surveillance Team |
| EPI_ISL_2995999, EPI_ISL_2996000 | Lighthouse Lab in Glasgow | Wellcome Sanger Institute for the COVID-19 Genomics UK (COG-UK) Consortium | Harper VanSteenhouse, Yumi Kasai, David Gray, Carol Clugston, Anna Dominiczak and Alex Alderton, Roberto Amato, Jeffrey Barrett, Sonia Goncalves, Ewan Harrison, David K. Jackson, Ian Johnston, Dominic Kwiatkowski, Cordelia Langford, John Sillitoe on behalf of the Wellcome Sanger Institute COVID-19 Surveillance Team |
| EPI_ISL_2996001 | Lighthouse Laboratory Plymouth | Wellcome Sanger Institute for the COVID-19 Genomics UK (COG-UK) Consortium | Lighthouse Laboratory Plymouth and Alex Alderton, Roberto Amato, Jeffrey Barrett, Sonia Goncalves, Ewan Harrison, David K. Jackson, Ian Johnston, Dominic Kwiatkowski, Cordelia Langford, John Sillitoe on behalf of the Wellcome Sanger Institute COVID-19 Surveillance Team |
| EPI_ISL_2996002, EPI_ISL_2996003 | Lighthouse Lab in Glasgow | Wellcome Sanger Institute for the COVID-19 Genomics UK (COG-UK) Consortium | Harper VanSteenhouse, Yumi Kasai, David Gray, Carol Clugston, Anna Dominiczak and Alex Alderton, Roberto Amato, Jeffrey Barrett, Sonia Goncalves, Ewan Harrison, David K. Jackson, Ian Johnston, Dominic Kwiatkowski, Cordelia Langford, John Sillitoe on behalf of the Wellcome Sanger Institute COVID-19 Surveillance Team |

| | | | |
|---|---|---|---|
| EPI_ISL_2996004, EPI_ISL_2996005 | Lighthouse Laboratory Plymouth | Wellcome Sanger Institute for the COVID-19 Genomics UK (COG-UK) Consortium | Lighthouse Laboratory Plymouth and Alex Alderton, Roberto Amato, Jeffrey Barrett, Sonia Goncalves, Ewan Harrison, David K. Jackson, Ian Johnston, Dominic Kwiatkowski, Cordelia Langford, John Sillitoe on behalf of the Wellcome Sanger Institute COVID-19 Surveillance Team |
| EPI_ISL_2996006, EPI_ISL_2996007, EPI_ISL_2996008, EPI_ISL_2996009, EPI_ISL_2996010, EPI_ISL_2996011 | Lighthouse Lab in Glasgow | Wellcome Sanger Institute for the COVID-19 Genomics UK (COG-UK) Consortium | Harper VanSteenhouse, Yumi Kasai, David Gray, Carol Clugston, Anna Dominiczak and Alex Alderton, Roberto Amato, Jeffrey Barrett, Sonia Goncalves, Ewan Harrison, David K. Jackson, Ian Johnston, Dominic Kwiatkowski, Cordelia Langford, John Sillitoe on behalf of the Wellcome Sanger Institute COVID-19 Surveillance Team |
| EPI_ISL_2996012 | Berkshire and Surrey Pathology Services Lighthouse Laboratory | Wellcome Sanger Institute for the COVID-19 Genomics UK (COG-UK) Consortium | Berkshire and Surrey Pathology Services Lighthouse Laboratory and Alex Alderton, Roberto Amato, Jeffrey Barrett, Sonia Goncalves, Ewan Harrison, David K. Jackson, Ian Johnston, Dominic Kwiatkowski, Cordelia Langford, John Sillitoe on behalf of the Wellcome Sanger Institute COVID-19 Surveillance Team |
| EPI_ISL_2996013, EPI_ISL_2996014 | Lighthouse Laboratory Plymouth | Wellcome Sanger Institute for the COVID-19 Genomics UK (COG-UK) Consortium | Lighthouse Laboratory Plymouth and Alex Alderton, Roberto Amato, Jeffrey Barrett, Sonia Goncalves, Ewan Harrison, David K. Jackson, Ian Johnston, Dominic Kwiatkowski, Cordelia Langford, John Sillitoe on behalf of the Wellcome Sanger Institute COVID-19 Surveillance Team |
| EPI_ISL_2996015, EPI_ISL_2996016, EPI_ISL_2996017, EPI_ISL_2996018, EPI_ISL_2996019, EPI_ISL_2996020, EPI_ISL_2996021, EPI_ISL_2996022, EPI_ISL_2996023 | Lighthouse Lab in Glasgow | Wellcome Sanger Institute for the COVID-19 Genomics UK (COG-UK) Consortium | Harper VanSteenhouse, Yumi Kasai, David Gray, Carol Clugston, Anna Dominiczak and Alex Alderton, Roberto Amato, Jeffrey Barrett, Sonia Goncalves, Ewan Harrison, David K. Jackson, Ian Johnston, Dominic Kwiatkowski, Cordelia Langford, John Sillitoe on behalf of the Wellcome Sanger Institute COVID-19 Surveillance Team |
| EPI_ISL_2996024 | Lighthouse Laboratory Plymouth | Wellcome Sanger Institute for the COVID-19 Genomics UK (COG-UK) Consortium | Lighthouse Laboratory Plymouth and Alex Alderton, Roberto Amato, Jeffrey Barrett, Sonia Goncalves, Ewan Harrison, David K. Jackson, Ian Johnston, Dominic Kwiatkowski, Cordelia Langford, John Sillitoe on behalf of the Wellcome Sanger Institute COVID-19 Surveillance Team |
| EPI_ISL_2996025 | Berkshire and Surrey Pathology Services Lighthouse Laboratory | Wellcome Sanger Institute for the COVID-19 Genomics UK (COG-UK) Consortium | Berkshire and Surrey Pathology Services Lighthouse Laboratory and Alex Alderton, Roberto Amato, Jeffrey Barrett, Sonia Goncalves, Ewan Harrison, David K. Jackson, Ian Johnston, Dominic Kwiatkowski, Cordelia Langford, John Sillitoe on behalf of the Wellcome Sanger Institute COVID-19 Surveillance Team |
| EPI_ISL_2996026, EPI_ISL_2996027, EPI_ISL_2996028 | Lighthouse Lab in Glasgow | Wellcome Sanger Institute for the COVID-19 Genomics UK (COG-UK) Consortium | Harper VanSteenhouse, Yumi Kasai, David Gray, Carol Clugston, Anna Dominiczak and Alex Alderton, Roberto Amato, Jeffrey Barrett, Sonia Goncalves, Ewan Harrison, David K. Jackson, Ian Johnston, Dominic Kwiatkowski, Cordelia Langford, John Sillitoe on behalf of the Wellcome Sanger Institute COVID-19 Surveillance Team |
| EPI_ISL_2996029, EPI_ISL_2996030 | Lighthouse Laboratory Plymouth | Wellcome Sanger Institute for the COVID-19 Genomics UK (COG-UK) Consortium | Lighthouse Laboratory Plymouth and Alex Alderton, Roberto Amato, Jeffrey Barrett, Sonia Goncalves, Ewan Harrison, David K. Jackson, Ian Johnston, Dominic Kwiatkowski, Cordelia Langford, John Sillitoe on behalf of the Wellcome Sanger Institute COVID-19 Surveillance Team |
| EPI_ISL_2996031 | Lighthouse Lab in Glasgow | Wellcome Sanger Institute for the COVID-19 Genomics UK (COG-UK) Consortium | Harper VanSteenhouse, Yumi Kasai, David Gray, Carol Clugston, Anna Dominiczak and Alex Alderton, Roberto Amato, Jeffrey Barrett, Sonia Goncalves, Ewan Harrison, David K. Jackson, Ian Johnston, Dominic Kwiatkowski, Cordelia Langford, John Sillitoe on behalf of the Wellcome Sanger Institute COVID-19 Surveillance Team |
| EPI_ISL_2996032 | Berkshire and Surrey Pathology Services Lighthouse Laboratory | Wellcome Sanger Institute for the COVID-19 Genomics UK (COG-UK) Consortium | Berkshire and Surrey Pathology Services Lighthouse Laboratory and Alex Alderton, Roberto Amato, Jeffrey Barrett, Sonia Goncalves, Ewan Harrison, David K. Jackson, Ian Johnston, Dominic Kwiatkowski, Cordelia Langford, John Sillitoe on behalf of the Wellcome Sanger Institute COVID-19 Surveillance Team |
| EPI_ISL_2996033 | Lighthouse Lab in Glasgow | Wellcome Sanger Institute for the COVID-19 Genomics UK (COG-UK) Consortium | Harper VanSteenhouse, Yumi Kasai, David Gray, Carol Clugston, Anna Dominiczak and Alex Alderton, Roberto Amato, Jeffrey Barrett, Sonia Goncalves, Ewan Harrison, David K. Jackson, Ian Johnston, Dominic Kwiatkowski, Cordelia Langford, John Sillitoe on behalf of the Wellcome Sanger Institute COVID-19 Surveillance Team |
| EPI_ISL_2996034 | Berkshire and Surrey Pathology Services Lighthouse Laboratory | Wellcome Sanger Institute for the COVID-19 Genomics UK (COG-UK) Consortium | Berkshire and Surrey Pathology Services Lighthouse Laboratory and Alex Alderton, Roberto Amato, Jeffrey Barrett, Sonia Goncalves, Ewan Harrison, David K. Jackson, Ian Johnston, Dominic Kwiatkowski, Cordelia Langford, John Sillitoe on behalf of the Wellcome Sanger Institute COVID-19 Surveillance Team |
| EPI_ISL_2996035 | Lighthouse Lab in Glasgow | Wellcome Sanger Institute for the COVID-19 Genomics UK (COG-UK) Consortium | Harper VanSteenhouse, Yumi Kasai, David Gray, Carol Clugston, Anna Dominiczak and Alex Alderton, Roberto Amato, Jeffrey Barrett, Sonia Goncalves, Ewan Harrison, David K. Jackson, Ian Johnston, Dominic Kwiatkowski, Cordelia Langford, John Sillitoe on behalf of the Wellcome Sanger Institute COVID-19 Surveillance Team |
| EPI_ISL_2996036 | Berkshire and Surrey Pathology Services Lighthouse Laboratory | Wellcome Sanger Institute for the COVID-19 Genomics UK (COG-UK) Consortium | Berkshire and Surrey Pathology Services Lighthouse Laboratory and Alex Alderton, Roberto Amato, Jeffrey Barrett, Sonia Goncalves, Ewan Harrison, David K. Jackson, Ian Johnston, Dominic Kwiatkowski, Cordelia Langford, John Sillitoe on behalf of the Wellcome Sanger Institute COVID-19 Surveillance Team |
| EPI_ISL_2996037, EPI_ISL_2996038 | Lighthouse Laboratory Plymouth | Wellcome Sanger Institute for the COVID-19 Genomics UK (COG-UK) Consortium | Lighthouse Laboratory Plymouth and Alex Alderton, Roberto Amato, Jeffrey Barrett, Sonia Goncalves, Ewan Harrison, David K. Jackson, Ian Johnston, Dominic Kwiatkowski, Cordelia Langford, John Sillitoe on behalf of the Wellcome Sanger Institute COVID-19 Surveillance Team |
| EPI_ISL_2996039 | Berkshire and Surrey Pathology Services Lighthouse Laboratory | Wellcome Sanger Institute for the COVID-19 Genomics UK (COG-UK) Consortium | Berkshire and Surrey Pathology Services Lighthouse Laboratory and Alex Alderton, Roberto Amato, Jeffrey Barrett, Sonia Goncalves, Ewan Harrison, David K. Jackson, Ian Johnston, Dominic Kwiatkowski, Cordelia Langford, John Sillitoe on behalf of the Wellcome Sanger Institute COVID-19 Surveillance Team |
| EPI_ISL_2996040 | Lighthouse Laboratory Plymouth | Wellcome Sanger Institute for the COVID-19 Genomics UK (COG-UK) Consortium | Lighthouse Laboratory Plymouth and Alex Alderton, Roberto Amato, Jeffrey Barrett, Sonia Goncalves, Ewan Harrison, David K. Jackson, Ian Johnston, Dominic Kwiatkowski, Cordelia Langford, John Sillitoe on behalf of the Wellcome Sanger Institute COVID-19 Surveillance Team |
| EPI_ISL_2996041, EPI_ISL_2996042, EPI_ISL_2996043, EPI_ISL_2996044 | Lighthouse Lab in Glasgow | Wellcome Sanger Institute for the COVID-19 Genomics UK (COG-UK) Consortium | Harper VanSteenhouse, Yumi Kasai, David Gray, Carol Clugston, Anna Dominiczak and Alex Alderton, Roberto Amato, Jeffrey Barrett, Sonia Goncalves, Ewan Harrison, David K. Jackson, Ian Johnston, Dominic Kwiatkowski, Cordelia Langford, John Sillitoe on behalf of the Wellcome Sanger Institute COVID-19 Surveillance Team |
| EPI_ISL_2996045 | Berkshire and Surrey Pathology Services Lighthouse Laboratory | Wellcome Sanger Institute for the COVID-19 Genomics UK (COG-UK) Consortium | Berkshire and Surrey Pathology Services Lighthouse Laboratory and Alex Alderton, Roberto Amato, Jeffrey Barrett, Sonia Goncalves, Ewan Harrison, David K. Jackson, Ian Johnston, Dominic Kwiatkowski, Cordelia Langford, John Sillitoe on behalf of the Wellcome Sanger Institute COVID-19 Surveillance Team |
| EPI_ISL_2996046, EPI_ISL_2996047 | Lighthouse Laboratory Plymouth | Wellcome Sanger Institute for the COVID-19 Genomics UK (COG-UK) Consortium | Lighthouse Laboratory Plymouth and Alex Alderton, Roberto Amato, Jeffrey Barrett, Sonia Goncalves, Ewan Harrison, David K. Jackson, Ian Johnston, Dominic Kwiatkowski, Cordelia Langford, John Sillitoe on behalf of the Wellcome Sanger Institute COVID-19 Surveillance Team |
| EPI_ISL_2996048 | Berkshire and Surrey Pathology Services Lighthouse Laboratory | Wellcome Sanger Institute for the COVID-19 Genomics UK (COG-UK) Consortium | Berkshire and Surrey Pathology Services Lighthouse Laboratory and Alex Alderton, Roberto Amato, Jeffrey Barrett, Sonia Goncalves, Ewan Harrison, David K. Jackson, Ian Johnston, Dominic Kwiatkowski, Cordelia Langford, John Sillitoe on behalf of the Wellcome Sanger Institute COVID-19 Surveillance Team |
| EPI_ISL_2996049, EPI_ISL_2996050 | Lighthouse Lab in Glasgow | Wellcome Sanger Institute for the COVID-19 Genomics UK (COG-UK) Consortium | Harper VanSteenhouse, Yumi Kasai, David Gray, Carol Clugston, Anna Dominiczak and Alex Alderton, Roberto Amato, Jeffrey Barrett, Sonia Goncalves, Ewan Harrison, David K. Jackson, Ian Johnston, Dominic Kwiatkowski, Cordelia Langford, John Sillitoe on behalf of the Wellcome Sanger Institute COVID-19 Surveillance Team |
| EPI_ISL_2996051 | Lighthouse Laboratory Plymouth | Wellcome Sanger Institute for the COVID-19 Genomics UK (COG-UK) Consortium | Lighthouse Laboratory Plymouth and Alex Alderton, Roberto Amato, Jeffrey Barrett, Sonia Goncalves, Ewan Harrison, David K. Jackson, Ian Johnston, Dominic Kwiatkowski, Cordelia Langford, John Sillitoe on behalf of the Wellcome Sanger Institute COVID-19 Surveillance Team |
| EPI_ISL_2996052, EPI_ISL_2996053 | Lighthouse Lab in Glasgow | Wellcome Sanger Institute for the COVID-19 Genomics UK (COG-UK) Consortium | Harper VanSteenhouse, Yumi Kasai, David Gray, Carol Clugston, Anna Dominiczak and Alex Alderton, Roberto Amato, Jeffrey Barrett, Sonia Goncalves, Ewan Harrison, David K. Jackson, Ian Johnston, Dominic Kwiatkowski, Cordelia Langford, John Sillitoe on behalf of the Wellcome Sanger Institute COVID-19 Surveillance Team |
| EPI_ISL_2996054 | Lighthouse Laboratory Plymouth | Wellcome Sanger Institute for the COVID-19 Genomics UK (COG-UK) Consortium | Lighthouse Laboratory Plymouth and Alex Alderton, Roberto Amato, Jeffrey Barrett, Sonia Goncalves, Ewan Harrison, David K. Jackson, Ian Johnston, Dominic Kwiatkowski, Cordelia Langford, John Sillitoe on behalf of the Wellcome Sanger Institute COVID-19 Surveillance Team |
| EPI_ISL_2996055 | Lighthouse Lab in Glasgow | Wellcome Sanger Institute for the COVID-19 Genomics UK (COG-UK) Consortium | Harper VanSteenhouse, Yumi Kasai, David Gray, Carol Clugston, Anna Dominiczak and Alex Alderton, Roberto Amato, Jeffrey Barrett, Sonia Goncalves, Ewan Harrison, David K. Jackson, Ian Johnston, Dominic Kwiatkowski, Cordelia Langford, John Sillitoe on behalf of the Wellcome Sanger Institute COVID-19 Surveillance Team |
| EPI_ISL_2996056 | Lighthouse Laboratory Plymouth | Wellcome Sanger Institute for the COVID-19 Genomics UK (COG-UK) Consortium | Lighthouse Laboratory Plymouth and Alex Alderton, Roberto Amato, Jeffrey Barrett, Sonia Goncalves, Ewan Harrison, David K. Jackson, Ian Johnston, Dominic Kwiatkowski, Cordelia Langford, John Sillitoe on behalf of the Wellcome Sanger Institute COVID-19 Surveillance Team |
| EPI_ISL_2996057, EPI_ISL_2996058 | Lighthouse Lab in Glasgow | Wellcome Sanger Institute for the COVID-19 Genomics UK (COG-UK) Consortium | Harper VanSteenhouse, Yumi Kasai, David Gray, Carol Clugston, Anna Dominiczak and Alex Alderton, Roberto Amato, Jeffrey Barrett, Sonia Goncalves, Ewan Harrison, David K. Jackson, Ian Johnston, Dominic Kwiatkowski, Cordelia Langford, John Sillitoe on behalf of the Wellcome Sanger Institute COVID-19 Surveillance Team |
| EPI_ISL_2996059 | Berkshire and Surrey Pathology Services Lighthouse Laboratory | Wellcome Sanger Institute for the COVID-19 Genomics UK (COG-UK) Consortium | Berkshire and Surrey Pathology Services Lighthouse Laboratory and Alex Alderton, Roberto Amato, Jeffrey Barrett, Sonia Goncalves, Ewan Harrison, David K. Jackson, Ian Johnston, Dominic Kwiatkowski, Cordelia Langford, John Sillitoe on behalf of the Wellcome Sanger Institute COVID-19 Surveillance Team |
| EPI_ISL_2996060, EPI_ISL_2996061 | Lighthouse Lab in Glasgow | Wellcome Sanger Institute for the COVID-19 Genomics UK (COG-UK) Consortium | Harper VanSteenhouse, Yumi Kasai, David Gray, Carol Clugston, Anna Dominiczak and Alex Alderton, Roberto Amato, Jeffrey Barrett, Sonia Goncalves, Ewan Harrison, David K. Jackson, Ian Johnston, Dominic Kwiatkowski, Cordelia Langford, John Sillitoe on behalf of the Wellcome Sanger Institute COVID-19 Surveillance Team |
| EPI_ISL_2996062 | Lighthouse Laboratory Plymouth | Wellcome Sanger Institute for the COVID-19 Genomics UK (COG-UK) Consortium | Lighthouse Laboratory Plymouth and Alex Alderton, Roberto Amato, Jeffrey Barrett, Sonia Goncalves, Ewan Harrison, David K. Jackson, Ian Johnston, Dominic Kwiatkowski, Cordelia Langford, John Sillitoe on behalf of the Wellcome Sanger Institute COVID-19 Surveillance Team |
| EPI_ISL_2996063 | Lighthouse Lab in Glasgow | Wellcome Sanger Institute for the COVID-19 Genomics UK (COG-UK) Consortium | Harper VanSteenhouse, Yumi Kasai, David Gray, Carol Clugston, Anna Dominiczak and Alex Alderton, Roberto Amato, Jeffrey Barrett, Sonia Goncalves, Ewan Harrison, David K. Jackson, Ian Johnston, Dominic Kwiatkowski, Cordelia Langford, John Sillitoe on behalf of the Wellcome Sanger Institute COVID-19 Surveillance Team |
| EPI_ISL_2996064 | Lighthouse Laboratory Plymouth | Wellcome Sanger Institute for the COVID-19 Genomics UK (COG-UK) Consortium | Lighthouse Laboratory Plymouth and Alex Alderton, Roberto Amato, Jeffrey Barrett, Sonia Goncalves, Ewan Harrison, David K. Jackson, Ian Johnston, Dominic Kwiatkowski, Cordelia Langford, John Sillitoe on behalf of the Wellcome Sanger Institute COVID-19 Surveillance Team |
| EPI_ISL_2996065, EPI_ISL_2996066 | Lighthouse Lab in Glasgow | Wellcome Sanger Institute for the COVID-19 Genomics UK (COG-UK) Consortium | Harper VanSteenhouse, Yumi Kasai, David Gray, Carol Clugston, Anna Dominiczak and Alex Alderton, Roberto Amato, Jeffrey Barrett, Sonia Goncalves, Ewan Harrison, David K. Jackson, Ian Johnston, Dominic Kwiatkowski, Cordelia Langford, John Sillitoe on behalf of the Wellcome Sanger Institute COVID-19 Surveillance Team |
| EPI_ISL_2996067, EPI_ISL_2996068 | Berkshire and Surrey Pathology Services Lighthouse Laboratory | Wellcome Sanger Institute for the COVID-19 Genomics UK (COG-UK) Consortium | Berkshire and Surrey Pathology Services Lighthouse Laboratory and Alex Alderton, Roberto Amato, Jeffrey Barrett, Sonia Goncalves, Ewan Harrison, David K. Jackson, Ian Johnston, Dominic Kwiatkowski, Cordelia Langford, John Sillitoe on behalf of the Wellcome Sanger Institute COVID-19 Surveillance Team |
| EPI_ISL_2996069 | Lighthouse Laboratory Plymouth | Wellcome Sanger Institute for the COVID-19 Genomics UK (COG-UK) Consortium | Lighthouse Laboratory Plymouth and Alex Alderton, Roberto Amato, Jeffrey Barrett, Sonia Goncalves, Ewan Harrison, David K. Jackson, Ian Johnston, Dominic Kwiatkowski, Cordelia Langford, John Sillitoe on behalf of the Wellcome Sanger Institute COVID-19 Surveillance Team |
| EPI_ISL_2996070 | Lighthouse Lab in Glasgow | Wellcome Sanger Institute for the COVID-19 Genomics UK (COG-UK) Consortium | Harper VanSteenhouse, Yumi Kasai, David Gray, Carol Clugston, Anna Dominiczak and Alex Alderton, Roberto Amato, Jeffrey Barrett, Sonia Goncalves, Ewan Harrison, David K. Jackson, Ian Johnston, Dominic Kwiatkowski, Cordelia Langford, John Sillitoe on behalf of the Wellcome Sanger Institute COVID-19 Surveillance Team |
| EPI_ISL_2996071, EPI_ISL_2996072 | Lighthouse Laboratory Plymouth | Wellcome Sanger Institute for the COVID-19 Genomics UK (COG-UK) Consortium | Lighthouse Laboratory Plymouth and Alex Alderton, Roberto Amato, Jeffrey Barrett, Sonia Goncalves, Ewan Harrison, David K. Jackson, Ian Johnston, Dominic Kwiatkowski, Cordelia Langford, John Sillitoe on behalf of the Wellcome Sanger Institute COVID-19 Surveillance Team |
| EPI_ISL_2996073 | Berkshire and Surrey Pathology Services Lighthouse Laboratory | Wellcome Sanger Institute for the COVID-19 Genomics UK (COG-UK) | Berkshire and Surrey Pathology Services Lighthouse Laboratory and Alex Alderton, Roberto Amato, Jeffrey Barrett, Sonia Goncalves, Ewan Harrison, David K. Jackson, Ian Johnston, Dominic Kwiatkowski, Cordelia Langford, John Sillitoe on behalf |

| | | | |
|---|---|---|---|
| | | Consortium | of the Wellcome Sanger Institute COVID-19 Surveillance Team |
| EPI_ISL_2996074 | Lighthouse Laboratory Plymouth | Wellcome Sanger Institute for the COVID-19 Genomics UK (COG-UK) Consortium | Lighthouse Laboratory Plymouth and Alex Alderton, Roberto Amato, Jeffrey Barrett, Sonia Goncalves, Ewan Harrison, David K. Jackson, Ian Johnston, Dominic Kwiatkowski, Cordelia Langford, John Sillitoe on behalf of the Wellcome Sanger Institute COVID-19 Surveillance Team |
| EPI_ISL_2996075, EPI_ISL_2996076, EPI_ISL_2996077, EPI_ISL_2996078, EPI_ISL_2996079, EPI_ISL_2996080, EPI_ISL_2996081 | Lighthouse Lab in Glasgow | Wellcome Sanger Institute for the COVID-19 Genomics UK (COG-UK) Consortium | Harper VanSteenhouse, Yumi Kasai, David Gray, Carol Clugston, Anna Dominiczak and Alex Alderton, Roberto Amato, Jeffrey Barrett, Sonia Goncalves, Ewan Harrison, David K. Jackson, Ian Johnston, Dominic Kwiatkowski, Cordelia Langford, John Sillitoe on behalf of the Wellcome Sanger Institute COVID-19 Surveillance Team |
| EPI_ISL_2996082 | Lighthouse Laboratory Plymouth | Wellcome Sanger Institute for the COVID-19 Genomics UK (COG-UK) Consortium | Lighthouse Laboratory Plymouth and Alex Alderton, Roberto Amato, Jeffrey Barrett, Sonia Goncalves, Ewan Harrison, David K. Jackson, Ian Johnston, Dominic Kwiatkowski, Cordelia Langford, John Sillitoe on behalf of the Wellcome Sanger Institute COVID-19 Surveillance Team |
| EPI_ISL_2996083, EPI_ISL_2996084, EPI_ISL_2996085 | Lighthouse Lab in Glasgow | Wellcome Sanger Institute for the COVID-19 Genomics UK (COG-UK) Consortium | Harper VanSteenhouse, Yumi Kasai, David Gray, Carol Clugston, Anna Dominiczak and Alex Alderton, Roberto Amato, Jeffrey Barrett, Sonia Goncalves, Ewan Harrison, David K. Jackson, Ian Johnston, Dominic Kwiatkowski, Cordelia Langford, John Sillitoe on behalf of the Wellcome Sanger Institute COVID-19 Surveillance Team |
| EPI_ISL_2996086 | Lighthouse Laboratory Plymouth | Wellcome Sanger Institute for the COVID-19 Genomics UK (COG-UK) Consortium | Lighthouse Laboratory Plymouth and Alex Alderton, Roberto Amato, Jeffrey Barrett, Sonia Goncalves, Ewan Harrison, David K. Jackson, Ian Johnston, Dominic Kwiatkowski, Cordelia Langford, John Sillitoe on behalf of the Wellcome Sanger Institute COVID-19 Surveillance Team |
| EPI_ISL_2996087 | Berkshire and Surrey Pathology Services Lighthouse Laboratory | Wellcome Sanger Institute for the COVID-19 Genomics UK (COG-UK) Consortium | Berkshire and Surrey Pathology Services Lighthouse Laboratory and Alex Alderton, Roberto Amato, Jeffrey Barrett, Sonia Goncalves, Ewan Harrison, David K. Jackson, Ian Johnston, Dominic Kwiatkowski, Cordelia Langford, John Sillitoe on behalf of the Wellcome Sanger Institute COVID-19 Surveillance Team |
| EPI_ISL_2996088, EPI_ISL_2996089 | Lighthouse Lab in Glasgow | Wellcome Sanger Institute for the COVID-19 Genomics UK (COG-UK) Consortium | Harper VanSteenhouse, Yumi Kasai, David Gray, Carol Clugston, Anna Dominiczak and Alex Alderton, Roberto Amato, Jeffrey Barrett, Sonia Goncalves, Ewan Harrison, David K. Jackson, Ian Johnston, Dominic Kwiatkowski, Cordelia Langford, John Sillitoe on behalf of the Wellcome Sanger Institute COVID-19 Surveillance Team |
| EPI_ISL_2996090 | Lighthouse Laboratory Plymouth | Wellcome Sanger Institute for the COVID-19 Genomics UK (COG-UK) Consortium | Lighthouse Laboratory Plymouth and Alex Alderton, Roberto Amato, Jeffrey Barrett, Sonia Goncalves, Ewan Harrison, David K. Jackson, Ian Johnston, Dominic Kwiatkowski, Cordelia Langford, John Sillitoe on behalf of the Wellcome Sanger Institute COVID-19 Surveillance Team |
| EPI_ISL_2996091 | Lighthouse Lab in Glasgow | Wellcome Sanger Institute for the COVID-19 Genomics UK (COG-UK) Consortium | Harper VanSteenhouse, Yumi Kasai, David Gray, Carol Clugston, Anna Dominiczak and Alex Alderton, Roberto Amato, Jeffrey Barrett, Sonia Goncalves, Ewan Harrison, David K. Jackson, Ian Johnston, Dominic Kwiatkowski, Cordelia Langford, John Sillitoe on behalf of the Wellcome Sanger Institute COVID-19 Surveillance Team |
| EPI_ISL_2996092 | Lighthouse Laboratory Plymouth | Wellcome Sanger Institute for the COVID-19 Genomics UK (COG-UK) Consortium | Lighthouse Laboratory Plymouth and Alex Alderton, Roberto Amato, Jeffrey Barrett, Sonia Goncalves, Ewan Harrison, David K. Jackson, Ian Johnston, Dominic Kwiatkowski, Cordelia Langford, John Sillitoe on behalf of the Wellcome Sanger Institute COVID-19 Surveillance Team |
| EPI_ISL_2996093 | Berkshire and Surrey Pathology Services Lighthouse Laboratory | Wellcome Sanger Institute for the COVID-19 Genomics UK (COG-UK) Consortium | Berkshire and Surrey Pathology Services Lighthouse Laboratory and Alex Alderton, Roberto Amato, Jeffrey Barrett, Sonia Goncalves, Ewan Harrison, David K. Jackson, Ian Johnston, Dominic Kwiatkowski, Cordelia Langford, John Sillitoe on behalf of the Wellcome Sanger Institute COVID-19 Surveillance Team |
| EPI_ISL_2996094, EPI_ISL_2996095, EPI_ISL_2996096 | Lighthouse Lab in Glasgow | Wellcome Sanger Institute for the COVID-19 Genomics UK (COG-UK) Consortium | Harper VanSteenhouse, Yumi Kasai, David Gray, Carol Clugston, Anna Dominiczak and Alex Alderton, Roberto Amato, Jeffrey Barrett, Sonia Goncalves, Ewan Harrison, David K. Jackson, Ian Johnston, Dominic Kwiatkowski, Cordelia Langford, John Sillitoe on behalf of the Wellcome Sanger Institute COVID-19 Surveillance Team |
| EPI_ISL_2996097, EPI_ISL_2996098 | Lighthouse Laboratory Plymouth | Wellcome Sanger Institute for the COVID-19 Genomics UK (COG-UK) Consortium | Lighthouse Laboratory Plymouth and Alex Alderton, Roberto Amato, Jeffrey Barrett, Sonia Goncalves, Ewan Harrison, David K. Jackson, Ian Johnston, Dominic Kwiatkowski, Cordelia Langford, John Sillitoe on behalf of the Wellcome Sanger Institute COVID-19 Surveillance Team |
| EPI_ISL_2996099 | Lighthouse Lab in Glasgow | Wellcome Sanger Institute for the COVID-19 Genomics UK (COG-UK) Consortium | Harper VanSteenhouse, Yumi Kasai, David Gray, Carol Clugston, Anna Dominiczak and Alex Alderton, Roberto Amato, Jeffrey Barrett, Sonia Goncalves, Ewan Harrison, David K. Jackson, Ian Johnston, Dominic Kwiatkowski, Cordelia Langford, John Sillitoe on behalf of the Wellcome Sanger Institute COVID-19 Surveillance Team |
| EPI_ISL_2996101 | Berkshire and Surrey Pathology Services Lighthouse Laboratory | Wellcome Sanger Institute for the COVID-19 Genomics UK (COG-UK) Consortium | Berkshire and Surrey Pathology Services Lighthouse Laboratory and Alex Alderton, Roberto Amato, Jeffrey Barrett, Sonia Goncalves, Ewan Harrison, David K. Jackson, Ian Johnston, Dominic Kwiatkowski, Cordelia Langford, John Sillitoe on behalf of the Wellcome Sanger Institute COVID-19 Surveillance Team |
| EPI_ISL_2996102 | Lighthouse Lab in Glasgow | Wellcome Sanger Institute for the COVID-19 Genomics UK (COG-UK) Consortium | Harper VanSteenhouse, Yumi Kasai, David Gray, Carol Clugston, Anna Dominiczak and Alex Alderton, Roberto Amato, Jeffrey Barrett, Sonia Goncalves, Ewan Harrison, David K. Jackson, Ian Johnston, Dominic Kwiatkowski, Cordelia Langford, John Sillitoe on behalf of the Wellcome Sanger Institute COVID-19 Surveillance Team |
| EPI_ISL_2996103, EPI_ISL_2996104 | Lighthouse Laboratory Plymouth | Wellcome Sanger Institute for the COVID-19 Genomics UK (COG-UK) Consortium | Lighthouse Laboratory Plymouth and Alex Alderton, Roberto Amato, Jeffrey Barrett, Sonia Goncalves, Ewan Harrison, David K. Jackson, Ian Johnston, Dominic Kwiatkowski, Cordelia Langford, John Sillitoe on behalf of the Wellcome Sanger Institute COVID-19 Surveillance Team |
| EPI_ISL_2996105 | Lighthouse Lab in Glasgow | Wellcome Sanger Institute for the COVID-19 Genomics UK (COG-UK) Consortium | Harper VanSteenhouse, Yumi Kasai, David Gray, Carol Clugston, Anna Dominiczak and Alex Alderton, Roberto Amato, Jeffrey Barrett, Sonia Goncalves, Ewan Harrison, David K. Jackson, Ian Johnston, Dominic Kwiatkowski, Cordelia Langford, John Sillitoe on behalf of the Wellcome Sanger Institute COVID-19 Surveillance Team |
| EPI_ISL_2996106 | Berkshire and Surrey Pathology Services Lighthouse Laboratory | Wellcome Sanger Institute for the COVID-19 Genomics UK (COG-UK) Consortium | Berkshire and Surrey Pathology Services Lighthouse Laboratory and Alex Alderton, Roberto Amato, Jeffrey Barrett, Sonia Goncalves, Ewan Harrison, David K. Jackson, Ian Johnston, Dominic Kwiatkowski, Cordelia Langford, John Sillitoe on behalf of the Wellcome Sanger Institute COVID-19 Surveillance Team |
| EPI_ISL_2996107, EPI_ISL_2996108, EPI_ISL_2996109 | Lighthouse Lab in Glasgow | Wellcome Sanger Institute for the COVID-19 Genomics UK (COG-UK) Consortium | Harper VanSteenhouse, Yumi Kasai, David Gray, Carol Clugston, Anna Dominiczak and Alex Alderton, Roberto Amato, Jeffrey Barrett, Sonia Goncalves, Ewan Harrison, David K. Jackson, Ian Johnston, Dominic Kwiatkowski, Cordelia Langford, John Sillitoe on behalf of the Wellcome Sanger Institute COVID-19 Surveillance Team |
| EPI_ISL_2996110 | Lighthouse Laboratory Plymouth | Wellcome Sanger Institute for the COVID-19 Genomics UK (COG-UK) Consortium | Lighthouse Laboratory Plymouth and Alex Alderton, Roberto Amato, Jeffrey Barrett, Sonia Goncalves, Ewan Harrison, David K. Jackson, Ian Johnston, Dominic Kwiatkowski, Cordelia Langford, John Sillitoe on behalf of the Wellcome Sanger Institute COVID-19 Surveillance Team |
| EPI_ISL_2996111 | Berkshire and Surrey Pathology Services Lighthouse Laboratory | Wellcome Sanger Institute for the COVID-19 Genomics UK (COG-UK) Consortium | Berkshire and Surrey Pathology Services Lighthouse Laboratory and Alex Alderton, Roberto Amato, Jeffrey Barrett, Sonia Goncalves, Ewan Harrison, David K. Jackson, Ian Johnston, Dominic Kwiatkowski, Cordelia Langford, John Sillitoe on behalf of the Wellcome Sanger Institute COVID-19 Surveillance Team |
| EPI_ISL_2996112 | Lighthouse Laboratory Plymouth | Wellcome Sanger Institute for the COVID-19 Genomics UK (COG-UK) Consortium | Lighthouse Laboratory Plymouth and Alex Alderton, Roberto Amato, Jeffrey Barrett, Sonia Goncalves, Ewan Harrison, David K. Jackson, Ian Johnston, Dominic Kwiatkowski, Cordelia Langford, John Sillitoe on behalf of the Wellcome Sanger Institute COVID-19 Surveillance Team |
| EPI_ISL_2996113 | Lighthouse Lab in Glasgow | Wellcome Sanger Institute for the COVID-19 Genomics UK (COG-UK) Consortium | Harper VanSteenhouse, Yumi Kasai, David Gray, Carol Clugston, Anna Dominiczak and Alex Alderton, Roberto Amato, Jeffrey Barrett, Sonia Goncalves, Ewan Harrison, David K. Jackson, Ian Johnston, Dominic Kwiatkowski, Cordelia Langford, John Sillitoe on behalf of the Wellcome Sanger Institute COVID-19 Surveillance Team |
| EPI_ISL_2996114 | Berkshire and Surrey Pathology Services Lighthouse Laboratory | Wellcome Sanger Institute for the COVID-19 Genomics UK (COG-UK) Consortium | Berkshire and Surrey Pathology Services Lighthouse Laboratory and Alex Alderton, Roberto Amato, Jeffrey Barrett, Sonia Goncalves, Ewan Harrison, David K. Jackson, Ian Johnston, Dominic Kwiatkowski, Cordelia Langford, John Sillitoe on behalf of the Wellcome Sanger Institute COVID-19 Surveillance Team |
| EPI_ISL_2996115 | Lighthouse Laboratory Plymouth | Wellcome Sanger Institute for the COVID-19 Genomics UK (COG-UK) Consortium | Lighthouse Laboratory Plymouth and Alex Alderton, Roberto Amato, Jeffrey Barrett, Sonia Goncalves, Ewan Harrison, David K. Jackson, Ian Johnston, Dominic Kwiatkowski, Cordelia Langford, John Sillitoe on behalf of the Wellcome Sanger Institute COVID-19 Surveillance Team |
| EPI_ISL_2996116 | Lighthouse Lab in Glasgow | Wellcome Sanger Institute for the COVID-19 Genomics UK (COG-UK) Consortium | Harper VanSteenhouse, Yumi Kasai, David Gray, Carol Clugston, Anna Dominiczak and Alex Alderton, Roberto Amato, Jeffrey Barrett, Sonia Goncalves, Ewan Harrison, David K. Jackson, Ian Johnston, Dominic Kwiatkowski, Cordelia Langford, John Sillitoe on behalf of the Wellcome Sanger Institute COVID-19 Surveillance Team |
| EPI_ISL_2996117 | Lighthouse Laboratory Plymouth | Wellcome Sanger Institute for the COVID-19 Genomics UK (COG-UK) Consortium | Lighthouse Laboratory Plymouth and Alex Alderton, Roberto Amato, Jeffrey Barrett, Sonia Goncalves, Ewan Harrison, David K. Jackson, Ian Johnston, Dominic Kwiatkowski, Cordelia Langford, John Sillitoe on behalf of the Wellcome Sanger Institute COVID-19 Surveillance Team |
| EPI_ISL_2996118 | Lighthouse Lab in Glasgow | Wellcome Sanger Institute for the COVID-19 Genomics UK (COG-UK) Consortium | Harper VanSteenhouse, Yumi Kasai, David Gray, Carol Clugston, Anna Dominiczak and Alex Alderton, Roberto Amato, Jeffrey Barrett, Sonia Goncalves, Ewan Harrison, David K. Jackson, Ian Johnston, Dominic Kwiatkowski, Cordelia Langford, John Sillitoe on behalf of the Wellcome Sanger Institute COVID-19 Surveillance Team |
| EPI_ISL_2996119 | Berkshire and Surrey Pathology Services Lighthouse Laboratory | Wellcome Sanger Institute for the COVID-19 Genomics UK (COG-UK) Consortium | Berkshire and Surrey Pathology Services Lighthouse Laboratory and Alex Alderton, Roberto Amato, Jeffrey Barrett, Sonia Goncalves, Ewan Harrison, David K. Jackson, Ian Johnston, Dominic Kwiatkowski, Cordelia Langford, John Sillitoe on behalf of the Wellcome Sanger Institute COVID-19 Surveillance Team |
| EPI_ISL_2996120 | Lighthouse Laboratory Plymouth | Wellcome Sanger Institute for the COVID-19 Genomics UK (COG-UK) Consortium | Lighthouse Laboratory Plymouth and Alex Alderton, Roberto Amato, Jeffrey Barrett, Sonia Goncalves, Ewan Harrison, David K. Jackson, Ian Johnston, Dominic Kwiatkowski, Cordelia Langford, John Sillitoe on behalf of the Wellcome Sanger Institute COVID-19 Surveillance Team |
| EPI_ISL_2996121, EPI_ISL_2996122 | Lighthouse Lab in Glasgow | Wellcome Sanger Institute for the COVID-19 Genomics UK (COG-UK) Consortium | Harper VanSteenhouse, Yumi Kasai, David Gray, Carol Clugston, Anna Dominiczak and Alex Alderton, Roberto Amato, Jeffrey Barrett, Sonia Goncalves, Ewan Harrison, David K. Jackson, Ian Johnston, Dominic Kwiatkowski, Cordelia Langford, John Sillitoe on behalf of the Wellcome Sanger Institute COVID-19 Surveillance Team |
| EPI_ISL_2996123 | Berkshire and Surrey Pathology Services Lighthouse Laboratory | Wellcome Sanger Institute for the COVID-19 Genomics UK (COG-UK) Consortium | Berkshire and Surrey Pathology Services Lighthouse Laboratory and Alex Alderton, Roberto Amato, Jeffrey Barrett, Sonia Goncalves, Ewan Harrison, David K. Jackson, Ian Johnston, Dominic Kwiatkowski, Cordelia Langford, John Sillitoe on behalf of the Wellcome Sanger Institute COVID-19 Surveillance Team |
| EPI_ISL_2996124 | Lighthouse Lab in Glasgow | Wellcome Sanger Institute for the COVID-19 Genomics UK (COG-UK) Consortium | Harper VanSteenhouse, Yumi Kasai, David Gray, Carol Clugston, Anna Dominiczak and Alex Alderton, Roberto Amato, Jeffrey Barrett, Sonia Goncalves, Ewan Harrison, David K. Jackson, Ian Johnston, Dominic Kwiatkowski, Cordelia Langford, John Sillitoe on behalf of the Wellcome Sanger Institute COVID-19 Surveillance Team |
| EPI_ISL_2996125 | Lighthouse Laboratory Plymouth | Wellcome Sanger Institute for the COVID-19 Genomics UK (COG-UK) Consortium | Lighthouse Laboratory Plymouth and Alex Alderton, Roberto Amato, Jeffrey Barrett, Sonia Goncalves, Ewan Harrison, David K. Jackson, Ian Johnston, Dominic Kwiatkowski, Cordelia Langford, John Sillitoe on behalf of the Wellcome Sanger Institute COVID-19 Surveillance Team |
| EPI_ISL_2996126 | Lighthouse Lab in Glasgow | Wellcome Sanger Institute for the COVID-19 Genomics UK (COG-UK) Consortium | Harper VanSteenhouse, Yumi Kasai, David Gray, Carol Clugston, Anna Dominiczak and Alex Alderton, Roberto Amato, Jeffrey Barrett, Sonia Goncalves, Ewan Harrison, David K. Jackson, Ian Johnston, Dominic Kwiatkowski, Cordelia Langford, John Sillitoe on behalf of the Wellcome Sanger Institute COVID-19 Surveillance Team |
| EPI_ISL_2996127 | Berkshire and Surrey Pathology Services Lighthouse Laboratory | Wellcome Sanger Institute for the COVID-19 Genomics UK (COG-UK) Consortium | Berkshire and Surrey Pathology Services Lighthouse Laboratory and Alex Alderton, Roberto Amato, Jeffrey Barrett, Sonia Goncalves, Ewan Harrison, David K. Jackson, Ian Johnston, Dominic Kwiatkowski, Cordelia Langford, John Sillitoe on behalf of the Wellcome Sanger Institute COVID-19 Surveillance Team |
| EPI_ISL_2996128, EPI_ISL_2996129, EPI_ISL_2996130, EPI_ISL_2996131, EPI_ISL_2996132, EPI_ISL_2996133, EPI_ISL_2996134 | Lighthouse Lab in Glasgow | Wellcome Sanger Institute for the COVID-19 Genomics UK (COG-UK) Consortium | Harper VanSteenhouse, Yumi Kasai, David Gray, Carol Clugston, Anna Dominiczak and Alex Alderton, Roberto Amato, Jeffrey Barrett, Sonia Goncalves, Ewan Harrison, David K. Jackson, Ian Johnston, Dominic Kwiatkowski, Cordelia Langford, John Sillitoe on behalf of the Wellcome Sanger Institute COVID-19 Surveillance Team |
| EPI_ISL_2996135 | Berkshire and Surrey Pathology Services Lighthouse Laboratory | Wellcome Sanger Institute for the COVID-19 Genomics UK (COG-UK) Consortium | Berkshire and Surrey Pathology Services Lighthouse Laboratory and Alex Alderton, Roberto Amato, Jeffrey Barrett, Sonia Goncalves, Ewan Harrison, David K. Jackson, Ian Johnston, Dominic Kwiatkowski, Cordelia Langford, John Sillitoe on behalf of the Wellcome Sanger Institute COVID-19 Surveillance Team |
| EPI_ISL_2996136 | Lighthouse Laboratory Plymouth | Wellcome Sanger Institute for the COVID-19 Genomics UK (COG-UK) | Lighthouse Laboratory Plymouth and Alex Alderton, Roberto Amato, Jeffrey Barrett, Sonia Goncalves, Ewan Harrison, David K. Jackson, Ian Johnston, Dominic Kwiatkowski, Cordelia Langford, John Sillitoe on behalf of the Wellcome Sanger |

| | | | |
|---|---|---|---|
| | | Consortium | Institute COVID-19 Surveillance Team |
| EPI_ISL_2996137 | Berkshire and Surrey Pathology Services Lighthouse Laboratory | Wellcome Sanger Institute for the COVID-19 Genomics UK (COG-UK) Consortium | Berkshire and Surrey Pathology Services Lighthouse Laboratory and Alex Alderton, Roberto Amato, Jeffrey Barrett, Sonia Goncalves, Ewan Harrison, David K. Jackson, Ian Johnston, Dominic Kwiatkowski, Cordelia Langford, John Sillitoe on behalf of the Wellcome Sanger Institute COVID-19 Surveillance Team |
| EPI_ISL_2996138 | Lighthouse Lab in Glasgow | Wellcome Sanger Institute for the COVID-19 Genomics UK (COG-UK) Consortium | Harper VanSteenhouse, Yumi Kasai, David Gray, Carol Clugston, Anna Dominiczak and Alex Alderton, Roberto Amato, Jeffrey Barrett, Sonia Goncalves, Ewan Harrison, David K. Jackson, Ian Johnston, Dominic Kwiatkowski, Cordelia Langford, John Sillitoe on behalf of the Wellcome Sanger Institute COVID-19 Surveillance Team |
| EPI_ISL_2996139, EPI_ISL_2996140 | Berkshire and Surrey Pathology Services Lighthouse Laboratory | Wellcome Sanger Institute for the COVID-19 Genomics UK (COG-UK) Consortium | Berkshire and Surrey Pathology Services Lighthouse Laboratory and Alex Alderton, Roberto Amato, Jeffrey Barrett, Sonia Goncalves, Ewan Harrison, David K. Jackson, Ian Johnston, Dominic Kwiatkowski, Cordelia Langford, John Sillitoe on behalf of the Wellcome Sanger Institute COVID-19 Surveillance Team |
| EPI_ISL_2996141 | Lighthouse Lab in Glasgow | Wellcome Sanger Institute for the COVID-19 Genomics UK (COG-UK) Consortium | Harper VanSteenhouse, Yumi Kasai, David Gray, Carol Clugston, Anna Dominiczak and Alex Alderton, Roberto Amato, Jeffrey Barrett, Sonia Goncalves, Ewan Harrison, David K. Jackson, Ian Johnston, Dominic Kwiatkowski, Cordelia Langford, John Sillitoe on behalf of the Wellcome Sanger Institute COVID-19 Surveillance Team |
| EPI_ISL_2996142 | Berkshire and Surrey Pathology Services Lighthouse Laboratory | Wellcome Sanger Institute for the COVID-19 Genomics UK (COG-UK) Consortium | Berkshire and Surrey Pathology Services Lighthouse Laboratory and Alex Alderton, Roberto Amato, Jeffrey Barrett, Sonia Goncalves, Ewan Harrison, David K. Jackson, Ian Johnston, Dominic Kwiatkowski, Cordelia Langford, John Sillitoe on behalf of the Wellcome Sanger Institute COVID-19 Surveillance Team |
| EPI_ISL_2996143, EPI_ISL_2996144, EPI_ISL_2996145, EPI_ISL_2996146, EPI_ISL_2996147, EPI_ISL_2996148 | Lighthouse Lab in Glasgow | Wellcome Sanger Institute for the COVID-19 Genomics UK (COG-UK) Consortium | Harper VanSteenhouse, Yumi Kasai, David Gray, Carol Clugston, Anna Dominiczak and Alex Alderton, Roberto Amato, Jeffrey Barrett, Sonia Goncalves, Ewan Harrison, David K. Jackson, Ian Johnston, Dominic Kwiatkowski, Cordelia Langford, John Sillitoe on behalf of the Wellcome Sanger Institute COVID-19 Surveillance Team |
| EPI_ISL_2996149 | Berkshire and Surrey Pathology Services Lighthouse Laboratory | Wellcome Sanger Institute for the COVID-19 Genomics UK (COG-UK) Consortium | Berkshire and Surrey Pathology Services Lighthouse Laboratory and Alex Alderton, Roberto Amato, Jeffrey Barrett, Sonia Goncalves, Ewan Harrison, David K. Jackson, Ian Johnston, Dominic Kwiatkowski, Cordelia Langford, John Sillitoe on behalf of the Wellcome Sanger Institute COVID-19 Surveillance Team |
| EPI_ISL_2996150 | Lighthouse Lab in Glasgow | Wellcome Sanger Institute for the COVID-19 Genomics UK (COG-UK) Consortium | Harper VanSteenhouse, Yumi Kasai, David Gray, Carol Clugston, Anna Dominiczak and Alex Alderton, Roberto Amato, Jeffrey Barrett, Sonia Goncalves, Ewan Harrison, David K. Jackson, Ian Johnston, Dominic Kwiatkowski, Cordelia Langford, John Sillitoe on behalf of the Wellcome Sanger Institute COVID-19 Surveillance Team |
| EPI_ISL_2996151 | Berkshire and Surrey Pathology Services Lighthouse Laboratory | Wellcome Sanger Institute for the COVID-19 Genomics UK (COG-UK) Consortium | Berkshire and Surrey Pathology Services Lighthouse Laboratory and Alex Alderton, Roberto Amato, Jeffrey Barrett, Sonia Goncalves, Ewan Harrison, David K. Jackson, Ian Johnston, Dominic Kwiatkowski, Cordelia Langford, John Sillitoe on behalf of the Wellcome Sanger Institute COVID-19 Surveillance Team |
| EPI_ISL_2996152 | Lighthouse Lab in Glasgow | Wellcome Sanger Institute for the COVID-19 Genomics UK (COG-UK) Consortium | Harper VanSteenhouse, Yumi Kasai, David Gray, Carol Clugston, Anna Dominiczak and Alex Alderton, Roberto Amato, Jeffrey Barrett, Sonia Goncalves, Ewan Harrison, David K. Jackson, Ian Johnston, Dominic Kwiatkowski, Cordelia Langford, John Sillitoe on behalf of the Wellcome Sanger Institute COVID-19 Surveillance Team |
| EPI_ISL_2996153, EPI_ISL_2996154 | Berkshire and Surrey Pathology Services Lighthouse Laboratory | Wellcome Sanger Institute for the COVID-19 Genomics UK (COG-UK) Consortium | Berkshire and Surrey Pathology Services Lighthouse Laboratory and Alex Alderton, Roberto Amato, Jeffrey Barrett, Sonia Goncalves, Ewan Harrison, David K. Jackson, Ian Johnston, Dominic Kwiatkowski, Cordelia Langford, John Sillitoe on behalf of the Wellcome Sanger Institute COVID-19 Surveillance Team |
| EPI_ISL_2996155, EPI_ISL_2996156 | Lighthouse Lab in Glasgow | Wellcome Sanger Institute for the COVID-19 Genomics UK (COG-UK) Consortium | Harper VanSteenhouse, Yumi Kasai, David Gray, Carol Clugston, Anna Dominiczak and Alex Alderton, Roberto Amato, Jeffrey Barrett, Sonia Goncalves, Ewan Harrison, David K. Jackson, Ian Johnston, Dominic Kwiatkowski, Cordelia Langford, John Sillitoe on behalf of the Wellcome Sanger Institute COVID-19 Surveillance Team |
| EPI_ISL_2996157 | Berkshire and Surrey Pathology Services Lighthouse Laboratory | Wellcome Sanger Institute for the COVID-19 Genomics UK (COG-UK) Consortium | Berkshire and Surrey Pathology Services Lighthouse Laboratory and Alex Alderton, Roberto Amato, Jeffrey Barrett, Sonia Goncalves, Ewan Harrison, David K. Jackson, Ian Johnston, Dominic Kwiatkowski, Cordelia Langford, John Sillitoe on behalf of the Wellcome Sanger Institute COVID-19 Surveillance Team |
| EPI_ISL_2996158 | Lighthouse Laboratory Plymouth | Wellcome Sanger Institute for the COVID-19 Genomics UK (COG-UK) Consortium | Lighthouse Laboratory Plymouth and Alex Alderton, Roberto Amato, Jeffrey Barrett, Sonia Goncalves, Ewan Harrison, David K. Jackson, Ian Johnston, Dominic Kwiatkowski, Cordelia Langford, John Sillitoe on behalf of the Wellcome Sanger Institute COVID-19 Surveillance Team |
| EPI_ISL_2996159 | Berkshire and Surrey Pathology Services Lighthouse Laboratory | Wellcome Sanger Institute for the COVID-19 Genomics UK (COG-UK) Consortium | Berkshire and Surrey Pathology Services Lighthouse Laboratory and Alex Alderton, Roberto Amato, Jeffrey Barrett, Sonia Goncalves, Ewan Harrison, David K. Jackson, Ian Johnston, Dominic Kwiatkowski, Cordelia Langford, John Sillitoe on behalf of the Wellcome Sanger Institute COVID-19 Surveillance Team |
| EPI_ISL_2996160 | Lighthouse Laboratory Plymouth | Wellcome Sanger Institute for the COVID-19 Genomics UK (COG-UK) Consortium | Lighthouse Laboratory Plymouth and Alex Alderton, Roberto Amato, Jeffrey Barrett, Sonia Goncalves, Ewan Harrison, David K. Jackson, Ian Johnston, Dominic Kwiatkowski, Cordelia Langford, John Sillitoe on behalf of the Wellcome Sanger Institute COVID-19 Surveillance Team |
| EPI_ISL_2996161 | Lighthouse Lab in Glasgow | Wellcome Sanger Institute for the COVID-19 Genomics UK (COG-UK) Consortium | Harper VanSteenhouse, Yumi Kasai, David Gray, Carol Clugston, Anna Dominiczak and Alex Alderton, Roberto Amato, Jeffrey Barrett, Sonia Goncalves, Ewan Harrison, David K. Jackson, Ian Johnston, Dominic Kwiatkowski, Cordelia Langford, John Sillitoe on behalf of the Wellcome Sanger Institute COVID-19 Surveillance Team |
| EPI_ISL_2996162 | Berkshire and Surrey Pathology Services Lighthouse Laboratory | Wellcome Sanger Institute for the COVID-19 Genomics UK (COG-UK) Consortium | Berkshire and Surrey Pathology Services Lighthouse Laboratory and Alex Alderton, Roberto Amato, Jeffrey Barrett, Sonia Goncalves, Ewan Harrison, David K. Jackson, Ian Johnston, Dominic Kwiatkowski, Cordelia Langford, John Sillitoe on behalf of the Wellcome Sanger Institute COVID-19 Surveillance Team |
| EPI_ISL_2996163 | Lighthouse Lab in Glasgow | Wellcome Sanger Institute for the COVID-19 Genomics UK (COG-UK) Consortium | Harper VanSteenhouse, Yumi Kasai, David Gray, Carol Clugston, Anna Dominiczak and Alex Alderton, Roberto Amato, Jeffrey Barrett, Sonia Goncalves, Ewan Harrison, David K. Jackson, Ian Johnston, Dominic Kwiatkowski, Cordelia Langford, John Sillitoe on behalf of the Wellcome Sanger Institute COVID-19 Surveillance Team |
| EPI_ISL_2996164 | Lighthouse Laboratory Plymouth | Wellcome Sanger Institute for the COVID-19 Genomics UK (COG-UK) Consortium | Lighthouse Laboratory Plymouth and Alex Alderton, Roberto Amato, Jeffrey Barrett, Sonia Goncalves, Ewan Harrison, David K. Jackson, Ian Johnston, Dominic Kwiatkowski, Cordelia Langford, John Sillitoe on behalf of the Wellcome Sanger Institute COVID-19 Surveillance Team |
| EPI_ISL_2996165, EPI_ISL_2996166 | Lighthouse Lab in Glasgow | Wellcome Sanger Institute for the COVID-19 Genomics UK (COG-UK) Consortium | Harper VanSteenhouse, Yumi Kasai, David Gray, Carol Clugston, Anna Dominiczak and Alex Alderton, Roberto Amato, Jeffrey Barrett, Sonia Goncalves, Ewan Harrison, David K. Jackson, Ian Johnston, Dominic Kwiatkowski, Cordelia Langford, John Sillitoe on behalf of the Wellcome Sanger Institute COVID-19 Surveillance Team |
| EPI_ISL_2996167 | Lighthouse Laboratory Plymouth | Wellcome Sanger Institute for the COVID-19 Genomics UK (COG-UK) Consortium | Lighthouse Laboratory Plymouth and Alex Alderton, Roberto Amato, Jeffrey Barrett, Sonia Goncalves, Ewan Harrison, David K. Jackson, Ian Johnston, Dominic Kwiatkowski, Cordelia Langford, John Sillitoe on behalf of the Wellcome Sanger Institute COVID-19 Surveillance Team |
| EPI_ISL_2996168 | Berkshire and Surrey Pathology Services Lighthouse Laboratory | Wellcome Sanger Institute for the COVID-19 Genomics UK (COG-UK) Consortium | Berkshire and Surrey Pathology Services Lighthouse Laboratory and Alex Alderton, Roberto Amato, Jeffrey Barrett, Sonia Goncalves, Ewan Harrison, David K. Jackson, Ian Johnston, Dominic Kwiatkowski, Cordelia Langford, John Sillitoe on behalf of the Wellcome Sanger Institute COVID-19 Surveillance Team |
| EPI_ISL_2996169 | Lighthouse Laboratory Plymouth | Wellcome Sanger Institute for the COVID-19 Genomics UK (COG-UK) Consortium | Lighthouse Laboratory Plymouth and Alex Alderton, Roberto Amato, Jeffrey Barrett, Sonia Goncalves, Ewan Harrison, David K. Jackson, Ian Johnston, Dominic Kwiatkowski, Cordelia Langford, John Sillitoe on behalf of the Wellcome Sanger Institute COVID-19 Surveillance Team |
| EPI_ISL_2996170, EPI_ISL_2996171 | Lighthouse Lab in Glasgow | Wellcome Sanger Institute for the COVID-19 Genomics UK (COG-UK) Consortium | Harper VanSteenhouse, Yumi Kasai, David Gray, Carol Clugston, Anna Dominiczak and Alex Alderton, Roberto Amato, Jeffrey Barrett, Sonia Goncalves, Ewan Harrison, David K. Jackson, Ian Johnston, Dominic Kwiatkowski, Cordelia Langford, John Sillitoe on behalf of the Wellcome Sanger Institute COVID-19 Surveillance Team |
| EPI_ISL_2996172 | Berkshire and Surrey Pathology Services Lighthouse Laboratory | Wellcome Sanger Institute for the COVID-19 Genomics UK (COG-UK) Consortium | Berkshire and Surrey Pathology Services Lighthouse Laboratory and Alex Alderton, Roberto Amato, Jeffrey Barrett, Sonia Goncalves, Ewan Harrison, David K. Jackson, Ian Johnston, Dominic Kwiatkowski, Cordelia Langford, John Sillitoe on behalf of the Wellcome Sanger Institute COVID-19 Surveillance Team |
| EPI_ISL_2996173 | Lighthouse Laboratory Plymouth | Wellcome Sanger Institute for the COVID-19 Genomics UK (COG-UK) Consortium | Lighthouse Laboratory Plymouth and Alex Alderton, Roberto Amato, Jeffrey Barrett, Sonia Goncalves, Ewan Harrison, David K. Jackson, Ian Johnston, Dominic Kwiatkowski, Cordelia Langford, John Sillitoe on behalf of the Wellcome Sanger Institute COVID-19 Surveillance Team |
| EPI_ISL_2996174 | Lighthouse Lab in Glasgow | Wellcome Sanger Institute for the COVID-19 Genomics UK (COG-UK) Consortium | Harper VanSteenhouse, Yumi Kasai, David Gray, Carol Clugston, Anna Dominiczak and Alex Alderton, Roberto Amato, Jeffrey Barrett, Sonia Goncalves, Ewan Harrison, David K. Jackson, Ian Johnston, Dominic Kwiatkowski, Cordelia Langford, John Sillitoe on behalf of the Wellcome Sanger Institute COVID-19 Surveillance Team |
| EPI_ISL_2996175, EPI_ISL_2996176 | Lighthouse Laboratory Plymouth | Wellcome Sanger Institute for the COVID-19 Genomics UK (COG-UK) Consortium | Lighthouse Laboratory Plymouth and Alex Alderton, Roberto Amato, Jeffrey Barrett, Sonia Goncalves, Ewan Harrison, David K. Jackson, Ian Johnston, Dominic Kwiatkowski, Cordelia Langford, John Sillitoe on behalf of the Wellcome Sanger Institute COVID-19 Surveillance Team |
| EPI_ISL_2996177, EPI_ISL_2996178 | Lighthouse Lab in Glasgow | Wellcome Sanger Institute for the COVID-19 Genomics UK (COG-UK) Consortium | Harper VanSteenhouse, Yumi Kasai, David Gray, Carol Clugston, Anna Dominiczak and Alex Alderton, Roberto Amato, Jeffrey Barrett, Sonia Goncalves, Ewan Harrison, David K. Jackson, Ian Johnston, Dominic Kwiatkowski, Cordelia Langford, John Sillitoe on behalf of the Wellcome Sanger Institute COVID-19 Surveillance Team |
| EPI_ISL_2996179 | Berkshire and Surrey Pathology Services Lighthouse Laboratory | Wellcome Sanger Institute for the COVID-19 Genomics UK (COG-UK) Consortium | Berkshire and Surrey Pathology Services Lighthouse Laboratory and Alex Alderton, Roberto Amato, Jeffrey Barrett, Sonia Goncalves, Ewan Harrison, David K. Jackson, Ian Johnston, Dominic Kwiatkowski, Cordelia Langford, John Sillitoe on behalf of the Wellcome Sanger Institute COVID-19 Surveillance Team |
| EPI_ISL_2996180 | Lighthouse Lab in Glasgow | Wellcome Sanger Institute for the COVID-19 Genomics UK (COG-UK) Consortium | Harper VanSteenhouse, Yumi Kasai, David Gray, Carol Clugston, Anna Dominiczak and Alex Alderton, Roberto Amato, Jeffrey Barrett, Sonia Goncalves, Ewan Harrison, David K. Jackson, Ian Johnston, Dominic Kwiatkowski, Cordelia Langford, John Sillitoe on behalf of the Wellcome Sanger Institute COVID-19 Surveillance Team |
| EPI_ISL_2996181 | Berkshire and Surrey Pathology Services Lighthouse Laboratory | Wellcome Sanger Institute for the COVID-19 Genomics UK (COG-UK) Consortium | Berkshire and Surrey Pathology Services Lighthouse Laboratory and Alex Alderton, Roberto Amato, Jeffrey Barrett, Sonia Goncalves, Ewan Harrison, David K. Jackson, Ian Johnston, Dominic Kwiatkowski, Cordelia Langford, John Sillitoe on behalf of the Wellcome Sanger Institute COVID-19 Surveillance Team |
| EPI_ISL_2996182 | Lighthouse Lab in Glasgow | Wellcome Sanger Institute for the COVID-19 Genomics UK (COG-UK) Consortium | Harper VanSteenhouse, Yumi Kasai, David Gray, Carol Clugston, Anna Dominiczak and Alex Alderton, Roberto Amato, Jeffrey Barrett, Sonia Goncalves, Ewan Harrison, David K. Jackson, Ian Johnston, Dominic Kwiatkowski, Cordelia Langford, John Sillitoe on behalf of the Wellcome Sanger Institute COVID-19 Surveillance Team |
| EPI_ISL_2996183 | Lighthouse Laboratory Plymouth | Wellcome Sanger Institute for the COVID-19 Genomics UK (COG-UK) Consortium | Lighthouse Laboratory Plymouth and Alex Alderton, Roberto Amato, Jeffrey Barrett, Sonia Goncalves, Ewan Harrison, David K. Jackson, Ian Johnston, Dominic Kwiatkowski, Cordelia Langford, John Sillitoe on behalf of the Wellcome Sanger Institute COVID-19 Surveillance Team |
| EPI_ISL_2996184, EPI_ISL_2996185 | Lighthouse Lab in Glasgow | Wellcome Sanger Institute for the COVID-19 Genomics UK (COG-UK) Consortium | Harper VanSteenhouse, Yumi Kasai, David Gray, Carol Clugston, Anna Dominiczak and Alex Alderton, Roberto Amato, Jeffrey Barrett, Sonia Goncalves, Ewan Harrison, David K. Jackson, Ian Johnston, Dominic Kwiatkowski, Cordelia Langford, John Sillitoe on behalf of the Wellcome Sanger Institute COVID-19 Surveillance Team |
| EPI_ISL_2996186, EPI_ISL_2996187, EPI_ISL_2996188, EPI_ISL_2996189 | Lighthouse Laboratory Plymouth | Wellcome Sanger Institute for the COVID-19 Genomics UK (COG-UK) Consortium | Lighthouse Laboratory Plymouth and Alex Alderton, Roberto Amato, Jeffrey Barrett, Sonia Goncalves, Ewan Harrison, David K. Jackson, Ian Johnston, Dominic Kwiatkowski, Cordelia Langford, John Sillitoe on behalf of the Wellcome Sanger Institute COVID-19 Surveillance Team |
| EPI_ISL_2996190 | Berkshire and Surrey Pathology Services Lighthouse Laboratory | Wellcome Sanger Institute for the COVID-19 Genomics UK (COG-UK) Consortium | Berkshire and Surrey Pathology Services Lighthouse Laboratory and Alex Alderton, Roberto Amato, Jeffrey Barrett, Sonia Goncalves, Ewan Harrison, David K. Jackson, Ian Johnston, Dominic Kwiatkowski, Cordelia Langford, John Sillitoe on behalf of the Wellcome Sanger Institute COVID-19 Surveillance Team |
| EPI_ISL_2996191 | Lighthouse Laboratory Plymouth | Wellcome Sanger Institute for the COVID-19 Genomics UK (COG-UK) Consortium | Lighthouse Laboratory Plymouth and Alex Alderton, Roberto Amato, Jeffrey Barrett, Sonia Goncalves, Ewan Harrison, David K. Jackson, Ian Johnston, Dominic Kwiatkowski, Cordelia Langford, John Sillitoe on behalf of the Wellcome Sanger Institute COVID-19 Surveillance Team |
| EPI_ISL_2996192 | Lighthouse Lab in Glasgow | Wellcome Sanger Institute for the COVID-19 Genomics UK (COG-UK) Consortium | Harper VanSteenhouse, Yumi Kasai, David Gray, Carol Clugston, Anna Dominiczak and Alex Alderton, Roberto Amato, Jeffrey Barrett, Sonia Goncalves, Ewan Harrison, David K. Jackson, Ian Johnston, Dominic Kwiatkowski, Cordelia Langford, John Sillitoe on behalf of the Wellcome Sanger Institute COVID-19 Surveillance Team |
| EPI_ISL_2996193 | Berkshire and Surrey Pathology Services Lighthouse Laboratory | Wellcome Sanger Institute for the COVID-19 Genomics UK (COG-UK) Consortium | Berkshire and Surrey Pathology Services Lighthouse Laboratory and Alex Alderton, Roberto Amato, Jeffrey Barrett, Sonia Goncalves, Ewan Harrison, David K. Jackson, Ian Johnston, Dominic Kwiatkowski, Cordelia Langford, John Sillitoe on behalf of the Wellcome Sanger Institute COVID-19 Surveillance Team |
| EPI_ISL_2996194, EPI_ISL_2996195, | Lighthouse Laboratory Plymouth | Wellcome Sanger Institute for the COVID-19 Genomics UK (COG-UK) Consortium | Lighthouse Laboratory Plymouth and Alex Alderton, Roberto Amato, Jeffrey Barrett, Sonia Goncalves, Ewan Harrison, David K. Jackson, Ian Johnston, Dominic Kwiatkowski, Cordelia Langford, John Sillitoe on behalf of the Wellcome Sanger Institute COVID-19 Surveillance Team |

| | | | |
|---|---|---|---|
| EPI_ISL_2996196, EPI_ISL_2996197 | | | |
| EPI_ISL_2996198, EPI_ISL_2996199, EPI_ISL_2996200, EPI_ISL_2996201, EPI_ISL_2996202, EPI_ISL_2996203 | Lighthouse Lab in Glasgow | Wellcome Sanger Institute for the COVID-19 Genomics UK (COG-UK) Consortium | Harper VanSteenhouse, Yumi Kasai, David Gray, Carol Clugston, Anna Dominiczak and Alex Alderton, Roberto Amato, Jeffrey Barrett, Sonia Goncalves, Ewan Harrison, David K. Jackson, Ian Johnston, Dominic Kwiatkowski, Cordelia Langford, John Sillitoe on behalf of the Wellcome Sanger Institute COVID-19 Surveillance Team |
| EPI_ISL_2996204 | Berkshire and Surrey Pathology Services Lighthouse Laboratory | Wellcome Sanger Institute for the COVID-19 Genomics UK (COG-UK) Consortium | Berkshire and Surrey Pathology Services Lighthouse Laboratory and Alex Alderton, Roberto Amato, Jeffrey Barrett, Sonia Goncalves, Ewan Harrison, David K. Jackson, Ian Johnston, Dominic Kwiatkowski, Cordelia Langford, John Sillitoe on behalf of the Wellcome Sanger Institute COVID-19 Surveillance Team |
| EPI_ISL_2996205, EPI_ISL_2996206, EPI_ISL_2996207, EPI_ISL_2996208 | Lighthouse Lab in Glasgow | Wellcome Sanger Institute for the COVID-19 Genomics UK (COG-UK) Consortium | Harper VanSteenhouse, Yumi Kasai, David Gray, Carol Clugston, Anna Dominiczak and Alex Alderton, Roberto Amato, Jeffrey Barrett, Sonia Goncalves, Ewan Harrison, David K. Jackson, Ian Johnston, Dominic Kwiatkowski, Cordelia Langford, John Sillitoe on behalf of the Wellcome Sanger Institute COVID-19 Surveillance Team |
| EPI_ISL_2996209 | Lighthouse Laboratory Plymouth | Wellcome Sanger Institute for the COVID-19 Genomics UK (COG-UK) Consortium | Lighthouse Laboratory Plymouth and Alex Alderton, Roberto Amato, Jeffrey Barrett, Sonia Goncalves, Ewan Harrison, David K. Jackson, Ian Johnston, Dominic Kwiatkowski, Cordelia Langford, John Sillitoe on behalf of the Wellcome Sanger Institute COVID-19 Surveillance Team |
| EPI_ISL_2996210 | Lighthouse Lab in Glasgow | Wellcome Sanger Institute for the COVID-19 Genomics UK (COG-UK) Consortium | Harper VanSteenhouse, Yumi Kasai, David Gray, Carol Clugston, Anna Dominiczak and Alex Alderton, Roberto Amato, Jeffrey Barrett, Sonia Goncalves, Ewan Harrison, David K. Jackson, Ian Johnston, Dominic Kwiatkowski, Cordelia Langford, John Sillitoe on behalf of the Wellcome Sanger Institute COVID-19 Surveillance Team |
| EPI_ISL_2996211, EPI_ISL_2996212 | Berkshire and Surrey Pathology Services Lighthouse Laboratory | Wellcome Sanger Institute for the COVID-19 Genomics UK (COG-UK) Consortium | Berkshire and Surrey Pathology Services Lighthouse Laboratory and Alex Alderton, Roberto Amato, Jeffrey Barrett, Sonia Goncalves, Ewan Harrison, David K. Jackson, Ian Johnston, Dominic Kwiatkowski, Cordelia Langford, John Sillitoe on behalf of the Wellcome Sanger Institute COVID-19 Surveillance Team |
| EPI_ISL_2996213, EPI_ISL_2996214, EPI_ISL_2996215, EPI_ISL_2996216, EPI_ISL_2996217 | Lighthouse Lab in Glasgow | Wellcome Sanger Institute for the COVID-19 Genomics UK (COG-UK) Consortium | Harper VanSteenhouse, Yumi Kasai, David Gray, Carol Clugston, Anna Dominiczak and Alex Alderton, Roberto Amato, Jeffrey Barrett, Sonia Goncalves, Ewan Harrison, David K. Jackson, Ian Johnston, Dominic Kwiatkowski, Cordelia Langford, John Sillitoe on behalf of the Wellcome Sanger Institute COVID-19 Surveillance Team |
| EPI_ISL_2996218, EPI_ISL_2996219 | Berkshire and Surrey Pathology Services Lighthouse Laboratory | Wellcome Sanger Institute for the COVID-19 Genomics UK (COG-UK) Consortium | Berkshire and Surrey Pathology Services Lighthouse Laboratory and Alex Alderton, Roberto Amato, Jeffrey Barrett, Sonia Goncalves, Ewan Harrison, David K. Jackson, Ian Johnston, Dominic Kwiatkowski, Cordelia Langford, John Sillitoe on behalf of the Wellcome Sanger Institute COVID-19 Surveillance Team |
| EPI_ISL_2996220, EPI_ISL_2996221 | Lighthouse Lab in Glasgow | Wellcome Sanger Institute for the COVID-19 Genomics UK (COG-UK) Consortium | Harper VanSteenhouse, Yumi Kasai, David Gray, Carol Clugston, Anna Dominiczak and Alex Alderton, Roberto Amato, Jeffrey Barrett, Sonia Goncalves, Ewan Harrison, David K. Jackson, Ian Johnston, Dominic Kwiatkowski, Cordelia Langford, John Sillitoe on behalf of the Wellcome Sanger Institute COVID-19 Surveillance Team |
| EPI_ISL_2996222 | Lighthouse Laboratory Plymouth | Wellcome Sanger Institute for the COVID-19 Genomics UK (COG-UK) Consortium | Lighthouse Laboratory Plymouth and Alex Alderton, Roberto Amato, Jeffrey Barrett, Sonia Goncalves, Ewan Harrison, David K. Jackson, Ian Johnston, Dominic Kwiatkowski, Cordelia Langford, John Sillitoe on behalf of the Wellcome Sanger Institute COVID-19 Surveillance Team |
| EPI_ISL_2996223, EPI_ISL_2996224, EPI_ISL_2996225, EPI_ISL_2996226, EPI_ISL_2996227, EPI_ISL_2996228 | Berkshire and Surrey Pathology Services Lighthouse Laboratory | Wellcome Sanger Institute for the COVID-19 Genomics UK (COG-UK) Consortium | Berkshire and Surrey Pathology Services Lighthouse Laboratory and Alex Alderton, Roberto Amato, Jeffrey Barrett, Sonia Goncalves, Ewan Harrison, David K. Jackson, Ian Johnston, Dominic Kwiatkowski, Cordelia Langford, John Sillitoe on behalf of the Wellcome Sanger Institute COVID-19 Surveillance Team |
| EPI_ISL_2996229 | Lighthouse Lab in Glasgow | Wellcome Sanger Institute for the COVID-19 Genomics UK (COG-UK) Consortium | Harper VanSteenhouse, Yumi Kasai, David Gray, Carol Clugston, Anna Dominiczak and Alex Alderton, Roberto Amato, Jeffrey Barrett, Sonia Goncalves, Ewan Harrison, David K. Jackson, Ian Johnston, Dominic Kwiatkowski, Cordelia Langford, John Sillitoe on behalf of the Wellcome Sanger Institute COVID-19 Surveillance Team |
| EPI_ISL_2996230 | Berkshire and Surrey Pathology Services Lighthouse Laboratory | Wellcome Sanger Institute for the COVID-19 Genomics UK (COG-UK) Consortium | Berkshire and Surrey Pathology Services Lighthouse Laboratory and Alex Alderton, Roberto Amato, Jeffrey Barrett, Sonia Goncalves, Ewan Harrison, David K. Jackson, Ian Johnston, Dominic Kwiatkowski, Cordelia Langford, John Sillitoe on behalf of the Wellcome Sanger Institute COVID-19 Surveillance Team |
| EPI_ISL_2996231, EPI_ISL_2996232 | Lighthouse Lab in Glasgow | Wellcome Sanger Institute for the COVID-19 Genomics UK (COG-UK) Consortium | Harper VanSteenhouse, Yumi Kasai, David Gray, Carol Clugston, Anna Dominiczak and Alex Alderton, Roberto Amato, Jeffrey Barrett, Sonia Goncalves, Ewan Harrison, David K. Jackson, Ian Johnston, Dominic Kwiatkowski, Cordelia Langford, John Sillitoe on behalf of the Wellcome Sanger Institute COVID-19 Surveillance Team |
| EPI_ISL_2996233 | Lighthouse Laboratory Plymouth | Wellcome Sanger Institute for the COVID-19 Genomics UK (COG-UK) Consortium | Lighthouse Laboratory Plymouth and Alex Alderton, Roberto Amato, Jeffrey Barrett, Sonia Goncalves, Ewan Harrison, David K. Jackson, Ian Johnston, Dominic Kwiatkowski, Cordelia Langford, John Sillitoe on behalf of the Wellcome Sanger Institute COVID-19 Surveillance Team |
| EPI_ISL_2996234, EPI_ISL_2996235, EPI_ISL_2996236, EPI_ISL_2996237, EPI_ISL_2996238, EPI_ISL_2996239, EPI_ISL_2996240 | Lighthouse Lab in Glasgow | Wellcome Sanger Institute for the COVID-19 Genomics UK (COG-UK) Consortium | Harper VanSteenhouse, Yumi Kasai, David Gray, Carol Clugston, Anna Dominiczak and Alex Alderton, Roberto Amato, Jeffrey Barrett, Sonia Goncalves, Ewan Harrison, David K. Jackson, Ian Johnston, Dominic Kwiatkowski, Cordelia Langford, John Sillitoe on behalf of the Wellcome Sanger Institute COVID-19 Surveillance Team |
| EPI_ISL_2996241 | Lighthouse Laboratory Plymouth | Wellcome Sanger Institute for the COVID-19 Genomics UK (COG-UK) Consortium | Lighthouse Laboratory Plymouth and Alex Alderton, Roberto Amato, Jeffrey Barrett, Sonia Goncalves, Ewan Harrison, David K. Jackson, Ian Johnston, Dominic Kwiatkowski, Cordelia Langford, John Sillitoe on behalf of the Wellcome Sanger Institute COVID-19 Surveillance Team |
| EPI_ISL_2996242 | Berkshire and Surrey Pathology Services Lighthouse Laboratory | Wellcome Sanger Institute for the COVID-19 Genomics UK (COG-UK) Consortium | Berkshire and Surrey Pathology Services Lighthouse Laboratory and Alex Alderton, Roberto Amato, Jeffrey Barrett, Sonia Goncalves, Ewan Harrison, David K. Jackson, Ian Johnston, Dominic Kwiatkowski, Cordelia Langford, John Sillitoe on behalf of the Wellcome Sanger Institute COVID-19 Surveillance Team |
| EPI_ISL_2996243 | Lighthouse Laboratory Plymouth | Wellcome Sanger Institute for the COVID-19 Genomics UK (COG-UK) Consortium | Lighthouse Laboratory Plymouth and Alex Alderton, Roberto Amato, Jeffrey Barrett, Sonia Goncalves, Ewan Harrison, David K. Jackson, Ian Johnston, Dominic Kwiatkowski, Cordelia Langford, John Sillitoe on behalf of the Wellcome Sanger Institute COVID-19 Surveillance Team |
| EPI_ISL_2996244 | Lighthouse Lab in Glasgow | Wellcome Sanger Institute for the COVID-19 Genomics UK (COG-UK) Consortium | Harper VanSteenhouse, Yumi Kasai, David Gray, Carol Clugston, Anna Dominiczak and Alex Alderton, Roberto Amato, Jeffrey Barrett, Sonia Goncalves, Ewan Harrison, David K. Jackson, Ian Johnston, Dominic Kwiatkowski, Cordelia Langford, John Sillitoe on behalf of the Wellcome Sanger Institute COVID-19 Surveillance Team |
| EPI_ISL_2996245, EPI_ISL_2996246 | Berkshire and Surrey Pathology Services Lighthouse Laboratory | Wellcome Sanger Institute for the COVID-19 Genomics UK (COG-UK) Consortium | Berkshire and Surrey Pathology Services Lighthouse Laboratory and Alex Alderton, Roberto Amato, Jeffrey Barrett, Sonia Goncalves, Ewan Harrison, David K. Jackson, Ian Johnston, Dominic Kwiatkowski, Cordelia Langford, John Sillitoe on behalf of the Wellcome Sanger Institute COVID-19 Surveillance Team |
| EPI_ISL_2996247 | Lighthouse Lab in Glasgow | Wellcome Sanger Institute for the COVID-19 Genomics UK (COG-UK) Consortium | Harper VanSteenhouse, Yumi Kasai, David Gray, Carol Clugston, Anna Dominiczak and Alex Alderton, Roberto Amato, Jeffrey Barrett, Sonia Goncalves, Ewan Harrison, David K. Jackson, Ian Johnston, Dominic Kwiatkowski, Cordelia Langford, John Sillitoe on behalf of the Wellcome Sanger Institute COVID-19 Surveillance Team |
| EPI_ISL_2996248, EPI_ISL_2996249, EPI_ISL_2996250 | Berkshire and Surrey Pathology Services Lighthouse Laboratory | Wellcome Sanger Institute for the COVID-19 Genomics UK (COG-UK) Consortium | Berkshire and Surrey Pathology Services Lighthouse Laboratory and Alex Alderton, Roberto Amato, Jeffrey Barrett, Sonia Goncalves, Ewan Harrison, David K. Jackson, Ian Johnston, Dominic Kwiatkowski, Cordelia Langford, John Sillitoe on behalf of the Wellcome Sanger Institute COVID-19 Surveillance Team |
| EPI_ISL_2996251, EPI_ISL_2996252, EPI_ISL_2996253 | Lighthouse Lab in Glasgow | Wellcome Sanger Institute for the COVID-19 Genomics UK (COG-UK) Consortium | Harper VanSteenhouse, Yumi Kasai, David Gray, Carol Clugston, Anna Dominiczak and Alex Alderton, Roberto Amato, Jeffrey Barrett, Sonia Goncalves, Ewan Harrison, David K. Jackson, Ian Johnston, Dominic Kwiatkowski, Cordelia Langford, John Sillitoe on behalf of the Wellcome Sanger Institute COVID-19 Surveillance Team |
| EPI_ISL_2996254 | Berkshire and Surrey Pathology Services Lighthouse Laboratory | Wellcome Sanger Institute for the COVID-19 Genomics UK (COG-UK) Consortium | Berkshire and Surrey Pathology Services Lighthouse Laboratory and Alex Alderton, Roberto Amato, Jeffrey Barrett, Sonia Goncalves, Ewan Harrison, David K. Jackson, Ian Johnston, Dominic Kwiatkowski, Cordelia Langford, John Sillitoe on behalf of the Wellcome Sanger Institute COVID-19 Surveillance Team |
| EPI_ISL_2996255 | Lighthouse Lab in Glasgow | Wellcome Sanger Institute for the COVID-19 Genomics UK (COG-UK) Consortium | Harper VanSteenhouse, Yumi Kasai, David Gray, Carol Clugston, Anna Dominiczak and Alex Alderton, Roberto Amato, Jeffrey Barrett, Sonia Goncalves, Ewan Harrison, David K. Jackson, Ian Johnston, Dominic Kwiatkowski, Cordelia Langford, John Sillitoe on behalf of the Wellcome Sanger Institute COVID-19 Surveillance Team |
| EPI_ISL_2996256 | Lighthouse Laboratory Plymouth | Wellcome Sanger Institute for the COVID-19 Genomics UK (COG-UK) Consortium | Lighthouse Laboratory Plymouth and Alex Alderton, Roberto Amato, Jeffrey Barrett, Sonia Goncalves, Ewan Harrison, David K. Jackson, Ian Johnston, Dominic Kwiatkowski, Cordelia Langford, John Sillitoe on behalf of the Wellcome Sanger Institute COVID-19 Surveillance Team |
| EPI_ISL_2996257, EPI_ISL_2996258 | Lighthouse Lab in Glasgow | Wellcome Sanger Institute for the COVID-19 Genomics UK (COG-UK) Consortium | Harper VanSteenhouse, Yumi Kasai, David Gray, Carol Clugston, Anna Dominiczak and Alex Alderton, Roberto Amato, Jeffrey Barrett, Sonia Goncalves, Ewan Harrison, David K. Jackson, Ian Johnston, Dominic Kwiatkowski, Cordelia Langford, John Sillitoe on behalf of the Wellcome Sanger Institute COVID-19 Surveillance Team |
| EPI_ISL_2996259, EPI_ISL_2996260 | Lighthouse Laboratory Plymouth | Wellcome Sanger Institute for the COVID-19 Genomics UK (COG-UK) Consortium | Lighthouse Laboratory Plymouth and Alex Alderton, Roberto Amato, Jeffrey Barrett, Sonia Goncalves, Ewan Harrison, David K. Jackson, Ian Johnston, Dominic Kwiatkowski, Cordelia Langford, John Sillitoe on behalf of the Wellcome Sanger Institute COVID-19 Surveillance Team |
| EPI_ISL_2996261 | Berkshire and Surrey Pathology Services Lighthouse Laboratory | Wellcome Sanger Institute for the COVID-19 Genomics UK (COG-UK) Consortium | Berkshire and Surrey Pathology Services Lighthouse Laboratory and Alex Alderton, Roberto Amato, Jeffrey Barrett, Sonia Goncalves, Ewan Harrison, David K. Jackson, Ian Johnston, Dominic Kwiatkowski, Cordelia Langford, John Sillitoe on behalf of the Wellcome Sanger Institute COVID-19 Surveillance Team |
| EPI_ISL_2996262, EPI_ISL_2996263, EPI_ISL_2996264 | Lighthouse Lab in Glasgow | Wellcome Sanger Institute for the COVID-19 Genomics UK (COG-UK) Consortium | Harper VanSteenhouse, Yumi Kasai, David Gray, Carol Clugston, Anna Dominiczak and Alex Alderton, Roberto Amato, Jeffrey Barrett, Sonia Goncalves, Ewan Harrison, David K. Jackson, Ian Johnston, Dominic Kwiatkowski, Cordelia Langford, John Sillitoe on behalf of the Wellcome Sanger Institute COVID-19 Surveillance Team |
| EPI_ISL_2996265, EPI_ISL_2996266, EPI_ISL_2996267 | Berkshire and Surrey Pathology Services Lighthouse Laboratory | Wellcome Sanger Institute for the COVID-19 Genomics UK (COG-UK) Consortium | Berkshire and Surrey Pathology Services Lighthouse Laboratory and Alex Alderton, Roberto Amato, Jeffrey Barrett, Sonia Goncalves, Ewan Harrison, David K. Jackson, Ian Johnston, Dominic Kwiatkowski, Cordelia Langford, John Sillitoe on behalf of the Wellcome Sanger Institute COVID-19 Surveillance Team |
| EPI_ISL_2996268 | Lighthouse Lab in Glasgow | Wellcome Sanger Institute for the COVID-19 Genomics UK (COG-UK) Consortium | Harper VanSteenhouse, Yumi Kasai, David Gray, Carol Clugston, Anna Dominiczak and Alex Alderton, Roberto Amato, Jeffrey Barrett, Sonia Goncalves, Ewan Harrison, David K. Jackson, Ian Johnston, Dominic Kwiatkowski, Cordelia Langford, John Sillitoe on behalf of the Wellcome Sanger Institute COVID-19 Surveillance Team |
| EPI_ISL_2996269 | Berkshire and Surrey Pathology Services Lighthouse Laboratory | Wellcome Sanger Institute for the COVID-19 Genomics UK (COG-UK) Consortium | Berkshire and Surrey Pathology Services Lighthouse Laboratory and Alex Alderton, Roberto Amato, Jeffrey Barrett, Sonia Goncalves, Ewan Harrison, David K. Jackson, Ian Johnston, Dominic Kwiatkowski, Cordelia Langford, John Sillitoe on behalf of the Wellcome Sanger Institute COVID-19 Surveillance Team |
| EPI_ISL_2996270 | Lighthouse Laboratory Plymouth | Wellcome Sanger Institute for the COVID-19 Genomics UK (COG-UK) Consortium | Lighthouse Laboratory Plymouth and Alex Alderton, Roberto Amato, Jeffrey Barrett, Sonia Goncalves, Ewan Harrison, David K. Jackson, Ian Johnston, Dominic Kwiatkowski, Cordelia Langford, John Sillitoe on behalf of the Wellcome Sanger Institute COVID-19 Surveillance Team |

| | | | |
|---|---|---|---|
| EPI_ISL_2996271, EPI_ISL_2996272 | Lighthouse Lab in Glasgow | Wellcome Sanger Institute for the COVID-19 Genomics UK (COG-UK) Consortium | Harper VanSteenhouse, Yumi Kasai, David Gray, Carol Clugston, Anna Dominiczak and Alex Alderton, Roberto Amato, Jeffrey Barrett, Sonia Goncalves, Ewan Harrison, David K. Jackson, Ian Johnston, Dominic Kwiatkowski, Cordelia Langford, John Sillitoe on behalf of the Wellcome Sanger Institute COVID-19 Surveillance Team |
| EPI_ISL_2996273 | Lighthouse Laboratory Plymouth | Wellcome Sanger Institute for the COVID-19 Genomics UK (COG-UK) Consortium | Lighthouse Laboratory Plymouth and Alex Alderton, Roberto Amato, Jeffrey Barrett, Sonia Goncalves, Ewan Harrison, David K. Jackson, Ian Johnston, Dominic Kwiatkowski, Cordelia Langford, John Sillitoe on behalf of the Wellcome Sanger Institute COVID-19 Surveillance Team |
| EPI_ISL_2996274 | Lighthouse Lab in Glasgow | Wellcome Sanger Institute for the COVID-19 Genomics UK (COG-UK) Consortium | Harper VanSteenhouse, Yumi Kasai, David Gray, Carol Clugston, Anna Dominiczak and Alex Alderton, Roberto Amato, Jeffrey Barrett, Sonia Goncalves, Ewan Harrison, David K. Jackson, Ian Johnston, Dominic Kwiatkowski, Cordelia Langford, John Sillitoe on behalf of the Wellcome Sanger Institute COVID-19 Surveillance Team |
| EPI_ISL_2996275 | Lighthouse Laboratory Plymouth | Wellcome Sanger Institute for the COVID-19 Genomics UK (COG-UK) Consortium | Lighthouse Laboratory Plymouth and Alex Alderton, Roberto Amato, Jeffrey Barrett, Sonia Goncalves, Ewan Harrison, David K. Jackson, Ian Johnston, Dominic Kwiatkowski, Cordelia Langford, John Sillitoe on behalf of the Wellcome Sanger Institute COVID-19 Surveillance Team |
| EPI_ISL_2996276, EPI_ISL_2996277 | Berkshire and Surrey Pathology Services Lighthouse Laboratory | Wellcome Sanger Institute for the COVID-19 Genomics UK (COG-UK) Consortium | Berkshire and Surrey Pathology Services Lighthouse Laboratory and Alex Alderton, Roberto Amato, Jeffrey Barrett, Sonia Goncalves, Ewan Harrison, David K. Jackson, Ian Johnston, Dominic Kwiatkowski, Cordelia Langford, John Sillitoe on behalf of the Wellcome Sanger Institute COVID-19 Surveillance Team |
| EPI_ISL_2996278, EPI_ISL_2996279 | Lighthouse Lab in Glasgow | Wellcome Sanger Institute for the COVID-19 Genomics UK (COG-UK) Consortium | Harper VanSteenhouse, Yumi Kasai, David Gray, Carol Clugston, Anna Dominiczak and Alex Alderton, Roberto Amato, Jeffrey Barrett, Sonia Goncalves, Ewan Harrison, David K. Jackson, Ian Johnston, Dominic Kwiatkowski, Cordelia Langford, John Sillitoe on behalf of the Wellcome Sanger Institute COVID-19 Surveillance Team |
| EPI_ISL_2996280 | Berkshire and Surrey Pathology Services Lighthouse Laboratory | Wellcome Sanger Institute for the COVID-19 Genomics UK (COG-UK) Consortium | Berkshire and Surrey Pathology Services Lighthouse Laboratory and Alex Alderton, Roberto Amato, Jeffrey Barrett, Sonia Goncalves, Ewan Harrison, David K. Jackson, Ian Johnston, Dominic Kwiatkowski, Cordelia Langford, John Sillitoe on behalf of the Wellcome Sanger Institute COVID-19 Surveillance Team |
| EPI_ISL_2996281 | Lighthouse Laboratory Plymouth | Wellcome Sanger Institute for the COVID-19 Genomics UK (COG-UK) Consortium | Lighthouse Laboratory Plymouth and Alex Alderton, Roberto Amato, Jeffrey Barrett, Sonia Goncalves, Ewan Harrison, David K. Jackson, Ian Johnston, Dominic Kwiatkowski, Cordelia Langford, John Sillitoe on behalf of the Wellcome Sanger Institute COVID-19 Surveillance Team |
| EPI_ISL_2996282 | Lighthouse Lab in Glasgow | Wellcome Sanger Institute for the COVID-19 Genomics UK (COG-UK) Consortium | Harper VanSteenhouse, Yumi Kasai, David Gray, Carol Clugston, Anna Dominiczak and Alex Alderton, Roberto Amato, Jeffrey Barrett, Sonia Goncalves, Ewan Harrison, David K. Jackson, Ian Johnston, Dominic Kwiatkowski, Cordelia Langford, John Sillitoe on behalf of the Wellcome Sanger Institute COVID-19 Surveillance Team |
| EPI_ISL_2996283 | Lighthouse Laboratory Plymouth | Wellcome Sanger Institute for the COVID-19 Genomics UK (COG-UK) Consortium | Lighthouse Laboratory Plymouth and Alex Alderton, Roberto Amato, Jeffrey Barrett, Sonia Goncalves, Ewan Harrison, David K. Jackson, Ian Johnston, Dominic Kwiatkowski, Cordelia Langford, John Sillitoe on behalf of the Wellcome Sanger Institute COVID-19 Surveillance Team |
| EPI_ISL_2996284 | Berkshire and Surrey Pathology Services Lighthouse Laboratory | Wellcome Sanger Institute for the COVID-19 Genomics UK (COG-UK) Consortium | Berkshire and Surrey Pathology Services Lighthouse Laboratory and Alex Alderton, Roberto Amato, Jeffrey Barrett, Sonia Goncalves, Ewan Harrison, David K. Jackson, Ian Johnston, Dominic Kwiatkowski, Cordelia Langford, John Sillitoe on behalf of the Wellcome Sanger Institute COVID-19 Surveillance Team |
| EPI_ISL_2996285, EPI_ISL_2996286 | Lighthouse Laboratory Plymouth | Wellcome Sanger Institute for the COVID-19 Genomics UK (COG-UK) Consortium | Lighthouse Laboratory Plymouth and Alex Alderton, Roberto Amato, Jeffrey Barrett, Sonia Goncalves, Ewan Harrison, David K. Jackson, Ian Johnston, Dominic Kwiatkowski, Cordelia Langford, John Sillitoe on behalf of the Wellcome Sanger Institute COVID-19 Surveillance Team |
| EPI_ISL_2996287, EPI_ISL_2996288, EPI_ISL_2996289 | Lighthouse Lab in Glasgow | Wellcome Sanger Institute for the COVID-19 Genomics UK (COG-UK) Consortium | Harper VanSteenhouse, Yumi Kasai, David Gray, Carol Clugston, Anna Dominiczak and Alex Alderton, Roberto Amato, Jeffrey Barrett, Sonia Goncalves, Ewan Harrison, David K. Jackson, Ian Johnston, Dominic Kwiatkowski, Cordelia Langford, John Sillitoe on behalf of the Wellcome Sanger Institute COVID-19 Surveillance Team |
| EPI_ISL_2996290 | Lighthouse Laboratory Plymouth | Wellcome Sanger Institute for the COVID-19 Genomics UK (COG-UK) Consortium | Lighthouse Laboratory Plymouth and Alex Alderton, Roberto Amato, Jeffrey Barrett, Sonia Goncalves, Ewan Harrison, David K. Jackson, Ian Johnston, Dominic Kwiatkowski, Cordelia Langford, John Sillitoe on behalf of the Wellcome Sanger Institute COVID-19 Surveillance Team |
| EPI_ISL_2996291, EPI_ISL_2996292, EPI_ISL_2996293 | Lighthouse Lab in Glasgow | Wellcome Sanger Institute for the COVID-19 Genomics UK (COG-UK) Consortium | Harper VanSteenhouse, Yumi Kasai, David Gray, Carol Clugston, Anna Dominiczak and Alex Alderton, Roberto Amato, Jeffrey Barrett, Sonia Goncalves, Ewan Harrison, David K. Jackson, Ian Johnston, Dominic Kwiatkowski, Cordelia Langford, John Sillitoe on behalf of the Wellcome Sanger Institute COVID-19 Surveillance Team |
| EPI_ISL_2996294 | Berkshire and Surrey Pathology Services Lighthouse Laboratory | Wellcome Sanger Institute for the COVID-19 Genomics UK (COG-UK) Consortium | Berkshire and Surrey Pathology Services Lighthouse Laboratory and Alex Alderton, Roberto Amato, Jeffrey Barrett, Sonia Goncalves, Ewan Harrison, David K. Jackson, Ian Johnston, Dominic Kwiatkowski, Cordelia Langford, John Sillitoe on behalf of the Wellcome Sanger Institute COVID-19 Surveillance Team |
| EPI_ISL_2996295 | Lighthouse Lab in Glasgow | Wellcome Sanger Institute for the COVID-19 Genomics UK (COG-UK) Consortium | Harper VanSteenhouse, Yumi Kasai, David Gray, Carol Clugston, Anna Dominiczak and Alex Alderton, Roberto Amato, Jeffrey Barrett, Sonia Goncalves, Ewan Harrison, David K. Jackson, Ian Johnston, Dominic Kwiatkowski, Cordelia Langford, John Sillitoe on behalf of the Wellcome Sanger Institute COVID-19 Surveillance Team |
| EPI_ISL_2996296 | Lighthouse Laboratory Plymouth | Wellcome Sanger Institute for the COVID-19 Genomics UK (COG-UK) Consortium | Lighthouse Laboratory Plymouth and Alex Alderton, Roberto Amato, Jeffrey Barrett, Sonia Goncalves, Ewan Harrison, David K. Jackson, Ian Johnston, Dominic Kwiatkowski, Cordelia Langford, John Sillitoe on behalf of the Wellcome Sanger Institute COVID-19 Surveillance Team |
| EPI_ISL_2996297, EPI_ISL_2996298, EPI_ISL_2996299, EPI_ISL_2996300, EPI_ISL_2996301, EPI_ISL_2996302 | Lighthouse Lab in Glasgow | Wellcome Sanger Institute for the COVID-19 Genomics UK (COG-UK) Consortium | Harper VanSteenhouse, Yumi Kasai, David Gray, Carol Clugston, Anna Dominiczak and Alex Alderton, Roberto Amato, Jeffrey Barrett, Sonia Goncalves, Ewan Harrison, David K. Jackson, Ian Johnston, Dominic Kwiatkowski, Cordelia Langford, John Sillitoe on behalf of the Wellcome Sanger Institute COVID-19 Surveillance Team |
| EPI_ISL_2996303 | Berkshire and Surrey Pathology Services Lighthouse Laboratory | Wellcome Sanger Institute for the COVID-19 Genomics UK (COG-UK) Consortium | Berkshire and Surrey Pathology Services Lighthouse Laboratory and Alex Alderton, Roberto Amato, Jeffrey Barrett, Sonia Goncalves, Ewan Harrison, David K. Jackson, Ian Johnston, Dominic Kwiatkowski, Cordelia Langford, John Sillitoe on behalf of the Wellcome Sanger Institute COVID-19 Surveillance Team |
| EPI_ISL_2996304, EPI_ISL_2996305 | Lighthouse Laboratory Plymouth | Wellcome Sanger Institute for the COVID-19 Genomics UK (COG-UK) Consortium | Lighthouse Laboratory Plymouth and Alex Alderton, Roberto Amato, Jeffrey Barrett, Sonia Goncalves, Ewan Harrison, David K. Jackson, Ian Johnston, Dominic Kwiatkowski, Cordelia Langford, John Sillitoe on behalf of the Wellcome Sanger Institute COVID-19 Surveillance Team |
| EPI_ISL_2996306, EPI_ISL_2996307, EPI_ISL_2996308 | Lighthouse Lab in Glasgow | Wellcome Sanger Institute for the COVID-19 Genomics UK (COG-UK) Consortium | Harper VanSteenhouse, Yumi Kasai, David Gray, Carol Clugston, Anna Dominiczak and Alex Alderton, Roberto Amato, Jeffrey Barrett, Sonia Goncalves, Ewan Harrison, David K. Jackson, Ian Johnston, Dominic Kwiatkowski, Cordelia Langford, John Sillitoe on behalf of the Wellcome Sanger Institute COVID-19 Surveillance Team |
| EPI_ISL_2996309 | Berkshire and Surrey Pathology Services Lighthouse Laboratory | Wellcome Sanger Institute for the COVID-19 Genomics UK (COG-UK) Consortium | Berkshire and Surrey Pathology Services Lighthouse Laboratory and Alex Alderton, Roberto Amato, Jeffrey Barrett, Sonia Goncalves, Ewan Harrison, David K. Jackson, Ian Johnston, Dominic Kwiatkowski, Cordelia Langford, John Sillitoe on behalf of the Wellcome Sanger Institute COVID-19 Surveillance Team |
| EPI_ISL_2996310 | Lighthouse Lab in Glasgow | Wellcome Sanger Institute for the COVID-19 Genomics UK (COG-UK) Consortium | Harper VanSteenhouse, Yumi Kasai, David Gray, Carol Clugston, Anna Dominiczak and Alex Alderton, Roberto Amato, Jeffrey Barrett, Sonia Goncalves, Ewan Harrison, David K. Jackson, Ian Johnston, Dominic Kwiatkowski, Cordelia Langford, John Sillitoe on behalf of the Wellcome Sanger Institute COVID-19 Surveillance Team |
| EPI_ISL_2996311, EPI_ISL_2996312, EPI_ISL_2996313 | Lighthouse Laboratory Plymouth | Wellcome Sanger Institute for the COVID-19 Genomics UK (COG-UK) Consortium | Lighthouse Laboratory Plymouth and Alex Alderton, Roberto Amato, Jeffrey Barrett, Sonia Goncalves, Ewan Harrison, David K. Jackson, Ian Johnston, Dominic Kwiatkowski, Cordelia Langford, John Sillitoe on behalf of the Wellcome Sanger Institute COVID-19 Surveillance Team |
| EPI_ISL_2996314, EPI_ISL_2996315 | Lighthouse Lab in Glasgow | Wellcome Sanger Institute for the COVID-19 Genomics UK (COG-UK) Consortium | Harper VanSteenhouse, Yumi Kasai, David Gray, Carol Clugston, Anna Dominiczak and Alex Alderton, Roberto Amato, Jeffrey Barrett, Sonia Goncalves, Ewan Harrison, David K. Jackson, Ian Johnston, Dominic Kwiatkowski, Cordelia Langford, John Sillitoe on behalf of the Wellcome Sanger Institute COVID-19 Surveillance Team |
| EPI_ISL_2996316 | Berkshire and Surrey Pathology Services Lighthouse Laboratory | Wellcome Sanger Institute for the COVID-19 Genomics UK (COG-UK) Consortium | Berkshire and Surrey Pathology Services Lighthouse Laboratory and Alex Alderton, Roberto Amato, Jeffrey Barrett, Sonia Goncalves, Ewan Harrison, David K. Jackson, Ian Johnston, Dominic Kwiatkowski, Cordelia Langford, John Sillitoe on behalf of the Wellcome Sanger Institute COVID-19 Surveillance Team |
| EPI_ISL_2996317 | Lighthouse Laboratory Plymouth | Wellcome Sanger Institute for the COVID-19 Genomics UK (COG-UK) Consortium | Lighthouse Laboratory Plymouth and Alex Alderton, Roberto Amato, Jeffrey Barrett, Sonia Goncalves, Ewan Harrison, David K. Jackson, Ian Johnston, Dominic Kwiatkowski, Cordelia Langford, John Sillitoe on behalf of the Wellcome Sanger Institute COVID-19 Surveillance Team |
| EPI_ISL_2996318, EPI_ISL_2996319 | Berkshire and Surrey Pathology Services Lighthouse Laboratory | Wellcome Sanger Institute for the COVID-19 Genomics UK (COG-UK) Consortium | Berkshire and Surrey Pathology Services Lighthouse Laboratory and Alex Alderton, Roberto Amato, Jeffrey Barrett, Sonia Goncalves, Ewan Harrison, David K. Jackson, Ian Johnston, Dominic Kwiatkowski, Cordelia Langford, John Sillitoe on behalf of the Wellcome Sanger Institute COVID-19 Surveillance Team |
| EPI_ISL_2996320 | Lighthouse Lab in Glasgow | Wellcome Sanger Institute for the COVID-19 Genomics UK (COG-UK) Consortium | Harper VanSteenhouse, Yumi Kasai, David Gray, Carol Clugston, Anna Dominiczak and Alex Alderton, Roberto Amato, Jeffrey Barrett, Sonia Goncalves, Ewan Harrison, David K. Jackson, Ian Johnston, Dominic Kwiatkowski, Cordelia Langford, John Sillitoe on behalf of the Wellcome Sanger Institute COVID-19 Surveillance Team |
| EPI_ISL_2996321 | Lighthouse Laboratory Plymouth | Wellcome Sanger Institute for the COVID-19 Genomics UK (COG-UK) Consortium | Lighthouse Laboratory Plymouth and Alex Alderton, Roberto Amato, Jeffrey Barrett, Sonia Goncalves, Ewan Harrison, David K. Jackson, Ian Johnston, Dominic Kwiatkowski, Cordelia Langford, John Sillitoe on behalf of the Wellcome Sanger Institute COVID-19 Surveillance Team |
| EPI_ISL_2996322 | Lighthouse Lab in Glasgow | Wellcome Sanger Institute for the COVID-19 Genomics UK (COG-UK) Consortium | Harper VanSteenhouse, Yumi Kasai, David Gray, Carol Clugston, Anna Dominiczak and Alex Alderton, Roberto Amato, Jeffrey Barrett, Sonia Goncalves, Ewan Harrison, David K. Jackson, Ian Johnston, Dominic Kwiatkowski, Cordelia Langford, John Sillitoe on behalf of the Wellcome Sanger Institute COVID-19 Surveillance Team |
| EPI_ISL_2996323 | Lighthouse Laboratory Plymouth | Wellcome Sanger Institute for the COVID-19 Genomics UK (COG-UK) Consortium | Lighthouse Laboratory Plymouth and Alex Alderton, Roberto Amato, Jeffrey Barrett, Sonia Goncalves, Ewan Harrison, David K. Jackson, Ian Johnston, Dominic Kwiatkowski, Cordelia Langford, John Sillitoe on behalf of the Wellcome Sanger Institute COVID-19 Surveillance Team |
| EPI_ISL_2996324 | Lighthouse Lab in Glasgow | Wellcome Sanger Institute for the COVID-19 Genomics UK (COG-UK) Consortium | Harper VanSteenhouse, Yumi Kasai, David Gray, Carol Clugston, Anna Dominiczak and Alex Alderton, Roberto Amato, Jeffrey Barrett, Sonia Goncalves, Ewan Harrison, David K. Jackson, Ian Johnston, Dominic Kwiatkowski, Cordelia Langford, John Sillitoe on behalf of the Wellcome Sanger Institute COVID-19 Surveillance Team |
| EPI_ISL_2996325 | Lighthouse Laboratory Plymouth | Wellcome Sanger Institute for the COVID-19 Genomics UK (COG-UK) Consortium | Lighthouse Laboratory Plymouth and Alex Alderton, Roberto Amato, Jeffrey Barrett, Sonia Goncalves, Ewan Harrison, David K. Jackson, Ian Johnston, Dominic Kwiatkowski, Cordelia Langford, John Sillitoe on behalf of the Wellcome Sanger Institute COVID-19 Surveillance Team |
| EPI_ISL_2996326, EPI_ISL_2996327 | Lighthouse Lab in Glasgow | Wellcome Sanger Institute for the COVID-19 Genomics UK (COG-UK) Consortium | Harper VanSteenhouse, Yumi Kasai, David Gray, Carol Clugston, Anna Dominiczak and Alex Alderton, Roberto Amato, Jeffrey Barrett, Sonia Goncalves, Ewan Harrison, David K. Jackson, Ian Johnston, Dominic Kwiatkowski, Cordelia Langford, John Sillitoe on behalf of the Wellcome Sanger Institute COVID-19 Surveillance Team |
| EPI_ISL_2996328 | Lighthouse Laboratory Plymouth | Wellcome Sanger Institute for the COVID-19 Genomics UK (COG-UK) Consortium | Lighthouse Laboratory Plymouth and Alex Alderton, Roberto Amato, Jeffrey Barrett, Sonia Goncalves, Ewan Harrison, David K. Jackson, Ian Johnston, Dominic Kwiatkowski, Cordelia Langford, John Sillitoe on behalf of the Wellcome Sanger Institute COVID-19 Surveillance Team |
| EPI_ISL_2996329, EPI_ISL_2996330 | Lighthouse Lab in Glasgow | Wellcome Sanger Institute for the COVID-19 Genomics UK (COG-UK) Consortium | Harper VanSteenhouse, Yumi Kasai, David Gray, Carol Clugston, Anna Dominiczak and Alex Alderton, Roberto Amato, Jeffrey Barrett, Sonia Goncalves, Ewan Harrison, David K. Jackson, Ian Johnston, Dominic Kwiatkowski, Cordelia Langford, John Sillitoe on behalf of the Wellcome Sanger Institute COVID-19 Surveillance Team |
| EPI_ISL_2996331, EPI_ISL_2996332, EPI_ISL_2996333, EPI_ISL_2996334, EPI_ISL_2996335, EPI_ISL_2996336, EPI_ISL_2996337 | Lighthouse Laboratory Plymouth | Wellcome Sanger Institute for the COVID-19 Genomics UK (COG-UK) Consortium | Lighthouse Laboratory Plymouth and Alex Alderton, Roberto Amato, Jeffrey Barrett, Sonia Goncalves, Ewan Harrison, David K. Jackson, Ian Johnston, Dominic Kwiatkowski, Cordelia Langford, John Sillitoe on behalf of the Wellcome Sanger Institute COVID-19 Surveillance Team |
| EPI_ISL_2996338, EPI_ISL_2996339 | Lighthouse Lab in Milton Keynes | Wellcome Sanger Institute for the COVID-19 Genomics UK (COG-UK) Consortium | The Lighthouse Lab in Milton Keynes and Alex Alderton, Roberto Amato, Jeffrey Barrett, Sonia Goncalves, Ewan Harrison, David K. Jackson, Ian Johnston, Dominic Kwiatkowski, Cordelia Langford, John Sillitoe on behalf of the Wellcome Sanger Institute COVID-19 Surveillance Team |

| | | | |
|---|---|---|---|
| EPI_ISL_2996340, EPI_ISL_2996341, EPI_ISL_2996342, EPI_ISL_2996343 | Lighthouse Laboratory Plymouth | Wellcome Sanger Institute for the COVID-19 Genomics UK (COG-UK) Consortium | Lighthouse Laboratory Plymouth and Alex Alderton, Roberto Amato, Jeffrey Barrett, Sonia Goncalves, Ewan Harrison, David K. Jackson, Ian Johnston, Dominic Kwiatkowski, Cordelia Langford, John Sillitoe on behalf of the Wellcome Sanger Institute COVID-19 Surveillance Team |
| EPI_ISL_2996344 | Lighthouse Lab in Milton Keynes | Wellcome Sanger Institute for the COVID-19 Genomics UK (COG-UK) Consortium | The Lighthouse Lab in Milton Keynes and Alex Alderton, Roberto Amato, Jeffrey Barrett, Sonia Goncalves, Ewan Harrison, David K. Jackson, Ian Johnston, Dominic Kwiatkowski, Cordelia Langford, John Sillitoe on behalf of the Wellcome Sanger Institute COVID-19 Surveillance Team |
| EPI_ISL_2996345 | Lighthouse Laboratory Plymouth | Wellcome Sanger Institute for the COVID-19 Genomics UK (COG-UK) Consortium | Lighthouse Laboratory Plymouth and Alex Alderton, Roberto Amato, Jeffrey Barrett, Sonia Goncalves, Ewan Harrison, David K. Jackson, Ian Johnston, Dominic Kwiatkowski, Cordelia Langford, John Sillitoe on behalf of the Wellcome Sanger Institute COVID-19 Surveillance Team |
| EPI_ISL_2996346 | Lighthouse Lab in Milton Keynes | Wellcome Sanger Institute for the COVID-19 Genomics UK (COG-UK) Consortium | The Lighthouse Lab in Milton Keynes and Alex Alderton, Roberto Amato, Jeffrey Barrett, Sonia Goncalves, Ewan Harrison, David K. Jackson, Ian Johnston, Dominic Kwiatkowski, Cordelia Langford, John Sillitoe on behalf of the Wellcome Sanger Institute COVID-19 Surveillance Team |
| EPI_ISL_2996347, EPI_ISL_2996348 | Lighthouse Laboratory Plymouth | Wellcome Sanger Institute for the COVID-19 Genomics UK (COG-UK) Consortium | Lighthouse Laboratory Plymouth and Alex Alderton, Roberto Amato, Jeffrey Barrett, Sonia Goncalves, Ewan Harrison, David K. Jackson, Ian Johnston, Dominic Kwiatkowski, Cordelia Langford, John Sillitoe on behalf of the Wellcome Sanger Institute COVID-19 Surveillance Team |
| EPI_ISL_2996349 | Lighthouse Lab in Milton Keynes | Wellcome Sanger Institute for the COVID-19 Genomics UK (COG-UK) Consortium | The Lighthouse Lab in Milton Keynes and Alex Alderton, Roberto Amato, Jeffrey Barrett, Sonia Goncalves, Ewan Harrison, David K. Jackson, Ian Johnston, Dominic Kwiatkowski, Cordelia Langford, John Sillitoe on behalf of the Wellcome Sanger Institute COVID-19 Surveillance Team |
| EPI_ISL_2996350 | Lighthouse Laboratory Plymouth | Wellcome Sanger Institute for the COVID-19 Genomics UK (COG-UK) Consortium | Lighthouse Laboratory Plymouth and Alex Alderton, Roberto Amato, Jeffrey Barrett, Sonia Goncalves, Ewan Harrison, David K. Jackson, Ian Johnston, Dominic Kwiatkowski, Cordelia Langford, John Sillitoe on behalf of the Wellcome Sanger Institute COVID-19 Surveillance Team |
| EPI_ISL_2996351, EPI_ISL_2996352 | Lighthouse Lab in Milton Keynes | Wellcome Sanger Institute for the COVID-19 Genomics UK (COG-UK) Consortium | The Lighthouse Lab in Milton Keynes and Alex Alderton, Roberto Amato, Jeffrey Barrett, Sonia Goncalves, Ewan Harrison, David K. Jackson, Ian Johnston, Dominic Kwiatkowski, Cordelia Langford, John Sillitoe on behalf of the Wellcome Sanger Institute COVID-19 Surveillance Team |
| EPI_ISL_2996353, EPI_ISL_2996354, EPI_ISL_2996355, EPI_ISL_2996356, EPI_ISL_2996357, EPI_ISL_2996358 | Lighthouse Laboratory Plymouth | Wellcome Sanger Institute for the COVID-19 Genomics UK (COG-UK) Consortium | Lighthouse Laboratory Plymouth and Alex Alderton, Roberto Amato, Jeffrey Barrett, Sonia Goncalves, Ewan Harrison, David K. Jackson, Ian Johnston, Dominic Kwiatkowski, Cordelia Langford, John Sillitoe on behalf of the Wellcome Sanger Institute COVID-19 Surveillance Team |
| EPI_ISL_2996359 | Lighthouse Lab in Milton Keynes | Wellcome Sanger Institute for the COVID-19 Genomics UK (COG-UK) Consortium | The Lighthouse Lab in Milton Keynes and Alex Alderton, Roberto Amato, Jeffrey Barrett, Sonia Goncalves, Ewan Harrison, David K. Jackson, Ian Johnston, Dominic Kwiatkowski, Cordelia Langford, John Sillitoe on behalf of the Wellcome Sanger Institute COVID-19 Surveillance Team |
| EPI_ISL_2996360, EPI_ISL_2996361, EPI_ISL_2996362 | Lighthouse Laboratory Plymouth | Wellcome Sanger Institute for the COVID-19 Genomics UK (COG-UK) Consortium | Lighthouse Laboratory Plymouth and Alex Alderton, Roberto Amato, Jeffrey Barrett, Sonia Goncalves, Ewan Harrison, David K. Jackson, Ian Johnston, Dominic Kwiatkowski, Cordelia Langford, John Sillitoe on behalf of the Wellcome Sanger Institute COVID-19 Surveillance Team |
| EPI_ISL_2996363 | Lighthouse Lab in Milton Keynes | Wellcome Sanger Institute for the COVID-19 Genomics UK (COG-UK) Consortium | The Lighthouse Lab in Milton Keynes and Alex Alderton, Roberto Amato, Jeffrey Barrett, Sonia Goncalves, Ewan Harrison, David K. Jackson, Ian Johnston, Dominic Kwiatkowski, Cordelia Langford, John Sillitoe on behalf of the Wellcome Sanger Institute COVID-19 Surveillance Team |
| EPI_ISL_2996364 | Lighthouse Laboratory Plymouth | Wellcome Sanger Institute for the COVID-19 Genomics UK (COG-UK) Consortium | Lighthouse Laboratory Plymouth and Alex Alderton, Roberto Amato, Jeffrey Barrett, Sonia Goncalves, Ewan Harrison, David K. Jackson, Ian Johnston, Dominic Kwiatkowski, Cordelia Langford, John Sillitoe on behalf of the Wellcome Sanger Institute COVID-19 Surveillance Team |
| EPI_ISL_2996365, EPI_ISL_2996366 | Lighthouse Lab in Milton Keynes | Wellcome Sanger Institute for the COVID-19 Genomics UK (COG-UK) Consortium | The Lighthouse Lab in Milton Keynes and Alex Alderton, Roberto Amato, Jeffrey Barrett, Sonia Goncalves, Ewan Harrison, David K. Jackson, Ian Johnston, Dominic Kwiatkowski, Cordelia Langford, John Sillitoe on behalf of the Wellcome Sanger Institute COVID-19 Surveillance Team |
| EPI_ISL_2996367 | Lighthouse Laboratory Plymouth | Wellcome Sanger Institute for the COVID-19 Genomics UK (COG-UK) Consortium | Lighthouse Laboratory Plymouth and Alex Alderton, Roberto Amato, Jeffrey Barrett, Sonia Goncalves, Ewan Harrison, David K. Jackson, Ian Johnston, Dominic Kwiatkowski, Cordelia Langford, John Sillitoe on behalf of the Wellcome Sanger Institute COVID-19 Surveillance Team |
| EPI_ISL_2996368 | Lighthouse Lab in Milton Keynes | Wellcome Sanger Institute for the COVID-19 Genomics UK (COG-UK) Consortium | The Lighthouse Lab in Milton Keynes and Alex Alderton, Roberto Amato, Jeffrey Barrett, Sonia Goncalves, Ewan Harrison, David K. Jackson, Ian Johnston, Dominic Kwiatkowski, Cordelia Langford, John Sillitoe on behalf of the Wellcome Sanger Institute COVID-19 Surveillance Team |
| EPI_ISL_2996369, EPI_ISL_2996370, EPI_ISL_2996371 | Lighthouse Laboratory Plymouth | Wellcome Sanger Institute for the COVID-19 Genomics UK (COG-UK) Consortium | Lighthouse Laboratory Plymouth and Alex Alderton, Roberto Amato, Jeffrey Barrett, Sonia Goncalves, Ewan Harrison, David K. Jackson, Ian Johnston, Dominic Kwiatkowski, Cordelia Langford, John Sillitoe on behalf of the Wellcome Sanger Institute COVID-19 Surveillance Team |
| EPI_ISL_2996372 | Lighthouse Lab in Milton Keynes | Wellcome Sanger Institute for the COVID-19 Genomics UK (COG-UK) Consortium | The Lighthouse Lab in Milton Keynes and Alex Alderton, Roberto Amato, Jeffrey Barrett, Sonia Goncalves, Ewan Harrison, David K. Jackson, Ian Johnston, Dominic Kwiatkowski, Cordelia Langford, John Sillitoe on behalf of the Wellcome Sanger Institute COVID-19 Surveillance Team |
| EPI_ISL_2996373, EPI_ISL_2996374, EPI_ISL_2996375 | Lighthouse Laboratory Plymouth | Wellcome Sanger Institute for the COVID-19 Genomics UK (COG-UK) Consortium | Lighthouse Laboratory Plymouth and Alex Alderton, Roberto Amato, Jeffrey Barrett, Sonia Goncalves, Ewan Harrison, David K. Jackson, Ian Johnston, Dominic Kwiatkowski, Cordelia Langford, John Sillitoe on behalf of the Wellcome Sanger Institute COVID-19 Surveillance Team |
| EPI_ISL_2996376 | Lighthouse Lab in Milton Keynes | Wellcome Sanger Institute for the COVID-19 Genomics UK (COG-UK) Consortium | The Lighthouse Lab in Milton Keynes and Alex Alderton, Roberto Amato, Jeffrey Barrett, Sonia Goncalves, Ewan Harrison, David K. Jackson, Ian Johnston, Dominic Kwiatkowski, Cordelia Langford, John Sillitoe on behalf of the Wellcome Sanger Institute COVID-19 Surveillance Team |
| EPI_ISL_2996377, EPI_ISL_2996378 | Lighthouse Laboratory Plymouth | Wellcome Sanger Institute for the COVID-19 Genomics UK (COG-UK) Consortium | Lighthouse Laboratory Plymouth and Alex Alderton, Roberto Amato, Jeffrey Barrett, Sonia Goncalves, Ewan Harrison, David K. Jackson, Ian Johnston, Dominic Kwiatkowski, Cordelia Langford, John Sillitoe on behalf of the Wellcome Sanger Institute COVID-19 Surveillance Team |
| EPI_ISL_2996379 | Lighthouse Lab in Milton Keynes | Wellcome Sanger Institute for the COVID-19 Genomics UK (COG-UK) Consortium | The Lighthouse Lab in Milton Keynes and Alex Alderton, Roberto Amato, Jeffrey Barrett, Sonia Goncalves, Ewan Harrison, David K. Jackson, Ian Johnston, Dominic Kwiatkowski, Cordelia Langford, John Sillitoe on behalf of the Wellcome Sanger Institute COVID-19 Surveillance Team |
| EPI_ISL_2996380, EPI_ISL_2996381, EPI_ISL_2996382, EPI_ISL_2996383, EPI_ISL_2996384, EPI_ISL_2996385, EPI_ISL_2996386, EPI_ISL_2996387, EPI_ISL_2996388, EPI_ISL_2996389 | Lighthouse Laboratory Plymouth | Wellcome Sanger Institute for the COVID-19 Genomics UK (COG-UK) Consortium | Lighthouse Laboratory Plymouth and Alex Alderton, Roberto Amato, Jeffrey Barrett, Sonia Goncalves, Ewan Harrison, David K. Jackson, Ian Johnston, Dominic Kwiatkowski, Cordelia Langford, John Sillitoe on behalf of the Wellcome Sanger Institute COVID-19 Surveillance Team |
| EPI_ISL_2996390, EPI_ISL_2996391 | Lighthouse Lab in Milton Keynes | Wellcome Sanger Institute for the COVID-19 Genomics UK (COG-UK) Consortium | The Lighthouse Lab in Milton Keynes and Alex Alderton, Roberto Amato, Jeffrey Barrett, Sonia Goncalves, Ewan Harrison, David K. Jackson, Ian Johnston, Dominic Kwiatkowski, Cordelia Langford, John Sillitoe on behalf of the Wellcome Sanger Institute COVID-19 Surveillance Team |
| EPI_ISL_2996392, EPI_ISL_2996393, EPI_ISL_2996394, EPI_ISL_2996395, EPI_ISL_2996396, EPI_ISL_2996397 | Lighthouse Laboratory Plymouth | Wellcome Sanger Institute for the COVID-19 Genomics UK (COG-UK) Consortium | Lighthouse Laboratory Plymouth and Alex Alderton, Roberto Amato, Jeffrey Barrett, Sonia Goncalves, Ewan Harrison, David K. Jackson, Ian Johnston, Dominic Kwiatkowski, Cordelia Langford, John Sillitoe on behalf of the Wellcome Sanger Institute COVID-19 Surveillance Team |
| EPI_ISL_2996398, EPI_ISL_2996399, EPI_ISL_2996400, EPI_ISL_2996401 | Lighthouse Lab in Milton Keynes | Wellcome Sanger Institute for the COVID-19 Genomics UK (COG-UK) Consortium | The Lighthouse Lab in Milton Keynes and Alex Alderton, Roberto Amato, Jeffrey Barrett, Sonia Goncalves, Ewan Harrison, David K. Jackson, Ian Johnston, Dominic Kwiatkowski, Cordelia Langford, John Sillitoe on behalf of the Wellcome Sanger Institute COVID-19 Surveillance Team |
| EPI_ISL_2996402 | Lighthouse Laboratory Plymouth | Wellcome Sanger Institute for the COVID-19 Genomics UK (COG-UK) Consortium | Lighthouse Laboratory Plymouth and Alex Alderton, Roberto Amato, Jeffrey Barrett, Sonia Goncalves, Ewan Harrison, David K. Jackson, Ian Johnston, Dominic Kwiatkowski, Cordelia Langford, John Sillitoe on behalf of the Wellcome Sanger Institute COVID-19 Surveillance Team |
| EPI_ISL_2996403 | Lighthouse Lab in Milton Keynes | Wellcome Sanger Institute for the COVID-19 Genomics UK (COG-UK) Consortium | The Lighthouse Lab in Milton Keynes and Alex Alderton, Roberto Amato, Jeffrey Barrett, Sonia Goncalves, Ewan Harrison, David K. Jackson, Ian Johnston, Dominic Kwiatkowski, Cordelia Langford, John Sillitoe on behalf of the Wellcome Sanger Institute COVID-19 Surveillance Team |
| EPI_ISL_2996404, EPI_ISL_2996405 | Lighthouse Laboratory Plymouth | Wellcome Sanger Institute for the COVID-19 Genomics UK (COG-UK) Consortium | Lighthouse Laboratory Plymouth and Alex Alderton, Roberto Amato, Jeffrey Barrett, Sonia Goncalves, Ewan Harrison, David K. Jackson, Ian Johnston, Dominic Kwiatkowski, Cordelia Langford, John Sillitoe on behalf of the Wellcome Sanger Institute COVID-19 Surveillance Team |
| EPI_ISL_2996406 | Lighthouse Lab in Milton Keynes | Wellcome Sanger Institute for the COVID-19 Genomics UK (COG-UK) Consortium | The Lighthouse Lab in Milton Keynes and Alex Alderton, Roberto Amato, Jeffrey Barrett, Sonia Goncalves, Ewan Harrison, David K. Jackson, Ian Johnston, Dominic Kwiatkowski, Cordelia Langford, John Sillitoe on behalf of the Wellcome Sanger Institute COVID-19 Surveillance Team |
| EPI_ISL_2996407, EPI_ISL_2996408 | Lighthouse Laboratory Plymouth | Wellcome Sanger Institute for the COVID-19 Genomics UK (COG-UK) Consortium | Lighthouse Laboratory Plymouth and Alex Alderton, Roberto Amato, Jeffrey Barrett, Sonia Goncalves, Ewan Harrison, David K. Jackson, Ian Johnston, Dominic Kwiatkowski, Cordelia Langford, John Sillitoe on behalf of the Wellcome Sanger Institute COVID-19 Surveillance Team |
| EPI_ISL_2996409, EPI_ISL_2996410, EPI_ISL_2996411 | Lighthouse Lab in Milton Keynes | Wellcome Sanger Institute for the COVID-19 Genomics UK (COG-UK) Consortium | The Lighthouse Lab in Milton Keynes and Alex Alderton, Roberto Amato, Jeffrey Barrett, Sonia Goncalves, Ewan Harrison, David K. Jackson, Ian Johnston, Dominic Kwiatkowski, Cordelia Langford, John Sillitoe on behalf of the Wellcome Sanger Institute COVID-19 Surveillance Team |
| EPI_ISL_2996412, EPI_ISL_2996413, EPI_ISL_2996414, EPI_ISL_2996415, EPI_ISL_2996416, EPI_ISL_2996417, EPI_ISL_2996418, EPI_ISL_2996419, EPI_ISL_2996420, EPI_ISL_2996421, EPI_ISL_2996422 | | | |
| see above | Lighthouse Laboratory Plymouth | Wellcome Sanger Institute for the COVID-19 Genomics UK (COG-UK) Consortium | Lighthouse Laboratory Plymouth and Alex Alderton, Roberto Amato, Jeffrey Barrett, Sonia Goncalves, Ewan Harrison, David K. Jackson, Ian Johnston, Dominic Kwiatkowski, Cordelia Langford, John Sillitoe on behalf of the Wellcome Sanger Institute COVID-19 Surveillance Team |
| EPI_ISL_2996423 | Lighthouse Lab in Milton Keynes | Wellcome Sanger Institute for the COVID-19 Genomics UK (COG-UK) Consortium | The Lighthouse Lab in Milton Keynes and Alex Alderton, Roberto Amato, Jeffrey Barrett, Sonia Goncalves, Ewan Harrison, David K. Jackson, Ian Johnston, Dominic Kwiatkowski, Cordelia Langford, John Sillitoe on behalf of the Wellcome Sanger Institute COVID-19 Surveillance Team |
| EPI_ISL_2996424, | Lighthouse Laboratory Plymouth | Wellcome Sanger Institute for the COVID-19 Genomics UK (COG-UK) | Lighthouse Laboratory Plymouth and Alex Alderton, Roberto Amato, Jeffrey Barrett, Sonia Goncalves, Ewan Harrison, David K. Jackson, Ian Johnston, Dominic Kwiatkowski, Cordelia Langford, John Sillitoe on behalf of the Wellcome Sanger |

| | | | |
|---|---|---|---|
| EPI_ISL_2996425, EPI_ISL_2996426 | | Consortium | Institute COVID-19 Surveillance Team |
| EPI_ISL_2996427, EPI_ISL_2996428 | Lighthouse Lab in Milton Keynes | Wellcome Sanger Institute for the COVID-19 Genomics UK (COG-UK) Consortium | The Lighthouse Lab in Milton Keynes and Alex Alderton, Roberto Amato, Jeffrey Barrett, Sonia Goncalves, Ewan Harrison, David K. Jackson, Ian Johnston, Dominic Kwiatkowski, Cordelia Langford, John Sillitoe on behalf of the Wellcome Sanger Institute COVID-19 Surveillance Team |
| EPI_ISL_2996429, EPI_ISL_2996430 | Lighthouse Laboratory Plymouth | Wellcome Sanger Institute for the COVID-19 Genomics UK (COG-UK) Consortium | Lighthouse Laboratory Plymouth and Alex Alderton, Roberto Amato, Jeffrey Barrett, Sonia Goncalves, Ewan Harrison, David K. Jackson, Ian Johnston, Dominic Kwiatkowski, Cordelia Langford, John Sillitoe on behalf of the Wellcome Sanger Institute COVID-19 Surveillance Team |
| EPI_ISL_2996431 | Lighthouse Lab in Milton Keynes | Wellcome Sanger Institute for the COVID-19 Genomics UK (COG-UK) Consortium | The Lighthouse Lab in Milton Keynes and Alex Alderton, Roberto Amato, Jeffrey Barrett, Sonia Goncalves, Ewan Harrison, David K. Jackson, Ian Johnston, Dominic Kwiatkowski, Cordelia Langford, John Sillitoe on behalf of the Wellcome Sanger Institute COVID-19 Surveillance Team |
| EPI_ISL_2996432 | Lighthouse Laboratory Plymouth | Wellcome Sanger Institute for the COVID-19 Genomics UK (COG-UK) Consortium | Lighthouse Laboratory Plymouth and Alex Alderton, Roberto Amato, Jeffrey Barrett, Sonia Goncalves, Ewan Harrison, David K. Jackson, Ian Johnston, Dominic Kwiatkowski, Cordelia Langford, John Sillitoe on behalf of the Wellcome Sanger Institute COVID-19 Surveillance Team |
| EPI_ISL_2996433 | Lighthouse Lab in Milton Keynes | Wellcome Sanger Institute for the COVID-19 Genomics UK (COG-UK) Consortium | The Lighthouse Lab in Milton Keynes and Alex Alderton, Roberto Amato, Jeffrey Barrett, Sonia Goncalves, Ewan Harrison, David K. Jackson, Ian Johnston, Dominic Kwiatkowski, Cordelia Langford, John Sillitoe on behalf of the Wellcome Sanger Institute COVID-19 Surveillance Team |
| EPI_ISL_2996434, EPI_ISL_2996435 | Lighthouse Laboratory Plymouth | Wellcome Sanger Institute for the COVID-19 Genomics UK (COG-UK) Consortium | Lighthouse Laboratory Plymouth and Alex Alderton, Roberto Amato, Jeffrey Barrett, Sonia Goncalves, Ewan Harrison, David K. Jackson, Ian Johnston, Dominic Kwiatkowski, Cordelia Langford, John Sillitoe on behalf of the Wellcome Sanger Institute COVID-19 Surveillance Team |
| EPI_ISL_2996436 | Lighthouse Lab in Milton Keynes | Wellcome Sanger Institute for the COVID-19 Genomics UK (COG-UK) Consortium | The Lighthouse Lab in Milton Keynes and Alex Alderton, Roberto Amato, Jeffrey Barrett, Sonia Goncalves, Ewan Harrison, David K. Jackson, Ian Johnston, Dominic Kwiatkowski, Cordelia Langford, John Sillitoe on behalf of the Wellcome Sanger Institute COVID-19 Surveillance Team |
| EPI_ISL_2996437, EPI_ISL_2996438, EPI_ISL_2996439 | Lighthouse Laboratory Plymouth | Wellcome Sanger Institute for the COVID-19 Genomics UK (COG-UK) Consortium | Lighthouse Laboratory Plymouth and Alex Alderton, Roberto Amato, Jeffrey Barrett, Sonia Goncalves, Ewan Harrison, David K. Jackson, Ian Johnston, Dominic Kwiatkowski, Cordelia Langford, John Sillitoe on behalf of the Wellcome Sanger Institute COVID-19 Surveillance Team |
| EPI_ISL_2996440, EPI_ISL_2996441 | Lighthouse Lab in Milton Keynes | Wellcome Sanger Institute for the COVID-19 Genomics UK (COG-UK) Consortium | The Lighthouse Lab in Milton Keynes and Alex Alderton, Roberto Amato, Jeffrey Barrett, Sonia Goncalves, Ewan Harrison, David K. Jackson, Ian Johnston, Dominic Kwiatkowski, Cordelia Langford, John Sillitoe on behalf of the Wellcome Sanger Institute COVID-19 Surveillance Team |
| EPI_ISL_2996442, EPI_ISL_2996443, EPI_ISL_2996444, EPI_ISL_2996445, EPI_ISL_2996446, EPI_ISL_2996447, EPI_ISL_2996448, EPI_ISL_2996449, EPI_ISL_2996450, EPI_ISL_2996451, EPI_ISL_2996452, EPI_ISL_2996453 | | | |
| see above | Lighthouse Laboratory Plymouth | Wellcome Sanger Institute for the COVID-19 Genomics UK (COG-UK) Consortium | Lighthouse Laboratory Plymouth and Alex Alderton, Roberto Amato, Jeffrey Barrett, Sonia Goncalves, Ewan Harrison, David K. Jackson, Ian Johnston, Dominic Kwiatkowski, Cordelia Langford, John Sillitoe on behalf of the Wellcome Sanger Institute COVID-19 Surveillance Team |
| EPI_ISL_2996454 | Lighthouse Lab in Milton Keynes | Wellcome Sanger Institute for the COVID-19 Genomics UK (COG-UK) Consortium | The Lighthouse Lab in Milton Keynes and Alex Alderton, Roberto Amato, Jeffrey Barrett, Sonia Goncalves, Ewan Harrison, David K. Jackson, Ian Johnston, Dominic Kwiatkowski, Cordelia Langford, John Sillitoe on behalf of the Wellcome Sanger Institute COVID-19 Surveillance Team |
| EPI_ISL_2996455, EPI_ISL_2996456, EPI_ISL_2996457 | Lighthouse Laboratory Plymouth | Wellcome Sanger Institute for the COVID-19 Genomics UK (COG-UK) Consortium | Lighthouse Laboratory Plymouth and Alex Alderton, Roberto Amato, Jeffrey Barrett, Sonia Goncalves, Ewan Harrison, David K. Jackson, Ian Johnston, Dominic Kwiatkowski, Cordelia Langford, John Sillitoe on behalf of the Wellcome Sanger Institute COVID-19 Surveillance Team |
| EPI_ISL_2996458 | Lighthouse Lab in Milton Keynes | Wellcome Sanger Institute for the COVID-19 Genomics UK (COG-UK) Consortium | The Lighthouse Lab in Milton Keynes and Alex Alderton, Roberto Amato, Jeffrey Barrett, Sonia Goncalves, Ewan Harrison, David K. Jackson, Ian Johnston, Dominic Kwiatkowski, Cordelia Langford, John Sillitoe on behalf of the Wellcome Sanger Institute COVID-19 Surveillance Team |
| EPI_ISL_2996459, EPI_ISL_2996460, EPI_ISL_2996461, EPI_ISL_2996462, EPI_ISL_2996463, EPI_ISL_2996464 | Lighthouse Laboratory Plymouth | Wellcome Sanger Institute for the COVID-19 Genomics UK (COG-UK) Consortium | Lighthouse Laboratory Plymouth and Alex Alderton, Roberto Amato, Jeffrey Barrett, Sonia Goncalves, Ewan Harrison, David K. Jackson, Ian Johnston, Dominic Kwiatkowski, Cordelia Langford, John Sillitoe on behalf of the Wellcome Sanger Institute COVID-19 Surveillance Team |
| EPI_ISL_2996465 | Lighthouse Lab in Milton Keynes | Wellcome Sanger Institute for the COVID-19 Genomics UK (COG-UK) Consortium | The Lighthouse Lab in Milton Keynes and Alex Alderton, Roberto Amato, Jeffrey Barrett, Sonia Goncalves, Ewan Harrison, David K. Jackson, Ian Johnston, Dominic Kwiatkowski, Cordelia Langford, John Sillitoe on behalf of the Wellcome Sanger Institute COVID-19 Surveillance Team |
| EPI_ISL_2996466 | Lighthouse Laboratory Plymouth | Wellcome Sanger Institute for the COVID-19 Genomics UK (COG-UK) Consortium | Lighthouse Laboratory Plymouth and Alex Alderton, Roberto Amato, Jeffrey Barrett, Sonia Goncalves, Ewan Harrison, David K. Jackson, Ian Johnston, Dominic Kwiatkowski, Cordelia Langford, John Sillitoe on behalf of the Wellcome Sanger Institute COVID-19 Surveillance Team |
| EPI_ISL_2996467 | Lighthouse Lab in Milton Keynes | Wellcome Sanger Institute for the COVID-19 Genomics UK (COG-UK) Consortium | The Lighthouse Lab in Milton Keynes and Alex Alderton, Roberto Amato, Jeffrey Barrett, Sonia Goncalves, Ewan Harrison, David K. Jackson, Ian Johnston, Dominic Kwiatkowski, Cordelia Langford, John Sillitoe on behalf of the Wellcome Sanger Institute COVID-19 Surveillance Team |
| EPI_ISL_2996468, EPI_ISL_2996469, EPI_ISL_2996470, EPI_ISL_2996471 | Lighthouse Laboratory Plymouth | Wellcome Sanger Institute for the COVID-19 Genomics UK (COG-UK) Consortium | Lighthouse Laboratory Plymouth and Alex Alderton, Roberto Amato, Jeffrey Barrett, Sonia Goncalves, Ewan Harrison, David K. Jackson, Ian Johnston, Dominic Kwiatkowski, Cordelia Langford, John Sillitoe on behalf of the Wellcome Sanger Institute COVID-19 Surveillance Team |
| EPI_ISL_2996472 | Lighthouse Lab in Milton Keynes | Wellcome Sanger Institute for the COVID-19 Genomics UK (COG-UK) Consortium | The Lighthouse Lab in Milton Keynes and Alex Alderton, Roberto Amato, Jeffrey Barrett, Sonia Goncalves, Ewan Harrison, David K. Jackson, Ian Johnston, Dominic Kwiatkowski, Cordelia Langford, John Sillitoe on behalf of the Wellcome Sanger Institute COVID-19 Surveillance Team |
| EPI_ISL_2996473, EPI_ISL_2996474, EPI_ISL_2996475, EPI_ISL_2996476, EPI_ISL_2996477 | Lighthouse Laboratory Plymouth | Wellcome Sanger Institute for the COVID-19 Genomics UK (COG-UK) Consortium | Lighthouse Laboratory Plymouth and Alex Alderton, Roberto Amato, Jeffrey Barrett, Sonia Goncalves, Ewan Harrison, David K. Jackson, Ian Johnston, Dominic Kwiatkowski, Cordelia Langford, John Sillitoe on behalf of the Wellcome Sanger Institute COVID-19 Surveillance Team |
| EPI_ISL_2996478 | Lighthouse Lab in Milton Keynes | Wellcome Sanger Institute for the COVID-19 Genomics UK (COG-UK) Consortium | The Lighthouse Lab in Milton Keynes and Alex Alderton, Roberto Amato, Jeffrey Barrett, Sonia Goncalves, Ewan Harrison, David K. Jackson, Ian Johnston, Dominic Kwiatkowski, Cordelia Langford, John Sillitoe on behalf of the Wellcome Sanger Institute COVID-19 Surveillance Team |
| EPI_ISL_2996479, EPI_ISL_2996480, EPI_ISL_2996481, EPI_ISL_2996482, EPI_ISL_2996483, EPI_ISL_2996484, EPI_ISL_2996485, EPI_ISL_2996486, EPI_ISL_2996487, EPI_ISL_2996488, EPI_ISL_2996489, EPI_ISL_2996490, EPI_ISL_2996491, EPI_ISL_2996492, EPI_ISL_2996493 | | | |
| see above | Lighthouse Laboratory Plymouth | Wellcome Sanger Institute for the COVID-19 Genomics UK (COG-UK) Consortium | Lighthouse Laboratory Plymouth and Alex Alderton, Roberto Amato, Jeffrey Barrett, Sonia Goncalves, Ewan Harrison, David K. Jackson, Ian Johnston, Dominic Kwiatkowski, Cordelia Langford, John Sillitoe on behalf of the Wellcome Sanger Institute COVID-19 Surveillance Team |
| EPI_ISL_2996494 | Lighthouse Lab in Milton Keynes | Wellcome Sanger Institute for the COVID-19 Genomics UK (COG-UK) Consortium | The Lighthouse Lab in Milton Keynes and Alex Alderton, Roberto Amato, Jeffrey Barrett, Sonia Goncalves, Ewan Harrison, David K. Jackson, Ian Johnston, Dominic Kwiatkowski, Cordelia Langford, John Sillitoe on behalf of the Wellcome Sanger Institute COVID-19 Surveillance Team |
| EPI_ISL_2996495 | Lighthouse Laboratory Plymouth | Wellcome Sanger Institute for the COVID-19 Genomics UK (COG-UK) Consortium | Lighthouse Laboratory Plymouth and Alex Alderton, Roberto Amato, Jeffrey Barrett, Sonia Goncalves, Ewan Harrison, David K. Jackson, Ian Johnston, Dominic Kwiatkowski, Cordelia Langford, John Sillitoe on behalf of the Wellcome Sanger Institute COVID-19 Surveillance Team |
| EPI_ISL_2996496 | Lighthouse Lab in Milton Keynes | Wellcome Sanger Institute for the COVID-19 Genomics UK (COG-UK) Consortium | The Lighthouse Lab in Milton Keynes and Alex Alderton, Roberto Amato, Jeffrey Barrett, Sonia Goncalves, Ewan Harrison, David K. Jackson, Ian Johnston, Dominic Kwiatkowski, Cordelia Langford, John Sillitoe on behalf of the Wellcome Sanger Institute COVID-19 Surveillance Team |
| EPI_ISL_2996497, EPI_ISL_2996498, EPI_ISL_2996499, EPI_ISL_2996500 | Lighthouse Laboratory Plymouth | Wellcome Sanger Institute for the COVID-19 Genomics UK (COG-UK) Consortium | Lighthouse Laboratory Plymouth and Alex Alderton, Roberto Amato, Jeffrey Barrett, Sonia Goncalves, Ewan Harrison, David K. Jackson, Ian Johnston, Dominic Kwiatkowski, Cordelia Langford, John Sillitoe on behalf of the Wellcome Sanger Institute COVID-19 Surveillance Team |
| EPI_ISL_2996501 | Lighthouse Lab in Milton Keynes | Wellcome Sanger Institute for the COVID-19 Genomics UK (COG-UK) Consortium | The Lighthouse Lab in Milton Keynes and Alex Alderton, Roberto Amato, Jeffrey Barrett, Sonia Goncalves, Ewan Harrison, David K. Jackson, Ian Johnston, Dominic Kwiatkowski, Cordelia Langford, John Sillitoe on behalf of the Wellcome Sanger Institute COVID-19 Surveillance Team |
| EPI_ISL_2996502, EPI_ISL_2996503, EPI_ISL_2996504, EPI_ISL_2996505 | Lighthouse Laboratory Plymouth | Wellcome Sanger Institute for the COVID-19 Genomics UK (COG-UK) Consortium | Lighthouse Laboratory Plymouth and Alex Alderton, Roberto Amato, Jeffrey Barrett, Sonia Goncalves, Ewan Harrison, David K. Jackson, Ian Johnston, Dominic Kwiatkowski, Cordelia Langford, John Sillitoe on behalf of the Wellcome Sanger Institute COVID-19 Surveillance Team |
| EPI_ISL_2996506 | Lighthouse Lab in Milton Keynes | Wellcome Sanger Institute for the COVID-19 Genomics UK (COG-UK) Consortium | The Lighthouse Lab in Milton Keynes and Alex Alderton, Roberto Amato, Jeffrey Barrett, Sonia Goncalves, Ewan Harrison, David K. Jackson, Ian Johnston, Dominic Kwiatkowski, Cordelia Langford, John Sillitoe on behalf of the Wellcome Sanger Institute COVID-19 Surveillance Team |
| EPI_ISL_2996507, EPI_ISL_2996508, EPI_ISL_2996509, EPI_ISL_2996510 | Lighthouse Laboratory Plymouth | Wellcome Sanger Institute for the COVID-19 Genomics UK (COG-UK) Consortium | Lighthouse Laboratory Plymouth and Alex Alderton, Roberto Amato, Jeffrey Barrett, Sonia Goncalves, Ewan Harrison, David K. Jackson, Ian Johnston, Dominic Kwiatkowski, Cordelia Langford, John Sillitoe on behalf of the Wellcome Sanger Institute COVID-19 Surveillance Team |
| EPI_ISL_2996511 | Lighthouse Lab in Milton Keynes | Wellcome Sanger Institute for the COVID-19 Genomics UK (COG-UK) Consortium | The Lighthouse Lab in Milton Keynes and Alex Alderton, Roberto Amato, Jeffrey Barrett, Sonia Goncalves, Ewan Harrison, David K. Jackson, Ian Johnston, Dominic Kwiatkowski, Cordelia Langford, John Sillitoe on behalf of the Wellcome Sanger Institute COVID-19 Surveillance Team |
| EPI_ISL_2996512 | Lighthouse Laboratory Plymouth | Wellcome Sanger Institute for the COVID-19 Genomics UK (COG-UK) Consortium | Lighthouse Laboratory Plymouth and Alex Alderton, Roberto Amato, Jeffrey Barrett, Sonia Goncalves, Ewan Harrison, David K. Jackson, Ian Johnston, Dominic Kwiatkowski, Cordelia Langford, John Sillitoe on behalf of the Wellcome Sanger Institute COVID-19 Surveillance Team |
| EPI_ISL_2996513 | Lighthouse Lab in Milton Keynes | Wellcome Sanger Institute for the COVID-19 Genomics UK (COG-UK) Consortium | The Lighthouse Lab in Milton Keynes and Alex Alderton, Roberto Amato, Jeffrey Barrett, Sonia Goncalves, Ewan Harrison, David K. Jackson, Ian Johnston, Dominic Kwiatkowski, Cordelia Langford, John Sillitoe on behalf of the Wellcome Sanger Institute COVID-19 Surveillance Team |
| EPI_ISL_2996514, EPI_ISL_2996515, EPI_ISL_2996516, EPI_ISL_2996517 | Lighthouse Laboratory Plymouth | Wellcome Sanger Institute for the COVID-19 Genomics UK (COG-UK) Consortium | Lighthouse Laboratory Plymouth and Alex Alderton, Roberto Amato, Jeffrey Barrett, Sonia Goncalves, Ewan Harrison, David K. Jackson, Ian Johnston, Dominic Kwiatkowski, Cordelia Langford, John Sillitoe on behalf of the Wellcome Sanger Institute COVID-19 Surveillance Team |
| EPI_ISL_2996518 | Lighthouse Lab in Milton Keynes | Wellcome Sanger Institute for the COVID-19 Genomics UK (COG-UK) Consortium | The Lighthouse Lab in Milton Keynes and Alex Alderton, Roberto Amato, Jeffrey Barrett, Sonia Goncalves, Ewan Harrison, David K. Jackson, Ian Johnston, Dominic Kwiatkowski, Cordelia Langford, John Sillitoe on behalf of the Wellcome Sanger Institute COVID-19 Surveillance Team |

| | | | |
|---|---|---|---|
| EPI_ISL_2996519 | Lighthouse Laboratory Plymouth | Wellcome Sanger Institute for the COVID-19 Genomics UK (COG-UK) Consortium | Lighthouse Laboratory Plymouth and Alex Alderton, Roberto Amato, Jeffrey Barrett, Sonia Goncalves, Ewan Harrison, David K. Jackson, Ian Johnston, Dominic Kwiatkowski, Cordelia Langford, John Sillitoe on behalf of the Wellcome Sanger Institute COVID-19 Surveillance Team |
| EPI_ISL_2996520 | Lighthouse Lab in Milton Keynes | Wellcome Sanger Institute for the COVID-19 Genomics UK (COG-UK) Consortium | The Lighthouse Lab in Milton Keynes and Alex Alderton, Roberto Amato, Jeffrey Barrett, Sonia Goncalves, Ewan Harrison, David K. Jackson, Ian Johnston, Dominic Kwiatkowski, Cordelia Langford, John Sillitoe on behalf of the Wellcome Sanger Institute COVID-19 Surveillance Team |
| EPI_ISL_2996521 | Lighthouse Laboratory Plymouth | Wellcome Sanger Institute for the COVID-19 Genomics UK (COG-UK) Consortium | Lighthouse Laboratory Plymouth and Alex Alderton, Roberto Amato, Jeffrey Barrett, Sonia Goncalves, Ewan Harrison, David K. Jackson, Ian Johnston, Dominic Kwiatkowski, Cordelia Langford, John Sillitoe on behalf of the Wellcome Sanger Institute COVID-19 Surveillance Team |
| EPI_ISL_2996522, EPI_ISL_2996523, EPI_ISL_2996524 | Lighthouse Lab in Milton Keynes | Wellcome Sanger Institute for the COVID-19 Genomics UK (COG-UK) Consortium | The Lighthouse Lab in Milton Keynes and Alex Alderton, Roberto Amato, Jeffrey Barrett, Sonia Goncalves, Ewan Harrison, David K. Jackson, Ian Johnston, Dominic Kwiatkowski, Cordelia Langford, John Sillitoe on behalf of the Wellcome Sanger Institute COVID-19 Surveillance Team |
| EPI_ISL_2996525, EPI_ISL_2996526, EPI_ISL_2996527, EPI_ISL_2996528, EPI_ISL_2996529 | Lighthouse Laboratory Plymouth | Wellcome Sanger Institute for the COVID-19 Genomics UK (COG-UK) Consortium | Lighthouse Laboratory Plymouth and Alex Alderton, Roberto Amato, Jeffrey Barrett, Sonia Goncalves, Ewan Harrison, David K. Jackson, Ian Johnston, Dominic Kwiatkowski, Cordelia Langford, John Sillitoe on behalf of the Wellcome Sanger Institute COVID-19 Surveillance Team |
| EPI_ISL_2996530, EPI_ISL_2996531 | Lighthouse Lab in Milton Keynes | Wellcome Sanger Institute for the COVID-19 Genomics UK (COG-UK) Consortium | The Lighthouse Lab in Milton Keynes and Alex Alderton, Roberto Amato, Jeffrey Barrett, Sonia Goncalves, Ewan Harrison, David K. Jackson, Ian Johnston, Dominic Kwiatkowski, Cordelia Langford, John Sillitoe on behalf of the Wellcome Sanger Institute COVID-19 Surveillance Team |
| EPI_ISL_2996532, EPI_ISL_2996533, EPI_ISL_2996534, EPI_ISL_2996535, EPI_ISL_2996536 | Lighthouse Laboratory Plymouth | Wellcome Sanger Institute for the COVID-19 Genomics UK (COG-UK) Consortium | Lighthouse Laboratory Plymouth and Alex Alderton, Roberto Amato, Jeffrey Barrett, Sonia Goncalves, Ewan Harrison, David K. Jackson, Ian Johnston, Dominic Kwiatkowski, Cordelia Langford, John Sillitoe on behalf of the Wellcome Sanger Institute COVID-19 Surveillance Team |
| EPI_ISL_2996537 | Lighthouse Lab in Milton Keynes | Wellcome Sanger Institute for the COVID-19 Genomics UK (COG-UK) Consortium | The Lighthouse Lab in Milton Keynes and Alex Alderton, Roberto Amato, Jeffrey Barrett, Sonia Goncalves, Ewan Harrison, David K. Jackson, Ian Johnston, Dominic Kwiatkowski, Cordelia Langford, John Sillitoe on behalf of the Wellcome Sanger Institute COVID-19 Surveillance Team |
| EPI_ISL_2996538, EPI_ISL_2996539, EPI_ISL_2996540 | Lighthouse Laboratory Plymouth | Wellcome Sanger Institute for the COVID-19 Genomics UK (COG-UK) Consortium | Lighthouse Laboratory Plymouth and Alex Alderton, Roberto Amato, Jeffrey Barrett, Sonia Goncalves, Ewan Harrison, David K. Jackson, Ian Johnston, Dominic Kwiatkowski, Cordelia Langford, John Sillitoe on behalf of the Wellcome Sanger Institute COVID-19 Surveillance Team |
| EPI_ISL_2996541, EPI_ISL_2996542 | Lighthouse Lab in Milton Keynes | Wellcome Sanger Institute for the COVID-19 Genomics UK (COG-UK) Consortium | The Lighthouse Lab in Milton Keynes and Alex Alderton, Roberto Amato, Jeffrey Barrett, Sonia Goncalves, Ewan Harrison, David K. Jackson, Ian Johnston, Dominic Kwiatkowski, Cordelia Langford, John Sillitoe on behalf of the Wellcome Sanger Institute COVID-19 Surveillance Team |
| EPI_ISL_2996543 | Lighthouse Laboratory Plymouth | Wellcome Sanger Institute for the COVID-19 Genomics UK (COG-UK) Consortium | Lighthouse Laboratory Plymouth and Alex Alderton, Roberto Amato, Jeffrey Barrett, Sonia Goncalves, Ewan Harrison, David K. Jackson, Ian Johnston, Dominic Kwiatkowski, Cordelia Langford, John Sillitoe on behalf of the Wellcome Sanger Institute COVID-19 Surveillance Team |
| EPI_ISL_2996544 | Lighthouse Lab in Milton Keynes | Wellcome Sanger Institute for the COVID-19 Genomics UK (COG-UK) Consortium | The Lighthouse Lab in Milton Keynes and Alex Alderton, Roberto Amato, Jeffrey Barrett, Sonia Goncalves, Ewan Harrison, David K. Jackson, Ian Johnston, Dominic Kwiatkowski, Cordelia Langford, John Sillitoe on behalf of the Wellcome Sanger Institute COVID-19 Surveillance Team |
| EPI_ISL_2996545 | Lighthouse Laboratory Plymouth | Wellcome Sanger Institute for the COVID-19 Genomics UK (COG-UK) Consortium | Lighthouse Laboratory Plymouth and Alex Alderton, Roberto Amato, Jeffrey Barrett, Sonia Goncalves, Ewan Harrison, David K. Jackson, Ian Johnston, Dominic Kwiatkowski, Cordelia Langford, John Sillitoe on behalf of the Wellcome Sanger Institute COVID-19 Surveillance Team |
| EPI_ISL_2996546 | Lighthouse Lab in Milton Keynes | Wellcome Sanger Institute for the COVID-19 Genomics UK (COG-UK) Consortium | The Lighthouse Lab in Milton Keynes and Alex Alderton, Roberto Amato, Jeffrey Barrett, Sonia Goncalves, Ewan Harrison, David K. Jackson, Ian Johnston, Dominic Kwiatkowski, Cordelia Langford, John Sillitoe on behalf of the Wellcome Sanger Institute COVID-19 Surveillance Team |
| EPI_ISL_2996547, EPI_ISL_2996548, EPI_ISL_2996549, EPI_ISL_2996550, EPI_ISL_2996551, EPI_ISL_2996552, EPI_ISL_2996553, EPI_ISL_2996554 | Lighthouse Laboratory Plymouth | Wellcome Sanger Institute for the COVID-19 Genomics UK (COG-UK) Consortium | Lighthouse Laboratory Plymouth and Alex Alderton, Roberto Amato, Jeffrey Barrett, Sonia Goncalves, Ewan Harrison, David K. Jackson, Ian Johnston, Dominic Kwiatkowski, Cordelia Langford, John Sillitoe on behalf of the Wellcome Sanger Institute COVID-19 Surveillance Team |
| EPI_ISL_2996555, EPI_ISL_2996556 | Lighthouse Lab in Milton Keynes | Wellcome Sanger Institute for the COVID-19 Genomics UK (COG-UK) Consortium | The Lighthouse Lab in Milton Keynes and Alex Alderton, Roberto Amato, Jeffrey Barrett, Sonia Goncalves, Ewan Harrison, David K. Jackson, Ian Johnston, Dominic Kwiatkowski, Cordelia Langford, John Sillitoe on behalf of the Wellcome Sanger Institute COVID-19 Surveillance Team |
| EPI_ISL_2996557 | Lighthouse Laboratory Plymouth | Wellcome Sanger Institute for the COVID-19 Genomics UK (COG-UK) Consortium | Lighthouse Laboratory Plymouth and Alex Alderton, Roberto Amato, Jeffrey Barrett, Sonia Goncalves, Ewan Harrison, David K. Jackson, Ian Johnston, Dominic Kwiatkowski, Cordelia Langford, John Sillitoe on behalf of the Wellcome Sanger Institute COVID-19 Surveillance Team |
| EPI_ISL_2996558, EPI_ISL_2996559 | Lighthouse Lab in Milton Keynes | Wellcome Sanger Institute for the COVID-19 Genomics UK (COG-UK) Consortium | The Lighthouse Lab in Milton Keynes and Alex Alderton, Roberto Amato, Jeffrey Barrett, Sonia Goncalves, Ewan Harrison, David K. Jackson, Ian Johnston, Dominic Kwiatkowski, Cordelia Langford, John Sillitoe on behalf of the Wellcome Sanger Institute COVID-19 Surveillance Team |
| EPI_ISL_2996560 | Lighthouse Laboratory Plymouth | Wellcome Sanger Institute for the COVID-19 Genomics UK (COG-UK) Consortium | Lighthouse Laboratory Plymouth and Alex Alderton, Roberto Amato, Jeffrey Barrett, Sonia Goncalves, Ewan Harrison, David K. Jackson, Ian Johnston, Dominic Kwiatkowski, Cordelia Langford, John Sillitoe on behalf of the Wellcome Sanger Institute COVID-19 Surveillance Team |
| EPI_ISL_2996561 | Lighthouse Lab in Milton Keynes | Wellcome Sanger Institute for the COVID-19 Genomics UK (COG-UK) Consortium | The Lighthouse Lab in Milton Keynes and Alex Alderton, Roberto Amato, Jeffrey Barrett, Sonia Goncalves, Ewan Harrison, David K. Jackson, Ian Johnston, Dominic Kwiatkowski, Cordelia Langford, John Sillitoe on behalf of the Wellcome Sanger Institute COVID-19 Surveillance Team |
| EPI_ISL_2996562 | Lighthouse Laboratory Plymouth | Wellcome Sanger Institute for the COVID-19 Genomics UK (COG-UK) Consortium | Lighthouse Laboratory Plymouth and Alex Alderton, Roberto Amato, Jeffrey Barrett, Sonia Goncalves, Ewan Harrison, David K. Jackson, Ian Johnston, Dominic Kwiatkowski, Cordelia Langford, John Sillitoe on behalf of the Wellcome Sanger Institute COVID-19 Surveillance Team |
| EPI_ISL_2996563 | Lighthouse Lab in Milton Keynes | Wellcome Sanger Institute for the COVID-19 Genomics UK (COG-UK) Consortium | The Lighthouse Lab in Milton Keynes and Alex Alderton, Roberto Amato, Jeffrey Barrett, Sonia Goncalves, Ewan Harrison, David K. Jackson, Ian Johnston, Dominic Kwiatkowski, Cordelia Langford, John Sillitoe on behalf of the Wellcome Sanger Institute COVID-19 Surveillance Team |
| EPI_ISL_2996564, EPI_ISL_2996565, EPI_ISL_2996566, EPI_ISL_2996567, EPI_ISL_2996568, EPI_ISL_2996569, EPI_ISL_2996570, EPI_ISL_2996571, EPI_ISL_2996572, EPI_ISL_2996573, EPI_ISL_2996574, EPI_ISL_2996575, EPI_ISL_2996576, EPI_ISL_2996577, EPI_ISL_2996578, EPI_ISL_2996579, EPI_ISL_2996580, EPI_ISL_2996581 | | | |
| see above | Lighthouse Laboratory Plymouth | Wellcome Sanger Institute for the COVID-19 Genomics UK (COG-UK) Consortium | Lighthouse Laboratory Plymouth and Alex Alderton, Roberto Amato, Jeffrey Barrett, Sonia Goncalves, Ewan Harrison, David K. Jackson, Ian Johnston, Dominic Kwiatkowski, Cordelia Langford, John Sillitoe on behalf of the Wellcome Sanger Institute COVID-19 Surveillance Team |
| EPI_ISL_2996582 | Lighthouse Lab in Milton Keynes | Wellcome Sanger Institute for the COVID-19 Genomics UK (COG-UK) Consortium | The Lighthouse Lab in Milton Keynes and Alex Alderton, Roberto Amato, Jeffrey Barrett, Sonia Goncalves, Ewan Harrison, David K. Jackson, Ian Johnston, Dominic Kwiatkowski, Cordelia Langford, John Sillitoe on behalf of the Wellcome Sanger Institute COVID-19 Surveillance Team |
| EPI_ISL_2996583, EPI_ISL_2996584, EPI_ISL_2996585, EPI_ISL_2996586, EPI_ISL_2996587, EPI_ISL_2996588, EPI_ISL_2996589 | Lighthouse Laboratory Plymouth | Wellcome Sanger Institute for the COVID-19 Genomics UK (COG-UK) Consortium | Lighthouse Laboratory Plymouth and Alex Alderton, Roberto Amato, Jeffrey Barrett, Sonia Goncalves, Ewan Harrison, David K. Jackson, Ian Johnston, Dominic Kwiatkowski, Cordelia Langford, John Sillitoe on behalf of the Wellcome Sanger Institute COVID-19 Surveillance Team |
| EPI_ISL_2996590, EPI_ISL_2996591, EPI_ISL_2996592 | Lighthouse Lab in Milton Keynes | Wellcome Sanger Institute for the COVID-19 Genomics UK (COG-UK) Consortium | The Lighthouse Lab in Milton Keynes and Alex Alderton, Roberto Amato, Jeffrey Barrett, Sonia Goncalves, Ewan Harrison, David K. Jackson, Ian Johnston, Dominic Kwiatkowski, Cordelia Langford, John Sillitoe on behalf of the Wellcome Sanger Institute COVID-19 Surveillance Team |
| EPI_ISL_2996593, EPI_ISL_2996594, EPI_ISL_2996595, EPI_ISL_2996596, EPI_ISL_2996597, EPI_ISL_2996598 | Lighthouse Laboratory Plymouth | Wellcome Sanger Institute for the COVID-19 Genomics UK (COG-UK) Consortium | Lighthouse Laboratory Plymouth and Alex Alderton, Roberto Amato, Jeffrey Barrett, Sonia Goncalves, Ewan Harrison, David K. Jackson, Ian Johnston, Dominic Kwiatkowski, Cordelia Langford, John Sillitoe on behalf of the Wellcome Sanger Institute COVID-19 Surveillance Team |
| EPI_ISL_2996599 | Lighthouse Lab in Milton Keynes | Wellcome Sanger Institute for the COVID-19 Genomics UK (COG-UK) Consortium | The Lighthouse Lab in Milton Keynes and Alex Alderton, Roberto Amato, Jeffrey Barrett, Sonia Goncalves, Ewan Harrison, David K. Jackson, Ian Johnston, Dominic Kwiatkowski, Cordelia Langford, John Sillitoe on behalf of the Wellcome Sanger Institute COVID-19 Surveillance Team |
| EPI_ISL_2996600, EPI_ISL_2996601 | Lighthouse Laboratory Plymouth | Wellcome Sanger Institute for the COVID-19 Genomics UK (COG-UK) Consortium | Lighthouse Laboratory Plymouth and Alex Alderton, Roberto Amato, Jeffrey Barrett, Sonia Goncalves, Ewan Harrison, David K. Jackson, Ian Johnston, Dominic Kwiatkowski, Cordelia Langford, John Sillitoe on behalf of the Wellcome Sanger Institute COVID-19 Surveillance Team |
| EPI_ISL_2996602 | Lighthouse Lab in Milton Keynes | Wellcome Sanger Institute for the COVID-19 Genomics UK (COG-UK) Consortium | The Lighthouse Lab in Milton Keynes and Alex Alderton, Roberto Amato, Jeffrey Barrett, Sonia Goncalves, Ewan Harrison, David K. Jackson, Ian Johnston, Dominic Kwiatkowski, Cordelia Langford, John Sillitoe on behalf of the Wellcome Sanger Institute COVID-19 Surveillance Team |
| EPI_ISL_2996603, EPI_ISL_2996604, EPI_ISL_2996605, EPI_ISL_2996606 | Lighthouse Laboratory Plymouth | Wellcome Sanger Institute for the COVID-19 Genomics UK (COG-UK) Consortium | Lighthouse Laboratory Plymouth and Alex Alderton, Roberto Amato, Jeffrey Barrett, Sonia Goncalves, Ewan Harrison, David K. Jackson, Ian Johnston, Dominic Kwiatkowski, Cordelia Langford, John Sillitoe on behalf of the Wellcome Sanger Institute COVID-19 Surveillance Team |
| EPI_ISL_2996607 | Lighthouse Lab in Milton Keynes | Wellcome Sanger Institute for the COVID-19 Genomics UK (COG-UK) Consortium | The Lighthouse Lab in Milton Keynes and Alex Alderton, Roberto Amato, Jeffrey Barrett, Sonia Goncalves, Ewan Harrison, David K. Jackson, Ian Johnston, Dominic Kwiatkowski, Cordelia Langford, John Sillitoe on behalf of the Wellcome Sanger Institute COVID-19 Surveillance Team |
| EPI_ISL_2996608 | Lighthouse Laboratory Plymouth | Wellcome Sanger Institute for the COVID-19 Genomics UK (COG-UK) Consortium | Lighthouse Laboratory Plymouth and Alex Alderton, Roberto Amato, Jeffrey Barrett, Sonia Goncalves, Ewan Harrison, David K. Jackson, Ian Johnston, Dominic Kwiatkowski, Cordelia Langford, John Sillitoe on behalf of the Wellcome Sanger Institute COVID-19 Surveillance Team |
| EPI_ISL_2996609 | Lighthouse Lab in Milton Keynes | Wellcome Sanger Institute for the COVID-19 Genomics UK (COG-UK) Consortium | The Lighthouse Lab in Milton Keynes and Alex Alderton, Roberto Amato, Jeffrey Barrett, Sonia Goncalves, Ewan Harrison, David K. Jackson, Ian Johnston, Dominic Kwiatkowski, Cordelia Langford, John Sillitoe on behalf of the Wellcome Sanger Institute COVID-19 Surveillance Team |

| | | | |
|---|---|---|---|
| EPI_ISL_2996610, EPI_ISL_2996611 | Lighthouse Laboratory Plymouth | Wellcome Sanger Institute for the COVID-19 Genomics UK (COG-UK) Consortium | Lighthouse Laboratory Plymouth and Alex Alderton, Roberto Amato, Jeffrey Barrett, Sonia Goncalves, Ewan Harrison, David K. Jackson, Ian Johnston, Dominic Kwiatkowski, Cordelia Langford, John Sillitoe on behalf of the Wellcome Sanger Institute COVID-19 Surveillance Team |
| EPI_ISL_2996612 | Lighthouse Lab in Milton Keynes | Wellcome Sanger Institute for the COVID-19 Genomics UK (COG-UK) Consortium | The Lighthouse Lab in Milton Keynes and Alex Alderton, Roberto Amato, Jeffrey Barrett, Sonia Goncalves, Ewan Harrison, David K. Jackson, Ian Johnston, Dominic Kwiatkowski, Cordelia Langford, John Sillitoe on behalf of the Wellcome Sanger Institute COVID-19 Surveillance Team |
| EPI_ISL_2996613 | Lighthouse Laboratory Plymouth | Wellcome Sanger Institute for the COVID-19 Genomics UK (COG-UK) Consortium | Lighthouse Laboratory Plymouth and Alex Alderton, Roberto Amato, Jeffrey Barrett, Sonia Goncalves, Ewan Harrison, David K. Jackson, Ian Johnston, Dominic Kwiatkowski, Cordelia Langford, John Sillitoe on behalf of the Wellcome Sanger Institute COVID-19 Surveillance Team |
| EPI_ISL_2996614 | Lighthouse Lab in Milton Keynes | Wellcome Sanger Institute for the COVID-19 Genomics UK (COG-UK) Consortium | The Lighthouse Lab in Milton Keynes and Alex Alderton, Roberto Amato, Jeffrey Barrett, Sonia Goncalves, Ewan Harrison, David K. Jackson, Ian Johnston, Dominic Kwiatkowski, Cordelia Langford, John Sillitoe on behalf of the Wellcome Sanger Institute COVID-19 Surveillance Team |
| EPI_ISL_2996615, EPI_ISL_2996616, EPI_ISL_2996617 | Lighthouse Laboratory Plymouth | Wellcome Sanger Institute for the COVID-19 Genomics UK (COG-UK) Consortium | Lighthouse Laboratory Plymouth and Alex Alderton, Roberto Amato, Jeffrey Barrett, Sonia Goncalves, Ewan Harrison, David K. Jackson, Ian Johnston, Dominic Kwiatkowski, Cordelia Langford, John Sillitoe on behalf of the Wellcome Sanger Institute COVID-19 Surveillance Team |
| EPI_ISL_2996618 | Lighthouse Lab in Milton Keynes | Wellcome Sanger Institute for the COVID-19 Genomics UK (COG-UK) Consortium | The Lighthouse Lab in Milton Keynes and Alex Alderton, Roberto Amato, Jeffrey Barrett, Sonia Goncalves, Ewan Harrison, David K. Jackson, Ian Johnston, Dominic Kwiatkowski, Cordelia Langford, John Sillitoe on behalf of the Wellcome Sanger Institute COVID-19 Surveillance Team |
| EPI_ISL_2996619, EPI_ISL_2996620, EPI_ISL_2996621, EPI_ISL_2996622, EPI_ISL_2996623 | Lighthouse Laboratory Plymouth | Wellcome Sanger Institute for the COVID-19 Genomics UK (COG-UK) Consortium | Lighthouse Laboratory Plymouth and Alex Alderton, Roberto Amato, Jeffrey Barrett, Sonia Goncalves, Ewan Harrison, David K. Jackson, Ian Johnston, Dominic Kwiatkowski, Cordelia Langford, John Sillitoe on behalf of the Wellcome Sanger Institute COVID-19 Surveillance Team |
| EPI_ISL_2996624 | Lighthouse Lab in Milton Keynes | Wellcome Sanger Institute for the COVID-19 Genomics UK (COG-UK) Consortium | The Lighthouse Lab in Milton Keynes and Alex Alderton, Roberto Amato, Jeffrey Barrett, Sonia Goncalves, Ewan Harrison, David K. Jackson, Ian Johnston, Dominic Kwiatkowski, Cordelia Langford, John Sillitoe on behalf of the Wellcome Sanger Institute COVID-19 Surveillance Team |
| EPI_ISL_2996625, EPI_ISL_2996626, EPI_ISL_2996627 | Lighthouse Laboratory Plymouth | Wellcome Sanger Institute for the COVID-19 Genomics UK (COG-UK) Consortium | Lighthouse Laboratory Plymouth and Alex Alderton, Roberto Amato, Jeffrey Barrett, Sonia Goncalves, Ewan Harrison, David K. Jackson, Ian Johnston, Dominic Kwiatkowski, Cordelia Langford, John Sillitoe on behalf of the Wellcome Sanger Institute COVID-19 Surveillance Team |
| EPI_ISL_2996628 | Lighthouse Lab in Milton Keynes | Wellcome Sanger Institute for the COVID-19 Genomics UK (COG-UK) Consortium | The Lighthouse Lab in Milton Keynes and Alex Alderton, Roberto Amato, Jeffrey Barrett, Sonia Goncalves, Ewan Harrison, David K. Jackson, Ian Johnston, Dominic Kwiatkowski, Cordelia Langford, John Sillitoe on behalf of the Wellcome Sanger Institute COVID-19 Surveillance Team |
| EPI_ISL_2996629 | Lighthouse Laboratory Plymouth | Wellcome Sanger Institute for the COVID-19 Genomics UK (COG-UK) Consortium | Lighthouse Laboratory Plymouth and Alex Alderton, Roberto Amato, Jeffrey Barrett, Sonia Goncalves, Ewan Harrison, David K. Jackson, Ian Johnston, Dominic Kwiatkowski, Cordelia Langford, John Sillitoe on behalf of the Wellcome Sanger Institute COVID-19 Surveillance Team |
| EPI_ISL_2996630 | Lighthouse Lab in Milton Keynes | Wellcome Sanger Institute for the COVID-19 Genomics UK (COG-UK) Consortium | The Lighthouse Lab in Milton Keynes and Alex Alderton, Roberto Amato, Jeffrey Barrett, Sonia Goncalves, Ewan Harrison, David K. Jackson, Ian Johnston, Dominic Kwiatkowski, Cordelia Langford, John Sillitoe on behalf of the Wellcome Sanger Institute COVID-19 Surveillance Team |
| EPI_ISL_2996631, EPI_ISL_2996632 | Lighthouse Laboratory Plymouth | Wellcome Sanger Institute for the COVID-19 Genomics UK (COG-UK) Consortium | Lighthouse Laboratory Plymouth and Alex Alderton, Roberto Amato, Jeffrey Barrett, Sonia Goncalves, Ewan Harrison, David K. Jackson, Ian Johnston, Dominic Kwiatkowski, Cordelia Langford, John Sillitoe on behalf of the Wellcome Sanger Institute COVID-19 Surveillance Team |
| EPI_ISL_2996633 | Lighthouse Lab in Milton Keynes | Wellcome Sanger Institute for the COVID-19 Genomics UK (COG-UK) Consortium | The Lighthouse Lab in Milton Keynes and Alex Alderton, Roberto Amato, Jeffrey Barrett, Sonia Goncalves, Ewan Harrison, David K. Jackson, Ian Johnston, Dominic Kwiatkowski, Cordelia Langford, John Sillitoe on behalf of the Wellcome Sanger Institute COVID-19 Surveillance Team |
| EPI_ISL_2996634, EPI_ISL_2996635, EPI_ISL_2996636 | Lighthouse Laboratory Plymouth | Wellcome Sanger Institute for the COVID-19 Genomics UK (COG-UK) Consortium | Lighthouse Laboratory Plymouth and Alex Alderton, Roberto Amato, Jeffrey Barrett, Sonia Goncalves, Ewan Harrison, David K. Jackson, Ian Johnston, Dominic Kwiatkowski, Cordelia Langford, John Sillitoe on behalf of the Wellcome Sanger Institute COVID-19 Surveillance Team |
| EPI_ISL_2996637 | Lighthouse Lab in Milton Keynes | Wellcome Sanger Institute for the COVID-19 Genomics UK (COG-UK) Consortium | The Lighthouse Lab in Milton Keynes and Alex Alderton, Roberto Amato, Jeffrey Barrett, Sonia Goncalves, Ewan Harrison, David K. Jackson, Ian Johnston, Dominic Kwiatkowski, Cordelia Langford, John Sillitoe on behalf of the Wellcome Sanger Institute COVID-19 Surveillance Team |
| EPI_ISL_2996638, EPI_ISL_2996639, EPI_ISL_2996640 | Lighthouse Laboratory Plymouth | Wellcome Sanger Institute for the COVID-19 Genomics UK (COG-UK) Consortium | Lighthouse Laboratory Plymouth and Alex Alderton, Roberto Amato, Jeffrey Barrett, Sonia Goncalves, Ewan Harrison, David K. Jackson, Ian Johnston, Dominic Kwiatkowski, Cordelia Langford, John Sillitoe on behalf of the Wellcome Sanger Institute COVID-19 Surveillance Team |
| EPI_ISL_2996641 | Lighthouse Lab in Milton Keynes | Wellcome Sanger Institute for the COVID-19 Genomics UK (COG-UK) Consortium | The Lighthouse Lab in Milton Keynes and Alex Alderton, Roberto Amato, Jeffrey Barrett, Sonia Goncalves, Ewan Harrison, David K. Jackson, Ian Johnston, Dominic Kwiatkowski, Cordelia Langford, John Sillitoe on behalf of the Wellcome Sanger Institute COVID-19 Surveillance Team |
| EPI_ISL_2996642, EPI_ISL_2996643, EPI_ISL_2996644, EPI_ISL_2996645, EPI_ISL_2996646, EPI_ISL_2996647, EPI_ISL_2996648 | Lighthouse Laboratory Plymouth | Wellcome Sanger Institute for the COVID-19 Genomics UK (COG-UK) Consortium | Lighthouse Laboratory Plymouth and Alex Alderton, Roberto Amato, Jeffrey Barrett, Sonia Goncalves, Ewan Harrison, David K. Jackson, Ian Johnston, Dominic Kwiatkowski, Cordelia Langford, John Sillitoe on behalf of the Wellcome Sanger Institute COVID-19 Surveillance Team |
| EPI_ISL_2996651, EPI_ISL_2996653, EPI_ISL_2996654, EPI_ISL_2996655, EPI_ISL_2996657, EPI_ISL_2996665 | Lighthouse Lab in Glasgow | Wellcome Sanger Institute for the COVID-19 Genomics UK (COG-UK) Consortium | Harper VanSteenhouse, Yumi Kasai, David Gray, Carol Clugston, Anna Dominiczak and Alex Alderton, Roberto Amato, Jeffrey Barrett, Sonia Goncalves, Ewan Harrison, David K. Jackson, Ian Johnston, Dominic Kwiatkowski, Cordelia Langford, John Sillitoe on behalf of the Wellcome Sanger Institute COVID-19 Surveillance Team |
| EPI_ISL_2996674 | Lighthouse Lab in Alderley Park | Wellcome Sanger Institute for the COVID-19 Genomics UK (COG-UK) Consortium | Jacquelyn Wynn, Mairead Hyland, The Lighthouse Lab in Alderley Park and Alex Alderton, Roberto Amato, Jeffrey Barrett, Sonia Goncalves, Ewan Harrison, David K. Jackson, Ian Johnston, Dominic Kwiatkowski, Cordelia Langford, John Sillitoe on behalf of the Wellcome Sanger Institute COVID-19 Surveillance Team |
| EPI_ISL_2996677, EPI_ISL_2996678 | Lighthouse Lab in Glasgow | Wellcome Sanger Institute for the COVID-19 Genomics UK (COG-UK) Consortium | Harper VanSteenhouse, Yumi Kasai, David Gray, Carol Clugston, Anna Dominiczak and Alex Alderton, Roberto Amato, Jeffrey Barrett, Sonia Goncalves, Ewan Harrison, David K. Jackson, Ian Johnston, Dominic Kwiatkowski, Cordelia Langford, John Sillitoe on behalf of the Wellcome Sanger Institute COVID-19 Surveillance Team |
| EPI_ISL_2996686 | Lighthouse Lab in Alderley Park | Wellcome Sanger Institute for the COVID-19 Genomics UK (COG-UK) Consortium | Jacquelyn Wynn, Mairead Hyland, The Lighthouse Lab in Alderley Park and Alex Alderton, Roberto Amato, Jeffrey Barrett, Sonia Goncalves, Ewan Harrison, David K. Jackson, Ian Johnston, Dominic Kwiatkowski, Cordelia Langford, John Sillitoe on behalf of the Wellcome Sanger Institute COVID-19 Surveillance Team |
| EPI_ISL_2996700, EPI_ISL_2996701, EPI_ISL_2996702, EPI_ISL_2996707, EPI_ISL_2996709 | Lighthouse Lab in Glasgow | Wellcome Sanger Institute for the COVID-19 Genomics UK (COG-UK) Consortium | Harper VanSteenhouse, Yumi Kasai, David Gray, Carol Clugston, Anna Dominiczak and Alex Alderton, Roberto Amato, Jeffrey Barrett, Sonia Goncalves, Ewan Harrison, David K. Jackson, Ian Johnston, Dominic Kwiatkowski, Cordelia Langford, John Sillitoe on behalf of the Wellcome Sanger Institute COVID-19 Surveillance Team |
| EPI_ISL_2996712 | Lighthouse Lab in Milton Keynes | Wellcome Sanger Institute for the COVID-19 Genomics UK (COG-UK) Consortium | The Lighthouse Lab in Milton Keynes and Alex Alderton, Roberto Amato, Jeffrey Barrett, Sonia Goncalves, Ewan Harrison, David K. Jackson, Ian Johnston, Dominic Kwiatkowski, Cordelia Langford, John Sillitoe on behalf of the Wellcome Sanger Institute COVID-19 Surveillance Team |
| EPI_ISL_2996715, EPI_ISL_2996716, EPI_ISL_2996720, EPI_ISL_2996721, EPI_ISL_2996723, EPI_ISL_2996732 | Lighthouse Lab in Glasgow | Wellcome Sanger Institute for the COVID-19 Genomics UK (COG-UK) Consortium | Harper VanSteenhouse, Yumi Kasai, David Gray, Carol Clugston, Anna Dominiczak and Alex Alderton, Roberto Amato, Jeffrey Barrett, Sonia Goncalves, Ewan Harrison, David K. Jackson, Ian Johnston, Dominic Kwiatkowski, Cordelia Langford, John Sillitoe on behalf of the Wellcome Sanger Institute COVID-19 Surveillance Team |
| EPI_ISL_2996733 | Lighthouse Lab in Milton Keynes | Wellcome Sanger Institute for the COVID-19 Genomics UK (COG-UK) Consortium | The Lighthouse Lab in Milton Keynes and Alex Alderton, Roberto Amato, Jeffrey Barrett, Sonia Goncalves, Ewan Harrison, David K. Jackson, Ian Johnston, Dominic Kwiatkowski, Cordelia Langford, John Sillitoe on behalf of the Wellcome Sanger Institute COVID-19 Surveillance Team |
| EPI_ISL_2996734 | Lighthouse Lab in Glasgow | Wellcome Sanger Institute for the COVID-19 Genomics UK (COG-UK) Consortium | Harper VanSteenhouse, Yumi Kasai, David Gray, Carol Clugston, Anna Dominiczak and Alex Alderton, Roberto Amato, Jeffrey Barrett, Sonia Goncalves, Ewan Harrison, David K. Jackson, Ian Johnston, Dominic Kwiatkowski, Cordelia Langford, John Sillitoe on behalf of the Wellcome Sanger Institute COVID-19 Surveillance Team |
| EPI_ISL_2996735 | Lighthouse Lab in Milton Keynes | Wellcome Sanger Institute for the COVID-19 Genomics UK (COG-UK) Consortium | The Lighthouse Lab in Milton Keynes and Alex Alderton, Roberto Amato, Jeffrey Barrett, Sonia Goncalves, Ewan Harrison, David K. Jackson, Ian Johnston, Dominic Kwiatkowski, Cordelia Langford, John Sillitoe on behalf of the Wellcome Sanger Institute COVID-19 Surveillance Team |
| EPI_ISL_2996736, EPI_ISL_2996737 | Berkshire and Surrey Pathology Services Lighthouse Laboratory | Wellcome Sanger Institute for the COVID-19 Genomics UK (COG-UK) Consortium | Berkshire and Surrey Pathology Services Lighthouse Laboratory and Alex Alderton, Roberto Amato, Jeffrey Barrett, Sonia Goncalves, Ewan Harrison, David K. Jackson, Ian Johnston, Dominic Kwiatkowski, Cordelia Langford, John Sillitoe on behalf of the Wellcome Sanger Institute COVID-19 Surveillance Team |
| EPI_ISL_2996738 | Lighthouse Lab in Milton Keynes | Wellcome Sanger Institute for the COVID-19 Genomics UK (COG-UK) Consortium | The Lighthouse Lab in Milton Keynes and Alex Alderton, Roberto Amato, Jeffrey Barrett, Sonia Goncalves, Ewan Harrison, David K. Jackson, Ian Johnston, Dominic Kwiatkowski, Cordelia Langford, John Sillitoe on behalf of the Wellcome Sanger Institute COVID-19 Surveillance Team |
| EPI_ISL_2996739 | Lighthouse Lab in Glasgow | Wellcome Sanger Institute for the COVID-19 Genomics UK (COG-UK) Consortium | Harper VanSteenhouse, Yumi Kasai, David Gray, Carol Clugston, Anna Dominiczak and Alex Alderton, Roberto Amato, Jeffrey Barrett, Sonia Goncalves, Ewan Harrison, David K. Jackson, Ian Johnston, Dominic Kwiatkowski, Cordelia Langford, John Sillitoe on behalf of the Wellcome Sanger Institute COVID-19 Surveillance Team |
| EPI_ISL_2996740 | Lighthouse Lab in Milton Keynes | Wellcome Sanger Institute for the COVID-19 Genomics UK (COG-UK) Consortium | The Lighthouse Lab in Milton Keynes and Alex Alderton, Roberto Amato, Jeffrey Barrett, Sonia Goncalves, Ewan Harrison, David K. Jackson, Ian Johnston, Dominic Kwiatkowski, Cordelia Langford, John Sillitoe on behalf of the Wellcome Sanger Institute COVID-19 Surveillance Team |
| EPI_ISL_2996741 | Lighthouse Laboratory Plymouth | Wellcome Sanger Institute for the COVID-19 Genomics UK (COG-UK) Consortium | Lighthouse Laboratory Plymouth and Alex Alderton, Roberto Amato, Jeffrey Barrett, Sonia Goncalves, Ewan Harrison, David K. Jackson, Ian Johnston, Dominic Kwiatkowski, Cordelia Langford, John Sillitoe on behalf of the Wellcome Sanger Institute COVID-19 Surveillance Team |
| EPI_ISL_2996742 | Lighthouse Lab in Milton Keynes | Wellcome Sanger Institute for the COVID-19 Genomics UK (COG-UK) Consortium | The Lighthouse Lab in Milton Keynes and Alex Alderton, Roberto Amato, Jeffrey Barrett, Sonia Goncalves, Ewan Harrison, David K. Jackson, Ian Johnston, Dominic Kwiatkowski, Cordelia Langford, John Sillitoe on behalf of the Wellcome Sanger Institute COVID-19 Surveillance Team |
| EPI_ISL_2996743 | Lighthouse Laboratory Plymouth | Wellcome Sanger Institute for the COVID-19 Genomics UK (COG-UK) Consortium | Lighthouse Laboratory Plymouth and Alex Alderton, Roberto Amato, Jeffrey Barrett, Sonia Goncalves, Ewan Harrison, David K. Jackson, Ian Johnston, Dominic Kwiatkowski, Cordelia Langford, John Sillitoe on behalf of the Wellcome Sanger Institute COVID-19 Surveillance Team |

| | | | |
|---|---|---|---|
| EPI_ISL_2996744 | Berkshire and Surrey Pathology Services Lighthouse Laboratory | Wellcome Sanger Institute for the COVID-19 Genomics UK (COG-UK) Consortium | Berkshire and Surrey Pathology Services Lighthouse Laboratory and Alex Alderton, Roberto Amato, Jeffrey Barrett, Sonia Goncalves, Ewan Harrison, David K. Jackson, Ian Johnston, Dominic Kwiatkowski, Cordelia Langford, John Sillitoe on behalf of the Wellcome Sanger Institute COVID-19 Surveillance Team |
| EPI_ISL_2996745 | Lighthouse Lab in Glasgow | Wellcome Sanger Institute for the COVID-19 Genomics UK (COG-UK) Consortium | Harper VanSteenhouse, Yumi Kasai, David Gray, Carol Clugston, Anna Dominiczak and Alex Alderton, Roberto Amato, Jeffrey Barrett, Sonia Goncalves, Ewan Harrison, David K. Jackson, Ian Johnston, Dominic Kwiatkowski, Cordelia Langford, John Sillitoe on behalf of the Wellcome Sanger Institute COVID-19 Surveillance Team |
| EPI_ISL_2996746 | Berkshire and Surrey Pathology Services Lighthouse Laboratory | Wellcome Sanger Institute for the COVID-19 Genomics UK (COG-UK) Consortium | Berkshire and Surrey Pathology Services Lighthouse Laboratory and Alex Alderton, Roberto Amato, Jeffrey Barrett, Sonia Goncalves, Ewan Harrison, David K. Jackson, Ian Johnston, Dominic Kwiatkowski, Cordelia Langford, John Sillitoe on behalf of the Wellcome Sanger Institute COVID-19 Surveillance Team |
| EPI_ISL_2996747 | Lighthouse Lab in Glasgow | Wellcome Sanger Institute for the COVID-19 Genomics UK (COG-UK) Consortium | Harper VanSteenhouse, Yumi Kasai, David Gray, Carol Clugston, Anna Dominiczak and Alex Alderton, Roberto Amato, Jeffrey Barrett, Sonia Goncalves, Ewan Harrison, David K. Jackson, Ian Johnston, Dominic Kwiatkowski, Cordelia Langford, John Sillitoe on behalf of the Wellcome Sanger Institute COVID-19 Surveillance Team |
| EPI_ISL_2996748 | Lighthouse Lab in Milton Keynes | Wellcome Sanger Institute for the COVID-19 Genomics UK (COG-UK) Consortium | The Lighthouse Lab in Milton Keynes and Alex Alderton, Roberto Amato, Jeffrey Barrett, Sonia Goncalves, Ewan Harrison, David K. Jackson, Ian Johnston, Dominic Kwiatkowski, Cordelia Langford, John Sillitoe on behalf of the Wellcome Sanger Institute COVID-19 Surveillance Team |
| EPI_ISL_2996749 | Berkshire and Surrey Pathology Services Lighthouse Laboratory | Wellcome Sanger Institute for the COVID-19 Genomics UK (COG-UK) Consortium | Berkshire and Surrey Pathology Services Lighthouse Laboratory and Alex Alderton, Roberto Amato, Jeffrey Barrett, Sonia Goncalves, Ewan Harrison, David K. Jackson, Ian Johnston, Dominic Kwiatkowski, Cordelia Langford, John Sillitoe on behalf of the Wellcome Sanger Institute COVID-19 Surveillance Team |
| EPI_ISL_2996750 | Lighthouse Laboratory Plymouth | Wellcome Sanger Institute for the COVID-19 Genomics UK (COG-UK) Consortium | Lighthouse Laboratory Plymouth and Alex Alderton, Roberto Amato, Jeffrey Barrett, Sonia Goncalves, Ewan Harrison, David K. Jackson, Ian Johnston, Dominic Kwiatkowski, Cordelia Langford, John Sillitoe on behalf of the Wellcome Sanger Institute COVID-19 Surveillance Team |
| EPI_ISL_2996751, EPI_ISL_2996752 | Lighthouse Lab in Glasgow | Wellcome Sanger Institute for the COVID-19 Genomics UK (COG-UK) Consortium | Harper VanSteenhouse, Yumi Kasai, David Gray, Carol Clugston, Anna Dominiczak and Alex Alderton, Roberto Amato, Jeffrey Barrett, Sonia Goncalves, Ewan Harrison, David K. Jackson, Ian Johnston, Dominic Kwiatkowski, Cordelia Langford, John Sillitoe on behalf of the Wellcome Sanger Institute COVID-19 Surveillance Team |
| EPI_ISL_2996753 | Berkshire and Surrey Pathology Services Lighthouse Laboratory | Wellcome Sanger Institute for the COVID-19 Genomics UK (COG-UK) Consortium | Berkshire and Surrey Pathology Services Lighthouse Laboratory and Alex Alderton, Roberto Amato, Jeffrey Barrett, Sonia Goncalves, Ewan Harrison, David K. Jackson, Ian Johnston, Dominic Kwiatkowski, Cordelia Langford, John Sillitoe on behalf of the Wellcome Sanger Institute COVID-19 Surveillance Team |
| EPI_ISL_2996754 | Lighthouse Laboratory Plymouth | Wellcome Sanger Institute for the COVID-19 Genomics UK (COG-UK) Consortium | Lighthouse Laboratory Plymouth and Alex Alderton, Roberto Amato, Jeffrey Barrett, Sonia Goncalves, Ewan Harrison, David K. Jackson, Ian Johnston, Dominic Kwiatkowski, Cordelia Langford, John Sillitoe on behalf of the Wellcome Sanger Institute COVID-19 Surveillance Team |
| EPI_ISL_2996755 | Lighthouse Lab in Milton Keynes | Wellcome Sanger Institute for the COVID-19 Genomics UK (COG-UK) Consortium | The Lighthouse Lab in Milton Keynes and Alex Alderton, Roberto Amato, Jeffrey Barrett, Sonia Goncalves, Ewan Harrison, David K. Jackson, Ian Johnston, Dominic Kwiatkowski, Cordelia Langford, John Sillitoe on behalf of the Wellcome Sanger Institute COVID-19 Surveillance Team |
| EPI_ISL_2996756 | Lighthouse Laboratory Plymouth | Wellcome Sanger Institute for the COVID-19 Genomics UK (COG-UK) Consortium | Lighthouse Laboratory Plymouth and Alex Alderton, Roberto Amato, Jeffrey Barrett, Sonia Goncalves, Ewan Harrison, David K. Jackson, Ian Johnston, Dominic Kwiatkowski, Cordelia Langford, John Sillitoe on behalf of the Wellcome Sanger Institute COVID-19 Surveillance Team |
| EPI_ISL_2996757 | Lighthouse Lab in Milton Keynes | Wellcome Sanger Institute for the COVID-19 Genomics UK (COG-UK) Consortium | The Lighthouse Lab in Milton Keynes and Alex Alderton, Roberto Amato, Jeffrey Barrett, Sonia Goncalves, Ewan Harrison, David K. Jackson, Ian Johnston, Dominic Kwiatkowski, Cordelia Langford, John Sillitoe on behalf of the Wellcome Sanger Institute COVID-19 Surveillance Team |
| EPI_ISL_2996758 | Lighthouse Laboratory Plymouth | Wellcome Sanger Institute for the COVID-19 Genomics UK (COG-UK) Consortium | Lighthouse Laboratory Plymouth and Alex Alderton, Roberto Amato, Jeffrey Barrett, Sonia Goncalves, Ewan Harrison, David K. Jackson, Ian Johnston, Dominic Kwiatkowski, Cordelia Langford, John Sillitoe on behalf of the Wellcome Sanger Institute COVID-19 Surveillance Team |
| EPI_ISL_2996759 | Lighthouse Lab in Glasgow | Wellcome Sanger Institute for the COVID-19 Genomics UK (COG-UK) Consortium | Harper VanSteenhouse, Yumi Kasai, David Gray, Carol Clugston, Anna Dominiczak and Alex Alderton, Roberto Amato, Jeffrey Barrett, Sonia Goncalves, Ewan Harrison, David K. Jackson, Ian Johnston, Dominic Kwiatkowski, Cordelia Langford, John Sillitoe on behalf of the Wellcome Sanger Institute COVID-19 Surveillance Team |
| EPI_ISL_2996760 | Lighthouse Laboratory Plymouth | Wellcome Sanger Institute for the COVID-19 Genomics UK (COG-UK) Consortium | Lighthouse Laboratory Plymouth and Alex Alderton, Roberto Amato, Jeffrey Barrett, Sonia Goncalves, Ewan Harrison, David K. Jackson, Ian Johnston, Dominic Kwiatkowski, Cordelia Langford, John Sillitoe on behalf of the Wellcome Sanger Institute COVID-19 Surveillance Team |
| EPI_ISL_2996761 | Berkshire and Surrey Pathology Services Lighthouse Laboratory | Wellcome Sanger Institute for the COVID-19 Genomics UK (COG-UK) Consortium | Berkshire and Surrey Pathology Services Lighthouse Laboratory and Alex Alderton, Roberto Amato, Jeffrey Barrett, Sonia Goncalves, Ewan Harrison, David K. Jackson, Ian Johnston, Dominic Kwiatkowski, Cordelia Langford, John Sillitoe on behalf of the Wellcome Sanger Institute COVID-19 Surveillance Team |
| EPI_ISL_2996762 | Lighthouse Lab in Glasgow | Wellcome Sanger Institute for the COVID-19 Genomics UK (COG-UK) Consortium | Harper VanSteenhouse, Yumi Kasai, David Gray, Carol Clugston, Anna Dominiczak and Alex Alderton, Roberto Amato, Jeffrey Barrett, Sonia Goncalves, Ewan Harrison, David K. Jackson, Ian Johnston, Dominic Kwiatkowski, Cordelia Langford, John Sillitoe on behalf of the Wellcome Sanger Institute COVID-19 Surveillance Team |
| EPI_ISL_2996763 | Berkshire and Surrey Pathology Services Lighthouse Laboratory | Wellcome Sanger Institute for the COVID-19 Genomics UK (COG-UK) Consortium | Berkshire and Surrey Pathology Services Lighthouse Laboratory and Alex Alderton, Roberto Amato, Jeffrey Barrett, Sonia Goncalves, Ewan Harrison, David K. Jackson, Ian Johnston, Dominic Kwiatkowski, Cordelia Langford, John Sillitoe on behalf of the Wellcome Sanger Institute COVID-19 Surveillance Team |
| EPI_ISL_2996764 | Lighthouse Lab in Glasgow | Wellcome Sanger Institute for the COVID-19 Genomics UK (COG-UK) Consortium | Harper VanSteenhouse, Yumi Kasai, David Gray, Carol Clugston, Anna Dominiczak and Alex Alderton, Roberto Amato, Jeffrey Barrett, Sonia Goncalves, Ewan Harrison, David K. Jackson, Ian Johnston, Dominic Kwiatkowski, Cordelia Langford, John Sillitoe on behalf of the Wellcome Sanger Institute COVID-19 Surveillance Team |
| EPI_ISL_2996765 | Lighthouse Laboratory Plymouth | Wellcome Sanger Institute for the COVID-19 Genomics UK (COG-UK) Consortium | Lighthouse Laboratory Plymouth and Alex Alderton, Roberto Amato, Jeffrey Barrett, Sonia Goncalves, Ewan Harrison, David K. Jackson, Ian Johnston, Dominic Kwiatkowski, Cordelia Langford, John Sillitoe on behalf of the Wellcome Sanger Institute COVID-19 Surveillance Team |
| EPI_ISL_2996766 | Berkshire and Surrey Pathology Services Lighthouse Laboratory | Wellcome Sanger Institute for the COVID-19 Genomics UK (COG-UK) Consortium | Berkshire and Surrey Pathology Services Lighthouse Laboratory and Alex Alderton, Roberto Amato, Jeffrey Barrett, Sonia Goncalves, Ewan Harrison, David K. Jackson, Ian Johnston, Dominic Kwiatkowski, Cordelia Langford, John Sillitoe on behalf of the Wellcome Sanger Institute COVID-19 Surveillance Team |
| EPI_ISL_2996767, EPI_ISL_2996768 | Lighthouse Laboratory Plymouth | Wellcome Sanger Institute for the COVID-19 Genomics UK (COG-UK) Consortium | Lighthouse Laboratory Plymouth and Alex Alderton, Roberto Amato, Jeffrey Barrett, Sonia Goncalves, Ewan Harrison, David K. Jackson, Ian Johnston, Dominic Kwiatkowski, Cordelia Langford, John Sillitoe on behalf of the Wellcome Sanger Institute COVID-19 Surveillance Team |
| EPI_ISL_2996769, EPI_ISL_2996770 | Lighthouse Lab in Glasgow | Wellcome Sanger Institute for the COVID-19 Genomics UK (COG-UK) Consortium | Harper VanSteenhouse, Yumi Kasai, David Gray, Carol Clugston, Anna Dominiczak and Alex Alderton, Roberto Amato, Jeffrey Barrett, Sonia Goncalves, Ewan Harrison, David K. Jackson, Ian Johnston, Dominic Kwiatkowski, Cordelia Langford, John Sillitoe on behalf of the Wellcome Sanger Institute COVID-19 Surveillance Team |
| EPI_ISL_2996771, EPI_ISL_2996772 | Berkshire and Surrey Pathology Services Lighthouse Laboratory | Wellcome Sanger Institute for the COVID-19 Genomics UK (COG-UK) Consortium | Berkshire and Surrey Pathology Services Lighthouse Laboratory and Alex Alderton, Roberto Amato, Jeffrey Barrett, Sonia Goncalves, Ewan Harrison, David K. Jackson, Ian Johnston, Dominic Kwiatkowski, Cordelia Langford, John Sillitoe on behalf of the Wellcome Sanger Institute COVID-19 Surveillance Team |
| EPI_ISL_2996773 | Lighthouse Lab in Milton Keynes | Wellcome Sanger Institute for the COVID-19 Genomics UK (COG-UK) Consortium | The Lighthouse Lab in Milton Keynes and Alex Alderton, Roberto Amato, Jeffrey Barrett, Sonia Goncalves, Ewan Harrison, David K. Jackson, Ian Johnston, Dominic Kwiatkowski, Cordelia Langford, John Sillitoe on behalf of the Wellcome Sanger Institute COVID-19 Surveillance Team |
| EPI_ISL_2996774, EPI_ISL_2996775, EPI_ISL_2996776 | Berkshire and Surrey Pathology Services Lighthouse Laboratory | Wellcome Sanger Institute for the COVID-19 Genomics UK (COG-UK) Consortium | Berkshire and Surrey Pathology Services Lighthouse Laboratory and Alex Alderton, Roberto Amato, Jeffrey Barrett, Sonia Goncalves, Ewan Harrison, David K. Jackson, Ian Johnston, Dominic Kwiatkowski, Cordelia Langford, John Sillitoe on behalf of the Wellcome Sanger Institute COVID-19 Surveillance Team |
| EPI_ISL_2996777 | Lighthouse Laboratory Plymouth | Wellcome Sanger Institute for the COVID-19 Genomics UK (COG-UK) Consortium | Lighthouse Laboratory Plymouth and Alex Alderton, Roberto Amato, Jeffrey Barrett, Sonia Goncalves, Ewan Harrison, David K. Jackson, Ian Johnston, Dominic Kwiatkowski, Cordelia Langford, John Sillitoe on behalf of the Wellcome Sanger Institute COVID-19 Surveillance Team |
| EPI_ISL_2996778 | Lighthouse Lab in Milton Keynes | Wellcome Sanger Institute for the COVID-19 Genomics UK (COG-UK) Consortium | The Lighthouse Lab in Milton Keynes and Alex Alderton, Roberto Amato, Jeffrey Barrett, Sonia Goncalves, Ewan Harrison, David K. Jackson, Ian Johnston, Dominic Kwiatkowski, Cordelia Langford, John Sillitoe on behalf of the Wellcome Sanger Institute COVID-19 Surveillance Team |
| EPI_ISL_2996779 | Berkshire and Surrey Pathology Services Lighthouse Laboratory | Wellcome Sanger Institute for the COVID-19 Genomics UK (COG-UK) Consortium | Berkshire and Surrey Pathology Services Lighthouse Laboratory and Alex Alderton, Roberto Amato, Jeffrey Barrett, Sonia Goncalves, Ewan Harrison, David K. Jackson, Ian Johnston, Dominic Kwiatkowski, Cordelia Langford, John Sillitoe on behalf of the Wellcome Sanger Institute COVID-19 Surveillance Team |
| EPI_ISL_2996780 | Lighthouse Lab in Glasgow | Wellcome Sanger Institute for the COVID-19 Genomics UK (COG-UK) Consortium | Harper VanSteenhouse, Yumi Kasai, David Gray, Carol Clugston, Anna Dominiczak and Alex Alderton, Roberto Amato, Jeffrey Barrett, Sonia Goncalves, Ewan Harrison, David K. Jackson, Ian Johnston, Dominic Kwiatkowski, Cordelia Langford, John Sillitoe on behalf of the Wellcome Sanger Institute COVID-19 Surveillance Team |
| EPI_ISL_2996781 | Lighthouse Lab in Milton Keynes | Wellcome Sanger Institute for the COVID-19 Genomics UK (COG-UK) Consortium | The Lighthouse Lab in Milton Keynes and Alex Alderton, Roberto Amato, Jeffrey Barrett, Sonia Goncalves, Ewan Harrison, David K. Jackson, Ian Johnston, Dominic Kwiatkowski, Cordelia Langford, John Sillitoe on behalf of the Wellcome Sanger Institute COVID-19 Surveillance Team |
| EPI_ISL_2996782 | Lighthouse Laboratory Plymouth | Wellcome Sanger Institute for the COVID-19 Genomics UK (COG-UK) Consortium | Lighthouse Laboratory Plymouth and Alex Alderton, Roberto Amato, Jeffrey Barrett, Sonia Goncalves, Ewan Harrison, David K. Jackson, Ian Johnston, Dominic Kwiatkowski, Cordelia Langford, John Sillitoe on behalf of the Wellcome Sanger Institute COVID-19 Surveillance Team |
| EPI_ISL_2996783 | Lighthouse Lab in Milton Keynes | Wellcome Sanger Institute for the COVID-19 Genomics UK (COG-UK) Consortium | The Lighthouse Lab in Milton Keynes and Alex Alderton, Roberto Amato, Jeffrey Barrett, Sonia Goncalves, Ewan Harrison, David K. Jackson, Ian Johnston, Dominic Kwiatkowski, Cordelia Langford, John Sillitoe on behalf of the Wellcome Sanger Institute COVID-19 Surveillance Team |
| EPI_ISL_2996784 | Lighthouse Laboratory Plymouth | Wellcome Sanger Institute for the COVID-19 Genomics UK (COG-UK) Consortium | Lighthouse Laboratory Plymouth and Alex Alderton, Roberto Amato, Jeffrey Barrett, Sonia Goncalves, Ewan Harrison, David K. Jackson, Ian Johnston, Dominic Kwiatkowski, Cordelia Langford, John Sillitoe on behalf of the Wellcome Sanger Institute COVID-19 Surveillance Team |
| EPI_ISL_2996785 | Lighthouse Lab in Glasgow | Wellcome Sanger Institute for the COVID-19 Genomics UK (COG-UK) Consortium | Harper VanSteenhouse, Yumi Kasai, David Gray, Carol Clugston, Anna Dominiczak and Alex Alderton, Roberto Amato, Jeffrey Barrett, Sonia Goncalves, Ewan Harrison, David K. Jackson, Ian Johnston, Dominic Kwiatkowski, Cordelia Langford, John Sillitoe on behalf of the Wellcome Sanger Institute COVID-19 Surveillance Team |
| EPI_ISL_2996786 | Lighthouse Laboratory Plymouth | Wellcome Sanger Institute for the COVID-19 Genomics UK (COG-UK) Consortium | Lighthouse Laboratory Plymouth and Alex Alderton, Roberto Amato, Jeffrey Barrett, Sonia Goncalves, Ewan Harrison, David K. Jackson, Ian Johnston, Dominic Kwiatkowski, Cordelia Langford, John Sillitoe on behalf of the Wellcome Sanger Institute COVID-19 Surveillance Team |
| EPI_ISL_2996787 | Lighthouse Lab in Milton Keynes | Wellcome Sanger Institute for the COVID-19 Genomics UK (COG-UK) Consortium | The Lighthouse Lab in Milton Keynes and Alex Alderton, Roberto Amato, Jeffrey Barrett, Sonia Goncalves, Ewan Harrison, David K. Jackson, Ian Johnston, Dominic Kwiatkowski, Cordelia Langford, John Sillitoe on behalf of the Wellcome Sanger Institute COVID-19 Surveillance Team |
| EPI_ISL_2996788 | Lighthouse Lab in Glasgow | Wellcome Sanger Institute for the COVID-19 Genomics UK (COG-UK) Consortium | Harper VanSteenhouse, Yumi Kasai, David Gray, Carol Clugston, Anna Dominiczak and Alex Alderton, Roberto Amato, Jeffrey Barrett, Sonia Goncalves, Ewan Harrison, David K. Jackson, Ian Johnston, Dominic Kwiatkowski, Cordelia Langford, John Sillitoe on behalf of the Wellcome Sanger Institute COVID-19 Surveillance Team |
| EPI_ISL_2996789 | Berkshire and Surrey Pathology Services Lighthouse Laboratory | Wellcome Sanger Institute for the COVID-19 Genomics UK (COG-UK) Consortium | Berkshire and Surrey Pathology Services Lighthouse Laboratory and Alex Alderton, Roberto Amato, Jeffrey Barrett, Sonia Goncalves, Ewan Harrison, David K. Jackson, Ian Johnston, Dominic Kwiatkowski, Cordelia Langford, John Sillitoe on behalf of the Wellcome Sanger Institute COVID-19 Surveillance Team |
| EPI_ISL_2996790, EPI_ISL_2996791, EPI_ISL_2996792 | Lighthouse Lab in Milton Keynes | Wellcome Sanger Institute for the COVID-19 Genomics UK (COG-UK) Consortium | The Lighthouse Lab in Milton Keynes and Alex Alderton, Roberto Amato, Jeffrey Barrett, Sonia Goncalves, Ewan Harrison, David K. Jackson, Ian Johnston, Dominic Kwiatkowski, Cordelia Langford, John Sillitoe on behalf of the Wellcome Sanger Institute COVID-19 Surveillance Team |
| EPI_ISL_2996793, EPI_ISL_2996794 | Lighthouse Laboratory Plymouth | Wellcome Sanger Institute for the COVID-19 Genomics UK (COG-UK) Consortium | Lighthouse Laboratory Plymouth and Alex Alderton, Roberto Amato, Jeffrey Barrett, Sonia Goncalves, Ewan Harrison, David K. Jackson, Ian Johnston, Dominic Kwiatkowski, Cordelia Langford, John Sillitoe on behalf of the Wellcome Sanger Institute COVID-19 Surveillance Team |
| EPI_ISL_2996795 | Lighthouse Lab in Glasgow | Wellcome Sanger Institute for the COVID-19 Genomics UK (COG-UK) Consortium | Harper VanSteenhouse, Yumi Kasai, David Gray, Carol Clugston, Anna Dominiczak and Alex Alderton, Roberto Amato, Jeffrey Barrett, Sonia Goncalves, Ewan Harrison, David K. Jackson, Ian Johnston, Dominic Kwiatkowski, Cordelia Langford, John Sillitoe on behalf of the Wellcome Sanger Institute COVID-19 Surveillance Team |
| EPI_ISL_2996796 | Berkshire and Surrey Pathology Services Lighthouse Laboratory | Wellcome Sanger Institute for the COVID-19 Genomics UK (COG-UK) Consortium | Berkshire and Surrey Pathology Services Lighthouse Laboratory and Alex Alderton, Roberto Amato, Jeffrey Barrett, Sonia Goncalves, Ewan Harrison, David K. Jackson, Ian Johnston, Dominic Kwiatkowski, Cordelia Langford, John Sillitoe on behalf of the Wellcome Sanger Institute COVID-19 Surveillance Team |

| | | | |
|---|---|---|---|
| EPI_ISL_2996797, EPI_ISL_2996798 | Lighthouse Lab in Glasgow | Wellcome Sanger Institute for the COVID-19 Genomics UK (COG-UK) Consortium | Harper VanSteenhouse, Yumi Kasai, David Gray, Carol Clugston, Anna Dominiczak and Alex Alderton, Roberto Amato, Jeffrey Barrett, Sonia Goncalves, Ewan Harrison, David K. Jackson, Ian Johnston, Dominic Kwiatkowski, Cordelia Langford, John Sillitoe on behalf of the Wellcome Sanger Institute COVID-19 Surveillance Team |
| EPI_ISL_2996799, EPI_ISL_2996800 | Lighthouse Laboratory Plymouth | Wellcome Sanger Institute for the COVID-19 Genomics UK (COG-UK) Consortium | Lighthouse Laboratory Plymouth and Alex Alderton, Roberto Amato, Jeffrey Barrett, Sonia Goncalves, Ewan Harrison, David K. Jackson, Ian Johnston, Dominic Kwiatkowski, Cordelia Langford, John Sillitoe on behalf of the Wellcome Sanger Institute COVID-19 Surveillance Team |
| EPI_ISL_2996801 | Lighthouse Lab in Glasgow | Wellcome Sanger Institute for the COVID-19 Genomics UK (COG-UK) Consortium | Harper VanSteenhouse, Yumi Kasai, David Gray, Carol Clugston, Anna Dominiczak and Alex Alderton, Roberto Amato, Jeffrey Barrett, Sonia Goncalves, Ewan Harrison, David K. Jackson, Ian Johnston, Dominic Kwiatkowski, Cordelia Langford, John Sillitoe on behalf of the Wellcome Sanger Institute COVID-19 Surveillance Team |
| EPI_ISL_2996802 | Lighthouse Lab in Milton Keynes | Wellcome Sanger Institute for the COVID-19 Genomics UK (COG-UK) Consortium | The Lighthouse Lab in Milton Keynes and Alex Alderton, Roberto Amato, Jeffrey Barrett, Sonia Goncalves, Ewan Harrison, David K. Jackson, Ian Johnston, Dominic Kwiatkowski, Cordelia Langford, John Sillitoe on behalf of the Wellcome Sanger Institute COVID-19 Surveillance Team |
| EPI_ISL_2996803, EPI_ISL_2996804 | Berkshire and Surrey Pathology Services Lighthouse Laboratory | Wellcome Sanger Institute for the COVID-19 Genomics UK (COG-UK) Consortium | Berkshire and Surrey Pathology Services Lighthouse Laboratory and Alex Alderton, Roberto Amato, Jeffrey Barrett, Sonia Goncalves, Ewan Harrison, David K. Jackson, Ian Johnston, Dominic Kwiatkowski, Cordelia Langford, John Sillitoe on behalf of the Wellcome Sanger Institute COVID-19 Surveillance Team |
| EPI_ISL_2996805 | Lighthouse Lab in Glasgow | Wellcome Sanger Institute for the COVID-19 Genomics UK (COG-UK) Consortium | Harper VanSteenhouse, Yumi Kasai, David Gray, Carol Clugston, Anna Dominiczak and Alex Alderton, Roberto Amato, Jeffrey Barrett, Sonia Goncalves, Ewan Harrison, David K. Jackson, Ian Johnston, Dominic Kwiatkowski, Cordelia Langford, John Sillitoe on behalf of the Wellcome Sanger Institute COVID-19 Surveillance Team |
| EPI_ISL_2996806 | Lighthouse Lab in Milton Keynes | Wellcome Sanger Institute for the COVID-19 Genomics UK (COG-UK) Consortium | The Lighthouse Lab in Milton Keynes and Alex Alderton, Roberto Amato, Jeffrey Barrett, Sonia Goncalves, Ewan Harrison, David K. Jackson, Ian Johnston, Dominic Kwiatkowski, Cordelia Langford, John Sillitoe on behalf of the Wellcome Sanger Institute COVID-19 Surveillance Team |
| EPI_ISL_2996807, EPI_ISL_2996808 | Lighthouse Laboratory Plymouth | Wellcome Sanger Institute for the COVID-19 Genomics UK (COG-UK) Consortium | Lighthouse Laboratory Plymouth and Alex Alderton, Roberto Amato, Jeffrey Barrett, Sonia Goncalves, Ewan Harrison, David K. Jackson, Ian Johnston, Dominic Kwiatkowski, Cordelia Langford, John Sillitoe on behalf of the Wellcome Sanger Institute COVID-19 Surveillance Team |
| EPI_ISL_2996809 | Lighthouse Lab in Milton Keynes | Wellcome Sanger Institute for the COVID-19 Genomics UK (COG-UK) Consortium | The Lighthouse Lab in Milton Keynes and Alex Alderton, Roberto Amato, Jeffrey Barrett, Sonia Goncalves, Ewan Harrison, David K. Jackson, Ian Johnston, Dominic Kwiatkowski, Cordelia Langford, John Sillitoe on behalf of the Wellcome Sanger Institute COVID-19 Surveillance Team |
| EPI_ISL_2996810 | Lighthouse Laboratory Plymouth | Wellcome Sanger Institute for the COVID-19 Genomics UK (COG-UK) Consortium | Lighthouse Laboratory Plymouth and Alex Alderton, Roberto Amato, Jeffrey Barrett, Sonia Goncalves, Ewan Harrison, David K. Jackson, Ian Johnston, Dominic Kwiatkowski, Cordelia Langford, John Sillitoe on behalf of the Wellcome Sanger Institute COVID-19 Surveillance Team |
| EPI_ISL_2996811 | Lighthouse Lab in Milton Keynes | Wellcome Sanger Institute for the COVID-19 Genomics UK (COG-UK) Consortium | The Lighthouse Lab in Milton Keynes and Alex Alderton, Roberto Amato, Jeffrey Barrett, Sonia Goncalves, Ewan Harrison, David K. Jackson, Ian Johnston, Dominic Kwiatkowski, Cordelia Langford, John Sillitoe on behalf of the Wellcome Sanger Institute COVID-19 Surveillance Team |
| EPI_ISL_2996812 | Lighthouse Lab in Glasgow | Wellcome Sanger Institute for the COVID-19 Genomics UK (COG-UK) Consortium | Harper VanSteenhouse, Yumi Kasai, David Gray, Carol Clugston, Anna Dominiczak and Alex Alderton, Roberto Amato, Jeffrey Barrett, Sonia Goncalves, Ewan Harrison, David K. Jackson, Ian Johnston, Dominic Kwiatkowski, Cordelia Langford, John Sillitoe on behalf of the Wellcome Sanger Institute COVID-19 Surveillance Team |
| EPI_ISL_2996813, EPI_ISL_2996814, EPI_ISL_2996815 | Lighthouse Laboratory Plymouth | Wellcome Sanger Institute for the COVID-19 Genomics UK (COG-UK) Consortium | Lighthouse Laboratory Plymouth and Alex Alderton, Roberto Amato, Jeffrey Barrett, Sonia Goncalves, Ewan Harrison, David K. Jackson, Ian Johnston, Dominic Kwiatkowski, Cordelia Langford, John Sillitoe on behalf of the Wellcome Sanger Institute COVID-19 Surveillance Team |
| EPI_ISL_2996816 | Lighthouse Lab in Milton Keynes | Wellcome Sanger Institute for the COVID-19 Genomics UK (COG-UK) Consortium | The Lighthouse Lab in Milton Keynes and Alex Alderton, Roberto Amato, Jeffrey Barrett, Sonia Goncalves, Ewan Harrison, David K. Jackson, Ian Johnston, Dominic Kwiatkowski, Cordelia Langford, John Sillitoe on behalf of the Wellcome Sanger Institute COVID-19 Surveillance Team |
| EPI_ISL_2996817 | Berkshire and Surrey Pathology Services Lighthouse Laboratory | Wellcome Sanger Institute for the COVID-19 Genomics UK (COG-UK) Consortium | Berkshire and Surrey Pathology Services Lighthouse Laboratory and Alex Alderton, Roberto Amato, Jeffrey Barrett, Sonia Goncalves, Ewan Harrison, David K. Jackson, Ian Johnston, Dominic Kwiatkowski, Cordelia Langford, John Sillitoe on behalf of the Wellcome Sanger Institute COVID-19 Surveillance Team |
| EPI_ISL_2996818, EPI_ISL_2996819 | Lighthouse Laboratory Plymouth | Wellcome Sanger Institute for the COVID-19 Genomics UK (COG-UK) Consortium | Lighthouse Laboratory Plymouth and Alex Alderton, Roberto Amato, Jeffrey Barrett, Sonia Goncalves, Ewan Harrison, David K. Jackson, Ian Johnston, Dominic Kwiatkowski, Cordelia Langford, John Sillitoe on behalf of the Wellcome Sanger Institute COVID-19 Surveillance Team |
| EPI_ISL_2996820 | Lighthouse Lab in Milton Keynes | Wellcome Sanger Institute for the COVID-19 Genomics UK (COG-UK) Consortium | The Lighthouse Lab in Milton Keynes and Alex Alderton, Roberto Amato, Jeffrey Barrett, Sonia Goncalves, Ewan Harrison, David K. Jackson, Ian Johnston, Dominic Kwiatkowski, Cordelia Langford, John Sillitoe on behalf of the Wellcome Sanger Institute COVID-19 Surveillance Team |
| EPI_ISL_2996821 | Lighthouse Laboratory Plymouth | Wellcome Sanger Institute for the COVID-19 Genomics UK (COG-UK) Consortium | Lighthouse Laboratory Plymouth and Alex Alderton, Roberto Amato, Jeffrey Barrett, Sonia Goncalves, Ewan Harrison, David K. Jackson, Ian Johnston, Dominic Kwiatkowski, Cordelia Langford, John Sillitoe on behalf of the Wellcome Sanger Institute COVID-19 Surveillance Team |
| EPI_ISL_2996822, EPI_ISL_2996823 | Lighthouse Lab in Glasgow | Wellcome Sanger Institute for the COVID-19 Genomics UK (COG-UK) Consortium | Harper VanSteenhouse, Yumi Kasai, David Gray, Carol Clugston, Anna Dominiczak and Alex Alderton, Roberto Amato, Jeffrey Barrett, Sonia Goncalves, Ewan Harrison, David K. Jackson, Ian Johnston, Dominic Kwiatkowski, Cordelia Langford, John Sillitoe on behalf of the Wellcome Sanger Institute COVID-19 Surveillance Team |
| EPI_ISL_2996824 | Lighthouse Lab in Milton Keynes | Wellcome Sanger Institute for the COVID-19 Genomics UK (COG-UK) Consortium | The Lighthouse Lab in Milton Keynes and Alex Alderton, Roberto Amato, Jeffrey Barrett, Sonia Goncalves, Ewan Harrison, David K. Jackson, Ian Johnston, Dominic Kwiatkowski, Cordelia Langford, John Sillitoe on behalf of the Wellcome Sanger Institute COVID-19 Surveillance Team |
| EPI_ISL_2996825 | Berkshire and Surrey Pathology Services Lighthouse Laboratory | Wellcome Sanger Institute for the COVID-19 Genomics UK (COG-UK) Consortium | Berkshire and Surrey Pathology Services Lighthouse Laboratory and Alex Alderton, Roberto Amato, Jeffrey Barrett, Sonia Goncalves, Ewan Harrison, David K. Jackson, Ian Johnston, Dominic Kwiatkowski, Cordelia Langford, John Sillitoe on behalf of the Wellcome Sanger Institute COVID-19 Surveillance Team |
| EPI_ISL_2996826 | Lighthouse Lab in Glasgow | Wellcome Sanger Institute for the COVID-19 Genomics UK (COG-UK) Consortium | Harper VanSteenhouse, Yumi Kasai, David Gray, Carol Clugston, Anna Dominiczak and Alex Alderton, Roberto Amato, Jeffrey Barrett, Sonia Goncalves, Ewan Harrison, David K. Jackson, Ian Johnston, Dominic Kwiatkowski, Cordelia Langford, John Sillitoe on behalf of the Wellcome Sanger Institute COVID-19 Surveillance Team |
| EPI_ISL_2996827, EPI_ISL_2996828, EPI_ISL_2996829 | Lighthouse Laboratory Plymouth | Wellcome Sanger Institute for the COVID-19 Genomics UK (COG-UK) Consortium | Lighthouse Laboratory Plymouth and Alex Alderton, Roberto Amato, Jeffrey Barrett, Sonia Goncalves, Ewan Harrison, David K. Jackson, Ian Johnston, Dominic Kwiatkowski, Cordelia Langford, John Sillitoe on behalf of the Wellcome Sanger Institute COVID-19 Surveillance Team |
| EPI_ISL_2996830 | Berkshire and Surrey Pathology Services Lighthouse Laboratory | Wellcome Sanger Institute for the COVID-19 Genomics UK (COG-UK) Consortium | Berkshire and Surrey Pathology Services Lighthouse Laboratory and Alex Alderton, Roberto Amato, Jeffrey Barrett, Sonia Goncalves, Ewan Harrison, David K. Jackson, Ian Johnston, Dominic Kwiatkowski, Cordelia Langford, John Sillitoe on behalf of the Wellcome Sanger Institute COVID-19 Surveillance Team |
| EPI_ISL_2996831, EPI_ISL_2996832, EPI_ISL_2996833 | Lighthouse Lab in Glasgow | Wellcome Sanger Institute for the COVID-19 Genomics UK (COG-UK) Consortium | Harper VanSteenhouse, Yumi Kasai, David Gray, Carol Clugston, Anna Dominiczak and Alex Alderton, Roberto Amato, Jeffrey Barrett, Sonia Goncalves, Ewan Harrison, David K. Jackson, Ian Johnston, Dominic Kwiatkowski, Cordelia Langford, John Sillitoe on behalf of the Wellcome Sanger Institute COVID-19 Surveillance Team |
| EPI_ISL_2996834 | Lighthouse Lab in Milton Keynes | Wellcome Sanger Institute for the COVID-19 Genomics UK (COG-UK) Consortium | The Lighthouse Lab in Milton Keynes and Alex Alderton, Roberto Amato, Jeffrey Barrett, Sonia Goncalves, Ewan Harrison, David K. Jackson, Ian Johnston, Dominic Kwiatkowski, Cordelia Langford, John Sillitoe on behalf of the Wellcome Sanger Institute COVID-19 Surveillance Team |
| EPI_ISL_2996835, EPI_ISL_2996836 | Lighthouse Laboratory Plymouth | Wellcome Sanger Institute for the COVID-19 Genomics UK (COG-UK) Consortium | Lighthouse Laboratory Plymouth and Alex Alderton, Roberto Amato, Jeffrey Barrett, Sonia Goncalves, Ewan Harrison, David K. Jackson, Ian Johnston, Dominic Kwiatkowski, Cordelia Langford, John Sillitoe on behalf of the Wellcome Sanger Institute COVID-19 Surveillance Team |
| EPI_ISL_2996837 | Lighthouse Lab in Milton Keynes | Wellcome Sanger Institute for the COVID-19 Genomics UK (COG-UK) Consortium | The Lighthouse Lab in Milton Keynes and Alex Alderton, Roberto Amato, Jeffrey Barrett, Sonia Goncalves, Ewan Harrison, David K. Jackson, Ian Johnston, Dominic Kwiatkowski, Cordelia Langford, John Sillitoe on behalf of the Wellcome Sanger Institute COVID-19 Surveillance Team |
| EPI_ISL_2996838, EPI_ISL_2996839 | Lighthouse Lab in Glasgow | Wellcome Sanger Institute for the COVID-19 Genomics UK (COG-UK) Consortium | Harper VanSteenhouse, Yumi Kasai, David Gray, Carol Clugston, Anna Dominiczak and Alex Alderton, Roberto Amato, Jeffrey Barrett, Sonia Goncalves, Ewan Harrison, David K. Jackson, Ian Johnston, Dominic Kwiatkowski, Cordelia Langford, John Sillitoe on behalf of the Wellcome Sanger Institute COVID-19 Surveillance Team |
| EPI_ISL_2996840 | Lighthouse Lab in Milton Keynes | Wellcome Sanger Institute for the COVID-19 Genomics UK (COG-UK) Consortium | The Lighthouse Lab in Milton Keynes and Alex Alderton, Roberto Amato, Jeffrey Barrett, Sonia Goncalves, Ewan Harrison, David K. Jackson, Ian Johnston, Dominic Kwiatkowski, Cordelia Langford, John Sillitoe on behalf of the Wellcome Sanger Institute COVID-19 Surveillance Team |
| EPI_ISL_2996841, EPI_ISL_2996842 | Berkshire and Surrey Pathology Services Lighthouse Laboratory | Wellcome Sanger Institute for the COVID-19 Genomics UK (COG-UK) Consortium | Berkshire and Surrey Pathology Services Lighthouse Laboratory and Alex Alderton, Roberto Amato, Jeffrey Barrett, Sonia Goncalves, Ewan Harrison, David K. Jackson, Ian Johnston, Dominic Kwiatkowski, Cordelia Langford, John Sillitoe on behalf of the Wellcome Sanger Institute COVID-19 Surveillance Team |
| EPI_ISL_2996843 | Lighthouse Lab in Milton Keynes | Wellcome Sanger Institute for the COVID-19 Genomics UK (COG-UK) Consortium | The Lighthouse Lab in Milton Keynes and Alex Alderton, Roberto Amato, Jeffrey Barrett, Sonia Goncalves, Ewan Harrison, David K. Jackson, Ian Johnston, Dominic Kwiatkowski, Cordelia Langford, John Sillitoe on behalf of the Wellcome Sanger Institute COVID-19 Surveillance Team |
| EPI_ISL_2996844 | Lighthouse Laboratory Plymouth | Wellcome Sanger Institute for the COVID-19 Genomics UK (COG-UK) Consortium | Lighthouse Laboratory Plymouth and Alex Alderton, Roberto Amato, Jeffrey Barrett, Sonia Goncalves, Ewan Harrison, David K. Jackson, Ian Johnston, Dominic Kwiatkowski, Cordelia Langford, John Sillitoe on behalf of the Wellcome Sanger Institute COVID-19 Surveillance Team |
| EPI_ISL_2996845 | Lighthouse Lab in Milton Keynes | Wellcome Sanger Institute for the COVID-19 Genomics UK (COG-UK) Consortium | The Lighthouse Lab in Milton Keynes and Alex Alderton, Roberto Amato, Jeffrey Barrett, Sonia Goncalves, Ewan Harrison, David K. Jackson, Ian Johnston, Dominic Kwiatkowski, Cordelia Langford, John Sillitoe on behalf of the Wellcome Sanger Institute COVID-19 Surveillance Team |
| EPI_ISL_2996846 | Lighthouse Lab in Glasgow | Wellcome Sanger Institute for the COVID-19 Genomics UK (COG-UK) Consortium | Harper VanSteenhouse, Yumi Kasai, David Gray, Carol Clugston, Anna Dominiczak and Alex Alderton, Roberto Amato, Jeffrey Barrett, Sonia Goncalves, Ewan Harrison, David K. Jackson, Ian Johnston, Dominic Kwiatkowski, Cordelia Langford, John Sillitoe on behalf of the Wellcome Sanger Institute COVID-19 Surveillance Team |
| EPI_ISL_2996847 | Berkshire and Surrey Pathology Services Lighthouse Laboratory | Wellcome Sanger Institute for the COVID-19 Genomics UK (COG-UK) Consortium | Berkshire and Surrey Pathology Services Lighthouse Laboratory and Alex Alderton, Roberto Amato, Jeffrey Barrett, Sonia Goncalves, Ewan Harrison, David K. Jackson, Ian Johnston, Dominic Kwiatkowski, Cordelia Langford, John Sillitoe on behalf of the Wellcome Sanger Institute COVID-19 Surveillance Team |
| EPI_ISL_2996848, EPI_ISL_2996849 | Lighthouse Laboratory Plymouth | Wellcome Sanger Institute for the COVID-19 Genomics UK (COG-UK) Consortium | Lighthouse Laboratory Plymouth and Alex Alderton, Roberto Amato, Jeffrey Barrett, Sonia Goncalves, Ewan Harrison, David K. Jackson, Ian Johnston, Dominic Kwiatkowski, Cordelia Langford, John Sillitoe on behalf of the Wellcome Sanger Institute COVID-19 Surveillance Team |
| EPI_ISL_2996850, EPI_ISL_2996851, EPI_ISL_2996852, EPI_ISL_2996853 | Lighthouse Lab in Milton Keynes | Wellcome Sanger Institute for the COVID-19 Genomics UK (COG-UK) Consortium | The Lighthouse Lab in Milton Keynes and Alex Alderton, Roberto Amato, Jeffrey Barrett, Sonia Goncalves, Ewan Harrison, David K. Jackson, Ian Johnston, Dominic Kwiatkowski, Cordelia Langford, John Sillitoe on behalf of the Wellcome Sanger Institute COVID-19 Surveillance Team |
| EPI_ISL_2996854, EPI_ISL_2996855, EPI_ISL_2996856, EPI_ISL_2996857 | Berkshire and Surrey Pathology Services Lighthouse Laboratory | Wellcome Sanger Institute for the COVID-19 Genomics UK (COG-UK) Consortium | Berkshire and Surrey Pathology Services Lighthouse Laboratory and Alex Alderton, Roberto Amato, Jeffrey Barrett, Sonia Goncalves, Ewan Harrison, David K. Jackson, Ian Johnston, Dominic Kwiatkowski, Cordelia Langford, John Sillitoe on behalf of the Wellcome Sanger Institute COVID-19 Surveillance Team |
| EPI_ISL_2996858 | Lighthouse Laboratory Plymouth | Wellcome Sanger Institute for the COVID-19 Genomics UK (COG-UK) Consortium | Lighthouse Laboratory Plymouth and Alex Alderton, Roberto Amato, Jeffrey Barrett, Sonia Goncalves, Ewan Harrison, David K. Jackson, Ian Johnston, Dominic Kwiatkowski, Cordelia Langford, John Sillitoe on behalf of the Wellcome Sanger Institute COVID-19 Surveillance Team |
| EPI_ISL_2996859 | Lighthouse Lab in Milton Keynes | Wellcome Sanger Institute for the COVID-19 Genomics UK (COG-UK) Consortium | The Lighthouse Lab in Milton Keynes and Alex Alderton, Roberto Amato, Jeffrey Barrett, Sonia Goncalves, Ewan Harrison, David K. Jackson, Ian Johnston, Dominic Kwiatkowski, Cordelia Langford, John Sillitoe on behalf of the Wellcome Sanger Institute COVID-19 Surveillance Team |
| EPI_ISL_2996860 | Lighthouse Laboratory Plymouth | Wellcome Sanger Institute for the COVID-19 Genomics UK (COG-UK) Consortium | Lighthouse Laboratory Plymouth and Alex Alderton, Roberto Amato, Jeffrey Barrett, Sonia Goncalves, Ewan Harrison, David K. Jackson, Ian Johnston, Dominic Kwiatkowski, Cordelia Langford, John Sillitoe on behalf of the Wellcome Sanger Institute COVID-19 Surveillance Team |

| | | | |
|---|---|---|---|
| EPI_ISL_2996861 | Berkshire and Surrey Pathology Services Lighthouse Laboratory | Wellcome Sanger Institute for the COVID-19 Genomics UK (COG-UK) Consortium | Berkshire and Surrey Pathology Services Lighthouse Laboratory and Alex Alderton, Roberto Amato, Jeffrey Barrett, Sonia Goncalves, Ewan Harrison, David K. Jackson, Ian Johnston, Dominic Kwiatkowski, Cordelia Langford, John Sillitoe on behalf of the Wellcome Sanger Institute COVID-19 Surveillance Team |
| EPI_ISL_2996862 | Lighthouse Lab in Glasgow | Wellcome Sanger Institute for the COVID-19 Genomics UK (COG-UK) Consortium | Harper VanSteenhouse, Yumi Kasai, David Gray, Carol Clugston, Anna Dominiczak and Alex Alderton, Roberto Amato, Jeffrey Barrett, Sonia Goncalves, Ewan Harrison, David K. Jackson, Ian Johnston, Dominic Kwiatkowski, Cordelia Langford, John Sillitoe on behalf of the Wellcome Sanger Institute COVID-19 Surveillance Team |
| EPI_ISL_2996863 | Lighthouse Lab in Milton Keynes | Wellcome Sanger Institute for the COVID-19 Genomics UK (COG-UK) Consortium | The Lighthouse Lab in Milton Keynes and Alex Alderton, Roberto Amato, Jeffrey Barrett, Sonia Goncalves, Ewan Harrison, David K. Jackson, Ian Johnston, Dominic Kwiatkowski, Cordelia Langford, John Sillitoe on behalf of the Wellcome Sanger Institute COVID-19 Surveillance Team |
| EPI_ISL_2996864 | Lighthouse Lab in Glasgow | Wellcome Sanger Institute for the COVID-19 Genomics UK (COG-UK) Consortium | Harper VanSteenhouse, Yumi Kasai, David Gray, Carol Clugston, Anna Dominiczak and Alex Alderton, Roberto Amato, Jeffrey Barrett, Sonia Goncalves, Ewan Harrison, David K. Jackson, Ian Johnston, Dominic Kwiatkowski, Cordelia Langford, John Sillitoe on behalf of the Wellcome Sanger Institute COVID-19 Surveillance Team |
| EPI_ISL_2996865 | Berkshire and Surrey Pathology Services Lighthouse Laboratory | Wellcome Sanger Institute for the COVID-19 Genomics UK (COG-UK) Consortium | Berkshire and Surrey Pathology Services Lighthouse Laboratory and Alex Alderton, Roberto Amato, Jeffrey Barrett, Sonia Goncalves, Ewan Harrison, David K. Jackson, Ian Johnston, Dominic Kwiatkowski, Cordelia Langford, John Sillitoe on behalf of the Wellcome Sanger Institute COVID-19 Surveillance Team |
| EPI_ISL_2996866, EPI_ISL_2996867 | Lighthouse Laboratory Plymouth | Wellcome Sanger Institute for the COVID-19 Genomics UK (COG-UK) Consortium | Lighthouse Laboratory Plymouth and Alex Alderton, Roberto Amato, Jeffrey Barrett, Sonia Goncalves, Ewan Harrison, David K. Jackson, Ian Johnston, Dominic Kwiatkowski, Cordelia Langford, John Sillitoe on behalf of the Wellcome Sanger Institute COVID-19 Surveillance Team |
| EPI_ISL_2996868 | Lighthouse Lab in Milton Keynes | Wellcome Sanger Institute for the COVID-19 Genomics UK (COG-UK) Consortium | The Lighthouse Lab in Milton Keynes and Alex Alderton, Roberto Amato, Jeffrey Barrett, Sonia Goncalves, Ewan Harrison, David K. Jackson, Ian Johnston, Dominic Kwiatkowski, Cordelia Langford, John Sillitoe on behalf of the Wellcome Sanger Institute COVID-19 Surveillance Team |
| EPI_ISL_2996869 | Lighthouse Lab in Glasgow | Wellcome Sanger Institute for the COVID-19 Genomics UK (COG-UK) Consortium | Harper VanSteenhouse, Yumi Kasai, David Gray, Carol Clugston, Anna Dominiczak and Alex Alderton, Roberto Amato, Jeffrey Barrett, Sonia Goncalves, Ewan Harrison, David K. Jackson, Ian Johnston, Dominic Kwiatkowski, Cordelia Langford, John Sillitoe on behalf of the Wellcome Sanger Institute COVID-19 Surveillance Team |
| EPI_ISL_2996870 | Berkshire and Surrey Pathology Services Lighthouse Laboratory | Wellcome Sanger Institute for the COVID-19 Genomics UK (COG-UK) Consortium | Berkshire and Surrey Pathology Services Lighthouse Laboratory and Alex Alderton, Roberto Amato, Jeffrey Barrett, Sonia Goncalves, Ewan Harrison, David K. Jackson, Ian Johnston, Dominic Kwiatkowski, Cordelia Langford, John Sillitoe on behalf of the Wellcome Sanger Institute COVID-19 Surveillance Team |
| EPI_ISL_2996871, EPI_ISL_2996872 | Lighthouse Laboratory Plymouth | Wellcome Sanger Institute for the COVID-19 Genomics UK (COG-UK) Consortium | Lighthouse Laboratory Plymouth and Alex Alderton, Roberto Amato, Jeffrey Barrett, Sonia Goncalves, Ewan Harrison, David K. Jackson, Ian Johnston, Dominic Kwiatkowski, Cordelia Langford, John Sillitoe on behalf of the Wellcome Sanger Institute COVID-19 Surveillance Team |
| EPI_ISL_2996873 | Lighthouse Lab in Milton Keynes | Wellcome Sanger Institute for the COVID-19 Genomics UK (COG-UK) Consortium | The Lighthouse Lab in Milton Keynes and Alex Alderton, Roberto Amato, Jeffrey Barrett, Sonia Goncalves, Ewan Harrison, David K. Jackson, Ian Johnston, Dominic Kwiatkowski, Cordelia Langford, John Sillitoe on behalf of the Wellcome Sanger Institute COVID-19 Surveillance Team |
| EPI_ISL_2996874, EPI_ISL_2996875, EPI_ISL_2996876 | Lighthouse Lab in Glasgow | Wellcome Sanger Institute for the COVID-19 Genomics UK (COG-UK) Consortium | Harper VanSteenhouse, Yumi Kasai, David Gray, Carol Clugston, Anna Dominiczak and Alex Alderton, Roberto Amato, Jeffrey Barrett, Sonia Goncalves, Ewan Harrison, David K. Jackson, Ian Johnston, Dominic Kwiatkowski, Cordelia Langford, John Sillitoe on behalf of the Wellcome Sanger Institute COVID-19 Surveillance Team |
| EPI_ISL_2996877 | Berkshire and Surrey Pathology Services Lighthouse Laboratory | Wellcome Sanger Institute for the COVID-19 Genomics UK (COG-UK) Consortium | Berkshire and Surrey Pathology Services Lighthouse Laboratory and Alex Alderton, Roberto Amato, Jeffrey Barrett, Sonia Goncalves, Ewan Harrison, David K. Jackson, Ian Johnston, Dominic Kwiatkowski, Cordelia Langford, John Sillitoe on behalf of the Wellcome Sanger Institute COVID-19 Surveillance Team |
| EPI_ISL_2996878 | Lighthouse Lab in Glasgow | Wellcome Sanger Institute for the COVID-19 Genomics UK (COG-UK) Consortium | Harper VanSteenhouse, Yumi Kasai, David Gray, Carol Clugston, Anna Dominiczak and Alex Alderton, Roberto Amato, Jeffrey Barrett, Sonia Goncalves, Ewan Harrison, David K. Jackson, Ian Johnston, Dominic Kwiatkowski, Cordelia Langford, John Sillitoe on behalf of the Wellcome Sanger Institute COVID-19 Surveillance Team |
| EPI_ISL_2996879 | Berkshire and Surrey Pathology Services Lighthouse Laboratory | Wellcome Sanger Institute for the COVID-19 Genomics UK (COG-UK) Consortium | Berkshire and Surrey Pathology Services Lighthouse Laboratory and Alex Alderton, Roberto Amato, Jeffrey Barrett, Sonia Goncalves, Ewan Harrison, David K. Jackson, Ian Johnston, Dominic Kwiatkowski, Cordelia Langford, John Sillitoe on behalf of the Wellcome Sanger Institute COVID-19 Surveillance Team |
| EPI_ISL_2996880 | Lighthouse Lab in Milton Keynes | Wellcome Sanger Institute for the COVID-19 Genomics UK (COG-UK) Consortium | The Lighthouse Lab in Milton Keynes and Alex Alderton, Roberto Amato, Jeffrey Barrett, Sonia Goncalves, Ewan Harrison, David K. Jackson, Ian Johnston, Dominic Kwiatkowski, Cordelia Langford, John Sillitoe on behalf of the Wellcome Sanger Institute COVID-19 Surveillance Team |
| EPI_ISL_2996881, EPI_ISL_2996882 | Berkshire and Surrey Pathology Services Lighthouse Laboratory | Wellcome Sanger Institute for the COVID-19 Genomics UK (COG-UK) Consortium | Berkshire and Surrey Pathology Services Lighthouse Laboratory and Alex Alderton, Roberto Amato, Jeffrey Barrett, Sonia Goncalves, Ewan Harrison, David K. Jackson, Ian Johnston, Dominic Kwiatkowski, Cordelia Langford, John Sillitoe on behalf of the Wellcome Sanger Institute COVID-19 Surveillance Team |
| EPI_ISL_2996883 | Lighthouse Lab in Milton Keynes | Wellcome Sanger Institute for the COVID-19 Genomics UK (COG-UK) Consortium | The Lighthouse Lab in Milton Keynes and Alex Alderton, Roberto Amato, Jeffrey Barrett, Sonia Goncalves, Ewan Harrison, David K. Jackson, Ian Johnston, Dominic Kwiatkowski, Cordelia Langford, John Sillitoe on behalf of the Wellcome Sanger Institute COVID-19 Surveillance Team |
| EPI_ISL_2996884 | Berkshire and Surrey Pathology Services Lighthouse Laboratory | Wellcome Sanger Institute for the COVID-19 Genomics UK (COG-UK) Consortium | Berkshire and Surrey Pathology Services Lighthouse Laboratory and Alex Alderton, Roberto Amato, Jeffrey Barrett, Sonia Goncalves, Ewan Harrison, David K. Jackson, Ian Johnston, Dominic Kwiatkowski, Cordelia Langford, John Sillitoe on behalf of the Wellcome Sanger Institute COVID-19 Surveillance Team |
| EPI_ISL_2996885 | Lighthouse Laboratory Plymouth | Wellcome Sanger Institute for the COVID-19 Genomics UK (COG-UK) Consortium | Lighthouse Laboratory Plymouth and Alex Alderton, Roberto Amato, Jeffrey Barrett, Sonia Goncalves, Ewan Harrison, David K. Jackson, Ian Johnston, Dominic Kwiatkowski, Cordelia Langford, John Sillitoe on behalf of the Wellcome Sanger Institute COVID-19 Surveillance Team |
| EPI_ISL_2996886 | Lighthouse Lab in Milton Keynes | Wellcome Sanger Institute for the COVID-19 Genomics UK (COG-UK) Consortium | The Lighthouse Lab in Milton Keynes and Alex Alderton, Roberto Amato, Jeffrey Barrett, Sonia Goncalves, Ewan Harrison, David K. Jackson, Ian Johnston, Dominic Kwiatkowski, Cordelia Langford, John Sillitoe on behalf of the Wellcome Sanger Institute COVID-19 Surveillance Team |
| EPI_ISL_2996887, EPI_ISL_2996888, EPI_ISL_2996889 | Lighthouse Laboratory Plymouth | Wellcome Sanger Institute for the COVID-19 Genomics UK (COG-UK) Consortium | Lighthouse Laboratory Plymouth and Alex Alderton, Roberto Amato, Jeffrey Barrett, Sonia Goncalves, Ewan Harrison, David K. Jackson, Ian Johnston, Dominic Kwiatkowski, Cordelia Langford, John Sillitoe on behalf of the Wellcome Sanger Institute COVID-19 Surveillance Team |
| EPI_ISL_2996890, EPI_ISL_2996891 | Lighthouse Lab in Milton Keynes | Wellcome Sanger Institute for the COVID-19 Genomics UK (COG-UK) Consortium | The Lighthouse Lab in Milton Keynes and Alex Alderton, Roberto Amato, Jeffrey Barrett, Sonia Goncalves, Ewan Harrison, David K. Jackson, Ian Johnston, Dominic Kwiatkowski, Cordelia Langford, John Sillitoe on behalf of the Wellcome Sanger Institute COVID-19 Surveillance Team |
| EPI_ISL_2996892, EPI_ISL_2996893, EPI_ISL_2996894 | Berkshire and Surrey Pathology Services Lighthouse Laboratory | Wellcome Sanger Institute for the COVID-19 Genomics UK (COG-UK) Consortium | Berkshire and Surrey Pathology Services Lighthouse Laboratory and Alex Alderton, Roberto Amato, Jeffrey Barrett, Sonia Goncalves, Ewan Harrison, David K. Jackson, Ian Johnston, Dominic Kwiatkowski, Cordelia Langford, John Sillitoe on behalf of the Wellcome Sanger Institute COVID-19 Surveillance Team |
| EPI_ISL_2996895, EPI_ISL_2996896 | Lighthouse Laboratory Plymouth | Wellcome Sanger Institute for the COVID-19 Genomics UK (COG-UK) Consortium | Lighthouse Laboratory Plymouth and Alex Alderton, Roberto Amato, Jeffrey Barrett, Sonia Goncalves, Ewan Harrison, David K. Jackson, Ian Johnston, Dominic Kwiatkowski, Cordelia Langford, John Sillitoe on behalf of the Wellcome Sanger Institute COVID-19 Surveillance Team |
| EPI_ISL_2996897 | Berkshire and Surrey Pathology Services Lighthouse Laboratory | Wellcome Sanger Institute for the COVID-19 Genomics UK (COG-UK) Consortium | Berkshire and Surrey Pathology Services Lighthouse Laboratory and Alex Alderton, Roberto Amato, Jeffrey Barrett, Sonia Goncalves, Ewan Harrison, David K. Jackson, Ian Johnston, Dominic Kwiatkowski, Cordelia Langford, John Sillitoe on behalf of the Wellcome Sanger Institute COVID-19 Surveillance Team |
| EPI_ISL_2996898, EPI_ISL_2996899 | Lighthouse Lab in Milton Keynes | Wellcome Sanger Institute for the COVID-19 Genomics UK (COG-UK) Consortium | The Lighthouse Lab in Milton Keynes and Alex Alderton, Roberto Amato, Jeffrey Barrett, Sonia Goncalves, Ewan Harrison, David K. Jackson, Ian Johnston, Dominic Kwiatkowski, Cordelia Langford, John Sillitoe on behalf of the Wellcome Sanger Institute COVID-19 Surveillance Team |
| EPI_ISL_2996900 | Berkshire and Surrey Pathology Services Lighthouse Laboratory | Wellcome Sanger Institute for the COVID-19 Genomics UK (COG-UK) Consortium | Berkshire and Surrey Pathology Services Lighthouse Laboratory and Alex Alderton, Roberto Amato, Jeffrey Barrett, Sonia Goncalves, Ewan Harrison, David K. Jackson, Ian Johnston, Dominic Kwiatkowski, Cordelia Langford, John Sillitoe on behalf of the Wellcome Sanger Institute COVID-19 Surveillance Team |
| EPI_ISL_2996901, EPI_ISL_2996902 | Lighthouse Lab in Milton Keynes | Wellcome Sanger Institute for the COVID-19 Genomics UK (COG-UK) Consortium | The Lighthouse Lab in Milton Keynes and Alex Alderton, Roberto Amato, Jeffrey Barrett, Sonia Goncalves, Ewan Harrison, David K. Jackson, Ian Johnston, Dominic Kwiatkowski, Cordelia Langford, John Sillitoe on behalf of the Wellcome Sanger Institute COVID-19 Surveillance Team |
| EPI_ISL_2996903 | Lighthouse Lab in Glasgow | Wellcome Sanger Institute for the COVID-19 Genomics UK (COG-UK) Consortium | Harper VanSteenhouse, Yumi Kasai, David Gray, Carol Clugston, Anna Dominiczak and Alex Alderton, Roberto Amato, Jeffrey Barrett, Sonia Goncalves, Ewan Harrison, David K. Jackson, Ian Johnston, Dominic Kwiatkowski, Cordelia Langford, John Sillitoe on behalf of the Wellcome Sanger Institute COVID-19 Surveillance Team |
| EPI_ISL_2996904 | Berkshire and Surrey Pathology Services Lighthouse Laboratory | Wellcome Sanger Institute for the COVID-19 Genomics UK (COG-UK) Consortium | Berkshire and Surrey Pathology Services Lighthouse Laboratory and Alex Alderton, Roberto Amato, Jeffrey Barrett, Sonia Goncalves, Ewan Harrison, David K. Jackson, Ian Johnston, Dominic Kwiatkowski, Cordelia Langford, John Sillitoe on behalf of the Wellcome Sanger Institute COVID-19 Surveillance Team |
| EPI_ISL_2996905 | Lighthouse Lab in Milton Keynes | Wellcome Sanger Institute for the COVID-19 Genomics UK (COG-UK) Consortium | The Lighthouse Lab in Milton Keynes and Alex Alderton, Roberto Amato, Jeffrey Barrett, Sonia Goncalves, Ewan Harrison, David K. Jackson, Ian Johnston, Dominic Kwiatkowski, Cordelia Langford, John Sillitoe on behalf of the Wellcome Sanger Institute COVID-19 Surveillance Team |
| EPI_ISL_2996906 | Berkshire and Surrey Pathology Services Lighthouse Laboratory | Wellcome Sanger Institute for the COVID-19 Genomics UK (COG-UK) Consortium | Berkshire and Surrey Pathology Services Lighthouse Laboratory and Alex Alderton, Roberto Amato, Jeffrey Barrett, Sonia Goncalves, Ewan Harrison, David K. Jackson, Ian Johnston, Dominic Kwiatkowski, Cordelia Langford, John Sillitoe on behalf of the Wellcome Sanger Institute COVID-19 Surveillance Team |
| EPI_ISL_2996907 | Lighthouse Laboratory Plymouth | Wellcome Sanger Institute for the COVID-19 Genomics UK (COG-UK) Consortium | Lighthouse Laboratory Plymouth and Alex Alderton, Roberto Amato, Jeffrey Barrett, Sonia Goncalves, Ewan Harrison, David K. Jackson, Ian Johnston, Dominic Kwiatkowski, Cordelia Langford, John Sillitoe on behalf of the Wellcome Sanger Institute COVID-19 Surveillance Team |
| EPI_ISL_2996908 | Lighthouse Lab in Glasgow | Wellcome Sanger Institute for the COVID-19 Genomics UK (COG-UK) Consortium | Harper VanSteenhouse, Yumi Kasai, David Gray, Carol Clugston, Anna Dominiczak and Alex Alderton, Roberto Amato, Jeffrey Barrett, Sonia Goncalves, Ewan Harrison, David K. Jackson, Ian Johnston, Dominic Kwiatkowski, Cordelia Langford, John Sillitoe on behalf of the Wellcome Sanger Institute COVID-19 Surveillance Team |
| EPI_ISL_2996909, EPI_ISL_2996910, EPI_ISL_2996911 | Lighthouse Lab in Milton Keynes | Wellcome Sanger Institute for the COVID-19 Genomics UK (COG-UK) Consortium | The Lighthouse Lab in Milton Keynes and Alex Alderton, Roberto Amato, Jeffrey Barrett, Sonia Goncalves, Ewan Harrison, David K. Jackson, Ian Johnston, Dominic Kwiatkowski, Cordelia Langford, John Sillitoe on behalf of the Wellcome Sanger Institute COVID-19 Surveillance Team |
| EPI_ISL_2996912, EPI_ISL_2996913 | Lighthouse Laboratory Plymouth | Wellcome Sanger Institute for the COVID-19 Genomics UK (COG-UK) Consortium | Lighthouse Laboratory Plymouth and Alex Alderton, Roberto Amato, Jeffrey Barrett, Sonia Goncalves, Ewan Harrison, David K. Jackson, Ian Johnston, Dominic Kwiatkowski, Cordelia Langford, John Sillitoe on behalf of the Wellcome Sanger Institute COVID-19 Surveillance Team |
| EPI_ISL_2996914 | Lighthouse Lab in Glasgow | Wellcome Sanger Institute for the COVID-19 Genomics UK (COG-UK) Consortium | Harper VanSteenhouse, Yumi Kasai, David Gray, Carol Clugston, Anna Dominiczak and Alex Alderton, Roberto Amato, Jeffrey Barrett, Sonia Goncalves, Ewan Harrison, David K. Jackson, Ian Johnston, Dominic Kwiatkowski, Cordelia Langford, John Sillitoe on behalf of the Wellcome Sanger Institute COVID-19 Surveillance Team |
| EPI_ISL_2996915, EPI_ISL_2996916 | Lighthouse Lab in Milton Keynes | Wellcome Sanger Institute for the COVID-19 Genomics UK (COG-UK) Consortium | The Lighthouse Lab in Milton Keynes and Alex Alderton, Roberto Amato, Jeffrey Barrett, Sonia Goncalves, Ewan Harrison, David K. Jackson, Ian Johnston, Dominic Kwiatkowski, Cordelia Langford, John Sillitoe on behalf of the Wellcome Sanger Institute COVID-19 Surveillance Team |
| EPI_ISL_2996917 | Berkshire and Surrey Pathology Services Lighthouse Laboratory | Wellcome Sanger Institute for the COVID-19 Genomics UK (COG-UK) Consortium | Berkshire and Surrey Pathology Services Lighthouse Laboratory and Alex Alderton, Roberto Amato, Jeffrey Barrett, Sonia Goncalves, Ewan Harrison, David K. Jackson, Ian Johnston, Dominic Kwiatkowski, Cordelia Langford, John Sillitoe on behalf of the Wellcome Sanger Institute COVID-19 Surveillance Team |
| EPI_ISL_2996918 | Lighthouse Lab in Glasgow | Wellcome Sanger Institute for the COVID-19 Genomics UK (COG-UK) Consortium | Harper VanSteenhouse, Yumi Kasai, David Gray, Carol Clugston, Anna Dominiczak and Alex Alderton, Roberto Amato, Jeffrey Barrett, Sonia Goncalves, Ewan Harrison, David K. Jackson, Ian Johnston, Dominic Kwiatkowski, Cordelia Langford, John Sillitoe on behalf of the Wellcome Sanger Institute COVID-19 Surveillance Team |
| EPI_ISL_2996919 | Lighthouse Lab in Milton Keynes | Wellcome Sanger Institute for the COVID-19 Genomics UK (COG-UK) Consortium | The Lighthouse Lab in Milton Keynes and Alex Alderton, Roberto Amato, Jeffrey Barrett, Sonia Goncalves, Ewan Harrison, David K. Jackson, Ian Johnston, Dominic Kwiatkowski, Cordelia Langford, John Sillitoe on behalf of the Wellcome Sanger Institute COVID-19 Surveillance Team |
| EPI_ISL_2996920 | Lighthouse Lab in Glasgow | Wellcome Sanger Institute for the COVID-19 Genomics UK (COG-UK) Consortium | Harper VanSteenhouse, Yumi Kasai, David Gray, Carol Clugston, Anna Dominiczak and Alex Alderton, Roberto Amato, Jeffrey Barrett, Sonia Goncalves, Ewan Harrison, David K. Jackson, Ian Johnston, Dominic Kwiatkowski, Cordelia Langford, John Sillitoe on behalf of the Wellcome Sanger Institute COVID-19 Surveillance Team |
| EPI_ISL_2996921 | Lighthouse Lab in Milton Keynes | Wellcome Sanger Institute for the COVID-19 Genomics UK (COG-UK) | The Lighthouse Lab in Milton Keynes and Alex Alderton, Roberto Amato, Jeffrey Barrett, Sonia Goncalves, Ewan Harrison, David K. Jackson, Ian Johnston, Dominic Kwiatkowski, Cordelia Langford, John Sillitoe on behalf of the Wellcome Sanger |

| | | | |
|---|---|---|---|
| | | Consortium | Institute COVID-19 Surveillance Team |
| EPI_ISL_2996922 | Berkshire and Surrey Pathology Services Lighthouse Laboratory | Wellcome Sanger Institute for the COVID-19 Genomics UK (COG-UK) Consortium | Berkshire and Surrey Pathology Services Lighthouse Laboratory and Alex Alderton, Roberto Amato, Jeffrey Barrett, Sonia Goncalves, Ewan Harrison, David K. Jackson, Ian Johnston, Dominic Kwiatkowski, Cordelia Langford, John Sillitoe on behalf of the Wellcome Sanger Institute COVID-19 Surveillance Team |
| EPI_ISL_2996923 | Lighthouse Lab in Milton Keynes | Wellcome Sanger Institute for the COVID-19 Genomics UK (COG-UK) Consortium | The Lighthouse Lab in Milton Keynes and Alex Alderton, Roberto Amato, Jeffrey Barrett, Sonia Goncalves, Ewan Harrison, David K. Jackson, Ian Johnston, Dominic Kwiatkowski, Cordelia Langford, John Sillitoe on behalf of the Wellcome Sanger Institute COVID-19 Surveillance Team |
| EPI_ISL_2996924 | Lighthouse Lab in Glasgow | Wellcome Sanger Institute for the COVID-19 Genomics UK (COG-UK) Consortium | Harper VanSteenhouse, Yumi Kasai, David Gray, Carol Clugston, Anna Dominiczak and Alex Alderton, Roberto Amato, Jeffrey Barrett, Sonia Goncalves, Ewan Harrison, David K. Jackson, Ian Johnston, Dominic Kwiatkowski, Cordelia Langford, John Sillitoe on behalf of the Wellcome Sanger Institute COVID-19 Surveillance Team |
| EPI_ISL_2996925 | Lighthouse Laboratory Plymouth | Wellcome Sanger Institute for the COVID-19 Genomics UK (COG-UK) Consortium | Lighthouse Laboratory Plymouth and Alex Alderton, Roberto Amato, Jeffrey Barrett, Sonia Goncalves, Ewan Harrison, David K. Jackson, Ian Johnston, Dominic Kwiatkowski, Cordelia Langford, John Sillitoe on behalf of the Wellcome Sanger Institute COVID-19 Surveillance Team |
| EPI_ISL_2996926, EPI_ISL_2996927 | Lighthouse Lab in Milton Keynes | Wellcome Sanger Institute for the COVID-19 Genomics UK (COG-UK) Consortium | The Lighthouse Lab in Milton Keynes and Alex Alderton, Roberto Amato, Jeffrey Barrett, Sonia Goncalves, Ewan Harrison, David K. Jackson, Ian Johnston, Dominic Kwiatkowski, Cordelia Langford, John Sillitoe on behalf of the Wellcome Sanger Institute COVID-19 Surveillance Team |
| EPI_ISL_2996928 | Lighthouse Laboratory Plymouth | Wellcome Sanger Institute for the COVID-19 Genomics UK (COG-UK) Consortium | Lighthouse Laboratory Plymouth and Alex Alderton, Roberto Amato, Jeffrey Barrett, Sonia Goncalves, Ewan Harrison, David K. Jackson, Ian Johnston, Dominic Kwiatkowski, Cordelia Langford, John Sillitoe on behalf of the Wellcome Sanger Institute COVID-19 Surveillance Team |
| EPI_ISL_2996929 | Berkshire and Surrey Pathology Services Lighthouse Laboratory | Wellcome Sanger Institute for the COVID-19 Genomics UK (COG-UK) Consortium | Berkshire and Surrey Pathology Services Lighthouse Laboratory and Alex Alderton, Roberto Amato, Jeffrey Barrett, Sonia Goncalves, Ewan Harrison, David K. Jackson, Ian Johnston, Dominic Kwiatkowski, Cordelia Langford, John Sillitoe on behalf of the Wellcome Sanger Institute COVID-19 Surveillance Team |
| EPI_ISL_2996930 | Lighthouse Lab in Glasgow | Wellcome Sanger Institute for the COVID-19 Genomics UK (COG-UK) Consortium | Harper VanSteenhouse, Yumi Kasai, David Gray, Carol Clugston, Anna Dominiczak and Alex Alderton, Roberto Amato, Jeffrey Barrett, Sonia Goncalves, Ewan Harrison, David K. Jackson, Ian Johnston, Dominic Kwiatkowski, Cordelia Langford, John Sillitoe on behalf of the Wellcome Sanger Institute COVID-19 Surveillance Team |
| EPI_ISL_2996931 | Lighthouse Laboratory Plymouth | Wellcome Sanger Institute for the COVID-19 Genomics UK (COG-UK) Consortium | Lighthouse Laboratory Plymouth and Alex Alderton, Roberto Amato, Jeffrey Barrett, Sonia Goncalves, Ewan Harrison, David K. Jackson, Ian Johnston, Dominic Kwiatkowski, Cordelia Langford, John Sillitoe on behalf of the Wellcome Sanger Institute COVID-19 Surveillance Team |
| EPI_ISL_2996932, EPI_ISL_2996933 | Lighthouse Lab in Milton Keynes | Wellcome Sanger Institute for the COVID-19 Genomics UK (COG-UK) Consortium | The Lighthouse Lab in Milton Keynes and Alex Alderton, Roberto Amato, Jeffrey Barrett, Sonia Goncalves, Ewan Harrison, David K. Jackson, Ian Johnston, Dominic Kwiatkowski, Cordelia Langford, John Sillitoe on behalf of the Wellcome Sanger Institute COVID-19 Surveillance Team |
| EPI_ISL_2996934 | Berkshire and Surrey Pathology Services Lighthouse Laboratory | Wellcome Sanger Institute for the COVID-19 Genomics UK (COG-UK) Consortium | Berkshire and Surrey Pathology Services Lighthouse Laboratory and Alex Alderton, Roberto Amato, Jeffrey Barrett, Sonia Goncalves, Ewan Harrison, David K. Jackson, Ian Johnston, Dominic Kwiatkowski, Cordelia Langford, John Sillitoe on behalf of the Wellcome Sanger Institute COVID-19 Surveillance Team |
| EPI_ISL_2996935 | Lighthouse Laboratory Plymouth | Wellcome Sanger Institute for the COVID-19 Genomics UK (COG-UK) Consortium | Lighthouse Laboratory Plymouth and Alex Alderton, Roberto Amato, Jeffrey Barrett, Sonia Goncalves, Ewan Harrison, David K. Jackson, Ian Johnston, Dominic Kwiatkowski, Cordelia Langford, John Sillitoe on behalf of the Wellcome Sanger Institute COVID-19 Surveillance Team |
| EPI_ISL_2996936, EPI_ISL_2996937 | Berkshire and Surrey Pathology Services Lighthouse Laboratory | Wellcome Sanger Institute for the COVID-19 Genomics UK (COG-UK) Consortium | Berkshire and Surrey Pathology Services Lighthouse Laboratory and Alex Alderton, Roberto Amato, Jeffrey Barrett, Sonia Goncalves, Ewan Harrison, David K. Jackson, Ian Johnston, Dominic Kwiatkowski, Cordelia Langford, John Sillitoe on behalf of the Wellcome Sanger Institute COVID-19 Surveillance Team |
| EPI_ISL_2996938 | Lighthouse Lab in Glasgow | Wellcome Sanger Institute for the COVID-19 Genomics UK (COG-UK) Consortium | Harper VanSteenhouse, Yumi Kasai, David Gray, Carol Clugston, Anna Dominiczak and Alex Alderton, Roberto Amato, Jeffrey Barrett, Sonia Goncalves, Ewan Harrison, David K. Jackson, Ian Johnston, Dominic Kwiatkowski, Cordelia Langford, John Sillitoe on behalf of the Wellcome Sanger Institute COVID-19 Surveillance Team |
| EPI_ISL_2996939 | Lighthouse Lab in Milton Keynes | Wellcome Sanger Institute for the COVID-19 Genomics UK (COG-UK) Consortium | The Lighthouse Lab in Milton Keynes and Alex Alderton, Roberto Amato, Jeffrey Barrett, Sonia Goncalves, Ewan Harrison, David K. Jackson, Ian Johnston, Dominic Kwiatkowski, Cordelia Langford, John Sillitoe on behalf of the Wellcome Sanger Institute COVID-19 Surveillance Team |
| EPI_ISL_2996940 | Lighthouse Lab in Glasgow | Wellcome Sanger Institute for the COVID-19 Genomics UK (COG-UK) Consortium | Harper VanSteenhouse, Yumi Kasai, David Gray, Carol Clugston, Anna Dominiczak and Alex Alderton, Roberto Amato, Jeffrey Barrett, Sonia Goncalves, Ewan Harrison, David K. Jackson, Ian Johnston, Dominic Kwiatkowski, Cordelia Langford, John Sillitoe on behalf of the Wellcome Sanger Institute COVID-19 Surveillance Team |
| EPI_ISL_2996941 | Berkshire and Surrey Pathology Services Lighthouse Laboratory | Wellcome Sanger Institute for the COVID-19 Genomics UK (COG-UK) Consortium | Berkshire and Surrey Pathology Services Lighthouse Laboratory and Alex Alderton, Roberto Amato, Jeffrey Barrett, Sonia Goncalves, Ewan Harrison, David K. Jackson, Ian Johnston, Dominic Kwiatkowski, Cordelia Langford, John Sillitoe on behalf of the Wellcome Sanger Institute COVID-19 Surveillance Team |
| EPI_ISL_2996942 | Lighthouse Laboratory Plymouth | Wellcome Sanger Institute for the COVID-19 Genomics UK (COG-UK) Consortium | Lighthouse Laboratory Plymouth and Alex Alderton, Roberto Amato, Jeffrey Barrett, Sonia Goncalves, Ewan Harrison, David K. Jackson, Ian Johnston, Dominic Kwiatkowski, Cordelia Langford, John Sillitoe on behalf of the Wellcome Sanger Institute COVID-19 Surveillance Team |
| EPI_ISL_2996943, EPI_ISL_2996944 | Lighthouse Lab in Glasgow | Wellcome Sanger Institute for the COVID-19 Genomics UK (COG-UK) Consortium | Harper VanSteenhouse, Yumi Kasai, David Gray, Carol Clugston, Anna Dominiczak and Alex Alderton, Roberto Amato, Jeffrey Barrett, Sonia Goncalves, Ewan Harrison, David K. Jackson, Ian Johnston, Dominic Kwiatkowski, Cordelia Langford, John Sillitoe on behalf of the Wellcome Sanger Institute COVID-19 Surveillance Team |
| EPI_ISL_2996945 | Lighthouse Lab in Milton Keynes | Wellcome Sanger Institute for the COVID-19 Genomics UK (COG-UK) Consortium | The Lighthouse Lab in Milton Keynes and Alex Alderton, Roberto Amato, Jeffrey Barrett, Sonia Goncalves, Ewan Harrison, David K. Jackson, Ian Johnston, Dominic Kwiatkowski, Cordelia Langford, John Sillitoe on behalf of the Wellcome Sanger Institute COVID-19 Surveillance Team |
| EPI_ISL_2996946 | Berkshire and Surrey Pathology Services Lighthouse Laboratory | Wellcome Sanger Institute for the COVID-19 Genomics UK (COG-UK) Consortium | Berkshire and Surrey Pathology Services Lighthouse Laboratory and Alex Alderton, Roberto Amato, Jeffrey Barrett, Sonia Goncalves, Ewan Harrison, David K. Jackson, Ian Johnston, Dominic Kwiatkowski, Cordelia Langford, John Sillitoe on behalf of the Wellcome Sanger Institute COVID-19 Surveillance Team |
| EPI_ISL_2996947 | Lighthouse Lab in Milton Keynes | Wellcome Sanger Institute for the COVID-19 Genomics UK (COG-UK) Consortium | The Lighthouse Lab in Milton Keynes and Alex Alderton, Roberto Amato, Jeffrey Barrett, Sonia Goncalves, Ewan Harrison, David K. Jackson, Ian Johnston, Dominic Kwiatkowski, Cordelia Langford, John Sillitoe on behalf of the Wellcome Sanger Institute COVID-19 Surveillance Team |
| EPI_ISL_2996948 | Lighthouse Laboratory Plymouth | Wellcome Sanger Institute for the COVID-19 Genomics UK (COG-UK) Consortium | Lighthouse Laboratory Plymouth and Alex Alderton, Roberto Amato, Jeffrey Barrett, Sonia Goncalves, Ewan Harrison, David K. Jackson, Ian Johnston, Dominic Kwiatkowski, Cordelia Langford, John Sillitoe on behalf of the Wellcome Sanger Institute COVID-19 Surveillance Team |
| EPI_ISL_2996949 | Berkshire and Surrey Pathology Services Lighthouse Laboratory | Wellcome Sanger Institute for the COVID-19 Genomics UK (COG-UK) Consortium | Berkshire and Surrey Pathology Services Lighthouse Laboratory and Alex Alderton, Roberto Amato, Jeffrey Barrett, Sonia Goncalves, Ewan Harrison, David K. Jackson, Ian Johnston, Dominic Kwiatkowski, Cordelia Langford, John Sillitoe on behalf of the Wellcome Sanger Institute COVID-19 Surveillance Team |
| EPI_ISL_2996950, EPI_ISL_2996951, EPI_ISL_2996952, EPI_ISL_2996953 | Lighthouse Lab in Glasgow | Wellcome Sanger Institute for the COVID-19 Genomics UK (COG-UK) Consortium | Harper VanSteenhouse, Yumi Kasai, David Gray, Carol Clugston, Anna Dominiczak and Alex Alderton, Roberto Amato, Jeffrey Barrett, Sonia Goncalves, Ewan Harrison, David K. Jackson, Ian Johnston, Dominic Kwiatkowski, Cordelia Langford, John Sillitoe on behalf of the Wellcome Sanger Institute COVID-19 Surveillance Team |
| EPI_ISL_2996954 | Berkshire and Surrey Pathology Services Lighthouse Laboratory | Wellcome Sanger Institute for the COVID-19 Genomics UK (COG-UK) Consortium | Berkshire and Surrey Pathology Services Lighthouse Laboratory and Alex Alderton, Roberto Amato, Jeffrey Barrett, Sonia Goncalves, Ewan Harrison, David K. Jackson, Ian Johnston, Dominic Kwiatkowski, Cordelia Langford, John Sillitoe on behalf of the Wellcome Sanger Institute COVID-19 Surveillance Team |
| EPI_ISL_2996955 | Lighthouse Laboratory Plymouth | Wellcome Sanger Institute for the COVID-19 Genomics UK (COG-UK) Consortium | Lighthouse Laboratory Plymouth and Alex Alderton, Roberto Amato, Jeffrey Barrett, Sonia Goncalves, Ewan Harrison, David K. Jackson, Ian Johnston, Dominic Kwiatkowski, Cordelia Langford, John Sillitoe on behalf of the Wellcome Sanger Institute COVID-19 Surveillance Team |
| EPI_ISL_2996956 | Berkshire and Surrey Pathology Services Lighthouse Laboratory | Wellcome Sanger Institute for the COVID-19 Genomics UK (COG-UK) Consortium | Berkshire and Surrey Pathology Services Lighthouse Laboratory and Alex Alderton, Roberto Amato, Jeffrey Barrett, Sonia Goncalves, Ewan Harrison, David K. Jackson, Ian Johnston, Dominic Kwiatkowski, Cordelia Langford, John Sillitoe on behalf of the Wellcome Sanger Institute COVID-19 Surveillance Team |
| EPI_ISL_2996957, EPI_ISL_2996958 | Lighthouse Laboratory Plymouth | Wellcome Sanger Institute for the COVID-19 Genomics UK (COG-UK) Consortium | Lighthouse Laboratory Plymouth and Alex Alderton, Roberto Amato, Jeffrey Barrett, Sonia Goncalves, Ewan Harrison, David K. Jackson, Ian Johnston, Dominic Kwiatkowski, Cordelia Langford, John Sillitoe on behalf of the Wellcome Sanger Institute COVID-19 Surveillance Team |
| EPI_ISL_2996959 | Lighthouse Lab in Glasgow | Wellcome Sanger Institute for the COVID-19 Genomics UK (COG-UK) Consortium | Harper VanSteenhouse, Yumi Kasai, David Gray, Carol Clugston, Anna Dominiczak and Alex Alderton, Roberto Amato, Jeffrey Barrett, Sonia Goncalves, Ewan Harrison, David K. Jackson, Ian Johnston, Dominic Kwiatkowski, Cordelia Langford, John Sillitoe on behalf of the Wellcome Sanger Institute COVID-19 Surveillance Team |
| EPI_ISL_2996960 | Lighthouse Lab in Milton Keynes | Wellcome Sanger Institute for the COVID-19 Genomics UK (COG-UK) Consortium | The Lighthouse Lab in Milton Keynes and Alex Alderton, Roberto Amato, Jeffrey Barrett, Sonia Goncalves, Ewan Harrison, David K. Jackson, Ian Johnston, Dominic Kwiatkowski, Cordelia Langford, John Sillitoe on behalf of the Wellcome Sanger Institute COVID-19 Surveillance Team |
| EPI_ISL_2996961, EPI_ISL_2996962 | Lighthouse Laboratory Plymouth | Wellcome Sanger Institute for the COVID-19 Genomics UK (COG-UK) Consortium | Lighthouse Laboratory Plymouth and Alex Alderton, Roberto Amato, Jeffrey Barrett, Sonia Goncalves, Ewan Harrison, David K. Jackson, Ian Johnston, Dominic Kwiatkowski, Cordelia Langford, John Sillitoe on behalf of the Wellcome Sanger Institute COVID-19 Surveillance Team |
| EPI_ISL_2996963 | Berkshire and Surrey Pathology Services Lighthouse Laboratory | Wellcome Sanger Institute for the COVID-19 Genomics UK (COG-UK) Consortium | Berkshire and Surrey Pathology Services Lighthouse Laboratory and Alex Alderton, Roberto Amato, Jeffrey Barrett, Sonia Goncalves, Ewan Harrison, David K. Jackson, Ian Johnston, Dominic Kwiatkowski, Cordelia Langford, John Sillitoe on behalf of the Wellcome Sanger Institute COVID-19 Surveillance Team |
| EPI_ISL_2996964 | Lighthouse Laboratory Plymouth | Wellcome Sanger Institute for the COVID-19 Genomics UK (COG-UK) Consortium | Lighthouse Laboratory Plymouth and Alex Alderton, Roberto Amato, Jeffrey Barrett, Sonia Goncalves, Ewan Harrison, David K. Jackson, Ian Johnston, Dominic Kwiatkowski, Cordelia Langford, John Sillitoe on behalf of the Wellcome Sanger Institute COVID-19 Surveillance Team |
| EPI_ISL_2996965 | Lighthouse Lab in Glasgow | Wellcome Sanger Institute for the COVID-19 Genomics UK (COG-UK) Consortium | Harper VanSteenhouse, Yumi Kasai, David Gray, Carol Clugston, Anna Dominiczak and Alex Alderton, Roberto Amato, Jeffrey Barrett, Sonia Goncalves, Ewan Harrison, David K. Jackson, Ian Johnston, Dominic Kwiatkowski, Cordelia Langford, John Sillitoe on behalf of the Wellcome Sanger Institute COVID-19 Surveillance Team |
| EPI_ISL_2996966 | Lighthouse Laboratory Plymouth | Wellcome Sanger Institute for the COVID-19 Genomics UK (COG-UK) Consortium | Lighthouse Laboratory Plymouth and Alex Alderton, Roberto Amato, Jeffrey Barrett, Sonia Goncalves, Ewan Harrison, David K. Jackson, Ian Johnston, Dominic Kwiatkowski, Cordelia Langford, John Sillitoe on behalf of the Wellcome Sanger Institute COVID-19 Surveillance Team |
| EPI_ISL_2996967 | Lighthouse Lab in Milton Keynes | Wellcome Sanger Institute for the COVID-19 Genomics UK (COG-UK) Consortium | The Lighthouse Lab in Milton Keynes and Alex Alderton, Roberto Amato, Jeffrey Barrett, Sonia Goncalves, Ewan Harrison, David K. Jackson, Ian Johnston, Dominic Kwiatkowski, Cordelia Langford, John Sillitoe on behalf of the Wellcome Sanger Institute COVID-19 Surveillance Team |
| EPI_ISL_2996968 | Lighthouse Laboratory Plymouth | Wellcome Sanger Institute for the COVID-19 Genomics UK (COG-UK) Consortium | Lighthouse Laboratory Plymouth and Alex Alderton, Roberto Amato, Jeffrey Barrett, Sonia Goncalves, Ewan Harrison, David K. Jackson, Ian Johnston, Dominic Kwiatkowski, Cordelia Langford, John Sillitoe on behalf of the Wellcome Sanger Institute COVID-19 Surveillance Team |
| EPI_ISL_2996969 | Lighthouse Lab in Glasgow | Wellcome Sanger Institute for the COVID-19 Genomics UK (COG-UK) Consortium | Harper VanSteenhouse, Yumi Kasai, David Gray, Carol Clugston, Anna Dominiczak and Alex Alderton, Roberto Amato, Jeffrey Barrett, Sonia Goncalves, Ewan Harrison, David K. Jackson, Ian Johnston, Dominic Kwiatkowski, Cordelia Langford, John Sillitoe on behalf of the Wellcome Sanger Institute COVID-19 Surveillance Team |
| EPI_ISL_2996970 | Lighthouse Laboratory Plymouth | Wellcome Sanger Institute for the COVID-19 Genomics UK (COG-UK) Consortium | Lighthouse Laboratory Plymouth and Alex Alderton, Roberto Amato, Jeffrey Barrett, Sonia Goncalves, Ewan Harrison, David K. Jackson, Ian Johnston, Dominic Kwiatkowski, Cordelia Langford, John Sillitoe on behalf of the Wellcome Sanger Institute COVID-19 Surveillance Team |
| EPI_ISL_2996971 | Berkshire and Surrey Pathology Services Lighthouse Laboratory | Wellcome Sanger Institute for the COVID-19 Genomics UK (COG-UK) Consortium | Berkshire and Surrey Pathology Services Lighthouse Laboratory and Alex Alderton, Roberto Amato, Jeffrey Barrett, Sonia Goncalves, Ewan Harrison, David K. Jackson, Ian Johnston, Dominic Kwiatkowski, Cordelia Langford, John Sillitoe on behalf of the Wellcome Sanger Institute COVID-19 Surveillance Team |
| EPI_ISL_2996972 | Lighthouse Lab in Milton Keynes | Wellcome Sanger Institute for the COVID-19 Genomics UK (COG-UK) Consortium | The Lighthouse Lab in Milton Keynes and Alex Alderton, Roberto Amato, Jeffrey Barrett, Sonia Goncalves, Ewan Harrison, David K. Jackson, Ian Johnston, Dominic Kwiatkowski, Cordelia Langford, John Sillitoe on behalf of the Wellcome Sanger Institute COVID-19 Surveillance Team |
| EPI_ISL_2996973 | Berkshire and Surrey Pathology Services Lighthouse Laboratory | Wellcome Sanger Institute for the COVID-19 Genomics UK (COG-UK) Consortium | Berkshire and Surrey Pathology Services Lighthouse Laboratory and Alex Alderton, Roberto Amato, Jeffrey Barrett, Sonia Goncalves, Ewan Harrison, David K. Jackson, Ian Johnston, Dominic Kwiatkowski, Cordelia Langford, John Sillitoe on behalf of the Wellcome Sanger Institute COVID-19 Surveillance Team |
| EPI_ISL_2996974 | Lighthouse Lab in Glasgow | Wellcome Sanger Institute for the COVID-19 Genomics UK (COG-UK) | Harper VanSteenhouse, Yumi Kasai, David Gray, Carol Clugston, Anna Dominiczak and Alex Alderton, Roberto Amato, Jeffrey Barrett, Sonia Goncalves, Ewan Harrison, David K. Jackson, Ian Johnston, Dominic Kwiatkowski, Cordelia Langford, |

| | | | |
|---|---|---|---|
| | | Consortium | John Sillitoe on behalf of the Wellcome Sanger Institute COVID-19 Surveillance Team |
| EPI_ISL_2996975 | Lighthouse Laboratory Plymouth | Wellcome Sanger Institute for the COVID-19 Genomics UK (COG-UK) Consortium | Lighthouse Laboratory Plymouth and Alex Alderton, Roberto Amato, Jeffrey Barrett, Sonia Goncalves, Ewan Harrison, David K. Jackson, Ian Johnston, Dominic Kwiatkowski, Cordelia Langford, John Sillitoe on behalf of the Wellcome Sanger Institute COVID-19 Surveillance Team |
| EPI_ISL_2996976 | Lighthouse Lab in Glasgow | Wellcome Sanger Institute for the COVID-19 Genomics UK (COG-UK) Consortium | Harper VanSteenhouse, Yumi Kasai, David Gray, Carol Clugston, Anna Dominiczak and Alex Alderton, Roberto Amato, Jeffrey Barrett, Sonia Goncalves, Ewan Harrison, David K. Jackson, Ian Johnston, Dominic Kwiatkowski, Cordelia Langford, John Sillitoe on behalf of the Wellcome Sanger Institute COVID-19 Surveillance Team |
| EPI_ISL_2996977 | Lighthouse Laboratory Plymouth | Wellcome Sanger Institute for the COVID-19 Genomics UK (COG-UK) Consortium | Lighthouse Laboratory Plymouth and Alex Alderton, Roberto Amato, Jeffrey Barrett, Sonia Goncalves, Ewan Harrison, David K. Jackson, Ian Johnston, Dominic Kwiatkowski, Cordelia Langford, John Sillitoe on behalf of the Wellcome Sanger Institute COVID-19 Surveillance Team |
| EPI_ISL_2996978 | Lighthouse Lab in Glasgow | Wellcome Sanger Institute for the COVID-19 Genomics UK (COG-UK) Consortium | Harper VanSteenhouse, Yumi Kasai, David Gray, Carol Clugston, Anna Dominiczak and Alex Alderton, Roberto Amato, Jeffrey Barrett, Sonia Goncalves, Ewan Harrison, David K. Jackson, Ian Johnston, Dominic Kwiatkowski, Cordelia Langford, John Sillitoe on behalf of the Wellcome Sanger Institute COVID-19 Surveillance Team |
| EPI_ISL_2996979 | Lighthouse Laboratory Plymouth | Wellcome Sanger Institute for the COVID-19 Genomics UK (COG-UK) Consortium | Lighthouse Laboratory Plymouth and Alex Alderton, Roberto Amato, Jeffrey Barrett, Sonia Goncalves, Ewan Harrison, David K. Jackson, Ian Johnston, Dominic Kwiatkowski, Cordelia Langford, John Sillitoe on behalf of the Wellcome Sanger Institute COVID-19 Surveillance Team |
| EPI_ISL_2996980 | Lighthouse Lab in Milton Keynes | Wellcome Sanger Institute for the COVID-19 Genomics UK (COG-UK) Consortium | The Lighthouse Lab in Milton Keynes and Alex Alderton, Roberto Amato, Jeffrey Barrett, Sonia Goncalves, Ewan Harrison, David K. Jackson, Ian Johnston, Dominic Kwiatkowski, Cordelia Langford, John Sillitoe on behalf of the Wellcome Sanger Institute COVID-19 Surveillance Team |
| EPI_ISL_2996981, EPI_ISL_2996982 | Lighthouse Laboratory Plymouth | Wellcome Sanger Institute for the COVID-19 Genomics UK (COG-UK) Consortium | Lighthouse Laboratory Plymouth and Alex Alderton, Roberto Amato, Jeffrey Barrett, Sonia Goncalves, Ewan Harrison, David K. Jackson, Ian Johnston, Dominic Kwiatkowski, Cordelia Langford, John Sillitoe on behalf of the Wellcome Sanger Institute COVID-19 Surveillance Team |
| EPI_ISL_2996983 | Lighthouse Lab in Glasgow | Wellcome Sanger Institute for the COVID-19 Genomics UK (COG-UK) Consortium | Harper VanSteenhouse, Yumi Kasai, David Gray, Carol Clugston, Anna Dominiczak and Alex Alderton, Roberto Amato, Jeffrey Barrett, Sonia Goncalves, Ewan Harrison, David K. Jackson, Ian Johnston, Dominic Kwiatkowski, Cordelia Langford, John Sillitoe on behalf of the Wellcome Sanger Institute COVID-19 Surveillance Team |
| EPI_ISL_2996984 | Berkshire and Surrey Pathology Services Lighthouse Laboratory | Wellcome Sanger Institute for the COVID-19 Genomics UK (COG-UK) Consortium | Berkshire and Surrey Pathology Services Lighthouse Laboratory and Alex Alderton, Roberto Amato, Jeffrey Barrett, Sonia Goncalves, Ewan Harrison, David K. Jackson, Ian Johnston, Dominic Kwiatkowski, Cordelia Langford, John Sillitoe on behalf of the Wellcome Sanger Institute COVID-19 Surveillance Team |
| EPI_ISL_2996985 | Lighthouse Lab in Glasgow | Wellcome Sanger Institute for the COVID-19 Genomics UK (COG-UK) Consortium | Harper VanSteenhouse, Yumi Kasai, David Gray, Carol Clugston, Anna Dominiczak and Alex Alderton, Roberto Amato, Jeffrey Barrett, Sonia Goncalves, Ewan Harrison, David K. Jackson, Ian Johnston, Dominic Kwiatkowski, Cordelia Langford, John Sillitoe on behalf of the Wellcome Sanger Institute COVID-19 Surveillance Team |
| EPI_ISL_2996986 | Lighthouse Lab in Milton Keynes | Wellcome Sanger Institute for the COVID-19 Genomics UK (COG-UK) Consortium | The Lighthouse Lab in Milton Keynes and Alex Alderton, Roberto Amato, Jeffrey Barrett, Sonia Goncalves, Ewan Harrison, David K. Jackson, Ian Johnston, Dominic Kwiatkowski, Cordelia Langford, John Sillitoe on behalf of the Wellcome Sanger Institute COVID-19 Surveillance Team |
| EPI_ISL_2996987 | Berkshire and Surrey Pathology Services Lighthouse Laboratory | Wellcome Sanger Institute for the COVID-19 Genomics UK (COG-UK) Consortium | Berkshire and Surrey Pathology Services Lighthouse Laboratory and Alex Alderton, Roberto Amato, Jeffrey Barrett, Sonia Goncalves, Ewan Harrison, David K. Jackson, Ian Johnston, Dominic Kwiatkowski, Cordelia Langford, John Sillitoe on behalf of the Wellcome Sanger Institute COVID-19 Surveillance Team |
| EPI_ISL_2996988, EPI_ISL_2996989, EPI_ISL_2996990 | Lighthouse Laboratory Plymouth | Wellcome Sanger Institute for the COVID-19 Genomics UK (COG-UK) Consortium | Lighthouse Laboratory Plymouth and Alex Alderton, Roberto Amato, Jeffrey Barrett, Sonia Goncalves, Ewan Harrison, David K. Jackson, Ian Johnston, Dominic Kwiatkowski, Cordelia Langford, John Sillitoe on behalf of the Wellcome Sanger Institute COVID-19 Surveillance Team |
| EPI_ISL_2996991 | Berkshire and Surrey Pathology Services Lighthouse Laboratory | Wellcome Sanger Institute for the COVID-19 Genomics UK (COG-UK) Consortium | Berkshire and Surrey Pathology Services Lighthouse Laboratory and Alex Alderton, Roberto Amato, Jeffrey Barrett, Sonia Goncalves, Ewan Harrison, David K. Jackson, Ian Johnston, Dominic Kwiatkowski, Cordelia Langford, John Sillitoe on behalf of the Wellcome Sanger Institute COVID-19 Surveillance Team |
| EPI_ISL_2996992 | Lighthouse Lab in Glasgow | Wellcome Sanger Institute for the COVID-19 Genomics UK (COG-UK) Consortium | Harper VanSteenhouse, Yumi Kasai, David Gray, Carol Clugston, Anna Dominiczak and Alex Alderton, Roberto Amato, Jeffrey Barrett, Sonia Goncalves, Ewan Harrison, David K. Jackson, Ian Johnston, Dominic Kwiatkowski, Cordelia Langford, John Sillitoe on behalf of the Wellcome Sanger Institute COVID-19 Surveillance Team |
| EPI_ISL_2996993 | Lighthouse Lab in Milton Keynes | Wellcome Sanger Institute for the COVID-19 Genomics UK (COG-UK) Consortium | The Lighthouse Lab in Milton Keynes and Alex Alderton, Roberto Amato, Jeffrey Barrett, Sonia Goncalves, Ewan Harrison, David K. Jackson, Ian Johnston, Dominic Kwiatkowski, Cordelia Langford, John Sillitoe on behalf of the Wellcome Sanger Institute COVID-19 Surveillance Team |
| EPI_ISL_2996994 | Lighthouse Lab in Glasgow | Wellcome Sanger Institute for the COVID-19 Genomics UK (COG-UK) Consortium | Harper VanSteenhouse, Yumi Kasai, David Gray, Carol Clugston, Anna Dominiczak and Alex Alderton, Roberto Amato, Jeffrey Barrett, Sonia Goncalves, Ewan Harrison, David K. Jackson, Ian Johnston, Dominic Kwiatkowski, Cordelia Langford, John Sillitoe on behalf of the Wellcome Sanger Institute COVID-19 Surveillance Team |
| EPI_ISL_2996995, EPI_ISL_2996996 | Berkshire and Surrey Pathology Services Lighthouse Laboratory | Wellcome Sanger Institute for the COVID-19 Genomics UK (COG-UK) Consortium | Berkshire and Surrey Pathology Services Lighthouse Laboratory and Alex Alderton, Roberto Amato, Jeffrey Barrett, Sonia Goncalves, Ewan Harrison, David K. Jackson, Ian Johnston, Dominic Kwiatkowski, Cordelia Langford, John Sillitoe on behalf of the Wellcome Sanger Institute COVID-19 Surveillance Team |
| EPI_ISL_2996997, EPI_ISL_2996998 | Lighthouse Lab in Milton Keynes | Wellcome Sanger Institute for the COVID-19 Genomics UK (COG-UK) Consortium | The Lighthouse Lab in Milton Keynes and Alex Alderton, Roberto Amato, Jeffrey Barrett, Sonia Goncalves, Ewan Harrison, David K. Jackson, Ian Johnston, Dominic Kwiatkowski, Cordelia Langford, John Sillitoe on behalf of the Wellcome Sanger Institute COVID-19 Surveillance Team |
| EPI_ISL_2996999 | Lighthouse Lab in Glasgow | Wellcome Sanger Institute for the COVID-19 Genomics UK (COG-UK) Consortium | Harper VanSteenhouse, Yumi Kasai, David Gray, Carol Clugston, Anna Dominiczak and Alex Alderton, Roberto Amato, Jeffrey Barrett, Sonia Goncalves, Ewan Harrison, David K. Jackson, Ian Johnston, Dominic Kwiatkowski, Cordelia Langford, John Sillitoe on behalf of the Wellcome Sanger Institute COVID-19 Surveillance Team |
| EPI_ISL_2997000 | Lighthouse Laboratory Plymouth | Wellcome Sanger Institute for the COVID-19 Genomics UK (COG-UK) Consortium | Lighthouse Laboratory Plymouth and Alex Alderton, Roberto Amato, Jeffrey Barrett, Sonia Goncalves, Ewan Harrison, David K. Jackson, Ian Johnston, Dominic Kwiatkowski, Cordelia Langford, John Sillitoe on behalf of the Wellcome Sanger Institute COVID-19 Surveillance Team |
| EPI_ISL_2997001 | Lighthouse Lab in Glasgow | Wellcome Sanger Institute for the COVID-19 Genomics UK (COG-UK) Consortium | Harper VanSteenhouse, Yumi Kasai, David Gray, Carol Clugston, Anna Dominiczak and Alex Alderton, Roberto Amato, Jeffrey Barrett, Sonia Goncalves, Ewan Harrison, David K. Jackson, Ian Johnston, Dominic Kwiatkowski, Cordelia Langford, John Sillitoe on behalf of the Wellcome Sanger Institute COVID-19 Surveillance Team |
| EPI_ISL_2997002 | Lighthouse Lab in Milton Keynes | Wellcome Sanger Institute for the COVID-19 Genomics UK (COG-UK) Consortium | The Lighthouse Lab in Milton Keynes and Alex Alderton, Roberto Amato, Jeffrey Barrett, Sonia Goncalves, Ewan Harrison, David K. Jackson, Ian Johnston, Dominic Kwiatkowski, Cordelia Langford, John Sillitoe on behalf of the Wellcome Sanger Institute COVID-19 Surveillance Team |
| EPI_ISL_2997003 | Lighthouse Lab in Glasgow | Wellcome Sanger Institute for the COVID-19 Genomics UK (COG-UK) Consortium | Harper VanSteenhouse, Yumi Kasai, David Gray, Carol Clugston, Anna Dominiczak and Alex Alderton, Roberto Amato, Jeffrey Barrett, Sonia Goncalves, Ewan Harrison, David K. Jackson, Ian Johnston, Dominic Kwiatkowski, Cordelia Langford, John Sillitoe on behalf of the Wellcome Sanger Institute COVID-19 Surveillance Team |
| EPI_ISL_2997004 | Lighthouse Lab in Milton Keynes | Wellcome Sanger Institute for the COVID-19 Genomics UK (COG-UK) Consortium | The Lighthouse Lab in Milton Keynes and Alex Alderton, Roberto Amato, Jeffrey Barrett, Sonia Goncalves, Ewan Harrison, David K. Jackson, Ian Johnston, Dominic Kwiatkowski, Cordelia Langford, John Sillitoe on behalf of the Wellcome Sanger Institute COVID-19 Surveillance Team |
| EPI_ISL_2997005 | Lighthouse Laboratory Plymouth | Wellcome Sanger Institute for the COVID-19 Genomics UK (COG-UK) Consortium | Lighthouse Laboratory Plymouth and Alex Alderton, Roberto Amato, Jeffrey Barrett, Sonia Goncalves, Ewan Harrison, David K. Jackson, Ian Johnston, Dominic Kwiatkowski, Cordelia Langford, John Sillitoe on behalf of the Wellcome Sanger Institute COVID-19 Surveillance Team |
| EPI_ISL_2997006 | Lighthouse Lab in Glasgow | Wellcome Sanger Institute for the COVID-19 Genomics UK (COG-UK) Consortium | Harper VanSteenhouse, Yumi Kasai, David Gray, Carol Clugston, Anna Dominiczak and Alex Alderton, Roberto Amato, Jeffrey Barrett, Sonia Goncalves, Ewan Harrison, David K. Jackson, Ian Johnston, Dominic Kwiatkowski, Cordelia Langford, John Sillitoe on behalf of the Wellcome Sanger Institute COVID-19 Surveillance Team |
| EPI_ISL_2997007 | Lighthouse Lab in Milton Keynes | Wellcome Sanger Institute for the COVID-19 Genomics UK (COG-UK) Consortium | The Lighthouse Lab in Milton Keynes and Alex Alderton, Roberto Amato, Jeffrey Barrett, Sonia Goncalves, Ewan Harrison, David K. Jackson, Ian Johnston, Dominic Kwiatkowski, Cordelia Langford, John Sillitoe on behalf of the Wellcome Sanger Institute COVID-19 Surveillance Team |
| EPI_ISL_2997008 | Berkshire and Surrey Pathology Services Lighthouse Laboratory | Wellcome Sanger Institute for the COVID-19 Genomics UK (COG-UK) Consortium | Berkshire and Surrey Pathology Services Lighthouse Laboratory and Alex Alderton, Roberto Amato, Jeffrey Barrett, Sonia Goncalves, Ewan Harrison, David K. Jackson, Ian Johnston, Dominic Kwiatkowski, Cordelia Langford, John Sillitoe on behalf of the Wellcome Sanger Institute COVID-19 Surveillance Team |
| EPI_ISL_2997009, EPI_ISL_2997010, EPI_ISL_2997011 | Lighthouse Lab in Glasgow | Wellcome Sanger Institute for the COVID-19 Genomics UK (COG-UK) Consortium | Harper VanSteenhouse, Yumi Kasai, David Gray, Carol Clugston, Anna Dominiczak and Alex Alderton, Roberto Amato, Jeffrey Barrett, Sonia Goncalves, Ewan Harrison, David K. Jackson, Ian Johnston, Dominic Kwiatkowski, Cordelia Langford, John Sillitoe on behalf of the Wellcome Sanger Institute COVID-19 Surveillance Team |
| EPI_ISL_2997012, EPI_ISL_2997013 | Berkshire and Surrey Pathology Services Lighthouse Laboratory | Wellcome Sanger Institute for the COVID-19 Genomics UK (COG-UK) Consortium | Berkshire and Surrey Pathology Services Lighthouse Laboratory and Alex Alderton, Roberto Amato, Jeffrey Barrett, Sonia Goncalves, Ewan Harrison, David K. Jackson, Ian Johnston, Dominic Kwiatkowski, Cordelia Langford, John Sillitoe on behalf of the Wellcome Sanger Institute COVID-19 Surveillance Team |
| EPI_ISL_2997014 | Lighthouse Lab in Milton Keynes | Wellcome Sanger Institute for the COVID-19 Genomics UK (COG-UK) Consortium | The Lighthouse Lab in Milton Keynes and Alex Alderton, Roberto Amato, Jeffrey Barrett, Sonia Goncalves, Ewan Harrison, David K. Jackson, Ian Johnston, Dominic Kwiatkowski, Cordelia Langford, John Sillitoe on behalf of the Wellcome Sanger Institute COVID-19 Surveillance Team |
| EPI_ISL_2997015 | Berkshire and Surrey Pathology Services Lighthouse Laboratory | Wellcome Sanger Institute for the COVID-19 Genomics UK (COG-UK) Consortium | Berkshire and Surrey Pathology Services Lighthouse Laboratory and Alex Alderton, Roberto Amato, Jeffrey Barrett, Sonia Goncalves, Ewan Harrison, David K. Jackson, Ian Johnston, Dominic Kwiatkowski, Cordelia Langford, John Sillitoe on behalf of the Wellcome Sanger Institute COVID-19 Surveillance Team |
| EPI_ISL_2997016 | Lighthouse Lab in Milton Keynes | Wellcome Sanger Institute for the COVID-19 Genomics UK (COG-UK) Consortium | The Lighthouse Lab in Milton Keynes and Alex Alderton, Roberto Amato, Jeffrey Barrett, Sonia Goncalves, Ewan Harrison, David K. Jackson, Ian Johnston, Dominic Kwiatkowski, Cordelia Langford, John Sillitoe on behalf of the Wellcome Sanger Institute COVID-19 Surveillance Team |
| EPI_ISL_2997017, EPI_ISL_2997018, EPI_ISL_2997019 | Lighthouse Lab in Glasgow | Wellcome Sanger Institute for the COVID-19 Genomics UK (COG-UK) Consortium | Harper VanSteenhouse, Yumi Kasai, David Gray, Carol Clugston, Anna Dominiczak and Alex Alderton, Roberto Amato, Jeffrey Barrett, Sonia Goncalves, Ewan Harrison, David K. Jackson, Ian Johnston, Dominic Kwiatkowski, Cordelia Langford, John Sillitoe on behalf of the Wellcome Sanger Institute COVID-19 Surveillance Team |
| EPI_ISL_2997020, EPI_ISL_2997021, EPI_ISL_2997022 | Berkshire and Surrey Pathology Services Lighthouse Laboratory | Wellcome Sanger Institute for the COVID-19 Genomics UK (COG-UK) Consortium | Berkshire and Surrey Pathology Services Lighthouse Laboratory and Alex Alderton, Roberto Amato, Jeffrey Barrett, Sonia Goncalves, Ewan Harrison, David K. Jackson, Ian Johnston, Dominic Kwiatkowski, Cordelia Langford, John Sillitoe on behalf of the Wellcome Sanger Institute COVID-19 Surveillance Team |
| EPI_ISL_2997023 | Lighthouse Lab in Glasgow | Wellcome Sanger Institute for the COVID-19 Genomics UK (COG-UK) Consortium | Harper VanSteenhouse, Yumi Kasai, David Gray, Carol Clugston, Anna Dominiczak and Alex Alderton, Roberto Amato, Jeffrey Barrett, Sonia Goncalves, Ewan Harrison, David K. Jackson, Ian Johnston, Dominic Kwiatkowski, Cordelia Langford, John Sillitoe on behalf of the Wellcome Sanger Institute COVID-19 Surveillance Team |
| EPI_ISL_2997024 | Lighthouse Lab in Milton Keynes | Wellcome Sanger Institute for the COVID-19 Genomics UK (COG-UK) Consortium | The Lighthouse Lab in Milton Keynes and Alex Alderton, Roberto Amato, Jeffrey Barrett, Sonia Goncalves, Ewan Harrison, David K. Jackson, Ian Johnston, Dominic Kwiatkowski, Cordelia Langford, John Sillitoe on behalf of the Wellcome Sanger Institute COVID-19 Surveillance Team |
| EPI_ISL_2997025 | Berkshire and Surrey Pathology Services Lighthouse Laboratory | Wellcome Sanger Institute for the COVID-19 Genomics UK (COG-UK) Consortium | Berkshire and Surrey Pathology Services Lighthouse Laboratory and Alex Alderton, Roberto Amato, Jeffrey Barrett, Sonia Goncalves, Ewan Harrison, David K. Jackson, Ian Johnston, Dominic Kwiatkowski, Cordelia Langford, John Sillitoe on behalf of the Wellcome Sanger Institute COVID-19 Surveillance Team |
| EPI_ISL_2997026 | Lighthouse Laboratory Plymouth | Wellcome Sanger Institute for the COVID-19 Genomics UK (COG-UK) Consortium | Lighthouse Laboratory Plymouth and Alex Alderton, Roberto Amato, Jeffrey Barrett, Sonia Goncalves, Ewan Harrison, David K. Jackson, Ian Johnston, Dominic Kwiatkowski, Cordelia Langford, John Sillitoe on behalf of the Wellcome Sanger Institute COVID-19 Surveillance Team |
| EPI_ISL_2997027, EPI_ISL_2997028 | Lighthouse Lab in Milton Keynes | Wellcome Sanger Institute for the COVID-19 Genomics UK (COG-UK) Consortium | The Lighthouse Lab in Milton Keynes and Alex Alderton, Roberto Amato, Jeffrey Barrett, Sonia Goncalves, Ewan Harrison, David K. Jackson, Ian Johnston, Dominic Kwiatkowski, Cordelia Langford, John Sillitoe on behalf of the Wellcome Sanger Institute COVID-19 Surveillance Team |
| EPI_ISL_2997029 | Berkshire and Surrey Pathology Services Lighthouse Laboratory | Wellcome Sanger Institute for the COVID-19 Genomics UK (COG-UK) Consortium | Berkshire and Surrey Pathology Services Lighthouse Laboratory and Alex Alderton, Roberto Amato, Jeffrey Barrett, Sonia Goncalves, Ewan Harrison, David K. Jackson, Ian Johnston, Dominic Kwiatkowski, Cordelia Langford, John Sillitoe on behalf of the Wellcome Sanger Institute COVID-19 Surveillance Team |
| EPI_ISL_2997030 | Lighthouse Laboratory Plymouth | Wellcome Sanger Institute for the COVID-19 Genomics UK (COG-UK) Consortium | Lighthouse Laboratory Plymouth and Alex Alderton, Roberto Amato, Jeffrey Barrett, Sonia Goncalves, Ewan Harrison, David K. Jackson, Ian Johnston, Dominic Kwiatkowski, Cordelia Langford, John Sillitoe on behalf of the Wellcome Sanger Institute COVID-19 Surveillance Team |

| | | | |
|---|---|---|---|
| EPI_ISL_2997031 | Lighthouse Lab in Milton Keynes | Wellcome Sanger Institute for the COVID-19 Genomics UK (COG-UK) Consortium | The Lighthouse Lab in Milton Keynes and Alex Alderton, Roberto Amato, Jeffrey Barrett, Sonia Goncalves, Ewan Harrison, David K. Jackson, Ian Johnston, Dominic Kwiatkowski, Cordelia Langford, John Sillitoe on behalf of the Wellcome Sanger Institute COVID-19 Surveillance Team |
| EPI_ISL_2997032 | Lighthouse Laboratory Plymouth | Wellcome Sanger Institute for the COVID-19 Genomics UK (COG-UK) Consortium | Lighthouse Laboratory Plymouth and Alex Alderton, Roberto Amato, Jeffrey Barrett, Sonia Goncalves, Ewan Harrison, David K. Jackson, Ian Johnston, Dominic Kwiatkowski, Cordelia Langford, John Sillitoe on behalf of the Wellcome Sanger Institute COVID-19 Surveillance Team |
| EPI_ISL_2997033 | Lighthouse Lab in Glasgow | Wellcome Sanger Institute for the COVID-19 Genomics UK (COG-UK) Consortium | Harper VanSteenhouse, Yumi Kasai, David Gray, Carol Clugston, Anna Dominiczak and Alex Alderton, Roberto Amato, Jeffrey Barrett, Sonia Goncalves, Ewan Harrison, David K. Jackson, Ian Johnston, Dominic Kwiatkowski, Cordelia Langford, John Sillitoe on behalf of the Wellcome Sanger Institute COVID-19 Surveillance Team |
| EPI_ISL_2997034 | Lighthouse Laboratory Plymouth | Wellcome Sanger Institute for the COVID-19 Genomics UK (COG-UK) Consortium | Lighthouse Laboratory Plymouth and Alex Alderton, Roberto Amato, Jeffrey Barrett, Sonia Goncalves, Ewan Harrison, David K. Jackson, Ian Johnston, Dominic Kwiatkowski, Cordelia Langford, John Sillitoe on behalf of the Wellcome Sanger Institute COVID-19 Surveillance Team |
| EPI_ISL_2997035, EPI_ISL_2997036 | Berkshire and Surrey Pathology Services Lighthouse Laboratory | Wellcome Sanger Institute for the COVID-19 Genomics UK (COG-UK) Consortium | Berkshire and Surrey Pathology Services Lighthouse Laboratory and Alex Alderton, Roberto Amato, Jeffrey Barrett, Sonia Goncalves, Ewan Harrison, David K. Jackson, Ian Johnston, Dominic Kwiatkowski, Cordelia Langford, John Sillitoe on behalf of the Wellcome Sanger Institute COVID-19 Surveillance Team |
| EPI_ISL_2997037, EPI_ISL_2997038 | Lighthouse Lab in Milton Keynes | Wellcome Sanger Institute for the COVID-19 Genomics UK (COG-UK) Consortium | The Lighthouse Lab in Milton Keynes and Alex Alderton, Roberto Amato, Jeffrey Barrett, Sonia Goncalves, Ewan Harrison, David K. Jackson, Ian Johnston, Dominic Kwiatkowski, Cordelia Langford, John Sillitoe on behalf of the Wellcome Sanger Institute COVID-19 Surveillance Team |
| EPI_ISL_2997039 | Lighthouse Lab in Glasgow | Wellcome Sanger Institute for the COVID-19 Genomics UK (COG-UK) Consortium | Harper VanSteenhouse, Yumi Kasai, David Gray, Carol Clugston, Anna Dominiczak and Alex Alderton, Roberto Amato, Jeffrey Barrett, Sonia Goncalves, Ewan Harrison, David K. Jackson, Ian Johnston, Dominic Kwiatkowski, Cordelia Langford, John Sillitoe on behalf of the Wellcome Sanger Institute COVID-19 Surveillance Team |
| EPI_ISL_2997040, EPI_ISL_2997041 | Berkshire and Surrey Pathology Services Lighthouse Laboratory | Wellcome Sanger Institute for the COVID-19 Genomics UK (COG-UK) Consortium | Berkshire and Surrey Pathology Services Lighthouse Laboratory and Alex Alderton, Roberto Amato, Jeffrey Barrett, Sonia Goncalves, Ewan Harrison, David K. Jackson, Ian Johnston, Dominic Kwiatkowski, Cordelia Langford, John Sillitoe on behalf of the Wellcome Sanger Institute COVID-19 Surveillance Team |
| EPI_ISL_2997042 | Lighthouse Lab in Glasgow | Wellcome Sanger Institute for the COVID-19 Genomics UK (COG-UK) Consortium | Harper VanSteenhouse, Yumi Kasai, David Gray, Carol Clugston, Anna Dominiczak and Alex Alderton, Roberto Amato, Jeffrey Barrett, Sonia Goncalves, Ewan Harrison, David K. Jackson, Ian Johnston, Dominic Kwiatkowski, Cordelia Langford, John Sillitoe on behalf of the Wellcome Sanger Institute COVID-19 Surveillance Team |
| EPI_ISL_2997043 | Lighthouse Laboratory Plymouth | Wellcome Sanger Institute for the COVID-19 Genomics UK (COG-UK) Consortium | Lighthouse Laboratory Plymouth and Alex Alderton, Roberto Amato, Jeffrey Barrett, Sonia Goncalves, Ewan Harrison, David K. Jackson, Ian Johnston, Dominic Kwiatkowski, Cordelia Langford, John Sillitoe on behalf of the Wellcome Sanger Institute COVID-19 Surveillance Team |
| EPI_ISL_2997044 | Berkshire and Surrey Pathology Services Lighthouse Laboratory | Wellcome Sanger Institute for the COVID-19 Genomics UK (COG-UK) Consortium | Berkshire and Surrey Pathology Services Lighthouse Laboratory and Alex Alderton, Roberto Amato, Jeffrey Barrett, Sonia Goncalves, Ewan Harrison, David K. Jackson, Ian Johnston, Dominic Kwiatkowski, Cordelia Langford, John Sillitoe on behalf of the Wellcome Sanger Institute COVID-19 Surveillance Team |
| EPI_ISL_2997045, EPI_ISL_2997046 | Lighthouse Laboratory Plymouth | Wellcome Sanger Institute for the COVID-19 Genomics UK (COG-UK) Consortium | Lighthouse Laboratory Plymouth and Alex Alderton, Roberto Amato, Jeffrey Barrett, Sonia Goncalves, Ewan Harrison, David K. Jackson, Ian Johnston, Dominic Kwiatkowski, Cordelia Langford, John Sillitoe on behalf of the Wellcome Sanger Institute COVID-19 Surveillance Team |
| EPI_ISL_2997047 | Berkshire and Surrey Pathology Services Lighthouse Laboratory | Wellcome Sanger Institute for the COVID-19 Genomics UK (COG-UK) Consortium | Berkshire and Surrey Pathology Services Lighthouse Laboratory and Alex Alderton, Roberto Amato, Jeffrey Barrett, Sonia Goncalves, Ewan Harrison, David K. Jackson, Ian Johnston, Dominic Kwiatkowski, Cordelia Langford, John Sillitoe on behalf of the Wellcome Sanger Institute COVID-19 Surveillance Team |
| EPI_ISL_2997048, EPI_ISL_2997049 | Lighthouse Lab in Milton Keynes | Wellcome Sanger Institute for the COVID-19 Genomics UK (COG-UK) Consortium | The Lighthouse Lab in Milton Keynes and Alex Alderton, Roberto Amato, Jeffrey Barrett, Sonia Goncalves, Ewan Harrison, David K. Jackson, Ian Johnston, Dominic Kwiatkowski, Cordelia Langford, John Sillitoe on behalf of the Wellcome Sanger Institute COVID-19 Surveillance Team |
| EPI_ISL_2997050 | Lighthouse Lab in Glasgow | Wellcome Sanger Institute for the COVID-19 Genomics UK (COG-UK) Consortium | Harper VanSteenhouse, Yumi Kasai, David Gray, Carol Clugston, Anna Dominiczak and Alex Alderton, Roberto Amato, Jeffrey Barrett, Sonia Goncalves, Ewan Harrison, David K. Jackson, Ian Johnston, Dominic Kwiatkowski, Cordelia Langford, John Sillitoe on behalf of the Wellcome Sanger Institute COVID-19 Surveillance Team |
| EPI_ISL_2997051 | Lighthouse Lab in Milton Keynes | Wellcome Sanger Institute for the COVID-19 Genomics UK (COG-UK) Consortium | The Lighthouse Lab in Milton Keynes and Alex Alderton, Roberto Amato, Jeffrey Barrett, Sonia Goncalves, Ewan Harrison, David K. Jackson, Ian Johnston, Dominic Kwiatkowski, Cordelia Langford, John Sillitoe on behalf of the Wellcome Sanger Institute COVID-19 Surveillance Team |
| EPI_ISL_2997052 | Lighthouse Lab in Glasgow | Wellcome Sanger Institute for the COVID-19 Genomics UK (COG-UK) Consortium | Harper VanSteenhouse, Yumi Kasai, David Gray, Carol Clugston, Anna Dominiczak and Alex Alderton, Roberto Amato, Jeffrey Barrett, Sonia Goncalves, Ewan Harrison, David K. Jackson, Ian Johnston, Dominic Kwiatkowski, Cordelia Langford, John Sillitoe on behalf of the Wellcome Sanger Institute COVID-19 Surveillance Team |
| EPI_ISL_2997053 | Lighthouse Laboratory Plymouth | Wellcome Sanger Institute for the COVID-19 Genomics UK (COG-UK) Consortium | Lighthouse Laboratory Plymouth and Alex Alderton, Roberto Amato, Jeffrey Barrett, Sonia Goncalves, Ewan Harrison, David K. Jackson, Ian Johnston, Dominic Kwiatkowski, Cordelia Langford, John Sillitoe on behalf of the Wellcome Sanger Institute COVID-19 Surveillance Team |
| EPI_ISL_2997054 | Lighthouse Lab in Milton Keynes | Wellcome Sanger Institute for the COVID-19 Genomics UK (COG-UK) Consortium | The Lighthouse Lab in Milton Keynes and Alex Alderton, Roberto Amato, Jeffrey Barrett, Sonia Goncalves, Ewan Harrison, David K. Jackson, Ian Johnston, Dominic Kwiatkowski, Cordelia Langford, John Sillitoe on behalf of the Wellcome Sanger Institute COVID-19 Surveillance Team |
| EPI_ISL_2997055 | Lighthouse Laboratory Plymouth | Wellcome Sanger Institute for the COVID-19 Genomics UK (COG-UK) Consortium | Lighthouse Laboratory Plymouth and Alex Alderton, Roberto Amato, Jeffrey Barrett, Sonia Goncalves, Ewan Harrison, David K. Jackson, Ian Johnston, Dominic Kwiatkowski, Cordelia Langford, John Sillitoe on behalf of the Wellcome Sanger Institute COVID-19 Surveillance Team |
| EPI_ISL_2997056 | Lighthouse Lab in Milton Keynes | Wellcome Sanger Institute for the COVID-19 Genomics UK (COG-UK) Consortium | The Lighthouse Lab in Milton Keynes and Alex Alderton, Roberto Amato, Jeffrey Barrett, Sonia Goncalves, Ewan Harrison, David K. Jackson, Ian Johnston, Dominic Kwiatkowski, Cordelia Langford, John Sillitoe on behalf of the Wellcome Sanger Institute COVID-19 Surveillance Team |
| EPI_ISL_2997057 | Berkshire and Surrey Pathology Services Lighthouse Laboratory | Wellcome Sanger Institute for the COVID-19 Genomics UK (COG-UK) Consortium | Berkshire and Surrey Pathology Services Lighthouse Laboratory and Alex Alderton, Roberto Amato, Jeffrey Barrett, Sonia Goncalves, Ewan Harrison, David K. Jackson, Ian Johnston, Dominic Kwiatkowski, Cordelia Langford, John Sillitoe on behalf of the Wellcome Sanger Institute COVID-19 Surveillance Team |
| EPI_ISL_2997058 | Lighthouse Lab in Glasgow | Wellcome Sanger Institute for the COVID-19 Genomics UK (COG-UK) Consortium | Harper VanSteenhouse, Yumi Kasai, David Gray, Carol Clugston, Anna Dominiczak and Alex Alderton, Roberto Amato, Jeffrey Barrett, Sonia Goncalves, Ewan Harrison, David K. Jackson, Ian Johnston, Dominic Kwiatkowski, Cordelia Langford, John Sillitoe on behalf of the Wellcome Sanger Institute COVID-19 Surveillance Team |
| EPI_ISL_2997059, EPI_ISL_2997060, EPI_ISL_2997061, EPI_ISL_2997062, EPI_ISL_2997063 | Lighthouse Laboratory Plymouth | Wellcome Sanger Institute for the COVID-19 Genomics UK (COG-UK) Consortium | Lighthouse Laboratory Plymouth and Alex Alderton, Roberto Amato, Jeffrey Barrett, Sonia Goncalves, Ewan Harrison, David K. Jackson, Ian Johnston, Dominic Kwiatkowski, Cordelia Langford, John Sillitoe on behalf of the Wellcome Sanger Institute COVID-19 Surveillance Team |
| EPI_ISL_2997064 | Berkshire and Surrey Pathology Services Lighthouse Laboratory | Wellcome Sanger Institute for the COVID-19 Genomics UK (COG-UK) Consortium | Berkshire and Surrey Pathology Services Lighthouse Laboratory and Alex Alderton, Roberto Amato, Jeffrey Barrett, Sonia Goncalves, Ewan Harrison, David K. Jackson, Ian Johnston, Dominic Kwiatkowski, Cordelia Langford, John Sillitoe on behalf of the Wellcome Sanger Institute COVID-19 Surveillance Team |
| EPI_ISL_2997065 | Lighthouse Laboratory Plymouth | Wellcome Sanger Institute for the COVID-19 Genomics UK (COG-UK) Consortium | Lighthouse Laboratory Plymouth and Alex Alderton, Roberto Amato, Jeffrey Barrett, Sonia Goncalves, Ewan Harrison, David K. Jackson, Ian Johnston, Dominic Kwiatkowski, Cordelia Langford, John Sillitoe on behalf of the Wellcome Sanger Institute COVID-19 Surveillance Team |
| EPI_ISL_2997066 | Lighthouse Lab in Glasgow | Wellcome Sanger Institute for the COVID-19 Genomics UK (COG-UK) Consortium | Harper VanSteenhouse, Yumi Kasai, David Gray, Carol Clugston, Anna Dominiczak and Alex Alderton, Roberto Amato, Jeffrey Barrett, Sonia Goncalves, Ewan Harrison, David K. Jackson, Ian Johnston, Dominic Kwiatkowski, Cordelia Langford, John Sillitoe on behalf of the Wellcome Sanger Institute COVID-19 Surveillance Team |
| EPI_ISL_2997067 | Lighthouse Laboratory Plymouth | Wellcome Sanger Institute for the COVID-19 Genomics UK (COG-UK) Consortium | Lighthouse Laboratory Plymouth and Alex Alderton, Roberto Amato, Jeffrey Barrett, Sonia Goncalves, Ewan Harrison, David K. Jackson, Ian Johnston, Dominic Kwiatkowski, Cordelia Langford, John Sillitoe on behalf of the Wellcome Sanger Institute COVID-19 Surveillance Team |
| EPI_ISL_2997068, EPI_ISL_2997069 | Lighthouse Lab in Glasgow | Wellcome Sanger Institute for the COVID-19 Genomics UK (COG-UK) Consortium | Harper VanSteenhouse, Yumi Kasai, David Gray, Carol Clugston, Anna Dominiczak and Alex Alderton, Roberto Amato, Jeffrey Barrett, Sonia Goncalves, Ewan Harrison, David K. Jackson, Ian Johnston, Dominic Kwiatkowski, Cordelia Langford, John Sillitoe on behalf of the Wellcome Sanger Institute COVID-19 Surveillance Team |
| EPI_ISL_2997070 | Lighthouse Lab in Milton Keynes | Wellcome Sanger Institute for the COVID-19 Genomics UK (COG-UK) Consortium | The Lighthouse Lab in Milton Keynes and Alex Alderton, Roberto Amato, Jeffrey Barrett, Sonia Goncalves, Ewan Harrison, David K. Jackson, Ian Johnston, Dominic Kwiatkowski, Cordelia Langford, John Sillitoe on behalf of the Wellcome Sanger Institute COVID-19 Surveillance Team |
| EPI_ISL_2997071 | Lighthouse Laboratory Plymouth | Wellcome Sanger Institute for the COVID-19 Genomics UK (COG-UK) Consortium | Lighthouse Laboratory Plymouth and Alex Alderton, Roberto Amato, Jeffrey Barrett, Sonia Goncalves, Ewan Harrison, David K. Jackson, Ian Johnston, Dominic Kwiatkowski, Cordelia Langford, John Sillitoe on behalf of the Wellcome Sanger Institute COVID-19 Surveillance Team |
| EPI_ISL_2997072 | Lighthouse Lab in Glasgow | Wellcome Sanger Institute for the COVID-19 Genomics UK (COG-UK) Consortium | Harper VanSteenhouse, Yumi Kasai, David Gray, Carol Clugston, Anna Dominiczak and Alex Alderton, Roberto Amato, Jeffrey Barrett, Sonia Goncalves, Ewan Harrison, David K. Jackson, Ian Johnston, Dominic Kwiatkowski, Cordelia Langford, John Sillitoe on behalf of the Wellcome Sanger Institute COVID-19 Surveillance Team |
| EPI_ISL_2997073, EPI_ISL_2997074, EPI_ISL_2997075 | Lighthouse Lab in Milton Keynes | Wellcome Sanger Institute for the COVID-19 Genomics UK (COG-UK) Consortium | The Lighthouse Lab in Milton Keynes and Alex Alderton, Roberto Amato, Jeffrey Barrett, Sonia Goncalves, Ewan Harrison, David K. Jackson, Ian Johnston, Dominic Kwiatkowski, Cordelia Langford, John Sillitoe on behalf of the Wellcome Sanger Institute COVID-19 Surveillance Team |
| EPI_ISL_2997076, EPI_ISL_2997077 | Lighthouse Lab in Glasgow | Wellcome Sanger Institute for the COVID-19 Genomics UK (COG-UK) Consortium | Harper VanSteenhouse, Yumi Kasai, David Gray, Carol Clugston, Anna Dominiczak and Alex Alderton, Roberto Amato, Jeffrey Barrett, Sonia Goncalves, Ewan Harrison, David K. Jackson, Ian Johnston, Dominic Kwiatkowski, Cordelia Langford, John Sillitoe on behalf of the Wellcome Sanger Institute COVID-19 Surveillance Team |
| EPI_ISL_2997078 | Lighthouse Lab in Milton Keynes | Wellcome Sanger Institute for the COVID-19 Genomics UK (COG-UK) Consortium | The Lighthouse Lab in Milton Keynes and Alex Alderton, Roberto Amato, Jeffrey Barrett, Sonia Goncalves, Ewan Harrison, David K. Jackson, Ian Johnston, Dominic Kwiatkowski, Cordelia Langford, John Sillitoe on behalf of the Wellcome Sanger Institute COVID-19 Surveillance Team |
| EPI_ISL_2997079 | Berkshire and Surrey Pathology Services Lighthouse Laboratory | Wellcome Sanger Institute for the COVID-19 Genomics UK (COG-UK) Consortium | Berkshire and Surrey Pathology Services Lighthouse Laboratory and Alex Alderton, Roberto Amato, Jeffrey Barrett, Sonia Goncalves, Ewan Harrison, David K. Jackson, Ian Johnston, Dominic Kwiatkowski, Cordelia Langford, John Sillitoe on behalf of the Wellcome Sanger Institute COVID-19 Surveillance Team |
| EPI_ISL_2997080 | Lighthouse Lab in Glasgow | Wellcome Sanger Institute for the COVID-19 Genomics UK (COG-UK) Consortium | Harper VanSteenhouse, Yumi Kasai, David Gray, Carol Clugston, Anna Dominiczak and Alex Alderton, Roberto Amato, Jeffrey Barrett, Sonia Goncalves, Ewan Harrison, David K. Jackson, Ian Johnston, Dominic Kwiatkowski, Cordelia Langford, John Sillitoe on behalf of the Wellcome Sanger Institute COVID-19 Surveillance Team |
| EPI_ISL_2997081 | Lighthouse Lab in Milton Keynes | Wellcome Sanger Institute for the COVID-19 Genomics UK (COG-UK) Consortium | The Lighthouse Lab in Milton Keynes and Alex Alderton, Roberto Amato, Jeffrey Barrett, Sonia Goncalves, Ewan Harrison, David K. Jackson, Ian Johnston, Dominic Kwiatkowski, Cordelia Langford, John Sillitoe on behalf of the Wellcome Sanger Institute COVID-19 Surveillance Team |
| EPI_ISL_2997082, EPI_ISL_2997083 | Lighthouse Laboratory Plymouth | Wellcome Sanger Institute for the COVID-19 Genomics UK (COG-UK) Consortium | Lighthouse Laboratory Plymouth and Alex Alderton, Roberto Amato, Jeffrey Barrett, Sonia Goncalves, Ewan Harrison, David K. Jackson, Ian Johnston, Dominic Kwiatkowski, Cordelia Langford, John Sillitoe on behalf of the Wellcome Sanger Institute COVID-19 Surveillance Team |
| EPI_ISL_2997084 | Berkshire and Surrey Pathology Services Lighthouse Laboratory | Wellcome Sanger Institute for the COVID-19 Genomics UK (COG-UK) Consortium | Berkshire and Surrey Pathology Services Lighthouse Laboratory and Alex Alderton, Roberto Amato, Jeffrey Barrett, Sonia Goncalves, Ewan Harrison, David K. Jackson, Ian Johnston, Dominic Kwiatkowski, Cordelia Langford, John Sillitoe on behalf of the Wellcome Sanger Institute COVID-19 Surveillance Team |
| EPI_ISL_2997085, EPI_ISL_2997086 | Lighthouse Lab in Milton Keynes | Wellcome Sanger Institute for the COVID-19 Genomics UK (COG-UK) Consortium | The Lighthouse Lab in Milton Keynes and Alex Alderton, Roberto Amato, Jeffrey Barrett, Sonia Goncalves, Ewan Harrison, David K. Jackson, Ian Johnston, Dominic Kwiatkowski, Cordelia Langford, John Sillitoe on behalf of the Wellcome Sanger Institute COVID-19 Surveillance Team |
| EPI_ISL_2997087 | Lighthouse Laboratory Plymouth | Wellcome Sanger Institute for the COVID-19 Genomics UK (COG-UK) Consortium | Lighthouse Laboratory Plymouth and Alex Alderton, Roberto Amato, Jeffrey Barrett, Sonia Goncalves, Ewan Harrison, David K. Jackson, Ian Johnston, Dominic Kwiatkowski, Cordelia Langford, John Sillitoe on behalf of the Wellcome Sanger Institute COVID-19 Surveillance Team |
| EPI_ISL_2997088 | Berkshire and Surrey Pathology Services Lighthouse Laboratory | Wellcome Sanger Institute for the COVID-19 Genomics UK (COG-UK) Consortium | Berkshire and Surrey Pathology Services Lighthouse Laboratory and Alex Alderton, Roberto Amato, Jeffrey Barrett, Sonia Goncalves, Ewan Harrison, David K. Jackson, Ian Johnston, Dominic Kwiatkowski, Cordelia Langford, John Sillitoe on behalf of the Wellcome Sanger Institute COVID-19 Surveillance Team |
| EPI_ISL_2997089 | Lighthouse Lab in Glasgow | Wellcome Sanger Institute for the COVID-19 Genomics UK (COG-UK) | Harper VanSteenhouse, Yumi Kasai, David Gray, Carol Clugston, Anna Dominiczak and Alex Alderton, Roberto Amato, Jeffrey Barrett, Sonia Goncalves, Ewan Harrison, David K. Jackson, Ian Johnston, Dominic Kwiatkowski, Cordelia Langford, |

| | | | |
|---|---|---|---|
| | | Consortium | John Sillitoe on behalf of the Wellcome Sanger Institute COVID-19 Surveillance Team |
| EPI_ISL_2997090 | Lighthouse Lab in Milton Keynes | Wellcome Sanger Institute for the COVID-19 Genomics UK (COG-UK) Consortium | The Lighthouse Lab in Milton Keynes and Alex Alderton, Roberto Amato, Jeffrey Barrett, Sonia Goncalves, Ewan Harrison, David K. Jackson, Ian Johnston, Dominic Kwiatkowski, Cordelia Langford, John Sillitoe on behalf of the Wellcome Sanger Institute COVID-19 Surveillance Team |
| EPI_ISL_2997091 | Lighthouse Laboratory Plymouth | Wellcome Sanger Institute for the COVID-19 Genomics UK (COG-UK) Consortium | Lighthouse Laboratory Plymouth and Alex Alderton, Roberto Amato, Jeffrey Barrett, Sonia Goncalves, Ewan Harrison, David K. Jackson, Ian Johnston, Dominic Kwiatkowski, Cordelia Langford, John Sillitoe on behalf of the Wellcome Sanger Institute COVID-19 Surveillance Team |
| EPI_ISL_2997092 | Berkshire and Surrey Pathology Services Lighthouse Laboratory | Wellcome Sanger Institute for the COVID-19 Genomics UK (COG-UK) Consortium | Berkshire and Surrey Pathology Services Lighthouse Laboratory and Alex Alderton, Roberto Amato, Jeffrey Barrett, Sonia Goncalves, Ewan Harrison, David K. Jackson, Ian Johnston, Dominic Kwiatkowski, Cordelia Langford, John Sillitoe on behalf of the Wellcome Sanger Institute COVID-19 Surveillance Team |
| EPI_ISL_2997093 | Lighthouse Laboratory Plymouth | Wellcome Sanger Institute for the COVID-19 Genomics UK (COG-UK) Consortium | Lighthouse Laboratory Plymouth and Alex Alderton, Roberto Amato, Jeffrey Barrett, Sonia Goncalves, Ewan Harrison, David K. Jackson, Ian Johnston, Dominic Kwiatkowski, Cordelia Langford, John Sillitoe on behalf of the Wellcome Sanger Institute COVID-19 Surveillance Team |
| EPI_ISL_2997094 | Lighthouse Lab in Milton Keynes | Wellcome Sanger Institute for the COVID-19 Genomics UK (COG-UK) Consortium | The Lighthouse Lab in Milton Keynes and Alex Alderton, Roberto Amato, Jeffrey Barrett, Sonia Goncalves, Ewan Harrison, David K. Jackson, Ian Johnston, Dominic Kwiatkowski, Cordelia Langford, John Sillitoe on behalf of the Wellcome Sanger Institute COVID-19 Surveillance Team |
| EPI_ISL_2997095, EPI_ISL_2997096 | Lighthouse Lab in Glasgow | Wellcome Sanger Institute for the COVID-19 Genomics UK (COG-UK) Consortium | Harper VanSteenhouse, Yumi Kasai, David Gray, Carol Clugston, Anna Dominiczak and Alex Alderton, Roberto Amato, Jeffrey Barrett, Sonia Goncalves, Ewan Harrison, David K. Jackson, Ian Johnston, Dominic Kwiatkowski, Cordelia Langford, John Sillitoe on behalf of the Wellcome Sanger Institute COVID-19 Surveillance Team |
| EPI_ISL_2997097, EPI_ISL_2997098, EPI_ISL_2997099 | Lighthouse Lab in Milton Keynes | Wellcome Sanger Institute for the COVID-19 Genomics UK (COG-UK) Consortium | The Lighthouse Lab in Milton Keynes and Alex Alderton, Roberto Amato, Jeffrey Barrett, Sonia Goncalves, Ewan Harrison, David K. Jackson, Ian Johnston, Dominic Kwiatkowski, Cordelia Langford, John Sillitoe on behalf of the Wellcome Sanger Institute COVID-19 Surveillance Team |
| EPI_ISL_2997100 | Lighthouse Laboratory Plymouth | Wellcome Sanger Institute for the COVID-19 Genomics UK (COG-UK) Consortium | Lighthouse Laboratory Plymouth and Alex Alderton, Roberto Amato, Jeffrey Barrett, Sonia Goncalves, Ewan Harrison, David K. Jackson, Ian Johnston, Dominic Kwiatkowski, Cordelia Langford, John Sillitoe on behalf of the Wellcome Sanger Institute COVID-19 Surveillance Team |
| EPI_ISL_2997101, EPI_ISL_2997102, EPI_ISL_2997103, EPI_ISL_2997104 | Lighthouse Lab in Glasgow | Wellcome Sanger Institute for the COVID-19 Genomics UK (COG-UK) Consortium | Harper VanSteenhouse, Yumi Kasai, David Gray, Carol Clugston, Anna Dominiczak and Alex Alderton, Roberto Amato, Jeffrey Barrett, Sonia Goncalves, Ewan Harrison, David K. Jackson, Ian Johnston, Dominic Kwiatkowski, Cordelia Langford, John Sillitoe on behalf of the Wellcome Sanger Institute COVID-19 Surveillance Team |
| EPI_ISL_2997105 | Berkshire and Surrey Pathology Services Lighthouse Laboratory | Wellcome Sanger Institute for the COVID-19 Genomics UK (COG-UK) Consortium | Berkshire and Surrey Pathology Services Lighthouse Laboratory and Alex Alderton, Roberto Amato, Jeffrey Barrett, Sonia Goncalves, Ewan Harrison, David K. Jackson, Ian Johnston, Dominic Kwiatkowski, Cordelia Langford, John Sillitoe on behalf of the Wellcome Sanger Institute COVID-19 Surveillance Team |
| EPI_ISL_2997106 | Lighthouse Lab in Glasgow | Wellcome Sanger Institute for the COVID-19 Genomics UK (COG-UK) Consortium | Harper VanSteenhouse, Yumi Kasai, David Gray, Carol Clugston, Anna Dominiczak and Alex Alderton, Roberto Amato, Jeffrey Barrett, Sonia Goncalves, Ewan Harrison, David K. Jackson, Ian Johnston, Dominic Kwiatkowski, Cordelia Langford, John Sillitoe on behalf of the Wellcome Sanger Institute COVID-19 Surveillance Team |
| EPI_ISL_2997107 | Lighthouse Lab in Milton Keynes | Wellcome Sanger Institute for the COVID-19 Genomics UK (COG-UK) Consortium | The Lighthouse Lab in Milton Keynes and Alex Alderton, Roberto Amato, Jeffrey Barrett, Sonia Goncalves, Ewan Harrison, David K. Jackson, Ian Johnston, Dominic Kwiatkowski, Cordelia Langford, John Sillitoe on behalf of the Wellcome Sanger Institute COVID-19 Surveillance Team |
| EPI_ISL_2997108, EPI_ISL_2997109, EPI_ISL_2997110 | Lighthouse Laboratory Plymouth | Wellcome Sanger Institute for the COVID-19 Genomics UK (COG-UK) Consortium | Lighthouse Laboratory Plymouth and Alex Alderton, Roberto Amato, Jeffrey Barrett, Sonia Goncalves, Ewan Harrison, David K. Jackson, Ian Johnston, Dominic Kwiatkowski, Cordelia Langford, John Sillitoe on behalf of the Wellcome Sanger Institute COVID-19 Surveillance Team |
| EPI_ISL_2997111, EPI_ISL_2997112 | Lighthouse Lab in Milton Keynes | Wellcome Sanger Institute for the COVID-19 Genomics UK (COG-UK) Consortium | The Lighthouse Lab in Milton Keynes and Alex Alderton, Roberto Amato, Jeffrey Barrett, Sonia Goncalves, Ewan Harrison, David K. Jackson, Ian Johnston, Dominic Kwiatkowski, Cordelia Langford, John Sillitoe on behalf of the Wellcome Sanger Institute COVID-19 Surveillance Team |
| EPI_ISL_2997113 | Berkshire and Surrey Pathology Services Lighthouse Laboratory | Wellcome Sanger Institute for the COVID-19 Genomics UK (COG-UK) Consortium | Berkshire and Surrey Pathology Services Lighthouse Laboratory and Alex Alderton, Roberto Amato, Jeffrey Barrett, Sonia Goncalves, Ewan Harrison, David K. Jackson, Ian Johnston, Dominic Kwiatkowski, Cordelia Langford, John Sillitoe on behalf of the Wellcome Sanger Institute COVID-19 Surveillance Team |
| EPI_ISL_2997114 | Lighthouse Lab in Glasgow | Wellcome Sanger Institute for the COVID-19 Genomics UK (COG-UK) Consortium | Harper VanSteenhouse, Yumi Kasai, David Gray, Carol Clugston, Anna Dominiczak and Alex Alderton, Roberto Amato, Jeffrey Barrett, Sonia Goncalves, Ewan Harrison, David K. Jackson, Ian Johnston, Dominic Kwiatkowski, Cordelia Langford, John Sillitoe on behalf of the Wellcome Sanger Institute COVID-19 Surveillance Team |
| EPI_ISL_2997115 | Lighthouse Laboratory Plymouth | Wellcome Sanger Institute for the COVID-19 Genomics UK (COG-UK) Consortium | Lighthouse Laboratory Plymouth and Alex Alderton, Roberto Amato, Jeffrey Barrett, Sonia Goncalves, Ewan Harrison, David K. Jackson, Ian Johnston, Dominic Kwiatkowski, Cordelia Langford, John Sillitoe on behalf of the Wellcome Sanger Institute COVID-19 Surveillance Team |
| EPI_ISL_2997116, EPI_ISL_2997117 | Lighthouse Lab in Glasgow | Wellcome Sanger Institute for the COVID-19 Genomics UK (COG-UK) Consortium | Harper VanSteenhouse, Yumi Kasai, David Gray, Carol Clugston, Anna Dominiczak and Alex Alderton, Roberto Amato, Jeffrey Barrett, Sonia Goncalves, Ewan Harrison, David K. Jackson, Ian Johnston, Dominic Kwiatkowski, Cordelia Langford, John Sillitoe on behalf of the Wellcome Sanger Institute COVID-19 Surveillance Team |
| EPI_ISL_2997118 | Lighthouse Laboratory Plymouth | Wellcome Sanger Institute for the COVID-19 Genomics UK (COG-UK) Consortium | Lighthouse Laboratory Plymouth and Alex Alderton, Roberto Amato, Jeffrey Barrett, Sonia Goncalves, Ewan Harrison, David K. Jackson, Ian Johnston, Dominic Kwiatkowski, Cordelia Langford, John Sillitoe on behalf of the Wellcome Sanger Institute COVID-19 Surveillance Team |
| EPI_ISL_2997119 | Lighthouse Lab in Milton Keynes | Wellcome Sanger Institute for the COVID-19 Genomics UK (COG-UK) Consortium | The Lighthouse Lab in Milton Keynes and Alex Alderton, Roberto Amato, Jeffrey Barrett, Sonia Goncalves, Ewan Harrison, David K. Jackson, Ian Johnston, Dominic Kwiatkowski, Cordelia Langford, John Sillitoe on behalf of the Wellcome Sanger Institute COVID-19 Surveillance Team |
| EPI_ISL_2997120 | Berkshire and Surrey Pathology Services Lighthouse Laboratory | Wellcome Sanger Institute for the COVID-19 Genomics UK (COG-UK) Consortium | Berkshire and Surrey Pathology Services Lighthouse Laboratory and Alex Alderton, Roberto Amato, Jeffrey Barrett, Sonia Goncalves, Ewan Harrison, David K. Jackson, Ian Johnston, Dominic Kwiatkowski, Cordelia Langford, John Sillitoe on behalf of the Wellcome Sanger Institute COVID-19 Surveillance Team |
| EPI_ISL_2997121 | Lighthouse Lab in Glasgow | Wellcome Sanger Institute for the COVID-19 Genomics UK (COG-UK) Consortium | Harper VanSteenhouse, Yumi Kasai, David Gray, Carol Clugston, Anna Dominiczak and Alex Alderton, Roberto Amato, Jeffrey Barrett, Sonia Goncalves, Ewan Harrison, David K. Jackson, Ian Johnston, Dominic Kwiatkowski, Cordelia Langford, John Sillitoe on behalf of the Wellcome Sanger Institute COVID-19 Surveillance Team |
| EPI_ISL_2997122 | Lighthouse Laboratory Plymouth | Wellcome Sanger Institute for the COVID-19 Genomics UK (COG-UK) Consortium | Lighthouse Laboratory Plymouth and Alex Alderton, Roberto Amato, Jeffrey Barrett, Sonia Goncalves, Ewan Harrison, David K. Jackson, Ian Johnston, Dominic Kwiatkowski, Cordelia Langford, John Sillitoe on behalf of the Wellcome Sanger Institute COVID-19 Surveillance Team |
| EPI_ISL_2997123 | Lighthouse Lab in Milton Keynes | Wellcome Sanger Institute for the COVID-19 Genomics UK (COG-UK) Consortium | The Lighthouse Lab in Milton Keynes and Alex Alderton, Roberto Amato, Jeffrey Barrett, Sonia Goncalves, Ewan Harrison, David K. Jackson, Ian Johnston, Dominic Kwiatkowski, Cordelia Langford, John Sillitoe on behalf of the Wellcome Sanger Institute COVID-19 Surveillance Team |
| EPI_ISL_2997124 | Lighthouse Laboratory Plymouth | Wellcome Sanger Institute for the COVID-19 Genomics UK (COG-UK) Consortium | Lighthouse Laboratory Plymouth and Alex Alderton, Roberto Amato, Jeffrey Barrett, Sonia Goncalves, Ewan Harrison, David K. Jackson, Ian Johnston, Dominic Kwiatkowski, Cordelia Langford, John Sillitoe on behalf of the Wellcome Sanger Institute COVID-19 Surveillance Team |
| EPI_ISL_2997125 | Berkshire and Surrey Pathology Services Lighthouse Laboratory | Wellcome Sanger Institute for the COVID-19 Genomics UK (COG-UK) Consortium | Berkshire and Surrey Pathology Services Lighthouse Laboratory and Alex Alderton, Roberto Amato, Jeffrey Barrett, Sonia Goncalves, Ewan Harrison, David K. Jackson, Ian Johnston, Dominic Kwiatkowski, Cordelia Langford, John Sillitoe on behalf of the Wellcome Sanger Institute COVID-19 Surveillance Team |
| EPI_ISL_2997126 | Lighthouse Lab in Glasgow | Wellcome Sanger Institute for the COVID-19 Genomics UK (COG-UK) Consortium | Harper VanSteenhouse, Yumi Kasai, David Gray, Carol Clugston, Anna Dominiczak and Alex Alderton, Roberto Amato, Jeffrey Barrett, Sonia Goncalves, Ewan Harrison, David K. Jackson, Ian Johnston, Dominic Kwiatkowski, Cordelia Langford, John Sillitoe on behalf of the Wellcome Sanger Institute COVID-19 Surveillance Team |
| EPI_ISL_2997127 | Lighthouse Lab in Milton Keynes | Wellcome Sanger Institute for the COVID-19 Genomics UK (COG-UK) Consortium | The Lighthouse Lab in Milton Keynes and Alex Alderton, Roberto Amato, Jeffrey Barrett, Sonia Goncalves, Ewan Harrison, David K. Jackson, Ian Johnston, Dominic Kwiatkowski, Cordelia Langford, John Sillitoe on behalf of the Wellcome Sanger Institute COVID-19 Surveillance Team |
| EPI_ISL_2997128 | Lighthouse Lab in Glasgow | Wellcome Sanger Institute for the COVID-19 Genomics UK (COG-UK) Consortium | Harper VanSteenhouse, Yumi Kasai, David Gray, Carol Clugston, Anna Dominiczak and Alex Alderton, Roberto Amato, Jeffrey Barrett, Sonia Goncalves, Ewan Harrison, David K. Jackson, Ian Johnston, Dominic Kwiatkowski, Cordelia Langford, John Sillitoe on behalf of the Wellcome Sanger Institute COVID-19 Surveillance Team |
| EPI_ISL_2997129 | Lighthouse Laboratory Plymouth | Wellcome Sanger Institute for the COVID-19 Genomics UK (COG-UK) Consortium | Lighthouse Laboratory Plymouth and Alex Alderton, Roberto Amato, Jeffrey Barrett, Sonia Goncalves, Ewan Harrison, David K. Jackson, Ian Johnston, Dominic Kwiatkowski, Cordelia Langford, John Sillitoe on behalf of the Wellcome Sanger Institute COVID-19 Surveillance Team |
| EPI_ISL_2997130 | Lighthouse Lab in Milton Keynes | Wellcome Sanger Institute for the COVID-19 Genomics UK (COG-UK) Consortium | The Lighthouse Lab in Milton Keynes and Alex Alderton, Roberto Amato, Jeffrey Barrett, Sonia Goncalves, Ewan Harrison, David K. Jackson, Ian Johnston, Dominic Kwiatkowski, Cordelia Langford, John Sillitoe on behalf of the Wellcome Sanger Institute COVID-19 Surveillance Team |
| EPI_ISL_2997131 | Lighthouse Lab in Glasgow | Wellcome Sanger Institute for the COVID-19 Genomics UK (COG-UK) Consortium | Harper VanSteenhouse, Yumi Kasai, David Gray, Carol Clugston, Anna Dominiczak and Alex Alderton, Roberto Amato, Jeffrey Barrett, Sonia Goncalves, Ewan Harrison, David K. Jackson, Ian Johnston, Dominic Kwiatkowski, Cordelia Langford, John Sillitoe on behalf of the Wellcome Sanger Institute COVID-19 Surveillance Team |
| EPI_ISL_2997132, EPI_ISL_2997133 | Berkshire and Surrey Pathology Services Lighthouse Laboratory | Wellcome Sanger Institute for the COVID-19 Genomics UK (COG-UK) Consortium | Berkshire and Surrey Pathology Services Lighthouse Laboratory and Alex Alderton, Roberto Amato, Jeffrey Barrett, Sonia Goncalves, Ewan Harrison, David K. Jackson, Ian Johnston, Dominic Kwiatkowski, Cordelia Langford, John Sillitoe on behalf of the Wellcome Sanger Institute COVID-19 Surveillance Team |
| EPI_ISL_2997134 | Lighthouse Laboratory Plymouth | Wellcome Sanger Institute for the COVID-19 Genomics UK (COG-UK) Consortium | Lighthouse Laboratory Plymouth and Alex Alderton, Roberto Amato, Jeffrey Barrett, Sonia Goncalves, Ewan Harrison, David K. Jackson, Ian Johnston, Dominic Kwiatkowski, Cordelia Langford, John Sillitoe on behalf of the Wellcome Sanger Institute COVID-19 Surveillance Team |
| EPI_ISL_2997135 | Lighthouse Lab in Glasgow | Wellcome Sanger Institute for the COVID-19 Genomics UK (COG-UK) Consortium | Harper VanSteenhouse, Yumi Kasai, David Gray, Carol Clugston, Anna Dominiczak and Alex Alderton, Roberto Amato, Jeffrey Barrett, Sonia Goncalves, Ewan Harrison, David K. Jackson, Ian Johnston, Dominic Kwiatkowski, Cordelia Langford, John Sillitoe on behalf of the Wellcome Sanger Institute COVID-19 Surveillance Team |
| EPI_ISL_2997136 | Lighthouse Laboratory Plymouth | Wellcome Sanger Institute for the COVID-19 Genomics UK (COG-UK) Consortium | Lighthouse Laboratory Plymouth and Alex Alderton, Roberto Amato, Jeffrey Barrett, Sonia Goncalves, Ewan Harrison, David K. Jackson, Ian Johnston, Dominic Kwiatkowski, Cordelia Langford, John Sillitoe on behalf of the Wellcome Sanger Institute COVID-19 Surveillance Team |
| EPI_ISL_2997137 | Lighthouse Lab in Glasgow | Wellcome Sanger Institute for the COVID-19 Genomics UK (COG-UK) Consortium | Harper VanSteenhouse, Yumi Kasai, David Gray, Carol Clugston, Anna Dominiczak and Alex Alderton, Roberto Amato, Jeffrey Barrett, Sonia Goncalves, Ewan Harrison, David K. Jackson, Ian Johnston, Dominic Kwiatkowski, Cordelia Langford, John Sillitoe on behalf of the Wellcome Sanger Institute COVID-19 Surveillance Team |
| EPI_ISL_2997138 | Lighthouse Laboratory Plymouth | Wellcome Sanger Institute for the COVID-19 Genomics UK (COG-UK) Consortium | Lighthouse Laboratory Plymouth and Alex Alderton, Roberto Amato, Jeffrey Barrett, Sonia Goncalves, Ewan Harrison, David K. Jackson, Ian Johnston, Dominic Kwiatkowski, Cordelia Langford, John Sillitoe on behalf of the Wellcome Sanger Institute COVID-19 Surveillance Team |
| EPI_ISL_2997139 | Lighthouse Lab in Glasgow | Wellcome Sanger Institute for the COVID-19 Genomics UK (COG-UK) Consortium | Harper VanSteenhouse, Yumi Kasai, David Gray, Carol Clugston, Anna Dominiczak and Alex Alderton, Roberto Amato, Jeffrey Barrett, Sonia Goncalves, Ewan Harrison, David K. Jackson, Ian Johnston, Dominic Kwiatkowski, Cordelia Langford, John Sillitoe on behalf of the Wellcome Sanger Institute COVID-19 Surveillance Team |
| EPI_ISL_2997140 | Lighthouse Laboratory Plymouth | Wellcome Sanger Institute for the COVID-19 Genomics UK (COG-UK) Consortium | Lighthouse Laboratory Plymouth and Alex Alderton, Roberto Amato, Jeffrey Barrett, Sonia Goncalves, Ewan Harrison, David K. Jackson, Ian Johnston, Dominic Kwiatkowski, Cordelia Langford, John Sillitoe on behalf of the Wellcome Sanger Institute COVID-19 Surveillance Team |
| EPI_ISL_2997141 | Lighthouse Lab in Glasgow | Wellcome Sanger Institute for the COVID-19 Genomics UK (COG-UK) Consortium | Harper VanSteenhouse, Yumi Kasai, David Gray, Carol Clugston, Anna Dominiczak and Alex Alderton, Roberto Amato, Jeffrey Barrett, Sonia Goncalves, Ewan Harrison, David K. Jackson, Ian Johnston, Dominic Kwiatkowski, Cordelia Langford, John Sillitoe on behalf of the Wellcome Sanger Institute COVID-19 Surveillance Team |
| EPI_ISL_2997142 | Lighthouse Lab in Milton Keynes | Wellcome Sanger Institute for the COVID-19 Genomics UK (COG-UK) Consortium | The Lighthouse Lab in Milton Keynes and Alex Alderton, Roberto Amato, Jeffrey Barrett, Sonia Goncalves, Ewan Harrison, David K. Jackson, Ian Johnston, Dominic Kwiatkowski, Cordelia Langford, John Sillitoe on behalf of the Wellcome Sanger Institute COVID-19 Surveillance Team |
| EPI_ISL_2997143, EPI_ISL_2997144 | Berkshire and Surrey Pathology Services Lighthouse Laboratory | Wellcome Sanger Institute for the COVID-19 Genomics UK (COG-UK) Consortium | Berkshire and Surrey Pathology Services Lighthouse Laboratory and Alex Alderton, Roberto Amato, Jeffrey Barrett, Sonia Goncalves, Ewan Harrison, David K. Jackson, Ian Johnston, Dominic Kwiatkowski, Cordelia Langford, John Sillitoe on behalf of the Wellcome Sanger Institute COVID-19 Surveillance Team |

| | | | |
|---|---|---|---|
| EPI_ISL_2997145 | Lighthouse Lab in Glasgow | Wellcome Sanger Institute for the COVID-19 Genomics UK (COG-UK) Consortium | Harper VanSteenhouse, Yumi Kasai, David Gray, Carol Clugston, Anna Dominiczak and Alex Alderton, Roberto Amato, Jeffrey Barrett, Sonia Goncalves, Ewan Harrison, David K. Jackson, Ian Johnston, Dominic Kwiatkowski, Cordelia Langford, John Sillitoe on behalf of the Wellcome Sanger Institute COVID-19 Surveillance Team |
| EPI_ISL_2997146 | Lighthouse Laboratory Plymouth | Wellcome Sanger Institute for the COVID-19 Genomics UK (COG-UK) Consortium | Lighthouse Laboratory Plymouth and Alex Alderton, Roberto Amato, Jeffrey Barrett, Sonia Goncalves, Ewan Harrison, David K. Jackson, Ian Johnston, Dominic Kwiatkowski, Cordelia Langford, John Sillitoe on behalf of the Wellcome Sanger Institute COVID-19 Surveillance Team |
| EPI_ISL_2997147 | Berkshire and Surrey Pathology Services Lighthouse Laboratory | Wellcome Sanger Institute for the COVID-19 Genomics UK (COG-UK) Consortium | Berkshire and Surrey Pathology Services Lighthouse Laboratory and Alex Alderton, Roberto Amato, Jeffrey Barrett, Sonia Goncalves, Ewan Harrison, David K. Jackson, Ian Johnston, Dominic Kwiatkowski, Cordelia Langford, John Sillitoe on behalf of the Wellcome Sanger Institute COVID-19 Surveillance Team |
| EPI_ISL_2997148 | Lighthouse Lab in Glasgow | Wellcome Sanger Institute for the COVID-19 Genomics UK (COG-UK) Consortium | Harper VanSteenhouse, Yumi Kasai, David Gray, Carol Clugston, Anna Dominiczak and Alex Alderton, Roberto Amato, Jeffrey Barrett, Sonia Goncalves, Ewan Harrison, David K. Jackson, Ian Johnston, Dominic Kwiatkowski, Cordelia Langford, John Sillitoe on behalf of the Wellcome Sanger Institute COVID-19 Surveillance Team |
| EPI_ISL_2997149 | Lighthouse Lab in Milton Keynes | Wellcome Sanger Institute for the COVID-19 Genomics UK (COG-UK) Consortium | The Lighthouse Lab in Milton Keynes and Alex Alderton, Roberto Amato, Jeffrey Barrett, Sonia Goncalves, Ewan Harrison, David K. Jackson, Ian Johnston, Dominic Kwiatkowski, Cordelia Langford, John Sillitoe on behalf of the Wellcome Sanger Institute COVID-19 Surveillance Team |
| EPI_ISL_2997150 | Berkshire and Surrey Pathology Services Lighthouse Laboratory | Wellcome Sanger Institute for the COVID-19 Genomics UK (COG-UK) Consortium | Berkshire and Surrey Pathology Services Lighthouse Laboratory and Alex Alderton, Roberto Amato, Jeffrey Barrett, Sonia Goncalves, Ewan Harrison, David K. Jackson, Ian Johnston, Dominic Kwiatkowski, Cordelia Langford, John Sillitoe on behalf of the Wellcome Sanger Institute COVID-19 Surveillance Team |
| EPI_ISL_2997151 | Lighthouse Laboratory Plymouth | Wellcome Sanger Institute for the COVID-19 Genomics UK (COG-UK) Consortium | Lighthouse Laboratory Plymouth and Alex Alderton, Roberto Amato, Jeffrey Barrett, Sonia Goncalves, Ewan Harrison, David K. Jackson, Ian Johnston, Dominic Kwiatkowski, Cordelia Langford, John Sillitoe on behalf of the Wellcome Sanger Institute COVID-19 Surveillance Team |
| EPI_ISL_2997152, EPI_ISL_2997153 | Lighthouse Lab in Milton Keynes | Wellcome Sanger Institute for the COVID-19 Genomics UK (COG-UK) Consortium | The Lighthouse Lab in Milton Keynes and Alex Alderton, Roberto Amato, Jeffrey Barrett, Sonia Goncalves, Ewan Harrison, David K. Jackson, Ian Johnston, Dominic Kwiatkowski, Cordelia Langford, John Sillitoe on behalf of the Wellcome Sanger Institute COVID-19 Surveillance Team |
| EPI_ISL_2997154 | Lighthouse Lab in Glasgow | Wellcome Sanger Institute for the COVID-19 Genomics UK (COG-UK) Consortium | Harper VanSteenhouse, Yumi Kasai, David Gray, Carol Clugston, Anna Dominiczak and Alex Alderton, Roberto Amato, Jeffrey Barrett, Sonia Goncalves, Ewan Harrison, David K. Jackson, Ian Johnston, Dominic Kwiatkowski, Cordelia Langford, John Sillitoe on behalf of the Wellcome Sanger Institute COVID-19 Surveillance Team |
| EPI_ISL_2997155, EPI_ISL_2997156 | Lighthouse Lab in Milton Keynes | Wellcome Sanger Institute for the COVID-19 Genomics UK (COG-UK) Consortium | The Lighthouse Lab in Milton Keynes and Alex Alderton, Roberto Amato, Jeffrey Barrett, Sonia Goncalves, Ewan Harrison, David K. Jackson, Ian Johnston, Dominic Kwiatkowski, Cordelia Langford, John Sillitoe on behalf of the Wellcome Sanger Institute COVID-19 Surveillance Team |
| EPI_ISL_2997157 | Lighthouse Laboratory Plymouth | Wellcome Sanger Institute for the COVID-19 Genomics UK (COG-UK) Consortium | Lighthouse Laboratory Plymouth and Alex Alderton, Roberto Amato, Jeffrey Barrett, Sonia Goncalves, Ewan Harrison, David K. Jackson, Ian Johnston, Dominic Kwiatkowski, Cordelia Langford, John Sillitoe on behalf of the Wellcome Sanger Institute COVID-19 Surveillance Team |
| EPI_ISL_2997158, EPI_ISL_2997159 | Berkshire and Surrey Pathology Services Lighthouse Laboratory | Wellcome Sanger Institute for the COVID-19 Genomics UK (COG-UK) Consortium | Berkshire and Surrey Pathology Services Lighthouse Laboratory and Alex Alderton, Roberto Amato, Jeffrey Barrett, Sonia Goncalves, Ewan Harrison, David K. Jackson, Ian Johnston, Dominic Kwiatkowski, Cordelia Langford, John Sillitoe on behalf of the Wellcome Sanger Institute COVID-19 Surveillance Team |
| EPI_ISL_2997160, EPI_ISL_2997161, EPI_ISL_2997162, EPI_ISL_2997163 | Lighthouse Laboratory Plymouth | Wellcome Sanger Institute for the COVID-19 Genomics UK (COG-UK) Consortium | Lighthouse Laboratory Plymouth and Alex Alderton, Roberto Amato, Jeffrey Barrett, Sonia Goncalves, Ewan Harrison, David K. Jackson, Ian Johnston, Dominic Kwiatkowski, Cordelia Langford, John Sillitoe on behalf of the Wellcome Sanger Institute COVID-19 Surveillance Team |
| EPI_ISL_2997164 | Lighthouse Lab in Milton Keynes | Wellcome Sanger Institute for the COVID-19 Genomics UK (COG-UK) Consortium | The Lighthouse Lab in Milton Keynes and Alex Alderton, Roberto Amato, Jeffrey Barrett, Sonia Goncalves, Ewan Harrison, David K. Jackson, Ian Johnston, Dominic Kwiatkowski, Cordelia Langford, John Sillitoe on behalf of the Wellcome Sanger Institute COVID-19 Surveillance Team |
| EPI_ISL_2997165 | Berkshire and Surrey Pathology Services Lighthouse Laboratory | Wellcome Sanger Institute for the COVID-19 Genomics UK (COG-UK) Consortium | Berkshire and Surrey Pathology Services Lighthouse Laboratory and Alex Alderton, Roberto Amato, Jeffrey Barrett, Sonia Goncalves, Ewan Harrison, David K. Jackson, Ian Johnston, Dominic Kwiatkowski, Cordelia Langford, John Sillitoe on behalf of the Wellcome Sanger Institute COVID-19 Surveillance Team |
| EPI_ISL_2997166 | Lighthouse Lab in Milton Keynes | Wellcome Sanger Institute for the COVID-19 Genomics UK (COG-UK) Consortium | The Lighthouse Lab in Milton Keynes and Alex Alderton, Roberto Amato, Jeffrey Barrett, Sonia Goncalves, Ewan Harrison, David K. Jackson, Ian Johnston, Dominic Kwiatkowski, Cordelia Langford, John Sillitoe on behalf of the Wellcome Sanger Institute COVID-19 Surveillance Team |
| EPI_ISL_2997167, EPI_ISL_2997168, EPI_ISL_2997169, EPI_ISL_2997170 | Lighthouse Laboratory Plymouth | Wellcome Sanger Institute for the COVID-19 Genomics UK (COG-UK) Consortium | Lighthouse Laboratory Plymouth and Alex Alderton, Roberto Amato, Jeffrey Barrett, Sonia Goncalves, Ewan Harrison, David K. Jackson, Ian Johnston, Dominic Kwiatkowski, Cordelia Langford, John Sillitoe on behalf of the Wellcome Sanger Institute COVID-19 Surveillance Team |
| EPI_ISL_2997171 | Lighthouse Lab in Milton Keynes | Wellcome Sanger Institute for the COVID-19 Genomics UK (COG-UK) Consortium | The Lighthouse Lab in Milton Keynes and Alex Alderton, Roberto Amato, Jeffrey Barrett, Sonia Goncalves, Ewan Harrison, David K. Jackson, Ian Johnston, Dominic Kwiatkowski, Cordelia Langford, John Sillitoe on behalf of the Wellcome Sanger Institute COVID-19 Surveillance Team |
| EPI_ISL_2997172, EPI_ISL_2997173 | Lighthouse Laboratory Plymouth | Wellcome Sanger Institute for the COVID-19 Genomics UK (COG-UK) Consortium | Lighthouse Laboratory Plymouth and Alex Alderton, Roberto Amato, Jeffrey Barrett, Sonia Goncalves, Ewan Harrison, David K. Jackson, Ian Johnston, Dominic Kwiatkowski, Cordelia Langford, John Sillitoe on behalf of the Wellcome Sanger Institute COVID-19 Surveillance Team |
| EPI_ISL_2997174, EPI_ISL_2997175 | Berkshire and Surrey Pathology Services Lighthouse Laboratory | Wellcome Sanger Institute for the COVID-19 Genomics UK (COG-UK) Consortium | Berkshire and Surrey Pathology Services Lighthouse Laboratory and Alex Alderton, Roberto Amato, Jeffrey Barrett, Sonia Goncalves, Ewan Harrison, David K. Jackson, Ian Johnston, Dominic Kwiatkowski, Cordelia Langford, John Sillitoe on behalf of the Wellcome Sanger Institute COVID-19 Surveillance Team |
| EPI_ISL_2997176 | Lighthouse Laboratory Plymouth | Wellcome Sanger Institute for the COVID-19 Genomics UK (COG-UK) Consortium | Lighthouse Laboratory Plymouth and Alex Alderton, Roberto Amato, Jeffrey Barrett, Sonia Goncalves, Ewan Harrison, David K. Jackson, Ian Johnston, Dominic Kwiatkowski, Cordelia Langford, John Sillitoe on behalf of the Wellcome Sanger Institute COVID-19 Surveillance Team |
| EPI_ISL_2997177, EPI_ISL_2997178 | Berkshire and Surrey Pathology Services Lighthouse Laboratory | Wellcome Sanger Institute for the COVID-19 Genomics UK (COG-UK) Consortium | Berkshire and Surrey Pathology Services Lighthouse Laboratory and Alex Alderton, Roberto Amato, Jeffrey Barrett, Sonia Goncalves, Ewan Harrison, David K. Jackson, Ian Johnston, Dominic Kwiatkowski, Cordelia Langford, John Sillitoe on behalf of the Wellcome Sanger Institute COVID-19 Surveillance Team |
| EPI_ISL_2997179 | Lighthouse Laboratory Plymouth | Wellcome Sanger Institute for the COVID-19 Genomics UK (COG-UK) Consortium | Lighthouse Laboratory Plymouth and Alex Alderton, Roberto Amato, Jeffrey Barrett, Sonia Goncalves, Ewan Harrison, David K. Jackson, Ian Johnston, Dominic Kwiatkowski, Cordelia Langford, John Sillitoe on behalf of the Wellcome Sanger Institute COVID-19 Surveillance Team |
| EPI_ISL_2997180 | Berkshire and Surrey Pathology Services Lighthouse Laboratory | Wellcome Sanger Institute for the COVID-19 Genomics UK (COG-UK) Consortium | Berkshire and Surrey Pathology Services Lighthouse Laboratory and Alex Alderton, Roberto Amato, Jeffrey Barrett, Sonia Goncalves, Ewan Harrison, David K. Jackson, Ian Johnston, Dominic Kwiatkowski, Cordelia Langford, John Sillitoe on behalf of the Wellcome Sanger Institute COVID-19 Surveillance Team |
| EPI_ISL_2997181 | Lighthouse Lab in Milton Keynes | Wellcome Sanger Institute for the COVID-19 Genomics UK (COG-UK) Consortium | The Lighthouse Lab in Milton Keynes and Alex Alderton, Roberto Amato, Jeffrey Barrett, Sonia Goncalves, Ewan Harrison, David K. Jackson, Ian Johnston, Dominic Kwiatkowski, Cordelia Langford, John Sillitoe on behalf of the Wellcome Sanger Institute COVID-19 Surveillance Team |
| EPI_ISL_2997182, EPI_ISL_2997183, EPI_ISL_2997184 | Lighthouse Lab in Glasgow | Wellcome Sanger Institute for the COVID-19 Genomics UK (COG-UK) Consortium | Harper VanSteenhouse, Yumi Kasai, David Gray, Carol Clugston, Anna Dominiczak and Alex Alderton, Roberto Amato, Jeffrey Barrett, Sonia Goncalves, Ewan Harrison, David K. Jackson, Ian Johnston, Dominic Kwiatkowski, Cordelia Langford, John Sillitoe on behalf of the Wellcome Sanger Institute COVID-19 Surveillance Team |
| EPI_ISL_2997185 | Lighthouse Lab in Milton Keynes | Wellcome Sanger Institute for the COVID-19 Genomics UK (COG-UK) Consortium | The Lighthouse Lab in Milton Keynes and Alex Alderton, Roberto Amato, Jeffrey Barrett, Sonia Goncalves, Ewan Harrison, David K. Jackson, Ian Johnston, Dominic Kwiatkowski, Cordelia Langford, John Sillitoe on behalf of the Wellcome Sanger Institute COVID-19 Surveillance Team |
| EPI_ISL_2997186 | Lighthouse Laboratory Plymouth | Wellcome Sanger Institute for the COVID-19 Genomics UK (COG-UK) Consortium | Lighthouse Laboratory Plymouth and Alex Alderton, Roberto Amato, Jeffrey Barrett, Sonia Goncalves, Ewan Harrison, David K. Jackson, Ian Johnston, Dominic Kwiatkowski, Cordelia Langford, John Sillitoe on behalf of the Wellcome Sanger Institute COVID-19 Surveillance Team |
| EPI_ISL_2997187 | Berkshire and Surrey Pathology Services Lighthouse Laboratory | Wellcome Sanger Institute for the COVID-19 Genomics UK (COG-UK) Consortium | Berkshire and Surrey Pathology Services Lighthouse Laboratory and Alex Alderton, Roberto Amato, Jeffrey Barrett, Sonia Goncalves, Ewan Harrison, David K. Jackson, Ian Johnston, Dominic Kwiatkowski, Cordelia Langford, John Sillitoe on behalf of the Wellcome Sanger Institute COVID-19 Surveillance Team |
| EPI_ISL_2997188 | Lighthouse Laboratory Plymouth | Wellcome Sanger Institute for the COVID-19 Genomics UK (COG-UK) Consortium | Lighthouse Laboratory Plymouth and Alex Alderton, Roberto Amato, Jeffrey Barrett, Sonia Goncalves, Ewan Harrison, David K. Jackson, Ian Johnston, Dominic Kwiatkowski, Cordelia Langford, John Sillitoe on behalf of the Wellcome Sanger Institute COVID-19 Surveillance Team |
| EPI_ISL_2997189 | Berkshire and Surrey Pathology Services Lighthouse Laboratory | Wellcome Sanger Institute for the COVID-19 Genomics UK (COG-UK) Consortium | Berkshire and Surrey Pathology Services Lighthouse Laboratory and Alex Alderton, Roberto Amato, Jeffrey Barrett, Sonia Goncalves, Ewan Harrison, David K. Jackson, Ian Johnston, Dominic Kwiatkowski, Cordelia Langford, John Sillitoe on behalf of the Wellcome Sanger Institute COVID-19 Surveillance Team |
| EPI_ISL_2997190, EPI_ISL_2997191 | Lighthouse Lab in Glasgow | Wellcome Sanger Institute for the COVID-19 Genomics UK (COG-UK) Consortium | Harper VanSteenhouse, Yumi Kasai, David Gray, Carol Clugston, Anna Dominiczak and Alex Alderton, Roberto Amato, Jeffrey Barrett, Sonia Goncalves, Ewan Harrison, David K. Jackson, Ian Johnston, Dominic Kwiatkowski, Cordelia Langford, John Sillitoe on behalf of the Wellcome Sanger Institute COVID-19 Surveillance Team |
| EPI_ISL_2997192 | Berkshire and Surrey Pathology Services Lighthouse Laboratory | Wellcome Sanger Institute for the COVID-19 Genomics UK (COG-UK) Consortium | Berkshire and Surrey Pathology Services Lighthouse Laboratory and Alex Alderton, Roberto Amato, Jeffrey Barrett, Sonia Goncalves, Ewan Harrison, David K. Jackson, Ian Johnston, Dominic Kwiatkowski, Cordelia Langford, John Sillitoe on behalf of the Wellcome Sanger Institute COVID-19 Surveillance Team |
| EPI_ISL_2997193, EPI_ISL_2997194 | Lighthouse Lab in Glasgow | Wellcome Sanger Institute for the COVID-19 Genomics UK (COG-UK) Consortium | Harper VanSteenhouse, Yumi Kasai, David Gray, Carol Clugston, Anna Dominiczak and Alex Alderton, Roberto Amato, Jeffrey Barrett, Sonia Goncalves, Ewan Harrison, David K. Jackson, Ian Johnston, Dominic Kwiatkowski, Cordelia Langford, John Sillitoe on behalf of the Wellcome Sanger Institute COVID-19 Surveillance Team |
| EPI_ISL_2997195 | Lighthouse Lab in Milton Keynes | Wellcome Sanger Institute for the COVID-19 Genomics UK (COG-UK) Consortium | The Lighthouse Lab in Milton Keynes and Alex Alderton, Roberto Amato, Jeffrey Barrett, Sonia Goncalves, Ewan Harrison, David K. Jackson, Ian Johnston, Dominic Kwiatkowski, Cordelia Langford, John Sillitoe on behalf of the Wellcome Sanger Institute COVID-19 Surveillance Team |
| EPI_ISL_2997196, EPI_ISL_2997197 | Lighthouse Laboratory Plymouth | Wellcome Sanger Institute for the COVID-19 Genomics UK (COG-UK) Consortium | Lighthouse Laboratory Plymouth and Alex Alderton, Roberto Amato, Jeffrey Barrett, Sonia Goncalves, Ewan Harrison, David K. Jackson, Ian Johnston, Dominic Kwiatkowski, Cordelia Langford, John Sillitoe on behalf of the Wellcome Sanger Institute COVID-19 Surveillance Team |
| EPI_ISL_2997198 | Berkshire and Surrey Pathology Services Lighthouse Laboratory | Wellcome Sanger Institute for the COVID-19 Genomics UK (COG-UK) Consortium | Berkshire and Surrey Pathology Services Lighthouse Laboratory and Alex Alderton, Roberto Amato, Jeffrey Barrett, Sonia Goncalves, Ewan Harrison, David K. Jackson, Ian Johnston, Dominic Kwiatkowski, Cordelia Langford, John Sillitoe on behalf of the Wellcome Sanger Institute COVID-19 Surveillance Team |
| EPI_ISL_2997199 | Lighthouse Lab in Milton Keynes | Wellcome Sanger Institute for the COVID-19 Genomics UK (COG-UK) Consortium | The Lighthouse Lab in Milton Keynes and Alex Alderton, Roberto Amato, Jeffrey Barrett, Sonia Goncalves, Ewan Harrison, David K. Jackson, Ian Johnston, Dominic Kwiatkowski, Cordelia Langford, John Sillitoe on behalf of the Wellcome Sanger Institute COVID-19 Surveillance Team |
| EPI_ISL_2997200, EPI_ISL_2997201 | Lighthouse Lab in Glasgow | Wellcome Sanger Institute for the COVID-19 Genomics UK (COG-UK) Consortium | Harper VanSteenhouse, Yumi Kasai, David Gray, Carol Clugston, Anna Dominiczak and Alex Alderton, Roberto Amato, Jeffrey Barrett, Sonia Goncalves, Ewan Harrison, David K. Jackson, Ian Johnston, Dominic Kwiatkowski, Cordelia Langford, John Sillitoe on behalf of the Wellcome Sanger Institute COVID-19 Surveillance Team |
| EPI_ISL_2997202 | Lighthouse Lab in Milton Keynes | Wellcome Sanger Institute for the COVID-19 Genomics UK (COG-UK) Consortium | The Lighthouse Lab in Milton Keynes and Alex Alderton, Roberto Amato, Jeffrey Barrett, Sonia Goncalves, Ewan Harrison, David K. Jackson, Ian Johnston, Dominic Kwiatkowski, Cordelia Langford, John Sillitoe on behalf of the Wellcome Sanger Institute COVID-19 Surveillance Team |
| EPI_ISL_2997203 | Berkshire and Surrey Pathology Services Lighthouse Laboratory | Wellcome Sanger Institute for the COVID-19 Genomics UK (COG-UK) Consortium | Berkshire and Surrey Pathology Services Lighthouse Laboratory and Alex Alderton, Roberto Amato, Jeffrey Barrett, Sonia Goncalves, Ewan Harrison, David K. Jackson, Ian Johnston, Dominic Kwiatkowski, Cordelia Langford, John Sillitoe on behalf of the Wellcome Sanger Institute COVID-19 Surveillance Team |
| EPI_ISL_2997204, EPI_ISL_2997205 | Lighthouse Laboratory Plymouth | Wellcome Sanger Institute for the COVID-19 Genomics UK (COG-UK) Consortium | Lighthouse Laboratory Plymouth and Alex Alderton, Roberto Amato, Jeffrey Barrett, Sonia Goncalves, Ewan Harrison, David K. Jackson, Ian Johnston, Dominic Kwiatkowski, Cordelia Langford, John Sillitoe on behalf of the Wellcome Sanger Institute COVID-19 Surveillance Team |
| EPI_ISL_2997206 | Lighthouse Lab in Glasgow | Wellcome Sanger Institute for the COVID-19 Genomics UK (COG-UK) Consortium | Harper VanSteenhouse, Yumi Kasai, David Gray, Carol Clugston, Anna Dominiczak and Alex Alderton, Roberto Amato, Jeffrey Barrett, Sonia Goncalves, Ewan Harrison, David K. Jackson, Ian Johnston, Dominic Kwiatkowski, Cordelia Langford, John Sillitoe on behalf of the Wellcome Sanger Institute COVID-19 Surveillance Team |

| | | | |
|---|---|---|---|
| EPI_ISL_2997207 | Lighthouse Laboratory Plymouth | Wellcome Sanger Institute for the COVID-19 Genomics UK (COG-UK) Consortium | Lighthouse Laboratory Plymouth and Alex Alderton, Roberto Amato, Jeffrey Barrett, Sonia Goncalves, Ewan Harrison, David K. Jackson, Ian Johnston, Dominic Kwiatkowski, Cordelia Langford, John Sillitoe on behalf of the Wellcome Sanger Institute COVID-19 Surveillance Team |
| EPI_ISL_2997208 | Lighthouse Lab in Milton Keynes | Wellcome Sanger Institute for the COVID-19 Genomics UK (COG-UK) Consortium | The Lighthouse Lab in Milton Keynes and Alex Alderton, Roberto Amato, Jeffrey Barrett, Sonia Goncalves, Ewan Harrison, David K. Jackson, Ian Johnston, Dominic Kwiatkowski, Cordelia Langford, John Sillitoe on behalf of the Wellcome Sanger Institute COVID-19 Surveillance Team |
| EPI_ISL_2997209 | Berkshire and Surrey Pathology Services Lighthouse Laboratory | Wellcome Sanger Institute for the COVID-19 Genomics UK (COG-UK) Consortium | Berkshire and Surrey Pathology Services Lighthouse Laboratory and Alex Alderton, Roberto Amato, Jeffrey Barrett, Sonia Goncalves, Ewan Harrison, David K. Jackson, Ian Johnston, Dominic Kwiatkowski, Cordelia Langford, John Sillitoe on behalf of the Wellcome Sanger Institute COVID-19 Surveillance Team |
| EPI_ISL_2997210 | Lighthouse Lab in Milton Keynes | Wellcome Sanger Institute for the COVID-19 Genomics UK (COG-UK) Consortium | The Lighthouse Lab in Milton Keynes and Alex Alderton, Roberto Amato, Jeffrey Barrett, Sonia Goncalves, Ewan Harrison, David K. Jackson, Ian Johnston, Dominic Kwiatkowski, Cordelia Langford, John Sillitoe on behalf of the Wellcome Sanger Institute COVID-19 Surveillance Team |
| EPI_ISL_2997211 | Berkshire and Surrey Pathology Services Lighthouse Laboratory | Wellcome Sanger Institute for the COVID-19 Genomics UK (COG-UK) Consortium | Berkshire and Surrey Pathology Services Lighthouse Laboratory and Alex Alderton, Roberto Amato, Jeffrey Barrett, Sonia Goncalves, Ewan Harrison, David K. Jackson, Ian Johnston, Dominic Kwiatkowski, Cordelia Langford, John Sillitoe on behalf of the Wellcome Sanger Institute COVID-19 Surveillance Team |
| EPI_ISL_2997212 | Lighthouse Laboratory Plymouth | Wellcome Sanger Institute for the COVID-19 Genomics UK (COG-UK) Consortium | Lighthouse Laboratory Plymouth and Alex Alderton, Roberto Amato, Jeffrey Barrett, Sonia Goncalves, Ewan Harrison, David K. Jackson, Ian Johnston, Dominic Kwiatkowski, Cordelia Langford, John Sillitoe on behalf of the Wellcome Sanger Institute COVID-19 Surveillance Team |
| EPI_ISL_2997213, EPI_ISL_2997214 | Berkshire and Surrey Pathology Services Lighthouse Laboratory | Wellcome Sanger Institute for the COVID-19 Genomics UK (COG-UK) Consortium | Berkshire and Surrey Pathology Services Lighthouse Laboratory and Alex Alderton, Roberto Amato, Jeffrey Barrett, Sonia Goncalves, Ewan Harrison, David K. Jackson, Ian Johnston, Dominic Kwiatkowski, Cordelia Langford, John Sillitoe on behalf of the Wellcome Sanger Institute COVID-19 Surveillance Team |
| EPI_ISL_2997215 | Lighthouse Lab in Milton Keynes | Wellcome Sanger Institute for the COVID-19 Genomics UK (COG-UK) Consortium | The Lighthouse Lab in Milton Keynes and Alex Alderton, Roberto Amato, Jeffrey Barrett, Sonia Goncalves, Ewan Harrison, David K. Jackson, Ian Johnston, Dominic Kwiatkowski, Cordelia Langford, John Sillitoe on behalf of the Wellcome Sanger Institute COVID-19 Surveillance Team |
| EPI_ISL_2997216 | Berkshire and Surrey Pathology Services Lighthouse Laboratory | Wellcome Sanger Institute for the COVID-19 Genomics UK (COG-UK) Consortium | Berkshire and Surrey Pathology Services Lighthouse Laboratory and Alex Alderton, Roberto Amato, Jeffrey Barrett, Sonia Goncalves, Ewan Harrison, David K. Jackson, Ian Johnston, Dominic Kwiatkowski, Cordelia Langford, John Sillitoe on behalf of the Wellcome Sanger Institute COVID-19 Surveillance Team |
| EPI_ISL_2997217 | Lighthouse Lab in Glasgow | Wellcome Sanger Institute for the COVID-19 Genomics UK (COG-UK) Consortium | Harper VanSteenhouse, Yumi Kasai, David Gray, Carol Clugston, Anna Dominiczak and Alex Alderton, Roberto Amato, Jeffrey Barrett, Sonia Goncalves, Ewan Harrison, David K. Jackson, Ian Johnston, Dominic Kwiatkowski, Cordelia Langford, John Sillitoe on behalf of the Wellcome Sanger Institute COVID-19 Surveillance Team |
| EPI_ISL_2997218 | Lighthouse Laboratory Plymouth | Wellcome Sanger Institute for the COVID-19 Genomics UK (COG-UK) Consortium | Lighthouse Laboratory Plymouth and Alex Alderton, Roberto Amato, Jeffrey Barrett, Sonia Goncalves, Ewan Harrison, David K. Jackson, Ian Johnston, Dominic Kwiatkowski, Cordelia Langford, John Sillitoe on behalf of the Wellcome Sanger Institute COVID-19 Surveillance Team |
| EPI_ISL_2997219, EPI_ISL_2997220 | Lighthouse Lab in Milton Keynes | Wellcome Sanger Institute for the COVID-19 Genomics UK (COG-UK) Consortium | The Lighthouse Lab in Milton Keynes and Alex Alderton, Roberto Amato, Jeffrey Barrett, Sonia Goncalves, Ewan Harrison, David K. Jackson, Ian Johnston, Dominic Kwiatkowski, Cordelia Langford, John Sillitoe on behalf of the Wellcome Sanger Institute COVID-19 Surveillance Team |
| EPI_ISL_2997221 | Lighthouse Lab in Glasgow | Wellcome Sanger Institute for the COVID-19 Genomics UK (COG-UK) Consortium | Harper VanSteenhouse, Yumi Kasai, David Gray, Carol Clugston, Anna Dominiczak and Alex Alderton, Roberto Amato, Jeffrey Barrett, Sonia Goncalves, Ewan Harrison, David K. Jackson, Ian Johnston, Dominic Kwiatkowski, Cordelia Langford, John Sillitoe on behalf of the Wellcome Sanger Institute COVID-19 Surveillance Team |
| EPI_ISL_2997222, EPI_ISL_2997223 | Lighthouse Laboratory Plymouth | Wellcome Sanger Institute for the COVID-19 Genomics UK (COG-UK) Consortium | Lighthouse Laboratory Plymouth and Alex Alderton, Roberto Amato, Jeffrey Barrett, Sonia Goncalves, Ewan Harrison, David K. Jackson, Ian Johnston, Dominic Kwiatkowski, Cordelia Langford, John Sillitoe on behalf of the Wellcome Sanger Institute COVID-19 Surveillance Team |
| EPI_ISL_2997224 | Berkshire and Surrey Pathology Services Lighthouse Laboratory | Wellcome Sanger Institute for the COVID-19 Genomics UK (COG-UK) Consortium | Berkshire and Surrey Pathology Services Lighthouse Laboratory and Alex Alderton, Roberto Amato, Jeffrey Barrett, Sonia Goncalves, Ewan Harrison, David K. Jackson, Ian Johnston, Dominic Kwiatkowski, Cordelia Langford, John Sillitoe on behalf of the Wellcome Sanger Institute COVID-19 Surveillance Team |
| EPI_ISL_2997225 | Lighthouse Lab in Glasgow | Wellcome Sanger Institute for the COVID-19 Genomics UK (COG-UK) Consortium | Harper VanSteenhouse, Yumi Kasai, David Gray, Carol Clugston, Anna Dominiczak and Alex Alderton, Roberto Amato, Jeffrey Barrett, Sonia Goncalves, Ewan Harrison, David K. Jackson, Ian Johnston, Dominic Kwiatkowski, Cordelia Langford, John Sillitoe on behalf of the Wellcome Sanger Institute COVID-19 Surveillance Team |
| EPI_ISL_2997226 | Lighthouse Lab in Milton Keynes | Wellcome Sanger Institute for the COVID-19 Genomics UK (COG-UK) Consortium | The Lighthouse Lab in Milton Keynes and Alex Alderton, Roberto Amato, Jeffrey Barrett, Sonia Goncalves, Ewan Harrison, David K. Jackson, Ian Johnston, Dominic Kwiatkowski, Cordelia Langford, John Sillitoe on behalf of the Wellcome Sanger Institute COVID-19 Surveillance Team |
| EPI_ISL_2997227 | Lighthouse Laboratory Plymouth | Wellcome Sanger Institute for the COVID-19 Genomics UK (COG-UK) Consortium | Lighthouse Laboratory Plymouth and Alex Alderton, Roberto Amato, Jeffrey Barrett, Sonia Goncalves, Ewan Harrison, David K. Jackson, Ian Johnston, Dominic Kwiatkowski, Cordelia Langford, John Sillitoe on behalf of the Wellcome Sanger Institute COVID-19 Surveillance Team |
| EPI_ISL_2997228 | Lighthouse Lab in Glasgow | Wellcome Sanger Institute for the COVID-19 Genomics UK (COG-UK) Consortium | Harper VanSteenhouse, Yumi Kasai, David Gray, Carol Clugston, Anna Dominiczak and Alex Alderton, Roberto Amato, Jeffrey Barrett, Sonia Goncalves, Ewan Harrison, David K. Jackson, Ian Johnston, Dominic Kwiatkowski, Cordelia Langford, John Sillitoe on behalf of the Wellcome Sanger Institute COVID-19 Surveillance Team |
| EPI_ISL_2997229, EPI_ISL_2997230 | Lighthouse Laboratory Plymouth | Wellcome Sanger Institute for the COVID-19 Genomics UK (COG-UK) Consortium | Lighthouse Laboratory Plymouth and Alex Alderton, Roberto Amato, Jeffrey Barrett, Sonia Goncalves, Ewan Harrison, David K. Jackson, Ian Johnston, Dominic Kwiatkowski, Cordelia Langford, John Sillitoe on behalf of the Wellcome Sanger Institute COVID-19 Surveillance Team |
| EPI_ISL_2997231 | Lighthouse Lab in Milton Keynes | Wellcome Sanger Institute for the COVID-19 Genomics UK (COG-UK) Consortium | The Lighthouse Lab in Milton Keynes and Alex Alderton, Roberto Amato, Jeffrey Barrett, Sonia Goncalves, Ewan Harrison, David K. Jackson, Ian Johnston, Dominic Kwiatkowski, Cordelia Langford, John Sillitoe on behalf of the Wellcome Sanger Institute COVID-19 Surveillance Team |
| EPI_ISL_2997232 | Lighthouse Laboratory Plymouth | Wellcome Sanger Institute for the COVID-19 Genomics UK (COG-UK) Consortium | Lighthouse Laboratory Plymouth and Alex Alderton, Roberto Amato, Jeffrey Barrett, Sonia Goncalves, Ewan Harrison, David K. Jackson, Ian Johnston, Dominic Kwiatkowski, Cordelia Langford, John Sillitoe on behalf of the Wellcome Sanger Institute COVID-19 Surveillance Team |
| EPI_ISL_2997233 | Berkshire and Surrey Pathology Services Lighthouse Laboratory | Wellcome Sanger Institute for the COVID-19 Genomics UK (COG-UK) Consortium | Berkshire and Surrey Pathology Services Lighthouse Laboratory and Alex Alderton, Roberto Amato, Jeffrey Barrett, Sonia Goncalves, Ewan Harrison, David K. Jackson, Ian Johnston, Dominic Kwiatkowski, Cordelia Langford, John Sillitoe on behalf of the Wellcome Sanger Institute COVID-19 Surveillance Team |
| EPI_ISL_2997234, EPI_ISL_2997235, EPI_ISL_2997236 | Lighthouse Lab in Glasgow | Wellcome Sanger Institute for the COVID-19 Genomics UK (COG-UK) Consortium | Harper VanSteenhouse, Yumi Kasai, David Gray, Carol Clugston, Anna Dominiczak and Alex Alderton, Roberto Amato, Jeffrey Barrett, Sonia Goncalves, Ewan Harrison, David K. Jackson, Ian Johnston, Dominic Kwiatkowski, Cordelia Langford, John Sillitoe on behalf of the Wellcome Sanger Institute COVID-19 Surveillance Team |
| EPI_ISL_2997237 | Lighthouse Laboratory Plymouth | Wellcome Sanger Institute for the COVID-19 Genomics UK (COG-UK) Consortium | Lighthouse Laboratory Plymouth and Alex Alderton, Roberto Amato, Jeffrey Barrett, Sonia Goncalves, Ewan Harrison, David K. Jackson, Ian Johnston, Dominic Kwiatkowski, Cordelia Langford, John Sillitoe on behalf of the Wellcome Sanger Institute COVID-19 Surveillance Team |
| EPI_ISL_2997238 | Lighthouse Lab in Glasgow | Wellcome Sanger Institute for the COVID-19 Genomics UK (COG-UK) Consortium | Harper VanSteenhouse, Yumi Kasai, David Gray, Carol Clugston, Anna Dominiczak and Alex Alderton, Roberto Amato, Jeffrey Barrett, Sonia Goncalves, Ewan Harrison, David K. Jackson, Ian Johnston, Dominic Kwiatkowski, Cordelia Langford, John Sillitoe on behalf of the Wellcome Sanger Institute COVID-19 Surveillance Team |
| EPI_ISL_2997239 | Lighthouse Lab in Milton Keynes | Wellcome Sanger Institute for the COVID-19 Genomics UK (COG-UK) Consortium | The Lighthouse Lab in Milton Keynes and Alex Alderton, Roberto Amato, Jeffrey Barrett, Sonia Goncalves, Ewan Harrison, David K. Jackson, Ian Johnston, Dominic Kwiatkowski, Cordelia Langford, John Sillitoe on behalf of the Wellcome Sanger Institute COVID-19 Surveillance Team |
| EPI_ISL_2997240, EPI_ISL_2997241 | Lighthouse Laboratory Plymouth | Wellcome Sanger Institute for the COVID-19 Genomics UK (COG-UK) Consortium | Lighthouse Laboratory Plymouth and Alex Alderton, Roberto Amato, Jeffrey Barrett, Sonia Goncalves, Ewan Harrison, David K. Jackson, Ian Johnston, Dominic Kwiatkowski, Cordelia Langford, John Sillitoe on behalf of the Wellcome Sanger Institute COVID-19 Surveillance Team |
| EPI_ISL_2997242 | Lighthouse Lab in Glasgow | Wellcome Sanger Institute for the COVID-19 Genomics UK (COG-UK) Consortium | Harper VanSteenhouse, Yumi Kasai, David Gray, Carol Clugston, Anna Dominiczak and Alex Alderton, Roberto Amato, Jeffrey Barrett, Sonia Goncalves, Ewan Harrison, David K. Jackson, Ian Johnston, Dominic Kwiatkowski, Cordelia Langford, John Sillitoe on behalf of the Wellcome Sanger Institute COVID-19 Surveillance Team |
| EPI_ISL_2997243 | Berkshire and Surrey Pathology Services Lighthouse Laboratory | Wellcome Sanger Institute for the COVID-19 Genomics UK (COG-UK) Consortium | Berkshire and Surrey Pathology Services Lighthouse Laboratory and Alex Alderton, Roberto Amato, Jeffrey Barrett, Sonia Goncalves, Ewan Harrison, David K. Jackson, Ian Johnston, Dominic Kwiatkowski, Cordelia Langford, John Sillitoe on behalf of the Wellcome Sanger Institute COVID-19 Surveillance Team |
| EPI_ISL_2997244 | Lighthouse Lab in Milton Keynes | Wellcome Sanger Institute for the COVID-19 Genomics UK (COG-UK) Consortium | The Lighthouse Lab in Milton Keynes and Alex Alderton, Roberto Amato, Jeffrey Barrett, Sonia Goncalves, Ewan Harrison, David K. Jackson, Ian Johnston, Dominic Kwiatkowski, Cordelia Langford, John Sillitoe on behalf of the Wellcome Sanger Institute COVID-19 Surveillance Team |
| EPI_ISL_2997245 | Berkshire and Surrey Pathology Services Lighthouse Laboratory | Wellcome Sanger Institute for the COVID-19 Genomics UK (COG-UK) Consortium | Berkshire and Surrey Pathology Services Lighthouse Laboratory and Alex Alderton, Roberto Amato, Jeffrey Barrett, Sonia Goncalves, Ewan Harrison, David K. Jackson, Ian Johnston, Dominic Kwiatkowski, Cordelia Langford, John Sillitoe on behalf of the Wellcome Sanger Institute COVID-19 Surveillance Team |
| EPI_ISL_2997246 | Lighthouse Laboratory Plymouth | Wellcome Sanger Institute for the COVID-19 Genomics UK (COG-UK) Consortium | Lighthouse Laboratory Plymouth and Alex Alderton, Roberto Amato, Jeffrey Barrett, Sonia Goncalves, Ewan Harrison, David K. Jackson, Ian Johnston, Dominic Kwiatkowski, Cordelia Langford, John Sillitoe on behalf of the Wellcome Sanger Institute COVID-19 Surveillance Team |
| EPI_ISL_2997247 | Lighthouse Lab in Milton Keynes | Wellcome Sanger Institute for the COVID-19 Genomics UK (COG-UK) Consortium | The Lighthouse Lab in Milton Keynes and Alex Alderton, Roberto Amato, Jeffrey Barrett, Sonia Goncalves, Ewan Harrison, David K. Jackson, Ian Johnston, Dominic Kwiatkowski, Cordelia Langford, John Sillitoe on behalf of the Wellcome Sanger Institute COVID-19 Surveillance Team |
| EPI_ISL_2997248 | Lighthouse Lab in Glasgow | Wellcome Sanger Institute for the COVID-19 Genomics UK (COG-UK) Consortium | Harper VanSteenhouse, Yumi Kasai, David Gray, Carol Clugston, Anna Dominiczak and Alex Alderton, Roberto Amato, Jeffrey Barrett, Sonia Goncalves, Ewan Harrison, David K. Jackson, Ian Johnston, Dominic Kwiatkowski, Cordelia Langford, John Sillitoe on behalf of the Wellcome Sanger Institute COVID-19 Surveillance Team |
| EPI_ISL_2997249 | Berkshire and Surrey Pathology Services Lighthouse Laboratory | Wellcome Sanger Institute for the COVID-19 Genomics UK (COG-UK) Consortium | Berkshire and Surrey Pathology Services Lighthouse Laboratory and Alex Alderton, Roberto Amato, Jeffrey Barrett, Sonia Goncalves, Ewan Harrison, David K. Jackson, Ian Johnston, Dominic Kwiatkowski, Cordelia Langford, John Sillitoe on behalf of the Wellcome Sanger Institute COVID-19 Surveillance Team |
| EPI_ISL_2997250 | Lighthouse Lab in Milton Keynes | Wellcome Sanger Institute for the COVID-19 Genomics UK (COG-UK) Consortium | The Lighthouse Lab in Milton Keynes and Alex Alderton, Roberto Amato, Jeffrey Barrett, Sonia Goncalves, Ewan Harrison, David K. Jackson, Ian Johnston, Dominic Kwiatkowski, Cordelia Langford, John Sillitoe on behalf of the Wellcome Sanger Institute COVID-19 Surveillance Team |
| EPI_ISL_2997251, EPI_ISL_2997252 | Berkshire and Surrey Pathology Services Lighthouse Laboratory | Wellcome Sanger Institute for the COVID-19 Genomics UK (COG-UK) Consortium | Berkshire and Surrey Pathology Services Lighthouse Laboratory and Alex Alderton, Roberto Amato, Jeffrey Barrett, Sonia Goncalves, Ewan Harrison, David K. Jackson, Ian Johnston, Dominic Kwiatkowski, Cordelia Langford, John Sillitoe on behalf of the Wellcome Sanger Institute COVID-19 Surveillance Team |
| EPI_ISL_2997253, EPI_ISL_2997254 | Lighthouse Lab in Glasgow | Wellcome Sanger Institute for the COVID-19 Genomics UK (COG-UK) Consortium | Harper VanSteenhouse, Yumi Kasai, David Gray, Carol Clugston, Anna Dominiczak and Alex Alderton, Roberto Amato, Jeffrey Barrett, Sonia Goncalves, Ewan Harrison, David K. Jackson, Ian Johnston, Dominic Kwiatkowski, Cordelia Langford, John Sillitoe on behalf of the Wellcome Sanger Institute COVID-19 Surveillance Team |
| EPI_ISL_2997255 | Berkshire and Surrey Pathology Services Lighthouse Laboratory | Wellcome Sanger Institute for the COVID-19 Genomics UK (COG-UK) Consortium | Berkshire and Surrey Pathology Services Lighthouse Laboratory and Alex Alderton, Roberto Amato, Jeffrey Barrett, Sonia Goncalves, Ewan Harrison, David K. Jackson, Ian Johnston, Dominic Kwiatkowski, Cordelia Langford, John Sillitoe on behalf of the Wellcome Sanger Institute COVID-19 Surveillance Team |
| EPI_ISL_2997256 | Lighthouse Laboratory Plymouth | Wellcome Sanger Institute for the COVID-19 Genomics UK (COG-UK) Consortium | Lighthouse Laboratory Plymouth and Alex Alderton, Roberto Amato, Jeffrey Barrett, Sonia Goncalves, Ewan Harrison, David K. Jackson, Ian Johnston, Dominic Kwiatkowski, Cordelia Langford, John Sillitoe on behalf of the Wellcome Sanger Institute COVID-19 Surveillance Team |
| EPI_ISL_2997257 | Lighthouse Lab in Glasgow | Wellcome Sanger Institute for the COVID-19 Genomics UK (COG-UK) Consortium | Harper VanSteenhouse, Yumi Kasai, David Gray, Carol Clugston, Anna Dominiczak and Alex Alderton, Roberto Amato, Jeffrey Barrett, Sonia Goncalves, Ewan Harrison, David K. Jackson, Ian Johnston, Dominic Kwiatkowski, Cordelia Langford, John Sillitoe on behalf of the Wellcome Sanger Institute COVID-19 Surveillance Team |
| EPI_ISL_2997258 | Lighthouse Lab in Milton Keynes | Wellcome Sanger Institute for the COVID-19 Genomics UK (COG-UK) Consortium | The Lighthouse Lab in Milton Keynes and Alex Alderton, Roberto Amato, Jeffrey Barrett, Sonia Goncalves, Ewan Harrison, David K. Jackson, Ian Johnston, Dominic Kwiatkowski, Cordelia Langford, John Sillitoe on behalf of the Wellcome Sanger Institute COVID-19 Surveillance Team |
| EPI_ISL_2997259 | Lighthouse Lab in Glasgow | Wellcome Sanger Institute for the COVID-19 Genomics UK (COG-UK) Consortium | Harper VanSteenhouse, Yumi Kasai, David Gray, Carol Clugston, Anna Dominiczak and Alex Alderton, Roberto Amato, Jeffrey Barrett, Sonia Goncalves, Ewan Harrison, David K. Jackson, Ian Johnston, Dominic Kwiatkowski, Cordelia Langford, John Sillitoe on behalf of the Wellcome Sanger Institute COVID-19 Surveillance Team |
| EPI_ISL_2997260, EPI_ISL_2997261 | Lighthouse Laboratory Plymouth | Wellcome Sanger Institute for the COVID-19 Genomics UK (COG-UK) Consortium | Lighthouse Laboratory Plymouth and Alex Alderton, Roberto Amato, Jeffrey Barrett, Sonia Goncalves, Ewan Harrison, David K. Jackson, Ian Johnston, Dominic Kwiatkowski, Cordelia Langford, John Sillitoe on behalf of the Wellcome Sanger Institute COVID-19 Surveillance Team |

| | | | |
|---|---|---|---|
| EPI_ISL_2997262, EPI_ISL_2997263 | Berkshire and Surrey Pathology Services Lighthouse Laboratory | Wellcome Sanger Institute for the COVID-19 Genomics UK (COG-UK) Consortium | Berkshire and Surrey Pathology Services Lighthouse Laboratory and Alex Alderton, Roberto Amato, Jeffrey Barrett, Sonia Goncalves, Ewan Harrison, David K. Jackson, Ian Johnston, Dominic Kwiatkowski, Cordelia Langford, John Sillitoe on behalf of the Wellcome Sanger Institute COVID-19 Surveillance Team |
| EPI_ISL_2997264, EPI_ISL_2997265, EPI_ISL_2997266 | Lighthouse Lab in Glasgow | Wellcome Sanger Institute for the COVID-19 Genomics UK (COG-UK) Consortium | Harper VanSteenhouse, Yumi Kasai, David Gray, Carol Clugston, Anna Dominiczak and Alex Alderton, Roberto Amato, Jeffrey Barrett, Sonia Goncalves, Ewan Harrison, David K. Jackson, Ian Johnston, Dominic Kwiatkowski, Cordelia Langford, John Sillitoe on behalf of the Wellcome Sanger Institute COVID-19 Surveillance Team |
| EPI_ISL_2997267 | Lighthouse Laboratory Plymouth | Wellcome Sanger Institute for the COVID-19 Genomics UK (COG-UK) Consortium | Lighthouse Laboratory Plymouth and Alex Alderton, Roberto Amato, Jeffrey Barrett, Sonia Goncalves, Ewan Harrison, David K. Jackson, Ian Johnston, Dominic Kwiatkowski, Cordelia Langford, John Sillitoe on behalf of the Wellcome Sanger Institute COVID-19 Surveillance Team |
| EPI_ISL_2997268, EPI_ISL_2997269 | Lighthouse Lab in Milton Keynes | Wellcome Sanger Institute for the COVID-19 Genomics UK (COG-UK) Consortium | The Lighthouse Lab in Milton Keynes and Alex Alderton, Roberto Amato, Jeffrey Barrett, Sonia Goncalves, Ewan Harrison, David K. Jackson, Ian Johnston, Dominic Kwiatkowski, Cordelia Langford, John Sillitoe on behalf of the Wellcome Sanger Institute COVID-19 Surveillance Team |
| EPI_ISL_2997270, EPI_ISL_2997271 | Lighthouse Laboratory Plymouth | Wellcome Sanger Institute for the COVID-19 Genomics UK (COG-UK) Consortium | Lighthouse Laboratory Plymouth and Alex Alderton, Roberto Amato, Jeffrey Barrett, Sonia Goncalves, Ewan Harrison, David K. Jackson, Ian Johnston, Dominic Kwiatkowski, Cordelia Langford, John Sillitoe on behalf of the Wellcome Sanger Institute COVID-19 Surveillance Team |
| EPI_ISL_2997272 | Lighthouse Lab in Glasgow | Wellcome Sanger Institute for the COVID-19 Genomics UK (COG-UK) Consortium | Harper VanSteenhouse, Yumi Kasai, David Gray, Carol Clugston, Anna Dominiczak and Alex Alderton, Roberto Amato, Jeffrey Barrett, Sonia Goncalves, Ewan Harrison, David K. Jackson, Ian Johnston, Dominic Kwiatkowski, Cordelia Langford, John Sillitoe on behalf of the Wellcome Sanger Institute COVID-19 Surveillance Team |
| EPI_ISL_2997273 | Lighthouse Lab in Milton Keynes | Wellcome Sanger Institute for the COVID-19 Genomics UK (COG-UK) Consortium | The Lighthouse Lab in Milton Keynes and Alex Alderton, Roberto Amato, Jeffrey Barrett, Sonia Goncalves, Ewan Harrison, David K. Jackson, Ian Johnston, Dominic Kwiatkowski, Cordelia Langford, John Sillitoe on behalf of the Wellcome Sanger Institute COVID-19 Surveillance Team |
| EPI_ISL_2997274 | Berkshire and Surrey Pathology Services Lighthouse Laboratory | Wellcome Sanger Institute for the COVID-19 Genomics UK (COG-UK) Consortium | Berkshire and Surrey Pathology Services Lighthouse Laboratory and Alex Alderton, Roberto Amato, Jeffrey Barrett, Sonia Goncalves, Ewan Harrison, David K. Jackson, Ian Johnston, Dominic Kwiatkowski, Cordelia Langford, John Sillitoe on behalf of the Wellcome Sanger Institute COVID-19 Surveillance Team |
| EPI_ISL_2997275, EPI_ISL_2997276 | Lighthouse Lab in Milton Keynes | Wellcome Sanger Institute for the COVID-19 Genomics UK (COG-UK) Consortium | The Lighthouse Lab in Milton Keynes and Alex Alderton, Roberto Amato, Jeffrey Barrett, Sonia Goncalves, Ewan Harrison, David K. Jackson, Ian Johnston, Dominic Kwiatkowski, Cordelia Langford, John Sillitoe on behalf of the Wellcome Sanger Institute COVID-19 Surveillance Team |
| EPI_ISL_2997277 | Lighthouse Laboratory Plymouth | Wellcome Sanger Institute for the COVID-19 Genomics UK (COG-UK) Consortium | Lighthouse Laboratory Plymouth and Alex Alderton, Roberto Amato, Jeffrey Barrett, Sonia Goncalves, Ewan Harrison, David K. Jackson, Ian Johnston, Dominic Kwiatkowski, Cordelia Langford, John Sillitoe on behalf of the Wellcome Sanger Institute COVID-19 Surveillance Team |
| EPI_ISL_2997278 | Berkshire and Surrey Pathology Services Lighthouse Laboratory | Wellcome Sanger Institute for the COVID-19 Genomics UK (COG-UK) Consortium | Berkshire and Surrey Pathology Services Lighthouse Laboratory and Alex Alderton, Roberto Amato, Jeffrey Barrett, Sonia Goncalves, Ewan Harrison, David K. Jackson, Ian Johnston, Dominic Kwiatkowski, Cordelia Langford, John Sillitoe on behalf of the Wellcome Sanger Institute COVID-19 Surveillance Team |
| EPI_ISL_2997279, EPI_ISL_2997280, EPI_ISL_2997281 | Lighthouse Lab in Glasgow | Wellcome Sanger Institute for the COVID-19 Genomics UK (COG-UK) Consortium | Harper VanSteenhouse, Yumi Kasai, David Gray, Carol Clugston, Anna Dominiczak and Alex Alderton, Roberto Amato, Jeffrey Barrett, Sonia Goncalves, Ewan Harrison, David K. Jackson, Ian Johnston, Dominic Kwiatkowski, Cordelia Langford, John Sillitoe on behalf of the Wellcome Sanger Institute COVID-19 Surveillance Team |
| EPI_ISL_2997282 | Lighthouse Lab in Milton Keynes | Wellcome Sanger Institute for the COVID-19 Genomics UK (COG-UK) Consortium | The Lighthouse Lab in Milton Keynes and Alex Alderton, Roberto Amato, Jeffrey Barrett, Sonia Goncalves, Ewan Harrison, David K. Jackson, Ian Johnston, Dominic Kwiatkowski, Cordelia Langford, John Sillitoe on behalf of the Wellcome Sanger Institute COVID-19 Surveillance Team |
| EPI_ISL_2997283, EPI_ISL_2997284 | Berkshire and Surrey Pathology Services Lighthouse Laboratory | Wellcome Sanger Institute for the COVID-19 Genomics UK (COG-UK) Consortium | Berkshire and Surrey Pathology Services Lighthouse Laboratory and Alex Alderton, Roberto Amato, Jeffrey Barrett, Sonia Goncalves, Ewan Harrison, David K. Jackson, Ian Johnston, Dominic Kwiatkowski, Cordelia Langford, John Sillitoe on behalf of the Wellcome Sanger Institute COVID-19 Surveillance Team |
| EPI_ISL_2997285 | Lighthouse Laboratory Plymouth | Wellcome Sanger Institute for the COVID-19 Genomics UK (COG-UK) Consortium | Lighthouse Laboratory Plymouth and Alex Alderton, Roberto Amato, Jeffrey Barrett, Sonia Goncalves, Ewan Harrison, David K. Jackson, Ian Johnston, Dominic Kwiatkowski, Cordelia Langford, John Sillitoe on behalf of the Wellcome Sanger Institute COVID-19 Surveillance Team |
| EPI_ISL_2997286, EPI_ISL_2997287 | Lighthouse Lab in Glasgow | Wellcome Sanger Institute for the COVID-19 Genomics UK (COG-UK) Consortium | Harper VanSteenhouse, Yumi Kasai, David Gray, Carol Clugston, Anna Dominiczak and Alex Alderton, Roberto Amato, Jeffrey Barrett, Sonia Goncalves, Ewan Harrison, David K. Jackson, Ian Johnston, Dominic Kwiatkowski, Cordelia Langford, John Sillitoe on behalf of the Wellcome Sanger Institute COVID-19 Surveillance Team |
| EPI_ISL_2997288 | Lighthouse Lab in Milton Keynes | Wellcome Sanger Institute for the COVID-19 Genomics UK (COG-UK) Consortium | The Lighthouse Lab in Milton Keynes and Alex Alderton, Roberto Amato, Jeffrey Barrett, Sonia Goncalves, Ewan Harrison, David K. Jackson, Ian Johnston, Dominic Kwiatkowski, Cordelia Langford, John Sillitoe on behalf of the Wellcome Sanger Institute COVID-19 Surveillance Team |
| EPI_ISL_2997289 | Lighthouse Lab in Glasgow | Wellcome Sanger Institute for the COVID-19 Genomics UK (COG-UK) Consortium | Harper VanSteenhouse, Yumi Kasai, David Gray, Carol Clugston, Anna Dominiczak and Alex Alderton, Roberto Amato, Jeffrey Barrett, Sonia Goncalves, Ewan Harrison, David K. Jackson, Ian Johnston, Dominic Kwiatkowski, Cordelia Langford, John Sillitoe on behalf of the Wellcome Sanger Institute COVID-19 Surveillance Team |
| EPI_ISL_2997290, EPI_ISL_2997291 | Lighthouse Lab in Milton Keynes | Wellcome Sanger Institute for the COVID-19 Genomics UK (COG-UK) Consortium | The Lighthouse Lab in Milton Keynes and Alex Alderton, Roberto Amato, Jeffrey Barrett, Sonia Goncalves, Ewan Harrison, David K. Jackson, Ian Johnston, Dominic Kwiatkowski, Cordelia Langford, John Sillitoe on behalf of the Wellcome Sanger Institute COVID-19 Surveillance Team |
| EPI_ISL_2997292 | Berkshire and Surrey Pathology Services Lighthouse Laboratory | Wellcome Sanger Institute for the COVID-19 Genomics UK (COG-UK) Consortium | Berkshire and Surrey Pathology Services Lighthouse Laboratory and Alex Alderton, Roberto Amato, Jeffrey Barrett, Sonia Goncalves, Ewan Harrison, David K. Jackson, Ian Johnston, Dominic Kwiatkowski, Cordelia Langford, John Sillitoe on behalf of the Wellcome Sanger Institute COVID-19 Surveillance Team |
| EPI_ISL_2997293, EPI_ISL_2997294, EPI_ISL_2997295 | Lighthouse Laboratory Plymouth | Wellcome Sanger Institute for the COVID-19 Genomics UK (COG-UK) Consortium | Lighthouse Laboratory Plymouth and Alex Alderton, Roberto Amato, Jeffrey Barrett, Sonia Goncalves, Ewan Harrison, David K. Jackson, Ian Johnston, Dominic Kwiatkowski, Cordelia Langford, John Sillitoe on behalf of the Wellcome Sanger Institute COVID-19 Surveillance Team |
| EPI_ISL_2997296 | Lighthouse Lab in Glasgow | Wellcome Sanger Institute for the COVID-19 Genomics UK (COG-UK) Consortium | Harper VanSteenhouse, Yumi Kasai, David Gray, Carol Clugston, Anna Dominiczak and Alex Alderton, Roberto Amato, Jeffrey Barrett, Sonia Goncalves, Ewan Harrison, David K. Jackson, Ian Johnston, Dominic Kwiatkowski, Cordelia Langford, John Sillitoe on behalf of the Wellcome Sanger Institute COVID-19 Surveillance Team |
| EPI_ISL_2997297 | Lighthouse Laboratory Plymouth | Wellcome Sanger Institute for the COVID-19 Genomics UK (COG-UK) Consortium | Lighthouse Laboratory Plymouth and Alex Alderton, Roberto Amato, Jeffrey Barrett, Sonia Goncalves, Ewan Harrison, David K. Jackson, Ian Johnston, Dominic Kwiatkowski, Cordelia Langford, John Sillitoe on behalf of the Wellcome Sanger Institute COVID-19 Surveillance Team |
| EPI_ISL_2997298, EPI_ISL_2997299 | Lighthouse Lab in Glasgow | Wellcome Sanger Institute for the COVID-19 Genomics UK (COG-UK) Consortium | Harper VanSteenhouse, Yumi Kasai, David Gray, Carol Clugston, Anna Dominiczak and Alex Alderton, Roberto Amato, Jeffrey Barrett, Sonia Goncalves, Ewan Harrison, David K. Jackson, Ian Johnston, Dominic Kwiatkowski, Cordelia Langford, John Sillitoe on behalf of the Wellcome Sanger Institute COVID-19 Surveillance Team |
| EPI_ISL_2997300 | Lighthouse Lab in Milton Keynes | Wellcome Sanger Institute for the COVID-19 Genomics UK (COG-UK) Consortium | The Lighthouse Lab in Milton Keynes and Alex Alderton, Roberto Amato, Jeffrey Barrett, Sonia Goncalves, Ewan Harrison, David K. Jackson, Ian Johnston, Dominic Kwiatkowski, Cordelia Langford, John Sillitoe on behalf of the Wellcome Sanger Institute COVID-19 Surveillance Team |
| EPI_ISL_2997301, EPI_ISL_2997302 | Lighthouse Laboratory Plymouth | Wellcome Sanger Institute for the COVID-19 Genomics UK (COG-UK) Consortium | Lighthouse Laboratory Plymouth and Alex Alderton, Roberto Amato, Jeffrey Barrett, Sonia Goncalves, Ewan Harrison, David K. Jackson, Ian Johnston, Dominic Kwiatkowski, Cordelia Langford, John Sillitoe on behalf of the Wellcome Sanger Institute COVID-19 Surveillance Team |
| EPI_ISL_2997303 | Lighthouse Lab in Milton Keynes | Wellcome Sanger Institute for the COVID-19 Genomics UK (COG-UK) Consortium | The Lighthouse Lab in Milton Keynes and Alex Alderton, Roberto Amato, Jeffrey Barrett, Sonia Goncalves, Ewan Harrison, David K. Jackson, Ian Johnston, Dominic Kwiatkowski, Cordelia Langford, John Sillitoe on behalf of the Wellcome Sanger Institute COVID-19 Surveillance Team |
| EPI_ISL_2997304 | Berkshire and Surrey Pathology Services Lighthouse Laboratory | Wellcome Sanger Institute for the COVID-19 Genomics UK (COG-UK) Consortium | Berkshire and Surrey Pathology Services Lighthouse Laboratory and Alex Alderton, Roberto Amato, Jeffrey Barrett, Sonia Goncalves, Ewan Harrison, David K. Jackson, Ian Johnston, Dominic Kwiatkowski, Cordelia Langford, John Sillitoe on behalf of the Wellcome Sanger Institute COVID-19 Surveillance Team |
| EPI_ISL_2997305, EPI_ISL_2997306, EPI_ISL_2997307, EPI_ISL_2997308 | Lighthouse Lab in Milton Keynes | Wellcome Sanger Institute for the COVID-19 Genomics UK (COG-UK) Consortium | The Lighthouse Lab in Milton Keynes and Alex Alderton, Roberto Amato, Jeffrey Barrett, Sonia Goncalves, Ewan Harrison, David K. Jackson, Ian Johnston, Dominic Kwiatkowski, Cordelia Langford, John Sillitoe on behalf of the Wellcome Sanger Institute COVID-19 Surveillance Team |
| EPI_ISL_2997309 | Lighthouse Lab in Glasgow | Wellcome Sanger Institute for the COVID-19 Genomics UK (COG-UK) Consortium | Harper VanSteenhouse, Yumi Kasai, David Gray, Carol Clugston, Anna Dominiczak and Alex Alderton, Roberto Amato, Jeffrey Barrett, Sonia Goncalves, Ewan Harrison, David K. Jackson, Ian Johnston, Dominic Kwiatkowski, Cordelia Langford, John Sillitoe on behalf of the Wellcome Sanger Institute COVID-19 Surveillance Team |
| EPI_ISL_2997310, EPI_ISL_2997311 | Lighthouse Lab in Milton Keynes | Wellcome Sanger Institute for the COVID-19 Genomics UK (COG-UK) Consortium | The Lighthouse Lab in Milton Keynes and Alex Alderton, Roberto Amato, Jeffrey Barrett, Sonia Goncalves, Ewan Harrison, David K. Jackson, Ian Johnston, Dominic Kwiatkowski, Cordelia Langford, John Sillitoe on behalf of the Wellcome Sanger Institute COVID-19 Surveillance Team |
| EPI_ISL_2997312 | Berkshire and Surrey Pathology Services Lighthouse Laboratory | Wellcome Sanger Institute for the COVID-19 Genomics UK (COG-UK) Consortium | Berkshire and Surrey Pathology Services Lighthouse Laboratory and Alex Alderton, Roberto Amato, Jeffrey Barrett, Sonia Goncalves, Ewan Harrison, David K. Jackson, Ian Johnston, Dominic Kwiatkowski, Cordelia Langford, John Sillitoe on behalf of the Wellcome Sanger Institute COVID-19 Surveillance Team |
| EPI_ISL_2997313, EPI_ISL_2997314, EPI_ISL_2997315, EPI_ISL_2997316 | Lighthouse Lab in Glasgow | Wellcome Sanger Institute for the COVID-19 Genomics UK (COG-UK) Consortium | Harper VanSteenhouse, Yumi Kasai, David Gray, Carol Clugston, Anna Dominiczak and Alex Alderton, Roberto Amato, Jeffrey Barrett, Sonia Goncalves, Ewan Harrison, David K. Jackson, Ian Johnston, Dominic Kwiatkowski, Cordelia Langford, John Sillitoe on behalf of the Wellcome Sanger Institute COVID-19 Surveillance Team |
| EPI_ISL_2997317 | Lighthouse Lab in Milton Keynes | Wellcome Sanger Institute for the COVID-19 Genomics UK (COG-UK) Consortium | The Lighthouse Lab in Milton Keynes and Alex Alderton, Roberto Amato, Jeffrey Barrett, Sonia Goncalves, Ewan Harrison, David K. Jackson, Ian Johnston, Dominic Kwiatkowski, Cordelia Langford, John Sillitoe on behalf of the Wellcome Sanger Institute COVID-19 Surveillance Team |
| EPI_ISL_2997318 | Lighthouse Lab in Glasgow | Wellcome Sanger Institute for the COVID-19 Genomics UK (COG-UK) Consortium | Harper VanSteenhouse, Yumi Kasai, David Gray, Carol Clugston, Anna Dominiczak and Alex Alderton, Roberto Amato, Jeffrey Barrett, Sonia Goncalves, Ewan Harrison, David K. Jackson, Ian Johnston, Dominic Kwiatkowski, Cordelia Langford, John Sillitoe on behalf of the Wellcome Sanger Institute COVID-19 Surveillance Team |
| EPI_ISL_2997319, EPI_ISL_2997320 | Lighthouse Laboratory Plymouth | Wellcome Sanger Institute for the COVID-19 Genomics UK (COG-UK) Consortium | Lighthouse Laboratory Plymouth and Alex Alderton, Roberto Amato, Jeffrey Barrett, Sonia Goncalves, Ewan Harrison, David K. Jackson, Ian Johnston, Dominic Kwiatkowski, Cordelia Langford, John Sillitoe on behalf of the Wellcome Sanger Institute COVID-19 Surveillance Team |
| EPI_ISL_2997321, EPI_ISL_2997322 | Lighthouse Lab in Glasgow | Wellcome Sanger Institute for the COVID-19 Genomics UK (COG-UK) Consortium | Harper VanSteenhouse, Yumi Kasai, David Gray, Carol Clugston, Anna Dominiczak and Alex Alderton, Roberto Amato, Jeffrey Barrett, Sonia Goncalves, Ewan Harrison, David K. Jackson, Ian Johnston, Dominic Kwiatkowski, Cordelia Langford, John Sillitoe on behalf of the Wellcome Sanger Institute COVID-19 Surveillance Team |
| EPI_ISL_2997323 | Lighthouse Laboratory Plymouth | Wellcome Sanger Institute for the COVID-19 Genomics UK (COG-UK) Consortium | Lighthouse Laboratory Plymouth and Alex Alderton, Roberto Amato, Jeffrey Barrett, Sonia Goncalves, Ewan Harrison, David K. Jackson, Ian Johnston, Dominic Kwiatkowski, Cordelia Langford, John Sillitoe on behalf of the Wellcome Sanger Institute COVID-19 Surveillance Team |
| EPI_ISL_2997324 | Lighthouse Lab in Glasgow | Wellcome Sanger Institute for the COVID-19 Genomics UK (COG-UK) Consortium | Harper VanSteenhouse, Yumi Kasai, David Gray, Carol Clugston, Anna Dominiczak and Alex Alderton, Roberto Amato, Jeffrey Barrett, Sonia Goncalves, Ewan Harrison, David K. Jackson, Ian Johnston, Dominic Kwiatkowski, Cordelia Langford, John Sillitoe on behalf of the Wellcome Sanger Institute COVID-19 Surveillance Team |
| EPI_ISL_2997325 | Lighthouse Laboratory Plymouth | Wellcome Sanger Institute for the COVID-19 Genomics UK (COG-UK) Consortium | Lighthouse Laboratory Plymouth and Alex Alderton, Roberto Amato, Jeffrey Barrett, Sonia Goncalves, Ewan Harrison, David K. Jackson, Ian Johnston, Dominic Kwiatkowski, Cordelia Langford, John Sillitoe on behalf of the Wellcome Sanger Institute COVID-19 Surveillance Team |
| EPI_ISL_2997326, EPI_ISL_2997327 | Lighthouse Lab in Milton Keynes | Wellcome Sanger Institute for the COVID-19 Genomics UK (COG-UK) Consortium | The Lighthouse Lab in Milton Keynes and Alex Alderton, Roberto Amato, Jeffrey Barrett, Sonia Goncalves, Ewan Harrison, David K. Jackson, Ian Johnston, Dominic Kwiatkowski, Cordelia Langford, John Sillitoe on behalf of the Wellcome Sanger Institute COVID-19 Surveillance Team |
| EPI_ISL_2997328, EPI_ISL_2997329, EPI_ISL_2997330 | Lighthouse Laboratory Plymouth | Wellcome Sanger Institute for the COVID-19 Genomics UK (COG-UK) Consortium | Lighthouse Laboratory Plymouth and Alex Alderton, Roberto Amato, Jeffrey Barrett, Sonia Goncalves, Ewan Harrison, David K. Jackson, Ian Johnston, Dominic Kwiatkowski, Cordelia Langford, John Sillitoe on behalf of the Wellcome Sanger Institute COVID-19 Surveillance Team |

| | | | |
|---|---|---|---|
| EPI_ISL_2997331, EPI_ISL_2997332 | Lighthouse Lab in Milton Keynes | Wellcome Sanger Institute for the COVID-19 Genomics UK (COG-UK) Consortium | The Lighthouse Lab in Milton Keynes and Alex Alderton, Roberto Amato, Jeffrey Barrett, Sonia Goncalves, Ewan Harrison, David K. Jackson, Ian Johnston, Dominic Kwiatkowski, Cordelia Langford, John Sillitoe on behalf of the Wellcome Sanger Institute COVID-19 Surveillance Team |
| EPI_ISL_2997333 | Berkshire and Surrey Pathology Services Lighthouse Laboratory | Wellcome Sanger Institute for the COVID-19 Genomics UK (COG-UK) Consortium | Berkshire and Surrey Pathology Services Lighthouse Laboratory and Alex Alderton, Roberto Amato, Jeffrey Barrett, Sonia Goncalves, Ewan Harrison, David K. Jackson, Ian Johnston, Dominic Kwiatkowski, Cordelia Langford, John Sillitoe on behalf of the Wellcome Sanger Institute COVID-19 Surveillance Team |
| EPI_ISL_2997334 | Lighthouse Lab in Milton Keynes | Wellcome Sanger Institute for the COVID-19 Genomics UK (COG-UK) Consortium | The Lighthouse Lab in Milton Keynes and Alex Alderton, Roberto Amato, Jeffrey Barrett, Sonia Goncalves, Ewan Harrison, David K. Jackson, Ian Johnston, Dominic Kwiatkowski, Cordelia Langford, John Sillitoe on behalf of the Wellcome Sanger Institute COVID-19 Surveillance Team |
| EPI_ISL_2997335 | Berkshire and Surrey Pathology Services Lighthouse Laboratory | Wellcome Sanger Institute for the COVID-19 Genomics UK (COG-UK) Consortium | Berkshire and Surrey Pathology Services Lighthouse Laboratory and Alex Alderton, Roberto Amato, Jeffrey Barrett, Sonia Goncalves, Ewan Harrison, David K. Jackson, Ian Johnston, Dominic Kwiatkowski, Cordelia Langford, John Sillitoe on behalf of the Wellcome Sanger Institute COVID-19 Surveillance Team |
| EPI_ISL_2997336 | Lighthouse Laboratory Plymouth | Wellcome Sanger Institute for the COVID-19 Genomics UK (COG-UK) Consortium | Lighthouse Laboratory Plymouth and Alex Alderton, Roberto Amato, Jeffrey Barrett, Sonia Goncalves, Ewan Harrison, David K. Jackson, Ian Johnston, Dominic Kwiatkowski, Cordelia Langford, John Sillitoe on behalf of the Wellcome Sanger Institute COVID-19 Surveillance Team |
| EPI_ISL_2997337, EPI_ISL_2997338 | Berkshire and Surrey Pathology Services Lighthouse Laboratory | Wellcome Sanger Institute for the COVID-19 Genomics UK (COG-UK) Consortium | Berkshire and Surrey Pathology Services Lighthouse Laboratory and Alex Alderton, Roberto Amato, Jeffrey Barrett, Sonia Goncalves, Ewan Harrison, David K. Jackson, Ian Johnston, Dominic Kwiatkowski, Cordelia Langford, John Sillitoe on behalf of the Wellcome Sanger Institute COVID-19 Surveillance Team |
| EPI_ISL_2997339 | Lighthouse Lab in Milton Keynes | Wellcome Sanger Institute for the COVID-19 Genomics UK (COG-UK) Consortium | The Lighthouse Lab in Milton Keynes and Alex Alderton, Roberto Amato, Jeffrey Barrett, Sonia Goncalves, Ewan Harrison, David K. Jackson, Ian Johnston, Dominic Kwiatkowski, Cordelia Langford, John Sillitoe on behalf of the Wellcome Sanger Institute COVID-19 Surveillance Team |
| EPI_ISL_2997340 | Lighthouse Laboratory Plymouth | Wellcome Sanger Institute for the COVID-19 Genomics UK (COG-UK) Consortium | Lighthouse Laboratory Plymouth and Alex Alderton, Roberto Amato, Jeffrey Barrett, Sonia Goncalves, Ewan Harrison, David K. Jackson, Ian Johnston, Dominic Kwiatkowski, Cordelia Langford, John Sillitoe on behalf of the Wellcome Sanger Institute COVID-19 Surveillance Team |
| EPI_ISL_2997341 | Lighthouse Lab in Glasgow | Wellcome Sanger Institute for the COVID-19 Genomics UK (COG-UK) Consortium | Harper VanSteenhouse, Yumi Kasai, David Gray, Carol Clugston, Anna Dominiczak and Alex Alderton, Roberto Amato, Jeffrey Barrett, Sonia Goncalves, Ewan Harrison, David K. Jackson, Ian Johnston, Dominic Kwiatkowski, Cordelia Langford, John Sillitoe on behalf of the Wellcome Sanger Institute COVID-19 Surveillance Team |
| EPI_ISL_2997342 | Lighthouse Lab in Milton Keynes | Wellcome Sanger Institute for the COVID-19 Genomics UK (COG-UK) Consortium | The Lighthouse Lab in Milton Keynes and Alex Alderton, Roberto Amato, Jeffrey Barrett, Sonia Goncalves, Ewan Harrison, David K. Jackson, Ian Johnston, Dominic Kwiatkowski, Cordelia Langford, John Sillitoe on behalf of the Wellcome Sanger Institute COVID-19 Surveillance Team |
| EPI_ISL_2997343 | Berkshire and Surrey Pathology Services Lighthouse Laboratory | Wellcome Sanger Institute for the COVID-19 Genomics UK (COG-UK) Consortium | Berkshire and Surrey Pathology Services Lighthouse Laboratory and Alex Alderton, Roberto Amato, Jeffrey Barrett, Sonia Goncalves, Ewan Harrison, David K. Jackson, Ian Johnston, Dominic Kwiatkowski, Cordelia Langford, John Sillitoe on behalf of the Wellcome Sanger Institute COVID-19 Surveillance Team |
| EPI_ISL_2997344, EPI_ISL_2997345 | Lighthouse Lab in Milton Keynes | Wellcome Sanger Institute for the COVID-19 Genomics UK (COG-UK) Consortium | The Lighthouse Lab in Milton Keynes and Alex Alderton, Roberto Amato, Jeffrey Barrett, Sonia Goncalves, Ewan Harrison, David K. Jackson, Ian Johnston, Dominic Kwiatkowski, Cordelia Langford, John Sillitoe on behalf of the Wellcome Sanger Institute COVID-19 Surveillance Team |
| EPI_ISL_2997346 | Berkshire and Surrey Pathology Services Lighthouse Laboratory | Wellcome Sanger Institute for the COVID-19 Genomics UK (COG-UK) Consortium | Berkshire and Surrey Pathology Services Lighthouse Laboratory and Alex Alderton, Roberto Amato, Jeffrey Barrett, Sonia Goncalves, Ewan Harrison, David K. Jackson, Ian Johnston, Dominic Kwiatkowski, Cordelia Langford, John Sillitoe on behalf of the Wellcome Sanger Institute COVID-19 Surveillance Team |
| EPI_ISL_2997347 | Lighthouse Lab in Glasgow | Wellcome Sanger Institute for the COVID-19 Genomics UK (COG-UK) Consortium | Harper VanSteenhouse, Yumi Kasai, David Gray, Carol Clugston, Anna Dominiczak and Alex Alderton, Roberto Amato, Jeffrey Barrett, Sonia Goncalves, Ewan Harrison, David K. Jackson, Ian Johnston, Dominic Kwiatkowski, Cordelia Langford, John Sillitoe on behalf of the Wellcome Sanger Institute COVID-19 Surveillance Team |
| EPI_ISL_2997348 | Berkshire and Surrey Pathology Services Lighthouse Laboratory | Wellcome Sanger Institute for the COVID-19 Genomics UK (COG-UK) Consortium | Berkshire and Surrey Pathology Services Lighthouse Laboratory and Alex Alderton, Roberto Amato, Jeffrey Barrett, Sonia Goncalves, Ewan Harrison, David K. Jackson, Ian Johnston, Dominic Kwiatkowski, Cordelia Langford, John Sillitoe on behalf of the Wellcome Sanger Institute COVID-19 Surveillance Team |
| EPI_ISL_2997349 | Lighthouse Lab in Glasgow | Wellcome Sanger Institute for the COVID-19 Genomics UK (COG-UK) Consortium | Harper VanSteenhouse, Yumi Kasai, David Gray, Carol Clugston, Anna Dominiczak and Alex Alderton, Roberto Amato, Jeffrey Barrett, Sonia Goncalves, Ewan Harrison, David K. Jackson, Ian Johnston, Dominic Kwiatkowski, Cordelia Langford, John Sillitoe on behalf of the Wellcome Sanger Institute COVID-19 Surveillance Team |
| EPI_ISL_2997350 | Lighthouse Lab in Milton Keynes | Wellcome Sanger Institute for the COVID-19 Genomics UK (COG-UK) Consortium | The Lighthouse Lab in Milton Keynes and Alex Alderton, Roberto Amato, Jeffrey Barrett, Sonia Goncalves, Ewan Harrison, David K. Jackson, Ian Johnston, Dominic Kwiatkowski, Cordelia Langford, John Sillitoe on behalf of the Wellcome Sanger Institute COVID-19 Surveillance Team |
| EPI_ISL_2997351, EPI_ISL_2997352 | Lighthouse Laboratory Plymouth | Wellcome Sanger Institute for the COVID-19 Genomics UK (COG-UK) Consortium | Lighthouse Laboratory Plymouth and Alex Alderton, Roberto Amato, Jeffrey Barrett, Sonia Goncalves, Ewan Harrison, David K. Jackson, Ian Johnston, Dominic Kwiatkowski, Cordelia Langford, John Sillitoe on behalf of the Wellcome Sanger Institute COVID-19 Surveillance Team |
| EPI_ISL_2997353, EPI_ISL_2997354, EPI_ISL_2997355 | Lighthouse Lab in Milton Keynes | Wellcome Sanger Institute for the COVID-19 Genomics UK (COG-UK) Consortium | The Lighthouse Lab in Milton Keynes and Alex Alderton, Roberto Amato, Jeffrey Barrett, Sonia Goncalves, Ewan Harrison, David K. Jackson, Ian Johnston, Dominic Kwiatkowski, Cordelia Langford, John Sillitoe on behalf of the Wellcome Sanger Institute COVID-19 Surveillance Team |
| EPI_ISL_2997356, EPI_ISL_2997357, EPI_ISL_2997358, EPI_ISL_2997359, EPI_ISL_2997360 | Berkshire and Surrey Pathology Services Lighthouse Laboratory | Wellcome Sanger Institute for the COVID-19 Genomics UK (COG-UK) Consortium | Berkshire and Surrey Pathology Services Lighthouse Laboratory and Alex Alderton, Roberto Amato, Jeffrey Barrett, Sonia Goncalves, Ewan Harrison, David K. Jackson, Ian Johnston, Dominic Kwiatkowski, Cordelia Langford, John Sillitoe on behalf of the Wellcome Sanger Institute COVID-19 Surveillance Team |
| EPI_ISL_2997361 | Lighthouse Lab in Glasgow | Wellcome Sanger Institute for the COVID-19 Genomics UK (COG-UK) Consortium | Harper VanSteenhouse, Yumi Kasai, David Gray, Carol Clugston, Anna Dominiczak and Alex Alderton, Roberto Amato, Jeffrey Barrett, Sonia Goncalves, Ewan Harrison, David K. Jackson, Ian Johnston, Dominic Kwiatkowski, Cordelia Langford, John Sillitoe on behalf of the Wellcome Sanger Institute COVID-19 Surveillance Team |
| EPI_ISL_2997362, EPI_ISL_2997363, EPI_ISL_2997364, EPI_ISL_2997365, EPI_ISL_2997366, EPI_ISL_2997367, EPI_ISL_2997368, EPI_ISL_2997369 | Berkshire and Surrey Pathology Services Lighthouse Laboratory | Wellcome Sanger Institute for the COVID-19 Genomics UK (COG-UK) Consortium | Berkshire and Surrey Pathology Services Lighthouse Laboratory and Alex Alderton, Roberto Amato, Jeffrey Barrett, Sonia Goncalves, Ewan Harrison, David K. Jackson, Ian Johnston, Dominic Kwiatkowski, Cordelia Langford, John Sillitoe on behalf of the Wellcome Sanger Institute COVID-19 Surveillance Team |
| EPI_ISL_2997370 | Lighthouse Lab in Glasgow | Wellcome Sanger Institute for the COVID-19 Genomics UK (COG-UK) Consortium | Harper VanSteenhouse, Yumi Kasai, David Gray, Carol Clugston, Anna Dominiczak and Alex Alderton, Roberto Amato, Jeffrey Barrett, Sonia Goncalves, Ewan Harrison, David K. Jackson, Ian Johnston, Dominic Kwiatkowski, Cordelia Langford, John Sillitoe on behalf of the Wellcome Sanger Institute COVID-19 Surveillance Team |
| EPI_ISL_2997371 | Berkshire and Surrey Pathology Services Lighthouse Laboratory | Wellcome Sanger Institute for the COVID-19 Genomics UK (COG-UK) Consortium | Berkshire and Surrey Pathology Services Lighthouse Laboratory and Alex Alderton, Roberto Amato, Jeffrey Barrett, Sonia Goncalves, Ewan Harrison, David K. Jackson, Ian Johnston, Dominic Kwiatkowski, Cordelia Langford, John Sillitoe on behalf of the Wellcome Sanger Institute COVID-19 Surveillance Team |
| EPI_ISL_2997372, EPI_ISL_2997373 | Lighthouse Lab in Glasgow | Wellcome Sanger Institute for the COVID-19 Genomics UK (COG-UK) Consortium | Harper VanSteenhouse, Yumi Kasai, David Gray, Carol Clugston, Anna Dominiczak and Alex Alderton, Roberto Amato, Jeffrey Barrett, Sonia Goncalves, Ewan Harrison, David K. Jackson, Ian Johnston, Dominic Kwiatkowski, Cordelia Langford, John Sillitoe on behalf of the Wellcome Sanger Institute COVID-19 Surveillance Team |
| EPI_ISL_2997374 | Berkshire and Surrey Pathology Services Lighthouse Laboratory | Wellcome Sanger Institute for the COVID-19 Genomics UK (COG-UK) Consortium | Berkshire and Surrey Pathology Services Lighthouse Laboratory and Alex Alderton, Roberto Amato, Jeffrey Barrett, Sonia Goncalves, Ewan Harrison, David K. Jackson, Ian Johnston, Dominic Kwiatkowski, Cordelia Langford, John Sillitoe on behalf of the Wellcome Sanger Institute COVID-19 Surveillance Team |
| EPI_ISL_2997375 | Lighthouse Lab in Glasgow | Wellcome Sanger Institute for the COVID-19 Genomics UK (COG-UK) Consortium | Harper VanSteenhouse, Yumi Kasai, David Gray, Carol Clugston, Anna Dominiczak and Alex Alderton, Roberto Amato, Jeffrey Barrett, Sonia Goncalves, Ewan Harrison, David K. Jackson, Ian Johnston, Dominic Kwiatkowski, Cordelia Langford, John Sillitoe on behalf of the Wellcome Sanger Institute COVID-19 Surveillance Team |
| EPI_ISL_2997376 | Berkshire and Surrey Pathology Services Lighthouse Laboratory | Wellcome Sanger Institute for the COVID-19 Genomics UK (COG-UK) Consortium | Berkshire and Surrey Pathology Services Lighthouse Laboratory and Alex Alderton, Roberto Amato, Jeffrey Barrett, Sonia Goncalves, Ewan Harrison, David K. Jackson, Ian Johnston, Dominic Kwiatkowski, Cordelia Langford, John Sillitoe on behalf of the Wellcome Sanger Institute COVID-19 Surveillance Team |
| EPI_ISL_2997377 | Lighthouse Lab in Glasgow | Wellcome Sanger Institute for the COVID-19 Genomics UK (COG-UK) Consortium | Harper VanSteenhouse, Yumi Kasai, David Gray, Carol Clugston, Anna Dominiczak and Alex Alderton, Roberto Amato, Jeffrey Barrett, Sonia Goncalves, Ewan Harrison, David K. Jackson, Ian Johnston, Dominic Kwiatkowski, Cordelia Langford, John Sillitoe on behalf of the Wellcome Sanger Institute COVID-19 Surveillance Team |
| EPI_ISL_2997378 | Berkshire and Surrey Pathology Services Lighthouse Laboratory | Wellcome Sanger Institute for the COVID-19 Genomics UK (COG-UK) Consortium | Berkshire and Surrey Pathology Services Lighthouse Laboratory and Alex Alderton, Roberto Amato, Jeffrey Barrett, Sonia Goncalves, Ewan Harrison, David K. Jackson, Ian Johnston, Dominic Kwiatkowski, Cordelia Langford, John Sillitoe on behalf of the Wellcome Sanger Institute COVID-19 Surveillance Team |
| EPI_ISL_2997379 | Lighthouse Lab in Glasgow | Wellcome Sanger Institute for the COVID-19 Genomics UK (COG-UK) Consortium | Harper VanSteenhouse, Yumi Kasai, David Gray, Carol Clugston, Anna Dominiczak and Alex Alderton, Roberto Amato, Jeffrey Barrett, Sonia Goncalves, Ewan Harrison, David K. Jackson, Ian Johnston, Dominic Kwiatkowski, Cordelia Langford, John Sillitoe on behalf of the Wellcome Sanger Institute COVID-19 Surveillance Team |
| EPI_ISL_2997380 | Berkshire and Surrey Pathology Services Lighthouse Laboratory | Wellcome Sanger Institute for the COVID-19 Genomics UK (COG-UK) Consortium | Berkshire and Surrey Pathology Services Lighthouse Laboratory and Alex Alderton, Roberto Amato, Jeffrey Barrett, Sonia Goncalves, Ewan Harrison, David K. Jackson, Ian Johnston, Dominic Kwiatkowski, Cordelia Langford, John Sillitoe on behalf of the Wellcome Sanger Institute COVID-19 Surveillance Team |
| EPI_ISL_2997381 | Lighthouse Lab in Glasgow | Wellcome Sanger Institute for the COVID-19 Genomics UK (COG-UK) Consortium | Harper VanSteenhouse, Yumi Kasai, David Gray, Carol Clugston, Anna Dominiczak and Alex Alderton, Roberto Amato, Jeffrey Barrett, Sonia Goncalves, Ewan Harrison, David K. Jackson, Ian Johnston, Dominic Kwiatkowski, Cordelia Langford, John Sillitoe on behalf of the Wellcome Sanger Institute COVID-19 Surveillance Team |
| EPI_ISL_2997382, EPI_ISL_2997383, EPI_ISL_2997384 | Berkshire and Surrey Pathology Services Lighthouse Laboratory | Wellcome Sanger Institute for the COVID-19 Genomics UK (COG-UK) Consortium | Berkshire and Surrey Pathology Services Lighthouse Laboratory and Alex Alderton, Roberto Amato, Jeffrey Barrett, Sonia Goncalves, Ewan Harrison, David K. Jackson, Ian Johnston, Dominic Kwiatkowski, Cordelia Langford, John Sillitoe on behalf of the Wellcome Sanger Institute COVID-19 Surveillance Team |
| EPI_ISL_2997385 | Lighthouse Lab in Glasgow | Wellcome Sanger Institute for the COVID-19 Genomics UK (COG-UK) Consortium | Harper VanSteenhouse, Yumi Kasai, David Gray, Carol Clugston, Anna Dominiczak and Alex Alderton, Roberto Amato, Jeffrey Barrett, Sonia Goncalves, Ewan Harrison, David K. Jackson, Ian Johnston, Dominic Kwiatkowski, Cordelia Langford, John Sillitoe on behalf of the Wellcome Sanger Institute COVID-19 Surveillance Team |
| EPI_ISL_2997386 | Berkshire and Surrey Pathology Services Lighthouse Laboratory | Wellcome Sanger Institute for the COVID-19 Genomics UK (COG-UK) Consortium | Berkshire and Surrey Pathology Services Lighthouse Laboratory and Alex Alderton, Roberto Amato, Jeffrey Barrett, Sonia Goncalves, Ewan Harrison, David K. Jackson, Ian Johnston, Dominic Kwiatkowski, Cordelia Langford, John Sillitoe on behalf of the Wellcome Sanger Institute COVID-19 Surveillance Team |
| EPI_ISL_2997387 | Lighthouse Lab in Glasgow | Wellcome Sanger Institute for the COVID-19 Genomics UK (COG-UK) Consortium | Harper VanSteenhouse, Yumi Kasai, David Gray, Carol Clugston, Anna Dominiczak and Alex Alderton, Roberto Amato, Jeffrey Barrett, Sonia Goncalves, Ewan Harrison, David K. Jackson, Ian Johnston, Dominic Kwiatkowski, Cordelia Langford, John Sillitoe on behalf of the Wellcome Sanger Institute COVID-19 Surveillance Team |
| EPI_ISL_2997388, EPI_ISL_2997389 | Berkshire and Surrey Pathology Services Lighthouse Laboratory | Wellcome Sanger Institute for the COVID-19 Genomics UK (COG-UK) Consortium | Berkshire and Surrey Pathology Services Lighthouse Laboratory and Alex Alderton, Roberto Amato, Jeffrey Barrett, Sonia Goncalves, Ewan Harrison, David K. Jackson, Ian Johnston, Dominic Kwiatkowski, Cordelia Langford, John Sillitoe on behalf of the Wellcome Sanger Institute COVID-19 Surveillance Team |
| EPI_ISL_2997390, EPI_ISL_2997391, EPI_ISL_2997392 | Lighthouse Lab in Glasgow | Wellcome Sanger Institute for the COVID-19 Genomics UK (COG-UK) Consortium | Harper VanSteenhouse, Yumi Kasai, David Gray, Carol Clugston, Anna Dominiczak and Alex Alderton, Roberto Amato, Jeffrey Barrett, Sonia Goncalves, Ewan Harrison, David K. Jackson, Ian Johnston, Dominic Kwiatkowski, Cordelia Langford, John Sillitoe on behalf of the Wellcome Sanger Institute COVID-19 Surveillance Team |
| EPI_ISL_2997393, EPI_ISL_2997394, | Berkshire and Surrey Pathology Services Lighthouse Laboratory | Wellcome Sanger Institute for the COVID-19 Genomics UK (COG-UK) Consortium | Berkshire and Surrey Pathology Services Lighthouse Laboratory and Alex Alderton, Roberto Amato, Jeffrey Barrett, Sonia Goncalves, Ewan Harrison, David K. Jackson, Ian Johnston, Dominic Kwiatkowski, Cordelia Langford, John Sillitoe on behalf of the Wellcome Sanger Institute COVID-19 Surveillance Team |

| | | | |
|---|---|---|---|
| EPI_ISL_2997395, EPI_ISL_2997396, EPI_ISL_2997397, EPI_ISL_2997398, EPI_ISL_2997399 | | | |
| EPI_ISL_2997400, EPI_ISL_2997401 | Lighthouse Lab in Glasgow | Wellcome Sanger Institute for the COVID-19 Genomics UK (COG-UK) Consortium | Harper VanSteenhouse, Yumi Kasai, David Gray, Carol Clugston, Anna Dominiczak and Alex Alderton, Roberto Amato, Jeffrey Barrett, Sonia Goncalves, Ewan Harrison, David K. Jackson, Ian Johnston, Dominic Kwiatkowski, Cordelia Langford, John Sillitoe on behalf of the Wellcome Sanger Institute COVID-19 Surveillance Team |
| EPI_ISL_2997402, EPI_ISL_2997403, EPI_ISL_2997404, EPI_ISL_2997405, EPI_ISL_2997406 | Berkshire and Surrey Pathology Services Lighthouse Laboratory | Wellcome Sanger Institute for the COVID-19 Genomics UK (COG-UK) Consortium | Berkshire and Surrey Pathology Services Lighthouse Laboratory and Alex Alderton, Roberto Amato, Jeffrey Barrett, Sonia Goncalves, Ewan Harrison, David K. Jackson, Ian Johnston, Dominic Kwiatkowski, Cordelia Langford, John Sillitoe on behalf of the Wellcome Sanger Institute COVID-19 Surveillance Team |
| EPI_ISL_2997407, EPI_ISL_2997408 | Lighthouse Lab in Glasgow | Wellcome Sanger Institute for the COVID-19 Genomics UK (COG-UK) Consortium | Harper VanSteenhouse, Yumi Kasai, David Gray, Carol Clugston, Anna Dominiczak and Alex Alderton, Roberto Amato, Jeffrey Barrett, Sonia Goncalves, Ewan Harrison, David K. Jackson, Ian Johnston, Dominic Kwiatkowski, Cordelia Langford, John Sillitoe on behalf of the Wellcome Sanger Institute COVID-19 Surveillance Team |
| EPI_ISL_2997409 | Berkshire and Surrey Pathology Services Lighthouse Laboratory | Wellcome Sanger Institute for the COVID-19 Genomics UK (COG-UK) Consortium | Berkshire and Surrey Pathology Services Lighthouse Laboratory and Alex Alderton, Roberto Amato, Jeffrey Barrett, Sonia Goncalves, Ewan Harrison, David K. Jackson, Ian Johnston, Dominic Kwiatkowski, Cordelia Langford, John Sillitoe on behalf of the Wellcome Sanger Institute COVID-19 Surveillance Team |
| EPI_ISL_2997410 | Lighthouse Lab in Glasgow | Wellcome Sanger Institute for the COVID-19 Genomics UK (COG-UK) Consortium | Harper VanSteenhouse, Yumi Kasai, David Gray, Carol Clugston, Anna Dominiczak and Alex Alderton, Roberto Amato, Jeffrey Barrett, Sonia Goncalves, Ewan Harrison, David K. Jackson, Ian Johnston, Dominic Kwiatkowski, Cordelia Langford, John Sillitoe on behalf of the Wellcome Sanger Institute COVID-19 Surveillance Team |
| EPI_ISL_2997411, EPI_ISL_2997412, EPI_ISL_2997413, EPI_ISL_2997414, EPI_ISL_2997415, EPI_ISL_2997416, EPI_ISL_2997417, EPI_ISL_2997418 | Berkshire and Surrey Pathology Services Lighthouse Laboratory | Wellcome Sanger Institute for the COVID-19 Genomics UK (COG-UK) Consortium | Berkshire and Surrey Pathology Services Lighthouse Laboratory and Alex Alderton, Roberto Amato, Jeffrey Barrett, Sonia Goncalves, Ewan Harrison, David K. Jackson, Ian Johnston, Dominic Kwiatkowski, Cordelia Langford, John Sillitoe on behalf of the Wellcome Sanger Institute COVID-19 Surveillance Team |
| EPI_ISL_2997419, EPI_ISL_2997420 | Lighthouse Lab in Glasgow | Wellcome Sanger Institute for the COVID-19 Genomics UK (COG-UK) Consortium | Harper VanSteenhouse, Yumi Kasai, David Gray, Carol Clugston, Anna Dominiczak and Alex Alderton, Roberto Amato, Jeffrey Barrett, Sonia Goncalves, Ewan Harrison, David K. Jackson, Ian Johnston, Dominic Kwiatkowski, Cordelia Langford, John Sillitoe on behalf of the Wellcome Sanger Institute COVID-19 Surveillance Team |
| EPI_ISL_2997421 | Berkshire and Surrey Pathology Services Lighthouse Laboratory | Wellcome Sanger Institute for the COVID-19 Genomics UK (COG-UK) Consortium | Berkshire and Surrey Pathology Services Lighthouse Laboratory and Alex Alderton, Roberto Amato, Jeffrey Barrett, Sonia Goncalves, Ewan Harrison, David K. Jackson, Ian Johnston, Dominic Kwiatkowski, Cordelia Langford, John Sillitoe on behalf of the Wellcome Sanger Institute COVID-19 Surveillance Team |
| EPI_ISL_2997422, EPI_ISL_2997423 | Lighthouse Lab in Glasgow | Wellcome Sanger Institute for the COVID-19 Genomics UK (COG-UK) Consortium | Harper VanSteenhouse, Yumi Kasai, David Gray, Carol Clugston, Anna Dominiczak and Alex Alderton, Roberto Amato, Jeffrey Barrett, Sonia Goncalves, Ewan Harrison, David K. Jackson, Ian Johnston, Dominic Kwiatkowski, Cordelia Langford, John Sillitoe on behalf of the Wellcome Sanger Institute COVID-19 Surveillance Team |
| EPI_ISL_2997424 | Berkshire and Surrey Pathology Services Lighthouse Laboratory | Wellcome Sanger Institute for the COVID-19 Genomics UK (COG-UK) Consortium | Berkshire and Surrey Pathology Services Lighthouse Laboratory and Alex Alderton, Roberto Amato, Jeffrey Barrett, Sonia Goncalves, Ewan Harrison, David K. Jackson, Ian Johnston, Dominic Kwiatkowski, Cordelia Langford, John Sillitoe on behalf of the Wellcome Sanger Institute COVID-19 Surveillance Team |
| EPI_ISL_2997425 | Lighthouse Lab in Glasgow | Wellcome Sanger Institute for the COVID-19 Genomics UK (COG-UK) Consortium | Harper VanSteenhouse, Yumi Kasai, David Gray, Carol Clugston, Anna Dominiczak and Alex Alderton, Roberto Amato, Jeffrey Barrett, Sonia Goncalves, Ewan Harrison, David K. Jackson, Ian Johnston, Dominic Kwiatkowski, Cordelia Langford, John Sillitoe on behalf of the Wellcome Sanger Institute COVID-19 Surveillance Team |
| EPI_ISL_2997426 | Berkshire and Surrey Pathology Services Lighthouse Laboratory | Wellcome Sanger Institute for the COVID-19 Genomics UK (COG-UK) Consortium | Berkshire and Surrey Pathology Services Lighthouse Laboratory and Alex Alderton, Roberto Amato, Jeffrey Barrett, Sonia Goncalves, Ewan Harrison, David K. Jackson, Ian Johnston, Dominic Kwiatkowski, Cordelia Langford, John Sillitoe on behalf of the Wellcome Sanger Institute COVID-19 Surveillance Team |
| EPI_ISL_2997427 | Lighthouse Lab in Glasgow | Wellcome Sanger Institute for the COVID-19 Genomics UK (COG-UK) Consortium | Harper VanSteenhouse, Yumi Kasai, David Gray, Carol Clugston, Anna Dominiczak and Alex Alderton, Roberto Amato, Jeffrey Barrett, Sonia Goncalves, Ewan Harrison, David K. Jackson, Ian Johnston, Dominic Kwiatkowski, Cordelia Langford, John Sillitoe on behalf of the Wellcome Sanger Institute COVID-19 Surveillance Team |
| EPI_ISL_2997428 | Berkshire and Surrey Pathology Services Lighthouse Laboratory | Wellcome Sanger Institute for the COVID-19 Genomics UK (COG-UK) Consortium | Berkshire and Surrey Pathology Services Lighthouse Laboratory and Alex Alderton, Roberto Amato, Jeffrey Barrett, Sonia Goncalves, Ewan Harrison, David K. Jackson, Ian Johnston, Dominic Kwiatkowski, Cordelia Langford, John Sillitoe on behalf of the Wellcome Sanger Institute COVID-19 Surveillance Team |
| EPI_ISL_2997429 | Lighthouse Lab in Glasgow | Wellcome Sanger Institute for the COVID-19 Genomics UK (COG-UK) Consortium | Harper VanSteenhouse, Yumi Kasai, David Gray, Carol Clugston, Anna Dominiczak and Alex Alderton, Roberto Amato, Jeffrey Barrett, Sonia Goncalves, Ewan Harrison, David K. Jackson, Ian Johnston, Dominic Kwiatkowski, Cordelia Langford, John Sillitoe on behalf of the Wellcome Sanger Institute COVID-19 Surveillance Team |
| EPI_ISL_2997430, EPI_ISL_2997431, EPI_ISL_2997432, EPI_ISL_2997433, EPI_ISL_2997434 | Berkshire and Surrey Pathology Services Lighthouse Laboratory | Wellcome Sanger Institute for the COVID-19 Genomics UK (COG-UK) Consortium | Berkshire and Surrey Pathology Services Lighthouse Laboratory and Alex Alderton, Roberto Amato, Jeffrey Barrett, Sonia Goncalves, Ewan Harrison, David K. Jackson, Ian Johnston, Dominic Kwiatkowski, Cordelia Langford, John Sillitoe on behalf of the Wellcome Sanger Institute COVID-19 Surveillance Team |
| EPI_ISL_2997435, EPI_ISL_2997436 | Lighthouse Lab in Glasgow | Wellcome Sanger Institute for the COVID-19 Genomics UK (COG-UK) Consortium | Harper VanSteenhouse, Yumi Kasai, David Gray, Carol Clugston, Anna Dominiczak and Alex Alderton, Roberto Amato, Jeffrey Barrett, Sonia Goncalves, Ewan Harrison, David K. Jackson, Ian Johnston, Dominic Kwiatkowski, Cordelia Langford, John Sillitoe on behalf of the Wellcome Sanger Institute COVID-19 Surveillance Team |
| EPI_ISL_2997437, EPI_ISL_2997438, EPI_ISL_2997439, EPI_ISL_2997440 | Berkshire and Surrey Pathology Services Lighthouse Laboratory | Wellcome Sanger Institute for the COVID-19 Genomics UK (COG-UK) Consortium | Berkshire and Surrey Pathology Services Lighthouse Laboratory and Alex Alderton, Roberto Amato, Jeffrey Barrett, Sonia Goncalves, Ewan Harrison, David K. Jackson, Ian Johnston, Dominic Kwiatkowski, Cordelia Langford, John Sillitoe on behalf of the Wellcome Sanger Institute COVID-19 Surveillance Team |
| EPI_ISL_2997441, EPI_ISL_2997442 | Lighthouse Lab in Glasgow | Wellcome Sanger Institute for the COVID-19 Genomics UK (COG-UK) Consortium | Harper VanSteenhouse, Yumi Kasai, David Gray, Carol Clugston, Anna Dominiczak and Alex Alderton, Roberto Amato, Jeffrey Barrett, Sonia Goncalves, Ewan Harrison, David K. Jackson, Ian Johnston, Dominic Kwiatkowski, Cordelia Langford, John Sillitoe on behalf of the Wellcome Sanger Institute COVID-19 Surveillance Team |
| EPI_ISL_2997443, EPI_ISL_2997444, EPI_ISL_2997445, EPI_ISL_2997446 | Berkshire and Surrey Pathology Services Lighthouse Laboratory | Wellcome Sanger Institute for the COVID-19 Genomics UK (COG-UK) Consortium | Berkshire and Surrey Pathology Services Lighthouse Laboratory and Alex Alderton, Roberto Amato, Jeffrey Barrett, Sonia Goncalves, Ewan Harrison, David K. Jackson, Ian Johnston, Dominic Kwiatkowski, Cordelia Langford, John Sillitoe on behalf of the Wellcome Sanger Institute COVID-19 Surveillance Team |
| EPI_ISL_2997447 | Lighthouse Lab in Glasgow | Wellcome Sanger Institute for the COVID-19 Genomics UK (COG-UK) Consortium | Harper VanSteenhouse, Yumi Kasai, David Gray, Carol Clugston, Anna Dominiczak and Alex Alderton, Roberto Amato, Jeffrey Barrett, Sonia Goncalves, Ewan Harrison, David K. Jackson, Ian Johnston, Dominic Kwiatkowski, Cordelia Langford, John Sillitoe on behalf of the Wellcome Sanger Institute COVID-19 Surveillance Team |
| EPI_ISL_2997448 | Berkshire and Surrey Pathology Services Lighthouse Laboratory | Wellcome Sanger Institute for the COVID-19 Genomics UK (COG-UK) Consortium | Berkshire and Surrey Pathology Services Lighthouse Laboratory and Alex Alderton, Roberto Amato, Jeffrey Barrett, Sonia Goncalves, Ewan Harrison, David K. Jackson, Ian Johnston, Dominic Kwiatkowski, Cordelia Langford, John Sillitoe on behalf of the Wellcome Sanger Institute COVID-19 Surveillance Team |
| EPI_ISL_2997449 | Lighthouse Lab in Glasgow | Wellcome Sanger Institute for the COVID-19 Genomics UK (COG-UK) Consortium | Harper VanSteenhouse, Yumi Kasai, David Gray, Carol Clugston, Anna Dominiczak and Alex Alderton, Roberto Amato, Jeffrey Barrett, Sonia Goncalves, Ewan Harrison, David K. Jackson, Ian Johnston, Dominic Kwiatkowski, Cordelia Langford, John Sillitoe on behalf of the Wellcome Sanger Institute COVID-19 Surveillance Team |
| EPI_ISL_2997450, EPI_ISL_2997451 | Berkshire and Surrey Pathology Services Lighthouse Laboratory | Wellcome Sanger Institute for the COVID-19 Genomics UK (COG-UK) Consortium | Berkshire and Surrey Pathology Services Lighthouse Laboratory and Alex Alderton, Roberto Amato, Jeffrey Barrett, Sonia Goncalves, Ewan Harrison, David K. Jackson, Ian Johnston, Dominic Kwiatkowski, Cordelia Langford, John Sillitoe on behalf of the Wellcome Sanger Institute COVID-19 Surveillance Team |
| EPI_ISL_2997452, EPI_ISL_2997453 | Lighthouse Lab in Glasgow | Wellcome Sanger Institute for the COVID-19 Genomics UK (COG-UK) Consortium | Harper VanSteenhouse, Yumi Kasai, David Gray, Carol Clugston, Anna Dominiczak and Alex Alderton, Roberto Amato, Jeffrey Barrett, Sonia Goncalves, Ewan Harrison, David K. Jackson, Ian Johnston, Dominic Kwiatkowski, Cordelia Langford, John Sillitoe on behalf of the Wellcome Sanger Institute COVID-19 Surveillance Team |
| EPI_ISL_2997454 | Berkshire and Surrey Pathology Services Lighthouse Laboratory | Wellcome Sanger Institute for the COVID-19 Genomics UK (COG-UK) Consortium | Berkshire and Surrey Pathology Services Lighthouse Laboratory and Alex Alderton, Roberto Amato, Jeffrey Barrett, Sonia Goncalves, Ewan Harrison, David K. Jackson, Ian Johnston, Dominic Kwiatkowski, Cordelia Langford, John Sillitoe on behalf of the Wellcome Sanger Institute COVID-19 Surveillance Team |
| EPI_ISL_2997455, EPI_ISL_2997456 | Lighthouse Lab in Glasgow | Wellcome Sanger Institute for the COVID-19 Genomics UK (COG-UK) Consortium | Harper VanSteenhouse, Yumi Kasai, David Gray, Carol Clugston, Anna Dominiczak and Alex Alderton, Roberto Amato, Jeffrey Barrett, Sonia Goncalves, Ewan Harrison, David K. Jackson, Ian Johnston, Dominic Kwiatkowski, Cordelia Langford, John Sillitoe on behalf of the Wellcome Sanger Institute COVID-19 Surveillance Team |
| EPI_ISL_2997457, EPI_ISL_2997458, EPI_ISL_2997459, EPI_ISL_2997460, EPI_ISL_2997461 | Berkshire and Surrey Pathology Services Lighthouse Laboratory | Wellcome Sanger Institute for the COVID-19 Genomics UK (COG-UK) Consortium | Berkshire and Surrey Pathology Services Lighthouse Laboratory and Alex Alderton, Roberto Amato, Jeffrey Barrett, Sonia Goncalves, Ewan Harrison, David K. Jackson, Ian Johnston, Dominic Kwiatkowski, Cordelia Langford, John Sillitoe on behalf of the Wellcome Sanger Institute COVID-19 Surveillance Team |
| EPI_ISL_2997462, EPI_ISL_2997463 | Lighthouse Lab in Glasgow | Wellcome Sanger Institute for the COVID-19 Genomics UK (COG-UK) Consortium | Harper VanSteenhouse, Yumi Kasai, David Gray, Carol Clugston, Anna Dominiczak and Alex Alderton, Roberto Amato, Jeffrey Barrett, Sonia Goncalves, Ewan Harrison, David K. Jackson, Ian Johnston, Dominic Kwiatkowski, Cordelia Langford, John Sillitoe on behalf of the Wellcome Sanger Institute COVID-19 Surveillance Team |
| EPI_ISL_2997464, EPI_ISL_2997465, EPI_ISL_2997466, EPI_ISL_2997467, EPI_ISL_2997468, EPI_ISL_2997469 | Berkshire and Surrey Pathology Services Lighthouse Laboratory | Wellcome Sanger Institute for the COVID-19 Genomics UK (COG-UK) Consortium | Berkshire and Surrey Pathology Services Lighthouse Laboratory and Alex Alderton, Roberto Amato, Jeffrey Barrett, Sonia Goncalves, Ewan Harrison, David K. Jackson, Ian Johnston, Dominic Kwiatkowski, Cordelia Langford, John Sillitoe on behalf of the Wellcome Sanger Institute COVID-19 Surveillance Team |
| EPI_ISL_2997470 | Lighthouse Lab in Glasgow | Wellcome Sanger Institute for the COVID-19 Genomics UK (COG-UK) Consortium | Harper VanSteenhouse, Yumi Kasai, David Gray, Carol Clugston, Anna Dominiczak and Alex Alderton, Roberto Amato, Jeffrey Barrett, Sonia Goncalves, Ewan Harrison, David K. Jackson, Ian Johnston, Dominic Kwiatkowski, Cordelia Langford, John Sillitoe on behalf of the Wellcome Sanger Institute COVID-19 Surveillance Team |
| EPI_ISL_2997471 | Berkshire and Surrey Pathology Services Lighthouse Laboratory | Wellcome Sanger Institute for the COVID-19 Genomics UK (COG-UK) Consortium | Berkshire and Surrey Pathology Services Lighthouse Laboratory and Alex Alderton, Roberto Amato, Jeffrey Barrett, Sonia Goncalves, Ewan Harrison, David K. Jackson, Ian Johnston, Dominic Kwiatkowski, Cordelia Langford, John Sillitoe on behalf of the Wellcome Sanger Institute COVID-19 Surveillance Team |
| EPI_ISL_2997472 | Lighthouse Lab in Glasgow | Wellcome Sanger Institute for the COVID-19 Genomics UK (COG-UK) Consortium | Harper VanSteenhouse, Yumi Kasai, David Gray, Carol Clugston, Anna Dominiczak and Alex Alderton, Roberto Amato, Jeffrey Barrett, Sonia Goncalves, Ewan Harrison, David K. Jackson, Ian Johnston, Dominic Kwiatkowski, Cordelia Langford, John Sillitoe on behalf of the Wellcome Sanger Institute COVID-19 Surveillance Team |

EPI_ISL_2997473, EPI_ISL_2997474, EPI_ISL_2997475, EPI_ISL_2997476, EPI_ISL_2997477, EPI_ISL_2997478, EPI_ISL_2997479, EPI_ISL_2997480, EPI_ISL_2997481, EPI_ISL_2997482, EPI_ISL_2997483

| | | | |
|---|---|---|---|
| see above | Berkshire and Surrey Pathology Services Lighthouse Laboratory | Wellcome Sanger Institute for the COVID-19 Genomics UK (COG-UK) Consortium | Berkshire and Surrey Pathology Services Lighthouse Laboratory and Alex Alderton, Roberto Amato, Jeffrey Barrett, Sonia Goncalves, Ewan Harrison, David K. Jackson, Ian Johnston, Dominic Kwiatkowski, Cordelia Langford, John Sillitoe on behalf of the Wellcome Sanger Institute COVID-19 Surveillance Team |
| EPI_ISL_2997484, EPI_ISL_2997485 | Lighthouse Lab in Glasgow | Wellcome Sanger Institute for the COVID-19 Genomics UK (COG-UK) Consortium | Harper VanSteenhouse, Yumi Kasai, David Gray, Carol Clugston, Anna Dominiczak and Alex Alderton, Roberto Amato, Jeffrey Barrett, Sonia Goncalves, Ewan Harrison, David K. Jackson, Ian Johnston, Dominic Kwiatkowski, Cordelia Langford, John Sillitoe on behalf of the Wellcome Sanger Institute COVID-19 Surveillance Team |
| EPI_ISL_2997486, EPI_ISL_2997487 | Berkshire and Surrey Pathology Services Lighthouse Laboratory | Wellcome Sanger Institute for the COVID-19 Genomics UK (COG-UK) Consortium | Berkshire and Surrey Pathology Services Lighthouse Laboratory and Alex Alderton, Roberto Amato, Jeffrey Barrett, Sonia Goncalves, Ewan Harrison, David K. Jackson, Ian Johnston, Dominic Kwiatkowski, Cordelia Langford, John Sillitoe on behalf of the Wellcome Sanger Institute COVID-19 Surveillance Team |
| EPI_ISL_2997488, EPI_ISL_2997489, EPI_ISL_2997490 | Lighthouse Lab in Glasgow | Wellcome Sanger Institute for the COVID-19 Genomics UK (COG-UK) Consortium | Harper VanSteenhouse, Yumi Kasai, David Gray, Carol Clugston, Anna Dominiczak and Alex Alderton, Roberto Amato, Jeffrey Barrett, Sonia Goncalves, Ewan Harrison, David K. Jackson, Ian Johnston, Dominic Kwiatkowski, Cordelia Langford, John Sillitoe on behalf of the Wellcome Sanger Institute COVID-19 Surveillance Team |
| EPI_ISL_2997491, EPI_ISL_2997492 | Berkshire and Surrey Pathology Services Lighthouse Laboratory | Wellcome Sanger Institute for the COVID-19 Genomics UK (COG-UK) Consortium | Berkshire and Surrey Pathology Services Lighthouse Laboratory and Alex Alderton, Roberto Amato, Jeffrey Barrett, Sonia Goncalves, Ewan Harrison, David K. Jackson, Ian Johnston, Dominic Kwiatkowski, Cordelia Langford, John Sillitoe on behalf of the Wellcome Sanger Institute COVID-19 Surveillance Team |
| EPI_ISL_2997493 | Lighthouse Lab in Glasgow | Wellcome Sanger Institute for the COVID-19 Genomics UK (COG-UK) Consortium | Harper VanSteenhouse, Yumi Kasai, David Gray, Carol Clugston, Anna Dominiczak and Alex Alderton, Roberto Amato, Jeffrey Barrett, Sonia Goncalves, Ewan Harrison, David K. Jackson, Ian Johnston, Dominic Kwiatkowski, Cordelia Langford, John Sillitoe on behalf of the Wellcome Sanger Institute COVID-19 Surveillance Team |
| EPI_ISL_2997494, EPI_ISL_2997495 | Berkshire and Surrey Pathology Services Lighthouse Laboratory | Wellcome Sanger Institute for the COVID-19 Genomics UK (COG-UK) Consortium | Berkshire and Surrey Pathology Services Lighthouse Laboratory and Alex Alderton, Roberto Amato, Jeffrey Barrett, Sonia Goncalves, Ewan Harrison, David K. Jackson, Ian Johnston, Dominic Kwiatkowski, Cordelia Langford, John Sillitoe on behalf of the Wellcome Sanger Institute COVID-19 Surveillance Team |
| EPI_ISL_2997496 | Lighthouse Lab in Glasgow | Wellcome Sanger Institute for the COVID-19 Genomics UK (COG-UK) Consortium | Harper VanSteenhouse, Yumi Kasai, David Gray, Carol Clugston, Anna Dominiczak and Alex Alderton, Roberto Amato, Jeffrey Barrett, Sonia Goncalves, Ewan Harrison, David K. Jackson, Ian Johnston, Dominic Kwiatkowski, Cordelia Langford, John Sillitoe on behalf of the Wellcome Sanger Institute COVID-19 Surveillance Team |
| EPI_ISL_2997497, EPI_ISL_2997498, EPI_ISL_2997499, EPI_ISL_2997500, EPI_ISL_2997501, EPI_ISL_2997502, EPI_ISL_2997503, EPI_ISL_2997504, EPI_ISL_2997505, EPI_ISL_2997506, EPI_ISL_2997507, EPI_ISL_2997508, EPI_ISL_2997509 | | | |
| see above | Berkshire and Surrey Pathology Services Lighthouse Laboratory | Wellcome Sanger Institute for the COVID-19 Genomics UK (COG-UK) Consortium | Berkshire and Surrey Pathology Services Lighthouse Laboratory and Alex Alderton, Roberto Amato, Jeffrey Barrett, Sonia Goncalves, Ewan Harrison, David K. Jackson, Ian Johnston, Dominic Kwiatkowski, Cordelia Langford, John Sillitoe on behalf of the Wellcome Sanger Institute COVID-19 Surveillance Team |
| EPI_ISL_2997510 | Lighthouse Lab in Glasgow | Wellcome Sanger Institute for the COVID-19 Genomics UK (COG-UK) Consortium | Harper VanSteenhouse, Yumi Kasai, David Gray, Carol Clugston, Anna Dominiczak and Alex Alderton, Roberto Amato, Jeffrey Barrett, Sonia Goncalves, Ewan Harrison, David K. Jackson, Ian Johnston, Dominic Kwiatkowski, Cordelia Langford, John Sillitoe on behalf of the Wellcome Sanger Institute COVID-19 Surveillance Team |
| EPI_ISL_2997511, EPI_ISL_2997512 | Berkshire and Surrey Pathology Services Lighthouse Laboratory | Wellcome Sanger Institute for the COVID-19 Genomics UK (COG-UK) Consortium | Berkshire and Surrey Pathology Services Lighthouse Laboratory and Alex Alderton, Roberto Amato, Jeffrey Barrett, Sonia Goncalves, Ewan Harrison, David K. Jackson, Ian Johnston, Dominic Kwiatkowski, Cordelia Langford, John Sillitoe on behalf of the Wellcome Sanger Institute COVID-19 Surveillance Team |
| EPI_ISL_2997513 | Lighthouse Lab in Glasgow | Wellcome Sanger Institute for the COVID-19 Genomics UK (COG-UK) Consortium | Harper VanSteenhouse, Yumi Kasai, David Gray, Carol Clugston, Anna Dominiczak and Alex Alderton, Roberto Amato, Jeffrey Barrett, Sonia Goncalves, Ewan Harrison, David K. Jackson, Ian Johnston, Dominic Kwiatkowski, Cordelia Langford, John Sillitoe on behalf of the Wellcome Sanger Institute COVID-19 Surveillance Team |
| EPI_ISL_2997514 | Berkshire and Surrey Pathology Services Lighthouse Laboratory | Wellcome Sanger Institute for the COVID-19 Genomics UK (COG-UK) Consortium | Berkshire and Surrey Pathology Services Lighthouse Laboratory and Alex Alderton, Roberto Amato, Jeffrey Barrett, Sonia Goncalves, Ewan Harrison, David K. Jackson, Ian Johnston, Dominic Kwiatkowski, Cordelia Langford, John Sillitoe on behalf of the Wellcome Sanger Institute COVID-19 Surveillance Team |
| EPI_ISL_2997515 | Lighthouse Lab in Glasgow | Wellcome Sanger Institute for the COVID-19 Genomics UK (COG-UK) Consortium | Harper VanSteenhouse, Yumi Kasai, David Gray, Carol Clugston, Anna Dominiczak and Alex Alderton, Roberto Amato, Jeffrey Barrett, Sonia Goncalves, Ewan Harrison, David K. Jackson, Ian Johnston, Dominic Kwiatkowski, Cordelia Langford, John Sillitoe on behalf of the Wellcome Sanger Institute COVID-19 Surveillance Team |
| EPI_ISL_2997516 | Berkshire and Surrey Pathology Services Lighthouse Laboratory | Wellcome Sanger Institute for the COVID-19 Genomics UK (COG-UK) Consortium | Berkshire and Surrey Pathology Services Lighthouse Laboratory and Alex Alderton, Roberto Amato, Jeffrey Barrett, Sonia Goncalves, Ewan Harrison, David K. Jackson, Ian Johnston, Dominic Kwiatkowski, Cordelia Langford, John Sillitoe on behalf of the Wellcome Sanger Institute COVID-19 Surveillance Team |
| EPI_ISL_2997517 | Lighthouse Lab in Glasgow | Wellcome Sanger Institute for the COVID-19 Genomics UK (COG-UK) Consortium | Harper VanSteenhouse, Yumi Kasai, David Gray, Carol Clugston, Anna Dominiczak and Alex Alderton, Roberto Amato, Jeffrey Barrett, Sonia Goncalves, Ewan Harrison, David K. Jackson, Ian Johnston, Dominic Kwiatkowski, Cordelia Langford, John Sillitoe on behalf of the Wellcome Sanger Institute COVID-19 Surveillance Team |
| EPI_ISL_2997518, EPI_ISL_2997519, EPI_ISL_2997520 | Berkshire and Surrey Pathology Services Lighthouse Laboratory | Wellcome Sanger Institute for the COVID-19 Genomics UK (COG-UK) Consortium | Berkshire and Surrey Pathology Services Lighthouse Laboratory and Alex Alderton, Roberto Amato, Jeffrey Barrett, Sonia Goncalves, Ewan Harrison, David K. Jackson, Ian Johnston, Dominic Kwiatkowski, Cordelia Langford, John Sillitoe on behalf of the Wellcome Sanger Institute COVID-19 Surveillance Team |
| EPI_ISL_2997521, EPI_ISL_2997522 | Lighthouse Lab in Glasgow | Wellcome Sanger Institute for the COVID-19 Genomics UK (COG-UK) Consortium | Harper VanSteenhouse, Yumi Kasai, David Gray, Carol Clugston, Anna Dominiczak and Alex Alderton, Roberto Amato, Jeffrey Barrett, Sonia Goncalves, Ewan Harrison, David K. Jackson, Ian Johnston, Dominic Kwiatkowski, Cordelia Langford, John Sillitoe on behalf of the Wellcome Sanger Institute COVID-19 Surveillance Team |
| EPI_ISL_2997523, EPI_ISL_2997524 | Berkshire and Surrey Pathology Services Lighthouse Laboratory | Wellcome Sanger Institute for the COVID-19 Genomics UK (COG-UK) Consortium | Berkshire and Surrey Pathology Services Lighthouse Laboratory and Alex Alderton, Roberto Amato, Jeffrey Barrett, Sonia Goncalves, Ewan Harrison, David K. Jackson, Ian Johnston, Dominic Kwiatkowski, Cordelia Langford, John Sillitoe on behalf of the Wellcome Sanger Institute COVID-19 Surveillance Team |
| EPI_ISL_2997525 | Lighthouse Lab in Glasgow | Wellcome Sanger Institute for the COVID-19 Genomics UK (COG-UK) Consortium | Harper VanSteenhouse, Yumi Kasai, David Gray, Carol Clugston, Anna Dominiczak and Alex Alderton, Roberto Amato, Jeffrey Barrett, Sonia Goncalves, Ewan Harrison, David K. Jackson, Ian Johnston, Dominic Kwiatkowski, Cordelia Langford, John Sillitoe on behalf of the Wellcome Sanger Institute COVID-19 Surveillance Team |
| EPI_ISL_2997526, EPI_ISL_2997527, EPI_ISL_2997528, EPI_ISL_2997529, EPI_ISL_2997530, EPI_ISL_2997531, EPI_ISL_2997532 | Berkshire and Surrey Pathology Services Lighthouse Laboratory | Wellcome Sanger Institute for the COVID-19 Genomics UK (COG-UK) Consortium | Berkshire and Surrey Pathology Services Lighthouse Laboratory and Alex Alderton, Roberto Amato, Jeffrey Barrett, Sonia Goncalves, Ewan Harrison, David K. Jackson, Ian Johnston, Dominic Kwiatkowski, Cordelia Langford, John Sillitoe on behalf of the Wellcome Sanger Institute COVID-19 Surveillance Team |
| EPI_ISL_2997533 | Lighthouse Lab in Glasgow | Wellcome Sanger Institute for the COVID-19 Genomics UK (COG-UK) Consortium | Harper VanSteenhouse, Yumi Kasai, David Gray, Carol Clugston, Anna Dominiczak and Alex Alderton, Roberto Amato, Jeffrey Barrett, Sonia Goncalves, Ewan Harrison, David K. Jackson, Ian Johnston, Dominic Kwiatkowski, Cordelia Langford, John Sillitoe on behalf of the Wellcome Sanger Institute COVID-19 Surveillance Team |
| EPI_ISL_2997534, EPI_ISL_2997535, EPI_ISL_2997536, EPI_ISL_2997537, EPI_ISL_2997538 | Berkshire and Surrey Pathology Services Lighthouse Laboratory | Wellcome Sanger Institute for the COVID-19 Genomics UK (COG-UK) Consortium | Berkshire and Surrey Pathology Services Lighthouse Laboratory and Alex Alderton, Roberto Amato, Jeffrey Barrett, Sonia Goncalves, Ewan Harrison, David K. Jackson, Ian Johnston, Dominic Kwiatkowski, Cordelia Langford, John Sillitoe on behalf of the Wellcome Sanger Institute COVID-19 Surveillance Team |
| EPI_ISL_2997539 | Lighthouse Lab in Glasgow | Wellcome Sanger Institute for the COVID-19 Genomics UK (COG-UK) Consortium | Harper VanSteenhouse, Yumi Kasai, David Gray, Carol Clugston, Anna Dominiczak and Alex Alderton, Roberto Amato, Jeffrey Barrett, Sonia Goncalves, Ewan Harrison, David K. Jackson, Ian Johnston, Dominic Kwiatkowski, Cordelia Langford, John Sillitoe on behalf of the Wellcome Sanger Institute COVID-19 Surveillance Team |
| EPI_ISL_2997540, EPI_ISL_2997541, EPI_ISL_2997542, EPI_ISL_2997543, EPI_ISL_2997544 | Berkshire and Surrey Pathology Services Lighthouse Laboratory | Wellcome Sanger Institute for the COVID-19 Genomics UK (COG-UK) Consortium | Berkshire and Surrey Pathology Services Lighthouse Laboratory and Alex Alderton, Roberto Amato, Jeffrey Barrett, Sonia Goncalves, Ewan Harrison, David K. Jackson, Ian Johnston, Dominic Kwiatkowski, Cordelia Langford, John Sillitoe on behalf of the Wellcome Sanger Institute COVID-19 Surveillance Team |
| EPI_ISL_2997545, EPI_ISL_2997546, EPI_ISL_2997547, EPI_ISL_2997548, EPI_ISL_2997549 | Lighthouse Lab in Glasgow | Wellcome Sanger Institute for the COVID-19 Genomics UK (COG-UK) Consortium | Harper VanSteenhouse, Yumi Kasai, David Gray, Carol Clugston, Anna Dominiczak and Alex Alderton, Roberto Amato, Jeffrey Barrett, Sonia Goncalves, Ewan Harrison, David K. Jackson, Ian Johnston, Dominic Kwiatkowski, Cordelia Langford, John Sillitoe on behalf of the Wellcome Sanger Institute COVID-19 Surveillance Team |
| EPI_ISL_2997550, EPI_ISL_2997551 | Berkshire and Surrey Pathology Services Lighthouse Laboratory | Wellcome Sanger Institute for the COVID-19 Genomics UK (COG-UK) Consortium | Berkshire and Surrey Pathology Services Lighthouse Laboratory and Alex Alderton, Roberto Amato, Jeffrey Barrett, Sonia Goncalves, Ewan Harrison, David K. Jackson, Ian Johnston, Dominic Kwiatkowski, Cordelia Langford, John Sillitoe on behalf of the Wellcome Sanger Institute COVID-19 Surveillance Team |
| EPI_ISL_2997552 | Lighthouse Lab in Glasgow | Wellcome Sanger Institute for the COVID-19 Genomics UK (COG-UK) Consortium | Harper VanSteenhouse, Yumi Kasai, David Gray, Carol Clugston, Anna Dominiczak and Alex Alderton, Roberto Amato, Jeffrey Barrett, Sonia Goncalves, Ewan Harrison, David K. Jackson, Ian Johnston, Dominic Kwiatkowski, Cordelia Langford, John Sillitoe on behalf of the Wellcome Sanger Institute COVID-19 Surveillance Team |
| EPI_ISL_2997553, EPI_ISL_2997554, EPI_ISL_2997555, EPI_ISL_2997556, EPI_ISL_2997557, EPI_ISL_2997558 | Berkshire and Surrey Pathology Services Lighthouse Laboratory | Wellcome Sanger Institute for the COVID-19 Genomics UK (COG-UK) Consortium | Berkshire and Surrey Pathology Services Lighthouse Laboratory and Alex Alderton, Roberto Amato, Jeffrey Barrett, Sonia Goncalves, Ewan Harrison, David K. Jackson, Ian Johnston, Dominic Kwiatkowski, Cordelia Langford, John Sillitoe on behalf of the Wellcome Sanger Institute COVID-19 Surveillance Team |
| EPI_ISL_2997559 | Lighthouse Lab in Glasgow | Wellcome Sanger Institute for the COVID-19 Genomics UK (COG-UK) Consortium | Harper VanSteenhouse, Yumi Kasai, David Gray, Carol Clugston, Anna Dominiczak and Alex Alderton, Roberto Amato, Jeffrey Barrett, Sonia Goncalves, Ewan Harrison, David K. Jackson, Ian Johnston, Dominic Kwiatkowski, Cordelia Langford, John Sillitoe on behalf of the Wellcome Sanger Institute COVID-19 Surveillance Team |
| EPI_ISL_2997560, EPI_ISL_2997561, EPI_ISL_2997562 | Berkshire and Surrey Pathology Services Lighthouse Laboratory | Wellcome Sanger Institute for the COVID-19 Genomics UK (COG-UK) Consortium | Berkshire and Surrey Pathology Services Lighthouse Laboratory and Alex Alderton, Roberto Amato, Jeffrey Barrett, Sonia Goncalves, Ewan Harrison, David K. Jackson, Ian Johnston, Dominic Kwiatkowski, Cordelia Langford, John Sillitoe on behalf of the Wellcome Sanger Institute COVID-19 Surveillance Team |
| EPI_ISL_2997563 | Lighthouse Lab in Glasgow | Wellcome Sanger Institute for the COVID-19 Genomics UK (COG-UK) Consortium | Harper VanSteenhouse, Yumi Kasai, David Gray, Carol Clugston, Anna Dominiczak and Alex Alderton, Roberto Amato, Jeffrey Barrett, Sonia Goncalves, Ewan Harrison, David K. Jackson, Ian Johnston, Dominic Kwiatkowski, Cordelia Langford, John Sillitoe on behalf of the Wellcome Sanger Institute COVID-19 Surveillance Team |
| EPI_ISL_2997564, EPI_ISL_2997565, EPI_ISL_2997566, EPI_ISL_2997567, EPI_ISL_2997568 | Berkshire and Surrey Pathology Services Lighthouse Laboratory | Wellcome Sanger Institute for the COVID-19 Genomics UK (COG-UK) Consortium | Berkshire and Surrey Pathology Services Lighthouse Laboratory and Alex Alderton, Roberto Amato, Jeffrey Barrett, Sonia Goncalves, Ewan Harrison, David K. Jackson, Ian Johnston, Dominic Kwiatkowski, Cordelia Langford, John Sillitoe on behalf of the Wellcome Sanger Institute COVID-19 Surveillance Team |
| EPI_ISL_2997569 | Lighthouse Lab in Glasgow | Wellcome Sanger Institute for the COVID-19 Genomics UK (COG-UK) Consortium | Harper VanSteenhouse, Yumi Kasai, David Gray, Carol Clugston, Anna Dominiczak and Alex Alderton, Roberto Amato, Jeffrey Barrett, Sonia Goncalves, Ewan Harrison, David K. Jackson, Ian Johnston, Dominic Kwiatkowski, Cordelia Langford, John Sillitoe on behalf of the Wellcome Sanger Institute COVID-19 Surveillance Team |
| EPI_ISL_2997570, | Berkshire and Surrey Pathology Services Lighthouse Laboratory | Wellcome Sanger Institute for the COVID-19 Genomics UK (COG-UK) | Berkshire and Surrey Pathology Services Lighthouse Laboratory and Alex Alderton, Roberto Amato, Jeffrey Barrett, Sonia Goncalves, Ewan Harrison, David K. Jackson, Ian Johnston, Dominic Kwiatkowski, Cordelia Langford, John Sillitoe on behalf |

| | | | |
|---|---|---|---|
| EPI_ISL_2997571, EPI_ISL_2997572, EPI_ISL_2997573 | | Consortium | of the Wellcome Sanger Institute COVID-19 Surveillance Team |
| EPI_ISL_2997574 | Lighthouse Lab in Glasgow | Wellcome Sanger Institute for the COVID-19 Genomics UK (COG-UK) Consortium | Harper VanSteenhouse, Yumi Kasai, David Gray, Carol Clugston, Anna Dominiczak and Alex Alderton, Roberto Amato, Jeffrey Barrett, Sonia Goncalves, Ewan Harrison, David K. Jackson, Ian Johnston, Dominic Kwiatkowski, Cordelia Langford, John Sillitoe on behalf of the Wellcome Sanger Institute COVID-19 Surveillance Team |
| EPI_ISL_2997575, EPI_ISL_2997576, EPI_ISL_2997577, EPI_ISL_2997578, EPI_ISL_2997579 | Berkshire and Surrey Pathology Services Lighthouse Laboratory | Wellcome Sanger Institute for the COVID-19 Genomics UK (COG-UK) Consortium | Berkshire and Surrey Pathology Services Lighthouse Laboratory and Alex Alderton, Roberto Amato, Jeffrey Barrett, Sonia Goncalves, Ewan Harrison, David K. Jackson, Ian Johnston, Dominic Kwiatkowski, Cordelia Langford, John Sillitoe on behalf of the Wellcome Sanger Institute COVID-19 Surveillance Team |
| EPI_ISL_2997580 | Lighthouse Lab in Glasgow | Wellcome Sanger Institute for the COVID-19 Genomics UK (COG-UK) Consortium | Harper VanSteenhouse, Yumi Kasai, David Gray, Carol Clugston, Anna Dominiczak and Alex Alderton, Roberto Amato, Jeffrey Barrett, Sonia Goncalves, Ewan Harrison, David K. Jackson, Ian Johnston, Dominic Kwiatkowski, Cordelia Langford, John Sillitoe on behalf of the Wellcome Sanger Institute COVID-19 Surveillance Team |
| EPI_ISL_2997581, EPI_ISL_2997582, EPI_ISL_2997583, EPI_ISL_2997584, EPI_ISL_2997585, EPI_ISL_2997586 | Berkshire and Surrey Pathology Services Lighthouse Laboratory | Wellcome Sanger Institute for the COVID-19 Genomics UK (COG-UK) Consortium | Berkshire and Surrey Pathology Services Lighthouse Laboratory and Alex Alderton, Roberto Amato, Jeffrey Barrett, Sonia Goncalves, Ewan Harrison, David K. Jackson, Ian Johnston, Dominic Kwiatkowski, Cordelia Langford, John Sillitoe on behalf of the Wellcome Sanger Institute COVID-19 Surveillance Team |
| EPI_ISL_2997587, EPI_ISL_2997588 | Lighthouse Lab in Glasgow | Wellcome Sanger Institute for the COVID-19 Genomics UK (COG-UK) Consortium | Harper VanSteenhouse, Yumi Kasai, David Gray, Carol Clugston, Anna Dominiczak and Alex Alderton, Roberto Amato, Jeffrey Barrett, Sonia Goncalves, Ewan Harrison, David K. Jackson, Ian Johnston, Dominic Kwiatkowski, Cordelia Langford, John Sillitoe on behalf of the Wellcome Sanger Institute COVID-19 Surveillance Team |
| EPI_ISL_2997589, EPI_ISL_2997590, EPI_ISL_2997591 | Berkshire and Surrey Pathology Services Lighthouse Laboratory | Wellcome Sanger Institute for the COVID-19 Genomics UK (COG-UK) Consortium | Berkshire and Surrey Pathology Services Lighthouse Laboratory and Alex Alderton, Roberto Amato, Jeffrey Barrett, Sonia Goncalves, Ewan Harrison, David K. Jackson, Ian Johnston, Dominic Kwiatkowski, Cordelia Langford, John Sillitoe on behalf of the Wellcome Sanger Institute COVID-19 Surveillance Team |
| EPI_ISL_2997592, EPI_ISL_2997593 | Lighthouse Lab in Glasgow | Wellcome Sanger Institute for the COVID-19 Genomics UK (COG-UK) Consortium | Harper VanSteenhouse, Yumi Kasai, David Gray, Carol Clugston, Anna Dominiczak and Alex Alderton, Roberto Amato, Jeffrey Barrett, Sonia Goncalves, Ewan Harrison, David K. Jackson, Ian Johnston, Dominic Kwiatkowski, Cordelia Langford, John Sillitoe on behalf of the Wellcome Sanger Institute COVID-19 Surveillance Team |
| EPI_ISL_2997594, EPI_ISL_2997595, EPI_ISL_2997596 | Berkshire and Surrey Pathology Services Lighthouse Laboratory | Wellcome Sanger Institute for the COVID-19 Genomics UK (COG-UK) Consortium | Berkshire and Surrey Pathology Services Lighthouse Laboratory and Alex Alderton, Roberto Amato, Jeffrey Barrett, Sonia Goncalves, Ewan Harrison, David K. Jackson, Ian Johnston, Dominic Kwiatkowski, Cordelia Langford, John Sillitoe on behalf of the Wellcome Sanger Institute COVID-19 Surveillance Team |
| EPI_ISL_2997597, EPI_ISL_2997598 | Lighthouse Lab in Glasgow | Wellcome Sanger Institute for the COVID-19 Genomics UK (COG-UK) Consortium | Harper VanSteenhouse, Yumi Kasai, David Gray, Carol Clugston, Anna Dominiczak and Alex Alderton, Roberto Amato, Jeffrey Barrett, Sonia Goncalves, Ewan Harrison, David K. Jackson, Ian Johnston, Dominic Kwiatkowski, Cordelia Langford, John Sillitoe on behalf of the Wellcome Sanger Institute COVID-19 Surveillance Team |
| EPI_ISL_2997599 | Berkshire and Surrey Pathology Services Lighthouse Laboratory | Wellcome Sanger Institute for the COVID-19 Genomics UK (COG-UK) Consortium | Berkshire and Surrey Pathology Services Lighthouse Laboratory and Alex Alderton, Roberto Amato, Jeffrey Barrett, Sonia Goncalves, Ewan Harrison, David K. Jackson, Ian Johnston, Dominic Kwiatkowski, Cordelia Langford, John Sillitoe on behalf of the Wellcome Sanger Institute COVID-19 Surveillance Team |
| EPI_ISL_2997600 | Lighthouse Lab in Glasgow | Wellcome Sanger Institute for the COVID-19 Genomics UK (COG-UK) Consortium | Harper VanSteenhouse, Yumi Kasai, David Gray, Carol Clugston, Anna Dominiczak and Alex Alderton, Roberto Amato, Jeffrey Barrett, Sonia Goncalves, Ewan Harrison, David K. Jackson, Ian Johnston, Dominic Kwiatkowski, Cordelia Langford, John Sillitoe on behalf of the Wellcome Sanger Institute COVID-19 Surveillance Team |
| EPI_ISL_2997601, EPI_ISL_2997602, EPI_ISL_2997603, EPI_ISL_2997604, EPI_ISL_2997605 | Berkshire and Surrey Pathology Services Lighthouse Laboratory | Wellcome Sanger Institute for the COVID-19 Genomics UK (COG-UK) Consortium | Berkshire and Surrey Pathology Services Lighthouse Laboratory and Alex Alderton, Roberto Amato, Jeffrey Barrett, Sonia Goncalves, Ewan Harrison, David K. Jackson, Ian Johnston, Dominic Kwiatkowski, Cordelia Langford, John Sillitoe on behalf of the Wellcome Sanger Institute COVID-19 Surveillance Team |
| EPI_ISL_2997606, EPI_ISL_2997607, EPI_ISL_2997608 | Lighthouse Lab in Glasgow | Wellcome Sanger Institute for the COVID-19 Genomics UK (COG-UK) Consortium | Harper VanSteenhouse, Yumi Kasai, David Gray, Carol Clugston, Anna Dominiczak and Alex Alderton, Roberto Amato, Jeffrey Barrett, Sonia Goncalves, Ewan Harrison, David K. Jackson, Ian Johnston, Dominic Kwiatkowski, Cordelia Langford, John Sillitoe on behalf of the Wellcome Sanger Institute COVID-19 Surveillance Team |
| EPI_ISL_2997609, EPI_ISL_2997610, EPI_ISL_2997611, EPI_ISL_2997612, EPI_ISL_2997613, EPI_ISL_2997614, EPI_ISL_2997615, EPI_ISL_2997616, EPI_ISL_2997617, EPI_ISL_2997618, EPI_ISL_2997619, EPI_ISL_2997620 | | | |
| see above | Berkshire and Surrey Pathology Services Lighthouse Laboratory | Wellcome Sanger Institute for the COVID-19 Genomics UK (COG-UK) Consortium | Berkshire and Surrey Pathology Services Lighthouse Laboratory and Alex Alderton, Roberto Amato, Jeffrey Barrett, Sonia Goncalves, Ewan Harrison, David K. Jackson, Ian Johnston, Dominic Kwiatkowski, Cordelia Langford, John Sillitoe on behalf of the Wellcome Sanger Institute COVID-19 Surveillance Team |
| EPI_ISL_2997621 | Lighthouse Lab in Glasgow | Wellcome Sanger Institute for the COVID-19 Genomics UK (COG-UK) Consortium | Harper VanSteenhouse, Yumi Kasai, David Gray, Carol Clugston, Anna Dominiczak and Alex Alderton, Roberto Amato, Jeffrey Barrett, Sonia Goncalves, Ewan Harrison, David K. Jackson, Ian Johnston, Dominic Kwiatkowski, Cordelia Langford, John Sillitoe on behalf of the Wellcome Sanger Institute COVID-19 Surveillance Team |
| EPI_ISL_2997622, EPI_ISL_2997623, EPI_ISL_2997624, EPI_ISL_2997625, EPI_ISL_2997626, EPI_ISL_2997627, EPI_ISL_2997628 | Berkshire and Surrey Pathology Services Lighthouse Laboratory | Wellcome Sanger Institute for the COVID-19 Genomics UK (COG-UK) Consortium | Berkshire and Surrey Pathology Services Lighthouse Laboratory and Alex Alderton, Roberto Amato, Jeffrey Barrett, Sonia Goncalves, Ewan Harrison, David K. Jackson, Ian Johnston, Dominic Kwiatkowski, Cordelia Langford, John Sillitoe on behalf of the Wellcome Sanger Institute COVID-19 Surveillance Team |
| EPI_ISL_2997629 | Lighthouse Lab in Glasgow | Wellcome Sanger Institute for the COVID-19 Genomics UK (COG-UK) Consortium | Harper VanSteenhouse, Yumi Kasai, David Gray, Carol Clugston, Anna Dominiczak and Alex Alderton, Roberto Amato, Jeffrey Barrett, Sonia Goncalves, Ewan Harrison, David K. Jackson, Ian Johnston, Dominic Kwiatkowski, Cordelia Langford, John Sillitoe on behalf of the Wellcome Sanger Institute COVID-19 Surveillance Team |
| EPI_ISL_2997630, EPI_ISL_2997631, EPI_ISL_2997632, EPI_ISL_2997633, EPI_ISL_2997634, EPI_ISL_2997635 | Berkshire and Surrey Pathology Services Lighthouse Laboratory | Wellcome Sanger Institute for the COVID-19 Genomics UK (COG-UK) Consortium | Berkshire and Surrey Pathology Services Lighthouse Laboratory and Alex Alderton, Roberto Amato, Jeffrey Barrett, Sonia Goncalves, Ewan Harrison, David K. Jackson, Ian Johnston, Dominic Kwiatkowski, Cordelia Langford, John Sillitoe on behalf of the Wellcome Sanger Institute COVID-19 Surveillance Team |
| EPI_ISL_2997636 | Lighthouse Lab in Glasgow | Wellcome Sanger Institute for the COVID-19 Genomics UK (COG-UK) Consortium | Harper VanSteenhouse, Yumi Kasai, David Gray, Carol Clugston, Anna Dominiczak and Alex Alderton, Roberto Amato, Jeffrey Barrett, Sonia Goncalves, Ewan Harrison, David K. Jackson, Ian Johnston, Dominic Kwiatkowski, Cordelia Langford, John Sillitoe on behalf of the Wellcome Sanger Institute COVID-19 Surveillance Team |
| EPI_ISL_2997637, EPI_ISL_2997638, EPI_ISL_2997639 | Berkshire and Surrey Pathology Services Lighthouse Laboratory | Wellcome Sanger Institute for the COVID-19 Genomics UK (COG-UK) Consortium | Berkshire and Surrey Pathology Services Lighthouse Laboratory and Alex Alderton, Roberto Amato, Jeffrey Barrett, Sonia Goncalves, Ewan Harrison, David K. Jackson, Ian Johnston, Dominic Kwiatkowski, Cordelia Langford, John Sillitoe on behalf of the Wellcome Sanger Institute COVID-19 Surveillance Team |
| EPI_ISL_2997640 | Lighthouse Lab in Glasgow | Wellcome Sanger Institute for the COVID-19 Genomics UK (COG-UK) Consortium | Harper VanSteenhouse, Yumi Kasai, David Gray, Carol Clugston, Anna Dominiczak and Alex Alderton, Roberto Amato, Jeffrey Barrett, Sonia Goncalves, Ewan Harrison, David K. Jackson, Ian Johnston, Dominic Kwiatkowski, Cordelia Langford, John Sillitoe on behalf of the Wellcome Sanger Institute COVID-19 Surveillance Team |
| EPI_ISL_2997641, EPI_ISL_2997642, EPI_ISL_2997643, EPI_ISL_2997644, EPI_ISL_2997645, EPI_ISL_2997646 | Berkshire and Surrey Pathology Services Lighthouse Laboratory | Wellcome Sanger Institute for the COVID-19 Genomics UK (COG-UK) Consortium | Berkshire and Surrey Pathology Services Lighthouse Laboratory and Alex Alderton, Roberto Amato, Jeffrey Barrett, Sonia Goncalves, Ewan Harrison, David K. Jackson, Ian Johnston, Dominic Kwiatkowski, Cordelia Langford, John Sillitoe on behalf of the Wellcome Sanger Institute COVID-19 Surveillance Team |
| EPI_ISL_2997647 | Lighthouse Lab in Glasgow | Wellcome Sanger Institute for the COVID-19 Genomics UK (COG-UK) Consortium | Harper VanSteenhouse, Yumi Kasai, David Gray, Carol Clugston, Anna Dominiczak and Alex Alderton, Roberto Amato, Jeffrey Barrett, Sonia Goncalves, Ewan Harrison, David K. Jackson, Ian Johnston, Dominic Kwiatkowski, Cordelia Langford, John Sillitoe on behalf of the Wellcome Sanger Institute COVID-19 Surveillance Team |
| EPI_ISL_2997648, EPI_ISL_2997649, EPI_ISL_2997650, EPI_ISL_2997651, EPI_ISL_2997652 | Berkshire and Surrey Pathology Services Lighthouse Laboratory | Wellcome Sanger Institute for the COVID-19 Genomics UK (COG-UK) Consortium | Berkshire and Surrey Pathology Services Lighthouse Laboratory and Alex Alderton, Roberto Amato, Jeffrey Barrett, Sonia Goncalves, Ewan Harrison, David K. Jackson, Ian Johnston, Dominic Kwiatkowski, Cordelia Langford, John Sillitoe on behalf of the Wellcome Sanger Institute COVID-19 Surveillance Team |
| EPI_ISL_2997653, EPI_ISL_2997654 | Lighthouse Lab in Glasgow | Wellcome Sanger Institute for the COVID-19 Genomics UK (COG-UK) Consortium | Harper VanSteenhouse, Yumi Kasai, David Gray, Carol Clugston, Anna Dominiczak and Alex Alderton, Roberto Amato, Jeffrey Barrett, Sonia Goncalves, Ewan Harrison, David K. Jackson, Ian Johnston, Dominic Kwiatkowski, Cordelia Langford, John Sillitoe on behalf of the Wellcome Sanger Institute COVID-19 Surveillance Team |
| EPI_ISL_2997655, EPI_ISL_2997656 | Berkshire and Surrey Pathology Services Lighthouse Laboratory | Wellcome Sanger Institute for the COVID-19 Genomics UK (COG-UK) Consortium | Berkshire and Surrey Pathology Services Lighthouse Laboratory and Alex Alderton, Roberto Amato, Jeffrey Barrett, Sonia Goncalves, Ewan Harrison, David K. Jackson, Ian Johnston, Dominic Kwiatkowski, Cordelia Langford, John Sillitoe on behalf of the Wellcome Sanger Institute COVID-19 Surveillance Team |
| EPI_ISL_2997657 | Lighthouse Lab in Glasgow | Wellcome Sanger Institute for the COVID-19 Genomics UK (COG-UK) Consortium | Harper VanSteenhouse, Yumi Kasai, David Gray, Carol Clugston, Anna Dominiczak and Alex Alderton, Roberto Amato, Jeffrey Barrett, Sonia Goncalves, Ewan Harrison, David K. Jackson, Ian Johnston, Dominic Kwiatkowski, Cordelia Langford, John Sillitoe on behalf of the Wellcome Sanger Institute COVID-19 Surveillance Team |
| EPI_ISL_2997658 | Berkshire and Surrey Pathology Services Lighthouse Laboratory | Wellcome Sanger Institute for the COVID-19 Genomics UK (COG-UK) Consortium | Berkshire and Surrey Pathology Services Lighthouse Laboratory and Alex Alderton, Roberto Amato, Jeffrey Barrett, Sonia Goncalves, Ewan Harrison, David K. Jackson, Ian Johnston, Dominic Kwiatkowski, Cordelia Langford, John Sillitoe on behalf of the Wellcome Sanger Institute COVID-19 Surveillance Team |
| EPI_ISL_2997659 | Lighthouse Lab in Glasgow | Wellcome Sanger Institute for the COVID-19 Genomics UK (COG-UK) Consortium | Harper VanSteenhouse, Yumi Kasai, David Gray, Carol Clugston, Anna Dominiczak and Alex Alderton, Roberto Amato, Jeffrey Barrett, Sonia Goncalves, Ewan Harrison, David K. Jackson, Ian Johnston, Dominic Kwiatkowski, Cordelia Langford, John Sillitoe on behalf of the Wellcome Sanger Institute COVID-19 Surveillance Team |
| EPI_ISL_2997660, EPI_ISL_2997661 | Berkshire and Surrey Pathology Services Lighthouse Laboratory | Wellcome Sanger Institute for the COVID-19 Genomics UK (COG-UK) Consortium | Berkshire and Surrey Pathology Services Lighthouse Laboratory and Alex Alderton, Roberto Amato, Jeffrey Barrett, Sonia Goncalves, Ewan Harrison, David K. Jackson, Ian Johnston, Dominic Kwiatkowski, Cordelia Langford, John Sillitoe on behalf of the Wellcome Sanger Institute COVID-19 Surveillance Team |
| EPI_ISL_2997662, | Lighthouse Lab in Glasgow | Wellcome Sanger Institute for the COVID-19 Genomics UK (COG-UK) | Harper VanSteenhouse, Yumi Kasai, David Gray, Carol Clugston, Anna Dominiczak and Alex Alderton, Roberto Amato, Jeffrey Barrett, Sonia Goncalves, Ewan Harrison, David K. Jackson, Ian Johnston, Dominic Kwiatkowski, Cordelia Langford, |

| | | | |
|---|---|---|---|
| EPI_ISL_2997663, EPI_ISL_2997664 | | Consortium | John Sillitoe on behalf of the Wellcome Sanger Institute COVID-19 Surveillance Team |
| EPI_ISL_2997665, EPI_ISL_2997666, EPI_ISL_2997667 | Berkshire and Surrey Pathology Services Lighthouse Laboratory | Wellcome Sanger Institute for the COVID-19 Genomics UK (COG-UK) Consortium | Berkshire and Surrey Pathology Services Lighthouse Laboratory and Alex Alderton, Roberto Amato, Jeffrey Barrett, Sonia Goncalves, Ewan Harrison, David K. Jackson, Ian Johnston, Dominic Kwiatkowski, Cordelia Langford, John Sillitoe on behalf of the Wellcome Sanger Institute COVID-19 Surveillance Team |
| EPI_ISL_2997668 | Lighthouse Lab in Glasgow | Wellcome Sanger Institute for the COVID-19 Genomics UK (COG-UK) Consortium | Harper VanSteenhouse, Yumi Kasai, David Gray, Carol Clugston, Anna Dominiczak and Alex Alderton, Roberto Amato, Jeffrey Barrett, Sonia Goncalves, Ewan Harrison, David K. Jackson, Ian Johnston, Dominic Kwiatkowski, Cordelia Langford, John Sillitoe on behalf of the Wellcome Sanger Institute COVID-19 Surveillance Team |
| EPI_ISL_2997669, EPI_ISL_2997670 | Lighthouse Lab in Milton Keynes | Wellcome Sanger Institute for the COVID-19 Genomics UK (COG-UK) Consortium | The Lighthouse Lab in Milton Keynes and Alex Alderton, Roberto Amato, Jeffrey Barrett, Sonia Goncalves, Ewan Harrison, David K. Jackson, Ian Johnston, Dominic Kwiatkowski, Cordelia Langford, John Sillitoe on behalf of the Wellcome Sanger Institute COVID-19 Surveillance Team |
| EPI_ISL_2997671, EPI_ISL_2997672 | Lighthouse Lab in Glasgow | Wellcome Sanger Institute for the COVID-19 Genomics UK (COG-UK) Consortium | Harper VanSteenhouse, Yumi Kasai, David Gray, Carol Clugston, Anna Dominiczak and Alex Alderton, Roberto Amato, Jeffrey Barrett, Sonia Goncalves, Ewan Harrison, David K. Jackson, Ian Johnston, Dominic Kwiatkowski, Cordelia Langford, John Sillitoe on behalf of the Wellcome Sanger Institute COVID-19 Surveillance Team |
| EPI_ISL_2997673, EPI_ISL_2997674 | Lighthouse Laboratory Plymouth | Wellcome Sanger Institute for the COVID-19 Genomics UK (COG-UK) Consortium | Lighthouse Laboratory Plymouth and Alex Alderton, Roberto Amato, Jeffrey Barrett, Sonia Goncalves, Ewan Harrison, David K. Jackson, Ian Johnston, Dominic Kwiatkowski, Cordelia Langford, John Sillitoe on behalf of the Wellcome Sanger Institute COVID-19 Surveillance Team |
| EPI_ISL_2997675 | Lighthouse Lab in Milton Keynes | Wellcome Sanger Institute for the COVID-19 Genomics UK (COG-UK) Consortium | The Lighthouse Lab in Milton Keynes and Alex Alderton, Roberto Amato, Jeffrey Barrett, Sonia Goncalves, Ewan Harrison, David K. Jackson, Ian Johnston, Dominic Kwiatkowski, Cordelia Langford, John Sillitoe on behalf of the Wellcome Sanger Institute COVID-19 Surveillance Team |
| EPI_ISL_2997676 | Lighthouse Lab in Glasgow | Wellcome Sanger Institute for the COVID-19 Genomics UK (COG-UK) Consortium | Harper VanSteenhouse, Yumi Kasai, David Gray, Carol Clugston, Anna Dominiczak and Alex Alderton, Roberto Amato, Jeffrey Barrett, Sonia Goncalves, Ewan Harrison, David K. Jackson, Ian Johnston, Dominic Kwiatkowski, Cordelia Langford, John Sillitoe on behalf of the Wellcome Sanger Institute COVID-19 Surveillance Team |
| EPI_ISL_2997677, EPI_ISL_2997678 | Lighthouse Laboratory Plymouth | Wellcome Sanger Institute for the COVID-19 Genomics UK (COG-UK) Consortium | Lighthouse Laboratory Plymouth and Alex Alderton, Roberto Amato, Jeffrey Barrett, Sonia Goncalves, Ewan Harrison, David K. Jackson, Ian Johnston, Dominic Kwiatkowski, Cordelia Langford, John Sillitoe on behalf of the Wellcome Sanger Institute COVID-19 Surveillance Team |
| EPI_ISL_2997679, EPI_ISL_2997680 | Lighthouse Lab in Milton Keynes | Wellcome Sanger Institute for the COVID-19 Genomics UK (COG-UK) Consortium | The Lighthouse Lab in Milton Keynes and Alex Alderton, Roberto Amato, Jeffrey Barrett, Sonia Goncalves, Ewan Harrison, David K. Jackson, Ian Johnston, Dominic Kwiatkowski, Cordelia Langford, John Sillitoe on behalf of the Wellcome Sanger Institute COVID-19 Surveillance Team |
| EPI_ISL_2997681, EPI_ISL_2997682, EPI_ISL_2997683, EPI_ISL_2997684 | Lighthouse Laboratory Plymouth | Wellcome Sanger Institute for the COVID-19 Genomics UK (COG-UK) Consortium | Lighthouse Laboratory Plymouth and Alex Alderton, Roberto Amato, Jeffrey Barrett, Sonia Goncalves, Ewan Harrison, David K. Jackson, Ian Johnston, Dominic Kwiatkowski, Cordelia Langford, John Sillitoe on behalf of the Wellcome Sanger Institute COVID-19 Surveillance Team |
| EPI_ISL_2997685, EPI_ISL_2997686 | Lighthouse Lab in Milton Keynes | Wellcome Sanger Institute for the COVID-19 Genomics UK (COG-UK) Consortium | The Lighthouse Lab in Milton Keynes and Alex Alderton, Roberto Amato, Jeffrey Barrett, Sonia Goncalves, Ewan Harrison, David K. Jackson, Ian Johnston, Dominic Kwiatkowski, Cordelia Langford, John Sillitoe on behalf of the Wellcome Sanger Institute COVID-19 Surveillance Team |
| EPI_ISL_2997687 | Lighthouse Laboratory Plymouth | Wellcome Sanger Institute for the COVID-19 Genomics UK (COG-UK) Consortium | Lighthouse Laboratory Plymouth and Alex Alderton, Roberto Amato, Jeffrey Barrett, Sonia Goncalves, Ewan Harrison, David K. Jackson, Ian Johnston, Dominic Kwiatkowski, Cordelia Langford, John Sillitoe on behalf of the Wellcome Sanger Institute COVID-19 Surveillance Team |
| EPI_ISL_2997688 | Lighthouse Lab in Glasgow | Wellcome Sanger Institute for the COVID-19 Genomics UK (COG-UK) Consortium | Harper VanSteenhouse, Yumi Kasai, David Gray, Carol Clugston, Anna Dominiczak and Alex Alderton, Roberto Amato, Jeffrey Barrett, Sonia Goncalves, Ewan Harrison, David K. Jackson, Ian Johnston, Dominic Kwiatkowski, Cordelia Langford, John Sillitoe on behalf of the Wellcome Sanger Institute COVID-19 Surveillance Team |
| EPI_ISL_2997689 | Lighthouse Laboratory Plymouth | Wellcome Sanger Institute for the COVID-19 Genomics UK (COG-UK) Consortium | Lighthouse Laboratory Plymouth and Alex Alderton, Roberto Amato, Jeffrey Barrett, Sonia Goncalves, Ewan Harrison, David K. Jackson, Ian Johnston, Dominic Kwiatkowski, Cordelia Langford, John Sillitoe on behalf of the Wellcome Sanger Institute COVID-19 Surveillance Team |
| EPI_ISL_2997690, EPI_ISL_2997691 | Lighthouse Lab in Glasgow | Wellcome Sanger Institute for the COVID-19 Genomics UK (COG-UK) Consortium | Harper VanSteenhouse, Yumi Kasai, David Gray, Carol Clugston, Anna Dominiczak and Alex Alderton, Roberto Amato, Jeffrey Barrett, Sonia Goncalves, Ewan Harrison, David K. Jackson, Ian Johnston, Dominic Kwiatkowski, Cordelia Langford, John Sillitoe on behalf of the Wellcome Sanger Institute COVID-19 Surveillance Team |
| EPI_ISL_2997692, EPI_ISL_2997693 | Lighthouse Laboratory Plymouth | Wellcome Sanger Institute for the COVID-19 Genomics UK (COG-UK) Consortium | Lighthouse Laboratory Plymouth and Alex Alderton, Roberto Amato, Jeffrey Barrett, Sonia Goncalves, Ewan Harrison, David K. Jackson, Ian Johnston, Dominic Kwiatkowski, Cordelia Langford, John Sillitoe on behalf of the Wellcome Sanger Institute COVID-19 Surveillance Team |
| EPI_ISL_2997694, EPI_ISL_2997695 | Lighthouse Lab in Glasgow | Wellcome Sanger Institute for the COVID-19 Genomics UK (COG-UK) Consortium | Harper VanSteenhouse, Yumi Kasai, David Gray, Carol Clugston, Anna Dominiczak and Alex Alderton, Roberto Amato, Jeffrey Barrett, Sonia Goncalves, Ewan Harrison, David K. Jackson, Ian Johnston, Dominic Kwiatkowski, Cordelia Langford, John Sillitoe on behalf of the Wellcome Sanger Institute COVID-19 Surveillance Team |
| EPI_ISL_2997696 | Lighthouse Laboratory Plymouth | Wellcome Sanger Institute for the COVID-19 Genomics UK (COG-UK) Consortium | Lighthouse Laboratory Plymouth and Alex Alderton, Roberto Amato, Jeffrey Barrett, Sonia Goncalves, Ewan Harrison, David K. Jackson, Ian Johnston, Dominic Kwiatkowski, Cordelia Langford, John Sillitoe on behalf of the Wellcome Sanger Institute COVID-19 Surveillance Team |
| EPI_ISL_2997697 | Lighthouse Lab in Glasgow | Wellcome Sanger Institute for the COVID-19 Genomics UK (COG-UK) Consortium | Harper VanSteenhouse, Yumi Kasai, David Gray, Carol Clugston, Anna Dominiczak and Alex Alderton, Roberto Amato, Jeffrey Barrett, Sonia Goncalves, Ewan Harrison, David K. Jackson, Ian Johnston, Dominic Kwiatkowski, Cordelia Langford, John Sillitoe on behalf of the Wellcome Sanger Institute COVID-19 Surveillance Team |
| EPI_ISL_2997698, EPI_ISL_2997699 | Lighthouse Laboratory Plymouth | Wellcome Sanger Institute for the COVID-19 Genomics UK (COG-UK) Consortium | Lighthouse Laboratory Plymouth and Alex Alderton, Roberto Amato, Jeffrey Barrett, Sonia Goncalves, Ewan Harrison, David K. Jackson, Ian Johnston, Dominic Kwiatkowski, Cordelia Langford, John Sillitoe on behalf of the Wellcome Sanger Institute COVID-19 Surveillance Team |
| EPI_ISL_2997700 | Lighthouse Lab in Milton Keynes | Wellcome Sanger Institute for the COVID-19 Genomics UK (COG-UK) Consortium | The Lighthouse Lab in Milton Keynes and Alex Alderton, Roberto Amato, Jeffrey Barrett, Sonia Goncalves, Ewan Harrison, David K. Jackson, Ian Johnston, Dominic Kwiatkowski, Cordelia Langford, John Sillitoe on behalf of the Wellcome Sanger Institute COVID-19 Surveillance Team |
| EPI_ISL_2997701, EPI_ISL_2997702 | Lighthouse Laboratory Plymouth | Wellcome Sanger Institute for the COVID-19 Genomics UK (COG-UK) Consortium | Lighthouse Laboratory Plymouth and Alex Alderton, Roberto Amato, Jeffrey Barrett, Sonia Goncalves, Ewan Harrison, David K. Jackson, Ian Johnston, Dominic Kwiatkowski, Cordelia Langford, John Sillitoe on behalf of the Wellcome Sanger Institute COVID-19 Surveillance Team |
| EPI_ISL_2997703, EPI_ISL_2997704 | Lighthouse Lab in Milton Keynes | Wellcome Sanger Institute for the COVID-19 Genomics UK (COG-UK) Consortium | The Lighthouse Lab in Milton Keynes and Alex Alderton, Roberto Amato, Jeffrey Barrett, Sonia Goncalves, Ewan Harrison, David K. Jackson, Ian Johnston, Dominic Kwiatkowski, Cordelia Langford, John Sillitoe on behalf of the Wellcome Sanger Institute COVID-19 Surveillance Team |
| EPI_ISL_2997705, EPI_ISL_2997706, EPI_ISL_2997707, EPI_ISL_2997708 | Lighthouse Laboratory Plymouth | Wellcome Sanger Institute for the COVID-19 Genomics UK (COG-UK) Consortium | Lighthouse Laboratory Plymouth and Alex Alderton, Roberto Amato, Jeffrey Barrett, Sonia Goncalves, Ewan Harrison, David K. Jackson, Ian Johnston, Dominic Kwiatkowski, Cordelia Langford, John Sillitoe on behalf of the Wellcome Sanger Institute COVID-19 Surveillance Team |
| EPI_ISL_2997709, EPI_ISL_2997710, EPI_ISL_2997711 | Lighthouse Lab in Milton Keynes | Wellcome Sanger Institute for the COVID-19 Genomics UK (COG-UK) Consortium | The Lighthouse Lab in Milton Keynes and Alex Alderton, Roberto Amato, Jeffrey Barrett, Sonia Goncalves, Ewan Harrison, David K. Jackson, Ian Johnston, Dominic Kwiatkowski, Cordelia Langford, John Sillitoe on behalf of the Wellcome Sanger Institute COVID-19 Surveillance Team |
| EPI_ISL_2997712 | Lighthouse Lab in Glasgow | Wellcome Sanger Institute for the COVID-19 Genomics UK (COG-UK) Consortium | Harper VanSteenhouse, Yumi Kasai, David Gray, Carol Clugston, Anna Dominiczak and Alex Alderton, Roberto Amato, Jeffrey Barrett, Sonia Goncalves, Ewan Harrison, David K. Jackson, Ian Johnston, Dominic Kwiatkowski, Cordelia Langford, John Sillitoe on behalf of the Wellcome Sanger Institute COVID-19 Surveillance Team |
| EPI_ISL_2997713, EPI_ISL_2997714, EPI_ISL_2997715 | Lighthouse Laboratory Plymouth | Wellcome Sanger Institute for the COVID-19 Genomics UK (COG-UK) Consortium | Lighthouse Laboratory Plymouth and Alex Alderton, Roberto Amato, Jeffrey Barrett, Sonia Goncalves, Ewan Harrison, David K. Jackson, Ian Johnston, Dominic Kwiatkowski, Cordelia Langford, John Sillitoe on behalf of the Wellcome Sanger Institute COVID-19 Surveillance Team |
| EPI_ISL_2997716 | Lighthouse Lab in Milton Keynes | Wellcome Sanger Institute for the COVID-19 Genomics UK (COG-UK) Consortium | The Lighthouse Lab in Milton Keynes and Alex Alderton, Roberto Amato, Jeffrey Barrett, Sonia Goncalves, Ewan Harrison, David K. Jackson, Ian Johnston, Dominic Kwiatkowski, Cordelia Langford, John Sillitoe on behalf of the Wellcome Sanger Institute COVID-19 Surveillance Team |
| EPI_ISL_2997717 | Lighthouse Lab in Glasgow | Wellcome Sanger Institute for the COVID-19 Genomics UK (COG-UK) Consortium | Harper VanSteenhouse, Yumi Kasai, David Gray, Carol Clugston, Anna Dominiczak and Alex Alderton, Roberto Amato, Jeffrey Barrett, Sonia Goncalves, Ewan Harrison, David K. Jackson, Ian Johnston, Dominic Kwiatkowski, Cordelia Langford, John Sillitoe on behalf of the Wellcome Sanger Institute COVID-19 Surveillance Team |
| EPI_ISL_2997718 | Lighthouse Laboratory Plymouth | Wellcome Sanger Institute for the COVID-19 Genomics UK (COG-UK) Consortium | Lighthouse Laboratory Plymouth and Alex Alderton, Roberto Amato, Jeffrey Barrett, Sonia Goncalves, Ewan Harrison, David K. Jackson, Ian Johnston, Dominic Kwiatkowski, Cordelia Langford, John Sillitoe on behalf of the Wellcome Sanger Institute COVID-19 Surveillance Team |
| EPI_ISL_2997719, EPI_ISL_2997720 | Lighthouse Lab in Glasgow | Wellcome Sanger Institute for the COVID-19 Genomics UK (COG-UK) Consortium | Harper VanSteenhouse, Yumi Kasai, David Gray, Carol Clugston, Anna Dominiczak and Alex Alderton, Roberto Amato, Jeffrey Barrett, Sonia Goncalves, Ewan Harrison, David K. Jackson, Ian Johnston, Dominic Kwiatkowski, Cordelia Langford, John Sillitoe on behalf of the Wellcome Sanger Institute COVID-19 Surveillance Team |
| EPI_ISL_2997721, EPI_ISL_2997722 | Lighthouse Lab in Milton Keynes | Wellcome Sanger Institute for the COVID-19 Genomics UK (COG-UK) Consortium | The Lighthouse Lab in Milton Keynes and Alex Alderton, Roberto Amato, Jeffrey Barrett, Sonia Goncalves, Ewan Harrison, David K. Jackson, Ian Johnston, Dominic Kwiatkowski, Cordelia Langford, John Sillitoe on behalf of the Wellcome Sanger Institute COVID-19 Surveillance Team |
| EPI_ISL_2997723 | Lighthouse Laboratory Plymouth | Wellcome Sanger Institute for the COVID-19 Genomics UK (COG-UK) Consortium | Lighthouse Laboratory Plymouth and Alex Alderton, Roberto Amato, Jeffrey Barrett, Sonia Goncalves, Ewan Harrison, David K. Jackson, Ian Johnston, Dominic Kwiatkowski, Cordelia Langford, John Sillitoe on behalf of the Wellcome Sanger Institute COVID-19 Surveillance Team |
| EPI_ISL_2997724 | Lighthouse Lab in Milton Keynes | Wellcome Sanger Institute for the COVID-19 Genomics UK (COG-UK) Consortium | The Lighthouse Lab in Milton Keynes and Alex Alderton, Roberto Amato, Jeffrey Barrett, Sonia Goncalves, Ewan Harrison, David K. Jackson, Ian Johnston, Dominic Kwiatkowski, Cordelia Langford, John Sillitoe on behalf of the Wellcome Sanger Institute COVID-19 Surveillance Team |
| EPI_ISL_2997725 | Lighthouse Laboratory Plymouth | Wellcome Sanger Institute for the COVID-19 Genomics UK (COG-UK) Consortium | Lighthouse Laboratory Plymouth and Alex Alderton, Roberto Amato, Jeffrey Barrett, Sonia Goncalves, Ewan Harrison, David K. Jackson, Ian Johnston, Dominic Kwiatkowski, Cordelia Langford, John Sillitoe on behalf of the Wellcome Sanger Institute COVID-19 Surveillance Team |
| EPI_ISL_2997726, EPI_ISL_2997727 | Lighthouse Lab in Glasgow | Wellcome Sanger Institute for the COVID-19 Genomics UK (COG-UK) Consortium | Harper VanSteenhouse, Yumi Kasai, David Gray, Carol Clugston, Anna Dominiczak and Alex Alderton, Roberto Amato, Jeffrey Barrett, Sonia Goncalves, Ewan Harrison, David K. Jackson, Ian Johnston, Dominic Kwiatkowski, Cordelia Langford, John Sillitoe on behalf of the Wellcome Sanger Institute COVID-19 Surveillance Team |
| EPI_ISL_2997728, EPI_ISL_2997729, EPI_ISL_2997730 | Lighthouse Lab in Milton Keynes | Wellcome Sanger Institute for the COVID-19 Genomics UK (COG-UK) Consortium | The Lighthouse Lab in Milton Keynes and Alex Alderton, Roberto Amato, Jeffrey Barrett, Sonia Goncalves, Ewan Harrison, David K. Jackson, Ian Johnston, Dominic Kwiatkowski, Cordelia Langford, John Sillitoe on behalf of the Wellcome Sanger Institute COVID-19 Surveillance Team |
| EPI_ISL_2997731 | Lighthouse Laboratory Plymouth | Wellcome Sanger Institute for the COVID-19 Genomics UK (COG-UK) Consortium | Lighthouse Laboratory Plymouth and Alex Alderton, Roberto Amato, Jeffrey Barrett, Sonia Goncalves, Ewan Harrison, David K. Jackson, Ian Johnston, Dominic Kwiatkowski, Cordelia Langford, John Sillitoe on behalf of the Wellcome Sanger Institute COVID-19 Surveillance Team |
| EPI_ISL_2997732 | Lighthouse Lab in Glasgow | Wellcome Sanger Institute for the COVID-19 Genomics UK (COG-UK) Consortium | Harper VanSteenhouse, Yumi Kasai, David Gray, Carol Clugston, Anna Dominiczak and Alex Alderton, Roberto Amato, Jeffrey Barrett, Sonia Goncalves, Ewan Harrison, David K. Jackson, Ian Johnston, Dominic Kwiatkowski, Cordelia Langford, John Sillitoe on behalf of the Wellcome Sanger Institute COVID-19 Surveillance Team |
| EPI_ISL_2997733 | Lighthouse Lab in Milton Keynes | Wellcome Sanger Institute for the COVID-19 Genomics UK (COG-UK) Consortium | The Lighthouse Lab in Milton Keynes and Alex Alderton, Roberto Amato, Jeffrey Barrett, Sonia Goncalves, Ewan Harrison, David K. Jackson, Ian Johnston, Dominic Kwiatkowski, Cordelia Langford, John Sillitoe on behalf of the Wellcome Sanger Institute COVID-19 Surveillance Team |
| EPI_ISL_2997734 | Lighthouse Laboratory Plymouth | Wellcome Sanger Institute for the COVID-19 Genomics UK (COG-UK) Consortium | Lighthouse Laboratory Plymouth and Alex Alderton, Roberto Amato, Jeffrey Barrett, Sonia Goncalves, Ewan Harrison, David K. Jackson, Ian Johnston, Dominic Kwiatkowski, Cordelia Langford, John Sillitoe on behalf of the Wellcome Sanger Institute COVID-19 Surveillance Team |

| | | | |
|---|---|---|---|
| EPI_ISL_2997735 | Lighthouse Lab in Glasgow | Wellcome Sanger Institute for the COVID-19 Genomics UK (COG-UK) Consortium | Harper VanSteenhouse, Yumi Kasai, David Gray, Carol Clugston, Anna Dominiczak and Alex Alderton, Roberto Amato, Jeffrey Barrett, Sonia Goncalves, Ewan Harrison, David K. Jackson, Ian Johnston, Dominic Kwiatkowski, Cordelia Langford, John Sillitoe on behalf of the Wellcome Sanger Institute COVID-19 Surveillance Team |
| EPI_ISL_2997736, EPI_ISL_2997737, EPI_ISL_2997738 | Lighthouse Laboratory Plymouth | Wellcome Sanger Institute for the COVID-19 Genomics UK (COG-UK) Consortium | Lighthouse Laboratory Plymouth and Alex Alderton, Roberto Amato, Jeffrey Barrett, Sonia Goncalves, Ewan Harrison, David K. Jackson, Ian Johnston, Dominic Kwiatkowski, Cordelia Langford, John Sillitoe on behalf of the Wellcome Sanger Institute COVID-19 Surveillance Team |
| EPI_ISL_2997739, EPI_ISL_2997740, EPI_ISL_2997741 | Lighthouse Lab in Glasgow | Wellcome Sanger Institute for the COVID-19 Genomics UK (COG-UK) Consortium | Harper VanSteenhouse, Yumi Kasai, David Gray, Carol Clugston, Anna Dominiczak and Alex Alderton, Roberto Amato, Jeffrey Barrett, Sonia Goncalves, Ewan Harrison, David K. Jackson, Ian Johnston, Dominic Kwiatkowski, Cordelia Langford, John Sillitoe on behalf of the Wellcome Sanger Institute COVID-19 Surveillance Team |
| EPI_ISL_2997742 | Lighthouse Laboratory Plymouth | Wellcome Sanger Institute for the COVID-19 Genomics UK (COG-UK) Consortium | Lighthouse Laboratory Plymouth and Alex Alderton, Roberto Amato, Jeffrey Barrett, Sonia Goncalves, Ewan Harrison, David K. Jackson, Ian Johnston, Dominic Kwiatkowski, Cordelia Langford, John Sillitoe on behalf of the Wellcome Sanger Institute COVID-19 Surveillance Team |
| EPI_ISL_2997743, EPI_ISL_2997744, EPI_ISL_2997745 | Lighthouse Lab in Milton Keynes | Wellcome Sanger Institute for the COVID-19 Genomics UK (COG-UK) Consortium | The Lighthouse Lab in Milton Keynes and Alex Alderton, Roberto Amato, Jeffrey Barrett, Sonia Goncalves, Ewan Harrison, David K. Jackson, Ian Johnston, Dominic Kwiatkowski, Cordelia Langford, John Sillitoe on behalf of the Wellcome Sanger Institute COVID-19 Surveillance Team |
| EPI_ISL_2997746, EPI_ISL_2997747, EPI_ISL_2997748, EPI_ISL_2997749 | Lighthouse Laboratory Plymouth | Wellcome Sanger Institute for the COVID-19 Genomics UK (COG-UK) Consortium | Lighthouse Laboratory Plymouth and Alex Alderton, Roberto Amato, Jeffrey Barrett, Sonia Goncalves, Ewan Harrison, David K. Jackson, Ian Johnston, Dominic Kwiatkowski, Cordelia Langford, John Sillitoe on behalf of the Wellcome Sanger Institute COVID-19 Surveillance Team |
| EPI_ISL_2997750, EPI_ISL_2997751 | Lighthouse Lab in Glasgow | Wellcome Sanger Institute for the COVID-19 Genomics UK (COG-UK) Consortium | Harper VanSteenhouse, Yumi Kasai, David Gray, Carol Clugston, Anna Dominiczak and Alex Alderton, Roberto Amato, Jeffrey Barrett, Sonia Goncalves, Ewan Harrison, David K. Jackson, Ian Johnston, Dominic Kwiatkowski, Cordelia Langford, John Sillitoe on behalf of the Wellcome Sanger Institute COVID-19 Surveillance Team |
| EPI_ISL_2997752 | Lighthouse Laboratory Plymouth | Wellcome Sanger Institute for the COVID-19 Genomics UK (COG-UK) Consortium | Lighthouse Laboratory Plymouth and Alex Alderton, Roberto Amato, Jeffrey Barrett, Sonia Goncalves, Ewan Harrison, David K. Jackson, Ian Johnston, Dominic Kwiatkowski, Cordelia Langford, John Sillitoe on behalf of the Wellcome Sanger Institute COVID-19 Surveillance Team |
| EPI_ISL_2997753 | Lighthouse Lab in Glasgow | Wellcome Sanger Institute for the COVID-19 Genomics UK (COG-UK) Consortium | Harper VanSteenhouse, Yumi Kasai, David Gray, Carol Clugston, Anna Dominiczak and Alex Alderton, Roberto Amato, Jeffrey Barrett, Sonia Goncalves, Ewan Harrison, David K. Jackson, Ian Johnston, Dominic Kwiatkowski, Cordelia Langford, John Sillitoe on behalf of the Wellcome Sanger Institute COVID-19 Surveillance Team |
| EPI_ISL_2997754 | Lighthouse Laboratory Plymouth | Wellcome Sanger Institute for the COVID-19 Genomics UK (COG-UK) Consortium | Lighthouse Laboratory Plymouth and Alex Alderton, Roberto Amato, Jeffrey Barrett, Sonia Goncalves, Ewan Harrison, David K. Jackson, Ian Johnston, Dominic Kwiatkowski, Cordelia Langford, John Sillitoe on behalf of the Wellcome Sanger Institute COVID-19 Surveillance Team |
| EPI_ISL_2997755 | Lighthouse Lab in Glasgow | Wellcome Sanger Institute for the COVID-19 Genomics UK (COG-UK) Consortium | Harper VanSteenhouse, Yumi Kasai, David Gray, Carol Clugston, Anna Dominiczak and Alex Alderton, Roberto Amato, Jeffrey Barrett, Sonia Goncalves, Ewan Harrison, David K. Jackson, Ian Johnston, Dominic Kwiatkowski, Cordelia Langford, John Sillitoe on behalf of the Wellcome Sanger Institute COVID-19 Surveillance Team |
| EPI_ISL_2997756 | Lighthouse Laboratory Plymouth | Wellcome Sanger Institute for the COVID-19 Genomics UK (COG-UK) Consortium | Lighthouse Laboratory Plymouth and Alex Alderton, Roberto Amato, Jeffrey Barrett, Sonia Goncalves, Ewan Harrison, David K. Jackson, Ian Johnston, Dominic Kwiatkowski, Cordelia Langford, John Sillitoe on behalf of the Wellcome Sanger Institute COVID-19 Surveillance Team |
| EPI_ISL_2997757 | Lighthouse Lab in Glasgow | Wellcome Sanger Institute for the COVID-19 Genomics UK (COG-UK) Consortium | Harper VanSteenhouse, Yumi Kasai, David Gray, Carol Clugston, Anna Dominiczak and Alex Alderton, Roberto Amato, Jeffrey Barrett, Sonia Goncalves, Ewan Harrison, David K. Jackson, Ian Johnston, Dominic Kwiatkowski, Cordelia Langford, John Sillitoe on behalf of the Wellcome Sanger Institute COVID-19 Surveillance Team |
| EPI_ISL_2997758 | Lighthouse Lab in Milton Keynes | Wellcome Sanger Institute for the COVID-19 Genomics UK (COG-UK) Consortium | The Lighthouse Lab in Milton Keynes and Alex Alderton, Roberto Amato, Jeffrey Barrett, Sonia Goncalves, Ewan Harrison, David K. Jackson, Ian Johnston, Dominic Kwiatkowski, Cordelia Langford, John Sillitoe on behalf of the Wellcome Sanger Institute COVID-19 Surveillance Team |
| EPI_ISL_2997759 | Lighthouse Lab in Glasgow | Wellcome Sanger Institute for the COVID-19 Genomics UK (COG-UK) Consortium | Harper VanSteenhouse, Yumi Kasai, David Gray, Carol Clugston, Anna Dominiczak and Alex Alderton, Roberto Amato, Jeffrey Barrett, Sonia Goncalves, Ewan Harrison, David K. Jackson, Ian Johnston, Dominic Kwiatkowski, Cordelia Langford, John Sillitoe on behalf of the Wellcome Sanger Institute COVID-19 Surveillance Team |
| EPI_ISL_2997760 | Lighthouse Laboratory Plymouth | Wellcome Sanger Institute for the COVID-19 Genomics UK (COG-UK) Consortium | Lighthouse Laboratory Plymouth and Alex Alderton, Roberto Amato, Jeffrey Barrett, Sonia Goncalves, Ewan Harrison, David K. Jackson, Ian Johnston, Dominic Kwiatkowski, Cordelia Langford, John Sillitoe on behalf of the Wellcome Sanger Institute COVID-19 Surveillance Team |
| EPI_ISL_2997761 | Lighthouse Lab in Glasgow | Wellcome Sanger Institute for the COVID-19 Genomics UK (COG-UK) Consortium | Harper VanSteenhouse, Yumi Kasai, David Gray, Carol Clugston, Anna Dominiczak and Alex Alderton, Roberto Amato, Jeffrey Barrett, Sonia Goncalves, Ewan Harrison, David K. Jackson, Ian Johnston, Dominic Kwiatkowski, Cordelia Langford, John Sillitoe on behalf of the Wellcome Sanger Institute COVID-19 Surveillance Team |
| EPI_ISL_2997762, EPI_ISL_2997763, EPI_ISL_2997764 | Lighthouse Laboratory Plymouth | Wellcome Sanger Institute for the COVID-19 Genomics UK (COG-UK) Consortium | Lighthouse Laboratory Plymouth and Alex Alderton, Roberto Amato, Jeffrey Barrett, Sonia Goncalves, Ewan Harrison, David K. Jackson, Ian Johnston, Dominic Kwiatkowski, Cordelia Langford, John Sillitoe on behalf of the Wellcome Sanger Institute COVID-19 Surveillance Team |
| EPI_ISL_2997765 | Lighthouse Lab in Milton Keynes | Wellcome Sanger Institute for the COVID-19 Genomics UK (COG-UK) Consortium | The Lighthouse Lab in Milton Keynes and Alex Alderton, Roberto Amato, Jeffrey Barrett, Sonia Goncalves, Ewan Harrison, David K. Jackson, Ian Johnston, Dominic Kwiatkowski, Cordelia Langford, John Sillitoe on behalf of the Wellcome Sanger Institute COVID-19 Surveillance Team |
| EPI_ISL_2997766 | Lighthouse Laboratory Plymouth | Wellcome Sanger Institute for the COVID-19 Genomics UK (COG-UK) Consortium | Lighthouse Laboratory Plymouth and Alex Alderton, Roberto Amato, Jeffrey Barrett, Sonia Goncalves, Ewan Harrison, David K. Jackson, Ian Johnston, Dominic Kwiatkowski, Cordelia Langford, John Sillitoe on behalf of the Wellcome Sanger Institute COVID-19 Surveillance Team |
| EPI_ISL_2997767, EPI_ISL_2997768, EPI_ISL_2997769 | Lighthouse Lab in Milton Keynes | Wellcome Sanger Institute for the COVID-19 Genomics UK (COG-UK) Consortium | The Lighthouse Lab in Milton Keynes and Alex Alderton, Roberto Amato, Jeffrey Barrett, Sonia Goncalves, Ewan Harrison, David K. Jackson, Ian Johnston, Dominic Kwiatkowski, Cordelia Langford, John Sillitoe on behalf of the Wellcome Sanger Institute COVID-19 Surveillance Team |
| EPI_ISL_2997770 | Lighthouse Lab in Glasgow | Wellcome Sanger Institute for the COVID-19 Genomics UK (COG-UK) Consortium | Harper VanSteenhouse, Yumi Kasai, David Gray, Carol Clugston, Anna Dominiczak and Alex Alderton, Roberto Amato, Jeffrey Barrett, Sonia Goncalves, Ewan Harrison, David K. Jackson, Ian Johnston, Dominic Kwiatkowski, Cordelia Langford, John Sillitoe on behalf of the Wellcome Sanger Institute COVID-19 Surveillance Team |
| EPI_ISL_2997771 | Lighthouse Laboratory Plymouth | Wellcome Sanger Institute for the COVID-19 Genomics UK (COG-UK) Consortium | Lighthouse Laboratory Plymouth and Alex Alderton, Roberto Amato, Jeffrey Barrett, Sonia Goncalves, Ewan Harrison, David K. Jackson, Ian Johnston, Dominic Kwiatkowski, Cordelia Langford, John Sillitoe on behalf of the Wellcome Sanger Institute COVID-19 Surveillance Team |
| EPI_ISL_2997772 | Lighthouse Lab in Glasgow | Wellcome Sanger Institute for the COVID-19 Genomics UK (COG-UK) Consortium | Harper VanSteenhouse, Yumi Kasai, David Gray, Carol Clugston, Anna Dominiczak and Alex Alderton, Roberto Amato, Jeffrey Barrett, Sonia Goncalves, Ewan Harrison, David K. Jackson, Ian Johnston, Dominic Kwiatkowski, Cordelia Langford, John Sillitoe on behalf of the Wellcome Sanger Institute COVID-19 Surveillance Team |
| EPI_ISL_2997773 | Lighthouse Laboratory Plymouth | Wellcome Sanger Institute for the COVID-19 Genomics UK (COG-UK) Consortium | Lighthouse Laboratory Plymouth and Alex Alderton, Roberto Amato, Jeffrey Barrett, Sonia Goncalves, Ewan Harrison, David K. Jackson, Ian Johnston, Dominic Kwiatkowski, Cordelia Langford, John Sillitoe on behalf of the Wellcome Sanger Institute COVID-19 Surveillance Team |
| EPI_ISL_2997774 | Lighthouse Lab in Glasgow | Wellcome Sanger Institute for the COVID-19 Genomics UK (COG-UK) Consortium | Harper VanSteenhouse, Yumi Kasai, David Gray, Carol Clugston, Anna Dominiczak and Alex Alderton, Roberto Amato, Jeffrey Barrett, Sonia Goncalves, Ewan Harrison, David K. Jackson, Ian Johnston, Dominic Kwiatkowski, Cordelia Langford, John Sillitoe on behalf of the Wellcome Sanger Institute COVID-19 Surveillance Team |
| EPI_ISL_2997775 | Lighthouse Laboratory Plymouth | Wellcome Sanger Institute for the COVID-19 Genomics UK (COG-UK) Consortium | Lighthouse Laboratory Plymouth and Alex Alderton, Roberto Amato, Jeffrey Barrett, Sonia Goncalves, Ewan Harrison, David K. Jackson, Ian Johnston, Dominic Kwiatkowski, Cordelia Langford, John Sillitoe on behalf of the Wellcome Sanger Institute COVID-19 Surveillance Team |
| EPI_ISL_2997776 | Lighthouse Lab in Glasgow | Wellcome Sanger Institute for the COVID-19 Genomics UK (COG-UK) Consortium | Harper VanSteenhouse, Yumi Kasai, David Gray, Carol Clugston, Anna Dominiczak and Alex Alderton, Roberto Amato, Jeffrey Barrett, Sonia Goncalves, Ewan Harrison, David K. Jackson, Ian Johnston, Dominic Kwiatkowski, Cordelia Langford, John Sillitoe on behalf of the Wellcome Sanger Institute COVID-19 Surveillance Team |
| EPI_ISL_2997777 | Lighthouse Laboratory Plymouth | Wellcome Sanger Institute for the COVID-19 Genomics UK (COG-UK) Consortium | Lighthouse Laboratory Plymouth and Alex Alderton, Roberto Amato, Jeffrey Barrett, Sonia Goncalves, Ewan Harrison, David K. Jackson, Ian Johnston, Dominic Kwiatkowski, Cordelia Langford, John Sillitoe on behalf of the Wellcome Sanger Institute COVID-19 Surveillance Team |
| EPI_ISL_2997778, EPI_ISL_2997779 | Lighthouse Lab in Glasgow | Wellcome Sanger Institute for the COVID-19 Genomics UK (COG-UK) Consortium | Harper VanSteenhouse, Yumi Kasai, David Gray, Carol Clugston, Anna Dominiczak and Alex Alderton, Roberto Amato, Jeffrey Barrett, Sonia Goncalves, Ewan Harrison, David K. Jackson, Ian Johnston, Dominic Kwiatkowski, Cordelia Langford, John Sillitoe on behalf of the Wellcome Sanger Institute COVID-19 Surveillance Team |
| EPI_ISL_2997780 | Lighthouse Laboratory Plymouth | Wellcome Sanger Institute for the COVID-19 Genomics UK (COG-UK) Consortium | Lighthouse Laboratory Plymouth and Alex Alderton, Roberto Amato, Jeffrey Barrett, Sonia Goncalves, Ewan Harrison, David K. Jackson, Ian Johnston, Dominic Kwiatkowski, Cordelia Langford, John Sillitoe on behalf of the Wellcome Sanger Institute COVID-19 Surveillance Team |
| EPI_ISL_2997781 | Lighthouse Lab in Glasgow | Wellcome Sanger Institute for the COVID-19 Genomics UK (COG-UK) Consortium | Harper VanSteenhouse, Yumi Kasai, David Gray, Carol Clugston, Anna Dominiczak and Alex Alderton, Roberto Amato, Jeffrey Barrett, Sonia Goncalves, Ewan Harrison, David K. Jackson, Ian Johnston, Dominic Kwiatkowski, Cordelia Langford, John Sillitoe on behalf of the Wellcome Sanger Institute COVID-19 Surveillance Team |
| EPI_ISL_2997782, EPI_ISL_2997783, EPI_ISL_2997784, EPI_ISL_2997785 | Lighthouse Lab in Milton Keynes | Wellcome Sanger Institute for the COVID-19 Genomics UK (COG-UK) Consortium | The Lighthouse Lab in Milton Keynes and Alex Alderton, Roberto Amato, Jeffrey Barrett, Sonia Goncalves, Ewan Harrison, David K. Jackson, Ian Johnston, Dominic Kwiatkowski, Cordelia Langford, John Sillitoe on behalf of the Wellcome Sanger Institute COVID-19 Surveillance Team |
| EPI_ISL_2997786 | Lighthouse Laboratory Plymouth | Wellcome Sanger Institute for the COVID-19 Genomics UK (COG-UK) Consortium | Lighthouse Laboratory Plymouth and Alex Alderton, Roberto Amato, Jeffrey Barrett, Sonia Goncalves, Ewan Harrison, David K. Jackson, Ian Johnston, Dominic Kwiatkowski, Cordelia Langford, John Sillitoe on behalf of the Wellcome Sanger Institute COVID-19 Surveillance Team |
| EPI_ISL_2997787 | Lighthouse Lab in Glasgow | Wellcome Sanger Institute for the COVID-19 Genomics UK (COG-UK) Consortium | Harper VanSteenhouse, Yumi Kasai, David Gray, Carol Clugston, Anna Dominiczak and Alex Alderton, Roberto Amato, Jeffrey Barrett, Sonia Goncalves, Ewan Harrison, David K. Jackson, Ian Johnston, Dominic Kwiatkowski, Cordelia Langford, John Sillitoe on behalf of the Wellcome Sanger Institute COVID-19 Surveillance Team |
| EPI_ISL_2997788, EPI_ISL_2997789 | Lighthouse Laboratory Plymouth | Wellcome Sanger Institute for the COVID-19 Genomics UK (COG-UK) Consortium | Lighthouse Laboratory Plymouth and Alex Alderton, Roberto Amato, Jeffrey Barrett, Sonia Goncalves, Ewan Harrison, David K. Jackson, Ian Johnston, Dominic Kwiatkowski, Cordelia Langford, John Sillitoe on behalf of the Wellcome Sanger Institute COVID-19 Surveillance Team |
| EPI_ISL_2997790 | Lighthouse Lab in Glasgow | Wellcome Sanger Institute for the COVID-19 Genomics UK (COG-UK) Consortium | Harper VanSteenhouse, Yumi Kasai, David Gray, Carol Clugston, Anna Dominiczak and Alex Alderton, Roberto Amato, Jeffrey Barrett, Sonia Goncalves, Ewan Harrison, David K. Jackson, Ian Johnston, Dominic Kwiatkowski, Cordelia Langford, John Sillitoe on behalf of the Wellcome Sanger Institute COVID-19 Surveillance Team |
| EPI_ISL_2997791, EPI_ISL_2997792 | Lighthouse Lab in Milton Keynes | Wellcome Sanger Institute for the COVID-19 Genomics UK (COG-UK) Consortium | The Lighthouse Lab in Milton Keynes and Alex Alderton, Roberto Amato, Jeffrey Barrett, Sonia Goncalves, Ewan Harrison, David K. Jackson, Ian Johnston, Dominic Kwiatkowski, Cordelia Langford, John Sillitoe on behalf of the Wellcome Sanger Institute COVID-19 Surveillance Team |
| EPI_ISL_2997793, EPI_ISL_2997794, EPI_ISL_2997795, EPI_ISL_2997796 | Lighthouse Laboratory Plymouth | Wellcome Sanger Institute for the COVID-19 Genomics UK (COG-UK) Consortium | Lighthouse Laboratory Plymouth and Alex Alderton, Roberto Amato, Jeffrey Barrett, Sonia Goncalves, Ewan Harrison, David K. Jackson, Ian Johnston, Dominic Kwiatkowski, Cordelia Langford, John Sillitoe on behalf of the Wellcome Sanger Institute COVID-19 Surveillance Team |
| EPI_ISL_2997797 | Lighthouse Lab in Milton Keynes | Wellcome Sanger Institute for the COVID-19 Genomics UK (COG-UK) Consortium | The Lighthouse Lab in Milton Keynes and Alex Alderton, Roberto Amato, Jeffrey Barrett, Sonia Goncalves, Ewan Harrison, David K. Jackson, Ian Johnston, Dominic Kwiatkowski, Cordelia Langford, John Sillitoe on behalf of the Wellcome Sanger Institute COVID-19 Surveillance Team |
| EPI_ISL_2997798 | Lighthouse Laboratory Plymouth | Wellcome Sanger Institute for the COVID-19 Genomics UK (COG-UK) | Lighthouse Laboratory Plymouth and Alex Alderton, Roberto Amato, Jeffrey Barrett, Sonia Goncalves, Ewan Harrison, David K. Jackson, Ian Johnston, Dominic Kwiatkowski, Cordelia Langford, John Sillitoe on behalf of the Wellcome Sanger |

| | | | |
|---|---|---|---|
| | | Consortium | Institute COVID-19 Surveillance Team |
| EPI_ISL_2997799 | Lighthouse Lab in Milton Keynes | Wellcome Sanger Institute for the COVID-19 Genomics UK (COG-UK) Consortium | The Lighthouse Lab in Milton Keynes and Alex Alderton, Roberto Amato, Jeffrey Barrett, Sonia Goncalves, Ewan Harrison, David K. Jackson, Ian Johnston, Dominic Kwiatkowski, Cordelia Langford, John Sillitoe on behalf of the Wellcome Sanger Institute COVID-19 Surveillance Team |
| EPI_ISL_2997800 | Lighthouse Lab in Glasgow | Wellcome Sanger Institute for the COVID-19 Genomics UK (COG-UK) Consortium | Harper VanSteenhouse, Yumi Kasai, David Gray, Carol Clugston, Anna Dominiczak and Alex Alderton, Roberto Amato, Jeffrey Barrett, Sonia Goncalves, Ewan Harrison, David K. Jackson, Ian Johnston, Dominic Kwiatkowski, Cordelia Langford, John Sillitoe on behalf of the Wellcome Sanger Institute COVID-19 Surveillance Team |
| EPI_ISL_2997801, EPI_ISL_2997802, EPI_ISL_2997803 | Lighthouse Laboratory Plymouth | Wellcome Sanger Institute for the COVID-19 Genomics UK (COG-UK) Consortium | Lighthouse Laboratory Plymouth and Alex Alderton, Roberto Amato, Jeffrey Barrett, Sonia Goncalves, Ewan Harrison, David K. Jackson, Ian Johnston, Dominic Kwiatkowski, Cordelia Langford, John Sillitoe on behalf of the Wellcome Sanger Institute COVID-19 Surveillance Team |
| EPI_ISL_2997804, EPI_ISL_2997805 | Lighthouse Lab in Glasgow | Wellcome Sanger Institute for the COVID-19 Genomics UK (COG-UK) Consortium | Harper VanSteenhouse, Yumi Kasai, David Gray, Carol Clugston, Anna Dominiczak and Alex Alderton, Roberto Amato, Jeffrey Barrett, Sonia Goncalves, Ewan Harrison, David K. Jackson, Ian Johnston, Dominic Kwiatkowski, Cordelia Langford, John Sillitoe on behalf of the Wellcome Sanger Institute COVID-19 Surveillance Team |
| EPI_ISL_2997806, EPI_ISL_2997807, EPI_ISL_2997808, EPI_ISL_2997809 | Lighthouse Laboratory Plymouth | Wellcome Sanger Institute for the COVID-19 Genomics UK (COG-UK) Consortium | Lighthouse Laboratory Plymouth and Alex Alderton, Roberto Amato, Jeffrey Barrett, Sonia Goncalves, Ewan Harrison, David K. Jackson, Ian Johnston, Dominic Kwiatkowski, Cordelia Langford, John Sillitoe on behalf of the Wellcome Sanger Institute COVID-19 Surveillance Team |
| EPI_ISL_2997810, EPI_ISL_2997811, EPI_ISL_2997812 | Lighthouse Lab in Milton Keynes | Wellcome Sanger Institute for the COVID-19 Genomics UK (COG-UK) Consortium | The Lighthouse Lab in Milton Keynes and Alex Alderton, Roberto Amato, Jeffrey Barrett, Sonia Goncalves, Ewan Harrison, David K. Jackson, Ian Johnston, Dominic Kwiatkowski, Cordelia Langford, John Sillitoe on behalf of the Wellcome Sanger Institute COVID-19 Surveillance Team |
| EPI_ISL_2997813, EPI_ISL_2997814 | Lighthouse Lab in Glasgow | Wellcome Sanger Institute for the COVID-19 Genomics UK (COG-UK) Consortium | Harper VanSteenhouse, Yumi Kasai, David Gray, Carol Clugston, Anna Dominiczak and Alex Alderton, Roberto Amato, Jeffrey Barrett, Sonia Goncalves, Ewan Harrison, David K. Jackson, Ian Johnston, Dominic Kwiatkowski, Cordelia Langford, John Sillitoe on behalf of the Wellcome Sanger Institute COVID-19 Surveillance Team |
| EPI_ISL_2997815, EPI_ISL_2997816 | Lighthouse Laboratory Plymouth | Wellcome Sanger Institute for the COVID-19 Genomics UK (COG-UK) Consortium | Lighthouse Laboratory Plymouth and Alex Alderton, Roberto Amato, Jeffrey Barrett, Sonia Goncalves, Ewan Harrison, David K. Jackson, Ian Johnston, Dominic Kwiatkowski, Cordelia Langford, John Sillitoe on behalf of the Wellcome Sanger Institute COVID-19 Surveillance Team |
| EPI_ISL_2997817 | Lighthouse Lab in Milton Keynes | Wellcome Sanger Institute for the COVID-19 Genomics UK (COG-UK) Consortium | The Lighthouse Lab in Milton Keynes and Alex Alderton, Roberto Amato, Jeffrey Barrett, Sonia Goncalves, Ewan Harrison, David K. Jackson, Ian Johnston, Dominic Kwiatkowski, Cordelia Langford, John Sillitoe on behalf of the Wellcome Sanger Institute COVID-19 Surveillance Team |
| EPI_ISL_2997818 | Lighthouse Laboratory Plymouth | Wellcome Sanger Institute for the COVID-19 Genomics UK (COG-UK) Consortium | Lighthouse Laboratory Plymouth and Alex Alderton, Roberto Amato, Jeffrey Barrett, Sonia Goncalves, Ewan Harrison, David K. Jackson, Ian Johnston, Dominic Kwiatkowski, Cordelia Langford, John Sillitoe on behalf of the Wellcome Sanger Institute COVID-19 Surveillance Team |
| EPI_ISL_2997819, EPI_ISL_2997820 | Lighthouse Lab in Milton Keynes | Wellcome Sanger Institute for the COVID-19 Genomics UK (COG-UK) Consortium | The Lighthouse Lab in Milton Keynes and Alex Alderton, Roberto Amato, Jeffrey Barrett, Sonia Goncalves, Ewan Harrison, David K. Jackson, Ian Johnston, Dominic Kwiatkowski, Cordelia Langford, John Sillitoe on behalf of the Wellcome Sanger Institute COVID-19 Surveillance Team |
| EPI_ISL_2997821 | Lighthouse Laboratory Plymouth | Wellcome Sanger Institute for the COVID-19 Genomics UK (COG-UK) Consortium | Lighthouse Laboratory Plymouth and Alex Alderton, Roberto Amato, Jeffrey Barrett, Sonia Goncalves, Ewan Harrison, David K. Jackson, Ian Johnston, Dominic Kwiatkowski, Cordelia Langford, John Sillitoe on behalf of the Wellcome Sanger Institute COVID-19 Surveillance Team |
| EPI_ISL_2997822, EPI_ISL_2997823 | Lighthouse Lab in Milton Keynes | Wellcome Sanger Institute for the COVID-19 Genomics UK (COG-UK) Consortium | The Lighthouse Lab in Milton Keynes and Alex Alderton, Roberto Amato, Jeffrey Barrett, Sonia Goncalves, Ewan Harrison, David K. Jackson, Ian Johnston, Dominic Kwiatkowski, Cordelia Langford, John Sillitoe on behalf of the Wellcome Sanger Institute COVID-19 Surveillance Team |
| EPI_ISL_2997824 | Lighthouse Laboratory Plymouth | Wellcome Sanger Institute for the COVID-19 Genomics UK (COG-UK) Consortium | Lighthouse Laboratory Plymouth and Alex Alderton, Roberto Amato, Jeffrey Barrett, Sonia Goncalves, Ewan Harrison, David K. Jackson, Ian Johnston, Dominic Kwiatkowski, Cordelia Langford, John Sillitoe on behalf of the Wellcome Sanger Institute COVID-19 Surveillance Team |
| EPI_ISL_2997825, EPI_ISL_2997826 | Lighthouse Lab in Glasgow | Wellcome Sanger Institute for the COVID-19 Genomics UK (COG-UK) Consortium | Harper VanSteenhouse, Yumi Kasai, David Gray, Carol Clugston, Anna Dominiczak and Alex Alderton, Roberto Amato, Jeffrey Barrett, Sonia Goncalves, Ewan Harrison, David K. Jackson, Ian Johnston, Dominic Kwiatkowski, Cordelia Langford, John Sillitoe on behalf of the Wellcome Sanger Institute COVID-19 Surveillance Team |
| EPI_ISL_2997827, EPI_ISL_2997828 | Lighthouse Laboratory Plymouth | Wellcome Sanger Institute for the COVID-19 Genomics UK (COG-UK) Consortium | Lighthouse Laboratory Plymouth and Alex Alderton, Roberto Amato, Jeffrey Barrett, Sonia Goncalves, Ewan Harrison, David K. Jackson, Ian Johnston, Dominic Kwiatkowski, Cordelia Langford, John Sillitoe on behalf of the Wellcome Sanger Institute COVID-19 Surveillance Team |
| EPI_ISL_2997829 | Lighthouse Lab in Milton Keynes | Wellcome Sanger Institute for the COVID-19 Genomics UK (COG-UK) Consortium | The Lighthouse Lab in Milton Keynes and Alex Alderton, Roberto Amato, Jeffrey Barrett, Sonia Goncalves, Ewan Harrison, David K. Jackson, Ian Johnston, Dominic Kwiatkowski, Cordelia Langford, John Sillitoe on behalf of the Wellcome Sanger Institute COVID-19 Surveillance Team |
| EPI_ISL_2997830 | Lighthouse Laboratory Plymouth | Wellcome Sanger Institute for the COVID-19 Genomics UK (COG-UK) Consortium | Lighthouse Laboratory Plymouth and Alex Alderton, Roberto Amato, Jeffrey Barrett, Sonia Goncalves, Ewan Harrison, David K. Jackson, Ian Johnston, Dominic Kwiatkowski, Cordelia Langford, John Sillitoe on behalf of the Wellcome Sanger Institute COVID-19 Surveillance Team |
| EPI_ISL_2997831 | Lighthouse Lab in Glasgow | Wellcome Sanger Institute for the COVID-19 Genomics UK (COG-UK) Consortium | Harper VanSteenhouse, Yumi Kasai, David Gray, Carol Clugston, Anna Dominiczak and Alex Alderton, Roberto Amato, Jeffrey Barrett, Sonia Goncalves, Ewan Harrison, David K. Jackson, Ian Johnston, Dominic Kwiatkowski, Cordelia Langford, John Sillitoe on behalf of the Wellcome Sanger Institute COVID-19 Surveillance Team |
| EPI_ISL_2997832 | Lighthouse Laboratory Plymouth | Wellcome Sanger Institute for the COVID-19 Genomics UK (COG-UK) Consortium | Lighthouse Laboratory Plymouth and Alex Alderton, Roberto Amato, Jeffrey Barrett, Sonia Goncalves, Ewan Harrison, David K. Jackson, Ian Johnston, Dominic Kwiatkowski, Cordelia Langford, John Sillitoe on behalf of the Wellcome Sanger Institute COVID-19 Surveillance Team |
| EPI_ISL_2997833 | Lighthouse Lab in Milton Keynes | Wellcome Sanger Institute for the COVID-19 Genomics UK (COG-UK) Consortium | The Lighthouse Lab in Milton Keynes and Alex Alderton, Roberto Amato, Jeffrey Barrett, Sonia Goncalves, Ewan Harrison, David K. Jackson, Ian Johnston, Dominic Kwiatkowski, Cordelia Langford, John Sillitoe on behalf of the Wellcome Sanger Institute COVID-19 Surveillance Team |
| EPI_ISL_2997834, EPI_ISL_2997835 | Lighthouse Lab in Glasgow | Wellcome Sanger Institute for the COVID-19 Genomics UK (COG-UK) Consortium | Harper VanSteenhouse, Yumi Kasai, David Gray, Carol Clugston, Anna Dominiczak and Alex Alderton, Roberto Amato, Jeffrey Barrett, Sonia Goncalves, Ewan Harrison, David K. Jackson, Ian Johnston, Dominic Kwiatkowski, Cordelia Langford, John Sillitoe on behalf of the Wellcome Sanger Institute COVID-19 Surveillance Team |
| EPI_ISL_2997836 | Lighthouse Lab in Milton Keynes | Wellcome Sanger Institute for the COVID-19 Genomics UK (COG-UK) Consortium | The Lighthouse Lab in Milton Keynes and Alex Alderton, Roberto Amato, Jeffrey Barrett, Sonia Goncalves, Ewan Harrison, David K. Jackson, Ian Johnston, Dominic Kwiatkowski, Cordelia Langford, John Sillitoe on behalf of the Wellcome Sanger Institute COVID-19 Surveillance Team |
| EPI_ISL_2997837, EPI_ISL_2997838, EPI_ISL_2997839, EPI_ISL_2997840, EPI_ISL_2997841 | Lighthouse Laboratory Plymouth | Wellcome Sanger Institute for the COVID-19 Genomics UK (COG-UK) Consortium | Lighthouse Laboratory Plymouth and Alex Alderton, Roberto Amato, Jeffrey Barrett, Sonia Goncalves, Ewan Harrison, David K. Jackson, Ian Johnston, Dominic Kwiatkowski, Cordelia Langford, John Sillitoe on behalf of the Wellcome Sanger Institute COVID-19 Surveillance Team |
| EPI_ISL_2997842 | Lighthouse Lab in Glasgow | Wellcome Sanger Institute for the COVID-19 Genomics UK (COG-UK) Consortium | Harper VanSteenhouse, Yumi Kasai, David Gray, Carol Clugston, Anna Dominiczak and Alex Alderton, Roberto Amato, Jeffrey Barrett, Sonia Goncalves, Ewan Harrison, David K. Jackson, Ian Johnston, Dominic Kwiatkowski, Cordelia Langford, John Sillitoe on behalf of the Wellcome Sanger Institute COVID-19 Surveillance Team |
| EPI_ISL_2997843, EPI_ISL_2997844, EPI_ISL_2997845 | Lighthouse Laboratory Plymouth | Wellcome Sanger Institute for the COVID-19 Genomics UK (COG-UK) Consortium | Lighthouse Laboratory Plymouth and Alex Alderton, Roberto Amato, Jeffrey Barrett, Sonia Goncalves, Ewan Harrison, David K. Jackson, Ian Johnston, Dominic Kwiatkowski, Cordelia Langford, John Sillitoe on behalf of the Wellcome Sanger Institute COVID-19 Surveillance Team |
| EPI_ISL_2997846 | Lighthouse Lab in Milton Keynes | Wellcome Sanger Institute for the COVID-19 Genomics UK (COG-UK) Consortium | The Lighthouse Lab in Milton Keynes and Alex Alderton, Roberto Amato, Jeffrey Barrett, Sonia Goncalves, Ewan Harrison, David K. Jackson, Ian Johnston, Dominic Kwiatkowski, Cordelia Langford, John Sillitoe on behalf of the Wellcome Sanger Institute COVID-19 Surveillance Team |
| EPI_ISL_2997847, EPI_ISL_2997848 | Lighthouse Lab in Glasgow | Wellcome Sanger Institute for the COVID-19 Genomics UK (COG-UK) Consortium | Harper VanSteenhouse, Yumi Kasai, David Gray, Carol Clugston, Anna Dominiczak and Alex Alderton, Roberto Amato, Jeffrey Barrett, Sonia Goncalves, Ewan Harrison, David K. Jackson, Ian Johnston, Dominic Kwiatkowski, Cordelia Langford, John Sillitoe on behalf of the Wellcome Sanger Institute COVID-19 Surveillance Team |
| EPI_ISL_2997849 | Lighthouse Laboratory Plymouth | Wellcome Sanger Institute for the COVID-19 Genomics UK (COG-UK) Consortium | Lighthouse Laboratory Plymouth and Alex Alderton, Roberto Amato, Jeffrey Barrett, Sonia Goncalves, Ewan Harrison, David K. Jackson, Ian Johnston, Dominic Kwiatkowski, Cordelia Langford, John Sillitoe on behalf of the Wellcome Sanger Institute COVID-19 Surveillance Team |
| EPI_ISL_2997850 | Lighthouse Lab in Glasgow | Wellcome Sanger Institute for the COVID-19 Genomics UK (COG-UK) Consortium | Harper VanSteenhouse, Yumi Kasai, David Gray, Carol Clugston, Anna Dominiczak and Alex Alderton, Roberto Amato, Jeffrey Barrett, Sonia Goncalves, Ewan Harrison, David K. Jackson, Ian Johnston, Dominic Kwiatkowski, Cordelia Langford, John Sillitoe on behalf of the Wellcome Sanger Institute COVID-19 Surveillance Team |
| EPI_ISL_2997851 | Lighthouse Laboratory Plymouth | Wellcome Sanger Institute for the COVID-19 Genomics UK (COG-UK) Consortium | Lighthouse Laboratory Plymouth and Alex Alderton, Roberto Amato, Jeffrey Barrett, Sonia Goncalves, Ewan Harrison, David K. Jackson, Ian Johnston, Dominic Kwiatkowski, Cordelia Langford, John Sillitoe on behalf of the Wellcome Sanger Institute COVID-19 Surveillance Team |
| EPI_ISL_2997852, EPI_ISL_2997853 | Lighthouse Lab in Glasgow | Wellcome Sanger Institute for the COVID-19 Genomics UK (COG-UK) Consortium | Harper VanSteenhouse, Yumi Kasai, David Gray, Carol Clugston, Anna Dominiczak and Alex Alderton, Roberto Amato, Jeffrey Barrett, Sonia Goncalves, Ewan Harrison, David K. Jackson, Ian Johnston, Dominic Kwiatkowski, Cordelia Langford, John Sillitoe on behalf of the Wellcome Sanger Institute COVID-19 Surveillance Team |
| EPI_ISL_2997854 | Lighthouse Lab in Milton Keynes | Wellcome Sanger Institute for the COVID-19 Genomics UK (COG-UK) Consortium | The Lighthouse Lab in Milton Keynes and Alex Alderton, Roberto Amato, Jeffrey Barrett, Sonia Goncalves, Ewan Harrison, David K. Jackson, Ian Johnston, Dominic Kwiatkowski, Cordelia Langford, John Sillitoe on behalf of the Wellcome Sanger Institute COVID-19 Surveillance Team |
| EPI_ISL_2997855, EPI_ISL_2997856 | Lighthouse Laboratory Plymouth | Wellcome Sanger Institute for the COVID-19 Genomics UK (COG-UK) Consortium | Lighthouse Laboratory Plymouth and Alex Alderton, Roberto Amato, Jeffrey Barrett, Sonia Goncalves, Ewan Harrison, David K. Jackson, Ian Johnston, Dominic Kwiatkowski, Cordelia Langford, John Sillitoe on behalf of the Wellcome Sanger Institute COVID-19 Surveillance Team |
| EPI_ISL_2997857 | Lighthouse Lab in Milton Keynes | Wellcome Sanger Institute for the COVID-19 Genomics UK (COG-UK) Consortium | The Lighthouse Lab in Milton Keynes and Alex Alderton, Roberto Amato, Jeffrey Barrett, Sonia Goncalves, Ewan Harrison, David K. Jackson, Ian Johnston, Dominic Kwiatkowski, Cordelia Langford, John Sillitoe on behalf of the Wellcome Sanger Institute COVID-19 Surveillance Team |
| EPI_ISL_2997858 | Lighthouse Lab in Glasgow | Wellcome Sanger Institute for the COVID-19 Genomics UK (COG-UK) Consortium | Harper VanSteenhouse, Yumi Kasai, David Gray, Carol Clugston, Anna Dominiczak and Alex Alderton, Roberto Amato, Jeffrey Barrett, Sonia Goncalves, Ewan Harrison, David K. Jackson, Ian Johnston, Dominic Kwiatkowski, Cordelia Langford, John Sillitoe on behalf of the Wellcome Sanger Institute COVID-19 Surveillance Team |
| EPI_ISL_2997859, EPI_ISL_2997860 | Lighthouse Laboratory Plymouth | Wellcome Sanger Institute for the COVID-19 Genomics UK (COG-UK) Consortium | Lighthouse Laboratory Plymouth and Alex Alderton, Roberto Amato, Jeffrey Barrett, Sonia Goncalves, Ewan Harrison, David K. Jackson, Ian Johnston, Dominic Kwiatkowski, Cordelia Langford, John Sillitoe on behalf of the Wellcome Sanger Institute COVID-19 Surveillance Team |
| EPI_ISL_2997861 | Lighthouse Lab in Glasgow | Wellcome Sanger Institute for the COVID-19 Genomics UK (COG-UK) Consortium | Harper VanSteenhouse, Yumi Kasai, David Gray, Carol Clugston, Anna Dominiczak and Alex Alderton, Roberto Amato, Jeffrey Barrett, Sonia Goncalves, Ewan Harrison, David K. Jackson, Ian Johnston, Dominic Kwiatkowski, Cordelia Langford, John Sillitoe on behalf of the Wellcome Sanger Institute COVID-19 Surveillance Team |
| EPI_ISL_2997862, EPI_ISL_2997863 | Lighthouse Laboratory Plymouth | Wellcome Sanger Institute for the COVID-19 Genomics UK (COG-UK) Consortium | Lighthouse Laboratory Plymouth and Alex Alderton, Roberto Amato, Jeffrey Barrett, Sonia Goncalves, Ewan Harrison, David K. Jackson, Ian Johnston, Dominic Kwiatkowski, Cordelia Langford, John Sillitoe on behalf of the Wellcome Sanger Institute COVID-19 Surveillance Team |
| EPI_ISL_2997864 | Lighthouse Lab in Glasgow | Wellcome Sanger Institute for the COVID-19 Genomics UK (COG-UK) Consortium | Harper VanSteenhouse, Yumi Kasai, David Gray, Carol Clugston, Anna Dominiczak and Alex Alderton, Roberto Amato, Jeffrey Barrett, Sonia Goncalves, Ewan Harrison, David K. Jackson, Ian Johnston, Dominic Kwiatkowski, Cordelia Langford, John Sillitoe on behalf of the Wellcome Sanger Institute COVID-19 Surveillance Team |
| EPI_ISL_2997865, EPI_ISL_2997866 | Lighthouse Lab in Milton Keynes | Wellcome Sanger Institute for the COVID-19 Genomics UK (COG-UK) Consortium | The Lighthouse Lab in Milton Keynes and Alex Alderton, Roberto Amato, Jeffrey Barrett, Sonia Goncalves, Ewan Harrison, David K. Jackson, Ian Johnston, Dominic Kwiatkowski, Cordelia Langford, John Sillitoe on behalf of the Wellcome Sanger Institute COVID-19 Surveillance Team |
| EPI_ISL_2997867 | Lighthouse Lab in Glasgow | Wellcome Sanger Institute for the COVID-19 Genomics UK (COG-UK) | Harper VanSteenhouse, Yumi Kasai, David Gray, Carol Clugston, Anna Dominiczak and Alex Alderton, Roberto Amato, Jeffrey Barrett, Sonia Goncalves, Ewan Harrison, David K. Jackson, Ian Johnston, Dominic Kwiatkowski, Cordelia Langford, |

| | | | |
|---|---|---|---|
| | | Consortium | John Sillitoe on behalf of the Wellcome Sanger Institute COVID-19 Surveillance Team |
| EPI_ISL_2997868, EPI_ISL_2997869, EPI_ISL_2997870, EPI_ISL_2997871, EPI_ISL_2997872 | Lighthouse Laboratory Plymouth | Wellcome Sanger Institute for the COVID-19 Genomics UK (COG-UK) Consortium | Lighthouse Laboratory Plymouth and Alex Alderton, Roberto Amato, Jeffrey Barrett, Sonia Goncalves, Ewan Harrison, David K. Jackson, Ian Johnston, Dominic Kwiatkowski, Cordelia Langford, John Sillitoe on behalf of the Wellcome Sanger Institute COVID-19 Surveillance Team |
| EPI_ISL_2997873 | Lighthouse Lab in Glasgow | Wellcome Sanger Institute for the COVID-19 Genomics UK (COG-UK) Consortium | Harper VanSteenhouse, Yumi Kasai, David Gray, Carol Clugston, Anna Dominiczak and Alex Alderton, Roberto Amato, Jeffrey Barrett, Sonia Goncalves, Ewan Harrison, David K. Jackson, Ian Johnston, Dominic Kwiatkowski, Cordelia Langford, John Sillitoe on behalf of the Wellcome Sanger Institute COVID-19 Surveillance Team |
| EPI_ISL_2997874 | Lighthouse Laboratory Plymouth | Wellcome Sanger Institute for the COVID-19 Genomics UK (COG-UK) Consortium | Lighthouse Laboratory Plymouth and Alex Alderton, Roberto Amato, Jeffrey Barrett, Sonia Goncalves, Ewan Harrison, David K. Jackson, Ian Johnston, Dominic Kwiatkowski, Cordelia Langford, John Sillitoe on behalf of the Wellcome Sanger Institute COVID-19 Surveillance Team |
| EPI_ISL_2997875 | Lighthouse Lab in Glasgow | Wellcome Sanger Institute for the COVID-19 Genomics UK (COG-UK) Consortium | Harper VanSteenhouse, Yumi Kasai, David Gray, Carol Clugston, Anna Dominiczak and Alex Alderton, Roberto Amato, Jeffrey Barrett, Sonia Goncalves, Ewan Harrison, David K. Jackson, Ian Johnston, Dominic Kwiatkowski, Cordelia Langford, John Sillitoe on behalf of the Wellcome Sanger Institute COVID-19 Surveillance Team |
| EPI_ISL_2997876 | Lighthouse Laboratory Plymouth | Wellcome Sanger Institute for the COVID-19 Genomics UK (COG-UK) Consortium | Lighthouse Laboratory Plymouth and Alex Alderton, Roberto Amato, Jeffrey Barrett, Sonia Goncalves, Ewan Harrison, David K. Jackson, Ian Johnston, Dominic Kwiatkowski, Cordelia Langford, John Sillitoe on behalf of the Wellcome Sanger Institute COVID-19 Surveillance Team |
| EPI_ISL_2997877 | Lighthouse Lab in Glasgow | Wellcome Sanger Institute for the COVID-19 Genomics UK (COG-UK) Consortium | Harper VanSteenhouse, Yumi Kasai, David Gray, Carol Clugston, Anna Dominiczak and Alex Alderton, Roberto Amato, Jeffrey Barrett, Sonia Goncalves, Ewan Harrison, David K. Jackson, Ian Johnston, Dominic Kwiatkowski, Cordelia Langford, John Sillitoe on behalf of the Wellcome Sanger Institute COVID-19 Surveillance Team |
| EPI_ISL_2997878 | Lighthouse Laboratory Plymouth | Wellcome Sanger Institute for the COVID-19 Genomics UK (COG-UK) Consortium | Lighthouse Laboratory Plymouth and Alex Alderton, Roberto Amato, Jeffrey Barrett, Sonia Goncalves, Ewan Harrison, David K. Jackson, Ian Johnston, Dominic Kwiatkowski, Cordelia Langford, John Sillitoe on behalf of the Wellcome Sanger Institute COVID-19 Surveillance Team |
| EPI_ISL_2997879 | Lighthouse Lab in Milton Keynes | Wellcome Sanger Institute for the COVID-19 Genomics UK (COG-UK) Consortium | The Lighthouse Lab in Milton Keynes and Alex Alderton, Roberto Amato, Jeffrey Barrett, Sonia Goncalves, Ewan Harrison, David K. Jackson, Ian Johnston, Dominic Kwiatkowski, Cordelia Langford, John Sillitoe on behalf of the Wellcome Sanger Institute COVID-19 Surveillance Team |
| EPI_ISL_2997880 | Lighthouse Laboratory Plymouth | Wellcome Sanger Institute for the COVID-19 Genomics UK (COG-UK) Consortium | Lighthouse Laboratory Plymouth and Alex Alderton, Roberto Amato, Jeffrey Barrett, Sonia Goncalves, Ewan Harrison, David K. Jackson, Ian Johnston, Dominic Kwiatkowski, Cordelia Langford, John Sillitoe on behalf of the Wellcome Sanger Institute COVID-19 Surveillance Team |
| EPI_ISL_2997881 | Lighthouse Lab in Milton Keynes | Wellcome Sanger Institute for the COVID-19 Genomics UK (COG-UK) Consortium | The Lighthouse Lab in Milton Keynes and Alex Alderton, Roberto Amato, Jeffrey Barrett, Sonia Goncalves, Ewan Harrison, David K. Jackson, Ian Johnston, Dominic Kwiatkowski, Cordelia Langford, John Sillitoe on behalf of the Wellcome Sanger Institute COVID-19 Surveillance Team |
| EPI_ISL_2997882 | Lighthouse Laboratory Plymouth | Wellcome Sanger Institute for the COVID-19 Genomics UK (COG-UK) Consortium | Lighthouse Laboratory Plymouth and Alex Alderton, Roberto Amato, Jeffrey Barrett, Sonia Goncalves, Ewan Harrison, David K. Jackson, Ian Johnston, Dominic Kwiatkowski, Cordelia Langford, John Sillitoe on behalf of the Wellcome Sanger Institute COVID-19 Surveillance Team |
| EPI_ISL_2997883 | Lighthouse Lab in Milton Keynes | Wellcome Sanger Institute for the COVID-19 Genomics UK (COG-UK) Consortium | The Lighthouse Lab in Milton Keynes and Alex Alderton, Roberto Amato, Jeffrey Barrett, Sonia Goncalves, Ewan Harrison, David K. Jackson, Ian Johnston, Dominic Kwiatkowski, Cordelia Langford, John Sillitoe on behalf of the Wellcome Sanger Institute COVID-19 Surveillance Team |
| EPI_ISL_2997884 | Lighthouse Laboratory Plymouth | Wellcome Sanger Institute for the COVID-19 Genomics UK (COG-UK) Consortium | Lighthouse Laboratory Plymouth and Alex Alderton, Roberto Amato, Jeffrey Barrett, Sonia Goncalves, Ewan Harrison, David K. Jackson, Ian Johnston, Dominic Kwiatkowski, Cordelia Langford, John Sillitoe on behalf of the Wellcome Sanger Institute COVID-19 Surveillance Team |
| EPI_ISL_2997885, EPI_ISL_2997886 | Lighthouse Lab in Milton Keynes | Wellcome Sanger Institute for the COVID-19 Genomics UK (COG-UK) Consortium | The Lighthouse Lab in Milton Keynes and Alex Alderton, Roberto Amato, Jeffrey Barrett, Sonia Goncalves, Ewan Harrison, David K. Jackson, Ian Johnston, Dominic Kwiatkowski, Cordelia Langford, John Sillitoe on behalf of the Wellcome Sanger Institute COVID-19 Surveillance Team |
| EPI_ISL_2997887 | Lighthouse Laboratory Plymouth | Wellcome Sanger Institute for the COVID-19 Genomics UK (COG-UK) Consortium | Lighthouse Laboratory Plymouth and Alex Alderton, Roberto Amato, Jeffrey Barrett, Sonia Goncalves, Ewan Harrison, David K. Jackson, Ian Johnston, Dominic Kwiatkowski, Cordelia Langford, John Sillitoe on behalf of the Wellcome Sanger Institute COVID-19 Surveillance Team |
| EPI_ISL_2997888 | Lighthouse Lab in Milton Keynes | Wellcome Sanger Institute for the COVID-19 Genomics UK (COG-UK) Consortium | The Lighthouse Lab in Milton Keynes and Alex Alderton, Roberto Amato, Jeffrey Barrett, Sonia Goncalves, Ewan Harrison, David K. Jackson, Ian Johnston, Dominic Kwiatkowski, Cordelia Langford, John Sillitoe on behalf of the Wellcome Sanger Institute COVID-19 Surveillance Team |
| EPI_ISL_2997889 | Lighthouse Lab in Glasgow | Wellcome Sanger Institute for the COVID-19 Genomics UK (COG-UK) Consortium | Harper VanSteenhouse, Yumi Kasai, David Gray, Carol Clugston, Anna Dominiczak and Alex Alderton, Roberto Amato, Jeffrey Barrett, Sonia Goncalves, Ewan Harrison, David K. Jackson, Ian Johnston, Dominic Kwiatkowski, Cordelia Langford, John Sillitoe on behalf of the Wellcome Sanger Institute COVID-19 Surveillance Team |
| EPI_ISL_2997890, EPI_ISL_2997891, EPI_ISL_2997892 | Lighthouse Laboratory Plymouth | Wellcome Sanger Institute for the COVID-19 Genomics UK (COG-UK) Consortium | Lighthouse Laboratory Plymouth and Alex Alderton, Roberto Amato, Jeffrey Barrett, Sonia Goncalves, Ewan Harrison, David K. Jackson, Ian Johnston, Dominic Kwiatkowski, Cordelia Langford, John Sillitoe on behalf of the Wellcome Sanger Institute COVID-19 Surveillance Team |
| EPI_ISL_2997893 | Lighthouse Lab in Glasgow | Wellcome Sanger Institute for the COVID-19 Genomics UK (COG-UK) Consortium | Harper VanSteenhouse, Yumi Kasai, David Gray, Carol Clugston, Anna Dominiczak and Alex Alderton, Roberto Amato, Jeffrey Barrett, Sonia Goncalves, Ewan Harrison, David K. Jackson, Ian Johnston, Dominic Kwiatkowski, Cordelia Langford, John Sillitoe on behalf of the Wellcome Sanger Institute COVID-19 Surveillance Team |
| EPI_ISL_2997894, EPI_ISL_2997895 | Lighthouse Laboratory Plymouth | Wellcome Sanger Institute for the COVID-19 Genomics UK (COG-UK) Consortium | Lighthouse Laboratory Plymouth and Alex Alderton, Roberto Amato, Jeffrey Barrett, Sonia Goncalves, Ewan Harrison, David K. Jackson, Ian Johnston, Dominic Kwiatkowski, Cordelia Langford, John Sillitoe on behalf of the Wellcome Sanger Institute COVID-19 Surveillance Team |
| EPI_ISL_2997896 | Lighthouse Lab in Milton Keynes | Wellcome Sanger Institute for the COVID-19 Genomics UK (COG-UK) Consortium | The Lighthouse Lab in Milton Keynes and Alex Alderton, Roberto Amato, Jeffrey Barrett, Sonia Goncalves, Ewan Harrison, David K. Jackson, Ian Johnston, Dominic Kwiatkowski, Cordelia Langford, John Sillitoe on behalf of the Wellcome Sanger Institute COVID-19 Surveillance Team |
| EPI_ISL_2997897 | Lighthouse Laboratory Plymouth | Wellcome Sanger Institute for the COVID-19 Genomics UK (COG-UK) Consortium | Lighthouse Laboratory Plymouth and Alex Alderton, Roberto Amato, Jeffrey Barrett, Sonia Goncalves, Ewan Harrison, David K. Jackson, Ian Johnston, Dominic Kwiatkowski, Cordelia Langford, John Sillitoe on behalf of the Wellcome Sanger Institute COVID-19 Surveillance Team |
| EPI_ISL_2997898 | Lighthouse Lab in Glasgow | Wellcome Sanger Institute for the COVID-19 Genomics UK (COG-UK) Consortium | Harper VanSteenhouse, Yumi Kasai, David Gray, Carol Clugston, Anna Dominiczak and Alex Alderton, Roberto Amato, Jeffrey Barrett, Sonia Goncalves, Ewan Harrison, David K. Jackson, Ian Johnston, Dominic Kwiatkowski, Cordelia Langford, John Sillitoe on behalf of the Wellcome Sanger Institute COVID-19 Surveillance Team |
| EPI_ISL_2997899, EPI_ISL_2997900 | Lighthouse Laboratory Plymouth | Wellcome Sanger Institute for the COVID-19 Genomics UK (COG-UK) Consortium | Lighthouse Laboratory Plymouth and Alex Alderton, Roberto Amato, Jeffrey Barrett, Sonia Goncalves, Ewan Harrison, David K. Jackson, Ian Johnston, Dominic Kwiatkowski, Cordelia Langford, John Sillitoe on behalf of the Wellcome Sanger Institute COVID-19 Surveillance Team |
| EPI_ISL_2997901 | Lighthouse Lab in Glasgow | Wellcome Sanger Institute for the COVID-19 Genomics UK (COG-UK) Consortium | Harper VanSteenhouse, Yumi Kasai, David Gray, Carol Clugston, Anna Dominiczak and Alex Alderton, Roberto Amato, Jeffrey Barrett, Sonia Goncalves, Ewan Harrison, David K. Jackson, Ian Johnston, Dominic Kwiatkowski, Cordelia Langford, John Sillitoe on behalf of the Wellcome Sanger Institute COVID-19 Surveillance Team |
| EPI_ISL_2997902, EPI_ISL_2997903 | Lighthouse Laboratory Plymouth | Wellcome Sanger Institute for the COVID-19 Genomics UK (COG-UK) Consortium | Lighthouse Laboratory Plymouth and Alex Alderton, Roberto Amato, Jeffrey Barrett, Sonia Goncalves, Ewan Harrison, David K. Jackson, Ian Johnston, Dominic Kwiatkowski, Cordelia Langford, John Sillitoe on behalf of the Wellcome Sanger Institute COVID-19 Surveillance Team |
| EPI_ISL_2997904, EPI_ISL_2997905 | Lighthouse Lab in Glasgow | Wellcome Sanger Institute for the COVID-19 Genomics UK (COG-UK) Consortium | Harper VanSteenhouse, Yumi Kasai, David Gray, Carol Clugston, Anna Dominiczak and Alex Alderton, Roberto Amato, Jeffrey Barrett, Sonia Goncalves, Ewan Harrison, David K. Jackson, Ian Johnston, Dominic Kwiatkowski, Cordelia Langford, John Sillitoe on behalf of the Wellcome Sanger Institute COVID-19 Surveillance Team |
| EPI_ISL_2997906, EPI_ISL_2997907 | Lighthouse Laboratory Plymouth | Wellcome Sanger Institute for the COVID-19 Genomics UK (COG-UK) Consortium | Lighthouse Laboratory Plymouth and Alex Alderton, Roberto Amato, Jeffrey Barrett, Sonia Goncalves, Ewan Harrison, David K. Jackson, Ian Johnston, Dominic Kwiatkowski, Cordelia Langford, John Sillitoe on behalf of the Wellcome Sanger Institute COVID-19 Surveillance Team |
| EPI_ISL_2997908 | Lighthouse Lab in Milton Keynes | Wellcome Sanger Institute for the COVID-19 Genomics UK (COG-UK) Consortium | The Lighthouse Lab in Milton Keynes and Alex Alderton, Roberto Amato, Jeffrey Barrett, Sonia Goncalves, Ewan Harrison, David K. Jackson, Ian Johnston, Dominic Kwiatkowski, Cordelia Langford, John Sillitoe on behalf of the Wellcome Sanger Institute COVID-19 Surveillance Team |
| EPI_ISL_2997909 | Lighthouse Laboratory Plymouth | Wellcome Sanger Institute for the COVID-19 Genomics UK (COG-UK) Consortium | Lighthouse Laboratory Plymouth and Alex Alderton, Roberto Amato, Jeffrey Barrett, Sonia Goncalves, Ewan Harrison, David K. Jackson, Ian Johnston, Dominic Kwiatkowski, Cordelia Langford, John Sillitoe on behalf of the Wellcome Sanger Institute COVID-19 Surveillance Team |
| EPI_ISL_2997910, EPI_ISL_2997911 | Lighthouse Lab in Milton Keynes | Wellcome Sanger Institute for the COVID-19 Genomics UK (COG-UK) Consortium | The Lighthouse Lab in Milton Keynes and Alex Alderton, Roberto Amato, Jeffrey Barrett, Sonia Goncalves, Ewan Harrison, David K. Jackson, Ian Johnston, Dominic Kwiatkowski, Cordelia Langford, John Sillitoe on behalf of the Wellcome Sanger Institute COVID-19 Surveillance Team |
| EPI_ISL_2997912 | Lighthouse Laboratory Plymouth | Wellcome Sanger Institute for the COVID-19 Genomics UK (COG-UK) Consortium | Lighthouse Laboratory Plymouth and Alex Alderton, Roberto Amato, Jeffrey Barrett, Sonia Goncalves, Ewan Harrison, David K. Jackson, Ian Johnston, Dominic Kwiatkowski, Cordelia Langford, John Sillitoe on behalf of the Wellcome Sanger Institute COVID-19 Surveillance Team |
| EPI_ISL_2997913 | Lighthouse Lab in Milton Keynes | Wellcome Sanger Institute for the COVID-19 Genomics UK (COG-UK) Consortium | The Lighthouse Lab in Milton Keynes and Alex Alderton, Roberto Amato, Jeffrey Barrett, Sonia Goncalves, Ewan Harrison, David K. Jackson, Ian Johnston, Dominic Kwiatkowski, Cordelia Langford, John Sillitoe on behalf of the Wellcome Sanger Institute COVID-19 Surveillance Team |
| EPI_ISL_2997914 | Lighthouse Lab in Glasgow | Wellcome Sanger Institute for the COVID-19 Genomics UK (COG-UK) Consortium | Harper VanSteenhouse, Yumi Kasai, David Gray, Carol Clugston, Anna Dominiczak and Alex Alderton, Roberto Amato, Jeffrey Barrett, Sonia Goncalves, Ewan Harrison, David K. Jackson, Ian Johnston, Dominic Kwiatkowski, Cordelia Langford, John Sillitoe on behalf of the Wellcome Sanger Institute COVID-19 Surveillance Team |
| EPI_ISL_2997915, EPI_ISL_2997916 | Lighthouse Laboratory Plymouth | Wellcome Sanger Institute for the COVID-19 Genomics UK (COG-UK) Consortium | Lighthouse Laboratory Plymouth and Alex Alderton, Roberto Amato, Jeffrey Barrett, Sonia Goncalves, Ewan Harrison, David K. Jackson, Ian Johnston, Dominic Kwiatkowski, Cordelia Langford, John Sillitoe on behalf of the Wellcome Sanger Institute COVID-19 Surveillance Team |
| EPI_ISL_2997917 | Lighthouse Lab in Glasgow | Wellcome Sanger Institute for the COVID-19 Genomics UK (COG-UK) Consortium | Harper VanSteenhouse, Yumi Kasai, David Gray, Carol Clugston, Anna Dominiczak and Alex Alderton, Roberto Amato, Jeffrey Barrett, Sonia Goncalves, Ewan Harrison, David K. Jackson, Ian Johnston, Dominic Kwiatkowski, Cordelia Langford, John Sillitoe on behalf of the Wellcome Sanger Institute COVID-19 Surveillance Team |
| EPI_ISL_2997918 | Lighthouse Laboratory Plymouth | Wellcome Sanger Institute for the COVID-19 Genomics UK (COG-UK) Consortium | Lighthouse Laboratory Plymouth and Alex Alderton, Roberto Amato, Jeffrey Barrett, Sonia Goncalves, Ewan Harrison, David K. Jackson, Ian Johnston, Dominic Kwiatkowski, Cordelia Langford, John Sillitoe on behalf of the Wellcome Sanger Institute COVID-19 Surveillance Team |
| EPI_ISL_2997919 | Lighthouse Lab in Milton Keynes | Wellcome Sanger Institute for the COVID-19 Genomics UK (COG-UK) Consortium | The Lighthouse Lab in Milton Keynes and Alex Alderton, Roberto Amato, Jeffrey Barrett, Sonia Goncalves, Ewan Harrison, David K. Jackson, Ian Johnston, Dominic Kwiatkowski, Cordelia Langford, John Sillitoe on behalf of the Wellcome Sanger Institute COVID-19 Surveillance Team |
| EPI_ISL_2997920 | Lighthouse Laboratory Plymouth | Wellcome Sanger Institute for the COVID-19 Genomics UK (COG-UK) Consortium | Lighthouse Laboratory Plymouth and Alex Alderton, Roberto Amato, Jeffrey Barrett, Sonia Goncalves, Ewan Harrison, David K. Jackson, Ian Johnston, Dominic Kwiatkowski, Cordelia Langford, John Sillitoe on behalf of the Wellcome Sanger Institute COVID-19 Surveillance Team |
| EPI_ISL_2997921, EPI_ISL_2997922, EPI_ISL_2997923, EPI_ISL_2997924 | Lighthouse Lab in Milton Keynes | Wellcome Sanger Institute for the COVID-19 Genomics UK (COG-UK) Consortium | The Lighthouse Lab in Milton Keynes and Alex Alderton, Roberto Amato, Jeffrey Barrett, Sonia Goncalves, Ewan Harrison, David K. Jackson, Ian Johnston, Dominic Kwiatkowski, Cordelia Langford, John Sillitoe on behalf of the Wellcome Sanger Institute COVID-19 Surveillance Team |
| EPI_ISL_2997925 | Lighthouse Laboratory Plymouth | Wellcome Sanger Institute for the COVID-19 Genomics UK (COG-UK) Consortium | Lighthouse Laboratory Plymouth and Alex Alderton, Roberto Amato, Jeffrey Barrett, Sonia Goncalves, Ewan Harrison, David K. Jackson, Ian Johnston, Dominic Kwiatkowski, Cordelia Langford, John Sillitoe on behalf of the Wellcome Sanger Institute COVID-19 Surveillance Team |
| EPI_ISL_2997926 | Lighthouse Lab in Milton Keynes | Wellcome Sanger Institute for the COVID-19 Genomics UK (COG-UK) Consortium | The Lighthouse Lab in Milton Keynes and Alex Alderton, Roberto Amato, Jeffrey Barrett, Sonia Goncalves, Ewan Harrison, David K. Jackson, Ian Johnston, Dominic Kwiatkowski, Cordelia Langford, John Sillitoe on behalf of the Wellcome Sanger Institute COVID-19 Surveillance Team |

| | | | |
|---|---|---|---|
| EPI_ISL_2997927 | Lighthouse Laboratory Plymouth | Wellcome Sanger Institute for the COVID-19 Genomics UK (COG-UK) Consortium | Lighthouse Laboratory Plymouth and Alex Alderton, Roberto Amato, Jeffrey Barrett, Sonia Goncalves, Ewan Harrison, David K. Jackson, Ian Johnston, Dominic Kwiatkowski, Cordelia Langford, John Sillitoe on behalf of the Wellcome Sanger Institute COVID-19 Surveillance Team |
| EPI_ISL_2997928 | Lighthouse Lab in Glasgow | Wellcome Sanger Institute for the COVID-19 Genomics UK (COG-UK) Consortium | Harper VanSteenhouse, Yumi Kasai, David Gray, Carol Clugston, Anna Dominiczak and Alex Alderton, Roberto Amato, Jeffrey Barrett, Sonia Goncalves, Ewan Harrison, David K. Jackson, Ian Johnston, Dominic Kwiatkowski, Cordelia Langford, John Sillitoe on behalf of the Wellcome Sanger Institute COVID-19 Surveillance Team |
| EPI_ISL_2997929 | Lighthouse Laboratory Plymouth | Wellcome Sanger Institute for the COVID-19 Genomics UK (COG-UK) Consortium | Lighthouse Laboratory Plymouth and Alex Alderton, Roberto Amato, Jeffrey Barrett, Sonia Goncalves, Ewan Harrison, David K. Jackson, Ian Johnston, Dominic Kwiatkowski, Cordelia Langford, John Sillitoe on behalf of the Wellcome Sanger Institute COVID-19 Surveillance Team |
| EPI_ISL_2997930, EPI_ISL_2997931, EPI_ISL_2997932, EPI_ISL_2997933 | Lighthouse Lab in Glasgow | Wellcome Sanger Institute for the COVID-19 Genomics UK (COG-UK) Consortium | Harper VanSteenhouse, Yumi Kasai, David Gray, Carol Clugston, Anna Dominiczak and Alex Alderton, Roberto Amato, Jeffrey Barrett, Sonia Goncalves, Ewan Harrison, David K. Jackson, Ian Johnston, Dominic Kwiatkowski, Cordelia Langford, John Sillitoe on behalf of the Wellcome Sanger Institute COVID-19 Surveillance Team |
| EPI_ISL_2997934 | Lighthouse Lab in Milton Keynes | Wellcome Sanger Institute for the COVID-19 Genomics UK (COG-UK) Consortium | The Lighthouse Lab in Milton Keynes and Alex Alderton, Roberto Amato, Jeffrey Barrett, Sonia Goncalves, Ewan Harrison, David K. Jackson, Ian Johnston, Dominic Kwiatkowski, Cordelia Langford, John Sillitoe on behalf of the Wellcome Sanger Institute COVID-19 Surveillance Team |
| EPI_ISL_2997935, EPI_ISL_2997936 | Lighthouse Laboratory Plymouth | Wellcome Sanger Institute for the COVID-19 Genomics UK (COG-UK) Consortium | Lighthouse Laboratory Plymouth and Alex Alderton, Roberto Amato, Jeffrey Barrett, Sonia Goncalves, Ewan Harrison, David K. Jackson, Ian Johnston, Dominic Kwiatkowski, Cordelia Langford, John Sillitoe on behalf of the Wellcome Sanger Institute COVID-19 Surveillance Team |
| EPI_ISL_2997937 | Lighthouse Lab in Glasgow | Wellcome Sanger Institute for the COVID-19 Genomics UK (COG-UK) Consortium | Harper VanSteenhouse, Yumi Kasai, David Gray, Carol Clugston, Anna Dominiczak and Alex Alderton, Roberto Amato, Jeffrey Barrett, Sonia Goncalves, Ewan Harrison, David K. Jackson, Ian Johnston, Dominic Kwiatkowski, Cordelia Langford, John Sillitoe on behalf of the Wellcome Sanger Institute COVID-19 Surveillance Team |
| EPI_ISL_2997938, EPI_ISL_2997939, EPI_ISL_2997940, EPI_ISL_2997941 | Lighthouse Laboratory Plymouth | Wellcome Sanger Institute for the COVID-19 Genomics UK (COG-UK) Consortium | Lighthouse Laboratory Plymouth and Alex Alderton, Roberto Amato, Jeffrey Barrett, Sonia Goncalves, Ewan Harrison, David K. Jackson, Ian Johnston, Dominic Kwiatkowski, Cordelia Langford, John Sillitoe on behalf of the Wellcome Sanger Institute COVID-19 Surveillance Team |
| EPI_ISL_2997942 | Lighthouse Lab in Milton Keynes | Wellcome Sanger Institute for the COVID-19 Genomics UK (COG-UK) Consortium | The Lighthouse Lab in Milton Keynes and Alex Alderton, Roberto Amato, Jeffrey Barrett, Sonia Goncalves, Ewan Harrison, David K. Jackson, Ian Johnston, Dominic Kwiatkowski, Cordelia Langford, John Sillitoe on behalf of the Wellcome Sanger Institute COVID-19 Surveillance Team |
| EPI_ISL_2997943 | Lighthouse Laboratory Plymouth | Wellcome Sanger Institute for the COVID-19 Genomics UK (COG-UK) Consortium | Lighthouse Laboratory Plymouth and Alex Alderton, Roberto Amato, Jeffrey Barrett, Sonia Goncalves, Ewan Harrison, David K. Jackson, Ian Johnston, Dominic Kwiatkowski, Cordelia Langford, John Sillitoe on behalf of the Wellcome Sanger Institute COVID-19 Surveillance Team |
| EPI_ISL_2997944 | Lighthouse Lab in Glasgow | Wellcome Sanger Institute for the COVID-19 Genomics UK (COG-UK) Consortium | Harper VanSteenhouse, Yumi Kasai, David Gray, Carol Clugston, Anna Dominiczak and Alex Alderton, Roberto Amato, Jeffrey Barrett, Sonia Goncalves, Ewan Harrison, David K. Jackson, Ian Johnston, Dominic Kwiatkowski, Cordelia Langford, John Sillitoe on behalf of the Wellcome Sanger Institute COVID-19 Surveillance Team |
| EPI_ISL_2997945, EPI_ISL_2997946 | Lighthouse Laboratory Plymouth | Wellcome Sanger Institute for the COVID-19 Genomics UK (COG-UK) Consortium | Lighthouse Laboratory Plymouth and Alex Alderton, Roberto Amato, Jeffrey Barrett, Sonia Goncalves, Ewan Harrison, David K. Jackson, Ian Johnston, Dominic Kwiatkowski, Cordelia Langford, John Sillitoe on behalf of the Wellcome Sanger Institute COVID-19 Surveillance Team |
| EPI_ISL_2997947, EPI_ISL_2997948, EPI_ISL_2997949, EPI_ISL_2997950, EPI_ISL_2997951, EPI_ISL_2997952 | Lighthouse Lab in Glasgow | Wellcome Sanger Institute for the COVID-19 Genomics UK (COG-UK) Consortium | Harper VanSteenhouse, Yumi Kasai, David Gray, Carol Clugston, Anna Dominiczak and Alex Alderton, Roberto Amato, Jeffrey Barrett, Sonia Goncalves, Ewan Harrison, David K. Jackson, Ian Johnston, Dominic Kwiatkowski, Cordelia Langford, John Sillitoe on behalf of the Wellcome Sanger Institute COVID-19 Surveillance Team |
| EPI_ISL_2997953 | Lighthouse Lab in Milton Keynes | Wellcome Sanger Institute for the COVID-19 Genomics UK (COG-UK) Consortium | The Lighthouse Lab in Milton Keynes and Alex Alderton, Roberto Amato, Jeffrey Barrett, Sonia Goncalves, Ewan Harrison, David K. Jackson, Ian Johnston, Dominic Kwiatkowski, Cordelia Langford, John Sillitoe on behalf of the Wellcome Sanger Institute COVID-19 Surveillance Team |
| EPI_ISL_2997954, EPI_ISL_2997955 | Lighthouse Laboratory Plymouth | Wellcome Sanger Institute for the COVID-19 Genomics UK (COG-UK) Consortium | Lighthouse Laboratory Plymouth and Alex Alderton, Roberto Amato, Jeffrey Barrett, Sonia Goncalves, Ewan Harrison, David K. Jackson, Ian Johnston, Dominic Kwiatkowski, Cordelia Langford, John Sillitoe on behalf of the Wellcome Sanger Institute COVID-19 Surveillance Team |
| EPI_ISL_2997956, EPI_ISL_2997957 | Lighthouse Lab in Milton Keynes | Wellcome Sanger Institute for the COVID-19 Genomics UK (COG-UK) Consortium | The Lighthouse Lab in Milton Keynes and Alex Alderton, Roberto Amato, Jeffrey Barrett, Sonia Goncalves, Ewan Harrison, David K. Jackson, Ian Johnston, Dominic Kwiatkowski, Cordelia Langford, John Sillitoe on behalf of the Wellcome Sanger Institute COVID-19 Surveillance Team |
| EPI_ISL_2997958, EPI_ISL_2997959, EPI_ISL_2997960 | Lighthouse Laboratory Plymouth | Wellcome Sanger Institute for the COVID-19 Genomics UK (COG-UK) Consortium | Lighthouse Laboratory Plymouth and Alex Alderton, Roberto Amato, Jeffrey Barrett, Sonia Goncalves, Ewan Harrison, David K. Jackson, Ian Johnston, Dominic Kwiatkowski, Cordelia Langford, John Sillitoe on behalf of the Wellcome Sanger Institute COVID-19 Surveillance Team |
| EPI_ISL_2997961 | Lighthouse Lab in Milton Keynes | Wellcome Sanger Institute for the COVID-19 Genomics UK (COG-UK) Consortium | The Lighthouse Lab in Milton Keynes and Alex Alderton, Roberto Amato, Jeffrey Barrett, Sonia Goncalves, Ewan Harrison, David K. Jackson, Ian Johnston, Dominic Kwiatkowski, Cordelia Langford, John Sillitoe on behalf of the Wellcome Sanger Institute COVID-19 Surveillance Team |
| EPI_ISL_2997962, EPI_ISL_2997963, EPI_ISL_2997964 | Lighthouse Laboratory Plymouth | Wellcome Sanger Institute for the COVID-19 Genomics UK (COG-UK) Consortium | Lighthouse Laboratory Plymouth and Alex Alderton, Roberto Amato, Jeffrey Barrett, Sonia Goncalves, Ewan Harrison, David K. Jackson, Ian Johnston, Dominic Kwiatkowski, Cordelia Langford, John Sillitoe on behalf of the Wellcome Sanger Institute COVID-19 Surveillance Team |
| EPI_ISL_2997965 | Lighthouse Lab in Milton Keynes | Wellcome Sanger Institute for the COVID-19 Genomics UK (COG-UK) Consortium | The Lighthouse Lab in Milton Keynes and Alex Alderton, Roberto Amato, Jeffrey Barrett, Sonia Goncalves, Ewan Harrison, David K. Jackson, Ian Johnston, Dominic Kwiatkowski, Cordelia Langford, John Sillitoe on behalf of the Wellcome Sanger Institute COVID-19 Surveillance Team |
| EPI_ISL_2997966 | Lighthouse Laboratory Plymouth | Wellcome Sanger Institute for the COVID-19 Genomics UK (COG-UK) Consortium | Lighthouse Laboratory Plymouth and Alex Alderton, Roberto Amato, Jeffrey Barrett, Sonia Goncalves, Ewan Harrison, David K. Jackson, Ian Johnston, Dominic Kwiatkowski, Cordelia Langford, John Sillitoe on behalf of the Wellcome Sanger Institute COVID-19 Surveillance Team |
| EPI_ISL_2997967 | Lighthouse Lab in Milton Keynes | Wellcome Sanger Institute for the COVID-19 Genomics UK (COG-UK) Consortium | The Lighthouse Lab in Milton Keynes and Alex Alderton, Roberto Amato, Jeffrey Barrett, Sonia Goncalves, Ewan Harrison, David K. Jackson, Ian Johnston, Dominic Kwiatkowski, Cordelia Langford, John Sillitoe on behalf of the Wellcome Sanger Institute COVID-19 Surveillance Team |
| EPI_ISL_2997968 | Lighthouse Laboratory Plymouth | Wellcome Sanger Institute for the COVID-19 Genomics UK (COG-UK) Consortium | Lighthouse Laboratory Plymouth and Alex Alderton, Roberto Amato, Jeffrey Barrett, Sonia Goncalves, Ewan Harrison, David K. Jackson, Ian Johnston, Dominic Kwiatkowski, Cordelia Langford, John Sillitoe on behalf of the Wellcome Sanger Institute COVID-19 Surveillance Team |
| EPI_ISL_2997969 | Lighthouse Lab in Milton Keynes | Wellcome Sanger Institute for the COVID-19 Genomics UK (COG-UK) Consortium | The Lighthouse Lab in Milton Keynes and Alex Alderton, Roberto Amato, Jeffrey Barrett, Sonia Goncalves, Ewan Harrison, David K. Jackson, Ian Johnston, Dominic Kwiatkowski, Cordelia Langford, John Sillitoe on behalf of the Wellcome Sanger Institute COVID-19 Surveillance Team |
| EPI_ISL_2997970 | Lighthouse Lab in Glasgow | Wellcome Sanger Institute for the COVID-19 Genomics UK (COG-UK) Consortium | Harper VanSteenhouse, Yumi Kasai, David Gray, Carol Clugston, Anna Dominiczak and Alex Alderton, Roberto Amato, Jeffrey Barrett, Sonia Goncalves, Ewan Harrison, David K. Jackson, Ian Johnston, Dominic Kwiatkowski, Cordelia Langford, John Sillitoe on behalf of the Wellcome Sanger Institute COVID-19 Surveillance Team |
| EPI_ISL_2997971 | Lighthouse Laboratory Plymouth | Wellcome Sanger Institute for the COVID-19 Genomics UK (COG-UK) Consortium | Lighthouse Laboratory Plymouth and Alex Alderton, Roberto Amato, Jeffrey Barrett, Sonia Goncalves, Ewan Harrison, David K. Jackson, Ian Johnston, Dominic Kwiatkowski, Cordelia Langford, John Sillitoe on behalf of the Wellcome Sanger Institute COVID-19 Surveillance Team |
| EPI_ISL_2997972 | Lighthouse Lab in Glasgow | Wellcome Sanger Institute for the COVID-19 Genomics UK (COG-UK) Consortium | Harper VanSteenhouse, Yumi Kasai, David Gray, Carol Clugston, Anna Dominiczak and Alex Alderton, Roberto Amato, Jeffrey Barrett, Sonia Goncalves, Ewan Harrison, David K. Jackson, Ian Johnston, Dominic Kwiatkowski, Cordelia Langford, John Sillitoe on behalf of the Wellcome Sanger Institute COVID-19 Surveillance Team |
| EPI_ISL_2997973 | Lighthouse Lab in Milton Keynes | Wellcome Sanger Institute for the COVID-19 Genomics UK (COG-UK) Consortium | The Lighthouse Lab in Milton Keynes and Alex Alderton, Roberto Amato, Jeffrey Barrett, Sonia Goncalves, Ewan Harrison, David K. Jackson, Ian Johnston, Dominic Kwiatkowski, Cordelia Langford, John Sillitoe on behalf of the Wellcome Sanger Institute COVID-19 Surveillance Team |
| EPI_ISL_2997974, EPI_ISL_2997975, EPI_ISL_2997976 | Lighthouse Lab in Glasgow | Wellcome Sanger Institute for the COVID-19 Genomics UK (COG-UK) Consortium | Harper VanSteenhouse, Yumi Kasai, David Gray, Carol Clugston, Anna Dominiczak and Alex Alderton, Roberto Amato, Jeffrey Barrett, Sonia Goncalves, Ewan Harrison, David K. Jackson, Ian Johnston, Dominic Kwiatkowski, Cordelia Langford, John Sillitoe on behalf of the Wellcome Sanger Institute COVID-19 Surveillance Team |
| EPI_ISL_2997977 | Lighthouse Lab in Milton Keynes | Wellcome Sanger Institute for the COVID-19 Genomics UK (COG-UK) Consortium | The Lighthouse Lab in Milton Keynes and Alex Alderton, Roberto Amato, Jeffrey Barrett, Sonia Goncalves, Ewan Harrison, David K. Jackson, Ian Johnston, Dominic Kwiatkowski, Cordelia Langford, John Sillitoe on behalf of the Wellcome Sanger Institute COVID-19 Surveillance Team |
| EPI_ISL_2997978 | Lighthouse Laboratory Plymouth | Wellcome Sanger Institute for the COVID-19 Genomics UK (COG-UK) Consortium | Lighthouse Laboratory Plymouth and Alex Alderton, Roberto Amato, Jeffrey Barrett, Sonia Goncalves, Ewan Harrison, David K. Jackson, Ian Johnston, Dominic Kwiatkowski, Cordelia Langford, John Sillitoe on behalf of the Wellcome Sanger Institute COVID-19 Surveillance Team |
| EPI_ISL_2997979, EPI_ISL_2997980 | Lighthouse Lab in Milton Keynes | Wellcome Sanger Institute for the COVID-19 Genomics UK (COG-UK) Consortium | The Lighthouse Lab in Milton Keynes and Alex Alderton, Roberto Amato, Jeffrey Barrett, Sonia Goncalves, Ewan Harrison, David K. Jackson, Ian Johnston, Dominic Kwiatkowski, Cordelia Langford, John Sillitoe on behalf of the Wellcome Sanger Institute COVID-19 Surveillance Team |
| EPI_ISL_2997981, EPI_ISL_2997982, EPI_ISL_2997983, EPI_ISL_2997984, EPI_ISL_2997985 | Lighthouse Laboratory Plymouth | Wellcome Sanger Institute for the COVID-19 Genomics UK (COG-UK) Consortium | Lighthouse Laboratory Plymouth and Alex Alderton, Roberto Amato, Jeffrey Barrett, Sonia Goncalves, Ewan Harrison, David K. Jackson, Ian Johnston, Dominic Kwiatkowski, Cordelia Langford, John Sillitoe on behalf of the Wellcome Sanger Institute COVID-19 Surveillance Team |
| EPI_ISL_2997986 | Lighthouse Lab in Milton Keynes | Wellcome Sanger Institute for the COVID-19 Genomics UK (COG-UK) Consortium | The Lighthouse Lab in Milton Keynes and Alex Alderton, Roberto Amato, Jeffrey Barrett, Sonia Goncalves, Ewan Harrison, David K. Jackson, Ian Johnston, Dominic Kwiatkowski, Cordelia Langford, John Sillitoe on behalf of the Wellcome Sanger Institute COVID-19 Surveillance Team |
| EPI_ISL_2997987, EPI_ISL_2997988, EPI_ISL_2997989 | Lighthouse Lab in Glasgow | Wellcome Sanger Institute for the COVID-19 Genomics UK (COG-UK) Consortium | Harper VanSteenhouse, Yumi Kasai, David Gray, Carol Clugston, Anna Dominiczak and Alex Alderton, Roberto Amato, Jeffrey Barrett, Sonia Goncalves, Ewan Harrison, David K. Jackson, Ian Johnston, Dominic Kwiatkowski, Cordelia Langford, John Sillitoe on behalf of the Wellcome Sanger Institute COVID-19 Surveillance Team |
| EPI_ISL_2997990 | Lighthouse Lab in Milton Keynes | Wellcome Sanger Institute for the COVID-19 Genomics UK (COG-UK) Consortium | The Lighthouse Lab in Milton Keynes and Alex Alderton, Roberto Amato, Jeffrey Barrett, Sonia Goncalves, Ewan Harrison, David K. Jackson, Ian Johnston, Dominic Kwiatkowski, Cordelia Langford, John Sillitoe on behalf of the Wellcome Sanger Institute COVID-19 Surveillance Team |
| EPI_ISL_2997991, EPI_ISL_2997992, EPI_ISL_2997993, EPI_ISL_2997994, EPI_ISL_2997995 | Lighthouse Laboratory Plymouth | Wellcome Sanger Institute for the COVID-19 Genomics UK (COG-UK) Consortium | Lighthouse Laboratory Plymouth and Alex Alderton, Roberto Amato, Jeffrey Barrett, Sonia Goncalves, Ewan Harrison, David K. Jackson, Ian Johnston, Dominic Kwiatkowski, Cordelia Langford, John Sillitoe on behalf of the Wellcome Sanger Institute COVID-19 Surveillance Team |
| EPI_ISL_2997996 | Lighthouse Lab in Glasgow | Wellcome Sanger Institute for the COVID-19 Genomics UK (COG-UK) Consortium | Harper VanSteenhouse, Yumi Kasai, David Gray, Carol Clugston, Anna Dominiczak and Alex Alderton, Roberto Amato, Jeffrey Barrett, Sonia Goncalves, Ewan Harrison, David K. Jackson, Ian Johnston, Dominic Kwiatkowski, Cordelia Langford, John Sillitoe on behalf of the Wellcome Sanger Institute COVID-19 Surveillance Team |

| | | | |
|---|---|---|---|
| EPI_ISL_2997997 | Lighthouse Lab in Milton Keynes | Wellcome Sanger Institute for the COVID-19 Genomics UK (COG-UK) Consortium | The Lighthouse Lab in Milton Keynes and Alex Alderton, Roberto Amato, Jeffrey Barrett, Sonia Goncalves, Ewan Harrison, David K. Jackson, Ian Johnston, Dominic Kwiatkowski, Cordelia Langford, John Sillitoe on behalf of the Wellcome Sanger Institute COVID-19 Surveillance Team |
| EPI_ISL_2997998, EPI_ISL_2997999 | Lighthouse Laboratory Plymouth | Wellcome Sanger Institute for the COVID-19 Genomics UK (COG-UK) Consortium | Lighthouse Laboratory Plymouth and Alex Alderton, Roberto Amato, Jeffrey Barrett, Sonia Goncalves, Ewan Harrison, David K. Jackson, Ian Johnston, Dominic Kwiatkowski, Cordelia Langford, John Sillitoe on behalf of the Wellcome Sanger Institute COVID-19 Surveillance Team |
| EPI_ISL_2998000, EPI_ISL_2998001 | Lighthouse Lab in Milton Keynes | Wellcome Sanger Institute for the COVID-19 Genomics UK (COG-UK) Consortium | The Lighthouse Lab in Milton Keynes and Alex Alderton, Roberto Amato, Jeffrey Barrett, Sonia Goncalves, Ewan Harrison, David K. Jackson, Ian Johnston, Dominic Kwiatkowski, Cordelia Langford, John Sillitoe on behalf of the Wellcome Sanger Institute COVID-19 Surveillance Team |
| EPI_ISL_2998002, EPI_ISL_2998003, EPI_ISL_2998004, EPI_ISL_2998005, EPI_ISL_2998006 | Lighthouse Laboratory Plymouth | Wellcome Sanger Institute for the COVID-19 Genomics UK (COG-UK) Consortium | Lighthouse Laboratory Plymouth and Alex Alderton, Roberto Amato, Jeffrey Barrett, Sonia Goncalves, Ewan Harrison, David K. Jackson, Ian Johnston, Dominic Kwiatkowski, Cordelia Langford, John Sillitoe on behalf of the Wellcome Sanger Institute COVID-19 Surveillance Team |
| EPI_ISL_2998007, EPI_ISL_2998008 | Lighthouse Lab in Milton Keynes | Wellcome Sanger Institute for the COVID-19 Genomics UK (COG-UK) Consortium | The Lighthouse Lab in Milton Keynes and Alex Alderton, Roberto Amato, Jeffrey Barrett, Sonia Goncalves, Ewan Harrison, David K. Jackson, Ian Johnston, Dominic Kwiatkowski, Cordelia Langford, John Sillitoe on behalf of the Wellcome Sanger Institute COVID-19 Surveillance Team |
| EPI_ISL_2998009 | Lighthouse Laboratory Plymouth | Wellcome Sanger Institute for the COVID-19 Genomics UK (COG-UK) Consortium | Lighthouse Laboratory Plymouth and Alex Alderton, Roberto Amato, Jeffrey Barrett, Sonia Goncalves, Ewan Harrison, David K. Jackson, Ian Johnston, Dominic Kwiatkowski, Cordelia Langford, John Sillitoe on behalf of the Wellcome Sanger Institute COVID-19 Surveillance Team |
| EPI_ISL_2998010 | Lighthouse Lab in Glasgow | Wellcome Sanger Institute for the COVID-19 Genomics UK (COG-UK) Consortium | Harper VanSteenhouse, Yumi Kasai, David Gray, Carol Clugston, Anna Dominiczak and Alex Alderton, Roberto Amato, Jeffrey Barrett, Sonia Goncalves, Ewan Harrison, David K. Jackson, Ian Johnston, Dominic Kwiatkowski, Cordelia Langford, John Sillitoe on behalf of the Wellcome Sanger Institute COVID-19 Surveillance Team |
| EPI_ISL_2998011, EPI_ISL_2998012 | Lighthouse Laboratory Plymouth | Wellcome Sanger Institute for the COVID-19 Genomics UK (COG-UK) Consortium | Lighthouse Laboratory Plymouth and Alex Alderton, Roberto Amato, Jeffrey Barrett, Sonia Goncalves, Ewan Harrison, David K. Jackson, Ian Johnston, Dominic Kwiatkowski, Cordelia Langford, John Sillitoe on behalf of the Wellcome Sanger Institute COVID-19 Surveillance Team |
| EPI_ISL_2998013 | Lighthouse Lab in Milton Keynes | Wellcome Sanger Institute for the COVID-19 Genomics UK (COG-UK) Consortium | The Lighthouse Lab in Milton Keynes and Alex Alderton, Roberto Amato, Jeffrey Barrett, Sonia Goncalves, Ewan Harrison, David K. Jackson, Ian Johnston, Dominic Kwiatkowski, Cordelia Langford, John Sillitoe on behalf of the Wellcome Sanger Institute COVID-19 Surveillance Team |
| EPI_ISL_2998014, EPI_ISL_2998015 | Lighthouse Laboratory Plymouth | Wellcome Sanger Institute for the COVID-19 Genomics UK (COG-UK) Consortium | Lighthouse Laboratory Plymouth and Alex Alderton, Roberto Amato, Jeffrey Barrett, Sonia Goncalves, Ewan Harrison, David K. Jackson, Ian Johnston, Dominic Kwiatkowski, Cordelia Langford, John Sillitoe on behalf of the Wellcome Sanger Institute COVID-19 Surveillance Team |
| EPI_ISL_2998016 | Lighthouse Lab in Milton Keynes | Wellcome Sanger Institute for the COVID-19 Genomics UK (COG-UK) Consortium | The Lighthouse Lab in Milton Keynes and Alex Alderton, Roberto Amato, Jeffrey Barrett, Sonia Goncalves, Ewan Harrison, David K. Jackson, Ian Johnston, Dominic Kwiatkowski, Cordelia Langford, John Sillitoe on behalf of the Wellcome Sanger Institute COVID-19 Surveillance Team |
| EPI_ISL_2998017 | Lighthouse Lab in Glasgow | Wellcome Sanger Institute for the COVID-19 Genomics UK (COG-UK) Consortium | Harper VanSteenhouse, Yumi Kasai, David Gray, Carol Clugston, Anna Dominiczak and Alex Alderton, Roberto Amato, Jeffrey Barrett, Sonia Goncalves, Ewan Harrison, David K. Jackson, Ian Johnston, Dominic Kwiatkowski, Cordelia Langford, John Sillitoe on behalf of the Wellcome Sanger Institute COVID-19 Surveillance Team |
| EPI_ISL_2998018 | Lighthouse Laboratory Plymouth | Wellcome Sanger Institute for the COVID-19 Genomics UK (COG-UK) Consortium | Lighthouse Laboratory Plymouth and Alex Alderton, Roberto Amato, Jeffrey Barrett, Sonia Goncalves, Ewan Harrison, David K. Jackson, Ian Johnston, Dominic Kwiatkowski, Cordelia Langford, John Sillitoe on behalf of the Wellcome Sanger Institute COVID-19 Surveillance Team |
| EPI_ISL_2998019, EPI_ISL_2998020, EPI_ISL_2998021, EPI_ISL_2998022 | Lighthouse Lab in Glasgow | Wellcome Sanger Institute for the COVID-19 Genomics UK (COG-UK) Consortium | Harper VanSteenhouse, Yumi Kasai, David Gray, Carol Clugston, Anna Dominiczak and Alex Alderton, Roberto Amato, Jeffrey Barrett, Sonia Goncalves, Ewan Harrison, David K. Jackson, Ian Johnston, Dominic Kwiatkowski, Cordelia Langford, John Sillitoe on behalf of the Wellcome Sanger Institute COVID-19 Surveillance Team |
| EPI_ISL_2998023, EPI_ISL_2998024, EPI_ISL_2998025 | Lighthouse Lab in Milton Keynes | Wellcome Sanger Institute for the COVID-19 Genomics UK (COG-UK) Consortium | The Lighthouse Lab in Milton Keynes and Alex Alderton, Roberto Amato, Jeffrey Barrett, Sonia Goncalves, Ewan Harrison, David K. Jackson, Ian Johnston, Dominic Kwiatkowski, Cordelia Langford, John Sillitoe on behalf of the Wellcome Sanger Institute COVID-19 Surveillance Team |
| EPI_ISL_2998026, EPI_ISL_2998027, EPI_ISL_2998028, EPI_ISL_2998029 | Lighthouse Laboratory Plymouth | Wellcome Sanger Institute for the COVID-19 Genomics UK (COG-UK) Consortium | Lighthouse Laboratory Plymouth and Alex Alderton, Roberto Amato, Jeffrey Barrett, Sonia Goncalves, Ewan Harrison, David K. Jackson, Ian Johnston, Dominic Kwiatkowski, Cordelia Langford, John Sillitoe on behalf of the Wellcome Sanger Institute COVID-19 Surveillance Team |
| EPI_ISL_2998030, EPI_ISL_2998031 | Lighthouse Lab in Glasgow | Wellcome Sanger Institute for the COVID-19 Genomics UK (COG-UK) Consortium | Harper VanSteenhouse, Yumi Kasai, David Gray, Carol Clugston, Anna Dominiczak and Alex Alderton, Roberto Amato, Jeffrey Barrett, Sonia Goncalves, Ewan Harrison, David K. Jackson, Ian Johnston, Dominic Kwiatkowski, Cordelia Langford, John Sillitoe on behalf of the Wellcome Sanger Institute COVID-19 Surveillance Team |
| EPI_ISL_2998032 | Lighthouse Laboratory Plymouth | Wellcome Sanger Institute for the COVID-19 Genomics UK (COG-UK) Consortium | Lighthouse Laboratory Plymouth and Alex Alderton, Roberto Amato, Jeffrey Barrett, Sonia Goncalves, Ewan Harrison, David K. Jackson, Ian Johnston, Dominic Kwiatkowski, Cordelia Langford, John Sillitoe on behalf of the Wellcome Sanger Institute COVID-19 Surveillance Team |
| EPI_ISL_2998033 | Lighthouse Lab in Milton Keynes | Wellcome Sanger Institute for the COVID-19 Genomics UK (COG-UK) Consortium | The Lighthouse Lab in Milton Keynes and Alex Alderton, Roberto Amato, Jeffrey Barrett, Sonia Goncalves, Ewan Harrison, David K. Jackson, Ian Johnston, Dominic Kwiatkowski, Cordelia Langford, John Sillitoe on behalf of the Wellcome Sanger Institute COVID-19 Surveillance Team |
| EPI_ISL_2998034, EPI_ISL_2998035 | Lighthouse Laboratory Plymouth | Wellcome Sanger Institute for the COVID-19 Genomics UK (COG-UK) Consortium | Lighthouse Laboratory Plymouth and Alex Alderton, Roberto Amato, Jeffrey Barrett, Sonia Goncalves, Ewan Harrison, David K. Jackson, Ian Johnston, Dominic Kwiatkowski, Cordelia Langford, John Sillitoe on behalf of the Wellcome Sanger Institute COVID-19 Surveillance Team |
| EPI_ISL_2998036 | Lighthouse Lab in Milton Keynes | Wellcome Sanger Institute for the COVID-19 Genomics UK (COG-UK) Consortium | The Lighthouse Lab in Milton Keynes and Alex Alderton, Roberto Amato, Jeffrey Barrett, Sonia Goncalves, Ewan Harrison, David K. Jackson, Ian Johnston, Dominic Kwiatkowski, Cordelia Langford, John Sillitoe on behalf of the Wellcome Sanger Institute COVID-19 Surveillance Team |
| EPI_ISL_2998037 | Lighthouse Laboratory Plymouth | Wellcome Sanger Institute for the COVID-19 Genomics UK (COG-UK) Consortium | Lighthouse Laboratory Plymouth and Alex Alderton, Roberto Amato, Jeffrey Barrett, Sonia Goncalves, Ewan Harrison, David K. Jackson, Ian Johnston, Dominic Kwiatkowski, Cordelia Langford, John Sillitoe on behalf of the Wellcome Sanger Institute COVID-19 Surveillance Team |
| EPI_ISL_2998038 | Lighthouse Lab in Milton Keynes | Wellcome Sanger Institute for the COVID-19 Genomics UK (COG-UK) Consortium | The Lighthouse Lab in Milton Keynes and Alex Alderton, Roberto Amato, Jeffrey Barrett, Sonia Goncalves, Ewan Harrison, David K. Jackson, Ian Johnston, Dominic Kwiatkowski, Cordelia Langford, John Sillitoe on behalf of the Wellcome Sanger Institute COVID-19 Surveillance Team |
| EPI_ISL_2998039, EPI_ISL_2998040, EPI_ISL_2998041, EPI_ISL_2998042, EPI_ISL_2998043, EPI_ISL_2998044, EPI_ISL_2998045, EPI_ISL_2998046, EPI_ISL_2998047, EPI_ISL_2998048, EPI_ISL_2998049, EPI_ISL_2998050, EPI_ISL_2998051 | | | |
| see above | Lighthouse Laboratory Plymouth | Wellcome Sanger Institute for the COVID-19 Genomics UK (COG-UK) Consortium | Lighthouse Laboratory Plymouth and Alex Alderton, Roberto Amato, Jeffrey Barrett, Sonia Goncalves, Ewan Harrison, David K. Jackson, Ian Johnston, Dominic Kwiatkowski, Cordelia Langford, John Sillitoe on behalf of the Wellcome Sanger Institute COVID-19 Surveillance Team |
| EPI_ISL_2998052 | Lighthouse Lab in Milton Keynes | Wellcome Sanger Institute for the COVID-19 Genomics UK (COG-UK) Consortium | The Lighthouse Lab in Milton Keynes and Alex Alderton, Roberto Amato, Jeffrey Barrett, Sonia Goncalves, Ewan Harrison, David K. Jackson, Ian Johnston, Dominic Kwiatkowski, Cordelia Langford, John Sillitoe on behalf of the Wellcome Sanger Institute COVID-19 Surveillance Team |
| EPI_ISL_2998053, EPI_ISL_2998054, EPI_ISL_2998055 | Lighthouse Laboratory Plymouth | Wellcome Sanger Institute for the COVID-19 Genomics UK (COG-UK) Consortium | Lighthouse Laboratory Plymouth and Alex Alderton, Roberto Amato, Jeffrey Barrett, Sonia Goncalves, Ewan Harrison, David K. Jackson, Ian Johnston, Dominic Kwiatkowski, Cordelia Langford, John Sillitoe on behalf of the Wellcome Sanger Institute COVID-19 Surveillance Team |
| EPI_ISL_2998056 | Lighthouse Lab in Glasgow | Wellcome Sanger Institute for the COVID-19 Genomics UK (COG-UK) Consortium | Harper VanSteenhouse, Yumi Kasai, David Gray, Carol Clugston, Anna Dominiczak and Alex Alderton, Roberto Amato, Jeffrey Barrett, Sonia Goncalves, Ewan Harrison, David K. Jackson, Ian Johnston, Dominic Kwiatkowski, Cordelia Langford, John Sillitoe on behalf of the Wellcome Sanger Institute COVID-19 Surveillance Team |
| EPI_ISL_2998057, EPI_ISL_2998058 | Lighthouse Lab in Milton Keynes | Wellcome Sanger Institute for the COVID-19 Genomics UK (COG-UK) Consortium | The Lighthouse Lab in Milton Keynes and Alex Alderton, Roberto Amato, Jeffrey Barrett, Sonia Goncalves, Ewan Harrison, David K. Jackson, Ian Johnston, Dominic Kwiatkowski, Cordelia Langford, John Sillitoe on behalf of the Wellcome Sanger Institute COVID-19 Surveillance Team |
| EPI_ISL_2998059 | Lighthouse Laboratory Plymouth | Wellcome Sanger Institute for the COVID-19 Genomics UK (COG-UK) Consortium | Lighthouse Laboratory Plymouth and Alex Alderton, Roberto Amato, Jeffrey Barrett, Sonia Goncalves, Ewan Harrison, David K. Jackson, Ian Johnston, Dominic Kwiatkowski, Cordelia Langford, John Sillitoe on behalf of the Wellcome Sanger Institute COVID-19 Surveillance Team |
| EPI_ISL_2998060, EPI_ISL_2998061 | Lighthouse Lab in Milton Keynes | Wellcome Sanger Institute for the COVID-19 Genomics UK (COG-UK) Consortium | The Lighthouse Lab in Milton Keynes and Alex Alderton, Roberto Amato, Jeffrey Barrett, Sonia Goncalves, Ewan Harrison, David K. Jackson, Ian Johnston, Dominic Kwiatkowski, Cordelia Langford, John Sillitoe on behalf of the Wellcome Sanger Institute COVID-19 Surveillance Team |
| EPI_ISL_2998062 | Lighthouse Lab in Glasgow | Wellcome Sanger Institute for the COVID-19 Genomics UK (COG-UK) Consortium | Harper VanSteenhouse, Yumi Kasai, David Gray, Carol Clugston, Anna Dominiczak and Alex Alderton, Roberto Amato, Jeffrey Barrett, Sonia Goncalves, Ewan Harrison, David K. Jackson, Ian Johnston, Dominic Kwiatkowski, Cordelia Langford, John Sillitoe on behalf of the Wellcome Sanger Institute COVID-19 Surveillance Team |
| EPI_ISL_2998063 | Lighthouse Laboratory Plymouth | Wellcome Sanger Institute for the COVID-19 Genomics UK (COG-UK) Consortium | Lighthouse Laboratory Plymouth and Alex Alderton, Roberto Amato, Jeffrey Barrett, Sonia Goncalves, Ewan Harrison, David K. Jackson, Ian Johnston, Dominic Kwiatkowski, Cordelia Langford, John Sillitoe on behalf of the Wellcome Sanger Institute COVID-19 Surveillance Team |
| EPI_ISL_2998064 | Lighthouse Lab in Glasgow | Wellcome Sanger Institute for the COVID-19 Genomics UK (COG-UK) Consortium | Harper VanSteenhouse, Yumi Kasai, David Gray, Carol Clugston, Anna Dominiczak and Alex Alderton, Roberto Amato, Jeffrey Barrett, Sonia Goncalves, Ewan Harrison, David K. Jackson, Ian Johnston, Dominic Kwiatkowski, Cordelia Langford, John Sillitoe on behalf of the Wellcome Sanger Institute COVID-19 Surveillance Team |
| EPI_ISL_2998065, EPI_ISL_2998066, EPI_ISL_2998067 | Lighthouse Laboratory Plymouth | Wellcome Sanger Institute for the COVID-19 Genomics UK (COG-UK) Consortium | Lighthouse Laboratory Plymouth and Alex Alderton, Roberto Amato, Jeffrey Barrett, Sonia Goncalves, Ewan Harrison, David K. Jackson, Ian Johnston, Dominic Kwiatkowski, Cordelia Langford, John Sillitoe on behalf of the Wellcome Sanger Institute COVID-19 Surveillance Team |
| EPI_ISL_2998068 | Lighthouse Lab in Glasgow | Wellcome Sanger Institute for the COVID-19 Genomics UK (COG-UK) Consortium | Harper VanSteenhouse, Yumi Kasai, David Gray, Carol Clugston, Anna Dominiczak and Alex Alderton, Roberto Amato, Jeffrey Barrett, Sonia Goncalves, Ewan Harrison, David K. Jackson, Ian Johnston, Dominic Kwiatkowski, Cordelia Langford, John Sillitoe on behalf of the Wellcome Sanger Institute COVID-19 Surveillance Team |
| EPI_ISL_2998069, EPI_ISL_2998070, EPI_ISL_2998071 | Lighthouse Laboratory Plymouth | Wellcome Sanger Institute for the COVID-19 Genomics UK (COG-UK) Consortium | Lighthouse Laboratory Plymouth and Alex Alderton, Roberto Amato, Jeffrey Barrett, Sonia Goncalves, Ewan Harrison, David K. Jackson, Ian Johnston, Dominic Kwiatkowski, Cordelia Langford, John Sillitoe on behalf of the Wellcome Sanger Institute COVID-19 Surveillance Team |
| EPI_ISL_2998072, EPI_ISL_2998073, EPI_ISL_2998074 | Lighthouse Lab in Glasgow | Wellcome Sanger Institute for the COVID-19 Genomics UK (COG-UK) Consortium | Harper VanSteenhouse, Yumi Kasai, David Gray, Carol Clugston, Anna Dominiczak and Alex Alderton, Roberto Amato, Jeffrey Barrett, Sonia Goncalves, Ewan Harrison, David K. Jackson, Ian Johnston, Dominic Kwiatkowski, Cordelia Langford, John Sillitoe on behalf of the Wellcome Sanger Institute COVID-19 Surveillance Team |
| EPI_ISL_2998075, EPI_ISL_2998076, EPI_ISL_2998077 | Lighthouse Lab in Milton Keynes | Wellcome Sanger Institute for the COVID-19 Genomics UK (COG-UK) Consortium | The Lighthouse Lab in Milton Keynes and Alex Alderton, Roberto Amato, Jeffrey Barrett, Sonia Goncalves, Ewan Harrison, David K. Jackson, Ian Johnston, Dominic Kwiatkowski, Cordelia Langford, John Sillitoe on behalf of the Wellcome Sanger Institute COVID-19 Surveillance Team |
| EPI_ISL_2998078 | Lighthouse Lab in Glasgow | Wellcome Sanger Institute for the COVID-19 Genomics UK (COG-UK) Consortium | Harper VanSteenhouse, Yumi Kasai, David Gray, Carol Clugston, Anna Dominiczak and Alex Alderton, Roberto Amato, Jeffrey Barrett, Sonia Goncalves, Ewan Harrison, David K. Jackson, Ian Johnston, Dominic Kwiatkowski, Cordelia Langford, John Sillitoe on behalf of the Wellcome Sanger Institute COVID-19 Surveillance Team |

| | | | |
|---|---|---|---|
| EPI_ISL_2998079, EPI_ISL_2998080 | Lighthouse Laboratory Plymouth | Wellcome Sanger Institute for the COVID-19 Genomics UK (COG-UK) Consortium | Lighthouse Laboratory Plymouth and Alex Alderton, Roberto Amato, Jeffrey Barrett, Sonia Goncalves, Ewan Harrison, David K. Jackson, Ian Johnston, Dominic Kwiatkowski, Cordelia Langford, John Sillitoe on behalf of the Wellcome Sanger Institute COVID-19 Surveillance Team |
| EPI_ISL_2998081 | Lighthouse Lab in Milton Keynes | Wellcome Sanger Institute for the COVID-19 Genomics UK (COG-UK) Consortium | The Lighthouse Lab in Milton Keynes and Alex Alderton, Roberto Amato, Jeffrey Barrett, Sonia Goncalves, Ewan Harrison, David K. Jackson, Ian Johnston, Dominic Kwiatkowski, Cordelia Langford, John Sillitoe on behalf of the Wellcome Sanger Institute COVID-19 Surveillance Team |
| EPI_ISL_2998082 | Lighthouse Laboratory Plymouth | Wellcome Sanger Institute for the COVID-19 Genomics UK (COG-UK) Consortium | Lighthouse Laboratory Plymouth and Alex Alderton, Roberto Amato, Jeffrey Barrett, Sonia Goncalves, Ewan Harrison, David K. Jackson, Ian Johnston, Dominic Kwiatkowski, Cordelia Langford, John Sillitoe on behalf of the Wellcome Sanger Institute COVID-19 Surveillance Team |
| EPI_ISL_2998083 | Lighthouse Lab in Milton Keynes | Wellcome Sanger Institute for the COVID-19 Genomics UK (COG-UK) Consortium | The Lighthouse Lab in Milton Keynes and Alex Alderton, Roberto Amato, Jeffrey Barrett, Sonia Goncalves, Ewan Harrison, David K. Jackson, Ian Johnston, Dominic Kwiatkowski, Cordelia Langford, John Sillitoe on behalf of the Wellcome Sanger Institute COVID-19 Surveillance Team |
| EPI_ISL_2998084, EPI_ISL_2998085, EPI_ISL_2998086, EPI_ISL_2998087 | Lighthouse Laboratory Plymouth | Wellcome Sanger Institute for the COVID-19 Genomics UK (COG-UK) Consortium | Lighthouse Laboratory Plymouth and Alex Alderton, Roberto Amato, Jeffrey Barrett, Sonia Goncalves, Ewan Harrison, David K. Jackson, Ian Johnston, Dominic Kwiatkowski, Cordelia Langford, John Sillitoe on behalf of the Wellcome Sanger Institute COVID-19 Surveillance Team |
| EPI_ISL_2998088, EPI_ISL_2998089 | Lighthouse Lab in Glasgow | Wellcome Sanger Institute for the COVID-19 Genomics UK (COG-UK) Consortium | Harper VanSteenhouse, Yumi Kasai, David Gray, Carol Clugston, Anna Dominiczak and Alex Alderton, Roberto Amato, Jeffrey Barrett, Sonia Goncalves, Ewan Harrison, David K. Jackson, Ian Johnston, Dominic Kwiatkowski, Cordelia Langford, John Sillitoe on behalf of the Wellcome Sanger Institute COVID-19 Surveillance Team |
| EPI_ISL_2998090 | Lighthouse Lab in Milton Keynes | Wellcome Sanger Institute for the COVID-19 Genomics UK (COG-UK) Consortium | The Lighthouse Lab in Milton Keynes and Alex Alderton, Roberto Amato, Jeffrey Barrett, Sonia Goncalves, Ewan Harrison, David K. Jackson, Ian Johnston, Dominic Kwiatkowski, Cordelia Langford, John Sillitoe on behalf of the Wellcome Sanger Institute COVID-19 Surveillance Team |
| EPI_ISL_2998091 | Lighthouse Laboratory Plymouth | Wellcome Sanger Institute for the COVID-19 Genomics UK (COG-UK) Consortium | Lighthouse Laboratory Plymouth and Alex Alderton, Roberto Amato, Jeffrey Barrett, Sonia Goncalves, Ewan Harrison, David K. Jackson, Ian Johnston, Dominic Kwiatkowski, Cordelia Langford, John Sillitoe on behalf of the Wellcome Sanger Institute COVID-19 Surveillance Team |
| EPI_ISL_2998092, EPI_ISL_2998093, EPI_ISL_2998094 | Lighthouse Lab in Glasgow | Wellcome Sanger Institute for the COVID-19 Genomics UK (COG-UK) Consortium | Harper VanSteenhouse, Yumi Kasai, David Gray, Carol Clugston, Anna Dominiczak and Alex Alderton, Roberto Amato, Jeffrey Barrett, Sonia Goncalves, Ewan Harrison, David K. Jackson, Ian Johnston, Dominic Kwiatkowski, Cordelia Langford, John Sillitoe on behalf of the Wellcome Sanger Institute COVID-19 Surveillance Team |
| EPI_ISL_2998095 | Lighthouse Laboratory Plymouth | Wellcome Sanger Institute for the COVID-19 Genomics UK (COG-UK) Consortium | Lighthouse Laboratory Plymouth and Alex Alderton, Roberto Amato, Jeffrey Barrett, Sonia Goncalves, Ewan Harrison, David K. Jackson, Ian Johnston, Dominic Kwiatkowski, Cordelia Langford, John Sillitoe on behalf of the Wellcome Sanger Institute COVID-19 Surveillance Team |
| EPI_ISL_2998096 | Lighthouse Lab in Glasgow | Wellcome Sanger Institute for the COVID-19 Genomics UK (COG-UK) Consortium | Harper VanSteenhouse, Yumi Kasai, David Gray, Carol Clugston, Anna Dominiczak and Alex Alderton, Roberto Amato, Jeffrey Barrett, Sonia Goncalves, Ewan Harrison, David K. Jackson, Ian Johnston, Dominic Kwiatkowski, Cordelia Langford, John Sillitoe on behalf of the Wellcome Sanger Institute COVID-19 Surveillance Team |
| EPI_ISL_2998097, EPI_ISL_2998098 | Lighthouse Lab in Milton Keynes | Wellcome Sanger Institute for the COVID-19 Genomics UK (COG-UK) Consortium | The Lighthouse Lab in Milton Keynes and Alex Alderton, Roberto Amato, Jeffrey Barrett, Sonia Goncalves, Ewan Harrison, David K. Jackson, Ian Johnston, Dominic Kwiatkowski, Cordelia Langford, John Sillitoe on behalf of the Wellcome Sanger Institute COVID-19 Surveillance Team |
| EPI_ISL_2998099 | Lighthouse Lab in Glasgow | Wellcome Sanger Institute for the COVID-19 Genomics UK (COG-UK) Consortium | Harper VanSteenhouse, Yumi Kasai, David Gray, Carol Clugston, Anna Dominiczak and Alex Alderton, Roberto Amato, Jeffrey Barrett, Sonia Goncalves, Ewan Harrison, David K. Jackson, Ian Johnston, Dominic Kwiatkowski, Cordelia Langford, John Sillitoe on behalf of the Wellcome Sanger Institute COVID-19 Surveillance Team |
| EPI_ISL_2998100 | Lighthouse Laboratory Plymouth | Wellcome Sanger Institute for the COVID-19 Genomics UK (COG-UK) Consortium | Lighthouse Laboratory Plymouth and Alex Alderton, Roberto Amato, Jeffrey Barrett, Sonia Goncalves, Ewan Harrison, David K. Jackson, Ian Johnston, Dominic Kwiatkowski, Cordelia Langford, John Sillitoe on behalf of the Wellcome Sanger Institute COVID-19 Surveillance Team |
| EPI_ISL_2998101, EPI_ISL_2998102 | Lighthouse Lab in Milton Keynes | Wellcome Sanger Institute for the COVID-19 Genomics UK (COG-UK) Consortium | The Lighthouse Lab in Milton Keynes and Alex Alderton, Roberto Amato, Jeffrey Barrett, Sonia Goncalves, Ewan Harrison, David K. Jackson, Ian Johnston, Dominic Kwiatkowski, Cordelia Langford, John Sillitoe on behalf of the Wellcome Sanger Institute COVID-19 Surveillance Team |
| EPI_ISL_2998103, EPI_ISL_2998104, EPI_ISL_2998105 | Lighthouse Laboratory Plymouth | Wellcome Sanger Institute for the COVID-19 Genomics UK (COG-UK) Consortium | Lighthouse Laboratory Plymouth and Alex Alderton, Roberto Amato, Jeffrey Barrett, Sonia Goncalves, Ewan Harrison, David K. Jackson, Ian Johnston, Dominic Kwiatkowski, Cordelia Langford, John Sillitoe on behalf of the Wellcome Sanger Institute COVID-19 Surveillance Team |
| EPI_ISL_2998106, EPI_ISL_2998107, EPI_ISL_2998108 | Lighthouse Lab in Milton Keynes | Wellcome Sanger Institute for the COVID-19 Genomics UK (COG-UK) Consortium | The Lighthouse Lab in Milton Keynes and Alex Alderton, Roberto Amato, Jeffrey Barrett, Sonia Goncalves, Ewan Harrison, David K. Jackson, Ian Johnston, Dominic Kwiatkowski, Cordelia Langford, John Sillitoe on behalf of the Wellcome Sanger Institute COVID-19 Surveillance Team |
| EPI_ISL_2998109, EPI_ISL_2998110, EPI_ISL_2998111, EPI_ISL_2998112, EPI_ISL_2998113, EPI_ISL_2998114 | Lighthouse Laboratory Plymouth | Wellcome Sanger Institute for the COVID-19 Genomics UK (COG-UK) Consortium | Lighthouse Laboratory Plymouth and Alex Alderton, Roberto Amato, Jeffrey Barrett, Sonia Goncalves, Ewan Harrison, David K. Jackson, Ian Johnston, Dominic Kwiatkowski, Cordelia Langford, John Sillitoe on behalf of the Wellcome Sanger Institute COVID-19 Surveillance Team |
| EPI_ISL_2998115 | Lighthouse Lab in Glasgow | Wellcome Sanger Institute for the COVID-19 Genomics UK (COG-UK) Consortium | Harper VanSteenhouse, Yumi Kasai, David Gray, Carol Clugston, Anna Dominiczak and Alex Alderton, Roberto Amato, Jeffrey Barrett, Sonia Goncalves, Ewan Harrison, David K. Jackson, Ian Johnston, Dominic Kwiatkowski, Cordelia Langford, John Sillitoe on behalf of the Wellcome Sanger Institute COVID-19 Surveillance Team |
| EPI_ISL_2998116 | Lighthouse Laboratory Plymouth | Wellcome Sanger Institute for the COVID-19 Genomics UK (COG-UK) Consortium | Lighthouse Laboratory Plymouth and Alex Alderton, Roberto Amato, Jeffrey Barrett, Sonia Goncalves, Ewan Harrison, David K. Jackson, Ian Johnston, Dominic Kwiatkowski, Cordelia Langford, John Sillitoe on behalf of the Wellcome Sanger Institute COVID-19 Surveillance Team |
| EPI_ISL_2998117 | Lighthouse Lab in Glasgow | Wellcome Sanger Institute for the COVID-19 Genomics UK (COG-UK) Consortium | Harper VanSteenhouse, Yumi Kasai, David Gray, Carol Clugston, Anna Dominiczak and Alex Alderton, Roberto Amato, Jeffrey Barrett, Sonia Goncalves, Ewan Harrison, David K. Jackson, Ian Johnston, Dominic Kwiatkowski, Cordelia Langford, John Sillitoe on behalf of the Wellcome Sanger Institute COVID-19 Surveillance Team |
| EPI_ISL_2998118, EPI_ISL_2998119, EPI_ISL_2998120 | Lighthouse Lab in Milton Keynes | Wellcome Sanger Institute for the COVID-19 Genomics UK (COG-UK) Consortium | The Lighthouse Lab in Milton Keynes and Alex Alderton, Roberto Amato, Jeffrey Barrett, Sonia Goncalves, Ewan Harrison, David K. Jackson, Ian Johnston, Dominic Kwiatkowski, Cordelia Langford, John Sillitoe on behalf of the Wellcome Sanger Institute COVID-19 Surveillance Team |
| EPI_ISL_2998121 | Lighthouse Lab in Glasgow | Wellcome Sanger Institute for the COVID-19 Genomics UK (COG-UK) Consortium | Harper VanSteenhouse, Yumi Kasai, David Gray, Carol Clugston, Anna Dominiczak and Alex Alderton, Roberto Amato, Jeffrey Barrett, Sonia Goncalves, Ewan Harrison, David K. Jackson, Ian Johnston, Dominic Kwiatkowski, Cordelia Langford, John Sillitoe on behalf of the Wellcome Sanger Institute COVID-19 Surveillance Team |
| EPI_ISL_2998122 | Lighthouse Lab in Milton Keynes | Wellcome Sanger Institute for the COVID-19 Genomics UK (COG-UK) Consortium | The Lighthouse Lab in Milton Keynes and Alex Alderton, Roberto Amato, Jeffrey Barrett, Sonia Goncalves, Ewan Harrison, David K. Jackson, Ian Johnston, Dominic Kwiatkowski, Cordelia Langford, John Sillitoe on behalf of the Wellcome Sanger Institute COVID-19 Surveillance Team |
| EPI_ISL_2998123 | Lighthouse Laboratory Plymouth | Wellcome Sanger Institute for the COVID-19 Genomics UK (COG-UK) Consortium | Lighthouse Laboratory Plymouth and Alex Alderton, Roberto Amato, Jeffrey Barrett, Sonia Goncalves, Ewan Harrison, David K. Jackson, Ian Johnston, Dominic Kwiatkowski, Cordelia Langford, John Sillitoe on behalf of the Wellcome Sanger Institute COVID-19 Surveillance Team |
| EPI_ISL_2998124 | Lighthouse Lab in Glasgow | Wellcome Sanger Institute for the COVID-19 Genomics UK (COG-UK) Consortium | Harper VanSteenhouse, Yumi Kasai, David Gray, Carol Clugston, Anna Dominiczak and Alex Alderton, Roberto Amato, Jeffrey Barrett, Sonia Goncalves, Ewan Harrison, David K. Jackson, Ian Johnston, Dominic Kwiatkowski, Cordelia Langford, John Sillitoe on behalf of the Wellcome Sanger Institute COVID-19 Surveillance Team |
| EPI_ISL_2998125 | Lighthouse Laboratory Plymouth | Wellcome Sanger Institute for the COVID-19 Genomics UK (COG-UK) Consortium | Lighthouse Laboratory Plymouth and Alex Alderton, Roberto Amato, Jeffrey Barrett, Sonia Goncalves, Ewan Harrison, David K. Jackson, Ian Johnston, Dominic Kwiatkowski, Cordelia Langford, John Sillitoe on behalf of the Wellcome Sanger Institute COVID-19 Surveillance Team |
| EPI_ISL_2998126 | Lighthouse Lab in Glasgow | Wellcome Sanger Institute for the COVID-19 Genomics UK (COG-UK) Consortium | Harper VanSteenhouse, Yumi Kasai, David Gray, Carol Clugston, Anna Dominiczak and Alex Alderton, Roberto Amato, Jeffrey Barrett, Sonia Goncalves, Ewan Harrison, David K. Jackson, Ian Johnston, Dominic Kwiatkowski, Cordelia Langford, John Sillitoe on behalf of the Wellcome Sanger Institute COVID-19 Surveillance Team |
| EPI_ISL_2998127, EPI_ISL_2998128 | Lighthouse Laboratory Plymouth | Wellcome Sanger Institute for the COVID-19 Genomics UK (COG-UK) Consortium | Lighthouse Laboratory Plymouth and Alex Alderton, Roberto Amato, Jeffrey Barrett, Sonia Goncalves, Ewan Harrison, David K. Jackson, Ian Johnston, Dominic Kwiatkowski, Cordelia Langford, John Sillitoe on behalf of the Wellcome Sanger Institute COVID-19 Surveillance Team |
| EPI_ISL_2998129 | Lighthouse Lab in Milton Keynes | Wellcome Sanger Institute for the COVID-19 Genomics UK (COG-UK) Consortium | The Lighthouse Lab in Milton Keynes and Alex Alderton, Roberto Amato, Jeffrey Barrett, Sonia Goncalves, Ewan Harrison, David K. Jackson, Ian Johnston, Dominic Kwiatkowski, Cordelia Langford, John Sillitoe on behalf of the Wellcome Sanger Institute COVID-19 Surveillance Team |
| EPI_ISL_2998130, EPI_ISL_2998131, EPI_ISL_2998132, EPI_ISL_2998133 | Lighthouse Laboratory Plymouth | Wellcome Sanger Institute for the COVID-19 Genomics UK (COG-UK) Consortium | Lighthouse Laboratory Plymouth and Alex Alderton, Roberto Amato, Jeffrey Barrett, Sonia Goncalves, Ewan Harrison, David K. Jackson, Ian Johnston, Dominic Kwiatkowski, Cordelia Langford, John Sillitoe on behalf of the Wellcome Sanger Institute COVID-19 Surveillance Team |
| EPI_ISL_2998134 | Lighthouse Lab in Milton Keynes | Wellcome Sanger Institute for the COVID-19 Genomics UK (COG-UK) Consortium | The Lighthouse Lab in Milton Keynes and Alex Alderton, Roberto Amato, Jeffrey Barrett, Sonia Goncalves, Ewan Harrison, David K. Jackson, Ian Johnston, Dominic Kwiatkowski, Cordelia Langford, John Sillitoe on behalf of the Wellcome Sanger Institute COVID-19 Surveillance Team |
| EPI_ISL_2998135 | Lighthouse Laboratory Plymouth | Wellcome Sanger Institute for the COVID-19 Genomics UK (COG-UK) Consortium | Lighthouse Laboratory Plymouth and Alex Alderton, Roberto Amato, Jeffrey Barrett, Sonia Goncalves, Ewan Harrison, David K. Jackson, Ian Johnston, Dominic Kwiatkowski, Cordelia Langford, John Sillitoe on behalf of the Wellcome Sanger Institute COVID-19 Surveillance Team |
| EPI_ISL_2998136 | Lighthouse Lab in Glasgow | Wellcome Sanger Institute for the COVID-19 Genomics UK (COG-UK) Consortium | Harper VanSteenhouse, Yumi Kasai, David Gray, Carol Clugston, Anna Dominiczak and Alex Alderton, Roberto Amato, Jeffrey Barrett, Sonia Goncalves, Ewan Harrison, David K. Jackson, Ian Johnston, Dominic Kwiatkowski, Cordelia Langford, John Sillitoe on behalf of the Wellcome Sanger Institute COVID-19 Surveillance Team |
| EPI_ISL_2998137, EPI_ISL_2998138 | Lighthouse Lab in Milton Keynes | Wellcome Sanger Institute for the COVID-19 Genomics UK (COG-UK) Consortium | The Lighthouse Lab in Milton Keynes and Alex Alderton, Roberto Amato, Jeffrey Barrett, Sonia Goncalves, Ewan Harrison, David K. Jackson, Ian Johnston, Dominic Kwiatkowski, Cordelia Langford, John Sillitoe on behalf of the Wellcome Sanger Institute COVID-19 Surveillance Team |
| EPI_ISL_2998139 | Lighthouse Lab in Glasgow | Wellcome Sanger Institute for the COVID-19 Genomics UK (COG-UK) Consortium | Harper VanSteenhouse, Yumi Kasai, David Gray, Carol Clugston, Anna Dominiczak and Alex Alderton, Roberto Amato, Jeffrey Barrett, Sonia Goncalves, Ewan Harrison, David K. Jackson, Ian Johnston, Dominic Kwiatkowski, Cordelia Langford, John Sillitoe on behalf of the Wellcome Sanger Institute COVID-19 Surveillance Team |
| EPI_ISL_2998140, EPI_ISL_2998141 | Lighthouse Lab in Milton Keynes | Wellcome Sanger Institute for the COVID-19 Genomics UK (COG-UK) Consortium | The Lighthouse Lab in Milton Keynes and Alex Alderton, Roberto Amato, Jeffrey Barrett, Sonia Goncalves, Ewan Harrison, David K. Jackson, Ian Johnston, Dominic Kwiatkowski, Cordelia Langford, John Sillitoe on behalf of the Wellcome Sanger Institute COVID-19 Surveillance Team |
| EPI_ISL_2998142 | Lighthouse Laboratory Plymouth | Wellcome Sanger Institute for the COVID-19 Genomics UK (COG-UK) Consortium | Lighthouse Laboratory Plymouth and Alex Alderton, Roberto Amato, Jeffrey Barrett, Sonia Goncalves, Ewan Harrison, David K. Jackson, Ian Johnston, Dominic Kwiatkowski, Cordelia Langford, John Sillitoe on behalf of the Wellcome Sanger Institute COVID-19 Surveillance Team |
| EPI_ISL_2998143, EPI_ISL_2998144, EPI_ISL_2998145 | Lighthouse Lab in Glasgow | Wellcome Sanger Institute for the COVID-19 Genomics UK (COG-UK) Consortium | Harper VanSteenhouse, Yumi Kasai, David Gray, Carol Clugston, Anna Dominiczak and Alex Alderton, Roberto Amato, Jeffrey Barrett, Sonia Goncalves, Ewan Harrison, David K. Jackson, Ian Johnston, Dominic Kwiatkowski, Cordelia Langford, John Sillitoe on behalf of the Wellcome Sanger Institute COVID-19 Surveillance Team |
| EPI_ISL_2998146 | Lighthouse Laboratory Plymouth | Wellcome Sanger Institute for the COVID-19 Genomics UK (COG-UK) Consortium | Lighthouse Laboratory Plymouth and Alex Alderton, Roberto Amato, Jeffrey Barrett, Sonia Goncalves, Ewan Harrison, David K. Jackson, Ian Johnston, Dominic Kwiatkowski, Cordelia Langford, John Sillitoe on behalf of the Wellcome Sanger Institute COVID-19 Surveillance Team |

| | | | |
|---|---|---|---|
| EPI_ISL_2998147 | Lighthouse Lab in Glasgow | Wellcome Sanger Institute for the COVID-19 Genomics UK (COG-UK) Consortium | Harper VanSteenhouse, Yumi Kasai, David Gray, Carol Clugston, Anna Dominiczak and Alex Alderton, Roberto Amato, Jeffrey Barrett, Sonia Goncalves, Ewan Harrison, David K. Jackson, Ian Johnston, Dominic Kwiatkowski, Cordelia Langford, John Sillitoe on behalf of the Wellcome Sanger Institute COVID-19 Surveillance Team |
| EPI_ISL_2998148, EPI_ISL_2998149 | Lighthouse Laboratory Plymouth | Wellcome Sanger Institute for the COVID-19 Genomics UK (COG-UK) Consortium | Lighthouse Laboratory Plymouth and Alex Alderton, Roberto Amato, Jeffrey Barrett, Sonia Goncalves, Ewan Harrison, David K. Jackson, Ian Johnston, Dominic Kwiatkowski, Cordelia Langford, John Sillitoe on behalf of the Wellcome Sanger Institute COVID-19 Surveillance Team |
| EPI_ISL_2998150 | Lighthouse Lab in Milton Keynes | Wellcome Sanger Institute for the COVID-19 Genomics UK (COG-UK) Consortium | The Lighthouse Lab in Milton Keynes and Alex Alderton, Roberto Amato, Jeffrey Barrett, Sonia Goncalves, Ewan Harrison, David K. Jackson, Ian Johnston, Dominic Kwiatkowski, Cordelia Langford, John Sillitoe on behalf of the Wellcome Sanger Institute COVID-19 Surveillance Team |
| EPI_ISL_2998151 | Lighthouse Lab in Glasgow | Wellcome Sanger Institute for the COVID-19 Genomics UK (COG-UK) Consortium | Harper VanSteenhouse, Yumi Kasai, David Gray, Carol Clugston, Anna Dominiczak and Alex Alderton, Roberto Amato, Jeffrey Barrett, Sonia Goncalves, Ewan Harrison, David K. Jackson, Ian Johnston, Dominic Kwiatkowski, Cordelia Langford, John Sillitoe on behalf of the Wellcome Sanger Institute COVID-19 Surveillance Team |
| EPI_ISL_2998152, EPI_ISL_2998153 | Lighthouse Laboratory Plymouth | Wellcome Sanger Institute for the COVID-19 Genomics UK (COG-UK) Consortium | Lighthouse Laboratory Plymouth and Alex Alderton, Roberto Amato, Jeffrey Barrett, Sonia Goncalves, Ewan Harrison, David K. Jackson, Ian Johnston, Dominic Kwiatkowski, Cordelia Langford, John Sillitoe on behalf of the Wellcome Sanger Institute COVID-19 Surveillance Team |
| EPI_ISL_2998154, EPI_ISL_2998155 | Lighthouse Lab in Milton Keynes | Wellcome Sanger Institute for the COVID-19 Genomics UK (COG-UK) Consortium | The Lighthouse Lab in Milton Keynes and Alex Alderton, Roberto Amato, Jeffrey Barrett, Sonia Goncalves, Ewan Harrison, David K. Jackson, Ian Johnston, Dominic Kwiatkowski, Cordelia Langford, John Sillitoe on behalf of the Wellcome Sanger Institute COVID-19 Surveillance Team |
| EPI_ISL_2998156 | Lighthouse Laboratory Plymouth | Wellcome Sanger Institute for the COVID-19 Genomics UK (COG-UK) Consortium | Lighthouse Laboratory Plymouth and Alex Alderton, Roberto Amato, Jeffrey Barrett, Sonia Goncalves, Ewan Harrison, David K. Jackson, Ian Johnston, Dominic Kwiatkowski, Cordelia Langford, John Sillitoe on behalf of the Wellcome Sanger Institute COVID-19 Surveillance Team |
| EPI_ISL_2998157, EPI_ISL_2998158, EPI_ISL_2998159 | Lighthouse Lab in Milton Keynes | Wellcome Sanger Institute for the COVID-19 Genomics UK (COG-UK) Consortium | The Lighthouse Lab in Milton Keynes and Alex Alderton, Roberto Amato, Jeffrey Barrett, Sonia Goncalves, Ewan Harrison, David K. Jackson, Ian Johnston, Dominic Kwiatkowski, Cordelia Langford, John Sillitoe on behalf of the Wellcome Sanger Institute COVID-19 Surveillance Team |
| EPI_ISL_2998160 | Lighthouse Lab in Glasgow | Wellcome Sanger Institute for the COVID-19 Genomics UK (COG-UK) Consortium | Harper VanSteenhouse, Yumi Kasai, David Gray, Carol Clugston, Anna Dominiczak and Alex Alderton, Roberto Amato, Jeffrey Barrett, Sonia Goncalves, Ewan Harrison, David K. Jackson, Ian Johnston, Dominic Kwiatkowski, Cordelia Langford, John Sillitoe on behalf of the Wellcome Sanger Institute COVID-19 Surveillance Team |
| EPI_ISL_2998161, EPI_ISL_2998162 | Lighthouse Laboratory Plymouth | Wellcome Sanger Institute for the COVID-19 Genomics UK (COG-UK) Consortium | Lighthouse Laboratory Plymouth and Alex Alderton, Roberto Amato, Jeffrey Barrett, Sonia Goncalves, Ewan Harrison, David K. Jackson, Ian Johnston, Dominic Kwiatkowski, Cordelia Langford, John Sillitoe on behalf of the Wellcome Sanger Institute COVID-19 Surveillance Team |
| EPI_ISL_2998163 | Lighthouse Lab in Milton Keynes | Wellcome Sanger Institute for the COVID-19 Genomics UK (COG-UK) Consortium | The Lighthouse Lab in Milton Keynes and Alex Alderton, Roberto Amato, Jeffrey Barrett, Sonia Goncalves, Ewan Harrison, David K. Jackson, Ian Johnston, Dominic Kwiatkowski, Cordelia Langford, John Sillitoe on behalf of the Wellcome Sanger Institute COVID-19 Surveillance Team |
| EPI_ISL_2998164, EPI_ISL_2998165, EPI_ISL_2998166 | Lighthouse Laboratory Plymouth | Wellcome Sanger Institute for the COVID-19 Genomics UK (COG-UK) Consortium | Lighthouse Laboratory Plymouth and Alex Alderton, Roberto Amato, Jeffrey Barrett, Sonia Goncalves, Ewan Harrison, David K. Jackson, Ian Johnston, Dominic Kwiatkowski, Cordelia Langford, John Sillitoe on behalf of the Wellcome Sanger Institute COVID-19 Surveillance Team |
| EPI_ISL_2998167 | Lighthouse Lab in Milton Keynes | Wellcome Sanger Institute for the COVID-19 Genomics UK (COG-UK) Consortium | The Lighthouse Lab in Milton Keynes and Alex Alderton, Roberto Amato, Jeffrey Barrett, Sonia Goncalves, Ewan Harrison, David K. Jackson, Ian Johnston, Dominic Kwiatkowski, Cordelia Langford, John Sillitoe on behalf of the Wellcome Sanger Institute COVID-19 Surveillance Team |
| EPI_ISL_2998168, EPI_ISL_2998169, EPI_ISL_2998170, EPI_ISL_2998171, EPI_ISL_2998172, EPI_ISL_2998173, EPI_ISL_2998174, EPI_ISL_2998175, EPI_ISL_2998176 | Lighthouse Laboratory Plymouth | Wellcome Sanger Institute for the COVID-19 Genomics UK (COG-UK) Consortium | Lighthouse Laboratory Plymouth and Alex Alderton, Roberto Amato, Jeffrey Barrett, Sonia Goncalves, Ewan Harrison, David K. Jackson, Ian Johnston, Dominic Kwiatkowski, Cordelia Langford, John Sillitoe on behalf of the Wellcome Sanger Institute COVID-19 Surveillance Team |
| EPI_ISL_2998177 | Lighthouse Lab in Glasgow | Wellcome Sanger Institute for the COVID-19 Genomics UK (COG-UK) Consortium | Harper VanSteenhouse, Yumi Kasai, David Gray, Carol Clugston, Anna Dominiczak and Alex Alderton, Roberto Amato, Jeffrey Barrett, Sonia Goncalves, Ewan Harrison, David K. Jackson, Ian Johnston, Dominic Kwiatkowski, Cordelia Langford, John Sillitoe on behalf of the Wellcome Sanger Institute COVID-19 Surveillance Team |
| EPI_ISL_2998178, EPI_ISL_2998179 | Lighthouse Laboratory Plymouth | Wellcome Sanger Institute for the COVID-19 Genomics UK (COG-UK) Consortium | Lighthouse Laboratory Plymouth and Alex Alderton, Roberto Amato, Jeffrey Barrett, Sonia Goncalves, Ewan Harrison, David K. Jackson, Ian Johnston, Dominic Kwiatkowski, Cordelia Langford, John Sillitoe on behalf of the Wellcome Sanger Institute COVID-19 Surveillance Team |
| EPI_ISL_2998180 | Lighthouse Lab in Milton Keynes | Wellcome Sanger Institute for the COVID-19 Genomics UK (COG-UK) Consortium | The Lighthouse Lab in Milton Keynes and Alex Alderton, Roberto Amato, Jeffrey Barrett, Sonia Goncalves, Ewan Harrison, David K. Jackson, Ian Johnston, Dominic Kwiatkowski, Cordelia Langford, John Sillitoe on behalf of the Wellcome Sanger Institute COVID-19 Surveillance Team |
| EPI_ISL_2998181, EPI_ISL_2998182 | Lighthouse Laboratory Plymouth | Wellcome Sanger Institute for the COVID-19 Genomics UK (COG-UK) Consortium | Lighthouse Laboratory Plymouth and Alex Alderton, Roberto Amato, Jeffrey Barrett, Sonia Goncalves, Ewan Harrison, David K. Jackson, Ian Johnston, Dominic Kwiatkowski, Cordelia Langford, John Sillitoe on behalf of the Wellcome Sanger Institute COVID-19 Surveillance Team |
| EPI_ISL_2998183 | Lighthouse Lab in Glasgow | Wellcome Sanger Institute for the COVID-19 Genomics UK (COG-UK) Consortium | Harper VanSteenhouse, Yumi Kasai, David Gray, Carol Clugston, Anna Dominiczak and Alex Alderton, Roberto Amato, Jeffrey Barrett, Sonia Goncalves, Ewan Harrison, David K. Jackson, Ian Johnston, Dominic Kwiatkowski, Cordelia Langford, John Sillitoe on behalf of the Wellcome Sanger Institute COVID-19 Surveillance Team |
| EPI_ISL_2998184, EPI_ISL_2998185, EPI_ISL_2998186 | Lighthouse Laboratory Plymouth | Wellcome Sanger Institute for the COVID-19 Genomics UK (COG-UK) Consortium | Lighthouse Laboratory Plymouth and Alex Alderton, Roberto Amato, Jeffrey Barrett, Sonia Goncalves, Ewan Harrison, David K. Jackson, Ian Johnston, Dominic Kwiatkowski, Cordelia Langford, John Sillitoe on behalf of the Wellcome Sanger Institute COVID-19 Surveillance Team |
| EPI_ISL_2998187 | Lighthouse Lab in Glasgow | Wellcome Sanger Institute for the COVID-19 Genomics UK (COG-UK) Consortium | Harper VanSteenhouse, Yumi Kasai, David Gray, Carol Clugston, Anna Dominiczak and Alex Alderton, Roberto Amato, Jeffrey Barrett, Sonia Goncalves, Ewan Harrison, David K. Jackson, Ian Johnston, Dominic Kwiatkowski, Cordelia Langford, John Sillitoe on behalf of the Wellcome Sanger Institute COVID-19 Surveillance Team |
| EPI_ISL_2998188 | Lighthouse Laboratory Plymouth | Wellcome Sanger Institute for the COVID-19 Genomics UK (COG-UK) Consortium | Lighthouse Laboratory Plymouth and Alex Alderton, Roberto Amato, Jeffrey Barrett, Sonia Goncalves, Ewan Harrison, David K. Jackson, Ian Johnston, Dominic Kwiatkowski, Cordelia Langford, John Sillitoe on behalf of the Wellcome Sanger Institute COVID-19 Surveillance Team |
| EPI_ISL_2998189, EPI_ISL_2998190, EPI_ISL_2998191 | Lighthouse Lab in Glasgow | Wellcome Sanger Institute for the COVID-19 Genomics UK (COG-UK) Consortium | Harper VanSteenhouse, Yumi Kasai, David Gray, Carol Clugston, Anna Dominiczak and Alex Alderton, Roberto Amato, Jeffrey Barrett, Sonia Goncalves, Ewan Harrison, David K. Jackson, Ian Johnston, Dominic Kwiatkowski, Cordelia Langford, John Sillitoe on behalf of the Wellcome Sanger Institute COVID-19 Surveillance Team |
| EPI_ISL_2998192 | Lighthouse Laboratory Plymouth | Wellcome Sanger Institute for the COVID-19 Genomics UK (COG-UK) Consortium | Lighthouse Laboratory Plymouth and Alex Alderton, Roberto Amato, Jeffrey Barrett, Sonia Goncalves, Ewan Harrison, David K. Jackson, Ian Johnston, Dominic Kwiatkowski, Cordelia Langford, John Sillitoe on behalf of the Wellcome Sanger Institute COVID-19 Surveillance Team |
| EPI_ISL_2998193 | Lighthouse Lab in Glasgow | Wellcome Sanger Institute for the COVID-19 Genomics UK (COG-UK) Consortium | Harper VanSteenhouse, Yumi Kasai, David Gray, Carol Clugston, Anna Dominiczak and Alex Alderton, Roberto Amato, Jeffrey Barrett, Sonia Goncalves, Ewan Harrison, David K. Jackson, Ian Johnston, Dominic Kwiatkowski, Cordelia Langford, John Sillitoe on behalf of the Wellcome Sanger Institute COVID-19 Surveillance Team |
| EPI_ISL_2998194 | Lighthouse Lab in Milton Keynes | Wellcome Sanger Institute for the COVID-19 Genomics UK (COG-UK) Consortium | The Lighthouse Lab in Milton Keynes and Alex Alderton, Roberto Amato, Jeffrey Barrett, Sonia Goncalves, Ewan Harrison, David K. Jackson, Ian Johnston, Dominic Kwiatkowski, Cordelia Langford, John Sillitoe on behalf of the Wellcome Sanger Institute COVID-19 Surveillance Team |
| EPI_ISL_2998195 | Lighthouse Lab in Glasgow | Wellcome Sanger Institute for the COVID-19 Genomics UK (COG-UK) Consortium | Harper VanSteenhouse, Yumi Kasai, David Gray, Carol Clugston, Anna Dominiczak and Alex Alderton, Roberto Amato, Jeffrey Barrett, Sonia Goncalves, Ewan Harrison, David K. Jackson, Ian Johnston, Dominic Kwiatkowski, Cordelia Langford, John Sillitoe on behalf of the Wellcome Sanger Institute COVID-19 Surveillance Team |
| EPI_ISL_2998196 | Lighthouse Laboratory Plymouth | Wellcome Sanger Institute for the COVID-19 Genomics UK (COG-UK) Consortium | Lighthouse Laboratory Plymouth and Alex Alderton, Roberto Amato, Jeffrey Barrett, Sonia Goncalves, Ewan Harrison, David K. Jackson, Ian Johnston, Dominic Kwiatkowski, Cordelia Langford, John Sillitoe on behalf of the Wellcome Sanger Institute COVID-19 Surveillance Team |
| EPI_ISL_2998197, EPI_ISL_2998198, EPI_ISL_2998199, EPI_ISL_2998200 | Lighthouse Lab in Milton Keynes | Wellcome Sanger Institute for the COVID-19 Genomics UK (COG-UK) Consortium | The Lighthouse Lab in Milton Keynes and Alex Alderton, Roberto Amato, Jeffrey Barrett, Sonia Goncalves, Ewan Harrison, David K. Jackson, Ian Johnston, Dominic Kwiatkowski, Cordelia Langford, John Sillitoe on behalf of the Wellcome Sanger Institute COVID-19 Surveillance Team |
| EPI_ISL_2998201 | Lighthouse Laboratory Plymouth | Wellcome Sanger Institute for the COVID-19 Genomics UK (COG-UK) Consortium | Lighthouse Laboratory Plymouth and Alex Alderton, Roberto Amato, Jeffrey Barrett, Sonia Goncalves, Ewan Harrison, David K. Jackson, Ian Johnston, Dominic Kwiatkowski, Cordelia Langford, John Sillitoe on behalf of the Wellcome Sanger Institute COVID-19 Surveillance Team |
| EPI_ISL_2998202 | Lighthouse Lab in Glasgow | Wellcome Sanger Institute for the COVID-19 Genomics UK (COG-UK) Consortium | Harper VanSteenhouse, Yumi Kasai, David Gray, Carol Clugston, Anna Dominiczak and Alex Alderton, Roberto Amato, Jeffrey Barrett, Sonia Goncalves, Ewan Harrison, David K. Jackson, Ian Johnston, Dominic Kwiatkowski, Cordelia Langford, John Sillitoe on behalf of the Wellcome Sanger Institute COVID-19 Surveillance Team |
| EPI_ISL_2998203 | Lighthouse Laboratory Plymouth | Wellcome Sanger Institute for the COVID-19 Genomics UK (COG-UK) Consortium | Lighthouse Laboratory Plymouth and Alex Alderton, Roberto Amato, Jeffrey Barrett, Sonia Goncalves, Ewan Harrison, David K. Jackson, Ian Johnston, Dominic Kwiatkowski, Cordelia Langford, John Sillitoe on behalf of the Wellcome Sanger Institute COVID-19 Surveillance Team |
| EPI_ISL_2998204 | Lighthouse Lab in Milton Keynes | Wellcome Sanger Institute for the COVID-19 Genomics UK (COG-UK) Consortium | The Lighthouse Lab in Milton Keynes and Alex Alderton, Roberto Amato, Jeffrey Barrett, Sonia Goncalves, Ewan Harrison, David K. Jackson, Ian Johnston, Dominic Kwiatkowski, Cordelia Langford, John Sillitoe on behalf of the Wellcome Sanger Institute COVID-19 Surveillance Team |
| EPI_ISL_2998205, EPI_ISL_2998206 | Lighthouse Laboratory Plymouth | Wellcome Sanger Institute for the COVID-19 Genomics UK (COG-UK) Consortium | Lighthouse Laboratory Plymouth and Alex Alderton, Roberto Amato, Jeffrey Barrett, Sonia Goncalves, Ewan Harrison, David K. Jackson, Ian Johnston, Dominic Kwiatkowski, Cordelia Langford, John Sillitoe on behalf of the Wellcome Sanger Institute COVID-19 Surveillance Team |
| EPI_ISL_2998207, EPI_ISL_2998208 | Lighthouse Lab in Glasgow | Wellcome Sanger Institute for the COVID-19 Genomics UK (COG-UK) Consortium | Harper VanSteenhouse, Yumi Kasai, David Gray, Carol Clugston, Anna Dominiczak and Alex Alderton, Roberto Amato, Jeffrey Barrett, Sonia Goncalves, Ewan Harrison, David K. Jackson, Ian Johnston, Dominic Kwiatkowski, Cordelia Langford, John Sillitoe on behalf of the Wellcome Sanger Institute COVID-19 Surveillance Team |
| EPI_ISL_2998209, EPI_ISL_2998210 | Lighthouse Laboratory Plymouth | Wellcome Sanger Institute for the COVID-19 Genomics UK (COG-UK) Consortium | Lighthouse Laboratory Plymouth and Alex Alderton, Roberto Amato, Jeffrey Barrett, Sonia Goncalves, Ewan Harrison, David K. Jackson, Ian Johnston, Dominic Kwiatkowski, Cordelia Langford, John Sillitoe on behalf of the Wellcome Sanger Institute COVID-19 Surveillance Team |
| EPI_ISL_2998211 | Lighthouse Lab in Milton Keynes | Wellcome Sanger Institute for the COVID-19 Genomics UK (COG-UK) Consortium | The Lighthouse Lab in Milton Keynes and Alex Alderton, Roberto Amato, Jeffrey Barrett, Sonia Goncalves, Ewan Harrison, David K. Jackson, Ian Johnston, Dominic Kwiatkowski, Cordelia Langford, John Sillitoe on behalf of the Wellcome Sanger Institute COVID-19 Surveillance Team |
| EPI_ISL_2998212 | Lighthouse Laboratory Plymouth | Wellcome Sanger Institute for the COVID-19 Genomics UK (COG-UK) Consortium | Lighthouse Laboratory Plymouth and Alex Alderton, Roberto Amato, Jeffrey Barrett, Sonia Goncalves, Ewan Harrison, David K. Jackson, Ian Johnston, Dominic Kwiatkowski, Cordelia Langford, John Sillitoe on behalf of the Wellcome Sanger Institute COVID-19 Surveillance Team |
| EPI_ISL_2998213 | Lighthouse Lab in Milton Keynes | Wellcome Sanger Institute for the COVID-19 Genomics UK (COG-UK) Consortium | The Lighthouse Lab in Milton Keynes and Alex Alderton, Roberto Amato, Jeffrey Barrett, Sonia Goncalves, Ewan Harrison, David K. Jackson, Ian Johnston, Dominic Kwiatkowski, Cordelia Langford, John Sillitoe on behalf of the Wellcome Sanger Institute COVID-19 Surveillance Team |
| EPI_ISL_2998214 | Lighthouse Laboratory Plymouth | Wellcome Sanger Institute for the COVID-19 Genomics UK (COG-UK) Consortium | Lighthouse Laboratory Plymouth and Alex Alderton, Roberto Amato, Jeffrey Barrett, Sonia Goncalves, Ewan Harrison, David K. Jackson, Ian Johnston, Dominic Kwiatkowski, Cordelia Langford, John Sillitoe on behalf of the Wellcome Sanger Institute COVID-19 Surveillance Team |

| | | | |
|---|---|---|---|
| EPI_ISL_2998215 | Lighthouse Lab in Glasgow | Wellcome Sanger Institute for the COVID-19 Genomics UK (COG-UK) Consortium | Harper VanSteenhouse, Yumi Kasai, David Gray, Carol Clugston, Anna Dominiczak and Alex Alderton, Roberto Amato, Jeffrey Barrett, Sonia Goncalves, Ewan Harrison, David K. Jackson, Ian Johnston, Dominic Kwiatkowski, Cordelia Langford, John Sillitoe on behalf of the Wellcome Sanger Institute COVID-19 Surveillance Team |
| EPI_ISL_2998216 | Lighthouse Laboratory Plymouth | Wellcome Sanger Institute for the COVID-19 Genomics UK (COG-UK) Consortium | Lighthouse Laboratory Plymouth and Alex Alderton, Roberto Amato, Jeffrey Barrett, Sonia Goncalves, Ewan Harrison, David K. Jackson, Ian Johnston, Dominic Kwiatkowski, Cordelia Langford, John Sillitoe on behalf of the Wellcome Sanger Institute COVID-19 Surveillance Team |
| EPI_ISL_2998217 | Lighthouse Lab in Milton Keynes | Wellcome Sanger Institute for the COVID-19 Genomics UK (COG-UK) Consortium | The Lighthouse Lab in Milton Keynes and Alex Alderton, Roberto Amato, Jeffrey Barrett, Sonia Goncalves, Ewan Harrison, David K. Jackson, Ian Johnston, Dominic Kwiatkowski, Cordelia Langford, John Sillitoe on behalf of the Wellcome Sanger Institute COVID-19 Surveillance Team |
| EPI_ISL_2998218 | Lighthouse Lab in Glasgow | Wellcome Sanger Institute for the COVID-19 Genomics UK (COG-UK) Consortium | Harper VanSteenhouse, Yumi Kasai, David Gray, Carol Clugston, Anna Dominiczak and Alex Alderton, Roberto Amato, Jeffrey Barrett, Sonia Goncalves, Ewan Harrison, David K. Jackson, Ian Johnston, Dominic Kwiatkowski, Cordelia Langford, John Sillitoe on behalf of the Wellcome Sanger Institute COVID-19 Surveillance Team |
| EPI_ISL_2998219, EPI_ISL_2998220 | Lighthouse Laboratory Plymouth | Wellcome Sanger Institute for the COVID-19 Genomics UK (COG-UK) Consortium | Lighthouse Laboratory Plymouth and Alex Alderton, Roberto Amato, Jeffrey Barrett, Sonia Goncalves, Ewan Harrison, David K. Jackson, Ian Johnston, Dominic Kwiatkowski, Cordelia Langford, John Sillitoe on behalf of the Wellcome Sanger Institute COVID-19 Surveillance Team |
| EPI_ISL_2998221 | Lighthouse Lab in Milton Keynes | Wellcome Sanger Institute for the COVID-19 Genomics UK (COG-UK) Consortium | The Lighthouse Lab in Milton Keynes and Alex Alderton, Roberto Amato, Jeffrey Barrett, Sonia Goncalves, Ewan Harrison, David K. Jackson, Ian Johnston, Dominic Kwiatkowski, Cordelia Langford, John Sillitoe on behalf of the Wellcome Sanger Institute COVID-19 Surveillance Team |
| EPI_ISL_2998222, EPI_ISL_2998223, EPI_ISL_2998224 | Lighthouse Laboratory Plymouth | Wellcome Sanger Institute for the COVID-19 Genomics UK (COG-UK) Consortium | Lighthouse Laboratory Plymouth and Alex Alderton, Roberto Amato, Jeffrey Barrett, Sonia Goncalves, Ewan Harrison, David K. Jackson, Ian Johnston, Dominic Kwiatkowski, Cordelia Langford, John Sillitoe on behalf of the Wellcome Sanger Institute COVID-19 Surveillance Team |
| EPI_ISL_2998225 | Lighthouse Lab in Glasgow | Wellcome Sanger Institute for the COVID-19 Genomics UK (COG-UK) Consortium | Harper VanSteenhouse, Yumi Kasai, David Gray, Carol Clugston, Anna Dominiczak and Alex Alderton, Roberto Amato, Jeffrey Barrett, Sonia Goncalves, Ewan Harrison, David K. Jackson, Ian Johnston, Dominic Kwiatkowski, Cordelia Langford, John Sillitoe on behalf of the Wellcome Sanger Institute COVID-19 Surveillance Team |
| EPI_ISL_2998226, EPI_ISL_2998227, EPI_ISL_2998228 | Lighthouse Laboratory Plymouth | Wellcome Sanger Institute for the COVID-19 Genomics UK (COG-UK) Consortium | Lighthouse Laboratory Plymouth and Alex Alderton, Roberto Amato, Jeffrey Barrett, Sonia Goncalves, Ewan Harrison, David K. Jackson, Ian Johnston, Dominic Kwiatkowski, Cordelia Langford, John Sillitoe on behalf of the Wellcome Sanger Institute COVID-19 Surveillance Team |
| EPI_ISL_2998229 | Lighthouse Lab in Milton Keynes | Wellcome Sanger Institute for the COVID-19 Genomics UK (COG-UK) Consortium | The Lighthouse Lab in Milton Keynes and Alex Alderton, Roberto Amato, Jeffrey Barrett, Sonia Goncalves, Ewan Harrison, David K. Jackson, Ian Johnston, Dominic Kwiatkowski, Cordelia Langford, John Sillitoe on behalf of the Wellcome Sanger Institute COVID-19 Surveillance Team |
| EPI_ISL_2998230 | Lighthouse Lab in Glasgow | Wellcome Sanger Institute for the COVID-19 Genomics UK (COG-UK) Consortium | Harper VanSteenhouse, Yumi Kasai, David Gray, Carol Clugston, Anna Dominiczak and Alex Alderton, Roberto Amato, Jeffrey Barrett, Sonia Goncalves, Ewan Harrison, David K. Jackson, Ian Johnston, Dominic Kwiatkowski, Cordelia Langford, John Sillitoe on behalf of the Wellcome Sanger Institute COVID-19 Surveillance Team |
| EPI_ISL_2998231, EPI_ISL_2998232, EPI_ISL_2998233 | Lighthouse Lab in Milton Keynes | Wellcome Sanger Institute for the COVID-19 Genomics UK (COG-UK) Consortium | The Lighthouse Lab in Milton Keynes and Alex Alderton, Roberto Amato, Jeffrey Barrett, Sonia Goncalves, Ewan Harrison, David K. Jackson, Ian Johnston, Dominic Kwiatkowski, Cordelia Langford, John Sillitoe on behalf of the Wellcome Sanger Institute COVID-19 Surveillance Team |
| EPI_ISL_2998234 | Lighthouse Laboratory Plymouth | Wellcome Sanger Institute for the COVID-19 Genomics UK (COG-UK) Consortium | Lighthouse Laboratory Plymouth and Alex Alderton, Roberto Amato, Jeffrey Barrett, Sonia Goncalves, Ewan Harrison, David K. Jackson, Ian Johnston, Dominic Kwiatkowski, Cordelia Langford, John Sillitoe on behalf of the Wellcome Sanger Institute COVID-19 Surveillance Team |
| EPI_ISL_2998235 | Lighthouse Lab in Glasgow | Wellcome Sanger Institute for the COVID-19 Genomics UK (COG-UK) Consortium | Harper VanSteenhouse, Yumi Kasai, David Gray, Carol Clugston, Anna Dominiczak and Alex Alderton, Roberto Amato, Jeffrey Barrett, Sonia Goncalves, Ewan Harrison, David K. Jackson, Ian Johnston, Dominic Kwiatkowski, Cordelia Langford, John Sillitoe on behalf of the Wellcome Sanger Institute COVID-19 Surveillance Team |
| EPI_ISL_2998236 | Lighthouse Laboratory Plymouth | Wellcome Sanger Institute for the COVID-19 Genomics UK (COG-UK) Consortium | Lighthouse Laboratory Plymouth and Alex Alderton, Roberto Amato, Jeffrey Barrett, Sonia Goncalves, Ewan Harrison, David K. Jackson, Ian Johnston, Dominic Kwiatkowski, Cordelia Langford, John Sillitoe on behalf of the Wellcome Sanger Institute COVID-19 Surveillance Team |
| EPI_ISL_2998237 | Lighthouse Lab in Milton Keynes | Wellcome Sanger Institute for the COVID-19 Genomics UK (COG-UK) Consortium | The Lighthouse Lab in Milton Keynes and Alex Alderton, Roberto Amato, Jeffrey Barrett, Sonia Goncalves, Ewan Harrison, David K. Jackson, Ian Johnston, Dominic Kwiatkowski, Cordelia Langford, John Sillitoe on behalf of the Wellcome Sanger Institute COVID-19 Surveillance Team |
| EPI_ISL_2998238, EPI_ISL_2998239 | Lighthouse Laboratory Plymouth | Wellcome Sanger Institute for the COVID-19 Genomics UK (COG-UK) Consortium | Lighthouse Laboratory Plymouth and Alex Alderton, Roberto Amato, Jeffrey Barrett, Sonia Goncalves, Ewan Harrison, David K. Jackson, Ian Johnston, Dominic Kwiatkowski, Cordelia Langford, John Sillitoe on behalf of the Wellcome Sanger Institute COVID-19 Surveillance Team |
| EPI_ISL_2998240 | Lighthouse Lab in Glasgow | Wellcome Sanger Institute for the COVID-19 Genomics UK (COG-UK) Consortium | Harper VanSteenhouse, Yumi Kasai, David Gray, Carol Clugston, Anna Dominiczak and Alex Alderton, Roberto Amato, Jeffrey Barrett, Sonia Goncalves, Ewan Harrison, David K. Jackson, Ian Johnston, Dominic Kwiatkowski, Cordelia Langford, John Sillitoe on behalf of the Wellcome Sanger Institute COVID-19 Surveillance Team |
| EPI_ISL_2998241, EPI_ISL_2998242, EPI_ISL_2998243, EPI_ISL_2998244, EPI_ISL_2998245, EPI_ISL_2998246, EPI_ISL_2998247 | Lighthouse Laboratory Plymouth | Wellcome Sanger Institute for the COVID-19 Genomics UK (COG-UK) Consortium | Lighthouse Laboratory Plymouth and Alex Alderton, Roberto Amato, Jeffrey Barrett, Sonia Goncalves, Ewan Harrison, David K. Jackson, Ian Johnston, Dominic Kwiatkowski, Cordelia Langford, John Sillitoe on behalf of the Wellcome Sanger Institute COVID-19 Surveillance Team |
| EPI_ISL_2998248 | Lighthouse Lab in Milton Keynes | Wellcome Sanger Institute for the COVID-19 Genomics UK (COG-UK) Consortium | The Lighthouse Lab in Milton Keynes and Alex Alderton, Roberto Amato, Jeffrey Barrett, Sonia Goncalves, Ewan Harrison, David K. Jackson, Ian Johnston, Dominic Kwiatkowski, Cordelia Langford, John Sillitoe on behalf of the Wellcome Sanger Institute COVID-19 Surveillance Team |
| EPI_ISL_2998249 | Lighthouse Lab in Glasgow | Wellcome Sanger Institute for the COVID-19 Genomics UK (COG-UK) Consortium | Harper VanSteenhouse, Yumi Kasai, David Gray, Carol Clugston, Anna Dominiczak and Alex Alderton, Roberto Amato, Jeffrey Barrett, Sonia Goncalves, Ewan Harrison, David K. Jackson, Ian Johnston, Dominic Kwiatkowski, Cordelia Langford, John Sillitoe on behalf of the Wellcome Sanger Institute COVID-19 Surveillance Team |
| EPI_ISL_2998250 | Lighthouse Lab in Milton Keynes | Wellcome Sanger Institute for the COVID-19 Genomics UK (COG-UK) Consortium | The Lighthouse Lab in Milton Keynes and Alex Alderton, Roberto Amato, Jeffrey Barrett, Sonia Goncalves, Ewan Harrison, David K. Jackson, Ian Johnston, Dominic Kwiatkowski, Cordelia Langford, John Sillitoe on behalf of the Wellcome Sanger Institute COVID-19 Surveillance Team |
| EPI_ISL_2998251 | Lighthouse Lab in Glasgow | Wellcome Sanger Institute for the COVID-19 Genomics UK (COG-UK) Consortium | Harper VanSteenhouse, Yumi Kasai, David Gray, Carol Clugston, Anna Dominiczak and Alex Alderton, Roberto Amato, Jeffrey Barrett, Sonia Goncalves, Ewan Harrison, David K. Jackson, Ian Johnston, Dominic Kwiatkowski, Cordelia Langford, John Sillitoe on behalf of the Wellcome Sanger Institute COVID-19 Surveillance Team |
| EPI_ISL_2998252 | Lighthouse Lab in Milton Keynes | Wellcome Sanger Institute for the COVID-19 Genomics UK (COG-UK) Consortium | The Lighthouse Lab in Milton Keynes and Alex Alderton, Roberto Amato, Jeffrey Barrett, Sonia Goncalves, Ewan Harrison, David K. Jackson, Ian Johnston, Dominic Kwiatkowski, Cordelia Langford, John Sillitoe on behalf of the Wellcome Sanger Institute COVID-19 Surveillance Team |
| EPI_ISL_2998253, EPI_ISL_2998254, EPI_ISL_2998255, EPI_ISL_2998256 | Lighthouse Laboratory Plymouth | Wellcome Sanger Institute for the COVID-19 Genomics UK (COG-UK) Consortium | Lighthouse Laboratory Plymouth and Alex Alderton, Roberto Amato, Jeffrey Barrett, Sonia Goncalves, Ewan Harrison, David K. Jackson, Ian Johnston, Dominic Kwiatkowski, Cordelia Langford, John Sillitoe on behalf of the Wellcome Sanger Institute COVID-19 Surveillance Team |
| EPI_ISL_2998257 | Lighthouse Lab in Glasgow | Wellcome Sanger Institute for the COVID-19 Genomics UK (COG-UK) Consortium | Harper VanSteenhouse, Yumi Kasai, David Gray, Carol Clugston, Anna Dominiczak and Alex Alderton, Roberto Amato, Jeffrey Barrett, Sonia Goncalves, Ewan Harrison, David K. Jackson, Ian Johnston, Dominic Kwiatkowski, Cordelia Langford, John Sillitoe on behalf of the Wellcome Sanger Institute COVID-19 Surveillance Team |
| EPI_ISL_2998258 | Lighthouse Lab in Milton Keynes | Wellcome Sanger Institute for the COVID-19 Genomics UK (COG-UK) Consortium | The Lighthouse Lab in Milton Keynes and Alex Alderton, Roberto Amato, Jeffrey Barrett, Sonia Goncalves, Ewan Harrison, David K. Jackson, Ian Johnston, Dominic Kwiatkowski, Cordelia Langford, John Sillitoe on behalf of the Wellcome Sanger Institute COVID-19 Surveillance Team |
| EPI_ISL_2998259, EPI_ISL_2998260 | Lighthouse Lab in Glasgow | Wellcome Sanger Institute for the COVID-19 Genomics UK (COG-UK) Consortium | Harper VanSteenhouse, Yumi Kasai, David Gray, Carol Clugston, Anna Dominiczak and Alex Alderton, Roberto Amato, Jeffrey Barrett, Sonia Goncalves, Ewan Harrison, David K. Jackson, Ian Johnston, Dominic Kwiatkowski, Cordelia Langford, John Sillitoe on behalf of the Wellcome Sanger Institute COVID-19 Surveillance Team |
| EPI_ISL_2998261 | Lighthouse Lab in Milton Keynes | Wellcome Sanger Institute for the COVID-19 Genomics UK (COG-UK) Consortium | The Lighthouse Lab in Milton Keynes and Alex Alderton, Roberto Amato, Jeffrey Barrett, Sonia Goncalves, Ewan Harrison, David K. Jackson, Ian Johnston, Dominic Kwiatkowski, Cordelia Langford, John Sillitoe on behalf of the Wellcome Sanger Institute COVID-19 Surveillance Team |
| EPI_ISL_2998262, EPI_ISL_2998263, EPI_ISL_2998264 | Lighthouse Lab in Glasgow | Wellcome Sanger Institute for the COVID-19 Genomics UK (COG-UK) Consortium | Harper VanSteenhouse, Yumi Kasai, David Gray, Carol Clugston, Anna Dominiczak and Alex Alderton, Roberto Amato, Jeffrey Barrett, Sonia Goncalves, Ewan Harrison, David K. Jackson, Ian Johnston, Dominic Kwiatkowski, Cordelia Langford, John Sillitoe on behalf of the Wellcome Sanger Institute COVID-19 Surveillance Team |
| EPI_ISL_2998265, EPI_ISL_2998266, EPI_ISL_2998267 | Lighthouse Laboratory Plymouth | Wellcome Sanger Institute for the COVID-19 Genomics UK (COG-UK) Consortium | Lighthouse Laboratory Plymouth and Alex Alderton, Roberto Amato, Jeffrey Barrett, Sonia Goncalves, Ewan Harrison, David K. Jackson, Ian Johnston, Dominic Kwiatkowski, Cordelia Langford, John Sillitoe on behalf of the Wellcome Sanger Institute COVID-19 Surveillance Team |
| EPI_ISL_2998268 | Lighthouse Lab in Glasgow | Wellcome Sanger Institute for the COVID-19 Genomics UK (COG-UK) Consortium | Harper VanSteenhouse, Yumi Kasai, David Gray, Carol Clugston, Anna Dominiczak and Alex Alderton, Roberto Amato, Jeffrey Barrett, Sonia Goncalves, Ewan Harrison, David K. Jackson, Ian Johnston, Dominic Kwiatkowski, Cordelia Langford, John Sillitoe on behalf of the Wellcome Sanger Institute COVID-19 Surveillance Team |
| EPI_ISL_2998269 | Lighthouse Lab in Milton Keynes | Wellcome Sanger Institute for the COVID-19 Genomics UK (COG-UK) Consortium | The Lighthouse Lab in Milton Keynes and Alex Alderton, Roberto Amato, Jeffrey Barrett, Sonia Goncalves, Ewan Harrison, David K. Jackson, Ian Johnston, Dominic Kwiatkowski, Cordelia Langford, John Sillitoe on behalf of the Wellcome Sanger Institute COVID-19 Surveillance Team |
| EPI_ISL_2998270 | Lighthouse Laboratory Plymouth | Wellcome Sanger Institute for the COVID-19 Genomics UK (COG-UK) Consortium | Lighthouse Laboratory Plymouth and Alex Alderton, Roberto Amato, Jeffrey Barrett, Sonia Goncalves, Ewan Harrison, David K. Jackson, Ian Johnston, Dominic Kwiatkowski, Cordelia Langford, John Sillitoe on behalf of the Wellcome Sanger Institute COVID-19 Surveillance Team |
| EPI_ISL_2998271 | Lighthouse Lab in Glasgow | Wellcome Sanger Institute for the COVID-19 Genomics UK (COG-UK) Consortium | Harper VanSteenhouse, Yumi Kasai, David Gray, Carol Clugston, Anna Dominiczak and Alex Alderton, Roberto Amato, Jeffrey Barrett, Sonia Goncalves, Ewan Harrison, David K. Jackson, Ian Johnston, Dominic Kwiatkowski, Cordelia Langford, John Sillitoe on behalf of the Wellcome Sanger Institute COVID-19 Surveillance Team |
| EPI_ISL_2998272, EPI_ISL_2998273 | Lighthouse Laboratory Plymouth | Wellcome Sanger Institute for the COVID-19 Genomics UK (COG-UK) Consortium | Lighthouse Laboratory Plymouth and Alex Alderton, Roberto Amato, Jeffrey Barrett, Sonia Goncalves, Ewan Harrison, David K. Jackson, Ian Johnston, Dominic Kwiatkowski, Cordelia Langford, John Sillitoe on behalf of the Wellcome Sanger Institute COVID-19 Surveillance Team |
| EPI_ISL_2998274 | Lighthouse Lab in Milton Keynes | Wellcome Sanger Institute for the COVID-19 Genomics UK (COG-UK) Consortium | The Lighthouse Lab in Milton Keynes and Alex Alderton, Roberto Amato, Jeffrey Barrett, Sonia Goncalves, Ewan Harrison, David K. Jackson, Ian Johnston, Dominic Kwiatkowski, Cordelia Langford, John Sillitoe on behalf of the Wellcome Sanger Institute COVID-19 Surveillance Team |
| EPI_ISL_2998275 | Lighthouse Lab in Glasgow | Wellcome Sanger Institute for the COVID-19 Genomics UK (COG-UK) Consortium | Harper VanSteenhouse, Yumi Kasai, David Gray, Carol Clugston, Anna Dominiczak and Alex Alderton, Roberto Amato, Jeffrey Barrett, Sonia Goncalves, Ewan Harrison, David K. Jackson, Ian Johnston, Dominic Kwiatkowski, Cordelia Langford, John Sillitoe on behalf of the Wellcome Sanger Institute COVID-19 Surveillance Team |
| EPI_ISL_2998276, EPI_ISL_2998277 | Lighthouse Lab in Milton Keynes | Wellcome Sanger Institute for the COVID-19 Genomics UK (COG-UK) Consortium | The Lighthouse Lab in Milton Keynes and Alex Alderton, Roberto Amato, Jeffrey Barrett, Sonia Goncalves, Ewan Harrison, David K. Jackson, Ian Johnston, Dominic Kwiatkowski, Cordelia Langford, John Sillitoe on behalf of the Wellcome Sanger Institute COVID-19 Surveillance Team |
| EPI_ISL_2998278 | Lighthouse Laboratory Plymouth | Wellcome Sanger Institute for the COVID-19 Genomics UK (COG-UK) Consortium | Lighthouse Laboratory Plymouth and Alex Alderton, Roberto Amato, Jeffrey Barrett, Sonia Goncalves, Ewan Harrison, David K. Jackson, Ian Johnston, Dominic Kwiatkowski, Cordelia Langford, John Sillitoe on behalf of the Wellcome Sanger Institute COVID-19 Surveillance Team |

| | | | |
|---|---|---|---|
| EPI_ISL_2998279 | Lighthouse Lab in Glasgow | Wellcome Sanger Institute for the COVID-19 Genomics UK (COG-UK) Consortium | Harper VanSteenhouse, Yumi Kasai, David Gray, Carol Clugston, Anna Dominiczak and Alex Alderton, Roberto Amato, Jeffrey Barrett, Sonia Goncalves, Ewan Harrison, David K. Jackson, Ian Johnston, Dominic Kwiatkowski, Cordelia Langford, John Sillitoe on behalf of the Wellcome Sanger Institute COVID-19 Surveillance Team |
| EPI_ISL_2998280, EPI_ISL_2998281 | Lighthouse Laboratory Plymouth | Wellcome Sanger Institute for the COVID-19 Genomics UK (COG-UK) Consortium | Lighthouse Laboratory Plymouth and Alex Alderton, Roberto Amato, Jeffrey Barrett, Sonia Goncalves, Ewan Harrison, David K. Jackson, Ian Johnston, Dominic Kwiatkowski, Cordelia Langford, John Sillitoe on behalf of the Wellcome Sanger Institute COVID-19 Surveillance Team |
| EPI_ISL_2998282, EPI_ISL_2998283, EPI_ISL_2998284, EPI_ISL_2998285 | Lighthouse Lab in Milton Keynes | Wellcome Sanger Institute for the COVID-19 Genomics UK (COG-UK) Consortium | The Lighthouse Lab in Milton Keynes and Alex Alderton, Roberto Amato, Jeffrey Barrett, Sonia Goncalves, Ewan Harrison, David K. Jackson, Ian Johnston, Dominic Kwiatkowski, Cordelia Langford, John Sillitoe on behalf of the Wellcome Sanger Institute COVID-19 Surveillance Team |
| EPI_ISL_2998286 | Lighthouse Laboratory Plymouth | Wellcome Sanger Institute for the COVID-19 Genomics UK (COG-UK) Consortium | Lighthouse Laboratory Plymouth and Alex Alderton, Roberto Amato, Jeffrey Barrett, Sonia Goncalves, Ewan Harrison, David K. Jackson, Ian Johnston, Dominic Kwiatkowski, Cordelia Langford, John Sillitoe on behalf of the Wellcome Sanger Institute COVID-19 Surveillance Team |
| EPI_ISL_2998287, EPI_ISL_2998288 | Lighthouse Lab in Glasgow | Wellcome Sanger Institute for the COVID-19 Genomics UK (COG-UK) Consortium | Harper VanSteenhouse, Yumi Kasai, David Gray, Carol Clugston, Anna Dominiczak and Alex Alderton, Roberto Amato, Jeffrey Barrett, Sonia Goncalves, Ewan Harrison, David K. Jackson, Ian Johnston, Dominic Kwiatkowski, Cordelia Langford, John Sillitoe on behalf of the Wellcome Sanger Institute COVID-19 Surveillance Team |
| EPI_ISL_2998289 | Lighthouse Laboratory Plymouth | Wellcome Sanger Institute for the COVID-19 Genomics UK (COG-UK) Consortium | Lighthouse Laboratory Plymouth and Alex Alderton, Roberto Amato, Jeffrey Barrett, Sonia Goncalves, Ewan Harrison, David K. Jackson, Ian Johnston, Dominic Kwiatkowski, Cordelia Langford, John Sillitoe on behalf of the Wellcome Sanger Institute COVID-19 Surveillance Team |
| EPI_ISL_2998290 | Lighthouse Lab in Glasgow | Wellcome Sanger Institute for the COVID-19 Genomics UK (COG-UK) Consortium | Harper VanSteenhouse, Yumi Kasai, David Gray, Carol Clugston, Anna Dominiczak and Alex Alderton, Roberto Amato, Jeffrey Barrett, Sonia Goncalves, Ewan Harrison, David K. Jackson, Ian Johnston, Dominic Kwiatkowski, Cordelia Langford, John Sillitoe on behalf of the Wellcome Sanger Institute COVID-19 Surveillance Team |
| EPI_ISL_2998291 | Lighthouse Laboratory Plymouth | Wellcome Sanger Institute for the COVID-19 Genomics UK (COG-UK) Consortium | Lighthouse Laboratory Plymouth and Alex Alderton, Roberto Amato, Jeffrey Barrett, Sonia Goncalves, Ewan Harrison, David K. Jackson, Ian Johnston, Dominic Kwiatkowski, Cordelia Langford, John Sillitoe on behalf of the Wellcome Sanger Institute COVID-19 Surveillance Team |
| EPI_ISL_2998292 | Lighthouse Lab in Glasgow | Wellcome Sanger Institute for the COVID-19 Genomics UK (COG-UK) Consortium | Harper VanSteenhouse, Yumi Kasai, David Gray, Carol Clugston, Anna Dominiczak and Alex Alderton, Roberto Amato, Jeffrey Barrett, Sonia Goncalves, Ewan Harrison, David K. Jackson, Ian Johnston, Dominic Kwiatkowski, Cordelia Langford, John Sillitoe on behalf of the Wellcome Sanger Institute COVID-19 Surveillance Team |
| EPI_ISL_2998293, EPI_ISL_2998294, EPI_ISL_2998295, EPI_ISL_2998296, EPI_ISL_2998297, EPI_ISL_2998298 | Lighthouse Laboratory Plymouth | Wellcome Sanger Institute for the COVID-19 Genomics UK (COG-UK) Consortium | Lighthouse Laboratory Plymouth and Alex Alderton, Roberto Amato, Jeffrey Barrett, Sonia Goncalves, Ewan Harrison, David K. Jackson, Ian Johnston, Dominic Kwiatkowski, Cordelia Langford, John Sillitoe on behalf of the Wellcome Sanger Institute COVID-19 Surveillance Team |
| EPI_ISL_2998299 | Lighthouse Lab in Milton Keynes | Wellcome Sanger Institute for the COVID-19 Genomics UK (COG-UK) Consortium | The Lighthouse Lab in Milton Keynes and Alex Alderton, Roberto Amato, Jeffrey Barrett, Sonia Goncalves, Ewan Harrison, David K. Jackson, Ian Johnston, Dominic Kwiatkowski, Cordelia Langford, John Sillitoe on behalf of the Wellcome Sanger Institute COVID-19 Surveillance Team |
| EPI_ISL_2998300 | Lighthouse Lab in Glasgow | Wellcome Sanger Institute for the COVID-19 Genomics UK (COG-UK) Consortium | Harper VanSteenhouse, Yumi Kasai, David Gray, Carol Clugston, Anna Dominiczak and Alex Alderton, Roberto Amato, Jeffrey Barrett, Sonia Goncalves, Ewan Harrison, David K. Jackson, Ian Johnston, Dominic Kwiatkowski, Cordelia Langford, John Sillitoe on behalf of the Wellcome Sanger Institute COVID-19 Surveillance Team |
| EPI_ISL_2998301 | Lighthouse Laboratory Plymouth | Wellcome Sanger Institute for the COVID-19 Genomics UK (COG-UK) Consortium | Lighthouse Laboratory Plymouth and Alex Alderton, Roberto Amato, Jeffrey Barrett, Sonia Goncalves, Ewan Harrison, David K. Jackson, Ian Johnston, Dominic Kwiatkowski, Cordelia Langford, John Sillitoe on behalf of the Wellcome Sanger Institute COVID-19 Surveillance Team |
| EPI_ISL_2998302, EPI_ISL_2998303, EPI_ISL_2998304 | Lighthouse Lab in Glasgow | Wellcome Sanger Institute for the COVID-19 Genomics UK (COG-UK) Consortium | Harper VanSteenhouse, Yumi Kasai, David Gray, Carol Clugston, Anna Dominiczak and Alex Alderton, Roberto Amato, Jeffrey Barrett, Sonia Goncalves, Ewan Harrison, David K. Jackson, Ian Johnston, Dominic Kwiatkowski, Cordelia Langford, John Sillitoe on behalf of the Wellcome Sanger Institute COVID-19 Surveillance Team |
| EPI_ISL_2998305 | Lighthouse Laboratory Plymouth | Wellcome Sanger Institute for the COVID-19 Genomics UK (COG-UK) Consortium | Lighthouse Laboratory Plymouth and Alex Alderton, Roberto Amato, Jeffrey Barrett, Sonia Goncalves, Ewan Harrison, David K. Jackson, Ian Johnston, Dominic Kwiatkowski, Cordelia Langford, John Sillitoe on behalf of the Wellcome Sanger Institute COVID-19 Surveillance Team |
| EPI_ISL_2998306, EPI_ISL_2998307 | Lighthouse Lab in Milton Keynes | Wellcome Sanger Institute for the COVID-19 Genomics UK (COG-UK) Consortium | The Lighthouse Lab in Milton Keynes and Alex Alderton, Roberto Amato, Jeffrey Barrett, Sonia Goncalves, Ewan Harrison, David K. Jackson, Ian Johnston, Dominic Kwiatkowski, Cordelia Langford, John Sillitoe on behalf of the Wellcome Sanger Institute COVID-19 Surveillance Team |
| EPI_ISL_2998308 | Lighthouse Lab in Glasgow | Wellcome Sanger Institute for the COVID-19 Genomics UK (COG-UK) Consortium | Harper VanSteenhouse, Yumi Kasai, David Gray, Carol Clugston, Anna Dominiczak and Alex Alderton, Roberto Amato, Jeffrey Barrett, Sonia Goncalves, Ewan Harrison, David K. Jackson, Ian Johnston, Dominic Kwiatkowski, Cordelia Langford, John Sillitoe on behalf of the Wellcome Sanger Institute COVID-19 Surveillance Team |
| EPI_ISL_2998309, EPI_ISL_2998310, EPI_ISL_2998311 | Lighthouse Laboratory Plymouth | Wellcome Sanger Institute for the COVID-19 Genomics UK (COG-UK) Consortium | Lighthouse Laboratory Plymouth and Alex Alderton, Roberto Amato, Jeffrey Barrett, Sonia Goncalves, Ewan Harrison, David K. Jackson, Ian Johnston, Dominic Kwiatkowski, Cordelia Langford, John Sillitoe on behalf of the Wellcome Sanger Institute COVID-19 Surveillance Team |
| EPI_ISL_2998312 | Lighthouse Lab in Glasgow | Wellcome Sanger Institute for the COVID-19 Genomics UK (COG-UK) Consortium | Harper VanSteenhouse, Yumi Kasai, David Gray, Carol Clugston, Anna Dominiczak and Alex Alderton, Roberto Amato, Jeffrey Barrett, Sonia Goncalves, Ewan Harrison, David K. Jackson, Ian Johnston, Dominic Kwiatkowski, Cordelia Langford, John Sillitoe on behalf of the Wellcome Sanger Institute COVID-19 Surveillance Team |
| EPI_ISL_2998313 | Lighthouse Laboratory Plymouth | Wellcome Sanger Institute for the COVID-19 Genomics UK (COG-UK) Consortium | Lighthouse Laboratory Plymouth and Alex Alderton, Roberto Amato, Jeffrey Barrett, Sonia Goncalves, Ewan Harrison, David K. Jackson, Ian Johnston, Dominic Kwiatkowski, Cordelia Langford, John Sillitoe on behalf of the Wellcome Sanger Institute COVID-19 Surveillance Team |
| EPI_ISL_2998314, EPI_ISL_2998315 | Lighthouse Lab in Glasgow | Wellcome Sanger Institute for the COVID-19 Genomics UK (COG-UK) Consortium | Harper VanSteenhouse, Yumi Kasai, David Gray, Carol Clugston, Anna Dominiczak and Alex Alderton, Roberto Amato, Jeffrey Barrett, Sonia Goncalves, Ewan Harrison, David K. Jackson, Ian Johnston, Dominic Kwiatkowski, Cordelia Langford, John Sillitoe on behalf of the Wellcome Sanger Institute COVID-19 Surveillance Team |
| EPI_ISL_2998316, EPI_ISL_2998317, EPI_ISL_2998318 | Lighthouse Lab in Milton Keynes | Wellcome Sanger Institute for the COVID-19 Genomics UK (COG-UK) Consortium | The Lighthouse Lab in Milton Keynes and Alex Alderton, Roberto Amato, Jeffrey Barrett, Sonia Goncalves, Ewan Harrison, David K. Jackson, Ian Johnston, Dominic Kwiatkowski, Cordelia Langford, John Sillitoe on behalf of the Wellcome Sanger Institute COVID-19 Surveillance Team |
| EPI_ISL_2998319, EPI_ISL_2998320, EPI_ISL_2998321 | Lighthouse Laboratory Plymouth | Wellcome Sanger Institute for the COVID-19 Genomics UK (COG-UK) Consortium | Lighthouse Laboratory Plymouth and Alex Alderton, Roberto Amato, Jeffrey Barrett, Sonia Goncalves, Ewan Harrison, David K. Jackson, Ian Johnston, Dominic Kwiatkowski, Cordelia Langford, John Sillitoe on behalf of the Wellcome Sanger Institute COVID-19 Surveillance Team |
| EPI_ISL_2998322 | Lighthouse Lab in Glasgow | Wellcome Sanger Institute for the COVID-19 Genomics UK (COG-UK) Consortium | Harper VanSteenhouse, Yumi Kasai, David Gray, Carol Clugston, Anna Dominiczak and Alex Alderton, Roberto Amato, Jeffrey Barrett, Sonia Goncalves, Ewan Harrison, David K. Jackson, Ian Johnston, Dominic Kwiatkowski, Cordelia Langford, John Sillitoe on behalf of the Wellcome Sanger Institute COVID-19 Surveillance Team |
| EPI_ISL_2998323 | Lighthouse Laboratory Plymouth | Wellcome Sanger Institute for the COVID-19 Genomics UK (COG-UK) Consortium | Lighthouse Laboratory Plymouth and Alex Alderton, Roberto Amato, Jeffrey Barrett, Sonia Goncalves, Ewan Harrison, David K. Jackson, Ian Johnston, Dominic Kwiatkowski, Cordelia Langford, John Sillitoe on behalf of the Wellcome Sanger Institute COVID-19 Surveillance Team |
| EPI_ISL_2998324, EPI_ISL_2998325, EPI_ISL_2998326 | Lighthouse Lab in Milton Keynes | Wellcome Sanger Institute for the COVID-19 Genomics UK (COG-UK) Consortium | The Lighthouse Lab in Milton Keynes and Alex Alderton, Roberto Amato, Jeffrey Barrett, Sonia Goncalves, Ewan Harrison, David K. Jackson, Ian Johnston, Dominic Kwiatkowski, Cordelia Langford, John Sillitoe on behalf of the Wellcome Sanger Institute COVID-19 Surveillance Team |
| EPI_ISL_2998327, EPI_ISL_2998328 | Lighthouse Lab in Glasgow | Wellcome Sanger Institute for the COVID-19 Genomics UK (COG-UK) Consortium | Harper VanSteenhouse, Yumi Kasai, David Gray, Carol Clugston, Anna Dominiczak and Alex Alderton, Roberto Amato, Jeffrey Barrett, Sonia Goncalves, Ewan Harrison, David K. Jackson, Ian Johnston, Dominic Kwiatkowski, Cordelia Langford, John Sillitoe on behalf of the Wellcome Sanger Institute COVID-19 Surveillance Team |
| EPI_ISL_2998329, EPI_ISL_2998330 | Lighthouse Laboratory Plymouth | Wellcome Sanger Institute for the COVID-19 Genomics UK (COG-UK) Consortium | Lighthouse Laboratory Plymouth and Alex Alderton, Roberto Amato, Jeffrey Barrett, Sonia Goncalves, Ewan Harrison, David K. Jackson, Ian Johnston, Dominic Kwiatkowski, Cordelia Langford, John Sillitoe on behalf of the Wellcome Sanger Institute COVID-19 Surveillance Team |
| EPI_ISL_2998331 | Lighthouse Lab in Glasgow | Wellcome Sanger Institute for the COVID-19 Genomics UK (COG-UK) Consortium | Harper VanSteenhouse, Yumi Kasai, David Gray, Carol Clugston, Anna Dominiczak and Alex Alderton, Roberto Amato, Jeffrey Barrett, Sonia Goncalves, Ewan Harrison, David K. Jackson, Ian Johnston, Dominic Kwiatkowski, Cordelia Langford, John Sillitoe on behalf of the Wellcome Sanger Institute COVID-19 Surveillance Team |
| EPI_ISL_2998332 | Lighthouse Laboratory Plymouth | Wellcome Sanger Institute for the COVID-19 Genomics UK (COG-UK) Consortium | Lighthouse Laboratory Plymouth and Alex Alderton, Roberto Amato, Jeffrey Barrett, Sonia Goncalves, Ewan Harrison, David K. Jackson, Ian Johnston, Dominic Kwiatkowski, Cordelia Langford, John Sillitoe on behalf of the Wellcome Sanger Institute COVID-19 Surveillance Team |
| EPI_ISL_2998333 | Lighthouse Lab in Glasgow | Wellcome Sanger Institute for the COVID-19 Genomics UK (COG-UK) Consortium | Harper VanSteenhouse, Yumi Kasai, David Gray, Carol Clugston, Anna Dominiczak and Alex Alderton, Roberto Amato, Jeffrey Barrett, Sonia Goncalves, Ewan Harrison, David K. Jackson, Ian Johnston, Dominic Kwiatkowski, Cordelia Langford, John Sillitoe on behalf of the Wellcome Sanger Institute COVID-19 Surveillance Team |
| EPI_ISL_2998334, EPI_ISL_2998335 | Lighthouse Laboratory Plymouth | Wellcome Sanger Institute for the COVID-19 Genomics UK (COG-UK) Consortium | Lighthouse Laboratory Plymouth and Alex Alderton, Roberto Amato, Jeffrey Barrett, Sonia Goncalves, Ewan Harrison, David K. Jackson, Ian Johnston, Dominic Kwiatkowski, Cordelia Langford, John Sillitoe on behalf of the Wellcome Sanger Institute COVID-19 Surveillance Team |
| EPI_ISL_2998336, EPI_ISL_2998337, EPI_ISL_2998338 | Lighthouse Lab in Milton Keynes | Wellcome Sanger Institute for the COVID-19 Genomics UK (COG-UK) Consortium | The Lighthouse Lab in Milton Keynes and Alex Alderton, Roberto Amato, Jeffrey Barrett, Sonia Goncalves, Ewan Harrison, David K. Jackson, Ian Johnston, Dominic Kwiatkowski, Cordelia Langford, John Sillitoe on behalf of the Wellcome Sanger Institute COVID-19 Surveillance Team |
| EPI_ISL_2998339 | Lighthouse Lab in Glasgow | Wellcome Sanger Institute for the COVID-19 Genomics UK (COG-UK) Consortium | Harper VanSteenhouse, Yumi Kasai, David Gray, Carol Clugston, Anna Dominiczak and Alex Alderton, Roberto Amato, Jeffrey Barrett, Sonia Goncalves, Ewan Harrison, David K. Jackson, Ian Johnston, Dominic Kwiatkowski, Cordelia Langford, John Sillitoe on behalf of the Wellcome Sanger Institute COVID-19 Surveillance Team |
| EPI_ISL_2998340, EPI_ISL_2998341 | Lighthouse Laboratory Plymouth | Wellcome Sanger Institute for the COVID-19 Genomics UK (COG-UK) Consortium | Lighthouse Laboratory Plymouth and Alex Alderton, Roberto Amato, Jeffrey Barrett, Sonia Goncalves, Ewan Harrison, David K. Jackson, Ian Johnston, Dominic Kwiatkowski, Cordelia Langford, John Sillitoe on behalf of the Wellcome Sanger Institute COVID-19 Surveillance Team |
| EPI_ISL_2998342 | Lighthouse Lab in Milton Keynes | Wellcome Sanger Institute for the COVID-19 Genomics UK (COG-UK) Consortium | The Lighthouse Lab in Milton Keynes and Alex Alderton, Roberto Amato, Jeffrey Barrett, Sonia Goncalves, Ewan Harrison, David K. Jackson, Ian Johnston, Dominic Kwiatkowski, Cordelia Langford, John Sillitoe on behalf of the Wellcome Sanger Institute COVID-19 Surveillance Team |
| EPI_ISL_2998343, EPI_ISL_2998344 | Lighthouse Laboratory Plymouth | Wellcome Sanger Institute for the COVID-19 Genomics UK (COG-UK) Consortium | Lighthouse Laboratory Plymouth and Alex Alderton, Roberto Amato, Jeffrey Barrett, Sonia Goncalves, Ewan Harrison, David K. Jackson, Ian Johnston, Dominic Kwiatkowski, Cordelia Langford, John Sillitoe on behalf of the Wellcome Sanger Institute COVID-19 Surveillance Team |
| EPI_ISL_2998345 | Lighthouse Lab in Glasgow | Wellcome Sanger Institute for the COVID-19 Genomics UK (COG-UK) Consortium | Harper VanSteenhouse, Yumi Kasai, David Gray, Carol Clugston, Anna Dominiczak and Alex Alderton, Roberto Amato, Jeffrey Barrett, Sonia Goncalves, Ewan Harrison, David K. Jackson, Ian Johnston, Dominic Kwiatkowski, Cordelia Langford, John Sillitoe on behalf of the Wellcome Sanger Institute COVID-19 Surveillance Team |
| EPI_ISL_2998346, EPI_ISL_2998347, EPI_ISL_2998348 | Lighthouse Laboratory Plymouth | Wellcome Sanger Institute for the COVID-19 Genomics UK (COG-UK) Consortium | Lighthouse Laboratory Plymouth and Alex Alderton, Roberto Amato, Jeffrey Barrett, Sonia Goncalves, Ewan Harrison, David K. Jackson, Ian Johnston, Dominic Kwiatkowski, Cordelia Langford, John Sillitoe on behalf of the Wellcome Sanger Institute COVID-19 Surveillance Team |
| EPI_ISL_2998349, EPI_ISL_2998350 | Lighthouse Lab in Glasgow | Wellcome Sanger Institute for the COVID-19 Genomics UK (COG-UK) Consortium | Harper VanSteenhouse, Yumi Kasai, David Gray, Carol Clugston, Anna Dominiczak and Alex Alderton, Roberto Amato, Jeffrey Barrett, Sonia Goncalves, Ewan Harrison, David K. Jackson, Ian Johnston, Dominic Kwiatkowski, Cordelia Langford, John Sillitoe on behalf of the Wellcome Sanger Institute COVID-19 Surveillance Team |

| | | | |
|---|---|---|---|
| EPI_ISL_2998351 | Lighthouse Laboratory Plymouth | Wellcome Sanger Institute for the COVID-19 Genomics UK (COG-UK) Consortium | Lighthouse Laboratory Plymouth and Alex Alderton, Roberto Amato, Jeffrey Barrett, Sonia Goncalves, Ewan Harrison, David K. Jackson, Ian Johnston, Dominic Kwiatkowski, Cordelia Langford, John Sillitoe on behalf of the Wellcome Sanger Institute COVID-19 Surveillance Team |
| EPI_ISL_2998352 | Lighthouse Lab in Glasgow | Wellcome Sanger Institute for the COVID-19 Genomics UK (COG-UK) Consortium | Harper VanSteenhouse, Yumi Kasai, David Gray, Carol Clugston, Anna Dominiczak and Alex Alderton, Roberto Amato, Jeffrey Barrett, Sonia Goncalves, Ewan Harrison, David K. Jackson, Ian Johnston, Dominic Kwiatkowski, Cordelia Langford, John Sillitoe on behalf of the Wellcome Sanger Institute COVID-19 Surveillance Team |
| EPI_ISL_2998353, EPI_ISL_2998354 | Lighthouse Laboratory Plymouth | Wellcome Sanger Institute for the COVID-19 Genomics UK (COG-UK) Consortium | Lighthouse Laboratory Plymouth and Alex Alderton, Roberto Amato, Jeffrey Barrett, Sonia Goncalves, Ewan Harrison, David K. Jackson, Ian Johnston, Dominic Kwiatkowski, Cordelia Langford, John Sillitoe on behalf of the Wellcome Sanger Institute COVID-19 Surveillance Team |
| EPI_ISL_2998355 | Lighthouse Lab in Milton Keynes | Wellcome Sanger Institute for the COVID-19 Genomics UK (COG-UK) Consortium | The Lighthouse Lab in Milton Keynes and Alex Alderton, Roberto Amato, Jeffrey Barrett, Sonia Goncalves, Ewan Harrison, David K. Jackson, Ian Johnston, Dominic Kwiatkowski, Cordelia Langford, John Sillitoe on behalf of the Wellcome Sanger Institute COVID-19 Surveillance Team |
| EPI_ISL_2998356, EPI_ISL_2998357 | Lighthouse Lab in Glasgow | Wellcome Sanger Institute for the COVID-19 Genomics UK (COG-UK) Consortium | Harper VanSteenhouse, Yumi Kasai, David Gray, Carol Clugston, Anna Dominiczak and Alex Alderton, Roberto Amato, Jeffrey Barrett, Sonia Goncalves, Ewan Harrison, David K. Jackson, Ian Johnston, Dominic Kwiatkowski, Cordelia Langford, John Sillitoe on behalf of the Wellcome Sanger Institute COVID-19 Surveillance Team |
| EPI_ISL_2998358 | Lighthouse Laboratory Plymouth | Wellcome Sanger Institute for the COVID-19 Genomics UK (COG-UK) Consortium | Lighthouse Laboratory Plymouth and Alex Alderton, Roberto Amato, Jeffrey Barrett, Sonia Goncalves, Ewan Harrison, David K. Jackson, Ian Johnston, Dominic Kwiatkowski, Cordelia Langford, John Sillitoe on behalf of the Wellcome Sanger Institute COVID-19 Surveillance Team |
| EPI_ISL_2998359 | Lighthouse Lab in Milton Keynes | Wellcome Sanger Institute for the COVID-19 Genomics UK (COG-UK) Consortium | The Lighthouse Lab in Milton Keynes and Alex Alderton, Roberto Amato, Jeffrey Barrett, Sonia Goncalves, Ewan Harrison, David K. Jackson, Ian Johnston, Dominic Kwiatkowski, Cordelia Langford, John Sillitoe on behalf of the Wellcome Sanger Institute COVID-19 Surveillance Team |
| EPI_ISL_2998360 | Lighthouse Laboratory Plymouth | Wellcome Sanger Institute for the COVID-19 Genomics UK (COG-UK) Consortium | Lighthouse Laboratory Plymouth and Alex Alderton, Roberto Amato, Jeffrey Barrett, Sonia Goncalves, Ewan Harrison, David K. Jackson, Ian Johnston, Dominic Kwiatkowski, Cordelia Langford, John Sillitoe on behalf of the Wellcome Sanger Institute COVID-19 Surveillance Team |
| EPI_ISL_2998361, EPI_ISL_2998362 | Lighthouse Lab in Glasgow | Wellcome Sanger Institute for the COVID-19 Genomics UK (COG-UK) Consortium | Harper VanSteenhouse, Yumi Kasai, David Gray, Carol Clugston, Anna Dominiczak and Alex Alderton, Roberto Amato, Jeffrey Barrett, Sonia Goncalves, Ewan Harrison, David K. Jackson, Ian Johnston, Dominic Kwiatkowski, Cordelia Langford, John Sillitoe on behalf of the Wellcome Sanger Institute COVID-19 Surveillance Team |
| EPI_ISL_2998363 | Lighthouse Laboratory Plymouth | Wellcome Sanger Institute for the COVID-19 Genomics UK (COG-UK) Consortium | Lighthouse Laboratory Plymouth and Alex Alderton, Roberto Amato, Jeffrey Barrett, Sonia Goncalves, Ewan Harrison, David K. Jackson, Ian Johnston, Dominic Kwiatkowski, Cordelia Langford, John Sillitoe on behalf of the Wellcome Sanger Institute COVID-19 Surveillance Team |
| EPI_ISL_2998364 | Microbiology Department, Laboratori Clínic Metropolitana Nord. Hospital Universitari Germans Trias i Pujol | Can Ruti SARS-CoV-2 Sequencing Hub (HUGTiP/IrsiCaixa/IGTP) | Marc Noguera-Julian, Pilar Armengol, Ignacio Blanco, Antoni E Bordoy, Francesc Catala-Moll, Pere-Joan Cardona, Maria Casadellà, Cristina Casañ, Gemma Clara, Bonaventura Clotet, Montserrat Giménez, Mercedes Guerrero, Anna Not, David Panisello, Roger Paredes, Mariona Parera, Alexia París, Verónica Saludes, Alba Sánchez, Laia Soler, Lauro Sumoy, and Elisa Martró on behalf of the Can Ruti SARS-CoV-2 Sequencing Hub |
| EPI_ISL_2998365 | Lighthouse Laboratory Plymouth | Wellcome Sanger Institute for the COVID-19 Genomics UK (COG-UK) Consortium | Lighthouse Laboratory Plymouth and Alex Alderton, Roberto Amato, Jeffrey Barrett, Sonia Goncalves, Ewan Harrison, David K. Jackson, Ian Johnston, Dominic Kwiatkowski, Cordelia Langford, John Sillitoe on behalf of the Wellcome Sanger Institute COVID-19 Surveillance Team |
| EPI_ISL_2998366, EPI_ISL_2998367 | Lighthouse Lab in Glasgow | Wellcome Sanger Institute for the COVID-19 Genomics UK (COG-UK) Consortium | Harper VanSteenhouse, Yumi Kasai, David Gray, Carol Clugston, Anna Dominiczak and Alex Alderton, Roberto Amato, Jeffrey Barrett, Sonia Goncalves, Ewan Harrison, David K. Jackson, Ian Johnston, Dominic Kwiatkowski, Cordelia Langford, John Sillitoe on behalf of the Wellcome Sanger Institute COVID-19 Surveillance Team |
| EPI_ISL_2998368 | Microbiology Department, Laboratori Clínic Metropolitana Nord. Hospital Universitari Germans Trias i Pujol | Can Ruti SARS-CoV-2 Sequencing Hub (HUGTiP/IrsiCaixa/IGTP) | Marc Noguera-Julian, Pilar Armengol, Ignacio Blanco, Antoni E Bordoy, Francesc Catala-Moll, Pere-Joan Cardona, Maria Casadellà, Cristina Casañ, Gemma Clara, Bonaventura Clotet, Montserrat Giménez, Mercedes Guerrero, Anna Not, David Panisello, Roger Paredes, Mariona Parera, Alexia París, Verónica Saludes, Alba Sánchez, Laia Soler, Lauro Sumoy, and Elisa Martró on behalf of the Can Ruti SARS-CoV-2 Sequencing Hub |
| EPI_ISL_2998369, EPI_ISL_2998370 | Lighthouse Laboratory Plymouth | Wellcome Sanger Institute for the COVID-19 Genomics UK (COG-UK) Consortium | Lighthouse Laboratory Plymouth and Alex Alderton, Roberto Amato, Jeffrey Barrett, Sonia Goncalves, Ewan Harrison, David K. Jackson, Ian Johnston, Dominic Kwiatkowski, Cordelia Langford, John Sillitoe on behalf of the Wellcome Sanger Institute COVID-19 Surveillance Team |
| EPI_ISL_2998371 | Microbiologia CATLAB | Can Ruti SARS-CoV-2 Sequencing Hub (HUGTiP/IrsiCaixa/IGTP) | Marc Noguera-Julian, Pilar Armengol, Ignacio Blanco, Antoni E Bordoy, Francesc Catala-Moll, Pere-Joan Cardona, Maria Casadellà, Cristina Casañ, Gemma Clara, Bonaventura Clotet, Montserrat Giménez, Mercedes Guerrero, Anna Not, David Panisello, Roger Paredes, Mariona Parera, Alexia París, Verónica Saludes, Alba Sánchez, Laia Soler, Lauro Sumoy, and Elisa Martró on behalf of the Can Ruti SARS-CoV-2 Sequencing Hub |
| EPI_ISL_2998372 | Lighthouse Laboratory Plymouth | Wellcome Sanger Institute for the COVID-19 Genomics UK (COG-UK) Consortium | Lighthouse Laboratory Plymouth and Alex Alderton, Roberto Amato, Jeffrey Barrett, Sonia Goncalves, Ewan Harrison, David K. Jackson, Ian Johnston, Dominic Kwiatkowski, Cordelia Langford, John Sillitoe on behalf of the Wellcome Sanger Institute COVID-19 Surveillance Team |
| EPI_ISL_2998373 | Lighthouse Lab in Glasgow | Wellcome Sanger Institute for the COVID-19 Genomics UK (COG-UK) Consortium | Harper VanSteenhouse, Yumi Kasai, David Gray, Carol Clugston, Anna Dominiczak and Alex Alderton, Roberto Amato, Jeffrey Barrett, Sonia Goncalves, Ewan Harrison, David K. Jackson, Ian Johnston, Dominic Kwiatkowski, Cordelia Langford, John Sillitoe on behalf of the Wellcome Sanger Institute COVID-19 Surveillance Team |
| EPI_ISL_2998375 | Lighthouse Lab in Milton Keynes | Wellcome Sanger Institute for the COVID-19 Genomics UK (COG-UK) Consortium | The Lighthouse Lab in Milton Keynes and Alex Alderton, Roberto Amato, Jeffrey Barrett, Sonia Goncalves, Ewan Harrison, David K. Jackson, Ian Johnston, Dominic Kwiatkowski, Cordelia Langford, John Sillitoe on behalf of the Wellcome Sanger Institute COVID-19 Surveillance Team |
| EPI_ISL_2998376 | Lighthouse Laboratory Plymouth | Wellcome Sanger Institute for the COVID-19 Genomics UK (COG-UK) Consortium | Lighthouse Laboratory Plymouth and Alex Alderton, Roberto Amato, Jeffrey Barrett, Sonia Goncalves, Ewan Harrison, David K. Jackson, Ian Johnston, Dominic Kwiatkowski, Cordelia Langford, John Sillitoe on behalf of the Wellcome Sanger Institute COVID-19 Surveillance Team |
| EPI_ISL_2998377 | Lighthouse Lab in Glasgow | Wellcome Sanger Institute for the COVID-19 Genomics UK (COG-UK) Consortium | Harper VanSteenhouse, Yumi Kasai, David Gray, Carol Clugston, Anna Dominiczak and Alex Alderton, Roberto Amato, Jeffrey Barrett, Sonia Goncalves, Ewan Harrison, David K. Jackson, Ian Johnston, Dominic Kwiatkowski, Cordelia Langford, John Sillitoe on behalf of the Wellcome Sanger Institute COVID-19 Surveillance Team |
| EPI_ISL_2998378 | Microbiology Department, Laboratori Clínic Metropolitana Nord. Hospital Universitari Germans Trias i Pujol | Can Ruti SARS-CoV-2 Sequencing Hub (HUGTiP/IrsiCaixa/IGTP) | Marc Noguera-Julian, Pilar Armengol, Ignacio Blanco, Antoni E Bordoy, Francesc Catala-Moll, Pere-Joan Cardona, Maria Casadellà, Cristina Casañ, Gemma Clara, Bonaventura Clotet, Montserrat Giménez, Mercedes Guerrero, Anna Not, David Panisello, Roger Paredes, Mariona Parera, Alexia París, Verónica Saludes, Alba Sánchez, Laia Soler, Lauro Sumoy, and Elisa Martró on behalf of the Can Ruti SARS-CoV-2 Sequencing Hub |
| EPI_ISL_2998379 | Lighthouse Lab in Milton Keynes | Wellcome Sanger Institute for the COVID-19 Genomics UK (COG-UK) Consortium | The Lighthouse Lab in Milton Keynes and Alex Alderton, Roberto Amato, Jeffrey Barrett, Sonia Goncalves, Ewan Harrison, David K. Jackson, Ian Johnston, Dominic Kwiatkowski, Cordelia Langford, John Sillitoe on behalf of the Wellcome Sanger Institute COVID-19 Surveillance Team |
| EPI_ISL_2998380 | Lighthouse Lab in Glasgow | Wellcome Sanger Institute for the COVID-19 Genomics UK (COG-UK) Consortium | Harper VanSteenhouse, Yumi Kasai, David Gray, Carol Clugston, Anna Dominiczak and Alex Alderton, Roberto Amato, Jeffrey Barrett, Sonia Goncalves, Ewan Harrison, David K. Jackson, Ian Johnston, Dominic Kwiatkowski, Cordelia Langford, John Sillitoe on behalf of the Wellcome Sanger Institute COVID-19 Surveillance Team |
| EPI_ISL_2998381 | Lighthouse Lab in Milton Keynes | Wellcome Sanger Institute for the COVID-19 Genomics UK (COG-UK) Consortium | The Lighthouse Lab in Milton Keynes and Alex Alderton, Roberto Amato, Jeffrey Barrett, Sonia Goncalves, Ewan Harrison, David K. Jackson, Ian Johnston, Dominic Kwiatkowski, Cordelia Langford, John Sillitoe on behalf of the Wellcome Sanger Institute COVID-19 Surveillance Team |
| EPI_ISL_2998382 | Microbiology Department, Laboratori Clínic Metropolitana Nord. Hospital Universitari Germans Trias i Pujol | Can Ruti SARS-CoV-2 Sequencing Hub (HUGTiP/IrsiCaixa/IGTP) | Marc Noguera-Julian, Pilar Armengol, Ignacio Blanco, Antoni E Bordoy, Francesc Catala-Moll, Pere-Joan Cardona, Maria Casadellà, Cristina Casañ, Gemma Clara, Bonaventura Clotet, Montserrat Giménez, Mercedes Guerrero, Anna Not, David Panisello, Roger Paredes, Mariona Parera, Alexia París, Verónica Saludes, Alba Sánchez, Laia Soler, Lauro Sumoy, and Elisa Martró on behalf of the Can Ruti SARS-CoV-2 Sequencing Hub |
| EPI_ISL_2998383 | Lighthouse Lab in Milton Keynes | Wellcome Sanger Institute for the COVID-19 Genomics UK (COG-UK) Consortium | The Lighthouse Lab in Milton Keynes and Alex Alderton, Roberto Amato, Jeffrey Barrett, Sonia Goncalves, Ewan Harrison, David K. Jackson, Ian Johnston, Dominic Kwiatkowski, Cordelia Langford, John Sillitoe on behalf of the Wellcome Sanger Institute COVID-19 Surveillance Team |
| EPI_ISL_2998384 | Lighthouse Lab in Glasgow | Wellcome Sanger Institute for the COVID-19 Genomics UK (COG-UK) Consortium | Harper VanSteenhouse, Yumi Kasai, David Gray, Carol Clugston, Anna Dominiczak and Alex Alderton, Roberto Amato, Jeffrey Barrett, Sonia Goncalves, Ewan Harrison, David K. Jackson, Ian Johnston, Dominic Kwiatkowski, Cordelia Langford, John Sillitoe on behalf of the Wellcome Sanger Institute COVID-19 Surveillance Team |
| EPI_ISL_2998385 | Lighthouse Lab in Milton Keynes | Wellcome Sanger Institute for the COVID-19 Genomics UK (COG-UK) Consortium | The Lighthouse Lab in Milton Keynes and Alex Alderton, Roberto Amato, Jeffrey Barrett, Sonia Goncalves, Ewan Harrison, David K. Jackson, Ian Johnston, Dominic Kwiatkowski, Cordelia Langford, John Sillitoe on behalf of the Wellcome Sanger Institute COVID-19 Surveillance Team |
| EPI_ISL_2998386 | Microbiology Department, Laboratori Clínic Metropolitana Nord. Hospital Universitari Germans Trias i Pujol | Can Ruti SARS-CoV-2 Sequencing Hub (HUGTiP/IrsiCaixa/IGTP) | Marc Noguera-Julian, Pilar Armengol, Ignacio Blanco, Antoni E Bordoy, Francesc Catala-Moll, Pere-Joan Cardona, Maria Casadellà, Cristina Casañ, Gemma Clara, Bonaventura Clotet, Montserrat Giménez, Mercedes Guerrero, Anna Not, David Panisello, Roger Paredes, Mariona Parera, Alexia París, Verónica Saludes, Alba Sánchez, Laia Soler, Lauro Sumoy, and Elisa Martró on behalf of the Can Ruti SARS-CoV-2 Sequencing Hub |
| EPI_ISL_2998387, EPI_ISL_2998388 | Lighthouse Laboratory Plymouth | Wellcome Sanger Institute for the COVID-19 Genomics UK (COG-UK) Consortium | Lighthouse Laboratory Plymouth and Alex Alderton, Roberto Amato, Jeffrey Barrett, Sonia Goncalves, Ewan Harrison, David K. Jackson, Ian Johnston, Dominic Kwiatkowski, Cordelia Langford, John Sillitoe on behalf of the Wellcome Sanger Institute COVID-19 Surveillance Team |
| EPI_ISL_2998389 | Microbiology Department, Laboratori Clínic Metropolitana Nord. Hospital Universitari Germans Trias i Pujol | Can Ruti SARS-CoV-2 Sequencing Hub (HUGTiP/IrsiCaixa/IGTP) | Marc Noguera-Julian, Pilar Armengol, Ignacio Blanco, Antoni E Bordoy, Francesc Catala-Moll, Pere-Joan Cardona, Maria Casadellà, Cristina Casañ, Gemma Clara, Bonaventura Clotet, Montserrat Giménez, Mercedes Guerrero, Anna Not, David Panisello, Roger Paredes, Mariona Parera, Alexia París, Verónica Saludes, Alba Sánchez, Laia Soler, Lauro Sumoy, and Elisa Martró on behalf of the Can Ruti SARS-CoV-2 Sequencing Hub |
| EPI_ISL_2998390, EPI_ISL_2998391 | Lighthouse Laboratory Plymouth | Wellcome Sanger Institute for the COVID-19 Genomics UK (COG-UK) Consortium | Lighthouse Laboratory Plymouth and Alex Alderton, Roberto Amato, Jeffrey Barrett, Sonia Goncalves, Ewan Harrison, David K. Jackson, Ian Johnston, Dominic Kwiatkowski, Cordelia Langford, John Sillitoe on behalf of the Wellcome Sanger Institute COVID-19 Surveillance Team |
| EPI_ISL_2998392 | Lighthouse Lab in Glasgow | Wellcome Sanger Institute for the COVID-19 Genomics UK (COG-UK) Consortium | Harper VanSteenhouse, Yumi Kasai, David Gray, Carol Clugston, Anna Dominiczak and Alex Alderton, Roberto Amato, Jeffrey Barrett, Sonia Goncalves, Ewan Harrison, David K. Jackson, Ian Johnston, Dominic Kwiatkowski, Cordelia Langford, John Sillitoe on behalf of the Wellcome Sanger Institute COVID-19 Surveillance Team |
| EPI_ISL_2998394 | Lighthouse Laboratory Plymouth | Wellcome Sanger Institute for the COVID-19 Genomics UK (COG-UK) Consortium | Lighthouse Laboratory Plymouth and Alex Alderton, Roberto Amato, Jeffrey Barrett, Sonia Goncalves, Ewan Harrison, David K. Jackson, Ian Johnston, Dominic Kwiatkowski, Cordelia Langford, John Sillitoe on behalf of the Wellcome Sanger Institute COVID-19 Surveillance Team |
| EPI_ISL_2998395 | Lighthouse Lab in Glasgow | Wellcome Sanger Institute for the COVID-19 Genomics UK (COG-UK) Consortium | Harper VanSteenhouse, Yumi Kasai, David Gray, Carol Clugston, Anna Dominiczak and Alex Alderton, Roberto Amato, Jeffrey Barrett, Sonia Goncalves, Ewan Harrison, David K. Jackson, Ian Johnston, Dominic Kwiatkowski, Cordelia Langford, John Sillitoe on behalf of the Wellcome Sanger Institute COVID-19 Surveillance Team |
| EPI_ISL_2998396 | Microbiology Department, Laboratori Clínic Metropolitana Nord. Hospital Universitari Germans Trias i Pujol | Can Ruti SARS-CoV-2 Sequencing Hub (HUGTiP/IrsiCaixa/IGTP) | Marc Noguera-Julian, Pilar Armengol, Ignacio Blanco, Antoni E Bordoy, Francesc Catala-Moll, Pere-Joan Cardona, Maria Casadellà, Cristina Casañ, Gemma Clara, Bonaventura Clotet, Montserrat Giménez, Mercedes Guerrero, Anna Not, David Panisello, Roger Paredes, Mariona Parera, Alexia París, Verónica Saludes, Alba Sánchez, Laia Soler, Lauro Sumoy, and Elisa Martró on behalf of the Can Ruti SARS-CoV-2 Sequencing Hub |
| EPI_ISL_2998397, EPI_ISL_2998398 | Lighthouse Laboratory Plymouth | Wellcome Sanger Institute for the COVID-19 Genomics UK (COG-UK) Consortium | Lighthouse Laboratory Plymouth and Alex Alderton, Roberto Amato, Jeffrey Barrett, Sonia Goncalves, Ewan Harrison, David K. Jackson, Ian Johnston, Dominic Kwiatkowski, Cordelia Langford, John Sillitoe on behalf of the Wellcome Sanger Institute COVID-19 Surveillance Team |
| EPI_ISL_2998399 | Microbiology Department, Laboratori Clínic Metropolitana Nord. Hospital Universitari Germans Trias i Pujol | Can Ruti SARS-CoV-2 Sequencing Hub (HUGTiP/IrsiCaixa/IGTP) | Marc Noguera-Julian, Pilar Armengol, Ignacio Blanco, Antoni E Bordoy, Francesc Catala-Moll, Pere-Joan Cardona, Maria Casadellà, Cristina Casañ, Gemma Clara, Bonaventura Clotet, Montserrat Giménez, Mercedes Guerrero, Anna Not, David Panisello, Roger Paredes, Mariona Parera, Alexia París, Verónica Saludes, Alba Sánchez, Laia Soler, Lauro Sumoy, and Elisa Martró on behalf of the Can Ruti SARS-CoV-2 Sequencing Hub |
| EPI_ISL_2998400, EPI_ISL_2998401 | Lighthouse Lab in Milton Keynes | Wellcome Sanger Institute for the COVID-19 Genomics UK (COG-UK) Consortium | The Lighthouse Lab in Milton Keynes and Alex Alderton, Roberto Amato, Jeffrey Barrett, Sonia Goncalves, Ewan Harrison, David K. Jackson, Ian Johnston, Dominic Kwiatkowski, Cordelia Langford, John Sillitoe on behalf of the Wellcome Sanger Institute COVID-19 Surveillance Team |
| EPI_ISL_2998402 | Lighthouse Laboratory Plymouth | Wellcome Sanger Institute for the COVID-19 Genomics UK (COG-UK) Consortium | Lighthouse Laboratory Plymouth and Alex Alderton, Roberto Amato, Jeffrey Barrett, Sonia Goncalves, Ewan Harrison, David K. Jackson, Ian Johnston, Dominic Kwiatkowski, Cordelia Langford, John Sillitoe on behalf of the Wellcome Sanger Institute COVID-19 Surveillance Team |
| EPI_ISL_2998403 | Microbiology Department, Laboratori Clínic Metropolitana Nord. Hospital Universitari Germans Trias i Pujol | Can Ruti SARS-CoV-2 Sequencing Hub (HUGTiP/IrsiCaixa/IGTP) | Marc Noguera-Julian, Pilar Armengol, Ignacio Blanco, Antoni E Bordoy, Francesc Catala-Moll, Pere-Joan Cardona, Maria Casadellà, Cristina Casañ, Gemma Clara, Bonaventura Clotet, Montserrat Giménez, Mercedes Guerrero, Anna Not, David Panisello, Roger Paredes, Mariona Parera, Alexia París, Verónica Saludes, Alba Sánchez, Laia Soler, Lauro Sumoy, and Elisa Martró on behalf of the Can Ruti SARS-CoV-2 Sequencing Hub |
| EPI_ISL_2998404 | Lighthouse Laboratory Plymouth | Wellcome Sanger Institute for the COVID-19 Genomics UK (COG-UK) Consortium | Lighthouse Laboratory Plymouth and Alex Alderton, Roberto Amato, Jeffrey Barrett, Sonia Goncalves, Ewan Harrison, David K. Jackson, Ian Johnston, Dominic Kwiatkowski, Cordelia Langford, John Sillitoe on behalf of the Wellcome Sanger Institute COVID-19 Surveillance Team |
| EPI_ISL_2998405, EPI_ISL_2998406 | Lighthouse Lab in Milton Keynes | Wellcome Sanger Institute for the COVID-19 Genomics UK (COG-UK) Consortium | The Lighthouse Lab in Milton Keynes and Alex Alderton, Roberto Amato, Jeffrey Barrett, Sonia Goncalves, Ewan Harrison, David K. Jackson, Ian Johnston, Dominic Kwiatkowski, Cordelia Langford, John Sillitoe on behalf of the Wellcome Sanger Institute COVID-19 Surveillance Team |
| EPI_ISL_2998407 | Microbiology Department, Laboratori Clínic Metropolitana Nord. Hospital Universitari Germans Trias i Pujol | Can Ruti SARS-CoV-2 Sequencing Hub (HUGTiP/IrsiCaixa/IGTP) | Marc Noguera-Julian, Pilar Armengol, Ignacio Blanco, Antoni E Bordoy, Francesc Catala-Moll, Pere-Joan Cardona, Maria Casadellà, Cristina Casañ, Gemma Clara, Bonaventura Clotet, Montserrat Giménez, Mercedes Guerrero, Anna Not, David Panisello, Roger Paredes, Mariona Parera, Alexia París, Verónica Saludes, Alba Sánchez, Laia Soler, Lauro Sumoy, and Elisa Martró on behalf of the Can Ruti SARS-CoV-2 Sequencing Hub |

| | | | |
|---|---|---|---|
| EPI_ISL_2998408, EPI_ISL_2998409 | Lighthouse Lab in Milton Keynes | Wellcome Sanger Institute for the COVID-19 Genomics UK (COG-UK) Consortium | The Lighthouse Lab in Milton Keynes and Alex Alderton, Roberto Amato, Jeffrey Barrett, Sonia Goncalves, Ewan Harrison, David K. Jackson, Ian Johnston, Dominic Kwiatkowski, Cordelia Langford, John Sillitoe on behalf of the Wellcome Sanger Institute COVID-19 Surveillance Team |
| EPI_ISL_2998410 | Microbiologia CATLAB | Can Ruti SARS-CoV-2 Sequencing Hub (HUGTiP/IrsiCaixa/IGTP) | Marc Noguera-Julian, Pilar Armengol, Ignacio Blanco, Antoni E Bordoy, Francesc Catala-Moll, Pere-Joan Cardona, Maria Casadellà, Cristina Casañ, Gemma Clara, Bonaventura Clotet, Montserrat Giménez, Mercedes Guerrero, Anna Not, David Panisello, Roger Paredes, Mariona Parera, Alexia París, Verónica Saludes, Alba Sánchez, Laia Soler, Lauro Sumoy, and Elisa Martró on behalf of the Can Ruti SARS-CoV-2 Sequencing Hub |
| EPI_ISL_2998411, EPI_ISL_2998412, EPI_ISL_2998413 | Lighthouse Laboratory Plymouth | Wellcome Sanger Institute for the COVID-19 Genomics UK (COG-UK) Consortium | Lighthouse Laboratory Plymouth and Alex Alderton, Roberto Amato, Jeffrey Barrett, Sonia Goncalves, Ewan Harrison, David K. Jackson, Ian Johnston, Dominic Kwiatkowski, Cordelia Langford, John Sillitoe on behalf of the Wellcome Sanger Institute COVID-19 Surveillance Team |
| EPI_ISL_2998414 | Microbiologia CATLAB | Can Ruti SARS-CoV-2 Sequencing Hub (HUGTiP/IrsiCaixa/IGTP) | Marc Noguera-Julian, Pilar Armengol, Ignacio Blanco, Antoni E Bordoy, Francesc Catala-Moll, Pere-Joan Cardona, Maria Casadellà, Cristina Casañ, Gemma Clara, Bonaventura Clotet, Montserrat Giménez, Mercedes Guerrero, Anna Not, David Panisello, Roger Paredes, Mariona Parera, Alexia París, Verónica Saludes, Alba Sánchez, Laia Soler, Lauro Sumoy, and Elisa Martró on behalf of the Can Ruti SARS-CoV-2 Sequencing Hub |
| EPI_ISL_2998415, EPI_ISL_2998416 | Lighthouse Laboratory Plymouth | Wellcome Sanger Institute for the COVID-19 Genomics UK (COG-UK) Consortium | Lighthouse Laboratory Plymouth and Alex Alderton, Roberto Amato, Jeffrey Barrett, Sonia Goncalves, Ewan Harrison, David K. Jackson, Ian Johnston, Dominic Kwiatkowski, Cordelia Langford, John Sillitoe on behalf of the Wellcome Sanger Institute COVID-19 Surveillance Team |
| EPI_ISL_2998417 | Microbiologia CATLAB | Can Ruti SARS-CoV-2 Sequencing Hub (HUGTiP/IrsiCaixa/IGTP) | Marc Noguera-Julian, Pilar Armengol, Ignacio Blanco, Antoni E Bordoy, Francesc Catala-Moll, Pere-Joan Cardona, Maria Casadellà, Cristina Casañ, Gemma Clara, Bonaventura Clotet, Montserrat Giménez, Mercedes Guerrero, Anna Not, David Panisello, Roger Paredes, Mariona Parera, Alexia París, Verónica Saludes, Alba Sánchez, Laia Soler, Lauro Sumoy, and Elisa Martró on behalf of the Can Ruti SARS-CoV-2 Sequencing Hub |
| EPI_ISL_2998418 | Lighthouse Lab in Milton Keynes | Wellcome Sanger Institute for the COVID-19 Genomics UK (COG-UK) Consortium | The Lighthouse Lab in Milton Keynes and Alex Alderton, Roberto Amato, Jeffrey Barrett, Sonia Goncalves, Ewan Harrison, David K. Jackson, Ian Johnston, Dominic Kwiatkowski, Cordelia Langford, John Sillitoe on behalf of the Wellcome Sanger Institute COVID-19 Surveillance Team |
| EPI_ISL_2998419 | Lighthouse Laboratory Plymouth | Wellcome Sanger Institute for the COVID-19 Genomics UK (COG-UK) Consortium | Lighthouse Laboratory Plymouth and Alex Alderton, Roberto Amato, Jeffrey Barrett, Sonia Goncalves, Ewan Harrison, David K. Jackson, Ian Johnston, Dominic Kwiatkowski, Cordelia Langford, John Sillitoe on behalf of the Wellcome Sanger Institute COVID-19 Surveillance Team |
| EPI_ISL_2998420 | Microbiology Department, Laboratori Clínic Metropolitana Nord. Hospital Universitari Germans Trias i Pujol | Can Ruti SARS-CoV-2 Sequencing Hub (HUGTiP/IrsiCaixa/IGTP) | Marc Noguera-Julian, Pilar Armengol, Ignacio Blanco, Antoni E Bordoy, Francesc Catala-Moll, Pere-Joan Cardona, Maria Casadellà, Cristina Casañ, Gemma Clara, Bonaventura Clotet, Montserrat Giménez, Mercedes Guerrero, Anna Not, David Panisello, Roger Paredes, Mariona Parera, Alexia París, Verónica Saludes, Alba Sánchez, Laia Soler, Lauro Sumoy, and Elisa Martró on behalf of the Can Ruti SARS-CoV-2 Sequencing Hub |
| EPI_ISL_2998421 | Lighthouse Lab in Milton Keynes | Wellcome Sanger Institute for the COVID-19 Genomics UK (COG-UK) Consortium | The Lighthouse Lab in Milton Keynes and Alex Alderton, Roberto Amato, Jeffrey Barrett, Sonia Goncalves, Ewan Harrison, David K. Jackson, Ian Johnston, Dominic Kwiatkowski, Cordelia Langford, John Sillitoe on behalf of the Wellcome Sanger Institute COVID-19 Surveillance Team |
| EPI_ISL_2998422 | Lighthouse Lab in Glasgow | Wellcome Sanger Institute for the COVID-19 Genomics UK (COG-UK) Consortium | Harper VanSteenhouse, Yumi Kasai, David Gray, Carol Clugston, Anna Dominiczak and Alex Alderton, Roberto Amato, Jeffrey Barrett, Sonia Goncalves, Ewan Harrison, David K. Jackson, Ian Johnston, Dominic Kwiatkowski, Cordelia Langford, John Sillitoe on behalf of the Wellcome Sanger Institute COVID-19 Surveillance Team |
| EPI_ISL_2998423, EPI_ISL_2998424 | Lighthouse Lab in Milton Keynes | Wellcome Sanger Institute for the COVID-19 Genomics UK (COG-UK) Consortium | The Lighthouse Lab in Milton Keynes and Alex Alderton, Roberto Amato, Jeffrey Barrett, Sonia Goncalves, Ewan Harrison, David K. Jackson, Ian Johnston, Dominic Kwiatkowski, Cordelia Langford, John Sillitoe on behalf of the Wellcome Sanger Institute COVID-19 Surveillance Team |
| EPI_ISL_2998425 | Microbiology Department, Laboratori Clínic Metropolitana Nord. Hospital Universitari Germans Trias i Pujol | Can Ruti SARS-CoV-2 Sequencing Hub (HUGTiP/IrsiCaixa/IGTP) | Marc Noguera-Julian, Pilar Armengol, Ignacio Blanco, Antoni E Bordoy, Francesc Catala-Moll, Pere-Joan Cardona, Maria Casadellà, Cristina Casañ, Gemma Clara, Bonaventura Clotet, Montserrat Giménez, Mercedes Guerrero, Anna Not, David Panisello, Roger Paredes, Mariona Parera, Alexia París, Verónica Saludes, Alba Sánchez, Laia Soler, Lauro Sumoy, and Elisa Martró on behalf of the Can Ruti SARS-CoV-2 Sequencing Hub |
| EPI_ISL_2998426 | Lighthouse Laboratory Plymouth | Wellcome Sanger Institute for the COVID-19 Genomics UK (COG-UK) Consortium | Lighthouse Laboratory Plymouth and Alex Alderton, Roberto Amato, Jeffrey Barrett, Sonia Goncalves, Ewan Harrison, David K. Jackson, Ian Johnston, Dominic Kwiatkowski, Cordelia Langford, John Sillitoe on behalf of the Wellcome Sanger Institute COVID-19 Surveillance Team |
| EPI_ISL_2998427, EPI_ISL_2998428 | Lighthouse Lab in Glasgow | Wellcome Sanger Institute for the COVID-19 Genomics UK (COG-UK) Consortium | Harper VanSteenhouse, Yumi Kasai, David Gray, Carol Clugston, Anna Dominiczak and Alex Alderton, Roberto Amato, Jeffrey Barrett, Sonia Goncalves, Ewan Harrison, David K. Jackson, Ian Johnston, Dominic Kwiatkowski, Cordelia Langford, John Sillitoe on behalf of the Wellcome Sanger Institute COVID-19 Surveillance Team |
| EPI_ISL_2998429 | Microbiology Department, Laboratori Clínic Metropolitana Nord. Hospital Universitari Germans Trias i Pujol | Can Ruti SARS-CoV-2 Sequencing Hub (HUGTiP/IrsiCaixa/IGTP) | Marc Noguera-Julian, Pilar Armengol, Ignacio Blanco, Antoni E Bordoy, Francesc Catala-Moll, Pere-Joan Cardona, Maria Casadellà, Cristina Casañ, Gemma Clara, Bonaventura Clotet, Montserrat Giménez, Mercedes Guerrero, Anna Not, David Panisello, Roger Paredes, Mariona Parera, Alexia París, Verónica Saludes, Alba Sánchez, Laia Soler, Lauro Sumoy, and Elisa Martró on behalf of the Can Ruti SARS-CoV-2 Sequencing Hub |
| EPI_ISL_2998430 | Lighthouse Lab in Glasgow | Wellcome Sanger Institute for the COVID-19 Genomics UK (COG-UK) Consortium | Harper VanSteenhouse, Yumi Kasai, David Gray, Carol Clugston, Anna Dominiczak and Alex Alderton, Roberto Amato, Jeffrey Barrett, Sonia Goncalves, Ewan Harrison, David K. Jackson, Ian Johnston, Dominic Kwiatkowski, Cordelia Langford, John Sillitoe on behalf of the Wellcome Sanger Institute COVID-19 Surveillance Team |
| EPI_ISL_2998431 | Lighthouse Laboratory Plymouth | Wellcome Sanger Institute for the COVID-19 Genomics UK (COG-UK) Consortium | Lighthouse Laboratory Plymouth and Alex Alderton, Roberto Amato, Jeffrey Barrett, Sonia Goncalves, Ewan Harrison, David K. Jackson, Ian Johnston, Dominic Kwiatkowski, Cordelia Langford, John Sillitoe on behalf of the Wellcome Sanger Institute COVID-19 Surveillance Team |
| EPI_ISL_2998432 | Lighthouse Lab in Glasgow | Wellcome Sanger Institute for the COVID-19 Genomics UK (COG-UK) Consortium | Harper VanSteenhouse, Yumi Kasai, David Gray, Carol Clugston, Anna Dominiczak and Alex Alderton, Roberto Amato, Jeffrey Barrett, Sonia Goncalves, Ewan Harrison, David K. Jackson, Ian Johnston, Dominic Kwiatkowski, Cordelia Langford, John Sillitoe on behalf of the Wellcome Sanger Institute COVID-19 Surveillance Team |
| EPI_ISL_2998433 | Microbiology Department, Laboratori Clínic Metropolitana Nord. Hospital Universitari Germans Trias i Pujol | Can Ruti SARS-CoV-2 Sequencing Hub (HUGTiP/IrsiCaixa/IGTP) | Marc Noguera-Julian, Pilar Armengol, Ignacio Blanco, Antoni E Bordoy, Francesc Catala-Moll, Pere-Joan Cardona, Maria Casadellà, Cristina Casañ, Gemma Clara, Bonaventura Clotet, Montserrat Giménez, Mercedes Guerrero, Anna Not, David Panisello, Roger Paredes, Mariona Parera, Alexia París, Verónica Saludes, Alba Sánchez, Laia Soler, Lauro Sumoy, and Elisa Martró on behalf of the Can Ruti SARS-CoV-2 Sequencing Hub |
| EPI_ISL_2998434, EPI_ISL_2998435 | Lighthouse Laboratory Plymouth | Wellcome Sanger Institute for the COVID-19 Genomics UK (COG-UK) Consortium | Lighthouse Laboratory Plymouth and Alex Alderton, Roberto Amato, Jeffrey Barrett, Sonia Goncalves, Ewan Harrison, David K. Jackson, Ian Johnston, Dominic Kwiatkowski, Cordelia Langford, John Sillitoe on behalf of the Wellcome Sanger Institute COVID-19 Surveillance Team |
| EPI_ISL_2998436 | Microbiology Department, Laboratori Clínic Metropolitana Nord. Hospital Universitari Germans Trias i Pujol | Can Ruti SARS-CoV-2 Sequencing Hub (HUGTiP/IrsiCaixa/IGTP) | Marc Noguera-Julian, Pilar Armengol, Ignacio Blanco, Antoni E Bordoy, Francesc Catala-Moll, Pere-Joan Cardona, Maria Casadellà, Cristina Casañ, Gemma Clara, Bonaventura Clotet, Montserrat Giménez, Mercedes Guerrero, Anna Not, David Panisello, Roger Paredes, Mariona Parera, Alexia París, Verónica Saludes, Alba Sánchez, Laia Soler, Lauro Sumoy, and Elisa Martró on behalf of the Can Ruti SARS-CoV-2 Sequencing Hub |
| EPI_ISL_2998437 | Lighthouse Laboratory Plymouth | Wellcome Sanger Institute for the COVID-19 Genomics UK (COG-UK) Consortium | Lighthouse Laboratory Plymouth and Alex Alderton, Roberto Amato, Jeffrey Barrett, Sonia Goncalves, Ewan Harrison, David K. Jackson, Ian Johnston, Dominic Kwiatkowski, Cordelia Langford, John Sillitoe on behalf of the Wellcome Sanger Institute COVID-19 Surveillance Team |
| EPI_ISL_2998438 | Lighthouse Lab in Milton Keynes | Wellcome Sanger Institute for the COVID-19 Genomics UK (COG-UK) Consortium | The Lighthouse Lab in Milton Keynes and Alex Alderton, Roberto Amato, Jeffrey Barrett, Sonia Goncalves, Ewan Harrison, David K. Jackson, Ian Johnston, Dominic Kwiatkowski, Cordelia Langford, John Sillitoe on behalf of the Wellcome Sanger Institute COVID-19 Surveillance Team |
| EPI_ISL_2998439 | Microbiology Department, Laboratori Clínic Metropolitana Nord. Hospital Universitari Germans Trias i Pujol | Can Ruti SARS-CoV-2 Sequencing Hub (HUGTiP/IrsiCaixa/IGTP) | Marc Noguera-Julian, Pilar Armengol, Ignacio Blanco, Antoni E Bordoy, Francesc Catala-Moll, Pere-Joan Cardona, Maria Casadellà, Cristina Casañ, Gemma Clara, Bonaventura Clotet, Montserrat Giménez, Mercedes Guerrero, Anna Not, David Panisello, Roger Paredes, Mariona Parera, Alexia París, Verónica Saludes, Alba Sánchez, Laia Soler, Lauro Sumoy, and Elisa Martró on behalf of the Can Ruti SARS-CoV-2 Sequencing Hub |
| EPI_ISL_2998440, EPI_ISL_2998441, EPI_ISL_2998442 | Lighthouse Laboratory Plymouth | Wellcome Sanger Institute for the COVID-19 Genomics UK (COG-UK) Consortium | Lighthouse Laboratory Plymouth and Alex Alderton, Roberto Amato, Jeffrey Barrett, Sonia Goncalves, Ewan Harrison, David K. Jackson, Ian Johnston, Dominic Kwiatkowski, Cordelia Langford, John Sillitoe on behalf of the Wellcome Sanger Institute COVID-19 Surveillance Team |
| EPI_ISL_2998443 | Microbiology Department, Laboratori Clínic Metropolitana Nord. Hospital Universitari Germans Trias i Pujol | Can Ruti SARS-CoV-2 Sequencing Hub (HUGTiP/IrsiCaixa/IGTP) | Marc Noguera-Julian, Pilar Armengol, Ignacio Blanco, Antoni E Bordoy, Francesc Catala-Moll, Pere-Joan Cardona, Maria Casadellà, Cristina Casañ, Gemma Clara, Bonaventura Clotet, Montserrat Giménez, Mercedes Guerrero, Anna Not, David Panisello, Roger Paredes, Mariona Parera, Alexia París, Verónica Saludes, Alba Sánchez, Laia Soler, Lauro Sumoy, and Elisa Martró on behalf of the Can Ruti SARS-CoV-2 Sequencing Hub |
| EPI_ISL_2998444, EPI_ISL_2998445 | Lighthouse Laboratory Plymouth | Wellcome Sanger Institute for the COVID-19 Genomics UK (COG-UK) Consortium | Lighthouse Laboratory Plymouth and Alex Alderton, Roberto Amato, Jeffrey Barrett, Sonia Goncalves, Ewan Harrison, David K. Jackson, Ian Johnston, Dominic Kwiatkowski, Cordelia Langford, John Sillitoe on behalf of the Wellcome Sanger Institute COVID-19 Surveillance Team |
| EPI_ISL_2998446, EPI_ISL_2998448, EPI_ISL_2998449 | Lighthouse Lab in Milton Keynes | Wellcome Sanger Institute for the COVID-19 Genomics UK (COG-UK) Consortium | The Lighthouse Lab in Milton Keynes and Alex Alderton, Roberto Amato, Jeffrey Barrett, Sonia Goncalves, Ewan Harrison, David K. Jackson, Ian Johnston, Dominic Kwiatkowski, Cordelia Langford, John Sillitoe on behalf of the Wellcome Sanger Institute COVID-19 Surveillance Team |
| EPI_ISL_2998450 | Microbiology Department, Laboratori Clínic Metropolitana Nord. Hospital Universitari Germans Trias i Pujol | Can Ruti SARS-CoV-2 Sequencing Hub (HUGTiP/IrsiCaixa/IGTP) | Marc Noguera-Julian, Pilar Armengol, Ignacio Blanco, Antoni E Bordoy, Francesc Catala-Moll, Pere-Joan Cardona, Maria Casadellà, Cristina Casañ, Gemma Clara, Bonaventura Clotet, Montserrat Giménez, Mercedes Guerrero, Anna Not, David Panisello, Roger Paredes, Mariona Parera, Alexia París, Verónica Saludes, Alba Sánchez, Laia Soler, Lauro Sumoy, and Elisa Martró on behalf of the Can Ruti SARS-CoV-2 Sequencing Hub |
| EPI_ISL_2998451 | Lighthouse Laboratory Plymouth | Wellcome Sanger Institute for the COVID-19 Genomics UK (COG-UK) Consortium | Lighthouse Laboratory Plymouth and Alex Alderton, Roberto Amato, Jeffrey Barrett, Sonia Goncalves, Ewan Harrison, David K. Jackson, Ian Johnston, Dominic Kwiatkowski, Cordelia Langford, John Sillitoe on behalf of the Wellcome Sanger Institute COVID-19 Surveillance Team |
| EPI_ISL_2998452 | Lighthouse Lab in Milton Keynes | Wellcome Sanger Institute for the COVID-19 Genomics UK (COG-UK) Consortium | The Lighthouse Lab in Milton Keynes and Alex Alderton, Roberto Amato, Jeffrey Barrett, Sonia Goncalves, Ewan Harrison, David K. Jackson, Ian Johnston, Dominic Kwiatkowski, Cordelia Langford, John Sillitoe on behalf of the Wellcome Sanger Institute COVID-19 Surveillance Team |
| EPI_ISL_2998453, EPI_ISL_2998455, EPI_ISL_2998456 | Lighthouse Laboratory Plymouth | Wellcome Sanger Institute for the COVID-19 Genomics UK (COG-UK) Consortium | Lighthouse Laboratory Plymouth and Alex Alderton, Roberto Amato, Jeffrey Barrett, Sonia Goncalves, Ewan Harrison, David K. Jackson, Ian Johnston, Dominic Kwiatkowski, Cordelia Langford, John Sillitoe on behalf of the Wellcome Sanger Institute COVID-19 Surveillance Team |
| EPI_ISL_2998457 | Microbiologia CATLAB | Can Ruti SARS-CoV-2 Sequencing Hub (HUGTiP/IrsiCaixa/IGTP) | Marc Noguera-Julian, Pilar Armengol, Ignacio Blanco, Antoni E Bordoy, Francesc Catala-Moll, Pere-Joan Cardona, Maria Casadellà, Cristina Casañ, Gemma Clara, Bonaventura Clotet, Montserrat Giménez, Mercedes Guerrero, Anna Not, David Panisello, Roger Paredes, Mariona Parera, Alexia París, Verónica Saludes, Alba Sánchez, Laia Soler, Lauro Sumoy, and Elisa Martró on behalf of the Can Ruti SARS-CoV-2 Sequencing Hub |
| EPI_ISL_2998458 | Lighthouse Laboratory Plymouth | Wellcome Sanger Institute for the COVID-19 Genomics UK (COG-UK) Consortium | Lighthouse Laboratory Plymouth and Alex Alderton, Roberto Amato, Jeffrey Barrett, Sonia Goncalves, Ewan Harrison, David K. Jackson, Ian Johnston, Dominic Kwiatkowski, Cordelia Langford, John Sillitoe on behalf of the Wellcome Sanger Institute COVID-19 Surveillance Team |
| EPI_ISL_2998459 | Lighthouse Lab in Glasgow | Wellcome Sanger Institute for the COVID-19 Genomics UK (COG-UK) Consortium | Harper VanSteenhouse, Yumi Kasai, David Gray, Carol Clugston, Anna Dominiczak and Alex Alderton, Roberto Amato, Jeffrey Barrett, Sonia Goncalves, Ewan Harrison, David K. Jackson, Ian Johnston, Dominic Kwiatkowski, Cordelia Langford, John Sillitoe on behalf of the Wellcome Sanger Institute COVID-19 Surveillance Team |
| EPI_ISL_2998460 | Lighthouse Laboratory Plymouth | Wellcome Sanger Institute for the COVID-19 Genomics UK (COG-UK) Consortium | Lighthouse Laboratory Plymouth and Alex Alderton, Roberto Amato, Jeffrey Barrett, Sonia Goncalves, Ewan Harrison, David K. Jackson, Ian Johnston, Dominic Kwiatkowski, Cordelia Langford, John Sillitoe on behalf of the Wellcome Sanger Institute COVID-19 Surveillance Team |
| EPI_ISL_2998461 | Microbiologia CATLAB | Can Ruti SARS-CoV-2 Sequencing Hub (HUGTiP/IrsiCaixa/IGTP) | Marc Noguera-Julian, Pilar Armengol, Ignacio Blanco, Antoni E Bordoy, Francesc Catala-Moll, Pere-Joan Cardona, Maria Casadellà, Cristina Casañ, Gemma Clara, Bonaventura Clotet, Montserrat Giménez, Mercedes Guerrero, Anna Not, David Panisello, Roger Paredes, Mariona Parera, Alexia París, Verónica Saludes, Alba Sánchez, Laia Soler, Lauro Sumoy, and Elisa Martró on behalf of the Can Ruti SARS-CoV-2 Sequencing Hub |
| EPI_ISL_2998462 | Lighthouse Lab in Milton Keynes | Wellcome Sanger Institute for the COVID-19 Genomics UK (COG-UK) Consortium | The Lighthouse Lab in Milton Keynes and Alex Alderton, Roberto Amato, Jeffrey Barrett, Sonia Goncalves, Ewan Harrison, David K. Jackson, Ian Johnston, Dominic Kwiatkowski, Cordelia Langford, John Sillitoe on behalf of the Wellcome Sanger Institute COVID-19 Surveillance Team |
| EPI_ISL_2998463 | Lighthouse Laboratory Plymouth | Wellcome Sanger Institute for the COVID-19 Genomics UK (COG-UK) Consortium | Lighthouse Laboratory Plymouth and Alex Alderton, Roberto Amato, Jeffrey Barrett, Sonia Goncalves, Ewan Harrison, David K. Jackson, Ian Johnston, Dominic Kwiatkowski, Cordelia Langford, John Sillitoe on behalf of the Wellcome Sanger Institute COVID-19 Surveillance Team |
| EPI_ISL_2998464 | Microbiology Department, Laboratori Clínic Metropolitana Nord. Hospital Universitari Germans Trias i Pujol | Can Ruti SARS-CoV-2 Sequencing Hub (HUGTiP/IrsiCaixa/IGTP) | Marc Noguera-Julian, Pilar Armengol, Ignacio Blanco, Antoni E Bordoy, Francesc Catala-Moll, Pere-Joan Cardona, Maria Casadellà, Cristina Casañ, Gemma Clara, Bonaventura Clotet, Montserrat Giménez, Mercedes Guerrero, Anna Not, David Panisello, Roger Paredes, Mariona Parera, Alexia París, Verónica Saludes, Alba Sánchez, Laia Soler, Lauro Sumoy, and Elisa Martró on behalf of the Can Ruti SARS-CoV-2 Sequencing Hub |
| EPI_ISL_2998465 | Lighthouse Lab in Glasgow | Wellcome Sanger Institute for the COVID-19 Genomics UK (COG-UK) Consortium | Harper VanSteenhouse, Yumi Kasai, David Gray, Carol Clugston, Anna Dominiczak and Alex Alderton, Roberto Amato, Jeffrey Barrett, Sonia Goncalves, Ewan Harrison, David K. Jackson, Ian Johnston, Dominic Kwiatkowski, Cordelia Langford, John Sillitoe on behalf of the Wellcome Sanger Institute COVID-19 Surveillance Team |
| EPI_ISL_2998466 | Lighthouse Laboratory Plymouth | Wellcome Sanger Institute for the COVID-19 Genomics UK (COG-UK) Consortium | Lighthouse Laboratory Plymouth and Alex Alderton, Roberto Amato, Jeffrey Barrett, Sonia Goncalves, Ewan Harrison, David K. Jackson, Ian Johnston, Dominic Kwiatkowski, Cordelia Langford, John Sillitoe on behalf of the Wellcome Sanger Institute COVID-19 Surveillance Team |

| | | | |
|---|---|---|---|
| EPI_ISL_2998467 | Lighthouse Lab in Milton Keynes | Wellcome Sanger Institute for the COVID-19 Genomics UK (COG-UK) Consortium | The Lighthouse Lab in Milton Keynes and Alex Alderton, Roberto Amato, Jeffrey Barrett, Sonia Goncalves, Ewan Harrison, David K. Jackson, Ian Johnston, Dominic Kwiatkowski, Cordelia Langford, John Sillitoe on behalf of the Wellcome Sanger Institute COVID-19 Surveillance Team |
| EPI_ISL_2998468 | Microbiology Department, Laboratori Clínic Metropolitana Nord. Hospital Universitari Germans Trias i Pujol | Can Ruti SARS-CoV-2 Sequencing Hub (HUGTiP/IrsiCaixa/IGTP) | Marc Noguera-Julian, Pilar Armengol, Ignacio Blanco, Antoni E Bordoy, Francesc Catala-Moll, Pere-Joan Cardona, Maria Casadellà, Cristina Casañ, Gemma Clara, Bonaventura Clotet, Montserrat Giménez, Mercedes Guerrero, Anna Not, David Panisello, Roger Paredes, Mariona Parera, Alexia París, Verónica Saludes, Alba Sánchez, Laia Soler, Lauro Sumoy, and Elisa Martró on behalf of the Can Ruti SARS-CoV-2 Sequencing Hub |
| EPI_ISL_2998469 | Lighthouse Laboratory Plymouth | Wellcome Sanger Institute for the COVID-19 Genomics UK (COG-UK) Consortium | Lighthouse Laboratory Plymouth and Alex Alderton, Roberto Amato, Jeffrey Barrett, Sonia Goncalves, Ewan Harrison, David K. Jackson, Ian Johnston, Dominic Kwiatkowski, Cordelia Langford, John Sillitoe on behalf of the Wellcome Sanger Institute COVID-19 Surveillance Team |
| EPI_ISL_2998470 | Lighthouse Lab in Milton Keynes | Wellcome Sanger Institute for the COVID-19 Genomics UK (COG-UK) Consortium | The Lighthouse Lab in Milton Keynes and Alex Alderton, Roberto Amato, Jeffrey Barrett, Sonia Goncalves, Ewan Harrison, David K. Jackson, Ian Johnston, Dominic Kwiatkowski, Cordelia Langford, John Sillitoe on behalf of the Wellcome Sanger Institute COVID-19 Surveillance Team |
| EPI_ISL_2998471 | Microbiology Department, Laboratori Clínic Metropolitana Nord. Hospital Universitari Germans Trias i Pujol | Can Ruti SARS-CoV-2 Sequencing Hub (HUGTiP/IrsiCaixa/IGTP) | Marc Noguera-Julian, Pilar Armengol, Ignacio Blanco, Antoni E Bordoy, Francesc Catala-Moll, Pere-Joan Cardona, Maria Casadellà, Cristina Casañ, Gemma Clara, Bonaventura Clotet, Montserrat Giménez, Mercedes Guerrero, Anna Not, David Panisello, Roger Paredes, Mariona Parera, Alexia París, Verónica Saludes, Alba Sánchez, Laia Soler, Lauro Sumoy, and Elisa Martró on behalf of the Can Ruti SARS-CoV-2 Sequencing Hub |
| EPI_ISL_2998472 | Lighthouse Laboratory Plymouth | Wellcome Sanger Institute for the COVID-19 Genomics UK (COG-UK) Consortium | Lighthouse Laboratory Plymouth and Alex Alderton, Roberto Amato, Jeffrey Barrett, Sonia Goncalves, Ewan Harrison, David K. Jackson, Ian Johnston, Dominic Kwiatkowski, Cordelia Langford, John Sillitoe on behalf of the Wellcome Sanger Institute COVID-19 Surveillance Team |
| EPI_ISL_2998473 | Lighthouse Lab in Milton Keynes | Wellcome Sanger Institute for the COVID-19 Genomics UK (COG-UK) Consortium | The Lighthouse Lab in Milton Keynes and Alex Alderton, Roberto Amato, Jeffrey Barrett, Sonia Goncalves, Ewan Harrison, David K. Jackson, Ian Johnston, Dominic Kwiatkowski, Cordelia Langford, John Sillitoe on behalf of the Wellcome Sanger Institute COVID-19 Surveillance Team |
| EPI_ISL_2998475 | Lighthouse Lab in Glasgow | Wellcome Sanger Institute for the COVID-19 Genomics UK (COG-UK) Consortium | Harper VanSteenhouse, Yumi Kasai, David Gray, Carol Clugston, Anna Dominiczak and Alex Alderton, Roberto Amato, Jeffrey Barrett, Sonia Goncalves, Ewan Harrison, David K. Jackson, Ian Johnston, Dominic Kwiatkowski, Cordelia Langford, John Sillitoe on behalf of the Wellcome Sanger Institute COVID-19 Surveillance Team |
| EPI_ISL_2998476, EPI_ISL_2998477 | Lighthouse Laboratory Plymouth | Wellcome Sanger Institute for the COVID-19 Genomics UK (COG-UK) Consortium | Lighthouse Laboratory Plymouth and Alex Alderton, Roberto Amato, Jeffrey Barrett, Sonia Goncalves, Ewan Harrison, David K. Jackson, Ian Johnston, Dominic Kwiatkowski, Cordelia Langford, John Sillitoe on behalf of the Wellcome Sanger Institute COVID-19 Surveillance Team |
| EPI_ISL_2998478 | Microbiology Department, Laboratori Clínic Metropolitana Nord. Hospital Universitari Germans Trias i Pujol | Can Ruti SARS-CoV-2 Sequencing Hub (HUGTiP/IrsiCaixa/IGTP) | Marc Noguera-Julian, Pilar Armengol, Ignacio Blanco, Antoni E Bordoy, Francesc Catala-Moll, Pere-Joan Cardona, Maria Casadellà, Cristina Casañ, Gemma Clara, Bonaventura Clotet, Montserrat Giménez, Mercedes Guerrero, Anna Not, David Panisello, Roger Paredes, Mariona Parera, Alexia París, Verónica Saludes, Alba Sánchez, Laia Soler, Lauro Sumoy, and Elisa Martró on behalf of the Can Ruti SARS-CoV-2 Sequencing Hub |
| EPI_ISL_2998479 | Lighthouse Lab in Milton Keynes | Wellcome Sanger Institute for the COVID-19 Genomics UK (COG-UK) Consortium | The Lighthouse Lab in Milton Keynes and Alex Alderton, Roberto Amato, Jeffrey Barrett, Sonia Goncalves, Ewan Harrison, David K. Jackson, Ian Johnston, Dominic Kwiatkowski, Cordelia Langford, John Sillitoe on behalf of the Wellcome Sanger Institute COVID-19 Surveillance Team |
| EPI_ISL_2998480 | Lighthouse Laboratory Plymouth | Wellcome Sanger Institute for the COVID-19 Genomics UK (COG-UK) Consortium | Lighthouse Laboratory Plymouth and Alex Alderton, Roberto Amato, Jeffrey Barrett, Sonia Goncalves, Ewan Harrison, David K. Jackson, Ian Johnston, Dominic Kwiatkowski, Cordelia Langford, John Sillitoe on behalf of the Wellcome Sanger Institute COVID-19 Surveillance Team |
| EPI_ISL_2998481 | Lighthouse Lab in Glasgow | Wellcome Sanger Institute for the COVID-19 Genomics UK (COG-UK) Consortium | Harper VanSteenhouse, Yumi Kasai, David Gray, Carol Clugston, Anna Dominiczak and Alex Alderton, Roberto Amato, Jeffrey Barrett, Sonia Goncalves, Ewan Harrison, David K. Jackson, Ian Johnston, Dominic Kwiatkowski, Cordelia Langford, John Sillitoe on behalf of the Wellcome Sanger Institute COVID-19 Surveillance Team |
| EPI_ISL_2998482 | Microbiology Department, Laboratori Clínic Metropolitana Nord. Hospital Universitari Germans Trias i Pujol | Can Ruti SARS-CoV-2 Sequencing Hub (HUGTiP/IrsiCaixa/IGTP) | Marc Noguera-Julian, Pilar Armengol, Ignacio Blanco, Antoni E Bordoy, Francesc Catala-Moll, Pere-Joan Cardona, Maria Casadellà, Cristina Casañ, Gemma Clara, Bonaventura Clotet, Montserrat Giménez, Mercedes Guerrero, Anna Not, David Panisello, Roger Paredes, Mariona Parera, Alexia París, Verónica Saludes, Alba Sánchez, Laia Soler, Lauro Sumoy, and Elisa Martró on behalf of the Can Ruti SARS-CoV-2 Sequencing Hub |
| EPI_ISL_2998483, EPI_ISL_2998484, EPI_ISL_2998486 | Lighthouse Laboratory Plymouth | Wellcome Sanger Institute for the COVID-19 Genomics UK (COG-UK) Consortium | Lighthouse Laboratory Plymouth and Alex Alderton, Roberto Amato, Jeffrey Barrett, Sonia Goncalves, Ewan Harrison, David K. Jackson, Ian Johnston, Dominic Kwiatkowski, Cordelia Langford, John Sillitoe on behalf of the Wellcome Sanger Institute COVID-19 Surveillance Team |
| EPI_ISL_2998487 | Lighthouse Lab in Glasgow | Wellcome Sanger Institute for the COVID-19 Genomics UK (COG-UK) Consortium | Harper VanSteenhouse, Yumi Kasai, David Gray, Carol Clugston, Anna Dominiczak and Alex Alderton, Roberto Amato, Jeffrey Barrett, Sonia Goncalves, Ewan Harrison, David K. Jackson, Ian Johnston, Dominic Kwiatkowski, Cordelia Langford, John Sillitoe on behalf of the Wellcome Sanger Institute COVID-19 Surveillance Team |
| EPI_ISL_2998488 | Microbiology Department, Laboratori Clínic Metropolitana Nord. Hospital Universitari Germans Trias i Pujol | Can Ruti SARS-CoV-2 Sequencing Hub (HUGTiP/IrsiCaixa/IGTP) | Marc Noguera-Julian, Pilar Armengol, Ignacio Blanco, Antoni E Bordoy, Francesc Catala-Moll, Pere-Joan Cardona, Maria Casadellà, Cristina Casañ, Gemma Clara, Bonaventura Clotet, Montserrat Giménez, Mercedes Guerrero, Anna Not, David Panisello, Roger Paredes, Mariona Parera, Alexia París, Verónica Saludes, Alba Sánchez, Laia Soler, Lauro Sumoy, and Elisa Martró on behalf of the Can Ruti SARS-CoV-2 Sequencing Hub |
| EPI_ISL_2998489 | Lighthouse Lab in Glasgow | Wellcome Sanger Institute for the COVID-19 Genomics UK (COG-UK) Consortium | Harper VanSteenhouse, Yumi Kasai, David Gray, Carol Clugston, Anna Dominiczak and Alex Alderton, Roberto Amato, Jeffrey Barrett, Sonia Goncalves, Ewan Harrison, David K. Jackson, Ian Johnston, Dominic Kwiatkowski, Cordelia Langford, John Sillitoe on behalf of the Wellcome Sanger Institute COVID-19 Surveillance Team |
| EPI_ISL_2998490 | Lighthouse Laboratory Plymouth | Wellcome Sanger Institute for the COVID-19 Genomics UK (COG-UK) Consortium | Lighthouse Laboratory Plymouth and Alex Alderton, Roberto Amato, Jeffrey Barrett, Sonia Goncalves, Ewan Harrison, David K. Jackson, Ian Johnston, Dominic Kwiatkowski, Cordelia Langford, John Sillitoe on behalf of the Wellcome Sanger Institute COVID-19 Surveillance Team |
| EPI_ISL_2998491 | Microbiology Department, Laboratori Clínic Metropolitana Nord. Hospital Universitari Germans Trias i Pujol | Can Ruti SARS-CoV-2 Sequencing Hub (HUGTiP/IrsiCaixa/IGTP) | Marc Noguera-Julian, Pilar Armengol, Ignacio Blanco, Antoni E Bordoy, Francesc Catala-Moll, Pere-Joan Cardona, Maria Casadellà, Cristina Casañ, Gemma Clara, Bonaventura Clotet, Montserrat Giménez, Mercedes Guerrero, Anna Not, David Panisello, Roger Paredes, Mariona Parera, Alexia París, Verónica Saludes, Alba Sánchez, Laia Soler, Lauro Sumoy, and Elisa Martró on behalf of the Can Ruti SARS-CoV-2 Sequencing Hub |
| EPI_ISL_2998492, EPI_ISL_2998493 | Lighthouse Lab in Glasgow | Wellcome Sanger Institute for the COVID-19 Genomics UK (COG-UK) Consortium | Harper VanSteenhouse, Yumi Kasai, David Gray, Carol Clugston, Anna Dominiczak and Alex Alderton, Roberto Amato, Jeffrey Barrett, Sonia Goncalves, Ewan Harrison, David K. Jackson, Ian Johnston, Dominic Kwiatkowski, Cordelia Langford, John Sillitoe on behalf of the Wellcome Sanger Institute COVID-19 Surveillance Team |
| EPI_ISL_2998494 | Lighthouse Laboratory Plymouth | Wellcome Sanger Institute for the COVID-19 Genomics UK (COG-UK) Consortium | Lighthouse Laboratory Plymouth and Alex Alderton, Roberto Amato, Jeffrey Barrett, Sonia Goncalves, Ewan Harrison, David K. Jackson, Ian Johnston, Dominic Kwiatkowski, Cordelia Langford, John Sillitoe on behalf of the Wellcome Sanger Institute COVID-19 Surveillance Team |
| EPI_ISL_2998495 | Microbiologia CATLAB | Can Ruti SARS-CoV-2 Sequencing Hub (HUGTiP/IrsiCaixa/IGTP) | Marc Noguera-Julian, Pilar Armengol, Ignacio Blanco, Antoni E Bordoy, Francesc Catala-Moll, Pere-Joan Cardona, Maria Casadellà, Cristina Casañ, Gemma Clara, Bonaventura Clotet, Montserrat Giménez, Mercedes Guerrero, Anna Not, David Panisello, Roger Paredes, Mariona Parera, Alexia París, Verónica Saludes, Alba Sánchez, Laia Soler, Lauro Sumoy, and Elisa Martró on behalf of the Can Ruti SARS-CoV-2 Sequencing Hub |
| EPI_ISL_2998496 | Lighthouse Laboratory Plymouth | Wellcome Sanger Institute for the COVID-19 Genomics UK (COG-UK) Consortium | Lighthouse Laboratory Plymouth and Alex Alderton, Roberto Amato, Jeffrey Barrett, Sonia Goncalves, Ewan Harrison, David K. Jackson, Ian Johnston, Dominic Kwiatkowski, Cordelia Langford, John Sillitoe on behalf of the Wellcome Sanger Institute COVID-19 Surveillance Team |
| EPI_ISL_2998497 | Lighthouse Lab in Milton Keynes | Wellcome Sanger Institute for the COVID-19 Genomics UK (COG-UK) Consortium | The Lighthouse Lab in Milton Keynes and Alex Alderton, Roberto Amato, Jeffrey Barrett, Sonia Goncalves, Ewan Harrison, David K. Jackson, Ian Johnston, Dominic Kwiatkowski, Cordelia Langford, John Sillitoe on behalf of the Wellcome Sanger Institute COVID-19 Surveillance Team |
| EPI_ISL_2998498 | Microbiologia CATLAB | Can Ruti SARS-CoV-2 Sequencing Hub (HUGTiP/IrsiCaixa/IGTP) | Marc Noguera-Julian, Pilar Armengol, Ignacio Blanco, Antoni E Bordoy, Francesc Catala-Moll, Pere-Joan Cardona, Maria Casadellà, Cristina Casañ, Gemma Clara, Bonaventura Clotet, Montserrat Giménez, Mercedes Guerrero, Anna Not, David Panisello, Roger Paredes, Mariona Parera, Alexia París, Verónica Saludes, Alba Sánchez, Laia Soler, Lauro Sumoy, and Elisa Martró on behalf of the Can Ruti SARS-CoV-2 Sequencing Hub |
| EPI_ISL_2998499 | Lighthouse Laboratory Plymouth | Wellcome Sanger Institute for the COVID-19 Genomics UK (COG-UK) Consortium | Lighthouse Laboratory Plymouth and Alex Alderton, Roberto Amato, Jeffrey Barrett, Sonia Goncalves, Ewan Harrison, David K. Jackson, Ian Johnston, Dominic Kwiatkowski, Cordelia Langford, John Sillitoe on behalf of the Wellcome Sanger Institute COVID-19 Surveillance Team |
| EPI_ISL_2998500 | Lighthouse Lab in Milton Keynes | Wellcome Sanger Institute for the COVID-19 Genomics UK (COG-UK) Consortium | The Lighthouse Lab in Milton Keynes and Alex Alderton, Roberto Amato, Jeffrey Barrett, Sonia Goncalves, Ewan Harrison, David K. Jackson, Ian Johnston, Dominic Kwiatkowski, Cordelia Langford, John Sillitoe on behalf of the Wellcome Sanger Institute COVID-19 Surveillance Team |
| EPI_ISL_2998501 | Microbiologia CATLAB | Can Ruti SARS-CoV-2 Sequencing Hub (HUGTiP/IrsiCaixa/IGTP) | Marc Noguera-Julian, Pilar Armengol, Ignacio Blanco, Antoni E Bordoy, Francesc Catala-Moll, Pere-Joan Cardona, Maria Casadellà, Cristina Casañ, Gemma Clara, Bonaventura Clotet, Montserrat Giménez, Mercedes Guerrero, Anna Not, David Panisello, Roger Paredes, Mariona Parera, Alexia París, Verónica Saludes, Alba Sánchez, Laia Soler, Lauro Sumoy, and Elisa Martró on behalf of the Can Ruti SARS-CoV-2 Sequencing Hub |
| EPI_ISL_2998502 | Lighthouse Lab in Glasgow | Wellcome Sanger Institute for the COVID-19 Genomics UK (COG-UK) Consortium | Harper VanSteenhouse, Yumi Kasai, David Gray, Carol Clugston, Anna Dominiczak and Alex Alderton, Roberto Amato, Jeffrey Barrett, Sonia Goncalves, Ewan Harrison, David K. Jackson, Ian Johnston, Dominic Kwiatkowski, Cordelia Langford, John Sillitoe on behalf of the Wellcome Sanger Institute COVID-19 Surveillance Team |
| EPI_ISL_2998503 | Lighthouse Laboratory Plymouth | Wellcome Sanger Institute for the COVID-19 Genomics UK (COG-UK) Consortium | Lighthouse Laboratory Plymouth and Alex Alderton, Roberto Amato, Jeffrey Barrett, Sonia Goncalves, Ewan Harrison, David K. Jackson, Ian Johnston, Dominic Kwiatkowski, Cordelia Langford, John Sillitoe on behalf of the Wellcome Sanger Institute COVID-19 Surveillance Team |
| EPI_ISL_2998504 | Lighthouse Lab in Glasgow | Wellcome Sanger Institute for the COVID-19 Genomics UK (COG-UK) Consortium | Harper VanSteenhouse, Yumi Kasai, David Gray, Carol Clugston, Anna Dominiczak and Alex Alderton, Roberto Amato, Jeffrey Barrett, Sonia Goncalves, Ewan Harrison, David K. Jackson, Ian Johnston, Dominic Kwiatkowski, Cordelia Langford, John Sillitoe on behalf of the Wellcome Sanger Institute COVID-19 Surveillance Team |
| EPI_ISL_2998505 | Microbiology Department, Laboratori Clínic Metropolitana Nord. Hospital Universitari Germans Trias i Pujol | Can Ruti SARS-CoV-2 Sequencing Hub (HUGTiP/IrsiCaixa/IGTP) | Marc Noguera-Julian, Pilar Armengol, Ignacio Blanco, Antoni E Bordoy, Francesc Catala-Moll, Pere-Joan Cardona, Maria Casadellà, Cristina Casañ, Gemma Clara, Bonaventura Clotet, Montserrat Giménez, Mercedes Guerrero, Anna Not, David Panisello, Roger Paredes, Mariona Parera, Alexia París, Verónica Saludes, Alba Sánchez, Laia Soler, Lauro Sumoy, and Elisa Martró on behalf of the Can Ruti SARS-CoV-2 Sequencing Hub |
| EPI_ISL_2998507 | Lighthouse Lab in Glasgow | Wellcome Sanger Institute for the COVID-19 Genomics UK (COG-UK) Consortium | Harper VanSteenhouse, Yumi Kasai, David Gray, Carol Clugston, Anna Dominiczak and Alex Alderton, Roberto Amato, Jeffrey Barrett, Sonia Goncalves, Ewan Harrison, David K. Jackson, Ian Johnston, Dominic Kwiatkowski, Cordelia Langford, John Sillitoe on behalf of the Wellcome Sanger Institute COVID-19 Surveillance Team |
| EPI_ISL_2998508 | Microbiology Department, Laboratori Clínic Metropolitana Nord. Hospital Universitari Germans Trias i Pujol | Can Ruti SARS-CoV-2 Sequencing Hub (HUGTiP/IrsiCaixa/IGTP) | Marc Noguera-Julian, Pilar Armengol, Ignacio Blanco, Antoni E Bordoy, Francesc Catala-Moll, Pere-Joan Cardona, Maria Casadellà, Cristina Casañ, Gemma Clara, Bonaventura Clotet, Montserrat Giménez, Mercedes Guerrero, Anna Not, David Panisello, Roger Paredes, Mariona Parera, Alexia París, Verónica Saludes, Alba Sánchez, Laia Soler, Lauro Sumoy, and Elisa Martró on behalf of the Can Ruti SARS-CoV-2 Sequencing Hub |
| EPI_ISL_2998509, EPI_ISL_2998510 | Lighthouse Laboratory Plymouth | Wellcome Sanger Institute for the COVID-19 Genomics UK (COG-UK) Consortium | Lighthouse Laboratory Plymouth and Alex Alderton, Roberto Amato, Jeffrey Barrett, Sonia Goncalves, Ewan Harrison, David K. Jackson, Ian Johnston, Dominic Kwiatkowski, Cordelia Langford, John Sillitoe on behalf of the Wellcome Sanger Institute COVID-19 Surveillance Team |
| EPI_ISL_2998511 | Lighthouse Lab in Glasgow | Wellcome Sanger Institute for the COVID-19 Genomics UK (COG-UK) Consortium | Harper VanSteenhouse, Yumi Kasai, David Gray, Carol Clugston, Anna Dominiczak and Alex Alderton, Roberto Amato, Jeffrey Barrett, Sonia Goncalves, Ewan Harrison, David K. Jackson, Ian Johnston, Dominic Kwiatkowski, Cordelia Langford, John Sillitoe on behalf of the Wellcome Sanger Institute COVID-19 Surveillance Team |
| EPI_ISL_2998512 | Microbiology Department, Laboratori Clínic Metropolitana Nord. Hospital Universitari Germans Trias i Pujol | Can Ruti SARS-CoV-2 Sequencing Hub (HUGTiP/IrsiCaixa/IGTP) | Marc Noguera-Julian, Pilar Armengol, Ignacio Blanco, Antoni E Bordoy, Francesc Catala-Moll, Pere-Joan Cardona, Maria Casadellà, Cristina Casañ, Gemma Clara, Bonaventura Clotet, Montserrat Giménez, Mercedes Guerrero, Anna Not, David Panisello, Roger Paredes, Mariona Parera, Alexia París, Verónica Saludes, Alba Sánchez, Laia Soler, Lauro Sumoy, and Elisa Martró on behalf of the Can Ruti SARS-CoV-2 Sequencing Hub |
| EPI_ISL_2998513 | Lighthouse Lab in Glasgow | Wellcome Sanger Institute for the COVID-19 Genomics UK (COG-UK) Consortium | Harper VanSteenhouse, Yumi Kasai, David Gray, Carol Clugston, Anna Dominiczak and Alex Alderton, Roberto Amato, Jeffrey Barrett, Sonia Goncalves, Ewan Harrison, David K. Jackson, Ian Johnston, Dominic Kwiatkowski, Cordelia Langford, John Sillitoe on behalf of the Wellcome Sanger Institute COVID-19 Surveillance Team |
| EPI_ISL_2998514 | Lighthouse Laboratory Plymouth | Wellcome Sanger Institute for the COVID-19 Genomics UK (COG-UK) Consortium | Lighthouse Laboratory Plymouth and Alex Alderton, Roberto Amato, Jeffrey Barrett, Sonia Goncalves, Ewan Harrison, David K. Jackson, Ian Johnston, Dominic Kwiatkowski, Cordelia Langford, John Sillitoe on behalf of the Wellcome Sanger Institute COVID-19 Surveillance Team |
| EPI_ISL_2998516 | Lighthouse Lab in Milton Keynes | Wellcome Sanger Institute for the COVID-19 Genomics UK (COG-UK) Consortium | The Lighthouse Lab in Milton Keynes and Alex Alderton, Roberto Amato, Jeffrey Barrett, Sonia Goncalves, Ewan Harrison, David K. Jackson, Ian Johnston, Dominic Kwiatkowski, Cordelia Langford, John Sillitoe on behalf of the Wellcome Sanger Institute COVID-19 Surveillance Team |
| EPI_ISL_2998517 | Lighthouse Laboratory Plymouth | Wellcome Sanger Institute for the COVID-19 Genomics UK (COG-UK) Consortium | Lighthouse Laboratory Plymouth and Alex Alderton, Roberto Amato, Jeffrey Barrett, Sonia Goncalves, Ewan Harrison, David K. Jackson, Ian Johnston, Dominic Kwiatkowski, Cordelia Langford, John Sillitoe on behalf of the Wellcome Sanger Institute COVID-19 Surveillance Team |
| EPI_ISL_2998518 | Lighthouse Lab in Milton Keynes | Wellcome Sanger Institute for the COVID-19 Genomics UK (COG-UK) Consortium | The Lighthouse Lab in Milton Keynes and Alex Alderton, Roberto Amato, Jeffrey Barrett, Sonia Goncalves, Ewan Harrison, David K. Jackson, Ian Johnston, Dominic Kwiatkowski, Cordelia Langford, John Sillitoe on behalf of the Wellcome Sanger Institute COVID-19 Surveillance Team |
| EPI_ISL_2998519 | Microbiology Department, Laboratori Clínic Metropolitana Nord. Hospital Universitari Germans Trias i Pujol | Can Ruti SARS-CoV-2 Sequencing Hub (HUGTiP/IrsiCaixa/IGTP) | Marc Noguera-Julian, Pilar Armengol, Ignacio Blanco, Antoni E Bordoy, Francesc Catala-Moll, Pere-Joan Cardona, Maria Casadellà, Cristina Casañ, Gemma Clara, Bonaventura Clotet, Montserrat Giménez, Mercedes Guerrero, Anna Not, David Panisello, Roger Paredes, Mariona Parera, Alexia París, Verónica Saludes, Alba Sánchez, Laia Soler, Lauro Sumoy, and Elisa Martró on behalf of the Can Ruti SARS-CoV-2 Sequencing Hub |
| EPI_ISL_2998520, | Lighthouse Laboratory Plymouth | Wellcome Sanger Institute for the COVID-19 Genomics UK (COG-UK) | Lighthouse Laboratory Plymouth and Alex Alderton, Roberto Amato, Jeffrey Barrett, Sonia Goncalves, Ewan Harrison, David K. Jackson, Ian Johnston, Dominic Kwiatkowski, Cordelia Langford, John Sillitoe on behalf of the Wellcome |

| | | | |
|---|---|---|---|
| EPI_ISL_2998521 | | Consortium | Institute COVID-19 Surveillance Team |
| EPI_ISL_2998522 | Microbiology Department, Laboratori Clínic Metropolitana Nord. Hospital Universitari Germans Trias i Pujol | Can Ruti SARS-CoV-2 Sequencing Hub (HUGTiP/IrsiCaixa/IGTP) | Marc Noguera-Julian, Pilar Armengol, Ignacio Blanco, Antoni E Bordoy, Francesc Catala-Moll, Pere-Joan Cardona, Maria Casadellà, Cristina Casañ, Gemma Clara, Bonaventura Clotet, Montserrat Giménez, Mercedes Guerrero, Anna Not, David Panisello, Roger Paredes, Mariona Parera, Alexia París, Verónica Saludes, Alba Sánchez, Laia Soler, Lauro Sumoy, and Elisa Martró on behalf of the Can Ruti SARS-CoV-2 Sequencing Hub |
| EPI_ISL_2998523 | Lighthouse Lab in Milton Keynes | Wellcome Sanger Institute for the COVID-19 Genomics UK (COG-UK) Consortium | The Lighthouse Lab in Milton Keynes and Alex Alderton, Roberto Amato, Jeffrey Barrett, Sonia Goncalves, Ewan Harrison, David K. Jackson, Ian Johnston, Dominic Kwiatkowski, Cordelia Langford, John Sillitoe on behalf of the Wellcome Sanger Institute COVID-19 Surveillance Team |
| EPI_ISL_2998524 | Lighthouse Laboratory Plymouth | Wellcome Sanger Institute for the COVID-19 Genomics UK (COG-UK) Consortium | Lighthouse Laboratory Plymouth and Alex Alderton, Roberto Amato, Jeffrey Barrett, Sonia Goncalves, Ewan Harrison, David K. Jackson, Ian Johnston, Dominic Kwiatkowski, Cordelia Langford, John Sillitoe on behalf of the Wellcome Sanger Institute COVID-19 Surveillance Team |
| EPI_ISL_2998526 | Lighthouse Lab in Glasgow | Wellcome Sanger Institute for the COVID-19 Genomics UK (COG-UK) Consortium | Harper VanSteenhouse, Yumi Kasai, David Gray, Carol Clugston, Anna Dominiczak and Alex Alderton, Roberto Amato, Jeffrey Barrett, Sonia Goncalves, Ewan Harrison, David K. Jackson, Ian Johnston, Dominic Kwiatkowski, Cordelia Langford, John Sillitoe on behalf of the Wellcome Sanger Institute COVID-19 Surveillance Team |
| EPI_ISL_2998527 | Lighthouse Laboratory Plymouth | Wellcome Sanger Institute for the COVID-19 Genomics UK (COG-UK) Consortium | Lighthouse Laboratory Plymouth and Alex Alderton, Roberto Amato, Jeffrey Barrett, Sonia Goncalves, Ewan Harrison, David K. Jackson, Ian Johnston, Dominic Kwiatkowski, Cordelia Langford, John Sillitoe on behalf of the Wellcome Sanger Institute COVID-19 Surveillance Team |
| EPI_ISL_2998528 | Lighthouse Lab in Milton Keynes | Wellcome Sanger Institute for the COVID-19 Genomics UK (COG-UK) Consortium | The Lighthouse Lab in Milton Keynes and Alex Alderton, Roberto Amato, Jeffrey Barrett, Sonia Goncalves, Ewan Harrison, David K. Jackson, Ian Johnston, Dominic Kwiatkowski, Cordelia Langford, John Sillitoe on behalf of the Wellcome Sanger Institute COVID-19 Surveillance Team |
| EPI_ISL_2998529 | Microbiology Department, Laboratori Clínic Metropolitana Nord. Hospital Universitari Germans Trias i Pujol | Can Ruti SARS-CoV-2 Sequencing Hub (HUGTiP/IrsiCaixa/IGTP) | Marc Noguera-Julian, Pilar Armengol, Ignacio Blanco, Antoni E Bordoy, Francesc Catala-Moll, Pere-Joan Cardona, Maria Casadellà, Cristina Casañ, Gemma Clara, Bonaventura Clotet, Montserrat Giménez, Mercedes Guerrero, Anna Not, David Panisello, Roger Paredes, Mariona Parera, Alexia París, Verónica Saludes, Alba Sánchez, Laia Soler, Lauro Sumoy, and Elisa Martró on behalf of the Can Ruti SARS-CoV-2 Sequencing Hub |
| EPI_ISL_2998530, EPI_ISL_2998531 | Lighthouse Laboratory Plymouth | Wellcome Sanger Institute for the COVID-19 Genomics UK (COG-UK) Consortium | Lighthouse Laboratory Plymouth and Alex Alderton, Roberto Amato, Jeffrey Barrett, Sonia Goncalves, Ewan Harrison, David K. Jackson, Ian Johnston, Dominic Kwiatkowski, Cordelia Langford, John Sillitoe on behalf of the Wellcome Sanger Institute COVID-19 Surveillance Team |
| EPI_ISL_2998532 | Lighthouse Lab in Milton Keynes | Wellcome Sanger Institute for the COVID-19 Genomics UK (COG-UK) Consortium | The Lighthouse Lab in Milton Keynes and Alex Alderton, Roberto Amato, Jeffrey Barrett, Sonia Goncalves, Ewan Harrison, David K. Jackson, Ian Johnston, Dominic Kwiatkowski, Cordelia Langford, John Sillitoe on behalf of the Wellcome Sanger Institute COVID-19 Surveillance Team |
| EPI_ISL_2998533 | Microbiologia CATLAB | Can Ruti SARS-CoV-2 Sequencing Hub (HUGTiP/IrsiCaixa/IGTP) | Marc Noguera-Julian, Pilar Armengol, Ignacio Blanco, Antoni E Bordoy, Francesc Catala-Moll, Pere-Joan Cardona, Maria Casadellà, Cristina Casañ, Gemma Clara, Bonaventura Clotet, Montserrat Giménez, Mercedes Guerrero, Anna Not, David Panisello, Roger Paredes, Mariona Parera, Alexia París, Verónica Saludes, Alba Sánchez, Laia Soler, Lauro Sumoy, and Elisa Martró on behalf of the Can Ruti SARS-CoV-2 Sequencing Hub |
| EPI_ISL_2998534, EPI_ISL_2998535 | Lighthouse Laboratory Plymouth | Wellcome Sanger Institute for the COVID-19 Genomics UK (COG-UK) Consortium | Lighthouse Laboratory Plymouth and Alex Alderton, Roberto Amato, Jeffrey Barrett, Sonia Goncalves, Ewan Harrison, David K. Jackson, Ian Johnston, Dominic Kwiatkowski, Cordelia Langford, John Sillitoe on behalf of the Wellcome Sanger Institute COVID-19 Surveillance Team |
| EPI_ISL_2998537 | Lighthouse Lab in Glasgow | Wellcome Sanger Institute for the COVID-19 Genomics UK (COG-UK) Consortium | Harper VanSteenhouse, Yumi Kasai, David Gray, Carol Clugston, Anna Dominiczak and Alex Alderton, Roberto Amato, Jeffrey Barrett, Sonia Goncalves, Ewan Harrison, David K. Jackson, Ian Johnston, Dominic Kwiatkowski, Cordelia Langford, John Sillitoe on behalf of the Wellcome Sanger Institute COVID-19 Surveillance Team |
| EPI_ISL_2998538 | Lighthouse Lab in Milton Keynes | Wellcome Sanger Institute for the COVID-19 Genomics UK (COG-UK) Consortium | The Lighthouse Lab in Milton Keynes and Alex Alderton, Roberto Amato, Jeffrey Barrett, Sonia Goncalves, Ewan Harrison, David K. Jackson, Ian Johnston, Dominic Kwiatkowski, Cordelia Langford, John Sillitoe on behalf of the Wellcome Sanger Institute COVID-19 Surveillance Team |
| EPI_ISL_2998539 | Microbiologia CATLAB | Can Ruti SARS-CoV-2 Sequencing Hub (HUGTiP/IrsiCaixa/IGTP) | Marc Noguera-Julian, Pilar Armengol, Ignacio Blanco, Antoni E Bordoy, Francesc Catala-Moll, Pere-Joan Cardona, Maria Casadellà, Cristina Casañ, Gemma Clara, Bonaventura Clotet, Montserrat Giménez, Mercedes Guerrero, Anna Not, David Panisello, Roger Paredes, Mariona Parera, Alexia París, Verónica Saludes, Alba Sánchez, Laia Soler, Lauro Sumoy, and Elisa Martró on behalf of the Can Ruti SARS-CoV-2 Sequencing Hub |
| EPI_ISL_2998540 | Lighthouse Laboratory Plymouth | Wellcome Sanger Institute for the COVID-19 Genomics UK (COG-UK) Consortium | Lighthouse Laboratory Plymouth and Alex Alderton, Roberto Amato, Jeffrey Barrett, Sonia Goncalves, Ewan Harrison, David K. Jackson, Ian Johnston, Dominic Kwiatkowski, Cordelia Langford, John Sillitoe on behalf of the Wellcome Sanger Institute COVID-19 Surveillance Team |
| EPI_ISL_2998541 | Lighthouse Lab in Milton Keynes | Wellcome Sanger Institute for the COVID-19 Genomics UK (COG-UK) Consortium | The Lighthouse Lab in Milton Keynes and Alex Alderton, Roberto Amato, Jeffrey Barrett, Sonia Goncalves, Ewan Harrison, David K. Jackson, Ian Johnston, Dominic Kwiatkowski, Cordelia Langford, John Sillitoe on behalf of the Wellcome Sanger Institute COVID-19 Surveillance Team |
| EPI_ISL_2998542 | Microbiology Department, Laboratori Clínic Metropolitana Nord. Hospital Universitari Germans Trias i Pujol | Can Ruti SARS-CoV-2 Sequencing Hub (HUGTiP/IrsiCaixa/IGTP) | Marc Noguera-Julian, Pilar Armengol, Ignacio Blanco, Antoni E Bordoy, Francesc Catala-Moll, Pere-Joan Cardona, Maria Casadellà, Cristina Casañ, Gemma Clara, Bonaventura Clotet, Montserrat Giménez, Mercedes Guerrero, Anna Not, David Panisello, Roger Paredes, Mariona Parera, Alexia París, Verónica Saludes, Alba Sánchez, Laia Soler, Lauro Sumoy, and Elisa Martró on behalf of the Can Ruti SARS-CoV-2 Sequencing Hub |
| EPI_ISL_2998543 | Lighthouse Lab in Milton Keynes | Wellcome Sanger Institute for the COVID-19 Genomics UK (COG-UK) Consortium | The Lighthouse Lab in Milton Keynes and Alex Alderton, Roberto Amato, Jeffrey Barrett, Sonia Goncalves, Ewan Harrison, David K. Jackson, Ian Johnston, Dominic Kwiatkowski, Cordelia Langford, John Sillitoe on behalf of the Wellcome Sanger Institute COVID-19 Surveillance Team |
| EPI_ISL_2998544 | Lighthouse Lab in Glasgow | Wellcome Sanger Institute for the COVID-19 Genomics UK (COG-UK) Consortium | Harper VanSteenhouse, Yumi Kasai, David Gray, Carol Clugston, Anna Dominiczak and Alex Alderton, Roberto Amato, Jeffrey Barrett, Sonia Goncalves, Ewan Harrison, David K. Jackson, Ian Johnston, Dominic Kwiatkowski, Cordelia Langford, John Sillitoe on behalf of the Wellcome Sanger Institute COVID-19 Surveillance Team |
| EPI_ISL_2998545 | Lighthouse Laboratory Plymouth | Wellcome Sanger Institute for the COVID-19 Genomics UK (COG-UK) Consortium | Lighthouse Laboratory Plymouth and Alex Alderton, Roberto Amato, Jeffrey Barrett, Sonia Goncalves, Ewan Harrison, David K. Jackson, Ian Johnston, Dominic Kwiatkowski, Cordelia Langford, John Sillitoe on behalf of the Wellcome Sanger Institute COVID-19 Surveillance Team |
| EPI_ISL_2998546 | Microbiology Department, Laboratori Clínic Metropolitana Nord. Hospital Universitari Germans Trias i Pujol | Can Ruti SARS-CoV-2 Sequencing Hub (HUGTiP/IrsiCaixa/IGTP) | Marc Noguera-Julian, Pilar Armengol, Ignacio Blanco, Antoni E Bordoy, Francesc Catala-Moll, Pere-Joan Cardona, Maria Casadellà, Cristina Casañ, Gemma Clara, Bonaventura Clotet, Montserrat Giménez, Mercedes Guerrero, Anna Not, David Panisello, Roger Paredes, Mariona Parera, Alexia París, Verónica Saludes, Alba Sánchez, Laia Soler, Lauro Sumoy, and Elisa Martró on behalf of the Can Ruti SARS-CoV-2 Sequencing Hub |
| EPI_ISL_2998547 | Lighthouse Lab in Glasgow | Wellcome Sanger Institute for the COVID-19 Genomics UK (COG-UK) Consortium | Harper VanSteenhouse, Yumi Kasai, David Gray, Carol Clugston, Anna Dominiczak and Alex Alderton, Roberto Amato, Jeffrey Barrett, Sonia Goncalves, Ewan Harrison, David K. Jackson, Ian Johnston, Dominic Kwiatkowski, Cordelia Langford, John Sillitoe on behalf of the Wellcome Sanger Institute COVID-19 Surveillance Team |
| EPI_ISL_2998548 | Lighthouse Laboratory Plymouth | Wellcome Sanger Institute for the COVID-19 Genomics UK (COG-UK) Consortium | Lighthouse Laboratory Plymouth and Alex Alderton, Roberto Amato, Jeffrey Barrett, Sonia Goncalves, Ewan Harrison, David K. Jackson, Ian Johnston, Dominic Kwiatkowski, Cordelia Langford, John Sillitoe on behalf of the Wellcome Sanger Institute COVID-19 Surveillance Team |
| EPI_ISL_2998549 | Microbiology Department, Laboratori Clínic Metropolitana Nord. Hospital Universitari Germans Trias i Pujol | Can Ruti SARS-CoV-2 Sequencing Hub (HUGTiP/IrsiCaixa/IGTP) | Marc Noguera-Julian, Pilar Armengol, Ignacio Blanco, Antoni E Bordoy, Francesc Catala-Moll, Pere-Joan Cardona, Maria Casadellà, Cristina Casañ, Gemma Clara, Bonaventura Clotet, Montserrat Giménez, Mercedes Guerrero, Anna Not, David Panisello, Roger Paredes, Mariona Parera, Alexia París, Verónica Saludes, Alba Sánchez, Laia Soler, Lauro Sumoy, and Elisa Martró on behalf of the Can Ruti SARS-CoV-2 Sequencing Hub |
| EPI_ISL_2998550, EPI_ISL_2998551 | Lighthouse Lab in Milton Keynes | Wellcome Sanger Institute for the COVID-19 Genomics UK (COG-UK) Consortium | The Lighthouse Lab in Milton Keynes and Alex Alderton, Roberto Amato, Jeffrey Barrett, Sonia Goncalves, Ewan Harrison, David K. Jackson, Ian Johnston, Dominic Kwiatkowski, Cordelia Langford, John Sillitoe on behalf of the Wellcome Sanger Institute COVID-19 Surveillance Team |
| EPI_ISL_2998552 | Lighthouse Laboratory Plymouth | Wellcome Sanger Institute for the COVID-19 Genomics UK (COG-UK) Consortium | Lighthouse Laboratory Plymouth and Alex Alderton, Roberto Amato, Jeffrey Barrett, Sonia Goncalves, Ewan Harrison, David K. Jackson, Ian Johnston, Dominic Kwiatkowski, Cordelia Langford, John Sillitoe on behalf of the Wellcome Sanger Institute COVID-19 Surveillance Team |
| EPI_ISL_2998553 | Microbiology Department, Laboratori Clínic Metropolitana Nord. Hospital Universitari Germans Trias i Pujol | Can Ruti SARS-CoV-2 Sequencing Hub (HUGTiP/IrsiCaixa/IGTP) | Marc Noguera-Julian, Pilar Armengol, Ignacio Blanco, Antoni E Bordoy, Francesc Catala-Moll, Pere-Joan Cardona, Maria Casadellà, Cristina Casañ, Gemma Clara, Bonaventura Clotet, Montserrat Giménez, Mercedes Guerrero, Anna Not, David Panisello, Roger Paredes, Mariona Parera, Alexia París, Verónica Saludes, Alba Sánchez, Laia Soler, Lauro Sumoy, and Elisa Martró on behalf of the Can Ruti SARS-CoV-2 Sequencing Hub |
| EPI_ISL_2998554 | Lighthouse Lab in Milton Keynes | Wellcome Sanger Institute for the COVID-19 Genomics UK (COG-UK) Consortium | The Lighthouse Lab in Milton Keynes and Alex Alderton, Roberto Amato, Jeffrey Barrett, Sonia Goncalves, Ewan Harrison, David K. Jackson, Ian Johnston, Dominic Kwiatkowski, Cordelia Langford, John Sillitoe on behalf of the Wellcome Sanger Institute COVID-19 Surveillance Team |
| EPI_ISL_2998555, EPI_ISL_2998557, EPI_ISL_2998558 | Lighthouse Laboratory Plymouth | Wellcome Sanger Institute for the COVID-19 Genomics UK (COG-UK) Consortium | Lighthouse Laboratory Plymouth and Alex Alderton, Roberto Amato, Jeffrey Barrett, Sonia Goncalves, Ewan Harrison, David K. Jackson, Ian Johnston, Dominic Kwiatkowski, Cordelia Langford, John Sillitoe on behalf of the Wellcome Sanger Institute COVID-19 Surveillance Team |
| EPI_ISL_2998559 | Microbiology Department, Laboratori Clínic Metropolitana Nord. Hospital Universitari Germans Trias i Pujol | Can Ruti SARS-CoV-2 Sequencing Hub (HUGTiP/IrsiCaixa/IGTP) | Marc Noguera-Julian, Pilar Armengol, Ignacio Blanco, Antoni E Bordoy, Francesc Catala-Moll, Pere-Joan Cardona, Maria Casadellà, Cristina Casañ, Gemma Clara, Bonaventura Clotet, Montserrat Giménez, Mercedes Guerrero, Anna Not, David Panisello, Roger Paredes, Mariona Parera, Alexia París, Verónica Saludes, Alba Sánchez, Laia Soler, Lauro Sumoy, and Elisa Martró on behalf of the Can Ruti SARS-CoV-2 Sequencing Hub |
| EPI_ISL_2998560, EPI_ISL_2998561, EPI_ISL_2998562 | Lighthouse Laboratory Plymouth | Wellcome Sanger Institute for the COVID-19 Genomics UK (COG-UK) Consortium | Lighthouse Laboratory Plymouth and Alex Alderton, Roberto Amato, Jeffrey Barrett, Sonia Goncalves, Ewan Harrison, David K. Jackson, Ian Johnston, Dominic Kwiatkowski, Cordelia Langford, John Sillitoe on behalf of the Wellcome Sanger Institute COVID-19 Surveillance Team |
| EPI_ISL_2998563 | Microbiology Department, Laboratori Clínic Metropolitana Nord. Hospital Universitari Germans Trias i Pujol | Can Ruti SARS-CoV-2 Sequencing Hub (HUGTiP/IrsiCaixa/IGTP) | Marc Noguera-Julian, Pilar Armengol, Ignacio Blanco, Antoni E Bordoy, Francesc Catala-Moll, Pere-Joan Cardona, Maria Casadellà, Cristina Casañ, Gemma Clara, Bonaventura Clotet, Montserrat Giménez, Mercedes Guerrero, Anna Not, David Panisello, Roger Paredes, Mariona Parera, Alexia París, Verónica Saludes, Alba Sánchez, Laia Soler, Lauro Sumoy, and Elisa Martró on behalf of the Can Ruti SARS-CoV-2 Sequencing Hub |
| EPI_ISL_2998564, EPI_ISL_2998565, EPI_ISL_2998566 | Lighthouse Lab in Glasgow | Wellcome Sanger Institute for the COVID-19 Genomics UK (COG-UK) Consortium | Harper VanSteenhouse, Yumi Kasai, David Gray, Carol Clugston, Anna Dominiczak and Alex Alderton, Roberto Amato, Jeffrey Barrett, Sonia Goncalves, Ewan Harrison, David K. Jackson, Ian Johnston, Dominic Kwiatkowski, Cordelia Langford, John Sillitoe on behalf of the Wellcome Sanger Institute COVID-19 Surveillance Team |
| EPI_ISL_2998567 | Microbiology Department, Laboratori Clínic Metropolitana Nord. Hospital Universitari Germans Trias i Pujol | Can Ruti SARS-CoV-2 Sequencing Hub (HUGTiP/IrsiCaixa/IGTP) | Marc Noguera-Julian, Pilar Armengol, Ignacio Blanco, Antoni E Bordoy, Francesc Catala-Moll, Pere-Joan Cardona, Maria Casadellà, Cristina Casañ, Gemma Clara, Bonaventura Clotet, Montserrat Giménez, Mercedes Guerrero, Anna Not, David Panisello, Roger Paredes, Mariona Parera, Alexia París, Verónica Saludes, Alba Sánchez, Laia Soler, Lauro Sumoy, and Elisa Martró on behalf of the Can Ruti SARS-CoV-2 Sequencing Hub |
| EPI_ISL_2998568 | Lighthouse Lab in Milton Keynes | Wellcome Sanger Institute for the COVID-19 Genomics UK (COG-UK) Consortium | The Lighthouse Lab in Milton Keynes and Alex Alderton, Roberto Amato, Jeffrey Barrett, Sonia Goncalves, Ewan Harrison, David K. Jackson, Ian Johnston, Dominic Kwiatkowski, Cordelia Langford, John Sillitoe on behalf of the Wellcome Sanger Institute COVID-19 Surveillance Team |
| EPI_ISL_2998569 | Lighthouse Laboratory Plymouth | Wellcome Sanger Institute for the COVID-19 Genomics UK (COG-UK) Consortium | Lighthouse Laboratory Plymouth and Alex Alderton, Roberto Amato, Jeffrey Barrett, Sonia Goncalves, Ewan Harrison, David K. Jackson, Ian Johnston, Dominic Kwiatkowski, Cordelia Langford, John Sillitoe on behalf of the Wellcome Sanger Institute COVID-19 Surveillance Team |
| EPI_ISL_2998570 | Lighthouse Lab in Milton Keynes | Wellcome Sanger Institute for the COVID-19 Genomics UK (COG-UK) Consortium | The Lighthouse Lab in Milton Keynes and Alex Alderton, Roberto Amato, Jeffrey Barrett, Sonia Goncalves, Ewan Harrison, David K. Jackson, Ian Johnston, Dominic Kwiatkowski, Cordelia Langford, John Sillitoe on behalf of the Wellcome Sanger Institute COVID-19 Surveillance Team |
| EPI_ISL_2998572 | Lighthouse Laboratory Plymouth | Wellcome Sanger Institute for the COVID-19 Genomics UK (COG-UK) Consortium | Lighthouse Laboratory Plymouth and Alex Alderton, Roberto Amato, Jeffrey Barrett, Sonia Goncalves, Ewan Harrison, David K. Jackson, Ian Johnston, Dominic Kwiatkowski, Cordelia Langford, John Sillitoe on behalf of the Wellcome Sanger Institute COVID-19 Surveillance Team |
| EPI_ISL_2998573 | Lighthouse Lab in Milton Keynes | Wellcome Sanger Institute for the COVID-19 Genomics UK (COG-UK) Consortium | The Lighthouse Lab in Milton Keynes and Alex Alderton, Roberto Amato, Jeffrey Barrett, Sonia Goncalves, Ewan Harrison, David K. Jackson, Ian Johnston, Dominic Kwiatkowski, Cordelia Langford, John Sillitoe on behalf of the Wellcome Sanger Institute COVID-19 Surveillance Team |
| EPI_ISL_2998574 | Lighthouse Lab in Glasgow | Wellcome Sanger Institute for the COVID-19 Genomics UK (COG-UK) Consortium | Harper VanSteenhouse, Yumi Kasai, David Gray, Carol Clugston, Anna Dominiczak and Alex Alderton, Roberto Amato, Jeffrey Barrett, Sonia Goncalves, Ewan Harrison, David K. Jackson, Ian Johnston, Dominic Kwiatkowski, Cordelia Langford, John Sillitoe on behalf of the Wellcome Sanger Institute COVID-19 Surveillance Team |
| EPI_ISL_2998575 | Microbiologia CATLAB | Can Ruti SARS-CoV-2 Sequencing Hub (HUGTiP/IrsiCaixa/IGTP) | Marc Noguera-Julian, Pilar Armengol, Ignacio Blanco, Antoni E Bordoy, Francesc Catala-Moll, Pere-Joan Cardona, Maria Casadellà, Cristina Casañ, Gemma Clara, Bonaventura Clotet, Montserrat Giménez, Mercedes Guerrero, Anna Not, David Panisello, Roger Paredes, Mariona Parera, Alexia París, Verónica Saludes, Alba Sánchez, Laia Soler, Lauro Sumoy, and Elisa Martró on behalf of the Can Ruti SARS-CoV-2 Sequencing Hub |
| EPI_ISL_2998576, EPI_ISL_2998577 | Lighthouse Laboratory Plymouth | Wellcome Sanger Institute for the COVID-19 Genomics UK (COG-UK) Consortium | Lighthouse Laboratory Plymouth and Alex Alderton, Roberto Amato, Jeffrey Barrett, Sonia Goncalves, Ewan Harrison, David K. Jackson, Ian Johnston, Dominic Kwiatkowski, Cordelia Langford, John Sillitoe on behalf of the Wellcome Sanger Institute COVID-19 Surveillance Team |
| EPI_ISL_2998578 | Microbiologia CATLAB | Can Ruti SARS-CoV-2 Sequencing Hub (HUGTiP/IrsiCaixa/IGTP) | Marc Noguera-Julian, Pilar Armengol, Ignacio Blanco, Antoni E Bordoy, Francesc Catala-Moll, Pere-Joan Cardona, Maria Casadellà, Cristina Casañ, Gemma Clara, Bonaventura Clotet, Montserrat Giménez, Mercedes Guerrero, Anna Not, David Panisello, Roger Paredes, Mariona Parera, Alexia París, Verónica Saludes, Alba Sánchez, Laia Soler, Lauro Sumoy, and Elisa Martró on behalf of the Can Ruti SARS-CoV-2 Sequencing Hub |

| | | | |
|---|---|---|---|
| EPI_ISL_2998579, EPI_ISL_2998580 | Lighthouse Lab in Milton Keynes | Wellcome Sanger Institute for the COVID-19 Genomics UK (COG-UK) Consortium | The Lighthouse Lab in Milton Keynes and Alex Alderton, Roberto Amato, Jeffrey Barrett, Sonia Goncalves, Ewan Harrison, David K. Jackson, Ian Johnston, Dominic Kwiatkowski, Cordelia Langford, John Sillitoe on behalf of the Wellcome Sanger Institute COVID-19 Surveillance Team |
| EPI_ISL_2998581 | Microbiology Department, Laboratori Clínic Metropolitana Nord. Hospital Universitari Germans Trias i Pujol | Can Ruti SARS-CoV-2 Sequencing Hub (HUGTiP/IrsiCaixa/IGTP) | Marc Noguera-Julian, Pilar Armengol, Ignacio Blanco, Antoni E Bordoy, Francesc Catala-Moll, Pere-Joan Cardona, Maria Casadellà, Cristina Casañ, Gemma Clara, Bonaventura Clotet, Montserrat Giménez, Mercedes Guerrero, Anna Not, David Panisello, Roger Paredes, Mariona Parera, Alexia París, Verónica Saludes, Alba Sánchez, Laia Soler, Lauro Sumoy, and Elisa Martró on behalf of the Can Ruti SARS-CoV-2 Sequencing Hub |
| EPI_ISL_2998582 | Lighthouse Lab in Milton Keynes | Wellcome Sanger Institute for the COVID-19 Genomics UK (COG-UK) Consortium | The Lighthouse Lab in Milton Keynes and Alex Alderton, Roberto Amato, Jeffrey Barrett, Sonia Goncalves, Ewan Harrison, David K. Jackson, Ian Johnston, Dominic Kwiatkowski, Cordelia Langford, John Sillitoe on behalf of the Wellcome Sanger Institute COVID-19 Surveillance Team |
| EPI_ISL_2998583 | Lighthouse Laboratory Plymouth | Wellcome Sanger Institute for the COVID-19 Genomics UK (COG-UK) Consortium | Lighthouse Laboratory Plymouth and Alex Alderton, Roberto Amato, Jeffrey Barrett, Sonia Goncalves, Ewan Harrison, David K. Jackson, Ian Johnston, Dominic Kwiatkowski, Cordelia Langford, John Sillitoe on behalf of the Wellcome Sanger Institute COVID-19 Surveillance Team |
| EPI_ISL_2998584 | Microbiology Department, Laboratori Clínic Metropolitana Nord. Hospital Universitari Germans Trias i Pujol | Can Ruti SARS-CoV-2 Sequencing Hub (HUGTiP/IrsiCaixa/IGTP) | Marc Noguera-Julian, Pilar Armengol, Ignacio Blanco, Antoni E Bordoy, Francesc Catala-Moll, Pere-Joan Cardona, Maria Casadellà, Cristina Casañ, Gemma Clara, Bonaventura Clotet, Montserrat Giménez, Mercedes Guerrero, Anna Not, David Panisello, Roger Paredes, Mariona Parera, Alexia París, Verónica Saludes, Alba Sánchez, Laia Soler, Lauro Sumoy, and Elisa Martró on behalf of the Can Ruti SARS-CoV-2 Sequencing Hub |
| EPI_ISL_2998585 | Lighthouse Laboratory Plymouth | Wellcome Sanger Institute for the COVID-19 Genomics UK (COG-UK) Consortium | Lighthouse Laboratory Plymouth and Alex Alderton, Roberto Amato, Jeffrey Barrett, Sonia Goncalves, Ewan Harrison, David K. Jackson, Ian Johnston, Dominic Kwiatkowski, Cordelia Langford, John Sillitoe on behalf of the Wellcome Sanger Institute COVID-19 Surveillance Team |
| EPI_ISL_2998586, EPI_ISL_2998587 | Lighthouse Lab in Glasgow | Wellcome Sanger Institute for the COVID-19 Genomics UK (COG-UK) Consortium | Harper VanSteenhouse, Yumi Kasai, David Gray, Carol Clugston, Anna Dominiczak and Alex Alderton, Roberto Amato, Jeffrey Barrett, Sonia Goncalves, Ewan Harrison, David K. Jackson, Ian Johnston, Dominic Kwiatkowski, Cordelia Langford, John Sillitoe on behalf of the Wellcome Sanger Institute COVID-19 Surveillance Team |
| EPI_ISL_2998588 | Microbiology Department, Laboratori Clínic Metropolitana Nord. Hospital Universitari Germans Trias i Pujol | Can Ruti SARS-CoV-2 Sequencing Hub (HUGTiP/IrsiCaixa/IGTP) | Marc Noguera-Julian, Pilar Armengol, Ignacio Blanco, Antoni E Bordoy, Francesc Catala-Moll, Pere-Joan Cardona, Maria Casadellà, Cristina Casañ, Gemma Clara, Bonaventura Clotet, Montserrat Giménez, Mercedes Guerrero, Anna Not, David Panisello, Roger Paredes, Mariona Parera, Alexia París, Verónica Saludes, Alba Sánchez, Laia Soler, Lauro Sumoy, and Elisa Martró on behalf of the Can Ruti SARS-CoV-2 Sequencing Hub |
| EPI_ISL_2998589, EPI_ISL_2998590, EPI_ISL_2998591 | Lighthouse Laboratory Plymouth | Wellcome Sanger Institute for the COVID-19 Genomics UK (COG-UK) Consortium | Lighthouse Laboratory Plymouth and Alex Alderton, Roberto Amato, Jeffrey Barrett, Sonia Goncalves, Ewan Harrison, David K. Jackson, Ian Johnston, Dominic Kwiatkowski, Cordelia Langford, John Sillitoe on behalf of the Wellcome Sanger Institute COVID-19 Surveillance Team |
| EPI_ISL_2998592 | Microbiology Department, Laboratori Clínic Metropolitana Nord. Hospital Universitari Germans Trias i Pujol | Can Ruti SARS-CoV-2 Sequencing Hub (HUGTiP/IrsiCaixa/IGTP) | Marc Noguera-Julian, Pilar Armengol, Ignacio Blanco, Antoni E Bordoy, Francesc Catala-Moll, Pere-Joan Cardona, Maria Casadellà, Cristina Casañ, Gemma Clara, Bonaventura Clotet, Montserrat Giménez, Mercedes Guerrero, Anna Not, David Panisello, Roger Paredes, Mariona Parera, Alexia París, Verónica Saludes, Alba Sánchez, Laia Soler, Lauro Sumoy, and Elisa Martró on behalf of the Can Ruti SARS-CoV-2 Sequencing Hub |
| EPI_ISL_2998593 | Lighthouse Lab in Milton Keynes | Wellcome Sanger Institute for the COVID-19 Genomics UK (COG-UK) Consortium | The Lighthouse Lab in Milton Keynes and Alex Alderton, Roberto Amato, Jeffrey Barrett, Sonia Goncalves, Ewan Harrison, David K. Jackson, Ian Johnston, Dominic Kwiatkowski, Cordelia Langford, John Sillitoe on behalf of the Wellcome Sanger Institute COVID-19 Surveillance Team |
| EPI_ISL_2998594 | Lighthouse Laboratory Plymouth | Wellcome Sanger Institute for the COVID-19 Genomics UK (COG-UK) Consortium | Lighthouse Laboratory Plymouth and Alex Alderton, Roberto Amato, Jeffrey Barrett, Sonia Goncalves, Ewan Harrison, David K. Jackson, Ian Johnston, Dominic Kwiatkowski, Cordelia Langford, John Sillitoe on behalf of the Wellcome Sanger Institute COVID-19 Surveillance Team |
| EPI_ISL_2998595 | Microbiology Department, Laboratori Clínic Metropolitana Nord. Hospital Universitari Germans Trias i Pujol | Can Ruti SARS-CoV-2 Sequencing Hub (HUGTiP/IrsiCaixa/IGTP) | Marc Noguera-Julian, Pilar Armengol, Ignacio Blanco, Antoni E Bordoy, Francesc Catala-Moll, Pere-Joan Cardona, Maria Casadellà, Cristina Casañ, Gemma Clara, Bonaventura Clotet, Montserrat Giménez, Mercedes Guerrero, Anna Not, David Panisello, Roger Paredes, Mariona Parera, Alexia París, Verónica Saludes, Alba Sánchez, Laia Soler, Lauro Sumoy, and Elisa Martró on behalf of the Can Ruti SARS-CoV-2 Sequencing Hub |
| EPI_ISL_2998596, EPI_ISL_2998597, EPI_ISL_2998598 | Lighthouse Laboratory Plymouth | Wellcome Sanger Institute for the COVID-19 Genomics UK (COG-UK) Consortium | Lighthouse Laboratory Plymouth and Alex Alderton, Roberto Amato, Jeffrey Barrett, Sonia Goncalves, Ewan Harrison, David K. Jackson, Ian Johnston, Dominic Kwiatkowski, Cordelia Langford, John Sillitoe on behalf of the Wellcome Sanger Institute COVID-19 Surveillance Team |
| EPI_ISL_2998599 | Microbiology Department, Laboratori Clínic Metropolitana Nord. Hospital Universitari Germans Trias i Pujol | Can Ruti SARS-CoV-2 Sequencing Hub (HUGTiP/IrsiCaixa/IGTP) | Marc Noguera-Julian, Pilar Armengol, Ignacio Blanco, Antoni E Bordoy, Francesc Catala-Moll, Pere-Joan Cardona, Maria Casadellà, Cristina Casañ, Gemma Clara, Bonaventura Clotet, Montserrat Giménez, Mercedes Guerrero, Anna Not, David Panisello, Roger Paredes, Mariona Parera, Alexia París, Verónica Saludes, Alba Sánchez, Laia Soler, Lauro Sumoy, and Elisa Martró on behalf of the Can Ruti SARS-CoV-2 Sequencing Hub |
| EPI_ISL_2998600, EPI_ISL_2998601, EPI_ISL_2998602, EPI_ISL_2998603, EPI_ISL_2998604 | Lighthouse Laboratory Plymouth | Wellcome Sanger Institute for the COVID-19 Genomics UK (COG-UK) Consortium | Lighthouse Laboratory Plymouth and Alex Alderton, Roberto Amato, Jeffrey Barrett, Sonia Goncalves, Ewan Harrison, David K. Jackson, Ian Johnston, Dominic Kwiatkowski, Cordelia Langford, John Sillitoe on behalf of the Wellcome Sanger Institute COVID-19 Surveillance Team |
| EPI_ISL_2998605 | Microbiology Department, Laboratori Clínic Metropolitana Nord. Hospital Universitari Germans Trias i Pujol | Can Ruti SARS-CoV-2 Sequencing Hub (HUGTiP/IrsiCaixa/IGTP) | Marc Noguera-Julian, Pilar Armengol, Ignacio Blanco, Antoni E Bordoy, Francesc Catala-Moll, Pere-Joan Cardona, Maria Casadellà, Cristina Casañ, Gemma Clara, Bonaventura Clotet, Montserrat Giménez, Mercedes Guerrero, Anna Not, David Panisello, Roger Paredes, Mariona Parera, Alexia París, Verónica Saludes, Alba Sánchez, Laia Soler, Lauro Sumoy, and Elisa Martró on behalf of the Can Ruti SARS-CoV-2 Sequencing Hub |
| EPI_ISL_2998606 | Lighthouse Lab in Glasgow | Wellcome Sanger Institute for the COVID-19 Genomics UK (COG-UK) Consortium | Harper VanSteenhouse, Yumi Kasai, David Gray, Carol Clugston, Anna Dominiczak and Alex Alderton, Roberto Amato, Jeffrey Barrett, Sonia Goncalves, Ewan Harrison, David K. Jackson, Ian Johnston, Dominic Kwiatkowski, Cordelia Langford, John Sillitoe on behalf of the Wellcome Sanger Institute COVID-19 Surveillance Team |
| EPI_ISL_2998607 | Lighthouse Laboratory Plymouth | Wellcome Sanger Institute for the COVID-19 Genomics UK (COG-UK) Consortium | Lighthouse Laboratory Plymouth and Alex Alderton, Roberto Amato, Jeffrey Barrett, Sonia Goncalves, Ewan Harrison, David K. Jackson, Ian Johnston, Dominic Kwiatkowski, Cordelia Langford, John Sillitoe on behalf of the Wellcome Sanger Institute COVID-19 Surveillance Team |
| EPI_ISL_2998608 | Microbiology Department, Laboratori Clínic Metropolitana Nord. Hospital Universitari Germans Trias i Pujol | Can Ruti SARS-CoV-2 Sequencing Hub (HUGTiP/IrsiCaixa/IGTP) | Marc Noguera-Julian, Pilar Armengol, Ignacio Blanco, Antoni E Bordoy, Francesc Catala-Moll, Pere-Joan Cardona, Maria Casadellà, Cristina Casañ, Gemma Clara, Bonaventura Clotet, Montserrat Giménez, Mercedes Guerrero, Anna Not, David Panisello, Roger Paredes, Mariona Parera, Alexia París, Verónica Saludes, Alba Sánchez, Laia Soler, Lauro Sumoy, and Elisa Martró on behalf of the Can Ruti SARS-CoV-2 Sequencing Hub |
| EPI_ISL_2998609, EPI_ISL_2998610, EPI_ISL_2998611 | Lighthouse Laboratory Plymouth | Wellcome Sanger Institute for the COVID-19 Genomics UK (COG-UK) Consortium | Lighthouse Laboratory Plymouth and Alex Alderton, Roberto Amato, Jeffrey Barrett, Sonia Goncalves, Ewan Harrison, David K. Jackson, Ian Johnston, Dominic Kwiatkowski, Cordelia Langford, John Sillitoe on behalf of the Wellcome Sanger Institute COVID-19 Surveillance Team |
| EPI_ISL_2998612 | Microbiology Department, Laboratori Clínic Metropolitana Nord. Hospital Universitari Germans Trias i Pujol | Can Ruti SARS-CoV-2 Sequencing Hub (HUGTiP/IrsiCaixa/IGTP) | Marc Noguera-Julian, Pilar Armengol, Ignacio Blanco, Antoni E Bordoy, Francesc Catala-Moll, Pere-Joan Cardona, Maria Casadellà, Cristina Casañ, Gemma Clara, Bonaventura Clotet, Montserrat Giménez, Mercedes Guerrero, Anna Not, David Panisello, Roger Paredes, Mariona Parera, Alexia París, Verónica Saludes, Alba Sánchez, Laia Soler, Lauro Sumoy, and Elisa Martró on behalf of the Can Ruti SARS-CoV-2 Sequencing Hub |
| EPI_ISL_2998613 | Lighthouse Lab in Milton Keynes | Wellcome Sanger Institute for the COVID-19 Genomics UK (COG-UK) Consortium | The Lighthouse Lab in Milton Keynes and Alex Alderton, Roberto Amato, Jeffrey Barrett, Sonia Goncalves, Ewan Harrison, David K. Jackson, Ian Johnston, Dominic Kwiatkowski, Cordelia Langford, John Sillitoe on behalf of the Wellcome Sanger Institute COVID-19 Surveillance Team |
| EPI_ISL_2998614 | Lighthouse Lab in Glasgow | Wellcome Sanger Institute for the COVID-19 Genomics UK (COG-UK) Consortium | Harper VanSteenhouse, Yumi Kasai, David Gray, Carol Clugston, Anna Dominiczak and Alex Alderton, Roberto Amato, Jeffrey Barrett, Sonia Goncalves, Ewan Harrison, David K. Jackson, Ian Johnston, Dominic Kwiatkowski, Cordelia Langford, John Sillitoe on behalf of the Wellcome Sanger Institute COVID-19 Surveillance Team |
| EPI_ISL_2998615 | Lighthouse Laboratory Plymouth | Wellcome Sanger Institute for the COVID-19 Genomics UK (COG-UK) Consortium | Lighthouse Laboratory Plymouth and Alex Alderton, Roberto Amato, Jeffrey Barrett, Sonia Goncalves, Ewan Harrison, David K. Jackson, Ian Johnston, Dominic Kwiatkowski, Cordelia Langford, John Sillitoe on behalf of the Wellcome Sanger Institute COVID-19 Surveillance Team |
| EPI_ISL_2998616 | Microbiologia CATLAB | Can Ruti SARS-CoV-2 Sequencing Hub (HUGTiP/IrsiCaixa/IGTP) | Marc Noguera-Julian, Pilar Armengol, Ignacio Blanco, Antoni E Bordoy, Francesc Catala-Moll, Pere-Joan Cardona, Maria Casadellà, Cristina Casañ, Gemma Clara, Bonaventura Clotet, Montserrat Giménez, Mercedes Guerrero, Anna Not, David Panisello, Roger Paredes, Mariona Parera, Alexia París, Verónica Saludes, Alba Sánchez, Laia Soler, Lauro Sumoy, and Elisa Martró on behalf of the Can Ruti SARS-CoV-2 Sequencing Hub |
| EPI_ISL_2998617 | Lighthouse Lab in Milton Keynes | Wellcome Sanger Institute for the COVID-19 Genomics UK (COG-UK) Consortium | The Lighthouse Lab in Milton Keynes and Alex Alderton, Roberto Amato, Jeffrey Barrett, Sonia Goncalves, Ewan Harrison, David K. Jackson, Ian Johnston, Dominic Kwiatkowski, Cordelia Langford, John Sillitoe on behalf of the Wellcome Sanger Institute COVID-19 Surveillance Team |
| EPI_ISL_2998618 | Lighthouse Laboratory Plymouth | Wellcome Sanger Institute for the COVID-19 Genomics UK (COG-UK) Consortium | Lighthouse Laboratory Plymouth and Alex Alderton, Roberto Amato, Jeffrey Barrett, Sonia Goncalves, Ewan Harrison, David K. Jackson, Ian Johnston, Dominic Kwiatkowski, Cordelia Langford, John Sillitoe on behalf of the Wellcome Sanger Institute COVID-19 Surveillance Team |
| EPI_ISL_2998620 | Lighthouse Lab in Glasgow | Wellcome Sanger Institute for the COVID-19 Genomics UK (COG-UK) Consortium | Harper VanSteenhouse, Yumi Kasai, David Gray, Carol Clugston, Anna Dominiczak and Alex Alderton, Roberto Amato, Jeffrey Barrett, Sonia Goncalves, Ewan Harrison, David K. Jackson, Ian Johnston, Dominic Kwiatkowski, Cordelia Langford, John Sillitoe on behalf of the Wellcome Sanger Institute COVID-19 Surveillance Team |
| EPI_ISL_2998621 | Lighthouse Lab in Milton Keynes | Wellcome Sanger Institute for the COVID-19 Genomics UK (COG-UK) Consortium | The Lighthouse Lab in Milton Keynes and Alex Alderton, Roberto Amato, Jeffrey Barrett, Sonia Goncalves, Ewan Harrison, David K. Jackson, Ian Johnston, Dominic Kwiatkowski, Cordelia Langford, John Sillitoe on behalf of the Wellcome Sanger Institute COVID-19 Surveillance Team |
| EPI_ISL_2998622 | Microbiology Department, Laboratori Clínic Metropolitana Nord. Hospital Universitari Germans Trias i Pujol | Can Ruti SARS-CoV-2 Sequencing Hub (HUGTiP/IrsiCaixa/IGTP) | Marc Noguera-Julian, Pilar Armengol, Ignacio Blanco, Antoni E Bordoy, Francesc Catala-Moll, Pere-Joan Cardona, Maria Casadellà, Cristina Casañ, Gemma Clara, Bonaventura Clotet, Montserrat Giménez, Mercedes Guerrero, Anna Not, David Panisello, Roger Paredes, Mariona Parera, Alexia París, Verónica Saludes, Alba Sánchez, Laia Soler, Lauro Sumoy, and Elisa Martró on behalf of the Can Ruti SARS-CoV-2 Sequencing Hub |
| EPI_ISL_2998623, EPI_ISL_2998624, EPI_ISL_2998625 | Lighthouse Laboratory Plymouth | Wellcome Sanger Institute for the COVID-19 Genomics UK (COG-UK) Consortium | Lighthouse Laboratory Plymouth and Alex Alderton, Roberto Amato, Jeffrey Barrett, Sonia Goncalves, Ewan Harrison, David K. Jackson, Ian Johnston, Dominic Kwiatkowski, Cordelia Langford, John Sillitoe on behalf of the Wellcome Sanger Institute COVID-19 Surveillance Team |
| EPI_ISL_2998626 | Microbiology Department, Laboratori Clínic Metropolitana Nord. Hospital Universitari Germans Trias i Pujol | Can Ruti SARS-CoV-2 Sequencing Hub (HUGTiP/IrsiCaixa/IGTP) | Marc Noguera-Julian, Pilar Armengol, Ignacio Blanco, Antoni E Bordoy, Francesc Catala-Moll, Pere-Joan Cardona, Maria Casadellà, Cristina Casañ, Gemma Clara, Bonaventura Clotet, Montserrat Giménez, Mercedes Guerrero, Anna Not, David Panisello, Roger Paredes, Mariona Parera, Alexia París, Verónica Saludes, Alba Sánchez, Laia Soler, Lauro Sumoy, and Elisa Martró on behalf of the Can Ruti SARS-CoV-2 Sequencing Hub |
| EPI_ISL_2998627 | Lighthouse Lab in Milton Keynes | Wellcome Sanger Institute for the COVID-19 Genomics UK (COG-UK) Consortium | The Lighthouse Lab in Milton Keynes and Alex Alderton, Roberto Amato, Jeffrey Barrett, Sonia Goncalves, Ewan Harrison, David K. Jackson, Ian Johnston, Dominic Kwiatkowski, Cordelia Langford, John Sillitoe on behalf of the Wellcome Sanger Institute COVID-19 Surveillance Team |
| EPI_ISL_2998628, EPI_ISL_2998629 | Lighthouse Laboratory Plymouth | Wellcome Sanger Institute for the COVID-19 Genomics UK (COG-UK) Consortium | Lighthouse Laboratory Plymouth and Alex Alderton, Roberto Amato, Jeffrey Barrett, Sonia Goncalves, Ewan Harrison, David K. Jackson, Ian Johnston, Dominic Kwiatkowski, Cordelia Langford, John Sillitoe on behalf of the Wellcome Sanger Institute COVID-19 Surveillance Team |
| EPI_ISL_2998630 | Microbiology Department, Laboratori Clínic Metropolitana Nord. Hospital Universitari Germans Trias i Pujol | Can Ruti SARS-CoV-2 Sequencing Hub (HUGTiP/IrsiCaixa/IGTP) | Marc Noguera-Julian, Pilar Armengol, Ignacio Blanco, Antoni E Bordoy, Francesc Catala-Moll, Pere-Joan Cardona, Maria Casadellà, Cristina Casañ, Gemma Clara, Bonaventura Clotet, Montserrat Giménez, Mercedes Guerrero, Anna Not, David Panisello, Roger Paredes, Mariona Parera, Alexia París, Verónica Saludes, Alba Sánchez, Laia Soler, Lauro Sumoy, and Elisa Martró on behalf of the Can Ruti SARS-CoV-2 Sequencing Hub |
| EPI_ISL_2998631 | Lighthouse Lab in Glasgow | Wellcome Sanger Institute for the COVID-19 Genomics UK (COG-UK) Consortium | Harper VanSteenhouse, Yumi Kasai, David Gray, Carol Clugston, Anna Dominiczak and Alex Alderton, Roberto Amato, Jeffrey Barrett, Sonia Goncalves, Ewan Harrison, David K. Jackson, Ian Johnston, Dominic Kwiatkowski, Cordelia Langford, John Sillitoe on behalf of the Wellcome Sanger Institute COVID-19 Surveillance Team |
| EPI_ISL_2998632 | Lighthouse Lab in Milton Keynes | Wellcome Sanger Institute for the COVID-19 Genomics UK (COG-UK) Consortium | The Lighthouse Lab in Milton Keynes and Alex Alderton, Roberto Amato, Jeffrey Barrett, Sonia Goncalves, Ewan Harrison, David K. Jackson, Ian Johnston, Dominic Kwiatkowski, Cordelia Langford, John Sillitoe on behalf of the Wellcome Sanger Institute COVID-19 Surveillance Team |
| EPI_ISL_2998633 | Microbiology Department, Laboratori Clínic Metropolitana Nord. Hospital Universitari Germans Trias i Pujol | Can Ruti SARS-CoV-2 Sequencing Hub (HUGTiP/IrsiCaixa/IGTP) | Marc Noguera-Julian, Pilar Armengol, Ignacio Blanco, Antoni E Bordoy, Francesc Catala-Moll, Pere-Joan Cardona, Maria Casadellà, Cristina Casañ, Gemma Clara, Bonaventura Clotet, Montserrat Giménez, Mercedes Guerrero, Anna Not, David Panisello, Roger Paredes, Mariona Parera, Alexia París, Verónica Saludes, Alba Sánchez, Laia Soler, Lauro Sumoy, and Elisa Martró on behalf of the Can Ruti SARS-CoV-2 Sequencing Hub |
| EPI_ISL_2998634, EPI_ISL_2998635, EPI_ISL_2998636 | Lighthouse Laboratory Plymouth | Wellcome Sanger Institute for the COVID-19 Genomics UK (COG-UK) Consortium | Lighthouse Laboratory Plymouth and Alex Alderton, Roberto Amato, Jeffrey Barrett, Sonia Goncalves, Ewan Harrison, David K. Jackson, Ian Johnston, Dominic Kwiatkowski, Cordelia Langford, John Sillitoe on behalf of the Wellcome Sanger Institute COVID-19 Surveillance Team |
| EPI_ISL_2998637 | Microbiology Department, Laboratori Clínic Metropolitana Nord. Hospital Universitari Germans Trias i Pujol | Can Ruti SARS-CoV-2 Sequencing Hub (HUGTiP/IrsiCaixa/IGTP) | Marc Noguera-Julian, Pilar Armengol, Ignacio Blanco, Antoni E Bordoy, Francesc Catala-Moll, Pere-Joan Cardona, Maria Casadellà, Cristina Casañ, Gemma Clara, Bonaventura Clotet, Montserrat Giménez, Mercedes Guerrero, Anna Not, David Panisello, Roger Paredes, Mariona Parera, Alexia París, Verónica Saludes, Alba Sánchez, Laia Soler, Lauro Sumoy, and Elisa Martró on behalf of the Can Ruti SARS-CoV-2 Sequencing Hub |
| EPI_ISL_2998638 | Lighthouse Lab in Glasgow | Wellcome Sanger Institute for the COVID-19 Genomics UK (COG-UK) Consortium | Harper VanSteenhouse, Yumi Kasai, David Gray, Carol Clugston, Anna Dominiczak and Alex Alderton, Roberto Amato, Jeffrey Barrett, Sonia Goncalves, Ewan Harrison, David K. Jackson, Ian Johnston, Dominic Kwiatkowski, Cordelia Langford, John Sillitoe on behalf of the Wellcome Sanger Institute COVID-19 Surveillance Team |

| | | | |
|---|---|---|---|
| EPI_ISL_2998639, EPI_ISL_2998640 | Lighthouse Laboratory Plymouth | Wellcome Sanger Institute for the COVID-19 Genomics UK (COG-UK) Consortium | Lighthouse Laboratory Plymouth and Alex Alderton, Roberto Amato, Jeffrey Barrett, Sonia Goncalves, Ewan Harrison, David K. Jackson, Ian Johnston, Dominic Kwiatkowski, Cordelia Langford, John Sillitoe on behalf of the Wellcome Sanger Institute COVID-19 Surveillance Team |
| EPI_ISL_2998641 | Microbiology Department, Laboratori Clínic Metropolitana Nord. Hospital Universitari Germans Trias i Pujol | Can Ruti SARS-CoV-2 Sequencing Hub (HUGTiP/IrsiCaixa/IGTP) | Marc Noguera-Julian, Pilar Armengol, Ignacio Blanco, Antoni E Bordoy, Francesc Catala-Moll, Pere-Joan Cardona, Maria Casadellà, Cristina Casañ, Gemma Clara, Bonaventura Clotet, Montserrat Giménez, Mercedes Guerrero, Anna Not, David Panisello, Roger Paredes, Mariona Parera, Alexia París, Verónica Saludes, Alba Sánchez, Laia Soler, Lauro Sumoy, and Elisa Martró on behalf of the Can Ruti SARS-CoV-2 Sequencing Hub |
| EPI_ISL_2998642, EPI_ISL_2998643 | Lighthouse Lab in Milton Keynes | Wellcome Sanger Institute for the COVID-19 Genomics UK (COG-UK) Consortium | The Lighthouse Lab in Milton Keynes and Alex Alderton, Roberto Amato, Jeffrey Barrett, Sonia Goncalves, Ewan Harrison, David K. Jackson, Ian Johnston, Dominic Kwiatkowski, Cordelia Langford, John Sillitoe on behalf of the Wellcome Sanger Institute COVID-19 Surveillance Team |
| EPI_ISL_2998644 | Microbiology Department, Laboratori Clínic Metropolitana Nord. Hospital Universitari Germans Trias i Pujol | Can Ruti SARS-CoV-2 Sequencing Hub (HUGTiP/IrsiCaixa/IGTP) | Marc Noguera-Julian, Pilar Armengol, Ignacio Blanco, Antoni E Bordoy, Francesc Catala-Moll, Pere-Joan Cardona, Maria Casadellà, Cristina Casañ, Gemma Clara, Bonaventura Clotet, Montserrat Giménez, Mercedes Guerrero, Anna Not, David Panisello, Roger Paredes, Mariona Parera, Alexia París, Verónica Saludes, Alba Sánchez, Laia Soler, Lauro Sumoy, and Elisa Martró on behalf of the Can Ruti SARS-CoV-2 Sequencing Hub |
| EPI_ISL_2998645 | Lighthouse Lab in Glasgow | Wellcome Sanger Institute for the COVID-19 Genomics UK (COG-UK) Consortium | Harper VanSteenhouse, Yumi Kasai, David Gray, Carol Clugston, Anna Dominiczak and Alex Alderton, Roberto Amato, Jeffrey Barrett, Sonia Goncalves, Ewan Harrison, David K. Jackson, Ian Johnston, Dominic Kwiatkowski, Cordelia Langford, John Sillitoe on behalf of the Wellcome Sanger Institute COVID-19 Surveillance Team |
| EPI_ISL_2998646 | Lighthouse Laboratory Plymouth | Wellcome Sanger Institute for the COVID-19 Genomics UK (COG-UK) Consortium | Lighthouse Laboratory Plymouth and Alex Alderton, Roberto Amato, Jeffrey Barrett, Sonia Goncalves, Ewan Harrison, David K. Jackson, Ian Johnston, Dominic Kwiatkowski, Cordelia Langford, John Sillitoe on behalf of the Wellcome Sanger Institute COVID-19 Surveillance Team |
| EPI_ISL_2998647 | Lighthouse Lab in Milton Keynes | Wellcome Sanger Institute for the COVID-19 Genomics UK (COG-UK) Consortium | The Lighthouse Lab in Milton Keynes and Alex Alderton, Roberto Amato, Jeffrey Barrett, Sonia Goncalves, Ewan Harrison, David K. Jackson, Ian Johnston, Dominic Kwiatkowski, Cordelia Langford, John Sillitoe on behalf of the Wellcome Sanger Institute COVID-19 Surveillance Team |
| EPI_ISL_2998648 | Microbiology Department, Laboratori Clínic Metropolitana Nord. Hospital Universitari Germans Trias i Pujol | Can Ruti SARS-CoV-2 Sequencing Hub (HUGTiP/IrsiCaixa/IGTP) | Marc Noguera-Julian, Pilar Armengol, Ignacio Blanco, Antoni E Bordoy, Francesc Catala-Moll, Pere-Joan Cardona, Maria Casadellà, Cristina Casañ, Gemma Clara, Bonaventura Clotet, Montserrat Giménez, Mercedes Guerrero, Anna Not, David Panisello, Roger Paredes, Mariona Parera, Alexia París, Verónica Saludes, Alba Sánchez, Laia Soler, Lauro Sumoy, and Elisa Martró on behalf of the Can Ruti SARS-CoV-2 Sequencing Hub |
| EPI_ISL_2998649, EPI_ISL_2998650 | Lighthouse Laboratory Plymouth | Wellcome Sanger Institute for the COVID-19 Genomics UK (COG-UK) Consortium | Lighthouse Laboratory Plymouth and Alex Alderton, Roberto Amato, Jeffrey Barrett, Sonia Goncalves, Ewan Harrison, David K. Jackson, Ian Johnston, Dominic Kwiatkowski, Cordelia Langford, John Sillitoe on behalf of the Wellcome Sanger Institute COVID-19 Surveillance Team |
| EPI_ISL_2998651 | Microbiologia CATLAB | Can Ruti SARS-CoV-2 Sequencing Hub (HUGTiP/IrsiCaixa/IGTP) | Marc Noguera-Julian, Pilar Armengol, Ignacio Blanco, Antoni E Bordoy, Francesc Catala-Moll, Pere-Joan Cardona, Maria Casadellà, Cristina Casañ, Gemma Clara, Bonaventura Clotet, Montserrat Giménez, Mercedes Guerrero, Anna Not, David Panisello, Roger Paredes, Mariona Parera, Alexia París, Verónica Saludes, Alba Sánchez, Laia Soler, Lauro Sumoy, and Elisa Martró on behalf of the Can Ruti SARS-CoV-2 Sequencing Hub |
| EPI_ISL_2998652, EPI_ISL_2998653 | Lighthouse Laboratory Plymouth | Wellcome Sanger Institute for the COVID-19 Genomics UK (COG-UK) Consortium | Lighthouse Laboratory Plymouth and Alex Alderton, Roberto Amato, Jeffrey Barrett, Sonia Goncalves, Ewan Harrison, David K. Jackson, Ian Johnston, Dominic Kwiatkowski, Cordelia Langford, John Sillitoe on behalf of the Wellcome Sanger Institute COVID-19 Surveillance Team |
| EPI_ISL_2998654 | Lighthouse Lab in Milton Keynes | Wellcome Sanger Institute for the COVID-19 Genomics UK (COG-UK) Consortium | The Lighthouse Lab in Milton Keynes and Alex Alderton, Roberto Amato, Jeffrey Barrett, Sonia Goncalves, Ewan Harrison, David K. Jackson, Ian Johnston, Dominic Kwiatkowski, Cordelia Langford, John Sillitoe on behalf of the Wellcome Sanger Institute COVID-19 Surveillance Team |
| EPI_ISL_2998655 | Microbiologia CATLAB | Can Ruti SARS-CoV-2 Sequencing Hub (HUGTiP/IrsiCaixa/IGTP) | Marc Noguera-Julian, Pilar Armengol, Ignacio Blanco, Antoni E Bordoy, Francesc Catala-Moll, Pere-Joan Cardona, Maria Casadellà, Cristina Casañ, Gemma Clara, Bonaventura Clotet, Montserrat Giménez, Mercedes Guerrero, Anna Not, David Panisello, Roger Paredes, Mariona Parera, Alexia París, Verónica Saludes, Alba Sánchez, Laia Soler, Lauro Sumoy, and Elisa Martró on behalf of the Can Ruti SARS-CoV-2 Sequencing Hub |
| EPI_ISL_2998656 | Lighthouse Laboratory Plymouth | Wellcome Sanger Institute for the COVID-19 Genomics UK (COG-UK) Consortium | Lighthouse Laboratory Plymouth and Alex Alderton, Roberto Amato, Jeffrey Barrett, Sonia Goncalves, Ewan Harrison, David K. Jackson, Ian Johnston, Dominic Kwiatkowski, Cordelia Langford, John Sillitoe on behalf of the Wellcome Sanger Institute COVID-19 Surveillance Team |
| EPI_ISL_2998657 | Lighthouse Lab in Milton Keynes | Wellcome Sanger Institute for the COVID-19 Genomics UK (COG-UK) Consortium | The Lighthouse Lab in Milton Keynes and Alex Alderton, Roberto Amato, Jeffrey Barrett, Sonia Goncalves, Ewan Harrison, David K. Jackson, Ian Johnston, Dominic Kwiatkowski, Cordelia Langford, John Sillitoe on behalf of the Wellcome Sanger Institute COVID-19 Surveillance Team |
| EPI_ISL_2998658 | Lighthouse Laboratory Plymouth | Wellcome Sanger Institute for the COVID-19 Genomics UK (COG-UK) Consortium | Lighthouse Laboratory Plymouth and Alex Alderton, Roberto Amato, Jeffrey Barrett, Sonia Goncalves, Ewan Harrison, David K. Jackson, Ian Johnston, Dominic Kwiatkowski, Cordelia Langford, John Sillitoe on behalf of the Wellcome Sanger Institute COVID-19 Surveillance Team |
| EPI_ISL_2998659 | Microbiologia CATLAB | Can Ruti SARS-CoV-2 Sequencing Hub (HUGTiP/IrsiCaixa/IGTP) | Marc Noguera-Julian, Pilar Armengol, Ignacio Blanco, Antoni E Bordoy, Francesc Catala-Moll, Pere-Joan Cardona, Maria Casadellà, Cristina Casañ, Gemma Clara, Bonaventura Clotet, Montserrat Giménez, Mercedes Guerrero, Anna Not, David Panisello, Roger Paredes, Mariona Parera, Alexia París, Verónica Saludes, Alba Sánchez, Laia Soler, Lauro Sumoy, and Elisa Martró on behalf of the Can Ruti SARS-CoV-2 Sequencing Hub |
| EPI_ISL_2998660, EPI_ISL_2998661 | Lighthouse Laboratory Plymouth | Wellcome Sanger Institute for the COVID-19 Genomics UK (COG-UK) Consortium | Lighthouse Laboratory Plymouth and Alex Alderton, Roberto Amato, Jeffrey Barrett, Sonia Goncalves, Ewan Harrison, David K. Jackson, Ian Johnston, Dominic Kwiatkowski, Cordelia Langford, John Sillitoe on behalf of the Wellcome Sanger Institute COVID-19 Surveillance Team |
| EPI_ISL_2998662 | Microbiology Department, Laboratori Clínic Metropolitana Nord. Hospital Universitari Germans Trias i Pujol | Can Ruti SARS-CoV-2 Sequencing Hub (HUGTiP/IrsiCaixa/IGTP) | Marc Noguera-Julian, Pilar Armengol, Ignacio Blanco, Antoni E Bordoy, Francesc Catala-Moll, Pere-Joan Cardona, Maria Casadellà, Cristina Casañ, Gemma Clara, Bonaventura Clotet, Montserrat Giménez, Mercedes Guerrero, Anna Not, David Panisello, Roger Paredes, Mariona Parera, Alexia París, Verónica Saludes, Alba Sánchez, Laia Soler, Lauro Sumoy, and Elisa Martró on behalf of the Can Ruti SARS-CoV-2 Sequencing Hub |
| EPI_ISL_2998663, EPI_ISL_2998664 | Lighthouse Laboratory Plymouth | Wellcome Sanger Institute for the COVID-19 Genomics UK (COG-UK) Consortium | Lighthouse Laboratory Plymouth and Alex Alderton, Roberto Amato, Jeffrey Barrett, Sonia Goncalves, Ewan Harrison, David K. Jackson, Ian Johnston, Dominic Kwiatkowski, Cordelia Langford, John Sillitoe on behalf of the Wellcome Sanger Institute COVID-19 Surveillance Team |
| EPI_ISL_2998665 | Lighthouse Lab in Glasgow | Wellcome Sanger Institute for the COVID-19 Genomics UK (COG-UK) Consortium | Harper VanSteenhouse, Yumi Kasai, David Gray, Carol Clugston, Anna Dominiczak and Alex Alderton, Roberto Amato, Jeffrey Barrett, Sonia Goncalves, Ewan Harrison, David K. Jackson, Ian Johnston, Dominic Kwiatkowski, Cordelia Langford, John Sillitoe on behalf of the Wellcome Sanger Institute COVID-19 Surveillance Team |
| EPI_ISL_2998666 | Microbiology Department, Laboratori Clínic Metropolitana Nord. Hospital Universitari Germans Trias i Pujol | Can Ruti SARS-CoV-2 Sequencing Hub (HUGTiP/IrsiCaixa/IGTP) | Marc Noguera-Julian, Pilar Armengol, Ignacio Blanco, Antoni E Bordoy, Francesc Catala-Moll, Pere-Joan Cardona, Maria Casadellà, Cristina Casañ, Gemma Clara, Bonaventura Clotet, Montserrat Giménez, Mercedes Guerrero, Anna Not, David Panisello, Roger Paredes, Mariona Parera, Alexia París, Verónica Saludes, Alba Sánchez, Laia Soler, Lauro Sumoy, and Elisa Martró on behalf of the Can Ruti SARS-CoV-2 Sequencing Hub |
| EPI_ISL_2998667 | Lighthouse Laboratory Plymouth | Wellcome Sanger Institute for the COVID-19 Genomics UK (COG-UK) Consortium | Lighthouse Laboratory Plymouth and Alex Alderton, Roberto Amato, Jeffrey Barrett, Sonia Goncalves, Ewan Harrison, David K. Jackson, Ian Johnston, Dominic Kwiatkowski, Cordelia Langford, John Sillitoe on behalf of the Wellcome Sanger Institute COVID-19 Surveillance Team |
| EPI_ISL_2998668 | Lighthouse Lab in Glasgow | Wellcome Sanger Institute for the COVID-19 Genomics UK (COG-UK) Consortium | Harper VanSteenhouse, Yumi Kasai, David Gray, Carol Clugston, Anna Dominiczak and Alex Alderton, Roberto Amato, Jeffrey Barrett, Sonia Goncalves, Ewan Harrison, David K. Jackson, Ian Johnston, Dominic Kwiatkowski, Cordelia Langford, John Sillitoe on behalf of the Wellcome Sanger Institute COVID-19 Surveillance Team |
| EPI_ISL_2998669 | Lighthouse Lab in Milton Keynes | Wellcome Sanger Institute for the COVID-19 Genomics UK (COG-UK) Consortium | The Lighthouse Lab in Milton Keynes and Alex Alderton, Roberto Amato, Jeffrey Barrett, Sonia Goncalves, Ewan Harrison, David K. Jackson, Ian Johnston, Dominic Kwiatkowski, Cordelia Langford, John Sillitoe on behalf of the Wellcome Sanger Institute COVID-19 Surveillance Team |
| EPI_ISL_2998670 | Microbiology Department, Laboratori Clínic Metropolitana Nord. Hospital Universitari Germans Trias i Pujol | Can Ruti SARS-CoV-2 Sequencing Hub (HUGTiP/IrsiCaixa/IGTP) | Marc Noguera-Julian, Pilar Armengol, Ignacio Blanco, Antoni E Bordoy, Francesc Catala-Moll, Pere-Joan Cardona, Maria Casadellà, Cristina Casañ, Gemma Clara, Bonaventura Clotet, Montserrat Giménez, Mercedes Guerrero, Anna Not, David Panisello, Roger Paredes, Mariona Parera, Alexia París, Verónica Saludes, Alba Sánchez, Laia Soler, Lauro Sumoy, and Elisa Martró on behalf of the Can Ruti SARS-CoV-2 Sequencing Hub |
| EPI_ISL_2998671 | Lighthouse Laboratory Plymouth | Wellcome Sanger Institute for the COVID-19 Genomics UK (COG-UK) Consortium | Lighthouse Laboratory Plymouth and Alex Alderton, Roberto Amato, Jeffrey Barrett, Sonia Goncalves, Ewan Harrison, David K. Jackson, Ian Johnston, Dominic Kwiatkowski, Cordelia Langford, John Sillitoe on behalf of the Wellcome Sanger Institute COVID-19 Surveillance Team |
| EPI_ISL_2998672 | Lighthouse Lab in Milton Keynes | Wellcome Sanger Institute for the COVID-19 Genomics UK (COG-UK) Consortium | The Lighthouse Lab in Milton Keynes and Alex Alderton, Roberto Amato, Jeffrey Barrett, Sonia Goncalves, Ewan Harrison, David K. Jackson, Ian Johnston, Dominic Kwiatkowski, Cordelia Langford, John Sillitoe on behalf of the Wellcome Sanger Institute COVID-19 Surveillance Team |
| EPI_ISL_2998674 | Lighthouse Laboratory Plymouth | Wellcome Sanger Institute for the COVID-19 Genomics UK (COG-UK) Consortium | Lighthouse Laboratory Plymouth and Alex Alderton, Roberto Amato, Jeffrey Barrett, Sonia Goncalves, Ewan Harrison, David K. Jackson, Ian Johnston, Dominic Kwiatkowski, Cordelia Langford, John Sillitoe on behalf of the Wellcome Sanger Institute COVID-19 Surveillance Team |
| EPI_ISL_2998675, EPI_ISL_2998676 | Lighthouse Lab in Glasgow | Wellcome Sanger Institute for the COVID-19 Genomics UK (COG-UK) Consortium | Harper VanSteenhouse, Yumi Kasai, David Gray, Carol Clugston, Anna Dominiczak and Alex Alderton, Roberto Amato, Jeffrey Barrett, Sonia Goncalves, Ewan Harrison, David K. Jackson, Ian Johnston, Dominic Kwiatkowski, Cordelia Langford, John Sillitoe on behalf of the Wellcome Sanger Institute COVID-19 Surveillance Team |
| EPI_ISL_2998677 | Microbiology Department, Laboratori Clínic Metropolitana Nord. Hospital Universitari Germans Trias i Pujol | Can Ruti SARS-CoV-2 Sequencing Hub (HUGTiP/IrsiCaixa/IGTP) | Marc Noguera-Julian, Pilar Armengol, Ignacio Blanco, Antoni E Bordoy, Francesc Catala-Moll, Pere-Joan Cardona, Maria Casadellà, Cristina Casañ, Gemma Clara, Bonaventura Clotet, Montserrat Giménez, Mercedes Guerrero, Anna Not, David Panisello, Roger Paredes, Mariona Parera, Alexia París, Verónica Saludes, Alba Sánchez, Laia Soler, Lauro Sumoy, and Elisa Martró on behalf of the Can Ruti SARS-CoV-2 Sequencing Hub |
| EPI_ISL_2998678, EPI_ISL_2998679 | Lighthouse Laboratory Plymouth | Wellcome Sanger Institute for the COVID-19 Genomics UK (COG-UK) Consortium | Lighthouse Laboratory Plymouth and Alex Alderton, Roberto Amato, Jeffrey Barrett, Sonia Goncalves, Ewan Harrison, David K. Jackson, Ian Johnston, Dominic Kwiatkowski, Cordelia Langford, John Sillitoe on behalf of the Wellcome Sanger Institute COVID-19 Surveillance Team |
| EPI_ISL_2998680 | Lighthouse Lab in Glasgow | Wellcome Sanger Institute for the COVID-19 Genomics UK (COG-UK) Consortium | Harper VanSteenhouse, Yumi Kasai, David Gray, Carol Clugston, Anna Dominiczak and Alex Alderton, Roberto Amato, Jeffrey Barrett, Sonia Goncalves, Ewan Harrison, David K. Jackson, Ian Johnston, Dominic Kwiatkowski, Cordelia Langford, John Sillitoe on behalf of the Wellcome Sanger Institute COVID-19 Surveillance Team |
| EPI_ISL_2998681 | Microbiology Department, Laboratori Clínic Metropolitana Nord. Hospital Universitari Germans Trias i Pujol | Can Ruti SARS-CoV-2 Sequencing Hub (HUGTiP/IrsiCaixa/IGTP) | Marc Noguera-Julian, Pilar Armengol, Ignacio Blanco, Antoni E Bordoy, Francesc Catala-Moll, Pere-Joan Cardona, Maria Casadellà, Cristina Casañ, Gemma Clara, Bonaventura Clotet, Montserrat Giménez, Mercedes Guerrero, Anna Not, David Panisello, Roger Paredes, Mariona Parera, Alexia París, Verónica Saludes, Alba Sánchez, Laia Soler, Lauro Sumoy, and Elisa Martró on behalf of the Can Ruti SARS-CoV-2 Sequencing Hub |
| EPI_ISL_2998682 | Lighthouse Lab in Milton Keynes | Wellcome Sanger Institute for the COVID-19 Genomics UK (COG-UK) Consortium | The Lighthouse Lab in Milton Keynes and Alex Alderton, Roberto Amato, Jeffrey Barrett, Sonia Goncalves, Ewan Harrison, David K. Jackson, Ian Johnston, Dominic Kwiatkowski, Cordelia Langford, John Sillitoe on behalf of the Wellcome Sanger Institute COVID-19 Surveillance Team |
| EPI_ISL_2998683 | Lighthouse Laboratory Plymouth | Wellcome Sanger Institute for the COVID-19 Genomics UK (COG-UK) Consortium | Lighthouse Laboratory Plymouth and Alex Alderton, Roberto Amato, Jeffrey Barrett, Sonia Goncalves, Ewan Harrison, David K. Jackson, Ian Johnston, Dominic Kwiatkowski, Cordelia Langford, John Sillitoe on behalf of the Wellcome Sanger Institute COVID-19 Surveillance Team |
| EPI_ISL_2998684 | Lighthouse Lab in Milton Keynes | Wellcome Sanger Institute for the COVID-19 Genomics UK (COG-UK) Consortium | The Lighthouse Lab in Milton Keynes and Alex Alderton, Roberto Amato, Jeffrey Barrett, Sonia Goncalves, Ewan Harrison, David K. Jackson, Ian Johnston, Dominic Kwiatkowski, Cordelia Langford, John Sillitoe on behalf of the Wellcome Sanger Institute COVID-19 Surveillance Team |
| EPI_ISL_2998685 | Microbiology Department, Laboratori Clínic Metropolitana Nord. Hospital Universitari Germans Trias i Pujol | Can Ruti SARS-CoV-2 Sequencing Hub (HUGTiP/IrsiCaixa/IGTP) | Marc Noguera-Julian, Pilar Armengol, Ignacio Blanco, Antoni E Bordoy, Francesc Catala-Moll, Pere-Joan Cardona, Maria Casadellà, Cristina Casañ, Gemma Clara, Bonaventura Clotet, Montserrat Giménez, Mercedes Guerrero, Anna Not, David Panisello, Roger Paredes, Mariona Parera, Alexia París, Verónica Saludes, Alba Sánchez, Laia Soler, Lauro Sumoy, and Elisa Martró on behalf of the Can Ruti SARS-CoV-2 Sequencing Hub |
| EPI_ISL_2998686 | Lighthouse Lab in Milton Keynes | Wellcome Sanger Institute for the COVID-19 Genomics UK (COG-UK) Consortium | The Lighthouse Lab in Milton Keynes and Alex Alderton, Roberto Amato, Jeffrey Barrett, Sonia Goncalves, Ewan Harrison, David K. Jackson, Ian Johnston, Dominic Kwiatkowski, Cordelia Langford, John Sillitoe on behalf of the Wellcome Sanger Institute COVID-19 Surveillance Team |
| EPI_ISL_2998687 | Lighthouse Laboratory Plymouth | Wellcome Sanger Institute for the COVID-19 Genomics UK (COG-UK) Consortium | Lighthouse Laboratory Plymouth and Alex Alderton, Roberto Amato, Jeffrey Barrett, Sonia Goncalves, Ewan Harrison, David K. Jackson, Ian Johnston, Dominic Kwiatkowski, Cordelia Langford, John Sillitoe on behalf of the Wellcome Sanger Institute COVID-19 Surveillance Team |
| EPI_ISL_2998688 | Lighthouse Lab in Milton Keynes | Wellcome Sanger Institute for the COVID-19 Genomics UK (COG-UK) Consortium | The Lighthouse Lab in Milton Keynes and Alex Alderton, Roberto Amato, Jeffrey Barrett, Sonia Goncalves, Ewan Harrison, David K. Jackson, Ian Johnston, Dominic Kwiatkowski, Cordelia Langford, John Sillitoe on behalf of the Wellcome Sanger Institute COVID-19 Surveillance Team |
| EPI_ISL_2998689 | Microbiology Department, Laboratori Clínic Metropolitana Nord. Hospital Universitari Germans Trias i Pujol | Can Ruti SARS-CoV-2 Sequencing Hub (HUGTiP/IrsiCaixa/IGTP) | Marc Noguera-Julian, Pilar Armengol, Ignacio Blanco, Antoni E Bordoy, Francesc Catala-Moll, Pere-Joan Cardona, Maria Casadellà, Cristina Casañ, Gemma Clara, Bonaventura Clotet, Montserrat Giménez, Mercedes Guerrero, Anna Not, David Panisello, Roger Paredes, Mariona Parera, Alexia París, Verónica Saludes, Alba Sánchez, Laia Soler, Lauro Sumoy, and Elisa Martró on behalf of the Can Ruti SARS-CoV-2 Sequencing Hub |
| EPI_ISL_2998690, EPI_ISL_2998691, EPI_ISL_2998692, EPI_ISL_2998693 | Lighthouse Lab in Glasgow | Wellcome Sanger Institute for the COVID-19 Genomics UK (COG-UK) Consortium | Harper VanSteenhouse, Yumi Kasai, David Gray, Carol Clugston, Anna Dominiczak and Alex Alderton, Roberto Amato, Jeffrey Barrett, Sonia Goncalves, Ewan Harrison, David K. Jackson, Ian Johnston, Dominic Kwiatkowski, Cordelia Langford, John Sillitoe on behalf of the Wellcome Sanger Institute COVID-19 Surveillance Team |
| EPI_ISL_2998694, EPI_ISL_2998695 | Lighthouse Lab in Milton Keynes | Wellcome Sanger Institute for the COVID-19 Genomics UK (COG-UK) Consortium | The Lighthouse Lab in Milton Keynes and Alex Alderton, Roberto Amato, Jeffrey Barrett, Sonia Goncalves, Ewan Harrison, David K. Jackson, Ian Johnston, Dominic Kwiatkowski, Cordelia Langford, John Sillitoe on behalf of the Wellcome Sanger Institute COVID-19 Surveillance Team |

| | | | |
|---|---|---|---|
| EPI_ISL_2998696 | Lighthouse Laboratory Plymouth | Wellcome Sanger Institute for the COVID-19 Genomics UK (COG-UK) Consortium | Lighthouse Laboratory Plymouth and Alex Alderton, Roberto Amato, Jeffrey Barrett, Sonia Goncalves, Ewan Harrison, David K. Jackson, Ian Johnston, Dominic Kwiatkowski, Cordelia Langford, John Sillitoe on behalf of the Wellcome Sanger Institute COVID-19 Surveillance Team |
| EPI_ISL_2998697 | Lighthouse Lab in Glasgow | Wellcome Sanger Institute for the COVID-19 Genomics UK (COG-UK) Consortium | Harper VanSteenhouse, Yumi Kasai, David Gray, Carol Clugston, Anna Dominiczak and Alex Alderton, Roberto Amato, Jeffrey Barrett, Sonia Goncalves, Ewan Harrison, David K. Jackson, Ian Johnston, Dominic Kwiatkowski, Cordelia Langford, John Sillitoe on behalf of the Wellcome Sanger Institute COVID-19 Surveillance Team |
| EPI_ISL_2998698, EPI_ISL_2998699, EPI_ISL_2998700 | Lighthouse Lab in Milton Keynes | Wellcome Sanger Institute for the COVID-19 Genomics UK (COG-UK) Consortium | The Lighthouse Lab in Milton Keynes and Alex Alderton, Roberto Amato, Jeffrey Barrett, Sonia Goncalves, Ewan Harrison, David K. Jackson, Ian Johnston, Dominic Kwiatkowski, Cordelia Langford, John Sillitoe on behalf of the Wellcome Sanger Institute COVID-19 Surveillance Team |
| EPI_ISL_2998701 | Lighthouse Lab in Glasgow | Wellcome Sanger Institute for the COVID-19 Genomics UK (COG-UK) Consortium | Harper VanSteenhouse, Yumi Kasai, David Gray, Carol Clugston, Anna Dominiczak and Alex Alderton, Roberto Amato, Jeffrey Barrett, Sonia Goncalves, Ewan Harrison, David K. Jackson, Ian Johnston, Dominic Kwiatkowski, Cordelia Langford, John Sillitoe on behalf of the Wellcome Sanger Institute COVID-19 Surveillance Team |
| EPI_ISL_2998702, EPI_ISL_2998703 | Lighthouse Laboratory Plymouth | Wellcome Sanger Institute for the COVID-19 Genomics UK (COG-UK) Consortium | Lighthouse Laboratory Plymouth and Alex Alderton, Roberto Amato, Jeffrey Barrett, Sonia Goncalves, Ewan Harrison, David K. Jackson, Ian Johnston, Dominic Kwiatkowski, Cordelia Langford, John Sillitoe on behalf of the Wellcome Sanger Institute COVID-19 Surveillance Team |
| EPI_ISL_2998704 | Lighthouse Lab in Glasgow | Wellcome Sanger Institute for the COVID-19 Genomics UK (COG-UK) Consortium | Harper VanSteenhouse, Yumi Kasai, David Gray, Carol Clugston, Anna Dominiczak and Alex Alderton, Roberto Amato, Jeffrey Barrett, Sonia Goncalves, Ewan Harrison, David K. Jackson, Ian Johnston, Dominic Kwiatkowski, Cordelia Langford, John Sillitoe on behalf of the Wellcome Sanger Institute COVID-19 Surveillance Team |
| EPI_ISL_2998705, EPI_ISL_2998706 | Lighthouse Laboratory Plymouth | Wellcome Sanger Institute for the COVID-19 Genomics UK (COG-UK) Consortium | Lighthouse Laboratory Plymouth and Alex Alderton, Roberto Amato, Jeffrey Barrett, Sonia Goncalves, Ewan Harrison, David K. Jackson, Ian Johnston, Dominic Kwiatkowski, Cordelia Langford, John Sillitoe on behalf of the Wellcome Sanger Institute COVID-19 Surveillance Team |
| EPI_ISL_2998707 | Lighthouse Lab in Milton Keynes | Wellcome Sanger Institute for the COVID-19 Genomics UK (COG-UK) Consortium | The Lighthouse Lab in Milton Keynes and Alex Alderton, Roberto Amato, Jeffrey Barrett, Sonia Goncalves, Ewan Harrison, David K. Jackson, Ian Johnston, Dominic Kwiatkowski, Cordelia Langford, John Sillitoe on behalf of the Wellcome Sanger Institute COVID-19 Surveillance Team |
| EPI_ISL_2998708 | Lighthouse Laboratory Plymouth | Wellcome Sanger Institute for the COVID-19 Genomics UK (COG-UK) Consortium | Lighthouse Laboratory Plymouth and Alex Alderton, Roberto Amato, Jeffrey Barrett, Sonia Goncalves, Ewan Harrison, David K. Jackson, Ian Johnston, Dominic Kwiatkowski, Cordelia Langford, John Sillitoe on behalf of the Wellcome Sanger Institute COVID-19 Surveillance Team |
| EPI_ISL_2998709 | Lighthouse Lab in Glasgow | Wellcome Sanger Institute for the COVID-19 Genomics UK (COG-UK) Consortium | Harper VanSteenhouse, Yumi Kasai, David Gray, Carol Clugston, Anna Dominiczak and Alex Alderton, Roberto Amato, Jeffrey Barrett, Sonia Goncalves, Ewan Harrison, David K. Jackson, Ian Johnston, Dominic Kwiatkowski, Cordelia Langford, John Sillitoe on behalf of the Wellcome Sanger Institute COVID-19 Surveillance Team |
| EPI_ISL_2998710 | Lighthouse Laboratory Plymouth | Wellcome Sanger Institute for the COVID-19 Genomics UK (COG-UK) Consortium | Lighthouse Laboratory Plymouth and Alex Alderton, Roberto Amato, Jeffrey Barrett, Sonia Goncalves, Ewan Harrison, David K. Jackson, Ian Johnston, Dominic Kwiatkowski, Cordelia Langford, John Sillitoe on behalf of the Wellcome Sanger Institute COVID-19 Surveillance Team |
| EPI_ISL_2998711 | Lighthouse Lab in Glasgow | Wellcome Sanger Institute for the COVID-19 Genomics UK (COG-UK) Consortium | Harper VanSteenhouse, Yumi Kasai, David Gray, Carol Clugston, Anna Dominiczak and Alex Alderton, Roberto Amato, Jeffrey Barrett, Sonia Goncalves, Ewan Harrison, David K. Jackson, Ian Johnston, Dominic Kwiatkowski, Cordelia Langford, John Sillitoe on behalf of the Wellcome Sanger Institute COVID-19 Surveillance Team |
| EPI_ISL_2998712, EPI_ISL_2998713 | Lighthouse Lab in Milton Keynes | Wellcome Sanger Institute for the COVID-19 Genomics UK (COG-UK) Consortium | The Lighthouse Lab in Milton Keynes and Alex Alderton, Roberto Amato, Jeffrey Barrett, Sonia Goncalves, Ewan Harrison, David K. Jackson, Ian Johnston, Dominic Kwiatkowski, Cordelia Langford, John Sillitoe on behalf of the Wellcome Sanger Institute COVID-19 Surveillance Team |
| EPI_ISL_2998714 | Lighthouse Lab in Glasgow | Wellcome Sanger Institute for the COVID-19 Genomics UK (COG-UK) Consortium | Harper VanSteenhouse, Yumi Kasai, David Gray, Carol Clugston, Anna Dominiczak and Alex Alderton, Roberto Amato, Jeffrey Barrett, Sonia Goncalves, Ewan Harrison, David K. Jackson, Ian Johnston, Dominic Kwiatkowski, Cordelia Langford, John Sillitoe on behalf of the Wellcome Sanger Institute COVID-19 Surveillance Team |
| EPI_ISL_2998715 | Lighthouse Laboratory Plymouth | Wellcome Sanger Institute for the COVID-19 Genomics UK (COG-UK) Consortium | Lighthouse Laboratory Plymouth and Alex Alderton, Roberto Amato, Jeffrey Barrett, Sonia Goncalves, Ewan Harrison, David K. Jackson, Ian Johnston, Dominic Kwiatkowski, Cordelia Langford, John Sillitoe on behalf of the Wellcome Sanger Institute COVID-19 Surveillance Team |
| EPI_ISL_2998716 | Lighthouse Lab in Glasgow | Wellcome Sanger Institute for the COVID-19 Genomics UK (COG-UK) Consortium | Harper VanSteenhouse, Yumi Kasai, David Gray, Carol Clugston, Anna Dominiczak and Alex Alderton, Roberto Amato, Jeffrey Barrett, Sonia Goncalves, Ewan Harrison, David K. Jackson, Ian Johnston, Dominic Kwiatkowski, Cordelia Langford, John Sillitoe on behalf of the Wellcome Sanger Institute COVID-19 Surveillance Team |
| EPI_ISL_2998717 | Lighthouse Laboratory Plymouth | Wellcome Sanger Institute for the COVID-19 Genomics UK (COG-UK) Consortium | Lighthouse Laboratory Plymouth and Alex Alderton, Roberto Amato, Jeffrey Barrett, Sonia Goncalves, Ewan Harrison, David K. Jackson, Ian Johnston, Dominic Kwiatkowski, Cordelia Langford, John Sillitoe on behalf of the Wellcome Sanger Institute COVID-19 Surveillance Team |
| EPI_ISL_2998718, EPI_ISL_2998719 | Lighthouse Lab in Milton Keynes | Wellcome Sanger Institute for the COVID-19 Genomics UK (COG-UK) Consortium | The Lighthouse Lab in Milton Keynes and Alex Alderton, Roberto Amato, Jeffrey Barrett, Sonia Goncalves, Ewan Harrison, David K. Jackson, Ian Johnston, Dominic Kwiatkowski, Cordelia Langford, John Sillitoe on behalf of the Wellcome Sanger Institute COVID-19 Surveillance Team |
| EPI_ISL_2998720 | Lighthouse Lab in Glasgow | Wellcome Sanger Institute for the COVID-19 Genomics UK (COG-UK) Consortium | Harper VanSteenhouse, Yumi Kasai, David Gray, Carol Clugston, Anna Dominiczak and Alex Alderton, Roberto Amato, Jeffrey Barrett, Sonia Goncalves, Ewan Harrison, David K. Jackson, Ian Johnston, Dominic Kwiatkowski, Cordelia Langford, John Sillitoe on behalf of the Wellcome Sanger Institute COVID-19 Surveillance Team |
| EPI_ISL_2998721 | Lighthouse Lab in Milton Keynes | Wellcome Sanger Institute for the COVID-19 Genomics UK (COG-UK) Consortium | The Lighthouse Lab in Milton Keynes and Alex Alderton, Roberto Amato, Jeffrey Barrett, Sonia Goncalves, Ewan Harrison, David K. Jackson, Ian Johnston, Dominic Kwiatkowski, Cordelia Langford, John Sillitoe on behalf of the Wellcome Sanger Institute COVID-19 Surveillance Team |
| EPI_ISL_2998722 | Lighthouse Lab in Glasgow | Wellcome Sanger Institute for the COVID-19 Genomics UK (COG-UK) Consortium | Harper VanSteenhouse, Yumi Kasai, David Gray, Carol Clugston, Anna Dominiczak and Alex Alderton, Roberto Amato, Jeffrey Barrett, Sonia Goncalves, Ewan Harrison, David K. Jackson, Ian Johnston, Dominic Kwiatkowski, Cordelia Langford, John Sillitoe on behalf of the Wellcome Sanger Institute COVID-19 Surveillance Team |
| EPI_ISL_2998723, EPI_ISL_2998724, EPI_ISL_2998725, EPI_ISL_2998726, EPI_ISL_2998727, EPI_ISL_2998728 | Lighthouse Laboratory Plymouth | Wellcome Sanger Institute for the COVID-19 Genomics UK (COG-UK) Consortium | Lighthouse Laboratory Plymouth and Alex Alderton, Roberto Amato, Jeffrey Barrett, Sonia Goncalves, Ewan Harrison, David K. Jackson, Ian Johnston, Dominic Kwiatkowski, Cordelia Langford, John Sillitoe on behalf of the Wellcome Sanger Institute COVID-19 Surveillance Team |
| EPI_ISL_2998729 | Lighthouse Lab in Glasgow | Wellcome Sanger Institute for the COVID-19 Genomics UK (COG-UK) Consortium | Harper VanSteenhouse, Yumi Kasai, David Gray, Carol Clugston, Anna Dominiczak and Alex Alderton, Roberto Amato, Jeffrey Barrett, Sonia Goncalves, Ewan Harrison, David K. Jackson, Ian Johnston, Dominic Kwiatkowski, Cordelia Langford, John Sillitoe on behalf of the Wellcome Sanger Institute COVID-19 Surveillance Team |
| EPI_ISL_2998730, EPI_ISL_2998731 | Lighthouse Laboratory Plymouth | Wellcome Sanger Institute for the COVID-19 Genomics UK (COG-UK) Consortium | Lighthouse Laboratory Plymouth and Alex Alderton, Roberto Amato, Jeffrey Barrett, Sonia Goncalves, Ewan Harrison, David K. Jackson, Ian Johnston, Dominic Kwiatkowski, Cordelia Langford, John Sillitoe on behalf of the Wellcome Sanger Institute COVID-19 Surveillance Team |
| EPI_ISL_2998732 | Lighthouse Lab in Glasgow | Wellcome Sanger Institute for the COVID-19 Genomics UK (COG-UK) Consortium | Harper VanSteenhouse, Yumi Kasai, David Gray, Carol Clugston, Anna Dominiczak and Alex Alderton, Roberto Amato, Jeffrey Barrett, Sonia Goncalves, Ewan Harrison, David K. Jackson, Ian Johnston, Dominic Kwiatkowski, Cordelia Langford, John Sillitoe on behalf of the Wellcome Sanger Institute COVID-19 Surveillance Team |
| EPI_ISL_2998733 | Lighthouse Laboratory Plymouth | Wellcome Sanger Institute for the COVID-19 Genomics UK (COG-UK) Consortium | Lighthouse Laboratory Plymouth and Alex Alderton, Roberto Amato, Jeffrey Barrett, Sonia Goncalves, Ewan Harrison, David K. Jackson, Ian Johnston, Dominic Kwiatkowski, Cordelia Langford, John Sillitoe on behalf of the Wellcome Sanger Institute COVID-19 Surveillance Team |
| EPI_ISL_2998734 | Lighthouse Lab in Milton Keynes | Wellcome Sanger Institute for the COVID-19 Genomics UK (COG-UK) Consortium | The Lighthouse Lab in Milton Keynes and Alex Alderton, Roberto Amato, Jeffrey Barrett, Sonia Goncalves, Ewan Harrison, David K. Jackson, Ian Johnston, Dominic Kwiatkowski, Cordelia Langford, John Sillitoe on behalf of the Wellcome Sanger Institute COVID-19 Surveillance Team |
| EPI_ISL_2998735 | Berkshire and Surrey Pathology Services Lighthouse Laboratory | Wellcome Sanger Institute for the COVID-19 Genomics UK (COG-UK) Consortium | Berkshire and Surrey Pathology Services Lighthouse Laboratory and Alex Alderton, Roberto Amato, Jeffrey Barrett, Sonia Goncalves, Ewan Harrison, David K. Jackson, Ian Johnston, Dominic Kwiatkowski, Cordelia Langford, John Sillitoe on behalf of the Wellcome Sanger Institute COVID-19 Surveillance Team |
| EPI_ISL_2998736 | Lighthouse Lab in Milton Keynes | Wellcome Sanger Institute for the COVID-19 Genomics UK (COG-UK) Consortium | The Lighthouse Lab in Milton Keynes and Alex Alderton, Roberto Amato, Jeffrey Barrett, Sonia Goncalves, Ewan Harrison, David K. Jackson, Ian Johnston, Dominic Kwiatkowski, Cordelia Langford, John Sillitoe on behalf of the Wellcome Sanger Institute COVID-19 Surveillance Team |
| EPI_ISL_2998737 | Berkshire and Surrey Pathology Services Lighthouse Laboratory | Wellcome Sanger Institute for the COVID-19 Genomics UK (COG-UK) Consortium | Berkshire and Surrey Pathology Services Lighthouse Laboratory and Alex Alderton, Roberto Amato, Jeffrey Barrett, Sonia Goncalves, Ewan Harrison, David K. Jackson, Ian Johnston, Dominic Kwiatkowski, Cordelia Langford, John Sillitoe on behalf of the Wellcome Sanger Institute COVID-19 Surveillance Team |
| EPI_ISL_2998738 | Lighthouse Lab in Glasgow | Wellcome Sanger Institute for the COVID-19 Genomics UK (COG-UK) Consortium | Harper VanSteenhouse, Yumi Kasai, David Gray, Carol Clugston, Anna Dominiczak and Alex Alderton, Roberto Amato, Jeffrey Barrett, Sonia Goncalves, Ewan Harrison, David K. Jackson, Ian Johnston, Dominic Kwiatkowski, Cordelia Langford, John Sillitoe on behalf of the Wellcome Sanger Institute COVID-19 Surveillance Team |
| EPI_ISL_2998739 | Berkshire and Surrey Pathology Services Lighthouse Laboratory | Wellcome Sanger Institute for the COVID-19 Genomics UK (COG-UK) Consortium | Berkshire and Surrey Pathology Services Lighthouse Laboratory and Alex Alderton, Roberto Amato, Jeffrey Barrett, Sonia Goncalves, Ewan Harrison, David K. Jackson, Ian Johnston, Dominic Kwiatkowski, Cordelia Langford, John Sillitoe on behalf of the Wellcome Sanger Institute COVID-19 Surveillance Team |
| EPI_ISL_2998740 | Lighthouse Lab in Milton Keynes | Wellcome Sanger Institute for the COVID-19 Genomics UK (COG-UK) Consortium | The Lighthouse Lab in Milton Keynes and Alex Alderton, Roberto Amato, Jeffrey Barrett, Sonia Goncalves, Ewan Harrison, David K. Jackson, Ian Johnston, Dominic Kwiatkowski, Cordelia Langford, John Sillitoe on behalf of the Wellcome Sanger Institute COVID-19 Surveillance Team |
| EPI_ISL_2998741 | Lighthouse Lab in Glasgow | Wellcome Sanger Institute for the COVID-19 Genomics UK (COG-UK) Consortium | Harper VanSteenhouse, Yumi Kasai, David Gray, Carol Clugston, Anna Dominiczak and Alex Alderton, Roberto Amato, Jeffrey Barrett, Sonia Goncalves, Ewan Harrison, David K. Jackson, Ian Johnston, Dominic Kwiatkowski, Cordelia Langford, John Sillitoe on behalf of the Wellcome Sanger Institute COVID-19 Surveillance Team |
| EPI_ISL_2998742 | Lighthouse Laboratory Plymouth | Wellcome Sanger Institute for the COVID-19 Genomics UK (COG-UK) Consortium | Lighthouse Laboratory Plymouth and Alex Alderton, Roberto Amato, Jeffrey Barrett, Sonia Goncalves, Ewan Harrison, David K. Jackson, Ian Johnston, Dominic Kwiatkowski, Cordelia Langford, John Sillitoe on behalf of the Wellcome Sanger Institute COVID-19 Surveillance Team |
| EPI_ISL_2998743 | Lighthouse Lab in Glasgow | Wellcome Sanger Institute for the COVID-19 Genomics UK (COG-UK) Consortium | Harper VanSteenhouse, Yumi Kasai, David Gray, Carol Clugston, Anna Dominiczak and Alex Alderton, Roberto Amato, Jeffrey Barrett, Sonia Goncalves, Ewan Harrison, David K. Jackson, Ian Johnston, Dominic Kwiatkowski, Cordelia Langford, John Sillitoe on behalf of the Wellcome Sanger Institute COVID-19 Surveillance Team |
| EPI_ISL_2998744 | Berkshire and Surrey Pathology Services Lighthouse Laboratory | Wellcome Sanger Institute for the COVID-19 Genomics UK (COG-UK) Consortium | Berkshire and Surrey Pathology Services Lighthouse Laboratory and Alex Alderton, Roberto Amato, Jeffrey Barrett, Sonia Goncalves, Ewan Harrison, David K. Jackson, Ian Johnston, Dominic Kwiatkowski, Cordelia Langford, John Sillitoe on behalf of the Wellcome Sanger Institute COVID-19 Surveillance Team |
| EPI_ISL_2998745 | Lighthouse Lab in Glasgow | Wellcome Sanger Institute for the COVID-19 Genomics UK (COG-UK) Consortium | Harper VanSteenhouse, Yumi Kasai, David Gray, Carol Clugston, Anna Dominiczak and Alex Alderton, Roberto Amato, Jeffrey Barrett, Sonia Goncalves, Ewan Harrison, David K. Jackson, Ian Johnston, Dominic Kwiatkowski, Cordelia Langford, John Sillitoe on behalf of the Wellcome Sanger Institute COVID-19 Surveillance Team |
| EPI_ISL_2998746 | Berkshire and Surrey Pathology Services Lighthouse Laboratory | Wellcome Sanger Institute for the COVID-19 Genomics UK (COG-UK) Consortium | Berkshire and Surrey Pathology Services Lighthouse Laboratory and Alex Alderton, Roberto Amato, Jeffrey Barrett, Sonia Goncalves, Ewan Harrison, David K. Jackson, Ian Johnston, Dominic Kwiatkowski, Cordelia Langford, John Sillitoe on behalf of the Wellcome Sanger Institute COVID-19 Surveillance Team |
| EPI_ISL_2998747 | Lighthouse Lab in Glasgow | Wellcome Sanger Institute for the COVID-19 Genomics UK (COG-UK) Consortium | Harper VanSteenhouse, Yumi Kasai, David Gray, Carol Clugston, Anna Dominiczak and Alex Alderton, Roberto Amato, Jeffrey Barrett, Sonia Goncalves, Ewan Harrison, David K. Jackson, Ian Johnston, Dominic Kwiatkowski, Cordelia Langford, John Sillitoe on behalf of the Wellcome Sanger Institute COVID-19 Surveillance Team |
| EPI_ISL_2998748, EPI_ISL_2998749 | Lighthouse Laboratory Plymouth | Wellcome Sanger Institute for the COVID-19 Genomics UK (COG-UK) Consortium | Lighthouse Laboratory Plymouth and Alex Alderton, Roberto Amato, Jeffrey Barrett, Sonia Goncalves, Ewan Harrison, David K. Jackson, Ian Johnston, Dominic Kwiatkowski, Cordelia Langford, John Sillitoe on behalf of the Wellcome Sanger Institute COVID-19 Surveillance Team |
| EPI_ISL_2998750 | Berkshire and Surrey Pathology Services Lighthouse Laboratory | Wellcome Sanger Institute for the COVID-19 Genomics UK (COG-UK) Consortium | Berkshire and Surrey Pathology Services Lighthouse Laboratory and Alex Alderton, Roberto Amato, Jeffrey Barrett, Sonia Goncalves, Ewan Harrison, David K. Jackson, Ian Johnston, Dominic Kwiatkowski, Cordelia Langford, John Sillitoe on behalf of the Wellcome Sanger Institute COVID-19 Surveillance Team |
| EPI_ISL_2998751 | Lighthouse Lab in Glasgow | Wellcome Sanger Institute for the COVID-19 Genomics UK (COG-UK) Consortium | Harper VanSteenhouse, Yumi Kasai, David Gray, Carol Clugston, Anna Dominiczak and Alex Alderton, Roberto Amato, Jeffrey Barrett, Sonia Goncalves, Ewan Harrison, David K. Jackson, Ian Johnston, Dominic Kwiatkowski, Cordelia Langford, John Sillitoe on behalf of the Wellcome Sanger Institute COVID-19 Surveillance Team |

| | | | |
|---|---|---|---|
| EPI_ISL_2998752 | Lighthouse Laboratory Plymouth | Wellcome Sanger Institute for the COVID-19 Genomics UK (COG-UK) Consortium | Lighthouse Laboratory Plymouth and Alex Alderton, Roberto Amato, Jeffrey Barrett, Sonia Goncalves, Ewan Harrison, David K. Jackson, Ian Johnston, Dominic Kwiatkowski, Cordelia Langford, John Sillitoe on behalf of the Wellcome Sanger Institute COVID-19 Surveillance Team |
| EPI_ISL_2998753 | Berkshire and Surrey Pathology Services Lighthouse Laboratory | Wellcome Sanger Institute for the COVID-19 Genomics UK (COG-UK) Consortium | Berkshire and Surrey Pathology Services Lighthouse Laboratory and Alex Alderton, Roberto Amato, Jeffrey Barrett, Sonia Goncalves, Ewan Harrison, David K. Jackson, Ian Johnston, Dominic Kwiatkowski, Cordelia Langford, John Sillitoe on behalf of the Wellcome Sanger Institute COVID-19 Surveillance Team |
| EPI_ISL_2998754 | Lighthouse Lab in Glasgow | Wellcome Sanger Institute for the COVID-19 Genomics UK (COG-UK) Consortium | Harper VanSteenhouse, Yumi Kasai, David Gray, Carol Clugston, Anna Dominiczak and Alex Alderton, Roberto Amato, Jeffrey Barrett, Sonia Goncalves, Ewan Harrison, David K. Jackson, Ian Johnston, Dominic Kwiatkowski, Cordelia Langford, John Sillitoe on behalf of the Wellcome Sanger Institute COVID-19 Surveillance Team |
| EPI_ISL_2998755, EPI_ISL_2998756 | Lighthouse Laboratory Plymouth | Wellcome Sanger Institute for the COVID-19 Genomics UK (COG-UK) Consortium | Lighthouse Laboratory Plymouth and Alex Alderton, Roberto Amato, Jeffrey Barrett, Sonia Goncalves, Ewan Harrison, David K. Jackson, Ian Johnston, Dominic Kwiatkowski, Cordelia Langford, John Sillitoe on behalf of the Wellcome Sanger Institute COVID-19 Surveillance Team |
| EPI_ISL_2998757 | Lighthouse Lab in Milton Keynes | Wellcome Sanger Institute for the COVID-19 Genomics UK (COG-UK) Consortium | The Lighthouse Lab in Milton Keynes and Alex Alderton, Roberto Amato, Jeffrey Barrett, Sonia Goncalves, Ewan Harrison, David K. Jackson, Ian Johnston, Dominic Kwiatkowski, Cordelia Langford, John Sillitoe on behalf of the Wellcome Sanger Institute COVID-19 Surveillance Team |
| EPI_ISL_2998758 | Lighthouse Laboratory Plymouth | Wellcome Sanger Institute for the COVID-19 Genomics UK (COG-UK) Consortium | Lighthouse Laboratory Plymouth and Alex Alderton, Roberto Amato, Jeffrey Barrett, Sonia Goncalves, Ewan Harrison, David K. Jackson, Ian Johnston, Dominic Kwiatkowski, Cordelia Langford, John Sillitoe on behalf of the Wellcome Sanger Institute COVID-19 Surveillance Team |
| EPI_ISL_2998759 | Berkshire and Surrey Pathology Services Lighthouse Laboratory | Wellcome Sanger Institute for the COVID-19 Genomics UK (COG-UK) Consortium | Berkshire and Surrey Pathology Services Lighthouse Laboratory and Alex Alderton, Roberto Amato, Jeffrey Barrett, Sonia Goncalves, Ewan Harrison, David K. Jackson, Ian Johnston, Dominic Kwiatkowski, Cordelia Langford, John Sillitoe on behalf of the Wellcome Sanger Institute COVID-19 Surveillance Team |
| EPI_ISL_2998760 | Lighthouse Laboratory Plymouth | Wellcome Sanger Institute for the COVID-19 Genomics UK (COG-UK) Consortium | Lighthouse Laboratory Plymouth and Alex Alderton, Roberto Amato, Jeffrey Barrett, Sonia Goncalves, Ewan Harrison, David K. Jackson, Ian Johnston, Dominic Kwiatkowski, Cordelia Langford, John Sillitoe on behalf of the Wellcome Sanger Institute COVID-19 Surveillance Team |
| EPI_ISL_2998761 | Lighthouse Lab in Glasgow | Wellcome Sanger Institute for the COVID-19 Genomics UK (COG-UK) Consortium | Harper VanSteenhouse, Yumi Kasai, David Gray, Carol Clugston, Anna Dominiczak and Alex Alderton, Roberto Amato, Jeffrey Barrett, Sonia Goncalves, Ewan Harrison, David K. Jackson, Ian Johnston, Dominic Kwiatkowski, Cordelia Langford, John Sillitoe on behalf of the Wellcome Sanger Institute COVID-19 Surveillance Team |
| EPI_ISL_2998762, EPI_ISL_2998763 | Lighthouse Laboratory Plymouth | Wellcome Sanger Institute for the COVID-19 Genomics UK (COG-UK) Consortium | Lighthouse Laboratory Plymouth and Alex Alderton, Roberto Amato, Jeffrey Barrett, Sonia Goncalves, Ewan Harrison, David K. Jackson, Ian Johnston, Dominic Kwiatkowski, Cordelia Langford, John Sillitoe on behalf of the Wellcome Sanger Institute COVID-19 Surveillance Team |
| EPI_ISL_2998764 | Lighthouse Lab in Glasgow | Wellcome Sanger Institute for the COVID-19 Genomics UK (COG-UK) Consortium | Harper VanSteenhouse, Yumi Kasai, David Gray, Carol Clugston, Anna Dominiczak and Alex Alderton, Roberto Amato, Jeffrey Barrett, Sonia Goncalves, Ewan Harrison, David K. Jackson, Ian Johnston, Dominic Kwiatkowski, Cordelia Langford, John Sillitoe on behalf of the Wellcome Sanger Institute COVID-19 Surveillance Team |
| EPI_ISL_2998765 | Lighthouse Lab in Milton Keynes | Wellcome Sanger Institute for the COVID-19 Genomics UK (COG-UK) Consortium | The Lighthouse Lab in Milton Keynes and Alex Alderton, Roberto Amato, Jeffrey Barrett, Sonia Goncalves, Ewan Harrison, David K. Jackson, Ian Johnston, Dominic Kwiatkowski, Cordelia Langford, John Sillitoe on behalf of the Wellcome Sanger Institute COVID-19 Surveillance Team |
| EPI_ISL_2998766, EPI_ISL_2998767 | Lighthouse Laboratory Plymouth | Wellcome Sanger Institute for the COVID-19 Genomics UK (COG-UK) Consortium | Lighthouse Laboratory Plymouth and Alex Alderton, Roberto Amato, Jeffrey Barrett, Sonia Goncalves, Ewan Harrison, David K. Jackson, Ian Johnston, Dominic Kwiatkowski, Cordelia Langford, John Sillitoe on behalf of the Wellcome Sanger Institute COVID-19 Surveillance Team |
| EPI_ISL_2998768 | Berkshire and Surrey Pathology Services Lighthouse Laboratory | Wellcome Sanger Institute for the COVID-19 Genomics UK (COG-UK) Consortium | Berkshire and Surrey Pathology Services Lighthouse Laboratory and Alex Alderton, Roberto Amato, Jeffrey Barrett, Sonia Goncalves, Ewan Harrison, David K. Jackson, Ian Johnston, Dominic Kwiatkowski, Cordelia Langford, John Sillitoe on behalf of the Wellcome Sanger Institute COVID-19 Surveillance Team |
| EPI_ISL_2998769 | Lighthouse Laboratory Plymouth | Wellcome Sanger Institute for the COVID-19 Genomics UK (COG-UK) Consortium | Lighthouse Laboratory Plymouth and Alex Alderton, Roberto Amato, Jeffrey Barrett, Sonia Goncalves, Ewan Harrison, David K. Jackson, Ian Johnston, Dominic Kwiatkowski, Cordelia Langford, John Sillitoe on behalf of the Wellcome Sanger Institute COVID-19 Surveillance Team |
| EPI_ISL_2998770 | Lighthouse Lab in Milton Keynes | Wellcome Sanger Institute for the COVID-19 Genomics UK (COG-UK) Consortium | The Lighthouse Lab in Milton Keynes and Alex Alderton, Roberto Amato, Jeffrey Barrett, Sonia Goncalves, Ewan Harrison, David K. Jackson, Ian Johnston, Dominic Kwiatkowski, Cordelia Langford, John Sillitoe on behalf of the Wellcome Sanger Institute COVID-19 Surveillance Team |
| EPI_ISL_2998771, EPI_ISL_2998772, EPI_ISL_2998773 | Lighthouse Lab in Glasgow | Wellcome Sanger Institute for the COVID-19 Genomics UK (COG-UK) Consortium | Harper VanSteenhouse, Yumi Kasai, David Gray, Carol Clugston, Anna Dominiczak and Alex Alderton, Roberto Amato, Jeffrey Barrett, Sonia Goncalves, Ewan Harrison, David K. Jackson, Ian Johnston, Dominic Kwiatkowski, Cordelia Langford, John Sillitoe on behalf of the Wellcome Sanger Institute COVID-19 Surveillance Team |
| EPI_ISL_2998774, EPI_ISL_2998775 | Lighthouse Lab in Milton Keynes | Wellcome Sanger Institute for the COVID-19 Genomics UK (COG-UK) Consortium | The Lighthouse Lab in Milton Keynes and Alex Alderton, Roberto Amato, Jeffrey Barrett, Sonia Goncalves, Ewan Harrison, David K. Jackson, Ian Johnston, Dominic Kwiatkowski, Cordelia Langford, John Sillitoe on behalf of the Wellcome Sanger Institute COVID-19 Surveillance Team |
| EPI_ISL_2998776 | Lighthouse Laboratory Plymouth | Wellcome Sanger Institute for the COVID-19 Genomics UK (COG-UK) Consortium | Lighthouse Laboratory Plymouth and Alex Alderton, Roberto Amato, Jeffrey Barrett, Sonia Goncalves, Ewan Harrison, David K. Jackson, Ian Johnston, Dominic Kwiatkowski, Cordelia Langford, John Sillitoe on behalf of the Wellcome Sanger Institute COVID-19 Surveillance Team |
| EPI_ISL_2998777 | Berkshire and Surrey Pathology Services Lighthouse Laboratory | Wellcome Sanger Institute for the COVID-19 Genomics UK (COG-UK) Consortium | Berkshire and Surrey Pathology Services Lighthouse Laboratory and Alex Alderton, Roberto Amato, Jeffrey Barrett, Sonia Goncalves, Ewan Harrison, David K. Jackson, Ian Johnston, Dominic Kwiatkowski, Cordelia Langford, John Sillitoe on behalf of the Wellcome Sanger Institute COVID-19 Surveillance Team |
| EPI_ISL_2998778, EPI_ISL_2998779 | Lighthouse Lab in Milton Keynes | Wellcome Sanger Institute for the COVID-19 Genomics UK (COG-UK) Consortium | The Lighthouse Lab in Milton Keynes and Alex Alderton, Roberto Amato, Jeffrey Barrett, Sonia Goncalves, Ewan Harrison, David K. Jackson, Ian Johnston, Dominic Kwiatkowski, Cordelia Langford, John Sillitoe on behalf of the Wellcome Sanger Institute COVID-19 Surveillance Team |
| EPI_ISL_2998780 | Lighthouse Lab in Glasgow | Wellcome Sanger Institute for the COVID-19 Genomics UK (COG-UK) Consortium | Harper VanSteenhouse, Yumi Kasai, David Gray, Carol Clugston, Anna Dominiczak and Alex Alderton, Roberto Amato, Jeffrey Barrett, Sonia Goncalves, Ewan Harrison, David K. Jackson, Ian Johnston, Dominic Kwiatkowski, Cordelia Langford, John Sillitoe on behalf of the Wellcome Sanger Institute COVID-19 Surveillance Team |
| EPI_ISL_2998781 | Berkshire and Surrey Pathology Services Lighthouse Laboratory | Wellcome Sanger Institute for the COVID-19 Genomics UK (COG-UK) Consortium | Berkshire and Surrey Pathology Services Lighthouse Laboratory and Alex Alderton, Roberto Amato, Jeffrey Barrett, Sonia Goncalves, Ewan Harrison, David K. Jackson, Ian Johnston, Dominic Kwiatkowski, Cordelia Langford, John Sillitoe on behalf of the Wellcome Sanger Institute COVID-19 Surveillance Team |
| EPI_ISL_2998782 | Lighthouse Lab in Milton Keynes | Wellcome Sanger Institute for the COVID-19 Genomics UK (COG-UK) Consortium | The Lighthouse Lab in Milton Keynes and Alex Alderton, Roberto Amato, Jeffrey Barrett, Sonia Goncalves, Ewan Harrison, David K. Jackson, Ian Johnston, Dominic Kwiatkowski, Cordelia Langford, John Sillitoe on behalf of the Wellcome Sanger Institute COVID-19 Surveillance Team |
| EPI_ISL_2998783, EPI_ISL_2998784 | Berkshire and Surrey Pathology Services Lighthouse Laboratory | Wellcome Sanger Institute for the COVID-19 Genomics UK (COG-UK) Consortium | Berkshire and Surrey Pathology Services Lighthouse Laboratory and Alex Alderton, Roberto Amato, Jeffrey Barrett, Sonia Goncalves, Ewan Harrison, David K. Jackson, Ian Johnston, Dominic Kwiatkowski, Cordelia Langford, John Sillitoe on behalf of the Wellcome Sanger Institute COVID-19 Surveillance Team |
| EPI_ISL_2998785 | Lighthouse Lab in Milton Keynes | Wellcome Sanger Institute for the COVID-19 Genomics UK (COG-UK) Consortium | The Lighthouse Lab in Milton Keynes and Alex Alderton, Roberto Amato, Jeffrey Barrett, Sonia Goncalves, Ewan Harrison, David K. Jackson, Ian Johnston, Dominic Kwiatkowski, Cordelia Langford, John Sillitoe on behalf of the Wellcome Sanger Institute COVID-19 Surveillance Team |
| EPI_ISL_2998786 | Berkshire and Surrey Pathology Services Lighthouse Laboratory | Wellcome Sanger Institute for the COVID-19 Genomics UK (COG-UK) Consortium | Berkshire and Surrey Pathology Services Lighthouse Laboratory and Alex Alderton, Roberto Amato, Jeffrey Barrett, Sonia Goncalves, Ewan Harrison, David K. Jackson, Ian Johnston, Dominic Kwiatkowski, Cordelia Langford, John Sillitoe on behalf of the Wellcome Sanger Institute COVID-19 Surveillance Team |
| EPI_ISL_2998787 | Lighthouse Lab in Glasgow | Wellcome Sanger Institute for the COVID-19 Genomics UK (COG-UK) Consortium | Harper VanSteenhouse, Yumi Kasai, David Gray, Carol Clugston, Anna Dominiczak and Alex Alderton, Roberto Amato, Jeffrey Barrett, Sonia Goncalves, Ewan Harrison, David K. Jackson, Ian Johnston, Dominic Kwiatkowski, Cordelia Langford, John Sillitoe on behalf of the Wellcome Sanger Institute COVID-19 Surveillance Team |
| EPI_ISL_2998788 | Lighthouse Lab in Milton Keynes | Wellcome Sanger Institute for the COVID-19 Genomics UK (COG-UK) Consortium | The Lighthouse Lab in Milton Keynes and Alex Alderton, Roberto Amato, Jeffrey Barrett, Sonia Goncalves, Ewan Harrison, David K. Jackson, Ian Johnston, Dominic Kwiatkowski, Cordelia Langford, John Sillitoe on behalf of the Wellcome Sanger Institute COVID-19 Surveillance Team |
| EPI_ISL_2998789 | Berkshire and Surrey Pathology Services Lighthouse Laboratory | Wellcome Sanger Institute for the COVID-19 Genomics UK (COG-UK) Consortium | Berkshire and Surrey Pathology Services Lighthouse Laboratory and Alex Alderton, Roberto Amato, Jeffrey Barrett, Sonia Goncalves, Ewan Harrison, David K. Jackson, Ian Johnston, Dominic Kwiatkowski, Cordelia Langford, John Sillitoe on behalf of the Wellcome Sanger Institute COVID-19 Surveillance Team |
| EPI_ISL_2998790 | Lighthouse Laboratory Plymouth | Wellcome Sanger Institute for the COVID-19 Genomics UK (COG-UK) Consortium | Lighthouse Laboratory Plymouth and Alex Alderton, Roberto Amato, Jeffrey Barrett, Sonia Goncalves, Ewan Harrison, David K. Jackson, Ian Johnston, Dominic Kwiatkowski, Cordelia Langford, John Sillitoe on behalf of the Wellcome Sanger Institute COVID-19 Surveillance Team |
| EPI_ISL_2998791, EPI_ISL_2998792 | Berkshire and Surrey Pathology Services Lighthouse Laboratory | Wellcome Sanger Institute for the COVID-19 Genomics UK (COG-UK) Consortium | Berkshire and Surrey Pathology Services Lighthouse Laboratory and Alex Alderton, Roberto Amato, Jeffrey Barrett, Sonia Goncalves, Ewan Harrison, David K. Jackson, Ian Johnston, Dominic Kwiatkowski, Cordelia Langford, John Sillitoe on behalf of the Wellcome Sanger Institute COVID-19 Surveillance Team |
| EPI_ISL_2998793 | Lighthouse Laboratory Plymouth | Wellcome Sanger Institute for the COVID-19 Genomics UK (COG-UK) Consortium | Lighthouse Laboratory Plymouth and Alex Alderton, Roberto Amato, Jeffrey Barrett, Sonia Goncalves, Ewan Harrison, David K. Jackson, Ian Johnston, Dominic Kwiatkowski, Cordelia Langford, John Sillitoe on behalf of the Wellcome Sanger Institute COVID-19 Surveillance Team |
| EPI_ISL_2998794 | Berkshire and Surrey Pathology Services Lighthouse Laboratory | Wellcome Sanger Institute for the COVID-19 Genomics UK (COG-UK) Consortium | Berkshire and Surrey Pathology Services Lighthouse Laboratory and Alex Alderton, Roberto Amato, Jeffrey Barrett, Sonia Goncalves, Ewan Harrison, David K. Jackson, Ian Johnston, Dominic Kwiatkowski, Cordelia Langford, John Sillitoe on behalf of the Wellcome Sanger Institute COVID-19 Surveillance Team |
| EPI_ISL_2998795 | Lighthouse Laboratory Plymouth | Wellcome Sanger Institute for the COVID-19 Genomics UK (COG-UK) Consortium | Lighthouse Laboratory Plymouth and Alex Alderton, Roberto Amato, Jeffrey Barrett, Sonia Goncalves, Ewan Harrison, David K. Jackson, Ian Johnston, Dominic Kwiatkowski, Cordelia Langford, John Sillitoe on behalf of the Wellcome Sanger Institute COVID-19 Surveillance Team |
| EPI_ISL_2998796 | Lighthouse Lab in Glasgow | Wellcome Sanger Institute for the COVID-19 Genomics UK (COG-UK) Consortium | Harper VanSteenhouse, Yumi Kasai, David Gray, Carol Clugston, Anna Dominiczak and Alex Alderton, Roberto Amato, Jeffrey Barrett, Sonia Goncalves, Ewan Harrison, David K. Jackson, Ian Johnston, Dominic Kwiatkowski, Cordelia Langford, John Sillitoe on behalf of the Wellcome Sanger Institute COVID-19 Surveillance Team |
| EPI_ISL_2998797 | Lighthouse Laboratory Plymouth | Wellcome Sanger Institute for the COVID-19 Genomics UK (COG-UK) Consortium | Lighthouse Laboratory Plymouth and Alex Alderton, Roberto Amato, Jeffrey Barrett, Sonia Goncalves, Ewan Harrison, David K. Jackson, Ian Johnston, Dominic Kwiatkowski, Cordelia Langford, John Sillitoe on behalf of the Wellcome Sanger Institute COVID-19 Surveillance Team |
| EPI_ISL_2998798 | Lighthouse Lab in Glasgow | Wellcome Sanger Institute for the COVID-19 Genomics UK (COG-UK) Consortium | Harper VanSteenhouse, Yumi Kasai, David Gray, Carol Clugston, Anna Dominiczak and Alex Alderton, Roberto Amato, Jeffrey Barrett, Sonia Goncalves, Ewan Harrison, David K. Jackson, Ian Johnston, Dominic Kwiatkowski, Cordelia Langford, John Sillitoe on behalf of the Wellcome Sanger Institute COVID-19 Surveillance Team |
| EPI_ISL_2998799 | Lighthouse Lab in Milton Keynes | Wellcome Sanger Institute for the COVID-19 Genomics UK (COG-UK) Consortium | The Lighthouse Lab in Milton Keynes and Alex Alderton, Roberto Amato, Jeffrey Barrett, Sonia Goncalves, Ewan Harrison, David K. Jackson, Ian Johnston, Dominic Kwiatkowski, Cordelia Langford, John Sillitoe on behalf of the Wellcome Sanger Institute COVID-19 Surveillance Team |
| EPI_ISL_2998800, EPI_ISL_2998801 | Lighthouse Laboratory Plymouth | Wellcome Sanger Institute for the COVID-19 Genomics UK (COG-UK) Consortium | Lighthouse Laboratory Plymouth and Alex Alderton, Roberto Amato, Jeffrey Barrett, Sonia Goncalves, Ewan Harrison, David K. Jackson, Ian Johnston, Dominic Kwiatkowski, Cordelia Langford, John Sillitoe on behalf of the Wellcome Sanger Institute COVID-19 Surveillance Team |
| EPI_ISL_2998802 | Lighthouse Lab in Milton Keynes | Wellcome Sanger Institute for the COVID-19 Genomics UK (COG-UK) Consortium | The Lighthouse Lab in Milton Keynes and Alex Alderton, Roberto Amato, Jeffrey Barrett, Sonia Goncalves, Ewan Harrison, David K. Jackson, Ian Johnston, Dominic Kwiatkowski, Cordelia Langford, John Sillitoe on behalf of the Wellcome Sanger Institute COVID-19 Surveillance Team |
| EPI_ISL_2998803 | Lighthouse Lab in Glasgow | Wellcome Sanger Institute for the COVID-19 Genomics UK (COG-UK) Consortium | Harper VanSteenhouse, Yumi Kasai, David Gray, Carol Clugston, Anna Dominiczak and Alex Alderton, Roberto Amato, Jeffrey Barrett, Sonia Goncalves, Ewan Harrison, David K. Jackson, Ian Johnston, Dominic Kwiatkowski, Cordelia Langford, John Sillitoe on behalf of the Wellcome Sanger Institute COVID-19 Surveillance Team |
| EPI_ISL_2998804 | Lighthouse Lab in Milton Keynes | Wellcome Sanger Institute for the COVID-19 Genomics UK (COG-UK) Consortium | The Lighthouse Lab in Milton Keynes and Alex Alderton, Roberto Amato, Jeffrey Barrett, Sonia Goncalves, Ewan Harrison, David K. Jackson, Ian Johnston, Dominic Kwiatkowski, Cordelia Langford, John Sillitoe on behalf of the Wellcome Sanger Institute COVID-19 Surveillance Team |
| EPI_ISL_2998805 | Lighthouse Laboratory Plymouth | Wellcome Sanger Institute for the COVID-19 Genomics UK (COG-UK) Consortium | Lighthouse Laboratory Plymouth and Alex Alderton, Roberto Amato, Jeffrey Barrett, Sonia Goncalves, Ewan Harrison, David K. Jackson, Ian Johnston, Dominic Kwiatkowski, Cordelia Langford, John Sillitoe on behalf of the Wellcome Sanger Institute COVID-19 Surveillance Team |
| EPI_ISL_2998806 | Lighthouse Lab in Milton Keynes | Wellcome Sanger Institute for the COVID-19 Genomics UK (COG-UK) Consortium | The Lighthouse Lab in Milton Keynes and Alex Alderton, Roberto Amato, Jeffrey Barrett, Sonia Goncalves, Ewan Harrison, David K. Jackson, Ian Johnston, Dominic Kwiatkowski, Cordelia Langford, John Sillitoe on behalf of the Wellcome Sanger Institute COVID-19 Surveillance Team |

| | | | |
|---|---|---|---|
| EPI_ISL_2998807 | Lighthouse Lab in Glasgow | Wellcome Sanger Institute for the COVID-19 Genomics UK (COG-UK) Consortium | Harper VanSteenhouse, Yumi Kasai, David Gray, Carol Clugston, Anna Dominiczak and Alex Alderton, Roberto Amato, Jeffrey Barrett, Sonia Goncalves, Ewan Harrison, David K. Jackson, Ian Johnston, Dominic Kwiatkowski, Cordelia Langford, John Sillitoe on behalf of the Wellcome Sanger Institute COVID-19 Surveillance Team |
| EPI_ISL_2998808, EPI_ISL_2998809, EPI_ISL_2998810 | Lighthouse Laboratory Plymouth | Wellcome Sanger Institute for the COVID-19 Genomics UK (COG-UK) Consortium | Lighthouse Laboratory Plymouth and Alex Alderton, Roberto Amato, Jeffrey Barrett, Sonia Goncalves, Ewan Harrison, David K. Jackson, Ian Johnston, Dominic Kwiatkowski, Cordelia Langford, John Sillitoe on behalf of the Wellcome Sanger Institute COVID-19 Surveillance Team |
| EPI_ISL_2998811, EPI_ISL_2998812, EPI_ISL_2998813 | Lighthouse Lab in Glasgow | Wellcome Sanger Institute for the COVID-19 Genomics UK (COG-UK) Consortium | Harper VanSteenhouse, Yumi Kasai, David Gray, Carol Clugston, Anna Dominiczak and Alex Alderton, Roberto Amato, Jeffrey Barrett, Sonia Goncalves, Ewan Harrison, David K. Jackson, Ian Johnston, Dominic Kwiatkowski, Cordelia Langford, John Sillitoe on behalf of the Wellcome Sanger Institute COVID-19 Surveillance Team |
| EPI_ISL_2998814 | Lighthouse Lab in Milton Keynes | Wellcome Sanger Institute for the COVID-19 Genomics UK (COG-UK) Consortium | The Lighthouse Lab in Milton Keynes and Alex Alderton, Roberto Amato, Jeffrey Barrett, Sonia Goncalves, Ewan Harrison, David K. Jackson, Ian Johnston, Dominic Kwiatkowski, Cordelia Langford, John Sillitoe on behalf of the Wellcome Sanger Institute COVID-19 Surveillance Team |
| EPI_ISL_2998815 | Berkshire and Surrey Pathology Services Lighthouse Laboratory | Wellcome Sanger Institute for the COVID-19 Genomics UK (COG-UK) Consortium | Berkshire and Surrey Pathology Services Lighthouse Laboratory and Alex Alderton, Roberto Amato, Jeffrey Barrett, Sonia Goncalves, Ewan Harrison, David K. Jackson, Ian Johnston, Dominic Kwiatkowski, Cordelia Langford, John Sillitoe on behalf of the Wellcome Sanger Institute COVID-19 Surveillance Team |
| EPI_ISL_2998816 | Lighthouse Laboratory Plymouth | Wellcome Sanger Institute for the COVID-19 Genomics UK (COG-UK) Consortium | Lighthouse Laboratory Plymouth and Alex Alderton, Roberto Amato, Jeffrey Barrett, Sonia Goncalves, Ewan Harrison, David K. Jackson, Ian Johnston, Dominic Kwiatkowski, Cordelia Langford, John Sillitoe on behalf of the Wellcome Sanger Institute COVID-19 Surveillance Team |
| EPI_ISL_2998817, EPI_ISL_2998818 | Berkshire and Surrey Pathology Services Lighthouse Laboratory | Wellcome Sanger Institute for the COVID-19 Genomics UK (COG-UK) Consortium | Berkshire and Surrey Pathology Services Lighthouse Laboratory and Alex Alderton, Roberto Amato, Jeffrey Barrett, Sonia Goncalves, Ewan Harrison, David K. Jackson, Ian Johnston, Dominic Kwiatkowski, Cordelia Langford, John Sillitoe on behalf of the Wellcome Sanger Institute COVID-19 Surveillance Team |
| EPI_ISL_2998819, EPI_ISL_2998820 | Lighthouse Laboratory Plymouth | Wellcome Sanger Institute for the COVID-19 Genomics UK (COG-UK) Consortium | Lighthouse Laboratory Plymouth and Alex Alderton, Roberto Amato, Jeffrey Barrett, Sonia Goncalves, Ewan Harrison, David K. Jackson, Ian Johnston, Dominic Kwiatkowski, Cordelia Langford, John Sillitoe on behalf of the Wellcome Sanger Institute COVID-19 Surveillance Team |
| EPI_ISL_2998821, EPI_ISL_2998822 | Lighthouse Lab in Milton Keynes | Wellcome Sanger Institute for the COVID-19 Genomics UK (COG-UK) Consortium | The Lighthouse Lab in Milton Keynes and Alex Alderton, Roberto Amato, Jeffrey Barrett, Sonia Goncalves, Ewan Harrison, David K. Jackson, Ian Johnston, Dominic Kwiatkowski, Cordelia Langford, John Sillitoe on behalf of the Wellcome Sanger Institute COVID-19 Surveillance Team |
| EPI_ISL_2998823, EPI_ISL_2998824, EPI_ISL_2998825 | Lighthouse Lab in Glasgow | Wellcome Sanger Institute for the COVID-19 Genomics UK (COG-UK) Consortium | Harper VanSteenhouse, Yumi Kasai, David Gray, Carol Clugston, Anna Dominiczak and Alex Alderton, Roberto Amato, Jeffrey Barrett, Sonia Goncalves, Ewan Harrison, David K. Jackson, Ian Johnston, Dominic Kwiatkowski, Cordelia Langford, John Sillitoe on behalf of the Wellcome Sanger Institute COVID-19 Surveillance Team |
| EPI_ISL_2998826, EPI_ISL_2998827 | Lighthouse Laboratory Plymouth | Wellcome Sanger Institute for the COVID-19 Genomics UK (COG-UK) Consortium | Lighthouse Laboratory Plymouth and Alex Alderton, Roberto Amato, Jeffrey Barrett, Sonia Goncalves, Ewan Harrison, David K. Jackson, Ian Johnston, Dominic Kwiatkowski, Cordelia Langford, John Sillitoe on behalf of the Wellcome Sanger Institute COVID-19 Surveillance Team |
| EPI_ISL_2998828 | Lighthouse Lab in Glasgow | Wellcome Sanger Institute for the COVID-19 Genomics UK (COG-UK) Consortium | Harper VanSteenhouse, Yumi Kasai, David Gray, Carol Clugston, Anna Dominiczak and Alex Alderton, Roberto Amato, Jeffrey Barrett, Sonia Goncalves, Ewan Harrison, David K. Jackson, Ian Johnston, Dominic Kwiatkowski, Cordelia Langford, John Sillitoe on behalf of the Wellcome Sanger Institute COVID-19 Surveillance Team |
| EPI_ISL_2998829 | Lighthouse Laboratory Plymouth | Wellcome Sanger Institute for the COVID-19 Genomics UK (COG-UK) Consortium | Lighthouse Laboratory Plymouth and Alex Alderton, Roberto Amato, Jeffrey Barrett, Sonia Goncalves, Ewan Harrison, David K. Jackson, Ian Johnston, Dominic Kwiatkowski, Cordelia Langford, John Sillitoe on behalf of the Wellcome Sanger Institute COVID-19 Surveillance Team |
| EPI_ISL_2998830 | Lighthouse Lab in Glasgow | Wellcome Sanger Institute for the COVID-19 Genomics UK (COG-UK) Consortium | Harper VanSteenhouse, Yumi Kasai, David Gray, Carol Clugston, Anna Dominiczak and Alex Alderton, Roberto Amato, Jeffrey Barrett, Sonia Goncalves, Ewan Harrison, David K. Jackson, Ian Johnston, Dominic Kwiatkowski, Cordelia Langford, John Sillitoe on behalf of the Wellcome Sanger Institute COVID-19 Surveillance Team |
| EPI_ISL_2998831 | Lighthouse Lab in Milton Keynes | Wellcome Sanger Institute for the COVID-19 Genomics UK (COG-UK) Consortium | The Lighthouse Lab in Milton Keynes and Alex Alderton, Roberto Amato, Jeffrey Barrett, Sonia Goncalves, Ewan Harrison, David K. Jackson, Ian Johnston, Dominic Kwiatkowski, Cordelia Langford, John Sillitoe on behalf of the Wellcome Sanger Institute COVID-19 Surveillance Team |
| EPI_ISL_2998832 | Lighthouse Laboratory Plymouth | Wellcome Sanger Institute for the COVID-19 Genomics UK (COG-UK) Consortium | Lighthouse Laboratory Plymouth and Alex Alderton, Roberto Amato, Jeffrey Barrett, Sonia Goncalves, Ewan Harrison, David K. Jackson, Ian Johnston, Dominic Kwiatkowski, Cordelia Langford, John Sillitoe on behalf of the Wellcome Sanger Institute COVID-19 Surveillance Team |
| EPI_ISL_2998833 | Lighthouse Lab in Glasgow | Wellcome Sanger Institute for the COVID-19 Genomics UK (COG-UK) Consortium | Harper VanSteenhouse, Yumi Kasai, David Gray, Carol Clugston, Anna Dominiczak and Alex Alderton, Roberto Amato, Jeffrey Barrett, Sonia Goncalves, Ewan Harrison, David K. Jackson, Ian Johnston, Dominic Kwiatkowski, Cordelia Langford, John Sillitoe on behalf of the Wellcome Sanger Institute COVID-19 Surveillance Team |
| EPI_ISL_2998834 | Lighthouse Lab in Milton Keynes | Wellcome Sanger Institute for the COVID-19 Genomics UK (COG-UK) Consortium | The Lighthouse Lab in Milton Keynes and Alex Alderton, Roberto Amato, Jeffrey Barrett, Sonia Goncalves, Ewan Harrison, David K. Jackson, Ian Johnston, Dominic Kwiatkowski, Cordelia Langford, John Sillitoe on behalf of the Wellcome Sanger Institute COVID-19 Surveillance Team |
| EPI_ISL_2998835 | Lighthouse Lab in Glasgow | Wellcome Sanger Institute for the COVID-19 Genomics UK (COG-UK) Consortium | Harper VanSteenhouse, Yumi Kasai, David Gray, Carol Clugston, Anna Dominiczak and Alex Alderton, Roberto Amato, Jeffrey Barrett, Sonia Goncalves, Ewan Harrison, David K. Jackson, Ian Johnston, Dominic Kwiatkowski, Cordelia Langford, John Sillitoe on behalf of the Wellcome Sanger Institute COVID-19 Surveillance Team |
| EPI_ISL_2998836 | Lighthouse Lab in Milton Keynes | Wellcome Sanger Institute for the COVID-19 Genomics UK (COG-UK) Consortium | The Lighthouse Lab in Milton Keynes and Alex Alderton, Roberto Amato, Jeffrey Barrett, Sonia Goncalves, Ewan Harrison, David K. Jackson, Ian Johnston, Dominic Kwiatkowski, Cordelia Langford, John Sillitoe on behalf of the Wellcome Sanger Institute COVID-19 Surveillance Team |
| EPI_ISL_2998837 | Lighthouse Lab in Glasgow | Wellcome Sanger Institute for the COVID-19 Genomics UK (COG-UK) Consortium | Harper VanSteenhouse, Yumi Kasai, David Gray, Carol Clugston, Anna Dominiczak and Alex Alderton, Roberto Amato, Jeffrey Barrett, Sonia Goncalves, Ewan Harrison, David K. Jackson, Ian Johnston, Dominic Kwiatkowski, Cordelia Langford, John Sillitoe on behalf of the Wellcome Sanger Institute COVID-19 Surveillance Team |
| EPI_ISL_2998838, EPI_ISL_2998839 | Lighthouse Lab in Milton Keynes | Wellcome Sanger Institute for the COVID-19 Genomics UK (COG-UK) Consortium | The Lighthouse Lab in Milton Keynes and Alex Alderton, Roberto Amato, Jeffrey Barrett, Sonia Goncalves, Ewan Harrison, David K. Jackson, Ian Johnston, Dominic Kwiatkowski, Cordelia Langford, John Sillitoe on behalf of the Wellcome Sanger Institute COVID-19 Surveillance Team |
| EPI_ISL_2998840 | Lighthouse Laboratory Plymouth | Wellcome Sanger Institute for the COVID-19 Genomics UK (COG-UK) Consortium | Lighthouse Laboratory Plymouth and Alex Alderton, Roberto Amato, Jeffrey Barrett, Sonia Goncalves, Ewan Harrison, David K. Jackson, Ian Johnston, Dominic Kwiatkowski, Cordelia Langford, John Sillitoe on behalf of the Wellcome Sanger Institute COVID-19 Surveillance Team |
| EPI_ISL_2998841 | Lighthouse Lab in Glasgow | Wellcome Sanger Institute for the COVID-19 Genomics UK (COG-UK) Consortium | Harper VanSteenhouse, Yumi Kasai, David Gray, Carol Clugston, Anna Dominiczak and Alex Alderton, Roberto Amato, Jeffrey Barrett, Sonia Goncalves, Ewan Harrison, David K. Jackson, Ian Johnston, Dominic Kwiatkowski, Cordelia Langford, John Sillitoe on behalf of the Wellcome Sanger Institute COVID-19 Surveillance Team |
| EPI_ISL_2998842, EPI_ISL_2998843 | Lighthouse Lab in Milton Keynes | Wellcome Sanger Institute for the COVID-19 Genomics UK (COG-UK) Consortium | The Lighthouse Lab in Milton Keynes and Alex Alderton, Roberto Amato, Jeffrey Barrett, Sonia Goncalves, Ewan Harrison, David K. Jackson, Ian Johnston, Dominic Kwiatkowski, Cordelia Langford, John Sillitoe on behalf of the Wellcome Sanger Institute COVID-19 Surveillance Team |
| EPI_ISL_2998844 | Berkshire and Surrey Pathology Services Lighthouse Laboratory | Wellcome Sanger Institute for the COVID-19 Genomics UK (COG-UK) Consortium | Berkshire and Surrey Pathology Services Lighthouse Laboratory and Alex Alderton, Roberto Amato, Jeffrey Barrett, Sonia Goncalves, Ewan Harrison, David K. Jackson, Ian Johnston, Dominic Kwiatkowski, Cordelia Langford, John Sillitoe on behalf of the Wellcome Sanger Institute COVID-19 Surveillance Team |
| EPI_ISL_2998845 | Lighthouse Lab in Milton Keynes | Wellcome Sanger Institute for the COVID-19 Genomics UK (COG-UK) Consortium | The Lighthouse Lab in Milton Keynes and Alex Alderton, Roberto Amato, Jeffrey Barrett, Sonia Goncalves, Ewan Harrison, David K. Jackson, Ian Johnston, Dominic Kwiatkowski, Cordelia Langford, John Sillitoe on behalf of the Wellcome Sanger Institute COVID-19 Surveillance Team |
| EPI_ISL_2998846, EPI_ISL_2998847, EPI_ISL_2998848 | Lighthouse Laboratory Plymouth | Wellcome Sanger Institute for the COVID-19 Genomics UK (COG-UK) Consortium | Lighthouse Laboratory Plymouth and Alex Alderton, Roberto Amato, Jeffrey Barrett, Sonia Goncalves, Ewan Harrison, David K. Jackson, Ian Johnston, Dominic Kwiatkowski, Cordelia Langford, John Sillitoe on behalf of the Wellcome Sanger Institute COVID-19 Surveillance Team |
| EPI_ISL_2998849, EPI_ISL_2998850 | Lighthouse Lab in Glasgow | Wellcome Sanger Institute for the COVID-19 Genomics UK (COG-UK) Consortium | Harper VanSteenhouse, Yumi Kasai, David Gray, Carol Clugston, Anna Dominiczak and Alex Alderton, Roberto Amato, Jeffrey Barrett, Sonia Goncalves, Ewan Harrison, David K. Jackson, Ian Johnston, Dominic Kwiatkowski, Cordelia Langford, John Sillitoe on behalf of the Wellcome Sanger Institute COVID-19 Surveillance Team |
| EPI_ISL_2998851 | Lighthouse Lab in Milton Keynes | Wellcome Sanger Institute for the COVID-19 Genomics UK (COG-UK) Consortium | The Lighthouse Lab in Milton Keynes and Alex Alderton, Roberto Amato, Jeffrey Barrett, Sonia Goncalves, Ewan Harrison, David K. Jackson, Ian Johnston, Dominic Kwiatkowski, Cordelia Langford, John Sillitoe on behalf of the Wellcome Sanger Institute COVID-19 Surveillance Team |
| EPI_ISL_2998852 | Lighthouse Laboratory Plymouth | Wellcome Sanger Institute for the COVID-19 Genomics UK (COG-UK) Consortium | Lighthouse Laboratory Plymouth and Alex Alderton, Roberto Amato, Jeffrey Barrett, Sonia Goncalves, Ewan Harrison, David K. Jackson, Ian Johnston, Dominic Kwiatkowski, Cordelia Langford, John Sillitoe on behalf of the Wellcome Sanger Institute COVID-19 Surveillance Team |
| EPI_ISL_2998853, EPI_ISL_2998854 | Lighthouse Lab in Milton Keynes | Wellcome Sanger Institute for the COVID-19 Genomics UK (COG-UK) Consortium | The Lighthouse Lab in Milton Keynes and Alex Alderton, Roberto Amato, Jeffrey Barrett, Sonia Goncalves, Ewan Harrison, David K. Jackson, Ian Johnston, Dominic Kwiatkowski, Cordelia Langford, John Sillitoe on behalf of the Wellcome Sanger Institute COVID-19 Surveillance Team |
| EPI_ISL_2998855 | Lighthouse Laboratory Plymouth | Wellcome Sanger Institute for the COVID-19 Genomics UK (COG-UK) Consortium | Lighthouse Laboratory Plymouth and Alex Alderton, Roberto Amato, Jeffrey Barrett, Sonia Goncalves, Ewan Harrison, David K. Jackson, Ian Johnston, Dominic Kwiatkowski, Cordelia Langford, John Sillitoe on behalf of the Wellcome Sanger Institute COVID-19 Surveillance Team |
| EPI_ISL_2998856, EPI_ISL_2998857 | Berkshire and Surrey Pathology Services Lighthouse Laboratory | Wellcome Sanger Institute for the COVID-19 Genomics UK (COG-UK) Consortium | Berkshire and Surrey Pathology Services Lighthouse Laboratory and Alex Alderton, Roberto Amato, Jeffrey Barrett, Sonia Goncalves, Ewan Harrison, David K. Jackson, Ian Johnston, Dominic Kwiatkowski, Cordelia Langford, John Sillitoe on behalf of the Wellcome Sanger Institute COVID-19 Surveillance Team |
| EPI_ISL_2998858 | Lighthouse Laboratory Plymouth | Wellcome Sanger Institute for the COVID-19 Genomics UK (COG-UK) Consortium | Lighthouse Laboratory Plymouth and Alex Alderton, Roberto Amato, Jeffrey Barrett, Sonia Goncalves, Ewan Harrison, David K. Jackson, Ian Johnston, Dominic Kwiatkowski, Cordelia Langford, John Sillitoe on behalf of the Wellcome Sanger Institute COVID-19 Surveillance Team |
| EPI_ISL_2998859 | Berkshire and Surrey Pathology Services Lighthouse Laboratory | Wellcome Sanger Institute for the COVID-19 Genomics UK (COG-UK) Consortium | Berkshire and Surrey Pathology Services Lighthouse Laboratory and Alex Alderton, Roberto Amato, Jeffrey Barrett, Sonia Goncalves, Ewan Harrison, David K. Jackson, Ian Johnston, Dominic Kwiatkowski, Cordelia Langford, John Sillitoe on behalf of the Wellcome Sanger Institute COVID-19 Surveillance Team |
| EPI_ISL_2998860, EPI_ISL_2998861 | Lighthouse Lab in Glasgow | Wellcome Sanger Institute for the COVID-19 Genomics UK (COG-UK) Consortium | Harper VanSteenhouse, Yumi Kasai, David Gray, Carol Clugston, Anna Dominiczak and Alex Alderton, Roberto Amato, Jeffrey Barrett, Sonia Goncalves, Ewan Harrison, David K. Jackson, Ian Johnston, Dominic Kwiatkowski, Cordelia Langford, John Sillitoe on behalf of the Wellcome Sanger Institute COVID-19 Surveillance Team |
| EPI_ISL_2998862 | Lighthouse Lab in Milton Keynes | Wellcome Sanger Institute for the COVID-19 Genomics UK (COG-UK) Consortium | The Lighthouse Lab in Milton Keynes and Alex Alderton, Roberto Amato, Jeffrey Barrett, Sonia Goncalves, Ewan Harrison, David K. Jackson, Ian Johnston, Dominic Kwiatkowski, Cordelia Langford, John Sillitoe on behalf of the Wellcome Sanger Institute COVID-19 Surveillance Team |
| EPI_ISL_2998863 | Berkshire and Surrey Pathology Services Lighthouse Laboratory | Wellcome Sanger Institute for the COVID-19 Genomics UK (COG-UK) Consortium | Berkshire and Surrey Pathology Services Lighthouse Laboratory and Alex Alderton, Roberto Amato, Jeffrey Barrett, Sonia Goncalves, Ewan Harrison, David K. Jackson, Ian Johnston, Dominic Kwiatkowski, Cordelia Langford, John Sillitoe on behalf of the Wellcome Sanger Institute COVID-19 Surveillance Team |
| EPI_ISL_2998864 | Lighthouse Lab in Glasgow | Wellcome Sanger Institute for the COVID-19 Genomics UK (COG-UK) Consortium | Harper VanSteenhouse, Yumi Kasai, David Gray, Carol Clugston, Anna Dominiczak and Alex Alderton, Roberto Amato, Jeffrey Barrett, Sonia Goncalves, Ewan Harrison, David K. Jackson, Ian Johnston, Dominic Kwiatkowski, Cordelia Langford, John Sillitoe on behalf of the Wellcome Sanger Institute COVID-19 Surveillance Team |
| EPI_ISL_2998865 | Berkshire and Surrey Pathology Services Lighthouse Laboratory | Wellcome Sanger Institute for the COVID-19 Genomics UK (COG-UK) Consortium | Berkshire and Surrey Pathology Services Lighthouse Laboratory and Alex Alderton, Roberto Amato, Jeffrey Barrett, Sonia Goncalves, Ewan Harrison, David K. Jackson, Ian Johnston, Dominic Kwiatkowski, Cordelia Langford, John Sillitoe on behalf of the Wellcome Sanger Institute COVID-19 Surveillance Team |
| EPI_ISL_2998866 | Lighthouse Lab in Milton Keynes | Wellcome Sanger Institute for the COVID-19 Genomics UK (COG-UK) Consortium | The Lighthouse Lab in Milton Keynes and Alex Alderton, Roberto Amato, Jeffrey Barrett, Sonia Goncalves, Ewan Harrison, David K. Jackson, Ian Johnston, Dominic Kwiatkowski, Cordelia Langford, John Sillitoe on behalf of the Wellcome Sanger Institute COVID-19 Surveillance Team |
| EPI_ISL_2998867 | Berkshire and Surrey Pathology Services Lighthouse Laboratory | Wellcome Sanger Institute for the COVID-19 Genomics UK (COG-UK) Consortium | Berkshire and Surrey Pathology Services Lighthouse Laboratory and Alex Alderton, Roberto Amato, Jeffrey Barrett, Sonia Goncalves, Ewan Harrison, David K. Jackson, Ian Johnston, Dominic Kwiatkowski, Cordelia Langford, John Sillitoe on behalf of the Wellcome Sanger Institute COVID-19 Surveillance Team |

| | | | |
|---|---|---|---|
| EPI_ISL_2998868 | Lighthouse Lab in Glasgow | Wellcome Sanger Institute for the COVID-19 Genomics UK (COG-UK) Consortium | Harper VanSteenhouse, Yumi Kasai, David Gray, Carol Clugston, Anna Dominiczak and Alex Alderton, Roberto Amato, Jeffrey Barrett, Sonia Goncalves, Ewan Harrison, David K. Jackson, Ian Johnston, Dominic Kwiatkowski, Cordelia Langford, John Sillitoe on behalf of the Wellcome Sanger Institute COVID-19 Surveillance Team |
| EPI_ISL_2998869 | Lighthouse Lab in Milton Keynes | Wellcome Sanger Institute for the COVID-19 Genomics UK (COG-UK) Consortium | The Lighthouse Lab in Milton Keynes and Alex Alderton, Roberto Amato, Jeffrey Barrett, Sonia Goncalves, Ewan Harrison, David K. Jackson, Ian Johnston, Dominic Kwiatkowski, Cordelia Langford, John Sillitoe on behalf of the Wellcome Sanger Institute COVID-19 Surveillance Team |
| EPI_ISL_2998870, EPI_ISL_2998871 | Lighthouse Laboratory Plymouth | Wellcome Sanger Institute for the COVID-19 Genomics UK (COG-UK) Consortium | Lighthouse Laboratory Plymouth and Alex Alderton, Roberto Amato, Jeffrey Barrett, Sonia Goncalves, Ewan Harrison, David K. Jackson, Ian Johnston, Dominic Kwiatkowski, Cordelia Langford, John Sillitoe on behalf of the Wellcome Sanger Institute COVID-19 Surveillance Team |
| EPI_ISL_2998872 | Lighthouse Lab in Glasgow | Wellcome Sanger Institute for the COVID-19 Genomics UK (COG-UK) Consortium | Harper VanSteenhouse, Yumi Kasai, David Gray, Carol Clugston, Anna Dominiczak and Alex Alderton, Roberto Amato, Jeffrey Barrett, Sonia Goncalves, Ewan Harrison, David K. Jackson, Ian Johnston, Dominic Kwiatkowski, Cordelia Langford, John Sillitoe on behalf of the Wellcome Sanger Institute COVID-19 Surveillance Team |
| EPI_ISL_2998873, EPI_ISL_2998874, EPI_ISL_2998875, EPI_ISL_2998876 | Berkshire and Surrey Pathology Services Lighthouse Laboratory | Wellcome Sanger Institute for the COVID-19 Genomics UK (COG-UK) Consortium | Berkshire and Surrey Pathology Services Lighthouse Laboratory and Alex Alderton, Roberto Amato, Jeffrey Barrett, Sonia Goncalves, Ewan Harrison, David K. Jackson, Ian Johnston, Dominic Kwiatkowski, Cordelia Langford, John Sillitoe on behalf of the Wellcome Sanger Institute COVID-19 Surveillance Team |
| EPI_ISL_2998877 | Lighthouse Lab in Milton Keynes | Wellcome Sanger Institute for the COVID-19 Genomics UK (COG-UK) Consortium | The Lighthouse Lab in Milton Keynes and Alex Alderton, Roberto Amato, Jeffrey Barrett, Sonia Goncalves, Ewan Harrison, David K. Jackson, Ian Johnston, Dominic Kwiatkowski, Cordelia Langford, John Sillitoe on behalf of the Wellcome Sanger Institute COVID-19 Surveillance Team |
| EPI_ISL_2998878 | Lighthouse Laboratory Plymouth | Wellcome Sanger Institute for the COVID-19 Genomics UK (COG-UK) Consortium | Lighthouse Laboratory Plymouth and Alex Alderton, Roberto Amato, Jeffrey Barrett, Sonia Goncalves, Ewan Harrison, David K. Jackson, Ian Johnston, Dominic Kwiatkowski, Cordelia Langford, John Sillitoe on behalf of the Wellcome Sanger Institute COVID-19 Surveillance Team |
| EPI_ISL_2998879 | Berkshire and Surrey Pathology Services Lighthouse Laboratory | Wellcome Sanger Institute for the COVID-19 Genomics UK (COG-UK) Consortium | Berkshire and Surrey Pathology Services Lighthouse Laboratory and Alex Alderton, Roberto Amato, Jeffrey Barrett, Sonia Goncalves, Ewan Harrison, David K. Jackson, Ian Johnston, Dominic Kwiatkowski, Cordelia Langford, John Sillitoe on behalf of the Wellcome Sanger Institute COVID-19 Surveillance Team |
| EPI_ISL_2998880 | Lighthouse Lab in Glasgow | Wellcome Sanger Institute for the COVID-19 Genomics UK (COG-UK) Consortium | Harper VanSteenhouse, Yumi Kasai, David Gray, Carol Clugston, Anna Dominiczak and Alex Alderton, Roberto Amato, Jeffrey Barrett, Sonia Goncalves, Ewan Harrison, David K. Jackson, Ian Johnston, Dominic Kwiatkowski, Cordelia Langford, John Sillitoe on behalf of the Wellcome Sanger Institute COVID-19 Surveillance Team |
| EPI_ISL_2998881 | Lighthouse Laboratory Plymouth | Wellcome Sanger Institute for the COVID-19 Genomics UK (COG-UK) Consortium | Lighthouse Laboratory Plymouth and Alex Alderton, Roberto Amato, Jeffrey Barrett, Sonia Goncalves, Ewan Harrison, David K. Jackson, Ian Johnston, Dominic Kwiatkowski, Cordelia Langford, John Sillitoe on behalf of the Wellcome Sanger Institute COVID-19 Surveillance Team |
| EPI_ISL_2998882 | Berkshire and Surrey Pathology Services Lighthouse Laboratory | Wellcome Sanger Institute for the COVID-19 Genomics UK (COG-UK) Consortium | Berkshire and Surrey Pathology Services Lighthouse Laboratory and Alex Alderton, Roberto Amato, Jeffrey Barrett, Sonia Goncalves, Ewan Harrison, David K. Jackson, Ian Johnston, Dominic Kwiatkowski, Cordelia Langford, John Sillitoe on behalf of the Wellcome Sanger Institute COVID-19 Surveillance Team |
| EPI_ISL_2998883 | Lighthouse Lab in Milton Keynes | Wellcome Sanger Institute for the COVID-19 Genomics UK (COG-UK) Consortium | The Lighthouse Lab in Milton Keynes and Alex Alderton, Roberto Amato, Jeffrey Barrett, Sonia Goncalves, Ewan Harrison, David K. Jackson, Ian Johnston, Dominic Kwiatkowski, Cordelia Langford, John Sillitoe on behalf of the Wellcome Sanger Institute COVID-19 Surveillance Team |
| EPI_ISL_2998884 | Lighthouse Lab in Glasgow | Wellcome Sanger Institute for the COVID-19 Genomics UK (COG-UK) Consortium | Harper VanSteenhouse, Yumi Kasai, David Gray, Carol Clugston, Anna Dominiczak and Alex Alderton, Roberto Amato, Jeffrey Barrett, Sonia Goncalves, Ewan Harrison, David K. Jackson, Ian Johnston, Dominic Kwiatkowski, Cordelia Langford, John Sillitoe on behalf of the Wellcome Sanger Institute COVID-19 Surveillance Team |
| EPI_ISL_2998885 | Lighthouse Laboratory Plymouth | Wellcome Sanger Institute for the COVID-19 Genomics UK (COG-UK) Consortium | Lighthouse Laboratory Plymouth and Alex Alderton, Roberto Amato, Jeffrey Barrett, Sonia Goncalves, Ewan Harrison, David K. Jackson, Ian Johnston, Dominic Kwiatkowski, Cordelia Langford, John Sillitoe on behalf of the Wellcome Sanger Institute COVID-19 Surveillance Team |
| EPI_ISL_2998886 | Berkshire and Surrey Pathology Services Lighthouse Laboratory | Wellcome Sanger Institute for the COVID-19 Genomics UK (COG-UK) Consortium | Berkshire and Surrey Pathology Services Lighthouse Laboratory and Alex Alderton, Roberto Amato, Jeffrey Barrett, Sonia Goncalves, Ewan Harrison, David K. Jackson, Ian Johnston, Dominic Kwiatkowski, Cordelia Langford, John Sillitoe on behalf of the Wellcome Sanger Institute COVID-19 Surveillance Team |
| EPI_ISL_2998887 | Lighthouse Lab in Glasgow | Wellcome Sanger Institute for the COVID-19 Genomics UK (COG-UK) Consortium | Harper VanSteenhouse, Yumi Kasai, David Gray, Carol Clugston, Anna Dominiczak and Alex Alderton, Roberto Amato, Jeffrey Barrett, Sonia Goncalves, Ewan Harrison, David K. Jackson, Ian Johnston, Dominic Kwiatkowski, Cordelia Langford, John Sillitoe on behalf of the Wellcome Sanger Institute COVID-19 Surveillance Team |
| EPI_ISL_2998888, EPI_ISL_2998889 | Lighthouse Lab in Milton Keynes | Wellcome Sanger Institute for the COVID-19 Genomics UK (COG-UK) Consortium | The Lighthouse Lab in Milton Keynes and Alex Alderton, Roberto Amato, Jeffrey Barrett, Sonia Goncalves, Ewan Harrison, David K. Jackson, Ian Johnston, Dominic Kwiatkowski, Cordelia Langford, John Sillitoe on behalf of the Wellcome Sanger Institute COVID-19 Surveillance Team |
| EPI_ISL_2998890 | Berkshire and Surrey Pathology Services Lighthouse Laboratory | Wellcome Sanger Institute for the COVID-19 Genomics UK (COG-UK) Consortium | Berkshire and Surrey Pathology Services Lighthouse Laboratory and Alex Alderton, Roberto Amato, Jeffrey Barrett, Sonia Goncalves, Ewan Harrison, David K. Jackson, Ian Johnston, Dominic Kwiatkowski, Cordelia Langford, John Sillitoe on behalf of the Wellcome Sanger Institute COVID-19 Surveillance Team |
| EPI_ISL_2998891 | Lighthouse Laboratory Plymouth | Wellcome Sanger Institute for the COVID-19 Genomics UK (COG-UK) Consortium | Lighthouse Laboratory Plymouth and Alex Alderton, Roberto Amato, Jeffrey Barrett, Sonia Goncalves, Ewan Harrison, David K. Jackson, Ian Johnston, Dominic Kwiatkowski, Cordelia Langford, John Sillitoe on behalf of the Wellcome Sanger Institute COVID-19 Surveillance Team |
| EPI_ISL_2998892 | Lighthouse Lab in Milton Keynes | Wellcome Sanger Institute for the COVID-19 Genomics UK (COG-UK) Consortium | The Lighthouse Lab in Milton Keynes and Alex Alderton, Roberto Amato, Jeffrey Barrett, Sonia Goncalves, Ewan Harrison, David K. Jackson, Ian Johnston, Dominic Kwiatkowski, Cordelia Langford, John Sillitoe on behalf of the Wellcome Sanger Institute COVID-19 Surveillance Team |
| EPI_ISL_2998893, EPI_ISL_2998894, EPI_ISL_2998895, EPI_ISL_2998896 | Lighthouse Lab in Glasgow | Wellcome Sanger Institute for the COVID-19 Genomics UK (COG-UK) Consortium | Harper VanSteenhouse, Yumi Kasai, David Gray, Carol Clugston, Anna Dominiczak and Alex Alderton, Roberto Amato, Jeffrey Barrett, Sonia Goncalves, Ewan Harrison, David K. Jackson, Ian Johnston, Dominic Kwiatkowski, Cordelia Langford, John Sillitoe on behalf of the Wellcome Sanger Institute COVID-19 Surveillance Team |
| EPI_ISL_2998897 | Lighthouse Lab in Milton Keynes | Wellcome Sanger Institute for the COVID-19 Genomics UK (COG-UK) Consortium | The Lighthouse Lab in Milton Keynes and Alex Alderton, Roberto Amato, Jeffrey Barrett, Sonia Goncalves, Ewan Harrison, David K. Jackson, Ian Johnston, Dominic Kwiatkowski, Cordelia Langford, John Sillitoe on behalf of the Wellcome Sanger Institute COVID-19 Surveillance Team |
| EPI_ISL_2998898, EPI_ISL_2998899, EPI_ISL_2998900 | Lighthouse Lab in Glasgow | Wellcome Sanger Institute for the COVID-19 Genomics UK (COG-UK) Consortium | Harper VanSteenhouse, Yumi Kasai, David Gray, Carol Clugston, Anna Dominiczak and Alex Alderton, Roberto Amato, Jeffrey Barrett, Sonia Goncalves, Ewan Harrison, David K. Jackson, Ian Johnston, Dominic Kwiatkowski, Cordelia Langford, John Sillitoe on behalf of the Wellcome Sanger Institute COVID-19 Surveillance Team |
| EPI_ISL_2998901 | Lighthouse Lab in Milton Keynes | Wellcome Sanger Institute for the COVID-19 Genomics UK (COG-UK) Consortium | The Lighthouse Lab in Milton Keynes and Alex Alderton, Roberto Amato, Jeffrey Barrett, Sonia Goncalves, Ewan Harrison, David K. Jackson, Ian Johnston, Dominic Kwiatkowski, Cordelia Langford, John Sillitoe on behalf of the Wellcome Sanger Institute COVID-19 Surveillance Team |
| EPI_ISL_2998902 | Lighthouse Laboratory Plymouth | Wellcome Sanger Institute for the COVID-19 Genomics UK (COG-UK) Consortium | Lighthouse Laboratory Plymouth and Alex Alderton, Roberto Amato, Jeffrey Barrett, Sonia Goncalves, Ewan Harrison, David K. Jackson, Ian Johnston, Dominic Kwiatkowski, Cordelia Langford, John Sillitoe on behalf of the Wellcome Sanger Institute COVID-19 Surveillance Team |
| EPI_ISL_2998903, EPI_ISL_2998904 | Lighthouse Lab in Glasgow | Wellcome Sanger Institute for the COVID-19 Genomics UK (COG-UK) Consortium | Harper VanSteenhouse, Yumi Kasai, David Gray, Carol Clugston, Anna Dominiczak and Alex Alderton, Roberto Amato, Jeffrey Barrett, Sonia Goncalves, Ewan Harrison, David K. Jackson, Ian Johnston, Dominic Kwiatkowski, Cordelia Langford, John Sillitoe on behalf of the Wellcome Sanger Institute COVID-19 Surveillance Team |
| EPI_ISL_2998905, EPI_ISL_2998906 | Lighthouse Laboratory Plymouth | Wellcome Sanger Institute for the COVID-19 Genomics UK (COG-UK) Consortium | Lighthouse Laboratory Plymouth and Alex Alderton, Roberto Amato, Jeffrey Barrett, Sonia Goncalves, Ewan Harrison, David K. Jackson, Ian Johnston, Dominic Kwiatkowski, Cordelia Langford, John Sillitoe on behalf of the Wellcome Sanger Institute COVID-19 Surveillance Team |
| EPI_ISL_2998907 | Berkshire and Surrey Pathology Services Lighthouse Laboratory | Wellcome Sanger Institute for the COVID-19 Genomics UK (COG-UK) Consortium | Berkshire and Surrey Pathology Services Lighthouse Laboratory and Alex Alderton, Roberto Amato, Jeffrey Barrett, Sonia Goncalves, Ewan Harrison, David K. Jackson, Ian Johnston, Dominic Kwiatkowski, Cordelia Langford, John Sillitoe on behalf of the Wellcome Sanger Institute COVID-19 Surveillance Team |
| EPI_ISL_2998908 | Lighthouse Lab in Glasgow | Wellcome Sanger Institute for the COVID-19 Genomics UK (COG-UK) Consortium | Harper VanSteenhouse, Yumi Kasai, David Gray, Carol Clugston, Anna Dominiczak and Alex Alderton, Roberto Amato, Jeffrey Barrett, Sonia Goncalves, Ewan Harrison, David K. Jackson, Ian Johnston, Dominic Kwiatkowski, Cordelia Langford, John Sillitoe on behalf of the Wellcome Sanger Institute COVID-19 Surveillance Team |
| EPI_ISL_2998909 | Lighthouse Laboratory Plymouth | Wellcome Sanger Institute for the COVID-19 Genomics UK (COG-UK) Consortium | Lighthouse Laboratory Plymouth and Alex Alderton, Roberto Amato, Jeffrey Barrett, Sonia Goncalves, Ewan Harrison, David K. Jackson, Ian Johnston, Dominic Kwiatkowski, Cordelia Langford, John Sillitoe on behalf of the Wellcome Sanger Institute COVID-19 Surveillance Team |
| EPI_ISL_2998910, EPI_ISL_2998911, EPI_ISL_2998912 | Lighthouse Lab in Glasgow | Wellcome Sanger Institute for the COVID-19 Genomics UK (COG-UK) Consortium | Harper VanSteenhouse, Yumi Kasai, David Gray, Carol Clugston, Anna Dominiczak and Alex Alderton, Roberto Amato, Jeffrey Barrett, Sonia Goncalves, Ewan Harrison, David K. Jackson, Ian Johnston, Dominic Kwiatkowski, Cordelia Langford, John Sillitoe on behalf of the Wellcome Sanger Institute COVID-19 Surveillance Team |
| EPI_ISL_2998913 | Lighthouse Lab in Milton Keynes | Wellcome Sanger Institute for the COVID-19 Genomics UK (COG-UK) Consortium | The Lighthouse Lab in Milton Keynes and Alex Alderton, Roberto Amato, Jeffrey Barrett, Sonia Goncalves, Ewan Harrison, David K. Jackson, Ian Johnston, Dominic Kwiatkowski, Cordelia Langford, John Sillitoe on behalf of the Wellcome Sanger Institute COVID-19 Surveillance Team |
| EPI_ISL_2998914 | Lighthouse Laboratory Plymouth | Wellcome Sanger Institute for the COVID-19 Genomics UK (COG-UK) Consortium | Lighthouse Laboratory Plymouth and Alex Alderton, Roberto Amato, Jeffrey Barrett, Sonia Goncalves, Ewan Harrison, David K. Jackson, Ian Johnston, Dominic Kwiatkowski, Cordelia Langford, John Sillitoe on behalf of the Wellcome Sanger Institute COVID-19 Surveillance Team |
| EPI_ISL_2998915 | Lighthouse Lab in Milton Keynes | Wellcome Sanger Institute for the COVID-19 Genomics UK (COG-UK) Consortium | The Lighthouse Lab in Milton Keynes and Alex Alderton, Roberto Amato, Jeffrey Barrett, Sonia Goncalves, Ewan Harrison, David K. Jackson, Ian Johnston, Dominic Kwiatkowski, Cordelia Langford, John Sillitoe on behalf of the Wellcome Sanger Institute COVID-19 Surveillance Team |
| EPI_ISL_2998916 | Berkshire and Surrey Pathology Services Lighthouse Laboratory | Wellcome Sanger Institute for the COVID-19 Genomics UK (COG-UK) Consortium | Berkshire and Surrey Pathology Services Lighthouse Laboratory and Alex Alderton, Roberto Amato, Jeffrey Barrett, Sonia Goncalves, Ewan Harrison, David K. Jackson, Ian Johnston, Dominic Kwiatkowski, Cordelia Langford, John Sillitoe on behalf of the Wellcome Sanger Institute COVID-19 Surveillance Team |
| EPI_ISL_2998917 | Lighthouse Laboratory Plymouth | Wellcome Sanger Institute for the COVID-19 Genomics UK (COG-UK) Consortium | Lighthouse Laboratory Plymouth and Alex Alderton, Roberto Amato, Jeffrey Barrett, Sonia Goncalves, Ewan Harrison, David K. Jackson, Ian Johnston, Dominic Kwiatkowski, Cordelia Langford, John Sillitoe on behalf of the Wellcome Sanger Institute COVID-19 Surveillance Team |
| EPI_ISL_2998918 | Lighthouse Lab in Glasgow | Wellcome Sanger Institute for the COVID-19 Genomics UK (COG-UK) Consortium | Harper VanSteenhouse, Yumi Kasai, David Gray, Carol Clugston, Anna Dominiczak and Alex Alderton, Roberto Amato, Jeffrey Barrett, Sonia Goncalves, Ewan Harrison, David K. Jackson, Ian Johnston, Dominic Kwiatkowski, Cordelia Langford, John Sillitoe on behalf of the Wellcome Sanger Institute COVID-19 Surveillance Team |
| EPI_ISL_2998919, EPI_ISL_2998920 | Lighthouse Lab in Milton Keynes | Wellcome Sanger Institute for the COVID-19 Genomics UK (COG-UK) Consortium | The Lighthouse Lab in Milton Keynes and Alex Alderton, Roberto Amato, Jeffrey Barrett, Sonia Goncalves, Ewan Harrison, David K. Jackson, Ian Johnston, Dominic Kwiatkowski, Cordelia Langford, John Sillitoe on behalf of the Wellcome Sanger Institute COVID-19 Surveillance Team |
| EPI_ISL_2998921 | Lighthouse Lab in Glasgow | Wellcome Sanger Institute for the COVID-19 Genomics UK (COG-UK) Consortium | Harper VanSteenhouse, Yumi Kasai, David Gray, Carol Clugston, Anna Dominiczak and Alex Alderton, Roberto Amato, Jeffrey Barrett, Sonia Goncalves, Ewan Harrison, David K. Jackson, Ian Johnston, Dominic Kwiatkowski, Cordelia Langford, John Sillitoe on behalf of the Wellcome Sanger Institute COVID-19 Surveillance Team |
| EPI_ISL_2998922 | Lighthouse Lab in Milton Keynes | Wellcome Sanger Institute for the COVID-19 Genomics UK (COG-UK) Consortium | The Lighthouse Lab in Milton Keynes and Alex Alderton, Roberto Amato, Jeffrey Barrett, Sonia Goncalves, Ewan Harrison, David K. Jackson, Ian Johnston, Dominic Kwiatkowski, Cordelia Langford, John Sillitoe on behalf of the Wellcome Sanger Institute COVID-19 Surveillance Team |
| EPI_ISL_2998923 | Berkshire and Surrey Pathology Services Lighthouse Laboratory | Wellcome Sanger Institute for the COVID-19 Genomics UK (COG-UK) Consortium | Berkshire and Surrey Pathology Services Lighthouse Laboratory and Alex Alderton, Roberto Amato, Jeffrey Barrett, Sonia Goncalves, Ewan Harrison, David K. Jackson, Ian Johnston, Dominic Kwiatkowski, Cordelia Langford, John Sillitoe on behalf of the Wellcome Sanger Institute COVID-19 Surveillance Team |
| EPI_ISL_2998924 | Lighthouse Laboratory Plymouth | Wellcome Sanger Institute for the COVID-19 Genomics UK (COG-UK) Consortium | Lighthouse Laboratory Plymouth and Alex Alderton, Roberto Amato, Jeffrey Barrett, Sonia Goncalves, Ewan Harrison, David K. Jackson, Ian Johnston, Dominic Kwiatkowski, Cordelia Langford, John Sillitoe on behalf of the Wellcome Sanger Institute COVID-19 Surveillance Team |
| EPI_ISL_2998925 | Lighthouse Lab in Glasgow | Wellcome Sanger Institute for the COVID-19 Genomics UK (COG-UK) Consortium | Harper VanSteenhouse, Yumi Kasai, David Gray, Carol Clugston, Anna Dominiczak and Alex Alderton, Roberto Amato, Jeffrey Barrett, Sonia Goncalves, Ewan Harrison, David K. Jackson, Ian Johnston, Dominic Kwiatkowski, Cordelia Langford, John Sillitoe on behalf of the Wellcome Sanger Institute COVID-19 Surveillance Team |

| | | | |
|---|---|---|---|
| EPI_ISL_2998926, EPI_ISL_2998927 | Lighthouse Laboratory Plymouth | Wellcome Sanger Institute for the COVID-19 Genomics UK (COG-UK) Consortium | Lighthouse Laboratory Plymouth and Alex Alderton, Roberto Amato, Jeffrey Barrett, Sonia Goncalves, Ewan Harrison, David K. Jackson, Ian Johnston, Dominic Kwiatkowski, Cordelia Langford, John Sillitoe on behalf of the Wellcome Sanger Institute COVID-19 Surveillance Team |
| EPI_ISL_2998928 | Lighthouse Lab in Glasgow | Wellcome Sanger Institute for the COVID-19 Genomics UK (COG-UK) Consortium | Harper VanSteenhouse, Yumi Kasai, David Gray, Carol Clugston, Anna Dominiczak and Alex Alderton, Roberto Amato, Jeffrey Barrett, Sonia Goncalves, Ewan Harrison, David K. Jackson, Ian Johnston, Dominic Kwiatkowski, Cordelia Langford, John Sillitoe on behalf of the Wellcome Sanger Institute COVID-19 Surveillance Team |
| EPI_ISL_2998929 | Berkshire and Surrey Pathology Services Lighthouse Laboratory | Wellcome Sanger Institute for the COVID-19 Genomics UK (COG-UK) Consortium | Berkshire and Surrey Pathology Services Lighthouse Laboratory and Alex Alderton, Roberto Amato, Jeffrey Barrett, Sonia Goncalves, Ewan Harrison, David K. Jackson, Ian Johnston, Dominic Kwiatkowski, Cordelia Langford, John Sillitoe on behalf of the Wellcome Sanger Institute COVID-19 Surveillance Team |
| EPI_ISL_2998930 | Lighthouse Lab in Milton Keynes | Wellcome Sanger Institute for the COVID-19 Genomics UK (COG-UK) Consortium | The Lighthouse Lab in Milton Keynes and Alex Alderton, Roberto Amato, Jeffrey Barrett, Sonia Goncalves, Ewan Harrison, David K. Jackson, Ian Johnston, Dominic Kwiatkowski, Cordelia Langford, John Sillitoe on behalf of the Wellcome Sanger Institute COVID-19 Surveillance Team |
| EPI_ISL_2998931 | Lighthouse Laboratory Plymouth | Wellcome Sanger Institute for the COVID-19 Genomics UK (COG-UK) Consortium | Lighthouse Laboratory Plymouth and Alex Alderton, Roberto Amato, Jeffrey Barrett, Sonia Goncalves, Ewan Harrison, David K. Jackson, Ian Johnston, Dominic Kwiatkowski, Cordelia Langford, John Sillitoe on behalf of the Wellcome Sanger Institute COVID-19 Surveillance Team |
| EPI_ISL_2998932, EPI_ISL_2998933 | Berkshire and Surrey Pathology Services Lighthouse Laboratory | Wellcome Sanger Institute for the COVID-19 Genomics UK (COG-UK) Consortium | Berkshire and Surrey Pathology Services Lighthouse Laboratory and Alex Alderton, Roberto Amato, Jeffrey Barrett, Sonia Goncalves, Ewan Harrison, David K. Jackson, Ian Johnston, Dominic Kwiatkowski, Cordelia Langford, John Sillitoe on behalf of the Wellcome Sanger Institute COVID-19 Surveillance Team |
| EPI_ISL_2998934 | Lighthouse Lab in Glasgow | Wellcome Sanger Institute for the COVID-19 Genomics UK (COG-UK) Consortium | Harper VanSteenhouse, Yumi Kasai, David Gray, Carol Clugston, Anna Dominiczak and Alex Alderton, Roberto Amato, Jeffrey Barrett, Sonia Goncalves, Ewan Harrison, David K. Jackson, Ian Johnston, Dominic Kwiatkowski, Cordelia Langford, John Sillitoe on behalf of the Wellcome Sanger Institute COVID-19 Surveillance Team |
| EPI_ISL_2998935, EPI_ISL_2998936 | Berkshire and Surrey Pathology Services Lighthouse Laboratory | Wellcome Sanger Institute for the COVID-19 Genomics UK (COG-UK) Consortium | Berkshire and Surrey Pathology Services Lighthouse Laboratory and Alex Alderton, Roberto Amato, Jeffrey Barrett, Sonia Goncalves, Ewan Harrison, David K. Jackson, Ian Johnston, Dominic Kwiatkowski, Cordelia Langford, John Sillitoe on behalf of the Wellcome Sanger Institute COVID-19 Surveillance Team |
| EPI_ISL_2998937, EPI_ISL_2998938 | Lighthouse Laboratory Plymouth | Wellcome Sanger Institute for the COVID-19 Genomics UK (COG-UK) Consortium | Lighthouse Laboratory Plymouth and Alex Alderton, Roberto Amato, Jeffrey Barrett, Sonia Goncalves, Ewan Harrison, David K. Jackson, Ian Johnston, Dominic Kwiatkowski, Cordelia Langford, John Sillitoe on behalf of the Wellcome Sanger Institute COVID-19 Surveillance Team |
| EPI_ISL_2998939 | Lighthouse Lab in Milton Keynes | Wellcome Sanger Institute for the COVID-19 Genomics UK (COG-UK) Consortium | The Lighthouse Lab in Milton Keynes and Alex Alderton, Roberto Amato, Jeffrey Barrett, Sonia Goncalves, Ewan Harrison, David K. Jackson, Ian Johnston, Dominic Kwiatkowski, Cordelia Langford, John Sillitoe on behalf of the Wellcome Sanger Institute COVID-19 Surveillance Team |
| EPI_ISL_2998940 | Lighthouse Laboratory Plymouth | Wellcome Sanger Institute for the COVID-19 Genomics UK (COG-UK) Consortium | Lighthouse Laboratory Plymouth and Alex Alderton, Roberto Amato, Jeffrey Barrett, Sonia Goncalves, Ewan Harrison, David K. Jackson, Ian Johnston, Dominic Kwiatkowski, Cordelia Langford, John Sillitoe on behalf of the Wellcome Sanger Institute COVID-19 Surveillance Team |
| EPI_ISL_2998941 | Lighthouse Lab in Glasgow | Wellcome Sanger Institute for the COVID-19 Genomics UK (COG-UK) Consortium | Harper VanSteenhouse, Yumi Kasai, David Gray, Carol Clugston, Anna Dominiczak and Alex Alderton, Roberto Amato, Jeffrey Barrett, Sonia Goncalves, Ewan Harrison, David K. Jackson, Ian Johnston, Dominic Kwiatkowski, Cordelia Langford, John Sillitoe on behalf of the Wellcome Sanger Institute COVID-19 Surveillance Team |
| EPI_ISL_2998942 | Lighthouse Lab in Milton Keynes | Wellcome Sanger Institute for the COVID-19 Genomics UK (COG-UK) Consortium | The Lighthouse Lab in Milton Keynes and Alex Alderton, Roberto Amato, Jeffrey Barrett, Sonia Goncalves, Ewan Harrison, David K. Jackson, Ian Johnston, Dominic Kwiatkowski, Cordelia Langford, John Sillitoe on behalf of the Wellcome Sanger Institute COVID-19 Surveillance Team |
| EPI_ISL_2998943 | Lighthouse Laboratory Plymouth | Wellcome Sanger Institute for the COVID-19 Genomics UK (COG-UK) Consortium | Lighthouse Laboratory Plymouth and Alex Alderton, Roberto Amato, Jeffrey Barrett, Sonia Goncalves, Ewan Harrison, David K. Jackson, Ian Johnston, Dominic Kwiatkowski, Cordelia Langford, John Sillitoe on behalf of the Wellcome Sanger Institute COVID-19 Surveillance Team |
| EPI_ISL_2998944 | Lighthouse Lab in Milton Keynes | Wellcome Sanger Institute for the COVID-19 Genomics UK (COG-UK) Consortium | The Lighthouse Lab in Milton Keynes and Alex Alderton, Roberto Amato, Jeffrey Barrett, Sonia Goncalves, Ewan Harrison, David K. Jackson, Ian Johnston, Dominic Kwiatkowski, Cordelia Langford, John Sillitoe on behalf of the Wellcome Sanger Institute COVID-19 Surveillance Team |
| EPI_ISL_2998945 | Lighthouse Laboratory Plymouth | Wellcome Sanger Institute for the COVID-19 Genomics UK (COG-UK) Consortium | Lighthouse Laboratory Plymouth and Alex Alderton, Roberto Amato, Jeffrey Barrett, Sonia Goncalves, Ewan Harrison, David K. Jackson, Ian Johnston, Dominic Kwiatkowski, Cordelia Langford, John Sillitoe on behalf of the Wellcome Sanger Institute COVID-19 Surveillance Team |
| EPI_ISL_2998946 | Lighthouse Lab in Glasgow | Wellcome Sanger Institute for the COVID-19 Genomics UK (COG-UK) Consortium | Harper VanSteenhouse, Yumi Kasai, David Gray, Carol Clugston, Anna Dominiczak and Alex Alderton, Roberto Amato, Jeffrey Barrett, Sonia Goncalves, Ewan Harrison, David K. Jackson, Ian Johnston, Dominic Kwiatkowski, Cordelia Langford, John Sillitoe on behalf of the Wellcome Sanger Institute COVID-19 Surveillance Team |
| EPI_ISL_2998947 | Lighthouse Lab in Milton Keynes | Wellcome Sanger Institute for the COVID-19 Genomics UK (COG-UK) Consortium | The Lighthouse Lab in Milton Keynes and Alex Alderton, Roberto Amato, Jeffrey Barrett, Sonia Goncalves, Ewan Harrison, David K. Jackson, Ian Johnston, Dominic Kwiatkowski, Cordelia Langford, John Sillitoe on behalf of the Wellcome Sanger Institute COVID-19 Surveillance Team |
| EPI_ISL_2998948 | Berkshire and Surrey Pathology Services Lighthouse Laboratory | Wellcome Sanger Institute for the COVID-19 Genomics UK (COG-UK) Consortium | Berkshire and Surrey Pathology Services Lighthouse Laboratory and Alex Alderton, Roberto Amato, Jeffrey Barrett, Sonia Goncalves, Ewan Harrison, David K. Jackson, Ian Johnston, Dominic Kwiatkowski, Cordelia Langford, John Sillitoe on behalf of the Wellcome Sanger Institute COVID-19 Surveillance Team |
| EPI_ISL_2998949, EPI_ISL_2998950 | Lighthouse Lab in Glasgow | Wellcome Sanger Institute for the COVID-19 Genomics UK (COG-UK) Consortium | Harper VanSteenhouse, Yumi Kasai, David Gray, Carol Clugston, Anna Dominiczak and Alex Alderton, Roberto Amato, Jeffrey Barrett, Sonia Goncalves, Ewan Harrison, David K. Jackson, Ian Johnston, Dominic Kwiatkowski, Cordelia Langford, John Sillitoe on behalf of the Wellcome Sanger Institute COVID-19 Surveillance Team |
| EPI_ISL_2998951 | Berkshire and Surrey Pathology Services Lighthouse Laboratory | Wellcome Sanger Institute for the COVID-19 Genomics UK (COG-UK) Consortium | Berkshire and Surrey Pathology Services Lighthouse Laboratory and Alex Alderton, Roberto Amato, Jeffrey Barrett, Sonia Goncalves, Ewan Harrison, David K. Jackson, Ian Johnston, Dominic Kwiatkowski, Cordelia Langford, John Sillitoe on behalf of the Wellcome Sanger Institute COVID-19 Surveillance Team |
| EPI_ISL_2998952 | Lighthouse Lab in Glasgow | Wellcome Sanger Institute for the COVID-19 Genomics UK (COG-UK) Consortium | Harper VanSteenhouse, Yumi Kasai, David Gray, Carol Clugston, Anna Dominiczak and Alex Alderton, Roberto Amato, Jeffrey Barrett, Sonia Goncalves, Ewan Harrison, David K. Jackson, Ian Johnston, Dominic Kwiatkowski, Cordelia Langford, John Sillitoe on behalf of the Wellcome Sanger Institute COVID-19 Surveillance Team |
| EPI_ISL_2998953 | Lighthouse Laboratory Plymouth | Wellcome Sanger Institute for the COVID-19 Genomics UK (COG-UK) Consortium | Lighthouse Laboratory Plymouth and Alex Alderton, Roberto Amato, Jeffrey Barrett, Sonia Goncalves, Ewan Harrison, David K. Jackson, Ian Johnston, Dominic Kwiatkowski, Cordelia Langford, John Sillitoe on behalf of the Wellcome Sanger Institute COVID-19 Surveillance Team |
| EPI_ISL_2998954, EPI_ISL_2998955 | Lighthouse Lab in Glasgow | Wellcome Sanger Institute for the COVID-19 Genomics UK (COG-UK) Consortium | Harper VanSteenhouse, Yumi Kasai, David Gray, Carol Clugston, Anna Dominiczak and Alex Alderton, Roberto Amato, Jeffrey Barrett, Sonia Goncalves, Ewan Harrison, David K. Jackson, Ian Johnston, Dominic Kwiatkowski, Cordelia Langford, John Sillitoe on behalf of the Wellcome Sanger Institute COVID-19 Surveillance Team |
| EPI_ISL_2998956 | Lighthouse Lab in Milton Keynes | Wellcome Sanger Institute for the COVID-19 Genomics UK (COG-UK) Consortium | The Lighthouse Lab in Milton Keynes and Alex Alderton, Roberto Amato, Jeffrey Barrett, Sonia Goncalves, Ewan Harrison, David K. Jackson, Ian Johnston, Dominic Kwiatkowski, Cordelia Langford, John Sillitoe on behalf of the Wellcome Sanger Institute COVID-19 Surveillance Team |
| EPI_ISL_2998957 | Berkshire and Surrey Pathology Services Lighthouse Laboratory | Wellcome Sanger Institute for the COVID-19 Genomics UK (COG-UK) Consortium | Berkshire and Surrey Pathology Services Lighthouse Laboratory and Alex Alderton, Roberto Amato, Jeffrey Barrett, Sonia Goncalves, Ewan Harrison, David K. Jackson, Ian Johnston, Dominic Kwiatkowski, Cordelia Langford, John Sillitoe on behalf of the Wellcome Sanger Institute COVID-19 Surveillance Team |
| EPI_ISL_2998958 | Lighthouse Lab in Glasgow | Wellcome Sanger Institute for the COVID-19 Genomics UK (COG-UK) Consortium | Harper VanSteenhouse, Yumi Kasai, David Gray, Carol Clugston, Anna Dominiczak and Alex Alderton, Roberto Amato, Jeffrey Barrett, Sonia Goncalves, Ewan Harrison, David K. Jackson, Ian Johnston, Dominic Kwiatkowski, Cordelia Langford, John Sillitoe on behalf of the Wellcome Sanger Institute COVID-19 Surveillance Team |
| EPI_ISL_2998959, EPI_ISL_2998960, EPI_ISL_2998961 | Berkshire and Surrey Pathology Services Lighthouse Laboratory | Wellcome Sanger Institute for the COVID-19 Genomics UK (COG-UK) Consortium | Berkshire and Surrey Pathology Services Lighthouse Laboratory and Alex Alderton, Roberto Amato, Jeffrey Barrett, Sonia Goncalves, Ewan Harrison, David K. Jackson, Ian Johnston, Dominic Kwiatkowski, Cordelia Langford, John Sillitoe on behalf of the Wellcome Sanger Institute COVID-19 Surveillance Team |
| EPI_ISL_2998962 | Lighthouse Lab in Milton Keynes | Wellcome Sanger Institute for the COVID-19 Genomics UK (COG-UK) Consortium | The Lighthouse Lab in Milton Keynes and Alex Alderton, Roberto Amato, Jeffrey Barrett, Sonia Goncalves, Ewan Harrison, David K. Jackson, Ian Johnston, Dominic Kwiatkowski, Cordelia Langford, John Sillitoe on behalf of the Wellcome Sanger Institute COVID-19 Surveillance Team |
| EPI_ISL_2998963 | Lighthouse Laboratory Plymouth | Wellcome Sanger Institute for the COVID-19 Genomics UK (COG-UK) Consortium | Lighthouse Laboratory Plymouth and Alex Alderton, Roberto Amato, Jeffrey Barrett, Sonia Goncalves, Ewan Harrison, David K. Jackson, Ian Johnston, Dominic Kwiatkowski, Cordelia Langford, John Sillitoe on behalf of the Wellcome Sanger Institute COVID-19 Surveillance Team |
| EPI_ISL_2998964 | Lighthouse Lab in Glasgow | Wellcome Sanger Institute for the COVID-19 Genomics UK (COG-UK) Consortium | Harper VanSteenhouse, Yumi Kasai, David Gray, Carol Clugston, Anna Dominiczak and Alex Alderton, Roberto Amato, Jeffrey Barrett, Sonia Goncalves, Ewan Harrison, David K. Jackson, Ian Johnston, Dominic Kwiatkowski, Cordelia Langford, John Sillitoe on behalf of the Wellcome Sanger Institute COVID-19 Surveillance Team |
| EPI_ISL_2998965 | Lighthouse Laboratory Plymouth | Wellcome Sanger Institute for the COVID-19 Genomics UK (COG-UK) Consortium | Lighthouse Laboratory Plymouth and Alex Alderton, Roberto Amato, Jeffrey Barrett, Sonia Goncalves, Ewan Harrison, David K. Jackson, Ian Johnston, Dominic Kwiatkowski, Cordelia Langford, John Sillitoe on behalf of the Wellcome Sanger Institute COVID-19 Surveillance Team |
| EPI_ISL_2998966 | Lighthouse Lab in Glasgow | Wellcome Sanger Institute for the COVID-19 Genomics UK (COG-UK) Consortium | Harper VanSteenhouse, Yumi Kasai, David Gray, Carol Clugston, Anna Dominiczak and Alex Alderton, Roberto Amato, Jeffrey Barrett, Sonia Goncalves, Ewan Harrison, David K. Jackson, Ian Johnston, Dominic Kwiatkowski, Cordelia Langford, John Sillitoe on behalf of the Wellcome Sanger Institute COVID-19 Surveillance Team |
| EPI_ISL_2998967 | Lighthouse Lab in Milton Keynes | Wellcome Sanger Institute for the COVID-19 Genomics UK (COG-UK) Consortium | The Lighthouse Lab in Milton Keynes and Alex Alderton, Roberto Amato, Jeffrey Barrett, Sonia Goncalves, Ewan Harrison, David K. Jackson, Ian Johnston, Dominic Kwiatkowski, Cordelia Langford, John Sillitoe on behalf of the Wellcome Sanger Institute COVID-19 Surveillance Team |
| EPI_ISL_2998968, EPI_ISL_2998969 | Lighthouse Laboratory Plymouth | Wellcome Sanger Institute for the COVID-19 Genomics UK (COG-UK) Consortium | Lighthouse Laboratory Plymouth and Alex Alderton, Roberto Amato, Jeffrey Barrett, Sonia Goncalves, Ewan Harrison, David K. Jackson, Ian Johnston, Dominic Kwiatkowski, Cordelia Langford, John Sillitoe on behalf of the Wellcome Sanger Institute COVID-19 Surveillance Team |
| EPI_ISL_2998970 | Lighthouse Lab in Milton Keynes | Wellcome Sanger Institute for the COVID-19 Genomics UK (COG-UK) Consortium | The Lighthouse Lab in Milton Keynes and Alex Alderton, Roberto Amato, Jeffrey Barrett, Sonia Goncalves, Ewan Harrison, David K. Jackson, Ian Johnston, Dominic Kwiatkowski, Cordelia Langford, John Sillitoe on behalf of the Wellcome Sanger Institute COVID-19 Surveillance Team |
| EPI_ISL_2998971 | Berkshire and Surrey Pathology Services Lighthouse Laboratory | Wellcome Sanger Institute for the COVID-19 Genomics UK (COG-UK) Consortium | Berkshire and Surrey Pathology Services Lighthouse Laboratory and Alex Alderton, Roberto Amato, Jeffrey Barrett, Sonia Goncalves, Ewan Harrison, David K. Jackson, Ian Johnston, Dominic Kwiatkowski, Cordelia Langford, John Sillitoe on behalf of the Wellcome Sanger Institute COVID-19 Surveillance Team |
| EPI_ISL_2998972 | Lighthouse Laboratory Plymouth | Wellcome Sanger Institute for the COVID-19 Genomics UK (COG-UK) Consortium | Lighthouse Laboratory Plymouth and Alex Alderton, Roberto Amato, Jeffrey Barrett, Sonia Goncalves, Ewan Harrison, David K. Jackson, Ian Johnston, Dominic Kwiatkowski, Cordelia Langford, John Sillitoe on behalf of the Wellcome Sanger Institute COVID-19 Surveillance Team |
| EPI_ISL_2998973, EPI_ISL_2998974, EPI_ISL_2998975 | Lighthouse Lab in Milton Keynes | Wellcome Sanger Institute for the COVID-19 Genomics UK (COG-UK) Consortium | The Lighthouse Lab in Milton Keynes and Alex Alderton, Roberto Amato, Jeffrey Barrett, Sonia Goncalves, Ewan Harrison, David K. Jackson, Ian Johnston, Dominic Kwiatkowski, Cordelia Langford, John Sillitoe on behalf of the Wellcome Sanger Institute COVID-19 Surveillance Team |
| EPI_ISL_2998976, EPI_ISL_2998977 | Berkshire and Surrey Pathology Services Lighthouse Laboratory | Wellcome Sanger Institute for the COVID-19 Genomics UK (COG-UK) Consortium | Berkshire and Surrey Pathology Services Lighthouse Laboratory and Alex Alderton, Roberto Amato, Jeffrey Barrett, Sonia Goncalves, Ewan Harrison, David K. Jackson, Ian Johnston, Dominic Kwiatkowski, Cordelia Langford, John Sillitoe on behalf of the Wellcome Sanger Institute COVID-19 Surveillance Team |
| EPI_ISL_2998978 | Lighthouse Laboratory Plymouth | Wellcome Sanger Institute for the COVID-19 Genomics UK (COG-UK) Consortium | Lighthouse Laboratory Plymouth and Alex Alderton, Roberto Amato, Jeffrey Barrett, Sonia Goncalves, Ewan Harrison, David K. Jackson, Ian Johnston, Dominic Kwiatkowski, Cordelia Langford, John Sillitoe on behalf of the Wellcome Sanger Institute COVID-19 Surveillance Team |
| EPI_ISL_2998979, EPI_ISL_2998980 | Berkshire and Surrey Pathology Services Lighthouse Laboratory | Wellcome Sanger Institute for the COVID-19 Genomics UK (COG-UK) Consortium | Berkshire and Surrey Pathology Services Lighthouse Laboratory and Alex Alderton, Roberto Amato, Jeffrey Barrett, Sonia Goncalves, Ewan Harrison, David K. Jackson, Ian Johnston, Dominic Kwiatkowski, Cordelia Langford, John Sillitoe on behalf of the Wellcome Sanger Institute COVID-19 Surveillance Team |
| EPI_ISL_2998981 | Lighthouse Lab in Milton Keynes | Wellcome Sanger Institute for the COVID-19 Genomics UK (COG-UK) Consortium | The Lighthouse Lab in Milton Keynes and Alex Alderton, Roberto Amato, Jeffrey Barrett, Sonia Goncalves, Ewan Harrison, David K. Jackson, Ian Johnston, Dominic Kwiatkowski, Cordelia Langford, John Sillitoe on behalf of the Wellcome Sanger Institute COVID-19 Surveillance Team |
| EPI_ISL_2998982 | Lighthouse Laboratory Plymouth | Wellcome Sanger Institute for the COVID-19 Genomics UK (COG-UK) Consortium | Lighthouse Laboratory Plymouth and Alex Alderton, Roberto Amato, Jeffrey Barrett, Sonia Goncalves, Ewan Harrison, David K. Jackson, Ian Johnston, Dominic Kwiatkowski, Cordelia Langford, John Sillitoe on behalf of the Wellcome Sanger Institute COVID-19 Surveillance Team |

| | | | |
|---|---|---|---|
| EPI_ISL_2998983 | Lighthouse Lab in Glasgow | Wellcome Sanger Institute for the COVID-19 Genomics UK (COG-UK) Consortium | Harper VanSteenhouse, Yumi Kasai, David Gray, Carol Clugston, Anna Dominiczak and Alex Alderton, Roberto Amato, Jeffrey Barrett, Sonia Goncalves, Ewan Harrison, David K. Jackson, Ian Johnston, Dominic Kwiatkowski, Cordelia Langford, John Sillitoe on behalf of the Wellcome Sanger Institute COVID-19 Surveillance Team |
| EPI_ISL_2998984 | Berkshire and Surrey Pathology Services Lighthouse Laboratory | Wellcome Sanger Institute for the COVID-19 Genomics UK (COG-UK) Consortium | Berkshire and Surrey Pathology Services Lighthouse Laboratory and Alex Alderton, Roberto Amato, Jeffrey Barrett, Sonia Goncalves, Ewan Harrison, David K. Jackson, Ian Johnston, Dominic Kwiatkowski, Cordelia Langford, John Sillitoe on behalf of the Wellcome Sanger Institute COVID-19 Surveillance Team |
| EPI_ISL_2998985, EPI_ISL_2998986 | Lighthouse Lab in Milton Keynes | Wellcome Sanger Institute for the COVID-19 Genomics UK (COG-UK) Consortium | The Lighthouse Lab in Milton Keynes and Alex Alderton, Roberto Amato, Jeffrey Barrett, Sonia Goncalves, Ewan Harrison, David K. Jackson, Ian Johnston, Dominic Kwiatkowski, Cordelia Langford, John Sillitoe on behalf of the Wellcome Sanger Institute COVID-19 Surveillance Team |
| EPI_ISL_2998987, EPI_ISL_2998988 | Lighthouse Laboratory Plymouth | Wellcome Sanger Institute for the COVID-19 Genomics UK (COG-UK) Consortium | Lighthouse Laboratory Plymouth and Alex Alderton, Roberto Amato, Jeffrey Barrett, Sonia Goncalves, Ewan Harrison, David K. Jackson, Ian Johnston, Dominic Kwiatkowski, Cordelia Langford, John Sillitoe on behalf of the Wellcome Sanger Institute COVID-19 Surveillance Team |
| EPI_ISL_2998989 | Berkshire and Surrey Pathology Services Lighthouse Laboratory | Wellcome Sanger Institute for the COVID-19 Genomics UK (COG-UK) Consortium | Berkshire and Surrey Pathology Services Lighthouse Laboratory and Alex Alderton, Roberto Amato, Jeffrey Barrett, Sonia Goncalves, Ewan Harrison, David K. Jackson, Ian Johnston, Dominic Kwiatkowski, Cordelia Langford, John Sillitoe on behalf of the Wellcome Sanger Institute COVID-19 Surveillance Team |
| EPI_ISL_2998990 | Lighthouse Lab in Glasgow | Wellcome Sanger Institute for the COVID-19 Genomics UK (COG-UK) Consortium | Harper VanSteenhouse, Yumi Kasai, David Gray, Carol Clugston, Anna Dominiczak and Alex Alderton, Roberto Amato, Jeffrey Barrett, Sonia Goncalves, Ewan Harrison, David K. Jackson, Ian Johnston, Dominic Kwiatkowski, Cordelia Langford, John Sillitoe on behalf of the Wellcome Sanger Institute COVID-19 Surveillance Team |
| EPI_ISL_2998991 | Lighthouse Laboratory Plymouth | Wellcome Sanger Institute for the COVID-19 Genomics UK (COG-UK) Consortium | Lighthouse Laboratory Plymouth and Alex Alderton, Roberto Amato, Jeffrey Barrett, Sonia Goncalves, Ewan Harrison, David K. Jackson, Ian Johnston, Dominic Kwiatkowski, Cordelia Langford, John Sillitoe on behalf of the Wellcome Sanger Institute COVID-19 Surveillance Team |
| EPI_ISL_2998992, EPI_ISL_2998993 | Berkshire and Surrey Pathology Services Lighthouse Laboratory | Wellcome Sanger Institute for the COVID-19 Genomics UK (COG-UK) Consortium | Berkshire and Surrey Pathology Services Lighthouse Laboratory and Alex Alderton, Roberto Amato, Jeffrey Barrett, Sonia Goncalves, Ewan Harrison, David K. Jackson, Ian Johnston, Dominic Kwiatkowski, Cordelia Langford, John Sillitoe on behalf of the Wellcome Sanger Institute COVID-19 Surveillance Team |
| EPI_ISL_2998994, EPI_ISL_2998995, EPI_ISL_2998996 | Lighthouse Lab in Milton Keynes | Wellcome Sanger Institute for the COVID-19 Genomics UK (COG-UK) Consortium | The Lighthouse Lab in Milton Keynes and Alex Alderton, Roberto Amato, Jeffrey Barrett, Sonia Goncalves, Ewan Harrison, David K. Jackson, Ian Johnston, Dominic Kwiatkowski, Cordelia Langford, John Sillitoe on behalf of the Wellcome Sanger Institute COVID-19 Surveillance Team |
| EPI_ISL_2998997 | Lighthouse Laboratory Plymouth | Wellcome Sanger Institute for the COVID-19 Genomics UK (COG-UK) Consortium | Lighthouse Laboratory Plymouth and Alex Alderton, Roberto Amato, Jeffrey Barrett, Sonia Goncalves, Ewan Harrison, David K. Jackson, Ian Johnston, Dominic Kwiatkowski, Cordelia Langford, John Sillitoe on behalf of the Wellcome Sanger Institute COVID-19 Surveillance Team |
| EPI_ISL_2998998 | Berkshire and Surrey Pathology Services Lighthouse Laboratory | Wellcome Sanger Institute for the COVID-19 Genomics UK (COG-UK) Consortium | Berkshire and Surrey Pathology Services Lighthouse Laboratory and Alex Alderton, Roberto Amato, Jeffrey Barrett, Sonia Goncalves, Ewan Harrison, David K. Jackson, Ian Johnston, Dominic Kwiatkowski, Cordelia Langford, John Sillitoe on behalf of the Wellcome Sanger Institute COVID-19 Surveillance Team |
| EPI_ISL_2998999 | Lighthouse Lab in Glasgow | Wellcome Sanger Institute for the COVID-19 Genomics UK (COG-UK) Consortium | Harper VanSteenhouse, Yumi Kasai, David Gray, Carol Clugston, Anna Dominiczak and Alex Alderton, Roberto Amato, Jeffrey Barrett, Sonia Goncalves, Ewan Harrison, David K. Jackson, Ian Johnston, Dominic Kwiatkowski, Cordelia Langford, John Sillitoe on behalf of the Wellcome Sanger Institute COVID-19 Surveillance Team |
| EPI_ISL_2999000 | Lighthouse Laboratory Plymouth | Wellcome Sanger Institute for the COVID-19 Genomics UK (COG-UK) Consortium | Lighthouse Laboratory Plymouth and Alex Alderton, Roberto Amato, Jeffrey Barrett, Sonia Goncalves, Ewan Harrison, David K. Jackson, Ian Johnston, Dominic Kwiatkowski, Cordelia Langford, John Sillitoe on behalf of the Wellcome Sanger Institute COVID-19 Surveillance Team |
| EPI_ISL_2999001 | Lighthouse Lab in Alderley Park | Wellcome Sanger Institute for the COVID-19 Genomics UK (COG-UK) Consortium | Jacquelyn Wynn, Mairead Hyland, The Lighthouse Lab in Alderley Park and Alex Alderton, Roberto Amato, Jeffrey Barrett, Sonia Goncalves, Ewan Harrison, David K. Jackson, Ian Johnston, Dominic Kwiatkowski, Cordelia Langford, John Sillitoe on behalf of the Wellcome Sanger Institute COVID-19 Surveillance Team |
| EPI_ISL_2999002, EPI_ISL_2999003, EPI_ISL_2999004 | Lighthouse Laboratory Plymouth | Wellcome Sanger Institute for the COVID-19 Genomics UK (COG-UK) Consortium | Lighthouse Laboratory Plymouth and Alex Alderton, Roberto Amato, Jeffrey Barrett, Sonia Goncalves, Ewan Harrison, David K. Jackson, Ian Johnston, Dominic Kwiatkowski, Cordelia Langford, John Sillitoe on behalf of the Wellcome Sanger Institute COVID-19 Surveillance Team |
| EPI_ISL_2999005 | Lighthouse Lab in Alderley Park | Wellcome Sanger Institute for the COVID-19 Genomics UK (COG-UK) Consortium | Jacquelyn Wynn, Mairead Hyland, The Lighthouse Lab in Alderley Park and Alex Alderton, Roberto Amato, Jeffrey Barrett, Sonia Goncalves, Ewan Harrison, David K. Jackson, Ian Johnston, Dominic Kwiatkowski, Cordelia Langford, John Sillitoe on behalf of the Wellcome Sanger Institute COVID-19 Surveillance Team |
| EPI_ISL_2999006 | Lighthouse Lab in Glasgow | Wellcome Sanger Institute for the COVID-19 Genomics UK (COG-UK) Consortium | Harper VanSteenhouse, Yumi Kasai, David Gray, Carol Clugston, Anna Dominiczak and Alex Alderton, Roberto Amato, Jeffrey Barrett, Sonia Goncalves, Ewan Harrison, David K. Jackson, Ian Johnston, Dominic Kwiatkowski, Cordelia Langford, John Sillitoe on behalf of the Wellcome Sanger Institute COVID-19 Surveillance Team |
| EPI_ISL_2999007 | Lighthouse Laboratory Plymouth | Wellcome Sanger Institute for the COVID-19 Genomics UK (COG-UK) Consortium | Lighthouse Laboratory Plymouth and Alex Alderton, Roberto Amato, Jeffrey Barrett, Sonia Goncalves, Ewan Harrison, David K. Jackson, Ian Johnston, Dominic Kwiatkowski, Cordelia Langford, John Sillitoe on behalf of the Wellcome Sanger Institute COVID-19 Surveillance Team |
| EPI_ISL_2999008, EPI_ISL_2999009 | Lighthouse Lab in Glasgow | Wellcome Sanger Institute for the COVID-19 Genomics UK (COG-UK) Consortium | Harper VanSteenhouse, Yumi Kasai, David Gray, Carol Clugston, Anna Dominiczak and Alex Alderton, Roberto Amato, Jeffrey Barrett, Sonia Goncalves, Ewan Harrison, David K. Jackson, Ian Johnston, Dominic Kwiatkowski, Cordelia Langford, John Sillitoe on behalf of the Wellcome Sanger Institute COVID-19 Surveillance Team |
| EPI_ISL_2999010, EPI_ISL_2999011 | Lighthouse Laboratory Plymouth | Wellcome Sanger Institute for the COVID-19 Genomics UK (COG-UK) Consortium | Lighthouse Laboratory Plymouth and Alex Alderton, Roberto Amato, Jeffrey Barrett, Sonia Goncalves, Ewan Harrison, David K. Jackson, Ian Johnston, Dominic Kwiatkowski, Cordelia Langford, John Sillitoe on behalf of the Wellcome Sanger Institute COVID-19 Surveillance Team |
| EPI_ISL_2999012, EPI_ISL_2999013, EPI_ISL_2999014 | Lighthouse Lab in Alderley Park | Wellcome Sanger Institute for the COVID-19 Genomics UK (COG-UK) Consortium | Jacquelyn Wynn, Mairead Hyland, The Lighthouse Lab in Alderley Park and Alex Alderton, Roberto Amato, Jeffrey Barrett, Sonia Goncalves, Ewan Harrison, David K. Jackson, Ian Johnston, Dominic Kwiatkowski, Cordelia Langford, John Sillitoe on behalf of the Wellcome Sanger Institute COVID-19 Surveillance Team |
| EPI_ISL_2999015 | Lighthouse Lab in Glasgow | Wellcome Sanger Institute for the COVID-19 Genomics UK (COG-UK) Consortium | Harper VanSteenhouse, Yumi Kasai, David Gray, Carol Clugston, Anna Dominiczak and Alex Alderton, Roberto Amato, Jeffrey Barrett, Sonia Goncalves, Ewan Harrison, David K. Jackson, Ian Johnston, Dominic Kwiatkowski, Cordelia Langford, John Sillitoe on behalf of the Wellcome Sanger Institute COVID-19 Surveillance Team |
| EPI_ISL_2999016 | Lighthouse Laboratory Plymouth | Wellcome Sanger Institute for the COVID-19 Genomics UK (COG-UK) Consortium | Lighthouse Laboratory Plymouth and Alex Alderton, Roberto Amato, Jeffrey Barrett, Sonia Goncalves, Ewan Harrison, David K. Jackson, Ian Johnston, Dominic Kwiatkowski, Cordelia Langford, John Sillitoe on behalf of the Wellcome Sanger Institute COVID-19 Surveillance Team |
| EPI_ISL_2999017 | Lighthouse Lab in Alderley Park | Wellcome Sanger Institute for the COVID-19 Genomics UK (COG-UK) Consortium | Jacquelyn Wynn, Mairead Hyland, The Lighthouse Lab in Alderley Park and Alex Alderton, Roberto Amato, Jeffrey Barrett, Sonia Goncalves, Ewan Harrison, David K. Jackson, Ian Johnston, Dominic Kwiatkowski, Cordelia Langford, John Sillitoe on behalf of the Wellcome Sanger Institute COVID-19 Surveillance Team |
| EPI_ISL_2999018 | Lighthouse Lab in Glasgow | Wellcome Sanger Institute for the COVID-19 Genomics UK (COG-UK) Consortium | Harper VanSteenhouse, Yumi Kasai, David Gray, Carol Clugston, Anna Dominiczak and Alex Alderton, Roberto Amato, Jeffrey Barrett, Sonia Goncalves, Ewan Harrison, David K. Jackson, Ian Johnston, Dominic Kwiatkowski, Cordelia Langford, John Sillitoe on behalf of the Wellcome Sanger Institute COVID-19 Surveillance Team |
| EPI_ISL_2999019 | Lighthouse Lab in Alderley Park | Wellcome Sanger Institute for the COVID-19 Genomics UK (COG-UK) Consortium | Jacquelyn Wynn, Mairead Hyland, The Lighthouse Lab in Alderley Park and Alex Alderton, Roberto Amato, Jeffrey Barrett, Sonia Goncalves, Ewan Harrison, David K. Jackson, Ian Johnston, Dominic Kwiatkowski, Cordelia Langford, John Sillitoe on behalf of the Wellcome Sanger Institute COVID-19 Surveillance Team |
| EPI_ISL_2999020, EPI_ISL_2999021, EPI_ISL_2999022 | Lighthouse Lab in Glasgow | Wellcome Sanger Institute for the COVID-19 Genomics UK (COG-UK) Consortium | Harper VanSteenhouse, Yumi Kasai, David Gray, Carol Clugston, Anna Dominiczak and Alex Alderton, Roberto Amato, Jeffrey Barrett, Sonia Goncalves, Ewan Harrison, David K. Jackson, Ian Johnston, Dominic Kwiatkowski, Cordelia Langford, John Sillitoe on behalf of the Wellcome Sanger Institute COVID-19 Surveillance Team |
| EPI_ISL_2999023 | Lighthouse Laboratory Plymouth | Wellcome Sanger Institute for the COVID-19 Genomics UK (COG-UK) Consortium | Lighthouse Laboratory Plymouth and Alex Alderton, Roberto Amato, Jeffrey Barrett, Sonia Goncalves, Ewan Harrison, David K. Jackson, Ian Johnston, Dominic Kwiatkowski, Cordelia Langford, John Sillitoe on behalf of the Wellcome Sanger Institute COVID-19 Surveillance Team |
| EPI_ISL_2999024 | Lighthouse Lab in Glasgow | Wellcome Sanger Institute for the COVID-19 Genomics UK (COG-UK) Consortium | Harper VanSteenhouse, Yumi Kasai, David Gray, Carol Clugston, Anna Dominiczak and Alex Alderton, Roberto Amato, Jeffrey Barrett, Sonia Goncalves, Ewan Harrison, David K. Jackson, Ian Johnston, Dominic Kwiatkowski, Cordelia Langford, John Sillitoe on behalf of the Wellcome Sanger Institute COVID-19 Surveillance Team |
| EPI_ISL_2999025 | Lighthouse Laboratory Plymouth | Wellcome Sanger Institute for the COVID-19 Genomics UK (COG-UK) Consortium | Lighthouse Laboratory Plymouth and Alex Alderton, Roberto Amato, Jeffrey Barrett, Sonia Goncalves, Ewan Harrison, David K. Jackson, Ian Johnston, Dominic Kwiatkowski, Cordelia Langford, John Sillitoe on behalf of the Wellcome Sanger Institute COVID-19 Surveillance Team |
| EPI_ISL_2999026, EPI_ISL_2999027 | Lighthouse Lab in Glasgow | Wellcome Sanger Institute for the COVID-19 Genomics UK (COG-UK) Consortium | Harper VanSteenhouse, Yumi Kasai, David Gray, Carol Clugston, Anna Dominiczak and Alex Alderton, Roberto Amato, Jeffrey Barrett, Sonia Goncalves, Ewan Harrison, David K. Jackson, Ian Johnston, Dominic Kwiatkowski, Cordelia Langford, John Sillitoe on behalf of the Wellcome Sanger Institute COVID-19 Surveillance Team |
| EPI_ISL_2999028 | Lighthouse Laboratory Plymouth | Wellcome Sanger Institute for the COVID-19 Genomics UK (COG-UK) Consortium | Lighthouse Laboratory Plymouth and Alex Alderton, Roberto Amato, Jeffrey Barrett, Sonia Goncalves, Ewan Harrison, David K. Jackson, Ian Johnston, Dominic Kwiatkowski, Cordelia Langford, John Sillitoe on behalf of the Wellcome Sanger Institute COVID-19 Surveillance Team |
| EPI_ISL_2999029, EPI_ISL_2999030 | Lighthouse Lab in Alderley Park | Wellcome Sanger Institute for the COVID-19 Genomics UK (COG-UK) Consortium | Jacquelyn Wynn, Mairead Hyland, The Lighthouse Lab in Alderley Park and Alex Alderton, Roberto Amato, Jeffrey Barrett, Sonia Goncalves, Ewan Harrison, David K. Jackson, Ian Johnston, Dominic Kwiatkowski, Cordelia Langford, John Sillitoe on behalf of the Wellcome Sanger Institute COVID-19 Surveillance Team |
| EPI_ISL_2999031 | Lighthouse Lab in Glasgow | Wellcome Sanger Institute for the COVID-19 Genomics UK (COG-UK) Consortium | Harper VanSteenhouse, Yumi Kasai, David Gray, Carol Clugston, Anna Dominiczak and Alex Alderton, Roberto Amato, Jeffrey Barrett, Sonia Goncalves, Ewan Harrison, David K. Jackson, Ian Johnston, Dominic Kwiatkowski, Cordelia Langford, John Sillitoe on behalf of the Wellcome Sanger Institute COVID-19 Surveillance Team |
| EPI_ISL_2999032 | Lighthouse Laboratory Plymouth | Wellcome Sanger Institute for the COVID-19 Genomics UK (COG-UK) Consortium | Lighthouse Laboratory Plymouth and Alex Alderton, Roberto Amato, Jeffrey Barrett, Sonia Goncalves, Ewan Harrison, David K. Jackson, Ian Johnston, Dominic Kwiatkowski, Cordelia Langford, John Sillitoe on behalf of the Wellcome Sanger Institute COVID-19 Surveillance Team |
| EPI_ISL_2999033 | Lighthouse Lab in Glasgow | Wellcome Sanger Institute for the COVID-19 Genomics UK (COG-UK) Consortium | Harper VanSteenhouse, Yumi Kasai, David Gray, Carol Clugston, Anna Dominiczak and Alex Alderton, Roberto Amato, Jeffrey Barrett, Sonia Goncalves, Ewan Harrison, David K. Jackson, Ian Johnston, Dominic Kwiatkowski, Cordelia Langford, John Sillitoe on behalf of the Wellcome Sanger Institute COVID-19 Surveillance Team |
| EPI_ISL_2999034 | Lighthouse Lab in Alderley Park | Wellcome Sanger Institute for the COVID-19 Genomics UK (COG-UK) Consortium | Jacquelyn Wynn, Mairead Hyland, The Lighthouse Lab in Alderley Park and Alex Alderton, Roberto Amato, Jeffrey Barrett, Sonia Goncalves, Ewan Harrison, David K. Jackson, Ian Johnston, Dominic Kwiatkowski, Cordelia Langford, John Sillitoe on behalf of the Wellcome Sanger Institute COVID-19 Surveillance Team |
| EPI_ISL_2999035 | Lighthouse Lab in Glasgow | Wellcome Sanger Institute for the COVID-19 Genomics UK (COG-UK) Consortium | Harper VanSteenhouse, Yumi Kasai, David Gray, Carol Clugston, Anna Dominiczak and Alex Alderton, Roberto Amato, Jeffrey Barrett, Sonia Goncalves, Ewan Harrison, David K. Jackson, Ian Johnston, Dominic Kwiatkowski, Cordelia Langford, John Sillitoe on behalf of the Wellcome Sanger Institute COVID-19 Surveillance Team |
| EPI_ISL_2999036 | Lighthouse Laboratory Plymouth | Wellcome Sanger Institute for the COVID-19 Genomics UK (COG-UK) Consortium | Lighthouse Laboratory Plymouth and Alex Alderton, Roberto Amato, Jeffrey Barrett, Sonia Goncalves, Ewan Harrison, David K. Jackson, Ian Johnston, Dominic Kwiatkowski, Cordelia Langford, John Sillitoe on behalf of the Wellcome Sanger Institute COVID-19 Surveillance Team |
| EPI_ISL_2999037, EPI_ISL_2999038 | Lighthouse Lab in Glasgow | Wellcome Sanger Institute for the COVID-19 Genomics UK (COG-UK) Consortium | Harper VanSteenhouse, Yumi Kasai, David Gray, Carol Clugston, Anna Dominiczak and Alex Alderton, Roberto Amato, Jeffrey Barrett, Sonia Goncalves, Ewan Harrison, David K. Jackson, Ian Johnston, Dominic Kwiatkowski, Cordelia Langford, John Sillitoe on behalf of the Wellcome Sanger Institute COVID-19 Surveillance Team |
| EPI_ISL_2999039 | Lighthouse Laboratory Plymouth | Wellcome Sanger Institute for the COVID-19 Genomics UK (COG-UK) Consortium | Lighthouse Laboratory Plymouth and Alex Alderton, Roberto Amato, Jeffrey Barrett, Sonia Goncalves, Ewan Harrison, David K. Jackson, Ian Johnston, Dominic Kwiatkowski, Cordelia Langford, John Sillitoe on behalf of the Wellcome Sanger Institute COVID-19 Surveillance Team |
| EPI_ISL_2999040, EPI_ISL_2999041 | Lighthouse Lab in Glasgow | Wellcome Sanger Institute for the COVID-19 Genomics UK (COG-UK) Consortium | Harper VanSteenhouse, Yumi Kasai, David Gray, Carol Clugston, Anna Dominiczak and Alex Alderton, Roberto Amato, Jeffrey Barrett, Sonia Goncalves, Ewan Harrison, David K. Jackson, Ian Johnston, Dominic Kwiatkowski, Cordelia Langford, John Sillitoe on behalf of the Wellcome Sanger Institute COVID-19 Surveillance Team |
| EPI_ISL_2999042, EPI_ISL_2999043 | Lighthouse Lab in Alderley Park | Wellcome Sanger Institute for the COVID-19 Genomics UK (COG-UK) Consortium | Jacquelyn Wynn, Mairead Hyland, The Lighthouse Lab in Alderley Park and Alex Alderton, Roberto Amato, Jeffrey Barrett, Sonia Goncalves, Ewan Harrison, David K. Jackson, Ian Johnston, Dominic Kwiatkowski, Cordelia Langford, John Sillitoe on behalf of the Wellcome Sanger Institute COVID-19 Surveillance Team |
| EPI_ISL_2999044 | Lighthouse Lab in Glasgow | Wellcome Sanger Institute for the COVID-19 Genomics UK (COG-UK) | Harper VanSteenhouse, Yumi Kasai, David Gray, Carol Clugston, Anna Dominiczak and Alex Alderton, Roberto Amato, Jeffrey Barrett, Sonia Goncalves, Ewan Harrison, David K. Jackson, Ian Johnston, Dominic Kwiatkowski, Cordelia Langford, |

| | | | |
|---|---|---|---|
| | | Consortium | John Sillitoe on behalf of the Wellcome Sanger Institute COVID-19 Surveillance Team |
| EPI_ISL_2999045 | Lighthouse Laboratory Plymouth | Wellcome Sanger Institute for the COVID-19 Genomics UK (COG-UK) Consortium | Lighthouse Laboratory Plymouth and Alex Alderton, Roberto Amato, Jeffrey Barrett, Sonia Goncalves, Ewan Harrison, David K. Jackson, Ian Johnston, Dominic Kwiatkowski, Cordelia Langford, John Sillitoe on behalf of the Wellcome Sanger Institute COVID-19 Surveillance Team |
| EPI_ISL_2999046 | Lighthouse Lab in Alderley Park | Wellcome Sanger Institute for the COVID-19 Genomics UK (COG-UK) Consortium | Jacquelyn Wynn, Mairead Hyland, The Lighthouse Lab in Alderley Park and Alex Alderton, Roberto Amato, Jeffrey Barrett, Sonia Goncalves, Ewan Harrison, David K. Jackson, Ian Johnston, Dominic Kwiatkowski, Cordelia Langford, John Sillitoe on behalf of the Wellcome Sanger Institute COVID-19 Surveillance Team |
| EPI_ISL_2999047, EPI_ISL_2999048 | Lighthouse Lab in Glasgow | Wellcome Sanger Institute for the COVID-19 Genomics UK (COG-UK) Consortium | Harper VanSteenhouse, Yumi Kasai, David Gray, Carol Clugston, Anna Dominiczak and Alex Alderton, Roberto Amato, Jeffrey Barrett, Sonia Goncalves, Ewan Harrison, David K. Jackson, Ian Johnston, Dominic Kwiatkowski, Cordelia Langford, John Sillitoe on behalf of the Wellcome Sanger Institute COVID-19 Surveillance Team |
| EPI_ISL_2999049, EPI_ISL_2999050 | Lighthouse Laboratory Plymouth | Wellcome Sanger Institute for the COVID-19 Genomics UK (COG-UK) Consortium | Lighthouse Laboratory Plymouth and Alex Alderton, Roberto Amato, Jeffrey Barrett, Sonia Goncalves, Ewan Harrison, David K. Jackson, Ian Johnston, Dominic Kwiatkowski, Cordelia Langford, John Sillitoe on behalf of the Wellcome Sanger Institute COVID-19 Surveillance Team |
| EPI_ISL_2999051, EPI_ISL_2999052 | Lighthouse Lab in Alderley Park | Wellcome Sanger Institute for the COVID-19 Genomics UK (COG-UK) Consortium | Jacquelyn Wynn, Mairead Hyland, The Lighthouse Lab in Alderley Park and Alex Alderton, Roberto Amato, Jeffrey Barrett, Sonia Goncalves, Ewan Harrison, David K. Jackson, Ian Johnston, Dominic Kwiatkowski, Cordelia Langford, John Sillitoe on behalf of the Wellcome Sanger Institute COVID-19 Surveillance Team |
| EPI_ISL_2999053 | Lighthouse Lab in Glasgow | Wellcome Sanger Institute for the COVID-19 Genomics UK (COG-UK) Consortium | Harper VanSteenhouse, Yumi Kasai, David Gray, Carol Clugston, Anna Dominiczak and Alex Alderton, Roberto Amato, Jeffrey Barrett, Sonia Goncalves, Ewan Harrison, David K. Jackson, Ian Johnston, Dominic Kwiatkowski, Cordelia Langford, John Sillitoe on behalf of the Wellcome Sanger Institute COVID-19 Surveillance Team |
| EPI_ISL_2999054 | Lighthouse Laboratory Plymouth | Wellcome Sanger Institute for the COVID-19 Genomics UK (COG-UK) Consortium | Lighthouse Laboratory Plymouth and Alex Alderton, Roberto Amato, Jeffrey Barrett, Sonia Goncalves, Ewan Harrison, David K. Jackson, Ian Johnston, Dominic Kwiatkowski, Cordelia Langford, John Sillitoe on behalf of the Wellcome Sanger Institute COVID-19 Surveillance Team |
| EPI_ISL_2999055 | Lighthouse Lab in Alderley Park | Wellcome Sanger Institute for the COVID-19 Genomics UK (COG-UK) Consortium | Jacquelyn Wynn, Mairead Hyland, The Lighthouse Lab in Alderley Park and Alex Alderton, Roberto Amato, Jeffrey Barrett, Sonia Goncalves, Ewan Harrison, David K. Jackson, Ian Johnston, Dominic Kwiatkowski, Cordelia Langford, John Sillitoe on behalf of the Wellcome Sanger Institute COVID-19 Surveillance Team |
| EPI_ISL_2999056 | Lighthouse Lab in Glasgow | Wellcome Sanger Institute for the COVID-19 Genomics UK (COG-UK) Consortium | Harper VanSteenhouse, Yumi Kasai, David Gray, Carol Clugston, Anna Dominiczak and Alex Alderton, Roberto Amato, Jeffrey Barrett, Sonia Goncalves, Ewan Harrison, David K. Jackson, Ian Johnston, Dominic Kwiatkowski, Cordelia Langford, John Sillitoe on behalf of the Wellcome Sanger Institute COVID-19 Surveillance Team |
| EPI_ISL_2999057 | Lighthouse Laboratory Plymouth | Wellcome Sanger Institute for the COVID-19 Genomics UK (COG-UK) Consortium | Lighthouse Laboratory Plymouth and Alex Alderton, Roberto Amato, Jeffrey Barrett, Sonia Goncalves, Ewan Harrison, David K. Jackson, Ian Johnston, Dominic Kwiatkowski, Cordelia Langford, John Sillitoe on behalf of the Wellcome Sanger Institute COVID-19 Surveillance Team |
| EPI_ISL_2999058, EPI_ISL_2999059 | Lighthouse Lab in Alderley Park | Wellcome Sanger Institute for the COVID-19 Genomics UK (COG-UK) Consortium | Jacquelyn Wynn, Mairead Hyland, The Lighthouse Lab in Alderley Park and Alex Alderton, Roberto Amato, Jeffrey Barrett, Sonia Goncalves, Ewan Harrison, David K. Jackson, Ian Johnston, Dominic Kwiatkowski, Cordelia Langford, John Sillitoe on behalf of the Wellcome Sanger Institute COVID-19 Surveillance Team |
| EPI_ISL_2999060 | Lighthouse Laboratory Plymouth | Wellcome Sanger Institute for the COVID-19 Genomics UK (COG-UK) Consortium | Lighthouse Laboratory Plymouth and Alex Alderton, Roberto Amato, Jeffrey Barrett, Sonia Goncalves, Ewan Harrison, David K. Jackson, Ian Johnston, Dominic Kwiatkowski, Cordelia Langford, John Sillitoe on behalf of the Wellcome Sanger Institute COVID-19 Surveillance Team |
| EPI_ISL_2999061 | Lighthouse Lab in Alderley Park | Wellcome Sanger Institute for the COVID-19 Genomics UK (COG-UK) Consortium | Jacquelyn Wynn, Mairead Hyland, The Lighthouse Lab in Alderley Park and Alex Alderton, Roberto Amato, Jeffrey Barrett, Sonia Goncalves, Ewan Harrison, David K. Jackson, Ian Johnston, Dominic Kwiatkowski, Cordelia Langford, John Sillitoe on behalf of the Wellcome Sanger Institute COVID-19 Surveillance Team |
| EPI_ISL_2999062, EPI_ISL_2999063, EPI_ISL_2999064 | Lighthouse Lab in Glasgow | Wellcome Sanger Institute for the COVID-19 Genomics UK (COG-UK) Consortium | Harper VanSteenhouse, Yumi Kasai, David Gray, Carol Clugston, Anna Dominiczak and Alex Alderton, Roberto Amato, Jeffrey Barrett, Sonia Goncalves, Ewan Harrison, David K. Jackson, Ian Johnston, Dominic Kwiatkowski, Cordelia Langford, John Sillitoe on behalf of the Wellcome Sanger Institute COVID-19 Surveillance Team |
| EPI_ISL_2999065 | Lighthouse Laboratory Plymouth | Wellcome Sanger Institute for the COVID-19 Genomics UK (COG-UK) Consortium | Lighthouse Laboratory Plymouth and Alex Alderton, Roberto Amato, Jeffrey Barrett, Sonia Goncalves, Ewan Harrison, David K. Jackson, Ian Johnston, Dominic Kwiatkowski, Cordelia Langford, John Sillitoe on behalf of the Wellcome Sanger Institute COVID-19 Surveillance Team |
| EPI_ISL_2999066, EPI_ISL_2999067, EPI_ISL_2999068 | Lighthouse Lab in Glasgow | Wellcome Sanger Institute for the COVID-19 Genomics UK (COG-UK) Consortium | Harper VanSteenhouse, Yumi Kasai, David Gray, Carol Clugston, Anna Dominiczak and Alex Alderton, Roberto Amato, Jeffrey Barrett, Sonia Goncalves, Ewan Harrison, David K. Jackson, Ian Johnston, Dominic Kwiatkowski, Cordelia Langford, John Sillitoe on behalf of the Wellcome Sanger Institute COVID-19 Surveillance Team |
| EPI_ISL_2999069 | Lighthouse Lab in Alderley Park | Wellcome Sanger Institute for the COVID-19 Genomics UK (COG-UK) Consortium | Jacquelyn Wynn, Mairead Hyland, The Lighthouse Lab in Alderley Park and Alex Alderton, Roberto Amato, Jeffrey Barrett, Sonia Goncalves, Ewan Harrison, David K. Jackson, Ian Johnston, Dominic Kwiatkowski, Cordelia Langford, John Sillitoe on behalf of the Wellcome Sanger Institute COVID-19 Surveillance Team |
| EPI_ISL_2999070 | Lighthouse Laboratory Plymouth | Wellcome Sanger Institute for the COVID-19 Genomics UK (COG-UK) Consortium | Lighthouse Laboratory Plymouth and Alex Alderton, Roberto Amato, Jeffrey Barrett, Sonia Goncalves, Ewan Harrison, David K. Jackson, Ian Johnston, Dominic Kwiatkowski, Cordelia Langford, John Sillitoe on behalf of the Wellcome Sanger Institute COVID-19 Surveillance Team |
| EPI_ISL_2999071 | Lighthouse Lab in Alderley Park | Wellcome Sanger Institute for the COVID-19 Genomics UK (COG-UK) Consortium | Jacquelyn Wynn, Mairead Hyland, The Lighthouse Lab in Alderley Park and Alex Alderton, Roberto Amato, Jeffrey Barrett, Sonia Goncalves, Ewan Harrison, David K. Jackson, Ian Johnston, Dominic Kwiatkowski, Cordelia Langford, John Sillitoe on behalf of the Wellcome Sanger Institute COVID-19 Surveillance Team |
| EPI_ISL_2999072, EPI_ISL_2999073, EPI_ISL_2999074 | Lighthouse Laboratory Plymouth | Wellcome Sanger Institute for the COVID-19 Genomics UK (COG-UK) Consortium | Lighthouse Laboratory Plymouth and Alex Alderton, Roberto Amato, Jeffrey Barrett, Sonia Goncalves, Ewan Harrison, David K. Jackson, Ian Johnston, Dominic Kwiatkowski, Cordelia Langford, John Sillitoe on behalf of the Wellcome Sanger Institute COVID-19 Surveillance Team |
| EPI_ISL_2999075, EPI_ISL_2999076 | Lighthouse Lab in Alderley Park | Wellcome Sanger Institute for the COVID-19 Genomics UK (COG-UK) Consortium | Jacquelyn Wynn, Mairead Hyland, The Lighthouse Lab in Alderley Park and Alex Alderton, Roberto Amato, Jeffrey Barrett, Sonia Goncalves, Ewan Harrison, David K. Jackson, Ian Johnston, Dominic Kwiatkowski, Cordelia Langford, John Sillitoe on behalf of the Wellcome Sanger Institute COVID-19 Surveillance Team |
| EPI_ISL_2999077 | Lighthouse Lab in Glasgow | Wellcome Sanger Institute for the COVID-19 Genomics UK (COG-UK) Consortium | Harper VanSteenhouse, Yumi Kasai, David Gray, Carol Clugston, Anna Dominiczak and Alex Alderton, Roberto Amato, Jeffrey Barrett, Sonia Goncalves, Ewan Harrison, David K. Jackson, Ian Johnston, Dominic Kwiatkowski, Cordelia Langford, John Sillitoe on behalf of the Wellcome Sanger Institute COVID-19 Surveillance Team |
| EPI_ISL_2999078, EPI_ISL_2999079 | Lighthouse Laboratory Plymouth | Wellcome Sanger Institute for the COVID-19 Genomics UK (COG-UK) Consortium | Lighthouse Laboratory Plymouth and Alex Alderton, Roberto Amato, Jeffrey Barrett, Sonia Goncalves, Ewan Harrison, David K. Jackson, Ian Johnston, Dominic Kwiatkowski, Cordelia Langford, John Sillitoe on behalf of the Wellcome Sanger Institute COVID-19 Surveillance Team |
| EPI_ISL_2999080 | Lighthouse Lab in Alderley Park | Wellcome Sanger Institute for the COVID-19 Genomics UK (COG-UK) Consortium | Jacquelyn Wynn, Mairead Hyland, The Lighthouse Lab in Alderley Park and Alex Alderton, Roberto Amato, Jeffrey Barrett, Sonia Goncalves, Ewan Harrison, David K. Jackson, Ian Johnston, Dominic Kwiatkowski, Cordelia Langford, John Sillitoe on behalf of the Wellcome Sanger Institute COVID-19 Surveillance Team |
| EPI_ISL_2999081 | Lighthouse Lab in Glasgow | Wellcome Sanger Institute for the COVID-19 Genomics UK (COG-UK) Consortium | Harper VanSteenhouse, Yumi Kasai, David Gray, Carol Clugston, Anna Dominiczak and Alex Alderton, Roberto Amato, Jeffrey Barrett, Sonia Goncalves, Ewan Harrison, David K. Jackson, Ian Johnston, Dominic Kwiatkowski, Cordelia Langford, John Sillitoe on behalf of the Wellcome Sanger Institute COVID-19 Surveillance Team |
| EPI_ISL_2999082 | Lighthouse Laboratory Plymouth | Wellcome Sanger Institute for the COVID-19 Genomics UK (COG-UK) Consortium | Lighthouse Laboratory Plymouth and Alex Alderton, Roberto Amato, Jeffrey Barrett, Sonia Goncalves, Ewan Harrison, David K. Jackson, Ian Johnston, Dominic Kwiatkowski, Cordelia Langford, John Sillitoe on behalf of the Wellcome Sanger Institute COVID-19 Surveillance Team |
| EPI_ISL_2999083 | Lighthouse Lab in Glasgow | Wellcome Sanger Institute for the COVID-19 Genomics UK (COG-UK) Consortium | Harper VanSteenhouse, Yumi Kasai, David Gray, Carol Clugston, Anna Dominiczak and Alex Alderton, Roberto Amato, Jeffrey Barrett, Sonia Goncalves, Ewan Harrison, David K. Jackson, Ian Johnston, Dominic Kwiatkowski, Cordelia Langford, John Sillitoe on behalf of the Wellcome Sanger Institute COVID-19 Surveillance Team |
| EPI_ISL_2999084, EPI_ISL_2999085 | Lighthouse Laboratory Plymouth | Wellcome Sanger Institute for the COVID-19 Genomics UK (COG-UK) Consortium | Lighthouse Laboratory Plymouth and Alex Alderton, Roberto Amato, Jeffrey Barrett, Sonia Goncalves, Ewan Harrison, David K. Jackson, Ian Johnston, Dominic Kwiatkowski, Cordelia Langford, John Sillitoe on behalf of the Wellcome Sanger Institute COVID-19 Surveillance Team |
| EPI_ISL_2999086 | Lighthouse Lab in Alderley Park | Wellcome Sanger Institute for the COVID-19 Genomics UK (COG-UK) Consortium | Jacquelyn Wynn, Mairead Hyland, The Lighthouse Lab in Alderley Park and Alex Alderton, Roberto Amato, Jeffrey Barrett, Sonia Goncalves, Ewan Harrison, David K. Jackson, Ian Johnston, Dominic Kwiatkowski, Cordelia Langford, John Sillitoe on behalf of the Wellcome Sanger Institute COVID-19 Surveillance Team |
| EPI_ISL_2999087 | Lighthouse Laboratory Plymouth | Wellcome Sanger Institute for the COVID-19 Genomics UK (COG-UK) Consortium | Lighthouse Laboratory Plymouth and Alex Alderton, Roberto Amato, Jeffrey Barrett, Sonia Goncalves, Ewan Harrison, David K. Jackson, Ian Johnston, Dominic Kwiatkowski, Cordelia Langford, John Sillitoe on behalf of the Wellcome Sanger Institute COVID-19 Surveillance Team |
| EPI_ISL_2999088, EPI_ISL_2999089 | Lighthouse Lab in Glasgow | Wellcome Sanger Institute for the COVID-19 Genomics UK (COG-UK) Consortium | Harper VanSteenhouse, Yumi Kasai, David Gray, Carol Clugston, Anna Dominiczak and Alex Alderton, Roberto Amato, Jeffrey Barrett, Sonia Goncalves, Ewan Harrison, David K. Jackson, Ian Johnston, Dominic Kwiatkowski, Cordelia Langford, John Sillitoe on behalf of the Wellcome Sanger Institute COVID-19 Surveillance Team |
| EPI_ISL_2999090 | Lighthouse Lab in Alderley Park | Wellcome Sanger Institute for the COVID-19 Genomics UK (COG-UK) Consortium | Jacquelyn Wynn, Mairead Hyland, The Lighthouse Lab in Alderley Park and Alex Alderton, Roberto Amato, Jeffrey Barrett, Sonia Goncalves, Ewan Harrison, David K. Jackson, Ian Johnston, Dominic Kwiatkowski, Cordelia Langford, John Sillitoe on behalf of the Wellcome Sanger Institute COVID-19 Surveillance Team |
| EPI_ISL_2999091 | Lighthouse Lab in Glasgow | Wellcome Sanger Institute for the COVID-19 Genomics UK (COG-UK) Consortium | Harper VanSteenhouse, Yumi Kasai, David Gray, Carol Clugston, Anna Dominiczak and Alex Alderton, Roberto Amato, Jeffrey Barrett, Sonia Goncalves, Ewan Harrison, David K. Jackson, Ian Johnston, Dominic Kwiatkowski, Cordelia Langford, John Sillitoe on behalf of the Wellcome Sanger Institute COVID-19 Surveillance Team |
| EPI_ISL_2999092 | Lighthouse Laboratory Plymouth | Wellcome Sanger Institute for the COVID-19 Genomics UK (COG-UK) Consortium | Lighthouse Laboratory Plymouth and Alex Alderton, Roberto Amato, Jeffrey Barrett, Sonia Goncalves, Ewan Harrison, David K. Jackson, Ian Johnston, Dominic Kwiatkowski, Cordelia Langford, John Sillitoe on behalf of the Wellcome Sanger Institute COVID-19 Surveillance Team |
| EPI_ISL_2999093, EPI_ISL_2999094, EPI_ISL_2999095 | Lighthouse Lab in Alderley Park | Wellcome Sanger Institute for the COVID-19 Genomics UK (COG-UK) Consortium | Jacquelyn Wynn, Mairead Hyland, The Lighthouse Lab in Alderley Park and Alex Alderton, Roberto Amato, Jeffrey Barrett, Sonia Goncalves, Ewan Harrison, David K. Jackson, Ian Johnston, Dominic Kwiatkowski, Cordelia Langford, John Sillitoe on behalf of the Wellcome Sanger Institute COVID-19 Surveillance Team |
| EPI_ISL_2999096 | Lighthouse Laboratory Plymouth | Wellcome Sanger Institute for the COVID-19 Genomics UK (COG-UK) Consortium | Lighthouse Laboratory Plymouth and Alex Alderton, Roberto Amato, Jeffrey Barrett, Sonia Goncalves, Ewan Harrison, David K. Jackson, Ian Johnston, Dominic Kwiatkowski, Cordelia Langford, John Sillitoe on behalf of the Wellcome Sanger Institute COVID-19 Surveillance Team |
| EPI_ISL_2999097 | Lighthouse Lab in Alderley Park | Wellcome Sanger Institute for the COVID-19 Genomics UK (COG-UK) Consortium | Jacquelyn Wynn, Mairead Hyland, The Lighthouse Lab in Alderley Park and Alex Alderton, Roberto Amato, Jeffrey Barrett, Sonia Goncalves, Ewan Harrison, David K. Jackson, Ian Johnston, Dominic Kwiatkowski, Cordelia Langford, John Sillitoe on behalf of the Wellcome Sanger Institute COVID-19 Surveillance Team |
| EPI_ISL_2999098 | Lighthouse Lab in Glasgow | Wellcome Sanger Institute for the COVID-19 Genomics UK (COG-UK) Consortium | Harper VanSteenhouse, Yumi Kasai, David Gray, Carol Clugston, Anna Dominiczak and Alex Alderton, Roberto Amato, Jeffrey Barrett, Sonia Goncalves, Ewan Harrison, David K. Jackson, Ian Johnston, Dominic Kwiatkowski, Cordelia Langford, John Sillitoe on behalf of the Wellcome Sanger Institute COVID-19 Surveillance Team |
| EPI_ISL_2999099 | Lighthouse Lab in Alderley Park | Wellcome Sanger Institute for the COVID-19 Genomics UK (COG-UK) Consortium | Jacquelyn Wynn, Mairead Hyland, The Lighthouse Lab in Alderley Park and Alex Alderton, Roberto Amato, Jeffrey Barrett, Sonia Goncalves, Ewan Harrison, David K. Jackson, Ian Johnston, Dominic Kwiatkowski, Cordelia Langford, John Sillitoe on behalf of the Wellcome Sanger Institute COVID-19 Surveillance Team |
| EPI_ISL_2999100, EPI_ISL_2999101 | Lighthouse Lab in Glasgow | Wellcome Sanger Institute for the COVID-19 Genomics UK (COG-UK) Consortium | Harper VanSteenhouse, Yumi Kasai, David Gray, Carol Clugston, Anna Dominiczak and Alex Alderton, Roberto Amato, Jeffrey Barrett, Sonia Goncalves, Ewan Harrison, David K. Jackson, Ian Johnston, Dominic Kwiatkowski, Cordelia Langford, John Sillitoe on behalf of the Wellcome Sanger Institute COVID-19 Surveillance Team |
| EPI_ISL_2999102 | Lighthouse Lab in Alderley Park | Wellcome Sanger Institute for the COVID-19 Genomics UK (COG-UK) Consortium | Jacquelyn Wynn, Mairead Hyland, The Lighthouse Lab in Alderley Park and Alex Alderton, Roberto Amato, Jeffrey Barrett, Sonia Goncalves, Ewan Harrison, David K. Jackson, Ian Johnston, Dominic Kwiatkowski, Cordelia Langford, John Sillitoe on behalf of the Wellcome Sanger Institute COVID-19 Surveillance Team |
| EPI_ISL_2999103, EPI_ISL_2999104 | Lighthouse Lab in Glasgow | Wellcome Sanger Institute for the COVID-19 Genomics UK (COG-UK) Consortium | Harper VanSteenhouse, Yumi Kasai, David Gray, Carol Clugston, Anna Dominiczak and Alex Alderton, Roberto Amato, Jeffrey Barrett, Sonia Goncalves, Ewan Harrison, David K. Jackson, Ian Johnston, Dominic Kwiatkowski, Cordelia Langford, John Sillitoe on behalf of the Wellcome Sanger Institute COVID-19 Surveillance Team |
| EPI_ISL_2999105, EPI_ISL_2999106 | Lighthouse Laboratory Plymouth | Wellcome Sanger Institute for the COVID-19 Genomics UK (COG-UK) Consortium | Lighthouse Laboratory Plymouth and Alex Alderton, Roberto Amato, Jeffrey Barrett, Sonia Goncalves, Ewan Harrison, David K. Jackson, Ian Johnston, Dominic Kwiatkowski, Cordelia Langford, John Sillitoe on behalf of the Wellcome Sanger Institute COVID-19 Surveillance Team |

| | | | |
|---|---|---|---|
| EPI_ISL_2999107 | Lighthouse Lab in Alderley Park | Wellcome Sanger Institute for the COVID-19 Genomics UK (COG-UK) Consortium | Jacquelyn Wynn, Mairead Hyland, The Lighthouse Lab in Alderley Park and Alex Alderton, Roberto Amato, Jeffrey Barrett, Sonia Goncalves, Ewan Harrison, David K. Jackson, Ian Johnston, Dominic Kwiatkowski, Cordelia Langford, John Sillitoe on behalf of the Wellcome Sanger Institute COVID-19 Surveillance Team |
| EPI_ISL_2999108 | Lighthouse Lab in Glasgow | Wellcome Sanger Institute for the COVID-19 Genomics UK (COG-UK) Consortium | Harper VanSteenhouse, Yumi Kasai, David Gray, Carol Clugston, Anna Dominiczak and Alex Alderton, Roberto Amato, Jeffrey Barrett, Sonia Goncalves, Ewan Harrison, David K. Jackson, Ian Johnston, Dominic Kwiatkowski, Cordelia Langford, John Sillitoe on behalf of the Wellcome Sanger Institute COVID-19 Surveillance Team |
| EPI_ISL_2999109, EPI_ISL_2999110 | Lighthouse Lab in Alderley Park | Wellcome Sanger Institute for the COVID-19 Genomics UK (COG-UK) Consortium | Jacquelyn Wynn, Mairead Hyland, The Lighthouse Lab in Alderley Park and Alex Alderton, Roberto Amato, Jeffrey Barrett, Sonia Goncalves, Ewan Harrison, David K. Jackson, Ian Johnston, Dominic Kwiatkowski, Cordelia Langford, John Sillitoe on behalf of the Wellcome Sanger Institute COVID-19 Surveillance Team |
| EPI_ISL_2999111 | Lighthouse Laboratory Plymouth | Wellcome Sanger Institute for the COVID-19 Genomics UK (COG-UK) Consortium | Lighthouse Laboratory Plymouth and Alex Alderton, Roberto Amato, Jeffrey Barrett, Sonia Goncalves, Ewan Harrison, David K. Jackson, Ian Johnston, Dominic Kwiatkowski, Cordelia Langford, John Sillitoe on behalf of the Wellcome Sanger Institute COVID-19 Surveillance Team |
| EPI_ISL_2999112 | Lighthouse Lab in Glasgow | Wellcome Sanger Institute for the COVID-19 Genomics UK (COG-UK) Consortium | Harper VanSteenhouse, Yumi Kasai, David Gray, Carol Clugston, Anna Dominiczak and Alex Alderton, Roberto Amato, Jeffrey Barrett, Sonia Goncalves, Ewan Harrison, David K. Jackson, Ian Johnston, Dominic Kwiatkowski, Cordelia Langford, John Sillitoe on behalf of the Wellcome Sanger Institute COVID-19 Surveillance Team |
| EPI_ISL_2999113, EPI_ISL_2999114 | Lighthouse Lab in Alderley Park | Wellcome Sanger Institute for the COVID-19 Genomics UK (COG-UK) Consortium | Jacquelyn Wynn, Mairead Hyland, The Lighthouse Lab in Alderley Park and Alex Alderton, Roberto Amato, Jeffrey Barrett, Sonia Goncalves, Ewan Harrison, David K. Jackson, Ian Johnston, Dominic Kwiatkowski, Cordelia Langford, John Sillitoe on behalf of the Wellcome Sanger Institute COVID-19 Surveillance Team |
| EPI_ISL_2999115 | Lighthouse Lab in Glasgow | Wellcome Sanger Institute for the COVID-19 Genomics UK (COG-UK) Consortium | Harper VanSteenhouse, Yumi Kasai, David Gray, Carol Clugston, Anna Dominiczak and Alex Alderton, Roberto Amato, Jeffrey Barrett, Sonia Goncalves, Ewan Harrison, David K. Jackson, Ian Johnston, Dominic Kwiatkowski, Cordelia Langford, John Sillitoe on behalf of the Wellcome Sanger Institute COVID-19 Surveillance Team |
| EPI_ISL_2999116 | Lighthouse Lab in Alderley Park | Wellcome Sanger Institute for the COVID-19 Genomics UK (COG-UK) Consortium | Jacquelyn Wynn, Mairead Hyland, The Lighthouse Lab in Alderley Park and Alex Alderton, Roberto Amato, Jeffrey Barrett, Sonia Goncalves, Ewan Harrison, David K. Jackson, Ian Johnston, Dominic Kwiatkowski, Cordelia Langford, John Sillitoe on behalf of the Wellcome Sanger Institute COVID-19 Surveillance Team |
| EPI_ISL_2999117, EPI_ISL_2999118 | Lighthouse Lab in Glasgow | Wellcome Sanger Institute for the COVID-19 Genomics UK (COG-UK) Consortium | Harper VanSteenhouse, Yumi Kasai, David Gray, Carol Clugston, Anna Dominiczak and Alex Alderton, Roberto Amato, Jeffrey Barrett, Sonia Goncalves, Ewan Harrison, David K. Jackson, Ian Johnston, Dominic Kwiatkowski, Cordelia Langford, John Sillitoe on behalf of the Wellcome Sanger Institute COVID-19 Surveillance Team |
| EPI_ISL_2999119 | Lighthouse Lab in Alderley Park | Wellcome Sanger Institute for the COVID-19 Genomics UK (COG-UK) Consortium | Jacquelyn Wynn, Mairead Hyland, The Lighthouse Lab in Alderley Park and Alex Alderton, Roberto Amato, Jeffrey Barrett, Sonia Goncalves, Ewan Harrison, David K. Jackson, Ian Johnston, Dominic Kwiatkowski, Cordelia Langford, John Sillitoe on behalf of the Wellcome Sanger Institute COVID-19 Surveillance Team |
| EPI_ISL_2999120 | Lighthouse Lab in Glasgow | Wellcome Sanger Institute for the COVID-19 Genomics UK (COG-UK) Consortium | Harper VanSteenhouse, Yumi Kasai, David Gray, Carol Clugston, Anna Dominiczak and Alex Alderton, Roberto Amato, Jeffrey Barrett, Sonia Goncalves, Ewan Harrison, David K. Jackson, Ian Johnston, Dominic Kwiatkowski, Cordelia Langford, John Sillitoe on behalf of the Wellcome Sanger Institute COVID-19 Surveillance Team |
| EPI_ISL_2999121 | Lighthouse Lab in Alderley Park | Wellcome Sanger Institute for the COVID-19 Genomics UK (COG-UK) Consortium | Jacquelyn Wynn, Mairead Hyland, The Lighthouse Lab in Alderley Park and Alex Alderton, Roberto Amato, Jeffrey Barrett, Sonia Goncalves, Ewan Harrison, David K. Jackson, Ian Johnston, Dominic Kwiatkowski, Cordelia Langford, John Sillitoe on behalf of the Wellcome Sanger Institute COVID-19 Surveillance Team |
| EPI_ISL_2999122 | Lighthouse Laboratory Plymouth | Wellcome Sanger Institute for the COVID-19 Genomics UK (COG-UK) Consortium | Lighthouse Laboratory Plymouth and Alex Alderton, Roberto Amato, Jeffrey Barrett, Sonia Goncalves, Ewan Harrison, David K. Jackson, Ian Johnston, Dominic Kwiatkowski, Cordelia Langford, John Sillitoe on behalf of the Wellcome Sanger Institute COVID-19 Surveillance Team |
| EPI_ISL_2999123 | Lighthouse Lab in Glasgow | Wellcome Sanger Institute for the COVID-19 Genomics UK (COG-UK) Consortium | Harper VanSteenhouse, Yumi Kasai, David Gray, Carol Clugston, Anna Dominiczak and Alex Alderton, Roberto Amato, Jeffrey Barrett, Sonia Goncalves, Ewan Harrison, David K. Jackson, Ian Johnston, Dominic Kwiatkowski, Cordelia Langford, John Sillitoe on behalf of the Wellcome Sanger Institute COVID-19 Surveillance Team |
| EPI_ISL_2999124 | Lighthouse Laboratory Plymouth | Wellcome Sanger Institute for the COVID-19 Genomics UK (COG-UK) Consortium | Lighthouse Laboratory Plymouth and Alex Alderton, Roberto Amato, Jeffrey Barrett, Sonia Goncalves, Ewan Harrison, David K. Jackson, Ian Johnston, Dominic Kwiatkowski, Cordelia Langford, John Sillitoe on behalf of the Wellcome Sanger Institute COVID-19 Surveillance Team |
| EPI_ISL_2999125 | Lighthouse Lab in Glasgow | Wellcome Sanger Institute for the COVID-19 Genomics UK (COG-UK) Consortium | Harper VanSteenhouse, Yumi Kasai, David Gray, Carol Clugston, Anna Dominiczak and Alex Alderton, Roberto Amato, Jeffrey Barrett, Sonia Goncalves, Ewan Harrison, David K. Jackson, Ian Johnston, Dominic Kwiatkowski, Cordelia Langford, John Sillitoe on behalf of the Wellcome Sanger Institute COVID-19 Surveillance Team |
| EPI_ISL_2999126 | Lighthouse Lab in Alderley Park | Wellcome Sanger Institute for the COVID-19 Genomics UK (COG-UK) Consortium | Jacquelyn Wynn, Mairead Hyland, The Lighthouse Lab in Alderley Park and Alex Alderton, Roberto Amato, Jeffrey Barrett, Sonia Goncalves, Ewan Harrison, David K. Jackson, Ian Johnston, Dominic Kwiatkowski, Cordelia Langford, John Sillitoe on behalf of the Wellcome Sanger Institute COVID-19 Surveillance Team |
| EPI_ISL_2999127 | Lighthouse Lab in Glasgow | Wellcome Sanger Institute for the COVID-19 Genomics UK (COG-UK) Consortium | Harper VanSteenhouse, Yumi Kasai, David Gray, Carol Clugston, Anna Dominiczak and Alex Alderton, Roberto Amato, Jeffrey Barrett, Sonia Goncalves, Ewan Harrison, David K. Jackson, Ian Johnston, Dominic Kwiatkowski, Cordelia Langford, John Sillitoe on behalf of the Wellcome Sanger Institute COVID-19 Surveillance Team |
| EPI_ISL_2999128 | Lighthouse Lab in Alderley Park | Wellcome Sanger Institute for the COVID-19 Genomics UK (COG-UK) Consortium | Jacquelyn Wynn, Mairead Hyland, The Lighthouse Lab in Alderley Park and Alex Alderton, Roberto Amato, Jeffrey Barrett, Sonia Goncalves, Ewan Harrison, David K. Jackson, Ian Johnston, Dominic Kwiatkowski, Cordelia Langford, John Sillitoe on behalf of the Wellcome Sanger Institute COVID-19 Surveillance Team |
| EPI_ISL_2999129 | Lighthouse Lab in Glasgow | Wellcome Sanger Institute for the COVID-19 Genomics UK (COG-UK) Consortium | Harper VanSteenhouse, Yumi Kasai, David Gray, Carol Clugston, Anna Dominiczak and Alex Alderton, Roberto Amato, Jeffrey Barrett, Sonia Goncalves, Ewan Harrison, David K. Jackson, Ian Johnston, Dominic Kwiatkowski, Cordelia Langford, John Sillitoe on behalf of the Wellcome Sanger Institute COVID-19 Surveillance Team |
| EPI_ISL_2999130 | Lighthouse Laboratory Plymouth | Wellcome Sanger Institute for the COVID-19 Genomics UK (COG-UK) Consortium | Lighthouse Laboratory Plymouth and Alex Alderton, Roberto Amato, Jeffrey Barrett, Sonia Goncalves, Ewan Harrison, David K. Jackson, Ian Johnston, Dominic Kwiatkowski, Cordelia Langford, John Sillitoe on behalf of the Wellcome Sanger Institute COVID-19 Surveillance Team |
| EPI_ISL_2999131, EPI_ISL_2999132 | Lighthouse Lab in Alderley Park | Wellcome Sanger Institute for the COVID-19 Genomics UK (COG-UK) Consortium | Jacquelyn Wynn, Mairead Hyland, The Lighthouse Lab in Alderley Park and Alex Alderton, Roberto Amato, Jeffrey Barrett, Sonia Goncalves, Ewan Harrison, David K. Jackson, Ian Johnston, Dominic Kwiatkowski, Cordelia Langford, John Sillitoe on behalf of the Wellcome Sanger Institute COVID-19 Surveillance Team |
| EPI_ISL_2999133 | Lighthouse Lab in Glasgow | Wellcome Sanger Institute for the COVID-19 Genomics UK (COG-UK) Consortium | Harper VanSteenhouse, Yumi Kasai, David Gray, Carol Clugston, Anna Dominiczak and Alex Alderton, Roberto Amato, Jeffrey Barrett, Sonia Goncalves, Ewan Harrison, David K. Jackson, Ian Johnston, Dominic Kwiatkowski, Cordelia Langford, John Sillitoe on behalf of the Wellcome Sanger Institute COVID-19 Surveillance Team |
| EPI_ISL_2999134 | Lighthouse Laboratory Plymouth | Wellcome Sanger Institute for the COVID-19 Genomics UK (COG-UK) Consortium | Lighthouse Laboratory Plymouth and Alex Alderton, Roberto Amato, Jeffrey Barrett, Sonia Goncalves, Ewan Harrison, David K. Jackson, Ian Johnston, Dominic Kwiatkowski, Cordelia Langford, John Sillitoe on behalf of the Wellcome Sanger Institute COVID-19 Surveillance Team |
| EPI_ISL_2999135 | Lighthouse Lab in Alderley Park | Wellcome Sanger Institute for the COVID-19 Genomics UK (COG-UK) Consortium | Jacquelyn Wynn, Mairead Hyland, The Lighthouse Lab in Alderley Park and Alex Alderton, Roberto Amato, Jeffrey Barrett, Sonia Goncalves, Ewan Harrison, David K. Jackson, Ian Johnston, Dominic Kwiatkowski, Cordelia Langford, John Sillitoe on behalf of the Wellcome Sanger Institute COVID-19 Surveillance Team |
| EPI_ISL_2999136, EPI_ISL_2999137, EPI_ISL_2999138, EPI_ISL_2999139 | Lighthouse Laboratory Plymouth | Wellcome Sanger Institute for the COVID-19 Genomics UK (COG-UK) Consortium | Lighthouse Laboratory Plymouth and Alex Alderton, Roberto Amato, Jeffrey Barrett, Sonia Goncalves, Ewan Harrison, David K. Jackson, Ian Johnston, Dominic Kwiatkowski, Cordelia Langford, John Sillitoe on behalf of the Wellcome Sanger Institute COVID-19 Surveillance Team |
| EPI_ISL_2999140 | Lighthouse Lab in Alderley Park | Wellcome Sanger Institute for the COVID-19 Genomics UK (COG-UK) Consortium | Jacquelyn Wynn, Mairead Hyland, The Lighthouse Lab in Alderley Park and Alex Alderton, Roberto Amato, Jeffrey Barrett, Sonia Goncalves, Ewan Harrison, David K. Jackson, Ian Johnston, Dominic Kwiatkowski, Cordelia Langford, John Sillitoe on behalf of the Wellcome Sanger Institute COVID-19 Surveillance Team |
| EPI_ISL_2999141 | Lighthouse Laboratory Plymouth | Wellcome Sanger Institute for the COVID-19 Genomics UK (COG-UK) Consortium | Lighthouse Laboratory Plymouth and Alex Alderton, Roberto Amato, Jeffrey Barrett, Sonia Goncalves, Ewan Harrison, David K. Jackson, Ian Johnston, Dominic Kwiatkowski, Cordelia Langford, John Sillitoe on behalf of the Wellcome Sanger Institute COVID-19 Surveillance Team |
| EPI_ISL_2999142 | Lighthouse Lab in Alderley Park | Wellcome Sanger Institute for the COVID-19 Genomics UK (COG-UK) Consortium | Jacquelyn Wynn, Mairead Hyland, The Lighthouse Lab in Alderley Park and Alex Alderton, Roberto Amato, Jeffrey Barrett, Sonia Goncalves, Ewan Harrison, David K. Jackson, Ian Johnston, Dominic Kwiatkowski, Cordelia Langford, John Sillitoe on behalf of the Wellcome Sanger Institute COVID-19 Surveillance Team |
| EPI_ISL_2999143 | Lighthouse Lab in Glasgow | Wellcome Sanger Institute for the COVID-19 Genomics UK (COG-UK) Consortium | Harper VanSteenhouse, Yumi Kasai, David Gray, Carol Clugston, Anna Dominiczak and Alex Alderton, Roberto Amato, Jeffrey Barrett, Sonia Goncalves, Ewan Harrison, David K. Jackson, Ian Johnston, Dominic Kwiatkowski, Cordelia Langford, John Sillitoe on behalf of the Wellcome Sanger Institute COVID-19 Surveillance Team |
| EPI_ISL_2999144 | Lighthouse Laboratory Plymouth | Wellcome Sanger Institute for the COVID-19 Genomics UK (COG-UK) Consortium | Lighthouse Laboratory Plymouth and Alex Alderton, Roberto Amato, Jeffrey Barrett, Sonia Goncalves, Ewan Harrison, David K. Jackson, Ian Johnston, Dominic Kwiatkowski, Cordelia Langford, John Sillitoe on behalf of the Wellcome Sanger Institute COVID-19 Surveillance Team |
| EPI_ISL_2999145 | Lighthouse Lab in Glasgow | Wellcome Sanger Institute for the COVID-19 Genomics UK (COG-UK) Consortium | Harper VanSteenhouse, Yumi Kasai, David Gray, Carol Clugston, Anna Dominiczak and Alex Alderton, Roberto Amato, Jeffrey Barrett, Sonia Goncalves, Ewan Harrison, David K. Jackson, Ian Johnston, Dominic Kwiatkowski, Cordelia Langford, John Sillitoe on behalf of the Wellcome Sanger Institute COVID-19 Surveillance Team |
| EPI_ISL_2999146 | Lighthouse Lab in Alderley Park | Wellcome Sanger Institute for the COVID-19 Genomics UK (COG-UK) Consortium | Jacquelyn Wynn, Mairead Hyland, The Lighthouse Lab in Alderley Park and Alex Alderton, Roberto Amato, Jeffrey Barrett, Sonia Goncalves, Ewan Harrison, David K. Jackson, Ian Johnston, Dominic Kwiatkowski, Cordelia Langford, John Sillitoe on behalf of the Wellcome Sanger Institute COVID-19 Surveillance Team |
| EPI_ISL_2999147, EPI_ISL_2999148 | Lighthouse Laboratory Plymouth | Wellcome Sanger Institute for the COVID-19 Genomics UK (COG-UK) Consortium | Lighthouse Laboratory Plymouth and Alex Alderton, Roberto Amato, Jeffrey Barrett, Sonia Goncalves, Ewan Harrison, David K. Jackson, Ian Johnston, Dominic Kwiatkowski, Cordelia Langford, John Sillitoe on behalf of the Wellcome Sanger Institute COVID-19 Surveillance Team |
| EPI_ISL_2999149, EPI_ISL_2999150 | Lighthouse Lab in Glasgow | Wellcome Sanger Institute for the COVID-19 Genomics UK (COG-UK) Consortium | Harper VanSteenhouse, Yumi Kasai, David Gray, Carol Clugston, Anna Dominiczak and Alex Alderton, Roberto Amato, Jeffrey Barrett, Sonia Goncalves, Ewan Harrison, David K. Jackson, Ian Johnston, Dominic Kwiatkowski, Cordelia Langford, John Sillitoe on behalf of the Wellcome Sanger Institute COVID-19 Surveillance Team |
| EPI_ISL_2999151, EPI_ISL_2999152 | Lighthouse Laboratory Plymouth | Wellcome Sanger Institute for the COVID-19 Genomics UK (COG-UK) Consortium | Lighthouse Laboratory Plymouth and Alex Alderton, Roberto Amato, Jeffrey Barrett, Sonia Goncalves, Ewan Harrison, David K. Jackson, Ian Johnston, Dominic Kwiatkowski, Cordelia Langford, John Sillitoe on behalf of the Wellcome Sanger Institute COVID-19 Surveillance Team |
| EPI_ISL_2999153 | Lighthouse Lab in Glasgow | Wellcome Sanger Institute for the COVID-19 Genomics UK (COG-UK) Consortium | Harper VanSteenhouse, Yumi Kasai, David Gray, Carol Clugston, Anna Dominiczak and Alex Alderton, Roberto Amato, Jeffrey Barrett, Sonia Goncalves, Ewan Harrison, David K. Jackson, Ian Johnston, Dominic Kwiatkowski, Cordelia Langford, John Sillitoe on behalf of the Wellcome Sanger Institute COVID-19 Surveillance Team |
| EPI_ISL_2999154, EPI_ISL_2999155 | Lighthouse Lab in Alderley Park | Wellcome Sanger Institute for the COVID-19 Genomics UK (COG-UK) Consortium | Jacquelyn Wynn, Mairead Hyland, The Lighthouse Lab in Alderley Park and Alex Alderton, Roberto Amato, Jeffrey Barrett, Sonia Goncalves, Ewan Harrison, David K. Jackson, Ian Johnston, Dominic Kwiatkowski, Cordelia Langford, John Sillitoe on behalf of the Wellcome Sanger Institute COVID-19 Surveillance Team |
| EPI_ISL_2999156 | Lighthouse Laboratory Plymouth | Wellcome Sanger Institute for the COVID-19 Genomics UK (COG-UK) Consortium | Lighthouse Laboratory Plymouth and Alex Alderton, Roberto Amato, Jeffrey Barrett, Sonia Goncalves, Ewan Harrison, David K. Jackson, Ian Johnston, Dominic Kwiatkowski, Cordelia Langford, John Sillitoe on behalf of the Wellcome Sanger Institute COVID-19 Surveillance Team |
| EPI_ISL_2999157, EPI_ISL_2999158 | Lighthouse Lab in Alderley Park | Wellcome Sanger Institute for the COVID-19 Genomics UK (COG-UK) Consortium | Jacquelyn Wynn, Mairead Hyland, The Lighthouse Lab in Alderley Park and Alex Alderton, Roberto Amato, Jeffrey Barrett, Sonia Goncalves, Ewan Harrison, David K. Jackson, Ian Johnston, Dominic Kwiatkowski, Cordelia Langford, John Sillitoe on behalf of the Wellcome Sanger Institute COVID-19 Surveillance Team |
| EPI_ISL_2999159 | Lighthouse Lab in Glasgow | Wellcome Sanger Institute for the COVID-19 Genomics UK (COG-UK) Consortium | Harper VanSteenhouse, Yumi Kasai, David Gray, Carol Clugston, Anna Dominiczak and Alex Alderton, Roberto Amato, Jeffrey Barrett, Sonia Goncalves, Ewan Harrison, David K. Jackson, Ian Johnston, Dominic Kwiatkowski, Cordelia Langford, John Sillitoe on behalf of the Wellcome Sanger Institute COVID-19 Surveillance Team |
| EPI_ISL_2999160 | Lighthouse Lab in Alderley Park | Wellcome Sanger Institute for the COVID-19 Genomics UK (COG-UK) Consortium | Jacquelyn Wynn, Mairead Hyland, The Lighthouse Lab in Alderley Park and Alex Alderton, Roberto Amato, Jeffrey Barrett, Sonia Goncalves, Ewan Harrison, David K. Jackson, Ian Johnston, Dominic Kwiatkowski, Cordelia Langford, John Sillitoe on behalf of the Wellcome Sanger Institute COVID-19 Surveillance Team |
| EPI_ISL_2999161 | Lighthouse Lab in Glasgow | Wellcome Sanger Institute for the COVID-19 Genomics UK (COG-UK) Consortium | Harper VanSteenhouse, Yumi Kasai, David Gray, Carol Clugston, Anna Dominiczak and Alex Alderton, Roberto Amato, Jeffrey Barrett, Sonia Goncalves, Ewan Harrison, David K. Jackson, Ian Johnston, Dominic Kwiatkowski, Cordelia Langford, John Sillitoe on behalf of the Wellcome Sanger Institute COVID-19 Surveillance Team |
| EPI_ISL_2999162 | Lighthouse Laboratory Plymouth | Wellcome Sanger Institute for the COVID-19 Genomics UK (COG-UK) Consortium | Lighthouse Laboratory Plymouth and Alex Alderton, Roberto Amato, Jeffrey Barrett, Sonia Goncalves, Ewan Harrison, David K. Jackson, Ian Johnston, Dominic Kwiatkowski, Cordelia Langford, John Sillitoe on behalf of the Wellcome Sanger Institute COVID-19 Surveillance Team |
| EPI_ISL_2999163 | Lighthouse Lab in Glasgow | Wellcome Sanger Institute for the COVID-19 Genomics UK (COG-UK) | Harper VanSteenhouse, Yumi Kasai, David Gray, Carol Clugston, Anna Dominiczak and Alex Alderton, Roberto Amato, Jeffrey Barrett, Sonia Goncalves, Ewan Harrison, David K. Jackson, Ian Johnston, Dominic Kwiatkowski, Cordelia Langford, |

| | | | |
|---|---|---|---|
| | | Consortium | John Sillitoe on behalf of the Wellcome Sanger Institute COVID-19 Surveillance Team |
| EPI_ISL_2999164 | Lighthouse Laboratory Plymouth | Wellcome Sanger Institute for the COVID-19 Genomics UK (COG-UK) Consortium | Lighthouse Laboratory Plymouth and Alex Alderton, Roberto Amato, Jeffrey Barrett, Sonia Goncalves, Ewan Harrison, David K. Jackson, Ian Johnston, Dominic Kwiatkowski, Cordelia Langford, John Sillitoe on behalf of the Wellcome Sanger Institute COVID-19 Surveillance Team |
| EPI_ISL_2999165 | Lighthouse Lab in Alderley Park | Wellcome Sanger Institute for the COVID-19 Genomics UK (COG-UK) Consortium | Jacquelyn Wynn, Mairead Hyland, The Lighthouse Lab in Alderley Park and Alex Alderton, Roberto Amato, Jeffrey Barrett, Sonia Goncalves, Ewan Harrison, David K. Jackson, Ian Johnston, Dominic Kwiatkowski, Cordelia Langford, John Sillitoe on behalf of the Wellcome Sanger Institute COVID-19 Surveillance Team |
| EPI_ISL_2999166 | Lighthouse Lab in Glasgow | Wellcome Sanger Institute for the COVID-19 Genomics UK (COG-UK) Consortium | Harper VanSteenhouse, Yumi Kasai, David Gray, Carol Clugston, Anna Dominiczak and Alex Alderton, Roberto Amato, Jeffrey Barrett, Sonia Goncalves, Ewan Harrison, David K. Jackson, Ian Johnston, Dominic Kwiatkowski, Cordelia Langford, John Sillitoe on behalf of the Wellcome Sanger Institute COVID-19 Surveillance Team |
| EPI_ISL_2999167 | Lighthouse Lab in Alderley Park | Wellcome Sanger Institute for the COVID-19 Genomics UK (COG-UK) Consortium | Jacquelyn Wynn, Mairead Hyland, The Lighthouse Lab in Alderley Park and Alex Alderton, Roberto Amato, Jeffrey Barrett, Sonia Goncalves, Ewan Harrison, David K. Jackson, Ian Johnston, Dominic Kwiatkowski, Cordelia Langford, John Sillitoe on behalf of the Wellcome Sanger Institute COVID-19 Surveillance Team |
| EPI_ISL_2999168 | Lighthouse Lab in Glasgow | Wellcome Sanger Institute for the COVID-19 Genomics UK (COG-UK) Consortium | Harper VanSteenhouse, Yumi Kasai, David Gray, Carol Clugston, Anna Dominiczak and Alex Alderton, Roberto Amato, Jeffrey Barrett, Sonia Goncalves, Ewan Harrison, David K. Jackson, Ian Johnston, Dominic Kwiatkowski, Cordelia Langford, John Sillitoe on behalf of the Wellcome Sanger Institute COVID-19 Surveillance Team |
| EPI_ISL_2999169, EPI_ISL_2999170 | Lighthouse Laboratory Plymouth | Wellcome Sanger Institute for the COVID-19 Genomics UK (COG-UK) Consortium | Lighthouse Laboratory Plymouth and Alex Alderton, Roberto Amato, Jeffrey Barrett, Sonia Goncalves, Ewan Harrison, David K. Jackson, Ian Johnston, Dominic Kwiatkowski, Cordelia Langford, John Sillitoe on behalf of the Wellcome Sanger Institute COVID-19 Surveillance Team |
| EPI_ISL_2999171, EPI_ISL_2999172, EPI_ISL_2999173, EPI_ISL_2999174 | Lighthouse Lab in Alderley Park | Wellcome Sanger Institute for the COVID-19 Genomics UK (COG-UK) Consortium | Jacquelyn Wynn, Mairead Hyland, The Lighthouse Lab in Alderley Park and Alex Alderton, Roberto Amato, Jeffrey Barrett, Sonia Goncalves, Ewan Harrison, David K. Jackson, Ian Johnston, Dominic Kwiatkowski, Cordelia Langford, John Sillitoe on behalf of the Wellcome Sanger Institute COVID-19 Surveillance Team |
| EPI_ISL_2999175 | Lighthouse Lab in Glasgow | Wellcome Sanger Institute for the COVID-19 Genomics UK (COG-UK) Consortium | Harper VanSteenhouse, Yumi Kasai, David Gray, Carol Clugston, Anna Dominiczak and Alex Alderton, Roberto Amato, Jeffrey Barrett, Sonia Goncalves, Ewan Harrison, David K. Jackson, Ian Johnston, Dominic Kwiatkowski, Cordelia Langford, John Sillitoe on behalf of the Wellcome Sanger Institute COVID-19 Surveillance Team |
| EPI_ISL_2999176 | Lighthouse Laboratory Plymouth | Wellcome Sanger Institute for the COVID-19 Genomics UK (COG-UK) Consortium | Lighthouse Laboratory Plymouth and Alex Alderton, Roberto Amato, Jeffrey Barrett, Sonia Goncalves, Ewan Harrison, David K. Jackson, Ian Johnston, Dominic Kwiatkowski, Cordelia Langford, John Sillitoe on behalf of the Wellcome Sanger Institute COVID-19 Surveillance Team |
| EPI_ISL_2999177 | Lighthouse Lab in Glasgow | Wellcome Sanger Institute for the COVID-19 Genomics UK (COG-UK) Consortium | Harper VanSteenhouse, Yumi Kasai, David Gray, Carol Clugston, Anna Dominiczak and Alex Alderton, Roberto Amato, Jeffrey Barrett, Sonia Goncalves, Ewan Harrison, David K. Jackson, Ian Johnston, Dominic Kwiatkowski, Cordelia Langford, John Sillitoe on behalf of the Wellcome Sanger Institute COVID-19 Surveillance Team |
| EPI_ISL_2999178 | Lighthouse Laboratory Plymouth | Wellcome Sanger Institute for the COVID-19 Genomics UK (COG-UK) Consortium | Lighthouse Laboratory Plymouth and Alex Alderton, Roberto Amato, Jeffrey Barrett, Sonia Goncalves, Ewan Harrison, David K. Jackson, Ian Johnston, Dominic Kwiatkowski, Cordelia Langford, John Sillitoe on behalf of the Wellcome Sanger Institute COVID-19 Surveillance Team |
| EPI_ISL_2999179 | Lighthouse Lab in Alderley Park | Wellcome Sanger Institute for the COVID-19 Genomics UK (COG-UK) Consortium | Jacquelyn Wynn, Mairead Hyland, The Lighthouse Lab in Alderley Park and Alex Alderton, Roberto Amato, Jeffrey Barrett, Sonia Goncalves, Ewan Harrison, David K. Jackson, Ian Johnston, Dominic Kwiatkowski, Cordelia Langford, John Sillitoe on behalf of the Wellcome Sanger Institute COVID-19 Surveillance Team |
| EPI_ISL_2999180 | Lighthouse Lab in Glasgow | Wellcome Sanger Institute for the COVID-19 Genomics UK (COG-UK) Consortium | Harper VanSteenhouse, Yumi Kasai, David Gray, Carol Clugston, Anna Dominiczak and Alex Alderton, Roberto Amato, Jeffrey Barrett, Sonia Goncalves, Ewan Harrison, David K. Jackson, Ian Johnston, Dominic Kwiatkowski, Cordelia Langford, John Sillitoe on behalf of the Wellcome Sanger Institute COVID-19 Surveillance Team |
| EPI_ISL_2999181, EPI_ISL_2999182, EPI_ISL_2999183 | Lighthouse Laboratory Plymouth | Wellcome Sanger Institute for the COVID-19 Genomics UK (COG-UK) Consortium | Lighthouse Laboratory Plymouth and Alex Alderton, Roberto Amato, Jeffrey Barrett, Sonia Goncalves, Ewan Harrison, David K. Jackson, Ian Johnston, Dominic Kwiatkowski, Cordelia Langford, John Sillitoe on behalf of the Wellcome Sanger Institute COVID-19 Surveillance Team |
| EPI_ISL_2999184, EPI_ISL_2999185 | Lighthouse Lab in Alderley Park | Wellcome Sanger Institute for the COVID-19 Genomics UK (COG-UK) Consortium | Jacquelyn Wynn, Mairead Hyland, The Lighthouse Lab in Alderley Park and Alex Alderton, Roberto Amato, Jeffrey Barrett, Sonia Goncalves, Ewan Harrison, David K. Jackson, Ian Johnston, Dominic Kwiatkowski, Cordelia Langford, John Sillitoe on behalf of the Wellcome Sanger Institute COVID-19 Surveillance Team |
| EPI_ISL_2999186, EPI_ISL_2999187 | Lighthouse Lab in Glasgow | Wellcome Sanger Institute for the COVID-19 Genomics UK (COG-UK) Consortium | Harper VanSteenhouse, Yumi Kasai, David Gray, Carol Clugston, Anna Dominiczak and Alex Alderton, Roberto Amato, Jeffrey Barrett, Sonia Goncalves, Ewan Harrison, David K. Jackson, Ian Johnston, Dominic Kwiatkowski, Cordelia Langford, John Sillitoe on behalf of the Wellcome Sanger Institute COVID-19 Surveillance Team |
| EPI_ISL_2999188 | Lighthouse Lab in Alderley Park | Wellcome Sanger Institute for the COVID-19 Genomics UK (COG-UK) Consortium | Jacquelyn Wynn, Mairead Hyland, The Lighthouse Lab in Alderley Park and Alex Alderton, Roberto Amato, Jeffrey Barrett, Sonia Goncalves, Ewan Harrison, David K. Jackson, Ian Johnston, Dominic Kwiatkowski, Cordelia Langford, John Sillitoe on behalf of the Wellcome Sanger Institute COVID-19 Surveillance Team |
| EPI_ISL_2999189, EPI_ISL_2999190, EPI_ISL_2999191, EPI_ISL_2999192, EPI_ISL_2999193 | Lighthouse Laboratory Plymouth | Wellcome Sanger Institute for the COVID-19 Genomics UK (COG-UK) Consortium | Lighthouse Laboratory Plymouth and Alex Alderton, Roberto Amato, Jeffrey Barrett, Sonia Goncalves, Ewan Harrison, David K. Jackson, Ian Johnston, Dominic Kwiatkowski, Cordelia Langford, John Sillitoe on behalf of the Wellcome Sanger Institute COVID-19 Surveillance Team |
| EPI_ISL_2999194 | Lighthouse Lab in Alderley Park | Wellcome Sanger Institute for the COVID-19 Genomics UK (COG-UK) Consortium | Jacquelyn Wynn, Mairead Hyland, The Lighthouse Lab in Alderley Park and Alex Alderton, Roberto Amato, Jeffrey Barrett, Sonia Goncalves, Ewan Harrison, David K. Jackson, Ian Johnston, Dominic Kwiatkowski, Cordelia Langford, John Sillitoe on behalf of the Wellcome Sanger Institute COVID-19 Surveillance Team |
| EPI_ISL_2999195 | Lighthouse Lab in Glasgow | Wellcome Sanger Institute for the COVID-19 Genomics UK (COG-UK) Consortium | Harper VanSteenhouse, Yumi Kasai, David Gray, Carol Clugston, Anna Dominiczak and Alex Alderton, Roberto Amato, Jeffrey Barrett, Sonia Goncalves, Ewan Harrison, David K. Jackson, Ian Johnston, Dominic Kwiatkowski, Cordelia Langford, John Sillitoe on behalf of the Wellcome Sanger Institute COVID-19 Surveillance Team |
| EPI_ISL_2999196 | Lighthouse Lab in Alderley Park | Wellcome Sanger Institute for the COVID-19 Genomics UK (COG-UK) Consortium | Jacquelyn Wynn, Mairead Hyland, The Lighthouse Lab in Alderley Park and Alex Alderton, Roberto Amato, Jeffrey Barrett, Sonia Goncalves, Ewan Harrison, David K. Jackson, Ian Johnston, Dominic Kwiatkowski, Cordelia Langford, John Sillitoe on behalf of the Wellcome Sanger Institute COVID-19 Surveillance Team |
| EPI_ISL_2999197 | Lighthouse Laboratory Plymouth | Wellcome Sanger Institute for the COVID-19 Genomics UK (COG-UK) Consortium | Lighthouse Laboratory Plymouth and Alex Alderton, Roberto Amato, Jeffrey Barrett, Sonia Goncalves, Ewan Harrison, David K. Jackson, Ian Johnston, Dominic Kwiatkowski, Cordelia Langford, John Sillitoe on behalf of the Wellcome Sanger Institute COVID-19 Surveillance Team |
| EPI_ISL_2999198 | Lighthouse Lab in Glasgow | Wellcome Sanger Institute for the COVID-19 Genomics UK (COG-UK) Consortium | Harper VanSteenhouse, Yumi Kasai, David Gray, Carol Clugston, Anna Dominiczak and Alex Alderton, Roberto Amato, Jeffrey Barrett, Sonia Goncalves, Ewan Harrison, David K. Jackson, Ian Johnston, Dominic Kwiatkowski, Cordelia Langford, John Sillitoe on behalf of the Wellcome Sanger Institute COVID-19 Surveillance Team |
| EPI_ISL_2999199 | Lighthouse Laboratory Plymouth | Wellcome Sanger Institute for the COVID-19 Genomics UK (COG-UK) Consortium | Lighthouse Laboratory Plymouth and Alex Alderton, Roberto Amato, Jeffrey Barrett, Sonia Goncalves, Ewan Harrison, David K. Jackson, Ian Johnston, Dominic Kwiatkowski, Cordelia Langford, John Sillitoe on behalf of the Wellcome Sanger Institute COVID-19 Surveillance Team |
| EPI_ISL_2999200 | Lighthouse Lab in Alderley Park | Wellcome Sanger Institute for the COVID-19 Genomics UK (COG-UK) Consortium | Jacquelyn Wynn, Mairead Hyland, The Lighthouse Lab in Alderley Park and Alex Alderton, Roberto Amato, Jeffrey Barrett, Sonia Goncalves, Ewan Harrison, David K. Jackson, Ian Johnston, Dominic Kwiatkowski, Cordelia Langford, John Sillitoe on behalf of the Wellcome Sanger Institute COVID-19 Surveillance Team |
| EPI_ISL_2999201, EPI_ISL_2999202 | Lighthouse Laboratory Plymouth | Wellcome Sanger Institute for the COVID-19 Genomics UK (COG-UK) Consortium | Lighthouse Laboratory Plymouth and Alex Alderton, Roberto Amato, Jeffrey Barrett, Sonia Goncalves, Ewan Harrison, David K. Jackson, Ian Johnston, Dominic Kwiatkowski, Cordelia Langford, John Sillitoe on behalf of the Wellcome Sanger Institute COVID-19 Surveillance Team |
| EPI_ISL_2999203 | Lighthouse Lab in Glasgow | Wellcome Sanger Institute for the COVID-19 Genomics UK (COG-UK) Consortium | Harper VanSteenhouse, Yumi Kasai, David Gray, Carol Clugston, Anna Dominiczak and Alex Alderton, Roberto Amato, Jeffrey Barrett, Sonia Goncalves, Ewan Harrison, David K. Jackson, Ian Johnston, Dominic Kwiatkowski, Cordelia Langford, John Sillitoe on behalf of the Wellcome Sanger Institute COVID-19 Surveillance Team |
| EPI_ISL_2999204, EPI_ISL_2999205, EPI_ISL_2999206 | Lighthouse Lab in Alderley Park | Wellcome Sanger Institute for the COVID-19 Genomics UK (COG-UK) Consortium | Jacquelyn Wynn, Mairead Hyland, The Lighthouse Lab in Alderley Park and Alex Alderton, Roberto Amato, Jeffrey Barrett, Sonia Goncalves, Ewan Harrison, David K. Jackson, Ian Johnston, Dominic Kwiatkowski, Cordelia Langford, John Sillitoe on behalf of the Wellcome Sanger Institute COVID-19 Surveillance Team |
| EPI_ISL_2999207, EPI_ISL_2999208, EPI_ISL_2999209, EPI_ISL_2999210, EPI_ISL_2999211 | Lighthouse Lab in Glasgow | Wellcome Sanger Institute for the COVID-19 Genomics UK (COG-UK) Consortium | Harper VanSteenhouse, Yumi Kasai, David Gray, Carol Clugston, Anna Dominiczak and Alex Alderton, Roberto Amato, Jeffrey Barrett, Sonia Goncalves, Ewan Harrison, David K. Jackson, Ian Johnston, Dominic Kwiatkowski, Cordelia Langford, John Sillitoe on behalf of the Wellcome Sanger Institute COVID-19 Surveillance Team |
| EPI_ISL_2999212, EPI_ISL_2999213, EPI_ISL_2999214 | Lighthouse Lab in Alderley Park | Wellcome Sanger Institute for the COVID-19 Genomics UK (COG-UK) Consortium | Jacquelyn Wynn, Mairead Hyland, The Lighthouse Lab in Alderley Park and Alex Alderton, Roberto Amato, Jeffrey Barrett, Sonia Goncalves, Ewan Harrison, David K. Jackson, Ian Johnston, Dominic Kwiatkowski, Cordelia Langford, John Sillitoe on behalf of the Wellcome Sanger Institute COVID-19 Surveillance Team |
| EPI_ISL_2999215, EPI_ISL_2999216 | Lighthouse Lab in Glasgow | Wellcome Sanger Institute for the COVID-19 Genomics UK (COG-UK) Consortium | Harper VanSteenhouse, Yumi Kasai, David Gray, Carol Clugston, Anna Dominiczak and Alex Alderton, Roberto Amato, Jeffrey Barrett, Sonia Goncalves, Ewan Harrison, David K. Jackson, Ian Johnston, Dominic Kwiatkowski, Cordelia Langford, John Sillitoe on behalf of the Wellcome Sanger Institute COVID-19 Surveillance Team |
| EPI_ISL_2999217 | Lighthouse Laboratory Plymouth | Wellcome Sanger Institute for the COVID-19 Genomics UK (COG-UK) Consortium | Lighthouse Laboratory Plymouth and Alex Alderton, Roberto Amato, Jeffrey Barrett, Sonia Goncalves, Ewan Harrison, David K. Jackson, Ian Johnston, Dominic Kwiatkowski, Cordelia Langford, John Sillitoe on behalf of the Wellcome Sanger Institute COVID-19 Surveillance Team |
| EPI_ISL_2999218, EPI_ISL_2999219 | Lighthouse Lab in Glasgow | Wellcome Sanger Institute for the COVID-19 Genomics UK (COG-UK) Consortium | Harper VanSteenhouse, Yumi Kasai, David Gray, Carol Clugston, Anna Dominiczak and Alex Alderton, Roberto Amato, Jeffrey Barrett, Sonia Goncalves, Ewan Harrison, David K. Jackson, Ian Johnston, Dominic Kwiatkowski, Cordelia Langford, John Sillitoe on behalf of the Wellcome Sanger Institute COVID-19 Surveillance Team |
| EPI_ISL_2999220, EPI_ISL_2999221 | Lighthouse Laboratory Plymouth | Wellcome Sanger Institute for the COVID-19 Genomics UK (COG-UK) Consortium | Lighthouse Laboratory Plymouth and Alex Alderton, Roberto Amato, Jeffrey Barrett, Sonia Goncalves, Ewan Harrison, David K. Jackson, Ian Johnston, Dominic Kwiatkowski, Cordelia Langford, John Sillitoe on behalf of the Wellcome Sanger Institute COVID-19 Surveillance Team |
| EPI_ISL_2999222, EPI_ISL_2999223, EPI_ISL_2999224, EPI_ISL_2999225, EPI_ISL_2999226 | Lighthouse Lab in Alderley Park | Wellcome Sanger Institute for the COVID-19 Genomics UK (COG-UK) Consortium | Jacquelyn Wynn, Mairead Hyland, The Lighthouse Lab in Alderley Park and Alex Alderton, Roberto Amato, Jeffrey Barrett, Sonia Goncalves, Ewan Harrison, David K. Jackson, Ian Johnston, Dominic Kwiatkowski, Cordelia Langford, John Sillitoe on behalf of the Wellcome Sanger Institute COVID-19 Surveillance Team |
| EPI_ISL_2999227 | Lighthouse Lab in Glasgow | Wellcome Sanger Institute for the COVID-19 Genomics UK (COG-UK) Consortium | Harper VanSteenhouse, Yumi Kasai, David Gray, Carol Clugston, Anna Dominiczak and Alex Alderton, Roberto Amato, Jeffrey Barrett, Sonia Goncalves, Ewan Harrison, David K. Jackson, Ian Johnston, Dominic Kwiatkowski, Cordelia Langford, John Sillitoe on behalf of the Wellcome Sanger Institute COVID-19 Surveillance Team |
| EPI_ISL_2999228 | Lighthouse Lab in Alderley Park | Wellcome Sanger Institute for the COVID-19 Genomics UK (COG-UK) Consortium | Jacquelyn Wynn, Mairead Hyland, The Lighthouse Lab in Alderley Park and Alex Alderton, Roberto Amato, Jeffrey Barrett, Sonia Goncalves, Ewan Harrison, David K. Jackson, Ian Johnston, Dominic Kwiatkowski, Cordelia Langford, John Sillitoe on behalf of the Wellcome Sanger Institute COVID-19 Surveillance Team |
| EPI_ISL_2999229 | Lighthouse Lab in Glasgow | Wellcome Sanger Institute for the COVID-19 Genomics UK (COG-UK) Consortium | Harper VanSteenhouse, Yumi Kasai, David Gray, Carol Clugston, Anna Dominiczak and Alex Alderton, Roberto Amato, Jeffrey Barrett, Sonia Goncalves, Ewan Harrison, David K. Jackson, Ian Johnston, Dominic Kwiatkowski, Cordelia Langford, John Sillitoe on behalf of the Wellcome Sanger Institute COVID-19 Surveillance Team |
| EPI_ISL_2999230 | Lighthouse Laboratory Plymouth | Wellcome Sanger Institute for the COVID-19 Genomics UK (COG-UK) Consortium | Lighthouse Laboratory Plymouth and Alex Alderton, Roberto Amato, Jeffrey Barrett, Sonia Goncalves, Ewan Harrison, David K. Jackson, Ian Johnston, Dominic Kwiatkowski, Cordelia Langford, John Sillitoe on behalf of the Wellcome Sanger Institute COVID-19 Surveillance Team |

| | | | |
|---|---|---|---|
| EPI_ISL_2999231, EPI_ISL_2999232 | Lighthouse Lab in Glasgow | Wellcome Sanger Institute for the COVID-19 Genomics UK (COG-UK) Consortium | Harper VanSteenhouse, Yumi Kasai, David Gray, Carol Clugston, Anna Dominiczak and Alex Alderton, Roberto Amato, Jeffrey Barrett, Sonia Goncalves, Ewan Harrison, David K. Jackson, Ian Johnston, Dominic Kwiatkowski, Cordelia Langford, John Sillitoe on behalf of the Wellcome Sanger Institute COVID-19 Surveillance Team |
| EPI_ISL_2999233 | Lighthouse Lab in Alderley Park | Wellcome Sanger Institute for the COVID-19 Genomics UK (COG-UK) Consortium | Jacquelyn Wynn, Mairead Hyland, The Lighthouse Lab in Alderley Park and Alex Alderton, Roberto Amato, Jeffrey Barrett, Sonia Goncalves, Ewan Harrison, David K. Jackson, Ian Johnston, Dominic Kwiatkowski, Cordelia Langford, John Sillitoe on behalf of the Wellcome Sanger Institute COVID-19 Surveillance Team |
| EPI_ISL_2999234 | Lighthouse Lab in Glasgow | Wellcome Sanger Institute for the COVID-19 Genomics UK (COG-UK) Consortium | Harper VanSteenhouse, Yumi Kasai, David Gray, Carol Clugston, Anna Dominiczak and Alex Alderton, Roberto Amato, Jeffrey Barrett, Sonia Goncalves, Ewan Harrison, David K. Jackson, Ian Johnston, Dominic Kwiatkowski, Cordelia Langford, John Sillitoe on behalf of the Wellcome Sanger Institute COVID-19 Surveillance Team |
| EPI_ISL_2999235 | Lighthouse Lab in Alderley Park | Wellcome Sanger Institute for the COVID-19 Genomics UK (COG-UK) Consortium | Jacquelyn Wynn, Mairead Hyland, The Lighthouse Lab in Alderley Park and Alex Alderton, Roberto Amato, Jeffrey Barrett, Sonia Goncalves, Ewan Harrison, David K. Jackson, Ian Johnston, Dominic Kwiatkowski, Cordelia Langford, John Sillitoe on behalf of the Wellcome Sanger Institute COVID-19 Surveillance Team |
| EPI_ISL_2999236, EPI_ISL_2999237, EPI_ISL_2999238 | Lighthouse Laboratory Plymouth | Wellcome Sanger Institute for the COVID-19 Genomics UK (COG-UK) Consortium | Lighthouse Laboratory Plymouth and Alex Alderton, Roberto Amato, Jeffrey Barrett, Sonia Goncalves, Ewan Harrison, David K. Jackson, Ian Johnston, Dominic Kwiatkowski, Cordelia Langford, John Sillitoe on behalf of the Wellcome Sanger Institute COVID-19 Surveillance Team |
| EPI_ISL_2999239, EPI_ISL_2999240 | Lighthouse Lab in Alderley Park | Wellcome Sanger Institute for the COVID-19 Genomics UK (COG-UK) Consortium | Jacquelyn Wynn, Mairead Hyland, The Lighthouse Lab in Alderley Park and Alex Alderton, Roberto Amato, Jeffrey Barrett, Sonia Goncalves, Ewan Harrison, David K. Jackson, Ian Johnston, Dominic Kwiatkowski, Cordelia Langford, John Sillitoe on behalf of the Wellcome Sanger Institute COVID-19 Surveillance Team |
| EPI_ISL_2999241 | Lighthouse Laboratory Plymouth | Wellcome Sanger Institute for the COVID-19 Genomics UK (COG-UK) Consortium | Lighthouse Laboratory Plymouth and Alex Alderton, Roberto Amato, Jeffrey Barrett, Sonia Goncalves, Ewan Harrison, David K. Jackson, Ian Johnston, Dominic Kwiatkowski, Cordelia Langford, John Sillitoe on behalf of the Wellcome Sanger Institute COVID-19 Surveillance Team |
| EPI_ISL_2999242 | Lighthouse Lab in Alderley Park | Wellcome Sanger Institute for the COVID-19 Genomics UK (COG-UK) Consortium | Jacquelyn Wynn, Mairead Hyland, The Lighthouse Lab in Alderley Park and Alex Alderton, Roberto Amato, Jeffrey Barrett, Sonia Goncalves, Ewan Harrison, David K. Jackson, Ian Johnston, Dominic Kwiatkowski, Cordelia Langford, John Sillitoe on behalf of the Wellcome Sanger Institute COVID-19 Surveillance Team |
| EPI_ISL_2999243 | Lighthouse Laboratory Plymouth | Wellcome Sanger Institute for the COVID-19 Genomics UK (COG-UK) Consortium | Lighthouse Laboratory Plymouth and Alex Alderton, Roberto Amato, Jeffrey Barrett, Sonia Goncalves, Ewan Harrison, David K. Jackson, Ian Johnston, Dominic Kwiatkowski, Cordelia Langford, John Sillitoe on behalf of the Wellcome Sanger Institute COVID-19 Surveillance Team |
| EPI_ISL_2999244, EPI_ISL_2999245 | Lighthouse Lab in Glasgow | Wellcome Sanger Institute for the COVID-19 Genomics UK (COG-UK) Consortium | Harper VanSteenhouse, Yumi Kasai, David Gray, Carol Clugston, Anna Dominiczak and Alex Alderton, Roberto Amato, Jeffrey Barrett, Sonia Goncalves, Ewan Harrison, David K. Jackson, Ian Johnston, Dominic Kwiatkowski, Cordelia Langford, John Sillitoe on behalf of the Wellcome Sanger Institute COVID-19 Surveillance Team |
| EPI_ISL_2999246 | Lighthouse Lab in Alderley Park | Wellcome Sanger Institute for the COVID-19 Genomics UK (COG-UK) Consortium | Jacquelyn Wynn, Mairead Hyland, The Lighthouse Lab in Alderley Park and Alex Alderton, Roberto Amato, Jeffrey Barrett, Sonia Goncalves, Ewan Harrison, David K. Jackson, Ian Johnston, Dominic Kwiatkowski, Cordelia Langford, John Sillitoe on behalf of the Wellcome Sanger Institute COVID-19 Surveillance Team |
| EPI_ISL_2999247 | Lighthouse Laboratory Plymouth | Wellcome Sanger Institute for the COVID-19 Genomics UK (COG-UK) Consortium | Lighthouse Laboratory Plymouth and Alex Alderton, Roberto Amato, Jeffrey Barrett, Sonia Goncalves, Ewan Harrison, David K. Jackson, Ian Johnston, Dominic Kwiatkowski, Cordelia Langford, John Sillitoe on behalf of the Wellcome Sanger Institute COVID-19 Surveillance Team |
| EPI_ISL_2999248, EPI_ISL_2999249, EPI_ISL_2999250 | Lighthouse Lab in Glasgow | Wellcome Sanger Institute for the COVID-19 Genomics UK (COG-UK) Consortium | Harper VanSteenhouse, Yumi Kasai, David Gray, Carol Clugston, Anna Dominiczak and Alex Alderton, Roberto Amato, Jeffrey Barrett, Sonia Goncalves, Ewan Harrison, David K. Jackson, Ian Johnston, Dominic Kwiatkowski, Cordelia Langford, John Sillitoe on behalf of the Wellcome Sanger Institute COVID-19 Surveillance Team |
| EPI_ISL_2999251 | Lighthouse Laboratory Plymouth | Wellcome Sanger Institute for the COVID-19 Genomics UK (COG-UK) Consortium | Lighthouse Laboratory Plymouth and Alex Alderton, Roberto Amato, Jeffrey Barrett, Sonia Goncalves, Ewan Harrison, David K. Jackson, Ian Johnston, Dominic Kwiatkowski, Cordelia Langford, John Sillitoe on behalf of the Wellcome Sanger Institute COVID-19 Surveillance Team |
| EPI_ISL_2999252 | Lighthouse Lab in Alderley Park | Wellcome Sanger Institute for the COVID-19 Genomics UK (COG-UK) Consortium | Jacquelyn Wynn, Mairead Hyland, The Lighthouse Lab in Alderley Park and Alex Alderton, Roberto Amato, Jeffrey Barrett, Sonia Goncalves, Ewan Harrison, David K. Jackson, Ian Johnston, Dominic Kwiatkowski, Cordelia Langford, John Sillitoe on behalf of the Wellcome Sanger Institute COVID-19 Surveillance Team |
| EPI_ISL_2999253, EPI_ISL_2999254 | Lighthouse Lab in Glasgow | Wellcome Sanger Institute for the COVID-19 Genomics UK (COG-UK) Consortium | Harper VanSteenhouse, Yumi Kasai, David Gray, Carol Clugston, Anna Dominiczak and Alex Alderton, Roberto Amato, Jeffrey Barrett, Sonia Goncalves, Ewan Harrison, David K. Jackson, Ian Johnston, Dominic Kwiatkowski, Cordelia Langford, John Sillitoe on behalf of the Wellcome Sanger Institute COVID-19 Surveillance Team |
| EPI_ISL_2999255, EPI_ISL_2999256, EPI_ISL_2999257 | Lighthouse Laboratory Plymouth | Wellcome Sanger Institute for the COVID-19 Genomics UK (COG-UK) Consortium | Lighthouse Laboratory Plymouth and Alex Alderton, Roberto Amato, Jeffrey Barrett, Sonia Goncalves, Ewan Harrison, David K. Jackson, Ian Johnston, Dominic Kwiatkowski, Cordelia Langford, John Sillitoe on behalf of the Wellcome Sanger Institute COVID-19 Surveillance Team |
| EPI_ISL_2999258, EPI_ISL_2999259 | Lighthouse Lab in Glasgow | Wellcome Sanger Institute for the COVID-19 Genomics UK (COG-UK) Consortium | Harper VanSteenhouse, Yumi Kasai, David Gray, Carol Clugston, Anna Dominiczak and Alex Alderton, Roberto Amato, Jeffrey Barrett, Sonia Goncalves, Ewan Harrison, David K. Jackson, Ian Johnston, Dominic Kwiatkowski, Cordelia Langford, John Sillitoe on behalf of the Wellcome Sanger Institute COVID-19 Surveillance Team |
| EPI_ISL_2999260 | Lighthouse Lab in Milton Keynes | Wellcome Sanger Institute for the COVID-19 Genomics UK (COG-UK) Consortium | The Lighthouse Lab in Milton Keynes and Alex Alderton, Roberto Amato, Jeffrey Barrett, Sonia Goncalves, Ewan Harrison, David K. Jackson, Ian Johnston, Dominic Kwiatkowski, Cordelia Langford, John Sillitoe on behalf of the Wellcome Sanger Institute COVID-19 Surveillance Team |
| EPI_ISL_2999261, EPI_ISL_2999262, EPI_ISL_2999263 | Lighthouse Lab in Alderley Park | Wellcome Sanger Institute for the COVID-19 Genomics UK (COG-UK) Consortium | Jacquelyn Wynn, Mairead Hyland, The Lighthouse Lab in Alderley Park and Alex Alderton, Roberto Amato, Jeffrey Barrett, Sonia Goncalves, Ewan Harrison, David K. Jackson, Ian Johnston, Dominic Kwiatkowski, Cordelia Langford, John Sillitoe on behalf of the Wellcome Sanger Institute COVID-19 Surveillance Team |
| EPI_ISL_2999264, EPI_ISL_2999265, EPI_ISL_2999266, EPI_ISL_2999267, EPI_ISL_2999268, EPI_ISL_2999269, EPI_ISL_2999270, EPI_ISL_2999271, EPI_ISL_2999272, EPI_ISL_2999273, EPI_ISL_2999274, EPI_ISL_2999275, EPI_ISL_2999276, EPI_ISL_2999277, EPI_ISL_2999278 | | | |
| see above | Lighthouse Lab in Glasgow | Wellcome Sanger Institute for the COVID-19 Genomics UK (COG-UK) Consortium | Harper VanSteenhouse, Yumi Kasai, David Gray, Carol Clugston, Anna Dominiczak and Alex Alderton, Roberto Amato, Jeffrey Barrett, Sonia Goncalves, Ewan Harrison, David K. Jackson, Ian Johnston, Dominic Kwiatkowski, Cordelia Langford, John Sillitoe on behalf of the Wellcome Sanger Institute COVID-19 Surveillance Team |
| EPI_ISL_2999279, EPI_ISL_2999280 | Lighthouse Lab in Alderley Park | Wellcome Sanger Institute for the COVID-19 Genomics UK (COG-UK) Consortium | Jacquelyn Wynn, Mairead Hyland, The Lighthouse Lab in Alderley Park and Alex Alderton, Roberto Amato, Jeffrey Barrett, Sonia Goncalves, Ewan Harrison, David K. Jackson, Ian Johnston, Dominic Kwiatkowski, Cordelia Langford, John Sillitoe on behalf of the Wellcome Sanger Institute COVID-19 Surveillance Team |
| EPI_ISL_2999281, EPI_ISL_2999282, EPI_ISL_2999283, EPI_ISL_2999285, EPI_ISL_2999286, EPI_ISL_2999287, EPI_ISL_2999288, EPI_ISL_2999289, EPI_ISL_2999290, EPI_ISL_2999291, EPI_ISL_2999292, EPI_ISL_2999293, EPI_ISL_2999294, EPI_ISL_2999295, EPI_ISL_2999296, EPI_ISL_2999297, EPI_ISL_2999298, EPI_ISL_2999299, EPI_ISL_2999300, EPI_ISL_2999301, EPI_ISL_2999302, EPI_ISL_2999303, EPI_ISL_2999304, EPI_ISL_2999305, EPI_ISL_2999306, EPI_ISL_2999307, EPI_ISL_2999308, EPI_ISL_2999309, EPI_ISL_2999310, EPI_ISL_2999311, EPI_ISL_2999312, EPI_ISL_2999313, EPI_ISL_2999314, EPI_ISL_2999315, EPI_ISL_2999316, EPI_ISL_2999317, EPI_ISL_2999318, EPI_ISL_2999319, EPI_ISL_2999320, EPI_ISL_2999321, EPI_ISL_2999322, EPI_ISL_2999323, EPI_ISL_2999324, EPI_ISL_2999325, EPI_ISL_2999326, EPI_ISL_2999327 | | | |
| see above | Lighthouse Lab in Glasgow | Wellcome Sanger Institute for the COVID-19 Genomics UK (COG-UK) Consortium | Harper VanSteenhouse, Yumi Kasai, David Gray, Carol Clugston, Anna Dominiczak and Alex Alderton, Roberto Amato, Jeffrey Barrett, Sonia Goncalves, Ewan Harrison, David K. Jackson, Ian Johnston, Dominic Kwiatkowski, Cordelia Langford, John Sillitoe on behalf of the Wellcome Sanger Institute COVID-19 Surveillance Team |
| EPI_ISL_2999328 | Lighthouse Lab in Alderley Park | Wellcome Sanger Institute for the COVID-19 Genomics UK (COG-UK) Consortium | Jacquelyn Wynn, Mairead Hyland, The Lighthouse Lab in Alderley Park and Alex Alderton, Roberto Amato, Jeffrey Barrett, Sonia Goncalves, Ewan Harrison, David K. Jackson, Ian Johnston, Dominic Kwiatkowski, Cordelia Langford, John Sillitoe on behalf of the Wellcome Sanger Institute COVID-19 Surveillance Team |
| EPI_ISL_2999329, EPI_ISL_2999330 | Lighthouse Lab in Glasgow | Wellcome Sanger Institute for the COVID-19 Genomics UK (COG-UK) Consortium | Harper VanSteenhouse, Yumi Kasai, David Gray, Carol Clugston, Anna Dominiczak and Alex Alderton, Roberto Amato, Jeffrey Barrett, Sonia Goncalves, Ewan Harrison, David K. Jackson, Ian Johnston, Dominic Kwiatkowski, Cordelia Langford, John Sillitoe on behalf of the Wellcome Sanger Institute COVID-19 Surveillance Team |
| EPI_ISL_2999331, EPI_ISL_2999332 | Lighthouse Lab in Alderley Park | Wellcome Sanger Institute for the COVID-19 Genomics UK (COG-UK) Consortium | Jacquelyn Wynn, Mairead Hyland, The Lighthouse Lab in Alderley Park and Alex Alderton, Roberto Amato, Jeffrey Barrett, Sonia Goncalves, Ewan Harrison, David K. Jackson, Ian Johnston, Dominic Kwiatkowski, Cordelia Langford, John Sillitoe on behalf of the Wellcome Sanger Institute COVID-19 Surveillance Team |
| EPI_ISL_2999333, EPI_ISL_2999334, EPI_ISL_2999335, EPI_ISL_2999336 | Lighthouse Lab in Glasgow | Wellcome Sanger Institute for the COVID-19 Genomics UK (COG-UK) Consortium | Harper VanSteenhouse, Yumi Kasai, David Gray, Carol Clugston, Anna Dominiczak and Alex Alderton, Roberto Amato, Jeffrey Barrett, Sonia Goncalves, Ewan Harrison, David K. Jackson, Ian Johnston, Dominic Kwiatkowski, Cordelia Langford, John Sillitoe on behalf of the Wellcome Sanger Institute COVID-19 Surveillance Team |
| EPI_ISL_2999337 | Lighthouse Lab in Alderley Park | Wellcome Sanger Institute for the COVID-19 Genomics UK (COG-UK) Consortium | Jacquelyn Wynn, Mairead Hyland, The Lighthouse Lab in Alderley Park and Alex Alderton, Roberto Amato, Jeffrey Barrett, Sonia Goncalves, Ewan Harrison, David K. Jackson, Ian Johnston, Dominic Kwiatkowski, Cordelia Langford, John Sillitoe on behalf of the Wellcome Sanger Institute COVID-19 Surveillance Team |
| EPI_ISL_2999338, EPI_ISL_2999339, EPI_ISL_2999340, EPI_ISL_2999341 | Lighthouse Lab in Glasgow | Wellcome Sanger Institute for the COVID-19 Genomics UK (COG-UK) Consortium | Harper VanSteenhouse, Yumi Kasai, David Gray, Carol Clugston, Anna Dominiczak and Alex Alderton, Roberto Amato, Jeffrey Barrett, Sonia Goncalves, Ewan Harrison, David K. Jackson, Ian Johnston, Dominic Kwiatkowski, Cordelia Langford, John Sillitoe on behalf of the Wellcome Sanger Institute COVID-19 Surveillance Team |
| EPI_ISL_2999342 | Lighthouse Lab in Alderley Park | Wellcome Sanger Institute for the COVID-19 Genomics UK (COG-UK) Consortium | Jacquelyn Wynn, Mairead Hyland, The Lighthouse Lab in Alderley Park and Alex Alderton, Roberto Amato, Jeffrey Barrett, Sonia Goncalves, Ewan Harrison, David K. Jackson, Ian Johnston, Dominic Kwiatkowski, Cordelia Langford, John Sillitoe on behalf of the Wellcome Sanger Institute COVID-19 Surveillance Team |
| EPI_ISL_2999343, EPI_ISL_2999344, EPI_ISL_2999345, EPI_ISL_2999346, EPI_ISL_2999347, EPI_ISL_2999348, EPI_ISL_2999349 | Lighthouse Lab in Glasgow | Wellcome Sanger Institute for the COVID-19 Genomics UK (COG-UK) Consortium | Harper VanSteenhouse, Yumi Kasai, David Gray, Carol Clugston, Anna Dominiczak and Alex Alderton, Roberto Amato, Jeffrey Barrett, Sonia Goncalves, Ewan Harrison, David K. Jackson, Ian Johnston, Dominic Kwiatkowski, Cordelia Langford, John Sillitoe on behalf of the Wellcome Sanger Institute COVID-19 Surveillance Team |
| EPI_ISL_2999350 | Lighthouse Lab in Alderley Park | Wellcome Sanger Institute for the COVID-19 Genomics UK (COG-UK) Consortium | Jacquelyn Wynn, Mairead Hyland, The Lighthouse Lab in Alderley Park and Alex Alderton, Roberto Amato, Jeffrey Barrett, Sonia Goncalves, Ewan Harrison, David K. Jackson, Ian Johnston, Dominic Kwiatkowski, Cordelia Langford, John Sillitoe on behalf of the Wellcome Sanger Institute COVID-19 Surveillance Team |
| EPI_ISL_2999351, EPI_ISL_2999352, EPI_ISL_2999353, EPI_ISL_2999354 | Lighthouse Lab in Glasgow | Wellcome Sanger Institute for the COVID-19 Genomics UK (COG-UK) Consortium | Harper VanSteenhouse, Yumi Kasai, David Gray, Carol Clugston, Anna Dominiczak and Alex Alderton, Roberto Amato, Jeffrey Barrett, Sonia Goncalves, Ewan Harrison, David K. Jackson, Ian Johnston, Dominic Kwiatkowski, Cordelia Langford, John Sillitoe on behalf of the Wellcome Sanger Institute COVID-19 Surveillance Team |
| EPI_ISL_2999355 | Lighthouse Lab in Alderley Park | Wellcome Sanger Institute for the COVID-19 Genomics UK (COG-UK) Consortium | Jacquelyn Wynn, Mairead Hyland, The Lighthouse Lab in Alderley Park and Alex Alderton, Roberto Amato, Jeffrey Barrett, Sonia Goncalves, Ewan Harrison, David K. Jackson, Ian Johnston, Dominic Kwiatkowski, Cordelia Langford, John Sillitoe on behalf of the Wellcome Sanger Institute COVID-19 Surveillance Team |
| EPI_ISL_2999356 | Lighthouse Lab in Glasgow | Wellcome Sanger Institute for the COVID-19 Genomics UK (COG-UK) Consortium | Harper VanSteenhouse, Yumi Kasai, David Gray, Carol Clugston, Anna Dominiczak and Alex Alderton, Roberto Amato, Jeffrey Barrett, Sonia Goncalves, Ewan Harrison, David K. Jackson, Ian Johnston, Dominic Kwiatkowski, Cordelia Langford, John Sillitoe on behalf of the Wellcome Sanger Institute COVID-19 Surveillance Team |
| EPI_ISL_2999357 | Lighthouse Lab in Alderley Park | Wellcome Sanger Institute for the COVID-19 Genomics UK (COG-UK) Consortium | Jacquelyn Wynn, Mairead Hyland, The Lighthouse Lab in Alderley Park and Alex Alderton, Roberto Amato, Jeffrey Barrett, Sonia Goncalves, Ewan Harrison, David K. Jackson, Ian Johnston, Dominic Kwiatkowski, Cordelia Langford, John Sillitoe on behalf of the Wellcome Sanger Institute COVID-19 Surveillance Team |
| EPI_ISL_2999358, EPI_ISL_2999359, | Lighthouse Lab in Glasgow | Wellcome Sanger Institute for the COVID-19 Genomics UK (COG-UK) Consortium | Harper VanSteenhouse, Yumi Kasai, David Gray, Carol Clugston, Anna Dominiczak and Alex Alderton, Roberto Amato, Jeffrey Barrett, Sonia Goncalves, Ewan Harrison, David K. Jackson, Ian Johnston, Dominic Kwiatkowski, Cordelia Langford, John Sillitoe on behalf of the Wellcome Sanger Institute COVID-19 Surveillance Team |

| | | | |
|---|---|---|---|
| EPI_ISL_2999360 | | | |
| EPI_ISL_2999361 | Lighthouse Lab in Alderley Park | Wellcome Sanger Institute for the COVID-19 Genomics UK (COG-UK) Consortium | Jacquelyn Wynn, Mairead Hyland, The Lighthouse Lab in Alderley Park and Alex Alderton, Roberto Amato, Jeffrey Barrett, Sonia Goncalves, Ewan Harrison, David K. Jackson, Ian Johnston, Dominic Kwiatkowski, Cordelia Langford, John Sillitoe on behalf of the Wellcome Sanger Institute COVID-19 Surveillance Team |
| EPI_ISL_2999362, EPI_ISL_2999363, EPI_ISL_2999364 | Lighthouse Lab in Glasgow | Wellcome Sanger Institute for the COVID-19 Genomics UK (COG-UK) Consortium | Harper VanSteenhouse, Yumi Kasai, David Gray, Carol Clugston, Anna Dominiczak and Alex Alderton, Roberto Amato, Jeffrey Barrett, Sonia Goncalves, Ewan Harrison, David K. Jackson, Ian Johnston, Dominic Kwiatkowski, Cordelia Langford, John Sillitoe on behalf of the Wellcome Sanger Institute COVID-19 Surveillance Team |
| EPI_ISL_2999365 | Lighthouse Lab in Alderley Park | Wellcome Sanger Institute for the COVID-19 Genomics UK (COG-UK) Consortium | Jacquelyn Wynn, Mairead Hyland, The Lighthouse Lab in Alderley Park and Alex Alderton, Roberto Amato, Jeffrey Barrett, Sonia Goncalves, Ewan Harrison, David K. Jackson, Ian Johnston, Dominic Kwiatkowski, Cordelia Langford, John Sillitoe on behalf of the Wellcome Sanger Institute COVID-19 Surveillance Team |
| EPI_ISL_2999366, EPI_ISL_2999367, EPI_ISL_2999368, EPI_ISL_2999369 | Lighthouse Lab in Glasgow | Wellcome Sanger Institute for the COVID-19 Genomics UK (COG-UK) Consortium | Harper VanSteenhouse, Yumi Kasai, David Gray, Carol Clugston, Anna Dominiczak and Alex Alderton, Roberto Amato, Jeffrey Barrett, Sonia Goncalves, Ewan Harrison, David K. Jackson, Ian Johnston, Dominic Kwiatkowski, Cordelia Langford, John Sillitoe on behalf of the Wellcome Sanger Institute COVID-19 Surveillance Team |
| EPI_ISL_2999370 | Lighthouse Lab in Alderley Park | Wellcome Sanger Institute for the COVID-19 Genomics UK (COG-UK) Consortium | Jacquelyn Wynn, Mairead Hyland, The Lighthouse Lab in Alderley Park and Alex Alderton, Roberto Amato, Jeffrey Barrett, Sonia Goncalves, Ewan Harrison, David K. Jackson, Ian Johnston, Dominic Kwiatkowski, Cordelia Langford, John Sillitoe on behalf of the Wellcome Sanger Institute COVID-19 Surveillance Team |
| EPI_ISL_2999371, EPI_ISL_2999372, EPI_ISL_2999373, EPI_ISL_2999374, EPI_ISL_2999375, EPI_ISL_2999376, EPI_ISL_2999377, EPI_ISL_2999378, EPI_ISL_2999379, EPI_ISL_2999380, EPI_ISL_2999381 | | | |
| see above | Lighthouse Lab in Glasgow | Wellcome Sanger Institute for the COVID-19 Genomics UK (COG-UK) Consortium | Harper VanSteenhouse, Yumi Kasai, David Gray, Carol Clugston, Anna Dominiczak and Alex Alderton, Roberto Amato, Jeffrey Barrett, Sonia Goncalves, Ewan Harrison, David K. Jackson, Ian Johnston, Dominic Kwiatkowski, Cordelia Langford, John Sillitoe on behalf of the Wellcome Sanger Institute COVID-19 Surveillance Team |
| EPI_ISL_2999382 | Lighthouse Lab in Alderley Park | Wellcome Sanger Institute for the COVID-19 Genomics UK (COG-UK) Consortium | Jacquelyn Wynn, Mairead Hyland, The Lighthouse Lab in Alderley Park and Alex Alderton, Roberto Amato, Jeffrey Barrett, Sonia Goncalves, Ewan Harrison, David K. Jackson, Ian Johnston, Dominic Kwiatkowski, Cordelia Langford, John Sillitoe on behalf of the Wellcome Sanger Institute COVID-19 Surveillance Team |
| EPI_ISL_2999383 | Lighthouse Lab in Glasgow | Wellcome Sanger Institute for the COVID-19 Genomics UK (COG-UK) Consortium | Harper VanSteenhouse, Yumi Kasai, David Gray, Carol Clugston, Anna Dominiczak and Alex Alderton, Roberto Amato, Jeffrey Barrett, Sonia Goncalves, Ewan Harrison, David K. Jackson, Ian Johnston, Dominic Kwiatkowski, Cordelia Langford, John Sillitoe on behalf of the Wellcome Sanger Institute COVID-19 Surveillance Team |
| EPI_ISL_2999384, EPI_ISL_2999385 | Lighthouse Lab in Alderley Park | Wellcome Sanger Institute for the COVID-19 Genomics UK (COG-UK) Consortium | Jacquelyn Wynn, Mairead Hyland, The Lighthouse Lab in Alderley Park and Alex Alderton, Roberto Amato, Jeffrey Barrett, Sonia Goncalves, Ewan Harrison, David K. Jackson, Ian Johnston, Dominic Kwiatkowski, Cordelia Langford, John Sillitoe on behalf of the Wellcome Sanger Institute COVID-19 Surveillance Team |
| EPI_ISL_2999386 | Lighthouse Lab in Glasgow | Wellcome Sanger Institute for the COVID-19 Genomics UK (COG-UK) Consortium | Harper VanSteenhouse, Yumi Kasai, David Gray, Carol Clugston, Anna Dominiczak and Alex Alderton, Roberto Amato, Jeffrey Barrett, Sonia Goncalves, Ewan Harrison, David K. Jackson, Ian Johnston, Dominic Kwiatkowski, Cordelia Langford, John Sillitoe on behalf of the Wellcome Sanger Institute COVID-19 Surveillance Team |
| EPI_ISL_2999387 | Lighthouse Lab in Alderley Park | Wellcome Sanger Institute for the COVID-19 Genomics UK (COG-UK) Consortium | Jacquelyn Wynn, Mairead Hyland, The Lighthouse Lab in Alderley Park and Alex Alderton, Roberto Amato, Jeffrey Barrett, Sonia Goncalves, Ewan Harrison, David K. Jackson, Ian Johnston, Dominic Kwiatkowski, Cordelia Langford, John Sillitoe on behalf of the Wellcome Sanger Institute COVID-19 Surveillance Team |
| EPI_ISL_2999388, EPI_ISL_2999389, EPI_ISL_2999390, EPI_ISL_2999391, EPI_ISL_2999392, EPI_ISL_2999393, EPI_ISL_2999394 | Lighthouse Lab in Milton Keynes | Wellcome Sanger Institute for the COVID-19 Genomics UK (COG-UK) Consortium | The Lighthouse Lab in Milton Keynes and Alex Alderton, Roberto Amato, Jeffrey Barrett, Sonia Goncalves, Ewan Harrison, David K. Jackson, Ian Johnston, Dominic Kwiatkowski, Cordelia Langford, John Sillitoe on behalf of the Wellcome Sanger Institute COVID-19 Surveillance Team |
| EPI_ISL_2999395 | Lighthouse Lab in Alderley Park | Wellcome Sanger Institute for the COVID-19 Genomics UK (COG-UK) Consortium | Jacquelyn Wynn, Mairead Hyland, The Lighthouse Lab in Alderley Park and Alex Alderton, Roberto Amato, Jeffrey Barrett, Sonia Goncalves, Ewan Harrison, David K. Jackson, Ian Johnston, Dominic Kwiatkowski, Cordelia Langford, John Sillitoe on behalf of the Wellcome Sanger Institute COVID-19 Surveillance Team |
| EPI_ISL_2999396 | Berkshire and Surrey Pathology Services Lighthouse Laboratory | Wellcome Sanger Institute for the COVID-19 Genomics UK (COG-UK) Consortium | Berkshire and Surrey Pathology Services Lighthouse Laboratory and Alex Alderton, Roberto Amato, Jeffrey Barrett, Sonia Goncalves, Ewan Harrison, David K. Jackson, Ian Johnston, Dominic Kwiatkowski, Cordelia Langford, John Sillitoe on behalf of the Wellcome Sanger Institute COVID-19 Surveillance Team |
| EPI_ISL_2999397, EPI_ISL_2999398, EPI_ISL_2999399, EPI_ISL_2999400, EPI_ISL_2999401 | Lighthouse Lab in Milton Keynes | Wellcome Sanger Institute for the COVID-19 Genomics UK (COG-UK) Consortium | The Lighthouse Lab in Milton Keynes and Alex Alderton, Roberto Amato, Jeffrey Barrett, Sonia Goncalves, Ewan Harrison, David K. Jackson, Ian Johnston, Dominic Kwiatkowski, Cordelia Langford, John Sillitoe on behalf of the Wellcome Sanger Institute COVID-19 Surveillance Team |
| EPI_ISL_2999402 | Lighthouse Lab in Alderley Park | Wellcome Sanger Institute for the COVID-19 Genomics UK (COG-UK) Consortium | Jacquelyn Wynn, Mairead Hyland, The Lighthouse Lab in Alderley Park and Alex Alderton, Roberto Amato, Jeffrey Barrett, Sonia Goncalves, Ewan Harrison, David K. Jackson, Ian Johnston, Dominic Kwiatkowski, Cordelia Langford, John Sillitoe on behalf of the Wellcome Sanger Institute COVID-19 Surveillance Team |
| EPI_ISL_2999403 | Berkshire and Surrey Pathology Services Lighthouse Laboratory | Wellcome Sanger Institute for the COVID-19 Genomics UK (COG-UK) Consortium | Berkshire and Surrey Pathology Services Lighthouse Laboratory and Alex Alderton, Roberto Amato, Jeffrey Barrett, Sonia Goncalves, Ewan Harrison, David K. Jackson, Ian Johnston, Dominic Kwiatkowski, Cordelia Langford, John Sillitoe on behalf of the Wellcome Sanger Institute COVID-19 Surveillance Team |
| EPI_ISL_2999404 | Lighthouse Laboratory Plymouth | Wellcome Sanger Institute for the COVID-19 Genomics UK (COG-UK) Consortium | Lighthouse Laboratory Plymouth and Alex Alderton, Roberto Amato, Jeffrey Barrett, Sonia Goncalves, Ewan Harrison, David K. Jackson, Ian Johnston, Dominic Kwiatkowski, Cordelia Langford, John Sillitoe on behalf of the Wellcome Sanger Institute COVID-19 Surveillance Team |
| EPI_ISL_2999405, EPI_ISL_2999406 | Lighthouse Lab in Milton Keynes | Wellcome Sanger Institute for the COVID-19 Genomics UK (COG-UK) Consortium | The Lighthouse Lab in Milton Keynes and Alex Alderton, Roberto Amato, Jeffrey Barrett, Sonia Goncalves, Ewan Harrison, David K. Jackson, Ian Johnston, Dominic Kwiatkowski, Cordelia Langford, John Sillitoe on behalf of the Wellcome Sanger Institute COVID-19 Surveillance Team |
| EPI_ISL_2999407 | Lighthouse Lab in Alderley Park | Wellcome Sanger Institute for the COVID-19 Genomics UK (COG-UK) Consortium | Jacquelyn Wynn, Mairead Hyland, The Lighthouse Lab in Alderley Park and Alex Alderton, Roberto Amato, Jeffrey Barrett, Sonia Goncalves, Ewan Harrison, David K. Jackson, Ian Johnston, Dominic Kwiatkowski, Cordelia Langford, John Sillitoe on behalf of the Wellcome Sanger Institute COVID-19 Surveillance Team |
| EPI_ISL_2999408 | Lighthouse Lab in Milton Keynes | Wellcome Sanger Institute for the COVID-19 Genomics UK (COG-UK) Consortium | The Lighthouse Lab in Milton Keynes and Alex Alderton, Roberto Amato, Jeffrey Barrett, Sonia Goncalves, Ewan Harrison, David K. Jackson, Ian Johnston, Dominic Kwiatkowski, Cordelia Langford, John Sillitoe on behalf of the Wellcome Sanger Institute COVID-19 Surveillance Team |
| EPI_ISL_2999409, EPI_ISL_2999410, EPI_ISL_2999411, EPI_ISL_2999412, EPI_ISL_2999413 | Lighthouse Lab in Alderley Park | Wellcome Sanger Institute for the COVID-19 Genomics UK (COG-UK) Consortium | Jacquelyn Wynn, Mairead Hyland, The Lighthouse Lab in Alderley Park and Alex Alderton, Roberto Amato, Jeffrey Barrett, Sonia Goncalves, Ewan Harrison, David K. Jackson, Ian Johnston, Dominic Kwiatkowski, Cordelia Langford, John Sillitoe on behalf of the Wellcome Sanger Institute COVID-19 Surveillance Team |
| EPI_ISL_2999414 | Lighthouse Lab in Milton Keynes | Wellcome Sanger Institute for the COVID-19 Genomics UK (COG-UK) Consortium | The Lighthouse Lab in Milton Keynes and Alex Alderton, Roberto Amato, Jeffrey Barrett, Sonia Goncalves, Ewan Harrison, David K. Jackson, Ian Johnston, Dominic Kwiatkowski, Cordelia Langford, John Sillitoe on behalf of the Wellcome Sanger Institute COVID-19 Surveillance Team |
| EPI_ISL_2999415 | Lighthouse Lab in Alderley Park | Wellcome Sanger Institute for the COVID-19 Genomics UK (COG-UK) Consortium | Jacquelyn Wynn, Mairead Hyland, The Lighthouse Lab in Alderley Park and Alex Alderton, Roberto Amato, Jeffrey Barrett, Sonia Goncalves, Ewan Harrison, David K. Jackson, Ian Johnston, Dominic Kwiatkowski, Cordelia Langford, John Sillitoe on behalf of the Wellcome Sanger Institute COVID-19 Surveillance Team |
| EPI_ISL_2999416, EPI_ISL_2999417 | Lighthouse Laboratory Plymouth | Wellcome Sanger Institute for the COVID-19 Genomics UK (COG-UK) Consortium | Lighthouse Laboratory Plymouth and Alex Alderton, Roberto Amato, Jeffrey Barrett, Sonia Goncalves, Ewan Harrison, David K. Jackson, Ian Johnston, Dominic Kwiatkowski, Cordelia Langford, John Sillitoe on behalf of the Wellcome Sanger Institute COVID-19 Surveillance Team |
| EPI_ISL_2999418, EPI_ISL_2999419 | Lighthouse Lab in Milton Keynes | Wellcome Sanger Institute for the COVID-19 Genomics UK (COG-UK) Consortium | The Lighthouse Lab in Milton Keynes and Alex Alderton, Roberto Amato, Jeffrey Barrett, Sonia Goncalves, Ewan Harrison, David K. Jackson, Ian Johnston, Dominic Kwiatkowski, Cordelia Langford, John Sillitoe on behalf of the Wellcome Sanger Institute COVID-19 Surveillance Team |
| EPI_ISL_2999420, EPI_ISL_2999421 | Lighthouse Lab in Alderley Park | Wellcome Sanger Institute for the COVID-19 Genomics UK (COG-UK) Consortium | Jacquelyn Wynn, Mairead Hyland, The Lighthouse Lab in Alderley Park and Alex Alderton, Roberto Amato, Jeffrey Barrett, Sonia Goncalves, Ewan Harrison, David K. Jackson, Ian Johnston, Dominic Kwiatkowski, Cordelia Langford, John Sillitoe on behalf of the Wellcome Sanger Institute COVID-19 Surveillance Team |
| EPI_ISL_2999422 | Lighthouse Lab in Milton Keynes | Wellcome Sanger Institute for the COVID-19 Genomics UK (COG-UK) Consortium | The Lighthouse Lab in Milton Keynes and Alex Alderton, Roberto Amato, Jeffrey Barrett, Sonia Goncalves, Ewan Harrison, David K. Jackson, Ian Johnston, Dominic Kwiatkowski, Cordelia Langford, John Sillitoe on behalf of the Wellcome Sanger Institute COVID-19 Surveillance Team |
| EPI_ISL_2999423, EPI_ISL_2999424 | Lighthouse Laboratory Plymouth | Wellcome Sanger Institute for the COVID-19 Genomics UK (COG-UK) Consortium | Lighthouse Laboratory Plymouth and Alex Alderton, Roberto Amato, Jeffrey Barrett, Sonia Goncalves, Ewan Harrison, David K. Jackson, Ian Johnston, Dominic Kwiatkowski, Cordelia Langford, John Sillitoe on behalf of the Wellcome Sanger Institute COVID-19 Surveillance Team |
| EPI_ISL_2999425 | Lighthouse Lab in Milton Keynes | Wellcome Sanger Institute for the COVID-19 Genomics UK (COG-UK) Consortium | The Lighthouse Lab in Milton Keynes and Alex Alderton, Roberto Amato, Jeffrey Barrett, Sonia Goncalves, Ewan Harrison, David K. Jackson, Ian Johnston, Dominic Kwiatkowski, Cordelia Langford, John Sillitoe on behalf of the Wellcome Sanger Institute COVID-19 Surveillance Team |
| EPI_ISL_2999426 | Lighthouse Lab in Alderley Park | Wellcome Sanger Institute for the COVID-19 Genomics UK (COG-UK) Consortium | Jacquelyn Wynn, Mairead Hyland, The Lighthouse Lab in Alderley Park and Alex Alderton, Roberto Amato, Jeffrey Barrett, Sonia Goncalves, Ewan Harrison, David K. Jackson, Ian Johnston, Dominic Kwiatkowski, Cordelia Langford, John Sillitoe on behalf of the Wellcome Sanger Institute COVID-19 Surveillance Team |
| EPI_ISL_2999427, EPI_ISL_2999428, EPI_ISL_2999429 | Lighthouse Laboratory Plymouth | Wellcome Sanger Institute for the COVID-19 Genomics UK (COG-UK) Consortium | Lighthouse Laboratory Plymouth and Alex Alderton, Roberto Amato, Jeffrey Barrett, Sonia Goncalves, Ewan Harrison, David K. Jackson, Ian Johnston, Dominic Kwiatkowski, Cordelia Langford, John Sillitoe on behalf of the Wellcome Sanger Institute COVID-19 Surveillance Team |
| EPI_ISL_2999430 | Lighthouse Lab in Milton Keynes | Wellcome Sanger Institute for the COVID-19 Genomics UK (COG-UK) Consortium | The Lighthouse Lab in Milton Keynes and Alex Alderton, Roberto Amato, Jeffrey Barrett, Sonia Goncalves, Ewan Harrison, David K. Jackson, Ian Johnston, Dominic Kwiatkowski, Cordelia Langford, John Sillitoe on behalf of the Wellcome Sanger Institute COVID-19 Surveillance Team |
| EPI_ISL_2999431, EPI_ISL_2999432 | Lighthouse Lab in Alderley Park | Wellcome Sanger Institute for the COVID-19 Genomics UK (COG-UK) Consortium | Jacquelyn Wynn, Mairead Hyland, The Lighthouse Lab in Alderley Park and Alex Alderton, Roberto Amato, Jeffrey Barrett, Sonia Goncalves, Ewan Harrison, David K. Jackson, Ian Johnston, Dominic Kwiatkowski, Cordelia Langford, John Sillitoe on behalf of the Wellcome Sanger Institute COVID-19 Surveillance Team |
| EPI_ISL_2999433 | Lighthouse Laboratory Plymouth | Wellcome Sanger Institute for the COVID-19 Genomics UK (COG-UK) Consortium | Lighthouse Laboratory Plymouth and Alex Alderton, Roberto Amato, Jeffrey Barrett, Sonia Goncalves, Ewan Harrison, David K. Jackson, Ian Johnston, Dominic Kwiatkowski, Cordelia Langford, John Sillitoe on behalf of the Wellcome Sanger Institute COVID-19 Surveillance Team |
| EPI_ISL_2999434 | Lighthouse Lab in Milton Keynes | Wellcome Sanger Institute for the COVID-19 Genomics UK (COG-UK) Consortium | The Lighthouse Lab in Milton Keynes and Alex Alderton, Roberto Amato, Jeffrey Barrett, Sonia Goncalves, Ewan Harrison, David K. Jackson, Ian Johnston, Dominic Kwiatkowski, Cordelia Langford, John Sillitoe on behalf of the Wellcome Sanger Institute COVID-19 Surveillance Team |
| EPI_ISL_2999435, EPI_ISL_2999436 | Lighthouse Laboratory Plymouth | Wellcome Sanger Institute for the COVID-19 Genomics UK (COG-UK) Consortium | Lighthouse Laboratory Plymouth and Alex Alderton, Roberto Amato, Jeffrey Barrett, Sonia Goncalves, Ewan Harrison, David K. Jackson, Ian Johnston, Dominic Kwiatkowski, Cordelia Langford, John Sillitoe on behalf of the Wellcome Sanger Institute COVID-19 Surveillance Team |
| EPI_ISL_2999437 | Lighthouse Lab in Milton Keynes | Wellcome Sanger Institute for the COVID-19 Genomics UK (COG-UK) Consortium | The Lighthouse Lab in Milton Keynes and Alex Alderton, Roberto Amato, Jeffrey Barrett, Sonia Goncalves, Ewan Harrison, David K. Jackson, Ian Johnston, Dominic Kwiatkowski, Cordelia Langford, John Sillitoe on behalf of the Wellcome Sanger Institute COVID-19 Surveillance Team |

| | | | |
|---|---|---|---|
| EPI_ISL_2999438 | Lighthouse Laboratory Plymouth | Wellcome Sanger Institute for the COVID-19 Genomics UK (COG-UK) Consortium | Lighthouse Laboratory Plymouth and Alex Alderton, Roberto Amato, Jeffrey Barrett, Sonia Goncalves, Ewan Harrison, David K. Jackson, Ian Johnston, Dominic Kwiatkowski, Cordelia Langford, John Sillitoe on behalf of the Wellcome Sanger Institute COVID-19 Surveillance Team |
| EPI_ISL_2999439, EPI_ISL_2999440 | Lighthouse Lab in Milton Keynes | Wellcome Sanger Institute for the COVID-19 Genomics UK (COG-UK) Consortium | The Lighthouse Lab in Milton Keynes and Alex Alderton, Roberto Amato, Jeffrey Barrett, Sonia Goncalves, Ewan Harrison, David K. Jackson, Ian Johnston, Dominic Kwiatkowski, Cordelia Langford, John Sillitoe on behalf of the Wellcome Sanger Institute COVID-19 Surveillance Team |
| EPI_ISL_2999441, EPI_ISL_2999442, EPI_ISL_2999443 | Lighthouse Lab in Alderley Park | Wellcome Sanger Institute for the COVID-19 Genomics UK (COG-UK) Consortium | Jacquelyn Wynn, Mairead Hyland, The Lighthouse Lab in Alderley Park and Alex Alderton, Roberto Amato, Jeffrey Barrett, Sonia Goncalves, Ewan Harrison, David K. Jackson, Ian Johnston, Dominic Kwiatkowski, Cordelia Langford, John Sillitoe on behalf of the Wellcome Sanger Institute COVID-19 Surveillance Team |
| EPI_ISL_2999444, EPI_ISL_2999445 | Lighthouse Laboratory Plymouth | Wellcome Sanger Institute for the COVID-19 Genomics UK (COG-UK) Consortium | Lighthouse Laboratory Plymouth and Alex Alderton, Roberto Amato, Jeffrey Barrett, Sonia Goncalves, Ewan Harrison, David K. Jackson, Ian Johnston, Dominic Kwiatkowski, Cordelia Langford, John Sillitoe on behalf of the Wellcome Sanger Institute COVID-19 Surveillance Team |
| EPI_ISL_2999446 | Lighthouse Lab in Alderley Park | Wellcome Sanger Institute for the COVID-19 Genomics UK (COG-UK) Consortium | Jacquelyn Wynn, Mairead Hyland, The Lighthouse Lab in Alderley Park and Alex Alderton, Roberto Amato, Jeffrey Barrett, Sonia Goncalves, Ewan Harrison, David K. Jackson, Ian Johnston, Dominic Kwiatkowski, Cordelia Langford, John Sillitoe on behalf of the Wellcome Sanger Institute COVID-19 Surveillance Team |
| EPI_ISL_2999447 | Lighthouse Laboratory Plymouth | Wellcome Sanger Institute for the COVID-19 Genomics UK (COG-UK) Consortium | Lighthouse Laboratory Plymouth and Alex Alderton, Roberto Amato, Jeffrey Barrett, Sonia Goncalves, Ewan Harrison, David K. Jackson, Ian Johnston, Dominic Kwiatkowski, Cordelia Langford, John Sillitoe on behalf of the Wellcome Sanger Institute COVID-19 Surveillance Team |
| EPI_ISL_2999448 | Lighthouse Lab in Alderley Park | Wellcome Sanger Institute for the COVID-19 Genomics UK (COG-UK) Consortium | Jacquelyn Wynn, Mairead Hyland, The Lighthouse Lab in Alderley Park and Alex Alderton, Roberto Amato, Jeffrey Barrett, Sonia Goncalves, Ewan Harrison, David K. Jackson, Ian Johnston, Dominic Kwiatkowski, Cordelia Langford, John Sillitoe on behalf of the Wellcome Sanger Institute COVID-19 Surveillance Team |
| EPI_ISL_2999449, EPI_ISL_2999450 | Lighthouse Lab in Milton Keynes | Wellcome Sanger Institute for the COVID-19 Genomics UK (COG-UK) Consortium | The Lighthouse Lab in Milton Keynes and Alex Alderton, Roberto Amato, Jeffrey Barrett, Sonia Goncalves, Ewan Harrison, David K. Jackson, Ian Johnston, Dominic Kwiatkowski, Cordelia Langford, John Sillitoe on behalf of the Wellcome Sanger Institute COVID-19 Surveillance Team |
| EPI_ISL_2999451, EPI_ISL_2999452 | Lighthouse Lab in Alderley Park | Wellcome Sanger Institute for the COVID-19 Genomics UK (COG-UK) Consortium | Jacquelyn Wynn, Mairead Hyland, The Lighthouse Lab in Alderley Park and Alex Alderton, Roberto Amato, Jeffrey Barrett, Sonia Goncalves, Ewan Harrison, David K. Jackson, Ian Johnston, Dominic Kwiatkowski, Cordelia Langford, John Sillitoe on behalf of the Wellcome Sanger Institute COVID-19 Surveillance Team |
| EPI_ISL_2999453 | Lighthouse Laboratory Plymouth | Wellcome Sanger Institute for the COVID-19 Genomics UK (COG-UK) Consortium | Lighthouse Laboratory Plymouth and Alex Alderton, Roberto Amato, Jeffrey Barrett, Sonia Goncalves, Ewan Harrison, David K. Jackson, Ian Johnston, Dominic Kwiatkowski, Cordelia Langford, John Sillitoe on behalf of the Wellcome Sanger Institute COVID-19 Surveillance Team |
| EPI_ISL_2999454, EPI_ISL_2999455, EPI_ISL_2999456, EPI_ISL_2999457, EPI_ISL_2999458 | Lighthouse Lab in Alderley Park | Wellcome Sanger Institute for the COVID-19 Genomics UK (COG-UK) Consortium | Jacquelyn Wynn, Mairead Hyland, The Lighthouse Lab in Alderley Park and Alex Alderton, Roberto Amato, Jeffrey Barrett, Sonia Goncalves, Ewan Harrison, David K. Jackson, Ian Johnston, Dominic Kwiatkowski, Cordelia Langford, John Sillitoe on behalf of the Wellcome Sanger Institute COVID-19 Surveillance Team |
| EPI_ISL_2999459, EPI_ISL_2999460 | Lighthouse Lab in Milton Keynes | Wellcome Sanger Institute for the COVID-19 Genomics UK (COG-UK) Consortium | The Lighthouse Lab in Milton Keynes and Alex Alderton, Roberto Amato, Jeffrey Barrett, Sonia Goncalves, Ewan Harrison, David K. Jackson, Ian Johnston, Dominic Kwiatkowski, Cordelia Langford, John Sillitoe on behalf of the Wellcome Sanger Institute COVID-19 Surveillance Team |
| EPI_ISL_2999461 | Lighthouse Lab in Alderley Park | Wellcome Sanger Institute for the COVID-19 Genomics UK (COG-UK) Consortium | Jacquelyn Wynn, Mairead Hyland, The Lighthouse Lab in Alderley Park and Alex Alderton, Roberto Amato, Jeffrey Barrett, Sonia Goncalves, Ewan Harrison, David K. Jackson, Ian Johnston, Dominic Kwiatkowski, Cordelia Langford, John Sillitoe on behalf of the Wellcome Sanger Institute COVID-19 Surveillance Team |
| EPI_ISL_2999462 | Lighthouse Laboratory Plymouth | Wellcome Sanger Institute for the COVID-19 Genomics UK (COG-UK) Consortium | Lighthouse Laboratory Plymouth and Alex Alderton, Roberto Amato, Jeffrey Barrett, Sonia Goncalves, Ewan Harrison, David K. Jackson, Ian Johnston, Dominic Kwiatkowski, Cordelia Langford, John Sillitoe on behalf of the Wellcome Sanger Institute COVID-19 Surveillance Team |
| EPI_ISL_2999463, EPI_ISL_2999464 | Lighthouse Lab in Alderley Park | Wellcome Sanger Institute for the COVID-19 Genomics UK (COG-UK) Consortium | Jacquelyn Wynn, Mairead Hyland, The Lighthouse Lab in Alderley Park and Alex Alderton, Roberto Amato, Jeffrey Barrett, Sonia Goncalves, Ewan Harrison, David K. Jackson, Ian Johnston, Dominic Kwiatkowski, Cordelia Langford, John Sillitoe on behalf of the Wellcome Sanger Institute COVID-19 Surveillance Team |
| EPI_ISL_2999465 | Lighthouse Lab in Milton Keynes | Wellcome Sanger Institute for the COVID-19 Genomics UK (COG-UK) Consortium | The Lighthouse Lab in Milton Keynes and Alex Alderton, Roberto Amato, Jeffrey Barrett, Sonia Goncalves, Ewan Harrison, David K. Jackson, Ian Johnston, Dominic Kwiatkowski, Cordelia Langford, John Sillitoe on behalf of the Wellcome Sanger Institute COVID-19 Surveillance Team |
| EPI_ISL_2999466, EPI_ISL_2999467 | Lighthouse Lab in Alderley Park | Wellcome Sanger Institute for the COVID-19 Genomics UK (COG-UK) Consortium | Jacquelyn Wynn, Mairead Hyland, The Lighthouse Lab in Alderley Park and Alex Alderton, Roberto Amato, Jeffrey Barrett, Sonia Goncalves, Ewan Harrison, David K. Jackson, Ian Johnston, Dominic Kwiatkowski, Cordelia Langford, John Sillitoe on behalf of the Wellcome Sanger Institute COVID-19 Surveillance Team |
| EPI_ISL_2999468 | Lighthouse Lab in Milton Keynes | Wellcome Sanger Institute for the COVID-19 Genomics UK (COG-UK) Consortium | The Lighthouse Lab in Milton Keynes and Alex Alderton, Roberto Amato, Jeffrey Barrett, Sonia Goncalves, Ewan Harrison, David K. Jackson, Ian Johnston, Dominic Kwiatkowski, Cordelia Langford, John Sillitoe on behalf of the Wellcome Sanger Institute COVID-19 Surveillance Team |
| EPI_ISL_2999469 | Lighthouse Laboratory Plymouth | Wellcome Sanger Institute for the COVID-19 Genomics UK (COG-UK) Consortium | Lighthouse Laboratory Plymouth and Alex Alderton, Roberto Amato, Jeffrey Barrett, Sonia Goncalves, Ewan Harrison, David K. Jackson, Ian Johnston, Dominic Kwiatkowski, Cordelia Langford, John Sillitoe on behalf of the Wellcome Sanger Institute COVID-19 Surveillance Team |
| EPI_ISL_2999470 | Lighthouse Lab in Milton Keynes | Wellcome Sanger Institute for the COVID-19 Genomics UK (COG-UK) Consortium | The Lighthouse Lab in Milton Keynes and Alex Alderton, Roberto Amato, Jeffrey Barrett, Sonia Goncalves, Ewan Harrison, David K. Jackson, Ian Johnston, Dominic Kwiatkowski, Cordelia Langford, John Sillitoe on behalf of the Wellcome Sanger Institute COVID-19 Surveillance Team |
| EPI_ISL_2999471, EPI_ISL_2999472 | Lighthouse Laboratory Plymouth | Wellcome Sanger Institute for the COVID-19 Genomics UK (COG-UK) Consortium | Lighthouse Laboratory Plymouth and Alex Alderton, Roberto Amato, Jeffrey Barrett, Sonia Goncalves, Ewan Harrison, David K. Jackson, Ian Johnston, Dominic Kwiatkowski, Cordelia Langford, John Sillitoe on behalf of the Wellcome Sanger Institute COVID-19 Surveillance Team |
| EPI_ISL_2999473 | Lighthouse Lab in Milton Keynes | Wellcome Sanger Institute for the COVID-19 Genomics UK (COG-UK) Consortium | The Lighthouse Lab in Milton Keynes and Alex Alderton, Roberto Amato, Jeffrey Barrett, Sonia Goncalves, Ewan Harrison, David K. Jackson, Ian Johnston, Dominic Kwiatkowski, Cordelia Langford, John Sillitoe on behalf of the Wellcome Sanger Institute COVID-19 Surveillance Team |
| EPI_ISL_2999474, EPI_ISL_2999475 | Lighthouse Laboratory Plymouth | Wellcome Sanger Institute for the COVID-19 Genomics UK (COG-UK) Consortium | Lighthouse Laboratory Plymouth and Alex Alderton, Roberto Amato, Jeffrey Barrett, Sonia Goncalves, Ewan Harrison, David K. Jackson, Ian Johnston, Dominic Kwiatkowski, Cordelia Langford, John Sillitoe on behalf of the Wellcome Sanger Institute COVID-19 Surveillance Team |
| EPI_ISL_2999476 | Lighthouse Lab in Milton Keynes | Wellcome Sanger Institute for the COVID-19 Genomics UK (COG-UK) Consortium | The Lighthouse Lab in Milton Keynes and Alex Alderton, Roberto Amato, Jeffrey Barrett, Sonia Goncalves, Ewan Harrison, David K. Jackson, Ian Johnston, Dominic Kwiatkowski, Cordelia Langford, John Sillitoe on behalf of the Wellcome Sanger Institute COVID-19 Surveillance Team |
| EPI_ISL_2999477 | Lighthouse Lab in Alderley Park | Wellcome Sanger Institute for the COVID-19 Genomics UK (COG-UK) Consortium | Jacquelyn Wynn, Mairead Hyland, The Lighthouse Lab in Alderley Park and Alex Alderton, Roberto Amato, Jeffrey Barrett, Sonia Goncalves, Ewan Harrison, David K. Jackson, Ian Johnston, Dominic Kwiatkowski, Cordelia Langford, John Sillitoe on behalf of the Wellcome Sanger Institute COVID-19 Surveillance Team |
| EPI_ISL_2999478 | Lighthouse Laboratory Plymouth | Wellcome Sanger Institute for the COVID-19 Genomics UK (COG-UK) Consortium | Lighthouse Laboratory Plymouth and Alex Alderton, Roberto Amato, Jeffrey Barrett, Sonia Goncalves, Ewan Harrison, David K. Jackson, Ian Johnston, Dominic Kwiatkowski, Cordelia Langford, John Sillitoe on behalf of the Wellcome Sanger Institute COVID-19 Surveillance Team |
| EPI_ISL_2999479 | Lighthouse Lab in Milton Keynes | Wellcome Sanger Institute for the COVID-19 Genomics UK (COG-UK) Consortium | The Lighthouse Lab in Milton Keynes and Alex Alderton, Roberto Amato, Jeffrey Barrett, Sonia Goncalves, Ewan Harrison, David K. Jackson, Ian Johnston, Dominic Kwiatkowski, Cordelia Langford, John Sillitoe on behalf of the Wellcome Sanger Institute COVID-19 Surveillance Team |
| EPI_ISL_2999480, EPI_ISL_2999481, EPI_ISL_2999482 | Lighthouse Laboratory Plymouth | Wellcome Sanger Institute for the COVID-19 Genomics UK (COG-UK) Consortium | Lighthouse Laboratory Plymouth and Alex Alderton, Roberto Amato, Jeffrey Barrett, Sonia Goncalves, Ewan Harrison, David K. Jackson, Ian Johnston, Dominic Kwiatkowski, Cordelia Langford, John Sillitoe on behalf of the Wellcome Sanger Institute COVID-19 Surveillance Team |
| EPI_ISL_2999483, EPI_ISL_2999484 | Lighthouse Lab in Milton Keynes | Wellcome Sanger Institute for the COVID-19 Genomics UK (COG-UK) Consortium | The Lighthouse Lab in Milton Keynes and Alex Alderton, Roberto Amato, Jeffrey Barrett, Sonia Goncalves, Ewan Harrison, David K. Jackson, Ian Johnston, Dominic Kwiatkowski, Cordelia Langford, John Sillitoe on behalf of the Wellcome Sanger Institute COVID-19 Surveillance Team |
| EPI_ISL_2999485 | Lighthouse Lab in Alderley Park | Wellcome Sanger Institute for the COVID-19 Genomics UK (COG-UK) Consortium | Jacquelyn Wynn, Mairead Hyland, The Lighthouse Lab in Alderley Park and Alex Alderton, Roberto Amato, Jeffrey Barrett, Sonia Goncalves, Ewan Harrison, David K. Jackson, Ian Johnston, Dominic Kwiatkowski, Cordelia Langford, John Sillitoe on behalf of the Wellcome Sanger Institute COVID-19 Surveillance Team |
| EPI_ISL_2999486 | Lighthouse Laboratory Plymouth | Wellcome Sanger Institute for the COVID-19 Genomics UK (COG-UK) Consortium | Lighthouse Laboratory Plymouth and Alex Alderton, Roberto Amato, Jeffrey Barrett, Sonia Goncalves, Ewan Harrison, David K. Jackson, Ian Johnston, Dominic Kwiatkowski, Cordelia Langford, John Sillitoe on behalf of the Wellcome Sanger Institute COVID-19 Surveillance Team |
| EPI_ISL_2999487 | Lighthouse Lab in Milton Keynes | Wellcome Sanger Institute for the COVID-19 Genomics UK (COG-UK) Consortium | The Lighthouse Lab in Milton Keynes and Alex Alderton, Roberto Amato, Jeffrey Barrett, Sonia Goncalves, Ewan Harrison, David K. Jackson, Ian Johnston, Dominic Kwiatkowski, Cordelia Langford, John Sillitoe on behalf of the Wellcome Sanger Institute COVID-19 Surveillance Team |
| EPI_ISL_2999488 | Lighthouse Lab in Alderley Park | Wellcome Sanger Institute for the COVID-19 Genomics UK (COG-UK) Consortium | Jacquelyn Wynn, Mairead Hyland, The Lighthouse Lab in Alderley Park and Alex Alderton, Roberto Amato, Jeffrey Barrett, Sonia Goncalves, Ewan Harrison, David K. Jackson, Ian Johnston, Dominic Kwiatkowski, Cordelia Langford, John Sillitoe on behalf of the Wellcome Sanger Institute COVID-19 Surveillance Team |
| EPI_ISL_2999489 | Lighthouse Lab in Milton Keynes | Wellcome Sanger Institute for the COVID-19 Genomics UK (COG-UK) Consortium | The Lighthouse Lab in Milton Keynes and Alex Alderton, Roberto Amato, Jeffrey Barrett, Sonia Goncalves, Ewan Harrison, David K. Jackson, Ian Johnston, Dominic Kwiatkowski, Cordelia Langford, John Sillitoe on behalf of the Wellcome Sanger Institute COVID-19 Surveillance Team |
| EPI_ISL_2999490, EPI_ISL_2999491, EPI_ISL_2999492 | Lighthouse Laboratory Plymouth | Wellcome Sanger Institute for the COVID-19 Genomics UK (COG-UK) Consortium | Lighthouse Laboratory Plymouth and Alex Alderton, Roberto Amato, Jeffrey Barrett, Sonia Goncalves, Ewan Harrison, David K. Jackson, Ian Johnston, Dominic Kwiatkowski, Cordelia Langford, John Sillitoe on behalf of the Wellcome Sanger Institute COVID-19 Surveillance Team |
| EPI_ISL_2999493 | Lighthouse Lab in Alderley Park | Wellcome Sanger Institute for the COVID-19 Genomics UK (COG-UK) Consortium | Jacquelyn Wynn, Mairead Hyland, The Lighthouse Lab in Alderley Park and Alex Alderton, Roberto Amato, Jeffrey Barrett, Sonia Goncalves, Ewan Harrison, David K. Jackson, Ian Johnston, Dominic Kwiatkowski, Cordelia Langford, John Sillitoe on behalf of the Wellcome Sanger Institute COVID-19 Surveillance Team |
| EPI_ISL_2999494 | Lighthouse Laboratory Plymouth | Wellcome Sanger Institute for the COVID-19 Genomics UK (COG-UK) Consortium | Lighthouse Laboratory Plymouth and Alex Alderton, Roberto Amato, Jeffrey Barrett, Sonia Goncalves, Ewan Harrison, David K. Jackson, Ian Johnston, Dominic Kwiatkowski, Cordelia Langford, John Sillitoe on behalf of the Wellcome Sanger Institute COVID-19 Surveillance Team |
| EPI_ISL_2999495, EPI_ISL_2999496 | Lighthouse Lab in Alderley Park | Wellcome Sanger Institute for the COVID-19 Genomics UK (COG-UK) Consortium | Jacquelyn Wynn, Mairead Hyland, The Lighthouse Lab in Alderley Park and Alex Alderton, Roberto Amato, Jeffrey Barrett, Sonia Goncalves, Ewan Harrison, David K. Jackson, Ian Johnston, Dominic Kwiatkowski, Cordelia Langford, John Sillitoe on behalf of the Wellcome Sanger Institute COVID-19 Surveillance Team |
| EPI_ISL_2999497 | Lighthouse Lab in Milton Keynes | Wellcome Sanger Institute for the COVID-19 Genomics UK (COG-UK) Consortium | The Lighthouse Lab in Milton Keynes and Alex Alderton, Roberto Amato, Jeffrey Barrett, Sonia Goncalves, Ewan Harrison, David K. Jackson, Ian Johnston, Dominic Kwiatkowski, Cordelia Langford, John Sillitoe on behalf of the Wellcome Sanger Institute COVID-19 Surveillance Team |
| EPI_ISL_2999498 | Lighthouse Laboratory Plymouth | Wellcome Sanger Institute for the COVID-19 Genomics UK (COG-UK) Consortium | Lighthouse Laboratory Plymouth and Alex Alderton, Roberto Amato, Jeffrey Barrett, Sonia Goncalves, Ewan Harrison, David K. Jackson, Ian Johnston, Dominic Kwiatkowski, Cordelia Langford, John Sillitoe on behalf of the Wellcome Sanger Institute COVID-19 Surveillance Team |
| EPI_ISL_2999499 | Lighthouse Lab in Alderley Park | Wellcome Sanger Institute for the COVID-19 Genomics UK (COG-UK) Consortium | Jacquelyn Wynn, Mairead Hyland, The Lighthouse Lab in Alderley Park and Alex Alderton, Roberto Amato, Jeffrey Barrett, Sonia Goncalves, Ewan Harrison, David K. Jackson, Ian Johnston, Dominic Kwiatkowski, Cordelia Langford, John Sillitoe on behalf of the Wellcome Sanger Institute COVID-19 Surveillance Team |
| EPI_ISL_2999500, EPI_ISL_2999501 | Lighthouse Laboratory Plymouth | Wellcome Sanger Institute for the COVID-19 Genomics UK (COG-UK) Consortium | Lighthouse Laboratory Plymouth and Alex Alderton, Roberto Amato, Jeffrey Barrett, Sonia Goncalves, Ewan Harrison, David K. Jackson, Ian Johnston, Dominic Kwiatkowski, Cordelia Langford, John Sillitoe on behalf of the Wellcome Sanger Institute COVID-19 Surveillance Team |
| EPI_ISL_2999502, EPI_ISL_2999503 | Lighthouse Lab in Alderley Park | Wellcome Sanger Institute for the COVID-19 Genomics UK (COG-UK) Consortium | Jacquelyn Wynn, Mairead Hyland, The Lighthouse Lab in Alderley Park and Alex Alderton, Roberto Amato, Jeffrey Barrett, Sonia Goncalves, Ewan Harrison, David K. Jackson, Ian Johnston, Dominic Kwiatkowski, Cordelia Langford, John Sillitoe on behalf of the Wellcome Sanger Institute COVID-19 Surveillance Team |

| | | | |
|---|---|---|---|
| EPI_ISL_2999504 | Lighthouse Lab in Milton Keynes | Wellcome Sanger Institute for the COVID-19 Genomics UK (COG-UK) Consortium | The Lighthouse Lab in Milton Keynes and Alex Alderton, Roberto Amato, Jeffrey Barrett, Sonia Goncalves, Ewan Harrison, David K. Jackson, Ian Johnston, Dominic Kwiatkowski, Cordelia Langford, John Sillitoe on behalf of the Wellcome Sanger Institute COVID-19 Surveillance Team |
| EPI_ISL_2999505, EPI_ISL_2999506 | Lighthouse Laboratory Plymouth | Wellcome Sanger Institute for the COVID-19 Genomics UK (COG-UK) Consortium | Lighthouse Laboratory Plymouth and Alex Alderton, Roberto Amato, Jeffrey Barrett, Sonia Goncalves, Ewan Harrison, David K. Jackson, Ian Johnston, Dominic Kwiatkowski, Cordelia Langford, John Sillitoe on behalf of the Wellcome Sanger Institute COVID-19 Surveillance Team |
| EPI_ISL_2999507 | Lighthouse Lab in Alderley Park | Wellcome Sanger Institute for the COVID-19 Genomics UK (COG-UK) Consortium | Jacquelyn Wynn, Mairead Hyland, The Lighthouse Lab in Alderley Park and Alex Alderton, Roberto Amato, Jeffrey Barrett, Sonia Goncalves, Ewan Harrison, David K. Jackson, Ian Johnston, Dominic Kwiatkowski, Cordelia Langford, John Sillitoe on behalf of the Wellcome Sanger Institute COVID-19 Surveillance Team |
| EPI_ISL_2999508 | Lighthouse Laboratory Plymouth | Wellcome Sanger Institute for the COVID-19 Genomics UK (COG-UK) Consortium | Lighthouse Laboratory Plymouth and Alex Alderton, Roberto Amato, Jeffrey Barrett, Sonia Goncalves, Ewan Harrison, David K. Jackson, Ian Johnston, Dominic Kwiatkowski, Cordelia Langford, John Sillitoe on behalf of the Wellcome Sanger Institute COVID-19 Surveillance Team |
| EPI_ISL_2999509 | Lighthouse Lab in Milton Keynes | Wellcome Sanger Institute for the COVID-19 Genomics UK (COG-UK) Consortium | The Lighthouse Lab in Milton Keynes and Alex Alderton, Roberto Amato, Jeffrey Barrett, Sonia Goncalves, Ewan Harrison, David K. Jackson, Ian Johnston, Dominic Kwiatkowski, Cordelia Langford, John Sillitoe on behalf of the Wellcome Sanger Institute COVID-19 Surveillance Team |
| EPI_ISL_2999510 | Lighthouse Lab in Alderley Park | Wellcome Sanger Institute for the COVID-19 Genomics UK (COG-UK) Consortium | Jacquelyn Wynn, Mairead Hyland, The Lighthouse Lab in Alderley Park and Alex Alderton, Roberto Amato, Jeffrey Barrett, Sonia Goncalves, Ewan Harrison, David K. Jackson, Ian Johnston, Dominic Kwiatkowski, Cordelia Langford, John Sillitoe on behalf of the Wellcome Sanger Institute COVID-19 Surveillance Team |
| EPI_ISL_2999511 | Lighthouse Laboratory Plymouth | Wellcome Sanger Institute for the COVID-19 Genomics UK (COG-UK) Consortium | Lighthouse Laboratory Plymouth and Alex Alderton, Roberto Amato, Jeffrey Barrett, Sonia Goncalves, Ewan Harrison, David K. Jackson, Ian Johnston, Dominic Kwiatkowski, Cordelia Langford, John Sillitoe on behalf of the Wellcome Sanger Institute COVID-19 Surveillance Team |
| EPI_ISL_2999512 | Lighthouse Lab in Alderley Park | Wellcome Sanger Institute for the COVID-19 Genomics UK (COG-UK) Consortium | Jacquelyn Wynn, Mairead Hyland, The Lighthouse Lab in Alderley Park and Alex Alderton, Roberto Amato, Jeffrey Barrett, Sonia Goncalves, Ewan Harrison, David K. Jackson, Ian Johnston, Dominic Kwiatkowski, Cordelia Langford, John Sillitoe on behalf of the Wellcome Sanger Institute COVID-19 Surveillance Team |
| EPI_ISL_2999513 | Lighthouse Lab in Milton Keynes | Wellcome Sanger Institute for the COVID-19 Genomics UK (COG-UK) Consortium | The Lighthouse Lab in Milton Keynes and Alex Alderton, Roberto Amato, Jeffrey Barrett, Sonia Goncalves, Ewan Harrison, David K. Jackson, Ian Johnston, Dominic Kwiatkowski, Cordelia Langford, John Sillitoe on behalf of the Wellcome Sanger Institute COVID-19 Surveillance Team |
| EPI_ISL_2999514 | Lighthouse Lab in Alderley Park | Wellcome Sanger Institute for the COVID-19 Genomics UK (COG-UK) Consortium | Jacquelyn Wynn, Mairead Hyland, The Lighthouse Lab in Alderley Park and Alex Alderton, Roberto Amato, Jeffrey Barrett, Sonia Goncalves, Ewan Harrison, David K. Jackson, Ian Johnston, Dominic Kwiatkowski, Cordelia Langford, John Sillitoe on behalf of the Wellcome Sanger Institute COVID-19 Surveillance Team |
| EPI_ISL_2999515 | Lighthouse Laboratory Plymouth | Wellcome Sanger Institute for the COVID-19 Genomics UK (COG-UK) Consortium | Lighthouse Laboratory Plymouth and Alex Alderton, Roberto Amato, Jeffrey Barrett, Sonia Goncalves, Ewan Harrison, David K. Jackson, Ian Johnston, Dominic Kwiatkowski, Cordelia Langford, John Sillitoe on behalf of the Wellcome Sanger Institute COVID-19 Surveillance Team |
| EPI_ISL_2999516 | Lighthouse Lab in Alderley Park | Wellcome Sanger Institute for the COVID-19 Genomics UK (COG-UK) Consortium | Jacquelyn Wynn, Mairead Hyland, The Lighthouse Lab in Alderley Park and Alex Alderton, Roberto Amato, Jeffrey Barrett, Sonia Goncalves, Ewan Harrison, David K. Jackson, Ian Johnston, Dominic Kwiatkowski, Cordelia Langford, John Sillitoe on behalf of the Wellcome Sanger Institute COVID-19 Surveillance Team |
| EPI_ISL_2999517, EPI_ISL_2999518 | Lighthouse Lab in Milton Keynes | Wellcome Sanger Institute for the COVID-19 Genomics UK (COG-UK) Consortium | The Lighthouse Lab in Milton Keynes and Alex Alderton, Roberto Amato, Jeffrey Barrett, Sonia Goncalves, Ewan Harrison, David K. Jackson, Ian Johnston, Dominic Kwiatkowski, Cordelia Langford, John Sillitoe on behalf of the Wellcome Sanger Institute COVID-19 Surveillance Team |
| EPI_ISL_2999519 | Lighthouse Lab in Alderley Park | Wellcome Sanger Institute for the COVID-19 Genomics UK (COG-UK) Consortium | Jacquelyn Wynn, Mairead Hyland, The Lighthouse Lab in Alderley Park and Alex Alderton, Roberto Amato, Jeffrey Barrett, Sonia Goncalves, Ewan Harrison, David K. Jackson, Ian Johnston, Dominic Kwiatkowski, Cordelia Langford, John Sillitoe on behalf of the Wellcome Sanger Institute COVID-19 Surveillance Team |
| EPI_ISL_2999520, EPI_ISL_2999521, EPI_ISL_2999522 | Lighthouse Lab in Milton Keynes | Wellcome Sanger Institute for the COVID-19 Genomics UK (COG-UK) Consortium | The Lighthouse Lab in Milton Keynes and Alex Alderton, Roberto Amato, Jeffrey Barrett, Sonia Goncalves, Ewan Harrison, David K. Jackson, Ian Johnston, Dominic Kwiatkowski, Cordelia Langford, John Sillitoe on behalf of the Wellcome Sanger Institute COVID-19 Surveillance Team |
| EPI_ISL_2999523 | Lighthouse Lab in Alderley Park | Wellcome Sanger Institute for the COVID-19 Genomics UK (COG-UK) Consortium | Jacquelyn Wynn, Mairead Hyland, The Lighthouse Lab in Alderley Park and Alex Alderton, Roberto Amato, Jeffrey Barrett, Sonia Goncalves, Ewan Harrison, David K. Jackson, Ian Johnston, Dominic Kwiatkowski, Cordelia Langford, John Sillitoe on behalf of the Wellcome Sanger Institute COVID-19 Surveillance Team |
| EPI_ISL_2999524 | Lighthouse Laboratory Plymouth | Wellcome Sanger Institute for the COVID-19 Genomics UK (COG-UK) Consortium | Lighthouse Laboratory Plymouth and Alex Alderton, Roberto Amato, Jeffrey Barrett, Sonia Goncalves, Ewan Harrison, David K. Jackson, Ian Johnston, Dominic Kwiatkowski, Cordelia Langford, John Sillitoe on behalf of the Wellcome Sanger Institute COVID-19 Surveillance Team |
| EPI_ISL_2999525, EPI_ISL_2999526 | Lighthouse Lab in Milton Keynes | Wellcome Sanger Institute for the COVID-19 Genomics UK (COG-UK) Consortium | The Lighthouse Lab in Milton Keynes and Alex Alderton, Roberto Amato, Jeffrey Barrett, Sonia Goncalves, Ewan Harrison, David K. Jackson, Ian Johnston, Dominic Kwiatkowski, Cordelia Langford, John Sillitoe on behalf of the Wellcome Sanger Institute COVID-19 Surveillance Team |
| EPI_ISL_2999527, EPI_ISL_2999528, EPI_ISL_2999529 | Lighthouse Lab in Alderley Park | Wellcome Sanger Institute for the COVID-19 Genomics UK (COG-UK) Consortium | Jacquelyn Wynn, Mairead Hyland, The Lighthouse Lab in Alderley Park and Alex Alderton, Roberto Amato, Jeffrey Barrett, Sonia Goncalves, Ewan Harrison, David K. Jackson, Ian Johnston, Dominic Kwiatkowski, Cordelia Langford, John Sillitoe on behalf of the Wellcome Sanger Institute COVID-19 Surveillance Team |
| EPI_ISL_2999530, EPI_ISL_2999531 | Lighthouse Lab in Milton Keynes | Wellcome Sanger Institute for the COVID-19 Genomics UK (COG-UK) Consortium | The Lighthouse Lab in Milton Keynes and Alex Alderton, Roberto Amato, Jeffrey Barrett, Sonia Goncalves, Ewan Harrison, David K. Jackson, Ian Johnston, Dominic Kwiatkowski, Cordelia Langford, John Sillitoe on behalf of the Wellcome Sanger Institute COVID-19 Surveillance Team |
| EPI_ISL_2999532 | Lighthouse Laboratory Plymouth | Wellcome Sanger Institute for the COVID-19 Genomics UK (COG-UK) Consortium | Lighthouse Laboratory Plymouth and Alex Alderton, Roberto Amato, Jeffrey Barrett, Sonia Goncalves, Ewan Harrison, David K. Jackson, Ian Johnston, Dominic Kwiatkowski, Cordelia Langford, John Sillitoe on behalf of the Wellcome Sanger Institute COVID-19 Surveillance Team |
| EPI_ISL_2999533, EPI_ISL_2999534 | Lighthouse Lab in Alderley Park | Wellcome Sanger Institute for the COVID-19 Genomics UK (COG-UK) Consortium | Jacquelyn Wynn, Mairead Hyland, The Lighthouse Lab in Alderley Park and Alex Alderton, Roberto Amato, Jeffrey Barrett, Sonia Goncalves, Ewan Harrison, David K. Jackson, Ian Johnston, Dominic Kwiatkowski, Cordelia Langford, John Sillitoe on behalf of the Wellcome Sanger Institute COVID-19 Surveillance Team |
| EPI_ISL_2999535 | Lighthouse Lab in Milton Keynes | Wellcome Sanger Institute for the COVID-19 Genomics UK (COG-UK) Consortium | The Lighthouse Lab in Milton Keynes and Alex Alderton, Roberto Amato, Jeffrey Barrett, Sonia Goncalves, Ewan Harrison, David K. Jackson, Ian Johnston, Dominic Kwiatkowski, Cordelia Langford, John Sillitoe on behalf of the Wellcome Sanger Institute COVID-19 Surveillance Team |
| EPI_ISL_2999536, EPI_ISL_2999537, EPI_ISL_2999538 | Lighthouse Laboratory Plymouth | Wellcome Sanger Institute for the COVID-19 Genomics UK (COG-UK) Consortium | Lighthouse Laboratory Plymouth and Alex Alderton, Roberto Amato, Jeffrey Barrett, Sonia Goncalves, Ewan Harrison, David K. Jackson, Ian Johnston, Dominic Kwiatkowski, Cordelia Langford, John Sillitoe on behalf of the Wellcome Sanger Institute COVID-19 Surveillance Team |
| EPI_ISL_2999539 | Lighthouse Lab in Alderley Park | Wellcome Sanger Institute for the COVID-19 Genomics UK (COG-UK) Consortium | Jacquelyn Wynn, Mairead Hyland, The Lighthouse Lab in Alderley Park and Alex Alderton, Roberto Amato, Jeffrey Barrett, Sonia Goncalves, Ewan Harrison, David K. Jackson, Ian Johnston, Dominic Kwiatkowski, Cordelia Langford, John Sillitoe on behalf of the Wellcome Sanger Institute COVID-19 Surveillance Team |
| EPI_ISL_2999540 | Lighthouse Lab in Milton Keynes | Wellcome Sanger Institute for the COVID-19 Genomics UK (COG-UK) Consortium | The Lighthouse Lab in Milton Keynes and Alex Alderton, Roberto Amato, Jeffrey Barrett, Sonia Goncalves, Ewan Harrison, David K. Jackson, Ian Johnston, Dominic Kwiatkowski, Cordelia Langford, John Sillitoe on behalf of the Wellcome Sanger Institute COVID-19 Surveillance Team |
| EPI_ISL_2999541, EPI_ISL_2999542, EPI_ISL_2999543, EPI_ISL_2999544 | Lighthouse Laboratory Plymouth | Wellcome Sanger Institute for the COVID-19 Genomics UK (COG-UK) Consortium | Lighthouse Laboratory Plymouth and Alex Alderton, Roberto Amato, Jeffrey Barrett, Sonia Goncalves, Ewan Harrison, David K. Jackson, Ian Johnston, Dominic Kwiatkowski, Cordelia Langford, John Sillitoe on behalf of the Wellcome Sanger Institute COVID-19 Surveillance Team |
| EPI_ISL_2999545, EPI_ISL_2999546 | Lighthouse Lab in Alderley Park | Wellcome Sanger Institute for the COVID-19 Genomics UK (COG-UK) Consortium | Jacquelyn Wynn, Mairead Hyland, The Lighthouse Lab in Alderley Park and Alex Alderton, Roberto Amato, Jeffrey Barrett, Sonia Goncalves, Ewan Harrison, David K. Jackson, Ian Johnston, Dominic Kwiatkowski, Cordelia Langford, John Sillitoe on behalf of the Wellcome Sanger Institute COVID-19 Surveillance Team |
| EPI_ISL_2999547 | Lighthouse Laboratory Plymouth | Wellcome Sanger Institute for the COVID-19 Genomics UK (COG-UK) Consortium | Lighthouse Laboratory Plymouth and Alex Alderton, Roberto Amato, Jeffrey Barrett, Sonia Goncalves, Ewan Harrison, David K. Jackson, Ian Johnston, Dominic Kwiatkowski, Cordelia Langford, John Sillitoe on behalf of the Wellcome Sanger Institute COVID-19 Surveillance Team |
| EPI_ISL_2999548 | Lighthouse Lab in Milton Keynes | Wellcome Sanger Institute for the COVID-19 Genomics UK (COG-UK) Consortium | The Lighthouse Lab in Milton Keynes and Alex Alderton, Roberto Amato, Jeffrey Barrett, Sonia Goncalves, Ewan Harrison, David K. Jackson, Ian Johnston, Dominic Kwiatkowski, Cordelia Langford, John Sillitoe on behalf of the Wellcome Sanger Institute COVID-19 Surveillance Team |
| EPI_ISL_2999549, EPI_ISL_2999550, EPI_ISL_2999551, EPI_ISL_2999552 | Lighthouse Laboratory Plymouth | Wellcome Sanger Institute for the COVID-19 Genomics UK (COG-UK) Consortium | Lighthouse Laboratory Plymouth and Alex Alderton, Roberto Amato, Jeffrey Barrett, Sonia Goncalves, Ewan Harrison, David K. Jackson, Ian Johnston, Dominic Kwiatkowski, Cordelia Langford, John Sillitoe on behalf of the Wellcome Sanger Institute COVID-19 Surveillance Team |
| EPI_ISL_2999553 | Lighthouse Lab in Milton Keynes | Wellcome Sanger Institute for the COVID-19 Genomics UK (COG-UK) Consortium | The Lighthouse Lab in Milton Keynes and Alex Alderton, Roberto Amato, Jeffrey Barrett, Sonia Goncalves, Ewan Harrison, David K. Jackson, Ian Johnston, Dominic Kwiatkowski, Cordelia Langford, John Sillitoe on behalf of the Wellcome Sanger Institute COVID-19 Surveillance Team |
| EPI_ISL_2999554, EPI_ISL_2999555, EPI_ISL_2999556 | Lighthouse Laboratory Plymouth | Wellcome Sanger Institute for the COVID-19 Genomics UK (COG-UK) Consortium | Lighthouse Laboratory Plymouth and Alex Alderton, Roberto Amato, Jeffrey Barrett, Sonia Goncalves, Ewan Harrison, David K. Jackson, Ian Johnston, Dominic Kwiatkowski, Cordelia Langford, John Sillitoe on behalf of the Wellcome Sanger Institute COVID-19 Surveillance Team |
| EPI_ISL_2999557 | Lighthouse Lab in Milton Keynes | Wellcome Sanger Institute for the COVID-19 Genomics UK (COG-UK) Consortium | The Lighthouse Lab in Milton Keynes and Alex Alderton, Roberto Amato, Jeffrey Barrett, Sonia Goncalves, Ewan Harrison, David K. Jackson, Ian Johnston, Dominic Kwiatkowski, Cordelia Langford, John Sillitoe on behalf of the Wellcome Sanger Institute COVID-19 Surveillance Team |
| EPI_ISL_2999558 | Lighthouse Lab in Alderley Park | Wellcome Sanger Institute for the COVID-19 Genomics UK (COG-UK) Consortium | Jacquelyn Wynn, Mairead Hyland, The Lighthouse Lab in Alderley Park and Alex Alderton, Roberto Amato, Jeffrey Barrett, Sonia Goncalves, Ewan Harrison, David K. Jackson, Ian Johnston, Dominic Kwiatkowski, Cordelia Langford, John Sillitoe on behalf of the Wellcome Sanger Institute COVID-19 Surveillance Team |
| EPI_ISL_2999559 | Lighthouse Lab in Milton Keynes | Wellcome Sanger Institute for the COVID-19 Genomics UK (COG-UK) Consortium | The Lighthouse Lab in Milton Keynes and Alex Alderton, Roberto Amato, Jeffrey Barrett, Sonia Goncalves, Ewan Harrison, David K. Jackson, Ian Johnston, Dominic Kwiatkowski, Cordelia Langford, John Sillitoe on behalf of the Wellcome Sanger Institute COVID-19 Surveillance Team |
| EPI_ISL_2999560 | Lighthouse Laboratory Plymouth | Wellcome Sanger Institute for the COVID-19 Genomics UK (COG-UK) Consortium | Lighthouse Laboratory Plymouth and Alex Alderton, Roberto Amato, Jeffrey Barrett, Sonia Goncalves, Ewan Harrison, David K. Jackson, Ian Johnston, Dominic Kwiatkowski, Cordelia Langford, John Sillitoe on behalf of the Wellcome Sanger Institute COVID-19 Surveillance Team |
| EPI_ISL_2999561 | Lighthouse Lab in Milton Keynes | Wellcome Sanger Institute for the COVID-19 Genomics UK (COG-UK) Consortium | The Lighthouse Lab in Milton Keynes and Alex Alderton, Roberto Amato, Jeffrey Barrett, Sonia Goncalves, Ewan Harrison, David K. Jackson, Ian Johnston, Dominic Kwiatkowski, Cordelia Langford, John Sillitoe on behalf of the Wellcome Sanger Institute COVID-19 Surveillance Team |
| EPI_ISL_2999562 | Lighthouse Lab in Alderley Park | Wellcome Sanger Institute for the COVID-19 Genomics UK (COG-UK) Consortium | Jacquelyn Wynn, Mairead Hyland, The Lighthouse Lab in Alderley Park and Alex Alderton, Roberto Amato, Jeffrey Barrett, Sonia Goncalves, Ewan Harrison, David K. Jackson, Ian Johnston, Dominic Kwiatkowski, Cordelia Langford, John Sillitoe on behalf of the Wellcome Sanger Institute COVID-19 Surveillance Team |
| EPI_ISL_2999563 | Lighthouse Lab in Milton Keynes | Wellcome Sanger Institute for the COVID-19 Genomics UK (COG-UK) Consortium | The Lighthouse Lab in Milton Keynes and Alex Alderton, Roberto Amato, Jeffrey Barrett, Sonia Goncalves, Ewan Harrison, David K. Jackson, Ian Johnston, Dominic Kwiatkowski, Cordelia Langford, John Sillitoe on behalf of the Wellcome Sanger Institute COVID-19 Surveillance Team |
| EPI_ISL_2999564, EPI_ISL_2999565 | Lighthouse Lab in Alderley Park | Wellcome Sanger Institute for the COVID-19 Genomics UK (COG-UK) Consortium | Jacquelyn Wynn, Mairead Hyland, The Lighthouse Lab in Alderley Park and Alex Alderton, Roberto Amato, Jeffrey Barrett, Sonia Goncalves, Ewan Harrison, David K. Jackson, Ian Johnston, Dominic Kwiatkowski, Cordelia Langford, John Sillitoe on behalf of the Wellcome Sanger Institute COVID-19 Surveillance Team |
| EPI_ISL_2999566 | Lighthouse Laboratory Plymouth | Wellcome Sanger Institute for the COVID-19 Genomics UK (COG-UK) Consortium | Lighthouse Laboratory Plymouth and Alex Alderton, Roberto Amato, Jeffrey Barrett, Sonia Goncalves, Ewan Harrison, David K. Jackson, Ian Johnston, Dominic Kwiatkowski, Cordelia Langford, John Sillitoe on behalf of the Wellcome Sanger Institute COVID-19 Surveillance Team |

| | | | |
|---|---|---|---|
| EPI_ISL_2999567 | Lighthouse Lab in Milton Keynes | Wellcome Sanger Institute for the COVID-19 Genomics UK (COG-UK) Consortium | The Lighthouse Lab in Milton Keynes and Alex Alderton, Roberto Amato, Jeffrey Barrett, Sonia Goncalves, Ewan Harrison, David K. Jackson, Ian Johnston, Dominic Kwiatkowski, Cordelia Langford, John Sillitoe on behalf of the Wellcome Sanger Institute COVID-19 Surveillance Team |
| EPI_ISL_2999568 | Lighthouse Lab in Alderley Park | Wellcome Sanger Institute for the COVID-19 Genomics UK (COG-UK) Consortium | Jacquelyn Wynn, Mairead Hyland, The Lighthouse Lab in Alderley Park and Alex Alderton, Roberto Amato, Jeffrey Barrett, Sonia Goncalves, Ewan Harrison, David K. Jackson, Ian Johnston, Dominic Kwiatkowski, Cordelia Langford, John Sillitoe on behalf of the Wellcome Sanger Institute COVID-19 Surveillance Team |
| EPI_ISL_2999569 | Lighthouse Laboratory Plymouth | Wellcome Sanger Institute for the COVID-19 Genomics UK (COG-UK) Consortium | Lighthouse Laboratory Plymouth and Alex Alderton, Roberto Amato, Jeffrey Barrett, Sonia Goncalves, Ewan Harrison, David K. Jackson, Ian Johnston, Dominic Kwiatkowski, Cordelia Langford, John Sillitoe on behalf of the Wellcome Sanger Institute COVID-19 Surveillance Team |
| EPI_ISL_2999570, EPI_ISL_2999571 | Lighthouse Lab in Milton Keynes | Wellcome Sanger Institute for the COVID-19 Genomics UK (COG-UK) Consortium | The Lighthouse Lab in Milton Keynes and Alex Alderton, Roberto Amato, Jeffrey Barrett, Sonia Goncalves, Ewan Harrison, David K. Jackson, Ian Johnston, Dominic Kwiatkowski, Cordelia Langford, John Sillitoe on behalf of the Wellcome Sanger Institute COVID-19 Surveillance Team |
| EPI_ISL_2999572 | Lighthouse Lab in Alderley Park | Wellcome Sanger Institute for the COVID-19 Genomics UK (COG-UK) Consortium | Jacquelyn Wynn, Mairead Hyland, The Lighthouse Lab in Alderley Park and Alex Alderton, Roberto Amato, Jeffrey Barrett, Sonia Goncalves, Ewan Harrison, David K. Jackson, Ian Johnston, Dominic Kwiatkowski, Cordelia Langford, John Sillitoe on behalf of the Wellcome Sanger Institute COVID-19 Surveillance Team |
| EPI_ISL_2999573 | Lighthouse Lab in Milton Keynes | Wellcome Sanger Institute for the COVID-19 Genomics UK (COG-UK) Consortium | The Lighthouse Lab in Milton Keynes and Alex Alderton, Roberto Amato, Jeffrey Barrett, Sonia Goncalves, Ewan Harrison, David K. Jackson, Ian Johnston, Dominic Kwiatkowski, Cordelia Langford, John Sillitoe on behalf of the Wellcome Sanger Institute COVID-19 Surveillance Team |
| EPI_ISL_2999574 | Lighthouse Laboratory Plymouth | Wellcome Sanger Institute for the COVID-19 Genomics UK (COG-UK) Consortium | Lighthouse Laboratory Plymouth and Alex Alderton, Roberto Amato, Jeffrey Barrett, Sonia Goncalves, Ewan Harrison, David K. Jackson, Ian Johnston, Dominic Kwiatkowski, Cordelia Langford, John Sillitoe on behalf of the Wellcome Sanger Institute COVID-19 Surveillance Team |
| EPI_ISL_2999575 | Lighthouse Lab in Milton Keynes | Wellcome Sanger Institute for the COVID-19 Genomics UK (COG-UK) Consortium | The Lighthouse Lab in Milton Keynes and Alex Alderton, Roberto Amato, Jeffrey Barrett, Sonia Goncalves, Ewan Harrison, David K. Jackson, Ian Johnston, Dominic Kwiatkowski, Cordelia Langford, John Sillitoe on behalf of the Wellcome Sanger Institute COVID-19 Surveillance Team |
| EPI_ISL_2999576 | Lighthouse Laboratory Plymouth | Wellcome Sanger Institute for the COVID-19 Genomics UK (COG-UK) Consortium | Lighthouse Laboratory Plymouth and Alex Alderton, Roberto Amato, Jeffrey Barrett, Sonia Goncalves, Ewan Harrison, David K. Jackson, Ian Johnston, Dominic Kwiatkowski, Cordelia Langford, John Sillitoe on behalf of the Wellcome Sanger Institute COVID-19 Surveillance Team |
| EPI_ISL_2999577 | Lighthouse Lab in Alderley Park | Wellcome Sanger Institute for the COVID-19 Genomics UK (COG-UK) Consortium | Jacquelyn Wynn, Mairead Hyland, The Lighthouse Lab in Alderley Park and Alex Alderton, Roberto Amato, Jeffrey Barrett, Sonia Goncalves, Ewan Harrison, David K. Jackson, Ian Johnston, Dominic Kwiatkowski, Cordelia Langford, John Sillitoe on behalf of the Wellcome Sanger Institute COVID-19 Surveillance Team |
| EPI_ISL_2999578, EPI_ISL_2999579 | Lighthouse Laboratory Plymouth | Wellcome Sanger Institute for the COVID-19 Genomics UK (COG-UK) Consortium | Lighthouse Laboratory Plymouth and Alex Alderton, Roberto Amato, Jeffrey Barrett, Sonia Goncalves, Ewan Harrison, David K. Jackson, Ian Johnston, Dominic Kwiatkowski, Cordelia Langford, John Sillitoe on behalf of the Wellcome Sanger Institute COVID-19 Surveillance Team |
| EPI_ISL_2999580, EPI_ISL_2999581 | Lighthouse Lab in Milton Keynes | Wellcome Sanger Institute for the COVID-19 Genomics UK (COG-UK) Consortium | The Lighthouse Lab in Milton Keynes and Alex Alderton, Roberto Amato, Jeffrey Barrett, Sonia Goncalves, Ewan Harrison, David K. Jackson, Ian Johnston, Dominic Kwiatkowski, Cordelia Langford, John Sillitoe on behalf of the Wellcome Sanger Institute COVID-19 Surveillance Team |
| EPI_ISL_2999582, EPI_ISL_2999583 | Lighthouse Lab in Alderley Park | Wellcome Sanger Institute for the COVID-19 Genomics UK (COG-UK) Consortium | Jacquelyn Wynn, Mairead Hyland, The Lighthouse Lab in Alderley Park and Alex Alderton, Roberto Amato, Jeffrey Barrett, Sonia Goncalves, Ewan Harrison, David K. Jackson, Ian Johnston, Dominic Kwiatkowski, Cordelia Langford, John Sillitoe on behalf of the Wellcome Sanger Institute COVID-19 Surveillance Team |
| EPI_ISL_2999584, EPI_ISL_2999585, EPI_ISL_2999586 | Lighthouse Laboratory Plymouth | Wellcome Sanger Institute for the COVID-19 Genomics UK (COG-UK) Consortium | Lighthouse Laboratory Plymouth and Alex Alderton, Roberto Amato, Jeffrey Barrett, Sonia Goncalves, Ewan Harrison, David K. Jackson, Ian Johnston, Dominic Kwiatkowski, Cordelia Langford, John Sillitoe on behalf of the Wellcome Sanger Institute COVID-19 Surveillance Team |
| EPI_ISL_2999587 | Lighthouse Lab in Milton Keynes | Wellcome Sanger Institute for the COVID-19 Genomics UK (COG-UK) Consortium | The Lighthouse Lab in Milton Keynes and Alex Alderton, Roberto Amato, Jeffrey Barrett, Sonia Goncalves, Ewan Harrison, David K. Jackson, Ian Johnston, Dominic Kwiatkowski, Cordelia Langford, John Sillitoe on behalf of the Wellcome Sanger Institute COVID-19 Surveillance Team |
| EPI_ISL_2999588 | Lighthouse Laboratory Plymouth | Wellcome Sanger Institute for the COVID-19 Genomics UK (COG-UK) Consortium | Lighthouse Laboratory Plymouth and Alex Alderton, Roberto Amato, Jeffrey Barrett, Sonia Goncalves, Ewan Harrison, David K. Jackson, Ian Johnston, Dominic Kwiatkowski, Cordelia Langford, John Sillitoe on behalf of the Wellcome Sanger Institute COVID-19 Surveillance Team |
| EPI_ISL_2999589 | Lighthouse Lab in Milton Keynes | Wellcome Sanger Institute for the COVID-19 Genomics UK (COG-UK) Consortium | The Lighthouse Lab in Milton Keynes and Alex Alderton, Roberto Amato, Jeffrey Barrett, Sonia Goncalves, Ewan Harrison, David K. Jackson, Ian Johnston, Dominic Kwiatkowski, Cordelia Langford, John Sillitoe on behalf of the Wellcome Sanger Institute COVID-19 Surveillance Team |
| EPI_ISL_2999590 | Lighthouse Laboratory Plymouth | Wellcome Sanger Institute for the COVID-19 Genomics UK (COG-UK) Consortium | Lighthouse Laboratory Plymouth and Alex Alderton, Roberto Amato, Jeffrey Barrett, Sonia Goncalves, Ewan Harrison, David K. Jackson, Ian Johnston, Dominic Kwiatkowski, Cordelia Langford, John Sillitoe on behalf of the Wellcome Sanger Institute COVID-19 Surveillance Team |
| EPI_ISL_2999591, EPI_ISL_2999592, EPI_ISL_2999593 | Lighthouse Lab in Alderley Park | Wellcome Sanger Institute for the COVID-19 Genomics UK (COG-UK) Consortium | Jacquelyn Wynn, Mairead Hyland, The Lighthouse Lab in Alderley Park and Alex Alderton, Roberto Amato, Jeffrey Barrett, Sonia Goncalves, Ewan Harrison, David K. Jackson, Ian Johnston, Dominic Kwiatkowski, Cordelia Langford, John Sillitoe on behalf of the Wellcome Sanger Institute COVID-19 Surveillance Team |
| EPI_ISL_2999594, EPI_ISL_2999595, EPI_ISL_2999596 | Lighthouse Laboratory Plymouth | Wellcome Sanger Institute for the COVID-19 Genomics UK (COG-UK) Consortium | Lighthouse Laboratory Plymouth and Alex Alderton, Roberto Amato, Jeffrey Barrett, Sonia Goncalves, Ewan Harrison, David K. Jackson, Ian Johnston, Dominic Kwiatkowski, Cordelia Langford, John Sillitoe on behalf of the Wellcome Sanger Institute COVID-19 Surveillance Team |
| EPI_ISL_2999597, EPI_ISL_2999598, EPI_ISL_2999599, EPI_ISL_2999600 | Lighthouse Lab in Alderley Park | Wellcome Sanger Institute for the COVID-19 Genomics UK (COG-UK) Consortium | Jacquelyn Wynn, Mairead Hyland, The Lighthouse Lab in Alderley Park and Alex Alderton, Roberto Amato, Jeffrey Barrett, Sonia Goncalves, Ewan Harrison, David K. Jackson, Ian Johnston, Dominic Kwiatkowski, Cordelia Langford, John Sillitoe on behalf of the Wellcome Sanger Institute COVID-19 Surveillance Team |
| EPI_ISL_2999601 | Lighthouse Laboratory Plymouth | Wellcome Sanger Institute for the COVID-19 Genomics UK (COG-UK) Consortium | Lighthouse Laboratory Plymouth and Alex Alderton, Roberto Amato, Jeffrey Barrett, Sonia Goncalves, Ewan Harrison, David K. Jackson, Ian Johnston, Dominic Kwiatkowski, Cordelia Langford, John Sillitoe on behalf of the Wellcome Sanger Institute COVID-19 Surveillance Team |
| EPI_ISL_2999602 | Lighthouse Lab in Alderley Park | Wellcome Sanger Institute for the COVID-19 Genomics UK (COG-UK) Consortium | Jacquelyn Wynn, Mairead Hyland, The Lighthouse Lab in Alderley Park and Alex Alderton, Roberto Amato, Jeffrey Barrett, Sonia Goncalves, Ewan Harrison, David K. Jackson, Ian Johnston, Dominic Kwiatkowski, Cordelia Langford, John Sillitoe on behalf of the Wellcome Sanger Institute COVID-19 Surveillance Team |
| EPI_ISL_2999603 | Lighthouse Lab in Milton Keynes | Wellcome Sanger Institute for the COVID-19 Genomics UK (COG-UK) Consortium | The Lighthouse Lab in Milton Keynes and Alex Alderton, Roberto Amato, Jeffrey Barrett, Sonia Goncalves, Ewan Harrison, David K. Jackson, Ian Johnston, Dominic Kwiatkowski, Cordelia Langford, John Sillitoe on behalf of the Wellcome Sanger Institute COVID-19 Surveillance Team |
| EPI_ISL_2999604, EPI_ISL_2999605 | Lighthouse Lab in Alderley Park | Wellcome Sanger Institute for the COVID-19 Genomics UK (COG-UK) Consortium | Jacquelyn Wynn, Mairead Hyland, The Lighthouse Lab in Alderley Park and Alex Alderton, Roberto Amato, Jeffrey Barrett, Sonia Goncalves, Ewan Harrison, David K. Jackson, Ian Johnston, Dominic Kwiatkowski, Cordelia Langford, John Sillitoe on behalf of the Wellcome Sanger Institute COVID-19 Surveillance Team |
| EPI_ISL_2999606 | Lighthouse Lab in Milton Keynes | Wellcome Sanger Institute for the COVID-19 Genomics UK (COG-UK) Consortium | The Lighthouse Lab in Milton Keynes and Alex Alderton, Roberto Amato, Jeffrey Barrett, Sonia Goncalves, Ewan Harrison, David K. Jackson, Ian Johnston, Dominic Kwiatkowski, Cordelia Langford, John Sillitoe on behalf of the Wellcome Sanger Institute COVID-19 Surveillance Team |
| EPI_ISL_2999607, EPI_ISL_2999608 | Lighthouse Lab in Alderley Park | Wellcome Sanger Institute for the COVID-19 Genomics UK (COG-UK) Consortium | Jacquelyn Wynn, Mairead Hyland, The Lighthouse Lab in Alderley Park and Alex Alderton, Roberto Amato, Jeffrey Barrett, Sonia Goncalves, Ewan Harrison, David K. Jackson, Ian Johnston, Dominic Kwiatkowski, Cordelia Langford, John Sillitoe on behalf of the Wellcome Sanger Institute COVID-19 Surveillance Team |
| EPI_ISL_2999609, EPI_ISL_2999610 | Lighthouse Laboratory Plymouth | Wellcome Sanger Institute for the COVID-19 Genomics UK (COG-UK) Consortium | Lighthouse Laboratory Plymouth and Alex Alderton, Roberto Amato, Jeffrey Barrett, Sonia Goncalves, Ewan Harrison, David K. Jackson, Ian Johnston, Dominic Kwiatkowski, Cordelia Langford, John Sillitoe on behalf of the Wellcome Sanger Institute COVID-19 Surveillance Team |
| EPI_ISL_2999611 | Lighthouse Lab in Milton Keynes | Wellcome Sanger Institute for the COVID-19 Genomics UK (COG-UK) Consortium | The Lighthouse Lab in Milton Keynes and Alex Alderton, Roberto Amato, Jeffrey Barrett, Sonia Goncalves, Ewan Harrison, David K. Jackson, Ian Johnston, Dominic Kwiatkowski, Cordelia Langford, John Sillitoe on behalf of the Wellcome Sanger Institute COVID-19 Surveillance Team |
| EPI_ISL_2999612 | Lighthouse Lab in Alderley Park | Wellcome Sanger Institute for the COVID-19 Genomics UK (COG-UK) Consortium | Jacquelyn Wynn, Mairead Hyland, The Lighthouse Lab in Alderley Park and Alex Alderton, Roberto Amato, Jeffrey Barrett, Sonia Goncalves, Ewan Harrison, David K. Jackson, Ian Johnston, Dominic Kwiatkowski, Cordelia Langford, John Sillitoe on behalf of the Wellcome Sanger Institute COVID-19 Surveillance Team |
| EPI_ISL_2999613, EPI_ISL_2999614 | Lighthouse Lab in Milton Keynes | Wellcome Sanger Institute for the COVID-19 Genomics UK (COG-UK) Consortium | The Lighthouse Lab in Milton Keynes and Alex Alderton, Roberto Amato, Jeffrey Barrett, Sonia Goncalves, Ewan Harrison, David K. Jackson, Ian Johnston, Dominic Kwiatkowski, Cordelia Langford, John Sillitoe on behalf of the Wellcome Sanger Institute COVID-19 Surveillance Team |
| EPI_ISL_2999615 | Lighthouse Laboratory Plymouth | Wellcome Sanger Institute for the COVID-19 Genomics UK (COG-UK) Consortium | Lighthouse Laboratory Plymouth and Alex Alderton, Roberto Amato, Jeffrey Barrett, Sonia Goncalves, Ewan Harrison, David K. Jackson, Ian Johnston, Dominic Kwiatkowski, Cordelia Langford, John Sillitoe on behalf of the Wellcome Sanger Institute COVID-19 Surveillance Team |
| EPI_ISL_2999616, EPI_ISL_2999617 | Lighthouse Lab in Milton Keynes | Wellcome Sanger Institute for the COVID-19 Genomics UK (COG-UK) Consortium | The Lighthouse Lab in Milton Keynes and Alex Alderton, Roberto Amato, Jeffrey Barrett, Sonia Goncalves, Ewan Harrison, David K. Jackson, Ian Johnston, Dominic Kwiatkowski, Cordelia Langford, John Sillitoe on behalf of the Wellcome Sanger Institute COVID-19 Surveillance Team |
| EPI_ISL_2999618, EPI_ISL_2999619, EPI_ISL_2999620, EPI_ISL_2999621 | Lighthouse Lab in Alderley Park | Wellcome Sanger Institute for the COVID-19 Genomics UK (COG-UK) Consortium | Jacquelyn Wynn, Mairead Hyland, The Lighthouse Lab in Alderley Park and Alex Alderton, Roberto Amato, Jeffrey Barrett, Sonia Goncalves, Ewan Harrison, David K. Jackson, Ian Johnston, Dominic Kwiatkowski, Cordelia Langford, John Sillitoe on behalf of the Wellcome Sanger Institute COVID-19 Surveillance Team |
| EPI_ISL_2999622, EPI_ISL_2999623 | Lighthouse Lab in Milton Keynes | Wellcome Sanger Institute for the COVID-19 Genomics UK (COG-UK) Consortium | The Lighthouse Lab in Milton Keynes and Alex Alderton, Roberto Amato, Jeffrey Barrett, Sonia Goncalves, Ewan Harrison, David K. Jackson, Ian Johnston, Dominic Kwiatkowski, Cordelia Langford, John Sillitoe on behalf of the Wellcome Sanger Institute COVID-19 Surveillance Team |
| EPI_ISL_2999624, EPI_ISL_2999625, EPI_ISL_2999626 | Lighthouse Laboratory Plymouth | Wellcome Sanger Institute for the COVID-19 Genomics UK (COG-UK) Consortium | Lighthouse Laboratory Plymouth and Alex Alderton, Roberto Amato, Jeffrey Barrett, Sonia Goncalves, Ewan Harrison, David K. Jackson, Ian Johnston, Dominic Kwiatkowski, Cordelia Langford, John Sillitoe on behalf of the Wellcome Sanger Institute COVID-19 Surveillance Team |
| EPI_ISL_2999627, EPI_ISL_2999628 | Lighthouse Lab in Milton Keynes | Wellcome Sanger Institute for the COVID-19 Genomics UK (COG-UK) Consortium | The Lighthouse Lab in Milton Keynes and Alex Alderton, Roberto Amato, Jeffrey Barrett, Sonia Goncalves, Ewan Harrison, David K. Jackson, Ian Johnston, Dominic Kwiatkowski, Cordelia Langford, John Sillitoe on behalf of the Wellcome Sanger Institute COVID-19 Surveillance Team |
| EPI_ISL_2999629 | Lighthouse Lab in Alderley Park | Wellcome Sanger Institute for the COVID-19 Genomics UK (COG-UK) Consortium | Jacquelyn Wynn, Mairead Hyland, The Lighthouse Lab in Alderley Park and Alex Alderton, Roberto Amato, Jeffrey Barrett, Sonia Goncalves, Ewan Harrison, David K. Jackson, Ian Johnston, Dominic Kwiatkowski, Cordelia Langford, John Sillitoe on behalf of the Wellcome Sanger Institute COVID-19 Surveillance Team |
| EPI_ISL_2999630 | Lighthouse Laboratory Plymouth | Wellcome Sanger Institute for the COVID-19 Genomics UK (COG-UK) Consortium | Lighthouse Laboratory Plymouth and Alex Alderton, Roberto Amato, Jeffrey Barrett, Sonia Goncalves, Ewan Harrison, David K. Jackson, Ian Johnston, Dominic Kwiatkowski, Cordelia Langford, John Sillitoe on behalf of the Wellcome Sanger Institute COVID-19 Surveillance Team |
| EPI_ISL_2999631 | Lighthouse Lab in Milton Keynes | Wellcome Sanger Institute for the COVID-19 Genomics UK (COG-UK) Consortium | The Lighthouse Lab in Milton Keynes and Alex Alderton, Roberto Amato, Jeffrey Barrett, Sonia Goncalves, Ewan Harrison, David K. Jackson, Ian Johnston, Dominic Kwiatkowski, Cordelia Langford, John Sillitoe on behalf of the Wellcome Sanger Institute COVID-19 Surveillance Team |
| EPI_ISL_2999632 | Lighthouse Laboratory Plymouth | Wellcome Sanger Institute for the COVID-19 Genomics UK (COG-UK) Consortium | Lighthouse Laboratory Plymouth and Alex Alderton, Roberto Amato, Jeffrey Barrett, Sonia Goncalves, Ewan Harrison, David K. Jackson, Ian Johnston, Dominic Kwiatkowski, Cordelia Langford, John Sillitoe on behalf of the Wellcome Sanger Institute COVID-19 Surveillance Team |
| EPI_ISL_2999633 | Lighthouse Lab in Alderley Park | Wellcome Sanger Institute for the COVID-19 Genomics UK (COG-UK) | Jacquelyn Wynn, Mairead Hyland, The Lighthouse Lab in Alderley Park and Alex Alderton, Roberto Amato, Jeffrey Barrett, Sonia Goncalves, Ewan Harrison, David K. Jackson, Ian Johnston, Dominic Kwiatkowski, Cordelia Langford, John Sillitoe on |

| | | | |
|---|---|---|---|
| | | Consortium | behalf of the Wellcome Sanger Institute COVID-19 Surveillance Team |
| EPI_ISL_2999634 | Lighthouse Lab in Milton Keynes | Wellcome Sanger Institute for the COVID-19 Genomics UK (COG-UK) Consortium | The Lighthouse Lab in Milton Keynes and Alex Alderton, Roberto Amato, Jeffrey Barrett, Sonia Goncalves, Ewan Harrison, David K. Jackson, Ian Johnston, Dominic Kwiatkowski, Cordelia Langford, John Sillitoe on behalf of the Wellcome Sanger Institute COVID-19 Surveillance Team |
| EPI_ISL_2999635, EPI_ISL_2999636 | Lighthouse Lab in Alderley Park | Wellcome Sanger Institute for the COVID-19 Genomics UK (COG-UK) Consortium | Jacquelyn Wynn, Mairead Hyland, The Lighthouse Lab in Alderley Park and Alex Alderton, Roberto Amato, Jeffrey Barrett, Sonia Goncalves, Ewan Harrison, David K. Jackson, Ian Johnston, Dominic Kwiatkowski, Cordelia Langford, John Sillitoe on behalf of the Wellcome Sanger Institute COVID-19 Surveillance Team |
| EPI_ISL_2999637 | Lighthouse Lab in Milton Keynes | Wellcome Sanger Institute for the COVID-19 Genomics UK (COG-UK) Consortium | The Lighthouse Lab in Milton Keynes and Alex Alderton, Roberto Amato, Jeffrey Barrett, Sonia Goncalves, Ewan Harrison, David K. Jackson, Ian Johnston, Dominic Kwiatkowski, Cordelia Langford, John Sillitoe on behalf of the Wellcome Sanger Institute COVID-19 Surveillance Team |
| EPI_ISL_2999638 | Lighthouse Laboratory Plymouth | Wellcome Sanger Institute for the COVID-19 Genomics UK (COG-UK) Consortium | Lighthouse Laboratory Plymouth and Alex Alderton, Roberto Amato, Jeffrey Barrett, Sonia Goncalves, Ewan Harrison, David K. Jackson, Ian Johnston, Dominic Kwiatkowski, Cordelia Langford, John Sillitoe on behalf of the Wellcome Sanger Institute COVID-19 Surveillance Team |
| EPI_ISL_2999639, EPI_ISL_2999640 | Lighthouse Lab in Milton Keynes | Wellcome Sanger Institute for the COVID-19 Genomics UK (COG-UK) Consortium | The Lighthouse Lab in Milton Keynes and Alex Alderton, Roberto Amato, Jeffrey Barrett, Sonia Goncalves, Ewan Harrison, David K. Jackson, Ian Johnston, Dominic Kwiatkowski, Cordelia Langford, John Sillitoe on behalf of the Wellcome Sanger Institute COVID-19 Surveillance Team |
| EPI_ISL_2999641, EPI_ISL_2999642, EPI_ISL_2999643 | Lighthouse Laboratory Plymouth | Wellcome Sanger Institute for the COVID-19 Genomics UK (COG-UK) Consortium | Lighthouse Laboratory Plymouth and Alex Alderton, Roberto Amato, Jeffrey Barrett, Sonia Goncalves, Ewan Harrison, David K. Jackson, Ian Johnston, Dominic Kwiatkowski, Cordelia Langford, John Sillitoe on behalf of the Wellcome Sanger Institute COVID-19 Surveillance Team |
| EPI_ISL_2999644 | Lighthouse Lab in Milton Keynes | Wellcome Sanger Institute for the COVID-19 Genomics UK (COG-UK) Consortium | The Lighthouse Lab in Milton Keynes and Alex Alderton, Roberto Amato, Jeffrey Barrett, Sonia Goncalves, Ewan Harrison, David K. Jackson, Ian Johnston, Dominic Kwiatkowski, Cordelia Langford, John Sillitoe on behalf of the Wellcome Sanger Institute COVID-19 Surveillance Team |
| EPI_ISL_2999645, EPI_ISL_2999646, EPI_ISL_2999647 | Lighthouse Lab in Alderley Park | Wellcome Sanger Institute for the COVID-19 Genomics UK (COG-UK) Consortium | Jacquelyn Wynn, Mairead Hyland, The Lighthouse Lab in Alderley Park and Alex Alderton, Roberto Amato, Jeffrey Barrett, Sonia Goncalves, Ewan Harrison, David K. Jackson, Ian Johnston, Dominic Kwiatkowski, Cordelia Langford, John Sillitoe on behalf of the Wellcome Sanger Institute COVID-19 Surveillance Team |
| EPI_ISL_2999648, EPI_ISL_2999649 | Lighthouse Lab in Milton Keynes | Wellcome Sanger Institute for the COVID-19 Genomics UK (COG-UK) Consortium | The Lighthouse Lab in Milton Keynes and Alex Alderton, Roberto Amato, Jeffrey Barrett, Sonia Goncalves, Ewan Harrison, David K. Jackson, Ian Johnston, Dominic Kwiatkowski, Cordelia Langford, John Sillitoe on behalf of the Wellcome Sanger Institute COVID-19 Surveillance Team |
| EPI_ISL_2999650, EPI_ISL_2999651, EPI_ISL_2999652, EPI_ISL_2999653, EPI_ISL_2999654, EPI_ISL_2999655, EPI_ISL_2999656, EPI_ISL_2999657, EPI_ISL_2999658, EPI_ISL_2999659, EPI_ISL_2999660, EPI_ISL_2999661, EPI_ISL_2999662, EPI_ISL_2999663, EPI_ISL_2999664, EPI_ISL_2999665, EPI_ISL_2999666, EPI_ISL_2999667, EPI_ISL_2999668, EPI_ISL_2999669, EPI_ISL_2999670, EPI_ISL_2999671, EPI_ISL_2999672, EPI_ISL_2999673, EPI_ISL_2999674, EPI_ISL_2999675, EPI_ISL_2999676, EPI_ISL_2999677, EPI_ISL_2999678, EPI_ISL_2999679 | | | |
| see above | Lighthouse Lab in Glasgow | Wellcome Sanger Institute for the COVID-19 Genomics UK (COG-UK) Consortium | Harper VanSteenhouse, Yumi Kasai, David Gray, Carol Clugston, Anna Dominiczak and Alex Alderton, Roberto Amato, Jeffrey Barrett, Sonia Goncalves, Ewan Harrison, David K. Jackson, Ian Johnston, Dominic Kwiatkowski, Cordelia Langford, John Sillitoe on behalf of the Wellcome Sanger Institute COVID-19 Surveillance Team |
| EPI_ISL_2999680 | Randox Laboratories | Wellcome Sanger Institute for the COVID-19 Genomics UK (COG-UK) Consortium | Randox Laboratories and Alex Alderton, Roberto Amato, Jeffrey Barrett, Sonia Goncalves, Ewan Harrison, David K. Jackson, Ian Johnston, Dominic Kwiatkowski, Cordelia Langford, John Sillitoe on behalf of the Wellcome Sanger Institute COVID-19 Surveillance Team |
| EPI_ISL_2999681, EPI_ISL_2999682, EPI_ISL_2999683, EPI_ISL_2999684, EPI_ISL_2999685, EPI_ISL_2999686, EPI_ISL_2999687, EPI_ISL_2999688, EPI_ISL_2999689, EPI_ISL_2999690, EPI_ISL_2999691, EPI_ISL_2999692, EPI_ISL_2999693, EPI_ISL_2999694, EPI_ISL_2999695, EPI_ISL_2999696, EPI_ISL_2999697, EPI_ISL_2999698, EPI_ISL_2999699, EPI_ISL_2999700 | | | |
| see above | Lighthouse Lab in Glasgow | Wellcome Sanger Institute for the COVID-19 Genomics UK (COG-UK) Consortium | Harper VanSteenhouse, Yumi Kasai, David Gray, Carol Clugston, Anna Dominiczak and Alex Alderton, Roberto Amato, Jeffrey Barrett, Sonia Goncalves, Ewan Harrison, David K. Jackson, Ian Johnston, Dominic Kwiatkowski, Cordelia Langford, John Sillitoe on behalf of the Wellcome Sanger Institute COVID-19 Surveillance Team |
| EPI_ISL_2999701 | Randox Laboratories | Wellcome Sanger Institute for the COVID-19 Genomics UK (COG-UK) Consortium | Randox Laboratories and Alex Alderton, Roberto Amato, Jeffrey Barrett, Sonia Goncalves, Ewan Harrison, David K. Jackson, Ian Johnston, Dominic Kwiatkowski, Cordelia Langford, John Sillitoe on behalf of the Wellcome Sanger Institute COVID-19 Surveillance Team |
| EPI_ISL_2999702, EPI_ISL_2999703 | Lighthouse Lab in Glasgow | Wellcome Sanger Institute for the COVID-19 Genomics UK (COG-UK) Consortium | Harper VanSteenhouse, Yumi Kasai, David Gray, Carol Clugston, Anna Dominiczak and Alex Alderton, Roberto Amato, Jeffrey Barrett, Sonia Goncalves, Ewan Harrison, David K. Jackson, Ian Johnston, Dominic Kwiatkowski, Cordelia Langford, John Sillitoe on behalf of the Wellcome Sanger Institute COVID-19 Surveillance Team |
| EPI_ISL_2999704, EPI_ISL_2999705, EPI_ISL_2999706 | Lighthouse Lab in Milton Keynes | Wellcome Sanger Institute for the COVID-19 Genomics UK (COG-UK) Consortium | The Lighthouse Lab in Milton Keynes and Alex Alderton, Roberto Amato, Jeffrey Barrett, Sonia Goncalves, Ewan Harrison, David K. Jackson, Ian Johnston, Dominic Kwiatkowski, Cordelia Langford, John Sillitoe on behalf of the Wellcome Sanger Institute COVID-19 Surveillance Team |
| EPI_ISL_2999707 | Lighthouse Laboratory Plymouth | Wellcome Sanger Institute for the COVID-19 Genomics UK (COG-UK) Consortium | Lighthouse Laboratory Plymouth and Alex Alderton, Roberto Amato, Jeffrey Barrett, Sonia Goncalves, Ewan Harrison, David K. Jackson, Ian Johnston, Dominic Kwiatkowski, Cordelia Langford, John Sillitoe on behalf of the Wellcome Sanger Institute COVID-19 Surveillance Team |
| EPI_ISL_2999708, EPI_ISL_2999709, EPI_ISL_2999710, EPI_ISL_2999711, EPI_ISL_2999712, EPI_ISL_2999713, EPI_ISL_2999714, EPI_ISL_2999715 | Lighthouse Lab in Milton Keynes | Wellcome Sanger Institute for the COVID-19 Genomics UK (COG-UK) Consortium | The Lighthouse Lab in Milton Keynes and Alex Alderton, Roberto Amato, Jeffrey Barrett, Sonia Goncalves, Ewan Harrison, David K. Jackson, Ian Johnston, Dominic Kwiatkowski, Cordelia Langford, John Sillitoe on behalf of the Wellcome Sanger Institute COVID-19 Surveillance Team |
| EPI_ISL_2999716 | Lighthouse Lab in Alderley Park | Wellcome Sanger Institute for the COVID-19 Genomics UK (COG-UK) Consortium | Jacquelyn Wynn, Mairead Hyland, The Lighthouse Lab in Alderley Park and Alex Alderton, Roberto Amato, Jeffrey Barrett, Sonia Goncalves, Ewan Harrison, David K. Jackson, Ian Johnston, Dominic Kwiatkowski, Cordelia Langford, John Sillitoe on behalf of the Wellcome Sanger Institute COVID-19 Surveillance Team |
| EPI_ISL_2999717, EPI_ISL_2999718, EPI_ISL_2999719, EPI_ISL_2999720, EPI_ISL_2999721, EPI_ISL_2999722, EPI_ISL_2999723, EPI_ISL_2999724, EPI_ISL_2999725, EPI_ISL_2999726 | Lighthouse Lab in Milton Keynes | Wellcome Sanger Institute for the COVID-19 Genomics UK (COG-UK) Consortium | The Lighthouse Lab in Milton Keynes and Alex Alderton, Roberto Amato, Jeffrey Barrett, Sonia Goncalves, Ewan Harrison, David K. Jackson, Ian Johnston, Dominic Kwiatkowski, Cordelia Langford, John Sillitoe on behalf of the Wellcome Sanger Institute COVID-19 Surveillance Team |
| EPI_ISL_2999727 | Lighthouse Lab in Alderley Park | Wellcome Sanger Institute for the COVID-19 Genomics UK (COG-UK) Consortium | Jacquelyn Wynn, Mairead Hyland, The Lighthouse Lab in Alderley Park and Alex Alderton, Roberto Amato, Jeffrey Barrett, Sonia Goncalves, Ewan Harrison, David K. Jackson, Ian Johnston, Dominic Kwiatkowski, Cordelia Langford, John Sillitoe on behalf of the Wellcome Sanger Institute COVID-19 Surveillance Team |
| EPI_ISL_2999728 | Berkshire and Surrey Pathology Services Lighthouse Laboratory | Wellcome Sanger Institute for the COVID-19 Genomics UK (COG-UK) Consortium | Berkshire and Surrey Pathology Services Lighthouse Laboratory and Alex Alderton, Roberto Amato, Jeffrey Barrett, Sonia Goncalves, Ewan Harrison, David K. Jackson, Ian Johnston, Dominic Kwiatkowski, Cordelia Langford, John Sillitoe on behalf of the Wellcome Sanger Institute COVID-19 Surveillance Team |
| EPI_ISL_2999729 | Lighthouse Lab in Alderley Park | Wellcome Sanger Institute for the COVID-19 Genomics UK (COG-UK) Consortium | Jacquelyn Wynn, Mairead Hyland, The Lighthouse Lab in Alderley Park and Alex Alderton, Roberto Amato, Jeffrey Barrett, Sonia Goncalves, Ewan Harrison, David K. Jackson, Ian Johnston, Dominic Kwiatkowski, Cordelia Langford, John Sillitoe on behalf of the Wellcome Sanger Institute COVID-19 Surveillance Team |
| EPI_ISL_2999730 | Berkshire and Surrey Pathology Services Lighthouse Laboratory | Wellcome Sanger Institute for the COVID-19 Genomics UK (COG-UK) Consortium | Berkshire and Surrey Pathology Services Lighthouse Laboratory and Alex Alderton, Roberto Amato, Jeffrey Barrett, Sonia Goncalves, Ewan Harrison, David K. Jackson, Ian Johnston, Dominic Kwiatkowski, Cordelia Langford, John Sillitoe on behalf of the Wellcome Sanger Institute COVID-19 Surveillance Team |
| EPI_ISL_2999731, EPI_ISL_2999732, EPI_ISL_2999733, EPI_ISL_2999734, EPI_ISL_2999735 | Lighthouse Lab in Alderley Park | Wellcome Sanger Institute for the COVID-19 Genomics UK (COG-UK) Consortium | Jacquelyn Wynn, Mairead Hyland, The Lighthouse Lab in Alderley Park and Alex Alderton, Roberto Amato, Jeffrey Barrett, Sonia Goncalves, Ewan Harrison, David K. Jackson, Ian Johnston, Dominic Kwiatkowski, Cordelia Langford, John Sillitoe on behalf of the Wellcome Sanger Institute COVID-19 Surveillance Team |
| EPI_ISL_2999736, EPI_ISL_2999737, EPI_ISL_2999738, EPI_ISL_2999739, EPI_ISL_2999740, EPI_ISL_2999741, EPI_ISL_2999742, EPI_ISL_2999743, EPI_ISL_2999744, EPI_ISL_2999745, EPI_ISL_2999746, EPI_ISL_2999747, EPI_ISL_2999748, EPI_ISL_2999749, EPI_ISL_2999750, EPI_ISL_2999751, EPI_ISL_2999752, EPI_ISL_2999753, EPI_ISL_2999754, EPI_ISL_2999755, EPI_ISL_2999756, EPI_ISL_2999757, EPI_ISL_2999758, EPI_ISL_2999759, EPI_ISL_2999760, EPI_ISL_2999761, EPI_ISL_2999762, EPI_ISL_2999763, EPI_ISL_2999764, EPI_ISL_2999765, EPI_ISL_2999766, EPI_ISL_2999767, EPI_ISL_2999768, EPI_ISL_2999769, EPI_ISL_2999770, EPI_ISL_2999771, EPI_ISL_2999772, EPI_ISL_2999773, EPI_ISL_2999774, EPI_ISL_2999775, EPI_ISL_2999776, EPI_ISL_2999777, EPI_ISL_2999778, EPI_ISL_2999779, EPI_ISL_2999780, EPI_ISL_2999781, EPI_ISL_2999782, EPI_ISL_2999783, EPI_ISL_2999784, EPI_ISL_2999786, EPI_ISL_2999787, EPI_ISL_2999788, EPI_ISL_2999789, EPI_ISL_2999790, EPI_ISL_2999791, EPI_ISL_2999792, EPI_ISL_2999793, EPI_ISL_2999794, EPI_ISL_2999795, EPI_ISL_2999796, EPI_ISL_2999797, EPI_ISL_2999798, EPI_ISL_2999799, EPI_ISL_2999800, EPI_ISL_2999801, EPI_ISL_2999802, EPI_ISL_2999803, EPI_ISL_2999804, EPI_ISL_2999805, EPI_ISL_2999806 | | | |
| see above | Lighthouse Lab in Glasgow | Wellcome Sanger Institute for the COVID-19 Genomics UK (COG-UK) Consortium | Harper VanSteenhouse, Yumi Kasai, David Gray, Carol Clugston, Anna Dominiczak and Alex Alderton, Roberto Amato, Jeffrey Barrett, Sonia Goncalves, Ewan Harrison, David K. Jackson, Ian Johnston, Dominic Kwiatkowski, Cordelia Langford, John Sillitoe on behalf of the Wellcome Sanger Institute COVID-19 Surveillance Team |
| EPI_ISL_2999808 | Berkshire and Surrey Pathology Services Lighthouse Laboratory | Wellcome Sanger Institute for the COVID-19 Genomics UK (COG-UK) Consortium | Berkshire and Surrey Pathology Services Lighthouse Laboratory and Alex Alderton, Roberto Amato, Jeffrey Barrett, Sonia Goncalves, Ewan Harrison, David K. Jackson, Ian Johnston, Dominic Kwiatkowski, Cordelia Langford, John Sillitoe on behalf of the Wellcome Sanger Institute COVID-19 Surveillance Team |
| EPI_ISL_2999809, EPI_ISL_2999810 | Health Services Laboratories | Wellcome Sanger Institute for the COVID-19 Genomics UK (COG-UK) Consortium | Health Services Laboratories and Alex Alderton, Roberto Amato, Jeffrey Barrett, Sonia Goncalves, Ewan Harrison, David K. Jackson, Ian Johnston, Dominic Kwiatkowski, Cordelia Langford, John Sillitoe on behalf of the Wellcome Sanger Institute COVID-19 Surveillance Team |
| EPI_ISL_2999811, EPI_ISL_2999812, EPI_ISL_2999813 | Berkshire and Surrey Pathology Services Lighthouse Laboratory | Wellcome Sanger Institute for the COVID-19 Genomics UK (COG-UK) Consortium | Berkshire and Surrey Pathology Services Lighthouse Laboratory and Alex Alderton, Roberto Amato, Jeffrey Barrett, Sonia Goncalves, Ewan Harrison, David K. Jackson, Ian Johnston, Dominic Kwiatkowski, Cordelia Langford, John Sillitoe on behalf of the Wellcome Sanger Institute COVID-19 Surveillance Team |
| EPI_ISL_2999814 | Health Services Laboratories | Wellcome Sanger Institute for the COVID-19 Genomics UK (COG-UK) Consortium | Health Services Laboratories and Alex Alderton, Roberto Amato, Jeffrey Barrett, Sonia Goncalves, Ewan Harrison, David K. Jackson, Ian Johnston, Dominic Kwiatkowski, Cordelia Langford, John Sillitoe on behalf of the Wellcome Sanger Institute COVID-19 Surveillance Team |
| EPI_ISL_2999815 | Berkshire and Surrey Pathology Services Lighthouse Laboratory | Wellcome Sanger Institute for the COVID-19 Genomics UK (COG-UK) Consortium | Berkshire and Surrey Pathology Services Lighthouse Laboratory and Alex Alderton, Roberto Amato, Jeffrey Barrett, Sonia Goncalves, Ewan Harrison, David K. Jackson, Ian Johnston, Dominic Kwiatkowski, Cordelia Langford, John Sillitoe on behalf of the Wellcome Sanger Institute COVID-19 Surveillance Team |
| EPI_ISL_2999816 | Lighthouse Lab in Alderley Park | Wellcome Sanger Institute for the COVID-19 Genomics UK (COG-UK) Consortium | Jacquelyn Wynn, Mairead Hyland, The Lighthouse Lab in Alderley Park and Alex Alderton, Roberto Amato, Jeffrey Barrett, Sonia Goncalves, Ewan Harrison, David K. Jackson, Ian Johnston, Dominic Kwiatkowski, Cordelia Langford, John Sillitoe on behalf of the Wellcome Sanger Institute COVID-19 Surveillance Team |
| EPI_ISL_2999817 | Berkshire and Surrey Pathology Services Lighthouse Laboratory | Wellcome Sanger Institute for the COVID-19 Genomics UK (COG-UK) Consortium | Berkshire and Surrey Pathology Services Lighthouse Laboratory and Alex Alderton, Roberto Amato, Jeffrey Barrett, Sonia Goncalves, Ewan Harrison, David K. Jackson, Ian Johnston, Dominic Kwiatkowski, Cordelia Langford, John Sillitoe on behalf of the Wellcome Sanger Institute COVID-19 Surveillance Team |
| EPI_ISL_2999818 | Lighthouse Lab in Alderley Park | Wellcome Sanger Institute for the COVID-19 Genomics UK (COG-UK) | Jacquelyn Wynn, Mairead Hyland, The Lighthouse Lab in Alderley Park and Alex Alderton, Roberto Amato, Jeffrey Barrett, Sonia Goncalves, Ewan Harrison, David K. Jackson, Ian Johnston, Dominic Kwiatkowski, Cordelia Langford, John Sillitoe on |

| | | | |
|---|---|---|---|
| | | Consortium | behalf of the Wellcome Sanger Institute COVID-19 Surveillance Team |
| EPI_ISL_2999819 | Berkshire and Surrey Pathology Services Lighthouse Laboratory | Wellcome Sanger Institute for the COVID-19 Genomics UK (COG-UK) Consortium | Berkshire and Surrey Pathology Services Lighthouse Laboratory and Alex Alderton, Roberto Amato, Jeffrey Barrett, Sonia Goncalves, Ewan Harrison, David K. Jackson, Ian Johnston, Dominic Kwiatkowski, Cordelia Langford, John Sillitoe on behalf of the Wellcome Sanger Institute COVID-19 Surveillance Team |
| EPI_ISL_2999820, EPI_ISL_2999821 | Health Services Laboratories | Wellcome Sanger Institute for the COVID-19 Genomics UK (COG-UK) Consortium | Health Services Laboratories and Alex Alderton, Roberto Amato, Jeffrey Barrett, Sonia Goncalves, Ewan Harrison, David K. Jackson, Ian Johnston, Dominic Kwiatkowski, Cordelia Langford, John Sillitoe on behalf of the Wellcome Sanger Institute COVID-19 Surveillance Team |
| EPI_ISL_2999822, EPI_ISL_2999823 | Berkshire and Surrey Pathology Services Lighthouse Laboratory | Wellcome Sanger Institute for the COVID-19 Genomics UK (COG-UK) Consortium | Berkshire and Surrey Pathology Services Lighthouse Laboratory and Alex Alderton, Roberto Amato, Jeffrey Barrett, Sonia Goncalves, Ewan Harrison, David K. Jackson, Ian Johnston, Dominic Kwiatkowski, Cordelia Langford, John Sillitoe on behalf of the Wellcome Sanger Institute COVID-19 Surveillance Team |
| EPI_ISL_2999824, EPI_ISL_2999825, EPI_ISL_2999826 | Lighthouse Lab in Alderley Park | Wellcome Sanger Institute for the COVID-19 Genomics UK (COG-UK) Consortium | Jacquelyn Wynn, Mairead Hyland, The Lighthouse Lab in Alderley Park and Alex Alderton, Roberto Amato, Jeffrey Barrett, Sonia Goncalves, Ewan Harrison, David K. Jackson, Ian Johnston, Dominic Kwiatkowski, Cordelia Langford, John Sillitoe on behalf of the Wellcome Sanger Institute COVID-19 Surveillance Team |
| EPI_ISL_2999827 | Health Services Laboratories | Wellcome Sanger Institute for the COVID-19 Genomics UK (COG-UK) Consortium | Health Services Laboratories and Alex Alderton, Roberto Amato, Jeffrey Barrett, Sonia Goncalves, Ewan Harrison, David K. Jackson, Ian Johnston, Dominic Kwiatkowski, Cordelia Langford, John Sillitoe on behalf of the Wellcome Sanger Institute COVID-19 Surveillance Team |
| EPI_ISL_2999828 | Lighthouse Lab in Alderley Park | Wellcome Sanger Institute for the COVID-19 Genomics UK (COG-UK) Consortium | Jacquelyn Wynn, Mairead Hyland, The Lighthouse Lab in Alderley Park and Alex Alderton, Roberto Amato, Jeffrey Barrett, Sonia Goncalves, Ewan Harrison, David K. Jackson, Ian Johnston, Dominic Kwiatkowski, Cordelia Langford, John Sillitoe on behalf of the Wellcome Sanger Institute COVID-19 Surveillance Team |
| EPI_ISL_2999829, EPI_ISL_2999830, EPI_ISL_2999831, EPI_ISL_2999832 | Berkshire and Surrey Pathology Services Lighthouse Laboratory | Wellcome Sanger Institute for the COVID-19 Genomics UK (COG-UK) Consortium | Berkshire and Surrey Pathology Services Lighthouse Laboratory and Alex Alderton, Roberto Amato, Jeffrey Barrett, Sonia Goncalves, Ewan Harrison, David K. Jackson, Ian Johnston, Dominic Kwiatkowski, Cordelia Langford, John Sillitoe on behalf of the Wellcome Sanger Institute COVID-19 Surveillance Team |
| EPI_ISL_2999833, EPI_ISL_2999834 | Health Services Laboratories | Wellcome Sanger Institute for the COVID-19 Genomics UK (COG-UK) Consortium | Health Services Laboratories and Alex Alderton, Roberto Amato, Jeffrey Barrett, Sonia Goncalves, Ewan Harrison, David K. Jackson, Ian Johnston, Dominic Kwiatkowski, Cordelia Langford, John Sillitoe on behalf of the Wellcome Sanger Institute COVID-19 Surveillance Team |
| EPI_ISL_2999835 | Lighthouse Lab in Alderley Park | Wellcome Sanger Institute for the COVID-19 Genomics UK (COG-UK) Consortium | Jacquelyn Wynn, Mairead Hyland, The Lighthouse Lab in Alderley Park and Alex Alderton, Roberto Amato, Jeffrey Barrett, Sonia Goncalves, Ewan Harrison, David K. Jackson, Ian Johnston, Dominic Kwiatkowski, Cordelia Langford, John Sillitoe on behalf of the Wellcome Sanger Institute COVID-19 Surveillance Team |
| EPI_ISL_2999836 | Berkshire and Surrey Pathology Services Lighthouse Laboratory | Wellcome Sanger Institute for the COVID-19 Genomics UK (COG-UK) Consortium | Berkshire and Surrey Pathology Services Lighthouse Laboratory and Alex Alderton, Roberto Amato, Jeffrey Barrett, Sonia Goncalves, Ewan Harrison, David K. Jackson, Ian Johnston, Dominic Kwiatkowski, Cordelia Langford, John Sillitoe on behalf of the Wellcome Sanger Institute COVID-19 Surveillance Team |
| EPI_ISL_2999837 | Health Services Laboratories | Wellcome Sanger Institute for the COVID-19 Genomics UK (COG-UK) Consortium | Health Services Laboratories and Alex Alderton, Roberto Amato, Jeffrey Barrett, Sonia Goncalves, Ewan Harrison, David K. Jackson, Ian Johnston, Dominic Kwiatkowski, Cordelia Langford, John Sillitoe on behalf of the Wellcome Sanger Institute COVID-19 Surveillance Team |
| EPI_ISL_2999838 | Berkshire and Surrey Pathology Services Lighthouse Laboratory | Wellcome Sanger Institute for the COVID-19 Genomics UK (COG-UK) Consortium | Berkshire and Surrey Pathology Services Lighthouse Laboratory and Alex Alderton, Roberto Amato, Jeffrey Barrett, Sonia Goncalves, Ewan Harrison, David K. Jackson, Ian Johnston, Dominic Kwiatkowski, Cordelia Langford, John Sillitoe on behalf of the Wellcome Sanger Institute COVID-19 Surveillance Team |
| EPI_ISL_2999839 | Lighthouse Lab in Alderley Park | Wellcome Sanger Institute for the COVID-19 Genomics UK (COG-UK) Consortium | Jacquelyn Wynn, Mairead Hyland, The Lighthouse Lab in Alderley Park and Alex Alderton, Roberto Amato, Jeffrey Barrett, Sonia Goncalves, Ewan Harrison, David K. Jackson, Ian Johnston, Dominic Kwiatkowski, Cordelia Langford, John Sillitoe on behalf of the Wellcome Sanger Institute COVID-19 Surveillance Team |
| EPI_ISL_2999840, EPI_ISL_2999841 | Berkshire and Surrey Pathology Services Lighthouse Laboratory | Wellcome Sanger Institute for the COVID-19 Genomics UK (COG-UK) Consortium | Berkshire and Surrey Pathology Services Lighthouse Laboratory and Alex Alderton, Roberto Amato, Jeffrey Barrett, Sonia Goncalves, Ewan Harrison, David K. Jackson, Ian Johnston, Dominic Kwiatkowski, Cordelia Langford, John Sillitoe on behalf of the Wellcome Sanger Institute COVID-19 Surveillance Team |
| EPI_ISL_2999842 | Lighthouse Lab in Alderley Park | Wellcome Sanger Institute for the COVID-19 Genomics UK (COG-UK) Consortium | Jacquelyn Wynn, Mairead Hyland, The Lighthouse Lab in Alderley Park and Alex Alderton, Roberto Amato, Jeffrey Barrett, Sonia Goncalves, Ewan Harrison, David K. Jackson, Ian Johnston, Dominic Kwiatkowski, Cordelia Langford, John Sillitoe on behalf of the Wellcome Sanger Institute COVID-19 Surveillance Team |
| EPI_ISL_2999843, EPI_ISL_2999844, EPI_ISL_2999845, EPI_ISL_2999846 | Health Services Laboratories | Wellcome Sanger Institute for the COVID-19 Genomics UK (COG-UK) Consortium | Health Services Laboratories and Alex Alderton, Roberto Amato, Jeffrey Barrett, Sonia Goncalves, Ewan Harrison, David K. Jackson, Ian Johnston, Dominic Kwiatkowski, Cordelia Langford, John Sillitoe on behalf of the Wellcome Sanger Institute COVID-19 Surveillance Team |
| EPI_ISL_2999847 | Lighthouse Lab in Alderley Park | Wellcome Sanger Institute for the COVID-19 Genomics UK (COG-UK) Consortium | Jacquelyn Wynn, Mairead Hyland, The Lighthouse Lab in Alderley Park and Alex Alderton, Roberto Amato, Jeffrey Barrett, Sonia Goncalves, Ewan Harrison, David K. Jackson, Ian Johnston, Dominic Kwiatkowski, Cordelia Langford, John Sillitoe on behalf of the Wellcome Sanger Institute COVID-19 Surveillance Team |
| EPI_ISL_2999848, EPI_ISL_2999849 | Health Services Laboratories | Wellcome Sanger Institute for the COVID-19 Genomics UK (COG-UK) Consortium | Health Services Laboratories and Alex Alderton, Roberto Amato, Jeffrey Barrett, Sonia Goncalves, Ewan Harrison, David K. Jackson, Ian Johnston, Dominic Kwiatkowski, Cordelia Langford, John Sillitoe on behalf of the Wellcome Sanger Institute COVID-19 Surveillance Team |
| EPI_ISL_2999850 | Berkshire and Surrey Pathology Services Lighthouse Laboratory | Wellcome Sanger Institute for the COVID-19 Genomics UK (COG-UK) Consortium | Berkshire and Surrey Pathology Services Lighthouse Laboratory and Alex Alderton, Roberto Amato, Jeffrey Barrett, Sonia Goncalves, Ewan Harrison, David K. Jackson, Ian Johnston, Dominic Kwiatkowski, Cordelia Langford, John Sillitoe on behalf of the Wellcome Sanger Institute COVID-19 Surveillance Team |
| EPI_ISL_2999851 | Lighthouse Lab in Alderley Park | Wellcome Sanger Institute for the COVID-19 Genomics UK (COG-UK) Consortium | Jacquelyn Wynn, Mairead Hyland, The Lighthouse Lab in Alderley Park and Alex Alderton, Roberto Amato, Jeffrey Barrett, Sonia Goncalves, Ewan Harrison, David K. Jackson, Ian Johnston, Dominic Kwiatkowski, Cordelia Langford, John Sillitoe on behalf of the Wellcome Sanger Institute COVID-19 Surveillance Team |
| EPI_ISL_2999852 | Berkshire and Surrey Pathology Services Lighthouse Laboratory | Wellcome Sanger Institute for the COVID-19 Genomics UK (COG-UK) Consortium | Berkshire and Surrey Pathology Services Lighthouse Laboratory and Alex Alderton, Roberto Amato, Jeffrey Barrett, Sonia Goncalves, Ewan Harrison, David K. Jackson, Ian Johnston, Dominic Kwiatkowski, Cordelia Langford, John Sillitoe on behalf of the Wellcome Sanger Institute COVID-19 Surveillance Team |
| EPI_ISL_2999853, EPI_ISL_2999854 | Health Services Laboratories | Wellcome Sanger Institute for the COVID-19 Genomics UK (COG-UK) Consortium | Health Services Laboratories and Alex Alderton, Roberto Amato, Jeffrey Barrett, Sonia Goncalves, Ewan Harrison, David K. Jackson, Ian Johnston, Dominic Kwiatkowski, Cordelia Langford, John Sillitoe on behalf of the Wellcome Sanger Institute COVID-19 Surveillance Team |
| EPI_ISL_2999855 | Berkshire and Surrey Pathology Services Lighthouse Laboratory | Wellcome Sanger Institute for the COVID-19 Genomics UK (COG-UK) Consortium | Berkshire and Surrey Pathology Services Lighthouse Laboratory and Alex Alderton, Roberto Amato, Jeffrey Barrett, Sonia Goncalves, Ewan Harrison, David K. Jackson, Ian Johnston, Dominic Kwiatkowski, Cordelia Langford, John Sillitoe on behalf of the Wellcome Sanger Institute COVID-19 Surveillance Team |
| EPI_ISL_2999856 | Health Services Laboratories | Wellcome Sanger Institute for the COVID-19 Genomics UK (COG-UK) Consortium | Health Services Laboratories and Alex Alderton, Roberto Amato, Jeffrey Barrett, Sonia Goncalves, Ewan Harrison, David K. Jackson, Ian Johnston, Dominic Kwiatkowski, Cordelia Langford, John Sillitoe on behalf of the Wellcome Sanger Institute COVID-19 Surveillance Team |
| EPI_ISL_2999857, EPI_ISL_2999858, EPI_ISL_2999859 | Berkshire and Surrey Pathology Services Lighthouse Laboratory | Wellcome Sanger Institute for the COVID-19 Genomics UK (COG-UK) Consortium | Berkshire and Surrey Pathology Services Lighthouse Laboratory and Alex Alderton, Roberto Amato, Jeffrey Barrett, Sonia Goncalves, Ewan Harrison, David K. Jackson, Ian Johnston, Dominic Kwiatkowski, Cordelia Langford, John Sillitoe on behalf of the Wellcome Sanger Institute COVID-19 Surveillance Team |
| EPI_ISL_2999860, EPI_ISL_2999861, EPI_ISL_2999862 | Lighthouse Lab in Alderley Park | Wellcome Sanger Institute for the COVID-19 Genomics UK (COG-UK) Consortium | Jacquelyn Wynn, Mairead Hyland, The Lighthouse Lab in Alderley Park and Alex Alderton, Roberto Amato, Jeffrey Barrett, Sonia Goncalves, Ewan Harrison, David K. Jackson, Ian Johnston, Dominic Kwiatkowski, Cordelia Langford, John Sillitoe on behalf of the Wellcome Sanger Institute COVID-19 Surveillance Team |
| EPI_ISL_2999863 | Health Services Laboratories | Wellcome Sanger Institute for the COVID-19 Genomics UK (COG-UK) Consortium | Health Services Laboratories and Alex Alderton, Roberto Amato, Jeffrey Barrett, Sonia Goncalves, Ewan Harrison, David K. Jackson, Ian Johnston, Dominic Kwiatkowski, Cordelia Langford, John Sillitoe on behalf of the Wellcome Sanger Institute COVID-19 Surveillance Team |
| EPI_ISL_2999864 | Berkshire and Surrey Pathology Services Lighthouse Laboratory | Wellcome Sanger Institute for the COVID-19 Genomics UK (COG-UK) Consortium | Berkshire and Surrey Pathology Services Lighthouse Laboratory and Alex Alderton, Roberto Amato, Jeffrey Barrett, Sonia Goncalves, Ewan Harrison, David K. Jackson, Ian Johnston, Dominic Kwiatkowski, Cordelia Langford, John Sillitoe on behalf of the Wellcome Sanger Institute COVID-19 Surveillance Team |
| EPI_ISL_2999865, EPI_ISL_2999866, EPI_ISL_2999867 | Lighthouse Lab in Alderley Park | Wellcome Sanger Institute for the COVID-19 Genomics UK (COG-UK) Consortium | Jacquelyn Wynn, Mairead Hyland, The Lighthouse Lab in Alderley Park and Alex Alderton, Roberto Amato, Jeffrey Barrett, Sonia Goncalves, Ewan Harrison, David K. Jackson, Ian Johnston, Dominic Kwiatkowski, Cordelia Langford, John Sillitoe on behalf of the Wellcome Sanger Institute COVID-19 Surveillance Team |
| EPI_ISL_2999868 | Health Services Laboratories | Wellcome Sanger Institute for the COVID-19 Genomics UK (COG-UK) Consortium | Health Services Laboratories and Alex Alderton, Roberto Amato, Jeffrey Barrett, Sonia Goncalves, Ewan Harrison, David K. Jackson, Ian Johnston, Dominic Kwiatkowski, Cordelia Langford, John Sillitoe on behalf of the Wellcome Sanger Institute COVID-19 Surveillance Team |
| EPI_ISL_2999869 | Lighthouse Lab in Alderley Park | Wellcome Sanger Institute for the COVID-19 Genomics UK (COG-UK) Consortium | Jacquelyn Wynn, Mairead Hyland, The Lighthouse Lab in Alderley Park and Alex Alderton, Roberto Amato, Jeffrey Barrett, Sonia Goncalves, Ewan Harrison, David K. Jackson, Ian Johnston, Dominic Kwiatkowski, Cordelia Langford, John Sillitoe on behalf of the Wellcome Sanger Institute COVID-19 Surveillance Team |
| EPI_ISL_2999870 | Health Services Laboratories | Wellcome Sanger Institute for the COVID-19 Genomics UK (COG-UK) Consortium | Health Services Laboratories and Alex Alderton, Roberto Amato, Jeffrey Barrett, Sonia Goncalves, Ewan Harrison, David K. Jackson, Ian Johnston, Dominic Kwiatkowski, Cordelia Langford, John Sillitoe on behalf of the Wellcome Sanger Institute COVID-19 Surveillance Team |
| EPI_ISL_2999871, EPI_ISL_2999872 | Berkshire and Surrey Pathology Services Lighthouse Laboratory | Wellcome Sanger Institute for the COVID-19 Genomics UK (COG-UK) Consortium | Berkshire and Surrey Pathology Services Lighthouse Laboratory and Alex Alderton, Roberto Amato, Jeffrey Barrett, Sonia Goncalves, Ewan Harrison, David K. Jackson, Ian Johnston, Dominic Kwiatkowski, Cordelia Langford, John Sillitoe on behalf of the Wellcome Sanger Institute COVID-19 Surveillance Team |
| EPI_ISL_2999873 | Health Services Laboratories | Wellcome Sanger Institute for the COVID-19 Genomics UK (COG-UK) Consortium | Health Services Laboratories and Alex Alderton, Roberto Amato, Jeffrey Barrett, Sonia Goncalves, Ewan Harrison, David K. Jackson, Ian Johnston, Dominic Kwiatkowski, Cordelia Langford, John Sillitoe on behalf of the Wellcome Sanger Institute COVID-19 Surveillance Team |
| EPI_ISL_2999874, EPI_ISL_2999875, EPI_ISL_2999876 | Berkshire and Surrey Pathology Services Lighthouse Laboratory | Wellcome Sanger Institute for the COVID-19 Genomics UK (COG-UK) Consortium | Berkshire and Surrey Pathology Services Lighthouse Laboratory and Alex Alderton, Roberto Amato, Jeffrey Barrett, Sonia Goncalves, Ewan Harrison, David K. Jackson, Ian Johnston, Dominic Kwiatkowski, Cordelia Langford, John Sillitoe on behalf of the Wellcome Sanger Institute COVID-19 Surveillance Team |
| EPI_ISL_2999877 | Lighthouse Lab in Alderley Park | Wellcome Sanger Institute for the COVID-19 Genomics UK (COG-UK) Consortium | Jacquelyn Wynn, Mairead Hyland, The Lighthouse Lab in Alderley Park and Alex Alderton, Roberto Amato, Jeffrey Barrett, Sonia Goncalves, Ewan Harrison, David K. Jackson, Ian Johnston, Dominic Kwiatkowski, Cordelia Langford, John Sillitoe on behalf of the Wellcome Sanger Institute COVID-19 Surveillance Team |
| EPI_ISL_2999878 | Berkshire and Surrey Pathology Services Lighthouse Laboratory | Wellcome Sanger Institute for the COVID-19 Genomics UK (COG-UK) Consortium | Berkshire and Surrey Pathology Services Lighthouse Laboratory and Alex Alderton, Roberto Amato, Jeffrey Barrett, Sonia Goncalves, Ewan Harrison, David K. Jackson, Ian Johnston, Dominic Kwiatkowski, Cordelia Langford, John Sillitoe on behalf of the Wellcome Sanger Institute COVID-19 Surveillance Team |
| EPI_ISL_2999879, EPI_ISL_2999880, EPI_ISL_2999881 | Lighthouse Lab in Alderley Park | Wellcome Sanger Institute for the COVID-19 Genomics UK (COG-UK) Consortium | Jacquelyn Wynn, Mairead Hyland, The Lighthouse Lab in Alderley Park and Alex Alderton, Roberto Amato, Jeffrey Barrett, Sonia Goncalves, Ewan Harrison, David K. Jackson, Ian Johnston, Dominic Kwiatkowski, Cordelia Langford, John Sillitoe on behalf of the Wellcome Sanger Institute COVID-19 Surveillance Team |
| EPI_ISL_2999882 | Berkshire and Surrey Pathology Services Lighthouse Laboratory | Wellcome Sanger Institute for the COVID-19 Genomics UK (COG-UK) Consortium | Berkshire and Surrey Pathology Services Lighthouse Laboratory and Alex Alderton, Roberto Amato, Jeffrey Barrett, Sonia Goncalves, Ewan Harrison, David K. Jackson, Ian Johnston, Dominic Kwiatkowski, Cordelia Langford, John Sillitoe on behalf of the Wellcome Sanger Institute COVID-19 Surveillance Team |
| EPI_ISL_2999883 | Health Services Laboratories | Wellcome Sanger Institute for the COVID-19 Genomics UK (COG-UK) Consortium | Health Services Laboratories and Alex Alderton, Roberto Amato, Jeffrey Barrett, Sonia Goncalves, Ewan Harrison, David K. Jackson, Ian Johnston, Dominic Kwiatkowski, Cordelia Langford, John Sillitoe on behalf of the Wellcome Sanger Institute COVID-19 Surveillance Team |
| EPI_ISL_2999884, | Berkshire and Surrey Pathology Services Lighthouse Laboratory | Wellcome Sanger Institute for the COVID-19 Genomics UK (COG-UK) | Berkshire and Surrey Pathology Services Lighthouse Laboratory and Alex Alderton, Roberto Amato, Jeffrey Barrett, Sonia Goncalves, Ewan Harrison, David K. Jackson, Ian Johnston, Dominic Kwiatkowski, Cordelia Langford, John Sillitoe on behalf |

| | | | |
|---|---|---|---|
| EPI_ISL_2999885, EPI_ISL_2999886, EPI_ISL_2999887, EPI_ISL_2999888 | | Consortium | of the Wellcome Sanger Institute COVID-19 Surveillance Team |
| EPI_ISL_2999889 | Lighthouse Lab in Alderley Park | Wellcome Sanger Institute for the COVID-19 Genomics UK (COG-UK) Consortium | Jacquelyn Wynn, Mairead Hyland, The Lighthouse Lab in Alderley Park and Alex Alderton, Roberto Amato, Jeffrey Barrett, Sonia Goncalves, Ewan Harrison, David K. Jackson, Ian Johnston, Dominic Kwiatkowski, Cordelia Langford, John Sillitoe on behalf of the Wellcome Sanger Institute COVID-19 Surveillance Team |
| EPI_ISL_2999890 | Berkshire and Surrey Pathology Services Lighthouse Laboratory | Wellcome Sanger Institute for the COVID-19 Genomics UK (COG-UK) Consortium | Berkshire and Surrey Pathology Services Lighthouse Laboratory and Alex Alderton, Roberto Amato, Jeffrey Barrett, Sonia Goncalves, Ewan Harrison, David K. Jackson, Ian Johnston, Dominic Kwiatkowski, Cordelia Langford, John Sillitoe on behalf of the Wellcome Sanger Institute COVID-19 Surveillance Team |
| EPI_ISL_2999891 | Lighthouse Lab in Alderley Park | Wellcome Sanger Institute for the COVID-19 Genomics UK (COG-UK) Consortium | Jacquelyn Wynn, Mairead Hyland, The Lighthouse Lab in Alderley Park and Alex Alderton, Roberto Amato, Jeffrey Barrett, Sonia Goncalves, Ewan Harrison, David K. Jackson, Ian Johnston, Dominic Kwiatkowski, Cordelia Langford, John Sillitoe on behalf of the Wellcome Sanger Institute COVID-19 Surveillance Team |
| EPI_ISL_2999892, EPI_ISL_2999893 | Berkshire and Surrey Pathology Services Lighthouse Laboratory | Wellcome Sanger Institute for the COVID-19 Genomics UK (COG-UK) Consortium | Berkshire and Surrey Pathology Services Lighthouse Laboratory and Alex Alderton, Roberto Amato, Jeffrey Barrett, Sonia Goncalves, Ewan Harrison, David K. Jackson, Ian Johnston, Dominic Kwiatkowski, Cordelia Langford, John Sillitoe on behalf of the Wellcome Sanger Institute COVID-19 Surveillance Team |
| EPI_ISL_2999894 | Lighthouse Lab in Alderley Park | Wellcome Sanger Institute for the COVID-19 Genomics UK (COG-UK) Consortium | Jacquelyn Wynn, Mairead Hyland, The Lighthouse Lab in Alderley Park and Alex Alderton, Roberto Amato, Jeffrey Barrett, Sonia Goncalves, Ewan Harrison, David K. Jackson, Ian Johnston, Dominic Kwiatkowski, Cordelia Langford, John Sillitoe on behalf of the Wellcome Sanger Institute COVID-19 Surveillance Team |
| EPI_ISL_2999895, EPI_ISL_2999896, EPI_ISL_2999897 | Berkshire and Surrey Pathology Services Lighthouse Laboratory | Wellcome Sanger Institute for the COVID-19 Genomics UK (COG-UK) Consortium | Berkshire and Surrey Pathology Services Lighthouse Laboratory and Alex Alderton, Roberto Amato, Jeffrey Barrett, Sonia Goncalves, Ewan Harrison, David K. Jackson, Ian Johnston, Dominic Kwiatkowski, Cordelia Langford, John Sillitoe on behalf of the Wellcome Sanger Institute COVID-19 Surveillance Team |
| EPI_ISL_2999898 | Health Services Laboratories | Wellcome Sanger Institute for the COVID-19 Genomics UK (COG-UK) Consortium | Health Services Laboratories and Alex Alderton, Roberto Amato, Jeffrey Barrett, Sonia Goncalves, Ewan Harrison, David K. Jackson, Ian Johnston, Dominic Kwiatkowski, Cordelia Langford, John Sillitoe on behalf of the Wellcome Sanger Institute COVID-19 Surveillance Team |
| EPI_ISL_2999899, EPI_ISL_2999900, EPI_ISL_2999901, EPI_ISL_2999902 | Berkshire and Surrey Pathology Services Lighthouse Laboratory | Wellcome Sanger Institute for the COVID-19 Genomics UK (COG-UK) Consortium | Berkshire and Surrey Pathology Services Lighthouse Laboratory and Alex Alderton, Roberto Amato, Jeffrey Barrett, Sonia Goncalves, Ewan Harrison, David K. Jackson, Ian Johnston, Dominic Kwiatkowski, Cordelia Langford, John Sillitoe on behalf of the Wellcome Sanger Institute COVID-19 Surveillance Team |
| EPI_ISL_2999903 | Health Services Laboratories | Wellcome Sanger Institute for the COVID-19 Genomics UK (COG-UK) Consortium | Health Services Laboratories and Alex Alderton, Roberto Amato, Jeffrey Barrett, Sonia Goncalves, Ewan Harrison, David K. Jackson, Ian Johnston, Dominic Kwiatkowski, Cordelia Langford, John Sillitoe on behalf of the Wellcome Sanger Institute COVID-19 Surveillance Team |
| EPI_ISL_2999904, EPI_ISL_2999905 | Berkshire and Surrey Pathology Services Lighthouse Laboratory | Wellcome Sanger Institute for the COVID-19 Genomics UK (COG-UK) Consortium | Berkshire and Surrey Pathology Services Lighthouse Laboratory and Alex Alderton, Roberto Amato, Jeffrey Barrett, Sonia Goncalves, Ewan Harrison, David K. Jackson, Ian Johnston, Dominic Kwiatkowski, Cordelia Langford, John Sillitoe on behalf of the Wellcome Sanger Institute COVID-19 Surveillance Team |
| EPI_ISL_2999906 | Health Services Laboratories | Wellcome Sanger Institute for the COVID-19 Genomics UK (COG-UK) Consortium | Health Services Laboratories and Alex Alderton, Roberto Amato, Jeffrey Barrett, Sonia Goncalves, Ewan Harrison, David K. Jackson, Ian Johnston, Dominic Kwiatkowski, Cordelia Langford, John Sillitoe on behalf of the Wellcome Sanger Institute COVID-19 Surveillance Team |
| EPI_ISL_2999907, EPI_ISL_2999908 | Berkshire and Surrey Pathology Services Lighthouse Laboratory | Wellcome Sanger Institute for the COVID-19 Genomics UK (COG-UK) Consortium | Berkshire and Surrey Pathology Services Lighthouse Laboratory and Alex Alderton, Roberto Amato, Jeffrey Barrett, Sonia Goncalves, Ewan Harrison, David K. Jackson, Ian Johnston, Dominic Kwiatkowski, Cordelia Langford, John Sillitoe on behalf of the Wellcome Sanger Institute COVID-19 Surveillance Team |
| EPI_ISL_2999909 | Lighthouse Lab in Alderley Park | Wellcome Sanger Institute for the COVID-19 Genomics UK (COG-UK) Consortium | Jacquelyn Wynn, Mairead Hyland, The Lighthouse Lab in Alderley Park and Alex Alderton, Roberto Amato, Jeffrey Barrett, Sonia Goncalves, Ewan Harrison, David K. Jackson, Ian Johnston, Dominic Kwiatkowski, Cordelia Langford, John Sillitoe on behalf of the Wellcome Sanger Institute COVID-19 Surveillance Team |
| EPI_ISL_2999910 | Berkshire and Surrey Pathology Services Lighthouse Laboratory | Wellcome Sanger Institute for the COVID-19 Genomics UK (COG-UK) Consortium | Berkshire and Surrey Pathology Services Lighthouse Laboratory and Alex Alderton, Roberto Amato, Jeffrey Barrett, Sonia Goncalves, Ewan Harrison, David K. Jackson, Ian Johnston, Dominic Kwiatkowski, Cordelia Langford, John Sillitoe on behalf of the Wellcome Sanger Institute COVID-19 Surveillance Team |
| EPI_ISL_2999911 | Lighthouse Lab in Alderley Park | Wellcome Sanger Institute for the COVID-19 Genomics UK (COG-UK) Consortium | Jacquelyn Wynn, Mairead Hyland, The Lighthouse Lab in Alderley Park and Alex Alderton, Roberto Amato, Jeffrey Barrett, Sonia Goncalves, Ewan Harrison, David K. Jackson, Ian Johnston, Dominic Kwiatkowski, Cordelia Langford, John Sillitoe on behalf of the Wellcome Sanger Institute COVID-19 Surveillance Team |
| EPI_ISL_2999912, EPI_ISL_2999913 | Berkshire and Surrey Pathology Services Lighthouse Laboratory | Wellcome Sanger Institute for the COVID-19 Genomics UK (COG-UK) Consortium | Berkshire and Surrey Pathology Services Lighthouse Laboratory and Alex Alderton, Roberto Amato, Jeffrey Barrett, Sonia Goncalves, Ewan Harrison, David K. Jackson, Ian Johnston, Dominic Kwiatkowski, Cordelia Langford, John Sillitoe on behalf of the Wellcome Sanger Institute COVID-19 Surveillance Team |
| EPI_ISL_2999914 | Health Services Laboratories | Wellcome Sanger Institute for the COVID-19 Genomics UK (COG-UK) Consortium | Health Services Laboratories and Alex Alderton, Roberto Amato, Jeffrey Barrett, Sonia Goncalves, Ewan Harrison, David K. Jackson, Ian Johnston, Dominic Kwiatkowski, Cordelia Langford, John Sillitoe on behalf of the Wellcome Sanger Institute COVID-19 Surveillance Team |
| EPI_ISL_2999915 | Berkshire and Surrey Pathology Services Lighthouse Laboratory | Wellcome Sanger Institute for the COVID-19 Genomics UK (COG-UK) Consortium | Berkshire and Surrey Pathology Services Lighthouse Laboratory and Alex Alderton, Roberto Amato, Jeffrey Barrett, Sonia Goncalves, Ewan Harrison, David K. Jackson, Ian Johnston, Dominic Kwiatkowski, Cordelia Langford, John Sillitoe on behalf of the Wellcome Sanger Institute COVID-19 Surveillance Team |
| EPI_ISL_2999916 | Lighthouse Lab in Alderley Park | Wellcome Sanger Institute for the COVID-19 Genomics UK (COG-UK) Consortium | Jacquelyn Wynn, Mairead Hyland, The Lighthouse Lab in Alderley Park and Alex Alderton, Roberto Amato, Jeffrey Barrett, Sonia Goncalves, Ewan Harrison, David K. Jackson, Ian Johnston, Dominic Kwiatkowski, Cordelia Langford, John Sillitoe on behalf of the Wellcome Sanger Institute COVID-19 Surveillance Team |
| EPI_ISL_2999917 | Berkshire and Surrey Pathology Services Lighthouse Laboratory | Wellcome Sanger Institute for the COVID-19 Genomics UK (COG-UK) Consortium | Berkshire and Surrey Pathology Services Lighthouse Laboratory and Alex Alderton, Roberto Amato, Jeffrey Barrett, Sonia Goncalves, Ewan Harrison, David K. Jackson, Ian Johnston, Dominic Kwiatkowski, Cordelia Langford, John Sillitoe on behalf of the Wellcome Sanger Institute COVID-19 Surveillance Team |
| EPI_ISL_2999918, EPI_ISL_2999919 | Health Services Laboratories | Wellcome Sanger Institute for the COVID-19 Genomics UK (COG-UK) Consortium | Health Services Laboratories and Alex Alderton, Roberto Amato, Jeffrey Barrett, Sonia Goncalves, Ewan Harrison, David K. Jackson, Ian Johnston, Dominic Kwiatkowski, Cordelia Langford, John Sillitoe on behalf of the Wellcome Sanger Institute COVID-19 Surveillance Team |
| EPI_ISL_2999920 | Lighthouse Lab in Alderley Park | Wellcome Sanger Institute for the COVID-19 Genomics UK (COG-UK) Consortium | Jacquelyn Wynn, Mairead Hyland, The Lighthouse Lab in Alderley Park and Alex Alderton, Roberto Amato, Jeffrey Barrett, Sonia Goncalves, Ewan Harrison, David K. Jackson, Ian Johnston, Dominic Kwiatkowski, Cordelia Langford, John Sillitoe on behalf of the Wellcome Sanger Institute COVID-19 Surveillance Team |
| EPI_ISL_2999921, EPI_ISL_2999922 | Health Services Laboratories | Wellcome Sanger Institute for the COVID-19 Genomics UK (COG-UK) Consortium | Health Services Laboratories and Alex Alderton, Roberto Amato, Jeffrey Barrett, Sonia Goncalves, Ewan Harrison, David K. Jackson, Ian Johnston, Dominic Kwiatkowski, Cordelia Langford, John Sillitoe on behalf of the Wellcome Sanger Institute COVID-19 Surveillance Team |
| EPI_ISL_2999923, EPI_ISL_2999924 | Berkshire and Surrey Pathology Services Lighthouse Laboratory | Wellcome Sanger Institute for the COVID-19 Genomics UK (COG-UK) Consortium | Berkshire and Surrey Pathology Services Lighthouse Laboratory and Alex Alderton, Roberto Amato, Jeffrey Barrett, Sonia Goncalves, Ewan Harrison, David K. Jackson, Ian Johnston, Dominic Kwiatkowski, Cordelia Langford, John Sillitoe on behalf of the Wellcome Sanger Institute COVID-19 Surveillance Team |
| EPI_ISL_2999925 | Health Services Laboratories | Wellcome Sanger Institute for the COVID-19 Genomics UK (COG-UK) Consortium | Health Services Laboratories and Alex Alderton, Roberto Amato, Jeffrey Barrett, Sonia Goncalves, Ewan Harrison, David K. Jackson, Ian Johnston, Dominic Kwiatkowski, Cordelia Langford, John Sillitoe on behalf of the Wellcome Sanger Institute COVID-19 Surveillance Team |
| EPI_ISL_2999926, EPI_ISL_2999928 | Berkshire and Surrey Pathology Services Lighthouse Laboratory | Wellcome Sanger Institute for the COVID-19 Genomics UK (COG-UK) Consortium | Berkshire and Surrey Pathology Services Lighthouse Laboratory and Alex Alderton, Roberto Amato, Jeffrey Barrett, Sonia Goncalves, Ewan Harrison, David K. Jackson, Ian Johnston, Dominic Kwiatkowski, Cordelia Langford, John Sillitoe on behalf of the Wellcome Sanger Institute COVID-19 Surveillance Team |
| EPI_ISL_2999929 | Health Services Laboratories | Wellcome Sanger Institute for the COVID-19 Genomics UK (COG-UK) Consortium | Health Services Laboratories and Alex Alderton, Roberto Amato, Jeffrey Barrett, Sonia Goncalves, Ewan Harrison, David K. Jackson, Ian Johnston, Dominic Kwiatkowski, Cordelia Langford, John Sillitoe on behalf of the Wellcome Sanger Institute COVID-19 Surveillance Team |
| EPI_ISL_2999930 | Berkshire and Surrey Pathology Services Lighthouse Laboratory | Wellcome Sanger Institute for the COVID-19 Genomics UK (COG-UK) Consortium | Berkshire and Surrey Pathology Services Lighthouse Laboratory and Alex Alderton, Roberto Amato, Jeffrey Barrett, Sonia Goncalves, Ewan Harrison, David K. Jackson, Ian Johnston, Dominic Kwiatkowski, Cordelia Langford, John Sillitoe on behalf of the Wellcome Sanger Institute COVID-19 Surveillance Team |
| EPI_ISL_2999931 | Lighthouse Lab in Alderley Park | Wellcome Sanger Institute for the COVID-19 Genomics UK (COG-UK) Consortium | Jacquelyn Wynn, Mairead Hyland, The Lighthouse Lab in Alderley Park and Alex Alderton, Roberto Amato, Jeffrey Barrett, Sonia Goncalves, Ewan Harrison, David K. Jackson, Ian Johnston, Dominic Kwiatkowski, Cordelia Langford, John Sillitoe on behalf of the Wellcome Sanger Institute COVID-19 Surveillance Team |
| EPI_ISL_2999932 | Berkshire and Surrey Pathology Services Lighthouse Laboratory | Wellcome Sanger Institute for the COVID-19 Genomics UK (COG-UK) Consortium | Berkshire and Surrey Pathology Services Lighthouse Laboratory and Alex Alderton, Roberto Amato, Jeffrey Barrett, Sonia Goncalves, Ewan Harrison, David K. Jackson, Ian Johnston, Dominic Kwiatkowski, Cordelia Langford, John Sillitoe on behalf of the Wellcome Sanger Institute COVID-19 Surveillance Team |
| EPI_ISL_2999933 | Lighthouse Lab in Alderley Park | Wellcome Sanger Institute for the COVID-19 Genomics UK (COG-UK) Consortium | Jacquelyn Wynn, Mairead Hyland, The Lighthouse Lab in Alderley Park and Alex Alderton, Roberto Amato, Jeffrey Barrett, Sonia Goncalves, Ewan Harrison, David K. Jackson, Ian Johnston, Dominic Kwiatkowski, Cordelia Langford, John Sillitoe on behalf of the Wellcome Sanger Institute COVID-19 Surveillance Team |
| EPI_ISL_2999934, EPI_ISL_2999935 | Berkshire and Surrey Pathology Services Lighthouse Laboratory | Wellcome Sanger Institute for the COVID-19 Genomics UK (COG-UK) Consortium | Berkshire and Surrey Pathology Services Lighthouse Laboratory and Alex Alderton, Roberto Amato, Jeffrey Barrett, Sonia Goncalves, Ewan Harrison, David K. Jackson, Ian Johnston, Dominic Kwiatkowski, Cordelia Langford, John Sillitoe on behalf of the Wellcome Sanger Institute COVID-19 Surveillance Team |
| EPI_ISL_2999936 | Health Services Laboratories | Wellcome Sanger Institute for the COVID-19 Genomics UK (COG-UK) Consortium | Health Services Laboratories and Alex Alderton, Roberto Amato, Jeffrey Barrett, Sonia Goncalves, Ewan Harrison, David K. Jackson, Ian Johnston, Dominic Kwiatkowski, Cordelia Langford, John Sillitoe on behalf of the Wellcome Sanger Institute COVID-19 Surveillance Team |
| EPI_ISL_2999937 | Berkshire and Surrey Pathology Services Lighthouse Laboratory | Wellcome Sanger Institute for the COVID-19 Genomics UK (COG-UK) Consortium | Berkshire and Surrey Pathology Services Lighthouse Laboratory and Alex Alderton, Roberto Amato, Jeffrey Barrett, Sonia Goncalves, Ewan Harrison, David K. Jackson, Ian Johnston, Dominic Kwiatkowski, Cordelia Langford, John Sillitoe on behalf of the Wellcome Sanger Institute COVID-19 Surveillance Team |
| EPI_ISL_2999938 | Lighthouse Lab in Alderley Park | Wellcome Sanger Institute for the COVID-19 Genomics UK (COG-UK) Consortium | Jacquelyn Wynn, Mairead Hyland, The Lighthouse Lab in Alderley Park and Alex Alderton, Roberto Amato, Jeffrey Barrett, Sonia Goncalves, Ewan Harrison, David K. Jackson, Ian Johnston, Dominic Kwiatkowski, Cordelia Langford, John Sillitoe on behalf of the Wellcome Sanger Institute COVID-19 Surveillance Team |
| EPI_ISL_2999939 | Berkshire and Surrey Pathology Services Lighthouse Laboratory | Wellcome Sanger Institute for the COVID-19 Genomics UK (COG-UK) Consortium | Berkshire and Surrey Pathology Services Lighthouse Laboratory and Alex Alderton, Roberto Amato, Jeffrey Barrett, Sonia Goncalves, Ewan Harrison, David K. Jackson, Ian Johnston, Dominic Kwiatkowski, Cordelia Langford, John Sillitoe on behalf of the Wellcome Sanger Institute COVID-19 Surveillance Team |
| EPI_ISL_2999940 | Lighthouse Lab in Alderley Park | Wellcome Sanger Institute for the COVID-19 Genomics UK (COG-UK) Consortium | Jacquelyn Wynn, Mairead Hyland, The Lighthouse Lab in Alderley Park and Alex Alderton, Roberto Amato, Jeffrey Barrett, Sonia Goncalves, Ewan Harrison, David K. Jackson, Ian Johnston, Dominic Kwiatkowski, Cordelia Langford, John Sillitoe on behalf of the Wellcome Sanger Institute COVID-19 Surveillance Team |
| EPI_ISL_2999941 | Berkshire and Surrey Pathology Services Lighthouse Laboratory | Wellcome Sanger Institute for the COVID-19 Genomics UK (COG-UK) Consortium | Berkshire and Surrey Pathology Services Lighthouse Laboratory and Alex Alderton, Roberto Amato, Jeffrey Barrett, Sonia Goncalves, Ewan Harrison, David K. Jackson, Ian Johnston, Dominic Kwiatkowski, Cordelia Langford, John Sillitoe on behalf of the Wellcome Sanger Institute COVID-19 Surveillance Team |
| EPI_ISL_2999942 | Lighthouse Lab in Alderley Park | Wellcome Sanger Institute for the COVID-19 Genomics UK (COG-UK) Consortium | Jacquelyn Wynn, Mairead Hyland, The Lighthouse Lab in Alderley Park and Alex Alderton, Roberto Amato, Jeffrey Barrett, Sonia Goncalves, Ewan Harrison, David K. Jackson, Ian Johnston, Dominic Kwiatkowski, Cordelia Langford, John Sillitoe on behalf of the Wellcome Sanger Institute COVID-19 Surveillance Team |
| EPI_ISL_2999943, EPI_ISL_2999944, EPI_ISL_2999945 | Berkshire and Surrey Pathology Services Lighthouse Laboratory | Wellcome Sanger Institute for the COVID-19 Genomics UK (COG-UK) Consortium | Berkshire and Surrey Pathology Services Lighthouse Laboratory and Alex Alderton, Roberto Amato, Jeffrey Barrett, Sonia Goncalves, Ewan Harrison, David K. Jackson, Ian Johnston, Dominic Kwiatkowski, Cordelia Langford, John Sillitoe on behalf of the Wellcome Sanger Institute COVID-19 Surveillance Team |
| EPI_ISL_2999946, EPI_ISL_2999947, EPI_ISL_2999948 | Lighthouse Lab in Alderley Park | Wellcome Sanger Institute for the COVID-19 Genomics UK (COG-UK) Consortium | Jacquelyn Wynn, Mairead Hyland, The Lighthouse Lab in Alderley Park and Alex Alderton, Roberto Amato, Jeffrey Barrett, Sonia Goncalves, Ewan Harrison, David K. Jackson, Ian Johnston, Dominic Kwiatkowski, Cordelia Langford, John Sillitoe on behalf of the Wellcome Sanger Institute COVID-19 Surveillance Team |

| | | | |
|---|---|---|---|
| EPI_ISL_2999949, EPI_ISL_2999950, EPI_ISL_2999951, EPI_ISL_2999952 | Berkshire and Surrey Pathology Services Lighthouse Laboratory | Wellcome Sanger Institute for the COVID-19 Genomics UK (COG-UK) Consortium | Berkshire and Surrey Pathology Services Lighthouse Laboratory and Alex Alderton, Roberto Amato, Jeffrey Barrett, Sonia Goncalves, Ewan Harrison, David K. Jackson, Ian Johnston, Dominic Kwiatkowski, Cordelia Langford, John Sillitoe on behalf of the Wellcome Sanger Institute COVID-19 Surveillance Team |
| EPI_ISL_2999953 | Lighthouse Lab in Alderley Park | Wellcome Sanger Institute for the COVID-19 Genomics UK (COG-UK) Consortium | Jacquelyn Wynn, Mairead Hyland, The Lighthouse Lab in Alderley Park and Alex Alderton, Roberto Amato, Jeffrey Barrett, Sonia Goncalves, Ewan Harrison, David K. Jackson, Ian Johnston, Dominic Kwiatkowski, Cordelia Langford, John Sillitoe on behalf of the Wellcome Sanger Institute COVID-19 Surveillance Team |
| EPI_ISL_2999954 | Health Services Laboratories | Wellcome Sanger Institute for the COVID-19 Genomics UK (COG-UK) Consortium | Health Services Laboratories and Alex Alderton, Roberto Amato, Jeffrey Barrett, Sonia Goncalves, Ewan Harrison, David K. Jackson, Ian Johnston, Dominic Kwiatkowski, Cordelia Langford, John Sillitoe on behalf of the Wellcome Sanger Institute COVID-19 Surveillance Team |
| EPI_ISL_2999955 | Berkshire and Surrey Pathology Services Lighthouse Laboratory | Wellcome Sanger Institute for the COVID-19 Genomics UK (COG-UK) Consortium | Berkshire and Surrey Pathology Services Lighthouse Laboratory and Alex Alderton, Roberto Amato, Jeffrey Barrett, Sonia Goncalves, Ewan Harrison, David K. Jackson, Ian Johnston, Dominic Kwiatkowski, Cordelia Langford, John Sillitoe on behalf of the Wellcome Sanger Institute COVID-19 Surveillance Team |
| EPI_ISL_2999956 | Lighthouse Lab in Alderley Park | Wellcome Sanger Institute for the COVID-19 Genomics UK (COG-UK) Consortium | Jacquelyn Wynn, Mairead Hyland, The Lighthouse Lab in Alderley Park and Alex Alderton, Roberto Amato, Jeffrey Barrett, Sonia Goncalves, Ewan Harrison, David K. Jackson, Ian Johnston, Dominic Kwiatkowski, Cordelia Langford, John Sillitoe on behalf of the Wellcome Sanger Institute COVID-19 Surveillance Team |
| EPI_ISL_2999957, EPI_ISL_2999958 | Health Services Laboratories | Wellcome Sanger Institute for the COVID-19 Genomics UK (COG-UK) Consortium | Health Services Laboratories and Alex Alderton, Roberto Amato, Jeffrey Barrett, Sonia Goncalves, Ewan Harrison, David K. Jackson, Ian Johnston, Dominic Kwiatkowski, Cordelia Langford, John Sillitoe on behalf of the Wellcome Sanger Institute COVID-19 Surveillance Team |
| EPI_ISL_2999959, EPI_ISL_2999960 | Berkshire and Surrey Pathology Services Lighthouse Laboratory | Wellcome Sanger Institute for the COVID-19 Genomics UK (COG-UK) Consortium | Berkshire and Surrey Pathology Services Lighthouse Laboratory and Alex Alderton, Roberto Amato, Jeffrey Barrett, Sonia Goncalves, Ewan Harrison, David K. Jackson, Ian Johnston, Dominic Kwiatkowski, Cordelia Langford, John Sillitoe on behalf of the Wellcome Sanger Institute COVID-19 Surveillance Team |
| EPI_ISL_2999961 | Health Services Laboratories | Wellcome Sanger Institute for the COVID-19 Genomics UK (COG-UK) Consortium | Health Services Laboratories and Alex Alderton, Roberto Amato, Jeffrey Barrett, Sonia Goncalves, Ewan Harrison, David K. Jackson, Ian Johnston, Dominic Kwiatkowski, Cordelia Langford, John Sillitoe on behalf of the Wellcome Sanger Institute COVID-19 Surveillance Team |
| EPI_ISL_2999962, EPI_ISL_2999963, EPI_ISL_2999964 | Berkshire and Surrey Pathology Services Lighthouse Laboratory | Wellcome Sanger Institute for the COVID-19 Genomics UK (COG-UK) Consortium | Berkshire and Surrey Pathology Services Lighthouse Laboratory and Alex Alderton, Roberto Amato, Jeffrey Barrett, Sonia Goncalves, Ewan Harrison, David K. Jackson, Ian Johnston, Dominic Kwiatkowski, Cordelia Langford, John Sillitoe on behalf of the Wellcome Sanger Institute COVID-19 Surveillance Team |
| EPI_ISL_2999965 | Health Services Laboratories | Wellcome Sanger Institute for the COVID-19 Genomics UK (COG-UK) Consortium | Health Services Laboratories and Alex Alderton, Roberto Amato, Jeffrey Barrett, Sonia Goncalves, Ewan Harrison, David K. Jackson, Ian Johnston, Dominic Kwiatkowski, Cordelia Langford, John Sillitoe on behalf of the Wellcome Sanger Institute COVID-19 Surveillance Team |
| EPI_ISL_2999966 | Lighthouse Lab in Alderley Park | Wellcome Sanger Institute for the COVID-19 Genomics UK (COG-UK) Consortium | Jacquelyn Wynn, Mairead Hyland, The Lighthouse Lab in Alderley Park and Alex Alderton, Roberto Amato, Jeffrey Barrett, Sonia Goncalves, Ewan Harrison, David K. Jackson, Ian Johnston, Dominic Kwiatkowski, Cordelia Langford, John Sillitoe on behalf of the Wellcome Sanger Institute COVID-19 Surveillance Team |
| EPI_ISL_2999967, EPI_ISL_2999968 | Berkshire and Surrey Pathology Services Lighthouse Laboratory | Wellcome Sanger Institute for the COVID-19 Genomics UK (COG-UK) Consortium | Berkshire and Surrey Pathology Services Lighthouse Laboratory and Alex Alderton, Roberto Amato, Jeffrey Barrett, Sonia Goncalves, Ewan Harrison, David K. Jackson, Ian Johnston, Dominic Kwiatkowski, Cordelia Langford, John Sillitoe on behalf of the Wellcome Sanger Institute COVID-19 Surveillance Team |
| EPI_ISL_2999969 | Health Services Laboratories | Wellcome Sanger Institute for the COVID-19 Genomics UK (COG-UK) Consortium | Health Services Laboratories and Alex Alderton, Roberto Amato, Jeffrey Barrett, Sonia Goncalves, Ewan Harrison, David K. Jackson, Ian Johnston, Dominic Kwiatkowski, Cordelia Langford, John Sillitoe on behalf of the Wellcome Sanger Institute COVID-19 Surveillance Team |
| EPI_ISL_2999970, EPI_ISL_2999971 | Berkshire and Surrey Pathology Services Lighthouse Laboratory | Wellcome Sanger Institute for the COVID-19 Genomics UK (COG-UK) Consortium | Berkshire and Surrey Pathology Services Lighthouse Laboratory and Alex Alderton, Roberto Amato, Jeffrey Barrett, Sonia Goncalves, Ewan Harrison, David K. Jackson, Ian Johnston, Dominic Kwiatkowski, Cordelia Langford, John Sillitoe on behalf of the Wellcome Sanger Institute COVID-19 Surveillance Team |
| EPI_ISL_2999972, EPI_ISL_2999973, EPI_ISL_2999974 | Lighthouse Lab in Alderley Park | Wellcome Sanger Institute for the COVID-19 Genomics UK (COG-UK) Consortium | Jacquelyn Wynn, Mairead Hyland, The Lighthouse Lab in Alderley Park and Alex Alderton, Roberto Amato, Jeffrey Barrett, Sonia Goncalves, Ewan Harrison, David K. Jackson, Ian Johnston, Dominic Kwiatkowski, Cordelia Langford, John Sillitoe on behalf of the Wellcome Sanger Institute COVID-19 Surveillance Team |
| EPI_ISL_2999975, EPI_ISL_2999976 | Berkshire and Surrey Pathology Services Lighthouse Laboratory | Wellcome Sanger Institute for the COVID-19 Genomics UK (COG-UK) Consortium | Berkshire and Surrey Pathology Services Lighthouse Laboratory and Alex Alderton, Roberto Amato, Jeffrey Barrett, Sonia Goncalves, Ewan Harrison, David K. Jackson, Ian Johnston, Dominic Kwiatkowski, Cordelia Langford, John Sillitoe on behalf of the Wellcome Sanger Institute COVID-19 Surveillance Team |
| EPI_ISL_2999977, EPI_ISL_2999978, EPI_ISL_2999979 | Health Services Laboratories | Wellcome Sanger Institute for the COVID-19 Genomics UK (COG-UK) Consortium | Health Services Laboratories and Alex Alderton, Roberto Amato, Jeffrey Barrett, Sonia Goncalves, Ewan Harrison, David K. Jackson, Ian Johnston, Dominic Kwiatkowski, Cordelia Langford, John Sillitoe on behalf of the Wellcome Sanger Institute COVID-19 Surveillance Team |
| EPI_ISL_2999980, EPI_ISL_2999981 | Lighthouse Lab in Alderley Park | Wellcome Sanger Institute for the COVID-19 Genomics UK (COG-UK) Consortium | Jacquelyn Wynn, Mairead Hyland, The Lighthouse Lab in Alderley Park and Alex Alderton, Roberto Amato, Jeffrey Barrett, Sonia Goncalves, Ewan Harrison, David K. Jackson, Ian Johnston, Dominic Kwiatkowski, Cordelia Langford, John Sillitoe on behalf of the Wellcome Sanger Institute COVID-19 Surveillance Team |
| EPI_ISL_2999982 | Health Services Laboratories | Wellcome Sanger Institute for the COVID-19 Genomics UK (COG-UK) Consortium | Health Services Laboratories and Alex Alderton, Roberto Amato, Jeffrey Barrett, Sonia Goncalves, Ewan Harrison, David K. Jackson, Ian Johnston, Dominic Kwiatkowski, Cordelia Langford, John Sillitoe on behalf of the Wellcome Sanger Institute COVID-19 Surveillance Team |
| EPI_ISL_2999983 | Berkshire and Surrey Pathology Services Lighthouse Laboratory | Wellcome Sanger Institute for the COVID-19 Genomics UK (COG-UK) Consortium | Berkshire and Surrey Pathology Services Lighthouse Laboratory and Alex Alderton, Roberto Amato, Jeffrey Barrett, Sonia Goncalves, Ewan Harrison, David K. Jackson, Ian Johnston, Dominic Kwiatkowski, Cordelia Langford, John Sillitoe on behalf of the Wellcome Sanger Institute COVID-19 Surveillance Team |
| EPI_ISL_2999984 | Health Services Laboratories | Wellcome Sanger Institute for the COVID-19 Genomics UK (COG-UK) Consortium | Health Services Laboratories and Alex Alderton, Roberto Amato, Jeffrey Barrett, Sonia Goncalves, Ewan Harrison, David K. Jackson, Ian Johnston, Dominic Kwiatkowski, Cordelia Langford, John Sillitoe on behalf of the Wellcome Sanger Institute COVID-19 Surveillance Team |
| EPI_ISL_2999985, EPI_ISL_2999986 | Berkshire and Surrey Pathology Services Lighthouse Laboratory | Wellcome Sanger Institute for the COVID-19 Genomics UK (COG-UK) Consortium | Berkshire and Surrey Pathology Services Lighthouse Laboratory and Alex Alderton, Roberto Amato, Jeffrey Barrett, Sonia Goncalves, Ewan Harrison, David K. Jackson, Ian Johnston, Dominic Kwiatkowski, Cordelia Langford, John Sillitoe on behalf of the Wellcome Sanger Institute COVID-19 Surveillance Team |
| EPI_ISL_2999987 | Health Services Laboratories | Wellcome Sanger Institute for the COVID-19 Genomics UK (COG-UK) Consortium | Health Services Laboratories and Alex Alderton, Roberto Amato, Jeffrey Barrett, Sonia Goncalves, Ewan Harrison, David K. Jackson, Ian Johnston, Dominic Kwiatkowski, Cordelia Langford, John Sillitoe on behalf of the Wellcome Sanger Institute COVID-19 Surveillance Team |
| EPI_ISL_2999988 | Lighthouse Lab in Alderley Park | Wellcome Sanger Institute for the COVID-19 Genomics UK (COG-UK) Consortium | Jacquelyn Wynn, Mairead Hyland, The Lighthouse Lab in Alderley Park and Alex Alderton, Roberto Amato, Jeffrey Barrett, Sonia Goncalves, Ewan Harrison, David K. Jackson, Ian Johnston, Dominic Kwiatkowski, Cordelia Langford, John Sillitoe on behalf of the Wellcome Sanger Institute COVID-19 Surveillance Team |
| EPI_ISL_2999989 | Health Services Laboratories | Wellcome Sanger Institute for the COVID-19 Genomics UK (COG-UK) Consortium | Health Services Laboratories and Alex Alderton, Roberto Amato, Jeffrey Barrett, Sonia Goncalves, Ewan Harrison, David K. Jackson, Ian Johnston, Dominic Kwiatkowski, Cordelia Langford, John Sillitoe on behalf of the Wellcome Sanger Institute COVID-19 Surveillance Team |
| EPI_ISL_2999990 | Berkshire and Surrey Pathology Services Lighthouse Laboratory | Wellcome Sanger Institute for the COVID-19 Genomics UK (COG-UK) Consortium | Berkshire and Surrey Pathology Services Lighthouse Laboratory and Alex Alderton, Roberto Amato, Jeffrey Barrett, Sonia Goncalves, Ewan Harrison, David K. Jackson, Ian Johnston, Dominic Kwiatkowski, Cordelia Langford, John Sillitoe on behalf of the Wellcome Sanger Institute COVID-19 Surveillance Team |
| EPI_ISL_2999991 | Lighthouse Lab in Alderley Park | Wellcome Sanger Institute for the COVID-19 Genomics UK (COG-UK) Consortium | Jacquelyn Wynn, Mairead Hyland, The Lighthouse Lab in Alderley Park and Alex Alderton, Roberto Amato, Jeffrey Barrett, Sonia Goncalves, Ewan Harrison, David K. Jackson, Ian Johnston, Dominic Kwiatkowski, Cordelia Langford, John Sillitoe on behalf of the Wellcome Sanger Institute COVID-19 Surveillance Team |
| EPI_ISL_2999992 | Berkshire and Surrey Pathology Services Lighthouse Laboratory | Wellcome Sanger Institute for the COVID-19 Genomics UK (COG-UK) Consortium | Berkshire and Surrey Pathology Services Lighthouse Laboratory and Alex Alderton, Roberto Amato, Jeffrey Barrett, Sonia Goncalves, Ewan Harrison, David K. Jackson, Ian Johnston, Dominic Kwiatkowski, Cordelia Langford, John Sillitoe on behalf of the Wellcome Sanger Institute COVID-19 Surveillance Team |
| EPI_ISL_2999993 | Lighthouse Lab in Alderley Park | Wellcome Sanger Institute for the COVID-19 Genomics UK (COG-UK) Consortium | Jacquelyn Wynn, Mairead Hyland, The Lighthouse Lab in Alderley Park and Alex Alderton, Roberto Amato, Jeffrey Barrett, Sonia Goncalves, Ewan Harrison, David K. Jackson, Ian Johnston, Dominic Kwiatkowski, Cordelia Langford, John Sillitoe on behalf of the Wellcome Sanger Institute COVID-19 Surveillance Team |
| EPI_ISL_2999994 | Berkshire and Surrey Pathology Services Lighthouse Laboratory | Wellcome Sanger Institute for the COVID-19 Genomics UK (COG-UK) Consortium | Berkshire and Surrey Pathology Services Lighthouse Laboratory and Alex Alderton, Roberto Amato, Jeffrey Barrett, Sonia Goncalves, Ewan Harrison, David K. Jackson, Ian Johnston, Dominic Kwiatkowski, Cordelia Langford, John Sillitoe on behalf of the Wellcome Sanger Institute COVID-19 Surveillance Team |
| EPI_ISL_2999995 | Health Services Laboratories | Wellcome Sanger Institute for the COVID-19 Genomics UK (COG-UK) Consortium | Health Services Laboratories and Alex Alderton, Roberto Amato, Jeffrey Barrett, Sonia Goncalves, Ewan Harrison, David K. Jackson, Ian Johnston, Dominic Kwiatkowski, Cordelia Langford, John Sillitoe on behalf of the Wellcome Sanger Institute COVID-19 Surveillance Team |
| EPI_ISL_2999996, EPI_ISL_2999997 | Berkshire and Surrey Pathology Services Lighthouse Laboratory | Wellcome Sanger Institute for the COVID-19 Genomics UK (COG-UK) Consortium | Berkshire and Surrey Pathology Services Lighthouse Laboratory and Alex Alderton, Roberto Amato, Jeffrey Barrett, Sonia Goncalves, Ewan Harrison, David K. Jackson, Ian Johnston, Dominic Kwiatkowski, Cordelia Langford, John Sillitoe on behalf of the Wellcome Sanger Institute COVID-19 Surveillance Team |
| EPI_ISL_2999998 | Lighthouse Lab in Alderley Park | Wellcome Sanger Institute for the COVID-19 Genomics UK (COG-UK) Consortium | Jacquelyn Wynn, Mairead Hyland, The Lighthouse Lab in Alderley Park and Alex Alderton, Roberto Amato, Jeffrey Barrett, Sonia Goncalves, Ewan Harrison, David K. Jackson, Ian Johnston, Dominic Kwiatkowski, Cordelia Langford, John Sillitoe on behalf of the Wellcome Sanger Institute COVID-19 Surveillance Team |
| EPI_ISL_2999999 | Berkshire and Surrey Pathology Services Lighthouse Laboratory | Wellcome Sanger Institute for the COVID-19 Genomics UK (COG-UK) Consortium | Berkshire and Surrey Pathology Services Lighthouse Laboratory and Alex Alderton, Roberto Amato, Jeffrey Barrett, Sonia Goncalves, Ewan Harrison, David K. Jackson, Ian Johnston, Dominic Kwiatkowski, Cordelia Langford, John Sillitoe on behalf of the Wellcome Sanger Institute COVID-19 Surveillance Team |
| EPI_ISL_3000000 | Lighthouse Lab in Alderley Park | Wellcome Sanger Institute for the COVID-19 Genomics UK (COG-UK) Consortium | Jacquelyn Wynn, Mairead Hyland, The Lighthouse Lab in Alderley Park and Alex Alderton, Roberto Amato, Jeffrey Barrett, Sonia Goncalves, Ewan Harrison, David K. Jackson, Ian Johnston, Dominic Kwiatkowski, Cordelia Langford, John Sillitoe on behalf of the Wellcome Sanger Institute COVID-19 Surveillance Team |
| EPI_ISL_3000001 | Health Services Laboratories | Wellcome Sanger Institute for the COVID-19 Genomics UK (COG-UK) Consortium | Health Services Laboratories and Alex Alderton, Roberto Amato, Jeffrey Barrett, Sonia Goncalves, Ewan Harrison, David K. Jackson, Ian Johnston, Dominic Kwiatkowski, Cordelia Langford, John Sillitoe on behalf of the Wellcome Sanger Institute COVID-19 Surveillance Team |
| EPI_ISL_3000002 | Lighthouse Lab in Alderley Park | Wellcome Sanger Institute for the COVID-19 Genomics UK (COG-UK) Consortium | Jacquelyn Wynn, Mairead Hyland, The Lighthouse Lab in Alderley Park and Alex Alderton, Roberto Amato, Jeffrey Barrett, Sonia Goncalves, Ewan Harrison, David K. Jackson, Ian Johnston, Dominic Kwiatkowski, Cordelia Langford, John Sillitoe on behalf of the Wellcome Sanger Institute COVID-19 Surveillance Team |
| EPI_ISL_3000003 | Berkshire and Surrey Pathology Services Lighthouse Laboratory | Wellcome Sanger Institute for the COVID-19 Genomics UK (COG-UK) Consortium | Berkshire and Surrey Pathology Services Lighthouse Laboratory and Alex Alderton, Roberto Amato, Jeffrey Barrett, Sonia Goncalves, Ewan Harrison, David K. Jackson, Ian Johnston, Dominic Kwiatkowski, Cordelia Langford, John Sillitoe on behalf of the Wellcome Sanger Institute COVID-19 Surveillance Team |
| EPI_ISL_3000004, EPI_ISL_3000005, EPI_ISL_3000006 | Lighthouse Lab in Alderley Park | Wellcome Sanger Institute for the COVID-19 Genomics UK (COG-UK) Consortium | Jacquelyn Wynn, Mairead Hyland, The Lighthouse Lab in Alderley Park and Alex Alderton, Roberto Amato, Jeffrey Barrett, Sonia Goncalves, Ewan Harrison, David K. Jackson, Ian Johnston, Dominic Kwiatkowski, Cordelia Langford, John Sillitoe on behalf of the Wellcome Sanger Institute COVID-19 Surveillance Team |
| EPI_ISL_3000007, EPI_ISL_3000008 | Health Services Laboratories | Wellcome Sanger Institute for the COVID-19 Genomics UK (COG-UK) Consortium | Health Services Laboratories and Alex Alderton, Roberto Amato, Jeffrey Barrett, Sonia Goncalves, Ewan Harrison, David K. Jackson, Ian Johnston, Dominic Kwiatkowski, Cordelia Langford, John Sillitoe on behalf of the Wellcome Sanger Institute COVID-19 Surveillance Team |
| EPI_ISL_3000009 | Lighthouse Lab in Alderley Park | Wellcome Sanger Institute for the COVID-19 Genomics UK (COG-UK) Consortium | Jacquelyn Wynn, Mairead Hyland, The Lighthouse Lab in Alderley Park and Alex Alderton, Roberto Amato, Jeffrey Barrett, Sonia Goncalves, Ewan Harrison, David K. Jackson, Ian Johnston, Dominic Kwiatkowski, Cordelia Langford, John Sillitoe on behalf of the Wellcome Sanger Institute COVID-19 Surveillance Team |
| EPI_ISL_3000010 | Berkshire and Surrey Pathology Services Lighthouse Laboratory | Wellcome Sanger Institute for the COVID-19 Genomics UK (COG-UK) Consortium | Berkshire and Surrey Pathology Services Lighthouse Laboratory and Alex Alderton, Roberto Amato, Jeffrey Barrett, Sonia Goncalves, Ewan Harrison, David K. Jackson, Ian Johnston, Dominic Kwiatkowski, Cordelia Langford, John Sillitoe on behalf of the Wellcome Sanger Institute COVID-19 Surveillance Team |
| EPI_ISL_3000011 | Health Services Laboratories | Wellcome Sanger Institute for the COVID-19 Genomics UK (COG-UK) Consortium | Health Services Laboratories and Alex Alderton, Roberto Amato, Jeffrey Barrett, Sonia Goncalves, Ewan Harrison, David K. Jackson, Ian Johnston, Dominic Kwiatkowski, Cordelia Langford, John Sillitoe on behalf of the Wellcome Sanger Institute COVID-19 Surveillance Team |

| | | | |
|---|---|---|---|
| EPI_ISL_3000012 | Berkshire and Surrey Pathology Services Lighthouse Laboratory | Wellcome Sanger Institute for the COVID-19 Genomics UK (COG-UK) Consortium | Berkshire and Surrey Pathology Services Lighthouse Laboratory and Alex Alderton, Roberto Amato, Jeffrey Barrett, Sonia Goncalves, Ewan Harrison, David K. Jackson, Ian Johnston, Dominic Kwiatkowski, Cordelia Langford, John Sillitoe on behalf of the Wellcome Sanger Institute COVID-19 Surveillance Team |
| EPI_ISL_3000013 | Health Services Laboratories | Wellcome Sanger Institute for the COVID-19 Genomics UK (COG-UK) Consortium | Health Services Laboratories and Alex Alderton, Roberto Amato, Jeffrey Barrett, Sonia Goncalves, Ewan Harrison, David K. Jackson, Ian Johnston, Dominic Kwiatkowski, Cordelia Langford, John Sillitoe on behalf of the Wellcome Sanger Institute COVID-19 Surveillance Team |
| EPI_ISL_3000014 | Berkshire and Surrey Pathology Services Lighthouse Laboratory | Wellcome Sanger Institute for the COVID-19 Genomics UK (COG-UK) Consortium | Berkshire and Surrey Pathology Services Lighthouse Laboratory and Alex Alderton, Roberto Amato, Jeffrey Barrett, Sonia Goncalves, Ewan Harrison, David K. Jackson, Ian Johnston, Dominic Kwiatkowski, Cordelia Langford, John Sillitoe on behalf of the Wellcome Sanger Institute COVID-19 Surveillance Team |
| EPI_ISL_3000015 | Lighthouse Lab in Alderley Park | Wellcome Sanger Institute for the COVID-19 Genomics UK (COG-UK) Consortium | Jacquelyn Wynn, Mairead Hyland, The Lighthouse Lab in Alderley Park and Alex Alderton, Roberto Amato, Jeffrey Barrett, Sonia Goncalves, Ewan Harrison, David K. Jackson, Ian Johnston, Dominic Kwiatkowski, Cordelia Langford, John Sillitoe on behalf of the Wellcome Sanger Institute COVID-19 Surveillance Team |
| EPI_ISL_3000016, EPI_ISL_3000017, EPI_ISL_3000018, EPI_ISL_3000019 | Berkshire and Surrey Pathology Services Lighthouse Laboratory | Wellcome Sanger Institute for the COVID-19 Genomics UK (COG-UK) Consortium | Berkshire and Surrey Pathology Services Lighthouse Laboratory and Alex Alderton, Roberto Amato, Jeffrey Barrett, Sonia Goncalves, Ewan Harrison, David K. Jackson, Ian Johnston, Dominic Kwiatkowski, Cordelia Langford, John Sillitoe on behalf of the Wellcome Sanger Institute COVID-19 Surveillance Team |
| EPI_ISL_3000020, EPI_ISL_3000021 | Lighthouse Lab in Alderley Park | Wellcome Sanger Institute for the COVID-19 Genomics UK (COG-UK) Consortium | Jacquelyn Wynn, Mairead Hyland, The Lighthouse Lab in Alderley Park and Alex Alderton, Roberto Amato, Jeffrey Barrett, Sonia Goncalves, Ewan Harrison, David K. Jackson, Ian Johnston, Dominic Kwiatkowski, Cordelia Langford, John Sillitoe on behalf of the Wellcome Sanger Institute COVID-19 Surveillance Team |
| EPI_ISL_3000022 | Berkshire and Surrey Pathology Services Lighthouse Laboratory | Wellcome Sanger Institute for the COVID-19 Genomics UK (COG-UK) Consortium | Berkshire and Surrey Pathology Services Lighthouse Laboratory and Alex Alderton, Roberto Amato, Jeffrey Barrett, Sonia Goncalves, Ewan Harrison, David K. Jackson, Ian Johnston, Dominic Kwiatkowski, Cordelia Langford, John Sillitoe on behalf of the Wellcome Sanger Institute COVID-19 Surveillance Team |
| EPI_ISL_3000023 | Lighthouse Lab in Alderley Park | Wellcome Sanger Institute for the COVID-19 Genomics UK (COG-UK) Consortium | Jacquelyn Wynn, Mairead Hyland, The Lighthouse Lab in Alderley Park and Alex Alderton, Roberto Amato, Jeffrey Barrett, Sonia Goncalves, Ewan Harrison, David K. Jackson, Ian Johnston, Dominic Kwiatkowski, Cordelia Langford, John Sillitoe on behalf of the Wellcome Sanger Institute COVID-19 Surveillance Team |
| EPI_ISL_3000024 | Health Services Laboratories | Wellcome Sanger Institute for the COVID-19 Genomics UK (COG-UK) Consortium | Health Services Laboratories and Alex Alderton, Roberto Amato, Jeffrey Barrett, Sonia Goncalves, Ewan Harrison, David K. Jackson, Ian Johnston, Dominic Kwiatkowski, Cordelia Langford, John Sillitoe on behalf of the Wellcome Sanger Institute COVID-19 Surveillance Team |
| EPI_ISL_3000025 | Berkshire and Surrey Pathology Services Lighthouse Laboratory | Wellcome Sanger Institute for the COVID-19 Genomics UK (COG-UK) Consortium | Berkshire and Surrey Pathology Services Lighthouse Laboratory and Alex Alderton, Roberto Amato, Jeffrey Barrett, Sonia Goncalves, Ewan Harrison, David K. Jackson, Ian Johnston, Dominic Kwiatkowski, Cordelia Langford, John Sillitoe on behalf of the Wellcome Sanger Institute COVID-19 Surveillance Team |
| EPI_ISL_3000026, EPI_ISL_3000027 | Lighthouse Lab in Alderley Park | Wellcome Sanger Institute for the COVID-19 Genomics UK (COG-UK) Consortium | Jacquelyn Wynn, Mairead Hyland, The Lighthouse Lab in Alderley Park and Alex Alderton, Roberto Amato, Jeffrey Barrett, Sonia Goncalves, Ewan Harrison, David K. Jackson, Ian Johnston, Dominic Kwiatkowski, Cordelia Langford, John Sillitoe on behalf of the Wellcome Sanger Institute COVID-19 Surveillance Team |
| EPI_ISL_3000028, EPI_ISL_3000029 | Berkshire and Surrey Pathology Services Lighthouse Laboratory | Wellcome Sanger Institute for the COVID-19 Genomics UK (COG-UK) Consortium | Berkshire and Surrey Pathology Services Lighthouse Laboratory and Alex Alderton, Roberto Amato, Jeffrey Barrett, Sonia Goncalves, Ewan Harrison, David K. Jackson, Ian Johnston, Dominic Kwiatkowski, Cordelia Langford, John Sillitoe on behalf of the Wellcome Sanger Institute COVID-19 Surveillance Team |
| EPI_ISL_3000030 | Health Services Laboratories | Wellcome Sanger Institute for the COVID-19 Genomics UK (COG-UK) Consortium | Health Services Laboratories and Alex Alderton, Roberto Amato, Jeffrey Barrett, Sonia Goncalves, Ewan Harrison, David K. Jackson, Ian Johnston, Dominic Kwiatkowski, Cordelia Langford, John Sillitoe on behalf of the Wellcome Sanger Institute COVID-19 Surveillance Team |
| EPI_ISL_3000031, EPI_ISL_3000032, EPI_ISL_3000033 | Berkshire and Surrey Pathology Services Lighthouse Laboratory | Wellcome Sanger Institute for the COVID-19 Genomics UK (COG-UK) Consortium | Berkshire and Surrey Pathology Services Lighthouse Laboratory and Alex Alderton, Roberto Amato, Jeffrey Barrett, Sonia Goncalves, Ewan Harrison, David K. Jackson, Ian Johnston, Dominic Kwiatkowski, Cordelia Langford, John Sillitoe on behalf of the Wellcome Sanger Institute COVID-19 Surveillance Team |
| EPI_ISL_3000034 | Lighthouse Lab in Alderley Park | Wellcome Sanger Institute for the COVID-19 Genomics UK (COG-UK) Consortium | Jacquelyn Wynn, Mairead Hyland, The Lighthouse Lab in Alderley Park and Alex Alderton, Roberto Amato, Jeffrey Barrett, Sonia Goncalves, Ewan Harrison, David K. Jackson, Ian Johnston, Dominic Kwiatkowski, Cordelia Langford, John Sillitoe on behalf of the Wellcome Sanger Institute COVID-19 Surveillance Team |
| EPI_ISL_3000035 | Health Services Laboratories | Wellcome Sanger Institute for the COVID-19 Genomics UK (COG-UK) Consortium | Health Services Laboratories and Alex Alderton, Roberto Amato, Jeffrey Barrett, Sonia Goncalves, Ewan Harrison, David K. Jackson, Ian Johnston, Dominic Kwiatkowski, Cordelia Langford, John Sillitoe on behalf of the Wellcome Sanger Institute COVID-19 Surveillance Team |
| EPI_ISL_3000036, EPI_ISL_3000037 | Berkshire and Surrey Pathology Services Lighthouse Laboratory | Wellcome Sanger Institute for the COVID-19 Genomics UK (COG-UK) Consortium | Berkshire and Surrey Pathology Services Lighthouse Laboratory and Alex Alderton, Roberto Amato, Jeffrey Barrett, Sonia Goncalves, Ewan Harrison, David K. Jackson, Ian Johnston, Dominic Kwiatkowski, Cordelia Langford, John Sillitoe on behalf of the Wellcome Sanger Institute COVID-19 Surveillance Team |
| EPI_ISL_3000038 | Health Services Laboratories | Wellcome Sanger Institute for the COVID-19 Genomics UK (COG-UK) Consortium | Health Services Laboratories and Alex Alderton, Roberto Amato, Jeffrey Barrett, Sonia Goncalves, Ewan Harrison, David K. Jackson, Ian Johnston, Dominic Kwiatkowski, Cordelia Langford, John Sillitoe on behalf of the Wellcome Sanger Institute COVID-19 Surveillance Team |
| EPI_ISL_3000039, EPI_ISL_3000040 | Berkshire and Surrey Pathology Services Lighthouse Laboratory | Wellcome Sanger Institute for the COVID-19 Genomics UK (COG-UK) Consortium | Berkshire and Surrey Pathology Services Lighthouse Laboratory and Alex Alderton, Roberto Amato, Jeffrey Barrett, Sonia Goncalves, Ewan Harrison, David K. Jackson, Ian Johnston, Dominic Kwiatkowski, Cordelia Langford, John Sillitoe on behalf of the Wellcome Sanger Institute COVID-19 Surveillance Team |
| EPI_ISL_3000041, EPI_ISL_3000042 | Health Services Laboratories | Wellcome Sanger Institute for the COVID-19 Genomics UK (COG-UK) Consortium | Health Services Laboratories and Alex Alderton, Roberto Amato, Jeffrey Barrett, Sonia Goncalves, Ewan Harrison, David K. Jackson, Ian Johnston, Dominic Kwiatkowski, Cordelia Langford, John Sillitoe on behalf of the Wellcome Sanger Institute COVID-19 Surveillance Team |
| EPI_ISL_3000043, EPI_ISL_3000044 | Berkshire and Surrey Pathology Services Lighthouse Laboratory | Wellcome Sanger Institute for the COVID-19 Genomics UK (COG-UK) Consortium | Berkshire and Surrey Pathology Services Lighthouse Laboratory and Alex Alderton, Roberto Amato, Jeffrey Barrett, Sonia Goncalves, Ewan Harrison, David K. Jackson, Ian Johnston, Dominic Kwiatkowski, Cordelia Langford, John Sillitoe on behalf of the Wellcome Sanger Institute COVID-19 Surveillance Team |
| EPI_ISL_3000045, EPI_ISL_3000046 | Lighthouse Lab in Alderley Park | Wellcome Sanger Institute for the COVID-19 Genomics UK (COG-UK) Consortium | Jacquelyn Wynn, Mairead Hyland, The Lighthouse Lab in Alderley Park and Alex Alderton, Roberto Amato, Jeffrey Barrett, Sonia Goncalves, Ewan Harrison, David K. Jackson, Ian Johnston, Dominic Kwiatkowski, Cordelia Langford, John Sillitoe on behalf of the Wellcome Sanger Institute COVID-19 Surveillance Team |
| EPI_ISL_3000047, EPI_ISL_3000048, EPI_ISL_3000049, EPI_ISL_3000050 | Berkshire and Surrey Pathology Services Lighthouse Laboratory | Wellcome Sanger Institute for the COVID-19 Genomics UK (COG-UK) Consortium | Berkshire and Surrey Pathology Services Lighthouse Laboratory and Alex Alderton, Roberto Amato, Jeffrey Barrett, Sonia Goncalves, Ewan Harrison, David K. Jackson, Ian Johnston, Dominic Kwiatkowski, Cordelia Langford, John Sillitoe on behalf of the Wellcome Sanger Institute COVID-19 Surveillance Team |
| EPI_ISL_3000051 | Lighthouse Lab in Alderley Park | Wellcome Sanger Institute for the COVID-19 Genomics UK (COG-UK) Consortium | Jacquelyn Wynn, Mairead Hyland, The Lighthouse Lab in Alderley Park and Alex Alderton, Roberto Amato, Jeffrey Barrett, Sonia Goncalves, Ewan Harrison, David K. Jackson, Ian Johnston, Dominic Kwiatkowski, Cordelia Langford, John Sillitoe on behalf of the Wellcome Sanger Institute COVID-19 Surveillance Team |
| EPI_ISL_3000052 | Health Services Laboratories | Wellcome Sanger Institute for the COVID-19 Genomics UK (COG-UK) Consortium | Health Services Laboratories and Alex Alderton, Roberto Amato, Jeffrey Barrett, Sonia Goncalves, Ewan Harrison, David K. Jackson, Ian Johnston, Dominic Kwiatkowski, Cordelia Langford, John Sillitoe on behalf of the Wellcome Sanger Institute COVID-19 Surveillance Team |
| EPI_ISL_3000053 | Berkshire and Surrey Pathology Services Lighthouse Laboratory | Wellcome Sanger Institute for the COVID-19 Genomics UK (COG-UK) Consortium | Berkshire and Surrey Pathology Services Lighthouse Laboratory and Alex Alderton, Roberto Amato, Jeffrey Barrett, Sonia Goncalves, Ewan Harrison, David K. Jackson, Ian Johnston, Dominic Kwiatkowski, Cordelia Langford, John Sillitoe on behalf of the Wellcome Sanger Institute COVID-19 Surveillance Team |
| EPI_ISL_3000054 | Lighthouse Lab in Alderley Park | Wellcome Sanger Institute for the COVID-19 Genomics UK (COG-UK) Consortium | Jacquelyn Wynn, Mairead Hyland, The Lighthouse Lab in Alderley Park and Alex Alderton, Roberto Amato, Jeffrey Barrett, Sonia Goncalves, Ewan Harrison, David K. Jackson, Ian Johnston, Dominic Kwiatkowski, Cordelia Langford, John Sillitoe on behalf of the Wellcome Sanger Institute COVID-19 Surveillance Team |
| EPI_ISL_3000055, EPI_ISL_3000056 | Health Services Laboratories | Wellcome Sanger Institute for the COVID-19 Genomics UK (COG-UK) Consortium | Health Services Laboratories and Alex Alderton, Roberto Amato, Jeffrey Barrett, Sonia Goncalves, Ewan Harrison, David K. Jackson, Ian Johnston, Dominic Kwiatkowski, Cordelia Langford, John Sillitoe on behalf of the Wellcome Sanger Institute COVID-19 Surveillance Team |
| EPI_ISL_3000057, EPI_ISL_3000058 | Berkshire and Surrey Pathology Services Lighthouse Laboratory | Wellcome Sanger Institute for the COVID-19 Genomics UK (COG-UK) Consortium | Berkshire and Surrey Pathology Services Lighthouse Laboratory and Alex Alderton, Roberto Amato, Jeffrey Barrett, Sonia Goncalves, Ewan Harrison, David K. Jackson, Ian Johnston, Dominic Kwiatkowski, Cordelia Langford, John Sillitoe on behalf of the Wellcome Sanger Institute COVID-19 Surveillance Team |
| EPI_ISL_3000059 | Lighthouse Lab in Alderley Park | Wellcome Sanger Institute for the COVID-19 Genomics UK (COG-UK) Consortium | Jacquelyn Wynn, Mairead Hyland, The Lighthouse Lab in Alderley Park and Alex Alderton, Roberto Amato, Jeffrey Barrett, Sonia Goncalves, Ewan Harrison, David K. Jackson, Ian Johnston, Dominic Kwiatkowski, Cordelia Langford, John Sillitoe on behalf of the Wellcome Sanger Institute COVID-19 Surveillance Team |
| EPI_ISL_3000060 | Berkshire and Surrey Pathology Services Lighthouse Laboratory | Wellcome Sanger Institute for the COVID-19 Genomics UK (COG-UK) Consortium | Berkshire and Surrey Pathology Services Lighthouse Laboratory and Alex Alderton, Roberto Amato, Jeffrey Barrett, Sonia Goncalves, Ewan Harrison, David K. Jackson, Ian Johnston, Dominic Kwiatkowski, Cordelia Langford, John Sillitoe on behalf of the Wellcome Sanger Institute COVID-19 Surveillance Team |
| EPI_ISL_3000061 | Health Services Laboratories | Wellcome Sanger Institute for the COVID-19 Genomics UK (COG-UK) Consortium | Health Services Laboratories and Alex Alderton, Roberto Amato, Jeffrey Barrett, Sonia Goncalves, Ewan Harrison, David K. Jackson, Ian Johnston, Dominic Kwiatkowski, Cordelia Langford, John Sillitoe on behalf of the Wellcome Sanger Institute COVID-19 Surveillance Team |
| EPI_ISL_3000062, EPI_ISL_3000063, EPI_ISL_3000064 | Berkshire and Surrey Pathology Services Lighthouse Laboratory | Wellcome Sanger Institute for the COVID-19 Genomics UK (COG-UK) Consortium | Berkshire and Surrey Pathology Services Lighthouse Laboratory and Alex Alderton, Roberto Amato, Jeffrey Barrett, Sonia Goncalves, Ewan Harrison, David K. Jackson, Ian Johnston, Dominic Kwiatkowski, Cordelia Langford, John Sillitoe on behalf of the Wellcome Sanger Institute COVID-19 Surveillance Team |
| EPI_ISL_3000065 | Health Services Laboratories | Wellcome Sanger Institute for the COVID-19 Genomics UK (COG-UK) Consortium | Health Services Laboratories and Alex Alderton, Roberto Amato, Jeffrey Barrett, Sonia Goncalves, Ewan Harrison, David K. Jackson, Ian Johnston, Dominic Kwiatkowski, Cordelia Langford, John Sillitoe on behalf of the Wellcome Sanger Institute COVID-19 Surveillance Team |
| EPI_ISL_3000066 | Lighthouse Lab in Alderley Park | Wellcome Sanger Institute for the COVID-19 Genomics UK (COG-UK) Consortium | Jacquelyn Wynn, Mairead Hyland, The Lighthouse Lab in Alderley Park and Alex Alderton, Roberto Amato, Jeffrey Barrett, Sonia Goncalves, Ewan Harrison, David K. Jackson, Ian Johnston, Dominic Kwiatkowski, Cordelia Langford, John Sillitoe on behalf of the Wellcome Sanger Institute COVID-19 Surveillance Team |
| EPI_ISL_3000067 | Health Services Laboratories | Wellcome Sanger Institute for the COVID-19 Genomics UK (COG-UK) Consortium | Health Services Laboratories and Alex Alderton, Roberto Amato, Jeffrey Barrett, Sonia Goncalves, Ewan Harrison, David K. Jackson, Ian Johnston, Dominic Kwiatkowski, Cordelia Langford, John Sillitoe on behalf of the Wellcome Sanger Institute COVID-19 Surveillance Team |
| EPI_ISL_3000068, EPI_ISL_3000069 | Berkshire and Surrey Pathology Services Lighthouse Laboratory | Wellcome Sanger Institute for the COVID-19 Genomics UK (COG-UK) Consortium | Berkshire and Surrey Pathology Services Lighthouse Laboratory and Alex Alderton, Roberto Amato, Jeffrey Barrett, Sonia Goncalves, Ewan Harrison, David K. Jackson, Ian Johnston, Dominic Kwiatkowski, Cordelia Langford, John Sillitoe on behalf of the Wellcome Sanger Institute COVID-19 Surveillance Team |
| EPI_ISL_3000070, EPI_ISL_3000071, EPI_ISL_3000072, EPI_ISL_3000073, EPI_ISL_3000074 | Lighthouse Lab in Alderley Park | Wellcome Sanger Institute for the COVID-19 Genomics UK (COG-UK) Consortium | Jacquelyn Wynn, Mairead Hyland, The Lighthouse Lab in Alderley Park and Alex Alderton, Roberto Amato, Jeffrey Barrett, Sonia Goncalves, Ewan Harrison, David K. Jackson, Ian Johnston, Dominic Kwiatkowski, Cordelia Langford, John Sillitoe on behalf of the Wellcome Sanger Institute COVID-19 Surveillance Team |
| EPI_ISL_3000075, EPI_ISL_3000076, EPI_ISL_3000077 | Berkshire and Surrey Pathology Services Lighthouse Laboratory | Wellcome Sanger Institute for the COVID-19 Genomics UK (COG-UK) Consortium | Berkshire and Surrey Pathology Services Lighthouse Laboratory and Alex Alderton, Roberto Amato, Jeffrey Barrett, Sonia Goncalves, Ewan Harrison, David K. Jackson, Ian Johnston, Dominic Kwiatkowski, Cordelia Langford, John Sillitoe on behalf of the Wellcome Sanger Institute COVID-19 Surveillance Team |
| EPI_ISL_3000078 | Health Services Laboratories | Wellcome Sanger Institute for the COVID-19 Genomics UK (COG-UK) Consortium | Health Services Laboratories and Alex Alderton, Roberto Amato, Jeffrey Barrett, Sonia Goncalves, Ewan Harrison, David K. Jackson, Ian Johnston, Dominic Kwiatkowski, Cordelia Langford, John Sillitoe on behalf of the Wellcome Sanger Institute COVID-19 Surveillance Team |
| EPI_ISL_3000079, EPI_ISL_3000080 | Berkshire and Surrey Pathology Services Lighthouse Laboratory | Wellcome Sanger Institute for the COVID-19 Genomics UK (COG-UK) Consortium | Berkshire and Surrey Pathology Services Lighthouse Laboratory and Alex Alderton, Roberto Amato, Jeffrey Barrett, Sonia Goncalves, Ewan Harrison, David K. Jackson, Ian Johnston, Dominic Kwiatkowski, Cordelia Langford, John Sillitoe on behalf of the Wellcome Sanger Institute COVID-19 Surveillance Team |

| | | | |
|---|---|---|---|
| EPI_ISL_3000081 | Health Services Laboratories | Wellcome Sanger Institute for the COVID-19 Genomics UK (COG-UK) Consortium | Health Services Laboratories and Alex Alderton, Roberto Amato, Jeffrey Barrett, Sonia Goncalves, Ewan Harrison, David K. Jackson, Ian Johnston, Dominic Kwiatkowski, Cordelia Langford, John Sillitoe on behalf of the Wellcome Sanger Institute COVID-19 Surveillance Team |
| EPI_ISL_3000082 | Lighthouse Lab in Alderley Park | Wellcome Sanger Institute for the COVID-19 Genomics UK (COG-UK) Consortium | Jacquelyn Wynn, Mairead Hyland, The Lighthouse Lab in Alderley Park and Alex Alderton, Roberto Amato, Jeffrey Barrett, Sonia Goncalves, Ewan Harrison, David K. Jackson, Ian Johnston, Dominic Kwiatkowski, Cordelia Langford, John Sillitoe on behalf of the Wellcome Sanger Institute COVID-19 Surveillance Team |
| EPI_ISL_3000083 | Berkshire and Surrey Pathology Services Lighthouse Laboratory | Wellcome Sanger Institute for the COVID-19 Genomics UK (COG-UK) Consortium | Berkshire and Surrey Pathology Services Lighthouse Laboratory and Alex Alderton, Roberto Amato, Jeffrey Barrett, Sonia Goncalves, Ewan Harrison, David K. Jackson, Ian Johnston, Dominic Kwiatkowski, Cordelia Langford, John Sillitoe on behalf of the Wellcome Sanger Institute COVID-19 Surveillance Team |
| EPI_ISL_3000084 | Lighthouse Lab in Alderley Park | Wellcome Sanger Institute for the COVID-19 Genomics UK (COG-UK) Consortium | Jacquelyn Wynn, Mairead Hyland, The Lighthouse Lab in Alderley Park and Alex Alderton, Roberto Amato, Jeffrey Barrett, Sonia Goncalves, Ewan Harrison, David K. Jackson, Ian Johnston, Dominic Kwiatkowski, Cordelia Langford, John Sillitoe on behalf of the Wellcome Sanger Institute COVID-19 Surveillance Team |
| EPI_ISL_3000085 | Health Services Laboratories | Wellcome Sanger Institute for the COVID-19 Genomics UK (COG-UK) Consortium | Health Services Laboratories and Alex Alderton, Roberto Amato, Jeffrey Barrett, Sonia Goncalves, Ewan Harrison, David K. Jackson, Ian Johnston, Dominic Kwiatkowski, Cordelia Langford, John Sillitoe on behalf of the Wellcome Sanger Institute COVID-19 Surveillance Team |
| EPI_ISL_3000086 | Lighthouse Lab in Alderley Park | Wellcome Sanger Institute for the COVID-19 Genomics UK (COG-UK) Consortium | Jacquelyn Wynn, Mairead Hyland, The Lighthouse Lab in Alderley Park and Alex Alderton, Roberto Amato, Jeffrey Barrett, Sonia Goncalves, Ewan Harrison, David K. Jackson, Ian Johnston, Dominic Kwiatkowski, Cordelia Langford, John Sillitoe on behalf of the Wellcome Sanger Institute COVID-19 Surveillance Team |
| EPI_ISL_3000087 | Health Services Laboratories | Wellcome Sanger Institute for the COVID-19 Genomics UK (COG-UK) Consortium | Health Services Laboratories and Alex Alderton, Roberto Amato, Jeffrey Barrett, Sonia Goncalves, Ewan Harrison, David K. Jackson, Ian Johnston, Dominic Kwiatkowski, Cordelia Langford, John Sillitoe on behalf of the Wellcome Sanger Institute COVID-19 Surveillance Team |
| EPI_ISL_3000088, EPI_ISL_3000089 | Berkshire and Surrey Pathology Services Lighthouse Laboratory | Wellcome Sanger Institute for the COVID-19 Genomics UK (COG-UK) Consortium | Berkshire and Surrey Pathology Services Lighthouse Laboratory and Alex Alderton, Roberto Amato, Jeffrey Barrett, Sonia Goncalves, Ewan Harrison, David K. Jackson, Ian Johnston, Dominic Kwiatkowski, Cordelia Langford, John Sillitoe on behalf of the Wellcome Sanger Institute COVID-19 Surveillance Team |
| EPI_ISL_3000090 | Health Services Laboratories | Wellcome Sanger Institute for the COVID-19 Genomics UK (COG-UK) Consortium | Health Services Laboratories and Alex Alderton, Roberto Amato, Jeffrey Barrett, Sonia Goncalves, Ewan Harrison, David K. Jackson, Ian Johnston, Dominic Kwiatkowski, Cordelia Langford, John Sillitoe on behalf of the Wellcome Sanger Institute COVID-19 Surveillance Team |
| EPI_ISL_3000091 | Berkshire and Surrey Pathology Services Lighthouse Laboratory | Wellcome Sanger Institute for the COVID-19 Genomics UK (COG-UK) Consortium | Berkshire and Surrey Pathology Services Lighthouse Laboratory and Alex Alderton, Roberto Amato, Jeffrey Barrett, Sonia Goncalves, Ewan Harrison, David K. Jackson, Ian Johnston, Dominic Kwiatkowski, Cordelia Langford, John Sillitoe on behalf of the Wellcome Sanger Institute COVID-19 Surveillance Team |
| EPI_ISL_3000092 | Health Services Laboratories | Wellcome Sanger Institute for the COVID-19 Genomics UK (COG-UK) Consortium | Health Services Laboratories and Alex Alderton, Roberto Amato, Jeffrey Barrett, Sonia Goncalves, Ewan Harrison, David K. Jackson, Ian Johnston, Dominic Kwiatkowski, Cordelia Langford, John Sillitoe on behalf of the Wellcome Sanger Institute COVID-19 Surveillance Team |
| EPI_ISL_3000093 | Lighthouse Lab in Alderley Park | Wellcome Sanger Institute for the COVID-19 Genomics UK (COG-UK) Consortium | Jacquelyn Wynn, Mairead Hyland, The Lighthouse Lab in Alderley Park and Alex Alderton, Roberto Amato, Jeffrey Barrett, Sonia Goncalves, Ewan Harrison, David K. Jackson, Ian Johnston, Dominic Kwiatkowski, Cordelia Langford, John Sillitoe on behalf of the Wellcome Sanger Institute COVID-19 Surveillance Team |
| EPI_ISL_3000094 | Health Services Laboratories | Wellcome Sanger Institute for the COVID-19 Genomics UK (COG-UK) Consortium | Health Services Laboratories and Alex Alderton, Roberto Amato, Jeffrey Barrett, Sonia Goncalves, Ewan Harrison, David K. Jackson, Ian Johnston, Dominic Kwiatkowski, Cordelia Langford, John Sillitoe on behalf of the Wellcome Sanger Institute COVID-19 Surveillance Team |
| EPI_ISL_3000095 | Berkshire and Surrey Pathology Services Lighthouse Laboratory | Wellcome Sanger Institute for the COVID-19 Genomics UK (COG-UK) Consortium | Berkshire and Surrey Pathology Services Lighthouse Laboratory and Alex Alderton, Roberto Amato, Jeffrey Barrett, Sonia Goncalves, Ewan Harrison, David K. Jackson, Ian Johnston, Dominic Kwiatkowski, Cordelia Langford, John Sillitoe on behalf of the Wellcome Sanger Institute COVID-19 Surveillance Team |
| EPI_ISL_3000096 | Lighthouse Lab in Alderley Park | Wellcome Sanger Institute for the COVID-19 Genomics UK (COG-UK) Consortium | Jacquelyn Wynn, Mairead Hyland, The Lighthouse Lab in Alderley Park and Alex Alderton, Roberto Amato, Jeffrey Barrett, Sonia Goncalves, Ewan Harrison, David K. Jackson, Ian Johnston, Dominic Kwiatkowski, Cordelia Langford, John Sillitoe on behalf of the Wellcome Sanger Institute COVID-19 Surveillance Team |
| EPI_ISL_3000097 | Berkshire and Surrey Pathology Services Lighthouse Laboratory | Wellcome Sanger Institute for the COVID-19 Genomics UK (COG-UK) Consortium | Berkshire and Surrey Pathology Services Lighthouse Laboratory and Alex Alderton, Roberto Amato, Jeffrey Barrett, Sonia Goncalves, Ewan Harrison, David K. Jackson, Ian Johnston, Dominic Kwiatkowski, Cordelia Langford, John Sillitoe on behalf of the Wellcome Sanger Institute COVID-19 Surveillance Team |
| EPI_ISL_3000098 | Lighthouse Lab in Alderley Park | Wellcome Sanger Institute for the COVID-19 Genomics UK (COG-UK) Consortium | Jacquelyn Wynn, Mairead Hyland, The Lighthouse Lab in Alderley Park and Alex Alderton, Roberto Amato, Jeffrey Barrett, Sonia Goncalves, Ewan Harrison, David K. Jackson, Ian Johnston, Dominic Kwiatkowski, Cordelia Langford, John Sillitoe on behalf of the Wellcome Sanger Institute COVID-19 Surveillance Team |
| EPI_ISL_3000099 | Berkshire and Surrey Pathology Services Lighthouse Laboratory | Wellcome Sanger Institute for the COVID-19 Genomics UK (COG-UK) Consortium | Berkshire and Surrey Pathology Services Lighthouse Laboratory and Alex Alderton, Roberto Amato, Jeffrey Barrett, Sonia Goncalves, Ewan Harrison, David K. Jackson, Ian Johnston, Dominic Kwiatkowski, Cordelia Langford, John Sillitoe on behalf of the Wellcome Sanger Institute COVID-19 Surveillance Team |
| EPI_ISL_3000100 | Lighthouse Lab in Alderley Park | Wellcome Sanger Institute for the COVID-19 Genomics UK (COG-UK) Consortium | Jacquelyn Wynn, Mairead Hyland, The Lighthouse Lab in Alderley Park and Alex Alderton, Roberto Amato, Jeffrey Barrett, Sonia Goncalves, Ewan Harrison, David K. Jackson, Ian Johnston, Dominic Kwiatkowski, Cordelia Langford, John Sillitoe on behalf of the Wellcome Sanger Institute COVID-19 Surveillance Team |
| EPI_ISL_3000101 | Berkshire and Surrey Pathology Services Lighthouse Laboratory | Wellcome Sanger Institute for the COVID-19 Genomics UK (COG-UK) Consortium | Berkshire and Surrey Pathology Services Lighthouse Laboratory and Alex Alderton, Roberto Amato, Jeffrey Barrett, Sonia Goncalves, Ewan Harrison, David K. Jackson, Ian Johnston, Dominic Kwiatkowski, Cordelia Langford, John Sillitoe on behalf of the Wellcome Sanger Institute COVID-19 Surveillance Team |
| EPI_ISL_3000102 | Health Services Laboratories | Wellcome Sanger Institute for the COVID-19 Genomics UK (COG-UK) Consortium | Health Services Laboratories and Alex Alderton, Roberto Amato, Jeffrey Barrett, Sonia Goncalves, Ewan Harrison, David K. Jackson, Ian Johnston, Dominic Kwiatkowski, Cordelia Langford, John Sillitoe on behalf of the Wellcome Sanger Institute COVID-19 Surveillance Team |
| EPI_ISL_3000103 | Lighthouse Lab in Alderley Park | Wellcome Sanger Institute for the COVID-19 Genomics UK (COG-UK) Consortium | Jacquelyn Wynn, Mairead Hyland, The Lighthouse Lab in Alderley Park and Alex Alderton, Roberto Amato, Jeffrey Barrett, Sonia Goncalves, Ewan Harrison, David K. Jackson, Ian Johnston, Dominic Kwiatkowski, Cordelia Langford, John Sillitoe on behalf of the Wellcome Sanger Institute COVID-19 Surveillance Team |
| EPI_ISL_3000104, EPI_ISL_3000105, EPI_ISL_3000106, EPI_ISL_3000107, EPI_ISL_3000108, EPI_ISL_3000109, EPI_ISL_3000110, EPI_ISL_3000111 | Berkshire and Surrey Pathology Services Lighthouse Laboratory | Wellcome Sanger Institute for the COVID-19 Genomics UK (COG-UK) Consortium | Berkshire and Surrey Pathology Services Lighthouse Laboratory and Alex Alderton, Roberto Amato, Jeffrey Barrett, Sonia Goncalves, Ewan Harrison, David K. Jackson, Ian Johnston, Dominic Kwiatkowski, Cordelia Langford, John Sillitoe on behalf of the Wellcome Sanger Institute COVID-19 Surveillance Team |
| EPI_ISL_3000112 | Health Services Laboratories | Wellcome Sanger Institute for the COVID-19 Genomics UK (COG-UK) Consortium | Health Services Laboratories and Alex Alderton, Roberto Amato, Jeffrey Barrett, Sonia Goncalves, Ewan Harrison, David K. Jackson, Ian Johnston, Dominic Kwiatkowski, Cordelia Langford, John Sillitoe on behalf of the Wellcome Sanger Institute COVID-19 Surveillance Team |
| EPI_ISL_3000113 | Berkshire and Surrey Pathology Services Lighthouse Laboratory | Wellcome Sanger Institute for the COVID-19 Genomics UK (COG-UK) Consortium | Berkshire and Surrey Pathology Services Lighthouse Laboratory and Alex Alderton, Roberto Amato, Jeffrey Barrett, Sonia Goncalves, Ewan Harrison, David K. Jackson, Ian Johnston, Dominic Kwiatkowski, Cordelia Langford, John Sillitoe on behalf of the Wellcome Sanger Institute COVID-19 Surveillance Team |
| EPI_ISL_3000114, EPI_ISL_3000115 | Health Services Laboratories | Wellcome Sanger Institute for the COVID-19 Genomics UK (COG-UK) Consortium | Health Services Laboratories and Alex Alderton, Roberto Amato, Jeffrey Barrett, Sonia Goncalves, Ewan Harrison, David K. Jackson, Ian Johnston, Dominic Kwiatkowski, Cordelia Langford, John Sillitoe on behalf of the Wellcome Sanger Institute COVID-19 Surveillance Team |
| EPI_ISL_3000116 | Berkshire and Surrey Pathology Services Lighthouse Laboratory | Wellcome Sanger Institute for the COVID-19 Genomics UK (COG-UK) Consortium | Berkshire and Surrey Pathology Services Lighthouse Laboratory and Alex Alderton, Roberto Amato, Jeffrey Barrett, Sonia Goncalves, Ewan Harrison, David K. Jackson, Ian Johnston, Dominic Kwiatkowski, Cordelia Langford, John Sillitoe on behalf of the Wellcome Sanger Institute COVID-19 Surveillance Team |
| EPI_ISL_3000117, EPI_ISL_3000118 | Health Services Laboratories | Wellcome Sanger Institute for the COVID-19 Genomics UK (COG-UK) Consortium | Health Services Laboratories and Alex Alderton, Roberto Amato, Jeffrey Barrett, Sonia Goncalves, Ewan Harrison, David K. Jackson, Ian Johnston, Dominic Kwiatkowski, Cordelia Langford, John Sillitoe on behalf of the Wellcome Sanger Institute COVID-19 Surveillance Team |
| EPI_ISL_3000119, EPI_ISL_3000120, EPI_ISL_3000121, EPI_ISL_3000122, EPI_ISL_3000123, EPI_ISL_3000124 | Berkshire and Surrey Pathology Services Lighthouse Laboratory | Wellcome Sanger Institute for the COVID-19 Genomics UK (COG-UK) Consortium | Berkshire and Surrey Pathology Services Lighthouse Laboratory and Alex Alderton, Roberto Amato, Jeffrey Barrett, Sonia Goncalves, Ewan Harrison, David K. Jackson, Ian Johnston, Dominic Kwiatkowski, Cordelia Langford, John Sillitoe on behalf of the Wellcome Sanger Institute COVID-19 Surveillance Team |
| EPI_ISL_3000125 | Lighthouse Lab in Alderley Park | Wellcome Sanger Institute for the COVID-19 Genomics UK (COG-UK) Consortium | Jacquelyn Wynn, Mairead Hyland, The Lighthouse Lab in Alderley Park and Alex Alderton, Roberto Amato, Jeffrey Barrett, Sonia Goncalves, Ewan Harrison, David K. Jackson, Ian Johnston, Dominic Kwiatkowski, Cordelia Langford, John Sillitoe on behalf of the Wellcome Sanger Institute COVID-19 Surveillance Team |
| EPI_ISL_3000126 | Berkshire and Surrey Pathology Services Lighthouse Laboratory | Wellcome Sanger Institute for the COVID-19 Genomics UK (COG-UK) Consortium | Berkshire and Surrey Pathology Services Lighthouse Laboratory and Alex Alderton, Roberto Amato, Jeffrey Barrett, Sonia Goncalves, Ewan Harrison, David K. Jackson, Ian Johnston, Dominic Kwiatkowski, Cordelia Langford, John Sillitoe on behalf of the Wellcome Sanger Institute COVID-19 Surveillance Team |
| EPI_ISL_3000127 | Lighthouse Lab in Alderley Park | Wellcome Sanger Institute for the COVID-19 Genomics UK (COG-UK) Consortium | Jacquelyn Wynn, Mairead Hyland, The Lighthouse Lab in Alderley Park and Alex Alderton, Roberto Amato, Jeffrey Barrett, Sonia Goncalves, Ewan Harrison, David K. Jackson, Ian Johnston, Dominic Kwiatkowski, Cordelia Langford, John Sillitoe on behalf of the Wellcome Sanger Institute COVID-19 Surveillance Team |
| EPI_ISL_3000128, EPI_ISL_3000129 | Berkshire and Surrey Pathology Services Lighthouse Laboratory | Wellcome Sanger Institute for the COVID-19 Genomics UK (COG-UK) Consortium | Berkshire and Surrey Pathology Services Lighthouse Laboratory and Alex Alderton, Roberto Amato, Jeffrey Barrett, Sonia Goncalves, Ewan Harrison, David K. Jackson, Ian Johnston, Dominic Kwiatkowski, Cordelia Langford, John Sillitoe on behalf of the Wellcome Sanger Institute COVID-19 Surveillance Team |
| EPI_ISL_3000130 | Lighthouse Lab in Alderley Park | Wellcome Sanger Institute for the COVID-19 Genomics UK (COG-UK) Consortium | Jacquelyn Wynn, Mairead Hyland, The Lighthouse Lab in Alderley Park and Alex Alderton, Roberto Amato, Jeffrey Barrett, Sonia Goncalves, Ewan Harrison, David K. Jackson, Ian Johnston, Dominic Kwiatkowski, Cordelia Langford, John Sillitoe on behalf of the Wellcome Sanger Institute COVID-19 Surveillance Team |
| EPI_ISL_3000131 | Berkshire and Surrey Pathology Services Lighthouse Laboratory | Wellcome Sanger Institute for the COVID-19 Genomics UK (COG-UK) Consortium | Berkshire and Surrey Pathology Services Lighthouse Laboratory and Alex Alderton, Roberto Amato, Jeffrey Barrett, Sonia Goncalves, Ewan Harrison, David K. Jackson, Ian Johnston, Dominic Kwiatkowski, Cordelia Langford, John Sillitoe on behalf of the Wellcome Sanger Institute COVID-19 Surveillance Team |
| EPI_ISL_3000132 | Health Services Laboratories | Wellcome Sanger Institute for the COVID-19 Genomics UK (COG-UK) Consortium | Health Services Laboratories and Alex Alderton, Roberto Amato, Jeffrey Barrett, Sonia Goncalves, Ewan Harrison, David K. Jackson, Ian Johnston, Dominic Kwiatkowski, Cordelia Langford, John Sillitoe on behalf of the Wellcome Sanger Institute COVID-19 Surveillance Team |
| EPI_ISL_3000133, EPI_ISL_3000134, EPI_ISL_3000135 | Berkshire and Surrey Pathology Services Lighthouse Laboratory | Wellcome Sanger Institute for the COVID-19 Genomics UK (COG-UK) Consortium | Berkshire and Surrey Pathology Services Lighthouse Laboratory and Alex Alderton, Roberto Amato, Jeffrey Barrett, Sonia Goncalves, Ewan Harrison, David K. Jackson, Ian Johnston, Dominic Kwiatkowski, Cordelia Langford, John Sillitoe on behalf of the Wellcome Sanger Institute COVID-19 Surveillance Team |
| EPI_ISL_3000136, EPI_ISL_3000137 | Health Services Laboratories | Wellcome Sanger Institute for the COVID-19 Genomics UK (COG-UK) Consortium | Health Services Laboratories and Alex Alderton, Roberto Amato, Jeffrey Barrett, Sonia Goncalves, Ewan Harrison, David K. Jackson, Ian Johnston, Dominic Kwiatkowski, Cordelia Langford, John Sillitoe on behalf of the Wellcome Sanger Institute COVID-19 Surveillance Team |
| EPI_ISL_3000138, EPI_ISL_3000139 | Berkshire and Surrey Pathology Services Lighthouse Laboratory | Wellcome Sanger Institute for the COVID-19 Genomics UK (COG-UK) Consortium | Berkshire and Surrey Pathology Services Lighthouse Laboratory and Alex Alderton, Roberto Amato, Jeffrey Barrett, Sonia Goncalves, Ewan Harrison, David K. Jackson, Ian Johnston, Dominic Kwiatkowski, Cordelia Langford, John Sillitoe on behalf of the Wellcome Sanger Institute COVID-19 Surveillance Team |
| EPI_ISL_3000140 | Lighthouse Lab in Alderley Park | Wellcome Sanger Institute for the COVID-19 Genomics UK (COG-UK) Consortium | Jacquelyn Wynn, Mairead Hyland, The Lighthouse Lab in Alderley Park and Alex Alderton, Roberto Amato, Jeffrey Barrett, Sonia Goncalves, Ewan Harrison, David K. Jackson, Ian Johnston, Dominic Kwiatkowski, Cordelia Langford, John Sillitoe on behalf of the Wellcome Sanger Institute COVID-19 Surveillance Team |
| EPI_ISL_3000141, EPI_ISL_3000142, | Berkshire and Surrey Pathology Services Lighthouse Laboratory | Wellcome Sanger Institute for the COVID-19 Genomics UK (COG-UK) Consortium | Berkshire and Surrey Pathology Services Lighthouse Laboratory and Alex Alderton, Roberto Amato, Jeffrey Barrett, Sonia Goncalves, Ewan Harrison, David K. Jackson, Ian Johnston, Dominic Kwiatkowski, Cordelia Langford, John Sillitoe on behalf of the Wellcome Sanger Institute COVID-19 Surveillance Team |

| | | | |
|---|---|---|---|
| EPI_ISL_3000143 | | | |
| EPI_ISL_3000144 | Lighthouse Lab in Alderley Park | Wellcome Sanger Institute for the COVID-19 Genomics UK (COG-UK) Consortium | Jacquelyn Wynn, Mairead Hyland, The Lighthouse Lab in Alderley Park and Alex Alderton, Roberto Amato, Jeffrey Barrett, Sonia Goncalves, Ewan Harrison, David K. Jackson, Ian Johnston, Dominic Kwiatkowski, Cordelia Langford, John Sillitoe on behalf of the Wellcome Sanger Institute COVID-19 Surveillance Team |
| EPI_ISL_3000145, EPI_ISL_3000146 | Berkshire and Surrey Pathology Services Lighthouse Laboratory | Wellcome Sanger Institute for the COVID-19 Genomics UK (COG-UK) Consortium | Berkshire and Surrey Pathology Services Lighthouse Laboratory and Alex Alderton, Roberto Amato, Jeffrey Barrett, Sonia Goncalves, Ewan Harrison, David K. Jackson, Ian Johnston, Dominic Kwiatkowski, Cordelia Langford, John Sillitoe on behalf of the Wellcome Sanger Institute COVID-19 Surveillance Team |
| EPI_ISL_3000147 | Health Services Laboratories | Wellcome Sanger Institute for the COVID-19 Genomics UK (COG-UK) Consortium | Health Services Laboratories and Alex Alderton, Roberto Amato, Jeffrey Barrett, Sonia Goncalves, Ewan Harrison, David K. Jackson, Ian Johnston, Dominic Kwiatkowski, Cordelia Langford, John Sillitoe on behalf of the Wellcome Sanger Institute COVID-19 Surveillance Team |
| EPI_ISL_3000148 | Lighthouse Lab in Alderley Park | Wellcome Sanger Institute for the COVID-19 Genomics UK (COG-UK) Consortium | Jacquelyn Wynn, Mairead Hyland, The Lighthouse Lab in Alderley Park and Alex Alderton, Roberto Amato, Jeffrey Barrett, Sonia Goncalves, Ewan Harrison, David K. Jackson, Ian Johnston, Dominic Kwiatkowski, Cordelia Langford, John Sillitoe on behalf of the Wellcome Sanger Institute COVID-19 Surveillance Team |
| EPI_ISL_3000149 | Berkshire and Surrey Pathology Services Lighthouse Laboratory | Wellcome Sanger Institute for the COVID-19 Genomics UK (COG-UK) Consortium | Berkshire and Surrey Pathology Services Lighthouse Laboratory and Alex Alderton, Roberto Amato, Jeffrey Barrett, Sonia Goncalves, Ewan Harrison, David K. Jackson, Ian Johnston, Dominic Kwiatkowski, Cordelia Langford, John Sillitoe on behalf of the Wellcome Sanger Institute COVID-19 Surveillance Team |
| EPI_ISL_3000150 | Health Services Laboratories | Wellcome Sanger Institute for the COVID-19 Genomics UK (COG-UK) Consortium | Health Services Laboratories and Alex Alderton, Roberto Amato, Jeffrey Barrett, Sonia Goncalves, Ewan Harrison, David K. Jackson, Ian Johnston, Dominic Kwiatkowski, Cordelia Langford, John Sillitoe on behalf of the Wellcome Sanger Institute COVID-19 Surveillance Team |
| EPI_ISL_3000151, EPI_ISL_3000152 | Berkshire and Surrey Pathology Services Lighthouse Laboratory | Wellcome Sanger Institute for the COVID-19 Genomics UK (COG-UK) Consortium | Berkshire and Surrey Pathology Services Lighthouse Laboratory and Alex Alderton, Roberto Amato, Jeffrey Barrett, Sonia Goncalves, Ewan Harrison, David K. Jackson, Ian Johnston, Dominic Kwiatkowski, Cordelia Langford, John Sillitoe on behalf of the Wellcome Sanger Institute COVID-19 Surveillance Team |
| EPI_ISL_3000153 | Health Services Laboratories | Wellcome Sanger Institute for the COVID-19 Genomics UK (COG-UK) Consortium | Health Services Laboratories and Alex Alderton, Roberto Amato, Jeffrey Barrett, Sonia Goncalves, Ewan Harrison, David K. Jackson, Ian Johnston, Dominic Kwiatkowski, Cordelia Langford, John Sillitoe on behalf of the Wellcome Sanger Institute COVID-19 Surveillance Team |
| EPI_ISL_3000154 | Berkshire and Surrey Pathology Services Lighthouse Laboratory | Wellcome Sanger Institute for the COVID-19 Genomics UK (COG-UK) Consortium | Berkshire and Surrey Pathology Services Lighthouse Laboratory and Alex Alderton, Roberto Amato, Jeffrey Barrett, Sonia Goncalves, Ewan Harrison, David K. Jackson, Ian Johnston, Dominic Kwiatkowski, Cordelia Langford, John Sillitoe on behalf of the Wellcome Sanger Institute COVID-19 Surveillance Team |
| EPI_ISL_3000155 | Clinical Microbiology Laboratory, Faculty of Medicine, Universitas Indonesia | Clinical Microbiology Laboratory, Faculty of Medicine, Universitas Indonesia | Ibrahim F, Natasha A, Yasmon A, Fitrhiyah, Saharman YR, Karuniawati A, Sudarmono P. |
| EPI_ISL_3000157, EPI_ISL_3000160 | Centers for Disease Control, R.O.C. (Taiwan) | Centers for Disease Control, R.O.C. (Taiwan) | Ji-Rong Yang, Yu-Chi-Lin, Jung-Jung Mu, Ming-Tsan-Liu |
| EPI_ISL_3000161, EPI_ISL_3000162, EPI_ISL_3000163, EPI_ISL_3000164, EPI_ISL_3000165, EPI_ISL_3000166, EPI_ISL_3000167, EPI_ISL_3000168, EPI_ISL_3000169, EPI_ISL_3000170, EPI_ISL_3000171, EPI_ISL_3000172, EPI_ISL_3000173, EPI_ISL_3000174, EPI_ISL_3000175, EPI_ISL_3000176, EPI_ISL_3000177, EPI_ISL_3000178, EPI_ISL_3000179, EPI_ISL_3000180, EPI_ISL_3000181, EPI_ISL_3000182, EPI_ISL_3000183, EPI_ISL_3000184, EPI_ISL_3000185, EPI_ISL_3000186, EPI_ISL_3000187, EPI_ISL_3000188, EPI_ISL_3000189, EPI_ISL_3000190, EPI_ISL_3000191, EPI_ISL_3000192, EPI_ISL_3000193, EPI_ISL_3000194, EPI_ISL_3000195, EPI_ISL_3000196, EPI_ISL_3000197, EPI_ISL_3000198, EPI_ISL_3000199, EPI_ISL_3000200, EPI_ISL_3000201, EPI_ISL_3000202, EPI_ISL_3000203, EPI_ISL_3000204, EPI_ISL_3000205, EPI_ISL_3000206, EPI_ISL_3000207, EPI_ISL_3000208, EPI_ISL_3000209, EPI_ISL_3000210, EPI_ISL_3000211, EPI_ISL_3000212, EPI_ISL_3000213, EPI_ISL_3000214, EPI_ISL_3000215, EPI_ISL_3000216, EPI_ISL_3000217, EPI_ISL_3000219, EPI_ISL_3000220, EPI_ISL_3000221, EPI_ISL_3000222, EPI_ISL_3000223, EPI_ISL_3000224, EPI_ISL_3000225, EPI_ISL_3000226, EPI_ISL_3000227, EPI_ISL_3000228, EPI_ISL_3000229, EPI_ISL_3000230, EPI_ISL_3000231, EPI_ISL_3000232, EPI_ISL_3000233, EPI_ISL_3000234, EPI_ISL_3000235, EPI_ISL_3000236, EPI_ISL_3000237, EPI_ISL_3000238, EPI_ISL_3000239, EPI_ISL_3000240, EPI_ISL_3000241, EPI_ISL_3000242, EPI_ISL_3000243, EPI_ISL_3000244, EPI_ISL_3000245, EPI_ISL_3000246, EPI_ISL_3000247, EPI_ISL_3000248, EPI_ISL_3000249, EPI_ISL_3000250, EPI_ISL_3000251, EPI_ISL_3000252, EPI_ISL_3000253, EPI_ISL_3000254, EPI_ISL_3000255, EPI_ISL_3000256, EPI_ISL_3000257, EPI_ISL_3000258, EPI_ISL_3000259, EPI_ISL_3000260, EPI_ISL_3000261, EPI_ISL_3000262, EPI_ISL_3000263, EPI_ISL_3000264, EPI_ISL_3000265, EPI_ISL_3000266, EPI_ISL_3000267, EPI_ISL_3000268, EPI_ISL_3000269, EPI_ISL_3000270, EPI_ISL_3000271, EPI_ISL_3000272, EPI_ISL_3000273, EPI_ISL_3000275, EPI_ISL_3000276, EPI_ISL_3000277, EPI_ISL_3000278, EPI_ISL_3000279, EPI_ISL_3000280, EPI_ISL_3000281, EPI_ISL_3000282, EPI_ISL_3000283, EPI_ISL_3000284, EPI_ISL_3000285, EPI_ISL_3000286, EPI_ISL_3000287, EPI_ISL_3000288, EPI_ISL_3000289, EPI_ISL_3000290, EPI_ISL_3000291, EPI_ISL_3000292, EPI_ISL_3000293, EPI_ISL_3000294, EPI_ISL_3000295, EPI_ISL_3000296, EPI_ISL_3000297, EPI_ISL_3000298, EPI_ISL_3000299, EPI_ISL_3000300, EPI_ISL_3000301, EPI_ISL_3000302, EPI_ISL_3000303, EPI_ISL_3000304, EPI_ISL_3000305, EPI_ISL_3000306, EPI_ISL_3000307, EPI_ISL_3000308, EPI_ISL_3000309, EPI_ISL_3000310, EPI_ISL_3000311, EPI_ISL_3000312, EPI_ISL_3000313, EPI_ISL_3000314, EPI_ISL_3000315, EPI_ISL_3000316, EPI_ISL_3000317, EPI_ISL_3000318, EPI_ISL_3000319, EPI_ISL_3000320, EPI_ISL_3000321, EPI_ISL_3000322, EPI_ISL_3000323, EPI_ISL_3000324 | | | |
| see above | Quadram Institute Bioscience | COVID-19 Genomics UK (COG-UK) Consortium | Dave J. Baker, Gemma L. Kay, Alp Aydin, Thanh Le-Viet, Steven Rudder, Ana P. Tedim, Anastasia Kolyva, Maria Diaz, Leonardo de Oliveira Martins, Nabil-Fareed Alikhan, Lizzie Meadows, Rachael Stanley, Rose K Davidson, Ngozi Elumogo, Muhammed Yasir, Sophie J. Prosolek, Ebenezer Foster-Nyarko, Martin Lott, Christopher Jeanes, Lewis G. Spurgin, Nicholas M. Thomson, Alexander J Trotter, Rachel Gilroy, Luke Bedford, Samuel Bloomfield, Claire Stuart, Andrew Bell, Reenesh Prakash, Lindsay Coupland, Samir Dervisevic, Alison E. Mather, John Wain, Mark Webber, Andrew J. Page, Justin O'Grady |
| EPI_ISL_3000325, EPI_ISL_3000326, EPI_ISL_3000327, EPI_ISL_3000328, EPI_ISL_3000329, EPI_ISL_3000330, EPI_ISL_3000331, EPI_ISL_3000332, EPI_ISL_3000333, EPI_ISL_3000334, EPI_ISL_3000335, EPI_ISL_3000336, EPI_ISL_3000337, EPI_ISL_3000338, EPI_ISL_3000339, EPI_ISL_3000340, EPI_ISL_3000341, EPI_ISL_3000342, EPI_ISL_3000343, EPI_ISL_3000344, EPI_ISL_3000345, EPI_ISL_3000346, EPI_ISL_3000347, EPI_ISL_3000348, EPI_ISL_3000349, EPI_ISL_3000350, EPI_ISL_3000351, EPI_ISL_3000352, EPI_ISL_3000353, EPI_ISL_3000354, EPI_ISL_3000355, EPI_ISL_3000356, EPI_ISL_3000357, EPI_ISL_3000358, EPI_ISL_3000359, EPI_ISL_3000360, EPI_ISL_3000361, EPI_ISL_3000362, EPI_ISL_3000363, EPI_ISL_3000364, EPI_ISL_3000365, EPI_ISL_3000366, EPI_ISL_3000367, EPI_ISL_3000369, EPI_ISL_3000370, EPI_ISL_3000371, EPI_ISL_3000372, EPI_ISL_3000373, EPI_ISL_3000374, EPI_ISL_3000375, EPI_ISL_3000376, EPI_ISL_3000377, EPI_ISL_3000379, EPI_ISL_3000380, EPI_ISL_3000381, EPI_ISL_3000382, EPI_ISL_3000383, EPI_ISL_3000384, EPI_ISL_3000385, EPI_ISL_3000386, EPI_ISL_3000387, EPI_ISL_3000388, EPI_ISL_3000389, EPI_ISL_3000390, EPI_ISL_3000391, EPI_ISL_3000392, EPI_ISL_3000393, EPI_ISL_3000394, EPI_ISL_3000395, EPI_ISL_3000396, EPI_ISL_3000397, EPI_ISL_3000398, EPI_ISL_3000399, EPI_ISL_3000400, EPI_ISL_3000401, EPI_ISL_3000402, EPI_ISL_3000403, EPI_ISL_3000404, EPI_ISL_3000405, EPI_ISL_3000406, EPI_ISL_3000407, EPI_ISL_3000408, EPI_ISL_3000410, EPI_ISL_3000411, EPI_ISL_3000412, EPI_ISL_3000413, EPI_ISL_3000414, EPI_ISL_3000415, EPI_ISL_3000416, EPI_ISL_3000417, EPI_ISL_3000418, EPI_ISL_3000419, EPI_ISL_3000420, EPI_ISL_3000421, EPI_ISL_3000422, EPI_ISL_3000423, EPI_ISL_3000424, EPI_ISL_3000425, EPI_ISL_3000426, EPI_ISL_3000427, EPI_ISL_3000428, EPI_ISL_3000429, EPI_ISL_3000430, EPI_ISL_3000431, EPI_ISL_3000432, EPI_ISL_3000433, EPI_ISL_3000434, EPI_ISL_3000435, EPI_ISL_3000436, EPI_ISL_3000437, EPI_ISL_3000438, EPI_ISL_3000439, EPI_ISL_3000440, EPI_ISL_3000441, EPI_ISL_3000442, EPI_ISL_3000443, EPI_ISL_3000444, EPI_ISL_3000445, EPI_ISL_3000446, EPI_ISL_3000447, EPI_ISL_3000448, EPI_ISL_3000449, EPI_ISL_3000450, EPI_ISL_3000451, EPI_ISL_3000452, EPI_ISL_3000453, EPI_ISL_3000454, EPI_ISL_3000455, EPI_ISL_3000456, EPI_ISL_3000457, EPI_ISL_3000458, EPI_ISL_3000459, EPI_ISL_3000460, EPI_ISL_3000461, EPI_ISL_3000462, EPI_ISL_3000463, EPI_ISL_3000464, EPI_ISL_3000465, EPI_ISL_3000466, EPI_ISL_3000467, EPI_ISL_3000468, EPI_ISL_3000469, EPI_ISL_3000470, EPI_ISL_3000471, EPI_ISL_3000472, EPI_ISL_3000473, EPI_ISL_3000474, EPI_ISL_3000475, EPI_ISL_3000476, EPI_ISL_3000477, EPI_ISL_3000478, EPI_ISL_3000479, EPI_ISL_3000480, EPI_ISL_3000481, EPI_ISL_3000482, EPI_ISL_3000483, EPI_ISL_3000484, EPI_ISL_3000485, EPI_ISL_3000486, EPI_ISL_3000487, EPI_ISL_3000488, EPI_ISL_3000489, EPI_ISL_3000490, EPI_ISL_3000492, EPI_ISL_3000493, EPI_ISL_3000494, EPI_ISL_3000495, EPI_ISL_3000496, EPI_ISL_3000497, EPI_ISL_3000498, EPI_ISL_3000499, EPI_ISL_3000500, EPI_ISL_3000501, EPI_ISL_3000502, EPI_ISL_3000503, EPI_ISL_3000504, EPI_ISL_3000505, EPI_ISL_3000506, EPI_ISL_3000507, EPI_ISL_3000508, EPI_ISL_3000509, EPI_ISL_3000510, EPI_ISL_3000511, EPI_ISL_3000512, EPI_ISL_3000513, EPI_ISL_3000514, EPI_ISL_3000515, EPI_ISL_3000516, EPI_ISL_3000517, EPI_ISL_3000518, EPI_ISL_3000519, EPI_ISL_3000520, EPI_ISL_3000521, EPI_ISL_3000522, EPI_ISL_3000523, EPI_ISL_3000524, EPI_ISL_3000525, EPI_ISL_3000526, EPI_ISL_3000527, EPI_ISL_3000528, EPI_ISL_3000529, EPI_ISL_3000530, EPI_ISL_3000531, EPI_ISL_3000532, EPI_ISL_3000533, EPI_ISL_3000534, EPI_ISL_3000535, EPI_ISL_3000536, EPI_ISL_3000537, EPI_ISL_3000538, EPI_ISL_3000539, EPI_ISL_3000540, EPI_ISL_3000541, EPI_ISL_3000542, EPI_ISL_3000543, EPI_ISL_3000544, EPI_ISL_3000545, EPI_ISL_3000546, EPI_ISL_3000547, EPI_ISL_3000548, EPI_ISL_3000549, EPI_ISL_3000550, EPI_ISL_3000551, EPI_ISL_3000552, EPI_ISL_3000553, EPI_ISL_3000554, EPI_ISL_3000555, EPI_ISL_3000556, EPI_ISL_3000557, EPI_ISL_3000558, EPI_ISL_3000559, EPI_ISL_3000560, EPI_ISL_3000561, EPI_ISL_3000562, EPI_ISL_3000563, EPI_ISL_3000564, EPI_ISL_3000565, EPI_ISL_3000566, EPI_ISL_3000567, EPI_ISL_3000568, EPI_ISL_3000569, EPI_ISL_3000570, EPI_ISL_3000571, EPI_ISL_3000572, EPI_ISL_3000573, EPI_ISL_3000574, EPI_ISL_3000575, EPI_ISL_3000576, EPI_ISL_3000577, EPI_ISL_3000578, EPI_ISL_3000579, EPI_ISL_3000580, EPI_ISL_3000581, EPI_ISL_3000582, EPI_ISL_3000583, EPI_ISL_3000584, EPI_ISL_3000585, EPI_ISL_3000586, EPI_ISL_3000587, EPI_ISL_3000588, EPI_ISL_3000589, EPI_ISL_3000590, EPI_ISL_3000591, EPI_ISL_3000592, EPI_ISL_3000593, EPI_ISL_3000594, EPI_ISL_3000595, EPI_ISL_3000596, EPI_ISL_3000597, EPI_ISL_3000598, EPI_ISL_3000599, EPI_ISL_3000600, EPI_ISL_3000601, EPI_ISL_3000602, EPI_ISL_3000603, EPI_ISL_3000604, EPI_ISL_3000605, EPI_ISL_3000606, EPI_ISL_3000607, EPI_ISL_3000608, EPI_ISL_3000609, EPI_ISL_3000610, EPI_ISL_3000611, EPI_ISL_3000612, EPI_ISL_3000613, EPI_ISL_3000614, EPI_ISL_3000615, EPI_ISL_3000616, EPI_ISL_3000617, EPI_ISL_3000619, EPI_ISL_3000620, EPI_ISL_3000621, EPI_ISL_3000622, EPI_ISL_3000623, EPI_ISL_3000624, EPI_ISL_3000625, EPI_ISL_3000626, EPI_ISL_3000627 | | | |
| see above | University of Birmingham | COVID-19 Genomics UK (COG-UK) Consortium | Institute of Microbiology, University of Birmingham: Claire McMurray, Joanne Stockton, Samuel Nicholls, Radoslaw Poplawski, Will Rowe, Josh Quick, Nicholas Loman. University of Birmingham Testing Laboratory: Celina M Whalley, Andrew Bosworth, Charlotte Poxon, Kasun Wanigasooriya, Oliver Pickles, Mike Kidd, Alex Richter, Andrew D Beggs PHE Heartlands Lab: Husam Osman, Andrew Bosworth. Queen Elizabeth Hospital: Anna Casey |
| EPI_ISL_3000628, EPI_ISL_3000629, EPI_ISL_3000630, EPI_ISL_3000631, EPI_ISL_3000632, EPI_ISL_3000633, EPI_ISL_3000634, EPI_ISL_3000635, EPI_ISL_3000636, EPI_ISL_3000637, EPI_ISL_3000638, EPI_ISL_3000639, EPI_ISL_3000640, EPI_ISL_3000641, EPI_ISL_3000642, EPI_ISL_3000643, EPI_ISL_3000644, EPI_ISL_3000645, EPI_ISL_3000646, EPI_ISL_3000647, EPI_ISL_3000648, EPI_ISL_3000649, EPI_ISL_3000650, EPI_ISL_3000651, EPI_ISL_3000652, EPI_ISL_3000653, EPI_ISL_3000654, EPI_ISL_3000655, EPI_ISL_3000656, EPI_ISL_3000657, EPI_ISL_3000658, EPI_ISL_3000659, EPI_ISL_3000661, EPI_ISL_3000663, EPI_ISL_3000664, EPI_ISL_3000665, EPI_ISL_3000666, EPI_ISL_3000668, EPI_ISL_3000669, EPI_ISL_3000670, EPI_ISL_3000671, EPI_ISL_3000672, EPI_ISL_3000673, EPI_ISL_3000674, EPI_ISL_3000675, EPI_ISL_3000676, EPI_ISL_3000677, EPI_ISL_3000678, EPI_ISL_3000679, EPI_ISL_3000680, EPI_ISL_3000681, EPI_ISL_3000682, EPI_ISL_3000683, EPI_ISL_3000684, EPI_ISL_3000685, EPI_ISL_3000686, EPI_ISL_3000687, EPI_ISL_3000688, EPI_ISL_3000689, EPI_ISL_3000690, EPI_ISL_3000691, EPI_ISL_3000692, EPI_ISL_3000693, EPI_ISL_3000694, EPI_ISL_3000695, EPI_ISL_3000696, EPI_ISL_3000697, EPI_ISL_3000698, EPI_ISL_3000699, EPI_ISL_3000700, EPI_ISL_3000701, EPI_ISL_3000702, EPI_ISL_3000703, EPI_ISL_3000704, EPI_ISL_3000705, EPI_ISL_3000706, EPI_ISL_3000707, EPI_ISL_3000708, EPI_ISL_3000709, EPI_ISL_3000710, EPI_ISL_3000711, EPI_ISL_3000712, EPI_ISL_3000713, EPI_ISL_3000714, EPI_ISL_3000715, EPI_ISL_3000716, EPI_ISL_3000717, EPI_ISL_3000718, EPI_ISL_3000719, EPI_ISL_3000720, EPI_ISL_3000721, EPI_ISL_3000722, EPI_ISL_3000723, EPI_ISL_3000724, EPI_ISL_3000725, EPI_ISL_3000726, EPI_ISL_3000727, EPI_ISL_3000728, EPI_ISL_3000729, EPI_ISL_3000730, EPI_ISL_3000731, EPI_ISL_3000732, EPI_ISL_3000733, EPI_ISL_3000734, EPI_ISL_3000735, EPI_ISL_3000736, EPI_ISL_3000737, EPI_ISL_3000738, EPI_ISL_3000739, EPI_ISL_3000740, EPI_ISL_3000741, EPI_ISL_3000742, EPI_ISL_3000743, EPI_ISL_3000744, EPI_ISL_3000745, EPI_ISL_3000746, EPI_ISL_3000747, EPI_ISL_3000748, EPI_ISL_3000749, EPI_ISL_3000750, EPI_ISL_3000751, EPI_ISL_3000752, EPI_ISL_3000753, EPI_ISL_3000754, EPI_ISL_3000755, EPI_ISL_3000756, EPI_ISL_3000757, EPI_ISL_3000758, EPI_ISL_3000759, EPI_ISL_3000760, EPI_ISL_3000761, EPI_ISL_3000762, EPI_ISL_3000763, EPI_ISL_3000764, EPI_ISL_3000765, EPI_ISL_3000766, EPI_ISL_3000767, EPI_ISL_3000768, EPI_ISL_3000769, EPI_ISL_3000770, EPI_ISL_3000771, EPI_ISL_3000772, EPI_ISL_3000773, EPI_ISL_3000774, EPI_ISL_3000775, EPI_ISL_3000776, EPI_ISL_3000777, EPI_ISL_3000778, EPI_ISL_3000779, EPI_ISL_3000780, EPI_ISL_3000781, EPI_ISL_3000782, EPI_ISL_3000783, EPI_ISL_3000784, EPI_ISL_3000785, EPI_ISL_3000786, EPI_ISL_3000787, EPI_ISL_3000788, EPI_ISL_3000789, EPI_ISL_3000791, EPI_ISL_3000792, EPI_ISL_3000793, EPI_ISL_3000794, EPI_ISL_3000795, EPI_ISL_3000796, EPI_ISL_3000797, EPI_ISL_3000798, EPI_ISL_3000799, EPI_ISL_3000800, EPI_ISL_3000801, EPI_ISL_3000802, EPI_ISL_3000803, EPI_ISL_3000804, EPI_ISL_3000805, EPI_ISL_3000806, EPI_ISL_3000807, EPI_ISL_3000808, EPI_ISL_3000809, EPI_ISL_3000810, EPI_ISL_3000811, EPI_ISL_3000812, EPI_ISL_3000813, EPI_ISL_3000814, EPI_ISL_3000815, EPI_ISL_3000816, EPI_ISL_3000817, EPI_ISL_3000818, EPI_ISL_3000819, EPI_ISL_3000820, EPI_ISL_3000821, EPI_ISL_3000822, EPI_ISL_3000823, EPI_ISL_3000824, EPI_ISL_3000825, EPI_ISL_3000826, EPI_ISL_3000827, EPI_ISL_3000828, EPI_ISL_3000829, EPI_ISL_3000830, EPI_ISL_3000831, EPI_ISL_3000832, EPI_ISL_3000833, EPI_ISL_3000834, EPI_ISL_3000835, EPI_ISL_3000836, EPI_ISL_3000837, EPI_ISL_3000838, EPI_ISL_3000839, EPI_ISL_3000840, EPI_ISL_3000841, EPI_ISL_3000842, EPI_ISL_3000843, EPI_ISL_3000844, EPI_ISL_3000845, EPI_ISL_3000846, EPI_ISL_3000847, EPI_ISL_3000848, EPI_ISL_3000849, EPI_ISL_3000850, EPI_ISL_3000851, EPI_ISL_3000852, EPI_ISL_3000853, EPI_ISL_3000854, EPI_ISL_3000855, EPI_ISL_3000856, EPI_ISL_3000857, EPI_ISL_3000858, EPI_ISL_3000859, EPI_ISL_3000860, EPI_ISL_3000861, EPI_ISL_3000862, EPI_ISL_3000863, EPI_ISL_3000864, EPI_ISL_3000865, EPI_ISL_3000866, EPI_ISL_3000867, EPI_ISL_3000868, EPI_ISL_3000869, EPI_ISL_3000870, EPI_ISL_3000871, EPI_ISL_3000872, EPI_ISL_3000873, EPI_ISL_3000874, EPI_ISL_3000875, EPI_ISL_3000876, EPI_ISL_3000877, EPI_ISL_3000878, EPI_ISL_3000879, EPI_ISL_3000880, EPI_ISL_3000881, EPI_ISL_3000882, EPI_ISL_3000883, EPI_ISL_3000884, EPI_ISL_3000885, EPI_ISL_3000886, EPI_ISL_3000887, EPI_ISL_3000888, EPI_ISL_3000889, EPI_ISL_3000890, EPI_ISL_3000891, EPI_ISL_3000892, EPI_ISL_3000893, EPI_ISL_3000894, EPI_ISL_3000895, EPI_ISL_3000896, EPI_ISL_3000897, EPI_ISL_3000898, EPI_ISL_3000899, EPI_ISL_3000900, EPI_ISL_3000901, EPI_ISL_3000902, EPI_ISL_3000903, EPI_ISL_3000904, EPI_ISL_3000905, EPI_ISL_3000906, EPI_ISL_3000907, EPI_ISL_3000908, EPI_ISL_3000909, EPI_ISL_3000910, EPI_ISL_3000911, EPI_ISL_3000912, EPI_ISL_3000913, EPI_ISL_3000914, EPI_ISL_3000915, EPI_ISL_3000916, EPI_ISL_3000917, EPI_ISL_3000918, EPI_ISL_3000919, EPI_ISL_3000920, EPI_ISL_3000921, EPI_ISL_3000922, EPI_ISL_3000923, EPI_ISL_3000924, EPI_ISL_3000925, EPI_ISL_3000926, EPI_ISL_3000927, EPI_ISL_3000928, EPI_ISL_3000929, EPI_ISL_3000930, EPI_ISL_3000931, EPI_ISL_3000932, EPI_ISL_3000933, EPI_ISL_3000934, EPI_ISL_3000935, EPI_ISL_3000936, EPI_ISL_3000937, EPI_ISL_3000938, EPI_ISL_3000939, EPI_ISL_3000940, EPI_ISL_3000941, EPI_ISL_3000942, EPI_ISL_3000943, EPI_ISL_3000944, EPI_ISL_3000945, EPI_ISL_3000946, EPI_ISL_3000947, EPI_ISL_3000948, EPI_ISL_3000949, EPI_ISL_3000950, EPI_ISL_3000951, EPI_ISL_3000952, EPI_ISL_3000953, EPI_ISL_3000954, EPI_ISL_3000955, EPI_ISL_3000956, EPI_ISL_3000957, EPI_ISL_3000958, EPI_ISL_3000959, EPI_ISL_3000960, EPI_ISL_3000961, EPI_ISL_3000962, EPI_ISL_3000963, EPI_ISL_3000964, EPI_ISL_3000965, EPI_ISL_3000966 | | | |
| see above | Department of Pathology, University of Cambridge | COVID-19 Genomics UK (COG-UK) Consortium | Aminu S. Jahun, Yasmin Chaudhry, Iliana Georgana, Myra Hosmillo, Rhys Izuagbe, William L. Hamilton, Martin D. Curran, Surendra Parmar, Ian Goodfellow |
| EPI_ISL_3000967, EPI_ISL_3000968, EPI_ISL_3000969, EPI_ISL_3000970, EPI_ISL_3000971, EPI_ISL_3000972, EPI_ISL_3000973, EPI_ISL_3000974, EPI_ISL_3000975, EPI_ISL_3000976, EPI_ISL_3000977, EPI_ISL_3000978, EPI_ISL_3000979, EPI_ISL_3000980, EPI_ISL_3000981, EPI_ISL_3000982, EPI_ISL_3000983, EPI_ISL_3000984, EPI_ISL_3000985, EPI_ISL_3000986, EPI_ISL_3000987, EPI_ISL_3000988, EPI_ISL_3000989, EPI_ISL_3000990, EPI_ISL_3000991, EPI_ISL_3000992, EPI_ISL_3000993, EPI_ISL_3000994, EPI_ISL_3000995, EPI_ISL_3000996, EPI_ISL_3000999, EPI_ISL_3001000, EPI_ISL_3001001, EPI_ISL_3001002, EPI_ISL_3001003, EPI_ISL_3001004, EPI_ISL_3001005, EPI_ISL_3001006, EPI_ISL_3001007, EPI_ISL_3001008, EPI_ISL_3001009, EPI_ISL_3001010, EPI_ISL_3001011, EPI_ISL_3001012, EPI_ISL_3001013, EPI_ISL_3001014, EPI_ISL_3001015, EPI_ISL_3001016, EPI_ISL_3001017, EPI_ISL_3001018, EPI_ISL_3001019, EPI_ISL_3001020, EPI_ISL_3001021, EPI_ISL_3001022, EPI_ISL_3001023, EPI_ISL_3001024, EPI_ISL_3001025, EPI_ISL_3001026, EPI_ISL_3001027, EPI_ISL_3001028, EPI_ISL_3001029, EPI_ISL_3001030, EPI_ISL_3001031, EPI_ISL_3001032, EPI_ISL_3001033, EPI_ISL_3001034, EPI_ISL_3001035, EPI_ISL_3001037, EPI_ISL_3001038, EPI_ISL_3001039, EPI_ISL_3001040, EPI_ISL_3001041, EPI_ISL_3001042, EPI_ISL_3001043, EPI_ISL_3001044, EPI_ISL_3001045, EPI_ISL_3001046, EPI_ISL_3001047, EPI_ISL_3001048, EPI_ISL_3001049, EPI_ISL_3001050, EPI_ISL_3001051, EPI_ISL_3001052, EPI_ISL_3001053, EPI_ISL_3001054, EPI_ISL_3001056, EPI_ISL_3001057, EPI_ISL_3001058, EPI_ISL_3001059, EPI_ISL_3001060, EPI_ISL_3001061, EPI_ISL_3001062, EPI_ISL_3001063, EPI_ISL_3001064, EPI_ISL_3001065, EPI_ISL_3001066, EPI_ISL_3001068, EPI_ISL_3001069, EPI_ISL_3001070, EPI_ISL_3001071, EPI_ISL_3001072, EPI_ISL_3001073, EPI_ISL_3001075, EPI_ISL_3001076, EPI_ISL_3001077, EPI_ISL_3001078, EPI_ISL_3001079, EPI_ISL_3001080, EPI_ISL_3001081, EPI_ISL_3001082, EPI_ISL_3001083, EPI_ISL_3001084, EPI_ISL_3001085, EPI_ISL_3001086, EPI_ISL_3001087, EPI_ISL_3001088, EPI_ISL_3001089, EPI_ISL_3001090, EPI_ISL_3001091, EPI_ISL_3001092, EPI_ISL_3001093, EPI_ISL_3001094, EPI_ISL_3001095, EPI_ISL_3001096, EPI_ISL_3001097, EPI_ISL_3001098, EPI_ISL_3001099, EPI_ISL_3001101, EPI_ISL_3001102, EPI_ISL_3001103, EPI_ISL_3001104, EPI_ISL_3001105, EPI_ISL_3001106, EPI_ISL_3001107, EPI_ISL_3001108, EPI_ISL_3001109, EPI_ISL_3001110, EPI_ISL_3001111, EPI_ISL_3001112, EPI_ISL_3001113, EPI_ISL_3001114, EPI_ISL_3001115, EPI_ISL_3001116, EPI_ISL_3001117, EPI_ISL_3001118, EPI_ISL_3001119, EPI_ISL_3001120, EPI_ISL_3001121, EPI_ISL_3001122, EPI_ISL_3001123, EPI_ISL_3001124, EPI_ISL_3001125, EPI_ISL_3001126, EPI_ISL_3001127, EPI_ISL_3001128, EPI_ISL_3001129, EPI_ISL_3001130, EPI_ISL_3001131, EPI_ISL_3001132, EPI_ISL_3001133, EPI_ISL_3001134, EPI_ISL_3001135, EPI_ISL_3001136, EPI_ISL_3001137, EPI_ISL_3001138, EPI_ISL_3001139, EPI_ISL_3001140, EPI_ISL_3001141, EPI_ISL_3001142, EPI_ISL_3001143, EPI_ISL_3001144, EPI_ISL_3001145, EPI_ISL_3001146, EPI_ISL_3001147, EPI_ISL_3001148, EPI_ISL_3001149, EPI_ISL_3001150, EPI_ISL_3001151, EPI_ISL_3001152, EPI_ISL_3001153 | | | |
| see above | West of Scotland Specialist Virology Centre, NHSGGC / MRC-University of Glasgow Centre for Virus Research | COVID-19 Genomics UK (COG-UK) Consortium | Ana da Silva Filipe, Natasha Johnson, Kathy Smollett, Daniel Mair, Stephen Carmichael, Alice Broos, Lily Tong, Jenna Nichols, Kyriaki Nomikou; Sarah McDonald; Ioulia Tsatsani, Richard Orton, Joseph Hughes, Sreenu Vattipally, David L Robertson; Alasdair MacLean, Rory Gunson; Sharif Shaaban, Matthew Holden; Rachel Blacow, Guy Mollett, Kathy Li, James Shepherd, Antonia Ho, Emma Thomson |
| EPI_ISL_3001154, EPI_ISL_3001155, EPI_ISL_3001156, EPI_ISL_3001157, EPI_ISL_3001159, EPI_ISL_3001160, EPI_ISL_3001161, EPI_ISL_3001162, EPI_ISL_3001163, EPI_ISL_3001164, EPI_ISL_3001165, EPI_ISL_3001166, EPI_ISL_3001167, EPI_ISL_3001168, EPI_ISL_3001169, EPI_ISL_3001170, EPI_ISL_3001171, EPI_ISL_3001172, EPI_ISL_3001173, EPI_ISL_3001174, EPI_ISL_3001175, EPI_ISL_3001176, EPI_ISL_3001177, EPI_ISL_3001178, EPI_ISL_3001179, EPI_ISL_3001180, EPI_ISL_3001181, EPI_ISL_3001182, EPI_ISL_3001183, EPI_ISL_3001184, EPI_ISL_3001185, EPI_ISL_3001186, EPI_ISL_3001187, EPI_ISL_3001188, EPI_ISL_3001190, EPI_ISL_3001191, EPI_ISL_3001192, EPI_ISL_3001193, EPI_ISL_3001194, EPI_ISL_3001195, EPI_ISL_3001196, EPI_ISL_3001197, EPI_ISL_3001198, EPI_ISL_3001199, EPI_ISL_3001200, EPI_ISL_3001201, EPI_ISL_3001202, EPI_ISL_3001203, EPI_ISL_3001204, EPI_ISL_3001205, EPI_ISL_3001206, EPI_ISL_3001207, EPI_ISL_3001208, EPI_ISL_3001209, EPI_ISL_3001210, EPI_ISL_3001211, EPI_ISL_3001212, EPI_ISL_3001213, EPI_ISL_3001214, EPI_ISL_3001215, EPI_ISL_3001216, EPI_ISL_3001217, EPI_ISL_3001218, EPI_ISL_3001219, EPI_ISL_3001220, EPI_ISL_3001221, EPI_ISL_3001222, EPI_ISL_3001223, EPI_ISL_3001224, EPI_ISL_3001225, EPI_ISL_3001226, EPI_ISL_3001227, EPI_ISL_3001228, EPI_ISL_3001229, EPI_ISL_3001230, EPI_ISL_3001231, EPI_ISL_3001232, EPI_ISL_3001233, EPI_ISL_3001234, EPI_ISL_3001235, EPI_ISL_3001236, EPI_ISL_3001237, EPI_ISL_3001238, EPI_ISL_3001239, EPI_ISL_3001240, EPI_ISL_3001241, EPI_ISL_3001242, EPI_ISL_3001243, EPI_ISL_3001244, EPI_ISL_3001245, EPI_ISL_3001246, EPI_ISL_3001247, EPI_ISL_3001248, EPI_ISL_3001249, EPI_ISL_3001250, EPI_ISL_3001251, EPI_ISL_3001252, EPI_ISL_3001253, EPI_ISL_3001254, EPI_ISL_3001255, EPI_ISL_3001256, EPI_ISL_3001258, EPI_ISL_3001259, EPI_ISL_3001260, EPI_ISL_3001261, EPI_ISL_3001262, EPI_ISL_3001263, EPI_ISL_3001264, EPI_ISL_3001266, EPI_ISL_3001267, EPI_ISL_3001269, EPI_ISL_3001270, EPI_ISL_3001271, EPI_ISL_3001272 | | | |
| see above | Virology Department, Royal Infirmary of Edinburgh, NHS Lothian / School of Biological Sciences, University of Edinburgh | COVID-19 Genomics UK (COG-UK) Consortium | McHugh M, Dewar R, Cotton S, Gallagher A, Maloney D, Fernandez G, O'Toole Á, Scher E, Hill V, McCrone JT, Colquhoun R, Yu X, Jackson B, Rambaut A, Templeton K |
| EPI_ISL_3001274, EPI_ISL_3001275, EPI_ISL_3001276, EPI_ISL_3001277, EPI_ISL_3001278, EPI_ISL_3001279, EPI_ISL_3001280, EPI_ISL_3001281, EPI_ISL_3001282, EPI_ISL_3001283, EPI_ISL_3001284, EPI_ISL_3001285, EPI_ISL_3001286, EPI_ISL_3001287, EPI_ISL_3001288, EPI_ISL_3001289, EPI_ISL_3001290, EPI_ISL_3001291, EPI_ISL_3001292, EPI_ISL_3001293, EPI_ISL_3001294, EPI_ISL_3001295, EPI_ISL_3001296, EPI_ISL_3001297, EPI_ISL_3001298, EPI_ISL_3001299, EPI_ISL_3001300, EPI_ISL_3001301, EPI_ISL_3001302, EPI_ISL_3001303, EPI_ISL_3001304, EPI_ISL_3001305, EPI_ISL_3001306, EPI_ISL_3001307, EPI_ISL_3001308, EPI_ISL_3001309, EPI_ISL_3001310, EPI_ISL_3001311, EPI_ISL_3001312, EPI_ISL_3001313, EPI_ISL_3001314, EPI_ISL_3001315, EPI_ISL_3001316, EPI_ISL_3001317, EPI_ISL_3001318, EPI_ISL_3001319, EPI_ISL_3001320, EPI_ISL_3001321, EPI_ISL_3001322, EPI_ISL_3001323, EPI_ISL_3001324, EPI_ISL_3001325, EPI_ISL_3001326, EPI_ISL_3001327, EPI_ISL_3001328, EPI_ISL_3001329, EPI_ISL_3001330, EPI_ISL_3001331, EPI_ISL_3001332, EPI_ISL_3001333, EPI_ISL_3001334, EPI_ISL_3001335 | | | |
| see above | University College London, Great Ormond Street Hospital for Children NHS Foundation Trust, Imperial College Healthcare NHS Trust | COVID-19 Genomics UK (COG-UK) Consortium | Sergi Castellano, Rachel Williams, Mark Kristiansen, Sunando Roy, Charlotte Williams, Nadua Bayzid, Marius Cotic, Helena Tutill, Patricia Dyal, Judith Breuer |
| EPI_ISL_3001336, EPI_ISL_3001337, EPI_ISL_3001338, EPI_ISL_3001339, EPI_ISL_3001340, EPI_ISL_3001341, EPI_ISL_3001342, EPI_ISL_3001343, EPI_ISL_3001344, EPI_ISL_3001345, EPI_ISL_3001346, EPI_ISL_3001347, EPI_ISL_3001348, EPI_ISL_3001349, EPI_ISL_3001350, EPI_ISL_3001351, EPI_ISL_3001352, EPI_ISL_3001353, EPI_ISL_3001354, EPI_ISL_3001355, EPI_ISL_3001356, EPI_ISL_3001357, EPI_ISL_3001358, EPI_ISL_3001359, EPI_ISL_3001360, EPI_ISL_3001361, EPI_ISL_3001362, EPI_ISL_3001363, EPI_ISL_3001364, EPI_ISL_3001365, EPI_ISL_3001366, EPI_ISL_3001367, EPI_ISL_3001369, EPI_ISL_3001370, EPI_ISL_3001371, EPI_ISL_3001372, EPI_ISL_3001373, EPI_ISL_3001374, EPI_ISL_3001375, EPI_ISL_3001376, EPI_ISL_3001377, EPI_ISL_3001378, EPI_ISL_3001379, EPI_ISL_3001380, EPI_ISL_3001381, EPI_ISL_3001382, EPI_ISL_3001383, EPI_ISL_3001384, EPI_ISL_3001385, EPI_ISL_3001386, EPI_ISL_3001387, EPI_ISL_3001388, EPI_ISL_3001389, EPI_ISL_3001390, EPI_ISL_3001391, EPI_ISL_3001392, EPI_ISL_3001393, EPI_ISL_3001394, EPI_ISL_3001395, EPI_ISL_3001396, EPI_ISL_3001397, EPI_ISL_3001398, EPI_ISL_3001399, EPI_ISL_3001400, EPI_ISL_3001401, EPI_ISL_3001402, EPI_ISL_3001403, EPI_ISL_3001404, EPI_ISL_3001405, EPI_ISL_3001406, EPI_ISL_3001407, EPI_ISL_3001408, EPI_ISL_3001409, EPI_ISL_3001410, EPI_ISL_3001411, EPI_ISL_3001412, EPI_ISL_3001413, EPI_ISL_3001414, EPI_ISL_3001415, EPI_ISL_3001416, EPI_ISL_3001417, EPI_ISL_3001418, EPI_ISL_3001419, EPI_ISL_3001420, EPI_ISL_3001421, EPI_ISL_3001422, EPI_ISL_3001423, EPI_ISL_3001424, EPI_ISL_3001425, EPI_ISL_3001426, EPI_ISL_3001427, EPI_ISL_3001428, EPI_ISL_3001429, EPI_ISL_3001430, EPI_ISL_3001431, EPI_ISL_3001432, EPI_ISL_3001433, EPI_ISL_3001434, EPI_ISL_3001435, EPI_ISL_3001436, EPI_ISL_3001437, EPI_ISL_3001438, EPI_ISL_3001439, EPI_ISL_3001440, EPI_ISL_3001441, EPI_ISL_3001442, EPI_ISL_3001443, EPI_ISL_3001444, EPI_ISL_3001445, EPI_ISL_3001446, EPI_ISL_3001447, EPI_ISL_3001448, EPI_ISL_3001449, EPI_ISL_3001450, EPI_ISL_3001451, EPI_ISL_3001452, EPI_ISL_3001453, EPI_ISL_3001454, EPI_ISL_3001455, EPI_ISL_3001456, EPI_ISL_3001457, EPI_ISL_3001458, EPI_ISL_3001459, EPI_ISL_3001460, EPI_ISL_3001461, EPI_ISL_3001462, EPI_ISL_3001463, EPI_ISL_3001464, EPI_ISL_3001465, EPI_ISL_3001466, EPI_ISL_3001467, EPI_ISL_3001468, EPI_ISL_3001469, EPI_ISL_3001470, EPI_ISL_3001471, EPI_ISL_3001472, EPI_ISL_3001473, EPI_ISL_3001474, EPI_ISL_3001475, EPI_ISL_3001476, EPI_ISL_3001477, EPI_ISL_3001478, EPI_ISL_3001479, EPI_ISL_3001480, EPI_ISL_3001481, EPI_ISL_3001482, EPI_ISL_3001483, EPI_ISL_3001484, EPI_ISL_3001485, EPI_ISL_3001486, EPI_ISL_3001487, EPI_ISL_3001488, EPI_ISL_3001489, EPI_ISL_3001490, | | | |

EPI_ISL_3001491, EPI_ISL_3001492, EPI_ISL_3001493, EPI_ISL_3001494, EPI_ISL_3001495, EPI_ISL_3001496, EPI_ISL_3001497, EPI_ISL_3001498, EPI_ISL_3001499, EPI_ISL_3001500, EPI_ISL_3001501, EPI_ISL_3001502, EPI_ISL_3001503, EPI_ISL_3001504, EPI_ISL_3001505, EPI_ISL_3001506, EPI_ISL_3001507, EPI_ISL_3001508, EPI_ISL_3001509, EPI_ISL_3001510, EPI_ISL_3001511, EPI_ISL_3001512,
EPI_ISL_3001513, EPI_ISL_3001514, EPI_ISL_3001515, EPI_ISL_3001516, EPI_ISL_3001517, EPI_ISL_3001518, EPI_ISL_3001519, EPI_ISL_3001520, EPI_ISL_3001521, EPI_ISL_3001522, EPI_ISL_3001523, EPI_ISL_3001524, EPI_ISL_3001525, EPI_ISL_3001526, EPI_ISL_3001527, EPI_ISL_3001528, EPI_ISL_3001529, EPI_ISL_3001530, EPI_ISL_3001531, EPI_ISL_3001532, EPI_ISL_3001533, EPI_ISL_3001534,
EPI_ISL_3001535, EPI_ISL_3001536, EPI_ISL_3001537, EPI_ISL_3001538, EPI_ISL_3001539, EPI_ISL_3001540, EPI_ISL_3001541, EPI_ISL_3001542, EPI_ISL_3001543, EPI_ISL_3001544, EPI_ISL_3001545, EPI_ISL_3001547, EPI_ISL_3001548, EPI_ISL_3001549, EPI_ISL_3001550, EPI_ISL_3001551, EPI_ISL_3001552, EPI_ISL_3001553, EPI_ISL_3001554, EPI_ISL_3001555, EPI_ISL_3001556, EPI_ISL_3001557,
EPI_ISL_3001558, EPI_ISL_3001559, EPI_ISL_3001560, EPI_ISL_3001561, EPI_ISL_3001562, EPI_ISL_3001563, EPI_ISL_3001564, EPI_ISL_3001565, EPI_ISL_3001566, EPI_ISL_3001567, EPI_ISL_3001568, EPI_ISL_3001569, EPI_ISL_3001570, EPI_ISL_3001571, EPI_ISL_3001572, EPI_ISL_3001573, EPI_ISL_3001574, EPI_ISL_3001575, EPI_ISL_3001576, EPI_ISL_3001577, EPI_ISL_3001578, EPI_ISL_3001579,
EPI_ISL_3001580, EPI_ISL_3001581, EPI_ISL_3001582, EPI_ISL_3001583, EPI_ISL_3001584, EPI_ISL_3001585, EPI_ISL_3001586, EPI_ISL_3001587, EPI_ISL_3001588, EPI_ISL_3001589, EPI_ISL_3001590, EPI_ISL_3001591, EPI_ISL_3001592, EPI_ISL_3001593, EPI_ISL_3001594, EPI_ISL_3001595, EPI_ISL_3001596, EPI_ISL_3001597, EPI_ISL_3001598, EPI_ISL_3001599, EPI_ISL_3001600, EPI_ISL_3001601,
EPI_ISL_3001602, EPI_ISL_3001603, EPI_ISL_3001604, EPI_ISL_3001605, EPI_ISL_3001606, EPI_ISL_3001607, EPI_ISL_3001608, EPI_ISL_3001609, EPI_ISL_3001610, EPI_ISL_3001611, EPI_ISL_3001612, EPI_ISL_3001613, EPI_ISL_3001614, EPI_ISL_3001615, EPI_ISL_3001616, EPI_ISL_3001617, EPI_ISL_3001618, EPI_ISL_3001619, EPI_ISL_3001620, EPI_ISL_3001621, EPI_ISL_3001622, EPI_ISL_3001623,
EPI_ISL_3001624, EPI_ISL_3001625, EPI_ISL_3001626, EPI_ISL_3001627, EPI_ISL_3001628, EPI_ISL_3001629, EPI_ISL_3001630, EPI_ISL_3001631, EPI_ISL_3001632, EPI_ISL_3001633, EPI_ISL_3001634, EPI_ISL_3001635, EPI_ISL_3001636, EPI_ISL_3001637, EPI_ISL_3001638, EPI_ISL_3001639, EPI_ISL_3001640, EPI_ISL_3001641, EPI_ISL_3001642, EPI_ISL_3001643, EPI_ISL_3001644, EPI_ISL_3001645,
EPI_ISL_3001646, EPI_ISL_3001647, EPI_ISL_3001648, EPI_ISL_3001649, EPI_ISL_3001650, EPI_ISL_3001651, EPI_ISL_3001652, EPI_ISL_3001653, EPI_ISL_3001654, EPI_ISL_3001655, EPI_ISL_3001656, EPI_ISL_3001657, EPI_ISL_3001658, EPI_ISL_3001659, EPI_ISL_3001660, EPI_ISL_3001661, EPI_ISL_3001662, EPI_ISL_3001663, EPI_ISL_3001664, EPI_ISL_3001665, EPI_ISL_3001666, EPI_ISL_3001667,
EPI_ISL_3001668, EPI_ISL_3001669, EPI_ISL_3001670, EPI_ISL_3001671, EPI_ISL_3001672, EPI_ISL_3001673, EPI_ISL_3001674, EPI_ISL_3001675, EPI_ISL_3001676, EPI_ISL_3001677, EPI_ISL_3001678, EPI_ISL_3001679, EPI_ISL_3001680, EPI_ISL_3001681, EPI_ISL_3001682, EPI_ISL_3001683, EPI_ISL_3001684, EPI_ISL_3001685, EPI_ISL_3001686, EPI_ISL_3001687, EPI_ISL_3001688, EPI_ISL_3001689,
EPI_ISL_3001690, EPI_ISL_3001691, EPI_ISL_3001692, EPI_ISL_3001693, EPI_ISL_3001694, EPI_ISL_3001695, EPI_ISL_3001696, EPI_ISL_3001697, EPI_ISL_3001698, EPI_ISL_3001699, EPI_ISL_3001700, EPI_ISL_3001701, EPI_ISL_3001702, EPI_ISL_3001703, EPI_ISL_3001704, EPI_ISL_3001705, EPI_ISL_3001706, EPI_ISL_3001707, EPI_ISL_3001708, EPI_ISL_3001709, EPI_ISL_3001710, EPI_ISL_3001711,
EPI_ISL_3001712, EPI_ISL_3001713, EPI_ISL_3001714, EPI_ISL_3001715, EPI_ISL_3001716, EPI_ISL_3001717, EPI_ISL_3001718, EPI_ISL_3001719, EPI_ISL_3001720, EPI_ISL_3001721, EPI_ISL_3001722, EPI_ISL_3001723, EPI_ISL_3001724, EPI_ISL_3001725, EPI_ISL_3001726, EPI_ISL_3001727, EPI_ISL_3001728, EPI_ISL_3001729, EPI_ISL_3001730, EPI_ISL_3001731, EPI_ISL_3001732, EPI_ISL_3001733,
EPI_ISL_3001734, EPI_ISL_3001735, EPI_ISL_3001736, EPI_ISL_3001737, EPI_ISL_3001738, EPI_ISL_3001739, EPI_ISL_3001740, EPI_ISL_3001741, EPI_ISL_3001742, EPI_ISL_3001743, EPI_ISL_3001744, EPI_ISL_3001745, EPI_ISL_3001746, EPI_ISL_3001747, EPI_ISL_3001748, EPI_ISL_3001749, EPI_ISL_3001750, EPI_ISL_3001751, EPI_ISL_3001752, EPI_ISL_3001753, EPI_ISL_3001754, EPI_ISL_3001755,
EPI_ISL_3001756, EPI_ISL_3001757, EPI_ISL_3001758, EPI_ISL_3001759, EPI_ISL_3001760, EPI_ISL_3001761, EPI_ISL_3001762, EPI_ISL_3001763, EPI_ISL_3001764, EPI_ISL_3001765, EPI_ISL_3001766, EPI_ISL_3001767, EPI_ISL_3001768, EPI_ISL_3001769, EPI_ISL_3001770, EPI_ISL_3001771, EPI_ISL_3001772, EPI_ISL_3001773, EPI_ISL_3001774, EPI_ISL_3001775, EPI_ISL_3001776, EPI_ISL_3001777,
EPI_ISL_3001778, EPI_ISL_3001779, EPI_ISL_3001780, EPI_ISL_3001781, EPI_ISL_3001782, EPI_ISL_3001783, EPI_ISL_3001784, EPI_ISL_3001785, EPI_ISL_3001786, EPI_ISL_3001787, EPI_ISL_3001788, EPI_ISL_3001789, EPI_ISL_3001790, EPI_ISL_3001791, EPI_ISL_3001792, EPI_ISL_3001793, EPI_ISL_3001794, EPI_ISL_3001795, EPI_ISL_3001796, EPI_ISL_3001797, EPI_ISL_3001798, EPI_ISL_3001799,
EPI_ISL_3001800, EPI_ISL_3001801, EPI_ISL_3001802, EPI_ISL_3001803, EPI_ISL_3001804, EPI_ISL_3001805, EPI_ISL_3001806, EPI_ISL_3001807, EPI_ISL_3001808, EPI_ISL_3001809, EPI_ISL_3001810, EPI_ISL_3001811, EPI_ISL_3001812, EPI_ISL_3001813, EPI_ISL_3001814, EPI_ISL_3001815, EPI_ISL_3001816, EPI_ISL_3001817, EPI_ISL_3001818, EPI_ISL_3001819, EPI_ISL_3001820, EPI_ISL_3001821,
EPI_ISL_3001822, EPI_ISL_3001823, EPI_ISL_3001824, EPI_ISL_3001825, EPI_ISL_3001826, EPI_ISL_3001827, EPI_ISL_3001828, EPI_ISL_3001829, EPI_ISL_3001830, EPI_ISL_3001831, EPI_ISL_3001832, EPI_ISL_3001833, EPI_ISL_3001834, EPI_ISL_3001835, EPI_ISL_3001836, EPI_ISL_3001837, EPI_ISL_3001838, EPI_ISL_3001839, EPI_ISL_3001840, EPI_ISL_3001842, EPI_ISL_3001843, EPI_ISL_3001844,
EPI_ISL_3001845, EPI_ISL_3001846, EPI_ISL_3001847, EPI_ISL_3001848, EPI_ISL_3001849, EPI_ISL_3001850, EPI_ISL_3001851, EPI_ISL_3001852, EPI_ISL_3001853, EPI_ISL_3001854, EPI_ISL_3001855, EPI_ISL_3001856, EPI_ISL_3001857, EPI_ISL_3001858, EPI_ISL_3001859, EPI_ISL_3001860, EPI_ISL_3001861, EPI_ISL_3001862, EPI_ISL_3001863, EPI_ISL_3001864, EPI_ISL_3001865, EPI_ISL_3001866,
EPI_ISL_3001867, EPI_ISL_3001868, EPI_ISL_3001869, EPI_ISL_3001870, EPI_ISL_3001871, EPI_ISL_3001872, EPI_ISL_3001873, EPI_ISL_3001874, EPI_ISL_3001875, EPI_ISL_3001876, EPI_ISL_3001877, EPI_ISL_3001878, EPI_ISL_3001879, EPI_ISL_3001880, EPI_ISL_3001881, EPI_ISL_3001882, EPI_ISL_3001883, EPI_ISL_3001884, EPI_ISL_3001885, EPI_ISL_3001886, EPI_ISL_3001887, EPI_ISL_3001888,
EPI_ISL_3001889, EPI_ISL_3001890, EPI_ISL_3001891, EPI_ISL_3001892, EPI_ISL_3001893, EPI_ISL_3001894, EPI_ISL_3001895, EPI_ISL_3001896, EPI_ISL_3001897, EPI_ISL_3001898, EPI_ISL_3001899, EPI_ISL_3001900, EPI_ISL_3001901, EPI_ISL_3001902, EPI_ISL_3001903, EPI_ISL_3001904, EPI_ISL_3001905, EPI_ISL_3001906, EPI_ISL_3001907, EPI_ISL_3001908, EPI_ISL_3001909, EPI_ISL_3001910,
EPI_ISL_3001911, EPI_ISL_3001912, EPI_ISL_3001913, EPI_ISL_3001914, EPI_ISL_3001915, EPI_ISL_3001916, EPI_ISL_3001917, EPI_ISL_3001918, EPI_ISL_3001919, EPI_ISL_3001920, EPI_ISL_3001921, EPI_ISL_3001922, EPI_ISL_3001923, EPI_ISL_3001924, EPI_ISL_3001925, EPI_ISL_3001926, EPI_ISL_3001927, EPI_ISL_3001928, EPI_ISL_3001929, EPI_ISL_3001930, EPI_ISL_3001931, EPI_ISL_3001932,
EPI_ISL_3001933, EPI_ISL_3001934, EPI_ISL_3001935, EPI_ISL_3001936, EPI_ISL_3001937, EPI_ISL_3001938, EPI_ISL_3001939, EPI_ISL_3001940, EPI_ISL_3001941, EPI_ISL_3001942, EPI_ISL_3001943, EPI_ISL_3001944, EPI_ISL_3001945, EPI_ISL_3001946, EPI_ISL_3001947, EPI_ISL_3001948, EPI_ISL_3001949, EPI_ISL_3001950, EPI_ISL_3001951, EPI_ISL_3001953, EPI_ISL_3001954, EPI_ISL_3001955,
EPI_ISL_3001956, EPI_ISL_3001957, EPI_ISL_3001958, EPI_ISL_3001959, EPI_ISL_3001960, EPI_ISL_3001961, EPI_ISL_3001962, EPI_ISL_3001963, EPI_ISL_3001964, EPI_ISL_3001965, EPI_ISL_3001966, EPI_ISL_3001967, EPI_ISL_3001968, EPI_ISL_3001969, EPI_ISL_3001970, EPI_ISL_3001971, EPI_ISL_3001972, EPI_ISL_3001973, EPI_ISL_3001974, EPI_ISL_3001975, EPI_ISL_3001976, EPI_ISL_3001977,
EPI_ISL_3001978, EPI_ISL_3001979, EPI_ISL_3001980, EPI_ISL_3001981, EPI_ISL_3001982, EPI_ISL_3001983, EPI_ISL_3001984, EPI_ISL_3001985, EPI_ISL_3001986, EPI_ISL_3001987, EPI_ISL_3001988, EPI_ISL_3001989, EPI_ISL_3001990, EPI_ISL_3001991, EPI_ISL_3001992, EPI_ISL_3001993, EPI_ISL_3001994, EPI_ISL_3001995, EPI_ISL_3001996, EPI_ISL_3001997, EPI_ISL_3001998, EPI_ISL_3001999,
EPI_ISL_3002000, EPI_ISL_3002001, EPI_ISL_3002002, EPI_ISL_3002003, EPI_ISL_3002004, EPI_ISL_3002005, EPI_ISL_3002006, EPI_ISL_3002007, EPI_ISL_3002008, EPI_ISL_3002009, EPI_ISL_3002010, EPI_ISL_3002011, EPI_ISL_3002012, EPI_ISL_3002013, EPI_ISL_3002014, EPI_ISL_3002015, EPI_ISL_3002016, EPI_ISL_3002017, EPI_ISL_3002018, EPI_ISL_3002019, EPI_ISL_3002020, EPI_ISL_3002021,
EPI_ISL_3002022, EPI_ISL_3002023, EPI_ISL_3002024, EPI_ISL_3002025, EPI_ISL_3002026, EPI_ISL_3002027, EPI_ISL_3002028, EPI_ISL_3002029, EPI_ISL_3002030, EPI_ISL_3002031, EPI_ISL_3002032, EPI_ISL_3002033, EPI_ISL_3002034, EPI_ISL_3002035, EPI_ISL_3002036, EPI_ISL_3002037, EPI_ISL_3002038, EPI_ISL_3002039, EPI_ISL_3002040, EPI_ISL_3002041, EPI_ISL_3002042, EPI_ISL_3002043,
EPI_ISL_3002044, EPI_ISL_3002045, EPI_ISL_3002046, EPI_ISL_3002047, EPI_ISL_3002048, EPI_ISL_3002049, EPI_ISL_3002050, EPI_ISL_3002051, EPI_ISL_3002052, EPI_ISL_3002053, EPI_ISL_3002054, EPI_ISL_3002055, EPI_ISL_3002056, EPI_ISL_3002057, EPI_ISL_3002058, EPI_ISL_3002059, EPI_ISL_3002060, EPI_ISL_3002061, EPI_ISL_3002062, EPI_ISL_3002063, EPI_ISL_3002064, EPI_ISL_3002065,
EPI_ISL_3002066, EPI_ISL_3002067, EPI_ISL_3002068, EPI_ISL_3002069, EPI_ISL_3002070, EPI_ISL_3002071, EPI_ISL_3002072, EPI_ISL_3002073, EPI_ISL_3002074, EPI_ISL_3002075, EPI_ISL_3002076, EPI_ISL_3002077, EPI_ISL_3002078, EPI_ISL_3002079, EPI_ISL_3002080, EPI_ISL_3002081, EPI_ISL_3002082, EPI_ISL_3002083, EPI_ISL_3002084, EPI_ISL_3002085, EPI_ISL_3002086, EPI_ISL_3002087,
EPI_ISL_3002088, EPI_ISL_3002089, EPI_ISL_3002090, EPI_ISL_3002091, EPI_ISL_3002092, EPI_ISL_3002093, EPI_ISL_3002094, EPI_ISL_3002095, EPI_ISL_3002096, EPI_ISL_3002097, EPI_ISL_3002098, EPI_ISL_3002099, EPI_ISL_3002100, EPI_ISL_3002101, EPI_ISL_3002102, EPI_ISL_3002103, EPI_ISL_3002104, EPI_ISL_3002105, EPI_ISL_3002106, EPI_ISL_3002107, EPI_ISL_3002108, EPI_ISL_3002109,
EPI_ISL_3002110, EPI_ISL_3002111, EPI_ISL_3002112, EPI_ISL_3002113, EPI_ISL_3002114, EPI_ISL_3002115, EPI_ISL_3002116, EPI_ISL_3002117, EPI_ISL_3002118, EPI_ISL_3002119, EPI_ISL_3002120, EPI_ISL_3002121, EPI_ISL_3002122, EPI_ISL_3002123, EPI_ISL_3002124, EPI_ISL_3002125, EPI_ISL_3002126, EPI_ISL_3002127, EPI_ISL_3002128, EPI_ISL_3002129, EPI_ISL_3002130, EPI_ISL_3002131,
EPI_ISL_3002132, EPI_ISL_3002133, EPI_ISL_3002134, EPI_ISL_3002135, EPI_ISL_3002136, EPI_ISL_3002137, EPI_ISL_3002138, EPI_ISL_3002139, EPI_ISL_3002140, EPI_ISL_3002141, EPI_ISL_3002142, EPI_ISL_3002143, EPI_ISL_3002144, EPI_ISL_3002145, EPI_ISL_3002146, EPI_ISL_3002147, EPI_ISL_3002148, EPI_ISL_3002149, EPI_ISL_3002150, EPI_ISL_3002151, EPI_ISL_3002152, EPI_ISL_3002153,
EPI_ISL_3002154, EPI_ISL_3002155, EPI_ISL_3002156, EPI_ISL_3002157, EPI_ISL_3002158, EPI_ISL_3002159, EPI_ISL_3002160, EPI_ISL_3002162, EPI_ISL_3002163, EPI_ISL_3002164, EPI_ISL_3002165, EPI_ISL_3002166, EPI_ISL_3002167, EPI_ISL_3002168, EPI_ISL_3002169, EPI_ISL_3002170, EPI_ISL_3002171, EPI_ISL_3002172, EPI_ISL_3002173, EPI_ISL_3002174, EPI_ISL_3002175, EPI_ISL_3002176,
EPI_ISL_3002177, EPI_ISL_3002179, EPI_ISL_3002180, EPI_ISL_3002181, EPI_ISL_3002182, EPI_ISL_3002183, EPI_ISL_3002184, EPI_ISL_3002185, EPI_ISL_3002186, EPI_ISL_3002187, EPI_ISL_3002188, EPI_ISL_3002189, EPI_ISL_3002190, EPI_ISL_3002191, EPI_ISL_3002192, EPI_ISL_3002193, EPI_ISL_3002194, EPI_ISL_3002195, EPI_ISL_3002196, EPI_ISL_3002197, EPI_ISL_3002198, EPI_ISL_3002199,
EPI_ISL_3002200, EPI_ISL_3002201, EPI_ISL_3002202, EPI_ISL_3002203, EPI_ISL_3002204, EPI_ISL_3002205, EPI_ISL_3002206, EPI_ISL_3002207, EPI_ISL_3002208, EPI_ISL_3002209, EPI_ISL_3002210, EPI_ISL_3002211, EPI_ISL_3002212, EPI_ISL_3002213, EPI_ISL_3002214, EPI_ISL_3002215, EPI_ISL_3002216, EPI_ISL_3002217, EPI_ISL_3002218, EPI_ISL_3002219, EPI_ISL_3002220, EPI_ISL_3002221,
EPI_ISL_3002222, EPI_ISL_3002223, EPI_ISL_3002224, EPI_ISL_3002225, EPI_ISL_3002226, EPI_ISL_3002227, EPI_ISL_3002228, EPI_ISL_3002229, EPI_ISL_3002230, EPI_ISL_3002231, EPI_ISL_3002232, EPI_ISL_3002233, EPI_ISL_3002234, EPI_ISL_3002235, EPI_ISL_3002236, EPI_ISL_3002237, EPI_ISL_3002238, EPI_ISL_3002239, EPI_ISL_3002240, EPI_ISL_3002241, EPI_ISL_3002242, EPI_ISL_3002243,
EPI_ISL_3002244, EPI_ISL_3002245, EPI_ISL_3002246, EPI_ISL_3002247, EPI_ISL_3002248, EPI_ISL_3002249, EPI_ISL_3002250, EPI_ISL_3002251, EPI_ISL_3002252, EPI_ISL_3002253, EPI_ISL_3002254, EPI_ISL_3002255, EPI_ISL_3002256, EPI_ISL_3002257, EPI_ISL_3002258, EPI_ISL_3002259, EPI_ISL_3002260, EPI_ISL_3002261, EPI_ISL_3002262, EPI_ISL_3002263, EPI_ISL_3002264, EPI_ISL_3002265,
EPI_ISL_3002266, EPI_ISL_3002267, EPI_ISL_3002268, EPI_ISL_3002269, EPI_ISL_3002270, EPI_ISL_3002271, EPI_ISL_3002272, EPI_ISL_3002273, EPI_ISL_3002274, EPI_ISL_3002275, EPI_ISL_3002276, EPI_ISL_3002277, EPI_ISL_3002278, EPI_ISL_3002279, EPI_ISL_3002280, EPI_ISL_3002281, EPI_ISL_3002282, EPI_ISL_3002283, EPI_ISL_3002284, EPI_ISL_3002285, EPI_ISL_3002286, EPI_ISL_3002287,
EPI_ISL_3002288, EPI_ISL_3002289, EPI_ISL_3002290, EPI_ISL_3002291, EPI_ISL_3002292, EPI_ISL_3002293, EPI_ISL_3002294, EPI_ISL_3002295

| | | | |
|---|---|---|---|
| see above | Northumbria University / South Tees Hospitals NHS Foundation Trust / North Cumbria Integrated Care NHS Foundation Trust / North Tees and Hartlepool NHS Foundation Trust / Newcastle Hospitals NHS Foundation Trust | COVID-19 Genomics UK (COG-UK) Consortium | Darren L Smith,Andrew Nelson,Matthew Bashton,Greg R Young,Joshua Loh,John Allan,Mohammad A Tariq,Giles S Holt,Gary Black,Wen C Yew,Lynn Dover,Paul Baker,Steve Liggett,Sarah Essex,Jane Greenaway,Debra Padgett,Clive Graham,Garren Scott,Edward Barton,Emma Swindells,Brendan Payne,Jennifer Collins,Yusri Taha,Gary Eltringham |

EPI_ISL_3002296, EPI_ISL_3002297, EPI_ISL_3002298, EPI_ISL_3002299, EPI_ISL_3002300, EPI_ISL_3002301, EPI_ISL_3002302, EPI_ISL_3002303, EPI_ISL_3002304, EPI_ISL_3002305, EPI_ISL_3002306, EPI_ISL_3002307, EPI_ISL_3002308, EPI_ISL_3002309, EPI_ISL_3002310, EPI_ISL_3002311, EPI_ISL_3002312, EPI_ISL_3002313, EPI_ISL_3002314, EPI_ISL_3002315, EPI_ISL_3002316, EPI_ISL_3002317,
EPI_ISL_3002318, EPI_ISL_3002319, EPI_ISL_3002320, EPI_ISL_3002321, EPI_ISL_3002322, EPI_ISL_3002323, EPI_ISL_3002324, EPI_ISL_3002325, EPI_ISL_3002326, EPI_ISL_3002327, EPI_ISL_3002328, EPI_ISL_3002329, EPI_ISL_3002330, EPI_ISL_3002331, EPI_ISL_3002332, EPI_ISL_3002333, EPI_ISL_3002334, EPI_ISL_3002335, EPI_ISL_3002336, EPI_ISL_3002337, EPI_ISL_3002338, EPI_ISL_3002339,
EPI_ISL_3002340, EPI_ISL_3002341, EPI_ISL_3002342, EPI_ISL_3002343, EPI_ISL_3002344, EPI_ISL_3002345, EPI_ISL_3002346, EPI_ISL_3002347, EPI_ISL_3002348, EPI_ISL_3002349, EPI_ISL_3002350, EPI_ISL_3002351, EPI_ISL_3002352, EPI_ISL_3002353, EPI_ISL_3002354, EPI_ISL_3002355, EPI_ISL_3002356, EPI_ISL_3002357, EPI_ISL_3002358, EPI_ISL_3002359, EPI_ISL_3002360, EPI_ISL_3002361,
EPI_ISL_3002362, EPI_ISL_3002363, EPI_ISL_3002364, EPI_ISL_3002365, EPI_ISL_3002366, EPI_ISL_3002367, EPI_ISL_3002368, EPI_ISL_3002369, EPI_ISL_3002370, EPI_ISL_3002371, EPI_ISL_3002372, EPI_ISL_3002373, EPI_ISL_3002374, EPI_ISL_3002375, EPI_ISL_3002376, EPI_ISL_3002377, EPI_ISL_3002378, EPI_ISL_3002379, EPI_ISL_3002380, EPI_ISL_3002381, EPI_ISL_3002382, EPI_ISL_3002383,
EPI_ISL_3002384, EPI_ISL_3002385, EPI_ISL_3002386, EPI_ISL_3002387, EPI_ISL_3002388, EPI_ISL_3002389, EPI_ISL_3002390, EPI_ISL_3002391, EPI_ISL_3002392, EPI_ISL_3002393, EPI_ISL_3002394, EPI_ISL_3002395, EPI_ISL_3002396, EPI_ISL_3002397, EPI_ISL_3002398, EPI_ISL_3002399, EPI_ISL_3002400, EPI_ISL_3002401, EPI_ISL_3002402, EPI_ISL_3002403, EPI_ISL_3002404, EPI_ISL_3002405,
EPI_ISL_3002406, EPI_ISL_3002407, EPI_ISL_3002408, EPI_ISL_3002409, EPI_ISL_3002410, EPI_ISL_3002411, EPI_ISL_3002412, EPI_ISL_3002413, EPI_ISL_3002414, EPI_ISL_3002415, EPI_ISL_3002416, EPI_ISL_3002417, EPI_ISL_3002418, EPI_ISL_3002419, EPI_ISL_3002420, EPI_ISL_3002421, EPI_ISL_3002422, EPI_ISL_3002423, EPI_ISL_3002424, EPI_ISL_3002425, EPI_ISL_3002426, EPI_ISL_3002427,
EPI_ISL_3002428, EPI_ISL_3002429, EPI_ISL_3002430, EPI_ISL_3002431, EPI_ISL_3002432, EPI_ISL_3002433, EPI_ISL_3002434, EPI_ISL_3002435, EPI_ISL_3002436, EPI_ISL_3002437, EPI_ISL_3002438, EPI_ISL_3002439, EPI_ISL_3002440, EPI_ISL_3002441, EPI_ISL_3002442, EPI_ISL_3002443, EPI_ISL_3002444, EPI_ISL_3002445, EPI_ISL_3002446, EPI_ISL_3002447, EPI_ISL_3002448, EPI_ISL_3002449,
EPI_ISL_3002450, EPI_ISL_3002451, EPI_ISL_3002452, EPI_ISL_3002453, EPI_ISL_3002454, EPI_ISL_3002455, EPI_ISL_3002456, EPI_ISL_3002457, EPI_ISL_3002458, EPI_ISL_3002459, EPI_ISL_3002460, EPI_ISL_3002461, EPI_ISL_3002462, EPI_ISL_3002463, EPI_ISL_3002464, EPI_ISL_3002465, EPI_ISL_3002466, EPI_ISL_3002467, EPI_ISL_3002468, EPI_ISL_3002469, EPI_ISL_3002470, EPI_ISL_3002471,
EPI_ISL_3002472, EPI_ISL_3002473, EPI_ISL_3002474, EPI_ISL_3002475, EPI_ISL_3002476, EPI_ISL_3002477, EPI_ISL_3002478, EPI_ISL_3002479, EPI_ISL_3002480, EPI_ISL_3002481, EPI_ISL_3002482, EPI_ISL_3002483, EPI_ISL_3002484, EPI_ISL_3002485, EPI_ISL_3002486, EPI_ISL_3002487, EPI_ISL_3002488, EPI_ISL_3002489, EPI_ISL_3002490, EPI_ISL_3002491, EPI_ISL_3002492, EPI_ISL_3002493,
EPI_ISL_3002495, EPI_ISL_3002496, EPI_ISL_3002497, EPI_ISL_3002498, EPI_ISL_3002499, EPI_ISL_3002500, EPI_ISL_3002501, EPI_ISL_3002502, EPI_ISL_3002503, EPI_ISL_3002504, EPI_ISL_3002505, EPI_ISL_3002506, EPI_ISL_3002507, EPI_ISL_3002508, EPI_ISL_3002509, EPI_ISL_3002510, EPI_ISL_3002511, EPI_ISL_3002512, EPI_ISL_3002513, EPI_ISL_3002514, EPI_ISL_3002515, EPI_ISL_3002516,
EPI_ISL_3002517, EPI_ISL_3002518, EPI_ISL_3002519, EPI_ISL_3002520, EPI_ISL_3002521, EPI_ISL_3002522, EPI_ISL_3002523, EPI_ISL_3002524, EPI_ISL_3002525, EPI_ISL_3002526, EPI_ISL_3002527, EPI_ISL_3002528, EPI_ISL_3002529, EPI_ISL_3002530, EPI_ISL_3002531, EPI_ISL_3002532, EPI_ISL_3002533, EPI_ISL_3002534, EPI_ISL_3002535, EPI_ISL_3002536, EPI_ISL_3002537, EPI_ISL_3002538,
EPI_ISL_3002539, EPI_ISL_3002540, EPI_ISL_3002541, EPI_ISL_3002542, EPI_ISL_3002543, EPI_ISL_3002544, EPI_ISL_3002545, EPI_ISL_3002546, EPI_ISL_3002547, EPI_ISL_3002548, EPI_ISL_3002549, EPI_ISL_3002550, EPI_ISL_3002551, EPI_ISL_3002552, EPI_ISL_3002553, EPI_ISL_3002554, EPI_ISL_3002555, EPI_ISL_3002556, EPI_ISL_3002557, EPI_ISL_3002558, EPI_ISL_3002559, EPI_ISL_3002560,
EPI_ISL_3002561, EPI_ISL_3002562, EPI_ISL_3002563, EPI_ISL_3002564, EPI_ISL_3002565, EPI_ISL_3002566, EPI_ISL_3002567, EPI_ISL_3002568, EPI_ISL_3002569, EPI_ISL_3002570, EPI_ISL_3002571, EPI_ISL_3002572, EPI_ISL_3002573, EPI_ISL_3002574, EPI_ISL_3002575, EPI_ISL_3002576, EPI_ISL_3002577, EPI_ISL_3002578, EPI_ISL_3002579, EPI_ISL_3002580, EPI_ISL_3002581, EPI_ISL_3002582,
EPI_ISL_3002583, EPI_ISL_3002584, EPI_ISL_3002585, EPI_ISL_3002586, EPI_ISL_3002587, EPI_ISL_3002588, EPI_ISL_3002589, EPI_ISL_3002590, EPI_ISL_3002591, EPI_ISL_3002592, EPI_ISL_3002593, EPI_ISL_3002594, EPI_ISL_3002595, EPI_ISL_3002596, EPI_ISL_3002597, EPI_ISL_3002598, EPI_ISL_3002599, EPI_ISL_3002600, EPI_ISL_3002601, EPI_ISL_3002602, EPI_ISL_3002603, EPI_ISL_3002604,
EPI_ISL_3002605, EPI_ISL_3002606, EPI_ISL_3002607, EPI_ISL_3002608, EPI_ISL_3002609, EPI_ISL_3002610, EPI_ISL_3002611, EPI_ISL_3002612, EPI_ISL_3002613, EPI_ISL_3002614, EPI_ISL_3002615, EPI_ISL_3002616, EPI_ISL_3002617, EPI_ISL_3002618, EPI_ISL_3002619, EPI_ISL_3002620, EPI_ISL_3002621, EPI_ISL_3002622, EPI_ISL_3002623, EPI_ISL_3002624, EPI_ISL_3002625, EPI_ISL_3002626,
EPI_ISL_3002627, EPI_ISL_3002628, EPI_ISL_3002629, EPI_ISL_3002630, EPI_ISL_3002631, EPI_ISL_3002632, EPI_ISL_3002633, EPI_ISL_3002634, EPI_ISL_3002635, EPI_ISL_3002636, EPI_ISL_3002637, EPI_ISL_3002638, EPI_ISL_3002639, EPI_ISL_3002640, EPI_ISL_3002641, EPI_ISL_3002642, EPI_ISL_3002643, EPI_ISL_3002644, EPI_ISL_3002645, EPI_ISL_3002646, EPI_ISL_3002647, EPI_ISL_3002648,
EPI_ISL_3002649, EPI_ISL_3002650, EPI_ISL_3002651, EPI_ISL_3002652, EPI_ISL_3002653, EPI_ISL_3002654, EPI_ISL_3002655, EPI_ISL_3002656, EPI_ISL_3002657, EPI_ISL_3002658, EPI_ISL_3002659, EPI_ISL_3002660, EPI_ISL_3002661, EPI_ISL_3002662, EPI_ISL_3002663, EPI_ISL_3002664, EPI_ISL_3002665, EPI_ISL_3002666, EPI_ISL_3002667, EPI_ISL_3002668, EPI_ISL_3002669, EPI_ISL_3002670,
EPI_ISL_3002671, EPI_ISL_3002672, EPI_ISL_3002673, EPI_ISL_3002674, EPI_ISL_3002675, EPI_ISL_3002676, EPI_ISL_3002677, EPI_ISL_3002678, EPI_ISL_3002679, EPI_ISL_3002680, EPI_ISL_3002681, EPI_ISL_3002682, EPI_ISL_3002683, EPI_ISL_3002684, EPI_ISL_3002685, EPI_ISL_3002686, EPI_ISL_3002687, EPI_ISL_3002688, EPI_ISL_3002689, EPI_ISL_3002690, EPI_ISL_3002691, EPI_ISL_3002692,
EPI_ISL_3002693, EPI_ISL_3002694, EPI_ISL_3002695, EPI_ISL_3002696, EPI_ISL_3002697, EPI_ISL_3002698, EPI_ISL_3002699, EPI_ISL_3002700, EPI_ISL_3002701, EPI_ISL_3002702, EPI_ISL_3002703, EPI_ISL_3002704, EPI_ISL_3002705, EPI_ISL_3002706, EPI_ISL_3002707, EPI_ISL_3002708, EPI_ISL_3002709, EPI_ISL_3002710, EPI_ISL_3002711, EPI_ISL_3002712, EPI_ISL_3002713, EPI_ISL_3002714,
EPI_ISL_3002715, EPI_ISL_3002716, EPI_ISL_3002717, EPI_ISL_3002718, EPI_ISL_3002719, EPI_ISL_3002720, EPI_ISL_3002721, EPI_ISL_3002722, EPI_ISL_3002723, EPI_ISL_3002724, EPI_ISL_3002725, EPI_ISL_3002727, EPI_ISL_3002728, EPI_ISL_3002729, EPI_ISL_3002730, EPI_ISL_3002731, EPI_ISL_3002732, EPI_ISL_3002733, EPI_ISL_3002734, EPI_ISL_3002735, EPI_ISL_3002736, EPI_ISL_3002737,
EPI_ISL_3002738, EPI_ISL_3002739, EPI_ISL_3002740, EPI_ISL_3002741, EPI_ISL_3002742, EPI_ISL_3002743, EPI_ISL_3002744, EPI_ISL_3002745, EPI_ISL_3002746, EPI_ISL_3002747, EPI_ISL_3002748, EPI_ISL_3002749, EPI_ISL_3002750, EPI_ISL_3002751, EPI_ISL_3002752, EPI_ISL_3002753, EPI_ISL_3002754, EPI_ISL_3002755, EPI_ISL_3002756, EPI_ISL_3002757, EPI_ISL_3002758, EPI_ISL_3002759,
EPI_ISL_3002760, EPI_ISL_3002761, EPI_ISL_3002762, EPI_ISL_3002763, EPI_ISL_3002764, EPI_ISL_3002765, EPI_ISL_3002766, EPI_ISL_3002767, EPI_ISL_3002768, EPI_ISL_3002769, EPI_ISL_3002770, EPI_ISL_3002771, EPI_ISL_3002772, EPI_ISL_3002773, EPI_ISL_3002774, EPI_ISL_3002775, EPI_ISL_3002776, EPI_ISL_3002777, EPI_ISL_3002778, EPI_ISL_3002779, EPI_ISL_3002780, EPI_ISL_3002781,
EPI_ISL_3002782, EPI_ISL_3002783, EPI_ISL_3002784, EPI_ISL_3002785, EPI_ISL_3002786, EPI_ISL_3002787, EPI_ISL_3002788, EPI_ISL_3002789, EPI_ISL_3002790, EPI_ISL_3002791, EPI_ISL_3002792, EPI_ISL_3002793, EPI_ISL_3002794, EPI_ISL_3002795, EPI_ISL_3002796, EPI_ISL_3002797, EPI_ISL_3002798, EPI_ISL_3002799, EPI_ISL_3002800, EPI_ISL_3002801, EPI_ISL_3002802, EPI_ISL_3002803,
EPI_ISL_3002804, EPI_ISL_3002805, EPI_ISL_3002806, EPI_ISL_3002807, EPI_ISL_3002808, EPI_ISL_3002809, EPI_ISL_3002810, EPI_ISL_3002811, EPI_ISL_3002812, EPI_ISL_3002813, EPI_ISL_3002814, EPI_ISL_3002815, EPI_ISL_3002816, EPI_ISL_3002817, EPI_ISL_3002818, EPI_ISL_3002819, EPI_ISL_3002820, EPI_ISL_3002821, EPI_ISL_3002822, EPI_ISL_3002823, EPI_ISL_3002824, EPI_ISL_3002825,
EPI_ISL_3002826, EPI_ISL_3002827, EPI_ISL_3002828, EPI_ISL_3002829, EPI_ISL_3002830, EPI_ISL_3002831, EPI_ISL_3002832, EPI_ISL_3002833, EPI_ISL_3002834, EPI_ISL_3002835, EPI_ISL_3002836, EPI_ISL_3002837, EPI_ISL_3002838, EPI_ISL_3002839, EPI_ISL_3002840, EPI_ISL_3002841, EPI_ISL_3002842, EPI_ISL_3002843, EPI_ISL_3002844, EPI_ISL_3002845, EPI_ISL_3002846, EPI_ISL_3002847,
EPI_ISL_3002848, EPI_ISL_3002849, EPI_ISL_3002850, EPI_ISL_3002851, EPI_ISL_3002852, EPI_ISL_3002853, EPI_ISL_3002854, EPI_ISL_3002855, EPI_ISL_3002856, EPI_ISL_3002857, EPI_ISL_3002858, EPI_ISL_3002859, EPI_ISL_3002860, EPI_ISL_3002861, EPI_ISL_3002862, EPI_ISL_3002863, EPI_ISL_3002864, EPI_ISL_3002865, EPI_ISL_3002866, EPI_ISL_3002867, EPI_ISL_3002868, EPI_ISL_3002869,
EPI_ISL_3002870, EPI_ISL_3002871, EPI_ISL_3002872, EPI_ISL_3002873, EPI_ISL_3002874, EPI_ISL_3002875, EPI_ISL_3002876, EPI_ISL_3002877, EPI_ISL_3002878, EPI_ISL_3002879, EPI_ISL_3002880, EPI_ISL_3002881, EPI_ISL_3002882, EPI_ISL_3002883, EPI_ISL_3002884, EPI_ISL_3002885, EPI_ISL_3002886, EPI_ISL_3002887, EPI_ISL_3002888, EPI_ISL_3002889, EPI_ISL_3002890, EPI_ISL_3002891,
EPI_ISL_3002892, EPI_ISL_3002893, EPI_ISL_3002894, EPI_ISL_3002895, EPI_ISL_3002896, EPI_ISL_3002897, EPI_ISL_3002898, EPI_ISL_3002899, EPI_ISL_3002900, EPI_ISL_3002901, EPI_ISL_3002902, EPI_ISL_3002903, EPI_ISL_3002904, EPI_ISL_3002905, EPI_ISL_3002906, EPI_ISL_3002907, EPI_ISL_3002908, EPI_ISL_3002909, EPI_ISL_3002910, EPI_ISL_3002911, EPI_ISL_3002912, EPI_ISL_3002913,
EPI_ISL_3002914, EPI_ISL_3002915, EPI_ISL_3002916, EPI_ISL_3002917, EPI_ISL_3002918, EPI_ISL_3002919, EPI_ISL_3002920, EPI_ISL_3002921, EPI_ISL_3002922, EPI_ISL_3002923, EPI_ISL_3002924, EPI_ISL_3002925, EPI_ISL_3002926, EPI_ISL_3002927, EPI_ISL_3002928, EPI_ISL_3002929, EPI_ISL_3002930, EPI_ISL_3002931, EPI_ISL_3002933, EPI_ISL_3002934, EPI_ISL_3002935, EPI_ISL_3002936,
EPI_ISL_3002937, EPI_ISL_3002938, EPI_ISL_3002939, EPI_ISL_3002940, EPI_ISL_3002941, EPI_ISL_3002942, EPI_ISL_3002943, EPI_ISL_3002944, EPI_ISL_3002945, EPI_ISL_3002946, EPI_ISL_3002947, EPI_ISL_3002948, EPI_ISL_3002949, EPI_ISL_3002950, EPI_ISL_3002951, EPI_ISL_3002952, EPI_ISL_3002953, EPI_ISL_3002954, EPI_ISL_3002955, EPI_ISL_3002956, EPI_ISL_3002957, EPI_ISL_3002958,
EPI_ISL_3002959, EPI_ISL_3002960, EPI_ISL_3002961, EPI_ISL_3002962, EPI_ISL_3002963, EPI_ISL_3002964, EPI_ISL_3002965, EPI_ISL_3002966, EPI_ISL_3002967, EPI_ISL_3002968, EPI_ISL_3002969, EPI_ISL_3002970, EPI_ISL_3002971, EPI_ISL_3002972, EPI_ISL_3002973, EPI_ISL_3002974, EPI_ISL_3002975, EPI_ISL_3002976, EPI_ISL_3002977, EPI_ISL_3002978, EPI_ISL_3002979, EPI_ISL_3002980,
EPI_ISL_3002981, EPI_ISL_3002982, EPI_ISL_3002983, EPI_ISL_3002984, EPI_ISL_3002985, EPI_ISL_3002986, EPI_ISL_3002987, EPI_ISL_3002988, EPI_ISL_3002989, EPI_ISL_3002990, EPI_ISL_3002991, EPI_ISL_3002992, EPI_ISL_3002993, EPI_ISL_3002994, EPI_ISL_3002995, EPI_ISL_3002996, EPI_ISL_3002997, EPI_ISL_3002998, EPI_ISL_3002999, EPI_ISL_3003000, EPI_ISL_3003001, EPI_ISL_3003002,
EPI_ISL_3003003, EPI_ISL_3003004, EPI_ISL_3003006, EPI_ISL_3003007, EPI_ISL_3003008, EPI_ISL_3003009, EPI_ISL_3003010, EPI_ISL_3003011, EPI_ISL_3003012, EPI_ISL_3003013, EPI_ISL_3003014, EPI_ISL_3003016, EPI_ISL_3003017, EPI_ISL_3003018, EPI_ISL_3003019, EPI_ISL_3003020, EPI_ISL_3003021, EPI_ISL_3003022, EPI_ISL_3003023, EPI_ISL_3003024, EPI_ISL_3003026, EPI_ISL_3003027,
EPI_ISL_3003028, EPI_ISL_3003029, EPI_ISL_3003030, EPI_ISL_3003031, EPI_ISL_3003032, EPI_ISL_3003033, EPI_ISL_3003034, EPI_ISL_3003035, EPI_ISL_3003036, EPI_ISL_3003037, EPI_ISL_3003038, EPI_ISL_3003039, EPI_ISL_3003040, EPI_ISL_3003041, EPI_ISL_3003042, EPI_ISL_3003043, EPI_ISL_3003044, EPI_ISL_3003045, EPI_ISL_3003046, EPI_ISL_3003047, EPI_ISL_3003048, EPI_ISL_3003049,
EPI_ISL_3003050, EPI_ISL_3003051, EPI_ISL_3003052, EPI_ISL_3003053, EPI_ISL_3003054, EPI_ISL_3003055, EPI_ISL_3003056, EPI_ISL_3003057, EPI_ISL_3003058, EPI_ISL_3003059, EPI_ISL_3003060, EPI_ISL_3003061, EPI_ISL_3003062, EPI_ISL_3003063, EPI_ISL_3003064, EPI_ISL_3003065, EPI_ISL_3003066, EPI_ISL_3003067, EPI_ISL_3003068, EPI_ISL_3003069, EPI_ISL_3003070, EPI_ISL_3003071,
EPI_ISL_3003072, EPI_ISL_3003073, EPI_ISL_3003074, EPI_ISL_3003075, EPI_ISL_3003076, EPI_ISL_3003077, EPI_ISL_3003078, EPI_ISL_3003079, EPI_ISL_3003080, EPI_ISL_3003081, EPI_ISL_3003082, EPI_ISL_3003083, EPI_ISL_3003084, EPI_ISL_3003085, EPI_ISL_3003086, EPI_ISL_3003087, EPI_ISL_3003088, EPI_ISL_3003089, EPI_ISL_3003090, EPI_ISL_3003091, EPI_ISL_3003092, EPI_ISL_3003093,
EPI_ISL_3003094, EPI_ISL_3003095, EPI_ISL_3003096, EPI_ISL_3003097, EPI_ISL_3003098, EPI_ISL_3003099, EPI_ISL_3003100, EPI_ISL_3003101, EPI_ISL_3003102, EPI_ISL_3003103, EPI_ISL_3003104, EPI_ISL_3003105, EPI_ISL_3003106, EPI_ISL_3003107, EPI_ISL_3003108, EPI_ISL_3003110, EPI_ISL_3003111, EPI_ISL_3003112, EPI_ISL_3003113, EPI_ISL_3003114, EPI_ISL_3003115, EPI_ISL_3003116,
EPI_ISL_3003117, EPI_ISL_3003118, EPI_ISL_3003119, EPI_ISL_3003120, EPI_ISL_3003121, EPI_ISL_3003122, EPI_ISL_3003123, EPI_ISL_3003124, EPI_ISL_3003125, EPI_ISL_3003126, EPI_ISL_3003127, EPI_ISL_3003128, EPI_ISL_3003129, EPI_ISL_3003130, EPI_ISL_3003131, EPI_ISL_3003132, EPI_ISL_3003133, EPI_ISL_3003134, EPI_ISL_3003135, EPI_ISL_3003136, EPI_ISL_3003137, EPI_ISL_3003138,
EPI_ISL_3003139, EPI_ISL_3003140, EPI_ISL_3003141, EPI_ISL_3003142, EPI_ISL_3003143, EPI_ISL_3003144, EPI_ISL_3003145, EPI_ISL_3003146, EPI_ISL_3003147, EPI_ISL_3003148, EPI_ISL_3003149, EPI_ISL_3003150, EPI_ISL_3003151, EPI_ISL_3003152, EPI_ISL_3003153, EPI_ISL_3003154, EPI_ISL_3003155, EPI_ISL_3003156, EPI_ISL_3003157, EPI_ISL_3003158, EPI_ISL_3003159, EPI_ISL_3003160,
EPI_ISL_3003161, EPI_ISL_3003162, EPI_ISL_3003163, EPI_ISL_3003164, EPI_ISL_3003165, EPI_ISL_3003166, EPI_ISL_3003167, EPI_ISL_3003168, EPI_ISL_3003169, EPI_ISL_3003170, EPI_ISL_3003171, EPI_ISL_3003172, EPI_ISL_3003173, EPI_ISL_3003174, EPI_ISL_3003175, EPI_ISL_3003176, EPI_ISL_3003177, EPI_ISL_3003178, EPI_ISL_3003179, EPI_ISL_3003180, EPI_ISL_3003181, EPI_ISL_3003182,
EPI_ISL_3003183, EPI_ISL_3003184, EPI_ISL_3003185, EPI_ISL_3003186, EPI_ISL_3003187, EPI_ISL_3003188, EPI_ISL_3003189, EPI_ISL_3003190, EPI_ISL_3003191, EPI_ISL_3003192, EPI_ISL_3003193, EPI_ISL_3003194, EPI_ISL_3003195, EPI_ISL_3003196, EPI_ISL_3003197, EPI_ISL_3003198

| | | | |
|---|---|---|---|
| see above | Quadram Institute Bioscience | COVID-19 Genomics UK (COG-UK) Consortium | Dave J. Baker, Gemma L. Kay, Alp Aydin, Thanh Le-Viet, Steven Rudder, Ana P. Tedim, Anastasia Kolyva, Maria Diaz, Leonardo de Oliveira Martins, Nabil-Fareed Alikhan, Lizzie Meadows, Rachael Stanley, Rose K Davidson, Ngozi Elumogo, Muhammed Yasir, Sophie J. Prosolek, Ebenezer Foster-Nyarko, Martin Lott, Christopher Jeanes, Lewis G. Spurgin, Nicholas M. Thomson, Alexander J Trotter, Rachel Gilroy, Luke Bedford, Samuel Bloomfield, Claire Stuart, Andrew Bell, Reenesh Prakash, Lindsay Coupland, Samir Dervisevic, Alison E. Mather, John Wain, Mark Webber, Andrew J. Page, Justin O'Grady |

EPI_ISL_3003200, EPI_ISL_3003201, EPI_ISL_3003202, EPI_ISL_3003203, EPI_ISL_3003204, EPI_ISL_3003205, EPI_ISL_3003206, EPI_ISL_3003207, EPI_ISL_3003208, EPI_ISL_3003209, EPI_ISL_3003210, EPI_ISL_3003211, EPI_ISL_3003212, EPI_ISL_3003213, EPI_ISL_3003214, EPI_ISL_3003215, EPI_ISL_3003216, EPI_ISL_3003217, EPI_ISL_3003218, EPI_ISL_3003219, EPI_ISL_3003220, EPI_ISL_3003221,
EPI_ISL_3003222, EPI_ISL_3003223, EPI_ISL_3003224, EPI_ISL_3003225, EPI_ISL_3003226, EPI_ISL_3003227, EPI_ISL_3003228, EPI_ISL_3003229, EPI_ISL_3003230, EPI_ISL_3003231, EPI_ISL_3003232, EPI_ISL_3003233, EPI_ISL_3003234, EPI_ISL_3003235, EPI_ISL_3003236, EPI_ISL_3003237, EPI_ISL_3003238, EPI_ISL_3003239, EPI_ISL_3003240, EPI_ISL_3003241, EPI_ISL_3003242, EPI_ISL_3003243,
EPI_ISL_3003244, EPI_ISL_3003245, EPI_ISL_3003246, EPI_ISL_3003247, EPI_ISL_3003248, EPI_ISL_3003249, EPI_ISL_3003250, EPI_ISL_3003251, EPI_ISL_3003252, EPI_ISL_3003253, EPI_ISL_3003254, EPI_ISL_3003255, EPI_ISL_3003256, EPI_ISL_3003257, EPI_ISL_3003258, EPI_ISL_3003259, EPI_ISL_3003260, EPI_ISL_3003261, EPI_ISL_3003262, EPI_ISL_3003263, EPI_ISL_3003264, EPI_ISL_3003265,
EPI_ISL_3003266

| | | | |
|---|---|---|---|
| see above | Queens Medical Centre, Clinical Microbiology Department / DeepSeq Nottingham | COVID-19 Genomics UK (COG-UK) Consortium | Gemma Clark, Wendy Smith, Manjinder Khakh, Vicki M Fleming, Michelle M Lister, Hannah Howson-Wells, Jonathan Ball, Timothy Byaruhanga, Jayasree Dey, Emily Park, Jack Hill, Patrick McClure, Joseph Chappell, Theocharis Tsoleridis, Nadine Holmes, Matthew Carlisle, Christopher Moore, Fei Sang, Johnny Debebe, Victoria Wright, Matthew Loose |

EPI_ISL_3003491, EPI_ISL_3003492, EPI_ISL_3003493, EPI_ISL_3003494, EPI_ISL_3003495, EPI_ISL_3003496, EPI_ISL_3003497, EPI_ISL_3003498, EPI_ISL_3003499, EPI_ISL_3003501, EPI_ISL_3003502, EPI_ISL_3003504, EPI_ISL_3003505, EPI_ISL_3003507, EPI_ISL_3003509, EPI_ISL_3003510, EPI_ISL_3003511, EPI_ISL_3003512, EPI_ISL_3003513, EPI_ISL_3003514, EPI_ISL_3003515, EPI_ISL_3003516,
EPI_ISL_3003518, EPI_ISL_3003519, EPI_ISL_3003521, EPI_ISL_3003522, EPI_ISL_3003524, EPI_ISL_3003525, EPI_ISL_3003526, EPI_ISL_3003527, EPI_ISL_3003528, EPI_ISL_3003529, EPI_ISL_3003530, EPI_ISL_3003531, EPI_ISL_3003532, EPI_ISL_3003533, EPI_ISL_3003534, EPI_ISL_3003535, EPI_ISL_3003536, EPI_ISL_3003537, EPI_ISL_3003539, EPI_ISL_3003540, EPI_ISL_3003541, EPI_ISL_3003542,
EPI_ISL_3003543, EPI_ISL_3003544, EPI_ISL_3003546, EPI_ISL_3003547, EPI_ISL_3003548, EPI_ISL_3003549, EPI_ISL_3003550, EPI_ISL_3003551, EPI_ISL_3003552, EPI_ISL_3003553, EPI_ISL_3003554, EPI_ISL_3003555, EPI_ISL_3003556, EPI_ISL_3003557, EPI_ISL_3003559, EPI_ISL_3003560, EPI_ISL_3003561, EPI_ISL_3003562, EPI_ISL_3003563, EPI_ISL_3003564, EPI_ISL_3003566, EPI_ISL_3003567,
EPI_ISL_3003568, EPI_ISL_3003569, EPI_ISL_3003570, EPI_ISL_3003571, EPI_ISL_3003572, EPI_ISL_3003573, EPI_ISL_3003574, EPI_ISL_3003575, EPI_ISL_3003576, EPI_ISL_3003577, EPI_ISL_3003578, EPI_ISL_3003580, EPI_ISL_3003581, EPI_ISL_3003582, EPI_ISL_3003583, EPI_ISL_3003584, EPI_ISL_3003585, EPI_ISL_3003586, EPI_ISL_3003587, EPI_ISL_3003588, EPI_ISL_3003589, EPI_ISL_3003590,
EPI_ISL_3003591, EPI_ISL_3003593, EPI_ISL_3003594, EPI_ISL_3003595, EPI_ISL_3003596, EPI_ISL_3003597, EPI_ISL_3003599, EPI_ISL_3003600, EPI_ISL_3003601, EPI_ISL_3003602, EPI_ISL_3003604, EPI_ISL_3003605, EPI_ISL_3003606, EPI_ISL_3003607, EPI_ISL_3003608, EPI_ISL_3003610, EPI_ISL_3003611, EPI_ISL_3003612, EPI_ISL_3003614, EPI_ISL_3003615, EPI_ISL_3003616, EPI_ISL_3003617,
EPI_ISL_3003619, EPI_ISL_3003620, EPI_ISL_3003621, EPI_ISL_3003622, EPI_ISL_3003623, EPI_ISL_3003624, EPI_ISL_3003625, EPI_ISL_3003626, EPI_ISL_3003627, EPI_ISL_3003628, EPI_ISL_3003629, EPI_ISL_3003630, EPI_ISL_3003631, EPI_ISL_3003632, EPI_ISL_3003633, EPI_ISL_3003634, EPI_ISL_3003635, EPI_ISL_3003636, EPI_ISL_3003637, EPI_ISL_3003638, EPI_ISL_3003639, EPI_ISL_3003640,
EPI_ISL_3003641, EPI_ISL_3003642, EPI_ISL_3003643, EPI_ISL_3003646, EPI_ISL_3003647, EPI_ISL_3003648, EPI_ISL_3003649, EPI_ISL_3003650, EPI_ISL_3003651, EPI_ISL_3003652, EPI_ISL_3003653, EPI_ISL_3003654, EPI_ISL_3003655, EPI_ISL_3003656, EPI_ISL_3003657, EPI_ISL_3003658, EPI_ISL_3003659, EPI_ISL_3003660, EPI_ISL_3003661, EPI_ISL_3003662, EPI_ISL_3003663, EPI_ISL_3003666,
EPI_ISL_3003667, EPI_ISL_3003668, EPI_ISL_3003670, EPI_ISL_3003671, EPI_ISL_3003673, EPI_ISL_3003674, EPI_ISL_3003675, EPI_ISL_3003676, EPI_ISL_3003677, EPI_ISL_3003678, EPI_ISL_3003680, EPI_ISL_3003681, EPI_ISL_3003682, EPI_ISL_3003683, EPI_ISL_3003684, EPI_ISL_3003685, EPI_ISL_3003686, EPI_ISL_3003687, EPI_ISL_3003688

| | | | |
|---|---|---|---|
| see above | Oxford Viromics, NDM, University of Oxford; Oxford University Hospitals; Basingstoke and North Hampshire Hospital | COVID-19 Genomics UK (COG-UK) Consortium | Tanya Golubchik, David Bonsall, George Macintyre, Amy Trebes, Mariateresa de Cesare, Catrin Moore, Alex Mobbs, Anita Justice, Robert Shaw, Monique Andersson, Timothy Peto, Emma Wise, Nathan Moore, Jessica Lynch, Nick Cortes, Matilde Mori, Stephen Kidd, David Buck, John Todd, Christophe Fraser |

EPI_ISL_3003729, EPI_ISL_3003730, EPI_ISL_3003731, EPI_ISL_3003732, EPI_ISL_3003733, EPI_ISL_3003734, EPI_ISL_3003735, EPI_ISL_3003736, EPI_ISL_3003737, EPI_ISL_3003738, EPI_ISL_3003739, EPI_ISL_3003740, EPI_ISL_3003741, EPI_ISL_3003742, EPI_ISL_3003743, EPI_ISL_3003744, EPI_ISL_3003745, EPI_ISL_3003746, EPI_ISL_3003747, EPI_ISL_3003748, EPI_ISL_3003749, EPI_ISL_3003750,
EPI_ISL_3003751, EPI_ISL_3003752, EPI_ISL_3003753, EPI_ISL_3003754, EPI_ISL_3003755, EPI_ISL_3003756, EPI_ISL_3003757, EPI_ISL_3003758, EPI_ISL_3003759, EPI_ISL_3003760, EPI_ISL_3003761, EPI_ISL_3003762, EPI_ISL_3003763, EPI_ISL_3003764, EPI_ISL_3003765, EPI_ISL_3003766, EPI_ISL_3003767, EPI_ISL_3003768, EPI_ISL_3003769, EPI_ISL_3003770, EPI_ISL_3003771, EPI_ISL_3003772,
EPI_ISL_3003773, EPI_ISL_3003774, EPI_ISL_3003775, EPI_ISL_3003776, EPI_ISL_3003777, EPI_ISL_3003778, EPI_ISL_3003779, EPI_ISL_3003780, EPI_ISL_3003781, EPI_ISL_3003782, EPI_ISL_3003783, EPI_ISL_3003784, EPI_ISL_3003785, EPI_ISL_3003786, EPI_ISL_3003787, EPI_ISL_3003788, EPI_ISL_3003789, EPI_ISL_3003790, EPI_ISL_3003791, EPI_ISL_3003792, EPI_ISL_3003793, EPI_ISL_3003794,
EPI_ISL_3003795, EPI_ISL_3003796, EPI_ISL_3003797, EPI_ISL_3003798, EPI_ISL_3003799, EPI_ISL_3003800, EPI_ISL_3003801, EPI_ISL_3003802, EPI_ISL_3003803, EPI_ISL_3003805, EPI_ISL_3003806, EPI_ISL_3003807, EPI_ISL_3003808, EPI_ISL_3003809, EPI_ISL_3003810, EPI_ISL_3003811, EPI_ISL_3003812, EPI_ISL_3003813, EPI_ISL_3003814, EPI_ISL_3003815, EPI_ISL_3003816, EPI_ISL_3003817,
EPI_ISL_3003818, EPI_ISL_3003819, EPI_ISL_3003820, EPI_ISL_3003821, EPI_ISL_3003822, EPI_ISL_3003823, EPI_ISL_3003824, EPI_ISL_3003825, EPI_ISL_3003826, EPI_ISL_3003827, EPI_ISL_3003828, EPI_ISL_3003829, EPI_ISL_3003830, EPI_ISL_3003831, EPI_ISL_3003832, EPI_ISL_3003833, EPI_ISL_3003834, EPI_ISL_3003835, EPI_ISL_3003836, EPI_ISL_3003837, EPI_ISL_3003838, EPI_ISL_3003839,
EPI_ISL_3003840, EPI_ISL_3003841, EPI_ISL_3003842, EPI_ISL_3003843, EPI_ISL_3003844, EPI_ISL_3003845, EPI_ISL_3003846, EPI_ISL_3003847, EPI_ISL_3003848, EPI_ISL_3003849, EPI_ISL_3003850, EPI_ISL_3003851, EPI_ISL_3003852, EPI_ISL_3003853, EPI_ISL_3003854, EPI_ISL_3003855, EPI_ISL_3003856, EPI_ISL_3003857, EPI_ISL_3003858, EPI_ISL_3003859, EPI_ISL_3003860, EPI_ISL_3003861,

EPI_ISL_3003862, EPI_ISL_3003863, EPI_ISL_3003864, EPI_ISL_3003865, EPI_ISL_3003866, EPI_ISL_3003867, EPI_ISL_3003868, EPI_ISL_3003869, EPI_ISL_3003870, EPI_ISL_3003871, EPI_ISL_3003872, EPI_ISL_3003873, EPI_ISL_3003874, EPI_ISL_3003875, EPI_ISL_3003876, EPI_ISL_3003877, EPI_ISL_3003878, EPI_ISL_3003879, EPI_ISL_3003880, EPI_ISL_3003881, EPI_ISL_3003882, EPI_ISL_3003883, EPI_ISL_3003884, EPI_ISL_3003885, EPI_ISL_3003886, EPI_ISL_3003887, EPI_ISL_3003888, EPI_ISL_3003889, EPI_ISL_3003890, EPI_ISL_3003891, EPI_ISL_3003892, EPI_ISL_3003893, EPI_ISL_3003894, EPI_ISL_3003895, EPI_ISL_3003896, EPI_ISL_3003897, EPI_ISL_3003898, EPI_ISL_3003899, EPI_ISL_3003900, EPI_ISL_3003901, EPI_ISL_3003902, EPI_ISL_3003903, EPI_ISL_3003904, EPI_ISL_3003905, EPI_ISL_3003906, EPI_ISL_3003907, EPI_ISL_3003908, EPI_ISL_3003909, EPI_ISL_3003910, EPI_ISL_3003911, EPI_ISL_3003912, EPI_ISL_3003914, EPI_ISL_3003915, EPI_ISL_3003916, EPI_ISL_3003917, EPI_ISL_3003918, EPI_ISL_3003919, EPI_ISL_3003920, EPI_ISL_3003921, EPI_ISL_3003923, EPI_ISL_3003924, EPI_ISL_3003925, EPI_ISL_3003926, EPI_ISL_3003927, EPI_ISL_3003928, EPI_ISL_3003929, EPI_ISL_3003930, EPI_ISL_3003931, EPI_ISL_3003932, EPI_ISL_3003933, EPI_ISL_3003934, EPI_ISL_3003935, EPI_ISL_3003936, EPI_ISL_3003937, EPI_ISL_3003938, EPI_ISL_3003939, EPI_ISL_3003940, EPI_ISL_3003941, EPI_ISL_3003942, EPI_ISL_3003943, EPI_ISL_3003944, EPI_ISL_3003945, EPI_ISL_3003946, EPI_ISL_3003947, EPI_ISL_3003948, EPI_ISL_3003949, EPI_ISL_3003950, EPI_ISL_3003951, EPI_ISL_3003952, EPI_ISL_3003953, EPI_ISL_3003954, EPI_ISL_3003955, EPI_ISL_3003956, EPI_ISL_3003957, EPI_ISL_3003958, EPI_ISL_3003959, EPI_ISL_3003960, EPI_ISL_3003961, EPI_ISL_3003962, EPI_ISL_3003963, EPI_ISL_3003964, EPI_ISL_3003965, EPI_ISL_3003966, EPI_ISL_3003967, EPI_ISL_3003968, EPI_ISL_3003969, EPI_ISL_3003970, EPI_ISL_3003971, EPI_ISL_3003972, EPI_ISL_3003973, EPI_ISL_3003974, EPI_ISL_3003975, EPI_ISL_3003976, EPI_ISL_3003977, EPI_ISL_3003978, EPI_ISL_3003979, EPI_ISL_3003980, EPI_ISL_3003982, EPI_ISL_3003983, EPI_ISL_3003984, EPI_ISL_3003985, EPI_ISL_3003986, EPI_ISL_3003987, EPI_ISL_3003988, EPI_ISL_3003989, EPI_ISL_3003990, EPI_ISL_3003991, EPI_ISL_3003992, EPI_ISL_3003993, EPI_ISL_3003994, EPI_ISL_3003995, EPI_ISL_3003996, EPI_ISL_3003997, EPI_ISL_3003998, EPI_ISL_3003999, EPI_ISL_3004000, EPI_ISL_3004001, EPI_ISL_3004002, EPI_ISL_3004003, EPI_ISL_3004004, EPI_ISL_3004005, EPI_ISL_3004006, EPI_ISL_3004007, EPI_ISL_3004008, EPI_ISL_3004009, EPI_ISL_3004010, EPI_ISL_3004011, EPI_ISL_3004012, EPI_ISL_3004013, EPI_ISL_3004014, EPI_ISL_3004015, EPI_ISL_3004016, EPI_ISL_3004017, EPI_ISL_3004018, EPI_ISL_3004019, EPI_ISL_3004020, EPI_ISL_3004021, EPI_ISL_3004022, EPI_ISL_3004023, EPI_ISL_3004024, EPI_ISL_3004025, EPI_ISL_3004026, EPI_ISL_3004027, EPI_ISL_3004028, EPI_ISL_3004029, EPI_ISL_3004030, EPI_ISL_3004031, EPI_ISL_3004032, EPI_ISL_3004033, EPI_ISL_3004034, EPI_ISL_3004035, EPI_ISL_3004036, EPI_ISL_3004037, EPI_ISL_3004038, EPI_ISL_3004039, EPI_ISL_3004040, EPI_ISL_3004041, EPI_ISL_3004042, EPI_ISL_3004043, EPI_ISL_3004044, EPI_ISL_3004045, EPI_ISL_3004046, EPI_ISL_3004047, EPI_ISL_3004048, EPI_ISL_3004049, EPI_ISL_3004050, EPI_ISL_3004051, EPI_ISL_3004052, EPI_ISL_3004053, EPI_ISL_3004054, EPI_ISL_3004055, EPI_ISL_3004056, EPI_ISL_3004057, EPI_ISL_3004058, EPI_ISL_3004059, EPI_ISL_3004060, EPI_ISL_3004061, EPI_ISL_3004062, EPI_ISL_3004063, EPI_ISL_3004064, EPI_ISL_3004065, EPI_ISL_3004066, EPI_ISL_3004067, EPI_ISL_3004068, EPI_ISL_3004069, EPI_ISL_3004070, EPI_ISL_3004071, EPI_ISL_3004072, EPI_ISL_3004073, EPI_ISL_3004074, EPI_ISL_3004075, EPI_ISL_3004076, EPI_ISL_3004077, EPI_ISL_3004078, EPI_ISL_3004079, EPI_ISL_3004080, EPI_ISL_3004081, EPI_ISL_3004082, EPI_ISL_3004083, EPI_ISL_3004084, EPI_ISL_3004085, EPI_ISL_3004086, EPI_ISL_3004087, EPI_ISL_3004088, EPI_ISL_3004089, EPI_ISL_3004090, EPI_ISL_3004091, EPI_ISL_3004092, EPI_ISL_3004093, EPI_ISL_3004094, EPI_ISL_3004095, EPI_ISL_3004096, EPI_ISL_3004097, EPI_ISL_3004098, EPI_ISL_3004099, EPI_ISL_3004100, EPI_ISL_3004101, EPI_ISL_3004102, EPI_ISL_3004103, EPI_ISL_3004104, EPI_ISL_3004105, EPI_ISL_3004106, EPI_ISL_3004107, EPI_ISL_3004108, EPI_ISL_3004109, EPI_ISL_3004110, EPI_ISL_3004111, EPI_ISL_3004112, EPI_ISL_3004113, EPI_ISL_3004114, EPI_ISL_3004115, EPI_ISL_3004116, EPI_ISL_3004117, EPI_ISL_3004118, EPI_ISL_3004119, EPI_ISL_3004120, EPI_ISL_3004121, EPI_ISL_3004123, EPI_ISL_3004124, EPI_ISL_3004125, EPI_ISL_3004126, EPI_ISL_3004127, EPI_ISL_3004128, EPI_ISL_3004130, EPI_ISL_3004132, EPI_ISL_3004133, EPI_ISL_3004134, EPI_ISL_3004135, EPI_ISL_3004136, EPI_ISL_3004138, EPI_ISL_3004139, EPI_ISL_3004142, EPI_ISL_3004143, EPI_ISL_3004144, EPI_ISL_3004145, EPI_ISL_3004146, EPI_ISL_3004148, EPI_ISL_3004149, EPI_ISL_3004150, EPI_ISL_3004151, EPI_ISL_3004152, EPI_ISL_3004153, EPI_ISL_3004154, EPI_ISL_3004156, EPI_ISL_3004157, EPI_ISL_3004158, EPI_ISL_3004159, EPI_ISL_3004160, EPI_ISL_3004161, EPI_ISL_3004162, EPI_ISL_3004163, EPI_ISL_3004164, EPI_ISL_3004166, EPI_ISL_3004167, EPI_ISL_3004168, EPI_ISL_3004169, EPI_ISL_3004170, EPI_ISL_3004171, EPI_ISL_3004172, EPI_ISL_3004173, EPI_ISL_3004174, EPI_ISL_3004175, EPI_ISL_3004176, EPI_ISL_3004177, EPI_ISL_3004178, EPI_ISL_3004179, EPI_ISL_3004180, EPI_ISL_3004181, EPI_ISL_3004183, EPI_ISL_3004184, EPI_ISL_3004185, EPI_ISL_3004186, EPI_ISL_3004187, EPI_ISL_3004188, EPI_ISL_3004189, EPI_ISL_3004190, EPI_ISL_3004191, EPI_ISL_3004192, EPI_ISL_3004193, EPI_ISL_3004194, EPI_ISL_3004195, EPI_ISL_3004196, EPI_ISL_3004197, EPI_ISL_3004198, EPI_ISL_3004199, EPI_ISL_3004200, EPI_ISL_3004201, EPI_ISL_3004202, EPI_ISL_3004203, EPI_ISL_3004204, EPI_ISL_3004205, EPI_ISL_3004206, EPI_ISL_3004207, EPI_ISL_3004208, EPI_ISL_3004209, EPI_ISL_3004210, EPI_ISL_3004211, EPI_ISL_3004212, EPI_ISL_3004213, EPI_ISL_3004214, EPI_ISL_3004215, EPI_ISL_3004216, EPI_ISL_3004217, EPI_ISL_3004218, EPI_ISL_3004220, EPI_ISL_3004221, EPI_ISL_3004223, EPI_ISL_3004224, EPI_ISL_3004226, EPI_ISL_3004227, EPI_ISL_3004228, EPI_ISL_3004229, EPI_ISL_3004230, EPI_ISL_3004231, EPI_ISL_3004232, EPI_ISL_3004233, EPI_ISL_3004235, EPI_ISL_3004236, EPI_ISL_3004237, EPI_ISL_3004238, EPI_ISL_3004239, EPI_ISL_3004240, EPI_ISL_3004241, EPI_ISL_3004242, EPI_ISL_3004243, EPI_ISL_3004244, EPI_ISL_3004245, EPI_ISL_3004246, EPI_ISL_3004248, EPI_ISL_3004249, EPI_ISL_3004250, EPI_ISL_3004251, EPI_ISL_3004252, EPI_ISL_3004253, EPI_ISL_3004255, EPI_ISL_3004257, EPI_ISL_3004258, EPI_ISL_3004259, EPI_ISL_3004261, EPI_ISL_3004262, EPI_ISL_3004263, EPI_ISL_3004264, EPI_ISL_3004265, EPI_ISL_3004266, EPI_ISL_3004267, EPI_ISL_3004270, EPI_ISL_3004272, EPI_ISL_3004273, EPI_ISL_3004274, EPI_ISL_3004275, EPI_ISL_3004276, EPI_ISL_3004277, EPI_ISL_3004278, EPI_ISL_3004280, EPI_ISL_3004281, EPI_ISL_3004283, EPI_ISL_3004284, EPI_ISL_3004285, EPI_ISL_3004286, EPI_ISL_3004287, EPI_ISL_3004288, EPI_ISL_3004290, EPI_ISL_3004291, EPI_ISL_3004292, EPI_ISL_3004293, EPI_ISL_3004294, EPI_ISL_3004295, EPI_ISL_3004296, EPI_ISL_3004297, EPI_ISL_3004298, EPI_ISL_3004299, EPI_ISL_3004300, EPI_ISL_3004301, EPI_ISL_3004302, EPI_ISL_3004303, EPI_ISL_3004304, EPI_ISL_3004305, EPI_ISL_3004306, EPI_ISL_3004307, EPI_ISL_3004308, EPI_ISL_3004309, EPI_ISL_3004310, EPI_ISL_3004311, EPI_ISL_3004312, EPI_ISL_3004313, EPI_ISL_3004314, EPI_ISL_3004315, EPI_ISL_3004316, EPI_ISL_3004317, EPI_ISL_3004318, EPI_ISL_3004319, EPI_ISL_3004320, EPI_ISL_3004321, EPI_ISL_3004322, EPI_ISL_3004323, EPI_ISL_3004324, EPI_ISL_3004325, EPI_ISL_3004326, EPI_ISL_3004327, EPI_ISL_3004328, EPI_ISL_3004329, EPI_ISL_3004330, EPI_ISL_3004331, EPI_ISL_3004332, EPI_ISL_3004334, EPI_ISL_3004335, EPI_ISL_3004336, EPI_ISL_3004337, EPI_ISL_3004338, EPI_ISL_3004339, EPI_ISL_3004340, EPI_ISL_3004342, EPI_ISL_3004343, EPI_ISL_3004344, EPI_ISL_3004345, EPI_ISL_3004347, EPI_ISL_3004348, EPI_ISL_3004349, EPI_ISL_3004350, EPI_ISL_3004351, EPI_ISL_3004353, EPI_ISL_3004354, EPI_ISL_3004355, EPI_ISL_3004356, EPI_ISL_3004357, EPI_ISL_3004358, EPI_ISL_3004359, EPI_ISL_3004360, EPI_ISL_3004361, EPI_ISL_3004364, EPI_ISL_3004365, EPI_ISL_3004366, EPI_ISL_3004367, EPI_ISL_3004368, EPI_ISL_3004369

| | | | |
|---|---|---|---|
| see above | Respiratory Virus Unit, Microbiology Services Colindale, Public Health England | COVID-19 Genomics UK (COG-UK) Consortium | PHE Covid Sequencing Team |

EPI_ISL_3004370, EPI_ISL_3004371, EPI_ISL_3004372, EPI_ISL_3004373, EPI_ISL_3004374, EPI_ISL_3004375, EPI_ISL_3004376, EPI_ISL_3004377, EPI_ISL_3004378, EPI_ISL_3004379, EPI_ISL_3004380, EPI_ISL_3004381, EPI_ISL_3004382, EPI_ISL_3004383, EPI_ISL_3004384, EPI_ISL_3004385, EPI_ISL_3004386, EPI_ISL_3004387, EPI_ISL_3004388, EPI_ISL_3004389, EPI_ISL_3004390, EPI_ISL_3004391, EPI_ISL_3004392, EPI_ISL_3004393, EPI_ISL_3004394, EPI_ISL_3004395, EPI_ISL_3004396, EPI_ISL_3004397, EPI_ISL_3004398, EPI_ISL_3004399, EPI_ISL_3004400, EPI_ISL_3004401, EPI_ISL_3004402, EPI_ISL_3004403, EPI_ISL_3004404, EPI_ISL_3004405, EPI_ISL_3004406, EPI_ISL_3004407, EPI_ISL_3004408, EPI_ISL_3004409, EPI_ISL_3004410, EPI_ISL_3004411, EPI_ISL_3004412, EPI_ISL_3004413, EPI_ISL_3004414, EPI_ISL_3004415, EPI_ISL_3004416, EPI_ISL_3004417, EPI_ISL_3004418, EPI_ISL_3004419, EPI_ISL_3004420, EPI_ISL_3004421, EPI_ISL_3004422, EPI_ISL_3004423, EPI_ISL_3004424, EPI_ISL_3004425, EPI_ISL_3004426, EPI_ISL_3004427, EPI_ISL_3004428, EPI_ISL_3004429, EPI_ISL_3004430, EPI_ISL_3004431, EPI_ISL_3004432, EPI_ISL_3004433, EPI_ISL_3004434, EPI_ISL_3004435, EPI_ISL_3004436, EPI_ISL_3004437, EPI_ISL_3004438, EPI_ISL_3004439, EPI_ISL_3004440, EPI_ISL_3004441, EPI_ISL_3004442, EPI_ISL_3004443, EPI_ISL_3004444, EPI_ISL_3004445, EPI_ISL_3004446, EPI_ISL_3004447, EPI_ISL_3004448, EPI_ISL_3004449, EPI_ISL_3004450, EPI_ISL_3004451, EPI_ISL_3004452, EPI_ISL_3004453, EPI_ISL_3004454, EPI_ISL_3004455, EPI_ISL_3004456, EPI_ISL_3004457, EPI_ISL_3004458, EPI_ISL_3004459, EPI_ISL_3004460, EPI_ISL_3004461, EPI_ISL_3004462, EPI_ISL_3004463, EPI_ISL_3004464, EPI_ISL_3004465, EPI_ISL_3004466, EPI_ISL_3004467, EPI_ISL_3004468, EPI_ISL_3004469, EPI_ISL_3004470, EPI_ISL_3004471, EPI_ISL_3004472, EPI_ISL_3004473, EPI_ISL_3004474, EPI_ISL_3004475, EPI_ISL_3004476, EPI_ISL_3004477, EPI_ISL_3004478, EPI_ISL_3004479, EPI_ISL_3004480, EPI_ISL_3004481, EPI_ISL_3004482, EPI_ISL_3004483, EPI_ISL_3004484, EPI_ISL_3004485, EPI_ISL_3004486, EPI_ISL_3004487, EPI_ISL_3004488, EPI_ISL_3004489, EPI_ISL_3004490, EPI_ISL_3004491, EPI_ISL_3004492, EPI_ISL_3004493, EPI_ISL_3004494, EPI_ISL_3004495, EPI_ISL_3004496, EPI_ISL_3004497, EPI_ISL_3004498, EPI_ISL_3004499, EPI_ISL_3004500, EPI_ISL_3004501, EPI_ISL_3004502, EPI_ISL_3004503, EPI_ISL_3004504, EPI_ISL_3004505, EPI_ISL_3004506, EPI_ISL_3004507, EPI_ISL_3004508, EPI_ISL_3004509, EPI_ISL_3004510, EPI_ISL_3004511, EPI_ISL_3004512, EPI_ISL_3004513, EPI_ISL_3004514, EPI_ISL_3004515, EPI_ISL_3004516, EPI_ISL_3004517, EPI_ISL_3004518, EPI_ISL_3004519, EPI_ISL_3004520, EPI_ISL_3004521, EPI_ISL_3004522, EPI_ISL_3004523, EPI_ISL_3004524, EPI_ISL_3004525, EPI_ISL_3004526, EPI_ISL_3004527, EPI_ISL_3004528, EPI_ISL_3004529, EPI_ISL_3004530, EPI_ISL_3004531, EPI_ISL_3004532, EPI_ISL_3004533, EPI_ISL_3004534, EPI_ISL_3004535, EPI_ISL_3004536, EPI_ISL_3004537, EPI_ISL_3004538, EPI_ISL_3004539, EPI_ISL_3004540, EPI_ISL_3004541, EPI_ISL_3004542, EPI_ISL_3004543, EPI_ISL_3004544, EPI_ISL_3004545, EPI_ISL_3004546, EPI_ISL_3004547, EPI_ISL_3004548, EPI_ISL_3004549, EPI_ISL_3004550, EPI_ISL_3004551, EPI_ISL_3004552, EPI_ISL_3004553, EPI_ISL_3004554, EPI_ISL_3004555, EPI_ISL_3004556, EPI_ISL_3004557, EPI_ISL_3004558, EPI_ISL_3004559, EPI_ISL_3004560, EPI_ISL_3004561, EPI_ISL_3004562, EPI_ISL_3004563, EPI_ISL_3004564, EPI_ISL_3004565, EPI_ISL_3004566, EPI_ISL_3004567, EPI_ISL_3004568, EPI_ISL_3004569, EPI_ISL_3004570, EPI_ISL_3004571, EPI_ISL_3004572, EPI_ISL_3004574, EPI_ISL_3004575, EPI_ISL_3004576, EPI_ISL_3004577, EPI_ISL_3004578, EPI_ISL_3004579, EPI_ISL_3004580, EPI_ISL_3004581, EPI_ISL_3004582, EPI_ISL_3004583, EPI_ISL_3004584, EPI_ISL_3004585, EPI_ISL_3004586, EPI_ISL_3004587, EPI_ISL_3004588, EPI_ISL_3004589, EPI_ISL_3004590, EPI_ISL_3004591, EPI_ISL_3004592, EPI_ISL_3004593, EPI_ISL_3004594, EPI_ISL_3004595, EPI_ISL_3004596, EPI_ISL_3004597, EPI_ISL_3004598, EPI_ISL_3004599, EPI_ISL_3004600, EPI_ISL_3004601, EPI_ISL_3004602, EPI_ISL_3004603, EPI_ISL_3004604, EPI_ISL_3004605, EPI_ISL_3004606, EPI_ISL_3004607, EPI_ISL_3004608, EPI_ISL_3004609, EPI_ISL_3004610, EPI_ISL_3004611, EPI_ISL_3004612, EPI_ISL_3004613, EPI_ISL_3004614, EPI_ISL_3004615, EPI_ISL_3004616, EPI_ISL_3004617, EPI_ISL_3004618, EPI_ISL_3004619, EPI_ISL_3004620, EPI_ISL_3004621, EPI_ISL_3004622, EPI_ISL_3004623, EPI_ISL_3004624, EPI_ISL_3004625, EPI_ISL_3004626, EPI_ISL_3004627, EPI_ISL_3004628, EPI_ISL_3004629, EPI_ISL_3004630, EPI_ISL_3004631, EPI_ISL_3004632, EPI_ISL_3004633, EPI_ISL_3004634, EPI_ISL_3004635, EPI_ISL_3004636, EPI_ISL_3004637, EPI_ISL_3004638, EPI_ISL_3004639, EPI_ISL_3004640, EPI_ISL_3004641, EPI_ISL_3004642, EPI_ISL_3004643, EPI_ISL_3004644, EPI_ISL_3004645, EPI_ISL_3004646, EPI_ISL_3004647, EPI_ISL_3004648, EPI_ISL_3004649, EPI_ISL_3004651, EPI_ISL_3004652, EPI_ISL_3004653, EPI_ISL_3004654, EPI_ISL_3004655, EPI_ISL_3004656, EPI_ISL_3004657, EPI_ISL_3004658, EPI_ISL_3004659, EPI_ISL_3004660, EPI_ISL_3004661, EPI_ISL_3004662, EPI_ISL_3004663, EPI_ISL_3004664, EPI_ISL_3004665, EPI_ISL_3004666, EPI_ISL_3004667, EPI_ISL_3004668, EPI_ISL_3004669, EPI_ISL_3004670, EPI_ISL_3004671, EPI_ISL_3004672, EPI_ISL_3004673, EPI_ISL_3004674, EPI_ISL_3004675, EPI_ISL_3004676, EPI_ISL_3004677, EPI_ISL_3004678, EPI_ISL_3004679, EPI_ISL_3004680, EPI_ISL_3004681, EPI_ISL_3004682, EPI_ISL_3004683, EPI_ISL_3004684, EPI_ISL_3004685, EPI_ISL_3004686, EPI_ISL_3004687, EPI_ISL_3004688, EPI_ISL_3004689, EPI_ISL_3004690, EPI_ISL_3004691, EPI_ISL_3004692, EPI_ISL_3004693, EPI_ISL_3004694, EPI_ISL_3004695, EPI_ISL_3004696, EPI_ISL_3004697, EPI_ISL_3004698, EPI_ISL_3004699, EPI_ISL_3004700, EPI_ISL_3004701, EPI_ISL_3004702, EPI_ISL_3004703, EPI_ISL_3004704, EPI_ISL_3004705, EPI_ISL_3004706, EPI_ISL_3004707, EPI_ISL_3004708, EPI_ISL_3004709, EPI_ISL_3004710, EPI_ISL_3004711, EPI_ISL_3004712, EPI_ISL_3004713, EPI_ISL_3004714, EPI_ISL_3004715, EPI_ISL_3004716, EPI_ISL_3004717, EPI_ISL_3004718, EPI_ISL_3004719, EPI_ISL_3004720, EPI_ISL_3004721, EPI_ISL_3004722, EPI_ISL_3004723, EPI_ISL_3004724, EPI_ISL_3004725, EPI_ISL_3004726, EPI_ISL_3004727, EPI_ISL_3004728, EPI_ISL_3004729, EPI_ISL_3004730, EPI_ISL_3004731, EPI_ISL_3004732, EPI_ISL_3004733, EPI_ISL_3004734, EPI_ISL_3004735, EPI_ISL_3004736, EPI_ISL_3004737, EPI_ISL_3004738, EPI_ISL_3004739, EPI_ISL_3004740, EPI_ISL_3004741, EPI_ISL_3004742, EPI_ISL_3004743, EPI_ISL_3004744, EPI_ISL_3004745, EPI_ISL_3004746, EPI_ISL_3004747, EPI_ISL_3004748, EPI_ISL_3004749, EPI_ISL_3004750, EPI_ISL_3004751, EPI_ISL_3004752, EPI_ISL_3004753, EPI_ISL_3004754, EPI_ISL_3004755, EPI_ISL_3004756, EPI_ISL_3004757, EPI_ISL_3004758, EPI_ISL_3004759, EPI_ISL_3004760, EPI_ISL_3004761, EPI_ISL_3004762, EPI_ISL_3004763, EPI_ISL_3004764, EPI_ISL_3004765, EPI_ISL_3004766, EPI_ISL_3004767, EPI_ISL_3004768, EPI_ISL_3004769, EPI_ISL_3004770, EPI_ISL_3004771, EPI_ISL_3004772, EPI_ISL_3004773, EPI_ISL_3004774, EPI_ISL_3004775, EPI_ISL_3004776, EPI_ISL_3004777, EPI_ISL_3004778, EPI_ISL_3004779, EPI_ISL_3004780, EPI_ISL_3004781, EPI_ISL_3004782, EPI_ISL_3004783, EPI_ISL_3004784, EPI_ISL_3004785, EPI_ISL_3004786, EPI_ISL_3004787, EPI_ISL_3004788, EPI_ISL_3004789, EPI_ISL_3004790, EPI_ISL_3004791, EPI_ISL_3004792, EPI_ISL_3004793, EPI_ISL_3004794, EPI_ISL_3004795, EPI_ISL_3004796, EPI_ISL_3004797, EPI_ISL_3004798, EPI_ISL_3004799, EPI_ISL_3004800, EPI_ISL_3004801, EPI_ISL_3004802, EPI_ISL_3004803, EPI_ISL_3004804, EPI_ISL_3004805, EPI_ISL_3004806, EPI_ISL_3004807, EPI_ISL_3004808, EPI_ISL_3004809, EPI_ISL_3004810, EPI_ISL_3004811, EPI_ISL_3004812, EPI_ISL_3004813, EPI_ISL_3004814, EPI_ISL_3004815, EPI_ISL_3004816, EPI_ISL_3004817, EPI_ISL_3004818, EPI_ISL_3004819, EPI_ISL_3004820, EPI_ISL_3004821, EPI_ISL_3004822, EPI_ISL_3004823, EPI_ISL_3004824, EPI_ISL_3004825, EPI_ISL_3004826, EPI_ISL_3004827, EPI_ISL_3004828, EPI_ISL_3004829, EPI_ISL_3004830, EPI_ISL_3004831, EPI_ISL_3004832, EPI_ISL_3004833, EPI_ISL_3004834, EPI_ISL_3004835, EPI_ISL_3004836, EPI_ISL_3004837, EPI_ISL_3004838, EPI_ISL_3004839, EPI_ISL_3004840, EPI_ISL_3004841, EPI_ISL_3004842, EPI_ISL_3004843, EPI_ISL_3004844, EPI_ISL_3004845, EPI_ISL_3004846, EPI_ISL_3004847, EPI_ISL_3004848, EPI_ISL_3004849, EPI_ISL_3004850, EPI_ISL_3004851, EPI_ISL_3004852, EPI_ISL_3004853, EPI_ISL_3004854, EPI_ISL_3004855, EPI_ISL_3004856, EPI_ISL_3004857, EPI_ISL_3004858, EPI_ISL_3004859, EPI_ISL_3004860, EPI_ISL_3004861, EPI_ISL_3004862, EPI_ISL_3004863, EPI_ISL_3004864, EPI_ISL_3004865, EPI_ISL_3004866, EPI_ISL_3004867, EPI_ISL_3004868, EPI_ISL_3004869, EPI_ISL_3004870, EPI_ISL_3004871, EPI_ISL_3004872, EPI_ISL_3004873, EPI_ISL_3004874, EPI_ISL_3004875, EPI_ISL_3004876, EPI_ISL_3004877, EPI_ISL_3004878, EPI_ISL_3004879, EPI_ISL_3004880, EPI_ISL_3004881, EPI_ISL_3004882, EPI_ISL_3004884, EPI_ISL_3004885, EPI_ISL_3004886, EPI_ISL_3004887, EPI_ISL_3004888, EPI_ISL_3004889, EPI_ISL_3004890, EPI_ISL_3004891, EPI_ISL_3004892, EPI_ISL_3004893, EPI_ISL_3004894, EPI_ISL_3004895, EPI_ISL_3004896, EPI_ISL_3004897, EPI_ISL_3004898, EPI_ISL_3004899, EPI_ISL_3004900, EPI_ISL_3004901, EPI_ISL_3004902, EPI_ISL_3004903, EPI_ISL_3004904, EPI_ISL_3004905, EPI_ISL_3004906, EPI_ISL_3004907, EPI_ISL_3004908, EPI_ISL_3004909, EPI_ISL_3004910, EPI_ISL_3004911, EPI_ISL_3004912, EPI_ISL_3004913, EPI_ISL_3004914, EPI_ISL_3004915, EPI_ISL_3004916, EPI_ISL_3004917, EPI_ISL_3004918, EPI_ISL_3004919, EPI_ISL_3004920, EPI_ISL_3004921, EPI_ISL_3004922, EPI_ISL_3004923, EPI_ISL_3004924, EPI_ISL_3004925, EPI_ISL_3004926, EPI_ISL_3004927, EPI_ISL_3004928, EPI_ISL_3004929, EPI_ISL_3004930, EPI_ISL_3004931, EPI_ISL_3004932, EPI_ISL_3004933, EPI_ISL_3004934, EPI_ISL_3004935, EPI_ISL_3004937, EPI_ISL_3004938, EPI_ISL_3004939, EPI_ISL_3004940, EPI_ISL_3004941, EPI_ISL_3004942, EPI_ISL_3004943, EPI_ISL_3004944, EPI_ISL_3004945, EPI_ISL_3004946, EPI_ISL_3004947, EPI_ISL_3004948, EPI_ISL_3004949, EPI_ISL_3004950, EPI_ISL_3004951, EPI_ISL_3004952, EPI_ISL_3004953, EPI_ISL_3004954, EPI_ISL_3004955, EPI_ISL_3004956, EPI_ISL_3004957, EPI_ISL_3004958, EPI_ISL_3004959, EPI_ISL_3004960, EPI_ISL_3004961, EPI_ISL_3004962, EPI_ISL_3004963, EPI_ISL_3004964, EPI_ISL_3004965, EPI_ISL_3004966, EPI_ISL_3004967, EPI_ISL_3004968, EPI_ISL_3004969, EPI_ISL_3004970, EPI_ISL_3004971, EPI_ISL_3004972, EPI_ISL_3004973, EPI_ISL_3004974, EPI_ISL_3004975, EPI_ISL_3004976, EPI_ISL_3004977, EPI_ISL_3004978, EPI_ISL_3004979, EPI_ISL_3004980, EPI_ISL_3004981, EPI_ISL_3004982, EPI_ISL_3004983, EPI_ISL_3004984, EPI_ISL_3004985, EPI_ISL_3004986, EPI_ISL_3004987, EPI_ISL_3004988, EPI_ISL_3004989, EPI_ISL_3004990, EPI_ISL_3004991, EPI_ISL_3004992, EPI_ISL_3004993, EPI_ISL_3004994, EPI_ISL_3004995, EPI_ISL_3004996, EPI_ISL_3004997, EPI_ISL_3004998, EPI_ISL_3004999, EPI_ISL_3005000, EPI_ISL_3005001, EPI_ISL_3005002, EPI_ISL_3005003, EPI_ISL_3005004, EPI_ISL_3005005, EPI_ISL_3005006, EPI_ISL_3005007, EPI_ISL_3005008, EPI_ISL_3005009, EPI_ISL_3005010, EPI_ISL_3005011, EPI_ISL_3005012, EPI_ISL_3005013, EPI_ISL_3005014, EPI_ISL_3005015, EPI_ISL_3005016, EPI_ISL_3005017, EPI_ISL_3005018, EPI_ISL_3005019, EPI_ISL_3005020, EPI_ISL_3005021, EPI_ISL_3005022, EPI_ISL_3005023, EPI_ISL_3005024, EPI_ISL_3005025, EPI_ISL_3005026, EPI_ISL_3005027, EPI_ISL_3005028, EPI_ISL_3005029, EPI_ISL_3005030, EPI_ISL_3005031, EPI_ISL_3005032, EPI_ISL_3005033, EPI_ISL_3005034, EPI_ISL_3005035, EPI_ISL_3005036, EPI_ISL_3005037, EPI_ISL_3005038, EPI_ISL_3005039, EPI_ISL_3005040, EPI_ISL_3005041, EPI_ISL_3005042, EPI_ISL_3005043, EPI_ISL_3005044, EPI_ISL_3005045, EPI_ISL_3005046, EPI_ISL_3005047, EPI_ISL_3005048, EPI_ISL_3005049, EPI_ISL_3005050, EPI_ISL_3005051, EPI_ISL_3005052, EPI_ISL_3005053, EPI_ISL_3005054, EPI_ISL_3005055, EPI_ISL_3005056, EPI_ISL_3005057, EPI_ISL_3005058, EPI_ISL_3005059, EPI_ISL_3005060, EPI_ISL_3005061, EPI_ISL_3005062, EPI_ISL_3005063, EPI_ISL_3005064, EPI_ISL_3005065, EPI_ISL_3005066, EPI_ISL_3005067, EPI_ISL_3005068, EPI_ISL_3005069, EPI_ISL_3005070, EPI_ISL_3005071, EPI_ISL_3005072, EPI_ISL_3005073, EPI_ISL_3005074, EPI_ISL_3005075, EPI_ISL_3005076, EPI_ISL_3005077, EPI_ISL_3005078, EPI_ISL_3005079, EPI_ISL_3005080, EPI_ISL_3005081, EPI_ISL_3005082, EPI_ISL_3005083, EPI_ISL_3005084, EPI_ISL_3005085, EPI_ISL_3005086, EPI_ISL_3005087, EPI_ISL_3005088, EPI_ISL_3005089, EPI_ISL_3005090, EPI_ISL_3005091, EPI_ISL_3005092, EPI_ISL_3005093, EPI_ISL_3005094, EPI_ISL_3005095, EPI_ISL_3005096, EPI_ISL_3005097, EPI_ISL_3005098, EPI_ISL_3005099, EPI_ISL_3005100, EPI_ISL_3005101, EPI_ISL_3005102, EPI_ISL_3005103, EPI_ISL_3005104, EPI_ISL_3005105, EPI_ISL_3005106, EPI_ISL_3005107, EPI_ISL_3005108, EPI_ISL_3005109, EPI_ISL_3005110, EPI_ISL_3005111, EPI_ISL_3005112, EPI_ISL_3005113, EPI_ISL_3005114, EPI_ISL_3005115, EPI_ISL_3005116, EPI_ISL_3005117, EPI_ISL_3005118, EPI_ISL_3005119, EPI_ISL_3005120, EPI_ISL_3005121, EPI_ISL_3005122, EPI_ISL_3005123, EPI_ISL_3005124, EPI_ISL_3005125, EPI_ISL_3005126, EPI_ISL_3005127, EPI_ISL_3005128, EPI_ISL_3005129, EPI_ISL_3005130, EPI_ISL_3005131, EPI_ISL_3005132, EPI_ISL_3005133, EPI_ISL_3005134, EPI_ISL_3005135, EPI_ISL_3005136, EPI_ISL_3005137, EPI_ISL_3005138, EPI_ISL_3005139, EPI_ISL_3005140, EPI_ISL_3005141, EPI_ISL_3005142, EPI_ISL_3005143, EPI_ISL_3005144, EPI_ISL_3005145, EPI_ISL_3005146, EPI_ISL_3005147, EPI_ISL_3005148, EPI_ISL_3005149, EPI_ISL_3005150, EPI_ISL_3005151, EPI_ISL_3005152, EPI_ISL_3005153, EPI_ISL_3005154, EPI_ISL_3005156, EPI_ISL_3005157, EPI_ISL_3005158, EPI_ISL_3005159, EPI_ISL_3005160, EPI_ISL_3005161, EPI_ISL_3005162, EPI_ISL_3005163, EPI_ISL_3005164, EPI_ISL_3005165, EPI_ISL_3005166, EPI_ISL_3005167, EPI_ISL_3005168, EPI_ISL_3005169, EPI_ISL_3005170, EPI_ISL_3005171, EPI_ISL_3005172, EPI_ISL_3005173, EPI_ISL_3005174, EPI_ISL_3005175, EPI_ISL_3005176, EPI_ISL_3005177, EPI_ISL_3005178, EPI_ISL_3005179, EPI_ISL_3005180, EPI_ISL_3005181, EPI_ISL_3005182, EPI_ISL_3005183, EPI_ISL_3005184, EPI_ISL_3005185, EPI_ISL_3005186, EPI_ISL_3005187, EPI_ISL_3005188, EPI_ISL_3005189, EPI_ISL_3005190, EPI_ISL_3005191, EPI_ISL_3005192, EPI_ISL_3005193, EPI_ISL_3005194, EPI_ISL_3005195, EPI_ISL_3005196, EPI_ISL_3005197, EPI_ISL_3005198, EPI_ISL_3005199, EPI_ISL_3005200, EPI_ISL_3005201, EPI_ISL_3005202, EPI_ISL_3005203, EPI_ISL_3005204, EPI_ISL_3005205, EPI_ISL_3005206, EPI_ISL_3005207, EPI_ISL_3005208, EPI_ISL_3005209, EPI_ISL_3005210, EPI_ISL_3005211, EPI_ISL_3005212, EPI_ISL_3005213, EPI_ISL_3005214, EPI_ISL_3005215, EPI_ISL_3005216, EPI_ISL_3005217, EPI_ISL_3005218, EPI_ISL_3005219, EPI_ISL_3005220, EPI_ISL_3005221, EPI_ISL_3005222, EPI_ISL_3005223, EPI_ISL_3005224, EPI_ISL_3005225, EPI_ISL_3005226, EPI_ISL_3005227, EPI_ISL_3005228, EPI_ISL_3005229, EPI_ISL_3005230, EPI_ISL_3005231, EPI_ISL_3005232, EPI_ISL_3005233, EPI_ISL_3005234, EPI_ISL_3005235, EPI_ISL_3005236, EPI_ISL_3005237, EPI_ISL_3005238, EPI_ISL_3005239, EPI_ISL_3005240, EPI_ISL_3005241, EPI_ISL_3005242, EPI_ISL_3005243, EPI_ISL_3005244, EPI_ISL_3005245, EPI_ISL_3005246, EPI_ISL_3005247, EPI_ISL_3005248, EPI_ISL_3005249, EPI_ISL_3005250, EPI_ISL_3005251, EPI_ISL_3005252, EPI_ISL_3005253, EPI_ISL_3005254, EPI_ISL_3005255, EPI_ISL_3005256, EPI_ISL_3005257, EPI_ISL_3005258, EPI_ISL_3005259, EPI_ISL_3005260, EPI_ISL_3005261, EPI_ISL_3005262, EPI_ISL_3005263, EPI_ISL_3005264, EPI_ISL_3005265, EPI_ISL_3005266, EPI_ISL_3005267, EPI_ISL_3005268, EPI_ISL_3005269, EPI_ISL_3005270, EPI_ISL_3005271, EPI_ISL_3005272, EPI_ISL_3005273, EPI_ISL_3005274, EPI_ISL_3005275, EPI_ISL_3005276, EPI_ISL_3005277, EPI_ISL_3005278, EPI_ISL_3005279, EPI_ISL_3005280, EPI_ISL_3005281, EPI_ISL_3005282, EPI_ISL_3005283, EPI_ISL_3005284, EPI_ISL_3005285, EPI_ISL_3005286, EPI_ISL_3005287, EPI_ISL_3005288, EPI_ISL_3005289, EPI_ISL_3005290, EPI_ISL_3005291, EPI_ISL_3005292, EPI_ISL_3005293, EPI_ISL_3005294, EPI_ISL_3005295, EPI_ISL_3005296, EPI_ISL_3005297, EPI_ISL_3005298, EPI_ISL_3005299, EPI_ISL_3005300, EPI_ISL_3005301, EPI_ISL_3005302, EPI_ISL_3005303, EPI_ISL_3005304, EPI_ISL_3005305, EPI_ISL_3005306, EPI_ISL_3005307, EPI_ISL_3005308, EPI_ISL_3005309, EPI_ISL_3005310, EPI_ISL_3005311, EPI_ISL_3005312, EPI_ISL_3005313, EPI_ISL_3005314, EPI_ISL_3005315, EPI_ISL_3005316, EPI_ISL_3005317, EPI_ISL_3005318, EPI_ISL_3005319, EPI_ISL_3005320, EPI_ISL_3005321, EPI_ISL_3005322, EPI_ISL_3005323, EPI_ISL_3005324, EPI_ISL_3005325, EPI_ISL_3005326, EPI_ISL_3005327, EPI_ISL_3005328, EPI_ISL_3005329, EPI_ISL_3005330, EPI_ISL_3005331, EPI_ISL_3005332, EPI_ISL_3005333, EPI_ISL_3005334, EPI_ISL_3005335, EPI_ISL_3005336, EPI_ISL_3005337, EPI_ISL_3005338, EPI_ISL_3005339, EPI_ISL_3005340, EPI_ISL_3005341, EPI_ISL_3005342, EPI_ISL_3005343, EPI_ISL_3005344, EPI_ISL_3005345, EPI_ISL_3005346, EPI_ISL_3005347, EPI_ISL_3005348, EPI_ISL_3005349, EPI_ISL_3005350, EPI_ISL_3005351, EPI_ISL_3005352, EPI_ISL_3005353, EPI_ISL_3005354, EPI_ISL_3005355, EPI_ISL_3005356, EPI_ISL_3005357, EPI_ISL_3005358, EPI_ISL_3005359, EPI_ISL_3005360, EPI_ISL_3005361, EPI_ISL_3005362, EPI_ISL_3005363, EPI_ISL_3005364, EPI_ISL_3005365, EPI_ISL_3005366, EPI_ISL_3005367, EPI_ISL_3005368, EPI_ISL_3005369, EPI_ISL_3005370, EPI_ISL_3005371, EPI_ISL_3005372, EPI_ISL_3005373, EPI_ISL_3005374, EPI_ISL_3005375, EPI_ISL_3005376, EPI_ISL_3005377, EPI_ISL_3005378, EPI_ISL_3005379, EPI_ISL_3005380, EPI_ISL_3005381, EPI_ISL_3005382, EPI_ISL_3005383, EPI_ISL_3005384, EPI_ISL_3005385, EPI_ISL_3005386, EPI_ISL_3005387, EPI_ISL_3005388, EPI_ISL_3005389, EPI_ISL_3005390, EPI_ISL_3005391, EPI_ISL_3005392, EPI_ISL_3005393, EPI_ISL_3005394, EPI_ISL_3005395, EPI_ISL_3005396, EPI_ISL_3005397, EPI_ISL_3005398, EPI_ISL_3005399, EPI_ISL_3005400, EPI_ISL_3005401, EPI_ISL_3005402, EPI_ISL_3005403, EPI_ISL_3005404, EPI_ISL_3005405, EPI_ISL_3005406, EPI_ISL_3005407, EPI_ISL_3005408, EPI_ISL_3005409, EPI_ISL_3005410, EPI_ISL_3005411, EPI_ISL_3005412, EPI_ISL_3005413, EPI_ISL_3005414, EPI_ISL_3005415, EPI_ISL_3005416, EPI_ISL_3005417, EPI_ISL_3005418, EPI_ISL_3005419, EPI_ISL_3005420, EPI_ISL_3005421, EPI_ISL_3005422, EPI_ISL_3005423, EPI_ISL_3005424, EPI_ISL_3005425, EPI_ISL_3005426, EPI_ISL_3005427, EPI_ISL_3005428, EPI_ISL_3005429, EPI_ISL_3005430, EPI_ISL_3005431, EPI_ISL_3005432, EPI_ISL_3005433, EPI_ISL_3005434, EPI_ISL_3005435, EPI_ISL_3005436, EPI_ISL_3005437, EPI_ISL_3005438, EPI_ISL_3005439, EPI_ISL_3005440, EPI_ISL_3005441, EPI_ISL_3005442, EPI_ISL_3005443, EPI_ISL_3005444, EPI_ISL_3005445, EPI_ISL_3005446, EPI_ISL_3005447, EPI_ISL_3005448, EPI_ISL_3005449, EPI_ISL_3005450, EPI_ISL_3005451, EPI_ISL_3005452, EPI_ISL_3005453, EPI_ISL_3005454, EPI_ISL_3005455, EPI_ISL_3005456, EPI_ISL_3005457, EPI_ISL_3005458, EPI_ISL_3005459, EPI_ISL_3005460, EPI_ISL_3005461, EPI_ISL_3005462, EPI_ISL_3005463, EPI_ISL_3005464, EPI_ISL_3005465, EPI_ISL_3005466, EPI_ISL_3005467, EPI_ISL_3005468, EPI_ISL_3005469, EPI_ISL_3005470, EPI_ISL_3005471, EPI_ISL_3005472, EPI_ISL_3005473, EPI_ISL_3005474, EPI_ISL_3005475, EPI_ISL_3005476, EPI_ISL_3005477, EPI_ISL_3005478, EPI_ISL_3005479, EPI_ISL_3005480, EPI_ISL_3005481, EPI_ISL_3005482, EPI_ISL_3005483, EPI_ISL_3005484, EPI_ISL_3005485, EPI_ISL_3005486, EPI_ISL_3005487, EPI_ISL_3005488, EPI_ISL_3005489, EPI_ISL_3005490, EPI_ISL_3005491, EPI_ISL_3005492, EPI_ISL_3005493, EPI_ISL_3005494, EPI_ISL_3005495, EPI_ISL_3005496, EPI_ISL_3005497, EPI_ISL_3005498, EPI_ISL_3005499, EPI_ISL_3005500, EPI_ISL_3005501, EPI_ISL_3005502, EPI_ISL_3005503, EPI_ISL_3005504, EPI_ISL_3005505, EPI_ISL_3005506, EPI_ISL_3005507, EPI_ISL_3005508, EPI_ISL_3005509, EPI_ISL_3005510, EPI_ISL_3005511, EPI_ISL_3005512, EPI_ISL_3005513, EPI_ISL_3005514, EPI_ISL_3005515, EPI_ISL_3005516, EPI_ISL_3005517, EPI_ISL_3005518, EPI_ISL_3005519, EPI_ISL_3005520, EPI_ISL_3005521, EPI_ISL_3005522, EPI_ISL_3005523, EPI_ISL_3005524, EPI_ISL_3005525, EPI_ISL_3005526, EPI_ISL_3005527, EPI_ISL_3005528, EPI_ISL_3005529, EPI_ISL_3005530, EPI_ISL_3005531, EPI_ISL_3005532, EPI_ISL_3005533, EPI_ISL_3005534, EPI_ISL_3005535, EPI_ISL_3005536, EPI_ISL_3005537, EPI_ISL_3005538, EPI_ISL_3005539, EPI_ISL_3005540, EPI_ISL_3005541, EPI_ISL_3005542, EPI_ISL_3005543, EPI_ISL_3005544, EPI_ISL_3005545, EPI_ISL_3005546, EPI_ISL_3005547, EPI_ISL_3005548, EPI_ISL_3005549, EPI_ISL_3005550, EPI_ISL_3005551, EPI_ISL_3005552, EPI_ISL_3005553, EPI_ISL_3005554, EPI_ISL_3005555, EPI_ISL_3005556, EPI_ISL_3005557, EPI_ISL_3005558, EPI_ISL_3005559, EPI_ISL_3005560, EPI_ISL_3005561, EPI_ISL_3005562, EPI_ISL_3005563, EPI_ISL_3005564, EPI_ISL_3005565, EPI_ISL_3005566, EPI_ISL_3005567, EPI_ISL_3005568, EPI_ISL_3005569, EPI_ISL_3005570, EPI_ISL_3005571, EPI_ISL_3005572, EPI_ISL_3005573, EPI_ISL_3005574, EPI_ISL_3005575, EPI_ISL_3005576, EPI_ISL_3005577, EPI_ISL_3005578, EPI_ISL_3005579, EPI_ISL_3005580, EPI_ISL_3005581, EPI_ISL_3005582, EPI_ISL_3005583, EPI_ISL_3005584, EPI_ISL_3005585, EPI_ISL_3005586, EPI_ISL_3005587, EPI_ISL_3005588, EPI_ISL_3005589, EPI_ISL_3005590, EPI_ISL_3005591, EPI_ISL_3005592, EPI_ISL_3005593, EPI_ISL_3005594, EPI_ISL_3005595, EPI_ISL_3005596, EPI_ISL_3005597, EPI_ISL_3005598, EPI_ISL_3005599, EPI_ISL_3005600, EPI_ISL_3005601, EPI_ISL_3005602, EPI_ISL_3005603, EPI_ISL_3005604, EPI_ISL_3005605, EPI_ISL_3005606, EPI_ISL_3005607, EPI_ISL_3005608, EPI_ISL_3005609, EPI_ISL_3005610, EPI_ISL_3005611, EPI_ISL_3005612, EPI_ISL_3005613, EPI_ISL_3005614, EPI_ISL_3005615, EPI_ISL_3005616, EPI_ISL_3005617, EPI_ISL_3005618, EPI_ISL_3005619, EPI_ISL_3005620, EPI_ISL_3005621, EPI_ISL_3005622, EPI_ISL_3005623, EPI_ISL_3005624, EPI_ISL_3005625, EPI_ISL_3005626, EPI_ISL_3005627, EPI_ISL_3005628, EPI_ISL_3005629, EPI_ISL_3005630, EPI_ISL_3005631, EPI_ISL_3005632, EPI_ISL_3005633, EPI_ISL_3005634, EPI_ISL_3005635, EPI_ISL_3005636, EPI_ISL_3005637, EPI_ISL_3005638, EPI_ISL_3005639, EPI_ISL_3005640, EPI_ISL_3005641, EPI_ISL_3005642, EPI_ISL_3005643, EPI_ISL_3005644, EPI_ISL_3005645, EPI_ISL_3005646, EPI_ISL_3005647, EPI_ISL_3005648, EPI_ISL_3005649, EPI_ISL_3005650, EPI_ISL_3005651, EPI_ISL_3005652, EPI_ISL_3005653, EPI_ISL_3005654, EPI_ISL_3005655, EPI_ISL_3005656, EPI_ISL_3005657, EPI_ISL_3005658, EPI_ISL_3005659, EPI_ISL_3005660, EPI_ISL_3005661, EPI_ISL_3005662, EPI_ISL_3005663, EPI_ISL_3005664, EPI_ISL_3005665, EPI_ISL_3005666, EPI_ISL_3005667, EPI_ISL_3005668, EPI_ISL_3005669, EPI_ISL_3005670, EPI_ISL_3005671, EPI_ISL_3005672, EPI_ISL_3005673, EPI_ISL_3005674, EPI_ISL_3005675, EPI_ISL_3005676, EPI_ISL_3005677, EPI_ISL_3005678, EPI_ISL_3005679, EPI_ISL_3005680, EPI_ISL_3005681, EPI_ISL_3005682, EPI_ISL_3005683, EPI_ISL_3005684, EPI_ISL_3005685, EPI_ISL_3005686, EPI_ISL_3005687, EPI_ISL_3005688, EPI_ISL_3005689, EPI_ISL_3005690, EPI_ISL_3005691, EPI_ISL_3005692, EPI_ISL_3005693, EPI_ISL_3005694, EPI_ISL_3005695, EPI_ISL_3005696, EPI_ISL_3005697, EPI_ISL_3005698, EPI_ISL_3005699, EPI_ISL_3005700, EPI_ISL_3005701, EPI_ISL_3005702, EPI_ISL_3005703, EPI_ISL_3005704, EPI_ISL_3005705, EPI_ISL_3005706, EPI_ISL_3005707, EPI_ISL_3005708, EPI_ISL_3005709, EPI_ISL_3005710, EPI_ISL_3005711, EPI_ISL_3005712, EPI_ISL_3005713, EPI_ISL_3005714, EPI_ISL_3005715, EPI_ISL_3005716, EPI_ISL_3005717, EPI_ISL_3005718, EPI_ISL_3005719, EPI_ISL_3005720, EPI_ISL_3005721, EPI_ISL_3005722, EPI_ISL_3005723, EPI_ISL_3005724, EPI_ISL_3005725, EPI_ISL_3005726, EPI_ISL_3005727, EPI_ISL_3005728, EPI_ISL_3005729, EPI_ISL_3005730, EPI_ISL_3005731, EPI_ISL_3005732, EPI_ISL_3005733, EPI_ISL_3005734, EPI_ISL_3005735, EPI_ISL_3005736, EPI_ISL_3005737, EPI_ISL_3005738, EPI_ISL_3005739, EPI_ISL_3005740, EPI_ISL_3005741, EPI_ISL_3005742, EPI_ISL_3005743, EPI_ISL_3005744, EPI_ISL_3005745, EPI_ISL_3005746, EPI_ISL_3005747, EPI_ISL_3005748, EPI_ISL_3005749, EPI_ISL_3005750, EPI_ISL_3005751, EPI_ISL_3005752, EPI_ISL_3005753, EPI_ISL_3005754, EPI_ISL_3005755, EPI_ISL_3005756, EPI_ISL_3005757, EPI_ISL_3005758, EPI_ISL_3005759, EPI_ISL_3005760, EPI_ISL_3005761, EPI_ISL_3005762, EPI_ISL_3005763, EPI_ISL_3005764, EPI_ISL_3005765, EPI_ISL_3005766, EPI_ISL_3005767, EPI_ISL_3005768, EPI_ISL_3005769, EPI_ISL_3005770, EPI_ISL_3005771, EPI_ISL_3005772, EPI_ISL_3005773, EPI_ISL_3005774, EPI_ISL_3005775, EPI_ISL_3005776, EPI_ISL_3005777, EPI_ISL_3005778, EPI_ISL_3005779, EPI_ISL_3005780, EPI_ISL_3005781, EPI_ISL_3005782, EPI_ISL_3005783, EPI_ISL_3005784, EPI_ISL_3005785, EPI_ISL_3005786, EPI_ISL_3005787, EPI_ISL_3005788, EPI_ISL_3005789, EPI_ISL_3005790, EPI_ISL_3005791, EPI_ISL_3005792, EPI_ISL_3005793, EPI_ISL_3005794, EPI_ISL_3005795, EPI_ISL_3005796, EPI_ISL_3005797, EPI_ISL_3005798, EPI_ISL_3005799, EPI_ISL_3005800, EPI_ISL_3005801, EPI_ISL_3005802, EPI_ISL_3005803, EPI_ISL_3005804, EPI_ISL_3005805, EPI_ISL_3005806, EPI_ISL_3005807, EPI_ISL_3005808, EPI_ISL_3005809, EPI_ISL_3005810, EPI_ISL_3005811, EPI_ISL_3005812, EPI_ISL_3005813, EPI_ISL_3005814, EPI_ISL_3005815, EPI_ISL_3005816, EPI_ISL_3005817, EPI_ISL_3005818, EPI_ISL_3005819, EPI_ISL_3005820, EPI_ISL_3005821, EPI_ISL_3005822, EPI_ISL_3005823, EPI_ISL_3005824, EPI_ISL_3005825, EPI_ISL_3005826, EPI_ISL_3005827, EPI_ISL_3005828, EPI_ISL_3005829, EPI_ISL_3005830, EPI_ISL_3005831, EPI_ISL_3005832, EPI_ISL_3005833, EPI_ISL_3005834, EPI_ISL_3005835, EPI_ISL_3005836, EPI_ISL_3005837, EPI_ISL_3005838, EPI_ISL_3005839, EPI_ISL_3005840, EPI_ISL_3005841, EPI_ISL_3005842, EPI_ISL_3005843, EPI_ISL_3005844, EPI_ISL_3005845, EPI_ISL_3005846, EPI_ISL_3005847, EPI_ISL_3005848, EPI_ISL_3005849, EPI_ISL_3005850, EPI_ISL_3005851, EPI_ISL_3005852, EPI_ISL_3005853, EPI_ISL_3005854, EPI_ISL_3005855, EPI_ISL_3005856, EPI_ISL_3005857, EPI_ISL_3005858, EPI_ISL_3005859, EPI_ISL_3005860, EPI_ISL_3005861, EPI_ISL_3005862, EPI_ISL_3005863, EPI_ISL_3005864, EPI_ISL_3005865, EPI_ISL_3005866, EPI_ISL_3005867, EPI_ISL_3005868, EPI_ISL_3005869, EPI_ISL_3005870, EPI_ISL_3005871, EPI_ISL_3005872, EPI_ISL_3005873, EPI_ISL_3005874, EPI_ISL_3005875, EPI_ISL_3005876, EPI_ISL_3005877, EPI_ISL_3005878, EPI_ISL_3005879, EPI_ISL_3005880, EPI_ISL_3005881, EPI_ISL_3005882, EPI_ISL_3005883, EPI_ISL_3005884, EPI_ISL_3005885, EPI_ISL_3005886, EPI_ISL_3005887, EPI_ISL_3005888, EPI_ISL_3005889, EPI_ISL_3005890, EPI_ISL_3005891, EPI_ISL_3005892, EPI_ISL_3005893, EPI_ISL_3005894, EPI_ISL_3005895, EPI_ISL_3005896, EPI_ISL_3005897, EPI_ISL_3005898, EPI_ISL_3005899, EPI_ISL_3005900, EPI_ISL_3005901, EPI_ISL_3005902, EPI_ISL_3005903, EPI_ISL_3005904, EPI_ISL_3005905, EPI_ISL_3005906, EPI_ISL_3005907, EPI_ISL_3005908, EPI_ISL_3005909, EPI_ISL_3005910, EPI_ISL_3005911, EPI_ISL_3005912, EPI_ISL_3005913, EPI_ISL_3005914, EPI_ISL_3005915, EPI_ISL_3005916, EPI_ISL_3005917, EPI_ISL_3005918, EPI_ISL_3005919, EPI_ISL_3005920, EPI_ISL_3005921, EPI_ISL_3005922, EPI_ISL_3005923, EPI_ISL_3005924, EPI_ISL_3005925, EPI_ISL_3005926, EPI_ISL_3005927, EPI_ISL_3005928, EPI_ISL_3005929, EPI_ISL_3005930, EPI_ISL_3005931, EPI_ISL_3005932, EPI_ISL_3005933, EPI_ISL_3005934, EPI_ISL_3005935, EPI_ISL_3005936, EPI_ISL_3005937, EPI_ISL_3005938, EPI_ISL_3005939, EPI_ISL_3005940, EPI_ISL_3005941, EPI_ISL_3005942, EPI_ISL_3005943, EPI_ISL_3005944, EPI_ISL_3005945, EPI_ISL_3005946, EPI_ISL_3005947, EPI_ISL_3005948, EPI_ISL_3005949, EPI_ISL_3005951, EPI_ISL_3005952, EPI_ISL_3005953, EPI_ISL_3005954, EPI_ISL_3005955, EPI_ISL_3005956, EPI_ISL_3005957, EPI_ISL_3005958, EPI_ISL_3005959, EPI_ISL_3005960, EPI_ISL_3005961, EPI_ISL_3005962, EPI_ISL_3005963, EPI_ISL_3005964, EPI_ISL_3005965, EPI_ISL_3005966, EPI_ISL_3005967, EPI_ISL_3005968, EPI_ISL_3005969, EPI_ISL_3005970, EPI_ISL_3005971, EPI_ISL_3005972, EPI_ISL_3005973, EPI_ISL_3005974, EPI_ISL_3005975, EPI_ISL_3005976, EPI_ISL_3005977, EPI_ISL_3005978, EPI_ISL_3005979, EPI_ISL_3005980, EPI_ISL_3005981, EPI_ISL_3005982, EPI_ISL_3005983, EPI_ISL_3005984, EPI_ISL_3005985, EPI_ISL_3005986, EPI_ISL_3005987, EPI_ISL_3005988, EPI_ISL_3005989, EPI_ISL_3005990, EPI_ISL_3005991, EPI_ISL_3005992, EPI_ISL_3005993, EPI_ISL_3005994, EPI_ISL_3005995, EPI_ISL_3005996, EPI_ISL_3005997, EPI_ISL_3005998, EPI_ISL_3005999, EPI_ISL_3006000, EPI_ISL_3006001, EPI_ISL_3006002, EPI_ISL_3006003, EPI_ISL_3006004, EPI_ISL_3006005, EPI_ISL_3006006, EPI_ISL_3006007, EPI_ISL_3006008, EPI_ISL_3006009, EPI_ISL_3006010, EPI_ISL_3006011, EPI_ISL_3006012, EPI_ISL_3006013, EPI_ISL_3006014, EPI_ISL_3006015, EPI_ISL_3006016, EPI_ISL_3006017, EPI_ISL_3006018, EPI_ISL_3006019, EPI_ISL_3006020, EPI_ISL_3006021, EPI_ISL_3006022, EPI_ISL_3006023, EPI_ISL_3006024, EPI_ISL_3006025, EPI_ISL_3006026, EPI_ISL_3006027, EPI_ISL_3006028, EPI_ISL_3006029, EPI_ISL_3006030, EPI_ISL_3006031, EPI_ISL_3006032, EPI_ISL_3006033, EPI_ISL_3006034, EPI_ISL_3006035, EPI_ISL_3006036, EPI_ISL_3006037, EPI_ISL_3006038, EPI_ISL_3006039, EPI_ISL_3006040, EPI_ISL_3006041, EPI_ISL_3006042, EPI_ISL_3006043, EPI_ISL_3006044, EPI_ISL_3006045, EPI_ISL_3006046, EPI_ISL_3006047, EPI_ISL_3006048, EPI_ISL_3006049, EPI_ISL_3006050, EPI_ISL_3006051, EPI_ISL_3006052, EPI_ISL_3006053, EPI_ISL_3006054, EPI_ISL_3006055, EPI_ISL_3006056, EPI_ISL_3006057, EPI_ISL_3006058, EPI_ISL_3006059, EPI_ISL_3006060, EPI_ISL_3006061, EPI_ISL_3006062, EPI_ISL_3006063, EPI_ISL_3006064, EPI_ISL_3006065, EPI_ISL_3006066, EPI_ISL_3006067, EPI_ISL_3006068, EPI_ISL_3006069, EPI_ISL_3006070, EPI_ISL_3006071, EPI_ISL_3006072, EPI_ISL_3006073, EPI_ISL_3006074, EPI_ISL_3006075, EPI_ISL_3006076, EPI_ISL_3006077, EPI_ISL_3006078, EPI_ISL_3006079, EPI_ISL_3006080, EPI_ISL_3006081, EPI_ISL_3006082, EPI_ISL_3006083, EPI_ISL_3006084, EPI_ISL_3006085, EPI_ISL_3006086, EPI_ISL_3006087, EPI_ISL_3006088, EPI_ISL_3006089, EPI_ISL_3006090, EPI_ISL_3006091, EPI_ISL_3006092, EPI_ISL_3006093, EPI_ISL_3006094, EPI_ISL_3006095

| | | | |
|---|---|---|---|
| see above | Originating lab: Wales Specialist Virology Centre Sequencing lab: Pathogen Genomics Unit | Public Health Wales Microbiology Cardiff Wales Specialist Virology Centre | Catherine Moore, Johnathan Evans, Laura Gifford, Malorie Perry, Simon Cottrell, Angela Marchbank, Alec Birchley, Alexander Adams, Amy Gaskin, Bree Gatica-Wilcox, Jason Coombes, Joel Southgate, Lauren Gilbert, Lee Graham, Nicole Pacchiarini, Sara Kumziene-Summerhayes, Sarah Taylor, Sophie Jones, Sara Rey, Matthew Bull, Joanne Watkins, Sally Corden, Tom Connor |

EPI_ISL_3006096, EPI_ISL_3006097, EPI_ISL_3006098, EPI_ISL_3006099, EPI_ISL_3006100, EPI_ISL_3006101, EPI_ISL_3006102, EPI_ISL_3006103, EPI_ISL_3006104, EPI_ISL_3006105, EPI_ISL_3006106, EPI_ISL_3006107, EPI_ISL_3006108, EPI_ISL_3006109, EPI_ISL_3006110, EPI_ISL_3006111, EPI_ISL_3006112, EPI_ISL_3006113, EPI_ISL_3006114, EPI_ISL_3006115, EPI_ISL_3006116, EPI_ISL_3006117, EPI_ISL_3006118, EPI_ISL_3006119, EPI_ISL_3006120, EPI_ISL_3006121, EPI_ISL_3006122, EPI_ISL_3006123, EPI_ISL_3006124, EPI_ISL_3006125, EPI_ISL_3006126, EPI_ISL_3006127, EPI_ISL_3006128, EPI_ISL_3006129, EPI_ISL_3006130, EPI_ISL_3006131, EPI_ISL_3006132, EPI_ISL_3006133, EPI_ISL_3006134, EPI_ISL_3006135, EPI_ISL_3006136, EPI_ISL_3006137, EPI_ISL_3006138, EPI_ISL_3006139, EPI_ISL_3006140, EPI_ISL_3006141, EPI_ISL_3006142, EPI_ISL_3006143, EPI_ISL_3006144, EPI_ISL_3006145, EPI_ISL_3006146, EPI_ISL_3006147, EPI_ISL_3006148, EPI_ISL_3006149, EPI_ISL_3006150, EPI_ISL_3006151, EPI_ISL_3006152, EPI_ISL_3006153, EPI_ISL_3006154, EPI_ISL_3006155, EPI_ISL_3006156, EPI_ISL_3006157, EPI_ISL_3006158, EPI_ISL_3006159, EPI_ISL_3006160, EPI_ISL_3006161,

| | | | |
|---|---|---|---|
| EPI_ISL_3006162, EPI_ISL_3006163, EPI_ISL_3006164, EPI_ISL_3006165, EPI_ISL_3006166, EPI_ISL_3006167, EPI_ISL_3006168, EPI_ISL_3006169, EPI_ISL_3006170, EPI_ISL_3006171, EPI_ISL_3006172, EPI_ISL_3006173, EPI_ISL_3006174, EPI_ISL_3006175, EPI_ISL_3006176, EPI_ISL_3006177, EPI_ISL_3006178, EPI_ISL_3006179, EPI_ISL_3006180, EPI_ISL_3006181, EPI_ISL_3006182, EPI_ISL_3006183, EPI_ISL_3006184, EPI_ISL_3006185, EPI_ISL_3006186, EPI_ISL_3006187, EPI_ISL_3006188, EPI_ISL_3006189, EPI_ISL_3006190, EPI_ISL_3006191, EPI_ISL_3006192, EPI_ISL_3006193, EPI_ISL_3006194, EPI_ISL_3006195, EPI_ISL_3006196, EPI_ISL_3006197, EPI_ISL_3006198, EPI_ISL_3006199, EPI_ISL_3006200, EPI_ISL_3006201, EPI_ISL_3006202, EPI_ISL_3006203, EPI_ISL_3006204, EPI_ISL_3006205, EPI_ISL_3006206, EPI_ISL_3006207, EPI_ISL_3006208, EPI_ISL_3006209, EPI_ISL_3006210, EPI_ISL_3006211, EPI_ISL_3006212, EPI_ISL_3006213, EPI_ISL_3006214, EPI_ISL_3006215, EPI_ISL_3006216, EPI_ISL_3006217, EPI_ISL_3006218, EPI_ISL_3006219, EPI_ISL_3006220, EPI_ISL_3006221, EPI_ISL_3006222, EPI_ISL_3006223, EPI_ISL_3006224, EPI_ISL_3006225, EPI_ISL_3006226, EPI_ISL_3006227, EPI_ISL_3006228, EPI_ISL_3006229, EPI_ISL_3006230, EPI_ISL_3006231, EPI_ISL_3006232, EPI_ISL_3006233, EPI_ISL_3006234, EPI_ISL_3006235, EPI_ISL_3006236, EPI_ISL_3006237, EPI_ISL_3006238, EPI_ISL_3006239, EPI_ISL_3006240, EPI_ISL_3006241, EPI_ISL_3006242, EPI_ISL_3006243, EPI_ISL_3006244, EPI_ISL_3006245, EPI_ISL_3006246, EPI_ISL_3006247, EPI_ISL_3006248, EPI_ISL_3006249, EPI_ISL_3006250, EPI_ISL_3006251, EPI_ISL_3006252, EPI_ISL_3006253, EPI_ISL_3006254, EPI_ISL_3006255, EPI_ISL_3006256, EPI_ISL_3006257, EPI_ISL_3006258, EPI_ISL_3006259, EPI_ISL_3006260, EPI_ISL_3006261, EPI_ISL_3006262, EPI_ISL_3006263, EPI_ISL_3006264, EPI_ISL_3006265, EPI_ISL_3006266, EPI_ISL_3006267, EPI_ISL_3006268, EPI_ISL_3006269, EPI_ISL_3006270, EPI_ISL_3006271, EPI_ISL_3006272, EPI_ISL_3006273, EPI_ISL_3006274, EPI_ISL_3006275, EPI_ISL_3006276, EPI_ISL_3006277, EPI_ISL_3006278, EPI_ISL_3006279, EPI_ISL_3006280, EPI_ISL_3006281, EPI_ISL_3006282, EPI_ISL_3006283, EPI_ISL_3006284, EPI_ISL_3006285, EPI_ISL_3006286, EPI_ISL_3006287, EPI_ISL_3006288, EPI_ISL_3006289, EPI_ISL_3006290, EPI_ISL_3006291, EPI_ISL_3006292, EPI_ISL_3006293, EPI_ISL_3006294, EPI_ISL_3006295, EPI_ISL_3006296, EPI_ISL_3006297, EPI_ISL_3006298, EPI_ISL_3006299, EPI_ISL_3006300, EPI_ISL_3006301, EPI_ISL_3006302, EPI_ISL_3006303, EPI_ISL_3006304, EPI_ISL_3006305, EPI_ISL_3006306, EPI_ISL_3006307, EPI_ISL_3006308, EPI_ISL_3006309, EPI_ISL_3006310, EPI_ISL_3006311, EPI_ISL_3006312, EPI_ISL_3006313, EPI_ISL_3006314, EPI_ISL_3006315, EPI_ISL_3006316, EPI_ISL_3006317, EPI_ISL_3006318, EPI_ISL_3006319, EPI_ISL_3006320, EPI_ISL_3006321, EPI_ISL_3006322, EPI_ISL_3006323, EPI_ISL_3006324, EPI_ISL_3006325, EPI_ISL_3006326, EPI_ISL_3006327, EPI_ISL_3006328, EPI_ISL_3006329, EPI_ISL_3006330, EPI_ISL_3006331, EPI_ISL_3006332, EPI_ISL_3006333, EPI_ISL_3006334, EPI_ISL_3006335, EPI_ISL_3006336, EPI_ISL_3006337, EPI_ISL_3006338, EPI_ISL_3006339, EPI_ISL_3006340, EPI_ISL_3006341, EPI_ISL_3006342, EPI_ISL_3006343, EPI_ISL_3006344, EPI_ISL_3006345, EPI_ISL_3006346, EPI_ISL_3006347, EPI_ISL_3006348, EPI_ISL_3006349, EPI_ISL_3006350, EPI_ISL_3006351, EPI_ISL_3006352, EPI_ISL_3006353, EPI_ISL_3006354, EPI_ISL_3006355, EPI_ISL_3006356, EPI_ISL_3006357, EPI_ISL_3006358, EPI_ISL_3006359, EPI_ISL_3006360, EPI_ISL_3006361, EPI_ISL_3006362, EPI_ISL_3006363, EPI_ISL_3006364, EPI_ISL_3006365, EPI_ISL_3006366, EPI_ISL_3006367, EPI_ISL_3006368, EPI_ISL_3006369, EPI_ISL_3006370, EPI_ISL_3006371, EPI_ISL_3006372, EPI_ISL_3006373, EPI_ISL_3006374, EPI_ISL_3006375, EPI_ISL_3006376, EPI_ISL_3006377, EPI_ISL_3006378, EPI_ISL_3006379, EPI_ISL_3006380, EPI_ISL_3006381, EPI_ISL_3006382, EPI_ISL_3006383, EPI_ISL_3006384, EPI_ISL_3006385, EPI_ISL_3006386, EPI_ISL_3006387, EPI_ISL_3006388, EPI_ISL_3006389, EPI_ISL_3006390, EPI_ISL_3006391, EPI_ISL_3006392, EPI_ISL_3006393, EPI_ISL_3006394, EPI_ISL_3006395, EPI_ISL_3006396, EPI_ISL_3006397, EPI_ISL_3006398, EPI_ISL_3006399, EPI_ISL_3006400, EPI_ISL_3006401 | | | |
| see above | Centre for Enzyme Innovation, University of Portsmouth / Translational Research Laboratory, Portsmouth Hospitals NHS Trust / Hampshire Hospitals NHS Foundation Trust / Bournemouth University / University Hospitals Dorset NHS Foundation Trust / Brighton and Sussex University Hospitals NHS Trust / Dartford and Gravesham NHS Trust / Isle of Wight NHS Trust / Maidstone and Tunbridge Wells NHS Trust / University Hospital Southampton NHS Foundation Trust | COVID-19 Genomics UK (COG-UK) Consortium | Samuel Robson, Angela Beckett, Salman Goudarzi, Christopher Fearn, Kate Cook, Katie Loveson, Sharon Glaysher, Scott Elliott, Kelly Bicknell, Sarah Wyllie, Allyson Lloyd, Robert Impey, Anoop Chauhan, Stephen Kidd, Nathan Moore, Nick Cortes, Claire Thomas, Anna Mantzaouratou, Sarah Buchan, Magdalena Barrow, Andrew Butt, Liz Sheridan, Jonnie Seymour, Dorian Crudgington, Ben Macklin, Mohammed Hassan-Ibrahim, Cassandra Malone, Benjamin Cogger, Kevin Tucker, Samirakhon Raupova, Rachael Jeremiah, Anibolina Castigador, Emily Macnaughton, Karen Withell, Kordo Saeed, Jacqui Prieto, Adhyana Mahanama, Buddhini Samaraweera, Siona Silvieri, Emanuela Pelosi, Eleri Wilson-Davies, Sarah Jeremiah, Helen Wheeler, Matthew Harvey, Thea Sass, Helen Umpleby, Stephen Aplin |
| EPI_ISL_3006402, EPI_ISL_3006403, EPI_ISL_3006405, EPI_ISL_3006406, EPI_ISL_3006407, EPI_ISL_3006408, EPI_ISL_3006409, EPI_ISL_3006410, EPI_ISL_3006411, EPI_ISL_3006412, EPI_ISL_3006413, EPI_ISL_3006414, EPI_ISL_3006415, EPI_ISL_3006416, EPI_ISL_3006417, EPI_ISL_3006418, EPI_ISL_3006419, EPI_ISL_3006420, EPI_ISL_3006421, EPI_ISL_3006422, EPI_ISL_3006423, EPI_ISL_3006424, EPI_ISL_3006425, EPI_ISL_3006426, EPI_ISL_3006427, EPI_ISL_3006428, EPI_ISL_3006429, EPI_ISL_3006430, EPI_ISL_3006431, EPI_ISL_3006432, EPI_ISL_3006433, EPI_ISL_3006434, EPI_ISL_3006435, EPI_ISL_3006436, EPI_ISL_3006437, EPI_ISL_3006438, EPI_ISL_3006439, EPI_ISL_3006440, EPI_ISL_3006441, EPI_ISL_3006442, EPI_ISL_3006443, EPI_ISL_3006444, EPI_ISL_3006445, EPI_ISL_3006446, EPI_ISL_3006447, EPI_ISL_3006448, EPI_ISL_3006449, EPI_ISL_3006450, EPI_ISL_3006451, EPI_ISL_3006452, EPI_ISL_3006453, EPI_ISL_3006454, EPI_ISL_3006455, EPI_ISL_3006456, EPI_ISL_3006457, EPI_ISL_3006458, EPI_ISL_3006459, EPI_ISL_3006460, EPI_ISL_3006461, EPI_ISL_3006462, EPI_ISL_3006463, EPI_ISL_3006464, EPI_ISL_3006465, EPI_ISL_3006466, EPI_ISL_3006467, EPI_ISL_3006468, EPI_ISL_3006469, EPI_ISL_3006470, EPI_ISL_3006471, EPI_ISL_3006472, EPI_ISL_3006473, EPI_ISL_3006474, EPI_ISL_3006475, EPI_ISL_3006476, EPI_ISL_3006477, EPI_ISL_3006478, EPI_ISL_3006479, EPI_ISL_3006480, EPI_ISL_3006481, EPI_ISL_3006482, EPI_ISL_3006483, EPI_ISL_3006484, EPI_ISL_3006485, EPI_ISL_3006486, EPI_ISL_3006487, EPI_ISL_3006488, EPI_ISL_3006489, EPI_ISL_3006490, EPI_ISL_3006491, EPI_ISL_3006492, EPI_ISL_3006493, EPI_ISL_3006494, EPI_ISL_3006495, EPI_ISL_3006496, EPI_ISL_3006497, EPI_ISL_3006498, EPI_ISL_3006499, EPI_ISL_3006500, EPI_ISL_3006501, EPI_ISL_3006502, EPI_ISL_3006503, EPI_ISL_3006504, EPI_ISL_3006505, EPI_ISL_3006506, EPI_ISL_3006507, EPI_ISL_3006508, EPI_ISL_3006509, EPI_ISL_3006510, EPI_ISL_3006511, EPI_ISL_3006512, EPI_ISL_3006513, EPI_ISL_3006514, EPI_ISL_3006515, EPI_ISL_3006516, EPI_ISL_3006517, EPI_ISL_3006518, EPI_ISL_3006519, EPI_ISL_3006520, EPI_ISL_3006521, EPI_ISL_3006522, EPI_ISL_3006523, EPI_ISL_3006524, EPI_ISL_3006525, EPI_ISL_3006526, EPI_ISL_3006527, EPI_ISL_3006528, EPI_ISL_3006529, EPI_ISL_3006530, EPI_ISL_3006531, EPI_ISL_3006532, EPI_ISL_3006533, EPI_ISL_3006534, EPI_ISL_3006535, EPI_ISL_3006536, EPI_ISL_3006537, EPI_ISL_3006538, EPI_ISL_3006539, EPI_ISL_3006540, EPI_ISL_3006541, EPI_ISL_3006542, EPI_ISL_3006543, EPI_ISL_3006544, EPI_ISL_3006545, EPI_ISL_3006546, EPI_ISL_3006547, EPI_ISL_3006548, EPI_ISL_3006549, EPI_ISL_3006550, EPI_ISL_3006551, EPI_ISL_3006552, EPI_ISL_3006553, EPI_ISL_3006554, EPI_ISL_3006555, EPI_ISL_3006556, EPI_ISL_3006557, EPI_ISL_3006558, EPI_ISL_3006559, EPI_ISL_3006560, EPI_ISL_3006561, EPI_ISL_3006562, EPI_ISL_3006563, EPI_ISL_3006564, EPI_ISL_3006565, EPI_ISL_3006566, EPI_ISL_3006567, EPI_ISL_3006568, EPI_ISL_3006569, EPI_ISL_3006570, EPI_ISL_3006571, EPI_ISL_3006572, EPI_ISL_3006573, EPI_ISL_3006574, EPI_ISL_3006575, EPI_ISL_3006576, EPI_ISL_3006577, EPI_ISL_3006578, EPI_ISL_3006579, EPI_ISL_3006580, EPI_ISL_3006581, EPI_ISL_3006582, EPI_ISL_3006583, EPI_ISL_3006584, EPI_ISL_3006585, EPI_ISL_3006586, EPI_ISL_3006587, EPI_ISL_3006588, EPI_ISL_3006589, EPI_ISL_3006590, EPI_ISL_3006591, EPI_ISL_3006592, EPI_ISL_3006593, EPI_ISL_3006594, EPI_ISL_3006595, EPI_ISL_3006596, EPI_ISL_3006597, EPI_ISL_3006598, EPI_ISL_3006599, EPI_ISL_3006600, EPI_ISL_3006601, EPI_ISL_3006602, EPI_ISL_3006603, EPI_ISL_3006604, EPI_ISL_3006605, EPI_ISL_3006606, EPI_ISL_3006607, EPI_ISL_3006608, EPI_ISL_3006609, EPI_ISL_3006610, EPI_ISL_3006611, EPI_ISL_3006612, EPI_ISL_3006613, EPI_ISL_3006614, EPI_ISL_3006615, EPI_ISL_3006616, EPI_ISL_3006617, EPI_ISL_3006618, EPI_ISL_3006619, EPI_ISL_3006620, EPI_ISL_3006621, EPI_ISL_3006622, EPI_ISL_3006623, EPI_ISL_3006624, EPI_ISL_3006625, EPI_ISL_3006626, EPI_ISL_3006627, EPI_ISL_3006628, EPI_ISL_3006629, EPI_ISL_3006630, EPI_ISL_3006631, EPI_ISL_3006632, EPI_ISL_3006633, EPI_ISL_3006634, EPI_ISL_3006635, EPI_ISL_3006636, EPI_ISL_3006637, EPI_ISL_3006638, EPI_ISL_3006639, EPI_ISL_3006640, EPI_ISL_3006641, EPI_ISL_3006642, EPI_ISL_3006643, EPI_ISL_3006644, EPI_ISL_3006645, EPI_ISL_3006646, EPI_ISL_3006647, EPI_ISL_3006648, EPI_ISL_3006649, EPI_ISL_3006650, EPI_ISL_3006651, EPI_ISL_3006652, EPI_ISL_3006653, EPI_ISL_3006654, EPI_ISL_3006655, EPI_ISL_3006656, EPI_ISL_3006657, EPI_ISL_3006658, EPI_ISL_3006659, EPI_ISL_3006660, EPI_ISL_3006661, EPI_ISL_3006662, EPI_ISL_3006663, EPI_ISL_3006664, EPI_ISL_3006665, EPI_ISL_3006666, EPI_ISL_3006667, EPI_ISL_3006668, EPI_ISL_3006669, EPI_ISL_3006670, EPI_ISL_3006671, EPI_ISL_3006672, EPI_ISL_3006673, EPI_ISL_3006674, EPI_ISL_3006675, EPI_ISL_3006676, EPI_ISL_3006677, EPI_ISL_3006678, EPI_ISL_3006679, EPI_ISL_3006680, EPI_ISL_3006681, EPI_ISL_3006682, EPI_ISL_3006683, EPI_ISL_3006684, EPI_ISL_3006685, EPI_ISL_3006686, EPI_ISL_3006687, EPI_ISL_3006688, EPI_ISL_3006689, EPI_ISL_3006690, EPI_ISL_3006691, EPI_ISL_3006692, EPI_ISL_3006693, EPI_ISL_3006694, EPI_ISL_3006695, EPI_ISL_3006696, EPI_ISL_3006697, EPI_ISL_3006698, EPI_ISL_3006699, EPI_ISL_3006700, EPI_ISL_3006701, EPI_ISL_3006702, EPI_ISL_3006703, EPI_ISL_3006704, EPI_ISL_3006705, EPI_ISL_3006706, EPI_ISL_3006707, EPI_ISL_3006708, EPI_ISL_3006709, EPI_ISL_3006710, EPI_ISL_3006711, EPI_ISL_3006712, EPI_ISL_3006713, EPI_ISL_3006714, EPI_ISL_3006715, EPI_ISL_3006716, EPI_ISL_3006717, EPI_ISL_3006718, EPI_ISL_3006719, EPI_ISL_3006720, EPI_ISL_3006721, EPI_ISL_3006722, EPI_ISL_3006723, EPI_ISL_3006724, EPI_ISL_3006725, EPI_ISL_3006726, EPI_ISL_3006727, EPI_ISL_3006728, EPI_ISL_3006729, EPI_ISL_3006730, EPI_ISL_3006731, EPI_ISL_3006732, EPI_ISL_3006733, EPI_ISL_3006734, EPI_ISL_3006735 | | | |
| see above | Virology Department, Sheffield Teaching Hospitals NHS Foundation Trust/Department of Infection, Immunity and Cardiovascular Disease, The Medical School, University of Sheffield | COVID-19 Genomics UK (COG-UK) Consortium | Thushan de Silva, Matthew Parker, Nikki Smith, Adri Angyal, Rebecca Brown, Luke Green, Rachel Tucker, Paul Parsons, Danielle Groves, Katie Johnson, Laura Carrilero, Alex Keeley, Dave Partridge, Matthew Wyles, Benjamin Lindsey, Mehmet Yavuz, Mohammad Raza, Cariad Evans |
| EPI_ISL_3006737, EPI_ISL_3006740, EPI_ISL_3006741, EPI_ISL_3006742, EPI_ISL_3006743, EPI_ISL_3006744, EPI_ISL_3006745, EPI_ISL_3006746, EPI_ISL_3006747, EPI_ISL_3006748, EPI_ISL_3006749, EPI_ISL_3006750, EPI_ISL_3006751, EPI_ISL_3006753, EPI_ISL_3006754 | | | |
| see above | Institute of Molecular and Translational Medicine / Laboratory of Experimental Medicine, Faculty of Medicine and Dentistry, Palacky University and University Hospital Olomouc | Institute of Molecular and Translational Medicine / Laboratory of Experimental Medicine, Faculty of Medicine and Dentistry, Palacky University | Rastislav Slavkovský, Hana Jaworek, Vladimíra Koudeláková, Barbora Blumová, Marián Hajdúch |
| EPI_ISL_3006755, EPI_ISL_3006756, EPI_ISL_3006757 | CGB laboratoř a.s., Kořenského 1210/10 703 00 Ostrava - Vítkovice, Czech Republic | Institute of Molecular and Translational Medicine / Laboratory of Experimental Medicine, Faculty of Medicine and Dentistry, Palacky University | Rastislav Slavkovský, Hana Jaworek, Vladimíra Koudeláková, Barbora Blumová, Marián Hajdúch |
| EPI_ISL_3006758 | Institute of Molecular and Translational Medicine / Laboratory of Experimental Medicine, Faculty of Medicine and Dentistry, Palacky University and University Hospital Olomouc | Institute of Molecular and Translational Medicine / Laboratory of Experimental Medicine, Faculty of Medicine and Dentistry, Palacky University | Rastislav Slavkovský, Hana Jaworek, Vladimíra Koudeláková, Barbora Blumová, Marián Hajdúch |
| EPI_ISL_3006759, EPI_ISL_3006761 | CGB laboratoř a.s., Kořenského 1210/10 703 00 Ostrava - Vítkovice, Czech Republic | Institute of Molecular and Translational Medicine / Laboratory of Experimental Medicine, Faculty of Medicine and Dentistry, Palacky University | Rastislav Slavkovský, Hana Jaworek, Vladimíra Koudeláková, Barbora Blumová, Marián Hajdúch |
| EPI_ISL_3006762, EPI_ISL_3006763, EPI_ISL_3006764 | Institute of Molecular and Translational Medicine / Laboratory of Experimental Medicine, Faculty of Medicine and Dentistry, Palacky University and University Hospital Olomouc | Institute of Molecular and Translational Medicine / Laboratory of Experimental Medicine, Faculty of Medicine and Dentistry, Palacky University | Rastislav Slavkovský, Hana Jaworek, Vladimíra Koudeláková, Barbora Blumová, Marián Hajdúch |
| EPI_ISL_3006794 | AULSS 2 Marca Trevigiana | Istituto Zooprofilattico Sperimentale delle Venezie | Adelaide Milani, Alessia Schivo, Annalisa Salviato, Elisa Palumbo, Erika Giorgia Quaranta, Luca Tassoni, Ambra Pastori, Edoardo Giussani, Alice Fusaro, Isabella Monne, Calogero Terregino, Antonia Ricci |
| EPI_ISL_3006796 | AULSS 7 Pedemontana | Istituto Zooprofilattico Sperimentale delle Venezie | Adelaide Milani, Alessia Schivo, Annalisa Salviato, Elisa Palumbo, Erika Giorgia Quaranta, Luca Tassoni, Ambra Pastori, Edoardo Giussani, Alice Fusaro, Isabella Monne, Calogero Terregino, Antonia Ricci |
| EPI_ISL_3006797, EPI_ISL_3006798, EPI_ISL_3006799 | Fujian Center for Disease Control and Prevention | Fujian Center for Disease Control and Prevention | Lin Qi, Huang Zhimiao, Weng Yuwei |
| EPI_ISL_3006803, EPI_ISL_3006804, EPI_ISL_3006807, EPI_ISL_3006812, EPI_ISL_3006815, EPI_ISL_3006816, EPI_ISL_3006817, EPI_ISL_3006818, EPI_ISL_3006820, EPI_ISL_3006821, EPI_ISL_3006823, EPI_ISL_3006824, EPI_ISL_3006825, EPI_ISL_3006826, EPI_ISL_3006827, EPI_ISL_3006828, EPI_ISL_3006829 | | | |
| see above | Medini Rai Medical College and Hospital, Medininagar, Palamu | Institute of Life Sciences - INSACOG | Sunil K. Raghav, Safal Walia, Arup Ghosh, Atimukta Jha, Amol M. Kanampalliwar, Omprakash Shiriwas, Sana Fatma, Shifu Aggarwal, Rupesh Dash, Rajeeb Swain, Punit Prasad, INSACOG Consortium, Ajay Parida |
| EPI_ISL_3006831, EPI_ISL_3006832, EPI_ISL_3006833, EPI_ISL_3006834, EPI_ISL_3006835, EPI_ISL_3006838, EPI_ISL_3006839, EPI_ISL_3006841, EPI_ISL_3006842, EPI_ISL_3006843, EPI_ISL_3006846, EPI_ISL_3006849, EPI_ISL_3006852, EPI_ISL_3006864, EPI_ISL_3006865, EPI_ISL_3006866, EPI_ISL_3006872, EPI_ISL_3006873, EPI_ISL_3006878 | | | |
| see above | Patliputra Medical College & Hospital (PMCH), Dhanbad | Institute of Life Sciences - INSACOG | Sunil K. Raghav, Safal Walia, Arup Ghosh, Atimukta Jha, Amol M. Kanampalliwar, Omprakash Shiriwas, Sana Fatma, Shifu Aggarwal, Rupesh Dash, Rajeeb Swain, Punit Prasad, INSACOG Consortium, Ajay Parida |
| EPI_ISL_3006894, EPI_ISL_3006917, EPI_ISL_3006958, EPI_ISL_3006961, EPI_ISL_3006962, EPI_ISL_3006965 | Chhattisgarh Institute of Medical Sciences (CIMS), Bilaspur | Institute of Life Sciences - INSACOG | Sunil K. Raghav, Safal Walia, Arup Ghosh, Atimukta Jha, Amol M. Kanampalliwar, Omprakash Shiriwas, Sana Fatma, Shifu Aggarwal, Rupesh Dash, Rajeeb Swain, Punit Prasad, INSACOG Consortium, Ajay Parida |
| EPI_ISL_3006967, EPI_ISL_3006969, EPI_ISL_3006973, EPI_ISL_3006975, EPI_ISL_3006977, EPI_ISL_3006979, EPI_ISL_3006980, EPI_ISL_3006981, EPI_ISL_3006982, EPI_ISL_3006983, EPI_ISL_3006984 | | | |
| see above | District Public Health Laboratory, Angul | Institute of Life Sciences - INSACOG | Sunil K. Raghav, Safal Walia, Arup Ghosh, Atimukta Jha, Amol M. Kanampalliwar, Omprakash Shiriwas, Sana Fatma, Shifu Aggarwal, Rupesh Dash, Rajeeb Swain, Punit Prasad, INSACOG Consortium, Ajay Parida |
| EPI_ISL_3006985, EPI_ISL_3006986, EPI_ISL_3006988, EPI_ISL_3006991 | Pandit Raghunath Murmu Medical College, Baripada | Institute of Life Sciences - INSACOG | Sunil K. Raghav, Safal Walia, Arup Ghosh, Atimukta Jha, Amol M. Kanampalliwar, Omprakash Shiriwas, Sana Fatma, Shifu Aggarwal, Rupesh Dash, Rajeeb Swain, Punit Prasad, INSACOG Consortium, Ajay Parida |
| EPI_ISL_3007000, EPI_ISL_3007001 | Kalinga Institute of Medical Sciences(KIMS), Bhubaneswar | Institute of Life Sciences - INSACOG | Sunil K. Raghav, Safal Walia, Arup Ghosh, Atimukta Jha, Amol M. Kanampalliwar, Omprakash Shiriwas, Sana Fatma, Shifu Aggarwal, Rupesh Dash, Rajeeb Swain, Punit Prasad, INSACOG Consortium, Ajay Parida |
| EPI_ISL_3007009, EPI_ISL_3007027 | District Public Health Laboratory, Angul | Institute of Life Sciences - INSACOG | Sunil K. Raghav, Safal Walia, Arup Ghosh, Atimukta Jha, Amol M. Kanampalliwar, Omprakash Shiriwas, Sana Fatma, Shifu Aggarwal, Rupesh Dash, Rajeeb Swain, Punit Prasad, INSACOG Consortium, Ajay Parida |
| EPI_ISL_3007040, EPI_ISL_3007043 | inDNA Life Sciecnces, Bhubaneswar | Institute of Life Sciences - INSACOG | Sunil K. Raghav, Safal Walia, Arup Ghosh, Atimukta Jha, Amol M. Kanampalliwar, Omprakash Shiriwas, Sana Fatma, Shifu Aggarwal, Rupesh Dash, Rajeeb Swain, Punit Prasad, INSACOG Consortium, Ajay Parida |
| EPI_ISL_3007048 | District Head Quarters Hospital, Nayagarh | Institute of Life Sciences - INSACOG | Sunil K. Raghav, Safal Walia, Arup Ghosh, Atimukta Jha, Amol M. Kanampalliwar, Omprakash Shiriwas, Sana Fatma, Shifu Aggarwal, Rupesh Dash, Rajeeb Swain, Punit Prasad, INSACOG Consortium, Ajay Parida |
| EPI_ISL_3007061, EPI_ISL_3007064, EPI_ISL_3007065, EPI_ISL_3007071, EPI_ISL_3007072, EPI_ISL_3007074, EPI_ISL_3007077, EPI_ISL_3007080, EPI_ISL_3007084, EPI_ISL_3007085, EPI_ISL_3007090, EPI_ISL_3007091, EPI_ISL_3007094 | | | |
| see above | Regional Medical Research Center, Bhubaneswar | Institute of Life Sciences - INSACOG | Sunil K. Raghav, Safal Walia, Arup Ghosh, Atimukta Jha, Amol M. Kanampalliwar, Omprakash Shiriwas, Sana Fatma, Shifu Aggarwal, Rupesh Dash, Rajeeb Swain, Punit Prasad, INSACOG Consortium, Ajay Parida |
| EPI_ISL_3007101, EPI_ISL_3007103, EPI_ISL_3007104, EPI_ISL_3007105, EPI_ISL_3007106, EPI_ISL_3007107, EPI_ISL_3007108, EPI_ISL_3007109, EPI_ISL_3007110, EPI_ISL_3007111, EPI_ISL_3007112, EPI_ISL_3007113, EPI_ISL_3007114, EPI_ISL_3007115, EPI_ISL_3007116, EPI_ISL_3007117, EPI_ISL_3007118, EPI_ISL_3007120, EPI_ISL_3007122, EPI_ISL_3007123, EPI_ISL_3007125, EPI_ISL_3007132, EPI_ISL_3007134, EPI_ISL_3007135, EPI_ISL_3007136, EPI_ISL_3007138 | | | |
| see above | District Public Health Laboratory, Angul | Institute of Life Sciences - INSACOG | Sunil K. Raghav, Safal Walia, Arup Ghosh, Atimukta Jha, Amol M. Kanampalliwar, Omprakash Shiriwas, Sana Fatma, Shifu Aggarwal, Rupesh Dash, Rajeeb Swain, Punit Prasad, INSACOG Consortium, Ajay Parida |
| EPI_ISL_3007139, EPI_ISL_3007140, EPI_ISL_3007144, EPI_ISL_3007145 | Pandit Raghunath Murmu Medical College, Baripada | Institute of Life Sciences - INSACOG | Sunil K. Raghav, Safal Walia, Arup Ghosh, Atimukta Jha, Amol M. Kanampalliwar, Omprakash Shiriwas, Sana Fatma, Shifu Aggarwal, Rupesh Dash, Rajeeb Swain, Punit Prasad, INSACOG Consortium, Ajay Parida |
| EPI_ISL_3007146 | District Public Health Laboratory, Angul | Institute of Life Sciences - INSACOG | Sunil K. Raghav, Safal Walia, Arup Ghosh, Atimukta Jha, Amol M. Kanampalliwar, Omprakash Shiriwas, Sana Fatma, Shifu Aggarwal, Rupesh Dash, Rajeeb Swain, Punit Prasad, INSACOG Consortium, Ajay Parida |
| EPI_ISL_3007147, EPI_ISL_3007148, EPI_ISL_3007149, EPI_ISL_3007151, EPI_ISL_3007152, EPI_ISL_3007153, EPI_ISL_3007154, EPI_ISL_3007155, EPI_ISL_3007156, EPI_ISL_3007157, EPI_ISL_3007158, EPI_ISL_3007160, EPI_ISL_3007161, EPI_ISL_3007163, EPI_ISL_3007164, EPI_ISL_3007165, EPI_ISL_3007166, EPI_ISL_3007167, EPI_ISL_3007169, EPI_ISL_3007170, EPI_ISL_3007171, EPI_ISL_3007173, EPI_ISL_3007174, EPI_ISL_3007175 | | | |
| see above | Institute of Life Sciences, Bhubaneswar | Institute of Life Sciences - INSACOG | Sunil K. Raghav, Safal Walia, Arup Ghosh, Atimukta Jha, Amol M. Kanampalliwar, Omprakash Shiriwas, Sana Fatma, Shifu Aggarwal, Rupesh Dash, Rajeeb Swain, Punit Prasad, INSACOG Consortium, Ajay Parida |

| | | | |
|---|---|---|---|
| EPI_ISL_3007177, EPI_ISL_3007178, EPI_ISL_3007179, EPI_ISL_3007180, EPI_ISL_3007181, EPI_ISL_3007182, EPI_ISL_3007183 | Karolinska University Hospital Solna | Karolinska University Hospital | Jan Albert, Tobias Allander, Annelie Bjerkner, Sandra Broddesson, Robert Dyrdak, Martin Ekman, Lynda Eneh, Lina Guerra Blomqvist, Karolina Ininbergs, Tanja Normark, Isak Sylvin, Zhibing Yun, Martina Wahlund, Valtteri Wirta |
| EPI_ISL_3007187 | Synlab Medilab | Karolinska University Hospital | Jan Albert, Tobias Allander, Annelie Bjerkner, Sandra Broddesson, Robert Dyrdak, Martin Ekman, Lynda Eneh, Lina Guerra Blomqvist, Karolina Ininbergs, Tanja Normark, Isak Sylvin, Zhibing Yun, Martina Wahlund, Valtteri Wirta |
| EPI_ISL_3007188 | Unilabs | Karolinska University Hospital | Jan Albert, Tobias Allander, Annelie Bjerkner, Sandra Broddesson, Robert Dyrdak, Martin Ekman, Lynda Eneh, Lina Guerra Blomqvist, Karolina Ininbergs, Tanja Normark, Isak Sylvin, Zhibing Yun, Martina Wahlund, Valtteri Wirta |
| EPI_ISL_3007189 | Karolinska University Hospital Solna | Karolinska University Hospital | Jan Albert, Tobias Allander, Annelie Bjerkner, Sandra Broddesson, Robert Dyrdak, Martin Ekman, Lynda Eneh, Lina Guerra Blomqvist, Karolina Ininbergs, Tanja Normark, Isak Sylvin, Zhibing Yun, Martina Wahlund, Valtteri Wirta |
| EPI_ISL_3007190, EPI_ISL_3007191, EPI_ISL_3007192, EPI_ISL_3007193, EPI_ISL_3007194, EPI_ISL_3007195 | Unilabs | Karolinska University Hospital | Jan Albert, Tobias Allander, Annelie Bjerkner, Sandra Broddesson, Robert Dyrdak, Martin Ekman, Lynda Eneh, Lina Guerra Blomqvist, Karolina Ininbergs, Tanja Normark, Isak Sylvin, Zhibing Yun, Martina Wahlund, Valtteri Wirta |
| EPI_ISL_3007196 | Synlab Medilab | Karolinska University Hospital | Jan Albert, Tobias Allander, Annelie Bjerkner, Sandra Broddesson, Robert Dyrdak, Martin Ekman, Lynda Eneh, Lina Guerra Blomqvist, Karolina Ininbergs, Tanja Normark, Isak Sylvin, Zhibing Yun, Martina Wahlund, Valtteri Wirta |
| EPI_ISL_3007197, EPI_ISL_3007198 | Unilabs | Karolinska University Hospital | Jan Albert, Tobias Allander, Annelie Bjerkner, Sandra Broddesson, Robert Dyrdak, Martin Ekman, Lynda Eneh, Lina Guerra Blomqvist, Karolina Ininbergs, Tanja Normark, Isak Sylvin, Zhibing Yun, Martina Wahlund, Valtteri Wirta |
| EPI_ISL_3007199 | Synlab Medilab | Karolinska University Hospital | Jan Albert, Tobias Allander, Annelie Bjerkner, Sandra Broddesson, Robert Dyrdak, Martin Ekman, Lynda Eneh, Lina Guerra Blomqvist, Karolina Ininbergs, Tanja Normark, Isak Sylvin, Zhibing Yun, Martina Wahlund, Valtteri Wirta |
| EPI_ISL_3007200, EPI_ISL_3007201, EPI_ISL_3007202, EPI_ISL_3007204, EPI_ISL_3007205, EPI_ISL_3007206, EPI_ISL_3007208 | Unilabs | Karolinska University Hospital | Jan Albert, Tobias Allander, Annelie Bjerkner, Sandra Broddesson, Robert Dyrdak, Martin Ekman, Lynda Eneh, Lina Guerra Blomqvist, Karolina Ininbergs, Tanja Normark, Isak Sylvin, Zhibing Yun, Martina Wahlund, Valtteri Wirta |
| EPI_ISL_3007209, EPI_ISL_3007211 | Synlab Medilab | Karolinska University Hospital | Jan Albert, Tobias Allander, Annelie Bjerkner, Sandra Broddesson, Robert Dyrdak, Martin Ekman, Lynda Eneh, Lina Guerra Blomqvist, Karolina Ininbergs, Tanja Normark, Isak Sylvin, Zhibing Yun, Martina Wahlund, Valtteri Wirta |
| EPI_ISL_3007212, EPI_ISL_3007213 | Unilabs | Karolinska University Hospital | Jan Albert, Tobias Allander, Annelie Bjerkner, Sandra Broddesson, Robert Dyrdak, Martin Ekman, Lynda Eneh, Lina Guerra Blomqvist, Karolina Ininbergs, Tanja Normark, Isak Sylvin, Zhibing Yun, Martina Wahlund, Valtteri Wirta |
| EPI_ISL_3007214, EPI_ISL_3007215, EPI_ISL_3007216 | Karolinska University Hospital Solna | Karolinska University Hospital | Jan Albert, Tobias Allander, Annelie Bjerkner, Sandra Broddesson, Robert Dyrdak, Martin Ekman, Lynda Eneh, Lina Guerra Blomqvist, Karolina Ininbergs, Tanja Normark, Isak Sylvin, Zhibing Yun, Martina Wahlund, Valtteri Wirta |
| EPI_ISL_3007217 | Unilabs | Karolinska University Hospital | Jan Albert, Tobias Allander, Annelie Bjerkner, Sandra Broddesson, Robert Dyrdak, Martin Ekman, Lynda Eneh, Lina Guerra Blomqvist, Karolina Ininbergs, Tanja Normark, Isak Sylvin, Zhibing Yun, Martina Wahlund, Valtteri Wirta |
| EPI_ISL_3007218 | Karolinska University Hospital Solna | Karolinska University Hospital | Jan Albert, Tobias Allander, Annelie Bjerkner, Sandra Broddesson, Robert Dyrdak, Martin Ekman, Lynda Eneh, Lina Guerra Blomqvist, Karolina Ininbergs, Tanja Normark, Isak Sylvin, Zhibing Yun, Martina Wahlund, Valtteri Wirta |
| EPI_ISL_3007219, EPI_ISL_3007220 | Unilabs | Karolinska University Hospital | Jan Albert, Tobias Allander, Annelie Bjerkner, Sandra Broddesson, Robert Dyrdak, Martin Ekman, Lynda Eneh, Lina Guerra Blomqvist, Karolina Ininbergs, Tanja Normark, Isak Sylvin, Zhibing Yun, Martina Wahlund, Valtteri Wirta |
| EPI_ISL_3007222 | Karolinska University Hospital Solna | Karolinska University Hospital | Jan Albert, Tobias Allander, Annelie Bjerkner, Sandra Broddesson, Robert Dyrdak, Martin Ekman, Lynda Eneh, Lina Guerra Blomqvist, Karolina Ininbergs, Tanja Normark, Isak Sylvin, Zhibing Yun, Martina Wahlund, Valtteri Wirta |
| EPI_ISL_3007223 | Unilabs | Karolinska University Hospital | Jan Albert, Tobias Allander, Annelie Bjerkner, Sandra Broddesson, Robert Dyrdak, Martin Ekman, Lynda Eneh, Lina Guerra Blomqvist, Karolina Ininbergs, Tanja Normark, Isak Sylvin, Zhibing Yun, Martina Wahlund, Valtteri Wirta |
| EPI_ISL_3007224 | Synlab Medilab | Karolinska University Hospital | Jan Albert, Tobias Allander, Annelie Bjerkner, Sandra Broddesson, Robert Dyrdak, Martin Ekman, Lynda Eneh, Lina Guerra Blomqvist, Karolina Ininbergs, Tanja Normark, Isak Sylvin, Zhibing Yun, Martina Wahlund, Valtteri Wirta |
| EPI_ISL_3007225 | Unilabs | Karolinska University Hospital | Jan Albert, Tobias Allander, Annelie Bjerkner, Sandra Broddesson, Robert Dyrdak, Martin Ekman, Lynda Eneh, Lina Guerra Blomqvist, Karolina Ininbergs, Tanja Normark, Isak Sylvin, Zhibing Yun, Martina Wahlund, Valtteri Wirta |
| EPI_ISL_3007226 | Synlab Medilab | Karolinska University Hospital | Jan Albert, Tobias Allander, Annelie Bjerkner, Sandra Broddesson, Robert Dyrdak, Martin Ekman, Lynda Eneh, Lina Guerra Blomqvist, Karolina Ininbergs, Tanja Normark, Isak Sylvin, Zhibing Yun, Martina Wahlund, Valtteri Wirta |
| EPI_ISL_3007227 | Unilabs | Karolinska University Hospital | Jan Albert, Tobias Allander, Annelie Bjerkner, Sandra Broddesson, Robert Dyrdak, Martin Ekman, Lynda Eneh, Lina Guerra Blomqvist, Karolina Ininbergs, Tanja Normark, Isak Sylvin, Zhibing Yun, Martina Wahlund, Valtteri Wirta |
| EPI_ISL_3007228 | Karolinska University Hospital Solna | Karolinska University Hospital | Jan Albert, Tobias Allander, Annelie Bjerkner, Sandra Broddesson, Robert Dyrdak, Martin Ekman, Lynda Eneh, Lina Guerra Blomqvist, Karolina Ininbergs, Tanja Normark, Isak Sylvin, Zhibing Yun, Martina Wahlund, Valtteri Wirta |
| EPI_ISL_3007229 | Unilabs | Karolinska University Hospital | Jan Albert, Tobias Allander, Annelie Bjerkner, Sandra Broddesson, Robert Dyrdak, Martin Ekman, Lynda Eneh, Lina Guerra Blomqvist, Karolina Ininbergs, Tanja Normark, Isak Sylvin, Zhibing Yun, Martina Wahlund, Valtteri Wirta |
| EPI_ISL_3007230 | Karolinska University Hospital Solna | Karolinska University Hospital | Jan Albert, Tobias Allander, Annelie Bjerkner, Sandra Broddesson, Robert Dyrdak, Martin Ekman, Lynda Eneh, Lina Guerra Blomqvist, Karolina Ininbergs, Tanja Normark, Isak Sylvin, Zhibing Yun, Martina Wahlund, Valtteri Wirta |
| EPI_ISL_3007231 | Synlab Medilab | Karolinska University Hospital | Jan Albert, Tobias Allander, Annelie Bjerkner, Sandra Broddesson, Robert Dyrdak, Martin Ekman, Lynda Eneh, Lina Guerra Blomqvist, Karolina Ininbergs, Tanja Normark, Isak Sylvin, Zhibing Yun, Martina Wahlund, Valtteri Wirta |
| EPI_ISL_3007232 | Unilabs | Karolinska University Hospital | Jan Albert, Tobias Allander, Annelie Bjerkner, Sandra Broddesson, Robert Dyrdak, Martin Ekman, Lynda Eneh, Lina Guerra Blomqvist, Karolina Ininbergs, Tanja Normark, Isak Sylvin, Zhibing Yun, Martina Wahlund, Valtteri Wirta |
| EPI_ISL_3007233 | Synlab Medilab | Karolinska University Hospital | Jan Albert, Tobias Allander, Annelie Bjerkner, Sandra Broddesson, Robert Dyrdak, Martin Ekman, Lynda Eneh, Lina Guerra Blomqvist, Karolina Ininbergs, Tanja Normark, Isak Sylvin, Zhibing Yun, Martina Wahlund, Valtteri Wirta |
| EPI_ISL_3007234 | Karolinska University Hospital Solna | Karolinska University Hospital | Jan Albert, Tobias Allander, Annelie Bjerkner, Sandra Broddesson, Robert Dyrdak, Martin Ekman, Lynda Eneh, Lina Guerra Blomqvist, Karolina Ininbergs, Tanja Normark, Isak Sylvin, Zhibing Yun, Martina Wahlund, Valtteri Wirta |
| EPI_ISL_3007235 | Synlab Medilab | Karolinska University Hospital | Jan Albert, Tobias Allander, Annelie Bjerkner, Sandra Broddesson, Robert Dyrdak, Martin Ekman, Lynda Eneh, Lina Guerra Blomqvist, Karolina Ininbergs, Tanja Normark, Isak Sylvin, Zhibing Yun, Martina Wahlund, Valtteri Wirta |
| EPI_ISL_3007236 | Unilabs | Karolinska University Hospital | Jan Albert, Tobias Allander, Annelie Bjerkner, Sandra Broddesson, Robert Dyrdak, Martin Ekman, Lynda Eneh, Lina Guerra Blomqvist, Karolina Ininbergs, Tanja Normark, Isak Sylvin, Zhibing Yun, Martina Wahlund, Valtteri Wirta |
| EPI_ISL_3007237, EPI_ISL_3007238 | Karolinska University Hospital Solna | Karolinska University Hospital | Jan Albert, Tobias Allander, Annelie Bjerkner, Sandra Broddesson, Robert Dyrdak, Martin Ekman, Lynda Eneh, Lina Guerra Blomqvist, Karolina Ininbergs, Tanja Normark, Isak Sylvin, Zhibing Yun, Martina Wahlund, Valtteri Wirta |
| EPI_ISL_3007239 | Unilabs | Karolinska University Hospital | Jan Albert, Tobias Allander, Annelie Bjerkner, Sandra Broddesson, Robert Dyrdak, Martin Ekman, Lynda Eneh, Lina Guerra Blomqvist, Karolina Ininbergs, Tanja Normark, Isak Sylvin, Zhibing Yun, Martina Wahlund, Valtteri Wirta |
| EPI_ISL_3007240 | Synlab Medilab | Karolinska University Hospital | Jan Albert, Tobias Allander, Annelie Bjerkner, Sandra Broddesson, Robert Dyrdak, Martin Ekman, Lynda Eneh, Lina Guerra Blomqvist, Karolina Ininbergs, Tanja Normark, Isak Sylvin, Zhibing Yun, Martina Wahlund, Valtteri Wirta |
| EPI_ISL_3007241, EPI_ISL_3007242 | Unilabs | Karolinska University Hospital | Jan Albert, Tobias Allander, Annelie Bjerkner, Sandra Broddesson, Robert Dyrdak, Martin Ekman, Lynda Eneh, Lina Guerra Blomqvist, Karolina Ininbergs, Tanja Normark, Isak Sylvin, Zhibing Yun, Martina Wahlund, Valtteri Wirta |
| EPI_ISL_3007243 | Synlab Medilab | Karolinska University Hospital | Jan Albert, Tobias Allander, Annelie Bjerkner, Sandra Broddesson, Robert Dyrdak, Martin Ekman, Lynda Eneh, Lina Guerra Blomqvist, Karolina Ininbergs, Tanja Normark, Isak Sylvin, Zhibing Yun, Martina Wahlund, Valtteri Wirta |
| EPI_ISL_3007244, EPI_ISL_3007245, EPI_ISL_3007246 | Unilabs | Karolinska University Hospital | Jan Albert, Tobias Allander, Annelie Bjerkner, Sandra Broddesson, Robert Dyrdak, Martin Ekman, Lynda Eneh, Lina Guerra Blomqvist, Karolina Ininbergs, Tanja Normark, Isak Sylvin, Zhibing Yun, Martina Wahlund, Valtteri Wirta |
| EPI_ISL_3007247 | Karolinska University Hospital Solna | Karolinska University Hospital | Jan Albert, Tobias Allander, Annelie Bjerkner, Sandra Broddesson, Robert Dyrdak, Martin Ekman, Lynda Eneh, Lina Guerra Blomqvist, Karolina Ininbergs, Tanja Normark, Isak Sylvin, Zhibing Yun, Martina Wahlund, Valtteri Wirta |
| EPI_ISL_3007248, EPI_ISL_3007249, EPI_ISL_3007250, EPI_ISL_3007253, EPI_ISL_3007254, EPI_ISL_3007255, EPI_ISL_3007256, EPI_ISL_3007257, EPI_ISL_3007258, EPI_ISL_3007259, EPI_ISL_3007260 | | | |
| see above | Karolinska University Hospital Huddinge | Karolinska University Hospital | Jan Albert, Tobias Allander, Annelie Bjerkner, Sandra Broddesson, Robert Dyrdak, Martin Ekman, Lynda Eneh, Lina Guerra Blomqvist, Karolina Ininbergs, Tanja Normark, Isak Sylvin, Zhibing Yun, Martina Wahlund, Valtteri Wirta |
| EPI_ISL_3007261, EPI_ISL_3007262, EPI_ISL_3007263, EPI_ISL_3007266, EPI_ISL_3007268, EPI_ISL_3007269, EPI_ISL_3007270, EPI_ISL_3007274, EPI_ISL_3007275, EPI_ISL_3007278 | AULSS 3 Venezia | UOSD Genetica e Citogenetica - Azienda ULSS 3 Serenissima; Istituto Zooprofilattico Sperimentale delle Venezie | Mosé Favarato, Elisa Squarcina, Laura Squarzon, Claudia Perini, Laura Bevilacqua, Adelaide Milani, Alessia Schivo, Annalisa Salviato, Elisa Palumbo, Erika Giorgia Quaranta, Luca Tassoni, Ambra Pastori, Edoardo Giussani, Alice Fusaro, Isabella Monne, Calogero Terregino, Antonia Ricci |
| EPI_ISL_3007280 | Histopath | NSW Health Pathology - Institute of Clinical Pathology and Medical Research; Westmead Hospital; University of Sydney | CIDM-PH et al. |
| EPI_ISL_3007281, EPI_ISL_3007282, EPI_ISL_3007283, EPI_ISL_3007284, EPI_ISL_3007285, EPI_ISL_3007286, EPI_ISL_3007287, EPI_ISL_3007288 | Sydney South West Pathology Service (SSWPS) - Liverpool Hospital - NSW Health Pathology | NSW Health Pathology - Institute of Clinical Pathology and Medical Research; Westmead Hospital; University of Sydney | CIDM-PH et al. |
| EPI_ISL_3007289, EPI_ISL_3007290, EPI_ISL_3007291 | Laverty Pathology | NSW Health Pathology - Institute of Clinical Pathology and Medical Research; Westmead Hospital; University of Sydney | CIDM-PH et al. |
| EPI_ISL_3007292, | Australian Clinical Labs | NSW Health Pathology - Institute of Clinical Pathology and Medical | CIDM-PH et al. |

| | | | |
|---|---|---|---|
| EPI_ISL_3007293 | | Research; Westmead Hospital; University of Sydney | |
| EPI_ISL_3007294 | Sydney South West Pathology Service (SSWPS) - Liverpool Hospital - NSW Health Pathology | NSW Health Pathology - Institute of Clinical Pathology and Medical Research; Westmead Hospital; University of Sydney | CIDM-PH et al. |
| EPI_ISL_3007295 | St Vincent's Pathology (SydPath) | NSW Health Pathology - Institute of Clinical Pathology and Medical Research; Westmead Hospital; University of Sydney | CIDM-PH et al. |
| EPI_ISL_3007296 | Histopath | NSW Health Pathology - Institute of Clinical Pathology and Medical Research; Westmead Hospital; University of Sydney | CIDM-PH et al. |
| EPI_ISL_3007297 | Pathology West - NSW Health Pathology | NSW Health Pathology - Institute of Clinical Pathology and Medical Research; Westmead Hospital; University of Sydney | CIDM-PH et al. |
| EPI_ISL_3007298 | 4Cyte Pathology | NSW Health Pathology - Institute of Clinical Pathology and Medical Research; Westmead Hospital; University of Sydney | CIDM-PH et al. |
| EPI_ISL_3007299, EPI_ISL_3007300 | Laverty Pathology | NSW Health Pathology - Institute of Clinical Pathology and Medical Research; Westmead Hospital; University of Sydney | CIDM-PH et al. |
| EPI_ISL_3007301 | Sydney South West Pathology Service (SSWPS) - Liverpool Hospital - NSW Health Pathology | NSW Health Pathology - Institute of Clinical Pathology and Medical Research; Westmead Hospital; University of Sydney | CIDM-PH et al. |
| EPI_ISL_3007302 | Histopath | NSW Health Pathology - Institute of Clinical Pathology and Medical Research; Westmead Hospital; University of Sydney | CIDM-PH et al. |
| EPI_ISL_3007303, EPI_ISL_3007304 | LKO | Jessa | Severine Berden et al. on behalf of the Jessa_cmdLab |
| EPI_ISL_3007305, EPI_ISL_3007306 | Ospedale Sandro Pertini ASL Roma2 | Department of General Diagnostics; Department of Virology; Istituto Zooprofilattico Sperimentale del Lazio e della Toscana (IZSLT) | Patricia Alba, Giuseppe Manna, Elena L. Diaconu, Fabiola Feltrin, Raffaela Conti, Teresa Scicluna, Virginia Carfora, Antonella Cersini, Alessia Franco, Antonio Battisti. |
| EPI_ISL_3007307, EPI_ISL_3007309, EPI_ISL_3007310, EPI_ISL_3007311, EPI_ISL_3007312, EPI_ISL_3007316, EPI_ISL_3007317, EPI_ISL_3007318, EPI_ISL_3007322, EPI_ISL_3007323, EPI_ISL_3007324, EPI_ISL_3007325, EPI_ISL_3007326, EPI_ISL_3007327, EPI_ISL_3007328, EPI_ISL_3007329, EPI_ISL_3007331, EPI_ISL_3007333, EPI_ISL_3007334, EPI_ISL_3007335, EPI_ISL_3007336, EPI_ISL_3007339, EPI_ISL_3007340, EPI_ISL_3007342, EPI_ISL_3007343, EPI_ISL_3007345, EPI_ISL_3007346, EPI_ISL_3007348, EPI_ISL_3007350, EPI_ISL_3007352, EPI_ISL_3007354, EPI_ISL_3007355, EPI_ISL_3007357, EPI_ISL_3007362, EPI_ISL_3007363, EPI_ISL_3007364, EPI_ISL_3007366, EPI_ISL_3007368, EPI_ISL_3007371, EPI_ISL_3007372, EPI_ISL_3007373, EPI_ISL_3007375, EPI_ISL_3007377, EPI_ISL_3007378, EPI_ISL_3007379, EPI_ISL_3007380, EPI_ISL_3007382, EPI_ISL_3007383, EPI_ISL_3007384, EPI_ISL_3007385, EPI_ISL_3007386, EPI_ISL_3007388, EPI_ISL_3007390, EPI_ISL_3007391, EPI_ISL_3007392, EPI_ISL_3007393, EPI_ISL_3007394, EPI_ISL_3007395, EPI_ISL_3007397, EPI_ISL_3007398, EPI_ISL_3007399, EPI_ISL_3007401, EPI_ISL_3007402, EPI_ISL_3007403, EPI_ISL_3007404, EPI_ISL_3007405, EPI_ISL_3007407, EPI_ISL_3007408, EPI_ISL_3007409, EPI_ISL_3007410, EPI_ISL_3007412, EPI_ISL_3007414, EPI_ISL_3007417, EPI_ISL_3007418, EPI_ISL_3007419, EPI_ISL_3007422, EPI_ISL_3007423, EPI_ISL_3007424, EPI_ISL_3007425, EPI_ISL_3007428, EPI_ISL_3007429, EPI_ISL_3007430, EPI_ISL_3007431, EPI_ISL_3007432, EPI_ISL_3007433, EPI_ISL_3007434, EPI_ISL_3007435, EPI_ISL_3007436, EPI_ISL_3007437, EPI_ISL_3007438, EPI_ISL_3007439, EPI_ISL_3007442, EPI_ISL_3007443, EPI_ISL_3007444, EPI_ISL_3007445, EPI_ISL_3007449, EPI_ISL_3007450, EPI_ISL_3007451, EPI_ISL_3007452, EPI_ISL_3007454, EPI_ISL_3007455, EPI_ISL_3007456, EPI_ISL_3007457, EPI_ISL_3007458, EPI_ISL_3007464, EPI_ISL_3007466, EPI_ISL_3007467, EPI_ISL_3007468, EPI_ISL_3007469, EPI_ISL_3007470, EPI_ISL_3007471, EPI_ISL_3007474, EPI_ISL_3007475, EPI_ISL_3007477, EPI_ISL_3007479, EPI_ISL_3007480, EPI_ISL_3007481, EPI_ISL_3007482, EPI_ISL_3007484, EPI_ISL_3007485 | | | |
| see above | Broad Institute Clinical Research Sequencing Platform | Infectious Disease Program, Broad Institute of Harvard and MIT | Siddle,K.J., Adams,G., Pearlman,L., Gladden-Young,A., Vicente,G., Blumenstiel,B., DeFelice,M., Lee,M., McGovern,S., Lagerborg,K., Rudy,M., DeRuff,K., Carter,A., Normandin,E., Bauer,M., Reilly,S., Tomkins-Tinch,C., Loreth,C., Chaluvadi,S., Meldrim,J., Granger,B., Lemieux,J.E., Birren,B.W., Sabeti,P.C., Larkin,K., Dodge,S., Lennon,N., Madoff,L., Brown,C., Gallagher,G., Smole,S., Park,D.J., Gabriel,S., and MacInnis,B.L. |
| EPI_ISL_3007490, EPI_ISL_3007492, EPI_ISL_3007493, EPI_ISL_3007494, EPI_ISL_3007495, EPI_ISL_3007496, EPI_ISL_3007497, EPI_ISL_3007498, EPI_ISL_3007499, EPI_ISL_3007500, EPI_ISL_3007501, EPI_ISL_3007503, EPI_ISL_3007504, EPI_ISL_3007505, EPI_ISL_3007506, EPI_ISL_3007507, EPI_ISL_3007508, EPI_ISL_3007509, EPI_ISL_3007510, EPI_ISL_3007513, EPI_ISL_3007514, EPI_ISL_3007515 | | | |
| see above | IN State Department of Health Laboratory Services | IN State Department of Health Laboratory Services | Cassandra Campion, Jamie Yeadon, Brian Pope, Lixia Liu, Kyle Brownlee, Melissa Hindenlang, Mark Glazier |
| EPI_ISL_3007523, EPI_ISL_3007524 | Viollier AG | Department of Biosystems Science and Engineering, ETH Zürich | Chaoran Chen, Sarah Nadeau, Catharine Aquino, Ivan Topolsky, Kim Philipp Jablonski, Lara Fuhrmann, Daniel Ehrsam, Isabel Stürmer, Andreia Cabral de Gouvea, Maria Domenica Moccia, Simon Grüter, Timothy Sykes, Lennart Opitz, Griffin White, Laura Neff, Doris Popovic, Andrea Patrignani, Jay Tracy, Ralph Schlapbach, Christiane Beckmann, Maurice Redondo, Olivier Kobel, Christoph Noppen, Niko Beerenwinkel, Tanja Stadler |
| EPI_ISL_3007526, EPI_ISL_3007527, EPI_ISL_3007529, EPI_ISL_3007530, EPI_ISL_3007532, EPI_ISL_3007533, EPI_ISL_3007534, EPI_ISL_3007535, EPI_ISL_3007537 | Viollier AG | Department of Biosystems Science and Engineering, ETH Zürich | Andrea Patrizia Salzmann, Henriette Kurth, Christiane Beckmann, Ivan Topolsky, Chaoran Chen, Sarah Nadeau, Kim Philipp Jablonski, Lara Fuhrmann, Niko Beerenwinkel, Christoph Noppen, Olivier Kobel, Maurice Redondo, Tanja Stadler |
| EPI_ISL_3007538 | National Public Health Laboratory, National Centre for Infectious Diseases | National Public Health Laboratory, National Centre for Infectious Diseases | Tze Minn Mak, Katherine Ching, Zhenyang Zhou, Royce Ang, Lin Cui, Raymond Tzer Pin Lin |
| EPI_ISL_3007539 | Viollier AG | Department of Biosystems Science and Engineering, ETH Zürich | Andrea Patrizia Salzmann, Henriette Kurth, Christiane Beckmann, Ivan Topolsky, Chaoran Chen, Sarah Nadeau, Kim Philipp Jablonski, Lara Fuhrmann, Niko Beerenwinkel, Christoph Noppen, Olivier Kobel, Maurice Redondo, Tanja Stadler |
| EPI_ISL_3007540, EPI_ISL_3007542, EPI_ISL_3007544, EPI_ISL_3007546, EPI_ISL_3007548 | National Public Health Laboratory, National Centre for Infectious Diseases | National Public Health Laboratory, National Centre for Infectious Diseases | Tze Minn Mak, Katherine Ching, Zhenyang Zhou, Royce Ang, Lin Cui, Raymond Tzer Pin Lin |
| EPI_ISL_3007549 | Viollier AG | Department of Biosystems Science and Engineering, ETH Zürich | Andrea Patrizia Salzmann, Henriette Kurth, Christiane Beckmann, Ivan Topolsky, Chaoran Chen, Sarah Nadeau, Kim Philipp Jablonski, Lara Fuhrmann, Niko Beerenwinkel, Christoph Noppen, Olivier Kobel, Maurice Redondo, Tanja Stadler |
| EPI_ISL_3007550 | National Public Health Laboratory, National Centre for Infectious Diseases | National Public Health Laboratory, National Centre for Infectious Diseases | Tze Minn Mak, Katherine Ching, Zhenyang Zhou, Royce Ang, Lin Cui, Raymond Tzer Pin Lin |
| EPI_ISL_3007551 | Viollier AG | Department of Biosystems Science and Engineering, ETH Zürich | Andrea Patrizia Salzmann, Henriette Kurth, Christiane Beckmann, Ivan Topolsky, Chaoran Chen, Sarah Nadeau, Kim Philipp Jablonski, Lara Fuhrmann, Niko Beerenwinkel, Christoph Noppen, Olivier Kobel, Maurice Redondo, Tanja Stadler |
| EPI_ISL_3007552, EPI_ISL_3007554, EPI_ISL_3007556, EPI_ISL_3007558 | National Public Health Laboratory, National Centre for Infectious Diseases | National Public Health Laboratory, National Centre for Infectious Diseases | Tze Minn Mak, Katherine Ching, Zhenyang Zhou, Royce Ang, Lin Cui, Raymond Tzer Pin Lin |
| EPI_ISL_3007559 | Viollier AG | Department of Biosystems Science and Engineering, ETH Zürich | Andrea Patrizia Salzmann, Henriette Kurth, Christiane Beckmann, Ivan Topolsky, Chaoran Chen, Sarah Nadeau, Kim Philipp Jablonski, Lara Fuhrmann, Niko Beerenwinkel, Christoph Noppen, Olivier Kobel, Maurice Redondo, Tanja Stadler |
| EPI_ISL_3007560 | National Public Health Laboratory, National Centre for Infectious Diseases | National Public Health Laboratory, National Centre for Infectious Diseases | Tze Minn Mak, Katherine Ching, Zhenyang Zhou, Royce Ang, Lin Cui, Raymond Tzer Pin Lin |
| EPI_ISL_3007561 | Viollier AG | Department of Biosystems Science and Engineering, ETH Zürich | Andrea Patrizia Salzmann, Henriette Kurth, Christiane Beckmann, Ivan Topolsky, Chaoran Chen, Sarah Nadeau, Kim Philipp Jablonski, Lara Fuhrmann, Niko Beerenwinkel, Christoph Noppen, Olivier Kobel, Maurice Redondo, Tanja Stadler |
| EPI_ISL_3007562 | National Public Health Laboratory, National Centre for Infectious Diseases | National Public Health Laboratory, National Centre for Infectious Diseases | Tze Minn Mak, Katherine Ching, Zhenyang Zhou, Royce Ang, Lin Cui, Raymond Tzer Pin Lin |
| EPI_ISL_3007563 | Viollier AG | Department of Biosystems Science and Engineering, ETH Zürich | Andrea Patrizia Salzmann, Henriette Kurth, Christiane Beckmann, Ivan Topolsky, Chaoran Chen, Sarah Nadeau, Kim Philipp Jablonski, Lara Fuhrmann, Niko Beerenwinkel, Christoph Noppen, Olivier Kobel, Maurice Redondo, Tanja Stadler |
| EPI_ISL_3007564 | National Public Health Laboratory, National Centre for Infectious Diseases | National Public Health Laboratory, National Centre for Infectious Diseases | Tze Minn Mak, Katherine Ching, Zhenyang Zhou, Royce Ang, Lin Cui, Raymond Tzer Pin Lin |
| EPI_ISL_3007565 | Viollier AG | Department of Biosystems Science and Engineering, ETH Zürich | Andrea Patrizia Salzmann, Henriette Kurth, Christiane Beckmann, Ivan Topolsky, Chaoran Chen, Sarah Nadeau, Kim Philipp Jablonski, Lara Fuhrmann, Niko Beerenwinkel, Christoph Noppen, Olivier Kobel, Maurice Redondo, Tanja Stadler |
| EPI_ISL_3007566 | National Public Health Laboratory, National Centre for Infectious Diseases | National Public Health Laboratory, National Centre for Infectious Diseases | Tze Minn Mak, Katherine Ching, Zhenyang Zhou, Royce Ang, Lin Cui, Raymond Tzer Pin Lin |
| EPI_ISL_3007567 | Viollier AG | Department of Biosystems Science and Engineering, ETH Zürich | Andrea Patrizia Salzmann, Henriette Kurth, Christiane Beckmann, Ivan Topolsky, Chaoran Chen, Sarah Nadeau, Kim Philipp Jablonski, Lara Fuhrmann, Niko Beerenwinkel, Christoph Noppen, Olivier Kobel, Maurice Redondo, Tanja Stadler |
| EPI_ISL_3007568 | National Public Health Laboratory, National Centre for Infectious Diseases | National Public Health Laboratory, National Centre for Infectious Diseases | Tze Minn Mak, Katherine Ching, Zhenyang Zhou, Royce Ang, Lin Cui, Raymond Tzer Pin Lin |
| EPI_ISL_3007569 | Viollier AG | Department of Biosystems Science and Engineering, ETH Zürich | Andrea Patrizia Salzmann, Henriette Kurth, Christiane Beckmann, Ivan Topolsky, Chaoran Chen, Sarah Nadeau, Kim Philipp Jablonski, Lara Fuhrmann, Niko Beerenwinkel, Christoph Noppen, Olivier Kobel, Maurice Redondo, Tanja Stadler |
| EPI_ISL_3007570 | National Public Health Laboratory, National Centre for Infectious Diseases | National Public Health Laboratory, National Centre for Infectious Diseases | Tze Minn Mak, Katherine Ching, Zhenyang Zhou, Royce Ang, Lin Cui, Raymond Tzer Pin Lin |
| EPI_ISL_3007571 | Viollier AG | Department of Biosystems Science and Engineering, ETH Zürich | Andrea Patrizia Salzmann, Henriette Kurth, Christiane Beckmann, Ivan Topolsky, Chaoran Chen, Sarah Nadeau, Kim Philipp Jablonski, Lara Fuhrmann, Niko Beerenwinkel, Christoph Noppen, Olivier Kobel, Maurice Redondo, Tanja Stadler |
| EPI_ISL_3007572 | National Public Health Laboratory, National Centre for Infectious Diseases | National Public Health Laboratory, National Centre for Infectious Diseases | Tze Minn Mak, Katherine Ching, Zhenyang Zhou, Royce Ang, Lin Cui, Raymond Tzer Pin Lin |
| EPI_ISL_3007573 | Viollier AG | Department of Biosystems Science and Engineering, ETH Zürich | Andrea Patrizia Salzmann, Henriette Kurth, Christiane Beckmann, Ivan Topolsky, Chaoran Chen, Sarah Nadeau, Kim Philipp Jablonski, Lara Fuhrmann, Niko Beerenwinkel, Christoph Noppen, Olivier Kobel, Maurice Redondo, Tanja Stadler |
| EPI_ISL_3007574 | National Public Health Laboratory, National Centre for Infectious Diseases | National Public Health Laboratory, National Centre for Infectious Diseases | Tze Minn Mak, Katherine Ching, Zhenyang Zhou, Royce Ang, Lin Cui, Raymond Tzer Pin Lin |
| EPI_ISL_3007575 | Viollier AG | Department of Biosystems Science and Engineering, ETH Zürich | Andrea Patrizia Salzmann, Henriette Kurth, Christiane Beckmann, Ivan Topolsky, Chaoran Chen, Sarah Nadeau, Kim Philipp Jablonski, Lara Fuhrmann, Niko Beerenwinkel, Christoph Noppen, Olivier Kobel, Maurice Redondo, Tanja Stadler |
| EPI_ISL_3007576 | National Public Health Laboratory, National Centre for Infectious Diseases | National Public Health Laboratory, National Centre for Infectious Diseases | Tze Minn Mak, Katherine Ching, Zhenyang Zhou, Royce Ang, Lin Cui, Raymond Tzer Pin Lin |
| EPI_ISL_3007577 | Viollier AG | Department of Biosystems Science and Engineering, ETH Zürich | Andrea Patrizia Salzmann, Henriette Kurth, Christiane Beckmann, Ivan Topolsky, Chaoran Chen, Sarah Nadeau, Kim Philipp Jablonski, Lara Fuhrmann, Niko Beerenwinkel, Christoph Noppen, Olivier Kobel, Maurice Redondo, Tanja Stadler |
| EPI_ISL_3007578 | National Public Health Laboratory, National Centre for Infectious Diseases | National Public Health Laboratory, National Centre for Infectious Diseases | Tze Minn Mak, Katherine Ching, Zhenyang Zhou, Royce Ang, Lin Cui, Raymond Tzer Pin Lin |
| EPI_ISL_3007579 | Viollier AG | Department of Biosystems Science and Engineering, ETH Zürich | Andrea Patrizia Salzmann, Henriette Kurth, Christiane Beckmann, Ivan Topolsky, Chaoran Chen, Sarah Nadeau, Kim Philipp Jablonski, Lara Fuhrmann, Niko Beerenwinkel, Christoph Noppen, Olivier Kobel, Maurice Redondo, Tanja Stadler |
| EPI_ISL_3007580 | National Public Health Laboratory, National Centre for Infectious | National Public Health Laboratory, National Centre for Infectious | Tze Minn Mak, Katherine Ching, Zhenyang Zhou, Royce Ang, Lin Cui, Raymond Tzer Pin Lin |

| | | | |
|---|---|---|---|
| | Diseases | Diseases | |
| EPI_ISL_3007581 | Viollier AG | Department of Biosystems Science and Engineering, ETH Zürich | Andrea Patrizia Salzmann, Henriette Kurth, Christiane Beckmann, Ivan Topolsky, Chaoran Chen, Sarah Nadeau, Kim Philipp Jablonski, Lara Fuhrmann, Niko Beerenwinkel, Christoph Noppen, Olivier Kobel, Maurice Redondo, Tanja Stadler |
| EPI_ISL_3007582 | National Public Health Laboratory, National Centre for Infectious Diseases | National Public Health Laboratory, National Centre for Infectious Diseases | Tze Minn Mak, Katherine Ching, Zhenyang Zhou, Royce Ang, Lin Cui, Raymond Tzer Pin Lin |
| EPI_ISL_3007583, EPI_ISL_3007584 | Viollier AG | Department of Biosystems Science and Engineering, ETH Zürich | Andrea Patrizia Salzmann, Henriette Kurth, Christiane Beckmann, Ivan Topolsky, Chaoran Chen, Sarah Nadeau, Kim Philipp Jablonski, Lara Fuhrmann, Niko Beerenwinkel, Christoph Noppen, Olivier Kobel, Maurice Redondo, Tanja Stadler |
| EPI_ISL_3007585 | National Public Health Laboratory, National Centre for Infectious Diseases | National Public Health Laboratory, National Centre for Infectious Diseases | Tze Minn Mak, Katherine Ching, Zhenyang Zhou, Royce Ang, Lin Cui, Raymond Tzer Pin Lin |
| EPI_ISL_3007586 | Viollier AG | Department of Biosystems Science and Engineering, ETH Zürich | Andrea Patrizia Salzmann, Henriette Kurth, Christiane Beckmann, Ivan Topolsky, Chaoran Chen, Sarah Nadeau, Kim Philipp Jablonski, Lara Fuhrmann, Niko Beerenwinkel, Christoph Noppen, Olivier Kobel, Maurice Redondo, Tanja Stadler |
| EPI_ISL_3007587 | National Public Health Laboratory, National Centre for Infectious Diseases | National Public Health Laboratory, National Centre for Infectious Diseases | Tze Minn Mak, Katherine Ching, Zhenyang Zhou, Royce Ang, Lin Cui, Raymond Tzer Pin Lin |
| EPI_ISL_3007594, EPI_ISL_3007595, EPI_ISL_3007596, EPI_ISL_3007597, EPI_ISL_3007599, EPI_ISL_3007602, EPI_ISL_3007604, EPI_ISL_3007605, EPI_ISL_3007607, EPI_ISL_3007608, EPI_ISL_3007609, EPI_ISL_3007611, EPI_ISL_3007613, EPI_ISL_3007614, EPI_ISL_3007615, EPI_ISL_3007616, EPI_ISL_3007617, EPI_ISL_3007618, EPI_ISL_3007619, EPI_ISL_3007620, EPI_ISL_3007623, EPI_ISL_3007624, EPI_ISL_3007626 | | | |
| see above | Viollier AG | Department of Biosystems Science and Engineering, ETH Zürich | Andrea Patrizia Salzmann, Henriette Kurth, Christiane Beckmann, Ivan Topolsky, Chaoran Chen, Sarah Nadeau, Kim Philipp Jablonski, Lara Fuhrmann, Niko Beerenwinkel, Christoph Noppen, Olivier Kobel, Maurice Redondo, Tanja Stadler |
| EPI_ISL_3007627, EPI_ISL_3007629, EPI_ISL_3007630, EPI_ISL_3007631, EPI_ISL_3007632, EPI_ISL_3007633, EPI_ISL_3007634, EPI_ISL_3007635, EPI_ISL_3007636, EPI_ISL_3007637, EPI_ISL_3007638, EPI_ISL_3007639, EPI_ISL_3007640, EPI_ISL_3007641, EPI_ISL_3007642, EPI_ISL_3007643, EPI_ISL_3007644, EPI_ISL_3007645, EPI_ISL_3007646, EPI_ISL_3007648 | | | |
| see above | Dianalabs SA | Genesupport | Tanguy ARAUD, Geraldine Jost, Katia Jaton, Nadia Liassine |
| EPI_ISL_3007650 | NCSLPH | NCSLPH | Chase K, Miller MC, Greene S, Glover W |
| EPI_ISL_3007652 | LABORATORIO DE BIOLOGIA MOLECULAR CENTRO MEDICO NAVAL SEMAR | LABORATORIO DE PATOLOGÍA MOLECULAR CENTRO MEDICO NAVAL SEMAR | Meixueiro Calderon Claudia, Morales Ramirez Omar, Baltierra Uribe Shantal, Zuno Coronado Teutli, Garcia Alegria Irma, Alvarez Jimenez Violeta Dejanira, Lopez Bustos Abraham, Cabrera Gonzalez Moises, Lievano Esquinca Jazmin, Figueroa Antonio Victor, |
| EPI_ISL_3007656 | HOSPITAL CAN MISSES | HOSPITAL UNIVERSITARIO SON ESPASES | Servicio de Microbiología, Hospital Universitario Son Espases |
| EPI_ISL_3007657 | HOSPITAL UNIVERSITARIO SON ESPASES | HOSPITAL UNIVERSITARIO SON ESPASES | Servicio de Microbiología, Hospital Universitario Son Espases |
| EPI_ISL_3007659 | HOSPITAL CAN MISSES | HOSPITAL UNIVERSITARIO SON ESPASES | Servicio de Microbiología, Hospital Universitario Son Espases |
| EPI_ISL_3007660, EPI_ISL_3007661, EPI_ISL_3007662, EPI_ISL_3007663, EPI_ISL_3007664, EPI_ISL_3007665 | HOSPITAL UNIVERSITARIO SON ESPASES | HOSPITAL UNIVERSITARIO SON ESPASES | Servicio de Microbiología, Hospital Universitario Son Espases |
| EPI_ISL_3007666, EPI_ISL_3007667 | HOSPITAL CAN MISSES | HOSPITAL UNIVERSITARIO SON ESPASES | Servicio de Microbiología, Hospital Universitario Son Espases |
| EPI_ISL_3007668, EPI_ISL_3007671, EPI_ISL_3007672, EPI_ISL_3007673, EPI_ISL_3007674, EPI_ISL_3007675 | HOSPITAL UNIVERSITARIO SON ESPASES | HOSPITAL UNIVERSITARIO SON ESPASES | Servicio de Microbiología, Hospital Universitario Son Espases |
| EPI_ISL_3007676 | HOSPITAL CAN MISSES | HOSPITAL UNIVERSITARIO SON ESPASES | Servicio de Microbiología, Hospital Universitario Son Espases |
| EPI_ISL_3007677 | HOSPITAL UNIVERSITARIO SON ESPASES | HOSPITAL UNIVERSITARIO SON ESPASES | Servicio de Microbiología, Hospital Universitario Son Espases |
| EPI_ISL_3007678, EPI_ISL_3007679 | HOSPITAL CAN MISSES | HOSPITAL UNIVERSITARIO SON ESPASES | Servicio de Microbiología, Hospital Universitario Son Espases |
| EPI_ISL_3007681 | LABORATORIO DE BIOLOGÍA MOLECULAR CENTRO MEDICO NAVAL SEMAR | LABORATORIO DE PATOLOGÍA MOLECULAR CENTRO MEDICO NAVAL SEMAR | Meixueiro Calderon Claudia, Morales Ramirez Omar, Baltierra Uribe Shantal, Zuno Coronado Teutli, Garcia Alegria Irma, Alvarez Jimenez Violeta Dejanira, Lopez Bustos Abraham, Cabrera Gonzalez Moises, Lievano Esquinca Jazmin, Figueroa Antonio Victor, |
| EPI_ISL_3007682 | HOSPITAL UNIVERSITARIO SON ESPASES | HOSPITAL UNIVERSITARIO SON ESPASES | Servicio de Microbiología, Hospital Universitario Son Espases |
| EPI_ISL_3007683 | HOSPITAL MATEU ORFILA | HOSPITAL UNIVERSITARIO SON ESPASES | Servicio de Microbiología, Hospital Universitario Son Espases |
| EPI_ISL_3007684, EPI_ISL_3007685, EPI_ISL_3007686 | HOSPITAL UNIVERSITARIO SON ESPASES | HOSPITAL UNIVERSITARIO SON ESPASES | Servicio de Microbiología, Hospital Universitario Son Espases |
| EPI_ISL_3007687 | HOSPITAL CAN MISSES | HOSPITAL UNIVERSITARIO SON ESPASES | Servicio de Microbiología, Hospital Universitario Son Espases |
| EPI_ISL_3007688 | HOSPITAL UNIVERSITARIO SON ESPASES | HOSPITAL UNIVERSITARIO SON ESPASES | Servicio de Microbiología, Hospital Universitario Son Espases |
| EPI_ISL_3007690, EPI_ISL_3007691 | HOSPITAL CAN MISSES | HOSPITAL UNIVERSITARIO SON ESPASES | Servicio de Microbiología, Hospital Universitario Son Espases |
| EPI_ISL_3007692, EPI_ISL_3007693, EPI_ISL_3007694, EPI_ISL_3007696, EPI_ISL_3007697, EPI_ISL_3007698 | HOSPITAL UNIVERSITARIO SON ESPASES | HOSPITAL UNIVERSITARIO SON ESPASES | Servicio de Microbiología, Hospital Universitario Son Espases |
| EPI_ISL_3007699 | HOSPITAL CAN MISSES | HOSPITAL UNIVERSITARIO SON ESPASES | Servicio de Microbiología, Hospital Universitario Son Espases |
| EPI_ISL_3007700, EPI_ISL_3007702, EPI_ISL_3007703, EPI_ISL_3007704, EPI_ISL_3007705 | HOSPITAL UNIVERSITARIO SON ESPASES | HOSPITAL UNIVERSITARIO SON ESPASES | Servicio de Microbiología, Hospital Universitario Son Espases |
| EPI_ISL_3007706 | HOSPITAL CAN MISSES | HOSPITAL UNIVERSITARIO SON ESPASES | Servicio de Microbiología, Hospital Universitario Son Espases |
| EPI_ISL_3007707 | HOSPITAL UNIVERSITARIO SON ESPASES | HOSPITAL UNIVERSITARIO SON ESPASES | Servicio de Microbiología, Hospital Universitario Son Espases |
| EPI_ISL_3007708 | HOSPITAL CAN MISSES | HOSPITAL UNIVERSITARIO SON ESPASES | Servicio de Microbiología, Hospital Universitario Son Espases |
| EPI_ISL_3007709, EPI_ISL_3007710 | HOSPITAL UNIVERSITARIO SON ESPASES | HOSPITAL UNIVERSITARIO SON ESPASES | Servicio de Microbiología, Hospital Universitario Son Espases |
| EPI_ISL_3007711 | HOSPITAL MATEU ORFILA | HOSPITAL UNIVERSITARIO SON ESPASES | Servicio de Microbiología, Hospital Universitario Son Espases |
| EPI_ISL_3007712, EPI_ISL_3007713, EPI_ISL_3007714, EPI_ISL_3007716, EPI_ISL_3007717 | HOSPITAL UNIVERSITARIO SON ESPASES | HOSPITAL UNIVERSITARIO SON ESPASES | Servicio de Microbiología, Hospital Universitario Son Espases |
| EPI_ISL_3007718 | HOSPITAL CAN MISSES | HOSPITAL UNIVERSITARIO SON ESPASES | Servicio de Microbiología, Hospital Universitario Son Espases |
| EPI_ISL_3007719, EPI_ISL_3007720, EPI_ISL_3007721, EPI_ISL_3007723, EPI_ISL_3007724, EPI_ISL_3007725, EPI_ISL_3007726, EPI_ISL_3007728, EPI_ISL_3007729, EPI_ISL_3007731, EPI_ISL_3007732, EPI_ISL_3007733, EPI_ISL_3007734, EPI_ISL_3007735, EPI_ISL_3007736 | | | |
| see above | HOSPITAL UNIVERSITARIO SON ESPASES | HOSPITAL UNIVERSITARIO SON ESPASES | Servicio de Microbiología, Hospital Universitario Son Espases |
| EPI_ISL_3007742, EPI_ISL_3007743, EPI_ISL_3007744 | NCSLPH | NCSLPH | Chase K, Miller MC, Greene S, Glover W |
| EPI_ISL_3007745 | IN State Department of Health Laboratory Services | IN State Department of Health Laboratory Services | Cassandra Campion, Jamie Yeadon, Brian Pope, Lixia Liu, Kyle Brownlee, Melissa Hindenlang, Mark Glazier |
| EPI_ISL_3007746 | NCSLPH | NCSLPH | Chase K, Miller MC, Greene S, Glover W |
| EPI_ISL_3007747, EPI_ISL_3007748, EPI_ISL_3007749, EPI_ISL_3007750, EPI_ISL_3007751, EPI_ISL_3007752, EPI_ISL_3007753, EPI_ISL_3007754, EPI_ISL_3007755, EPI_ISL_3007756, EPI_ISL_3007757, EPI_ISL_3007758, EPI_ISL_3007759, EPI_ISL_3007760, EPI_ISL_3007761, EPI_ISL_3007762, EPI_ISL_3007763, EPI_ISL_3007764, EPI_ISL_3007765, EPI_ISL_3007766, EPI_ISL_3007767, EPI_ISL_3007768, EPI_ISL_3007769, EPI_ISL_3007770, EPI_ISL_3007771, EPI_ISL_3007772, EPI_ISL_3007773, EPI_ISL_3007774, EPI_ISL_3007775, EPI_ISL_3007776, EPI_ISL_3007777, EPI_ISL_3007778, EPI_ISL_3007779, EPI_ISL_3007780, EPI_ISL_3007781, EPI_ISL_3007782, EPI_ISL_3007783, EPI_ISL_3007784, EPI_ISL_3007785, EPI_ISL_3007786, EPI_ISL_3007787, EPI_ISL_3007788, EPI_ISL_3007789, EPI_ISL_3007790, EPI_ISL_3007791, EPI_ISL_3007792, EPI_ISL_3007793, EPI_ISL_3007794, EPI_ISL_3007795, EPI_ISL_3007796, EPI_ISL_3007797, EPI_ISL_3007798, EPI_ISL_3007799, EPI_ISL_3007800, EPI_ISL_3007801, EPI_ISL_3007802, EPI_ISL_3007803, EPI_ISL_3007804, EPI_ISL_3007805, EPI_ISL_3007806, EPI_ISL_3007807, EPI_ISL_3007808, EPI_ISL_3007809, EPI_ISL_3007810, EPI_ISL_3007811 | | | |
| see above | N.F. Gamaleya Research Center for Epidemiology and Microbiology | WHO National Influenza Centre Russian Federation | Andrey Komissarov, Artem Fadeev, Kseniya Komissarova, Oula Mansour, Kirill Varchenko, Mikhail Bakaev, Tamila Musaeva, Veronika Eder, Maria Pisareva, Nikita Yolshin, Daria Danilenko, Elizaveta Divisenko, Alina Odintsova, Elena Shidlovskaya, Ludmila Vasilchenko, Nadezhda Kuznetsova, Maria Nikiforova, Alexey Shchetinin, Artem Tkachuk, Andrei Pochtovyi, Andrei Siniavin, Denis Kleymenov, Evgeniia Bykonia, Liubov Popova, Elena Mazunina, Inna Dolzhikova, Anna Kovyrshina, Andrei Botikov, Dmitry Shcheblyakov, Vladimir Gushchin, Denis Logunov, Alexander Gintsburg, Ksenia Safina, Elena Nabieva, Georgii Bazykin, Dmitry Lioznov |
| EPI_ISL_3007812 | Magadan Regional Dispensary of Phthisiology and Infectious | WHO National Influenza Centre Russian Federation | Andrey Komissarov, Artem Fadeev, Kseniya Komissarova, Oula Mansour, Kirill Varchenko, Mikhail Bakaev, Tamila Musaeva, Veronika Eder, Maria Pisareva, Nikita Yolshin, Daria Danilenko, Ksenia Safina, Elena Nabieva, Georgii Bazykin, Dmitry |

| | | | |
|---|---|---|---|
| | Diseases | | Lioznov |
| EPI_ISL_3007813, EPI_ISL_3007814, EPI_ISL_3007815, EPI_ISL_3007816 | Chelyabinsk Regional Dermatovenerologic Dispensary | WHO National Influenza Centre Russian Federation | Andrey Komissarov, Artem Fadeev, Kseniya Komissarova, Oula Mansour, Kirill Varchenko, Mikhail Bakaev, Tamila Musaeva, Veronika Eder, Maria Pisareva, Nikita Yolshin, Daria Danilenko, Ksenia Safina, Elena Nabieva, Georgii Bazykin, Dmitry Lioznov |
| EPI_ISL_3007817, EPI_ISL_3007818, EPI_ISL_3007819 | N. A. Semashko Republican Clinical Hospital | WHO National Influenza Centre Russian Federation | Andrey Komissarov, Artem Fadeev, Kseniya Komissarova, Oula Mansour, Kirill Varchenko, Mikhail Bakaev, Tamila Musaeva, Veronika Eder, Maria Pisareva, Nikita Yolshin, Daria Danilenko, Ksenia Safina, Elena Nabieva, Georgii Bazykin, Dmitry Lioznov |
| EPI_ISL_3007821, EPI_ISL_3007822, EPI_ISL_3007823, EPI_ISL_3007824, EPI_ISL_3007825, EPI_ISL_3007832, EPI_ISL_3007835, EPI_ISL_3007836, EPI_ISL_3007837 | Srirama Chandra Bhanja Medical College and Hospital, Cuttack | Institute of Life Sciences - INSACOG | Sunil K. Raghav, Safal Walia, Arup Ghosh, Atimukta Jha, Amol M. Kanampalliwar, Omprakash Shiriwas, Sana Fatma, Shifu Aggarwal, Rupesh Dash, Rajeeb Swain, Punit Prasad, INSACOG Consortium, Ajay Parida |
| EPI_ISL_3007845, EPI_ISL_3007850, EPI_ISL_3007854, EPI_ISL_3007856, EPI_ISL_3007857, EPI_ISL_3007860, EPI_ISL_3007861, EPI_ISL_3007866, EPI_ISL_3007867, EPI_ISL_3007868, EPI_ISL_3007870, EPI_ISL_3007871, EPI_ISL_3007872 | | | |
| see above | Fakir Mohan Medical College & Hospital, Balasore | Institute of Life Sciences - INSACOG | Sunil K. Raghav, Safal Walia, Arup Ghosh, Atimukta Jha, Amol M. Kanampalliwar, Omprakash Shiriwas, Sana Fatma, Shifu Aggarwal, Rupesh Dash, Rajeeb Swain, Punit Prasad, INSACOG Consortium, Ajay Parida |
| EPI_ISL_3007873, EPI_ISL_3007874, EPI_ISL_3007875, EPI_ISL_3007877, EPI_ISL_3007878, EPI_ISL_3007879, EPI_ISL_3007880, EPI_ISL_3007882, EPI_ISL_3007883, EPI_ISL_3007885 | IISER Berhampur | Institute of Life Sciences - INSACOG | Sunil K. Raghav, Safal Walia, Arup Ghosh, Atimukta Jha, Amol M. Kanampalliwar, Omprakash Shiriwas, Sana Fatma, Shifu Aggarwal, Rupesh Dash, Rajeeb Swain, Punit Prasad, INSACOG Consortium, Ajay Parida |
| EPI_ISL_3007887, EPI_ISL_3007888, EPI_ISL_3007890, EPI_ISL_3007893, EPI_ISL_3007894, EPI_ISL_3007895, EPI_ISL_3007898, EPI_ISL_3007900, EPI_ISL_3007902, EPI_ISL_3007903, EPI_ISL_3007904, EPI_ISL_3007905, EPI_ISL_3007906, EPI_ISL_3007909, EPI_ISL_3007910, EPI_ISL_3007911, EPI_ISL_3007913, EPI_ISL_3007914, EPI_ISL_3007916 | | | |
| see above | Government Medical College, Bettiah | Institute of Life Sciences - INSACOG | Sunil K. Raghav, Safal Walia, Arup Ghosh, Atimukta Jha, Amol M. Kanampalliwar, Omprakash Shiriwas, Sana Fatma, Shifu Aggarwal, Rupesh Dash, Rajeeb Swain, Punit Prasad, INSACOG Consortium, Ajay Parida |
| EPI_ISL_3007926, EPI_ISL_3007927, EPI_ISL_3007928, EPI_ISL_3007929, EPI_ISL_3007930 | District Headquarter Hospital, Puri | Institute of Life Sciences - INSACOG | Sunil K. Raghav, Safal Walia, Arup Ghosh, Atimukta Jha, Amol M. Kanampalliwar, Omprakash Shiriwas, Sana Fatma, Shifu Aggarwal, Rupesh Dash, Rajeeb Swain, Punit Prasad, INSACOG Consortium, Ajay Parida |
| EPI_ISL_3007935, EPI_ISL_3007940 | District Head Quarters Hospital, Nayagarh | Institute of Life Sciences - INSACOG | Sunil K. Raghav, Safal Walia, Arup Ghosh, Atimukta Jha, Amol M. Kanampalliwar, Omprakash Shiriwas, Sana Fatma, Shifu Aggarwal, Rupesh Dash, Rajeeb Swain, Punit Prasad, INSACOG Consortium, Ajay Parida |
| EPI_ISL_3007945, EPI_ISL_3007947, EPI_ISL_3007948, EPI_ISL_3007949, EPI_ISL_3007950, EPI_ISL_3007951, EPI_ISL_3007952 | Late Baliram Kashyap Memorial Government Medical College, Jagdalpur | Institute of Life Sciences - INSACOG | Sunil K. Raghav, Safal Walia, Arup Ghosh, Atimukta Jha, Amol M. Kanampalliwar, Omprakash Shiriwas, Sana Fatma, Shifu Aggarwal, Rupesh Dash, Rajeeb Swain, Punit Prasad, INSACOG Consortium, Ajay Parida |
| EPI_ISL_3007963 | All India Institute of Medical Sciences, Bhubaneswar | Institute of Life Sciences - INSACOG | Sunil K. Raghav, Safal Walia, Arup Ghosh, Atimukta Jha, Amol M. Kanampalliwar, Omprakash Shiriwas, Sana Fatma, Shifu Aggarwal, Rupesh Dash, Rajeeb Swain, Punit Prasad, INSACOG Consortium, Ajay Parida |
| EPI_ISL_3007971 | Srirama Chandra Bhanja Medical College and Hospital, Cuttack | Institute of Life Sciences - INSACOG | Sunil K. Raghav, Safal Walia, Arup Ghosh, Atimukta Jha, Amol M. Kanampalliwar, Omprakash Shiriwas, Sana Fatma, Shifu Aggarwal, Rupesh Dash, Rajeeb Swain, Punit Prasad, INSACOG Consortium, Ajay Parida |
| EPI_ISL_3007981, EPI_ISL_3007982, EPI_ISL_3007985, EPI_ISL_3007986, EPI_ISL_3007988, EPI_ISL_3007989, EPI_ISL_3007990, EPI_ISL_3007992 | Kalinga Institute of Medical Sciences (KIMS), Bhubaneswar | Institute of Life Sciences - INSACOG | Sunil K. Raghav, Safal Walia, Arup Ghosh, Atimukta Jha, Amol M. Kanampalliwar, Omprakash Shiriwas, Sana Fatma, Shifu Aggarwal, Rupesh Dash, Rajeeb Swain, Punit Prasad, INSACOG Consortium, Ajay Parida |
| EPI_ISL_3007997, EPI_ISL_3007998, EPI_ISL_3007999 | Indira Gandhi Institute of Medical Sciences, Patna | Institute of Life Sciences - INSACOG | Sunil K. Raghav, Safal Walia, Arup Ghosh, Atimukta Jha, Amol M. Kanampalliwar, Omprakash Shiriwas, Sana Fatma, Shifu Aggarwal, Rupesh Dash, Rajeeb Swain, Punit Prasad, INSACOG Consortium, Ajay Parida |
| EPI_ISL_3008002, EPI_ISL_3008003, EPI_ISL_3008007, EPI_ISL_3008008, EPI_ISL_3008009, EPI_ISL_3008011, EPI_ISL_3008012, EPI_ISL_3008013, EPI_ISL_3008016, EPI_ISL_3008017, EPI_ISL_3008020, EPI_ISL_3008025, EPI_ISL_3008028, EPI_ISL_3008029, EPI_ISL_3008041, EPI_ISL_3008042, EPI_ISL_3008044, EPI_ISL_3008045, EPI_ISL_3008051, EPI_ISL_3008052, EPI_ISL_3008055, EPI_ISL_3008056, EPI_ISL_3008071, EPI_ISL_3008078 | | | |
| see above | District Public Health Laboratory, Angul | Institute of Life Sciences - INSACOG | Sunil K. Raghav, Safal Walia, Arup Ghosh, Atimukta Jha, Amol M. Kanampalliwar, Omprakash Shiriwas, Sana Fatma, Shifu Aggarwal, Rupesh Dash, Rajeeb Swain, Punit Prasad, INSACOG Consortium, Ajay Parida |
| EPI_ISL_3008079, EPI_ISL_3008080, EPI_ISL_3008081, EPI_ISL_3008082, EPI_ISL_3008083, EPI_ISL_3008084, EPI_ISL_3008085, EPI_ISL_3008086, EPI_ISL_3008087, EPI_ISL_3008091, EPI_ISL_3008092, EPI_ISL_3008093, EPI_ISL_3008094, EPI_ISL_3008097, EPI_ISL_3008098, EPI_ISL_3008099, EPI_ISL_3008102, EPI_ISL_3008104, EPI_ISL_3008107, EPI_ISL_3008108, EPI_ISL_3008111, EPI_ISL_3008114, EPI_ISL_3008115, EPI_ISL_3008116, EPI_ISL_3008117, EPI_ISL_3008118 | | | |
| see above | Regional Medical Research Center, Bhubaneswar | Institute of Life Sciences - INSACOG | Sunil K. Raghav, Safal Walia, Arup Ghosh, Atimukta Jha, Amol M. Kanampalliwar, Omprakash Shiriwas, Sana Fatma, Shifu Aggarwal, Rupesh Dash, Rajeeb Swain, Punit Prasad, INSACOG Consortium, Ajay Parida |
| EPI_ISL_3008119, EPI_ISL_3008123, EPI_ISL_3008124, EPI_ISL_3008125 | District Headquarter Hospital, Baripada | Institute of Life Sciences - INSACOG | Sunil K. Raghav, Safal Walia, Arup Ghosh, Atimukta Jha, Amol M. Kanampalliwar, Omprakash Shiriwas, Sana Fatma, Shifu Aggarwal, Rupesh Dash, Rajeeb Swain, Punit Prasad, INSACOG Consortium, Ajay Parida |
| EPI_ISL_3008128, EPI_ISL_3008129, EPI_ISL_3008130, EPI_ISL_3008131, EPI_ISL_3008134, EPI_ISL_3008136, EPI_ISL_3008137, EPI_ISL_3008138, EPI_ISL_3008139, EPI_ISL_3008140, EPI_ISL_3008141, EPI_ISL_3008142, EPI_ISL_3008143, EPI_ISL_3008144, EPI_ISL_3008146, EPI_ISL_3008148, EPI_ISL_3008152, EPI_ISL_3008154, EPI_ISL_3008156, EPI_ISL_3008158, EPI_ISL_3008160 | | | |
| see above | Institute of Life Sciences, Bhubaneswar | Institute of Life Sciences - INSACOG | Sunil K. Raghav, Safal Walia, Arup Ghosh, Atimukta Jha, Amol M. Kanampalliwar, Omprakash Shiriwas, Sana Fatma, Shifu Aggarwal, Rupesh Dash, Rajeeb Swain, Punit Prasad, INSACOG Consortium, Ajay Parida |
| EPI_ISL_3008213, EPI_ISL_3008214, EPI_ISL_3008215, EPI_ISL_3008216, EPI_ISL_3008217, EPI_ISL_3008219, EPI_ISL_3008220, EPI_ISL_3008222, EPI_ISL_3008223, EPI_ISL_3008225, EPI_ISL_3008226, EPI_ISL_3008227, EPI_ISL_3008228, EPI_ISL_3008231, EPI_ISL_3008233, EPI_ISL_3008234, EPI_ISL_3008235, EPI_ISL_3008236, EPI_ISL_3008238, EPI_ISL_3008239, EPI_ISL_3008240, EPI_ISL_3008241, EPI_ISL_3008242, EPI_ISL_3008243, EPI_ISL_3008244, EPI_ISL_3008245, EPI_ISL_3008246, EPI_ISL_3008247, EPI_ISL_3008248, EPI_ISL_3008249, EPI_ISL_3008251, EPI_ISL_3008253, EPI_ISL_3008257, EPI_ISL_3008258, EPI_ISL_3008259, EPI_ISL_3008260, EPI_ISL_3008261 | | | |
| see above | Hospital General Universitario Gregorio Marañón | Hospital General Universitario Gregorio Marañón | Sergio Buenestado Serrano, Pedro Sola Campoy, Laura Pérez-Lago, Cristina Rodriguez-Grande, Marta Herranz Martin, Victor Manuel de la Cueva, Julia Suárez, Pilar Catalán, Patricia Muñoz, Darío García de Viedma |
| EPI_ISL_3008262, EPI_ISL_3008263, EPI_ISL_3008264, EPI_ISL_3008265, EPI_ISL_3008266, EPI_ISL_3008267, EPI_ISL_3008268, EPI_ISL_3008269, EPI_ISL_3008270, EPI_ISL_3008271, EPI_ISL_3008272, EPI_ISL_3008273, EPI_ISL_3008274, EPI_ISL_3008275, EPI_ISL_3008276, EPI_ISL_3008277, EPI_ISL_3008278, EPI_ISL_3008279, EPI_ISL_3008280, EPI_ISL_3008281, EPI_ISL_3008282, EPI_ISL_3008283, EPI_ISL_3008284, EPI_ISL_3008285 | | | |
| see above | Azienda Sanitaria dell'Alto Adige - Laboratorio Aziendale di Microbiologia e Virologia | Azienda Sanitaria dell'Alto Adige | Irene Bianconi |
| EPI_ISL_3008287, EPI_ISL_3008289, EPI_ISL_3008291 | Centre Hospitalier Universitaire Clermont-Ferrand | CHU Clermont-Ferrand, service de virologie | Bisseux Maxime, Mirand Audrey, Combes Patricia, Henquell Cécile |
| EPI_ISL_3008292 | HOPITAL DU MONT DORE | CHU Clermont-Ferrand, service de virologie | Bisseux Maxime, Mirand Audrey, Combes Patricia, Henquell Cécile |
| EPI_ISL_3008293, EPI_ISL_3008300 | Furst Medical Laboratory | Norwegian Institute of Public Health, Department of Virology | Kathrine Stene-Johansen, Kamilla Heddeland Instefjord, Hilde Elshaug, Garcia Llorente Ignacio, Jon Bråte, Engebretsen Serina Beate,Pedersen Benedikte Nevjen, Line Victoria Moen, Debech Nadia, Atiya R Ali,Marie Paulsen Madsen, Rasmus Riis Kopperud, Hilde Vollan, Karoline Bragstad, Olav Hungnes |
| EPI_ISL_3008311, EPI_ISL_3008313 | Unilabs Laboratory Medicine | Norwegian Institute of Public Health, Department of Virology | Kathrine Stene-Johansen, Kamilla Heddeland Instefjord, Hilde Elshaug, Garcia Llorente Ignacio, Jon Bråte, Engebretsen Serina Beate,Pedersen Benedikte Nevjen, Line Victoria Moen, Debech Nadia, Atiya R Ali,Marie Paulsen Madsen, Rasmus Riis Kopperud, Hilde Vollan, Karoline Bragstad, Olav Hungnes |
| EPI_ISL_3008314, EPI_ISL_3008315 | Furst Medical Laboratory | Norwegian Institute of Public Health, Department of Virology | Kathrine Stene-Johansen, Kamilla Heddeland Instefjord, Hilde Elshaug, Garcia Llorente Ignacio, Jon Bråte, Engebretsen Serina Beate,Pedersen Benedikte Nevjen, Line Victoria Moen, Debech Nadia, Atiya R Ali,Marie Paulsen Madsen, Rasmus Riis Kopperud, Hilde Vollan, Karoline Bragstad, Olav Hungnes |
| EPI_ISL_3008316 | Hospital of Southern Norway - Kristiansand, Department of Medical Microbiology | Norwegian Institute of Public Health, Department of Virology | Kathrine Stene-Johansen, Kamilla Heddeland Instefjord, Hilde Elshaug, Garcia Llorente Ignacio, Jon Bråte, Engebretsen Serina Beate,Pedersen Benedikte Nevjen, Line Victoria Moen, Debech Nadia, Atiya R Ali,Marie Paulsen Madsen, Rasmus Riis Kopperud, Hilde Vollan, Karoline Bragstad, Olav Hungnes |
| EPI_ISL_3008318, EPI_ISL_3008319, EPI_ISL_3008320, EPI_ISL_3008322, EPI_ISL_3008324, EPI_ISL_3008327, | Furst Medical Laboratory | Norwegian Institute of Public Health, Department of Virology | Kathrine Stene-Johansen, Kamilla Heddeland Instefjord, Hilde Elshaug, Garcia Llorente Ignacio, Jon Bråte, Engebretsen Serina Beate,Pedersen Benedikte Nevjen, Line Victoria Moen, Debech Nadia, Atiya R Ali,Marie Paulsen Madsen, Rasmus Riis Kopperud, Hilde Vollan, Karoline Bragstad, Olav Hungnes |

| | | | |
|---|---|---|---|
| EPI_ISL_3008330 | | | |
| EPI_ISL_3008332 | Hospital of Southern Norway - Kristiansand, Department of Medical Microbiology | Norwegian Institute of Public Health, Department of Virology | Kathrine Stene-Johansen, Kamilla Heddeland Instefjord, Hilde Elshaug, Garcia Llorente Ignacio, Jon Bråte, Engebretsen Serina Beate,Pedersen Benedikte Nevjen, Line Victoria Moen, Debech Nadia, Atiya R Ali,Marie Paulsen Madsen, Rasmus Riis Kopperud, Hilde Vollan, Karoline Bragstad, Olav Hungnes |
| EPI_ISL_3008350 | Unilabs Laboratory Medicine | Norwegian Institute of Public Health, Department of Virology | Kathrine Stene-Johansen, Kamilla Heddeland Instefjord, Hilde Elshaug, Garcia Llorente Ignacio, Jon Bråte, Engebretsen Serina Beate,Pedersen Benedikte Nevjen, Line Victoria Moen, Debech Nadia, Atiya R Ali,Marie Paulsen Madsen, Rasmus Riis Kopperud, Hilde Vollan, Karoline Bragstad, Olav Hungnes |
| EPI_ISL_3008352, EPI_ISL_3008356 | Furst Medical Laboratory | Norwegian Institute of Public Health, Department of Virology | Kathrine Stene-Johansen, Kamilla Heddeland Instefjord, Hilde Elshaug, Garcia Llorente Ignacio, Jon Bråte, Engebretsen Serina Beate,Pedersen Benedikte Nevjen, Line Victoria Moen, Debech Nadia, Atiya R Ali,Marie Paulsen Madsen, Rasmus Riis Kopperud, Hilde Vollan, Karoline Bragstad, Olav Hungnes |
| EPI_ISL_3008364, EPI_ISL_3008374 | Vestfold Hospital, Toensberg Department of Microbiology | Norwegian Institute of Public Health, Department of Virology | Kathrine Stene-Johansen, Kamilla Heddeland Instefjord, Hilde Elshaug, Garcia Llorente Ignacio, Jon Bråte, Engebretsen Serina Beate,Pedersen Benedikte Nevjen, Line Victoria Moen, Debech Nadia, Atiya R Ali,Marie Paulsen Madsen, Rasmus Riis Kopperud, Hilde Vollan, Karoline Bragstad, Olav Hungnes |
| EPI_ISL_3008377 | Hospital of Southern Norway - Kristiansand, Department of Medical Microbiology | Norwegian Institute of Public Health, Department of Virology | Kathrine Stene-Johansen, Kamilla Heddeland Instefjord, Hilde Elshaug, Garcia Llorente Ignacio, Jon Bråte, Engebretsen Serina Beate,Pedersen Benedikte Nevjen, Line Victoria Moen, Debech Nadia, Atiya R Ali,Marie Paulsen Madsen, Rasmus Riis Kopperud, Hilde Vollan, Karoline Bragstad, Olav Hungnes |
| EPI_ISL_3008379 | City of Milwaukee Health Department Laboratory | City of Milwaukee Health Department Laboratory | Manjeet Khubbar, Amy Bauer, Jennifer Lentz, Samantha Scott, Sanjib Bhattacharyya |
| EPI_ISL_3008380 | Hospital of Southern Norway - Kristiansand, Department of Medical Microbiology | Norwegian Institute of Public Health, Department of Virology | Kathrine Stene-Johansen, Kamilla Heddeland Instefjord, Hilde Elshaug, Garcia Llorente Ignacio, Jon Bråte, Engebretsen Serina Beate,Pedersen Benedikte Nevjen, Line Victoria Moen, Debech Nadia, Atiya R Ali,Marie Paulsen Madsen, Rasmus Riis Kopperud, Hilde Vollan, Karoline Bragstad, Olav Hungnes |
| EPI_ISL_3008459, EPI_ISL_3008460, EPI_ISL_3008462, EPI_ISL_3008464, EPI_ISL_3008465, EPI_ISL_3008466, EPI_ISL_3008467, EPI_ISL_3008468, EPI_ISL_3008469, EPI_ISL_3008472, EPI_ISL_3008473, EPI_ISL_3008474, EPI_ISL_3008475, EPI_ISL_3008476, EPI_ISL_3008477, EPI_ISL_3008479, EPI_ISL_3008480, EPI_ISL_3008481, EPI_ISL_3008482, EPI_ISL_3008485, EPI_ISL_3008486, EPI_ISL_3008487, EPI_ISL_3008488, EPI_ISL_3008489, EPI_ISL_3008490, EPI_ISL_3008491, EPI_ISL_3008492, EPI_ISL_3008493, EPI_ISL_3008494, EPI_ISL_3008495, EPI_ISL_3008496, EPI_ISL_3008498, EPI_ISL_3008499, EPI_ISL_3008500, EPI_ISL_3008501, EPI_ISL_3008502, EPI_ISL_3008503, EPI_ISL_3008504, EPI_ISL_3008505, EPI_ISL_3008506, EPI_ISL_3008507, EPI_ISL_3008508, EPI_ISL_3008509, EPI_ISL_3008510, EPI_ISL_3008511, EPI_ISL_3008512, EPI_ISL_3008513, EPI_ISL_3008514, EPI_ISL_3008515, EPI_ISL_3008516, EPI_ISL_3008517, EPI_ISL_3008518, EPI_ISL_3008520, EPI_ISL_3008521, EPI_ISL_3008522, EPI_ISL_3008523, EPI_ISL_3008524, EPI_ISL_3008525, EPI_ISL_3008526 | | | |
| see above | Hospital Universitari i Politècnic La Fe de València | SeqCOVID-SPAIN consortium/IBV(CSIC) | María Dolores Gómez Ruiz, Eva González Barberá, Ana Gil Brusola, Salvador Giner Almaraz, José Luis López Hontangas and SeqCOVID-SPAIN consortium |
| EPI_ISL_3008530, EPI_ISL_3008534, EPI_ISL_3008537, EPI_ISL_3008543, EPI_ISL_3008545, EPI_ISL_3008546, EPI_ISL_3008547 | Servicio de Microbiología. Hospital General Universitario de Castellón | SeqCOVID-SPAIN consortium/IBV(CSIC) | Rosario Moreno Muñoz, María Dolores Tirado Balaguer and SeqCOVID-SPAIN consortium |
| EPI_ISL_3008548, EPI_ISL_3008550, EPI_ISL_3008551, EPI_ISL_3008552, EPI_ISL_3008553, EPI_ISL_3008555, EPI_ISL_3008556, EPI_ISL_3008558, EPI_ISL_3008559, EPI_ISL_3008560, EPI_ISL_3008561, EPI_ISL_3008562, EPI_ISL_3008564, EPI_ISL_3008566, EPI_ISL_3008568, EPI_ISL_3008569, EPI_ISL_3008570 | | | |
| see above | Klinisch Laboratorium ZNA | Klinisch Laboratorium ZNA | Verstrepen et al. |
| EPI_ISL_3008572 | Hospital General de Requena | SeqCOVID-SPAIN consortium/IBV(CSIC) | Martín Yago López, María Ángeles Goberna Bravo, Carolina Pla Cortes and SeqCOVID-SPAIN consortium |
| EPI_ISL_3008574, EPI_ISL_3008579, EPI_ISL_3008591 | Servicio de Microbiología. Hospital General Universitario de Castellón | SeqCOVID-SPAIN consortium/IBV(CSIC) | Rosario Moreno Muñoz, María Dolores Tirado Balaguer and SeqCOVID-SPAIN consortium |
| EPI_ISL_3008593 | Epimol – Molecular Epidemiology Group (UV-FISABIO) | SeqCOVID-SPAIN consortium/IBV(CSIC) | María Alma Bracho and SeqCOVID-SPAIN consortium |
| EPI_ISL_3008594 | Synlab - Hospital de Manises | SeqCOVID-SPAIN consortium/IBV(CSIC) | Amparo Broseta Tamarit, Carlos Gulin Blanco and SeqCOVID-SPAIN consortium |
| EPI_ISL_3008595, EPI_ISL_3008596, EPI_ISL_3008597 | Servicio de Microbiología. Hospital Arnau de Vilanova | SeqCOVID-SPAIN consortium/IBV(CSIC) | Victoria Domínguez, Rocío Falcón, Amparo Farga and SeqCOVID-SPAIN consortium |
| EPI_ISL_3008600 | Epimol – Molecular Epidemiology Group (UV-FISABIO) | SeqCOVID-SPAIN consortium/IBV(CSIC) | María Alma Bracho and SeqCOVID-SPAIN consortium |
| EPI_ISL_3008601, EPI_ISL_3008602, EPI_ISL_3008603, EPI_ISL_3008604, EPI_ISL_3008605, EPI_ISL_3008606, EPI_ISL_3008607, EPI_ISL_3008608, EPI_ISL_3008609, EPI_ISL_3008610, EPI_ISL_3008611, EPI_ISL_3008612, EPI_ISL_3008613, EPI_ISL_3008614, EPI_ISL_3008615, EPI_ISL_3008617, EPI_ISL_3008618, EPI_ISL_3008619, EPI_ISL_3008620, EPI_ISL_3008621, EPI_ISL_3008622, EPI_ISL_3008623, EPI_ISL_3008624, EPI_ISL_3008625, EPI_ISL_3008626, EPI_ISL_3008627, EPI_ISL_3008628, EPI_ISL_3008629, EPI_ISL_3008630, EPI_ISL_3008631, EPI_ISL_3008632, EPI_ISL_3008633, EPI_ISL_3008634, EPI_ISL_3008635, EPI_ISL_3008636, EPI_ISL_3008637, EPI_ISL_3008638, EPI_ISL_3008639, EPI_ISL_3008641, EPI_ISL_3008642, EPI_ISL_3008643, EPI_ISL_3008644, EPI_ISL_3008645, EPI_ISL_3008646, EPI_ISL_3008647, EPI_ISL_3008648, EPI_ISL_3008649, EPI_ISL_3008650, EPI_ISL_3008651, EPI_ISL_3008652, EPI_ISL_3008653, EPI_ISL_3008654, EPI_ISL_3008655, EPI_ISL_3008656 | | | |
| see above | Servicio de Microbiología. Hospital Clínico Universitario de Valencia | SeqCOVID-SPAIN consortium/IBV(CSIC) | David Navarro Ortega, Eliseo Albert Vicent, Ignacio Torres and SeqCOVID-SPAIN consortium |
| EPI_ISL_3008657, EPI_ISL_3008659, EPI_ISL_3008661, EPI_ISL_3008663, EPI_ISL_3008664, EPI_ISL_3008665, EPI_ISL_3008666, EPI_ISL_3008667, EPI_ISL_3008668, EPI_ISL_3008669, EPI_ISL_3008670, EPI_ISL_3008673, EPI_ISL_3008676, EPI_ISL_3008677, EPI_ISL_3008678, EPI_ISL_3008679, EPI_ISL_3008680, EPI_ISL_3008681, EPI_ISL_3008682, EPI_ISL_3008683, EPI_ISL_3008684, EPI_ISL_3008685, EPI_ISL_3008686, EPI_ISL_3008687, EPI_ISL_3008688, EPI_ISL_3008689, EPI_ISL_3008690, EPI_ISL_3008691, EPI_ISL_3008692, EPI_ISL_3008693, EPI_ISL_3008694, EPI_ISL_3008695, EPI_ISL_3008696, EPI_ISL_3008697, EPI_ISL_3008698, EPI_ISL_3008700, EPI_ISL_3008701, EPI_ISL_3008703, EPI_ISL_3008704, EPI_ISL_3008705, EPI_ISL_3008706, EPI_ISL_3008707, EPI_ISL_3008708, EPI_ISL_3008709, EPI_ISL_3008710, EPI_ISL_3008711, EPI_ISL_3008712, EPI_ISL_3008713, EPI_ISL_3008714, EPI_ISL_3008715, EPI_ISL_3008716, EPI_ISL_3008717, EPI_ISL_3008718, EPI_ISL_3008719, EPI_ISL_3008720, EPI_ISL_3008721, EPI_ISL_3008722, EPI_ISL_3008723, EPI_ISL_3008724, EPI_ISL_3008725, EPI_ISL_3008726, EPI_ISL_3008727, EPI_ISL_3008728, EPI_ISL_3008729, EPI_ISL_3008730, EPI_ISL_3008732, EPI_ISL_3008733, EPI_ISL_3008734, EPI_ISL_3008735, EPI_ISL_3008736, EPI_ISL_3008737, EPI_ISL_3008738, EPI_ISL_3008739, EPI_ISL_3008741, EPI_ISL_3008742, EPI_ISL_3008743, EPI_ISL_3008744, EPI_ISL_3008745, EPI_ISL_3008746, EPI_ISL_3008747, EPI_ISL_3008749, EPI_ISL_3008750, EPI_ISL_3008751, EPI_ISL_3008752, EPI_ISL_3008753, EPI_ISL_3008754, EPI_ISL_3008755, EPI_ISL_3008756 | | | |
| see above | Platform BIS UZA/UAntwerpen | Labo Klinische Biologie, UZA | Marie Le Mercier, Jasmine Coppens, Kathleen Holemans, Ines Verbesselt, Basil Britto Xavier, Christine Lammens, Veerle Matheeussen, Herman Goossens |
| EPI_ISL_3008757, EPI_ISL_3008758 | Algemeen Medisch Labo | Labo Klinische Biologie, UZA | Marie Le Mercier, Jasmine Coppens, Kathleen Holemans, Ines Verbesselt, Basil Britto Xavier, Christine Lammens, Veerle Matheeussen, Herman Goossens |
| EPI_ISL_3008759, EPI_ISL_3008760, EPI_ISL_3008761, EPI_ISL_3008762, EPI_ISL_3008763, EPI_ISL_3008764, EPI_ISL_3008766, EPI_ISL_3008767, EPI_ISL_3008768 | CENTRUM VOOR MEDISCHE ANALYSE | Labo Klinische Biologie, UZA | Marie Le Mercier, Jasmine Coppens, Kathleen Holemans, Ines Verbesselt, Basil Britto Xavier, Christine Lammens, Veerle Matheeussen, Herman Goossens |
| EPI_ISL_3008771, EPI_ISL_3008772, EPI_ISL_3008773, EPI_ISL_3008774, EPI_ISL_3008775, EPI_ISL_3008777, EPI_ISL_3008778, EPI_ISL_3008779, EPI_ISL_3008780, EPI_ISL_3008781, EPI_ISL_3008782, EPI_ISL_3008783, EPI_ISL_3008785, EPI_ISL_3008786, EPI_ISL_3008787, EPI_ISL_3008788, EPI_ISL_3008789, EPI_ISL_3008790, EPI_ISL_3008791, EPI_ISL_3008792, EPI_ISL_3008793, EPI_ISL_3008795, EPI_ISL_3008796, EPI_ISL_3008797, EPI_ISL_3008798, EPI_ISL_3008799, EPI_ISL_3008800, EPI_ISL_3008801, EPI_ISL_3008805, EPI_ISL_3008806, EPI_ISL_3008807, EPI_ISL_3008808, EPI_ISL_3008809, EPI_ISL_3008810, EPI_ISL_3008812 | | | |
| see above | Platform BIS UZA/UAntwerpen | Labo Klinische Biologie, UZA | Marie Le Mercier, Jasmine Coppens, Kathleen Holemans, Ines Verbesselt, Basil Britto Xavier, Christine Lammens, Veerle Matheeussen, Herman Goossens |
| EPI_ISL_3008814, EPI_ISL_3008815, EPI_ISL_3008816, EPI_ISL_3008818, EPI_ISL_3008819, EPI_ISL_3008820, EPI_ISL_3008821, EPI_ISL_3008822, EPI_ISL_3008823, EPI_ISL_3008824, EPI_ISL_3008825, EPI_ISL_3008826, EPI_ISL_3008827, EPI_ISL_3008828, EPI_ISL_3008829 | | | |
| see above | LHUB-ULB | Labo Klinische Biologie, UZA | Marie Le Mercier, Jasmine Coppens, Kathleen Holemans, Ines Verbesselt, Basil Britto Xavier, Christine Lammens, Veerle Matheeussen, Herman Goossens |
| EPI_ISL_3008831 | Center of Advanced Studies and Technology, Molecular Genetics Laboratory | Center of Advanced Studies and Technology, Molecular Genetics Laboratory | Anaclerio Federico, De Fabritiis Simone |
| EPI_ISL_3008832, EPI_ISL_3008833, EPI_ISL_3008834, EPI_ISL_3008835, EPI_ISL_3008836, EPI_ISL_3008837, EPI_ISL_3008838, EPI_ISL_3008839, EPI_ISL_3008840, EPI_ISL_3008841, EPI_ISL_3008842, EPI_ISL_3008843, EPI_ISL_3008844, EPI_ISL_3008845, EPI_ISL_3008846, EPI_ISL_3008847, EPI_ISL_3008848, EPI_ISL_3008849, EPI_ISL_3008850, EPI_ISL_3008851, EPI_ISL_3008852, EPI_ISL_3008853, EPI_ISL_3008854, EPI_ISL_3008855, EPI_ISL_3008856, EPI_ISL_3008857, EPI_ISL_3008858, EPI_ISL_3008859, EPI_ISL_3008860, EPI_ISL_3008861, EPI_ISL_3008862, EPI_ISL_3008863 | | | |
| see above | Kansas Health and Environmental Lab | Kansas Health and Environmental Lab | Mike Grose, Katherine Wiggins, Jonathan Barnell, Ben Olsen, and Phil Adam |
| EPI_ISL_3008864, EPI_ISL_3008865, EPI_ISL_3008867, EPI_ISL_3008868 | University of Liège COVID-19 testing center | GIGA Medical Genomics | Keith Durkin, Maria Artesi, Bouchra Boujemla, Nathalie Renotte, Nadine Cambisano, Sébastien Bontems, Cécile Meex, Claire Gourzonès, Olivier Ek, Laurent Gillet, Marie-Pierre Hayette, Vincent Bours |
| EPI_ISL_3008872, EPI_ISL_3008874, EPI_ISL_3008875, EPI_ISL_3008879, EPI_ISL_3008880, EPI_ISL_3008881, EPI_ISL_3008882 | SYNLAB | GIGA Medical Genomics | Keith Durkin, Maria Artesi, Bouchra Boujemla, Nathalie Renotte, Nadine Cambisano, Sébastien Bontems, Cécile Meex, Claire Gourzonès, Olivier Ek, Laurent Gillet, Marie-Pierre Hayette, Vincent Bours |
| EPI_ISL_3008884, EPI_ISL_3008889, EPI_ISL_3008890 | Department of Clinical Microbiology | GIGA Medical Genomics | Keith Durkin, Maria Artesi, Bouchra Boujemla, Nathalie Renotte, Nadine Cambisano, Sébastien Bontems, Cécile Meex, Claire Gourzonès, Olivier Ek, Laurent Gillet, Marie-Pierre Hayette, Vincent Bours |
| EPI_ISL_3008896, EPI_ISL_3008897, EPI_ISL_3008898, EPI_ISL_3008899, EPI_ISL_3008900, EPI_ISL_3008901, EPI_ISL_3008902, EPI_ISL_3008903, EPI_ISL_3008904, EPI_ISL_3008905, EPI_ISL_3008906, EPI_ISL_3008907, EPI_ISL_3008908, EPI_ISL_3008909, EPI_ISL_3008910, EPI_ISL_3008911, EPI_ISL_3008913, EPI_ISL_3008914, EPI_ISL_3008915, EPI_ISL_3008916, EPI_ISL_3008917, EPI_ISL_3008918, EPI_ISL_3008919, EPI_ISL_3008920, EPI_ISL_3008921, EPI_ISL_3008922, EPI_ISL_3008923, EPI_ISL_3008924, EPI_ISL_3008925, EPI_ISL_3008926 | | | |
| see above | Kansas Health and Environmental Lab | Kansas Health and Environmental Lab | Mike Grose, Katherine Wiggins, Jonathan Barnell, Ben Olsen, and Phil Adam |
| EPI_ISL_3008927, EPI_ISL_3008928, EPI_ISL_3008929, EPI_ISL_3008930, EPI_ISL_3008931, EPI_ISL_3008932, EPI_ISL_3008933, EPI_ISL_3008934, EPI_ISL_3008935, EPI_ISL_3008936, EPI_ISL_3008937, EPI_ISL_3008938, EPI_ISL_3008939, EPI_ISL_3008940, EPI_ISL_3008941, EPI_ISL_3008942, EPI_ISL_3008943, EPI_ISL_3008944, EPI_ISL_3008945, EPI_ISL_3008946, EPI_ISL_3008947, EPI_ISL_3008948, EPI_ISL_3008949, EPI_ISL_3008950, EPI_ISL_3008951, EPI_ISL_3008952, EPI_ISL_3008953, EPI_ISL_3008954, EPI_ISL_3008955, EPI_ISL_3008956, EPI_ISL_3008957, EPI_ISL_3008958, EPI_ISL_3008959, EPI_ISL_3008960, EPI_ISL_3008961, EPI_ISL_3008962, EPI_ISL_3008963, EPI_ISL_3008964, EPI_ISL_3008965, EPI_ISL_3008966, EPI_ISL_3008967, EPI_ISL_3008968, EPI_ISL_3008969, EPI_ISL_3008970, EPI_ISL_3008971, EPI_ISL_3008972, EPI_ISL_3008973, EPI_ISL_3008974, EPI_ISL_3008975, EPI_ISL_3008976, EPI_ISL_3008977, EPI_ISL_3008978, EPI_ISL_3008979, EPI_ISL_3008980, EPI_ISL_3008981, EPI_ISL_3008982, EPI_ISL_3008983, EPI_ISL_3008984, EPI_ISL_3008985, EPI_ISL_3008986, EPI_ISL_3008987, EPI_ISL_3008988, EPI_ISL_3008989, EPI_ISL_3008990, EPI_ISL_3008991, EPI_ISL_3008992, EPI_ISL_3008993, EPI_ISL_3008994, EPI_ISL_3008995, EPI_ISL_3008996, EPI_ISL_3008997, EPI_ISL_3008998, EPI_ISL_3008999, EPI_ISL_3009000, EPI_ISL_3009001, EPI_ISL_3009002, EPI_ISL_3009003, EPI_ISL_3009004, EPI_ISL_3009005, EPI_ISL_3009006, EPI_ISL_3009007, EPI_ISL_3009008, EPI_ISL_3009009, EPI_ISL_3009010, EPI_ISL_3009011, EPI_ISL_3009012, EPI_ISL_3009013, EPI_ISL_3009014, EPI_ISL_3009015, EPI_ISL_3009016, EPI_ISL_3009017, EPI_ISL_3009018, EPI_ISL_3009019, EPI_ISL_3009020, EPI_ISL_3009021, EPI_ISL_3009022, EPI_ISL_3009023, EPI_ISL_3009024, EPI_ISL_3009025, EPI_ISL_3009026, EPI_ISL_3009027, EPI_ISL_3009028, EPI_ISL_3009029, EPI_ISL_3009030, EPI_ISL_3009031, EPI_ISL_3009032, EPI_ISL_3009033, EPI_ISL_3009034, EPI_ISL_3009035, EPI_ISL_3009036, EPI_ISL_3009037, EPI_ISL_3009038, EPI_ISL_3009039, EPI_ISL_3009040, EPI_ISL_3009041, EPI_ISL_3009042, EPI_ISL_3009043, EPI_ISL_3009044, EPI_ISL_3009045, EPI_ISL_3009046 | | | |
| see above | National Public Health Laboratory, National Centre for Infectious Diseases | National Public Health Laboratory, National Centre for Infectious Diseases | Tze Minn Mak, Katherine Ching, Zhenyang Zhou, Royce Ang, Lin Cui, Raymond Tzer Pin Lin |

| | | | |
|---|---|---|---|
| EPI_ISL_3009051 | SIESP CHIETI - DRIVE IN ORTONA | Istituto Zooprofilattico Sperimentale dell'Abruzzo e Molise "G. Caporale" | Lorusso A, Marcacci M, Di Domenico M, Ancora M, Curini V, Di Lollo Valeria, Mangone I, Rinaldi A, Delli Compagni E, Scialabba S, Caporale M, Di Pasquale A, Cammà C, Puglia I, Calistri P, Savini G |
| EPI_ISL_3009055, EPI_ISL_3009056, EPI_ISL_3009057, EPI_ISL_3009058, EPI_ISL_3009059, EPI_ISL_3009060, EPI_ISL_3009061, EPI_ISL_3009062, EPI_ISL_3009063, EPI_ISL_3009064, EPI_ISL_3009065, EPI_ISL_3009066, EPI_ISL_3009067, EPI_ISL_3009068, EPI_ISL_3009069, EPI_ISL_3009070, EPI_ISL_3009071, EPI_ISL_3009072, EPI_ISL_3009073, EPI_ISL_3009074, EPI_ISL_3009075, EPI_ISL_3009076 | | | |
| see above | OSPEDALE CIVILE TERAMO - CENTRO TRASFUSIONALE | Istituto Zooprofilattico Sperimentale dell'Abruzzo e Molise "G. Caporale" | Lorusso A, Marcacci M, Di Domenico M, Ancora M, Curini V, Di Lollo Valeria, Mangone I, Rinaldi A, Delli Compagni E, Scialabba S, Caporale M, Di Pasquale A, Cammà C, Puglia I, Calistri P, Savini G |
| EPI_ISL_3009080, EPI_ISL_3009081, EPI_ISL_3009082, EPI_ISL_3009083, EPI_ISL_3009084, EPI_ISL_3009085, EPI_ISL_3009086, EPI_ISL_3009087, EPI_ISL_3009088, EPI_ISL_3009089, EPI_ISL_3009090, EPI_ISL_3009091, EPI_ISL_3009092, EPI_ISL_3009093, EPI_ISL_3009094, EPI_ISL_3009095, EPI_ISL_3009096, EPI_ISL_3009097, EPI_ISL_3009098, EPI_ISL_3009099, EPI_ISL_3009100, EPI_ISL_3009101, EPI_ISL_3009102 | | | |
| see above | Clinical Reference Laboratory | Kansas Health and Environmental Lab | Mike Grose, Katherine Wiggins, Jonathan Barnell, Ben Olsen, and Phil Adam |
| EPI_ISL_3009103, EPI_ISL_3009104, EPI_ISL_3009105, EPI_ISL_3009106, EPI_ISL_3009107, EPI_ISL_3009108 | Kansas Health and Environmental Lab | Kansas Health and Environmental Lab | Mike Grose, Katherine Wiggins, Jonathan Barnell, Ben Olsen, and Phil Adam |
| EPI_ISL_3009109, EPI_ISL_3009110, EPI_ISL_3009111, EPI_ISL_3009112, EPI_ISL_3009113, EPI_ISL_3009114, EPI_ISL_3009115, EPI_ISL_3009117, EPI_ISL_3009118, EPI_ISL_3009119, EPI_ISL_3009120, EPI_ISL_3009121, EPI_ISL_3009122, EPI_ISL_3009124, EPI_ISL_3009125, EPI_ISL_3009126 | | | |
| see above | Arizona State Public Health Laboratory | Arizona State Public Health Laboratory | Trung Huynh, Jessica Escobar, Katherine Fullerton, Nobuko Fukushima, Matthew Contursi, Stacy White, Linda Getsinger, Victor Waddell |
| EPI_ISL_3009149, EPI_ISL_3009169 | Maryland Genomics, Institute for Genome Sciences, University of Maryland School of Medicine | Maryland Genomics, Institute for Genome Sciences, University of Maryland School of Medicine | Tallon, Luke J; Sadzewicz, Lisa D; Humphrys, Mike; Ott, Sandra; Roussey, Holly; Santana-Cruz, Ivette; Vavikolanu, Kranthi; Lim, Jonathan;Hazen, Tracy;Fraser, Claire M; Ravel, Jacques |
| EPI_ISL_3009176 | Wojewódzki Szpital Specjalistyczny Laboratorium Mikrobiologiczne | 1. Academic Center for Pathomorphological and Genetic-Molecular Diagnostics ltd, Bialystok, Poland 2. National Institute of Public Health - National Institute of Hygiene, Warsaw, Poland | Radosław Charkiewicz, Jacek Nikliński, Przemysław Biecek. Joanna Reszeć, Piotr Majewski, Anetta Sulewska, Piotr Karabowicz, Joanna Kiśluk, Janusz Dzięcioł, Konrad Raczkowski, Katarzyna Zacharczuk, Magdalena Nowakowska, Małgorzata Sadkowska-Todys, Tomasz Wołkowicz |
| EPI_ISL_3009180 | NZOZ Meditest Diagnostyka Medyczna | 1. Academic Center for Pathomorphological and Genetic-Molecular Diagnostics ltd, Bialystok, Poland 2. National Institute of Public Health - National Institute of Hygiene, Warsaw, Poland | Radosław Charkiewicz, Jacek Nikliński, Przemysław Biecek. Joanna Reszeć, Piotr Majewski, Anetta Sulewska, Piotr Karabowicz, Joanna Kiśluk, Janusz Dzięcioł, Konrad Raczkowski, Katarzyna Zacharczuk, Magdalena Nowakowska, Małgorzata Sadkowska-Todys, Tomasz Wołkowicz |
| EPI_ISL_3009181 | Laboratorium Epidemiologii WSSE w Szczecinie | 1. Academic Center for Pathomorphological and Genetic-Molecular Diagnostics ltd, Bialystok, Poland 2. National Institute of Public Health - National Institute of Hygiene, Warsaw, Poland | Radosław Charkiewicz, Jacek Nikliński, Przemysław Biecek. Joanna Reszeć, Piotr Majewski, Anetta Sulewska, Piotr Karabowicz, Joanna Kiśluk, Janusz Dzięcioł, Konrad Raczkowski, Katarzyna Zacharczuk, Magdalena Nowakowska, Małgorzata Sadkowska-Todys, Tomasz Wołkowicz |
| EPI_ISL_3009182 | LABORATORIUM BADAŃ KLINICZNYCH WSSE w OPOLU | 1. Academic Center for Pathomorphological and Genetic-Molecular Diagnostics ltd, Bialystok, Poland 2. National Institute of Public Health - National Institute of Hygiene, Warsaw, Poland | Radosław Charkiewicz, Jacek Nikliński, Przemysław Biecek. Joanna Reszeć, Piotr Majewski, Anetta Sulewska, Piotr Karabowicz, Joanna Kiśluk, Janusz Dzięcioł, Konrad Raczkowski, Katarzyna Zacharczuk, Magdalena Nowakowska, Małgorzata Sadkowska-Todys, Tomasz Wołkowicz |
| EPI_ISL_3009193, EPI_ISL_3009234, EPI_ISL_3009245, EPI_ISL_3009284, EPI_ISL_3009323, EPI_ISL_3009330, EPI_ISL_3009340, EPI_ISL_3009366, EPI_ISL_3009396, EPI_ISL_3009429 | TGen North | TGen North | Jolene Bowers, Heather Centner, Brett Van Tassel, Chris French, Hayley Yaglom, Darrin Lemmer, Dave Engelthaler |
| EPI_ISL_3009447 | TGen North | TGen North | "Jolene Bowers, Heather Centner, Brett Van Tassel, Chris French, Hayley Yaglom, Darrin Lemmer, Dave Engelthaler" |
| EPI_ISL_3009460, EPI_ISL_3009464 | TGen North | TGen North | Jolene Bowers, Heather Centner, Brett Van Tassel, Chris French, Hayley Yaglom, Darrin Lemmer, Dave Engelthaler |
| EPI_ISL_3009467, EPI_ISL_3009471, EPI_ISL_3009472, EPI_ISL_3009473, EPI_ISL_3009474, EPI_ISL_3009476, EPI_ISL_3009477, EPI_ISL_3009478, EPI_ISL_3009479, EPI_ISL_3009480, EPI_ISL_3009481, EPI_ISL_3009482, EPI_ISL_3009483, EPI_ISL_3009485, EPI_ISL_3009486, EPI_ISL_3009488, EPI_ISL_3009489, EPI_ISL_3009490, EPI_ISL_3009491, EPI_ISL_3009492, EPI_ISL_3009493, EPI_ISL_3009494, EPI_ISL_3009496, EPI_ISL_3009498, EPI_ISL_3009500, EPI_ISL_3009502, EPI_ISL_3009503, EPI_ISL_3009504, EPI_ISL_3009506, EPI_ISL_3009507, EPI_ISL_3009508, EPI_ISL_3009512, EPI_ISL_3009513, EPI_ISL_3009515, EPI_ISL_3009516, EPI_ISL_3009517, EPI_ISL_3009518, EPI_ISL_3009519, EPI_ISL_3009520, EPI_ISL_3009521, EPI_ISL_3009522, EPI_ISL_3009523, EPI_ISL_3009524, EPI_ISL_3009525, EPI_ISL_3009526, EPI_ISL_3009527, EPI_ISL_3009528, EPI_ISL_3009529, EPI_ISL_3009530, EPI_ISL_3009531, EPI_ISL_3009532, EPI_ISL_3009533, EPI_ISL_3009534, EPI_ISL_3009535, EPI_ISL_3009536, EPI_ISL_3009537, EPI_ISL_3009538, EPI_ISL_3009539, EPI_ISL_3009541, EPI_ISL_3009542, EPI_ISL_3009543, EPI_ISL_3009544, EPI_ISL_3009546, EPI_ISL_3009548, EPI_ISL_3009549, EPI_ISL_3009550, EPI_ISL_3009551, EPI_ISL_3009552, EPI_ISL_3009553, EPI_ISL_3009554, EPI_ISL_3009555, EPI_ISL_3009556, EPI_ISL_3009557, EPI_ISL_3009560, EPI_ISL_3009561, EPI_ISL_3009562, EPI_ISL_3009563, EPI_ISL_3009564, EPI_ISL_3009565, EPI_ISL_3009566, EPI_ISL_3009568, EPI_ISL_3009569, EPI_ISL_3009570, EPI_ISL_3009571, EPI_ISL_3009573, EPI_ISL_3009574, EPI_ISL_3009575, EPI_ISL_3009576, EPI_ISL_3009577, EPI_ISL_3009579, EPI_ISL_3009580, EPI_ISL_3009582, EPI_ISL_3009583, EPI_ISL_3009584, EPI_ISL_3009586, EPI_ISL_3009587, EPI_ISL_3009590, EPI_ISL_3009591, EPI_ISL_3009592, EPI_ISL_3009593, EPI_ISL_3009594, EPI_ISL_3009595, EPI_ISL_3009596, EPI_ISL_3009597, EPI_ISL_3009598, EPI_ISL_3009600, EPI_ISL_3009601, EPI_ISL_3009602, EPI_ISL_3009603, EPI_ISL_3009604, EPI_ISL_3009605, EPI_ISL_3009606, EPI_ISL_3009607, EPI_ISL_3009609, EPI_ISL_3009610, EPI_ISL_3009611, EPI_ISL_3009612, EPI_ISL_3009614, EPI_ISL_3009615, EPI_ISL_3009617, EPI_ISL_3009619, EPI_ISL_3009620, EPI_ISL_3009621, EPI_ISL_3009622, EPI_ISL_3009623, EPI_ISL_3009624, EPI_ISL_3009625, EPI_ISL_3009626, EPI_ISL_3009627, EPI_ISL_3009628, EPI_ISL_3009631, EPI_ISL_3009633, EPI_ISL_3009634, EPI_ISL_3009635, EPI_ISL_3009636, EPI_ISL_3009637, EPI_ISL_3009638, EPI_ISL_3009639, EPI_ISL_3009642, EPI_ISL_3009643, EPI_ISL_3009645, EPI_ISL_3009647, EPI_ISL_3009648, EPI_ISL_3009650, EPI_ISL_3009651, EPI_ISL_3009654, EPI_ISL_3009655, EPI_ISL_3009656, EPI_ISL_3009657, EPI_ISL_3009658, EPI_ISL_3009659, EPI_ISL_3009660, EPI_ISL_3009663, EPI_ISL_3009664, EPI_ISL_3009665, EPI_ISL_3009666, EPI_ISL_3009667, EPI_ISL_3009668, EPI_ISL_3009669, EPI_ISL_3009670, EPI_ISL_3009672, EPI_ISL_3009673, EPI_ISL_3009675, EPI_ISL_3009676, EPI_ISL_3009677, EPI_ISL_3009678, EPI_ISL_3009679, EPI_ISL_3009680, EPI_ISL_3009681, EPI_ISL_3009682, EPI_ISL_3009683, EPI_ISL_3009684, EPI_ISL_3009685, EPI_ISL_3009686, EPI_ISL_3009687, EPI_ISL_3009688, EPI_ISL_3009689, EPI_ISL_3009690, EPI_ISL_3009691, EPI_ISL_3009693, EPI_ISL_3009695, EPI_ISL_3009696, EPI_ISL_3009698, EPI_ISL_3009700, EPI_ISL_3009701, EPI_ISL_3009702, EPI_ISL_3009703, EPI_ISL_3009705, EPI_ISL_3009706, EPI_ISL_3009707, EPI_ISL_3009708, EPI_ISL_3009710, EPI_ISL_3009711, EPI_ISL_3009712, EPI_ISL_3009713, EPI_ISL_3009714, EPI_ISL_3009715, EPI_ISL_3009716, EPI_ISL_3009717, EPI_ISL_3009719, EPI_ISL_3009720, EPI_ISL_3009721, EPI_ISL_3009722, EPI_ISL_3009723, EPI_ISL_3009724, EPI_ISL_3009725, EPI_ISL_3009727, EPI_ISL_3009728, EPI_ISL_3009729, EPI_ISL_3009730, EPI_ISL_3009731, EPI_ISL_3009733, EPI_ISL_3009735, EPI_ISL_3009736, EPI_ISL_3009737, EPI_ISL_3009738, EPI_ISL_3009739, EPI_ISL_3009740, EPI_ISL_3009743, EPI_ISL_3009745, EPI_ISL_3009746, EPI_ISL_3009747, EPI_ISL_3009748, EPI_ISL_3009749, EPI_ISL_3009750, EPI_ISL_3009754, EPI_ISL_3009756, EPI_ISL_3009758, EPI_ISL_3009759, EPI_ISL_3009764, EPI_ISL_3009765, EPI_ISL_3009769, EPI_ISL_3009770, EPI_ISL_3009771, EPI_ISL_3009774, EPI_ISL_3009775, EPI_ISL_3009776, EPI_ISL_3009781, EPI_ISL_3009783, EPI_ISL_3009784, EPI_ISL_3009785, EPI_ISL_3009786, EPI_ISL_3009787, EPI_ISL_3009788, EPI_ISL_3009790, EPI_ISL_3009792, EPI_ISL_3009793, EPI_ISL_3009794, EPI_ISL_3009796, EPI_ISL_3009797, EPI_ISL_3009798, EPI_ISL_3009799, EPI_ISL_3009800, EPI_ISL_3009801, EPI_ISL_3009802, EPI_ISL_3009803, EPI_ISL_3009804, EPI_ISL_3009805, EPI_ISL_3009806, EPI_ISL_3009807, EPI_ISL_3009808, EPI_ISL_3009809, EPI_ISL_3009810, EPI_ISL_3009812 | | | |
| see above | Pt. Jawahar Lal Nehru Memorial Medical College, Raipur | Institute of Life Sciences - INSACOG | Sunil K. Raghav, Safal Walia, Arup Ghosh, Atimukta Jha, Amol M. Kanampalliwar, Omprakash Shiriwas, Sana Fatma, Shifu Aggarwal, Rupesh Dash, Rajeeb Swain, Punit Prasad, INSACOG Consortium, Ajay Parida |
| EPI_ISL_3009813, EPI_ISL_3009814, EPI_ISL_3009815, EPI_ISL_3009816, EPI_ISL_3009817, EPI_ISL_3009818, EPI_ISL_3009819, EPI_ISL_3009820, EPI_ISL_3009822, EPI_ISL_3009824, EPI_ISL_3009825, EPI_ISL_3009827, EPI_ISL_3009828, EPI_ISL_3009829, EPI_ISL_3009830, EPI_ISL_3009831, EPI_ISL_3009832, EPI_ISL_3009833, EPI_ISL_3009835, EPI_ISL_3009836, EPI_ISL_3009837, EPI_ISL_3009838, EPI_ISL_3009839, EPI_ISL_3009840, EPI_ISL_3009841, EPI_ISL_3009842, EPI_ISL_3009843, EPI_ISL_3009844, EPI_ISL_3009846, EPI_ISL_3009847, EPI_ISL_3009848 | | | |
| see above | Mahatma Gandhi Memorial Medical College & Hospital, Jamshedpur | Institute of Life Sciences - INSACOG | Sunil K. Raghav, Safal Walia, Arup Ghosh, Atimukta Jha, Amol M. Kanampalliwar, Omprakash Shiriwas, Sana Fatma, Shifu Aggarwal, Rupesh Dash, Rajeeb Swain, Punit Prasad, INSACOG Consortium, Ajay Parida |
| EPI_ISL_3009849, EPI_ISL_3009850, EPI_ISL_3009852, EPI_ISL_3009853, EPI_ISL_3009854, EPI_ISL_3009855, EPI_ISL_3009856, EPI_ISL_3009858, EPI_ISL_3009859, EPI_ISL_3009860, EPI_ISL_3009861, EPI_ISL_3009862, EPI_ISL_3009863, EPI_ISL_3009864, EPI_ISL_3009865, EPI_ISL_3009866, EPI_ISL_3009867, EPI_ISL_3009868, EPI_ISL_3009869, EPI_ISL_3009870, EPI_ISL_3009871, EPI_ISL_3009872, EPI_ISL_3009873, EPI_ISL_3009874, EPI_ISL_3009875, EPI_ISL_3009876, EPI_ISL_3009877, EPI_ISL_3009878, EPI_ISL_3009879, EPI_ISL_3009880, EPI_ISL_3009881, EPI_ISL_3009882, EPI_ISL_3009883, EPI_ISL_3009884, EPI_ISL_3009885, EPI_ISL_3009886, EPI_ISL_3009888, EPI_ISL_3009889, EPI_ISL_3009890, EPI_ISL_3009892, EPI_ISL_3009893, EPI_ISL_3009894, EPI_ISL_3009895, EPI_ISL_3009896, EPI_ISL_3009897, EPI_ISL_3009898, EPI_ISL_3009899, EPI_ISL_3009900, EPI_ISL_3009902, EPI_ISL_3009903, EPI_ISL_3009904, EPI_ISL_3009906, EPI_ISL_3009907, EPI_ISL_3009908, EPI_ISL_3009912, EPI_ISL_3009913, EPI_ISL_3009914, EPI_ISL_3009915, EPI_ISL_3009916, EPI_ISL_3009917, EPI_ISL_3009918, EPI_ISL_3009919, EPI_ISL_3009920, EPI_ISL_3009921, EPI_ISL_3009923, EPI_ISL_3009925, EPI_ISL_3009926, EPI_ISL_3009927, EPI_ISL_3009928, EPI_ISL_3009929, EPI_ISL_3009930, EPI_ISL_3009931, EPI_ISL_3009932, EPI_ISL_3009933, EPI_ISL_3009934, EPI_ISL_3009936, EPI_ISL_3009937, EPI_ISL_3009938, EPI_ISL_3009939, EPI_ISL_3009940, EPI_ISL_3009941, EPI_ISL_3009944, EPI_ISL_3009945, EPI_ISL_3009947, EPI_ISL_3009948, EPI_ISL_3009949, EPI_ISL_3009950, EPI_ISL_3009951, EPI_ISL_3009952, EPI_ISL_3009953, EPI_ISL_3009954, EPI_ISL_3009955, EPI_ISL_3009956, EPI_ISL_3009957, EPI_ISL_3009958, EPI_ISL_3009959, EPI_ISL_3009960, EPI_ISL_3009961, EPI_ISL_3009962, EPI_ISL_3009963, EPI_ISL_3009964, EPI_ISL_3009965, EPI_ISL_3009966, EPI_ISL_3009968, EPI_ISL_3009969, EPI_ISL_3009970, EPI_ISL_3009971, EPI_ISL_3009972, EPI_ISL_3009973, EPI_ISL_3009974, EPI_ISL_3009975, EPI_ISL_3009976, EPI_ISL_3009977, EPI_ISL_3009978, EPI_ISL_3009979, EPI_ISL_3009980, EPI_ISL_3009981, EPI_ISL_3009982, EPI_ISL_3009984, EPI_ISL_3009985, EPI_ISL_3009986, EPI_ISL_3009987, EPI_ISL_3009988, EPI_ISL_3009989, EPI_ISL_3009990, EPI_ISL_3009991, EPI_ISL_3009992, EPI_ISL_3009993, EPI_ISL_3009994, EPI_ISL_3009995 | | | |
| see above | Hospital Universitari Vall d'Hebron - Vall d'Hebron Institut de Recerca | Hospital Universitari Vall d'Hebron - Vall d'Hebron Institut de Recerca | Cristina Andrés, Maria Piñana, Alejandra González-Sánchez, Damir Garcia-Cehic, Ariadna Rando, Juliana Esperalba, Maria Gema Codina, Carla Castillo, Karen García, Rodrigo Vásquez, Maria Carmen Martin, Tomàs Pumarola, Josep Quer, Andrés Antón |
| EPI_ISL_3009996, EPI_ISL_3009997, EPI_ISL_3009998, EPI_ISL_3009999, EPI_ISL_3010000, EPI_ISL_3010001, EPI_ISL_3010002, EPI_ISL_3010003, EPI_ISL_3010004, EPI_ISL_3010005, EPI_ISL_3010006, EPI_ISL_3010007, EPI_ISL_3010008, EPI_ISL_3010009, EPI_ISL_3010010, EPI_ISL_3010011, EPI_ISL_3010012, EPI_ISL_3010013, EPI_ISL_3010014, EPI_ISL_3010015, EPI_ISL_3010022, EPI_ISL_3010023, EPI_ISL_3010024, EPI_ISL_3010025, EPI_ISL_3010026, EPI_ISL_3010027, EPI_ISL_3010028, EPI_ISL_3010029, EPI_ISL_3010030, EPI_ISL_3010031, EPI_ISL_3010032, EPI_ISL_3010033, EPI_ISL_3010034, EPI_ISL_3010035, EPI_ISL_3010036, EPI_ISL_3010037, EPI_ISL_3010038, EPI_ISL_3010039, EPI_ISL_3010040, EPI_ISL_3010041, EPI_ISL_3010042, EPI_ISL_3010043, EPI_ISL_3010044, EPI_ISL_3010045, EPI_ISL_3010046, EPI_ISL_3010047, EPI_ISL_3010048, EPI_ISL_3010049 | | | |
| see above | Hospital Universitari Arnau de Vilanova | Hospital Universitari Vall d'Hebron - Vall d'Hebron Institut de Recerca | Cristina Andrés, Maria Piñana, Alejandra González-Sánchez, Damir Garcia-Cehic, Ariadna Rando, Juliana Esperalba, Maria Gema Codina, Carla Castillo, Karen García, Rodrigo Vásquez, Maria Carmen Martin, Tomàs Pumarola, Josep Quer, Andrés Antón |
| EPI_ISL_3010056, EPI_ISL_3010057, EPI_ISL_3010058, EPI_ISL_3010059, EPI_ISL_3010060, EPI_ISL_3010061, EPI_ISL_3010062, EPI_ISL_3010063, EPI_ISL_3010064, EPI_ISL_3010065, EPI_ISL_3010067, EPI_ISL_3010068, EPI_ISL_3010069, EPI_ISL_3010071 | | | |
| see above | Hospital Universitari Joan XXIII | Hospital Universitari Joan XXIII | Gemma Recio, Cristina Gutiérrez, Clara Benavent, Carla Martín, Margarida Terrón, Ester Picó, Natalia Bastón |
| EPI_ISL_3010072, EPI_ISL_3010073, EPI_ISL_3010075, EPI_ISL_3010076, EPI_ISL_3010077, EPI_ISL_3010078, EPI_ISL_3010079, EPI_ISL_3010081, EPI_ISL_3010082, EPI_ISL_3010083 | Wyoming Public Health Laboratory | Wyoming Public Health Laboratory | Jim Mildenberger, Taylor Fearing, Channing Weber, Ashley Norberg, Chayse Rowley, Marley Goetz, Brian Dominguez, Elliot Thomasson, Sam Britz, Cari Sloma, Robert Petit, and Rob Christensen |
| EPI_ISL_3010107, EPI_ISL_3010113, EPI_ISL_3010115 | DC Public Health Lab/ Dept. of Forensic Sciences | DC Public Health Lab/ Dept. of Forensic Sciences | Janis Doss, Scott Nguyen, Elizabeth Zelaya, Sarah Scott, Connie Maza, Monica Mann, Brittany Hamilton, David Payne, Jocelyn Hauser |
| EPI_ISL_3010138 | University of Michigan Clinical Microbiology Laboratory | Lauring Lab, University of Michigan, Department of Microbiology and Immunology | Valesano |
| EPI_ISL_3010140, EPI_ISL_3010141, EPI_ISL_3010147, EPI_ISL_3010163, EPI_ISL_3010174 | IVY3 Central Lab, Vanderbilt University Medical Center | Lauring Lab, University of Michigan, Department of Microbiology and Immunology | Valesano |
| EPI_ISL_3010182, EPI_ISL_3010183, EPI_ISL_3010184, EPI_ISL_3010185, EPI_ISL_3010186, EPI_ISL_3010187, EPI_ISL_3010188, EPI_ISL_3010190, EPI_ISL_3010191, EPI_ISL_3010192, EPI_ISL_3010193, EPI_ISL_3010194, EPI_ISL_3010195, EPI_ISL_3010196, EPI_ISL_3010197, EPI_ISL_3010198, EPI_ISL_3010199, EPI_ISL_3010200, EPI_ISL_3010202, EPI_ISL_3010204, EPI_ISL_3010205, EPI_ISL_3010206, EPI_ISL_3010207, EPI_ISL_3010208, EPI_ISL_3010209, EPI_ISL_3010210, EPI_ISL_3010211, EPI_ISL_3010212, EPI_ISL_3010213, EPI_ISL_3010214, EPI_ISL_3010215, EPI_ISL_3010216, EPI_ISL_3010218, EPI_ISL_3010219, EPI_ISL_3010220, EPI_ISL_3010221, EPI_ISL_3010222, EPI_ISL_3010223, EPI_ISL_3010224, EPI_ISL_3010225, EPI_ISL_3010226, EPI_ISL_3010227, EPI_ISL_3010228, EPI_ISL_3010229, | | | |

| | | | |
|---|---|---|---|
| EPI_ISL_3010230, EPI_ISL_3010231, EPI_ISL_3010233, EPI_ISL_3010234, EPI_ISL_3010235, EPI_ISL_3010236, EPI_ISL_3010237, EPI_ISL_3010238, EPI_ISL_3010239, EPI_ISL_3010240, EPI_ISL_3010241, EPI_ISL_3010242, EPI_ISL_3010243, EPI_ISL_3010244, EPI_ISL_3010245, EPI_ISL_3010246, EPI_ISL_3010248, EPI_ISL_3010249, EPI_ISL_3010250, EPI_ISL_3010251, EPI_ISL_3010252, EPI_ISL_3010253, EPI_ISL_3010254 | | | |
| see above | Ochsner Health | Bioinfoexperts, LLC | Judy Crabtree, Amy Feehan, Rebecca Rose, David J. Nolan, Tessa LaFleur, Julia-Garcia-Diaz, Lucio Miele, Susanna L. Lamers |
| EPI_ISL_3010282 | Humboldt County Public Health Laboratory | Humboldt County Public Health Laboratory | Jeremy Corrigan |
| EPI_ISL_3010298, EPI_ISL_3010300, EPI_ISL_3010301, EPI_ISL_3010302, EPI_ISL_3010303, EPI_ISL_3010305, EPI_ISL_3010306, EPI_ISL_3010307, EPI_ISL_3010308, EPI_ISL_3010312, EPI_ISL_3010313, EPI_ISL_3010314, EPI_ISL_3010315, EPI_ISL_3010317, EPI_ISL_3010318, EPI_ISL_3010319, EPI_ISL_3010322, EPI_ISL_3010323, EPI_ISL_3010324, EPI_ISL_3010325, EPI_ISL_3010326, EPI_ISL_3010328, EPI_ISL_3010329, EPI_ISL_3010330, EPI_ISL_3010331 | | | |
| see above | Orange County Public Health Laboratory | Orange County Public Health Laboratory | Angelica Torres, Julia Wolfe |
| EPI_ISL_3010333, EPI_ISL_3010334, EPI_ISL_3010335, EPI_ISL_3010336, EPI_ISL_3010337, EPI_ISL_3010338, EPI_ISL_3010340, EPI_ISL_3010341, EPI_ISL_3010342, EPI_ISL_3010343, EPI_ISL_3010344, EPI_ISL_3010345, EPI_ISL_3010346, EPI_ISL_3010347, EPI_ISL_3010348, EPI_ISL_3010349, EPI_ISL_3010351, EPI_ISL_3010352, EPI_ISL_3010357, EPI_ISL_3010358, EPI_ISL_3010359, EPI_ISL_3010360, EPI_ISL_3010362, EPI_ISL_3010363, EPI_ISL_3010364, EPI_ISL_3010365, EPI_ISL_3010366, EPI_ISL_3010367, EPI_ISL_3010368, EPI_ISL_3010369, EPI_ISL_3010370, EPI_ISL_3010371, EPI_ISL_3010372, EPI_ISL_3010373, EPI_ISL_3010374, EPI_ISL_3010375, EPI_ISL_3010376, EPI_ISL_3010377, EPI_ISL_3010378, EPI_ISL_3010379, EPI_ISL_3010380, EPI_ISL_3010381, EPI_ISL_3010382, EPI_ISL_3010383, EPI_ISL_3010384, EPI_ISL_3010385, EPI_ISL_3010386, EPI_ISL_3010387, EPI_ISL_3010389, EPI_ISL_3010390, EPI_ISL_3010391, EPI_ISL_3010393, EPI_ISL_3010394, EPI_ISL_3010396, EPI_ISL_3010397 | | | |
| see above | Ochsner Health | Bioinfoexperts, LLC | Judy Crabtree, Amy Feehan, Rebecca Rose, David J. Nolan, Tessa LaFleur, Julia-Garcia-Diaz, Lucio Miele, Susanna L. Lamers |
| EPI_ISL_3010464, EPI_ISL_3010465, EPI_ISL_3010467, EPI_ISL_3010468, EPI_ISL_3010470, EPI_ISL_3010471, EPI_ISL_3010472 | GA Department of Public Health | GA Department of Public Health | Stacy Reeves, Jonathan Edwards, Cynthia Dixey, Tonia Parrott, Aliyah Fields, Taylor Smith |
| EPI_ISL_3010584, EPI_ISL_3010585, EPI_ISL_3010587, EPI_ISL_3010588, EPI_ISL_3010592, EPI_ISL_3010593, EPI_ISL_3010595, EPI_ISL_3010596, EPI_ISL_3010597, EPI_ISL_3010600, EPI_ISL_3010601, EPI_ISL_3010603, EPI_ISL_3010605 | | | |
| see above | Oregon State Public Health Laboratory | Oregon State Public Health Laboratory | Rafia Razzaque, Eugene Yeboah, Vanda Makris, Laura Tsaknaridis, John Fontana and Shane Sevey |
| EPI_ISL_3010620, EPI_ISL_3010621, EPI_ISL_3010622, EPI_ISL_3010623, EPI_ISL_3010626, EPI_ISL_3010627, EPI_ISL_3010628, EPI_ISL_3010629, EPI_ISL_3010630, EPI_ISL_3010633, EPI_ISL_3010634, EPI_ISL_3010635, EPI_ISL_3010636, EPI_ISL_3010637, EPI_ISL_3010638, EPI_ISL_3010639, EPI_ISL_3010640, EPI_ISL_3010641, EPI_ISL_3010642, EPI_ISL_3010643, EPI_ISL_3010644, EPI_ISL_3010647, EPI_ISL_3010650, EPI_ISL_3010651, EPI_ISL_3010652, EPI_ISL_3010654, EPI_ISL_3010655, EPI_ISL_3010656, EPI_ISL_3010657, EPI_ISL_3010658, EPI_ISL_3010659, EPI_ISL_3010660, EPI_ISL_3010661, EPI_ISL_3010662, EPI_ISL_3010663, EPI_ISL_3010664, EPI_ISL_3010665, EPI_ISL_3010666, EPI_ISL_3010667, EPI_ISL_3010669, EPI_ISL_3010671, EPI_ISL_3010673, EPI_ISL_3010674, EPI_ISL_3010675, EPI_ISL_3010676, EPI_ISL_3010677, EPI_ISL_3010679, EPI_ISL_3010681, EPI_ISL_3010682, EPI_ISL_3010683, EPI_ISL_3010684, EPI_ISL_3010685, EPI_ISL_3010686, EPI_ISL_3010687, EPI_ISL_3010688, EPI_ISL_3010690, EPI_ISL_3010691, EPI_ISL_3010692, EPI_ISL_3010694, EPI_ISL_3010695, EPI_ISL_3010697, EPI_ISL_3010698, EPI_ISL_3010699, EPI_ISL_3010700, EPI_ISL_3010701, EPI_ISL_3010702, EPI_ISL_3010703, EPI_ISL_3010704, EPI_ISL_3010705, EPI_ISL_3010706, EPI_ISL_3010707, EPI_ISL_3010708, EPI_ISL_3010709, EPI_ISL_3010710, EPI_ISL_3010711, EPI_ISL_3010712, EPI_ISL_3010713, EPI_ISL_3010716, EPI_ISL_3010717, EPI_ISL_3010718, EPI_ISL_3010719, EPI_ISL_3010720, EPI_ISL_3010721, EPI_ISL_3010723, EPI_ISL_3010725, EPI_ISL_3010726, EPI_ISL_3010728, EPI_ISL_3010729, EPI_ISL_3010730, EPI_ISL_3010731, EPI_ISL_3010732, EPI_ISL_3010733, EPI_ISL_3010734, EPI_ISL_3010735, EPI_ISL_3010736, EPI_ISL_3010737, EPI_ISL_3010738, EPI_ISL_3010739, EPI_ISL_3010740, EPI_ISL_3010743, EPI_ISL_3010744, EPI_ISL_3010745, EPI_ISL_3010746, EPI_ISL_3010750, EPI_ISL_3010751, EPI_ISL_3010752, EPI_ISL_3010753, EPI_ISL_3010755, EPI_ISL_3010756, EPI_ISL_3010758 | | | |
| see above | Pandemic Response Lab - NYC | Pandemic Response Lab, R&D | Henry Lee, Michael Hammerling, Melissa Hopkins, Cybill del Castillo, Shinyoung Clair Kang, William Ward, Pradeep Bugga, Sol Rey, Dylan Law, Isabel Fernandez Escapa, Haiping Hao, Jon Laurent |
| EPI_ISL_3010968, EPI_ISL_3010969, EPI_ISL_3010970, EPI_ISL_3010971, EPI_ISL_3010972, EPI_ISL_3010973, EPI_ISL_3010974, EPI_ISL_3010975, EPI_ISL_3010976, EPI_ISL_3010977, EPI_ISL_3010978, EPI_ISL_3010980, EPI_ISL_3010982, EPI_ISL_3010983, EPI_ISL_3010984, EPI_ISL_3010985, EPI_ISL_3010986, EPI_ISL_3010987, EPI_ISL_3010988, EPI_ISL_3010989, EPI_ISL_3010990, EPI_ISL_3010991, EPI_ISL_3010992, EPI_ISL_3010994, EPI_ISL_3010995, EPI_ISL_3010996, EPI_ISL_3010997, EPI_ISL_3010998, EPI_ISL_3010999, EPI_ISL_3011000, EPI_ISL_3011001, EPI_ISL_3011002, EPI_ISL_3011003, EPI_ISL_3011004, EPI_ISL_3011005, EPI_ISL_3011006, EPI_ISL_3011007, EPI_ISL_3011008, EPI_ISL_3011009, EPI_ISL_3011010, EPI_ISL_3011011, EPI_ISL_3011012, EPI_ISL_3011013, EPI_ISL_3011014, EPI_ISL_3011015, EPI_ISL_3011016, EPI_ISL_3011017, EPI_ISL_3011019, EPI_ISL_3011020, EPI_ISL_3011021, EPI_ISL_3011022, EPI_ISL_3011023, EPI_ISL_3011024, EPI_ISL_3011025, EPI_ISL_3011026, EPI_ISL_3011027, EPI_ISL_3011028, EPI_ISL_3011029, EPI_ISL_3011030, EPI_ISL_3011031, EPI_ISL_3011032, EPI_ISL_3011033, EPI_ISL_3011034, EPI_ISL_3011035, EPI_ISL_3011036, EPI_ISL_3011041, EPI_ISL_3011042, EPI_ISL_3011043, EPI_ISL_3011044, EPI_ISL_3011045, EPI_ISL_3011046, EPI_ISL_3011047, EPI_ISL_3011048, EPI_ISL_3011049, EPI_ISL_3011050, EPI_ISL_3011051, EPI_ISL_3011052, EPI_ISL_3011053, EPI_ISL_3011054, EPI_ISL_3011055, EPI_ISL_3011056, EPI_ISL_3011057, EPI_ISL_3011058, EPI_ISL_3011059, EPI_ISL_3011060, EPI_ISL_3011061, EPI_ISL_3011062, EPI_ISL_3011063, EPI_ISL_3011064, EPI_ISL_3011065, EPI_ISL_3011066, EPI_ISL_3011067, EPI_ISL_3011068, EPI_ISL_3011070, EPI_ISL_3011071, EPI_ISL_3011072, EPI_ISL_3011073, EPI_ISL_3011074, EPI_ISL_3011075, EPI_ISL_3011076, EPI_ISL_3011077, EPI_ISL_3011078, EPI_ISL_3011079, EPI_ISL_3011080, EPI_ISL_3011081, EPI_ISL_3011082, EPI_ISL_3011083, EPI_ISL_3011084, EPI_ISL_3011085, EPI_ISL_3011086, EPI_ISL_3011087, EPI_ISL_3011088, EPI_ISL_3011089, EPI_ISL_3011090, EPI_ISL_3011091, EPI_ISL_3011092, EPI_ISL_3011093, EPI_ISL_3011094, EPI_ISL_3011096, EPI_ISL_3011097, EPI_ISL_3011098, EPI_ISL_3011099, EPI_ISL_3011100, EPI_ISL_3011101, EPI_ISL_3011102, EPI_ISL_3011103, EPI_ISL_3011104, EPI_ISL_3011105, EPI_ISL_3011106, EPI_ISL_3011107, EPI_ISL_3011108, EPI_ISL_3011109, EPI_ISL_3011110, EPI_ISL_3011111, EPI_ISL_3011112, EPI_ISL_3011113, EPI_ISL_3011114, EPI_ISL_3011115, EPI_ISL_3011116, EPI_ISL_3011117, EPI_ISL_3011118, EPI_ISL_3011119, EPI_ISL_3011120, EPI_ISL_3011121, EPI_ISL_3011122, EPI_ISL_3011123, EPI_ISL_3011124, EPI_ISL_3011125, EPI_ISL_3011126, EPI_ISL_3011127, EPI_ISL_3011128, EPI_ISL_3011129, EPI_ISL_3011130, EPI_ISL_3011131, EPI_ISL_3011132, EPI_ISL_3011133, EPI_ISL_3011134, EPI_ISL_3011135, EPI_ISL_3011136, EPI_ISL_3011137, EPI_ISL_3011138, EPI_ISL_3011139, EPI_ISL_3011140, EPI_ISL_3011141, EPI_ISL_3011142, EPI_ISL_3011144, EPI_ISL_3011145, EPI_ISL_3011146, EPI_ISL_3011148, EPI_ISL_3011149, EPI_ISL_3011150, EPI_ISL_3011151, EPI_ISL_3011152, EPI_ISL_3011154, EPI_ISL_3011155, EPI_ISL_3011156, EPI_ISL_3011157, EPI_ISL_3011158, EPI_ISL_3011159, EPI_ISL_3011160, EPI_ISL_3011161, EPI_ISL_3011162, EPI_ISL_3011163, EPI_ISL_3011164, EPI_ISL_3011165, EPI_ISL_3011166, EPI_ISL_3011167, EPI_ISL_3011168, EPI_ISL_3011169, EPI_ISL_3011170, EPI_ISL_3011171, EPI_ISL_3011172, EPI_ISL_3011173, EPI_ISL_3011174, EPI_ISL_3011175, EPI_ISL_3011176 | | | |
| see above | Missouri State Public Health Laboratory | Missouri State Public Health Laboratory | Matthew Sinn, Joshua Barry, Ashley New |
| EPI_ISL_3011177, EPI_ISL_3011178, EPI_ISL_3011180, EPI_ISL_3011181, EPI_ISL_3011182 | Queensland Health Forensic and Scientific Services | Queensland Health Forensic and Scientific Services | Son Nguyen |
| EPI_ISL_3011186, EPI_ISL_3011187, EPI_ISL_3011240, EPI_ISL_3011255, EPI_ISL_3011260, EPI_ISL_3011270, EPI_ISL_3011271, EPI_ISL_3011272, EPI_ISL_3011273, EPI_ISL_3011274, EPI_ISL_3011275, EPI_ISL_3011276, EPI_ISL_3011277, EPI_ISL_3011278, EPI_ISL_3011279, EPI_ISL_3011281, EPI_ISL_3011282, EPI_ISL_3011283, EPI_ISL_3011286, EPI_ISL_3011289, EPI_ISL_3011295, EPI_ISL_3011296, EPI_ISL_3011298, EPI_ISL_3011301, EPI_ISL_3011302, EPI_ISL_3011303, EPI_ISL_3011304, EPI_ISL_3011305, EPI_ISL_3011306, EPI_ISL_3011307, EPI_ISL_3011308, EPI_ISL_3011309, EPI_ISL_3011310, EPI_ISL_3011311, EPI_ISL_3011312, EPI_ISL_3011313, EPI_ISL_3011314, EPI_ISL_3011315, EPI_ISL_3011316, EPI_ISL_3011318, EPI_ISL_3011319, EPI_ISL_3011320, EPI_ISL_3011321, EPI_ISL_3011322, EPI_ISL_3011323, EPI_ISL_3011324, EPI_ISL_3011325, EPI_ISL_3011327, EPI_ISL_3011328, EPI_ISL_3011329, EPI_ISL_3011330, EPI_ISL_3011335, EPI_ISL_3011353, EPI_ISL_3011365, EPI_ISL_3011366, EPI_ISL_3011368, EPI_ISL_3011369, EPI_ISL_3011370, EPI_ISL_3011371, EPI_ISL_3011372, EPI_ISL_3011374, EPI_ISL_3011375, EPI_ISL_3011376, EPI_ISL_3011377, EPI_ISL_3011378, EPI_ISL_3011379, EPI_ISL_3011381, EPI_ISL_3011382, EPI_ISL_3011383, EPI_ISL_3011384, EPI_ISL_3011385, EPI_ISL_3011386, EPI_ISL_3011387, EPI_ISL_3011388, EPI_ISL_3011389, EPI_ISL_3011390, EPI_ISL_3011391, EPI_ISL_3011392, EPI_ISL_3011393, EPI_ISL_3011395, EPI_ISL_3011396, EPI_ISL_3011397, EPI_ISL_3011398, EPI_ISL_3011399, EPI_ISL_3011400, EPI_ISL_3011401, EPI_ISL_3011402, EPI_ISL_3011403, EPI_ISL_3011404, EPI_ISL_3011405, EPI_ISL_3011406, EPI_ISL_3011407, EPI_ISL_3011408, EPI_ISL_3011409, EPI_ISL_3011410, EPI_ISL_3011411, EPI_ISL_3011412, EPI_ISL_3011413, EPI_ISL_3011414, EPI_ISL_3011415, EPI_ISL_3011416, EPI_ISL_3011417, EPI_ISL_3011418, EPI_ISL_3011419, EPI_ISL_3011420, EPI_ISL_3011421, EPI_ISL_3011422, EPI_ISL_3011423, EPI_ISL_3011424, EPI_ISL_3011425, EPI_ISL_3011426, EPI_ISL_3011427, EPI_ISL_3011428, EPI_ISL_3011429, EPI_ISL_3011430, EPI_ISL_3011431, EPI_ISL_3011432, EPI_ISL_3011433, EPI_ISL_3011434, EPI_ISL_3011436, EPI_ISL_3011437, EPI_ISL_3011438, EPI_ISL_3011439, EPI_ISL_3011440, EPI_ISL_3011441, EPI_ISL_3011443, EPI_ISL_3011444, EPI_ISL_3011446, EPI_ISL_3011447, EPI_ISL_3011448, EPI_ISL_3011449, EPI_ISL_3011450, EPI_ISL_3011454, EPI_ISL_3011456, EPI_ISL_3011457, EPI_ISL_3011458, EPI_ISL_3011460, EPI_ISL_3011461, EPI_ISL_3011462, EPI_ISL_3011463, EPI_ISL_3011465, EPI_ISL_3011466, EPI_ISL_3011467, EPI_ISL_3011469, EPI_ISL_3011471, EPI_ISL_3011472, EPI_ISL_3011473, EPI_ISL_3011474, EPI_ISL_3011475, EPI_ISL_3011476, EPI_ISL_3011477, EPI_ISL_3011478, EPI_ISL_3011479, EPI_ISL_3011480, EPI_ISL_3011482, EPI_ISL_3011483, EPI_ISL_3011487, EPI_ISL_3011490, EPI_ISL_3011491, EPI_ISL_3011492, EPI_ISL_3011495, EPI_ISL_3011496, EPI_ISL_3011498, EPI_ISL_3011500, EPI_ISL_3011504, EPI_ISL_3011507, EPI_ISL_3011509, EPI_ISL_3011510, EPI_ISL_3011511, EPI_ISL_3011512 | | | |
| see above | National Virus Reference Laboratory | National Virus Reference Laboratory | Zoe Yandle, Charlene Bennett, Gabriel Gonzalez, Michael Carr, Jonathan Dean, Cillian F De Gascun |
| EPI_ISL_3011524, EPI_ISL_3011525, EPI_ISL_3011526, EPI_ISL_3011527, EPI_ISL_3011528, EPI_ISL_3011529, EPI_ISL_3011535, EPI_ISL_3011536, EPI_ISL_3011537, EPI_ISL_3011539, EPI_ISL_3011540, EPI_ISL_3011541, EPI_ISL_3011542, EPI_ISL_3011543, EPI_ISL_3011544, EPI_ISL_3011545, EPI_ISL_3011546, EPI_ISL_3011547, EPI_ISL_3011548, EPI_ISL_3011549, EPI_ISL_3011550, EPI_ISL_3011551, EPI_ISL_3011552, EPI_ISL_3011553, EPI_ISL_3011554, EPI_ISL_3011556, EPI_ISL_3011557, EPI_ISL_3011580, EPI_ISL_3011585, EPI_ISL_3011588, EPI_ISL_3011591, EPI_ISL_3011593, EPI_ISL_3011596, EPI_ISL_3011598 | | | |
| see above | National Virus Reference Laboratory | National Virus Reference Laboratory | Guerrino Macori, Gabriel Gonzalez, Michael Carr, Zoe Yandle, Charlene Bennett, Jonathan Dean, Seamus Fanning, Cillian F De Gascun |
| EPI_ISL_3011719, EPI_ISL_3011973, EPI_ISL_3012019, EPI_ISL_3012091, EPI_ISL_3012126 | Maryland Genomics, Institute for Genome Sciences, University of Maryland School of Medicine | Maryland Genomics, Institute for Genome Sciences, University of Maryland School of Medicine | Tallon, Luke J; Sadzewicz, Lisa D; Humphrys, Mike; Ott, Sandra; Roussey, Holly; Santana-Cruz, Ivette; Vavikolanu, Kranthi; Lim, Jonathan;Hazen, Tracy;Fraser, Claire M; Ravel, Jacques |
| EPI_ISL_3012181, EPI_ISL_3012182, EPI_ISL_3012184, EPI_ISL_3012186, EPI_ISL_3012187, EPI_ISL_3012190, EPI_ISL_3012191, EPI_ISL_3012192, EPI_ISL_3012193, EPI_ISL_3012194, EPI_ISL_3012195, EPI_ISL_3012196, EPI_ISL_3012197, EPI_ISL_3012198, EPI_ISL_3012199, EPI_ISL_3012200, EPI_ISL_3012201, EPI_ISL_3012202, EPI_ISL_3012203, EPI_ISL_3012204, EPI_ISL_3012205, EPI_ISL_3012206, EPI_ISL_3012207, EPI_ISL_3012208, EPI_ISL_3012211, EPI_ISL_3012212, EPI_ISL_3012213, EPI_ISL_3012214, EPI_ISL_3012215, EPI_ISL_3012216, EPI_ISL_3012217, EPI_ISL_3012218, EPI_ISL_3012219, EPI_ISL_3012220, EPI_ISL_3012221, EPI_ISL_3012222, EPI_ISL_3012223, EPI_ISL_3012224, EPI_ISL_3012225, EPI_ISL_3012226, EPI_ISL_3012227, EPI_ISL_3012228, EPI_ISL_3012229, EPI_ISL_3012230, EPI_ISL_3012231, EPI_ISL_3012232, EPI_ISL_3012233, EPI_ISL_3012235, EPI_ISL_3012237, EPI_ISL_3012238 | | | |
| see above | Nucleic Acid Testing, National Reference Laboratory | GIGA Medical Genomics | Yvan Butera, Keith Durkin, Maria Artesi, Bouchra Boujemla, Robert Rutayisire, Reuben Sindayiheba, Patrick Tuyisenge, Esperence Umumararungu, Sébastien Bontems, Marie-Pierre Hayette, Nathalie Renotte, Swaibu Gatare, Jacob Souopgui, Sabin Nsanzimana, Vincent Bours, Léon Mutesa |
| EPI_ISL_3012245, EPI_ISL_3012246, EPI_ISL_3012247, EPI_ISL_3012248, EPI_ISL_3012249, EPI_ISL_3012250, EPI_ISL_3012251, EPI_ISL_3012255, EPI_ISL_3012257, EPI_ISL_3012258, EPI_ISL_3012260, EPI_ISL_3012261, EPI_ISL_3012262, EPI_ISL_3012263, EPI_ISL_3012264, EPI_ISL_3012265, EPI_ISL_3012266, EPI_ISL_3012267, EPI_ISL_3012268, EPI_ISL_3012269, EPI_ISL_3012270, EPI_ISL_3012271, EPI_ISL_3012272, EPI_ISL_3012274, EPI_ISL_3012275, EPI_ISL_3012276, EPI_ISL_3012277, EPI_ISL_3012279, EPI_ISL_3012280, EPI_ISL_3012281, EPI_ISL_3012282, EPI_ISL_3012283, EPI_ISL_3012285, EPI_ISL_3012299, EPI_ISL_3012324, EPI_ISL_3012325, EPI_ISL_3012326, EPI_ISL_3012327, EPI_ISL_3012328, EPI_ISL_3012332, EPI_ISL_3012335, EPI_ISL_3012336, EPI_ISL_3012338, EPI_ISL_3012339, EPI_ISL_3012340, EPI_ISL_3012341, EPI_ISL_3012342, EPI_ISL_3012343, EPI_ISL_3012344, EPI_ISL_3012345, EPI_ISL_3012346, EPI_ISL_3012347, EPI_ISL_3012349, EPI_ISL_3012350, EPI_ISL_3012351, EPI_ISL_3012352, EPI_ISL_3012353, EPI_ISL_3012355, EPI_ISL_3012356, EPI_ISL_3012357, EPI_ISL_3012358, EPI_ISL_3012360, EPI_ISL_3012364, EPI_ISL_3012365, EPI_ISL_3012366, EPI_ISL_3012368, EPI_ISL_3012369, EPI_ISL_3012370, EPI_ISL_3012371, EPI_ISL_3012373, EPI_ISL_3012376, EPI_ISL_3012377, EPI_ISL_3012378, EPI_ISL_3012379, EPI_ISL_3012380, EPI_ISL_3012381, EPI_ISL_3012382, EPI_ISL_3012383, EPI_ISL_3012385, EPI_ISL_3012386, EPI_ISL_3012387, EPI_ISL_3012388, EPI_ISL_3012389, EPI_ISL_3012390, EPI_ISL_3012391, EPI_ISL_3012392, EPI_ISL_3012393, EPI_ISL_3012394, EPI_ISL_3012395, EPI_ISL_3012396, EPI_ISL_3012397, EPI_ISL_3012398, EPI_ISL_3012399, EPI_ISL_3012400, EPI_ISL_3012401, EPI_ISL_3012402, EPI_ISL_3012404, EPI_ISL_3012415, EPI_ISL_3012449, EPI_ISL_3012450, EPI_ISL_3012452, EPI_ISL_3012453, EPI_ISL_3012455, EPI_ISL_3012458, EPI_ISL_3012459, EPI_ISL_3012461, EPI_ISL_3012462, EPI_ISL_3012463, EPI_ISL_3012464, EPI_ISL_3012465, EPI_ISL_3012466, EPI_ISL_3012467, EPI_ISL_3012468, EPI_ISL_3012470, EPI_ISL_3012471, EPI_ISL_3012475, EPI_ISL_3012476, EPI_ISL_3012477, EPI_ISL_3012479, EPI_ISL_3012481, EPI_ISL_3012482, EPI_ISL_3012483, EPI_ISL_3012484, EPI_ISL_3012485, EPI_ISL_3012486, EPI_ISL_3012487, EPI_ISL_3012488, EPI_ISL_3012489, EPI_ISL_3012490, EPI_ISL_3012491, EPI_ISL_3012492, EPI_ISL_3012493, EPI_ISL_3012494, EPI_ISL_3012495, EPI_ISL_3012496, EPI_ISL_3012497, EPI_ISL_3012498, EPI_ISL_3012499, EPI_ISL_3012500, EPI_ISL_3012501, EPI_ISL_3012502, EPI_ISL_3012503, EPI_ISL_3012504, EPI_ISL_3012505, EPI_ISL_3012506, EPI_ISL_3012507, EPI_ISL_3012508, EPI_ISL_3012509, EPI_ISL_3012510, EPI_ISL_3012511, EPI_ISL_3012512, EPI_ISL_3012513, EPI_ISL_3012514, EPI_ISL_3012515, EPI_ISL_3012516, EPI_ISL_3012517, EPI_ISL_3012519, EPI_ISL_3012520, EPI_ISL_3012521, EPI_ISL_3012522, EPI_ISL_3012523, EPI_ISL_3012524, EPI_ISL_3012525, EPI_ISL_3012526, EPI_ISL_3012527, EPI_ISL_3012528, EPI_ISL_3012529, EPI_ISL_3012531, EPI_ISL_3012533, EPI_ISL_3012534, EPI_ISL_3012535, EPI_ISL_3012536, EPI_ISL_3012537, EPI_ISL_3012540, EPI_ISL_3012541, EPI_ISL_3012543, EPI_ISL_3012544, EPI_ISL_3012545, EPI_ISL_3012546, EPI_ISL_3012547, EPI_ISL_3012548, EPI_ISL_3012549, EPI_ISL_3012550, EPI_ISL_3012551, EPI_ISL_3012552, EPI_ISL_3012553, EPI_ISL_3012554, EPI_ISL_3012563, EPI_ISL_3012564, EPI_ISL_3012570, EPI_ISL_3012571, EPI_ISL_3012572, EPI_ISL_3012573, EPI_ISL_3012574, EPI_ISL_3012575, EPI_ISL_3012576, EPI_ISL_3012577, EPI_ISL_3012578, EPI_ISL_3012579, EPI_ISL_3012580, EPI_ISL_3012581, EPI_ISL_3012589, EPI_ISL_3012590, EPI_ISL_3012591, EPI_ISL_3012592, EPI_ISL_3012593, EPI_ISL_3012594, EPI_ISL_3012595, EPI_ISL_3012599, EPI_ISL_3012601, EPI_ISL_3012602, EPI_ISL_3012603, EPI_ISL_3012604, EPI_ISL_3012605, EPI_ISL_3012606, EPI_ISL_3012607, EPI_ISL_3012608, EPI_ISL_3012609, EPI_ISL_3012610, EPI_ISL_3012611, EPI_ISL_3012612, EPI_ISL_3012613, EPI_ISL_3012614, EPI_ISL_3012616, EPI_ISL_3012617, EPI_ISL_3012619, EPI_ISL_3012620, EPI_ISL_3012621, EPI_ISL_3012622, EPI_ISL_3012623, EPI_ISL_3012624, EPI_ISL_3012625, EPI_ISL_3012627, EPI_ISL_3012628, EPI_ISL_3012629, EPI_ISL_3012636, EPI_ISL_3012637, EPI_ISL_3012638, EPI_ISL_3012639, EPI_ISL_3012640, EPI_ISL_3012641, EPI_ISL_3012642, EPI_ISL_3012646, EPI_ISL_3012649, EPI_ISL_3012650, EPI_ISL_3012651, EPI_ISL_3012655, EPI_ISL_3012656, EPI_ISL_3012657, EPI_ISL_3012658, EPI_ISL_3012659, EPI_ISL_3012660, EPI_ISL_3012661, EPI_ISL_3012662, EPI_ISL_3012663, EPI_ISL_3012664, EPI_ISL_3012666, EPI_ISL_3012667, EPI_ISL_3012668, EPI_ISL_3012669, EPI_ISL_3012670, EPI_ISL_3012671, EPI_ISL_3012672, EPI_ISL_3012673, EPI_ISL_3012674, EPI_ISL_3012675, EPI_ISL_3012676, EPI_ISL_3012677, EPI_ISL_3012678, EPI_ISL_3012679, EPI_ISL_3012680, EPI_ISL_3012681, EPI_ISL_3012682, EPI_ISL_3012683, EPI_ISL_3012684, EPI_ISL_3012685, EPI_ISL_3012686, EPI_ISL_3012687, EPI_ISL_3012688, EPI_ISL_3012689, EPI_ISL_3012691, EPI_ISL_3012692, EPI_ISL_3012693, EPI_ISL_3012694, EPI_ISL_3012695, EPI_ISL_3012696, EPI_ISL_3012697, EPI_ISL_3012698, EPI_ISL_3012699, EPI_ISL_3012700, EPI_ISL_3012701, EPI_ISL_3012702, EPI_ISL_3012703, EPI_ISL_3012704, EPI_ISL_3012705, EPI_ISL_3012706, EPI_ISL_3012707, EPI_ISL_3012708, EPI_ISL_3012709, EPI_ISL_3012710, EPI_ISL_3012711, EPI_ISL_3012713, EPI_ISL_3012714, EPI_ISL_3012715, EPI_ISL_3012716, EPI_ISL_3012717, EPI_ISL_3012718, EPI_ISL_3012719, EPI_ISL_3012720, EPI_ISL_3012721, EPI_ISL_3012722, EPI_ISL_3012723, EPI_ISL_3012724, EPI_ISL_3012725, EPI_ISL_3012726, EPI_ISL_3012727, EPI_ISL_3012728, EPI_ISL_3012729, EPI_ISL_3012730, EPI_ISL_3012731, EPI_ISL_3012732, EPI_ISL_3012734, EPI_ISL_3012735, EPI_ISL_3012738, EPI_ISL_3012739, EPI_ISL_3012740, EPI_ISL_3012741, EPI_ISL_3012742, EPI_ISL_3012743, EPI_ISL_3012744, EPI_ISL_3012745, EPI_ISL_3012747, EPI_ISL_3012748, EPI_ISL_3012749, EPI_ISL_3012751, EPI_ISL_3012752, EPI_ISL_3012753, EPI_ISL_3012754, EPI_ISL_3012755, EPI_ISL_3012756, EPI_ISL_3012757, EPI_ISL_3012758, EPI_ISL_3012759, EPI_ISL_3012760, EPI_ISL_3012761, EPI_ISL_3012762, EPI_ISL_3012763, EPI_ISL_3012764, EPI_ISL_3012765, EPI_ISL_3012766, EPI_ISL_3012767, EPI_ISL_3012768, EPI_ISL_3012769, EPI_ISL_3012773, EPI_ISL_3012775, EPI_ISL_3012776, EPI_ISL_3012777, EPI_ISL_3012779, EPI_ISL_3012780, EPI_ISL_3012781, EPI_ISL_3012782, EPI_ISL_3012783, EPI_ISL_3012784, EPI_ISL_3012788, EPI_ISL_3012789, EPI_ISL_3012790, EPI_ISL_3012792, EPI_ISL_3012794, EPI_ISL_3012796, EPI_ISL_3012797, EPI_ISL_3012798, EPI_ISL_3012799, EPI_ISL_3012802, EPI_ISL_3012803, EPI_ISL_3012804, EPI_ISL_3012805, EPI_ISL_3012806, EPI_ISL_3012807, EPI_ISL_3012808, EPI_ISL_3012809, EPI_ISL_3012812, EPI_ISL_3012813, EPI_ISL_3012814, EPI_ISL_3012817, EPI_ISL_3012818, EPI_ISL_3012819, EPI_ISL_3012820, EPI_ISL_3012821, EPI_ISL_3012822, EPI_ISL_3012823, EPI_ISL_3012824, EPI_ISL_3012825, EPI_ISL_3012826, EPI_ISL_3012827, EPI_ISL_3012828, EPI_ISL_3012829, EPI_ISL_3012830, EPI_ISL_3012831, EPI_ISL_3012832, EPI_ISL_3012833, EPI_ISL_3012834, EPI_ISL_3012835, EPI_ISL_3012836, EPI_ISL_3012840, EPI_ISL_3012841, EPI_ISL_3012842, EPI_ISL_3012845, EPI_ISL_3012846, EPI_ISL_3012847, EPI_ISL_3012849, EPI_ISL_3012850, EPI_ISL_3012851, EPI_ISL_3012857, EPI_ISL_3012858, EPI_ISL_3012859, EPI_ISL_3012864, EPI_ISL_3012865, EPI_ISL_3012866, EPI_ISL_3012868, EPI_ISL_3012869, EPI_ISL_3012870, EPI_ISL_3012871, EPI_ISL_3012872, EPI_ISL_3012874, EPI_ISL_3012875, EPI_ISL_3012877, EPI_ISL_3012878, EPI_ISL_3012879, EPI_ISL_3012880, EPI_ISL_3012881, EPI_ISL_3012882, EPI_ISL_3012883, EPI_ISL_3012884, EPI_ISL_3012885, EPI_ISL_3012886, EPI_ISL_3012887, EPI_ISL_3012888, EPI_ISL_3012889, EPI_ISL_3012890, EPI_ISL_3012891, EPI_ISL_3012892, EPI_ISL_3012893, EPI_ISL_3012894, EPI_ISL_3012895, EPI_ISL_3012896, EPI_ISL_3012897, EPI_ISL_3012898, EPI_ISL_3012899, EPI_ISL_3012900, EPI_ISL_3012901, EPI_ISL_3012902, EPI_ISL_3012903, EPI_ISL_3012904, EPI_ISL_3012905, EPI_ISL_3012906, EPI_ISL_3012907, EPI_ISL_3012908, EPI_ISL_3012909, EPI_ISL_3012910, EPI_ISL_3012911, EPI_ISL_3012912, EPI_ISL_3012913, EPI_ISL_3012914, EPI_ISL_3012916, EPI_ISL_3012917, EPI_ISL_3012918, EPI_ISL_3012919, EPI_ISL_3012920, EPI_ISL_3012923, EPI_ISL_3012925, EPI_ISL_3012926, EPI_ISL_3012927, EPI_ISL_3012928, EPI_ISL_3012929, EPI_ISL_3012930, EPI_ISL_3012931, EPI_ISL_3012932, EPI_ISL_3012933, EPI_ISL_3012934, EPI_ISL_3012935, EPI_ISL_3012936, EPI_ISL_3012937, EPI_ISL_3012938, EPI_ISL_3012939, EPI_ISL_3012940, EPI_ISL_3012941, EPI_ISL_3012942, EPI_ISL_3012943, EPI_ISL_3012944, EPI_ISL_3012946, EPI_ISL_3012947, EPI_ISL_3012957, EPI_ISL_3012958, EPI_ISL_3012959, EPI_ISL_3012960, EPI_ISL_3012961, EPI_ISL_3012962, EPI_ISL_3012963, EPI_ISL_3012964, EPI_ISL_3012965, EPI_ISL_3012966, EPI_ISL_3012967, EPI_ISL_3012969, EPI_ISL_3012970, EPI_ISL_3012971, EPI_ISL_3012972, EPI_ISL_3012973, EPI_ISL_3012974, EPI_ISL_3012975, EPI_ISL_3012976, EPI_ISL_3012977, EPI_ISL_3012978, EPI_ISL_3012979, EPI_ISL_3012980, EPI_ISL_3012981, EPI_ISL_3012982, EPI_ISL_3012983, EPI_ISL_3012984, EPI_ISL_3012985, EPI_ISL_3012986, EPI_ISL_3012987, EPI_ISL_3012988, EPI_ISL_3012991, EPI_ISL_3012992, EPI_ISL_3012993, EPI_ISL_3012995, EPI_ISL_3012998, EPI_ISL_3013000, EPI_ISL_3013002, EPI_ISL_3013004, EPI_ISL_3013008, EPI_ISL_3013010, EPI_ISL_3013011, EPI_ISL_3013012, EPI_ISL_3013013, EPI_ISL_3013014, EPI_ISL_3013015, EPI_ISL_3013016, EPI_ISL_3013018, EPI_ISL_3013021, EPI_ISL_3013022, EPI_ISL_3013024, EPI_ISL_3013025, EPI_ISL_3013026, EPI_ISL_3013027, EPI_ISL_3013028, EPI_ISL_3013029, EPI_ISL_3013030, EPI_ISL_3013031, EPI_ISL_3013032 | | | |
| see above | Public Health Ontario Laboratory | Public Health Ontario Laboratory | Vanessa G Allen, Philip Banh, Yao Chen, Richard de Borja, Alireza Eshaghi, Nahuel Fittipaldi, Christine Frantz, Jonathan B Gubbay, Jennifer L Guthrie, Lawrence Heisler, Esha Joshi, Michael Laszloffy, Aimin Li, Michael CY Li, Dean Maxwell, Sandeep Nagra, Samir N Patel, Jared Simpson, Karthikeyan Sivaraman, Ashleigh Sullivan, Yogi Sundaravadanam, Sarah Teatero, Andre Villegas, Matthew Watson, Sandra Zittermann |
| EPI_ISL_3013034, EPI_ISL_3013035, EPI_ISL_3013037, EPI_ISL_3013038, EPI_ISL_3013039 | AULSS 3 Venezia | UOSD Genetica e Citogenetica - Azienda ULSS 3 Serenissima; Istituto Zooprofilattico Sperimentale delle Venezie | Mosé Favarato, Elisa Squarcina, Laura Squarzon, Claudia Perini, Laura Bevilacqua, Adelaide Milani, Alessia Schivo, Annalisa Salviato, Elisa Palumbo, Erika Giorgia Quaranta, Luca Tassoni, Ambra Pastori, Edoardo Giussani, Alice Fusaro, Isabella Monne, Calogero Terregino, Antonia Ricci |
| EPI_ISL_3013042 | Lighthouse Lab in Alderley Park | Wellcome Sanger Institute for the COVID-19 Genomics UK (COG-UK) Consortium | Jacquelyn Wynn, Mairead Hyland, The Lighthouse Lab in Alderley Park and Alex Alderton, Roberto Amato, Jeffrey Barrett, Sonia Goncalves, Ewan Harrison, David K. Jackson, Ian Johnston, Dominic Kwiatkowski, Cordelia Langford, John Sillitoe on behalf of the Wellcome Sanger Institute COVID-19 Surveillance Team |
| EPI_ISL_3013043, | Lighthouse Laboratory Plymouth | Wellcome Sanger Institute for the COVID-19 Genomics UK (COG-UK) | Lighthouse Laboratory Plymouth and Alex Alderton, Roberto Amato, Jeffrey Barrett, Sonia Goncalves, Ewan Harrison, David K. Jackson, Ian Johnston, Dominic Kwiatkowski, Cordelia Langford, John Sillitoe on behalf of the Wellcome Sanger |

| | | | |
|---|---|---|---|
| EPI_ISL_3013044 | | Consortium | Institute COVID-19 Surveillance Team |
| EPI_ISL_3013045 | Lighthouse Lab in Milton Keynes | Wellcome Sanger Institute for the COVID-19 Genomics UK (COG-UK) Consortium | The Lighthouse Lab in Milton Keynes and Alex Alderton, Roberto Amato, Jeffrey Barrett, Sonia Goncalves, Ewan Harrison, David K. Jackson, Ian Johnston, Dominic Kwiatkowski, Cordelia Langford, John Sillitoe on behalf of the Wellcome Sanger Institute COVID-19 Surveillance Team |
| EPI_ISL_3013046 | Lighthouse Laboratory Plymouth | Wellcome Sanger Institute for the COVID-19 Genomics UK (COG-UK) Consortium | Lighthouse Laboratory Plymouth and Alex Alderton, Roberto Amato, Jeffrey Barrett, Sonia Goncalves, Ewan Harrison, David K. Jackson, Ian Johnston, Dominic Kwiatkowski, Cordelia Langford, John Sillitoe on behalf of the Wellcome Sanger Institute COVID-19 Surveillance Team |
| EPI_ISL_3013047 | Lighthouse Lab in Milton Keynes | Wellcome Sanger Institute for the COVID-19 Genomics UK (COG-UK) Consortium | The Lighthouse Lab in Milton Keynes and Alex Alderton, Roberto Amato, Jeffrey Barrett, Sonia Goncalves, Ewan Harrison, David K. Jackson, Ian Johnston, Dominic Kwiatkowski, Cordelia Langford, John Sillitoe on behalf of the Wellcome Sanger Institute COVID-19 Surveillance Team |
| EPI_ISL_3013048 | Lighthouse Laboratory Plymouth | Wellcome Sanger Institute for the COVID-19 Genomics UK (COG-UK) Consortium | Lighthouse Laboratory Plymouth and Alex Alderton, Roberto Amato, Jeffrey Barrett, Sonia Goncalves, Ewan Harrison, David K. Jackson, Ian Johnston, Dominic Kwiatkowski, Cordelia Langford, John Sillitoe on behalf of the Wellcome Sanger Institute COVID-19 Surveillance Team |
| EPI_ISL_3013049 | Lighthouse Lab in Alderley Park | Wellcome Sanger Institute for the COVID-19 Genomics UK (COG-UK) Consortium | Jacquelyn Wynn, Mairead Hyland, The Lighthouse Lab in Alderley Park and Alex Alderton, Roberto Amato, Jeffrey Barrett, Sonia Goncalves, Ewan Harrison, David K. Jackson, Ian Johnston, Dominic Kwiatkowski, Cordelia Langford, John Sillitoe on behalf of the Wellcome Sanger Institute COVID-19 Surveillance Team |
| EPI_ISL_3013050 | Lighthouse Lab in Milton Keynes | Wellcome Sanger Institute for the COVID-19 Genomics UK (COG-UK) Consortium | The Lighthouse Lab in Milton Keynes and Alex Alderton, Roberto Amato, Jeffrey Barrett, Sonia Goncalves, Ewan Harrison, David K. Jackson, Ian Johnston, Dominic Kwiatkowski, Cordelia Langford, John Sillitoe on behalf of the Wellcome Sanger Institute COVID-19 Surveillance Team |
| EPI_ISL_3013051, EPI_ISL_3013052 | Lighthouse Lab in Alderley Park | Wellcome Sanger Institute for the COVID-19 Genomics UK (COG-UK) Consortium | Jacquelyn Wynn, Mairead Hyland, The Lighthouse Lab in Alderley Park and Alex Alderton, Roberto Amato, Jeffrey Barrett, Sonia Goncalves, Ewan Harrison, David K. Jackson, Ian Johnston, Dominic Kwiatkowski, Cordelia Langford, John Sillitoe on behalf of the Wellcome Sanger Institute COVID-19 Surveillance Team |
| EPI_ISL_3013053 | Lighthouse Lab in Milton Keynes | Wellcome Sanger Institute for the COVID-19 Genomics UK (COG-UK) Consortium | The Lighthouse Lab in Milton Keynes and Alex Alderton, Roberto Amato, Jeffrey Barrett, Sonia Goncalves, Ewan Harrison, David K. Jackson, Ian Johnston, Dominic Kwiatkowski, Cordelia Langford, John Sillitoe on behalf of the Wellcome Sanger Institute COVID-19 Surveillance Team |
| EPI_ISL_3013054 | Lighthouse Lab in Glasgow | Wellcome Sanger Institute for the COVID-19 Genomics UK (COG-UK) Consortium | Harper VanSteenhouse, Yumi Kasai, David Gray, Carol Clugston, Anna Dominiczak and Alex Alderton, Roberto Amato, Jeffrey Barrett, Sonia Goncalves, Ewan Harrison, David K. Jackson, Ian Johnston, Dominic Kwiatkowski, Cordelia Langford, John Sillitoe on behalf of the Wellcome Sanger Institute COVID-19 Surveillance Team |
| EPI_ISL_3013055, EPI_ISL_3013056, EPI_ISL_3013057 | Lighthouse Lab in Alderley Park | Wellcome Sanger Institute for the COVID-19 Genomics UK (COG-UK) Consortium | Jacquelyn Wynn, Mairead Hyland, The Lighthouse Lab in Alderley Park and Alex Alderton, Roberto Amato, Jeffrey Barrett, Sonia Goncalves, Ewan Harrison, David K. Jackson, Ian Johnston, Dominic Kwiatkowski, Cordelia Langford, John Sillitoe on behalf of the Wellcome Sanger Institute COVID-19 Surveillance Team |
| EPI_ISL_3013058, EPI_ISL_3013059, EPI_ISL_3013060 | Lighthouse Laboratory Plymouth | Wellcome Sanger Institute for the COVID-19 Genomics UK (COG-UK) Consortium | Lighthouse Laboratory Plymouth and Alex Alderton, Roberto Amato, Jeffrey Barrett, Sonia Goncalves, Ewan Harrison, David K. Jackson, Ian Johnston, Dominic Kwiatkowski, Cordelia Langford, John Sillitoe on behalf of the Wellcome Sanger Institute COVID-19 Surveillance Team |
| EPI_ISL_3013061 | Lighthouse Lab in Alderley Park | Wellcome Sanger Institute for the COVID-19 Genomics UK (COG-UK) Consortium | Jacquelyn Wynn, Mairead Hyland, The Lighthouse Lab in Alderley Park and Alex Alderton, Roberto Amato, Jeffrey Barrett, Sonia Goncalves, Ewan Harrison, David K. Jackson, Ian Johnston, Dominic Kwiatkowski, Cordelia Langford, John Sillitoe on behalf of the Wellcome Sanger Institute COVID-19 Surveillance Team |
| EPI_ISL_3013062, EPI_ISL_3013063, EPI_ISL_3013064, EPI_ISL_3013065 | Lighthouse Lab in Milton Keynes | Wellcome Sanger Institute for the COVID-19 Genomics UK (COG-UK) Consortium | The Lighthouse Lab in Milton Keynes and Alex Alderton, Roberto Amato, Jeffrey Barrett, Sonia Goncalves, Ewan Harrison, David K. Jackson, Ian Johnston, Dominic Kwiatkowski, Cordelia Langford, John Sillitoe on behalf of the Wellcome Sanger Institute COVID-19 Surveillance Team |
| EPI_ISL_3013066 | Lighthouse Lab in Glasgow | Wellcome Sanger Institute for the COVID-19 Genomics UK (COG-UK) Consortium | Harper VanSteenhouse, Yumi Kasai, David Gray, Carol Clugston, Anna Dominiczak and Alex Alderton, Roberto Amato, Jeffrey Barrett, Sonia Goncalves, Ewan Harrison, David K. Jackson, Ian Johnston, Dominic Kwiatkowski, Cordelia Langford, John Sillitoe on behalf of the Wellcome Sanger Institute COVID-19 Surveillance Team |
| EPI_ISL_3013067, EPI_ISL_3013068 | Lighthouse Laboratory Plymouth | Wellcome Sanger Institute for the COVID-19 Genomics UK (COG-UK) Consortium | Lighthouse Laboratory Plymouth and Alex Alderton, Roberto Amato, Jeffrey Barrett, Sonia Goncalves, Ewan Harrison, David K. Jackson, Ian Johnston, Dominic Kwiatkowski, Cordelia Langford, John Sillitoe on behalf of the Wellcome Sanger Institute COVID-19 Surveillance Team |
| EPI_ISL_3013069, EPI_ISL_3013070 | Lighthouse Lab in Glasgow | Wellcome Sanger Institute for the COVID-19 Genomics UK (COG-UK) Consortium | Harper VanSteenhouse, Yumi Kasai, David Gray, Carol Clugston, Anna Dominiczak and Alex Alderton, Roberto Amato, Jeffrey Barrett, Sonia Goncalves, Ewan Harrison, David K. Jackson, Ian Johnston, Dominic Kwiatkowski, Cordelia Langford, John Sillitoe on behalf of the Wellcome Sanger Institute COVID-19 Surveillance Team |
| EPI_ISL_3013071 | Lighthouse Laboratory Plymouth | Wellcome Sanger Institute for the COVID-19 Genomics UK (COG-UK) Consortium | Lighthouse Laboratory Plymouth and Alex Alderton, Roberto Amato, Jeffrey Barrett, Sonia Goncalves, Ewan Harrison, David K. Jackson, Ian Johnston, Dominic Kwiatkowski, Cordelia Langford, John Sillitoe on behalf of the Wellcome Sanger Institute COVID-19 Surveillance Team |
| EPI_ISL_3013072, EPI_ISL_3013073, EPI_ISL_3013074 | Lighthouse Lab in Glasgow | Wellcome Sanger Institute for the COVID-19 Genomics UK (COG-UK) Consortium | Harper VanSteenhouse, Yumi Kasai, David Gray, Carol Clugston, Anna Dominiczak and Alex Alderton, Roberto Amato, Jeffrey Barrett, Sonia Goncalves, Ewan Harrison, David K. Jackson, Ian Johnston, Dominic Kwiatkowski, Cordelia Langford, John Sillitoe on behalf of the Wellcome Sanger Institute COVID-19 Surveillance Team |
| EPI_ISL_3013075 | Lighthouse Lab in Milton Keynes | Wellcome Sanger Institute for the COVID-19 Genomics UK (COG-UK) Consortium | The Lighthouse Lab in Milton Keynes and Alex Alderton, Roberto Amato, Jeffrey Barrett, Sonia Goncalves, Ewan Harrison, David K. Jackson, Ian Johnston, Dominic Kwiatkowski, Cordelia Langford, John Sillitoe on behalf of the Wellcome Sanger Institute COVID-19 Surveillance Team |
| EPI_ISL_3013076 | Lighthouse Laboratory Plymouth | Wellcome Sanger Institute for the COVID-19 Genomics UK (COG-UK) Consortium | Lighthouse Laboratory Plymouth and Alex Alderton, Roberto Amato, Jeffrey Barrett, Sonia Goncalves, Ewan Harrison, David K. Jackson, Ian Johnston, Dominic Kwiatkowski, Cordelia Langford, John Sillitoe on behalf of the Wellcome Sanger Institute COVID-19 Surveillance Team |
| EPI_ISL_3013077 | Lighthouse Lab in Milton Keynes | Wellcome Sanger Institute for the COVID-19 Genomics UK (COG-UK) Consortium | The Lighthouse Lab in Milton Keynes and Alex Alderton, Roberto Amato, Jeffrey Barrett, Sonia Goncalves, Ewan Harrison, David K. Jackson, Ian Johnston, Dominic Kwiatkowski, Cordelia Langford, John Sillitoe on behalf of the Wellcome Sanger Institute COVID-19 Surveillance Team |
| EPI_ISL_3013078 | Lighthouse Lab in Alderley Park | Wellcome Sanger Institute for the COVID-19 Genomics UK (COG-UK) Consortium | Jacquelyn Wynn, Mairead Hyland, The Lighthouse Lab in Alderley Park and Alex Alderton, Roberto Amato, Jeffrey Barrett, Sonia Goncalves, Ewan Harrison, David K. Jackson, Ian Johnston, Dominic Kwiatkowski, Cordelia Langford, John Sillitoe on behalf of the Wellcome Sanger Institute COVID-19 Surveillance Team |
| EPI_ISL_3013079 | Lighthouse Laboratory Plymouth | Wellcome Sanger Institute for the COVID-19 Genomics UK (COG-UK) Consortium | Lighthouse Laboratory Plymouth and Alex Alderton, Roberto Amato, Jeffrey Barrett, Sonia Goncalves, Ewan Harrison, David K. Jackson, Ian Johnston, Dominic Kwiatkowski, Cordelia Langford, John Sillitoe on behalf of the Wellcome Sanger Institute COVID-19 Surveillance Team |
| EPI_ISL_3013080 | Lighthouse Lab in Milton Keynes | Wellcome Sanger Institute for the COVID-19 Genomics UK (COG-UK) Consortium | The Lighthouse Lab in Milton Keynes and Alex Alderton, Roberto Amato, Jeffrey Barrett, Sonia Goncalves, Ewan Harrison, David K. Jackson, Ian Johnston, Dominic Kwiatkowski, Cordelia Langford, John Sillitoe on behalf of the Wellcome Sanger Institute COVID-19 Surveillance Team |
| EPI_ISL_3013081 | Lighthouse Laboratory Plymouth | Wellcome Sanger Institute for the COVID-19 Genomics UK (COG-UK) Consortium | Lighthouse Laboratory Plymouth and Alex Alderton, Roberto Amato, Jeffrey Barrett, Sonia Goncalves, Ewan Harrison, David K. Jackson, Ian Johnston, Dominic Kwiatkowski, Cordelia Langford, John Sillitoe on behalf of the Wellcome Sanger Institute COVID-19 Surveillance Team |
| EPI_ISL_3013082, EPI_ISL_3013083, EPI_ISL_3013084, EPI_ISL_3013085 | Lighthouse Lab in Milton Keynes | Wellcome Sanger Institute for the COVID-19 Genomics UK (COG-UK) Consortium | The Lighthouse Lab in Milton Keynes and Alex Alderton, Roberto Amato, Jeffrey Barrett, Sonia Goncalves, Ewan Harrison, David K. Jackson, Ian Johnston, Dominic Kwiatkowski, Cordelia Langford, John Sillitoe on behalf of the Wellcome Sanger Institute COVID-19 Surveillance Team |
| EPI_ISL_3013086, EPI_ISL_3013087 | Lighthouse Laboratory Plymouth | Wellcome Sanger Institute for the COVID-19 Genomics UK (COG-UK) Consortium | Lighthouse Laboratory Plymouth and Alex Alderton, Roberto Amato, Jeffrey Barrett, Sonia Goncalves, Ewan Harrison, David K. Jackson, Ian Johnston, Dominic Kwiatkowski, Cordelia Langford, John Sillitoe on behalf of the Wellcome Sanger Institute COVID-19 Surveillance Team |
| EPI_ISL_3013088, EPI_ISL_3013089 | Lighthouse Lab in Milton Keynes | Wellcome Sanger Institute for the COVID-19 Genomics UK (COG-UK) Consortium | The Lighthouse Lab in Milton Keynes and Alex Alderton, Roberto Amato, Jeffrey Barrett, Sonia Goncalves, Ewan Harrison, David K. Jackson, Ian Johnston, Dominic Kwiatkowski, Cordelia Langford, John Sillitoe on behalf of the Wellcome Sanger Institute COVID-19 Surveillance Team |
| EPI_ISL_3013090, EPI_ISL_3013091 | Lighthouse Lab in Alderley Park | Wellcome Sanger Institute for the COVID-19 Genomics UK (COG-UK) Consortium | Jacquelyn Wynn, Mairead Hyland, The Lighthouse Lab in Alderley Park and Alex Alderton, Roberto Amato, Jeffrey Barrett, Sonia Goncalves, Ewan Harrison, David K. Jackson, Ian Johnston, Dominic Kwiatkowski, Cordelia Langford, John Sillitoe on behalf of the Wellcome Sanger Institute COVID-19 Surveillance Team |
| EPI_ISL_3013092, EPI_ISL_3013093 | Lighthouse Laboratory Plymouth | Wellcome Sanger Institute for the COVID-19 Genomics UK (COG-UK) Consortium | Lighthouse Laboratory Plymouth and Alex Alderton, Roberto Amato, Jeffrey Barrett, Sonia Goncalves, Ewan Harrison, David K. Jackson, Ian Johnston, Dominic Kwiatkowski, Cordelia Langford, John Sillitoe on behalf of the Wellcome Sanger Institute COVID-19 Surveillance Team |
| EPI_ISL_3013094 | Lighthouse Lab in Milton Keynes | Wellcome Sanger Institute for the COVID-19 Genomics UK (COG-UK) Consortium | The Lighthouse Lab in Milton Keynes and Alex Alderton, Roberto Amato, Jeffrey Barrett, Sonia Goncalves, Ewan Harrison, David K. Jackson, Ian Johnston, Dominic Kwiatkowski, Cordelia Langford, John Sillitoe on behalf of the Wellcome Sanger Institute COVID-19 Surveillance Team |
| EPI_ISL_3013095, EPI_ISL_3013096 | Lighthouse Lab in Alderley Park | Wellcome Sanger Institute for the COVID-19 Genomics UK (COG-UK) Consortium | Jacquelyn Wynn, Mairead Hyland, The Lighthouse Lab in Alderley Park and Alex Alderton, Roberto Amato, Jeffrey Barrett, Sonia Goncalves, Ewan Harrison, David K. Jackson, Ian Johnston, Dominic Kwiatkowski, Cordelia Langford, John Sillitoe on behalf of the Wellcome Sanger Institute COVID-19 Surveillance Team |
| EPI_ISL_3013097 | Lighthouse Lab in Milton Keynes | Wellcome Sanger Institute for the COVID-19 Genomics UK (COG-UK) Consortium | The Lighthouse Lab in Milton Keynes and Alex Alderton, Roberto Amato, Jeffrey Barrett, Sonia Goncalves, Ewan Harrison, David K. Jackson, Ian Johnston, Dominic Kwiatkowski, Cordelia Langford, John Sillitoe on behalf of the Wellcome Sanger Institute COVID-19 Surveillance Team |
| EPI_ISL_3013098, EPI_ISL_3013099 | Lighthouse Lab in Alderley Park | Wellcome Sanger Institute for the COVID-19 Genomics UK (COG-UK) Consortium | Jacquelyn Wynn, Mairead Hyland, The Lighthouse Lab in Alderley Park and Alex Alderton, Roberto Amato, Jeffrey Barrett, Sonia Goncalves, Ewan Harrison, David K. Jackson, Ian Johnston, Dominic Kwiatkowski, Cordelia Langford, John Sillitoe on behalf of the Wellcome Sanger Institute COVID-19 Surveillance Team |
| EPI_ISL_3013100 | Lighthouse Lab in Milton Keynes | Wellcome Sanger Institute for the COVID-19 Genomics UK (COG-UK) Consortium | The Lighthouse Lab in Milton Keynes and Alex Alderton, Roberto Amato, Jeffrey Barrett, Sonia Goncalves, Ewan Harrison, David K. Jackson, Ian Johnston, Dominic Kwiatkowski, Cordelia Langford, John Sillitoe on behalf of the Wellcome Sanger Institute COVID-19 Surveillance Team |
| EPI_ISL_3013101 | Lighthouse Laboratory Plymouth | Wellcome Sanger Institute for the COVID-19 Genomics UK (COG-UK) Consortium | Lighthouse Laboratory Plymouth and Alex Alderton, Roberto Amato, Jeffrey Barrett, Sonia Goncalves, Ewan Harrison, David K. Jackson, Ian Johnston, Dominic Kwiatkowski, Cordelia Langford, John Sillitoe on behalf of the Wellcome Sanger Institute COVID-19 Surveillance Team |
| EPI_ISL_3013102 | Lighthouse Lab in Alderley Park | Wellcome Sanger Institute for the COVID-19 Genomics UK (COG-UK) Consortium | Jacquelyn Wynn, Mairead Hyland, The Lighthouse Lab in Alderley Park and Alex Alderton, Roberto Amato, Jeffrey Barrett, Sonia Goncalves, Ewan Harrison, David K. Jackson, Ian Johnston, Dominic Kwiatkowski, Cordelia Langford, John Sillitoe on behalf of the Wellcome Sanger Institute COVID-19 Surveillance Team |
| EPI_ISL_3013103, EPI_ISL_3013104, EPI_ISL_3013105 | Lighthouse Laboratory Plymouth | Wellcome Sanger Institute for the COVID-19 Genomics UK (COG-UK) Consortium | Lighthouse Laboratory Plymouth and Alex Alderton, Roberto Amato, Jeffrey Barrett, Sonia Goncalves, Ewan Harrison, David K. Jackson, Ian Johnston, Dominic Kwiatkowski, Cordelia Langford, John Sillitoe on behalf of the Wellcome Sanger Institute COVID-19 Surveillance Team |
| EPI_ISL_3013106 | Lighthouse Lab in Glasgow | Wellcome Sanger Institute for the COVID-19 Genomics UK (COG-UK) Consortium | Harper VanSteenhouse, Yumi Kasai, David Gray, Carol Clugston, Anna Dominiczak and Alex Alderton, Roberto Amato, Jeffrey Barrett, Sonia Goncalves, Ewan Harrison, David K. Jackson, Ian Johnston, Dominic Kwiatkowski, Cordelia Langford, John Sillitoe on behalf of the Wellcome Sanger Institute COVID-19 Surveillance Team |
| EPI_ISL_3013107 | Lighthouse Laboratory Plymouth | Wellcome Sanger Institute for the COVID-19 Genomics UK (COG-UK) Consortium | Lighthouse Laboratory Plymouth and Alex Alderton, Roberto Amato, Jeffrey Barrett, Sonia Goncalves, Ewan Harrison, David K. Jackson, Ian Johnston, Dominic Kwiatkowski, Cordelia Langford, John Sillitoe on behalf of the Wellcome Sanger Institute COVID-19 Surveillance Team |
| EPI_ISL_3013108, EPI_ISL_3013109, EPI_ISL_3013110 | Lighthouse Lab in Milton Keynes | Wellcome Sanger Institute for the COVID-19 Genomics UK (COG-UK) Consortium | The Lighthouse Lab in Milton Keynes and Alex Alderton, Roberto Amato, Jeffrey Barrett, Sonia Goncalves, Ewan Harrison, David K. Jackson, Ian Johnston, Dominic Kwiatkowski, Cordelia Langford, John Sillitoe on behalf of the Wellcome Sanger Institute COVID-19 Surveillance Team |

| | | | |
|---|---|---|---|
| EPI_ISL_3013111 | Lighthouse Laboratory Plymouth | Wellcome Sanger Institute for the COVID-19 Genomics UK (COG-UK) Consortium | Lighthouse Laboratory Plymouth and Alex Alderton, Roberto Amato, Jeffrey Barrett, Sonia Goncalves, Ewan Harrison, David K. Jackson, Ian Johnston, Dominic Kwiatkowski, Cordelia Langford, John Sillitoe on behalf of the Wellcome Sanger Institute COVID-19 Surveillance Team |
| EPI_ISL_3013112 | Lighthouse Lab in Milton Keynes | Wellcome Sanger Institute for the COVID-19 Genomics UK (COG-UK) Consortium | The Lighthouse Lab in Milton Keynes and Alex Alderton, Roberto Amato, Jeffrey Barrett, Sonia Goncalves, Ewan Harrison, David K. Jackson, Ian Johnston, Dominic Kwiatkowski, Cordelia Langford, John Sillitoe on behalf of the Wellcome Sanger Institute COVID-19 Surveillance Team |
| EPI_ISL_3013113 | Lighthouse Lab in Alderley Park | Wellcome Sanger Institute for the COVID-19 Genomics UK (COG-UK) Consortium | Jacquelyn Wynn, Mairead Hyland, The Lighthouse Lab in Alderley Park and Alex Alderton, Roberto Amato, Jeffrey Barrett, Sonia Goncalves, Ewan Harrison, David K. Jackson, Ian Johnston, Dominic Kwiatkowski, Cordelia Langford, John Sillitoe on behalf of the Wellcome Sanger Institute COVID-19 Surveillance Team |
| EPI_ISL_3013114 | Lighthouse Lab in Milton Keynes | Wellcome Sanger Institute for the COVID-19 Genomics UK (COG-UK) Consortium | The Lighthouse Lab in Milton Keynes and Alex Alderton, Roberto Amato, Jeffrey Barrett, Sonia Goncalves, Ewan Harrison, David K. Jackson, Ian Johnston, Dominic Kwiatkowski, Cordelia Langford, John Sillitoe on behalf of the Wellcome Sanger Institute COVID-19 Surveillance Team |
| EPI_ISL_3013115 | Lighthouse Lab in Glasgow | Wellcome Sanger Institute for the COVID-19 Genomics UK (COG-UK) Consortium | Harper VanSteenhouse, Yumi Kasai, David Gray, Carol Clugston, Anna Dominiczak and Alex Alderton, Roberto Amato, Jeffrey Barrett, Sonia Goncalves, Ewan Harrison, David K. Jackson, Ian Johnston, Dominic Kwiatkowski, Cordelia Langford, John Sillitoe on behalf of the Wellcome Sanger Institute COVID-19 Surveillance Team |
| EPI_ISL_3013116, EPI_ISL_3013117 | Lighthouse Lab in Alderley Park | Wellcome Sanger Institute for the COVID-19 Genomics UK (COG-UK) Consortium | Jacquelyn Wynn, Mairead Hyland, The Lighthouse Lab in Alderley Park and Alex Alderton, Roberto Amato, Jeffrey Barrett, Sonia Goncalves, Ewan Harrison, David K. Jackson, Ian Johnston, Dominic Kwiatkowski, Cordelia Langford, John Sillitoe on behalf of the Wellcome Sanger Institute COVID-19 Surveillance Team |
| EPI_ISL_3013118 | Lighthouse Lab in Glasgow | Wellcome Sanger Institute for the COVID-19 Genomics UK (COG-UK) Consortium | Harper VanSteenhouse, Yumi Kasai, David Gray, Carol Clugston, Anna Dominiczak and Alex Alderton, Roberto Amato, Jeffrey Barrett, Sonia Goncalves, Ewan Harrison, David K. Jackson, Ian Johnston, Dominic Kwiatkowski, Cordelia Langford, John Sillitoe on behalf of the Wellcome Sanger Institute COVID-19 Surveillance Team |
| EPI_ISL_3013119 | Lighthouse Lab in Milton Keynes | Wellcome Sanger Institute for the COVID-19 Genomics UK (COG-UK) Consortium | The Lighthouse Lab in Milton Keynes and Alex Alderton, Roberto Amato, Jeffrey Barrett, Sonia Goncalves, Ewan Harrison, David K. Jackson, Ian Johnston, Dominic Kwiatkowski, Cordelia Langford, John Sillitoe on behalf of the Wellcome Sanger Institute COVID-19 Surveillance Team |
| EPI_ISL_3013120 | Lighthouse Laboratory Plymouth | Wellcome Sanger Institute for the COVID-19 Genomics UK (COG-UK) Consortium | Lighthouse Laboratory Plymouth and Alex Alderton, Roberto Amato, Jeffrey Barrett, Sonia Goncalves, Ewan Harrison, David K. Jackson, Ian Johnston, Dominic Kwiatkowski, Cordelia Langford, John Sillitoe on behalf of the Wellcome Sanger Institute COVID-19 Surveillance Team |
| EPI_ISL_3013121 | Lighthouse Lab in Milton Keynes | Wellcome Sanger Institute for the COVID-19 Genomics UK (COG-UK) Consortium | The Lighthouse Lab in Milton Keynes and Alex Alderton, Roberto Amato, Jeffrey Barrett, Sonia Goncalves, Ewan Harrison, David K. Jackson, Ian Johnston, Dominic Kwiatkowski, Cordelia Langford, John Sillitoe on behalf of the Wellcome Sanger Institute COVID-19 Surveillance Team |
| EPI_ISL_3013122 | Lighthouse Laboratory Plymouth | Wellcome Sanger Institute for the COVID-19 Genomics UK (COG-UK) Consortium | Lighthouse Laboratory Plymouth and Alex Alderton, Roberto Amato, Jeffrey Barrett, Sonia Goncalves, Ewan Harrison, David K. Jackson, Ian Johnston, Dominic Kwiatkowski, Cordelia Langford, John Sillitoe on behalf of the Wellcome Sanger Institute COVID-19 Surveillance Team |
| EPI_ISL_3013123 | Lighthouse Lab in Milton Keynes | Wellcome Sanger Institute for the COVID-19 Genomics UK (COG-UK) Consortium | The Lighthouse Lab in Milton Keynes and Alex Alderton, Roberto Amato, Jeffrey Barrett, Sonia Goncalves, Ewan Harrison, David K. Jackson, Ian Johnston, Dominic Kwiatkowski, Cordelia Langford, John Sillitoe on behalf of the Wellcome Sanger Institute COVID-19 Surveillance Team |
| EPI_ISL_3013124, EPI_ISL_3013125 | Lighthouse Laboratory Plymouth | Wellcome Sanger Institute for the COVID-19 Genomics UK (COG-UK) Consortium | Lighthouse Laboratory Plymouth and Alex Alderton, Roberto Amato, Jeffrey Barrett, Sonia Goncalves, Ewan Harrison, David K. Jackson, Ian Johnston, Dominic Kwiatkowski, Cordelia Langford, John Sillitoe on behalf of the Wellcome Sanger Institute COVID-19 Surveillance Team |
| EPI_ISL_3013126, EPI_ISL_3013127 | Lighthouse Lab in Milton Keynes | Wellcome Sanger Institute for the COVID-19 Genomics UK (COG-UK) Consortium | The Lighthouse Lab in Milton Keynes and Alex Alderton, Roberto Amato, Jeffrey Barrett, Sonia Goncalves, Ewan Harrison, David K. Jackson, Ian Johnston, Dominic Kwiatkowski, Cordelia Langford, John Sillitoe on behalf of the Wellcome Sanger Institute COVID-19 Surveillance Team |
| EPI_ISL_3013128 | Lighthouse Lab in Alderley Park | Wellcome Sanger Institute for the COVID-19 Genomics UK (COG-UK) Consortium | Jacquelyn Wynn, Mairead Hyland, The Lighthouse Lab in Alderley Park and Alex Alderton, Roberto Amato, Jeffrey Barrett, Sonia Goncalves, Ewan Harrison, David K. Jackson, Ian Johnston, Dominic Kwiatkowski, Cordelia Langford, John Sillitoe on behalf of the Wellcome Sanger Institute COVID-19 Surveillance Team |
| EPI_ISL_3013129, EPI_ISL_3013130 | Lighthouse Lab in Glasgow | Wellcome Sanger Institute for the COVID-19 Genomics UK (COG-UK) Consortium | Harper VanSteenhouse, Yumi Kasai, David Gray, Carol Clugston, Anna Dominiczak and Alex Alderton, Roberto Amato, Jeffrey Barrett, Sonia Goncalves, Ewan Harrison, David K. Jackson, Ian Johnston, Dominic Kwiatkowski, Cordelia Langford, John Sillitoe on behalf of the Wellcome Sanger Institute COVID-19 Surveillance Team |
| EPI_ISL_3013131, EPI_ISL_3013132 | Lighthouse Lab in Milton Keynes | Wellcome Sanger Institute for the COVID-19 Genomics UK (COG-UK) Consortium | The Lighthouse Lab in Milton Keynes and Alex Alderton, Roberto Amato, Jeffrey Barrett, Sonia Goncalves, Ewan Harrison, David K. Jackson, Ian Johnston, Dominic Kwiatkowski, Cordelia Langford, John Sillitoe on behalf of the Wellcome Sanger Institute COVID-19 Surveillance Team |
| EPI_ISL_3013133 | Lighthouse Lab in Alderley Park | Wellcome Sanger Institute for the COVID-19 Genomics UK (COG-UK) Consortium | Jacquelyn Wynn, Mairead Hyland, The Lighthouse Lab in Alderley Park and Alex Alderton, Roberto Amato, Jeffrey Barrett, Sonia Goncalves, Ewan Harrison, David K. Jackson, Ian Johnston, Dominic Kwiatkowski, Cordelia Langford, John Sillitoe on behalf of the Wellcome Sanger Institute COVID-19 Surveillance Team |
| EPI_ISL_3013134 | Lighthouse Lab in Glasgow | Wellcome Sanger Institute for the COVID-19 Genomics UK (COG-UK) Consortium | Harper VanSteenhouse, Yumi Kasai, David Gray, Carol Clugston, Anna Dominiczak and Alex Alderton, Roberto Amato, Jeffrey Barrett, Sonia Goncalves, Ewan Harrison, David K. Jackson, Ian Johnston, Dominic Kwiatkowski, Cordelia Langford, John Sillitoe on behalf of the Wellcome Sanger Institute COVID-19 Surveillance Team |
| EPI_ISL_3013135 | Lighthouse Lab in Alderley Park | Wellcome Sanger Institute for the COVID-19 Genomics UK (COG-UK) Consortium | Jacquelyn Wynn, Mairead Hyland, The Lighthouse Lab in Alderley Park and Alex Alderton, Roberto Amato, Jeffrey Barrett, Sonia Goncalves, Ewan Harrison, David K. Jackson, Ian Johnston, Dominic Kwiatkowski, Cordelia Langford, John Sillitoe on behalf of the Wellcome Sanger Institute COVID-19 Surveillance Team |
| EPI_ISL_3013136 | Lighthouse Lab in Milton Keynes | Wellcome Sanger Institute for the COVID-19 Genomics UK (COG-UK) Consortium | The Lighthouse Lab in Milton Keynes and Alex Alderton, Roberto Amato, Jeffrey Barrett, Sonia Goncalves, Ewan Harrison, David K. Jackson, Ian Johnston, Dominic Kwiatkowski, Cordelia Langford, John Sillitoe on behalf of the Wellcome Sanger Institute COVID-19 Surveillance Team |
| EPI_ISL_3013137, EPI_ISL_3013138 | Lighthouse Lab in Alderley Park | Wellcome Sanger Institute for the COVID-19 Genomics UK (COG-UK) Consortium | Jacquelyn Wynn, Mairead Hyland, The Lighthouse Lab in Alderley Park and Alex Alderton, Roberto Amato, Jeffrey Barrett, Sonia Goncalves, Ewan Harrison, David K. Jackson, Ian Johnston, Dominic Kwiatkowski, Cordelia Langford, John Sillitoe on behalf of the Wellcome Sanger Institute COVID-19 Surveillance Team |
| EPI_ISL_3013139 | Lighthouse Laboratory Plymouth | Wellcome Sanger Institute for the COVID-19 Genomics UK (COG-UK) Consortium | Lighthouse Laboratory Plymouth and Alex Alderton, Roberto Amato, Jeffrey Barrett, Sonia Goncalves, Ewan Harrison, David K. Jackson, Ian Johnston, Dominic Kwiatkowski, Cordelia Langford, John Sillitoe on behalf of the Wellcome Sanger Institute COVID-19 Surveillance Team |
| EPI_ISL_3013140 | Lighthouse Lab in Alderley Park | Wellcome Sanger Institute for the COVID-19 Genomics UK (COG-UK) Consortium | Jacquelyn Wynn, Mairead Hyland, The Lighthouse Lab in Alderley Park and Alex Alderton, Roberto Amato, Jeffrey Barrett, Sonia Goncalves, Ewan Harrison, David K. Jackson, Ian Johnston, Dominic Kwiatkowski, Cordelia Langford, John Sillitoe on behalf of the Wellcome Sanger Institute COVID-19 Surveillance Team |
| EPI_ISL_3013141 | Lighthouse Laboratory Plymouth | Wellcome Sanger Institute for the COVID-19 Genomics UK (COG-UK) Consortium | Lighthouse Laboratory Plymouth and Alex Alderton, Roberto Amato, Jeffrey Barrett, Sonia Goncalves, Ewan Harrison, David K. Jackson, Ian Johnston, Dominic Kwiatkowski, Cordelia Langford, John Sillitoe on behalf of the Wellcome Sanger Institute COVID-19 Surveillance Team |
| EPI_ISL_3013142 | Lighthouse Lab in Milton Keynes | Wellcome Sanger Institute for the COVID-19 Genomics UK (COG-UK) Consortium | The Lighthouse Lab in Milton Keynes and Alex Alderton, Roberto Amato, Jeffrey Barrett, Sonia Goncalves, Ewan Harrison, David K. Jackson, Ian Johnston, Dominic Kwiatkowski, Cordelia Langford, John Sillitoe on behalf of the Wellcome Sanger Institute COVID-19 Surveillance Team |
| EPI_ISL_3013143 | Lighthouse Lab in Glasgow | Wellcome Sanger Institute for the COVID-19 Genomics UK (COG-UK) Consortium | Harper VanSteenhouse, Yumi Kasai, David Gray, Carol Clugston, Anna Dominiczak and Alex Alderton, Roberto Amato, Jeffrey Barrett, Sonia Goncalves, Ewan Harrison, David K. Jackson, Ian Johnston, Dominic Kwiatkowski, Cordelia Langford, John Sillitoe on behalf of the Wellcome Sanger Institute COVID-19 Surveillance Team |
| EPI_ISL_3013144 | Lighthouse Lab in Alderley Park | Wellcome Sanger Institute for the COVID-19 Genomics UK (COG-UK) Consortium | Jacquelyn Wynn, Mairead Hyland, The Lighthouse Lab in Alderley Park and Alex Alderton, Roberto Amato, Jeffrey Barrett, Sonia Goncalves, Ewan Harrison, David K. Jackson, Ian Johnston, Dominic Kwiatkowski, Cordelia Langford, John Sillitoe on behalf of the Wellcome Sanger Institute COVID-19 Surveillance Team |
| EPI_ISL_3013145 | Lighthouse Lab in Milton Keynes | Wellcome Sanger Institute for the COVID-19 Genomics UK (COG-UK) Consortium | The Lighthouse Lab in Milton Keynes and Alex Alderton, Roberto Amato, Jeffrey Barrett, Sonia Goncalves, Ewan Harrison, David K. Jackson, Ian Johnston, Dominic Kwiatkowski, Cordelia Langford, John Sillitoe on behalf of the Wellcome Sanger Institute COVID-19 Surveillance Team |
| EPI_ISL_3013146 | Lighthouse Lab in Alderley Park | Wellcome Sanger Institute for the COVID-19 Genomics UK (COG-UK) Consortium | Jacquelyn Wynn, Mairead Hyland, The Lighthouse Lab in Alderley Park and Alex Alderton, Roberto Amato, Jeffrey Barrett, Sonia Goncalves, Ewan Harrison, David K. Jackson, Ian Johnston, Dominic Kwiatkowski, Cordelia Langford, John Sillitoe on behalf of the Wellcome Sanger Institute COVID-19 Surveillance Team |
| EPI_ISL_3013147 | Lighthouse Lab in Glasgow | Wellcome Sanger Institute for the COVID-19 Genomics UK (COG-UK) Consortium | Harper VanSteenhouse, Yumi Kasai, David Gray, Carol Clugston, Anna Dominiczak and Alex Alderton, Roberto Amato, Jeffrey Barrett, Sonia Goncalves, Ewan Harrison, David K. Jackson, Ian Johnston, Dominic Kwiatkowski, Cordelia Langford, John Sillitoe on behalf of the Wellcome Sanger Institute COVID-19 Surveillance Team |
| EPI_ISL_3013148 | Lighthouse Lab in Alderley Park | Wellcome Sanger Institute for the COVID-19 Genomics UK (COG-UK) Consortium | Jacquelyn Wynn, Mairead Hyland, The Lighthouse Lab in Alderley Park and Alex Alderton, Roberto Amato, Jeffrey Barrett, Sonia Goncalves, Ewan Harrison, David K. Jackson, Ian Johnston, Dominic Kwiatkowski, Cordelia Langford, John Sillitoe on behalf of the Wellcome Sanger Institute COVID-19 Surveillance Team |
| EPI_ISL_3013149 | Lighthouse Laboratory Plymouth | Wellcome Sanger Institute for the COVID-19 Genomics UK (COG-UK) Consortium | Lighthouse Laboratory Plymouth and Alex Alderton, Roberto Amato, Jeffrey Barrett, Sonia Goncalves, Ewan Harrison, David K. Jackson, Ian Johnston, Dominic Kwiatkowski, Cordelia Langford, John Sillitoe on behalf of the Wellcome Sanger Institute COVID-19 Surveillance Team |
| EPI_ISL_3013150 | Lighthouse Lab in Glasgow | Wellcome Sanger Institute for the COVID-19 Genomics UK (COG-UK) Consortium | Harper VanSteenhouse, Yumi Kasai, David Gray, Carol Clugston, Anna Dominiczak and Alex Alderton, Roberto Amato, Jeffrey Barrett, Sonia Goncalves, Ewan Harrison, David K. Jackson, Ian Johnston, Dominic Kwiatkowski, Cordelia Langford, John Sillitoe on behalf of the Wellcome Sanger Institute COVID-19 Surveillance Team |
| EPI_ISL_3013151 | Lighthouse Lab in Milton Keynes | Wellcome Sanger Institute for the COVID-19 Genomics UK (COG-UK) Consortium | The Lighthouse Lab in Milton Keynes and Alex Alderton, Roberto Amato, Jeffrey Barrett, Sonia Goncalves, Ewan Harrison, David K. Jackson, Ian Johnston, Dominic Kwiatkowski, Cordelia Langford, John Sillitoe on behalf of the Wellcome Sanger Institute COVID-19 Surveillance Team |
| EPI_ISL_3013152, EPI_ISL_3013153 | Lighthouse Laboratory Plymouth | Wellcome Sanger Institute for the COVID-19 Genomics UK (COG-UK) Consortium | Lighthouse Laboratory Plymouth and Alex Alderton, Roberto Amato, Jeffrey Barrett, Sonia Goncalves, Ewan Harrison, David K. Jackson, Ian Johnston, Dominic Kwiatkowski, Cordelia Langford, John Sillitoe on behalf of the Wellcome Sanger Institute COVID-19 Surveillance Team |
| EPI_ISL_3013154, EPI_ISL_3013155 | Lighthouse Lab in Alderley Park | Wellcome Sanger Institute for the COVID-19 Genomics UK (COG-UK) Consortium | Jacquelyn Wynn, Mairead Hyland, The Lighthouse Lab in Alderley Park and Alex Alderton, Roberto Amato, Jeffrey Barrett, Sonia Goncalves, Ewan Harrison, David K. Jackson, Ian Johnston, Dominic Kwiatkowski, Cordelia Langford, John Sillitoe on behalf of the Wellcome Sanger Institute COVID-19 Surveillance Team |
| EPI_ISL_3013156, EPI_ISL_3013157 | Lighthouse Laboratory Plymouth | Wellcome Sanger Institute for the COVID-19 Genomics UK (COG-UK) Consortium | Lighthouse Laboratory Plymouth and Alex Alderton, Roberto Amato, Jeffrey Barrett, Sonia Goncalves, Ewan Harrison, David K. Jackson, Ian Johnston, Dominic Kwiatkowski, Cordelia Langford, John Sillitoe on behalf of the Wellcome Sanger Institute COVID-19 Surveillance Team |
| EPI_ISL_3013158 | Lighthouse Lab in Milton Keynes | Wellcome Sanger Institute for the COVID-19 Genomics UK (COG-UK) Consortium | The Lighthouse Lab in Milton Keynes and Alex Alderton, Roberto Amato, Jeffrey Barrett, Sonia Goncalves, Ewan Harrison, David K. Jackson, Ian Johnston, Dominic Kwiatkowski, Cordelia Langford, John Sillitoe on behalf of the Wellcome Sanger Institute COVID-19 Surveillance Team |
| EPI_ISL_3013159 | Lighthouse Laboratory Plymouth | Wellcome Sanger Institute for the COVID-19 Genomics UK (COG-UK) Consortium | Lighthouse Laboratory Plymouth and Alex Alderton, Roberto Amato, Jeffrey Barrett, Sonia Goncalves, Ewan Harrison, David K. Jackson, Ian Johnston, Dominic Kwiatkowski, Cordelia Langford, John Sillitoe on behalf of the Wellcome Sanger Institute COVID-19 Surveillance Team |
| EPI_ISL_3013160 | Lighthouse Lab in Milton Keynes | Wellcome Sanger Institute for the COVID-19 Genomics UK (COG-UK) Consortium | The Lighthouse Lab in Milton Keynes and Alex Alderton, Roberto Amato, Jeffrey Barrett, Sonia Goncalves, Ewan Harrison, David K. Jackson, Ian Johnston, Dominic Kwiatkowski, Cordelia Langford, John Sillitoe on behalf of the Wellcome Sanger Institute COVID-19 Surveillance Team |
| EPI_ISL_3013161 | Lighthouse Lab in Alderley Park | Wellcome Sanger Institute for the COVID-19 Genomics UK (COG-UK) Consortium | Jacquelyn Wynn, Mairead Hyland, The Lighthouse Lab in Alderley Park and Alex Alderton, Roberto Amato, Jeffrey Barrett, Sonia Goncalves, Ewan Harrison, David K. Jackson, Ian Johnston, Dominic Kwiatkowski, Cordelia Langford, John Sillitoe on behalf of the Wellcome Sanger Institute COVID-19 Surveillance Team |
| EPI_ISL_3013162, EPI_ISL_3013163 | Lighthouse Lab in Milton Keynes | Wellcome Sanger Institute for the COVID-19 Genomics UK (COG-UK) Consortium | The Lighthouse Lab in Milton Keynes and Alex Alderton, Roberto Amato, Jeffrey Barrett, Sonia Goncalves, Ewan Harrison, David K. Jackson, Ian Johnston, Dominic Kwiatkowski, Cordelia Langford, John Sillitoe on behalf of the Wellcome Sanger Institute COVID-19 Surveillance Team |
| EPI_ISL_3013164 | Lighthouse Laboratory Plymouth | Wellcome Sanger Institute for the COVID-19 Genomics UK (COG-UK) Consortium | Lighthouse Laboratory Plymouth and Alex Alderton, Roberto Amato, Jeffrey Barrett, Sonia Goncalves, Ewan Harrison, David K. Jackson, Ian Johnston, Dominic Kwiatkowski, Cordelia Langford, John Sillitoe on behalf of the Wellcome Sanger Institute COVID-19 Surveillance Team |
| EPI_ISL_3013165 | Lighthouse Lab in Alderley Park | Wellcome Sanger Institute for the COVID-19 Genomics UK (COG-UK) Consortium | Jacquelyn Wynn, Mairead Hyland, The Lighthouse Lab in Alderley Park and Alex Alderton, Roberto Amato, Jeffrey Barrett, Sonia Goncalves, Ewan Harrison, David K. Jackson, Ian Johnston, Dominic Kwiatkowski, Cordelia Langford, John Sillitoe on behalf of the Wellcome Sanger Institute COVID-19 Surveillance Team |

| | | | |
|---|---|---|---|
| EPI_ISL_3013166 | Lighthouse Lab in Milton Keynes | Wellcome Sanger Institute for the COVID-19 Genomics UK (COG-UK) Consortium | The Lighthouse Lab in Milton Keynes and Alex Alderton, Roberto Amato, Jeffrey Barrett, Sonia Goncalves, Ewan Harrison, David K. Jackson, Ian Johnston, Dominic Kwiatkowski, Cordelia Langford, John Sillitoe on behalf of the Wellcome Sanger Institute COVID-19 Surveillance Team |
| EPI_ISL_3013167 | Lighthouse Lab in Alderley Park | Wellcome Sanger Institute for the COVID-19 Genomics UK (COG-UK) Consortium | Jacquelyn Wynn, Mairead Hyland, The Lighthouse Lab in Alderley Park and Alex Alderton, Roberto Amato, Jeffrey Barrett, Sonia Goncalves, Ewan Harrison, David K. Jackson, Ian Johnston, Dominic Kwiatkowski, Cordelia Langford, John Sillitoe on behalf of the Wellcome Sanger Institute COVID-19 Surveillance Team |
| EPI_ISL_3013168 | Lighthouse Lab in Milton Keynes | Wellcome Sanger Institute for the COVID-19 Genomics UK (COG-UK) Consortium | The Lighthouse Lab in Milton Keynes and Alex Alderton, Roberto Amato, Jeffrey Barrett, Sonia Goncalves, Ewan Harrison, David K. Jackson, Ian Johnston, Dominic Kwiatkowski, Cordelia Langford, John Sillitoe on behalf of the Wellcome Sanger Institute COVID-19 Surveillance Team |
| EPI_ISL_3013169 | Lighthouse Laboratory Plymouth | Wellcome Sanger Institute for the COVID-19 Genomics UK (COG-UK) Consortium | Lighthouse Laboratory Plymouth and Alex Alderton, Roberto Amato, Jeffrey Barrett, Sonia Goncalves, Ewan Harrison, David K. Jackson, Ian Johnston, Dominic Kwiatkowski, Cordelia Langford, John Sillitoe on behalf of the Wellcome Sanger Institute COVID-19 Surveillance Team |
| EPI_ISL_3013170 | Lighthouse Lab in Glasgow | Wellcome Sanger Institute for the COVID-19 Genomics UK (COG-UK) Consortium | Harper VanSteenhouse, Yumi Kasai, David Gray, Carol Clugston, Anna Dominiczak and Alex Alderton, Roberto Amato, Jeffrey Barrett, Sonia Goncalves, Ewan Harrison, David K. Jackson, Ian Johnston, Dominic Kwiatkowski, Cordelia Langford, John Sillitoe on behalf of the Wellcome Sanger Institute COVID-19 Surveillance Team |
| EPI_ISL_3013171 | Lighthouse Lab in Milton Keynes | Wellcome Sanger Institute for the COVID-19 Genomics UK (COG-UK) Consortium | The Lighthouse Lab in Milton Keynes and Alex Alderton, Roberto Amato, Jeffrey Barrett, Sonia Goncalves, Ewan Harrison, David K. Jackson, Ian Johnston, Dominic Kwiatkowski, Cordelia Langford, John Sillitoe on behalf of the Wellcome Sanger Institute COVID-19 Surveillance Team |
| EPI_ISL_3013172 | Lighthouse Laboratory Plymouth | Wellcome Sanger Institute for the COVID-19 Genomics UK (COG-UK) Consortium | Lighthouse Laboratory Plymouth and Alex Alderton, Roberto Amato, Jeffrey Barrett, Sonia Goncalves, Ewan Harrison, David K. Jackson, Ian Johnston, Dominic Kwiatkowski, Cordelia Langford, John Sillitoe on behalf of the Wellcome Sanger Institute COVID-19 Surveillance Team |
| EPI_ISL_3013173 | Lighthouse Lab in Milton Keynes | Wellcome Sanger Institute for the COVID-19 Genomics UK (COG-UK) Consortium | The Lighthouse Lab in Milton Keynes and Alex Alderton, Roberto Amato, Jeffrey Barrett, Sonia Goncalves, Ewan Harrison, David K. Jackson, Ian Johnston, Dominic Kwiatkowski, Cordelia Langford, John Sillitoe on behalf of the Wellcome Sanger Institute COVID-19 Surveillance Team |
| EPI_ISL_3013174 | Lighthouse Lab in Alderley Park | Wellcome Sanger Institute for the COVID-19 Genomics UK (COG-UK) Consortium | Jacquelyn Wynn, Mairead Hyland, The Lighthouse Lab in Alderley Park and Alex Alderton, Roberto Amato, Jeffrey Barrett, Sonia Goncalves, Ewan Harrison, David K. Jackson, Ian Johnston, Dominic Kwiatkowski, Cordelia Langford, John Sillitoe on behalf of the Wellcome Sanger Institute COVID-19 Surveillance Team |
| EPI_ISL_3013175 | Lighthouse Laboratory Plymouth | Wellcome Sanger Institute for the COVID-19 Genomics UK (COG-UK) Consortium | Lighthouse Laboratory Plymouth and Alex Alderton, Roberto Amato, Jeffrey Barrett, Sonia Goncalves, Ewan Harrison, David K. Jackson, Ian Johnston, Dominic Kwiatkowski, Cordelia Langford, John Sillitoe on behalf of the Wellcome Sanger Institute COVID-19 Surveillance Team |
| EPI_ISL_3013176 | Lighthouse Lab in Milton Keynes | Wellcome Sanger Institute for the COVID-19 Genomics UK (COG-UK) Consortium | The Lighthouse Lab in Milton Keynes and Alex Alderton, Roberto Amato, Jeffrey Barrett, Sonia Goncalves, Ewan Harrison, David K. Jackson, Ian Johnston, Dominic Kwiatkowski, Cordelia Langford, John Sillitoe on behalf of the Wellcome Sanger Institute COVID-19 Surveillance Team |
| EPI_ISL_3013177 | Lighthouse Lab in Glasgow | Wellcome Sanger Institute for the COVID-19 Genomics UK (COG-UK) Consortium | Harper VanSteenhouse, Yumi Kasai, David Gray, Carol Clugston, Anna Dominiczak and Alex Alderton, Roberto Amato, Jeffrey Barrett, Sonia Goncalves, Ewan Harrison, David K. Jackson, Ian Johnston, Dominic Kwiatkowski, Cordelia Langford, John Sillitoe on behalf of the Wellcome Sanger Institute COVID-19 Surveillance Team |
| EPI_ISL_3013178 | Lighthouse Lab in Milton Keynes | Wellcome Sanger Institute for the COVID-19 Genomics UK (COG-UK) Consortium | The Lighthouse Lab in Milton Keynes and Alex Alderton, Roberto Amato, Jeffrey Barrett, Sonia Goncalves, Ewan Harrison, David K. Jackson, Ian Johnston, Dominic Kwiatkowski, Cordelia Langford, John Sillitoe on behalf of the Wellcome Sanger Institute COVID-19 Surveillance Team |
| EPI_ISL_3013179, EPI_ISL_3013180, EPI_ISL_3013181, EPI_ISL_3013182 | Lighthouse Lab in Alderley Park | Wellcome Sanger Institute for the COVID-19 Genomics UK (COG-UK) Consortium | Jacquelyn Wynn, Mairead Hyland, The Lighthouse Lab in Alderley Park and Alex Alderton, Roberto Amato, Jeffrey Barrett, Sonia Goncalves, Ewan Harrison, David K. Jackson, Ian Johnston, Dominic Kwiatkowski, Cordelia Langford, John Sillitoe on behalf of the Wellcome Sanger Institute COVID-19 Surveillance Team |
| EPI_ISL_3013183, EPI_ISL_3013184, EPI_ISL_3013185 | Lighthouse Lab in Milton Keynes | Wellcome Sanger Institute for the COVID-19 Genomics UK (COG-UK) Consortium | The Lighthouse Lab in Milton Keynes and Alex Alderton, Roberto Amato, Jeffrey Barrett, Sonia Goncalves, Ewan Harrison, David K. Jackson, Ian Johnston, Dominic Kwiatkowski, Cordelia Langford, John Sillitoe on behalf of the Wellcome Sanger Institute COVID-19 Surveillance Team |
| EPI_ISL_3013186, EPI_ISL_3013187, EPI_ISL_3013188 | Lighthouse Laboratory Plymouth | Wellcome Sanger Institute for the COVID-19 Genomics UK (COG-UK) Consortium | Lighthouse Laboratory Plymouth and Alex Alderton, Roberto Amato, Jeffrey Barrett, Sonia Goncalves, Ewan Harrison, David K. Jackson, Ian Johnston, Dominic Kwiatkowski, Cordelia Langford, John Sillitoe on behalf of the Wellcome Sanger Institute COVID-19 Surveillance Team |
| EPI_ISL_3013189 | Lighthouse Lab in Alderley Park | Wellcome Sanger Institute for the COVID-19 Genomics UK (COG-UK) Consortium | Jacquelyn Wynn, Mairead Hyland, The Lighthouse Lab in Alderley Park and Alex Alderton, Roberto Amato, Jeffrey Barrett, Sonia Goncalves, Ewan Harrison, David K. Jackson, Ian Johnston, Dominic Kwiatkowski, Cordelia Langford, John Sillitoe on behalf of the Wellcome Sanger Institute COVID-19 Surveillance Team |
| EPI_ISL_3013190 | Lighthouse Lab in Milton Keynes | Wellcome Sanger Institute for the COVID-19 Genomics UK (COG-UK) Consortium | The Lighthouse Lab in Milton Keynes and Alex Alderton, Roberto Amato, Jeffrey Barrett, Sonia Goncalves, Ewan Harrison, David K. Jackson, Ian Johnston, Dominic Kwiatkowski, Cordelia Langford, John Sillitoe on behalf of the Wellcome Sanger Institute COVID-19 Surveillance Team |
| EPI_ISL_3013191 | Lighthouse Laboratory Plymouth | Wellcome Sanger Institute for the COVID-19 Genomics UK (COG-UK) Consortium | Lighthouse Laboratory Plymouth and Alex Alderton, Roberto Amato, Jeffrey Barrett, Sonia Goncalves, Ewan Harrison, David K. Jackson, Ian Johnston, Dominic Kwiatkowski, Cordelia Langford, John Sillitoe on behalf of the Wellcome Sanger Institute COVID-19 Surveillance Team |
| EPI_ISL_3013192 | Lighthouse Lab in Alderley Park | Wellcome Sanger Institute for the COVID-19 Genomics UK (COG-UK) Consortium | Jacquelyn Wynn, Mairead Hyland, The Lighthouse Lab in Alderley Park and Alex Alderton, Roberto Amato, Jeffrey Barrett, Sonia Goncalves, Ewan Harrison, David K. Jackson, Ian Johnston, Dominic Kwiatkowski, Cordelia Langford, John Sillitoe on behalf of the Wellcome Sanger Institute COVID-19 Surveillance Team |
| EPI_ISL_3013193 | Lighthouse Lab in Milton Keynes | Wellcome Sanger Institute for the COVID-19 Genomics UK (COG-UK) Consortium | The Lighthouse Lab in Milton Keynes and Alex Alderton, Roberto Amato, Jeffrey Barrett, Sonia Goncalves, Ewan Harrison, David K. Jackson, Ian Johnston, Dominic Kwiatkowski, Cordelia Langford, John Sillitoe on behalf of the Wellcome Sanger Institute COVID-19 Surveillance Team |
| EPI_ISL_3013194 | Lighthouse Laboratory Plymouth | Wellcome Sanger Institute for the COVID-19 Genomics UK (COG-UK) Consortium | Lighthouse Laboratory Plymouth and Alex Alderton, Roberto Amato, Jeffrey Barrett, Sonia Goncalves, Ewan Harrison, David K. Jackson, Ian Johnston, Dominic Kwiatkowski, Cordelia Langford, John Sillitoe on behalf of the Wellcome Sanger Institute COVID-19 Surveillance Team |
| EPI_ISL_3013195 | Lighthouse Lab in Alderley Park | Wellcome Sanger Institute for the COVID-19 Genomics UK (COG-UK) Consortium | Jacquelyn Wynn, Mairead Hyland, The Lighthouse Lab in Alderley Park and Alex Alderton, Roberto Amato, Jeffrey Barrett, Sonia Goncalves, Ewan Harrison, David K. Jackson, Ian Johnston, Dominic Kwiatkowski, Cordelia Langford, John Sillitoe on behalf of the Wellcome Sanger Institute COVID-19 Surveillance Team |
| EPI_ISL_3013196 | Lighthouse Lab in Milton Keynes | Wellcome Sanger Institute for the COVID-19 Genomics UK (COG-UK) Consortium | The Lighthouse Lab in Milton Keynes and Alex Alderton, Roberto Amato, Jeffrey Barrett, Sonia Goncalves, Ewan Harrison, David K. Jackson, Ian Johnston, Dominic Kwiatkowski, Cordelia Langford, John Sillitoe on behalf of the Wellcome Sanger Institute COVID-19 Surveillance Team |
| EPI_ISL_3013197 | Lighthouse Laboratory Plymouth | Wellcome Sanger Institute for the COVID-19 Genomics UK (COG-UK) Consortium | Lighthouse Laboratory Plymouth and Alex Alderton, Roberto Amato, Jeffrey Barrett, Sonia Goncalves, Ewan Harrison, David K. Jackson, Ian Johnston, Dominic Kwiatkowski, Cordelia Langford, John Sillitoe on behalf of the Wellcome Sanger Institute COVID-19 Surveillance Team |
| EPI_ISL_3013198 | Lighthouse Lab in Glasgow | Wellcome Sanger Institute for the COVID-19 Genomics UK (COG-UK) Consortium | Harper VanSteenhouse, Yumi Kasai, David Gray, Carol Clugston, Anna Dominiczak and Alex Alderton, Roberto Amato, Jeffrey Barrett, Sonia Goncalves, Ewan Harrison, David K. Jackson, Ian Johnston, Dominic Kwiatkowski, Cordelia Langford, John Sillitoe on behalf of the Wellcome Sanger Institute COVID-19 Surveillance Team |
| EPI_ISL_3013199 | Lighthouse Laboratory Plymouth | Wellcome Sanger Institute for the COVID-19 Genomics UK (COG-UK) Consortium | Lighthouse Laboratory Plymouth and Alex Alderton, Roberto Amato, Jeffrey Barrett, Sonia Goncalves, Ewan Harrison, David K. Jackson, Ian Johnston, Dominic Kwiatkowski, Cordelia Langford, John Sillitoe on behalf of the Wellcome Sanger Institute COVID-19 Surveillance Team |
| EPI_ISL_3013200 | Lighthouse Lab in Glasgow | Wellcome Sanger Institute for the COVID-19 Genomics UK (COG-UK) Consortium | Harper VanSteenhouse, Yumi Kasai, David Gray, Carol Clugston, Anna Dominiczak and Alex Alderton, Roberto Amato, Jeffrey Barrett, Sonia Goncalves, Ewan Harrison, David K. Jackson, Ian Johnston, Dominic Kwiatkowski, Cordelia Langford, John Sillitoe on behalf of the Wellcome Sanger Institute COVID-19 Surveillance Team |
| EPI_ISL_3013201 | Lighthouse Lab in Milton Keynes | Wellcome Sanger Institute for the COVID-19 Genomics UK (COG-UK) Consortium | The Lighthouse Lab in Milton Keynes and Alex Alderton, Roberto Amato, Jeffrey Barrett, Sonia Goncalves, Ewan Harrison, David K. Jackson, Ian Johnston, Dominic Kwiatkowski, Cordelia Langford, John Sillitoe on behalf of the Wellcome Sanger Institute COVID-19 Surveillance Team |
| EPI_ISL_3013202, EPI_ISL_3013203, EPI_ISL_3013204 | Lighthouse Lab in Glasgow | Wellcome Sanger Institute for the COVID-19 Genomics UK (COG-UK) Consortium | Harper VanSteenhouse, Yumi Kasai, David Gray, Carol Clugston, Anna Dominiczak and Alex Alderton, Roberto Amato, Jeffrey Barrett, Sonia Goncalves, Ewan Harrison, David K. Jackson, Ian Johnston, Dominic Kwiatkowski, Cordelia Langford, John Sillitoe on behalf of the Wellcome Sanger Institute COVID-19 Surveillance Team |
| EPI_ISL_3013205, EPI_ISL_3013206, EPI_ISL_3013207, EPI_ISL_3013208, EPI_ISL_3013209 | Lighthouse Lab in Milton Keynes | Wellcome Sanger Institute for the COVID-19 Genomics UK (COG-UK) Consortium | The Lighthouse Lab in Milton Keynes and Alex Alderton, Roberto Amato, Jeffrey Barrett, Sonia Goncalves, Ewan Harrison, David K. Jackson, Ian Johnston, Dominic Kwiatkowski, Cordelia Langford, John Sillitoe on behalf of the Wellcome Sanger Institute COVID-19 Surveillance Team |
| EPI_ISL_3013210 | Lighthouse Lab in Alderley Park | Wellcome Sanger Institute for the COVID-19 Genomics UK (COG-UK) Consortium | Jacquelyn Wynn, Mairead Hyland, The Lighthouse Lab in Alderley Park and Alex Alderton, Roberto Amato, Jeffrey Barrett, Sonia Goncalves, Ewan Harrison, David K. Jackson, Ian Johnston, Dominic Kwiatkowski, Cordelia Langford, John Sillitoe on behalf of the Wellcome Sanger Institute COVID-19 Surveillance Team |
| EPI_ISL_3013211, EPI_ISL_3013212 | Lighthouse Laboratory Plymouth | Wellcome Sanger Institute for the COVID-19 Genomics UK (COG-UK) Consortium | Lighthouse Laboratory Plymouth and Alex Alderton, Roberto Amato, Jeffrey Barrett, Sonia Goncalves, Ewan Harrison, David K. Jackson, Ian Johnston, Dominic Kwiatkowski, Cordelia Langford, John Sillitoe on behalf of the Wellcome Sanger Institute COVID-19 Surveillance Team |
| EPI_ISL_3013213 | Lighthouse Lab in Alderley Park | Wellcome Sanger Institute for the COVID-19 Genomics UK (COG-UK) Consortium | Jacquelyn Wynn, Mairead Hyland, The Lighthouse Lab in Alderley Park and Alex Alderton, Roberto Amato, Jeffrey Barrett, Sonia Goncalves, Ewan Harrison, David K. Jackson, Ian Johnston, Dominic Kwiatkowski, Cordelia Langford, John Sillitoe on behalf of the Wellcome Sanger Institute COVID-19 Surveillance Team |
| EPI_ISL_3013214 | Lighthouse Laboratory Plymouth | Wellcome Sanger Institute for the COVID-19 Genomics UK (COG-UK) Consortium | Lighthouse Laboratory Plymouth and Alex Alderton, Roberto Amato, Jeffrey Barrett, Sonia Goncalves, Ewan Harrison, David K. Jackson, Ian Johnston, Dominic Kwiatkowski, Cordelia Langford, John Sillitoe on behalf of the Wellcome Sanger Institute COVID-19 Surveillance Team |
| EPI_ISL_3013215, EPI_ISL_3013216 | Lighthouse Lab in Alderley Park | Wellcome Sanger Institute for the COVID-19 Genomics UK (COG-UK) Consortium | Jacquelyn Wynn, Mairead Hyland, The Lighthouse Lab in Alderley Park and Alex Alderton, Roberto Amato, Jeffrey Barrett, Sonia Goncalves, Ewan Harrison, David K. Jackson, Ian Johnston, Dominic Kwiatkowski, Cordelia Langford, John Sillitoe on behalf of the Wellcome Sanger Institute COVID-19 Surveillance Team |
| EPI_ISL_3013217, EPI_ISL_3013218 | Lighthouse Laboratory Plymouth | Wellcome Sanger Institute for the COVID-19 Genomics UK (COG-UK) Consortium | Lighthouse Laboratory Plymouth and Alex Alderton, Roberto Amato, Jeffrey Barrett, Sonia Goncalves, Ewan Harrison, David K. Jackson, Ian Johnston, Dominic Kwiatkowski, Cordelia Langford, John Sillitoe on behalf of the Wellcome Sanger Institute COVID-19 Surveillance Team |
| EPI_ISL_3013219 | Lighthouse Lab in Glasgow | Wellcome Sanger Institute for the COVID-19 Genomics UK (COG-UK) Consortium | Harper VanSteenhouse, Yumi Kasai, David Gray, Carol Clugston, Anna Dominiczak and Alex Alderton, Roberto Amato, Jeffrey Barrett, Sonia Goncalves, Ewan Harrison, David K. Jackson, Ian Johnston, Dominic Kwiatkowski, Cordelia Langford, John Sillitoe on behalf of the Wellcome Sanger Institute COVID-19 Surveillance Team |
| EPI_ISL_3013220 | Lighthouse Lab in Milton Keynes | Wellcome Sanger Institute for the COVID-19 Genomics UK (COG-UK) Consortium | The Lighthouse Lab in Milton Keynes and Alex Alderton, Roberto Amato, Jeffrey Barrett, Sonia Goncalves, Ewan Harrison, David K. Jackson, Ian Johnston, Dominic Kwiatkowski, Cordelia Langford, John Sillitoe on behalf of the Wellcome Sanger Institute COVID-19 Surveillance Team |
| EPI_ISL_3013221 | Lighthouse Lab in Glasgow | Wellcome Sanger Institute for the COVID-19 Genomics UK (COG-UK) Consortium | Harper VanSteenhouse, Yumi Kasai, David Gray, Carol Clugston, Anna Dominiczak and Alex Alderton, Roberto Amato, Jeffrey Barrett, Sonia Goncalves, Ewan Harrison, David K. Jackson, Ian Johnston, Dominic Kwiatkowski, Cordelia Langford, John Sillitoe on behalf of the Wellcome Sanger Institute COVID-19 Surveillance Team |
| EPI_ISL_3013222 | Lighthouse Lab in Milton Keynes | Wellcome Sanger Institute for the COVID-19 Genomics UK (COG-UK) Consortium | The Lighthouse Lab in Milton Keynes and Alex Alderton, Roberto Amato, Jeffrey Barrett, Sonia Goncalves, Ewan Harrison, David K. Jackson, Ian Johnston, Dominic Kwiatkowski, Cordelia Langford, John Sillitoe on behalf of the Wellcome Sanger Institute COVID-19 Surveillance Team |
| EPI_ISL_3013223 | Lighthouse Lab in Glasgow | Wellcome Sanger Institute for the COVID-19 Genomics UK (COG-UK) Consortium | Harper VanSteenhouse, Yumi Kasai, David Gray, Carol Clugston, Anna Dominiczak and Alex Alderton, Roberto Amato, Jeffrey Barrett, Sonia Goncalves, Ewan Harrison, David K. Jackson, Ian Johnston, Dominic Kwiatkowski, Cordelia Langford, John Sillitoe on behalf of the Wellcome Sanger Institute COVID-19 Surveillance Team |

| | | | |
|---|---|---|---|
| EPI_ISL_3013224 | Lighthouse Laboratory Plymouth | Wellcome Sanger Institute for the COVID-19 Genomics UK (COG-UK) Consortium | Lighthouse Laboratory Plymouth and Alex Alderton, Roberto Amato, Jeffrey Barrett, Sonia Goncalves, Ewan Harrison, David K. Jackson, Ian Johnston, Dominic Kwiatkowski, Cordelia Langford, John Sillitoe on behalf of the Wellcome Sanger Institute COVID-19 Surveillance Team |
| EPI_ISL_3013225 | Lighthouse Lab in Alderley Park | Wellcome Sanger Institute for the COVID-19 Genomics UK (COG-UK) Consortium | Jacquelyn Wynn, Mairead Hyland, The Lighthouse Lab in Alderley Park and Alex Alderton, Roberto Amato, Jeffrey Barrett, Sonia Goncalves, Ewan Harrison, David K. Jackson, Ian Johnston, Dominic Kwiatkowski, Cordelia Langford, John Sillitoe on behalf of the Wellcome Sanger Institute COVID-19 Surveillance Team |
| EPI_ISL_3013226 | Lighthouse Laboratory Plymouth | Wellcome Sanger Institute for the COVID-19 Genomics UK (COG-UK) Consortium | Lighthouse Laboratory Plymouth and Alex Alderton, Roberto Amato, Jeffrey Barrett, Sonia Goncalves, Ewan Harrison, David K. Jackson, Ian Johnston, Dominic Kwiatkowski, Cordelia Langford, John Sillitoe on behalf of the Wellcome Sanger Institute COVID-19 Surveillance Team |
| EPI_ISL_3013227 | Lighthouse Lab in Milton Keynes | Wellcome Sanger Institute for the COVID-19 Genomics UK (COG-UK) Consortium | The Lighthouse Lab in Milton Keynes and Alex Alderton, Roberto Amato, Jeffrey Barrett, Sonia Goncalves, Ewan Harrison, David K. Jackson, Ian Johnston, Dominic Kwiatkowski, Cordelia Langford, John Sillitoe on behalf of the Wellcome Sanger Institute COVID-19 Surveillance Team |
| EPI_ISL_3013228, EPI_ISL_3013229 | Lighthouse Lab in Glasgow | Wellcome Sanger Institute for the COVID-19 Genomics UK (COG-UK) Consortium | Harper VanSteenhouse, Yumi Kasai, David Gray, Carol Clugston, Anna Dominiczak and Alex Alderton, Roberto Amato, Jeffrey Barrett, Sonia Goncalves, Ewan Harrison, David K. Jackson, Ian Johnston, Dominic Kwiatkowski, Cordelia Langford, John Sillitoe on behalf of the Wellcome Sanger Institute COVID-19 Surveillance Team |
| EPI_ISL_3013230 | Lighthouse Lab in Alderley Park | Wellcome Sanger Institute for the COVID-19 Genomics UK (COG-UK) Consortium | Jacquelyn Wynn, Mairead Hyland, The Lighthouse Lab in Alderley Park and Alex Alderton, Roberto Amato, Jeffrey Barrett, Sonia Goncalves, Ewan Harrison, David K. Jackson, Ian Johnston, Dominic Kwiatkowski, Cordelia Langford, John Sillitoe on behalf of the Wellcome Sanger Institute COVID-19 Surveillance Team |
| EPI_ISL_3013231, EPI_ISL_3013232 | Lighthouse Lab in Glasgow | Wellcome Sanger Institute for the COVID-19 Genomics UK (COG-UK) Consortium | Harper VanSteenhouse, Yumi Kasai, David Gray, Carol Clugston, Anna Dominiczak and Alex Alderton, Roberto Amato, Jeffrey Barrett, Sonia Goncalves, Ewan Harrison, David K. Jackson, Ian Johnston, Dominic Kwiatkowski, Cordelia Langford, John Sillitoe on behalf of the Wellcome Sanger Institute COVID-19 Surveillance Team |
| EPI_ISL_3013233 | Lighthouse Lab in Milton Keynes | Wellcome Sanger Institute for the COVID-19 Genomics UK (COG-UK) Consortium | The Lighthouse Lab in Milton Keynes and Alex Alderton, Roberto Amato, Jeffrey Barrett, Sonia Goncalves, Ewan Harrison, David K. Jackson, Ian Johnston, Dominic Kwiatkowski, Cordelia Langford, John Sillitoe on behalf of the Wellcome Sanger Institute COVID-19 Surveillance Team |
| EPI_ISL_3013234 | Lighthouse Lab in Glasgow | Wellcome Sanger Institute for the COVID-19 Genomics UK (COG-UK) Consortium | Harper VanSteenhouse, Yumi Kasai, David Gray, Carol Clugston, Anna Dominiczak and Alex Alderton, Roberto Amato, Jeffrey Barrett, Sonia Goncalves, Ewan Harrison, David K. Jackson, Ian Johnston, Dominic Kwiatkowski, Cordelia Langford, John Sillitoe on behalf of the Wellcome Sanger Institute COVID-19 Surveillance Team |
| EPI_ISL_3013235 | Lighthouse Laboratory Plymouth | Wellcome Sanger Institute for the COVID-19 Genomics UK (COG-UK) Consortium | Lighthouse Laboratory Plymouth and Alex Alderton, Roberto Amato, Jeffrey Barrett, Sonia Goncalves, Ewan Harrison, David K. Jackson, Ian Johnston, Dominic Kwiatkowski, Cordelia Langford, John Sillitoe on behalf of the Wellcome Sanger Institute COVID-19 Surveillance Team |
| EPI_ISL_3013236, EPI_ISL_3013237, EPI_ISL_3013238 | Lighthouse Lab in Alderley Park | Wellcome Sanger Institute for the COVID-19 Genomics UK (COG-UK) Consortium | Jacquelyn Wynn, Mairead Hyland, The Lighthouse Lab in Alderley Park and Alex Alderton, Roberto Amato, Jeffrey Barrett, Sonia Goncalves, Ewan Harrison, David K. Jackson, Ian Johnston, Dominic Kwiatkowski, Cordelia Langford, John Sillitoe on behalf of the Wellcome Sanger Institute COVID-19 Surveillance Team |
| EPI_ISL_3013239 | Lighthouse Lab in Milton Keynes | Wellcome Sanger Institute for the COVID-19 Genomics UK (COG-UK) Consortium | The Lighthouse Lab in Milton Keynes and Alex Alderton, Roberto Amato, Jeffrey Barrett, Sonia Goncalves, Ewan Harrison, David K. Jackson, Ian Johnston, Dominic Kwiatkowski, Cordelia Langford, John Sillitoe on behalf of the Wellcome Sanger Institute COVID-19 Surveillance Team |
| EPI_ISL_3013240 | Lighthouse Laboratory Plymouth | Wellcome Sanger Institute for the COVID-19 Genomics UK (COG-UK) Consortium | Lighthouse Laboratory Plymouth and Alex Alderton, Roberto Amato, Jeffrey Barrett, Sonia Goncalves, Ewan Harrison, David K. Jackson, Ian Johnston, Dominic Kwiatkowski, Cordelia Langford, John Sillitoe on behalf of the Wellcome Sanger Institute COVID-19 Surveillance Team |
| EPI_ISL_3013241 | Lighthouse Lab in Glasgow | Wellcome Sanger Institute for the COVID-19 Genomics UK (COG-UK) Consortium | Harper VanSteenhouse, Yumi Kasai, David Gray, Carol Clugston, Anna Dominiczak and Alex Alderton, Roberto Amato, Jeffrey Barrett, Sonia Goncalves, Ewan Harrison, David K. Jackson, Ian Johnston, Dominic Kwiatkowski, Cordelia Langford, John Sillitoe on behalf of the Wellcome Sanger Institute COVID-19 Surveillance Team |
| EPI_ISL_3013242 | Lighthouse Laboratory Plymouth | Wellcome Sanger Institute for the COVID-19 Genomics UK (COG-UK) Consortium | Lighthouse Laboratory Plymouth and Alex Alderton, Roberto Amato, Jeffrey Barrett, Sonia Goncalves, Ewan Harrison, David K. Jackson, Ian Johnston, Dominic Kwiatkowski, Cordelia Langford, John Sillitoe on behalf of the Wellcome Sanger Institute COVID-19 Surveillance Team |
| EPI_ISL_3013243 | Lighthouse Lab in Milton Keynes | Wellcome Sanger Institute for the COVID-19 Genomics UK (COG-UK) Consortium | The Lighthouse Lab in Milton Keynes and Alex Alderton, Roberto Amato, Jeffrey Barrett, Sonia Goncalves, Ewan Harrison, David K. Jackson, Ian Johnston, Dominic Kwiatkowski, Cordelia Langford, John Sillitoe on behalf of the Wellcome Sanger Institute COVID-19 Surveillance Team |
| EPI_ISL_3013244, EPI_ISL_3013245 | Lighthouse Lab in Alderley Park | Wellcome Sanger Institute for the COVID-19 Genomics UK (COG-UK) Consortium | Jacquelyn Wynn, Mairead Hyland, The Lighthouse Lab in Alderley Park and Alex Alderton, Roberto Amato, Jeffrey Barrett, Sonia Goncalves, Ewan Harrison, David K. Jackson, Ian Johnston, Dominic Kwiatkowski, Cordelia Langford, John Sillitoe on behalf of the Wellcome Sanger Institute COVID-19 Surveillance Team |
| EPI_ISL_3013246 | Lighthouse Lab in Milton Keynes | Wellcome Sanger Institute for the COVID-19 Genomics UK (COG-UK) Consortium | The Lighthouse Lab in Milton Keynes and Alex Alderton, Roberto Amato, Jeffrey Barrett, Sonia Goncalves, Ewan Harrison, David K. Jackson, Ian Johnston, Dominic Kwiatkowski, Cordelia Langford, John Sillitoe on behalf of the Wellcome Sanger Institute COVID-19 Surveillance Team |
| EPI_ISL_3013247, EPI_ISL_3013248 | Lighthouse Laboratory Plymouth | Wellcome Sanger Institute for the COVID-19 Genomics UK (COG-UK) Consortium | Lighthouse Laboratory Plymouth and Alex Alderton, Roberto Amato, Jeffrey Barrett, Sonia Goncalves, Ewan Harrison, David K. Jackson, Ian Johnston, Dominic Kwiatkowski, Cordelia Langford, John Sillitoe on behalf of the Wellcome Sanger Institute COVID-19 Surveillance Team |
| EPI_ISL_3013249, EPI_ISL_3013250 | Lighthouse Lab in Glasgow | Wellcome Sanger Institute for the COVID-19 Genomics UK (COG-UK) Consortium | Harper VanSteenhouse, Yumi Kasai, David Gray, Carol Clugston, Anna Dominiczak and Alex Alderton, Roberto Amato, Jeffrey Barrett, Sonia Goncalves, Ewan Harrison, David K. Jackson, Ian Johnston, Dominic Kwiatkowski, Cordelia Langford, John Sillitoe on behalf of the Wellcome Sanger Institute COVID-19 Surveillance Team |
| EPI_ISL_3013251 | Lighthouse Lab in Alderley Park | Wellcome Sanger Institute for the COVID-19 Genomics UK (COG-UK) Consortium | Jacquelyn Wynn, Mairead Hyland, The Lighthouse Lab in Alderley Park and Alex Alderton, Roberto Amato, Jeffrey Barrett, Sonia Goncalves, Ewan Harrison, David K. Jackson, Ian Johnston, Dominic Kwiatkowski, Cordelia Langford, John Sillitoe on behalf of the Wellcome Sanger Institute COVID-19 Surveillance Team |
| EPI_ISL_3013252 | Lighthouse Laboratory Plymouth | Wellcome Sanger Institute for the COVID-19 Genomics UK (COG-UK) Consortium | Lighthouse Laboratory Plymouth and Alex Alderton, Roberto Amato, Jeffrey Barrett, Sonia Goncalves, Ewan Harrison, David K. Jackson, Ian Johnston, Dominic Kwiatkowski, Cordelia Langford, John Sillitoe on behalf of the Wellcome Sanger Institute COVID-19 Surveillance Team |
| EPI_ISL_3013253, EPI_ISL_3013254 | Lighthouse Lab in Milton Keynes | Wellcome Sanger Institute for the COVID-19 Genomics UK (COG-UK) Consortium | The Lighthouse Lab in Milton Keynes and Alex Alderton, Roberto Amato, Jeffrey Barrett, Sonia Goncalves, Ewan Harrison, David K. Jackson, Ian Johnston, Dominic Kwiatkowski, Cordelia Langford, John Sillitoe on behalf of the Wellcome Sanger Institute COVID-19 Surveillance Team |
| EPI_ISL_3013255, EPI_ISL_3013256 | Lighthouse Laboratory Plymouth | Wellcome Sanger Institute for the COVID-19 Genomics UK (COG-UK) Consortium | Lighthouse Laboratory Plymouth and Alex Alderton, Roberto Amato, Jeffrey Barrett, Sonia Goncalves, Ewan Harrison, David K. Jackson, Ian Johnston, Dominic Kwiatkowski, Cordelia Langford, John Sillitoe on behalf of the Wellcome Sanger Institute COVID-19 Surveillance Team |
| EPI_ISL_3013257, EPI_ISL_3013258 | Lighthouse Lab in Alderley Park | Wellcome Sanger Institute for the COVID-19 Genomics UK (COG-UK) Consortium | Jacquelyn Wynn, Mairead Hyland, The Lighthouse Lab in Alderley Park and Alex Alderton, Roberto Amato, Jeffrey Barrett, Sonia Goncalves, Ewan Harrison, David K. Jackson, Ian Johnston, Dominic Kwiatkowski, Cordelia Langford, John Sillitoe on behalf of the Wellcome Sanger Institute COVID-19 Surveillance Team |
| EPI_ISL_3013259 | Lighthouse Lab in Milton Keynes | Wellcome Sanger Institute for the COVID-19 Genomics UK (COG-UK) Consortium | The Lighthouse Lab in Milton Keynes and Alex Alderton, Roberto Amato, Jeffrey Barrett, Sonia Goncalves, Ewan Harrison, David K. Jackson, Ian Johnston, Dominic Kwiatkowski, Cordelia Langford, John Sillitoe on behalf of the Wellcome Sanger Institute COVID-19 Surveillance Team |
| EPI_ISL_3013260 | Lighthouse Lab in Glasgow | Wellcome Sanger Institute for the COVID-19 Genomics UK (COG-UK) Consortium | Harper VanSteenhouse, Yumi Kasai, David Gray, Carol Clugston, Anna Dominiczak and Alex Alderton, Roberto Amato, Jeffrey Barrett, Sonia Goncalves, Ewan Harrison, David K. Jackson, Ian Johnston, Dominic Kwiatkowski, Cordelia Langford, John Sillitoe on behalf of the Wellcome Sanger Institute COVID-19 Surveillance Team |
| EPI_ISL_3013261, EPI_ISL_3013262 | Lighthouse Lab in Alderley Park | Wellcome Sanger Institute for the COVID-19 Genomics UK (COG-UK) Consortium | Jacquelyn Wynn, Mairead Hyland, The Lighthouse Lab in Alderley Park and Alex Alderton, Roberto Amato, Jeffrey Barrett, Sonia Goncalves, Ewan Harrison, David K. Jackson, Ian Johnston, Dominic Kwiatkowski, Cordelia Langford, John Sillitoe on behalf of the Wellcome Sanger Institute COVID-19 Surveillance Team |
| EPI_ISL_3013263 | Lighthouse Lab in Glasgow | Wellcome Sanger Institute for the COVID-19 Genomics UK (COG-UK) Consortium | Harper VanSteenhouse, Yumi Kasai, David Gray, Carol Clugston, Anna Dominiczak and Alex Alderton, Roberto Amato, Jeffrey Barrett, Sonia Goncalves, Ewan Harrison, David K. Jackson, Ian Johnston, Dominic Kwiatkowski, Cordelia Langford, John Sillitoe on behalf of the Wellcome Sanger Institute COVID-19 Surveillance Team |
| EPI_ISL_3013264 | Lighthouse Laboratory Plymouth | Wellcome Sanger Institute for the COVID-19 Genomics UK (COG-UK) Consortium | Lighthouse Laboratory Plymouth and Alex Alderton, Roberto Amato, Jeffrey Barrett, Sonia Goncalves, Ewan Harrison, David K. Jackson, Ian Johnston, Dominic Kwiatkowski, Cordelia Langford, John Sillitoe on behalf of the Wellcome Sanger Institute COVID-19 Surveillance Team |
| EPI_ISL_3013265, EPI_ISL_3013266 | Lighthouse Lab in Milton Keynes | Wellcome Sanger Institute for the COVID-19 Genomics UK (COG-UK) Consortium | The Lighthouse Lab in Milton Keynes and Alex Alderton, Roberto Amato, Jeffrey Barrett, Sonia Goncalves, Ewan Harrison, David K. Jackson, Ian Johnston, Dominic Kwiatkowski, Cordelia Langford, John Sillitoe on behalf of the Wellcome Sanger Institute COVID-19 Surveillance Team |
| EPI_ISL_3013267 | Lighthouse Laboratory Plymouth | Wellcome Sanger Institute for the COVID-19 Genomics UK (COG-UK) Consortium | Lighthouse Laboratory Plymouth and Alex Alderton, Roberto Amato, Jeffrey Barrett, Sonia Goncalves, Ewan Harrison, David K. Jackson, Ian Johnston, Dominic Kwiatkowski, Cordelia Langford, John Sillitoe on behalf of the Wellcome Sanger Institute COVID-19 Surveillance Team |
| EPI_ISL_3013268 | Lighthouse Lab in Milton Keynes | Wellcome Sanger Institute for the COVID-19 Genomics UK (COG-UK) Consortium | The Lighthouse Lab in Milton Keynes and Alex Alderton, Roberto Amato, Jeffrey Barrett, Sonia Goncalves, Ewan Harrison, David K. Jackson, Ian Johnston, Dominic Kwiatkowski, Cordelia Langford, John Sillitoe on behalf of the Wellcome Sanger Institute COVID-19 Surveillance Team |
| EPI_ISL_3013269, EPI_ISL_3013270, EPI_ISL_3013271 | Lighthouse Lab in Alderley Park | Wellcome Sanger Institute for the COVID-19 Genomics UK (COG-UK) Consortium | Jacquelyn Wynn, Mairead Hyland, The Lighthouse Lab in Alderley Park and Alex Alderton, Roberto Amato, Jeffrey Barrett, Sonia Goncalves, Ewan Harrison, David K. Jackson, Ian Johnston, Dominic Kwiatkowski, Cordelia Langford, John Sillitoe on behalf of the Wellcome Sanger Institute COVID-19 Surveillance Team |
| EPI_ISL_3013272 | Lighthouse Lab in Milton Keynes | Wellcome Sanger Institute for the COVID-19 Genomics UK (COG-UK) Consortium | The Lighthouse Lab in Milton Keynes and Alex Alderton, Roberto Amato, Jeffrey Barrett, Sonia Goncalves, Ewan Harrison, David K. Jackson, Ian Johnston, Dominic Kwiatkowski, Cordelia Langford, John Sillitoe on behalf of the Wellcome Sanger Institute COVID-19 Surveillance Team |
| EPI_ISL_3013273 | Lighthouse Laboratory Plymouth | Wellcome Sanger Institute for the COVID-19 Genomics UK (COG-UK) Consortium | Lighthouse Laboratory Plymouth and Alex Alderton, Roberto Amato, Jeffrey Barrett, Sonia Goncalves, Ewan Harrison, David K. Jackson, Ian Johnston, Dominic Kwiatkowski, Cordelia Langford, John Sillitoe on behalf of the Wellcome Sanger Institute COVID-19 Surveillance Team |
| EPI_ISL_3013274 | Lighthouse Lab in Glasgow | Wellcome Sanger Institute for the COVID-19 Genomics UK (COG-UK) Consortium | Harper VanSteenhouse, Yumi Kasai, David Gray, Carol Clugston, Anna Dominiczak and Alex Alderton, Roberto Amato, Jeffrey Barrett, Sonia Goncalves, Ewan Harrison, David K. Jackson, Ian Johnston, Dominic Kwiatkowski, Cordelia Langford, John Sillitoe on behalf of the Wellcome Sanger Institute COVID-19 Surveillance Team |
| EPI_ISL_3013275 | Lighthouse Lab in Milton Keynes | Wellcome Sanger Institute for the COVID-19 Genomics UK (COG-UK) Consortium | The Lighthouse Lab in Milton Keynes and Alex Alderton, Roberto Amato, Jeffrey Barrett, Sonia Goncalves, Ewan Harrison, David K. Jackson, Ian Johnston, Dominic Kwiatkowski, Cordelia Langford, John Sillitoe on behalf of the Wellcome Sanger Institute COVID-19 Surveillance Team |
| EPI_ISL_3013276 | Lighthouse Lab in Alderley Park | Wellcome Sanger Institute for the COVID-19 Genomics UK (COG-UK) Consortium | Jacquelyn Wynn, Mairead Hyland, The Lighthouse Lab in Alderley Park and Alex Alderton, Roberto Amato, Jeffrey Barrett, Sonia Goncalves, Ewan Harrison, David K. Jackson, Ian Johnston, Dominic Kwiatkowski, Cordelia Langford, John Sillitoe on behalf of the Wellcome Sanger Institute COVID-19 Surveillance Team |
| EPI_ISL_3013277 | Lighthouse Laboratory Plymouth | Wellcome Sanger Institute for the COVID-19 Genomics UK (COG-UK) Consortium | Lighthouse Laboratory Plymouth and Alex Alderton, Roberto Amato, Jeffrey Barrett, Sonia Goncalves, Ewan Harrison, David K. Jackson, Ian Johnston, Dominic Kwiatkowski, Cordelia Langford, John Sillitoe on behalf of the Wellcome Sanger Institute COVID-19 Surveillance Team |
| EPI_ISL_3013278 | Lighthouse Lab in Milton Keynes | Wellcome Sanger Institute for the COVID-19 Genomics UK (COG-UK) Consortium | The Lighthouse Lab in Milton Keynes and Alex Alderton, Roberto Amato, Jeffrey Barrett, Sonia Goncalves, Ewan Harrison, David K. Jackson, Ian Johnston, Dominic Kwiatkowski, Cordelia Langford, John Sillitoe on behalf of the Wellcome Sanger Institute COVID-19 Surveillance Team |
| EPI_ISL_3013279 | Lighthouse Laboratory Plymouth | Wellcome Sanger Institute for the COVID-19 Genomics UK (COG-UK) Consortium | Lighthouse Laboratory Plymouth and Alex Alderton, Roberto Amato, Jeffrey Barrett, Sonia Goncalves, Ewan Harrison, David K. Jackson, Ian Johnston, Dominic Kwiatkowski, Cordelia Langford, John Sillitoe on behalf of the Wellcome Sanger Institute COVID-19 Surveillance Team |
| EPI_ISL_3013280 | Lighthouse Lab in Alderley Park | Wellcome Sanger Institute for the COVID-19 Genomics UK (COG-UK) Consortium | Jacquelyn Wynn, Mairead Hyland, The Lighthouse Lab in Alderley Park and Alex Alderton, Roberto Amato, Jeffrey Barrett, Sonia Goncalves, Ewan Harrison, David K. Jackson, Ian Johnston, Dominic Kwiatkowski, Cordelia Langford, John Sillitoe on behalf of the Wellcome Sanger Institute COVID-19 Surveillance Team |
| EPI_ISL_3013281 | Lighthouse Laboratory Plymouth | Wellcome Sanger Institute for the COVID-19 Genomics UK (COG-UK) Consortium | Lighthouse Laboratory Plymouth and Alex Alderton, Roberto Amato, Jeffrey Barrett, Sonia Goncalves, Ewan Harrison, David K. Jackson, Ian Johnston, Dominic Kwiatkowski, Cordelia Langford, John Sillitoe on behalf of the Wellcome Sanger Institute COVID-19 Surveillance Team |

| | | | |
|---|---|---|---|
| EPI_ISL_3013282, EPI_ISL_3013283, EPI_ISL_3013284 | Lighthouse Lab in Glasgow | Wellcome Sanger Institute for the COVID-19 Genomics UK (COG-UK) Consortium | Harper VanSteenhouse, Yumi Kasai, David Gray, Carol Clugston, Anna Dominiczak and Alex Alderton, Roberto Amato, Jeffrey Barrett, Sonia Goncalves, Ewan Harrison, David K. Jackson, Ian Johnston, Dominic Kwiatkowski, Cordelia Langford, John Sillitoe on behalf of the Wellcome Sanger Institute COVID-19 Surveillance Team |
| EPI_ISL_3013285 | Lighthouse Lab in Milton Keynes | Wellcome Sanger Institute for the COVID-19 Genomics UK (COG-UK) Consortium | The Lighthouse Lab in Milton Keynes and Alex Alderton, Roberto Amato, Jeffrey Barrett, Sonia Goncalves, Ewan Harrison, David K. Jackson, Ian Johnston, Dominic Kwiatkowski, Cordelia Langford, John Sillitoe on behalf of the Wellcome Sanger Institute COVID-19 Surveillance Team |
| EPI_ISL_3013286 | Lighthouse Laboratory Plymouth | Wellcome Sanger Institute for the COVID-19 Genomics UK (COG-UK) Consortium | Lighthouse Laboratory Plymouth and Alex Alderton, Roberto Amato, Jeffrey Barrett, Sonia Goncalves, Ewan Harrison, David K. Jackson, Ian Johnston, Dominic Kwiatkowski, Cordelia Langford, John Sillitoe on behalf of the Wellcome Sanger Institute COVID-19 Surveillance Team |
| EPI_ISL_3013287 | Lighthouse Lab in Milton Keynes | Wellcome Sanger Institute for the COVID-19 Genomics UK (COG-UK) Consortium | The Lighthouse Lab in Milton Keynes and Alex Alderton, Roberto Amato, Jeffrey Barrett, Sonia Goncalves, Ewan Harrison, David K. Jackson, Ian Johnston, Dominic Kwiatkowski, Cordelia Langford, John Sillitoe on behalf of the Wellcome Sanger Institute COVID-19 Surveillance Team |
| EPI_ISL_3013288 | Lighthouse Lab in Glasgow | Wellcome Sanger Institute for the COVID-19 Genomics UK (COG-UK) Consortium | Harper VanSteenhouse, Yumi Kasai, David Gray, Carol Clugston, Anna Dominiczak and Alex Alderton, Roberto Amato, Jeffrey Barrett, Sonia Goncalves, Ewan Harrison, David K. Jackson, Ian Johnston, Dominic Kwiatkowski, Cordelia Langford, John Sillitoe on behalf of the Wellcome Sanger Institute COVID-19 Surveillance Team |
| EPI_ISL_3013289 | Lighthouse Lab in Milton Keynes | Wellcome Sanger Institute for the COVID-19 Genomics UK (COG-UK) Consortium | The Lighthouse Lab in Milton Keynes and Alex Alderton, Roberto Amato, Jeffrey Barrett, Sonia Goncalves, Ewan Harrison, David K. Jackson, Ian Johnston, Dominic Kwiatkowski, Cordelia Langford, John Sillitoe on behalf of the Wellcome Sanger Institute COVID-19 Surveillance Team |
| EPI_ISL_3013290, EPI_ISL_3013291 | Lighthouse Laboratory Plymouth | Wellcome Sanger Institute for the COVID-19 Genomics UK (COG-UK) Consortium | Lighthouse Laboratory Plymouth and Alex Alderton, Roberto Amato, Jeffrey Barrett, Sonia Goncalves, Ewan Harrison, David K. Jackson, Ian Johnston, Dominic Kwiatkowski, Cordelia Langford, John Sillitoe on behalf of the Wellcome Sanger Institute COVID-19 Surveillance Team |
| EPI_ISL_3013292 | Lighthouse Lab in Alderley Park | Wellcome Sanger Institute for the COVID-19 Genomics UK (COG-UK) Consortium | Jacquelyn Wynn, Mairead Hyland, The Lighthouse Lab in Alderley Park and Alex Alderton, Roberto Amato, Jeffrey Barrett, Sonia Goncalves, Ewan Harrison, David K. Jackson, Ian Johnston, Dominic Kwiatkowski, Cordelia Langford, John Sillitoe on behalf of the Wellcome Sanger Institute COVID-19 Surveillance Team |
| EPI_ISL_3013293 | Lighthouse Laboratory Plymouth | Wellcome Sanger Institute for the COVID-19 Genomics UK (COG-UK) Consortium | Lighthouse Laboratory Plymouth and Alex Alderton, Roberto Amato, Jeffrey Barrett, Sonia Goncalves, Ewan Harrison, David K. Jackson, Ian Johnston, Dominic Kwiatkowski, Cordelia Langford, John Sillitoe on behalf of the Wellcome Sanger Institute COVID-19 Surveillance Team |
| EPI_ISL_3013294, EPI_ISL_3013295 | Lighthouse Lab in Milton Keynes | Wellcome Sanger Institute for the COVID-19 Genomics UK (COG-UK) Consortium | The Lighthouse Lab in Milton Keynes and Alex Alderton, Roberto Amato, Jeffrey Barrett, Sonia Goncalves, Ewan Harrison, David K. Jackson, Ian Johnston, Dominic Kwiatkowski, Cordelia Langford, John Sillitoe on behalf of the Wellcome Sanger Institute COVID-19 Surveillance Team |
| EPI_ISL_3013296, EPI_ISL_3013297, EPI_ISL_3013298, EPI_ISL_3013299 | Lighthouse Lab in Alderley Park | Wellcome Sanger Institute for the COVID-19 Genomics UK (COG-UK) Consortium | Jacquelyn Wynn, Mairead Hyland, The Lighthouse Lab in Alderley Park and Alex Alderton, Roberto Amato, Jeffrey Barrett, Sonia Goncalves, Ewan Harrison, David K. Jackson, Ian Johnston, Dominic Kwiatkowski, Cordelia Langford, John Sillitoe on behalf of the Wellcome Sanger Institute COVID-19 Surveillance Team |
| EPI_ISL_3013300, EPI_ISL_3013301 | Lighthouse Lab in Glasgow | Wellcome Sanger Institute for the COVID-19 Genomics UK (COG-UK) Consortium | Harper VanSteenhouse, Yumi Kasai, David Gray, Carol Clugston, Anna Dominiczak and Alex Alderton, Roberto Amato, Jeffrey Barrett, Sonia Goncalves, Ewan Harrison, David K. Jackson, Ian Johnston, Dominic Kwiatkowski, Cordelia Langford, John Sillitoe on behalf of the Wellcome Sanger Institute COVID-19 Surveillance Team |
| EPI_ISL_3013302 | Lighthouse Lab in Alderley Park | Wellcome Sanger Institute for the COVID-19 Genomics UK (COG-UK) Consortium | Jacquelyn Wynn, Mairead Hyland, The Lighthouse Lab in Alderley Park and Alex Alderton, Roberto Amato, Jeffrey Barrett, Sonia Goncalves, Ewan Harrison, David K. Jackson, Ian Johnston, Dominic Kwiatkowski, Cordelia Langford, John Sillitoe on behalf of the Wellcome Sanger Institute COVID-19 Surveillance Team |
| EPI_ISL_3013303 | Lighthouse Laboratory Plymouth | Wellcome Sanger Institute for the COVID-19 Genomics UK (COG-UK) Consortium | Lighthouse Laboratory Plymouth and Alex Alderton, Roberto Amato, Jeffrey Barrett, Sonia Goncalves, Ewan Harrison, David K. Jackson, Ian Johnston, Dominic Kwiatkowski, Cordelia Langford, John Sillitoe on behalf of the Wellcome Sanger Institute COVID-19 Surveillance Team |
| EPI_ISL_3013304 | Lighthouse Lab in Alderley Park | Wellcome Sanger Institute for the COVID-19 Genomics UK (COG-UK) Consortium | Jacquelyn Wynn, Mairead Hyland, The Lighthouse Lab in Alderley Park and Alex Alderton, Roberto Amato, Jeffrey Barrett, Sonia Goncalves, Ewan Harrison, David K. Jackson, Ian Johnston, Dominic Kwiatkowski, Cordelia Langford, John Sillitoe on behalf of the Wellcome Sanger Institute COVID-19 Surveillance Team |
| EPI_ISL_3013305 | Lighthouse Laboratory Plymouth | Wellcome Sanger Institute for the COVID-19 Genomics UK (COG-UK) Consortium | Lighthouse Laboratory Plymouth and Alex Alderton, Roberto Amato, Jeffrey Barrett, Sonia Goncalves, Ewan Harrison, David K. Jackson, Ian Johnston, Dominic Kwiatkowski, Cordelia Langford, John Sillitoe on behalf of the Wellcome Sanger Institute COVID-19 Surveillance Team |
| EPI_ISL_3013306, EPI_ISL_3013307, EPI_ISL_3013308 | Lighthouse Lab in Glasgow | Wellcome Sanger Institute for the COVID-19 Genomics UK (COG-UK) Consortium | Harper VanSteenhouse, Yumi Kasai, David Gray, Carol Clugston, Anna Dominiczak and Alex Alderton, Roberto Amato, Jeffrey Barrett, Sonia Goncalves, Ewan Harrison, David K. Jackson, Ian Johnston, Dominic Kwiatkowski, Cordelia Langford, John Sillitoe on behalf of the Wellcome Sanger Institute COVID-19 Surveillance Team |
| EPI_ISL_3013309 | Lighthouse Laboratory Plymouth | Wellcome Sanger Institute for the COVID-19 Genomics UK (COG-UK) Consortium | Lighthouse Laboratory Plymouth and Alex Alderton, Roberto Amato, Jeffrey Barrett, Sonia Goncalves, Ewan Harrison, David K. Jackson, Ian Johnston, Dominic Kwiatkowski, Cordelia Langford, John Sillitoe on behalf of the Wellcome Sanger Institute COVID-19 Surveillance Team |
| EPI_ISL_3013310 | Lighthouse Lab in Milton Keynes | Wellcome Sanger Institute for the COVID-19 Genomics UK (COG-UK) Consortium | The Lighthouse Lab in Milton Keynes and Alex Alderton, Roberto Amato, Jeffrey Barrett, Sonia Goncalves, Ewan Harrison, David K. Jackson, Ian Johnston, Dominic Kwiatkowski, Cordelia Langford, John Sillitoe on behalf of the Wellcome Sanger Institute COVID-19 Surveillance Team |
| EPI_ISL_3013311 | Lighthouse Lab in Alderley Park | Wellcome Sanger Institute for the COVID-19 Genomics UK (COG-UK) Consortium | Jacquelyn Wynn, Mairead Hyland, The Lighthouse Lab in Alderley Park and Alex Alderton, Roberto Amato, Jeffrey Barrett, Sonia Goncalves, Ewan Harrison, David K. Jackson, Ian Johnston, Dominic Kwiatkowski, Cordelia Langford, John Sillitoe on behalf of the Wellcome Sanger Institute COVID-19 Surveillance Team |
| EPI_ISL_3013312 | Lighthouse Lab in Glasgow | Wellcome Sanger Institute for the COVID-19 Genomics UK (COG-UK) Consortium | Harper VanSteenhouse, Yumi Kasai, David Gray, Carol Clugston, Anna Dominiczak and Alex Alderton, Roberto Amato, Jeffrey Barrett, Sonia Goncalves, Ewan Harrison, David K. Jackson, Ian Johnston, Dominic Kwiatkowski, Cordelia Langford, John Sillitoe on behalf of the Wellcome Sanger Institute COVID-19 Surveillance Team |
| EPI_ISL_3013313 | Lighthouse Lab in Milton Keynes | Wellcome Sanger Institute for the COVID-19 Genomics UK (COG-UK) Consortium | The Lighthouse Lab in Milton Keynes and Alex Alderton, Roberto Amato, Jeffrey Barrett, Sonia Goncalves, Ewan Harrison, David K. Jackson, Ian Johnston, Dominic Kwiatkowski, Cordelia Langford, John Sillitoe on behalf of the Wellcome Sanger Institute COVID-19 Surveillance Team |
| EPI_ISL_3013314 | Lighthouse Lab in Alderley Park | Wellcome Sanger Institute for the COVID-19 Genomics UK (COG-UK) Consortium | Jacquelyn Wynn, Mairead Hyland, The Lighthouse Lab in Alderley Park and Alex Alderton, Roberto Amato, Jeffrey Barrett, Sonia Goncalves, Ewan Harrison, David K. Jackson, Ian Johnston, Dominic Kwiatkowski, Cordelia Langford, John Sillitoe on behalf of the Wellcome Sanger Institute COVID-19 Surveillance Team |
| EPI_ISL_3013315 | Lighthouse Lab in Milton Keynes | Wellcome Sanger Institute for the COVID-19 Genomics UK (COG-UK) Consortium | The Lighthouse Lab in Milton Keynes and Alex Alderton, Roberto Amato, Jeffrey Barrett, Sonia Goncalves, Ewan Harrison, David K. Jackson, Ian Johnston, Dominic Kwiatkowski, Cordelia Langford, John Sillitoe on behalf of the Wellcome Sanger Institute COVID-19 Surveillance Team |
| EPI_ISL_3013316 | Lighthouse Lab in Glasgow | Wellcome Sanger Institute for the COVID-19 Genomics UK (COG-UK) Consortium | Harper VanSteenhouse, Yumi Kasai, David Gray, Carol Clugston, Anna Dominiczak and Alex Alderton, Roberto Amato, Jeffrey Barrett, Sonia Goncalves, Ewan Harrison, David K. Jackson, Ian Johnston, Dominic Kwiatkowski, Cordelia Langford, John Sillitoe on behalf of the Wellcome Sanger Institute COVID-19 Surveillance Team |
| EPI_ISL_3013317 | Lighthouse Lab in Alderley Park | Wellcome Sanger Institute for the COVID-19 Genomics UK (COG-UK) Consortium | Jacquelyn Wynn, Mairead Hyland, The Lighthouse Lab in Alderley Park and Alex Alderton, Roberto Amato, Jeffrey Barrett, Sonia Goncalves, Ewan Harrison, David K. Jackson, Ian Johnston, Dominic Kwiatkowski, Cordelia Langford, John Sillitoe on behalf of the Wellcome Sanger Institute COVID-19 Surveillance Team |
| EPI_ISL_3013318 | Lighthouse Lab in Milton Keynes | Wellcome Sanger Institute for the COVID-19 Genomics UK (COG-UK) Consortium | The Lighthouse Lab in Milton Keynes and Alex Alderton, Roberto Amato, Jeffrey Barrett, Sonia Goncalves, Ewan Harrison, David K. Jackson, Ian Johnston, Dominic Kwiatkowski, Cordelia Langford, John Sillitoe on behalf of the Wellcome Sanger Institute COVID-19 Surveillance Team |
| EPI_ISL_3013319, EPI_ISL_3013320 | Lighthouse Lab in Glasgow | Wellcome Sanger Institute for the COVID-19 Genomics UK (COG-UK) Consortium | Harper VanSteenhouse, Yumi Kasai, David Gray, Carol Clugston, Anna Dominiczak and Alex Alderton, Roberto Amato, Jeffrey Barrett, Sonia Goncalves, Ewan Harrison, David K. Jackson, Ian Johnston, Dominic Kwiatkowski, Cordelia Langford, John Sillitoe on behalf of the Wellcome Sanger Institute COVID-19 Surveillance Team |
| EPI_ISL_3013321 | Lighthouse Lab in Milton Keynes | Wellcome Sanger Institute for the COVID-19 Genomics UK (COG-UK) Consortium | The Lighthouse Lab in Milton Keynes and Alex Alderton, Roberto Amato, Jeffrey Barrett, Sonia Goncalves, Ewan Harrison, David K. Jackson, Ian Johnston, Dominic Kwiatkowski, Cordelia Langford, John Sillitoe on behalf of the Wellcome Sanger Institute COVID-19 Surveillance Team |
| EPI_ISL_3013322 | Lighthouse Laboratory Plymouth | Wellcome Sanger Institute for the COVID-19 Genomics UK (COG-UK) Consortium | Lighthouse Laboratory Plymouth and Alex Alderton, Roberto Amato, Jeffrey Barrett, Sonia Goncalves, Ewan Harrison, David K. Jackson, Ian Johnston, Dominic Kwiatkowski, Cordelia Langford, John Sillitoe on behalf of the Wellcome Sanger Institute COVID-19 Surveillance Team |
| EPI_ISL_3013323 | Lighthouse Lab in Glasgow | Wellcome Sanger Institute for the COVID-19 Genomics UK (COG-UK) Consortium | Harper VanSteenhouse, Yumi Kasai, David Gray, Carol Clugston, Anna Dominiczak and Alex Alderton, Roberto Amato, Jeffrey Barrett, Sonia Goncalves, Ewan Harrison, David K. Jackson, Ian Johnston, Dominic Kwiatkowski, Cordelia Langford, John Sillitoe on behalf of the Wellcome Sanger Institute COVID-19 Surveillance Team |
| EPI_ISL_3013324 | Lighthouse Lab in Alderley Park | Wellcome Sanger Institute for the COVID-19 Genomics UK (COG-UK) Consortium | Jacquelyn Wynn, Mairead Hyland, The Lighthouse Lab in Alderley Park and Alex Alderton, Roberto Amato, Jeffrey Barrett, Sonia Goncalves, Ewan Harrison, David K. Jackson, Ian Johnston, Dominic Kwiatkowski, Cordelia Langford, John Sillitoe on behalf of the Wellcome Sanger Institute COVID-19 Surveillance Team |
| EPI_ISL_3013325 | Lighthouse Lab in Milton Keynes | Wellcome Sanger Institute for the COVID-19 Genomics UK (COG-UK) Consortium | The Lighthouse Lab in Milton Keynes and Alex Alderton, Roberto Amato, Jeffrey Barrett, Sonia Goncalves, Ewan Harrison, David K. Jackson, Ian Johnston, Dominic Kwiatkowski, Cordelia Langford, John Sillitoe on behalf of the Wellcome Sanger Institute COVID-19 Surveillance Team |
| EPI_ISL_3013326, EPI_ISL_3013327 | Lighthouse Lab in Glasgow | Wellcome Sanger Institute for the COVID-19 Genomics UK (COG-UK) Consortium | Harper VanSteenhouse, Yumi Kasai, David Gray, Carol Clugston, Anna Dominiczak and Alex Alderton, Roberto Amato, Jeffrey Barrett, Sonia Goncalves, Ewan Harrison, David K. Jackson, Ian Johnston, Dominic Kwiatkowski, Cordelia Langford, John Sillitoe on behalf of the Wellcome Sanger Institute COVID-19 Surveillance Team |
| EPI_ISL_3013328 | Lighthouse Lab in Milton Keynes | Wellcome Sanger Institute for the COVID-19 Genomics UK (COG-UK) Consortium | The Lighthouse Lab in Milton Keynes and Alex Alderton, Roberto Amato, Jeffrey Barrett, Sonia Goncalves, Ewan Harrison, David K. Jackson, Ian Johnston, Dominic Kwiatkowski, Cordelia Langford, John Sillitoe on behalf of the Wellcome Sanger Institute COVID-19 Surveillance Team |
| EPI_ISL_3013329 | Lighthouse Lab in Glasgow | Wellcome Sanger Institute for the COVID-19 Genomics UK (COG-UK) Consortium | Harper VanSteenhouse, Yumi Kasai, David Gray, Carol Clugston, Anna Dominiczak and Alex Alderton, Roberto Amato, Jeffrey Barrett, Sonia Goncalves, Ewan Harrison, David K. Jackson, Ian Johnston, Dominic Kwiatkowski, Cordelia Langford, John Sillitoe on behalf of the Wellcome Sanger Institute COVID-19 Surveillance Team |
| EPI_ISL_3013330 | Lighthouse Lab in Milton Keynes | Wellcome Sanger Institute for the COVID-19 Genomics UK (COG-UK) Consortium | The Lighthouse Lab in Milton Keynes and Alex Alderton, Roberto Amato, Jeffrey Barrett, Sonia Goncalves, Ewan Harrison, David K. Jackson, Ian Johnston, Dominic Kwiatkowski, Cordelia Langford, John Sillitoe on behalf of the Wellcome Sanger Institute COVID-19 Surveillance Team |
| EPI_ISL_3013331 | Lighthouse Lab in Alderley Park | Wellcome Sanger Institute for the COVID-19 Genomics UK (COG-UK) Consortium | Jacquelyn Wynn, Mairead Hyland, The Lighthouse Lab in Alderley Park and Alex Alderton, Roberto Amato, Jeffrey Barrett, Sonia Goncalves, Ewan Harrison, David K. Jackson, Ian Johnston, Dominic Kwiatkowski, Cordelia Langford, John Sillitoe on behalf of the Wellcome Sanger Institute COVID-19 Surveillance Team |
| EPI_ISL_3013332, EPI_ISL_3013333 | Lighthouse Lab in Glasgow | Wellcome Sanger Institute for the COVID-19 Genomics UK (COG-UK) Consortium | Harper VanSteenhouse, Yumi Kasai, David Gray, Carol Clugston, Anna Dominiczak and Alex Alderton, Roberto Amato, Jeffrey Barrett, Sonia Goncalves, Ewan Harrison, David K. Jackson, Ian Johnston, Dominic Kwiatkowski, Cordelia Langford, John Sillitoe on behalf of the Wellcome Sanger Institute COVID-19 Surveillance Team |
| EPI_ISL_3013334 | Lighthouse Lab in Milton Keynes | Wellcome Sanger Institute for the COVID-19 Genomics UK (COG-UK) Consortium | The Lighthouse Lab in Milton Keynes and Alex Alderton, Roberto Amato, Jeffrey Barrett, Sonia Goncalves, Ewan Harrison, David K. Jackson, Ian Johnston, Dominic Kwiatkowski, Cordelia Langford, John Sillitoe on behalf of the Wellcome Sanger Institute COVID-19 Surveillance Team |
| EPI_ISL_3013335 | Lighthouse Lab in Alderley Park | Wellcome Sanger Institute for the COVID-19 Genomics UK (COG-UK) Consortium | Jacquelyn Wynn, Mairead Hyland, The Lighthouse Lab in Alderley Park and Alex Alderton, Roberto Amato, Jeffrey Barrett, Sonia Goncalves, Ewan Harrison, David K. Jackson, Ian Johnston, Dominic Kwiatkowski, Cordelia Langford, John Sillitoe on behalf of the Wellcome Sanger Institute COVID-19 Surveillance Team |
| EPI_ISL_3013336 | Lighthouse Lab in Glasgow | Wellcome Sanger Institute for the COVID-19 Genomics UK (COG-UK) Consortium | Harper VanSteenhouse, Yumi Kasai, David Gray, Carol Clugston, Anna Dominiczak and Alex Alderton, Roberto Amato, Jeffrey Barrett, Sonia Goncalves, Ewan Harrison, David K. Jackson, Ian Johnston, Dominic Kwiatkowski, Cordelia Langford, John Sillitoe on behalf of the Wellcome Sanger Institute COVID-19 Surveillance Team |
| EPI_ISL_3013337 | Lighthouse Lab in Alderley Park | Wellcome Sanger Institute for the COVID-19 Genomics UK (COG-UK) Consortium | Jacquelyn Wynn, Mairead Hyland, The Lighthouse Lab in Alderley Park and Alex Alderton, Roberto Amato, Jeffrey Barrett, Sonia Goncalves, Ewan Harrison, David K. Jackson, Ian Johnston, Dominic Kwiatkowski, Cordelia Langford, John Sillitoe on behalf of the Wellcome Sanger Institute COVID-19 Surveillance Team |
| EPI_ISL_3013338, | Lighthouse Lab in Glasgow | Wellcome Sanger Institute for the COVID-19 Genomics UK (COG-UK) | Harper VanSteenhouse, Yumi Kasai, David Gray, Carol Clugston, Anna Dominiczak and Alex Alderton, Roberto Amato, Jeffrey Barrett, Sonia Goncalves, Ewan Harrison, David K. Jackson, Ian Johnston, Dominic Kwiatkowski, Cordelia Langford, |

| | | | |
|---|---|---|---|
| EPI_ISL_3013339 | | Consortium | John Sillitoe on behalf of the Wellcome Sanger Institute COVID-19 Surveillance Team |
| EPI_ISL_3013340 | Lighthouse Lab in Milton Keynes | Wellcome Sanger Institute for the COVID-19 Genomics UK (COG-UK) Consortium | The Lighthouse Lab in Milton Keynes and Alex Alderton, Roberto Amato, Jeffrey Barrett, Sonia Goncalves, Ewan Harrison, David K. Jackson, Ian Johnston, Dominic Kwiatkowski, Cordelia Langford, John Sillitoe on behalf of the Wellcome Sanger Institute COVID-19 Surveillance Team |
| EPI_ISL_3013341 | Lighthouse Lab in Glasgow | Wellcome Sanger Institute for the COVID-19 Genomics UK (COG-UK) Consortium | Harper VanSteenhouse, Yumi Kasai, David Gray, Carol Clugston, Anna Dominiczak and Alex Alderton, Roberto Amato, Jeffrey Barrett, Sonia Goncalves, Ewan Harrison, David K. Jackson, Ian Johnston, Dominic Kwiatkowski, Cordelia Langford, John Sillitoe on behalf of the Wellcome Sanger Institute COVID-19 Surveillance Team |
| EPI_ISL_3013342 | Lighthouse Lab in Milton Keynes | Wellcome Sanger Institute for the COVID-19 Genomics UK (COG-UK) Consortium | The Lighthouse Lab in Milton Keynes and Alex Alderton, Roberto Amato, Jeffrey Barrett, Sonia Goncalves, Ewan Harrison, David K. Jackson, Ian Johnston, Dominic Kwiatkowski, Cordelia Langford, John Sillitoe on behalf of the Wellcome Sanger Institute COVID-19 Surveillance Team |
| EPI_ISL_3013343 | Lighthouse Lab in Glasgow | Wellcome Sanger Institute for the COVID-19 Genomics UK (COG-UK) Consortium | Harper VanSteenhouse, Yumi Kasai, David Gray, Carol Clugston, Anna Dominiczak and Alex Alderton, Roberto Amato, Jeffrey Barrett, Sonia Goncalves, Ewan Harrison, David K. Jackson, Ian Johnston, Dominic Kwiatkowski, Cordelia Langford, John Sillitoe on behalf of the Wellcome Sanger Institute COVID-19 Surveillance Team |
| EPI_ISL_3013344 | Lighthouse Lab in Milton Keynes | Wellcome Sanger Institute for the COVID-19 Genomics UK (COG-UK) Consortium | The Lighthouse Lab in Milton Keynes and Alex Alderton, Roberto Amato, Jeffrey Barrett, Sonia Goncalves, Ewan Harrison, David K. Jackson, Ian Johnston, Dominic Kwiatkowski, Cordelia Langford, John Sillitoe on behalf of the Wellcome Sanger Institute COVID-19 Surveillance Team |
| EPI_ISL_3013345 | Lighthouse Lab in Alderley Park | Wellcome Sanger Institute for the COVID-19 Genomics UK (COG-UK) Consortium | Jacquelyn Wynn, Mairead Hyland, The Lighthouse Lab in Alderley Park and Alex Alderton, Roberto Amato, Jeffrey Barrett, Sonia Goncalves, Ewan Harrison, David K. Jackson, Ian Johnston, Dominic Kwiatkowski, Cordelia Langford, John Sillitoe on behalf of the Wellcome Sanger Institute COVID-19 Surveillance Team |
| EPI_ISL_3013346 | Lighthouse Lab in Milton Keynes | Wellcome Sanger Institute for the COVID-19 Genomics UK (COG-UK) Consortium | The Lighthouse Lab in Milton Keynes and Alex Alderton, Roberto Amato, Jeffrey Barrett, Sonia Goncalves, Ewan Harrison, David K. Jackson, Ian Johnston, Dominic Kwiatkowski, Cordelia Langford, John Sillitoe on behalf of the Wellcome Sanger Institute COVID-19 Surveillance Team |
| EPI_ISL_3013347 | Lighthouse Lab in Alderley Park | Wellcome Sanger Institute for the COVID-19 Genomics UK (COG-UK) Consortium | Jacquelyn Wynn, Mairead Hyland, The Lighthouse Lab in Alderley Park and Alex Alderton, Roberto Amato, Jeffrey Barrett, Sonia Goncalves, Ewan Harrison, David K. Jackson, Ian Johnston, Dominic Kwiatkowski, Cordelia Langford, John Sillitoe on behalf of the Wellcome Sanger Institute COVID-19 Surveillance Team |
| EPI_ISL_3013348 | Lighthouse Lab in Glasgow | Wellcome Sanger Institute for the COVID-19 Genomics UK (COG-UK) Consortium | Harper VanSteenhouse, Yumi Kasai, David Gray, Carol Clugston, Anna Dominiczak and Alex Alderton, Roberto Amato, Jeffrey Barrett, Sonia Goncalves, Ewan Harrison, David K. Jackson, Ian Johnston, Dominic Kwiatkowski, Cordelia Langford, John Sillitoe on behalf of the Wellcome Sanger Institute COVID-19 Surveillance Team |
| EPI_ISL_3013349 | Lighthouse Lab in Alderley Park | Wellcome Sanger Institute for the COVID-19 Genomics UK (COG-UK) Consortium | Jacquelyn Wynn, Mairead Hyland, The Lighthouse Lab in Alderley Park and Alex Alderton, Roberto Amato, Jeffrey Barrett, Sonia Goncalves, Ewan Harrison, David K. Jackson, Ian Johnston, Dominic Kwiatkowski, Cordelia Langford, John Sillitoe on behalf of the Wellcome Sanger Institute COVID-19 Surveillance Team |
| EPI_ISL_3013350 | Lighthouse Lab in Glasgow | Wellcome Sanger Institute for the COVID-19 Genomics UK (COG-UK) Consortium | Harper VanSteenhouse, Yumi Kasai, David Gray, Carol Clugston, Anna Dominiczak and Alex Alderton, Roberto Amato, Jeffrey Barrett, Sonia Goncalves, Ewan Harrison, David K. Jackson, Ian Johnston, Dominic Kwiatkowski, Cordelia Langford, John Sillitoe on behalf of the Wellcome Sanger Institute COVID-19 Surveillance Team |
| EPI_ISL_3013351 | Lighthouse Lab in Alderley Park | Wellcome Sanger Institute for the COVID-19 Genomics UK (COG-UK) Consortium | Jacquelyn Wynn, Mairead Hyland, The Lighthouse Lab in Alderley Park and Alex Alderton, Roberto Amato, Jeffrey Barrett, Sonia Goncalves, Ewan Harrison, David K. Jackson, Ian Johnston, Dominic Kwiatkowski, Cordelia Langford, John Sillitoe on behalf of the Wellcome Sanger Institute COVID-19 Surveillance Team |
| EPI_ISL_3013352, EPI_ISL_3013353 | Lighthouse Lab in Glasgow | Wellcome Sanger Institute for the COVID-19 Genomics UK (COG-UK) Consortium | Harper VanSteenhouse, Yumi Kasai, David Gray, Carol Clugston, Anna Dominiczak and Alex Alderton, Roberto Amato, Jeffrey Barrett, Sonia Goncalves, Ewan Harrison, David K. Jackson, Ian Johnston, Dominic Kwiatkowski, Cordelia Langford, John Sillitoe on behalf of the Wellcome Sanger Institute COVID-19 Surveillance Team |
| EPI_ISL_3013354 | Lighthouse Lab in Alderley Park | Wellcome Sanger Institute for the COVID-19 Genomics UK (COG-UK) Consortium | Jacquelyn Wynn, Mairead Hyland, The Lighthouse Lab in Alderley Park and Alex Alderton, Roberto Amato, Jeffrey Barrett, Sonia Goncalves, Ewan Harrison, David K. Jackson, Ian Johnston, Dominic Kwiatkowski, Cordelia Langford, John Sillitoe on behalf of the Wellcome Sanger Institute COVID-19 Surveillance Team |
| EPI_ISL_3013355, EPI_ISL_3013356 | Lighthouse Lab in Milton Keynes | Wellcome Sanger Institute for the COVID-19 Genomics UK (COG-UK) Consortium | The Lighthouse Lab in Milton Keynes and Alex Alderton, Roberto Amato, Jeffrey Barrett, Sonia Goncalves, Ewan Harrison, David K. Jackson, Ian Johnston, Dominic Kwiatkowski, Cordelia Langford, John Sillitoe on behalf of the Wellcome Sanger Institute COVID-19 Surveillance Team |
| EPI_ISL_3013357 | Lighthouse Lab in Alderley Park | Wellcome Sanger Institute for the COVID-19 Genomics UK (COG-UK) Consortium | Jacquelyn Wynn, Mairead Hyland, The Lighthouse Lab in Alderley Park and Alex Alderton, Roberto Amato, Jeffrey Barrett, Sonia Goncalves, Ewan Harrison, David K. Jackson, Ian Johnston, Dominic Kwiatkowski, Cordelia Langford, John Sillitoe on behalf of the Wellcome Sanger Institute COVID-19 Surveillance Team |
| EPI_ISL_3013358, EPI_ISL_3013359, EPI_ISL_3013360, EPI_ISL_3013361 | Lighthouse Lab in Glasgow | Wellcome Sanger Institute for the COVID-19 Genomics UK (COG-UK) Consortium | Harper VanSteenhouse, Yumi Kasai, David Gray, Carol Clugston, Anna Dominiczak and Alex Alderton, Roberto Amato, Jeffrey Barrett, Sonia Goncalves, Ewan Harrison, David K. Jackson, Ian Johnston, Dominic Kwiatkowski, Cordelia Langford, John Sillitoe on behalf of the Wellcome Sanger Institute COVID-19 Surveillance Team |
| EPI_ISL_3013362 | Lighthouse Lab in Alderley Park | Wellcome Sanger Institute for the COVID-19 Genomics UK (COG-UK) Consortium | Jacquelyn Wynn, Mairead Hyland, The Lighthouse Lab in Alderley Park and Alex Alderton, Roberto Amato, Jeffrey Barrett, Sonia Goncalves, Ewan Harrison, David K. Jackson, Ian Johnston, Dominic Kwiatkowski, Cordelia Langford, John Sillitoe on behalf of the Wellcome Sanger Institute COVID-19 Surveillance Team |
| EPI_ISL_3013363 | Lighthouse Lab in Milton Keynes | Wellcome Sanger Institute for the COVID-19 Genomics UK (COG-UK) Consortium | The Lighthouse Lab in Milton Keynes and Alex Alderton, Roberto Amato, Jeffrey Barrett, Sonia Goncalves, Ewan Harrison, David K. Jackson, Ian Johnston, Dominic Kwiatkowski, Cordelia Langford, John Sillitoe on behalf of the Wellcome Sanger Institute COVID-19 Surveillance Team |
| EPI_ISL_3013364, EPI_ISL_3013365, EPI_ISL_3013366, EPI_ISL_3013367 | Lighthouse Lab in Glasgow | Wellcome Sanger Institute for the COVID-19 Genomics UK (COG-UK) Consortium | Harper VanSteenhouse, Yumi Kasai, David Gray, Carol Clugston, Anna Dominiczak and Alex Alderton, Roberto Amato, Jeffrey Barrett, Sonia Goncalves, Ewan Harrison, David K. Jackson, Ian Johnston, Dominic Kwiatkowski, Cordelia Langford, John Sillitoe on behalf of the Wellcome Sanger Institute COVID-19 Surveillance Team |
| EPI_ISL_3013368 | Lighthouse Lab in Milton Keynes | Wellcome Sanger Institute for the COVID-19 Genomics UK (COG-UK) Consortium | The Lighthouse Lab in Milton Keynes and Alex Alderton, Roberto Amato, Jeffrey Barrett, Sonia Goncalves, Ewan Harrison, David K. Jackson, Ian Johnston, Dominic Kwiatkowski, Cordelia Langford, John Sillitoe on behalf of the Wellcome Sanger Institute COVID-19 Surveillance Team |
| EPI_ISL_3013369 | Lighthouse Lab in Alderley Park | Wellcome Sanger Institute for the COVID-19 Genomics UK (COG-UK) Consortium | Jacquelyn Wynn, Mairead Hyland, The Lighthouse Lab in Alderley Park and Alex Alderton, Roberto Amato, Jeffrey Barrett, Sonia Goncalves, Ewan Harrison, David K. Jackson, Ian Johnston, Dominic Kwiatkowski, Cordelia Langford, John Sillitoe on behalf of the Wellcome Sanger Institute COVID-19 Surveillance Team |
| EPI_ISL_3013370 | Lighthouse Lab in Glasgow | Wellcome Sanger Institute for the COVID-19 Genomics UK (COG-UK) Consortium | Harper VanSteenhouse, Yumi Kasai, David Gray, Carol Clugston, Anna Dominiczak and Alex Alderton, Roberto Amato, Jeffrey Barrett, Sonia Goncalves, Ewan Harrison, David K. Jackson, Ian Johnston, Dominic Kwiatkowski, Cordelia Langford, John Sillitoe on behalf of the Wellcome Sanger Institute COVID-19 Surveillance Team |
| EPI_ISL_3013371, EPI_ISL_3013372 | Lighthouse Lab in Milton Keynes | Wellcome Sanger Institute for the COVID-19 Genomics UK (COG-UK) Consortium | The Lighthouse Lab in Milton Keynes and Alex Alderton, Roberto Amato, Jeffrey Barrett, Sonia Goncalves, Ewan Harrison, David K. Jackson, Ian Johnston, Dominic Kwiatkowski, Cordelia Langford, John Sillitoe on behalf of the Wellcome Sanger Institute COVID-19 Surveillance Team |
| EPI_ISL_3013373 | Lighthouse Lab in Alderley Park | Wellcome Sanger Institute for the COVID-19 Genomics UK (COG-UK) Consortium | Jacquelyn Wynn, Mairead Hyland, The Lighthouse Lab in Alderley Park and Alex Alderton, Roberto Amato, Jeffrey Barrett, Sonia Goncalves, Ewan Harrison, David K. Jackson, Ian Johnston, Dominic Kwiatkowski, Cordelia Langford, John Sillitoe on behalf of the Wellcome Sanger Institute COVID-19 Surveillance Team |
| EPI_ISL_3013374 | Lighthouse Lab in Milton Keynes | Wellcome Sanger Institute for the COVID-19 Genomics UK (COG-UK) Consortium | The Lighthouse Lab in Milton Keynes and Alex Alderton, Roberto Amato, Jeffrey Barrett, Sonia Goncalves, Ewan Harrison, David K. Jackson, Ian Johnston, Dominic Kwiatkowski, Cordelia Langford, John Sillitoe on behalf of the Wellcome Sanger Institute COVID-19 Surveillance Team |
| EPI_ISL_3013375, EPI_ISL_3013376 | Lighthouse Lab in Glasgow | Wellcome Sanger Institute for the COVID-19 Genomics UK (COG-UK) Consortium | Harper VanSteenhouse, Yumi Kasai, David Gray, Carol Clugston, Anna Dominiczak and Alex Alderton, Roberto Amato, Jeffrey Barrett, Sonia Goncalves, Ewan Harrison, David K. Jackson, Ian Johnston, Dominic Kwiatkowski, Cordelia Langford, John Sillitoe on behalf of the Wellcome Sanger Institute COVID-19 Surveillance Team |
| EPI_ISL_3013377 | Lighthouse Lab in Milton Keynes | Wellcome Sanger Institute for the COVID-19 Genomics UK (COG-UK) Consortium | The Lighthouse Lab in Milton Keynes and Alex Alderton, Roberto Amato, Jeffrey Barrett, Sonia Goncalves, Ewan Harrison, David K. Jackson, Ian Johnston, Dominic Kwiatkowski, Cordelia Langford, John Sillitoe on behalf of the Wellcome Sanger Institute COVID-19 Surveillance Team |
| EPI_ISL_3013378 | Lighthouse Lab in Alderley Park | Wellcome Sanger Institute for the COVID-19 Genomics UK (COG-UK) Consortium | Jacquelyn Wynn, Mairead Hyland, The Lighthouse Lab in Alderley Park and Alex Alderton, Roberto Amato, Jeffrey Barrett, Sonia Goncalves, Ewan Harrison, David K. Jackson, Ian Johnston, Dominic Kwiatkowski, Cordelia Langford, John Sillitoe on behalf of the Wellcome Sanger Institute COVID-19 Surveillance Team |
| EPI_ISL_3013379, EPI_ISL_3013380, EPI_ISL_3013381 | Lighthouse Lab in Milton Keynes | Wellcome Sanger Institute for the COVID-19 Genomics UK (COG-UK) Consortium | The Lighthouse Lab in Milton Keynes and Alex Alderton, Roberto Amato, Jeffrey Barrett, Sonia Goncalves, Ewan Harrison, David K. Jackson, Ian Johnston, Dominic Kwiatkowski, Cordelia Langford, John Sillitoe on behalf of the Wellcome Sanger Institute COVID-19 Surveillance Team |
| EPI_ISL_3013382 | Lighthouse Lab in Glasgow | Wellcome Sanger Institute for the COVID-19 Genomics UK (COG-UK) Consortium | Harper VanSteenhouse, Yumi Kasai, David Gray, Carol Clugston, Anna Dominiczak and Alex Alderton, Roberto Amato, Jeffrey Barrett, Sonia Goncalves, Ewan Harrison, David K. Jackson, Ian Johnston, Dominic Kwiatkowski, Cordelia Langford, John Sillitoe on behalf of the Wellcome Sanger Institute COVID-19 Surveillance Team |
| EPI_ISL_3013383, EPI_ISL_3013384 | Lighthouse Lab in Milton Keynes | Wellcome Sanger Institute for the COVID-19 Genomics UK (COG-UK) Consortium | The Lighthouse Lab in Milton Keynes and Alex Alderton, Roberto Amato, Jeffrey Barrett, Sonia Goncalves, Ewan Harrison, David K. Jackson, Ian Johnston, Dominic Kwiatkowski, Cordelia Langford, John Sillitoe on behalf of the Wellcome Sanger Institute COVID-19 Surveillance Team |
| EPI_ISL_3013385, EPI_ISL_3013386 | Lighthouse Lab in Glasgow | Wellcome Sanger Institute for the COVID-19 Genomics UK (COG-UK) Consortium | Harper VanSteenhouse, Yumi Kasai, David Gray, Carol Clugston, Anna Dominiczak and Alex Alderton, Roberto Amato, Jeffrey Barrett, Sonia Goncalves, Ewan Harrison, David K. Jackson, Ian Johnston, Dominic Kwiatkowski, Cordelia Langford, John Sillitoe on behalf of the Wellcome Sanger Institute COVID-19 Surveillance Team |
| EPI_ISL_3013387 | Lighthouse Lab in Alderley Park | Wellcome Sanger Institute for the COVID-19 Genomics UK (COG-UK) Consortium | Jacquelyn Wynn, Mairead Hyland, The Lighthouse Lab in Alderley Park and Alex Alderton, Roberto Amato, Jeffrey Barrett, Sonia Goncalves, Ewan Harrison, David K. Jackson, Ian Johnston, Dominic Kwiatkowski, Cordelia Langford, John Sillitoe on behalf of the Wellcome Sanger Institute COVID-19 Surveillance Team |
| EPI_ISL_3013388, EPI_ISL_3013389, EPI_ISL_3013390 | Lighthouse Lab in Glasgow | Wellcome Sanger Institute for the COVID-19 Genomics UK (COG-UK) Consortium | Harper VanSteenhouse, Yumi Kasai, David Gray, Carol Clugston, Anna Dominiczak and Alex Alderton, Roberto Amato, Jeffrey Barrett, Sonia Goncalves, Ewan Harrison, David K. Jackson, Ian Johnston, Dominic Kwiatkowski, Cordelia Langford, John Sillitoe on behalf of the Wellcome Sanger Institute COVID-19 Surveillance Team |
| EPI_ISL_3013391, EPI_ISL_3013392 | Lighthouse Lab in Milton Keynes | Wellcome Sanger Institute for the COVID-19 Genomics UK (COG-UK) Consortium | The Lighthouse Lab in Milton Keynes and Alex Alderton, Roberto Amato, Jeffrey Barrett, Sonia Goncalves, Ewan Harrison, David K. Jackson, Ian Johnston, Dominic Kwiatkowski, Cordelia Langford, John Sillitoe on behalf of the Wellcome Sanger Institute COVID-19 Surveillance Team |
| EPI_ISL_3013393 | Lighthouse Lab in Glasgow | Wellcome Sanger Institute for the COVID-19 Genomics UK (COG-UK) Consortium | Harper VanSteenhouse, Yumi Kasai, David Gray, Carol Clugston, Anna Dominiczak and Alex Alderton, Roberto Amato, Jeffrey Barrett, Sonia Goncalves, Ewan Harrison, David K. Jackson, Ian Johnston, Dominic Kwiatkowski, Cordelia Langford, John Sillitoe on behalf of the Wellcome Sanger Institute COVID-19 Surveillance Team |
| EPI_ISL_3013394 | Lighthouse Lab in Milton Keynes | Wellcome Sanger Institute for the COVID-19 Genomics UK (COG-UK) Consortium | The Lighthouse Lab in Milton Keynes and Alex Alderton, Roberto Amato, Jeffrey Barrett, Sonia Goncalves, Ewan Harrison, David K. Jackson, Ian Johnston, Dominic Kwiatkowski, Cordelia Langford, John Sillitoe on behalf of the Wellcome Sanger Institute COVID-19 Surveillance Team |
| EPI_ISL_3013395 | Lighthouse Lab in Alderley Park | Wellcome Sanger Institute for the COVID-19 Genomics UK (COG-UK) Consortium | Jacquelyn Wynn, Mairead Hyland, The Lighthouse Lab in Alderley Park and Alex Alderton, Roberto Amato, Jeffrey Barrett, Sonia Goncalves, Ewan Harrison, David K. Jackson, Ian Johnston, Dominic Kwiatkowski, Cordelia Langford, John Sillitoe on behalf of the Wellcome Sanger Institute COVID-19 Surveillance Team |
| EPI_ISL_3013396 | Lighthouse Lab in Glasgow | Wellcome Sanger Institute for the COVID-19 Genomics UK (COG-UK) Consortium | Harper VanSteenhouse, Yumi Kasai, David Gray, Carol Clugston, Anna Dominiczak and Alex Alderton, Roberto Amato, Jeffrey Barrett, Sonia Goncalves, Ewan Harrison, David K. Jackson, Ian Johnston, Dominic Kwiatkowski, Cordelia Langford, John Sillitoe on behalf of the Wellcome Sanger Institute COVID-19 Surveillance Team |
| EPI_ISL_3013397 | Lighthouse Lab in Milton Keynes | Wellcome Sanger Institute for the COVID-19 Genomics UK (COG-UK) Consortium | The Lighthouse Lab in Milton Keynes and Alex Alderton, Roberto Amato, Jeffrey Barrett, Sonia Goncalves, Ewan Harrison, David K. Jackson, Ian Johnston, Dominic Kwiatkowski, Cordelia Langford, John Sillitoe on behalf of the Wellcome Sanger Institute COVID-19 Surveillance Team |
| EPI_ISL_3013398 | Lighthouse Lab in Alderley Park | Wellcome Sanger Institute for the COVID-19 Genomics UK (COG-UK) Consortium | Jacquelyn Wynn, Mairead Hyland, The Lighthouse Lab in Alderley Park and Alex Alderton, Roberto Amato, Jeffrey Barrett, Sonia Goncalves, Ewan Harrison, David K. Jackson, Ian Johnston, Dominic Kwiatkowski, Cordelia Langford, John Sillitoe on behalf of the Wellcome Sanger Institute COVID-19 Surveillance Team |

| | | | |
|---|---|---|---|
| EPI_ISL_3013399 | Lighthouse Lab in Glasgow | Wellcome Sanger Institute for the COVID-19 Genomics UK (COG-UK) Consortium | Harper VanSteenhouse, Yumi Kasai, David Gray, Carol Clugston, Anna Dominiczak and Alex Alderton, Roberto Amato, Jeffrey Barrett, Sonia Goncalves, Ewan Harrison, David K. Jackson, Ian Johnston, Dominic Kwiatkowski, Cordelia Langford, John Sillitoe on behalf of the Wellcome Sanger Institute COVID-19 Surveillance Team |
| EPI_ISL_3013400 | Lighthouse Lab in Milton Keynes | Wellcome Sanger Institute for the COVID-19 Genomics UK (COG-UK) Consortium | The Lighthouse Lab in Milton Keynes and Alex Alderton, Roberto Amato, Jeffrey Barrett, Sonia Goncalves, Ewan Harrison, David K. Jackson, Ian Johnston, Dominic Kwiatkowski, Cordelia Langford, John Sillitoe on behalf of the Wellcome Sanger Institute COVID-19 Surveillance Team |
| EPI_ISL_3013401 | Lighthouse Lab in Alderley Park | Wellcome Sanger Institute for the COVID-19 Genomics UK (COG-UK) Consortium | Jacquelyn Wynn, Mairead Hyland, The Lighthouse Lab in Alderley Park and Alex Alderton, Roberto Amato, Jeffrey Barrett, Sonia Goncalves, Ewan Harrison, David K. Jackson, Ian Johnston, Dominic Kwiatkowski, Cordelia Langford, John Sillitoe on behalf of the Wellcome Sanger Institute COVID-19 Surveillance Team |
| EPI_ISL_3013402 | Lighthouse Lab in Glasgow | Wellcome Sanger Institute for the COVID-19 Genomics UK (COG-UK) Consortium | Harper VanSteenhouse, Yumi Kasai, David Gray, Carol Clugston, Anna Dominiczak and Alex Alderton, Roberto Amato, Jeffrey Barrett, Sonia Goncalves, Ewan Harrison, David K. Jackson, Ian Johnston, Dominic Kwiatkowski, Cordelia Langford, John Sillitoe on behalf of the Wellcome Sanger Institute COVID-19 Surveillance Team |
| EPI_ISL_3013403 | Lighthouse Lab in Milton Keynes | Wellcome Sanger Institute for the COVID-19 Genomics UK (COG-UK) Consortium | The Lighthouse Lab in Milton Keynes and Alex Alderton, Roberto Amato, Jeffrey Barrett, Sonia Goncalves, Ewan Harrison, David K. Jackson, Ian Johnston, Dominic Kwiatkowski, Cordelia Langford, John Sillitoe on behalf of the Wellcome Sanger Institute COVID-19 Surveillance Team |
| EPI_ISL_3013404 | Lighthouse Lab in Alderley Park | Wellcome Sanger Institute for the COVID-19 Genomics UK (COG-UK) Consortium | Jacquelyn Wynn, Mairead Hyland, The Lighthouse Lab in Alderley Park and Alex Alderton, Roberto Amato, Jeffrey Barrett, Sonia Goncalves, Ewan Harrison, David K. Jackson, Ian Johnston, Dominic Kwiatkowski, Cordelia Langford, John Sillitoe on behalf of the Wellcome Sanger Institute COVID-19 Surveillance Team |
| EPI_ISL_3013405 | Lighthouse Lab in Glasgow | Wellcome Sanger Institute for the COVID-19 Genomics UK (COG-UK) Consortium | Harper VanSteenhouse, Yumi Kasai, David Gray, Carol Clugston, Anna Dominiczak and Alex Alderton, Roberto Amato, Jeffrey Barrett, Sonia Goncalves, Ewan Harrison, David K. Jackson, Ian Johnston, Dominic Kwiatkowski, Cordelia Langford, John Sillitoe on behalf of the Wellcome Sanger Institute COVID-19 Surveillance Team |
| EPI_ISL_3013406 | Lighthouse Lab in Milton Keynes | Wellcome Sanger Institute for the COVID-19 Genomics UK (COG-UK) Consortium | The Lighthouse Lab in Milton Keynes and Alex Alderton, Roberto Amato, Jeffrey Barrett, Sonia Goncalves, Ewan Harrison, David K. Jackson, Ian Johnston, Dominic Kwiatkowski, Cordelia Langford, John Sillitoe on behalf of the Wellcome Sanger Institute COVID-19 Surveillance Team |
| EPI_ISL_3013407 | Lighthouse Lab in Glasgow | Wellcome Sanger Institute for the COVID-19 Genomics UK (COG-UK) Consortium | Harper VanSteenhouse, Yumi Kasai, David Gray, Carol Clugston, Anna Dominiczak and Alex Alderton, Roberto Amato, Jeffrey Barrett, Sonia Goncalves, Ewan Harrison, David K. Jackson, Ian Johnston, Dominic Kwiatkowski, Cordelia Langford, John Sillitoe on behalf of the Wellcome Sanger Institute COVID-19 Surveillance Team |
| EPI_ISL_3013408 | Lighthouse Lab in Milton Keynes | Wellcome Sanger Institute for the COVID-19 Genomics UK (COG-UK) Consortium | The Lighthouse Lab in Milton Keynes and Alex Alderton, Roberto Amato, Jeffrey Barrett, Sonia Goncalves, Ewan Harrison, David K. Jackson, Ian Johnston, Dominic Kwiatkowski, Cordelia Langford, John Sillitoe on behalf of the Wellcome Sanger Institute COVID-19 Surveillance Team |
| EPI_ISL_3013409 | Lighthouse Lab in Glasgow | Wellcome Sanger Institute for the COVID-19 Genomics UK (COG-UK) Consortium | Harper VanSteenhouse, Yumi Kasai, David Gray, Carol Clugston, Anna Dominiczak and Alex Alderton, Roberto Amato, Jeffrey Barrett, Sonia Goncalves, Ewan Harrison, David K. Jackson, Ian Johnston, Dominic Kwiatkowski, Cordelia Langford, John Sillitoe on behalf of the Wellcome Sanger Institute COVID-19 Surveillance Team |
| EPI_ISL_3013410 | Lighthouse Lab in Alderley Park | Wellcome Sanger Institute for the COVID-19 Genomics UK (COG-UK) Consortium | Jacquelyn Wynn, Mairead Hyland, The Lighthouse Lab in Alderley Park and Alex Alderton, Roberto Amato, Jeffrey Barrett, Sonia Goncalves, Ewan Harrison, David K. Jackson, Ian Johnston, Dominic Kwiatkowski, Cordelia Langford, John Sillitoe on behalf of the Wellcome Sanger Institute COVID-19 Surveillance Team |
| EPI_ISL_3013411 | Lighthouse Lab in Glasgow | Wellcome Sanger Institute for the COVID-19 Genomics UK (COG-UK) Consortium | Harper VanSteenhouse, Yumi Kasai, David Gray, Carol Clugston, Anna Dominiczak and Alex Alderton, Roberto Amato, Jeffrey Barrett, Sonia Goncalves, Ewan Harrison, David K. Jackson, Ian Johnston, Dominic Kwiatkowski, Cordelia Langford, John Sillitoe on behalf of the Wellcome Sanger Institute COVID-19 Surveillance Team |
| EPI_ISL_3013412 | Lighthouse Lab in Alderley Park | Wellcome Sanger Institute for the COVID-19 Genomics UK (COG-UK) Consortium | Jacquelyn Wynn, Mairead Hyland, The Lighthouse Lab in Alderley Park and Alex Alderton, Roberto Amato, Jeffrey Barrett, Sonia Goncalves, Ewan Harrison, David K. Jackson, Ian Johnston, Dominic Kwiatkowski, Cordelia Langford, John Sillitoe on behalf of the Wellcome Sanger Institute COVID-19 Surveillance Team |
| EPI_ISL_3013413, EPI_ISL_3013414, EPI_ISL_3013415 | Lighthouse Lab in Milton Keynes | Wellcome Sanger Institute for the COVID-19 Genomics UK (COG-UK) Consortium | The Lighthouse Lab in Milton Keynes and Alex Alderton, Roberto Amato, Jeffrey Barrett, Sonia Goncalves, Ewan Harrison, David K. Jackson, Ian Johnston, Dominic Kwiatkowski, Cordelia Langford, John Sillitoe on behalf of the Wellcome Sanger Institute COVID-19 Surveillance Team |
| EPI_ISL_3013416, EPI_ISL_3013417, EPI_ISL_3013418, EPI_ISL_3013419 | Lighthouse Lab in Alderley Park | Wellcome Sanger Institute for the COVID-19 Genomics UK (COG-UK) Consortium | Jacquelyn Wynn, Mairead Hyland, The Lighthouse Lab in Alderley Park and Alex Alderton, Roberto Amato, Jeffrey Barrett, Sonia Goncalves, Ewan Harrison, David K. Jackson, Ian Johnston, Dominic Kwiatkowski, Cordelia Langford, John Sillitoe on behalf of the Wellcome Sanger Institute COVID-19 Surveillance Team |
| EPI_ISL_3013420 | Lighthouse Lab in Milton Keynes | Wellcome Sanger Institute for the COVID-19 Genomics UK (COG-UK) Consortium | The Lighthouse Lab in Milton Keynes and Alex Alderton, Roberto Amato, Jeffrey Barrett, Sonia Goncalves, Ewan Harrison, David K. Jackson, Ian Johnston, Dominic Kwiatkowski, Cordelia Langford, John Sillitoe on behalf of the Wellcome Sanger Institute COVID-19 Surveillance Team |
| EPI_ISL_3013421 | Lighthouse Lab in Glasgow | Wellcome Sanger Institute for the COVID-19 Genomics UK (COG-UK) Consortium | Harper VanSteenhouse, Yumi Kasai, David Gray, Carol Clugston, Anna Dominiczak and Alex Alderton, Roberto Amato, Jeffrey Barrett, Sonia Goncalves, Ewan Harrison, David K. Jackson, Ian Johnston, Dominic Kwiatkowski, Cordelia Langford, John Sillitoe on behalf of the Wellcome Sanger Institute COVID-19 Surveillance Team |
| EPI_ISL_3013422, EPI_ISL_3013423 | Lighthouse Lab in Alderley Park | Wellcome Sanger Institute for the COVID-19 Genomics UK (COG-UK) Consortium | Jacquelyn Wynn, Mairead Hyland, The Lighthouse Lab in Alderley Park and Alex Alderton, Roberto Amato, Jeffrey Barrett, Sonia Goncalves, Ewan Harrison, David K. Jackson, Ian Johnston, Dominic Kwiatkowski, Cordelia Langford, John Sillitoe on behalf of the Wellcome Sanger Institute COVID-19 Surveillance Team |
| EPI_ISL_3013424 | Lighthouse Lab in Milton Keynes | Wellcome Sanger Institute for the COVID-19 Genomics UK (COG-UK) Consortium | The Lighthouse Lab in Milton Keynes and Alex Alderton, Roberto Amato, Jeffrey Barrett, Sonia Goncalves, Ewan Harrison, David K. Jackson, Ian Johnston, Dominic Kwiatkowski, Cordelia Langford, John Sillitoe on behalf of the Wellcome Sanger Institute COVID-19 Surveillance Team |
| EPI_ISL_3013425 | Lighthouse Lab in Alderley Park | Wellcome Sanger Institute for the COVID-19 Genomics UK (COG-UK) Consortium | Jacquelyn Wynn, Mairead Hyland, The Lighthouse Lab in Alderley Park and Alex Alderton, Roberto Amato, Jeffrey Barrett, Sonia Goncalves, Ewan Harrison, David K. Jackson, Ian Johnston, Dominic Kwiatkowski, Cordelia Langford, John Sillitoe on behalf of the Wellcome Sanger Institute COVID-19 Surveillance Team |
| EPI_ISL_3013426 | Lighthouse Lab in Milton Keynes | Wellcome Sanger Institute for the COVID-19 Genomics UK (COG-UK) Consortium | The Lighthouse Lab in Milton Keynes and Alex Alderton, Roberto Amato, Jeffrey Barrett, Sonia Goncalves, Ewan Harrison, David K. Jackson, Ian Johnston, Dominic Kwiatkowski, Cordelia Langford, John Sillitoe on behalf of the Wellcome Sanger Institute COVID-19 Surveillance Team |
| EPI_ISL_3013427 | Lighthouse Lab in Glasgow | Wellcome Sanger Institute for the COVID-19 Genomics UK (COG-UK) Consortium | Harper VanSteenhouse, Yumi Kasai, David Gray, Carol Clugston, Anna Dominiczak and Alex Alderton, Roberto Amato, Jeffrey Barrett, Sonia Goncalves, Ewan Harrison, David K. Jackson, Ian Johnston, Dominic Kwiatkowski, Cordelia Langford, John Sillitoe on behalf of the Wellcome Sanger Institute COVID-19 Surveillance Team |
| EPI_ISL_3013428 | Lighthouse Lab in Alderley Park | Wellcome Sanger Institute for the COVID-19 Genomics UK (COG-UK) Consortium | Jacquelyn Wynn, Mairead Hyland, The Lighthouse Lab in Alderley Park and Alex Alderton, Roberto Amato, Jeffrey Barrett, Sonia Goncalves, Ewan Harrison, David K. Jackson, Ian Johnston, Dominic Kwiatkowski, Cordelia Langford, John Sillitoe on behalf of the Wellcome Sanger Institute COVID-19 Surveillance Team |
| EPI_ISL_3013429, EPI_ISL_3013430, EPI_ISL_3013431, EPI_ISL_3013432 | Lighthouse Lab in Glasgow | Wellcome Sanger Institute for the COVID-19 Genomics UK (COG-UK) Consortium | Harper VanSteenhouse, Yumi Kasai, David Gray, Carol Clugston, Anna Dominiczak and Alex Alderton, Roberto Amato, Jeffrey Barrett, Sonia Goncalves, Ewan Harrison, David K. Jackson, Ian Johnston, Dominic Kwiatkowski, Cordelia Langford, John Sillitoe on behalf of the Wellcome Sanger Institute COVID-19 Surveillance Team |
| EPI_ISL_3013433, EPI_ISL_3013434, EPI_ISL_3013435, EPI_ISL_3013436, EPI_ISL_3013437, EPI_ISL_3013438 | Lighthouse Lab in Milton Keynes | Wellcome Sanger Institute for the COVID-19 Genomics UK (COG-UK) Consortium | The Lighthouse Lab in Milton Keynes and Alex Alderton, Roberto Amato, Jeffrey Barrett, Sonia Goncalves, Ewan Harrison, David K. Jackson, Ian Johnston, Dominic Kwiatkowski, Cordelia Langford, John Sillitoe on behalf of the Wellcome Sanger Institute COVID-19 Surveillance Team |
| EPI_ISL_3013439 | Lighthouse Lab in Alderley Park | Wellcome Sanger Institute for the COVID-19 Genomics UK (COG-UK) Consortium | Jacquelyn Wynn, Mairead Hyland, The Lighthouse Lab in Alderley Park and Alex Alderton, Roberto Amato, Jeffrey Barrett, Sonia Goncalves, Ewan Harrison, David K. Jackson, Ian Johnston, Dominic Kwiatkowski, Cordelia Langford, John Sillitoe on behalf of the Wellcome Sanger Institute COVID-19 Surveillance Team |
| EPI_ISL_3013440 | Lighthouse Lab in Milton Keynes | Wellcome Sanger Institute for the COVID-19 Genomics UK (COG-UK) Consortium | The Lighthouse Lab in Milton Keynes and Alex Alderton, Roberto Amato, Jeffrey Barrett, Sonia Goncalves, Ewan Harrison, David K. Jackson, Ian Johnston, Dominic Kwiatkowski, Cordelia Langford, John Sillitoe on behalf of the Wellcome Sanger Institute COVID-19 Surveillance Team |
| EPI_ISL_3013441, EPI_ISL_3013442, EPI_ISL_3013443 | Lighthouse Lab in Alderley Park | Wellcome Sanger Institute for the COVID-19 Genomics UK (COG-UK) Consortium | Jacquelyn Wynn, Mairead Hyland, The Lighthouse Lab in Alderley Park and Alex Alderton, Roberto Amato, Jeffrey Barrett, Sonia Goncalves, Ewan Harrison, David K. Jackson, Ian Johnston, Dominic Kwiatkowski, Cordelia Langford, John Sillitoe on behalf of the Wellcome Sanger Institute COVID-19 Surveillance Team |
| EPI_ISL_3013444 | Lighthouse Lab in Glasgow | Wellcome Sanger Institute for the COVID-19 Genomics UK (COG-UK) Consortium | Harper VanSteenhouse, Yumi Kasai, David Gray, Carol Clugston, Anna Dominiczak and Alex Alderton, Roberto Amato, Jeffrey Barrett, Sonia Goncalves, Ewan Harrison, David K. Jackson, Ian Johnston, Dominic Kwiatkowski, Cordelia Langford, John Sillitoe on behalf of the Wellcome Sanger Institute COVID-19 Surveillance Team |
| EPI_ISL_3013445, EPI_ISL_3013446, EPI_ISL_3013447 | Lighthouse Lab in Milton Keynes | Wellcome Sanger Institute for the COVID-19 Genomics UK (COG-UK) Consortium | The Lighthouse Lab in Milton Keynes and Alex Alderton, Roberto Amato, Jeffrey Barrett, Sonia Goncalves, Ewan Harrison, David K. Jackson, Ian Johnston, Dominic Kwiatkowski, Cordelia Langford, John Sillitoe on behalf of the Wellcome Sanger Institute COVID-19 Surveillance Team |
| EPI_ISL_3013448 | Lighthouse Lab in Glasgow | Wellcome Sanger Institute for the COVID-19 Genomics UK (COG-UK) Consortium | Harper VanSteenhouse, Yumi Kasai, David Gray, Carol Clugston, Anna Dominiczak and Alex Alderton, Roberto Amato, Jeffrey Barrett, Sonia Goncalves, Ewan Harrison, David K. Jackson, Ian Johnston, Dominic Kwiatkowski, Cordelia Langford, John Sillitoe on behalf of the Wellcome Sanger Institute COVID-19 Surveillance Team |
| EPI_ISL_3013449, EPI_ISL_3013450 | Lighthouse Lab in Milton Keynes | Wellcome Sanger Institute for the COVID-19 Genomics UK (COG-UK) Consortium | The Lighthouse Lab in Milton Keynes and Alex Alderton, Roberto Amato, Jeffrey Barrett, Sonia Goncalves, Ewan Harrison, David K. Jackson, Ian Johnston, Dominic Kwiatkowski, Cordelia Langford, John Sillitoe on behalf of the Wellcome Sanger Institute COVID-19 Surveillance Team |
| EPI_ISL_3013451 | Lighthouse Lab in Alderley Park | Wellcome Sanger Institute for the COVID-19 Genomics UK (COG-UK) Consortium | Jacquelyn Wynn, Mairead Hyland, The Lighthouse Lab in Alderley Park and Alex Alderton, Roberto Amato, Jeffrey Barrett, Sonia Goncalves, Ewan Harrison, David K. Jackson, Ian Johnston, Dominic Kwiatkowski, Cordelia Langford, John Sillitoe on behalf of the Wellcome Sanger Institute COVID-19 Surveillance Team |
| EPI_ISL_3013452 | Lighthouse Lab in Glasgow | Wellcome Sanger Institute for the COVID-19 Genomics UK (COG-UK) Consortium | Harper VanSteenhouse, Yumi Kasai, David Gray, Carol Clugston, Anna Dominiczak and Alex Alderton, Roberto Amato, Jeffrey Barrett, Sonia Goncalves, Ewan Harrison, David K. Jackson, Ian Johnston, Dominic Kwiatkowski, Cordelia Langford, John Sillitoe on behalf of the Wellcome Sanger Institute COVID-19 Surveillance Team |
| EPI_ISL_3013453, EPI_ISL_3013454 | Lighthouse Lab in Alderley Park | Wellcome Sanger Institute for the COVID-19 Genomics UK (COG-UK) Consortium | Jacquelyn Wynn, Mairead Hyland, The Lighthouse Lab in Alderley Park and Alex Alderton, Roberto Amato, Jeffrey Barrett, Sonia Goncalves, Ewan Harrison, David K. Jackson, Ian Johnston, Dominic Kwiatkowski, Cordelia Langford, John Sillitoe on behalf of the Wellcome Sanger Institute COVID-19 Surveillance Team |
| EPI_ISL_3013455, EPI_ISL_3013456, EPI_ISL_3013457 | Lighthouse Lab in Milton Keynes | Wellcome Sanger Institute for the COVID-19 Genomics UK (COG-UK) Consortium | The Lighthouse Lab in Milton Keynes and Alex Alderton, Roberto Amato, Jeffrey Barrett, Sonia Goncalves, Ewan Harrison, David K. Jackson, Ian Johnston, Dominic Kwiatkowski, Cordelia Langford, John Sillitoe on behalf of the Wellcome Sanger Institute COVID-19 Surveillance Team |
| EPI_ISL_3013458 | Lighthouse Lab in Glasgow | Wellcome Sanger Institute for the COVID-19 Genomics UK (COG-UK) Consortium | Harper VanSteenhouse, Yumi Kasai, David Gray, Carol Clugston, Anna Dominiczak and Alex Alderton, Roberto Amato, Jeffrey Barrett, Sonia Goncalves, Ewan Harrison, David K. Jackson, Ian Johnston, Dominic Kwiatkowski, Cordelia Langford, John Sillitoe on behalf of the Wellcome Sanger Institute COVID-19 Surveillance Team |
| EPI_ISL_3013459, EPI_ISL_3013460 | Lighthouse Lab in Alderley Park | Wellcome Sanger Institute for the COVID-19 Genomics UK (COG-UK) Consortium | Jacquelyn Wynn, Mairead Hyland, The Lighthouse Lab in Alderley Park and Alex Alderton, Roberto Amato, Jeffrey Barrett, Sonia Goncalves, Ewan Harrison, David K. Jackson, Ian Johnston, Dominic Kwiatkowski, Cordelia Langford, John Sillitoe on behalf of the Wellcome Sanger Institute COVID-19 Surveillance Team |
| EPI_ISL_3013461 | Lighthouse Lab in Milton Keynes | Wellcome Sanger Institute for the COVID-19 Genomics UK (COG-UK) Consortium | The Lighthouse Lab in Milton Keynes and Alex Alderton, Roberto Amato, Jeffrey Barrett, Sonia Goncalves, Ewan Harrison, David K. Jackson, Ian Johnston, Dominic Kwiatkowski, Cordelia Langford, John Sillitoe on behalf of the Wellcome Sanger Institute COVID-19 Surveillance Team |

| | | | |
|---|---|---|---|
| EPI_ISL_3013462, EPI_ISL_3013463 | Lighthouse Lab in Alderley Park | Wellcome Sanger Institute for the COVID-19 Genomics UK (COG-UK) Consortium | Jacquelyn Wynn, Mairead Hyland, The Lighthouse Lab in Alderley Park and Alex Alderton, Roberto Amato, Jeffrey Barrett, Sonia Goncalves, Ewan Harrison, David K. Jackson, Ian Johnston, Dominic Kwiatkowski, Cordelia Langford, John Sillitoe on behalf of the Wellcome Sanger Institute COVID-19 Surveillance Team |
| EPI_ISL_3013464 | Lighthouse Lab in Glasgow | Wellcome Sanger Institute for the COVID-19 Genomics UK (COG-UK) Consortium | Harper VanSteenhouse, Yumi Kasai, David Gray, Carol Clugston, Anna Dominiczak and Alex Alderton, Roberto Amato, Jeffrey Barrett, Sonia Goncalves, Ewan Harrison, David K. Jackson, Ian Johnston, Dominic Kwiatkowski, Cordelia Langford, John Sillitoe on behalf of the Wellcome Sanger Institute COVID-19 Surveillance Team |
| EPI_ISL_3013465 | Lighthouse Lab in Alderley Park | Wellcome Sanger Institute for the COVID-19 Genomics UK (COG-UK) Consortium | Jacquelyn Wynn, Mairead Hyland, The Lighthouse Lab in Alderley Park and Alex Alderton, Roberto Amato, Jeffrey Barrett, Sonia Goncalves, Ewan Harrison, David K. Jackson, Ian Johnston, Dominic Kwiatkowski, Cordelia Langford, John Sillitoe on behalf of the Wellcome Sanger Institute COVID-19 Surveillance Team |
| EPI_ISL_3013466, EPI_ISL_3013467, EPI_ISL_3013468, EPI_ISL_3013469, EPI_ISL_3013470 | Lighthouse Lab in Glasgow | Wellcome Sanger Institute for the COVID-19 Genomics UK (COG-UK) Consortium | Harper VanSteenhouse, Yumi Kasai, David Gray, Carol Clugston, Anna Dominiczak and Alex Alderton, Roberto Amato, Jeffrey Barrett, Sonia Goncalves, Ewan Harrison, David K. Jackson, Ian Johnston, Dominic Kwiatkowski, Cordelia Langford, John Sillitoe on behalf of the Wellcome Sanger Institute COVID-19 Surveillance Team |
| EPI_ISL_3013471 | Lighthouse Lab in Alderley Park | Wellcome Sanger Institute for the COVID-19 Genomics UK (COG-UK) Consortium | Jacquelyn Wynn, Mairead Hyland, The Lighthouse Lab in Alderley Park and Alex Alderton, Roberto Amato, Jeffrey Barrett, Sonia Goncalves, Ewan Harrison, David K. Jackson, Ian Johnston, Dominic Kwiatkowski, Cordelia Langford, John Sillitoe on behalf of the Wellcome Sanger Institute COVID-19 Surveillance Team |
| EPI_ISL_3013472 | Lighthouse Lab in Milton Keynes | Wellcome Sanger Institute for the COVID-19 Genomics UK (COG-UK) Consortium | The Lighthouse Lab in Milton Keynes and Alex Alderton, Roberto Amato, Jeffrey Barrett, Sonia Goncalves, Ewan Harrison, David K. Jackson, Ian Johnston, Dominic Kwiatkowski, Cordelia Langford, John Sillitoe on behalf of the Wellcome Sanger Institute COVID-19 Surveillance Team |
| EPI_ISL_3013473, EPI_ISL_3013474 | Lighthouse Lab in Alderley Park | Wellcome Sanger Institute for the COVID-19 Genomics UK (COG-UK) Consortium | Jacquelyn Wynn, Mairead Hyland, The Lighthouse Lab in Alderley Park and Alex Alderton, Roberto Amato, Jeffrey Barrett, Sonia Goncalves, Ewan Harrison, David K. Jackson, Ian Johnston, Dominic Kwiatkowski, Cordelia Langford, John Sillitoe on behalf of the Wellcome Sanger Institute COVID-19 Surveillance Team |
| EPI_ISL_3013475 | Lighthouse Lab in Glasgow | Wellcome Sanger Institute for the COVID-19 Genomics UK (COG-UK) Consortium | Harper VanSteenhouse, Yumi Kasai, David Gray, Carol Clugston, Anna Dominiczak and Alex Alderton, Roberto Amato, Jeffrey Barrett, Sonia Goncalves, Ewan Harrison, David K. Jackson, Ian Johnston, Dominic Kwiatkowski, Cordelia Langford, John Sillitoe on behalf of the Wellcome Sanger Institute COVID-19 Surveillance Team |
| EPI_ISL_3013476 | Lighthouse Lab in Alderley Park | Wellcome Sanger Institute for the COVID-19 Genomics UK (COG-UK) Consortium | Jacquelyn Wynn, Mairead Hyland, The Lighthouse Lab in Alderley Park and Alex Alderton, Roberto Amato, Jeffrey Barrett, Sonia Goncalves, Ewan Harrison, David K. Jackson, Ian Johnston, Dominic Kwiatkowski, Cordelia Langford, John Sillitoe on behalf of the Wellcome Sanger Institute COVID-19 Surveillance Team |
| EPI_ISL_3013477 | Lighthouse Lab in Glasgow | Wellcome Sanger Institute for the COVID-19 Genomics UK (COG-UK) Consortium | Harper VanSteenhouse, Yumi Kasai, David Gray, Carol Clugston, Anna Dominiczak and Alex Alderton, Roberto Amato, Jeffrey Barrett, Sonia Goncalves, Ewan Harrison, David K. Jackson, Ian Johnston, Dominic Kwiatkowski, Cordelia Langford, John Sillitoe on behalf of the Wellcome Sanger Institute COVID-19 Surveillance Team |
| EPI_ISL_3013478 | Lighthouse Lab in Alderley Park | Wellcome Sanger Institute for the COVID-19 Genomics UK (COG-UK) Consortium | Jacquelyn Wynn, Mairead Hyland, The Lighthouse Lab in Alderley Park and Alex Alderton, Roberto Amato, Jeffrey Barrett, Sonia Goncalves, Ewan Harrison, David K. Jackson, Ian Johnston, Dominic Kwiatkowski, Cordelia Langford, John Sillitoe on behalf of the Wellcome Sanger Institute COVID-19 Surveillance Team |
| EPI_ISL_3013479 | Lighthouse Lab in Glasgow | Wellcome Sanger Institute for the COVID-19 Genomics UK (COG-UK) Consortium | Harper VanSteenhouse, Yumi Kasai, David Gray, Carol Clugston, Anna Dominiczak and Alex Alderton, Roberto Amato, Jeffrey Barrett, Sonia Goncalves, Ewan Harrison, David K. Jackson, Ian Johnston, Dominic Kwiatkowski, Cordelia Langford, John Sillitoe on behalf of the Wellcome Sanger Institute COVID-19 Surveillance Team |
| EPI_ISL_3013480 | Lighthouse Lab in Milton Keynes | Wellcome Sanger Institute for the COVID-19 Genomics UK (COG-UK) Consortium | The Lighthouse Lab in Milton Keynes and Alex Alderton, Roberto Amato, Jeffrey Barrett, Sonia Goncalves, Ewan Harrison, David K. Jackson, Ian Johnston, Dominic Kwiatkowski, Cordelia Langford, John Sillitoe on behalf of the Wellcome Sanger Institute COVID-19 Surveillance Team |
| EPI_ISL_3013481 | Lighthouse Lab in Glasgow | Wellcome Sanger Institute for the COVID-19 Genomics UK (COG-UK) Consortium | Harper VanSteenhouse, Yumi Kasai, David Gray, Carol Clugston, Anna Dominiczak and Alex Alderton, Roberto Amato, Jeffrey Barrett, Sonia Goncalves, Ewan Harrison, David K. Jackson, Ian Johnston, Dominic Kwiatkowski, Cordelia Langford, John Sillitoe on behalf of the Wellcome Sanger Institute COVID-19 Surveillance Team |
| EPI_ISL_3013482, EPI_ISL_3013483 | Lighthouse Lab in Milton Keynes | Wellcome Sanger Institute for the COVID-19 Genomics UK (COG-UK) Consortium | The Lighthouse Lab in Milton Keynes and Alex Alderton, Roberto Amato, Jeffrey Barrett, Sonia Goncalves, Ewan Harrison, David K. Jackson, Ian Johnston, Dominic Kwiatkowski, Cordelia Langford, John Sillitoe on behalf of the Wellcome Sanger Institute COVID-19 Surveillance Team |
| EPI_ISL_3013484, EPI_ISL_3013485 | Lighthouse Lab in Glasgow | Wellcome Sanger Institute for the COVID-19 Genomics UK (COG-UK) Consortium | Harper VanSteenhouse, Yumi Kasai, David Gray, Carol Clugston, Anna Dominiczak and Alex Alderton, Roberto Amato, Jeffrey Barrett, Sonia Goncalves, Ewan Harrison, David K. Jackson, Ian Johnston, Dominic Kwiatkowski, Cordelia Langford, John Sillitoe on behalf of the Wellcome Sanger Institute COVID-19 Surveillance Team |
| EPI_ISL_3013486, EPI_ISL_3013487 | Lighthouse Lab in Milton Keynes | Wellcome Sanger Institute for the COVID-19 Genomics UK (COG-UK) Consortium | The Lighthouse Lab in Milton Keynes and Alex Alderton, Roberto Amato, Jeffrey Barrett, Sonia Goncalves, Ewan Harrison, David K. Jackson, Ian Johnston, Dominic Kwiatkowski, Cordelia Langford, John Sillitoe on behalf of the Wellcome Sanger Institute COVID-19 Surveillance Team |
| EPI_ISL_3013488, EPI_ISL_3013489, EPI_ISL_3013490 | Lighthouse Lab in Alderley Park | Wellcome Sanger Institute for the COVID-19 Genomics UK (COG-UK) Consortium | Jacquelyn Wynn, Mairead Hyland, The Lighthouse Lab in Alderley Park and Alex Alderton, Roberto Amato, Jeffrey Barrett, Sonia Goncalves, Ewan Harrison, David K. Jackson, Ian Johnston, Dominic Kwiatkowski, Cordelia Langford, John Sillitoe on behalf of the Wellcome Sanger Institute COVID-19 Surveillance Team |
| EPI_ISL_3013491 | Lighthouse Lab in Milton Keynes | Wellcome Sanger Institute for the COVID-19 Genomics UK (COG-UK) Consortium | The Lighthouse Lab in Milton Keynes and Alex Alderton, Roberto Amato, Jeffrey Barrett, Sonia Goncalves, Ewan Harrison, David K. Jackson, Ian Johnston, Dominic Kwiatkowski, Cordelia Langford, John Sillitoe on behalf of the Wellcome Sanger Institute COVID-19 Surveillance Team |
| EPI_ISL_3013492 | Lighthouse Lab in Alderley Park | Wellcome Sanger Institute for the COVID-19 Genomics UK (COG-UK) Consortium | Jacquelyn Wynn, Mairead Hyland, The Lighthouse Lab in Alderley Park and Alex Alderton, Roberto Amato, Jeffrey Barrett, Sonia Goncalves, Ewan Harrison, David K. Jackson, Ian Johnston, Dominic Kwiatkowski, Cordelia Langford, John Sillitoe on behalf of the Wellcome Sanger Institute COVID-19 Surveillance Team |
| EPI_ISL_3013493, EPI_ISL_3013494, EPI_ISL_3013495, EPI_ISL_3013496, EPI_ISL_3013497 | Lighthouse Lab in Glasgow | Wellcome Sanger Institute for the COVID-19 Genomics UK (COG-UK) Consortium | Harper VanSteenhouse, Yumi Kasai, David Gray, Carol Clugston, Anna Dominiczak and Alex Alderton, Roberto Amato, Jeffrey Barrett, Sonia Goncalves, Ewan Harrison, David K. Jackson, Ian Johnston, Dominic Kwiatkowski, Cordelia Langford, John Sillitoe on behalf of the Wellcome Sanger Institute COVID-19 Surveillance Team |
| EPI_ISL_3013498 | Lighthouse Lab in Alderley Park | Wellcome Sanger Institute for the COVID-19 Genomics UK (COG-UK) Consortium | Jacquelyn Wynn, Mairead Hyland, The Lighthouse Lab in Alderley Park and Alex Alderton, Roberto Amato, Jeffrey Barrett, Sonia Goncalves, Ewan Harrison, David K. Jackson, Ian Johnston, Dominic Kwiatkowski, Cordelia Langford, John Sillitoe on behalf of the Wellcome Sanger Institute COVID-19 Surveillance Team |
| EPI_ISL_3013499 | Lighthouse Lab in Glasgow | Wellcome Sanger Institute for the COVID-19 Genomics UK (COG-UK) Consortium | Harper VanSteenhouse, Yumi Kasai, David Gray, Carol Clugston, Anna Dominiczak and Alex Alderton, Roberto Amato, Jeffrey Barrett, Sonia Goncalves, Ewan Harrison, David K. Jackson, Ian Johnston, Dominic Kwiatkowski, Cordelia Langford, John Sillitoe on behalf of the Wellcome Sanger Institute COVID-19 Surveillance Team |
| EPI_ISL_3013500, EPI_ISL_3013501 | Lighthouse Lab in Alderley Park | Wellcome Sanger Institute for the COVID-19 Genomics UK (COG-UK) Consortium | Jacquelyn Wynn, Mairead Hyland, The Lighthouse Lab in Alderley Park and Alex Alderton, Roberto Amato, Jeffrey Barrett, Sonia Goncalves, Ewan Harrison, David K. Jackson, Ian Johnston, Dominic Kwiatkowski, Cordelia Langford, John Sillitoe on behalf of the Wellcome Sanger Institute COVID-19 Surveillance Team |
| EPI_ISL_3013502 | Lighthouse Lab in Glasgow | Wellcome Sanger Institute for the COVID-19 Genomics UK (COG-UK) Consortium | Harper VanSteenhouse, Yumi Kasai, David Gray, Carol Clugston, Anna Dominiczak and Alex Alderton, Roberto Amato, Jeffrey Barrett, Sonia Goncalves, Ewan Harrison, David K. Jackson, Ian Johnston, Dominic Kwiatkowski, Cordelia Langford, John Sillitoe on behalf of the Wellcome Sanger Institute COVID-19 Surveillance Team |
| EPI_ISL_3013503 | Lighthouse Lab in Milton Keynes | Wellcome Sanger Institute for the COVID-19 Genomics UK (COG-UK) Consortium | The Lighthouse Lab in Milton Keynes and Alex Alderton, Roberto Amato, Jeffrey Barrett, Sonia Goncalves, Ewan Harrison, David K. Jackson, Ian Johnston, Dominic Kwiatkowski, Cordelia Langford, John Sillitoe on behalf of the Wellcome Sanger Institute COVID-19 Surveillance Team |
| EPI_ISL_3013504, EPI_ISL_3013505 | Lighthouse Lab in Glasgow | Wellcome Sanger Institute for the COVID-19 Genomics UK (COG-UK) Consortium | Harper VanSteenhouse, Yumi Kasai, David Gray, Carol Clugston, Anna Dominiczak and Alex Alderton, Roberto Amato, Jeffrey Barrett, Sonia Goncalves, Ewan Harrison, David K. Jackson, Ian Johnston, Dominic Kwiatkowski, Cordelia Langford, John Sillitoe on behalf of the Wellcome Sanger Institute COVID-19 Surveillance Team |
| EPI_ISL_3013506 | Lighthouse Lab in Alderley Park | Wellcome Sanger Institute for the COVID-19 Genomics UK (COG-UK) Consortium | Jacquelyn Wynn, Mairead Hyland, The Lighthouse Lab in Alderley Park and Alex Alderton, Roberto Amato, Jeffrey Barrett, Sonia Goncalves, Ewan Harrison, David K. Jackson, Ian Johnston, Dominic Kwiatkowski, Cordelia Langford, John Sillitoe on behalf of the Wellcome Sanger Institute COVID-19 Surveillance Team |
| EPI_ISL_3013507 | Lighthouse Lab in Milton Keynes | Wellcome Sanger Institute for the COVID-19 Genomics UK (COG-UK) Consortium | The Lighthouse Lab in Milton Keynes and Alex Alderton, Roberto Amato, Jeffrey Barrett, Sonia Goncalves, Ewan Harrison, David K. Jackson, Ian Johnston, Dominic Kwiatkowski, Cordelia Langford, John Sillitoe on behalf of the Wellcome Sanger Institute COVID-19 Surveillance Team |
| EPI_ISL_3013508 | Lighthouse Lab in Glasgow | Wellcome Sanger Institute for the COVID-19 Genomics UK (COG-UK) Consortium | Harper VanSteenhouse, Yumi Kasai, David Gray, Carol Clugston, Anna Dominiczak and Alex Alderton, Roberto Amato, Jeffrey Barrett, Sonia Goncalves, Ewan Harrison, David K. Jackson, Ian Johnston, Dominic Kwiatkowski, Cordelia Langford, John Sillitoe on behalf of the Wellcome Sanger Institute COVID-19 Surveillance Team |
| EPI_ISL_3013509 | Lighthouse Lab in Milton Keynes | Wellcome Sanger Institute for the COVID-19 Genomics UK (COG-UK) Consortium | The Lighthouse Lab in Milton Keynes and Alex Alderton, Roberto Amato, Jeffrey Barrett, Sonia Goncalves, Ewan Harrison, David K. Jackson, Ian Johnston, Dominic Kwiatkowski, Cordelia Langford, John Sillitoe on behalf of the Wellcome Sanger Institute COVID-19 Surveillance Team |
| EPI_ISL_3013510 | Lighthouse Lab in Glasgow | Wellcome Sanger Institute for the COVID-19 Genomics UK (COG-UK) Consortium | Harper VanSteenhouse, Yumi Kasai, David Gray, Carol Clugston, Anna Dominiczak and Alex Alderton, Roberto Amato, Jeffrey Barrett, Sonia Goncalves, Ewan Harrison, David K. Jackson, Ian Johnston, Dominic Kwiatkowski, Cordelia Langford, John Sillitoe on behalf of the Wellcome Sanger Institute COVID-19 Surveillance Team |
| EPI_ISL_3013511, EPI_ISL_3013512, EPI_ISL_3013513 | Lighthouse Lab in Alderley Park | Wellcome Sanger Institute for the COVID-19 Genomics UK (COG-UK) Consortium | Jacquelyn Wynn, Mairead Hyland, The Lighthouse Lab in Alderley Park and Alex Alderton, Roberto Amato, Jeffrey Barrett, Sonia Goncalves, Ewan Harrison, David K. Jackson, Ian Johnston, Dominic Kwiatkowski, Cordelia Langford, John Sillitoe on behalf of the Wellcome Sanger Institute COVID-19 Surveillance Team |
| EPI_ISL_3013514 | Lighthouse Lab in Glasgow | Wellcome Sanger Institute for the COVID-19 Genomics UK (COG-UK) Consortium | Harper VanSteenhouse, Yumi Kasai, David Gray, Carol Clugston, Anna Dominiczak and Alex Alderton, Roberto Amato, Jeffrey Barrett, Sonia Goncalves, Ewan Harrison, David K. Jackson, Ian Johnston, Dominic Kwiatkowski, Cordelia Langford, John Sillitoe on behalf of the Wellcome Sanger Institute COVID-19 Surveillance Team |
| EPI_ISL_3013515 | Lighthouse Lab in Milton Keynes | Wellcome Sanger Institute for the COVID-19 Genomics UK (COG-UK) Consortium | The Lighthouse Lab in Milton Keynes and Alex Alderton, Roberto Amato, Jeffrey Barrett, Sonia Goncalves, Ewan Harrison, David K. Jackson, Ian Johnston, Dominic Kwiatkowski, Cordelia Langford, John Sillitoe on behalf of the Wellcome Sanger Institute COVID-19 Surveillance Team |
| EPI_ISL_3013516 | Lighthouse Lab in Glasgow | Wellcome Sanger Institute for the COVID-19 Genomics UK (COG-UK) Consortium | Harper VanSteenhouse, Yumi Kasai, David Gray, Carol Clugston, Anna Dominiczak and Alex Alderton, Roberto Amato, Jeffrey Barrett, Sonia Goncalves, Ewan Harrison, David K. Jackson, Ian Johnston, Dominic Kwiatkowski, Cordelia Langford, John Sillitoe on behalf of the Wellcome Sanger Institute COVID-19 Surveillance Team |
| EPI_ISL_3013517 | Lighthouse Lab in Alderley Park | Wellcome Sanger Institute for the COVID-19 Genomics UK (COG-UK) Consortium | Jacquelyn Wynn, Mairead Hyland, The Lighthouse Lab in Alderley Park and Alex Alderton, Roberto Amato, Jeffrey Barrett, Sonia Goncalves, Ewan Harrison, David K. Jackson, Ian Johnston, Dominic Kwiatkowski, Cordelia Langford, John Sillitoe on behalf of the Wellcome Sanger Institute COVID-19 Surveillance Team |
| EPI_ISL_3013518 | Lighthouse Lab in Milton Keynes | Wellcome Sanger Institute for the COVID-19 Genomics UK (COG-UK) Consortium | The Lighthouse Lab in Milton Keynes and Alex Alderton, Roberto Amato, Jeffrey Barrett, Sonia Goncalves, Ewan Harrison, David K. Jackson, Ian Johnston, Dominic Kwiatkowski, Cordelia Langford, John Sillitoe on behalf of the Wellcome Sanger Institute COVID-19 Surveillance Team |
| EPI_ISL_3013519 | Lighthouse Lab in Glasgow | Wellcome Sanger Institute for the COVID-19 Genomics UK (COG-UK) Consortium | Harper VanSteenhouse, Yumi Kasai, David Gray, Carol Clugston, Anna Dominiczak and Alex Alderton, Roberto Amato, Jeffrey Barrett, Sonia Goncalves, Ewan Harrison, David K. Jackson, Ian Johnston, Dominic Kwiatkowski, Cordelia Langford, John Sillitoe on behalf of the Wellcome Sanger Institute COVID-19 Surveillance Team |
| EPI_ISL_3013520 | Lighthouse Lab in Milton Keynes | Wellcome Sanger Institute for the COVID-19 Genomics UK (COG-UK) Consortium | The Lighthouse Lab in Milton Keynes and Alex Alderton, Roberto Amato, Jeffrey Barrett, Sonia Goncalves, Ewan Harrison, David K. Jackson, Ian Johnston, Dominic Kwiatkowski, Cordelia Langford, John Sillitoe on behalf of the Wellcome Sanger Institute COVID-19 Surveillance Team |
| EPI_ISL_3013521 | Lighthouse Lab in Glasgow | Wellcome Sanger Institute for the COVID-19 Genomics UK (COG-UK) Consortium | Harper VanSteenhouse, Yumi Kasai, David Gray, Carol Clugston, Anna Dominiczak and Alex Alderton, Roberto Amato, Jeffrey Barrett, Sonia Goncalves, Ewan Harrison, David K. Jackson, Ian Johnston, Dominic Kwiatkowski, Cordelia Langford, John Sillitoe on behalf of the Wellcome Sanger Institute COVID-19 Surveillance Team |
| EPI_ISL_3013522, EPI_ISL_3013523 | Lighthouse Lab in Milton Keynes | Wellcome Sanger Institute for the COVID-19 Genomics UK (COG-UK) Consortium | The Lighthouse Lab in Milton Keynes and Alex Alderton, Roberto Amato, Jeffrey Barrett, Sonia Goncalves, Ewan Harrison, David K. Jackson, Ian Johnston, Dominic Kwiatkowski, Cordelia Langford, John Sillitoe on behalf of the Wellcome Sanger Institute COVID-19 Surveillance Team |

| | | | |
|---|---|---|---|
| EPI_ISL_3013524 | Lighthouse Lab in Glasgow | Wellcome Sanger Institute for the COVID-19 Genomics UK (COG-UK) Consortium | Harper VanSteenhouse, Yumi Kasai, David Gray, Carol Clugston, Anna Dominiczak and Alex Alderton, Roberto Amato, Jeffrey Barrett, Sonia Goncalves, Ewan Harrison, David K. Jackson, Ian Johnston, Dominic Kwiatkowski, Cordelia Langford, John Sillitoe on behalf of the Wellcome Sanger Institute COVID-19 Surveillance Team |
| EPI_ISL_3013525 | Lighthouse Lab in Milton Keynes | Wellcome Sanger Institute for the COVID-19 Genomics UK (COG-UK) Consortium | The Lighthouse Lab in Milton Keynes and Alex Alderton, Roberto Amato, Jeffrey Barrett, Sonia Goncalves, Ewan Harrison, David K. Jackson, Ian Johnston, Dominic Kwiatkowski, Cordelia Langford, John Sillitoe on behalf of the Wellcome Sanger Institute COVID-19 Surveillance Team |
| EPI_ISL_3013526 | Lighthouse Lab in Glasgow | Wellcome Sanger Institute for the COVID-19 Genomics UK (COG-UK) Consortium | Harper VanSteenhouse, Yumi Kasai, David Gray, Carol Clugston, Anna Dominiczak and Alex Alderton, Roberto Amato, Jeffrey Barrett, Sonia Goncalves, Ewan Harrison, David K. Jackson, Ian Johnston, Dominic Kwiatkowski, Cordelia Langford, John Sillitoe on behalf of the Wellcome Sanger Institute COVID-19 Surveillance Team |
| EPI_ISL_3013527, EPI_ISL_3013528 | Lighthouse Lab in Alderley Park | Wellcome Sanger Institute for the COVID-19 Genomics UK (COG-UK) Consortium | Jacquelyn Wynn, Mairead Hyland, The Lighthouse Lab in Alderley Park and Alex Alderton, Roberto Amato, Jeffrey Barrett, Sonia Goncalves, Ewan Harrison, David K. Jackson, Ian Johnston, Dominic Kwiatkowski, Cordelia Langford, John Sillitoe on behalf of the Wellcome Sanger Institute COVID-19 Surveillance Team |
| EPI_ISL_3013529 | Lighthouse Lab in Milton Keynes | Wellcome Sanger Institute for the COVID-19 Genomics UK (COG-UK) Consortium | The Lighthouse Lab in Milton Keynes and Alex Alderton, Roberto Amato, Jeffrey Barrett, Sonia Goncalves, Ewan Harrison, David K. Jackson, Ian Johnston, Dominic Kwiatkowski, Cordelia Langford, John Sillitoe on behalf of the Wellcome Sanger Institute COVID-19 Surveillance Team |
| EPI_ISL_3013530, EPI_ISL_3013531 | Lighthouse Lab in Alderley Park | Wellcome Sanger Institute for the COVID-19 Genomics UK (COG-UK) Consortium | Jacquelyn Wynn, Mairead Hyland, The Lighthouse Lab in Alderley Park and Alex Alderton, Roberto Amato, Jeffrey Barrett, Sonia Goncalves, Ewan Harrison, David K. Jackson, Ian Johnston, Dominic Kwiatkowski, Cordelia Langford, John Sillitoe on behalf of the Wellcome Sanger Institute COVID-19 Surveillance Team |
| EPI_ISL_3013532 | Lighthouse Lab in Milton Keynes | Wellcome Sanger Institute for the COVID-19 Genomics UK (COG-UK) Consortium | The Lighthouse Lab in Milton Keynes and Alex Alderton, Roberto Amato, Jeffrey Barrett, Sonia Goncalves, Ewan Harrison, David K. Jackson, Ian Johnston, Dominic Kwiatkowski, Cordelia Langford, John Sillitoe on behalf of the Wellcome Sanger Institute COVID-19 Surveillance Team |
| EPI_ISL_3013533, EPI_ISL_3013534 | Lighthouse Lab in Glasgow | Wellcome Sanger Institute for the COVID-19 Genomics UK (COG-UK) Consortium | Harper VanSteenhouse, Yumi Kasai, David Gray, Carol Clugston, Anna Dominiczak and Alex Alderton, Roberto Amato, Jeffrey Barrett, Sonia Goncalves, Ewan Harrison, David K. Jackson, Ian Johnston, Dominic Kwiatkowski, Cordelia Langford, John Sillitoe on behalf of the Wellcome Sanger Institute COVID-19 Surveillance Team |
| EPI_ISL_3013535 | Lighthouse Lab in Alderley Park | Wellcome Sanger Institute for the COVID-19 Genomics UK (COG-UK) Consortium | Jacquelyn Wynn, Mairead Hyland, The Lighthouse Lab in Alderley Park and Alex Alderton, Roberto Amato, Jeffrey Barrett, Sonia Goncalves, Ewan Harrison, David K. Jackson, Ian Johnston, Dominic Kwiatkowski, Cordelia Langford, John Sillitoe on behalf of the Wellcome Sanger Institute COVID-19 Surveillance Team |
| EPI_ISL_3013536 | Lighthouse Lab in Glasgow | Wellcome Sanger Institute for the COVID-19 Genomics UK (COG-UK) Consortium | Harper VanSteenhouse, Yumi Kasai, David Gray, Carol Clugston, Anna Dominiczak and Alex Alderton, Roberto Amato, Jeffrey Barrett, Sonia Goncalves, Ewan Harrison, David K. Jackson, Ian Johnston, Dominic Kwiatkowski, Cordelia Langford, John Sillitoe on behalf of the Wellcome Sanger Institute COVID-19 Surveillance Team |
| EPI_ISL_3013537 | Lighthouse Lab in Milton Keynes | Wellcome Sanger Institute for the COVID-19 Genomics UK (COG-UK) Consortium | The Lighthouse Lab in Milton Keynes and Alex Alderton, Roberto Amato, Jeffrey Barrett, Sonia Goncalves, Ewan Harrison, David K. Jackson, Ian Johnston, Dominic Kwiatkowski, Cordelia Langford, John Sillitoe on behalf of the Wellcome Sanger Institute COVID-19 Surveillance Team |
| EPI_ISL_3013538 | Lighthouse Lab in Alderley Park | Wellcome Sanger Institute for the COVID-19 Genomics UK (COG-UK) Consortium | Jacquelyn Wynn, Mairead Hyland, The Lighthouse Lab in Alderley Park and Alex Alderton, Roberto Amato, Jeffrey Barrett, Sonia Goncalves, Ewan Harrison, David K. Jackson, Ian Johnston, Dominic Kwiatkowski, Cordelia Langford, John Sillitoe on behalf of the Wellcome Sanger Institute COVID-19 Surveillance Team |
| EPI_ISL_3013539, EPI_ISL_3013540 | Lighthouse Lab in Glasgow | Wellcome Sanger Institute for the COVID-19 Genomics UK (COG-UK) Consortium | Harper VanSteenhouse, Yumi Kasai, David Gray, Carol Clugston, Anna Dominiczak and Alex Alderton, Roberto Amato, Jeffrey Barrett, Sonia Goncalves, Ewan Harrison, David K. Jackson, Ian Johnston, Dominic Kwiatkowski, Cordelia Langford, John Sillitoe on behalf of the Wellcome Sanger Institute COVID-19 Surveillance Team |
| EPI_ISL_3013541 | Lighthouse Lab in Alderley Park | Wellcome Sanger Institute for the COVID-19 Genomics UK (COG-UK) Consortium | Jacquelyn Wynn, Mairead Hyland, The Lighthouse Lab in Alderley Park and Alex Alderton, Roberto Amato, Jeffrey Barrett, Sonia Goncalves, Ewan Harrison, David K. Jackson, Ian Johnston, Dominic Kwiatkowski, Cordelia Langford, John Sillitoe on behalf of the Wellcome Sanger Institute COVID-19 Surveillance Team |
| EPI_ISL_3013542 | Lighthouse Lab in Glasgow | Wellcome Sanger Institute for the COVID-19 Genomics UK (COG-UK) Consortium | Harper VanSteenhouse, Yumi Kasai, David Gray, Carol Clugston, Anna Dominiczak and Alex Alderton, Roberto Amato, Jeffrey Barrett, Sonia Goncalves, Ewan Harrison, David K. Jackson, Ian Johnston, Dominic Kwiatkowski, Cordelia Langford, John Sillitoe on behalf of the Wellcome Sanger Institute COVID-19 Surveillance Team |
| EPI_ISL_3013543 | Lighthouse Lab in Alderley Park | Wellcome Sanger Institute for the COVID-19 Genomics UK (COG-UK) Consortium | Jacquelyn Wynn, Mairead Hyland, The Lighthouse Lab in Alderley Park and Alex Alderton, Roberto Amato, Jeffrey Barrett, Sonia Goncalves, Ewan Harrison, David K. Jackson, Ian Johnston, Dominic Kwiatkowski, Cordelia Langford, John Sillitoe on behalf of the Wellcome Sanger Institute COVID-19 Surveillance Team |
| EPI_ISL_3013544 | Lighthouse Lab in Milton Keynes | Wellcome Sanger Institute for the COVID-19 Genomics UK (COG-UK) Consortium | The Lighthouse Lab in Milton Keynes and Alex Alderton, Roberto Amato, Jeffrey Barrett, Sonia Goncalves, Ewan Harrison, David K. Jackson, Ian Johnston, Dominic Kwiatkowski, Cordelia Langford, John Sillitoe on behalf of the Wellcome Sanger Institute COVID-19 Surveillance Team |
| EPI_ISL_3013545 | Lighthouse Lab in Alderley Park | Wellcome Sanger Institute for the COVID-19 Genomics UK (COG-UK) Consortium | Jacquelyn Wynn, Mairead Hyland, The Lighthouse Lab in Alderley Park and Alex Alderton, Roberto Amato, Jeffrey Barrett, Sonia Goncalves, Ewan Harrison, David K. Jackson, Ian Johnston, Dominic Kwiatkowski, Cordelia Langford, John Sillitoe on behalf of the Wellcome Sanger Institute COVID-19 Surveillance Team |
| EPI_ISL_3013546 | Lighthouse Lab in Milton Keynes | Wellcome Sanger Institute for the COVID-19 Genomics UK (COG-UK) Consortium | The Lighthouse Lab in Milton Keynes and Alex Alderton, Roberto Amato, Jeffrey Barrett, Sonia Goncalves, Ewan Harrison, David K. Jackson, Ian Johnston, Dominic Kwiatkowski, Cordelia Langford, John Sillitoe on behalf of the Wellcome Sanger Institute COVID-19 Surveillance Team |
| EPI_ISL_3013547 | Lighthouse Lab in Alderley Park | Wellcome Sanger Institute for the COVID-19 Genomics UK (COG-UK) Consortium | Jacquelyn Wynn, Mairead Hyland, The Lighthouse Lab in Alderley Park and Alex Alderton, Roberto Amato, Jeffrey Barrett, Sonia Goncalves, Ewan Harrison, David K. Jackson, Ian Johnston, Dominic Kwiatkowski, Cordelia Langford, John Sillitoe on behalf of the Wellcome Sanger Institute COVID-19 Surveillance Team |
| EPI_ISL_3013548 | Lighthouse Lab in Glasgow | Wellcome Sanger Institute for the COVID-19 Genomics UK (COG-UK) Consortium | Harper VanSteenhouse, Yumi Kasai, David Gray, Carol Clugston, Anna Dominiczak and Alex Alderton, Roberto Amato, Jeffrey Barrett, Sonia Goncalves, Ewan Harrison, David K. Jackson, Ian Johnston, Dominic Kwiatkowski, Cordelia Langford, John Sillitoe on behalf of the Wellcome Sanger Institute COVID-19 Surveillance Team |
| EPI_ISL_3013549, EPI_ISL_3013550 | Lighthouse Lab in Milton Keynes | Wellcome Sanger Institute for the COVID-19 Genomics UK (COG-UK) Consortium | The Lighthouse Lab in Milton Keynes and Alex Alderton, Roberto Amato, Jeffrey Barrett, Sonia Goncalves, Ewan Harrison, David K. Jackson, Ian Johnston, Dominic Kwiatkowski, Cordelia Langford, John Sillitoe on behalf of the Wellcome Sanger Institute COVID-19 Surveillance Team |
| EPI_ISL_3013551 | Lighthouse Lab in Glasgow | Wellcome Sanger Institute for the COVID-19 Genomics UK (COG-UK) Consortium | Harper VanSteenhouse, Yumi Kasai, David Gray, Carol Clugston, Anna Dominiczak and Alex Alderton, Roberto Amato, Jeffrey Barrett, Sonia Goncalves, Ewan Harrison, David K. Jackson, Ian Johnston, Dominic Kwiatkowski, Cordelia Langford, John Sillitoe on behalf of the Wellcome Sanger Institute COVID-19 Surveillance Team |
| EPI_ISL_3013552 | Lighthouse Lab in Alderley Park | Wellcome Sanger Institute for the COVID-19 Genomics UK (COG-UK) Consortium | Jacquelyn Wynn, Mairead Hyland, The Lighthouse Lab in Alderley Park and Alex Alderton, Roberto Amato, Jeffrey Barrett, Sonia Goncalves, Ewan Harrison, David K. Jackson, Ian Johnston, Dominic Kwiatkowski, Cordelia Langford, John Sillitoe on behalf of the Wellcome Sanger Institute COVID-19 Surveillance Team |
| EPI_ISL_3013553, EPI_ISL_3013554, EPI_ISL_3013555 | Lighthouse Lab in Glasgow | Wellcome Sanger Institute for the COVID-19 Genomics UK (COG-UK) Consortium | Harper VanSteenhouse, Yumi Kasai, David Gray, Carol Clugston, Anna Dominiczak and Alex Alderton, Roberto Amato, Jeffrey Barrett, Sonia Goncalves, Ewan Harrison, David K. Jackson, Ian Johnston, Dominic Kwiatkowski, Cordelia Langford, John Sillitoe on behalf of the Wellcome Sanger Institute COVID-19 Surveillance Team |
| EPI_ISL_3013556 | Lighthouse Lab in Alderley Park | Wellcome Sanger Institute for the COVID-19 Genomics UK (COG-UK) Consortium | Jacquelyn Wynn, Mairead Hyland, The Lighthouse Lab in Alderley Park and Alex Alderton, Roberto Amato, Jeffrey Barrett, Sonia Goncalves, Ewan Harrison, David K. Jackson, Ian Johnston, Dominic Kwiatkowski, Cordelia Langford, John Sillitoe on behalf of the Wellcome Sanger Institute COVID-19 Surveillance Team |
| EPI_ISL_3013557 | Lighthouse Lab in Glasgow | Wellcome Sanger Institute for the COVID-19 Genomics UK (COG-UK) Consortium | Harper VanSteenhouse, Yumi Kasai, David Gray, Carol Clugston, Anna Dominiczak and Alex Alderton, Roberto Amato, Jeffrey Barrett, Sonia Goncalves, Ewan Harrison, David K. Jackson, Ian Johnston, Dominic Kwiatkowski, Cordelia Langford, John Sillitoe on behalf of the Wellcome Sanger Institute COVID-19 Surveillance Team |
| EPI_ISL_3013558 | Lighthouse Lab in Alderley Park | Wellcome Sanger Institute for the COVID-19 Genomics UK (COG-UK) Consortium | Jacquelyn Wynn, Mairead Hyland, The Lighthouse Lab in Alderley Park and Alex Alderton, Roberto Amato, Jeffrey Barrett, Sonia Goncalves, Ewan Harrison, David K. Jackson, Ian Johnston, Dominic Kwiatkowski, Cordelia Langford, John Sillitoe on behalf of the Wellcome Sanger Institute COVID-19 Surveillance Team |
| EPI_ISL_3013559 | Lighthouse Lab in Glasgow | Wellcome Sanger Institute for the COVID-19 Genomics UK (COG-UK) Consortium | Harper VanSteenhouse, Yumi Kasai, David Gray, Carol Clugston, Anna Dominiczak and Alex Alderton, Roberto Amato, Jeffrey Barrett, Sonia Goncalves, Ewan Harrison, David K. Jackson, Ian Johnston, Dominic Kwiatkowski, Cordelia Langford, John Sillitoe on behalf of the Wellcome Sanger Institute COVID-19 Surveillance Team |
| EPI_ISL_3013560 | Lighthouse Lab in Milton Keynes | Wellcome Sanger Institute for the COVID-19 Genomics UK (COG-UK) Consortium | The Lighthouse Lab in Milton Keynes and Alex Alderton, Roberto Amato, Jeffrey Barrett, Sonia Goncalves, Ewan Harrison, David K. Jackson, Ian Johnston, Dominic Kwiatkowski, Cordelia Langford, John Sillitoe on behalf of the Wellcome Sanger Institute COVID-19 Surveillance Team |
| EPI_ISL_3013561, EPI_ISL_3013562 | Lighthouse Lab in Alderley Park | Wellcome Sanger Institute for the COVID-19 Genomics UK (COG-UK) Consortium | Jacquelyn Wynn, Mairead Hyland, The Lighthouse Lab in Alderley Park and Alex Alderton, Roberto Amato, Jeffrey Barrett, Sonia Goncalves, Ewan Harrison, David K. Jackson, Ian Johnston, Dominic Kwiatkowski, Cordelia Langford, John Sillitoe on behalf of the Wellcome Sanger Institute COVID-19 Surveillance Team |
| EPI_ISL_3013563 | Lighthouse Lab in Glasgow | Wellcome Sanger Institute for the COVID-19 Genomics UK (COG-UK) Consortium | Harper VanSteenhouse, Yumi Kasai, David Gray, Carol Clugston, Anna Dominiczak and Alex Alderton, Roberto Amato, Jeffrey Barrett, Sonia Goncalves, Ewan Harrison, David K. Jackson, Ian Johnston, Dominic Kwiatkowski, Cordelia Langford, John Sillitoe on behalf of the Wellcome Sanger Institute COVID-19 Surveillance Team |
| EPI_ISL_3013564, EPI_ISL_3013565 | Lighthouse Lab in Milton Keynes | Wellcome Sanger Institute for the COVID-19 Genomics UK (COG-UK) Consortium | The Lighthouse Lab in Milton Keynes and Alex Alderton, Roberto Amato, Jeffrey Barrett, Sonia Goncalves, Ewan Harrison, David K. Jackson, Ian Johnston, Dominic Kwiatkowski, Cordelia Langford, John Sillitoe on behalf of the Wellcome Sanger Institute COVID-19 Surveillance Team |
| EPI_ISL_3013566 | Lighthouse Lab in Alderley Park | Wellcome Sanger Institute for the COVID-19 Genomics UK (COG-UK) Consortium | Jacquelyn Wynn, Mairead Hyland, The Lighthouse Lab in Alderley Park and Alex Alderton, Roberto Amato, Jeffrey Barrett, Sonia Goncalves, Ewan Harrison, David K. Jackson, Ian Johnston, Dominic Kwiatkowski, Cordelia Langford, John Sillitoe on behalf of the Wellcome Sanger Institute COVID-19 Surveillance Team |
| EPI_ISL_3013567 | Lighthouse Lab in Glasgow | Wellcome Sanger Institute for the COVID-19 Genomics UK (COG-UK) Consortium | Harper VanSteenhouse, Yumi Kasai, David Gray, Carol Clugston, Anna Dominiczak and Alex Alderton, Roberto Amato, Jeffrey Barrett, Sonia Goncalves, Ewan Harrison, David K. Jackson, Ian Johnston, Dominic Kwiatkowski, Cordelia Langford, John Sillitoe on behalf of the Wellcome Sanger Institute COVID-19 Surveillance Team |
| EPI_ISL_3013568 | Lighthouse Lab in Alderley Park | Wellcome Sanger Institute for the COVID-19 Genomics UK (COG-UK) Consortium | Jacquelyn Wynn, Mairead Hyland, The Lighthouse Lab in Alderley Park and Alex Alderton, Roberto Amato, Jeffrey Barrett, Sonia Goncalves, Ewan Harrison, David K. Jackson, Ian Johnston, Dominic Kwiatkowski, Cordelia Langford, John Sillitoe on behalf of the Wellcome Sanger Institute COVID-19 Surveillance Team |
| EPI_ISL_3013569 | Lighthouse Lab in Milton Keynes | Wellcome Sanger Institute for the COVID-19 Genomics UK (COG-UK) Consortium | The Lighthouse Lab in Milton Keynes and Alex Alderton, Roberto Amato, Jeffrey Barrett, Sonia Goncalves, Ewan Harrison, David K. Jackson, Ian Johnston, Dominic Kwiatkowski, Cordelia Langford, John Sillitoe on behalf of the Wellcome Sanger Institute COVID-19 Surveillance Team |
| EPI_ISL_3013570 | Lighthouse Lab in Alderley Park | Wellcome Sanger Institute for the COVID-19 Genomics UK (COG-UK) Consortium | Jacquelyn Wynn, Mairead Hyland, The Lighthouse Lab in Alderley Park and Alex Alderton, Roberto Amato, Jeffrey Barrett, Sonia Goncalves, Ewan Harrison, David K. Jackson, Ian Johnston, Dominic Kwiatkowski, Cordelia Langford, John Sillitoe on behalf of the Wellcome Sanger Institute COVID-19 Surveillance Team |
| EPI_ISL_3013571 | Lighthouse Lab in Glasgow | Wellcome Sanger Institute for the COVID-19 Genomics UK (COG-UK) Consortium | Harper VanSteenhouse, Yumi Kasai, David Gray, Carol Clugston, Anna Dominiczak and Alex Alderton, Roberto Amato, Jeffrey Barrett, Sonia Goncalves, Ewan Harrison, David K. Jackson, Ian Johnston, Dominic Kwiatkowski, Cordelia Langford, John Sillitoe on behalf of the Wellcome Sanger Institute COVID-19 Surveillance Team |
| EPI_ISL_3013572 | Lighthouse Lab in Milton Keynes | Wellcome Sanger Institute for the COVID-19 Genomics UK (COG-UK) Consortium | The Lighthouse Lab in Milton Keynes and Alex Alderton, Roberto Amato, Jeffrey Barrett, Sonia Goncalves, Ewan Harrison, David K. Jackson, Ian Johnston, Dominic Kwiatkowski, Cordelia Langford, John Sillitoe on behalf of the Wellcome Sanger Institute COVID-19 Surveillance Team |
| EPI_ISL_3013573, EPI_ISL_3013574 | Lighthouse Lab in Alderley Park | Wellcome Sanger Institute for the COVID-19 Genomics UK (COG-UK) Consortium | Jacquelyn Wynn, Mairead Hyland, The Lighthouse Lab in Alderley Park and Alex Alderton, Roberto Amato, Jeffrey Barrett, Sonia Goncalves, Ewan Harrison, David K. Jackson, Ian Johnston, Dominic Kwiatkowski, Cordelia Langford, John Sillitoe on behalf of the Wellcome Sanger Institute COVID-19 Surveillance Team |
| EPI_ISL_3013575 | Lighthouse Lab in Glasgow | Wellcome Sanger Institute for the COVID-19 Genomics UK (COG-UK) Consortium | Harper VanSteenhouse, Yumi Kasai, David Gray, Carol Clugston, Anna Dominiczak and Alex Alderton, Roberto Amato, Jeffrey Barrett, Sonia Goncalves, Ewan Harrison, David K. Jackson, Ian Johnston, Dominic Kwiatkowski, Cordelia Langford, John Sillitoe on behalf of the Wellcome Sanger Institute COVID-19 Surveillance Team |
| EPI_ISL_3013576 | Lighthouse Lab in Alderley Park | Wellcome Sanger Institute for the COVID-19 Genomics UK (COG-UK) Consortium | Jacquelyn Wynn, Mairead Hyland, The Lighthouse Lab in Alderley Park and Alex Alderton, Roberto Amato, Jeffrey Barrett, Sonia Goncalves, Ewan Harrison, David K. Jackson, Ian Johnston, Dominic Kwiatkowski, Cordelia Langford, John Sillitoe on behalf of the Wellcome Sanger Institute COVID-19 Surveillance Team |
| EPI_ISL_3013577 | Lighthouse Lab in Milton Keynes | Wellcome Sanger Institute for the COVID-19 Genomics UK (COG-UK) Consortium | The Lighthouse Lab in Milton Keynes and Alex Alderton, Roberto Amato, Jeffrey Barrett, Sonia Goncalves, Ewan Harrison, David K. Jackson, Ian Johnston, Dominic Kwiatkowski, Cordelia Langford, John Sillitoe on behalf of the Wellcome Sanger Institute COVID-19 Surveillance Team |
| EPI_ISL_3013578, EPI_ISL_3013579 | Lighthouse Lab in Alderley Park | Wellcome Sanger Institute for the COVID-19 Genomics UK (COG-UK) Consortium | Jacquelyn Wynn, Mairead Hyland, The Lighthouse Lab in Alderley Park and Alex Alderton, Roberto Amato, Jeffrey Barrett, Sonia Goncalves, Ewan Harrison, David K. Jackson, Ian Johnston, Dominic Kwiatkowski, Cordelia Langford, John Sillitoe on behalf of the Wellcome Sanger Institute COVID-19 Surveillance Team |

| | | | |
|---|---|---|---|
| EPI_ISL_3013580 | Lighthouse Lab in Milton Keynes | Wellcome Sanger Institute for the COVID-19 Genomics UK (COG-UK) Consortium | The Lighthouse Lab in Milton Keynes and Alex Alderton, Roberto Amato, Jeffrey Barrett, Sonia Goncalves, Ewan Harrison, David K. Jackson, Ian Johnston, Dominic Kwiatkowski, Cordelia Langford, John Sillitoe on behalf of the Wellcome Sanger Institute COVID-19 Surveillance Team |
| EPI_ISL_3013581 | Lighthouse Lab in Alderley Park | Wellcome Sanger Institute for the COVID-19 Genomics UK (COG-UK) Consortium | Jacquelyn Wynn, Mairead Hyland, The Lighthouse Lab in Alderley Park and Alex Alderton, Roberto Amato, Jeffrey Barrett, Sonia Goncalves, Ewan Harrison, David K. Jackson, Ian Johnston, Dominic Kwiatkowski, Cordelia Langford, John Sillitoe on behalf of the Wellcome Sanger Institute COVID-19 Surveillance Team |
| EPI_ISL_3013582 | Lighthouse Lab in Glasgow | Wellcome Sanger Institute for the COVID-19 Genomics UK (COG-UK) Consortium | Harper VanSteenhouse, Yumi Kasai, David Gray, Carol Clugston, Anna Dominiczak and Alex Alderton, Roberto Amato, Jeffrey Barrett, Sonia Goncalves, Ewan Harrison, David K. Jackson, Ian Johnston, Dominic Kwiatkowski, Cordelia Langford, John Sillitoe on behalf of the Wellcome Sanger Institute COVID-19 Surveillance Team |
| EPI_ISL_3013583 | Lighthouse Lab in Milton Keynes | Wellcome Sanger Institute for the COVID-19 Genomics UK (COG-UK) Consortium | The Lighthouse Lab in Milton Keynes and Alex Alderton, Roberto Amato, Jeffrey Barrett, Sonia Goncalves, Ewan Harrison, David K. Jackson, Ian Johnston, Dominic Kwiatkowski, Cordelia Langford, John Sillitoe on behalf of the Wellcome Sanger Institute COVID-19 Surveillance Team |
| EPI_ISL_3013584 | Lighthouse Lab in Alderley Park | Wellcome Sanger Institute for the COVID-19 Genomics UK (COG-UK) Consortium | Jacquelyn Wynn, Mairead Hyland, The Lighthouse Lab in Alderley Park and Alex Alderton, Roberto Amato, Jeffrey Barrett, Sonia Goncalves, Ewan Harrison, David K. Jackson, Ian Johnston, Dominic Kwiatkowski, Cordelia Langford, John Sillitoe on behalf of the Wellcome Sanger Institute COVID-19 Surveillance Team |
| EPI_ISL_3013585 | Lighthouse Lab in Milton Keynes | Wellcome Sanger Institute for the COVID-19 Genomics UK (COG-UK) Consortium | The Lighthouse Lab in Milton Keynes and Alex Alderton, Roberto Amato, Jeffrey Barrett, Sonia Goncalves, Ewan Harrison, David K. Jackson, Ian Johnston, Dominic Kwiatkowski, Cordelia Langford, John Sillitoe on behalf of the Wellcome Sanger Institute COVID-19 Surveillance Team |
| EPI_ISL_3013586, EPI_ISL_3013587, EPI_ISL_3013588, EPI_ISL_3013589 | Lighthouse Lab in Glasgow | Wellcome Sanger Institute for the COVID-19 Genomics UK (COG-UK) Consortium | Harper VanSteenhouse, Yumi Kasai, David Gray, Carol Clugston, Anna Dominiczak and Alex Alderton, Roberto Amato, Jeffrey Barrett, Sonia Goncalves, Ewan Harrison, David K. Jackson, Ian Johnston, Dominic Kwiatkowski, Cordelia Langford, John Sillitoe on behalf of the Wellcome Sanger Institute COVID-19 Surveillance Team |
| EPI_ISL_3013590 | Lighthouse Lab in Alderley Park | Wellcome Sanger Institute for the COVID-19 Genomics UK (COG-UK) Consortium | Jacquelyn Wynn, Mairead Hyland, The Lighthouse Lab in Alderley Park and Alex Alderton, Roberto Amato, Jeffrey Barrett, Sonia Goncalves, Ewan Harrison, David K. Jackson, Ian Johnston, Dominic Kwiatkowski, Cordelia Langford, John Sillitoe on behalf of the Wellcome Sanger Institute COVID-19 Surveillance Team |
| EPI_ISL_3013591, EPI_ISL_3013592 | Health Services Laboratories | Wellcome Sanger Institute for the COVID-19 Genomics UK (COG-UK) Consortium | Health Services Laboratories and Alex Alderton, Roberto Amato, Jeffrey Barrett, Sonia Goncalves, Ewan Harrison, David K. Jackson, Ian Johnston, Dominic Kwiatkowski, Cordelia Langford, John Sillitoe on behalf of the Wellcome Sanger Institute COVID-19 Surveillance Team |
| EPI_ISL_3013593 | Lighthouse Lab in Glasgow | Wellcome Sanger Institute for the COVID-19 Genomics UK (COG-UK) Consortium | Harper VanSteenhouse, Yumi Kasai, David Gray, Carol Clugston, Anna Dominiczak and Alex Alderton, Roberto Amato, Jeffrey Barrett, Sonia Goncalves, Ewan Harrison, David K. Jackson, Ian Johnston, Dominic Kwiatkowski, Cordelia Langford, John Sillitoe on behalf of the Wellcome Sanger Institute COVID-19 Surveillance Team |
| EPI_ISL_3013594, EPI_ISL_3013595 | Lighthouse Lab in Milton Keynes | Wellcome Sanger Institute for the COVID-19 Genomics UK (COG-UK) Consortium | The Lighthouse Lab in Milton Keynes and Alex Alderton, Roberto Amato, Jeffrey Barrett, Sonia Goncalves, Ewan Harrison, David K. Jackson, Ian Johnston, Dominic Kwiatkowski, Cordelia Langford, John Sillitoe on behalf of the Wellcome Sanger Institute COVID-19 Surveillance Team |
| EPI_ISL_3013596 | Lighthouse Lab in Alderley Park | Wellcome Sanger Institute for the COVID-19 Genomics UK (COG-UK) Consortium | Jacquelyn Wynn, Mairead Hyland, The Lighthouse Lab in Alderley Park and Alex Alderton, Roberto Amato, Jeffrey Barrett, Sonia Goncalves, Ewan Harrison, David K. Jackson, Ian Johnston, Dominic Kwiatkowski, Cordelia Langford, John Sillitoe on behalf of the Wellcome Sanger Institute COVID-19 Surveillance Team |
| EPI_ISL_3013597, EPI_ISL_3013598, EPI_ISL_3013599 | Lighthouse Lab in Glasgow | Wellcome Sanger Institute for the COVID-19 Genomics UK (COG-UK) Consortium | Harper VanSteenhouse, Yumi Kasai, David Gray, Carol Clugston, Anna Dominiczak and Alex Alderton, Roberto Amato, Jeffrey Barrett, Sonia Goncalves, Ewan Harrison, David K. Jackson, Ian Johnston, Dominic Kwiatkowski, Cordelia Langford, John Sillitoe on behalf of the Wellcome Sanger Institute COVID-19 Surveillance Team |
| EPI_ISL_3013600, EPI_ISL_3013601 | Lighthouse Lab in Milton Keynes | Wellcome Sanger Institute for the COVID-19 Genomics UK (COG-UK) Consortium | The Lighthouse Lab in Milton Keynes and Alex Alderton, Roberto Amato, Jeffrey Barrett, Sonia Goncalves, Ewan Harrison, David K. Jackson, Ian Johnston, Dominic Kwiatkowski, Cordelia Langford, John Sillitoe on behalf of the Wellcome Sanger Institute COVID-19 Surveillance Team |
| EPI_ISL_3013602, EPI_ISL_3013603 | Lighthouse Lab in Alderley Park | Wellcome Sanger Institute for the COVID-19 Genomics UK (COG-UK) Consortium | Jacquelyn Wynn, Mairead Hyland, The Lighthouse Lab in Alderley Park and Alex Alderton, Roberto Amato, Jeffrey Barrett, Sonia Goncalves, Ewan Harrison, David K. Jackson, Ian Johnston, Dominic Kwiatkowski, Cordelia Langford, John Sillitoe on behalf of the Wellcome Sanger Institute COVID-19 Surveillance Team |
| EPI_ISL_3013604, EPI_ISL_3013605, EPI_ISL_3013606 | Lighthouse Lab in Milton Keynes | Wellcome Sanger Institute for the COVID-19 Genomics UK (COG-UK) Consortium | The Lighthouse Lab in Milton Keynes and Alex Alderton, Roberto Amato, Jeffrey Barrett, Sonia Goncalves, Ewan Harrison, David K. Jackson, Ian Johnston, Dominic Kwiatkowski, Cordelia Langford, John Sillitoe on behalf of the Wellcome Sanger Institute COVID-19 Surveillance Team |
| EPI_ISL_3013607 | Lighthouse Lab in Alderley Park | Wellcome Sanger Institute for the COVID-19 Genomics UK (COG-UK) Consortium | Jacquelyn Wynn, Mairead Hyland, The Lighthouse Lab in Alderley Park and Alex Alderton, Roberto Amato, Jeffrey Barrett, Sonia Goncalves, Ewan Harrison, David K. Jackson, Ian Johnston, Dominic Kwiatkowski, Cordelia Langford, John Sillitoe on behalf of the Wellcome Sanger Institute COVID-19 Surveillance Team |
| EPI_ISL_3013608 | Lighthouse Lab in Milton Keynes | Wellcome Sanger Institute for the COVID-19 Genomics UK (COG-UK) Consortium | The Lighthouse Lab in Milton Keynes and Alex Alderton, Roberto Amato, Jeffrey Barrett, Sonia Goncalves, Ewan Harrison, David K. Jackson, Ian Johnston, Dominic Kwiatkowski, Cordelia Langford, John Sillitoe on behalf of the Wellcome Sanger Institute COVID-19 Surveillance Team |
| EPI_ISL_3013609 | Health Services Laboratories | Wellcome Sanger Institute for the COVID-19 Genomics UK (COG-UK) Consortium | Health Services Laboratories and Alex Alderton, Roberto Amato, Jeffrey Barrett, Sonia Goncalves, Ewan Harrison, David K. Jackson, Ian Johnston, Dominic Kwiatkowski, Cordelia Langford, John Sillitoe on behalf of the Wellcome Sanger Institute COVID-19 Surveillance Team |
| EPI_ISL_3013610 | Lighthouse Lab in Milton Keynes | Wellcome Sanger Institute for the COVID-19 Genomics UK (COG-UK) Consortium | The Lighthouse Lab in Milton Keynes and Alex Alderton, Roberto Amato, Jeffrey Barrett, Sonia Goncalves, Ewan Harrison, David K. Jackson, Ian Johnston, Dominic Kwiatkowski, Cordelia Langford, John Sillitoe on behalf of the Wellcome Sanger Institute COVID-19 Surveillance Team |
| EPI_ISL_3013611 | Lighthouse Lab in Alderley Park | Wellcome Sanger Institute for the COVID-19 Genomics UK (COG-UK) Consortium | Jacquelyn Wynn, Mairead Hyland, The Lighthouse Lab in Alderley Park and Alex Alderton, Roberto Amato, Jeffrey Barrett, Sonia Goncalves, Ewan Harrison, David K. Jackson, Ian Johnston, Dominic Kwiatkowski, Cordelia Langford, John Sillitoe on behalf of the Wellcome Sanger Institute COVID-19 Surveillance Team |
| EPI_ISL_3013612 | Lighthouse Lab in Glasgow | Wellcome Sanger Institute for the COVID-19 Genomics UK (COG-UK) Consortium | Harper VanSteenhouse, Yumi Kasai, David Gray, Carol Clugston, Anna Dominiczak and Alex Alderton, Roberto Amato, Jeffrey Barrett, Sonia Goncalves, Ewan Harrison, David K. Jackson, Ian Johnston, Dominic Kwiatkowski, Cordelia Langford, John Sillitoe on behalf of the Wellcome Sanger Institute COVID-19 Surveillance Team |
| EPI_ISL_3013613 | Lighthouse Lab in Milton Keynes | Wellcome Sanger Institute for the COVID-19 Genomics UK (COG-UK) Consortium | The Lighthouse Lab in Milton Keynes and Alex Alderton, Roberto Amato, Jeffrey Barrett, Sonia Goncalves, Ewan Harrison, David K. Jackson, Ian Johnston, Dominic Kwiatkowski, Cordelia Langford, John Sillitoe on behalf of the Wellcome Sanger Institute COVID-19 Surveillance Team |
| EPI_ISL_3013614, EPI_ISL_3013615, EPI_ISL_3013616, EPI_ISL_3013617 | Lighthouse Lab in Alderley Park | Wellcome Sanger Institute for the COVID-19 Genomics UK (COG-UK) Consortium | Jacquelyn Wynn, Mairead Hyland, The Lighthouse Lab in Alderley Park and Alex Alderton, Roberto Amato, Jeffrey Barrett, Sonia Goncalves, Ewan Harrison, David K. Jackson, Ian Johnston, Dominic Kwiatkowski, Cordelia Langford, John Sillitoe on behalf of the Wellcome Sanger Institute COVID-19 Surveillance Team |
| EPI_ISL_3013618 | Lighthouse Lab in Glasgow | Wellcome Sanger Institute for the COVID-19 Genomics UK (COG-UK) Consortium | Harper VanSteenhouse, Yumi Kasai, David Gray, Carol Clugston, Anna Dominiczak and Alex Alderton, Roberto Amato, Jeffrey Barrett, Sonia Goncalves, Ewan Harrison, David K. Jackson, Ian Johnston, Dominic Kwiatkowski, Cordelia Langford, John Sillitoe on behalf of the Wellcome Sanger Institute COVID-19 Surveillance Team |
| EPI_ISL_3013619, EPI_ISL_3013620 | Lighthouse Lab in Alderley Park | Wellcome Sanger Institute for the COVID-19 Genomics UK (COG-UK) Consortium | Jacquelyn Wynn, Mairead Hyland, The Lighthouse Lab in Alderley Park and Alex Alderton, Roberto Amato, Jeffrey Barrett, Sonia Goncalves, Ewan Harrison, David K. Jackson, Ian Johnston, Dominic Kwiatkowski, Cordelia Langford, John Sillitoe on behalf of the Wellcome Sanger Institute COVID-19 Surveillance Team |
| EPI_ISL_3013621 | Health Services Laboratories | Wellcome Sanger Institute for the COVID-19 Genomics UK (COG-UK) Consortium | Health Services Laboratories and Alex Alderton, Roberto Amato, Jeffrey Barrett, Sonia Goncalves, Ewan Harrison, David K. Jackson, Ian Johnston, Dominic Kwiatkowski, Cordelia Langford, John Sillitoe on behalf of the Wellcome Sanger Institute COVID-19 Surveillance Team |
| EPI_ISL_3013622 | Lighthouse Lab in Glasgow | Wellcome Sanger Institute for the COVID-19 Genomics UK (COG-UK) Consortium | Harper VanSteenhouse, Yumi Kasai, David Gray, Carol Clugston, Anna Dominiczak and Alex Alderton, Roberto Amato, Jeffrey Barrett, Sonia Goncalves, Ewan Harrison, David K. Jackson, Ian Johnston, Dominic Kwiatkowski, Cordelia Langford, John Sillitoe on behalf of the Wellcome Sanger Institute COVID-19 Surveillance Team |
| EPI_ISL_3013623 | Health Services Laboratories | Wellcome Sanger Institute for the COVID-19 Genomics UK (COG-UK) Consortium | Health Services Laboratories and Alex Alderton, Roberto Amato, Jeffrey Barrett, Sonia Goncalves, Ewan Harrison, David K. Jackson, Ian Johnston, Dominic Kwiatkowski, Cordelia Langford, John Sillitoe on behalf of the Wellcome Sanger Institute COVID-19 Surveillance Team |
| EPI_ISL_3013624 | Lighthouse Lab in Glasgow | Wellcome Sanger Institute for the COVID-19 Genomics UK (COG-UK) Consortium | Harper VanSteenhouse, Yumi Kasai, David Gray, Carol Clugston, Anna Dominiczak and Alex Alderton, Roberto Amato, Jeffrey Barrett, Sonia Goncalves, Ewan Harrison, David K. Jackson, Ian Johnston, Dominic Kwiatkowski, Cordelia Langford, John Sillitoe on behalf of the Wellcome Sanger Institute COVID-19 Surveillance Team |
| EPI_ISL_3013625 | Lighthouse Lab in Alderley Park | Wellcome Sanger Institute for the COVID-19 Genomics UK (COG-UK) Consortium | Jacquelyn Wynn, Mairead Hyland, The Lighthouse Lab in Alderley Park and Alex Alderton, Roberto Amato, Jeffrey Barrett, Sonia Goncalves, Ewan Harrison, David K. Jackson, Ian Johnston, Dominic Kwiatkowski, Cordelia Langford, John Sillitoe on behalf of the Wellcome Sanger Institute COVID-19 Surveillance Team |
| EPI_ISL_3013626 | Lighthouse Lab in Milton Keynes | Wellcome Sanger Institute for the COVID-19 Genomics UK (COG-UK) Consortium | The Lighthouse Lab in Milton Keynes and Alex Alderton, Roberto Amato, Jeffrey Barrett, Sonia Goncalves, Ewan Harrison, David K. Jackson, Ian Johnston, Dominic Kwiatkowski, Cordelia Langford, John Sillitoe on behalf of the Wellcome Sanger Institute COVID-19 Surveillance Team |
| EPI_ISL_3013627 | Lighthouse Lab in Alderley Park | Wellcome Sanger Institute for the COVID-19 Genomics UK (COG-UK) Consortium | Jacquelyn Wynn, Mairead Hyland, The Lighthouse Lab in Alderley Park and Alex Alderton, Roberto Amato, Jeffrey Barrett, Sonia Goncalves, Ewan Harrison, David K. Jackson, Ian Johnston, Dominic Kwiatkowski, Cordelia Langford, John Sillitoe on behalf of the Wellcome Sanger Institute COVID-19 Surveillance Team |
| EPI_ISL_3013628, EPI_ISL_3013629, EPI_ISL_3013630 | Lighthouse Lab in Milton Keynes | Wellcome Sanger Institute for the COVID-19 Genomics UK (COG-UK) Consortium | The Lighthouse Lab in Milton Keynes and Alex Alderton, Roberto Amato, Jeffrey Barrett, Sonia Goncalves, Ewan Harrison, David K. Jackson, Ian Johnston, Dominic Kwiatkowski, Cordelia Langford, John Sillitoe on behalf of the Wellcome Sanger Institute COVID-19 Surveillance Team |
| EPI_ISL_3013631 | Lighthouse Lab in Alderley Park | Wellcome Sanger Institute for the COVID-19 Genomics UK (COG-UK) Consortium | Jacquelyn Wynn, Mairead Hyland, The Lighthouse Lab in Alderley Park and Alex Alderton, Roberto Amato, Jeffrey Barrett, Sonia Goncalves, Ewan Harrison, David K. Jackson, Ian Johnston, Dominic Kwiatkowski, Cordelia Langford, John Sillitoe on behalf of the Wellcome Sanger Institute COVID-19 Surveillance Team |
| EPI_ISL_3013632 | Health Services Laboratories | Wellcome Sanger Institute for the COVID-19 Genomics UK (COG-UK) Consortium | Health Services Laboratories and Alex Alderton, Roberto Amato, Jeffrey Barrett, Sonia Goncalves, Ewan Harrison, David K. Jackson, Ian Johnston, Dominic Kwiatkowski, Cordelia Langford, John Sillitoe on behalf of the Wellcome Sanger Institute COVID-19 Surveillance Team |
| EPI_ISL_3013633, EPI_ISL_3013634 | Lighthouse Lab in Glasgow | Wellcome Sanger Institute for the COVID-19 Genomics UK (COG-UK) Consortium | Harper VanSteenhouse, Yumi Kasai, David Gray, Carol Clugston, Anna Dominiczak and Alex Alderton, Roberto Amato, Jeffrey Barrett, Sonia Goncalves, Ewan Harrison, David K. Jackson, Ian Johnston, Dominic Kwiatkowski, Cordelia Langford, John Sillitoe on behalf of the Wellcome Sanger Institute COVID-19 Surveillance Team |
| EPI_ISL_3013635 | Lighthouse Laboratory Plymouth | Wellcome Sanger Institute for the COVID-19 Genomics UK (COG-UK) Consortium | Lighthouse Laboratory Plymouth and Alex Alderton, Roberto Amato, Jeffrey Barrett, Sonia Goncalves, Ewan Harrison, David K. Jackson, Ian Johnston, Dominic Kwiatkowski, Cordelia Langford, John Sillitoe on behalf of the Wellcome Sanger Institute COVID-19 Surveillance Team |
| EPI_ISL_3013636 | Lighthouse Lab in Milton Keynes | Wellcome Sanger Institute for the COVID-19 Genomics UK (COG-UK) Consortium | The Lighthouse Lab in Milton Keynes and Alex Alderton, Roberto Amato, Jeffrey Barrett, Sonia Goncalves, Ewan Harrison, David K. Jackson, Ian Johnston, Dominic Kwiatkowski, Cordelia Langford, John Sillitoe on behalf of the Wellcome Sanger Institute COVID-19 Surveillance Team |
| EPI_ISL_3013637 | Health Services Laboratories | Wellcome Sanger Institute for the COVID-19 Genomics UK (COG-UK) Consortium | Health Services Laboratories and Alex Alderton, Roberto Amato, Jeffrey Barrett, Sonia Goncalves, Ewan Harrison, David K. Jackson, Ian Johnston, Dominic Kwiatkowski, Cordelia Langford, John Sillitoe on behalf of the Wellcome Sanger Institute COVID-19 Surveillance Team |
| EPI_ISL_3013638, EPI_ISL_3013639 | Lighthouse Lab in Alderley Park | Wellcome Sanger Institute for the COVID-19 Genomics UK (COG-UK) Consortium | Jacquelyn Wynn, Mairead Hyland, The Lighthouse Lab in Alderley Park and Alex Alderton, Roberto Amato, Jeffrey Barrett, Sonia Goncalves, Ewan Harrison, David K. Jackson, Ian Johnston, Dominic Kwiatkowski, Cordelia Langford, John Sillitoe on behalf of the Wellcome Sanger Institute COVID-19 Surveillance Team |
| EPI_ISL_3013640 | Lighthouse Lab in Milton Keynes | Wellcome Sanger Institute for the COVID-19 Genomics UK (COG-UK) Consortium | The Lighthouse Lab in Milton Keynes and Alex Alderton, Roberto Amato, Jeffrey Barrett, Sonia Goncalves, Ewan Harrison, David K. Jackson, Ian Johnston, Dominic Kwiatkowski, Cordelia Langford, John Sillitoe on behalf of the Wellcome Sanger Institute COVID-19 Surveillance Team |

| | | | |
|---|---|---|---|
| EPI_ISL_3013641 | Lighthouse Lab in Glasgow | Wellcome Sanger Institute for the COVID-19 Genomics UK (COG-UK) Consortium | Harper VanSteenhouse, Yumi Kasai, David Gray, Carol Clugston, Anna Dominiczak and Alex Alderton, Roberto Amato, Jeffrey Barrett, Sonia Goncalves, Ewan Harrison, David K. Jackson, Ian Johnston, Dominic Kwiatkowski, Cordelia Langford, John Sillitoe on behalf of the Wellcome Sanger Institute COVID-19 Surveillance Team |
| EPI_ISL_3013642 | Lighthouse Lab in Alderley Park | Wellcome Sanger Institute for the COVID-19 Genomics UK (COG-UK) Consortium | Jacquelyn Wynn, Mairead Hyland, The Lighthouse Lab in Alderley Park and Alex Alderton, Roberto Amato, Jeffrey Barrett, Sonia Goncalves, Ewan Harrison, David K. Jackson, Ian Johnston, Dominic Kwiatkowski, Cordelia Langford, John Sillitoe on behalf of the Wellcome Sanger Institute COVID-19 Surveillance Team |
| EPI_ISL_3013643 | Lighthouse Lab in Glasgow | Wellcome Sanger Institute for the COVID-19 Genomics UK (COG-UK) Consortium | Harper VanSteenhouse, Yumi Kasai, David Gray, Carol Clugston, Anna Dominiczak and Alex Alderton, Roberto Amato, Jeffrey Barrett, Sonia Goncalves, Ewan Harrison, David K. Jackson, Ian Johnston, Dominic Kwiatkowski, Cordelia Langford, John Sillitoe on behalf of the Wellcome Sanger Institute COVID-19 Surveillance Team |
| EPI_ISL_3013644, EPI_ISL_3013645 | Lighthouse Lab in Milton Keynes | Wellcome Sanger Institute for the COVID-19 Genomics UK (COG-UK) Consortium | The Lighthouse Lab in Milton Keynes and Alex Alderton, Roberto Amato, Jeffrey Barrett, Sonia Goncalves, Ewan Harrison, David K. Jackson, Ian Johnston, Dominic Kwiatkowski, Cordelia Langford, John Sillitoe on behalf of the Wellcome Sanger Institute COVID-19 Surveillance Team |
| EPI_ISL_3013646 | Lighthouse Lab in Alderley Park | Wellcome Sanger Institute for the COVID-19 Genomics UK (COG-UK) Consortium | Jacquelyn Wynn, Mairead Hyland, The Lighthouse Lab in Alderley Park and Alex Alderton, Roberto Amato, Jeffrey Barrett, Sonia Goncalves, Ewan Harrison, David K. Jackson, Ian Johnston, Dominic Kwiatkowski, Cordelia Langford, John Sillitoe on behalf of the Wellcome Sanger Institute COVID-19 Surveillance Team |
| EPI_ISL_3013647, EPI_ISL_3013648 | Lighthouse Lab in Glasgow | Wellcome Sanger Institute for the COVID-19 Genomics UK (COG-UK) Consortium | Harper VanSteenhouse, Yumi Kasai, David Gray, Carol Clugston, Anna Dominiczak and Alex Alderton, Roberto Amato, Jeffrey Barrett, Sonia Goncalves, Ewan Harrison, David K. Jackson, Ian Johnston, Dominic Kwiatkowski, Cordelia Langford, John Sillitoe on behalf of the Wellcome Sanger Institute COVID-19 Surveillance Team |
| EPI_ISL_3013649 | Lighthouse Lab in Alderley Park | Wellcome Sanger Institute for the COVID-19 Genomics UK (COG-UK) Consortium | Jacquelyn Wynn, Mairead Hyland, The Lighthouse Lab in Alderley Park and Alex Alderton, Roberto Amato, Jeffrey Barrett, Sonia Goncalves, Ewan Harrison, David K. Jackson, Ian Johnston, Dominic Kwiatkowski, Cordelia Langford, John Sillitoe on behalf of the Wellcome Sanger Institute COVID-19 Surveillance Team |
| EPI_ISL_3013650 | Lighthouse Lab in Milton Keynes | Wellcome Sanger Institute for the COVID-19 Genomics UK (COG-UK) Consortium | The Lighthouse Lab in Milton Keynes and Alex Alderton, Roberto Amato, Jeffrey Barrett, Sonia Goncalves, Ewan Harrison, David K. Jackson, Ian Johnston, Dominic Kwiatkowski, Cordelia Langford, John Sillitoe on behalf of the Wellcome Sanger Institute COVID-19 Surveillance Team |
| EPI_ISL_3013651, EPI_ISL_3013652, EPI_ISL_3013653 | Lighthouse Lab in Alderley Park | Wellcome Sanger Institute for the COVID-19 Genomics UK (COG-UK) Consortium | Jacquelyn Wynn, Mairead Hyland, The Lighthouse Lab in Alderley Park and Alex Alderton, Roberto Amato, Jeffrey Barrett, Sonia Goncalves, Ewan Harrison, David K. Jackson, Ian Johnston, Dominic Kwiatkowski, Cordelia Langford, John Sillitoe on behalf of the Wellcome Sanger Institute COVID-19 Surveillance Team |
| EPI_ISL_3013654 | Lighthouse Lab in Milton Keynes | Wellcome Sanger Institute for the COVID-19 Genomics UK (COG-UK) Consortium | The Lighthouse Lab in Milton Keynes and Alex Alderton, Roberto Amato, Jeffrey Barrett, Sonia Goncalves, Ewan Harrison, David K. Jackson, Ian Johnston, Dominic Kwiatkowski, Cordelia Langford, John Sillitoe on behalf of the Wellcome Sanger Institute COVID-19 Surveillance Team |
| EPI_ISL_3013655 | Lighthouse Lab in Alderley Park | Wellcome Sanger Institute for the COVID-19 Genomics UK (COG-UK) Consortium | Jacquelyn Wynn, Mairead Hyland, The Lighthouse Lab in Alderley Park and Alex Alderton, Roberto Amato, Jeffrey Barrett, Sonia Goncalves, Ewan Harrison, David K. Jackson, Ian Johnston, Dominic Kwiatkowski, Cordelia Langford, John Sillitoe on behalf of the Wellcome Sanger Institute COVID-19 Surveillance Team |
| EPI_ISL_3013656, EPI_ISL_3013657 | Lighthouse Lab in Glasgow | Wellcome Sanger Institute for the COVID-19 Genomics UK (COG-UK) Consortium | Harper VanSteenhouse, Yumi Kasai, David Gray, Carol Clugston, Anna Dominiczak and Alex Alderton, Roberto Amato, Jeffrey Barrett, Sonia Goncalves, Ewan Harrison, David K. Jackson, Ian Johnston, Dominic Kwiatkowski, Cordelia Langford, John Sillitoe on behalf of the Wellcome Sanger Institute COVID-19 Surveillance Team |
| EPI_ISL_3013658 | Lighthouse Lab in Milton Keynes | Wellcome Sanger Institute for the COVID-19 Genomics UK (COG-UK) Consortium | The Lighthouse Lab in Milton Keynes and Alex Alderton, Roberto Amato, Jeffrey Barrett, Sonia Goncalves, Ewan Harrison, David K. Jackson, Ian Johnston, Dominic Kwiatkowski, Cordelia Langford, John Sillitoe on behalf of the Wellcome Sanger Institute COVID-19 Surveillance Team |
| EPI_ISL_3013659 | Lighthouse Lab in Alderley Park | Wellcome Sanger Institute for the COVID-19 Genomics UK (COG-UK) Consortium | Jacquelyn Wynn, Mairead Hyland, The Lighthouse Lab in Alderley Park and Alex Alderton, Roberto Amato, Jeffrey Barrett, Sonia Goncalves, Ewan Harrison, David K. Jackson, Ian Johnston, Dominic Kwiatkowski, Cordelia Langford, John Sillitoe on behalf of the Wellcome Sanger Institute COVID-19 Surveillance Team |
| EPI_ISL_3013660 | Health Services Laboratories | Wellcome Sanger Institute for the COVID-19 Genomics UK (COG-UK) Consortium | Health Services Laboratories and Alex Alderton, Roberto Amato, Jeffrey Barrett, Sonia Goncalves, Ewan Harrison, David K. Jackson, Ian Johnston, Dominic Kwiatkowski, Cordelia Langford, John Sillitoe on behalf of the Wellcome Sanger Institute COVID-19 Surveillance Team |
| EPI_ISL_3013661 | Lighthouse Lab in Milton Keynes | Wellcome Sanger Institute for the COVID-19 Genomics UK (COG-UK) Consortium | The Lighthouse Lab in Milton Keynes and Alex Alderton, Roberto Amato, Jeffrey Barrett, Sonia Goncalves, Ewan Harrison, David K. Jackson, Ian Johnston, Dominic Kwiatkowski, Cordelia Langford, John Sillitoe on behalf of the Wellcome Sanger Institute COVID-19 Surveillance Team |
| EPI_ISL_3013662 | Lighthouse Lab in Alderley Park | Wellcome Sanger Institute for the COVID-19 Genomics UK (COG-UK) Consortium | Jacquelyn Wynn, Mairead Hyland, The Lighthouse Lab in Alderley Park and Alex Alderton, Roberto Amato, Jeffrey Barrett, Sonia Goncalves, Ewan Harrison, David K. Jackson, Ian Johnston, Dominic Kwiatkowski, Cordelia Langford, John Sillitoe on behalf of the Wellcome Sanger Institute COVID-19 Surveillance Team |
| EPI_ISL_3013663 | Health Services Laboratories | Wellcome Sanger Institute for the COVID-19 Genomics UK (COG-UK) Consortium | Health Services Laboratories and Alex Alderton, Roberto Amato, Jeffrey Barrett, Sonia Goncalves, Ewan Harrison, David K. Jackson, Ian Johnston, Dominic Kwiatkowski, Cordelia Langford, John Sillitoe on behalf of the Wellcome Sanger Institute COVID-19 Surveillance Team |
| EPI_ISL_3013664 | Lighthouse Lab in Milton Keynes | Wellcome Sanger Institute for the COVID-19 Genomics UK (COG-UK) Consortium | The Lighthouse Lab in Milton Keynes and Alex Alderton, Roberto Amato, Jeffrey Barrett, Sonia Goncalves, Ewan Harrison, David K. Jackson, Ian Johnston, Dominic Kwiatkowski, Cordelia Langford, John Sillitoe on behalf of the Wellcome Sanger Institute COVID-19 Surveillance Team |
| EPI_ISL_3013665 | Lighthouse Lab in Glasgow | Wellcome Sanger Institute for the COVID-19 Genomics UK (COG-UK) Consortium | Harper VanSteenhouse, Yumi Kasai, David Gray, Carol Clugston, Anna Dominiczak and Alex Alderton, Roberto Amato, Jeffrey Barrett, Sonia Goncalves, Ewan Harrison, David K. Jackson, Ian Johnston, Dominic Kwiatkowski, Cordelia Langford, John Sillitoe on behalf of the Wellcome Sanger Institute COVID-19 Surveillance Team |
| EPI_ISL_3013666 | Lighthouse Lab in Alderley Park | Wellcome Sanger Institute for the COVID-19 Genomics UK (COG-UK) Consortium | Jacquelyn Wynn, Mairead Hyland, The Lighthouse Lab in Alderley Park and Alex Alderton, Roberto Amato, Jeffrey Barrett, Sonia Goncalves, Ewan Harrison, David K. Jackson, Ian Johnston, Dominic Kwiatkowski, Cordelia Langford, John Sillitoe on behalf of the Wellcome Sanger Institute COVID-19 Surveillance Team |
| EPI_ISL_3013667 | Lighthouse Lab in Glasgow | Wellcome Sanger Institute for the COVID-19 Genomics UK (COG-UK) Consortium | Harper VanSteenhouse, Yumi Kasai, David Gray, Carol Clugston, Anna Dominiczak and Alex Alderton, Roberto Amato, Jeffrey Barrett, Sonia Goncalves, Ewan Harrison, David K. Jackson, Ian Johnston, Dominic Kwiatkowski, Cordelia Langford, John Sillitoe on behalf of the Wellcome Sanger Institute COVID-19 Surveillance Team |
| EPI_ISL_3013668 | Lighthouse Lab in Alderley Park | Wellcome Sanger Institute for the COVID-19 Genomics UK (COG-UK) Consortium | Jacquelyn Wynn, Mairead Hyland, The Lighthouse Lab in Alderley Park and Alex Alderton, Roberto Amato, Jeffrey Barrett, Sonia Goncalves, Ewan Harrison, David K. Jackson, Ian Johnston, Dominic Kwiatkowski, Cordelia Langford, John Sillitoe on behalf of the Wellcome Sanger Institute COVID-19 Surveillance Team |
| EPI_ISL_3013669 | Lighthouse Lab in Glasgow | Wellcome Sanger Institute for the COVID-19 Genomics UK (COG-UK) Consortium | Harper VanSteenhouse, Yumi Kasai, David Gray, Carol Clugston, Anna Dominiczak and Alex Alderton, Roberto Amato, Jeffrey Barrett, Sonia Goncalves, Ewan Harrison, David K. Jackson, Ian Johnston, Dominic Kwiatkowski, Cordelia Langford, John Sillitoe on behalf of the Wellcome Sanger Institute COVID-19 Surveillance Team |
| EPI_ISL_3013670 | Lighthouse Lab in Milton Keynes | Wellcome Sanger Institute for the COVID-19 Genomics UK (COG-UK) Consortium | The Lighthouse Lab in Milton Keynes and Alex Alderton, Roberto Amato, Jeffrey Barrett, Sonia Goncalves, Ewan Harrison, David K. Jackson, Ian Johnston, Dominic Kwiatkowski, Cordelia Langford, John Sillitoe on behalf of the Wellcome Sanger Institute COVID-19 Surveillance Team |
| EPI_ISL_3013671, EPI_ISL_3013672 | Health Services Laboratories | Wellcome Sanger Institute for the COVID-19 Genomics UK (COG-UK) Consortium | Health Services Laboratories and Alex Alderton, Roberto Amato, Jeffrey Barrett, Sonia Goncalves, Ewan Harrison, David K. Jackson, Ian Johnston, Dominic Kwiatkowski, Cordelia Langford, John Sillitoe on behalf of the Wellcome Sanger Institute COVID-19 Surveillance Team |
| EPI_ISL_3013673 | Lighthouse Lab in Glasgow | Wellcome Sanger Institute for the COVID-19 Genomics UK (COG-UK) Consortium | Harper VanSteenhouse, Yumi Kasai, David Gray, Carol Clugston, Anna Dominiczak and Alex Alderton, Roberto Amato, Jeffrey Barrett, Sonia Goncalves, Ewan Harrison, David K. Jackson, Ian Johnston, Dominic Kwiatkowski, Cordelia Langford, John Sillitoe on behalf of the Wellcome Sanger Institute COVID-19 Surveillance Team |
| EPI_ISL_3013674 | Lighthouse Lab in Milton Keynes | Wellcome Sanger Institute for the COVID-19 Genomics UK (COG-UK) Consortium | The Lighthouse Lab in Milton Keynes and Alex Alderton, Roberto Amato, Jeffrey Barrett, Sonia Goncalves, Ewan Harrison, David K. Jackson, Ian Johnston, Dominic Kwiatkowski, Cordelia Langford, John Sillitoe on behalf of the Wellcome Sanger Institute COVID-19 Surveillance Team |
| EPI_ISL_3013675, EPI_ISL_3013676 | Lighthouse Lab in Glasgow | Wellcome Sanger Institute for the COVID-19 Genomics UK (COG-UK) Consortium | Harper VanSteenhouse, Yumi Kasai, David Gray, Carol Clugston, Anna Dominiczak and Alex Alderton, Roberto Amato, Jeffrey Barrett, Sonia Goncalves, Ewan Harrison, David K. Jackson, Ian Johnston, Dominic Kwiatkowski, Cordelia Langford, John Sillitoe on behalf of the Wellcome Sanger Institute COVID-19 Surveillance Team |
| EPI_ISL_3013677 | Lighthouse Lab in Milton Keynes | Wellcome Sanger Institute for the COVID-19 Genomics UK (COG-UK) Consortium | The Lighthouse Lab in Milton Keynes and Alex Alderton, Roberto Amato, Jeffrey Barrett, Sonia Goncalves, Ewan Harrison, David K. Jackson, Ian Johnston, Dominic Kwiatkowski, Cordelia Langford, John Sillitoe on behalf of the Wellcome Sanger Institute COVID-19 Surveillance Team |
| EPI_ISL_3013678 | Lighthouse Lab in Alderley Park | Wellcome Sanger Institute for the COVID-19 Genomics UK (COG-UK) Consortium | Jacquelyn Wynn, Mairead Hyland, The Lighthouse Lab in Alderley Park and Alex Alderton, Roberto Amato, Jeffrey Barrett, Sonia Goncalves, Ewan Harrison, David K. Jackson, Ian Johnston, Dominic Kwiatkowski, Cordelia Langford, John Sillitoe on behalf of the Wellcome Sanger Institute COVID-19 Surveillance Team |
| EPI_ISL_3013679 | Lighthouse Lab in Milton Keynes | Wellcome Sanger Institute for the COVID-19 Genomics UK (COG-UK) Consortium | The Lighthouse Lab in Milton Keynes and Alex Alderton, Roberto Amato, Jeffrey Barrett, Sonia Goncalves, Ewan Harrison, David K. Jackson, Ian Johnston, Dominic Kwiatkowski, Cordelia Langford, John Sillitoe on behalf of the Wellcome Sanger Institute COVID-19 Surveillance Team |
| EPI_ISL_3013680 | Health Services Laboratories | Wellcome Sanger Institute for the COVID-19 Genomics UK (COG-UK) Consortium | Health Services Laboratories and Alex Alderton, Roberto Amato, Jeffrey Barrett, Sonia Goncalves, Ewan Harrison, David K. Jackson, Ian Johnston, Dominic Kwiatkowski, Cordelia Langford, John Sillitoe on behalf of the Wellcome Sanger Institute COVID-19 Surveillance Team |
| EPI_ISL_3013681, EPI_ISL_3013682, EPI_ISL_3013683, EPI_ISL_3013684 | Lighthouse Lab in Alderley Park | Wellcome Sanger Institute for the COVID-19 Genomics UK (COG-UK) Consortium | Jacquelyn Wynn, Mairead Hyland, The Lighthouse Lab in Alderley Park and Alex Alderton, Roberto Amato, Jeffrey Barrett, Sonia Goncalves, Ewan Harrison, David K. Jackson, Ian Johnston, Dominic Kwiatkowski, Cordelia Langford, John Sillitoe on behalf of the Wellcome Sanger Institute COVID-19 Surveillance Team |
| EPI_ISL_3013685, EPI_ISL_3013686, EPI_ISL_3013687 | Lighthouse Lab in Glasgow | Wellcome Sanger Institute for the COVID-19 Genomics UK (COG-UK) Consortium | Harper VanSteenhouse, Yumi Kasai, David Gray, Carol Clugston, Anna Dominiczak and Alex Alderton, Roberto Amato, Jeffrey Barrett, Sonia Goncalves, Ewan Harrison, David K. Jackson, Ian Johnston, Dominic Kwiatkowski, Cordelia Langford, John Sillitoe on behalf of the Wellcome Sanger Institute COVID-19 Surveillance Team |
| EPI_ISL_3013688 | Lighthouse Lab in Milton Keynes | Wellcome Sanger Institute for the COVID-19 Genomics UK (COG-UK) Consortium | The Lighthouse Lab in Milton Keynes and Alex Alderton, Roberto Amato, Jeffrey Barrett, Sonia Goncalves, Ewan Harrison, David K. Jackson, Ian Johnston, Dominic Kwiatkowski, Cordelia Langford, John Sillitoe on behalf of the Wellcome Sanger Institute COVID-19 Surveillance Team |
| EPI_ISL_3013689, EPI_ISL_3013690 | Health Services Laboratories | Wellcome Sanger Institute for the COVID-19 Genomics UK (COG-UK) Consortium | Health Services Laboratories and Alex Alderton, Roberto Amato, Jeffrey Barrett, Sonia Goncalves, Ewan Harrison, David K. Jackson, Ian Johnston, Dominic Kwiatkowski, Cordelia Langford, John Sillitoe on behalf of the Wellcome Sanger Institute COVID-19 Surveillance Team |
| EPI_ISL_3013691 | Lighthouse Lab in Milton Keynes | Wellcome Sanger Institute for the COVID-19 Genomics UK (COG-UK) Consortium | The Lighthouse Lab in Milton Keynes and Alex Alderton, Roberto Amato, Jeffrey Barrett, Sonia Goncalves, Ewan Harrison, David K. Jackson, Ian Johnston, Dominic Kwiatkowski, Cordelia Langford, John Sillitoe on behalf of the Wellcome Sanger Institute COVID-19 Surveillance Team |
| EPI_ISL_3013692, EPI_ISL_3013693, EPI_ISL_3013694 | Lighthouse Lab in Alderley Park | Wellcome Sanger Institute for the COVID-19 Genomics UK (COG-UK) Consortium | Jacquelyn Wynn, Mairead Hyland, The Lighthouse Lab in Alderley Park and Alex Alderton, Roberto Amato, Jeffrey Barrett, Sonia Goncalves, Ewan Harrison, David K. Jackson, Ian Johnston, Dominic Kwiatkowski, Cordelia Langford, John Sillitoe on behalf of the Wellcome Sanger Institute COVID-19 Surveillance Team |
| EPI_ISL_3013695 | Lighthouse Lab in Glasgow | Wellcome Sanger Institute for the COVID-19 Genomics UK (COG-UK) Consortium | Harper VanSteenhouse, Yumi Kasai, David Gray, Carol Clugston, Anna Dominiczak and Alex Alderton, Roberto Amato, Jeffrey Barrett, Sonia Goncalves, Ewan Harrison, David K. Jackson, Ian Johnston, Dominic Kwiatkowski, Cordelia Langford, John Sillitoe on behalf of the Wellcome Sanger Institute COVID-19 Surveillance Team |
| EPI_ISL_3013696 | Lighthouse Lab in Alderley Park | Wellcome Sanger Institute for the COVID-19 Genomics UK (COG-UK) Consortium | Jacquelyn Wynn, Mairead Hyland, The Lighthouse Lab in Alderley Park and Alex Alderton, Roberto Amato, Jeffrey Barrett, Sonia Goncalves, Ewan Harrison, David K. Jackson, Ian Johnston, Dominic Kwiatkowski, Cordelia Langford, John Sillitoe on behalf of the Wellcome Sanger Institute COVID-19 Surveillance Team |
| EPI_ISL_3013697 | Health Services Laboratories | Wellcome Sanger Institute for the COVID-19 Genomics UK (COG-UK) Consortium | Health Services Laboratories and Alex Alderton, Roberto Amato, Jeffrey Barrett, Sonia Goncalves, Ewan Harrison, David K. Jackson, Ian Johnston, Dominic Kwiatkowski, Cordelia Langford, John Sillitoe on behalf of the Wellcome Sanger Institute COVID-19 Surveillance Team |
| EPI_ISL_3013698 | Lighthouse Lab in Glasgow | Wellcome Sanger Institute for the COVID-19 Genomics UK (COG-UK) Consortium | Harper VanSteenhouse, Yumi Kasai, David Gray, Carol Clugston, Anna Dominiczak and Alex Alderton, Roberto Amato, Jeffrey Barrett, Sonia Goncalves, Ewan Harrison, David K. Jackson, Ian Johnston, Dominic Kwiatkowski, Cordelia Langford, John Sillitoe on behalf of the Wellcome Sanger Institute COVID-19 Surveillance Team |

| | | | |
|---|---|---|---|
| EPI_ISL_3013699 | Lighthouse Lab in Milton Keynes | Wellcome Sanger Institute for the COVID-19 Genomics UK (COG-UK) Consortium | The Lighthouse Lab in Milton Keynes and Alex Alderton, Roberto Amato, Jeffrey Barrett, Sonia Goncalves, Ewan Harrison, David K. Jackson, Ian Johnston, Dominic Kwiatkowski, Cordelia Langford, John Sillitoe on behalf of the Wellcome Sanger Institute COVID-19 Surveillance Team |
| EPI_ISL_3013700 | Lighthouse Lab in Alderley Park | Wellcome Sanger Institute for the COVID-19 Genomics UK (COG-UK) Consortium | Jacquelyn Wynn, Mairead Hyland, The Lighthouse Lab in Alderley Park and Alex Alderton, Roberto Amato, Jeffrey Barrett, Sonia Goncalves, Ewan Harrison, David K. Jackson, Ian Johnston, Dominic Kwiatkowski, Cordelia Langford, John Sillitoe on behalf of the Wellcome Sanger Institute COVID-19 Surveillance Team |
| EPI_ISL_3013701, EPI_ISL_3013702 | Lighthouse Lab in Milton Keynes | Wellcome Sanger Institute for the COVID-19 Genomics UK (COG-UK) Consortium | The Lighthouse Lab in Milton Keynes and Alex Alderton, Roberto Amato, Jeffrey Barrett, Sonia Goncalves, Ewan Harrison, David K. Jackson, Ian Johnston, Dominic Kwiatkowski, Cordelia Langford, John Sillitoe on behalf of the Wellcome Sanger Institute COVID-19 Surveillance Team |
| EPI_ISL_3013703, EPI_ISL_3013704, EPI_ISL_3013705 | Lighthouse Lab in Glasgow | Wellcome Sanger Institute for the COVID-19 Genomics UK (COG-UK) Consortium | Harper VanSteenhouse, Yumi Kasai, David Gray, Carol Clugston, Anna Dominiczak and Alex Alderton, Roberto Amato, Jeffrey Barrett, Sonia Goncalves, Ewan Harrison, David K. Jackson, Ian Johnston, Dominic Kwiatkowski, Cordelia Langford, John Sillitoe on behalf of the Wellcome Sanger Institute COVID-19 Surveillance Team |
| EPI_ISL_3013706, EPI_ISL_3013707 | Lighthouse Lab in Alderley Park | Wellcome Sanger Institute for the COVID-19 Genomics UK (COG-UK) Consortium | Jacquelyn Wynn, Mairead Hyland, The Lighthouse Lab in Alderley Park and Alex Alderton, Roberto Amato, Jeffrey Barrett, Sonia Goncalves, Ewan Harrison, David K. Jackson, Ian Johnston, Dominic Kwiatkowski, Cordelia Langford, John Sillitoe on behalf of the Wellcome Sanger Institute COVID-19 Surveillance Team |
| EPI_ISL_3013708 | Health Services Laboratories | Wellcome Sanger Institute for the COVID-19 Genomics UK (COG-UK) Consortium | Health Services Laboratories and Alex Alderton, Roberto Amato, Jeffrey Barrett, Sonia Goncalves, Ewan Harrison, David K. Jackson, Ian Johnston, Dominic Kwiatkowski, Cordelia Langford, John Sillitoe on behalf of the Wellcome Sanger Institute COVID-19 Surveillance Team |
| EPI_ISL_3013709 | Lighthouse Lab in Alderley Park | Wellcome Sanger Institute for the COVID-19 Genomics UK (COG-UK) Consortium | Jacquelyn Wynn, Mairead Hyland, The Lighthouse Lab in Alderley Park and Alex Alderton, Roberto Amato, Jeffrey Barrett, Sonia Goncalves, Ewan Harrison, David K. Jackson, Ian Johnston, Dominic Kwiatkowski, Cordelia Langford, John Sillitoe on behalf of the Wellcome Sanger Institute COVID-19 Surveillance Team |
| EPI_ISL_3013710, EPI_ISL_3013711 | Lighthouse Lab in Milton Keynes | Wellcome Sanger Institute for the COVID-19 Genomics UK (COG-UK) Consortium | The Lighthouse Lab in Milton Keynes and Alex Alderton, Roberto Amato, Jeffrey Barrett, Sonia Goncalves, Ewan Harrison, David K. Jackson, Ian Johnston, Dominic Kwiatkowski, Cordelia Langford, John Sillitoe on behalf of the Wellcome Sanger Institute COVID-19 Surveillance Team |
| EPI_ISL_3013712 | Lighthouse Lab in Glasgow | Wellcome Sanger Institute for the COVID-19 Genomics UK (COG-UK) Consortium | Harper VanSteenhouse, Yumi Kasai, David Gray, Carol Clugston, Anna Dominiczak and Alex Alderton, Roberto Amato, Jeffrey Barrett, Sonia Goncalves, Ewan Harrison, David K. Jackson, Ian Johnston, Dominic Kwiatkowski, Cordelia Langford, John Sillitoe on behalf of the Wellcome Sanger Institute COVID-19 Surveillance Team |
| EPI_ISL_3013713 | Lighthouse Lab in Milton Keynes | Wellcome Sanger Institute for the COVID-19 Genomics UK (COG-UK) Consortium | The Lighthouse Lab in Milton Keynes and Alex Alderton, Roberto Amato, Jeffrey Barrett, Sonia Goncalves, Ewan Harrison, David K. Jackson, Ian Johnston, Dominic Kwiatkowski, Cordelia Langford, John Sillitoe on behalf of the Wellcome Sanger Institute COVID-19 Surveillance Team |
| EPI_ISL_3013714 | Health Services Laboratories | Wellcome Sanger Institute for the COVID-19 Genomics UK (COG-UK) Consortium | Health Services Laboratories and Alex Alderton, Roberto Amato, Jeffrey Barrett, Sonia Goncalves, Ewan Harrison, David K. Jackson, Ian Johnston, Dominic Kwiatkowski, Cordelia Langford, John Sillitoe on behalf of the Wellcome Sanger Institute COVID-19 Surveillance Team |
| EPI_ISL_3013715, EPI_ISL_3013716 | Lighthouse Lab in Milton Keynes | Wellcome Sanger Institute for the COVID-19 Genomics UK (COG-UK) Consortium | The Lighthouse Lab in Milton Keynes and Alex Alderton, Roberto Amato, Jeffrey Barrett, Sonia Goncalves, Ewan Harrison, David K. Jackson, Ian Johnston, Dominic Kwiatkowski, Cordelia Langford, John Sillitoe on behalf of the Wellcome Sanger Institute COVID-19 Surveillance Team |
| EPI_ISL_3013717 | Lighthouse Lab in Alderley Park | Wellcome Sanger Institute for the COVID-19 Genomics UK (COG-UK) Consortium | Jacquelyn Wynn, Mairead Hyland, The Lighthouse Lab in Alderley Park and Alex Alderton, Roberto Amato, Jeffrey Barrett, Sonia Goncalves, Ewan Harrison, David K. Jackson, Ian Johnston, Dominic Kwiatkowski, Cordelia Langford, John Sillitoe on behalf of the Wellcome Sanger Institute COVID-19 Surveillance Team |
| EPI_ISL_3013718 | Lighthouse Lab in Milton Keynes | Wellcome Sanger Institute for the COVID-19 Genomics UK (COG-UK) Consortium | The Lighthouse Lab in Milton Keynes and Alex Alderton, Roberto Amato, Jeffrey Barrett, Sonia Goncalves, Ewan Harrison, David K. Jackson, Ian Johnston, Dominic Kwiatkowski, Cordelia Langford, John Sillitoe on behalf of the Wellcome Sanger Institute COVID-19 Surveillance Team |
| EPI_ISL_3013719 | Lighthouse Lab in Alderley Park | Wellcome Sanger Institute for the COVID-19 Genomics UK (COG-UK) Consortium | Jacquelyn Wynn, Mairead Hyland, The Lighthouse Lab in Alderley Park and Alex Alderton, Roberto Amato, Jeffrey Barrett, Sonia Goncalves, Ewan Harrison, David K. Jackson, Ian Johnston, Dominic Kwiatkowski, Cordelia Langford, John Sillitoe on behalf of the Wellcome Sanger Institute COVID-19 Surveillance Team |
| EPI_ISL_3013720, EPI_ISL_3013721 | Health Services Laboratories | Wellcome Sanger Institute for the COVID-19 Genomics UK (COG-UK) Consortium | Health Services Laboratories and Alex Alderton, Roberto Amato, Jeffrey Barrett, Sonia Goncalves, Ewan Harrison, David K. Jackson, Ian Johnston, Dominic Kwiatkowski, Cordelia Langford, John Sillitoe on behalf of the Wellcome Sanger Institute COVID-19 Surveillance Team |
| EPI_ISL_3013722 | Lighthouse Laboratory Plymouth | Wellcome Sanger Institute for the COVID-19 Genomics UK (COG-UK) Consortium | Lighthouse Laboratory Plymouth and Alex Alderton, Roberto Amato, Jeffrey Barrett, Sonia Goncalves, Ewan Harrison, David K. Jackson, Ian Johnston, Dominic Kwiatkowski, Cordelia Langford, John Sillitoe on behalf of the Wellcome Sanger Institute COVID-19 Surveillance Team |
| EPI_ISL_3013723 | Lighthouse Lab in Alderley Park | Wellcome Sanger Institute for the COVID-19 Genomics UK (COG-UK) Consortium | Jacquelyn Wynn, Mairead Hyland, The Lighthouse Lab in Alderley Park and Alex Alderton, Roberto Amato, Jeffrey Barrett, Sonia Goncalves, Ewan Harrison, David K. Jackson, Ian Johnston, Dominic Kwiatkowski, Cordelia Langford, John Sillitoe on behalf of the Wellcome Sanger Institute COVID-19 Surveillance Team |
| EPI_ISL_3013724 | Lighthouse Lab in Glasgow | Wellcome Sanger Institute for the COVID-19 Genomics UK (COG-UK) Consortium | Harper VanSteenhouse, Yumi Kasai, David Gray, Carol Clugston, Anna Dominiczak and Alex Alderton, Roberto Amato, Jeffrey Barrett, Sonia Goncalves, Ewan Harrison, David K. Jackson, Ian Johnston, Dominic Kwiatkowski, Cordelia Langford, John Sillitoe on behalf of the Wellcome Sanger Institute COVID-19 Surveillance Team |
| EPI_ISL_3013725 | Lighthouse Lab in Alderley Park | Wellcome Sanger Institute for the COVID-19 Genomics UK (COG-UK) Consortium | Jacquelyn Wynn, Mairead Hyland, The Lighthouse Lab in Alderley Park and Alex Alderton, Roberto Amato, Jeffrey Barrett, Sonia Goncalves, Ewan Harrison, David K. Jackson, Ian Johnston, Dominic Kwiatkowski, Cordelia Langford, John Sillitoe on behalf of the Wellcome Sanger Institute COVID-19 Surveillance Team |
| EPI_ISL_3013726 | Lighthouse Lab in Milton Keynes | Wellcome Sanger Institute for the COVID-19 Genomics UK (COG-UK) Consortium | The Lighthouse Lab in Milton Keynes and Alex Alderton, Roberto Amato, Jeffrey Barrett, Sonia Goncalves, Ewan Harrison, David K. Jackson, Ian Johnston, Dominic Kwiatkowski, Cordelia Langford, John Sillitoe on behalf of the Wellcome Sanger Institute COVID-19 Surveillance Team |
| EPI_ISL_3013727 | Lighthouse Lab in Glasgow | Wellcome Sanger Institute for the COVID-19 Genomics UK (COG-UK) Consortium | Harper VanSteenhouse, Yumi Kasai, David Gray, Carol Clugston, Anna Dominiczak and Alex Alderton, Roberto Amato, Jeffrey Barrett, Sonia Goncalves, Ewan Harrison, David K. Jackson, Ian Johnston, Dominic Kwiatkowski, Cordelia Langford, John Sillitoe on behalf of the Wellcome Sanger Institute COVID-19 Surveillance Team |
| EPI_ISL_3013728 | Lighthouse Lab in Milton Keynes | Wellcome Sanger Institute for the COVID-19 Genomics UK (COG-UK) Consortium | The Lighthouse Lab in Milton Keynes and Alex Alderton, Roberto Amato, Jeffrey Barrett, Sonia Goncalves, Ewan Harrison, David K. Jackson, Ian Johnston, Dominic Kwiatkowski, Cordelia Langford, John Sillitoe on behalf of the Wellcome Sanger Institute COVID-19 Surveillance Team |
| EPI_ISL_3013729 | Lighthouse Lab in Glasgow | Wellcome Sanger Institute for the COVID-19 Genomics UK (COG-UK) Consortium | Harper VanSteenhouse, Yumi Kasai, David Gray, Carol Clugston, Anna Dominiczak and Alex Alderton, Roberto Amato, Jeffrey Barrett, Sonia Goncalves, Ewan Harrison, David K. Jackson, Ian Johnston, Dominic Kwiatkowski, Cordelia Langford, John Sillitoe on behalf of the Wellcome Sanger Institute COVID-19 Surveillance Team |
| EPI_ISL_3013730 | Lighthouse Lab in Milton Keynes | Wellcome Sanger Institute for the COVID-19 Genomics UK (COG-UK) Consortium | The Lighthouse Lab in Milton Keynes and Alex Alderton, Roberto Amato, Jeffrey Barrett, Sonia Goncalves, Ewan Harrison, David K. Jackson, Ian Johnston, Dominic Kwiatkowski, Cordelia Langford, John Sillitoe on behalf of the Wellcome Sanger Institute COVID-19 Surveillance Team |
| EPI_ISL_3013731, EPI_ISL_3013732, EPI_ISL_3013733 | Lighthouse Lab in Glasgow | Wellcome Sanger Institute for the COVID-19 Genomics UK (COG-UK) Consortium | Harper VanSteenhouse, Yumi Kasai, David Gray, Carol Clugston, Anna Dominiczak and Alex Alderton, Roberto Amato, Jeffrey Barrett, Sonia Goncalves, Ewan Harrison, David K. Jackson, Ian Johnston, Dominic Kwiatkowski, Cordelia Langford, John Sillitoe on behalf of the Wellcome Sanger Institute COVID-19 Surveillance Team |
| EPI_ISL_3013734, EPI_ISL_3013735, EPI_ISL_3013736 | Lighthouse Lab in Alderley Park | Wellcome Sanger Institute for the COVID-19 Genomics UK (COG-UK) Consortium | Jacquelyn Wynn, Mairead Hyland, The Lighthouse Lab in Alderley Park and Alex Alderton, Roberto Amato, Jeffrey Barrett, Sonia Goncalves, Ewan Harrison, David K. Jackson, Ian Johnston, Dominic Kwiatkowski, Cordelia Langford, John Sillitoe on behalf of the Wellcome Sanger Institute COVID-19 Surveillance Team |
| EPI_ISL_3013737 | Health Services Laboratories | Wellcome Sanger Institute for the COVID-19 Genomics UK (COG-UK) Consortium | Health Services Laboratories and Alex Alderton, Roberto Amato, Jeffrey Barrett, Sonia Goncalves, Ewan Harrison, David K. Jackson, Ian Johnston, Dominic Kwiatkowski, Cordelia Langford, John Sillitoe on behalf of the Wellcome Sanger Institute COVID-19 Surveillance Team |
| EPI_ISL_3013738, EPI_ISL_3013739 | Lighthouse Lab in Milton Keynes | Wellcome Sanger Institute for the COVID-19 Genomics UK (COG-UK) Consortium | The Lighthouse Lab in Milton Keynes and Alex Alderton, Roberto Amato, Jeffrey Barrett, Sonia Goncalves, Ewan Harrison, David K. Jackson, Ian Johnston, Dominic Kwiatkowski, Cordelia Langford, John Sillitoe on behalf of the Wellcome Sanger Institute COVID-19 Surveillance Team |
| EPI_ISL_3013740, EPI_ISL_3013741 | Lighthouse Lab in Glasgow | Wellcome Sanger Institute for the COVID-19 Genomics UK (COG-UK) Consortium | Harper VanSteenhouse, Yumi Kasai, David Gray, Carol Clugston, Anna Dominiczak and Alex Alderton, Roberto Amato, Jeffrey Barrett, Sonia Goncalves, Ewan Harrison, David K. Jackson, Ian Johnston, Dominic Kwiatkowski, Cordelia Langford, John Sillitoe on behalf of the Wellcome Sanger Institute COVID-19 Surveillance Team |
| EPI_ISL_3013742, EPI_ISL_3013743 | Lighthouse Lab in Milton Keynes | Wellcome Sanger Institute for the COVID-19 Genomics UK (COG-UK) Consortium | The Lighthouse Lab in Milton Keynes and Alex Alderton, Roberto Amato, Jeffrey Barrett, Sonia Goncalves, Ewan Harrison, David K. Jackson, Ian Johnston, Dominic Kwiatkowski, Cordelia Langford, John Sillitoe on behalf of the Wellcome Sanger Institute COVID-19 Surveillance Team |
| EPI_ISL_3013744 | Lighthouse Lab in Alderley Park | Wellcome Sanger Institute for the COVID-19 Genomics UK (COG-UK) Consortium | Jacquelyn Wynn, Mairead Hyland, The Lighthouse Lab in Alderley Park and Alex Alderton, Roberto Amato, Jeffrey Barrett, Sonia Goncalves, Ewan Harrison, David K. Jackson, Ian Johnston, Dominic Kwiatkowski, Cordelia Langford, John Sillitoe on behalf of the Wellcome Sanger Institute COVID-19 Surveillance Team |
| EPI_ISL_3013745 | Lighthouse Lab in Glasgow | Wellcome Sanger Institute for the COVID-19 Genomics UK (COG-UK) Consortium | Harper VanSteenhouse, Yumi Kasai, David Gray, Carol Clugston, Anna Dominiczak and Alex Alderton, Roberto Amato, Jeffrey Barrett, Sonia Goncalves, Ewan Harrison, David K. Jackson, Ian Johnston, Dominic Kwiatkowski, Cordelia Langford, John Sillitoe on behalf of the Wellcome Sanger Institute COVID-19 Surveillance Team |
| EPI_ISL_3013746 | Lighthouse Lab in Milton Keynes | Wellcome Sanger Institute for the COVID-19 Genomics UK (COG-UK) Consortium | The Lighthouse Lab in Milton Keynes and Alex Alderton, Roberto Amato, Jeffrey Barrett, Sonia Goncalves, Ewan Harrison, David K. Jackson, Ian Johnston, Dominic Kwiatkowski, Cordelia Langford, John Sillitoe on behalf of the Wellcome Sanger Institute COVID-19 Surveillance Team |
| EPI_ISL_3013747 | Lighthouse Lab in Glasgow | Wellcome Sanger Institute for the COVID-19 Genomics UK (COG-UK) Consortium | Harper VanSteenhouse, Yumi Kasai, David Gray, Carol Clugston, Anna Dominiczak and Alex Alderton, Roberto Amato, Jeffrey Barrett, Sonia Goncalves, Ewan Harrison, David K. Jackson, Ian Johnston, Dominic Kwiatkowski, Cordelia Langford, John Sillitoe on behalf of the Wellcome Sanger Institute COVID-19 Surveillance Team |
| EPI_ISL_3013748, EPI_ISL_3013749 | Lighthouse Lab in Milton Keynes | Wellcome Sanger Institute for the COVID-19 Genomics UK (COG-UK) Consortium | The Lighthouse Lab in Milton Keynes and Alex Alderton, Roberto Amato, Jeffrey Barrett, Sonia Goncalves, Ewan Harrison, David K. Jackson, Ian Johnston, Dominic Kwiatkowski, Cordelia Langford, John Sillitoe on behalf of the Wellcome Sanger Institute COVID-19 Surveillance Team |
| EPI_ISL_3013750 | Lighthouse Lab in Glasgow | Wellcome Sanger Institute for the COVID-19 Genomics UK (COG-UK) Consortium | Harper VanSteenhouse, Yumi Kasai, David Gray, Carol Clugston, Anna Dominiczak and Alex Alderton, Roberto Amato, Jeffrey Barrett, Sonia Goncalves, Ewan Harrison, David K. Jackson, Ian Johnston, Dominic Kwiatkowski, Cordelia Langford, John Sillitoe on behalf of the Wellcome Sanger Institute COVID-19 Surveillance Team |
| EPI_ISL_3013751 | Lighthouse Lab in Alderley Park | Wellcome Sanger Institute for the COVID-19 Genomics UK (COG-UK) Consortium | Jacquelyn Wynn, Mairead Hyland, The Lighthouse Lab in Alderley Park and Alex Alderton, Roberto Amato, Jeffrey Barrett, Sonia Goncalves, Ewan Harrison, David K. Jackson, Ian Johnston, Dominic Kwiatkowski, Cordelia Langford, John Sillitoe on behalf of the Wellcome Sanger Institute COVID-19 Surveillance Team |
| EPI_ISL_3013752 | Lighthouse Lab in Milton Keynes | Wellcome Sanger Institute for the COVID-19 Genomics UK (COG-UK) Consortium | The Lighthouse Lab in Milton Keynes and Alex Alderton, Roberto Amato, Jeffrey Barrett, Sonia Goncalves, Ewan Harrison, David K. Jackson, Ian Johnston, Dominic Kwiatkowski, Cordelia Langford, John Sillitoe on behalf of the Wellcome Sanger Institute COVID-19 Surveillance Team |
| EPI_ISL_3013753 | Lighthouse Lab in Glasgow | Wellcome Sanger Institute for the COVID-19 Genomics UK (COG-UK) Consortium | Harper VanSteenhouse, Yumi Kasai, David Gray, Carol Clugston, Anna Dominiczak and Alex Alderton, Roberto Amato, Jeffrey Barrett, Sonia Goncalves, Ewan Harrison, David K. Jackson, Ian Johnston, Dominic Kwiatkowski, Cordelia Langford, John Sillitoe on behalf of the Wellcome Sanger Institute COVID-19 Surveillance Team |
| EPI_ISL_3013754 | Health Services Laboratories | Wellcome Sanger Institute for the COVID-19 Genomics UK (COG-UK) Consortium | Health Services Laboratories and Alex Alderton, Roberto Amato, Jeffrey Barrett, Sonia Goncalves, Ewan Harrison, David K. Jackson, Ian Johnston, Dominic Kwiatkowski, Cordelia Langford, John Sillitoe on behalf of the Wellcome Sanger Institute COVID-19 Surveillance Team |
| EPI_ISL_3013755, EPI_ISL_3013756 | Lighthouse Lab in Milton Keynes | Wellcome Sanger Institute for the COVID-19 Genomics UK (COG-UK) Consortium | The Lighthouse Lab in Milton Keynes and Alex Alderton, Roberto Amato, Jeffrey Barrett, Sonia Goncalves, Ewan Harrison, David K. Jackson, Ian Johnston, Dominic Kwiatkowski, Cordelia Langford, John Sillitoe on behalf of the Wellcome Sanger Institute COVID-19 Surveillance Team |
| EPI_ISL_3013757, EPI_ISL_3013758 | Health Services Laboratories | Wellcome Sanger Institute for the COVID-19 Genomics UK (COG-UK) Consortium | Health Services Laboratories and Alex Alderton, Roberto Amato, Jeffrey Barrett, Sonia Goncalves, Ewan Harrison, David K. Jackson, Ian Johnston, Dominic Kwiatkowski, Cordelia Langford, John Sillitoe on behalf of the Wellcome Sanger Institute COVID-19 Surveillance Team |
| EPI_ISL_3013759 | Lighthouse Lab in Milton Keynes | Wellcome Sanger Institute for the COVID-19 Genomics UK (COG-UK) | The Lighthouse Lab in Milton Keynes and Alex Alderton, Roberto Amato, Jeffrey Barrett, Sonia Goncalves, Ewan Harrison, David K. Jackson, Ian Johnston, Dominic Kwiatkowski, Cordelia Langford, John Sillitoe on behalf of the Wellcome Sanger |

| | | | |
|---|---|---|---|
| | | Consortium | Institute COVID-19 Surveillance Team |
| EPI_ISL_3013760, EPI_ISL_3013761, EPI_ISL_3013762 | Lighthouse Lab in Alderley Park | Wellcome Sanger Institute for the COVID-19 Genomics UK (COG-UK) Consortium | Jacquelyn Wynn, Mairead Hyland, The Lighthouse Lab in Alderley Park and Alex Alderton, Roberto Amato, Jeffrey Barrett, Sonia Goncalves, Ewan Harrison, David K. Jackson, Ian Johnston, Dominic Kwiatkowski, Cordelia Langford, John Sillitoe on behalf of the Wellcome Sanger Institute COVID-19 Surveillance Team |
| EPI_ISL_3013763, EPI_ISL_3013764, EPI_ISL_3013765 | Lighthouse Lab in Glasgow | Wellcome Sanger Institute for the COVID-19 Genomics UK (COG-UK) Consortium | Harper VanSteenhouse, Yumi Kasai, David Gray, Carol Clugston, Anna Dominiczak and Alex Alderton, Roberto Amato, Jeffrey Barrett, Sonia Goncalves, Ewan Harrison, David K. Jackson, Ian Johnston, Dominic Kwiatkowski, Cordelia Langford, John Sillitoe on behalf of the Wellcome Sanger Institute COVID-19 Surveillance Team |
| EPI_ISL_3013766 | Health Services Laboratories | Wellcome Sanger Institute for the COVID-19 Genomics UK (COG-UK) Consortium | Health Services Laboratories and Alex Alderton, Roberto Amato, Jeffrey Barrett, Sonia Goncalves, Ewan Harrison, David K. Jackson, Ian Johnston, Dominic Kwiatkowski, Cordelia Langford, John Sillitoe on behalf of the Wellcome Sanger Institute COVID-19 Surveillance Team |
| EPI_ISL_3013767, EPI_ISL_3013768 | Lighthouse Lab in Milton Keynes | Wellcome Sanger Institute for the COVID-19 Genomics UK (COG-UK) Consortium | The Lighthouse Lab in Milton Keynes and Alex Alderton, Roberto Amato, Jeffrey Barrett, Sonia Goncalves, Ewan Harrison, David K. Jackson, Ian Johnston, Dominic Kwiatkowski, Cordelia Langford, John Sillitoe on behalf of the Wellcome Sanger Institute COVID-19 Surveillance Team |
| EPI_ISL_3013769, EPI_ISL_3013770 | Lighthouse Lab in Glasgow | Wellcome Sanger Institute for the COVID-19 Genomics UK (COG-UK) Consortium | Harper VanSteenhouse, Yumi Kasai, David Gray, Carol Clugston, Anna Dominiczak and Alex Alderton, Roberto Amato, Jeffrey Barrett, Sonia Goncalves, Ewan Harrison, David K. Jackson, Ian Johnston, Dominic Kwiatkowski, Cordelia Langford, John Sillitoe on behalf of the Wellcome Sanger Institute COVID-19 Surveillance Team |
| EPI_ISL_3013771 | Health Services Laboratories | Wellcome Sanger Institute for the COVID-19 Genomics UK (COG-UK) Consortium | Health Services Laboratories and Alex Alderton, Roberto Amato, Jeffrey Barrett, Sonia Goncalves, Ewan Harrison, David K. Jackson, Ian Johnston, Dominic Kwiatkowski, Cordelia Langford, John Sillitoe on behalf of the Wellcome Sanger Institute COVID-19 Surveillance Team |
| EPI_ISL_3013772 | Lighthouse Lab in Alderley Park | Wellcome Sanger Institute for the COVID-19 Genomics UK (COG-UK) Consortium | Jacquelyn Wynn, Mairead Hyland, The Lighthouse Lab in Alderley Park and Alex Alderton, Roberto Amato, Jeffrey Barrett, Sonia Goncalves, Ewan Harrison, David K. Jackson, Ian Johnston, Dominic Kwiatkowski, Cordelia Langford, John Sillitoe on behalf of the Wellcome Sanger Institute COVID-19 Surveillance Team |
| EPI_ISL_3013773 | Health Services Laboratories | Wellcome Sanger Institute for the COVID-19 Genomics UK (COG-UK) Consortium | Health Services Laboratories and Alex Alderton, Roberto Amato, Jeffrey Barrett, Sonia Goncalves, Ewan Harrison, David K. Jackson, Ian Johnston, Dominic Kwiatkowski, Cordelia Langford, John Sillitoe on behalf of the Wellcome Sanger Institute COVID-19 Surveillance Team |
| EPI_ISL_3013774 | Lighthouse Lab in Glasgow | Wellcome Sanger Institute for the COVID-19 Genomics UK (COG-UK) Consortium | Harper VanSteenhouse, Yumi Kasai, David Gray, Carol Clugston, Anna Dominiczak and Alex Alderton, Roberto Amato, Jeffrey Barrett, Sonia Goncalves, Ewan Harrison, David K. Jackson, Ian Johnston, Dominic Kwiatkowski, Cordelia Langford, John Sillitoe on behalf of the Wellcome Sanger Institute COVID-19 Surveillance Team |
| EPI_ISL_3013775 | Lighthouse Lab in Milton Keynes | Wellcome Sanger Institute for the COVID-19 Genomics UK (COG-UK) Consortium | The Lighthouse Lab in Milton Keynes and Alex Alderton, Roberto Amato, Jeffrey Barrett, Sonia Goncalves, Ewan Harrison, David K. Jackson, Ian Johnston, Dominic Kwiatkowski, Cordelia Langford, John Sillitoe on behalf of the Wellcome Sanger Institute COVID-19 Surveillance Team |
| EPI_ISL_3013776 | Lighthouse Lab in Alderley Park | Wellcome Sanger Institute for the COVID-19 Genomics UK (COG-UK) Consortium | Jacquelyn Wynn, Mairead Hyland, The Lighthouse Lab in Alderley Park and Alex Alderton, Roberto Amato, Jeffrey Barrett, Sonia Goncalves, Ewan Harrison, David K. Jackson, Ian Johnston, Dominic Kwiatkowski, Cordelia Langford, John Sillitoe on behalf of the Wellcome Sanger Institute COVID-19 Surveillance Team |
| EPI_ISL_3013777 | Health Services Laboratories | Wellcome Sanger Institute for the COVID-19 Genomics UK (COG-UK) Consortium | Health Services Laboratories and Alex Alderton, Roberto Amato, Jeffrey Barrett, Sonia Goncalves, Ewan Harrison, David K. Jackson, Ian Johnston, Dominic Kwiatkowski, Cordelia Langford, John Sillitoe on behalf of the Wellcome Sanger Institute COVID-19 Surveillance Team |
| EPI_ISL_3013778 | Lighthouse Lab in Glasgow | Wellcome Sanger Institute for the COVID-19 Genomics UK (COG-UK) Consortium | Harper VanSteenhouse, Yumi Kasai, David Gray, Carol Clugston, Anna Dominiczak and Alex Alderton, Roberto Amato, Jeffrey Barrett, Sonia Goncalves, Ewan Harrison, David K. Jackson, Ian Johnston, Dominic Kwiatkowski, Cordelia Langford, John Sillitoe on behalf of the Wellcome Sanger Institute COVID-19 Surveillance Team |
| EPI_ISL_3013779, EPI_ISL_3013780, EPI_ISL_3013781, EPI_ISL_3013782 | Lighthouse Lab in Alderley Park | Wellcome Sanger Institute for the COVID-19 Genomics UK (COG-UK) Consortium | Jacquelyn Wynn, Mairead Hyland, The Lighthouse Lab in Alderley Park and Alex Alderton, Roberto Amato, Jeffrey Barrett, Sonia Goncalves, Ewan Harrison, David K. Jackson, Ian Johnston, Dominic Kwiatkowski, Cordelia Langford, John Sillitoe on behalf of the Wellcome Sanger Institute COVID-19 Surveillance Team |
| EPI_ISL_3013783, EPI_ISL_3013784 | Lighthouse Lab in Glasgow | Wellcome Sanger Institute for the COVID-19 Genomics UK (COG-UK) Consortium | Harper VanSteenhouse, Yumi Kasai, David Gray, Carol Clugston, Anna Dominiczak and Alex Alderton, Roberto Amato, Jeffrey Barrett, Sonia Goncalves, Ewan Harrison, David K. Jackson, Ian Johnston, Dominic Kwiatkowski, Cordelia Langford, John Sillitoe on behalf of the Wellcome Sanger Institute COVID-19 Surveillance Team |
| EPI_ISL_3013785 | Lighthouse Lab in Milton Keynes | Wellcome Sanger Institute for the COVID-19 Genomics UK (COG-UK) Consortium | The Lighthouse Lab in Milton Keynes and Alex Alderton, Roberto Amato, Jeffrey Barrett, Sonia Goncalves, Ewan Harrison, David K. Jackson, Ian Johnston, Dominic Kwiatkowski, Cordelia Langford, John Sillitoe on behalf of the Wellcome Sanger Institute COVID-19 Surveillance Team |
| EPI_ISL_3013786, EPI_ISL_3013787 | Lighthouse Lab in Glasgow | Wellcome Sanger Institute for the COVID-19 Genomics UK (COG-UK) Consortium | Harper VanSteenhouse, Yumi Kasai, David Gray, Carol Clugston, Anna Dominiczak and Alex Alderton, Roberto Amato, Jeffrey Barrett, Sonia Goncalves, Ewan Harrison, David K. Jackson, Ian Johnston, Dominic Kwiatkowski, Cordelia Langford, John Sillitoe on behalf of the Wellcome Sanger Institute COVID-19 Surveillance Team |
| EPI_ISL_3013788 | Lighthouse Lab in Alderley Park | Wellcome Sanger Institute for the COVID-19 Genomics UK (COG-UK) Consortium | Jacquelyn Wynn, Mairead Hyland, The Lighthouse Lab in Alderley Park and Alex Alderton, Roberto Amato, Jeffrey Barrett, Sonia Goncalves, Ewan Harrison, David K. Jackson, Ian Johnston, Dominic Kwiatkowski, Cordelia Langford, John Sillitoe on behalf of the Wellcome Sanger Institute COVID-19 Surveillance Team |
| EPI_ISL_3013789 | Health Services Laboratories | Wellcome Sanger Institute for the COVID-19 Genomics UK (COG-UK) Consortium | Health Services Laboratories and Alex Alderton, Roberto Amato, Jeffrey Barrett, Sonia Goncalves, Ewan Harrison, David K. Jackson, Ian Johnston, Dominic Kwiatkowski, Cordelia Langford, John Sillitoe on behalf of the Wellcome Sanger Institute COVID-19 Surveillance Team |
| EPI_ISL_3013790 | Lighthouse Lab in Milton Keynes | Wellcome Sanger Institute for the COVID-19 Genomics UK (COG-UK) Consortium | The Lighthouse Lab in Milton Keynes and Alex Alderton, Roberto Amato, Jeffrey Barrett, Sonia Goncalves, Ewan Harrison, David K. Jackson, Ian Johnston, Dominic Kwiatkowski, Cordelia Langford, John Sillitoe on behalf of the Wellcome Sanger Institute COVID-19 Surveillance Team |
| EPI_ISL_3013791 | Health Services Laboratories | Wellcome Sanger Institute for the COVID-19 Genomics UK (COG-UK) Consortium | Health Services Laboratories and Alex Alderton, Roberto Amato, Jeffrey Barrett, Sonia Goncalves, Ewan Harrison, David K. Jackson, Ian Johnston, Dominic Kwiatkowski, Cordelia Langford, John Sillitoe on behalf of the Wellcome Sanger Institute COVID-19 Surveillance Team |
| EPI_ISL_3013792 | Lighthouse Lab in Alderley Park | Wellcome Sanger Institute for the COVID-19 Genomics UK (COG-UK) Consortium | Jacquelyn Wynn, Mairead Hyland, The Lighthouse Lab in Alderley Park and Alex Alderton, Roberto Amato, Jeffrey Barrett, Sonia Goncalves, Ewan Harrison, David K. Jackson, Ian Johnston, Dominic Kwiatkowski, Cordelia Langford, John Sillitoe on behalf of the Wellcome Sanger Institute COVID-19 Surveillance Team |
| EPI_ISL_3013793 | Lighthouse Lab in Glasgow | Wellcome Sanger Institute for the COVID-19 Genomics UK (COG-UK) Consortium | Harper VanSteenhouse, Yumi Kasai, David Gray, Carol Clugston, Anna Dominiczak and Alex Alderton, Roberto Amato, Jeffrey Barrett, Sonia Goncalves, Ewan Harrison, David K. Jackson, Ian Johnston, Dominic Kwiatkowski, Cordelia Langford, John Sillitoe on behalf of the Wellcome Sanger Institute COVID-19 Surveillance Team |
| EPI_ISL_3013794 | Lighthouse Lab in Milton Keynes | Wellcome Sanger Institute for the COVID-19 Genomics UK (COG-UK) Consortium | The Lighthouse Lab in Milton Keynes and Alex Alderton, Roberto Amato, Jeffrey Barrett, Sonia Goncalves, Ewan Harrison, David K. Jackson, Ian Johnston, Dominic Kwiatkowski, Cordelia Langford, John Sillitoe on behalf of the Wellcome Sanger Institute COVID-19 Surveillance Team |
| EPI_ISL_3013795 | Lighthouse Lab in Glasgow | Wellcome Sanger Institute for the COVID-19 Genomics UK (COG-UK) Consortium | Harper VanSteenhouse, Yumi Kasai, David Gray, Carol Clugston, Anna Dominiczak and Alex Alderton, Roberto Amato, Jeffrey Barrett, Sonia Goncalves, Ewan Harrison, David K. Jackson, Ian Johnston, Dominic Kwiatkowski, Cordelia Langford, John Sillitoe on behalf of the Wellcome Sanger Institute COVID-19 Surveillance Team |
| EPI_ISL_3013796, EPI_ISL_3013797 | Lighthouse Lab in Alderley Park | Wellcome Sanger Institute for the COVID-19 Genomics UK (COG-UK) Consortium | Jacquelyn Wynn, Mairead Hyland, The Lighthouse Lab in Alderley Park and Alex Alderton, Roberto Amato, Jeffrey Barrett, Sonia Goncalves, Ewan Harrison, David K. Jackson, Ian Johnston, Dominic Kwiatkowski, Cordelia Langford, John Sillitoe on behalf of the Wellcome Sanger Institute COVID-19 Surveillance Team |
| EPI_ISL_3013798, EPI_ISL_3013799 | Lighthouse Lab in Milton Keynes | Wellcome Sanger Institute for the COVID-19 Genomics UK (COG-UK) Consortium | The Lighthouse Lab in Milton Keynes and Alex Alderton, Roberto Amato, Jeffrey Barrett, Sonia Goncalves, Ewan Harrison, David K. Jackson, Ian Johnston, Dominic Kwiatkowski, Cordelia Langford, John Sillitoe on behalf of the Wellcome Sanger Institute COVID-19 Surveillance Team |
| EPI_ISL_3013800, EPI_ISL_3013801 | Lighthouse Lab in Glasgow | Wellcome Sanger Institute for the COVID-19 Genomics UK (COG-UK) Consortium | Harper VanSteenhouse, Yumi Kasai, David Gray, Carol Clugston, Anna Dominiczak and Alex Alderton, Roberto Amato, Jeffrey Barrett, Sonia Goncalves, Ewan Harrison, David K. Jackson, Ian Johnston, Dominic Kwiatkowski, Cordelia Langford, John Sillitoe on behalf of the Wellcome Sanger Institute COVID-19 Surveillance Team |
| EPI_ISL_3013802 | Lighthouse Lab in Milton Keynes | Wellcome Sanger Institute for the COVID-19 Genomics UK (COG-UK) Consortium | The Lighthouse Lab in Milton Keynes and Alex Alderton, Roberto Amato, Jeffrey Barrett, Sonia Goncalves, Ewan Harrison, David K. Jackson, Ian Johnston, Dominic Kwiatkowski, Cordelia Langford, John Sillitoe on behalf of the Wellcome Sanger Institute COVID-19 Surveillance Team |
| EPI_ISL_3013803, EPI_ISL_3013804, EPI_ISL_3013805 | Health Services Laboratories | Wellcome Sanger Institute for the COVID-19 Genomics UK (COG-UK) Consortium | Health Services Laboratories and Alex Alderton, Roberto Amato, Jeffrey Barrett, Sonia Goncalves, Ewan Harrison, David K. Jackson, Ian Johnston, Dominic Kwiatkowski, Cordelia Langford, John Sillitoe on behalf of the Wellcome Sanger Institute COVID-19 Surveillance Team |
| EPI_ISL_3013806 | Lighthouse Lab in Milton Keynes | Wellcome Sanger Institute for the COVID-19 Genomics UK (COG-UK) Consortium | The Lighthouse Lab in Milton Keynes and Alex Alderton, Roberto Amato, Jeffrey Barrett, Sonia Goncalves, Ewan Harrison, David K. Jackson, Ian Johnston, Dominic Kwiatkowski, Cordelia Langford, John Sillitoe on behalf of the Wellcome Sanger Institute COVID-19 Surveillance Team |
| EPI_ISL_3013807 | Lighthouse Lab in Glasgow | Wellcome Sanger Institute for the COVID-19 Genomics UK (COG-UK) Consortium | Harper VanSteenhouse, Yumi Kasai, David Gray, Carol Clugston, Anna Dominiczak and Alex Alderton, Roberto Amato, Jeffrey Barrett, Sonia Goncalves, Ewan Harrison, David K. Jackson, Ian Johnston, Dominic Kwiatkowski, Cordelia Langford, John Sillitoe on behalf of the Wellcome Sanger Institute COVID-19 Surveillance Team |
| EPI_ISL_3013808 | Lighthouse Lab in Alderley Park | Wellcome Sanger Institute for the COVID-19 Genomics UK (COG-UK) Consortium | Jacquelyn Wynn, Mairead Hyland, The Lighthouse Lab in Alderley Park and Alex Alderton, Roberto Amato, Jeffrey Barrett, Sonia Goncalves, Ewan Harrison, David K. Jackson, Ian Johnston, Dominic Kwiatkowski, Cordelia Langford, John Sillitoe on behalf of the Wellcome Sanger Institute COVID-19 Surveillance Team |
| EPI_ISL_3013809 | Health Services Laboratories | Wellcome Sanger Institute for the COVID-19 Genomics UK (COG-UK) Consortium | Health Services Laboratories and Alex Alderton, Roberto Amato, Jeffrey Barrett, Sonia Goncalves, Ewan Harrison, David K. Jackson, Ian Johnston, Dominic Kwiatkowski, Cordelia Langford, John Sillitoe on behalf of the Wellcome Sanger Institute COVID-19 Surveillance Team |
| EPI_ISL_3013810 | Lighthouse Lab in Milton Keynes | Wellcome Sanger Institute for the COVID-19 Genomics UK (COG-UK) Consortium | The Lighthouse Lab in Milton Keynes and Alex Alderton, Roberto Amato, Jeffrey Barrett, Sonia Goncalves, Ewan Harrison, David K. Jackson, Ian Johnston, Dominic Kwiatkowski, Cordelia Langford, John Sillitoe on behalf of the Wellcome Sanger Institute COVID-19 Surveillance Team |
| EPI_ISL_3013811 | Lighthouse Lab in Alderley Park | Wellcome Sanger Institute for the COVID-19 Genomics UK (COG-UK) Consortium | Jacquelyn Wynn, Mairead Hyland, The Lighthouse Lab in Alderley Park and Alex Alderton, Roberto Amato, Jeffrey Barrett, Sonia Goncalves, Ewan Harrison, David K. Jackson, Ian Johnston, Dominic Kwiatkowski, Cordelia Langford, John Sillitoe on behalf of the Wellcome Sanger Institute COVID-19 Surveillance Team |
| EPI_ISL_3013812 | Lighthouse Lab in Milton Keynes | Wellcome Sanger Institute for the COVID-19 Genomics UK (COG-UK) Consortium | The Lighthouse Lab in Milton Keynes and Alex Alderton, Roberto Amato, Jeffrey Barrett, Sonia Goncalves, Ewan Harrison, David K. Jackson, Ian Johnston, Dominic Kwiatkowski, Cordelia Langford, John Sillitoe on behalf of the Wellcome Sanger Institute COVID-19 Surveillance Team |
| EPI_ISL_3013813 | Lighthouse Lab in Alderley Park | Wellcome Sanger Institute for the COVID-19 Genomics UK (COG-UK) Consortium | Jacquelyn Wynn, Mairead Hyland, The Lighthouse Lab in Alderley Park and Alex Alderton, Roberto Amato, Jeffrey Barrett, Sonia Goncalves, Ewan Harrison, David K. Jackson, Ian Johnston, Dominic Kwiatkowski, Cordelia Langford, John Sillitoe on behalf of the Wellcome Sanger Institute COVID-19 Surveillance Team |
| EPI_ISL_3013814 | Lighthouse Lab in Milton Keynes | Wellcome Sanger Institute for the COVID-19 Genomics UK (COG-UK) Consortium | The Lighthouse Lab in Milton Keynes and Alex Alderton, Roberto Amato, Jeffrey Barrett, Sonia Goncalves, Ewan Harrison, David K. Jackson, Ian Johnston, Dominic Kwiatkowski, Cordelia Langford, John Sillitoe on behalf of the Wellcome Sanger Institute COVID-19 Surveillance Team |
| EPI_ISL_3013815 | Lighthouse Lab in Glasgow | Wellcome Sanger Institute for the COVID-19 Genomics UK (COG-UK) Consortium | Harper VanSteenhouse, Yumi Kasai, David Gray, Carol Clugston, Anna Dominiczak and Alex Alderton, Roberto Amato, Jeffrey Barrett, Sonia Goncalves, Ewan Harrison, David K. Jackson, Ian Johnston, Dominic Kwiatkowski, Cordelia Langford, John Sillitoe on behalf of the Wellcome Sanger Institute COVID-19 Surveillance Team |
| EPI_ISL_3013816 | Health Services Laboratories | Wellcome Sanger Institute for the COVID-19 Genomics UK (COG-UK) Consortium | Health Services Laboratories and Alex Alderton, Roberto Amato, Jeffrey Barrett, Sonia Goncalves, Ewan Harrison, David K. Jackson, Ian Johnston, Dominic Kwiatkowski, Cordelia Langford, John Sillitoe on behalf of the Wellcome Sanger Institute COVID-19 Surveillance Team |
| EPI_ISL_3013817 | Lighthouse Lab in Glasgow | Wellcome Sanger Institute for the COVID-19 Genomics UK (COG-UK) Consortium | Harper VanSteenhouse, Yumi Kasai, David Gray, Carol Clugston, Anna Dominiczak and Alex Alderton, Roberto Amato, Jeffrey Barrett, Sonia Goncalves, Ewan Harrison, David K. Jackson, Ian Johnston, Dominic Kwiatkowski, Cordelia Langford, John Sillitoe on behalf of the Wellcome Sanger Institute COVID-19 Surveillance Team |
| EPI_ISL_3013818 | Lighthouse Lab in Milton Keynes | Wellcome Sanger Institute for the COVID-19 Genomics UK (COG-UK) | The Lighthouse Lab in Milton Keynes and Alex Alderton, Roberto Amato, Jeffrey Barrett, Sonia Goncalves, Ewan Harrison, David K. Jackson, Ian Johnston, Dominic Kwiatkowski, Cordelia Langford, John Sillitoe on behalf of the Wellcome Sanger |

| | | | |
|---|---|---|---|
| | | Consortium | Institute COVID-19 Surveillance Team |
| EPI_ISL_3013819, EPI_ISL_3013820 | Lighthouse Lab in Alderley Park | Wellcome Sanger Institute for the COVID-19 Genomics UK (COG-UK) Consortium | Jacquelyn Wynn, Mairead Hyland, The Lighthouse Lab in Alderley Park and Alex Alderton, Roberto Amato, Jeffrey Barrett, Sonia Goncalves, Ewan Harrison, David K. Jackson, Ian Johnston, Dominic Kwiatkowski, Cordelia Langford, John Sillitoe on behalf of the Wellcome Sanger Institute COVID-19 Surveillance Team |
| EPI_ISL_3013821 | Health Services Laboratories | Wellcome Sanger Institute for the COVID-19 Genomics UK (COG-UK) Consortium | Health Services Laboratories and Alex Alderton, Roberto Amato, Jeffrey Barrett, Sonia Goncalves, Ewan Harrison, David K. Jackson, Ian Johnston, Dominic Kwiatkowski, Cordelia Langford, John Sillitoe on behalf of the Wellcome Sanger Institute COVID-19 Surveillance Team |
| EPI_ISL_3013822, EPI_ISL_3013823, EPI_ISL_3013824 | Lighthouse Lab in Glasgow | Wellcome Sanger Institute for the COVID-19 Genomics UK (COG-UK) Consortium | Harper VanSteenhouse, Yumi Kasai, David Gray, Carol Clugston, Anna Dominiczak and Alex Alderton, Roberto Amato, Jeffrey Barrett, Sonia Goncalves, Ewan Harrison, David K. Jackson, Ian Johnston, Dominic Kwiatkowski, Cordelia Langford, John Sillitoe on behalf of the Wellcome Sanger Institute COVID-19 Surveillance Team |
| EPI_ISL_3013825 | Lighthouse Lab in Alderley Park | Wellcome Sanger Institute for the COVID-19 Genomics UK (COG-UK) Consortium | Jacquelyn Wynn, Mairead Hyland, The Lighthouse Lab in Alderley Park and Alex Alderton, Roberto Amato, Jeffrey Barrett, Sonia Goncalves, Ewan Harrison, David K. Jackson, Ian Johnston, Dominic Kwiatkowski, Cordelia Langford, John Sillitoe on behalf of the Wellcome Sanger Institute COVID-19 Surveillance Team |
| EPI_ISL_3013827 | Lighthouse Lab in Milton Keynes | Wellcome Sanger Institute for the COVID-19 Genomics UK (COG-UK) Consortium | The Lighthouse Lab in Milton Keynes and Alex Alderton, Roberto Amato, Jeffrey Barrett, Sonia Goncalves, Ewan Harrison, David K. Jackson, Ian Johnston, Dominic Kwiatkowski, Cordelia Langford, John Sillitoe on behalf of the Wellcome Sanger Institute COVID-19 Surveillance Team |
| EPI_ISL_3013828 | Lighthouse Lab in Glasgow | Wellcome Sanger Institute for the COVID-19 Genomics UK (COG-UK) Consortium | Harper VanSteenhouse, Yumi Kasai, David Gray, Carol Clugston, Anna Dominiczak and Alex Alderton, Roberto Amato, Jeffrey Barrett, Sonia Goncalves, Ewan Harrison, David K. Jackson, Ian Johnston, Dominic Kwiatkowski, Cordelia Langford, John Sillitoe on behalf of the Wellcome Sanger Institute COVID-19 Surveillance Team |
| EPI_ISL_3013829 | Lighthouse Lab in Alderley Park | Wellcome Sanger Institute for the COVID-19 Genomics UK (COG-UK) Consortium | Jacquelyn Wynn, Mairead Hyland, The Lighthouse Lab in Alderley Park and Alex Alderton, Roberto Amato, Jeffrey Barrett, Sonia Goncalves, Ewan Harrison, David K. Jackson, Ian Johnston, Dominic Kwiatkowski, Cordelia Langford, John Sillitoe on behalf of the Wellcome Sanger Institute COVID-19 Surveillance Team |
| EPI_ISL_3013830 | Health Services Laboratories | Wellcome Sanger Institute for the COVID-19 Genomics UK (COG-UK) Consortium | Health Services Laboratories and Alex Alderton, Roberto Amato, Jeffrey Barrett, Sonia Goncalves, Ewan Harrison, David K. Jackson, Ian Johnston, Dominic Kwiatkowski, Cordelia Langford, John Sillitoe on behalf of the Wellcome Sanger Institute COVID-19 Surveillance Team |
| EPI_ISL_3013831 | Lighthouse Lab in Milton Keynes | Wellcome Sanger Institute for the COVID-19 Genomics UK (COG-UK) Consortium | The Lighthouse Lab in Milton Keynes and Alex Alderton, Roberto Amato, Jeffrey Barrett, Sonia Goncalves, Ewan Harrison, David K. Jackson, Ian Johnston, Dominic Kwiatkowski, Cordelia Langford, John Sillitoe on behalf of the Wellcome Sanger Institute COVID-19 Surveillance Team |
| EPI_ISL_3013832 | Health Services Laboratories | Wellcome Sanger Institute for the COVID-19 Genomics UK (COG-UK) Consortium | Health Services Laboratories and Alex Alderton, Roberto Amato, Jeffrey Barrett, Sonia Goncalves, Ewan Harrison, David K. Jackson, Ian Johnston, Dominic Kwiatkowski, Cordelia Langford, John Sillitoe on behalf of the Wellcome Sanger Institute COVID-19 Surveillance Team |
| EPI_ISL_3013833 | Lighthouse Lab in Alderley Park | Wellcome Sanger Institute for the COVID-19 Genomics UK (COG-UK) Consortium | Jacquelyn Wynn, Mairead Hyland, The Lighthouse Lab in Alderley Park and Alex Alderton, Roberto Amato, Jeffrey Barrett, Sonia Goncalves, Ewan Harrison, David K. Jackson, Ian Johnston, Dominic Kwiatkowski, Cordelia Langford, John Sillitoe on behalf of the Wellcome Sanger Institute COVID-19 Surveillance Team |
| EPI_ISL_3013834 | Lighthouse Lab in Glasgow | Wellcome Sanger Institute for the COVID-19 Genomics UK (COG-UK) Consortium | Harper VanSteenhouse, Yumi Kasai, David Gray, Carol Clugston, Anna Dominiczak and Alex Alderton, Roberto Amato, Jeffrey Barrett, Sonia Goncalves, Ewan Harrison, David K. Jackson, Ian Johnston, Dominic Kwiatkowski, Cordelia Langford, John Sillitoe on behalf of the Wellcome Sanger Institute COVID-19 Surveillance Team |
| EPI_ISL_3013835, EPI_ISL_3013836 | Lighthouse Lab in Alderley Park | Wellcome Sanger Institute for the COVID-19 Genomics UK (COG-UK) Consortium | Jacquelyn Wynn, Mairead Hyland, The Lighthouse Lab in Alderley Park and Alex Alderton, Roberto Amato, Jeffrey Barrett, Sonia Goncalves, Ewan Harrison, David K. Jackson, Ian Johnston, Dominic Kwiatkowski, Cordelia Langford, John Sillitoe on behalf of the Wellcome Sanger Institute COVID-19 Surveillance Team |
| EPI_ISL_3013837, EPI_ISL_3013838 | Lighthouse Lab in Milton Keynes | Wellcome Sanger Institute for the COVID-19 Genomics UK (COG-UK) Consortium | The Lighthouse Lab in Milton Keynes and Alex Alderton, Roberto Amato, Jeffrey Barrett, Sonia Goncalves, Ewan Harrison, David K. Jackson, Ian Johnston, Dominic Kwiatkowski, Cordelia Langford, John Sillitoe on behalf of the Wellcome Sanger Institute COVID-19 Surveillance Team |
| EPI_ISL_3013839 | Lighthouse Lab in Glasgow | Wellcome Sanger Institute for the COVID-19 Genomics UK (COG-UK) Consortium | Harper VanSteenhouse, Yumi Kasai, David Gray, Carol Clugston, Anna Dominiczak and Alex Alderton, Roberto Amato, Jeffrey Barrett, Sonia Goncalves, Ewan Harrison, David K. Jackson, Ian Johnston, Dominic Kwiatkowski, Cordelia Langford, John Sillitoe on behalf of the Wellcome Sanger Institute COVID-19 Surveillance Team |
| EPI_ISL_3013840 | Health Services Laboratories | Wellcome Sanger Institute for the COVID-19 Genomics UK (COG-UK) Consortium | Health Services Laboratories and Alex Alderton, Roberto Amato, Jeffrey Barrett, Sonia Goncalves, Ewan Harrison, David K. Jackson, Ian Johnston, Dominic Kwiatkowski, Cordelia Langford, John Sillitoe on behalf of the Wellcome Sanger Institute COVID-19 Surveillance Team |
| EPI_ISL_3013841, EPI_ISL_3013842 | Lighthouse Lab in Milton Keynes | Wellcome Sanger Institute for the COVID-19 Genomics UK (COG-UK) Consortium | The Lighthouse Lab in Milton Keynes and Alex Alderton, Roberto Amato, Jeffrey Barrett, Sonia Goncalves, Ewan Harrison, David K. Jackson, Ian Johnston, Dominic Kwiatkowski, Cordelia Langford, John Sillitoe on behalf of the Wellcome Sanger Institute COVID-19 Surveillance Team |
| EPI_ISL_3013843 | Lighthouse Lab in Glasgow | Wellcome Sanger Institute for the COVID-19 Genomics UK (COG-UK) Consortium | Harper VanSteenhouse, Yumi Kasai, David Gray, Carol Clugston, Anna Dominiczak and Alex Alderton, Roberto Amato, Jeffrey Barrett, Sonia Goncalves, Ewan Harrison, David K. Jackson, Ian Johnston, Dominic Kwiatkowski, Cordelia Langford, John Sillitoe on behalf of the Wellcome Sanger Institute COVID-19 Surveillance Team |
| EPI_ISL_3013844 | Lighthouse Lab in Milton Keynes | Wellcome Sanger Institute for the COVID-19 Genomics UK (COG-UK) Consortium | The Lighthouse Lab in Milton Keynes and Alex Alderton, Roberto Amato, Jeffrey Barrett, Sonia Goncalves, Ewan Harrison, David K. Jackson, Ian Johnston, Dominic Kwiatkowski, Cordelia Langford, John Sillitoe on behalf of the Wellcome Sanger Institute COVID-19 Surveillance Team |
| EPI_ISL_3013845 | Lighthouse Lab in Glasgow | Wellcome Sanger Institute for the COVID-19 Genomics UK (COG-UK) Consortium | Harper VanSteenhouse, Yumi Kasai, David Gray, Carol Clugston, Anna Dominiczak and Alex Alderton, Roberto Amato, Jeffrey Barrett, Sonia Goncalves, Ewan Harrison, David K. Jackson, Ian Johnston, Dominic Kwiatkowski, Cordelia Langford, John Sillitoe on behalf of the Wellcome Sanger Institute COVID-19 Surveillance Team |
| EPI_ISL_3013846, EPI_ISL_3013847, EPI_ISL_3013848 | Lighthouse Lab in Alderley Park | Wellcome Sanger Institute for the COVID-19 Genomics UK (COG-UK) Consortium | Jacquelyn Wynn, Mairead Hyland, The Lighthouse Lab in Alderley Park and Alex Alderton, Roberto Amato, Jeffrey Barrett, Sonia Goncalves, Ewan Harrison, David K. Jackson, Ian Johnston, Dominic Kwiatkowski, Cordelia Langford, John Sillitoe on behalf of the Wellcome Sanger Institute COVID-19 Surveillance Team |
| EPI_ISL_3013849 | Lighthouse Lab in Milton Keynes | Wellcome Sanger Institute for the COVID-19 Genomics UK (COG-UK) Consortium | The Lighthouse Lab in Milton Keynes and Alex Alderton, Roberto Amato, Jeffrey Barrett, Sonia Goncalves, Ewan Harrison, David K. Jackson, Ian Johnston, Dominic Kwiatkowski, Cordelia Langford, John Sillitoe on behalf of the Wellcome Sanger Institute COVID-19 Surveillance Team |
| EPI_ISL_3013850 | Lighthouse Lab in Alderley Park | Wellcome Sanger Institute for the COVID-19 Genomics UK (COG-UK) Consortium | Jacquelyn Wynn, Mairead Hyland, The Lighthouse Lab in Alderley Park and Alex Alderton, Roberto Amato, Jeffrey Barrett, Sonia Goncalves, Ewan Harrison, David K. Jackson, Ian Johnston, Dominic Kwiatkowski, Cordelia Langford, John Sillitoe on behalf of the Wellcome Sanger Institute COVID-19 Surveillance Team |
| EPI_ISL_3013851 | Health Services Laboratories | Wellcome Sanger Institute for the COVID-19 Genomics UK (COG-UK) Consortium | Health Services Laboratories and Alex Alderton, Roberto Amato, Jeffrey Barrett, Sonia Goncalves, Ewan Harrison, David K. Jackson, Ian Johnston, Dominic Kwiatkowski, Cordelia Langford, John Sillitoe on behalf of the Wellcome Sanger Institute COVID-19 Surveillance Team |
| EPI_ISL_3013852, EPI_ISL_3013853 | Lighthouse Lab in Alderley Park | Wellcome Sanger Institute for the COVID-19 Genomics UK (COG-UK) Consortium | Jacquelyn Wynn, Mairead Hyland, The Lighthouse Lab in Alderley Park and Alex Alderton, Roberto Amato, Jeffrey Barrett, Sonia Goncalves, Ewan Harrison, David K. Jackson, Ian Johnston, Dominic Kwiatkowski, Cordelia Langford, John Sillitoe on behalf of the Wellcome Sanger Institute COVID-19 Surveillance Team |
| EPI_ISL_3013854, EPI_ISL_3013855 | Lighthouse Lab in Glasgow | Wellcome Sanger Institute for the COVID-19 Genomics UK (COG-UK) Consortium | Harper VanSteenhouse, Yumi Kasai, David Gray, Carol Clugston, Anna Dominiczak and Alex Alderton, Roberto Amato, Jeffrey Barrett, Sonia Goncalves, Ewan Harrison, David K. Jackson, Ian Johnston, Dominic Kwiatkowski, Cordelia Langford, John Sillitoe on behalf of the Wellcome Sanger Institute COVID-19 Surveillance Team |
| EPI_ISL_3013856 | Lighthouse Lab in Alderley Park | Wellcome Sanger Institute for the COVID-19 Genomics UK (COG-UK) Consortium | Jacquelyn Wynn, Mairead Hyland, The Lighthouse Lab in Alderley Park and Alex Alderton, Roberto Amato, Jeffrey Barrett, Sonia Goncalves, Ewan Harrison, David K. Jackson, Ian Johnston, Dominic Kwiatkowski, Cordelia Langford, John Sillitoe on behalf of the Wellcome Sanger Institute COVID-19 Surveillance Team |
| EPI_ISL_3013857, EPI_ISL_3013858 | Lighthouse Lab in Glasgow | Wellcome Sanger Institute for the COVID-19 Genomics UK (COG-UK) Consortium | Harper VanSteenhouse, Yumi Kasai, David Gray, Carol Clugston, Anna Dominiczak and Alex Alderton, Roberto Amato, Jeffrey Barrett, Sonia Goncalves, Ewan Harrison, David K. Jackson, Ian Johnston, Dominic Kwiatkowski, Cordelia Langford, John Sillitoe on behalf of the Wellcome Sanger Institute COVID-19 Surveillance Team |
| EPI_ISL_3013859 | Lighthouse Lab in Alderley Park | Wellcome Sanger Institute for the COVID-19 Genomics UK (COG-UK) Consortium | Jacquelyn Wynn, Mairead Hyland, The Lighthouse Lab in Alderley Park and Alex Alderton, Roberto Amato, Jeffrey Barrett, Sonia Goncalves, Ewan Harrison, David K. Jackson, Ian Johnston, Dominic Kwiatkowski, Cordelia Langford, John Sillitoe on behalf of the Wellcome Sanger Institute COVID-19 Surveillance Team |
| EPI_ISL_3013860, EPI_ISL_3013861 | Health Services Laboratories | Wellcome Sanger Institute for the COVID-19 Genomics UK (COG-UK) Consortium | Health Services Laboratories and Alex Alderton, Roberto Amato, Jeffrey Barrett, Sonia Goncalves, Ewan Harrison, David K. Jackson, Ian Johnston, Dominic Kwiatkowski, Cordelia Langford, John Sillitoe on behalf of the Wellcome Sanger Institute COVID-19 Surveillance Team |
| EPI_ISL_3013862 | Lighthouse Lab in Milton Keynes | Wellcome Sanger Institute for the COVID-19 Genomics UK (COG-UK) Consortium | The Lighthouse Lab in Milton Keynes and Alex Alderton, Roberto Amato, Jeffrey Barrett, Sonia Goncalves, Ewan Harrison, David K. Jackson, Ian Johnston, Dominic Kwiatkowski, Cordelia Langford, John Sillitoe on behalf of the Wellcome Sanger Institute COVID-19 Surveillance Team |
| EPI_ISL_3013863, EPI_ISL_3013864 | Lighthouse Lab in Alderley Park | Wellcome Sanger Institute for the COVID-19 Genomics UK (COG-UK) Consortium | Jacquelyn Wynn, Mairead Hyland, The Lighthouse Lab in Alderley Park and Alex Alderton, Roberto Amato, Jeffrey Barrett, Sonia Goncalves, Ewan Harrison, David K. Jackson, Ian Johnston, Dominic Kwiatkowski, Cordelia Langford, John Sillitoe on behalf of the Wellcome Sanger Institute COVID-19 Surveillance Team |
| EPI_ISL_3013865 | Lighthouse Lab in Glasgow | Wellcome Sanger Institute for the COVID-19 Genomics UK (COG-UK) Consortium | Harper VanSteenhouse, Yumi Kasai, David Gray, Carol Clugston, Anna Dominiczak and Alex Alderton, Roberto Amato, Jeffrey Barrett, Sonia Goncalves, Ewan Harrison, David K. Jackson, Ian Johnston, Dominic Kwiatkowski, Cordelia Langford, John Sillitoe on behalf of the Wellcome Sanger Institute COVID-19 Surveillance Team |
| EPI_ISL_3013866 | Lighthouse Lab in Alderley Park | Wellcome Sanger Institute for the COVID-19 Genomics UK (COG-UK) Consortium | Jacquelyn Wynn, Mairead Hyland, The Lighthouse Lab in Alderley Park and Alex Alderton, Roberto Amato, Jeffrey Barrett, Sonia Goncalves, Ewan Harrison, David K. Jackson, Ian Johnston, Dominic Kwiatkowski, Cordelia Langford, John Sillitoe on behalf of the Wellcome Sanger Institute COVID-19 Surveillance Team |
| EPI_ISL_3013867 | Lighthouse Lab in Milton Keynes | Wellcome Sanger Institute for the COVID-19 Genomics UK (COG-UK) Consortium | The Lighthouse Lab in Milton Keynes and Alex Alderton, Roberto Amato, Jeffrey Barrett, Sonia Goncalves, Ewan Harrison, David K. Jackson, Ian Johnston, Dominic Kwiatkowski, Cordelia Langford, John Sillitoe on behalf of the Wellcome Sanger Institute COVID-19 Surveillance Team |
| EPI_ISL_3013868 | Health Services Laboratories | Wellcome Sanger Institute for the COVID-19 Genomics UK (COG-UK) Consortium | Health Services Laboratories and Alex Alderton, Roberto Amato, Jeffrey Barrett, Sonia Goncalves, Ewan Harrison, David K. Jackson, Ian Johnston, Dominic Kwiatkowski, Cordelia Langford, John Sillitoe on behalf of the Wellcome Sanger Institute COVID-19 Surveillance Team |
| EPI_ISL_3013869 | Lighthouse Lab in Alderley Park | Wellcome Sanger Institute for the COVID-19 Genomics UK (COG-UK) Consortium | Jacquelyn Wynn, Mairead Hyland, The Lighthouse Lab in Alderley Park and Alex Alderton, Roberto Amato, Jeffrey Barrett, Sonia Goncalves, Ewan Harrison, David K. Jackson, Ian Johnston, Dominic Kwiatkowski, Cordelia Langford, John Sillitoe on behalf of the Wellcome Sanger Institute COVID-19 Surveillance Team |
| EPI_ISL_3013870 | Health Services Laboratories | Wellcome Sanger Institute for the COVID-19 Genomics UK (COG-UK) Consortium | Health Services Laboratories and Alex Alderton, Roberto Amato, Jeffrey Barrett, Sonia Goncalves, Ewan Harrison, David K. Jackson, Ian Johnston, Dominic Kwiatkowski, Cordelia Langford, John Sillitoe on behalf of the Wellcome Sanger Institute COVID-19 Surveillance Team |
| EPI_ISL_3013871 | Lighthouse Lab in Milton Keynes | Wellcome Sanger Institute for the COVID-19 Genomics UK (COG-UK) Consortium | The Lighthouse Lab in Milton Keynes and Alex Alderton, Roberto Amato, Jeffrey Barrett, Sonia Goncalves, Ewan Harrison, David K. Jackson, Ian Johnston, Dominic Kwiatkowski, Cordelia Langford, John Sillitoe on behalf of the Wellcome Sanger Institute COVID-19 Surveillance Team |
| EPI_ISL_3013872 | Lighthouse Lab in Alderley Park | Wellcome Sanger Institute for the COVID-19 Genomics UK (COG-UK) Consortium | Jacquelyn Wynn, Mairead Hyland, The Lighthouse Lab in Alderley Park and Alex Alderton, Roberto Amato, Jeffrey Barrett, Sonia Goncalves, Ewan Harrison, David K. Jackson, Ian Johnston, Dominic Kwiatkowski, Cordelia Langford, John Sillitoe on behalf of the Wellcome Sanger Institute COVID-19 Surveillance Team |
| EPI_ISL_3013873 | Lighthouse Lab in Glasgow | Wellcome Sanger Institute for the COVID-19 Genomics UK (COG-UK) Consortium | Harper VanSteenhouse, Yumi Kasai, David Gray, Carol Clugston, Anna Dominiczak and Alex Alderton, Roberto Amato, Jeffrey Barrett, Sonia Goncalves, Ewan Harrison, David K. Jackson, Ian Johnston, Dominic Kwiatkowski, Cordelia Langford, John Sillitoe on behalf of the Wellcome Sanger Institute COVID-19 Surveillance Team |
| EPI_ISL_3013874 | Lighthouse Lab in Milton Keynes | Wellcome Sanger Institute for the COVID-19 Genomics UK (COG-UK) Consortium | The Lighthouse Lab in Milton Keynes and Alex Alderton, Roberto Amato, Jeffrey Barrett, Sonia Goncalves, Ewan Harrison, David K. Jackson, Ian Johnston, Dominic Kwiatkowski, Cordelia Langford, John Sillitoe on behalf of the Wellcome Sanger Institute COVID-19 Surveillance Team |
| EPI_ISL_3013875 | Lighthouse Lab in Alderley Park | Wellcome Sanger Institute for the COVID-19 Genomics UK (COG-UK) Consortium | Jacquelyn Wynn, Mairead Hyland, The Lighthouse Lab in Alderley Park and Alex Alderton, Roberto Amato, Jeffrey Barrett, Sonia Goncalves, Ewan Harrison, David K. Jackson, Ian Johnston, Dominic Kwiatkowski, Cordelia Langford, John Sillitoe on behalf of the Wellcome Sanger Institute COVID-19 Surveillance Team |
| EPI_ISL_3013876, | Lighthouse Lab in Milton Keynes | Wellcome Sanger Institute for the COVID-19 Genomics UK (COG-UK) | The Lighthouse Lab in Milton Keynes and Alex Alderton, Roberto Amato, Jeffrey Barrett, Sonia Goncalves, Ewan Harrison, David K. Jackson, Ian Johnston, Dominic Kwiatkowski, Cordelia Langford, John Sillitoe on behalf of the Wellcome Sanger |

| | | | |
|---|---|---|---|
| EPI_ISL_3013877 | | Consortium | Institute COVID-19 Surveillance Team |
| EPI_ISL_3013878 | Lighthouse Lab in Glasgow | Wellcome Sanger Institute for the COVID-19 Genomics UK (COG-UK) Consortium | Harper VanSteenhouse, Yumi Kasai, David Gray, Carol Clugston, Anna Dominiczak and Alex Alderton, Roberto Amato, Jeffrey Barrett, Sonia Goncalves, Ewan Harrison, David K. Jackson, Ian Johnston, Dominic Kwiatkowski, Cordelia Langford, John Sillitoe on behalf of the Wellcome Sanger Institute COVID-19 Surveillance Team |
| EPI_ISL_3013879 | Lighthouse Lab in Milton Keynes | Wellcome Sanger Institute for the COVID-19 Genomics UK (COG-UK) Consortium | The Lighthouse Lab in Milton Keynes and Alex Alderton, Roberto Amato, Jeffrey Barrett, Sonia Goncalves, Ewan Harrison, David K. Jackson, Ian Johnston, Dominic Kwiatkowski, Cordelia Langford, John Sillitoe on behalf of the Wellcome Sanger Institute COVID-19 Surveillance Team |
| EPI_ISL_3013880 | Health Services Laboratories | Wellcome Sanger Institute for the COVID-19 Genomics UK (COG-UK) Consortium | Health Services Laboratories and Alex Alderton, Roberto Amato, Jeffrey Barrett, Sonia Goncalves, Ewan Harrison, David K. Jackson, Ian Johnston, Dominic Kwiatkowski, Cordelia Langford, John Sillitoe on behalf of the Wellcome Sanger Institute COVID-19 Surveillance Team |
| EPI_ISL_3013881 | Lighthouse Lab in Glasgow | Wellcome Sanger Institute for the COVID-19 Genomics UK (COG-UK) Consortium | Harper VanSteenhouse, Yumi Kasai, David Gray, Carol Clugston, Anna Dominiczak and Alex Alderton, Roberto Amato, Jeffrey Barrett, Sonia Goncalves, Ewan Harrison, David K. Jackson, Ian Johnston, Dominic Kwiatkowski, Cordelia Langford, John Sillitoe on behalf of the Wellcome Sanger Institute COVID-19 Surveillance Team |
| EPI_ISL_3013882 | Lighthouse Lab in Alderley Park | Wellcome Sanger Institute for the COVID-19 Genomics UK (COG-UK) Consortium | Jacquelyn Wynn, Mairead Hyland, The Lighthouse Lab in Alderley Park and Alex Alderton, Roberto Amato, Jeffrey Barrett, Sonia Goncalves, Ewan Harrison, David K. Jackson, Ian Johnston, Dominic Kwiatkowski, Cordelia Langford, John Sillitoe on behalf of the Wellcome Sanger Institute COVID-19 Surveillance Team |
| EPI_ISL_3013883, EPI_ISL_3013884 | Lighthouse Lab in Glasgow | Wellcome Sanger Institute for the COVID-19 Genomics UK (COG-UK) Consortium | Harper VanSteenhouse, Yumi Kasai, David Gray, Carol Clugston, Anna Dominiczak and Alex Alderton, Roberto Amato, Jeffrey Barrett, Sonia Goncalves, Ewan Harrison, David K. Jackson, Ian Johnston, Dominic Kwiatkowski, Cordelia Langford, John Sillitoe on behalf of the Wellcome Sanger Institute COVID-19 Surveillance Team |
| EPI_ISL_3013885 | Lighthouse Lab in Milton Keynes | Wellcome Sanger Institute for the COVID-19 Genomics UK (COG-UK) Consortium | The Lighthouse Lab in Milton Keynes and Alex Alderton, Roberto Amato, Jeffrey Barrett, Sonia Goncalves, Ewan Harrison, David K. Jackson, Ian Johnston, Dominic Kwiatkowski, Cordelia Langford, John Sillitoe on behalf of the Wellcome Sanger Institute COVID-19 Surveillance Team |
| EPI_ISL_3013886 | Lighthouse Lab in Glasgow | Wellcome Sanger Institute for the COVID-19 Genomics UK (COG-UK) Consortium | Harper VanSteenhouse, Yumi Kasai, David Gray, Carol Clugston, Anna Dominiczak and Alex Alderton, Roberto Amato, Jeffrey Barrett, Sonia Goncalves, Ewan Harrison, David K. Jackson, Ian Johnston, Dominic Kwiatkowski, Cordelia Langford, John Sillitoe on behalf of the Wellcome Sanger Institute COVID-19 Surveillance Team |
| EPI_ISL_3013887 | Lighthouse Lab in Milton Keynes | Wellcome Sanger Institute for the COVID-19 Genomics UK (COG-UK) Consortium | The Lighthouse Lab in Milton Keynes and Alex Alderton, Roberto Amato, Jeffrey Barrett, Sonia Goncalves, Ewan Harrison, David K. Jackson, Ian Johnston, Dominic Kwiatkowski, Cordelia Langford, John Sillitoe on behalf of the Wellcome Sanger Institute COVID-19 Surveillance Team |
| EPI_ISL_3013888 | Lighthouse Lab in Glasgow | Wellcome Sanger Institute for the COVID-19 Genomics UK (COG-UK) Consortium | Harper VanSteenhouse, Yumi Kasai, David Gray, Carol Clugston, Anna Dominiczak and Alex Alderton, Roberto Amato, Jeffrey Barrett, Sonia Goncalves, Ewan Harrison, David K. Jackson, Ian Johnston, Dominic Kwiatkowski, Cordelia Langford, John Sillitoe on behalf of the Wellcome Sanger Institute COVID-19 Surveillance Team |
| EPI_ISL_3013889 | Health Services Laboratories | Wellcome Sanger Institute for the COVID-19 Genomics UK (COG-UK) Consortium | Health Services Laboratories and Alex Alderton, Roberto Amato, Jeffrey Barrett, Sonia Goncalves, Ewan Harrison, David K. Jackson, Ian Johnston, Dominic Kwiatkowski, Cordelia Langford, John Sillitoe on behalf of the Wellcome Sanger Institute COVID-19 Surveillance Team |
| EPI_ISL_3013890 | Lighthouse Lab in Glasgow | Wellcome Sanger Institute for the COVID-19 Genomics UK (COG-UK) Consortium | Harper VanSteenhouse, Yumi Kasai, David Gray, Carol Clugston, Anna Dominiczak and Alex Alderton, Roberto Amato, Jeffrey Barrett, Sonia Goncalves, Ewan Harrison, David K. Jackson, Ian Johnston, Dominic Kwiatkowski, Cordelia Langford, John Sillitoe on behalf of the Wellcome Sanger Institute COVID-19 Surveillance Team |
| EPI_ISL_3013891 | Lighthouse Lab in Milton Keynes | Wellcome Sanger Institute for the COVID-19 Genomics UK (COG-UK) Consortium | The Lighthouse Lab in Milton Keynes and Alex Alderton, Roberto Amato, Jeffrey Barrett, Sonia Goncalves, Ewan Harrison, David K. Jackson, Ian Johnston, Dominic Kwiatkowski, Cordelia Langford, John Sillitoe on behalf of the Wellcome Sanger Institute COVID-19 Surveillance Team |
| EPI_ISL_3013892 | Lighthouse Lab in Alderley Park | Wellcome Sanger Institute for the COVID-19 Genomics UK (COG-UK) Consortium | Jacquelyn Wynn, Mairead Hyland, The Lighthouse Lab in Alderley Park and Alex Alderton, Roberto Amato, Jeffrey Barrett, Sonia Goncalves, Ewan Harrison, David K. Jackson, Ian Johnston, Dominic Kwiatkowski, Cordelia Langford, John Sillitoe on behalf of the Wellcome Sanger Institute COVID-19 Surveillance Team |
| EPI_ISL_3013893 | Lighthouse Lab in Glasgow | Wellcome Sanger Institute for the COVID-19 Genomics UK (COG-UK) Consortium | Harper VanSteenhouse, Yumi Kasai, David Gray, Carol Clugston, Anna Dominiczak and Alex Alderton, Roberto Amato, Jeffrey Barrett, Sonia Goncalves, Ewan Harrison, David K. Jackson, Ian Johnston, Dominic Kwiatkowski, Cordelia Langford, John Sillitoe on behalf of the Wellcome Sanger Institute COVID-19 Surveillance Team |
| EPI_ISL_3013894 | Health Services Laboratories | Wellcome Sanger Institute for the COVID-19 Genomics UK (COG-UK) Consortium | Health Services Laboratories and Alex Alderton, Roberto Amato, Jeffrey Barrett, Sonia Goncalves, Ewan Harrison, David K. Jackson, Ian Johnston, Dominic Kwiatkowski, Cordelia Langford, John Sillitoe on behalf of the Wellcome Sanger Institute COVID-19 Surveillance Team |
| EPI_ISL_3013895 | Lighthouse Lab in Alderley Park | Wellcome Sanger Institute for the COVID-19 Genomics UK (COG-UK) Consortium | Jacquelyn Wynn, Mairead Hyland, The Lighthouse Lab in Alderley Park and Alex Alderton, Roberto Amato, Jeffrey Barrett, Sonia Goncalves, Ewan Harrison, David K. Jackson, Ian Johnston, Dominic Kwiatkowski, Cordelia Langford, John Sillitoe on behalf of the Wellcome Sanger Institute COVID-19 Surveillance Team |
| EPI_ISL_3013896 | Lighthouse Lab in Glasgow | Wellcome Sanger Institute for the COVID-19 Genomics UK (COG-UK) Consortium | Harper VanSteenhouse, Yumi Kasai, David Gray, Carol Clugston, Anna Dominiczak and Alex Alderton, Roberto Amato, Jeffrey Barrett, Sonia Goncalves, Ewan Harrison, David K. Jackson, Ian Johnston, Dominic Kwiatkowski, Cordelia Langford, John Sillitoe on behalf of the Wellcome Sanger Institute COVID-19 Surveillance Team |
| EPI_ISL_3013897 | Lighthouse Lab in Alderley Park | Wellcome Sanger Institute for the COVID-19 Genomics UK (COG-UK) Consortium | Jacquelyn Wynn, Mairead Hyland, The Lighthouse Lab in Alderley Park and Alex Alderton, Roberto Amato, Jeffrey Barrett, Sonia Goncalves, Ewan Harrison, David K. Jackson, Ian Johnston, Dominic Kwiatkowski, Cordelia Langford, John Sillitoe on behalf of the Wellcome Sanger Institute COVID-19 Surveillance Team |
| EPI_ISL_3013898, EPI_ISL_3013899, EPI_ISL_3013900 | Lighthouse Lab in Glasgow | Wellcome Sanger Institute for the COVID-19 Genomics UK (COG-UK) Consortium | Harper VanSteenhouse, Yumi Kasai, David Gray, Carol Clugston, Anna Dominiczak and Alex Alderton, Roberto Amato, Jeffrey Barrett, Sonia Goncalves, Ewan Harrison, David K. Jackson, Ian Johnston, Dominic Kwiatkowski, Cordelia Langford, John Sillitoe on behalf of the Wellcome Sanger Institute COVID-19 Surveillance Team |
| EPI_ISL_3013901, EPI_ISL_3013902, EPI_ISL_3013905, EPI_ISL_3013906, EPI_ISL_3013907, EPI_ISL_3013908, EPI_ISL_3013909, EPI_ISL_3013911, EPI_ISL_3013912, EPI_ISL_3013913, EPI_ISL_3013914, EPI_ISL_3013915, EPI_ISL_3013916, EPI_ISL_3013917, EPI_ISL_3013918, EPI_ISL_3013919, EPI_ISL_3013920, EPI_ISL_3013921, EPI_ISL_3013922, EPI_ISL_3013923, EPI_ISL_3013924, EPI_ISL_3013925, EPI_ISL_3013926, EPI_ISL_3013927, EPI_ISL_3013928, EPI_ISL_3013929, EPI_ISL_3013930, EPI_ISL_3013931, EPI_ISL_3013932, EPI_ISL_3013933, EPI_ISL_3013934, EPI_ISL_3013935, EPI_ISL_3013936, EPI_ISL_3013937, EPI_ISL_3013939, EPI_ISL_3013940, EPI_ISL_3013942, EPI_ISL_3013943, EPI_ISL_3013944, EPI_ISL_3013945, EPI_ISL_3013946 | | | |
| see above | Randox Laboratories | Wellcome Sanger Institute for the COVID-19 Genomics UK (COG-UK) Consortium | Randox Laboratories and Alex Alderton, Roberto Amato, Jeffrey Barrett, Sonia Goncalves, Ewan Harrison, David K. Jackson, Ian Johnston, Dominic Kwiatkowski, Cordelia Langford, John Sillitoe on behalf of the Wellcome Sanger Institute COVID-19 Surveillance Team |
| EPI_ISL_3013947 | Lighthouse Laboratory Plymouth | Wellcome Sanger Institute for the COVID-19 Genomics UK (COG-UK) Consortium | Lighthouse Laboratory Plymouth and Alex Alderton, Roberto Amato, Jeffrey Barrett, Sonia Goncalves, Ewan Harrison, David K. Jackson, Ian Johnston, Dominic Kwiatkowski, Cordelia Langford, John Sillitoe on behalf of the Wellcome Sanger Institute COVID-19 Surveillance Team |
| EPI_ISL_3013948 | Berkshire and Surrey Pathology Services Lighthouse Laboratory | Wellcome Sanger Institute for the COVID-19 Genomics UK (COG-UK) Consortium | Berkshire and Surrey Pathology Services Lighthouse Laboratory and Alex Alderton, Roberto Amato, Jeffrey Barrett, Sonia Goncalves, Ewan Harrison, David K. Jackson, Ian Johnston, Dominic Kwiatkowski, Cordelia Langford, John Sillitoe on behalf of the Wellcome Sanger Institute COVID-19 Surveillance Team |
| EPI_ISL_3013949, EPI_ISL_3013950, EPI_ISL_3013951, EPI_ISL_3013952 | Lighthouse Lab in Alderley Park | Wellcome Sanger Institute for the COVID-19 Genomics UK (COG-UK) Consortium | Jacquelyn Wynn, Mairead Hyland, The Lighthouse Lab in Alderley Park and Alex Alderton, Roberto Amato, Jeffrey Barrett, Sonia Goncalves, Ewan Harrison, David K. Jackson, Ian Johnston, Dominic Kwiatkowski, Cordelia Langford, John Sillitoe on behalf of the Wellcome Sanger Institute COVID-19 Surveillance Team |
| EPI_ISL_3013953 | Lighthouse Laboratory Plymouth | Wellcome Sanger Institute for the COVID-19 Genomics UK (COG-UK) Consortium | Lighthouse Laboratory Plymouth and Alex Alderton, Roberto Amato, Jeffrey Barrett, Sonia Goncalves, Ewan Harrison, David K. Jackson, Ian Johnston, Dominic Kwiatkowski, Cordelia Langford, John Sillitoe on behalf of the Wellcome Sanger Institute COVID-19 Surveillance Team |
| EPI_ISL_3013954, EPI_ISL_3013955, EPI_ISL_3013956, EPI_ISL_3013957, EPI_ISL_3013958, EPI_ISL_3013959, EPI_ISL_3013960, EPI_ISL_3013961, EPI_ISL_3013962, EPI_ISL_3013963, EPI_ISL_3013964, EPI_ISL_3013965, EPI_ISL_3013966, EPI_ISL_3013967, EPI_ISL_3013968, EPI_ISL_3013970, EPI_ISL_3013971, EPI_ISL_3013972 | | | |
| see above | Randox Laboratories | Wellcome Sanger Institute for the COVID-19 Genomics UK (COG-UK) Consortium | Randox Laboratories and Alex Alderton, Roberto Amato, Jeffrey Barrett, Sonia Goncalves, Ewan Harrison, David K. Jackson, Ian Johnston, Dominic Kwiatkowski, Cordelia Langford, John Sillitoe on behalf of the Wellcome Sanger Institute COVID-19 Surveillance Team |
| EPI_ISL_3013973 | Lighthouse Lab in Alderley Park | Wellcome Sanger Institute for the COVID-19 Genomics UK (COG-UK) Consortium | Jacquelyn Wynn, Mairead Hyland, The Lighthouse Lab in Alderley Park and Alex Alderton, Roberto Amato, Jeffrey Barrett, Sonia Goncalves, Ewan Harrison, David K. Jackson, Ian Johnston, Dominic Kwiatkowski, Cordelia Langford, John Sillitoe on behalf of the Wellcome Sanger Institute COVID-19 Surveillance Team |
| EPI_ISL_3013974 | Health Services Laboratories | Wellcome Sanger Institute for the COVID-19 Genomics UK (COG-UK) Consortium | Health Services Laboratories and Alex Alderton, Roberto Amato, Jeffrey Barrett, Sonia Goncalves, Ewan Harrison, David K. Jackson, Ian Johnston, Dominic Kwiatkowski, Cordelia Langford, John Sillitoe on behalf of the Wellcome Sanger Institute COVID-19 Surveillance Team |
| EPI_ISL_3013975 | IRCCS S.Lucia | National Institute for Infectious Diseases (INMI) L. Spallanzani I.R.C.C.S | CEM Gruber, F Messina, F Santini, M Rueca, G Bonfiglio, O Butera, B Bartolini, E Giombini, MR Capobianchi, A Di Caro |
| EPI_ISL_3013977 | IRCCS I.D.I. | National Institute for Infectious Diseases (INMI) L. Spallanzani I.R.C.C.S | CEM Gruber, F Messina, F Santini, M Rueca, G Bonfiglio, O Butera, B Bartolini, E Giombini, MR Capobianchi, A Di Caro |
| EPI_ISL_3013979 | IRCCS INMI Lazzaro Spallanzani | National Institute for Infectious Diseases (INMI) L. Spallanzani I.R.C.C.S | CEM Gruber, F Messina, F Santini, M Rueca, G Bonfiglio, O Butera, B Bartolini, E Giombini, MR Capobianchi, A Di Caro |
| EPI_ISL_3013980 | RM3 | National Institute for Infectious Diseases (INMI) L. Spallanzani I.R.C.C.S | CEM Gruber, F Messina, F Santini, M Rueca, G Bonfiglio, O Butera, B Bartolini, E Giombini, MR Capobianchi, A Di Caro |
| EPI_ISL_3013982 | Osp. S.Pertini | National Institute for Infectious Diseases (INMI) L. Spallanzani I.R.C.C.S | CEM Gruber, F Messina, F Santini, M Rueca, G Bonfiglio, O Butera, B Bartolini, E Giombini, MR Capobianchi, A Di Caro |
| EPI_ISL_3013983 | IRCCS INMI Lazzaro Spallanzani | National Institute for Infectious Diseases (INMI) L. Spallanzani I.R.C.C.S | CEM Gruber, F Messina, F Santini, M Rueca, G Bonfiglio, O Butera, B Bartolini, E Giombini, MR Capobianchi, A Di Caro |
| EPI_ISL_3013984 | Osp. S.Pertini | National Institute for Infectious Diseases (INMI) L. Spallanzani I.R.C.C.S | CEM Gruber, F Messina, F Santini, M Rueca, G Bonfiglio, O Butera, B Bartolini, E Giombini, MR Capobianchi, A Di Caro |
| EPI_ISL_3013985, EPI_ISL_3013986 | A.O. S.Camillo | National Institute for Infectious Diseases (INMI) L. Spallanzani I.R.C.C.S | CEM Gruber, F Messina, F Santini, M Rueca, G Bonfiglio, O Butera, B Bartolini, E Giombini, MR Capobianchi, A Di Caro |
| EPI_ISL_3013987 | Osp. S.Pertini | National Institute for Infectious Diseases (INMI) L. Spallanzani I.R.C.C.S | CEM Gruber, F Messina, F Santini, M Rueca, G Bonfiglio, O Butera, B Bartolini, E Giombini, MR Capobianchi, A Di Caro |
| EPI_ISL_3013989 | A.O. S.Camillo | National Institute for Infectious Diseases (INMI) L. Spallanzani I.R.C.C.S | CEM Gruber, F Messina, F Santini, M Rueca, G Bonfiglio, O Butera, B Bartolini, E Giombini, MR Capobianchi, A Di Caro |
| EPI_ISL_3013990 | Osp. S.Pertini | National Institute for Infectious Diseases (INMI) L. Spallanzani I.R.C.C.S | CEM Gruber, F Messina, F Santini, M Rueca, G Bonfiglio, O Butera, B Bartolini, E Giombini, MR Capobianchi, A Di Caro |
| EPI_ISL_3013991 | A.O. S.Camillo | National Institute for Infectious Diseases (INMI) L. Spallanzani I.R.C.C.S | CEM Gruber, F Messina, F Santini, M Rueca, G Bonfiglio, O Butera, B Bartolini, E Giombini, MR Capobianchi, A Di Caro |
| EPI_ISL_3013992 | IRCCS S.Lucia | National Institute for Infectious Diseases (INMI) L. Spallanzani I.R.C.C.S | CEM Gruber, F Messina, F Santini, M Rueca, G Bonfiglio, O Butera, B Bartolini, E Giombini, MR Capobianchi, A Di Caro |
| EPI_ISL_3013993 | Osp. S.Pertini | National Institute for Infectious Diseases (INMI) L. Spallanzani I.R.C.C.S | CEM Gruber, F Messina, F Santini, M Rueca, G Bonfiglio, O Butera, B Bartolini, E Giombini, MR Capobianchi, A Di Caro |

| | | | |
|---|---|---|---|
| EPI_ISL_3013994 | IRCCS I.D.I. | National Institute for Infectious Diseases (INMI) L. Spallanzani I.R.C.C.S | CEM Gruber, F Messina, F Santini, M Rueca, G Bonfiglio, O Butera, B Bartolini, E Giombini, MR Capobianchi, A Di Caro |
| EPI_ISL_3013995 | Osp. S.Pertini | National Institute for Infectious Diseases (INMI) L. Spallanzani I.R.C.C.S | CEM Gruber, F Messina, F Santini, M Rueca, G Bonfiglio, O Butera, B Bartolini, E Giombini, MR Capobianchi, A Di Caro |
| EPI_ISL_3013996 | IRCCS I.D.I. | National Institute for Infectious Diseases (INMI) L. Spallanzani I.R.C.C.S | CEM Gruber, F Messina, F Santini, M Rueca, G Bonfiglio, O Butera, B Bartolini, E Giombini, MR Capobianchi, A Di Caro |
| EPI_ISL_3013997 | IRCCS S.Lucia | National Institute for Infectious Diseases (INMI) L. Spallanzani I.R.C.C.S | CEM Gruber, F Messina, F Santini, M Rueca, G Bonfiglio, O Butera, B Bartolini, E Giombini, MR Capobianchi, A Di Caro |
| EPI_ISL_3013998 | Osp. S.Pertini | National Institute for Infectious Diseases (INMI) L. Spallanzani I.R.C.C.S | CEM Gruber, F Messina, F Santini, M Rueca, G Bonfiglio, O Butera, B Bartolini, E Giombini, MR Capobianchi, A Di Caro |
| EPI_ISL_3013999 | IRCCS S.Lucia | National Institute for Infectious Diseases (INMI) L. Spallanzani I.R.C.C.S | CEM Gruber, F Messina, F Santini, M Rueca, G Bonfiglio, O Butera, B Bartolini, E Giombini, MR Capobianchi, A Di Caro |
| EPI_ISL_3014001 | IRCCS I.D.I. | National Institute for Infectious Diseases (INMI) L. Spallanzani I.R.C.C.S | CEM Gruber, F Messina, F Santini, M Rueca, G Bonfiglio, O Butera, B Bartolini, E Giombini, MR Capobianchi, A Di Caro |
| EPI_ISL_3014002 | Centro Diagnostico Buonarroti | National Institute for Infectious Diseases (INMI) L. Spallanzani I.R.C.C.S | CEM Gruber, F Messina, F Santini, M Rueca, G Bonfiglio, O Butera, B Bartolini, E Giombini, MR Capobianchi, A Di Caro |
| EPI_ISL_3014003, EPI_ISL_3014004 | IRCCS I.D.I. | National Institute for Infectious Diseases (INMI) L. Spallanzani I.R.C.C.S | CEM Gruber, F Messina, F Santini, M Rueca, G Bonfiglio, O Butera, B Bartolini, E Giombini, MR Capobianchi, A Di Caro |
| EPI_ISL_3014005 | IRCCS Bambino Gesù | National Institute for Infectious Diseases (INMI) L. Spallanzani I.R.C.C.S | CEM Gruber, F Messina, F Santini, M Rueca, G Bonfiglio, O Butera, B Bartolini, E Giombini, MR Capobianchi, A Di Caro |
| EPI_ISL_3014006 | Centro Diagnostico Buonarroti | National Institute for Infectious Diseases (INMI) L. Spallanzani I.R.C.C.S | CEM Gruber, F Messina, F Santini, M Rueca, G Bonfiglio, O Butera, B Bartolini, E Giombini, MR Capobianchi, A Di Caro |
| EPI_ISL_3014007, EPI_ISL_3014008 | IRCCS Bambino Gesù | National Institute for Infectious Diseases (INMI) L. Spallanzani I.R.C.C.S | CEM Gruber, F Messina, F Santini, M Rueca, G Bonfiglio, O Butera, B Bartolini, E Giombini, MR Capobianchi, A Di Caro |
| EPI_ISL_3014009 | Centro Diagnostico Buonarroti | National Institute for Infectious Diseases (INMI) L. Spallanzani I.R.C.C.S | CEM Gruber, F Messina, F Santini, M Rueca, G Bonfiglio, O Butera, B Bartolini, E Giombini, MR Capobianchi, A Di Caro |
| EPI_ISL_3014010 | IRCCS Bambino Gesù | National Institute for Infectious Diseases (INMI) L. Spallanzani I.R.C.C.S | CEM Gruber, F Messina, F Santini, M Rueca, G Bonfiglio, O Butera, B Bartolini, E Giombini, MR Capobianchi, A Di Caro |
| EPI_ISL_3014011 | Lab. Micro. FPG | National Institute for Infectious Diseases (INMI) L. Spallanzani I.R.C.C.S | CEM Gruber, F Messina, F Santini, M Rueca, G Bonfiglio, O Butera, B Bartolini, E Giombini, MR Capobianchi, A Di Caro |
| EPI_ISL_3014013 | Osp. S.Maria Goretti | National Institute for Infectious Diseases (INMI) L. Spallanzani I.R.C.C.S | CEM Gruber, F Messina, F Santini, M Rueca, G Bonfiglio, O Butera, B Bartolini, E Giombini, MR Capobianchi, A Di Caro |
| EPI_ISL_3014015, EPI_ISL_3014016, EPI_ISL_3014017 | Osp. S.Giovanni Evangelista | National Institute for Infectious Diseases (INMI) L. Spallanzani I.R.C.C.S | CEM Gruber, F Messina, F Santini, M Rueca, G Bonfiglio, O Butera, B Bartolini, E Giombini, MR Capobianchi, A Di Caro |
| EPI_ISL_3014018 | GENZANO | National Institute for Infectious Diseases (INMI) L. Spallanzani I.R.C.C.S | CEM Gruber, F Messina, F Santini, M Rueca, G Bonfiglio, O Butera, B Bartolini, E Giombini, MR Capobianchi, A Di Caro |
| EPI_ISL_3014019 | UOC Patologia clinica AO San Giovanni Addolorata | National Institute for Infectious Diseases (INMI) L. Spallanzani I.R.C.C.S | CEM Gruber, F Messina, F Santini, M Rueca, G Bonfiglio, O Butera, B Bartolini, E Giombini, MR Capobianchi, A Di Caro |
| EPI_ISL_3014020 | GENZANO | National Institute for Infectious Diseases (INMI) L. Spallanzani I.R.C.C.S | CEM Gruber, F Messina, F Santini, M Rueca, G Bonfiglio, O Butera, B Bartolini, E Giombini, MR Capobianchi, A Di Caro |
| EPI_ISL_3014021, EPI_ISL_3014022 | Pol.U. Campus Biomedico | National Institute for Infectious Diseases (INMI) L. Spallanzani I.R.C.C.S | CEM Gruber, F Messina, F Santini, M Rueca, G Bonfiglio, O Butera, B Bartolini, E Giombini, MR Capobianchi, A Di Caro |
| EPI_ISL_3014023, EPI_ISL_3014024, EPI_ISL_3014025 | A.U. Tor Vergata | National Institute for Infectious Diseases (INMI) L. Spallanzani I.R.C.C.S | CEM Gruber, F Messina, F Santini, M Rueca, G Bonfiglio, O Butera, B Bartolini, E Giombini, MR Capobianchi, A Di Caro |
| EPI_ISL_3014026 | Osp. S.Eugenio | National Institute for Infectious Diseases (INMI) L. Spallanzani I.R.C.C.S | CEM Gruber, F Messina, F Santini, M Rueca, G Bonfiglio, O Butera, B Bartolini, E Giombini, MR Capobianchi, A Di Caro |
| EPI_ISL_3014027 | A.O. S.Camillo | National Institute for Infectious Diseases (INMI) L. Spallanzani I.R.C.C.S | CEM Gruber, F Messina, F Santini, M Rueca, G Bonfiglio, O Butera, B Bartolini, E Giombini, MR Capobianchi, A Di Caro |
| EPI_ISL_3014028 | Osp S. Paolo Civitavecchia/ RM4 | National Institute for Infectious Diseases (INMI) L. Spallanzani I.R.C.C.S | CEM Gruber, F Messina, F Santini, M Rueca, G Bonfiglio, O Butera, B Bartolini, E Giombini, MR Capobianchi, A Di Caro |
| EPI_ISL_3014029 | Osp.C. S.Carlo Di Nancy | National Institute for Infectious Diseases (INMI) L. Spallanzani I.R.C.C.S | CEM Gruber, F Messina, F Santini, M Rueca, G Bonfiglio, O Butera, B Bartolini, E Giombini, MR Capobianchi, A Di Caro |
| EPI_ISL_3014030 | RM3 (positivo ID abbott ADR) | National Institute for Infectious Diseases (INMI) L. Spallanzani I.R.C.C.S | CEM Gruber, F Messina, F Santini, M Rueca, G Bonfiglio, O Butera, B Bartolini, E Giombini, MR Capobianchi, A Di Caro |
| EPI_ISL_3014031 | ACISMOM | National Institute for Infectious Diseases (INMI) L. Spallanzani I.R.C.C.S | CEM Gruber, F Messina, F Santini, M Rueca, G Bonfiglio, O Butera, B Bartolini, E Giombini, MR Capobianchi, A Di Caro |
| EPI_ISL_3014032 | RM2 | National Institute for Infectious Diseases (INMI) L. Spallanzani I.R.C.C.S | CEM Gruber, F Messina, F Santini, M Rueca, G Bonfiglio, O Butera, B Bartolini, E Giombini, MR Capobianchi, A Di Caro |
| EPI_ISL_3014033 | C.C.A. Aurelia Hospital | National Institute for Infectious Diseases (INMI) L. Spallanzani I.R.C.C.S | CEM Gruber, F Messina, F Santini, M Rueca, G Bonfiglio, O Butera, B Bartolini, E Giombini, MR Capobianchi, A Di Caro |
| EPI_ISL_3014034 | A.O. S.Camillo | National Institute for Infectious Diseases (INMI) L. Spallanzani I.R.C.C.S | CEM Gruber, F Messina, F Santini, M Rueca, G Bonfiglio, O Butera, B Bartolini, E Giombini, MR Capobianchi, A Di Caro |
| EPI_ISL_3014035, EPI_ISL_3014037 | IRCCS INMI Lazzaro Spallanzani | National Institute for Infectious Diseases (INMI) L. Spallanzani I.R.C.C.S | CEM Gruber, F Messina, F Santini, M Rueca, G Bonfiglio, O Butera, B Bartolini, E Giombini, MR Capobianchi, A Di Caro |
| EPI_ISL_3014038, EPI_ISL_3014039, EPI_ISL_3014040, EPI_ISL_3014041, EPI_ISL_3014042, EPI_ISL_3014043, EPI_ISL_3014044, EPI_ISL_3014045 | Osp. S.Pertini | National Institute for Infectious Diseases (INMI) L. Spallanzani I.R.C.C.S | CEM Gruber, F Messina, F Santini, M Rueca, G Bonfiglio, O Butera, B Bartolini, E Giombini, MR Capobianchi, A Di Caro |
| EPI_ISL_3014046 | IRCCS S.Lucia | National Institute for Infectious Diseases (INMI) L. Spallanzani I.R.C.C.S | CEM Gruber, F Messina, F Santini, M Rueca, G Bonfiglio, O Butera, B Bartolini, E Giombini, MR Capobianchi, A Di Caro |
| EPI_ISL_3014047, EPI_ISL_3014048, EPI_ISL_3014050, EPI_ISL_3014051 | Osp. S.Pertini | National Institute for Infectious Diseases (INMI) L. Spallanzani I.R.C.C.S | CEM Gruber, F Messina, F Santini, M Rueca, G Bonfiglio, O Butera, B Bartolini, E Giombini, MR Capobianchi, A Di Caro |
| EPI_ISL_3014052, EPI_ISL_3014053 | IRCCS S.Lucia | National Institute for Infectious Diseases (INMI) L. Spallanzani I.R.C.C.S | CEM Gruber, F Messina, F Santini, M Rueca, G Bonfiglio, O Butera, B Bartolini, E Giombini, MR Capobianchi, A Di Caro |
| EPI_ISL_3014054 | Osp. S.Pertini | National Institute for Infectious Diseases (INMI) L. Spallanzani I.R.C.C.S | CEM Gruber, F Messina, F Santini, M Rueca, G Bonfiglio, O Butera, B Bartolini, E Giombini, MR Capobianchi, A Di Caro |
| EPI_ISL_3014055 | A.U. Umberto I | National Institute for Infectious Diseases (INMI) L. Spallanzani I.R.C.C.S | CEM Gruber, F Messina, F Santini, M Rueca, G Bonfiglio, O Butera, B Bartolini, E Giombini, MR Capobianchi, A Di Caro |
| EPI_ISL_3014056 | IRCCS S.Lucia | National Institute for Infectious Diseases (INMI) L. Spallanzani I.R.C.C.S | CEM Gruber, F Messina, F Santini, M Rueca, G Bonfiglio, O Butera, B Bartolini, E Giombini, MR Capobianchi, A Di Caro |
| EPI_ISL_3014057, EPI_ISL_3014058 | A.U. Umberto I | National Institute for Infectious Diseases (INMI) L. Spallanzani I.R.C.C.S | CEM Gruber, F Messina, F Santini, M Rueca, G Bonfiglio, O Butera, B Bartolini, E Giombini, MR Capobianchi, A Di Caro |
| EPI_ISL_3014059 | IRCCS S.Lucia | National Institute for Infectious Diseases (INMI) L. Spallanzani I.R.C.C.S | CEM Gruber, F Messina, F Santini, M Rueca, G Bonfiglio, O Butera, B Bartolini, E Giombini, MR Capobianchi, A Di Caro |

| | | | |
|---|---|---|---|
| EPI_ISL_3014060 | IRCCS I.D.I. | National Institute for Infectious Diseases (INMI) L. Spallanzani I.R.C.C.S | CEM Gruber, F Messina, F Santini, M Rueca, G Bonfiglio, O Butera, B Bartolini, E Giombini, MR Capobianchi, A Di Caro |
| EPI_ISL_3014061 | A.U. Umberto I | National Institute for Infectious Diseases (INMI) L. Spallanzani I.R.C.C.S | CEM Gruber, F Messina, F Santini, M Rueca, G Bonfiglio, O Butera, B Bartolini, E Giombini, MR Capobianchi, A Di Caro |
| EPI_ISL_3014063 | IRCCS I.D.I. | National Institute for Infectious Diseases (INMI) L. Spallanzani I.R.C.C.S | CEM Gruber, F Messina, F Santini, M Rueca, G Bonfiglio, O Butera, B Bartolini, E Giombini, MR Capobianchi, A Di Caro |
| EPI_ISL_3014064 | Centro Diagnostico Buonarroti | National Institute for Infectious Diseases (INMI) L. Spallanzani I.R.C.C.S | CEM Gruber, F Messina, F Santini, M Rueca, G Bonfiglio, O Butera, B Bartolini, E Giombini, MR Capobianchi, A Di Caro |
| EPI_ISL_3014065 | RM1 | National Institute for Infectious Diseases (INMI) L. Spallanzani I.R.C.C.S | CEM Gruber, F Messina, F Santini, M Rueca, G Bonfiglio, O Butera, B Bartolini, E Giombini, MR Capobianchi, A Di Caro |
| EPI_ISL_3014066 | IRCCS Bambino Gesù | National Institute for Infectious Diseases (INMI) L. Spallanzani I.R.C.C.S | CEM Gruber, F Messina, F Santini, M Rueca, G Bonfiglio, O Butera, B Bartolini, E Giombini, MR Capobianchi, A Di Caro |
| EPI_ISL_3014067 | Centro Diagnostico Buonarroti | National Institute for Infectious Diseases (INMI) L. Spallanzani I.R.C.C.S | CEM Gruber, F Messina, F Santini, M Rueca, G Bonfiglio, O Butera, B Bartolini, E Giombini, MR Capobianchi, A Di Caro |
| EPI_ISL_3014068 | IRCCS Bambino Gesù | National Institute for Infectious Diseases (INMI) L. Spallanzani I.R.C.C.S | CEM Gruber, F Messina, F Santini, M Rueca, G Bonfiglio, O Butera, B Bartolini, E Giombini, MR Capobianchi, A Di Caro |
| EPI_ISL_3014069, EPI_ISL_3014070, EPI_ISL_3014071, EPI_ISL_3014072, EPI_ISL_3014073, EPI_ISL_3014074, EPI_ISL_3014075 | A.U. Tor Vergata | National Institute for Infectious Diseases (INMI) L. Spallanzani I.R.C.C.S | CEM Gruber, F Messina, F Santini, M Rueca, G Bonfiglio, O Butera, B Bartolini, E Giombini, MR Capobianchi, A Di Caro |
| EPI_ISL_3014076, EPI_ISL_3014077, EPI_ISL_3014078 | IRCCS INMI Lazzaro Spallanzani | National Institute for Infectious Diseases (INMI) L. Spallanzani I.R.C.C.S | CEM Gruber, F Messina, F Santini, M Rueca, G Bonfiglio, O Butera, B Bartolini, E Giombini, MR Capobianchi, A Di Caro |
| EPI_ISL_3014079 | Osp. S.Eugenio | National Institute for Infectious Diseases (INMI) L. Spallanzani I.R.C.C.S | CEM Gruber, F Messina, F Santini, M Rueca, G Bonfiglio, O Butera, B Bartolini, E Giombini, MR Capobianchi, A Di Caro |
| EPI_ISL_3014080 | IRCCS INMI Lazzaro Spallanzani | National Institute for Infectious Diseases (INMI) L. Spallanzani I.R.C.C.S | CEM Gruber, F Messina, F Santini, M Rueca, G Bonfiglio, O Butera, B Bartolini, E Giombini, MR Capobianchi, A Di Caro |
| EPI_ISL_3014081, EPI_ISL_3014082, EPI_ISL_3014083 | Osp. S.Eugenio | National Institute for Infectious Diseases (INMI) L. Spallanzani I.R.C.C.S | CEM Gruber, F Messina, F Santini, M Rueca, G Bonfiglio, O Butera, B Bartolini, E Giombini, MR Capobianchi, A Di Caro |
| EPI_ISL_3014084, EPI_ISL_3014085, EPI_ISL_3014086, EPI_ISL_3014087, EPI_ISL_3014088, EPI_ISL_3014089, EPI_ISL_3014091, EPI_ISL_3014092, EPI_ISL_3014093, EPI_ISL_3014095, EPI_ISL_3014096, EPI_ISL_3014098, EPI_ISL_3014099, EPI_ISL_3014100, EPI_ISL_3014101, EPI_ISL_3014102, EPI_ISL_3014103, EPI_ISL_3014104, EPI_ISL_3014105, EPI_ISL_3014106, EPI_ISL_3014107, EPI_ISL_3014108, EPI_ISL_3014110, EPI_ISL_3014111, EPI_ISL_3014112, EPI_ISL_3014113, EPI_ISL_3014114, EPI_ISL_3014115, EPI_ISL_3014117, EPI_ISL_3014118, EPI_ISL_3014119, EPI_ISL_3014121, EPI_ISL_3014122, EPI_ISL_3014124, EPI_ISL_3014125, EPI_ISL_3014126, EPI_ISL_3014127, EPI_ISL_3014128, EPI_ISL_3014129, EPI_ISL_3014130, EPI_ISL_3014131, EPI_ISL_3014133, EPI_ISL_3014139, EPI_ISL_3014140, EPI_ISL_3014141, EPI_ISL_3014143, EPI_ISL_3014144, EPI_ISL_3014145, EPI_ISL_3014146, EPI_ISL_3014147, EPI_ISL_3014148, EPI_ISL_3014149, EPI_ISL_3014150, EPI_ISL_3014151, EPI_ISL_3014153, EPI_ISL_3014154, EPI_ISL_3014155, EPI_ISL_3014172 | | | |
| see above | Cotugno | TIGEM | Antonio Grimaldi Patrizia Annunziata Francesco Panariello Biancamaria Pierri Claudia Tiberio Teresa Giuliano Valentina Bouche Chiara Colantuono Maria Concetta Cuomo Denise Di Concilio Lucio Di Filippo Anna Manfredi Marcello Salvi Antonio Limone Luigi Atripaldi Pellegrino Cerino Andrea Ballabio Davide Cacchiarelli |
| EPI_ISL_3014176, EPI_ISL_3014183, EPI_ISL_3014229, EPI_ISL_3014287, EPI_ISL_3014292, EPI_ISL_3014306, EPI_ISL_3014307, EPI_ISL_3014308, EPI_ISL_3014396 | Università Federico II - Dipartimento di scienze mediche traslazionali - Napoli | TIGEM | Antonio Grimaldi Patrizia Annunziata Francesco Panariello Teresa Giuliano Michele Cennamo Valentina Bouche Chiara Colantuono Lucio Di Filippo Mariano Fiorenza Anna Manfredi Marcello Salvi Giuseppe Portella Andrea Ballabio Davide Cacchiarelli |
| EPI_ISL_3014408, EPI_ISL_3014409, EPI_ISL_3014410, EPI_ISL_3014411, EPI_ISL_3014412, EPI_ISL_3014413, EPI_ISL_3014414, EPI_ISL_3014415, EPI_ISL_3014416, EPI_ISL_3014417, EPI_ISL_3014418, EPI_ISL_3014419, EPI_ISL_3014420, EPI_ISL_3014421, EPI_ISL_3014422, EPI_ISL_3014423, EPI_ISL_3014424, EPI_ISL_3014425, EPI_ISL_3014426, EPI_ISL_3014427, EPI_ISL_3014428, EPI_ISL_3014429, EPI_ISL_3014430, EPI_ISL_3014431, EPI_ISL_3014432, EPI_ISL_3014433, EPI_ISL_3014434, EPI_ISL_3014435, EPI_ISL_3014436, EPI_ISL_3014437, EPI_ISL_3014438, EPI_ISL_3014439, EPI_ISL_3014440, EPI_ISL_3014441, EPI_ISL_3014442, EPI_ISL_3014443, EPI_ISL_3014444, EPI_ISL_3014445, EPI_ISL_3014446, EPI_ISL_3014447, EPI_ISL_3014448, EPI_ISL_3014449, EPI_ISL_3014450, EPI_ISL_3014451, EPI_ISL_3014452, EPI_ISL_3014453, EPI_ISL_3014454, EPI_ISL_3014455, EPI_ISL_3014456, EPI_ISL_3014457, EPI_ISL_3014458, EPI_ISL_3014459, EPI_ISL_3014460, EPI_ISL_3014461, EPI_ISL_3014462, EPI_ISL_3014463, EPI_ISL_3014464, EPI_ISL_3014465, EPI_ISL_3014466, EPI_ISL_3014467, EPI_ISL_3014468, EPI_ISL_3014469, EPI_ISL_3014470 | | | |
| see above | Unidade de apoio ao diagnostico da COVID - UNADIG | Bioinformatics Laboratory / LNCC | Luiz G P de Almeida, Alessandra P Lamarca, Ronaldo da Silva F Jr, Liliane Cavalcante, Alexandra L Gerber, Ana Paula de C Guimaraes, Douglas Terra Machado, Cassia Alves, Diana Mariani, Cintia Policarpo, Gleidson da Silva de Oliveira, Mario Sergio Ribeiro, Silvia Carvalho, Flavio Dias da Silva, Marcio Henrique de Oliveira Garcia, Leandro Magalhaes de Souza, Cristiane Gomes da Silva, Caio Luiz Pereira Ribeiro, Andrea Cony Cavalcanti, Claudia Maria Braga de Mello, Amilcar Tanuri, Ana Tereza R Vasconcelos |
| EPI_ISL_3014486, EPI_ISL_3014505, EPI_ISL_3014508, EPI_ISL_3014868, EPI_ISL_3014885, EPI_ISL_3014887, EPI_ISL_3014891, EPI_ISL_3014892, EPI_ISL_3014894, EPI_ISL_3014895, EPI_ISL_3014896, EPI_ISL_3014897, EPI_ISL_3014898, EPI_ISL_3014904, EPI_ISL_3014905, EPI_ISL_3014949, EPI_ISL_3014952, EPI_ISL_3014953, EPI_ISL_3014954, EPI_ISL_3014955, EPI_ISL_3014956, EPI_ISL_3014958, EPI_ISL_3014959, EPI_ISL_3014960, EPI_ISL_3014961, EPI_ISL_3014962, EPI_ISL_3014963, EPI_ISL_3014964, EPI_ISL_3014966, EPI_ISL_3014967, EPI_ISL_3014970, EPI_ISL_3014971, EPI_ISL_3014972, EPI_ISL_3014973, EPI_ISL_3014974, EPI_ISL_3014975, EPI_ISL_3014977, EPI_ISL_3014978, EPI_ISL_3014979, EPI_ISL_3014980, EPI_ISL_3014981, EPI_ISL_3014982, EPI_ISL_3014983, EPI_ISL_3014984, EPI_ISL_3014985, EPI_ISL_3014986, EPI_ISL_3014987, EPI_ISL_3014988, EPI_ISL_3014990, EPI_ISL_3014992, EPI_ISL_3014993, EPI_ISL_3014994, EPI_ISL_3014995, EPI_ISL_3014996, EPI_ISL_3014997, EPI_ISL_3014998, EPI_ISL_3014999, EPI_ISL_3015000, EPI_ISL_3015001, EPI_ISL_3015004, EPI_ISL_3015005, EPI_ISL_3015006, EPI_ISL_3015007, EPI_ISL_3015008, EPI_ISL_3015009, EPI_ISL_3015012, EPI_ISL_3015015, EPI_ISL_3015016, EPI_ISL_3015017, EPI_ISL_3015018, EPI_ISL_3015020, EPI_ISL_3015021, EPI_ISL_3015022, EPI_ISL_3015024, EPI_ISL_3015025, EPI_ISL_3015026, EPI_ISL_3015027, EPI_ISL_3015030, EPI_ISL_3015031, EPI_ISL_3015032, EPI_ISL_3015035, EPI_ISL_3015036, EPI_ISL_3015039 | | | |
| see above | Department of Virology and Immunology, University of Helsinki and Helsinki University Hospital, Huslab Finland | Department of Virology, Faculty of Medicine, University of Helsinki, Helsinki, Finland | Teemu Smura, Ravi Kant, Phuoc Truong, Hussein Alburkat, Hannimari Kallio-Kokko, Jenni Virtanen, Maija Suvanto, Essi Korhonen, Sari Hannula, Harri Kangas, Hanna Liimatainen, Satu Kurkela, Hanna Jarva, Maija Lappalainen, Pekka Ellonen, Olli Vapalahti |
| EPI_ISL_3015053, EPI_ISL_3015054, EPI_ISL_3015055, EPI_ISL_3015056, EPI_ISL_3015057, EPI_ISL_3015058, EPI_ISL_3015059, EPI_ISL_3015060, EPI_ISL_3015061, EPI_ISL_3015062, EPI_ISL_3015063, EPI_ISL_3015064, EPI_ISL_3015065, EPI_ISL_3015066, EPI_ISL_3015067, EPI_ISL_3015068, EPI_ISL_3015069, EPI_ISL_3015070, EPI_ISL_3015071, EPI_ISL_3015072, EPI_ISL_3015073, EPI_ISL_3015074, EPI_ISL_3015075, EPI_ISL_3015076, EPI_ISL_3015077, EPI_ISL_3015080, EPI_ISL_3015081, EPI_ISL_3015082, EPI_ISL_3015083, EPI_ISL_3015084, EPI_ISL_3015085 | | | |
| see above | Labor Berlin Charité Vivantes GmbH / Institut für Virologie | Charité Universitätsmedizin Berlin, Institut für Virologie/Labor Berlin | Peter Menzel, Christine Stephan, Rolf Schwarzer, Victor M Corman, Barbara Mühlemann, Terry Jones, Christian Drosten |
| EPI_ISL_3015086, EPI_ISL_3015087, EPI_ISL_3015088, EPI_ISL_3015089, EPI_ISL_3015090, EPI_ISL_3015091, EPI_ISL_3015092, EPI_ISL_3015093, EPI_ISL_3015094, EPI_ISL_3015095 | AULSS 2 Marca Trevigiana | Istituto Zooprofilattico Sperimentale delle Venezie | Adelaide Milani, Alessia Schivo, Annalisa Salviato, Elisa Palumbo, Erika Giorgia Quaranta, Luca Tassoni, Ambra Pastori, Edoardo Giussani, Alice Fusaro, Isabella Monne, Calogero Terregino, Antonia Ricci |
| EPI_ISL_3015097, EPI_ISL_3015098, EPI_ISL_3015100, EPI_ISL_3015101, EPI_ISL_3015102, EPI_ISL_3015103, EPI_ISL_3015104, EPI_ISL_3015107, EPI_ISL_3015109, EPI_ISL_3015110, EPI_ISL_3015111, EPI_ISL_3015112, EPI_ISL_3015113, EPI_ISL_3015114 | | | |
| see above | Orebro University Hospital, Dept Laboratory Medicine, Clinical Microbiology | Orebro University Hospital | Sundqvist M et al |
| EPI_ISL_3015115, EPI_ISL_3015116, EPI_ISL_3015117, EPI_ISL_3015118, EPI_ISL_3015119, EPI_ISL_3015120, EPI_ISL_3015121, EPI_ISL_3015122, EPI_ISL_3015123 | Histopath | NSW Health Pathology - Institute of Clinical Pathology and Medical Research; Westmead Hospital; University of Sydney | CIDM-PH et al. |
| EPI_ISL_3015124 | Pathology West - NSW Health Pathology | NSW Health Pathology - Institute of Clinical Pathology and Medical Research; Westmead Hospital; University of Sydney | CIDM-PH et al. |
| EPI_ISL_3015125, EPI_ISL_3015126 | Laverty Pathology | NSW Health Pathology - Institute of Clinical Pathology and Medical Research; Westmead Hospital; University of Sydney | CIDM-PH et al. |
| EPI_ISL_3015127 | Pathology West - NSW Health Pathology | NSW Health Pathology - Institute of Clinical Pathology and Medical Research; Westmead Hospital; University of Sydney | CIDM-PH et al. |
| EPI_ISL_3015128 | 4Cyte Pathology | NSW Health Pathology - Institute of Clinical Pathology and Medical Research; Westmead Hospital; University of Sydney | CIDM-PH et al. |
| EPI_ISL_3015129 | Douglass Hanly Moir Pathology | NSW Health Pathology - Institute of Clinical Pathology and Medical Research; Westmead Hospital; University of Sydney | CIDM-PH et al. |
| EPI_ISL_3015130, EPI_ISL_3015131, EPI_ISL_3015132, EPI_ISL_3015133, EPI_ISL_3015134, EPI_ISL_3015135, EPI_ISL_3015136, EPI_ISL_3015137, EPI_ISL_3015138, EPI_ISL_3015139, EPI_ISL_3015140 | | | |

| | | | |
|---|---|---|---|
| see above | St Vincent's Pathology (SydPath) | NSW Health Pathology - Institute of Clinical Pathology and Medical Research; Westmead Hospital; University of Sydney | CIDM-PH et al. |
| EPI_ISL_3015141, EPI_ISL_3015142, EPI_ISL_3015143, EPI_ISL_3015144, EPI_ISL_3015145, EPI_ISL_3015146 | Histopath | NSW Health Pathology - Institute of Clinical Pathology and Medical Research; Westmead Hospital; University of Sydney | CIDM-PH et al. |
| EPI_ISL_3015147 | Pathology West - NSW Health Pathology | NSW Health Pathology - Institute of Clinical Pathology and Medical Research; Westmead Hospital; University of Sydney | CIDM-PH et al. |
| EPI_ISL_3015148 | 4Cyte Pathology | NSW Health Pathology - Institute of Clinical Pathology and Medical Research; Westmead Hospital; University of Sydney | CIDM-PH et al. |
| EPI_ISL_3015149 | Laverty Pathology | NSW Health Pathology - Institute of Clinical Pathology and Medical Research; Westmead Hospital; University of Sydney | CIDM-PH et al. |
| EPI_ISL_3015150 | 4Cyte Pathology | NSW Health Pathology - Institute of Clinical Pathology and Medical Research; Westmead Hospital; University of Sydney | CIDM-PH et al. |
| EPI_ISL_3015151 | Australian Clinical Labs (formerly Healthscope Pathology) | NSW Health Pathology - Institute of Clinical Pathology and Medical Research; Westmead Hospital; University of Sydney | CIDM-PH et al. |
| EPI_ISL_3015152, EPI_ISL_3015153, EPI_ISL_3015154, EPI_ISL_3015155, EPI_ISL_3015156, EPI_ISL_3015157 | Laverty Pathology | NSW Health Pathology - Institute of Clinical Pathology and Medical Research; Westmead Hospital; University of Sydney | CIDM-PH et al. |
| EPI_ISL_3015158, EPI_ISL_3015159, EPI_ISL_3015160, EPI_ISL_3015161, EPI_ISL_3015162 | Sydney South West Pathology Service (SSWPS) - Liverpool Hospital - NSW Health Pathology | NSW Health Pathology - Institute of Clinical Pathology and Medical Research; Westmead Hospital; University of Sydney | CIDM-PH et al. |
| EPI_ISL_3015163, EPI_ISL_3015164, EPI_ISL_3015165, EPI_ISL_3015166, EPI_ISL_3015167, EPI_ISL_3015168, EPI_ISL_3015169 | St Vincent's Pathology (SydPath) | NSW Health Pathology - Institute of Clinical Pathology and Medical Research; Westmead Hospital; University of Sydney | CIDM-PH et al. |
| EPI_ISL_3015170, EPI_ISL_3015171, EPI_ISL_3015172, EPI_ISL_3015173, EPI_ISL_3015174, EPI_ISL_3015175, EPI_ISL_3015176 | Histopath | NSW Health Pathology - Institute of Clinical Pathology and Medical Research; Westmead Hospital; University of Sydney | CIDM-PH et al. |
| EPI_ISL_3015200, EPI_ISL_3015201, EPI_ISL_3015202, EPI_ISL_3015203 | Wexner Medical Center | The Ohio State University College of Medicine | Koenig,S.,Aidja,Z. |
| EPI_ISL_3015205, EPI_ISL_3015206 | UCSC Genomics Institute | UCSC Genomics Institute | Mark Akeson, Ikenna Anigbogu, Eric Beraut, Isabel Bjork, Molly Cassatt-Johnstone, Terren Chang, Russell Corbett-Detig, Namrita Dhillon, Maximilian Haeussler, David Haussler, Angie Hinrichs, Miten Jain, Joshua Kapp, A. Marm Kilpatrick, Jakob McBroome, Hugh Olsen, Jeremy Sanford, Beth Shapiro, Michael Stone, Bryan Thornlow, Yatish Turakhia, Ciara Wanket |
| EPI_ISL_3015207, EPI_ISL_3015208, EPI_ISL_3015209, EPI_ISL_3015210, EPI_ISL_3015211, EPI_ISL_3015212, EPI_ISL_3015213, EPI_ISL_3015214, EPI_ISL_3015215, EPI_ISL_3015216, EPI_ISL_3015217, EPI_ISL_3015218, EPI_ISL_3015219, EPI_ISL_3015220, EPI_ISL_3015222, EPI_ISL_3015223 | | | |
| see above | Mahatma Gandhi Memorial Medical College & Hospital, Jamshedpur | Institute of Life Sciences - INSACOG | Sunil K. Raghav, Safal Walia, Arup Ghosh, Atimukta Jha, Amol M. Kanampalliwar, Omprakash Shiriwas, Sana Fatma, Shifu Aggarwal, Rupesh Dash, Rajeeb Swain, Punit Prasad, INSACOG Consortium, Ajay Parida |
| EPI_ISL_3015224, EPI_ISL_3015225, EPI_ISL_3015226, EPI_ISL_3015227, EPI_ISL_3015228, EPI_ISL_3015229, EPI_ISL_3015230, EPI_ISL_3015231, EPI_ISL_3015232 | District Head Quarters Hospital, Nayagarh | Institute of Life Sciences - INSACOG | Sunil K. Raghav, Safal Walia, Arup Ghosh, Atimukta Jha, Amol M. Kanampalliwar, Omprakash Shiriwas, Sana Fatma, Shifu Aggarwal, Rupesh Dash, Rajeeb Swain, Punit Prasad, INSACOG Consortium, Ajay Parida |
| EPI_ISL_3015233, EPI_ISL_3015234, EPI_ISL_3015235, EPI_ISL_3015236, EPI_ISL_3015237, EPI_ISL_3015238, EPI_ISL_3015240, EPI_ISL_3015241, EPI_ISL_3015243, EPI_ISL_3015245, EPI_ISL_3015246, EPI_ISL_3015247, EPI_ISL_3015248, EPI_ISL_3015249, EPI_ISL_3015250, EPI_ISL_3015251, EPI_ISL_3015252, EPI_ISL_3015253, EPI_ISL_3015254, EPI_ISL_3015255, EPI_ISL_3015256, EPI_ISL_3015257, EPI_ISL_3015259, EPI_ISL_3015262, EPI_ISL_3015265 | | | |
| see above | S.C.B. Medical College and Hospital, Cuttack | Institute of Life Sciences - INSACOG | Sunil K. Raghav, Safal Walia, Arup Ghosh, Atimukta Jha, Amol M. Kanampalliwar, Omprakash Shiriwas, Sana Fatma, Shifu Aggarwal, Rupesh Dash, Rajeeb Swain, Punit Prasad, INSACOG Consortium, Ajay Parida |
| EPI_ISL_3015268, EPI_ISL_3015269 | District Headquarter Hospital, Baripada | Institute of Life Sciences - INSACOG | Sunil K. Raghav, Safal Walia, Arup Ghosh, Atimukta Jha, Amol M. Kanampalliwar, Omprakash Shiriwas, Sana Fatma, Shifu Aggarwal, Rupesh Dash, Rajeeb Swain, Punit Prasad, INSACOG Consortium, Ajay Parida |
| EPI_ISL_3015271, EPI_ISL_3015272, EPI_ISL_3015273, EPI_ISL_3015274, EPI_ISL_3015275, EPI_ISL_3015276, EPI_ISL_3015277, EPI_ISL_3015278, EPI_ISL_3015279, EPI_ISL_3015281, EPI_ISL_3015282, EPI_ISL_3015283, EPI_ISL_3015284, EPI_ISL_3015287, EPI_ISL_3015288 | | | |
| see above | Old District Headquarter Hospital, Jharsuguda | Institute of Life Sciences - INSACOG | Sunil K. Raghav, Safal Walia, Arup Ghosh, Atimukta Jha, Amol M. Kanampalliwar, Omprakash Shiriwas, Sana Fatma, Shifu Aggarwal, Rupesh Dash, Rajeeb Swain, Punit Prasad, INSACOG Consortium, Ajay Parida |
| EPI_ISL_3015290 | Veer Surendra Sai Institute of Medical Sciences and Research, Burla, Sambalpur | Institute of Life Sciences - INSACOG | Sunil K. Raghav, Safal Walia, Arup Ghosh, Atimukta Jha, Amol M. Kanampalliwar, Omprakash Shiriwas, Sana Fatma, Shifu Aggarwal, Rupesh Dash, Rajeeb Swain, Punit Prasad, INSACOG Consortium, Ajay Parida |
| EPI_ISL_3015294, EPI_ISL_3015295, EPI_ISL_3015296, EPI_ISL_3015297, EPI_ISL_3015298, EPI_ISL_3015299, EPI_ISL_3015300, EPI_ISL_3015302, EPI_ISL_3015303, EPI_ISL_3015305 | Rajendra Memorial Research Institute of Medical Sciences, Patna | Institute of Life Sciences - INSACOG | Sunil K. Raghav, Safal Walia, Arup Ghosh, Atimukta Jha, Amol M. Kanampalliwar, Omprakash Shiriwas, Sana Fatma, Shifu Aggarwal, Rupesh Dash, Rajeeb Swain, Punit Prasad, INSACOG Consortium, Ajay Parida |
| EPI_ISL_3015308, EPI_ISL_3015309, EPI_ISL_3015311, EPI_ISL_3015313, EPI_ISL_3015314 | Patna Medical College, Patna | Institute of Life Sciences - INSACOG | Sunil K. Raghav, Safal Walia, Arup Ghosh, Atimukta Jha, Amol M. Kanampalliwar, Omprakash Shiriwas, Sana Fatma, Shifu Aggarwal, Rupesh Dash, Rajeeb Swain, Punit Prasad, INSACOG Consortium, Ajay Parida |
| EPI_ISL_3015318, EPI_ISL_3015319, EPI_ISL_3015320, EPI_ISL_3015321, EPI_ISL_3015322, EPI_ISL_3015323, EPI_ISL_3015325, EPI_ISL_3015326, EPI_ISL_3015327, EPI_ISL_3015328, EPI_ISL_3015330, EPI_ISL_3015331, EPI_ISL_3015332, EPI_ISL_3015333, EPI_ISL_3015335, EPI_ISL_3015336, EPI_ISL_3015337, EPI_ISL_3015338, EPI_ISL_3015339, EPI_ISL_3015340, EPI_ISL_3015341, EPI_ISL_3015342, EPI_ISL_3015343, EPI_ISL_3015344, EPI_ISL_3015345, EPI_ISL_3015347, EPI_ISL_3015348, EPI_ISL_3015349, EPI_ISL_3015350, EPI_ISL_3015351, EPI_ISL_3015352, EPI_ISL_3015353, EPI_ISL_3015354, EPI_ISL_3015355, EPI_ISL_3015356, EPI_ISL_3015357, EPI_ISL_3015358, EPI_ISL_3015359, EPI_ISL_3015360, EPI_ISL_3015361, EPI_ISL_3015362, EPI_ISL_3015363, EPI_ISL_3015365, EPI_ISL_3015366, EPI_ISL_3015367, EPI_ISL_3015368, EPI_ISL_3015369, EPI_ISL_3015370, EPI_ISL_3015371, EPI_ISL_3015372, EPI_ISL_3015373, EPI_ISL_3015374, EPI_ISL_3015375, EPI_ISL_3015376, EPI_ISL_3015377, EPI_ISL_3015378 | | | |
| see above | Govt. Medical College Rajnandgaon | Institute of Life Sciences - INSACOG | Sunil K. Raghav, Safal Walia, Arup Ghosh, Atimukta Jha, Amol M. Kanampalliwar, Omprakash Shiriwas, Sana Fatma, Shifu Aggarwal, Rupesh Dash, Rajeeb Swain, Punit Prasad, INSACOG Consortium, Ajay Parida |
| EPI_ISL_3015379, EPI_ISL_3015380, EPI_ISL_3015381, EPI_ISL_3015382, EPI_ISL_3015383, EPI_ISL_3015384, EPI_ISL_3015385, EPI_ISL_3015386, EPI_ISL_3015387, EPI_ISL_3015388, EPI_ISL_3015389, EPI_ISL_3015390, EPI_ISL_3015391, EPI_ISL_3015392, EPI_ISL_3015393, EPI_ISL_3015395, EPI_ISL_3015398, EPI_ISL_3015399, EPI_ISL_3015400, EPI_ISL_3015401, EPI_ISL_3015402, EPI_ISL_3015403, EPI_ISL_3015404, EPI_ISL_3015406, EPI_ISL_3015407, EPI_ISL_3015408, EPI_ISL_3015409, EPI_ISL_3015410, EPI_ISL_3015411, EPI_ISL_3015413, EPI_ISL_3015414, EPI_ISL_3015415, EPI_ISL_3015416, EPI_ISL_3015417, EPI_ISL_3015419, EPI_ISL_3015421, EPI_ISL_3015422, EPI_ISL_3015423, EPI_ISL_3015424, EPI_ISL_3015425, EPI_ISL_3015426, EPI_ISL_3015427, EPI_ISL_3015428 | | | |
| see above | Pt. Jawahar Lal Nehru Memorial Medical College, Raipur | Institute of Life Sciences - INSACOG | Sunil K. Raghav, Safal Walia, Arup Ghosh, Atimukta Jha, Amol M. Kanampalliwar, Omprakash Shiriwas, Sana Fatma, Shifu Aggarwal, Rupesh Dash, Rajeeb Swain, Punit Prasad, INSACOG Consortium, Ajay Parida |
| EPI_ISL_3015432, EPI_ISL_3015434, EPI_ISL_3015440 | S.C.B. Medical College and Hospital, Cuttack | Institute of Life Sciences - INSACOG | Sunil K. Raghav, Safal Walia, Arup Ghosh, Atimukta Jha, Amol M. Kanampalliwar, Omprakash Shiriwas, Sana Fatma, Shifu Aggarwal, Rupesh Dash, Rajeeb Swain, Punit Prasad, INSACOG Consortium, Ajay Parida |
| EPI_ISL_3015442, EPI_ISL_3015447, | District Headquarter Hospital, Baripada | Institute of Life Sciences - INSACOG | Sunil K. Raghav, Safal Walia, Arup Ghosh, Atimukta Jha, Amol M. Kanampalliwar, Omprakash Shiriwas, Sana Fatma, Shifu Aggarwal, Rupesh Dash, Rajeeb Swain, Punit Prasad, INSACOG Consortium, Ajay Parida |

| | | | |
|---|---|---|---|
| EPI_ISL_3015448, EPI_ISL_3015455, EPI_ISL_3015456 | | | |
| EPI_ISL_3015461, EPI_ISL_3015462, EPI_ISL_3015463, EPI_ISL_3015464, EPI_ISL_3015465, EPI_ISL_3015466, EPI_ISL_3015467, EPI_ISL_3015468, EPI_ISL_3015469, EPI_ISL_3015470, EPI_ISL_3015471, EPI_ISL_3015472, EPI_ISL_3015473, EPI_ISL_3015474, EPI_ISL_3015476, EPI_ISL_3015477, EPI_ISL_3015478, EPI_ISL_3015479, EPI_ISL_3015480, EPI_ISL_3015481, EPI_ISL_3015482, EPI_ISL_3015483, EPI_ISL_3015484, EPI_ISL_3015485, EPI_ISL_3015486, EPI_ISL_3015487, EPI_ISL_3015488, EPI_ISL_3015489, EPI_ISL_3015490, EPI_ISL_3015491, EPI_ISL_3015492 | | | |
| see above | Regional Medical Research Center, Bhubaneswar | Institute of Life Sciences - INSACOG | Sunil K. Raghav, Safal Walia, Arup Ghosh, Atimukta Jha, Amol M. Kanampalliwar, Omprakash Shiriwas, Sana Fatma, Shifu Aggarwal, Rupesh Dash, Rajeeb Swain, Punit Prasad, INSACOG Consortium, Ajay Parida |
| EPI_ISL_3015493, EPI_ISL_3015495, EPI_ISL_3015497, EPI_ISL_3015498, EPI_ISL_3015499, EPI_ISL_3015501, EPI_ISL_3015502 | Shrinivas Center for Scientific Research and Development (SCSRD), Hazaribag | Institute of Life Sciences - INSACOG | Sunil K. Raghav, Safal Walia, Arup Ghosh, Atimukta Jha, Amol M. Kanampalliwar, Omprakash Shiriwas, Sana Fatma, Shifu Aggarwal, Rupesh Dash, Rajeeb Swain, Punit Prasad, INSACOG Consortium, Ajay Parida |
| EPI_ISL_3015503, EPI_ISL_3015505, EPI_ISL_3015506, EPI_ISL_3015507, EPI_ISL_3015508, EPI_ISL_3015509, EPI_ISL_3015510, EPI_ISL_3015512, EPI_ISL_3015513, EPI_ISL_3015514, EPI_ISL_3015515, EPI_ISL_3015516, EPI_ISL_3015517, EPI_ISL_3015518, EPI_ISL_3015519, EPI_ISL_3015520, EPI_ISL_3015521, EPI_ISL_3015524, EPI_ISL_3015525, EPI_ISL_3015526, EPI_ISL_3015527, EPI_ISL_3015528, EPI_ISL_3015529, EPI_ISL_3015530, EPI_ISL_3015532, EPI_ISL_3015534, EPI_ISL_3015535, EPI_ISL_3015536, EPI_ISL_3015537, EPI_ISL_3015538, EPI_ISL_3015539, EPI_ISL_3015540, EPI_ISL_3015541, EPI_ISL_3015542, EPI_ISL_3015543, EPI_ISL_3015544, EPI_ISL_3015545, EPI_ISL_3015546, EPI_ISL_3015547, EPI_ISL_3015548, EPI_ISL_3015549, EPI_ISL_3015550, EPI_ISL_3015551, EPI_ISL_3015553, EPI_ISL_3015554, EPI_ISL_3015555, EPI_ISL_3015556, EPI_ISL_3015557, EPI_ISL_3015558, EPI_ISL_3015559, EPI_ISL_3015560, EPI_ISL_3015561, EPI_ISL_3015562, EPI_ISL_3015563, EPI_ISL_3015564, EPI_ISL_3015565, EPI_ISL_3015566, EPI_ISL_3015567, EPI_ISL_3015568, EPI_ISL_3015569, EPI_ISL_3015570, EPI_ISL_3015571, EPI_ISL_3015573, EPI_ISL_3015574, EPI_ISL_3015575, EPI_ISL_3015576, EPI_ISL_3015577, EPI_ISL_3015578, EPI_ISL_3015579, EPI_ISL_3015580, EPI_ISL_3015581, EPI_ISL_3015582, EPI_ISL_3015583 | | | |
| see above | Pt. Jawahar Lal Nehru Memorial Medical College, Raipur | Institute of Life Sciences - INSACOG | Sunil K. Raghav, Safal Walia, Arup Ghosh, Atimukta Jha, Amol M. Kanampalliwar, Omprakash Shiriwas, Sana Fatma, Shifu Aggarwal, Rupesh Dash, Rajeeb Swain, Punit Prasad, INSACOG Consortium, Ajay Parida |
| EPI_ISL_3015584, EPI_ISL_3015586, EPI_ISL_3015587, EPI_ISL_3015588 | Institute of Life Sciences, Bhubaneswar | Institute of Life Sciences - INSACOG | Sunil K. Raghav, Safal Walia, Arup Ghosh, Atimukta Jha, Amol M. Kanampalliwar, Omprakash Shiriwas, Sana Fatma, Shifu Aggarwal, Rupesh Dash, Rajeeb Swain, Punit Prasad, INSACOG Consortium, Ajay Parida |
| EPI_ISL_3015593, EPI_ISL_3015594, EPI_ISL_3015595, EPI_ISL_3015596, EPI_ISL_3015597, EPI_ISL_3015599, EPI_ISL_3015600, EPI_ISL_3015601, EPI_ISL_3015603, EPI_ISL_3015604 | Seattle Flu Study | Seattle Flu Study | Deborah A. Nickerson, Chris D. Frazar, Jover Lee, Benjamin Pelle, Erica Ryke, Matthew Richardson, Amanda Adler, Elisabeth Brandstetter, Peter D. Han, Kairsten Fay, Misja Ilcisin, Kirsten Lacombe, Thomas R. Sibley, Melissa Truong, Caitlin R. Wolf, Karen Cowgill, Stephanie Schrag, Jeff Duchin, Michael Boeckh, Janet A. Englund, Michael Famulare, Barry R. Lutz, Mark J. Rieder, Lea M. Starita, Matthew Thompson, Helen Y. Chu, Trevor Bedford, Jay Shendure |
| EPI_ISL_3015617 | Incyte Diagnostics Spokane | Seattle Flu Study | Deborah A. Nickerson, Chris D. Frazar, Jover Lee, Benjamin Pelle, Erica Ryke, Matthew Richardson, Amanda Adler, Elisabeth Brandstetter, Peter D. Han, Kairsten Fay, Misja Ilcisin, Kirsten Lacombe, Thomas R. Sibley, Melissa Truong, Caitlin R. Wolf, Romesh Gautom, Geoff Melly, Brian Hiatt, Philip Dykema, Scott Lindquist, Michael Boeckh, Janet A. Englund, Michael Famulare, Barry R. Lutz, Mark J. Rieder, Lea M. Starita, Matthew Thompson, Helen Y. Chu, Jay Shendure, Trevor Bedford |
| EPI_ISL_3015657, EPI_ISL_3015659 | Northwest Laboratory | Seattle Flu Study | Deborah A. Nickerson, Chris D. Frazar, Jover Lee, Benjamin Pelle, Erica Ryke, Matthew Richardson, Amanda Adler, Elisabeth Brandstetter, Peter D. Han, Kairsten Fay, Misja Ilcisin, Kirsten Lacombe, Thomas R. Sibley, Melissa Truong, Caitlin R. Wolf, Romesh Gautom, Geoff Melly, Brian Hiatt, Philip Dykema, Scott Lindquist, Michael Boeckh, Janet A. Englund, Michael Famulare, Barry R. Lutz, Mark J. Rieder, Lea M. Starita, Matthew Thompson, Helen Y. Chu, Jay Shendure, Trevor Bedford |
| EPI_ISL_3015717, EPI_ISL_3015719, EPI_ISL_3015720, EPI_ISL_3015731, EPI_ISL_3015738, EPI_ISL_3015742 | Atlas Genomics | Seattle Flu Study | Deborah A. Nickerson, Chris D. Frazar, Jover Lee, Benjamin Pelle, Erica Ryke, Matthew Richardson, Amanda Adler, Elisabeth Brandstetter, Peter D. Han, Kairsten Fay, Misja Ilcisin, Kirsten Lacombe, Thomas R. Sibley, Melissa Truong, Caitlin R. Wolf, Romesh Gautom, Geoff Melly, Brian Hiatt, Philip Dykema, Scott Lindquist, Michael Boeckh, Janet A. Englund, Michael Famulare, Barry R. Lutz, Mark J. Rieder, Lea M. Starita, Matthew Thompson, Helen Y. Chu, Jay Shendure, Trevor Bedford |
| EPI_ISL_3015750 | Seattle Flu Study | Seattle Flu Study | Deborah A. Nickerson, Chris D. Frazar, Jover Lee, Benjamin Pelle, Erica Ryke, Matthew Richardson, Amanda Adler, Elisabeth Brandstetter, Peter D. Han, Kairsten Fay, Misja Ilcisin, Kirsten Lacombe, Thomas R. Sibley, Melissa Truong, Caitlin R. Wolf, Karen Cowgill, Stephanie Schrag, Jeff Duchin, Michael Boeckh, Janet A. Englund, Michael Famulare, Barry R. Lutz, Mark J. Rieder, Lea M. Starita, Matthew Thompson, Helen Y. Chu, Trevor Bedford, Jay Shendure |
| EPI_ISL_3015766, EPI_ISL_3015769, EPI_ISL_3015773, EPI_ISL_3015777 | Atlas Genomics | Seattle Flu Study | Deborah A. Nickerson, Chris D. Frazar, Jover Lee, Benjamin Pelle, Erica Ryke, Matthew Richardson, Amanda Adler, Elisabeth Brandstetter, Peter D. Han, Kairsten Fay, Misja Ilcisin, Kirsten Lacombe, Thomas R. Sibley, Melissa Truong, Caitlin R. Wolf, Romesh Gautom, Geoff Melly, Brian Hiatt, Philip Dykema, Scott Lindquist, Michael Boeckh, Janet A. Englund, Michael Famulare, Barry R. Lutz, Mark J. Rieder, Lea M. Starita, Matthew Thompson, Helen Y. Chu, Jay Shendure, Trevor Bedford |
| EPI_ISL_3015782 | Seattle Flu Study | Seattle Flu Study | Deborah A. Nickerson, Chris D. Frazar, Jover Lee, Benjamin Pelle, Erica Ryke, Matthew Richardson, Amanda Adler, Elisabeth Brandstetter, Peter D. Han, Kairsten Fay, Misja Ilcisin, Kirsten Lacombe, Thomas R. Sibley, Melissa Truong, Caitlin R. Wolf, Michael Boeckh, Janet A. Englund, Michael Famulare, Barry R. Lutz, Mark J. Rieder, Lea M. Starita, Matthew Thompson, Jay Shendure, Trevor Bedford, Helen Y. Chu |
| EPI_ISL_3015787, EPI_ISL_3015793, EPI_ISL_3015794, EPI_ISL_3015799, EPI_ISL_3015807, EPI_ISL_3015811, EPI_ISL_3015815, EPI_ISL_3015818, EPI_ISL_3015820, EPI_ISL_3015823, EPI_ISL_3015824, EPI_ISL_3015826, EPI_ISL_3015830, EPI_ISL_3015831, EPI_ISL_3015833, EPI_ISL_3015835, EPI_ISL_3015836, EPI_ISL_3015837, EPI_ISL_3015839, EPI_ISL_3015842 | | | |
| see above | Northwest Laboratory | Seattle Flu Study | Deborah A. Nickerson, Chris D. Frazar, Jover Lee, Benjamin Pelle, Erica Ryke, Matthew Richardson, Amanda Adler, Elisabeth Brandstetter, Peter D. Han, Kairsten Fay, Misja Ilcisin, Kirsten Lacombe, Thomas R. Sibley, Melissa Truong, Caitlin R. Wolf, Romesh Gautom, Geoff Melly, Brian Hiatt, Philip Dykema, Scott Lindquist, Michael Boeckh, Janet A. Englund, Michael Famulare, Barry R. Lutz, Mark J. Rieder, Lea M. Starita, Matthew Thompson, Helen Y. Chu, Jay Shendure, Trevor Bedford |
| EPI_ISL_3015939, EPI_ISL_3015941, EPI_ISL_3015942, EPI_ISL_3015944, EPI_ISL_3015945, EPI_ISL_3015946, EPI_ISL_3015947, EPI_ISL_3015948, EPI_ISL_3015949, EPI_ISL_3015950, EPI_ISL_3015951, EPI_ISL_3015952, EPI_ISL_3015953, EPI_ISL_3015954, EPI_ISL_3015956, EPI_ISL_3015957, EPI_ISL_3015958, EPI_ISL_3015959, EPI_ISL_3015960, EPI_ISL_3015961, EPI_ISL_3015962, EPI_ISL_3015963, EPI_ISL_3015964, EPI_ISL_3015965, EPI_ISL_3015967, EPI_ISL_3015968, EPI_ISL_3015969, EPI_ISL_3015972, EPI_ISL_3015973, EPI_ISL_3015974, EPI_ISL_3015975, EPI_ISL_3015976, EPI_ISL_3015977, EPI_ISL_3015978, EPI_ISL_3015979, EPI_ISL_3015981, EPI_ISL_3015982, EPI_ISL_3015983, EPI_ISL_3015984, EPI_ISL_3015985, EPI_ISL_3015986, EPI_ISL_3015987, EPI_ISL_3015988, EPI_ISL_3015989, EPI_ISL_3015990, EPI_ISL_3015991, EPI_ISL_3015992, EPI_ISL_3015993, EPI_ISL_3015994, EPI_ISL_3015995, EPI_ISL_3015996, EPI_ISL_3015997, EPI_ISL_3015998, EPI_ISL_3015999, EPI_ISL_3016000, EPI_ISL_3016001, EPI_ISL_3016002, EPI_ISL_3016003, EPI_ISL_3016004, EPI_ISL_3016005, EPI_ISL_3016006, EPI_ISL_3016007, EPI_ISL_3016008, EPI_ISL_3016010 | | | |
| see above | District Headquarter Hospital, Keonjhar | Institute of Life Sciences - INSACOG | Sunil K. Raghav, Safal Walia, Arup Ghosh, Atimukta Jha, Amol M. Kanampalliwar, Omprakash Shiriwas, Sana Fatma, Shifu Aggarwal, Rupesh Dash, Rajeeb Swain, Punit Prasad, INSACOG Consortium, Ajay Parida |
| EPI_ISL_3016011, EPI_ISL_3016012, EPI_ISL_3016013, EPI_ISL_3016015, EPI_ISL_3016016, EPI_ISL_3016017, EPI_ISL_3016018, EPI_ISL_3016019, EPI_ISL_3016021, EPI_ISL_3016022, EPI_ISL_3016023, EPI_ISL_3016024, EPI_ISL_3016025, EPI_ISL_3016026, EPI_ISL_3016027, EPI_ISL_3016028, EPI_ISL_3016029, EPI_ISL_3016030, EPI_ISL_3016031, EPI_ISL_3016032 | | | |
| see above | ZeusNjove Diagnostic LLP, Bhubaneswar | Institute of Life Sciences - INSACOG | Sunil K. Raghav, Safal Walia, Arup Ghosh, Atimukta Jha, Amol M. Kanampalliwar, Omprakash Shiriwas, Sana Fatma, Shifu Aggarwal, Rupesh Dash, Rajeeb Swain, Punit Prasad, INSACOG Consortium, Ajay Parida |
| EPI_ISL_3016033, EPI_ISL_3016034, EPI_ISL_3016036, EPI_ISL_3016038, EPI_ISL_3016039, EPI_ISL_3016040, EPI_ISL_3016042, EPI_ISL_3016043, EPI_ISL_3016044, EPI_ISL_3016045, EPI_ISL_3016046, EPI_ISL_3016047, EPI_ISL_3016048, EPI_ISL_3016049, EPI_ISL_3016050, EPI_ISL_3016051, EPI_ISL_3016052, EPI_ISL_3016053, EPI_ISL_3016054, EPI_ISL_3016055, EPI_ISL_3016056, EPI_ISL_3016057, EPI_ISL_3016058, EPI_ISL_3016059, EPI_ISL_3016060, EPI_ISL_3016061, EPI_ISL_3016062, EPI_ISL_3016063, EPI_ISL_3016064, EPI_ISL_3016065, EPI_ISL_3016066, EPI_ISL_3016067, EPI_ISL_3016068, EPI_ISL_3016069, EPI_ISL_3016070, EPI_ISL_3016071, EPI_ISL_3016072, EPI_ISL_3016074, EPI_ISL_3016075, EPI_ISL_3016076 | | | |
| see above | IISER Berhampur | Institute of Life Sciences - INSACOG | Sunil K. Raghav, Safal Walia, Arup Ghosh, Atimukta Jha, Amol M. Kanampalliwar, Omprakash Shiriwas, Sana Fatma, Shifu Aggarwal, Rupesh Dash, Rajeeb Swain, Punit Prasad, INSACOG Consortium, Ajay Parida |
| EPI_ISL_3016077, EPI_ISL_3016078, EPI_ISL_3016079, EPI_ISL_3016080, EPI_ISL_3016081, EPI_ISL_3016082, EPI_ISL_3016083, EPI_ISL_3016084, EPI_ISL_3016085, EPI_ISL_3016086, EPI_ISL_3016087, EPI_ISL_3016088, EPI_ISL_3016089, EPI_ISL_3016090, EPI_ISL_3016091, EPI_ISL_3016093, EPI_ISL_3016094, EPI_ISL_3016096, EPI_ISL_3016098, EPI_ISL_3016100, EPI_ISL_3016101, EPI_ISL_3016102, EPI_ISL_3016103, EPI_ISL_3016104, EPI_ISL_3016106, EPI_ISL_3016107, EPI_ISL_3016108, EPI_ISL_3016109, EPI_ISL_3016110, EPI_ISL_3016111, EPI_ISL_3016112, EPI_ISL_3016113, EPI_ISL_3016114 | | | |
| see above | Pt. Jawahar Lal Nehru Memorial Medical College, Raipur | Institute of Life Sciences - INSACOG | Sunil K. Raghav, Safal Walia, Arup Ghosh, Atimukta Jha, Amol M. Kanampalliwar, Omprakash Shiriwas, Sana Fatma, Shifu Aggarwal, Rupesh Dash, Rajeeb Swain, Punit Prasad, INSACOG Consortium, Ajay Parida |
| EPI_ISL_3016115, EPI_ISL_3016116, EPI_ISL_3016117, EPI_ISL_3016118, EPI_ISL_3016120, EPI_ISL_3016121, EPI_ISL_3016122, EPI_ISL_3016124, EPI_ISL_3016125 | MKCG Medical College And Hospital, Brahmapur | Institute of Life Sciences - INSACOG | Sunil K. Raghav, Safal Walia, Arup Ghosh, Atimukta Jha, Amol M. Kanampalliwar, Omprakash Shiriwas, Sana Fatma, Shifu Aggarwal, Rupesh Dash, Rajeeb Swain, Punit Prasad, INSACOG Consortium, Ajay Parida |
| EPI_ISL_3016126, EPI_ISL_3016127, EPI_ISL_3016128, EPI_ISL_3016129, EPI_ISL_3016130, EPI_ISL_3016131, EPI_ISL_3016133, EPI_ISL_3016134, EPI_ISL_3016137, EPI_ISL_3016141, EPI_ISL_3016142, EPI_ISL_3016143, EPI_ISL_3016149, EPI_ISL_3016151, EPI_ISL_3016152, EPI_ISL_3016153, EPI_ISL_3016154, EPI_ISL_3016155, EPI_ISL_3016156, EPI_ISL_3016157, EPI_ISL_3016158, EPI_ISL_3016159, EPI_ISL_3016160, EPI_ISL_3016161, EPI_ISL_3016162, EPI_ISL_3016163, EPI_ISL_3016164 | | | |
| see above | S.C.B. Medical College and Hospital, Cuttack | Institute of Life Sciences - INSACOG | Sunil K. Raghav, Safal Walia, Arup Ghosh, Atimukta Jha, Amol M. Kanampalliwar, Omprakash Shiriwas, Sana Fatma, Shifu Aggarwal, Rupesh Dash, Rajeeb Swain, Punit Prasad, INSACOG Consortium, Ajay Parida |
| EPI_ISL_3016165, EPI_ISL_3016166, EPI_ISL_3016167, EPI_ISL_3016168, EPI_ISL_3016169, EPI_ISL_3016170, EPI_ISL_3016171, EPI_ISL_3016173, EPI_ISL_3016174, EPI_ISL_3016175, EPI_ISL_3016176, EPI_ISL_3016177, EPI_ISL_3016178, EPI_ISL_3016179, EPI_ISL_3016180, EPI_ISL_3016181, EPI_ISL_3016182, EPI_ISL_3016183, EPI_ISL_3016184 | | | |
| see above | Vikash Multispeciality Hospital, Bargarh | Institute of Life Sciences - INSACOG | Sunil K. Raghav, Safal Walia, Arup Ghosh, Atimukta Jha, Amol M. Kanampalliwar, Omprakash Shiriwas, Sana Fatma, Shifu Aggarwal, Rupesh Dash, Rajeeb Swain, Punit Prasad, INSACOG Consortium, Ajay Parida |
| EPI_ISL_3016185, EPI_ISL_3016186, EPI_ISL_3016187, EPI_ISL_3016188, EPI_ISL_3016189, EPI_ISL_3016190, EPI_ISL_3016191, EPI_ISL_3016192, EPI_ISL_3016193, EPI_ISL_3016194, EPI_ISL_3016195, EPI_ISL_3016196, EPI_ISL_3016197, EPI_ISL_3016198 | | | |
| see above | Government Medical College, Bettiah | Institute of Life Sciences - INSACOG | Sunil K. Raghav, Safal Walia, Arup Ghosh, Atimukta Jha, Amol M. Kanampalliwar, Omprakash Shiriwas, Sana Fatma, Shifu Aggarwal, Rupesh Dash, Rajeeb Swain, Punit Prasad, INSACOG Consortium, Ajay Parida |
| EPI_ISL_3016199, EPI_ISL_3016200, EPI_ISL_3016201, EPI_ISL_3016202, EPI_ISL_3016203, EPI_ISL_3016204, EPI_ISL_3016205, EPI_ISL_3016206, EPI_ISL_3016207, EPI_ISL_3016208, EPI_ISL_3016209, EPI_ISL_3016210, EPI_ISL_3016211, EPI_ISL_3016212, EPI_ISL_3016213, EPI_ISL_3016214, EPI_ISL_3016215, EPI_ISL_3016216, EPI_ISL_3016217, EPI_ISL_3016218, EPI_ISL_3016219, EPI_ISL_3016220, EPI_ISL_3016221, EPI_ISL_3016222, EPI_ISL_3016223, EPI_ISL_3016224, EPI_ISL_3016225, EPI_ISL_3016226, EPI_ISL_3016227, EPI_ISL_3016228, EPI_ISL_3016229, EPI_ISL_3016230, EPI_ISL_3016231, EPI_ISL_3016232, EPI_ISL_3016233, EPI_ISL_3016234, EPI_ISL_3016236, EPI_ISL_3016237, EPI_ISL_3016238, EPI_ISL_3016240, EPI_ISL_3016242, EPI_ISL_3016243, EPI_ISL_3016244, EPI_ISL_3016245, EPI_ISL_3016246, EPI_ISL_3016247, EPI_ISL_3016249, EPI_ISL_3016250, EPI_ISL_3016251, EPI_ISL_3016252, EPI_ISL_3016253, EPI_ISL_3016254, EPI_ISL_3016255, EPI_ISL_3016256, EPI_ISL_3016257, EPI_ISL_3016258, EPI_ISL_3016259, EPI_ISL_3016260, EPI_ISL_3016261, EPI_ISL_3016262, EPI_ISL_3016263, EPI_ISL_3016264, EPI_ISL_3016266, EPI_ISL_3016267, EPI_ISL_3016268, EPI_ISL_3016269, EPI_ISL_3016270, EPI_ISL_3016271, EPI_ISL_3016273, EPI_ISL_3016274, EPI_ISL_3016275, EPI_ISL_3016276, EPI_ISL_3016278, EPI_ISL_3016279, EPI_ISL_3016280, EPI_ISL_3016281, EPI_ISL_3016282, EPI_ISL_3016283 | | | |
| see above | Pt. Jawahar Lal Nehru Memorial Medical College, Raipur | Institute of Life Sciences - INSACOG | Sunil K. Raghav, Safal Walia, Arup Ghosh, Atimukta Jha, Amol M. Kanampalliwar, Omprakash Shiriwas, Sana Fatma, Shifu Aggarwal, Rupesh Dash, Rajeeb Swain, Punit Prasad, INSACOG Consortium, Ajay Parida |
| EPI_ISL_3016285, EPI_ISL_3016286, EPI_ISL_3016287, EPI_ISL_3016288, EPI_ISL_3016290, EPI_ISL_3016291, EPI_ISL_3016294, EPI_ISL_3016295, EPI_ISL_3016296, EPI_ISL_3016297, EPI_ISL_3016298, EPI_ISL_3016299, EPI_ISL_3016300, EPI_ISL_3016301, EPI_ISL_3016302, EPI_ISL_3016303, EPI_ISL_3016304, EPI_ISL_3016305, EPI_ISL_3016308, EPI_ISL_3016309, EPI_ISL_3016310, EPI_ISL_3016311, EPI_ISL_3016312, EPI_ISL_3016313, EPI_ISL_3016315, EPI_ISL_3016316 | | | |
| see above | Institute of Life Sciences, Bhubaneswar | Institute of Life Sciences - INSACOG | Sunil K. Raghav, Safal Walia, Arup Ghosh, Atimukta Jha, Amol M. Kanampalliwar, Omprakash Shiriwas, Sana Fatma, Shifu Aggarwal, Rupesh Dash, Rajeeb Swain, Punit Prasad, INSACOG Consortium, Ajay Parida |
| EPI_ISL_3016332, EPI_ISL_3016334, EPI_ISL_3016335, EPI_ISL_3016340, EPI_ISL_3016341, EPI_ISL_3016342, EPI_ISL_3016343, EPI_ISL_3016344, EPI_ISL_3016345, EPI_ISL_3016346, EPI_ISL_3016348, EPI_ISL_3016349, EPI_ISL_3016351, EPI_ISL_3016352, EPI_ISL_3016354, EPI_ISL_3016358, EPI_ISL_3016359, EPI_ISL_3016362, EPI_ISL_3016363, EPI_ISL_3016365, EPI_ISL_3016366, EPI_ISL_3016367, EPI_ISL_3016368, EPI_ISL_3016369, EPI_ISL_3016370, EPI_ISL_3016371, EPI_ISL_3016372, EPI_ISL_3016373, EPI_ISL_3016375, EPI_ISL_3016376, EPI_ISL_3016377, EPI_ISL_3016378, EPI_ISL_3016379, EPI_ISL_3016382, EPI_ISL_3016388, EPI_ISL_3016390, EPI_ISL_3016391, EPI_ISL_3016392, EPI_ISL_3016394, EPI_ISL_3016395, EPI_ISL_3016396, EPI_ISL_3016398, EPI_ISL_3016399, EPI_ISL_3016400, EPI_ISL_3016401, EPI_ISL_3016402, EPI_ISL_3016403, EPI_ISL_3016404, EPI_ISL_3016405, EPI_ISL_3016406, EPI_ISL_3016407, EPI_ISL_3016408, EPI_ISL_3016409, EPI_ISL_3016410, EPI_ISL_3016411, EPI_ISL_3016412, EPI_ISL_3016415, EPI_ISL_3016416, EPI_ISL_3016417, EPI_ISL_3016418, EPI_ISL_3016420, EPI_ISL_3016421, EPI_ISL_3016422, EPI_ISL_3016423, EPI_ISL_3016425 | | | |
| see above | Washington State Department of Health Public Health Laboratories | Washington State Department of Health Public Health Laboratories | Drew MacKellar, Philip Dykema, Denny Russell, Joenice Gonzalez, Hannah Gray, Geoff Melly, Vanessa De Los Santos, Darren Lucas, JohnAric Peterson, Avi Singh, Rebecca Cao |

| | | | |
|---|---|---|---|
| EPI_ISL_3016434, EPI_ISL_3016436, EPI_ISL_3016437, EPI_ISL_3016439, EPI_ISL_3016440, EPI_ISL_3016442, EPI_ISL_3016443, EPI_ISL_3016445, EPI_ISL_3016447, EPI_ISL_3016448, EPI_ISL_3016449, EPI_ISL_3016450, EPI_ISL_3016451, EPI_ISL_3016453, EPI_ISL_3016456, EPI_ISL_3016457, EPI_ISL_3016458, EPI_ISL_3016460, EPI_ISL_3016463, EPI_ISL_3016466, EPI_ISL_3016467, EPI_ISL_3016468, EPI_ISL_3016469, EPI_ISL_3016470, EPI_ISL_3016472 | | | |
| see above | Valais Hospital, Central Institute | Valais Hospital, Central Institute | Alexis Dumoulin, Lorenzo Cerutti, Henri Pegeot, Melyssa Elies, Deborah Penet, Keith Harshman, Ioannis Xenarios, Emmanouil Dermitzakis |
| EPI_ISL_3016478, EPI_ISL_3016479, EPI_ISL_3016480, EPI_ISL_3016481, EPI_ISL_3016482, EPI_ISL_3016483, EPI_ISL_3016484, EPI_ISL_3016485, EPI_ISL_3016486, EPI_ISL_3016487, EPI_ISL_3016489, EPI_ISL_3016491, EPI_ISL_3016492, EPI_ISL_3016493, EPI_ISL_3016494, EPI_ISL_3016495, EPI_ISL_3016497, EPI_ISL_3016499, EPI_ISL_3016500, EPI_ISL_3016501, EPI_ISL_3016504, EPI_ISL_3016505, EPI_ISL_3016506, EPI_ISL_3016507, EPI_ISL_3016508, EPI_ISL_3016509, EPI_ISL_3016510, EPI_ISL_3016512, EPI_ISL_3016513, EPI_ISL_3016514, EPI_ISL_3016515, EPI_ISL_3016516, EPI_ISL_3016517, EPI_ISL_3016518, EPI_ISL_3016521, EPI_ISL_3016522, EPI_ISL_3016523, EPI_ISL_3016524, EPI_ISL_3016525, EPI_ISL_3016526, EPI_ISL_3016528, EPI_ISL_3016529, EPI_ISL_3016530, EPI_ISL_3016531, EPI_ISL_3016532, EPI_ISL_3016533, EPI_ISL_3016534, EPI_ISL_3016535, EPI_ISL_3016536, EPI_ISL_3016537, EPI_ISL_3016538, EPI_ISL_3016539, EPI_ISL_3016540, EPI_ISL_3016541, EPI_ISL_3016542, EPI_ISL_3016543, EPI_ISL_3016545, EPI_ISL_3016546, EPI_ISL_3016547, EPI_ISL_3016549, EPI_ISL_3016550, EPI_ISL_3016553, EPI_ISL_3016554, EPI_ISL_3016555, EPI_ISL_3016556, EPI_ISL_3016557, EPI_ISL_3016558, EPI_ISL_3016560, EPI_ISL_3016561, EPI_ISL_3016562, EPI_ISL_3016563, EPI_ISL_3016564, EPI_ISL_3016565, EPI_ISL_3016566, EPI_ISL_3016567, EPI_ISL_3016571, EPI_ISL_3016573, EPI_ISL_3016574, EPI_ISL_3016575, EPI_ISL_3016576, EPI_ISL_3016577, EPI_ISL_3016578, EPI_ISL_3016579, EPI_ISL_3016582, EPI_ISL_3016583, EPI_ISL_3016584, EPI_ISL_3016586, EPI_ISL_3016587, EPI_ISL_3016588, EPI_ISL_3016589, EPI_ISL_3016590, EPI_ISL_3016591, EPI_ISL_3016592, EPI_ISL_3016593, EPI_ISL_3016594, EPI_ISL_3016595, EPI_ISL_3016596, EPI_ISL_3016597, EPI_ISL_3016598, EPI_ISL_3016599, EPI_ISL_3016600, EPI_ISL_3016601, EPI_ISL_3016602, EPI_ISL_3016603, EPI_ISL_3016604, EPI_ISL_3016605, EPI_ISL_3016607, EPI_ISL_3016608, EPI_ISL_3016609, EPI_ISL_3016610, EPI_ISL_3016611, EPI_ISL_3016612, EPI_ISL_3016613, EPI_ISL_3016614, EPI_ISL_3016616, EPI_ISL_3016617, EPI_ISL_3016618, EPI_ISL_3016621, EPI_ISL_3016622, EPI_ISL_3016624, EPI_ISL_3016625, EPI_ISL_3016626, EPI_ISL_3016627, EPI_ISL_3016628, EPI_ISL_3016630, EPI_ISL_3016632, EPI_ISL_3016634, EPI_ISL_3016635, EPI_ISL_3016636, EPI_ISL_3016637, EPI_ISL_3016638, EPI_ISL_3016639, EPI_ISL_3016641, EPI_ISL_3016642, EPI_ISL_3016643, EPI_ISL_3016644, EPI_ISL_3016645, EPI_ISL_3016646, EPI_ISL_3016649, EPI_ISL_3016650, EPI_ISL_3016652, EPI_ISL_3016653, EPI_ISL_3016656, EPI_ISL_3016657, EPI_ISL_3016658, EPI_ISL_3016659, EPI_ISL_3016662, EPI_ISL_3016663, EPI_ISL_3016664, EPI_ISL_3016665, EPI_ISL_3016666, EPI_ISL_3016667, EPI_ISL_3016668, EPI_ISL_3016670, EPI_ISL_3016672, EPI_ISL_3016673, EPI_ISL_3016674, EPI_ISL_3016675, EPI_ISL_3016676, EPI_ISL_3016677, EPI_ISL_3016678, EPI_ISL_3016679, EPI_ISL_3016680, EPI_ISL_3016681, EPI_ISL_3016682, EPI_ISL_3016683, EPI_ISL_3016684, EPI_ISL_3016685, EPI_ISL_3016686, EPI_ISL_3016687, EPI_ISL_3016688, EPI_ISL_3016689, EPI_ISL_3016690, EPI_ISL_3016691, EPI_ISL_3016692, EPI_ISL_3016693, EPI_ISL_3016694, EPI_ISL_3016696, EPI_ISL_3016697, EPI_ISL_3016698, EPI_ISL_3016699, EPI_ISL_3016700, EPI_ISL_3016701, EPI_ISL_3016703, EPI_ISL_3016705, EPI_ISL_3016706, EPI_ISL_3016707, EPI_ISL_3016710, EPI_ISL_3016711, EPI_ISL_3016712, EPI_ISL_3016713, EPI_ISL_3016714, EPI_ISL_3016715, EPI_ISL_3016716, EPI_ISL_3016717, EPI_ISL_3016718, EPI_ISL_3016719, EPI_ISL_3016720, EPI_ISL_3016722, EPI_ISL_3016723, EPI_ISL_3016724, EPI_ISL_3016725 | | | |
| see above | Fulgent Genetics | Centers for Disease Control and Prevention Division of Viral Diseases, Pathogen Discovery | Dakota Howard, Dhwani Batra, Peter W. Cook, Kara Moser, Adrian Paskey, Jason Caravas, Benjamin Rambo-Martin, Shatavia Morrison, Christopher Gulvick, Scott Sammons, Yvette Unoarumhi, Darlene Wagner, Matthew Schmerer, Harry Gao, Mickey Li, John Gao, Joseph Fierro, Benafsh Sapra, Becky Tsai, Yan Meng, Doreen Ng, James Xie, Clinton R. Paden, Duncan MacCannell |
| EPI_ISL_3016726, EPI_ISL_3016727, EPI_ISL_3016728, EPI_ISL_3016729, EPI_ISL_3016730, EPI_ISL_3016731, EPI_ISL_3016732, EPI_ISL_3016733, EPI_ISL_3016734, EPI_ISL_3016735, EPI_ISL_3016736, EPI_ISL_3016737, EPI_ISL_3016738, EPI_ISL_3016739, EPI_ISL_3016740, EPI_ISL_3016741, EPI_ISL_3016742, EPI_ISL_3016743, EPI_ISL_3016744, EPI_ISL_3016745, EPI_ISL_3016746, EPI_ISL_3016747, EPI_ISL_3016748, EPI_ISL_3016749, EPI_ISL_3016750, EPI_ISL_3016751, EPI_ISL_3016752, EPI_ISL_3016753, EPI_ISL_3016754, EPI_ISL_3016755, EPI_ISL_3016756, EPI_ISL_3016757, EPI_ISL_3016758, EPI_ISL_3016759, EPI_ISL_3016760, EPI_ISL_3016761, EPI_ISL_3016762, EPI_ISL_3016763, EPI_ISL_3016764, EPI_ISL_3016765, EPI_ISL_3016766, EPI_ISL_3016767, EPI_ISL_3016768, EPI_ISL_3016769, EPI_ISL_3016770, EPI_ISL_3016771, EPI_ISL_3016772, EPI_ISL_3016773, EPI_ISL_3016774, EPI_ISL_3016775, EPI_ISL_3016776, EPI_ISL_3016777, EPI_ISL_3016778, EPI_ISL_3016779, EPI_ISL_3016780, EPI_ISL_3016781, EPI_ISL_3016782, EPI_ISL_3016783, EPI_ISL_3016784, EPI_ISL_3016785, EPI_ISL_3016786, EPI_ISL_3016787, EPI_ISL_3016788, EPI_ISL_3016789, EPI_ISL_3016790, EPI_ISL_3016791, EPI_ISL_3016792, EPI_ISL_3016793, EPI_ISL_3016794, EPI_ISL_3016795, EPI_ISL_3016796, EPI_ISL_3016797, EPI_ISL_3016798, EPI_ISL_3016799, EPI_ISL_3016800, EPI_ISL_3016803, EPI_ISL_3016804, EPI_ISL_3016805, EPI_ISL_3016806, EPI_ISL_3016807, EPI_ISL_3016808, EPI_ISL_3016811, EPI_ISL_3016812, EPI_ISL_3016813, EPI_ISL_3016814, EPI_ISL_3016815, EPI_ISL_3016816, EPI_ISL_3016817, EPI_ISL_3016818, EPI_ISL_3016819, EPI_ISL_3016820, EPI_ISL_3016821, EPI_ISL_3016823, EPI_ISL_3016826, EPI_ISL_3016827, EPI_ISL_3016831, EPI_ISL_3016832, EPI_ISL_3016836, EPI_ISL_3016837, EPI_ISL_3016838, EPI_ISL_3016839, EPI_ISL_3016843, EPI_ISL_3016844, EPI_ISL_3016845, EPI_ISL_3016846, EPI_ISL_3016847, EPI_ISL_3016848, EPI_ISL_3016850, EPI_ISL_3016851, EPI_ISL_3016852, EPI_ISL_3016853, EPI_ISL_3016854, EPI_ISL_3016855, EPI_ISL_3016856, EPI_ISL_3016857, EPI_ISL_3016858, EPI_ISL_3016859, EPI_ISL_3016860, EPI_ISL_3016861, EPI_ISL_3016862, EPI_ISL_3016863, EPI_ISL_3016864, EPI_ISL_3016866, EPI_ISL_3016867, EPI_ISL_3016868, EPI_ISL_3016870, EPI_ISL_3016871, EPI_ISL_3016872, EPI_ISL_3016873, EPI_ISL_3016874, EPI_ISL_3016875, EPI_ISL_3016876, EPI_ISL_3016877, EPI_ISL_3016878, EPI_ISL_3016880, EPI_ISL_3016883, EPI_ISL_3016885, EPI_ISL_3016887, EPI_ISL_3016888, EPI_ISL_3016890, EPI_ISL_3016892, EPI_ISL_3016893, EPI_ISL_3016894, EPI_ISL_3016895, EPI_ISL_3016896, EPI_ISL_3016897, EPI_ISL_3016898, EPI_ISL_3016899, EPI_ISL_3016900, EPI_ISL_3016902, EPI_ISL_3016903, EPI_ISL_3016904, EPI_ISL_3016905, EPI_ISL_3016906, EPI_ISL_3016907, EPI_ISL_3016909, EPI_ISL_3016910, EPI_ISL_3016911, EPI_ISL_3016912, EPI_ISL_3016914, EPI_ISL_3016916, EPI_ISL_3016917, EPI_ISL_3016919, EPI_ISL_3016920, EPI_ISL_3016921, EPI_ISL_3016922, EPI_ISL_3016924, EPI_ISL_3016925, EPI_ISL_3016926, EPI_ISL_3016927, EPI_ISL_3016928, EPI_ISL_3016929, EPI_ISL_3016932, EPI_ISL_3016933, EPI_ISL_3016934, EPI_ISL_3016935, EPI_ISL_3016936, EPI_ISL_3016937, EPI_ISL_3016938, EPI_ISL_3016939, EPI_ISL_3016942, EPI_ISL_3016943, EPI_ISL_3016944, EPI_ISL_3016946, EPI_ISL_3016947, EPI_ISL_3016952, EPI_ISL_3016953, EPI_ISL_3016954, EPI_ISL_3016955, EPI_ISL_3016956, EPI_ISL_3016957, EPI_ISL_3016958, EPI_ISL_3016959, EPI_ISL_3016960, EPI_ISL_3016961, EPI_ISL_3016962, EPI_ISL_3016963, EPI_ISL_3016964, EPI_ISL_3016965, EPI_ISL_3016968, EPI_ISL_3016970, EPI_ISL_3016971, EPI_ISL_3016972, EPI_ISL_3016973, EPI_ISL_3016974, EPI_ISL_3016977, EPI_ISL_3016978, EPI_ISL_3016979, EPI_ISL_3016980, EPI_ISL_3016981, EPI_ISL_3016983, EPI_ISL_3016984, EPI_ISL_3016985, EPI_ISL_3016986, EPI_ISL_3016987, EPI_ISL_3016988, EPI_ISL_3016989, EPI_ISL_3016990, EPI_ISL_3016991, EPI_ISL_3016993, EPI_ISL_3016995, EPI_ISL_3016997, EPI_ISL_3016999, EPI_ISL_3017000, EPI_ISL_3017003, EPI_ISL_3017004, EPI_ISL_3017006, EPI_ISL_3017007, EPI_ISL_3017008, EPI_ISL_3017009, EPI_ISL_3017010, EPI_ISL_3017012, EPI_ISL_3017013, EPI_ISL_3017015, EPI_ISL_3017016, EPI_ISL_3017018, EPI_ISL_3017020, EPI_ISL_3017021, EPI_ISL_3017022, EPI_ISL_3017023, EPI_ISL_3017024, EPI_ISL_3017027, EPI_ISL_3017029, EPI_ISL_3017030, EPI_ISL_3017031, EPI_ISL_3017032, EPI_ISL_3017033, EPI_ISL_3017034, EPI_ISL_3017035, EPI_ISL_3017036, EPI_ISL_3017037, EPI_ISL_3017038, EPI_ISL_3017039, EPI_ISL_3017041, EPI_ISL_3017042, EPI_ISL_3017043, EPI_ISL_3017044, EPI_ISL_3017045, EPI_ISL_3017046, EPI_ISL_3017047, EPI_ISL_3017048, EPI_ISL_3017049, EPI_ISL_3017050, EPI_ISL_3017051, EPI_ISL_3017052, EPI_ISL_3017053, EPI_ISL_3017054, EPI_ISL_3017055, EPI_ISL_3017058, EPI_ISL_3017059, EPI_ISL_3017060, EPI_ISL_3017061, EPI_ISL_3017062, EPI_ISL_3017063, EPI_ISL_3017064, EPI_ISL_3017065, EPI_ISL_3017066, EPI_ISL_3017068, EPI_ISL_3017069, EPI_ISL_3017071, EPI_ISL_3017072, EPI_ISL_3017073, EPI_ISL_3017074, EPI_ISL_3017075, EPI_ISL_3017076, EPI_ISL_3017077, EPI_ISL_3017078, EPI_ISL_3017079, EPI_ISL_3017080, EPI_ISL_3017081, EPI_ISL_3017082, EPI_ISL_3017083, EPI_ISL_3017084, EPI_ISL_3017085, EPI_ISL_3017086, EPI_ISL_3017087, EPI_ISL_3017088, EPI_ISL_3017089, EPI_ISL_3017090, EPI_ISL_3017091, EPI_ISL_3017092, EPI_ISL_3017093, EPI_ISL_3017095, EPI_ISL_3017096, EPI_ISL_3017097, EPI_ISL_3017098, EPI_ISL_3017104, EPI_ISL_3017106, EPI_ISL_3017107, EPI_ISL_3017109, EPI_ISL_3017111, EPI_ISL_3017112, EPI_ISL_3017113, EPI_ISL_3017114, EPI_ISL_3017115, EPI_ISL_3017117, EPI_ISL_3017119, EPI_ISL_3017120, EPI_ISL_3017121, EPI_ISL_3017122, EPI_ISL_3017124, EPI_ISL_3017125, EPI_ISL_3017126, EPI_ISL_3017127, EPI_ISL_3017128, EPI_ISL_3017129, EPI_ISL_3017130, EPI_ISL_3017131, EPI_ISL_3017132, EPI_ISL_3017133, EPI_ISL_3017134, EPI_ISL_3017135, EPI_ISL_3017136, EPI_ISL_3017137, EPI_ISL_3017138, EPI_ISL_3017139, EPI_ISL_3017142, EPI_ISL_3017143, EPI_ISL_3017144, EPI_ISL_3017145, EPI_ISL_3017146, EPI_ISL_3017148, EPI_ISL_3017149, EPI_ISL_3017150, EPI_ISL_3017151, EPI_ISL_3017152, EPI_ISL_3017153, EPI_ISL_3017154, EPI_ISL_3017155, EPI_ISL_3017156, EPI_ISL_3017157, EPI_ISL_3017158, EPI_ISL_3017159, EPI_ISL_3017160, EPI_ISL_3017161, EPI_ISL_3017162, EPI_ISL_3017163, EPI_ISL_3017164, EPI_ISL_3017166, EPI_ISL_3017167, EPI_ISL_3017169, EPI_ISL_3017171, EPI_ISL_3017172, EPI_ISL_3017173, EPI_ISL_3017175, EPI_ISL_3017177, EPI_ISL_3017178, EPI_ISL_3017179, EPI_ISL_3017181, EPI_ISL_3017182, EPI_ISL_3017183, EPI_ISL_3017184, EPI_ISL_3017186, EPI_ISL_3017187, EPI_ISL_3017188, EPI_ISL_3017189, EPI_ISL_3017190, EPI_ISL_3017191, EPI_ISL_3017192, EPI_ISL_3017193, EPI_ISL_3017194, EPI_ISL_3017197, EPI_ISL_3017198, EPI_ISL_3017199, EPI_ISL_3017200, EPI_ISL_3017201, EPI_ISL_3017202, EPI_ISL_3017203, EPI_ISL_3017204, EPI_ISL_3017205, EPI_ISL_3017206, EPI_ISL_3017207, EPI_ISL_3017209, EPI_ISL_3017211, EPI_ISL_3017212, EPI_ISL_3017213, EPI_ISL_3017214, EPI_ISL_3017215, EPI_ISL_3017216, EPI_ISL_3017218, EPI_ISL_3017219, EPI_ISL_3017220, EPI_ISL_3017222, EPI_ISL_3017223, EPI_ISL_3017225, EPI_ISL_3017227, EPI_ISL_3017228, EPI_ISL_3017229, EPI_ISL_3017231, EPI_ISL_3017233, EPI_ISL_3017234, EPI_ISL_3017236, EPI_ISL_3017237, EPI_ISL_3017238, EPI_ISL_3017239, EPI_ISL_3017240, EPI_ISL_3017241, EPI_ISL_3017243, EPI_ISL_3017248, EPI_ISL_3017250, EPI_ISL_3017251, EPI_ISL_3017252, EPI_ISL_3017253, EPI_ISL_3017254, EPI_ISL_3017257, EPI_ISL_3017258, EPI_ISL_3017259, EPI_ISL_3017260, EPI_ISL_3017261, EPI_ISL_3017262, EPI_ISL_3017263, EPI_ISL_3017265, EPI_ISL_3017266, EPI_ISL_3017267, EPI_ISL_3017269, EPI_ISL_3017270, EPI_ISL_3017271, EPI_ISL_3017272, EPI_ISL_3017273, EPI_ISL_3017274, EPI_ISL_3017275, EPI_ISL_3017276, EPI_ISL_3017277, EPI_ISL_3017278, EPI_ISL_3017280, EPI_ISL_3017281, EPI_ISL_3017282, EPI_ISL_3017283, EPI_ISL_3017284, EPI_ISL_3017286, EPI_ISL_3017287, EPI_ISL_3017288, EPI_ISL_3017289, EPI_ISL_3017290, EPI_ISL_3017291, EPI_ISL_3017292, EPI_ISL_3017293, EPI_ISL_3017294, EPI_ISL_3017296, EPI_ISL_3017298, EPI_ISL_3017299, EPI_ISL_3017300, EPI_ISL_3017302, EPI_ISL_3017303, EPI_ISL_3017305, EPI_ISL_3017306, EPI_ISL_3017307, EPI_ISL_3017308, EPI_ISL_3017309, EPI_ISL_3017310, EPI_ISL_3017311, EPI_ISL_3017315, EPI_ISL_3017321, EPI_ISL_3017323, EPI_ISL_3017324, EPI_ISL_3017325, EPI_ISL_3017326, EPI_ISL_3017328, EPI_ISL_3017330, EPI_ISL_3017331, EPI_ISL_3017332, EPI_ISL_3017335, EPI_ISL_3017336, EPI_ISL_3017337, EPI_ISL_3017338, EPI_ISL_3017339, EPI_ISL_3017341, EPI_ISL_3017342, EPI_ISL_3017344, EPI_ISL_3017345, EPI_ISL_3017346, EPI_ISL_3017347, EPI_ISL_3017350, EPI_ISL_3017351, EPI_ISL_3017352, EPI_ISL_3017353, EPI_ISL_3017355, EPI_ISL_3017356, EPI_ISL_3017357, EPI_ISL_3017358, EPI_ISL_3017359, EPI_ISL_3017360, EPI_ISL_3017361, EPI_ISL_3017362, EPI_ISL_3017363, EPI_ISL_3017364, EPI_ISL_3017365, EPI_ISL_3017366, EPI_ISL_3017367, EPI_ISL_3017368, EPI_ISL_3017370, EPI_ISL_3017371, EPI_ISL_3017372, EPI_ISL_3017375, EPI_ISL_3017376, EPI_ISL_3017377, EPI_ISL_3017378, EPI_ISL_3017384, EPI_ISL_3017385, EPI_ISL_3017386, EPI_ISL_3017387, EPI_ISL_3017389, EPI_ISL_3017390, EPI_ISL_3017392, EPI_ISL_3017393, EPI_ISL_3017394, EPI_ISL_3017395, EPI_ISL_3017396, EPI_ISL_3017397, EPI_ISL_3017399, EPI_ISL_3017401, EPI_ISL_3017402, EPI_ISL_3017403, EPI_ISL_3017404, EPI_ISL_3017405, EPI_ISL_3017406, EPI_ISL_3017407, EPI_ISL_3017408, EPI_ISL_3017409, EPI_ISL_3017410, EPI_ISL_3017412, EPI_ISL_3017413, EPI_ISL_3017414, EPI_ISL_3017415, EPI_ISL_3017416, EPI_ISL_3017417, EPI_ISL_3017418, EPI_ISL_3017419, EPI_ISL_3017420, EPI_ISL_3017421, EPI_ISL_3017422, EPI_ISL_3017423, EPI_ISL_3017425, EPI_ISL_3017427, EPI_ISL_3017428, EPI_ISL_3017429, EPI_ISL_3017431, EPI_ISL_3017434, EPI_ISL_3017435, EPI_ISL_3017437, EPI_ISL_3017438, EPI_ISL_3017441, EPI_ISL_3017442, EPI_ISL_3017443, EPI_ISL_3017444, EPI_ISL_3017445, EPI_ISL_3017446, EPI_ISL_3017447, EPI_ISL_3017448, EPI_ISL_3017449, EPI_ISL_3017450, EPI_ISL_3017453, EPI_ISL_3017454, EPI_ISL_3017455, EPI_ISL_3017456, EPI_ISL_3017457, EPI_ISL_3017459, EPI_ISL_3017461, EPI_ISL_3017462, EPI_ISL_3017463, EPI_ISL_3017464, EPI_ISL_3017465, EPI_ISL_3017466, EPI_ISL_3017467, EPI_ISL_3017468, EPI_ISL_3017469, EPI_ISL_3017470, EPI_ISL_3017471, EPI_ISL_3017472, EPI_ISL_3017473, EPI_ISL_3017476, EPI_ISL_3017477, EPI_ISL_3017478, EPI_ISL_3017479, EPI_ISL_3017482, EPI_ISL_3017484, EPI_ISL_3017485, EPI_ISL_3017486, EPI_ISL_3017487, EPI_ISL_3017488, EPI_ISL_3017489, EPI_ISL_3017490, EPI_ISL_3017491, EPI_ISL_3017493, EPI_ISL_3017497, EPI_ISL_3017498, EPI_ISL_3017499, EPI_ISL_3017500, EPI_ISL_3017501, EPI_ISL_3017502, EPI_ISL_3017503, EPI_ISL_3017504, EPI_ISL_3017505, EPI_ISL_3017506, EPI_ISL_3017507, EPI_ISL_3017510, EPI_ISL_3017511, EPI_ISL_3017516, EPI_ISL_3017517, EPI_ISL_3017518, EPI_ISL_3017522, EPI_ISL_3017523, EPI_ISL_3017526, EPI_ISL_3017527, EPI_ISL_3017528, EPI_ISL_3017529, EPI_ISL_3017530, EPI_ISL_3017531, EPI_ISL_3017532, EPI_ISL_3017534, EPI_ISL_3017535, EPI_ISL_3017536, EPI_ISL_3017537, EPI_ISL_3017538, EPI_ISL_3017539, EPI_ISL_3017540, EPI_ISL_3017541, EPI_ISL_3017542, EPI_ISL_3017543, EPI_ISL_3017544, EPI_ISL_3017546, EPI_ISL_3017547, EPI_ISL_3017549, EPI_ISL_3017551, EPI_ISL_3017553, EPI_ISL_3017554, EPI_ISL_3017556, EPI_ISL_3017559, EPI_ISL_3017560, EPI_ISL_3017561, EPI_ISL_3017562, EPI_ISL_3017563, EPI_ISL_3017564, EPI_ISL_3017565, EPI_ISL_3017566, EPI_ISL_3017569, EPI_ISL_3017572, EPI_ISL_3017584, EPI_ISL_3017590, EPI_ISL_3017595, EPI_ISL_3017596, EPI_ISL_3017600, EPI_ISL_3017601, EPI_ISL_3017603, EPI_ISL_3017605, EPI_ISL_3017606, EPI_ISL_3017611, EPI_ISL_3017613, EPI_ISL_3017616, EPI_ISL_3017617, EPI_ISL_3017620, EPI_ISL_3017622, EPI_ISL_3017623, EPI_ISL_3017624, EPI_ISL_3017625, EPI_ISL_3017626, EPI_ISL_3017627, EPI_ISL_3017630, EPI_ISL_3017631, EPI_ISL_3017634, EPI_ISL_3017636, EPI_ISL_3017639, EPI_ISL_3017643, EPI_ISL_3017644, EPI_ISL_3017646, EPI_ISL_3017648, EPI_ISL_3017649, EPI_ISL_3017650, EPI_ISL_3017651, EPI_ISL_3017653, EPI_ISL_3017655, EPI_ISL_3017656, EPI_ISL_3017657, EPI_ISL_3017658, EPI_ISL_3017659, EPI_ISL_3017660, EPI_ISL_3017667, EPI_ISL_3017669, EPI_ISL_3017670, EPI_ISL_3017671, EPI_ISL_3017672, EPI_ISL_3017673, EPI_ISL_3017674, EPI_ISL_3017675, EPI_ISL_3017676, EPI_ISL_3017677, EPI_ISL_3017678, EPI_ISL_3017679, EPI_ISL_3017680, EPI_ISL_3017682, EPI_ISL_3017683, EPI_ISL_3017685, EPI_ISL_3017686, EPI_ISL_3017687, EPI_ISL_3017688, EPI_ISL_3017689, EPI_ISL_3017690, EPI_ISL_3017691, EPI_ISL_3017692, EPI_ISL_3017693, EPI_ISL_3017694, EPI_ISL_3017695, EPI_ISL_3017698, EPI_ISL_3017699, EPI_ISL_3017700, EPI_ISL_3017701, EPI_ISL_3017702, EPI_ISL_3017703, EPI_ISL_3017704, EPI_ISL_3017707, EPI_ISL_3017709, EPI_ISL_3017710, EPI_ISL_3017713, EPI_ISL_3017714, EPI_ISL_3017715, EPI_ISL_3017716, EPI_ISL_3017718, EPI_ISL_3017720, EPI_ISL_3017722, EPI_ISL_3017724, EPI_ISL_3017725, EPI_ISL_3017726, EPI_ISL_3017727, EPI_ISL_3017729, EPI_ISL_3017732 | | | |
| see above | Laboratory Corporation of America | Centers for Disease Control and Prevention Division of Viral Diseases, Pathogen Discovery | Dakota Howard, Dhwani Batra, Peter W. Cook, Kara Moser, Adrian Paskey, Jason Caravas, Benjamin Rambo-Martin, Shatavia Morrison, Christopher Gulvick, Scott Sammons, Yvette Unoarumhi, Darlene Wagner, Matthew Schmerer, Minoo Agarwal, Eyad Almasri, Debbie Boles, Ayla Burns, Nuthawin Charoensri, Oren Cohen, Susan Countryman, Mary Ann Cristobal, Bobbi Croy, Suzanne Dale, Hrushikesh Deshmukh, Amanda Douglas, Vincent Drouillon, Marcia Eisenberg, Howard Engler, Rama Ghatti, Prashant Gupta, Susan Hicks, Jake Humphrey, Lax Iyer, Lisa Pfefferle, Manoj Jain, Matthew Robinson, Mohan Kolli, Brian Krueger, Tim Kuphal, Stanley Letovsky, Michael Levandoski, Craig Lukasik, Jonathan Meltzer, Brian Norvell, Mindy Nye, Scott Parker, Christos Petropoulos, John Pruitt, Steven Ragan, Scott Ryan, Mike Sapeta, Jana Schroth, Suresh Babu Selvaraju, Goran Stevovic, Amanda Suchanek, Andrea Throop, Lyndon Tilson, Thomas Urban, Joe Voshell, Kimberly Wagner, Jonathan Williams, Mary Williamson, Qian Zeng, Tricia Zwiefelhofer, Clinton R. Paden, Duncan MacCannell |
| EPI_ISL_3017733 | Polyanalytic SA | Genesupport | Tanguy ARAUD, Geraldine Jost, Katia Jaton, Nadia Liassine |
| EPI_ISL_3017734 | Laboratory Corporation of America | Centers for Disease Control and Prevention Division of Viral Diseases, Pathogen Discovery | Dakota Howard, Dhwani Batra, Peter W. Cook, Kara Moser, Adrian Paskey, Jason Caravas, Benjamin Rambo-Martin, Shatavia Morrison, Christopher Gulvick, Scott Sammons, Yvette Unoarumhi, Darlene Wagner, Matthew Schmerer, Minoo Agarwal, Eyad Almasri, Debbie Boles, Ayla Burns, Nuthawin Charoensri, Oren Cohen, Susan Countryman, Mary Ann Cristobal, Bobbi Croy, Suzanne Dale, Hrushikesh Deshmukh, Amanda Douglas, Vincent Drouillon, Marcia Eisenberg, Howard Engler, Rama Ghatti, Prashant Gupta, Susan Hicks, Jake Humphrey, Lax Iyer, Lisa Pfefferle, Manoj Jain, Matthew Robinson, Mohan Kolli, Brian Krueger, Tim Kuphal, Stanley Letovsky, Michael Levandoski, Craig Lukasik, Jonathan Meltzer, Brian Norvell, Mindy Nye, Scott Parker, Christos Petropoulos, John Pruitt, Steven Ragan, Scott Ryan, Mike Sapeta, Jana Schroth, Suresh Babu Selvaraju, Goran Stevovic, Amanda Suchanek, Andrea Throop, Lyndon Tilson, Thomas Urban, Joe Voshell, Kimberly Wagner, Jonathan Williams, Mary Williamson, Qian Zeng, Tricia Zwiefelhofer, Clinton R. Paden, Duncan MacCannell |
| EPI_ISL_3017735, EPI_ISL_3017736 | Dianalabs-Valais SA | Genesupport | Tanguy ARAUD, Geraldine Jost, Katia Jaton, Nadia Liassine |
| EPI_ISL_3017737 | Laboratory Corporation of America | Centers for Disease Control and Prevention Division of Viral Diseases, Pathogen Discovery | Dakota Howard, Dhwani Batra, Peter W. Cook, Kara Moser, Adrian Paskey, Jason Caravas, Benjamin Rambo-Martin, Shatavia Morrison, Christopher Gulvick, Scott Sammons, Yvette Unoarumhi, Darlene Wagner, Matthew Schmerer, Minoo Agarwal, Eyad Almasri, Debbie Boles, Ayla Burns, Nuthawin Charoensri, Oren Cohen, Susan Countryman, Mary Ann Cristobal, Bobbi Croy, Suzanne Dale, Hrushikesh Deshmukh, Amanda Douglas, Vincent Drouillon, Marcia Eisenberg, Howard Engler, Rama Ghatti, Prashant Gupta, Susan Hicks, Jake Humphrey, Lax Iyer, Lisa Pfefferle, Manoj Jain, Matthew Robinson, Mohan Kolli, Brian Krueger, Tim Kuphal, Stanley Letovsky, Michael Levandoski, Craig Lukasik, Jonathan Meltzer, Brian Norvell, Mindy Nye, Scott Parker, Christos Petropoulos, John Pruitt, Steven Ragan, Scott Ryan, Mike Sapeta, Jana Schroth, Suresh Babu Selvaraju, Goran Stevovic, Amanda Suchanek, Andrea Throop, Lyndon Tilson, Thomas Urban, Joe Voshell, Kimberly Wagner, Jonathan Williams, Mary Williamson, Qian Zeng, Tricia Zwiefelhofer, Clinton R. Paden, Duncan MacCannell |
| EPI_ISL_3017738, EPI_ISL_3017739 | Dianalabs-Valais SA | Genesupport | Tanguy ARAUD, Geraldine Jost, Katia Jaton, Nadia Liassine |
| EPI_ISL_3017740 | Laboratory Corporation of America | Centers for Disease Control and Prevention Division of Viral Diseases, Pathogen Discovery | Dakota Howard, Dhwani Batra, Peter W. Cook, Kara Moser, Adrian Paskey, Jason Caravas, Benjamin Rambo-Martin, Shatavia Morrison, Christopher Gulvick, Scott Sammons, Yvette Unoarumhi, Darlene Wagner, Matthew Schmerer, Minoo Agarwal, Eyad Almasri, Debbie Boles, Ayla Burns, Nuthawin Charoensri, Oren Cohen, Susan Countryman, Mary Ann Cristobal, Bobbi Croy, Suzanne Dale, Hrushikesh Deshmukh, Amanda Douglas, Vincent Drouillon, Marcia Eisenberg, Howard Engler, Rama Ghatti, Prashant Gupta, Susan Hicks, Jake Humphrey, Lax Iyer, Lisa Pfefferle, Manoj Jain, Matthew Robinson, Mohan Kolli, Brian Krueger, Tim Kuphal, Stanley Letovsky, Michael Levandoski, Craig Lukasik, Jonathan Meltzer, Brian Norvell, Mindy Nye, Scott Parker, Christos Petropoulos, John Pruitt, Steven Ragan, Scott Ryan, Mike Sapeta, Jana Schroth, Suresh Babu Selvaraju, Goran Stevovic, Amanda Suchanek, Andrea Throop, Lyndon Tilson, Thomas Urban, Joe Voshell, Kimberly Wagner, Jonathan Williams, Mary Williamson, Qian Zeng, Tricia Zwiefelhofer, Clinton R. Paden, Duncan MacCannell |
| EPI_ISL_3017741 | Polyanalytic SA | Genesupport | Tanguy ARAUD, Geraldine Jost, Katia Jaton, Nadia Liassine |
| EPI_ISL_3017742 | Dianalabs-Valais SA | Genesupport | Tanguy ARAUD, Geraldine Jost, Katia Jaton, Nadia Liassine |
| EPI_ISL_3017745, EPI_ISL_3017746, EPI_ISL_3017747, EPI_ISL_3017748, EPI_ISL_3017749, EPI_ISL_3017750, EPI_ISL_3017751, EPI_ISL_3017752, EPI_ISL_3017753, EPI_ISL_3017754, EPI_ISL_3017755, EPI_ISL_3017756, EPI_ISL_3017757, EPI_ISL_3017758, EPI_ISL_3017759, EPI_ISL_3017760, EPI_ISL_3017761, EPI_ISL_3017762, EPI_ISL_3017763, EPI_ISL_3017765, EPI_ISL_3017766, EPI_ISL_3017767, EPI_ISL_3017768, EPI_ISL_3017770, EPI_ISL_3017772, EPI_ISL_3017773, EPI_ISL_3017775, EPI_ISL_3017777, EPI_ISL_3017778, EPI_ISL_3017779, EPI_ISL_3017780, EPI_ISL_3017782, EPI_ISL_3017783, EPI_ISL_3017785, EPI_ISL_3017786, EPI_ISL_3017787, EPI_ISL_3017788, EPI_ISL_3017789, EPI_ISL_3017790, EPI_ISL_3017791, EPI_ISL_3017792, EPI_ISL_3017794, EPI_ISL_3017795, EPI_ISL_3017796, EPI_ISL_3017797, EPI_ISL_3017798, EPI_ISL_3017799, EPI_ISL_3017804, EPI_ISL_3017805, EPI_ISL_3017806, EPI_ISL_3017807, EPI_ISL_3017808, EPI_ISL_3017809, EPI_ISL_3017814, EPI_ISL_3017815, EPI_ISL_3017817, EPI_ISL_3017818, EPI_ISL_3017819, EPI_ISL_3017820, EPI_ISL_3017821, EPI_ISL_3017822, EPI_ISL_3017823, EPI_ISL_3017824, EPI_ISL_3017825, EPI_ISL_3017826, EPI_ISL_3017827, EPI_ISL_3017828, EPI_ISL_3017829, EPI_ISL_3017830, EPI_ISL_3017831, EPI_ISL_3017832, EPI_ISL_3017834, EPI_ISL_3017835, EPI_ISL_3017836, EPI_ISL_3017837, EPI_ISL_3017841, EPI_ISL_3017844, EPI_ISL_3017845, EPI_ISL_3017850, EPI_ISL_3017851, EPI_ISL_3017852, EPI_ISL_3017856, EPI_ISL_3017859, EPI_ISL_3017860, EPI_ISL_3017861, EPI_ISL_3017862, EPI_ISL_3017863, EPI_ISL_3017864, EPI_ISL_3017865, EPI_ISL_3017868, EPI_ISL_3017869, EPI_ISL_3017871, EPI_ISL_3017874, EPI_ISL_3017875, EPI_ISL_3017876, EPI_ISL_3017877, EPI_ISL_3017878, EPI_ISL_3017879, EPI_ISL_3017880, EPI_ISL_3017881, EPI_ISL_3017882, EPI_ISL_3017883, EPI_ISL_3017884, EPI_ISL_3017886, EPI_ISL_3017888, EPI_ISL_3017890, EPI_ISL_3017893, EPI_ISL_3017894, EPI_ISL_3017895, EPI_ISL_3017896, EPI_ISL_3017897, EPI_ISL_3017898, EPI_ISL_3017899, EPI_ISL_3017900, EPI_ISL_3017901, EPI_ISL_3017902, EPI_ISL_3017903, EPI_ISL_3017904, EPI_ISL_3017906, EPI_ISL_3017907, EPI_ISL_3017908, EPI_ISL_3017909, EPI_ISL_3017912, EPI_ISL_3017913, EPI_ISL_3017914, EPI_ISL_3017915, EPI_ISL_3017916, EPI_ISL_3017917, EPI_ISL_3017918, EPI_ISL_3017919, EPI_ISL_3017920, EPI_ISL_3017921, EPI_ISL_3017922, EPI_ISL_3017923, EPI_ISL_3017924, EPI_ISL_3017925, EPI_ISL_3017926, EPI_ISL_3017927, EPI_ISL_3017928, EPI_ISL_3017929, EPI_ISL_3017931, EPI_ISL_3017932, EPI_ISL_3017933, EPI_ISL_3017935, EPI_ISL_3017936, EPI_ISL_3017937, EPI_ISL_3017939, EPI_ISL_3017940, EPI_ISL_3017941, EPI_ISL_3017942, EPI_ISL_3017943, EPI_ISL_3017944, EPI_ISL_3017945, EPI_ISL_3017946, EPI_ISL_3017947, EPI_ISL_3017949, EPI_ISL_3017951, EPI_ISL_3017952, EPI_ISL_3017953, EPI_ISL_3017954, EPI_ISL_3017956, EPI_ISL_3017957, EPI_ISL_3017958, EPI_ISL_3017959, EPI_ISL_3017960, EPI_ISL_3017961, EPI_ISL_3017962, EPI_ISL_3017963, EPI_ISL_3017964, EPI_ISL_3017966, EPI_ISL_3017967, EPI_ISL_3017970, EPI_ISL_3017971, EPI_ISL_3017972, EPI_ISL_3017974, EPI_ISL_3017975, EPI_ISL_3017976, EPI_ISL_3017977, EPI_ISL_3017978, EPI_ISL_3017980, EPI_ISL_3017981, EPI_ISL_3017984, EPI_ISL_3017985, EPI_ISL_3017986, EPI_ISL_3017987, EPI_ISL_3017988, EPI_ISL_3017990, EPI_ISL_3017991, EPI_ISL_3017993, EPI_ISL_3017996, EPI_ISL_3017997, EPI_ISL_3017998, EPI_ISL_3017999, EPI_ISL_3018000, EPI_ISL_3018001, EPI_ISL_3018002, EPI_ISL_3018003, EPI_ISL_3018004, EPI_ISL_3018007, EPI_ISL_3018008, EPI_ISL_3018009, EPI_ISL_3018010, EPI_ISL_3018011, EPI_ISL_3018012, EPI_ISL_3018013, EPI_ISL_3018014, EPI_ISL_3018015, EPI_ISL_3018016, EPI_ISL_3018017, EPI_ISL_3018019, EPI_ISL_3018020, EPI_ISL_3018023, EPI_ISL_3018024, EPI_ISL_3018025, EPI_ISL_3018027, EPI_ISL_3018030, EPI_ISL_3018033 | | | |
| see above | Laboratory Corporation of America | Centers for Disease Control and Prevention Division of Viral Diseases, Pathogen Discovery | Dakota Howard, Dhwani Batra, Peter W. Cook, Kara Moser, Adrian Paskey, Jason Caravas, Benjamin Rambo-Martin, Shatavia Morrison, Christopher Gulvick, Scott Sammons, Yvette Unoarumhi, Darlene Wagner, Matthew Schmerer, Minoo Agarwal, Eyad Almasri, Debbie Boles, Ayla Burns, Nuthawin Charoensri, Oren Cohen, Susan Countryman, Mary Ann Cristobal, Bobbi Croy, Suzanne Dale, Hrushikesh Deshmukh, Amanda Douglas, Vincent Drouillon, Marcia Eisenberg, Howard Engler, Rama Ghatti, Prashant Gupta, Susan Hicks, Jake Humphrey, Lax Iyer, Lisa Pfefferle, Manoj Jain, Matthew Robinson, Mohan Kolli, Brian Krueger, Tim Kuphal, Stanley Letovsky, Michael Levandoski, Craig Lukasik, Jonathan Meltzer, Brian Norvell, Mindy Nye, Scott Parker, Christos Petropoulos, John Pruitt, Steven Ragan, Scott Ryan, Mike Sapeta, Jana Schroth, Suresh Babu Selvaraju, Goran Stevovic, Amanda Suchanek, Andrea Throop, Lyndon Tilson, Thomas Urban, Joe Voshell, Kimberly Wagner, Jonathan Williams, Mary Williamson, Qian Zeng, Tricia Zwiefelhofer, Clinton R. Paden, Duncan MacCannell |
| EPI_ISL_3018034 | National Center of Infectious and Parasitic Diseases | National Center of Infectious and Parasitic Diseases | Alexiev et al |
| EPI_ISL_3018035, EPI_ISL_3018036, EPI_ISL_3018037, EPI_ISL_3018038, EPI_ISL_3018039, EPI_ISL_3018040, EPI_ISL_3018041, EPI_ISL_3018042, EPI_ISL_3018043, EPI_ISL_3018044, EPI_ISL_3018045, EPI_ISL_3018046, EPI_ISL_3018047, EPI_ISL_3018048, EPI_ISL_3018050, EPI_ISL_3018051, EPI_ISL_3018053, EPI_ISL_3018054, EPI_ISL_3018055, EPI_ISL_3018056, EPI_ISL_3018057, EPI_ISL_3018058, EPI_ISL_3018060, EPI_ISL_3018063, EPI_ISL_3018065, EPI_ISL_3018066, EPI_ISL_3018067, EPI_ISL_3018068, EPI_ISL_3018069, EPI_ISL_3018070, EPI_ISL_3018071, EPI_ISL_3018072, EPI_ISL_3018073, EPI_ISL_3018074, EPI_ISL_3018075, EPI_ISL_3018076, EPI_ISL_3018077, EPI_ISL_3018078, EPI_ISL_3018079, EPI_ISL_3018080, EPI_ISL_3018081, EPI_ISL_3018082, EPI_ISL_3018083, EPI_ISL_3018084, EPI_ISL_3018085, EPI_ISL_3018086, EPI_ISL_3018087, EPI_ISL_3018088, EPI_ISL_3018089, EPI_ISL_3018090, EPI_ISL_3018091, EPI_ISL_3018092, EPI_ISL_3018093, EPI_ISL_3018094, EPI_ISL_3018096, EPI_ISL_3018097, EPI_ISL_3018098, EPI_ISL_3018099, EPI_ISL_3018100, EPI_ISL_3018101, EPI_ISL_3018102, EPI_ISL_3018103, EPI_ISL_3018104, EPI_ISL_3018105, EPI_ISL_3018107, EPI_ISL_3018108, EPI_ISL_3018109, EPI_ISL_3018110, EPI_ISL_3018111, EPI_ISL_3018112, EPI_ISL_3018113, EPI_ISL_3018114, EPI_ISL_3018115, EPI_ISL_3018116, EPI_ISL_3018117, EPI_ISL_3018118, EPI_ISL_3018119, EPI_ISL_3018120, EPI_ISL_3018121, EPI_ISL_3018122, EPI_ISL_3018123, EPI_ISL_3018124, EPI_ISL_3018125, EPI_ISL_3018126, EPI_ISL_3018128, EPI_ISL_3018129, EPI_ISL_3018130, EPI_ISL_3018131, EPI_ISL_3018132, EPI_ISL_3018133, EPI_ISL_3018134, EPI_ISL_3018135, EPI_ISL_3018136, EPI_ISL_3018137, EPI_ISL_3018138, EPI_ISL_3018139, EPI_ISL_3018142, EPI_ISL_3018143, EPI_ISL_3018144, EPI_ISL_3018145, EPI_ISL_3018146, EPI_ISL_3018147, EPI_ISL_3018148, EPI_ISL_3018149, EPI_ISL_3018152, EPI_ISL_3018153, EPI_ISL_3018155, EPI_ISL_3018158, EPI_ISL_3018163, EPI_ISL_3018165 | | | |
| see above | Laboratory Corporation of America | Centers for Disease Control and Prevention Division of Viral Diseases, Pathogen Discovery | Dakota Howard, Dhwani Batra, Peter W. Cook, Kara Moser, Adrian Paskey, Jason Caravas, Benjamin Rambo-Martin, Shatavia Morrison, Christopher Gulvick, Scott Sammons, Yvette Unoarumhi, Darlene Wagner, Matthew Schmerer, Minoo Agarwal, Eyad Almasri, Debbie Boles, Ayla Burns, Nuthawin Charoensri, Oren Cohen, Susan Countryman, Mary Ann Cristobal, Bobbi Croy, Suzanne Dale, Hrushikesh Deshmukh, Amanda Douglas, Vincent Drouillon, Marcia Eisenberg, Howard Engler, Rama Ghatti, Prashant Gupta, Susan Hicks, Jake Humphrey, Lax Iyer, Lisa Pfefferle, Manoj Jain, Matthew Robinson, Mohan Kolli, Brian Krueger, Tim Kuphal, Stanley Letovsky, Michael Levandoski, Craig Lukasik, Jonathan Meltzer, Brian Norvell, Mindy Nye, Scott Parker, Christos Petropoulos, John Pruitt, Steven Ragan, Scott Ryan, Mike Sapeta, Jana Schroth, Suresh Babu Selvaraju, Goran Stevovic, Amanda Suchanek, Andrea Throop, Lyndon Tilson, Thomas Urban, Joe Voshell, Kimberly Wagner, Jonathan Williams, Mary Williamson, Qian Zeng, Tricia Zwiefelhofer, Clinton R. Paden, Duncan MacCannell |

| | | | |
|---|---|---|---|
| EPI_ISL_3018166, EPI_ISL_3018167, EPI_ISL_3018168, EPI_ISL_3018169, EPI_ISL_3018170, EPI_ISL_3018171, EPI_ISL_3018172, EPI_ISL_3018173, EPI_ISL_3018174, EPI_ISL_3018175, EPI_ISL_3018176, EPI_ISL_3018177, EPI_ISL_3018178, EPI_ISL_3018179, EPI_ISL_3018180, EPI_ISL_3018181, EPI_ISL_3018183, EPI_ISL_3018184, EPI_ISL_3018185, EPI_ISL_3018186, EPI_ISL_3018187, EPI_ISL_3018188, EPI_ISL_3018189, EPI_ISL_3018190, EPI_ISL_3018191, EPI_ISL_3018192, EPI_ISL_3018193, EPI_ISL_3018194, EPI_ISL_3018195, EPI_ISL_3018196, EPI_ISL_3018197, EPI_ISL_3018198, EPI_ISL_3018199, EPI_ISL_3018200, EPI_ISL_3018201, EPI_ISL_3018202, EPI_ISL_3018203, EPI_ISL_3018204, EPI_ISL_3018205, EPI_ISL_3018206, EPI_ISL_3018207, EPI_ISL_3018208, EPI_ISL_3018209, EPI_ISL_3018210, EPI_ISL_3018211, EPI_ISL_3018212, EPI_ISL_3018213, EPI_ISL_3018214, EPI_ISL_3018215, EPI_ISL_3018216, EPI_ISL_3018217, EPI_ISL_3018218, EPI_ISL_3018220, EPI_ISL_3018221, EPI_ISL_3018222, EPI_ISL_3018223, EPI_ISL_3018224, EPI_ISL_3018225, EPI_ISL_3018226, EPI_ISL_3018227, EPI_ISL_3018228, EPI_ISL_3018229, EPI_ISL_3018230, EPI_ISL_3018231, EPI_ISL_3018233, EPI_ISL_3018235, EPI_ISL_3018237, EPI_ISL_3018238, EPI_ISL_3018239, EPI_ISL_3018240, EPI_ISL_3018242, EPI_ISL_3018243, EPI_ISL_3018244, EPI_ISL_3018245, EPI_ISL_3018246, EPI_ISL_3018247, EPI_ISL_3018248, EPI_ISL_3018249, EPI_ISL_3018251, EPI_ISL_3018253, EPI_ISL_3018254, EPI_ISL_3018256, EPI_ISL_3018257, EPI_ISL_3018258, EPI_ISL_3018259, EPI_ISL_3018260, EPI_ISL_3018261, EPI_ISL_3018262, EPI_ISL_3018263, EPI_ISL_3018264, EPI_ISL_3018265, EPI_ISL_3018267, EPI_ISL_3018268, EPI_ISL_3018269, EPI_ISL_3018270, EPI_ISL_3018271, EPI_ISL_3018272, EPI_ISL_3018273, EPI_ISL_3018276, EPI_ISL_3018277, EPI_ISL_3018278, EPI_ISL_3018279, EPI_ISL_3018280, EPI_ISL_3018281, EPI_ISL_3018282, EPI_ISL_3018283, EPI_ISL_3018284, EPI_ISL_3018285, EPI_ISL_3018286, EPI_ISL_3018287, EPI_ISL_3018288, EPI_ISL_3018291, EPI_ISL_3018292, EPI_ISL_3018294, EPI_ISL_3018295, EPI_ISL_3018296, EPI_ISL_3018297, EPI_ISL_3018298, EPI_ISL_3018299, EPI_ISL_3018300, EPI_ISL_3018301, EPI_ISL_3018302, EPI_ISL_3018303, EPI_ISL_3018304, EPI_ISL_3018305, EPI_ISL_3018306, EPI_ISL_3018308, EPI_ISL_3018309, EPI_ISL_3018310, EPI_ISL_3018311, EPI_ISL_3018312, EPI_ISL_3018313, EPI_ISL_3018314, EPI_ISL_3018315, EPI_ISL_3018316, EPI_ISL_3018317, EPI_ISL_3018319, EPI_ISL_3018320, EPI_ISL_3018321, EPI_ISL_3018322, EPI_ISL_3018323, EPI_ISL_3018325, EPI_ISL_3018328, EPI_ISL_3018329, EPI_ISL_3018330, EPI_ISL_3018331, EPI_ISL_3018332, EPI_ISL_3018333, EPI_ISL_3018334, EPI_ISL_3018335, EPI_ISL_3018336, EPI_ISL_3018337, EPI_ISL_3018339, EPI_ISL_3018340, EPI_ISL_3018341, EPI_ISL_3018342, EPI_ISL_3018343, EPI_ISL_3018345, EPI_ISL_3018346, EPI_ISL_3018347, EPI_ISL_3018348, EPI_ISL_3018349, EPI_ISL_3018350, EPI_ISL_3018351, EPI_ISL_3018352, EPI_ISL_3018353, EPI_ISL_3018354, EPI_ISL_3018355, EPI_ISL_3018357, EPI_ISL_3018358, EPI_ISL_3018359, EPI_ISL_3018360, EPI_ISL_3018362, EPI_ISL_3018363, EPI_ISL_3018364, EPI_ISL_3018365, EPI_ISL_3018366, EPI_ISL_3018367, EPI_ISL_3018368, EPI_ISL_3018369, EPI_ISL_3018370, EPI_ISL_3018372, EPI_ISL_3018373, EPI_ISL_3018375, EPI_ISL_3018376, EPI_ISL_3018378, EPI_ISL_3018379, EPI_ISL_3018381, EPI_ISL_3018383, EPI_ISL_3018384, EPI_ISL_3018385, EPI_ISL_3018387, EPI_ISL_3018389, EPI_ISL_3018390, EPI_ISL_3018392, EPI_ISL_3018393, EPI_ISL_3018394, EPI_ISL_3018395, EPI_ISL_3018396, EPI_ISL_3018397, EPI_ISL_3018398, EPI_ISL_3018400, EPI_ISL_3018401, EPI_ISL_3018402, EPI_ISL_3018403, EPI_ISL_3018405, EPI_ISL_3018406, EPI_ISL_3018407, EPI_ISL_3018409, EPI_ISL_3018412, EPI_ISL_3018413, EPI_ISL_3018415, EPI_ISL_3018416, EPI_ISL_3018418, EPI_ISL_3018419, EPI_ISL_3018420, EPI_ISL_3018421, EPI_ISL_3018422, EPI_ISL_3018423, EPI_ISL_3018424, EPI_ISL_3018425, EPI_ISL_3018426, EPI_ISL_3018427, EPI_ISL_3018429, EPI_ISL_3018430, EPI_ISL_3018431, EPI_ISL_3018432, EPI_ISL_3018433, EPI_ISL_3018435, EPI_ISL_3018436, EPI_ISL_3018438, EPI_ISL_3018439, EPI_ISL_3018441, EPI_ISL_3018442, EPI_ISL_3018443, EPI_ISL_3018444, EPI_ISL_3018445, EPI_ISL_3018446, EPI_ISL_3018447, EPI_ISL_3018448, EPI_ISL_3018450, EPI_ISL_3018451, EPI_ISL_3018452, EPI_ISL_3018453, EPI_ISL_3018454, EPI_ISL_3018456, EPI_ISL_3018457, EPI_ISL_3018459, EPI_ISL_3018460, EPI_ISL_3018461, EPI_ISL_3018462, EPI_ISL_3018463, EPI_ISL_3018464, EPI_ISL_3018465, EPI_ISL_3018466, EPI_ISL_3018467, EPI_ISL_3018468, EPI_ISL_3018469, EPI_ISL_3018471, EPI_ISL_3018472, EPI_ISL_3018473, EPI_ISL_3018474, EPI_ISL_3018475, EPI_ISL_3018476, EPI_ISL_3018477, EPI_ISL_3018478, EPI_ISL_3018479, EPI_ISL_3018480, EPI_ISL_3018482, EPI_ISL_3018484, EPI_ISL_3018485, EPI_ISL_3018486, EPI_ISL_3018487, EPI_ISL_3018488, EPI_ISL_3018489, EPI_ISL_3018490, EPI_ISL_3018491, EPI_ISL_3018492, EPI_ISL_3018495, EPI_ISL_3018496, EPI_ISL_3018497, EPI_ISL_3018498, EPI_ISL_3018499, EPI_ISL_3018502, EPI_ISL_3018504, EPI_ISL_3018505, EPI_ISL_3018506, EPI_ISL_3018507, EPI_ISL_3018511, EPI_ISL_3018513, EPI_ISL_3018514, EPI_ISL_3018515, EPI_ISL_3018516, EPI_ISL_3018517, EPI_ISL_3018518, EPI_ISL_3018519, EPI_ISL_3018520, EPI_ISL_3018521, EPI_ISL_3018523, EPI_ISL_3018524, EPI_ISL_3018525, EPI_ISL_3018526, EPI_ISL_3018527, EPI_ISL_3018528, EPI_ISL_3018529, EPI_ISL_3018530, EPI_ISL_3018531, EPI_ISL_3018532, EPI_ISL_3018533, EPI_ISL_3018534, EPI_ISL_3018535, EPI_ISL_3018536, EPI_ISL_3018538, EPI_ISL_3018539, EPI_ISL_3018540, EPI_ISL_3018541, EPI_ISL_3018542, EPI_ISL_3018543, EPI_ISL_3018544, EPI_ISL_3018545, EPI_ISL_3018546, EPI_ISL_3018547, EPI_ISL_3018548, EPI_ISL_3018549, EPI_ISL_3018552, EPI_ISL_3018554, EPI_ISL_3018555, EPI_ISL_3018556, EPI_ISL_3018557, EPI_ISL_3018558, EPI_ISL_3018561, EPI_ISL_3018562, EPI_ISL_3018564, EPI_ISL_3018565, EPI_ISL_3018566, EPI_ISL_3018567, EPI_ISL_3018568, EPI_ISL_3018569, EPI_ISL_3018570, EPI_ISL_3018571, EPI_ISL_3018572, EPI_ISL_3018573, EPI_ISL_3018574, EPI_ISL_3018576, EPI_ISL_3018577, EPI_ISL_3018578, EPI_ISL_3018579, EPI_ISL_3018580, EPI_ISL_3018581, EPI_ISL_3018582, EPI_ISL_3018584, EPI_ISL_3018585, EPI_ISL_3018586, EPI_ISL_3018587, EPI_ISL_3018588, EPI_ISL_3018589, EPI_ISL_3018590, EPI_ISL_3018591, EPI_ISL_3018593, EPI_ISL_3018594, EPI_ISL_3018595, EPI_ISL_3018596, EPI_ISL_3018598, EPI_ISL_3018601, EPI_ISL_3018602, EPI_ISL_3018603, EPI_ISL_3018604, EPI_ISL_3018606, EPI_ISL_3018607, EPI_ISL_3018608, EPI_ISL_3018609, EPI_ISL_3018610, EPI_ISL_3018611, EPI_ISL_3018612, EPI_ISL_3018613, EPI_ISL_3018614, EPI_ISL_3018615, EPI_ISL_3018616, EPI_ISL_3018617, EPI_ISL_3018618, EPI_ISL_3018619, EPI_ISL_3018620, EPI_ISL_3018621, EPI_ISL_3018622, EPI_ISL_3018623, EPI_ISL_3018624, EPI_ISL_3018625, EPI_ISL_3018626, EPI_ISL_3018627, EPI_ISL_3018628, EPI_ISL_3018629, EPI_ISL_3018631, EPI_ISL_3018632, EPI_ISL_3018633, EPI_ISL_3018634, EPI_ISL_3018635, EPI_ISL_3018636, EPI_ISL_3018638, EPI_ISL_3018639, EPI_ISL_3018640, EPI_ISL_3018641, EPI_ISL_3018642, EPI_ISL_3018643, EPI_ISL_3018644, EPI_ISL_3018645, EPI_ISL_3018647, EPI_ISL_3018648, EPI_ISL_3018649, EPI_ISL_3018652, EPI_ISL_3018654, EPI_ISL_3018655, EPI_ISL_3018656, EPI_ISL_3018657, EPI_ISL_3018662, EPI_ISL_3018663, EPI_ISL_3018665, EPI_ISL_3018666, EPI_ISL_3018667, EPI_ISL_3018668, EPI_ISL_3018669, EPI_ISL_3018671, EPI_ISL_3018672, EPI_ISL_3018674, EPI_ISL_3018676, EPI_ISL_3018677, EPI_ISL_3018679, EPI_ISL_3018680, EPI_ISL_3018681, EPI_ISL_3018682, EPI_ISL_3018684, EPI_ISL_3018685, EPI_ISL_3018686, EPI_ISL_3018687, EPI_ISL_3018688, EPI_ISL_3018689, EPI_ISL_3018690, EPI_ISL_3018691, EPI_ISL_3018692, EPI_ISL_3018694, EPI_ISL_3018695, EPI_ISL_3018697, EPI_ISL_3018698, EPI_ISL_3018699, EPI_ISL_3018700, EPI_ISL_3018701, EPI_ISL_3018702, EPI_ISL_3018704, EPI_ISL_3018706, EPI_ISL_3018707, EPI_ISL_3018708, EPI_ISL_3018709, EPI_ISL_3018710, EPI_ISL_3018711, EPI_ISL_3018714, EPI_ISL_3018715, EPI_ISL_3018716, EPI_ISL_3018717, EPI_ISL_3018718, EPI_ISL_3018719, EPI_ISL_3018720, EPI_ISL_3018721, EPI_ISL_3018723, EPI_ISL_3018724, EPI_ISL_3018725, EPI_ISL_3018726, EPI_ISL_3018727, EPI_ISL_3018728, EPI_ISL_3018729, EPI_ISL_3018730, EPI_ISL_3018731, EPI_ISL_3018732, EPI_ISL_3018734, EPI_ISL_3018735, EPI_ISL_3018736, EPI_ISL_3018738, EPI_ISL_3018739, EPI_ISL_3018740, EPI_ISL_3018741, EPI_ISL_3018742, EPI_ISL_3018743, EPI_ISL_3018744, EPI_ISL_3018746, EPI_ISL_3018748, EPI_ISL_3018749, EPI_ISL_3018750, EPI_ISL_3018751, EPI_ISL_3018752, EPI_ISL_3018753, EPI_ISL_3018755, EPI_ISL_3018756, EPI_ISL_3018758, EPI_ISL_3018759, EPI_ISL_3018760, EPI_ISL_3018761, EPI_ISL_3018762, EPI_ISL_3018763, EPI_ISL_3018764, EPI_ISL_3018765, EPI_ISL_3018766, EPI_ISL_3018767, EPI_ISL_3018768, EPI_ISL_3018769, EPI_ISL_3018770, EPI_ISL_3018771, EPI_ISL_3018772, EPI_ISL_3018773, EPI_ISL_3018774, EPI_ISL_3018775, EPI_ISL_3018776, EPI_ISL_3018777, EPI_ISL_3018779, EPI_ISL_3018780, EPI_ISL_3018781, EPI_ISL_3018782, EPI_ISL_3018783, EPI_ISL_3018784, EPI_ISL_3018785, EPI_ISL_3018786, EPI_ISL_3018787, EPI_ISL_3018788, EPI_ISL_3018789, EPI_ISL_3018791, EPI_ISL_3018793, EPI_ISL_3018794, EPI_ISL_3018795, EPI_ISL_3018796, EPI_ISL_3018797, EPI_ISL_3018798, EPI_ISL_3018799, EPI_ISL_3018800, EPI_ISL_3018801, EPI_ISL_3018802, EPI_ISL_3018803, EPI_ISL_3018804, EPI_ISL_3018805, EPI_ISL_3018807, EPI_ISL_3018808, EPI_ISL_3018809, EPI_ISL_3018810, EPI_ISL_3018811, EPI_ISL_3018812, EPI_ISL_3018813, EPI_ISL_3018814, EPI_ISL_3018817, EPI_ISL_3018818, EPI_ISL_3018819, EPI_ISL_3018821, EPI_ISL_3018822, EPI_ISL_3018823, EPI_ISL_3018825, EPI_ISL_3018826, EPI_ISL_3018827, EPI_ISL_3018828, EPI_ISL_3018829, EPI_ISL_3018830, EPI_ISL_3018831, EPI_ISL_3018832, EPI_ISL_3018833, EPI_ISL_3018834, EPI_ISL_3018835, EPI_ISL_3018836, EPI_ISL_3018837, EPI_ISL_3018839, EPI_ISL_3018841, EPI_ISL_3018842, EPI_ISL_3018843, EPI_ISL_3018845, EPI_ISL_3018846, EPI_ISL_3018847, EPI_ISL_3018848, EPI_ISL_3018850, EPI_ISL_3018853, EPI_ISL_3018854, EPI_ISL_3018855, EPI_ISL_3018856, EPI_ISL_3018857, EPI_ISL_3018858, EPI_ISL_3018862, EPI_ISL_3018863, EPI_ISL_3018864, EPI_ISL_3018865, EPI_ISL_3018866, EPI_ISL_3018867, EPI_ISL_3018868, EPI_ISL_3018869, EPI_ISL_3018870, EPI_ISL_3018871, EPI_ISL_3018873, EPI_ISL_3018874, EPI_ISL_3018875, EPI_ISL_3018877, EPI_ISL_3018878, EPI_ISL_3018880, EPI_ISL_3018881, EPI_ISL_3018882, EPI_ISL_3018884, EPI_ISL_3018885, EPI_ISL_3018886, EPI_ISL_3018887, EPI_ISL_3018888, EPI_ISL_3018889, EPI_ISL_3018890, EPI_ISL_3018891, EPI_ISL_3018892, EPI_ISL_3018893, EPI_ISL_3018894, EPI_ISL_3018895, EPI_ISL_3018896, EPI_ISL_3018897, EPI_ISL_3018898, EPI_ISL_3018900, EPI_ISL_3018901, EPI_ISL_3018902, EPI_ISL_3018903, EPI_ISL_3018905, EPI_ISL_3018906, EPI_ISL_3018907, EPI_ISL_3018908, EPI_ISL_3018909, EPI_ISL_3018911, EPI_ISL_3018912, EPI_ISL_3018913, EPI_ISL_3018914, EPI_ISL_3018915, EPI_ISL_3018916, EPI_ISL_3018917, EPI_ISL_3018918, EPI_ISL_3018919, EPI_ISL_3018920, EPI_ISL_3018921, EPI_ISL_3018922, EPI_ISL_3018923, EPI_ISL_3018924, EPI_ISL_3018925, EPI_ISL_3018927, EPI_ISL_3018928, EPI_ISL_3018929 | | | |
| see above | Quest Diagnostics Incorporated | Centers for Disease Control and Prevention Division of Viral Diseases, Pathogen Discovery | Dakota Howard, Dhwani Batra, Peter W. Cook, Kara Moser, Adrian Paskey, Jason Caravas, Benjamin Rambo-Martin, Shatavia Morrison, Christopher Gulvick, Scott Sammons, Yvette Unoarumhi, Darlene Wagner, Matthew Schmerer, S. H. Rosenthal, A. Gerasimova, R. M. Kagan, B. Anderson, M. Hua, Y. Liu, L.E. Bernstein, K.E. Livingston, A. Perez, I. A. Shlyakhter, R. V. Rolando, R. Owen, P. Tanpaiboon, F. Lacbawan, Clinton R. Paden, Duncan MacCannell |
| EPI_ISL_3018930, EPI_ISL_3018931, EPI_ISL_3018932, EPI_ISL_3018933, EPI_ISL_3018934, EPI_ISL_3018936, EPI_ISL_3018937, EPI_ISL_3018938, EPI_ISL_3018939, EPI_ISL_3018940, EPI_ISL_3018942, EPI_ISL_3018943, EPI_ISL_3018944, EPI_ISL_3018946, EPI_ISL_3018947, EPI_ISL_3018948, EPI_ISL_3018949, EPI_ISL_3018950, EPI_ISL_3018951, EPI_ISL_3018952, EPI_ISL_3018953, EPI_ISL_3018954, EPI_ISL_3018955, EPI_ISL_3018956, EPI_ISL_3018957, EPI_ISL_3018958, EPI_ISL_3018959, EPI_ISL_3018960, EPI_ISL_3018961, EPI_ISL_3018962, EPI_ISL_3018963, EPI_ISL_3018964 | | | |
| see above | Laboratorio di Microbiologia | Laboratorio di Microbiologia | Martinetti Lucchini Gladys, Valeria Spina |
| EPI_ISL_3018965, EPI_ISL_3018966, EPI_ISL_3018967, EPI_ISL_3018968, EPI_ISL_3018969, EPI_ISL_3018970, EPI_ISL_3018971, EPI_ISL_3018972, EPI_ISL_3018973, EPI_ISL_3018974, EPI_ISL_3018975, EPI_ISL_3018976, EPI_ISL_3018977, EPI_ISL_3018978, EPI_ISL_3018979, EPI_ISL_3018981, EPI_ISL_3018982, EPI_ISL_3018983, EPI_ISL_3018984, EPI_ISL_3018985, EPI_ISL_3018986, EPI_ISL_3018987, EPI_ISL_3018988, EPI_ISL_3018989, EPI_ISL_3018990, EPI_ISL_3018991, EPI_ISL_3018992, EPI_ISL_3018993, EPI_ISL_3018994, EPI_ISL_3018995, EPI_ISL_3018996, EPI_ISL_3018997, EPI_ISL_3018998, EPI_ISL_3018999, EPI_ISL_3019000, EPI_ISL_3019001, EPI_ISL_3019002, EPI_ISL_3019003, EPI_ISL_3019004, EPI_ISL_3019005, EPI_ISL_3019006, EPI_ISL_3019007, EPI_ISL_3019008, EPI_ISL_3019009, EPI_ISL_3019010, EPI_ISL_3019011, EPI_ISL_3019012, EPI_ISL_3019013, EPI_ISL_3019014, EPI_ISL_3019015, EPI_ISL_3019016, EPI_ISL_3019017, EPI_ISL_3019018, EPI_ISL_3019019, EPI_ISL_3019020, EPI_ISL_3019021, EPI_ISL_3019022, EPI_ISL_3019023, EPI_ISL_3019024, EPI_ISL_3019025, EPI_ISL_3019026, EPI_ISL_3019027, EPI_ISL_3019028, EPI_ISL_3019029, EPI_ISL_3019030, EPI_ISL_3019031, EPI_ISL_3019032, EPI_ISL_3019033, EPI_ISL_3019034, EPI_ISL_3019035, EPI_ISL_3019036, EPI_ISL_3019037, EPI_ISL_3019038, EPI_ISL_3019039, EPI_ISL_3019040, EPI_ISL_3019041, EPI_ISL_3019042, EPI_ISL_3019043, EPI_ISL_3019044, EPI_ISL_3019045, EPI_ISL_3019046, EPI_ISL_3019047, EPI_ISL_3019048, EPI_ISL_3019049, EPI_ISL_3019050, EPI_ISL_3019051, EPI_ISL_3019052, EPI_ISL_3019053, EPI_ISL_3019054, EPI_ISL_3019055, EPI_ISL_3019056, EPI_ISL_3019057, EPI_ISL_3019058, EPI_ISL_3019059, EPI_ISL_3019060 | | | |
| see above | INSACOG-Sikkim | National Institute of Biomedical Genomics – INSACOG | Arindam Maitra, Tara D Sharma, Nidhan Kumar Biswas, Kaden Zangmu Bhutia, Sreedhar Chinnaswamy Tshering Ongmu Bhutia, Saumitra Das |
| EPI_ISL_3019061, EPI_ISL_3019062, EPI_ISL_3019063, EPI_ISL_3019064, EPI_ISL_3019065, EPI_ISL_3019066, EPI_ISL_3019067, EPI_ISL_3019068, EPI_ISL_3019069, EPI_ISL_3019070, EPI_ISL_3019071, EPI_ISL_3019074, EPI_ISL_3019075, EPI_ISL_3019076, EPI_ISL_3019077, EPI_ISL_3019078, EPI_ISL_3019079, EPI_ISL_3019081, EPI_ISL_3019082, EPI_ISL_3019083, EPI_ISL_3019084, EPI_ISL_3019086, EPI_ISL_3019087, EPI_ISL_3019088, EPI_ISL_3019089, EPI_ISL_3019090, EPI_ISL_3019091, EPI_ISL_3019092, EPI_ISL_3019093, EPI_ISL_3019094, EPI_ISL_3019095, EPI_ISL_3019096, EPI_ISL_3019098, EPI_ISL_3019099, EPI_ISL_3019100, EPI_ISL_3019101, EPI_ISL_3019102, EPI_ISL_3019103, EPI_ISL_3019104, EPI_ISL_3019105, EPI_ISL_3019107, EPI_ISL_3019108, EPI_ISL_3019109, EPI_ISL_3019110, EPI_ISL_3019112, EPI_ISL_3019113, EPI_ISL_3019114, EPI_ISL_3019115 | | | |
| see above | INSACOG-Mizoram | National Institute of Biomedical Genomics – INSACOG | Arindam Maitra, Swagnik Roy, Nidhan Kumar Biswas, Gracy Laldinmawii, Sreedhar Chinnaswamy, N Senthil Kumar, Saumitra Das |
| EPI_ISL_3019116, EPI_ISL_3019117, EPI_ISL_3019118, EPI_ISL_3019119, EPI_ISL_3019120, EPI_ISL_3019121, EPI_ISL_3019122, EPI_ISL_3019123, EPI_ISL_3019124, EPI_ISL_3019125, EPI_ISL_3019126, EPI_ISL_3019127, EPI_ISL_3019128, EPI_ISL_3019129, EPI_ISL_3019130, EPI_ISL_3019131, EPI_ISL_3019132, EPI_ISL_3019133, EPI_ISL_3019134, EPI_ISL_3019135, EPI_ISL_3019136, EPI_ISL_3019137, EPI_ISL_3019138, EPI_ISL_3019139, EPI_ISL_3019140, EPI_ISL_3019141, EPI_ISL_3019142, EPI_ISL_3019143, EPI_ISL_3019144, EPI_ISL_3019145, EPI_ISL_3019146, EPI_ISL_3019147, EPI_ISL_3019148, EPI_ISL_3019149, EPI_ISL_3019150, EPI_ISL_3019151, EPI_ISL_3019152, EPI_ISL_3019153, EPI_ISL_3019154, EPI_ISL_3019155, EPI_ISL_3019156, EPI_ISL_3019157, EPI_ISL_3019158, EPI_ISL_3019159, EPI_ISL_3019160, EPI_ISL_3019161, EPI_ISL_3019162, EPI_ISL_3019163, EPI_ISL_3019164, EPI_ISL_3019165, EPI_ISL_3019166, EPI_ISL_3019167, EPI_ISL_3019168, EPI_ISL_3019169, EPI_ISL_3019170, EPI_ISL_3019171, EPI_ISL_3019172, EPI_ISL_3019173, EPI_ISL_3019174, EPI_ISL_3019175, EPI_ISL_3019176, EPI_ISL_3019177, EPI_ISL_3019178, EPI_ISL_3019179, EPI_ISL_3019180, EPI_ISL_3019181, EPI_ISL_3019182, EPI_ISL_3019183 | | | |
| see above | INSACOG-MEGHALAYA | National Institute of Biomedical Genomics – INSACOG | Arindam Maitra, R. Lyngdoh, Nidhan Kumar Biswas, E Shadap, Sreedhar Chinnaswamy, E Marak, Saumitra Das |
| EPI_ISL_3019184, EPI_ISL_3019185, EPI_ISL_3019186, EPI_ISL_3019187, EPI_ISL_3019188, EPI_ISL_3019189, EPI_ISL_3019190, EPI_ISL_3019191, EPI_ISL_3019192, EPI_ISL_3019193, EPI_ISL_3019194, EPI_ISL_3019195, EPI_ISL_3019196, EPI_ISL_3019197, EPI_ISL_3019198, EPI_ISL_3019199, EPI_ISL_3019200, EPI_ISL_3019201, EPI_ISL_3019202, EPI_ISL_3019203, EPI_ISL_3019204, EPI_ISL_3019205, EPI_ISL_3019206, EPI_ISL_3019207, EPI_ISL_3019208, EPI_ISL_3019209, EPI_ISL_3019210, EPI_ISL_3019211, EPI_ISL_3019212, EPI_ISL_3019213, EPI_ISL_3019214, EPI_ISL_3019215, EPI_ISL_3019216, EPI_ISL_3019217, EPI_ISL_3019218, EPI_ISL_3019219, EPI_ISL_3019220, EPI_ISL_3019221, EPI_ISL_3019222, EPI_ISL_3019223, EPI_ISL_3019224, EPI_ISL_3019225, EPI_ISL_3019226, EPI_ISL_3019227, EPI_ISL_3019228, EPI_ISL_3019229, EPI_ISL_3019230, EPI_ISL_3019231, EPI_ISL_3019232, EPI_ISL_3019233, EPI_ISL_3019234, EPI_ISL_3019235, EPI_ISL_3019236, EPI_ISL_3019237, EPI_ISL_3019238, EPI_ISL_3019239, EPI_ISL_3019240, EPI_ISL_3019241, EPI_ISL_3019242, EPI_ISL_3019243, EPI_ISL_3019244, EPI_ISL_3019245, EPI_ISL_3019246, EPI_ISL_3019247, EPI_ISL_3019248, EPI_ISL_3019249, EPI_ISL_3019250, EPI_ISL_3019251, EPI_ISL_3019252, EPI_ISL_3019253, EPI_ISL_3019254, EPI_ISL_3019255, EPI_ISL_3019256, EPI_ISL_3019257, EPI_ISL_3019258, EPI_ISL_3019259, EPI_ISL_3019260, EPI_ISL_3019261, EPI_ISL_3019262, EPI_ISL_3019263, EPI_ISL_3019264, EPI_ISL_3019265, EPI_ISL_3019266, EPI_ISL_3019267, EPI_ISL_3019268, EPI_ISL_3019269, EPI_ISL_3019270, EPI_ISL_3019271, EPI_ISL_3019273, EPI_ISL_3019274, EPI_ISL_3019275, EPI_ISL_3019276, EPI_ISL_3019277, EPI_ISL_3019279, EPI_ISL_3019280, EPI_ISL_3019281, EPI_ISL_3019282, EPI_ISL_3019283, EPI_ISL_3019284, EPI_ISL_3019285, EPI_ISL_3019286, EPI_ISL_3019287, EPI_ISL_3019288, EPI_ISL_3019289, EPI_ISL_3019290, EPI_ISL_3019291, EPI_ISL_3019292, EPI_ISL_3019293, EPI_ISL_3019294, EPI_ISL_3019295, EPI_ISL_3019296, EPI_ISL_3019297, EPI_ISL_3019298, EPI_ISL_3019299, EPI_ISL_3019300, EPI_ISL_3019301, EPI_ISL_3019302, EPI_ISL_3019303, EPI_ISL_3019305, EPI_ISL_3019306, EPI_ISL_3019307, EPI_ISL_3019308, EPI_ISL_3019309, EPI_ISL_3019310, EPI_ISL_3019311, EPI_ISL_3019312, EPI_ISL_3019313, EPI_ISL_3019314, EPI_ISL_3019316, EPI_ISL_3019317, EPI_ISL_3019318, EPI_ISL_3019319, EPI_ISL_3019320, EPI_ISL_3019321, EPI_ISL_3019322, EPI_ISL_3019323, EPI_ISL_3019324, EPI_ISL_3019325, EPI_ISL_3019326, EPI_ISL_3019327, EPI_ISL_3019328, EPI_ISL_3019329, EPI_ISL_3019330, EPI_ISL_3019331, EPI_ISL_3019332, EPI_ISL_3019333, EPI_ISL_3019334, EPI_ISL_3019335, EPI_ISL_3019336, EPI_ISL_3019337, EPI_ISL_3019338, EPI_ISL_3019339, EPI_ISL_3019340, EPI_ISL_3019341, EPI_ISL_3019342, EPI_ISL_3019343, EPI_ISL_3019344, EPI_ISL_3019345, EPI_ISL_3019346, EPI_ISL_3019347, EPI_ISL_3019348, EPI_ISL_3019349, EPI_ISL_3019350, EPI_ISL_3019351, EPI_ISL_3019352, EPI_ISL_3019353, EPI_ISL_3019354, EPI_ISL_3019355, EPI_ISL_3019356, EPI_ISL_3019357, EPI_ISL_3019358, EPI_ISL_3019359, EPI_ISL_3019360, EPI_ISL_3019361, EPI_ISL_3019362, EPI_ISL_3019363, EPI_ISL_3019364, EPI_ISL_3019365, EPI_ISL_3019366, EPI_ISL_3019367, EPI_ISL_3019368, EPI_ISL_3019369, EPI_ISL_3019370, EPI_ISL_3019371, EPI_ISL_3019372, EPI_ISL_3019373, EPI_ISL_3019374, EPI_ISL_3019375, EPI_ISL_3019376, EPI_ISL_3019377, EPI_ISL_3019378, EPI_ISL_3019379, EPI_ISL_3019380, EPI_ISL_3019381, EPI_ISL_3019382, EPI_ISL_3019383, EPI_ISL_3019384, EPI_ISL_3019385, EPI_ISL_3019386, EPI_ISL_3019387, EPI_ISL_3019388, EPI_ISL_3019389, EPI_ISL_3019390, EPI_ISL_3019391, EPI_ISL_3019392, EPI_ISL_3019393, EPI_ISL_3019394, EPI_ISL_3019395, EPI_ISL_3019396, EPI_ISL_3019397, EPI_ISL_3019398, EPI_ISL_3019399, EPI_ISL_3019400, EPI_ISL_3019401, EPI_ISL_3019402, EPI_ISL_3019403, EPI_ISL_3019404, EPI_ISL_3019405, EPI_ISL_3019406, EPI_ISL_3019407, EPI_ISL_3019408, EPI_ISL_3019409, EPI_ISL_3019410, EPI_ISL_3019411, EPI_ISL_3019412, EPI_ISL_3019413, EPI_ISL_3019414, EPI_ISL_3019415, EPI_ISL_3019416, EPI_ISL_3019417, EPI_ISL_3019418, EPI_ISL_3019419, EPI_ISL_3019420, EPI_ISL_3019421, EPI_ISL_3019422, EPI_ISL_3019423, EPI_ISL_3019424, EPI_ISL_3019425, EPI_ISL_3019426, EPI_ISL_3019427, EPI_ISL_3019428, EPI_ISL_3019429, EPI_ISL_3019430, EPI_ISL_3019431, EPI_ISL_3019432, EPI_ISL_3019433, EPI_ISL_3019434, EPI_ISL_3019435, EPI_ISL_3019436, EPI_ISL_3019437, EPI_ISL_3019438, EPI_ISL_3019439, EPI_ISL_3019440, EPI_ISL_3019441, EPI_ISL_3019442, EPI_ISL_3019443, EPI_ISL_3019444, EPI_ISL_3019445, EPI_ISL_3019446, EPI_ISL_3019447, EPI_ISL_3019448, EPI_ISL_3019449, EPI_ISL_3019450, EPI_ISL_3019451, EPI_ISL_3019453, EPI_ISL_3019455, EPI_ISL_3019456, EPI_ISL_3019457, EPI_ISL_3019459, EPI_ISL_3019460, EPI_ISL_3019461, EPI_ISL_3019462, EPI_ISL_3019463, EPI_ISL_3019464, EPI_ISL_3019465, EPI_ISL_3019466, EPI_ISL_3019467, EPI_ISL_3019468, EPI_ISL_3019469, EPI_ISL_3019470, EPI_ISL_3019471, EPI_ISL_3019472, EPI_ISL_3019473 | | | |
| see above | INSACOG-WB | National Institute of Biomedical Genomics – INSACOG | Arindam Maitra, Bhaswati Bandyopadhyay, Nidhan Kumar Biswas, Tamal Ghosh, Sreedhar Chinnaswamy, Ajay Chakraborti, Saumitra Das |
| EPI_ISL_3019474 | QUARANTINE CAMP | Hong Kong Department of Health | Alan K.L. Tsang, Peter C.W. Yip, Ken H.L. Ng, Edman T.K. Lam, Rickjason C.W. Chan |
| EPI_ISL_3019475, EPI_ISL_3019476, EPI_ISL_3019477, EPI_ISL_3019478, EPI_ISL_3019479, EPI_ISL_3019480, EPI_ISL_3019481, EPI_ISL_3019482, EPI_ISL_3019483, EPI_ISL_3019484, EPI_ISL_3019485, EPI_ISL_3019486 | | | |
| see above | Temporary Specimen Collection Centre at the AsiaWorld-Expo | Hong Kong Department of Health | Alan K.L. Tsang, Peter C.W. Yip, Ken H.L. Ng, Edman T.K. Lam, Rickjason C.W. Chan |
| EPI_ISL_3019487 | HOME QUARANTINE TASKFORCE | Hong Kong Department of Health | Alan K.L. Tsang, Peter C.W. Yip, Ken H.L. Ng, Edman T.K. Lam, Rickjason C.W. Chan |
| EPI_ISL_3019488 | TUEN MUN HOSPITAL | Hong Kong Department of Health | Alan K.L. Tsang, Peter C.W. Yip, Ken H.L. Ng, Edman T.K. Lam, Rickjason C.W. Chan |
| EPI_ISL_3019489 | North Lantau Hospital | Hong Kong Department of Health | Alan K.L. Tsang, Peter C.W. Yip, Ken H.L. Ng, Edman T.K. Lam, Rickjason C.W. Chan |
| EPI_ISL_3019490, EPI_ISL_3019492 | HOME QUARANTINE TASKFORCE | Hong Kong Department of Health | Alan K.L. Tsang, Peter C.W. Yip, Ken H.L. Ng, Edman T.K. Lam, Rickjason C.W. Chan |
| EPI_ISL_3019493 | Queen Mary Hospital | Hong Kong Department of Health | Alan K.L. Tsang, Peter C.W. Yip, Ken H.L. Ng, Edman T.K. Lam, Rickjason C.W. Chan |
| EPI_ISL_3019494 | North Lantau Hospital | Hong Kong Department of Health | Alan K.L. Tsang, Peter C.W. Yip, Ken H.L. Ng, Edman T.K. Lam, Rickjason C.W. Chan |
| EPI_ISL_3019495 | COMMUNITY TESTING CENTRE | Hong Kong Department of Health | Alan K.L. Tsang, Peter C.W. Yip, Ken H.L. Ng, Edman T.K. Lam, Rickjason C.W. Chan |
| EPI_ISL_3019496 | HOME QUARANTINE TASKFORCE | Hong Kong Department of Health | Alan K.L. Tsang, Peter C.W. Yip, Ken H.L. Ng, Edman T.K. Lam, Rickjason C.W. Chan |
| EPI_ISL_3019501, EPI_ISL_3019502, EPI_ISL_3019503, EPI_ISL_3019508, EPI_ISL_3019510, EPI_ISL_3019511, EPI_ISL_3019517, EPI_ISL_3019521, EPI_ISL_3019524, EPI_ISL_3019526, EPI_ISL_3019528, EPI_ISL_3019532, EPI_ISL_3019534, EPI_ISL_3019536, EPI_ISL_3019537, EPI_ISL_3019540, EPI_ISL_3019541, EPI_ISL_3019542, EPI_ISL_3019543, EPI_ISL_3019544 | | | |
| see above | Centro de Investigación Biomédica de La Rioja - Hospital San Pedro Logroño | SeqCOVID-SPAIN consortium/IBV(CSIC) | María de Toro, José Manuel Azcona Gutiérrez, María Pilar Bea Escudero, Miriam Blasco Alberdi and SeqCOVID-SPAIN consortium |
| EPI_ISL_3019546, EPI_ISL_3019547, EPI_ISL_3019549, EPI_ISL_3019550, EPI_ISL_3019551, EPI_ISL_3019552, EPI_ISL_3019553, EPI_ISL_3019554, EPI_ISL_3019555, EPI_ISL_3019557, EPI_ISL_3019558, EPI_ISL_3019559, EPI_ISL_3019560, EPI_ISL_3019561, EPI_ISL_3019562, EPI_ISL_3019564, EPI_ISL_3019566, EPI_ISL_3019567, EPI_ISL_3019568, EPI_ISL_3019569, EPI_ISL_3019570, EPI_ISL_3019571, EPI_ISL_3019572, EPI_ISL_3019573, EPI_ISL_3019574, EPI_ISL_3019575, EPI_ISL_3019576, EPI_ISL_3019577, EPI_ISL_3019578, EPI_ISL_3019579, EPI_ISL_3019580, EPI_ISL_3019581, EPI_ISL_3019582, EPI_ISL_3019583, EPI_ISL_3019584, EPI_ISL_3019585, EPI_ISL_3019586, EPI_ISL_3019587, EPI_ISL_3019588, EPI_ISL_3019589, EPI_ISL_3019590, EPI_ISL_3019591, EPI_ISL_3019592, EPI_ISL_3019593, EPI_ISL_3019594, EPI_ISL_3019595, EPI_ISL_3019596, EPI_ISL_3019597, EPI_ISL_3019599, EPI_ISL_3019600, EPI_ISL_3019601, EPI_ISL_3019602, EPI_ISL_3019603, EPI_ISL_3019605, EPI_ISL_3019606, EPI_ISL_3019607, EPI_ISL_3019608, EPI_ISL_3019609, EPI_ISL_3019611, EPI_ISL_3019612, EPI_ISL_3019614, EPI_ISL_3019615, EPI_ISL_3019616, EPI_ISL_3019617, EPI_ISL_3019618, EPI_ISL_3019619, EPI_ISL_3019620, EPI_ISL_3019621, EPI_ISL_3019622, EPI_ISL_3019623, EPI_ISL_3019624, EPI_ISL_3019625, EPI_ISL_3019626, EPI_ISL_3019627, EPI_ISL_3019628, EPI_ISL_3019629, EPI_ISL_3019630, EPI_ISL_3019632, EPI_ISL_3019633, EPI_ISL_3019634, EPI_ISL_3019635, EPI_ISL_3019636, EPI_ISL_3019637, EPI_ISL_3019638, EPI_ISL_3019639, EPI_ISL_3019640, EPI_ISL_3019641, EPI_ISL_3019643, EPI_ISL_3019644, EPI_ISL_3019645, EPI_ISL_3019646, EPI_ISL_3019647, EPI_ISL_3019648, EPI_ISL_3019649, EPI_ISL_3019650, EPI_ISL_3019651, EPI_ISL_3019652, EPI_ISL_3019653, EPI_ISL_3019654, EPI_ISL_3019655, EPI_ISL_3019658, EPI_ISL_3019659, EPI_ISL_3019660, EPI_ISL_3019661, EPI_ISL_3019662, EPI_ISL_3019663, EPI_ISL_3019664, EPI_ISL_3019665, EPI_ISL_3019666, EPI_ISL_3019667, EPI_ISL_3019668, EPI_ISL_3019669, EPI_ISL_3019670, EPI_ISL_3019671, EPI_ISL_3019672, EPI_ISL_3019673, EPI_ISL_3019674, EPI_ISL_3019675, EPI_ISL_3019676, EPI_ISL_3019677, EPI_ISL_3019678, EPI_ISL_3019681, EPI_ISL_3019683, EPI_ISL_3019685, EPI_ISL_3019686, EPI_ISL_3019687, EPI_ISL_3019688, EPI_ISL_3019689, EPI_ISL_3019690, EPI_ISL_3019691, EPI_ISL_3019692, EPI_ISL_3019693, EPI_ISL_3019695, EPI_ISL_3019696, EPI_ISL_3019697, EPI_ISL_3019698, EPI_ISL_3019699, EPI_ISL_3019701, EPI_ISL_3019702, EPI_ISL_3019704, EPI_ISL_3019705, EPI_ISL_3019706, EPI_ISL_3019707, EPI_ISL_3019708, EPI_ISL_3019709, EPI_ISL_3019710, EPI_ISL_3019712, EPI_ISL_3019713, EPI_ISL_3019714, EPI_ISL_3019715, EPI_ISL_3019716, EPI_ISL_3019717, EPI_ISL_3019720, EPI_ISL_3019721, EPI_ISL_3019722, EPI_ISL_3019723, EPI_ISL_3019724, EPI_ISL_3019725, EPI_ISL_3019727, EPI_ISL_3019728, EPI_ISL_3019729, EPI_ISL_3019731, EPI_ISL_3019732, EPI_ISL_3019733, EPI_ISL_3019734, EPI_ISL_3019735, EPI_ISL_3019736, EPI_ISL_3019737, EPI_ISL_3019738, EPI_ISL_3019739, EPI_ISL_3019740, EPI_ISL_3019741, EPI_ISL_3019743, EPI_ISL_3019744, EPI_ISL_3019745, EPI_ISL_3019746, EPI_ISL_3019747, EPI_ISL_3019748, EPI_ISL_3019750, EPI_ISL_3019751, EPI_ISL_3019752, EPI_ISL_3019753, EPI_ISL_3019754, EPI_ISL_3019755, EPI_ISL_3019756, EPI_ISL_3019757, EPI_ISL_3019758, EPI_ISL_3019759, EPI_ISL_3019760, EPI_ISL_3019762, EPI_ISL_3019764, EPI_ISL_3019765, EPI_ISL_3019766, EPI_ISL_3019767, EPI_ISL_3019768, EPI_ISL_3019769, EPI_ISL_3019770, EPI_ISL_3019771, EPI_ISL_3019772, EPI_ISL_3019773, EPI_ISL_3019774, EPI_ISL_3019775, EPI_ISL_3019776, EPI_ISL_3019777, EPI_ISL_3019778, EPI_ISL_3019779, EPI_ISL_3019780, EPI_ISL_3019782, EPI_ISL_3019783, EPI_ISL_3019784, EPI_ISL_3019788, EPI_ISL_3019789, EPI_ISL_3019792, EPI_ISL_3019793, EPI_ISL_3019794, EPI_ISL_3019798, EPI_ISL_3019799, EPI_ISL_3019800, EPI_ISL_3019801, EPI_ISL_3019802, EPI_ISL_3019803, EPI_ISL_3019804, EPI_ISL_3019805, EPI_ISL_3019806, EPI_ISL_3019807, EPI_ISL_3019808, EPI_ISL_3019809, EPI_ISL_3019810, EPI_ISL_3019812, EPI_ISL_3019813, EPI_ISL_3019814, EPI_ISL_3019815, EPI_ISL_3019816, EPI_ISL_3019817, EPI_ISL_3019818, EPI_ISL_3019819, EPI_ISL_3019820, EPI_ISL_3019821, EPI_ISL_3019822, EPI_ISL_3019823, EPI_ISL_3019824, EPI_ISL_3019825, EPI_ISL_3019826, EPI_ISL_3019827, EPI_ISL_3019828, EPI_ISL_3019829, EPI_ISL_3019832, EPI_ISL_3019833, EPI_ISL_3019834, EPI_ISL_3019836, EPI_ISL_3019837, EPI_ISL_3019838, EPI_ISL_3019839, EPI_ISL_3019840, EPI_ISL_3019842, EPI_ISL_3019843, EPI_ISL_3019844, EPI_ISL_3019845, EPI_ISL_3019847, EPI_ISL_3019848, EPI_ISL_3019850, EPI_ISL_3019852, EPI_ISL_3019853, EPI_ISL_3019854, EPI_ISL_3019856, EPI_ISL_3019857, EPI_ISL_3019858, EPI_ISL_3019861, EPI_ISL_3019862, EPI_ISL_3019863, EPI_ISL_3019864, EPI_ISL_3019865, EPI_ISL_3019867, EPI_ISL_3019868, EPI_ISL_3019869, EPI_ISL_3019870, EPI_ISL_3019871, EPI_ISL_3019872, EPI_ISL_3019874, EPI_ISL_3019876, EPI_ISL_3019877, EPI_ISL_3019878, EPI_ISL_3019879, EPI_ISL_3019881, EPI_ISL_3019882, EPI_ISL_3019883, EPI_ISL_3019884, EPI_ISL_3019885, EPI_ISL_3019886, EPI_ISL_3019889, EPI_ISL_3019890, EPI_ISL_3019891, EPI_ISL_3019892, EPI_ISL_3019893, EPI_ISL_3019894, EPI_ISL_3019895, EPI_ISL_3019896, EPI_ISL_3019897, EPI_ISL_3019898, EPI_ISL_3019899, EPI_ISL_3019900, EPI_ISL_3019901, EPI_ISL_3019902, EPI_ISL_3019904, EPI_ISL_3019905, EPI_ISL_3019906, EPI_ISL_3019907, EPI_ISL_3019908, EPI_ISL_3019910, EPI_ISL_3019911 | | | |
| see above | Shamir Medical Center (Asaf Harofe) | Shamir Medical Center (Asaf Harofe) | Nir Rainy,Chen Weiner,Reut Sorek Abramovich,Yevgeni Yegorov,Anna Vishnevsky,Patricia Benveniste-Lekovitz,Abu Hamad Ramzia,Adina Bar Chaim |
| EPI_ISL_3019913 | Centracare Laboratory Services | Minnesota Department of Health, Public Health Laboratory | Alexandra Lorentz, Jacob Garfin, Matt Plumb, and Xiong Wang |
| EPI_ISL_3019914, EPI_ISL_3019915, | Minnesota Department of Health, Public Health Laboratory | Minnesota Department of Health, Public Health Laboratory | Alexandra Lorentz, Jacob Garfin, Matt Plumb, and Xiong Wang |

| | | | |
|---|---|---|---|
| EPI_ISL_3019917, EPI_ISL_3019918, EPI_ISL_3019919, EPI_ISL_3019920, EPI_ISL_3019921, EPI_ISL_3019922, EPI_ISL_3019923, EPI_ISL_3019925 | | | |
| EPI_ISL_3019926 | Centracare Laboratory Services | Minnesota Department of Health, Public Health Laboratory | Alexandra Lorentz, Jacob Garfin, Matt Plumb, and Xiong Wang |
| EPI_ISL_3019927, EPI_ISL_3019928, EPI_ISL_3019929, EPI_ISL_3019930, EPI_ISL_3019932, EPI_ISL_3019933, EPI_ISL_3019934, EPI_ISL_3019935, EPI_ISL_3019936, EPI_ISL_3019937 | Minnesota Department of Health, Public Health Laboratory | Minnesota Department of Health, Public Health Laboratory | Alexandra Lorentz, Jacob Garfin, Matt Plumb, and Xiong Wang |
| EPI_ISL_3019945 | RSUD Dr. W.Z. Johanes kupang | National Institute of Health Research and Development | Hana Apsari Pawestri, Subangkit, Arie Ardiansyah Nugraha, Kartika Dewi Puspa, Hartanti Dian Ikawati, Krisna Pangesti, Triyani Soekarso, Fajar Nur Sulistiyohadi, Nelly Puspandari, Vivi Setiawaty |
| EPI_ISL_3019972, EPI_ISL_3019973 | Mayo Clinic & Mayo Clinic Laboratories | Minnesota Department of Health, Public Health Laboratory | Alexandra Lorentz, Jacob Garfin, Matt Plumb, and Xiong Wang |
| EPI_ISL_3019974 | Minnesota Department of Health, Public Health Laboratory | Minnesota Department of Health, Public Health Laboratory | Alexandra Lorentz, Jacob Garfin, Matt Plumb, and Xiong Wang |
| EPI_ISL_3019977, EPI_ISL_3019981, EPI_ISL_3019986, EPI_ISL_3019988 | Mayo Clinic & Mayo Clinic Laboratories | Minnesota Department of Health, Public Health Laboratory | Alexandra Lorentz, Jacob Garfin, Matt Plumb, and Xiong Wang |
| EPI_ISL_3020010, EPI_ISL_3020011, EPI_ISL_3020015, EPI_ISL_3020019, EPI_ISL_3020020, EPI_ISL_3020023, EPI_ISL_3020024, EPI_ISL_3020025, EPI_ISL_3020026, EPI_ISL_3020029, EPI_ISL_3020033, EPI_ISL_3020034, EPI_ISL_3020037, EPI_ISL_3020040, EPI_ISL_3020042, EPI_ISL_3020045, EPI_ISL_3020046, EPI_ISL_3020047, EPI_ISL_3020049, EPI_ISL_3020050, EPI_ISL_3020051, EPI_ISL_3020052, EPI_ISL_3020057, EPI_ISL_3020058, EPI_ISL_3020059, EPI_ISL_3020061, EPI_ISL_3020062, EPI_ISL_3020063, EPI_ISL_3020064 | | | |
| see above | Infinity Biologix | Centers for Disease Control and Prevention Division of Viral Diseases, Pathogen Discovery | Dakota Howard, Dhwani Batra, Peter W. Cook, Kara Moser, Adrian Paskey, Jason Caravas, Benjamin Rambo-Martin, Shatavia Morrison, Christopher Gulvick, Scott Sammons, Yvette Unoarumhi, Darlene Wagner, Matthew Schmerer, Christian Bixby, Yihe Wang, Jonathan Schultz, Chirayu Goswami, Russ Hager, Robin Grimwood, Clinton R. Paden, Duncan MacCannell |
| EPI_ISL_3020070, EPI_ISL_3020071, EPI_ISL_3020072, EPI_ISL_3020073 | TXDSHS | TXDSHS | Rashmi Tuladhar, Bonnie Oh, Jenny Zhang, Maliha Rahman, Mayela Pedrueza, Anita Pokharel, Karen Bobier, Lorraine Rodriguez, Myong Koag, Chun Wang, Rachel Lee, Grace Kubin |
| EPI_ISL_3020074, EPI_ISL_3020075, EPI_ISL_3020076, EPI_ISL_3020077, EPI_ISL_3020078, EPI_ISL_3020079, EPI_ISL_3020080, EPI_ISL_3020081, EPI_ISL_3020082, EPI_ISL_3020083, EPI_ISL_3020084, EPI_ISL_3020085, EPI_ISL_3020086, EPI_ISL_3020087, EPI_ISL_3020088, EPI_ISL_3020089, EPI_ISL_3020090, EPI_ISL_3020091, EPI_ISL_3020092, EPI_ISL_3020093, EPI_ISL_3020094, EPI_ISL_3020095, EPI_ISL_3020096, EPI_ISL_3020097, EPI_ISL_3020098, EPI_ISL_3020099, EPI_ISL_3020100, EPI_ISL_3020101, EPI_ISL_3020102, EPI_ISL_3020103, EPI_ISL_3020104 | | | |
| see above | Nebraska Public Health Laboratory | NPHL COVID-19 Response Team | NPHL COVID-19 Response Team |
| EPI_ISL_3020105, EPI_ISL_3020107, EPI_ISL_3020108, EPI_ISL_3020109, EPI_ISL_3020110, EPI_ISL_3020111, EPI_ISL_3020112, EPI_ISL_3020118, EPI_ISL_3020123, EPI_ISL_3020124, EPI_ISL_3020125, EPI_ISL_3020126, EPI_ISL_3020132, EPI_ISL_3020133, EPI_ISL_3020136, EPI_ISL_3020137, EPI_ISL_3020138, EPI_ISL_3020150, EPI_ISL_3020151, EPI_ISL_3020152 | | | |
| see above | MD PHL | MD PHL | Maryland Department of Health Laboratories Administration |
| EPI_ISL_3020155, EPI_ISL_3020156, EPI_ISL_3020157, EPI_ISL_3020158, EPI_ISL_3020159 | Clinical Hospital of Infectious Diseases „Sfanta Parascheva" Iasi | National Institute of Infectious Diseases-Prof. Dr. Matei Bals Molecular Diagnostics Laboratory | Corina Casangiu, Leontina Banica, Marius Surleac, Petre Milu, Robert Hohan, Simona Paraschiv, Dan Otelea |
| EPI_ISL_3020162, EPI_ISL_3020163 | National Institute of Infectious Diseases-Prof. Dr. Matei Bals Molecular Diagnostics Laboratory | National Institute of Infectious Diseases-Prof. Dr. Matei Bals Molecular Diagnostics Laboratory | Corina Casangiu, Leontina Banica, Marius Surleac, Petre Milu, Robert Hohan, Simona Paraschiv, Dan Otelea |
| EPI_ISL_3020279, EPI_ISL_3020280, EPI_ISL_3020281 | Hospital Universitari Vall d'Hebron - Vall d'Hebron Institut de Recerca | Hospital Universitari Vall d'Hebron - Vall d'Hebron Institut de Recerca | Cristina Andrés, Maria Piñana, Alejandra González-Sánchez, Damir Garcia-Cehic, Ariadna Rando, Juliana Esperalba, Maria Gema Codina, Carla Castillo, Karen García, Rodrigo Vásquez, Maria Carmen Martin, Tomàs Pumarola, Josep Quer, Andrés Antón |
| EPI_ISL_3020283 | Hospital Universitari Arnau de Vilanova | Hospital Universitari Vall d'Hebron - Vall d'Hebron Institut de Recerca | Cristina Andrés, Maria Piñana, Alejandra González-Sánchez, Damir Garcia-Cehic, Ariadna Rando, Juliana Esperalba, Maria Gema Codina, Carla Castillo, Karen García, Rodrigo Vásquez, Maria Carmen Martin, Tomàs Pumarola, Josep Quer, Andrés Antón |
| EPI_ISL_3020287, EPI_ISL_3020288, EPI_ISL_3020289, EPI_ISL_3020290, EPI_ISL_3020291, EPI_ISL_3020292, EPI_ISL_3020294, EPI_ISL_3020295, EPI_ISL_3020296, EPI_ISL_3020297, EPI_ISL_3020298, EPI_ISL_3020299, EPI_ISL_3020300, EPI_ISL_3020301, EPI_ISL_3020302, EPI_ISL_3020303, EPI_ISL_3020304, EPI_ISL_3020305, EPI_ISL_3020306, EPI_ISL_3020307, EPI_ISL_3020308, EPI_ISL_3020309, EPI_ISL_3020310, EPI_ISL_3020311, EPI_ISL_3020312, EPI_ISL_3020315, EPI_ISL_3020316, EPI_ISL_3020317, EPI_ISL_3020320, EPI_ISL_3020322, EPI_ISL_3020323, EPI_ISL_3020324, EPI_ISL_3020325, EPI_ISL_3020326, EPI_ISL_3020327, EPI_ISL_3020328 | | | |
| see above | Tampa General Hospital Esoteric Lab | Tampa General Hospital Esoteric Research & Development Lab | Jon Faughn, Deanna Becker, Dominic Uy, Jorge Mecias-Frias, Elaine Vendrone, Vicki Healer, Amorce Lima, Suzane Silbert |
| EPI_ISL_3020340 | Laboratory of virology and molecular diagnostics, Institute of Public Health | Laboratory of virology and molecular diagnostics, Institute of Public Health | Kuzmanovska M, Boshevska G, Janchevska E. |
| EPI_ISL_3020341 | University Clinic of Cardiology | Laboratory of virology and molecular diagnostics, Institute of Public Health | Kuzmanovska M, Boshevska G, Janchevska E. |
| EPI_ISL_3020342, EPI_ISL_3020343 | General Hospital Tetovo | Laboratory of virology and molecular diagnostics, Institute of Public Health | Kuzmanovska M, Boshevska G, Janchevska E. |
| EPI_ISL_3020347, EPI_ISL_3020348, EPI_ISL_3020349, EPI_ISL_3020350, EPI_ISL_3020351, EPI_ISL_3020352, EPI_ISL_3020353, EPI_ISL_3020354, EPI_ISL_3020355, EPI_ISL_3020356, EPI_ISL_3020357 | | | |
| see above | Tokyo Metropolitan Institute of Public Health | Tokyo Metropolitan Institute of Public Health | Masaki Hayashi, Takako Yamazaki, Akane Negishi,Hirofumi Miyake, Kenshirou Kuroki, Maya Isogai, Takushi Fujiwara, Mami Nagashima, Jun Suzuki, Kenji Sadamasu |
| EPI_ISL_3020361, EPI_ISL_3020362, EPI_ISL_3020364, EPI_ISL_3020365, EPI_ISL_3020367, EPI_ISL_3020368, EPI_ISL_3020370, EPI_ISL_3020371, EPI_ISL_3020372, EPI_ISL_3020373, EPI_ISL_3020374, EPI_ISL_3020375, EPI_ISL_3020376, EPI_ISL_3020378, EPI_ISL_3020379, EPI_ISL_3020380, EPI_ISL_3020383, EPI_ISL_3020384, EPI_ISL_3020385, EPI_ISL_3020387, EPI_ISL_3020388, EPI_ISL_3020389, EPI_ISL_3020390, EPI_ISL_3020392 | | | |
| see above | MA15 Lieferdienst Veloce | AGES, Institute for Medical Microbiology and Hygiene | Kathrin Lippert, Justine Schaeffer, Alena Chalupka, Norbert Handra, Julia Klikovits, Philipp Wanka, Alexander Indra |
| EPI_ISL_3020395 | Bezirkshauptmannschaft Neusiedl/See Gesundheitsabteilung | AGES, Institute for Medical Microbiology and Hygiene | Kathrin Lippert, Justine Schaeffer, Alena Chalupka, Norbert Handra, Julia Klikovits, Philipp Wanka, Alexander Indra |
| EPI_ISL_3020400, EPI_ISL_3020401, EPI_ISL_3020402, EPI_ISL_3020403, EPI_ISL_3020404, EPI_ISL_3020405, EPI_ISL_3020406, EPI_ISL_3020407, EPI_ISL_3020408, EPI_ISL_3020409, EPI_ISL_3020410, EPI_ISL_3020411, EPI_ISL_3020412 | | | |
| see above | MA15 Lieferdienst Veloce | AGES, Institute for Medical Microbiology and Hygiene | Kathrin Lippert, Justine Schaeffer, Alena Chalupka, Norbert Handra, Julia Klikovits, Philipp Wanka, Alexander Indra |
| EPI_ISL_3020413 | Bezirkshauptmannschaft Neusiedl/See Gesundheitsabteilung | AGES, Institute for Medical Microbiology and Hygiene | Kathrin Lippert, Justine Schaeffer, Alena Chalupka, Norbert Handra, Julia Klikovits, Philipp Wanka, Alexander Indra |
| EPI_ISL_3020414 | Landessanitätsdirektion Burgenland Amt der Bgld Landesregierung | AGES, Institute for Medical Microbiology and Hygiene | Kathrin Lippert, Justine Schaeffer, Alena Chalupka, Norbert Handra, Julia Klikovits, Philipp Wanka, Alexander Indra |
| EPI_ISL_3020416 | Bezirkshauptmannschaft Neusiedl/See Gesundheitsabteilung | AGES, Institute for Medical Microbiology and Hygiene | Kathrin Lippert, Justine Schaeffer, Alena Chalupka, Norbert Handra, Julia Klikovits, Philipp Wanka, Alexander Indra |
| EPI_ISL_3020417, EPI_ISL_3020419, EPI_ISL_3020420, EPI_ISL_3020421, EPI_ISL_3020422, EPI_ISL_3020423, EPI_ISL_3020424, EPI_ISL_3020425, EPI_ISL_3020426 | MA15 Lieferdienst Veloce | AGES, Institute for Medical Microbiology and Hygiene | Kathrin Lippert, Justine Schaeffer, Alena Chalupka, Norbert Handra, Julia Klikovits, Philipp Wanka, Alexander Indra |
| EPI_ISL_3020427 | Erstaufnahmestelle Ost Traiskirchen | AGES, Institute for Medical Microbiology and Hygiene | Kathrin Lippert, Justine Schaeffer, Alena Chalupka, Norbert Handra, Julia Klikovits, Philipp Wanka, Alexander Indra |
| EPI_ISL_3020428, EPI_ISL_3020429, EPI_ISL_3020430 | Bezirkshauptmannschaft Eisenstadt-Umgebung Gesundheitsamt | AGES, Institute for Medical Microbiology and Hygiene | Kathrin Lippert, Justine Schaeffer, Alena Chalupka, Norbert Handra, Julia Klikovits, Philipp Wanka, Alexander Indra |
| EPI_ISL_3020431, EPI_ISL_3020432, EPI_ISL_3020433, EPI_ISL_3020434, EPI_ISL_3020436, EPI_ISL_3020437, EPI_ISL_3020438, EPI_ISL_3020440 | MA15 Lieferdienst Veloce | AGES, Institute for Medical Microbiology and Hygiene | Kathrin Lippert, Justine Schaeffer, Alena Chalupka, Norbert Handra, Julia Klikovits, Philipp Wanka, Alexander Indra |

| | | | |
|---|---|---|---|
| EPI_ISL_3020445 | Bezirkshauptmannschaft Mattersburg Gesundheitsabteilung | AGES, Institute for Medical Microbiology and Hygiene | Kathrin Lippert, Justine Schaeffer, Alena Chalupka, Norbert Handra, Julia Klikovits, Philipp Wanka, Alexander Indra |
| EPI_ISL_3020447, EPI_ISL_3020448, EPI_ISL_3020449, EPI_ISL_3020450, EPI_ISL_3020451 | MA15 Lieferdienst Veloce | AGES, Institute for Medical Microbiology and Hygiene | Kathrin Lippert, Justine Schaeffer, Alena Chalupka, Norbert Handra, Julia Klikovits, Philipp Wanka, Alexander Indra |
| EPI_ISL_3020452 | Bezirkshauptmannschaft Eisenstadt-Umgebung Gesundheitsamt | AGES, Institute for Medical Microbiology and Hygiene | Kathrin Lippert, Justine Schaeffer, Alena Chalupka, Norbert Handra, Julia Klikovits, Philipp Wanka, Alexander Indra |
| EPI_ISL_3020455, EPI_ISL_3020457 | Landessanitätsdirektion Burgenland Amt der Bgld Landesregierung | AGES, Institute for Medical Microbiology and Hygiene | Kathrin Lippert, Justine Schaeffer, Alena Chalupka, Norbert Handra, Julia Klikovits, Philipp Wanka, Alexander Indra |
| EPI_ISL_3020458 | Bezirkshauptmannschaft Eisenstadt-Umgebung Gesundheitsamt | AGES, Institute for Medical Microbiology and Hygiene | Kathrin Lippert, Justine Schaeffer, Alena Chalupka, Norbert Handra, Julia Klikovits, Philipp Wanka, Alexander Indra |
| EPI_ISL_3020460, EPI_ISL_3020461 | MA15 Lieferdienst Veloce | AGES, Institute for Medical Microbiology and Hygiene | Kathrin Lippert, Justine Schaeffer, Alena Chalupka, Norbert Handra, Julia Klikovits, Philipp Wanka, Alexander Indra |
| EPI_ISL_3020506 | Wyoming Public Health Laboratory | Wyoming Public Health Laboratory | Jim Mildenberger, Taylor Fearing, Channing Weber, Ashley Norberg, Chayse Rowley, Marley Goetz, Brian Dominguez, Elliot Thomasson, Sam Britz, Cari Sloma, Robert Petit, and Rob Christensen |
| EPI_ISL_3020516, EPI_ISL_3020518, EPI_ISL_3020523 | SYNLAB Suomi | Expert Microbiology, National Institute for Health and Welfare | Soile Blomqvist, Jani Halkilahti, Kirsi Liitsola, Haider al-Hello, Päivi Laurila, Erika Lindh, Teemu Smura, Ravi Kant, Phuoc Truong, Olli Vapalahti, Sari Hannula, Pekka Ellonen, Niina Ikonen, Carita Savolainen-Kopra |
| EPI_ISL_3020527 | Vita Laboratoriot Oy | Expert Microbiology, National Institute for Health and Welfare | Soile Blomqvist, Jani Halkilahti, Kirsi Liitsola, Haider al-Hello, Päivi Laurila, Erika Lindh, Teemu Smura, Ravi Kant, Phuoc Truong, Olli Vapalahti, Sari Hannula, Pekka Ellonen, Niina Ikonen, Carita Savolainen-Kopra |
| EPI_ISL_3020539 | SataDiag | Expert Microbiology, National Institute for Health and Welfare | Soile Blomqvist, Jani Halkilahti, Kirsi Liitsola, Haider al-Hello, Päivi Laurila, Erika Lindh, Teemu Smura, Ravi Kant, Phuoc Truong, Olli Vapalahti, Sari Hannula, Pekka Ellonen, Niina Ikonen, Carita Savolainen-Kopra |
| EPI_ISL_3020542, EPI_ISL_3020543, EPI_ISL_3020544, EPI_ISL_3020545, EPI_ISL_3020546 | NordLab Kokkola | Expert Microbiology, National Institute for Health and Welfare | Soile Blomqvist, Jani Halkilahti, Kirsi Liitsola, Haider al-Hello, Päivi Laurila, Erika Lindh, Teemu Smura, Ravi Kant, Phuoc Truong, Olli Vapalahti, Sari Hannula, Pekka Ellonen, Niina Ikonen, Carita Savolainen-Kopra |
| EPI_ISL_3020584, EPI_ISL_3020586, EPI_ISL_3020589, EPI_ISL_3020590, EPI_ISL_3020593, EPI_ISL_3020597, EPI_ISL_3020603, EPI_ISL_3020604, EPI_ISL_3020608, EPI_ISL_3020609, EPI_ISL_3020612, EPI_ISL_3020619, EPI_ISL_3020622, EPI_ISL_3020625, EPI_ISL_3020629, EPI_ISL_3020632, EPI_ISL_3020636, EPI_ISL_3020638, EPI_ISL_3020644 | | | |
| see above | Fimlab Laboratoriot Oy Tampere | Expert Microbiology, National Institute for Health and Welfare | Soile Blomqvist, Jani Halkilahti, Kirsi Liitsola, Haider al-Hello, Päivi Laurila, Erika Lindh, Teemu Smura, Ravi Kant, Phuoc Truong, Olli Vapalahti, Sari Hannula, Pekka Ellonen, Niina Ikonen, Carita Savolainen-Kopra |
| EPI_ISL_3020650, EPI_ISL_3020651, EPI_ISL_3020652, EPI_ISL_3020653, EPI_ISL_3020656, EPI_ISL_3020657, EPI_ISL_3020660, EPI_ISL_3020665, EPI_ISL_3020669, EPI_ISL_3020670, EPI_ISL_3020671, EPI_ISL_3020672, EPI_ISL_3020673 | | | |
| see above | NordLab Oulu | Expert Microbiology, National Institute for Health and Welfare | Soile Blomqvist, Jani Halkilahti, Kirsi Liitsola, Haider al-Hello, Päivi Laurila, Erika Lindh, Teemu Smura, Ravi Kant, Phuoc Truong, Olli Vapalahti, Sari Hannula, Pekka Ellonen, Niina Ikonen, Carita Savolainen-Kopra |
| EPI_ISL_3020688 | Vita Laboratoriot Oy | Expert Microbiology, National Institute for Health and Welfare | Soile Blomqvist, Jani Halkilahti, Kirsi Liitsola, Haider al-Hello, Päivi Laurila, Erika Lindh, Teemu Smura, Ravi Kant, Phuoc Truong, Olli Vapalahti, Sari Hannula, Pekka Ellonen, Niina Ikonen, Carita Savolainen-Kopra |
| EPI_ISL_3020690 | SYNLAB Suomi | Expert Microbiology, National Institute for Health and Welfare | Soile Blomqvist, Jani Halkilahti, Kirsi Liitsola, Haider al-Hello, Päivi Laurila, Erika Lindh, Teemu Smura, Ravi Kant, Phuoc Truong, Olli Vapalahti, Sari Hannula, Pekka Ellonen, Niina Ikonen, Carita Savolainen-Kopra |
| EPI_ISL_3020700, EPI_ISL_3020706, EPI_ISL_3020708, EPI_ISL_3020711, EPI_ISL_3020712, EPI_ISL_3020713, EPI_ISL_3020717, EPI_ISL_3020732, EPI_ISL_3020734, EPI_ISL_3020745, EPI_ISL_3020747, EPI_ISL_3020750, EPI_ISL_3020751, EPI_ISL_3020753, EPI_ISL_3020757 | | | |
| see above | Fimlab Laboratoriot Oy Tampere | Expert Microbiology, National Institute for Health and Welfare | Soile Blomqvist, Jani Halkilahti, Kirsi Liitsola, Haider al-Hello, Päivi Laurila, Erika Lindh, Teemu Smura, Ravi Kant, Phuoc Truong, Olli Vapalahti, Sari Hannula, Pekka Ellonen, Niina Ikonen, Carita Savolainen-Kopra |
| EPI_ISL_3020768 | Vita Laboratoriot Oy | Expert Microbiology, National Institute for Health and Welfare | Soile Blomqvist, Jani Halkilahti, Kirsi Liitsola, Haider al-Hello, Päivi Laurila, Erika Lindh, Teemu Smura, Ravi Kant, Phuoc Truong, Olli Vapalahti, Sari Hannula, Pekka Ellonen, Niina Ikonen, Carita Savolainen-Kopra |
| EPI_ISL_3020783, EPI_ISL_3020784 | Islab, Pohjois-Savon aluelaboratorio | Expert Microbiology, National Institute for Health and Welfare | Soile Blomqvist, Jani Halkilahti, Kirsi Liitsola, Haider al-Hello, Päivi Laurila, Erika Lindh, Teemu Smura, Ravi Kant, Phuoc Truong, Olli Vapalahti, Sari Hannula, Pekka Ellonen, Niina Ikonen, Carita Savolainen-Kopra |
| EPI_ISL_3020810, EPI_ISL_3020823, EPI_ISL_3020829, EPI_ISL_3020838 | Fimlab Laboratoriot Oy Tampere | Expert Microbiology, National Institute for Health and Welfare | Soile Blomqvist, Jani Halkilahti, Kirsi Liitsola, Haider al-Hello, Päivi Laurila, Erika Lindh, Teemu Smura, Ravi Kant, Phuoc Truong, Olli Vapalahti, Sari Hannula, Pekka Ellonen, Niina Ikonen, Carita Savolainen-Kopra |
| EPI_ISL_3020856, EPI_ISL_3020857, EPI_ISL_3020858, EPI_ISL_3020859 | NordLab Oulu | Expert Microbiology, National Institute for Health and Welfare | Soile Blomqvist, Jani Halkilahti, Kirsi Liitsola, Haider al-Hello, Päivi Laurila, Erika Lindh, Teemu Smura, Ravi Kant, Phuoc Truong, Olli Vapalahti, Sari Hannula, Pekka Ellonen, Niina Ikonen, Carita Savolainen-Kopra |
| EPI_ISL_3020860, EPI_ISL_3020861, EPI_ISL_3020862, EPI_ISL_3020863, EPI_ISL_3020864 | Islab, Pohjois-Karjalan aluelaboratorio | Expert Microbiology, National Institute for Health and Welfare | Soile Blomqvist, Jani Halkilahti, Kirsi Liitsola, Haider al-Hello, Päivi Laurila, Erika Lindh, Teemu Smura, Ravi Kant, Phuoc Truong, Olli Vapalahti, Sari Hannula, Pekka Ellonen, Niina Ikonen, Carita Savolainen-Kopra |
| EPI_ISL_3020866 | Islab, Etelä-Savon aluelaboratorio | Expert Microbiology, National Institute for Health and Welfare | Soile Blomqvist, Jani Halkilahti, Kirsi Liitsola, Haider al-Hello, Päivi Laurila, Erika Lindh, Teemu Smura, Ravi Kant, Phuoc Truong, Olli Vapalahti, Sari Hannula, Pekka Ellonen, Niina Ikonen, Carita Savolainen-Kopra |
| EPI_ISL_3020873 | Vita Laboratoriot Oy | Expert Microbiology, National Institute for Health and Welfare | Soile Blomqvist, Jani Halkilahti, Kirsi Liitsola, Haider al-Hello, Päivi Laurila, Erika Lindh, Teemu Smura, Ravi Kant, Phuoc Truong, Olli Vapalahti, Sari Hannula, Pekka Ellonen, Niina Ikonen, Carita Savolainen-Kopra |
| EPI_ISL_3020874, EPI_ISL_3020875, EPI_ISL_3020876, EPI_ISL_3020877 | NordLab Kajaani | Expert Microbiology, National Institute for Health and Welfare | Soile Blomqvist, Jani Halkilahti, Kirsi Liitsola, Haider al-Hello, Päivi Laurila, Erika Lindh, Teemu Smura, Ravi Kant, Phuoc Truong, Olli Vapalahti, Sari Hannula, Pekka Ellonen, Niina Ikonen, Carita Savolainen-Kopra |
| EPI_ISL_3020878 | Vita Laboratoriot Oy | Expert Microbiology, National Institute for Health and Welfare | Soile Blomqvist, Jani Halkilahti, Kirsi Liitsola, Haider al-Hello, Päivi Laurila, Erika Lindh, Teemu Smura, Ravi Kant, Phuoc Truong, Olli Vapalahti, Sari Hannula, Pekka Ellonen, Niina Ikonen, Carita Savolainen-Kopra |
| EPI_ISL_3020882, EPI_ISL_3020884, EPI_ISL_3020885 | SYNLAB Suomi | Expert Microbiology, National Institute for Health and Welfare | Soile Blomqvist, Jani Halkilahti, Kirsi Liitsola, Haider al-Hello, Päivi Laurila, Erika Lindh, Teemu Smura, Ravi Kant, Phuoc Truong, Olli Vapalahti, Sari Hannula, Pekka Ellonen, Niina Ikonen, Carita Savolainen-Kopra |
| EPI_ISL_3020888, EPI_ISL_3020890 | NordLab Kajaani | Expert Microbiology, National Institute for Health and Welfare | Soile Blomqvist, Jani Halkilahti, Kirsi Liitsola, Haider al-Hello, Päivi Laurila, Erika Lindh, Teemu Smura, Ravi Kant, Phuoc Truong, Olli Vapalahti, Sari Hannula, Pekka Ellonen, Niina Ikonen, Carita Savolainen-Kopra |
| EPI_ISL_3020892, EPI_ISL_3020893 | Islab, Pohjois-Savon aluelaboratorio | Expert Microbiology, National Institute for Health and Welfare | Soile Blomqvist, Jani Halkilahti, Kirsi Liitsola, Haider al-Hello, Päivi Laurila, Erika Lindh, Teemu Smura, Ravi Kant, Phuoc Truong, Olli Vapalahti, Sari Hannula, Pekka Ellonen, Niina Ikonen, Carita Savolainen-Kopra |
| EPI_ISL_3020894 | NordLab Rovaniemi | Expert Microbiology, National Institute for Health and Welfare | Soile Blomqvist, Jani Halkilahti, Kirsi Liitsola, Haider al-Hello, Päivi Laurila, Erika Lindh, Teemu Smura, Ravi Kant, Phuoc Truong, Olli Vapalahti, Sari Hannula, Pekka Ellonen, Niina Ikonen, Carita Savolainen-Kopra |
| EPI_ISL_3020897, EPI_ISL_3020898, EPI_ISL_3020899 | NordLab Kajaani | Expert Microbiology, National Institute for Health and Welfare | Soile Blomqvist, Jani Halkilahti, Kirsi Liitsola, Haider al-Hello, Päivi Laurila, Erika Lindh, Teemu Smura, Ravi Kant, Phuoc Truong, Olli Vapalahti, Sari Hannula, Pekka Ellonen, Niina Ikonen, Carita Savolainen-Kopra |
| EPI_ISL_3020900, EPI_ISL_3020901 | Islab, Pohjois-Karjalan aluelaboratorio | Expert Microbiology, National Institute for Health and Welfare | Soile Blomqvist, Jani Halkilahti, Kirsi Liitsola, Haider al-Hello, Päivi Laurila, Erika Lindh, Teemu Smura, Ravi Kant, Phuoc Truong, Olli Vapalahti, Sari Hannula, Pekka Ellonen, Niina Ikonen, Carita Savolainen-Kopra |
| EPI_ISL_3020904 | Islab, Etelä-Savon aluelaboratorio | Expert Microbiology, National Institute for Health and Welfare | Soile Blomqvist, Jani Halkilahti, Kirsi Liitsola, Haider al-Hello, Päivi Laurila, Erika Lindh, Teemu Smura, Ravi Kant, Phuoc Truong, Olli Vapalahti, Sari Hannula, Pekka Ellonen, Niina Ikonen, Carita Savolainen-Kopra |
| EPI_ISL_3020905, EPI_ISL_3020906, EPI_ISL_3020907, EPI_ISL_3020912, EPI_ISL_3020913, EPI_ISL_3020916, EPI_ISL_3020917, EPI_ISL_3020920, EPI_ISL_3020923, EPI_ISL_3020926, EPI_ISL_3020929, EPI_ISL_3020930, EPI_ISL_3020933, EPI_ISL_3020934, EPI_ISL_3020935, EPI_ISL_3020936, EPI_ISL_3020940, EPI_ISL_3020943, EPI_ISL_3020944, EPI_ISL_3020949 | | | |
| see above | Fimlab Laboratoriot Oy Tampere | Expert Microbiology, National Institute for Health and Welfare | Soile Blomqvist, Jani Halkilahti, Kirsi Liitsola, Haider al-Hello, Päivi Laurila, Erika Lindh, Teemu Smura, Ravi Kant, Phuoc Truong, Olli Vapalahti, Sari Hannula, Pekka Ellonen, Niina Ikonen, Carita Savolainen-Kopra |
| EPI_ISL_3020950 | NordLab Oulu | Expert Microbiology, National Institute for Health and Welfare | Soile Blomqvist, Jani Halkilahti, Kirsi Liitsola, Haider al-Hello, Päivi Laurila, Erika Lindh, Teemu Smura, Ravi Kant, Phuoc Truong, Olli Vapalahti, Sari Hannula, Pekka Ellonen, Niina Ikonen, Carita Savolainen-Kopra |
| EPI_ISL_3020951, EPI_ISL_3020952, EPI_ISL_3020953, EPI_ISL_3020954, EPI_ISL_3020955 | Islab, Pohjois-Savon aluelaboratorio | Expert Microbiology, National Institute for Health and Welfare | Soile Blomqvist, Jani Halkilahti, Kirsi Liitsola, Haider al-Hello, Päivi Laurila, Erika Lindh, Teemu Smura, Ravi Kant, Phuoc Truong, Olli Vapalahti, Sari Hannula, Pekka Ellonen, Niina Ikonen, Carita Savolainen-Kopra |
| EPI_ISL_3020956 | Vita Laboratoriot Oy | Expert Microbiology, National Institute for Health and Welfare | Soile Blomqvist, Jani Halkilahti, Kirsi Liitsola, Haider al-Hello, Päivi Laurila, Erika Lindh, Teemu Smura, Ravi Kant, Phuoc Truong, Olli Vapalahti, Sari Hannula, Pekka Ellonen, Niina Ikonen, Carita Savolainen-Kopra |
| EPI_ISL_3020957, EPI_ISL_3020958, EPI_ISL_3020959, EPI_ISL_3020960, EPI_ISL_3020961, EPI_ISL_3020962, EPI_ISL_3020964, EPI_ISL_3020965, EPI_ISL_3020966, EPI_ISL_3020967, EPI_ISL_3020968, EPI_ISL_3020969, EPI_ISL_3020970, EPI_ISL_3020971, EPI_ISL_3020972 | | | |
| see above | SYNLAB Suomi | Expert Microbiology, National Institute for Health and Welfare | Soile Blomqvist, Jani Halkilahti, Kirsi Liitsola, Haider al-Hello, Päivi Laurila, Erika Lindh, Teemu Smura, Ravi Kant, Phuoc Truong, Olli Vapalahti, Sari Hannula, Pekka Ellonen, Niina Ikonen, Carita Savolainen-Kopra |
| EPI_ISL_3020973, EPI_ISL_3020974, EPI_ISL_3020975, EPI_ISL_3020976, EPI_ISL_3020977 | Vita Laboratoriot Oy | Expert Microbiology, National Institute for Health and Welfare | Soile Blomqvist, Jani Halkilahti, Kirsi Liitsola, Haider al-Hello, Päivi Laurila, Erika Lindh, Teemu Smura, Ravi Kant, Phuoc Truong, Olli Vapalahti, Sari Hannula, Pekka Ellonen, Niina Ikonen, Carita Savolainen-Kopra |
| EPI_ISL_3020978 | Fimlab Laboratoriot Oy Tampere | Expert Microbiology, National Institute for Health and Welfare | Soile Blomqvist, Jani Halkilahti, Kirsi Liitsola, Haider al-Hello, Päivi Laurila, Erika Lindh, Teemu Smura, Ravi Kant, Phuoc Truong, Olli Vapalahti, Sari Hannula, Pekka Ellonen, Niina Ikonen, Carita Savolainen-Kopra |
| EPI_ISL_3020979, EPI_ISL_3020980 | Seinäjoen keskussairaala | Expert Microbiology, National Institute for Health and Welfare | Soile Blomqvist, Jani Halkilahti, Kirsi Liitsola, Haider al-Hello, Päivi Laurila, Erika Lindh, Teemu Smura, Ravi Kant, Phuoc Truong, Olli Vapalahti, Sari Hannula, Pekka Ellonen, Niina Ikonen, Carita Savolainen-Kopra |
| EPI_ISL_3020981 | Islab, Pohjois-Savon aluelaboratorio | Expert Microbiology, National Institute for Health and Welfare | Soile Blomqvist, Jani Halkilahti, Kirsi Liitsola, Haider al-Hello, Päivi Laurila, Erika Lindh, Teemu Smura, Ravi Kant, Phuoc Truong, Olli Vapalahti, Sari Hannula, Pekka Ellonen, Niina Ikonen, Carita Savolainen-Kopra |
| EPI_ISL_3020982, EPI_ISL_3020983, | TYKS, Kliininen mikrobiologia | Expert Microbiology, National Institute for Health and Welfare | Soile Blomqvist, Jani Halkilahti, Kirsi Liitsola, Haider al-Hello, Päivi Laurila, Erika Lindh, Teemu Smura, Ravi Kant, Phuoc Truong, Olli Vapalahti, Sari Hannula, Pekka Ellonen, Niina Ikonen, Carita Savolainen-Kopra |

| | | | |
|---|---|---|---|
| EPI_ISL_3020984, EPI_ISL_3020985 | | | |
| EPI_ISL_3020986, EPI_ISL_3020987 | SataDiag | Expert Microbiology, National Institute for Health and Welfare | Soile Blomqvist, Jani Halkilahti, Kirsi Liitsola, Haider al-Hello, Päivi Laurila, Erika Lindh, Teemu Smura, Ravi Kant, Phuoc Truong, Olli Vapalahti, Sari Hannula, Pekka Ellonen, Niina Ikonen, Carita Savolainen-Kopra |
| EPI_ISL_3020994, EPI_ISL_3020995, EPI_ISL_3020996, EPI_ISL_3020997, EPI_ISL_3020998, EPI_ISL_3020999, EPI_ISL_3021000, EPI_ISL_3021001, EPI_ISL_3021004, EPI_ISL_3021005, EPI_ISL_3021006, EPI_ISL_3021007, EPI_ISL_3021008, EPI_ISL_3021009, EPI_ISL_3021010, EPI_ISL_3021011, EPI_ISL_3021012, EPI_ISL_3021013, EPI_ISL_3021014, EPI_ISL_3021015, EPI_ISL_3021016, EPI_ISL_3021017, EPI_ISL_3021018, EPI_ISL_3021019, EPI_ISL_3021020, EPI_ISL_3021021, EPI_ISL_3021022, EPI_ISL_3021023 | | | |
| see above | Nebraska Public Health Laboratory | NPHL COVID-19 Response Team | NPHL COVID-19 Response Team |
| EPI_ISL_3021024, EPI_ISL_3021025, EPI_ISL_3021029, EPI_ISL_3021032, EPI_ISL_3021036 | Virginia Division of Consolidated Laboratory Services | Virginia Division of Consolidated Laboratory Services | Virginia Division of Consolidated Laboratory Services |
| EPI_ISL_3021038, EPI_ISL_3021041, EPI_ISL_3021042, EPI_ISL_3021044, EPI_ISL_3021045, EPI_ISL_3021046, EPI_ISL_3021048, EPI_ISL_3021049, EPI_ISL_3021050, EPI_ISL_3021051, EPI_ISL_3021052, EPI_ISL_3021053, EPI_ISL_3021054, EPI_ISL_3021056, EPI_ISL_3021057, EPI_ISL_3021059, EPI_ISL_3021060, EPI_ISL_3021061, EPI_ISL_3021062, EPI_ISL_3021063, EPI_ISL_3021064, EPI_ISL_3021066, EPI_ISL_3021067, EPI_ISL_3021070, EPI_ISL_3021071, EPI_ISL_3021072, EPI_ISL_3021073, EPI_ISL_3021074, EPI_ISL_3021075, EPI_ISL_3021076, EPI_ISL_3021077, EPI_ISL_3021078, EPI_ISL_3021079, EPI_ISL_3021082, EPI_ISL_3021083, EPI_ISL_3021084, EPI_ISL_3021085, EPI_ISL_3021086, EPI_ISL_3021087, EPI_ISL_3021088, EPI_ISL_3021089, EPI_ISL_3021090, EPI_ISL_3021091, EPI_ISL_3021092, EPI_ISL_3021093, EPI_ISL_3021094, EPI_ISL_3021095, EPI_ISL_3021096, EPI_ISL_3021097, EPI_ISL_3021098, EPI_ISL_3021099, EPI_ISL_3021100, EPI_ISL_3021101, EPI_ISL_3021102, EPI_ISL_3021103, EPI_ISL_3021105, EPI_ISL_3021106, EPI_ISL_3021107, EPI_ISL_3021109, EPI_ISL_3021110, EPI_ISL_3021111, EPI_ISL_3021112, EPI_ISL_3021113, EPI_ISL_3021114, EPI_ISL_3021115, EPI_ISL_3021118, EPI_ISL_3021119, EPI_ISL_3021120, EPI_ISL_3021121, EPI_ISL_3021122, EPI_ISL_3021123, EPI_ISL_3021124, EPI_ISL_3021125, EPI_ISL_3021126, EPI_ISL_3021127, EPI_ISL_3021128, EPI_ISL_3021129, EPI_ISL_3021130, EPI_ISL_3021131, EPI_ISL_3021132, EPI_ISL_3021133, EPI_ISL_3021134, EPI_ISL_3021135, EPI_ISL_3021136, EPI_ISL_3021137, EPI_ISL_3021138, EPI_ISL_3021139, EPI_ISL_3021140, EPI_ISL_3021141, EPI_ISL_3021142, EPI_ISL_3021143, EPI_ISL_3021144, EPI_ISL_3021145, EPI_ISL_3021146, EPI_ISL_3021147, EPI_ISL_3021148, EPI_ISL_3021149, EPI_ISL_3021150, EPI_ISL_3021151, EPI_ISL_3021152, EPI_ISL_3021153, EPI_ISL_3021154, EPI_ISL_3021155, EPI_ISL_3021156, EPI_ISL_3021157, EPI_ISL_3021158, EPI_ISL_3021159, EPI_ISL_3021160, EPI_ISL_3021161, EPI_ISL_3021162, EPI_ISL_3021163, EPI_ISL_3021164, EPI_ISL_3021165, EPI_ISL_3021166, EPI_ISL_3021168, EPI_ISL_3021169, EPI_ISL_3021170, EPI_ISL_3021171, EPI_ISL_3021172, EPI_ISL_3021173, EPI_ISL_3021174, EPI_ISL_3021175, EPI_ISL_3021176, EPI_ISL_3021177, EPI_ISL_3021178, EPI_ISL_3021179, EPI_ISL_3021180, EPI_ISL_3021181, EPI_ISL_3021182, EPI_ISL_3021183, EPI_ISL_3021184, EPI_ISL_3021185, EPI_ISL_3021186, EPI_ISL_3021188, EPI_ISL_3021189, EPI_ISL_3021190, EPI_ISL_3021191, EPI_ISL_3021192, EPI_ISL_3021193, EPI_ISL_3021194, EPI_ISL_3021195, EPI_ISL_3021196, EPI_ISL_3021197, EPI_ISL_3021198, EPI_ISL_3021199, EPI_ISL_3021200, EPI_ISL_3021201, EPI_ISL_3021202, EPI_ISL_3021203, EPI_ISL_3021204, EPI_ISL_3021205, EPI_ISL_3021206, EPI_ISL_3021208, EPI_ISL_3021209, EPI_ISL_3021210, EPI_ISL_3021211, EPI_ISL_3021212, EPI_ISL_3021213, EPI_ISL_3021214, EPI_ISL_3021215, EPI_ISL_3021216, EPI_ISL_3021217, EPI_ISL_3021218, EPI_ISL_3021219, EPI_ISL_3021220, EPI_ISL_3021221, EPI_ISL_3021222, EPI_ISL_3021223, EPI_ISL_3021224, EPI_ISL_3021225, EPI_ISL_3021226, EPI_ISL_3021227, EPI_ISL_3021228, EPI_ISL_3021229, EPI_ISL_3021230, EPI_ISL_3021231, EPI_ISL_3021232, EPI_ISL_3021234, EPI_ISL_3021235, EPI_ISL_3021236, EPI_ISL_3021237, EPI_ISL_3021238, EPI_ISL_3021239, EPI_ISL_3021240, EPI_ISL_3021241, EPI_ISL_3021242, EPI_ISL_3021243, EPI_ISL_3021244, EPI_ISL_3021245, EPI_ISL_3021246, EPI_ISL_3021247, EPI_ISL_3021248, EPI_ISL_3021249, EPI_ISL_3021250, EPI_ISL_3021251, EPI_ISL_3021252, EPI_ISL_3021253, EPI_ISL_3021254, EPI_ISL_3021255, EPI_ISL_3021256, EPI_ISL_3021257, EPI_ISL_3021258, EPI_ISL_3021259, EPI_ISL_3021260, EPI_ISL_3021261, EPI_ISL_3021262, EPI_ISL_3021263, EPI_ISL_3021264, EPI_ISL_3021265, EPI_ISL_3021266, EPI_ISL_3021267, EPI_ISL_3021268, EPI_ISL_3021269, EPI_ISL_3021270, EPI_ISL_3021271 | | | |
| see above | Nevada State Public Health Laboratory | Nevada State Public Health Laboratory | Andrew Gorzalski, Mark Pandori |
| EPI_ISL_3021272, EPI_ISL_3021273, EPI_ISL_3021274, EPI_ISL_3021275, EPI_ISL_3021276, EPI_ISL_3021277, EPI_ISL_3021278, EPI_ISL_3021279, EPI_ISL_3021280, EPI_ISL_3021281 | Kansas Health and Environmental Lab | Kansas Health and Environmental Lab | Katherine Wiggins, Mike Grose, Jonathan Barnell, Ben Olsen, and Phil Adam |
| EPI_ISL_3021282, EPI_ISL_3021283, EPI_ISL_3021284, EPI_ISL_3021285, EPI_ISL_3021286, EPI_ISL_3021287, EPI_ISL_3021288, EPI_ISL_3021289, EPI_ISL_3021290, EPI_ISL_3021291, EPI_ISL_3021292, EPI_ISL_3021293, EPI_ISL_3021294, EPI_ISL_3021295, EPI_ISL_3021296 | | | |
| see above | Wichita State University - Molecular Diagnostics Lab | Kansas Health and Environmental Lab | Katherine Wiggins, Mike Grose, Jonathan Barnell, Ben Olsen, and Phil Adam |
| EPI_ISL_3021297, EPI_ISL_3021298, EPI_ISL_3021299, EPI_ISL_3021300, EPI_ISL_3021301, EPI_ISL_3021302, EPI_ISL_3021303, EPI_ISL_3021304, EPI_ISL_3021305, EPI_ISL_3021306, EPI_ISL_3021307, EPI_ISL_3021308, EPI_ISL_3021309, EPI_ISL_3021310, EPI_ISL_3021312, EPI_ISL_3021313, EPI_ISL_3021314, EPI_ISL_3021315, EPI_ISL_3021316, EPI_ISL_3021317, EPI_ISL_3021318, EPI_ISL_3021319, EPI_ISL_3021320, EPI_ISL_3021321, EPI_ISL_3021322, EPI_ISL_3021323, EPI_ISL_3021324, EPI_ISL_3021325, EPI_ISL_3021326, EPI_ISL_3021327, EPI_ISL_3021328, EPI_ISL_3021329 | | | |
| see above | Kansas Health and Environmental Lab | Kansas Health and Environmental Lab | Katherine Wiggins, Mike Grose, Jonathan Barnell, Ben Olsen, and Phil Adam |
| EPI_ISL_3021332, EPI_ISL_3021334, EPI_ISL_3021340 | Cantacuzino National Military-Medical Institute, Viral Respiratory Infections Laboratory | Cantacuzino Institute Virology | Carmen Cherciu, Luiza Ustea, Nicoleta Paraschiv, Sorin Dinu, Mihaela Oprea, Mihaela Lazar |
| EPI_ISL_3021364 | Clinical Emergency County Hospital, Craiova | Cantacuzino Institute Virology | Carmen Cherciu, Luiza Ustea, Nicoleta Paraschiv, Sorin Dinu, Mihaela Oprea, Mihaela Lazar |
| EPI_ISL_3021406 | Cantacuzino National Military-Medical Institute, Viral Respiratory Infections Laboratory | Cantacuzino Institute Virology | Carmen Cherciu,Luiza Ustea, Nicoleta Paraschiv, Sorin Dinu, Mihaela Oprea, Mihaela Lazar |
| EPI_ISL_3021408, EPI_ISL_3021409, EPI_ISL_3021412, EPI_ISL_3021413 | Clinical Emergency County Hospital, Craiova | Cantacuzino Institute Virology | Carmen Cherciu,Luiza Ustea, Nicoleta Paraschiv, Sorin Dinu, Mihaela Oprea, Mihaela Lazar |
| EPI_ISL_3021414, EPI_ISL_3021415 | Cantacuzino National Military-Medical Institute, Viral Respiratory Infections Laboratory | Cantacuzino Institute Virology | Carmen Cherciu,Luiza Ustea, Nicoleta Paraschiv, Sorin Dinu, Mihaela Oprea, Mihaela Lazar |
| EPI_ISL_3021417, EPI_ISL_3021419, EPI_ISL_3021420, EPI_ISL_3021421 | Clinical Emergency County Hospital, Craiova | Cantacuzino Institute Virology | Carmen Cherciu,Luiza Ustea, Nicoleta Paraschiv, Sorin Dinu, Mihaela Oprea, Mihaela Lazar |
| EPI_ISL_3021427 | Cantacuzino National Military-Medical Institute, Viral Respiratory Infections Laboratory | Cantacuzino Institute Virology | Carmen Cherciu,Luiza Ustea, Nicoleta Paraschiv, Sorin Dinu, Mihaela Oprea, Mihaela Lazar |
| EPI_ISL_3021428 | Clinical Emergency County Hospital, Craiova | Cantacuzino Institute Virology | Carmen Cherciu,Luiza Ustea, Nicoleta Paraschiv, Sorin Dinu, Mihaela Oprea, Mihaela Lazar |
| EPI_ISL_3021429, EPI_ISL_3021430, EPI_ISL_3021431, EPI_ISL_3021432, EPI_ISL_3021433, EPI_ISL_3021435, EPI_ISL_3021436, EPI_ISL_3021437, EPI_ISL_3021438, EPI_ISL_3021439, EPI_ISL_3021440, EPI_ISL_3021441, EPI_ISL_3021442, EPI_ISL_3021443, EPI_ISL_3021444, EPI_ISL_3021445, EPI_ISL_3021446, EPI_ISL_3021447, EPI_ISL_3021448, EPI_ISL_3021449, EPI_ISL_3021450, EPI_ISL_3021451, EPI_ISL_3021452 | | | |
| see above | Nebraska Public Health Laboratory | NPHL COVID-19 Response Team | NPHL COVID-19 Response Team |
| EPI_ISL_3021460 | Twin Cities Community Hospital Lab | County of San Luis Obispo Public Health Laboratory | Katrina Erwin, Frances Sidhu |
| EPI_ISL_3021462, EPI_ISL_3021465, EPI_ISL_3021466, EPI_ISL_3021468, EPI_ISL_3021469, EPI_ISL_3021470, EPI_ISL_3021471, EPI_ISL_3021472, EPI_ISL_3021473, EPI_ISL_3021475 | County of San Luis Obispo Public Health Laboratory | County of San Luis Obispo Public Health Laboratory | Katrina Erwin, Frances Sidhu |
| EPI_ISL_3021476 | Nucleic Acid Testing, National Reference Laboratory | GIGA Medical Genomics | Yvan Butera, Keith Durkin, Maria Artesi, Bouchra Boujemla, Robert Rutayisire, Reuben Sindayiheba, Patrick Tuyisenge, Esperence Umumararungu, Sébastien Bontems, Marie-Pierre Hayette, Nathalie Renotte, Swaibu Gatare, Jacob Souopgui, Sabin Nsanzimana, Vincent Bours, Léon Mutesa |
| EPI_ISL_3021549, EPI_ISL_3021550, EPI_ISL_3021551, EPI_ISL_3021552, EPI_ISL_3021553, EPI_ISL_3021554, EPI_ISL_3021555, EPI_ISL_3021559, EPI_ISL_3021560, EPI_ISL_3021561, EPI_ISL_3021562, EPI_ISL_3021563, EPI_ISL_3021564, EPI_ISL_3021565, EPI_ISL_3021566, EPI_ISL_3021567, EPI_ISL_3021568, EPI_ISL_3021570, EPI_ISL_3021571, EPI_ISL_3021572, EPI_ISL_3021573, EPI_ISL_3021574, EPI_ISL_3021575, EPI_ISL_3021576, EPI_ISL_3021577, EPI_ISL_3021578, EPI_ISL_3021579, EPI_ISL_3021581, EPI_ISL_3021584, EPI_ISL_3021585, EPI_ISL_3021587, EPI_ISL_3021588, EPI_ISL_3021589, EPI_ISL_3021590, EPI_ISL_3021591, EPI_ISL_3021593, EPI_ISL_3021594, EPI_ISL_3021599, EPI_ISL_3021600, EPI_ISL_3021601, EPI_ISL_3021602, EPI_ISL_3021603, EPI_ISL_3021604, EPI_ISL_3021605, EPI_ISL_3021606, EPI_ISL_3021607, EPI_ISL_3021609, EPI_ISL_3021610, EPI_ISL_3021611, EPI_ISL_3021612, EPI_ISL_3021613, EPI_ISL_3021614, EPI_ISL_3021615, EPI_ISL_3021617, EPI_ISL_3021618, EPI_ISL_3021619, EPI_ISL_3021620, EPI_ISL_3021621, EPI_ISL_3021622, EPI_ISL_3021623, EPI_ISL_3021624, EPI_ISL_3021625, EPI_ISL_3021626, EPI_ISL_3021628, EPI_ISL_3021629, EPI_ISL_3021630, EPI_ISL_3021631, EPI_ISL_3021632, EPI_ISL_3021633, EPI_ISL_3021634, EPI_ISL_3021635, EPI_ISL_3021636, EPI_ISL_3021638, EPI_ISL_3021641, EPI_ISL_3021644, EPI_ISL_3021645, EPI_ISL_3021648, EPI_ISL_3021649, EPI_ISL_3021650, EPI_ISL_3021651, EPI_ISL_3021652, EPI_ISL_3021653, EPI_ISL_3021654, EPI_ISL_3021657, EPI_ISL_3021661, EPI_ISL_3021663, EPI_ISL_3021664, EPI_ISL_3021667, EPI_ISL_3021668, EPI_ISL_3021670, EPI_ISL_3021671, EPI_ISL_3021673, EPI_ISL_3021675, EPI_ISL_3021676, EPI_ISL_3021677, EPI_ISL_3021679, EPI_ISL_3021681, EPI_ISL_3021683, EPI_ISL_3021684, EPI_ISL_3021685, EPI_ISL_3021686, EPI_ISL_3021688, EPI_ISL_3021689, EPI_ISL_3021690, EPI_ISL_3021691, EPI_ISL_3021692, EPI_ISL_3021693, EPI_ISL_3021694, EPI_ISL_3021695, EPI_ISL_3021696, EPI_ISL_3021697, EPI_ISL_3021698, EPI_ISL_3021699, EPI_ISL_3021700, EPI_ISL_3021701, EPI_ISL_3021703, EPI_ISL_3021705, EPI_ISL_3021708, EPI_ISL_3021709, EPI_ISL_3021710, EPI_ISL_3021711, EPI_ISL_3021712, EPI_ISL_3021713, EPI_ISL_3021714, EPI_ISL_3021716, EPI_ISL_3021717, EPI_ISL_3021718, EPI_ISL_3021719, EPI_ISL_3021720, EPI_ISL_3021721, EPI_ISL_3021722, EPI_ISL_3021723, EPI_ISL_3021724, EPI_ISL_3021725, EPI_ISL_3021726, EPI_ISL_3021728, EPI_ISL_3021729, EPI_ISL_3021731, EPI_ISL_3021732, EPI_ISL_3021733, EPI_ISL_3021735, EPI_ISL_3021736, EPI_ISL_3021737, EPI_ISL_3021738, EPI_ISL_3021739, EPI_ISL_3021740, EPI_ISL_3021741, EPI_ISL_3021742, EPI_ISL_3021744, EPI_ISL_3021745, EPI_ISL_3021746, EPI_ISL_3021747, EPI_ISL_3021748, EPI_ISL_3021749, EPI_ISL_3021750, EPI_ISL_3021751, EPI_ISL_3021752, EPI_ISL_3021754, EPI_ISL_3021757, EPI_ISL_3021758, EPI_ISL_3021759, EPI_ISL_3021760, EPI_ISL_3021761, EPI_ISL_3021762, EPI_ISL_3021763, EPI_ISL_3021764, EPI_ISL_3021766, EPI_ISL_3021767, EPI_ISL_3021768, EPI_ISL_3021769, EPI_ISL_3021770, EPI_ISL_3021771, EPI_ISL_3021772, EPI_ISL_3021773, EPI_ISL_3021774, EPI_ISL_3021775, EPI_ISL_3021776, EPI_ISL_3021777, EPI_ISL_3021778, EPI_ISL_3021780, EPI_ISL_3021781, EPI_ISL_3021782, EPI_ISL_3021783, EPI_ISL_3021784, EPI_ISL_3021785, EPI_ISL_3021786, EPI_ISL_3021787, EPI_ISL_3021790, EPI_ISL_3021792, EPI_ISL_3021793, EPI_ISL_3021794, EPI_ISL_3021795, EPI_ISL_3021796, EPI_ISL_3021797, EPI_ISL_3021798, EPI_ISL_3021799, EPI_ISL_3021800, EPI_ISL_3021801, EPI_ISL_3021802, EPI_ISL_3021803, EPI_ISL_3021804, EPI_ISL_3021805, EPI_ISL_3021806, EPI_ISL_3021807, EPI_ISL_3021808, EPI_ISL_3021809, EPI_ISL_3021811, EPI_ISL_3021812, EPI_ISL_3021813, EPI_ISL_3021814, EPI_ISL_3021815, EPI_ISL_3021818, EPI_ISL_3021819, EPI_ISL_3021820, EPI_ISL_3021821, EPI_ISL_3021823, EPI_ISL_3021825, EPI_ISL_3021826, EPI_ISL_3021827, EPI_ISL_3021830, EPI_ISL_3021831, EPI_ISL_3021832, EPI_ISL_3021833, EPI_ISL_3021834, EPI_ISL_3021835, EPI_ISL_3021836, EPI_ISL_3021837, EPI_ISL_3021838, EPI_ISL_3021839, EPI_ISL_3021840, EPI_ISL_3021842, EPI_ISL_3021843, EPI_ISL_3021844, EPI_ISL_3021845, EPI_ISL_3021846, EPI_ISL_3021847, EPI_ISL_3021848, EPI_ISL_3021849, EPI_ISL_3021851, EPI_ISL_3021852, EPI_ISL_3021853, EPI_ISL_3021854, EPI_ISL_3021855, EPI_ISL_3021856, EPI_ISL_3021858, EPI_ISL_3021859, EPI_ISL_3021860, EPI_ISL_3021861, EPI_ISL_3021862, EPI_ISL_3021863, EPI_ISL_3021864, EPI_ISL_3021865, EPI_ISL_3021866, EPI_ISL_3021867, EPI_ISL_3021869, EPI_ISL_3021871, EPI_ISL_3021872, EPI_ISL_3021873, EPI_ISL_3021874, EPI_ISL_3021875, EPI_ISL_3021876, EPI_ISL_3021877, EPI_ISL_3021878, EPI_ISL_3021879, EPI_ISL_3021880, EPI_ISL_3021881, EPI_ISL_3021882, EPI_ISL_3021883 | | | |
| see above | Shamir Medical Center (Asaf Harofe) | Shamir Medical Center (Asaf Harofe) | Nir Rainy,Chen Weiner,Reut Sorek Abramovich,Yevgeni Yegorov,Anna Vishnevsky,Patricia Benveniste-Lekovitz,Abu Hamad Ramzia,Adina Bar Chaim |
| EPI_ISL_3021884, EPI_ISL_3021885, EPI_ISL_3021886, EPI_ISL_3021887, EPI_ISL_3021888, EPI_ISL_3021889, EPI_ISL_3021890, EPI_ISL_3021891, EPI_ISL_3021892, EPI_ISL_3021893, EPI_ISL_3021894, EPI_ISL_3021895, EPI_ISL_3021897, EPI_ISL_3021898, EPI_ISL_3021899, EPI_ISL_3021900, EPI_ISL_3021901, EPI_ISL_3021902, EPI_ISL_3021903, EPI_ISL_3021904, EPI_ISL_3021905, EPI_ISL_3021906, EPI_ISL_3021907, EPI_ISL_3021908, EPI_ISL_3021909, EPI_ISL_3021910, EPI_ISL_3021911, EPI_ISL_3021912, EPI_ISL_3021913, EPI_ISL_3021914, EPI_ISL_3021915, EPI_ISL_3021916, EPI_ISL_3021917, EPI_ISL_3021918, EPI_ISL_3021920, EPI_ISL_3021921, EPI_ISL_3021922, EPI_ISL_3021923, EPI_ISL_3021924, EPI_ISL_3021925, EPI_ISL_3021926, EPI_ISL_3021927, EPI_ISL_3021928, EPI_ISL_3021929, EPI_ISL_3021930, EPI_ISL_3021931, EPI_ISL_3021932, EPI_ISL_3021933, EPI_ISL_3021934, EPI_ISL_3021935, EPI_ISL_3021936, EPI_ISL_3021937, EPI_ISL_3021938, EPI_ISL_3021939, EPI_ISL_3021940, EPI_ISL_3021941, EPI_ISL_3021942, EPI_ISL_3021943, EPI_ISL_3021944, EPI_ISL_3021945, EPI_ISL_3021946, EPI_ISL_3021947, EPI_ISL_3021948, EPI_ISL_3021949, EPI_ISL_3021950, EPI_ISL_3021951, EPI_ISL_3021952, EPI_ISL_3021953, EPI_ISL_3021954, EPI_ISL_3021955, EPI_ISL_3021956, EPI_ISL_3021957, EPI_ISL_3021958, EPI_ISL_3021959, EPI_ISL_3021960, EPI_ISL_3021961, EPI_ISL_3021962, EPI_ISL_3021963, EPI_ISL_3021964, EPI_ISL_3021965, EPI_ISL_3021966, EPI_ISL_3021967, EPI_ISL_3021968, EPI_ISL_3021969, EPI_ISL_3021970, EPI_ISL_3021971, EPI_ISL_3021972, EPI_ISL_3021973, EPI_ISL_3021974, EPI_ISL_3021975, EPI_ISL_3021976, EPI_ISL_3021977, EPI_ISL_3021978, EPI_ISL_3021979, EPI_ISL_3021980, EPI_ISL_3021981, EPI_ISL_3021982, EPI_ISL_3021983, EPI_ISL_3021984, EPI_ISL_3021985, EPI_ISL_3021986, EPI_ISL_3021987, EPI_ISL_3021988, EPI_ISL_3021989, EPI_ISL_3021990, EPI_ISL_3021991, EPI_ISL_3021992, EPI_ISL_3021993, EPI_ISL_3021994, EPI_ISL_3021995, EPI_ISL_3021996, EPI_ISL_3021997 | | | |
| see above | INSACOG-MANIPUR | National Institute of Biomedical Genomics – INSACOG | Arindam Maitra, R.K.Manojkumar Singh, Nidhan Kumar Biswas, Kh. Ranjana Devi, Sreedhar Chinnaswamy, L. Shivadutta Singh, Saumitra Das |
| EPI_ISL_3022014, EPI_ISL_3022015, EPI_ISL_3022016, EPI_ISL_3022018, EPI_ISL_3022019, EPI_ISL_3022020, EPI_ISL_3022021, EPI_ISL_3022022, EPI_ISL_3022023, EPI_ISL_3022024, EPI_ISL_3022025, EPI_ISL_3022026, EPI_ISL_3022027, EPI_ISL_3022029, EPI_ISL_3022030, EPI_ISL_3022031, EPI_ISL_3022032, EPI_ISL_3022033, EPI_ISL_3022034, EPI_ISL_3022035, EPI_ISL_3022037, EPI_ISL_3022038, EPI_ISL_3022039, EPI_ISL_3022040, EPI_ISL_3022042, EPI_ISL_3022043, EPI_ISL_3022044, EPI_ISL_3022045, EPI_ISL_3022046, EPI_ISL_3022047, EPI_ISL_3022048, EPI_ISL_3022049, EPI_ISL_3022050, EPI_ISL_3022051, EPI_ISL_3022052, EPI_ISL_3022053, EPI_ISL_3022054, EPI_ISL_3022055, EPI_ISL_3022057, EPI_ISL_3022058, EPI_ISL_3022059, EPI_ISL_3022060, EPI_ISL_3022062, EPI_ISL_3022064, EPI_ISL_3022065, EPI_ISL_3022068, EPI_ISL_3022069, EPI_ISL_3022070, EPI_ISL_3022071, EPI_ISL_3022072, EPI_ISL_3022073, EPI_ISL_3022074, EPI_ISL_3022075, EPI_ISL_3022076, EPI_ISL_3022077, EPI_ISL_3022078, EPI_ISL_3022079, EPI_ISL_3022080, EPI_ISL_3022081, EPI_ISL_3022082, EPI_ISL_3022083, EPI_ISL_3022085, EPI_ISL_3022086, EPI_ISL_3022087, EPI_ISL_3022088, EPI_ISL_3022089, | | | |

| | | | |
|---|---|---|---|
| EPI_ISL_3022090, EPI_ISL_3022091, EPI_ISL_3022092, EPI_ISL_3022093, EPI_ISL_3022094, EPI_ISL_3022095, EPI_ISL_3022096, EPI_ISL_3022098, EPI_ISL_3022099, EPI_ISL_3022100, EPI_ISL_3022101, EPI_ISL_3022102, EPI_ISL_3022103, EPI_ISL_3022104, EPI_ISL_3022105, EPI_ISL_3022106, EPI_ISL_3022107, EPI_ISL_3022108, EPI_ISL_3022109, EPI_ISL_3022110, EPI_ISL_3022111, EPI_ISL_3022112, EPI_ISL_3022113, EPI_ISL_3022114, EPI_ISL_3022115, EPI_ISL_3022116, EPI_ISL_3022117, EPI_ISL_3022118, EPI_ISL_3022120, EPI_ISL_3022121, EPI_ISL_3022123, EPI_ISL_3022124, EPI_ISL_3022125, EPI_ISL_3022126, EPI_ISL_3022129, EPI_ISL_3022130, EPI_ISL_3022131, EPI_ISL_3022132, EPI_ISL_3022133, EPI_ISL_3022134, EPI_ISL_3022135, EPI_ISL_3022137, EPI_ISL_3022138, EPI_ISL_3022139, EPI_ISL_3022140, EPI_ISL_3022141, EPI_ISL_3022142, EPI_ISL_3022143, EPI_ISL_3022144, EPI_ISL_3022145, EPI_ISL_3022146, EPI_ISL_3022147, EPI_ISL_3022148, EPI_ISL_3022149, EPI_ISL_3022150, EPI_ISL_3022151, EPI_ISL_3022152, EPI_ISL_3022153, EPI_ISL_3022155, EPI_ISL_3022156, EPI_ISL_3022160, EPI_ISL_3022161, EPI_ISL_3022163, EPI_ISL_3022164, EPI_ISL_3022165, EPI_ISL_3022166, EPI_ISL_3022168, EPI_ISL_3022169, EPI_ISL_3022171, EPI_ISL_3022172, EPI_ISL_3022173, EPI_ISL_3022174, EPI_ISL_3022175, EPI_ISL_3022176, EPI_ISL_3022177, EPI_ISL_3022178, EPI_ISL_3022179, EPI_ISL_3022180, EPI_ISL_3022181, EPI_ISL_3022182, EPI_ISL_3022183, EPI_ISL_3022184, EPI_ISL_3022185, EPI_ISL_3022186, EPI_ISL_3022187, EPI_ISL_3022189, EPI_ISL_3022191, EPI_ISL_3022192, EPI_ISL_3022193, EPI_ISL_3022194, EPI_ISL_3022195, EPI_ISL_3022196, EPI_ISL_3022198, EPI_ISL_3022200, EPI_ISL_3022201, EPI_ISL_3022203, EPI_ISL_3022205, EPI_ISL_3022206, EPI_ISL_3022208, EPI_ISL_3022209, EPI_ISL_3022210, EPI_ISL_3022211, EPI_ISL_3022212, EPI_ISL_3022213, EPI_ISL_3022214, EPI_ISL_3022216, EPI_ISL_3022217, EPI_ISL_3022218, EPI_ISL_3022219, EPI_ISL_3022220, EPI_ISL_3022221, EPI_ISL_3022223, EPI_ISL_3022224, EPI_ISL_3022225, EPI_ISL_3022226, EPI_ISL_3022227, EPI_ISL_3022228, EPI_ISL_3022229, EPI_ISL_3022232, EPI_ISL_3022233, EPI_ISL_3022234, EPI_ISL_3022236, EPI_ISL_3022237, EPI_ISL_3022238, EPI_ISL_3022239, EPI_ISL_3022240, EPI_ISL_3022241, EPI_ISL_3022242, EPI_ISL_3022245, EPI_ISL_3022246, EPI_ISL_3022247, EPI_ISL_3022249, EPI_ISL_3022252, EPI_ISL_3022253, EPI_ISL_3022254, EPI_ISL_3022255, EPI_ISL_3022257, EPI_ISL_3022258, EPI_ISL_3022259, EPI_ISL_3022260, EPI_ISL_3022261, EPI_ISL_3022262, EPI_ISL_3022264, EPI_ISL_3022266, EPI_ISL_3022267, EPI_ISL_3022268, EPI_ISL_3022271, EPI_ISL_3022273, EPI_ISL_3022274, EPI_ISL_3022275, EPI_ISL_3022277, EPI_ISL_3022279, EPI_ISL_3022280, EPI_ISL_3022281, EPI_ISL_3022282, EPI_ISL_3022283, EPI_ISL_3022284, EPI_ISL_3022285, EPI_ISL_3022286, EPI_ISL_3022288, EPI_ISL_3022289, EPI_ISL_3022290, EPI_ISL_3022291, EPI_ISL_3022292, EPI_ISL_3022293, EPI_ISL_3022294, EPI_ISL_3022295, EPI_ISL_3022296, EPI_ISL_3022297, EPI_ISL_3022298, EPI_ISL_3022299, EPI_ISL_3022300, EPI_ISL_3022301, EPI_ISL_3022302, EPI_ISL_3022303, EPI_ISL_3022304, EPI_ISL_3022305, EPI_ISL_3022307, EPI_ISL_3022308, EPI_ISL_3022309, EPI_ISL_3022310, EPI_ISL_3022311, EPI_ISL_3022312, EPI_ISL_3022313, EPI_ISL_3022314, EPI_ISL_3022315, EPI_ISL_3022316, EPI_ISL_3022317, EPI_ISL_3022318, EPI_ISL_3022319, EPI_ISL_3022321, EPI_ISL_3022322, EPI_ISL_3022323, EPI_ISL_3022324, EPI_ISL_3022325, EPI_ISL_3022326, EPI_ISL_3022327, EPI_ISL_3022329, EPI_ISL_3022330, EPI_ISL_3022331, EPI_ISL_3022332, EPI_ISL_3022334, EPI_ISL_3022335, EPI_ISL_3022336, EPI_ISL_3022337, EPI_ISL_3022338, EPI_ISL_3022339, EPI_ISL_3022340, EPI_ISL_3022341, EPI_ISL_3022343, EPI_ISL_3022345, EPI_ISL_3022346, EPI_ISL_3022347, EPI_ISL_3022348, EPI_ISL_3022349, EPI_ISL_3022350, EPI_ISL_3022351, EPI_ISL_3022352, EPI_ISL_3022353, EPI_ISL_3022354, EPI_ISL_3022355, EPI_ISL_3022356 | | | |
| see above | Shamir Medical Center (Asaf Harofe) | Shamir Medical Center (Asaf Harofe) | Nir Rainy,Chen Weiner,Reut Sorek Abramovich,Yevgeni Yegorov,Anna Vishnevsky,Patricia Benveniste-Lekovitz,Abu Hamad Ramzia,Adina Bar Chaim |
| EPI_ISL_3022894, EPI_ISL_3023027, EPI_ISL_3023151, EPI_ISL_3023332, EPI_ISL_3023353, EPI_ISL_3023357, EPI_ISL_3023358, EPI_ISL_3023370, EPI_ISL_3023371, EPI_ISL_3023376 | Hospital of the University of Pennsylvania Molecular Pathology Lab | Bushman Lab - University of Pennsylvania | John K. Everett, Kyle Rodino, Shantan Reddy, Aoife M. Roche, Young Hwang, Scott Sherrill-Mix, Samantha A. Whiteside, Jevon Graham-Wooten, Layla A. Khatib, Ayannah S. Fitzgerald, Arupa Ganguly, Mike Feldman, Brendan Kelly, Ronald G. Collman and Frederic Bushan |
| EPI_ISL_3023712, EPI_ISL_3023713, EPI_ISL_3023714, EPI_ISL_3023715, EPI_ISL_3023716, EPI_ISL_3023718, EPI_ISL_3023720, EPI_ISL_3023721, EPI_ISL_3023722, EPI_ISL_3023723, EPI_ISL_3023724, EPI_ISL_3023725, EPI_ISL_3023726, EPI_ISL_3023727, EPI_ISL_3023728, EPI_ISL_3023729, EPI_ISL_3023730, EPI_ISL_3023731, EPI_ISL_3023732, EPI_ISL_3023733, EPI_ISL_3023734, EPI_ISL_3023735, EPI_ISL_3023736, EPI_ISL_3023737, EPI_ISL_3023738, EPI_ISL_3023740, EPI_ISL_3023741, EPI_ISL_3023742, EPI_ISL_3023743, EPI_ISL_3023745, EPI_ISL_3023746, EPI_ISL_3023747, EPI_ISL_3023752, EPI_ISL_3023756, EPI_ISL_3023758, EPI_ISL_3023760, EPI_ISL_3023762, EPI_ISL_3023763 | | | |
| see above | NJDOH, Public Health and Environmental Laboratories | NJ_PHEL | Lindsey Bodnar, Shiv K. Verma, Ryan Pachucki, Chelsea San Filippo, Jacquelyn Deverell, Dana Woell, Allison Roder, Byeong Jeong |
| EPI_ISL_3023765, EPI_ISL_3023766, EPI_ISL_3023768, EPI_ISL_3023770, EPI_ISL_3023771, EPI_ISL_3023774, EPI_ISL_3023777, EPI_ISL_3023779, EPI_ISL_3023780, EPI_ISL_3023781, EPI_ISL_3023783, EPI_ISL_3023784, EPI_ISL_3023786, EPI_ISL_3023787, EPI_ISL_3023788, EPI_ISL_3023789 | | | |
| see above | CT Department of Public Health | CT Department of Public Health | Claire_Pearson,Tu_N_Nguyen |
| EPI_ISL_3024083, EPI_ISL_3024084, EPI_ISL_3024085, EPI_ISL_3024087, EPI_ISL_3024088, EPI_ISL_3024089 | University Hospital Limerick | University Hospital Limerick | Carolyn Meaney, Colm McDonnell, Patrick Stapleton, Daniel Hare |
| EPI_ISL_3024102, EPI_ISL_3024103, EPI_ISL_3024106, EPI_ISL_3024107, EPI_ISL_3024108, EPI_ISL_3024110, EPI_ISL_3024111, EPI_ISL_3024112, EPI_ISL_3024118, EPI_ISL_3024119, EPI_ISL_3024120, EPI_ISL_3024121, EPI_ISL_3024122, EPI_ISL_3024123, EPI_ISL_3024124, EPI_ISL_3024125, EPI_ISL_3024126, EPI_ISL_3024127, EPI_ISL_3024128, EPI_ISL_3024129, EPI_ISL_3024130, EPI_ISL_3024131, EPI_ISL_3024132 | | | |
| see above | Rhode Island State Health Laboratory | Rhode Island State Health Laboratory | Ewa King, Richard C. Huard, Kristin Carpenter-Azevedo |
| EPI_ISL_3024135, EPI_ISL_3024136, EPI_ISL_3024137, EPI_ISL_3024138, EPI_ISL_3024139, EPI_ISL_3024140, EPI_ISL_3024141, EPI_ISL_3024142, EPI_ISL_3024143, EPI_ISL_3024144, EPI_ISL_3024146, EPI_ISL_3024147, EPI_ISL_3024148, EPI_ISL_3024149, EPI_ISL_3024150, EPI_ISL_3024151, EPI_ISL_3024152, EPI_ISL_3024153, EPI_ISL_3024154, EPI_ISL_3024155, EPI_ISL_3024156, EPI_ISL_3024157, EPI_ISL_3024158, EPI_ISL_3024159, EPI_ISL_3024161, EPI_ISL_3024162, EPI_ISL_3024163, EPI_ISL_3024164, EPI_ISL_3024165, EPI_ISL_3024166, EPI_ISL_3024167, EPI_ISL_3024168, EPI_ISL_3024169, EPI_ISL_3024171, EPI_ISL_3024172, EPI_ISL_3024173, EPI_ISL_3024174, EPI_ISL_3024175, EPI_ISL_3024176, EPI_ISL_3024177, EPI_ISL_3024178, EPI_ISL_3024179, EPI_ISL_3024181, EPI_ISL_3024182, EPI_ISL_3024183, EPI_ISL_3024185, EPI_ISL_3024186, EPI_ISL_3024187, EPI_ISL_3024188, EPI_ISL_3024189, EPI_ISL_3024190, EPI_ISL_3024191, EPI_ISL_3024195, EPI_ISL_3024197, EPI_ISL_3024198, EPI_ISL_3024199, EPI_ISL_3024200, EPI_ISL_3024201, EPI_ISL_3024202, EPI_ISL_3024203, EPI_ISL_3024204, EPI_ISL_3024206, EPI_ISL_3024207, EPI_ISL_3024208, EPI_ISL_3024209, EPI_ISL_3024210, EPI_ISL_3024211, EPI_ISL_3024212, EPI_ISL_3024213, EPI_ISL_3024214, EPI_ISL_3024216, EPI_ISL_3024217, EPI_ISL_3024219, EPI_ISL_3024221, EPI_ISL_3024222, EPI_ISL_3024224, EPI_ISL_3024225, EPI_ISL_3024226, EPI_ISL_3024227, EPI_ISL_3024228, EPI_ISL_3024229, EPI_ISL_3024231, EPI_ISL_3024232, EPI_ISL_3024233, EPI_ISL_3024234, EPI_ISL_3024235, EPI_ISL_3024236, EPI_ISL_3024237, EPI_ISL_3024238, EPI_ISL_3024239, EPI_ISL_3024243, EPI_ISL_3024244, EPI_ISL_3024245, EPI_ISL_3024246, EPI_ISL_3024247, EPI_ISL_3024248, EPI_ISL_3024250, EPI_ISL_3024251, EPI_ISL_3024252, EPI_ISL_3024254, EPI_ISL_3024255, EPI_ISL_3024257, EPI_ISL_3024258, EPI_ISL_3024260, EPI_ISL_3024261, EPI_ISL_3024262, EPI_ISL_3024263, EPI_ISL_3024264, EPI_ISL_3024265, EPI_ISL_3024266, EPI_ISL_3024267, EPI_ISL_3024268, EPI_ISL_3024269, EPI_ISL_3024270, EPI_ISL_3024271, EPI_ISL_3024272, EPI_ISL_3024273, EPI_ISL_3024275, EPI_ISL_3024276, EPI_ISL_3024278, EPI_ISL_3024279, EPI_ISL_3024280, EPI_ISL_3024283, EPI_ISL_3024284, EPI_ISL_3024285, EPI_ISL_3024286, EPI_ISL_3024287, EPI_ISL_3024288, EPI_ISL_3024289, EPI_ISL_3024290, EPI_ISL_3024291, EPI_ISL_3024292, EPI_ISL_3024293, EPI_ISL_3024294, EPI_ISL_3024295, EPI_ISL_3024296, EPI_ISL_3024297, EPI_ISL_3024299, EPI_ISL_3024300, EPI_ISL_3024301, EPI_ISL_3024302, EPI_ISL_3024303, EPI_ISL_3024305, EPI_ISL_3024306, EPI_ISL_3024307, EPI_ISL_3024308, EPI_ISL_3024309, EPI_ISL_3024310, EPI_ISL_3024312, EPI_ISL_3024313, EPI_ISL_3024314, EPI_ISL_3024315, EPI_ISL_3024317, EPI_ISL_3024318, EPI_ISL_3024319, EPI_ISL_3024320, EPI_ISL_3024321, EPI_ISL_3024323, EPI_ISL_3024324, EPI_ISL_3024325, EPI_ISL_3024326, EPI_ISL_3024329, EPI_ISL_3024330, EPI_ISL_3024331, EPI_ISL_3024332, EPI_ISL_3024333, EPI_ISL_3024334, EPI_ISL_3024335, EPI_ISL_3024336, EPI_ISL_3024337, EPI_ISL_3024338, EPI_ISL_3024339, EPI_ISL_3024340, EPI_ISL_3024341, EPI_ISL_3024342, EPI_ISL_3024343, EPI_ISL_3024344, EPI_ISL_3024345, EPI_ISL_3024346, EPI_ISL_3024349, EPI_ISL_3024350, EPI_ISL_3024351, EPI_ISL_3024352, EPI_ISL_3024354, EPI_ISL_3024356, EPI_ISL_3024357, EPI_ISL_3024358, EPI_ISL_3024359, EPI_ISL_3024360, EPI_ISL_3024361, EPI_ISL_3024363, EPI_ISL_3024364, EPI_ISL_3024366, EPI_ISL_3024368, EPI_ISL_3024370, EPI_ISL_3024371, EPI_ISL_3024372, EPI_ISL_3024373, EPI_ISL_3024374, EPI_ISL_3024375, EPI_ISL_3024376, EPI_ISL_3024377, EPI_ISL_3024378, EPI_ISL_3024382, EPI_ISL_3024383, EPI_ISL_3024384, EPI_ISL_3024385, EPI_ISL_3024386, EPI_ISL_3024387, EPI_ISL_3024388, EPI_ISL_3024389, EPI_ISL_3024390, EPI_ISL_3024391, EPI_ISL_3024392, EPI_ISL_3024393, EPI_ISL_3024394, EPI_ISL_3024395, EPI_ISL_3024396, EPI_ISL_3024397, EPI_ISL_3024399, EPI_ISL_3024400, EPI_ISL_3024401, EPI_ISL_3024402, EPI_ISL_3024403, EPI_ISL_3024404, EPI_ISL_3024405, EPI_ISL_3024406, EPI_ISL_3024408, EPI_ISL_3024409, EPI_ISL_3024410, EPI_ISL_3024411, EPI_ISL_3024412, EPI_ISL_3024413, EPI_ISL_3024414, EPI_ISL_3024415, EPI_ISL_3024416, EPI_ISL_3024418, EPI_ISL_3024420, EPI_ISL_3024421, EPI_ISL_3024422, EPI_ISL_3024423, EPI_ISL_3024425, EPI_ISL_3024426, EPI_ISL_3024427, EPI_ISL_3024428, EPI_ISL_3024429, EPI_ISL_3024430, EPI_ISL_3024431, EPI_ISL_3024432, EPI_ISL_3024433, EPI_ISL_3024434, EPI_ISL_3024435, EPI_ISL_3024436, EPI_ISL_3024437, EPI_ISL_3024438, EPI_ISL_3024439, EPI_ISL_3024440, EPI_ISL_3024441, EPI_ISL_3024442, EPI_ISL_3024443, EPI_ISL_3024444, EPI_ISL_3024445, EPI_ISL_3024446, EPI_ISL_3024449, EPI_ISL_3024450, EPI_ISL_3024451, EPI_ISL_3024452, EPI_ISL_3024453, EPI_ISL_3024454, EPI_ISL_3024455, EPI_ISL_3024456, EPI_ISL_3024457, EPI_ISL_3024458, EPI_ISL_3024459, EPI_ISL_3024460, EPI_ISL_3024461, EPI_ISL_3024462, EPI_ISL_3024463, EPI_ISL_3024464, EPI_ISL_3024465, EPI_ISL_3024466, EPI_ISL_3024467, EPI_ISL_3024468, EPI_ISL_3024469, EPI_ISL_3024472, EPI_ISL_3024473, EPI_ISL_3024475, EPI_ISL_3024477, EPI_ISL_3024478, EPI_ISL_3024479, EPI_ISL_3024480, EPI_ISL_3024481, EPI_ISL_3024483, EPI_ISL_3024484, EPI_ISL_3024485, EPI_ISL_3024487, EPI_ISL_3024488, EPI_ISL_3024489, EPI_ISL_3024490, EPI_ISL_3024491, EPI_ISL_3024492, EPI_ISL_3024493, EPI_ISL_3024494, EPI_ISL_3024496, EPI_ISL_3024498 | | | |
| see above | Shamir Medical Center (Asaf Harofe) | Shamir Medical Center (Asaf Harofe) | Nir Rainy,Chen Weiner,Reut Sorek Abramovich,Yevgeni Yegorov,Anna Vishnevsky,Patricia Benveniste-Lekovitz,Abu Hamad Ramzia,Adina Bar Chaim |
| EPI_ISL_3024500, EPI_ISL_3024501, EPI_ISL_3024503, EPI_ISL_3024504, EPI_ISL_3024506, EPI_ISL_3024508, EPI_ISL_3024509, EPI_ISL_3024510, EPI_ISL_3024512, EPI_ISL_3024513 | POLOKWANE LABORATORY | National Institute for Communicable Diseases of the National Health Laboratory Service | Amoako DG, Everatt J, Scheepers C, Mohale T, Ntuli N, Mahlangu B, Mnguni A, Ismail A, Bhiman JN |
| EPI_ISL_3024514, EPI_ISL_3024516, EPI_ISL_3024517, EPI_ISL_3024518, EPI_ISL_3024519, EPI_ISL_3024520, EPI_ISL_3024522, EPI_ISL_3024523, EPI_ISL_3024525, EPI_ISL_3024526 | LEBOWAKGOMO LABORATORY | National Institute for Communicable Diseases of the National Health Laboratory Service | Amoako DG, Everatt J, Scheepers C, Mohale T, Ntuli N, Mahlangu B, Mnguni A, Ismail A, Bhiman JN |
| EPI_ISL_3024527, EPI_ISL_3024529, EPI_ISL_3024530, EPI_ISL_3024531, EPI_ISL_3024533, EPI_ISL_3024535, EPI_ISL_3024536, EPI_ISL_3024537, EPI_ISL_3024539, EPI_ISL_3024541, EPI_ISL_3024542, EPI_ISL_3024544, EPI_ISL_3024545, EPI_ISL_3024547, EPI_ISL_3024548, EPI_ISL_3024549, EPI_ISL_3024550, EPI_ISL_3024551, EPI_ISL_3024552, EPI_ISL_3024553, EPI_ISL_3024555, EPI_ISL_3024556, EPI_ISL_3024557, EPI_ISL_3024559, EPI_ISL_3024560, EPI_ISL_3024565, EPI_ISL_3024566, EPI_ISL_3024567, EPI_ISL_3024568, EPI_ISL_3024571, EPI_ISL_3024572 | | | |
| see above | MANKWENG LABORATORY | National Institute for Communicable Diseases of the National Health Laboratory Service | Amoako DG, Everatt J, Scheepers C, Mohale T, Ntuli N, Mahlangu B, Mnguni A, Ismail A, Bhiman JN |
| EPI_ISL_3024575, EPI_ISL_3024577, EPI_ISL_3024578, EPI_ISL_3024579, EPI_ISL_3024580, EPI_ISL_3024581, EPI_ISL_3024584, EPI_ISL_3024585, EPI_ISL_3024586, EPI_ISL_3024587, EPI_ISL_3024588, EPI_ISL_3024589, EPI_ISL_3024590, EPI_ISL_3024591, EPI_ISL_3024592, EPI_ISL_3024593, EPI_ISL_3024594, EPI_ISL_3024595, EPI_ISL_3024596, EPI_ISL_3024597, EPI_ISL_3024598, EPI_ISL_3024599, EPI_ISL_3024601, EPI_ISL_3024602, EPI_ISL_3024604, EPI_ISL_3024605, EPI_ISL_3024606, EPI_ISL_3024607, EPI_ISL_3024608, EPI_ISL_3024609, EPI_ISL_3024610, EPI_ISL_3024611, EPI_ISL_3024614, EPI_ISL_3024615, EPI_ISL_3024616, EPI_ISL_3024620, EPI_ISL_3024621, EPI_ISL_3024623, EPI_ISL_3024624, EPI_ISL_3024625 | | | |
| see above | HELEN JOSEPH LABORATORY | National Institute for Communicable Diseases of the National Health Laboratory Service | Amoako DG, Everatt J, Scheepers C, Mohale T, Ntuli N, Mahlangu B, Mnguni A, Ismail A, Bhiman JN |
| EPI_ISL_3024631, EPI_ISL_3024632, EPI_ISL_3024633, EPI_ISL_3024641 | TAMBO MEMORIAL LABORATORY | National Institute for Communicable Diseases of the National Health Laboratory Service | Amoako DG, Everatt J, Scheepers C, Mohale T, Ntuli N, Mahlangu B, Mnguni A, Ismail A, Bhiman JN |
| EPI_ISL_3024645 | KOPANONG LABORATORY | National Institute for Communicable Diseases of the National Health Laboratory Service | Amoako DG, Everatt J, Scheepers C, Mohale T, Ntuli N, Mahlangu B, Mnguni A, Ismail A, Bhiman JN |
| EPI_ISL_3024659, EPI_ISL_3024680 | CHRIS HANI BARAGWANATH LABORATORY | National Institute for Communicable Diseases of the National Health Laboratory Service | Amoako DG, Everatt J, Scheepers C, Mohale T, Ntuli N, Mahlangu B, Mnguni A, Ismail A, Bhiman JN |
| EPI_ISL_3024682 | MILPARK HOSPITAL | National Institute for Communicable Diseases of the National Health Laboratory Service | Sisonke Team, Amoako DG, Everatt J, Scheepers C, Mohale T, Ntuli N, Mahlangu B, Mnguni A, Ismail A, Bhiman JN |
| EPI_ISL_3024728, EPI_ISL_3024729 | INSIDE DIAGNÓSTICOS | Instituto Butantan | Dimas Tadeu Covas, Antonio Jorge Martins, Claudia Renata dos Santos Barros, David Schlesinger, Debora Botequio Moretti, Elaine Cristina Marqueze, Elaine Vieira Santos, Evandra Strazza Rodrigues, Heidge Fukumasu, Jayme Augusto de Souza-Neto, José Salvatore Leister Patané, Luiz Alcantara, Luiz Lehmann Coutinho, Maria Carolina Elias, Maurício Lacerda Nogueira, Rafael dos Santos Bezerra, Raul Machado Neto, Rejane Maria Tommasini Grotto, Ricardo Haddad, Sandra Coccuzzo Sampaio Vessoni, Simone Kashima, Svetoslav Nanev Slavov, Vincent Louis Viala |
| EPI_ISL_3024730, EPI_ISL_3024731, EPI_ISL_3024732, EPI_ISL_3024733 | INSIDE DIAGNÓSTICOS - UPA TATUAPÉ | Instituto Butantan | Dimas Tadeu Covas, Antonio Jorge Martins, Claudia Renata dos Santos Barros, David Schlesinger, Debora Botequio Moretti, Elaine Cristina Marqueze, Elaine Vieira Santos, Evandra Strazza Rodrigues, Heidge Fukumasu, Jayme Augusto de Souza-Neto, José Salvatore Leister Patané, Luiz Alcantara, Luiz Lehmann Coutinho, Maria Carolina Elias, Maurício Lacerda Nogueira, Rafael dos Santos Bezerra, Raul Machado Neto, Rejane Maria Tommasini Grotto, Ricardo Haddad, Sandra Coccuzzo Sampaio Vessoni, Simone Kashima, Svetoslav Nanev Slavov, Vincent Louis Viala |
| EPI_ISL_3024782, EPI_ISL_3024784, EPI_ISL_3024785, | Maryland Public Health Laboratory (MD PHL) | Maryland Public Health Laboratory (MD PHL) | Maryland Department of Health Laboratories Administration |

| | | | |
|---|---|---|---|
| EPI_ISL_3024787, EPI_ISL_3024790, EPI_ISL_3024791, EPI_ISL_3024792, EPI_ISL_3024794, EPI_ISL_3024795, EPI_ISL_3024796 | | | |
| EPI_ISL_3024798, EPI_ISL_3024799, EPI_ISL_3024800, EPI_ISL_3024801, EPI_ISL_3024802, EPI_ISL_3024803, EPI_ISL_3024804, EPI_ISL_3024805, EPI_ISL_3024806, EPI_ISL_3024807, EPI_ISL_3024808 | | | |
| see above | Genome Information Research Center, Research Institute for Microbial Diseases, Osaka University | Genome Information Research Center, Research Institute for Microbial Diseases, Osaka University | Daisuke Motooka, Kentaro Tanaka, Hiroya Oki, and Shota Nakamura |
| EPI_ISL_3024812 | Division of Emerging Infectious Diseases, Bureau of Infectious Diseases Diagnosis Control, Korea Disease Control and Prevention Agency | Division of Emerging Infectious Diseases, Bureau of Infectious Diseases Diagnosis Control, Korea Disease Control and Prevention Agency | Jeong-Ah Kim, Ae Kyung Park, Il-Hwan Kim, Heui Man Kim, Jeong-Min Kim, Chae Young Lee, Eun-Jin Kim |
| EPI_ISL_3025207, EPI_ISL_3025248, EPI_ISL_3025277, EPI_ISL_3025283, EPI_ISL_3025294, EPI_ISL_3025295, EPI_ISL_3025297, EPI_ISL_3025298, EPI_ISL_3025299, EPI_ISL_3025302, EPI_ISL_3025310 | | | |
| see above | UVA Clinical Microbiology Lab | Mathers Lab | Hardik I. Parikh, Katie E. Barry, Amy J. Mathers |
| EPI_ISL_3025340, EPI_ISL_3025341, EPI_ISL_3025342, EPI_ISL_3025343, EPI_ISL_3025344, EPI_ISL_3025346, EPI_ISL_3025347, EPI_ISL_3025349, EPI_ISL_3025350, EPI_ISL_3025351, EPI_ISL_3025352, EPI_ISL_3025353, EPI_ISL_3025354, EPI_ISL_3025355, EPI_ISL_3025356, EPI_ISL_3025358, EPI_ISL_3025359, EPI_ISL_3025360, EPI_ISL_3025361, EPI_ISL_3025362, EPI_ISL_3025364, EPI_ISL_3025365, EPI_ISL_3025366, EPI_ISL_3025367, EPI_ISL_3025368, EPI_ISL_3025369, EPI_ISL_3025370, EPI_ISL_3025371, EPI_ISL_3025372, EPI_ISL_3025373, EPI_ISL_3025374, EPI_ISL_3025375 | | | |
| see above | Alaska State Virology Laboratory | Alaska State Virology Laboratory | Stephanie DeRonde, Elva House, Jacob Zidek, Lisa Smith, Ph.D., Jack Chen, Ph.D. |
| EPI_ISL_3025386, EPI_ISL_3025396, EPI_ISL_3025397, EPI_ISL_3025398, EPI_ISL_3025399, EPI_ISL_3025400, EPI_ISL_3025401, EPI_ISL_3025402, EPI_ISL_3025403, EPI_ISL_3025404, EPI_ISL_3025405, EPI_ISL_3025407, EPI_ISL_3025408, EPI_ISL_3025409, EPI_ISL_3025410, EPI_ISL_3025411, EPI_ISL_3025412 | | | |
| see above | UCC (Universidad Central del Caribe) | Grubaugh Lab - Yale School of Public Health | Joseph Fauver, Mallery Breban, Isabel Ott, Tara Alpert, Mary Petrone, Anderson Brito, Chantal Vogels, Annie Watkins, Chaney Kalinich, Jessica Rothman, Kendall Billig, Tobias Koch, Fabiola Cruz-Lopez, David Rivera Aponte, Fabiola Fontanet, Natalie Machargo, Andrea Firpo, Vianca Aponte, Ernesto Vazquez, Patricia Serrano, Tania Mitwalli, Karolane Gonzalez, Nathan Grubaugh |
| EPI_ISL_3025422, EPI_ISL_3025424, EPI_ISL_3025426, EPI_ISL_3025429, EPI_ISL_3025430, EPI_ISL_3025431, EPI_ISL_3025432, EPI_ISL_3025435, EPI_ISL_3025436, EPI_ISL_3025437, EPI_ISL_3025438, EPI_ISL_3025439, EPI_ISL_3025440, EPI_ISL_3025441, EPI_ISL_3025442, EPI_ISL_3025443 | | | |
| see above | Yale Clinical Virology Lab | Grubaugh Lab - Yale School of Public Health | Joseph Fauver, Mallery Breban, Isabel Ott, Tara Alpert, Mary Petrone, Anderson Brito, Chantal Vogels, Annie Watkins, Chaney Kalinich, Jessica Rothman, Kendall Billig, Tobias Koch, Marie L. Landry, Nathan Grubaugh |
| EPI_ISL_3025444, EPI_ISL_3025445 | Yale Pathology Lab | Grubaugh Lab - Yale School of Public Health | Joseph Fauver, Mallery Breban, Isabel Ott, Tara Alpert, Mary Petrone, Anderson Brito, Chantal Vogels, Annie Watkins, Chaney Kalinich, Jessica Rothman, Kendall Billig, Tobias Koch, Jianhui Wang, Chen Liu, Pei Hui, Nathan Grubaugh |
| EPI_ISL_3025446, EPI_ISL_3025447, EPI_ISL_3025448, EPI_ISL_3025449, EPI_ISL_3025451, EPI_ISL_3025452, EPI_ISL_3025453, EPI_ISL_3025454, EPI_ISL_3025455, EPI_ISL_3025456, EPI_ISL_3025457, EPI_ISL_3025458, EPI_ISL_3025460 | | | |
| see above | NJDOH, Public Health and Environmental Laboratories | New Jersey Public Health and Environmental Laboratories (NJ PHEL) | Lindsey Bodnar, Shiv K. Verma, Ryan Pachucki, Chelsea San Filippo, Jacquelyn Deverell, Dana Woell, Allison Roder, Byeong Jeong |
| EPI_ISL_3025547, EPI_ISL_3025558, EPI_ISL_3025591, EPI_ISL_3025612, EPI_ISL_3025616 | Laboratory, Gravity Diagnostics | Laboratory, Gravity Diagnostics | Diagnostics,G. |
| EPI_ISL_3025648, EPI_ISL_3025658, EPI_ISL_3025669, EPI_ISL_3025670, EPI_ISL_3025674, EPI_ISL_3025675, EPI_ISL_3025676, EPI_ISL_3025687, EPI_ISL_3025688, EPI_ISL_3025691, EPI_ISL_3025776, EPI_ISL_3025777, EPI_ISL_3025778, EPI_ISL_3025779, EPI_ISL_3025780, EPI_ISL_3025781, EPI_ISL_3025790, EPI_ISL_3025791, EPI_ISL_3025792, EPI_ISL_3025795, EPI_ISL_3025796, EPI_ISL_3025826, EPI_ISL_3025827, EPI_ISL_3025828, EPI_ISL_3025830, EPI_ISL_3025833, EPI_ISL_3025835, EPI_ISL_3025837, EPI_ISL_3025838, EPI_ISL_3025842, EPI_ISL_3025843 | | | |
| see above | Clinical Microbiology, Infection Prevention and Control | Section for Molecular Diagnostics | Björn Hallström, Jonas Björkman |
| EPI_ISL_3025845, EPI_ISL_3025846, EPI_ISL_3025848, EPI_ISL_3025849, EPI_ISL_3025851, EPI_ISL_3025857, EPI_ISL_3025860, EPI_ISL_3025861, EPI_ISL_3025862, EPI_ISL_3025863, EPI_ISL_3025864, EPI_ISL_3025866, EPI_ISL_3025867, EPI_ISL_3025869, EPI_ISL_3025871, EPI_ISL_3025872, EPI_ISL_3025873, EPI_ISL_3025874, EPI_ISL_3025876, EPI_ISL_3025878, EPI_ISL_3025880, EPI_ISL_3025881, EPI_ISL_3025882, EPI_ISL_3025883 | | | |
| see above | Virginia Division of Consolidated Laboratory Services | Virginia Division of Consolidated Laboratory Services | Virginia Division of Consolidated Laboratory Services |
| EPI_ISL_3025900, EPI_ISL_3025901, EPI_ISL_3025902, EPI_ISL_3025903, EPI_ISL_3025904, EPI_ISL_3025905, EPI_ISL_3025906, EPI_ISL_3025907, EPI_ISL_3025908, EPI_ISL_3025909, EPI_ISL_3025910, EPI_ISL_3025911 | | | |
| see above | UHTL, University Hospitals | UHTL, University Hospitals | Sadri,N., Alouani,D., Song,X. |
| EPI_ISL_3025912, EPI_ISL_3025913, EPI_ISL_3025914, EPI_ISL_3025915, EPI_ISL_3025916, EPI_ISL_3025917, EPI_ISL_3025918, EPI_ISL_3025919, EPI_ISL_3025920, EPI_ISL_3025921, EPI_ISL_3025923, EPI_ISL_3025924, EPI_ISL_3025925, EPI_ISL_3025926, EPI_ISL_3025928, EPI_ISL_3025930, EPI_ISL_3025932, EPI_ISL_3025933, EPI_ISL_3025935, EPI_ISL_3025936, EPI_ISL_3025937, EPI_ISL_3025938, EPI_ISL_3025940, EPI_ISL_3025941, EPI_ISL_3025942, EPI_ISL_3025943, EPI_ISL_3025944, EPI_ISL_3025945, EPI_ISL_3025946, EPI_ISL_3025948, EPI_ISL_3025949, EPI_ISL_3025950, EPI_ISL_3025951, EPI_ISL_3025952, EPI_ISL_3025953, EPI_ISL_3025954, EPI_ISL_3025956, EPI_ISL_3025957, EPI_ISL_3025961, EPI_ISL_3025962, EPI_ISL_3025963, EPI_ISL_3025964, EPI_ISL_3025965, EPI_ISL_3025966, EPI_ISL_3025968, EPI_ISL_3025969, EPI_ISL_3025971, EPI_ISL_3025972, EPI_ISL_3025973, EPI_ISL_3025974, EPI_ISL_3025975, EPI_ISL_3025976, EPI_ISL_3025977, EPI_ISL_3025978, EPI_ISL_3025979, EPI_ISL_3025980, EPI_ISL_3025981, EPI_ISL_3025984, EPI_ISL_3025985, EPI_ISL_3025989, EPI_ISL_3025990, EPI_ISL_3025991, EPI_ISL_3025992, EPI_ISL_3025996, EPI_ISL_3025997, EPI_ISL_3025998, EPI_ISL_3026000, EPI_ISL_3026001 | | | |
| see above | UW Virology Lab | UW Virology Lab | Pavitra Roychoudhury, Hong Xie, Lasata Shrestha, Shah Mohamed Bakhash, Tien V. Nguyen, Noah R. Baker, Sean Ellis, Meei-Li Huang, Keith R Jerome, Alexander Greninger |
| EPI_ISL_3026002, EPI_ISL_3026003, EPI_ISL_3026004, EPI_ISL_3026005, EPI_ISL_3026006, EPI_ISL_3026007, EPI_ISL_3026011, EPI_ISL_3026012, EPI_ISL_3026014 | Laboratorio di Riferimento Regionale della Sicilia Occidentale per l'Emergenza COVID-19 | Laboratorio di Riferimento Regionale della Sicilia Occidentale per l’Emergenza COVID-19 | Fabio Tramuto, Carmelo Massimo Maida, Daniela Di Naro, Giulia Randazzo, Walter Mazzucco, Giorgio Graziano, Vincenzo Restivo, Claudio Costantino, Francesco Vitale |
| EPI_ISL_3026021 | HOSPITAL SAN RAFAEL DE ALAJUELA [ALAJUELA/ALAJUELA] | Inciensa, Instituto Costarricense de Investigación y Enseñanza en Nutrición y Salud | Francisco Duarte, Hebleen Porras, Claudio Soto-Garita, Estela Cordero, Adriana Godínez, Melany Calderón, José Luis Vargas, Mariela Gutiérrez, Joselyn Prado, Caterina Guzmán, Nazareth Ruíz & Christian Sanchez |
| EPI_ISL_3026022 | AREA DE SALUD EL GUARCO [EL GUARCO/CARTAGO] | Inciensa, Instituto Costarricense de Investigación y Enseñanza en Nutrición y Salud | Francisco Duarte, Hebleen Porras, Claudio Soto-Garita, Estela Cordero, Adriana Godínez, Melany Calderón, José Luis Vargas, Mariela Gutiérrez, Joselyn Prado, Caterina Guzmán, Nazareth Ruíz & Mónica Charpentier |
| EPI_ISL_3026023 | AREA DE SALUD FORTUNA [San Carlos/ALAJUELA] | Inciensa, Instituto Costarricense de Investigación y Enseñanza en Nutrición y Salud | Francisco Duarte, Hebleen Porras, Claudio Soto-Garita, Estela Cordero, Adriana Godínez, Melany Calderón, José Luis Vargas, Mariela Gutiérrez, Joselyn Prado, Caterina Guzmán, Nazareth Ruíz & Carolina Arrieta |
| EPI_ISL_3026024 | CLINICA CATOLICA | Inciensa, Instituto Costarricense de Investigación y Enseñanza en Nutrición y Salud | Francisco Duarte, Hebleen Porras, Claudio Soto-Garita, Estela Cordero, Adriana Godínez, Melany Calderón, José Luis Vargas, Mariela Gutiérrez, Joselyn Prado, Caterina Guzmán, Nazareth Ruíz & Claudia Escobar |
| EPI_ISL_3026025 | AREA DE SALUD Heredia-VIRILLA [Heredia/Heredia] | Inciensa, Instituto Costarricense de Investigación y Enseñanza en Nutrición y Salud | Francisco Duarte, Hebleen Porras, Claudio Soto-Garita, Estela Cordero, Adriana Godínez, Melany Calderón, José Luis Vargas, Mariela Gutiérrez, Joselyn Prado, Caterina Guzmán, Nazareth Ruíz & Laura Garro |
| EPI_ISL_3026026 | AREA DE SALUD CORREDORES [CORREDORES/PUNTARENAS] | Inciensa, Instituto Costarricense de Investigación y Enseñanza en Nutrición y Salud | Francisco Duarte, Hebleen Porras, Claudio Soto-Garita, Estela Cordero, Adriana Godínez, Melany Calderón, José Luis Vargas, Mariela Gutiérrez, Joselyn Prado, Caterina Guzmán, Nazareth Ruíz & María Victoria |
| EPI_ISL_3026027, EPI_ISL_3026028 | AREA DE SALUD FORTUNA [San Carlos/ALAJUELA] | Inciensa, Instituto Costarricense de Investigación y Enseñanza en Nutrición y Salud | Francisco Duarte, Hebleen Porras, Claudio Soto-Garita, Estela Cordero, Adriana Godínez, Melany Calderón, José Luis Vargas, Mariela Gutiérrez, Joselyn Prado, Caterina Guzmán, Nazareth Ruíz & Carolina Arrieta |
| EPI_ISL_3026029 | AREA DE SALUD PAVAS (COOPESALUD) [PAVAS/San Jose] | Inciensa, Instituto Costarricense de Investigación y Enseñanza en Nutrición y Salud | Francisco Duarte, Hebleen Porras, Claudio Soto-Garita, Estela Cordero, Adriana Godínez, Melany Calderón, José Luis Vargas, Mariela Gutiérrez, Joselyn Prado, Caterina Guzmán, Nazareth Ruíz |
| EPI_ISL_3026030 | AREA DE SALUD BARVA (COOPESIBA) [BARVA/Heredia] | Inciensa, Instituto Costarricense de Investigación y Enseñanza en Nutrición y Salud | Francisco Duarte, Hebleen Porras, Claudio Soto-Garita, Estela Cordero, Adriana Godínez, Melany Calderón, José Luis Vargas, Mariela Gutiérrez, Joselyn Prado, Caterina Guzmán, Nazareth Ruíz & Valeria Peralta |
| EPI_ISL_3026031 | AREA DE SALUD Guatuso [Guatuso/ALAJUELA] | Inciensa, Instituto Costarricense de Investigación y Enseñanza en Nutrición y Salud | Francisco Duarte, Hebleen Porras, Claudio Soto-Garita, Estela Cordero, Adriana Godínez, Melany Calderón, José Luis Vargas, Mariela Gutiérrez, Joselyn Prado, Caterina Guzmán, Nazareth Ruíz & Francisco Chacón |
| EPI_ISL_3026032 | AREA DE SALUD LIMON [LIMON/LIMON] | Inciensa, Instituto Costarricense de Investigación y Enseñanza en Nutrición y Salud | Francisco Duarte, Hebleen Porras, Claudio Soto-Garita, Estela Cordero, Adriana Godínez, Melany Calderón, José Luis Vargas, Mariela Gutiérrez, Joselyn Prado, Caterina Guzmán, Nazareth Ruíz & Karolina Hall |
| EPI_ISL_3026033 | AREA DE SALUD Desamparados 1 - CLINICA DR. MARCIAL FALLAS [GRIFO ALTO/Desamparados/San Jose] | Inciensa, Instituto Costarricense de Investigación y Enseñanza en Nutrición y Salud | Francisco Duarte, Hebleen Porras, Claudio Soto-Garita, Estela Cordero, Adriana Godínez, Melany Calderón, José Luis Vargas, Mariela Gutiérrez, Joselyn Prado, Caterina Guzmán, Nazareth Ruíz |
| EPI_ISL_3026034, EPI_ISL_3026035, EPI_ISL_3026036, EPI_ISL_3026037, EPI_ISL_3026038, EPI_ISL_3026039 | National Public Health Laboratory, National Centre for Infectious Diseases | National Public Health Laboratory, National Centre for Infectious Diseases | Tze Minn Mak, Zhenyang Zhou, Royce Ang, Lin Cui, Raymond Tzer Pin Lin |
| EPI_ISL_3026064, EPI_ISL_3026066, EPI_ISL_3026067, EPI_ISL_3026068, EPI_ISL_3026069, EPI_ISL_3026073, EPI_ISL_3026074, EPI_ISL_3026075, EPI_ISL_3026082, EPI_ISL_3026083, EPI_ISL_3026088, EPI_ISL_3026090, EPI_ISL_3026091, EPI_ISL_3026092, EPI_ISL_3026093, EPI_ISL_3026094, EPI_ISL_3026096, EPI_ISL_3026112, EPI_ISL_3026118, EPI_ISL_3026119, EPI_ISL_3026120, EPI_ISL_3026123, EPI_ISL_3026127, EPI_ISL_3026130, EPI_ISL_3026132, EPI_ISL_3026133, EPI_ISL_3026134, EPI_ISL_3026142, EPI_ISL_3026143, EPI_ISL_3026144, EPI_ISL_3026145, EPI_ISL_3026149, EPI_ISL_3026151, EPI_ISL_3026157, EPI_ISL_3026159, EPI_ISL_3026163, EPI_ISL_3026177, EPI_ISL_3026178, EPI_ISL_3026184, EPI_ISL_3026186, EPI_ISL_3026197, EPI_ISL_3026199, EPI_ISL_3026200, EPI_ISL_3026210, EPI_ISL_3026211, EPI_ISL_3026214, EPI_ISL_3026215, EPI_ISL_3026216, EPI_ISL_3026217, EPI_ISL_3026218, EPI_ISL_3026219, EPI_ISL_3026221, EPI_ISL_3026226, EPI_ISL_3026228, EPI_ISL_3026232, EPI_ISL_3026233, EPI_ISL_3026241, EPI_ISL_3026247, EPI_ISL_3026251, EPI_ISL_3026259, EPI_ISL_3026266, EPI_ISL_3026279, EPI_ISL_3026288, EPI_ISL_3026289, EPI_ISL_3026294, EPI_ISL_3026302, EPI_ISL_3026303, EPI_ISL_3026311, EPI_ISL_3026312, EPI_ISL_3026313, EPI_ISL_3026314, EPI_ISL_3026317, EPI_ISL_3026318, EPI_ISL_3026319, EPI_ISL_3026321, EPI_ISL_3026327, EPI_ISL_3026328, EPI_ISL_3026332, EPI_ISL_3026333, EPI_ISL_3026336, EPI_ISL_3026339, EPI_ISL_3026343, EPI_ISL_3026344, EPI_ISL_3026345, EPI_ISL_3026346, EPI_ISL_3026347, EPI_ISL_3026355, EPI_ISL_3026356, EPI_ISL_3026357, EPI_ISL_3026363, EPI_ISL_3026364, EPI_ISL_3026366, EPI_ISL_3026367, EPI_ISL_3026369, EPI_ISL_3026371, EPI_ISL_3026372, EPI_ISL_3026373, EPI_ISL_3026374, EPI_ISL_3026375, EPI_ISL_3026377, EPI_ISL_3026378, EPI_ISL_3026379, EPI_ISL_3026380, EPI_ISL_3026381, EPI_ISL_3026384, EPI_ISL_3026385, EPI_ISL_3026386, EPI_ISL_3026388, EPI_ISL_3026389, EPI_ISL_3026390, EPI_ISL_3026391, EPI_ISL_3026392, EPI_ISL_3026393, EPI_ISL_3026396, EPI_ISL_3026397, EPI_ISL_3026398, EPI_ISL_3026399, EPI_ISL_3026401, EPI_ISL_3026402, EPI_ISL_3026403, EPI_ISL_3026406, EPI_ISL_3026407, EPI_ISL_3026408, EPI_ISL_3026409, EPI_ISL_3026410, EPI_ISL_3026411, EPI_ISL_3026412, EPI_ISL_3026413, EPI_ISL_3026414, EPI_ISL_3026415, EPI_ISL_3026416, EPI_ISL_3026417, EPI_ISL_3026418, EPI_ISL_3026419, EPI_ISL_3026421, EPI_ISL_3026422, EPI_ISL_3026424, EPI_ISL_3026425, EPI_ISL_3026426, EPI_ISL_3026427, EPI_ISL_3026428, EPI_ISL_3026429, EPI_ISL_3026430, EPI_ISL_3026431, EPI_ISL_3026432, EPI_ISL_3026433, EPI_ISL_3026435, EPI_ISL_3026437, EPI_ISL_3026438, EPI_ISL_3026441, EPI_ISL_3026443, EPI_ISL_3026445, EPI_ISL_3026446, EPI_ISL_3026447, EPI_ISL_3026448, EPI_ISL_3026449, EPI_ISL_3026450, EPI_ISL_3026451, EPI_ISL_3026452, EPI_ISL_3026453, EPI_ISL_3026455, EPI_ISL_3026456, EPI_ISL_3026458, EPI_ISL_3026461, EPI_ISL_3026463, EPI_ISL_3026464, EPI_ISL_3026465, EPI_ISL_3026466, EPI_ISL_3026467, EPI_ISL_3026469, EPI_ISL_3026470, EPI_ISL_3026471, EPI_ISL_3026473, EPI_ISL_3026474, EPI_ISL_3026476, EPI_ISL_3026477, EPI_ISL_3026478, EPI_ISL_3026480, EPI_ISL_3026481, EPI_ISL_3026485, EPI_ISL_3026488, EPI_ISL_3026490, EPI_ISL_3026491, EPI_ISL_3026496, EPI_ISL_3026497, EPI_ISL_3026498, EPI_ISL_3026501, EPI_ISL_3026503, EPI_ISL_3026507, EPI_ISL_3026511, EPI_ISL_3026514, EPI_ISL_3026515, EPI_ISL_3026517, EPI_ISL_3026518, EPI_ISL_3026519, EPI_ISL_3026521, EPI_ISL_3026522, EPI_ISL_3026523, | | | |

| | | | |
|---|---|---|---|
| EPI_ISL_3026524, EPI_ISL_3026525, EPI_ISL_3026526, EPI_ISL_3026529, EPI_ISL_3026530, EPI_ISL_3026531, EPI_ISL_3026533, EPI_ISL_3026535, EPI_ISL_3026536, EPI_ISL_3026537, EPI_ISL_3026538, EPI_ISL_3026539, EPI_ISL_3026540, EPI_ISL_3026541, EPI_ISL_3026543, EPI_ISL_3026544, EPI_ISL_3026545, EPI_ISL_3026546, EPI_ISL_3026548, EPI_ISL_3026549, EPI_ISL_3026550, EPI_ISL_3026551, EPI_ISL_3026552, EPI_ISL_3026553, EPI_ISL_3026554, EPI_ISL_3026555, EPI_ISL_3026557, EPI_ISL_3026558, EPI_ISL_3026559, EPI_ISL_3026561, EPI_ISL_3026562, EPI_ISL_3026563, EPI_ISL_3026565, EPI_ISL_3026566, EPI_ISL_3026568, EPI_ISL_3026569, EPI_ISL_3026570, EPI_ISL_3026571, EPI_ISL_3026572, EPI_ISL_3026573, EPI_ISL_3026574, EPI_ISL_3026575, EPI_ISL_3026576, EPI_ISL_3026577, EPI_ISL_3026578, EPI_ISL_3026579, EPI_ISL_3026580, EPI_ISL_3026582, EPI_ISL_3026584, EPI_ISL_3026585, EPI_ISL_3026587, EPI_ISL_3026589, EPI_ISL_3026590, EPI_ISL_3026591, EPI_ISL_3026592, EPI_ISL_3026593, EPI_ISL_3026594, EPI_ISL_3026596, EPI_ISL_3026597, EPI_ISL_3026598, EPI_ISL_3026599, EPI_ISL_3026600, EPI_ISL_3026602, EPI_ISL_3026607, EPI_ISL_3026609, EPI_ISL_3026610, EPI_ISL_3026612, EPI_ISL_3026613, EPI_ISL_3026614, EPI_ISL_3026615, EPI_ISL_3026616, EPI_ISL_3026617, EPI_ISL_3026618, EPI_ISL_3026623, EPI_ISL_3026625, EPI_ISL_3026627, EPI_ISL_3026628, EPI_ISL_3026630, EPI_ISL_3026632, EPI_ISL_3026633, EPI_ISL_3026634, EPI_ISL_3026636, EPI_ISL_3026637, EPI_ISL_3026638, EPI_ISL_3026639, EPI_ISL_3026641, EPI_ISL_3026642, EPI_ISL_3026643, EPI_ISL_3026644, EPI_ISL_3026646, EPI_ISL_3026649, EPI_ISL_3026652, EPI_ISL_3026653, EPI_ISL_3026654, EPI_ISL_3026655, EPI_ISL_3026656, EPI_ISL_3026658, EPI_ISL_3026660, EPI_ISL_3026661, EPI_ISL_3026662, EPI_ISL_3026663, EPI_ISL_3026664, EPI_ISL_3026665, EPI_ISL_3026670, EPI_ISL_3026671, EPI_ISL_3026672, EPI_ISL_3026675, EPI_ISL_3026677, EPI_ISL_3026679, EPI_ISL_3026680, EPI_ISL_3026682, EPI_ISL_3026684, EPI_ISL_3026690, EPI_ISL_3026692, EPI_ISL_3026693, EPI_ISL_3026696, EPI_ISL_3026700, EPI_ISL_3026701, EPI_ISL_3026702, EPI_ISL_3026703, EPI_ISL_3026704, EPI_ISL_3026705, EPI_ISL_3026706, EPI_ISL_3026708, EPI_ISL_3026711, EPI_ISL_3026712, EPI_ISL_3026715, EPI_ISL_3026717, EPI_ISL_3026720, EPI_ISL_3026722, EPI_ISL_3026723, EPI_ISL_3026728, EPI_ISL_3026729, EPI_ISL_3026730, EPI_ISL_3026731, EPI_ISL_3026733, EPI_ISL_3026736, EPI_ISL_3026737, EPI_ISL_3026738, EPI_ISL_3026741, EPI_ISL_3026745, EPI_ISL_3026747, EPI_ISL_3026755, EPI_ISL_3026756, EPI_ISL_3026757, EPI_ISL_3026758, EPI_ISL_3026759, EPI_ISL_3026760, EPI_ISL_3026761, EPI_ISL_3026762, EPI_ISL_3026763, EPI_ISL_3026764, EPI_ISL_3026765, EPI_ISL_3026767, EPI_ISL_3026773, EPI_ISL_3026774, EPI_ISL_3026778, EPI_ISL_3026779, EPI_ISL_3026781, EPI_ISL_3026784, EPI_ISL_3026785, EPI_ISL_3026787, EPI_ISL_3026788, EPI_ISL_3026789, EPI_ISL_3026792, EPI_ISL_3026793, EPI_ISL_3026795, EPI_ISL_3026797, EPI_ISL_3026798, EPI_ISL_3026800, EPI_ISL_3026801, EPI_ISL_3026803, EPI_ISL_3026807, EPI_ISL_3026808, EPI_ISL_3026809, EPI_ISL_3026810, EPI_ISL_3026811, EPI_ISL_3026812, EPI_ISL_3026813, EPI_ISL_3026815, EPI_ISL_3026817, EPI_ISL_3026818, EPI_ISL_3026822, EPI_ISL_3026828, EPI_ISL_3026829, EPI_ISL_3026830, EPI_ISL_3026832, EPI_ISL_3026835, EPI_ISL_3026836, EPI_ISL_3026839, EPI_ISL_3026840, EPI_ISL_3026841, EPI_ISL_3026842, EPI_ISL_3026843, EPI_ISL_3026847, EPI_ISL_3026849, EPI_ISL_3026850, EPI_ISL_3026852, EPI_ISL_3026853 | | | |
| see above | Division of Emerging Infectious Diseases, Bureau of Infectious Diseases Diagnosis Control, Korea Disease Control and Prevention Agency | Division of Emerging Infectious Diseases, Bureau of Infectious Diseases Diagnosis Control, Korea Disease Control and Prevention Agency | Jeong-Ah Kim, Ae Kyung Park, Il-Hwan Kim, Heui Man Kim, Jeong-Min Kim, Chae Young Lee, Eun-Jin Kim |
| EPI_ISL_3026861, EPI_ISL_3026862 | Genome Information Research Center, Research Institute for Microbial Diseases, Osaka University | Genome Information Research Center, Research Institute for Microbial Diseases, Osaka University | Daisuke Motooka, Kentaro Tanaka, Hiroya Oki, and Shota Nakamura |
| EPI_ISL_3026866, EPI_ISL_3026875, EPI_ISL_3026876, EPI_ISL_3026879, EPI_ISL_3026880, EPI_ISL_3026885, EPI_ISL_3026886, EPI_ISL_3026888, EPI_ISL_3026889, EPI_ISL_3026897, EPI_ISL_3026898, EPI_ISL_3026899, EPI_ISL_3026900, EPI_ISL_3026901, EPI_ISL_3026902, EPI_ISL_3026903, EPI_ISL_3026904, EPI_ISL_3026906, EPI_ISL_3026907, EPI_ISL_3026910 | | | |
| see above | Microbiology Division, SC DHEC | Microbiology Division, SC DHEC | Freeman,J.S. |
| EPI_ISL_3026912, EPI_ISL_3026913, EPI_ISL_3026920, EPI_ISL_3026927, EPI_ISL_3026928, EPI_ISL_3026929, EPI_ISL_3026930, EPI_ISL_3026931, EPI_ISL_3026932, EPI_ISL_3026933, EPI_ISL_3026934 | | | |
| see above | ASL BARI | University of Bari Biomedical Sciences and Human Oncology | Maria Chironna |
| EPI_ISL_3026935 | Lighthouse Lab in Alderley Park | Wellcome Sanger Institute for the COVID-19 Genomics UK (COG-UK) Consortium | Jacquelyn Wynn, Mairead Hyland, The Lighthouse Lab in Alderley Park and Alex Alderton, Roberto Amato, Jeffrey Barrett, Sonia Goncalves, Ewan Harrison, David K. Jackson, Ian Johnston, Dominic Kwiatkowski, Cordelia Langford, John Sillitoe on behalf of the Wellcome Sanger Institute COVID-19 Surveillance Team |
| EPI_ISL_3026936 | Lighthouse Lab in Glasgow | Wellcome Sanger Institute for the COVID-19 Genomics UK (COG-UK) Consortium | Harper VanSteenhouse, Yumi Kasai, David Gray, Carol Clugston, Anna Dominiczak and Alex Alderton, Roberto Amato, Jeffrey Barrett, Sonia Goncalves, Ewan Harrison, David K. Jackson, Ian Johnston, Dominic Kwiatkowski, Cordelia Langford, John Sillitoe on behalf of the Wellcome Sanger Institute COVID-19 Surveillance Team |
| EPI_ISL_3026937 | Lighthouse Lab in Milton Keynes | Wellcome Sanger Institute for the COVID-19 Genomics UK (COG-UK) Consortium | The Lighthouse Lab in Milton Keynes and Alex Alderton, Roberto Amato, Jeffrey Barrett, Sonia Goncalves, Ewan Harrison, David K. Jackson, Ian Johnston, Dominic Kwiatkowski, Cordelia Langford, John Sillitoe on behalf of the Wellcome Sanger Institute COVID-19 Surveillance Team |
| EPI_ISL_3026938 | Lighthouse Lab in Glasgow | Wellcome Sanger Institute for the COVID-19 Genomics UK (COG-UK) Consortium | Harper VanSteenhouse, Yumi Kasai, David Gray, Carol Clugston, Anna Dominiczak and Alex Alderton, Roberto Amato, Jeffrey Barrett, Sonia Goncalves, Ewan Harrison, David K. Jackson, Ian Johnston, Dominic Kwiatkowski, Cordelia Langford, John Sillitoe on behalf of the Wellcome Sanger Institute COVID-19 Surveillance Team |
| EPI_ISL_3026939 | Lighthouse Lab in Milton Keynes | Wellcome Sanger Institute for the COVID-19 Genomics UK (COG-UK) Consortium | The Lighthouse Lab in Milton Keynes and Alex Alderton, Roberto Amato, Jeffrey Barrett, Sonia Goncalves, Ewan Harrison, David K. Jackson, Ian Johnston, Dominic Kwiatkowski, Cordelia Langford, John Sillitoe on behalf of the Wellcome Sanger Institute COVID-19 Surveillance Team |
| EPI_ISL_3026940 | Lighthouse Lab in Glasgow | Wellcome Sanger Institute for the COVID-19 Genomics UK (COG-UK) Consortium | Harper VanSteenhouse, Yumi Kasai, David Gray, Carol Clugston, Anna Dominiczak and Alex Alderton, Roberto Amato, Jeffrey Barrett, Sonia Goncalves, Ewan Harrison, David K. Jackson, Ian Johnston, Dominic Kwiatkowski, Cordelia Langford, John Sillitoe on behalf of the Wellcome Sanger Institute COVID-19 Surveillance Team |
| EPI_ISL_3026941 | Lighthouse Lab in Alderley Park | Wellcome Sanger Institute for the COVID-19 Genomics UK (COG-UK) Consortium | Jacquelyn Wynn, Mairead Hyland, The Lighthouse Lab in Alderley Park and Alex Alderton, Roberto Amato, Jeffrey Barrett, Sonia Goncalves, Ewan Harrison, David K. Jackson, Ian Johnston, Dominic Kwiatkowski, Cordelia Langford, John Sillitoe on behalf of the Wellcome Sanger Institute COVID-19 Surveillance Team |
| EPI_ISL_3026942 | Lighthouse Lab in Milton Keynes | Wellcome Sanger Institute for the COVID-19 Genomics UK (COG-UK) Consortium | The Lighthouse Lab in Milton Keynes and Alex Alderton, Roberto Amato, Jeffrey Barrett, Sonia Goncalves, Ewan Harrison, David K. Jackson, Ian Johnston, Dominic Kwiatkowski, Cordelia Langford, John Sillitoe on behalf of the Wellcome Sanger Institute COVID-19 Surveillance Team |
| EPI_ISL_3026943 | Lighthouse Lab in Alderley Park | Wellcome Sanger Institute for the COVID-19 Genomics UK (COG-UK) Consortium | Jacquelyn Wynn, Mairead Hyland, The Lighthouse Lab in Alderley Park and Alex Alderton, Roberto Amato, Jeffrey Barrett, Sonia Goncalves, Ewan Harrison, David K. Jackson, Ian Johnston, Dominic Kwiatkowski, Cordelia Langford, John Sillitoe on behalf of the Wellcome Sanger Institute COVID-19 Surveillance Team |
| EPI_ISL_3026944 | Lighthouse Lab in Glasgow | Wellcome Sanger Institute for the COVID-19 Genomics UK (COG-UK) Consortium | Harper VanSteenhouse, Yumi Kasai, David Gray, Carol Clugston, Anna Dominiczak and Alex Alderton, Roberto Amato, Jeffrey Barrett, Sonia Goncalves, Ewan Harrison, David K. Jackson, Ian Johnston, Dominic Kwiatkowski, Cordelia Langford, John Sillitoe on behalf of the Wellcome Sanger Institute COVID-19 Surveillance Team |
| EPI_ISL_3026945 | Lighthouse Lab in Alderley Park | Wellcome Sanger Institute for the COVID-19 Genomics UK (COG-UK) Consortium | Jacquelyn Wynn, Mairead Hyland, The Lighthouse Lab in Alderley Park and Alex Alderton, Roberto Amato, Jeffrey Barrett, Sonia Goncalves, Ewan Harrison, David K. Jackson, Ian Johnston, Dominic Kwiatkowski, Cordelia Langford, John Sillitoe on behalf of the Wellcome Sanger Institute COVID-19 Surveillance Team |
| EPI_ISL_3026946, EPI_ISL_3026947 | Lighthouse Lab in Glasgow | Wellcome Sanger Institute for the COVID-19 Genomics UK (COG-UK) Consortium | Harper VanSteenhouse, Yumi Kasai, David Gray, Carol Clugston, Anna Dominiczak and Alex Alderton, Roberto Amato, Jeffrey Barrett, Sonia Goncalves, Ewan Harrison, David K. Jackson, Ian Johnston, Dominic Kwiatkowski, Cordelia Langford, John Sillitoe on behalf of the Wellcome Sanger Institute COVID-19 Surveillance Team |
| EPI_ISL_3026948, EPI_ISL_3026949 | Lighthouse Lab in Milton Keynes | Wellcome Sanger Institute for the COVID-19 Genomics UK (COG-UK) Consortium | The Lighthouse Lab in Milton Keynes and Alex Alderton, Roberto Amato, Jeffrey Barrett, Sonia Goncalves, Ewan Harrison, David K. Jackson, Ian Johnston, Dominic Kwiatkowski, Cordelia Langford, John Sillitoe on behalf of the Wellcome Sanger Institute COVID-19 Surveillance Team |
| EPI_ISL_3026950, EPI_ISL_3026951 | Lighthouse Lab in Alderley Park | Wellcome Sanger Institute for the COVID-19 Genomics UK (COG-UK) Consortium | Jacquelyn Wynn, Mairead Hyland, The Lighthouse Lab in Alderley Park and Alex Alderton, Roberto Amato, Jeffrey Barrett, Sonia Goncalves, Ewan Harrison, David K. Jackson, Ian Johnston, Dominic Kwiatkowski, Cordelia Langford, John Sillitoe on behalf of the Wellcome Sanger Institute COVID-19 Surveillance Team |
| EPI_ISL_3026952 | Lighthouse Lab in Milton Keynes | Wellcome Sanger Institute for the COVID-19 Genomics UK (COG-UK) Consortium | The Lighthouse Lab in Milton Keynes and Alex Alderton, Roberto Amato, Jeffrey Barrett, Sonia Goncalves, Ewan Harrison, David K. Jackson, Ian Johnston, Dominic Kwiatkowski, Cordelia Langford, John Sillitoe on behalf of the Wellcome Sanger Institute COVID-19 Surveillance Team |
| EPI_ISL_3026953 | Lighthouse Lab in Glasgow | Wellcome Sanger Institute for the COVID-19 Genomics UK (COG-UK) Consortium | Harper VanSteenhouse, Yumi Kasai, David Gray, Carol Clugston, Anna Dominiczak and Alex Alderton, Roberto Amato, Jeffrey Barrett, Sonia Goncalves, Ewan Harrison, David K. Jackson, Ian Johnston, Dominic Kwiatkowski, Cordelia Langford, John Sillitoe on behalf of the Wellcome Sanger Institute COVID-19 Surveillance Team |
| EPI_ISL_3026954, EPI_ISL_3026955 | Lighthouse Lab in Milton Keynes | Wellcome Sanger Institute for the COVID-19 Genomics UK (COG-UK) Consortium | The Lighthouse Lab in Milton Keynes and Alex Alderton, Roberto Amato, Jeffrey Barrett, Sonia Goncalves, Ewan Harrison, David K. Jackson, Ian Johnston, Dominic Kwiatkowski, Cordelia Langford, John Sillitoe on behalf of the Wellcome Sanger Institute COVID-19 Surveillance Team |
| EPI_ISL_3026956, EPI_ISL_3026957 | Lighthouse Lab in Glasgow | Wellcome Sanger Institute for the COVID-19 Genomics UK (COG-UK) Consortium | Harper VanSteenhouse, Yumi Kasai, David Gray, Carol Clugston, Anna Dominiczak and Alex Alderton, Roberto Amato, Jeffrey Barrett, Sonia Goncalves, Ewan Harrison, David K. Jackson, Ian Johnston, Dominic Kwiatkowski, Cordelia Langford, John Sillitoe on behalf of the Wellcome Sanger Institute COVID-19 Surveillance Team |
| EPI_ISL_3026958, EPI_ISL_3026959, EPI_ISL_3026960, EPI_ISL_3026961 | Lighthouse Lab in Alderley Park | Wellcome Sanger Institute for the COVID-19 Genomics UK (COG-UK) Consortium | Jacquelyn Wynn, Mairead Hyland, The Lighthouse Lab in Alderley Park and Alex Alderton, Roberto Amato, Jeffrey Barrett, Sonia Goncalves, Ewan Harrison, David K. Jackson, Ian Johnston, Dominic Kwiatkowski, Cordelia Langford, John Sillitoe on behalf of the Wellcome Sanger Institute COVID-19 Surveillance Team |
| EPI_ISL_3026962 | Lighthouse Lab in Glasgow | Wellcome Sanger Institute for the COVID-19 Genomics UK (COG-UK) Consortium | Harper VanSteenhouse, Yumi Kasai, David Gray, Carol Clugston, Anna Dominiczak and Alex Alderton, Roberto Amato, Jeffrey Barrett, Sonia Goncalves, Ewan Harrison, David K. Jackson, Ian Johnston, Dominic Kwiatkowski, Cordelia Langford, John Sillitoe on behalf of the Wellcome Sanger Institute COVID-19 Surveillance Team |
| EPI_ISL_3026963, EPI_ISL_3026964 | Lighthouse Lab in Alderley Park | Wellcome Sanger Institute for the COVID-19 Genomics UK (COG-UK) Consortium | Jacquelyn Wynn, Mairead Hyland, The Lighthouse Lab in Alderley Park and Alex Alderton, Roberto Amato, Jeffrey Barrett, Sonia Goncalves, Ewan Harrison, David K. Jackson, Ian Johnston, Dominic Kwiatkowski, Cordelia Langford, John Sillitoe on behalf of the Wellcome Sanger Institute COVID-19 Surveillance Team |
| EPI_ISL_3026965 | Lighthouse Lab in Glasgow | Wellcome Sanger Institute for the COVID-19 Genomics UK (COG-UK) Consortium | Harper VanSteenhouse, Yumi Kasai, David Gray, Carol Clugston, Anna Dominiczak and Alex Alderton, Roberto Amato, Jeffrey Barrett, Sonia Goncalves, Ewan Harrison, David K. Jackson, Ian Johnston, Dominic Kwiatkowski, Cordelia Langford, John Sillitoe on behalf of the Wellcome Sanger Institute COVID-19 Surveillance Team |
| EPI_ISL_3026966, EPI_ISL_3026967, EPI_ISL_3026968 | Lighthouse Lab in Alderley Park | Wellcome Sanger Institute for the COVID-19 Genomics UK (COG-UK) Consortium | Jacquelyn Wynn, Mairead Hyland, The Lighthouse Lab in Alderley Park and Alex Alderton, Roberto Amato, Jeffrey Barrett, Sonia Goncalves, Ewan Harrison, David K. Jackson, Ian Johnston, Dominic Kwiatkowski, Cordelia Langford, John Sillitoe on behalf of the Wellcome Sanger Institute COVID-19 Surveillance Team |
| EPI_ISL_3026969 | Lighthouse Lab in Milton Keynes | Wellcome Sanger Institute for the COVID-19 Genomics UK (COG-UK) Consortium | The Lighthouse Lab in Milton Keynes and Alex Alderton, Roberto Amato, Jeffrey Barrett, Sonia Goncalves, Ewan Harrison, David K. Jackson, Ian Johnston, Dominic Kwiatkowski, Cordelia Langford, John Sillitoe on behalf of the Wellcome Sanger Institute COVID-19 Surveillance Team |
| EPI_ISL_3026970, EPI_ISL_3026971 | Lighthouse Lab in Alderley Park | Wellcome Sanger Institute for the COVID-19 Genomics UK (COG-UK) Consortium | Jacquelyn Wynn, Mairead Hyland, The Lighthouse Lab in Alderley Park and Alex Alderton, Roberto Amato, Jeffrey Barrett, Sonia Goncalves, Ewan Harrison, David K. Jackson, Ian Johnston, Dominic Kwiatkowski, Cordelia Langford, John Sillitoe on behalf of the Wellcome Sanger Institute COVID-19 Surveillance Team |
| EPI_ISL_3026972 | Lighthouse Lab in Glasgow | Wellcome Sanger Institute for the COVID-19 Genomics UK (COG-UK) Consortium | Harper VanSteenhouse, Yumi Kasai, David Gray, Carol Clugston, Anna Dominiczak and Alex Alderton, Roberto Amato, Jeffrey Barrett, Sonia Goncalves, Ewan Harrison, David K. Jackson, Ian Johnston, Dominic Kwiatkowski, Cordelia Langford, John Sillitoe on behalf of the Wellcome Sanger Institute COVID-19 Surveillance Team |
| EPI_ISL_3026973 | Lighthouse Lab in Alderley Park | Wellcome Sanger Institute for the COVID-19 Genomics UK (COG-UK) Consortium | Jacquelyn Wynn, Mairead Hyland, The Lighthouse Lab in Alderley Park and Alex Alderton, Roberto Amato, Jeffrey Barrett, Sonia Goncalves, Ewan Harrison, David K. Jackson, Ian Johnston, Dominic Kwiatkowski, Cordelia Langford, John Sillitoe on behalf of the Wellcome Sanger Institute COVID-19 Surveillance Team |
| EPI_ISL_3026974, EPI_ISL_3026975 | Lighthouse Lab in Milton Keynes | Wellcome Sanger Institute for the COVID-19 Genomics UK (COG-UK) Consortium | The Lighthouse Lab in Milton Keynes and Alex Alderton, Roberto Amato, Jeffrey Barrett, Sonia Goncalves, Ewan Harrison, David K. Jackson, Ian Johnston, Dominic Kwiatkowski, Cordelia Langford, John Sillitoe on behalf of the Wellcome Sanger Institute COVID-19 Surveillance Team |
| EPI_ISL_3026976 | Lighthouse Lab in Glasgow | Wellcome Sanger Institute for the COVID-19 Genomics UK (COG-UK) Consortium | Harper VanSteenhouse, Yumi Kasai, David Gray, Carol Clugston, Anna Dominiczak and Alex Alderton, Roberto Amato, Jeffrey Barrett, Sonia Goncalves, Ewan Harrison, David K. Jackson, Ian Johnston, Dominic Kwiatkowski, Cordelia Langford, John Sillitoe on behalf of the Wellcome Sanger Institute COVID-19 Surveillance Team |
| EPI_ISL_3026977 | Lighthouse Lab in Milton Keynes | Wellcome Sanger Institute for the COVID-19 Genomics UK (COG-UK) Consortium | The Lighthouse Lab in Milton Keynes and Alex Alderton, Roberto Amato, Jeffrey Barrett, Sonia Goncalves, Ewan Harrison, David K. Jackson, Ian Johnston, Dominic Kwiatkowski, Cordelia Langford, John Sillitoe on behalf of the Wellcome Sanger Institute COVID-19 Surveillance Team |
| EPI_ISL_3026978, EPI_ISL_3026979 | Lighthouse Lab in Glasgow | Wellcome Sanger Institute for the COVID-19 Genomics UK (COG-UK) Consortium | Harper VanSteenhouse, Yumi Kasai, David Gray, Carol Clugston, Anna Dominiczak and Alex Alderton, Roberto Amato, Jeffrey Barrett, Sonia Goncalves, Ewan Harrison, David K. Jackson, Ian Johnston, Dominic Kwiatkowski, Cordelia Langford, John Sillitoe on behalf of the Wellcome Sanger Institute COVID-19 Surveillance Team |
| EPI_ISL_3026980, EPI_ISL_3026981, EPI_ISL_3026982 | Lighthouse Lab in Milton Keynes | Wellcome Sanger Institute for the COVID-19 Genomics UK (COG-UK) Consortium | The Lighthouse Lab in Milton Keynes and Alex Alderton, Roberto Amato, Jeffrey Barrett, Sonia Goncalves, Ewan Harrison, David K. Jackson, Ian Johnston, Dominic Kwiatkowski, Cordelia Langford, John Sillitoe on behalf of the Wellcome Sanger Institute COVID-19 Surveillance Team |
| EPI_ISL_3026983, EPI_ISL_3026984, EPI_ISL_3026985 | Lighthouse Lab in Glasgow | Wellcome Sanger Institute for the COVID-19 Genomics UK (COG-UK) Consortium | Harper VanSteenhouse, Yumi Kasai, David Gray, Carol Clugston, Anna Dominiczak and Alex Alderton, Roberto Amato, Jeffrey Barrett, Sonia Goncalves, Ewan Harrison, David K. Jackson, Ian Johnston, Dominic Kwiatkowski, Cordelia Langford, John Sillitoe on behalf of the Wellcome Sanger Institute COVID-19 Surveillance Team |
| EPI_ISL_3026986, EPI_ISL_3026987 | Lighthouse Lab in Milton Keynes | Wellcome Sanger Institute for the COVID-19 Genomics UK (COG-UK) Consortium | The Lighthouse Lab in Milton Keynes and Alex Alderton, Roberto Amato, Jeffrey Barrett, Sonia Goncalves, Ewan Harrison, David K. Jackson, Ian Johnston, Dominic Kwiatkowski, Cordelia Langford, John Sillitoe on behalf of the Wellcome Sanger Institute COVID-19 Surveillance Team |

| | | | |
|---|---|---|---|
| EPI_ISL_3026988, EPI_ISL_3026989 | Lighthouse Lab in Glasgow | Wellcome Sanger Institute for the COVID-19 Genomics UK (COG-UK) Consortium | Harper VanSteenhouse, Yumi Kasai, David Gray, Carol Clugston, Anna Dominiczak and Alex Alderton, Roberto Amato, Jeffrey Barrett, Sonia Goncalves, Ewan Harrison, David K. Jackson, Ian Johnston, Dominic Kwiatkowski, Cordelia Langford, John Sillitoe on behalf of the Wellcome Sanger Institute COVID-19 Surveillance Team |
| EPI_ISL_3026990, EPI_ISL_3026991 | Lighthouse Lab in Milton Keynes | Wellcome Sanger Institute for the COVID-19 Genomics UK (COG-UK) Consortium | The Lighthouse Lab in Milton Keynes and Alex Alderton, Roberto Amato, Jeffrey Barrett, Sonia Goncalves, Ewan Harrison, David K. Jackson, Ian Johnston, Dominic Kwiatkowski, Cordelia Langford, John Sillitoe on behalf of the Wellcome Sanger Institute COVID-19 Surveillance Team |
| EPI_ISL_3026992, EPI_ISL_3026993 | Lighthouse Lab in Alderley Park | Wellcome Sanger Institute for the COVID-19 Genomics UK (COG-UK) Consortium | Jacquelyn Wynn, Mairead Hyland, The Lighthouse Lab in Alderley Park and Alex Alderton, Roberto Amato, Jeffrey Barrett, Sonia Goncalves, Ewan Harrison, David K. Jackson, Ian Johnston, Dominic Kwiatkowski, Cordelia Langford, John Sillitoe on behalf of the Wellcome Sanger Institute COVID-19 Surveillance Team |
| EPI_ISL_3026994 | Lighthouse Lab in Milton Keynes | Wellcome Sanger Institute for the COVID-19 Genomics UK (COG-UK) Consortium | The Lighthouse Lab in Milton Keynes and Alex Alderton, Roberto Amato, Jeffrey Barrett, Sonia Goncalves, Ewan Harrison, David K. Jackson, Ian Johnston, Dominic Kwiatkowski, Cordelia Langford, John Sillitoe on behalf of the Wellcome Sanger Institute COVID-19 Surveillance Team |
| EPI_ISL_3026995 | Lighthouse Lab in Glasgow | Wellcome Sanger Institute for the COVID-19 Genomics UK (COG-UK) Consortium | Harper VanSteenhouse, Yumi Kasai, David Gray, Carol Clugston, Anna Dominiczak and Alex Alderton, Roberto Amato, Jeffrey Barrett, Sonia Goncalves, Ewan Harrison, David K. Jackson, Ian Johnston, Dominic Kwiatkowski, Cordelia Langford, John Sillitoe on behalf of the Wellcome Sanger Institute COVID-19 Surveillance Team |
| EPI_ISL_3026996, EPI_ISL_3026997, EPI_ISL_3026998, EPI_ISL_3026999, EPI_ISL_3027000 | Lighthouse Lab in Milton Keynes | Wellcome Sanger Institute for the COVID-19 Genomics UK (COG-UK) Consortium | The Lighthouse Lab in Milton Keynes and Alex Alderton, Roberto Amato, Jeffrey Barrett, Sonia Goncalves, Ewan Harrison, David K. Jackson, Ian Johnston, Dominic Kwiatkowski, Cordelia Langford, John Sillitoe on behalf of the Wellcome Sanger Institute COVID-19 Surveillance Team |
| EPI_ISL_3027001 | Lighthouse Lab in Glasgow | Wellcome Sanger Institute for the COVID-19 Genomics UK (COG-UK) Consortium | Harper VanSteenhouse, Yumi Kasai, David Gray, Carol Clugston, Anna Dominiczak and Alex Alderton, Roberto Amato, Jeffrey Barrett, Sonia Goncalves, Ewan Harrison, David K. Jackson, Ian Johnston, Dominic Kwiatkowski, Cordelia Langford, John Sillitoe on behalf of the Wellcome Sanger Institute COVID-19 Surveillance Team |
| EPI_ISL_3027002 | Lighthouse Lab in Alderley Park | Wellcome Sanger Institute for the COVID-19 Genomics UK (COG-UK) Consortium | Jacquelyn Wynn, Mairead Hyland, The Lighthouse Lab in Alderley Park and Alex Alderton, Roberto Amato, Jeffrey Barrett, Sonia Goncalves, Ewan Harrison, David K. Jackson, Ian Johnston, Dominic Kwiatkowski, Cordelia Langford, John Sillitoe on behalf of the Wellcome Sanger Institute COVID-19 Surveillance Team |
| EPI_ISL_3027003, EPI_ISL_3027004 | Lighthouse Lab in Glasgow | Wellcome Sanger Institute for the COVID-19 Genomics UK (COG-UK) Consortium | Harper VanSteenhouse, Yumi Kasai, David Gray, Carol Clugston, Anna Dominiczak and Alex Alderton, Roberto Amato, Jeffrey Barrett, Sonia Goncalves, Ewan Harrison, David K. Jackson, Ian Johnston, Dominic Kwiatkowski, Cordelia Langford, John Sillitoe on behalf of the Wellcome Sanger Institute COVID-19 Surveillance Team |
| EPI_ISL_3027005, EPI_ISL_3027006, EPI_ISL_3027007 | Lighthouse Lab in Milton Keynes | Wellcome Sanger Institute for the COVID-19 Genomics UK (COG-UK) Consortium | The Lighthouse Lab in Milton Keynes and Alex Alderton, Roberto Amato, Jeffrey Barrett, Sonia Goncalves, Ewan Harrison, David K. Jackson, Ian Johnston, Dominic Kwiatkowski, Cordelia Langford, John Sillitoe on behalf of the Wellcome Sanger Institute COVID-19 Surveillance Team |
| EPI_ISL_3027008 | Lighthouse Lab in Glasgow | Wellcome Sanger Institute for the COVID-19 Genomics UK (COG-UK) Consortium | Harper VanSteenhouse, Yumi Kasai, David Gray, Carol Clugston, Anna Dominiczak and Alex Alderton, Roberto Amato, Jeffrey Barrett, Sonia Goncalves, Ewan Harrison, David K. Jackson, Ian Johnston, Dominic Kwiatkowski, Cordelia Langford, John Sillitoe on behalf of the Wellcome Sanger Institute COVID-19 Surveillance Team |
| EPI_ISL_3027009 | Lighthouse Lab in Alderley Park | Wellcome Sanger Institute for the COVID-19 Genomics UK (COG-UK) Consortium | Jacquelyn Wynn, Mairead Hyland, The Lighthouse Lab in Alderley Park and Alex Alderton, Roberto Amato, Jeffrey Barrett, Sonia Goncalves, Ewan Harrison, David K. Jackson, Ian Johnston, Dominic Kwiatkowski, Cordelia Langford, John Sillitoe on behalf of the Wellcome Sanger Institute COVID-19 Surveillance Team |
| EPI_ISL_3027010 | Lighthouse Lab in Glasgow | Wellcome Sanger Institute for the COVID-19 Genomics UK (COG-UK) Consortium | Harper VanSteenhouse, Yumi Kasai, David Gray, Carol Clugston, Anna Dominiczak and Alex Alderton, Roberto Amato, Jeffrey Barrett, Sonia Goncalves, Ewan Harrison, David K. Jackson, Ian Johnston, Dominic Kwiatkowski, Cordelia Langford, John Sillitoe on behalf of the Wellcome Sanger Institute COVID-19 Surveillance Team |
| EPI_ISL_3027011 | Lighthouse Lab in Alderley Park | Wellcome Sanger Institute for the COVID-19 Genomics UK (COG-UK) Consortium | Jacquelyn Wynn, Mairead Hyland, The Lighthouse Lab in Alderley Park and Alex Alderton, Roberto Amato, Jeffrey Barrett, Sonia Goncalves, Ewan Harrison, David K. Jackson, Ian Johnston, Dominic Kwiatkowski, Cordelia Langford, John Sillitoe on behalf of the Wellcome Sanger Institute COVID-19 Surveillance Team |
| EPI_ISL_3027012, EPI_ISL_3027013, EPI_ISL_3027014 | Lighthouse Lab in Milton Keynes | Wellcome Sanger Institute for the COVID-19 Genomics UK (COG-UK) Consortium | The Lighthouse Lab in Milton Keynes and Alex Alderton, Roberto Amato, Jeffrey Barrett, Sonia Goncalves, Ewan Harrison, David K. Jackson, Ian Johnston, Dominic Kwiatkowski, Cordelia Langford, John Sillitoe on behalf of the Wellcome Sanger Institute COVID-19 Surveillance Team |
| EPI_ISL_3027015 | Lighthouse Lab in Glasgow | Wellcome Sanger Institute for the COVID-19 Genomics UK (COG-UK) Consortium | Harper VanSteenhouse, Yumi Kasai, David Gray, Carol Clugston, Anna Dominiczak and Alex Alderton, Roberto Amato, Jeffrey Barrett, Sonia Goncalves, Ewan Harrison, David K. Jackson, Ian Johnston, Dominic Kwiatkowski, Cordelia Langford, John Sillitoe on behalf of the Wellcome Sanger Institute COVID-19 Surveillance Team |
| EPI_ISL_3027016, EPI_ISL_3027017 | Lighthouse Lab in Milton Keynes | Wellcome Sanger Institute for the COVID-19 Genomics UK (COG-UK) Consortium | The Lighthouse Lab in Milton Keynes and Alex Alderton, Roberto Amato, Jeffrey Barrett, Sonia Goncalves, Ewan Harrison, David K. Jackson, Ian Johnston, Dominic Kwiatkowski, Cordelia Langford, John Sillitoe on behalf of the Wellcome Sanger Institute COVID-19 Surveillance Team |
| EPI_ISL_3027018 | Lighthouse Lab in Glasgow | Wellcome Sanger Institute for the COVID-19 Genomics UK (COG-UK) Consortium | Harper VanSteenhouse, Yumi Kasai, David Gray, Carol Clugston, Anna Dominiczak and Alex Alderton, Roberto Amato, Jeffrey Barrett, Sonia Goncalves, Ewan Harrison, David K. Jackson, Ian Johnston, Dominic Kwiatkowski, Cordelia Langford, John Sillitoe on behalf of the Wellcome Sanger Institute COVID-19 Surveillance Team |
| EPI_ISL_3027019, EPI_ISL_3027020 | Lighthouse Lab in Alderley Park | Wellcome Sanger Institute for the COVID-19 Genomics UK (COG-UK) Consortium | Jacquelyn Wynn, Mairead Hyland, The Lighthouse Lab in Alderley Park and Alex Alderton, Roberto Amato, Jeffrey Barrett, Sonia Goncalves, Ewan Harrison, David K. Jackson, Ian Johnston, Dominic Kwiatkowski, Cordelia Langford, John Sillitoe on behalf of the Wellcome Sanger Institute COVID-19 Surveillance Team |
| EPI_ISL_3027021 | Lighthouse Lab in Glasgow | Wellcome Sanger Institute for the COVID-19 Genomics UK (COG-UK) Consortium | Harper VanSteenhouse, Yumi Kasai, David Gray, Carol Clugston, Anna Dominiczak and Alex Alderton, Roberto Amato, Jeffrey Barrett, Sonia Goncalves, Ewan Harrison, David K. Jackson, Ian Johnston, Dominic Kwiatkowski, Cordelia Langford, John Sillitoe on behalf of the Wellcome Sanger Institute COVID-19 Surveillance Team |
| EPI_ISL_3027022, EPI_ISL_3027023 | Lighthouse Lab in Milton Keynes | Wellcome Sanger Institute for the COVID-19 Genomics UK (COG-UK) Consortium | The Lighthouse Lab in Milton Keynes and Alex Alderton, Roberto Amato, Jeffrey Barrett, Sonia Goncalves, Ewan Harrison, David K. Jackson, Ian Johnston, Dominic Kwiatkowski, Cordelia Langford, John Sillitoe on behalf of the Wellcome Sanger Institute COVID-19 Surveillance Team |
| EPI_ISL_3027024, EPI_ISL_3027025 | Lighthouse Lab in Alderley Park | Wellcome Sanger Institute for the COVID-19 Genomics UK (COG-UK) Consortium | Jacquelyn Wynn, Mairead Hyland, The Lighthouse Lab in Alderley Park and Alex Alderton, Roberto Amato, Jeffrey Barrett, Sonia Goncalves, Ewan Harrison, David K. Jackson, Ian Johnston, Dominic Kwiatkowski, Cordelia Langford, John Sillitoe on behalf of the Wellcome Sanger Institute COVID-19 Surveillance Team |
| EPI_ISL_3027026 | Lighthouse Lab in Milton Keynes | Wellcome Sanger Institute for the COVID-19 Genomics UK (COG-UK) Consortium | The Lighthouse Lab in Milton Keynes and Alex Alderton, Roberto Amato, Jeffrey Barrett, Sonia Goncalves, Ewan Harrison, David K. Jackson, Ian Johnston, Dominic Kwiatkowski, Cordelia Langford, John Sillitoe on behalf of the Wellcome Sanger Institute COVID-19 Surveillance Team |
| EPI_ISL_3027027 | Lighthouse Lab in Alderley Park | Wellcome Sanger Institute for the COVID-19 Genomics UK (COG-UK) Consortium | Jacquelyn Wynn, Mairead Hyland, The Lighthouse Lab in Alderley Park and Alex Alderton, Roberto Amato, Jeffrey Barrett, Sonia Goncalves, Ewan Harrison, David K. Jackson, Ian Johnston, Dominic Kwiatkowski, Cordelia Langford, John Sillitoe on behalf of the Wellcome Sanger Institute COVID-19 Surveillance Team |
| EPI_ISL_3027028 | Lighthouse Lab in Milton Keynes | Wellcome Sanger Institute for the COVID-19 Genomics UK (COG-UK) Consortium | The Lighthouse Lab in Milton Keynes and Alex Alderton, Roberto Amato, Jeffrey Barrett, Sonia Goncalves, Ewan Harrison, David K. Jackson, Ian Johnston, Dominic Kwiatkowski, Cordelia Langford, John Sillitoe on behalf of the Wellcome Sanger Institute COVID-19 Surveillance Team |
| EPI_ISL_3027029 | Lighthouse Lab in Glasgow | Wellcome Sanger Institute for the COVID-19 Genomics UK (COG-UK) Consortium | Harper VanSteenhouse, Yumi Kasai, David Gray, Carol Clugston, Anna Dominiczak and Alex Alderton, Roberto Amato, Jeffrey Barrett, Sonia Goncalves, Ewan Harrison, David K. Jackson, Ian Johnston, Dominic Kwiatkowski, Cordelia Langford, John Sillitoe on behalf of the Wellcome Sanger Institute COVID-19 Surveillance Team |
| EPI_ISL_3027030 | Lighthouse Lab in Alderley Park | Wellcome Sanger Institute for the COVID-19 Genomics UK (COG-UK) Consortium | Jacquelyn Wynn, Mairead Hyland, The Lighthouse Lab in Alderley Park and Alex Alderton, Roberto Amato, Jeffrey Barrett, Sonia Goncalves, Ewan Harrison, David K. Jackson, Ian Johnston, Dominic Kwiatkowski, Cordelia Langford, John Sillitoe on behalf of the Wellcome Sanger Institute COVID-19 Surveillance Team |
| EPI_ISL_3027031 | Lighthouse Lab in Glasgow | Wellcome Sanger Institute for the COVID-19 Genomics UK (COG-UK) Consortium | Harper VanSteenhouse, Yumi Kasai, David Gray, Carol Clugston, Anna Dominiczak and Alex Alderton, Roberto Amato, Jeffrey Barrett, Sonia Goncalves, Ewan Harrison, David K. Jackson, Ian Johnston, Dominic Kwiatkowski, Cordelia Langford, John Sillitoe on behalf of the Wellcome Sanger Institute COVID-19 Surveillance Team |
| EPI_ISL_3027032 | Lighthouse Lab in Alderley Park | Wellcome Sanger Institute for the COVID-19 Genomics UK (COG-UK) Consortium | Jacquelyn Wynn, Mairead Hyland, The Lighthouse Lab in Alderley Park and Alex Alderton, Roberto Amato, Jeffrey Barrett, Sonia Goncalves, Ewan Harrison, David K. Jackson, Ian Johnston, Dominic Kwiatkowski, Cordelia Langford, John Sillitoe on behalf of the Wellcome Sanger Institute COVID-19 Surveillance Team |
| EPI_ISL_3027033 | Lighthouse Lab in Milton Keynes | Wellcome Sanger Institute for the COVID-19 Genomics UK (COG-UK) Consortium | The Lighthouse Lab in Milton Keynes and Alex Alderton, Roberto Amato, Jeffrey Barrett, Sonia Goncalves, Ewan Harrison, David K. Jackson, Ian Johnston, Dominic Kwiatkowski, Cordelia Langford, John Sillitoe on behalf of the Wellcome Sanger Institute COVID-19 Surveillance Team |
| EPI_ISL_3027034, EPI_ISL_3027035 | Lighthouse Lab in Alderley Park | Wellcome Sanger Institute for the COVID-19 Genomics UK (COG-UK) Consortium | Jacquelyn Wynn, Mairead Hyland, The Lighthouse Lab in Alderley Park and Alex Alderton, Roberto Amato, Jeffrey Barrett, Sonia Goncalves, Ewan Harrison, David K. Jackson, Ian Johnston, Dominic Kwiatkowski, Cordelia Langford, John Sillitoe on behalf of the Wellcome Sanger Institute COVID-19 Surveillance Team |
| EPI_ISL_3027036 | Lighthouse Lab in Glasgow | Wellcome Sanger Institute for the COVID-19 Genomics UK (COG-UK) Consortium | Harper VanSteenhouse, Yumi Kasai, David Gray, Carol Clugston, Anna Dominiczak and Alex Alderton, Roberto Amato, Jeffrey Barrett, Sonia Goncalves, Ewan Harrison, David K. Jackson, Ian Johnston, Dominic Kwiatkowski, Cordelia Langford, John Sillitoe on behalf of the Wellcome Sanger Institute COVID-19 Surveillance Team |
| EPI_ISL_3027037 | Lighthouse Lab in Milton Keynes | Wellcome Sanger Institute for the COVID-19 Genomics UK (COG-UK) Consortium | The Lighthouse Lab in Milton Keynes and Alex Alderton, Roberto Amato, Jeffrey Barrett, Sonia Goncalves, Ewan Harrison, David K. Jackson, Ian Johnston, Dominic Kwiatkowski, Cordelia Langford, John Sillitoe on behalf of the Wellcome Sanger Institute COVID-19 Surveillance Team |
| EPI_ISL_3027038 | Lighthouse Lab in Alderley Park | Wellcome Sanger Institute for the COVID-19 Genomics UK (COG-UK) Consortium | Jacquelyn Wynn, Mairead Hyland, The Lighthouse Lab in Alderley Park and Alex Alderton, Roberto Amato, Jeffrey Barrett, Sonia Goncalves, Ewan Harrison, David K. Jackson, Ian Johnston, Dominic Kwiatkowski, Cordelia Langford, John Sillitoe on behalf of the Wellcome Sanger Institute COVID-19 Surveillance Team |
| EPI_ISL_3027039 | Lighthouse Lab in Milton Keynes | Wellcome Sanger Institute for the COVID-19 Genomics UK (COG-UK) Consortium | The Lighthouse Lab in Milton Keynes and Alex Alderton, Roberto Amato, Jeffrey Barrett, Sonia Goncalves, Ewan Harrison, David K. Jackson, Ian Johnston, Dominic Kwiatkowski, Cordelia Langford, John Sillitoe on behalf of the Wellcome Sanger Institute COVID-19 Surveillance Team |
| EPI_ISL_3027040 | Lighthouse Lab in Alderley Park | Wellcome Sanger Institute for the COVID-19 Genomics UK (COG-UK) Consortium | Jacquelyn Wynn, Mairead Hyland, The Lighthouse Lab in Alderley Park and Alex Alderton, Roberto Amato, Jeffrey Barrett, Sonia Goncalves, Ewan Harrison, David K. Jackson, Ian Johnston, Dominic Kwiatkowski, Cordelia Langford, John Sillitoe on behalf of the Wellcome Sanger Institute COVID-19 Surveillance Team |
| EPI_ISL_3027041 | Lighthouse Lab in Milton Keynes | Wellcome Sanger Institute for the COVID-19 Genomics UK (COG-UK) Consortium | The Lighthouse Lab in Milton Keynes and Alex Alderton, Roberto Amato, Jeffrey Barrett, Sonia Goncalves, Ewan Harrison, David K. Jackson, Ian Johnston, Dominic Kwiatkowski, Cordelia Langford, John Sillitoe on behalf of the Wellcome Sanger Institute COVID-19 Surveillance Team |
| EPI_ISL_3027042 | Lighthouse Lab in Alderley Park | Wellcome Sanger Institute for the COVID-19 Genomics UK (COG-UK) Consortium | Jacquelyn Wynn, Mairead Hyland, The Lighthouse Lab in Alderley Park and Alex Alderton, Roberto Amato, Jeffrey Barrett, Sonia Goncalves, Ewan Harrison, David K. Jackson, Ian Johnston, Dominic Kwiatkowski, Cordelia Langford, John Sillitoe on behalf of the Wellcome Sanger Institute COVID-19 Surveillance Team |
| EPI_ISL_3027043 | Lighthouse Lab in Milton Keynes | Wellcome Sanger Institute for the COVID-19 Genomics UK (COG-UK) Consortium | The Lighthouse Lab in Milton Keynes and Alex Alderton, Roberto Amato, Jeffrey Barrett, Sonia Goncalves, Ewan Harrison, David K. Jackson, Ian Johnston, Dominic Kwiatkowski, Cordelia Langford, John Sillitoe on behalf of the Wellcome Sanger Institute COVID-19 Surveillance Team |
| EPI_ISL_3027044 | Lighthouse Lab in Glasgow | Wellcome Sanger Institute for the COVID-19 Genomics UK (COG-UK) Consortium | Harper VanSteenhouse, Yumi Kasai, David Gray, Carol Clugston, Anna Dominiczak and Alex Alderton, Roberto Amato, Jeffrey Barrett, Sonia Goncalves, Ewan Harrison, David K. Jackson, Ian Johnston, Dominic Kwiatkowski, Cordelia Langford, John Sillitoe on behalf of the Wellcome Sanger Institute COVID-19 Surveillance Team |
| EPI_ISL_3027045, EPI_ISL_3027046, EPI_ISL_3027047 | Lighthouse Lab in Milton Keynes | Wellcome Sanger Institute for the COVID-19 Genomics UK (COG-UK) Consortium | The Lighthouse Lab in Milton Keynes and Alex Alderton, Roberto Amato, Jeffrey Barrett, Sonia Goncalves, Ewan Harrison, David K. Jackson, Ian Johnston, Dominic Kwiatkowski, Cordelia Langford, John Sillitoe on behalf of the Wellcome Sanger Institute COVID-19 Surveillance Team |
| EPI_ISL_3027048 | Lighthouse Lab in Glasgow | Wellcome Sanger Institute for the COVID-19 Genomics UK (COG-UK) Consortium | Harper VanSteenhouse, Yumi Kasai, David Gray, Carol Clugston, Anna Dominiczak and Alex Alderton, Roberto Amato, Jeffrey Barrett, Sonia Goncalves, Ewan Harrison, David K. Jackson, Ian Johnston, Dominic Kwiatkowski, Cordelia Langford, John Sillitoe on behalf of the Wellcome Sanger Institute COVID-19 Surveillance Team |
| EPI_ISL_3027049, EPI_ISL_3027050 | Lighthouse Lab in Milton Keynes | Wellcome Sanger Institute for the COVID-19 Genomics UK (COG-UK) Consortium | The Lighthouse Lab in Milton Keynes and Alex Alderton, Roberto Amato, Jeffrey Barrett, Sonia Goncalves, Ewan Harrison, David K. Jackson, Ian Johnston, Dominic Kwiatkowski, Cordelia Langford, John Sillitoe on behalf of the Wellcome Sanger Institute COVID-19 Surveillance Team |

| | | | |
|---|---|---|---|
| EPI_ISL_3027051, EPI_ISL_3027052 | Lighthouse Lab in Alderley Park | Wellcome Sanger Institute for the COVID-19 Genomics UK (COG-UK) Consortium | Jacquelyn Wynn, Mairead Hyland, The Lighthouse Lab in Alderley Park and Alex Alderton, Roberto Amato, Jeffrey Barrett, Sonia Goncalves, Ewan Harrison, David K. Jackson, Ian Johnston, Dominic Kwiatkowski, Cordelia Langford, John Sillitoe on behalf of the Wellcome Sanger Institute COVID-19 Surveillance Team |
| EPI_ISL_3027053 | Lighthouse Lab in Milton Keynes | Wellcome Sanger Institute for the COVID-19 Genomics UK (COG-UK) Consortium | The Lighthouse Lab in Milton Keynes and Alex Alderton, Roberto Amato, Jeffrey Barrett, Sonia Goncalves, Ewan Harrison, David K. Jackson, Ian Johnston, Dominic Kwiatkowski, Cordelia Langford, John Sillitoe on behalf of the Wellcome Sanger Institute COVID-19 Surveillance Team |
| EPI_ISL_3027054 | Lighthouse Lab in Glasgow | Wellcome Sanger Institute for the COVID-19 Genomics UK (COG-UK) Consortium | Harper VanSteenhouse, Yumi Kasai, David Gray, Carol Clugston, Anna Dominiczak and Alex Alderton, Roberto Amato, Jeffrey Barrett, Sonia Goncalves, Ewan Harrison, David K. Jackson, Ian Johnston, Dominic Kwiatkowski, Cordelia Langford, John Sillitoe on behalf of the Wellcome Sanger Institute COVID-19 Surveillance Team |
| EPI_ISL_3027055, EPI_ISL_3027056, EPI_ISL_3027057, EPI_ISL_3027058 | Lighthouse Lab in Milton Keynes | Wellcome Sanger Institute for the COVID-19 Genomics UK (COG-UK) Consortium | The Lighthouse Lab in Milton Keynes and Alex Alderton, Roberto Amato, Jeffrey Barrett, Sonia Goncalves, Ewan Harrison, David K. Jackson, Ian Johnston, Dominic Kwiatkowski, Cordelia Langford, John Sillitoe on behalf of the Wellcome Sanger Institute COVID-19 Surveillance Team |
| EPI_ISL_3027059, EPI_ISL_3027060 | Lighthouse Lab in Glasgow | Wellcome Sanger Institute for the COVID-19 Genomics UK (COG-UK) Consortium | Harper VanSteenhouse, Yumi Kasai, David Gray, Carol Clugston, Anna Dominiczak and Alex Alderton, Roberto Amato, Jeffrey Barrett, Sonia Goncalves, Ewan Harrison, David K. Jackson, Ian Johnston, Dominic Kwiatkowski, Cordelia Langford, John Sillitoe on behalf of the Wellcome Sanger Institute COVID-19 Surveillance Team |
| EPI_ISL_3027061, EPI_ISL_3027062, EPI_ISL_3027063 | Lighthouse Lab in Alderley Park | Wellcome Sanger Institute for the COVID-19 Genomics UK (COG-UK) Consortium | Jacquelyn Wynn, Mairead Hyland, The Lighthouse Lab in Alderley Park and Alex Alderton, Roberto Amato, Jeffrey Barrett, Sonia Goncalves, Ewan Harrison, David K. Jackson, Ian Johnston, Dominic Kwiatkowski, Cordelia Langford, John Sillitoe on behalf of the Wellcome Sanger Institute COVID-19 Surveillance Team |
| EPI_ISL_3027064, EPI_ISL_3027065 | Lighthouse Lab in Glasgow | Wellcome Sanger Institute for the COVID-19 Genomics UK (COG-UK) Consortium | Harper VanSteenhouse, Yumi Kasai, David Gray, Carol Clugston, Anna Dominiczak and Alex Alderton, Roberto Amato, Jeffrey Barrett, Sonia Goncalves, Ewan Harrison, David K. Jackson, Ian Johnston, Dominic Kwiatkowski, Cordelia Langford, John Sillitoe on behalf of the Wellcome Sanger Institute COVID-19 Surveillance Team |
| EPI_ISL_3027066 | Lighthouse Lab in Milton Keynes | Wellcome Sanger Institute for the COVID-19 Genomics UK (COG-UK) Consortium | The Lighthouse Lab in Milton Keynes and Alex Alderton, Roberto Amato, Jeffrey Barrett, Sonia Goncalves, Ewan Harrison, David K. Jackson, Ian Johnston, Dominic Kwiatkowski, Cordelia Langford, John Sillitoe on behalf of the Wellcome Sanger Institute COVID-19 Surveillance Team |
| EPI_ISL_3027067 | Lighthouse Lab in Alderley Park | Wellcome Sanger Institute for the COVID-19 Genomics UK (COG-UK) Consortium | Jacquelyn Wynn, Mairead Hyland, The Lighthouse Lab in Alderley Park and Alex Alderton, Roberto Amato, Jeffrey Barrett, Sonia Goncalves, Ewan Harrison, David K. Jackson, Ian Johnston, Dominic Kwiatkowski, Cordelia Langford, John Sillitoe on behalf of the Wellcome Sanger Institute COVID-19 Surveillance Team |
| EPI_ISL_3027068 | Lighthouse Lab in Milton Keynes | Wellcome Sanger Institute for the COVID-19 Genomics UK (COG-UK) Consortium | The Lighthouse Lab in Milton Keynes and Alex Alderton, Roberto Amato, Jeffrey Barrett, Sonia Goncalves, Ewan Harrison, David K. Jackson, Ian Johnston, Dominic Kwiatkowski, Cordelia Langford, John Sillitoe on behalf of the Wellcome Sanger Institute COVID-19 Surveillance Team |
| EPI_ISL_3027069, EPI_ISL_3027070 | Lighthouse Lab in Glasgow | Wellcome Sanger Institute for the COVID-19 Genomics UK (COG-UK) Consortium | Harper VanSteenhouse, Yumi Kasai, David Gray, Carol Clugston, Anna Dominiczak and Alex Alderton, Roberto Amato, Jeffrey Barrett, Sonia Goncalves, Ewan Harrison, David K. Jackson, Ian Johnston, Dominic Kwiatkowski, Cordelia Langford, John Sillitoe on behalf of the Wellcome Sanger Institute COVID-19 Surveillance Team |
| EPI_ISL_3027071 | Lighthouse Lab in Alderley Park | Wellcome Sanger Institute for the COVID-19 Genomics UK (COG-UK) Consortium | Jacquelyn Wynn, Mairead Hyland, The Lighthouse Lab in Alderley Park and Alex Alderton, Roberto Amato, Jeffrey Barrett, Sonia Goncalves, Ewan Harrison, David K. Jackson, Ian Johnston, Dominic Kwiatkowski, Cordelia Langford, John Sillitoe on behalf of the Wellcome Sanger Institute COVID-19 Surveillance Team |
| EPI_ISL_3027072 | Lighthouse Lab in Milton Keynes | Wellcome Sanger Institute for the COVID-19 Genomics UK (COG-UK) Consortium | The Lighthouse Lab in Milton Keynes and Alex Alderton, Roberto Amato, Jeffrey Barrett, Sonia Goncalves, Ewan Harrison, David K. Jackson, Ian Johnston, Dominic Kwiatkowski, Cordelia Langford, John Sillitoe on behalf of the Wellcome Sanger Institute COVID-19 Surveillance Team |
| EPI_ISL_3027073 | Lighthouse Lab in Glasgow | Wellcome Sanger Institute for the COVID-19 Genomics UK (COG-UK) Consortium | Harper VanSteenhouse, Yumi Kasai, David Gray, Carol Clugston, Anna Dominiczak and Alex Alderton, Roberto Amato, Jeffrey Barrett, Sonia Goncalves, Ewan Harrison, David K. Jackson, Ian Johnston, Dominic Kwiatkowski, Cordelia Langford, John Sillitoe on behalf of the Wellcome Sanger Institute COVID-19 Surveillance Team |
| EPI_ISL_3027074 | Lighthouse Lab in Milton Keynes | Wellcome Sanger Institute for the COVID-19 Genomics UK (COG-UK) Consortium | The Lighthouse Lab in Milton Keynes and Alex Alderton, Roberto Amato, Jeffrey Barrett, Sonia Goncalves, Ewan Harrison, David K. Jackson, Ian Johnston, Dominic Kwiatkowski, Cordelia Langford, John Sillitoe on behalf of the Wellcome Sanger Institute COVID-19 Surveillance Team |
| EPI_ISL_3027075 | Lighthouse Lab in Alderley Park | Wellcome Sanger Institute for the COVID-19 Genomics UK (COG-UK) Consortium | Jacquelyn Wynn, Mairead Hyland, The Lighthouse Lab in Alderley Park and Alex Alderton, Roberto Amato, Jeffrey Barrett, Sonia Goncalves, Ewan Harrison, David K. Jackson, Ian Johnston, Dominic Kwiatkowski, Cordelia Langford, John Sillitoe on behalf of the Wellcome Sanger Institute COVID-19 Surveillance Team |
| EPI_ISL_3027076 | Lighthouse Lab in Glasgow | Wellcome Sanger Institute for the COVID-19 Genomics UK (COG-UK) Consortium | Harper VanSteenhouse, Yumi Kasai, David Gray, Carol Clugston, Anna Dominiczak and Alex Alderton, Roberto Amato, Jeffrey Barrett, Sonia Goncalves, Ewan Harrison, David K. Jackson, Ian Johnston, Dominic Kwiatkowski, Cordelia Langford, John Sillitoe on behalf of the Wellcome Sanger Institute COVID-19 Surveillance Team |
| EPI_ISL_3027077, EPI_ISL_3027078 | Lighthouse Lab in Alderley Park | Wellcome Sanger Institute for the COVID-19 Genomics UK (COG-UK) Consortium | Jacquelyn Wynn, Mairead Hyland, The Lighthouse Lab in Alderley Park and Alex Alderton, Roberto Amato, Jeffrey Barrett, Sonia Goncalves, Ewan Harrison, David K. Jackson, Ian Johnston, Dominic Kwiatkowski, Cordelia Langford, John Sillitoe on behalf of the Wellcome Sanger Institute COVID-19 Surveillance Team |
| EPI_ISL_3027079, EPI_ISL_3027080 | Lighthouse Lab in Glasgow | Wellcome Sanger Institute for the COVID-19 Genomics UK (COG-UK) Consortium | Harper VanSteenhouse, Yumi Kasai, David Gray, Carol Clugston, Anna Dominiczak and Alex Alderton, Roberto Amato, Jeffrey Barrett, Sonia Goncalves, Ewan Harrison, David K. Jackson, Ian Johnston, Dominic Kwiatkowski, Cordelia Langford, John Sillitoe on behalf of the Wellcome Sanger Institute COVID-19 Surveillance Team |
| EPI_ISL_3027081 | Lighthouse Lab in Alderley Park | Wellcome Sanger Institute for the COVID-19 Genomics UK (COG-UK) Consortium | Jacquelyn Wynn, Mairead Hyland, The Lighthouse Lab in Alderley Park and Alex Alderton, Roberto Amato, Jeffrey Barrett, Sonia Goncalves, Ewan Harrison, David K. Jackson, Ian Johnston, Dominic Kwiatkowski, Cordelia Langford, John Sillitoe on behalf of the Wellcome Sanger Institute COVID-19 Surveillance Team |
| EPI_ISL_3027082 | Lighthouse Lab in Milton Keynes | Wellcome Sanger Institute for the COVID-19 Genomics UK (COG-UK) Consortium | The Lighthouse Lab in Milton Keynes and Alex Alderton, Roberto Amato, Jeffrey Barrett, Sonia Goncalves, Ewan Harrison, David K. Jackson, Ian Johnston, Dominic Kwiatkowski, Cordelia Langford, John Sillitoe on behalf of the Wellcome Sanger Institute COVID-19 Surveillance Team |
| EPI_ISL_3027083, EPI_ISL_3027084 | Lighthouse Lab in Alderley Park | Wellcome Sanger Institute for the COVID-19 Genomics UK (COG-UK) Consortium | Jacquelyn Wynn, Mairead Hyland, The Lighthouse Lab in Alderley Park and Alex Alderton, Roberto Amato, Jeffrey Barrett, Sonia Goncalves, Ewan Harrison, David K. Jackson, Ian Johnston, Dominic Kwiatkowski, Cordelia Langford, John Sillitoe on behalf of the Wellcome Sanger Institute COVID-19 Surveillance Team |
| EPI_ISL_3027085 | Lighthouse Lab in Glasgow | Wellcome Sanger Institute for the COVID-19 Genomics UK (COG-UK) Consortium | Harper VanSteenhouse, Yumi Kasai, David Gray, Carol Clugston, Anna Dominiczak and Alex Alderton, Roberto Amato, Jeffrey Barrett, Sonia Goncalves, Ewan Harrison, David K. Jackson, Ian Johnston, Dominic Kwiatkowski, Cordelia Langford, John Sillitoe on behalf of the Wellcome Sanger Institute COVID-19 Surveillance Team |
| EPI_ISL_3027086, EPI_ISL_3027087 | Lighthouse Lab in Alderley Park | Wellcome Sanger Institute for the COVID-19 Genomics UK (COG-UK) Consortium | Jacquelyn Wynn, Mairead Hyland, The Lighthouse Lab in Alderley Park and Alex Alderton, Roberto Amato, Jeffrey Barrett, Sonia Goncalves, Ewan Harrison, David K. Jackson, Ian Johnston, Dominic Kwiatkowski, Cordelia Langford, John Sillitoe on behalf of the Wellcome Sanger Institute COVID-19 Surveillance Team |
| EPI_ISL_3027088, EPI_ISL_3027089 | Lighthouse Lab in Milton Keynes | Wellcome Sanger Institute for the COVID-19 Genomics UK (COG-UK) Consortium | The Lighthouse Lab in Milton Keynes and Alex Alderton, Roberto Amato, Jeffrey Barrett, Sonia Goncalves, Ewan Harrison, David K. Jackson, Ian Johnston, Dominic Kwiatkowski, Cordelia Langford, John Sillitoe on behalf of the Wellcome Sanger Institute COVID-19 Surveillance Team |
| EPI_ISL_3027090 | Lighthouse Lab in Glasgow | Wellcome Sanger Institute for the COVID-19 Genomics UK (COG-UK) Consortium | Harper VanSteenhouse, Yumi Kasai, David Gray, Carol Clugston, Anna Dominiczak and Alex Alderton, Roberto Amato, Jeffrey Barrett, Sonia Goncalves, Ewan Harrison, David K. Jackson, Ian Johnston, Dominic Kwiatkowski, Cordelia Langford, John Sillitoe on behalf of the Wellcome Sanger Institute COVID-19 Surveillance Team |
| EPI_ISL_3027091 | Lighthouse Lab in Milton Keynes | Wellcome Sanger Institute for the COVID-19 Genomics UK (COG-UK) Consortium | The Lighthouse Lab in Milton Keynes and Alex Alderton, Roberto Amato, Jeffrey Barrett, Sonia Goncalves, Ewan Harrison, David K. Jackson, Ian Johnston, Dominic Kwiatkowski, Cordelia Langford, John Sillitoe on behalf of the Wellcome Sanger Institute COVID-19 Surveillance Team |
| EPI_ISL_3027092 | Lighthouse Lab in Alderley Park | Wellcome Sanger Institute for the COVID-19 Genomics UK (COG-UK) Consortium | Jacquelyn Wynn, Mairead Hyland, The Lighthouse Lab in Alderley Park and Alex Alderton, Roberto Amato, Jeffrey Barrett, Sonia Goncalves, Ewan Harrison, David K. Jackson, Ian Johnston, Dominic Kwiatkowski, Cordelia Langford, John Sillitoe on behalf of the Wellcome Sanger Institute COVID-19 Surveillance Team |
| EPI_ISL_3027093 | Lighthouse Lab in Glasgow | Wellcome Sanger Institute for the COVID-19 Genomics UK (COG-UK) Consortium | Harper VanSteenhouse, Yumi Kasai, David Gray, Carol Clugston, Anna Dominiczak and Alex Alderton, Roberto Amato, Jeffrey Barrett, Sonia Goncalves, Ewan Harrison, David K. Jackson, Ian Johnston, Dominic Kwiatkowski, Cordelia Langford, John Sillitoe on behalf of the Wellcome Sanger Institute COVID-19 Surveillance Team |
| EPI_ISL_3027094, EPI_ISL_3027095 | Lighthouse Lab in Milton Keynes | Wellcome Sanger Institute for the COVID-19 Genomics UK (COG-UK) Consortium | The Lighthouse Lab in Milton Keynes and Alex Alderton, Roberto Amato, Jeffrey Barrett, Sonia Goncalves, Ewan Harrison, David K. Jackson, Ian Johnston, Dominic Kwiatkowski, Cordelia Langford, John Sillitoe on behalf of the Wellcome Sanger Institute COVID-19 Surveillance Team |
| EPI_ISL_3027096, EPI_ISL_3027097 | Lighthouse Lab in Alderley Park | Wellcome Sanger Institute for the COVID-19 Genomics UK (COG-UK) Consortium | Jacquelyn Wynn, Mairead Hyland, The Lighthouse Lab in Alderley Park and Alex Alderton, Roberto Amato, Jeffrey Barrett, Sonia Goncalves, Ewan Harrison, David K. Jackson, Ian Johnston, Dominic Kwiatkowski, Cordelia Langford, John Sillitoe on behalf of the Wellcome Sanger Institute COVID-19 Surveillance Team |
| EPI_ISL_3027098, EPI_ISL_3027099 | Lighthouse Lab in Milton Keynes | Wellcome Sanger Institute for the COVID-19 Genomics UK (COG-UK) Consortium | The Lighthouse Lab in Milton Keynes and Alex Alderton, Roberto Amato, Jeffrey Barrett, Sonia Goncalves, Ewan Harrison, David K. Jackson, Ian Johnston, Dominic Kwiatkowski, Cordelia Langford, John Sillitoe on behalf of the Wellcome Sanger Institute COVID-19 Surveillance Team |
| EPI_ISL_3027100, EPI_ISL_3027101 | Lighthouse Lab in Glasgow | Wellcome Sanger Institute for the COVID-19 Genomics UK (COG-UK) Consortium | Harper VanSteenhouse, Yumi Kasai, David Gray, Carol Clugston, Anna Dominiczak and Alex Alderton, Roberto Amato, Jeffrey Barrett, Sonia Goncalves, Ewan Harrison, David K. Jackson, Ian Johnston, Dominic Kwiatkowski, Cordelia Langford, John Sillitoe on behalf of the Wellcome Sanger Institute COVID-19 Surveillance Team |
| EPI_ISL_3027102, EPI_ISL_3027103 | Lighthouse Lab in Milton Keynes | Wellcome Sanger Institute for the COVID-19 Genomics UK (COG-UK) Consortium | The Lighthouse Lab in Milton Keynes and Alex Alderton, Roberto Amato, Jeffrey Barrett, Sonia Goncalves, Ewan Harrison, David K. Jackson, Ian Johnston, Dominic Kwiatkowski, Cordelia Langford, John Sillitoe on behalf of the Wellcome Sanger Institute COVID-19 Surveillance Team |
| EPI_ISL_3027104 | Lighthouse Lab in Alderley Park | Wellcome Sanger Institute for the COVID-19 Genomics UK (COG-UK) Consortium | Jacquelyn Wynn, Mairead Hyland, The Lighthouse Lab in Alderley Park and Alex Alderton, Roberto Amato, Jeffrey Barrett, Sonia Goncalves, Ewan Harrison, David K. Jackson, Ian Johnston, Dominic Kwiatkowski, Cordelia Langford, John Sillitoe on behalf of the Wellcome Sanger Institute COVID-19 Surveillance Team |
| EPI_ISL_3027105, EPI_ISL_3027106, EPI_ISL_3027107, EPI_ISL_3027108 | Lighthouse Lab in Milton Keynes | Wellcome Sanger Institute for the COVID-19 Genomics UK (COG-UK) Consortium | The Lighthouse Lab in Milton Keynes and Alex Alderton, Roberto Amato, Jeffrey Barrett, Sonia Goncalves, Ewan Harrison, David K. Jackson, Ian Johnston, Dominic Kwiatkowski, Cordelia Langford, John Sillitoe on behalf of the Wellcome Sanger Institute COVID-19 Surveillance Team |
| EPI_ISL_3027109 | Lighthouse Lab in Alderley Park | Wellcome Sanger Institute for the COVID-19 Genomics UK (COG-UK) Consortium | Jacquelyn Wynn, Mairead Hyland, The Lighthouse Lab in Alderley Park and Alex Alderton, Roberto Amato, Jeffrey Barrett, Sonia Goncalves, Ewan Harrison, David K. Jackson, Ian Johnston, Dominic Kwiatkowski, Cordelia Langford, John Sillitoe on behalf of the Wellcome Sanger Institute COVID-19 Surveillance Team |
| EPI_ISL_3027110, EPI_ISL_3027111 | Lighthouse Lab in Milton Keynes | Wellcome Sanger Institute for the COVID-19 Genomics UK (COG-UK) Consortium | The Lighthouse Lab in Milton Keynes and Alex Alderton, Roberto Amato, Jeffrey Barrett, Sonia Goncalves, Ewan Harrison, David K. Jackson, Ian Johnston, Dominic Kwiatkowski, Cordelia Langford, John Sillitoe on behalf of the Wellcome Sanger Institute COVID-19 Surveillance Team |
| EPI_ISL_3027112 | Lighthouse Lab in Alderley Park | Wellcome Sanger Institute for the COVID-19 Genomics UK (COG-UK) Consortium | Jacquelyn Wynn, Mairead Hyland, The Lighthouse Lab in Alderley Park and Alex Alderton, Roberto Amato, Jeffrey Barrett, Sonia Goncalves, Ewan Harrison, David K. Jackson, Ian Johnston, Dominic Kwiatkowski, Cordelia Langford, John Sillitoe on behalf of the Wellcome Sanger Institute COVID-19 Surveillance Team |
| EPI_ISL_3027113, EPI_ISL_3027114 | Lighthouse Lab in Glasgow | Wellcome Sanger Institute for the COVID-19 Genomics UK (COG-UK) Consortium | Harper VanSteenhouse, Yumi Kasai, David Gray, Carol Clugston, Anna Dominiczak and Alex Alderton, Roberto Amato, Jeffrey Barrett, Sonia Goncalves, Ewan Harrison, David K. Jackson, Ian Johnston, Dominic Kwiatkowski, Cordelia Langford, John Sillitoe on behalf of the Wellcome Sanger Institute COVID-19 Surveillance Team |
| EPI_ISL_3027115 | Lighthouse Lab in Alderley Park | Wellcome Sanger Institute for the COVID-19 Genomics UK (COG-UK) Consortium | Jacquelyn Wynn, Mairead Hyland, The Lighthouse Lab in Alderley Park and Alex Alderton, Roberto Amato, Jeffrey Barrett, Sonia Goncalves, Ewan Harrison, David K. Jackson, Ian Johnston, Dominic Kwiatkowski, Cordelia Langford, John Sillitoe on behalf of the Wellcome Sanger Institute COVID-19 Surveillance Team |
| EPI_ISL_3027116, EPI_ISL_3027117 | Lighthouse Lab in Glasgow | Wellcome Sanger Institute for the COVID-19 Genomics UK (COG-UK) Consortium | Harper VanSteenhouse, Yumi Kasai, David Gray, Carol Clugston, Anna Dominiczak and Alex Alderton, Roberto Amato, Jeffrey Barrett, Sonia Goncalves, Ewan Harrison, David K. Jackson, Ian Johnston, Dominic Kwiatkowski, Cordelia Langford, John Sillitoe on behalf of the Wellcome Sanger Institute COVID-19 Surveillance Team |
| EPI_ISL_3027118 | Lighthouse Lab in Alderley Park | Wellcome Sanger Institute for the COVID-19 Genomics UK (COG-UK) Consortium | Jacquelyn Wynn, Mairead Hyland, The Lighthouse Lab in Alderley Park and Alex Alderton, Roberto Amato, Jeffrey Barrett, Sonia Goncalves, Ewan Harrison, David K. Jackson, Ian Johnston, Dominic Kwiatkowski, Cordelia Langford, John Sillitoe on behalf of the Wellcome Sanger Institute COVID-19 Surveillance Team |

| | | | |
|---|---|---|---|
| EPI_ISL_3027119 | Lighthouse Lab in Glasgow | Wellcome Sanger Institute for the COVID-19 Genomics UK (COG-UK) Consortium | Harper VanSteenhouse, Yumi Kasai, David Gray, Carol Clugston, Anna Dominiczak and Alex Alderton, Roberto Amato, Jeffrey Barrett, Sonia Goncalves, Ewan Harrison, David K. Jackson, Ian Johnston, Dominic Kwiatkowski, Cordelia Langford, John Sillitoe on behalf of the Wellcome Sanger Institute COVID-19 Surveillance Team |
| EPI_ISL_3027120 | Lighthouse Lab in Milton Keynes | Wellcome Sanger Institute for the COVID-19 Genomics UK (COG-UK) Consortium | The Lighthouse Lab in Milton Keynes and Alex Alderton, Roberto Amato, Jeffrey Barrett, Sonia Goncalves, Ewan Harrison, David K. Jackson, Ian Johnston, Dominic Kwiatkowski, Cordelia Langford, John Sillitoe on behalf of the Wellcome Sanger Institute COVID-19 Surveillance Team |
| EPI_ISL_3027121, EPI_ISL_3027122 | Lighthouse Lab in Alderley Park | Wellcome Sanger Institute for the COVID-19 Genomics UK (COG-UK) Consortium | Jacquelyn Wynn, Mairead Hyland, The Lighthouse Lab in Alderley Park and Alex Alderton, Roberto Amato, Jeffrey Barrett, Sonia Goncalves, Ewan Harrison, David K. Jackson, Ian Johnston, Dominic Kwiatkowski, Cordelia Langford, John Sillitoe on behalf of the Wellcome Sanger Institute COVID-19 Surveillance Team |
| EPI_ISL_3027123 | Lighthouse Lab in Glasgow | Wellcome Sanger Institute for the COVID-19 Genomics UK (COG-UK) Consortium | Harper VanSteenhouse, Yumi Kasai, David Gray, Carol Clugston, Anna Dominiczak and Alex Alderton, Roberto Amato, Jeffrey Barrett, Sonia Goncalves, Ewan Harrison, David K. Jackson, Ian Johnston, Dominic Kwiatkowski, Cordelia Langford, John Sillitoe on behalf of the Wellcome Sanger Institute COVID-19 Surveillance Team |
| EPI_ISL_3027124 | Lighthouse Lab in Alderley Park | Wellcome Sanger Institute for the COVID-19 Genomics UK (COG-UK) Consortium | Jacquelyn Wynn, Mairead Hyland, The Lighthouse Lab in Alderley Park and Alex Alderton, Roberto Amato, Jeffrey Barrett, Sonia Goncalves, Ewan Harrison, David K. Jackson, Ian Johnston, Dominic Kwiatkowski, Cordelia Langford, John Sillitoe on behalf of the Wellcome Sanger Institute COVID-19 Surveillance Team |
| EPI_ISL_3027125, EPI_ISL_3027126, EPI_ISL_3027127 | Lighthouse Lab in Milton Keynes | Wellcome Sanger Institute for the COVID-19 Genomics UK (COG-UK) Consortium | The Lighthouse Lab in Milton Keynes and Alex Alderton, Roberto Amato, Jeffrey Barrett, Sonia Goncalves, Ewan Harrison, David K. Jackson, Ian Johnston, Dominic Kwiatkowski, Cordelia Langford, John Sillitoe on behalf of the Wellcome Sanger Institute COVID-19 Surveillance Team |
| EPI_ISL_3027128, EPI_ISL_3027129, EPI_ISL_3027130 | Lighthouse Lab in Alderley Park | Wellcome Sanger Institute for the COVID-19 Genomics UK (COG-UK) Consortium | Jacquelyn Wynn, Mairead Hyland, The Lighthouse Lab in Alderley Park and Alex Alderton, Roberto Amato, Jeffrey Barrett, Sonia Goncalves, Ewan Harrison, David K. Jackson, Ian Johnston, Dominic Kwiatkowski, Cordelia Langford, John Sillitoe on behalf of the Wellcome Sanger Institute COVID-19 Surveillance Team |
| EPI_ISL_3027131 | Lighthouse Lab in Milton Keynes | Wellcome Sanger Institute for the COVID-19 Genomics UK (COG-UK) Consortium | The Lighthouse Lab in Milton Keynes and Alex Alderton, Roberto Amato, Jeffrey Barrett, Sonia Goncalves, Ewan Harrison, David K. Jackson, Ian Johnston, Dominic Kwiatkowski, Cordelia Langford, John Sillitoe on behalf of the Wellcome Sanger Institute COVID-19 Surveillance Team |
| EPI_ISL_3027132, EPI_ISL_3027133, EPI_ISL_3027134 | Lighthouse Lab in Alderley Park | Wellcome Sanger Institute for the COVID-19 Genomics UK (COG-UK) Consortium | Jacquelyn Wynn, Mairead Hyland, The Lighthouse Lab in Alderley Park and Alex Alderton, Roberto Amato, Jeffrey Barrett, Sonia Goncalves, Ewan Harrison, David K. Jackson, Ian Johnston, Dominic Kwiatkowski, Cordelia Langford, John Sillitoe on behalf of the Wellcome Sanger Institute COVID-19 Surveillance Team |
| EPI_ISL_3027135 | Lighthouse Lab in Milton Keynes | Wellcome Sanger Institute for the COVID-19 Genomics UK (COG-UK) Consortium | The Lighthouse Lab in Milton Keynes and Alex Alderton, Roberto Amato, Jeffrey Barrett, Sonia Goncalves, Ewan Harrison, David K. Jackson, Ian Johnston, Dominic Kwiatkowski, Cordelia Langford, John Sillitoe on behalf of the Wellcome Sanger Institute COVID-19 Surveillance Team |
| EPI_ISL_3027136 | Lighthouse Lab in Alderley Park | Wellcome Sanger Institute for the COVID-19 Genomics UK (COG-UK) Consortium | Jacquelyn Wynn, Mairead Hyland, The Lighthouse Lab in Alderley Park and Alex Alderton, Roberto Amato, Jeffrey Barrett, Sonia Goncalves, Ewan Harrison, David K. Jackson, Ian Johnston, Dominic Kwiatkowski, Cordelia Langford, John Sillitoe on behalf of the Wellcome Sanger Institute COVID-19 Surveillance Team |
| EPI_ISL_3027137, EPI_ISL_3027138, EPI_ISL_3027139 | Lighthouse Lab in Milton Keynes | Wellcome Sanger Institute for the COVID-19 Genomics UK (COG-UK) Consortium | The Lighthouse Lab in Milton Keynes and Alex Alderton, Roberto Amato, Jeffrey Barrett, Sonia Goncalves, Ewan Harrison, David K. Jackson, Ian Johnston, Dominic Kwiatkowski, Cordelia Langford, John Sillitoe on behalf of the Wellcome Sanger Institute COVID-19 Surveillance Team |
| EPI_ISL_3027140, EPI_ISL_3027141 | Lighthouse Lab in Glasgow | Wellcome Sanger Institute for the COVID-19 Genomics UK (COG-UK) Consortium | Harper VanSteenhouse, Yumi Kasai, David Gray, Carol Clugston, Anna Dominiczak and Alex Alderton, Roberto Amato, Jeffrey Barrett, Sonia Goncalves, Ewan Harrison, David K. Jackson, Ian Johnston, Dominic Kwiatkowski, Cordelia Langford, John Sillitoe on behalf of the Wellcome Sanger Institute COVID-19 Surveillance Team |
| EPI_ISL_3027142 | Lighthouse Lab in Alderley Park | Wellcome Sanger Institute for the COVID-19 Genomics UK (COG-UK) Consortium | Jacquelyn Wynn, Mairead Hyland, The Lighthouse Lab in Alderley Park and Alex Alderton, Roberto Amato, Jeffrey Barrett, Sonia Goncalves, Ewan Harrison, David K. Jackson, Ian Johnston, Dominic Kwiatkowski, Cordelia Langford, John Sillitoe on behalf of the Wellcome Sanger Institute COVID-19 Surveillance Team |
| EPI_ISL_3027143, EPI_ISL_3027144, EPI_ISL_3027145, EPI_ISL_3027146 | Lighthouse Lab in Milton Keynes | Wellcome Sanger Institute for the COVID-19 Genomics UK (COG-UK) Consortium | The Lighthouse Lab in Milton Keynes and Alex Alderton, Roberto Amato, Jeffrey Barrett, Sonia Goncalves, Ewan Harrison, David K. Jackson, Ian Johnston, Dominic Kwiatkowski, Cordelia Langford, John Sillitoe on behalf of the Wellcome Sanger Institute COVID-19 Surveillance Team |
| EPI_ISL_3027147 | Lighthouse Lab in Glasgow | Wellcome Sanger Institute for the COVID-19 Genomics UK (COG-UK) Consortium | Harper VanSteenhouse, Yumi Kasai, David Gray, Carol Clugston, Anna Dominiczak and Alex Alderton, Roberto Amato, Jeffrey Barrett, Sonia Goncalves, Ewan Harrison, David K. Jackson, Ian Johnston, Dominic Kwiatkowski, Cordelia Langford, John Sillitoe on behalf of the Wellcome Sanger Institute COVID-19 Surveillance Team |
| EPI_ISL_3027148 | Lighthouse Lab in Alderley Park | Wellcome Sanger Institute for the COVID-19 Genomics UK (COG-UK) Consortium | Jacquelyn Wynn, Mairead Hyland, The Lighthouse Lab in Alderley Park and Alex Alderton, Roberto Amato, Jeffrey Barrett, Sonia Goncalves, Ewan Harrison, David K. Jackson, Ian Johnston, Dominic Kwiatkowski, Cordelia Langford, John Sillitoe on behalf of the Wellcome Sanger Institute COVID-19 Surveillance Team |
| EPI_ISL_3027149 | Lighthouse Lab in Glasgow | Wellcome Sanger Institute for the COVID-19 Genomics UK (COG-UK) Consortium | Harper VanSteenhouse, Yumi Kasai, David Gray, Carol Clugston, Anna Dominiczak and Alex Alderton, Roberto Amato, Jeffrey Barrett, Sonia Goncalves, Ewan Harrison, David K. Jackson, Ian Johnston, Dominic Kwiatkowski, Cordelia Langford, John Sillitoe on behalf of the Wellcome Sanger Institute COVID-19 Surveillance Team |
| EPI_ISL_3027150 | Lighthouse Lab in Milton Keynes | Wellcome Sanger Institute for the COVID-19 Genomics UK (COG-UK) Consortium | The Lighthouse Lab in Milton Keynes and Alex Alderton, Roberto Amato, Jeffrey Barrett, Sonia Goncalves, Ewan Harrison, David K. Jackson, Ian Johnston, Dominic Kwiatkowski, Cordelia Langford, John Sillitoe on behalf of the Wellcome Sanger Institute COVID-19 Surveillance Team |
| EPI_ISL_3027151 | Lighthouse Lab in Alderley Park | Wellcome Sanger Institute for the COVID-19 Genomics UK (COG-UK) Consortium | Jacquelyn Wynn, Mairead Hyland, The Lighthouse Lab in Alderley Park and Alex Alderton, Roberto Amato, Jeffrey Barrett, Sonia Goncalves, Ewan Harrison, David K. Jackson, Ian Johnston, Dominic Kwiatkowski, Cordelia Langford, John Sillitoe on behalf of the Wellcome Sanger Institute COVID-19 Surveillance Team |
| EPI_ISL_3027152 | Lighthouse Lab in Milton Keynes | Wellcome Sanger Institute for the COVID-19 Genomics UK (COG-UK) Consortium | The Lighthouse Lab in Milton Keynes and Alex Alderton, Roberto Amato, Jeffrey Barrett, Sonia Goncalves, Ewan Harrison, David K. Jackson, Ian Johnston, Dominic Kwiatkowski, Cordelia Langford, John Sillitoe on behalf of the Wellcome Sanger Institute COVID-19 Surveillance Team |
| EPI_ISL_3027153 | Lighthouse Lab in Alderley Park | Wellcome Sanger Institute for the COVID-19 Genomics UK (COG-UK) Consortium | Jacquelyn Wynn, Mairead Hyland, The Lighthouse Lab in Alderley Park and Alex Alderton, Roberto Amato, Jeffrey Barrett, Sonia Goncalves, Ewan Harrison, David K. Jackson, Ian Johnston, Dominic Kwiatkowski, Cordelia Langford, John Sillitoe on behalf of the Wellcome Sanger Institute COVID-19 Surveillance Team |
| EPI_ISL_3027154 | Lighthouse Lab in Glasgow | Wellcome Sanger Institute for the COVID-19 Genomics UK (COG-UK) Consortium | Harper VanSteenhouse, Yumi Kasai, David Gray, Carol Clugston, Anna Dominiczak and Alex Alderton, Roberto Amato, Jeffrey Barrett, Sonia Goncalves, Ewan Harrison, David K. Jackson, Ian Johnston, Dominic Kwiatkowski, Cordelia Langford, John Sillitoe on behalf of the Wellcome Sanger Institute COVID-19 Surveillance Team |
| EPI_ISL_3027155, EPI_ISL_3027156 | Lighthouse Lab in Alderley Park | Wellcome Sanger Institute for the COVID-19 Genomics UK (COG-UK) Consortium | Jacquelyn Wynn, Mairead Hyland, The Lighthouse Lab in Alderley Park and Alex Alderton, Roberto Amato, Jeffrey Barrett, Sonia Goncalves, Ewan Harrison, David K. Jackson, Ian Johnston, Dominic Kwiatkowski, Cordelia Langford, John Sillitoe on behalf of the Wellcome Sanger Institute COVID-19 Surveillance Team |
| EPI_ISL_3027157 | Lighthouse Lab in Milton Keynes | Wellcome Sanger Institute for the COVID-19 Genomics UK (COG-UK) Consortium | The Lighthouse Lab in Milton Keynes and Alex Alderton, Roberto Amato, Jeffrey Barrett, Sonia Goncalves, Ewan Harrison, David K. Jackson, Ian Johnston, Dominic Kwiatkowski, Cordelia Langford, John Sillitoe on behalf of the Wellcome Sanger Institute COVID-19 Surveillance Team |
| EPI_ISL_3027158, EPI_ISL_3027159 | Lighthouse Lab in Alderley Park | Wellcome Sanger Institute for the COVID-19 Genomics UK (COG-UK) Consortium | Jacquelyn Wynn, Mairead Hyland, The Lighthouse Lab in Alderley Park and Alex Alderton, Roberto Amato, Jeffrey Barrett, Sonia Goncalves, Ewan Harrison, David K. Jackson, Ian Johnston, Dominic Kwiatkowski, Cordelia Langford, John Sillitoe on behalf of the Wellcome Sanger Institute COVID-19 Surveillance Team |
| EPI_ISL_3027160 | Lighthouse Lab in Milton Keynes | Wellcome Sanger Institute for the COVID-19 Genomics UK (COG-UK) Consortium | The Lighthouse Lab in Milton Keynes and Alex Alderton, Roberto Amato, Jeffrey Barrett, Sonia Goncalves, Ewan Harrison, David K. Jackson, Ian Johnston, Dominic Kwiatkowski, Cordelia Langford, John Sillitoe on behalf of the Wellcome Sanger Institute COVID-19 Surveillance Team |
| EPI_ISL_3027161, EPI_ISL_3027162, EPI_ISL_3027163, EPI_ISL_3027164, EPI_ISL_3027165, EPI_ISL_3027166 | Lighthouse Lab in Alderley Park | Wellcome Sanger Institute for the COVID-19 Genomics UK (COG-UK) Consortium | Jacquelyn Wynn, Mairead Hyland, The Lighthouse Lab in Alderley Park and Alex Alderton, Roberto Amato, Jeffrey Barrett, Sonia Goncalves, Ewan Harrison, David K. Jackson, Ian Johnston, Dominic Kwiatkowski, Cordelia Langford, John Sillitoe on behalf of the Wellcome Sanger Institute COVID-19 Surveillance Team |
| EPI_ISL_3027167 | Lighthouse Lab in Milton Keynes | Wellcome Sanger Institute for the COVID-19 Genomics UK (COG-UK) Consortium | The Lighthouse Lab in Milton Keynes and Alex Alderton, Roberto Amato, Jeffrey Barrett, Sonia Goncalves, Ewan Harrison, David K. Jackson, Ian Johnston, Dominic Kwiatkowski, Cordelia Langford, John Sillitoe on behalf of the Wellcome Sanger Institute COVID-19 Surveillance Team |
| EPI_ISL_3027168, EPI_ISL_3027169, EPI_ISL_3027170, EPI_ISL_3027171, EPI_ISL_3027172, EPI_ISL_3027173, EPI_ISL_3027174, EPI_ISL_3027175, EPI_ISL_3027176 | Lighthouse Lab in Alderley Park | Wellcome Sanger Institute for the COVID-19 Genomics UK (COG-UK) Consortium | Jacquelyn Wynn, Mairead Hyland, The Lighthouse Lab in Alderley Park and Alex Alderton, Roberto Amato, Jeffrey Barrett, Sonia Goncalves, Ewan Harrison, David K. Jackson, Ian Johnston, Dominic Kwiatkowski, Cordelia Langford, John Sillitoe on behalf of the Wellcome Sanger Institute COVID-19 Surveillance Team |
| EPI_ISL_3027177 | Lighthouse Lab in Milton Keynes | Wellcome Sanger Institute for the COVID-19 Genomics UK (COG-UK) Consortium | The Lighthouse Lab in Milton Keynes and Alex Alderton, Roberto Amato, Jeffrey Barrett, Sonia Goncalves, Ewan Harrison, David K. Jackson, Ian Johnston, Dominic Kwiatkowski, Cordelia Langford, John Sillitoe on behalf of the Wellcome Sanger Institute COVID-19 Surveillance Team |
| EPI_ISL_3027178, EPI_ISL_3027179 | Lighthouse Lab in Glasgow | Wellcome Sanger Institute for the COVID-19 Genomics UK (COG-UK) Consortium | Harper VanSteenhouse, Yumi Kasai, David Gray, Carol Clugston, Anna Dominiczak and Alex Alderton, Roberto Amato, Jeffrey Barrett, Sonia Goncalves, Ewan Harrison, David K. Jackson, Ian Johnston, Dominic Kwiatkowski, Cordelia Langford, John Sillitoe on behalf of the Wellcome Sanger Institute COVID-19 Surveillance Team |
| EPI_ISL_3027180 | Lighthouse Lab in Alderley Park | Wellcome Sanger Institute for the COVID-19 Genomics UK (COG-UK) Consortium | Jacquelyn Wynn, Mairead Hyland, The Lighthouse Lab in Alderley Park and Alex Alderton, Roberto Amato, Jeffrey Barrett, Sonia Goncalves, Ewan Harrison, David K. Jackson, Ian Johnston, Dominic Kwiatkowski, Cordelia Langford, John Sillitoe on behalf of the Wellcome Sanger Institute COVID-19 Surveillance Team |
| EPI_ISL_3027181, EPI_ISL_3027182 | Lighthouse Lab in Milton Keynes | Wellcome Sanger Institute for the COVID-19 Genomics UK (COG-UK) Consortium | The Lighthouse Lab in Milton Keynes and Alex Alderton, Roberto Amato, Jeffrey Barrett, Sonia Goncalves, Ewan Harrison, David K. Jackson, Ian Johnston, Dominic Kwiatkowski, Cordelia Langford, John Sillitoe on behalf of the Wellcome Sanger Institute COVID-19 Surveillance Team |
| EPI_ISL_3027183, EPI_ISL_3027184, EPI_ISL_3027185 | Lighthouse Lab in Alderley Park | Wellcome Sanger Institute for the COVID-19 Genomics UK (COG-UK) Consortium | Jacquelyn Wynn, Mairead Hyland, The Lighthouse Lab in Alderley Park and Alex Alderton, Roberto Amato, Jeffrey Barrett, Sonia Goncalves, Ewan Harrison, David K. Jackson, Ian Johnston, Dominic Kwiatkowski, Cordelia Langford, John Sillitoe on behalf of the Wellcome Sanger Institute COVID-19 Surveillance Team |
| EPI_ISL_3027186 | Lighthouse Lab in Glasgow | Wellcome Sanger Institute for the COVID-19 Genomics UK (COG-UK) Consortium | Harper VanSteenhouse, Yumi Kasai, David Gray, Carol Clugston, Anna Dominiczak and Alex Alderton, Roberto Amato, Jeffrey Barrett, Sonia Goncalves, Ewan Harrison, David K. Jackson, Ian Johnston, Dominic Kwiatkowski, Cordelia Langford, John Sillitoe on behalf of the Wellcome Sanger Institute COVID-19 Surveillance Team |
| EPI_ISL_3027187, EPI_ISL_3027188 | Lighthouse Lab in Milton Keynes | Wellcome Sanger Institute for the COVID-19 Genomics UK (COG-UK) Consortium | The Lighthouse Lab in Milton Keynes and Alex Alderton, Roberto Amato, Jeffrey Barrett, Sonia Goncalves, Ewan Harrison, David K. Jackson, Ian Johnston, Dominic Kwiatkowski, Cordelia Langford, John Sillitoe on behalf of the Wellcome Sanger Institute COVID-19 Surveillance Team |
| EPI_ISL_3027189, EPI_ISL_3027190 | Lighthouse Lab in Alderley Park | Wellcome Sanger Institute for the COVID-19 Genomics UK (COG-UK) Consortium | Jacquelyn Wynn, Mairead Hyland, The Lighthouse Lab in Alderley Park and Alex Alderton, Roberto Amato, Jeffrey Barrett, Sonia Goncalves, Ewan Harrison, David K. Jackson, Ian Johnston, Dominic Kwiatkowski, Cordelia Langford, John Sillitoe on behalf of the Wellcome Sanger Institute COVID-19 Surveillance Team |

| | | | |
|---|---|---|---|
| EPI_ISL_3027191 | Lighthouse Lab in Milton Keynes | Wellcome Sanger Institute for the COVID-19 Genomics UK (COG-UK) Consortium | The Lighthouse Lab in Milton Keynes and Alex Alderton, Roberto Amato, Jeffrey Barrett, Sonia Goncalves, Ewan Harrison, David K. Jackson, Ian Johnston, Dominic Kwiatkowski, Cordelia Langford, John Sillitoe on behalf of the Wellcome Sanger Institute COVID-19 Surveillance Team |
| EPI_ISL_3027192, EPI_ISL_3027193, EPI_ISL_3027194, EPI_ISL_3027195, EPI_ISL_3027196, EPI_ISL_3027197, EPI_ISL_3027198 | Lighthouse Lab in Alderley Park | Wellcome Sanger Institute for the COVID-19 Genomics UK (COG-UK) Consortium | Jacquelyn Wynn, Mairead Hyland, The Lighthouse Lab in Alderley Park and Alex Alderton, Roberto Amato, Jeffrey Barrett, Sonia Goncalves, Ewan Harrison, David K. Jackson, Ian Johnston, Dominic Kwiatkowski, Cordelia Langford, John Sillitoe on behalf of the Wellcome Sanger Institute COVID-19 Surveillance Team |
| EPI_ISL_3027199 | Lighthouse Lab in Glasgow | Wellcome Sanger Institute for the COVID-19 Genomics UK (COG-UK) Consortium | Harper VanSteenhouse, Yumi Kasai, David Gray, Carol Clugston, Anna Dominiczak and Alex Alderton, Roberto Amato, Jeffrey Barrett, Sonia Goncalves, Ewan Harrison, David K. Jackson, Ian Johnston, Dominic Kwiatkowski, Cordelia Langford, John Sillitoe on behalf of the Wellcome Sanger Institute COVID-19 Surveillance Team |
| EPI_ISL_3027200 | Lighthouse Lab in Milton Keynes | Wellcome Sanger Institute for the COVID-19 Genomics UK (COG-UK) Consortium | The Lighthouse Lab in Milton Keynes and Alex Alderton, Roberto Amato, Jeffrey Barrett, Sonia Goncalves, Ewan Harrison, David K. Jackson, Ian Johnston, Dominic Kwiatkowski, Cordelia Langford, John Sillitoe on behalf of the Wellcome Sanger Institute COVID-19 Surveillance Team |
| EPI_ISL_3027201, EPI_ISL_3027202 | Lighthouse Lab in Alderley Park | Wellcome Sanger Institute for the COVID-19 Genomics UK (COG-UK) Consortium | Jacquelyn Wynn, Mairead Hyland, The Lighthouse Lab in Alderley Park and Alex Alderton, Roberto Amato, Jeffrey Barrett, Sonia Goncalves, Ewan Harrison, David K. Jackson, Ian Johnston, Dominic Kwiatkowski, Cordelia Langford, John Sillitoe on behalf of the Wellcome Sanger Institute COVID-19 Surveillance Team |
| EPI_ISL_3027203 | Lighthouse Lab in Milton Keynes | Wellcome Sanger Institute for the COVID-19 Genomics UK (COG-UK) Consortium | The Lighthouse Lab in Milton Keynes and Alex Alderton, Roberto Amato, Jeffrey Barrett, Sonia Goncalves, Ewan Harrison, David K. Jackson, Ian Johnston, Dominic Kwiatkowski, Cordelia Langford, John Sillitoe on behalf of the Wellcome Sanger Institute COVID-19 Surveillance Team |
| EPI_ISL_3027204 | Lighthouse Lab in Alderley Park | Wellcome Sanger Institute for the COVID-19 Genomics UK (COG-UK) Consortium | Jacquelyn Wynn, Mairead Hyland, The Lighthouse Lab in Alderley Park and Alex Alderton, Roberto Amato, Jeffrey Barrett, Sonia Goncalves, Ewan Harrison, David K. Jackson, Ian Johnston, Dominic Kwiatkowski, Cordelia Langford, John Sillitoe on behalf of the Wellcome Sanger Institute COVID-19 Surveillance Team |
| EPI_ISL_3027205 | Lighthouse Lab in Milton Keynes | Wellcome Sanger Institute for the COVID-19 Genomics UK (COG-UK) Consortium | The Lighthouse Lab in Milton Keynes and Alex Alderton, Roberto Amato, Jeffrey Barrett, Sonia Goncalves, Ewan Harrison, David K. Jackson, Ian Johnston, Dominic Kwiatkowski, Cordelia Langford, John Sillitoe on behalf of the Wellcome Sanger Institute COVID-19 Surveillance Team |
| EPI_ISL_3027206 | Lighthouse Lab in Alderley Park | Wellcome Sanger Institute for the COVID-19 Genomics UK (COG-UK) Consortium | Jacquelyn Wynn, Mairead Hyland, The Lighthouse Lab in Alderley Park and Alex Alderton, Roberto Amato, Jeffrey Barrett, Sonia Goncalves, Ewan Harrison, David K. Jackson, Ian Johnston, Dominic Kwiatkowski, Cordelia Langford, John Sillitoe on behalf of the Wellcome Sanger Institute COVID-19 Surveillance Team |
| EPI_ISL_3027207 | Lighthouse Lab in Milton Keynes | Wellcome Sanger Institute for the COVID-19 Genomics UK (COG-UK) Consortium | The Lighthouse Lab in Milton Keynes and Alex Alderton, Roberto Amato, Jeffrey Barrett, Sonia Goncalves, Ewan Harrison, David K. Jackson, Ian Johnston, Dominic Kwiatkowski, Cordelia Langford, John Sillitoe on behalf of the Wellcome Sanger Institute COVID-19 Surveillance Team |
| EPI_ISL_3027208 | Lighthouse Lab in Glasgow | Wellcome Sanger Institute for the COVID-19 Genomics UK (COG-UK) Consortium | Harper VanSteenhouse, Yumi Kasai, David Gray, Carol Clugston, Anna Dominiczak and Alex Alderton, Roberto Amato, Jeffrey Barrett, Sonia Goncalves, Ewan Harrison, David K. Jackson, Ian Johnston, Dominic Kwiatkowski, Cordelia Langford, John Sillitoe on behalf of the Wellcome Sanger Institute COVID-19 Surveillance Team |
| EPI_ISL_3027209, EPI_ISL_3027210 | Lighthouse Lab in Alderley Park | Wellcome Sanger Institute for the COVID-19 Genomics UK (COG-UK) Consortium | Jacquelyn Wynn, Mairead Hyland, The Lighthouse Lab in Alderley Park and Alex Alderton, Roberto Amato, Jeffrey Barrett, Sonia Goncalves, Ewan Harrison, David K. Jackson, Ian Johnston, Dominic Kwiatkowski, Cordelia Langford, John Sillitoe on behalf of the Wellcome Sanger Institute COVID-19 Surveillance Team |
| EPI_ISL_3027211 | Lighthouse Lab in Glasgow | Wellcome Sanger Institute for the COVID-19 Genomics UK (COG-UK) Consortium | Harper VanSteenhouse, Yumi Kasai, David Gray, Carol Clugston, Anna Dominiczak and Alex Alderton, Roberto Amato, Jeffrey Barrett, Sonia Goncalves, Ewan Harrison, David K. Jackson, Ian Johnston, Dominic Kwiatkowski, Cordelia Langford, John Sillitoe on behalf of the Wellcome Sanger Institute COVID-19 Surveillance Team |
| EPI_ISL_3027212, EPI_ISL_3027213, EPI_ISL_3027214 | Lighthouse Lab in Milton Keynes | Wellcome Sanger Institute for the COVID-19 Genomics UK (COG-UK) Consortium | The Lighthouse Lab in Milton Keynes and Alex Alderton, Roberto Amato, Jeffrey Barrett, Sonia Goncalves, Ewan Harrison, David K. Jackson, Ian Johnston, Dominic Kwiatkowski, Cordelia Langford, John Sillitoe on behalf of the Wellcome Sanger Institute COVID-19 Surveillance Team |
| EPI_ISL_3027215, EPI_ISL_3027216 | Lighthouse Lab in Alderley Park | Wellcome Sanger Institute for the COVID-19 Genomics UK (COG-UK) Consortium | Jacquelyn Wynn, Mairead Hyland, The Lighthouse Lab in Alderley Park and Alex Alderton, Roberto Amato, Jeffrey Barrett, Sonia Goncalves, Ewan Harrison, David K. Jackson, Ian Johnston, Dominic Kwiatkowski, Cordelia Langford, John Sillitoe on behalf of the Wellcome Sanger Institute COVID-19 Surveillance Team |
| EPI_ISL_3027217 | Lighthouse Lab in Glasgow | Wellcome Sanger Institute for the COVID-19 Genomics UK (COG-UK) Consortium | Harper VanSteenhouse, Yumi Kasai, David Gray, Carol Clugston, Anna Dominiczak and Alex Alderton, Roberto Amato, Jeffrey Barrett, Sonia Goncalves, Ewan Harrison, David K. Jackson, Ian Johnston, Dominic Kwiatkowski, Cordelia Langford, John Sillitoe on behalf of the Wellcome Sanger Institute COVID-19 Surveillance Team |
| EPI_ISL_3027218 | Lighthouse Lab in Milton Keynes | Wellcome Sanger Institute for the COVID-19 Genomics UK (COG-UK) Consortium | The Lighthouse Lab in Milton Keynes and Alex Alderton, Roberto Amato, Jeffrey Barrett, Sonia Goncalves, Ewan Harrison, David K. Jackson, Ian Johnston, Dominic Kwiatkowski, Cordelia Langford, John Sillitoe on behalf of the Wellcome Sanger Institute COVID-19 Surveillance Team |
| EPI_ISL_3027219 | Lighthouse Lab in Glasgow | Wellcome Sanger Institute for the COVID-19 Genomics UK (COG-UK) Consortium | Harper VanSteenhouse, Yumi Kasai, David Gray, Carol Clugston, Anna Dominiczak and Alex Alderton, Roberto Amato, Jeffrey Barrett, Sonia Goncalves, Ewan Harrison, David K. Jackson, Ian Johnston, Dominic Kwiatkowski, Cordelia Langford, John Sillitoe on behalf of the Wellcome Sanger Institute COVID-19 Surveillance Team |
| EPI_ISL_3027220 | Lighthouse Lab in Alderley Park | Wellcome Sanger Institute for the COVID-19 Genomics UK (COG-UK) Consortium | Jacquelyn Wynn, Mairead Hyland, The Lighthouse Lab in Alderley Park and Alex Alderton, Roberto Amato, Jeffrey Barrett, Sonia Goncalves, Ewan Harrison, David K. Jackson, Ian Johnston, Dominic Kwiatkowski, Cordelia Langford, John Sillitoe on behalf of the Wellcome Sanger Institute COVID-19 Surveillance Team |
| EPI_ISL_3027221 | Lighthouse Lab in Milton Keynes | Wellcome Sanger Institute for the COVID-19 Genomics UK (COG-UK) Consortium | The Lighthouse Lab in Milton Keynes and Alex Alderton, Roberto Amato, Jeffrey Barrett, Sonia Goncalves, Ewan Harrison, David K. Jackson, Ian Johnston, Dominic Kwiatkowski, Cordelia Langford, John Sillitoe on behalf of the Wellcome Sanger Institute COVID-19 Surveillance Team |
| EPI_ISL_3027222, EPI_ISL_3027223 | Lighthouse Lab in Alderley Park | Wellcome Sanger Institute for the COVID-19 Genomics UK (COG-UK) Consortium | Jacquelyn Wynn, Mairead Hyland, The Lighthouse Lab in Alderley Park and Alex Alderton, Roberto Amato, Jeffrey Barrett, Sonia Goncalves, Ewan Harrison, David K. Jackson, Ian Johnston, Dominic Kwiatkowski, Cordelia Langford, John Sillitoe on behalf of the Wellcome Sanger Institute COVID-19 Surveillance Team |
| EPI_ISL_3027224 | Lighthouse Lab in Glasgow | Wellcome Sanger Institute for the COVID-19 Genomics UK (COG-UK) Consortium | Harper VanSteenhouse, Yumi Kasai, David Gray, Carol Clugston, Anna Dominiczak and Alex Alderton, Roberto Amato, Jeffrey Barrett, Sonia Goncalves, Ewan Harrison, David K. Jackson, Ian Johnston, Dominic Kwiatkowski, Cordelia Langford, John Sillitoe on behalf of the Wellcome Sanger Institute COVID-19 Surveillance Team |
| EPI_ISL_3027225 | Lighthouse Lab in Alderley Park | Wellcome Sanger Institute for the COVID-19 Genomics UK (COG-UK) Consortium | Jacquelyn Wynn, Mairead Hyland, The Lighthouse Lab in Alderley Park and Alex Alderton, Roberto Amato, Jeffrey Barrett, Sonia Goncalves, Ewan Harrison, David K. Jackson, Ian Johnston, Dominic Kwiatkowski, Cordelia Langford, John Sillitoe on behalf of the Wellcome Sanger Institute COVID-19 Surveillance Team |
| EPI_ISL_3027226, EPI_ISL_3027227 | Lighthouse Lab in Glasgow | Wellcome Sanger Institute for the COVID-19 Genomics UK (COG-UK) Consortium | Harper VanSteenhouse, Yumi Kasai, David Gray, Carol Clugston, Anna Dominiczak and Alex Alderton, Roberto Amato, Jeffrey Barrett, Sonia Goncalves, Ewan Harrison, David K. Jackson, Ian Johnston, Dominic Kwiatkowski, Cordelia Langford, John Sillitoe on behalf of the Wellcome Sanger Institute COVID-19 Surveillance Team |
| EPI_ISL_3027228 | Lighthouse Lab in Alderley Park | Wellcome Sanger Institute for the COVID-19 Genomics UK (COG-UK) Consortium | Jacquelyn Wynn, Mairead Hyland, The Lighthouse Lab in Alderley Park and Alex Alderton, Roberto Amato, Jeffrey Barrett, Sonia Goncalves, Ewan Harrison, David K. Jackson, Ian Johnston, Dominic Kwiatkowski, Cordelia Langford, John Sillitoe on behalf of the Wellcome Sanger Institute COVID-19 Surveillance Team |
| EPI_ISL_3027229 | Lighthouse Lab in Glasgow | Wellcome Sanger Institute for the COVID-19 Genomics UK (COG-UK) Consortium | Harper VanSteenhouse, Yumi Kasai, David Gray, Carol Clugston, Anna Dominiczak and Alex Alderton, Roberto Amato, Jeffrey Barrett, Sonia Goncalves, Ewan Harrison, David K. Jackson, Ian Johnston, Dominic Kwiatkowski, Cordelia Langford, John Sillitoe on behalf of the Wellcome Sanger Institute COVID-19 Surveillance Team |
| EPI_ISL_3027230 | Lighthouse Lab in Alderley Park | Wellcome Sanger Institute for the COVID-19 Genomics UK (COG-UK) Consortium | Jacquelyn Wynn, Mairead Hyland, The Lighthouse Lab in Alderley Park and Alex Alderton, Roberto Amato, Jeffrey Barrett, Sonia Goncalves, Ewan Harrison, David K. Jackson, Ian Johnston, Dominic Kwiatkowski, Cordelia Langford, John Sillitoe on behalf of the Wellcome Sanger Institute COVID-19 Surveillance Team |
| EPI_ISL_3027231 | Lighthouse Lab in Glasgow | Wellcome Sanger Institute for the COVID-19 Genomics UK (COG-UK) Consortium | Harper VanSteenhouse, Yumi Kasai, David Gray, Carol Clugston, Anna Dominiczak and Alex Alderton, Roberto Amato, Jeffrey Barrett, Sonia Goncalves, Ewan Harrison, David K. Jackson, Ian Johnston, Dominic Kwiatkowski, Cordelia Langford, John Sillitoe on behalf of the Wellcome Sanger Institute COVID-19 Surveillance Team |
| EPI_ISL_3027232, EPI_ISL_3027233, EPI_ISL_3027234, EPI_ISL_3027235, EPI_ISL_3027236, EPI_ISL_3027237 | Lighthouse Lab in Alderley Park | Wellcome Sanger Institute for the COVID-19 Genomics UK (COG-UK) Consortium | Jacquelyn Wynn, Mairead Hyland, The Lighthouse Lab in Alderley Park and Alex Alderton, Roberto Amato, Jeffrey Barrett, Sonia Goncalves, Ewan Harrison, David K. Jackson, Ian Johnston, Dominic Kwiatkowski, Cordelia Langford, John Sillitoe on behalf of the Wellcome Sanger Institute COVID-19 Surveillance Team |
| EPI_ISL_3027238 | Lighthouse Lab in Glasgow | Wellcome Sanger Institute for the COVID-19 Genomics UK (COG-UK) Consortium | Harper VanSteenhouse, Yumi Kasai, David Gray, Carol Clugston, Anna Dominiczak and Alex Alderton, Roberto Amato, Jeffrey Barrett, Sonia Goncalves, Ewan Harrison, David K. Jackson, Ian Johnston, Dominic Kwiatkowski, Cordelia Langford, John Sillitoe on behalf of the Wellcome Sanger Institute COVID-19 Surveillance Team |
| EPI_ISL_3027239 | Lighthouse Lab in Milton Keynes | Wellcome Sanger Institute for the COVID-19 Genomics UK (COG-UK) Consortium | The Lighthouse Lab in Milton Keynes and Alex Alderton, Roberto Amato, Jeffrey Barrett, Sonia Goncalves, Ewan Harrison, David K. Jackson, Ian Johnston, Dominic Kwiatkowski, Cordelia Langford, John Sillitoe on behalf of the Wellcome Sanger Institute COVID-19 Surveillance Team |
| EPI_ISL_3027240 | Lighthouse Lab in Alderley Park | Wellcome Sanger Institute for the COVID-19 Genomics UK (COG-UK) Consortium | Jacquelyn Wynn, Mairead Hyland, The Lighthouse Lab in Alderley Park and Alex Alderton, Roberto Amato, Jeffrey Barrett, Sonia Goncalves, Ewan Harrison, David K. Jackson, Ian Johnston, Dominic Kwiatkowski, Cordelia Langford, John Sillitoe on behalf of the Wellcome Sanger Institute COVID-19 Surveillance Team |
| EPI_ISL_3027241 | Lighthouse Lab in Milton Keynes | Wellcome Sanger Institute for the COVID-19 Genomics UK (COG-UK) Consortium | The Lighthouse Lab in Milton Keynes and Alex Alderton, Roberto Amato, Jeffrey Barrett, Sonia Goncalves, Ewan Harrison, David K. Jackson, Ian Johnston, Dominic Kwiatkowski, Cordelia Langford, John Sillitoe on behalf of the Wellcome Sanger Institute COVID-19 Surveillance Team |
| EPI_ISL_3027242 | Lighthouse Lab in Glasgow | Wellcome Sanger Institute for the COVID-19 Genomics UK (COG-UK) Consortium | Harper VanSteenhouse, Yumi Kasai, David Gray, Carol Clugston, Anna Dominiczak and Alex Alderton, Roberto Amato, Jeffrey Barrett, Sonia Goncalves, Ewan Harrison, David K. Jackson, Ian Johnston, Dominic Kwiatkowski, Cordelia Langford, John Sillitoe on behalf of the Wellcome Sanger Institute COVID-19 Surveillance Team |
| EPI_ISL_3027243, EPI_ISL_3027244 | Lighthouse Lab in Alderley Park | Wellcome Sanger Institute for the COVID-19 Genomics UK (COG-UK) Consortium | Jacquelyn Wynn, Mairead Hyland, The Lighthouse Lab in Alderley Park and Alex Alderton, Roberto Amato, Jeffrey Barrett, Sonia Goncalves, Ewan Harrison, David K. Jackson, Ian Johnston, Dominic Kwiatkowski, Cordelia Langford, John Sillitoe on behalf of the Wellcome Sanger Institute COVID-19 Surveillance Team |
| EPI_ISL_3027245, EPI_ISL_3027246 | Lighthouse Lab in Glasgow | Wellcome Sanger Institute for the COVID-19 Genomics UK (COG-UK) Consortium | Harper VanSteenhouse, Yumi Kasai, David Gray, Carol Clugston, Anna Dominiczak and Alex Alderton, Roberto Amato, Jeffrey Barrett, Sonia Goncalves, Ewan Harrison, David K. Jackson, Ian Johnston, Dominic Kwiatkowski, Cordelia Langford, John Sillitoe on behalf of the Wellcome Sanger Institute COVID-19 Surveillance Team |
| EPI_ISL_3027247, EPI_ISL_3027248, EPI_ISL_3027249, EPI_ISL_3027250 | Lighthouse Lab in Alderley Park | Wellcome Sanger Institute for the COVID-19 Genomics UK (COG-UK) Consortium | Jacquelyn Wynn, Mairead Hyland, The Lighthouse Lab in Alderley Park and Alex Alderton, Roberto Amato, Jeffrey Barrett, Sonia Goncalves, Ewan Harrison, David K. Jackson, Ian Johnston, Dominic Kwiatkowski, Cordelia Langford, John Sillitoe on behalf of the Wellcome Sanger Institute COVID-19 Surveillance Team |
| EPI_ISL_3027251 | Lighthouse Lab in Glasgow | Wellcome Sanger Institute for the COVID-19 Genomics UK (COG-UK) Consortium | Harper VanSteenhouse, Yumi Kasai, David Gray, Carol Clugston, Anna Dominiczak and Alex Alderton, Roberto Amato, Jeffrey Barrett, Sonia Goncalves, Ewan Harrison, David K. Jackson, Ian Johnston, Dominic Kwiatkowski, Cordelia Langford, John Sillitoe on behalf of the Wellcome Sanger Institute COVID-19 Surveillance Team |
| EPI_ISL_3027252 | Lighthouse Lab in Milton Keynes | Wellcome Sanger Institute for the COVID-19 Genomics UK (COG-UK) Consortium | The Lighthouse Lab in Milton Keynes and Alex Alderton, Roberto Amato, Jeffrey Barrett, Sonia Goncalves, Ewan Harrison, David K. Jackson, Ian Johnston, Dominic Kwiatkowski, Cordelia Langford, John Sillitoe on behalf of the Wellcome Sanger Institute COVID-19 Surveillance Team |
| EPI_ISL_3027253, EPI_ISL_3027254 | Lighthouse Lab in Alderley Park | Wellcome Sanger Institute for the COVID-19 Genomics UK (COG-UK) Consortium | Jacquelyn Wynn, Mairead Hyland, The Lighthouse Lab in Alderley Park and Alex Alderton, Roberto Amato, Jeffrey Barrett, Sonia Goncalves, Ewan Harrison, David K. Jackson, Ian Johnston, Dominic Kwiatkowski, Cordelia Langford, John Sillitoe on behalf of the Wellcome Sanger Institute COVID-19 Surveillance Team |

| | | | |
|---|---|---|---|
| EPI_ISL_3027255, EPI_ISL_3027256 | Lighthouse Lab in Glasgow | Wellcome Sanger Institute for the COVID-19 Genomics UK (COG-UK) Consortium | Harper VanSteenhouse, Yumi Kasai, David Gray, Carol Clugston, Anna Dominiczak and Alex Alderton, Roberto Amato, Jeffrey Barrett, Sonia Goncalves, Ewan Harrison, David K. Jackson, Ian Johnston, Dominic Kwiatkowski, Cordelia Langford, John Sillitoe on behalf of the Wellcome Sanger Institute COVID-19 Surveillance Team |
| EPI_ISL_3027257, EPI_ISL_3027258 | Lighthouse Lab in Alderley Park | Wellcome Sanger Institute for the COVID-19 Genomics UK (COG-UK) Consortium | Jacquelyn Wynn, Mairead Hyland, The Lighthouse Lab in Alderley Park and Alex Alderton, Roberto Amato, Jeffrey Barrett, Sonia Goncalves, Ewan Harrison, David K. Jackson, Ian Johnston, Dominic Kwiatkowski, Cordelia Langford, John Sillitoe on behalf of the Wellcome Sanger Institute COVID-19 Surveillance Team |
| EPI_ISL_3027259 | Lighthouse Lab in Glasgow | Wellcome Sanger Institute for the COVID-19 Genomics UK (COG-UK) Consortium | Harper VanSteenhouse, Yumi Kasai, David Gray, Carol Clugston, Anna Dominiczak and Alex Alderton, Roberto Amato, Jeffrey Barrett, Sonia Goncalves, Ewan Harrison, David K. Jackson, Ian Johnston, Dominic Kwiatkowski, Cordelia Langford, John Sillitoe on behalf of the Wellcome Sanger Institute COVID-19 Surveillance Team |
| EPI_ISL_3027260 | Lighthouse Lab in Alderley Park | Wellcome Sanger Institute for the COVID-19 Genomics UK (COG-UK) Consortium | Jacquelyn Wynn, Mairead Hyland, The Lighthouse Lab in Alderley Park and Alex Alderton, Roberto Amato, Jeffrey Barrett, Sonia Goncalves, Ewan Harrison, David K. Jackson, Ian Johnston, Dominic Kwiatkowski, Cordelia Langford, John Sillitoe on behalf of the Wellcome Sanger Institute COVID-19 Surveillance Team |
| EPI_ISL_3027261, EPI_ISL_3027262 | Lighthouse Lab in Glasgow | Wellcome Sanger Institute for the COVID-19 Genomics UK (COG-UK) Consortium | Harper VanSteenhouse, Yumi Kasai, David Gray, Carol Clugston, Anna Dominiczak and Alex Alderton, Roberto Amato, Jeffrey Barrett, Sonia Goncalves, Ewan Harrison, David K. Jackson, Ian Johnston, Dominic Kwiatkowski, Cordelia Langford, John Sillitoe on behalf of the Wellcome Sanger Institute COVID-19 Surveillance Team |
| EPI_ISL_3027263 | Lighthouse Lab in Alderley Park | Wellcome Sanger Institute for the COVID-19 Genomics UK (COG-UK) Consortium | Jacquelyn Wynn, Mairead Hyland, The Lighthouse Lab in Alderley Park and Alex Alderton, Roberto Amato, Jeffrey Barrett, Sonia Goncalves, Ewan Harrison, David K. Jackson, Ian Johnston, Dominic Kwiatkowski, Cordelia Langford, John Sillitoe on behalf of the Wellcome Sanger Institute COVID-19 Surveillance Team |
| EPI_ISL_3027264 | Lighthouse Lab in Milton Keynes | Wellcome Sanger Institute for the COVID-19 Genomics UK (COG-UK) Consortium | The Lighthouse Lab in Milton Keynes and Alex Alderton, Roberto Amato, Jeffrey Barrett, Sonia Goncalves, Ewan Harrison, David K. Jackson, Ian Johnston, Dominic Kwiatkowski, Cordelia Langford, John Sillitoe on behalf of the Wellcome Sanger Institute COVID-19 Surveillance Team |
| EPI_ISL_3027265 | Lighthouse Lab in Alderley Park | Wellcome Sanger Institute for the COVID-19 Genomics UK (COG-UK) Consortium | Jacquelyn Wynn, Mairead Hyland, The Lighthouse Lab in Alderley Park and Alex Alderton, Roberto Amato, Jeffrey Barrett, Sonia Goncalves, Ewan Harrison, David K. Jackson, Ian Johnston, Dominic Kwiatkowski, Cordelia Langford, John Sillitoe on behalf of the Wellcome Sanger Institute COVID-19 Surveillance Team |
| EPI_ISL_3027266 | Lighthouse Lab in Milton Keynes | Wellcome Sanger Institute for the COVID-19 Genomics UK (COG-UK) Consortium | The Lighthouse Lab in Milton Keynes and Alex Alderton, Roberto Amato, Jeffrey Barrett, Sonia Goncalves, Ewan Harrison, David K. Jackson, Ian Johnston, Dominic Kwiatkowski, Cordelia Langford, John Sillitoe on behalf of the Wellcome Sanger Institute COVID-19 Surveillance Team |
| EPI_ISL_3027267, EPI_ISL_3027268, EPI_ISL_3027269, EPI_ISL_3027270 | Lighthouse Lab in Alderley Park | Wellcome Sanger Institute for the COVID-19 Genomics UK (COG-UK) Consortium | Jacquelyn Wynn, Mairead Hyland, The Lighthouse Lab in Alderley Park and Alex Alderton, Roberto Amato, Jeffrey Barrett, Sonia Goncalves, Ewan Harrison, David K. Jackson, Ian Johnston, Dominic Kwiatkowski, Cordelia Langford, John Sillitoe on behalf of the Wellcome Sanger Institute COVID-19 Surveillance Team |
| EPI_ISL_3027271 | Lighthouse Lab in Glasgow | Wellcome Sanger Institute for the COVID-19 Genomics UK (COG-UK) Consortium | Harper VanSteenhouse, Yumi Kasai, David Gray, Carol Clugston, Anna Dominiczak and Alex Alderton, Roberto Amato, Jeffrey Barrett, Sonia Goncalves, Ewan Harrison, David K. Jackson, Ian Johnston, Dominic Kwiatkowski, Cordelia Langford, John Sillitoe on behalf of the Wellcome Sanger Institute COVID-19 Surveillance Team |
| EPI_ISL_3027272 | Lighthouse Lab in Alderley Park | Wellcome Sanger Institute for the COVID-19 Genomics UK (COG-UK) Consortium | Jacquelyn Wynn, Mairead Hyland, The Lighthouse Lab in Alderley Park and Alex Alderton, Roberto Amato, Jeffrey Barrett, Sonia Goncalves, Ewan Harrison, David K. Jackson, Ian Johnston, Dominic Kwiatkowski, Cordelia Langford, John Sillitoe on behalf of the Wellcome Sanger Institute COVID-19 Surveillance Team |
| EPI_ISL_3027273 | Lighthouse Lab in Milton Keynes | Wellcome Sanger Institute for the COVID-19 Genomics UK (COG-UK) Consortium | The Lighthouse Lab in Milton Keynes and Alex Alderton, Roberto Amato, Jeffrey Barrett, Sonia Goncalves, Ewan Harrison, David K. Jackson, Ian Johnston, Dominic Kwiatkowski, Cordelia Langford, John Sillitoe on behalf of the Wellcome Sanger Institute COVID-19 Surveillance Team |
| EPI_ISL_3027274, EPI_ISL_3027275 | Lighthouse Lab in Glasgow | Wellcome Sanger Institute for the COVID-19 Genomics UK (COG-UK) Consortium | Harper VanSteenhouse, Yumi Kasai, David Gray, Carol Clugston, Anna Dominiczak and Alex Alderton, Roberto Amato, Jeffrey Barrett, Sonia Goncalves, Ewan Harrison, David K. Jackson, Ian Johnston, Dominic Kwiatkowski, Cordelia Langford, John Sillitoe on behalf of the Wellcome Sanger Institute COVID-19 Surveillance Team |
| EPI_ISL_3027276 | Lighthouse Lab in Milton Keynes | Wellcome Sanger Institute for the COVID-19 Genomics UK (COG-UK) Consortium | The Lighthouse Lab in Milton Keynes and Alex Alderton, Roberto Amato, Jeffrey Barrett, Sonia Goncalves, Ewan Harrison, David K. Jackson, Ian Johnston, Dominic Kwiatkowski, Cordelia Langford, John Sillitoe on behalf of the Wellcome Sanger Institute COVID-19 Surveillance Team |
| EPI_ISL_3027277, EPI_ISL_3027278 | Lighthouse Lab in Glasgow | Wellcome Sanger Institute for the COVID-19 Genomics UK (COG-UK) Consortium | Harper VanSteenhouse, Yumi Kasai, David Gray, Carol Clugston, Anna Dominiczak and Alex Alderton, Roberto Amato, Jeffrey Barrett, Sonia Goncalves, Ewan Harrison, David K. Jackson, Ian Johnston, Dominic Kwiatkowski, Cordelia Langford, John Sillitoe on behalf of the Wellcome Sanger Institute COVID-19 Surveillance Team |
| EPI_ISL_3027279, EPI_ISL_3027280 | Lighthouse Lab in Alderley Park | Wellcome Sanger Institute for the COVID-19 Genomics UK (COG-UK) Consortium | Jacquelyn Wynn, Mairead Hyland, The Lighthouse Lab in Alderley Park and Alex Alderton, Roberto Amato, Jeffrey Barrett, Sonia Goncalves, Ewan Harrison, David K. Jackson, Ian Johnston, Dominic Kwiatkowski, Cordelia Langford, John Sillitoe on behalf of the Wellcome Sanger Institute COVID-19 Surveillance Team |
| EPI_ISL_3027281 | Lighthouse Lab in Milton Keynes | Wellcome Sanger Institute for the COVID-19 Genomics UK (COG-UK) Consortium | The Lighthouse Lab in Milton Keynes and Alex Alderton, Roberto Amato, Jeffrey Barrett, Sonia Goncalves, Ewan Harrison, David K. Jackson, Ian Johnston, Dominic Kwiatkowski, Cordelia Langford, John Sillitoe on behalf of the Wellcome Sanger Institute COVID-19 Surveillance Team |
| EPI_ISL_3027282 | Lighthouse Lab in Alderley Park | Wellcome Sanger Institute for the COVID-19 Genomics UK (COG-UK) Consortium | Jacquelyn Wynn, Mairead Hyland, The Lighthouse Lab in Alderley Park and Alex Alderton, Roberto Amato, Jeffrey Barrett, Sonia Goncalves, Ewan Harrison, David K. Jackson, Ian Johnston, Dominic Kwiatkowski, Cordelia Langford, John Sillitoe on behalf of the Wellcome Sanger Institute COVID-19 Surveillance Team |
| EPI_ISL_3027283 | Lighthouse Lab in Milton Keynes | Wellcome Sanger Institute for the COVID-19 Genomics UK (COG-UK) Consortium | The Lighthouse Lab in Milton Keynes and Alex Alderton, Roberto Amato, Jeffrey Barrett, Sonia Goncalves, Ewan Harrison, David K. Jackson, Ian Johnston, Dominic Kwiatkowski, Cordelia Langford, John Sillitoe on behalf of the Wellcome Sanger Institute COVID-19 Surveillance Team |
| EPI_ISL_3027284, EPI_ISL_3027285, EPI_ISL_3027286 | Lighthouse Lab in Alderley Park | Wellcome Sanger Institute for the COVID-19 Genomics UK (COG-UK) Consortium | Jacquelyn Wynn, Mairead Hyland, The Lighthouse Lab in Alderley Park and Alex Alderton, Roberto Amato, Jeffrey Barrett, Sonia Goncalves, Ewan Harrison, David K. Jackson, Ian Johnston, Dominic Kwiatkowski, Cordelia Langford, John Sillitoe on behalf of the Wellcome Sanger Institute COVID-19 Surveillance Team |
| EPI_ISL_3027287 | Lighthouse Lab in Glasgow | Wellcome Sanger Institute for the COVID-19 Genomics UK (COG-UK) Consortium | Harper VanSteenhouse, Yumi Kasai, David Gray, Carol Clugston, Anna Dominiczak and Alex Alderton, Roberto Amato, Jeffrey Barrett, Sonia Goncalves, Ewan Harrison, David K. Jackson, Ian Johnston, Dominic Kwiatkowski, Cordelia Langford, John Sillitoe on behalf of the Wellcome Sanger Institute COVID-19 Surveillance Team |
| EPI_ISL_3027288, EPI_ISL_3027289 | Lighthouse Lab in Alderley Park | Wellcome Sanger Institute for the COVID-19 Genomics UK (COG-UK) Consortium | Jacquelyn Wynn, Mairead Hyland, The Lighthouse Lab in Alderley Park and Alex Alderton, Roberto Amato, Jeffrey Barrett, Sonia Goncalves, Ewan Harrison, David K. Jackson, Ian Johnston, Dominic Kwiatkowski, Cordelia Langford, John Sillitoe on behalf of the Wellcome Sanger Institute COVID-19 Surveillance Team |
| EPI_ISL_3027290 | Lighthouse Lab in Milton Keynes | Wellcome Sanger Institute for the COVID-19 Genomics UK (COG-UK) Consortium | The Lighthouse Lab in Milton Keynes and Alex Alderton, Roberto Amato, Jeffrey Barrett, Sonia Goncalves, Ewan Harrison, David K. Jackson, Ian Johnston, Dominic Kwiatkowski, Cordelia Langford, John Sillitoe on behalf of the Wellcome Sanger Institute COVID-19 Surveillance Team |
| EPI_ISL_3027291 | Lighthouse Lab in Alderley Park | Wellcome Sanger Institute for the COVID-19 Genomics UK (COG-UK) Consortium | Jacquelyn Wynn, Mairead Hyland, The Lighthouse Lab in Alderley Park and Alex Alderton, Roberto Amato, Jeffrey Barrett, Sonia Goncalves, Ewan Harrison, David K. Jackson, Ian Johnston, Dominic Kwiatkowski, Cordelia Langford, John Sillitoe on behalf of the Wellcome Sanger Institute COVID-19 Surveillance Team |
| EPI_ISL_3027292 | Lighthouse Lab in Milton Keynes | Wellcome Sanger Institute for the COVID-19 Genomics UK (COG-UK) Consortium | The Lighthouse Lab in Milton Keynes and Alex Alderton, Roberto Amato, Jeffrey Barrett, Sonia Goncalves, Ewan Harrison, David K. Jackson, Ian Johnston, Dominic Kwiatkowski, Cordelia Langford, John Sillitoe on behalf of the Wellcome Sanger Institute COVID-19 Surveillance Team |
| EPI_ISL_3027293, EPI_ISL_3027294 | Lighthouse Lab in Glasgow | Wellcome Sanger Institute for the COVID-19 Genomics UK (COG-UK) Consortium | Harper VanSteenhouse, Yumi Kasai, David Gray, Carol Clugston, Anna Dominiczak and Alex Alderton, Roberto Amato, Jeffrey Barrett, Sonia Goncalves, Ewan Harrison, David K. Jackson, Ian Johnston, Dominic Kwiatkowski, Cordelia Langford, John Sillitoe on behalf of the Wellcome Sanger Institute COVID-19 Surveillance Team |
| EPI_ISL_3027295 | Lighthouse Lab in Milton Keynes | Wellcome Sanger Institute for the COVID-19 Genomics UK (COG-UK) Consortium | The Lighthouse Lab in Milton Keynes and Alex Alderton, Roberto Amato, Jeffrey Barrett, Sonia Goncalves, Ewan Harrison, David K. Jackson, Ian Johnston, Dominic Kwiatkowski, Cordelia Langford, John Sillitoe on behalf of the Wellcome Sanger Institute COVID-19 Surveillance Team |
| EPI_ISL_3027296 | Lighthouse Lab in Alderley Park | Wellcome Sanger Institute for the COVID-19 Genomics UK (COG-UK) Consortium | Jacquelyn Wynn, Mairead Hyland, The Lighthouse Lab in Alderley Park and Alex Alderton, Roberto Amato, Jeffrey Barrett, Sonia Goncalves, Ewan Harrison, David K. Jackson, Ian Johnston, Dominic Kwiatkowski, Cordelia Langford, John Sillitoe on behalf of the Wellcome Sanger Institute COVID-19 Surveillance Team |
| EPI_ISL_3027297 | Lighthouse Lab in Glasgow | Wellcome Sanger Institute for the COVID-19 Genomics UK (COG-UK) Consortium | Harper VanSteenhouse, Yumi Kasai, David Gray, Carol Clugston, Anna Dominiczak and Alex Alderton, Roberto Amato, Jeffrey Barrett, Sonia Goncalves, Ewan Harrison, David K. Jackson, Ian Johnston, Dominic Kwiatkowski, Cordelia Langford, John Sillitoe on behalf of the Wellcome Sanger Institute COVID-19 Surveillance Team |
| EPI_ISL_3027298 | Lighthouse Lab in Alderley Park | Wellcome Sanger Institute for the COVID-19 Genomics UK (COG-UK) Consortium | Jacquelyn Wynn, Mairead Hyland, The Lighthouse Lab in Alderley Park and Alex Alderton, Roberto Amato, Jeffrey Barrett, Sonia Goncalves, Ewan Harrison, David K. Jackson, Ian Johnston, Dominic Kwiatkowski, Cordelia Langford, John Sillitoe on behalf of the Wellcome Sanger Institute COVID-19 Surveillance Team |
| EPI_ISL_3027299 | Lighthouse Lab in Glasgow | Wellcome Sanger Institute for the COVID-19 Genomics UK (COG-UK) Consortium | Harper VanSteenhouse, Yumi Kasai, David Gray, Carol Clugston, Anna Dominiczak and Alex Alderton, Roberto Amato, Jeffrey Barrett, Sonia Goncalves, Ewan Harrison, David K. Jackson, Ian Johnston, Dominic Kwiatkowski, Cordelia Langford, John Sillitoe on behalf of the Wellcome Sanger Institute COVID-19 Surveillance Team |
| EPI_ISL_3027300 | Lighthouse Lab in Alderley Park | Wellcome Sanger Institute for the COVID-19 Genomics UK (COG-UK) Consortium | Jacquelyn Wynn, Mairead Hyland, The Lighthouse Lab in Alderley Park and Alex Alderton, Roberto Amato, Jeffrey Barrett, Sonia Goncalves, Ewan Harrison, David K. Jackson, Ian Johnston, Dominic Kwiatkowski, Cordelia Langford, John Sillitoe on behalf of the Wellcome Sanger Institute COVID-19 Surveillance Team |
| EPI_ISL_3027301, EPI_ISL_3027302 | Lighthouse Lab in Glasgow | Wellcome Sanger Institute for the COVID-19 Genomics UK (COG-UK) Consortium | Harper VanSteenhouse, Yumi Kasai, David Gray, Carol Clugston, Anna Dominiczak and Alex Alderton, Roberto Amato, Jeffrey Barrett, Sonia Goncalves, Ewan Harrison, David K. Jackson, Ian Johnston, Dominic Kwiatkowski, Cordelia Langford, John Sillitoe on behalf of the Wellcome Sanger Institute COVID-19 Surveillance Team |
| EPI_ISL_3027303 | Lighthouse Lab in Alderley Park | Wellcome Sanger Institute for the COVID-19 Genomics UK (COG-UK) Consortium | Jacquelyn Wynn, Mairead Hyland, The Lighthouse Lab in Alderley Park and Alex Alderton, Roberto Amato, Jeffrey Barrett, Sonia Goncalves, Ewan Harrison, David K. Jackson, Ian Johnston, Dominic Kwiatkowski, Cordelia Langford, John Sillitoe on behalf of the Wellcome Sanger Institute COVID-19 Surveillance Team |
| EPI_ISL_3027304 | Lighthouse Lab in Glasgow | Wellcome Sanger Institute for the COVID-19 Genomics UK (COG-UK) Consortium | Harper VanSteenhouse, Yumi Kasai, David Gray, Carol Clugston, Anna Dominiczak and Alex Alderton, Roberto Amato, Jeffrey Barrett, Sonia Goncalves, Ewan Harrison, David K. Jackson, Ian Johnston, Dominic Kwiatkowski, Cordelia Langford, John Sillitoe on behalf of the Wellcome Sanger Institute COVID-19 Surveillance Team |
| EPI_ISL_3027305 | Lighthouse Lab in Alderley Park | Wellcome Sanger Institute for the COVID-19 Genomics UK (COG-UK) Consortium | Jacquelyn Wynn, Mairead Hyland, The Lighthouse Lab in Alderley Park and Alex Alderton, Roberto Amato, Jeffrey Barrett, Sonia Goncalves, Ewan Harrison, David K. Jackson, Ian Johnston, Dominic Kwiatkowski, Cordelia Langford, John Sillitoe on behalf of the Wellcome Sanger Institute COVID-19 Surveillance Team |
| EPI_ISL_3027306 | Lighthouse Lab in Milton Keynes | Wellcome Sanger Institute for the COVID-19 Genomics UK (COG-UK) Consortium | The Lighthouse Lab in Milton Keynes and Alex Alderton, Roberto Amato, Jeffrey Barrett, Sonia Goncalves, Ewan Harrison, David K. Jackson, Ian Johnston, Dominic Kwiatkowski, Cordelia Langford, John Sillitoe on behalf of the Wellcome Sanger Institute COVID-19 Surveillance Team |
| EPI_ISL_3027307 | Lighthouse Lab in Glasgow | Wellcome Sanger Institute for the COVID-19 Genomics UK (COG-UK) Consortium | Harper VanSteenhouse, Yumi Kasai, David Gray, Carol Clugston, Anna Dominiczak and Alex Alderton, Roberto Amato, Jeffrey Barrett, Sonia Goncalves, Ewan Harrison, David K. Jackson, Ian Johnston, Dominic Kwiatkowski, Cordelia Langford, John Sillitoe on behalf of the Wellcome Sanger Institute COVID-19 Surveillance Team |
| EPI_ISL_3027308, EPI_ISL_3027309 | Lighthouse Lab in Alderley Park | Wellcome Sanger Institute for the COVID-19 Genomics UK (COG-UK) Consortium | Jacquelyn Wynn, Mairead Hyland, The Lighthouse Lab in Alderley Park and Alex Alderton, Roberto Amato, Jeffrey Barrett, Sonia Goncalves, Ewan Harrison, David K. Jackson, Ian Johnston, Dominic Kwiatkowski, Cordelia Langford, John Sillitoe on behalf of the Wellcome Sanger Institute COVID-19 Surveillance Team |
| EPI_ISL_3027310, EPI_ISL_3027311 | Lighthouse Lab in Milton Keynes | Wellcome Sanger Institute for the COVID-19 Genomics UK (COG-UK) Consortium | The Lighthouse Lab in Milton Keynes and Alex Alderton, Roberto Amato, Jeffrey Barrett, Sonia Goncalves, Ewan Harrison, David K. Jackson, Ian Johnston, Dominic Kwiatkowski, Cordelia Langford, John Sillitoe on behalf of the Wellcome Sanger Institute COVID-19 Surveillance Team |
| EPI_ISL_3027312 | Lighthouse Lab in Alderley Park | Wellcome Sanger Institute for the COVID-19 Genomics UK (COG-UK) Consortium | Jacquelyn Wynn, Mairead Hyland, The Lighthouse Lab in Alderley Park and Alex Alderton, Roberto Amato, Jeffrey Barrett, Sonia Goncalves, Ewan Harrison, David K. Jackson, Ian Johnston, Dominic Kwiatkowski, Cordelia Langford, John Sillitoe on behalf of the Wellcome Sanger Institute COVID-19 Surveillance Team |
| EPI_ISL_3027313 | Lighthouse Lab in Milton Keynes | Wellcome Sanger Institute for the COVID-19 Genomics UK (COG-UK) Consortium | The Lighthouse Lab in Milton Keynes and Alex Alderton, Roberto Amato, Jeffrey Barrett, Sonia Goncalves, Ewan Harrison, David K. Jackson, Ian Johnston, Dominic Kwiatkowski, Cordelia Langford, John Sillitoe on behalf of the Wellcome Sanger Institute COVID-19 Surveillance Team |
| EPI_ISL_3027314 | Lighthouse Lab in Alderley Park | Wellcome Sanger Institute for the COVID-19 Genomics UK (COG-UK) Consortium | Jacquelyn Wynn, Mairead Hyland, The Lighthouse Lab in Alderley Park and Alex Alderton, Roberto Amato, Jeffrey Barrett, Sonia Goncalves, Ewan Harrison, David K. Jackson, Ian Johnston, Dominic Kwiatkowski, Cordelia Langford, John Sillitoe on behalf of the Wellcome Sanger Institute COVID-19 Surveillance Team |

| | | | |
|---|---|---|---|
| EPI_ISL_3027315 | Lighthouse Lab in Milton Keynes | Wellcome Sanger Institute for the COVID-19 Genomics UK (COG-UK) Consortium | The Lighthouse Lab in Milton Keynes and Alex Alderton, Roberto Amato, Jeffrey Barrett, Sonia Goncalves, Ewan Harrison, David K. Jackson, Ian Johnston, Dominic Kwiatkowski, Cordelia Langford, John Sillitoe on behalf of the Wellcome Sanger Institute COVID-19 Surveillance Team |
| EPI_ISL_3027316 | Lighthouse Lab in Alderley Park | Wellcome Sanger Institute for the COVID-19 Genomics UK (COG-UK) Consortium | Jacquelyn Wynn, Mairead Hyland, The Lighthouse Lab in Alderley Park and Alex Alderton, Roberto Amato, Jeffrey Barrett, Sonia Goncalves, Ewan Harrison, David K. Jackson, Ian Johnston, Dominic Kwiatkowski, Cordelia Langford, John Sillitoe on behalf of the Wellcome Sanger Institute COVID-19 Surveillance Team |
| EPI_ISL_3027317 | Lighthouse Lab in Glasgow | Wellcome Sanger Institute for the COVID-19 Genomics UK (COG-UK) Consortium | Harper VanSteenhouse, Yumi Kasai, David Gray, Carol Clugston, Anna Dominiczak and Alex Alderton, Roberto Amato, Jeffrey Barrett, Sonia Goncalves, Ewan Harrison, David K. Jackson, Ian Johnston, Dominic Kwiatkowski, Cordelia Langford, John Sillitoe on behalf of the Wellcome Sanger Institute COVID-19 Surveillance Team |
| EPI_ISL_3027318 | Lighthouse Lab in Milton Keynes | Wellcome Sanger Institute for the COVID-19 Genomics UK (COG-UK) Consortium | The Lighthouse Lab in Milton Keynes and Alex Alderton, Roberto Amato, Jeffrey Barrett, Sonia Goncalves, Ewan Harrison, David K. Jackson, Ian Johnston, Dominic Kwiatkowski, Cordelia Langford, John Sillitoe on behalf of the Wellcome Sanger Institute COVID-19 Surveillance Team |
| EPI_ISL_3027319 | Lighthouse Lab in Glasgow | Wellcome Sanger Institute for the COVID-19 Genomics UK (COG-UK) Consortium | Harper VanSteenhouse, Yumi Kasai, David Gray, Carol Clugston, Anna Dominiczak and Alex Alderton, Roberto Amato, Jeffrey Barrett, Sonia Goncalves, Ewan Harrison, David K. Jackson, Ian Johnston, Dominic Kwiatkowski, Cordelia Langford, John Sillitoe on behalf of the Wellcome Sanger Institute COVID-19 Surveillance Team |
| EPI_ISL_3027320 | Lighthouse Lab in Alderley Park | Wellcome Sanger Institute for the COVID-19 Genomics UK (COG-UK) Consortium | Jacquelyn Wynn, Mairead Hyland, The Lighthouse Lab in Alderley Park and Alex Alderton, Roberto Amato, Jeffrey Barrett, Sonia Goncalves, Ewan Harrison, David K. Jackson, Ian Johnston, Dominic Kwiatkowski, Cordelia Langford, John Sillitoe on behalf of the Wellcome Sanger Institute COVID-19 Surveillance Team |
| EPI_ISL_3027321 | Lighthouse Lab in Glasgow | Wellcome Sanger Institute for the COVID-19 Genomics UK (COG-UK) Consortium | Harper VanSteenhouse, Yumi Kasai, David Gray, Carol Clugston, Anna Dominiczak and Alex Alderton, Roberto Amato, Jeffrey Barrett, Sonia Goncalves, Ewan Harrison, David K. Jackson, Ian Johnston, Dominic Kwiatkowski, Cordelia Langford, John Sillitoe on behalf of the Wellcome Sanger Institute COVID-19 Surveillance Team |
| EPI_ISL_3027322, EPI_ISL_3027323, EPI_ISL_3027324 | Lighthouse Lab in Alderley Park | Wellcome Sanger Institute for the COVID-19 Genomics UK (COG-UK) Consortium | Jacquelyn Wynn, Mairead Hyland, The Lighthouse Lab in Alderley Park and Alex Alderton, Roberto Amato, Jeffrey Barrett, Sonia Goncalves, Ewan Harrison, David K. Jackson, Ian Johnston, Dominic Kwiatkowski, Cordelia Langford, John Sillitoe on behalf of the Wellcome Sanger Institute COVID-19 Surveillance Team |
| EPI_ISL_3027325 | Lighthouse Lab in Glasgow | Wellcome Sanger Institute for the COVID-19 Genomics UK (COG-UK) Consortium | Harper VanSteenhouse, Yumi Kasai, David Gray, Carol Clugston, Anna Dominiczak and Alex Alderton, Roberto Amato, Jeffrey Barrett, Sonia Goncalves, Ewan Harrison, David K. Jackson, Ian Johnston, Dominic Kwiatkowski, Cordelia Langford, John Sillitoe on behalf of the Wellcome Sanger Institute COVID-19 Surveillance Team |
| EPI_ISL_3027326 | Lighthouse Lab in Alderley Park | Wellcome Sanger Institute for the COVID-19 Genomics UK (COG-UK) Consortium | Jacquelyn Wynn, Mairead Hyland, The Lighthouse Lab in Alderley Park and Alex Alderton, Roberto Amato, Jeffrey Barrett, Sonia Goncalves, Ewan Harrison, David K. Jackson, Ian Johnston, Dominic Kwiatkowski, Cordelia Langford, John Sillitoe on behalf of the Wellcome Sanger Institute COVID-19 Surveillance Team |
| EPI_ISL_3027327 | Lighthouse Lab in Glasgow | Wellcome Sanger Institute for the COVID-19 Genomics UK (COG-UK) Consortium | Harper VanSteenhouse, Yumi Kasai, David Gray, Carol Clugston, Anna Dominiczak and Alex Alderton, Roberto Amato, Jeffrey Barrett, Sonia Goncalves, Ewan Harrison, David K. Jackson, Ian Johnston, Dominic Kwiatkowski, Cordelia Langford, John Sillitoe on behalf of the Wellcome Sanger Institute COVID-19 Surveillance Team |
| EPI_ISL_3027328, EPI_ISL_3027329, EPI_ISL_3027330, EPI_ISL_3027331, EPI_ISL_3027332 | Lighthouse Lab in Alderley Park | Wellcome Sanger Institute for the COVID-19 Genomics UK (COG-UK) Consortium | Jacquelyn Wynn, Mairead Hyland, The Lighthouse Lab in Alderley Park and Alex Alderton, Roberto Amato, Jeffrey Barrett, Sonia Goncalves, Ewan Harrison, David K. Jackson, Ian Johnston, Dominic Kwiatkowski, Cordelia Langford, John Sillitoe on behalf of the Wellcome Sanger Institute COVID-19 Surveillance Team |
| EPI_ISL_3027333 | Lighthouse Lab in Milton Keynes | Wellcome Sanger Institute for the COVID-19 Genomics UK (COG-UK) Consortium | The Lighthouse Lab in Milton Keynes and Alex Alderton, Roberto Amato, Jeffrey Barrett, Sonia Goncalves, Ewan Harrison, David K. Jackson, Ian Johnston, Dominic Kwiatkowski, Cordelia Langford, John Sillitoe on behalf of the Wellcome Sanger Institute COVID-19 Surveillance Team |
| EPI_ISL_3027334 | Lighthouse Lab in Alderley Park | Wellcome Sanger Institute for the COVID-19 Genomics UK (COG-UK) Consortium | Jacquelyn Wynn, Mairead Hyland, The Lighthouse Lab in Alderley Park and Alex Alderton, Roberto Amato, Jeffrey Barrett, Sonia Goncalves, Ewan Harrison, David K. Jackson, Ian Johnston, Dominic Kwiatkowski, Cordelia Langford, John Sillitoe on behalf of the Wellcome Sanger Institute COVID-19 Surveillance Team |
| EPI_ISL_3027335 | Lighthouse Lab in Glasgow | Wellcome Sanger Institute for the COVID-19 Genomics UK (COG-UK) Consortium | Harper VanSteenhouse, Yumi Kasai, David Gray, Carol Clugston, Anna Dominiczak and Alex Alderton, Roberto Amato, Jeffrey Barrett, Sonia Goncalves, Ewan Harrison, David K. Jackson, Ian Johnston, Dominic Kwiatkowski, Cordelia Langford, John Sillitoe on behalf of the Wellcome Sanger Institute COVID-19 Surveillance Team |
| EPI_ISL_3027336, EPI_ISL_3027337 | Lighthouse Lab in Milton Keynes | Wellcome Sanger Institute for the COVID-19 Genomics UK (COG-UK) Consortium | The Lighthouse Lab in Milton Keynes and Alex Alderton, Roberto Amato, Jeffrey Barrett, Sonia Goncalves, Ewan Harrison, David K. Jackson, Ian Johnston, Dominic Kwiatkowski, Cordelia Langford, John Sillitoe on behalf of the Wellcome Sanger Institute COVID-19 Surveillance Team |
| EPI_ISL_3027338, EPI_ISL_3027339, EPI_ISL_3027340 | Lighthouse Lab in Alderley Park | Wellcome Sanger Institute for the COVID-19 Genomics UK (COG-UK) Consortium | Jacquelyn Wynn, Mairead Hyland, The Lighthouse Lab in Alderley Park and Alex Alderton, Roberto Amato, Jeffrey Barrett, Sonia Goncalves, Ewan Harrison, David K. Jackson, Ian Johnston, Dominic Kwiatkowski, Cordelia Langford, John Sillitoe on behalf of the Wellcome Sanger Institute COVID-19 Surveillance Team |
| EPI_ISL_3027341, EPI_ISL_3027342 | Lighthouse Lab in Milton Keynes | Wellcome Sanger Institute for the COVID-19 Genomics UK (COG-UK) Consortium | The Lighthouse Lab in Milton Keynes and Alex Alderton, Roberto Amato, Jeffrey Barrett, Sonia Goncalves, Ewan Harrison, David K. Jackson, Ian Johnston, Dominic Kwiatkowski, Cordelia Langford, John Sillitoe on behalf of the Wellcome Sanger Institute COVID-19 Surveillance Team |
| EPI_ISL_3027343, EPI_ISL_3027344, EPI_ISL_3027345, EPI_ISL_3027346, EPI_ISL_3027347 | Lighthouse Lab in Alderley Park | Wellcome Sanger Institute for the COVID-19 Genomics UK (COG-UK) Consortium | Jacquelyn Wynn, Mairead Hyland, The Lighthouse Lab in Alderley Park and Alex Alderton, Roberto Amato, Jeffrey Barrett, Sonia Goncalves, Ewan Harrison, David K. Jackson, Ian Johnston, Dominic Kwiatkowski, Cordelia Langford, John Sillitoe on behalf of the Wellcome Sanger Institute COVID-19 Surveillance Team |
| EPI_ISL_3027348 | Lighthouse Lab in Milton Keynes | Wellcome Sanger Institute for the COVID-19 Genomics UK (COG-UK) Consortium | The Lighthouse Lab in Milton Keynes and Alex Alderton, Roberto Amato, Jeffrey Barrett, Sonia Goncalves, Ewan Harrison, David K. Jackson, Ian Johnston, Dominic Kwiatkowski, Cordelia Langford, John Sillitoe on behalf of the Wellcome Sanger Institute COVID-19 Surveillance Team |
| EPI_ISL_3027349, EPI_ISL_3027350, EPI_ISL_3027351, EPI_ISL_3027352, EPI_ISL_3027353, EPI_ISL_3027354, EPI_ISL_3027355 | Lighthouse Lab in Alderley Park | Wellcome Sanger Institute for the COVID-19 Genomics UK (COG-UK) Consortium | Jacquelyn Wynn, Mairead Hyland, The Lighthouse Lab in Alderley Park and Alex Alderton, Roberto Amato, Jeffrey Barrett, Sonia Goncalves, Ewan Harrison, David K. Jackson, Ian Johnston, Dominic Kwiatkowski, Cordelia Langford, John Sillitoe on behalf of the Wellcome Sanger Institute COVID-19 Surveillance Team |
| EPI_ISL_3027356 | Lighthouse Lab in Milton Keynes | Wellcome Sanger Institute for the COVID-19 Genomics UK (COG-UK) Consortium | The Lighthouse Lab in Milton Keynes and Alex Alderton, Roberto Amato, Jeffrey Barrett, Sonia Goncalves, Ewan Harrison, David K. Jackson, Ian Johnston, Dominic Kwiatkowski, Cordelia Langford, John Sillitoe on behalf of the Wellcome Sanger Institute COVID-19 Surveillance Team |
| EPI_ISL_3027357 | Lighthouse Lab in Glasgow | Wellcome Sanger Institute for the COVID-19 Genomics UK (COG-UK) Consortium | Harper VanSteenhouse, Yumi Kasai, David Gray, Carol Clugston, Anna Dominiczak and Alex Alderton, Roberto Amato, Jeffrey Barrett, Sonia Goncalves, Ewan Harrison, David K. Jackson, Ian Johnston, Dominic Kwiatkowski, Cordelia Langford, John Sillitoe on behalf of the Wellcome Sanger Institute COVID-19 Surveillance Team |
| EPI_ISL_3027358, EPI_ISL_3027359, EPI_ISL_3027360 | Lighthouse Lab in Alderley Park | Wellcome Sanger Institute for the COVID-19 Genomics UK (COG-UK) Consortium | Jacquelyn Wynn, Mairead Hyland, The Lighthouse Lab in Alderley Park and Alex Alderton, Roberto Amato, Jeffrey Barrett, Sonia Goncalves, Ewan Harrison, David K. Jackson, Ian Johnston, Dominic Kwiatkowski, Cordelia Langford, John Sillitoe on behalf of the Wellcome Sanger Institute COVID-19 Surveillance Team |
| EPI_ISL_3027361 | Lighthouse Lab in Glasgow | Wellcome Sanger Institute for the COVID-19 Genomics UK (COG-UK) Consortium | Harper VanSteenhouse, Yumi Kasai, David Gray, Carol Clugston, Anna Dominiczak and Alex Alderton, Roberto Amato, Jeffrey Barrett, Sonia Goncalves, Ewan Harrison, David K. Jackson, Ian Johnston, Dominic Kwiatkowski, Cordelia Langford, John Sillitoe on behalf of the Wellcome Sanger Institute COVID-19 Surveillance Team |
| EPI_ISL_3027362, EPI_ISL_3027363 | Lighthouse Lab in Milton Keynes | Wellcome Sanger Institute for the COVID-19 Genomics UK (COG-UK) Consortium | The Lighthouse Lab in Milton Keynes and Alex Alderton, Roberto Amato, Jeffrey Barrett, Sonia Goncalves, Ewan Harrison, David K. Jackson, Ian Johnston, Dominic Kwiatkowski, Cordelia Langford, John Sillitoe on behalf of the Wellcome Sanger Institute COVID-19 Surveillance Team |
| EPI_ISL_3027364, EPI_ISL_3027365, EPI_ISL_3027366, EPI_ISL_3027367, EPI_ISL_3027368 | Lighthouse Lab in Alderley Park | Wellcome Sanger Institute for the COVID-19 Genomics UK (COG-UK) Consortium | Jacquelyn Wynn, Mairead Hyland, The Lighthouse Lab in Alderley Park and Alex Alderton, Roberto Amato, Jeffrey Barrett, Sonia Goncalves, Ewan Harrison, David K. Jackson, Ian Johnston, Dominic Kwiatkowski, Cordelia Langford, John Sillitoe on behalf of the Wellcome Sanger Institute COVID-19 Surveillance Team |
| EPI_ISL_3027369 | Lighthouse Lab in Glasgow | Wellcome Sanger Institute for the COVID-19 Genomics UK (COG-UK) Consortium | Harper VanSteenhouse, Yumi Kasai, David Gray, Carol Clugston, Anna Dominiczak and Alex Alderton, Roberto Amato, Jeffrey Barrett, Sonia Goncalves, Ewan Harrison, David K. Jackson, Ian Johnston, Dominic Kwiatkowski, Cordelia Langford, John Sillitoe on behalf of the Wellcome Sanger Institute COVID-19 Surveillance Team |
| EPI_ISL_3027370, EPI_ISL_3027371 | Lighthouse Lab in Milton Keynes | Wellcome Sanger Institute for the COVID-19 Genomics UK (COG-UK) Consortium | The Lighthouse Lab in Milton Keynes and Alex Alderton, Roberto Amato, Jeffrey Barrett, Sonia Goncalves, Ewan Harrison, David K. Jackson, Ian Johnston, Dominic Kwiatkowski, Cordelia Langford, John Sillitoe on behalf of the Wellcome Sanger Institute COVID-19 Surveillance Team |
| EPI_ISL_3027372 | Lighthouse Lab in Alderley Park | Wellcome Sanger Institute for the COVID-19 Genomics UK (COG-UK) Consortium | Jacquelyn Wynn, Mairead Hyland, The Lighthouse Lab in Alderley Park and Alex Alderton, Roberto Amato, Jeffrey Barrett, Sonia Goncalves, Ewan Harrison, David K. Jackson, Ian Johnston, Dominic Kwiatkowski, Cordelia Langford, John Sillitoe on behalf of the Wellcome Sanger Institute COVID-19 Surveillance Team |
| EPI_ISL_3027373, EPI_ISL_3027374, EPI_ISL_3027375, EPI_ISL_3027376 | Lighthouse Lab in Milton Keynes | Wellcome Sanger Institute for the COVID-19 Genomics UK (COG-UK) Consortium | The Lighthouse Lab in Milton Keynes and Alex Alderton, Roberto Amato, Jeffrey Barrett, Sonia Goncalves, Ewan Harrison, David K. Jackson, Ian Johnston, Dominic Kwiatkowski, Cordelia Langford, John Sillitoe on behalf of the Wellcome Sanger Institute COVID-19 Surveillance Team |
| EPI_ISL_3027377, EPI_ISL_3027378, EPI_ISL_3027379 | Lighthouse Lab in Alderley Park | Wellcome Sanger Institute for the COVID-19 Genomics UK (COG-UK) Consortium | Jacquelyn Wynn, Mairead Hyland, The Lighthouse Lab in Alderley Park and Alex Alderton, Roberto Amato, Jeffrey Barrett, Sonia Goncalves, Ewan Harrison, David K. Jackson, Ian Johnston, Dominic Kwiatkowski, Cordelia Langford, John Sillitoe on behalf of the Wellcome Sanger Institute COVID-19 Surveillance Team |
| EPI_ISL_3027380 | Lighthouse Lab in Milton Keynes | Wellcome Sanger Institute for the COVID-19 Genomics UK (COG-UK) Consortium | The Lighthouse Lab in Milton Keynes and Alex Alderton, Roberto Amato, Jeffrey Barrett, Sonia Goncalves, Ewan Harrison, David K. Jackson, Ian Johnston, Dominic Kwiatkowski, Cordelia Langford, John Sillitoe on behalf of the Wellcome Sanger Institute COVID-19 Surveillance Team |
| EPI_ISL_3027381, EPI_ISL_3027382 | Lighthouse Lab in Alderley Park | Wellcome Sanger Institute for the COVID-19 Genomics UK (COG-UK) Consortium | Jacquelyn Wynn, Mairead Hyland, The Lighthouse Lab in Alderley Park and Alex Alderton, Roberto Amato, Jeffrey Barrett, Sonia Goncalves, Ewan Harrison, David K. Jackson, Ian Johnston, Dominic Kwiatkowski, Cordelia Langford, John Sillitoe on behalf of the Wellcome Sanger Institute COVID-19 Surveillance Team |
| EPI_ISL_3027383, EPI_ISL_3027384, EPI_ISL_3027385, EPI_ISL_3027386, EPI_ISL_3027387 | Lighthouse Lab in Milton Keynes | Wellcome Sanger Institute for the COVID-19 Genomics UK (COG-UK) Consortium | The Lighthouse Lab in Milton Keynes and Alex Alderton, Roberto Amato, Jeffrey Barrett, Sonia Goncalves, Ewan Harrison, David K. Jackson, Ian Johnston, Dominic Kwiatkowski, Cordelia Langford, John Sillitoe on behalf of the Wellcome Sanger Institute COVID-19 Surveillance Team |
| EPI_ISL_3027388, | Lighthouse Lab in Glasgow | Wellcome Sanger Institute for the COVID-19 Genomics UK (COG-UK) | Harper VanSteenhouse, Yumi Kasai, David Gray, Carol Clugston, Anna Dominiczak and Alex Alderton, Roberto Amato, Jeffrey Barrett, Sonia Goncalves, Ewan Harrison, David K. Jackson, Ian Johnston, Dominic Kwiatkowski, Cordelia Langford, |

| | | | |
|---|---|---|---|
| EPI_ISL_3027389 | | Consortium | John Sillitoe on behalf of the Wellcome Sanger Institute COVID-19 Surveillance Team |
| EPI_ISL_3027390, EPI_ISL_3027391, EPI_ISL_3027392, EPI_ISL_3027393 | Lighthouse Lab in Alderley Park | Wellcome Sanger Institute for the COVID-19 Genomics UK (COG-UK) Consortium | Jacquelyn Wynn, Mairead Hyland, The Lighthouse Lab in Alderley Park and Alex Alderton, Roberto Amato, Jeffrey Barrett, Sonia Goncalves, Ewan Harrison, David K. Jackson, Ian Johnston, Dominic Kwiatkowski, Cordelia Langford, John Sillitoe on behalf of the Wellcome Sanger Institute COVID-19 Surveillance Team |
| EPI_ISL_3027394, EPI_ISL_3027395, EPI_ISL_3027396 | Lighthouse Lab in Milton Keynes | Wellcome Sanger Institute for the COVID-19 Genomics UK (COG-UK) Consortium | The Lighthouse Lab in Milton Keynes and Alex Alderton, Roberto Amato, Jeffrey Barrett, Sonia Goncalves, Ewan Harrison, David K. Jackson, Ian Johnston, Dominic Kwiatkowski, Cordelia Langford, John Sillitoe on behalf of the Wellcome Sanger Institute COVID-19 Surveillance Team |
| EPI_ISL_3027397 | Lighthouse Lab in Alderley Park | Wellcome Sanger Institute for the COVID-19 Genomics UK (COG-UK) Consortium | Jacquelyn Wynn, Mairead Hyland, The Lighthouse Lab in Alderley Park and Alex Alderton, Roberto Amato, Jeffrey Barrett, Sonia Goncalves, Ewan Harrison, David K. Jackson, Ian Johnston, Dominic Kwiatkowski, Cordelia Langford, John Sillitoe on behalf of the Wellcome Sanger Institute COVID-19 Surveillance Team |
| EPI_ISL_3027398 | Lighthouse Lab in Milton Keynes | Wellcome Sanger Institute for the COVID-19 Genomics UK (COG-UK) Consortium | The Lighthouse Lab in Milton Keynes and Alex Alderton, Roberto Amato, Jeffrey Barrett, Sonia Goncalves, Ewan Harrison, David K. Jackson, Ian Johnston, Dominic Kwiatkowski, Cordelia Langford, John Sillitoe on behalf of the Wellcome Sanger Institute COVID-19 Surveillance Team |
| EPI_ISL_3027399 | Lighthouse Lab in Alderley Park | Wellcome Sanger Institute for the COVID-19 Genomics UK (COG-UK) Consortium | Jacquelyn Wynn, Mairead Hyland, The Lighthouse Lab in Alderley Park and Alex Alderton, Roberto Amato, Jeffrey Barrett, Sonia Goncalves, Ewan Harrison, David K. Jackson, Ian Johnston, Dominic Kwiatkowski, Cordelia Langford, John Sillitoe on behalf of the Wellcome Sanger Institute COVID-19 Surveillance Team |
| EPI_ISL_3027400 | Lighthouse Lab in Milton Keynes | Wellcome Sanger Institute for the COVID-19 Genomics UK (COG-UK) Consortium | The Lighthouse Lab in Milton Keynes and Alex Alderton, Roberto Amato, Jeffrey Barrett, Sonia Goncalves, Ewan Harrison, David K. Jackson, Ian Johnston, Dominic Kwiatkowski, Cordelia Langford, John Sillitoe on behalf of the Wellcome Sanger Institute COVID-19 Surveillance Team |
| EPI_ISL_3027401, EPI_ISL_3027402, EPI_ISL_3027403, EPI_ISL_3027404, EPI_ISL_3027405 | Lighthouse Lab in Alderley Park | Wellcome Sanger Institute for the COVID-19 Genomics UK (COG-UK) Consortium | Jacquelyn Wynn, Mairead Hyland, The Lighthouse Lab in Alderley Park and Alex Alderton, Roberto Amato, Jeffrey Barrett, Sonia Goncalves, Ewan Harrison, David K. Jackson, Ian Johnston, Dominic Kwiatkowski, Cordelia Langford, John Sillitoe on behalf of the Wellcome Sanger Institute COVID-19 Surveillance Team |
| EPI_ISL_3027406, EPI_ISL_3027407, EPI_ISL_3027408 | Lighthouse Lab in Milton Keynes | Wellcome Sanger Institute for the COVID-19 Genomics UK (COG-UK) Consortium | The Lighthouse Lab in Milton Keynes and Alex Alderton, Roberto Amato, Jeffrey Barrett, Sonia Goncalves, Ewan Harrison, David K. Jackson, Ian Johnston, Dominic Kwiatkowski, Cordelia Langford, John Sillitoe on behalf of the Wellcome Sanger Institute COVID-19 Surveillance Team |
| EPI_ISL_3027409, EPI_ISL_3027410, EPI_ISL_3027411 | Lighthouse Lab in Alderley Park | Wellcome Sanger Institute for the COVID-19 Genomics UK (COG-UK) Consortium | Jacquelyn Wynn, Mairead Hyland, The Lighthouse Lab in Alderley Park and Alex Alderton, Roberto Amato, Jeffrey Barrett, Sonia Goncalves, Ewan Harrison, David K. Jackson, Ian Johnston, Dominic Kwiatkowski, Cordelia Langford, John Sillitoe on behalf of the Wellcome Sanger Institute COVID-19 Surveillance Team |
| EPI_ISL_3027412 | Lighthouse Lab in Glasgow | Wellcome Sanger Institute for the COVID-19 Genomics UK (COG-UK) Consortium | Harper VanSteenhouse, Yumi Kasai, David Gray, Carol Clugston, Anna Dominiczak and Alex Alderton, Roberto Amato, Jeffrey Barrett, Sonia Goncalves, Ewan Harrison, David K. Jackson, Ian Johnston, Dominic Kwiatkowski, Cordelia Langford, John Sillitoe on behalf of the Wellcome Sanger Institute COVID-19 Surveillance Team |
| EPI_ISL_3027413 | Lighthouse Lab in Milton Keynes | Wellcome Sanger Institute for the COVID-19 Genomics UK (COG-UK) Consortium | The Lighthouse Lab in Milton Keynes and Alex Alderton, Roberto Amato, Jeffrey Barrett, Sonia Goncalves, Ewan Harrison, David K. Jackson, Ian Johnston, Dominic Kwiatkowski, Cordelia Langford, John Sillitoe on behalf of the Wellcome Sanger Institute COVID-19 Surveillance Team |
| EPI_ISL_3027414, EPI_ISL_3027415, EPI_ISL_3027416 | Lighthouse Lab in Alderley Park | Wellcome Sanger Institute for the COVID-19 Genomics UK (COG-UK) Consortium | Jacquelyn Wynn, Mairead Hyland, The Lighthouse Lab in Alderley Park and Alex Alderton, Roberto Amato, Jeffrey Barrett, Sonia Goncalves, Ewan Harrison, David K. Jackson, Ian Johnston, Dominic Kwiatkowski, Cordelia Langford, John Sillitoe on behalf of the Wellcome Sanger Institute COVID-19 Surveillance Team |
| EPI_ISL_3027417 | Lighthouse Lab in Milton Keynes | Wellcome Sanger Institute for the COVID-19 Genomics UK (COG-UK) Consortium | The Lighthouse Lab in Milton Keynes and Alex Alderton, Roberto Amato, Jeffrey Barrett, Sonia Goncalves, Ewan Harrison, David K. Jackson, Ian Johnston, Dominic Kwiatkowski, Cordelia Langford, John Sillitoe on behalf of the Wellcome Sanger Institute COVID-19 Surveillance Team |
| EPI_ISL_3027418 | Lighthouse Lab in Glasgow | Wellcome Sanger Institute for the COVID-19 Genomics UK (COG-UK) Consortium | Harper VanSteenhouse, Yumi Kasai, David Gray, Carol Clugston, Anna Dominiczak and Alex Alderton, Roberto Amato, Jeffrey Barrett, Sonia Goncalves, Ewan Harrison, David K. Jackson, Ian Johnston, Dominic Kwiatkowski, Cordelia Langford, John Sillitoe on behalf of the Wellcome Sanger Institute COVID-19 Surveillance Team |
| EPI_ISL_3027419, EPI_ISL_3027420 | Lighthouse Lab in Milton Keynes | Wellcome Sanger Institute for the COVID-19 Genomics UK (COG-UK) Consortium | The Lighthouse Lab in Milton Keynes and Alex Alderton, Roberto Amato, Jeffrey Barrett, Sonia Goncalves, Ewan Harrison, David K. Jackson, Ian Johnston, Dominic Kwiatkowski, Cordelia Langford, John Sillitoe on behalf of the Wellcome Sanger Institute COVID-19 Surveillance Team |
| EPI_ISL_3027421, EPI_ISL_3027422 | Lighthouse Lab in Alderley Park | Wellcome Sanger Institute for the COVID-19 Genomics UK (COG-UK) Consortium | Jacquelyn Wynn, Mairead Hyland, The Lighthouse Lab in Alderley Park and Alex Alderton, Roberto Amato, Jeffrey Barrett, Sonia Goncalves, Ewan Harrison, David K. Jackson, Ian Johnston, Dominic Kwiatkowski, Cordelia Langford, John Sillitoe on behalf of the Wellcome Sanger Institute COVID-19 Surveillance Team |
| EPI_ISL_3027423, EPI_ISL_3027424 | Lighthouse Lab in Milton Keynes | Wellcome Sanger Institute for the COVID-19 Genomics UK (COG-UK) Consortium | The Lighthouse Lab in Milton Keynes and Alex Alderton, Roberto Amato, Jeffrey Barrett, Sonia Goncalves, Ewan Harrison, David K. Jackson, Ian Johnston, Dominic Kwiatkowski, Cordelia Langford, John Sillitoe on behalf of the Wellcome Sanger Institute COVID-19 Surveillance Team |
| EPI_ISL_3027425 | Lighthouse Lab in Glasgow | Wellcome Sanger Institute for the COVID-19 Genomics UK (COG-UK) Consortium | Harper VanSteenhouse, Yumi Kasai, David Gray, Carol Clugston, Anna Dominiczak and Alex Alderton, Roberto Amato, Jeffrey Barrett, Sonia Goncalves, Ewan Harrison, David K. Jackson, Ian Johnston, Dominic Kwiatkowski, Cordelia Langford, John Sillitoe on behalf of the Wellcome Sanger Institute COVID-19 Surveillance Team |
| EPI_ISL_3027426 | Lighthouse Lab in Alderley Park | Wellcome Sanger Institute for the COVID-19 Genomics UK (COG-UK) Consortium | Jacquelyn Wynn, Mairead Hyland, The Lighthouse Lab in Alderley Park and Alex Alderton, Roberto Amato, Jeffrey Barrett, Sonia Goncalves, Ewan Harrison, David K. Jackson, Ian Johnston, Dominic Kwiatkowski, Cordelia Langford, John Sillitoe on behalf of the Wellcome Sanger Institute COVID-19 Surveillance Team |
| EPI_ISL_3027427 | Lighthouse Lab in Glasgow | Wellcome Sanger Institute for the COVID-19 Genomics UK (COG-UK) Consortium | Harper VanSteenhouse, Yumi Kasai, David Gray, Carol Clugston, Anna Dominiczak and Alex Alderton, Roberto Amato, Jeffrey Barrett, Sonia Goncalves, Ewan Harrison, David K. Jackson, Ian Johnston, Dominic Kwiatkowski, Cordelia Langford, John Sillitoe on behalf of the Wellcome Sanger Institute COVID-19 Surveillance Team |
| EPI_ISL_3027428, EPI_ISL_3027429, EPI_ISL_3027430, EPI_ISL_3027431 | Lighthouse Lab in Alderley Park | Wellcome Sanger Institute for the COVID-19 Genomics UK (COG-UK) Consortium | Jacquelyn Wynn, Mairead Hyland, The Lighthouse Lab in Alderley Park and Alex Alderton, Roberto Amato, Jeffrey Barrett, Sonia Goncalves, Ewan Harrison, David K. Jackson, Ian Johnston, Dominic Kwiatkowski, Cordelia Langford, John Sillitoe on behalf of the Wellcome Sanger Institute COVID-19 Surveillance Team |
| EPI_ISL_3027432 | Lighthouse Lab in Glasgow | Wellcome Sanger Institute for the COVID-19 Genomics UK (COG-UK) Consortium | Harper VanSteenhouse, Yumi Kasai, David Gray, Carol Clugston, Anna Dominiczak and Alex Alderton, Roberto Amato, Jeffrey Barrett, Sonia Goncalves, Ewan Harrison, David K. Jackson, Ian Johnston, Dominic Kwiatkowski, Cordelia Langford, John Sillitoe on behalf of the Wellcome Sanger Institute COVID-19 Surveillance Team |
| EPI_ISL_3027433 | Lighthouse Lab in Alderley Park | Wellcome Sanger Institute for the COVID-19 Genomics UK (COG-UK) Consortium | Jacquelyn Wynn, Mairead Hyland, The Lighthouse Lab in Alderley Park and Alex Alderton, Roberto Amato, Jeffrey Barrett, Sonia Goncalves, Ewan Harrison, David K. Jackson, Ian Johnston, Dominic Kwiatkowski, Cordelia Langford, John Sillitoe on behalf of the Wellcome Sanger Institute COVID-19 Surveillance Team |
| EPI_ISL_3027434, EPI_ISL_3027435 | Lighthouse Lab in Glasgow | Wellcome Sanger Institute for the COVID-19 Genomics UK (COG-UK) Consortium | Harper VanSteenhouse, Yumi Kasai, David Gray, Carol Clugston, Anna Dominiczak and Alex Alderton, Roberto Amato, Jeffrey Barrett, Sonia Goncalves, Ewan Harrison, David K. Jackson, Ian Johnston, Dominic Kwiatkowski, Cordelia Langford, John Sillitoe on behalf of the Wellcome Sanger Institute COVID-19 Surveillance Team |
| EPI_ISL_3027436 | Lighthouse Lab in Alderley Park | Wellcome Sanger Institute for the COVID-19 Genomics UK (COG-UK) Consortium | Jacquelyn Wynn, Mairead Hyland, The Lighthouse Lab in Alderley Park and Alex Alderton, Roberto Amato, Jeffrey Barrett, Sonia Goncalves, Ewan Harrison, David K. Jackson, Ian Johnston, Dominic Kwiatkowski, Cordelia Langford, John Sillitoe on behalf of the Wellcome Sanger Institute COVID-19 Surveillance Team |
| EPI_ISL_3027437 | Lighthouse Lab in Milton Keynes | Wellcome Sanger Institute for the COVID-19 Genomics UK (COG-UK) Consortium | The Lighthouse Lab in Milton Keynes and Alex Alderton, Roberto Amato, Jeffrey Barrett, Sonia Goncalves, Ewan Harrison, David K. Jackson, Ian Johnston, Dominic Kwiatkowski, Cordelia Langford, John Sillitoe on behalf of the Wellcome Sanger Institute COVID-19 Surveillance Team |
| EPI_ISL_3027438, EPI_ISL_3027439, EPI_ISL_3027440, EPI_ISL_3027441, EPI_ISL_3027442, EPI_ISL_3027443, EPI_ISL_3027444, EPI_ISL_3027445, EPI_ISL_3027446, EPI_ISL_3027447 | Lighthouse Lab in Alderley Park | Wellcome Sanger Institute for the COVID-19 Genomics UK (COG-UK) Consortium | Jacquelyn Wynn, Mairead Hyland, The Lighthouse Lab in Alderley Park and Alex Alderton, Roberto Amato, Jeffrey Barrett, Sonia Goncalves, Ewan Harrison, David K. Jackson, Ian Johnston, Dominic Kwiatkowski, Cordelia Langford, John Sillitoe on behalf of the Wellcome Sanger Institute COVID-19 Surveillance Team |
| EPI_ISL_3027448 | Lighthouse Lab in Milton Keynes | Wellcome Sanger Institute for the COVID-19 Genomics UK (COG-UK) Consortium | The Lighthouse Lab in Milton Keynes and Alex Alderton, Roberto Amato, Jeffrey Barrett, Sonia Goncalves, Ewan Harrison, David K. Jackson, Ian Johnston, Dominic Kwiatkowski, Cordelia Langford, John Sillitoe on behalf of the Wellcome Sanger Institute COVID-19 Surveillance Team |
| EPI_ISL_3027449 | Lighthouse Lab in Alderley Park | Wellcome Sanger Institute for the COVID-19 Genomics UK (COG-UK) Consortium | Jacquelyn Wynn, Mairead Hyland, The Lighthouse Lab in Alderley Park and Alex Alderton, Roberto Amato, Jeffrey Barrett, Sonia Goncalves, Ewan Harrison, David K. Jackson, Ian Johnston, Dominic Kwiatkowski, Cordelia Langford, John Sillitoe on behalf of the Wellcome Sanger Institute COVID-19 Surveillance Team |
| EPI_ISL_3027450 | Lighthouse Lab in Milton Keynes | Wellcome Sanger Institute for the COVID-19 Genomics UK (COG-UK) Consortium | The Lighthouse Lab in Milton Keynes and Alex Alderton, Roberto Amato, Jeffrey Barrett, Sonia Goncalves, Ewan Harrison, David K. Jackson, Ian Johnston, Dominic Kwiatkowski, Cordelia Langford, John Sillitoe on behalf of the Wellcome Sanger Institute COVID-19 Surveillance Team |
| EPI_ISL_3027451 | Lighthouse Lab in Alderley Park | Wellcome Sanger Institute for the COVID-19 Genomics UK (COG-UK) Consortium | Jacquelyn Wynn, Mairead Hyland, The Lighthouse Lab in Alderley Park and Alex Alderton, Roberto Amato, Jeffrey Barrett, Sonia Goncalves, Ewan Harrison, David K. Jackson, Ian Johnston, Dominic Kwiatkowski, Cordelia Langford, John Sillitoe on behalf of the Wellcome Sanger Institute COVID-19 Surveillance Team |
| EPI_ISL_3027452 | Berkshire and Surrey Pathology Services Lighthouse Laboratory | Wellcome Sanger Institute for the COVID-19 Genomics UK (COG-UK) Consortium | Berkshire and Surrey Pathology Services Lighthouse Laboratory and Alex Alderton, Roberto Amato, Jeffrey Barrett, Sonia Goncalves, Ewan Harrison, David K. Jackson, Ian Johnston, Dominic Kwiatkowski, Cordelia Langford, John Sillitoe on behalf of the Wellcome Sanger Institute COVID-19 Surveillance Team |
| EPI_ISL_3027453 | Lighthouse Lab in Glasgow | Wellcome Sanger Institute for the COVID-19 Genomics UK (COG-UK) Consortium | Harper VanSteenhouse, Yumi Kasai, David Gray, Carol Clugston, Anna Dominiczak and Alex Alderton, Roberto Amato, Jeffrey Barrett, Sonia Goncalves, Ewan Harrison, David K. Jackson, Ian Johnston, Dominic Kwiatkowski, Cordelia Langford, John Sillitoe on behalf of the Wellcome Sanger Institute COVID-19 Surveillance Team |
| EPI_ISL_3027454 | Lighthouse Lab in Milton Keynes | Wellcome Sanger Institute for the COVID-19 Genomics UK (COG-UK) Consortium | The Lighthouse Lab in Milton Keynes and Alex Alderton, Roberto Amato, Jeffrey Barrett, Sonia Goncalves, Ewan Harrison, David K. Jackson, Ian Johnston, Dominic Kwiatkowski, Cordelia Langford, John Sillitoe on behalf of the Wellcome Sanger Institute COVID-19 Surveillance Team |
| EPI_ISL_3027455 | Berkshire and Surrey Pathology Services Lighthouse Laboratory | Wellcome Sanger Institute for the COVID-19 Genomics UK (COG-UK) Consortium | Berkshire and Surrey Pathology Services Lighthouse Laboratory and Alex Alderton, Roberto Amato, Jeffrey Barrett, Sonia Goncalves, Ewan Harrison, David K. Jackson, Ian Johnston, Dominic Kwiatkowski, Cordelia Langford, John Sillitoe on behalf of the Wellcome Sanger Institute COVID-19 Surveillance Team |
| EPI_ISL_3027456 | Lighthouse Lab in Milton Keynes | Wellcome Sanger Institute for the COVID-19 Genomics UK (COG-UK) Consortium | The Lighthouse Lab in Milton Keynes and Alex Alderton, Roberto Amato, Jeffrey Barrett, Sonia Goncalves, Ewan Harrison, David K. Jackson, Ian Johnston, Dominic Kwiatkowski, Cordelia Langford, John Sillitoe on behalf of the Wellcome Sanger Institute COVID-19 Surveillance Team |
| EPI_ISL_3027457 | Lighthouse Lab in Alderley Park | Wellcome Sanger Institute for the COVID-19 Genomics UK (COG-UK) | Jacquelyn Wynn, Mairead Hyland, The Lighthouse Lab in Alderley Park and Alex Alderton, Roberto Amato, Jeffrey Barrett, Sonia Goncalves, Ewan Harrison, David K. Jackson, Ian Johnston, Dominic Kwiatkowski, Cordelia Langford, John Sillitoe on |

| | | | |
|---|---|---|---|
| | | Consortium | behalf of the Wellcome Sanger Institute COVID-19 Surveillance Team |
| EPI_ISL_3027458 | Lighthouse Lab in Milton Keynes | Wellcome Sanger Institute for the COVID-19 Genomics UK (COG-UK) Consortium | The Lighthouse Lab in Milton Keynes and Alex Alderton, Roberto Amato, Jeffrey Barrett, Sonia Goncalves, Ewan Harrison, David K. Jackson, Ian Johnston, Dominic Kwiatkowski, Cordelia Langford, John Sillitoe on behalf of the Wellcome Sanger Institute COVID-19 Surveillance Team |
| EPI_ISL_3027459 | Lighthouse Lab in Alderley Park | Wellcome Sanger Institute for the COVID-19 Genomics UK (COG-UK) Consortium | Jacquelyn Wynn, Mairead Hyland, The Lighthouse Lab in Alderley Park and Alex Alderton, Roberto Amato, Jeffrey Barrett, Sonia Goncalves, Ewan Harrison, David K. Jackson, Ian Johnston, Dominic Kwiatkowski, Cordelia Langford, John Sillitoe on behalf of the Wellcome Sanger Institute COVID-19 Surveillance Team |
| EPI_ISL_3027460, EPI_ISL_3027461 | Lighthouse Lab in Milton Keynes | Wellcome Sanger Institute for the COVID-19 Genomics UK (COG-UK) Consortium | The Lighthouse Lab in Milton Keynes and Alex Alderton, Roberto Amato, Jeffrey Barrett, Sonia Goncalves, Ewan Harrison, David K. Jackson, Ian Johnston, Dominic Kwiatkowski, Cordelia Langford, John Sillitoe on behalf of the Wellcome Sanger Institute COVID-19 Surveillance Team |
| EPI_ISL_3027462, EPI_ISL_3027463 | Berkshire and Surrey Pathology Services Lighthouse Laboratory | Wellcome Sanger Institute for the COVID-19 Genomics UK (COG-UK) Consortium | Berkshire and Surrey Pathology Services Lighthouse Laboratory and Alex Alderton, Roberto Amato, Jeffrey Barrett, Sonia Goncalves, Ewan Harrison, David K. Jackson, Ian Johnston, Dominic Kwiatkowski, Cordelia Langford, John Sillitoe on behalf of the Wellcome Sanger Institute COVID-19 Surveillance Team |
| EPI_ISL_3027464 | Lighthouse Lab in Alderley Park | Wellcome Sanger Institute for the COVID-19 Genomics UK (COG-UK) Consortium | Jacquelyn Wynn, Mairead Hyland, The Lighthouse Lab in Alderley Park and Alex Alderton, Roberto Amato, Jeffrey Barrett, Sonia Goncalves, Ewan Harrison, David K. Jackson, Ian Johnston, Dominic Kwiatkowski, Cordelia Langford, John Sillitoe on behalf of the Wellcome Sanger Institute COVID-19 Surveillance Team |
| EPI_ISL_3027465 | Lighthouse Lab in Glasgow | Wellcome Sanger Institute for the COVID-19 Genomics UK (COG-UK) Consortium | Harper VanSteenhouse, Yumi Kasai, David Gray, Carol Clugston, Anna Dominiczak and Alex Alderton, Roberto Amato, Jeffrey Barrett, Sonia Goncalves, Ewan Harrison, David K. Jackson, Ian Johnston, Dominic Kwiatkowski, Cordelia Langford, John Sillitoe on behalf of the Wellcome Sanger Institute COVID-19 Surveillance Team |
| EPI_ISL_3027466 | Lighthouse Lab in Alderley Park | Wellcome Sanger Institute for the COVID-19 Genomics UK (COG-UK) Consortium | Jacquelyn Wynn, Mairead Hyland, The Lighthouse Lab in Alderley Park and Alex Alderton, Roberto Amato, Jeffrey Barrett, Sonia Goncalves, Ewan Harrison, David K. Jackson, Ian Johnston, Dominic Kwiatkowski, Cordelia Langford, John Sillitoe on behalf of the Wellcome Sanger Institute COVID-19 Surveillance Team |
| EPI_ISL_3027467 | Lighthouse Lab in Glasgow | Wellcome Sanger Institute for the COVID-19 Genomics UK (COG-UK) Consortium | Harper VanSteenhouse, Yumi Kasai, David Gray, Carol Clugston, Anna Dominiczak and Alex Alderton, Roberto Amato, Jeffrey Barrett, Sonia Goncalves, Ewan Harrison, David K. Jackson, Ian Johnston, Dominic Kwiatkowski, Cordelia Langford, John Sillitoe on behalf of the Wellcome Sanger Institute COVID-19 Surveillance Team |
| EPI_ISL_3027468 | Lighthouse Lab in Alderley Park | Wellcome Sanger Institute for the COVID-19 Genomics UK (COG-UK) Consortium | Jacquelyn Wynn, Mairead Hyland, The Lighthouse Lab in Alderley Park and Alex Alderton, Roberto Amato, Jeffrey Barrett, Sonia Goncalves, Ewan Harrison, David K. Jackson, Ian Johnston, Dominic Kwiatkowski, Cordelia Langford, John Sillitoe on behalf of the Wellcome Sanger Institute COVID-19 Surveillance Team |
| EPI_ISL_3027469 | Lighthouse Lab in Glasgow | Wellcome Sanger Institute for the COVID-19 Genomics UK (COG-UK) Consortium | Harper VanSteenhouse, Yumi Kasai, David Gray, Carol Clugston, Anna Dominiczak and Alex Alderton, Roberto Amato, Jeffrey Barrett, Sonia Goncalves, Ewan Harrison, David K. Jackson, Ian Johnston, Dominic Kwiatkowski, Cordelia Langford, John Sillitoe on behalf of the Wellcome Sanger Institute COVID-19 Surveillance Team |
| EPI_ISL_3027470 | Lighthouse Lab in Milton Keynes | Wellcome Sanger Institute for the COVID-19 Genomics UK (COG-UK) Consortium | The Lighthouse Lab in Milton Keynes and Alex Alderton, Roberto Amato, Jeffrey Barrett, Sonia Goncalves, Ewan Harrison, David K. Jackson, Ian Johnston, Dominic Kwiatkowski, Cordelia Langford, John Sillitoe on behalf of the Wellcome Sanger Institute COVID-19 Surveillance Team |
| EPI_ISL_3027471 | Lighthouse Lab in Glasgow | Wellcome Sanger Institute for the COVID-19 Genomics UK (COG-UK) Consortium | Harper VanSteenhouse, Yumi Kasai, David Gray, Carol Clugston, Anna Dominiczak and Alex Alderton, Roberto Amato, Jeffrey Barrett, Sonia Goncalves, Ewan Harrison, David K. Jackson, Ian Johnston, Dominic Kwiatkowski, Cordelia Langford, John Sillitoe on behalf of the Wellcome Sanger Institute COVID-19 Surveillance Team |
| EPI_ISL_3027472 | Lighthouse Lab in Alderley Park | Wellcome Sanger Institute for the COVID-19 Genomics UK (COG-UK) Consortium | Jacquelyn Wynn, Mairead Hyland, The Lighthouse Lab in Alderley Park and Alex Alderton, Roberto Amato, Jeffrey Barrett, Sonia Goncalves, Ewan Harrison, David K. Jackson, Ian Johnston, Dominic Kwiatkowski, Cordelia Langford, John Sillitoe on behalf of the Wellcome Sanger Institute COVID-19 Surveillance Team |
| EPI_ISL_3027473 | Lighthouse Lab in Glasgow | Wellcome Sanger Institute for the COVID-19 Genomics UK (COG-UK) Consortium | Harper VanSteenhouse, Yumi Kasai, David Gray, Carol Clugston, Anna Dominiczak and Alex Alderton, Roberto Amato, Jeffrey Barrett, Sonia Goncalves, Ewan Harrison, David K. Jackson, Ian Johnston, Dominic Kwiatkowski, Cordelia Langford, John Sillitoe on behalf of the Wellcome Sanger Institute COVID-19 Surveillance Team |
| EPI_ISL_3027474, EPI_ISL_3027475, EPI_ISL_3027476 | Lighthouse Lab in Alderley Park | Wellcome Sanger Institute for the COVID-19 Genomics UK (COG-UK) Consortium | Jacquelyn Wynn, Mairead Hyland, The Lighthouse Lab in Alderley Park and Alex Alderton, Roberto Amato, Jeffrey Barrett, Sonia Goncalves, Ewan Harrison, David K. Jackson, Ian Johnston, Dominic Kwiatkowski, Cordelia Langford, John Sillitoe on behalf of the Wellcome Sanger Institute COVID-19 Surveillance Team |
| EPI_ISL_3027477 | Lighthouse Lab in Glasgow | Wellcome Sanger Institute for the COVID-19 Genomics UK (COG-UK) Consortium | Harper VanSteenhouse, Yumi Kasai, David Gray, Carol Clugston, Anna Dominiczak and Alex Alderton, Roberto Amato, Jeffrey Barrett, Sonia Goncalves, Ewan Harrison, David K. Jackson, Ian Johnston, Dominic Kwiatkowski, Cordelia Langford, John Sillitoe on behalf of the Wellcome Sanger Institute COVID-19 Surveillance Team |
| EPI_ISL_3027478 | Berkshire and Surrey Pathology Services Lighthouse Laboratory | Wellcome Sanger Institute for the COVID-19 Genomics UK (COG-UK) Consortium | Berkshire and Surrey Pathology Services Lighthouse Laboratory and Alex Alderton, Roberto Amato, Jeffrey Barrett, Sonia Goncalves, Ewan Harrison, David K. Jackson, Ian Johnston, Dominic Kwiatkowski, Cordelia Langford, John Sillitoe on behalf of the Wellcome Sanger Institute COVID-19 Surveillance Team |
| EPI_ISL_3027479 | Lighthouse Lab in Milton Keynes | Wellcome Sanger Institute for the COVID-19 Genomics UK (COG-UK) Consortium | The Lighthouse Lab in Milton Keynes and Alex Alderton, Roberto Amato, Jeffrey Barrett, Sonia Goncalves, Ewan Harrison, David K. Jackson, Ian Johnston, Dominic Kwiatkowski, Cordelia Langford, John Sillitoe on behalf of the Wellcome Sanger Institute COVID-19 Surveillance Team |
| EPI_ISL_3027480, EPI_ISL_3027481 | Lighthouse Lab in Alderley Park | Wellcome Sanger Institute for the COVID-19 Genomics UK (COG-UK) Consortium | Jacquelyn Wynn, Mairead Hyland, The Lighthouse Lab in Alderley Park and Alex Alderton, Roberto Amato, Jeffrey Barrett, Sonia Goncalves, Ewan Harrison, David K. Jackson, Ian Johnston, Dominic Kwiatkowski, Cordelia Langford, John Sillitoe on behalf of the Wellcome Sanger Institute COVID-19 Surveillance Team |
| EPI_ISL_3027482, EPI_ISL_3027483 | Lighthouse Lab in Glasgow | Wellcome Sanger Institute for the COVID-19 Genomics UK (COG-UK) Consortium | Harper VanSteenhouse, Yumi Kasai, David Gray, Carol Clugston, Anna Dominiczak and Alex Alderton, Roberto Amato, Jeffrey Barrett, Sonia Goncalves, Ewan Harrison, David K. Jackson, Ian Johnston, Dominic Kwiatkowski, Cordelia Langford, John Sillitoe on behalf of the Wellcome Sanger Institute COVID-19 Surveillance Team |
| EPI_ISL_3027484 | Lighthouse Lab in Milton Keynes | Wellcome Sanger Institute for the COVID-19 Genomics UK (COG-UK) Consortium | The Lighthouse Lab in Milton Keynes and Alex Alderton, Roberto Amato, Jeffrey Barrett, Sonia Goncalves, Ewan Harrison, David K. Jackson, Ian Johnston, Dominic Kwiatkowski, Cordelia Langford, John Sillitoe on behalf of the Wellcome Sanger Institute COVID-19 Surveillance Team |
| EPI_ISL_3027485 | Lighthouse Lab in Alderley Park | Wellcome Sanger Institute for the COVID-19 Genomics UK (COG-UK) Consortium | Jacquelyn Wynn, Mairead Hyland, The Lighthouse Lab in Alderley Park and Alex Alderton, Roberto Amato, Jeffrey Barrett, Sonia Goncalves, Ewan Harrison, David K. Jackson, Ian Johnston, Dominic Kwiatkowski, Cordelia Langford, John Sillitoe on behalf of the Wellcome Sanger Institute COVID-19 Surveillance Team |
| EPI_ISL_3027486 | Lighthouse Lab in Glasgow | Wellcome Sanger Institute for the COVID-19 Genomics UK (COG-UK) Consortium | Harper VanSteenhouse, Yumi Kasai, David Gray, Carol Clugston, Anna Dominiczak and Alex Alderton, Roberto Amato, Jeffrey Barrett, Sonia Goncalves, Ewan Harrison, David K. Jackson, Ian Johnston, Dominic Kwiatkowski, Cordelia Langford, John Sillitoe on behalf of the Wellcome Sanger Institute COVID-19 Surveillance Team |
| EPI_ISL_3027487, EPI_ISL_3027488 | Lighthouse Lab in Milton Keynes | Wellcome Sanger Institute for the COVID-19 Genomics UK (COG-UK) Consortium | The Lighthouse Lab in Milton Keynes and Alex Alderton, Roberto Amato, Jeffrey Barrett, Sonia Goncalves, Ewan Harrison, David K. Jackson, Ian Johnston, Dominic Kwiatkowski, Cordelia Langford, John Sillitoe on behalf of the Wellcome Sanger Institute COVID-19 Surveillance Team |
| EPI_ISL_3027489, EPI_ISL_3027490, EPI_ISL_3027491, EPI_ISL_3027492, EPI_ISL_3027493 | Lighthouse Lab in Alderley Park | Wellcome Sanger Institute for the COVID-19 Genomics UK (COG-UK) Consortium | Jacquelyn Wynn, Mairead Hyland, The Lighthouse Lab in Alderley Park and Alex Alderton, Roberto Amato, Jeffrey Barrett, Sonia Goncalves, Ewan Harrison, David K. Jackson, Ian Johnston, Dominic Kwiatkowski, Cordelia Langford, John Sillitoe on behalf of the Wellcome Sanger Institute COVID-19 Surveillance Team |
| EPI_ISL_3027494 | Berkshire and Surrey Pathology Services Lighthouse Laboratory | Wellcome Sanger Institute for the COVID-19 Genomics UK (COG-UK) Consortium | Berkshire and Surrey Pathology Services Lighthouse Laboratory and Alex Alderton, Roberto Amato, Jeffrey Barrett, Sonia Goncalves, Ewan Harrison, David K. Jackson, Ian Johnston, Dominic Kwiatkowski, Cordelia Langford, John Sillitoe on behalf of the Wellcome Sanger Institute COVID-19 Surveillance Team |
| EPI_ISL_3027495 | Lighthouse Lab in Alderley Park | Wellcome Sanger Institute for the COVID-19 Genomics UK (COG-UK) Consortium | Jacquelyn Wynn, Mairead Hyland, The Lighthouse Lab in Alderley Park and Alex Alderton, Roberto Amato, Jeffrey Barrett, Sonia Goncalves, Ewan Harrison, David K. Jackson, Ian Johnston, Dominic Kwiatkowski, Cordelia Langford, John Sillitoe on behalf of the Wellcome Sanger Institute COVID-19 Surveillance Team |
| EPI_ISL_3027496, EPI_ISL_3027497 | Berkshire and Surrey Pathology Services Lighthouse Laboratory | Wellcome Sanger Institute for the COVID-19 Genomics UK (COG-UK) Consortium | Berkshire and Surrey Pathology Services Lighthouse Laboratory and Alex Alderton, Roberto Amato, Jeffrey Barrett, Sonia Goncalves, Ewan Harrison, David K. Jackson, Ian Johnston, Dominic Kwiatkowski, Cordelia Langford, John Sillitoe on behalf of the Wellcome Sanger Institute COVID-19 Surveillance Team |
| EPI_ISL_3027498 | Lighthouse Lab in Glasgow | Wellcome Sanger Institute for the COVID-19 Genomics UK (COG-UK) Consortium | Harper VanSteenhouse, Yumi Kasai, David Gray, Carol Clugston, Anna Dominiczak and Alex Alderton, Roberto Amato, Jeffrey Barrett, Sonia Goncalves, Ewan Harrison, David K. Jackson, Ian Johnston, Dominic Kwiatkowski, Cordelia Langford, John Sillitoe on behalf of the Wellcome Sanger Institute COVID-19 Surveillance Team |
| EPI_ISL_3027499 | Lighthouse Lab in Milton Keynes | Wellcome Sanger Institute for the COVID-19 Genomics UK (COG-UK) Consortium | The Lighthouse Lab in Milton Keynes and Alex Alderton, Roberto Amato, Jeffrey Barrett, Sonia Goncalves, Ewan Harrison, David K. Jackson, Ian Johnston, Dominic Kwiatkowski, Cordelia Langford, John Sillitoe on behalf of the Wellcome Sanger Institute COVID-19 Surveillance Team |
| EPI_ISL_3027500 | Lighthouse Lab in Glasgow | Wellcome Sanger Institute for the COVID-19 Genomics UK (COG-UK) Consortium | Harper VanSteenhouse, Yumi Kasai, David Gray, Carol Clugston, Anna Dominiczak and Alex Alderton, Roberto Amato, Jeffrey Barrett, Sonia Goncalves, Ewan Harrison, David K. Jackson, Ian Johnston, Dominic Kwiatkowski, Cordelia Langford, John Sillitoe on behalf of the Wellcome Sanger Institute COVID-19 Surveillance Team |
| EPI_ISL_3027501, EPI_ISL_3027502 | Berkshire and Surrey Pathology Services Lighthouse Laboratory | Wellcome Sanger Institute for the COVID-19 Genomics UK (COG-UK) Consortium | Berkshire and Surrey Pathology Services Lighthouse Laboratory and Alex Alderton, Roberto Amato, Jeffrey Barrett, Sonia Goncalves, Ewan Harrison, David K. Jackson, Ian Johnston, Dominic Kwiatkowski, Cordelia Langford, John Sillitoe on behalf of the Wellcome Sanger Institute COVID-19 Surveillance Team |
| EPI_ISL_3027503, EPI_ISL_3027504 | Lighthouse Lab in Alderley Park | Wellcome Sanger Institute for the COVID-19 Genomics UK (COG-UK) Consortium | Jacquelyn Wynn, Mairead Hyland, The Lighthouse Lab in Alderley Park and Alex Alderton, Roberto Amato, Jeffrey Barrett, Sonia Goncalves, Ewan Harrison, David K. Jackson, Ian Johnston, Dominic Kwiatkowski, Cordelia Langford, John Sillitoe on behalf of the Wellcome Sanger Institute COVID-19 Surveillance Team |
| EPI_ISL_3027505, EPI_ISL_3027506 | Lighthouse Lab in Glasgow | Wellcome Sanger Institute for the COVID-19 Genomics UK (COG-UK) Consortium | Harper VanSteenhouse, Yumi Kasai, David Gray, Carol Clugston, Anna Dominiczak and Alex Alderton, Roberto Amato, Jeffrey Barrett, Sonia Goncalves, Ewan Harrison, David K. Jackson, Ian Johnston, Dominic Kwiatkowski, Cordelia Langford, John Sillitoe on behalf of the Wellcome Sanger Institute COVID-19 Surveillance Team |
| EPI_ISL_3027507, EPI_ISL_3027508 | Lighthouse Lab in Alderley Park | Wellcome Sanger Institute for the COVID-19 Genomics UK (COG-UK) Consortium | Jacquelyn Wynn, Mairead Hyland, The Lighthouse Lab in Alderley Park and Alex Alderton, Roberto Amato, Jeffrey Barrett, Sonia Goncalves, Ewan Harrison, David K. Jackson, Ian Johnston, Dominic Kwiatkowski, Cordelia Langford, John Sillitoe on behalf of the Wellcome Sanger Institute COVID-19 Surveillance Team |
| EPI_ISL_3027509 | Lighthouse Lab in Glasgow | Wellcome Sanger Institute for the COVID-19 Genomics UK (COG-UK) Consortium | Harper VanSteenhouse, Yumi Kasai, David Gray, Carol Clugston, Anna Dominiczak and Alex Alderton, Roberto Amato, Jeffrey Barrett, Sonia Goncalves, Ewan Harrison, David K. Jackson, Ian Johnston, Dominic Kwiatkowski, Cordelia Langford, John Sillitoe on behalf of the Wellcome Sanger Institute COVID-19 Surveillance Team |
| EPI_ISL_3027510 | Berkshire and Surrey Pathology Services Lighthouse Laboratory | Wellcome Sanger Institute for the COVID-19 Genomics UK (COG-UK) Consortium | Berkshire and Surrey Pathology Services Lighthouse Laboratory and Alex Alderton, Roberto Amato, Jeffrey Barrett, Sonia Goncalves, Ewan Harrison, David K. Jackson, Ian Johnston, Dominic Kwiatkowski, Cordelia Langford, John Sillitoe on behalf of the Wellcome Sanger Institute COVID-19 Surveillance Team |
| EPI_ISL_3027511, EPI_ISL_3027512 | Lighthouse Lab in Alderley Park | Wellcome Sanger Institute for the COVID-19 Genomics UK (COG-UK) Consortium | Jacquelyn Wynn, Mairead Hyland, The Lighthouse Lab in Alderley Park and Alex Alderton, Roberto Amato, Jeffrey Barrett, Sonia Goncalves, Ewan Harrison, David K. Jackson, Ian Johnston, Dominic Kwiatkowski, Cordelia Langford, John Sillitoe on behalf of the Wellcome Sanger Institute COVID-19 Surveillance Team |
| EPI_ISL_3027513 | Lighthouse Lab in Glasgow | Wellcome Sanger Institute for the COVID-19 Genomics UK (COG-UK) Consortium | Harper VanSteenhouse, Yumi Kasai, David Gray, Carol Clugston, Anna Dominiczak and Alex Alderton, Roberto Amato, Jeffrey Barrett, Sonia Goncalves, Ewan Harrison, David K. Jackson, Ian Johnston, Dominic Kwiatkowski, Cordelia Langford, John Sillitoe on behalf of the Wellcome Sanger Institute COVID-19 Surveillance Team |
| EPI_ISL_3027514 | Lighthouse Lab in Alderley Park | Wellcome Sanger Institute for the COVID-19 Genomics UK (COG-UK) Consortium | Jacquelyn Wynn, Mairead Hyland, The Lighthouse Lab in Alderley Park and Alex Alderton, Roberto Amato, Jeffrey Barrett, Sonia Goncalves, Ewan Harrison, David K. Jackson, Ian Johnston, Dominic Kwiatkowski, Cordelia Langford, John Sillitoe on behalf of the Wellcome Sanger Institute COVID-19 Surveillance Team |
| EPI_ISL_3027515 | Lighthouse Lab in Glasgow | Wellcome Sanger Institute for the COVID-19 Genomics UK (COG-UK) Consortium | Harper VanSteenhouse, Yumi Kasai, David Gray, Carol Clugston, Anna Dominiczak and Alex Alderton, Roberto Amato, Jeffrey Barrett, Sonia Goncalves, Ewan Harrison, David K. Jackson, Ian Johnston, Dominic Kwiatkowski, Cordelia Langford, John Sillitoe on behalf of the Wellcome Sanger Institute COVID-19 Surveillance Team |
| EPI_ISL_3027516 | Lighthouse Lab in Milton Keynes | Wellcome Sanger Institute for the COVID-19 Genomics UK (COG-UK) Consortium | The Lighthouse Lab in Milton Keynes and Alex Alderton, Roberto Amato, Jeffrey Barrett, Sonia Goncalves, Ewan Harrison, David K. Jackson, Ian Johnston, Dominic Kwiatkowski, Cordelia Langford, John Sillitoe on behalf of the Wellcome Sanger Institute COVID-19 Surveillance Team |
| EPI_ISL_3027517 | Lighthouse Lab in Alderley Park | Wellcome Sanger Institute for the COVID-19 Genomics UK (COG-UK) Consortium | Jacquelyn Wynn, Mairead Hyland, The Lighthouse Lab in Alderley Park and Alex Alderton, Roberto Amato, Jeffrey Barrett, Sonia Goncalves, Ewan Harrison, David K. Jackson, Ian Johnston, Dominic Kwiatkowski, Cordelia Langford, John Sillitoe on behalf of the Wellcome Sanger Institute COVID-19 Surveillance Team |

| | | | |
|---|---|---|---|
| EPI_ISL_3027518 | Lighthouse Lab in Glasgow | Wellcome Sanger Institute for the COVID-19 Genomics UK (COG-UK) Consortium | Harper VanSteenhouse, Yumi Kasai, David Gray, Carol Clugston, Anna Dominiczak and Alex Alderton, Roberto Amato, Jeffrey Barrett, Sonia Goncalves, Ewan Harrison, David K. Jackson, Ian Johnston, Dominic Kwiatkowski, Cordelia Langford, John Sillitoe on behalf of the Wellcome Sanger Institute COVID-19 Surveillance Team |
| EPI_ISL_3027519, EPI_ISL_3027520, EPI_ISL_3027521, EPI_ISL_3027522 | Lighthouse Lab in Milton Keynes | Wellcome Sanger Institute for the COVID-19 Genomics UK (COG-UK) Consortium | The Lighthouse Lab in Milton Keynes and Alex Alderton, Roberto Amato, Jeffrey Barrett, Sonia Goncalves, Ewan Harrison, David K. Jackson, Ian Johnston, Dominic Kwiatkowski, Cordelia Langford, John Sillitoe on behalf of the Wellcome Sanger Institute COVID-19 Surveillance Team |
| EPI_ISL_3027523 | Lighthouse Lab in Glasgow | Wellcome Sanger Institute for the COVID-19 Genomics UK (COG-UK) Consortium | Harper VanSteenhouse, Yumi Kasai, David Gray, Carol Clugston, Anna Dominiczak and Alex Alderton, Roberto Amato, Jeffrey Barrett, Sonia Goncalves, Ewan Harrison, David K. Jackson, Ian Johnston, Dominic Kwiatkowski, Cordelia Langford, John Sillitoe on behalf of the Wellcome Sanger Institute COVID-19 Surveillance Team |
| EPI_ISL_3027524, EPI_ISL_3027525, EPI_ISL_3027526 | Berkshire and Surrey Pathology Services Lighthouse Laboratory | Wellcome Sanger Institute for the COVID-19 Genomics UK (COG-UK) Consortium | Berkshire and Surrey Pathology Services Lighthouse Laboratory and Alex Alderton, Roberto Amato, Jeffrey Barrett, Sonia Goncalves, Ewan Harrison, David K. Jackson, Ian Johnston, Dominic Kwiatkowski, Cordelia Langford, John Sillitoe on behalf of the Wellcome Sanger Institute COVID-19 Surveillance Team |
| EPI_ISL_3027527 | Lighthouse Lab in Milton Keynes | Wellcome Sanger Institute for the COVID-19 Genomics UK (COG-UK) Consortium | The Lighthouse Lab in Milton Keynes and Alex Alderton, Roberto Amato, Jeffrey Barrett, Sonia Goncalves, Ewan Harrison, David K. Jackson, Ian Johnston, Dominic Kwiatkowski, Cordelia Langford, John Sillitoe on behalf of the Wellcome Sanger Institute COVID-19 Surveillance Team |
| EPI_ISL_3027528 | Lighthouse Lab in Glasgow | Wellcome Sanger Institute for the COVID-19 Genomics UK (COG-UK) Consortium | Harper VanSteenhouse, Yumi Kasai, David Gray, Carol Clugston, Anna Dominiczak and Alex Alderton, Roberto Amato, Jeffrey Barrett, Sonia Goncalves, Ewan Harrison, David K. Jackson, Ian Johnston, Dominic Kwiatkowski, Cordelia Langford, John Sillitoe on behalf of the Wellcome Sanger Institute COVID-19 Surveillance Team |
| EPI_ISL_3027529, EPI_ISL_3027530 | Lighthouse Lab in Milton Keynes | Wellcome Sanger Institute for the COVID-19 Genomics UK (COG-UK) Consortium | The Lighthouse Lab in Milton Keynes and Alex Alderton, Roberto Amato, Jeffrey Barrett, Sonia Goncalves, Ewan Harrison, David K. Jackson, Ian Johnston, Dominic Kwiatkowski, Cordelia Langford, John Sillitoe on behalf of the Wellcome Sanger Institute COVID-19 Surveillance Team |
| EPI_ISL_3027531 | Berkshire and Surrey Pathology Services Lighthouse Laboratory | Wellcome Sanger Institute for the COVID-19 Genomics UK (COG-UK) Consortium | Berkshire and Surrey Pathology Services Lighthouse Laboratory and Alex Alderton, Roberto Amato, Jeffrey Barrett, Sonia Goncalves, Ewan Harrison, David K. Jackson, Ian Johnston, Dominic Kwiatkowski, Cordelia Langford, John Sillitoe on behalf of the Wellcome Sanger Institute COVID-19 Surveillance Team |
| EPI_ISL_3027532 | Lighthouse Lab in Glasgow | Wellcome Sanger Institute for the COVID-19 Genomics UK (COG-UK) Consortium | Harper VanSteenhouse, Yumi Kasai, David Gray, Carol Clugston, Anna Dominiczak and Alex Alderton, Roberto Amato, Jeffrey Barrett, Sonia Goncalves, Ewan Harrison, David K. Jackson, Ian Johnston, Dominic Kwiatkowski, Cordelia Langford, John Sillitoe on behalf of the Wellcome Sanger Institute COVID-19 Surveillance Team |
| EPI_ISL_3027533 | Lighthouse Lab in Milton Keynes | Wellcome Sanger Institute for the COVID-19 Genomics UK (COG-UK) Consortium | The Lighthouse Lab in Milton Keynes and Alex Alderton, Roberto Amato, Jeffrey Barrett, Sonia Goncalves, Ewan Harrison, David K. Jackson, Ian Johnston, Dominic Kwiatkowski, Cordelia Langford, John Sillitoe on behalf of the Wellcome Sanger Institute COVID-19 Surveillance Team |
| EPI_ISL_3027534 | Berkshire and Surrey Pathology Services Lighthouse Laboratory | Wellcome Sanger Institute for the COVID-19 Genomics UK (COG-UK) Consortium | Berkshire and Surrey Pathology Services Lighthouse Laboratory and Alex Alderton, Roberto Amato, Jeffrey Barrett, Sonia Goncalves, Ewan Harrison, David K. Jackson, Ian Johnston, Dominic Kwiatkowski, Cordelia Langford, John Sillitoe on behalf of the Wellcome Sanger Institute COVID-19 Surveillance Team |
| EPI_ISL_3027535 | Lighthouse Lab in Alderley Park | Wellcome Sanger Institute for the COVID-19 Genomics UK (COG-UK) Consortium | Jacquelyn Wynn, Mairead Hyland, The Lighthouse Lab in Alderley Park and Alex Alderton, Roberto Amato, Jeffrey Barrett, Sonia Goncalves, Ewan Harrison, David K. Jackson, Ian Johnston, Dominic Kwiatkowski, Cordelia Langford, John Sillitoe on behalf of the Wellcome Sanger Institute COVID-19 Surveillance Team |
| EPI_ISL_3027536, EPI_ISL_3027537 | Berkshire and Surrey Pathology Services Lighthouse Laboratory | Wellcome Sanger Institute for the COVID-19 Genomics UK (COG-UK) Consortium | Berkshire and Surrey Pathology Services Lighthouse Laboratory and Alex Alderton, Roberto Amato, Jeffrey Barrett, Sonia Goncalves, Ewan Harrison, David K. Jackson, Ian Johnston, Dominic Kwiatkowski, Cordelia Langford, John Sillitoe on behalf of the Wellcome Sanger Institute COVID-19 Surveillance Team |
| EPI_ISL_3027538 | Lighthouse Lab in Glasgow | Wellcome Sanger Institute for the COVID-19 Genomics UK (COG-UK) Consortium | Harper VanSteenhouse, Yumi Kasai, David Gray, Carol Clugston, Anna Dominiczak and Alex Alderton, Roberto Amato, Jeffrey Barrett, Sonia Goncalves, Ewan Harrison, David K. Jackson, Ian Johnston, Dominic Kwiatkowski, Cordelia Langford, John Sillitoe on behalf of the Wellcome Sanger Institute COVID-19 Surveillance Team |
| EPI_ISL_3027539, EPI_ISL_3027540, EPI_ISL_3027541, EPI_ISL_3027542 | Lighthouse Lab in Milton Keynes | Wellcome Sanger Institute for the COVID-19 Genomics UK (COG-UK) Consortium | The Lighthouse Lab in Milton Keynes and Alex Alderton, Roberto Amato, Jeffrey Barrett, Sonia Goncalves, Ewan Harrison, David K. Jackson, Ian Johnston, Dominic Kwiatkowski, Cordelia Langford, John Sillitoe on behalf of the Wellcome Sanger Institute COVID-19 Surveillance Team |
| EPI_ISL_3027543 | Lighthouse Lab in Glasgow | Wellcome Sanger Institute for the COVID-19 Genomics UK (COG-UK) Consortium | Harper VanSteenhouse, Yumi Kasai, David Gray, Carol Clugston, Anna Dominiczak and Alex Alderton, Roberto Amato, Jeffrey Barrett, Sonia Goncalves, Ewan Harrison, David K. Jackson, Ian Johnston, Dominic Kwiatkowski, Cordelia Langford, John Sillitoe on behalf of the Wellcome Sanger Institute COVID-19 Surveillance Team |
| EPI_ISL_3027544 | Lighthouse Lab in Milton Keynes | Wellcome Sanger Institute for the COVID-19 Genomics UK (COG-UK) Consortium | The Lighthouse Lab in Milton Keynes and Alex Alderton, Roberto Amato, Jeffrey Barrett, Sonia Goncalves, Ewan Harrison, David K. Jackson, Ian Johnston, Dominic Kwiatkowski, Cordelia Langford, John Sillitoe on behalf of the Wellcome Sanger Institute COVID-19 Surveillance Team |
| EPI_ISL_3027545 | Berkshire and Surrey Pathology Services Lighthouse Laboratory | Wellcome Sanger Institute for the COVID-19 Genomics UK (COG-UK) Consortium | Berkshire and Surrey Pathology Services Lighthouse Laboratory and Alex Alderton, Roberto Amato, Jeffrey Barrett, Sonia Goncalves, Ewan Harrison, David K. Jackson, Ian Johnston, Dominic Kwiatkowski, Cordelia Langford, John Sillitoe on behalf of the Wellcome Sanger Institute COVID-19 Surveillance Team |
| EPI_ISL_3027546 | Lighthouse Lab in Milton Keynes | Wellcome Sanger Institute for the COVID-19 Genomics UK (COG-UK) Consortium | The Lighthouse Lab in Milton Keynes and Alex Alderton, Roberto Amato, Jeffrey Barrett, Sonia Goncalves, Ewan Harrison, David K. Jackson, Ian Johnston, Dominic Kwiatkowski, Cordelia Langford, John Sillitoe on behalf of the Wellcome Sanger Institute COVID-19 Surveillance Team |
| EPI_ISL_3027547 | Lighthouse Lab in Glasgow | Wellcome Sanger Institute for the COVID-19 Genomics UK (COG-UK) Consortium | Harper VanSteenhouse, Yumi Kasai, David Gray, Carol Clugston, Anna Dominiczak and Alex Alderton, Roberto Amato, Jeffrey Barrett, Sonia Goncalves, Ewan Harrison, David K. Jackson, Ian Johnston, Dominic Kwiatkowski, Cordelia Langford, John Sillitoe on behalf of the Wellcome Sanger Institute COVID-19 Surveillance Team |
| EPI_ISL_3027548 | Lighthouse Lab in Alderley Park | Wellcome Sanger Institute for the COVID-19 Genomics UK (COG-UK) Consortium | Jacquelyn Wynn, Mairead Hyland, The Lighthouse Lab in Alderley Park and Alex Alderton, Roberto Amato, Jeffrey Barrett, Sonia Goncalves, Ewan Harrison, David K. Jackson, Ian Johnston, Dominic Kwiatkowski, Cordelia Langford, John Sillitoe on behalf of the Wellcome Sanger Institute COVID-19 Surveillance Team |
| EPI_ISL_3027549 | Berkshire and Surrey Pathology Services Lighthouse Laboratory | Wellcome Sanger Institute for the COVID-19 Genomics UK (COG-UK) Consortium | Berkshire and Surrey Pathology Services Lighthouse Laboratory and Alex Alderton, Roberto Amato, Jeffrey Barrett, Sonia Goncalves, Ewan Harrison, David K. Jackson, Ian Johnston, Dominic Kwiatkowski, Cordelia Langford, John Sillitoe on behalf of the Wellcome Sanger Institute COVID-19 Surveillance Team |
| EPI_ISL_3027550, EPI_ISL_3027551 | Lighthouse Lab in Alderley Park | Wellcome Sanger Institute for the COVID-19 Genomics UK (COG-UK) Consortium | Jacquelyn Wynn, Mairead Hyland, The Lighthouse Lab in Alderley Park and Alex Alderton, Roberto Amato, Jeffrey Barrett, Sonia Goncalves, Ewan Harrison, David K. Jackson, Ian Johnston, Dominic Kwiatkowski, Cordelia Langford, John Sillitoe on behalf of the Wellcome Sanger Institute COVID-19 Surveillance Team |
| EPI_ISL_3027552 | Lighthouse Lab in Milton Keynes | Wellcome Sanger Institute for the COVID-19 Genomics UK (COG-UK) Consortium | The Lighthouse Lab in Milton Keynes and Alex Alderton, Roberto Amato, Jeffrey Barrett, Sonia Goncalves, Ewan Harrison, David K. Jackson, Ian Johnston, Dominic Kwiatkowski, Cordelia Langford, John Sillitoe on behalf of the Wellcome Sanger Institute COVID-19 Surveillance Team |
| EPI_ISL_3027553 | Lighthouse Lab in Alderley Park | Wellcome Sanger Institute for the COVID-19 Genomics UK (COG-UK) Consortium | Jacquelyn Wynn, Mairead Hyland, The Lighthouse Lab in Alderley Park and Alex Alderton, Roberto Amato, Jeffrey Barrett, Sonia Goncalves, Ewan Harrison, David K. Jackson, Ian Johnston, Dominic Kwiatkowski, Cordelia Langford, John Sillitoe on behalf of the Wellcome Sanger Institute COVID-19 Surveillance Team |
| EPI_ISL_3027554 | Lighthouse Lab in Glasgow | Wellcome Sanger Institute for the COVID-19 Genomics UK (COG-UK) Consortium | Harper VanSteenhouse, Yumi Kasai, David Gray, Carol Clugston, Anna Dominiczak and Alex Alderton, Roberto Amato, Jeffrey Barrett, Sonia Goncalves, Ewan Harrison, David K. Jackson, Ian Johnston, Dominic Kwiatkowski, Cordelia Langford, John Sillitoe on behalf of the Wellcome Sanger Institute COVID-19 Surveillance Team |
| EPI_ISL_3027555 | Lighthouse Lab in Milton Keynes | Wellcome Sanger Institute for the COVID-19 Genomics UK (COG-UK) Consortium | The Lighthouse Lab in Milton Keynes and Alex Alderton, Roberto Amato, Jeffrey Barrett, Sonia Goncalves, Ewan Harrison, David K. Jackson, Ian Johnston, Dominic Kwiatkowski, Cordelia Langford, John Sillitoe on behalf of the Wellcome Sanger Institute COVID-19 Surveillance Team |
| EPI_ISL_3027556, EPI_ISL_3027557 | Berkshire and Surrey Pathology Services Lighthouse Laboratory | Wellcome Sanger Institute for the COVID-19 Genomics UK (COG-UK) Consortium | Berkshire and Surrey Pathology Services Lighthouse Laboratory and Alex Alderton, Roberto Amato, Jeffrey Barrett, Sonia Goncalves, Ewan Harrison, David K. Jackson, Ian Johnston, Dominic Kwiatkowski, Cordelia Langford, John Sillitoe on behalf of the Wellcome Sanger Institute COVID-19 Surveillance Team |
| EPI_ISL_3027558 | Lighthouse Lab in Milton Keynes | Wellcome Sanger Institute for the COVID-19 Genomics UK (COG-UK) Consortium | The Lighthouse Lab in Milton Keynes and Alex Alderton, Roberto Amato, Jeffrey Barrett, Sonia Goncalves, Ewan Harrison, David K. Jackson, Ian Johnston, Dominic Kwiatkowski, Cordelia Langford, John Sillitoe on behalf of the Wellcome Sanger Institute COVID-19 Surveillance Team |
| EPI_ISL_3027559 | Lighthouse Lab in Alderley Park | Wellcome Sanger Institute for the COVID-19 Genomics UK (COG-UK) Consortium | Jacquelyn Wynn, Mairead Hyland, The Lighthouse Lab in Alderley Park and Alex Alderton, Roberto Amato, Jeffrey Barrett, Sonia Goncalves, Ewan Harrison, David K. Jackson, Ian Johnston, Dominic Kwiatkowski, Cordelia Langford, John Sillitoe on behalf of the Wellcome Sanger Institute COVID-19 Surveillance Team |
| EPI_ISL_3027560 | Berkshire and Surrey Pathology Services Lighthouse Laboratory | Wellcome Sanger Institute for the COVID-19 Genomics UK (COG-UK) Consortium | Berkshire and Surrey Pathology Services Lighthouse Laboratory and Alex Alderton, Roberto Amato, Jeffrey Barrett, Sonia Goncalves, Ewan Harrison, David K. Jackson, Ian Johnston, Dominic Kwiatkowski, Cordelia Langford, John Sillitoe on behalf of the Wellcome Sanger Institute COVID-19 Surveillance Team |
| EPI_ISL_3027561, EPI_ISL_3027562 | Lighthouse Lab in Glasgow | Wellcome Sanger Institute for the COVID-19 Genomics UK (COG-UK) Consortium | Harper VanSteenhouse, Yumi Kasai, David Gray, Carol Clugston, Anna Dominiczak and Alex Alderton, Roberto Amato, Jeffrey Barrett, Sonia Goncalves, Ewan Harrison, David K. Jackson, Ian Johnston, Dominic Kwiatkowski, Cordelia Langford, John Sillitoe on behalf of the Wellcome Sanger Institute COVID-19 Surveillance Team |
| EPI_ISL_3027563, EPI_ISL_3027564 | Berkshire and Surrey Pathology Services Lighthouse Laboratory | Wellcome Sanger Institute for the COVID-19 Genomics UK (COG-UK) Consortium | Berkshire and Surrey Pathology Services Lighthouse Laboratory and Alex Alderton, Roberto Amato, Jeffrey Barrett, Sonia Goncalves, Ewan Harrison, David K. Jackson, Ian Johnston, Dominic Kwiatkowski, Cordelia Langford, John Sillitoe on behalf of the Wellcome Sanger Institute COVID-19 Surveillance Team |
| EPI_ISL_3027565 | Lighthouse Lab in Milton Keynes | Wellcome Sanger Institute for the COVID-19 Genomics UK (COG-UK) Consortium | The Lighthouse Lab in Milton Keynes and Alex Alderton, Roberto Amato, Jeffrey Barrett, Sonia Goncalves, Ewan Harrison, David K. Jackson, Ian Johnston, Dominic Kwiatkowski, Cordelia Langford, John Sillitoe on behalf of the Wellcome Sanger Institute COVID-19 Surveillance Team |
| EPI_ISL_3027566 | Lighthouse Lab in Glasgow | Wellcome Sanger Institute for the COVID-19 Genomics UK (COG-UK) Consortium | Harper VanSteenhouse, Yumi Kasai, David Gray, Carol Clugston, Anna Dominiczak and Alex Alderton, Roberto Amato, Jeffrey Barrett, Sonia Goncalves, Ewan Harrison, David K. Jackson, Ian Johnston, Dominic Kwiatkowski, Cordelia Langford, John Sillitoe on behalf of the Wellcome Sanger Institute COVID-19 Surveillance Team |
| EPI_ISL_3027567, EPI_ISL_3027568 | Lighthouse Lab in Milton Keynes | Wellcome Sanger Institute for the COVID-19 Genomics UK (COG-UK) Consortium | The Lighthouse Lab in Milton Keynes and Alex Alderton, Roberto Amato, Jeffrey Barrett, Sonia Goncalves, Ewan Harrison, David K. Jackson, Ian Johnston, Dominic Kwiatkowski, Cordelia Langford, John Sillitoe on behalf of the Wellcome Sanger Institute COVID-19 Surveillance Team |
| EPI_ISL_3027569, EPI_ISL_3027570 | Lighthouse Lab in Alderley Park | Wellcome Sanger Institute for the COVID-19 Genomics UK (COG-UK) Consortium | Jacquelyn Wynn, Mairead Hyland, The Lighthouse Lab in Alderley Park and Alex Alderton, Roberto Amato, Jeffrey Barrett, Sonia Goncalves, Ewan Harrison, David K. Jackson, Ian Johnston, Dominic Kwiatkowski, Cordelia Langford, John Sillitoe on behalf of the Wellcome Sanger Institute COVID-19 Surveillance Team |
| EPI_ISL_3027571 | Lighthouse Lab in Glasgow | Wellcome Sanger Institute for the COVID-19 Genomics UK (COG-UK) Consortium | Harper VanSteenhouse, Yumi Kasai, David Gray, Carol Clugston, Anna Dominiczak and Alex Alderton, Roberto Amato, Jeffrey Barrett, Sonia Goncalves, Ewan Harrison, David K. Jackson, Ian Johnston, Dominic Kwiatkowski, Cordelia Langford, John Sillitoe on behalf of the Wellcome Sanger Institute COVID-19 Surveillance Team |
| EPI_ISL_3027572, EPI_ISL_3027573 | Berkshire and Surrey Pathology Services Lighthouse Laboratory | Wellcome Sanger Institute for the COVID-19 Genomics UK (COG-UK) Consortium | Berkshire and Surrey Pathology Services Lighthouse Laboratory and Alex Alderton, Roberto Amato, Jeffrey Barrett, Sonia Goncalves, Ewan Harrison, David K. Jackson, Ian Johnston, Dominic Kwiatkowski, Cordelia Langford, John Sillitoe on behalf of the Wellcome Sanger Institute COVID-19 Surveillance Team |
| EPI_ISL_3027574 | Lighthouse Lab in Milton Keynes | Wellcome Sanger Institute for the COVID-19 Genomics UK (COG-UK) Consortium | The Lighthouse Lab in Milton Keynes and Alex Alderton, Roberto Amato, Jeffrey Barrett, Sonia Goncalves, Ewan Harrison, David K. Jackson, Ian Johnston, Dominic Kwiatkowski, Cordelia Langford, John Sillitoe on behalf of the Wellcome Sanger Institute COVID-19 Surveillance Team |
| EPI_ISL_3027575 | Lighthouse Lab in Glasgow | Wellcome Sanger Institute for the COVID-19 Genomics UK (COG-UK) Consortium | Harper VanSteenhouse, Yumi Kasai, David Gray, Carol Clugston, Anna Dominiczak and Alex Alderton, Roberto Amato, Jeffrey Barrett, Sonia Goncalves, Ewan Harrison, David K. Jackson, Ian Johnston, Dominic Kwiatkowski, Cordelia Langford, John Sillitoe on behalf of the Wellcome Sanger Institute COVID-19 Surveillance Team |
| EPI_ISL_3027576 | Lighthouse Lab in Alderley Park | Wellcome Sanger Institute for the COVID-19 Genomics UK (COG-UK) Consortium | Jacquelyn Wynn, Mairead Hyland, The Lighthouse Lab in Alderley Park and Alex Alderton, Roberto Amato, Jeffrey Barrett, Sonia Goncalves, Ewan Harrison, David K. Jackson, Ian Johnston, Dominic Kwiatkowski, Cordelia Langford, John Sillitoe on behalf of the Wellcome Sanger Institute COVID-19 Surveillance Team |
| EPI_ISL_3027577, EPI_ISL_3027578, | Lighthouse Lab in Glasgow | Wellcome Sanger Institute for the COVID-19 Genomics UK (COG-UK) Consortium | Harper VanSteenhouse, Yumi Kasai, David Gray, Carol Clugston, Anna Dominiczak and Alex Alderton, Roberto Amato, Jeffrey Barrett, Sonia Goncalves, Ewan Harrison, David K. Jackson, Ian Johnston, Dominic Kwiatkowski, Cordelia Langford, John Sillitoe on behalf of the Wellcome Sanger Institute COVID-19 Surveillance Team |

| | | | |
|---|---|---|---|
| EPI_ISL_3027579 | | | |
| EPI_ISL_3027580 | Lighthouse Lab in Milton Keynes | Wellcome Sanger Institute for the COVID-19 Genomics UK (COG-UK) Consortium | The Lighthouse Lab in Milton Keynes and Alex Alderton, Roberto Amato, Jeffrey Barrett, Sonia Goncalves, Ewan Harrison, David K. Jackson, Ian Johnston, Dominic Kwiatkowski, Cordelia Langford, John Sillitoe on behalf of the Wellcome Sanger Institute COVID-19 Surveillance Team |
| EPI_ISL_3027581 | Berkshire and Surrey Pathology Services Lighthouse Laboratory | Wellcome Sanger Institute for the COVID-19 Genomics UK (COG-UK) Consortium | Berkshire and Surrey Pathology Services Lighthouse Laboratory and Alex Alderton, Roberto Amato, Jeffrey Barrett, Sonia Goncalves, Ewan Harrison, David K. Jackson, Ian Johnston, Dominic Kwiatkowski, Cordelia Langford, John Sillitoe on behalf of the Wellcome Sanger Institute COVID-19 Surveillance Team |
| EPI_ISL_3027582 | Lighthouse Lab in Glasgow | Wellcome Sanger Institute for the COVID-19 Genomics UK (COG-UK) Consortium | Harper VanSteenhouse, Yumi Kasai, David Gray, Carol Clugston, Anna Dominiczak and Alex Alderton, Roberto Amato, Jeffrey Barrett, Sonia Goncalves, Ewan Harrison, David K. Jackson, Ian Johnston, Dominic Kwiatkowski, Cordelia Langford, John Sillitoe on behalf of the Wellcome Sanger Institute COVID-19 Surveillance Team |
| EPI_ISL_3027583 | Berkshire and Surrey Pathology Services Lighthouse Laboratory | Wellcome Sanger Institute for the COVID-19 Genomics UK (COG-UK) Consortium | Berkshire and Surrey Pathology Services Lighthouse Laboratory and Alex Alderton, Roberto Amato, Jeffrey Barrett, Sonia Goncalves, Ewan Harrison, David K. Jackson, Ian Johnston, Dominic Kwiatkowski, Cordelia Langford, John Sillitoe on behalf of the Wellcome Sanger Institute COVID-19 Surveillance Team |
| EPI_ISL_3027584 | Lighthouse Lab in Glasgow | Wellcome Sanger Institute for the COVID-19 Genomics UK (COG-UK) Consortium | Harper VanSteenhouse, Yumi Kasai, David Gray, Carol Clugston, Anna Dominiczak and Alex Alderton, Roberto Amato, Jeffrey Barrett, Sonia Goncalves, Ewan Harrison, David K. Jackson, Ian Johnston, Dominic Kwiatkowski, Cordelia Langford, John Sillitoe on behalf of the Wellcome Sanger Institute COVID-19 Surveillance Team |
| EPI_ISL_3027585 | Lighthouse Lab in Milton Keynes | Wellcome Sanger Institute for the COVID-19 Genomics UK (COG-UK) Consortium | The Lighthouse Lab in Milton Keynes and Alex Alderton, Roberto Amato, Jeffrey Barrett, Sonia Goncalves, Ewan Harrison, David K. Jackson, Ian Johnston, Dominic Kwiatkowski, Cordelia Langford, John Sillitoe on behalf of the Wellcome Sanger Institute COVID-19 Surveillance Team |
| EPI_ISL_3027586, EPI_ISL_3027587 | Berkshire and Surrey Pathology Services Lighthouse Laboratory | Wellcome Sanger Institute for the COVID-19 Genomics UK (COG-UK) Consortium | Berkshire and Surrey Pathology Services Lighthouse Laboratory and Alex Alderton, Roberto Amato, Jeffrey Barrett, Sonia Goncalves, Ewan Harrison, David K. Jackson, Ian Johnston, Dominic Kwiatkowski, Cordelia Langford, John Sillitoe on behalf of the Wellcome Sanger Institute COVID-19 Surveillance Team |
| EPI_ISL_3027588 | Lighthouse Lab in Milton Keynes | Wellcome Sanger Institute for the COVID-19 Genomics UK (COG-UK) Consortium | The Lighthouse Lab in Milton Keynes and Alex Alderton, Roberto Amato, Jeffrey Barrett, Sonia Goncalves, Ewan Harrison, David K. Jackson, Ian Johnston, Dominic Kwiatkowski, Cordelia Langford, John Sillitoe on behalf of the Wellcome Sanger Institute COVID-19 Surveillance Team |
| EPI_ISL_3027589 | Lighthouse Lab in Glasgow | Wellcome Sanger Institute for the COVID-19 Genomics UK (COG-UK) Consortium | Harper VanSteenhouse, Yumi Kasai, David Gray, Carol Clugston, Anna Dominiczak and Alex Alderton, Roberto Amato, Jeffrey Barrett, Sonia Goncalves, Ewan Harrison, David K. Jackson, Ian Johnston, Dominic Kwiatkowski, Cordelia Langford, John Sillitoe on behalf of the Wellcome Sanger Institute COVID-19 Surveillance Team |
| EPI_ISL_3027590 | Berkshire and Surrey Pathology Services Lighthouse Laboratory | Wellcome Sanger Institute for the COVID-19 Genomics UK (COG-UK) Consortium | Berkshire and Surrey Pathology Services Lighthouse Laboratory and Alex Alderton, Roberto Amato, Jeffrey Barrett, Sonia Goncalves, Ewan Harrison, David K. Jackson, Ian Johnston, Dominic Kwiatkowski, Cordelia Langford, John Sillitoe on behalf of the Wellcome Sanger Institute COVID-19 Surveillance Team |
| EPI_ISL_3027591 | Lighthouse Lab in Glasgow | Wellcome Sanger Institute for the COVID-19 Genomics UK (COG-UK) Consortium | Harper VanSteenhouse, Yumi Kasai, David Gray, Carol Clugston, Anna Dominiczak and Alex Alderton, Roberto Amato, Jeffrey Barrett, Sonia Goncalves, Ewan Harrison, David K. Jackson, Ian Johnston, Dominic Kwiatkowski, Cordelia Langford, John Sillitoe on behalf of the Wellcome Sanger Institute COVID-19 Surveillance Team |
| EPI_ISL_3027592 | Lighthouse Lab in Milton Keynes | Wellcome Sanger Institute for the COVID-19 Genomics UK (COG-UK) Consortium | The Lighthouse Lab in Milton Keynes and Alex Alderton, Roberto Amato, Jeffrey Barrett, Sonia Goncalves, Ewan Harrison, David K. Jackson, Ian Johnston, Dominic Kwiatkowski, Cordelia Langford, John Sillitoe on behalf of the Wellcome Sanger Institute COVID-19 Surveillance Team |
| EPI_ISL_3027593 | Berkshire and Surrey Pathology Services Lighthouse Laboratory | Wellcome Sanger Institute for the COVID-19 Genomics UK (COG-UK) Consortium | Berkshire and Surrey Pathology Services Lighthouse Laboratory and Alex Alderton, Roberto Amato, Jeffrey Barrett, Sonia Goncalves, Ewan Harrison, David K. Jackson, Ian Johnston, Dominic Kwiatkowski, Cordelia Langford, John Sillitoe on behalf of the Wellcome Sanger Institute COVID-19 Surveillance Team |
| EPI_ISL_3027594 | Lighthouse Lab in Alderley Park | Wellcome Sanger Institute for the COVID-19 Genomics UK (COG-UK) Consortium | Jacquelyn Wynn, Mairead Hyland, The Lighthouse Lab in Alderley Park and Alex Alderton, Roberto Amato, Jeffrey Barrett, Sonia Goncalves, Ewan Harrison, David K. Jackson, Ian Johnston, Dominic Kwiatkowski, Cordelia Langford, John Sillitoe on behalf of the Wellcome Sanger Institute COVID-19 Surveillance Team |
| EPI_ISL_3027595 | Berkshire and Surrey Pathology Services Lighthouse Laboratory | Wellcome Sanger Institute for the COVID-19 Genomics UK (COG-UK) Consortium | Berkshire and Surrey Pathology Services Lighthouse Laboratory and Alex Alderton, Roberto Amato, Jeffrey Barrett, Sonia Goncalves, Ewan Harrison, David K. Jackson, Ian Johnston, Dominic Kwiatkowski, Cordelia Langford, John Sillitoe on behalf of the Wellcome Sanger Institute COVID-19 Surveillance Team |
| EPI_ISL_3027596 | Lighthouse Lab in Alderley Park | Wellcome Sanger Institute for the COVID-19 Genomics UK (COG-UK) Consortium | Jacquelyn Wynn, Mairead Hyland, The Lighthouse Lab in Alderley Park and Alex Alderton, Roberto Amato, Jeffrey Barrett, Sonia Goncalves, Ewan Harrison, David K. Jackson, Ian Johnston, Dominic Kwiatkowski, Cordelia Langford, John Sillitoe on behalf of the Wellcome Sanger Institute COVID-19 Surveillance Team |
| EPI_ISL_3027597 | Lighthouse Lab in Milton Keynes | Wellcome Sanger Institute for the COVID-19 Genomics UK (COG-UK) Consortium | The Lighthouse Lab in Milton Keynes and Alex Alderton, Roberto Amato, Jeffrey Barrett, Sonia Goncalves, Ewan Harrison, David K. Jackson, Ian Johnston, Dominic Kwiatkowski, Cordelia Langford, John Sillitoe on behalf of the Wellcome Sanger Institute COVID-19 Surveillance Team |
| EPI_ISL_3027598, EPI_ISL_3027599, EPI_ISL_3027600 | Berkshire and Surrey Pathology Services Lighthouse Laboratory | Wellcome Sanger Institute for the COVID-19 Genomics UK (COG-UK) Consortium | Berkshire and Surrey Pathology Services Lighthouse Laboratory and Alex Alderton, Roberto Amato, Jeffrey Barrett, Sonia Goncalves, Ewan Harrison, David K. Jackson, Ian Johnston, Dominic Kwiatkowski, Cordelia Langford, John Sillitoe on behalf of the Wellcome Sanger Institute COVID-19 Surveillance Team |
| EPI_ISL_3027601 | Lighthouse Lab in Glasgow | Wellcome Sanger Institute for the COVID-19 Genomics UK (COG-UK) Consortium | Harper VanSteenhouse, Yumi Kasai, David Gray, Carol Clugston, Anna Dominiczak and Alex Alderton, Roberto Amato, Jeffrey Barrett, Sonia Goncalves, Ewan Harrison, David K. Jackson, Ian Johnston, Dominic Kwiatkowski, Cordelia Langford, John Sillitoe on behalf of the Wellcome Sanger Institute COVID-19 Surveillance Team |
| EPI_ISL_3027602, EPI_ISL_3027603, EPI_ISL_3027604 | Lighthouse Lab in Milton Keynes | Wellcome Sanger Institute for the COVID-19 Genomics UK (COG-UK) Consortium | The Lighthouse Lab in Milton Keynes and Alex Alderton, Roberto Amato, Jeffrey Barrett, Sonia Goncalves, Ewan Harrison, David K. Jackson, Ian Johnston, Dominic Kwiatkowski, Cordelia Langford, John Sillitoe on behalf of the Wellcome Sanger Institute COVID-19 Surveillance Team |
| EPI_ISL_3027605, EPI_ISL_3027606 | Lighthouse Lab in Alderley Park | Wellcome Sanger Institute for the COVID-19 Genomics UK (COG-UK) Consortium | Jacquelyn Wynn, Mairead Hyland, The Lighthouse Lab in Alderley Park and Alex Alderton, Roberto Amato, Jeffrey Barrett, Sonia Goncalves, Ewan Harrison, David K. Jackson, Ian Johnston, Dominic Kwiatkowski, Cordelia Langford, John Sillitoe on behalf of the Wellcome Sanger Institute COVID-19 Surveillance Team |
| EPI_ISL_3027607 | Lighthouse Lab in Milton Keynes | Wellcome Sanger Institute for the COVID-19 Genomics UK (COG-UK) Consortium | The Lighthouse Lab in Milton Keynes and Alex Alderton, Roberto Amato, Jeffrey Barrett, Sonia Goncalves, Ewan Harrison, David K. Jackson, Ian Johnston, Dominic Kwiatkowski, Cordelia Langford, John Sillitoe on behalf of the Wellcome Sanger Institute COVID-19 Surveillance Team |
| EPI_ISL_3027608, EPI_ISL_3027609 | Berkshire and Surrey Pathology Services Lighthouse Laboratory | Wellcome Sanger Institute for the COVID-19 Genomics UK (COG-UK) Consortium | Berkshire and Surrey Pathology Services Lighthouse Laboratory and Alex Alderton, Roberto Amato, Jeffrey Barrett, Sonia Goncalves, Ewan Harrison, David K. Jackson, Ian Johnston, Dominic Kwiatkowski, Cordelia Langford, John Sillitoe on behalf of the Wellcome Sanger Institute COVID-19 Surveillance Team |
| EPI_ISL_3027610 | Lighthouse Lab in Milton Keynes | Wellcome Sanger Institute for the COVID-19 Genomics UK (COG-UK) Consortium | The Lighthouse Lab in Milton Keynes and Alex Alderton, Roberto Amato, Jeffrey Barrett, Sonia Goncalves, Ewan Harrison, David K. Jackson, Ian Johnston, Dominic Kwiatkowski, Cordelia Langford, John Sillitoe on behalf of the Wellcome Sanger Institute COVID-19 Surveillance Team |
| EPI_ISL_3027611 | Lighthouse Lab in Alderley Park | Wellcome Sanger Institute for the COVID-19 Genomics UK (COG-UK) Consortium | Jacquelyn Wynn, Mairead Hyland, The Lighthouse Lab in Alderley Park and Alex Alderton, Roberto Amato, Jeffrey Barrett, Sonia Goncalves, Ewan Harrison, David K. Jackson, Ian Johnston, Dominic Kwiatkowski, Cordelia Langford, John Sillitoe on behalf of the Wellcome Sanger Institute COVID-19 Surveillance Team |
| EPI_ISL_3027612, EPI_ISL_3027613, EPI_ISL_3027614 | Berkshire and Surrey Pathology Services Lighthouse Laboratory | Wellcome Sanger Institute for the COVID-19 Genomics UK (COG-UK) Consortium | Berkshire and Surrey Pathology Services Lighthouse Laboratory and Alex Alderton, Roberto Amato, Jeffrey Barrett, Sonia Goncalves, Ewan Harrison, David K. Jackson, Ian Johnston, Dominic Kwiatkowski, Cordelia Langford, John Sillitoe on behalf of the Wellcome Sanger Institute COVID-19 Surveillance Team |
| EPI_ISL_3027615 | Lighthouse Lab in Milton Keynes | Wellcome Sanger Institute for the COVID-19 Genomics UK (COG-UK) Consortium | The Lighthouse Lab in Milton Keynes and Alex Alderton, Roberto Amato, Jeffrey Barrett, Sonia Goncalves, Ewan Harrison, David K. Jackson, Ian Johnston, Dominic Kwiatkowski, Cordelia Langford, John Sillitoe on behalf of the Wellcome Sanger Institute COVID-19 Surveillance Team |
| EPI_ISL_3027616, EPI_ISL_3027617, EPI_ISL_3027618, EPI_ISL_3027619, EPI_ISL_3027620 | Lighthouse Lab in Alderley Park | Wellcome Sanger Institute for the COVID-19 Genomics UK (COG-UK) Consortium | Jacquelyn Wynn, Mairead Hyland, The Lighthouse Lab in Alderley Park and Alex Alderton, Roberto Amato, Jeffrey Barrett, Sonia Goncalves, Ewan Harrison, David K. Jackson, Ian Johnston, Dominic Kwiatkowski, Cordelia Langford, John Sillitoe on behalf of the Wellcome Sanger Institute COVID-19 Surveillance Team |
| EPI_ISL_3027621 | Lighthouse Lab in Glasgow | Wellcome Sanger Institute for the COVID-19 Genomics UK (COG-UK) Consortium | Harper VanSteenhouse, Yumi Kasai, David Gray, Carol Clugston, Anna Dominiczak and Alex Alderton, Roberto Amato, Jeffrey Barrett, Sonia Goncalves, Ewan Harrison, David K. Jackson, Ian Johnston, Dominic Kwiatkowski, Cordelia Langford, John Sillitoe on behalf of the Wellcome Sanger Institute COVID-19 Surveillance Team |
| EPI_ISL_3027622 | Lighthouse Lab in Alderley Park | Wellcome Sanger Institute for the COVID-19 Genomics UK (COG-UK) Consortium | Jacquelyn Wynn, Mairead Hyland, The Lighthouse Lab in Alderley Park and Alex Alderton, Roberto Amato, Jeffrey Barrett, Sonia Goncalves, Ewan Harrison, David K. Jackson, Ian Johnston, Dominic Kwiatkowski, Cordelia Langford, John Sillitoe on behalf of the Wellcome Sanger Institute COVID-19 Surveillance Team |
| EPI_ISL_3027623 | Lighthouse Lab in Milton Keynes | Wellcome Sanger Institute for the COVID-19 Genomics UK (COG-UK) Consortium | The Lighthouse Lab in Milton Keynes and Alex Alderton, Roberto Amato, Jeffrey Barrett, Sonia Goncalves, Ewan Harrison, David K. Jackson, Ian Johnston, Dominic Kwiatkowski, Cordelia Langford, John Sillitoe on behalf of the Wellcome Sanger Institute COVID-19 Surveillance Team |
| EPI_ISL_3027624, EPI_ISL_3027625 | Lighthouse Lab in Alderley Park | Wellcome Sanger Institute for the COVID-19 Genomics UK (COG-UK) Consortium | Jacquelyn Wynn, Mairead Hyland, The Lighthouse Lab in Alderley Park and Alex Alderton, Roberto Amato, Jeffrey Barrett, Sonia Goncalves, Ewan Harrison, David K. Jackson, Ian Johnston, Dominic Kwiatkowski, Cordelia Langford, John Sillitoe on behalf of the Wellcome Sanger Institute COVID-19 Surveillance Team |
| EPI_ISL_3027626 | Lighthouse Lab in Milton Keynes | Wellcome Sanger Institute for the COVID-19 Genomics UK (COG-UK) Consortium | The Lighthouse Lab in Milton Keynes and Alex Alderton, Roberto Amato, Jeffrey Barrett, Sonia Goncalves, Ewan Harrison, David K. Jackson, Ian Johnston, Dominic Kwiatkowski, Cordelia Langford, John Sillitoe on behalf of the Wellcome Sanger Institute COVID-19 Surveillance Team |
| EPI_ISL_3027627, EPI_ISL_3027628 | Lighthouse Lab in Alderley Park | Wellcome Sanger Institute for the COVID-19 Genomics UK (COG-UK) Consortium | Jacquelyn Wynn, Mairead Hyland, The Lighthouse Lab in Alderley Park and Alex Alderton, Roberto Amato, Jeffrey Barrett, Sonia Goncalves, Ewan Harrison, David K. Jackson, Ian Johnston, Dominic Kwiatkowski, Cordelia Langford, John Sillitoe on behalf of the Wellcome Sanger Institute COVID-19 Surveillance Team |
| EPI_ISL_3027629 | Lighthouse Lab in Glasgow | Wellcome Sanger Institute for the COVID-19 Genomics UK (COG-UK) Consortium | Harper VanSteenhouse, Yumi Kasai, David Gray, Carol Clugston, Anna Dominiczak and Alex Alderton, Roberto Amato, Jeffrey Barrett, Sonia Goncalves, Ewan Harrison, David K. Jackson, Ian Johnston, Dominic Kwiatkowski, Cordelia Langford, John Sillitoe on behalf of the Wellcome Sanger Institute COVID-19 Surveillance Team |
| EPI_ISL_3027630, EPI_ISL_3027631 | Lighthouse Lab in Alderley Park | Wellcome Sanger Institute for the COVID-19 Genomics UK (COG-UK) Consortium | Jacquelyn Wynn, Mairead Hyland, The Lighthouse Lab in Alderley Park and Alex Alderton, Roberto Amato, Jeffrey Barrett, Sonia Goncalves, Ewan Harrison, David K. Jackson, Ian Johnston, Dominic Kwiatkowski, Cordelia Langford, John Sillitoe on behalf of the Wellcome Sanger Institute COVID-19 Surveillance Team |
| EPI_ISL_3027632 | Lighthouse Lab in Glasgow | Wellcome Sanger Institute for the COVID-19 Genomics UK (COG-UK) Consortium | Harper VanSteenhouse, Yumi Kasai, David Gray, Carol Clugston, Anna Dominiczak and Alex Alderton, Roberto Amato, Jeffrey Barrett, Sonia Goncalves, Ewan Harrison, David K. Jackson, Ian Johnston, Dominic Kwiatkowski, Cordelia Langford, John Sillitoe on behalf of the Wellcome Sanger Institute COVID-19 Surveillance Team |
| EPI_ISL_3027633 | Lighthouse Lab in Alderley Park | Wellcome Sanger Institute for the COVID-19 Genomics UK (COG-UK) Consortium | Jacquelyn Wynn, Mairead Hyland, The Lighthouse Lab in Alderley Park and Alex Alderton, Roberto Amato, Jeffrey Barrett, Sonia Goncalves, Ewan Harrison, David K. Jackson, Ian Johnston, Dominic Kwiatkowski, Cordelia Langford, John Sillitoe on behalf of the Wellcome Sanger Institute COVID-19 Surveillance Team |
| EPI_ISL_3027634 | Berkshire and Surrey Pathology Services Lighthouse Laboratory | Wellcome Sanger Institute for the COVID-19 Genomics UK (COG-UK) Consortium | Berkshire and Surrey Pathology Services Lighthouse Laboratory and Alex Alderton, Roberto Amato, Jeffrey Barrett, Sonia Goncalves, Ewan Harrison, David K. Jackson, Ian Johnston, Dominic Kwiatkowski, Cordelia Langford, John Sillitoe on behalf of the Wellcome Sanger Institute COVID-19 Surveillance Team |
| EPI_ISL_3027635 | Lighthouse Lab in Milton Keynes | Wellcome Sanger Institute for the COVID-19 Genomics UK (COG-UK) Consortium | The Lighthouse Lab in Milton Keynes and Alex Alderton, Roberto Amato, Jeffrey Barrett, Sonia Goncalves, Ewan Harrison, David K. Jackson, Ian Johnston, Dominic Kwiatkowski, Cordelia Langford, John Sillitoe on behalf of the Wellcome Sanger Institute COVID-19 Surveillance Team |
| EPI_ISL_3027636 | Lighthouse Lab in Glasgow | Wellcome Sanger Institute for the COVID-19 Genomics UK (COG-UK) Consortium | Harper VanSteenhouse, Yumi Kasai, David Gray, Carol Clugston, Anna Dominiczak and Alex Alderton, Roberto Amato, Jeffrey Barrett, Sonia Goncalves, Ewan Harrison, David K. Jackson, Ian Johnston, Dominic Kwiatkowski, Cordelia Langford, John Sillitoe on behalf of the Wellcome Sanger Institute COVID-19 Surveillance Team |
| EPI_ISL_3027637 | Lighthouse Lab in Alderley Park | Wellcome Sanger Institute for the COVID-19 Genomics UK (COG-UK) Consortium | Jacquelyn Wynn, Mairead Hyland, The Lighthouse Lab in Alderley Park and Alex Alderton, Roberto Amato, Jeffrey Barrett, Sonia Goncalves, Ewan Harrison, David K. Jackson, Ian Johnston, Dominic Kwiatkowski, Cordelia Langford, John Sillitoe on behalf of the Wellcome Sanger Institute COVID-19 Surveillance Team |

| | | | |
|---|---|---|---|
| EPI_ISL_3027638 | Berkshire and Surrey Pathology Services Lighthouse Laboratory | Wellcome Sanger Institute for the COVID-19 Genomics UK (COG-UK) Consortium | Berkshire and Surrey Pathology Services Lighthouse Laboratory and Alex Alderton, Roberto Amato, Jeffrey Barrett, Sonia Goncalves, Ewan Harrison, David K. Jackson, Ian Johnston, Dominic Kwiatkowski, Cordelia Langford, John Sillitoe on behalf of the Wellcome Sanger Institute COVID-19 Surveillance Team |
| EPI_ISL_3027639 | Lighthouse Lab in Alderley Park | Wellcome Sanger Institute for the COVID-19 Genomics UK (COG-UK) Consortium | Jacquelyn Wynn, Mairead Hyland, The Lighthouse Lab in Alderley Park and Alex Alderton, Roberto Amato, Jeffrey Barrett, Sonia Goncalves, Ewan Harrison, David K. Jackson, Ian Johnston, Dominic Kwiatkowski, Cordelia Langford, John Sillitoe on behalf of the Wellcome Sanger Institute COVID-19 Surveillance Team |
| EPI_ISL_3027640 | Lighthouse Lab in Glasgow | Wellcome Sanger Institute for the COVID-19 Genomics UK (COG-UK) Consortium | Harper VanSteenhouse, Yumi Kasai, David Gray, Carol Clugston, Anna Dominiczak and Alex Alderton, Roberto Amato, Jeffrey Barrett, Sonia Goncalves, Ewan Harrison, David K. Jackson, Ian Johnston, Dominic Kwiatkowski, Cordelia Langford, John Sillitoe on behalf of the Wellcome Sanger Institute COVID-19 Surveillance Team |
| EPI_ISL_3027641 | Lighthouse Lab in Alderley Park | Wellcome Sanger Institute for the COVID-19 Genomics UK (COG-UK) Consortium | Jacquelyn Wynn, Mairead Hyland, The Lighthouse Lab in Alderley Park and Alex Alderton, Roberto Amato, Jeffrey Barrett, Sonia Goncalves, Ewan Harrison, David K. Jackson, Ian Johnston, Dominic Kwiatkowski, Cordelia Langford, John Sillitoe on behalf of the Wellcome Sanger Institute COVID-19 Surveillance Team |
| EPI_ISL_3027642 | Lighthouse Lab in Milton Keynes | Wellcome Sanger Institute for the COVID-19 Genomics UK (COG-UK) Consortium | The Lighthouse Lab in Milton Keynes and Alex Alderton, Roberto Amato, Jeffrey Barrett, Sonia Goncalves, Ewan Harrison, David K. Jackson, Ian Johnston, Dominic Kwiatkowski, Cordelia Langford, John Sillitoe on behalf of the Wellcome Sanger Institute COVID-19 Surveillance Team |
| EPI_ISL_3027643 | Berkshire and Surrey Pathology Services Lighthouse Laboratory | Wellcome Sanger Institute for the COVID-19 Genomics UK (COG-UK) Consortium | Berkshire and Surrey Pathology Services Lighthouse Laboratory and Alex Alderton, Roberto Amato, Jeffrey Barrett, Sonia Goncalves, Ewan Harrison, David K. Jackson, Ian Johnston, Dominic Kwiatkowski, Cordelia Langford, John Sillitoe on behalf of the Wellcome Sanger Institute COVID-19 Surveillance Team |
| EPI_ISL_3027644, EPI_ISL_3027645 | Lighthouse Lab in Alderley Park | Wellcome Sanger Institute for the COVID-19 Genomics UK (COG-UK) Consortium | Jacquelyn Wynn, Mairead Hyland, The Lighthouse Lab in Alderley Park and Alex Alderton, Roberto Amato, Jeffrey Barrett, Sonia Goncalves, Ewan Harrison, David K. Jackson, Ian Johnston, Dominic Kwiatkowski, Cordelia Langford, John Sillitoe on behalf of the Wellcome Sanger Institute COVID-19 Surveillance Team |
| EPI_ISL_3027646 | Berkshire and Surrey Pathology Services Lighthouse Laboratory | Wellcome Sanger Institute for the COVID-19 Genomics UK (COG-UK) Consortium | Berkshire and Surrey Pathology Services Lighthouse Laboratory and Alex Alderton, Roberto Amato, Jeffrey Barrett, Sonia Goncalves, Ewan Harrison, David K. Jackson, Ian Johnston, Dominic Kwiatkowski, Cordelia Langford, John Sillitoe on behalf of the Wellcome Sanger Institute COVID-19 Surveillance Team |
| EPI_ISL_3027647, EPI_ISL_3027648 | Lighthouse Lab in Milton Keynes | Wellcome Sanger Institute for the COVID-19 Genomics UK (COG-UK) Consortium | The Lighthouse Lab in Milton Keynes and Alex Alderton, Roberto Amato, Jeffrey Barrett, Sonia Goncalves, Ewan Harrison, David K. Jackson, Ian Johnston, Dominic Kwiatkowski, Cordelia Langford, John Sillitoe on behalf of the Wellcome Sanger Institute COVID-19 Surveillance Team |
| EPI_ISL_3027649 | Berkshire and Surrey Pathology Services Lighthouse Laboratory | Wellcome Sanger Institute for the COVID-19 Genomics UK (COG-UK) Consortium | Berkshire and Surrey Pathology Services Lighthouse Laboratory and Alex Alderton, Roberto Amato, Jeffrey Barrett, Sonia Goncalves, Ewan Harrison, David K. Jackson, Ian Johnston, Dominic Kwiatkowski, Cordelia Langford, John Sillitoe on behalf of the Wellcome Sanger Institute COVID-19 Surveillance Team |
| EPI_ISL_3027650 | Lighthouse Lab in Glasgow | Wellcome Sanger Institute for the COVID-19 Genomics UK (COG-UK) Consortium | Harper VanSteenhouse, Yumi Kasai, David Gray, Carol Clugston, Anna Dominiczak and Alex Alderton, Roberto Amato, Jeffrey Barrett, Sonia Goncalves, Ewan Harrison, David K. Jackson, Ian Johnston, Dominic Kwiatkowski, Cordelia Langford, John Sillitoe on behalf of the Wellcome Sanger Institute COVID-19 Surveillance Team |
| EPI_ISL_3027651, EPI_ISL_3027652, EPI_ISL_3027653 | Lighthouse Lab in Milton Keynes | Wellcome Sanger Institute for the COVID-19 Genomics UK (COG-UK) Consortium | The Lighthouse Lab in Milton Keynes and Alex Alderton, Roberto Amato, Jeffrey Barrett, Sonia Goncalves, Ewan Harrison, David K. Jackson, Ian Johnston, Dominic Kwiatkowski, Cordelia Langford, John Sillitoe on behalf of the Wellcome Sanger Institute COVID-19 Surveillance Team |
| EPI_ISL_3027654, EPI_ISL_3027655 | Lighthouse Lab in Alderley Park | Wellcome Sanger Institute for the COVID-19 Genomics UK (COG-UK) Consortium | Jacquelyn Wynn, Mairead Hyland, The Lighthouse Lab in Alderley Park and Alex Alderton, Roberto Amato, Jeffrey Barrett, Sonia Goncalves, Ewan Harrison, David K. Jackson, Ian Johnston, Dominic Kwiatkowski, Cordelia Langford, John Sillitoe on behalf of the Wellcome Sanger Institute COVID-19 Surveillance Team |
| EPI_ISL_3027656 | Lighthouse Lab in Milton Keynes | Wellcome Sanger Institute for the COVID-19 Genomics UK (COG-UK) Consortium | The Lighthouse Lab in Milton Keynes and Alex Alderton, Roberto Amato, Jeffrey Barrett, Sonia Goncalves, Ewan Harrison, David K. Jackson, Ian Johnston, Dominic Kwiatkowski, Cordelia Langford, John Sillitoe on behalf of the Wellcome Sanger Institute COVID-19 Surveillance Team |
| EPI_ISL_3027657 | Lighthouse Lab in Alderley Park | Wellcome Sanger Institute for the COVID-19 Genomics UK (COG-UK) Consortium | Jacquelyn Wynn, Mairead Hyland, The Lighthouse Lab in Alderley Park and Alex Alderton, Roberto Amato, Jeffrey Barrett, Sonia Goncalves, Ewan Harrison, David K. Jackson, Ian Johnston, Dominic Kwiatkowski, Cordelia Langford, John Sillitoe on behalf of the Wellcome Sanger Institute COVID-19 Surveillance Team |
| EPI_ISL_3027658, EPI_ISL_3027659 | Berkshire and Surrey Pathology Services Lighthouse Laboratory | Wellcome Sanger Institute for the COVID-19 Genomics UK (COG-UK) Consortium | Berkshire and Surrey Pathology Services Lighthouse Laboratory and Alex Alderton, Roberto Amato, Jeffrey Barrett, Sonia Goncalves, Ewan Harrison, David K. Jackson, Ian Johnston, Dominic Kwiatkowski, Cordelia Langford, John Sillitoe on behalf of the Wellcome Sanger Institute COVID-19 Surveillance Team |
| EPI_ISL_3027660 | Lighthouse Lab in Glasgow | Wellcome Sanger Institute for the COVID-19 Genomics UK (COG-UK) Consortium | Harper VanSteenhouse, Yumi Kasai, David Gray, Carol Clugston, Anna Dominiczak and Alex Alderton, Roberto Amato, Jeffrey Barrett, Sonia Goncalves, Ewan Harrison, David K. Jackson, Ian Johnston, Dominic Kwiatkowski, Cordelia Langford, John Sillitoe on behalf of the Wellcome Sanger Institute COVID-19 Surveillance Team |
| EPI_ISL_3027661 | Lighthouse Lab in Milton Keynes | Wellcome Sanger Institute for the COVID-19 Genomics UK (COG-UK) Consortium | The Lighthouse Lab in Milton Keynes and Alex Alderton, Roberto Amato, Jeffrey Barrett, Sonia Goncalves, Ewan Harrison, David K. Jackson, Ian Johnston, Dominic Kwiatkowski, Cordelia Langford, John Sillitoe on behalf of the Wellcome Sanger Institute COVID-19 Surveillance Team |
| EPI_ISL_3027662 | Berkshire and Surrey Pathology Services Lighthouse Laboratory | Wellcome Sanger Institute for the COVID-19 Genomics UK (COG-UK) Consortium | Berkshire and Surrey Pathology Services Lighthouse Laboratory and Alex Alderton, Roberto Amato, Jeffrey Barrett, Sonia Goncalves, Ewan Harrison, David K. Jackson, Ian Johnston, Dominic Kwiatkowski, Cordelia Langford, John Sillitoe on behalf of the Wellcome Sanger Institute COVID-19 Surveillance Team |
| EPI_ISL_3027663, EPI_ISL_3027664 | Lighthouse Lab in Alderley Park | Wellcome Sanger Institute for the COVID-19 Genomics UK (COG-UK) Consortium | Jacquelyn Wynn, Mairead Hyland, The Lighthouse Lab in Alderley Park and Alex Alderton, Roberto Amato, Jeffrey Barrett, Sonia Goncalves, Ewan Harrison, David K. Jackson, Ian Johnston, Dominic Kwiatkowski, Cordelia Langford, John Sillitoe on behalf of the Wellcome Sanger Institute COVID-19 Surveillance Team |
| EPI_ISL_3027665, EPI_ISL_3027666 | Lighthouse Lab in Milton Keynes | Wellcome Sanger Institute for the COVID-19 Genomics UK (COG-UK) Consortium | The Lighthouse Lab in Milton Keynes and Alex Alderton, Roberto Amato, Jeffrey Barrett, Sonia Goncalves, Ewan Harrison, David K. Jackson, Ian Johnston, Dominic Kwiatkowski, Cordelia Langford, John Sillitoe on behalf of the Wellcome Sanger Institute COVID-19 Surveillance Team |
| EPI_ISL_3027667 | Berkshire and Surrey Pathology Services Lighthouse Laboratory | Wellcome Sanger Institute for the COVID-19 Genomics UK (COG-UK) Consortium | Berkshire and Surrey Pathology Services Lighthouse Laboratory and Alex Alderton, Roberto Amato, Jeffrey Barrett, Sonia Goncalves, Ewan Harrison, David K. Jackson, Ian Johnston, Dominic Kwiatkowski, Cordelia Langford, John Sillitoe on behalf of the Wellcome Sanger Institute COVID-19 Surveillance Team |
| EPI_ISL_3027668, EPI_ISL_3027669 | Lighthouse Lab in Milton Keynes | Wellcome Sanger Institute for the COVID-19 Genomics UK (COG-UK) Consortium | The Lighthouse Lab in Milton Keynes and Alex Alderton, Roberto Amato, Jeffrey Barrett, Sonia Goncalves, Ewan Harrison, David K. Jackson, Ian Johnston, Dominic Kwiatkowski, Cordelia Langford, John Sillitoe on behalf of the Wellcome Sanger Institute COVID-19 Surveillance Team |
| EPI_ISL_3027670 | Lighthouse Lab in Glasgow | Wellcome Sanger Institute for the COVID-19 Genomics UK (COG-UK) Consortium | Harper VanSteenhouse, Yumi Kasai, David Gray, Carol Clugston, Anna Dominiczak and Alex Alderton, Roberto Amato, Jeffrey Barrett, Sonia Goncalves, Ewan Harrison, David K. Jackson, Ian Johnston, Dominic Kwiatkowski, Cordelia Langford, John Sillitoe on behalf of the Wellcome Sanger Institute COVID-19 Surveillance Team |
| EPI_ISL_3027671, EPI_ISL_3027672, EPI_ISL_3027673, EPI_ISL_3027674, EPI_ISL_3027675 | Lighthouse Lab in Milton Keynes | Wellcome Sanger Institute for the COVID-19 Genomics UK (COG-UK) Consortium | The Lighthouse Lab in Milton Keynes and Alex Alderton, Roberto Amato, Jeffrey Barrett, Sonia Goncalves, Ewan Harrison, David K. Jackson, Ian Johnston, Dominic Kwiatkowski, Cordelia Langford, John Sillitoe on behalf of the Wellcome Sanger Institute COVID-19 Surveillance Team |
| EPI_ISL_3027676 | Lighthouse Lab in Glasgow | Wellcome Sanger Institute for the COVID-19 Genomics UK (COG-UK) Consortium | Harper VanSteenhouse, Yumi Kasai, David Gray, Carol Clugston, Anna Dominiczak and Alex Alderton, Roberto Amato, Jeffrey Barrett, Sonia Goncalves, Ewan Harrison, David K. Jackson, Ian Johnston, Dominic Kwiatkowski, Cordelia Langford, John Sillitoe on behalf of the Wellcome Sanger Institute COVID-19 Surveillance Team |
| EPI_ISL_3027677, EPI_ISL_3027678, EPI_ISL_3027679, EPI_ISL_3027680 | Lighthouse Lab in Milton Keynes | Wellcome Sanger Institute for the COVID-19 Genomics UK (COG-UK) Consortium | The Lighthouse Lab in Milton Keynes and Alex Alderton, Roberto Amato, Jeffrey Barrett, Sonia Goncalves, Ewan Harrison, David K. Jackson, Ian Johnston, Dominic Kwiatkowski, Cordelia Langford, John Sillitoe on behalf of the Wellcome Sanger Institute COVID-19 Surveillance Team |
| EPI_ISL_3027681, EPI_ISL_3027682 | Lighthouse Lab in Alderley Park | Wellcome Sanger Institute for the COVID-19 Genomics UK (COG-UK) Consortium | Jacquelyn Wynn, Mairead Hyland, The Lighthouse Lab in Alderley Park and Alex Alderton, Roberto Amato, Jeffrey Barrett, Sonia Goncalves, Ewan Harrison, David K. Jackson, Ian Johnston, Dominic Kwiatkowski, Cordelia Langford, John Sillitoe on behalf of the Wellcome Sanger Institute COVID-19 Surveillance Team |
| EPI_ISL_3027683 | Berkshire and Surrey Pathology Services Lighthouse Laboratory | Wellcome Sanger Institute for the COVID-19 Genomics UK (COG-UK) Consortium | Berkshire and Surrey Pathology Services Lighthouse Laboratory and Alex Alderton, Roberto Amato, Jeffrey Barrett, Sonia Goncalves, Ewan Harrison, David K. Jackson, Ian Johnston, Dominic Kwiatkowski, Cordelia Langford, John Sillitoe on behalf of the Wellcome Sanger Institute COVID-19 Surveillance Team |
| EPI_ISL_3027684 | Lighthouse Lab in Alderley Park | Wellcome Sanger Institute for the COVID-19 Genomics UK (COG-UK) Consortium | Jacquelyn Wynn, Mairead Hyland, The Lighthouse Lab in Alderley Park and Alex Alderton, Roberto Amato, Jeffrey Barrett, Sonia Goncalves, Ewan Harrison, David K. Jackson, Ian Johnston, Dominic Kwiatkowski, Cordelia Langford, John Sillitoe on behalf of the Wellcome Sanger Institute COVID-19 Surveillance Team |
| EPI_ISL_3027685 | Lighthouse Lab in Glasgow | Wellcome Sanger Institute for the COVID-19 Genomics UK (COG-UK) Consortium | Harper VanSteenhouse, Yumi Kasai, David Gray, Carol Clugston, Anna Dominiczak and Alex Alderton, Roberto Amato, Jeffrey Barrett, Sonia Goncalves, Ewan Harrison, David K. Jackson, Ian Johnston, Dominic Kwiatkowski, Cordelia Langford, John Sillitoe on behalf of the Wellcome Sanger Institute COVID-19 Surveillance Team |
| EPI_ISL_3027686 | Lighthouse Lab in Alderley Park | Wellcome Sanger Institute for the COVID-19 Genomics UK (COG-UK) Consortium | Jacquelyn Wynn, Mairead Hyland, The Lighthouse Lab in Alderley Park and Alex Alderton, Roberto Amato, Jeffrey Barrett, Sonia Goncalves, Ewan Harrison, David K. Jackson, Ian Johnston, Dominic Kwiatkowski, Cordelia Langford, John Sillitoe on behalf of the Wellcome Sanger Institute COVID-19 Surveillance Team |
| EPI_ISL_3027687 | Lighthouse Lab in Glasgow | Wellcome Sanger Institute for the COVID-19 Genomics UK (COG-UK) Consortium | Harper VanSteenhouse, Yumi Kasai, David Gray, Carol Clugston, Anna Dominiczak and Alex Alderton, Roberto Amato, Jeffrey Barrett, Sonia Goncalves, Ewan Harrison, David K. Jackson, Ian Johnston, Dominic Kwiatkowski, Cordelia Langford, John Sillitoe on behalf of the Wellcome Sanger Institute COVID-19 Surveillance Team |
| EPI_ISL_3027688 | Lighthouse Lab in Milton Keynes | Wellcome Sanger Institute for the COVID-19 Genomics UK (COG-UK) Consortium | The Lighthouse Lab in Milton Keynes and Alex Alderton, Roberto Amato, Jeffrey Barrett, Sonia Goncalves, Ewan Harrison, David K. Jackson, Ian Johnston, Dominic Kwiatkowski, Cordelia Langford, John Sillitoe on behalf of the Wellcome Sanger Institute COVID-19 Surveillance Team |
| EPI_ISL_3027689 | Berkshire and Surrey Pathology Services Lighthouse Laboratory | Wellcome Sanger Institute for the COVID-19 Genomics UK (COG-UK) Consortium | Berkshire and Surrey Pathology Services Lighthouse Laboratory and Alex Alderton, Roberto Amato, Jeffrey Barrett, Sonia Goncalves, Ewan Harrison, David K. Jackson, Ian Johnston, Dominic Kwiatkowski, Cordelia Langford, John Sillitoe on behalf of the Wellcome Sanger Institute COVID-19 Surveillance Team |
| EPI_ISL_3027690 | Lighthouse Lab in Glasgow | Wellcome Sanger Institute for the COVID-19 Genomics UK (COG-UK) Consortium | Harper VanSteenhouse, Yumi Kasai, David Gray, Carol Clugston, Anna Dominiczak and Alex Alderton, Roberto Amato, Jeffrey Barrett, Sonia Goncalves, Ewan Harrison, David K. Jackson, Ian Johnston, Dominic Kwiatkowski, Cordelia Langford, John Sillitoe on behalf of the Wellcome Sanger Institute COVID-19 Surveillance Team |
| EPI_ISL_3027691 | Berkshire and Surrey Pathology Services Lighthouse Laboratory | Wellcome Sanger Institute for the COVID-19 Genomics UK (COG-UK) Consortium | Berkshire and Surrey Pathology Services Lighthouse Laboratory and Alex Alderton, Roberto Amato, Jeffrey Barrett, Sonia Goncalves, Ewan Harrison, David K. Jackson, Ian Johnston, Dominic Kwiatkowski, Cordelia Langford, John Sillitoe on behalf of the Wellcome Sanger Institute COVID-19 Surveillance Team |
| EPI_ISL_3027692 | Lighthouse Lab in Alderley Park | Wellcome Sanger Institute for the COVID-19 Genomics UK (COG-UK) Consortium | Jacquelyn Wynn, Mairead Hyland, The Lighthouse Lab in Alderley Park and Alex Alderton, Roberto Amato, Jeffrey Barrett, Sonia Goncalves, Ewan Harrison, David K. Jackson, Ian Johnston, Dominic Kwiatkowski, Cordelia Langford, John Sillitoe on behalf of the Wellcome Sanger Institute COVID-19 Surveillance Team |
| EPI_ISL_3027693 | Berkshire and Surrey Pathology Services Lighthouse Laboratory | Wellcome Sanger Institute for the COVID-19 Genomics UK (COG-UK) Consortium | Berkshire and Surrey Pathology Services Lighthouse Laboratory and Alex Alderton, Roberto Amato, Jeffrey Barrett, Sonia Goncalves, Ewan Harrison, David K. Jackson, Ian Johnston, Dominic Kwiatkowski, Cordelia Langford, John Sillitoe on behalf of the Wellcome Sanger Institute COVID-19 Surveillance Team |
| EPI_ISL_3027694 | Lighthouse Lab in Alderley Park | Wellcome Sanger Institute for the COVID-19 Genomics UK (COG-UK) Consortium | Jacquelyn Wynn, Mairead Hyland, The Lighthouse Lab in Alderley Park and Alex Alderton, Roberto Amato, Jeffrey Barrett, Sonia Goncalves, Ewan Harrison, David K. Jackson, Ian Johnston, Dominic Kwiatkowski, Cordelia Langford, John Sillitoe on behalf of the Wellcome Sanger Institute COVID-19 Surveillance Team |
| EPI_ISL_3027695 | Lighthouse Lab in Glasgow | Wellcome Sanger Institute for the COVID-19 Genomics UK (COG-UK) Consortium | Harper VanSteenhouse, Yumi Kasai, David Gray, Carol Clugston, Anna Dominiczak and Alex Alderton, Roberto Amato, Jeffrey Barrett, Sonia Goncalves, Ewan Harrison, David K. Jackson, Ian Johnston, Dominic Kwiatkowski, Cordelia Langford, John Sillitoe on behalf of the Wellcome Sanger Institute COVID-19 Surveillance Team |
| EPI_ISL_3027696 | Lighthouse Lab in Alderley Park | Wellcome Sanger Institute for the COVID-19 Genomics UK (COG-UK) Consortium | Jacquelyn Wynn, Mairead Hyland, The Lighthouse Lab in Alderley Park and Alex Alderton, Roberto Amato, Jeffrey Barrett, Sonia Goncalves, Ewan Harrison, David K. Jackson, Ian Johnston, Dominic Kwiatkowski, Cordelia Langford, John Sillitoe on behalf of the Wellcome Sanger Institute COVID-19 Surveillance Team |
| EPI_ISL_3027697 | Lighthouse Lab in Milton Keynes | Wellcome Sanger Institute for the COVID-19 Genomics UK (COG-UK) Consortium | The Lighthouse Lab in Milton Keynes and Alex Alderton, Roberto Amato, Jeffrey Barrett, Sonia Goncalves, Ewan Harrison, David K. Jackson, Ian Johnston, Dominic Kwiatkowski, Cordelia Langford, John Sillitoe on behalf of the Wellcome Sanger Institute COVID-19 Surveillance Team |

| | | | |
|---|---|---|---|
| EPI_ISL_3027698 | Lighthouse Lab in Alderley Park | Wellcome Sanger Institute for the COVID-19 Genomics UK (COG-UK) Consortium | Jacquelyn Wynn, Mairead Hyland, The Lighthouse Lab in Alderley Park and Alex Alderton, Roberto Amato, Jeffrey Barrett, Sonia Goncalves, Ewan Harrison, David K. Jackson, Ian Johnston, Dominic Kwiatkowski, Cordelia Langford, John Sillitoe on behalf of the Wellcome Sanger Institute COVID-19 Surveillance Team |
| EPI_ISL_3027699 | Lighthouse Lab in Milton Keynes | Wellcome Sanger Institute for the COVID-19 Genomics UK (COG-UK) Consortium | The Lighthouse Lab in Milton Keynes and Alex Alderton, Roberto Amato, Jeffrey Barrett, Sonia Goncalves, Ewan Harrison, David K. Jackson, Ian Johnston, Dominic Kwiatkowski, Cordelia Langford, John Sillitoe on behalf of the Wellcome Sanger Institute COVID-19 Surveillance Team |
| EPI_ISL_3027700 | Lighthouse Lab in Alderley Park | Wellcome Sanger Institute for the COVID-19 Genomics UK (COG-UK) Consortium | Jacquelyn Wynn, Mairead Hyland, The Lighthouse Lab in Alderley Park and Alex Alderton, Roberto Amato, Jeffrey Barrett, Sonia Goncalves, Ewan Harrison, David K. Jackson, Ian Johnston, Dominic Kwiatkowski, Cordelia Langford, John Sillitoe on behalf of the Wellcome Sanger Institute COVID-19 Surveillance Team |
| EPI_ISL_3027701, EPI_ISL_3027702 | Berkshire and Surrey Pathology Services Lighthouse Laboratory | Wellcome Sanger Institute for the COVID-19 Genomics UK (COG-UK) Consortium | Berkshire and Surrey Pathology Services Lighthouse Laboratory and Alex Alderton, Roberto Amato, Jeffrey Barrett, Sonia Goncalves, Ewan Harrison, David K. Jackson, Ian Johnston, Dominic Kwiatkowski, Cordelia Langford, John Sillitoe on behalf of the Wellcome Sanger Institute COVID-19 Surveillance Team |
| EPI_ISL_3027703, EPI_ISL_3027704, EPI_ISL_3027705 | Lighthouse Lab in Milton Keynes | Wellcome Sanger Institute for the COVID-19 Genomics UK (COG-UK) Consortium | The Lighthouse Lab in Milton Keynes and Alex Alderton, Roberto Amato, Jeffrey Barrett, Sonia Goncalves, Ewan Harrison, David K. Jackson, Ian Johnston, Dominic Kwiatkowski, Cordelia Langford, John Sillitoe on behalf of the Wellcome Sanger Institute COVID-19 Surveillance Team |
| EPI_ISL_3027706 | Lighthouse Lab in Glasgow | Wellcome Sanger Institute for the COVID-19 Genomics UK (COG-UK) Consortium | Harper VanSteenhouse, Yumi Kasai, David Gray, Carol Clugston, Anna Dominiczak and Alex Alderton, Roberto Amato, Jeffrey Barrett, Sonia Goncalves, Ewan Harrison, David K. Jackson, Ian Johnston, Dominic Kwiatkowski, Cordelia Langford, John Sillitoe on behalf of the Wellcome Sanger Institute COVID-19 Surveillance Team |
| EPI_ISL_3027707, EPI_ISL_3027708 | Lighthouse Lab in Milton Keynes | Wellcome Sanger Institute for the COVID-19 Genomics UK (COG-UK) Consortium | The Lighthouse Lab in Milton Keynes and Alex Alderton, Roberto Amato, Jeffrey Barrett, Sonia Goncalves, Ewan Harrison, David K. Jackson, Ian Johnston, Dominic Kwiatkowski, Cordelia Langford, John Sillitoe on behalf of the Wellcome Sanger Institute COVID-19 Surveillance Team |
| EPI_ISL_3027709 | Lighthouse Lab in Alderley Park | Wellcome Sanger Institute for the COVID-19 Genomics UK (COG-UK) Consortium | Jacquelyn Wynn, Mairead Hyland, The Lighthouse Lab in Alderley Park and Alex Alderton, Roberto Amato, Jeffrey Barrett, Sonia Goncalves, Ewan Harrison, David K. Jackson, Ian Johnston, Dominic Kwiatkowski, Cordelia Langford, John Sillitoe on behalf of the Wellcome Sanger Institute COVID-19 Surveillance Team |
| EPI_ISL_3027710 | Lighthouse Lab in Glasgow | Wellcome Sanger Institute for the COVID-19 Genomics UK (COG-UK) Consortium | Harper VanSteenhouse, Yumi Kasai, David Gray, Carol Clugston, Anna Dominiczak and Alex Alderton, Roberto Amato, Jeffrey Barrett, Sonia Goncalves, Ewan Harrison, David K. Jackson, Ian Johnston, Dominic Kwiatkowski, Cordelia Langford, John Sillitoe on behalf of the Wellcome Sanger Institute COVID-19 Surveillance Team |
| EPI_ISL_3027711 | Berkshire and Surrey Pathology Services Lighthouse Laboratory | Wellcome Sanger Institute for the COVID-19 Genomics UK (COG-UK) Consortium | Berkshire and Surrey Pathology Services Lighthouse Laboratory and Alex Alderton, Roberto Amato, Jeffrey Barrett, Sonia Goncalves, Ewan Harrison, David K. Jackson, Ian Johnston, Dominic Kwiatkowski, Cordelia Langford, John Sillitoe on behalf of the Wellcome Sanger Institute COVID-19 Surveillance Team |
| EPI_ISL_3027712, EPI_ISL_3027713, EPI_ISL_3027714 | Lighthouse Lab in Alderley Park | Wellcome Sanger Institute for the COVID-19 Genomics UK (COG-UK) Consortium | Jacquelyn Wynn, Mairead Hyland, The Lighthouse Lab in Alderley Park and Alex Alderton, Roberto Amato, Jeffrey Barrett, Sonia Goncalves, Ewan Harrison, David K. Jackson, Ian Johnston, Dominic Kwiatkowski, Cordelia Langford, John Sillitoe on behalf of the Wellcome Sanger Institute COVID-19 Surveillance Team |
| EPI_ISL_3027715 | Lighthouse Lab in Glasgow | Wellcome Sanger Institute for the COVID-19 Genomics UK (COG-UK) Consortium | Harper VanSteenhouse, Yumi Kasai, David Gray, Carol Clugston, Anna Dominiczak and Alex Alderton, Roberto Amato, Jeffrey Barrett, Sonia Goncalves, Ewan Harrison, David K. Jackson, Ian Johnston, Dominic Kwiatkowski, Cordelia Langford, John Sillitoe on behalf of the Wellcome Sanger Institute COVID-19 Surveillance Team |
| EPI_ISL_3027716 | Lighthouse Lab in Alderley Park | Wellcome Sanger Institute for the COVID-19 Genomics UK (COG-UK) Consortium | Jacquelyn Wynn, Mairead Hyland, The Lighthouse Lab in Alderley Park and Alex Alderton, Roberto Amato, Jeffrey Barrett, Sonia Goncalves, Ewan Harrison, David K. Jackson, Ian Johnston, Dominic Kwiatkowski, Cordelia Langford, John Sillitoe on behalf of the Wellcome Sanger Institute COVID-19 Surveillance Team |
| EPI_ISL_3027717 | Lighthouse Lab in Milton Keynes | Wellcome Sanger Institute for the COVID-19 Genomics UK (COG-UK) Consortium | The Lighthouse Lab in Milton Keynes and Alex Alderton, Roberto Amato, Jeffrey Barrett, Sonia Goncalves, Ewan Harrison, David K. Jackson, Ian Johnston, Dominic Kwiatkowski, Cordelia Langford, John Sillitoe on behalf of the Wellcome Sanger Institute COVID-19 Surveillance Team |
| EPI_ISL_3027718 | Lighthouse Lab in Glasgow | Wellcome Sanger Institute for the COVID-19 Genomics UK (COG-UK) Consortium | Harper VanSteenhouse, Yumi Kasai, David Gray, Carol Clugston, Anna Dominiczak and Alex Alderton, Roberto Amato, Jeffrey Barrett, Sonia Goncalves, Ewan Harrison, David K. Jackson, Ian Johnston, Dominic Kwiatkowski, Cordelia Langford, John Sillitoe on behalf of the Wellcome Sanger Institute COVID-19 Surveillance Team |
| EPI_ISL_3027719, EPI_ISL_3027720, EPI_ISL_3027721 | Lighthouse Lab in Alderley Park | Wellcome Sanger Institute for the COVID-19 Genomics UK (COG-UK) Consortium | Jacquelyn Wynn, Mairead Hyland, The Lighthouse Lab in Alderley Park and Alex Alderton, Roberto Amato, Jeffrey Barrett, Sonia Goncalves, Ewan Harrison, David K. Jackson, Ian Johnston, Dominic Kwiatkowski, Cordelia Langford, John Sillitoe on behalf of the Wellcome Sanger Institute COVID-19 Surveillance Team |
| EPI_ISL_3027722 | Lighthouse Lab in Glasgow | Wellcome Sanger Institute for the COVID-19 Genomics UK (COG-UK) Consortium | Harper VanSteenhouse, Yumi Kasai, David Gray, Carol Clugston, Anna Dominiczak and Alex Alderton, Roberto Amato, Jeffrey Barrett, Sonia Goncalves, Ewan Harrison, David K. Jackson, Ian Johnston, Dominic Kwiatkowski, Cordelia Langford, John Sillitoe on behalf of the Wellcome Sanger Institute COVID-19 Surveillance Team |
| EPI_ISL_3027723 | Lighthouse Lab in Milton Keynes | Wellcome Sanger Institute for the COVID-19 Genomics UK (COG-UK) Consortium | The Lighthouse Lab in Milton Keynes and Alex Alderton, Roberto Amato, Jeffrey Barrett, Sonia Goncalves, Ewan Harrison, David K. Jackson, Ian Johnston, Dominic Kwiatkowski, Cordelia Langford, John Sillitoe on behalf of the Wellcome Sanger Institute COVID-19 Surveillance Team |
| EPI_ISL_3027724 | Lighthouse Lab in Alderley Park | Wellcome Sanger Institute for the COVID-19 Genomics UK (COG-UK) Consortium | Jacquelyn Wynn, Mairead Hyland, The Lighthouse Lab in Alderley Park and Alex Alderton, Roberto Amato, Jeffrey Barrett, Sonia Goncalves, Ewan Harrison, David K. Jackson, Ian Johnston, Dominic Kwiatkowski, Cordelia Langford, John Sillitoe on behalf of the Wellcome Sanger Institute COVID-19 Surveillance Team |
| EPI_ISL_3027725, EPI_ISL_3027726 | Berkshire and Surrey Pathology Services Lighthouse Laboratory | Wellcome Sanger Institute for the COVID-19 Genomics UK (COG-UK) Consortium | Berkshire and Surrey Pathology Services Lighthouse Laboratory and Alex Alderton, Roberto Amato, Jeffrey Barrett, Sonia Goncalves, Ewan Harrison, David K. Jackson, Ian Johnston, Dominic Kwiatkowski, Cordelia Langford, John Sillitoe on behalf of the Wellcome Sanger Institute COVID-19 Surveillance Team |
| EPI_ISL_3027727 | Lighthouse Lab in Glasgow | Wellcome Sanger Institute for the COVID-19 Genomics UK (COG-UK) Consortium | Harper VanSteenhouse, Yumi Kasai, David Gray, Carol Clugston, Anna Dominiczak and Alex Alderton, Roberto Amato, Jeffrey Barrett, Sonia Goncalves, Ewan Harrison, David K. Jackson, Ian Johnston, Dominic Kwiatkowski, Cordelia Langford, John Sillitoe on behalf of the Wellcome Sanger Institute COVID-19 Surveillance Team |
| EPI_ISL_3027728 | Berkshire and Surrey Pathology Services Lighthouse Laboratory | Wellcome Sanger Institute for the COVID-19 Genomics UK (COG-UK) Consortium | Berkshire and Surrey Pathology Services Lighthouse Laboratory and Alex Alderton, Roberto Amato, Jeffrey Barrett, Sonia Goncalves, Ewan Harrison, David K. Jackson, Ian Johnston, Dominic Kwiatkowski, Cordelia Langford, John Sillitoe on behalf of the Wellcome Sanger Institute COVID-19 Surveillance Team |
| EPI_ISL_3027729 | Lighthouse Lab in Milton Keynes | Wellcome Sanger Institute for the COVID-19 Genomics UK (COG-UK) Consortium | The Lighthouse Lab in Milton Keynes and Alex Alderton, Roberto Amato, Jeffrey Barrett, Sonia Goncalves, Ewan Harrison, David K. Jackson, Ian Johnston, Dominic Kwiatkowski, Cordelia Langford, John Sillitoe on behalf of the Wellcome Sanger Institute COVID-19 Surveillance Team |
| EPI_ISL_3027730 | Lighthouse Lab in Glasgow | Wellcome Sanger Institute for the COVID-19 Genomics UK (COG-UK) Consortium | Harper VanSteenhouse, Yumi Kasai, David Gray, Carol Clugston, Anna Dominiczak and Alex Alderton, Roberto Amato, Jeffrey Barrett, Sonia Goncalves, Ewan Harrison, David K. Jackson, Ian Johnston, Dominic Kwiatkowski, Cordelia Langford, John Sillitoe on behalf of the Wellcome Sanger Institute COVID-19 Surveillance Team |
| EPI_ISL_3027731 | Berkshire and Surrey Pathology Services Lighthouse Laboratory | Wellcome Sanger Institute for the COVID-19 Genomics UK (COG-UK) Consortium | Berkshire and Surrey Pathology Services Lighthouse Laboratory and Alex Alderton, Roberto Amato, Jeffrey Barrett, Sonia Goncalves, Ewan Harrison, David K. Jackson, Ian Johnston, Dominic Kwiatkowski, Cordelia Langford, John Sillitoe on behalf of the Wellcome Sanger Institute COVID-19 Surveillance Team |
| EPI_ISL_3027732 | Lighthouse Lab in Milton Keynes | Wellcome Sanger Institute for the COVID-19 Genomics UK (COG-UK) Consortium | The Lighthouse Lab in Milton Keynes and Alex Alderton, Roberto Amato, Jeffrey Barrett, Sonia Goncalves, Ewan Harrison, David K. Jackson, Ian Johnston, Dominic Kwiatkowski, Cordelia Langford, John Sillitoe on behalf of the Wellcome Sanger Institute COVID-19 Surveillance Team |
| EPI_ISL_3027733 | Lighthouse Lab in Alderley Park | Wellcome Sanger Institute for the COVID-19 Genomics UK (COG-UK) Consortium | Jacquelyn Wynn, Mairead Hyland, The Lighthouse Lab in Alderley Park and Alex Alderton, Roberto Amato, Jeffrey Barrett, Sonia Goncalves, Ewan Harrison, David K. Jackson, Ian Johnston, Dominic Kwiatkowski, Cordelia Langford, John Sillitoe on behalf of the Wellcome Sanger Institute COVID-19 Surveillance Team |
| EPI_ISL_3027734 | Lighthouse Lab in Milton Keynes | Wellcome Sanger Institute for the COVID-19 Genomics UK (COG-UK) Consortium | The Lighthouse Lab in Milton Keynes and Alex Alderton, Roberto Amato, Jeffrey Barrett, Sonia Goncalves, Ewan Harrison, David K. Jackson, Ian Johnston, Dominic Kwiatkowski, Cordelia Langford, John Sillitoe on behalf of the Wellcome Sanger Institute COVID-19 Surveillance Team |
| EPI_ISL_3027735 | Berkshire and Surrey Pathology Services Lighthouse Laboratory | Wellcome Sanger Institute for the COVID-19 Genomics UK (COG-UK) Consortium | Berkshire and Surrey Pathology Services Lighthouse Laboratory and Alex Alderton, Roberto Amato, Jeffrey Barrett, Sonia Goncalves, Ewan Harrison, David K. Jackson, Ian Johnston, Dominic Kwiatkowski, Cordelia Langford, John Sillitoe on behalf of the Wellcome Sanger Institute COVID-19 Surveillance Team |
| EPI_ISL_3027736 | Lighthouse Lab in Alderley Park | Wellcome Sanger Institute for the COVID-19 Genomics UK (COG-UK) Consortium | Jacquelyn Wynn, Mairead Hyland, The Lighthouse Lab in Alderley Park and Alex Alderton, Roberto Amato, Jeffrey Barrett, Sonia Goncalves, Ewan Harrison, David K. Jackson, Ian Johnston, Dominic Kwiatkowski, Cordelia Langford, John Sillitoe on behalf of the Wellcome Sanger Institute COVID-19 Surveillance Team |
| EPI_ISL_3027737, EPI_ISL_3027738 | Lighthouse Lab in Glasgow | Wellcome Sanger Institute for the COVID-19 Genomics UK (COG-UK) Consortium | Harper VanSteenhouse, Yumi Kasai, David Gray, Carol Clugston, Anna Dominiczak and Alex Alderton, Roberto Amato, Jeffrey Barrett, Sonia Goncalves, Ewan Harrison, David K. Jackson, Ian Johnston, Dominic Kwiatkowski, Cordelia Langford, John Sillitoe on behalf of the Wellcome Sanger Institute COVID-19 Surveillance Team |
| EPI_ISL_3027739, EPI_ISL_3027740 | Lighthouse Lab in Milton Keynes | Wellcome Sanger Institute for the COVID-19 Genomics UK (COG-UK) Consortium | The Lighthouse Lab in Milton Keynes and Alex Alderton, Roberto Amato, Jeffrey Barrett, Sonia Goncalves, Ewan Harrison, David K. Jackson, Ian Johnston, Dominic Kwiatkowski, Cordelia Langford, John Sillitoe on behalf of the Wellcome Sanger Institute COVID-19 Surveillance Team |
| EPI_ISL_3027741, EPI_ISL_3027742 | Lighthouse Lab in Alderley Park | Wellcome Sanger Institute for the COVID-19 Genomics UK (COG-UK) Consortium | Jacquelyn Wynn, Mairead Hyland, The Lighthouse Lab in Alderley Park and Alex Alderton, Roberto Amato, Jeffrey Barrett, Sonia Goncalves, Ewan Harrison, David K. Jackson, Ian Johnston, Dominic Kwiatkowski, Cordelia Langford, John Sillitoe on behalf of the Wellcome Sanger Institute COVID-19 Surveillance Team |
| EPI_ISL_3027743 | Berkshire and Surrey Pathology Services Lighthouse Laboratory | Wellcome Sanger Institute for the COVID-19 Genomics UK (COG-UK) Consortium | Berkshire and Surrey Pathology Services Lighthouse Laboratory and Alex Alderton, Roberto Amato, Jeffrey Barrett, Sonia Goncalves, Ewan Harrison, David K. Jackson, Ian Johnston, Dominic Kwiatkowski, Cordelia Langford, John Sillitoe on behalf of the Wellcome Sanger Institute COVID-19 Surveillance Team |
| EPI_ISL_3027744 | Lighthouse Lab in Glasgow | Wellcome Sanger Institute for the COVID-19 Genomics UK (COG-UK) Consortium | Harper VanSteenhouse, Yumi Kasai, David Gray, Carol Clugston, Anna Dominiczak and Alex Alderton, Roberto Amato, Jeffrey Barrett, Sonia Goncalves, Ewan Harrison, David K. Jackson, Ian Johnston, Dominic Kwiatkowski, Cordelia Langford, John Sillitoe on behalf of the Wellcome Sanger Institute COVID-19 Surveillance Team |
| EPI_ISL_3027745 | Berkshire and Surrey Pathology Services Lighthouse Laboratory | Wellcome Sanger Institute for the COVID-19 Genomics UK (COG-UK) Consortium | Berkshire and Surrey Pathology Services Lighthouse Laboratory and Alex Alderton, Roberto Amato, Jeffrey Barrett, Sonia Goncalves, Ewan Harrison, David K. Jackson, Ian Johnston, Dominic Kwiatkowski, Cordelia Langford, John Sillitoe on behalf of the Wellcome Sanger Institute COVID-19 Surveillance Team |
| EPI_ISL_3027746 | Lighthouse Lab in Milton Keynes | Wellcome Sanger Institute for the COVID-19 Genomics UK (COG-UK) Consortium | The Lighthouse Lab in Milton Keynes and Alex Alderton, Roberto Amato, Jeffrey Barrett, Sonia Goncalves, Ewan Harrison, David K. Jackson, Ian Johnston, Dominic Kwiatkowski, Cordelia Langford, John Sillitoe on behalf of the Wellcome Sanger Institute COVID-19 Surveillance Team |
| EPI_ISL_3027747, EPI_ISL_3027748, EPI_ISL_3027749, EPI_ISL_3027750, EPI_ISL_3027751, EPI_ISL_3027752, EPI_ISL_3027753, EPI_ISL_3027754, EPI_ISL_3027755, EPI_ISL_3027756, EPI_ISL_3027757, EPI_ISL_3027758, EPI_ISL_3027759, EPI_ISL_3027760, EPI_ISL_3027761, EPI_ISL_3027762, EPI_ISL_3027763, EPI_ISL_3027764, EPI_ISL_3027765, EPI_ISL_3027766, EPI_ISL_3027767, EPI_ISL_3027768, EPI_ISL_3027769, EPI_ISL_3027770, EPI_ISL_3027771, EPI_ISL_3027772, EPI_ISL_3027773, EPI_ISL_3027774, EPI_ISL_3027775, EPI_ISL_3027776, EPI_ISL_3027777, EPI_ISL_3027778, EPI_ISL_3027779, EPI_ISL_3027780, EPI_ISL_3027781, EPI_ISL_3027782, EPI_ISL_3027783, EPI_ISL_3027784, EPI_ISL_3027785, EPI_ISL_3027786, EPI_ISL_3027787, EPI_ISL_3027788, EPI_ISL_3027789, EPI_ISL_3027790, EPI_ISL_3027791, EPI_ISL_3027792, EPI_ISL_3027793, EPI_ISL_3027794, EPI_ISL_3027795, EPI_ISL_3027796, EPI_ISL_3027797, EPI_ISL_3027798, EPI_ISL_3027799, EPI_ISL_3027800, EPI_ISL_3027801, EPI_ISL_3027802, EPI_ISL_3027803, EPI_ISL_3027804, EPI_ISL_3027805, EPI_ISL_3027806, EPI_ISL_3027807, EPI_ISL_3027808, EPI_ISL_3027809, EPI_ISL_3027810, EPI_ISL_3027811, EPI_ISL_3027812, EPI_ISL_3027813, EPI_ISL_3027814, EPI_ISL_3027815, EPI_ISL_3027816, EPI_ISL_3027817, EPI_ISL_3027818, EPI_ISL_3027819, EPI_ISL_3027820, EPI_ISL_3027821, EPI_ISL_3027822, EPI_ISL_3027823, EPI_ISL_3027824, EPI_ISL_3027825, EPI_ISL_3027826, EPI_ISL_3027827, EPI_ISL_3027828, EPI_ISL_3027829, EPI_ISL_3027830, EPI_ISL_3027831, EPI_ISL_3027832, EPI_ISL_3027833, EPI_ISL_3027834, EPI_ISL_3027835, EPI_ISL_3027836, EPI_ISL_3027837, EPI_ISL_3027838, EPI_ISL_3027839, EPI_ISL_3027840, EPI_ISL_3027841, EPI_ISL_3027842, EPI_ISL_3027843, EPI_ISL_3027844, EPI_ISL_3027845, EPI_ISL_3027846, EPI_ISL_3027847, EPI_ISL_3027848, EPI_ISL_3027849, EPI_ISL_3027850, EPI_ISL_3027851, EPI_ISL_3027852, EPI_ISL_3027853, EPI_ISL_3027854, EPI_ISL_3027855, EPI_ISL_3027857, EPI_ISL_3027858, EPI_ISL_3027859, EPI_ISL_3027860, EPI_ISL_3027861, EPI_ISL_3027862, EPI_ISL_3027863, EPI_ISL_3027865, EPI_ISL_3027866, EPI_ISL_3027867, EPI_ISL_3027868, EPI_ISL_3027869, EPI_ISL_3027870, EPI_ISL_3027871, EPI_ISL_3027872, EPI_ISL_3027873, EPI_ISL_3027874, EPI_ISL_3027875, EPI_ISL_3027876, EPI_ISL_3027877, EPI_ISL_3027878, EPI_ISL_3027879, EPI_ISL_3027880, EPI_ISL_3027881, EPI_ISL_3027882, EPI_ISL_3027883, EPI_ISL_3027884, EPI_ISL_3027885, EPI_ISL_3027886, EPI_ISL_3027887, EPI_ISL_3027888, EPI_ISL_3027889, EPI_ISL_3027890, EPI_ISL_3027891, EPI_ISL_3027892, EPI_ISL_3027893, EPI_ISL_3027894 | | | |
| see above | Lighthouse Lab in Glasgow | Wellcome Sanger Institute for the COVID-19 Genomics UK (COG-UK) Consortium | Harper VanSteenhouse, Yumi Kasai, David Gray, Carol Clugston, Anna Dominiczak and Alex Alderton, Roberto Amato, Jeffrey Barrett, Sonia Goncalves, Ewan Harrison, David K. Jackson, Ian Johnston, Dominic Kwiatkowski, Cordelia Langford, John Sillitoe on behalf of the Wellcome Sanger Institute COVID-19 Surveillance Team |
| EPI_ISL_3027895, EPI_ISL_3027896, EPI_ISL_3027897, EPI_ISL_3027898, EPI_ISL_3027899, EPI_ISL_3027900 | Berkshire and Surrey Pathology Services Lighthouse Laboratory | Wellcome Sanger Institute for the COVID-19 Genomics UK (COG-UK) Consortium | Berkshire and Surrey Pathology Services Lighthouse Laboratory and Alex Alderton, Roberto Amato, Jeffrey Barrett, Sonia Goncalves, Ewan Harrison, David K. Jackson, Ian Johnston, Dominic Kwiatkowski, Cordelia Langford, John Sillitoe on behalf of the Wellcome Sanger Institute COVID-19 Surveillance Team |

| | | | |
|---|---|---|---|
| EPI_ISL_3027901, EPI_ISL_3027902, EPI_ISL_3027903, EPI_ISL_3027904, EPI_ISL_3027905, EPI_ISL_3027906, EPI_ISL_3027907 | Lighthouse Lab in Glasgow | Wellcome Sanger Institute for the COVID-19 Genomics UK (COG-UK) Consortium | Harper VanSteenhouse, Yumi Kasai, David Gray, Carol Clugston, Anna Dominiczak and Alex Alderton, Roberto Amato, Jeffrey Barrett, Sonia Goncalves, Ewan Harrison, David K. Jackson, Ian Johnston, Dominic Kwiatkowski, Cordelia Langford, John Sillitoe on behalf of the Wellcome Sanger Institute COVID-19 Surveillance Team |
| EPI_ISL_3027908, EPI_ISL_3027909 | Lighthouse Lab in Alderley Park | Wellcome Sanger Institute for the COVID-19 Genomics UK (COG-UK) Consortium | Jacquelyn Wynn, Mairead Hyland, The Lighthouse Lab in Alderley Park and Alex Alderton, Roberto Amato, Jeffrey Barrett, Sonia Goncalves, Ewan Harrison, David K. Jackson, Ian Johnston, Dominic Kwiatkowski, Cordelia Langford, John Sillitoe on behalf of the Wellcome Sanger Institute COVID-19 Surveillance Team |
| EPI_ISL_3027910, EPI_ISL_3027911, EPI_ISL_3027912, EPI_ISL_3027913 | Lighthouse Lab in Glasgow | Wellcome Sanger Institute for the COVID-19 Genomics UK (COG-UK) Consortium | Harper VanSteenhouse, Yumi Kasai, David Gray, Carol Clugston, Anna Dominiczak and Alex Alderton, Roberto Amato, Jeffrey Barrett, Sonia Goncalves, Ewan Harrison, David K. Jackson, Ian Johnston, Dominic Kwiatkowski, Cordelia Langford, John Sillitoe on behalf of the Wellcome Sanger Institute COVID-19 Surveillance Team |
| EPI_ISL_3027914 | Berkshire and Surrey Pathology Services Lighthouse Laboratory | Wellcome Sanger Institute for the COVID-19 Genomics UK (COG-UK) Consortium | Berkshire and Surrey Pathology Services Lighthouse Laboratory and Alex Alderton, Roberto Amato, Jeffrey Barrett, Sonia Goncalves, Ewan Harrison, David K. Jackson, Ian Johnston, Dominic Kwiatkowski, Cordelia Langford, John Sillitoe on behalf of the Wellcome Sanger Institute COVID-19 Surveillance Team |
| EPI_ISL_3027915 | Lighthouse Lab in Glasgow | Wellcome Sanger Institute for the COVID-19 Genomics UK (COG-UK) Consortium | Harper VanSteenhouse, Yumi Kasai, David Gray, Carol Clugston, Anna Dominiczak and Alex Alderton, Roberto Amato, Jeffrey Barrett, Sonia Goncalves, Ewan Harrison, David K. Jackson, Ian Johnston, Dominic Kwiatkowski, Cordelia Langford, John Sillitoe on behalf of the Wellcome Sanger Institute COVID-19 Surveillance Team |
| EPI_ISL_3027917, EPI_ISL_3027918, EPI_ISL_3027919, EPI_ISL_3027920, EPI_ISL_3027921, EPI_ISL_3027922, EPI_ISL_3027923, EPI_ISL_3027924, EPI_ISL_3027926, EPI_ISL_3027927, EPI_ISL_3027928, EPI_ISL_3027929, EPI_ISL_3027930, EPI_ISL_3027931, EPI_ISL_3027932, EPI_ISL_3027933, EPI_ISL_3027934, EPI_ISL_3027935, EPI_ISL_3027936, EPI_ISL_3027937, EPI_ISL_3027938, EPI_ISL_3027939, EPI_ISL_3027941, EPI_ISL_3027942, EPI_ISL_3027943, EPI_ISL_3027945, EPI_ISL_3027946, EPI_ISL_3027947, EPI_ISL_3027948, EPI_ISL_3027949, EPI_ISL_3027950, EPI_ISL_3027951, EPI_ISL_3027952, EPI_ISL_3027953, EPI_ISL_3027954, EPI_ISL_3027955, EPI_ISL_3027956, EPI_ISL_3027957, EPI_ISL_3027959, EPI_ISL_3027960, EPI_ISL_3027961, EPI_ISL_3027962, EPI_ISL_3027964, EPI_ISL_3027966, EPI_ISL_3027967, EPI_ISL_3027968, EPI_ISL_3027969, EPI_ISL_3027970, EPI_ISL_3027971, EPI_ISL_3027972, EPI_ISL_3027973, EPI_ISL_3027974, EPI_ISL_3027976, EPI_ISL_3027977, EPI_ISL_3027978, EPI_ISL_3027979, EPI_ISL_3027980, EPI_ISL_3027981, EPI_ISL_3027982, EPI_ISL_3027983, EPI_ISL_3027984, EPI_ISL_3027985, EPI_ISL_3027987, EPI_ISL_3027988, EPI_ISL_3027989, EPI_ISL_3027990, EPI_ISL_3027991, EPI_ISL_3027992, EPI_ISL_3027993, EPI_ISL_3027994, EPI_ISL_3027995, EPI_ISL_3027996, EPI_ISL_3027997, EPI_ISL_3027998, EPI_ISL_3027999, EPI_ISL_3028000, EPI_ISL_3028001, EPI_ISL_3028002 | | | |
| see above | Microbiology Department, Laboratori Clínic Metropolitana Nord. Hospital Universitari Germans Trias i Pujol | Can Ruti SARS-CoV-2 Sequencing Hub (HUGTiP/IrsiCaixa/IGTP) | Marc Noguera-Julian, Pilar Armengol, Ignacio Blanco, Antoni E Bordoy, Francesc Catala-Moll, Pere-Joan Cardona, Maria Casadellà, Cristina Casañ, Gemma Clara, Bonaventura Clotet, Montserrat Giménez, Mercedes Guerrero, Anna Not, David Panisello, Roger Paredes, Mariona Parera, Alexia París, Verónica Saludes, Alba Sánchez, Laia Soler, Lauro Sumoy, and Elisa Martró on behalf of the Can Ruti SARS-CoV-2 Sequencing Hub |
| EPI_ISL_3028004 | Microbiologia CATLAB | Can Ruti SARS-CoV-2 Sequencing Hub (HUGTiP/IrsiCaixa/IGTP) | Marc Noguera-Julian, Pilar Armengol, Ignacio Blanco, Antoni E Bordoy, Francesc Catala-Moll, Pere-Joan Cardona, Maria Casadellà, Cristina Casañ, Gemma Clara, Bonaventura Clotet, Montserrat Giménez, Mercedes Guerrero, Anna Not, David Panisello, Roger Paredes, Mariona Parera, Alexia París, Verónica Saludes, Alba Sánchez, Laia Soler, Lauro Sumoy, and Elisa Martró on behalf of the Can Ruti SARS-CoV-2 Sequencing Hub |
| EPI_ISL_3028005, EPI_ISL_3028006, EPI_ISL_3028007 | Microbiology Department, Laboratori Clínic Metropolitana Nord. Hospital Universitari Germans Trias i Pujol | Can Ruti SARS-CoV-2 Sequencing Hub (HUGTiP/IrsiCaixa/IGTP) | Marc Noguera-Julian, Pilar Armengol, Ignacio Blanco, Antoni E Bordoy, Francesc Catala-Moll, Pere-Joan Cardona, Maria Casadellà, Cristina Casañ, Gemma Clara, Bonaventura Clotet, Montserrat Giménez, Mercedes Guerrero, Anna Not, David Panisello, Roger Paredes, Mariona Parera, Alexia París, Verónica Saludes, Alba Sánchez, Laia Soler, Lauro Sumoy, and Elisa Martró on behalf of the Can Ruti SARS-CoV-2 Sequencing Hub |
| EPI_ISL_3028008, EPI_ISL_3028009 | Microbiologia CATLAB | Can Ruti SARS-CoV-2 Sequencing Hub (HUGTiP/IrsiCaixa/IGTP) | Marc Noguera-Julian, Pilar Armengol, Ignacio Blanco, Antoni E Bordoy, Francesc Catala-Moll, Pere-Joan Cardona, Maria Casadellà, Cristina Casañ, Gemma Clara, Bonaventura Clotet, Montserrat Giménez, Mercedes Guerrero, Anna Not, David Panisello, Roger Paredes, Mariona Parera, Alexia París, Verónica Saludes, Alba Sánchez, Laia Soler, Lauro Sumoy, and Elisa Martró on behalf of the Can Ruti SARS-CoV-2 Sequencing Hub |
| EPI_ISL_3028010, EPI_ISL_3028011, EPI_ISL_3028012 | Microbiology Department, Laboratori Clínic Metropolitana Nord. Hospital Universitari Germans Trias i Pujol | Can Ruti SARS-CoV-2 Sequencing Hub (HUGTiP/IrsiCaixa/IGTP) | Marc Noguera-Julian, Pilar Armengol, Ignacio Blanco, Antoni E Bordoy, Francesc Catala-Moll, Pere-Joan Cardona, Maria Casadellà, Cristina Casañ, Gemma Clara, Bonaventura Clotet, Montserrat Giménez, Mercedes Guerrero, Anna Not, David Panisello, Roger Paredes, Mariona Parera, Alexia París, Verónica Saludes, Alba Sánchez, Laia Soler, Lauro Sumoy, and Elisa Martró on behalf of the Can Ruti SARS-CoV-2 Sequencing Hub |
| EPI_ISL_3028013 | Microbiologia CATLAB | Can Ruti SARS-CoV-2 Sequencing Hub (HUGTiP/IrsiCaixa/IGTP) | Marc Noguera-Julian, Pilar Armengol, Ignacio Blanco, Antoni E Bordoy, Francesc Catala-Moll, Pere-Joan Cardona, Maria Casadellà, Cristina Casañ, Gemma Clara, Bonaventura Clotet, Montserrat Giménez, Mercedes Guerrero, Anna Not, David Panisello, Roger Paredes, Mariona Parera, Alexia París, Verónica Saludes, Alba Sánchez, Laia Soler, Lauro Sumoy, and Elisa Martró on behalf of the Can Ruti SARS-CoV-2 Sequencing Hub |
| EPI_ISL_3028014, EPI_ISL_3028015, EPI_ISL_3028016 | Microbiology Department, Laboratori Clínic Metropolitana Nord. Hospital Universitari Germans Trias i Pujol | Can Ruti SARS-CoV-2 Sequencing Hub (HUGTiP/IrsiCaixa/IGTP) | Marc Noguera-Julian, Pilar Armengol, Ignacio Blanco, Antoni E Bordoy, Francesc Catala-Moll, Pere-Joan Cardona, Maria Casadellà, Cristina Casañ, Gemma Clara, Bonaventura Clotet, Montserrat Giménez, Mercedes Guerrero, Anna Not, David Panisello, Roger Paredes, Mariona Parera, Alexia París, Verónica Saludes, Alba Sánchez, Laia Soler, Lauro Sumoy, and Elisa Martró on behalf of the Can Ruti SARS-CoV-2 Sequencing Hub |
| EPI_ISL_3028017, EPI_ISL_3028018 | Microbiologia CATLAB | Can Ruti SARS-CoV-2 Sequencing Hub (HUGTiP/IrsiCaixa/IGTP) | Marc Noguera-Julian, Pilar Armengol, Ignacio Blanco, Antoni E Bordoy, Francesc Catala-Moll, Pere-Joan Cardona, Maria Casadellà, Cristina Casañ, Gemma Clara, Bonaventura Clotet, Montserrat Giménez, Mercedes Guerrero, Anna Not, David Panisello, Roger Paredes, Mariona Parera, Alexia París, Verónica Saludes, Alba Sánchez, Laia Soler, Lauro Sumoy, and Elisa Martró on behalf of the Can Ruti SARS-CoV-2 Sequencing Hub |
| EPI_ISL_3028019, EPI_ISL_3028020, EPI_ISL_3028021 | Microbiology Department, Laboratori Clínic Metropolitana Nord. Hospital Universitari Germans Trias i Pujol | Can Ruti SARS-CoV-2 Sequencing Hub (HUGTiP/IrsiCaixa/IGTP) | Marc Noguera-Julian, Pilar Armengol, Ignacio Blanco, Antoni E Bordoy, Francesc Catala-Moll, Pere-Joan Cardona, Maria Casadellà, Cristina Casañ, Gemma Clara, Bonaventura Clotet, Montserrat Giménez, Mercedes Guerrero, Anna Not, David Panisello, Roger Paredes, Mariona Parera, Alexia París, Verónica Saludes, Alba Sánchez, Laia Soler, Lauro Sumoy, and Elisa Martró on behalf of the Can Ruti SARS-CoV-2 Sequencing Hub |
| EPI_ISL_3028022 | Microbiologia CATLAB | Can Ruti SARS-CoV-2 Sequencing Hub (HUGTiP/IrsiCaixa/IGTP) | Marc Noguera-Julian, Pilar Armengol, Ignacio Blanco, Antoni E Bordoy, Francesc Catala-Moll, Pere-Joan Cardona, Maria Casadellà, Cristina Casañ, Gemma Clara, Bonaventura Clotet, Montserrat Giménez, Mercedes Guerrero, Anna Not, David Panisello, Roger Paredes, Mariona Parera, Alexia París, Verónica Saludes, Alba Sánchez, Laia Soler, Lauro Sumoy, and Elisa Martró on behalf of the Can Ruti SARS-CoV-2 Sequencing Hub |
| EPI_ISL_3028023, EPI_ISL_3028024, EPI_ISL_3028025 | Microbiology Department, Laboratori Clínic Metropolitana Nord. Hospital Universitari Germans Trias i Pujol | Can Ruti SARS-CoV-2 Sequencing Hub (HUGTiP/IrsiCaixa/IGTP) | Marc Noguera-Julian, Pilar Armengol, Ignacio Blanco, Antoni E Bordoy, Francesc Catala-Moll, Pere-Joan Cardona, Maria Casadellà, Cristina Casañ, Gemma Clara, Bonaventura Clotet, Montserrat Giménez, Mercedes Guerrero, Anna Not, David Panisello, Roger Paredes, Mariona Parera, Alexia París, Verónica Saludes, Alba Sánchez, Laia Soler, Lauro Sumoy, and Elisa Martró on behalf of the Can Ruti SARS-CoV-2 Sequencing Hub |
| EPI_ISL_3028026, EPI_ISL_3028027 | Microbiologia CATLAB | Can Ruti SARS-CoV-2 Sequencing Hub (HUGTiP/IrsiCaixa/IGTP) | Marc Noguera-Julian, Pilar Armengol, Ignacio Blanco, Antoni E Bordoy, Francesc Catala-Moll, Pere-Joan Cardona, Maria Casadellà, Cristina Casañ, Gemma Clara, Bonaventura Clotet, Montserrat Giménez, Mercedes Guerrero, Anna Not, David Panisello, Roger Paredes, Mariona Parera, Alexia París, Verónica Saludes, Alba Sánchez, Laia Soler, Lauro Sumoy, and Elisa Martró on behalf of the Can Ruti SARS-CoV-2 Sequencing Hub |
| EPI_ISL_3028028, EPI_ISL_3028029, EPI_ISL_3028030 | Microbiology Department, Laboratori Clínic Metropolitana Nord. Hospital Universitari Germans Trias i Pujol | Can Ruti SARS-CoV-2 Sequencing Hub (HUGTiP/IrsiCaixa/IGTP) | Marc Noguera-Julian, Pilar Armengol, Ignacio Blanco, Antoni E Bordoy, Francesc Catala-Moll, Pere-Joan Cardona, Maria Casadellà, Cristina Casañ, Gemma Clara, Bonaventura Clotet, Montserrat Giménez, Mercedes Guerrero, Anna Not, David Panisello, Roger Paredes, Mariona Parera, Alexia París, Verónica Saludes, Alba Sánchez, Laia Soler, Lauro Sumoy, and Elisa Martró on behalf of the Can Ruti SARS-CoV-2 Sequencing Hub |
| EPI_ISL_3028031, EPI_ISL_3028032 | Microbiologia CATLAB | Can Ruti SARS-CoV-2 Sequencing Hub (HUGTiP/IrsiCaixa/IGTP) | Marc Noguera-Julian, Pilar Armengol, Ignacio Blanco, Antoni E Bordoy, Francesc Catala-Moll, Pere-Joan Cardona, Maria Casadellà, Cristina Casañ, Gemma Clara, Bonaventura Clotet, Montserrat Giménez, Mercedes Guerrero, Anna Not, David Panisello, Roger Paredes, Mariona Parera, Alexia París, Verónica Saludes, Alba Sánchez, Laia Soler, Lauro Sumoy, and Elisa Martró on behalf of the Can Ruti SARS-CoV-2 Sequencing Hub |
| EPI_ISL_3028033, EPI_ISL_3028034, EPI_ISL_3028035 | Microbiology Department, Laboratori Clínic Metropolitana Nord. Hospital Universitari Germans Trias i Pujol | Can Ruti SARS-CoV-2 Sequencing Hub (HUGTiP/IrsiCaixa/IGTP) | Marc Noguera-Julian, Pilar Armengol, Ignacio Blanco, Antoni E Bordoy, Francesc Catala-Moll, Pere-Joan Cardona, Maria Casadellà, Cristina Casañ, Gemma Clara, Bonaventura Clotet, Montserrat Giménez, Mercedes Guerrero, Anna Not, David Panisello, Roger Paredes, Mariona Parera, Alexia París, Verónica Saludes, Alba Sánchez, Laia Soler, Lauro Sumoy, and Elisa Martró on behalf of the Can Ruti SARS-CoV-2 Sequencing Hub |
| EPI_ISL_3028036, EPI_ISL_3028037 | Microbiologia CATLAB | Can Ruti SARS-CoV-2 Sequencing Hub (HUGTiP/IrsiCaixa/IGTP) | Marc Noguera-Julian, Pilar Armengol, Ignacio Blanco, Antoni E Bordoy, Francesc Catala-Moll, Pere-Joan Cardona, Maria Casadellà, Cristina Casañ, Gemma Clara, Bonaventura Clotet, Montserrat Giménez, Mercedes Guerrero, Anna Not, David Panisello, Roger Paredes, Mariona Parera, Alexia París, Verónica Saludes, Alba Sánchez, Laia Soler, Lauro Sumoy, and Elisa Martró on behalf of the Can Ruti SARS-CoV-2 Sequencing Hub |
| EPI_ISL_3028038, EPI_ISL_3028039, EPI_ISL_3028040 | Microbiology Department, Laboratori Clínic Metropolitana Nord. Hospital Universitari Germans Trias i Pujol | Can Ruti SARS-CoV-2 Sequencing Hub (HUGTiP/IrsiCaixa/IGTP) | Marc Noguera-Julian, Pilar Armengol, Ignacio Blanco, Antoni E Bordoy, Francesc Catala-Moll, Pere-Joan Cardona, Maria Casadellà, Cristina Casañ, Gemma Clara, Bonaventura Clotet, Montserrat Giménez, Mercedes Guerrero, Anna Not, David Panisello, Roger Paredes, Mariona Parera, Alexia París, Verónica Saludes, Alba Sánchez, Laia Soler, Lauro Sumoy, and Elisa Martró on behalf of the Can Ruti SARS-CoV-2 Sequencing Hub |
| EPI_ISL_3028165, EPI_ISL_3028166, EPI_ISL_3028167, EPI_ISL_3028168, EPI_ISL_3028169, EPI_ISL_3028170, EPI_ISL_3028171, EPI_ISL_3028172 | AULSS 1 Dolomiti | Istituto Zooprofilattico Sperimentale delle Venezie | Adelaide Milani, Alessia Schivo, Annalisa Salviato, Elisa Palumbo, Erika Giorgia Quaranta, Luca Tassoni, Ambra Pastori, Edoardo Giussani, Alice Fusaro, Isabella Monne, Calogero Terregino, Antonia Ricci |
| EPI_ISL_3028173, EPI_ISL_3028174, EPI_ISL_3028175, EPI_ISL_3028176, EPI_ISL_3028177, EPI_ISL_3028178, EPI_ISL_3028179, EPI_ISL_3028180, EPI_ISL_3028181, EPI_ISL_3028182, EPI_ISL_3028183, EPI_ISL_3028184, EPI_ISL_3028185, EPI_ISL_3028186, EPI_ISL_3028187, EPI_ISL_3028188, EPI_ISL_3028189, EPI_ISL_3028190, EPI_ISL_3028191, EPI_ISL_3028192, EPI_ISL_3028193, EPI_ISL_3028194, EPI_ISL_3028195, EPI_ISL_3028196, EPI_ISL_3028197, EPI_ISL_3028198, EPI_ISL_3028199, EPI_ISL_3028200, EPI_ISL_3028201, EPI_ISL_3028202, EPI_ISL_3028203, EPI_ISL_3028204, EPI_ISL_3028205, EPI_ISL_3028206, EPI_ISL_3028207, EPI_ISL_3028208, EPI_ISL_3028209, EPI_ISL_3028211, EPI_ISL_3028212, EPI_ISL_3028213, EPI_ISL_3028214, EPI_ISL_3028215, EPI_ISL_3028216 | | | |
| see above | AULSS 9 Scaligera | Istituto Zooprofilattico Sperimentale delle Venezie | Adelaide Milani, Alessia Schivo, Annalisa Salviato, Elisa Palumbo, Erika Giorgia Quaranta, Luca Tassoni, Ambra Pastori, Edoardo Giussani, Alice Fusaro, Isabella Monne, Calogero Terregino, Antonia Ricci |
| EPI_ISL_3028218 | AULSS 7 Pedemontana | Istituto Zooprofilattico Sperimentale delle Venezie | Adelaide Milani, Alessia Schivo, Annalisa Salviato, Elisa Palumbo, Erika Giorgia Quaranta, Luca Tassoni, Ambra Pastori, Edoardo Giussani, Alice Fusaro, Isabella Monne, Calogero Terregino, Antonia Ricci |
| EPI_ISL_3028219 | AULSS 9 Scaligera | Istituto Zooprofilattico Sperimentale delle Venezie | Adelaide Milani, Alessia Schivo, Annalisa Salviato, Elisa Palumbo, Erika Giorgia Quaranta, Luca Tassoni, Ambra Pastori, Edoardo Giussani, Alice Fusaro, Isabella Monne, Calogero Terregino, Antonia Ricci |
| EPI_ISL_3028220, EPI_ISL_3028221, EPI_ISL_3028222, EPI_ISL_3028223, EPI_ISL_3028224, EPI_ISL_3028226, EPI_ISL_3028227, EPI_ISL_3028228, EPI_ISL_3028229 | AULSS 7 Pedemontana | Istituto Zooprofilattico Sperimentale delle Venezie | Adelaide Milani, Alessia Schivo, Annalisa Salviato, Elisa Palumbo, Erika Giorgia Quaranta, Luca Tassoni, Ambra Pastori, Edoardo Giussani, Alice Fusaro, Isabella Monne, Calogero Terregino, Antonia Ricci |
| EPI_ISL_3028231, EPI_ISL_3028232, EPI_ISL_3028233, EPI_ISL_3028234, EPI_ISL_3028235, EPI_ISL_3028237 | AULSS 6 Euganea | Istituto Zooprofilattico Sperimentale delle Venezie | Adelaide Milani, Alessia Schivo, Annalisa Salviato, Elisa Palumbo, Erika Giorgia Quaranta, Luca Tassoni, Ambra Pastori, Edoardo Giussani, Alice Fusaro, Isabella Monne, Calogero Terregino, Antonia Ricci |
| EPI_ISL_3028239, EPI_ISL_3028241, EPI_ISL_3028243, EPI_ISL_3028244, EPI_ISL_3028246, EPI_ISL_3028247 | AULSS 2 Marca Trevigiana | Istituto Zooprofilattico Sperimentale delle Venezie | Adelaide Milani, Alessia Schivo, Annalisa Salviato, Elisa Palumbo, Erika Giorgia Quaranta, Luca Tassoni, Ambra Pastori, Edoardo Giussani, Alice Fusaro, Isabella Monne, Calogero Terregino, Antonia Ricci |

| | | | |
|---|---|---|---|
| EPI_ISL_3028261 | Molekylær Medicinsk Afdeling, Aarhus University Hospital, Aarhus, Denmark | Statens Serum Institut Bioinformatics and Microbial Genomics | Danish Covid-19 Genome Consortium |
| EPI_ISL_3028262, EPI_ISL_3028263, EPI_ISL_3028264, EPI_ISL_3028265, EPI_ISL_3028266, EPI_ISL_3028267, EPI_ISL_3028268, EPI_ISL_3028270, EPI_ISL_3028271, EPI_ISL_3028272, EPI_ISL_3028273 | | | |
| see above | Department of Bacteria, Parasites and Fungi, Statens Serum Institut, Copenhagen, Denmark | Statens Serum Institut Bioinformatics and Microbial Genomics | Danish Covid-19 Genome Consortium |
| EPI_ISL_3028274 | Molekylær Medicinsk Afdeling, Aarhus University Hospital, Aarhus, Denmark | Statens Serum Institut Bioinformatics and Microbial Genomics | Danish Covid-19 Genome Consortium |
| EPI_ISL_3028275, EPI_ISL_3028276, EPI_ISL_3028277, EPI_ISL_3028278, EPI_ISL_3028279, EPI_ISL_3028280, EPI_ISL_3028281, EPI_ISL_3028282, EPI_ISL_3028283, EPI_ISL_3028284, EPI_ISL_3028285, EPI_ISL_3028286, EPI_ISL_3028287, EPI_ISL_3028288, EPI_ISL_3028289, EPI_ISL_3028290, EPI_ISL_3028291, EPI_ISL_3028292, EPI_ISL_3028293, EPI_ISL_3028294, EPI_ISL_3028295, EPI_ISL_3028296, EPI_ISL_3028297, EPI_ISL_3028298, EPI_ISL_3028299, EPI_ISL_3028300, EPI_ISL_3028301, EPI_ISL_3028302, EPI_ISL_3028303, EPI_ISL_3028304, EPI_ISL_3028305, EPI_ISL_3028306, EPI_ISL_3028307, EPI_ISL_3028308, EPI_ISL_3028309, EPI_ISL_3028310, EPI_ISL_3028311, EPI_ISL_3028312, EPI_ISL_3028313, EPI_ISL_3028314, EPI_ISL_3028315, EPI_ISL_3028316, EPI_ISL_3028317, EPI_ISL_3028318, EPI_ISL_3028319, EPI_ISL_3028320, EPI_ISL_3028321, EPI_ISL_3028322, EPI_ISL_3028323, EPI_ISL_3028324, EPI_ISL_3028325, EPI_ISL_3028326, EPI_ISL_3028327, EPI_ISL_3028328, EPI_ISL_3028329, EPI_ISL_3028330, EPI_ISL_3028331, EPI_ISL_3028332, EPI_ISL_3028333, EPI_ISL_3028334, EPI_ISL_3028335, EPI_ISL_3028336, EPI_ISL_3028337, EPI_ISL_3028338, EPI_ISL_3028339, EPI_ISL_3028340, EPI_ISL_3028341, EPI_ISL_3028342, EPI_ISL_3028343, EPI_ISL_3028344, EPI_ISL_3028345, EPI_ISL_3028346, EPI_ISL_3028347, EPI_ISL_3028348, EPI_ISL_3028349, EPI_ISL_3028350, EPI_ISL_3028351, EPI_ISL_3028352, EPI_ISL_3028353, EPI_ISL_3028354, EPI_ISL_3028355, EPI_ISL_3028356, EPI_ISL_3028357, EPI_ISL_3028358, EPI_ISL_3028359, EPI_ISL_3028360, EPI_ISL_3028361, EPI_ISL_3028362, EPI_ISL_3028363, EPI_ISL_3028364, EPI_ISL_3028365, EPI_ISL_3028366, EPI_ISL_3028367, EPI_ISL_3028368, EPI_ISL_3028369, EPI_ISL_3028370, EPI_ISL_3028371, EPI_ISL_3028372, EPI_ISL_3028373, EPI_ISL_3028374, EPI_ISL_3028375, EPI_ISL_3028376, EPI_ISL_3028377, EPI_ISL_3028378, EPI_ISL_3028379, EPI_ISL_3028380, EPI_ISL_3028381, EPI_ISL_3028382, EPI_ISL_3028383, EPI_ISL_3028384, EPI_ISL_3028385, EPI_ISL_3028386, EPI_ISL_3028387, EPI_ISL_3028388, EPI_ISL_3028389, EPI_ISL_3028390, EPI_ISL_3028391, EPI_ISL_3028392, EPI_ISL_3028393, EPI_ISL_3028394, EPI_ISL_3028395, EPI_ISL_3028396, EPI_ISL_3028397, EPI_ISL_3028398, EPI_ISL_3028399, EPI_ISL_3028400, EPI_ISL_3028401, EPI_ISL_3028402, EPI_ISL_3028403, EPI_ISL_3028404, EPI_ISL_3028405, EPI_ISL_3028406, EPI_ISL_3028407, EPI_ISL_3028408 | | | |
| see above | Department of Bacteria, Parasites and Fungi, Statens Serum Institut, Copenhagen, Denmark | Statens Serum Institut Bioinformatics and Microbial Genomics | Danish Covid-19 Genome Consortium |
| EPI_ISL_3028409 | Department of Clinical Microbiology, Odense University Hospital, Odense, Denmark | Statens Serum Institut Bioinformatics and Microbial Genomics | Danish Covid-19 Genome Consortium |
| EPI_ISL_3028410, EPI_ISL_3028411, EPI_ISL_3028412, EPI_ISL_3028413, EPI_ISL_3028414, EPI_ISL_3028415, EPI_ISL_3028416, EPI_ISL_3028417, EPI_ISL_3028418, EPI_ISL_3028419, EPI_ISL_3028420, EPI_ISL_3028421, EPI_ISL_3028422, EPI_ISL_3028423, EPI_ISL_3028424, EPI_ISL_3028425, EPI_ISL_3028426, EPI_ISL_3028427, EPI_ISL_3028428, EPI_ISL_3028429, EPI_ISL_3028430, EPI_ISL_3028431, EPI_ISL_3028432, EPI_ISL_3028433, EPI_ISL_3028434, EPI_ISL_3028435, EPI_ISL_3028436, EPI_ISL_3028437, EPI_ISL_3028438, EPI_ISL_3028439, EPI_ISL_3028440, EPI_ISL_3028441, EPI_ISL_3028442, EPI_ISL_3028443, EPI_ISL_3028444, EPI_ISL_3028445, EPI_ISL_3028446, EPI_ISL_3028447, EPI_ISL_3028448, EPI_ISL_3028449, EPI_ISL_3028450, EPI_ISL_3028451, EPI_ISL_3028452, EPI_ISL_3028453, EPI_ISL_3028454, EPI_ISL_3028455, EPI_ISL_3028456, EPI_ISL_3028457, EPI_ISL_3028458, EPI_ISL_3028459, EPI_ISL_3028460, EPI_ISL_3028461, EPI_ISL_3028462, EPI_ISL_3028463, EPI_ISL_3028464 | | | |
| see above | Department of Bacteria, Parasites and Fungi, Statens Serum Institut, Copenhagen, Denmark | Statens Serum Institut Bioinformatics and Microbial Genomics | Danish Covid-19 Genome Consortium |
| EPI_ISL_3028465 | Molekylær Medicinsk Afdeling, Aarhus University Hospital, Aarhus, Denmark | Statens Serum Institut Bioinformatics and Microbial Genomics | Danish Covid-19 Genome Consortium |
| EPI_ISL_3028466, EPI_ISL_3028467, EPI_ISL_3028468, EPI_ISL_3028469, EPI_ISL_3028470 | Department of Bacteria, Parasites and Fungi, Statens Serum Institut, Copenhagen, Denmark | Statens Serum Institut Bioinformatics and Microbial Genomics | Danish Covid-19 Genome Consortium |
| EPI_ISL_3028471 | Molekylær Medicinsk Afdeling, Aarhus University Hospital, Aarhus, Denmark | Statens Serum Institut Bioinformatics and Microbial Genomics | Danish Covid-19 Genome Consortium |
| EPI_ISL_3028472, EPI_ISL_3028473, EPI_ISL_3028474, EPI_ISL_3028475, EPI_ISL_3028476, EPI_ISL_3028477, EPI_ISL_3028478, EPI_ISL_3028480, EPI_ISL_3028481, EPI_ISL_3028482, EPI_ISL_3028483 | | | |
| see above | Department of Bacteria, Parasites and Fungi, Statens Serum Institut, Copenhagen, Denmark | Statens Serum Institut Bioinformatics and Microbial Genomics | Danish Covid-19 Genome Consortium |
| EPI_ISL_3028484 | Molekylær Medicinsk Afdeling, Aarhus University Hospital, Aarhus, Denmark | Statens Serum Institut Bioinformatics and Microbial Genomics | Danish Covid-19 Genome Consortium |
| EPI_ISL_3028485, EPI_ISL_3028486, EPI_ISL_3028487, EPI_ISL_3028488, EPI_ISL_3028489, EPI_ISL_3028490, EPI_ISL_3028491, EPI_ISL_3028492, EPI_ISL_3028493, EPI_ISL_3028494, EPI_ISL_3028495, EPI_ISL_3028496, EPI_ISL_3028497, EPI_ISL_3028498, EPI_ISL_3028499, EPI_ISL_3028500, EPI_ISL_3028501, EPI_ISL_3028502, EPI_ISL_3028503, EPI_ISL_3028504, EPI_ISL_3028505, EPI_ISL_3028506, EPI_ISL_3028507, EPI_ISL_3028508, EPI_ISL_3028509, EPI_ISL_3028511, EPI_ISL_3028512, EPI_ISL_3028513, EPI_ISL_3028514, EPI_ISL_3028515, EPI_ISL_3028516, EPI_ISL_3028517, EPI_ISL_3028518, EPI_ISL_3028519 | | | |
| see above | Department of Bacteria, Parasites and Fungi, Statens Serum Institut, Copenhagen, Denmark | Statens Serum Institut Bioinformatics and Microbial Genomics | Danish Covid-19 Genome Consortium |
| EPI_ISL_3028520 | Molekylær Medicinsk Afdeling, Aarhus University Hospital, Aarhus, Denmark | Statens Serum Institut Bioinformatics and Microbial Genomics | Danish Covid-19 Genome Consortium |
| EPI_ISL_3028521, EPI_ISL_3028522, EPI_ISL_3028523, EPI_ISL_3028524, EPI_ISL_3028525, EPI_ISL_3028526, EPI_ISL_3028527, EPI_ISL_3028528, EPI_ISL_3028529, EPI_ISL_3028530, EPI_ISL_3028531, EPI_ISL_3028532, EPI_ISL_3028533, EPI_ISL_3028534, EPI_ISL_3028535, EPI_ISL_3028536, EPI_ISL_3028537, EPI_ISL_3028538, EPI_ISL_3028539, EPI_ISL_3028540, EPI_ISL_3028541, EPI_ISL_3028542, EPI_ISL_3028543, EPI_ISL_3028544, EPI_ISL_3028545, EPI_ISL_3028546, EPI_ISL_3028548, EPI_ISL_3028549, EPI_ISL_3028550, EPI_ISL_3028551, EPI_ISL_3028552, EPI_ISL_3028553, EPI_ISL_3028554, EPI_ISL_3028555, EPI_ISL_3028556, EPI_ISL_3028557, EPI_ISL_3028559, EPI_ISL_3028560, EPI_ISL_3028561, EPI_ISL_3028562, EPI_ISL_3028563, EPI_ISL_3028564, EPI_ISL_3028565, EPI_ISL_3028566, EPI_ISL_3028567, EPI_ISL_3028568, EPI_ISL_3028569, EPI_ISL_3028570, EPI_ISL_3028571, EPI_ISL_3028572, EPI_ISL_3028573, EPI_ISL_3028574, EPI_ISL_3028575, EPI_ISL_3028576, EPI_ISL_3028577, EPI_ISL_3028578 | | | |
| see above | Department of Bacteria, Parasites and Fungi, Statens Serum Institut, Copenhagen, Denmark | Statens Serum Institut Bioinformatics and Microbial Genomics | Danish Covid-19 Genome Consortium |
| EPI_ISL_3028579 | Department of Clinical Microbiology, Odense University Hospital, Odense, Denmark | Statens Serum Institut Bioinformatics and Microbial Genomics | Danish Covid-19 Genome Consortium |
| EPI_ISL_3028580, EPI_ISL_3028581, EPI_ISL_3028582, EPI_ISL_3028583, EPI_ISL_3028584, EPI_ISL_3028585, EPI_ISL_3028586, EPI_ISL_3028587, EPI_ISL_3028588, EPI_ISL_3028589, EPI_ISL_3028590, EPI_ISL_3028591, EPI_ISL_3028592, EPI_ISL_3028593 | | | |
| see above | Department of Bacteria, Parasites and Fungi, Statens Serum Institut, Copenhagen, Denmark | Statens Serum Institut Bioinformatics and Microbial Genomics | Danish Covid-19 Genome Consortium |
| EPI_ISL_3028594, EPI_ISL_3028595 | Department of Clinical Microbiology, Odense University Hospital, Odense, Denmark | Statens Serum Institut Bioinformatics and Microbial Genomics | Danish Covid-19 Genome Consortium |
| EPI_ISL_3028596, EPI_ISL_3028597, EPI_ISL_3028598, EPI_ISL_3028599, EPI_ISL_3028600, EPI_ISL_3028601, EPI_ISL_3028602, EPI_ISL_3028603, EPI_ISL_3028604, EPI_ISL_3028605, EPI_ISL_3028606, EPI_ISL_3028607, EPI_ISL_3028608 | | | |
| see above | Department of Bacteria, Parasites and Fungi, Statens Serum Institut, Copenhagen, Denmark | Statens Serum Institut Bioinformatics and Microbial Genomics | Danish Covid-19 Genome Consortium |
| EPI_ISL_3028609 | Molekylær Medicinsk Afdeling, Aarhus University Hospital, Aarhus, Denmark | Statens Serum Institut Bioinformatics and Microbial Genomics | Danish Covid-19 Genome Consortium |
| EPI_ISL_3028610, EPI_ISL_3028611, EPI_ISL_3028612, EPI_ISL_3028613, EPI_ISL_3028614, EPI_ISL_3028615, EPI_ISL_3028616, EPI_ISL_3028617, EPI_ISL_3028619, EPI_ISL_3028620, EPI_ISL_3028621, EPI_ISL_3028622, EPI_ISL_3028623, EPI_ISL_3028624, EPI_ISL_3028625, EPI_ISL_3028626, EPI_ISL_3028627, EPI_ISL_3028628, EPI_ISL_3028629, EPI_ISL_3028630, EPI_ISL_3028631, EPI_ISL_3028632, EPI_ISL_3028633, EPI_ISL_3028634, EPI_ISL_3028635, EPI_ISL_3028636, EPI_ISL_3028637, EPI_ISL_3028638, EPI_ISL_3028640, EPI_ISL_3028641, EPI_ISL_3028642, EPI_ISL_3028643, EPI_ISL_3028644, EPI_ISL_3028645, EPI_ISL_3028646, EPI_ISL_3028647, EPI_ISL_3028648, EPI_ISL_3028649, EPI_ISL_3028650, EPI_ISL_3028651, EPI_ISL_3028652, EPI_ISL_3028653, EPI_ISL_3028654 | | | |
| see above | Department of Bacteria, Parasites and Fungi, Statens Serum Institut, Copenhagen, Denmark | Statens Serum Institut Bioinformatics and Microbial Genomics | Danish Covid-19 Genome Consortium |
| EPI_ISL_3028655 | Department of Clinical Microbiology, Odense University Hospital, Odense, Denmark | Statens Serum Institut Bioinformatics and Microbial Genomics | Danish Covid-19 Genome Consortium |
| EPI_ISL_3028656, EPI_ISL_3028657, EPI_ISL_3028658, EPI_ISL_3028659, EPI_ISL_3028660, EPI_ISL_3028661, EPI_ISL_3028662, EPI_ISL_3028663, EPI_ISL_3028664, EPI_ISL_3028665, EPI_ISL_3028666, EPI_ISL_3028667, EPI_ISL_3028668, EPI_ISL_3028669, EPI_ISL_3028670, EPI_ISL_3028671, EPI_ISL_3028672 | | | |
| see above | Department of Bacteria, Parasites and Fungi, Statens Serum Institut, Copenhagen, Denmark | Statens Serum Institut Bioinformatics and Microbial Genomics | Danish Covid-19 Genome Consortium |
| EPI_ISL_3028673 | Molekylær Medicinsk Afdeling, Aarhus University Hospital, Aarhus, Denmark | Statens Serum Institut Bioinformatics and Microbial Genomics | Danish Covid-19 Genome Consortium |
| EPI_ISL_3028674, EPI_ISL_3028675, EPI_ISL_3028676, EPI_ISL_3028677, EPI_ISL_3028678, EPI_ISL_3028679, EPI_ISL_3028680, EPI_ISL_3028681, EPI_ISL_3028682, EPI_ISL_3028683, EPI_ISL_3028684, EPI_ISL_3028685, EPI_ISL_3028686, EPI_ISL_3028687, EPI_ISL_3028688, EPI_ISL_3028689, EPI_ISL_3028690, EPI_ISL_3028691, EPI_ISL_3028692, EPI_ISL_3028693, EPI_ISL_3028694, EPI_ISL_3028695, EPI_ISL_3028696, EPI_ISL_3028697, EPI_ISL_3028698, EPI_ISL_3028699, EPI_ISL_3028700, EPI_ISL_3028701, EPI_ISL_3028702, EPI_ISL_3028703, EPI_ISL_3028704, EPI_ISL_3028705, EPI_ISL_3028706, EPI_ISL_3028707, EPI_ISL_3028708, EPI_ISL_3028709, EPI_ISL_3028710, EPI_ISL_3028711, EPI_ISL_3028712, EPI_ISL_3028713, EPI_ISL_3028714, EPI_ISL_3028715, EPI_ISL_3028716, EPI_ISL_3028717, EPI_ISL_3028718, EPI_ISL_3028719, EPI_ISL_3028720, EPI_ISL_3028721, EPI_ISL_3028722, EPI_ISL_3028723, EPI_ISL_3028724, EPI_ISL_3028725 | | | |
| see above | Department of Bacteria, Parasites and Fungi, Statens Serum Institut, Copenhagen, Denmark | Statens Serum Institut Bioinformatics and Microbial Genomics | Danish Covid-19 Genome Consortium |
| EPI_ISL_3028726 | Molekylær Medicinsk Afdeling, Aarhus University Hospital, Aarhus, Denmark | Statens Serum Institut Bioinformatics and Microbial Genomics | Danish Covid-19 Genome Consortium |
| EPI_ISL_3028727, EPI_ISL_3028728, EPI_ISL_3028729, EPI_ISL_3028730, EPI_ISL_3028731, EPI_ISL_3028732, EPI_ISL_3028733, EPI_ISL_3028734, EPI_ISL_3028735, EPI_ISL_3028736, EPI_ISL_3028737, EPI_ISL_3028738, EPI_ISL_3028739, EPI_ISL_3028740, EPI_ISL_3028741, EPI_ISL_3028742, EPI_ISL_3028743, EPI_ISL_3028744, EPI_ISL_3028745, EPI_ISL_3028746, EPI_ISL_3028748, EPI_ISL_3028749, EPI_ISL_3028750, EPI_ISL_3028751, EPI_ISL_3028752, EPI_ISL_3028753, EPI_ISL_3028754, EPI_ISL_3028755, EPI_ISL_3028756, EPI_ISL_3028757, EPI_ISL_3028758, EPI_ISL_3028759, EPI_ISL_3028760, EPI_ISL_3028761, EPI_ISL_3028762, EPI_ISL_3028763, EPI_ISL_3028764, EPI_ISL_3028765, EPI_ISL_3028766, EPI_ISL_3028767, EPI_ISL_3028768, EPI_ISL_3028769, EPI_ISL_3028770, EPI_ISL_3028771, EPI_ISL_3028772, EPI_ISL_3028773, EPI_ISL_3028774, EPI_ISL_3028775, EPI_ISL_3028776, EPI_ISL_3028777, EPI_ISL_3028778, EPI_ISL_3028779, EPI_ISL_3028780, EPI_ISL_3028781, EPI_ISL_3028782, EPI_ISL_3028783, EPI_ISL_3028784, EPI_ISL_3028785, EPI_ISL_3028786, EPI_ISL_3028787, EPI_ISL_3028788, EPI_ISL_3028789, EPI_ISL_3028790, EPI_ISL_3028791, EPI_ISL_3028792, EPI_ISL_3028793, EPI_ISL_3028794, EPI_ISL_3028795, EPI_ISL_3028796, EPI_ISL_3028797, EPI_ISL_3028798, EPI_ISL_3028799, EPI_ISL_3028800 | | | |
| see above | Department of Bacteria, Parasites and Fungi, Statens Serum Institut, Copenhagen, Denmark | Statens Serum Institut Bioinformatics and Microbial Genomics | Danish Covid-19 Genome Consortium |
| EPI_ISL_3028801 | Department of Clinical Microbiology, Odense University Hospital, Odense, Denmark | Statens Serum Institut Bioinformatics and Microbial Genomics | Danish Covid-19 Genome Consortium |
| EPI_ISL_3028802, EPI_ISL_3028803, EPI_ISL_3028804, EPI_ISL_3028805, EPI_ISL_3028806, EPI_ISL_3028807, EPI_ISL_3028808, EPI_ISL_3028809, EPI_ISL_3028810, EPI_ISL_3028811, EPI_ISL_3028812, EPI_ISL_3028813, EPI_ISL_3028814, EPI_ISL_3028815, EPI_ISL_3028816, EPI_ISL_3028817, EPI_ISL_3028818, EPI_ISL_3028819, EPI_ISL_3028820, EPI_ISL_3028821, EPI_ISL_3028822, EPI_ISL_3028823, EPI_ISL_3028824, EPI_ISL_3028826, EPI_ISL_3028827 | | | |
| see above | Department of Bacteria, Parasites and Fungi, Statens Serum Institut, Copenhagen, Denmark | Statens Serum Institut Bioinformatics and Microbial Genomics | Danish Covid-19 Genome Consortium |
| EPI_ISL_3028828 | Molekylær Medicinsk Afdeling, Aarhus University Hospital, | Statens Serum Institut Bioinformatics and Microbial Genomics | Danish Covid-19 Genome Consortium |

| | | | |
|---|---|---|---|
| | Aarhus, Denmark | | |
| EPI_ISL_3028829, EPI_ISL_3028830, EPI_ISL_3028831, EPI_ISL_3028832, EPI_ISL_3028833, EPI_ISL_3028834, EPI_ISL_3028835, EPI_ISL_3028836, EPI_ISL_3028837, EPI_ISL_3028838, EPI_ISL_3028839, EPI_ISL_3028840, EPI_ISL_3028841, EPI_ISL_3028842, EPI_ISL_3028843, EPI_ISL_3028844, EPI_ISL_3028845, EPI_ISL_3028846, EPI_ISL_3028847 | | | |
| see above | Department of Bacteria, Parasites and Fungi, Statens Serum Institut, Copenhagen, Denmark | Statens Serum Institut Bioinformatics and Microbial Genomics | Danish Covid-19 Genome Consortium |
| EPI_ISL_3028848 | Department of Clinical Microbiology, Odense University Hospital, Odense, Denmark | Statens Serum Institut Bioinformatics and Microbial Genomics | Danish Covid-19 Genome Consortium |
| EPI_ISL_3028849, EPI_ISL_3028850, EPI_ISL_3028851, EPI_ISL_3028852, EPI_ISL_3028853, EPI_ISL_3028854, EPI_ISL_3028855, EPI_ISL_3028856, EPI_ISL_3028857, EPI_ISL_3028858, EPI_ISL_3028859, EPI_ISL_3028860, EPI_ISL_3028861, EPI_ISL_3028862, EPI_ISL_3028863, EPI_ISL_3028864, EPI_ISL_3028866, EPI_ISL_3028867, EPI_ISL_3028868 | | | |
| see above | Department of Bacteria, Parasites and Fungi, Statens Serum Institut, Copenhagen, Denmark | Statens Serum Institut Bioinformatics and Microbial Genomics | Danish Covid-19 Genome Consortium |
| EPI_ISL_3028869 | Department of Clinical Microbiology, Odense University Hospital, Odense, Denmark | Statens Serum Institut Bioinformatics and Microbial Genomics | Danish Covid-19 Genome Consortium |
| EPI_ISL_3028870, EPI_ISL_3028871, EPI_ISL_3028872, EPI_ISL_3028873, EPI_ISL_3028874, EPI_ISL_3028875, EPI_ISL_3028876, EPI_ISL_3028877, EPI_ISL_3028878, EPI_ISL_3028879, EPI_ISL_3028880, EPI_ISL_3028882, EPI_ISL_3028883, EPI_ISL_3028884, EPI_ISL_3028885, EPI_ISL_3028886, EPI_ISL_3028887, EPI_ISL_3028888, EPI_ISL_3028889, EPI_ISL_3028890, EPI_ISL_3028891, EPI_ISL_3028892, EPI_ISL_3028893, EPI_ISL_3028894, EPI_ISL_3028895, EPI_ISL_3028896, EPI_ISL_3028897, EPI_ISL_3028898, EPI_ISL_3028899, EPI_ISL_3028900, EPI_ISL_3028901, EPI_ISL_3028902 | | | |
| see above | Department of Bacteria, Parasites and Fungi, Statens Serum Institut, Copenhagen, Denmark | Statens Serum Institut Bioinformatics and Microbial Genomics | Danish Covid-19 Genome Consortium |
| EPI_ISL_3028903 | Department of Clinical Microbiology, Odense University Hospital, Odense, Denmark | Statens Serum Institut Bioinformatics and Microbial Genomics | Danish Covid-19 Genome Consortium |
| EPI_ISL_3028904, EPI_ISL_3028905, EPI_ISL_3028906, EPI_ISL_3028907, EPI_ISL_3028908, EPI_ISL_3028909, EPI_ISL_3028910, EPI_ISL_3028911, EPI_ISL_3028912, EPI_ISL_3028913, EPI_ISL_3028914, EPI_ISL_3028915, EPI_ISL_3028916, EPI_ISL_3028917, EPI_ISL_3028918, EPI_ISL_3028920, EPI_ISL_3028921, EPI_ISL_3028922, EPI_ISL_3028923, EPI_ISL_3028924, EPI_ISL_3028925, EPI_ISL_3028926, EPI_ISL_3028927, EPI_ISL_3028928, EPI_ISL_3028929, EPI_ISL_3028930, EPI_ISL_3028931, EPI_ISL_3028932, EPI_ISL_3028933, EPI_ISL_3028934, EPI_ISL_3028936, EPI_ISL_3028937, EPI_ISL_3028938, EPI_ISL_3028939, EPI_ISL_3028940, EPI_ISL_3028941, EPI_ISL_3028942, EPI_ISL_3028943, EPI_ISL_3028944, EPI_ISL_3028945, EPI_ISL_3028946, EPI_ISL_3028947, EPI_ISL_3028948, EPI_ISL_3028949, EPI_ISL_3028950, EPI_ISL_3028951, EPI_ISL_3028952, EPI_ISL_3028953, EPI_ISL_3028954, EPI_ISL_3028955, EPI_ISL_3028957, EPI_ISL_3028959, EPI_ISL_3028960, EPI_ISL_3028961, EPI_ISL_3028962, EPI_ISL_3028963, EPI_ISL_3028964, EPI_ISL_3028965, EPI_ISL_3028966, EPI_ISL_3028967, EPI_ISL_3028968, EPI_ISL_3028969, EPI_ISL_3028970, EPI_ISL_3028971, EPI_ISL_3028972, EPI_ISL_3028973, EPI_ISL_3028974, EPI_ISL_3028975, EPI_ISL_3028976, EPI_ISL_3028977, EPI_ISL_3028978, EPI_ISL_3028979, EPI_ISL_3028980, EPI_ISL_3028981, EPI_ISL_3028982, EPI_ISL_3028983, EPI_ISL_3028984, EPI_ISL_3028985, EPI_ISL_3028986, EPI_ISL_3028987, EPI_ISL_3028988, EPI_ISL_3028989, EPI_ISL_3028990, EPI_ISL_3028991, EPI_ISL_3028992, EPI_ISL_3028993, EPI_ISL_3028994, EPI_ISL_3028995, EPI_ISL_3028996, EPI_ISL_3028997, EPI_ISL_3028998, EPI_ISL_3028999, EPI_ISL_3029000, EPI_ISL_3029001, EPI_ISL_3029002, EPI_ISL_3029003, EPI_ISL_3029004, EPI_ISL_3029005, EPI_ISL_3029007, EPI_ISL_3029008, EPI_ISL_3029009, EPI_ISL_3029010, EPI_ISL_3029011, EPI_ISL_3029012, EPI_ISL_3029013, EPI_ISL_3029015, EPI_ISL_3029016, EPI_ISL_3029017, EPI_ISL_3029018, EPI_ISL_3029019, EPI_ISL_3029020, EPI_ISL_3029021, EPI_ISL_3029022, EPI_ISL_3029023, EPI_ISL_3029024, EPI_ISL_3029025, EPI_ISL_3029026, EPI_ISL_3029027, EPI_ISL_3029028, EPI_ISL_3029029, EPI_ISL_3029030, EPI_ISL_3029031, EPI_ISL_3029032, EPI_ISL_3029033, EPI_ISL_3029034, EPI_ISL_3029035, EPI_ISL_3029036, EPI_ISL_3029037, EPI_ISL_3029038, EPI_ISL_3029039, EPI_ISL_3029040, EPI_ISL_3029041, EPI_ISL_3029042, EPI_ISL_3029043, EPI_ISL_3029044, EPI_ISL_3029045, EPI_ISL_3029046, EPI_ISL_3029047, EPI_ISL_3029048, EPI_ISL_3029049, EPI_ISL_3029050, EPI_ISL_3029051, EPI_ISL_3029052, EPI_ISL_3029053, EPI_ISL_3029054, EPI_ISL_3029055, EPI_ISL_3029056, EPI_ISL_3029057, EPI_ISL_3029058, EPI_ISL_3029059, EPI_ISL_3029060, EPI_ISL_3029061, EPI_ISL_3029062, EPI_ISL_3029063, EPI_ISL_3029064, EPI_ISL_3029065, EPI_ISL_3029066, EPI_ISL_3029067, EPI_ISL_3029068, EPI_ISL_3029069, EPI_ISL_3029070, EPI_ISL_3029071, EPI_ISL_3029072, EPI_ISL_3029073, EPI_ISL_3029074, EPI_ISL_3029075, EPI_ISL_3029076, EPI_ISL_3029077, EPI_ISL_3029078, EPI_ISL_3029079, EPI_ISL_3029080, EPI_ISL_3029081, EPI_ISL_3029082, EPI_ISL_3029083, EPI_ISL_3029084, EPI_ISL_3029085, EPI_ISL_3029086, EPI_ISL_3029087, EPI_ISL_3029088, EPI_ISL_3029089, EPI_ISL_3029090, EPI_ISL_3029091, EPI_ISL_3029092, EPI_ISL_3029093, EPI_ISL_3029094, EPI_ISL_3029095, EPI_ISL_3029096, EPI_ISL_3029097, EPI_ISL_3029098, EPI_ISL_3029099, EPI_ISL_3029100, EPI_ISL_3029101, EPI_ISL_3029102, EPI_ISL_3029103, EPI_ISL_3029104, EPI_ISL_3029105, EPI_ISL_3029106, EPI_ISL_3029107, EPI_ISL_3029108, EPI_ISL_3029109, EPI_ISL_3029110, EPI_ISL_3029111, EPI_ISL_3029112, EPI_ISL_3029113, EPI_ISL_3029114, EPI_ISL_3029115, EPI_ISL_3029116, EPI_ISL_3029117, EPI_ISL_3029118, EPI_ISL_3029119, EPI_ISL_3029120, EPI_ISL_3029121, EPI_ISL_3029122, EPI_ISL_3029123, EPI_ISL_3029124, EPI_ISL_3029125, EPI_ISL_3029126, EPI_ISL_3029127, EPI_ISL_3029128, EPI_ISL_3029129, EPI_ISL_3029130, EPI_ISL_3029132, EPI_ISL_3029133, EPI_ISL_3029134, EPI_ISL_3029135, EPI_ISL_3029136, EPI_ISL_3029138, EPI_ISL_3029139, EPI_ISL_3029140, EPI_ISL_3029141, EPI_ISL_3029142, EPI_ISL_3029143, EPI_ISL_3029144, EPI_ISL_3029145, EPI_ISL_3029146, EPI_ISL_3029147, EPI_ISL_3029148, EPI_ISL_3029149, EPI_ISL_3029150, EPI_ISL_3029151, EPI_ISL_3029152, EPI_ISL_3029153, EPI_ISL_3029154, EPI_ISL_3029155, EPI_ISL_3029156, EPI_ISL_3029157, EPI_ISL_3029158, EPI_ISL_3029159, EPI_ISL_3029160, EPI_ISL_3029161, EPI_ISL_3029162, EPI_ISL_3029163, EPI_ISL_3029164, EPI_ISL_3029165, EPI_ISL_3029166, EPI_ISL_3029167, EPI_ISL_3029168, EPI_ISL_3029169, EPI_ISL_3029170, EPI_ISL_3029171, EPI_ISL_3029172, EPI_ISL_3029173, EPI_ISL_3029174, EPI_ISL_3029175, EPI_ISL_3029176, EPI_ISL_3029177, EPI_ISL_3029178, EPI_ISL_3029179, EPI_ISL_3029180, EPI_ISL_3029181, EPI_ISL_3029182, EPI_ISL_3029183, EPI_ISL_3029184, EPI_ISL_3029185, EPI_ISL_3029186, EPI_ISL_3029187, EPI_ISL_3029188, EPI_ISL_3029189, EPI_ISL_3029190, EPI_ISL_3029191, EPI_ISL_3029192, EPI_ISL_3029193, EPI_ISL_3029194, EPI_ISL_3029195, EPI_ISL_3029196, EPI_ISL_3029197, EPI_ISL_3029198, EPI_ISL_3029199, EPI_ISL_3029200, EPI_ISL_3029201, EPI_ISL_3029202, EPI_ISL_3029203, EPI_ISL_3029204, EPI_ISL_3029205, EPI_ISL_3029206, EPI_ISL_3029207, EPI_ISL_3029208, EPI_ISL_3029209, EPI_ISL_3029210, EPI_ISL_3029211, EPI_ISL_3029212, EPI_ISL_3029213, EPI_ISL_3029214, EPI_ISL_3029215, EPI_ISL_3029216, EPI_ISL_3029217, EPI_ISL_3029218, EPI_ISL_3029219, EPI_ISL_3029220, EPI_ISL_3029221, EPI_ISL_3029222, EPI_ISL_3029223, EPI_ISL_3029224, EPI_ISL_3029225, EPI_ISL_3029226, EPI_ISL_3029227, EPI_ISL_3029228, EPI_ISL_3029229, EPI_ISL_3029230, EPI_ISL_3029231, EPI_ISL_3029232, EPI_ISL_3029234, EPI_ISL_3029235, EPI_ISL_3029236, EPI_ISL_3029237, EPI_ISL_3029238, EPI_ISL_3029239, EPI_ISL_3029240, EPI_ISL_3029241, EPI_ISL_3029242, EPI_ISL_3029244, EPI_ISL_3029245, EPI_ISL_3029246, EPI_ISL_3029247 | | | |
| see above | Department of Bacteria, Parasites and Fungi, Statens Serum Institut, Copenhagen, Denmark | Statens Serum Institut Bioinformatics and Microbial Genomics | Danish Covid-19 Genome Consortium |
| EPI_ISL_3029248 | Molekylær Medicinsk Afdeling, Aarhus University Hospital, Aarhus, Denmark | Statens Serum Institut Bioinformatics and Microbial Genomics | Danish Covid-19 Genome Consortium |
| EPI_ISL_3029249, EPI_ISL_3029250, EPI_ISL_3029251, EPI_ISL_3029252, EPI_ISL_3029253, EPI_ISL_3029254, EPI_ISL_3029255, EPI_ISL_3029256, EPI_ISL_3029257, EPI_ISL_3029258, EPI_ISL_3029259, EPI_ISL_3029260, EPI_ISL_3029261, EPI_ISL_3029262, EPI_ISL_3029263, EPI_ISL_3029264, EPI_ISL_3029265, EPI_ISL_3029266, EPI_ISL_3029267, EPI_ISL_3029268, EPI_ISL_3029269, EPI_ISL_3029270, EPI_ISL_3029271, EPI_ISL_3029272, EPI_ISL_3029273, EPI_ISL_3029274, EPI_ISL_3029275, EPI_ISL_3029276, EPI_ISL_3029277, EPI_ISL_3029278, EPI_ISL_3029279, EPI_ISL_3029280, EPI_ISL_3029281, EPI_ISL_3029282, EPI_ISL_3029283, EPI_ISL_3029284, EPI_ISL_3029285, EPI_ISL_3029286, EPI_ISL_3029287, EPI_ISL_3029288, EPI_ISL_3029289, EPI_ISL_3029290, EPI_ISL_3029291, EPI_ISL_3029292, EPI_ISL_3029293, EPI_ISL_3029294, EPI_ISL_3029295, EPI_ISL_3029296 | | | |
| see above | Department of Bacteria, Parasites and Fungi, Statens Serum Institut, Copenhagen, Denmark | Statens Serum Institut Bioinformatics and Microbial Genomics | Danish Covid-19 Genome Consortium |
| EPI_ISL_3029297 | Department of Clinical Microbiology, Odense University Hospital, Odense, Denmark | Statens Serum Institut Bioinformatics and Microbial Genomics | Danish Covid-19 Genome Consortium |
| EPI_ISL_3029298 | Molekylær Medicinsk Afdeling, Aarhus University Hospital, Aarhus, Denmark | Statens Serum Institut Bioinformatics and Microbial Genomics | Danish Covid-19 Genome Consortium |
| EPI_ISL_3029299, EPI_ISL_3029300, EPI_ISL_3029301, EPI_ISL_3029302, EPI_ISL_3029303, EPI_ISL_3029304, EPI_ISL_3029305, EPI_ISL_3029306, EPI_ISL_3029307, EPI_ISL_3029308, EPI_ISL_3029309, EPI_ISL_3029310, EPI_ISL_3029311, EPI_ISL_3029312, EPI_ISL_3029313, EPI_ISL_3029314, EPI_ISL_3029315, EPI_ISL_3029316, EPI_ISL_3029317, EPI_ISL_3029318, EPI_ISL_3029319, EPI_ISL_3029320, EPI_ISL_3029321, EPI_ISL_3029322, EPI_ISL_3029323, EPI_ISL_3029324, EPI_ISL_3029325, EPI_ISL_3029326, EPI_ISL_3029328, EPI_ISL_3029329, EPI_ISL_3029330, EPI_ISL_3029331, EPI_ISL_3029332, EPI_ISL_3029333, EPI_ISL_3029335, EPI_ISL_3029336, EPI_ISL_3029337, EPI_ISL_3029338, EPI_ISL_3029339, EPI_ISL_3029340, EPI_ISL_3029341 | | | |
| see above | Department of Bacteria, Parasites and Fungi, Statens Serum Institut, Copenhagen, Denmark | Statens Serum Institut Bioinformatics and Microbial Genomics | Danish Covid-19 Genome Consortium |
| EPI_ISL_3029342 | Molekylær Medicinsk Afdeling, Aarhus University Hospital, Aarhus, Denmark | Statens Serum Institut Bioinformatics and Microbial Genomics | Danish Covid-19 Genome Consortium |
| EPI_ISL_3029343, EPI_ISL_3029344, EPI_ISL_3029345, EPI_ISL_3029346, EPI_ISL_3029347, EPI_ISL_3029348, EPI_ISL_3029349, EPI_ISL_3029350, EPI_ISL_3029351, EPI_ISL_3029352, EPI_ISL_3029353, EPI_ISL_3029354, EPI_ISL_3029356, EPI_ISL_3029357, EPI_ISL_3029358, EPI_ISL_3029359, EPI_ISL_3029360, EPI_ISL_3029361, EPI_ISL_3029362, EPI_ISL_3029363, EPI_ISL_3029364, EPI_ISL_3029365, EPI_ISL_3029366, EPI_ISL_3029368, EPI_ISL_3029369, EPI_ISL_3029370, EPI_ISL_3029371, EPI_ISL_3029372, EPI_ISL_3029373, EPI_ISL_3029374, EPI_ISL_3029375, EPI_ISL_3029376, EPI_ISL_3029377, EPI_ISL_3029378, EPI_ISL_3029379, EPI_ISL_3029380, EPI_ISL_3029381, EPI_ISL_3029382, EPI_ISL_3029383 | | | |
| see above | Department of Bacteria, Parasites and Fungi, Statens Serum Institut, Copenhagen, Denmark | Statens Serum Institut Bioinformatics and Microbial Genomics | Danish Covid-19 Genome Consortium |
| EPI_ISL_3029384 | Department of Clinical Microbiology, Odense University Hospital, Odense, Denmark | Statens Serum Institut Bioinformatics and Microbial Genomics | Danish Covid-19 Genome Consortium |
| EPI_ISL_3029385, EPI_ISL_3029386, EPI_ISL_3029387, EPI_ISL_3029388, EPI_ISL_3029389, EPI_ISL_3029390, EPI_ISL_3029391 | Department of Bacteria, Parasites and Fungi, Statens Serum Institut, Copenhagen, Denmark | Statens Serum Institut Bioinformatics and Microbial Genomics | Danish Covid-19 Genome Consortium |
| EPI_ISL_3029392 | Molekylær Medicinsk Afdeling, Aarhus University Hospital, Aarhus, Denmark | Statens Serum Institut Bioinformatics and Microbial Genomics | Danish Covid-19 Genome Consortium |
| EPI_ISL_3029393, EPI_ISL_3029394, EPI_ISL_3029395, EPI_ISL_3029396, EPI_ISL_3029397, EPI_ISL_3029398, EPI_ISL_3029399, EPI_ISL_3029400, EPI_ISL_3029401, EPI_ISL_3029402, EPI_ISL_3029403, EPI_ISL_3029404, EPI_ISL_3029405, EPI_ISL_3029406, EPI_ISL_3029407, EPI_ISL_3029408, EPI_ISL_3029409, EPI_ISL_3029410, EPI_ISL_3029411, EPI_ISL_3029413, EPI_ISL_3029414, EPI_ISL_3029415, EPI_ISL_3029416, EPI_ISL_3029417, EPI_ISL_3029418, EPI_ISL_3029419, EPI_ISL_3029420, EPI_ISL_3029421, EPI_ISL_3029422, EPI_ISL_3029423, EPI_ISL_3029424, EPI_ISL_3029425, EPI_ISL_3029426, EPI_ISL_3029427, EPI_ISL_3029428, EPI_ISL_3029429, EPI_ISL_3029430, EPI_ISL_3029431, EPI_ISL_3029432, EPI_ISL_3029433, EPI_ISL_3029434, EPI_ISL_3029435, EPI_ISL_3029436, EPI_ISL_3029438, EPI_ISL_3029439, EPI_ISL_3029440, EPI_ISL_3029441, EPI_ISL_3029442, EPI_ISL_3029444, EPI_ISL_3029445, EPI_ISL_3029446, EPI_ISL_3029447, EPI_ISL_3029448 | | | |
| see above | Department of Bacteria, Parasites and Fungi, Statens Serum Institut, Copenhagen, Denmark | Statens Serum Institut Bioinformatics and Microbial Genomics | Danish Covid-19 Genome Consortium |
| EPI_ISL_3029449 | Molekylær Medicinsk Afdeling, Aarhus University Hospital, Aarhus, Denmark | Statens Serum Institut Bioinformatics and Microbial Genomics | Danish Covid-19 Genome Consortium |
| EPI_ISL_3029450, EPI_ISL_3029451, EPI_ISL_3029452, EPI_ISL_3029453, EPI_ISL_3029454, EPI_ISL_3029455, EPI_ISL_3029456, EPI_ISL_3029457, EPI_ISL_3029458, EPI_ISL_3029459, EPI_ISL_3029460, EPI_ISL_3029461, EPI_ISL_3029462, EPI_ISL_3029463, EPI_ISL_3029464, EPI_ISL_3029465, EPI_ISL_3029466, EPI_ISL_3029467, EPI_ISL_3029468, EPI_ISL_3029469, EPI_ISL_3029470, EPI_ISL_3029471, EPI_ISL_3029472, EPI_ISL_3029473, EPI_ISL_3029474, EPI_ISL_3029475, EPI_ISL_3029476, EPI_ISL_3029477, EPI_ISL_3029478, EPI_ISL_3029479, EPI_ISL_3029481, EPI_ISL_3029482, EPI_ISL_3029483, EPI_ISL_3029484, EPI_ISL_3029485, EPI_ISL_3029486, EPI_ISL_3029487, EPI_ISL_3029488, EPI_ISL_3029489, EPI_ISL_3029490, EPI_ISL_3029491, EPI_ISL_3029492, EPI_ISL_3029493, EPI_ISL_3029494, EPI_ISL_3029495, EPI_ISL_3029496, EPI_ISL_3029497, EPI_ISL_3029498, EPI_ISL_3029499, EPI_ISL_3029500, EPI_ISL_3029501, EPI_ISL_3029502, EPI_ISL_3029504, EPI_ISL_3029505, EPI_ISL_3029506, EPI_ISL_3029507, EPI_ISL_3029508, EPI_ISL_3029509, EPI_ISL_3029510, EPI_ISL_3029511, EPI_ISL_3029512, EPI_ISL_3029513, EPI_ISL_3029514, EPI_ISL_3029515, EPI_ISL_3029516, EPI_ISL_3029517, EPI_ISL_3029518, EPI_ISL_3029519, EPI_ISL_3029520, EPI_ISL_3029521, EPI_ISL_3029522, EPI_ISL_3029523, EPI_ISL_3029524, EPI_ISL_3029525, EPI_ISL_3029526, EPI_ISL_3029527, EPI_ISL_3029528, EPI_ISL_3029530, EPI_ISL_3029531, EPI_ISL_3029532, EPI_ISL_3029533, EPI_ISL_3029534, EPI_ISL_3029535, EPI_ISL_3029536, EPI_ISL_3029537, EPI_ISL_3029538, EPI_ISL_3029539, EPI_ISL_3029541, EPI_ISL_3029542, EPI_ISL_3029543, EPI_ISL_3029544, EPI_ISL_3029545, EPI_ISL_3029546, EPI_ISL_3029547, EPI_ISL_3029548, EPI_ISL_3029549, EPI_ISL_3029550, EPI_ISL_3029551 | | | |
| see above | Department of Bacteria, Parasites and Fungi, Statens Serum Institut, Copenhagen, Denmark | Statens Serum Institut Bioinformatics and Microbial Genomics | Danish Covid-19 Genome Consortium |
| EPI_ISL_3029552 | Molekylær Medicinsk Afdeling, Aarhus University Hospital, Aarhus, Denmark | Statens Serum Institut Bioinformatics and Microbial Genomics | Danish Covid-19 Genome Consortium |
| EPI_ISL_3029553, EPI_ISL_3029554, EPI_ISL_3029555, EPI_ISL_3029556, EPI_ISL_3029557, EPI_ISL_3029558, EPI_ISL_3029559, EPI_ISL_3029560, EPI_ISL_3029561, EPI_ISL_3029562, EPI_ISL_3029563, EPI_ISL_3029564, EPI_ISL_3029565, EPI_ISL_3029566, EPI_ISL_3029567, EPI_ISL_3029568, EPI_ISL_3029569, EPI_ISL_3029570, EPI_ISL_3029571, EPI_ISL_3029572, EPI_ISL_3029573, EPI_ISL_3029574, EPI_ISL_3029575, EPI_ISL_3029576, EPI_ISL_3029577, EPI_ISL_3029578, EPI_ISL_3029579, EPI_ISL_3029580, EPI_ISL_3029581, EPI_ISL_3029582, EPI_ISL_3029583, EPI_ISL_3029584, EPI_ISL_3029585, EPI_ISL_3029586, EPI_ISL_3029587, EPI_ISL_3029588, EPI_ISL_3029589, EPI_ISL_3029590, EPI_ISL_3029591, EPI_ISL_3029592, EPI_ISL_3029593, EPI_ISL_3029594, EPI_ISL_3029595, EPI_ISL_3029596, EPI_ISL_3029597, EPI_ISL_3029598, EPI_ISL_3029599, EPI_ISL_3029600, EPI_ISL_3029601, EPI_ISL_3029602, EPI_ISL_3029603, EPI_ISL_3029604, EPI_ISL_3029605, EPI_ISL_3029606, EPI_ISL_3029607, EPI_ISL_3029608, EPI_ISL_3029609, EPI_ISL_3029610, EPI_ISL_3029611, EPI_ISL_3029612, EPI_ISL_3029613, EPI_ISL_3029614, EPI_ISL_3029615, EPI_ISL_3029616, EPI_ISL_3029617, EPI_ISL_3029618, EPI_ISL_3029619, EPI_ISL_3029620, EPI_ISL_3029621, EPI_ISL_3029622, EPI_ISL_3029623, EPI_ISL_3029624, EPI_ISL_3029625, EPI_ISL_3029626, EPI_ISL_3029627, EPI_ISL_3029628, EPI_ISL_3029629, EPI_ISL_3029630, EPI_ISL_3029631, EPI_ISL_3029632, EPI_ISL_3029633, EPI_ISL_3029634, EPI_ISL_3029635, EPI_ISL_3029636, EPI_ISL_3029637, EPI_ISL_3029638, EPI_ISL_3029639, EPI_ISL_3029640, EPI_ISL_3029641, EPI_ISL_3029642, EPI_ISL_3029643, EPI_ISL_3029645, EPI_ISL_3029646, EPI_ISL_3029647, EPI_ISL_3029648, EPI_ISL_3029649, EPI_ISL_3029650, EPI_ISL_3029651, EPI_ISL_3029652, EPI_ISL_3029653, EPI_ISL_3029654, EPI_ISL_3029655, EPI_ISL_3029656, EPI_ISL_3029657, EPI_ISL_3029658, EPI_ISL_3029659, EPI_ISL_3029660, EPI_ISL_3029661, EPI_ISL_3029662, EPI_ISL_3029663, EPI_ISL_3029664, EPI_ISL_3029665, EPI_ISL_3029666, EPI_ISL_3029667, EPI_ISL_3029668, EPI_ISL_3029669, EPI_ISL_3029670, EPI_ISL_3029671, EPI_ISL_3029672, EPI_ISL_3029673, EPI_ISL_3029674, EPI_ISL_3029675, EPI_ISL_3029676, EPI_ISL_3029677, EPI_ISL_3029678, EPI_ISL_3029679, EPI_ISL_3029680, EPI_ISL_3029681, EPI_ISL_3029682, EPI_ISL_3029683, EPI_ISL_3029684, EPI_ISL_3029685, EPI_ISL_3029686, EPI_ISL_3029687, EPI_ISL_3029688, EPI_ISL_3029689, EPI_ISL_3029690, EPI_ISL_3029691, EPI_ISL_3029692, EPI_ISL_3029693, EPI_ISL_3029694, EPI_ISL_3029695, EPI_ISL_3029696, EPI_ISL_3029697, EPI_ISL_3029698, EPI_ISL_3029699, EPI_ISL_3029700, EPI_ISL_3029701, EPI_ISL_3029702, EPI_ISL_3029703, EPI_ISL_3029704, EPI_ISL_3029705, EPI_ISL_3029706, EPI_ISL_3029707, EPI_ISL_3029708, EPI_ISL_3029709, EPI_ISL_3029710, EPI_ISL_3029711, EPI_ISL_3029712, EPI_ISL_3029713, EPI_ISL_3029714, EPI_ISL_3029715, EPI_ISL_3029716, EPI_ISL_3029717, EPI_ISL_3029718, EPI_ISL_3029719 | | | |
| see above | Department of Bacteria, Parasites and Fungi, Statens Serum Institut, Copenhagen, Denmark | Statens Serum Institut Bioinformatics and Microbial Genomics | Danish Covid-19 Genome Consortium |
| EPI_ISL_3029720 | Molekylær Medicinsk Afdeling, Aarhus University Hospital, Aarhus, Denmark | Statens Serum Institut Bioinformatics and Microbial Genomics | Danish Covid-19 Genome Consortium |

EPI_ISL_3029721, EPI_ISL_3029722, EPI_ISL_3029723, EPI_ISL_3029724, EPI_ISL_3029725, EPI_ISL_3029726, EPI_ISL_3029727, EPI_ISL_3029728, EPI_ISL_3029729, EPI_ISL_3029730, EPI_ISL_3029731, EPI_ISL_3029732, EPI_ISL_3029733, EPI_ISL_3029734

| | | | |
|---|---|---|---|
| see above | Department of Bacteria, Parasites and Fungi, Statens Serum Institut, Copenhagen, Denmark | Statens Serum Institut Bioinformatics and Microbial Genomics | Danish Covid-19 Genome Consortium |
| EPI_ISL_3029735 | Molekylær Medicinsk Afdeling, Aarhus University Hospital, Aarhus, Denmark | Statens Serum Institut Bioinformatics and Microbial Genomics | Danish Covid-19 Genome Consortium |

EPI_ISL_3029736, EPI_ISL_3029737, EPI_ISL_3029738, EPI_ISL_3029739, EPI_ISL_3029740, EPI_ISL_3029741, EPI_ISL_3029742, EPI_ISL_3029743, EPI_ISL_3029744, EPI_ISL_3029745, EPI_ISL_3029746, EPI_ISL_3029748, EPI_ISL_3029749, EPI_ISL_3029750, EPI_ISL_3029751, EPI_ISL_3029752, EPI_ISL_3029753, EPI_ISL_3029754, EPI_ISL_3029755, EPI_ISL_3029756, EPI_ISL_3029757, EPI_ISL_3029758, EPI_ISL_3029759, EPI_ISL_3029760, EPI_ISL_3029761, EPI_ISL_3029762, EPI_ISL_3029763, EPI_ISL_3029764, EPI_ISL_3029765, EPI_ISL_3029766, EPI_ISL_3029767, EPI_ISL_3029768, EPI_ISL_3029769, EPI_ISL_3029770, EPI_ISL_3029771, EPI_ISL_3029772, EPI_ISL_3029773, EPI_ISL_3029774, EPI_ISL_3029775, EPI_ISL_3029776, EPI_ISL_3029777, EPI_ISL_3029778, EPI_ISL_3029779, EPI_ISL_3029780, EPI_ISL_3029781, EPI_ISL_3029782, EPI_ISL_3029783, EPI_ISL_3029784, EPI_ISL_3029785, EPI_ISL_3029786, EPI_ISL_3029787, EPI_ISL_3029788, EPI_ISL_3029789, EPI_ISL_3029790, EPI_ISL_3029791, EPI_ISL_3029792, EPI_ISL_3029793, EPI_ISL_3029794, EPI_ISL_3029795, EPI_ISL_3029796

| | | | |
|---|---|---|---|
| see above | Department of Bacteria, Parasites and Fungi, Statens Serum Institut, Copenhagen, Denmark | Statens Serum Institut Bioinformatics and Microbial Genomics | Danish Covid-19 Genome Consortium |
| EPI_ISL_3029797 | Department of Clinical Microbiology, Odense University Hospital, Odense, Denmark | Statens Serum Institut Bioinformatics and Microbial Genomics | Danish Covid-19 Genome Consortium |

EPI_ISL_3029798, EPI_ISL_3029799, EPI_ISL_3029800, EPI_ISL_3029801, EPI_ISL_3029802, EPI_ISL_3029803, EPI_ISL_3029804, EPI_ISL_3029805, EPI_ISL_3029807, EPI_ISL_3029808, EPI_ISL_3029809, EPI_ISL_3029810, EPI_ISL_3029811, EPI_ISL_3029812, EPI_ISL_3029813, EPI_ISL_3029814, EPI_ISL_3029815, EPI_ISL_3029816

| | | | |
|---|---|---|---|
| see above | Department of Bacteria, Parasites and Fungi, Statens Serum Institut, Copenhagen, Denmark | Statens Serum Institut Bioinformatics and Microbial Genomics | Danish Covid-19 Genome Consortium |
| EPI_ISL_3029817 | AULSS 6 Euganea | Istituto Zooprofilattico Sperimentale delle Venezie | Adelaide Milani, Alessia Schivo, Annalisa Salviato, Elisa Palumbo, Erika Giorgia Quaranta, Luca Tassoni, Ambra Pastori, Edoardo Giussani, Alice Fusaro, Isabella Monne, Calogero Terregino, Antonia Ricci |
| EPI_ISL_3029818, EPI_ISL_3029819, EPI_ISL_3029820, EPI_ISL_3029821 | AULSS 5 Polesana | Istituto Zooprofilattico Sperimentale delle Venezie | Adelaide Milani, Alessia Schivo, Annalisa Salviato, Elisa Palumbo, Erika Giorgia Quaranta, Luca Tassoni, Ambra Pastori, Edoardo Giussani, Alice Fusaro, Isabella Monne, Calogero Terregino, Antonia Ricci |
| EPI_ISL_3029822, EPI_ISL_3029823 | AULSS 1 Dolomiti | Istituto Zooprofilattico Sperimentale delle Venezie | Adelaide Milani, Alessia Schivo, Annalisa Salviato, Elisa Palumbo, Erika Giorgia Quaranta, Luca Tassoni, Ambra Pastori, Edoardo Giussani, Alice Fusaro, Isabella Monne, Calogero Terregino, Antonia Ricci |
| EPI_ISL_3029824 | AULSS 5 Polesana | Istituto Zooprofilattico Sperimentale delle Venezie | Adelaide Milani, Alessia Schivo, Annalisa Salviato, Elisa Palumbo, Erika Giorgia Quaranta, Luca Tassoni, Ambra Pastori, Edoardo Giussani, Alice Fusaro, Isabella Monne, Calogero Terregino, Antonia Ricci |
| EPI_ISL_3029825, EPI_ISL_3029826, EPI_ISL_3029827, EPI_ISL_3029828 | AULSS 1 Dolomiti | Istituto Zooprofilattico Sperimentale delle Venezie | Adelaide Milani, Alessia Schivo, Annalisa Salviato, Elisa Palumbo, Erika Giorgia Quaranta, Luca Tassoni, Ambra Pastori, Edoardo Giussani, Alice Fusaro, Isabella Monne, Calogero Terregino, Antonia Ricci |
| EPI_ISL_3029835, EPI_ISL_3029836 | Genome Information Research Center, Research Institute for Microbial Diseases, Osaka University | Genome Information Research Center, Research Institute for Microbial Diseases, Osaka University | Daisuke Motooka, Kentaro Tanaka, Hiroya Oki, and Shota Nakamura |
| EPI_ISL_3029848, EPI_ISL_3029849 | Ospedale Giovanni Paolo II Ragusa RG | AOU Polclinico "G Rodolico-S.Marco" PO Rodolico and BIOMETEC UNICT CATANIA | STEFANI AND SCALIA |
| EPI_ISL_3029855, EPI_ISL_3029856, EPI_ISL_3029857, EPI_ISL_3029858, EPI_ISL_3029859, EPI_ISL_3029860, EPI_ISL_3029861 | Olive View Medical Center | Los Angeles County Public Health Laboratories | P. Hemarajata et al. |
| EPI_ISL_3029863, EPI_ISL_3029864 | Huntington Memorial Hospital | Los Angeles County Public Health Laboratories | P. Hemarajata et al. |
| EPI_ISL_3029870, EPI_ISL_3029871 | Martin Luther King, Jr. Outpatient Center | Los Angeles County Public Health Laboratories | P. Hemarajata et al. |
| EPI_ISL_3029872 | Henry Mayo Newhall Hospital | Los Angeles County Public Health Laboratories | P. Hemarajata et al. |
| EPI_ISL_3029898 | Ospedale Giovanni Paolo II Ragusa RG | AOU Polclinico "G Rodolico-S.Marco" PO Rodolico and BIOMETEC UNICT CATANIA | STEFANI AND SCALIA |
| EPI_ISL_3029904 | General Hospital Tetovo | Laboratory of Virology and Molecular Diagnostics, Institute of Public Health of Republic of North Macedonia | Kuzmanovska et al. |
| EPI_ISL_3029905 | RSUP Dr. Kariadi | Eijkman Institute for Molecular Biology, National Research and Innovation Agency | Lydia V. Panggalo, Iskandar Adnan, Sukma Oktavianthi, Lidwina Priliani, Edison Johar, Frilasita A Yudhaputri, Muhammad Rezki Rasyak, Willy Agustine, Hidayat Trimarsanto, Safarina G Malik, Khin Saw Myint, Amin Soebandrio |
| EPI_ISL_3029906 | RSUP Dr. Kariadi | Eijkman Institute for Molecular Biology, National Research and Innovation Agency | Sukma Oktavianthi, Lidwina Priliani, Edison Johar, Frilasita A Yudhaputri, Muhammad Rezki Rasyak, Willy Agustine, Hidayat Trimarsanto, Lydia V. Panggalo, Iskandar Adnan, Safarina G Malik, Khin Saw Myint, Amin Soebandrio |
| EPI_ISL_3029907 | RSUP Dr. Kariadi | Eijkman Institute for Molecular Biology, National Research and Innovation Agency | Lidwina Priliani, Edison Johar, Frilasita A Yudhaputri, Muhammad Rezki Rasyak, Willy Agustine, Hidayat Trimarsanto, Lydia V. Panggalo, Iskandar Adnan, Sukma Oktavianthi, Safarina G Malik, Khin Saw Myint, Amin Soebandrio |
| EPI_ISL_3029908 | RSUP Dr. Kariadi | Eijkman Institute for Molecular Biology, National Research and Innovation Agency | Edison Johar, Frilasita A Yudhaputri, Muhammad Rezki Rasyak, Willy Agustine, Hidayat Trimarsanto, Lydia V. Panggalo, Iskandar Adnan, Sukma Oktavianthi, Lidwina Priliani, Safarina G Malik, Khin Saw Myint, Amin Soebandrio |
| EPI_ISL_3029910 | RSUP Dr. Kariadi | Eijkman Institute for Molecular Biology, National Research and Innovation Agency | Muhammad Rezki Rasyak, Willy Agustine, Hidayat Trimarsanto, Lydia V. Panggalo, Iskandar Adnan, Sukma Oktavianthi, Lidwina Priliani, Edison Johar, Frilasita A Yudhaputri, Safarina G Malik, Khin Saw Myint, Amin Soebandrio |
| EPI_ISL_3029911 | RSUP Dr. Kariadi | Eijkman Institute for Molecular Biology, National Research and Innovation Agency | Willy Agustine, Hidayat Trimarsanto, Lydia V. Panggalo, Iskandar Adnan, Sukma Oktavianthi, Lidwina Priliani, Edison Johar, Frilasita A Yudhaputri, Muhammad Rezki Rasyak, Safarina G Malik, Khin Saw Myint, Amin Soebandrio |
| EPI_ISL_3029912 | RSUP Dr. Kariadi | Eijkman Institute for Molecular Biology, National Research and Innovation Agency | Hidayat Trimarsanto, Lydia V. Panggalo, Iskandar Adnan, Sukma Oktavianthi, Lidwina Priliani, Edison Johar, Frilasita A Yudhaputri, Muhammad Rezki Rasyak, Willy Agustine, Safarina G Malik, Khin Saw Myint, Amin Soebandrio |
| EPI_ISL_3029913 | RSUP Dr. Kariadi | Eijkman Institute for Molecular Biology, National Research and Innovation Agency | Lydia V. Panggalo, Iskandar Adnan, Sukma Oktavianthi, Lidwina Priliani, Edison Johar, Frilasita A Yudhaputri, Muhammad Rezki Rasyak, Willy Agustine, Hidayat Trimarsanto, Safarina G Malik, Khin Saw Myint, Amin Soebandrio |
| EPI_ISL_3029914 | RSUP Dr. Kariadi | Eijkman Institute for Molecular Biology, National Research and Innovation Agency | Iskandar Adnan, Sukma Oktavianthi, Lidwina Priliani, Edison Johar, Frilasita A Yudhaputri, Muhammad Rezki Rasyak, Willy Agustine, Hidayat Trimarsanto, Lydia V. Panggalo, Safarina G Malik, Khin Saw Myint, Amin Soebandrio |
| EPI_ISL_3029915 | RSUP Dr. Kariadi | Eijkman Institute for Molecular Biology, National Research and Innovation Agency | Lidwina Priliani, Edison Johar, Frilasita A Yudhaputri, Muhammad Rezki Rasyak, Willy Agustine, Hidayat Trimarsanto, Lydia V. Panggalo, Iskandar Adnan, Sukma Oktavianthi, Safarina G Malik, Khin Saw Myint, Amin Soebandrio |
| EPI_ISL_3029916 | RSUP Dr. Kariadi | Eijkman Institute for Molecular Biology, National Research and Innovation Agency | Edison Johar, Frilasita A Yudhaputri, Muhammad Rezki Rasyak, Willy Agustine, Hidayat Trimarsanto, Lydia V. Panggalo, Iskandar Adnan, Sukma Oktavianthi, Lidwina Priliani, Safarina G Malik, Khin Saw Myint, Amin Soebandrio |
| EPI_ISL_3029917 | RSUP Dr. Kariadi | Eijkman Institute for Molecular Biology, National Research and Innovation Agency | Frilasita A Yudhaputri, Muhammad Rezki Rasyak, Willy Agustine, Hidayat Trimarsanto, Lydia V. Panggalo, Iskandar Adnan, Sukma Oktavianthi, Lidwina Priliani, Edison Johar, Safarina G Malik, Khin Saw Myint, Amin Soebandrio |
| EPI_ISL_3029918 | RSUP Dr. Kariadi | Eijkman Institute for Molecular Biology, National Research and Innovation Agency | Muhammad Rezki Rasyak, Willy Agustine, Hidayat Trimarsanto, Lydia V. Panggalo, Iskandar Adnan, Sukma Oktavianthi, Lidwina Priliani, Edison Johar, Frilasita A Yudhaputri, Safarina G Malik, Khin Saw Myint, Amin Soebandrio |
| EPI_ISL_3029919 | RSUP Dr. Kariadi | Eijkman Institute for Molecular Biology, National Research and Innovation Agency | Hidayat Trimarsanto, Lydia V. Panggalo, Iskandar Adnan, Sukma Oktavianthi, Lidwina Priliani, Edison Johar, Frilasita A Yudhaputri, Muhammad Rezki Rasyak, Willy Agustine, Safarina G Malik, Khin Saw Myint, Amin Soebandrio |
| EPI_ISL_3029920 | RSUP Dr. Kariadi | Eijkman Institute for Molecular Biology, National Research and Innovation Agency | Lydia V. Panggalo, Iskandar Adnan, Sukma Oktavianthi, Lidwina Priliani, Edison Johar, Frilasita A Yudhaputri, Muhammad Rezki Rasyak, Willy Agustine, Hidayat Trimarsanto, Safarina G Malik, Khin Saw Myint, Amin Soebandrio |
| EPI_ISL_3029921 | RSUP Dr. Kariadi | Eijkman Institute for Molecular Biology, National Research and Innovation Agency | Iskandar Adnan, Sukma Oktavianthi, Lidwina Priliani, Edison Johar, Frilasita A Yudhaputri, Muhammad Rezki Rasyak, Willy Agustine, Hidayat Trimarsanto, Lydia V. Panggalo, Safarina G Malik, Khin Saw Myint, Amin Soebandrio |
| EPI_ISL_3029922 | RSUP Dr. Kariadi | Eijkman Institute for Molecular Biology, National Research and Innovation Agency | Sukma Oktavianthi, Lidwina Priliani, Edison Johar, Frilasita A Yudhaputri, Muhammad Rezki Rasyak, Willy Agustine, Hidayat Trimarsanto, Lydia V. Panggalo, Iskandar Adnan, Safarina G Malik, Khin Saw Myint, Amin Soebandrio |
| EPI_ISL_3029923 | RSUP Dr. Kariadi | Eijkman Institute for Molecular Biology, National Research and Innovation Agency | Edison Johar, Frilasita A Yudhaputri, Muhammad Rezki Rasyak, Willy Agustine, Hidayat Trimarsanto, Lydia V. Panggalo, Iskandar Adnan, Sukma Oktavianthi, Lidwina Priliani, Safarina G Malik, Khin Saw Myint, Amin Soebandrio |
| EPI_ISL_3029924 | RSUP Dr. Kariadi | Eijkman Institute for Molecular Biology, National Research and Innovation Agency | Frilasita A Yudhaputri, Muhammad Rezki Rasyak, Willy Agustine, Hidayat Trimarsanto, Lydia V. Panggalo, Iskandar Adnan, Sukma Oktavianthi, Lidwina Priliani, Edison Johar, Safarina G Malik, Khin Saw Myint, Amin Soebandrio |
| EPI_ISL_3029925 | RSUP Dr. Kariadi | Eijkman Institute for Molecular Biology, National Research and Innovation Agency | Muhammad Rezki Rasyak, Willy Agustine, Hidayat Trimarsanto, Lydia V. Panggalo, Iskandar Adnan, Sukma Oktavianthi, Lidwina Priliani, Edison Johar, Frilasita A Yudhaputri, Safarina G Malik, Khin Saw Myint, Amin Soebandrio |
| EPI_ISL_3029926 | RSUP Dr. Kariadi | Eijkman Institute for Molecular Biology, National Research and Innovation Agency | Hidayat Trimarsanto, Lydia V. Panggalo, Iskandar Adnan, Sukma Oktavianthi, Lidwina Priliani, Edison Johar, Frilasita A Yudhaputri, Muhammad Rezki Rasyak, Willy Agustine, Safarina G Malik, Khin Saw Myint, Amin Soebandrio |
| EPI_ISL_3029930 | RSUP Dr. Kariadi | Eijkman Institute for Molecular Biology, National Research and Innovation Agency | Edison Johar, Frilasita A Yudhaputri, Muhammad Rezki Rasyak, Willy Agustine, Hidayat Trimarsanto, Lydia V. Panggalo, Iskandar Adnan, Sukma Oktavianthi, Lidwina Priliani, Safarina G Malik, Khin Saw Myint, Amin Soebandrio |

| | | | |
|---|---|---|---|
| EPI_ISL_3029931 | RS Pondok Indah Pondok Indah | Eijkman Institute for Molecular Biology, National Research and Innovation Agency | Willy Agustine, Hidayat Trimarsanto, Lydia V. Panggalo, Iskandar Adnan, Sukma Oktavianthi, Lidwina Priliani, Edison Johar, Frilasita A Yudhaputri, Muhammad Rezki Rasyak, Safarina G Malik, Khin Saw Myint, Amin Soebandrio |
| EPI_ISL_3029932 | RS Pondok Indah Pondok Indah | Eijkman Institute for Molecular Biology, National Research and Innovation Agency | Lydia V. Panggalo, Iskandar Adnan, Sukma Oktavianthi, Lidwina Priliani, Edison Johar, Frilasita A Yudhaputri, Muhammad Rezki Rasyak, Willy Agustine, Hidayat Trimarsanto, Safarina G Malik, Khin Saw Myint, Amin Soebandrio |
| EPI_ISL_3029933 | RSU Dinda Tangerang | Eijkman Institute for Molecular Biology, National Research and Innovation Agency | Iskandar Adnan, Sukma Oktavianthi, Lidwina Priliani, Edison Johar, Frilasita A Yudhaputri, Muhammad Rezki Rasyak, Willy Agustine, Hidayat Trimarsanto, Lydia V. Panggalo, Safarina G Malik, Khin Saw Myint, Amin Soebandrio |
| EPI_ISL_3029934 | RS Hermina Depok | Eijkman Institute for Molecular Biology, National Research and Innovation Agency | Lidwina Priliani, Edison Johar, Frilasita A Yudhaputri, Muhammad Rezki Rasyak, Willy Agustine, Hidayat Trimarsanto, Lydia V. Panggalo, Iskandar Adnan, Sukma Oktavianthi, Safarina G Malik, Khin Saw Myint, Amin Soebandrio |
| EPI_ISL_3029935 | RS Hermina Depok | Eijkman Institute for Molecular Biology, National Research and Innovation Agency | Edison Johar, Frilasita A Yudhaputri, Muhammad Rezki Rasyak, Willy Agustine, Hidayat Trimarsanto, Lydia V. Panggalo, Iskandar Adnan, Sukma Oktavianthi, Lidwina Priliani, Safarina G Malik, Khin Saw Myint, Amin Soebandrio |
| EPI_ISL_3029937 | RS Hermina Periuk | Eijkman Institute for Molecular Biology, National Research and Innovation Agency | Hidayat Trimarsanto, Lydia V. Panggalo, Iskandar Adnan, Sukma Oktavianthi, Lidwina Priliani, Edison Johar, Frilasita A Yudhaputri, Muhammad Rezki Rasyak, Willy Agustine, Safarina G Malik, Khin Saw Myint, Amin Soebandrio |
| EPI_ISL_3029938 | RS Premier Bintaro | Eijkman Institute for Molecular Biology, National Research and Innovation Agency | Lydia V. Panggalo, Iskandar Adnan, Sukma Oktavianthi, Lidwina Priliani, Edison Johar, Frilasita A Yudhaputri, Muhammad Rezki Rasyak, Willy Agustine, Hidayat Trimarsanto, Safarina G Malik, Khin Saw Myint, Amin Soebandrio |
| EPI_ISL_3029939 | RSU HGA Depok | Eijkman Institute for Molecular Biology, National Research and Innovation Agency | Iskandar Adnan, Sukma Oktavianthi, Lidwina Priliani, Edison Johar, Frilasita A Yudhaputri, Muhammad Rezki Rasyak, Willy Agustine, Hidayat Trimarsanto, Lydia V. Panggalo, Safarina G Malik, Khin Saw Myint, Amin Soebandrio |
| EPI_ISL_3029940 | RS Pelni | Eijkman Institute for Molecular Biology, National Research and Innovation Agency | Sukma Oktavianthi, Lidwina Priliani, Edison Johar, Frilasita A Yudhaputri, Muhammad Rezki Rasyak, Willy Agustine, Hidayat Trimarsanto, Lydia V. Panggalo, Iskandar Adnan, Safarina G Malik, Khin Saw Myint, Amin Soebandrio |
| EPI_ISL_3029941 | RS Mitra Keluarga Cibubur | Eijkman Institute for Molecular Biology, National Research and Innovation Agency | Edison Johar, Frilasita A Yudhaputri, Muhammad Rezki Rasyak, Willy Agustine, Hidayat Trimarsanto, Lydia V. Panggalo, Iskandar Adnan, Sukma Oktavianthi, Lidwina Priliani, Safarina G Malik, Khin Saw Myint, Amin Soebandrio |
| EPI_ISL_3029942 | RS Mitra Keluarga Cibubur | Eijkman Institute for Molecular Biology, National Research and Innovation Agency | Frilasita A Yudhaputri, Muhammad Rezki Rasyak, Willy Agustine, Hidayat Trimarsanto, Lydia V. Panggalo, Iskandar Adnan, Sukma Oktavianthi, Lidwina Priliani, Edison Johar, Safarina G Malik, Khin Saw Myint, Amin Soebandrio |
| EPI_ISL_3029943 | Dinas Kesehatan Provinsi Nusa Tenggara Timur | Eijkman Institute for Molecular Biology, National Research and Innovation Agency | Muhammad Rezki Rasyak, Willy Agustine, Hidayat Trimarsanto, Lydia V. Panggalo, Iskandar Adnan, Sukma Oktavianthi, Lidwina Priliani, Edison Johar, Frilasita A Yudhaputri, Safarina G Malik, Khin Saw Myint, Amin Soebandrio |
| EPI_ISL_3029946 | Dinas Kesehatan Provinsi Nusa Tenggara Timur | Eijkman Institute for Molecular Biology, National Research and Innovation Agency | Sukma Oktavianthi, Lidwina Priliani, Edison Johar, Frilasita A Yudhaputri, Muhammad Rezki Rasyak, Willy Agustine, Hidayat Trimarsanto, Lydia V. Panggalo, Iskandar Adnan, Safarina G Malik, Khin Saw Myint, Amin Soebandrio |
| EPI_ISL_3029950 | Dinas Kesehatan Provinsi Nusa Tenggara Timur | Eijkman Institute for Molecular Biology, National Research and Innovation Agency | Lidwina Priliani, Edison Johar, Frilasita A Yudhaputri, Muhammad Rezki Rasyak, Willy Agustine, Hidayat Trimarsanto, Lydia V. Panggalo, Iskandar Adnan, Sukma Oktavianthi, Safarina G Malik, Khin Saw Myint, Amin Soebandrio |
| EPI_ISL_3029953 | Dinas Kesehatan Provinsi Nusa Tenggara Timur | Eijkman Institute for Molecular Biology, National Research and Innovation Agency | Lydia V. Panggalo, Iskandar Adnan, Sukma Oktavianthi, Lidwina Priliani, Edison Johar, Frilasita A Yudhaputri, Muhammad Rezki Rasyak, Willy Agustine, Hidayat Trimarsanto, Safarina G Malik, Khin Saw Myint, Amin Soebandrio |
| EPI_ISL_3029954 | Dinas Kesehatan Provinsi Nusa Tenggara Timur | Eijkman Institute for Molecular Biology, National Research and Innovation Agency | Iskandar Adnan, Sukma Oktavianthi, Lidwina Priliani, Edison Johar, Frilasita A Yudhaputri, Muhammad Rezki Rasyak, Willy Agustine, Hidayat Trimarsanto, Lydia V. Panggalo, Safarina G Malik, Khin Saw Myint, Amin Soebandrio |
| EPI_ISL_3029955 | Dinas Kesehatan Provinsi Nusa Tenggara Timur | Eijkman Institute for Molecular Biology, National Research and Innovation Agency | Edison Johar, Frilasita A Yudhaputri, Muhammad Rezki Rasyak, Willy Agustine, Hidayat Trimarsanto, Lydia V. Panggalo, Iskandar Adnan, Sukma Oktavianthi, Lidwina Priliani, Safarina G Malik, Khin Saw Myint, Amin Soebandrio |
| EPI_ISL_3029956 | Dinas Kesehatan Provinsi Nusa Tenggara Timur | Eijkman Institute for Molecular Biology, National Research and Innovation Agency | Muhammad Rezki Rasyak, Willy Agustine, Hidayat Trimarsanto, Lydia V. Panggalo, Iskandar Adnan, Sukma Oktavianthi, Lidwina Priliani, Edison Johar, Frilasita A Yudhaputri, Safarina G Malik, Khin Saw Myint, Amin Soebandrio |
| EPI_ISL_3029961 | Dinas Kesehatan Provinsi Nusa Tenggara Timur | Eijkman Institute for Molecular Biology, National Research and Innovation Agency | Sukma Oktavianthi, Lidwina Priliani, Edison Johar, Frilasita A Yudhaputri, Muhammad Rezki Rasyak, Willy Agustine, Hidayat Trimarsanto, Lydia V. Panggalo, Iskandar Adnan, Safarina G Malik, Khin Saw Myint, Amin Soebandrio |
| EPI_ISL_3029962 | Dinas Kesehatan Provinsi Nusa Tenggara Timur | Eijkman Institute for Molecular Biology, National Research and Innovation Agency | Lidwina Priliani, Edison Johar, Frilasita A Yudhaputri, Muhammad Rezki Rasyak, Willy Agustine, Hidayat Trimarsanto, Lydia V. Panggalo, Iskandar Adnan, Sukma Oktavianthi, Safarina G Malik, Khin Saw Myint, Amin Soebandrio |
| EPI_ISL_3029963 | Dinas Kesehatan Provinsi Nusa Tenggara Timur | Eijkman Institute for Molecular Biology, National Research and Innovation Agency | Edison Johar, Frilasita A Yudhaputri, Muhammad Rezki Rasyak, Willy Agustine, Hidayat Trimarsanto, Lydia V. Panggalo, Iskandar Adnan, Sukma Oktavianthi, Lidwina Priliani, Safarina G Malik, Khin Saw Myint, Amin Soebandrio |
| EPI_ISL_3029964 | Dinas Kesehatan Provinsi Nusa Tenggara Timur | Eijkman Institute for Molecular Biology, National Research and Innovation Agency | Frilasita A Yudhaputri, Muhammad Rezki Rasyak, Willy Agustine, Hidayat Trimarsanto, Lydia V. Panggalo, Iskandar Adnan, Sukma Oktavianthi, Lidwina Priliani, Edison Johar, Safarina G Malik, Khin Saw Myint, Amin Soebandrio |
| EPI_ISL_3029965 | Dinas Kesehatan Provinsi Nusa Tenggara Timur | Eijkman Institute for Molecular Biology, National Research and Innovation Agency | Willy Agustine, Hidayat Trimarsanto, Lydia V. Panggalo, Iskandar Adnan, Sukma Oktavianthi, Lidwina Priliani, Edison Johar, Frilasita A Yudhaputri, Muhammad Rezki Rasyak, Safarina G Malik, Khin Saw Myint, Amin Soebandrio |
| EPI_ISL_3029966 | Dinas Kesehatan Provinsi Nusa Tenggara Timur | Eijkman Institute for Molecular Biology, National Research and Innovation Agency | Lydia V. Panggalo, Iskandar Adnan, Sukma Oktavianthi, Lidwina Priliani, Edison Johar, Frilasita A Yudhaputri, Muhammad Rezki Rasyak, Willy Agustine, Hidayat Trimarsanto, Safarina G Malik, Khin Saw Myint, Amin Soebandrio |
| EPI_ISL_3029967 | Dinas Kesehatan Provinsi Nusa Tenggara Timur | Eijkman Institute for Molecular Biology, National Research and Innovation Agency | Iskandar Adnan, Sukma Oktavianthi, Lidwina Priliani, Edison Johar, Frilasita A Yudhaputri, Muhammad Rezki Rasyak, Willy Agustine, Hidayat Trimarsanto, Lydia V. Panggalo, Safarina G Malik, Khin Saw Myint, Amin Soebandrio |
| EPI_ISL_3029971 | Dinas Kesehatan Provinsi Nusa Tenggara Timur | Eijkman Institute for Molecular Biology, National Research and Innovation Agency | Muhammad Rezki Rasyak, Willy Agustine, Hidayat Trimarsanto, Lydia V. Panggalo, Iskandar Adnan, Sukma Oktavianthi, Lidwina Priliani, Edison Johar, Frilasita A Yudhaputri, Safarina G Malik, Khin Saw Myint, Amin Soebandrio |
| EPI_ISL_3029972 | Dinas Kesehatan Provinsi Nusa Tenggara Timur | Eijkman Institute for Molecular Biology, National Research and Innovation Agency | Hidayat Trimarsanto, Lydia V. Panggalo, Iskandar Adnan, Sukma Oktavianthi, Lidwina Priliani, Edison Johar, Frilasita A Yudhaputri, Muhammad Rezki Rasyak, Willy Agustine, Safarina G Malik, Khin Saw Myint, Amin Soebandrio |
| EPI_ISL_3029973 | Dinas Kesehatan Provinsi Nusa Tenggara Timur | Eijkman Institute for Molecular Biology, National Research and Innovation Agency | Lydia V. Panggalo, Iskandar Adnan, Sukma Oktavianthi, Lidwina Priliani, Edison Johar, Frilasita A Yudhaputri, Muhammad Rezki Rasyak, Willy Agustine, Hidayat Trimarsanto, Safarina G Malik, Khin Saw Myint, Amin Soebandrio |
| EPI_ISL_3029974 | Dinas Kesehatan Provinsi Nusa Tenggara Timur | Eijkman Institute for Molecular Biology, National Research and Innovation Agency | Iskandar Adnan, Sukma Oktavianthi, Lidwina Priliani, Edison Johar, Frilasita A Yudhaputri, Muhammad Rezki Rasyak, Willy Agustine, Hidayat Trimarsanto, Lydia V. Panggalo, Safarina G Malik, Khin Saw Myint, Amin Soebandrio |
| EPI_ISL_3029976 | Dinas Kesehatan Provinsi Nusa Tenggara Timur | Eijkman Institute for Molecular Biology, National Research and Innovation Agency | Lidwina Priliani, Edison Johar, Frilasita A Yudhaputri, Muhammad Rezki Rasyak, Willy Agustine, Hidayat Trimarsanto, Lydia V. Panggalo, Iskandar Adnan, Sukma Oktavianthi, Safarina G Malik, Khin Saw Myint, Amin Soebandrio |
| EPI_ISL_3029977 | Dinas Kesehatan Provinsi Nusa Tenggara Timur | Eijkman Institute for Molecular Biology, National Research and Innovation Agency | Edison Johar, Frilasita A Yudhaputri, Muhammad Rezki Rasyak, Willy Agustine, Hidayat Trimarsanto, Lydia V. Panggalo, Iskandar Adnan, Sukma Oktavianthi, Lidwina Priliani, Safarina G Malik, Khin Saw Myint, Amin Soebandrio |
| EPI_ISL_3029978 | Dinas Kesehatan Provinsi Nusa Tenggara Timur | Eijkman Institute for Molecular Biology, National Research and Innovation Agency | Muhammad Rezki Rasyak, Willy Agustine, Hidayat Trimarsanto, Lydia V. Panggalo, Iskandar Adnan, Sukma Oktavianthi, Lidwina Priliani, Edison Johar, Frilasita A Yudhaputri, Safarina G Malik, Khin Saw Myint, Amin Soebandrio |
| EPI_ISL_3029979 | Dinas Kesehatan Provinsi Nusa Tenggara Timur | Eijkman Institute for Molecular Biology, National Research and Innovation Agency | Willy Agustine, Hidayat Trimarsanto, Lydia V. Panggalo, Iskandar Adnan, Sukma Oktavianthi, Lidwina Priliani, Edison Johar, Frilasita A Yudhaputri, Muhammad Rezki Rasyak, Safarina G Malik, Khin Saw Myint, Amin Soebandrio |
| EPI_ISL_3029980 | Dinas Kesehatan Provinsi Nusa Tenggara Timur | Eijkman Institute for Molecular Biology, National Research and Innovation Agency | Hidayat Trimarsanto, Lydia V. Panggalo, Iskandar Adnan, Sukma Oktavianthi, Lidwina Priliani, Edison Johar, Frilasita A Yudhaputri, Muhammad Rezki Rasyak, Willy Agustine, Safarina G Malik, Khin Saw Myint, Amin Soebandrio |
| EPI_ISL_3029981 | Dinas Kesehatan Provinsi Nusa Tenggara Timur | Eijkman Institute for Molecular Biology, National Research and Innovation Agency | Lydia V. Panggalo, Iskandar Adnan, Sukma Oktavianthi, Lidwina Priliani, Edison Johar, Frilasita A Yudhaputri, Muhammad Rezki Rasyak, Willy Agustine, Hidayat Trimarsanto, Safarina G Malik, Khin Saw Myint, Amin Soebandrio |
| EPI_ISL_3029982 | Dinas Kesehatan Provinsi Nusa Tenggara Timur | Eijkman Institute for Molecular Biology, National Research and Innovation Agency | Iskandar Adnan, Sukma Oktavianthi, Lidwina Priliani, Edison Johar, Frilasita A Yudhaputri, Muhammad Rezki Rasyak, Willy Agustine, Hidayat Trimarsanto, Lydia V. Panggalo, Safarina G Malik, Khin Saw Myint, Amin Soebandrio |
| EPI_ISL_3029983 | Dinas Kesehatan Provinsi Nusa Tenggara Timur | Eijkman Institute for Molecular Biology, National Research and Innovation Agency | Sukma Oktavianthi, Lidwina Priliani, Edison Johar, Frilasita A Yudhaputri, Muhammad Rezki Rasyak, Willy Agustine, Hidayat Trimarsanto, Lydia V. Panggalo, Iskandar Adnan, Safarina G Malik, Khin Saw Myint, Amin Soebandrio |
| EPI_ISL_3029984 | Dinas Kesehatan Provinsi Nusa Tenggara Timur | Eijkman Institute for Molecular Biology, National Research and Innovation Agency | Lidwina Priliani, Edison Johar, Frilasita A Yudhaputri, Muhammad Rezki Rasyak, Willy Agustine, Hidayat Trimarsanto, Lydia V. Panggalo, Iskandar Adnan, Sukma Oktavianthi, Safarina G Malik, Khin Saw Myint, Amin Soebandrio |
| EPI_ISL_3029985 | Dinas Kesehatan Provinsi Nusa Tenggara Timur | Eijkman Institute for Molecular Biology, National Research and Innovation Agency | Edison Johar, Frilasita A Yudhaputri, Muhammad Rezki Rasyak, Willy Agustine, Hidayat Trimarsanto, Lydia V. Panggalo, Iskandar Adnan, Sukma Oktavianthi, Lidwina Priliani, Safarina G Malik, Khin Saw Myint, Amin Soebandrio |
| EPI_ISL_3029986 | Dinas Kesehatan Provinsi Nusa Tenggara Timur | Eijkman Institute for Molecular Biology, National Research and Innovation Agency | Frilasita A Yudhaputri, Muhammad Rezki Rasyak, Willy Agustine, Hidayat Trimarsanto, Lydia V. Panggalo, Iskandar Adnan, Sukma Oktavianthi, Lidwina Priliani, Edison Johar, Safarina G Malik, Khin Saw Myint, Amin Soebandrio |
| EPI_ISL_3029987 | Dinas Kesehatan Provinsi Nusa Tenggara Timur | Eijkman Institute for Molecular Biology, National Research and Innovation Agency | Muhammad Rezki Rasyak, Willy Agustine, Hidayat Trimarsanto, Lydia V. Panggalo, Iskandar Adnan, Sukma Oktavianthi, Lidwina Priliani, Edison Johar, Frilasita A Yudhaputri, Safarina G Malik, Khin Saw Myint, Amin Soebandrio |
| EPI_ISL_3029988 | Dinas Kesehatan Provinsi Nusa Tenggara Timur | Eijkman Institute for Molecular Biology, National Research and Innovation Agency | Willy Agustine, Hidayat Trimarsanto, Lydia V. Panggalo, Iskandar Adnan, Sukma Oktavianthi, Lidwina Priliani, Edison Johar, Frilasita A Yudhaputri, Muhammad Rezki Rasyak, Safarina G Malik, Khin Saw Myint, Amin Soebandrio |
| EPI_ISL_3029989 | Dinas Kesehatan Provinsi Nusa Tenggara Timur | Eijkman Institute for Molecular Biology, National Research and Innovation Agency | Hidayat Trimarsanto, Lydia V. Panggalo, Iskandar Adnan, Sukma Oktavianthi, Lidwina Priliani, Edison Johar, Frilasita A Yudhaputri, Muhammad Rezki Rasyak, Willy Agustine, Safarina G Malik, Khin Saw Myint, Amin Soebandrio |
| EPI_ISL_3029990 | Dinas Kesehatan Provinsi Nusa Tenggara Timur | Eijkman Institute for Molecular Biology, National Research and Innovation Agency | Sukma Oktavianthi, Lidwina Priliani, Edison Johar, Frilasita A Yudhaputri, Muhammad Rezki Rasyak, Willy Agustine, Hidayat Trimarsanto, Lydia V. Panggalo, Iskandar Adnan, Safarina G Malik, Khin Saw Myint, Amin Soebandrio |
| EPI_ISL_3029991 | Dinas Kesehatan Provinsi Nusa Tenggara Timur | Eijkman Institute for Molecular Biology, National Research and Innovation Agency | Lidwina Priliani, Edison Johar, Frilasita A Yudhaputri, Muhammad Rezki Rasyak, Willy Agustine, Hidayat Trimarsanto, Lydia V. Panggalo, Iskandar Adnan, Sukma Oktavianthi, Safarina G Malik, Khin Saw Myint, Amin Soebandrio |

| | | | |
|---|---|---|---|
| EPI_ISL_3029992 | Dinas Kesehatan Provinsi Nusa Tenggara Timur | Eijkman Institute for Molecular Biology, National Research and Innovation Agency | Edison Johar, Frilasita A Yudhaputri, Muhammad Rezki Rasyak, Willy Agustine, Hidayat Trimarsanto, Lydia V. Panggalo, Iskandar Adnan, Sukma Oktavianthi, Lidwina Priliani, Safarina G Malik, Khin Saw Myint, Amin Soebandrio |
| EPI_ISL_3029993 | Dinas Kesehatan Provinsi Nusa Tenggara Timur | Eijkman Institute for Molecular Biology, National Research and Innovation Agency | Frilasita A Yudhaputri, Muhammad Rezki Rasyak, Willy Agustine, Hidayat Trimarsanto, Lydia V. Panggalo, Iskandar Adnan, Sukma Oktavianthi, Lidwina Priliani, Edison Johar, Safarina G Malik, Khin Saw Myint, Amin Soebandrio |
| EPI_ISL_3029994 | RS Sentra Medika Cibinong | Eijkman Institute for Molecular Biology, National Research and Innovation Agency | Muhammad Rezki Rasyak, Willy Agustine, Hidayat Trimarsanto, Lydia V. Panggalo, Iskandar Adnan, Sukma Oktavianthi, Lidwina Priliani, Edison Johar, Frilasita A Yudhaputri, Safarina G Malik, Khin Saw Myint, Amin Soebandrio |
| EPI_ISL_3029995 | RS Mitra Keluarga Cibubur | Eijkman Institute for Molecular Biology, National Research and Innovation Agency | Willy Agustine, Hidayat Trimarsanto, Lydia V. Panggalo, Iskandar Adnan, Sukma Oktavianthi, Lidwina Priliani, Edison Johar, Frilasita A Yudhaputri, Muhammad Rezki Rasyak, Safarina G Malik, Khin Saw Myint, Amin Soebandrio |
| EPI_ISL_3029996 | RSCM | Eijkman Institute for Molecular Biology, National Research and Innovation Agency | Lydia V. Panggalo, Iskandar Adnan, Sukma Oktavianthi, Lidwina Priliani, Edison Johar, Frilasita A Yudhaputri, Muhammad Rezki Rasyak, Willy Agustine, Hidayat Trimarsanto, Safarina G Malik, Khin Saw Myint, Amin Soebandrio |
| EPI_ISL_3029997 | RS Pondok Indah Pondok Indah | Eijkman Institute for Molecular Biology, National Research and Innovation Agency | Iskandar Adnan, Sukma Oktavianthi, Lidwina Priliani, Edison Johar, Frilasita A Yudhaputri, Muhammad Rezki Rasyak, Willy Agustine, Hidayat Trimarsanto, Lydia V. Panggalo, Safarina G Malik, Khin Saw Myint, Amin Soebandrio |
| EPI_ISL_3029998 | RS Pondok Indah Pondok Indah | Eijkman Institute for Molecular Biology, National Research and Innovation Agency | Edison Johar, Frilasita A Yudhaputri, Muhammad Rezki Rasyak, Willy Agustine, Hidayat Trimarsanto, Lydia V. Panggalo, Iskandar Adnan, Sukma Oktavianthi, Lidwina Priliani, Safarina G Malik, Khin Saw Myint, Amin Soebandrio |
| EPI_ISL_3029999 | RS Mitra Keluarga Cibubur | Eijkman Institute for Molecular Biology, National Research and Innovation Agency | Frilasita A Yudhaputri, Muhammad Rezki Rasyak, Willy Agustine, Hidayat Trimarsanto, Lydia V. Panggalo, Iskandar Adnan, Sukma Oktavianthi, Lidwina Priliani, Edison Johar, Safarina G Malik, Khin Saw Myint, Amin Soebandrio |
| EPI_ISL_3030000 | RS Grha MM2100 | Eijkman Institute for Molecular Biology, National Research and Innovation Agency | Muhammad Rezki Rasyak, Willy Agustine, Hidayat Trimarsanto, Lydia V. Panggalo, Iskandar Adnan, Sukma Oktavianthi, Lidwina Priliani, Edison Johar, Frilasita A Yudhaputri, Safarina G Malik, Khin Saw Myint, Amin Soebandrio |
| EPI_ISL_3030001 | RS Mitra Keluarga Kemayoran | Eijkman Institute for Molecular Biology, National Research and Innovation Agency | Edison Johar, Frilasita A Yudhaputri, Muhammad Rezki Rasyak, Willy Agustine, Hidayat Trimarsanto, Lydia V. Panggalo, Iskandar Adnan, Sukma Oktavianthi, Lidwina Priliani, Safarina G Malik, Khin Saw Myint, Amin Soebandrio |
| EPI_ISL_3030040 | Puskesmas Pakutandang | Eijkman Institute for Molecular Biology, National Research and Innovation Agency; West Java Health Laboratory | Frilasita A Yudhaputri, Muhammad Rezki Rasyak, Willy Agustine, Hidayat Trimarsanto, Lydia V. Panggalo, Iskandar Adnan, Sukma Oktavianthi, Lidwina Priliani, Edison Johar, Azzania Fibriani, Ema Rahmawati, Ryan Bayusantika Ristandi, Rifky Waluyajati Rachman, Cut Nur Cinthia Alamanda, Safarina G Malik, Khin Saw Myint, Amin Soebandrio |
| EPI_ISL_3030042 | Puskesmas Luragung | Eijkman Institute for Molecular Biology, National Research and Innovation Agency; West Java Health Laboratory | Hidayat Trimarsanto, Lydia V. Panggalo, Iskandar Adnan, Sukma Oktavianthi, Lidwina Priliani, Edison Johar, Frilasita A Yudhaputri, Muhammad Rezki Rasyak, Willy Agustine, Azzania Fibriani, Ema Rahmawati, Ryan Bayusantika Ristandi, Rifky Waluyajati Rachman, Cut Nur Cinthia Alamanda, Safarina G Malik, Khin Saw Myint, Amin Soebandrio |
| EPI_ISL_3030043 | Puskesmas Hantara | Eijkman Institute for Molecular Biology, National Research and Innovation Agency; West Java Health Laboratory | Lydia V. Panggalo, Iskandar Adnan, Sukma Oktavianthi, Lidwina Priliani, Edison Johar, Frilasita A Yudhaputri, Muhammad Rezki Rasyak, Willy Agustine, Hidayat Trimarsanto, Azzania Fibriani, Ema Rahmawati, Ryan Bayusantika Ristandi, Rifky Waluyajati Rachman, Cut Nur Cinthia Alamanda, Safarina G Malik, Khin Saw Myint, Amin Soebandrio |
| EPI_ISL_3030045 | Puskesmas Jatihandap, Dinas Kesehatan Kota Bandung | Eijkman Institute for Molecular Biology, National Research and Innovation Agency; West Java Health Laboratory | Edison Johar, Frilasita A Yudhaputri, Muhammad Rezki Rasyak, Willy Agustine, Hidayat Trimarsanto, Lydia V. Panggalo, Iskandar Adnan, Sukma Oktavianthi, Lidwina Priliani, Azzania Fibriani, Ema Rahmawati, Ryan Bayusantika Ristandi, Rifky Waluyajati Rachman, Cut Nur Cinthia Alamanda, Ade Fany Safitri, Ranza Fadhlil Pratama, Tati Sutarti Sastrowiyoto, Husna Nugrahapraja, Safarina G Malik, Khin Saw Myint, Amin Soebandrio |
| EPI_ISL_3030046 | Puskesmas Jatihandap, Dinas Kesehatan Kota Bandung | Eijkman Institute for Molecular Biology, National Research and Innovation Agency; West Java Health Laboratory | Frilasita A Yudhaputri, Muhammad Rezki Rasyak, Willy Agustine, Hidayat Trimarsanto, Lydia V. Panggalo, Iskandar Adnan, Sukma Oktavianthi, Lidwina Priliani, Edison Johar, Azzania Fibriani, Ema Rahmawati, Ryan Bayusantika Ristandi, Rifky Waluyajati Rachman, Cut Nur Cinthia Alamanda, Ade Fany Safitri, Ranza Fadhlil Pratama, Tati Sutarti Sastrowiyoto, Husna Nugrahapraja, Safarina G Malik, Khin Saw Myint, Amin Soebandrio |
| EPI_ISL_3030048 | Puskesmas Sukawarna, Dinas Kesehatan Kota Bandung | Eijkman Institute for Molecular Biology, National Research and Innovation Agency; West Java Health Laboratory | Willy Agustine, Hidayat Trimarsanto, Lydia V. Panggalo, Iskandar Adnan, Sukma Oktavianthi, Lidwina Priliani, Edison Johar, Frilasita A Yudhaputri, Muhammad Rezki Rasyak, Azzania Fibriani, Ema Rahmawati, Ryan Bayusantika Ristandi, Rifky Waluyajati Rachman, Cut Nur Cinthia Alamanda, Ade Fany Safitri, Ranza Fadhlil Pratama, Tati Sutarti Sastrowiyoto, Husna Nugrahapraja, Safarina G Malik, Khin Saw Myint, Amin Soebandrio |
| EPI_ISL_3030049 | Puskesmas Arcamanik, Dinas Kesehatan Kota Bandung | Eijkman Institute for Molecular Biology, National Research and Innovation Agency; West Java Health Laboratory | Hidayat Trimarsanto, Lydia V. Panggalo, Iskandar Adnan, Sukma Oktavianthi, Lidwina Priliani, Edison Johar, Frilasita A Yudhaputri, Muhammad Rezki Rasyak, Willy Agustine, Azzania Fibriani, Ema Rahmawati, Ryan Bayusantika Ristandi, Rifky Waluyajati Rachman, Cut Nur Cinthia Alamanda, Usman Abdul Jabar, Sheila Widia Trisakti, Tati Sutarti Sastrowiyoto, Husna Nugrahapraja, Safarina G Malik, Khin Saw Myint, Amin Soebandrio |
| EPI_ISL_3030050 | RS Muhammadiyah Bandung, Dinas Kesehatan Kota Bandung | Eijkman Institute for Molecular Biology, National Research and Innovation Agency; West Java Health Laboratory | Lydia V. Panggalo, Iskandar Adnan, Sukma Oktavianthi, Lidwina Priliani, Edison Johar, Frilasita A Yudhaputri, Muhammad Rezki Rasyak, Willy Agustine, Hidayat Trimarsanto, Azzania Fibriani, Ema Rahmawati, Ryan Bayusantika Ristandi, Rifky Waluyajati Rachman, Cut Nur Cinthia Alamanda, Ade Fany Safitri, Ranza Fadhlil Pratama, Tati Sutarti Sastrowiyoto, Husna Nugrahapraja, Safarina G Malik, Khin Saw Myint, Amin Soebandrio |
| EPI_ISL_3030051 | RS Muhammadiyah Bandung, Dinas Kesehatan Kota Bandung | Eijkman Institute for Molecular Biology, National Research and Innovation Agency; West Java Health Laboratory | Iskandar Adnan, Sukma Oktavianthi, Lidwina Priliani, Edison Johar, Frilasita A Yudhaputri, Muhammad Rezki Rasyak, Willy Agustine, Hidayat Trimarsanto, Lydia V. Panggalo, Azzania Fibriani, Ema Rahmawati, Ryan Bayusantika Ristandi, Rifky Waluyajati Rachman, Cut Nur Cinthia Alamanda, Ade Fany Safitri, Ranza Fadhlil Pratama, Tati Sutarti Sastrowiyoto, Husna Nugrahapraja, Safarina G Malik, Khin Saw Myint, Amin Soebandrio |
| EPI_ISL_3030052 | RS Khusus Gigi dan Mulut Kota Bandung, Dinas Kesehatan Kota Bandung | Eijkman Institute for Molecular Biology, National Research and Innovation Agency; West Java Health Laboratory | Sukma Oktavianthi, Lidwina Priliani, Edison Johar, Frilasita A Yudhaputri, Muhammad Rezki Rasyak, Willy Agustine, Hidayat Trimarsanto, Lydia V. Panggalo, Iskandar Adnan, Azzania Fibriani, Ema Rahmawati, Ryan Bayusantika Ristandi, Rifky Waluyajati Rachman, Cut Nur Cinthia Alamanda, Ade Fany Safitri, Ranza Fadhlil Pratama, Tati Sutarti Sastrowiyoto, Husna Nugrahapraja, Safarina G Malik, Khin Saw Myint, Amin Soebandrio |
| EPI_ISL_3030053 | Puskesmas Sarijadi, Dinas Kesehatan Kota Bandung | Eijkman Institute for Molecular Biology, National Research and Innovation Agency; West Java Health Laboratory | Lidwina Priliani, Edison Johar, Frilasita A Yudhaputri, Muhammad Rezki Rasyak, Willy Agustine, Hidayat Trimarsanto, Lydia V. Panggalo, Iskandar Adnan, Sukma Oktavianthi, Azzania Fibriani, Ema Rahmawati, Ryan Bayusantika Ristandi, Rifky Waluyajati Rachman, Cut Nur Cinthia Alamanda, Ade Fany Safitri, Ranza Fadhlil Pratama, Tati Sutarti Sastrowiyoto, Husna Nugrahapraja, Safarina G Malik, Khin Saw Myint, Amin Soebandrio |
| EPI_ISL_3030054 | RSIA Grha Bunda, Dinas Kesehatan Kota Bandung | Eijkman Institute for Molecular Biology, National Research and Innovation Agency; West Java Health Laboratory | Edison Johar, Frilasita A Yudhaputri, Muhammad Rezki Rasyak, Willy Agustine, Hidayat Trimarsanto, Lydia V. Panggalo, Iskandar Adnan, Sukma Oktavianthi, Lidwina Priliani, Azzania Fibriani, Ema Rahmawati, Ryan Bayusantika Ristandi, Rifky Waluyajati Rachman, Cut Nur Cinthia Alamanda, Ade Fany Safitri, Ranza Fadhlil Pratama, Tati Sutarti Sastrowiyoto, Husna Nugrahapraja, Safarina G Malik, Khin Saw Myint, Amin Soebandrio |
| EPI_ISL_3030055 | Laboratorium Kesehatan Daerah Kota Bandung, Dinas Kesehatan Kota Bandung | Eijkman Institute for Molecular Biology, National Research and Innovation Agency; West Java Health Laboratory | Frilasita A Yudhaputri, Muhammad Rezki Rasyak, Willy Agustine, Hidayat Trimarsanto, Lydia V. Panggalo, Iskandar Adnan, Sukma Oktavianthi, Lidwina Priliani, Edison Johar, Azzania Fibriani, Ema Rahmawati, Ryan Bayusantika Ristandi, Rifky Waluyajati Rachman, Cut Nur Cinthia Alamanda, Usman Abdul Jabar, Sheila Widia Trisakti, Tati Sutarti Sastrowiyoto, Husna Nugrahapraja, Safarina G Malik, Khin Saw Myint, Amin Soebandrio |
| EPI_ISL_3030056 | RS Khusus Gigi dan Mulut Kota Bandung, Dinas Kesehatan Kota Bandung | Eijkman Institute for Molecular Biology, National Research and Innovation Agency; West Java Health Laboratory | Muhammad Rezki Rasyak, Willy Agustine, Hidayat Trimarsanto, Lydia V. Panggalo, Iskandar Adnan, Sukma Oktavianthi, Lidwina Priliani, Edison Johar, Frilasita A Yudhaputri, Azzania Fibriani, Ema Rahmawati, Ryan Bayusantika Ristandi, Rifky Waluyajati Rachman, Cut Nur Cinthia Alamanda, Ade Fany Safitri, Ranza Fadhlil Pratama, Tati Sutarti Sastrowiyoto, Husna Nugrahapraja, Safarina G Malik, Khin Saw Myint, Amin Soebandrio |
| EPI_ISL_3030057 | RS Khusus Gigi dan Mulut Kota Bandung, Dinas Kesehatan Kota Bandung | Eijkman Institute for Molecular Biology, National Research and Innovation Agency; West Java Health Laboratory | Willy Agustine, Hidayat Trimarsanto, Lydia V. Panggalo, Iskandar Adnan, Sukma Oktavianthi, Lidwina Priliani, Edison Johar, Frilasita A Yudhaputri, Muhammad Rezki Rasyak, Azzania Fibriani, Ema Rahmawati, Ryan Bayusantika Ristandi, Rifky Waluyajati Rachman, Cut Nur Cinthia Alamanda, Ade Fany Safitri, Ranza Fadhlil Pratama, Tati Sutarti Sastrowiyoto, Husna Nugrahapraja, Safarina G Malik, Khin Saw Myint, Amin Soebandrio |
| EPI_ISL_3030058 | Puskesmas Jatihandap, Dinas Kesehatan Kota Bandung | Eijkman Institute for Molecular Biology, National Research and Innovation Agency; West Java Health Laboratory | Hidayat Trimarsanto, Lydia V. Panggalo, Iskandar Adnan, Sukma Oktavianthi, Lidwina Priliani, Edison Johar, Frilasita A Yudhaputri, Muhammad Rezki Rasyak, Willy Agustine, Azzania Fibriani, Ema Rahmawati, Ryan Bayusantika Ristandi, Rifky Waluyajati Rachman, Cut Nur Cinthia Alamanda, Ade Fany Safitri, Ranza Fadhlil Pratama, Tati Sutarti Sastrowiyoto, Husna Nugrahapraja, Safarina G Malik, Khin Saw Myint, Amin Soebandrio |
| EPI_ISL_3030059 | Puskesmas Sarijadi, Dinas Kesehatan Kota Bandung | Eijkman Institute for Molecular Biology, National Research and Innovation Agency; West Java Health Laboratory | Lydia V. Panggalo, Iskandar Adnan, Sukma Oktavianthi, Lidwina Priliani, Edison Johar, Frilasita A Yudhaputri, Muhammad Rezki Rasyak, Willy Agustine, Hidayat Trimarsanto, Azzania Fibriani, Ema Rahmawati, Ryan Bayusantika Ristandi, Rifky Waluyajati Rachman, Cut Nur Cinthia Alamanda, Ade Fany Safitri, Ranza Fadhlil Pratama, Tati Sutarti Sastrowiyoto, Husna Nugrahapraja, Safarina G Malik, Khin Saw Myint, Amin Soebandrio |
| EPI_ISL_3030060 | CH Simone Veil | Institut Pasteur | Marion Barbet, Sylvie Behillil, Méline Bizard, Angela Brisebarre, Camille Capel, Vincent Enouf, Louise Lefrançois, Frédéric Lemoine, Christophe Malabat, Corinne Maufrais, Emmanuelle Permal, Etienne Simon-Lorière, Maud Vanpeene, Sylvie Van der Werf, Guiheneuf Raphaël |
| EPI_ISL_3030062, EPI_ISL_3030063, EPI_ISL_3030064, EPI_ISL_3030066, EPI_ISL_3030067, EPI_ISL_3030068, EPI_ISL_3030069, EPI_ISL_3030070, EPI_ISL_3030071, EPI_ISL_3030072, EPI_ISL_3030073, EPI_ISL_3030074, EPI_ISL_3030075, EPI_ISL_3030077, EPI_ISL_3030078, EPI_ISL_3030079, EPI_ISL_3030080, EPI_ISL_3030081 | | | |
| see above | IN State Department of Health Laboratory Services | IN State Department of Health Laboratory Services | Cassandra Campion, Jamie Yeadon, Brian Pope, Lixia Liu, Kyle Brownlee, Melissa Hindenlang, Mark Glazier |
| EPI_ISL_3030083 | Orebro University Hospital, Dept Laboratory Medicine, Clinical Microbiology | Orebro University Hospital | Sundqvist M et al |
| EPI_ISL_3030085 | Labormedizinisches Zentrum Dr Risch | Clinical Bacteriology | Tim Roloff, Fanny Wegner, Helena MB Seth-Smith, Alfredo Mari, Karoline Leuzinger, Julia Bielicki, Nadia Wohlwend,Martin Risch, Lorenz Risch, Manuel Battegay, Hans Hirsch, Adrian Egli |
| EPI_ISL_3030102 | Clinical Virology | Clinical Bacteriology | Tim Roloff, Fanny Wegner, Helena MB Seth-Smith, Alfredo Mari, Karoline Leuzinger, Julia Bielicki, Manuel Battegay, Hans Hirsch, Adrian Egli |
| EPI_ISL_3030103 | Institute of Microbiology and Immunology, Faculty of Medicine, University of Ljubljana | Institute of Microbiology and Immunology, Faculty of Medicine, University of Ljubljana | Alen Suljič, Samo Zakotnik, Tomaž Mark Zorec, Matic Brvar, Doroteja Vlaj, Andraž Celar, Dominika Šturm, Patricija Pozvek, Špela Pleh, Miša Korva, Mario Poljak, Tatjana Avšič – Županc |
| EPI_ISL_3030104 | Kantonsspital Baselland | Clinical Bacteriology | Tim Roloff, Fanny Wegner, Helena MB Seth-Smith, Alfredo Mari, Karoline Leuzinger, Julia Bielicki, Manuel Battegay, Hans Hirsch, Adrian Egli |
| EPI_ISL_3030105 | Institute of Microbiology and Immunology, Faculty of Medicine, University of Ljubljana | Institute of Microbiology and Immunology, Faculty of Medicine, University of Ljubljana | Alen Suljič, Samo Zakotnik, Tomaž Mark Zorec, Matic Brvar, Doroteja Vlaj, Andraž Celar, Dominika Šturm, Patricija Pozvek, Špela Pleh, Miša Korva, Mario Poljak, Tatjana Avšič – Županc |
| EPI_ISL_3030106 | Clinical Virology | Clinical Bacteriology | Tim Roloff, Fanny Wegner, Helena MB Seth-Smith, Alfredo Mari, Karoline Leuzinger, Julia Bielicki, Manuel Battegay, Hans Hirsch, Adrian Egli |
| EPI_ISL_3030107 | Institute of Microbiology and Immunology, Faculty of Medicine, University of Ljubljana | Institute of Microbiology and Immunology, Faculty of Medicine, University of Ljubljana | Alen Suljič, Samo Zakotnik, Tomaž Mark Zorec, Matic Brvar, Doroteja Vlaj, Andraž Celar, Dominika Šturm, Patricija Pozvek, Špela Pleh, Miša Korva, Mario Poljak, Tatjana Avšič – Županc |
| EPI_ISL_3030109 | Clinical Virology | Clinical Bacteriology | Tim Roloff, Fanny Wegner, Helena MB Seth-Smith, Alfredo Mari, Karoline Leuzinger, Julia Bielicki, Manuel Battegay, Hans Hirsch, Adrian Egli |
| EPI_ISL_3030110, EPI_ISL_3030112 | Institute of Microbiology and Immunology, Faculty of Medicine, University of Ljubljana | Institute of Microbiology and Immunology, Faculty of Medicine, University of Ljubljana | Alen Suljič, Samo Zakotnik, Tomaž Mark Zorec, Matic Brvar, Doroteja Vlaj, Andraž Celar, Dominika Šturm, Patricija Pozvek, Špela Pleh, Miša Korva, Mario Poljak, Tatjana Avšič – Županc |
| EPI_ISL_3030113 | Rothen Medizinische Laboratorien AG | Clinical Bacteriology | Tim Roloff, Fanny Wegner, Helena MB Seth-Smith, Alfredo Mari, Karoline Leuzinger, Julia Bielicki, Ingrid Steffen, Manuel Battegay, Hans Hirsch, Adrian Egli |
| EPI_ISL_3030115 | Synlab Suisse SA | Clinical Bacteriology | Tim Roloff, Fanny Wegner, Helena MB Seth-Smith, Alfredo Mari, Karoline Leuzinger, Julia Bielicki, Manuel Battegay, Hans Hirsch, Adrian Egli |
| EPI_ISL_3030116 | Institute of Microbiology and Immunology, Faculty of Medicine, University of Ljubljana | Institute of Microbiology and Immunology, Faculty of Medicine, University of Ljubljana | Alen Suljič, Samo Zakotnik, Tomaž Mark Zorec, Matic Brvar, Doroteja Vlaj, Andraž Celar, Dominika Šturm, Patricija Pozvek, Špela Pleh, Miša Korva, Mario Poljak, Tatjana Avšič – Županc |
| EPI_ISL_3030117, EPI_ISL_3030119 | Clinical Virology | Clinical Bacteriology | Tim Roloff, Fanny Wegner, Helena MB Seth-Smith, Alfredo Mari, Karoline Leuzinger, Julia Bielicki, Manuel Battegay, Hans Hirsch, Adrian Egli |
| EPI_ISL_3030120 | Institute of Microbiology and Immunology, Faculty of Medicine, University of Ljubljana | Institute of Microbiology and Immunology, Faculty of Medicine, University of Ljubljana | Alen Suljič, Samo Zakotnik, Tomaž Mark Zorec, Matic Brvar, Doroteja Vlaj, Andraž Celar, Dominika Šturm, Patricija Pozvek, Špela Pleh, Miša Korva, Mario Poljak, Tatjana Avšič – Županc |
| EPI_ISL_3030123 | Clinical Virology | Clinical Bacteriology | Tim Roloff, Fanny Wegner, Helena MB Seth-Smith, Alfredo Mari, Karoline Leuzinger, Julia Bielicki, Manuel Battegay, Hans Hirsch, Adrian Egli |
| EPI_ISL_3030124 | Institute of Microbiology and Immunology, Faculty of Medicine, University of Ljubljana | Institute of Microbiology and Immunology, Faculty of Medicine, University of Ljubljana | Alen Suljič, Samo Zakotnik, Tomaž Mark Zorec, Matic Brvar, Doroteja Vlaj, Andraž Celar, Dominika Šturm, Patricija Pozvek, Špela Pleh, Miša Korva, Mario Poljak, Tatjana Avšič – Županc |
| EPI_ISL_3030125 | Clinical Virology | Clinical Bacteriology | Tim Roloff, Fanny Wegner, Helena MB Seth-Smith, Alfredo Mari, Karoline Leuzinger, Julia Bielicki, Manuel Battegay, Hans Hirsch, Adrian Egli |
| EPI_ISL_3030126 | Institute of Microbiology and Immunology, Faculty of Medicine, University of Ljubljana | Institute of Microbiology and Immunology, Faculty of Medicine, University of Ljubljana | Alen Suljič, Samo Zakotnik, Tomaž Mark Zorec, Matic Brvar, Doroteja Vlaj, Andraž Celar, Dominika Šturm, Patricija Pozvek, Špela Pleh, Miša Korva, Mario Poljak, Tatjana Avšič – Županc |
| EPI_ISL_3030127 | Clinical Virology | Clinical Bacteriology | Tim Roloff, Fanny Wegner, Helena MB Seth-Smith, Alfredo Mari, Karoline Leuzinger, Julia Bielicki, Manuel Battegay, Hans Hirsch, Adrian Egli |

| | | | |
|---|---|---|---|
| EPI_ISL_3030128 | Institute of Microbiology and Immunology, Faculty of Medicine, University of Ljubljana | Institute of Microbiology and Immunology, Faculty of Medicine, University of Ljubljana | Alen Suljič, Samo Zakotnik, Tomaž Mark Zorec, Matic Brvar, Doroteja Vlaj, Andraž Celar, Dominika Šturm, Patricija Pozvek, Špela Pleh, Miša Korva, Mario Poljak, Tatjana Avšič – Županc |
| EPI_ISL_3030129 | Clinical Virology | Clinical Bacteriology | Tim Roloff, Fanny Wegner, Helena MB Seth-Smith, Alfredo Mari, Karoline Leuzinger, Julia Bielicki, Manuel Battegay, Hans Hirsch, Adrian Egli |
| EPI_ISL_3030130 | Institute of Microbiology and Immunology, Faculty of Medicine, University of Ljubljana | Institute of Microbiology and Immunology, Faculty of Medicine, University of Ljubljana | Alen Suljič, Samo Zakotnik, Tomaž Mark Zorec, Matic Brvar, Doroteja Vlaj, Andraž Celar, Dominika Šturm, Patricija Pozvek, Špela Pleh, Miša Korva, Mario Poljak, Tatjana Avšič – Županc |
| EPI_ISL_3030131 | Clinical Virology | Clinical Bacteriology | Tim Roloff, Fanny Wegner, Helena MB Seth-Smith, Alfredo Mari, Karoline Leuzinger, Julia Bielicki, Manuel Battegay, Hans Hirsch, Adrian Egli |
| EPI_ISL_3030132 | Institute of Microbiology and Immunology, Faculty of Medicine, University of Ljubljana | Institute of Microbiology and Immunology, Faculty of Medicine, University of Ljubljana | Alen Suljič, Samo Zakotnik, Tomaž Mark Zorec, Matic Brvar, Doroteja Vlaj, Andraž Celar, Dominika Šturm, Patricija Pozvek, Špela Pleh, Miša Korva, Mario Poljak, Tatjana Avšič – Županc |
| EPI_ISL_3030133 | Clinical Virology | Clinical Bacteriology | Tim Roloff, Fanny Wegner, Helena MB Seth-Smith, Alfredo Mari, Karoline Leuzinger, Julia Bielicki, Manuel Battegay, Hans Hirsch, Adrian Egli |
| EPI_ISL_3030134 | Institute of Microbiology and Immunology, Faculty of Medicine, University of Ljubljana | Institute of Microbiology and Immunology, Faculty of Medicine, University of Ljubljana | Alen Suljič, Samo Zakotnik, Tomaž Mark Zorec, Matic Brvar, Doroteja Vlaj, Andraž Celar, Dominika Šturm, Patricija Pozvek, Špela Pleh, Miša Korva, Mario Poljak, Tatjana Avšič – Županc |
| EPI_ISL_3030135 | Clinical Virology | Clinical Bacteriology | Tim Roloff, Fanny Wegner, Helena MB Seth-Smith, Alfredo Mari, Karoline Leuzinger, Julia Bielicki, Manuel Battegay, Hans Hirsch, Adrian Egli |
| EPI_ISL_3030136 | Institute of Microbiology and Immunology, Faculty of Medicine, University of Ljubljana | Institute of Microbiology and Immunology, Faculty of Medicine, University of Ljubljana | Alen Suljič, Samo Zakotnik, Tomaž Mark Zorec, Matic Brvar, Doroteja Vlaj, Andraž Celar, Dominika Šturm, Patricija Pozvek, Špela Pleh, Miša Korva, Mario Poljak, Tatjana Avšič – Županc |
| EPI_ISL_3030137 | Synlab Suisse SA | Clinical Bacteriology | Tim Roloff, Fanny Wegner, Helena MB Seth-Smith, Alfredo Mari, Karoline Leuzinger, Julia Bielicki, Manuel Battegay, Hans Hirsch, Adrian Egli |
| EPI_ISL_3030138 | Institute of Microbiology and Immunology, Faculty of Medicine, University of Ljubljana | Institute of Microbiology and Immunology, Faculty of Medicine, University of Ljubljana | Alen Suljič, Samo Zakotnik, Tomaž Mark Zorec, Matic Brvar, Doroteja Vlaj, Andraž Celar, Dominika Šturm, Patricija Pozvek, Špela Pleh, Miša Korva, Mario Poljak, Tatjana Avšič – Županc |
| EPI_ISL_3030139 | Synlab Suisse SA | Clinical Bacteriology | Tim Roloff, Fanny Wegner, Helena MB Seth-Smith, Alfredo Mari, Karoline Leuzinger, Julia Bielicki, Manuel Battegay, Hans Hirsch, Adrian Egli |
| EPI_ISL_3030140, EPI_ISL_3030141 | Institute of Microbiology and Immunology, Faculty of Medicine, University of Ljubljana | Institute of Microbiology and Immunology, Faculty of Medicine, University of Ljubljana | Alen Suljič, Samo Zakotnik, Tomaž Mark Zorec, Matic Brvar, Doroteja Vlaj, Andraž Celar, Dominika Šturm, Patricija Pozvek, Špela Pleh, Miša Korva, Mario Poljak, Tatjana Avšič – Županc |
| EPI_ISL_3030142, EPI_ISL_3030143 | Clinical Virology | Clinical Bacteriology | Tim Roloff, Fanny Wegner, Helena MB Seth-Smith, Alfredo Mari, Karoline Leuzinger, Julia Bielicki, Manuel Battegay, Hans Hirsch, Adrian Egli |
| EPI_ISL_3030144, EPI_ISL_3030146, EPI_ISL_3030147, EPI_ISL_3030149, EPI_ISL_3030150, EPI_ISL_3030151, EPI_ISL_3030152, EPI_ISL_3030153, EPI_ISL_3030154, EPI_ISL_3030155, EPI_ISL_3030156, EPI_ISL_3030157 | | | |
| see above | Institute of Microbiology and Immunology, Faculty of Medicine, University of Ljubljana | Institute of Microbiology and Immunology, Faculty of Medicine, University of Ljubljana | Alen Suljič, Samo Zakotnik, Tomaž Mark Zorec, Matic Brvar, Doroteja Vlaj, Andraž Celar, Dominika Šturm, Patricija Pozvek, Špela Pleh, Miša Korva, Mario Poljak, Tatjana Avšič – Županc |
| EPI_ISL_3030411 | Victorian Infectious Diseases Reference Laboratory (VIDRL) | VIDRL and MDU-PHL | Caly L., Seemann T., Sait, M.L., Druce J., Sherry, N.L. |
| EPI_ISL_3030412 | Royal Darwin Hospital Pathology | MDU-PHL | Meumann, E., Caly L., Seemann T., Sait, M.L., Druce J., Sherry, N.L. |
| EPI_ISL_3030413 | Microbiological Diagnostic Unit - Public Health Laboratory (MDU-PHL) | MDU-PHL | Seemann T., Sait, M.L., Sherry, N.L. |
| EPI_ISL_3030414, EPI_ISL_3030415, EPI_ISL_3030416 | Victorian Infectious Diseases Reference Laboratory (VIDRL) | VIDRL and MDU-PHL | Caly L., Seemann T., Sait, M.L., Druce J., Sherry, N.L. |
| EPI_ISL_3030419, EPI_ISL_3030421, EPI_ISL_3030423, EPI_ISL_3030424, EPI_ISL_3030425 | Microbiological Diagnostic Unit - Public Health Laboratory (MDU-PHL) | MDU-PHL | Seemann T., Sait, M.L., Sherry, N.L. |
| EPI_ISL_3030431, EPI_ISL_3030432, EPI_ISL_3030433 | MANKWENG LABORATORY | National Institute for Communicable Diseases of the National Health Laboratory Service | Amoako DG, Everatt J, Scheepers C, Mohale T, Ntuli N, Mahlangu B, Mnguni A, Ismail A, Bhiman JN |
| EPI_ISL_3030436 | HELEN JOSEPH LABORATORY | National Institute for Communicable Diseases of the National Health Laboratory Service | Amoako DG, Everatt J, Scheepers C, Mohale T, Ntuli N, Mahlangu B, Mnguni A, Ismail A, Bhiman JN |
| EPI_ISL_3030442, EPI_ISL_3030443, EPI_ISL_3030444, EPI_ISL_3030445, EPI_ISL_3030447, EPI_ISL_3030448, EPI_ISL_3030450, EPI_ISL_3030451, EPI_ISL_3030452, EPI_ISL_3030453, EPI_ISL_3030454, EPI_ISL_3030455, EPI_ISL_3030456, EPI_ISL_3030458, EPI_ISL_3030460, EPI_ISL_3030461, EPI_ISL_3030462, EPI_ISL_3030463, EPI_ISL_3030464 | | | |
| see above | Università degli Studi di Perugia | Istituto Zooprofilattico Sperimentale dell'Abruzzo e Molise "G. Caporale" | Mencacci A, Camilloni B, Biagetti M, Giammarioli M, Lorusso A, Marcacci M, Di Domenico M, Ancora M, Curini V, Mangone I, Rinaldi A, Delli Compagni E, Scialabba S, Di Pasquale A, Cammà C, Puglia I, Calistri P, Savini G |
| EPI_ISL_3030465, EPI_ISL_3030466, EPI_ISL_3030467 | Laboratorio Analisi Osp. Città di Castello - Azienda USL Umbria1 | Istituto Zooprofilattico Sperimentale dell'Abruzzo e Molise "G. Caporale" | Malagigi V, Tacconi P,Lorusso A, Marcacci M, Di Domenico M, Ancora M, Curini V, Mangone I, Rinaldi A, Delli Compagni E, Scialabba S, Di Pasquale A, Cammà C, Puglia I, Calistri P, Savini G |
| EPI_ISL_3030469 | Azienda Ospedaliera Terni | Istituto Zooprofilattico Sperimentale dell'Abruzzo e Molise "G. Caporale" | Palumbo M, Scaccetti A, Lorusso A, Marcacci M, Di Domenico M, Ancora M, Curini V, Mangone I, Rinaldi A, Delli Compagni E, Scialabba S, Di Pasquale A, Cammà C, Puglia I, Calistri P, Savini G |
| EPI_ISL_3030470, EPI_ISL_3030471, EPI_ISL_3030472, EPI_ISL_3030474, EPI_ISL_3030475, EPI_ISL_3030480, EPI_ISL_3030481, EPI_ISL_3030482, EPI_ISL_3030484, EPI_ISL_3030485, EPI_ISL_3030486 | | | |
| see above | Azienda USL Umbria 2 | Istituto Zooprofilattico Sperimentale dell'Abruzzo e Molise "G. Caporale" | Proietti A, Pistoni E, Lorusso A, Marcacci M, Di Domenico M, Ancora M, Curini V, Mangone I, Rinaldi A, Delli Compagni E, Scialabba S, Di Pasquale A, Cammà C, Puglia I, Calistri P, Savini G |
| EPI_ISL_3030490, EPI_ISL_3030491, EPI_ISL_3030492, EPI_ISL_3030493, EPI_ISL_3030494, EPI_ISL_3030495, EPI_ISL_3030496, EPI_ISL_3030497, EPI_ISL_3030498, EPI_ISL_3030499, EPI_ISL_3030500, EPI_ISL_3030501, EPI_ISL_3030502, EPI_ISL_3030503, EPI_ISL_3030504, EPI_ISL_3030505, EPI_ISL_3030506, EPI_ISL_3030507, EPI_ISL_3030508, EPI_ISL_3030509, EPI_ISL_3030510, EPI_ISL_3030511, EPI_ISL_3030512, EPI_ISL_3030513, EPI_ISL_3030514, EPI_ISL_3030515, EPI_ISL_3030516, EPI_ISL_3030517, EPI_ISL_3030518, EPI_ISL_3030519, EPI_ISL_3030520, EPI_ISL_3030521, EPI_ISL_3030522, EPI_ISL_3030523, EPI_ISL_3030524, EPI_ISL_3030525 | | | |
| see above | Dept. of Microbiology and Infection Control, Akershus University Hospital HF | Dept. of Microbiology and Infection Control, Akershus University Hospital HF | Hege Vangstein Aamot, Alexander Hesselberg Løvestad |
| EPI_ISL_3030526 | AMPATH LABORATORY | National Institute for Communicable Diseases of the National Health Laboratory Service | Amoako DG, Everatt J, Scheepers C, Mohale T, Ntuli N, Mahlangu B, Mnguni A, Ismail A, Bhiman JN |
| EPI_ISL_3030531 | Ampath Greenacres | National Institute for Communicable Diseases of the National Health Laboratory Service | Amoako DG, Everatt J, Scheepers C, Mohale T, Ntuli N, Mahlangu B, Mnguni A, Ismail A, Bhiman JN |
| EPI_ISL_3030533 | Toga LABORATORY | National Institute for Communicable Diseases of the National Health Laboratory Service | Amoako DG, Everatt J, Scheepers C, Mohale T, Ntuli N, Mahlangu B, Mnguni A, Ismail A, Bhiman JN |
| EPI_ISL_3030534, EPI_ISL_3030535, EPI_ISL_3030536, EPI_ISL_3030537, EPI_ISL_3030538, EPI_ISL_3030539, EPI_ISL_3030540 | Hospital Universitari Bellvitge | Microbiology Department | Sara Marti, Aida Gonzalez-Diaz, Laura Calatayud, Jordi Niubó, Carmen Ardanuy, Jordi Camara, M Angeles Domínguez |
| EPI_ISL_3030541, EPI_ISL_3030543, EPI_ISL_3030544, EPI_ISL_3030545, EPI_ISL_3030547, EPI_ISL_3030549 | CLILAB | Microbiology Department | Sara Marti, Aida Gonzalez-Diaz, Laura Calatayud, Jordi Niubó, Carmen Ardanuy, Jordi Camara, M Angeles Domínguez |
| EPI_ISL_3030550, EPI_ISL_3030551, EPI_ISL_3030552, EPI_ISL_3030553, EPI_ISL_3030554 | Hospital Universitari Bellvitge | Microbiology Department | Sara Marti, Aida Gonzalez-Diaz, Laura Calatayud, Jordi Niubó, Carmen Ardanuy, Jordi Camara, M Angeles Domínguez |
| EPI_ISL_3030556, EPI_ISL_3030558 | CLILAB | Microbiology Department | Sara Marti, Aida Gonzalez-Diaz, Laura Calatayud, Jordi Niubó, Carmen Ardanuy, Jordi Camara, M Angeles Domínguez |
| EPI_ISL_3030559, EPI_ISL_3030560, EPI_ISL_3030561, EPI_ISL_3030562, EPI_ISL_3030563, EPI_ISL_3030564, EPI_ISL_3030565, EPI_ISL_3030566, EPI_ISL_3030567, EPI_ISL_3030568, EPI_ISL_3030569, EPI_ISL_3030570, EPI_ISL_3030571, EPI_ISL_3030572, EPI_ISL_3030573, EPI_ISL_3030574, EPI_ISL_3030575, EPI_ISL_3030576, EPI_ISL_3030577, EPI_ISL_3030578, EPI_ISL_3030579, EPI_ISL_3030580, EPI_ISL_3030581, EPI_ISL_3030582, EPI_ISL_3030583, EPI_ISL_3030584, EPI_ISL_3030585, EPI_ISL_3030586, EPI_ISL_3030587, EPI_ISL_3030588, EPI_ISL_3030589, EPI_ISL_3030590, EPI_ISL_3030591, EPI_ISL_3030592, EPI_ISL_3030593, EPI_ISL_3030594, EPI_ISL_3030595, EPI_ISL_3030596, EPI_ISL_3030598, EPI_ISL_3030599, EPI_ISL_3030600, EPI_ISL_3030601, EPI_ISL_3030602, EPI_ISL_3030603, EPI_ISL_3030604, EPI_ISL_3030605, EPI_ISL_3030606, EPI_ISL_3030607, EPI_ISL_3030608, EPI_ISL_3030609, EPI_ISL_3030610, EPI_ISL_3030611, EPI_ISL_3030612, EPI_ISL_3030613, EPI_ISL_3030614 | | | |
| see above | Hospital Universitari Bellvitge | Microbiology Department | Sara Marti, Aida Gonzalez-Diaz, Laura Calatayud, Jordi Niubó, Carmen Ardanuy, Jordi Camara, M Angeles Domínguez |
| EPI_ISL_3030618, EPI_ISL_3030627, EPI_ISL_3030628, EPI_ISL_3030629, EPI_ISL_3030630, EPI_ISL_3030631, EPI_ISL_3030633, EPI_ISL_3030634, EPI_ISL_3030636, EPI_ISL_3030637, EPI_ISL_3030639, EPI_ISL_3030643, EPI_ISL_3030644, EPI_ISL_3030646, EPI_ISL_3030649, EPI_ISL_3030651, EPI_ISL_3030652, EPI_ISL_3030653, EPI_ISL_3030654, EPI_ISL_3030655, EPI_ISL_3030656, EPI_ISL_3030657, EPI_ISL_3030658, EPI_ISL_3030660, EPI_ISL_3030662, EPI_ISL_3030664, EPI_ISL_3030665 | | | |
| see above | Microbiology Department & Molecular Biology CORE \| CDB \| Hospital Clinic \| Barcelona | Microbiology Department & Molecular Biology CORE \| CDB \| Hospital Clinic \| Barcelona | Andrea Vergara Gomez, Jose Luis Villanueva Cañas, Miguel Julian Martínez Yoldi, Mª del Mar Mosquera Gutierrez, Mª Angeles Marcos Maeso |

| | | | |
|---|---|---|---|
| EPI_ISL_3030667, EPI_ISL_3030668, EPI_ISL_3030669, EPI_ISL_3030670, EPI_ISL_3030671, EPI_ISL_3030672, EPI_ISL_3030673, EPI_ISL_3030674, EPI_ISL_3030675, EPI_ISL_3030676, EPI_ISL_3030677, EPI_ISL_3030678, EPI_ISL_3030679, EPI_ISL_3030680, EPI_ISL_3030681, EPI_ISL_3030682, EPI_ISL_3030683, EPI_ISL_3030684, EPI_ISL_3030685, EPI_ISL_3030686, EPI_ISL_3030687, EPI_ISL_3030688, EPI_ISL_3030689, EPI_ISL_3030690, EPI_ISL_3030691, EPI_ISL_3030692, EPI_ISL_3030693, EPI_ISL_3030694, EPI_ISL_3030695, EPI_ISL_3030696 | | | |
| see above | Kansas Health and Environmental Lab | Kansas Health and Environmental Lab | Mike Grose, Jonathan Barnell, Ben Olsen, and Phil Adam |
| EPI_ISL_3030697, EPI_ISL_3030699, EPI_ISL_3030700, EPI_ISL_3030702, EPI_ISL_3030703, EPI_ISL_3030708, EPI_ISL_3030709, EPI_ISL_3030710, EPI_ISL_3030711, EPI_ISL_3030712, EPI_ISL_3030713, EPI_ISL_3030714, EPI_ISL_3030715, EPI_ISL_3030716, EPI_ISL_3030717, EPI_ISL_3030719, EPI_ISL_3030722, EPI_ISL_3030723, EPI_ISL_3030724, EPI_ISL_3030725, EPI_ISL_3030726, EPI_ISL_3030727, EPI_ISL_3030730, EPI_ISL_3030732, EPI_ISL_3030734, EPI_ISL_3030735, EPI_ISL_3030736, EPI_ISL_3030737, EPI_ISL_3030739, EPI_ISL_3030740, EPI_ISL_3030741 | | | |
| see above | MD PHL | MD PHL | Maryland Department of Health Laboratories Administration |
| EPI_ISL_3030749, EPI_ISL_3030751, EPI_ISL_3030753, EPI_ISL_3030754, EPI_ISL_3030755, EPI_ISL_3030756, EPI_ISL_3030758, EPI_ISL_3030761, EPI_ISL_3030762, EPI_ISL_3030765, EPI_ISL_3030768, EPI_ISL_3030772, EPI_ISL_3030773, EPI_ISL_3030775, EPI_ISL_3030776, EPI_ISL_3030777, EPI_ISL_3030784, EPI_ISL_3030789, EPI_ISL_3030792, EPI_ISL_3030796, EPI_ISL_3030797, EPI_ISL_3030799, EPI_ISL_3030801, EPI_ISL_3030802, EPI_ISL_3030805, EPI_ISL_3030806, EPI_ISL_3030808, EPI_ISL_3030809, EPI_ISL_3030810, EPI_ISL_3030811, EPI_ISL_3030814, EPI_ISL_3030815, EPI_ISL_3030817, EPI_ISL_3030819, EPI_ISL_3030824, EPI_ISL_3030826, EPI_ISL_3030828, EPI_ISL_3030829, EPI_ISL_3030832, EPI_ISL_3030833, EPI_ISL_3030903, EPI_ISL_3030905, EPI_ISL_3030912, EPI_ISL_3030913, EPI_ISL_3030915, EPI_ISL_3030917, EPI_ISL_3030918 | | | |
| see above | National Virus Reference Laboratory | National Virus Reference Laboratory | Zoe Yandle, Charlene Bennett, Gabriel Gonzalez, Michael Carr, Jonathan Dean, Cillian F De Gascun |
| EPI_ISL_3030921, EPI_ISL_3030922 | OSPEDALE CIVILE TERAMO - CENTRO TRASFUSIONALE | Istituto Zooprofilattico Sperimentale dell'Abruzzo e Molise "G. Caporale" | Lorusso A, Marcacci M, Di Domenico M, Ancora M, Curini V, Di Lollo Valeria, Mangone I, Rinaldi A, Delli Compagni E, Scialabba S, Caporale M, Di Pasquale A, Cammà C, Puglia I, Calistri P, Savini G |
| EPI_ISL_3030954 | URMC LABS | Wadsworth Center, New York State Department of Health | Kirsten St. George, Daryl M. Lamson, Alexis Russell, Matthew Shudt, Melissa A Leisner, Jonathan Plitnick, Catharine Prussing, Navjot Singh, John Kelly, Erasmus Schneider, Erica Lasek-Nesselquist |
| EPI_ISL_3031015 | Department of Medical Microbiology, St. Olavs hospital | Norwegian Institute of Public Health, Department of Virology | Kathrine Stene-Johansen, Kamilla Heddeland Instefjord, Hilde Elshaug, Garcia Llorente Ignacio, Jon Bråte, Engebretsen Serina Beate,Pedersen Benedikte Nevjen, Line Victoria Moen, Debech Nadia, Atiya R Ali,Marie Paulsen Madsen, Rasmus Riis Kopperud, Hilde Vollan, Karoline Bragstad, Olav Hungnes |
| EPI_ISL_3031016 | University Hospital of Northern Norway, Department for Microbiology and Infectious Disease Control | Norwegian Institute of Public Health, Department of Virology | Kathrine Stene-Johansen, Kamilla Heddeland Instefjord, Hilde Elshaug, Garcia Llorente Ignacio, Jon Bråte, Engebretsen Serina Beate,Pedersen Benedikte Nevjen, Line Victoria Moen, Debech Nadia, Atiya R Ali,Marie Paulsen Madsen, Rasmus Riis Kopperud, Hilde Vollan, Karoline Bragstad, Olav Hungnes |
| EPI_ISL_3031024, EPI_ISL_3031025 | Unilabs Laboratory Medicine | Norwegian Institute of Public Health, Department of Virology | Kathrine Stene-Johansen, Kamilla Heddeland Instefjord, Hilde Elshaug, Garcia Llorente Ignacio, Jon Bråte, Engebretsen Serina Beate,Pedersen Benedikte Nevjen, Line Victoria Moen, Debech Nadia, Atiya R Ali,Marie Paulsen Madsen, Rasmus Riis Kopperud, Hilde Vollan, Karoline Bragstad, Olav Hungnes |
| EPI_ISL_3031031, EPI_ISL_3031032 | Haukeland University Hospital, Dept. of Microbiology | Norwegian Institute of Public Health, Department of Virology | Kathrine Stene-Johansen, Kamilla Heddeland Instefjord, Hilde Elshaug, Garcia Llorente Ignacio, Jon Bråte, Engebretsen Serina Beate,Pedersen Benedikte Nevjen, Line Victoria Moen, Debech Nadia, Atiya R Ali,Marie Paulsen Madsen, Rasmus Riis Kopperud, Hilde Vollan, Karoline Bragstad, Olav Hungnes |
| EPI_ISL_3031034, EPI_ISL_3031035 | Ostfold Hospital Trust - Kalnes, Centre for Laboratory Medicine, Section for gene technology and infection serology | Norwegian Institute of Public Health, Department of Virology | Kathrine Stene-Johansen, Kamilla Heddeland Instefjord, Hilde Elshaug, Garcia Llorente Ignacio, Jon Bråte, Engebretsen Serina Beate,Pedersen Benedikte Nevjen, Line Victoria Moen, Debech Nadia, Atiya R Ali,Marie Paulsen Madsen, Rasmus Riis Kopperud, Hilde Vollan, Karoline Bragstad, Olav Hungnes |
| EPI_ISL_3031036 | Hospital of Southern Norway - Kristiansand, Department of Medical Microbiology | Norwegian Institute of Public Health, Department of Virology | Kathrine Stene-Johansen, Kamilla Heddeland Instefjord, Hilde Elshaug, Garcia Llorente Ignacio, Jon Bråte, Engebretsen Serina Beate,Pedersen Benedikte Nevjen, Line Victoria Moen, Debech Nadia, Atiya R Ali,Marie Paulsen Madsen, Rasmus Riis Kopperud, Hilde Vollan, Karoline Bragstad, Olav Hungnes |
| EPI_ISL_3031047, EPI_ISL_3031048, EPI_ISL_3031049 | Haukeland University Hospital, Dept. of Microbiology | Norwegian Institute of Public Health, Department of Virology | Kathrine Stene-Johansen, Kamilla Heddeland Instefjord, Hilde Elshaug, Garcia Llorente Ignacio, Jon Bråte, Engebretsen Serina Beate,Pedersen Benedikte Nevjen, Line Victoria Moen, Debech Nadia, Atiya R Ali,Marie Paulsen Madsen, Rasmus Riis Kopperud, Hilde Vollan, Karoline Bragstad, Olav Hungnes |
| EPI_ISL_3031052, EPI_ISL_3031053, EPI_ISL_3031054, EPI_ISL_3031055 | BOSTON HEART DIAGNOSTICS CORP | Wadsworth Center, New York State Department of Health | Kirsten St. George, Daryl M. Lamson, Alexis Russell, Matthew Shudt, Melissa A Leisner, Jonathan Plitnick, Catharine Prussing, Navjot Singh, John Kelly, Erasmus Schneider, Erica Lasek-Nesselquist |
| EPI_ISL_3031064, EPI_ISL_3031065, EPI_ISL_3031068, EPI_ISL_3031070, EPI_ISL_3031071, EPI_ISL_3031072, EPI_ISL_3031073, EPI_ISL_3031075, EPI_ISL_3031076, EPI_ISL_3031077 | NORTHWELL HEALTH LABORATORIES | Wadsworth Center, New York State Department of Health | Kirsten St. George, Daryl M. Lamson, Alexis Russell, Matthew Shudt, Melissa A Leisner, Jonathan Plitnick, Catharine Prussing, Navjot Singh, John Kelly, Erasmus Schneider, Erica Lasek-Nesselquist |
| EPI_ISL_3031078 | STONY BROOK UNIVERSITY HOSPITAL | Wadsworth Center, New York State Department of Health | Kirsten St. George, Daryl M. Lamson, Alexis Russell, Matthew Shudt, Melissa A Leisner, Jonathan Plitnick, Catharine Prussing, Navjot Singh, John Kelly, Erasmus Schneider, Erica Lasek-Nesselquist |
| EPI_ISL_3031132 | Vassar Brothers Medical Center Laboratory | Wadsworth Center, New York State Department of Health | Kirsten St. George, Daryl M. Lamson, Alexis Russell, Matthew Shudt, Melissa A Leisner, Jonathan Plitnick, Catharine Prussing, Navjot Singh, John Kelly, Erasmus Schneider, Erica Lasek-Nesselquist |
| EPI_ISL_3031140 | Vestfold Hospital, Toensberg Department of Microbiology | Norwegian Institute of Public Health, Department of Virology | Kathrine Stene-Johansen, Kamilla Heddeland Instefjord, Hilde Elshaug, Garcia Llorente Ignacio, Jon Bråte, Engebretsen Serina Beate,Pedersen Benedikte Nevjen, Line Victoria Moen, Debech Nadia, Atiya R Ali,Marie Paulsen Madsen, Rasmus Riis Kopperud, Hilde Vollan, Karoline Bragstad, Olav Hungnes |
| EPI_ISL_3031142, EPI_ISL_3031144 | Innlandet Hospital Trust, Division Lillehammer, Department for Medical Microbiology | Norwegian Institute of Public Health, Department of Virology | Kathrine Stene-Johansen, Kamilla Heddeland Instefjord, Hilde Elshaug, Garcia Llorente Ignacio, Jon Bråte, Engebretsen Serina Beate,Pedersen Benedikte Nevjen, Line Victoria Moen, Debech Nadia, Atiya R Ali,Marie Paulsen Madsen, Rasmus Riis Kopperud, Hilde Vollan, Karoline Bragstad, Olav Hungnes |
| EPI_ISL_3031158 | Department of Medical Microbiology, St. Olavs hospital | Norwegian Institute of Public Health, Department of Virology | Kathrine Stene-Johansen, Kamilla Heddeland Instefjord, Hilde Elshaug, Garcia Llorente Ignacio, Jon Bråte, Engebretsen Serina Beate,Pedersen Benedikte Nevjen, Line Victoria Moen, Debech Nadia, Atiya R Ali,Marie Paulsen Madsen, Rasmus Riis Kopperud, Hilde Vollan, Karoline Bragstad, Olav Hungnes |
| EPI_ISL_3031208, EPI_ISL_3031212 | STONY BROOK UNIVERSITY HOSPITAL | Wadsworth Center, New York State Department of Health | Kirsten St. George, Daryl M. Lamson, Alexis Russell, Matthew Shudt, Melissa A Leisner, Jonathan Plitnick, Catharine Prussing, Navjot Singh, John Kelly, Erasmus Schneider, Erica Lasek-Nesselquist |
| EPI_ISL_3031224, EPI_ISL_3031227, EPI_ISL_3031228, EPI_ISL_3031230, EPI_ISL_3031233, EPI_ISL_3031234, EPI_ISL_3031237, EPI_ISL_3031243, EPI_ISL_3031246, EPI_ISL_3031248, EPI_ISL_3031250, EPI_ISL_3031251, EPI_ISL_3031252, EPI_ISL_3031253, EPI_ISL_3031254, EPI_ISL_3031260, EPI_ISL_3031261, EPI_ISL_3031262, EPI_ISL_3031264, EPI_ISL_3031265, EPI_ISL_3031266, EPI_ISL_3031267, EPI_ISL_3031268, EPI_ISL_3031269, EPI_ISL_3031270 | | | |
| see above | URMC LABS | Wadsworth Center, New York State Department of Health | Kirsten St. George, Daryl M. Lamson, Alexis Russell, Matthew Shudt, Melissa A Leisner, Jonathan Plitnick, Catharine Prussing, Navjot Singh, John Kelly, Erasmus Schneider, Erica Lasek-Nesselquist |
| EPI_ISL_3031275, EPI_ISL_3031276 | Azarian Lab at Burnett School of Biomedical Sciences | Azarian Lab at Burnett School of Biomedical Sciences | Catherine Johnston, Eleonora Cella, Taj Azarian |
| EPI_ISL_3031347, EPI_ISL_3031348, EPI_ISL_3031349, EPI_ISL_3031350, EPI_ISL_3031351, EPI_ISL_3031352, EPI_ISL_3031353, EPI_ISL_3031354, EPI_ISL_3031355, EPI_ISL_3031356, EPI_ISL_3031357, EPI_ISL_3031358, EPI_ISL_3031359, EPI_ISL_3031360, EPI_ISL_3031361, EPI_ISL_3031362, EPI_ISL_3031363, EPI_ISL_3031364, EPI_ISL_3031365, EPI_ISL_3031366, EPI_ISL_3031368, EPI_ISL_3031369 | | | |
| see above | Arizona State Public Health Laboratory | Arizona State Public Health Laboratory | Trung Huynh, Jessica Escobar, Katherine Fullerton, Nobuko Fukushima, Matthew Contursi, Stacy White, Linda Getsinger, Victor Waddell |
| EPI_ISL_3031370, EPI_ISL_3031372 | Centre Hospitalier Universitaire Clermont-Ferrand | CHU Clermont-Ferrand, service de virologie | Bisseux Maxime, Mirand Audrey, Combes Patricia, Henquell Cécile |
| EPI_ISL_3031383 | KCRH KEMRI/DEID Surveillance Site | USAMRD-A, Basic Science Laboratory | Gathii Kimita, Josphat Nyataya, Esther Omuseni, Faith Sigei, Alan Lemtudo, Eric Muthanje, Brian Andika, Rehema Liyai, Rachel Githii, Clement Masakwe, Stephen Ochola, George Awinda, Carol Kifude, Beth Mutai, John Waitumbi |
| EPI_ISL_3031384 | KEMRI/DEID Surveillance Site (Kombewa Sub County Hospital) | USAMRD-A, Basic Science Laboratory | Gathii Kimita, Josphat Nyataya, Esther Omuseni, Faith Sigei, Alan Lemtudo, Eric Muthanje, Brian Andika, Rehema Liyai, Rachel Githii, Clement Masakwe, Stephen Ochola, George Awinda, Carol Kifude, Beth Mutai, John Waitumbi |
| EPI_ISL_3031389, EPI_ISL_3031390 | Port Health Clinic, Busia border point | USAMRD-A, Basic Science Laboratory | Gathii Kimita, Josphat Nyataya, Esther Omuseni, Faith Sigei, Alan Lemtudo, Eric Muthanje, Brian Andika, Rehema Liyai, Rachel Githii, Clement Masakwe, Stephen Ochola, George Awinda, Carol Kifude, Beth Mutai, John Waitumbi |
| EPI_ISL_3031391 | Othaya Sub-County Hospital | USAMRD-A, Basic Science Laboratory | Gathii Kimita, Josphat Nyataya, Esther Omuseni, Faith Sigei, Alan Lemtudo, Eric Muthanje, Brian Andika, Rehema Liyai, Rachel Githii, Clement Masakwe, Stephen Ochola, George Awinda, Carol Kifude, Beth Mutai, John Waitumbi |
| EPI_ISL_3031395 | Jaramogi Oginga Odinga Teaching and Referral Hospital | USAMRD-A, Basic Science Laboratory | Gathii Kimita, Josphat Nyataya, Esther Omuseni, Faith Sigei, Alan Lemtudo, Eric Muthanje, Brian Andika, Rehema Liyai, Rachel Githii, Clement Masakwe, Stephen Ochola, George Awinda, Carol Kifude, Beth Mutai, John Waitumbi |
| EPI_ISL_3031396 | Othaya Sub-County Hospital | USAMRD-A, Basic Science Laboratory | Gathii Kimita, Josphat Nyataya, Esther Omuseni, Faith Sigei, Alan Lemtudo, Eric Muthanje, Brian Andika, Rehema Liyai, Rachel Githii, Clement Masakwe, Stephen Ochola, George Awinda, Carol Kifude, Beth Mutai, John Waitumbi |
| EPI_ISL_3031398 | King'ong'o Dispensary | USAMRD-A, Basic Science Laboratory | Gathii Kimita, Josphat Nyataya, Esther Omuseni, Faith Sigei, Alan Lemtudo, Eric Muthanje, Brian Andika, Rehema Liyai, Rachel Githii, Clement Masakwe, Stephen Ochola, George Awinda, Carol Kifude, Beth Mutai, John Waitumbi |
| EPI_ISL_3031399 | Othaya Sub-County Hospital | USAMRD-A, Basic Science Laboratory | Gathii Kimita, Josphat Nyataya, Esther Omuseni, Faith Sigei, Alan Lemtudo, Eric Muthanje, Brian Andika, Rehema Liyai, Rachel Githii, Clement Masakwe, Stephen Ochola, George Awinda, Carol Kifude, Beth Mutai, John Waitumbi |
| EPI_ISL_3031400, EPI_ISL_3031401 | KCRH KEMRI/DEID Surveillance Site | USAMRD-A, Basic Science Laboratory | Gathii Kimita, Josphat Nyataya, Esther Omuseni, Faith Sigei, Alan Lemtudo, Eric Muthanje, Brian Andika, Rehema Liyai, Rachel Githii, Clement Masakwe, Stephen Ochola, George Awinda, Carol Kifude, Beth Mutai, John Waitumbi |
| EPI_ISL_3031403, EPI_ISL_3031404, EPI_ISL_3031406 | KEMRI/DEID Surveillance Site (Kombewa Sub County Hospital) | USAMRD-A, Basic Science Laboratory | Gathii Kimita, Josphat Nyataya, Esther Omuseni, Faith Sigei, Alan Lemtudo, Eric Muthanje, Brian Andika, Rehema Liyai, Rachel Githii, Clement Masakwe, Stephen Ochola, George Awinda, Carol Kifude, Beth Mutai, John Waitumbi |
| EPI_ISL_3031407, EPI_ISL_3031408, EPI_ISL_3031409, EPI_ISL_3031410, EPI_ISL_3031411, EPI_ISL_3031413, EPI_ISL_3031414, EPI_ISL_3031415, EPI_ISL_3031416, EPI_ISL_3031417 | KCRH KEMRI/DEID Surveillance Site | USAMRD-A, Basic Science Laboratory | Gathii Kimita, Josphat Nyataya, Esther Omuseni, Faith Sigei, Alan Lemtudo, Eric Muthanje, Brian Andika, Rehema Liyai, Rachel Githii, Clement Masakwe, Stephen Ochola, George Awinda, Carol Kifude, Beth Mutai, John Waitumbi |
| EPI_ISL_3031418, EPI_ISL_3031419 | Port Health Clinic, Busia border point | USAMRD-A, Basic Science Laboratory | Gathii Kimita, Josphat Nyataya, Esther Omuseni, Faith Sigei, Alan Lemtudo, Eric Muthanje, Brian Andika, Rehema Liyai, Rachel Githii, Clement Masakwe, Stephen Ochola, George Awinda, Carol Kifude, Beth Mutai, John Waitumbi |
| EPI_ISL_3031421, EPI_ISL_3031423, EPI_ISL_3031424, EPI_ISL_3031425, EPI_ISL_3031426, EPI_ISL_3031427, EPI_ISL_3031428, EPI_ISL_3031429, EPI_ISL_3031430, EPI_ISL_3031431, EPI_ISL_3031432, EPI_ISL_3031433 | | | |
| see above | Jaramogi Oginga Odinga Teaching and Referral Hospital | USAMRD-A, Basic Science Laboratory | Gathii Kimita, Josphat Nyataya, Esther Omuseni, Faith Sigei, Alan Lemtudo, Eric Muthanje, Brian Andika, Rehema Liyai, Rachel Githii, Clement Masakwe, Stephen Ochola, George Awinda, Carol Kifude, Beth Mutai, John Waitumbi |
| EPI_ISL_3031465, | Maryland Genomics, Institute for Genome Sciences, University | Maryland Genomics, Institute for Genome Sciences, University of | Tallon, Luke J; Sadzewicz, Lisa D; Humphrys, Mike; Ott, Sandra; Roussey, Holly; Santana-Cruz, Ivette; Vavikolanu, Kranthi; Lim, Jonathan;Hazen, Tracy;Fraser, Claire M; Ravel, Jacques |

| | | | |
|---|---|---|---|
| EPI_ISL_3031471, EPI_ISL_3031472, EPI_ISL_3031473, EPI_ISL_3031474 | of Maryland School of Medicine | Maryland School of Medicine | |
| EPI_ISL_3031475, EPI_ISL_3031476, EPI_ISL_3031479, EPI_ISL_3031480, EPI_ISL_3031481, EPI_ISL_3031482 | Banteay Meanchey Regional Laboratory | Virology Unit, Institut Pasteur du Cambodge | Leakhena Pum, Sophoannadedh Rath, Cecile Troupin, Ly Sovann, Kraing Sidonn, Yi Sengdoeurn, Chin Savuth, Chau Darapheak, Veasna Duong, Erik A Karlsson |
| EPI_ISL_3031485, EPI_ISL_3031486, EPI_ISL_3031487, EPI_ISL_3031488 | Kampong Cham Rapid Response Team | Virology Unit, Institut Pasteur du Cambodge | Leakhena Pum, Sophoannadedh Rath, Cecile Troupin, Ly Sovann, Kraing Sidonn, Yi Sengdoeurn, Chin Savuth, Chau Darapheak, Veasna Duong, Erik A Karlsson |
| EPI_ISL_3031489 | Kampong Thom Rapid Response Team | Virology Unit, Institut Pasteur du Cambodge | Leakhena Pum, Sophoannadedh Rath, Cecile Troupin, Ly Sovann, Kraing Sidonn, Yi Sengdoeurn, Chin Savuth, Chau Darapheak, Veasna Duong, Erik A Karlsson |
| EPI_ISL_3031490, EPI_ISL_3031491, EPI_ISL_3031494, EPI_ISL_3031496, EPI_ISL_3031497, EPI_ISL_3031498, EPI_ISL_3031499, EPI_ISL_3031500, EPI_ISL_3031501, EPI_ISL_3031502, EPI_ISL_3031503 | | | |
| see above | Oddar Meanchey Rapid Response Team | Virology Unit, Institut Pasteur du Cambodge | Leakhena Pum, Sophoannadedh Rath, Cecile Troupin, Ly Sovann, Kraing Sidonn, Yi Sengdoeurn, Chin Savuth, Chau Darapheak, Veasna Duong, Erik A Karlsson |
| EPI_ISL_3031504, EPI_ISL_3031505, EPI_ISL_3031516, EPI_ISL_3031525, EPI_ISL_3031527, EPI_ISL_3031528, EPI_ISL_3031529, EPI_ISL_3031530, EPI_ISL_3031534 | Virology Unit, Institut Pasteur du Cambodge | Virology Unit, Institut Pasteur du Cambodge | Leakhena Pum, Sophoannadedh Rath, Cecile Troupin, Ly Sovann, Kraing Sidonn, Yi Sengdoeurn, Chin Savuth, Chau Darapheak, Veasna Duong, Erik A Karlsson |
| EPI_ISL_3031559 | Pailin Rapid Response Team | Virology Unit, Institut Pasteur du Cambodge | Leakhena Pum, Sophoannadedh Rath, Cecile Troupin, Ly Sovann, Kraing Sidonn, Yi Sengdoeurn, Chin Savuth, Chau Darapheak, Veasna Duong, Erik A Karlsson |
| EPI_ISL_3031565, EPI_ISL_3031566, EPI_ISL_3031568, EPI_ISL_3031569, EPI_ISL_3031570, EPI_ISL_3031571, EPI_ISL_3031572, EPI_ISL_3031573, EPI_ISL_3031574, EPI_ISL_3031575, EPI_ISL_3031576, EPI_ISL_3031577, EPI_ISL_3031578, EPI_ISL_3031579, EPI_ISL_3031580, EPI_ISL_3031581, EPI_ISL_3031582, EPI_ISL_3031583, EPI_ISL_3031586, EPI_ISL_3031588, EPI_ISL_3031589, EPI_ISL_3031590, EPI_ISL_3031591, EPI_ISL_3031592, EPI_ISL_3031593, EPI_ISL_3031594, EPI_ISL_3031595, EPI_ISL_3031596, EPI_ISL_3031597, EPI_ISL_3031598, EPI_ISL_3031600, EPI_ISL_3031601, EPI_ISL_3031602 | | | |
| see above | Baylor Scott & White-Temple | Baylor Scott & White-Temple | Ari Rao, Linden Morales, Kimberly Walker, Marcus Volz, Shelby Johnson |
| EPI_ISL_3031603 | Baylor Scott & White-Waxahachie | Baylor Scott & White-Temple | Ari Rao, Linden Morales, Kimberly Walker, Marcus Volz, Shelby Johnson |
| EPI_ISL_3031604, EPI_ISL_3031605, EPI_ISL_3031606, EPI_ISL_3031607, EPI_ISL_3031608, EPI_ISL_3031609, EPI_ISL_3031612, EPI_ISL_3031613, EPI_ISL_3031614, EPI_ISL_3031615, EPI_ISL_3031616, EPI_ISL_3031617, EPI_ISL_3031619, EPI_ISL_3031620, EPI_ISL_3031621, EPI_ISL_3031622, EPI_ISL_3031623, EPI_ISL_3031624, EPI_ISL_3031625, EPI_ISL_3031626, EPI_ISL_3031627, EPI_ISL_3031628, EPI_ISL_3031629, EPI_ISL_3031630, EPI_ISL_3031631, EPI_ISL_3031632, EPI_ISL_3031633, EPI_ISL_3031634, EPI_ISL_3031635, EPI_ISL_3031636, EPI_ISL_3031637, EPI_ISL_3031638, EPI_ISL_3031639, EPI_ISL_3031640, EPI_ISL_3031641, EPI_ISL_3031642, EPI_ISL_3031643, EPI_ISL_3031644, EPI_ISL_3031646, EPI_ISL_3031647 | | | |
| see above | Baylor Scott & White-Temple | Baylor Scott & White-Temple | Ari Rao, Linden Morales, Kimberly Walker, Marcus Volz, Shelby Johnson |
| EPI_ISL_3031648 | Labo Analyses Med | National Reference Center for Viruses of Respiratory Infections, Institut Pasteur, Paris | Marion Barbet, Sylvie Behillil, Méline Bizard, Angela Brisebarre, Camille Capel, Vincent Enouf, Louise Lefrançois, Frédéric Lemoine, Christophe Malabat, Corinne Maufrais, Adrein Pain, Etienne Simon-Lorière, Maud Vanpeene, Sylvie Van der Werf ,Alexandra Ducancelle |
| EPI_ISL_3031649 | Labo Analyses Med | National Reference Center for Viruses of Respiratory Infections, Institut Pasteur, Paris | Marion Barbet, Sylvie Behillil, Méline Bizard, Angela Brisebarre, Camille Capel, Vincent Enouf, Louise Lefrançois, Frédéric Lemoine, Christophe Malabat, Corinne Maufrais, Victoire Baillet, Etienne Simon-Lorière, Maud Vanpeene, Sylvie Van der Werf ,Alexandra Ducancelle |
| EPI_ISL_3031657 | Hospital | National Reference Center for Viruses of Respiratory Infections, Institut Pasteur, Paris | Marion Barbet, Sylvie Behillil, Méline Bizard, Angela Brisebarre, Camille Capel, Vincent Enouf, Louise Lefrançois, Frédéric Lemoine, Christophe Malabat, Corinne Maufrais, Hub de Bioinformatique et Biostatistique, Etienne Simon-Lorière, Maud Vanpeene, Sylvie Van der Werf ,Alexandra Ducancelle |
| EPI_ISL_3031658 | Hospital | National Reference Center for Viruses of Respiratory Infections, Institut Pasteur, Paris | Marion Barbet, Sylvie Behillil, Méline Bizard, Angela Brisebarre, Camille Capel, Vincent Enouf, Louise Lefrançois, Frédéric Lemoine, Christophe Malabat, Corinne Maufrais, Adrien Pain, Etienne Simon-Lorière, Maud Vanpeene, Sylvie Van der Werf ,Alexandra Ducancelle |
| EPI_ISL_3031659 | Hospital | National Reference Center for Viruses of Respiratory Infections, Institut Pasteur, Paris | Marion Barbet, Sylvie Behillil, Méline Bizard, Angela Brisebarre, Camille Capel, Vincent Enouf, Louise Lefrançois, Frédéric Lemoine, Christophe Malabat, Corinne Maufrais, Hub de Bioinformatique et Biostatistique, Etienne Simon-Lorière, Maud Vanpeene, Sylvie Van der Werf ,Alexandra Ducancelle |
| EPI_ISL_3031660, EPI_ISL_3031661, EPI_ISL_3031663, EPI_ISL_3031664, EPI_ISL_3031665 | Labo Analyses Med | National Reference Center for Viruses of Respiratory Infections, Institut Pasteur, Paris | Marion Barbet, Sylvie Behillil, Méline Bizard, Angela Brisebarre, Camille Capel, Vincent Enouf, Louise Lefrançois, Frédéric Lemoine, Christophe Malabat, Corinne Maufrais, Hub de Bioinformatique et Biostatistique, Etienne Simon-Lorière, Maud Vanpeene, Sylvie Van der Werf ,Alexandra Ducancelle |
| EPI_ISL_3031666, EPI_ISL_3031667 | Labo Analyses Med | National Reference Center for Viruses of Respiratory Infections, Institut Pasteur, Paris | Marion Barbet, Sylvie Behillil, Méline Bizard, Angela Brisebarre, Camille Capel, Vincent Enouf, Louise Lefrançois, Frédéric Lemoine, Christophe Malabat, Corinne Maufrais, Adrien Pain, Etienne Simon-Lorière, Maud Vanpeene, Sylvie Van der Werf ,Alexandra Ducancelle |
| EPI_ISL_3031668, EPI_ISL_3031669 | Labo Analyses Med | National Reference Center for Viruses of Respiratory Infections, Institut Pasteur, Paris | Marion Barbet, Sylvie Behillil, Méline Bizard, Angela Brisebarre, Camille Capel, Vincent Enouf, Louise Lefrançois, Frédéric Lemoine, Christophe Malabat, Corinne Maufrais, Hub de Bioinformatique et Biostatistique, Etienne Simon-Lorière, Maud Vanpeene, Sylvie Van der Werf ,Alexandra Ducancelle |
| EPI_ISL_3031670 | Labo Analyses Med | National Reference Center for Viruses of Respiratory Infections, Institut Pasteur, Paris | Marion Barbet, Sylvie Behillil, Méline Bizard, Angela Brisebarre, Camille Capel, Vincent Enouf, Louise Lefrançois, Frédéric Lemoine, Christophe Malabat, Corinne Maufrais, Adrien Pain, Etienne Simon-Lorière, Maud Vanpeene, Sylvie Van der Werf ,Alexandra Ducancelle |
| EPI_ISL_3031671, EPI_ISL_3031672, EPI_ISL_3031673 | Labo Analyses Med | National Reference Center for Viruses of Respiratory Infections, Institut Pasteur, Paris | Marion Barbet, Sylvie Behillil, Méline Bizard, Angela Brisebarre, Camille Capel, Vincent Enouf, Louise Lefrançois, Frédéric Lemoine, Christophe Malabat, Corinne Maufrais, Hub de Bioinformatique et Biostatistique, Etienne Simon-Lorière, Maud Vanpeene, Sylvie Van der Werf ,Alexandra Ducancelle |
| EPI_ISL_3031674 | Labo Analyses Med | National Reference Center for Viruses of Respiratory Infections, Institut Pasteur, Paris | Marion Barbet, Sylvie Behillil, Méline Bizard, Angela Brisebarre, Camille Capel, Vincent Enouf, Louise Lefrançois, Frédéric Lemoine, Christophe Malabat, Corinne Maufrais, Adrien Pain, Etienne Simon-Lorière, Maud Vanpeene, Sylvie Van der Werf ,Alexandra Ducancelle |
| EPI_ISL_3031675 | Labo Analyses Med | National Reference Center for Viruses of Respiratory Infections, Institut Pasteur, Paris | Marion Barbet, Sylvie Behillil, Méline Bizard, Angela Brisebarre, Camille Capel, Vincent Enouf, Louise Lefrançois, Frédéric Lemoine, Christophe Malabat, Corinne Maufrais, Hub de Bioinformatique et Biostatistique, Etienne Simon-Lorière, Maud Vanpeene, Sylvie Van der Werf ,Alexandra Ducancelle |
| EPI_ISL_3031676 | Labo Analyses Med | National Reference Center for Viruses of Respiratory Infections, Institut Pasteur, Paris | Marion Barbet, Sylvie Behillil, Méline Bizard, Angela Brisebarre, Camille Capel, Vincent Enouf, Louise Lefrançois, Frédéric Lemoine, Christophe Malabat, Corinne Maufrais, Adrien Pain, Etienne Simon-Lorière, Maud Vanpeene, Sylvie Van der Werf ,Alexandra Ducancelle |
| EPI_ISL_3031677 | Labo Analyses Med | National Reference Center for Viruses of Respiratory Infections, Institut Pasteur, Paris | Marion Barbet, Sylvie Behillil, Méline Bizard, Angela Brisebarre, Camille Capel, Vincent Enouf, Louise Lefrançois, Frédéric Lemoine, Christophe Malabat, Corinne Maufrais, Hub de Bioinformatique et Biostatistique, Etienne Simon-Lorière, Maud Vanpeene, Sylvie Van der Werf ,Alexandra Ducancelle |
| EPI_ISL_3031678 | Labo Analyses Med | National Reference Center for Viruses of Respiratory Infections, Institut Pasteur, Paris | Marion Barbet, Sylvie Behillil, Méline Bizard, Angela Brisebarre, Camille Capel, Vincent Enouf, Louise Lefrançois, Frédéric Lemoine, Christophe Malabat, Corinne Maufrais, Adrien Pain, Etienne Simon-Lorière, Maud Vanpeene, Sylvie Van der Werf ,Alexandra Ducancelle |
| EPI_ISL_3031679, EPI_ISL_3031680 | Labo Analyses Med | National Reference Center for Viruses of Respiratory Infections, Institut Pasteur, Paris | Marion Barbet, Sylvie Behillil, Méline Bizard, Angela Brisebarre, Camille Capel, Vincent Enouf, Louise Lefrançois, Frédéric Lemoine, Christophe Malabat, Corinne Maufrais, Hub de Bioinformatique et Biostatistique, Etienne Simon-Lorière, Maud Vanpeene, Sylvie Van der Werf ,Alexandra Ducancelle |
| EPI_ISL_3031681 | Hospital | National Reference Center for Viruses of Respiratory Infections, Institut Pasteur, Paris | Marion Barbet, Sylvie Behillil, Méline Bizard, Angela Brisebarre, Camille Capel, Vincent Enouf, Louise Lefrançois, Frédéric Lemoine, Christophe Malabat, Corinne Maufrais, Victoire Baillet, Etienne Simon-Lorière, Maud Vanpeene, Sylvie Van der Werf ,Alexandra Ducancelle |
| EPI_ISL_3031682 | Hospital | National Reference Center for Viruses of Respiratory Infections, Institut Pasteur, Paris | Marion Barbet, Sylvie Behillil, Méline Bizard, Angela Brisebarre, Camille Capel, Vincent Enouf, Louise Lefrançois, Frédéric Lemoine, Christophe Malabat, Corinne Maufrais, Hub de Bioinformatique et Biostatistique, Etienne Simon-Lorière, Maud Vanpeene, Sylvie Van der Werf ,Alexandra Ducancelle |
| EPI_ISL_3031683 | Hospital | National Reference Center for Viruses of Respiratory Infections, Institut Pasteur, Paris | Marion Barbet, Sylvie Behillil, Méline Bizard, Angela Brisebarre, Camille Capel, Vincent Enouf, Louise Lefrançois, Frédéric Lemoine, Christophe Malabat, Corinne Maufrais, Adrien Pain, Etienne Simon-Lorière, Maud Vanpeene, Sylvie Van der Werf ,Alexandra Ducancelle |
| EPI_ISL_3031685 | Hospital | National Reference Center for Viruses of Respiratory Infections, Institut Pasteur, Paris | Marion Barbet, Sylvie Behillil, Méline Bizard, Angela Brisebarre, Camille Capel, Vincent Enouf, Louise Lefrançois, Frédéric Lemoine, Christophe Malabat, Corinne Maufrais, Victoire Baillet, Etienne Simon-Lorière, Maud Vanpeene, Sylvie Van der Werf ,Alexandra Ducancelle |
| EPI_ISL_3031686, EPI_ISL_3031687 | Armies | National Reference Center for Viruses of Respiratory Infections, Institut Pasteur, Paris | Marion Barbet, Sylvie Behillil, Méline Bizard, Angela Brisebarre, Camille Capel, Vincent Enouf, Louise Lefrançois, Frédéric Lemoine, Christophe Malabat, Corinne Maufrais, Hub de Bioinformatique et Biostatistique, Etienne Simon-Lorière, Maud Vanpeene, Sylvie Van der Werf ,Alexandra Ducancelle |
| EPI_ISL_3031688 | Hospital | National Reference Center for Viruses of Respiratory Infections, Institut Pasteur, Paris | Marion Barbet, Sylvie Behillil, Méline Bizard, Angela Brisebarre, Camille Capel, Vincent Enouf, Louise Lefrançois, Frédéric Lemoine, Christophe Malabat, Corinne Maufrais, Victoire Baillet, Etienne Simon-Lorière, Maud Vanpeene, Sylvie Van der Werf ,Alexandra Ducancelle |
| EPI_ISL_3031689 | Hospital | National Reference Center for Viruses of Respiratory Infections, Institut Pasteur, Paris | Marion Barbet, Sylvie Behillil, Méline Bizard, Angela Brisebarre, Camille Capel, Vincent Enouf, Louise Lefrançois, Frédéric Lemoine, Christophe Malabat, Corinne Maufrais, Hub de Bioinformatique et Biostatistique, Etienne Simon-Lorière, Maud Vanpeene, Sylvie Van der Werf ,Alexandra Ducancelle |
| EPI_ISL_3031690, EPI_ISL_3031691, EPI_ISL_3031692, EPI_ISL_3031693, EPI_ISL_3031694, EPI_ISL_3031695, EPI_ISL_3031696 | Labo Analyses Med | National Reference Center for Viruses of Respiratory Infections, Institut Pasteur, Paris | Marion Barbet, Sylvie Behillil, Méline Bizard, Angela Brisebarre, Camille Capel, Vincent Enouf, Louise Lefrançois, Frédéric Lemoine, Christophe Malabat, Corinne Maufrais, Hub de Bioinformatique et Biostatistique, Etienne Simon-Lorière, Maud Vanpeene, Sylvie Van der Werf ,Alexandra Ducancelle |
| EPI_ISL_3031697, EPI_ISL_3031698, EPI_ISL_3031699, | Hospital | National Reference Center for Viruses of Respiratory Infections, Institut Pasteur, Paris | Marion Barbet, Sylvie Behillil, Méline Bizard, Angela Brisebarre, Camille Capel, Vincent Enouf, Louise Lefrançois, Frédéric Lemoine, Christophe Malabat, Corinne Maufrais, Hub de Bioinformatique et Biostatistique, Etienne Simon-Lorière, Maud Vanpeene, Sylvie Van der Werf ,Alexandra Ducancelle |

| | | | |
|---|---|---|---|
| EPI_ISL_3031700 | | | |
| EPI_ISL_3031701, EPI_ISL_3031702, EPI_ISL_3031705, EPI_ISL_3031706, EPI_ISL_3031710, EPI_ISL_3031711, EPI_ISL_3031712, EPI_ISL_3031713, EPI_ISL_3031714, EPI_ISL_3031715, EPI_ISL_3031716, EPI_ISL_3031718, EPI_ISL_3031719, EPI_ISL_3031720, EPI_ISL_3031721, EPI_ISL_3031722, EPI_ISL_3031723, EPI_ISL_3031724, EPI_ISL_3031725, EPI_ISL_3031726, EPI_ISL_3031727, EPI_ISL_3031728, EPI_ISL_3031730, EPI_ISL_3031731, EPI_ISL_3031732, EPI_ISL_3031733, EPI_ISL_3031734, EPI_ISL_3031735, EPI_ISL_3031736, EPI_ISL_3031737, EPI_ISL_3031738, EPI_ISL_3031739, EPI_ISL_3031740, EPI_ISL_3031741, EPI_ISL_3031742, EPI_ISL_3031743, EPI_ISL_3031744, EPI_ISL_3031745, EPI_ISL_3031746, EPI_ISL_3031747, EPI_ISL_3031748, EPI_ISL_3031749, EPI_ISL_3031750, EPI_ISL_3031751 | | | |
| see above | Labo Analyses Med | National Reference Center for Viruses of Respiratory Infections, Institut Pasteur, Paris | Marion Barbet, Sylvie Behillil, Méline Bizard, Angela Brisebarre, Camille Capel, Vincent Enouf, Louise Lefrançois, Frédéric Lemoine, Christophe Malabat, Corinne Maufrais, Hub de Bioinformatique et Biostatistique, Etienne Simon-Lorière, Maud Vanpeene, Sylvie Van der Werf ,Alexandra Ducancelle |
| EPI_ISL_3031757, EPI_ISL_3031758, EPI_ISL_3031759, EPI_ISL_3031760, EPI_ISL_3031761, EPI_ISL_3031762, EPI_ISL_3031763, EPI_ISL_3031764, EPI_ISL_3031765, EPI_ISL_3031766, EPI_ISL_3031767, EPI_ISL_3031768, EPI_ISL_3031769, EPI_ISL_3031770, EPI_ISL_3031771, EPI_ISL_3031772, EPI_ISL_3031773, EPI_ISL_3031774, EPI_ISL_3031775, EPI_ISL_3031776, EPI_ISL_3031777 | | | |
| see above | Kansas Health and Environmental Lab | Kansas Health and Environmental Lab | Mike Grose, Katherine Wiggins, Jonathan Barnell, Ben Olsen, and Phil Adam |
| EPI_ISL_3031779 | Clinical Reference Laboratory | Kansas Health and Environmental Lab | Mike Grose, Katherine Wiggins, Jonathan Barnell, Ben Olsen, and Phil Adam |
| EPI_ISL_3031781, EPI_ISL_3031782, EPI_ISL_3031783 | Labo Analyses Med | National Reference Center for Viruses of Respiratory Infections, Institut Pasteur, Paris | Marion Barbet, Sylvie Behillil, Méline Bizard, Angela Brisebarre, Camille Capel, Vincent Enouf, Louise Lefrançois, Frédéric Lemoine, Christophe Malabat, Corinne Maufrais, Hub de Bioinformatique et Statistique, Etienne Simon-Lorière, Maud Vanpeene, Sylvie Van der Werf |
| EPI_ISL_3031784, EPI_ISL_3031785 | Labo Analyses Med | National Reference Center for Viruses of Respiratory Infections, Institut Pasteur, Paris | Marion Barbet, Sylvie Behillil, Méline Bizard, Angela Brisebarre, Camille Capel, Vincent Enouf, Louise Lefrançois, Frédéric Lemoine, Christophe Malabat, Corinne Maufrais, Emmanuelle Permal, Etienne Simon-Lorière, Maud Vanpeene, Sylvie Van der Werf |
| EPI_ISL_3031786, EPI_ISL_3031787, EPI_ISL_3031788 | Labo Analyses Med | National Reference Center for Viruses of Respiratory Infections, Institut Pasteur, Paris | Marion Barbet, Sylvie Behillil, Méline Bizard, Angela Brisebarre, Camille Capel, Vincent Enouf, Louise Lefrançois, Frédéric Lemoine, Christophe Malabat, Corinne Maufrais, Hub de Bioinformatique et Statistique, Etienne Simon-Lorière, Maud Vanpeene, Sylvie Van der Werf |
| EPI_ISL_3031789 | Labo Analyses Med | National Reference Center for Viruses of Respiratory Infections, Institut Pasteur, Paris | Marion Barbet, Sylvie Behillil, Méline Bizard, Angela Brisebarre, Camille Capel, Vincent Enouf, Louise Lefrançois, Frédéric Lemoine, Christophe Malabat, Corinne Maufrais, Emmanuelle Permal, Etienne Simon-Lorière, Maud Vanpeene, Sylvie Van der Werf |
| EPI_ISL_3031790, EPI_ISL_3031792, EPI_ISL_3031793 | Hospital | National Reference Center for Viruses of Respiratory Infections, Institut Pasteur, Paris | Marion Barbet, Sylvie Behillil, Méline Bizard, Angela Brisebarre, Camille Capel, Vincent Enouf, Louise Lefrançois, Frédéric Lemoine, Christophe Malabat, Corinne Maufrais, Hub de Bioinformatique et Statistique, Etienne Simon-Lorière, Maud Vanpeene, Sylvie Van der Werf |
| EPI_ISL_3031796 | Labo Analyses Med | National Reference Center for Viruses of Respiratory Infections, Institut Pasteur, Paris | Marion Barbet, Sylvie Behillil, Méline Bizard, Angela Brisebarre, Camille Capel, Vincent Enouf, Louise Lefrançois, Frédéric Lemoine, Christophe Malabat, Corinne Maufrais, Victoire Baillet, Etienne Simon-Lorière, Maud Vanpeene, Sylvie Van der Werf |
| EPI_ISL_3031798, EPI_ISL_3031799, EPI_ISL_3031800, EPI_ISL_3031801, EPI_ISL_3031802, EPI_ISL_3031803, EPI_ISL_3031804 | Labo Analyses Med | National Reference Center for Viruses of Respiratory Infections, Institut Pasteur, Paris | Marion Barbet, Sylvie Behillil, Méline Bizard, Angela Brisebarre, Camille Capel, Vincent Enouf, Louise Lefrançois, Frédéric Lemoine, Christophe Malabat, Corinne Maufrais, Hub de Bioinformatique et Statistique, Etienne Simon-Lorière, Maud Vanpeene, Sylvie Van der Werf |
| EPI_ISL_3031805 | Labo Analyses Med | National Reference Center for Viruses of Respiratory Infections, Institut Pasteur, Paris | Marion Barbet, Sylvie Behillil, Méline Bizard, Angela Brisebarre, Camille Capel, Vincent Enouf, Louise Lefrançois, Frédéric Lemoine, Christophe Malabat, Corinne Maufrais, Victoire Baillet, Etienne Simon-Lorière, Maud Vanpeene, Sylvie Van der Werf |
| EPI_ISL_3031806 | Labo Analyses Med | National Reference Center for Viruses of Respiratory Infections, Institut Pasteur, Paris | Marion Barbet, Sylvie Behillil, Méline Bizard, Angela Brisebarre, Camille Capel, Vincent Enouf, Louise Lefrançois, Frédéric Lemoine, Christophe Malabat, Corinne Maufrais, Hub de Bioinformatique et Statistique, Etienne Simon-Lorière, Maud Vanpeene, Sylvie Van der Werf |
| EPI_ISL_3031807 | Labo Analyses Med | National Reference Center for Viruses of Respiratory Infections, Institut Pasteur, Paris | Marion Barbet, Sylvie Behillil, Méline Bizard, Angela Brisebarre, Camille Capel, Vincent Enouf, Louise Lefrançois, Frédéric Lemoine, Christophe Malabat, Corinne Maufrais, Victoire Baillet, Etienne Simon-Lorière, Maud Vanpeene, Sylvie Van der Werf |
| EPI_ISL_3031808, EPI_ISL_3031809, EPI_ISL_3031811, EPI_ISL_3031812, EPI_ISL_3031813, EPI_ISL_3031816, EPI_ISL_3031817, EPI_ISL_3031818, EPI_ISL_3031819, EPI_ISL_3031820, EPI_ISL_3031821 | | | |
| see above | Labo Analyses Med | National Reference Center for Viruses of Respiratory Infections, Institut Pasteur, Paris | Marion Barbet, Sylvie Behillil, Méline Bizard, Angela Brisebarre, Camille Capel, Vincent Enouf, Louise Lefrançois, Frédéric Lemoine, Christophe Malabat, Corinne Maufrais, Hub de Bioinformatique et Statistique, Etienne Simon-Lorière, Maud Vanpeene, Sylvie Van der Werf |
| EPI_ISL_3031822, EPI_ISL_3031823 | Labo Analyses Med | National Reference Center for Viruses of Respiratory Infections, Institut Pasteur, Paris | Marion Barbet, Sylvie Behillil, Méline Bizard, Angela Brisebarre, Camille Capel, Vincent Enouf, Louise Lefrançois, Frédéric Lemoine, Christophe Malabat, Corinne Maufrais, Victoire Baillet, Etienne Simon-Lorière, Maud Vanpeene, Sylvie Van der Werf |
| EPI_ISL_3031824, EPI_ISL_3031825 | Labo Analyses Med | National Reference Center for Viruses of Respiratory Infections, Institut Pasteur, Paris | Marion Barbet, Sylvie Behillil, Méline Bizard, Angela Brisebarre, Camille Capel, Vincent Enouf, Louise Lefrançois, Frédéric Lemoine, Christophe Malabat, Corinne Maufrais, Hub de Bioinformatique et Statistique, Etienne Simon-Lorière, Maud Vanpeene, Sylvie Van der Werf |
| EPI_ISL_3031826 | Labo Analyses Med | National Reference Center for Viruses of Respiratory Infections, Institut Pasteur, Paris | Marion Barbet, Sylvie Behillil, Méline Bizard, Angela Brisebarre, Camille Capel, Vincent Enouf, Louise Lefrançois, Frédéric Lemoine, Christophe Malabat, Corinne Maufrais, Victoire Baillet, Etienne Simon-Lorière, Maud Vanpeene, Sylvie Van der Werf |
| EPI_ISL_3031827, EPI_ISL_3031828, EPI_ISL_3031829, EPI_ISL_3031830, EPI_ISL_3031831 | Labo Analyses Med | National Reference Center for Viruses of Respiratory Infections, Institut Pasteur, Paris | Marion Barbet, Sylvie Behillil, Méline Bizard, Angela Brisebarre, Camille Capel, Vincent Enouf, Louise Lefrançois, Frédéric Lemoine, Christophe Malabat, Corinne Maufrais, Hub de Bioinformatique et Statistique, Etienne Simon-Lorière, Maud Vanpeene, Sylvie Van der Werf |
| EPI_ISL_3031832, EPI_ISL_3031833 | Labo Analyses Med | National Reference Center for Viruses of Respiratory Infections, Institut Pasteur, Paris | Marion Barbet, Sylvie Behillil, Méline Bizard, Angela Brisebarre, Camille Capel, Vincent Enouf, Louise Lefrançois, Frédéric Lemoine, Christophe Malabat, Corinne Maufrais, Victoire Baillet, Etienne Simon-Lorière, Maud Vanpeene, Sylvie Van der Werf |
| EPI_ISL_3031834, EPI_ISL_3031835, EPI_ISL_3031836 | Labo Analyses Med | National Reference Center for Viruses of Respiratory Infections, Institut Pasteur, Paris | Marion Barbet, Sylvie Behillil, Méline Bizard, Angela Brisebarre, Camille Capel, Vincent Enouf, Louise Lefrançois, Frédéric Lemoine, Christophe Malabat, Corinne Maufrais, Hub de Bioinformatique et Statistique, Etienne Simon-Lorière, Maud Vanpeene, Sylvie Van der Werf |
| EPI_ISL_3031837 | Labo Analyses Med | National Reference Center for Viruses of Respiratory Infections, Institut Pasteur, Paris | Marion Barbet, Sylvie Behillil, Méline Bizard, Angela Brisebarre, Camille Capel, Vincent Enouf, Louise Lefrançois, Frédéric Lemoine, Christophe Malabat, Corinne Maufrais, Victoire Baillet, Etienne Simon-Lorière, Maud Vanpeene, Sylvie Van der Werf |
| EPI_ISL_3031839 | Labo Analyses Med | National Reference Center for Viruses of Respiratory Infections, Institut Pasteur, Paris | Marion Barbet, Sylvie Behillil, Méline Bizard, Angela Brisebarre, Camille Capel, Vincent Enouf, Louise Lefrançois, Frédéric Lemoine, Christophe Malabat, Corinne Maufrais, Hub de Bioinformatique et Statistique, Etienne Simon-Lorière, Maud Vanpeene, Sylvie Van der Werf |
| EPI_ISL_3031840 | Labo Analyses Med | National Reference Center for Viruses of Respiratory Infections, Institut Pasteur, Paris | Marion Barbet, Sylvie Behillil, Méline Bizard, Angela Brisebarre, Camille Capel, Vincent Enouf, Louise Lefrançois, Frédéric Lemoine, Christophe Malabat, Corinne Maufrais, Victoire Baillet, Etienne Simon-Lorière, Maud Vanpeene, Sylvie Van der Werf |
| EPI_ISL_3031842, EPI_ISL_3031843, EPI_ISL_3031844, EPI_ISL_3031845, EPI_ISL_3031846 | Labo Analyses Med | National Reference Center for Viruses of Respiratory Infections, Institut Pasteur, Paris | Marion Barbet, Sylvie Behillil, Méline Bizard, Angela Brisebarre, Camille Capel, Vincent Enouf, Louise Lefrançois, Frédéric Lemoine, Christophe Malabat, Corinne Maufrais, Hub de Bioinformatique et Statistique, Etienne Simon-Lorière, Maud Vanpeene, Sylvie Van der Werf |
| EPI_ISL_3031847 | Labo Analyses Med | National Reference Center for Viruses of Respiratory Infections, Institut Pasteur, Paris | Marion Barbet, Sylvie Behillil, Méline Bizard, Angela Brisebarre, Camille Capel, Vincent Enouf, Louise Lefrançois, Frédéric Lemoine, Christophe Malabat, Corinne Maufrais, Victoire Baillet, Etienne Simon-Lorière, Maud Vanpeene, Sylvie Van der Werf |
| EPI_ISL_3031848, EPI_ISL_3031849 | Labo Analyses Med | National Reference Center for Viruses of Respiratory Infections, Institut Pasteur, Paris | Marion Barbet, Sylvie Behillil, Méline Bizard, Angela Brisebarre, Camille Capel, Vincent Enouf, Louise Lefrançois, Frédéric Lemoine, Christophe Malabat, Corinne Maufrais, Hub de Bioinformatique et Statistique, Etienne Simon-Lorière, Maud Vanpeene, Sylvie Van der Werf |
| EPI_ISL_3031850 | Labo Analyses Med | National Reference Center for Viruses of Respiratory Infections, Institut Pasteur, Paris | Marion Barbet, Sylvie Behillil, Méline Bizard, Angela Brisebarre, Camille Capel, Vincent Enouf, Louise Lefrançois, Frédéric Lemoine, Christophe Malabat, Corinne Maufrais, Victoire Baillet, Etienne Simon-Lorière, Maud Vanpeene, Sylvie Van der Werf |
| EPI_ISL_3031852 | Labo Analyses Med | National Reference Center for Viruses of Respiratory Infections, Institut Pasteur, Paris | Marion Barbet, Sylvie Behillil, Méline Bizard, Angela Brisebarre, Camille Capel, Vincent Enouf, Louise Lefrançois, Frédéric Lemoine, Christophe Malabat, Corinne Maufrais, Hub de Bioinformatique et Statistique, Etienne Simon-Lorière, Maud Vanpeene, Sylvie Van der Werf |
| EPI_ISL_3031853 | Labo Analyses Med | National Reference Center for Viruses of Respiratory Infections, Institut Pasteur, Paris | Marion Barbet, Sylvie Behillil, Méline Bizard, Angela Brisebarre, Camille Capel, Vincent Enouf, Louise Lefrançois, Frédéric Lemoine, Christophe Malabat, Corinne Maufrais, Victoire Baillet, Etienne Simon-Lorière, Maud Vanpeene, Sylvie Van der Werf |
| EPI_ISL_3031854 | Labo Analyses Med | National Reference Center for Viruses of Respiratory Infections, Institut Pasteur, Paris | Marion Barbet, Sylvie Behillil, Méline Bizard, Angela Brisebarre, Camille Capel, Vincent Enouf, Louise Lefrançois, Frédéric Lemoine, Christophe Malabat, Corinne Maufrais, Hub de Bioinformatique et Statistique, Etienne Simon-Lorière, Maud Vanpeene, Sylvie Van der Werf |
| EPI_ISL_3031856 | Labo Analyses Med | National Reference Center for Viruses of Respiratory Infections, Institut Pasteur, Paris | Marion Barbet, Sylvie Behillil, Méline Bizard, Angela Brisebarre, Camille Capel, Vincent Enouf, Louise Lefrançois, Frédéric Lemoine, Christophe Malabat, Corinne Maufrais, Victoire Baillet, Etienne Simon-Lorière, Maud Vanpeene, Sylvie Van der Werf |
| EPI_ISL_3031857, EPI_ISL_3031858, EPI_ISL_3031861 | Labo Analyses Med | National Reference Center for Viruses of Respiratory Infections, Institut Pasteur, Paris | Marion Barbet, Sylvie Behillil, Méline Bizard, Angela Brisebarre, Camille Capel, Vincent Enouf, Louise Lefrançois, Frédéric Lemoine, Christophe Malabat, Corinne Maufrais, Hub de Bioinformatique et Statistique, Etienne Simon-Lorière, Maud Vanpeene, Sylvie Van der Werf |
| EPI_ISL_3031862 | Labo Analyses Med | National Reference Center for Viruses of Respiratory Infections, Institut Pasteur, Paris | Marion Barbet, Sylvie Behillil, Méline Bizard, Angela Brisebarre, Camille Capel, Vincent Enouf, Louise Lefrançois, Frédéric Lemoine, Christophe Malabat, Corinne Maufrais, Victoire Baillet, Etienne Simon-Lorière, Maud Vanpeene, Sylvie Van der Werf |
| EPI_ISL_3031866, EPI_ISL_3031867, EPI_ISL_3031868, EPI_ISL_3031870, EPI_ISL_3031871, EPI_ISL_3031872, EPI_ISL_3031873, EPI_ISL_3031874, EPI_ISL_3031875, EPI_ISL_3031876, EPI_ISL_3031877, EPI_ISL_3031878, EPI_ISL_3031880, EPI_ISL_3031881, EPI_ISL_3031882, EPI_ISL_3031884, EPI_ISL_3031885, EPI_ISL_3031886, EPI_ISL_3031887, EPI_ISL_3031890, EPI_ISL_3031892, EPI_ISL_3031893, EPI_ISL_3031896, EPI_ISL_3031897, EPI_ISL_3031898, EPI_ISL_3031900, EPI_ISL_3031902, EPI_ISL_3031905, EPI_ISL_3031906, EPI_ISL_3031907, EPI_ISL_3031908, EPI_ISL_3031910, EPI_ISL_3031911, EPI_ISL_3031912 | | | |
| see above | Labo Analyses Med | National Reference Center for Viruses of Respiratory Infections, Institut Pasteur, Paris | Marion Barbet, Sylvie Behillil, Méline Bizard, Angela Brisebarre, Camille Capel, Vincent Enouf, Louise Lefrançois, Frédéric Lemoine, Christophe Malabat, Corinne Maufrais, Hub de Bioinformatique et Statistique, Etienne Simon-Lorière, Maud Vanpeene, Sylvie Van der Werf |
| EPI_ISL_3031918 | Hospital | National Reference Center for Viruses of Respiratory Infections, Institut Pasteur, Paris | Marion Barbet, Sylvie Behillil, Méline Bizard, Angela Brisebarre, Camille Capel, Vincent Enouf, Louise Lefrançois, Frédéric Lemoine, Christophe Malabat, Corinne Maufrais, Hub de Bioinformatique et Statistique, Etienne Simon-Lorière, Maud Vanpeene, Sylvie Van der Werf |
| EPI_ISL_3031920, EPI_ISL_3031922 | Labo Analyses Med | National Reference Center for Viruses of Respiratory Infections, Institut Pasteur, Paris | Marion Barbet, Sylvie Behillil, Méline Bizard, Angela Brisebarre, Camille Capel, Vincent Enouf, Louise Lefrançois, Frédéric Lemoine, Christophe Malabat, Corinne Maufrais, Hub de Bioinformatique et Statistique, Etienne Simon-Lorière, Maud Vanpeene, Sylvie Van der Werf |

| | | | |
|---|---|---|---|
| EPI_ISL_3031923, EPI_ISL_3031924 | Hospital | National Reference Center for Viruses of Respiratory Infections, Institut Pasteur, Paris | Marion Barbet, Sylvie Behillil, Méline Bizard, Angela Brisebarre, Camille Capel, Vincent Enouf, Louise Lefrançois, Frédéric Lemoine, Christophe Malabat, Corinne Maufrais, Hub de Bioinformatique et Statistique, Etienne Simon-Lorière, Maud Vanpeene, Sylvie Van der Werf |
| EPI_ISL_3031925, EPI_ISL_3031926, EPI_ISL_3031927, EPI_ISL_3031928, EPI_ISL_3031929, EPI_ISL_3031930, EPI_ISL_3031931, EPI_ISL_3031932, EPI_ISL_3031933 | Labo Analyses Med | National Reference Center for Viruses of Respiratory Infections, Institut Pasteur, Paris | Marion Barbet, Sylvie Behillil, Méline Bizard, Angela Brisebarre, Camille Capel, Vincent Enouf, Louise Lefrançois, Frédéric Lemoine, Christophe Malabat, Corinne Maufrais, Hub de Bioinformatique et Statistique, Etienne Simon-Lorière, Maud Vanpeene, Sylvie Van der Werf |
| EPI_ISL_3031939, EPI_ISL_3031940, EPI_ISL_3031941, EPI_ISL_3031942, EPI_ISL_3031943, EPI_ISL_3031944, EPI_ISL_3031966, EPI_ISL_3031988, EPI_ISL_3031998, EPI_ISL_3032004, EPI_ISL_3032028, EPI_ISL_3032029, EPI_ISL_3032030, EPI_ISL_3032033, EPI_ISL_3032034, EPI_ISL_3032035, EPI_ISL_3032036, EPI_ISL_3032037 | | | |
| see above | Institute of Virology, Medical Center, University of Freiburg, Freiburg, Germany | Institute of Virology, Clinial Virus Genomics, Medical Center, University of Freiburg, Freiburg, Germany | Jonas Fuchs, Lisa Kern, Lena Jaki, Sandra Reuter, Hajo Grundmann, Marcus Panning |
| EPI_ISL_3032039, EPI_ISL_3032042 | Willis-Knighton Medical Center Hospital Laboratory | LSUHS Emerging Viral Threat Laboratory | Gregory L. Ware, Alexander Mijalis, Jeremy P. Kamil, Jennifer L. Carroll, Maarten Van Diest, Rona S. Scott, Andrew D. Yurochko, Christopher G. Kevil, John A. Vanchiere |
| EPI_ISL_3032043, EPI_ISL_3032044, EPI_ISL_3032047, EPI_ISL_3032048 | LSUHS Emerging Viral Threat Laboratory | LSUHS Emerging Viral Threat Laboratory | Gregory L. Ware, Alexander Mijalis, Jeremy P. Kamil, Jennifer L. Carroll, Maarten Van Diest, Rona S. Scott, Andrew D. Yurochko, Christopher G. Kevil, John A. Vanchiere |
| EPI_ISL_3032101, EPI_ISL_3032102, EPI_ISL_3032103, EPI_ISL_3032104, EPI_ISL_3032105, EPI_ISL_3032106, EPI_ISL_3032107, EPI_ISL_3032108, EPI_ISL_3032109, EPI_ISL_3032110, EPI_ISL_3032111, EPI_ISL_3032112, EPI_ISL_3032113, EPI_ISL_3032114, EPI_ISL_3032115, EPI_ISL_3032116, EPI_ISL_3032117, EPI_ISL_3032118, EPI_ISL_3032119, EPI_ISL_3032120, EPI_ISL_3032121, EPI_ISL_3032122, EPI_ISL_3032123, EPI_ISL_3032124, EPI_ISL_3032125, EPI_ISL_3032126, EPI_ISL_3032127 | | | |
| see above | Kansas Health and Environmental Lab | Kansas Health and Environmental Lab | Mike Grose, Katherine Wiggins, Jonathan Barnell, Ben Olsen, and Phil Adam |
| EPI_ISL_3032153, EPI_ISL_3032154, EPI_ISL_3032155, EPI_ISL_3032159, EPI_ISL_3032162, EPI_ISL_3032163, EPI_ISL_3032179, EPI_ISL_3032180, EPI_ISL_3032183 | UW Virology Lab | UW Virology Lab | Pavitra Roychoudhury, Hong Xie, Lasata Shrestha, Shah Mohamed Bakhash, Tien V. Nguyen, Noah R. Baker, Sean Ellis, Meei-Li Huang, Keith R Jerome, Alexander Greninger |
| EPI_ISL_3032188, EPI_ISL_3032190, EPI_ISL_3032192, EPI_ISL_3032193, EPI_ISL_3032195, EPI_ISL_3032197, EPI_ISL_3032198, EPI_ISL_3032200, EPI_ISL_3032202 | SYNLAB | GIGA Medical Genomics | Keith Durkin, Maria Artesi, Bouchra Boujemla, Nathalie Renotte, Nadine Cambisano, Sébastien Bontems, Cécile Meex, Claire Gourzonès, Olivier Ek, Laurent Gillet, Marie-Pierre Hayette, Vincent Bours |
| EPI_ISL_3032203, EPI_ISL_3032204, EPI_ISL_3032205, EPI_ISL_3032206, EPI_ISL_3032207, EPI_ISL_3032208, EPI_ISL_3032209, EPI_ISL_3032210, EPI_ISL_3032211, EPI_ISL_3032212, EPI_ISL_3032216, EPI_ISL_3032217, EPI_ISL_3032218, EPI_ISL_3032219, EPI_ISL_3032220, EPI_ISL_3032221, EPI_ISL_3032222, EPI_ISL_3032223, EPI_ISL_3032224, EPI_ISL_3032225, EPI_ISL_3032226, EPI_ISL_3032227, EPI_ISL_3032228, EPI_ISL_3032229, EPI_ISL_3032230, EPI_ISL_3032231, EPI_ISL_3032232, EPI_ISL_3032233, EPI_ISL_3032234, EPI_ISL_3032235, EPI_ISL_3032236, EPI_ISL_3032238, EPI_ISL_3032240, EPI_ISL_3032241, EPI_ISL_3032242, EPI_ISL_3032243, EPI_ISL_3032244, EPI_ISL_3032245, EPI_ISL_3032247, EPI_ISL_3032248, EPI_ISL_3032249, EPI_ISL_3032250, EPI_ISL_3032252, EPI_ISL_3032255, EPI_ISL_3032256, EPI_ISL_3032257, EPI_ISL_3032258, EPI_ISL_3032259, EPI_ISL_3032260, EPI_ISL_3032261, EPI_ISL_3032263, EPI_ISL_3032264, EPI_ISL_3032266, EPI_ISL_3032267, EPI_ISL_3032268, EPI_ISL_3032269, EPI_ISL_3032270 | | | |
| see above | University of Liège COVID-19 testing center | GIGA Medical Genomics | Keith Durkin, Maria Artesi, Bouchra Boujemla, Nathalie Renotte, Nadine Cambisano, Sébastien Bontems, Cécile Meex, Claire Gourzonès, Olivier Ek, Laurent Gillet, Marie-Pierre Hayette, Vincent Bours |
| EPI_ISL_3032271, EPI_ISL_3032272, EPI_ISL_3032273, EPI_ISL_3032274, EPI_ISL_3032275, EPI_ISL_3032276, EPI_ISL_3032277, EPI_ISL_3032278, EPI_ISL_3032279, EPI_ISL_3032280, EPI_ISL_3032281, EPI_ISL_3032282, EPI_ISL_3032283, EPI_ISL_3032285, EPI_ISL_3032286, EPI_ISL_3032287, EPI_ISL_3032288, EPI_ISL_3032289, EPI_ISL_3032290, EPI_ISL_3032291, EPI_ISL_3032292, EPI_ISL_3032293 | | | |
| see above | Ospedale Santissima Annunziata | Istituto Zooprofilattico Sperimentale della Puglia e della Basilicata | Parisi A., Bianco A., Capozzi L., Del Sambro L., Simone D., Difato L., Ridolfi D., Giannico A, Tampoia M. |
| EPI_ISL_3032294, EPI_ISL_3032295, EPI_ISL_3032296, EPI_ISL_3032298, EPI_ISL_3032299, EPI_ISL_3032300 | Ospedale Vito Fazzi | Istituto Zooprofilattico Sperimentale della Puglia e della Basilicata | Parisi A., Bianco A., Capozzi L., Del Sambro L., Simone D., Difato L., Lobreglio G. |
| EPI_ISL_3032304, EPI_ISL_3032306, EPI_ISL_3032307, EPI_ISL_3032308, EPI_ISL_3032309 | Ospedale Santa Caterina Novella | Istituto Zooprofilattico Sperimentale della Puglia e della Basilicata | Parisi A., Bianco A., Capozzi L., Del Sambro L., Simone D., Difato L., Bruno A. R. |
| EPI_ISL_3032312, EPI_ISL_3032313 | I.R.C.C.S. "S. De Bellis" - Ente Ospedaliero | Istituto Zooprofilattico Sperimentale della Puglia e della Basilicata | Parisi A., Bianco A., Capozzi L., Del Sambro L., Simone D., Difato L., Notarnicola M., Lippolis A. |
| EPI_ISL_3032314, EPI_ISL_3032315 | Istituto Zooprofilattico Sperimentale della Puglia e della Basilicata | Istituto Zooprofilattico Sperimentale della Puglia e della Basilicata | Parisi A., Bianco A., Capozzi L., Del Sambro L., Simone D., Difato L., Pennuzzi G., Pace L., Galante D. |
| EPI_ISL_3032316, EPI_ISL_3032318, EPI_ISL_3032319, EPI_ISL_3032321, EPI_ISL_3032322, EPI_ISL_3032324, EPI_ISL_3032325, EPI_ISL_3032326, EPI_ISL_3032327, EPI_ISL_3032328, EPI_ISL_3032329, EPI_ISL_3032330 | | | |
| see above | Presidio di Brindisi Di Summa - Perrino | Istituto Zooprofilattico Sperimentale della Puglia e della Basilicata | Parisi A., Bianco A., Capozzi L., Del Sambro L., Simone D., Difato L., Pennuzzi G., Santoro A. |
| EPI_ISL_3032331, EPI_ISL_3032332, EPI_ISL_3032333 | BIOFAST | Instituto Butantan | Dimas Tadeu Covas, Antonio Jorge Martins, Claudia Renata dos Santos Barros, David Schlesinger, Debora Botequio Moretti, Elaine Cristina Marqueze, Elaine Vieira Santos, Evandra Strazza Rodrigues, Heidge Fukumasu, Jayme Augusto de Souza-Neto, José Salvatore Leister Patané, Luiz Alcantara, Luiz Lehmann Coutinho, Maria Carolina Elias, Maurício Lacerda Nogueira, Rafael dos Santos Bezerra, Raul Machado Neto, Rejane Maria Tommasini Grotto, Ricardo Haddad, Sandra Coccuzzo Sampaio Vessoni, Simone Kashima, Svetoslav Nanev Slavov, Vincent Louis Viala |
| EPI_ISL_3032334 | INSIDE DIAGNÓSTICOS | Instituto Butantan | Dimas Tadeu Covas, Antonio Jorge Martins, Claudia Renata dos Santos Barros, David Schlesinger, Debora Botequio Moretti, Elaine Cristina Marqueze, Elaine Vieira Santos, Evandra Strazza Rodrigues, Heidge Fukumasu, Jayme Augusto de Souza-Neto, José Salvatore Leister Patané, Luiz Alcantara, Luiz Lehmann Coutinho, Maria Carolina Elias, Maurício Lacerda Nogueira, Rafael dos Santos Bezerra, Raul Machado Neto, Rejane Maria Tommasini Grotto, Ricardo Haddad, Sandra Coccuzzo Sampaio Vessoni, Simone Kashima, Svetoslav Nanev Slavov, Vincent Louis Viala |
| EPI_ISL_3032335, EPI_ISL_3032336, EPI_ISL_3032337, EPI_ISL_3032338, EPI_ISL_3032339, EPI_ISL_3032340, EPI_ISL_3032341, EPI_ISL_3032342, EPI_ISL_3032343, EPI_ISL_3032344, EPI_ISL_3032345, EPI_ISL_3032346, EPI_ISL_3032347, EPI_ISL_3032348, EPI_ISL_3032349, EPI_ISL_3032350, EPI_ISL_3032352, EPI_ISL_3032353, EPI_ISL_3032354, EPI_ISL_3032356, EPI_ISL_3032357, EPI_ISL_3032358, EPI_ISL_3032359, EPI_ISL_3032360, EPI_ISL_3032361, EPI_ISL_3032362, EPI_ISL_3032363 | | | |
| see above | TXDSHS | TXDSHS | Rashmi Tuladhar, Bonnie Oh, Jenny Zhang, Maliha Rahman, Mayela Pedrueza, Anita Pokharel, Karen Bobier, Lorraine Rodriguez, Myong Koag, Chun Wang, Rachel Lee, Grace Kubin |
| EPI_ISL_3032364 | Wojewódzki Specjalistyczny Szpital im. Biegańskiego | National Institute of Public Health NIH - National Research Institute | Wołkowicz Tomasz, Zacharczuk Katarzyna, Sadkowska-Todys Małgorzata, Gierczyńki Rafał |
| EPI_ISL_3032365 | ALAB Laboratorium | National Institute of Public Health NIH - National Research Institute | Wołkowicz Tomasz, Zacharczuk Katarzyna, Sadkowska-Todys Małgorzata, Gierczyńki Rafał |
| EPI_ISL_3032366, EPI_ISL_3032367, EPI_ISL_3032368 | Wojewódzki Szpital Specjalistyczny Laboratorium Mikrobiologiczne | National Institute of Public Health NIH - National Research Institute | Wołkowicz Tomasz, Zacharczuk Katarzyna, Sadkowska-Todys Małgorzata, Gierczyńki Rafał |
| EPI_ISL_3032370 | Vitagenum | National Institute of Public Health NIH - National Research Institute | Wołkowicz Tomasz, Zacharczuk Katarzyna, Sadkowska-Todys Małgorzata, Gierczyńki Rafał |
| EPI_ISL_3032371 | Laboratorium Badawczo-Rozwojowe Vitagenum | National Institute of Public Health NIH - National Research Institute | Wołkowicz Tomasz, Zacharczuk Katarzyna, Sadkowska-Todys Małgorzata, Gierczyńki Rafał |
| EPI_ISL_3032372 | DL OBM WSSE Lublin | National Institute of Public Health NIH - National Research Institute | Wołkowicz Tomasz, Zacharczuk Katarzyna, Sadkowska-Todys Małgorzata, Gierczyńki Rafał |
| EPI_ISL_3032374 | Zakład Diagnostyki Laboratoryjnej MSWiA w Poznaniu | National Institute of Public Health NIH - National Research Institute | Wołkowicz Tomasz, Zacharczuk Katarzyna, Sadkowska-Todys Małgorzata, Gierczyńki Rafał |
| EPI_ISL_3032375, EPI_ISL_3032376 | GenXOne SA | National Institute of Public Health NIH - National Research Institute | Wołkowicz Tomasz, Zacharczuk Katarzyna, Sadkowska-Todys Małgorzata, Gierczyńki Rafał |
| EPI_ISL_3032377 | Instytut Mikroekologii Sp. z o.o. & Co. Sp.k. | National Institute of Public Health NIH - National Research Institute | Wołkowicz Tomasz, Zacharczuk Katarzyna, Sadkowska-Todys Małgorzata, Gierczyńki Rafał |
| EPI_ISL_3032378 | Diagnostyka Sp. z o.o. | National Institute of Public Health NIH - National Research Institute | Wołkowicz Tomasz, Zacharczuk Katarzyna, Sadkowska-Todys Małgorzata, Gierczyńki Rafał |
| EPI_ISL_3032379, EPI_ISL_3032381 | GenXOne SA | National Institute of Public Health NIH - National Research Institute | Wołkowicz Tomasz, Zacharczuk Katarzyna, Sadkowska-Todys Małgorzata, Gierczyńki Rafał |
| EPI_ISL_3032382 | Zakład Diagnostyki Laboratoryjnej SPWSZ w Szczecinie | National Institute of Public Health NIH - National Research Institute | Wołkowicz Tomasz, Zacharczuk Katarzyna, Sadkowska-Todys Małgorzata, Gierczyńki Rafał |
| EPI_ISL_3032383 | Laboratorium Analiz Lekarskich ALAB | National Institute of Public Health NIH - National Research Institute | Wołkowicz Tomasz, Zacharczuk Katarzyna, Sadkowska-Todys Małgorzata, Gierczyńki Rafał |
| EPI_ISL_3032384 | NZOZ Meditest Diagnostyka Medyczna | National Institute of Public Health NIH - National Research Institute | Wołkowicz Tomasz, Zacharczuk Katarzyna, Sadkowska-Todys Małgorzata, Gierczyńki Rafał |
| EPI_ISL_3032387 | NZOZ Medigen Diagnostyka Molekularna | National Institute of Public Health NIH - National Research Institute | Wołkowicz Tomasz, Zacharczuk Katarzyna, Sadkowska-Todys Małgorzata, Gierczyńki Rafał |

| | | | |
|---|---|---|---|
| EPI_ISL_3032388 | Medyczne Laboratorium Diagnostyka | National Institute of Public Health NIH - National Research Institute | Wołkowicz Tomasz, Zacharczuk Katarzyna, Sadkowska-Todys Małgorzata, Gierczyński Rafał |
| EPI_ISL_3032389 | National Institute of Public Health NIH - National Research Institute | National Institute of Public Health NIH - National Research Institute | Wołkowicz Tomasz, Zacharczuk Katarzyna, Sadkowska-Todys Małgorzata, Gierczyński Rafał |
| EPI_ISL_3032438, EPI_ISL_3032439, EPI_ISL_3032441, EPI_ISL_3032442, EPI_ISL_3032443, EPI_ISL_3032444, EPI_ISL_3032445, EPI_ISL_3032446, EPI_ISL_3032447, EPI_ISL_3032449, EPI_ISL_3032450, EPI_ISL_3032451, EPI_ISL_3032452, EPI_ISL_3032459, EPI_ISL_3032460 | | | |
| see above | CHU Pontchaillou | CHU Pontchaillou | GROLHIER Claire, DENOUAL Florent, ETCHEVERRY Amandine, SASSI Mohamed, JAGLINE Steven, FEBREAU Christine, PRONIER Charlotte, GALIBERT Marie Dominique, DE TAYRAC Marie, THIBAULT Vincent |
| EPI_ISL_3032461, EPI_ISL_3032462, EPI_ISL_3032463, EPI_ISL_3032464, EPI_ISL_3032465, EPI_ISL_3032467, EPI_ISL_3032468, EPI_ISL_3032469, EPI_ISL_3032470, EPI_ISL_3032471, EPI_ISL_3032472, EPI_ISL_3032473, EPI_ISL_3032474, EPI_ISL_3032476, EPI_ISL_3032477, EPI_ISL_3032478, EPI_ISL_3032479, EPI_ISL_3032480, EPI_ISL_3032481, EPI_ISL_3032482, EPI_ISL_3032483, EPI_ISL_3032484, EPI_ISL_3032485, EPI_ISL_3032486, EPI_ISL_3032487, EPI_ISL_3032488, EPI_ISL_3032489, EPI_ISL_3032490, EPI_ISL_3032491, EPI_ISL_3032492, EPI_ISL_3032493, EPI_ISL_3032494, EPI_ISL_3032495, EPI_ISL_3032496, EPI_ISL_3032497, EPI_ISL_3032498, EPI_ISL_3032500, EPI_ISL_3032501, EPI_ISL_3032502, EPI_ISL_3032503, EPI_ISL_3032504, EPI_ISL_3032506, EPI_ISL_3032507, EPI_ISL_3032508, EPI_ISL_3032509, EPI_ISL_3032510, EPI_ISL_3032511, EPI_ISL_3032513, EPI_ISL_3032514, EPI_ISL_3032515, EPI_ISL_3032516, EPI_ISL_3032517, EPI_ISL_3032518, EPI_ISL_3032519, EPI_ISL_3032520, EPI_ISL_3032521, EPI_ISL_3032522, EPI_ISL_3032525, EPI_ISL_3032526, EPI_ISL_3032527 | | | |
| see above | Division of Medical Virology, National Health Laboratory Service (NHLS), Tygerberg Hospital / Stellenbosch University | Division of Medical Virology, National Health Laboratory Service (NHLS), Tygerberg Hospital / Stellenbosch University | Susan Engelbrecht, Tongai Maponga, Kamela Mahlakwane, Gert van Zyl, Wolfgang Preiser |
| EPI_ISL_3032530, EPI_ISL_3032531, EPI_ISL_3032533, EPI_ISL_3032534, EPI_ISL_3032535, EPI_ISL_3032536 | PathCare, Cape Town | Division of Medical Virology, National Health Laboratory Service (NHLS), Tygerberg Hospital / Stellenbosch University | Jean Maritz, Petra Raimond, Susan Engelbrecht, Tongai Maponga, Bronwyn Kleinhans, Kamela Mahlakwane, Gert van Zyl, Wolfgang Preiser |
| EPI_ISL_3032537 | PathCare, Cape Town | Division of Medical Virology, National Health Laboratory Service (NHLS), Tygerberg Hospital / Stellenbosch University | Susan Engelbrecht, Tongai Maponga, Kamela Mahlakwane, Gert van Zyl, Wolfgang Preiser |
| EPI_ISL_3032538 | Division of Medical Virology, National Health Laboratory Service (NHLS), Tygerberg Hospital / Stellenbosch University | Division of Medical Virology, National Health Laboratory Service (NHLS), Tygerberg Hospital / Stellenbosch University | Susan Engelbrecht, Tongai Maponga, Kamela Mahlakwane, Gert van Zyl, Wolfgang Preiser |
| EPI_ISL_3032555, EPI_ISL_3032575, EPI_ISL_3032576, EPI_ISL_3032577, EPI_ISL_3032578, EPI_ISL_3032583, EPI_ISL_3032585, EPI_ISL_3032588, EPI_ISL_3032589, EPI_ISL_3032590, EPI_ISL_3032593, EPI_ISL_3032594, EPI_ISL_3032598, EPI_ISL_3032602, EPI_ISL_3032604, EPI_ISL_3032605, EPI_ISL_3032621, EPI_ISL_3032622, EPI_ISL_3032623, EPI_ISL_3032624, EPI_ISL_3032643, EPI_ISL_3032646, EPI_ISL_3032650, EPI_ISL_3032651, EPI_ISL_3032652, EPI_ISL_3032653, EPI_ISL_3032659, EPI_ISL_3032660, EPI_ISL_3032661, EPI_ISL_3032663, EPI_ISL_3032664, EPI_ISL_3032666, EPI_ISL_3032667, EPI_ISL_3032668, EPI_ISL_3032669, EPI_ISL_3032671, EPI_ISL_3032672, EPI_ISL_3032673, EPI_ISL_3032674, EPI_ISL_3032675, EPI_ISL_3032676, EPI_ISL_3032677, EPI_ISL_3032680, EPI_ISL_3032682, EPI_ISL_3032683, EPI_ISL_3032684, EPI_ISL_3032686, EPI_ISL_3032687, EPI_ISL_3032688, EPI_ISL_3032690, EPI_ISL_3032692, EPI_ISL_3032693, EPI_ISL_3032694, EPI_ISL_3032695, EPI_ISL_3032696, EPI_ISL_3032698, EPI_ISL_3032699, EPI_ISL_3032700, EPI_ISL_3032701, EPI_ISL_3032702, EPI_ISL_3032703, EPI_ISL_3032704, EPI_ISL_3032705, EPI_ISL_3032707, EPI_ISL_3032708, EPI_ISL_3032709, EPI_ISL_3032710, EPI_ISL_3032833, EPI_ISL_3032840 | | | |
| see above | Ministry of Health Turkey | Ministry of Health Turkey | Fatma Bayrakdar, Yasemin Cosgun, Suleyman Yalcin, Gulay Korukluoglu |
| EPI_ISL_3033091, EPI_ISL_3033096, EPI_ISL_3033118, EPI_ISL_3033126, EPI_ISL_3033131, EPI_ISL_3033132, EPI_ISL_3033134, EPI_ISL_3033135, EPI_ISL_3033137, EPI_ISL_3033138, EPI_ISL_3033139, EPI_ISL_3033140, EPI_ISL_3033142, EPI_ISL_3033143, EPI_ISL_3033148, EPI_ISL_3033149, EPI_ISL_3033151, EPI_ISL_3033152, EPI_ISL_3033153, EPI_ISL_3033160, EPI_ISL_3033162, EPI_ISL_3033164, EPI_ISL_3033165, EPI_ISL_3033172, EPI_ISL_3033173, EPI_ISL_3033177, EPI_ISL_3033181, EPI_ISL_3033182, EPI_ISL_3033183, EPI_ISL_3033186, EPI_ISL_3033192, EPI_ISL_3033194, EPI_ISL_3033195, EPI_ISL_3033199, EPI_ISL_3033200, EPI_ISL_3033201, EPI_ISL_3033203, EPI_ISL_3033211, EPI_ISL_3033214, EPI_ISL_3033219, EPI_ISL_3033225, EPI_ISL_3033226, EPI_ISL_3033231, EPI_ISL_3033239, EPI_ISL_3033241, EPI_ISL_3033242, EPI_ISL_3033243, EPI_ISL_3033244, EPI_ISL_3033246, EPI_ISL_3033247, EPI_ISL_3033248, EPI_ISL_3033255, EPI_ISL_3033259, EPI_ISL_3033260, EPI_ISL_3033261, EPI_ISL_3033262, EPI_ISL_3033263, EPI_ISL_3033267, EPI_ISL_3033269, EPI_ISL_3033272, EPI_ISL_3033274, EPI_ISL_3033275, EPI_ISL_3033277, EPI_ISL_3033278, EPI_ISL_3033279, EPI_ISL_3033280, EPI_ISL_3033281, EPI_ISL_3033282, EPI_ISL_3033283, EPI_ISL_3033285, EPI_ISL_3033287, EPI_ISL_3033290, EPI_ISL_3033293, EPI_ISL_3033294, EPI_ISL_3033296, EPI_ISL_3033301, EPI_ISL_3033303, EPI_ISL_3033304, EPI_ISL_3033306, EPI_ISL_3033307, EPI_ISL_3033308, EPI_ISL_3033309, EPI_ISL_3033310, EPI_ISL_3033311, EPI_ISL_3033313, EPI_ISL_3033316, EPI_ISL_3033317, EPI_ISL_3033318, EPI_ISL_3033319, EPI_ISL_3033322 | | | |
| see above | State Laboratories Division, Hawaii State Department of Health | State Laboratories Division, Hawaii State Department of Health | Pamela O'Brien, Drew Kuwazaki, Ayana Garnet, Razvan Sultana, Edward Desmond |
| EPI_ISL_3033386, EPI_ISL_3033387, EPI_ISL_3033388, EPI_ISL_3033389, EPI_ISL_3033390, EPI_ISL_3033391 | LESP Yucatan | Instituto de Diagnostico y Referencia Epidemiologicos (INDRE) | Claudia Wong-Arambula, Abril Rodriguez-Maldonado, Vanessa Rivero-Arredondo, Ariadna Medina-Benitez, Joaquin Quiroz-Mercado, Sergio Rangel-Guerrero, Maribel Gonzalez-Villa, Natividad Cruz-Ortiz, Tatiana Nunez-Garcia, Gisela Barrera-Badillo, Lucia Hernandez-Rivas, Irma Lopez-Martinez, Ernesto Ramirez-Gonzalez. |
| EPI_ISL_3033392 | LESP Tabasco | Instituto de Diagnostico y Referencia Epidemiologicos (INDRE) | Claudia Wong-Arambula, Abril Rodriguez-Maldonado, Vanessa Rivero-Arredondo, Ariadna Medina-Benitez, Joaquin Quiroz-Mercado, Sergio Rangel-Guerrero, Maribel Gonzalez-Villa, Natividad Cruz-Ortiz, Tatiana Nunez-Garcia, Gisela Barrera-Badillo, Lucia Hernandez-Rivas, Irma Lopez-Martinez, Ernesto Ramirez-Gonzalez. |
| EPI_ISL_3033393, EPI_ISL_3033394, EPI_ISL_3033395, EPI_ISL_3033396, EPI_ISL_3033397, EPI_ISL_3033398, EPI_ISL_3033399, EPI_ISL_3033400, EPI_ISL_3033401, EPI_ISL_3033402, EPI_ISL_3033403 | | | |
| see above | LESP Hidalgo | Instituto de Diagnostico y Referencia Epidemiologicos (INDRE) | Claudia Wong-Arambula, Abril Rodriguez-Maldonado, Vanessa Rivero-Arredondo, Ariadna Medina-Benitez, Joaquin Quiroz-Mercado, Sergio Rangel-Guerrero, Maribel Gonzalez-Villa, Natividad Cruz-Ortiz, Tatiana Nunez-Garcia, Gisela Barrera-Badillo, Lucia Hernandez-Rivas, Irma Lopez-Martinez, Ernesto Ramirez-Gonzalez. |
| EPI_ISL_3033404 | LESP Baja California | Instituto de Diagnostico y Referencia Epidemiologicos (INDRE) | Claudia Wong-Arambula, Abril Rodriguez-Maldonado, Vanessa Rivero-Arredondo, Ariadna Medina-Benitez, Joaquin Quiroz-Mercado, Sergio Rangel-Guerrero, Maribel Gonzalez-Villa, Natividad Cruz-Ortiz, Tatiana Nunez-Garcia, Gisela Barrera-Badillo, Lucia Hernandez-Rivas, Irma Lopez-Martinez, Ernesto Ramirez-Gonzalez. |
| EPI_ISL_3033405, EPI_ISL_3033406, EPI_ISL_3033407, EPI_ISL_3033408 | LESP Sinaloa | Instituto de Diagnostico y Referencia Epidemiologicos (INDRE) | Claudia Wong-Arambula, Abril Rodriguez-Maldonado, Vanessa Rivero-Arredondo, Ariadna Medina-Benitez, Joaquin Quiroz-Mercado, Sergio Rangel-Guerrero, Maribel Gonzalez-Villa, Natividad Cruz-Ortiz, Tatiana Nunez-Garcia, Gisela Barrera-Badillo, Lucia Hernandez-Rivas, Irma Lopez-Martinez, Ernesto Ramirez-Gonzalez. |
| EPI_ISL_3033409 | LESP Quintana Roo | Instituto de Diagnostico y Referencia Epidemiologicos (INDRE) | Claudia Wong-Arambula, Abril Rodriguez-Maldonado, Vanessa Rivero-Arredondo, Ariadna Medina-Benitez, Joaquin Quiroz-Mercado, Sergio Rangel-Guerrero, Maribel Gonzalez-Villa, Natividad Cruz-Ortiz, Tatiana Nunez-Garcia, Gisela Barrera-Badillo, Lucia Hernandez-Rivas, Irma Lopez-Martinez, Ernesto Ramirez-Gonzalez. |
| EPI_ISL_3033410, EPI_ISL_3033411, EPI_ISL_3033412, EPI_ISL_3033413, EPI_ISL_3033414, EPI_ISL_3033415, EPI_ISL_3033416, EPI_ISL_3033417, EPI_ISL_3033418, EPI_ISL_3033419, EPI_ISL_3033420, EPI_ISL_3033421, EPI_ISL_3033422, EPI_ISL_3033423, EPI_ISL_3033424, EPI_ISL_3033425, EPI_ISL_3033426, EPI_ISL_3033427, EPI_ISL_3033428, EPI_ISL_3033429 | | | |
| see above | LESP Baja California Sur | Instituto de Diagnostico y Referencia Epidemiologicos (INDRE) | Claudia Wong-Arambula, Abril Rodriguez-Maldonado, Vanessa Rivero-Arredondo, Ariadna Medina-Benitez, Joaquin Quiroz-Mercado, Sergio Rangel-Guerrero, Maribel Gonzalez-Villa, Natividad Cruz-Ortiz, Tatiana Nunez-Garcia, Gisela Barrera-Badillo, Lucia Hernandez-Rivas, Irma Lopez-Martinez, Ernesto Ramirez-Gonzalez. |
| EPI_ISL_3033430, EPI_ISL_3033431, EPI_ISL_3033432 | LESP Sonora | Instituto de Diagnostico y Referencia Epidemiologicos (INDRE) | Claudia Wong-Arambula, Abril Rodriguez-Maldonado, Vanessa Rivero-Arredondo, Ariadna Medina-Benitez, Joaquin Quiroz-Mercado, Sergio Rangel-Guerrero, Maribel Gonzalez-Villa, Natividad Cruz-Ortiz, Tatiana Nunez-Garcia, Gisela Barrera-Badillo, Lucia Hernandez-Rivas, Irma Lopez-Martinez, Ernesto Ramirez-Gonzalez. |
| EPI_ISL_3033433 | LESP Colima | Instituto de Diagnostico y Referencia Epidemiologicos (INDRE) | Claudia Wong-Arambula, Abril Rodriguez-Maldonado, Vanessa Rivero-Arredondo, Ariadna Medina-Benitez, Joaquin Quiroz-Mercado, Sergio Rangel-Guerrero, Maribel Gonzalez-Villa, Natividad Cruz-Ortiz, Tatiana Nunez-Garcia, Gisela Barrera-Badillo, Lucia Hernandez-Rivas, Irma Lopez-Martinez, Ernesto Ramirez-Gonzalez. |
| EPI_ISL_3033434, EPI_ISL_3033435 | LESP Tabasco | Instituto de Diagnostico y Referencia Epidemiologicos (INDRE) | Claudia Wong-Arambula, Abril Rodriguez-Maldonado, Vanessa Rivero-Arredondo, Ariadna Medina-Benitez, Joaquin Quiroz-Mercado, Sergio Rangel-Guerrero, Maribel Gonzalez-Villa, Natividad Cruz-Ortiz, Tatiana Nunez-Garcia, Gisela Barrera-Badillo, Lucia Hernandez-Rivas, Irma Lopez-Martinez, Ernesto Ramirez-Gonzalez. |
| EPI_ISL_3033436 | LESP Zacatecas | Instituto de Diagnostico y Referencia Epidemiologicos (INDRE) | Claudia Wong-Arambula, Abril Rodriguez-Maldonado, Vanessa Rivero-Arredondo, Ariadna Medina-Benitez, Joaquin Quiroz-Mercado, Sergio Rangel-Guerrero, Maribel Gonzalez-Villa, Natividad Cruz-Ortiz, Tatiana Nunez-Garcia, Gisela Barrera-Badillo, Lucia Hernandez-Rivas, Irma Lopez-Martinez, Ernesto Ramirez-Gonzalez. |
| EPI_ISL_3033437 | LESP Baja California Sur | Instituto de Diagnostico y Referencia Epidemiologicos (INDRE) | Claudia Wong-Arambula, Abril Rodriguez-Maldonado, Vanessa Rivero-Arredondo, Ariadna Medina-Benitez, Joaquin Quiroz-Mercado, Sergio Rangel-Guerrero, Maribel Gonzalez-Villa, Natividad Cruz-Ortiz, Tatiana Nunez-Garcia, Gisela Barrera-Badillo, Lucia Hernandez-Rivas, Irma Lopez-Martinez, Ernesto Ramirez-Gonzalez. |
| EPI_ISL_3033438, EPI_ISL_3033439, EPI_ISL_3033440 | LESP Jalisco | Instituto de Diagnostico y Referencia Epidemiologicos (INDRE) | Claudia Wong-Arambula, Abril Rodriguez-Maldonado, Vanessa Rivero-Arredondo, Ariadna Medina-Benitez, Joaquin Quiroz-Mercado, Sergio Rangel-Guerrero, Maribel Gonzalez-Villa, Natividad Cruz-Ortiz, Tatiana Nunez-Garcia, Gisela Barrera-Badillo, Lucia Hernandez-Rivas, Irma Lopez-Martinez, Ernesto Ramirez-Gonzalez. |
| EPI_ISL_3033441, EPI_ISL_3033442, EPI_ISL_3033443, EPI_ISL_3033444, EPI_ISL_3033445, EPI_ISL_3033446, EPI_ISL_3033447 | LESP Michoacan | Instituto de Diagnostico y Referencia Epidemiologicos (INDRE) | Claudia Wong-Arambula, Abril Rodriguez-Maldonado, Vanessa Rivero-Arredondo, Ariadna Medina-Benitez, Joaquin Quiroz-Mercado, Sergio Rangel-Guerrero, Maribel Gonzalez-Villa, Natividad Cruz-Ortiz, Tatiana Nunez-Garcia, Gisela Barrera-Badillo, Lucia Hernandez-Rivas, Irma Lopez-Martinez, Ernesto Ramirez-Gonzalez. |
| EPI_ISL_3033448, EPI_ISL_3033449 | LESP Sinaloa | Instituto de Diagnostico y Referencia Epidemiologicos (INDRE) | Claudia Wong-Arambula, Abril Rodriguez-Maldonado, Vanessa Rivero-Arredondo, Ariadna Medina-Benitez, Joaquin Quiroz-Mercado, Sergio Rangel-Guerrero, Maribel Gonzalez-Villa, Natividad Cruz-Ortiz, Tatiana Nunez-Garcia, Gisela Barrera-Badillo, Lucia Hernandez-Rivas, Irma Lopez-Martinez, Ernesto Ramirez-Gonzalez. |
| EPI_ISL_3033450 | LESP Ciudad de Mexico | Instituto de Diagnostico y Referencia Epidemiologicos (INDRE) | Claudia Wong-Arambula, Abril Rodriguez-Maldonado, Vanessa Rivero-Arredondo, Ariadna Medina-Benitez, Joaquin Quiroz-Mercado, Sergio Rangel-Guerrero, Maribel Gonzalez-Villa, Natividad Cruz-Ortiz, Tatiana Nunez-Garcia, Gisela Barrera-Badillo, Lucia Hernandez-Rivas, Irma Lopez-Martinez, Ernesto Ramirez-Gonzalez. |
| EPI_ISL_3033451 | LESP Queretaro | Instituto de Diagnostico y Referencia Epidemiologicos (INDRE) | Claudia Wong-Arambula, Abril Rodriguez-Maldonado, Vanessa Rivero-Arredondo, Ariadna Medina-Benitez, Joaquin Quiroz-Mercado, Sergio Rangel-Guerrero, Maribel Gonzalez-Villa, Natividad Cruz-Ortiz, Tatiana Nunez-Garcia, Gisela Barrera-Badillo, Lucia Hernandez-Rivas, Irma Lopez-Martinez, Ernesto Ramirez-Gonzalez. |
| EPI_ISL_3033452 | LESP Nuevo Leon | Instituto de Diagnostico y Referencia Epidemiologicos (INDRE) | Claudia Wong-Arambula, Abril Rodriguez-Maldonado, Vanessa Rivero-Arredondo, Ariadna Medina-Benitez, Joaquin Quiroz-Mercado, Sergio Rangel-Guerrero, Maribel Gonzalez-Villa, Natividad Cruz-Ortiz, Tatiana Nunez-Garcia, Gisela Barrera-Badillo, Lucia Hernandez-Rivas, Irma Lopez-Martinez, Ernesto Ramirez-Gonzalez. |
| EPI_ISL_3033453, EPI_ISL_3033454 | LESP Tabasco | Instituto de Diagnostico y Referencia Epidemiologicos (INDRE) | Claudia Wong-Arambula, Abril Rodriguez-Maldonado, Vanessa Rivero-Arredondo, Ariadna Medina-Benitez, Joaquin Quiroz-Mercado, Sergio Rangel-Guerrero, Maribel Gonzalez-Villa, Natividad Cruz-Ortiz, Tatiana Nunez-Garcia, Gisela Barrera-Badillo, Lucia Hernandez-Rivas, Irma Lopez-Martinez, Ernesto Ramirez-Gonzalez. |
| EPI_ISL_3033455 | LESP Guanajuato | Instituto de Diagnostico y Referencia Epidemiologicos (INDRE) | Claudia Wong-Arambula, Abril Rodriguez-Maldonado, Vanessa Rivero-Arredondo, Ariadna Medina-Benitez, Joaquin Quiroz-Mercado, Sergio Rangel-Guerrero, Maribel Gonzalez-Villa, Natividad Cruz-Ortiz, Tatiana Nunez-Garcia, Gisela Barrera-Badillo, Lucia Hernandez-Rivas, Irma Lopez-Martinez, Ernesto Ramirez-Gonzalez. |
| EPI_ISL_3033456, EPI_ISL_3033457, EPI_ISL_3033458, EPI_ISL_3033459 | LESP Puebla | Instituto de Diagnostico y Referencia Epidemiologicos (INDRE) | Claudia Wong-Arambula, Abril Rodriguez-Maldonado, Vanessa Rivero-Arredondo, Ariadna Medina-Benitez, Joaquin Quiroz-Mercado, Sergio Rangel-Guerrero, Maribel Gonzalez-Villa, Natividad Cruz-Ortiz, Tatiana Nunez-Garcia, Gisela Barrera-Badillo, Lucia Hernandez-Rivas, Irma Lopez-Martinez, Ernesto Ramirez-Gonzalez. |
| EPI_ISL_3033460, EPI_ISL_3033461 | LESP Tabasco | Instituto de Diagnostico y Referencia Epidemiologicos (INDRE) | Claudia Wong-Arambula, Abril Rodriguez-Maldonado, Vanessa Rivero-Arredondo, Ariadna Medina-Benitez, Joaquin Quiroz-Mercado, Sergio Rangel-Guerrero, Maribel Gonzalez-Villa, Natividad Cruz-Ortiz, Tatiana Nunez-Garcia, Gisela Barrera-Badillo, Lucia Hernandez-Rivas, Irma Lopez-Martinez, Ernesto Ramirez-Gonzalez. |
| EPI_ISL_3037245, EPI_ISL_3037247, | Monterey County Public Health Laboratory | Monterey County Public Health Laboratory | Monterey County Public Health Laboratory |

| | | | |
|---|---|---|---|
| EPI_ISL_3037248, EPI_ISL_3037249 | | | |
| EPI_ISL_3038875, EPI_ISL_3038876, EPI_ISL_3038877, EPI_ISL_3038878, EPI_ISL_3038879, EPI_ISL_3038880, EPI_ISL_3038881, EPI_ISL_3038882, EPI_ISL_3038883, EPI_ISL_3038884, EPI_ISL_3038885, EPI_ISL_3038886, EPI_ISL_3038887, EPI_ISL_3038888, EPI_ISL_3038889, EPI_ISL_3038895, EPI_ISL_3038896 | | | |
| see above | Wyoming Public Health Laboratory | Wyoming Public Health Laboratory | Jim Mildenberger, Taylor Fearing, Channing Weber, Ashley Norberg, Chayse Rowley, Marley Goetz, Brian Dominguez, Elliot Thomasson, Sam Britz, Cari Sloma, Robert Petit, and Rob Christensen |
| EPI_ISL_3038898, EPI_ISL_3038899 | PathWest Laboratory Medicine WA | PathWest Laboratory Medicine WA Microbial Surveillance Unit | PathWest Laboratory Medicine WA Microbial Surveillance Unit |
| EPI_ISL_3038903 | Regional Medical Sciences Center 8 Udonthani | National Institute of Health, Department of Medical Sciences, Ministry of Public Health, Thailand | Pilailuk Okada; Siripaporn Phuygun; Nuttida Thongpramul; Sittiporn Parnmen; Ratana Tacharoenmuang; Pakorn Piromtong; Natchaya Khiadsang; Thanutsapa Thanadachakul; Warawan Wongboot; sirikanda wimol; Sunthareeya Waicharoen; |
| EPI_ISL_3038904 | Siriraj Hospital | National Institute of Health, Department of Medical Sciences, Ministry of Public Health, Thailand | Pilailuk Okada; Siripaporn Phuygun; Nuttida Thongpramul; Sittiporn Parnmen; Ratana Tacharoenmuang; Pakorn Piromtong; Natchaya Khiadsang; Thanutsapa Thanadachakul; Warawan Wongboot; sirikanda wimol; Sunthareeya Waicharoen; |
| EPI_ISL_3038905, EPI_ISL_3038906, EPI_ISL_3038907 | Karunvej Pathumthani Hospital | National Institute of Health, Department of Medical Sciences, Ministry of Public Health, Thailand | Pilailuk Okada; Siripaporn Phuygun; Nuttida Thongpramul; Sittiporn Parnmen; Ratana Tacharoenmuang; Pakorn Piromtong; Natchaya Khiadsang; Thanutsapa Thanadachakul; Warawan Wongboot; sirikanda wimol; Sunthareeya Waicharoen; |
| EPI_ISL_3038908, EPI_ISL_3038909 | Chulabhorn Hospital | National Institute of Health, Department of Medical Sciences, Ministry of Public Health, Thailand | Pilailuk Okada; Siripaporn Phuygun; Nuttida Thongpramul; Sittiporn Parnmen; Ratana Tacharoenmuang; Pakorn Piromtong; Natchaya Khiadsang; Thanutsapa Thanadachakul; Warawan Wongboot; sirikanda wimol; Sunthareeya Waicharoen; |
| EPI_ISL_3038910 | Regional Medical Sciences Center 8 Udonthani | National Institute of Health, Department of Medical Sciences, Ministry of Public Health, Thailand | Pilailuk Okada; Siripaporn Phuygun; Nuttida Thongpramul; Sittiporn Parnmen; Ratana Tacharoenmuang; Pakorn Piromtong; Natchaya Khiadsang; Thanutsapa Thanadachakul; Warawan Wongboot; sirikanda wimol; Sunthareeya Waicharoen; |
| EPI_ISL_3038911 | Regional Medical Sciences Center 10 Ubonratchathani | National Institute of Health, Department of Medical Sciences, Ministry of Public Health, Thailand | Pilailuk Okada; Siripaporn Phuygun; Nuttida Thongpramul; Sittiporn Parnmen; Ratana Tacharoenmuang; Pakorn Piromtong; Natchaya Khiadsang; Thanutsapa Thanadachakul; Warawan Wongboot; sirikanda wimol; Sunthareeya Waicharoen; |
| EPI_ISL_3038912 | Regional Medical Science Center 2 Phitsanulok | National Institute of Health, Department of Medical Sciences, Ministry of Public Health, Thailand | Pilailuk Okada; Siripaporn Phuygun; Nuttida Thongpramul; Sittiporn Parnmen; Ratana Tacharoenmuang; Pakorn Piromtong; Natchaya Khiadsang; Thanutsapa Thanadachakul; Warawan Wongboot; sirikanda wimol; Sunthareeya Waicharoen; |
| EPI_ISL_3039347, EPI_ISL_3039349, EPI_ISL_3039350, EPI_ISL_3039353, EPI_ISL_3039355, EPI_ISL_3039356, EPI_ISL_3039357, EPI_ISL_3039358, EPI_ISL_3039359, EPI_ISL_3039361, EPI_ISL_3039362, EPI_ISL_3039363, EPI_ISL_3039366, EPI_ISL_3039367, EPI_ISL_3039368, EPI_ISL_3039370, EPI_ISL_3039372, EPI_ISL_3039373, EPI_ISL_3039374, EPI_ISL_3039376, EPI_ISL_3039377, EPI_ISL_3039379, EPI_ISL_3039380, EPI_ISL_3039382, EPI_ISL_3039383, EPI_ISL_3039384, EPI_ISL_3039386, EPI_ISL_3039387, EPI_ISL_3039388, EPI_ISL_3039391, EPI_ISL_3039396, EPI_ISL_3039397, EPI_ISL_3039398, EPI_ISL_3039399, EPI_ISL_3039400, EPI_ISL_3039401, EPI_ISL_3039404, EPI_ISL_3039405, EPI_ISL_3039406, EPI_ISL_3039407, EPI_ISL_3039412, EPI_ISL_3039413, EPI_ISL_3039414, EPI_ISL_3039415, EPI_ISL_3039417, EPI_ISL_3039419, EPI_ISL_3039420, EPI_ISL_3039421, EPI_ISL_3039422, EPI_ISL_3039426, EPI_ISL_3039427, EPI_ISL_3039428, EPI_ISL_3039429, EPI_ISL_3039430, EPI_ISL_3039431, EPI_ISL_3039434, EPI_ISL_3039435, EPI_ISL_3039436, EPI_ISL_3039438 | | | |
| see above | Institute of Microbiology and Immunology, Faculty of Medicine, University of Ljubljana | Institute of Microbiology and Immunology, Faculty of Medicine, University of Ljubljana | Alen Suljič, Samo Zakotnik, Tomaž Mark Zorec, Matic Brvar, Doroteja Vlaj, Andraž Celar, Dominika Šturm, Patricija Pozvek, Špela Pleh, Miša Korva, Mario Poljak, Tatjana Avšič – Županc |
| EPI_ISL_3039508, EPI_ISL_3039509, EPI_ISL_3039511, EPI_ISL_3039512, EPI_ISL_3039513, EPI_ISL_3039514, EPI_ISL_3039515, EPI_ISL_3039516, EPI_ISL_3039518, EPI_ISL_3039519, EPI_ISL_3039520, EPI_ISL_3039521, EPI_ISL_3039522, EPI_ISL_3039523, EPI_ISL_3039524, EPI_ISL_3039526, EPI_ISL_3039527, EPI_ISL_3039528, EPI_ISL_3039529, EPI_ISL_3039530, EPI_ISL_3039531, EPI_ISL_3039532, EPI_ISL_3039533, EPI_ISL_3039534, EPI_ISL_3039535, EPI_ISL_3039536 | | | |
| see above | CHU Pontchaillou | CHU Pontchaillou | GROLHIER Claire, DENOUAL Florent, ETCHEVERRY Amandine, SASSI Mohamed, JAGLINE Steven, FEBREAU Christine, PRONIER Charlotte, GALIBERT Marie Dominique, DE TAYRAC Marie, THIBAULT Vincent |
| EPI_ISL_3039537, EPI_ISL_3039538, EPI_ISL_3039539, EPI_ISL_3039540, EPI_ISL_3039541, EPI_ISL_3039542, EPI_ISL_3039544, EPI_ISL_3039545, EPI_ISL_3039547, EPI_ISL_3039548, EPI_ISL_3039549, EPI_ISL_3039550, EPI_ISL_3039551, EPI_ISL_3039552, EPI_ISL_3039553, EPI_ISL_3039554, EPI_ISL_3039555, EPI_ISL_3039556, EPI_ISL_3039557, EPI_ISL_3039558, EPI_ISL_3039559, EPI_ISL_3039560, EPI_ISL_3039561, EPI_ISL_3039562, EPI_ISL_3039563, EPI_ISL_3039564, EPI_ISL_3039565, EPI_ISL_3039566, EPI_ISL_3039567, EPI_ISL_3039568, EPI_ISL_3039570, EPI_ISL_3039571, EPI_ISL_3039573, EPI_ISL_3039574, EPI_ISL_3039575, EPI_ISL_3039576, EPI_ISL_3039577, EPI_ISL_3039578, EPI_ISL_3039579, EPI_ISL_3039581 | | | |
| see above | ZOL | Jessa | Berden et al. on behalf of the Jessa_cmdLab |
| EPI_ISL_3039582 | LKO | Jessa | Berden et al. on behalf of the Jessa_cmdLab |
| EPI_ISL_3039583, EPI_ISL_3039584, EPI_ISL_3039585, EPI_ISL_3039586, EPI_ISL_3039587, EPI_ISL_3039588, EPI_ISL_3039589, EPI_ISL_3039590, EPI_ISL_3039591, EPI_ISL_3039592, EPI_ISL_3039593, EPI_ISL_3039594, EPI_ISL_3039595, EPI_ISL_3039596, EPI_ISL_3039597, EPI_ISL_3039598, EPI_ISL_3039599, EPI_ISL_3039600 | | | |
| see above | Jessa | Jessa | Berden et al. on behalf of the Jessa_cmdLab |
| EPI_ISL_3039602, EPI_ISL_3039603, EPI_ISL_3039604, EPI_ISL_3039605, EPI_ISL_3039606, EPI_ISL_3039607, EPI_ISL_3039608, EPI_ISL_3039609, EPI_ISL_3039610, EPI_ISL_3039611, EPI_ISL_3039612 | | | |
| see above | Department of Health Technology and Informatics, The Hong Kong Polytechnic University | Department of Health Technology and Informatics, The Hong Kong Polytechnic University | Gilman Kit-Hang Siu, Lam-Kwong Lee, Kenneth Siu-Sing Leung, Jake Siu-Lun Leung, Wing-Hei Lo, Timothy Ting-Leung Ng, Chloe Toi-Mei Chan, Hoi-Ching Jim, Ivan Tak-Fai Wong, Kingsley King-Gee Tam, Hiu-Yin Lao, Denise Sze-Hang Wong, Alan Ka-Lun Wu, Miranda Chong-Yee Yau, Yvette Wai-Man Lai, Kitty Sau-Chun Fung, Sandy Ka-Yee Chau, Barry Kin-Chung Wong, Wing-Kin To, Kristine Luk, Alex Yat-Man Ho, Tak-Lun Que, Kam-Tong Yip, Wing Cheong Yam, David Ho-Keung Shum, Shea Ping Yip |
| EPI_ISL_3039619 | Institute of medical Microbiology and Hospital Hygiene, Otto-von-Guericke University Magdeburg | Institute of medical Microbiology and Hospital Hygiene, Otto-von-Guericke University Magdeburg | Prof. Dr. Achim Kaasch, Aljoscha Tersteegen |
| EPI_ISL_3039622, EPI_ISL_3039623, EPI_ISL_3039624, EPI_ISL_3039625, EPI_ISL_3039627, EPI_ISL_3039629 | AULSS 9 Scaligera - Legnago | Istituto Zooprofilattico Sperimentale delle Venezie | Adelaide Milani, Alessia Schivo, Annalisa Salviato, Elisa Palumbo, Erika Giorgia Quaranta, Luca Tassoni, Ambra Pastori, Edoardo Giussani, Alice Fusaro, Isabella Monne, Calogero Terregino, Antonia Ricci |
| EPI_ISL_3039630, EPI_ISL_3039631, EPI_ISL_3039632, EPI_ISL_3039633, EPI_ISL_3039634, EPI_ISL_3039635, EPI_ISL_3039636, EPI_ISL_3039637, EPI_ISL_3039638, EPI_ISL_3039639, EPI_ISL_3039640, EPI_ISL_3039641, EPI_ISL_3039642, EPI_ISL_3039643, EPI_ISL_3039644, EPI_ISL_3039645, EPI_ISL_3039646, EPI_ISL_3039647, EPI_ISL_3039648, EPI_ISL_3039649, EPI_ISL_3039650, EPI_ISL_3039651, EPI_ISL_3039652, EPI_ISL_3039653 | | | |
| see above | AULSS 9 Scaligera - Villafranca | Istituto Zooprofilattico Sperimentale delle Venezie | Adelaide Milani, Alessia Schivo, Annalisa Salviato, Elisa Palumbo, Erika Giorgia Quaranta, Luca Tassoni, Ambra Pastori, Edoardo Giussani, Alice Fusaro, Isabella Monne, Calogero Terregino, Antonia Ricci |
| EPI_ISL_3039654, EPI_ISL_3039655, EPI_ISL_3039656, EPI_ISL_3039657, EPI_ISL_3039658, EPI_ISL_3039659, EPI_ISL_3039660, EPI_ISL_3039661, EPI_ISL_3039662, EPI_ISL_3039663, EPI_ISL_3039664, EPI_ISL_3039665, EPI_ISL_3039666, EPI_ISL_3039667, EPI_ISL_3039668, EPI_ISL_3039669, EPI_ISL_3039670, EPI_ISL_3039671, EPI_ISL_3039673, EPI_ISL_3039674, EPI_ISL_3039675, EPI_ISL_3039676, EPI_ISL_3039677 | | | |
| see above | AULSS 6 Euganea | Istituto Zooprofilattico Sperimentale delle Venezie | Adelaide Milani, Alessia Schivo, Annalisa Salviato, Elisa Palumbo, Erika Giorgia Quaranta, Luca Tassoni, Ambra Pastori, Edoardo Giussani, Alice Fusaro, Isabella Monne, Calogero Terregino, Antonia Ricci |
| EPI_ISL_3039679 | AULSS 7 Pedemontana - Bassano | Istituto Zooprofilattico Sperimentale delle Venezie | Adelaide Milani, Alessia Schivo, Annalisa Salviato, Elisa Palumbo, Erika Giorgia Quaranta, Luca Tassoni, Ambra Pastori, Edoardo Giussani, Alice Fusaro, Isabella Monne, Calogero Terregino, Antonia Ricci |
| EPI_ISL_3039680, EPI_ISL_3039681, EPI_ISL_3039682, EPI_ISL_3039683 | AULSS 6 Euganea | Istituto Zooprofilattico Sperimentale delle Venezie | Adelaide Milani, Alessia Schivo, Annalisa Salviato, Elisa Palumbo, Erika Giorgia Quaranta, Luca Tassoni, Ambra Pastori, Edoardo Giussani, Alice Fusaro, Isabella Monne, Calogero Terregino, Antonia Ricci |
| EPI_ISL_3039684, EPI_ISL_3039685 | AULSS 7 Pedemontana - Bassano | Istituto Zooprofilattico Sperimentale delle Venezie | Adelaide Milani, Alessia Schivo, Annalisa Salviato, Elisa Palumbo, Erika Giorgia Quaranta, Luca Tassoni, Ambra Pastori, Edoardo Giussani, Alice Fusaro, Isabella Monne, Calogero Terregino, Antonia Ricci |
| EPI_ISL_3039686, EPI_ISL_3039687, EPI_ISL_3039688, EPI_ISL_3039689 | AULSS 7 Pedemontana - Santorso | Istituto Zooprofilattico Sperimentale delle Venezie | Adelaide Milani, Alessia Schivo, Annalisa Salviato, Elisa Palumbo, Erika Giorgia Quaranta, Luca Tassoni, Ambra Pastori, Edoardo Giussani, Alice Fusaro, Isabella Monne, Calogero Terregino, Antonia Ricci |
| EPI_ISL_3039691, EPI_ISL_3039692, EPI_ISL_3039693, EPI_ISL_3039694, EPI_ISL_3039695, EPI_ISL_3039696, EPI_ISL_3039699, EPI_ISL_3039700, EPI_ISL_3039701 | AULSS 8 Berica | Istituto Zooprofilattico Sperimentale delle Venezie | Adelaide Milani, Alessia Schivo, Annalisa Salviato, Elisa Palumbo, Erika Giorgia Quaranta, Luca Tassoni, Ambra Pastori, Edoardo Giussani, Alice Fusaro, Isabella Monne, Calogero Terregino, Antonia Ricci |
| EPI_ISL_3039704, EPI_ISL_3039706 | AOUIVR | Istituto Zooprofilattico Sperimentale delle Venezie | Adelaide Milani, Alessia Schivo, Annalisa Salviato, Elisa Palumbo, Erika Giorgia Quaranta, Luca Tassoni, Ambra Pastori, Edoardo Giussani, Alice Fusaro, Isabella Monne, Calogero Terregino, Antonia Ricci |
| EPI_ISL_3039707 | AULSS 2 Marca Trevigiana | Istituto Zooprofilattico Sperimentale delle Venezie | Adelaide Milani, Alessia Schivo, Annalisa Salviato, Elisa Palumbo, Erika Giorgia Quaranta, Luca Tassoni, Ambra Pastori, Edoardo Giussani, Alice Fusaro, Isabella Monne, Calogero Terregino, Antonia Ricci |
| EPI_ISL_3039708, EPI_ISL_3039709, EPI_ISL_3039710, EPI_ISL_3039711, EPI_ISL_3039712, EPI_ISL_3039713, EPI_ISL_3039714 | AOUIVR | Istituto Zooprofilattico Sperimentale delle Venezie | Adelaide Milani, Alessia Schivo, Annalisa Salviato, Elisa Palumbo, Erika Giorgia Quaranta, Luca Tassoni, Ambra Pastori, Edoardo Giussani, Alice Fusaro, Isabella Monne, Calogero Terregino, Antonia Ricci |
| EPI_ISL_3039716, EPI_ISL_3039722 | CHU Pontchaillou | CHU Pontchaillou | GROLHIER Claire, DENOUAL Florent, ETCHEVERRY Amandine, SASSI Mohamed, JAGLINE Steven, FEBREAU Christine, PRONIER Charlotte, GALIBERT Marie Dominique, DE TAYRAC Marie, THIBAULT Vincent |
| EPI_ISL_3039725 | A.S.L.TO5 | Fondazione del Piemonte per l'Oncologia IRCCS | Antonino Sottile, Silvia Brossa, Paola Marino, Giorgio Giardina |
| EPI_ISL_3039726 | A.S.L.CN1 | Fondazione del Piemonte per l'Oncologia IRCCS | Antonino Sottile, Silvia Brossa, Paola Marino, Giorgio Giardina |
| EPI_ISL_3039727 | A.S.L.ALESSANDRIA | Fondazione del Piemonte per l'Oncologia IRCCS | Antonino Sottile, Silvia Brossa, Paola Marino, Giorgio Giardina |
| EPI_ISL_3039728, EPI_ISL_3039729 | A.S.L.BIELLA | Fondazione del Piemonte per l'Oncologia IRCCS | Antonino Sottile, Silvia Brossa, Paola Marino, Giorgio Giardina |
| EPI_ISL_3039730 | A.S.L.ALESSANDRIA | Fondazione del Piemonte per l'Oncologia IRCCS | Antonino Sottile, Silvia Brossa, Paola Marino, Giorgio Giardina |

| | | | |
|---|---|---|---|
| EPI_ISL_3039732, EPI_ISL_3039734, EPI_ISL_3039735, EPI_ISL_3039737, EPI_ISL_3039738, EPI_ISL_3039739, EPI_ISL_3039740, EPI_ISL_3039741, EPI_ISL_3039743, EPI_ISL_3039744, EPI_ISL_3039746, EPI_ISL_3039747, EPI_ISL_3039748, EPI_ISL_3039749, EPI_ISL_3039751, EPI_ISL_3039752, EPI_ISL_3039753, EPI_ISL_3039754, EPI_ISL_3039755 | | | |
| see above | A.S.L.CN1 | Fondazione del Piemonte per l'Oncologia IRCCS | Antonino Sottile, Silvia Brossa, Paola Marino, Giorgio Giardina |
| EPI_ISL_3039759 | A.S.L.ASTI | Fondazione del Piemonte per l'Oncologia IRCCS | Antonino Sottile, Silvia Brossa, Paola Marino, Giorgio Giardina |
| EPI_ISL_3039760, EPI_ISL_3039761 | A.S.L.CN1 | Fondazione del Piemonte per l'Oncologia IRCCS | Antonino Sottile, Silvia Brossa, Paola Marino, Giorgio Giardina |
| EPI_ISL_3039762 | A.S.L.TO4 | Fondazione del Piemonte per l'Oncologia IRCCS | Antonino Sottile, Silvia Brossa, Paola Marino, Giorgio Giardina |
| EPI_ISL_3039764 | A.S.L.ASTI | Fondazione del Piemonte per l'Oncologia IRCCS | Antonino Sottile, Silvia Brossa, Paola Marino, Giorgio Giardina |
| EPI_ISL_3039765 | A.S.L.TO4 | Fondazione del Piemonte per l'Oncologia IRCCS | Antonino Sottile, Silvia Brossa, Paola Marino, Giorgio Giardina |
| EPI_ISL_3039766 | A.S.L.ASTI | Fondazione del Piemonte per l'Oncologia IRCCS | Antonino Sottile, Silvia Brossa, Paola Marino, Giorgio Giardina |
| EPI_ISL_3039768, EPI_ISL_3039769, EPI_ISL_3039770, EPI_ISL_3039771, EPI_ISL_3039772, EPI_ISL_3039773, EPI_ISL_3039774, EPI_ISL_3039775, EPI_ISL_3039776, EPI_ISL_3039777, EPI_ISL_3039778, EPI_ISL_3039779, EPI_ISL_3039780, EPI_ISL_3039781, EPI_ISL_3039782 | | | |
| see above | A.S.L.NOVARA-BORGOMANERO | Fondazione del Piemonte per l'Oncologia IRCCS | Antonino Sottile, Silvia Brossa, Paola Marino, Giorgio Giardina |
| EPI_ISL_3039783, EPI_ISL_3039784, EPI_ISL_3039785, EPI_ISL_3039786, EPI_ISL_3039787, EPI_ISL_3039788, EPI_ISL_3039789, EPI_ISL_3039791, EPI_ISL_3039792, EPI_ISL_3039794, EPI_ISL_3039795 | | | |
| see above | A.S.L.VCO | Fondazione del Piemonte per l'Oncologia IRCCS | Antonino Sottile, Silvia Brossa, Paola Marino, Giorgio Giardina |
| EPI_ISL_3039796 | A.S.L.VERCELLI | Fondazione del Piemonte per l'Oncologia IRCCS | Antonino Sottile, Silvia Brossa, Paola Marino, Giorgio Giardina |
| EPI_ISL_3039797 | A.S.L.VCO | Fondazione del Piemonte per l'Oncologia IRCCS | Antonino Sottile, Silvia Brossa, Paola Marino, Giorgio Giardina |
| EPI_ISL_3039798 | A.S.L.VERCELLI | Fondazione del Piemonte per l'Oncologia IRCCS | Antonino Sottile, Silvia Brossa, Paola Marino, Giorgio Giardina |
| EPI_ISL_3039799 | A.S.L.CN1 | Fondazione del Piemonte per l'Oncologia IRCCS | Antonino Sottile, Silvia Brossa, Paola Marino, Giorgio Giardina |
| EPI_ISL_3039800 | A.S.L.NOVARA-BORGOMANERO | Fondazione del Piemonte per l'Oncologia IRCCS | Antonino Sottile, Silvia Brossa, Paola Marino, Giorgio Giardina |
| EPI_ISL_3039803 | A.S.L.VERCELLI | Fondazione del Piemonte per l'Oncologia IRCCS | Antonino Sottile, Silvia Brossa, Paola Marino, Giorgio Giardina |
| EPI_ISL_3039804 | A.S.L.NOVARA-BORGOMANERO | Fondazione del Piemonte per l'Oncologia IRCCS | Antonino Sottile, Silvia Brossa, Paola Marino, Giorgio Giardina |
| EPI_ISL_3039806 | A.S.L.VERCELLI | Fondazione del Piemonte per l'Oncologia IRCCS | Antonino Sottile, Silvia Brossa, Paola Marino, Giorgio Giardina |
| EPI_ISL_3039807 | A.S.L.NOVARA-BORGOMANERO | Fondazione del Piemonte per l'Oncologia IRCCS | Antonino Sottile, Silvia Brossa, Paola Marino, Giorgio Giardina |
| EPI_ISL_3039808 | A.S.L.VERCELLI | Fondazione del Piemonte per l'Oncologia IRCCS | Antonino Sottile, Silvia Brossa, Paola Marino, Giorgio Giardina |
| EPI_ISL_3039809, EPI_ISL_3039810, EPI_ISL_3039811, EPI_ISL_3039812, EPI_ISL_3039813, EPI_ISL_3039814, EPI_ISL_3039815, EPI_ISL_3039816, EPI_ISL_3039818, EPI_ISL_3039819, EPI_ISL_3039821, EPI_ISL_3039822 | | | |
| see above | A.S.L.NOVARA-BORGOMANERO | Fondazione del Piemonte per l'Oncologia IRCCS | Antonino Sottile, Silvia Brossa, Paola Marino, Giorgio Giardina |
| EPI_ISL_3039823, EPI_ISL_3039825 | A.S.L.ALESSANDRIA | Fondazione del Piemonte per l'Oncologia IRCCS | Antonino Sottile, Silvia Brossa, Paola Marino, Giorgio Giardina |
| EPI_ISL_3039826, EPI_ISL_3039827 | A.S.L.NOVARA-BORGOMANERO | Fondazione del Piemonte per l'Oncologia IRCCS | Antonino Sottile, Silvia Brossa, Paola Marino, Giorgio Giardina |
| EPI_ISL_3039828, EPI_ISL_3039829 | A.S.L.ALESSANDRIA | Fondazione del Piemonte per l'Oncologia IRCCS | Antonino Sottile, Silvia Brossa, Paola Marino, Giorgio Giardina |
| EPI_ISL_3039830, EPI_ISL_3039831, EPI_ISL_3039832, EPI_ISL_3039833, EPI_ISL_3039834, EPI_ISL_3039835, EPI_ISL_3039836, EPI_ISL_3039838, EPI_ISL_3039839, EPI_ISL_3039840, EPI_ISL_3039841, EPI_ISL_3039842, EPI_ISL_3039843, EPI_ISL_3039844, EPI_ISL_3039845, EPI_ISL_3039846, EPI_ISL_3039847, EPI_ISL_3039848, EPI_ISL_3039849, EPI_ISL_3039850, EPI_ISL_3039851, EPI_ISL_3039852, EPI_ISL_3039853, EPI_ISL_3039854, EPI_ISL_3039855, EPI_ISL_3039856, EPI_ISL_3039857, EPI_ISL_3039858, EPI_ISL_3039859, EPI_ISL_3039860, EPI_ISL_3039861, EPI_ISL_3039862, EPI_ISL_3039863, EPI_ISL_3039864, EPI_ISL_3039865, EPI_ISL_3039866, EPI_ISL_3039867, EPI_ISL_3039868, EPI_ISL_3039869, EPI_ISL_3039870, EPI_ISL_3039871, EPI_ISL_3039872, EPI_ISL_3039873, EPI_ISL_3039874, EPI_ISL_3039875, EPI_ISL_3039876, EPI_ISL_3039877, EPI_ISL_3039878, EPI_ISL_3039879, EPI_ISL_3039880, EPI_ISL_3039881, EPI_ISL_3039882, EPI_ISL_3039883, EPI_ISL_3039884, EPI_ISL_3039886, EPI_ISL_3039887, EPI_ISL_3039888, EPI_ISL_3039889, EPI_ISL_3039890, EPI_ISL_3039891, EPI_ISL_3039892, EPI_ISL_3039896, EPI_ISL_3039897, EPI_ISL_3039898, EPI_ISL_3039899, EPI_ISL_3039900, EPI_ISL_3039901, EPI_ISL_3039902, EPI_ISL_3039903, EPI_ISL_3039904, EPI_ISL_3039907, EPI_ISL_3039908, EPI_ISL_3039909, EPI_ISL_3039910, EPI_ISL_3039912, EPI_ISL_3039913, EPI_ISL_3039914, EPI_ISL_3039917, EPI_ISL_3039919, EPI_ISL_3039920, EPI_ISL_3039921, EPI_ISL_3039922, EPI_ISL_3039923, EPI_ISL_3039924, EPI_ISL_3039925, EPI_ISL_3039926, EPI_ISL_3039928, EPI_ISL_3039929, EPI_ISL_3039930, EPI_ISL_3039931, EPI_ISL_3039932, EPI_ISL_3039933, EPI_ISL_3039934, EPI_ISL_3039935, EPI_ISL_3039936, EPI_ISL_3039937, EPI_ISL_3039939, EPI_ISL_3039940, EPI_ISL_3039941, EPI_ISL_3039942, EPI_ISL_3039943, EPI_ISL_3039944, EPI_ISL_3039945, EPI_ISL_3039946, EPI_ISL_3039948, EPI_ISL_3039949, EPI_ISL_3039950, EPI_ISL_3039951, EPI_ISL_3039952, EPI_ISL_3039953, EPI_ISL_3039954, EPI_ISL_3039955, EPI_ISL_3039956, EPI_ISL_3039958, EPI_ISL_3039960, EPI_ISL_3039961, EPI_ISL_3039962, EPI_ISL_3039966, EPI_ISL_3039967, EPI_ISL_3039968, EPI_ISL_3039969, EPI_ISL_3039970, EPI_ISL_3039971, EPI_ISL_3039972, EPI_ISL_3039977, EPI_ISL_3039979, EPI_ISL_3039980, EPI_ISL_3039981, EPI_ISL_3039982, EPI_ISL_3039983, EPI_ISL_3039984, EPI_ISL_3039990, EPI_ISL_3039991, EPI_ISL_3039992, EPI_ISL_3039993, EPI_ISL_3039994, EPI_ISL_3039995, EPI_ISL_3039996, EPI_ISL_3040001, EPI_ISL_3040002, EPI_ISL_3040003, EPI_ISL_3040004, EPI_ISL_3040005, EPI_ISL_3040006, EPI_ISL_3040007, EPI_ISL_3040011, EPI_ISL_3040012, EPI_ISL_3040013, EPI_ISL_3040014, EPI_ISL_3040015, EPI_ISL_3040016, EPI_ISL_3040019, EPI_ISL_3040021, EPI_ISL_3040022, EPI_ISL_3040023, EPI_ISL_3040024, EPI_ISL_3040026, EPI_ISL_3040031, EPI_ISL_3040032, EPI_ISL_3040033, EPI_ISL_3040034, EPI_ISL_3040035, EPI_ISL_3040037, EPI_ISL_3040042, EPI_ISL_3040046 | | | |
| see above | Austrian Agency for Health and Food Safety (AGES) | Bergthaler laboratory, CeMM Research Center for Molecular Medicine of the Austrian Academy of Sciences | Lukas Endler, Anna Schedl, Fabian Amman, Petr Triska, Matthew Thornton, Thomas Penz, Benedikt Agerer, Maelle Le Moing, Michael Schuster, Bekir Erguner, Jan Laine, Martin Senekowitsch, Christoph Bock, Andreas Bergthaler |
| EPI_ISL_3040105 | Universität Innsbruck, Institut für Mikrobiologie | Bergthaler laboratory, CeMM Research Center for Molecular Medicine of the Austrian Academy of Sciences | Lukas Endler, Anna Schedl, Fabian Amman, Petr Triska, Matthew Thornton, Thomas Penz, Benedikt Agerer, Maelle Le Moing, Michael Schuster, Bekir Erguner, Jan Laine, Martin Senekowitsch, Christoph Bock, Andreas Bergthaler |
| EPI_ISL_3040111, EPI_ISL_3040112, EPI_ISL_3040115 | Institute for Medical and Chemical Laboratory Diagnostics, Kepler UniversitÃ¤tsklinikum | Bergthaler laboratory, CeMM Research Center for Molecular Medicine of the Austrian Academy of Sciences | Lukas Endler, Anna Schedl, Fabian Amman, Petr Triska, Matthew Thornton, Thomas Penz, Benedikt Agerer, Maelle Le Moing, Michael Schuster, Bekir Erguner, Jan Laine, Martin Senekowitsch, Christoph Bock, Andreas Bergthaler |
| EPI_ISL_3040119 | Vienna COVID-19 Detection Initiative (VCDI) | Bergthaler laboratory, CeMM Research Center for Molecular Medicine of the Austrian Academy of Sciences | Lukas Endler, Anna Schedl, Fabian Amman, Petr Triska, Matthew Thornton, Thomas Penz, Benedikt Agerer, Maelle Le Moing, Michael Schuster, Bekir Erguner, Jan Laine, Martin Senekowitsch, Christoph Bock, Andreas Bergthaler |
| EPI_ISL_3040129, EPI_ISL_3040130, EPI_ISL_3040133, EPI_ISL_3040134 | Fondation Congolaise pour la recherche medicale (FCRM), Francine Ntoumi | Fondation Congolaise pour la Recherche Médicale | Mfoutou Mapanguy Claujens Chastel; Batchi-Bouyou Armel Landry; Dr. Jean Claude Djontu; Dr. Abel Lissom; Prof. Dr. Thirumalaisamy P. Velavan; Prof. Francine Ntoumi |
| EPI_ISL_3040154, EPI_ISL_3040155, EPI_ISL_3040156, EPI_ISL_3040157, EPI_ISL_3040158, EPI_ISL_3040159, EPI_ISL_3040160, EPI_ISL_3040161, EPI_ISL_3040162, EPI_ISL_3040163, EPI_ISL_3040164, EPI_ISL_3040165, EPI_ISL_3040166, EPI_ISL_3040167, EPI_ISL_3040168, EPI_ISL_3040169, EPI_ISL_3040170, EPI_ISL_3040171, EPI_ISL_3040172, EPI_ISL_3040173, EPI_ISL_3040174, EPI_ISL_3040175, EPI_ISL_3040176, EPI_ISL_3040177, EPI_ISL_3040178, EPI_ISL_3040179, EPI_ISL_3040180, EPI_ISL_3040181, EPI_ISL_3040182, EPI_ISL_3040183, EPI_ISL_3040184, EPI_ISL_3040185, EPI_ISL_3040186, EPI_ISL_3040187, EPI_ISL_3040188, EPI_ISL_3040190, EPI_ISL_3040191, EPI_ISL_3040192, EPI_ISL_3040193, EPI_ISL_3040194, EPI_ISL_3040195, EPI_ISL_3040196, EPI_ISL_3040197, EPI_ISL_3040198, EPI_ISL_3040199, EPI_ISL_3040200, EPI_ISL_3040201, EPI_ISL_3040202, EPI_ISL_3040204, EPI_ISL_3040205, EPI_ISL_3040206, EPI_ISL_3040207, EPI_ISL_3040208, EPI_ISL_3040209, EPI_ISL_3040210, EPI_ISL_3040211, EPI_ISL_3040212, EPI_ISL_3040213, EPI_ISL_3040214, EPI_ISL_3040215, EPI_ISL_3040216, EPI_ISL_3040217, EPI_ISL_3040218, EPI_ISL_3040219, EPI_ISL_3040220, EPI_ISL_3040221, EPI_ISL_3040222, EPI_ISL_3040223, EPI_ISL_3040224, EPI_ISL_3040225, EPI_ISL_3040226, EPI_ISL_3040227, EPI_ISL_3040228, EPI_ISL_3040229, EPI_ISL_3040230, EPI_ISL_3040231, EPI_ISL_3040232, EPI_ISL_3040233, EPI_ISL_3040234, EPI_ISL_3040235, EPI_ISL_3040236, EPI_ISL_3040237, EPI_ISL_3040238, EPI_ISL_3040239, EPI_ISL_3040240, EPI_ISL_3040242, EPI_ISL_3040243, EPI_ISL_3040244, EPI_ISL_3040245, EPI_ISL_3040246, EPI_ISL_3040247, EPI_ISL_3040248, EPI_ISL_3040249, EPI_ISL_3040250, EPI_ISL_3040251, EPI_ISL_3040252, EPI_ISL_3040253, EPI_ISL_3040254, EPI_ISL_3040255, EPI_ISL_3040256, EPI_ISL_3040257, EPI_ISL_3040258, EPI_ISL_3040259, EPI_ISL_3040260, EPI_ISL_3040261, EPI_ISL_3040262, EPI_ISL_3040263, EPI_ISL_3040264, EPI_ISL_3040265, EPI_ISL_3040266, EPI_ISL_3040267, EPI_ISL_3040268, EPI_ISL_3040269, EPI_ISL_3040270, EPI_ISL_3040271, EPI_ISL_3040272, EPI_ISL_3040273, EPI_ISL_3040274, EPI_ISL_3040275, EPI_ISL_3040276, EPI_ISL_3040277, EPI_ISL_3040278, EPI_ISL_3040279, EPI_ISL_3040280, EPI_ISL_3040281, EPI_ISL_3040282, EPI_ISL_3040283, EPI_ISL_3040284, EPI_ISL_3040285, EPI_ISL_3040286, EPI_ISL_3040287, EPI_ISL_3040288, EPI_ISL_3040289, EPI_ISL_3040290, EPI_ISL_3040291, EPI_ISL_3040292, EPI_ISL_3040293, EPI_ISL_3040294, EPI_ISL_3040295, EPI_ISL_3040296, EPI_ISL_3040297, EPI_ISL_3040298, EPI_ISL_3040299, EPI_ISL_3040300, EPI_ISL_3040301, EPI_ISL_3040302, EPI_ISL_3040303, EPI_ISL_3040304, EPI_ISL_3040305, EPI_ISL_3040306, EPI_ISL_3040307, EPI_ISL_3040308, EPI_ISL_3040309, EPI_ISL_3040310, EPI_ISL_3040311, EPI_ISL_3040313, EPI_ISL_3040314, EPI_ISL_3040315, EPI_ISL_3040316, EPI_ISL_3040317, EPI_ISL_3040318, EPI_ISL_3040319, EPI_ISL_3040320, EPI_ISL_3040321, EPI_ISL_3040322, EPI_ISL_3040323, EPI_ISL_3040324, EPI_ISL_3040325, EPI_ISL_3040326, EPI_ISL_3040327, EPI_ISL_3040328, EPI_ISL_3040329, EPI_ISL_3040330, EPI_ISL_3040331, EPI_ISL_3040332, EPI_ISL_3040333, EPI_ISL_3040334, EPI_ISL_3040335, EPI_ISL_3040336, EPI_ISL_3040337, EPI_ISL_3040338, EPI_ISL_3040339, EPI_ISL_3040340, EPI_ISL_3040341, EPI_ISL_3040342, EPI_ISL_3040343, EPI_ISL_3040344, EPI_ISL_3040345, EPI_ISL_3040346, EPI_ISL_3040347, EPI_ISL_3040348, EPI_ISL_3040349, EPI_ISL_3040350, EPI_ISL_3040351, EPI_ISL_3040352, EPI_ISL_3040353, EPI_ISL_3040354, EPI_ISL_3040355, EPI_ISL_3040356, EPI_ISL_3040357, EPI_ISL_3040358, EPI_ISL_3040359, EPI_ISL_3040360, EPI_ISL_3040361, EPI_ISL_3040362, EPI_ISL_3040363, EPI_ISL_3040364, EPI_ISL_3040365, EPI_ISL_3040366, EPI_ISL_3040367, EPI_ISL_3040368, EPI_ISL_3040369, EPI_ISL_3040370, EPI_ISL_3040371, EPI_ISL_3040372, EPI_ISL_3040373, EPI_ISL_3040374, EPI_ISL_3040375, EPI_ISL_3040376, EPI_ISL_3040378, EPI_ISL_3040379, EPI_ISL_3040380, EPI_ISL_3040381, EPI_ISL_3040382, EPI_ISL_3040383, EPI_ISL_3040384, EPI_ISL_3040385, EPI_ISL_3040386, EPI_ISL_3040387, EPI_ISL_3040388, EPI_ISL_3040389, EPI_ISL_3040390, EPI_ISL_3040391, EPI_ISL_3040392, EPI_ISL_3040393, EPI_ISL_3040394, EPI_ISL_3040395, EPI_ISL_3040396, EPI_ISL_3040397, EPI_ISL_3040398, EPI_ISL_3040399, EPI_ISL_3040401, EPI_ISL_3040402, EPI_ISL_3040403, EPI_ISL_3040404, EPI_ISL_3040405, EPI_ISL_3040406, EPI_ISL_3040407, EPI_ISL_3040408, EPI_ISL_3040409, EPI_ISL_3040410, EPI_ISL_3040411, EPI_ISL_3040412, EPI_ISL_3040413, EPI_ISL_3040414, EPI_ISL_3040415, EPI_ISL_3040416, EPI_ISL_3040417, EPI_ISL_3040418, EPI_ISL_3040419, EPI_ISL_3040420, EPI_ISL_3040421, EPI_ISL_3040422, EPI_ISL_3040423, EPI_ISL_3040425, EPI_ISL_3040426, EPI_ISL_3040427, EPI_ISL_3040428, EPI_ISL_3040429, EPI_ISL_3040430, EPI_ISL_3040431, EPI_ISL_3040432, EPI_ISL_3040433, EPI_ISL_3040434, EPI_ISL_3040435, EPI_ISL_3040436, EPI_ISL_3040437, EPI_ISL_3040438, EPI_ISL_3040439, EPI_ISL_3040440, EPI_ISL_3040441, EPI_ISL_3040442, EPI_ISL_3040443, EPI_ISL_3040444, EPI_ISL_3040445, EPI_ISL_3040446, EPI_ISL_3040447, EPI_ISL_3040448, EPI_ISL_3040449, EPI_ISL_3040450, EPI_ISL_3040451, EPI_ISL_3040452, EPI_ISL_3040453, EPI_ISL_3040454, EPI_ISL_3040455, EPI_ISL_3040456, EPI_ISL_3040457, EPI_ISL_3040458, EPI_ISL_3040459, EPI_ISL_3040460, EPI_ISL_3040462, EPI_ISL_3040463, EPI_ISL_3040464, EPI_ISL_3040465, EPI_ISL_3040466, EPI_ISL_3040467, EPI_ISL_3040468, EPI_ISL_3040469, EPI_ISL_3040470, EPI_ISL_3040471, EPI_ISL_3040472, EPI_ISL_3040473, EPI_ISL_3040474, EPI_ISL_3040475, EPI_ISL_3040476, EPI_ISL_3040477, EPI_ISL_3040478, EPI_ISL_3040479, EPI_ISL_3040480, EPI_ISL_3040481, EPI_ISL_3040482, EPI_ISL_3040483, EPI_ISL_3040484, EPI_ISL_3040485, EPI_ISL_3040486, EPI_ISL_3040487, EPI_ISL_3040488, EPI_ISL_3040489, EPI_ISL_3040490, EPI_ISL_3040491, EPI_ISL_3040492, EPI_ISL_3040494, EPI_ISL_3040495, EPI_ISL_3040496, EPI_ISL_3040497, EPI_ISL_3040498, EPI_ISL_3040500, EPI_ISL_3040501, EPI_ISL_3040502, EPI_ISL_3040503, EPI_ISL_3040504, EPI_ISL_3040505, EPI_ISL_3040506, EPI_ISL_3040507, EPI_ISL_3040509, EPI_ISL_3040510, EPI_ISL_3040511, EPI_ISL_3040512, EPI_ISL_3040513, EPI_ISL_3040514, EPI_ISL_3040515, EPI_ISL_3040516, EPI_ISL_3040517, EPI_ISL_3040518, EPI_ISL_3040519, EPI_ISL_3040520, EPI_ISL_3040521, EPI_ISL_3040522, EPI_ISL_3040523, EPI_ISL_3040524, EPI_ISL_3040525, EPI_ISL_3040526, EPI_ISL_3040527, EPI_ISL_3040528, EPI_ISL_3040529, EPI_ISL_3040530, EPI_ISL_3040531, EPI_ISL_3040532, EPI_ISL_3040534, EPI_ISL_3040535, EPI_ISL_3040536, EPI_ISL_3040537, EPI_ISL_3040538, EPI_ISL_3040539, EPI_ISL_3040541, EPI_ISL_3040542, EPI_ISL_3040543, EPI_ISL_3040544, EPI_ISL_3040545, EPI_ISL_3040546, EPI_ISL_3040547, EPI_ISL_3040548, EPI_ISL_3040549, EPI_ISL_3040550, EPI_ISL_3040551, EPI_ISL_3040552, EPI_ISL_3040553, EPI_ISL_3040554, EPI_ISL_3040555, EPI_ISL_3040556, EPI_ISL_3040557, EPI_ISL_3040558, EPI_ISL_3040559, EPI_ISL_3040560, EPI_ISL_3040561, EPI_ISL_3040562, EPI_ISL_3040563, EPI_ISL_3040564, EPI_ISL_3040565, EPI_ISL_3040566, EPI_ISL_3040567, EPI_ISL_3040568, EPI_ISL_3040569, EPI_ISL_3040570, EPI_ISL_3040571, EPI_ISL_3040572, EPI_ISL_3040573, EPI_ISL_3040574, EPI_ISL_3040575, EPI_ISL_3040576, EPI_ISL_3040577, EPI_ISL_3040578, EPI_ISL_3040579, EPI_ISL_3040580, EPI_ISL_3040581, EPI_ISL_3040582, EPI_ISL_3040583, EPI_ISL_3040584, EPI_ISL_3040585, EPI_ISL_3040586, EPI_ISL_3040587, EPI_ISL_3040588, EPI_ISL_3040589, EPI_ISL_3040590, EPI_ISL_3040591, EPI_ISL_3040592, EPI_ISL_3040593, EPI_ISL_3040594, EPI_ISL_3040595, EPI_ISL_3040596, EPI_ISL_3040597, EPI_ISL_3040598, EPI_ISL_3040599, EPI_ISL_3040600, EPI_ISL_3040601, EPI_ISL_3040602, EPI_ISL_3040603, EPI_ISL_3040604, EPI_ISL_3040605, EPI_ISL_3040606, EPI_ISL_3040607, EPI_ISL_3040608, EPI_ISL_3040609, EPI_ISL_3040610, EPI_ISL_3040611, EPI_ISL_3040612, EPI_ISL_3040613, EPI_ISL_3040614, EPI_ISL_3040615, EPI_ISL_3040616, EPI_ISL_3040617, EPI_ISL_3040618, EPI_ISL_3040619, EPI_ISL_3040620, EPI_ISL_3040621, EPI_ISL_3040622, EPI_ISL_3040623, EPI_ISL_3040624, EPI_ISL_3040625, EPI_ISL_3040626, EPI_ISL_3040627, EPI_ISL_3040628, EPI_ISL_3040629, EPI_ISL_3040630, EPI_ISL_3040631, EPI_ISL_3040632, EPI_ISL_3040633, EPI_ISL_3040634, EPI_ISL_3040635, EPI_ISL_3040636, EPI_ISL_3040637, EPI_ISL_3040638, EPI_ISL_3040639, EPI_ISL_3040640, EPI_ISL_3040641, EPI_ISL_3040642, EPI_ISL_3040643, EPI_ISL_3040644, EPI_ISL_3040645, EPI_ISL_3040647, EPI_ISL_3040648, EPI_ISL_3040649, EPI_ISL_3040650, EPI_ISL_3040651, EPI_ISL_3040652, EPI_ISL_3040653, EPI_ISL_3040654, EPI_ISL_3040655, EPI_ISL_3040656, EPI_ISL_3040657, EPI_ISL_3040658, EPI_ISL_3040659, EPI_ISL_3040660, EPI_ISL_3040661, EPI_ISL_3040663, EPI_ISL_3040664, EPI_ISL_3040665, EPI_ISL_3040666, EPI_ISL_3040667, EPI_ISL_3040668, EPI_ISL_3040669, EPI_ISL_3040670, EPI_ISL_3040671, EPI_ISL_3040672, EPI_ISL_3040673, EPI_ISL_3040674, EPI_ISL_3040676, EPI_ISL_3040677, EPI_ISL_3040678, EPI_ISL_3040679, EPI_ISL_3040680, EPI_ISL_3040681, EPI_ISL_3040682, EPI_ISL_3040683, EPI_ISL_3040684, EPI_ISL_3040686, EPI_ISL_3040687, EPI_ISL_3040688, EPI_ISL_3040689, EPI_ISL_3040690, EPI_ISL_3040691, EPI_ISL_3040692, EPI_ISL_3040693, EPI_ISL_3040694, EPI_ISL_3040695, EPI_ISL_3040696, EPI_ISL_3040697, EPI_ISL_3040698, EPI_ISL_3040699, EPI_ISL_3040700, EPI_ISL_3040701, EPI_ISL_3040702, EPI_ISL_3040703, EPI_ISL_3040704, EPI_ISL_3040705, EPI_ISL_3040706, EPI_ISL_3040707, EPI_ISL_3040708, EPI_ISL_3040709, EPI_ISL_3040710, EPI_ISL_3040711, EPI_ISL_3040712, EPI_ISL_3040713, EPI_ISL_3040714, EPI_ISL_3040715, EPI_ISL_3040716, EPI_ISL_3040717, EPI_ISL_3040718, EPI_ISL_3040719, EPI_ISL_3040720, EPI_ISL_3040721, EPI_ISL_3040722, EPI_ISL_3040723, EPI_ISL_3040724, EPI_ISL_3040725, EPI_ISL_3040726, EPI_ISL_3040727, EPI_ISL_3040728, EPI_ISL_3040729, EPI_ISL_3040730, EPI_ISL_3040731, EPI_ISL_3040733, EPI_ISL_3040734, EPI_ISL_3040735, EPI_ISL_3040736, EPI_ISL_3040737, EPI_ISL_3040738, EPI_ISL_3040739, EPI_ISL_3040740, EPI_ISL_3040741, EPI_ISL_3040742, EPI_ISL_3040743, EPI_ISL_3040744, EPI_ISL_3040745, EPI_ISL_3040746, EPI_ISL_3040747, EPI_ISL_3040748, EPI_ISL_3040749, EPI_ISL_3040750, EPI_ISL_3040751, EPI_ISL_3040752, EPI_ISL_3040753, EPI_ISL_3040754, EPI_ISL_3040755, EPI_ISL_3040756, EPI_ISL_3040757, EPI_ISL_3040758, EPI_ISL_3040759, EPI_ISL_3040760, EPI_ISL_3040761, EPI_ISL_3040762, EPI_ISL_3040763, EPI_ISL_3040764, EPI_ISL_3040765, EPI_ISL_3040766, EPI_ISL_3040767, EPI_ISL_3040768, EPI_ISL_3040769, EPI_ISL_3040770, EPI_ISL_3040771, EPI_ISL_3040772, EPI_ISL_3040773, EPI_ISL_3040774, EPI_ISL_3040776, EPI_ISL_3040777, EPI_ISL_3040778, EPI_ISL_3040779, EPI_ISL_3040780, EPI_ISL_3040781, EPI_ISL_3040782, EPI_ISL_3040783, EPI_ISL_3040784, EPI_ISL_3040785, EPI_ISL_3040786, EPI_ISL_3040787, EPI_ISL_3040788, EPI_ISL_3040789, EPI_ISL_3040790, EPI_ISL_3040791, EPI_ISL_3040793, EPI_ISL_3040794, EPI_ISL_3040795, EPI_ISL_3040796, EPI_ISL_3040797, EPI_ISL_3040798, EPI_ISL_3040799, EPI_ISL_3040800, EPI_ISL_3040801, EPI_ISL_3040802, EPI_ISL_3040803, EPI_ISL_3040804, EPI_ISL_3040805, EPI_ISL_3040806, EPI_ISL_3040807, EPI_ISL_3040808, EPI_ISL_3040809, EPI_ISL_3040810, EPI_ISL_3040811, EPI_ISL_3040812, EPI_ISL_3040813, EPI_ISL_3040814, EPI_ISL_3040815, EPI_ISL_3040816, EPI_ISL_3040817, EPI_ISL_3040818, EPI_ISL_3040819, EPI_ISL_3040820, EPI_ISL_3040821, EPI_ISL_3040822, EPI_ISL_3040823, EPI_ISL_3040824, EPI_ISL_3040825, EPI_ISL_3040826, EPI_ISL_3040827, EPI_ISL_3040828, EPI_ISL_3040829, EPI_ISL_3040830, EPI_ISL_3040831, EPI_ISL_3040832, EPI_ISL_3040833, EPI_ISL_3040834, EPI_ISL_3040835, EPI_ISL_3040836, EPI_ISL_3040837, EPI_ISL_3040838, EPI_ISL_3040839, EPI_ISL_3040840, EPI_ISL_3040841, EPI_ISL_3040842, EPI_ISL_3040843, EPI_ISL_3040844, EPI_ISL_3040845, EPI_ISL_3040846, EPI_ISL_3040847, EPI_ISL_3040848, EPI_ISL_3040849, EPI_ISL_3040850, EPI_ISL_3040852, EPI_ISL_3040853, EPI_ISL_3040854, EPI_ISL_3040855, EPI_ISL_3040856, EPI_ISL_3040857, EPI_ISL_3040858, EPI_ISL_3040859, EPI_ISL_3040860, EPI_ISL_3040861, EPI_ISL_3040862, EPI_ISL_3040863, EPI_ISL_3040864, EPI_ISL_3040865, EPI_ISL_3040866, EPI_ISL_3040867, EPI_ISL_3040868, EPI_ISL_3040869, EPI_ISL_3040870, EPI_ISL_3040871, EPI_ISL_3040872, EPI_ISL_3040873, EPI_ISL_3040874, EPI_ISL_3040875, EPI_ISL_3040876, EPI_ISL_3040878, EPI_ISL_3040879, EPI_ISL_3040880, EPI_ISL_3040881, EPI_ISL_3040882, EPI_ISL_3040883, EPI_ISL_3040884, EPI_ISL_3040885, EPI_ISL_3040886, EPI_ISL_3040887, EPI_ISL_3040888, EPI_ISL_3040889, EPI_ISL_3040890, EPI_ISL_3040891, EPI_ISL_3040892, EPI_ISL_3040893, EPI_ISL_3040894, EPI_ISL_3040895, EPI_ISL_3040896, EPI_ISL_3040897, EPI_ISL_3040898, EPI_ISL_3040899, EPI_ISL_3040900, EPI_ISL_3040901, EPI_ISL_3040902, EPI_ISL_3040903, EPI_ISL_3040904, EPI_ISL_3040905, EPI_ISL_3040906, EPI_ISL_3040907, EPI_ISL_3040908, EPI_ISL_3040909, EPI_ISL_3040910, EPI_ISL_3040911, EPI_ISL_3040912, EPI_ISL_3040914, EPI_ISL_3040915, EPI_ISL_3040916, EPI_ISL_3040917, EPI_ISL_3040918, EPI_ISL_3040919, EPI_ISL_3040920, EPI_ISL_3040921, EPI_ISL_3040922, EPI_ISL_3040923, EPI_ISL_3040924, EPI_ISL_3040925, EPI_ISL_3040926, EPI_ISL_3040927, EPI_ISL_3040928, EPI_ISL_3040929, EPI_ISL_3040930, EPI_ISL_3040931, EPI_ISL_3040932, EPI_ISL_3040933, EPI_ISL_3040934, EPI_ISL_3040935, EPI_ISL_3040936, EPI_ISL_3040937, EPI_ISL_3040938, EPI_ISL_3040939, EPI_ISL_3040940, EPI_ISL_3040941, EPI_ISL_3040942, EPI_ISL_3040943, EPI_ISL_3040944, EPI_ISL_3040945, EPI_ISL_3040946, EPI_ISL_3040947, EPI_ISL_3040948, EPI_ISL_3040949, EPI_ISL_3040950, EPI_ISL_3040951, EPI_ISL_3040952, EPI_ISL_3040953, EPI_ISL_3040954, EPI_ISL_3040955, EPI_ISL_3040956, EPI_ISL_3040957, EPI_ISL_3040959, EPI_ISL_3040960, EPI_ISL_3040961, EPI_ISL_3040962, EPI_ISL_3040963, EPI_ISL_3040964, EPI_ISL_3040965, EPI_ISL_3040966, EPI_ISL_3040967, EPI_ISL_3040968, EPI_ISL_3040969, EPI_ISL_3040970, EPI_ISL_3040971, EPI_ISL_3040972, EPI_ISL_3040973, EPI_ISL_3040974, EPI_ISL_3040975, EPI_ISL_3040976, EPI_ISL_3040977, EPI_ISL_3040978, EPI_ISL_3040979, EPI_ISL_3040980, EPI_ISL_3040981, EPI_ISL_3040982, EPI_ISL_3040983, EPI_ISL_3040984, EPI_ISL_3040985, EPI_ISL_3040986, EPI_ISL_3040987, EPI_ISL_3040988, EPI_ISL_3040989, EPI_ISL_3040990, EPI_ISL_3040991, EPI_ISL_3040992, EPI_ISL_3040993, EPI_ISL_3040994, EPI_ISL_3040995, EPI_ISL_3040996, EPI_ISL_3040997, EPI_ISL_3040998, EPI_ISL_3040999, EPI_ISL_3041000, EPI_ISL_3041001, EPI_ISL_3041002, EPI_ISL_3041003, EPI_ISL_3041004, EPI_ISL_3041005, EPI_ISL_3041007, EPI_ISL_3041008, EPI_ISL_3041009, EPI_ISL_3041010, EPI_ISL_3041011, EPI_ISL_3041012, EPI_ISL_3041013, EPI_ISL_3041014, EPI_ISL_3041015, EPI_ISL_3041016, EPI_ISL_3041017, EPI_ISL_3041018, EPI_ISL_3041019, EPI_ISL_3041020, EPI_ISL_3041021, EPI_ISL_3041022, EPI_ISL_3041023, EPI_ISL_3041024, EPI_ISL_3041025, EPI_ISL_3041026, EPI_ISL_3041027, EPI_ISL_3041028, EPI_ISL_3041029, EPI_ISL_3041030, EPI_ISL_3041031, EPI_ISL_3041032, EPI_ISL_3041033, EPI_ISL_3041034, EPI_ISL_3041035, EPI_ISL_3041036, EPI_ISL_3041037, EPI_ISL_3041038, EPI_ISL_3041039, EPI_ISL_3041040, EPI_ISL_3041041, EPI_ISL_3041042, EPI_ISL_3041043, EPI_ISL_3041044, EPI_ISL_3041045, EPI_ISL_3041046, EPI_ISL_3041048, EPI_ISL_3041049, EPI_ISL_3041050, EPI_ISL_3041051, EPI_ISL_3041052, EPI_ISL_3041053, EPI_ISL_3041054, EPI_ISL_3041055, EPI_ISL_3041056, EPI_ISL_3041057, EPI_ISL_3041058, EPI_ISL_3041059, EPI_ISL_3041060, EPI_ISL_3041061, EPI_ISL_3041062, EPI_ISL_3041063, EPI_ISL_3041064, EPI_ISL_3041065, EPI_ISL_3041066, EPI_ISL_3041067, EPI_ISL_3041068, EPI_ISL_3041069, EPI_ISL_3041070, EPI_ISL_3041071, EPI_ISL_3041072, EPI_ISL_3041073, EPI_ISL_3041074, EPI_ISL_3041075, EPI_ISL_3041076, EPI_ISL_3041077, EPI_ISL_3041078, EPI_ISL_3041079, EPI_ISL_3041080, EPI_ISL_3041081, EPI_ISL_3041082, EPI_ISL_3041083, EPI_ISL_3041084, EPI_ISL_3041085, EPI_ISL_3041086, EPI_ISL_3041087, EPI_ISL_3041088, EPI_ISL_3041089, EPI_ISL_3041090, EPI_ISL_3041091, EPI_ISL_3041092, EPI_ISL_3041093, EPI_ISL_3041094, EPI_ISL_3041095, EPI_ISL_3041096, EPI_ISL_3041097, EPI_ISL_3041098, EPI_ISL_3041099, EPI_ISL_3041100, EPI_ISL_3041101, EPI_ISL_3041102, EPI_ISL_3041103, EPI_ISL_3041104, EPI_ISL_3041105, EPI_ISL_3041106, EPI_ISL_3041107, EPI_ISL_3041108, EPI_ISL_3041109, EPI_ISL_3041110, EPI_ISL_3041111, EPI_ISL_3041112, EPI_ISL_3041113, EPI_ISL_3041114, EPI_ISL_3041115, EPI_ISL_3041116, EPI_ISL_3041117, EPI_ISL_3041118, EPI_ISL_3041119, EPI_ISL_3041120, EPI_ISL_3041121, EPI_ISL_3041122, EPI_ISL_3041123, EPI_ISL_3041124, EPI_ISL_3041125, EPI_ISL_3041126, EPI_ISL_3041127, EPI_ISL_3041128, EPI_ISL_3041129, EPI_ISL_3041130, EPI_ISL_3041131, EPI_ISL_3041132, EPI_ISL_3041133, EPI_ISL_3041134, EPI_ISL_3041135, EPI_ISL_3041136, EPI_ISL_3041137, EPI_ISL_3041138, EPI_ISL_3041139, EPI_ISL_3041140, EPI_ISL_3041141, EPI_ISL_3041142, EPI_ISL_3041143, EPI_ISL_3041144, EPI_ISL_3041145, EPI_ISL_3041146, EPI_ISL_3041147, | | | |

| | | | |
|---|---|---|---|
| EPI_ISL_3041148, EPI_ISL_3041149, EPI_ISL_3041150, EPI_ISL_3041151, EPI_ISL_3041152, EPI_ISL_3041153, EPI_ISL_3041154, EPI_ISL_3041155, EPI_ISL_3041156, EPI_ISL_3041157, EPI_ISL_3041158, EPI_ISL_3041159, EPI_ISL_3041160, EPI_ISL_3041162, EPI_ISL_3041163, EPI_ISL_3041164, EPI_ISL_3041165, EPI_ISL_3041166, EPI_ISL_3041167, EPI_ISL_3041168, EPI_ISL_3041169, EPI_ISL_3041170, EPI_ISL_3041171, EPI_ISL_3041172, EPI_ISL_3041173, EPI_ISL_3041174, EPI_ISL_3041175, EPI_ISL_3041176, EPI_ISL_3041177, EPI_ISL_3041178, EPI_ISL_3041179, EPI_ISL_3041180, EPI_ISL_3041181, EPI_ISL_3041182, EPI_ISL_3041183, EPI_ISL_3041184, EPI_ISL_3041185, EPI_ISL_3041186, EPI_ISL_3041187, EPI_ISL_3041188, EPI_ISL_3041189, EPI_ISL_3041190, EPI_ISL_3041191, EPI_ISL_3041192, EPI_ISL_3041193, EPI_ISL_3041194, EPI_ISL_3041195, EPI_ISL_3041196, EPI_ISL_3041197, EPI_ISL_3041198, EPI_ISL_3041199, EPI_ISL_3041200, EPI_ISL_3041201, EPI_ISL_3041202, EPI_ISL_3041203, EPI_ISL_3041204, EPI_ISL_3041205, EPI_ISL_3041206, EPI_ISL_3041207, EPI_ISL_3041208, EPI_ISL_3041209, EPI_ISL_3041210, EPI_ISL_3041211, EPI_ISL_3041212, EPI_ISL_3041213, EPI_ISL_3041214, EPI_ISL_3041215, EPI_ISL_3041216, EPI_ISL_3041217, EPI_ISL_3041218, EPI_ISL_3041219, EPI_ISL_3041220, EPI_ISL_3041221, EPI_ISL_3041222, EPI_ISL_3041223, EPI_ISL_3041224, EPI_ISL_3041225, EPI_ISL_3041226, EPI_ISL_3041227, EPI_ISL_3041228, EPI_ISL_3041229, EPI_ISL_3041230, EPI_ISL_3041231, EPI_ISL_3041232, EPI_ISL_3041233, EPI_ISL_3041234, EPI_ISL_3041235, EPI_ISL_3041236, EPI_ISL_3041237, EPI_ISL_3041238, EPI_ISL_3041239, EPI_ISL_3041240, EPI_ISL_3041241, EPI_ISL_3041242, EPI_ISL_3041243, EPI_ISL_3041244, EPI_ISL_3041245, EPI_ISL_3041246, EPI_ISL_3041247, EPI_ISL_3041248, EPI_ISL_3041249, EPI_ISL_3041250, EPI_ISL_3041251, EPI_ISL_3041252, EPI_ISL_3041253, EPI_ISL_3041254, EPI_ISL_3041255, EPI_ISL_3041256, EPI_ISL_3041257, EPI_ISL_3041258, EPI_ISL_3041259, EPI_ISL_3041260, EPI_ISL_3041261, EPI_ISL_3041262, EPI_ISL_3041263, EPI_ISL_3041264, EPI_ISL_3041265, EPI_ISL_3041266, EPI_ISL_3041267, EPI_ISL_3041268, EPI_ISL_3041269, EPI_ISL_3041270, EPI_ISL_3041271, EPI_ISL_3041272, EPI_ISL_3041273, EPI_ISL_3041274, EPI_ISL_3041275, EPI_ISL_3041276, EPI_ISL_3041277, EPI_ISL_3041278, EPI_ISL_3041279, EPI_ISL_3041280, EPI_ISL_3041281, EPI_ISL_3041282, EPI_ISL_3041283, EPI_ISL_3041284, EPI_ISL_3041285, EPI_ISL_3041286, EPI_ISL_3041287, EPI_ISL_3041288, EPI_ISL_3041289, EPI_ISL_3041290, EPI_ISL_3041291, EPI_ISL_3041292, EPI_ISL_3041293, EPI_ISL_3041295, EPI_ISL_3041296, EPI_ISL_3041297, EPI_ISL_3041298, EPI_ISL_3041299, EPI_ISL_3041300, EPI_ISL_3041301, EPI_ISL_3041302, EPI_ISL_3041303, EPI_ISL_3041304, EPI_ISL_3041305, EPI_ISL_3041307, EPI_ISL_3041308, EPI_ISL_3041309, EPI_ISL_3041310, EPI_ISL_3041311, EPI_ISL_3041312, EPI_ISL_3041313, EPI_ISL_3041314, EPI_ISL_3041315, EPI_ISL_3041316, EPI_ISL_3041317, EPI_ISL_3041318, EPI_ISL_3041319, EPI_ISL_3041320, EPI_ISL_3041321, EPI_ISL_3041322, EPI_ISL_3041323, EPI_ISL_3041324, EPI_ISL_3041325, EPI_ISL_3041326, EPI_ISL_3041327, EPI_ISL_3041328, EPI_ISL_3041329, EPI_ISL_3041330, EPI_ISL_3041331, EPI_ISL_3041332, EPI_ISL_3041334, EPI_ISL_3041335, EPI_ISL_3041336, EPI_ISL_3041337, EPI_ISL_3041339, EPI_ISL_3041340, EPI_ISL_3041341, EPI_ISL_3041342, EPI_ISL_3041343, EPI_ISL_3041344, EPI_ISL_3041345, EPI_ISL_3041346, EPI_ISL_3041347, EPI_ISL_3041348, EPI_ISL_3041349, EPI_ISL_3041350, EPI_ISL_3041351, EPI_ISL_3041352, EPI_ISL_3041353, EPI_ISL_3041354, EPI_ISL_3041355, EPI_ISL_3041356, EPI_ISL_3041357, EPI_ISL_3041358, EPI_ISL_3041359, EPI_ISL_3041360, EPI_ISL_3041361, EPI_ISL_3041362, EPI_ISL_3041363, EPI_ISL_3041364, EPI_ISL_3041365, EPI_ISL_3041366, EPI_ISL_3041367, EPI_ISL_3041368, EPI_ISL_3041369, EPI_ISL_3041370, EPI_ISL_3041371, EPI_ISL_3041372, EPI_ISL_3041373, EPI_ISL_3041374, EPI_ISL_3041375, EPI_ISL_3041376, EPI_ISL_3041377, EPI_ISL_3041378, EPI_ISL_3041380, EPI_ISL_3041382, EPI_ISL_3041383, EPI_ISL_3041384, EPI_ISL_3041385, EPI_ISL_3041386, EPI_ISL_3041387, EPI_ISL_3041388, EPI_ISL_3041389, EPI_ISL_3041390, EPI_ISL_3041391, EPI_ISL_3041392, EPI_ISL_3041393, EPI_ISL_3041394, EPI_ISL_3041395, EPI_ISL_3041396, EPI_ISL_3041397, EPI_ISL_3041398, EPI_ISL_3041399, EPI_ISL_3041400, EPI_ISL_3041401, EPI_ISL_3041402, EPI_ISL_3041403, EPI_ISL_3041404, EPI_ISL_3041405, EPI_ISL_3041406, EPI_ISL_3041407, EPI_ISL_3041408, EPI_ISL_3041409, EPI_ISL_3041410, EPI_ISL_3041411, EPI_ISL_3041412, EPI_ISL_3041413, EPI_ISL_3041414, EPI_ISL_3041415, EPI_ISL_3041416, EPI_ISL_3041417, EPI_ISL_3041418, EPI_ISL_3041419, EPI_ISL_3041420, EPI_ISL_3041421, EPI_ISL_3041422, EPI_ISL_3041423, EPI_ISL_3041424, EPI_ISL_3041425, EPI_ISL_3041426, EPI_ISL_3041427, EPI_ISL_3041428, EPI_ISL_3041429, EPI_ISL_3041430, EPI_ISL_3041431, EPI_ISL_3041432, EPI_ISL_3041433, EPI_ISL_3041434, EPI_ISL_3041435, EPI_ISL_3041436, EPI_ISL_3041437, EPI_ISL_3041438, EPI_ISL_3041439, EPI_ISL_3041440, EPI_ISL_3041441, EPI_ISL_3041442, EPI_ISL_3041443, EPI_ISL_3041444, EPI_ISL_3041445, EPI_ISL_3041446, EPI_ISL_3041447, EPI_ISL_3041448, EPI_ISL_3041449, EPI_ISL_3041450, EPI_ISL_3041451, EPI_ISL_3041452, EPI_ISL_3041453, EPI_ISL_3041454, EPI_ISL_3041455, EPI_ISL_3041456, EPI_ISL_3041457, EPI_ISL_3041458, EPI_ISL_3041459 | | | |
| see above | Department of Bacteria, Parasites and Fungi, Statens Serum Institut, Copenhagen, Denmark | Statens Serum Institut Bioinformatics and Microbial Genomics | Danish Covid-19 Genome Consortium |
| EPI_ISL_3041460, EPI_ISL_3041461 | Lighthouse Lab in Glasgow | Wellcome Sanger Institute for the COVID-19 Genomics UK (COG-UK) Consortium | Harper VanSteenhouse, Yumi Kasai, David Gray, Carol Clugston, Anna Dominiczak and Alex Alderton, Roberto Amato, Jeffrey Barrett, Sonia Goncalves, Ewan Harrison, David K. Jackson, Ian Johnston, Dominic Kwiatkowski, Cordelia Langford, John Sillitoe on behalf of the Wellcome Sanger Institute COVID-19 Surveillance Team |
| EPI_ISL_3041462 | Lighthouse Lab in Milton Keynes | Wellcome Sanger Institute for the COVID-19 Genomics UK (COG-UK) Consortium | The Lighthouse Lab in Milton Keynes and Alex Alderton, Roberto Amato, Jeffrey Barrett, Sonia Goncalves, Ewan Harrison, David K. Jackson, Ian Johnston, Dominic Kwiatkowski, Cordelia Langford, John Sillitoe on behalf of the Wellcome Sanger Institute COVID-19 Surveillance Team |
| EPI_ISL_3041463 | Lighthouse Lab in Glasgow | Wellcome Sanger Institute for the COVID-19 Genomics UK (COG-UK) Consortium | Harper VanSteenhouse, Yumi Kasai, David Gray, Carol Clugston, Anna Dominiczak and Alex Alderton, Roberto Amato, Jeffrey Barrett, Sonia Goncalves, Ewan Harrison, David K. Jackson, Ian Johnston, Dominic Kwiatkowski, Cordelia Langford, John Sillitoe on behalf of the Wellcome Sanger Institute COVID-19 Surveillance Team |
| EPI_ISL_3041464 | Lighthouse Lab in Milton Keynes | Wellcome Sanger Institute for the COVID-19 Genomics UK (COG-UK) Consortium | The Lighthouse Lab in Milton Keynes and Alex Alderton, Roberto Amato, Jeffrey Barrett, Sonia Goncalves, Ewan Harrison, David K. Jackson, Ian Johnston, Dominic Kwiatkowski, Cordelia Langford, John Sillitoe on behalf of the Wellcome Sanger Institute COVID-19 Surveillance Team |
| EPI_ISL_3041465 | Lighthouse Lab in Glasgow | Wellcome Sanger Institute for the COVID-19 Genomics UK (COG-UK) Consortium | Harper VanSteenhouse, Yumi Kasai, David Gray, Carol Clugston, Anna Dominiczak and Alex Alderton, Roberto Amato, Jeffrey Barrett, Sonia Goncalves, Ewan Harrison, David K. Jackson, Ian Johnston, Dominic Kwiatkowski, Cordelia Langford, John Sillitoe on behalf of the Wellcome Sanger Institute COVID-19 Surveillance Team |
| EPI_ISL_3041466 | Lighthouse Lab in Milton Keynes | Wellcome Sanger Institute for the COVID-19 Genomics UK (COG-UK) Consortium | The Lighthouse Lab in Milton Keynes and Alex Alderton, Roberto Amato, Jeffrey Barrett, Sonia Goncalves, Ewan Harrison, David K. Jackson, Ian Johnston, Dominic Kwiatkowski, Cordelia Langford, John Sillitoe on behalf of the Wellcome Sanger Institute COVID-19 Surveillance Team |
| EPI_ISL_3041467, EPI_ISL_3041468, EPI_ISL_3041469, EPI_ISL_3041470, EPI_ISL_3041471, EPI_ISL_3041472 | Lighthouse Lab in Glasgow | Wellcome Sanger Institute for the COVID-19 Genomics UK (COG-UK) Consortium | Harper VanSteenhouse, Yumi Kasai, David Gray, Carol Clugston, Anna Dominiczak and Alex Alderton, Roberto Amato, Jeffrey Barrett, Sonia Goncalves, Ewan Harrison, David K. Jackson, Ian Johnston, Dominic Kwiatkowski, Cordelia Langford, John Sillitoe on behalf of the Wellcome Sanger Institute COVID-19 Surveillance Team |
| EPI_ISL_3041473 | Lighthouse Lab in Milton Keynes | Wellcome Sanger Institute for the COVID-19 Genomics UK (COG-UK) Consortium | The Lighthouse Lab in Milton Keynes and Alex Alderton, Roberto Amato, Jeffrey Barrett, Sonia Goncalves, Ewan Harrison, David K. Jackson, Ian Johnston, Dominic Kwiatkowski, Cordelia Langford, John Sillitoe on behalf of the Wellcome Sanger Institute COVID-19 Surveillance Team |
| EPI_ISL_3041474, EPI_ISL_3041475, EPI_ISL_3041476 | Lighthouse Lab in Glasgow | Wellcome Sanger Institute for the COVID-19 Genomics UK (COG-UK) Consortium | Harper VanSteenhouse, Yumi Kasai, David Gray, Carol Clugston, Anna Dominiczak and Alex Alderton, Roberto Amato, Jeffrey Barrett, Sonia Goncalves, Ewan Harrison, David K. Jackson, Ian Johnston, Dominic Kwiatkowski, Cordelia Langford, John Sillitoe on behalf of the Wellcome Sanger Institute COVID-19 Surveillance Team |
| EPI_ISL_3041477 | Lighthouse Lab in Milton Keynes | Wellcome Sanger Institute for the COVID-19 Genomics UK (COG-UK) Consortium | The Lighthouse Lab in Milton Keynes and Alex Alderton, Roberto Amato, Jeffrey Barrett, Sonia Goncalves, Ewan Harrison, David K. Jackson, Ian Johnston, Dominic Kwiatkowski, Cordelia Langford, John Sillitoe on behalf of the Wellcome Sanger Institute COVID-19 Surveillance Team |
| EPI_ISL_3041478, EPI_ISL_3041479 | Lighthouse Lab in Glasgow | Wellcome Sanger Institute for the COVID-19 Genomics UK (COG-UK) Consortium | Harper VanSteenhouse, Yumi Kasai, David Gray, Carol Clugston, Anna Dominiczak and Alex Alderton, Roberto Amato, Jeffrey Barrett, Sonia Goncalves, Ewan Harrison, David K. Jackson, Ian Johnston, Dominic Kwiatkowski, Cordelia Langford, John Sillitoe on behalf of the Wellcome Sanger Institute COVID-19 Surveillance Team |
| EPI_ISL_3041480 | Lighthouse Lab in Milton Keynes | Wellcome Sanger Institute for the COVID-19 Genomics UK (COG-UK) Consortium | The Lighthouse Lab in Milton Keynes and Alex Alderton, Roberto Amato, Jeffrey Barrett, Sonia Goncalves, Ewan Harrison, David K. Jackson, Ian Johnston, Dominic Kwiatkowski, Cordelia Langford, John Sillitoe on behalf of the Wellcome Sanger Institute COVID-19 Surveillance Team |
| EPI_ISL_3041481 | Lighthouse Lab in Glasgow | Wellcome Sanger Institute for the COVID-19 Genomics UK (COG-UK) Consortium | Harper VanSteenhouse, Yumi Kasai, David Gray, Carol Clugston, Anna Dominiczak and Alex Alderton, Roberto Amato, Jeffrey Barrett, Sonia Goncalves, Ewan Harrison, David K. Jackson, Ian Johnston, Dominic Kwiatkowski, Cordelia Langford, John Sillitoe on behalf of the Wellcome Sanger Institute COVID-19 Surveillance Team |
| EPI_ISL_3041482 | Lighthouse Lab in Milton Keynes | Wellcome Sanger Institute for the COVID-19 Genomics UK (COG-UK) Consortium | The Lighthouse Lab in Milton Keynes and Alex Alderton, Roberto Amato, Jeffrey Barrett, Sonia Goncalves, Ewan Harrison, David K. Jackson, Ian Johnston, Dominic Kwiatkowski, Cordelia Langford, John Sillitoe on behalf of the Wellcome Sanger Institute COVID-19 Surveillance Team |
| EPI_ISL_3041483, EPI_ISL_3041484, EPI_ISL_3041485, EPI_ISL_3041486, EPI_ISL_3041487, EPI_ISL_3041488, EPI_ISL_3041489, EPI_ISL_3041490, EPI_ISL_3041491, EPI_ISL_3041492, EPI_ISL_3041493 | | | |
| see above | Lighthouse Lab in Glasgow | Wellcome Sanger Institute for the COVID-19 Genomics UK (COG-UK) Consortium | Harper VanSteenhouse, Yumi Kasai, David Gray, Carol Clugston, Anna Dominiczak and Alex Alderton, Roberto Amato, Jeffrey Barrett, Sonia Goncalves, Ewan Harrison, David K. Jackson, Ian Johnston, Dominic Kwiatkowski, Cordelia Langford, John Sillitoe on behalf of the Wellcome Sanger Institute COVID-19 Surveillance Team |
| EPI_ISL_3041494 | Lighthouse Lab in Milton Keynes | Wellcome Sanger Institute for the COVID-19 Genomics UK (COG-UK) Consortium | The Lighthouse Lab in Milton Keynes and Alex Alderton, Roberto Amato, Jeffrey Barrett, Sonia Goncalves, Ewan Harrison, David K. Jackson, Ian Johnston, Dominic Kwiatkowski, Cordelia Langford, John Sillitoe on behalf of the Wellcome Sanger Institute COVID-19 Surveillance Team |
| EPI_ISL_3041495, EPI_ISL_3041496, EPI_ISL_3041497, EPI_ISL_3041498, EPI_ISL_3041499, EPI_ISL_3041500, EPI_ISL_3041501, EPI_ISL_3041502, EPI_ISL_3041503 | Lighthouse Lab in Glasgow | Wellcome Sanger Institute for the COVID-19 Genomics UK (COG-UK) Consortium | Harper VanSteenhouse, Yumi Kasai, David Gray, Carol Clugston, Anna Dominiczak and Alex Alderton, Roberto Amato, Jeffrey Barrett, Sonia Goncalves, Ewan Harrison, David K. Jackson, Ian Johnston, Dominic Kwiatkowski, Cordelia Langford, John Sillitoe on behalf of the Wellcome Sanger Institute COVID-19 Surveillance Team |
| EPI_ISL_3041504 | Lighthouse Lab in Milton Keynes | Wellcome Sanger Institute for the COVID-19 Genomics UK (COG-UK) Consortium | The Lighthouse Lab in Milton Keynes and Alex Alderton, Roberto Amato, Jeffrey Barrett, Sonia Goncalves, Ewan Harrison, David K. Jackson, Ian Johnston, Dominic Kwiatkowski, Cordelia Langford, John Sillitoe on behalf of the Wellcome Sanger Institute COVID-19 Surveillance Team |
| EPI_ISL_3041505 | Lighthouse Lab in Glasgow | Wellcome Sanger Institute for the COVID-19 Genomics UK (COG-UK) Consortium | Harper VanSteenhouse, Yumi Kasai, David Gray, Carol Clugston, Anna Dominiczak and Alex Alderton, Roberto Amato, Jeffrey Barrett, Sonia Goncalves, Ewan Harrison, David K. Jackson, Ian Johnston, Dominic Kwiatkowski, Cordelia Langford, John Sillitoe on behalf of the Wellcome Sanger Institute COVID-19 Surveillance Team |
| EPI_ISL_3041506 | Lighthouse Lab in Milton Keynes | Wellcome Sanger Institute for the COVID-19 Genomics UK (COG-UK) Consortium | The Lighthouse Lab in Milton Keynes and Alex Alderton, Roberto Amato, Jeffrey Barrett, Sonia Goncalves, Ewan Harrison, David K. Jackson, Ian Johnston, Dominic Kwiatkowski, Cordelia Langford, John Sillitoe on behalf of the Wellcome Sanger Institute COVID-19 Surveillance Team |
| EPI_ISL_3041507 | Lighthouse Lab in Glasgow | Wellcome Sanger Institute for the COVID-19 Genomics UK (COG-UK) Consortium | Harper VanSteenhouse, Yumi Kasai, David Gray, Carol Clugston, Anna Dominiczak and Alex Alderton, Roberto Amato, Jeffrey Barrett, Sonia Goncalves, Ewan Harrison, David K. Jackson, Ian Johnston, Dominic Kwiatkowski, Cordelia Langford, John Sillitoe on behalf of the Wellcome Sanger Institute COVID-19 Surveillance Team |
| EPI_ISL_3041508 | Lighthouse Lab in Milton Keynes | Wellcome Sanger Institute for the COVID-19 Genomics UK (COG-UK) Consortium | The Lighthouse Lab in Milton Keynes and Alex Alderton, Roberto Amato, Jeffrey Barrett, Sonia Goncalves, Ewan Harrison, David K. Jackson, Ian Johnston, Dominic Kwiatkowski, Cordelia Langford, John Sillitoe on behalf of the Wellcome Sanger Institute COVID-19 Surveillance Team |
| EPI_ISL_3041509, EPI_ISL_3041510, EPI_ISL_3041511 | Lighthouse Lab in Glasgow | Wellcome Sanger Institute for the COVID-19 Genomics UK (COG-UK) Consortium | Harper VanSteenhouse, Yumi Kasai, David Gray, Carol Clugston, Anna Dominiczak and Alex Alderton, Roberto Amato, Jeffrey Barrett, Sonia Goncalves, Ewan Harrison, David K. Jackson, Ian Johnston, Dominic Kwiatkowski, Cordelia Langford, John Sillitoe on behalf of the Wellcome Sanger Institute COVID-19 Surveillance Team |
| EPI_ISL_3041512, EPI_ISL_3041513 | Lighthouse Lab in Milton Keynes | Wellcome Sanger Institute for the COVID-19 Genomics UK (COG-UK) Consortium | The Lighthouse Lab in Milton Keynes and Alex Alderton, Roberto Amato, Jeffrey Barrett, Sonia Goncalves, Ewan Harrison, David K. Jackson, Ian Johnston, Dominic Kwiatkowski, Cordelia Langford, John Sillitoe on behalf of the Wellcome Sanger Institute COVID-19 Surveillance Team |
| EPI_ISL_3041514, EPI_ISL_3041515, EPI_ISL_3041516 | Lighthouse Lab in Glasgow | Wellcome Sanger Institute for the COVID-19 Genomics UK (COG-UK) Consortium | Harper VanSteenhouse, Yumi Kasai, David Gray, Carol Clugston, Anna Dominiczak and Alex Alderton, Roberto Amato, Jeffrey Barrett, Sonia Goncalves, Ewan Harrison, David K. Jackson, Ian Johnston, Dominic Kwiatkowski, Cordelia Langford, John Sillitoe on behalf of the Wellcome Sanger Institute COVID-19 Surveillance Team |
| EPI_ISL_3041517 | Lighthouse Lab in Milton Keynes | Wellcome Sanger Institute for the COVID-19 Genomics UK (COG-UK) Consortium | The Lighthouse Lab in Milton Keynes and Alex Alderton, Roberto Amato, Jeffrey Barrett, Sonia Goncalves, Ewan Harrison, David K. Jackson, Ian Johnston, Dominic Kwiatkowski, Cordelia Langford, John Sillitoe on behalf of the Wellcome Sanger Institute COVID-19 Surveillance Team |
| EPI_ISL_3041518 | Lighthouse Lab in Glasgow | Wellcome Sanger Institute for the COVID-19 Genomics UK (COG-UK) Consortium | Harper VanSteenhouse, Yumi Kasai, David Gray, Carol Clugston, Anna Dominiczak and Alex Alderton, Roberto Amato, Jeffrey Barrett, Sonia Goncalves, Ewan Harrison, David K. Jackson, Ian Johnston, Dominic Kwiatkowski, Cordelia Langford, John Sillitoe on behalf of the Wellcome Sanger Institute COVID-19 Surveillance Team |
| EPI_ISL_3041519 | Lighthouse Lab in Milton Keynes | Wellcome Sanger Institute for the COVID-19 Genomics UK (COG-UK) Consortium | The Lighthouse Lab in Milton Keynes and Alex Alderton, Roberto Amato, Jeffrey Barrett, Sonia Goncalves, Ewan Harrison, David K. Jackson, Ian Johnston, Dominic Kwiatkowski, Cordelia Langford, John Sillitoe on behalf of the Wellcome Sanger Institute COVID-19 Surveillance Team |
| EPI_ISL_3041520 | Lighthouse Lab in Glasgow | Wellcome Sanger Institute for the COVID-19 Genomics UK (COG-UK) Consortium | Harper VanSteenhouse, Yumi Kasai, David Gray, Carol Clugston, Anna Dominiczak and Alex Alderton, Roberto Amato, Jeffrey Barrett, Sonia Goncalves, Ewan Harrison, David K. Jackson, Ian Johnston, Dominic Kwiatkowski, Cordelia Langford, John Sillitoe on behalf of the Wellcome Sanger Institute COVID-19 Surveillance Team |
| EPI_ISL_3041521 | Lighthouse Lab in Milton Keynes | Wellcome Sanger Institute for the COVID-19 Genomics UK (COG-UK) Consortium | The Lighthouse Lab in Milton Keynes and Alex Alderton, Roberto Amato, Jeffrey Barrett, Sonia Goncalves, Ewan Harrison, David K. Jackson, Ian Johnston, Dominic Kwiatkowski, Cordelia Langford, John Sillitoe on behalf of the Wellcome Sanger Institute COVID-19 Surveillance Team |
| EPI_ISL_3041522 | Lighthouse Lab in Glasgow | Wellcome Sanger Institute for the COVID-19 Genomics UK (COG-UK) Consortium | Harper VanSteenhouse, Yumi Kasai, David Gray, Carol Clugston, Anna Dominiczak and Alex Alderton, Roberto Amato, Jeffrey Barrett, Sonia Goncalves, Ewan Harrison, David K. Jackson, Ian Johnston, Dominic Kwiatkowski, Cordelia Langford, John Sillitoe on behalf of the Wellcome Sanger Institute COVID-19 Surveillance Team |
| EPI_ISL_3041523 | Lighthouse Lab in Milton Keynes | Wellcome Sanger Institute for the COVID-19 Genomics UK (COG-UK) Consortium | The Lighthouse Lab in Milton Keynes and Alex Alderton, Roberto Amato, Jeffrey Barrett, Sonia Goncalves, Ewan Harrison, David K. Jackson, Ian Johnston, Dominic Kwiatkowski, Cordelia Langford, John Sillitoe on behalf of the Wellcome Sanger Institute COVID-19 Surveillance Team |

| | | | |
|---|---|---|---|
| EPI_ISL_3041524, EPI_ISL_3041525, EPI_ISL_3041526 | Lighthouse Lab in Glasgow | Wellcome Sanger Institute for the COVID-19 Genomics UK (COG-UK) Consortium | Harper VanSteenhouse, Yumi Kasai, David Gray, Carol Clugston, Anna Dominiczak and Alex Alderton, Roberto Amato, Jeffrey Barrett, Sonia Goncalves, Ewan Harrison, David K. Jackson, Ian Johnston, Dominic Kwiatkowski, Cordelia Langford, John Sillitoe on behalf of the Wellcome Sanger Institute COVID-19 Surveillance Team |
| EPI_ISL_3041527 | Lighthouse Lab in Milton Keynes | Wellcome Sanger Institute for the COVID-19 Genomics UK (COG-UK) Consortium | The Lighthouse Lab in Milton Keynes and Alex Alderton, Roberto Amato, Jeffrey Barrett, Sonia Goncalves, Ewan Harrison, David K. Jackson, Ian Johnston, Dominic Kwiatkowski, Cordelia Langford, John Sillitoe on behalf of the Wellcome Sanger Institute COVID-19 Surveillance Team |
| EPI_ISL_3041528, EPI_ISL_3041529 | Lighthouse Lab in Glasgow | Wellcome Sanger Institute for the COVID-19 Genomics UK (COG-UK) Consortium | Harper VanSteenhouse, Yumi Kasai, David Gray, Carol Clugston, Anna Dominiczak and Alex Alderton, Roberto Amato, Jeffrey Barrett, Sonia Goncalves, Ewan Harrison, David K. Jackson, Ian Johnston, Dominic Kwiatkowski, Cordelia Langford, John Sillitoe on behalf of the Wellcome Sanger Institute COVID-19 Surveillance Team |
| EPI_ISL_3041530, EPI_ISL_3041531 | Lighthouse Lab in Milton Keynes | Wellcome Sanger Institute for the COVID-19 Genomics UK (COG-UK) Consortium | The Lighthouse Lab in Milton Keynes and Alex Alderton, Roberto Amato, Jeffrey Barrett, Sonia Goncalves, Ewan Harrison, David K. Jackson, Ian Johnston, Dominic Kwiatkowski, Cordelia Langford, John Sillitoe on behalf of the Wellcome Sanger Institute COVID-19 Surveillance Team |
| EPI_ISL_3041532, EPI_ISL_3041533, EPI_ISL_3041534 | Lighthouse Lab in Glasgow | Wellcome Sanger Institute for the COVID-19 Genomics UK (COG-UK) Consortium | Harper VanSteenhouse, Yumi Kasai, David Gray, Carol Clugston, Anna Dominiczak and Alex Alderton, Roberto Amato, Jeffrey Barrett, Sonia Goncalves, Ewan Harrison, David K. Jackson, Ian Johnston, Dominic Kwiatkowski, Cordelia Langford, John Sillitoe on behalf of the Wellcome Sanger Institute COVID-19 Surveillance Team |
| EPI_ISL_3041535 | Lighthouse Lab in Milton Keynes | Wellcome Sanger Institute for the COVID-19 Genomics UK (COG-UK) Consortium | The Lighthouse Lab in Milton Keynes and Alex Alderton, Roberto Amato, Jeffrey Barrett, Sonia Goncalves, Ewan Harrison, David K. Jackson, Ian Johnston, Dominic Kwiatkowski, Cordelia Langford, John Sillitoe on behalf of the Wellcome Sanger Institute COVID-19 Surveillance Team |
| EPI_ISL_3041536, EPI_ISL_3041537, EPI_ISL_3041538, EPI_ISL_3041539 | Lighthouse Lab in Glasgow | Wellcome Sanger Institute for the COVID-19 Genomics UK (COG-UK) Consortium | Harper VanSteenhouse, Yumi Kasai, David Gray, Carol Clugston, Anna Dominiczak and Alex Alderton, Roberto Amato, Jeffrey Barrett, Sonia Goncalves, Ewan Harrison, David K. Jackson, Ian Johnston, Dominic Kwiatkowski, Cordelia Langford, John Sillitoe on behalf of the Wellcome Sanger Institute COVID-19 Surveillance Team |
| EPI_ISL_3041540 | Lighthouse Lab in Milton Keynes | Wellcome Sanger Institute for the COVID-19 Genomics UK (COG-UK) Consortium | The Lighthouse Lab in Milton Keynes and Alex Alderton, Roberto Amato, Jeffrey Barrett, Sonia Goncalves, Ewan Harrison, David K. Jackson, Ian Johnston, Dominic Kwiatkowski, Cordelia Langford, John Sillitoe on behalf of the Wellcome Sanger Institute COVID-19 Surveillance Team |
| EPI_ISL_3041541, EPI_ISL_3041542 | Lighthouse Lab in Glasgow | Wellcome Sanger Institute for the COVID-19 Genomics UK (COG-UK) Consortium | Harper VanSteenhouse, Yumi Kasai, David Gray, Carol Clugston, Anna Dominiczak and Alex Alderton, Roberto Amato, Jeffrey Barrett, Sonia Goncalves, Ewan Harrison, David K. Jackson, Ian Johnston, Dominic Kwiatkowski, Cordelia Langford, John Sillitoe on behalf of the Wellcome Sanger Institute COVID-19 Surveillance Team |
| EPI_ISL_3041543 | Lighthouse Lab in Milton Keynes | Wellcome Sanger Institute for the COVID-19 Genomics UK (COG-UK) Consortium | The Lighthouse Lab in Milton Keynes and Alex Alderton, Roberto Amato, Jeffrey Barrett, Sonia Goncalves, Ewan Harrison, David K. Jackson, Ian Johnston, Dominic Kwiatkowski, Cordelia Langford, John Sillitoe on behalf of the Wellcome Sanger Institute COVID-19 Surveillance Team |
| EPI_ISL_3041544, EPI_ISL_3041545 | Lighthouse Lab in Glasgow | Wellcome Sanger Institute for the COVID-19 Genomics UK (COG-UK) Consortium | Harper VanSteenhouse, Yumi Kasai, David Gray, Carol Clugston, Anna Dominiczak and Alex Alderton, Roberto Amato, Jeffrey Barrett, Sonia Goncalves, Ewan Harrison, David K. Jackson, Ian Johnston, Dominic Kwiatkowski, Cordelia Langford, John Sillitoe on behalf of the Wellcome Sanger Institute COVID-19 Surveillance Team |
| EPI_ISL_3041546 | Lighthouse Lab in Milton Keynes | Wellcome Sanger Institute for the COVID-19 Genomics UK (COG-UK) Consortium | The Lighthouse Lab in Milton Keynes and Alex Alderton, Roberto Amato, Jeffrey Barrett, Sonia Goncalves, Ewan Harrison, David K. Jackson, Ian Johnston, Dominic Kwiatkowski, Cordelia Langford, John Sillitoe on behalf of the Wellcome Sanger Institute COVID-19 Surveillance Team |
| EPI_ISL_3041547, EPI_ISL_3041548 | Lighthouse Lab in Glasgow | Wellcome Sanger Institute for the COVID-19 Genomics UK (COG-UK) Consortium | Harper VanSteenhouse, Yumi Kasai, David Gray, Carol Clugston, Anna Dominiczak and Alex Alderton, Roberto Amato, Jeffrey Barrett, Sonia Goncalves, Ewan Harrison, David K. Jackson, Ian Johnston, Dominic Kwiatkowski, Cordelia Langford, John Sillitoe on behalf of the Wellcome Sanger Institute COVID-19 Surveillance Team |
| EPI_ISL_3041549 | Lighthouse Lab in Milton Keynes | Wellcome Sanger Institute for the COVID-19 Genomics UK (COG-UK) Consortium | The Lighthouse Lab in Milton Keynes and Alex Alderton, Roberto Amato, Jeffrey Barrett, Sonia Goncalves, Ewan Harrison, David K. Jackson, Ian Johnston, Dominic Kwiatkowski, Cordelia Langford, John Sillitoe on behalf of the Wellcome Sanger Institute COVID-19 Surveillance Team |
| EPI_ISL_3041550 | Lighthouse Lab in Glasgow | Wellcome Sanger Institute for the COVID-19 Genomics UK (COG-UK) Consortium | Harper VanSteenhouse, Yumi Kasai, David Gray, Carol Clugston, Anna Dominiczak and Alex Alderton, Roberto Amato, Jeffrey Barrett, Sonia Goncalves, Ewan Harrison, David K. Jackson, Ian Johnston, Dominic Kwiatkowski, Cordelia Langford, John Sillitoe on behalf of the Wellcome Sanger Institute COVID-19 Surveillance Team |
| EPI_ISL_3041551 | Lighthouse Lab in Milton Keynes | Wellcome Sanger Institute for the COVID-19 Genomics UK (COG-UK) Consortium | The Lighthouse Lab in Milton Keynes and Alex Alderton, Roberto Amato, Jeffrey Barrett, Sonia Goncalves, Ewan Harrison, David K. Jackson, Ian Johnston, Dominic Kwiatkowski, Cordelia Langford, John Sillitoe on behalf of the Wellcome Sanger Institute COVID-19 Surveillance Team |
| EPI_ISL_3041552, EPI_ISL_3041553 | Lighthouse Lab in Glasgow | Wellcome Sanger Institute for the COVID-19 Genomics UK (COG-UK) Consortium | Harper VanSteenhouse, Yumi Kasai, David Gray, Carol Clugston, Anna Dominiczak and Alex Alderton, Roberto Amato, Jeffrey Barrett, Sonia Goncalves, Ewan Harrison, David K. Jackson, Ian Johnston, Dominic Kwiatkowski, Cordelia Langford, John Sillitoe on behalf of the Wellcome Sanger Institute COVID-19 Surveillance Team |
| EPI_ISL_3041554 | Lighthouse Lab in Milton Keynes | Wellcome Sanger Institute for the COVID-19 Genomics UK (COG-UK) Consortium | The Lighthouse Lab in Milton Keynes and Alex Alderton, Roberto Amato, Jeffrey Barrett, Sonia Goncalves, Ewan Harrison, David K. Jackson, Ian Johnston, Dominic Kwiatkowski, Cordelia Langford, John Sillitoe on behalf of the Wellcome Sanger Institute COVID-19 Surveillance Team |
| EPI_ISL_3041555, EPI_ISL_3041556, EPI_ISL_3041557, EPI_ISL_3041558, EPI_ISL_3041559, EPI_ISL_3041560 | Lighthouse Lab in Glasgow | Wellcome Sanger Institute for the COVID-19 Genomics UK (COG-UK) Consortium | Harper VanSteenhouse, Yumi Kasai, David Gray, Carol Clugston, Anna Dominiczak and Alex Alderton, Roberto Amato, Jeffrey Barrett, Sonia Goncalves, Ewan Harrison, David K. Jackson, Ian Johnston, Dominic Kwiatkowski, Cordelia Langford, John Sillitoe on behalf of the Wellcome Sanger Institute COVID-19 Surveillance Team |
| EPI_ISL_3041561, EPI_ISL_3041562 | Lighthouse Lab in Milton Keynes | Wellcome Sanger Institute for the COVID-19 Genomics UK (COG-UK) Consortium | The Lighthouse Lab in Milton Keynes and Alex Alderton, Roberto Amato, Jeffrey Barrett, Sonia Goncalves, Ewan Harrison, David K. Jackson, Ian Johnston, Dominic Kwiatkowski, Cordelia Langford, John Sillitoe on behalf of the Wellcome Sanger Institute COVID-19 Surveillance Team |
| EPI_ISL_3041563 | Lighthouse Lab in Glasgow | Wellcome Sanger Institute for the COVID-19 Genomics UK (COG-UK) Consortium | Harper VanSteenhouse, Yumi Kasai, David Gray, Carol Clugston, Anna Dominiczak and Alex Alderton, Roberto Amato, Jeffrey Barrett, Sonia Goncalves, Ewan Harrison, David K. Jackson, Ian Johnston, Dominic Kwiatkowski, Cordelia Langford, John Sillitoe on behalf of the Wellcome Sanger Institute COVID-19 Surveillance Team |
| EPI_ISL_3041564 | Lighthouse Lab in Milton Keynes | Wellcome Sanger Institute for the COVID-19 Genomics UK (COG-UK) Consortium | The Lighthouse Lab in Milton Keynes and Alex Alderton, Roberto Amato, Jeffrey Barrett, Sonia Goncalves, Ewan Harrison, David K. Jackson, Ian Johnston, Dominic Kwiatkowski, Cordelia Langford, John Sillitoe on behalf of the Wellcome Sanger Institute COVID-19 Surveillance Team |
| EPI_ISL_3041565, EPI_ISL_3041566 | Lighthouse Lab in Glasgow | Wellcome Sanger Institute for the COVID-19 Genomics UK (COG-UK) Consortium | Harper VanSteenhouse, Yumi Kasai, David Gray, Carol Clugston, Anna Dominiczak and Alex Alderton, Roberto Amato, Jeffrey Barrett, Sonia Goncalves, Ewan Harrison, David K. Jackson, Ian Johnston, Dominic Kwiatkowski, Cordelia Langford, John Sillitoe on behalf of the Wellcome Sanger Institute COVID-19 Surveillance Team |
| EPI_ISL_3041567 | Lighthouse Lab in Milton Keynes | Wellcome Sanger Institute for the COVID-19 Genomics UK (COG-UK) Consortium | The Lighthouse Lab in Milton Keynes and Alex Alderton, Roberto Amato, Jeffrey Barrett, Sonia Goncalves, Ewan Harrison, David K. Jackson, Ian Johnston, Dominic Kwiatkowski, Cordelia Langford, John Sillitoe on behalf of the Wellcome Sanger Institute COVID-19 Surveillance Team |
| EPI_ISL_3041568 | Lighthouse Lab in Glasgow | Wellcome Sanger Institute for the COVID-19 Genomics UK (COG-UK) Consortium | Harper VanSteenhouse, Yumi Kasai, David Gray, Carol Clugston, Anna Dominiczak and Alex Alderton, Roberto Amato, Jeffrey Barrett, Sonia Goncalves, Ewan Harrison, David K. Jackson, Ian Johnston, Dominic Kwiatkowski, Cordelia Langford, John Sillitoe on behalf of the Wellcome Sanger Institute COVID-19 Surveillance Team |
| EPI_ISL_3041569 | Lighthouse Lab in Milton Keynes | Wellcome Sanger Institute for the COVID-19 Genomics UK (COG-UK) Consortium | The Lighthouse Lab in Milton Keynes and Alex Alderton, Roberto Amato, Jeffrey Barrett, Sonia Goncalves, Ewan Harrison, David K. Jackson, Ian Johnston, Dominic Kwiatkowski, Cordelia Langford, John Sillitoe on behalf of the Wellcome Sanger Institute COVID-19 Surveillance Team |
| EPI_ISL_3041570, EPI_ISL_3041571, EPI_ISL_3041572, EPI_ISL_3041573, EPI_ISL_3041574, EPI_ISL_3041575, EPI_ISL_3041576 | Lighthouse Lab in Glasgow | Wellcome Sanger Institute for the COVID-19 Genomics UK (COG-UK) Consortium | Harper VanSteenhouse, Yumi Kasai, David Gray, Carol Clugston, Anna Dominiczak and Alex Alderton, Roberto Amato, Jeffrey Barrett, Sonia Goncalves, Ewan Harrison, David K. Jackson, Ian Johnston, Dominic Kwiatkowski, Cordelia Langford, John Sillitoe on behalf of the Wellcome Sanger Institute COVID-19 Surveillance Team |
| EPI_ISL_3041577 | Lighthouse Lab in Milton Keynes | Wellcome Sanger Institute for the COVID-19 Genomics UK (COG-UK) Consortium | The Lighthouse Lab in Milton Keynes and Alex Alderton, Roberto Amato, Jeffrey Barrett, Sonia Goncalves, Ewan Harrison, David K. Jackson, Ian Johnston, Dominic Kwiatkowski, Cordelia Langford, John Sillitoe on behalf of the Wellcome Sanger Institute COVID-19 Surveillance Team |
| EPI_ISL_3041578, EPI_ISL_3041579, EPI_ISL_3041580 | Lighthouse Lab in Glasgow | Wellcome Sanger Institute for the COVID-19 Genomics UK (COG-UK) Consortium | Harper VanSteenhouse, Yumi Kasai, David Gray, Carol Clugston, Anna Dominiczak and Alex Alderton, Roberto Amato, Jeffrey Barrett, Sonia Goncalves, Ewan Harrison, David K. Jackson, Ian Johnston, Dominic Kwiatkowski, Cordelia Langford, John Sillitoe on behalf of the Wellcome Sanger Institute COVID-19 Surveillance Team |
| EPI_ISL_3041581 | Lighthouse Lab in Milton Keynes | Wellcome Sanger Institute for the COVID-19 Genomics UK (COG-UK) Consortium | The Lighthouse Lab in Milton Keynes and Alex Alderton, Roberto Amato, Jeffrey Barrett, Sonia Goncalves, Ewan Harrison, David K. Jackson, Ian Johnston, Dominic Kwiatkowski, Cordelia Langford, John Sillitoe on behalf of the Wellcome Sanger Institute COVID-19 Surveillance Team |
| EPI_ISL_3041582, EPI_ISL_3041583, EPI_ISL_3041584 | Lighthouse Lab in Glasgow | Wellcome Sanger Institute for the COVID-19 Genomics UK (COG-UK) Consortium | Harper VanSteenhouse, Yumi Kasai, David Gray, Carol Clugston, Anna Dominiczak and Alex Alderton, Roberto Amato, Jeffrey Barrett, Sonia Goncalves, Ewan Harrison, David K. Jackson, Ian Johnston, Dominic Kwiatkowski, Cordelia Langford, John Sillitoe on behalf of the Wellcome Sanger Institute COVID-19 Surveillance Team |
| EPI_ISL_3041585, EPI_ISL_3041586 | Lighthouse Lab in Milton Keynes | Wellcome Sanger Institute for the COVID-19 Genomics UK (COG-UK) Consortium | The Lighthouse Lab in Milton Keynes and Alex Alderton, Roberto Amato, Jeffrey Barrett, Sonia Goncalves, Ewan Harrison, David K. Jackson, Ian Johnston, Dominic Kwiatkowski, Cordelia Langford, John Sillitoe on behalf of the Wellcome Sanger Institute COVID-19 Surveillance Team |
| EPI_ISL_3041587 | Lighthouse Lab in Glasgow | Wellcome Sanger Institute for the COVID-19 Genomics UK (COG-UK) Consortium | Harper VanSteenhouse, Yumi Kasai, David Gray, Carol Clugston, Anna Dominiczak and Alex Alderton, Roberto Amato, Jeffrey Barrett, Sonia Goncalves, Ewan Harrison, David K. Jackson, Ian Johnston, Dominic Kwiatkowski, Cordelia Langford, John Sillitoe on behalf of the Wellcome Sanger Institute COVID-19 Surveillance Team |
| EPI_ISL_3041588, EPI_ISL_3041589 | Lighthouse Lab in Milton Keynes | Wellcome Sanger Institute for the COVID-19 Genomics UK (COG-UK) Consortium | The Lighthouse Lab in Milton Keynes and Alex Alderton, Roberto Amato, Jeffrey Barrett, Sonia Goncalves, Ewan Harrison, David K. Jackson, Ian Johnston, Dominic Kwiatkowski, Cordelia Langford, John Sillitoe on behalf of the Wellcome Sanger Institute COVID-19 Surveillance Team |
| EPI_ISL_3041590 | Lighthouse Lab in Glasgow | Wellcome Sanger Institute for the COVID-19 Genomics UK (COG-UK) Consortium | Harper VanSteenhouse, Yumi Kasai, David Gray, Carol Clugston, Anna Dominiczak and Alex Alderton, Roberto Amato, Jeffrey Barrett, Sonia Goncalves, Ewan Harrison, David K. Jackson, Ian Johnston, Dominic Kwiatkowski, Cordelia Langford, John Sillitoe on behalf of the Wellcome Sanger Institute COVID-19 Surveillance Team |
| EPI_ISL_3041591 | Lighthouse Lab in Milton Keynes | Wellcome Sanger Institute for the COVID-19 Genomics UK (COG-UK) Consortium | The Lighthouse Lab in Milton Keynes and Alex Alderton, Roberto Amato, Jeffrey Barrett, Sonia Goncalves, Ewan Harrison, David K. Jackson, Ian Johnston, Dominic Kwiatkowski, Cordelia Langford, John Sillitoe on behalf of the Wellcome Sanger Institute COVID-19 Surveillance Team |
| EPI_ISL_3041592, EPI_ISL_3041593, EPI_ISL_3041594, EPI_ISL_3041595 | Lighthouse Lab in Glasgow | Wellcome Sanger Institute for the COVID-19 Genomics UK (COG-UK) Consortium | Harper VanSteenhouse, Yumi Kasai, David Gray, Carol Clugston, Anna Dominiczak and Alex Alderton, Roberto Amato, Jeffrey Barrett, Sonia Goncalves, Ewan Harrison, David K. Jackson, Ian Johnston, Dominic Kwiatkowski, Cordelia Langford, John Sillitoe on behalf of the Wellcome Sanger Institute COVID-19 Surveillance Team |
| EPI_ISL_3041596, EPI_ISL_3041597 | Lighthouse Lab in Milton Keynes | Wellcome Sanger Institute for the COVID-19 Genomics UK (COG-UK) Consortium | The Lighthouse Lab in Milton Keynes and Alex Alderton, Roberto Amato, Jeffrey Barrett, Sonia Goncalves, Ewan Harrison, David K. Jackson, Ian Johnston, Dominic Kwiatkowski, Cordelia Langford, John Sillitoe on behalf of the Wellcome Sanger Institute COVID-19 Surveillance Team |

| | | | |
|---|---|---|---|
| EPI_ISL_3041598, EPI_ISL_3041599, EPI_ISL_3041600, EPI_ISL_3041601, EPI_ISL_3041602, EPI_ISL_3041603 | Lighthouse Lab in Glasgow | Wellcome Sanger Institute for the COVID-19 Genomics UK (COG-UK) Consortium | Harper VanSteenhouse, Yumi Kasai, David Gray, Carol Clugston, Anna Dominiczak and Alex Alderton, Roberto Amato, Jeffrey Barrett, Sonia Goncalves, Ewan Harrison, David K. Jackson, Ian Johnston, Dominic Kwiatkowski, Cordelia Langford, John Sillitoe on behalf of the Wellcome Sanger Institute COVID-19 Surveillance Team |
| EPI_ISL_3041604 | Lighthouse Lab in Milton Keynes | Wellcome Sanger Institute for the COVID-19 Genomics UK (COG-UK) Consortium | The Lighthouse Lab in Milton Keynes and Alex Alderton, Roberto Amato, Jeffrey Barrett, Sonia Goncalves, Ewan Harrison, David K. Jackson, Ian Johnston, Dominic Kwiatkowski, Cordelia Langford, John Sillitoe on behalf of the Wellcome Sanger Institute COVID-19 Surveillance Team |
| EPI_ISL_3041605, EPI_ISL_3041606, EPI_ISL_3041607, EPI_ISL_3041608, EPI_ISL_3041609, EPI_ISL_3041610 | Lighthouse Lab in Glasgow | Wellcome Sanger Institute for the COVID-19 Genomics UK (COG-UK) Consortium | Harper VanSteenhouse, Yumi Kasai, David Gray, Carol Clugston, Anna Dominiczak and Alex Alderton, Roberto Amato, Jeffrey Barrett, Sonia Goncalves, Ewan Harrison, David K. Jackson, Ian Johnston, Dominic Kwiatkowski, Cordelia Langford, John Sillitoe on behalf of the Wellcome Sanger Institute COVID-19 Surveillance Team |
| EPI_ISL_3041611 | Lighthouse Lab in Milton Keynes | Wellcome Sanger Institute for the COVID-19 Genomics UK (COG-UK) Consortium | The Lighthouse Lab in Milton Keynes and Alex Alderton, Roberto Amato, Jeffrey Barrett, Sonia Goncalves, Ewan Harrison, David K. Jackson, Ian Johnston, Dominic Kwiatkowski, Cordelia Langford, John Sillitoe on behalf of the Wellcome Sanger Institute COVID-19 Surveillance Team |
| EPI_ISL_3041612, EPI_ISL_3041613, EPI_ISL_3041614 | Lighthouse Lab in Glasgow | Wellcome Sanger Institute for the COVID-19 Genomics UK (COG-UK) Consortium | Harper VanSteenhouse, Yumi Kasai, David Gray, Carol Clugston, Anna Dominiczak and Alex Alderton, Roberto Amato, Jeffrey Barrett, Sonia Goncalves, Ewan Harrison, David K. Jackson, Ian Johnston, Dominic Kwiatkowski, Cordelia Langford, John Sillitoe on behalf of the Wellcome Sanger Institute COVID-19 Surveillance Team |
| EPI_ISL_3041615 | Lighthouse Lab in Milton Keynes | Wellcome Sanger Institute for the COVID-19 Genomics UK (COG-UK) Consortium | The Lighthouse Lab in Milton Keynes and Alex Alderton, Roberto Amato, Jeffrey Barrett, Sonia Goncalves, Ewan Harrison, David K. Jackson, Ian Johnston, Dominic Kwiatkowski, Cordelia Langford, John Sillitoe on behalf of the Wellcome Sanger Institute COVID-19 Surveillance Team |
| EPI_ISL_3041616, EPI_ISL_3041617, EPI_ISL_3041618, EPI_ISL_3041619, EPI_ISL_3041620, EPI_ISL_3041621, EPI_ISL_3041622, EPI_ISL_3041623, EPI_ISL_3041624 | Lighthouse Lab in Glasgow | Wellcome Sanger Institute for the COVID-19 Genomics UK (COG-UK) Consortium | Harper VanSteenhouse, Yumi Kasai, David Gray, Carol Clugston, Anna Dominiczak and Alex Alderton, Roberto Amato, Jeffrey Barrett, Sonia Goncalves, Ewan Harrison, David K. Jackson, Ian Johnston, Dominic Kwiatkowski, Cordelia Langford, John Sillitoe on behalf of the Wellcome Sanger Institute COVID-19 Surveillance Team |
| EPI_ISL_3041625 | Lighthouse Lab in Milton Keynes | Wellcome Sanger Institute for the COVID-19 Genomics UK (COG-UK) Consortium | The Lighthouse Lab in Milton Keynes and Alex Alderton, Roberto Amato, Jeffrey Barrett, Sonia Goncalves, Ewan Harrison, David K. Jackson, Ian Johnston, Dominic Kwiatkowski, Cordelia Langford, John Sillitoe on behalf of the Wellcome Sanger Institute COVID-19 Surveillance Team |
| EPI_ISL_3041626, EPI_ISL_3041627, EPI_ISL_3041628, EPI_ISL_3041629, EPI_ISL_3041630 | Lighthouse Lab in Glasgow | Wellcome Sanger Institute for the COVID-19 Genomics UK (COG-UK) Consortium | Harper VanSteenhouse, Yumi Kasai, David Gray, Carol Clugston, Anna Dominiczak and Alex Alderton, Roberto Amato, Jeffrey Barrett, Sonia Goncalves, Ewan Harrison, David K. Jackson, Ian Johnston, Dominic Kwiatkowski, Cordelia Langford, John Sillitoe on behalf of the Wellcome Sanger Institute COVID-19 Surveillance Team |
| EPI_ISL_3041631 | Lighthouse Lab in Milton Keynes | Wellcome Sanger Institute for the COVID-19 Genomics UK (COG-UK) Consortium | The Lighthouse Lab in Milton Keynes and Alex Alderton, Roberto Amato, Jeffrey Barrett, Sonia Goncalves, Ewan Harrison, David K. Jackson, Ian Johnston, Dominic Kwiatkowski, Cordelia Langford, John Sillitoe on behalf of the Wellcome Sanger Institute COVID-19 Surveillance Team |
| EPI_ISL_3041632, EPI_ISL_3041633, EPI_ISL_3041634 | Lighthouse Lab in Glasgow | Wellcome Sanger Institute for the COVID-19 Genomics UK (COG-UK) Consortium | Harper VanSteenhouse, Yumi Kasai, David Gray, Carol Clugston, Anna Dominiczak and Alex Alderton, Roberto Amato, Jeffrey Barrett, Sonia Goncalves, Ewan Harrison, David K. Jackson, Ian Johnston, Dominic Kwiatkowski, Cordelia Langford, John Sillitoe on behalf of the Wellcome Sanger Institute COVID-19 Surveillance Team |
| EPI_ISL_3041635 | Lighthouse Lab in Milton Keynes | Wellcome Sanger Institute for the COVID-19 Genomics UK (COG-UK) Consortium | The Lighthouse Lab in Milton Keynes and Alex Alderton, Roberto Amato, Jeffrey Barrett, Sonia Goncalves, Ewan Harrison, David K. Jackson, Ian Johnston, Dominic Kwiatkowski, Cordelia Langford, John Sillitoe on behalf of the Wellcome Sanger Institute COVID-19 Surveillance Team |
| EPI_ISL_3041636, EPI_ISL_3041637, EPI_ISL_3041638 | Lighthouse Lab in Glasgow | Wellcome Sanger Institute for the COVID-19 Genomics UK (COG-UK) Consortium | Harper VanSteenhouse, Yumi Kasai, David Gray, Carol Clugston, Anna Dominiczak and Alex Alderton, Roberto Amato, Jeffrey Barrett, Sonia Goncalves, Ewan Harrison, David K. Jackson, Ian Johnston, Dominic Kwiatkowski, Cordelia Langford, John Sillitoe on behalf of the Wellcome Sanger Institute COVID-19 Surveillance Team |
| EPI_ISL_3041639 | Lighthouse Lab in Milton Keynes | Wellcome Sanger Institute for the COVID-19 Genomics UK (COG-UK) Consortium | The Lighthouse Lab in Milton Keynes and Alex Alderton, Roberto Amato, Jeffrey Barrett, Sonia Goncalves, Ewan Harrison, David K. Jackson, Ian Johnston, Dominic Kwiatkowski, Cordelia Langford, John Sillitoe on behalf of the Wellcome Sanger Institute COVID-19 Surveillance Team |
| EPI_ISL_3041640, EPI_ISL_3041641, EPI_ISL_3041642, EPI_ISL_3041643, EPI_ISL_3041644, EPI_ISL_3041645, EPI_ISL_3041646 | Lighthouse Lab in Glasgow | Wellcome Sanger Institute for the COVID-19 Genomics UK (COG-UK) Consortium | Harper VanSteenhouse, Yumi Kasai, David Gray, Carol Clugston, Anna Dominiczak and Alex Alderton, Roberto Amato, Jeffrey Barrett, Sonia Goncalves, Ewan Harrison, David K. Jackson, Ian Johnston, Dominic Kwiatkowski, Cordelia Langford, John Sillitoe on behalf of the Wellcome Sanger Institute COVID-19 Surveillance Team |
| EPI_ISL_3041647 | Lighthouse Lab in Milton Keynes | Wellcome Sanger Institute for the COVID-19 Genomics UK (COG-UK) Consortium | The Lighthouse Lab in Milton Keynes and Alex Alderton, Roberto Amato, Jeffrey Barrett, Sonia Goncalves, Ewan Harrison, David K. Jackson, Ian Johnston, Dominic Kwiatkowski, Cordelia Langford, John Sillitoe on behalf of the Wellcome Sanger Institute COVID-19 Surveillance Team |
| EPI_ISL_3041648, EPI_ISL_3041649, EPI_ISL_3041650 | Lighthouse Lab in Glasgow | Wellcome Sanger Institute for the COVID-19 Genomics UK (COG-UK) Consortium | Harper VanSteenhouse, Yumi Kasai, David Gray, Carol Clugston, Anna Dominiczak and Alex Alderton, Roberto Amato, Jeffrey Barrett, Sonia Goncalves, Ewan Harrison, David K. Jackson, Ian Johnston, Dominic Kwiatkowski, Cordelia Langford, John Sillitoe on behalf of the Wellcome Sanger Institute COVID-19 Surveillance Team |
| EPI_ISL_3041651 | Lighthouse Lab in Milton Keynes | Wellcome Sanger Institute for the COVID-19 Genomics UK (COG-UK) Consortium | The Lighthouse Lab in Milton Keynes and Alex Alderton, Roberto Amato, Jeffrey Barrett, Sonia Goncalves, Ewan Harrison, David K. Jackson, Ian Johnston, Dominic Kwiatkowski, Cordelia Langford, John Sillitoe on behalf of the Wellcome Sanger Institute COVID-19 Surveillance Team |
| EPI_ISL_3041652, EPI_ISL_3041653, EPI_ISL_3041654 | Lighthouse Lab in Glasgow | Wellcome Sanger Institute for the COVID-19 Genomics UK (COG-UK) Consortium | Harper VanSteenhouse, Yumi Kasai, David Gray, Carol Clugston, Anna Dominiczak and Alex Alderton, Roberto Amato, Jeffrey Barrett, Sonia Goncalves, Ewan Harrison, David K. Jackson, Ian Johnston, Dominic Kwiatkowski, Cordelia Langford, John Sillitoe on behalf of the Wellcome Sanger Institute COVID-19 Surveillance Team |
| EPI_ISL_3041655 | Lighthouse Lab in Milton Keynes | Wellcome Sanger Institute for the COVID-19 Genomics UK (COG-UK) Consortium | The Lighthouse Lab in Milton Keynes and Alex Alderton, Roberto Amato, Jeffrey Barrett, Sonia Goncalves, Ewan Harrison, David K. Jackson, Ian Johnston, Dominic Kwiatkowski, Cordelia Langford, John Sillitoe on behalf of the Wellcome Sanger Institute COVID-19 Surveillance Team |
| EPI_ISL_3041656, EPI_ISL_3041657, EPI_ISL_3041658, EPI_ISL_3041659 | Lighthouse Lab in Glasgow | Wellcome Sanger Institute for the COVID-19 Genomics UK (COG-UK) Consortium | Harper VanSteenhouse, Yumi Kasai, David Gray, Carol Clugston, Anna Dominiczak and Alex Alderton, Roberto Amato, Jeffrey Barrett, Sonia Goncalves, Ewan Harrison, David K. Jackson, Ian Johnston, Dominic Kwiatkowski, Cordelia Langford, John Sillitoe on behalf of the Wellcome Sanger Institute COVID-19 Surveillance Team |
| EPI_ISL_3041660 | Lighthouse Lab in Milton Keynes | Wellcome Sanger Institute for the COVID-19 Genomics UK (COG-UK) Consortium | The Lighthouse Lab in Milton Keynes and Alex Alderton, Roberto Amato, Jeffrey Barrett, Sonia Goncalves, Ewan Harrison, David K. Jackson, Ian Johnston, Dominic Kwiatkowski, Cordelia Langford, John Sillitoe on behalf of the Wellcome Sanger Institute COVID-19 Surveillance Team |
| EPI_ISL_3041661, EPI_ISL_3041662 | Lighthouse Lab in Glasgow | Wellcome Sanger Institute for the COVID-19 Genomics UK (COG-UK) Consortium | Harper VanSteenhouse, Yumi Kasai, David Gray, Carol Clugston, Anna Dominiczak and Alex Alderton, Roberto Amato, Jeffrey Barrett, Sonia Goncalves, Ewan Harrison, David K. Jackson, Ian Johnston, Dominic Kwiatkowski, Cordelia Langford, John Sillitoe on behalf of the Wellcome Sanger Institute COVID-19 Surveillance Team |
| EPI_ISL_3041663 | Lighthouse Lab in Milton Keynes | Wellcome Sanger Institute for the COVID-19 Genomics UK (COG-UK) Consortium | The Lighthouse Lab in Milton Keynes and Alex Alderton, Roberto Amato, Jeffrey Barrett, Sonia Goncalves, Ewan Harrison, David K. Jackson, Ian Johnston, Dominic Kwiatkowski, Cordelia Langford, John Sillitoe on behalf of the Wellcome Sanger Institute COVID-19 Surveillance Team |
| EPI_ISL_3041664, EPI_ISL_3041665 | Lighthouse Lab in Glasgow | Wellcome Sanger Institute for the COVID-19 Genomics UK (COG-UK) Consortium | Harper VanSteenhouse, Yumi Kasai, David Gray, Carol Clugston, Anna Dominiczak and Alex Alderton, Roberto Amato, Jeffrey Barrett, Sonia Goncalves, Ewan Harrison, David K. Jackson, Ian Johnston, Dominic Kwiatkowski, Cordelia Langford, John Sillitoe on behalf of the Wellcome Sanger Institute COVID-19 Surveillance Team |
| EPI_ISL_3041666 | Lighthouse Lab in Milton Keynes | Wellcome Sanger Institute for the COVID-19 Genomics UK (COG-UK) Consortium | The Lighthouse Lab in Milton Keynes and Alex Alderton, Roberto Amato, Jeffrey Barrett, Sonia Goncalves, Ewan Harrison, David K. Jackson, Ian Johnston, Dominic Kwiatkowski, Cordelia Langford, John Sillitoe on behalf of the Wellcome Sanger Institute COVID-19 Surveillance Team |
| EPI_ISL_3041667, EPI_ISL_3041668, EPI_ISL_3041669 | Lighthouse Lab in Glasgow | Wellcome Sanger Institute for the COVID-19 Genomics UK (COG-UK) Consortium | Harper VanSteenhouse, Yumi Kasai, David Gray, Carol Clugston, Anna Dominiczak and Alex Alderton, Roberto Amato, Jeffrey Barrett, Sonia Goncalves, Ewan Harrison, David K. Jackson, Ian Johnston, Dominic Kwiatkowski, Cordelia Langford, John Sillitoe on behalf of the Wellcome Sanger Institute COVID-19 Surveillance Team |
| EPI_ISL_3041670 | Lighthouse Lab in Milton Keynes | Wellcome Sanger Institute for the COVID-19 Genomics UK (COG-UK) Consortium | The Lighthouse Lab in Milton Keynes and Alex Alderton, Roberto Amato, Jeffrey Barrett, Sonia Goncalves, Ewan Harrison, David K. Jackson, Ian Johnston, Dominic Kwiatkowski, Cordelia Langford, John Sillitoe on behalf of the Wellcome Sanger Institute COVID-19 Surveillance Team |
| EPI_ISL_3041671 | Lighthouse Lab in Glasgow | Wellcome Sanger Institute for the COVID-19 Genomics UK (COG-UK) Consortium | Harper VanSteenhouse, Yumi Kasai, David Gray, Carol Clugston, Anna Dominiczak and Alex Alderton, Roberto Amato, Jeffrey Barrett, Sonia Goncalves, Ewan Harrison, David K. Jackson, Ian Johnston, Dominic Kwiatkowski, Cordelia Langford, John Sillitoe on behalf of the Wellcome Sanger Institute COVID-19 Surveillance Team |
| EPI_ISL_3041672 | Lighthouse Lab in Milton Keynes | Wellcome Sanger Institute for the COVID-19 Genomics UK (COG-UK) Consortium | The Lighthouse Lab in Milton Keynes and Alex Alderton, Roberto Amato, Jeffrey Barrett, Sonia Goncalves, Ewan Harrison, David K. Jackson, Ian Johnston, Dominic Kwiatkowski, Cordelia Langford, John Sillitoe on behalf of the Wellcome Sanger Institute COVID-19 Surveillance Team |
| EPI_ISL_3041673, EPI_ISL_3041674, EPI_ISL_3041675, EPI_ISL_3041676, EPI_ISL_3041677, EPI_ISL_3041678, | Lighthouse Lab in Glasgow | Wellcome Sanger Institute for the COVID-19 Genomics UK (COG-UK) Consortium | Harper VanSteenhouse, Yumi Kasai, David Gray, Carol Clugston, Anna Dominiczak and Alex Alderton, Roberto Amato, Jeffrey Barrett, Sonia Goncalves, Ewan Harrison, David K. Jackson, Ian Johnston, Dominic Kwiatkowski, Cordelia Langford, John Sillitoe on behalf of the Wellcome Sanger Institute COVID-19 Surveillance Team |

| | | | |
|---|---|---|---|
| EPI_ISL_3041679 | | | |
| EPI_ISL_3041680 | Lighthouse Lab in Milton Keynes | Wellcome Sanger Institute for the COVID-19 Genomics UK (COG-UK) Consortium | The Lighthouse Lab in Milton Keynes and Alex Alderton, Roberto Amato, Jeffrey Barrett, Sonia Goncalves, Ewan Harrison, David K. Jackson, Ian Johnston, Dominic Kwiatkowski, Cordelia Langford, John Sillitoe on behalf of the Wellcome Sanger Institute COVID-19 Surveillance Team |
| EPI_ISL_3041681, EPI_ISL_3041682, EPI_ISL_3041683, EPI_ISL_3041684, EPI_ISL_3041685, EPI_ISL_3041686 | Lighthouse Lab in Glasgow | Wellcome Sanger Institute for the COVID-19 Genomics UK (COG-UK) Consortium | Harper VanSteenhouse, Yumi Kasai, David Gray, Carol Clugston, Anna Dominiczak and Alex Alderton, Roberto Amato, Jeffrey Barrett, Sonia Goncalves, Ewan Harrison, David K. Jackson, Ian Johnston, Dominic Kwiatkowski, Cordelia Langford, John Sillitoe on behalf of the Wellcome Sanger Institute COVID-19 Surveillance Team |
| EPI_ISL_3041687 | Lighthouse Lab in Milton Keynes | Wellcome Sanger Institute for the COVID-19 Genomics UK (COG-UK) Consortium | The Lighthouse Lab in Milton Keynes and Alex Alderton, Roberto Amato, Jeffrey Barrett, Sonia Goncalves, Ewan Harrison, David K. Jackson, Ian Johnston, Dominic Kwiatkowski, Cordelia Langford, John Sillitoe on behalf of the Wellcome Sanger Institute COVID-19 Surveillance Team |
| EPI_ISL_3041688 | Lighthouse Lab in Glasgow | Wellcome Sanger Institute for the COVID-19 Genomics UK (COG-UK) Consortium | Harper VanSteenhouse, Yumi Kasai, David Gray, Carol Clugston, Anna Dominiczak and Alex Alderton, Roberto Amato, Jeffrey Barrett, Sonia Goncalves, Ewan Harrison, David K. Jackson, Ian Johnston, Dominic Kwiatkowski, Cordelia Langford, John Sillitoe on behalf of the Wellcome Sanger Institute COVID-19 Surveillance Team |
| EPI_ISL_3041689, EPI_ISL_3041690, EPI_ISL_3041691, EPI_ISL_3041692, EPI_ISL_3041693, EPI_ISL_3041694 | Lighthouse Lab in Milton Keynes | Wellcome Sanger Institute for the COVID-19 Genomics UK (COG-UK) Consortium | The Lighthouse Lab in Milton Keynes and Alex Alderton, Roberto Amato, Jeffrey Barrett, Sonia Goncalves, Ewan Harrison, David K. Jackson, Ian Johnston, Dominic Kwiatkowski, Cordelia Langford, John Sillitoe on behalf of the Wellcome Sanger Institute COVID-19 Surveillance Team |
| EPI_ISL_3041695 | Lighthouse Lab in Glasgow | Wellcome Sanger Institute for the COVID-19 Genomics UK (COG-UK) Consortium | Harper VanSteenhouse, Yumi Kasai, David Gray, Carol Clugston, Anna Dominiczak and Alex Alderton, Roberto Amato, Jeffrey Barrett, Sonia Goncalves, Ewan Harrison, David K. Jackson, Ian Johnston, Dominic Kwiatkowski, Cordelia Langford, John Sillitoe on behalf of the Wellcome Sanger Institute COVID-19 Surveillance Team |
| EPI_ISL_3041696, EPI_ISL_3041697 | Lighthouse Lab in Milton Keynes | Wellcome Sanger Institute for the COVID-19 Genomics UK (COG-UK) Consortium | The Lighthouse Lab in Milton Keynes and Alex Alderton, Roberto Amato, Jeffrey Barrett, Sonia Goncalves, Ewan Harrison, David K. Jackson, Ian Johnston, Dominic Kwiatkowski, Cordelia Langford, John Sillitoe on behalf of the Wellcome Sanger Institute COVID-19 Surveillance Team |
| EPI_ISL_3041698, EPI_ISL_3041699, EPI_ISL_3041700, EPI_ISL_3041701, EPI_ISL_3041702, EPI_ISL_3041703, EPI_ISL_3041704, EPI_ISL_3041705, EPI_ISL_3041706, EPI_ISL_3041707, EPI_ISL_3041708, EPI_ISL_3041709 | | | |
| see above | Lighthouse Lab in Glasgow | Wellcome Sanger Institute for the COVID-19 Genomics UK (COG-UK) Consortium | Harper VanSteenhouse, Yumi Kasai, David Gray, Carol Clugston, Anna Dominiczak and Alex Alderton, Roberto Amato, Jeffrey Barrett, Sonia Goncalves, Ewan Harrison, David K. Jackson, Ian Johnston, Dominic Kwiatkowski, Cordelia Langford, John Sillitoe on behalf of the Wellcome Sanger Institute COVID-19 Surveillance Team |
| EPI_ISL_3041710, EPI_ISL_3041711, EPI_ISL_3041712 | Lighthouse Lab in Milton Keynes | Wellcome Sanger Institute for the COVID-19 Genomics UK (COG-UK) Consortium | The Lighthouse Lab in Milton Keynes and Alex Alderton, Roberto Amato, Jeffrey Barrett, Sonia Goncalves, Ewan Harrison, David K. Jackson, Ian Johnston, Dominic Kwiatkowski, Cordelia Langford, John Sillitoe on behalf of the Wellcome Sanger Institute COVID-19 Surveillance Team |
| EPI_ISL_3041713, EPI_ISL_3041714 | Lighthouse Lab in Glasgow | Wellcome Sanger Institute for the COVID-19 Genomics UK (COG-UK) Consortium | Harper VanSteenhouse, Yumi Kasai, David Gray, Carol Clugston, Anna Dominiczak and Alex Alderton, Roberto Amato, Jeffrey Barrett, Sonia Goncalves, Ewan Harrison, David K. Jackson, Ian Johnston, Dominic Kwiatkowski, Cordelia Langford, John Sillitoe on behalf of the Wellcome Sanger Institute COVID-19 Surveillance Team |
| EPI_ISL_3041715 | Lighthouse Lab in Milton Keynes | Wellcome Sanger Institute for the COVID-19 Genomics UK (COG-UK) Consortium | The Lighthouse Lab in Milton Keynes and Alex Alderton, Roberto Amato, Jeffrey Barrett, Sonia Goncalves, Ewan Harrison, David K. Jackson, Ian Johnston, Dominic Kwiatkowski, Cordelia Langford, John Sillitoe on behalf of the Wellcome Sanger Institute COVID-19 Surveillance Team |
| EPI_ISL_3041716 | Lighthouse Lab in Glasgow | Wellcome Sanger Institute for the COVID-19 Genomics UK (COG-UK) Consortium | Harper VanSteenhouse, Yumi Kasai, David Gray, Carol Clugston, Anna Dominiczak and Alex Alderton, Roberto Amato, Jeffrey Barrett, Sonia Goncalves, Ewan Harrison, David K. Jackson, Ian Johnston, Dominic Kwiatkowski, Cordelia Langford, John Sillitoe on behalf of the Wellcome Sanger Institute COVID-19 Surveillance Team |
| EPI_ISL_3041717, EPI_ISL_3041718 | Lighthouse Lab in Milton Keynes | Wellcome Sanger Institute for the COVID-19 Genomics UK (COG-UK) Consortium | The Lighthouse Lab in Milton Keynes and Alex Alderton, Roberto Amato, Jeffrey Barrett, Sonia Goncalves, Ewan Harrison, David K. Jackson, Ian Johnston, Dominic Kwiatkowski, Cordelia Langford, John Sillitoe on behalf of the Wellcome Sanger Institute COVID-19 Surveillance Team |
| EPI_ISL_3041719, EPI_ISL_3041720, EPI_ISL_3041721 | Lighthouse Lab in Glasgow | Wellcome Sanger Institute for the COVID-19 Genomics UK (COG-UK) Consortium | Harper VanSteenhouse, Yumi Kasai, David Gray, Carol Clugston, Anna Dominiczak and Alex Alderton, Roberto Amato, Jeffrey Barrett, Sonia Goncalves, Ewan Harrison, David K. Jackson, Ian Johnston, Dominic Kwiatkowski, Cordelia Langford, John Sillitoe on behalf of the Wellcome Sanger Institute COVID-19 Surveillance Team |
| EPI_ISL_3041722, EPI_ISL_3041723 | Lighthouse Lab in Milton Keynes | Wellcome Sanger Institute for the COVID-19 Genomics UK (COG-UK) Consortium | The Lighthouse Lab in Milton Keynes and Alex Alderton, Roberto Amato, Jeffrey Barrett, Sonia Goncalves, Ewan Harrison, David K. Jackson, Ian Johnston, Dominic Kwiatkowski, Cordelia Langford, John Sillitoe on behalf of the Wellcome Sanger Institute COVID-19 Surveillance Team |
| EPI_ISL_3041724, EPI_ISL_3041725, EPI_ISL_3041726 | Lighthouse Lab in Glasgow | Wellcome Sanger Institute for the COVID-19 Genomics UK (COG-UK) Consortium | Harper VanSteenhouse, Yumi Kasai, David Gray, Carol Clugston, Anna Dominiczak and Alex Alderton, Roberto Amato, Jeffrey Barrett, Sonia Goncalves, Ewan Harrison, David K. Jackson, Ian Johnston, Dominic Kwiatkowski, Cordelia Langford, John Sillitoe on behalf of the Wellcome Sanger Institute COVID-19 Surveillance Team |
| EPI_ISL_3041727 | Lighthouse Lab in Milton Keynes | Wellcome Sanger Institute for the COVID-19 Genomics UK (COG-UK) Consortium | The Lighthouse Lab in Milton Keynes and Alex Alderton, Roberto Amato, Jeffrey Barrett, Sonia Goncalves, Ewan Harrison, David K. Jackson, Ian Johnston, Dominic Kwiatkowski, Cordelia Langford, John Sillitoe on behalf of the Wellcome Sanger Institute COVID-19 Surveillance Team |
| EPI_ISL_3041728 | Lighthouse Lab in Glasgow | Wellcome Sanger Institute for the COVID-19 Genomics UK (COG-UK) Consortium | Harper VanSteenhouse, Yumi Kasai, David Gray, Carol Clugston, Anna Dominiczak and Alex Alderton, Roberto Amato, Jeffrey Barrett, Sonia Goncalves, Ewan Harrison, David K. Jackson, Ian Johnston, Dominic Kwiatkowski, Cordelia Langford, John Sillitoe on behalf of the Wellcome Sanger Institute COVID-19 Surveillance Team |
| EPI_ISL_3041729 | Lighthouse Lab in Milton Keynes | Wellcome Sanger Institute for the COVID-19 Genomics UK (COG-UK) Consortium | The Lighthouse Lab in Milton Keynes and Alex Alderton, Roberto Amato, Jeffrey Barrett, Sonia Goncalves, Ewan Harrison, David K. Jackson, Ian Johnston, Dominic Kwiatkowski, Cordelia Langford, John Sillitoe on behalf of the Wellcome Sanger Institute COVID-19 Surveillance Team |
| EPI_ISL_3041730, EPI_ISL_3041731, EPI_ISL_3041732, EPI_ISL_3041733, EPI_ISL_3041734, EPI_ISL_3041735, EPI_ISL_3041736 | Lighthouse Lab in Glasgow | Wellcome Sanger Institute for the COVID-19 Genomics UK (COG-UK) Consortium | Harper VanSteenhouse, Yumi Kasai, David Gray, Carol Clugston, Anna Dominiczak and Alex Alderton, Roberto Amato, Jeffrey Barrett, Sonia Goncalves, Ewan Harrison, David K. Jackson, Ian Johnston, Dominic Kwiatkowski, Cordelia Langford, John Sillitoe on behalf of the Wellcome Sanger Institute COVID-19 Surveillance Team |
| EPI_ISL_3041737 | Lighthouse Lab in Milton Keynes | Wellcome Sanger Institute for the COVID-19 Genomics UK (COG-UK) Consortium | The Lighthouse Lab in Milton Keynes and Alex Alderton, Roberto Amato, Jeffrey Barrett, Sonia Goncalves, Ewan Harrison, David K. Jackson, Ian Johnston, Dominic Kwiatkowski, Cordelia Langford, John Sillitoe on behalf of the Wellcome Sanger Institute COVID-19 Surveillance Team |
| EPI_ISL_3041738 | Lighthouse Lab in Glasgow | Wellcome Sanger Institute for the COVID-19 Genomics UK (COG-UK) Consortium | Harper VanSteenhouse, Yumi Kasai, David Gray, Carol Clugston, Anna Dominiczak and Alex Alderton, Roberto Amato, Jeffrey Barrett, Sonia Goncalves, Ewan Harrison, David K. Jackson, Ian Johnston, Dominic Kwiatkowski, Cordelia Langford, John Sillitoe on behalf of the Wellcome Sanger Institute COVID-19 Surveillance Team |
| EPI_ISL_3041739 | Lighthouse Lab in Milton Keynes | Wellcome Sanger Institute for the COVID-19 Genomics UK (COG-UK) Consortium | The Lighthouse Lab in Milton Keynes and Alex Alderton, Roberto Amato, Jeffrey Barrett, Sonia Goncalves, Ewan Harrison, David K. Jackson, Ian Johnston, Dominic Kwiatkowski, Cordelia Langford, John Sillitoe on behalf of the Wellcome Sanger Institute COVID-19 Surveillance Team |
| EPI_ISL_3041740, EPI_ISL_3041741, EPI_ISL_3041742, EPI_ISL_3041743, EPI_ISL_3041744, EPI_ISL_3041745, EPI_ISL_3041746, EPI_ISL_3041747 | Lighthouse Lab in Glasgow | Wellcome Sanger Institute for the COVID-19 Genomics UK (COG-UK) Consortium | Harper VanSteenhouse, Yumi Kasai, David Gray, Carol Clugston, Anna Dominiczak and Alex Alderton, Roberto Amato, Jeffrey Barrett, Sonia Goncalves, Ewan Harrison, David K. Jackson, Ian Johnston, Dominic Kwiatkowski, Cordelia Langford, John Sillitoe on behalf of the Wellcome Sanger Institute COVID-19 Surveillance Team |
| EPI_ISL_3041748 | Lighthouse Lab in Milton Keynes | Wellcome Sanger Institute for the COVID-19 Genomics UK (COG-UK) Consortium | The Lighthouse Lab in Milton Keynes and Alex Alderton, Roberto Amato, Jeffrey Barrett, Sonia Goncalves, Ewan Harrison, David K. Jackson, Ian Johnston, Dominic Kwiatkowski, Cordelia Langford, John Sillitoe on behalf of the Wellcome Sanger Institute COVID-19 Surveillance Team |
| EPI_ISL_3041749, EPI_ISL_3041750 | Lighthouse Lab in Glasgow | Wellcome Sanger Institute for the COVID-19 Genomics UK (COG-UK) Consortium | Harper VanSteenhouse, Yumi Kasai, David Gray, Carol Clugston, Anna Dominiczak and Alex Alderton, Roberto Amato, Jeffrey Barrett, Sonia Goncalves, Ewan Harrison, David K. Jackson, Ian Johnston, Dominic Kwiatkowski, Cordelia Langford, John Sillitoe on behalf of the Wellcome Sanger Institute COVID-19 Surveillance Team |
| EPI_ISL_3041751 | Lighthouse Lab in Milton Keynes | Wellcome Sanger Institute for the COVID-19 Genomics UK (COG-UK) Consortium | The Lighthouse Lab in Milton Keynes and Alex Alderton, Roberto Amato, Jeffrey Barrett, Sonia Goncalves, Ewan Harrison, David K. Jackson, Ian Johnston, Dominic Kwiatkowski, Cordelia Langford, John Sillitoe on behalf of the Wellcome Sanger Institute COVID-19 Surveillance Team |
| EPI_ISL_3041752 | Lighthouse Lab in Glasgow | Wellcome Sanger Institute for the COVID-19 Genomics UK (COG-UK) Consortium | Harper VanSteenhouse, Yumi Kasai, David Gray, Carol Clugston, Anna Dominiczak and Alex Alderton, Roberto Amato, Jeffrey Barrett, Sonia Goncalves, Ewan Harrison, David K. Jackson, Ian Johnston, Dominic Kwiatkowski, Cordelia Langford, John Sillitoe on behalf of the Wellcome Sanger Institute COVID-19 Surveillance Team |
| EPI_ISL_3041753 | Lighthouse Lab in Milton Keynes | Wellcome Sanger Institute for the COVID-19 Genomics UK (COG-UK) Consortium | The Lighthouse Lab in Milton Keynes and Alex Alderton, Roberto Amato, Jeffrey Barrett, Sonia Goncalves, Ewan Harrison, David K. Jackson, Ian Johnston, Dominic Kwiatkowski, Cordelia Langford, John Sillitoe on behalf of the Wellcome Sanger Institute COVID-19 Surveillance Team |
| EPI_ISL_3041754, EPI_ISL_3041755, EPI_ISL_3041756 | Lighthouse Lab in Glasgow | Wellcome Sanger Institute for the COVID-19 Genomics UK (COG-UK) Consortium | Harper VanSteenhouse, Yumi Kasai, David Gray, Carol Clugston, Anna Dominiczak and Alex Alderton, Roberto Amato, Jeffrey Barrett, Sonia Goncalves, Ewan Harrison, David K. Jackson, Ian Johnston, Dominic Kwiatkowski, Cordelia Langford, John Sillitoe on behalf of the Wellcome Sanger Institute COVID-19 Surveillance Team |
| EPI_ISL_3041757, EPI_ISL_3041758 | Lighthouse Lab in Milton Keynes | Wellcome Sanger Institute for the COVID-19 Genomics UK (COG-UK) Consortium | The Lighthouse Lab in Milton Keynes and Alex Alderton, Roberto Amato, Jeffrey Barrett, Sonia Goncalves, Ewan Harrison, David K. Jackson, Ian Johnston, Dominic Kwiatkowski, Cordelia Langford, John Sillitoe on behalf of the Wellcome Sanger Institute COVID-19 Surveillance Team |
| EPI_ISL_3041759, EPI_ISL_3041760 | Lighthouse Lab in Glasgow | Wellcome Sanger Institute for the COVID-19 Genomics UK (COG-UK) Consortium | Harper VanSteenhouse, Yumi Kasai, David Gray, Carol Clugston, Anna Dominiczak and Alex Alderton, Roberto Amato, Jeffrey Barrett, Sonia Goncalves, Ewan Harrison, David K. Jackson, Ian Johnston, Dominic Kwiatkowski, Cordelia Langford, John Sillitoe on behalf of the Wellcome Sanger Institute COVID-19 Surveillance Team |
| EPI_ISL_3041761 | Lighthouse Lab in Milton Keynes | Wellcome Sanger Institute for the COVID-19 Genomics UK (COG-UK) Consortium | The Lighthouse Lab in Milton Keynes and Alex Alderton, Roberto Amato, Jeffrey Barrett, Sonia Goncalves, Ewan Harrison, David K. Jackson, Ian Johnston, Dominic Kwiatkowski, Cordelia Langford, John Sillitoe on behalf of the Wellcome Sanger Institute COVID-19 Surveillance Team |
| EPI_ISL_3041762 | Lighthouse Lab in Glasgow | Wellcome Sanger Institute for the COVID-19 Genomics UK (COG-UK) Consortium | Harper VanSteenhouse, Yumi Kasai, David Gray, Carol Clugston, Anna Dominiczak and Alex Alderton, Roberto Amato, Jeffrey Barrett, Sonia Goncalves, Ewan Harrison, David K. Jackson, Ian Johnston, Dominic Kwiatkowski, Cordelia Langford, John Sillitoe on behalf of the Wellcome Sanger Institute COVID-19 Surveillance Team |
| EPI_ISL_3041763, | Lighthouse Lab in Milton Keynes | Wellcome Sanger Institute for the COVID-19 Genomics UK (COG-UK) | The Lighthouse Lab in Milton Keynes and Alex Alderton, Roberto Amato, Jeffrey Barrett, Sonia Goncalves, Ewan Harrison, David K. Jackson, Ian Johnston, Dominic Kwiatkowski, Cordelia Langford, John Sillitoe on behalf of the Wellcome Sanger |

| | | | |
|---|---|---|---|
| EPI_ISL_3041764 | | Consortium | Institute COVID-19 Surveillance Team |
| EPI_ISL_3041765, EPI_ISL_3041766, EPI_ISL_3041767, EPI_ISL_3041768 | Lighthouse Lab in Glasgow | Wellcome Sanger Institute for the COVID-19 Genomics UK (COG-UK) Consortium | Harper VanSteenhouse, Yumi Kasai, David Gray, Carol Clugston, Anna Dominiczak and Alex Alderton, Roberto Amato, Jeffrey Barrett, Sonia Goncalves, Ewan Harrison, David K. Jackson, Ian Johnston, Dominic Kwiatkowski, Cordelia Langford, John Sillitoe on behalf of the Wellcome Sanger Institute COVID-19 Surveillance Team |
| EPI_ISL_3041769, EPI_ISL_3041770 | Lighthouse Lab in Milton Keynes | Wellcome Sanger Institute for the COVID-19 Genomics UK (COG-UK) Consortium | The Lighthouse Lab in Milton Keynes and Alex Alderton, Roberto Amato, Jeffrey Barrett, Sonia Goncalves, Ewan Harrison, David K. Jackson, Ian Johnston, Dominic Kwiatkowski, Cordelia Langford, John Sillitoe on behalf of the Wellcome Sanger Institute COVID-19 Surveillance Team |
| EPI_ISL_3041771, EPI_ISL_3041772, EPI_ISL_3041773 | Lighthouse Lab in Glasgow | Wellcome Sanger Institute for the COVID-19 Genomics UK (COG-UK) Consortium | Harper VanSteenhouse, Yumi Kasai, David Gray, Carol Clugston, Anna Dominiczak and Alex Alderton, Roberto Amato, Jeffrey Barrett, Sonia Goncalves, Ewan Harrison, David K. Jackson, Ian Johnston, Dominic Kwiatkowski, Cordelia Langford, John Sillitoe on behalf of the Wellcome Sanger Institute COVID-19 Surveillance Team |
| EPI_ISL_3041774 | Lighthouse Lab in Milton Keynes | Wellcome Sanger Institute for the COVID-19 Genomics UK (COG-UK) Consortium | The Lighthouse Lab in Milton Keynes and Alex Alderton, Roberto Amato, Jeffrey Barrett, Sonia Goncalves, Ewan Harrison, David K. Jackson, Ian Johnston, Dominic Kwiatkowski, Cordelia Langford, John Sillitoe on behalf of the Wellcome Sanger Institute COVID-19 Surveillance Team |
| EPI_ISL_3041775, EPI_ISL_3041776 | Randox Laboratories | Wellcome Sanger Institute for the COVID-19 Genomics UK (COG-UK) Consortium | Randox Laboratories and Alex Alderton, Roberto Amato, Jeffrey Barrett, Sonia Goncalves, Ewan Harrison, David K. Jackson, Ian Johnston, Dominic Kwiatkowski, Cordelia Langford, John Sillitoe on behalf of the Wellcome Sanger Institute COVID-19 Surveillance Team |
| EPI_ISL_3041777, EPI_ISL_3041778 | Lighthouse Lab in Milton Keynes | Wellcome Sanger Institute for the COVID-19 Genomics UK (COG-UK) Consortium | The Lighthouse Lab in Milton Keynes and Alex Alderton, Roberto Amato, Jeffrey Barrett, Sonia Goncalves, Ewan Harrison, David K. Jackson, Ian Johnston, Dominic Kwiatkowski, Cordelia Langford, John Sillitoe on behalf of the Wellcome Sanger Institute COVID-19 Surveillance Team |
| EPI_ISL_3041779 | Randox Laboratories | Wellcome Sanger Institute for the COVID-19 Genomics UK (COG-UK) Consortium | Randox Laboratories and Alex Alderton, Roberto Amato, Jeffrey Barrett, Sonia Goncalves, Ewan Harrison, David K. Jackson, Ian Johnston, Dominic Kwiatkowski, Cordelia Langford, John Sillitoe on behalf of the Wellcome Sanger Institute COVID-19 Surveillance Team |
| EPI_ISL_3041780, EPI_ISL_3041781 | Lighthouse Lab in Milton Keynes | Wellcome Sanger Institute for the COVID-19 Genomics UK (COG-UK) Consortium | The Lighthouse Lab in Milton Keynes and Alex Alderton, Roberto Amato, Jeffrey Barrett, Sonia Goncalves, Ewan Harrison, David K. Jackson, Ian Johnston, Dominic Kwiatkowski, Cordelia Langford, John Sillitoe on behalf of the Wellcome Sanger Institute COVID-19 Surveillance Team |
| EPI_ISL_3041782 | Lighthouse Lab in Alderley Park | Wellcome Sanger Institute for the COVID-19 Genomics UK (COG-UK) Consortium | Jacquelyn Wynn, Mairead Hyland, The Lighthouse Lab in Alderley Park and Alex Alderton, Roberto Amato, Jeffrey Barrett, Sonia Goncalves, Ewan Harrison, David K. Jackson, Ian Johnston, Dominic Kwiatkowski, Cordelia Langford, John Sillitoe on behalf of the Wellcome Sanger Institute COVID-19 Surveillance Team |
| EPI_ISL_3041783, EPI_ISL_3041784 | Randox Laboratories | Wellcome Sanger Institute for the COVID-19 Genomics UK (COG-UK) Consortium | Randox Laboratories and Alex Alderton, Roberto Amato, Jeffrey Barrett, Sonia Goncalves, Ewan Harrison, David K. Jackson, Ian Johnston, Dominic Kwiatkowski, Cordelia Langford, John Sillitoe on behalf of the Wellcome Sanger Institute COVID-19 Surveillance Team |
| EPI_ISL_3041785, EPI_ISL_3041786 | Lighthouse Lab in Alderley Park | Wellcome Sanger Institute for the COVID-19 Genomics UK (COG-UK) Consortium | Jacquelyn Wynn, Mairead Hyland, The Lighthouse Lab in Alderley Park and Alex Alderton, Roberto Amato, Jeffrey Barrett, Sonia Goncalves, Ewan Harrison, David K. Jackson, Ian Johnston, Dominic Kwiatkowski, Cordelia Langford, John Sillitoe on behalf of the Wellcome Sanger Institute COVID-19 Surveillance Team |
| EPI_ISL_3041787 | Randox Laboratories | Wellcome Sanger Institute for the COVID-19 Genomics UK (COG-UK) Consortium | Randox Laboratories and Alex Alderton, Roberto Amato, Jeffrey Barrett, Sonia Goncalves, Ewan Harrison, David K. Jackson, Ian Johnston, Dominic Kwiatkowski, Cordelia Langford, John Sillitoe on behalf of the Wellcome Sanger Institute COVID-19 Surveillance Team |
| EPI_ISL_3041788, EPI_ISL_3041789 | Lighthouse Lab in Milton Keynes | Wellcome Sanger Institute for the COVID-19 Genomics UK (COG-UK) Consortium | The Lighthouse Lab in Milton Keynes and Alex Alderton, Roberto Amato, Jeffrey Barrett, Sonia Goncalves, Ewan Harrison, David K. Jackson, Ian Johnston, Dominic Kwiatkowski, Cordelia Langford, John Sillitoe on behalf of the Wellcome Sanger Institute COVID-19 Surveillance Team |
| EPI_ISL_3041790 | Lighthouse Lab in Alderley Park | Wellcome Sanger Institute for the COVID-19 Genomics UK (COG-UK) Consortium | Jacquelyn Wynn, Mairead Hyland, The Lighthouse Lab in Alderley Park and Alex Alderton, Roberto Amato, Jeffrey Barrett, Sonia Goncalves, Ewan Harrison, David K. Jackson, Ian Johnston, Dominic Kwiatkowski, Cordelia Langford, John Sillitoe on behalf of the Wellcome Sanger Institute COVID-19 Surveillance Team |
| EPI_ISL_3041791 | Lighthouse Lab in Milton Keynes | Wellcome Sanger Institute for the COVID-19 Genomics UK (COG-UK) Consortium | The Lighthouse Lab in Milton Keynes and Alex Alderton, Roberto Amato, Jeffrey Barrett, Sonia Goncalves, Ewan Harrison, David K. Jackson, Ian Johnston, Dominic Kwiatkowski, Cordelia Langford, John Sillitoe on behalf of the Wellcome Sanger Institute COVID-19 Surveillance Team |
| EPI_ISL_3041792 | Randox Laboratories | Wellcome Sanger Institute for the COVID-19 Genomics UK (COG-UK) Consortium | Randox Laboratories and Alex Alderton, Roberto Amato, Jeffrey Barrett, Sonia Goncalves, Ewan Harrison, David K. Jackson, Ian Johnston, Dominic Kwiatkowski, Cordelia Langford, John Sillitoe on behalf of the Wellcome Sanger Institute COVID-19 Surveillance Team |
| EPI_ISL_3041793 | Berkshire and Surrey Pathology Services Lighthouse Laboratory | Wellcome Sanger Institute for the COVID-19 Genomics UK (COG-UK) Consortium | Berkshire and Surrey Pathology Services Lighthouse Laboratory and Alex Alderton, Roberto Amato, Jeffrey Barrett, Sonia Goncalves, Ewan Harrison, David K. Jackson, Ian Johnston, Dominic Kwiatkowski, Cordelia Langford, John Sillitoe on behalf of the Wellcome Sanger Institute COVID-19 Surveillance Team |
| EPI_ISL_3041794 | Randox Laboratories | Wellcome Sanger Institute for the COVID-19 Genomics UK (COG-UK) Consortium | Randox Laboratories and Alex Alderton, Roberto Amato, Jeffrey Barrett, Sonia Goncalves, Ewan Harrison, David K. Jackson, Ian Johnston, Dominic Kwiatkowski, Cordelia Langford, John Sillitoe on behalf of the Wellcome Sanger Institute COVID-19 Surveillance Team |
| EPI_ISL_3041795 | Lighthouse Lab in Alderley Park | Wellcome Sanger Institute for the COVID-19 Genomics UK (COG-UK) Consortium | Jacquelyn Wynn, Mairead Hyland, The Lighthouse Lab in Alderley Park and Alex Alderton, Roberto Amato, Jeffrey Barrett, Sonia Goncalves, Ewan Harrison, David K. Jackson, Ian Johnston, Dominic Kwiatkowski, Cordelia Langford, John Sillitoe on behalf of the Wellcome Sanger Institute COVID-19 Surveillance Team |
| EPI_ISL_3041796 | Randox Laboratories | Wellcome Sanger Institute for the COVID-19 Genomics UK (COG-UK) Consortium | Randox Laboratories and Alex Alderton, Roberto Amato, Jeffrey Barrett, Sonia Goncalves, Ewan Harrison, David K. Jackson, Ian Johnston, Dominic Kwiatkowski, Cordelia Langford, John Sillitoe on behalf of the Wellcome Sanger Institute COVID-19 Surveillance Team |
| EPI_ISL_3041797 | Berkshire and Surrey Pathology Services Lighthouse Laboratory | Wellcome Sanger Institute for the COVID-19 Genomics UK (COG-UK) Consortium | Berkshire and Surrey Pathology Services Lighthouse Laboratory and Alex Alderton, Roberto Amato, Jeffrey Barrett, Sonia Goncalves, Ewan Harrison, David K. Jackson, Ian Johnston, Dominic Kwiatkowski, Cordelia Langford, John Sillitoe on behalf of the Wellcome Sanger Institute COVID-19 Surveillance Team |
| EPI_ISL_3041798 | Lighthouse Lab in Milton Keynes | Wellcome Sanger Institute for the COVID-19 Genomics UK (COG-UK) Consortium | The Lighthouse Lab in Milton Keynes and Alex Alderton, Roberto Amato, Jeffrey Barrett, Sonia Goncalves, Ewan Harrison, David K. Jackson, Ian Johnston, Dominic Kwiatkowski, Cordelia Langford, John Sillitoe on behalf of the Wellcome Sanger Institute COVID-19 Surveillance Team |
| EPI_ISL_3041799, EPI_ISL_3041800, EPI_ISL_3041801 | Randox Laboratories | Wellcome Sanger Institute for the COVID-19 Genomics UK (COG-UK) Consortium | Randox Laboratories and Alex Alderton, Roberto Amato, Jeffrey Barrett, Sonia Goncalves, Ewan Harrison, David K. Jackson, Ian Johnston, Dominic Kwiatkowski, Cordelia Langford, John Sillitoe on behalf of the Wellcome Sanger Institute COVID-19 Surveillance Team |
| EPI_ISL_3041802 | Lighthouse Lab in Milton Keynes | Wellcome Sanger Institute for the COVID-19 Genomics UK (COG-UK) Consortium | The Lighthouse Lab in Milton Keynes and Alex Alderton, Roberto Amato, Jeffrey Barrett, Sonia Goncalves, Ewan Harrison, David K. Jackson, Ian Johnston, Dominic Kwiatkowski, Cordelia Langford, John Sillitoe on behalf of the Wellcome Sanger Institute COVID-19 Surveillance Team |
| EPI_ISL_3041803 | Berkshire and Surrey Pathology Services Lighthouse Laboratory | Wellcome Sanger Institute for the COVID-19 Genomics UK (COG-UK) Consortium | Berkshire and Surrey Pathology Services Lighthouse Laboratory and Alex Alderton, Roberto Amato, Jeffrey Barrett, Sonia Goncalves, Ewan Harrison, David K. Jackson, Ian Johnston, Dominic Kwiatkowski, Cordelia Langford, John Sillitoe on behalf of the Wellcome Sanger Institute COVID-19 Surveillance Team |
| EPI_ISL_3041804 | Lighthouse Lab in Alderley Park | Wellcome Sanger Institute for the COVID-19 Genomics UK (COG-UK) Consortium | Jacquelyn Wynn, Mairead Hyland, The Lighthouse Lab in Alderley Park and Alex Alderton, Roberto Amato, Jeffrey Barrett, Sonia Goncalves, Ewan Harrison, David K. Jackson, Ian Johnston, Dominic Kwiatkowski, Cordelia Langford, John Sillitoe on behalf of the Wellcome Sanger Institute COVID-19 Surveillance Team |
| EPI_ISL_3041805, EPI_ISL_3041806 | Randox Laboratories | Wellcome Sanger Institute for the COVID-19 Genomics UK (COG-UK) Consortium | Randox Laboratories and Alex Alderton, Roberto Amato, Jeffrey Barrett, Sonia Goncalves, Ewan Harrison, David K. Jackson, Ian Johnston, Dominic Kwiatkowski, Cordelia Langford, John Sillitoe on behalf of the Wellcome Sanger Institute COVID-19 Surveillance Team |
| EPI_ISL_3041807, EPI_ISL_3041808 | Lighthouse Lab in Milton Keynes | Wellcome Sanger Institute for the COVID-19 Genomics UK (COG-UK) Consortium | The Lighthouse Lab in Milton Keynes and Alex Alderton, Roberto Amato, Jeffrey Barrett, Sonia Goncalves, Ewan Harrison, David K. Jackson, Ian Johnston, Dominic Kwiatkowski, Cordelia Langford, John Sillitoe on behalf of the Wellcome Sanger Institute COVID-19 Surveillance Team |
| EPI_ISL_3041809, EPI_ISL_3041810, EPI_ISL_3041811 | Lighthouse Lab in Alderley Park | Wellcome Sanger Institute for the COVID-19 Genomics UK (COG-UK) Consortium | Jacquelyn Wynn, Mairead Hyland, The Lighthouse Lab in Alderley Park and Alex Alderton, Roberto Amato, Jeffrey Barrett, Sonia Goncalves, Ewan Harrison, David K. Jackson, Ian Johnston, Dominic Kwiatkowski, Cordelia Langford, John Sillitoe on behalf of the Wellcome Sanger Institute COVID-19 Surveillance Team |
| EPI_ISL_3041812 | Lighthouse Lab in Milton Keynes | Wellcome Sanger Institute for the COVID-19 Genomics UK (COG-UK) Consortium | The Lighthouse Lab in Milton Keynes and Alex Alderton, Roberto Amato, Jeffrey Barrett, Sonia Goncalves, Ewan Harrison, David K. Jackson, Ian Johnston, Dominic Kwiatkowski, Cordelia Langford, John Sillitoe on behalf of the Wellcome Sanger Institute COVID-19 Surveillance Team |
| EPI_ISL_3041813 | Berkshire and Surrey Pathology Services Lighthouse Laboratory | Wellcome Sanger Institute for the COVID-19 Genomics UK (COG-UK) Consortium | Berkshire and Surrey Pathology Services Lighthouse Laboratory and Alex Alderton, Roberto Amato, Jeffrey Barrett, Sonia Goncalves, Ewan Harrison, David K. Jackson, Ian Johnston, Dominic Kwiatkowski, Cordelia Langford, John Sillitoe on behalf of the Wellcome Sanger Institute COVID-19 Surveillance Team |
| EPI_ISL_3041814, EPI_ISL_3041815 | Lighthouse Lab in Alderley Park | Wellcome Sanger Institute for the COVID-19 Genomics UK (COG-UK) Consortium | Jacquelyn Wynn, Mairead Hyland, The Lighthouse Lab in Alderley Park and Alex Alderton, Roberto Amato, Jeffrey Barrett, Sonia Goncalves, Ewan Harrison, David K. Jackson, Ian Johnston, Dominic Kwiatkowski, Cordelia Langford, John Sillitoe on behalf of the Wellcome Sanger Institute COVID-19 Surveillance Team |
| EPI_ISL_3041816 | Lighthouse Lab in Milton Keynes | Wellcome Sanger Institute for the COVID-19 Genomics UK (COG-UK) Consortium | The Lighthouse Lab in Milton Keynes and Alex Alderton, Roberto Amato, Jeffrey Barrett, Sonia Goncalves, Ewan Harrison, David K. Jackson, Ian Johnston, Dominic Kwiatkowski, Cordelia Langford, John Sillitoe on behalf of the Wellcome Sanger Institute COVID-19 Surveillance Team |
| EPI_ISL_3041817, EPI_ISL_3041818, EPI_ISL_3041819 | Lighthouse Lab in Alderley Park | Wellcome Sanger Institute for the COVID-19 Genomics UK (COG-UK) Consortium | Jacquelyn Wynn, Mairead Hyland, The Lighthouse Lab in Alderley Park and Alex Alderton, Roberto Amato, Jeffrey Barrett, Sonia Goncalves, Ewan Harrison, David K. Jackson, Ian Johnston, Dominic Kwiatkowski, Cordelia Langford, John Sillitoe on behalf of the Wellcome Sanger Institute COVID-19 Surveillance Team |
| EPI_ISL_3041820, EPI_ISL_3041821, EPI_ISL_3041822 | Lighthouse Lab in Milton Keynes | Wellcome Sanger Institute for the COVID-19 Genomics UK (COG-UK) Consortium | The Lighthouse Lab in Milton Keynes and Alex Alderton, Roberto Amato, Jeffrey Barrett, Sonia Goncalves, Ewan Harrison, David K. Jackson, Ian Johnston, Dominic Kwiatkowski, Cordelia Langford, John Sillitoe on behalf of the Wellcome Sanger Institute COVID-19 Surveillance Team |
| EPI_ISL_3041823 | Lighthouse Lab in Alderley Park | Wellcome Sanger Institute for the COVID-19 Genomics UK (COG-UK) Consortium | Jacquelyn Wynn, Mairead Hyland, The Lighthouse Lab in Alderley Park and Alex Alderton, Roberto Amato, Jeffrey Barrett, Sonia Goncalves, Ewan Harrison, David K. Jackson, Ian Johnston, Dominic Kwiatkowski, Cordelia Langford, John Sillitoe on behalf of the Wellcome Sanger Institute COVID-19 Surveillance Team |
| EPI_ISL_3041824 | Lighthouse Lab in Milton Keynes | Wellcome Sanger Institute for the COVID-19 Genomics UK (COG-UK) Consortium | The Lighthouse Lab in Milton Keynes and Alex Alderton, Roberto Amato, Jeffrey Barrett, Sonia Goncalves, Ewan Harrison, David K. Jackson, Ian Johnston, Dominic Kwiatkowski, Cordelia Langford, John Sillitoe on behalf of the Wellcome Sanger Institute COVID-19 Surveillance Team |
| EPI_ISL_3041825 | Lighthouse Lab in Alderley Park | Wellcome Sanger Institute for the COVID-19 Genomics UK (COG-UK) Consortium | Jacquelyn Wynn, Mairead Hyland, The Lighthouse Lab in Alderley Park and Alex Alderton, Roberto Amato, Jeffrey Barrett, Sonia Goncalves, Ewan Harrison, David K. Jackson, Ian Johnston, Dominic Kwiatkowski, Cordelia Langford, John Sillitoe on behalf of the Wellcome Sanger Institute COVID-19 Surveillance Team |
| EPI_ISL_3041826 | Lighthouse Lab in Milton Keynes | Wellcome Sanger Institute for the COVID-19 Genomics UK (COG-UK) Consortium | The Lighthouse Lab in Milton Keynes and Alex Alderton, Roberto Amato, Jeffrey Barrett, Sonia Goncalves, Ewan Harrison, David K. Jackson, Ian Johnston, Dominic Kwiatkowski, Cordelia Langford, John Sillitoe on behalf of the Wellcome Sanger Institute COVID-19 Surveillance Team |
| EPI_ISL_3041827 | Berkshire and Surrey Pathology Services Lighthouse Laboratory | Wellcome Sanger Institute for the COVID-19 Genomics UK (COG-UK) Consortium | Berkshire and Surrey Pathology Services Lighthouse Laboratory and Alex Alderton, Roberto Amato, Jeffrey Barrett, Sonia Goncalves, Ewan Harrison, David K. Jackson, Ian Johnston, Dominic Kwiatkowski, Cordelia Langford, John Sillitoe on behalf of the Wellcome Sanger Institute COVID-19 Surveillance Team |
| EPI_ISL_3041828 | Randox Laboratories | Wellcome Sanger Institute for the COVID-19 Genomics UK (COG-UK) Consortium | Randox Laboratories and Alex Alderton, Roberto Amato, Jeffrey Barrett, Sonia Goncalves, Ewan Harrison, David K. Jackson, Ian Johnston, Dominic Kwiatkowski, Cordelia Langford, John Sillitoe on behalf of the Wellcome Sanger Institute COVID-19 Surveillance Team |
| EPI_ISL_3041829 | Berkshire and Surrey Pathology Services Lighthouse Laboratory | Wellcome Sanger Institute for the COVID-19 Genomics UK (COG-UK) Consortium | Berkshire and Surrey Pathology Services Lighthouse Laboratory and Alex Alderton, Roberto Amato, Jeffrey Barrett, Sonia Goncalves, Ewan Harrison, David K. Jackson, Ian Johnston, Dominic Kwiatkowski, Cordelia Langford, John Sillitoe on behalf of the Wellcome Sanger Institute COVID-19 Surveillance Team |

| | | | |
|---|---|---|---|
| EPI_ISL_3041830 | Lighthouse Lab in Milton Keynes | Wellcome Sanger Institute for the COVID-19 Genomics UK (COG-UK) Consortium | The Lighthouse Lab in Milton Keynes and Alex Alderton, Roberto Amato, Jeffrey Barrett, Sonia Goncalves, Ewan Harrison, David K. Jackson, Ian Johnston, Dominic Kwiatkowski, Cordelia Langford, John Sillitoe on behalf of the Wellcome Sanger Institute COVID-19 Surveillance Team |
| EPI_ISL_3041831 | Randox Laboratories | Wellcome Sanger Institute for the COVID-19 Genomics UK (COG-UK) Consortium | Randox Laboratories and Alex Alderton, Roberto Amato, Jeffrey Barrett, Sonia Goncalves, Ewan Harrison, David K. Jackson, Ian Johnston, Dominic Kwiatkowski, Cordelia Langford, John Sillitoe on behalf of the Wellcome Sanger Institute COVID-19 Surveillance Team |
| EPI_ISL_3041832 | Lighthouse Lab in Milton Keynes | Wellcome Sanger Institute for the COVID-19 Genomics UK (COG-UK) Consortium | The Lighthouse Lab in Milton Keynes and Alex Alderton, Roberto Amato, Jeffrey Barrett, Sonia Goncalves, Ewan Harrison, David K. Jackson, Ian Johnston, Dominic Kwiatkowski, Cordelia Langford, John Sillitoe on behalf of the Wellcome Sanger Institute COVID-19 Surveillance Team |
| EPI_ISL_3041833 | Lighthouse Lab in Alderley Park | Wellcome Sanger Institute for the COVID-19 Genomics UK (COG-UK) Consortium | Jacquelyn Wynn, Mairead Hyland, The Lighthouse Lab in Alderley Park and Alex Alderton, Roberto Amato, Jeffrey Barrett, Sonia Goncalves, Ewan Harrison, David K. Jackson, Ian Johnston, Dominic Kwiatkowski, Cordelia Langford, John Sillitoe on behalf of the Wellcome Sanger Institute COVID-19 Surveillance Team |
| EPI_ISL_3041834 | Berkshire and Surrey Pathology Services Lighthouse Laboratory | Wellcome Sanger Institute for the COVID-19 Genomics UK (COG-UK) Consortium | Berkshire and Surrey Pathology Services Lighthouse Laboratory and Alex Alderton, Roberto Amato, Jeffrey Barrett, Sonia Goncalves, Ewan Harrison, David K. Jackson, Ian Johnston, Dominic Kwiatkowski, Cordelia Langford, John Sillitoe on behalf of the Wellcome Sanger Institute COVID-19 Surveillance Team |
| EPI_ISL_3041835 | Randox Laboratories | Wellcome Sanger Institute for the COVID-19 Genomics UK (COG-UK) Consortium | Randox Laboratories and Alex Alderton, Roberto Amato, Jeffrey Barrett, Sonia Goncalves, Ewan Harrison, David K. Jackson, Ian Johnston, Dominic Kwiatkowski, Cordelia Langford, John Sillitoe on behalf of the Wellcome Sanger Institute COVID-19 Surveillance Team |
| EPI_ISL_3041836 | Lighthouse Lab in Milton Keynes | Wellcome Sanger Institute for the COVID-19 Genomics UK (COG-UK) Consortium | The Lighthouse Lab in Milton Keynes and Alex Alderton, Roberto Amato, Jeffrey Barrett, Sonia Goncalves, Ewan Harrison, David K. Jackson, Ian Johnston, Dominic Kwiatkowski, Cordelia Langford, John Sillitoe on behalf of the Wellcome Sanger Institute COVID-19 Surveillance Team |
| EPI_ISL_3041837 | Lighthouse Lab in Alderley Park | Wellcome Sanger Institute for the COVID-19 Genomics UK (COG-UK) Consortium | Jacquelyn Wynn, Mairead Hyland, The Lighthouse Lab in Alderley Park and Alex Alderton, Roberto Amato, Jeffrey Barrett, Sonia Goncalves, Ewan Harrison, David K. Jackson, Ian Johnston, Dominic Kwiatkowski, Cordelia Langford, John Sillitoe on behalf of the Wellcome Sanger Institute COVID-19 Surveillance Team |
| EPI_ISL_3041838, EPI_ISL_3041839 | Randox Laboratories | Wellcome Sanger Institute for the COVID-19 Genomics UK (COG-UK) Consortium | Randox Laboratories and Alex Alderton, Roberto Amato, Jeffrey Barrett, Sonia Goncalves, Ewan Harrison, David K. Jackson, Ian Johnston, Dominic Kwiatkowski, Cordelia Langford, John Sillitoe on behalf of the Wellcome Sanger Institute COVID-19 Surveillance Team |
| EPI_ISL_3041840 | Lighthouse Lab in Milton Keynes | Wellcome Sanger Institute for the COVID-19 Genomics UK (COG-UK) Consortium | The Lighthouse Lab in Milton Keynes and Alex Alderton, Roberto Amato, Jeffrey Barrett, Sonia Goncalves, Ewan Harrison, David K. Jackson, Ian Johnston, Dominic Kwiatkowski, Cordelia Langford, John Sillitoe on behalf of the Wellcome Sanger Institute COVID-19 Surveillance Team |
| EPI_ISL_3041841 | Randox Laboratories | Wellcome Sanger Institute for the COVID-19 Genomics UK (COG-UK) Consortium | Randox Laboratories and Alex Alderton, Roberto Amato, Jeffrey Barrett, Sonia Goncalves, Ewan Harrison, David K. Jackson, Ian Johnston, Dominic Kwiatkowski, Cordelia Langford, John Sillitoe on behalf of the Wellcome Sanger Institute COVID-19 Surveillance Team |
| EPI_ISL_3041842 | Lighthouse Lab in Alderley Park | Wellcome Sanger Institute for the COVID-19 Genomics UK (COG-UK) Consortium | Jacquelyn Wynn, Mairead Hyland, The Lighthouse Lab in Alderley Park and Alex Alderton, Roberto Amato, Jeffrey Barrett, Sonia Goncalves, Ewan Harrison, David K. Jackson, Ian Johnston, Dominic Kwiatkowski, Cordelia Langford, John Sillitoe on behalf of the Wellcome Sanger Institute COVID-19 Surveillance Team |
| EPI_ISL_3041843 | Randox Laboratories | Wellcome Sanger Institute for the COVID-19 Genomics UK (COG-UK) Consortium | Randox Laboratories and Alex Alderton, Roberto Amato, Jeffrey Barrett, Sonia Goncalves, Ewan Harrison, David K. Jackson, Ian Johnston, Dominic Kwiatkowski, Cordelia Langford, John Sillitoe on behalf of the Wellcome Sanger Institute COVID-19 Surveillance Team |
| EPI_ISL_3041844 | Lighthouse Lab in Alderley Park | Wellcome Sanger Institute for the COVID-19 Genomics UK (COG-UK) Consortium | Jacquelyn Wynn, Mairead Hyland, The Lighthouse Lab in Alderley Park and Alex Alderton, Roberto Amato, Jeffrey Barrett, Sonia Goncalves, Ewan Harrison, David K. Jackson, Ian Johnston, Dominic Kwiatkowski, Cordelia Langford, John Sillitoe on behalf of the Wellcome Sanger Institute COVID-19 Surveillance Team |
| EPI_ISL_3041845 | Lighthouse Lab in Milton Keynes | Wellcome Sanger Institute for the COVID-19 Genomics UK (COG-UK) Consortium | The Lighthouse Lab in Milton Keynes and Alex Alderton, Roberto Amato, Jeffrey Barrett, Sonia Goncalves, Ewan Harrison, David K. Jackson, Ian Johnston, Dominic Kwiatkowski, Cordelia Langford, John Sillitoe on behalf of the Wellcome Sanger Institute COVID-19 Surveillance Team |
| EPI_ISL_3041846 | Lighthouse Lab in Alderley Park | Wellcome Sanger Institute for the COVID-19 Genomics UK (COG-UK) Consortium | Jacquelyn Wynn, Mairead Hyland, The Lighthouse Lab in Alderley Park and Alex Alderton, Roberto Amato, Jeffrey Barrett, Sonia Goncalves, Ewan Harrison, David K. Jackson, Ian Johnston, Dominic Kwiatkowski, Cordelia Langford, John Sillitoe on behalf of the Wellcome Sanger Institute COVID-19 Surveillance Team |
| EPI_ISL_3041847, EPI_ISL_3041848 | Lighthouse Lab in Milton Keynes | Wellcome Sanger Institute for the COVID-19 Genomics UK (COG-UK) Consortium | The Lighthouse Lab in Milton Keynes and Alex Alderton, Roberto Amato, Jeffrey Barrett, Sonia Goncalves, Ewan Harrison, David K. Jackson, Ian Johnston, Dominic Kwiatkowski, Cordelia Langford, John Sillitoe on behalf of the Wellcome Sanger Institute COVID-19 Surveillance Team |
| EPI_ISL_3041849 | Berkshire and Surrey Pathology Services Lighthouse Laboratory | Wellcome Sanger Institute for the COVID-19 Genomics UK (COG-UK) Consortium | Berkshire and Surrey Pathology Services Lighthouse Laboratory and Alex Alderton, Roberto Amato, Jeffrey Barrett, Sonia Goncalves, Ewan Harrison, David K. Jackson, Ian Johnston, Dominic Kwiatkowski, Cordelia Langford, John Sillitoe on behalf of the Wellcome Sanger Institute COVID-19 Surveillance Team |
| EPI_ISL_3041850 | Randox Laboratories | Wellcome Sanger Institute for the COVID-19 Genomics UK (COG-UK) Consortium | Randox Laboratories and Alex Alderton, Roberto Amato, Jeffrey Barrett, Sonia Goncalves, Ewan Harrison, David K. Jackson, Ian Johnston, Dominic Kwiatkowski, Cordelia Langford, John Sillitoe on behalf of the Wellcome Sanger Institute COVID-19 Surveillance Team |
| EPI_ISL_3041851 | Lighthouse Lab in Alderley Park | Wellcome Sanger Institute for the COVID-19 Genomics UK (COG-UK) Consortium | Jacquelyn Wynn, Mairead Hyland, The Lighthouse Lab in Alderley Park and Alex Alderton, Roberto Amato, Jeffrey Barrett, Sonia Goncalves, Ewan Harrison, David K. Jackson, Ian Johnston, Dominic Kwiatkowski, Cordelia Langford, John Sillitoe on behalf of the Wellcome Sanger Institute COVID-19 Surveillance Team |
| EPI_ISL_3041852, EPI_ISL_3041853 | Randox Laboratories | Wellcome Sanger Institute for the COVID-19 Genomics UK (COG-UK) Consortium | Randox Laboratories and Alex Alderton, Roberto Amato, Jeffrey Barrett, Sonia Goncalves, Ewan Harrison, David K. Jackson, Ian Johnston, Dominic Kwiatkowski, Cordelia Langford, John Sillitoe on behalf of the Wellcome Sanger Institute COVID-19 Surveillance Team |
| EPI_ISL_3041854 | Lighthouse Lab in Milton Keynes | Wellcome Sanger Institute for the COVID-19 Genomics UK (COG-UK) Consortium | The Lighthouse Lab in Milton Keynes and Alex Alderton, Roberto Amato, Jeffrey Barrett, Sonia Goncalves, Ewan Harrison, David K. Jackson, Ian Johnston, Dominic Kwiatkowski, Cordelia Langford, John Sillitoe on behalf of the Wellcome Sanger Institute COVID-19 Surveillance Team |
| EPI_ISL_3041855, EPI_ISL_3041856 | Lighthouse Lab in Alderley Park | Wellcome Sanger Institute for the COVID-19 Genomics UK (COG-UK) Consortium | Jacquelyn Wynn, Mairead Hyland, The Lighthouse Lab in Alderley Park and Alex Alderton, Roberto Amato, Jeffrey Barrett, Sonia Goncalves, Ewan Harrison, David K. Jackson, Ian Johnston, Dominic Kwiatkowski, Cordelia Langford, John Sillitoe on behalf of the Wellcome Sanger Institute COVID-19 Surveillance Team |
| EPI_ISL_3041857 | Lighthouse Lab in Milton Keynes | Wellcome Sanger Institute for the COVID-19 Genomics UK (COG-UK) Consortium | The Lighthouse Lab in Milton Keynes and Alex Alderton, Roberto Amato, Jeffrey Barrett, Sonia Goncalves, Ewan Harrison, David K. Jackson, Ian Johnston, Dominic Kwiatkowski, Cordelia Langford, John Sillitoe on behalf of the Wellcome Sanger Institute COVID-19 Surveillance Team |
| EPI_ISL_3041858, EPI_ISL_3041859 | Lighthouse Lab in Alderley Park | Wellcome Sanger Institute for the COVID-19 Genomics UK (COG-UK) Consortium | Jacquelyn Wynn, Mairead Hyland, The Lighthouse Lab in Alderley Park and Alex Alderton, Roberto Amato, Jeffrey Barrett, Sonia Goncalves, Ewan Harrison, David K. Jackson, Ian Johnston, Dominic Kwiatkowski, Cordelia Langford, John Sillitoe on behalf of the Wellcome Sanger Institute COVID-19 Surveillance Team |
| EPI_ISL_3041860, EPI_ISL_3041861 | Berkshire and Surrey Pathology Services Lighthouse Laboratory | Wellcome Sanger Institute for the COVID-19 Genomics UK (COG-UK) Consortium | Berkshire and Surrey Pathology Services Lighthouse Laboratory and Alex Alderton, Roberto Amato, Jeffrey Barrett, Sonia Goncalves, Ewan Harrison, David K. Jackson, Ian Johnston, Dominic Kwiatkowski, Cordelia Langford, John Sillitoe on behalf of the Wellcome Sanger Institute COVID-19 Surveillance Team |
| EPI_ISL_3041862, EPI_ISL_3041863 | Lighthouse Lab in Milton Keynes | Wellcome Sanger Institute for the COVID-19 Genomics UK (COG-UK) Consortium | The Lighthouse Lab in Milton Keynes and Alex Alderton, Roberto Amato, Jeffrey Barrett, Sonia Goncalves, Ewan Harrison, David K. Jackson, Ian Johnston, Dominic Kwiatkowski, Cordelia Langford, John Sillitoe on behalf of the Wellcome Sanger Institute COVID-19 Surveillance Team |
| EPI_ISL_3041864 | Berkshire and Surrey Pathology Services Lighthouse Laboratory | Wellcome Sanger Institute for the COVID-19 Genomics UK (COG-UK) Consortium | Berkshire and Surrey Pathology Services Lighthouse Laboratory and Alex Alderton, Roberto Amato, Jeffrey Barrett, Sonia Goncalves, Ewan Harrison, David K. Jackson, Ian Johnston, Dominic Kwiatkowski, Cordelia Langford, John Sillitoe on behalf of the Wellcome Sanger Institute COVID-19 Surveillance Team |
| EPI_ISL_3041865 | Randox Laboratories | Wellcome Sanger Institute for the COVID-19 Genomics UK (COG-UK) Consortium | Randox Laboratories and Alex Alderton, Roberto Amato, Jeffrey Barrett, Sonia Goncalves, Ewan Harrison, David K. Jackson, Ian Johnston, Dominic Kwiatkowski, Cordelia Langford, John Sillitoe on behalf of the Wellcome Sanger Institute COVID-19 Surveillance Team |
| EPI_ISL_3041866 | Berkshire and Surrey Pathology Services Lighthouse Laboratory | Wellcome Sanger Institute for the COVID-19 Genomics UK (COG-UK) Consortium | Berkshire and Surrey Pathology Services Lighthouse Laboratory and Alex Alderton, Roberto Amato, Jeffrey Barrett, Sonia Goncalves, Ewan Harrison, David K. Jackson, Ian Johnston, Dominic Kwiatkowski, Cordelia Langford, John Sillitoe on behalf of the Wellcome Sanger Institute COVID-19 Surveillance Team |
| EPI_ISL_3041867 | Lighthouse Lab in Milton Keynes | Wellcome Sanger Institute for the COVID-19 Genomics UK (COG-UK) Consortium | The Lighthouse Lab in Milton Keynes and Alex Alderton, Roberto Amato, Jeffrey Barrett, Sonia Goncalves, Ewan Harrison, David K. Jackson, Ian Johnston, Dominic Kwiatkowski, Cordelia Langford, John Sillitoe on behalf of the Wellcome Sanger Institute COVID-19 Surveillance Team |
| EPI_ISL_3041868 | Randox Laboratories | Wellcome Sanger Institute for the COVID-19 Genomics UK (COG-UK) Consortium | Randox Laboratories and Alex Alderton, Roberto Amato, Jeffrey Barrett, Sonia Goncalves, Ewan Harrison, David K. Jackson, Ian Johnston, Dominic Kwiatkowski, Cordelia Langford, John Sillitoe on behalf of the Wellcome Sanger Institute COVID-19 Surveillance Team |
| EPI_ISL_3041869, EPI_ISL_3041870 | Lighthouse Lab in Alderley Park | Wellcome Sanger Institute for the COVID-19 Genomics UK (COG-UK) Consortium | Jacquelyn Wynn, Mairead Hyland, The Lighthouse Lab in Alderley Park and Alex Alderton, Roberto Amato, Jeffrey Barrett, Sonia Goncalves, Ewan Harrison, David K. Jackson, Ian Johnston, Dominic Kwiatkowski, Cordelia Langford, John Sillitoe on behalf of the Wellcome Sanger Institute COVID-19 Surveillance Team |
| EPI_ISL_3041871 | Lighthouse Lab in Milton Keynes | Wellcome Sanger Institute for the COVID-19 Genomics UK (COG-UK) Consortium | The Lighthouse Lab in Milton Keynes and Alex Alderton, Roberto Amato, Jeffrey Barrett, Sonia Goncalves, Ewan Harrison, David K. Jackson, Ian Johnston, Dominic Kwiatkowski, Cordelia Langford, John Sillitoe on behalf of the Wellcome Sanger Institute COVID-19 Surveillance Team |
| EPI_ISL_3041872 | Berkshire and Surrey Pathology Services Lighthouse Laboratory | Wellcome Sanger Institute for the COVID-19 Genomics UK (COG-UK) Consortium | Berkshire and Surrey Pathology Services Lighthouse Laboratory and Alex Alderton, Roberto Amato, Jeffrey Barrett, Sonia Goncalves, Ewan Harrison, David K. Jackson, Ian Johnston, Dominic Kwiatkowski, Cordelia Langford, John Sillitoe on behalf of the Wellcome Sanger Institute COVID-19 Surveillance Team |
| EPI_ISL_3041873 | Randox Laboratories | Wellcome Sanger Institute for the COVID-19 Genomics UK (COG-UK) Consortium | Randox Laboratories and Alex Alderton, Roberto Amato, Jeffrey Barrett, Sonia Goncalves, Ewan Harrison, David K. Jackson, Ian Johnston, Dominic Kwiatkowski, Cordelia Langford, John Sillitoe on behalf of the Wellcome Sanger Institute COVID-19 Surveillance Team |
| EPI_ISL_3041874, EPI_ISL_3041875 | Lighthouse Lab in Alderley Park | Wellcome Sanger Institute for the COVID-19 Genomics UK (COG-UK) Consortium | Jacquelyn Wynn, Mairead Hyland, The Lighthouse Lab in Alderley Park and Alex Alderton, Roberto Amato, Jeffrey Barrett, Sonia Goncalves, Ewan Harrison, David K. Jackson, Ian Johnston, Dominic Kwiatkowski, Cordelia Langford, John Sillitoe on behalf of the Wellcome Sanger Institute COVID-19 Surveillance Team |
| EPI_ISL_3041876, EPI_ISL_3041877 | Randox Laboratories | Wellcome Sanger Institute for the COVID-19 Genomics UK (COG-UK) Consortium | Randox Laboratories and Alex Alderton, Roberto Amato, Jeffrey Barrett, Sonia Goncalves, Ewan Harrison, David K. Jackson, Ian Johnston, Dominic Kwiatkowski, Cordelia Langford, John Sillitoe on behalf of the Wellcome Sanger Institute COVID-19 Surveillance Team |
| EPI_ISL_3041878 | Lighthouse Lab in Milton Keynes | Wellcome Sanger Institute for the COVID-19 Genomics UK (COG-UK) Consortium | The Lighthouse Lab in Milton Keynes and Alex Alderton, Roberto Amato, Jeffrey Barrett, Sonia Goncalves, Ewan Harrison, David K. Jackson, Ian Johnston, Dominic Kwiatkowski, Cordelia Langford, John Sillitoe on behalf of the Wellcome Sanger Institute COVID-19 Surveillance Team |
| EPI_ISL_3041879 | Lighthouse Lab in Alderley Park | Wellcome Sanger Institute for the COVID-19 Genomics UK (COG-UK) Consortium | Jacquelyn Wynn, Mairead Hyland, The Lighthouse Lab in Alderley Park and Alex Alderton, Roberto Amato, Jeffrey Barrett, Sonia Goncalves, Ewan Harrison, David K. Jackson, Ian Johnston, Dominic Kwiatkowski, Cordelia Langford, John Sillitoe on behalf of the Wellcome Sanger Institute COVID-19 Surveillance Team |
| EPI_ISL_3041880, EPI_ISL_3041881 | Lighthouse Lab in Milton Keynes | Wellcome Sanger Institute for the COVID-19 Genomics UK (COG-UK) Consortium | The Lighthouse Lab in Milton Keynes and Alex Alderton, Roberto Amato, Jeffrey Barrett, Sonia Goncalves, Ewan Harrison, David K. Jackson, Ian Johnston, Dominic Kwiatkowski, Cordelia Langford, John Sillitoe on behalf of the Wellcome Sanger Institute COVID-19 Surveillance Team |
| EPI_ISL_3041882 | Randox Laboratories | Wellcome Sanger Institute for the COVID-19 Genomics UK (COG-UK) Consortium | Randox Laboratories and Alex Alderton, Roberto Amato, Jeffrey Barrett, Sonia Goncalves, Ewan Harrison, David K. Jackson, Ian Johnston, Dominic Kwiatkowski, Cordelia Langford, John Sillitoe on behalf of the Wellcome Sanger Institute COVID-19 Surveillance Team |
| EPI_ISL_3041883 | Lighthouse Lab in Milton Keynes | Wellcome Sanger Institute for the COVID-19 Genomics UK (COG-UK) Consortium | The Lighthouse Lab in Milton Keynes and Alex Alderton, Roberto Amato, Jeffrey Barrett, Sonia Goncalves, Ewan Harrison, David K. Jackson, Ian Johnston, Dominic Kwiatkowski, Cordelia Langford, John Sillitoe on behalf of the Wellcome Sanger Institute COVID-19 Surveillance Team |
| EPI_ISL_3041884 | Randox Laboratories | Wellcome Sanger Institute for the COVID-19 Genomics UK (COG-UK) Consortium | Randox Laboratories and Alex Alderton, Roberto Amato, Jeffrey Barrett, Sonia Goncalves, Ewan Harrison, David K. Jackson, Ian Johnston, Dominic Kwiatkowski, Cordelia Langford, John Sillitoe on behalf of the Wellcome Sanger Institute COVID-19 Surveillance Team |
| EPI_ISL_3041885 | Lighthouse Lab in Milton Keynes | Wellcome Sanger Institute for the COVID-19 Genomics UK (COG-UK) Consortium | The Lighthouse Lab in Milton Keynes and Alex Alderton, Roberto Amato, Jeffrey Barrett, Sonia Goncalves, Ewan Harrison, David K. Jackson, Ian Johnston, Dominic Kwiatkowski, Cordelia Langford, John Sillitoe on behalf of the Wellcome Sanger Institute COVID-19 Surveillance Team |

| | | | |
|---|---|---|---|
| EPI_ISL_3041886 | Lighthouse Lab in Alderley Park | Wellcome Sanger Institute for the COVID-19 Genomics UK (COG-UK) Consortium | Jacquelyn Wynn, Mairead Hyland, The Lighthouse Lab in Alderley Park and Alex Alderton, Roberto Amato, Jeffrey Barrett, Sonia Goncalves, Ewan Harrison, David K. Jackson, Ian Johnston, Dominic Kwiatkowski, Cordelia Langford, John Sillitoe on behalf of the Wellcome Sanger Institute COVID-19 Surveillance Team |
| EPI_ISL_3041887 | Randox Laboratories | Wellcome Sanger Institute for the COVID-19 Genomics UK (COG-UK) Consortium | Randox Laboratories and Alex Alderton, Roberto Amato, Jeffrey Barrett, Sonia Goncalves, Ewan Harrison, David K. Jackson, Ian Johnston, Dominic Kwiatkowski, Cordelia Langford, John Sillitoe on behalf of the Wellcome Sanger Institute COVID-19 Surveillance Team |
| EPI_ISL_3041888, EPI_ISL_3041889 | Lighthouse Lab in Milton Keynes | Wellcome Sanger Institute for the COVID-19 Genomics UK (COG-UK) Consortium | The Lighthouse Lab in Milton Keynes and Alex Alderton, Roberto Amato, Jeffrey Barrett, Sonia Goncalves, Ewan Harrison, David K. Jackson, Ian Johnston, Dominic Kwiatkowski, Cordelia Langford, John Sillitoe on behalf of the Wellcome Sanger Institute COVID-19 Surveillance Team |
| EPI_ISL_3041890 | Randox Laboratories | Wellcome Sanger Institute for the COVID-19 Genomics UK (COG-UK) Consortium | Randox Laboratories and Alex Alderton, Roberto Amato, Jeffrey Barrett, Sonia Goncalves, Ewan Harrison, David K. Jackson, Ian Johnston, Dominic Kwiatkowski, Cordelia Langford, John Sillitoe on behalf of the Wellcome Sanger Institute COVID-19 Surveillance Team |
| EPI_ISL_3041891 | Lighthouse Lab in Milton Keynes | Wellcome Sanger Institute for the COVID-19 Genomics UK (COG-UK) Consortium | The Lighthouse Lab in Milton Keynes and Alex Alderton, Roberto Amato, Jeffrey Barrett, Sonia Goncalves, Ewan Harrison, David K. Jackson, Ian Johnston, Dominic Kwiatkowski, Cordelia Langford, John Sillitoe on behalf of the Wellcome Sanger Institute COVID-19 Surveillance Team |
| EPI_ISL_3041892 | Lighthouse Lab in Alderley Park | Wellcome Sanger Institute for the COVID-19 Genomics UK (COG-UK) Consortium | Jacquelyn Wynn, Mairead Hyland, The Lighthouse Lab in Alderley Park and Alex Alderton, Roberto Amato, Jeffrey Barrett, Sonia Goncalves, Ewan Harrison, David K. Jackson, Ian Johnston, Dominic Kwiatkowski, Cordelia Langford, John Sillitoe on behalf of the Wellcome Sanger Institute COVID-19 Surveillance Team |
| EPI_ISL_3041893 | Randox Laboratories | Wellcome Sanger Institute for the COVID-19 Genomics UK (COG-UK) Consortium | Randox Laboratories and Alex Alderton, Roberto Amato, Jeffrey Barrett, Sonia Goncalves, Ewan Harrison, David K. Jackson, Ian Johnston, Dominic Kwiatkowski, Cordelia Langford, John Sillitoe on behalf of the Wellcome Sanger Institute COVID-19 Surveillance Team |
| EPI_ISL_3041894 | Lighthouse Lab in Milton Keynes | Wellcome Sanger Institute for the COVID-19 Genomics UK (COG-UK) Consortium | The Lighthouse Lab in Milton Keynes and Alex Alderton, Roberto Amato, Jeffrey Barrett, Sonia Goncalves, Ewan Harrison, David K. Jackson, Ian Johnston, Dominic Kwiatkowski, Cordelia Langford, John Sillitoe on behalf of the Wellcome Sanger Institute COVID-19 Surveillance Team |
| EPI_ISL_3041895 | Berkshire and Surrey Pathology Services Lighthouse Laboratory | Wellcome Sanger Institute for the COVID-19 Genomics UK (COG-UK) Consortium | Berkshire and Surrey Pathology Services Lighthouse Laboratory and Alex Alderton, Roberto Amato, Jeffrey Barrett, Sonia Goncalves, Ewan Harrison, David K. Jackson, Ian Johnston, Dominic Kwiatkowski, Cordelia Langford, John Sillitoe on behalf of the Wellcome Sanger Institute COVID-19 Surveillance Team |
| EPI_ISL_3041896 | Lighthouse Lab in Milton Keynes | Wellcome Sanger Institute for the COVID-19 Genomics UK (COG-UK) Consortium | The Lighthouse Lab in Milton Keynes and Alex Alderton, Roberto Amato, Jeffrey Barrett, Sonia Goncalves, Ewan Harrison, David K. Jackson, Ian Johnston, Dominic Kwiatkowski, Cordelia Langford, John Sillitoe on behalf of the Wellcome Sanger Institute COVID-19 Surveillance Team |
| EPI_ISL_3041897, EPI_ISL_3041898 | Randox Laboratories | Wellcome Sanger Institute for the COVID-19 Genomics UK (COG-UK) Consortium | Randox Laboratories and Alex Alderton, Roberto Amato, Jeffrey Barrett, Sonia Goncalves, Ewan Harrison, David K. Jackson, Ian Johnston, Dominic Kwiatkowski, Cordelia Langford, John Sillitoe on behalf of the Wellcome Sanger Institute COVID-19 Surveillance Team |
| EPI_ISL_3041899 | Lighthouse Lab in Milton Keynes | Wellcome Sanger Institute for the COVID-19 Genomics UK (COG-UK) Consortium | The Lighthouse Lab in Milton Keynes and Alex Alderton, Roberto Amato, Jeffrey Barrett, Sonia Goncalves, Ewan Harrison, David K. Jackson, Ian Johnston, Dominic Kwiatkowski, Cordelia Langford, John Sillitoe on behalf of the Wellcome Sanger Institute COVID-19 Surveillance Team |
| EPI_ISL_3041900 | Randox Laboratories | Wellcome Sanger Institute for the COVID-19 Genomics UK (COG-UK) Consortium | Randox Laboratories and Alex Alderton, Roberto Amato, Jeffrey Barrett, Sonia Goncalves, Ewan Harrison, David K. Jackson, Ian Johnston, Dominic Kwiatkowski, Cordelia Langford, John Sillitoe on behalf of the Wellcome Sanger Institute COVID-19 Surveillance Team |
| EPI_ISL_3041901, EPI_ISL_3041902 | Lighthouse Lab in Alderley Park | Wellcome Sanger Institute for the COVID-19 Genomics UK (COG-UK) Consortium | Jacquelyn Wynn, Mairead Hyland, The Lighthouse Lab in Alderley Park and Alex Alderton, Roberto Amato, Jeffrey Barrett, Sonia Goncalves, Ewan Harrison, David K. Jackson, Ian Johnston, Dominic Kwiatkowski, Cordelia Langford, John Sillitoe on behalf of the Wellcome Sanger Institute COVID-19 Surveillance Team |
| EPI_ISL_3041903 | Lighthouse Lab in Milton Keynes | Wellcome Sanger Institute for the COVID-19 Genomics UK (COG-UK) Consortium | The Lighthouse Lab in Milton Keynes and Alex Alderton, Roberto Amato, Jeffrey Barrett, Sonia Goncalves, Ewan Harrison, David K. Jackson, Ian Johnston, Dominic Kwiatkowski, Cordelia Langford, John Sillitoe on behalf of the Wellcome Sanger Institute COVID-19 Surveillance Team |
| EPI_ISL_3041904 | Lighthouse Lab in Alderley Park | Wellcome Sanger Institute for the COVID-19 Genomics UK (COG-UK) Consortium | Jacquelyn Wynn, Mairead Hyland, The Lighthouse Lab in Alderley Park and Alex Alderton, Roberto Amato, Jeffrey Barrett, Sonia Goncalves, Ewan Harrison, David K. Jackson, Ian Johnston, Dominic Kwiatkowski, Cordelia Langford, John Sillitoe on behalf of the Wellcome Sanger Institute COVID-19 Surveillance Team |
| EPI_ISL_3041905, EPI_ISL_3041906 | Lighthouse Lab in Milton Keynes | Wellcome Sanger Institute for the COVID-19 Genomics UK (COG-UK) Consortium | The Lighthouse Lab in Milton Keynes and Alex Alderton, Roberto Amato, Jeffrey Barrett, Sonia Goncalves, Ewan Harrison, David K. Jackson, Ian Johnston, Dominic Kwiatkowski, Cordelia Langford, John Sillitoe on behalf of the Wellcome Sanger Institute COVID-19 Surveillance Team |
| EPI_ISL_3041907 | Randox Laboratories | Wellcome Sanger Institute for the COVID-19 Genomics UK (COG-UK) Consortium | Randox Laboratories and Alex Alderton, Roberto Amato, Jeffrey Barrett, Sonia Goncalves, Ewan Harrison, David K. Jackson, Ian Johnston, Dominic Kwiatkowski, Cordelia Langford, John Sillitoe on behalf of the Wellcome Sanger Institute COVID-19 Surveillance Team |
| EPI_ISL_3041908 | Lighthouse Lab in Milton Keynes | Wellcome Sanger Institute for the COVID-19 Genomics UK (COG-UK) Consortium | The Lighthouse Lab in Milton Keynes and Alex Alderton, Roberto Amato, Jeffrey Barrett, Sonia Goncalves, Ewan Harrison, David K. Jackson, Ian Johnston, Dominic Kwiatkowski, Cordelia Langford, John Sillitoe on behalf of the Wellcome Sanger Institute COVID-19 Surveillance Team |
| EPI_ISL_3041909 | Randox Laboratories | Wellcome Sanger Institute for the COVID-19 Genomics UK (COG-UK) Consortium | Randox Laboratories and Alex Alderton, Roberto Amato, Jeffrey Barrett, Sonia Goncalves, Ewan Harrison, David K. Jackson, Ian Johnston, Dominic Kwiatkowski, Cordelia Langford, John Sillitoe on behalf of the Wellcome Sanger Institute COVID-19 Surveillance Team |
| EPI_ISL_3041910, EPI_ISL_3041911 | Lighthouse Lab in Milton Keynes | Wellcome Sanger Institute for the COVID-19 Genomics UK (COG-UK) Consortium | The Lighthouse Lab in Milton Keynes and Alex Alderton, Roberto Amato, Jeffrey Barrett, Sonia Goncalves, Ewan Harrison, David K. Jackson, Ian Johnston, Dominic Kwiatkowski, Cordelia Langford, John Sillitoe on behalf of the Wellcome Sanger Institute COVID-19 Surveillance Team |
| EPI_ISL_3041912 | Randox Laboratories | Wellcome Sanger Institute for the COVID-19 Genomics UK (COG-UK) Consortium | Randox Laboratories and Alex Alderton, Roberto Amato, Jeffrey Barrett, Sonia Goncalves, Ewan Harrison, David K. Jackson, Ian Johnston, Dominic Kwiatkowski, Cordelia Langford, John Sillitoe on behalf of the Wellcome Sanger Institute COVID-19 Surveillance Team |
| EPI_ISL_3041913, EPI_ISL_3041914 | Berkshire and Surrey Pathology Services Lighthouse Laboratory | Wellcome Sanger Institute for the COVID-19 Genomics UK (COG-UK) Consortium | Berkshire and Surrey Pathology Services Lighthouse Laboratory and Alex Alderton, Roberto Amato, Jeffrey Barrett, Sonia Goncalves, Ewan Harrison, David K. Jackson, Ian Johnston, Dominic Kwiatkowski, Cordelia Langford, John Sillitoe on behalf of the Wellcome Sanger Institute COVID-19 Surveillance Team |
| EPI_ISL_3041915 | Lighthouse Lab in Alderley Park | Wellcome Sanger Institute for the COVID-19 Genomics UK (COG-UK) Consortium | Jacquelyn Wynn, Mairead Hyland, The Lighthouse Lab in Alderley Park and Alex Alderton, Roberto Amato, Jeffrey Barrett, Sonia Goncalves, Ewan Harrison, David K. Jackson, Ian Johnston, Dominic Kwiatkowski, Cordelia Langford, John Sillitoe on behalf of the Wellcome Sanger Institute COVID-19 Surveillance Team |
| EPI_ISL_3041916 | Lighthouse Lab in Milton Keynes | Wellcome Sanger Institute for the COVID-19 Genomics UK (COG-UK) Consortium | The Lighthouse Lab in Milton Keynes and Alex Alderton, Roberto Amato, Jeffrey Barrett, Sonia Goncalves, Ewan Harrison, David K. Jackson, Ian Johnston, Dominic Kwiatkowski, Cordelia Langford, John Sillitoe on behalf of the Wellcome Sanger Institute COVID-19 Surveillance Team |
| EPI_ISL_3041917 | Lighthouse Lab in Alderley Park | Wellcome Sanger Institute for the COVID-19 Genomics UK (COG-UK) Consortium | Jacquelyn Wynn, Mairead Hyland, The Lighthouse Lab in Alderley Park and Alex Alderton, Roberto Amato, Jeffrey Barrett, Sonia Goncalves, Ewan Harrison, David K. Jackson, Ian Johnston, Dominic Kwiatkowski, Cordelia Langford, John Sillitoe on behalf of the Wellcome Sanger Institute COVID-19 Surveillance Team |
| EPI_ISL_3041918 | Lighthouse Lab in Milton Keynes | Wellcome Sanger Institute for the COVID-19 Genomics UK (COG-UK) Consortium | The Lighthouse Lab in Milton Keynes and Alex Alderton, Roberto Amato, Jeffrey Barrett, Sonia Goncalves, Ewan Harrison, David K. Jackson, Ian Johnston, Dominic Kwiatkowski, Cordelia Langford, John Sillitoe on behalf of the Wellcome Sanger Institute COVID-19 Surveillance Team |
| EPI_ISL_3041919, EPI_ISL_3041920 | Randox Laboratories | Wellcome Sanger Institute for the COVID-19 Genomics UK (COG-UK) Consortium | Randox Laboratories and Alex Alderton, Roberto Amato, Jeffrey Barrett, Sonia Goncalves, Ewan Harrison, David K. Jackson, Ian Johnston, Dominic Kwiatkowski, Cordelia Langford, John Sillitoe on behalf of the Wellcome Sanger Institute COVID-19 Surveillance Team |
| EPI_ISL_3041921 | Lighthouse Lab in Milton Keynes | Wellcome Sanger Institute for the COVID-19 Genomics UK (COG-UK) Consortium | The Lighthouse Lab in Milton Keynes and Alex Alderton, Roberto Amato, Jeffrey Barrett, Sonia Goncalves, Ewan Harrison, David K. Jackson, Ian Johnston, Dominic Kwiatkowski, Cordelia Langford, John Sillitoe on behalf of the Wellcome Sanger Institute COVID-19 Surveillance Team |
| EPI_ISL_3041922 | Lighthouse Lab in Alderley Park | Wellcome Sanger Institute for the COVID-19 Genomics UK (COG-UK) Consortium | Jacquelyn Wynn, Mairead Hyland, The Lighthouse Lab in Alderley Park and Alex Alderton, Roberto Amato, Jeffrey Barrett, Sonia Goncalves, Ewan Harrison, David K. Jackson, Ian Johnston, Dominic Kwiatkowski, Cordelia Langford, John Sillitoe on behalf of the Wellcome Sanger Institute COVID-19 Surveillance Team |
| EPI_ISL_3041923 | Randox Laboratories | Wellcome Sanger Institute for the COVID-19 Genomics UK (COG-UK) Consortium | Randox Laboratories and Alex Alderton, Roberto Amato, Jeffrey Barrett, Sonia Goncalves, Ewan Harrison, David K. Jackson, Ian Johnston, Dominic Kwiatkowski, Cordelia Langford, John Sillitoe on behalf of the Wellcome Sanger Institute COVID-19 Surveillance Team |
| EPI_ISL_3041924, EPI_ISL_3041925 | Lighthouse Lab in Alderley Park | Wellcome Sanger Institute for the COVID-19 Genomics UK (COG-UK) Consortium | Jacquelyn Wynn, Mairead Hyland, The Lighthouse Lab in Alderley Park and Alex Alderton, Roberto Amato, Jeffrey Barrett, Sonia Goncalves, Ewan Harrison, David K. Jackson, Ian Johnston, Dominic Kwiatkowski, Cordelia Langford, John Sillitoe on behalf of the Wellcome Sanger Institute COVID-19 Surveillance Team |
| EPI_ISL_3041926, EPI_ISL_3041927 | Randox Laboratories | Wellcome Sanger Institute for the COVID-19 Genomics UK (COG-UK) Consortium | Randox Laboratories and Alex Alderton, Roberto Amato, Jeffrey Barrett, Sonia Goncalves, Ewan Harrison, David K. Jackson, Ian Johnston, Dominic Kwiatkowski, Cordelia Langford, John Sillitoe on behalf of the Wellcome Sanger Institute COVID-19 Surveillance Team |
| EPI_ISL_3041928 | Lighthouse Lab in Alderley Park | Wellcome Sanger Institute for the COVID-19 Genomics UK (COG-UK) Consortium | Jacquelyn Wynn, Mairead Hyland, The Lighthouse Lab in Alderley Park and Alex Alderton, Roberto Amato, Jeffrey Barrett, Sonia Goncalves, Ewan Harrison, David K. Jackson, Ian Johnston, Dominic Kwiatkowski, Cordelia Langford, John Sillitoe on behalf of the Wellcome Sanger Institute COVID-19 Surveillance Team |
| EPI_ISL_3041929 | Berkshire and Surrey Pathology Services Lighthouse Laboratory | Wellcome Sanger Institute for the COVID-19 Genomics UK (COG-UK) Consortium | Berkshire and Surrey Pathology Services Lighthouse Laboratory and Alex Alderton, Roberto Amato, Jeffrey Barrett, Sonia Goncalves, Ewan Harrison, David K. Jackson, Ian Johnston, Dominic Kwiatkowski, Cordelia Langford, John Sillitoe on behalf of the Wellcome Sanger Institute COVID-19 Surveillance Team |
| EPI_ISL_3041930, EPI_ISL_3041931, EPI_ISL_3041932 | Lighthouse Lab in Alderley Park | Wellcome Sanger Institute for the COVID-19 Genomics UK (COG-UK) Consortium | Jacquelyn Wynn, Mairead Hyland, The Lighthouse Lab in Alderley Park and Alex Alderton, Roberto Amato, Jeffrey Barrett, Sonia Goncalves, Ewan Harrison, David K. Jackson, Ian Johnston, Dominic Kwiatkowski, Cordelia Langford, John Sillitoe on behalf of the Wellcome Sanger Institute COVID-19 Surveillance Team |
| EPI_ISL_3041933 | Randox Laboratories | Wellcome Sanger Institute for the COVID-19 Genomics UK (COG-UK) Consortium | Randox Laboratories and Alex Alderton, Roberto Amato, Jeffrey Barrett, Sonia Goncalves, Ewan Harrison, David K. Jackson, Ian Johnston, Dominic Kwiatkowski, Cordelia Langford, John Sillitoe on behalf of the Wellcome Sanger Institute COVID-19 Surveillance Team |
| EPI_ISL_3041934 | Lighthouse Lab in Alderley Park | Wellcome Sanger Institute for the COVID-19 Genomics UK (COG-UK) Consortium | Jacquelyn Wynn, Mairead Hyland, The Lighthouse Lab in Alderley Park and Alex Alderton, Roberto Amato, Jeffrey Barrett, Sonia Goncalves, Ewan Harrison, David K. Jackson, Ian Johnston, Dominic Kwiatkowski, Cordelia Langford, John Sillitoe on behalf of the Wellcome Sanger Institute COVID-19 Surveillance Team |
| EPI_ISL_3041935 | Randox Laboratories | Wellcome Sanger Institute for the COVID-19 Genomics UK (COG-UK) Consortium | Randox Laboratories and Alex Alderton, Roberto Amato, Jeffrey Barrett, Sonia Goncalves, Ewan Harrison, David K. Jackson, Ian Johnston, Dominic Kwiatkowski, Cordelia Langford, John Sillitoe on behalf of the Wellcome Sanger Institute COVID-19 Surveillance Team |
| EPI_ISL_3041936 | Lighthouse Lab in Alderley Park | Wellcome Sanger Institute for the COVID-19 Genomics UK (COG-UK) Consortium | Jacquelyn Wynn, Mairead Hyland, The Lighthouse Lab in Alderley Park and Alex Alderton, Roberto Amato, Jeffrey Barrett, Sonia Goncalves, Ewan Harrison, David K. Jackson, Ian Johnston, Dominic Kwiatkowski, Cordelia Langford, John Sillitoe on behalf of the Wellcome Sanger Institute COVID-19 Surveillance Team |
| EPI_ISL_3041937 | Randox Laboratories | Wellcome Sanger Institute for the COVID-19 Genomics UK (COG-UK) Consortium | Randox Laboratories and Alex Alderton, Roberto Amato, Jeffrey Barrett, Sonia Goncalves, Ewan Harrison, David K. Jackson, Ian Johnston, Dominic Kwiatkowski, Cordelia Langford, John Sillitoe on behalf of the Wellcome Sanger Institute COVID-19 Surveillance Team |
| EPI_ISL_3041938, EPI_ISL_3041939, EPI_ISL_3041940, EPI_ISL_3041941, EPI_ISL_3041942, EPI_ISL_3041943 | Lighthouse Lab in Milton Keynes | Wellcome Sanger Institute for the COVID-19 Genomics UK (COG-UK) Consortium | The Lighthouse Lab in Milton Keynes and Alex Alderton, Roberto Amato, Jeffrey Barrett, Sonia Goncalves, Ewan Harrison, David K. Jackson, Ian Johnston, Dominic Kwiatkowski, Cordelia Langford, John Sillitoe on behalf of the Wellcome Sanger Institute COVID-19 Surveillance Team |
| EPI_ISL_3041944 | Randox Laboratories | Wellcome Sanger Institute for the COVID-19 Genomics UK (COG-UK) Consortium | Randox Laboratories and Alex Alderton, Roberto Amato, Jeffrey Barrett, Sonia Goncalves, Ewan Harrison, David K. Jackson, Ian Johnston, Dominic Kwiatkowski, Cordelia Langford, John Sillitoe on behalf of the Wellcome Sanger Institute COVID-19 Surveillance Team |

| | | | |
|---|---|---|---|
| EPI_ISL_3041945 | Lighthouse Lab in Milton Keynes | Wellcome Sanger Institute for the COVID-19 Genomics UK (COG-UK) Consortium | The Lighthouse Lab in Milton Keynes and Alex Alderton, Roberto Amato, Jeffrey Barrett, Sonia Goncalves, Ewan Harrison, David K. Jackson, Ian Johnston, Dominic Kwiatkowski, Cordelia Langford, John Sillitoe on behalf of the Wellcome Sanger Institute COVID-19 Surveillance Team |
| EPI_ISL_3041946 | Randox Laboratories | Wellcome Sanger Institute for the COVID-19 Genomics UK (COG-UK) Consortium | Randox Laboratories and Alex Alderton, Roberto Amato, Jeffrey Barrett, Sonia Goncalves, Ewan Harrison, David K. Jackson, Ian Johnston, Dominic Kwiatkowski, Cordelia Langford, John Sillitoe on behalf of the Wellcome Sanger Institute COVID-19 Surveillance Team |
| EPI_ISL_3041947 | Lighthouse Lab in Milton Keynes | Wellcome Sanger Institute for the COVID-19 Genomics UK (COG-UK) Consortium | The Lighthouse Lab in Milton Keynes and Alex Alderton, Roberto Amato, Jeffrey Barrett, Sonia Goncalves, Ewan Harrison, David K. Jackson, Ian Johnston, Dominic Kwiatkowski, Cordelia Langford, John Sillitoe on behalf of the Wellcome Sanger Institute COVID-19 Surveillance Team |
| EPI_ISL_3041948 | Lighthouse Lab in Alderley Park | Wellcome Sanger Institute for the COVID-19 Genomics UK (COG-UK) Consortium | Jacquelyn Wynn, Mairead Hyland, The Lighthouse Lab in Alderley Park and Alex Alderton, Roberto Amato, Jeffrey Barrett, Sonia Goncalves, Ewan Harrison, David K. Jackson, Ian Johnston, Dominic Kwiatkowski, Cordelia Langford, John Sillitoe on behalf of the Wellcome Sanger Institute COVID-19 Surveillance Team |
| EPI_ISL_3041949 | Lighthouse Lab in Milton Keynes | Wellcome Sanger Institute for the COVID-19 Genomics UK (COG-UK) Consortium | The Lighthouse Lab in Milton Keynes and Alex Alderton, Roberto Amato, Jeffrey Barrett, Sonia Goncalves, Ewan Harrison, David K. Jackson, Ian Johnston, Dominic Kwiatkowski, Cordelia Langford, John Sillitoe on behalf of the Wellcome Sanger Institute COVID-19 Surveillance Team |
| EPI_ISL_3041950 | Lighthouse Lab in Alderley Park | Wellcome Sanger Institute for the COVID-19 Genomics UK (COG-UK) Consortium | Jacquelyn Wynn, Mairead Hyland, The Lighthouse Lab in Alderley Park and Alex Alderton, Roberto Amato, Jeffrey Barrett, Sonia Goncalves, Ewan Harrison, David K. Jackson, Ian Johnston, Dominic Kwiatkowski, Cordelia Langford, John Sillitoe on behalf of the Wellcome Sanger Institute COVID-19 Surveillance Team |
| EPI_ISL_3041951 | Berkshire and Surrey Pathology Services Lighthouse Laboratory | Wellcome Sanger Institute for the COVID-19 Genomics UK (COG-UK) Consortium | Berkshire and Surrey Pathology Services Lighthouse Laboratory and Alex Alderton, Roberto Amato, Jeffrey Barrett, Sonia Goncalves, Ewan Harrison, David K. Jackson, Ian Johnston, Dominic Kwiatkowski, Cordelia Langford, John Sillitoe on behalf of the Wellcome Sanger Institute COVID-19 Surveillance Team |
| EPI_ISL_3041952 | Lighthouse Lab in Alderley Park | Wellcome Sanger Institute for the COVID-19 Genomics UK (COG-UK) Consortium | Jacquelyn Wynn, Mairead Hyland, The Lighthouse Lab in Alderley Park and Alex Alderton, Roberto Amato, Jeffrey Barrett, Sonia Goncalves, Ewan Harrison, David K. Jackson, Ian Johnston, Dominic Kwiatkowski, Cordelia Langford, John Sillitoe on behalf of the Wellcome Sanger Institute COVID-19 Surveillance Team |
| EPI_ISL_3041953, EPI_ISL_3041954, EPI_ISL_3041955 | Lighthouse Lab in Milton Keynes | Wellcome Sanger Institute for the COVID-19 Genomics UK (COG-UK) Consortium | The Lighthouse Lab in Milton Keynes and Alex Alderton, Roberto Amato, Jeffrey Barrett, Sonia Goncalves, Ewan Harrison, David K. Jackson, Ian Johnston, Dominic Kwiatkowski, Cordelia Langford, John Sillitoe on behalf of the Wellcome Sanger Institute COVID-19 Surveillance Team |
| EPI_ISL_3041956 | Randox Laboratories | Wellcome Sanger Institute for the COVID-19 Genomics UK (COG-UK) Consortium | Randox Laboratories and Alex Alderton, Roberto Amato, Jeffrey Barrett, Sonia Goncalves, Ewan Harrison, David K. Jackson, Ian Johnston, Dominic Kwiatkowski, Cordelia Langford, John Sillitoe on behalf of the Wellcome Sanger Institute COVID-19 Surveillance Team |
| EPI_ISL_3041957 | Berkshire and Surrey Pathology Services Lighthouse Laboratory | Wellcome Sanger Institute for the COVID-19 Genomics UK (COG-UK) Consortium | Berkshire and Surrey Pathology Services Lighthouse Laboratory and Alex Alderton, Roberto Amato, Jeffrey Barrett, Sonia Goncalves, Ewan Harrison, David K. Jackson, Ian Johnston, Dominic Kwiatkowski, Cordelia Langford, John Sillitoe on behalf of the Wellcome Sanger Institute COVID-19 Surveillance Team |
| EPI_ISL_3041958 | Randox Laboratories | Wellcome Sanger Institute for the COVID-19 Genomics UK (COG-UK) Consortium | Randox Laboratories and Alex Alderton, Roberto Amato, Jeffrey Barrett, Sonia Goncalves, Ewan Harrison, David K. Jackson, Ian Johnston, Dominic Kwiatkowski, Cordelia Langford, John Sillitoe on behalf of the Wellcome Sanger Institute COVID-19 Surveillance Team |
| EPI_ISL_3041959 | Lighthouse Lab in Milton Keynes | Wellcome Sanger Institute for the COVID-19 Genomics UK (COG-UK) Consortium | The Lighthouse Lab in Milton Keynes and Alex Alderton, Roberto Amato, Jeffrey Barrett, Sonia Goncalves, Ewan Harrison, David K. Jackson, Ian Johnston, Dominic Kwiatkowski, Cordelia Langford, John Sillitoe on behalf of the Wellcome Sanger Institute COVID-19 Surveillance Team |
| EPI_ISL_3041960 | Lighthouse Lab in Alderley Park | Wellcome Sanger Institute for the COVID-19 Genomics UK (COG-UK) Consortium | Jacquelyn Wynn, Mairead Hyland, The Lighthouse Lab in Alderley Park and Alex Alderton, Roberto Amato, Jeffrey Barrett, Sonia Goncalves, Ewan Harrison, David K. Jackson, Ian Johnston, Dominic Kwiatkowski, Cordelia Langford, John Sillitoe on behalf of the Wellcome Sanger Institute COVID-19 Surveillance Team |
| EPI_ISL_3041961 | Randox Laboratories | Wellcome Sanger Institute for the COVID-19 Genomics UK (COG-UK) Consortium | Randox Laboratories and Alex Alderton, Roberto Amato, Jeffrey Barrett, Sonia Goncalves, Ewan Harrison, David K. Jackson, Ian Johnston, Dominic Kwiatkowski, Cordelia Langford, John Sillitoe on behalf of the Wellcome Sanger Institute COVID-19 Surveillance Team |
| EPI_ISL_3041962 | Lighthouse Lab in Milton Keynes | Wellcome Sanger Institute for the COVID-19 Genomics UK (COG-UK) Consortium | The Lighthouse Lab in Milton Keynes and Alex Alderton, Roberto Amato, Jeffrey Barrett, Sonia Goncalves, Ewan Harrison, David K. Jackson, Ian Johnston, Dominic Kwiatkowski, Cordelia Langford, John Sillitoe on behalf of the Wellcome Sanger Institute COVID-19 Surveillance Team |
| EPI_ISL_3041963 | Lighthouse Lab in Alderley Park | Wellcome Sanger Institute for the COVID-19 Genomics UK (COG-UK) Consortium | Jacquelyn Wynn, Mairead Hyland, The Lighthouse Lab in Alderley Park and Alex Alderton, Roberto Amato, Jeffrey Barrett, Sonia Goncalves, Ewan Harrison, David K. Jackson, Ian Johnston, Dominic Kwiatkowski, Cordelia Langford, John Sillitoe on behalf of the Wellcome Sanger Institute COVID-19 Surveillance Team |
| EPI_ISL_3041964 | Randox Laboratories | Wellcome Sanger Institute for the COVID-19 Genomics UK (COG-UK) Consortium | Randox Laboratories and Alex Alderton, Roberto Amato, Jeffrey Barrett, Sonia Goncalves, Ewan Harrison, David K. Jackson, Ian Johnston, Dominic Kwiatkowski, Cordelia Langford, John Sillitoe on behalf of the Wellcome Sanger Institute COVID-19 Surveillance Team |
| EPI_ISL_3041965 | Berkshire and Surrey Pathology Services Lighthouse Laboratory | Wellcome Sanger Institute for the COVID-19 Genomics UK (COG-UK) Consortium | Berkshire and Surrey Pathology Services Lighthouse Laboratory and Alex Alderton, Roberto Amato, Jeffrey Barrett, Sonia Goncalves, Ewan Harrison, David K. Jackson, Ian Johnston, Dominic Kwiatkowski, Cordelia Langford, John Sillitoe on behalf of the Wellcome Sanger Institute COVID-19 Surveillance Team |
| EPI_ISL_3041966 | Lighthouse Lab in Milton Keynes | Wellcome Sanger Institute for the COVID-19 Genomics UK (COG-UK) Consortium | The Lighthouse Lab in Milton Keynes and Alex Alderton, Roberto Amato, Jeffrey Barrett, Sonia Goncalves, Ewan Harrison, David K. Jackson, Ian Johnston, Dominic Kwiatkowski, Cordelia Langford, John Sillitoe on behalf of the Wellcome Sanger Institute COVID-19 Surveillance Team |
| EPI_ISL_3041967 | Randox Laboratories | Wellcome Sanger Institute for the COVID-19 Genomics UK (COG-UK) Consortium | Randox Laboratories and Alex Alderton, Roberto Amato, Jeffrey Barrett, Sonia Goncalves, Ewan Harrison, David K. Jackson, Ian Johnston, Dominic Kwiatkowski, Cordelia Langford, John Sillitoe on behalf of the Wellcome Sanger Institute COVID-19 Surveillance Team |
| EPI_ISL_3041968 | Lighthouse Lab in Milton Keynes | Wellcome Sanger Institute for the COVID-19 Genomics UK (COG-UK) Consortium | The Lighthouse Lab in Milton Keynes and Alex Alderton, Roberto Amato, Jeffrey Barrett, Sonia Goncalves, Ewan Harrison, David K. Jackson, Ian Johnston, Dominic Kwiatkowski, Cordelia Langford, John Sillitoe on behalf of the Wellcome Sanger Institute COVID-19 Surveillance Team |
| EPI_ISL_3041969 | Randox Laboratories | Wellcome Sanger Institute for the COVID-19 Genomics UK (COG-UK) Consortium | Randox Laboratories and Alex Alderton, Roberto Amato, Jeffrey Barrett, Sonia Goncalves, Ewan Harrison, David K. Jackson, Ian Johnston, Dominic Kwiatkowski, Cordelia Langford, John Sillitoe on behalf of the Wellcome Sanger Institute COVID-19 Surveillance Team |
| EPI_ISL_3041970 | Lighthouse Lab in Milton Keynes | Wellcome Sanger Institute for the COVID-19 Genomics UK (COG-UK) Consortium | The Lighthouse Lab in Milton Keynes and Alex Alderton, Roberto Amato, Jeffrey Barrett, Sonia Goncalves, Ewan Harrison, David K. Jackson, Ian Johnston, Dominic Kwiatkowski, Cordelia Langford, John Sillitoe on behalf of the Wellcome Sanger Institute COVID-19 Surveillance Team |
| EPI_ISL_3041971 | Lighthouse Lab in Alderley Park | Wellcome Sanger Institute for the COVID-19 Genomics UK (COG-UK) Consortium | Jacquelyn Wynn, Mairead Hyland, The Lighthouse Lab in Alderley Park and Alex Alderton, Roberto Amato, Jeffrey Barrett, Sonia Goncalves, Ewan Harrison, David K. Jackson, Ian Johnston, Dominic Kwiatkowski, Cordelia Langford, John Sillitoe on behalf of the Wellcome Sanger Institute COVID-19 Surveillance Team |
| EPI_ISL_3041972 | Berkshire and Surrey Pathology Services Lighthouse Laboratory | Wellcome Sanger Institute for the COVID-19 Genomics UK (COG-UK) Consortium | Berkshire and Surrey Pathology Services Lighthouse Laboratory and Alex Alderton, Roberto Amato, Jeffrey Barrett, Sonia Goncalves, Ewan Harrison, David K. Jackson, Ian Johnston, Dominic Kwiatkowski, Cordelia Langford, John Sillitoe on behalf of the Wellcome Sanger Institute COVID-19 Surveillance Team |
| EPI_ISL_3041973 | Lighthouse Lab in Milton Keynes | Wellcome Sanger Institute for the COVID-19 Genomics UK (COG-UK) Consortium | The Lighthouse Lab in Milton Keynes and Alex Alderton, Roberto Amato, Jeffrey Barrett, Sonia Goncalves, Ewan Harrison, David K. Jackson, Ian Johnston, Dominic Kwiatkowski, Cordelia Langford, John Sillitoe on behalf of the Wellcome Sanger Institute COVID-19 Surveillance Team |
| EPI_ISL_3041974 | Lighthouse Lab in Alderley Park | Wellcome Sanger Institute for the COVID-19 Genomics UK (COG-UK) Consortium | Jacquelyn Wynn, Mairead Hyland, The Lighthouse Lab in Alderley Park and Alex Alderton, Roberto Amato, Jeffrey Barrett, Sonia Goncalves, Ewan Harrison, David K. Jackson, Ian Johnston, Dominic Kwiatkowski, Cordelia Langford, John Sillitoe on behalf of the Wellcome Sanger Institute COVID-19 Surveillance Team |
| EPI_ISL_3041975, EPI_ISL_3041976, EPI_ISL_3041977, EPI_ISL_3041978 | Randox Laboratories | Wellcome Sanger Institute for the COVID-19 Genomics UK (COG-UK) Consortium | Randox Laboratories and Alex Alderton, Roberto Amato, Jeffrey Barrett, Sonia Goncalves, Ewan Harrison, David K. Jackson, Ian Johnston, Dominic Kwiatkowski, Cordelia Langford, John Sillitoe on behalf of the Wellcome Sanger Institute COVID-19 Surveillance Team |
| EPI_ISL_3041979 | Lighthouse Lab in Milton Keynes | Wellcome Sanger Institute for the COVID-19 Genomics UK (COG-UK) Consortium | The Lighthouse Lab in Milton Keynes and Alex Alderton, Roberto Amato, Jeffrey Barrett, Sonia Goncalves, Ewan Harrison, David K. Jackson, Ian Johnston, Dominic Kwiatkowski, Cordelia Langford, John Sillitoe on behalf of the Wellcome Sanger Institute COVID-19 Surveillance Team |
| EPI_ISL_3041980 | Randox Laboratories | Wellcome Sanger Institute for the COVID-19 Genomics UK (COG-UK) Consortium | Randox Laboratories and Alex Alderton, Roberto Amato, Jeffrey Barrett, Sonia Goncalves, Ewan Harrison, David K. Jackson, Ian Johnston, Dominic Kwiatkowski, Cordelia Langford, John Sillitoe on behalf of the Wellcome Sanger Institute COVID-19 Surveillance Team |
| EPI_ISL_3041981, EPI_ISL_3041982, EPI_ISL_3041983, EPI_ISL_3041984, EPI_ISL_3041985 | Lighthouse Lab in Alderley Park | Wellcome Sanger Institute for the COVID-19 Genomics UK (COG-UK) Consortium | Jacquelyn Wynn, Mairead Hyland, The Lighthouse Lab in Alderley Park and Alex Alderton, Roberto Amato, Jeffrey Barrett, Sonia Goncalves, Ewan Harrison, David K. Jackson, Ian Johnston, Dominic Kwiatkowski, Cordelia Langford, John Sillitoe on behalf of the Wellcome Sanger Institute COVID-19 Surveillance Team |
| EPI_ISL_3041986, EPI_ISL_3041987 | Lighthouse Lab in Milton Keynes | Wellcome Sanger Institute for the COVID-19 Genomics UK (COG-UK) Consortium | The Lighthouse Lab in Milton Keynes and Alex Alderton, Roberto Amato, Jeffrey Barrett, Sonia Goncalves, Ewan Harrison, David K. Jackson, Ian Johnston, Dominic Kwiatkowski, Cordelia Langford, John Sillitoe on behalf of the Wellcome Sanger Institute COVID-19 Surveillance Team |
| EPI_ISL_3041988, EPI_ISL_3041989 | Lighthouse Lab in Alderley Park | Wellcome Sanger Institute for the COVID-19 Genomics UK (COG-UK) Consortium | Jacquelyn Wynn, Mairead Hyland, The Lighthouse Lab in Alderley Park and Alex Alderton, Roberto Amato, Jeffrey Barrett, Sonia Goncalves, Ewan Harrison, David K. Jackson, Ian Johnston, Dominic Kwiatkowski, Cordelia Langford, John Sillitoe on behalf of the Wellcome Sanger Institute COVID-19 Surveillance Team |
| EPI_ISL_3041990 | Lighthouse Lab in Milton Keynes | Wellcome Sanger Institute for the COVID-19 Genomics UK (COG-UK) Consortium | The Lighthouse Lab in Milton Keynes and Alex Alderton, Roberto Amato, Jeffrey Barrett, Sonia Goncalves, Ewan Harrison, David K. Jackson, Ian Johnston, Dominic Kwiatkowski, Cordelia Langford, John Sillitoe on behalf of the Wellcome Sanger Institute COVID-19 Surveillance Team |
| EPI_ISL_3041991 | Lighthouse Lab in Alderley Park | Wellcome Sanger Institute for the COVID-19 Genomics UK (COG-UK) Consortium | Jacquelyn Wynn, Mairead Hyland, The Lighthouse Lab in Alderley Park and Alex Alderton, Roberto Amato, Jeffrey Barrett, Sonia Goncalves, Ewan Harrison, David K. Jackson, Ian Johnston, Dominic Kwiatkowski, Cordelia Langford, John Sillitoe on behalf of the Wellcome Sanger Institute COVID-19 Surveillance Team |
| EPI_ISL_3041992 | Randox Laboratories | Wellcome Sanger Institute for the COVID-19 Genomics UK (COG-UK) Consortium | Randox Laboratories and Alex Alderton, Roberto Amato, Jeffrey Barrett, Sonia Goncalves, Ewan Harrison, David K. Jackson, Ian Johnston, Dominic Kwiatkowski, Cordelia Langford, John Sillitoe on behalf of the Wellcome Sanger Institute COVID-19 Surveillance Team |
| EPI_ISL_3041993 | Lighthouse Lab in Alderley Park | Wellcome Sanger Institute for the COVID-19 Genomics UK (COG-UK) Consortium | Jacquelyn Wynn, Mairead Hyland, The Lighthouse Lab in Alderley Park and Alex Alderton, Roberto Amato, Jeffrey Barrett, Sonia Goncalves, Ewan Harrison, David K. Jackson, Ian Johnston, Dominic Kwiatkowski, Cordelia Langford, John Sillitoe on behalf of the Wellcome Sanger Institute COVID-19 Surveillance Team |
| EPI_ISL_3041994, EPI_ISL_3041995 | Randox Laboratories | Wellcome Sanger Institute for the COVID-19 Genomics UK (COG-UK) Consortium | Randox Laboratories and Alex Alderton, Roberto Amato, Jeffrey Barrett, Sonia Goncalves, Ewan Harrison, David K. Jackson, Ian Johnston, Dominic Kwiatkowski, Cordelia Langford, John Sillitoe on behalf of the Wellcome Sanger Institute COVID-19 Surveillance Team |
| EPI_ISL_3041996, EPI_ISL_3041997, EPI_ISL_3041998 | Lighthouse Lab in Alderley Park | Wellcome Sanger Institute for the COVID-19 Genomics UK (COG-UK) Consortium | Jacquelyn Wynn, Mairead Hyland, The Lighthouse Lab in Alderley Park and Alex Alderton, Roberto Amato, Jeffrey Barrett, Sonia Goncalves, Ewan Harrison, David K. Jackson, Ian Johnston, Dominic Kwiatkowski, Cordelia Langford, John Sillitoe on behalf of the Wellcome Sanger Institute COVID-19 Surveillance Team |
| EPI_ISL_3041999 | Randox Laboratories | Wellcome Sanger Institute for the COVID-19 Genomics UK (COG-UK) Consortium | Randox Laboratories and Alex Alderton, Roberto Amato, Jeffrey Barrett, Sonia Goncalves, Ewan Harrison, David K. Jackson, Ian Johnston, Dominic Kwiatkowski, Cordelia Langford, John Sillitoe on behalf of the Wellcome Sanger Institute COVID-19 Surveillance Team |
| EPI_ISL_3042000 | Lighthouse Lab in Alderley Park | Wellcome Sanger Institute for the COVID-19 Genomics UK (COG-UK) Consortium | Jacquelyn Wynn, Mairead Hyland, The Lighthouse Lab in Alderley Park and Alex Alderton, Roberto Amato, Jeffrey Barrett, Sonia Goncalves, Ewan Harrison, David K. Jackson, Ian Johnston, Dominic Kwiatkowski, Cordelia Langford, John Sillitoe on behalf of the Wellcome Sanger Institute COVID-19 Surveillance Team |
| EPI_ISL_3042001 | Lighthouse Lab in Milton Keynes | Wellcome Sanger Institute for the COVID-19 Genomics UK (COG-UK) | The Lighthouse Lab in Milton Keynes and Alex Alderton, Roberto Amato, Jeffrey Barrett, Sonia Goncalves, Ewan Harrison, David K. Jackson, Ian Johnston, Dominic Kwiatkowski, Cordelia Langford, John Sillitoe on behalf of the Wellcome Sanger |

| | | | |
|---|---|---|---|
| | | Consortium | Institute COVID-19 Surveillance Team |
| EPI_ISL_3042002 | Randox Laboratories | Wellcome Sanger Institute for the COVID-19 Genomics UK (COG-UK) Consortium | Randox Laboratories and Alex Alderton, Roberto Amato, Jeffrey Barrett, Sonia Goncalves, Ewan Harrison, David K. Jackson, Ian Johnston, Dominic Kwiatkowski, Cordelia Langford, John Sillitoe on behalf of the Wellcome Sanger Institute COVID-19 Surveillance Team |
| EPI_ISL_3042003 | Berkshire and Surrey Pathology Services Lighthouse Laboratory | Wellcome Sanger Institute for the COVID-19 Genomics UK (COG-UK) Consortium | Berkshire and Surrey Pathology Services Lighthouse Laboratory and Alex Alderton, Roberto Amato, Jeffrey Barrett, Sonia Goncalves, Ewan Harrison, David K. Jackson, Ian Johnston, Dominic Kwiatkowski, Cordelia Langford, John Sillitoe on behalf of the Wellcome Sanger Institute COVID-19 Surveillance Team |
| EPI_ISL_3042004 | Randox Laboratories | Wellcome Sanger Institute for the COVID-19 Genomics UK (COG-UK) Consortium | Randox Laboratories and Alex Alderton, Roberto Amato, Jeffrey Barrett, Sonia Goncalves, Ewan Harrison, David K. Jackson, Ian Johnston, Dominic Kwiatkowski, Cordelia Langford, John Sillitoe on behalf of the Wellcome Sanger Institute COVID-19 Surveillance Team |
| EPI_ISL_3042005, EPI_ISL_3042006 | Lighthouse Lab in Milton Keynes | Wellcome Sanger Institute for the COVID-19 Genomics UK (COG-UK) Consortium | The Lighthouse Lab in Milton Keynes and Alex Alderton, Roberto Amato, Jeffrey Barrett, Sonia Goncalves, Ewan Harrison, David K. Jackson, Ian Johnston, Dominic Kwiatkowski, Cordelia Langford, John Sillitoe on behalf of the Wellcome Sanger Institute COVID-19 Surveillance Team |
| EPI_ISL_3042007 | Randox Laboratories | Wellcome Sanger Institute for the COVID-19 Genomics UK (COG-UK) Consortium | Randox Laboratories and Alex Alderton, Roberto Amato, Jeffrey Barrett, Sonia Goncalves, Ewan Harrison, David K. Jackson, Ian Johnston, Dominic Kwiatkowski, Cordelia Langford, John Sillitoe on behalf of the Wellcome Sanger Institute COVID-19 Surveillance Team |
| EPI_ISL_3042008 | Lighthouse Lab in Alderley Park | Wellcome Sanger Institute for the COVID-19 Genomics UK (COG-UK) Consortium | Jacquelyn Wynn, Mairead Hyland, The Lighthouse Lab in Alderley Park and Alex Alderton, Roberto Amato, Jeffrey Barrett, Sonia Goncalves, Ewan Harrison, David K. Jackson, Ian Johnston, Dominic Kwiatkowski, Cordelia Langford, John Sillitoe on behalf of the Wellcome Sanger Institute COVID-19 Surveillance Team |
| EPI_ISL_3042009 | Randox Laboratories | Wellcome Sanger Institute for the COVID-19 Genomics UK (COG-UK) Consortium | Randox Laboratories and Alex Alderton, Roberto Amato, Jeffrey Barrett, Sonia Goncalves, Ewan Harrison, David K. Jackson, Ian Johnston, Dominic Kwiatkowski, Cordelia Langford, John Sillitoe on behalf of the Wellcome Sanger Institute COVID-19 Surveillance Team |
| EPI_ISL_3042010 | Lighthouse Lab in Alderley Park | Wellcome Sanger Institute for the COVID-19 Genomics UK (COG-UK) Consortium | Jacquelyn Wynn, Mairead Hyland, The Lighthouse Lab in Alderley Park and Alex Alderton, Roberto Amato, Jeffrey Barrett, Sonia Goncalves, Ewan Harrison, David K. Jackson, Ian Johnston, Dominic Kwiatkowski, Cordelia Langford, John Sillitoe on behalf of the Wellcome Sanger Institute COVID-19 Surveillance Team |
| EPI_ISL_3042011 | Randox Laboratories | Wellcome Sanger Institute for the COVID-19 Genomics UK (COG-UK) Consortium | Randox Laboratories and Alex Alderton, Roberto Amato, Jeffrey Barrett, Sonia Goncalves, Ewan Harrison, David K. Jackson, Ian Johnston, Dominic Kwiatkowski, Cordelia Langford, John Sillitoe on behalf of the Wellcome Sanger Institute COVID-19 Surveillance Team |
| EPI_ISL_3042012 | Lighthouse Lab in Alderley Park | Wellcome Sanger Institute for the COVID-19 Genomics UK (COG-UK) Consortium | Jacquelyn Wynn, Mairead Hyland, The Lighthouse Lab in Alderley Park and Alex Alderton, Roberto Amato, Jeffrey Barrett, Sonia Goncalves, Ewan Harrison, David K. Jackson, Ian Johnston, Dominic Kwiatkowski, Cordelia Langford, John Sillitoe on behalf of the Wellcome Sanger Institute COVID-19 Surveillance Team |
| EPI_ISL_3042013 | Lighthouse Lab in Milton Keynes | Wellcome Sanger Institute for the COVID-19 Genomics UK (COG-UK) Consortium | The Lighthouse Lab in Milton Keynes and Alex Alderton, Roberto Amato, Jeffrey Barrett, Sonia Goncalves, Ewan Harrison, David K. Jackson, Ian Johnston, Dominic Kwiatkowski, Cordelia Langford, John Sillitoe on behalf of the Wellcome Sanger Institute COVID-19 Surveillance Team |
| EPI_ISL_3042014, EPI_ISL_3042015 | Lighthouse Lab in Alderley Park | Wellcome Sanger Institute for the COVID-19 Genomics UK (COG-UK) Consortium | Jacquelyn Wynn, Mairead Hyland, The Lighthouse Lab in Alderley Park and Alex Alderton, Roberto Amato, Jeffrey Barrett, Sonia Goncalves, Ewan Harrison, David K. Jackson, Ian Johnston, Dominic Kwiatkowski, Cordelia Langford, John Sillitoe on behalf of the Wellcome Sanger Institute COVID-19 Surveillance Team |
| EPI_ISL_3042016 | Berkshire and Surrey Pathology Services Lighthouse Laboratory | Wellcome Sanger Institute for the COVID-19 Genomics UK (COG-UK) Consortium | Berkshire and Surrey Pathology Services Lighthouse Laboratory and Alex Alderton, Roberto Amato, Jeffrey Barrett, Sonia Goncalves, Ewan Harrison, David K. Jackson, Ian Johnston, Dominic Kwiatkowski, Cordelia Langford, John Sillitoe on behalf of the Wellcome Sanger Institute COVID-19 Surveillance Team |
| EPI_ISL_3042017 | Lighthouse Lab in Milton Keynes | Wellcome Sanger Institute for the COVID-19 Genomics UK (COG-UK) Consortium | The Lighthouse Lab in Milton Keynes and Alex Alderton, Roberto Amato, Jeffrey Barrett, Sonia Goncalves, Ewan Harrison, David K. Jackson, Ian Johnston, Dominic Kwiatkowski, Cordelia Langford, John Sillitoe on behalf of the Wellcome Sanger Institute COVID-19 Surveillance Team |
| EPI_ISL_3042018 | Lighthouse Lab in Alderley Park | Wellcome Sanger Institute for the COVID-19 Genomics UK (COG-UK) Consortium | Jacquelyn Wynn, Mairead Hyland, The Lighthouse Lab in Alderley Park and Alex Alderton, Roberto Amato, Jeffrey Barrett, Sonia Goncalves, Ewan Harrison, David K. Jackson, Ian Johnston, Dominic Kwiatkowski, Cordelia Langford, John Sillitoe on behalf of the Wellcome Sanger Institute COVID-19 Surveillance Team |
| EPI_ISL_3042019 | Randox Laboratories | Wellcome Sanger Institute for the COVID-19 Genomics UK (COG-UK) Consortium | Randox Laboratories and Alex Alderton, Roberto Amato, Jeffrey Barrett, Sonia Goncalves, Ewan Harrison, David K. Jackson, Ian Johnston, Dominic Kwiatkowski, Cordelia Langford, John Sillitoe on behalf of the Wellcome Sanger Institute COVID-19 Surveillance Team |
| EPI_ISL_3042020, EPI_ISL_3042021 | Lighthouse Lab in Alderley Park | Wellcome Sanger Institute for the COVID-19 Genomics UK (COG-UK) Consortium | Jacquelyn Wynn, Mairead Hyland, The Lighthouse Lab in Alderley Park and Alex Alderton, Roberto Amato, Jeffrey Barrett, Sonia Goncalves, Ewan Harrison, David K. Jackson, Ian Johnston, Dominic Kwiatkowski, Cordelia Langford, John Sillitoe on behalf of the Wellcome Sanger Institute COVID-19 Surveillance Team |
| EPI_ISL_3042022 | Lighthouse Lab in Milton Keynes | Wellcome Sanger Institute for the COVID-19 Genomics UK (COG-UK) Consortium | The Lighthouse Lab in Milton Keynes and Alex Alderton, Roberto Amato, Jeffrey Barrett, Sonia Goncalves, Ewan Harrison, David K. Jackson, Ian Johnston, Dominic Kwiatkowski, Cordelia Langford, John Sillitoe on behalf of the Wellcome Sanger Institute COVID-19 Surveillance Team |
| EPI_ISL_3042023 | Berkshire and Surrey Pathology Services Lighthouse Laboratory | Wellcome Sanger Institute for the COVID-19 Genomics UK (COG-UK) Consortium | Berkshire and Surrey Pathology Services Lighthouse Laboratory and Alex Alderton, Roberto Amato, Jeffrey Barrett, Sonia Goncalves, Ewan Harrison, David K. Jackson, Ian Johnston, Dominic Kwiatkowski, Cordelia Langford, John Sillitoe on behalf of the Wellcome Sanger Institute COVID-19 Surveillance Team |
| EPI_ISL_3042024, EPI_ISL_3042025 | Randox Laboratories | Wellcome Sanger Institute for the COVID-19 Genomics UK (COG-UK) Consortium | Randox Laboratories and Alex Alderton, Roberto Amato, Jeffrey Barrett, Sonia Goncalves, Ewan Harrison, David K. Jackson, Ian Johnston, Dominic Kwiatkowski, Cordelia Langford, John Sillitoe on behalf of the Wellcome Sanger Institute COVID-19 Surveillance Team |
| EPI_ISL_3042026 | Lighthouse Lab in Alderley Park | Wellcome Sanger Institute for the COVID-19 Genomics UK (COG-UK) Consortium | Jacquelyn Wynn, Mairead Hyland, The Lighthouse Lab in Alderley Park and Alex Alderton, Roberto Amato, Jeffrey Barrett, Sonia Goncalves, Ewan Harrison, David K. Jackson, Ian Johnston, Dominic Kwiatkowski, Cordelia Langford, John Sillitoe on behalf of the Wellcome Sanger Institute COVID-19 Surveillance Team |
| EPI_ISL_3042027 | Lighthouse Lab in Milton Keynes | Wellcome Sanger Institute for the COVID-19 Genomics UK (COG-UK) Consortium | The Lighthouse Lab in Milton Keynes and Alex Alderton, Roberto Amato, Jeffrey Barrett, Sonia Goncalves, Ewan Harrison, David K. Jackson, Ian Johnston, Dominic Kwiatkowski, Cordelia Langford, John Sillitoe on behalf of the Wellcome Sanger Institute COVID-19 Surveillance Team |
| EPI_ISL_3042028 | Berkshire and Surrey Pathology Services Lighthouse Laboratory | Wellcome Sanger Institute for the COVID-19 Genomics UK (COG-UK) Consortium | Berkshire and Surrey Pathology Services Lighthouse Laboratory and Alex Alderton, Roberto Amato, Jeffrey Barrett, Sonia Goncalves, Ewan Harrison, David K. Jackson, Ian Johnston, Dominic Kwiatkowski, Cordelia Langford, John Sillitoe on behalf of the Wellcome Sanger Institute COVID-19 Surveillance Team |
| EPI_ISL_3042029 | Randox Laboratories | Wellcome Sanger Institute for the COVID-19 Genomics UK (COG-UK) Consortium | Randox Laboratories and Alex Alderton, Roberto Amato, Jeffrey Barrett, Sonia Goncalves, Ewan Harrison, David K. Jackson, Ian Johnston, Dominic Kwiatkowski, Cordelia Langford, John Sillitoe on behalf of the Wellcome Sanger Institute COVID-19 Surveillance Team |
| EPI_ISL_3042030, EPI_ISL_3042031, EPI_ISL_3042032, EPI_ISL_3042033 | Lighthouse Lab in Alderley Park | Wellcome Sanger Institute for the COVID-19 Genomics UK (COG-UK) Consortium | Jacquelyn Wynn, Mairead Hyland, The Lighthouse Lab in Alderley Park and Alex Alderton, Roberto Amato, Jeffrey Barrett, Sonia Goncalves, Ewan Harrison, David K. Jackson, Ian Johnston, Dominic Kwiatkowski, Cordelia Langford, John Sillitoe on behalf of the Wellcome Sanger Institute COVID-19 Surveillance Team |
| EPI_ISL_3042034 | Lighthouse Lab in Milton Keynes | Wellcome Sanger Institute for the COVID-19 Genomics UK (COG-UK) Consortium | The Lighthouse Lab in Milton Keynes and Alex Alderton, Roberto Amato, Jeffrey Barrett, Sonia Goncalves, Ewan Harrison, David K. Jackson, Ian Johnston, Dominic Kwiatkowski, Cordelia Langford, John Sillitoe on behalf of the Wellcome Sanger Institute COVID-19 Surveillance Team |
| EPI_ISL_3042035 | Lighthouse Lab in Alderley Park | Wellcome Sanger Institute for the COVID-19 Genomics UK (COG-UK) Consortium | Jacquelyn Wynn, Mairead Hyland, The Lighthouse Lab in Alderley Park and Alex Alderton, Roberto Amato, Jeffrey Barrett, Sonia Goncalves, Ewan Harrison, David K. Jackson, Ian Johnston, Dominic Kwiatkowski, Cordelia Langford, John Sillitoe on behalf of the Wellcome Sanger Institute COVID-19 Surveillance Team |
| EPI_ISL_3042036, EPI_ISL_3042037 | Randox Laboratories | Wellcome Sanger Institute for the COVID-19 Genomics UK (COG-UK) Consortium | Randox Laboratories and Alex Alderton, Roberto Amato, Jeffrey Barrett, Sonia Goncalves, Ewan Harrison, David K. Jackson, Ian Johnston, Dominic Kwiatkowski, Cordelia Langford, John Sillitoe on behalf of the Wellcome Sanger Institute COVID-19 Surveillance Team |
| EPI_ISL_3042038, EPI_ISL_3042039 | Lighthouse Lab in Milton Keynes | Wellcome Sanger Institute for the COVID-19 Genomics UK (COG-UK) Consortium | The Lighthouse Lab in Milton Keynes and Alex Alderton, Roberto Amato, Jeffrey Barrett, Sonia Goncalves, Ewan Harrison, David K. Jackson, Ian Johnston, Dominic Kwiatkowski, Cordelia Langford, John Sillitoe on behalf of the Wellcome Sanger Institute COVID-19 Surveillance Team |
| EPI_ISL_3042040, EPI_ISL_3042041 | Lighthouse Lab in Alderley Park | Wellcome Sanger Institute for the COVID-19 Genomics UK (COG-UK) Consortium | Jacquelyn Wynn, Mairead Hyland, The Lighthouse Lab in Alderley Park and Alex Alderton, Roberto Amato, Jeffrey Barrett, Sonia Goncalves, Ewan Harrison, David K. Jackson, Ian Johnston, Dominic Kwiatkowski, Cordelia Langford, John Sillitoe on behalf of the Wellcome Sanger Institute COVID-19 Surveillance Team |
| EPI_ISL_3042042, EPI_ISL_3042043 | Randox Laboratories | Wellcome Sanger Institute for the COVID-19 Genomics UK (COG-UK) Consortium | Randox Laboratories and Alex Alderton, Roberto Amato, Jeffrey Barrett, Sonia Goncalves, Ewan Harrison, David K. Jackson, Ian Johnston, Dominic Kwiatkowski, Cordelia Langford, John Sillitoe on behalf of the Wellcome Sanger Institute COVID-19 Surveillance Team |
| EPI_ISL_3042044 | Lighthouse Lab in Alderley Park | Wellcome Sanger Institute for the COVID-19 Genomics UK (COG-UK) Consortium | Jacquelyn Wynn, Mairead Hyland, The Lighthouse Lab in Alderley Park and Alex Alderton, Roberto Amato, Jeffrey Barrett, Sonia Goncalves, Ewan Harrison, David K. Jackson, Ian Johnston, Dominic Kwiatkowski, Cordelia Langford, John Sillitoe on behalf of the Wellcome Sanger Institute COVID-19 Surveillance Team |
| EPI_ISL_3042045 | Lighthouse Lab in Milton Keynes | Wellcome Sanger Institute for the COVID-19 Genomics UK (COG-UK) Consortium | The Lighthouse Lab in Milton Keynes and Alex Alderton, Roberto Amato, Jeffrey Barrett, Sonia Goncalves, Ewan Harrison, David K. Jackson, Ian Johnston, Dominic Kwiatkowski, Cordelia Langford, John Sillitoe on behalf of the Wellcome Sanger Institute COVID-19 Surveillance Team |
| EPI_ISL_3042046 | Randox Laboratories | Wellcome Sanger Institute for the COVID-19 Genomics UK (COG-UK) Consortium | Randox Laboratories and Alex Alderton, Roberto Amato, Jeffrey Barrett, Sonia Goncalves, Ewan Harrison, David K. Jackson, Ian Johnston, Dominic Kwiatkowski, Cordelia Langford, John Sillitoe on behalf of the Wellcome Sanger Institute COVID-19 Surveillance Team |
| EPI_ISL_3042047 | Lighthouse Lab in Alderley Park | Wellcome Sanger Institute for the COVID-19 Genomics UK (COG-UK) Consortium | Jacquelyn Wynn, Mairead Hyland, The Lighthouse Lab in Alderley Park and Alex Alderton, Roberto Amato, Jeffrey Barrett, Sonia Goncalves, Ewan Harrison, David K. Jackson, Ian Johnston, Dominic Kwiatkowski, Cordelia Langford, John Sillitoe on behalf of the Wellcome Sanger Institute COVID-19 Surveillance Team |
| EPI_ISL_3042048 | Lighthouse Laboratory Plymouth | Wellcome Sanger Institute for the COVID-19 Genomics UK (COG-UK) Consortium | Lighthouse Laboratory Plymouth and Alex Alderton, Roberto Amato, Jeffrey Barrett, Sonia Goncalves, Ewan Harrison, David K. Jackson, Ian Johnston, Dominic Kwiatkowski, Cordelia Langford, John Sillitoe on behalf of the Wellcome Sanger Institute COVID-19 Surveillance Team |
| EPI_ISL_3042049 | Lighthouse Lab in Milton Keynes | Wellcome Sanger Institute for the COVID-19 Genomics UK (COG-UK) Consortium | The Lighthouse Lab in Milton Keynes and Alex Alderton, Roberto Amato, Jeffrey Barrett, Sonia Goncalves, Ewan Harrison, David K. Jackson, Ian Johnston, Dominic Kwiatkowski, Cordelia Langford, John Sillitoe on behalf of the Wellcome Sanger Institute COVID-19 Surveillance Team |
| EPI_ISL_3042050, EPI_ISL_3042051 | Lighthouse Laboratory Plymouth | Wellcome Sanger Institute for the COVID-19 Genomics UK (COG-UK) Consortium | Lighthouse Laboratory Plymouth and Alex Alderton, Roberto Amato, Jeffrey Barrett, Sonia Goncalves, Ewan Harrison, David K. Jackson, Ian Johnston, Dominic Kwiatkowski, Cordelia Langford, John Sillitoe on behalf of the Wellcome Sanger Institute COVID-19 Surveillance Team |
| EPI_ISL_3042052 | Lighthouse Lab in Alderley Park | Wellcome Sanger Institute for the COVID-19 Genomics UK (COG-UK) Consortium | Jacquelyn Wynn, Mairead Hyland, The Lighthouse Lab in Alderley Park and Alex Alderton, Roberto Amato, Jeffrey Barrett, Sonia Goncalves, Ewan Harrison, David K. Jackson, Ian Johnston, Dominic Kwiatkowski, Cordelia Langford, John Sillitoe on behalf of the Wellcome Sanger Institute COVID-19 Surveillance Team |
| EPI_ISL_3042053 | Lighthouse Laboratory Plymouth | Wellcome Sanger Institute for the COVID-19 Genomics UK (COG-UK) Consortium | Lighthouse Laboratory Plymouth and Alex Alderton, Roberto Amato, Jeffrey Barrett, Sonia Goncalves, Ewan Harrison, David K. Jackson, Ian Johnston, Dominic Kwiatkowski, Cordelia Langford, John Sillitoe on behalf of the Wellcome Sanger Institute COVID-19 Surveillance Team |
| EPI_ISL_3042054 | Lighthouse Lab in Alderley Park | Wellcome Sanger Institute for the COVID-19 Genomics UK (COG-UK) Consortium | Jacquelyn Wynn, Mairead Hyland, The Lighthouse Lab in Alderley Park and Alex Alderton, Roberto Amato, Jeffrey Barrett, Sonia Goncalves, Ewan Harrison, David K. Jackson, Ian Johnston, Dominic Kwiatkowski, Cordelia Langford, John Sillitoe on behalf of the Wellcome Sanger Institute COVID-19 Surveillance Team |
| EPI_ISL_3042055 | Lighthouse Laboratory Plymouth | Wellcome Sanger Institute for the COVID-19 Genomics UK (COG-UK) Consortium | Lighthouse Laboratory Plymouth and Alex Alderton, Roberto Amato, Jeffrey Barrett, Sonia Goncalves, Ewan Harrison, David K. Jackson, Ian Johnston, Dominic Kwiatkowski, Cordelia Langford, John Sillitoe on behalf of the Wellcome Sanger Institute COVID-19 Surveillance Team |
| EPI_ISL_3042056 | Lighthouse Lab in Alderley Park | Wellcome Sanger Institute for the COVID-19 Genomics UK (COG-UK) Consortium | Jacquelyn Wynn, Mairead Hyland, The Lighthouse Lab in Alderley Park and Alex Alderton, Roberto Amato, Jeffrey Barrett, Sonia Goncalves, Ewan Harrison, David K. Jackson, Ian Johnston, Dominic Kwiatkowski, Cordelia Langford, John Sillitoe on behalf of the Wellcome Sanger Institute COVID-19 Surveillance Team |
| EPI_ISL_3042057, | Lighthouse Lab in Milton Keynes | Wellcome Sanger Institute for the COVID-19 Genomics UK (COG-UK) | The Lighthouse Lab in Milton Keynes and Alex Alderton, Roberto Amato, Jeffrey Barrett, Sonia Goncalves, Ewan Harrison, David K. Jackson, Ian Johnston, Dominic Kwiatkowski, Cordelia Langford, John Sillitoe on behalf of the Wellcome Sanger |

| | | | |
|---|---|---|---|
| EPI_ISL_3042058 | | Consortium | Institute COVID-19 Surveillance Team |
| EPI_ISL_3042059 | Lighthouse Laboratory Plymouth | Wellcome Sanger Institute for the COVID-19 Genomics UK (COG-UK) Consortium | Lighthouse Laboratory Plymouth and Alex Alderton, Roberto Amato, Jeffrey Barrett, Sonia Goncalves, Ewan Harrison, David K. Jackson, Ian Johnston, Dominic Kwiatkowski, Cordelia Langford, John Sillitoe on behalf of the Wellcome Sanger Institute COVID-19 Surveillance Team |
| EPI_ISL_3042060 | Berkshire and Surrey Pathology Services Lighthouse Laboratory | Wellcome Sanger Institute for the COVID-19 Genomics UK (COG-UK) Consortium | Berkshire and Surrey Pathology Services Lighthouse Laboratory and Alex Alderton, Roberto Amato, Jeffrey Barrett, Sonia Goncalves, Ewan Harrison, David K. Jackson, Ian Johnston, Dominic Kwiatkowski, Cordelia Langford, John Sillitoe on behalf of the Wellcome Sanger Institute COVID-19 Surveillance Team |
| EPI_ISL_3042061 | Lighthouse Lab in Milton Keynes | Wellcome Sanger Institute for the COVID-19 Genomics UK (COG-UK) Consortium | The Lighthouse Lab in Milton Keynes and Alex Alderton, Roberto Amato, Jeffrey Barrett, Sonia Goncalves, Ewan Harrison, David K. Jackson, Ian Johnston, Dominic Kwiatkowski, Cordelia Langford, John Sillitoe on behalf of the Wellcome Sanger Institute COVID-19 Surveillance Team |
| EPI_ISL_3042062, EPI_ISL_3042063 | Berkshire and Surrey Pathology Services Lighthouse Laboratory | Wellcome Sanger Institute for the COVID-19 Genomics UK (COG-UK) Consortium | Berkshire and Surrey Pathology Services Lighthouse Laboratory and Alex Alderton, Roberto Amato, Jeffrey Barrett, Sonia Goncalves, Ewan Harrison, David K. Jackson, Ian Johnston, Dominic Kwiatkowski, Cordelia Langford, John Sillitoe on behalf of the Wellcome Sanger Institute COVID-19 Surveillance Team |
| EPI_ISL_3042064, EPI_ISL_3042065 | Lighthouse Lab in Alderley Park | Wellcome Sanger Institute for the COVID-19 Genomics UK (COG-UK) Consortium | Jacquelyn Wynn, Mairead Hyland, The Lighthouse Lab in Alderley Park and Alex Alderton, Roberto Amato, Jeffrey Barrett, Sonia Goncalves, Ewan Harrison, David K. Jackson, Ian Johnston, Dominic Kwiatkowski, Cordelia Langford, John Sillitoe on behalf of the Wellcome Sanger Institute COVID-19 Surveillance Team |
| EPI_ISL_3042066 | Berkshire and Surrey Pathology Services Lighthouse Laboratory | Wellcome Sanger Institute for the COVID-19 Genomics UK (COG-UK) Consortium | Berkshire and Surrey Pathology Services Lighthouse Laboratory and Alex Alderton, Roberto Amato, Jeffrey Barrett, Sonia Goncalves, Ewan Harrison, David K. Jackson, Ian Johnston, Dominic Kwiatkowski, Cordelia Langford, John Sillitoe on behalf of the Wellcome Sanger Institute COVID-19 Surveillance Team |
| EPI_ISL_3042067, EPI_ISL_3042068, EPI_ISL_3042069 | Lighthouse Laboratory Plymouth | Wellcome Sanger Institute for the COVID-19 Genomics UK (COG-UK) Consortium | Lighthouse Laboratory Plymouth and Alex Alderton, Roberto Amato, Jeffrey Barrett, Sonia Goncalves, Ewan Harrison, David K. Jackson, Ian Johnston, Dominic Kwiatkowski, Cordelia Langford, John Sillitoe on behalf of the Wellcome Sanger Institute COVID-19 Surveillance Team |
| EPI_ISL_3042070, EPI_ISL_3042071 | Lighthouse Lab in Alderley Park | Wellcome Sanger Institute for the COVID-19 Genomics UK (COG-UK) Consortium | Jacquelyn Wynn, Mairead Hyland, The Lighthouse Lab in Alderley Park and Alex Alderton, Roberto Amato, Jeffrey Barrett, Sonia Goncalves, Ewan Harrison, David K. Jackson, Ian Johnston, Dominic Kwiatkowski, Cordelia Langford, John Sillitoe on behalf of the Wellcome Sanger Institute COVID-19 Surveillance Team |
| EPI_ISL_3042072 | Berkshire and Surrey Pathology Services Lighthouse Laboratory | Wellcome Sanger Institute for the COVID-19 Genomics UK (COG-UK) Consortium | Berkshire and Surrey Pathology Services Lighthouse Laboratory and Alex Alderton, Roberto Amato, Jeffrey Barrett, Sonia Goncalves, Ewan Harrison, David K. Jackson, Ian Johnston, Dominic Kwiatkowski, Cordelia Langford, John Sillitoe on behalf of the Wellcome Sanger Institute COVID-19 Surveillance Team |
| EPI_ISL_3042073 | Lighthouse Lab in Alderley Park | Wellcome Sanger Institute for the COVID-19 Genomics UK (COG-UK) Consortium | Jacquelyn Wynn, Mairead Hyland, The Lighthouse Lab in Alderley Park and Alex Alderton, Roberto Amato, Jeffrey Barrett, Sonia Goncalves, Ewan Harrison, David K. Jackson, Ian Johnston, Dominic Kwiatkowski, Cordelia Langford, John Sillitoe on behalf of the Wellcome Sanger Institute COVID-19 Surveillance Team |
| EPI_ISL_3042074, EPI_ISL_3042075 | Berkshire and Surrey Pathology Services Lighthouse Laboratory | Wellcome Sanger Institute for the COVID-19 Genomics UK (COG-UK) Consortium | Berkshire and Surrey Pathology Services Lighthouse Laboratory and Alex Alderton, Roberto Amato, Jeffrey Barrett, Sonia Goncalves, Ewan Harrison, David K. Jackson, Ian Johnston, Dominic Kwiatkowski, Cordelia Langford, John Sillitoe on behalf of the Wellcome Sanger Institute COVID-19 Surveillance Team |
| EPI_ISL_3042077 | Lighthouse Lab in Alderley Park | Wellcome Sanger Institute for the COVID-19 Genomics UK (COG-UK) Consortium | Jacquelyn Wynn, Mairead Hyland, The Lighthouse Lab in Alderley Park and Alex Alderton, Roberto Amato, Jeffrey Barrett, Sonia Goncalves, Ewan Harrison, David K. Jackson, Ian Johnston, Dominic Kwiatkowski, Cordelia Langford, John Sillitoe on behalf of the Wellcome Sanger Institute COVID-19 Surveillance Team |
| EPI_ISL_3042078 | Berkshire and Surrey Pathology Services Lighthouse Laboratory | Wellcome Sanger Institute for the COVID-19 Genomics UK (COG-UK) Consortium | Berkshire and Surrey Pathology Services Lighthouse Laboratory and Alex Alderton, Roberto Amato, Jeffrey Barrett, Sonia Goncalves, Ewan Harrison, David K. Jackson, Ian Johnston, Dominic Kwiatkowski, Cordelia Langford, John Sillitoe on behalf of the Wellcome Sanger Institute COVID-19 Surveillance Team |
| EPI_ISL_3042079 | Lighthouse Laboratory Plymouth | Wellcome Sanger Institute for the COVID-19 Genomics UK (COG-UK) Consortium | Lighthouse Laboratory Plymouth and Alex Alderton, Roberto Amato, Jeffrey Barrett, Sonia Goncalves, Ewan Harrison, David K. Jackson, Ian Johnston, Dominic Kwiatkowski, Cordelia Langford, John Sillitoe on behalf of the Wellcome Sanger Institute COVID-19 Surveillance Team |
| EPI_ISL_3042080, EPI_ISL_3042081 | Lighthouse Lab in Milton Keynes | Wellcome Sanger Institute for the COVID-19 Genomics UK (COG-UK) Consortium | The Lighthouse Lab in Milton Keynes and Alex Alderton, Roberto Amato, Jeffrey Barrett, Sonia Goncalves, Ewan Harrison, David K. Jackson, Ian Johnston, Dominic Kwiatkowski, Cordelia Langford, John Sillitoe on behalf of the Wellcome Sanger Institute COVID-19 Surveillance Team |
| EPI_ISL_3042082, EPI_ISL_3042083 | Lighthouse Laboratory Plymouth | Wellcome Sanger Institute for the COVID-19 Genomics UK (COG-UK) Consortium | Lighthouse Laboratory Plymouth and Alex Alderton, Roberto Amato, Jeffrey Barrett, Sonia Goncalves, Ewan Harrison, David K. Jackson, Ian Johnston, Dominic Kwiatkowski, Cordelia Langford, John Sillitoe on behalf of the Wellcome Sanger Institute COVID-19 Surveillance Team |
| EPI_ISL_3042084 | Lighthouse Lab in Milton Keynes | Wellcome Sanger Institute for the COVID-19 Genomics UK (COG-UK) Consortium | The Lighthouse Lab in Milton Keynes and Alex Alderton, Roberto Amato, Jeffrey Barrett, Sonia Goncalves, Ewan Harrison, David K. Jackson, Ian Johnston, Dominic Kwiatkowski, Cordelia Langford, John Sillitoe on behalf of the Wellcome Sanger Institute COVID-19 Surveillance Team |
| EPI_ISL_3042085 | Lighthouse Laboratory Plymouth | Wellcome Sanger Institute for the COVID-19 Genomics UK (COG-UK) Consortium | Lighthouse Laboratory Plymouth and Alex Alderton, Roberto Amato, Jeffrey Barrett, Sonia Goncalves, Ewan Harrison, David K. Jackson, Ian Johnston, Dominic Kwiatkowski, Cordelia Langford, John Sillitoe on behalf of the Wellcome Sanger Institute COVID-19 Surveillance Team |
| EPI_ISL_3042086 | Lighthouse Lab in Milton Keynes | Wellcome Sanger Institute for the COVID-19 Genomics UK (COG-UK) Consortium | The Lighthouse Lab in Milton Keynes and Alex Alderton, Roberto Amato, Jeffrey Barrett, Sonia Goncalves, Ewan Harrison, David K. Jackson, Ian Johnston, Dominic Kwiatkowski, Cordelia Langford, John Sillitoe on behalf of the Wellcome Sanger Institute COVID-19 Surveillance Team |
| EPI_ISL_3042087, EPI_ISL_3042088, EPI_ISL_3042089, EPI_ISL_3042090 | Lighthouse Laboratory Plymouth | Wellcome Sanger Institute for the COVID-19 Genomics UK (COG-UK) Consortium | Lighthouse Laboratory Plymouth and Alex Alderton, Roberto Amato, Jeffrey Barrett, Sonia Goncalves, Ewan Harrison, David K. Jackson, Ian Johnston, Dominic Kwiatkowski, Cordelia Langford, John Sillitoe on behalf of the Wellcome Sanger Institute COVID-19 Surveillance Team |
| EPI_ISL_3042091 | Lighthouse Lab in Alderley Park | Wellcome Sanger Institute for the COVID-19 Genomics UK (COG-UK) Consortium | Jacquelyn Wynn, Mairead Hyland, The Lighthouse Lab in Alderley Park and Alex Alderton, Roberto Amato, Jeffrey Barrett, Sonia Goncalves, Ewan Harrison, David K. Jackson, Ian Johnston, Dominic Kwiatkowski, Cordelia Langford, John Sillitoe on behalf of the Wellcome Sanger Institute COVID-19 Surveillance Team |
| EPI_ISL_3042092 | Berkshire and Surrey Pathology Services Lighthouse Laboratory | Wellcome Sanger Institute for the COVID-19 Genomics UK (COG-UK) Consortium | Berkshire and Surrey Pathology Services Lighthouse Laboratory and Alex Alderton, Roberto Amato, Jeffrey Barrett, Sonia Goncalves, Ewan Harrison, David K. Jackson, Ian Johnston, Dominic Kwiatkowski, Cordelia Langford, John Sillitoe on behalf of the Wellcome Sanger Institute COVID-19 Surveillance Team |
| EPI_ISL_3042093 | Lighthouse Laboratory Plymouth | Wellcome Sanger Institute for the COVID-19 Genomics UK (COG-UK) Consortium | Lighthouse Laboratory Plymouth and Alex Alderton, Roberto Amato, Jeffrey Barrett, Sonia Goncalves, Ewan Harrison, David K. Jackson, Ian Johnston, Dominic Kwiatkowski, Cordelia Langford, John Sillitoe on behalf of the Wellcome Sanger Institute COVID-19 Surveillance Team |
| EPI_ISL_3042094 | Lighthouse Lab in Alderley Park | Wellcome Sanger Institute for the COVID-19 Genomics UK (COG-UK) Consortium | Jacquelyn Wynn, Mairead Hyland, The Lighthouse Lab in Alderley Park and Alex Alderton, Roberto Amato, Jeffrey Barrett, Sonia Goncalves, Ewan Harrison, David K. Jackson, Ian Johnston, Dominic Kwiatkowski, Cordelia Langford, John Sillitoe on behalf of the Wellcome Sanger Institute COVID-19 Surveillance Team |
| EPI_ISL_3042095, EPI_ISL_3042096 | Berkshire and Surrey Pathology Services Lighthouse Laboratory | Wellcome Sanger Institute for the COVID-19 Genomics UK (COG-UK) Consortium | Berkshire and Surrey Pathology Services Lighthouse Laboratory and Alex Alderton, Roberto Amato, Jeffrey Barrett, Sonia Goncalves, Ewan Harrison, David K. Jackson, Ian Johnston, Dominic Kwiatkowski, Cordelia Langford, John Sillitoe on behalf of the Wellcome Sanger Institute COVID-19 Surveillance Team |
| EPI_ISL_3042097, EPI_ISL_3042098 | Lighthouse Lab in Milton Keynes | Wellcome Sanger Institute for the COVID-19 Genomics UK (COG-UK) Consortium | The Lighthouse Lab in Milton Keynes and Alex Alderton, Roberto Amato, Jeffrey Barrett, Sonia Goncalves, Ewan Harrison, David K. Jackson, Ian Johnston, Dominic Kwiatkowski, Cordelia Langford, John Sillitoe on behalf of the Wellcome Sanger Institute COVID-19 Surveillance Team |
| EPI_ISL_3042099, EPI_ISL_3042100 | Lighthouse Lab in Alderley Park | Wellcome Sanger Institute for the COVID-19 Genomics UK (COG-UK) Consortium | Jacquelyn Wynn, Mairead Hyland, The Lighthouse Lab in Alderley Park and Alex Alderton, Roberto Amato, Jeffrey Barrett, Sonia Goncalves, Ewan Harrison, David K. Jackson, Ian Johnston, Dominic Kwiatkowski, Cordelia Langford, John Sillitoe on behalf of the Wellcome Sanger Institute COVID-19 Surveillance Team |
| EPI_ISL_3042101 | Lighthouse Lab in Milton Keynes | Wellcome Sanger Institute for the COVID-19 Genomics UK (COG-UK) Consortium | The Lighthouse Lab in Milton Keynes and Alex Alderton, Roberto Amato, Jeffrey Barrett, Sonia Goncalves, Ewan Harrison, David K. Jackson, Ian Johnston, Dominic Kwiatkowski, Cordelia Langford, John Sillitoe on behalf of the Wellcome Sanger Institute COVID-19 Surveillance Team |
| EPI_ISL_3042102 | Lighthouse Lab in Alderley Park | Wellcome Sanger Institute for the COVID-19 Genomics UK (COG-UK) Consortium | Jacquelyn Wynn, Mairead Hyland, The Lighthouse Lab in Alderley Park and Alex Alderton, Roberto Amato, Jeffrey Barrett, Sonia Goncalves, Ewan Harrison, David K. Jackson, Ian Johnston, Dominic Kwiatkowski, Cordelia Langford, John Sillitoe on behalf of the Wellcome Sanger Institute COVID-19 Surveillance Team |
| EPI_ISL_3042103 | Lighthouse Lab in Milton Keynes | Wellcome Sanger Institute for the COVID-19 Genomics UK (COG-UK) Consortium | The Lighthouse Lab in Milton Keynes and Alex Alderton, Roberto Amato, Jeffrey Barrett, Sonia Goncalves, Ewan Harrison, David K. Jackson, Ian Johnston, Dominic Kwiatkowski, Cordelia Langford, John Sillitoe on behalf of the Wellcome Sanger Institute COVID-19 Surveillance Team |
| EPI_ISL_3042104 | Lighthouse Lab in Alderley Park | Wellcome Sanger Institute for the COVID-19 Genomics UK (COG-UK) Consortium | Jacquelyn Wynn, Mairead Hyland, The Lighthouse Lab in Alderley Park and Alex Alderton, Roberto Amato, Jeffrey Barrett, Sonia Goncalves, Ewan Harrison, David K. Jackson, Ian Johnston, Dominic Kwiatkowski, Cordelia Langford, John Sillitoe on behalf of the Wellcome Sanger Institute COVID-19 Surveillance Team |
| EPI_ISL_3042105 | Lighthouse Lab in Milton Keynes | Wellcome Sanger Institute for the COVID-19 Genomics UK (COG-UK) Consortium | The Lighthouse Lab in Milton Keynes and Alex Alderton, Roberto Amato, Jeffrey Barrett, Sonia Goncalves, Ewan Harrison, David K. Jackson, Ian Johnston, Dominic Kwiatkowski, Cordelia Langford, John Sillitoe on behalf of the Wellcome Sanger Institute COVID-19 Surveillance Team |
| EPI_ISL_3042106 | Lighthouse Laboratory Plymouth | Wellcome Sanger Institute for the COVID-19 Genomics UK (COG-UK) Consortium | Lighthouse Laboratory Plymouth and Alex Alderton, Roberto Amato, Jeffrey Barrett, Sonia Goncalves, Ewan Harrison, David K. Jackson, Ian Johnston, Dominic Kwiatkowski, Cordelia Langford, John Sillitoe on behalf of the Wellcome Sanger Institute COVID-19 Surveillance Team |
| EPI_ISL_3042107, EPI_ISL_3042108 | Lighthouse Lab in Milton Keynes | Wellcome Sanger Institute for the COVID-19 Genomics UK (COG-UK) Consortium | The Lighthouse Lab in Milton Keynes and Alex Alderton, Roberto Amato, Jeffrey Barrett, Sonia Goncalves, Ewan Harrison, David K. Jackson, Ian Johnston, Dominic Kwiatkowski, Cordelia Langford, John Sillitoe on behalf of the Wellcome Sanger Institute COVID-19 Surveillance Team |
| EPI_ISL_3042109, EPI_ISL_3042110 | Lighthouse Laboratory Plymouth | Wellcome Sanger Institute for the COVID-19 Genomics UK (COG-UK) Consortium | Lighthouse Laboratory Plymouth and Alex Alderton, Roberto Amato, Jeffrey Barrett, Sonia Goncalves, Ewan Harrison, David K. Jackson, Ian Johnston, Dominic Kwiatkowski, Cordelia Langford, John Sillitoe on behalf of the Wellcome Sanger Institute COVID-19 Surveillance Team |
| EPI_ISL_3042111 | Berkshire and Surrey Pathology Services Lighthouse Laboratory | Wellcome Sanger Institute for the COVID-19 Genomics UK (COG-UK) Consortium | Berkshire and Surrey Pathology Services Lighthouse Laboratory and Alex Alderton, Roberto Amato, Jeffrey Barrett, Sonia Goncalves, Ewan Harrison, David K. Jackson, Ian Johnston, Dominic Kwiatkowski, Cordelia Langford, John Sillitoe on behalf of the Wellcome Sanger Institute COVID-19 Surveillance Team |
| EPI_ISL_3042112 | Lighthouse Laboratory Plymouth | Wellcome Sanger Institute for the COVID-19 Genomics UK (COG-UK) Consortium | Lighthouse Laboratory Plymouth and Alex Alderton, Roberto Amato, Jeffrey Barrett, Sonia Goncalves, Ewan Harrison, David K. Jackson, Ian Johnston, Dominic Kwiatkowski, Cordelia Langford, John Sillitoe on behalf of the Wellcome Sanger Institute COVID-19 Surveillance Team |
| EPI_ISL_3042113, EPI_ISL_3042114 | Berkshire and Surrey Pathology Services Lighthouse Laboratory | Wellcome Sanger Institute for the COVID-19 Genomics UK (COG-UK) Consortium | Berkshire and Surrey Pathology Services Lighthouse Laboratory and Alex Alderton, Roberto Amato, Jeffrey Barrett, Sonia Goncalves, Ewan Harrison, David K. Jackson, Ian Johnston, Dominic Kwiatkowski, Cordelia Langford, John Sillitoe on behalf of the Wellcome Sanger Institute COVID-19 Surveillance Team |
| EPI_ISL_3042115 | Lighthouse Lab in Milton Keynes | Wellcome Sanger Institute for the COVID-19 Genomics UK (COG-UK) Consortium | The Lighthouse Lab in Milton Keynes and Alex Alderton, Roberto Amato, Jeffrey Barrett, Sonia Goncalves, Ewan Harrison, David K. Jackson, Ian Johnston, Dominic Kwiatkowski, Cordelia Langford, John Sillitoe on behalf of the Wellcome Sanger Institute COVID-19 Surveillance Team |
| EPI_ISL_3042116, EPI_ISL_3042117 | Lighthouse Lab in Alderley Park | Wellcome Sanger Institute for the COVID-19 Genomics UK (COG-UK) Consortium | Jacquelyn Wynn, Mairead Hyland, The Lighthouse Lab in Alderley Park and Alex Alderton, Roberto Amato, Jeffrey Barrett, Sonia Goncalves, Ewan Harrison, David K. Jackson, Ian Johnston, Dominic Kwiatkowski, Cordelia Langford, John Sillitoe on behalf of the Wellcome Sanger Institute COVID-19 Surveillance Team |
| EPI_ISL_3042118 | Lighthouse Lab in Milton Keynes | Wellcome Sanger Institute for the COVID-19 Genomics UK (COG-UK) Consortium | The Lighthouse Lab in Milton Keynes and Alex Alderton, Roberto Amato, Jeffrey Barrett, Sonia Goncalves, Ewan Harrison, David K. Jackson, Ian Johnston, Dominic Kwiatkowski, Cordelia Langford, John Sillitoe on behalf of the Wellcome Sanger Institute COVID-19 Surveillance Team |
| EPI_ISL_3042119 | Berkshire and Surrey Pathology Services Lighthouse Laboratory | Wellcome Sanger Institute for the COVID-19 Genomics UK (COG-UK) Consortium | Berkshire and Surrey Pathology Services Lighthouse Laboratory and Alex Alderton, Roberto Amato, Jeffrey Barrett, Sonia Goncalves, Ewan Harrison, David K. Jackson, Ian Johnston, Dominic Kwiatkowski, Cordelia Langford, John Sillitoe on behalf of the Wellcome Sanger Institute COVID-19 Surveillance Team |
| EPI_ISL_3042120 | Lighthouse Lab in Milton Keynes | Wellcome Sanger Institute for the COVID-19 Genomics UK (COG-UK) Consortium | The Lighthouse Lab in Milton Keynes and Alex Alderton, Roberto Amato, Jeffrey Barrett, Sonia Goncalves, Ewan Harrison, David K. Jackson, Ian Johnston, Dominic Kwiatkowski, Cordelia Langford, John Sillitoe on behalf of the Wellcome Sanger Institute COVID-19 Surveillance Team |

| | | | |
|---|---|---|---|
| EPI_ISL_3042121 | Lighthouse Laboratory Plymouth | Wellcome Sanger Institute for the COVID-19 Genomics UK (COG-UK) Consortium | Lighthouse Laboratory Plymouth and Alex Alderton, Roberto Amato, Jeffrey Barrett, Sonia Goncalves, Ewan Harrison, David K. Jackson, Ian Johnston, Dominic Kwiatkowski, Cordelia Langford, John Sillitoe on behalf of the Wellcome Sanger Institute COVID-19 Surveillance Team |
| EPI_ISL_3042122 | Berkshire and Surrey Pathology Services Lighthouse Laboratory | Wellcome Sanger Institute for the COVID-19 Genomics UK (COG-UK) Consortium | Berkshire and Surrey Pathology Services Lighthouse Laboratory and Alex Alderton, Roberto Amato, Jeffrey Barrett, Sonia Goncalves, Ewan Harrison, David K. Jackson, Ian Johnston, Dominic Kwiatkowski, Cordelia Langford, John Sillitoe on behalf of the Wellcome Sanger Institute COVID-19 Surveillance Team |
| EPI_ISL_3042123 | Lighthouse Lab in Milton Keynes | Wellcome Sanger Institute for the COVID-19 Genomics UK (COG-UK) Consortium | The Lighthouse Lab in Milton Keynes and Alex Alderton, Roberto Amato, Jeffrey Barrett, Sonia Goncalves, Ewan Harrison, David K. Jackson, Ian Johnston, Dominic Kwiatkowski, Cordelia Langford, John Sillitoe on behalf of the Wellcome Sanger Institute COVID-19 Surveillance Team |
| EPI_ISL_3042124 | Berkshire and Surrey Pathology Services Lighthouse Laboratory | Wellcome Sanger Institute for the COVID-19 Genomics UK (COG-UK) Consortium | Berkshire and Surrey Pathology Services Lighthouse Laboratory and Alex Alderton, Roberto Amato, Jeffrey Barrett, Sonia Goncalves, Ewan Harrison, David K. Jackson, Ian Johnston, Dominic Kwiatkowski, Cordelia Langford, John Sillitoe on behalf of the Wellcome Sanger Institute COVID-19 Surveillance Team |
| EPI_ISL_3042125 | Lighthouse Lab in Milton Keynes | Wellcome Sanger Institute for the COVID-19 Genomics UK (COG-UK) Consortium | The Lighthouse Lab in Milton Keynes and Alex Alderton, Roberto Amato, Jeffrey Barrett, Sonia Goncalves, Ewan Harrison, David K. Jackson, Ian Johnston, Dominic Kwiatkowski, Cordelia Langford, John Sillitoe on behalf of the Wellcome Sanger Institute COVID-19 Surveillance Team |
| EPI_ISL_3042126 | Lighthouse Lab in Alderley Park | Wellcome Sanger Institute for the COVID-19 Genomics UK (COG-UK) Consortium | Jacquelyn Wynn, Mairead Hyland, The Lighthouse Lab in Alderley Park and Alex Alderton, Roberto Amato, Jeffrey Barrett, Sonia Goncalves, Ewan Harrison, David K. Jackson, Ian Johnston, Dominic Kwiatkowski, Cordelia Langford, John Sillitoe on behalf of the Wellcome Sanger Institute COVID-19 Surveillance Team |
| EPI_ISL_3042127 | Lighthouse Lab in Milton Keynes | Wellcome Sanger Institute for the COVID-19 Genomics UK (COG-UK) Consortium | The Lighthouse Lab in Milton Keynes and Alex Alderton, Roberto Amato, Jeffrey Barrett, Sonia Goncalves, Ewan Harrison, David K. Jackson, Ian Johnston, Dominic Kwiatkowski, Cordelia Langford, John Sillitoe on behalf of the Wellcome Sanger Institute COVID-19 Surveillance Team |
| EPI_ISL_3042128 | Berkshire and Surrey Pathology Services Lighthouse Laboratory | Wellcome Sanger Institute for the COVID-19 Genomics UK (COG-UK) Consortium | Berkshire and Surrey Pathology Services Lighthouse Laboratory and Alex Alderton, Roberto Amato, Jeffrey Barrett, Sonia Goncalves, Ewan Harrison, David K. Jackson, Ian Johnston, Dominic Kwiatkowski, Cordelia Langford, John Sillitoe on behalf of the Wellcome Sanger Institute COVID-19 Surveillance Team |
| EPI_ISL_3042129 | Lighthouse Laboratory Plymouth | Wellcome Sanger Institute for the COVID-19 Genomics UK (COG-UK) Consortium | Lighthouse Laboratory Plymouth and Alex Alderton, Roberto Amato, Jeffrey Barrett, Sonia Goncalves, Ewan Harrison, David K. Jackson, Ian Johnston, Dominic Kwiatkowski, Cordelia Langford, John Sillitoe on behalf of the Wellcome Sanger Institute COVID-19 Surveillance Team |
| EPI_ISL_3042130 | Lighthouse Lab in Alderley Park | Wellcome Sanger Institute for the COVID-19 Genomics UK (COG-UK) Consortium | Jacquelyn Wynn, Mairead Hyland, The Lighthouse Lab in Alderley Park and Alex Alderton, Roberto Amato, Jeffrey Barrett, Sonia Goncalves, Ewan Harrison, David K. Jackson, Ian Johnston, Dominic Kwiatkowski, Cordelia Langford, John Sillitoe on behalf of the Wellcome Sanger Institute COVID-19 Surveillance Team |
| EPI_ISL_3042131 | Lighthouse Lab in Milton Keynes | Wellcome Sanger Institute for the COVID-19 Genomics UK (COG-UK) Consortium | The Lighthouse Lab in Milton Keynes and Alex Alderton, Roberto Amato, Jeffrey Barrett, Sonia Goncalves, Ewan Harrison, David K. Jackson, Ian Johnston, Dominic Kwiatkowski, Cordelia Langford, John Sillitoe on behalf of the Wellcome Sanger Institute COVID-19 Surveillance Team |
| EPI_ISL_3042132 | Berkshire and Surrey Pathology Services Lighthouse Laboratory | Wellcome Sanger Institute for the COVID-19 Genomics UK (COG-UK) Consortium | Berkshire and Surrey Pathology Services Lighthouse Laboratory and Alex Alderton, Roberto Amato, Jeffrey Barrett, Sonia Goncalves, Ewan Harrison, David K. Jackson, Ian Johnston, Dominic Kwiatkowski, Cordelia Langford, John Sillitoe on behalf of the Wellcome Sanger Institute COVID-19 Surveillance Team |
| EPI_ISL_3042133 | Lighthouse Lab in Milton Keynes | Wellcome Sanger Institute for the COVID-19 Genomics UK (COG-UK) Consortium | The Lighthouse Lab in Milton Keynes and Alex Alderton, Roberto Amato, Jeffrey Barrett, Sonia Goncalves, Ewan Harrison, David K. Jackson, Ian Johnston, Dominic Kwiatkowski, Cordelia Langford, John Sillitoe on behalf of the Wellcome Sanger Institute COVID-19 Surveillance Team |
| EPI_ISL_3042134, EPI_ISL_3042135, EPI_ISL_3042136 | Lighthouse Lab in Alderley Park | Wellcome Sanger Institute for the COVID-19 Genomics UK (COG-UK) Consortium | Jacquelyn Wynn, Mairead Hyland, The Lighthouse Lab in Alderley Park and Alex Alderton, Roberto Amato, Jeffrey Barrett, Sonia Goncalves, Ewan Harrison, David K. Jackson, Ian Johnston, Dominic Kwiatkowski, Cordelia Langford, John Sillitoe on behalf of the Wellcome Sanger Institute COVID-19 Surveillance Team |
| EPI_ISL_3042137, EPI_ISL_3042138, EPI_ISL_3042139 | Lighthouse Lab in Milton Keynes | Wellcome Sanger Institute for the COVID-19 Genomics UK (COG-UK) Consortium | The Lighthouse Lab in Milton Keynes and Alex Alderton, Roberto Amato, Jeffrey Barrett, Sonia Goncalves, Ewan Harrison, David K. Jackson, Ian Johnston, Dominic Kwiatkowski, Cordelia Langford, John Sillitoe on behalf of the Wellcome Sanger Institute COVID-19 Surveillance Team |
| EPI_ISL_3042140 | Lighthouse Laboratory Plymouth | Wellcome Sanger Institute for the COVID-19 Genomics UK (COG-UK) Consortium | Lighthouse Laboratory Plymouth and Alex Alderton, Roberto Amato, Jeffrey Barrett, Sonia Goncalves, Ewan Harrison, David K. Jackson, Ian Johnston, Dominic Kwiatkowski, Cordelia Langford, John Sillitoe on behalf of the Wellcome Sanger Institute COVID-19 Surveillance Team |
| EPI_ISL_3042141 | Lighthouse Lab in Alderley Park | Wellcome Sanger Institute for the COVID-19 Genomics UK (COG-UK) Consortium | Jacquelyn Wynn, Mairead Hyland, The Lighthouse Lab in Alderley Park and Alex Alderton, Roberto Amato, Jeffrey Barrett, Sonia Goncalves, Ewan Harrison, David K. Jackson, Ian Johnston, Dominic Kwiatkowski, Cordelia Langford, John Sillitoe on behalf of the Wellcome Sanger Institute COVID-19 Surveillance Team |
| EPI_ISL_3042142 | Lighthouse Laboratory Plymouth | Wellcome Sanger Institute for the COVID-19 Genomics UK (COG-UK) Consortium | Lighthouse Laboratory Plymouth and Alex Alderton, Roberto Amato, Jeffrey Barrett, Sonia Goncalves, Ewan Harrison, David K. Jackson, Ian Johnston, Dominic Kwiatkowski, Cordelia Langford, John Sillitoe on behalf of the Wellcome Sanger Institute COVID-19 Surveillance Team |
| EPI_ISL_3042143 | Lighthouse Lab in Alderley Park | Wellcome Sanger Institute for the COVID-19 Genomics UK (COG-UK) Consortium | Jacquelyn Wynn, Mairead Hyland, The Lighthouse Lab in Alderley Park and Alex Alderton, Roberto Amato, Jeffrey Barrett, Sonia Goncalves, Ewan Harrison, David K. Jackson, Ian Johnston, Dominic Kwiatkowski, Cordelia Langford, John Sillitoe on behalf of the Wellcome Sanger Institute COVID-19 Surveillance Team |
| EPI_ISL_3042144 | Berkshire and Surrey Pathology Services Lighthouse Laboratory | Wellcome Sanger Institute for the COVID-19 Genomics UK (COG-UK) Consortium | Berkshire and Surrey Pathology Services Lighthouse Laboratory and Alex Alderton, Roberto Amato, Jeffrey Barrett, Sonia Goncalves, Ewan Harrison, David K. Jackson, Ian Johnston, Dominic Kwiatkowski, Cordelia Langford, John Sillitoe on behalf of the Wellcome Sanger Institute COVID-19 Surveillance Team |
| EPI_ISL_3042145 | Lighthouse Lab in Milton Keynes | Wellcome Sanger Institute for the COVID-19 Genomics UK (COG-UK) Consortium | The Lighthouse Lab in Milton Keynes and Alex Alderton, Roberto Amato, Jeffrey Barrett, Sonia Goncalves, Ewan Harrison, David K. Jackson, Ian Johnston, Dominic Kwiatkowski, Cordelia Langford, John Sillitoe on behalf of the Wellcome Sanger Institute COVID-19 Surveillance Team |
| EPI_ISL_3042146 | Lighthouse Laboratory Plymouth | Wellcome Sanger Institute for the COVID-19 Genomics UK (COG-UK) Consortium | Lighthouse Laboratory Plymouth and Alex Alderton, Roberto Amato, Jeffrey Barrett, Sonia Goncalves, Ewan Harrison, David K. Jackson, Ian Johnston, Dominic Kwiatkowski, Cordelia Langford, John Sillitoe on behalf of the Wellcome Sanger Institute COVID-19 Surveillance Team |
| EPI_ISL_3042147 | Lighthouse Lab in Milton Keynes | Wellcome Sanger Institute for the COVID-19 Genomics UK (COG-UK) Consortium | The Lighthouse Lab in Milton Keynes and Alex Alderton, Roberto Amato, Jeffrey Barrett, Sonia Goncalves, Ewan Harrison, David K. Jackson, Ian Johnston, Dominic Kwiatkowski, Cordelia Langford, John Sillitoe on behalf of the Wellcome Sanger Institute COVID-19 Surveillance Team |
| EPI_ISL_3042148 | Berkshire and Surrey Pathology Services Lighthouse Laboratory | Wellcome Sanger Institute for the COVID-19 Genomics UK (COG-UK) Consortium | Berkshire and Surrey Pathology Services Lighthouse Laboratory and Alex Alderton, Roberto Amato, Jeffrey Barrett, Sonia Goncalves, Ewan Harrison, David K. Jackson, Ian Johnston, Dominic Kwiatkowski, Cordelia Langford, John Sillitoe on behalf of the Wellcome Sanger Institute COVID-19 Surveillance Team |
| EPI_ISL_3042149, EPI_ISL_3042150 | Lighthouse Lab in Milton Keynes | Wellcome Sanger Institute for the COVID-19 Genomics UK (COG-UK) Consortium | The Lighthouse Lab in Milton Keynes and Alex Alderton, Roberto Amato, Jeffrey Barrett, Sonia Goncalves, Ewan Harrison, David K. Jackson, Ian Johnston, Dominic Kwiatkowski, Cordelia Langford, John Sillitoe on behalf of the Wellcome Sanger Institute COVID-19 Surveillance Team |
| EPI_ISL_3042151 | Lighthouse Lab in Alderley Park | Wellcome Sanger Institute for the COVID-19 Genomics UK (COG-UK) Consortium | Jacquelyn Wynn, Mairead Hyland, The Lighthouse Lab in Alderley Park and Alex Alderton, Roberto Amato, Jeffrey Barrett, Sonia Goncalves, Ewan Harrison, David K. Jackson, Ian Johnston, Dominic Kwiatkowski, Cordelia Langford, John Sillitoe on behalf of the Wellcome Sanger Institute COVID-19 Surveillance Team |
| EPI_ISL_3042152 | Berkshire and Surrey Pathology Services Lighthouse Laboratory | Wellcome Sanger Institute for the COVID-19 Genomics UK (COG-UK) Consortium | Berkshire and Surrey Pathology Services Lighthouse Laboratory and Alex Alderton, Roberto Amato, Jeffrey Barrett, Sonia Goncalves, Ewan Harrison, David K. Jackson, Ian Johnston, Dominic Kwiatkowski, Cordelia Langford, John Sillitoe on behalf of the Wellcome Sanger Institute COVID-19 Surveillance Team |
| EPI_ISL_3042153 | Lighthouse Lab in Milton Keynes | Wellcome Sanger Institute for the COVID-19 Genomics UK (COG-UK) Consortium | The Lighthouse Lab in Milton Keynes and Alex Alderton, Roberto Amato, Jeffrey Barrett, Sonia Goncalves, Ewan Harrison, David K. Jackson, Ian Johnston, Dominic Kwiatkowski, Cordelia Langford, John Sillitoe on behalf of the Wellcome Sanger Institute COVID-19 Surveillance Team |
| EPI_ISL_3042154 | Lighthouse Lab in Alderley Park | Wellcome Sanger Institute for the COVID-19 Genomics UK (COG-UK) Consortium | Jacquelyn Wynn, Mairead Hyland, The Lighthouse Lab in Alderley Park and Alex Alderton, Roberto Amato, Jeffrey Barrett, Sonia Goncalves, Ewan Harrison, David K. Jackson, Ian Johnston, Dominic Kwiatkowski, Cordelia Langford, John Sillitoe on behalf of the Wellcome Sanger Institute COVID-19 Surveillance Team |
| EPI_ISL_3042155 | Lighthouse Laboratory Plymouth | Wellcome Sanger Institute for the COVID-19 Genomics UK (COG-UK) Consortium | Lighthouse Laboratory Plymouth and Alex Alderton, Roberto Amato, Jeffrey Barrett, Sonia Goncalves, Ewan Harrison, David K. Jackson, Ian Johnston, Dominic Kwiatkowski, Cordelia Langford, John Sillitoe on behalf of the Wellcome Sanger Institute COVID-19 Surveillance Team |
| EPI_ISL_3042156 | Lighthouse Lab in Alderley Park | Wellcome Sanger Institute for the COVID-19 Genomics UK (COG-UK) Consortium | Jacquelyn Wynn, Mairead Hyland, The Lighthouse Lab in Alderley Park and Alex Alderton, Roberto Amato, Jeffrey Barrett, Sonia Goncalves, Ewan Harrison, David K. Jackson, Ian Johnston, Dominic Kwiatkowski, Cordelia Langford, John Sillitoe on behalf of the Wellcome Sanger Institute COVID-19 Surveillance Team |
| EPI_ISL_3042157 | Lighthouse Lab in Milton Keynes | Wellcome Sanger Institute for the COVID-19 Genomics UK (COG-UK) Consortium | The Lighthouse Lab in Milton Keynes and Alex Alderton, Roberto Amato, Jeffrey Barrett, Sonia Goncalves, Ewan Harrison, David K. Jackson, Ian Johnston, Dominic Kwiatkowski, Cordelia Langford, John Sillitoe on behalf of the Wellcome Sanger Institute COVID-19 Surveillance Team |
| EPI_ISL_3042158 | Berkshire and Surrey Pathology Services Lighthouse Laboratory | Wellcome Sanger Institute for the COVID-19 Genomics UK (COG-UK) Consortium | Berkshire and Surrey Pathology Services Lighthouse Laboratory and Alex Alderton, Roberto Amato, Jeffrey Barrett, Sonia Goncalves, Ewan Harrison, David K. Jackson, Ian Johnston, Dominic Kwiatkowski, Cordelia Langford, John Sillitoe on behalf of the Wellcome Sanger Institute COVID-19 Surveillance Team |
| EPI_ISL_3042159 | Lighthouse Lab in Alderley Park | Wellcome Sanger Institute for the COVID-19 Genomics UK (COG-UK) Consortium | Jacquelyn Wynn, Mairead Hyland, The Lighthouse Lab in Alderley Park and Alex Alderton, Roberto Amato, Jeffrey Barrett, Sonia Goncalves, Ewan Harrison, David K. Jackson, Ian Johnston, Dominic Kwiatkowski, Cordelia Langford, John Sillitoe on behalf of the Wellcome Sanger Institute COVID-19 Surveillance Team |
| EPI_ISL_3042160 | Berkshire and Surrey Pathology Services Lighthouse Laboratory | Wellcome Sanger Institute for the COVID-19 Genomics UK (COG-UK) Consortium | Berkshire and Surrey Pathology Services Lighthouse Laboratory and Alex Alderton, Roberto Amato, Jeffrey Barrett, Sonia Goncalves, Ewan Harrison, David K. Jackson, Ian Johnston, Dominic Kwiatkowski, Cordelia Langford, John Sillitoe on behalf of the Wellcome Sanger Institute COVID-19 Surveillance Team |
| EPI_ISL_3042161, EPI_ISL_3042162 | Lighthouse Lab in Alderley Park | Wellcome Sanger Institute for the COVID-19 Genomics UK (COG-UK) Consortium | Jacquelyn Wynn, Mairead Hyland, The Lighthouse Lab in Alderley Park and Alex Alderton, Roberto Amato, Jeffrey Barrett, Sonia Goncalves, Ewan Harrison, David K. Jackson, Ian Johnston, Dominic Kwiatkowski, Cordelia Langford, John Sillitoe on behalf of the Wellcome Sanger Institute COVID-19 Surveillance Team |
| EPI_ISL_3042163 | Lighthouse Laboratory Plymouth | Wellcome Sanger Institute for the COVID-19 Genomics UK (COG-UK) Consortium | Lighthouse Laboratory Plymouth and Alex Alderton, Roberto Amato, Jeffrey Barrett, Sonia Goncalves, Ewan Harrison, David K. Jackson, Ian Johnston, Dominic Kwiatkowski, Cordelia Langford, John Sillitoe on behalf of the Wellcome Sanger Institute COVID-19 Surveillance Team |
| EPI_ISL_3042164 | Berkshire and Surrey Pathology Services Lighthouse Laboratory | Wellcome Sanger Institute for the COVID-19 Genomics UK (COG-UK) Consortium | Berkshire and Surrey Pathology Services Lighthouse Laboratory and Alex Alderton, Roberto Amato, Jeffrey Barrett, Sonia Goncalves, Ewan Harrison, David K. Jackson, Ian Johnston, Dominic Kwiatkowski, Cordelia Langford, John Sillitoe on behalf of the Wellcome Sanger Institute COVID-19 Surveillance Team |
| EPI_ISL_3042165 | Lighthouse Lab in Alderley Park | Wellcome Sanger Institute for the COVID-19 Genomics UK (COG-UK) Consortium | Jacquelyn Wynn, Mairead Hyland, The Lighthouse Lab in Alderley Park and Alex Alderton, Roberto Amato, Jeffrey Barrett, Sonia Goncalves, Ewan Harrison, David K. Jackson, Ian Johnston, Dominic Kwiatkowski, Cordelia Langford, John Sillitoe on behalf of the Wellcome Sanger Institute COVID-19 Surveillance Team |
| EPI_ISL_3042166 | Lighthouse Laboratory Plymouth | Wellcome Sanger Institute for the COVID-19 Genomics UK (COG-UK) Consortium | Lighthouse Laboratory Plymouth and Alex Alderton, Roberto Amato, Jeffrey Barrett, Sonia Goncalves, Ewan Harrison, David K. Jackson, Ian Johnston, Dominic Kwiatkowski, Cordelia Langford, John Sillitoe on behalf of the Wellcome Sanger Institute COVID-19 Surveillance Team |
| EPI_ISL_3042167, EPI_ISL_3042168 | Lighthouse Lab in Milton Keynes | Wellcome Sanger Institute for the COVID-19 Genomics UK (COG-UK) Consortium | The Lighthouse Lab in Milton Keynes and Alex Alderton, Roberto Amato, Jeffrey Barrett, Sonia Goncalves, Ewan Harrison, David K. Jackson, Ian Johnston, Dominic Kwiatkowski, Cordelia Langford, John Sillitoe on behalf of the Wellcome Sanger Institute COVID-19 Surveillance Team |
| EPI_ISL_3042169 | Lighthouse Laboratory Plymouth | Wellcome Sanger Institute for the COVID-19 Genomics UK (COG-UK) Consortium | Lighthouse Laboratory Plymouth and Alex Alderton, Roberto Amato, Jeffrey Barrett, Sonia Goncalves, Ewan Harrison, David K. Jackson, Ian Johnston, Dominic Kwiatkowski, Cordelia Langford, John Sillitoe on behalf of the Wellcome Sanger Institute COVID-19 Surveillance Team |
| EPI_ISL_3042170 | Lighthouse Lab in Alderley Park | Wellcome Sanger Institute for the COVID-19 Genomics UK (COG-UK) Consortium | Jacquelyn Wynn, Mairead Hyland, The Lighthouse Lab in Alderley Park and Alex Alderton, Roberto Amato, Jeffrey Barrett, Sonia Goncalves, Ewan Harrison, David K. Jackson, Ian Johnston, Dominic Kwiatkowski, Cordelia Langford, John Sillitoe on behalf of the Wellcome Sanger Institute COVID-19 Surveillance Team |
| EPI_ISL_3042171, EPI_ISL_3042172 | Lighthouse Laboratory Plymouth | Wellcome Sanger Institute for the COVID-19 Genomics UK (COG-UK) Consortium | Lighthouse Laboratory Plymouth and Alex Alderton, Roberto Amato, Jeffrey Barrett, Sonia Goncalves, Ewan Harrison, David K. Jackson, Ian Johnston, Dominic Kwiatkowski, Cordelia Langford, John Sillitoe on behalf of the Wellcome Sanger Institute COVID-19 Surveillance Team |
| EPI_ISL_3042173 | Lighthouse Lab in Milton Keynes | Wellcome Sanger Institute for the COVID-19 Genomics UK (COG-UK) | The Lighthouse Lab in Milton Keynes and Alex Alderton, Roberto Amato, Jeffrey Barrett, Sonia Goncalves, Ewan Harrison, David K. Jackson, Ian Johnston, Dominic Kwiatkowski, Cordelia Langford, John Sillitoe on behalf of the Wellcome Sanger |

| | | | |
|---|---|---|---|
| | | Consortium | Institute COVID-19 Surveillance Team |
| EPI_ISL_3042174 | Berkshire and Surrey Pathology Services Lighthouse Laboratory | Wellcome Sanger Institute for the COVID-19 Genomics UK (COG-UK) Consortium | Berkshire and Surrey Pathology Services Lighthouse Laboratory and Alex Alderton, Roberto Amato, Jeffrey Barrett, Sonia Goncalves, Ewan Harrison, David K. Jackson, Ian Johnston, Dominic Kwiatkowski, Cordelia Langford, John Sillitoe on behalf of the Wellcome Sanger Institute COVID-19 Surveillance Team |
| EPI_ISL_3042175 | Lighthouse Lab in Alderley Park | Wellcome Sanger Institute for the COVID-19 Genomics UK (COG-UK) Consortium | Jacquelyn Wynn, Mairead Hyland, The Lighthouse Lab in Alderley Park and Alex Alderton, Roberto Amato, Jeffrey Barrett, Sonia Goncalves, Ewan Harrison, David K. Jackson, Ian Johnston, Dominic Kwiatkowski, Cordelia Langford, John Sillitoe on behalf of the Wellcome Sanger Institute COVID-19 Surveillance Team |
| EPI_ISL_3042176 | Berkshire and Surrey Pathology Services Lighthouse Laboratory | Wellcome Sanger Institute for the COVID-19 Genomics UK (COG-UK) Consortium | Berkshire and Surrey Pathology Services Lighthouse Laboratory and Alex Alderton, Roberto Amato, Jeffrey Barrett, Sonia Goncalves, Ewan Harrison, David K. Jackson, Ian Johnston, Dominic Kwiatkowski, Cordelia Langford, John Sillitoe on behalf of the Wellcome Sanger Institute COVID-19 Surveillance Team |
| EPI_ISL_3042177, EPI_ISL_3042178 | Lighthouse Lab in Milton Keynes | Wellcome Sanger Institute for the COVID-19 Genomics UK (COG-UK) Consortium | The Lighthouse Lab in Milton Keynes and Alex Alderton, Roberto Amato, Jeffrey Barrett, Sonia Goncalves, Ewan Harrison, David K. Jackson, Ian Johnston, Dominic Kwiatkowski, Cordelia Langford, John Sillitoe on behalf of the Wellcome Sanger Institute COVID-19 Surveillance Team |
| EPI_ISL_3042179 | Lighthouse Laboratory Plymouth | Wellcome Sanger Institute for the COVID-19 Genomics UK (COG-UK) Consortium | Lighthouse Laboratory Plymouth and Alex Alderton, Roberto Amato, Jeffrey Barrett, Sonia Goncalves, Ewan Harrison, David K. Jackson, Ian Johnston, Dominic Kwiatkowski, Cordelia Langford, John Sillitoe on behalf of the Wellcome Sanger Institute COVID-19 Surveillance Team |
| EPI_ISL_3042180 | Berkshire and Surrey Pathology Services Lighthouse Laboratory | Wellcome Sanger Institute for the COVID-19 Genomics UK (COG-UK) Consortium | Berkshire and Surrey Pathology Services Lighthouse Laboratory and Alex Alderton, Roberto Amato, Jeffrey Barrett, Sonia Goncalves, Ewan Harrison, David K. Jackson, Ian Johnston, Dominic Kwiatkowski, Cordelia Langford, John Sillitoe on behalf of the Wellcome Sanger Institute COVID-19 Surveillance Team |
| EPI_ISL_3042181 | Lighthouse Lab in Milton Keynes | Wellcome Sanger Institute for the COVID-19 Genomics UK (COG-UK) Consortium | The Lighthouse Lab in Milton Keynes and Alex Alderton, Roberto Amato, Jeffrey Barrett, Sonia Goncalves, Ewan Harrison, David K. Jackson, Ian Johnston, Dominic Kwiatkowski, Cordelia Langford, John Sillitoe on behalf of the Wellcome Sanger Institute COVID-19 Surveillance Team |
| EPI_ISL_3042182 | Berkshire and Surrey Pathology Services Lighthouse Laboratory | Wellcome Sanger Institute for the COVID-19 Genomics UK (COG-UK) Consortium | Berkshire and Surrey Pathology Services Lighthouse Laboratory and Alex Alderton, Roberto Amato, Jeffrey Barrett, Sonia Goncalves, Ewan Harrison, David K. Jackson, Ian Johnston, Dominic Kwiatkowski, Cordelia Langford, John Sillitoe on behalf of the Wellcome Sanger Institute COVID-19 Surveillance Team |
| EPI_ISL_3042183 | Lighthouse Lab in Milton Keynes | Wellcome Sanger Institute for the COVID-19 Genomics UK (COG-UK) Consortium | The Lighthouse Lab in Milton Keynes and Alex Alderton, Roberto Amato, Jeffrey Barrett, Sonia Goncalves, Ewan Harrison, David K. Jackson, Ian Johnston, Dominic Kwiatkowski, Cordelia Langford, John Sillitoe on behalf of the Wellcome Sanger Institute COVID-19 Surveillance Team |
| EPI_ISL_3042184 | Berkshire and Surrey Pathology Services Lighthouse Laboratory | Wellcome Sanger Institute for the COVID-19 Genomics UK (COG-UK) Consortium | Berkshire and Surrey Pathology Services Lighthouse Laboratory and Alex Alderton, Roberto Amato, Jeffrey Barrett, Sonia Goncalves, Ewan Harrison, David K. Jackson, Ian Johnston, Dominic Kwiatkowski, Cordelia Langford, John Sillitoe on behalf of the Wellcome Sanger Institute COVID-19 Surveillance Team |
| EPI_ISL_3042185, EPI_ISL_3042186, EPI_ISL_3042187 | Lighthouse Lab in Alderley Park | Wellcome Sanger Institute for the COVID-19 Genomics UK (COG-UK) Consortium | Jacquelyn Wynn, Mairead Hyland, The Lighthouse Lab in Alderley Park and Alex Alderton, Roberto Amato, Jeffrey Barrett, Sonia Goncalves, Ewan Harrison, David K. Jackson, Ian Johnston, Dominic Kwiatkowski, Cordelia Langford, John Sillitoe on behalf of the Wellcome Sanger Institute COVID-19 Surveillance Team |
| EPI_ISL_3042188 | Lighthouse Laboratory Plymouth | Wellcome Sanger Institute for the COVID-19 Genomics UK (COG-UK) Consortium | Lighthouse Laboratory Plymouth and Alex Alderton, Roberto Amato, Jeffrey Barrett, Sonia Goncalves, Ewan Harrison, David K. Jackson, Ian Johnston, Dominic Kwiatkowski, Cordelia Langford, John Sillitoe on behalf of the Wellcome Sanger Institute COVID-19 Surveillance Team |
| EPI_ISL_3042189 | Lighthouse Lab in Milton Keynes | Wellcome Sanger Institute for the COVID-19 Genomics UK (COG-UK) Consortium | The Lighthouse Lab in Milton Keynes and Alex Alderton, Roberto Amato, Jeffrey Barrett, Sonia Goncalves, Ewan Harrison, David K. Jackson, Ian Johnston, Dominic Kwiatkowski, Cordelia Langford, John Sillitoe on behalf of the Wellcome Sanger Institute COVID-19 Surveillance Team |
| EPI_ISL_3042190 | Berkshire and Surrey Pathology Services Lighthouse Laboratory | Wellcome Sanger Institute for the COVID-19 Genomics UK (COG-UK) Consortium | Berkshire and Surrey Pathology Services Lighthouse Laboratory and Alex Alderton, Roberto Amato, Jeffrey Barrett, Sonia Goncalves, Ewan Harrison, David K. Jackson, Ian Johnston, Dominic Kwiatkowski, Cordelia Langford, John Sillitoe on behalf of the Wellcome Sanger Institute COVID-19 Surveillance Team |
| EPI_ISL_3042191 | Lighthouse Lab in Milton Keynes | Wellcome Sanger Institute for the COVID-19 Genomics UK (COG-UK) Consortium | The Lighthouse Lab in Milton Keynes and Alex Alderton, Roberto Amato, Jeffrey Barrett, Sonia Goncalves, Ewan Harrison, David K. Jackson, Ian Johnston, Dominic Kwiatkowski, Cordelia Langford, John Sillitoe on behalf of the Wellcome Sanger Institute COVID-19 Surveillance Team |
| EPI_ISL_3042192, EPI_ISL_3042193 | Lighthouse Laboratory Plymouth | Wellcome Sanger Institute for the COVID-19 Genomics UK (COG-UK) Consortium | Lighthouse Laboratory Plymouth and Alex Alderton, Roberto Amato, Jeffrey Barrett, Sonia Goncalves, Ewan Harrison, David K. Jackson, Ian Johnston, Dominic Kwiatkowski, Cordelia Langford, John Sillitoe on behalf of the Wellcome Sanger Institute COVID-19 Surveillance Team |
| EPI_ISL_3042194, EPI_ISL_3042195 | Lighthouse Lab in Alderley Park | Wellcome Sanger Institute for the COVID-19 Genomics UK (COG-UK) Consortium | Jacquelyn Wynn, Mairead Hyland, The Lighthouse Lab in Alderley Park and Alex Alderton, Roberto Amato, Jeffrey Barrett, Sonia Goncalves, Ewan Harrison, David K. Jackson, Ian Johnston, Dominic Kwiatkowski, Cordelia Langford, John Sillitoe on behalf of the Wellcome Sanger Institute COVID-19 Surveillance Team |
| EPI_ISL_3042196, EPI_ISL_3042197 | Berkshire and Surrey Pathology Services Lighthouse Laboratory | Wellcome Sanger Institute for the COVID-19 Genomics UK (COG-UK) Consortium | Berkshire and Surrey Pathology Services Lighthouse Laboratory and Alex Alderton, Roberto Amato, Jeffrey Barrett, Sonia Goncalves, Ewan Harrison, David K. Jackson, Ian Johnston, Dominic Kwiatkowski, Cordelia Langford, John Sillitoe on behalf of the Wellcome Sanger Institute COVID-19 Surveillance Team |
| EPI_ISL_3042198, EPI_ISL_3042199, EPI_ISL_3042200 | Lighthouse Laboratory Plymouth | Wellcome Sanger Institute for the COVID-19 Genomics UK (COG-UK) Consortium | Lighthouse Laboratory Plymouth and Alex Alderton, Roberto Amato, Jeffrey Barrett, Sonia Goncalves, Ewan Harrison, David K. Jackson, Ian Johnston, Dominic Kwiatkowski, Cordelia Langford, John Sillitoe on behalf of the Wellcome Sanger Institute COVID-19 Surveillance Team |
| EPI_ISL_3042201 | Lighthouse Lab in Milton Keynes | Wellcome Sanger Institute for the COVID-19 Genomics UK (COG-UK) Consortium | The Lighthouse Lab in Milton Keynes and Alex Alderton, Roberto Amato, Jeffrey Barrett, Sonia Goncalves, Ewan Harrison, David K. Jackson, Ian Johnston, Dominic Kwiatkowski, Cordelia Langford, John Sillitoe on behalf of the Wellcome Sanger Institute COVID-19 Surveillance Team |
| EPI_ISL_3042202 | Berkshire and Surrey Pathology Services Lighthouse Laboratory | Wellcome Sanger Institute for the COVID-19 Genomics UK (COG-UK) Consortium | Berkshire and Surrey Pathology Services Lighthouse Laboratory and Alex Alderton, Roberto Amato, Jeffrey Barrett, Sonia Goncalves, Ewan Harrison, David K. Jackson, Ian Johnston, Dominic Kwiatkowski, Cordelia Langford, John Sillitoe on behalf of the Wellcome Sanger Institute COVID-19 Surveillance Team |
| EPI_ISL_3042203 | Lighthouse Laboratory Plymouth | Wellcome Sanger Institute for the COVID-19 Genomics UK (COG-UK) Consortium | Lighthouse Laboratory Plymouth and Alex Alderton, Roberto Amato, Jeffrey Barrett, Sonia Goncalves, Ewan Harrison, David K. Jackson, Ian Johnston, Dominic Kwiatkowski, Cordelia Langford, John Sillitoe on behalf of the Wellcome Sanger Institute COVID-19 Surveillance Team |
| EPI_ISL_3042204 | Berkshire and Surrey Pathology Services Lighthouse Laboratory | Wellcome Sanger Institute for the COVID-19 Genomics UK (COG-UK) Consortium | Berkshire and Surrey Pathology Services Lighthouse Laboratory and Alex Alderton, Roberto Amato, Jeffrey Barrett, Sonia Goncalves, Ewan Harrison, David K. Jackson, Ian Johnston, Dominic Kwiatkowski, Cordelia Langford, John Sillitoe on behalf of the Wellcome Sanger Institute COVID-19 Surveillance Team |
| EPI_ISL_3042205, EPI_ISL_3042206 | Lighthouse Lab in Alderley Park | Wellcome Sanger Institute for the COVID-19 Genomics UK (COG-UK) Consortium | Jacquelyn Wynn, Mairead Hyland, The Lighthouse Lab in Alderley Park and Alex Alderton, Roberto Amato, Jeffrey Barrett, Sonia Goncalves, Ewan Harrison, David K. Jackson, Ian Johnston, Dominic Kwiatkowski, Cordelia Langford, John Sillitoe on behalf of the Wellcome Sanger Institute COVID-19 Surveillance Team |
| EPI_ISL_3042207, EPI_ISL_3042208, EPI_ISL_3042209 | Lighthouse Laboratory Plymouth | Wellcome Sanger Institute for the COVID-19 Genomics UK (COG-UK) Consortium | Lighthouse Laboratory Plymouth and Alex Alderton, Roberto Amato, Jeffrey Barrett, Sonia Goncalves, Ewan Harrison, David K. Jackson, Ian Johnston, Dominic Kwiatkowski, Cordelia Langford, John Sillitoe on behalf of the Wellcome Sanger Institute COVID-19 Surveillance Team |
| EPI_ISL_3042210, EPI_ISL_3042211 | Lighthouse Lab in Milton Keynes | Wellcome Sanger Institute for the COVID-19 Genomics UK (COG-UK) Consortium | The Lighthouse Lab in Milton Keynes and Alex Alderton, Roberto Amato, Jeffrey Barrett, Sonia Goncalves, Ewan Harrison, David K. Jackson, Ian Johnston, Dominic Kwiatkowski, Cordelia Langford, John Sillitoe on behalf of the Wellcome Sanger Institute COVID-19 Surveillance Team |
| EPI_ISL_3042212 | Berkshire and Surrey Pathology Services Lighthouse Laboratory | Wellcome Sanger Institute for the COVID-19 Genomics UK (COG-UK) Consortium | Berkshire and Surrey Pathology Services Lighthouse Laboratory and Alex Alderton, Roberto Amato, Jeffrey Barrett, Sonia Goncalves, Ewan Harrison, David K. Jackson, Ian Johnston, Dominic Kwiatkowski, Cordelia Langford, John Sillitoe on behalf of the Wellcome Sanger Institute COVID-19 Surveillance Team |
| EPI_ISL_3042213 | Lighthouse Lab in Milton Keynes | Wellcome Sanger Institute for the COVID-19 Genomics UK (COG-UK) Consortium | The Lighthouse Lab in Milton Keynes and Alex Alderton, Roberto Amato, Jeffrey Barrett, Sonia Goncalves, Ewan Harrison, David K. Jackson, Ian Johnston, Dominic Kwiatkowski, Cordelia Langford, John Sillitoe on behalf of the Wellcome Sanger Institute COVID-19 Surveillance Team |
| EPI_ISL_3042214, EPI_ISL_3042215 | Lighthouse Laboratory Plymouth | Wellcome Sanger Institute for the COVID-19 Genomics UK (COG-UK) Consortium | Lighthouse Laboratory Plymouth and Alex Alderton, Roberto Amato, Jeffrey Barrett, Sonia Goncalves, Ewan Harrison, David K. Jackson, Ian Johnston, Dominic Kwiatkowski, Cordelia Langford, John Sillitoe on behalf of the Wellcome Sanger Institute COVID-19 Surveillance Team |
| EPI_ISL_3042216 | Berkshire and Surrey Pathology Services Lighthouse Laboratory | Wellcome Sanger Institute for the COVID-19 Genomics UK (COG-UK) Consortium | Berkshire and Surrey Pathology Services Lighthouse Laboratory and Alex Alderton, Roberto Amato, Jeffrey Barrett, Sonia Goncalves, Ewan Harrison, David K. Jackson, Ian Johnston, Dominic Kwiatkowski, Cordelia Langford, John Sillitoe on behalf of the Wellcome Sanger Institute COVID-19 Surveillance Team |
| EPI_ISL_3042217, EPI_ISL_3042218 | Lighthouse Laboratory Plymouth | Wellcome Sanger Institute for the COVID-19 Genomics UK (COG-UK) Consortium | Lighthouse Laboratory Plymouth and Alex Alderton, Roberto Amato, Jeffrey Barrett, Sonia Goncalves, Ewan Harrison, David K. Jackson, Ian Johnston, Dominic Kwiatkowski, Cordelia Langford, John Sillitoe on behalf of the Wellcome Sanger Institute COVID-19 Surveillance Team |
| EPI_ISL_3042219 | Lighthouse Lab in Milton Keynes | Wellcome Sanger Institute for the COVID-19 Genomics UK (COG-UK) Consortium | The Lighthouse Lab in Milton Keynes and Alex Alderton, Roberto Amato, Jeffrey Barrett, Sonia Goncalves, Ewan Harrison, David K. Jackson, Ian Johnston, Dominic Kwiatkowski, Cordelia Langford, John Sillitoe on behalf of the Wellcome Sanger Institute COVID-19 Surveillance Team |
| EPI_ISL_3042220, EPI_ISL_3042221 | Berkshire and Surrey Pathology Services Lighthouse Laboratory | Wellcome Sanger Institute for the COVID-19 Genomics UK (COG-UK) Consortium | Berkshire and Surrey Pathology Services Lighthouse Laboratory and Alex Alderton, Roberto Amato, Jeffrey Barrett, Sonia Goncalves, Ewan Harrison, David K. Jackson, Ian Johnston, Dominic Kwiatkowski, Cordelia Langford, John Sillitoe on behalf of the Wellcome Sanger Institute COVID-19 Surveillance Team |
| EPI_ISL_3042222, EPI_ISL_3042223, EPI_ISL_3042224 | Lighthouse Laboratory Plymouth | Wellcome Sanger Institute for the COVID-19 Genomics UK (COG-UK) Consortium | Lighthouse Laboratory Plymouth and Alex Alderton, Roberto Amato, Jeffrey Barrett, Sonia Goncalves, Ewan Harrison, David K. Jackson, Ian Johnston, Dominic Kwiatkowski, Cordelia Langford, John Sillitoe on behalf of the Wellcome Sanger Institute COVID-19 Surveillance Team |
| EPI_ISL_3042225 | Berkshire and Surrey Pathology Services Lighthouse Laboratory | Wellcome Sanger Institute for the COVID-19 Genomics UK (COG-UK) Consortium | Berkshire and Surrey Pathology Services Lighthouse Laboratory and Alex Alderton, Roberto Amato, Jeffrey Barrett, Sonia Goncalves, Ewan Harrison, David K. Jackson, Ian Johnston, Dominic Kwiatkowski, Cordelia Langford, John Sillitoe on behalf of the Wellcome Sanger Institute COVID-19 Surveillance Team |
| EPI_ISL_3042226 | Lighthouse Laboratory Plymouth | Wellcome Sanger Institute for the COVID-19 Genomics UK (COG-UK) Consortium | Lighthouse Laboratory Plymouth and Alex Alderton, Roberto Amato, Jeffrey Barrett, Sonia Goncalves, Ewan Harrison, David K. Jackson, Ian Johnston, Dominic Kwiatkowski, Cordelia Langford, John Sillitoe on behalf of the Wellcome Sanger Institute COVID-19 Surveillance Team |
| EPI_ISL_3042227 | Lighthouse Lab in Milton Keynes | Wellcome Sanger Institute for the COVID-19 Genomics UK (COG-UK) Consortium | The Lighthouse Lab in Milton Keynes and Alex Alderton, Roberto Amato, Jeffrey Barrett, Sonia Goncalves, Ewan Harrison, David K. Jackson, Ian Johnston, Dominic Kwiatkowski, Cordelia Langford, John Sillitoe on behalf of the Wellcome Sanger Institute COVID-19 Surveillance Team |
| EPI_ISL_3042228 | Lighthouse Laboratory Plymouth | Wellcome Sanger Institute for the COVID-19 Genomics UK (COG-UK) Consortium | Lighthouse Laboratory Plymouth and Alex Alderton, Roberto Amato, Jeffrey Barrett, Sonia Goncalves, Ewan Harrison, David K. Jackson, Ian Johnston, Dominic Kwiatkowski, Cordelia Langford, John Sillitoe on behalf of the Wellcome Sanger Institute COVID-19 Surveillance Team |
| EPI_ISL_3042229, EPI_ISL_3042230 | Lighthouse Lab in Milton Keynes | Wellcome Sanger Institute for the COVID-19 Genomics UK (COG-UK) Consortium | The Lighthouse Lab in Milton Keynes and Alex Alderton, Roberto Amato, Jeffrey Barrett, Sonia Goncalves, Ewan Harrison, David K. Jackson, Ian Johnston, Dominic Kwiatkowski, Cordelia Langford, John Sillitoe on behalf of the Wellcome Sanger Institute COVID-19 Surveillance Team |
| EPI_ISL_3042231 | Lighthouse Laboratory Plymouth | Wellcome Sanger Institute for the COVID-19 Genomics UK (COG-UK) Consortium | Lighthouse Laboratory Plymouth and Alex Alderton, Roberto Amato, Jeffrey Barrett, Sonia Goncalves, Ewan Harrison, David K. Jackson, Ian Johnston, Dominic Kwiatkowski, Cordelia Langford, John Sillitoe on behalf of the Wellcome Sanger Institute COVID-19 Surveillance Team |
| EPI_ISL_3042232 | Lighthouse Lab in Milton Keynes | Wellcome Sanger Institute for the COVID-19 Genomics UK (COG-UK) Consortium | The Lighthouse Lab in Milton Keynes and Alex Alderton, Roberto Amato, Jeffrey Barrett, Sonia Goncalves, Ewan Harrison, David K. Jackson, Ian Johnston, Dominic Kwiatkowski, Cordelia Langford, John Sillitoe on behalf of the Wellcome Sanger Institute COVID-19 Surveillance Team |
| EPI_ISL_3042233, EPI_ISL_3042234 | Lighthouse Lab in Alderley Park | Wellcome Sanger Institute for the COVID-19 Genomics UK (COG-UK) Consortium | Jacquelyn Wynn, Mairead Hyland, The Lighthouse Lab in Alderley Park and Alex Alderton, Roberto Amato, Jeffrey Barrett, Sonia Goncalves, Ewan Harrison, David K. Jackson, Ian Johnston, Dominic Kwiatkowski, Cordelia Langford, John Sillitoe on behalf of the Wellcome Sanger Institute COVID-19 Surveillance Team |
| EPI_ISL_3042235 | Lighthouse Lab in Milton Keynes | Wellcome Sanger Institute for the COVID-19 Genomics UK (COG-UK) Consortium | The Lighthouse Lab in Milton Keynes and Alex Alderton, Roberto Amato, Jeffrey Barrett, Sonia Goncalves, Ewan Harrison, David K. Jackson, Ian Johnston, Dominic Kwiatkowski, Cordelia Langford, John Sillitoe on behalf of the Wellcome Sanger Institute COVID-19 Surveillance Team |

| | | | |
|---|---|---|---|
| EPI_ISL_3042236 | Berkshire and Surrey Pathology Services Lighthouse Laboratory | Wellcome Sanger Institute for the COVID-19 Genomics UK (COG-UK) Consortium | Berkshire and Surrey Pathology Services Lighthouse Laboratory and Alex Alderton, Roberto Amato, Jeffrey Barrett, Sonia Goncalves, Ewan Harrison, David K. Jackson, Ian Johnston, Dominic Kwiatkowski, Cordelia Langford, John Sillitoe on behalf of the Wellcome Sanger Institute COVID-19 Surveillance Team |
| EPI_ISL_3042237 | Lighthouse Lab in Alderley Park | Wellcome Sanger Institute for the COVID-19 Genomics UK (COG-UK) Consortium | Jacquelyn Wynn, Mairead Hyland, The Lighthouse Lab in Alderley Park and Alex Alderton, Roberto Amato, Jeffrey Barrett, Sonia Goncalves, Ewan Harrison, David K. Jackson, Ian Johnston, Dominic Kwiatkowski, Cordelia Langford, John Sillitoe on behalf of the Wellcome Sanger Institute COVID-19 Surveillance Team |
| EPI_ISL_3042238 | Berkshire and Surrey Pathology Services Lighthouse Laboratory | Wellcome Sanger Institute for the COVID-19 Genomics UK (COG-UK) Consortium | Berkshire and Surrey Pathology Services Lighthouse Laboratory and Alex Alderton, Roberto Amato, Jeffrey Barrett, Sonia Goncalves, Ewan Harrison, David K. Jackson, Ian Johnston, Dominic Kwiatkowski, Cordelia Langford, John Sillitoe on behalf of the Wellcome Sanger Institute COVID-19 Surveillance Team |
| EPI_ISL_3042239 | Lighthouse Laboratory Plymouth | Wellcome Sanger Institute for the COVID-19 Genomics UK (COG-UK) Consortium | Lighthouse Laboratory Plymouth and Alex Alderton, Roberto Amato, Jeffrey Barrett, Sonia Goncalves, Ewan Harrison, David K. Jackson, Ian Johnston, Dominic Kwiatkowski, Cordelia Langford, John Sillitoe on behalf of the Wellcome Sanger Institute COVID-19 Surveillance Team |
| EPI_ISL_3042240, EPI_ISL_3042241 | Lighthouse Lab in Milton Keynes | Wellcome Sanger Institute for the COVID-19 Genomics UK (COG-UK) Consortium | The Lighthouse Lab in Milton Keynes and Alex Alderton, Roberto Amato, Jeffrey Barrett, Sonia Goncalves, Ewan Harrison, David K. Jackson, Ian Johnston, Dominic Kwiatkowski, Cordelia Langford, John Sillitoe on behalf of the Wellcome Sanger Institute COVID-19 Surveillance Team |
| EPI_ISL_3042242 | Lighthouse Laboratory Plymouth | Wellcome Sanger Institute for the COVID-19 Genomics UK (COG-UK) Consortium | Lighthouse Laboratory Plymouth and Alex Alderton, Roberto Amato, Jeffrey Barrett, Sonia Goncalves, Ewan Harrison, David K. Jackson, Ian Johnston, Dominic Kwiatkowski, Cordelia Langford, John Sillitoe on behalf of the Wellcome Sanger Institute COVID-19 Surveillance Team |
| EPI_ISL_3042243, EPI_ISL_3042244 | Lighthouse Lab in Milton Keynes | Wellcome Sanger Institute for the COVID-19 Genomics UK (COG-UK) Consortium | The Lighthouse Lab in Milton Keynes and Alex Alderton, Roberto Amato, Jeffrey Barrett, Sonia Goncalves, Ewan Harrison, David K. Jackson, Ian Johnston, Dominic Kwiatkowski, Cordelia Langford, John Sillitoe on behalf of the Wellcome Sanger Institute COVID-19 Surveillance Team |
| EPI_ISL_3042245 | Lighthouse Laboratory Plymouth | Wellcome Sanger Institute for the COVID-19 Genomics UK (COG-UK) Consortium | Lighthouse Laboratory Plymouth and Alex Alderton, Roberto Amato, Jeffrey Barrett, Sonia Goncalves, Ewan Harrison, David K. Jackson, Ian Johnston, Dominic Kwiatkowski, Cordelia Langford, John Sillitoe on behalf of the Wellcome Sanger Institute COVID-19 Surveillance Team |
| EPI_ISL_3042246 | Berkshire and Surrey Pathology Services Lighthouse Laboratory | Wellcome Sanger Institute for the COVID-19 Genomics UK (COG-UK) Consortium | Berkshire and Surrey Pathology Services Lighthouse Laboratory and Alex Alderton, Roberto Amato, Jeffrey Barrett, Sonia Goncalves, Ewan Harrison, David K. Jackson, Ian Johnston, Dominic Kwiatkowski, Cordelia Langford, John Sillitoe on behalf of the Wellcome Sanger Institute COVID-19 Surveillance Team |
| EPI_ISL_3042247 | Lighthouse Lab in Milton Keynes | Wellcome Sanger Institute for the COVID-19 Genomics UK (COG-UK) Consortium | The Lighthouse Lab in Milton Keynes and Alex Alderton, Roberto Amato, Jeffrey Barrett, Sonia Goncalves, Ewan Harrison, David K. Jackson, Ian Johnston, Dominic Kwiatkowski, Cordelia Langford, John Sillitoe on behalf of the Wellcome Sanger Institute COVID-19 Surveillance Team |
| EPI_ISL_3042248, EPI_ISL_3042249, EPI_ISL_3042250 | Lighthouse Lab in Alderley Park | Wellcome Sanger Institute for the COVID-19 Genomics UK (COG-UK) Consortium | Jacquelyn Wynn, Mairead Hyland, The Lighthouse Lab in Alderley Park and Alex Alderton, Roberto Amato, Jeffrey Barrett, Sonia Goncalves, Ewan Harrison, David K. Jackson, Ian Johnston, Dominic Kwiatkowski, Cordelia Langford, John Sillitoe on behalf of the Wellcome Sanger Institute COVID-19 Surveillance Team |
| EPI_ISL_3042251 | Berkshire and Surrey Pathology Services Lighthouse Laboratory | Wellcome Sanger Institute for the COVID-19 Genomics UK (COG-UK) Consortium | Berkshire and Surrey Pathology Services Lighthouse Laboratory and Alex Alderton, Roberto Amato, Jeffrey Barrett, Sonia Goncalves, Ewan Harrison, David K. Jackson, Ian Johnston, Dominic Kwiatkowski, Cordelia Langford, John Sillitoe on behalf of the Wellcome Sanger Institute COVID-19 Surveillance Team |
| EPI_ISL_3042252 | Lighthouse Laboratory Plymouth | Wellcome Sanger Institute for the COVID-19 Genomics UK (COG-UK) Consortium | Lighthouse Laboratory Plymouth and Alex Alderton, Roberto Amato, Jeffrey Barrett, Sonia Goncalves, Ewan Harrison, David K. Jackson, Ian Johnston, Dominic Kwiatkowski, Cordelia Langford, John Sillitoe on behalf of the Wellcome Sanger Institute COVID-19 Surveillance Team |
| EPI_ISL_3042253, EPI_ISL_3042254 | Lighthouse Lab in Milton Keynes | Wellcome Sanger Institute for the COVID-19 Genomics UK (COG-UK) Consortium | The Lighthouse Lab in Milton Keynes and Alex Alderton, Roberto Amato, Jeffrey Barrett, Sonia Goncalves, Ewan Harrison, David K. Jackson, Ian Johnston, Dominic Kwiatkowski, Cordelia Langford, John Sillitoe on behalf of the Wellcome Sanger Institute COVID-19 Surveillance Team |
| EPI_ISL_3042255 | Berkshire and Surrey Pathology Services Lighthouse Laboratory | Wellcome Sanger Institute for the COVID-19 Genomics UK (COG-UK) Consortium | Berkshire and Surrey Pathology Services Lighthouse Laboratory and Alex Alderton, Roberto Amato, Jeffrey Barrett, Sonia Goncalves, Ewan Harrison, David K. Jackson, Ian Johnston, Dominic Kwiatkowski, Cordelia Langford, John Sillitoe on behalf of the Wellcome Sanger Institute COVID-19 Surveillance Team |
| EPI_ISL_3042256 | Lighthouse Lab in Milton Keynes | Wellcome Sanger Institute for the COVID-19 Genomics UK (COG-UK) Consortium | The Lighthouse Lab in Milton Keynes and Alex Alderton, Roberto Amato, Jeffrey Barrett, Sonia Goncalves, Ewan Harrison, David K. Jackson, Ian Johnston, Dominic Kwiatkowski, Cordelia Langford, John Sillitoe on behalf of the Wellcome Sanger Institute COVID-19 Surveillance Team |
| EPI_ISL_3042257 | Berkshire and Surrey Pathology Services Lighthouse Laboratory | Wellcome Sanger Institute for the COVID-19 Genomics UK (COG-UK) Consortium | Berkshire and Surrey Pathology Services Lighthouse Laboratory and Alex Alderton, Roberto Amato, Jeffrey Barrett, Sonia Goncalves, Ewan Harrison, David K. Jackson, Ian Johnston, Dominic Kwiatkowski, Cordelia Langford, John Sillitoe on behalf of the Wellcome Sanger Institute COVID-19 Surveillance Team |
| EPI_ISL_3042258, EPI_ISL_3042259 | Lighthouse Lab in Alderley Park | Wellcome Sanger Institute for the COVID-19 Genomics UK (COG-UK) Consortium | Jacquelyn Wynn, Mairead Hyland, The Lighthouse Lab in Alderley Park and Alex Alderton, Roberto Amato, Jeffrey Barrett, Sonia Goncalves, Ewan Harrison, David K. Jackson, Ian Johnston, Dominic Kwiatkowski, Cordelia Langford, John Sillitoe on behalf of the Wellcome Sanger Institute COVID-19 Surveillance Team |
| EPI_ISL_3042260 | Lighthouse Laboratory Plymouth | Wellcome Sanger Institute for the COVID-19 Genomics UK (COG-UK) Consortium | Lighthouse Laboratory Plymouth and Alex Alderton, Roberto Amato, Jeffrey Barrett, Sonia Goncalves, Ewan Harrison, David K. Jackson, Ian Johnston, Dominic Kwiatkowski, Cordelia Langford, John Sillitoe on behalf of the Wellcome Sanger Institute COVID-19 Surveillance Team |
| EPI_ISL_3042261 | Berkshire and Surrey Pathology Services Lighthouse Laboratory | Wellcome Sanger Institute for the COVID-19 Genomics UK (COG-UK) Consortium | Berkshire and Surrey Pathology Services Lighthouse Laboratory and Alex Alderton, Roberto Amato, Jeffrey Barrett, Sonia Goncalves, Ewan Harrison, David K. Jackson, Ian Johnston, Dominic Kwiatkowski, Cordelia Langford, John Sillitoe on behalf of the Wellcome Sanger Institute COVID-19 Surveillance Team |
| EPI_ISL_3042262 | Lighthouse Laboratory Plymouth | Wellcome Sanger Institute for the COVID-19 Genomics UK (COG-UK) Consortium | Lighthouse Laboratory Plymouth and Alex Alderton, Roberto Amato, Jeffrey Barrett, Sonia Goncalves, Ewan Harrison, David K. Jackson, Ian Johnston, Dominic Kwiatkowski, Cordelia Langford, John Sillitoe on behalf of the Wellcome Sanger Institute COVID-19 Surveillance Team |
| EPI_ISL_3042263 | Lighthouse Lab in Alderley Park | Wellcome Sanger Institute for the COVID-19 Genomics UK (COG-UK) Consortium | Jacquelyn Wynn, Mairead Hyland, The Lighthouse Lab in Alderley Park and Alex Alderton, Roberto Amato, Jeffrey Barrett, Sonia Goncalves, Ewan Harrison, David K. Jackson, Ian Johnston, Dominic Kwiatkowski, Cordelia Langford, John Sillitoe on behalf of the Wellcome Sanger Institute COVID-19 Surveillance Team |
| EPI_ISL_3042264 | Lighthouse Laboratory Plymouth | Wellcome Sanger Institute for the COVID-19 Genomics UK (COG-UK) Consortium | Lighthouse Laboratory Plymouth and Alex Alderton, Roberto Amato, Jeffrey Barrett, Sonia Goncalves, Ewan Harrison, David K. Jackson, Ian Johnston, Dominic Kwiatkowski, Cordelia Langford, John Sillitoe on behalf of the Wellcome Sanger Institute COVID-19 Surveillance Team |
| EPI_ISL_3042265 | Lighthouse Lab in Milton Keynes | Wellcome Sanger Institute for the COVID-19 Genomics UK (COG-UK) Consortium | The Lighthouse Lab in Milton Keynes and Alex Alderton, Roberto Amato, Jeffrey Barrett, Sonia Goncalves, Ewan Harrison, David K. Jackson, Ian Johnston, Dominic Kwiatkowski, Cordelia Langford, John Sillitoe on behalf of the Wellcome Sanger Institute COVID-19 Surveillance Team |
| EPI_ISL_3042266 | Lighthouse Lab in Alderley Park | Wellcome Sanger Institute for the COVID-19 Genomics UK (COG-UK) Consortium | Jacquelyn Wynn, Mairead Hyland, The Lighthouse Lab in Alderley Park and Alex Alderton, Roberto Amato, Jeffrey Barrett, Sonia Goncalves, Ewan Harrison, David K. Jackson, Ian Johnston, Dominic Kwiatkowski, Cordelia Langford, John Sillitoe on behalf of the Wellcome Sanger Institute COVID-19 Surveillance Team |
| EPI_ISL_3042267 | Berkshire and Surrey Pathology Services Lighthouse Laboratory | Wellcome Sanger Institute for the COVID-19 Genomics UK (COG-UK) Consortium | Berkshire and Surrey Pathology Services Lighthouse Laboratory and Alex Alderton, Roberto Amato, Jeffrey Barrett, Sonia Goncalves, Ewan Harrison, David K. Jackson, Ian Johnston, Dominic Kwiatkowski, Cordelia Langford, John Sillitoe on behalf of the Wellcome Sanger Institute COVID-19 Surveillance Team |
| EPI_ISL_3042268 | Lighthouse Lab in Milton Keynes | Wellcome Sanger Institute for the COVID-19 Genomics UK (COG-UK) Consortium | The Lighthouse Lab in Milton Keynes and Alex Alderton, Roberto Amato, Jeffrey Barrett, Sonia Goncalves, Ewan Harrison, David K. Jackson, Ian Johnston, Dominic Kwiatkowski, Cordelia Langford, John Sillitoe on behalf of the Wellcome Sanger Institute COVID-19 Surveillance Team |
| EPI_ISL_3042269 | Lighthouse Lab in Alderley Park | Wellcome Sanger Institute for the COVID-19 Genomics UK (COG-UK) Consortium | Jacquelyn Wynn, Mairead Hyland, The Lighthouse Lab in Alderley Park and Alex Alderton, Roberto Amato, Jeffrey Barrett, Sonia Goncalves, Ewan Harrison, David K. Jackson, Ian Johnston, Dominic Kwiatkowski, Cordelia Langford, John Sillitoe on behalf of the Wellcome Sanger Institute COVID-19 Surveillance Team |
| EPI_ISL_3042270 | Berkshire and Surrey Pathology Services Lighthouse Laboratory | Wellcome Sanger Institute for the COVID-19 Genomics UK (COG-UK) Consortium | Berkshire and Surrey Pathology Services Lighthouse Laboratory and Alex Alderton, Roberto Amato, Jeffrey Barrett, Sonia Goncalves, Ewan Harrison, David K. Jackson, Ian Johnston, Dominic Kwiatkowski, Cordelia Langford, John Sillitoe on behalf of the Wellcome Sanger Institute COVID-19 Surveillance Team |
| EPI_ISL_3042271 | Lighthouse Lab in Milton Keynes | Wellcome Sanger Institute for the COVID-19 Genomics UK (COG-UK) Consortium | The Lighthouse Lab in Milton Keynes and Alex Alderton, Roberto Amato, Jeffrey Barrett, Sonia Goncalves, Ewan Harrison, David K. Jackson, Ian Johnston, Dominic Kwiatkowski, Cordelia Langford, John Sillitoe on behalf of the Wellcome Sanger Institute COVID-19 Surveillance Team |
| EPI_ISL_3042272 | Berkshire and Surrey Pathology Services Lighthouse Laboratory | Wellcome Sanger Institute for the COVID-19 Genomics UK (COG-UK) Consortium | Berkshire and Surrey Pathology Services Lighthouse Laboratory and Alex Alderton, Roberto Amato, Jeffrey Barrett, Sonia Goncalves, Ewan Harrison, David K. Jackson, Ian Johnston, Dominic Kwiatkowski, Cordelia Langford, John Sillitoe on behalf of the Wellcome Sanger Institute COVID-19 Surveillance Team |
| EPI_ISL_3042273 | Lighthouse Laboratory Plymouth | Wellcome Sanger Institute for the COVID-19 Genomics UK (COG-UK) Consortium | Lighthouse Laboratory Plymouth and Alex Alderton, Roberto Amato, Jeffrey Barrett, Sonia Goncalves, Ewan Harrison, David K. Jackson, Ian Johnston, Dominic Kwiatkowski, Cordelia Langford, John Sillitoe on behalf of the Wellcome Sanger Institute COVID-19 Surveillance Team |
| EPI_ISL_3042274 | Lighthouse Lab in Milton Keynes | Wellcome Sanger Institute for the COVID-19 Genomics UK (COG-UK) Consortium | The Lighthouse Lab in Milton Keynes and Alex Alderton, Roberto Amato, Jeffrey Barrett, Sonia Goncalves, Ewan Harrison, David K. Jackson, Ian Johnston, Dominic Kwiatkowski, Cordelia Langford, John Sillitoe on behalf of the Wellcome Sanger Institute COVID-19 Surveillance Team |
| EPI_ISL_3042275 | Lighthouse Lab in Alderley Park | Wellcome Sanger Institute for the COVID-19 Genomics UK (COG-UK) Consortium | Jacquelyn Wynn, Mairead Hyland, The Lighthouse Lab in Alderley Park and Alex Alderton, Roberto Amato, Jeffrey Barrett, Sonia Goncalves, Ewan Harrison, David K. Jackson, Ian Johnston, Dominic Kwiatkowski, Cordelia Langford, John Sillitoe on behalf of the Wellcome Sanger Institute COVID-19 Surveillance Team |
| EPI_ISL_3042276 | Berkshire and Surrey Pathology Services Lighthouse Laboratory | Wellcome Sanger Institute for the COVID-19 Genomics UK (COG-UK) Consortium | Berkshire and Surrey Pathology Services Lighthouse Laboratory and Alex Alderton, Roberto Amato, Jeffrey Barrett, Sonia Goncalves, Ewan Harrison, David K. Jackson, Ian Johnston, Dominic Kwiatkowski, Cordelia Langford, John Sillitoe on behalf of the Wellcome Sanger Institute COVID-19 Surveillance Team |
| EPI_ISL_3042277, EPI_ISL_3042278 | Lighthouse Lab in Milton Keynes | Wellcome Sanger Institute for the COVID-19 Genomics UK (COG-UK) Consortium | The Lighthouse Lab in Milton Keynes and Alex Alderton, Roberto Amato, Jeffrey Barrett, Sonia Goncalves, Ewan Harrison, David K. Jackson, Ian Johnston, Dominic Kwiatkowski, Cordelia Langford, John Sillitoe on behalf of the Wellcome Sanger Institute COVID-19 Surveillance Team |
| EPI_ISL_3042279 | Lighthouse Laboratory Plymouth | Wellcome Sanger Institute for the COVID-19 Genomics UK (COG-UK) Consortium | Lighthouse Laboratory Plymouth and Alex Alderton, Roberto Amato, Jeffrey Barrett, Sonia Goncalves, Ewan Harrison, David K. Jackson, Ian Johnston, Dominic Kwiatkowski, Cordelia Langford, John Sillitoe on behalf of the Wellcome Sanger Institute COVID-19 Surveillance Team |
| EPI_ISL_3042280 | Berkshire and Surrey Pathology Services Lighthouse Laboratory | Wellcome Sanger Institute for the COVID-19 Genomics UK (COG-UK) Consortium | Berkshire and Surrey Pathology Services Lighthouse Laboratory and Alex Alderton, Roberto Amato, Jeffrey Barrett, Sonia Goncalves, Ewan Harrison, David K. Jackson, Ian Johnston, Dominic Kwiatkowski, Cordelia Langford, John Sillitoe on behalf of the Wellcome Sanger Institute COVID-19 Surveillance Team |
| EPI_ISL_3042281 | Lighthouse Lab in Milton Keynes | Wellcome Sanger Institute for the COVID-19 Genomics UK (COG-UK) Consortium | The Lighthouse Lab in Milton Keynes and Alex Alderton, Roberto Amato, Jeffrey Barrett, Sonia Goncalves, Ewan Harrison, David K. Jackson, Ian Johnston, Dominic Kwiatkowski, Cordelia Langford, John Sillitoe on behalf of the Wellcome Sanger Institute COVID-19 Surveillance Team |
| EPI_ISL_3042282 | Lighthouse Laboratory Plymouth | Wellcome Sanger Institute for the COVID-19 Genomics UK (COG-UK) Consortium | Lighthouse Laboratory Plymouth and Alex Alderton, Roberto Amato, Jeffrey Barrett, Sonia Goncalves, Ewan Harrison, David K. Jackson, Ian Johnston, Dominic Kwiatkowski, Cordelia Langford, John Sillitoe on behalf of the Wellcome Sanger Institute COVID-19 Surveillance Team |
| EPI_ISL_3042283 | Lighthouse Lab in Milton Keynes | Wellcome Sanger Institute for the COVID-19 Genomics UK (COG-UK) Consortium | The Lighthouse Lab in Milton Keynes and Alex Alderton, Roberto Amato, Jeffrey Barrett, Sonia Goncalves, Ewan Harrison, David K. Jackson, Ian Johnston, Dominic Kwiatkowski, Cordelia Langford, John Sillitoe on behalf of the Wellcome Sanger Institute COVID-19 Surveillance Team |
| EPI_ISL_3042284, EPI_ISL_3042285 | Berkshire and Surrey Pathology Services Lighthouse Laboratory | Wellcome Sanger Institute for the COVID-19 Genomics UK (COG-UK) Consortium | Berkshire and Surrey Pathology Services Lighthouse Laboratory and Alex Alderton, Roberto Amato, Jeffrey Barrett, Sonia Goncalves, Ewan Harrison, David K. Jackson, Ian Johnston, Dominic Kwiatkowski, Cordelia Langford, John Sillitoe on behalf of the Wellcome Sanger Institute COVID-19 Surveillance Team |
| EPI_ISL_3042286 | Lighthouse Laboratory Plymouth | Wellcome Sanger Institute for the COVID-19 Genomics UK (COG-UK) Consortium | Lighthouse Laboratory Plymouth and Alex Alderton, Roberto Amato, Jeffrey Barrett, Sonia Goncalves, Ewan Harrison, David K. Jackson, Ian Johnston, Dominic Kwiatkowski, Cordelia Langford, John Sillitoe on behalf of the Wellcome Sanger Institute COVID-19 Surveillance Team |
| EPI_ISL_3042287 | Lighthouse Lab in Alderley Park | Wellcome Sanger Institute for the COVID-19 Genomics UK (COG-UK) Consortium | Jacquelyn Wynn, Mairead Hyland, The Lighthouse Lab in Alderley Park and Alex Alderton, Roberto Amato, Jeffrey Barrett, Sonia Goncalves, Ewan Harrison, David K. Jackson, Ian Johnston, Dominic Kwiatkowski, Cordelia Langford, John Sillitoe on behalf of the Wellcome Sanger Institute COVID-19 Surveillance Team |
| EPI_ISL_3042288 | Berkshire and Surrey Pathology Services Lighthouse Laboratory | Wellcome Sanger Institute for the COVID-19 Genomics UK (COG-UK) Consortium | Berkshire and Surrey Pathology Services Lighthouse Laboratory and Alex Alderton, Roberto Amato, Jeffrey Barrett, Sonia Goncalves, Ewan Harrison, David K. Jackson, Ian Johnston, Dominic Kwiatkowski, Cordelia Langford, John Sillitoe on behalf of the Wellcome Sanger Institute COVID-19 Surveillance Team |

| | | | |
|---|---|---|---|
| EPI_ISL_3042289, EPI_ISL_3042290 | Lighthouse Laboratory Plymouth | Wellcome Sanger Institute for the COVID-19 Genomics UK (COG-UK) Consortium | Lighthouse Laboratory Plymouth and Alex Alderton, Roberto Amato, Jeffrey Barrett, Sonia Goncalves, Ewan Harrison, David K. Jackson, Ian Johnston, Dominic Kwiatkowski, Cordelia Langford, John Sillitoe on behalf of the Wellcome Sanger Institute COVID-19 Surveillance Team |
| EPI_ISL_3042291 | Lighthouse Lab in Milton Keynes | Wellcome Sanger Institute for the COVID-19 Genomics UK (COG-UK) Consortium | The Lighthouse Lab in Milton Keynes and Alex Alderton, Roberto Amato, Jeffrey Barrett, Sonia Goncalves, Ewan Harrison, David K. Jackson, Ian Johnston, Dominic Kwiatkowski, Cordelia Langford, John Sillitoe on behalf of the Wellcome Sanger Institute COVID-19 Surveillance Team |
| EPI_ISL_3042292 | Berkshire and Surrey Pathology Services Lighthouse Laboratory | Wellcome Sanger Institute for the COVID-19 Genomics UK (COG-UK) Consortium | Berkshire and Surrey Pathology Services Lighthouse Laboratory and Alex Alderton, Roberto Amato, Jeffrey Barrett, Sonia Goncalves, Ewan Harrison, David K. Jackson, Ian Johnston, Dominic Kwiatkowski, Cordelia Langford, John Sillitoe on behalf of the Wellcome Sanger Institute COVID-19 Surveillance Team |
| EPI_ISL_3042293 | Lighthouse Lab in Alderley Park | Wellcome Sanger Institute for the COVID-19 Genomics UK (COG-UK) Consortium | Jacquelyn Wynn, Mairead Hyland, The Lighthouse Lab in Alderley Park and Alex Alderton, Roberto Amato, Jeffrey Barrett, Sonia Goncalves, Ewan Harrison, David K. Jackson, Ian Johnston, Dominic Kwiatkowski, Cordelia Langford, John Sillitoe on behalf of the Wellcome Sanger Institute COVID-19 Surveillance Team |
| EPI_ISL_3042295 | Lighthouse Lab in Milton Keynes | Wellcome Sanger Institute for the COVID-19 Genomics UK (COG-UK) Consortium | The Lighthouse Lab in Milton Keynes and Alex Alderton, Roberto Amato, Jeffrey Barrett, Sonia Goncalves, Ewan Harrison, David K. Jackson, Ian Johnston, Dominic Kwiatkowski, Cordelia Langford, John Sillitoe on behalf of the Wellcome Sanger Institute COVID-19 Surveillance Team |
| EPI_ISL_3042296 | Lighthouse Laboratory Plymouth | Wellcome Sanger Institute for the COVID-19 Genomics UK (COG-UK) Consortium | Lighthouse Laboratory Plymouth and Alex Alderton, Roberto Amato, Jeffrey Barrett, Sonia Goncalves, Ewan Harrison, David K. Jackson, Ian Johnston, Dominic Kwiatkowski, Cordelia Langford, John Sillitoe on behalf of the Wellcome Sanger Institute COVID-19 Surveillance Team |
| EPI_ISL_3042297 | Lighthouse Lab in Milton Keynes | Wellcome Sanger Institute for the COVID-19 Genomics UK (COG-UK) Consortium | The Lighthouse Lab in Milton Keynes and Alex Alderton, Roberto Amato, Jeffrey Barrett, Sonia Goncalves, Ewan Harrison, David K. Jackson, Ian Johnston, Dominic Kwiatkowski, Cordelia Langford, John Sillitoe on behalf of the Wellcome Sanger Institute COVID-19 Surveillance Team |
| EPI_ISL_3042298 | Berkshire and Surrey Pathology Services Lighthouse Laboratory | Wellcome Sanger Institute for the COVID-19 Genomics UK (COG-UK) Consortium | Berkshire and Surrey Pathology Services Lighthouse Laboratory and Alex Alderton, Roberto Amato, Jeffrey Barrett, Sonia Goncalves, Ewan Harrison, David K. Jackson, Ian Johnston, Dominic Kwiatkowski, Cordelia Langford, John Sillitoe on behalf of the Wellcome Sanger Institute COVID-19 Surveillance Team |
| EPI_ISL_3042299 | Lighthouse Laboratory Plymouth | Wellcome Sanger Institute for the COVID-19 Genomics UK (COG-UK) Consortium | Lighthouse Laboratory Plymouth and Alex Alderton, Roberto Amato, Jeffrey Barrett, Sonia Goncalves, Ewan Harrison, David K. Jackson, Ian Johnston, Dominic Kwiatkowski, Cordelia Langford, John Sillitoe on behalf of the Wellcome Sanger Institute COVID-19 Surveillance Team |
| EPI_ISL_3042300 | Lighthouse Lab in Milton Keynes | Wellcome Sanger Institute for the COVID-19 Genomics UK (COG-UK) Consortium | The Lighthouse Lab in Milton Keynes and Alex Alderton, Roberto Amato, Jeffrey Barrett, Sonia Goncalves, Ewan Harrison, David K. Jackson, Ian Johnston, Dominic Kwiatkowski, Cordelia Langford, John Sillitoe on behalf of the Wellcome Sanger Institute COVID-19 Surveillance Team |
| EPI_ISL_3042301 | Lighthouse Lab in Alderley Park | Wellcome Sanger Institute for the COVID-19 Genomics UK (COG-UK) Consortium | Jacquelyn Wynn, Mairead Hyland, The Lighthouse Lab in Alderley Park and Alex Alderton, Roberto Amato, Jeffrey Barrett, Sonia Goncalves, Ewan Harrison, David K. Jackson, Ian Johnston, Dominic Kwiatkowski, Cordelia Langford, John Sillitoe on behalf of the Wellcome Sanger Institute COVID-19 Surveillance Team |
| EPI_ISL_3042302 | Lighthouse Lab in Milton Keynes | Wellcome Sanger Institute for the COVID-19 Genomics UK (COG-UK) Consortium | The Lighthouse Lab in Milton Keynes and Alex Alderton, Roberto Amato, Jeffrey Barrett, Sonia Goncalves, Ewan Harrison, David K. Jackson, Ian Johnston, Dominic Kwiatkowski, Cordelia Langford, John Sillitoe on behalf of the Wellcome Sanger Institute COVID-19 Surveillance Team |
| EPI_ISL_3042303 | Lighthouse Laboratory Plymouth | Wellcome Sanger Institute for the COVID-19 Genomics UK (COG-UK) Consortium | Lighthouse Laboratory Plymouth and Alex Alderton, Roberto Amato, Jeffrey Barrett, Sonia Goncalves, Ewan Harrison, David K. Jackson, Ian Johnston, Dominic Kwiatkowski, Cordelia Langford, John Sillitoe on behalf of the Wellcome Sanger Institute COVID-19 Surveillance Team |
| EPI_ISL_3042304, EPI_ISL_3042305 | Lighthouse Lab in Alderley Park | Wellcome Sanger Institute for the COVID-19 Genomics UK (COG-UK) Consortium | Jacquelyn Wynn, Mairead Hyland, The Lighthouse Lab in Alderley Park and Alex Alderton, Roberto Amato, Jeffrey Barrett, Sonia Goncalves, Ewan Harrison, David K. Jackson, Ian Johnston, Dominic Kwiatkowski, Cordelia Langford, John Sillitoe on behalf of the Wellcome Sanger Institute COVID-19 Surveillance Team |
| EPI_ISL_3042306 | Lighthouse Laboratory Plymouth | Wellcome Sanger Institute for the COVID-19 Genomics UK (COG-UK) Consortium | Lighthouse Laboratory Plymouth and Alex Alderton, Roberto Amato, Jeffrey Barrett, Sonia Goncalves, Ewan Harrison, David K. Jackson, Ian Johnston, Dominic Kwiatkowski, Cordelia Langford, John Sillitoe on behalf of the Wellcome Sanger Institute COVID-19 Surveillance Team |
| EPI_ISL_3042307 | Lighthouse Lab in Milton Keynes | Wellcome Sanger Institute for the COVID-19 Genomics UK (COG-UK) Consortium | The Lighthouse Lab in Milton Keynes and Alex Alderton, Roberto Amato, Jeffrey Barrett, Sonia Goncalves, Ewan Harrison, David K. Jackson, Ian Johnston, Dominic Kwiatkowski, Cordelia Langford, John Sillitoe on behalf of the Wellcome Sanger Institute COVID-19 Surveillance Team |
| EPI_ISL_3042308 | Lighthouse Laboratory Plymouth | Wellcome Sanger Institute for the COVID-19 Genomics UK (COG-UK) Consortium | Lighthouse Laboratory Plymouth and Alex Alderton, Roberto Amato, Jeffrey Barrett, Sonia Goncalves, Ewan Harrison, David K. Jackson, Ian Johnston, Dominic Kwiatkowski, Cordelia Langford, John Sillitoe on behalf of the Wellcome Sanger Institute COVID-19 Surveillance Team |
| EPI_ISL_3042309, EPI_ISL_3042310 | Berkshire and Surrey Pathology Services Lighthouse Laboratory | Wellcome Sanger Institute for the COVID-19 Genomics UK (COG-UK) Consortium | Berkshire and Surrey Pathology Services Lighthouse Laboratory and Alex Alderton, Roberto Amato, Jeffrey Barrett, Sonia Goncalves, Ewan Harrison, David K. Jackson, Ian Johnston, Dominic Kwiatkowski, Cordelia Langford, John Sillitoe on behalf of the Wellcome Sanger Institute COVID-19 Surveillance Team |
| EPI_ISL_3042311 | Lighthouse Lab in Milton Keynes | Wellcome Sanger Institute for the COVID-19 Genomics UK (COG-UK) Consortium | The Lighthouse Lab in Milton Keynes and Alex Alderton, Roberto Amato, Jeffrey Barrett, Sonia Goncalves, Ewan Harrison, David K. Jackson, Ian Johnston, Dominic Kwiatkowski, Cordelia Langford, John Sillitoe on behalf of the Wellcome Sanger Institute COVID-19 Surveillance Team |
| EPI_ISL_3042312 | Berkshire and Surrey Pathology Services Lighthouse Laboratory | Wellcome Sanger Institute for the COVID-19 Genomics UK (COG-UK) Consortium | Berkshire and Surrey Pathology Services Lighthouse Laboratory and Alex Alderton, Roberto Amato, Jeffrey Barrett, Sonia Goncalves, Ewan Harrison, David K. Jackson, Ian Johnston, Dominic Kwiatkowski, Cordelia Langford, John Sillitoe on behalf of the Wellcome Sanger Institute COVID-19 Surveillance Team |
| EPI_ISL_3042313 | Lighthouse Laboratory Plymouth | Wellcome Sanger Institute for the COVID-19 Genomics UK (COG-UK) Consortium | Lighthouse Laboratory Plymouth and Alex Alderton, Roberto Amato, Jeffrey Barrett, Sonia Goncalves, Ewan Harrison, David K. Jackson, Ian Johnston, Dominic Kwiatkowski, Cordelia Langford, John Sillitoe on behalf of the Wellcome Sanger Institute COVID-19 Surveillance Team |
| EPI_ISL_3042314 | Lighthouse Lab in Alderley Park | Wellcome Sanger Institute for the COVID-19 Genomics UK (COG-UK) Consortium | Jacquelyn Wynn, Mairead Hyland, The Lighthouse Lab in Alderley Park and Alex Alderton, Roberto Amato, Jeffrey Barrett, Sonia Goncalves, Ewan Harrison, David K. Jackson, Ian Johnston, Dominic Kwiatkowski, Cordelia Langford, John Sillitoe on behalf of the Wellcome Sanger Institute COVID-19 Surveillance Team |
| EPI_ISL_3042315 | Lighthouse Laboratory Plymouth | Wellcome Sanger Institute for the COVID-19 Genomics UK (COG-UK) Consortium | Lighthouse Laboratory Plymouth and Alex Alderton, Roberto Amato, Jeffrey Barrett, Sonia Goncalves, Ewan Harrison, David K. Jackson, Ian Johnston, Dominic Kwiatkowski, Cordelia Langford, John Sillitoe on behalf of the Wellcome Sanger Institute COVID-19 Surveillance Team |
| EPI_ISL_3042316 | Lighthouse Lab in Alderley Park | Wellcome Sanger Institute for the COVID-19 Genomics UK (COG-UK) Consortium | Jacquelyn Wynn, Mairead Hyland, The Lighthouse Lab in Alderley Park and Alex Alderton, Roberto Amato, Jeffrey Barrett, Sonia Goncalves, Ewan Harrison, David K. Jackson, Ian Johnston, Dominic Kwiatkowski, Cordelia Langford, John Sillitoe on behalf of the Wellcome Sanger Institute COVID-19 Surveillance Team |
| EPI_ISL_3042317, EPI_ISL_3042318, EPI_ISL_3042319 | Lighthouse Laboratory Plymouth | Wellcome Sanger Institute for the COVID-19 Genomics UK (COG-UK) Consortium | Lighthouse Laboratory Plymouth and Alex Alderton, Roberto Amato, Jeffrey Barrett, Sonia Goncalves, Ewan Harrison, David K. Jackson, Ian Johnston, Dominic Kwiatkowski, Cordelia Langford, John Sillitoe on behalf of the Wellcome Sanger Institute COVID-19 Surveillance Team |
| EPI_ISL_3042320, EPI_ISL_3042321 | Berkshire and Surrey Pathology Services Lighthouse Laboratory | Wellcome Sanger Institute for the COVID-19 Genomics UK (COG-UK) Consortium | Berkshire and Surrey Pathology Services Lighthouse Laboratory and Alex Alderton, Roberto Amato, Jeffrey Barrett, Sonia Goncalves, Ewan Harrison, David K. Jackson, Ian Johnston, Dominic Kwiatkowski, Cordelia Langford, John Sillitoe on behalf of the Wellcome Sanger Institute COVID-19 Surveillance Team |
| EPI_ISL_3042322, EPI_ISL_3042323 | Lighthouse Lab in Alderley Park | Wellcome Sanger Institute for the COVID-19 Genomics UK (COG-UK) Consortium | Jacquelyn Wynn, Mairead Hyland, The Lighthouse Lab in Alderley Park and Alex Alderton, Roberto Amato, Jeffrey Barrett, Sonia Goncalves, Ewan Harrison, David K. Jackson, Ian Johnston, Dominic Kwiatkowski, Cordelia Langford, John Sillitoe on behalf of the Wellcome Sanger Institute COVID-19 Surveillance Team |
| EPI_ISL_3042324 | Lighthouse Laboratory Plymouth | Wellcome Sanger Institute for the COVID-19 Genomics UK (COG-UK) Consortium | Lighthouse Laboratory Plymouth and Alex Alderton, Roberto Amato, Jeffrey Barrett, Sonia Goncalves, Ewan Harrison, David K. Jackson, Ian Johnston, Dominic Kwiatkowski, Cordelia Langford, John Sillitoe on behalf of the Wellcome Sanger Institute COVID-19 Surveillance Team |
| EPI_ISL_3042325 | Lighthouse Lab in Milton Keynes | Wellcome Sanger Institute for the COVID-19 Genomics UK (COG-UK) Consortium | The Lighthouse Lab in Milton Keynes and Alex Alderton, Roberto Amato, Jeffrey Barrett, Sonia Goncalves, Ewan Harrison, David K. Jackson, Ian Johnston, Dominic Kwiatkowski, Cordelia Langford, John Sillitoe on behalf of the Wellcome Sanger Institute COVID-19 Surveillance Team |
| EPI_ISL_3042326 | Lighthouse Lab in Alderley Park | Wellcome Sanger Institute for the COVID-19 Genomics UK (COG-UK) Consortium | Jacquelyn Wynn, Mairead Hyland, The Lighthouse Lab in Alderley Park and Alex Alderton, Roberto Amato, Jeffrey Barrett, Sonia Goncalves, Ewan Harrison, David K. Jackson, Ian Johnston, Dominic Kwiatkowski, Cordelia Langford, John Sillitoe on behalf of the Wellcome Sanger Institute COVID-19 Surveillance Team |
| EPI_ISL_3042327, EPI_ISL_3042328 | Lighthouse Laboratory Plymouth | Wellcome Sanger Institute for the COVID-19 Genomics UK (COG-UK) Consortium | Lighthouse Laboratory Plymouth and Alex Alderton, Roberto Amato, Jeffrey Barrett, Sonia Goncalves, Ewan Harrison, David K. Jackson, Ian Johnston, Dominic Kwiatkowski, Cordelia Langford, John Sillitoe on behalf of the Wellcome Sanger Institute COVID-19 Surveillance Team |
| EPI_ISL_3042329, EPI_ISL_3042330 | Lighthouse Lab in Alderley Park | Wellcome Sanger Institute for the COVID-19 Genomics UK (COG-UK) Consortium | Jacquelyn Wynn, Mairead Hyland, The Lighthouse Lab in Alderley Park and Alex Alderton, Roberto Amato, Jeffrey Barrett, Sonia Goncalves, Ewan Harrison, David K. Jackson, Ian Johnston, Dominic Kwiatkowski, Cordelia Langford, John Sillitoe on behalf of the Wellcome Sanger Institute COVID-19 Surveillance Team |
| EPI_ISL_3042331, EPI_ISL_3042332 | Lighthouse Lab in Milton Keynes | Wellcome Sanger Institute for the COVID-19 Genomics UK (COG-UK) Consortium | The Lighthouse Lab in Milton Keynes and Alex Alderton, Roberto Amato, Jeffrey Barrett, Sonia Goncalves, Ewan Harrison, David K. Jackson, Ian Johnston, Dominic Kwiatkowski, Cordelia Langford, John Sillitoe on behalf of the Wellcome Sanger Institute COVID-19 Surveillance Team |
| EPI_ISL_3042333 | Lighthouse Lab in Alderley Park | Wellcome Sanger Institute for the COVID-19 Genomics UK (COG-UK) Consortium | Jacquelyn Wynn, Mairead Hyland, The Lighthouse Lab in Alderley Park and Alex Alderton, Roberto Amato, Jeffrey Barrett, Sonia Goncalves, Ewan Harrison, David K. Jackson, Ian Johnston, Dominic Kwiatkowski, Cordelia Langford, John Sillitoe on behalf of the Wellcome Sanger Institute COVID-19 Surveillance Team |
| EPI_ISL_3042334 | Lighthouse Lab in Milton Keynes | Wellcome Sanger Institute for the COVID-19 Genomics UK (COG-UK) Consortium | The Lighthouse Lab in Milton Keynes and Alex Alderton, Roberto Amato, Jeffrey Barrett, Sonia Goncalves, Ewan Harrison, David K. Jackson, Ian Johnston, Dominic Kwiatkowski, Cordelia Langford, John Sillitoe on behalf of the Wellcome Sanger Institute COVID-19 Surveillance Team |
| EPI_ISL_3042335 | Berkshire and Surrey Pathology Services Lighthouse Laboratory | Wellcome Sanger Institute for the COVID-19 Genomics UK (COG-UK) Consortium | Berkshire and Surrey Pathology Services Lighthouse Laboratory and Alex Alderton, Roberto Amato, Jeffrey Barrett, Sonia Goncalves, Ewan Harrison, David K. Jackson, Ian Johnston, Dominic Kwiatkowski, Cordelia Langford, John Sillitoe on behalf of the Wellcome Sanger Institute COVID-19 Surveillance Team |
| EPI_ISL_3042336 | Lighthouse Lab in Milton Keynes | Wellcome Sanger Institute for the COVID-19 Genomics UK (COG-UK) Consortium | The Lighthouse Lab in Milton Keynes and Alex Alderton, Roberto Amato, Jeffrey Barrett, Sonia Goncalves, Ewan Harrison, David K. Jackson, Ian Johnston, Dominic Kwiatkowski, Cordelia Langford, John Sillitoe on behalf of the Wellcome Sanger Institute COVID-19 Surveillance Team |
| EPI_ISL_3042337 | Lighthouse Laboratory Plymouth | Wellcome Sanger Institute for the COVID-19 Genomics UK (COG-UK) Consortium | Lighthouse Laboratory Plymouth and Alex Alderton, Roberto Amato, Jeffrey Barrett, Sonia Goncalves, Ewan Harrison, David K. Jackson, Ian Johnston, Dominic Kwiatkowski, Cordelia Langford, John Sillitoe on behalf of the Wellcome Sanger Institute COVID-19 Surveillance Team |
| EPI_ISL_3042338 | Berkshire and Surrey Pathology Services Lighthouse Laboratory | Wellcome Sanger Institute for the COVID-19 Genomics UK (COG-UK) Consortium | Berkshire and Surrey Pathology Services Lighthouse Laboratory and Alex Alderton, Roberto Amato, Jeffrey Barrett, Sonia Goncalves, Ewan Harrison, David K. Jackson, Ian Johnston, Dominic Kwiatkowski, Cordelia Langford, John Sillitoe on behalf of the Wellcome Sanger Institute COVID-19 Surveillance Team |
| EPI_ISL_3042339 | Lighthouse Lab in Alderley Park | Wellcome Sanger Institute for the COVID-19 Genomics UK (COG-UK) Consortium | Jacquelyn Wynn, Mairead Hyland, The Lighthouse Lab in Alderley Park and Alex Alderton, Roberto Amato, Jeffrey Barrett, Sonia Goncalves, Ewan Harrison, David K. Jackson, Ian Johnston, Dominic Kwiatkowski, Cordelia Langford, John Sillitoe on behalf of the Wellcome Sanger Institute COVID-19 Surveillance Team |
| EPI_ISL_3042340 | Berkshire and Surrey Pathology Services Lighthouse Laboratory | Wellcome Sanger Institute for the COVID-19 Genomics UK (COG-UK) Consortium | Berkshire and Surrey Pathology Services Lighthouse Laboratory and Alex Alderton, Roberto Amato, Jeffrey Barrett, Sonia Goncalves, Ewan Harrison, David K. Jackson, Ian Johnston, Dominic Kwiatkowski, Cordelia Langford, John Sillitoe on behalf of the Wellcome Sanger Institute COVID-19 Surveillance Team |
| EPI_ISL_3042341 | Lighthouse Laboratory Plymouth | Wellcome Sanger Institute for the COVID-19 Genomics UK (COG-UK) Consortium | Lighthouse Laboratory Plymouth and Alex Alderton, Roberto Amato, Jeffrey Barrett, Sonia Goncalves, Ewan Harrison, David K. Jackson, Ian Johnston, Dominic Kwiatkowski, Cordelia Langford, John Sillitoe on behalf of the Wellcome Sanger Institute COVID-19 Surveillance Team |
| EPI_ISL_3042342 | Berkshire and Surrey Pathology Services Lighthouse Laboratory | Wellcome Sanger Institute for the COVID-19 Genomics UK (COG-UK) Consortium | Berkshire and Surrey Pathology Services Lighthouse Laboratory and Alex Alderton, Roberto Amato, Jeffrey Barrett, Sonia Goncalves, Ewan Harrison, David K. Jackson, Ian Johnston, Dominic Kwiatkowski, Cordelia Langford, John Sillitoe on behalf of the Wellcome Sanger Institute COVID-19 Surveillance Team |
| EPI_ISL_3042343 | Lighthouse Lab in Alderley Park | Wellcome Sanger Institute for the COVID-19 Genomics UK (COG-UK) Consortium | Jacquelyn Wynn, Mairead Hyland, The Lighthouse Lab in Alderley Park and Alex Alderton, Roberto Amato, Jeffrey Barrett, Sonia Goncalves, Ewan Harrison, David K. Jackson, Ian Johnston, Dominic Kwiatkowski, Cordelia Langford, John Sillitoe on behalf of the Wellcome Sanger Institute COVID-19 Surveillance Team |
| EPI_ISL_3042344 | Lighthouse Lab in Milton Keynes | Wellcome Sanger Institute for the COVID-19 Genomics UK (COG-UK) Consortium | The Lighthouse Lab in Milton Keynes and Alex Alderton, Roberto Amato, Jeffrey Barrett, Sonia Goncalves, Ewan Harrison, David K. Jackson, Ian Johnston, Dominic Kwiatkowski, Cordelia Langford, John Sillitoe on behalf of the Wellcome Sanger Institute COVID-19 Surveillance Team |

| | | | |
|---|---|---|---|
| EPI_ISL_3042345 | Lighthouse Laboratory Plymouth | Wellcome Sanger Institute for the COVID-19 Genomics UK (COG-UK) Consortium | Lighthouse Laboratory Plymouth and Alex Alderton, Roberto Amato, Jeffrey Barrett, Sonia Goncalves, Ewan Harrison, David K. Jackson, Ian Johnston, Dominic Kwiatkowski, Cordelia Langford, John Sillitoe on behalf of the Wellcome Sanger Institute COVID-19 Surveillance Team |
| EPI_ISL_3042346 | Berkshire and Surrey Pathology Services Lighthouse Laboratory | Wellcome Sanger Institute for the COVID-19 Genomics UK (COG-UK) Consortium | Berkshire and Surrey Pathology Services Lighthouse Laboratory and Alex Alderton, Roberto Amato, Jeffrey Barrett, Sonia Goncalves, Ewan Harrison, David K. Jackson, Ian Johnston, Dominic Kwiatkowski, Cordelia Langford, John Sillitoe on behalf of the Wellcome Sanger Institute COVID-19 Surveillance Team |
| EPI_ISL_3042347 | Lighthouse Laboratory Plymouth | Wellcome Sanger Institute for the COVID-19 Genomics UK (COG-UK) Consortium | Lighthouse Laboratory Plymouth and Alex Alderton, Roberto Amato, Jeffrey Barrett, Sonia Goncalves, Ewan Harrison, David K. Jackson, Ian Johnston, Dominic Kwiatkowski, Cordelia Langford, John Sillitoe on behalf of the Wellcome Sanger Institute COVID-19 Surveillance Team |
| EPI_ISL_3042348 | Lighthouse Lab in Alderley Park | Wellcome Sanger Institute for the COVID-19 Genomics UK (COG-UK) Consortium | Jacquelyn Wynn, Mairead Hyland, The Lighthouse Lab in Alderley Park and Alex Alderton, Roberto Amato, Jeffrey Barrett, Sonia Goncalves, Ewan Harrison, David K. Jackson, Ian Johnston, Dominic Kwiatkowski, Cordelia Langford, John Sillitoe on behalf of the Wellcome Sanger Institute COVID-19 Surveillance Team |
| EPI_ISL_3042349 | Lighthouse Lab in Milton Keynes | Wellcome Sanger Institute for the COVID-19 Genomics UK (COG-UK) Consortium | The Lighthouse Lab in Milton Keynes and Alex Alderton, Roberto Amato, Jeffrey Barrett, Sonia Goncalves, Ewan Harrison, David K. Jackson, Ian Johnston, Dominic Kwiatkowski, Cordelia Langford, John Sillitoe on behalf of the Wellcome Sanger Institute COVID-19 Surveillance Team |
| EPI_ISL_3042350 | Berkshire and Surrey Pathology Services Lighthouse Laboratory | Wellcome Sanger Institute for the COVID-19 Genomics UK (COG-UK) Consortium | Berkshire and Surrey Pathology Services Lighthouse Laboratory and Alex Alderton, Roberto Amato, Jeffrey Barrett, Sonia Goncalves, Ewan Harrison, David K. Jackson, Ian Johnston, Dominic Kwiatkowski, Cordelia Langford, John Sillitoe on behalf of the Wellcome Sanger Institute COVID-19 Surveillance Team |
| EPI_ISL_3042351 | Lighthouse Lab in Milton Keynes | Wellcome Sanger Institute for the COVID-19 Genomics UK (COG-UK) Consortium | The Lighthouse Lab in Milton Keynes and Alex Alderton, Roberto Amato, Jeffrey Barrett, Sonia Goncalves, Ewan Harrison, David K. Jackson, Ian Johnston, Dominic Kwiatkowski, Cordelia Langford, John Sillitoe on behalf of the Wellcome Sanger Institute COVID-19 Surveillance Team |
| EPI_ISL_3042352, EPI_ISL_3042353 | Berkshire and Surrey Pathology Services Lighthouse Laboratory | Wellcome Sanger Institute for the COVID-19 Genomics UK (COG-UK) Consortium | Berkshire and Surrey Pathology Services Lighthouse Laboratory and Alex Alderton, Roberto Amato, Jeffrey Barrett, Sonia Goncalves, Ewan Harrison, David K. Jackson, Ian Johnston, Dominic Kwiatkowski, Cordelia Langford, John Sillitoe on behalf of the Wellcome Sanger Institute COVID-19 Surveillance Team |
| EPI_ISL_3042354 | Lighthouse Laboratory Plymouth | Wellcome Sanger Institute for the COVID-19 Genomics UK (COG-UK) Consortium | Lighthouse Laboratory Plymouth and Alex Alderton, Roberto Amato, Jeffrey Barrett, Sonia Goncalves, Ewan Harrison, David K. Jackson, Ian Johnston, Dominic Kwiatkowski, Cordelia Langford, John Sillitoe on behalf of the Wellcome Sanger Institute COVID-19 Surveillance Team |
| EPI_ISL_3042355 | Lighthouse Lab in Alderley Park | Wellcome Sanger Institute for the COVID-19 Genomics UK (COG-UK) Consortium | Jacquelyn Wynn, Mairead Hyland, The Lighthouse Lab in Alderley Park and Alex Alderton, Roberto Amato, Jeffrey Barrett, Sonia Goncalves, Ewan Harrison, David K. Jackson, Ian Johnston, Dominic Kwiatkowski, Cordelia Langford, John Sillitoe on behalf of the Wellcome Sanger Institute COVID-19 Surveillance Team |
| EPI_ISL_3042356 | Lighthouse Lab in Milton Keynes | Wellcome Sanger Institute for the COVID-19 Genomics UK (COG-UK) Consortium | The Lighthouse Lab in Milton Keynes and Alex Alderton, Roberto Amato, Jeffrey Barrett, Sonia Goncalves, Ewan Harrison, David K. Jackson, Ian Johnston, Dominic Kwiatkowski, Cordelia Langford, John Sillitoe on behalf of the Wellcome Sanger Institute COVID-19 Surveillance Team |
| EPI_ISL_3042357 | Lighthouse Lab in Alderley Park | Wellcome Sanger Institute for the COVID-19 Genomics UK (COG-UK) Consortium | Jacquelyn Wynn, Mairead Hyland, The Lighthouse Lab in Alderley Park and Alex Alderton, Roberto Amato, Jeffrey Barrett, Sonia Goncalves, Ewan Harrison, David K. Jackson, Ian Johnston, Dominic Kwiatkowski, Cordelia Langford, John Sillitoe on behalf of the Wellcome Sanger Institute COVID-19 Surveillance Team |
| EPI_ISL_3042358 | Lighthouse Lab in Milton Keynes | Wellcome Sanger Institute for the COVID-19 Genomics UK (COG-UK) Consortium | The Lighthouse Lab in Milton Keynes and Alex Alderton, Roberto Amato, Jeffrey Barrett, Sonia Goncalves, Ewan Harrison, David K. Jackson, Ian Johnston, Dominic Kwiatkowski, Cordelia Langford, John Sillitoe on behalf of the Wellcome Sanger Institute COVID-19 Surveillance Team |
| EPI_ISL_3042359, EPI_ISL_3042360 | Lighthouse Laboratory Plymouth | Wellcome Sanger Institute for the COVID-19 Genomics UK (COG-UK) Consortium | Lighthouse Laboratory Plymouth and Alex Alderton, Roberto Amato, Jeffrey Barrett, Sonia Goncalves, Ewan Harrison, David K. Jackson, Ian Johnston, Dominic Kwiatkowski, Cordelia Langford, John Sillitoe on behalf of the Wellcome Sanger Institute COVID-19 Surveillance Team |
| EPI_ISL_3042361 | Lighthouse Lab in Milton Keynes | Wellcome Sanger Institute for the COVID-19 Genomics UK (COG-UK) Consortium | The Lighthouse Lab in Milton Keynes and Alex Alderton, Roberto Amato, Jeffrey Barrett, Sonia Goncalves, Ewan Harrison, David K. Jackson, Ian Johnston, Dominic Kwiatkowski, Cordelia Langford, John Sillitoe on behalf of the Wellcome Sanger Institute COVID-19 Surveillance Team |
| EPI_ISL_3042362, EPI_ISL_3042363, EPI_ISL_3042364, EPI_ISL_3042365 | Berkshire and Surrey Pathology Services Lighthouse Laboratory | Wellcome Sanger Institute for the COVID-19 Genomics UK (COG-UK) Consortium | Berkshire and Surrey Pathology Services Lighthouse Laboratory and Alex Alderton, Roberto Amato, Jeffrey Barrett, Sonia Goncalves, Ewan Harrison, David K. Jackson, Ian Johnston, Dominic Kwiatkowski, Cordelia Langford, John Sillitoe on behalf of the Wellcome Sanger Institute COVID-19 Surveillance Team |
| EPI_ISL_3042366, EPI_ISL_3042367 | Lighthouse Laboratory Plymouth | Wellcome Sanger Institute for the COVID-19 Genomics UK (COG-UK) Consortium | Lighthouse Laboratory Plymouth and Alex Alderton, Roberto Amato, Jeffrey Barrett, Sonia Goncalves, Ewan Harrison, David K. Jackson, Ian Johnston, Dominic Kwiatkowski, Cordelia Langford, John Sillitoe on behalf of the Wellcome Sanger Institute COVID-19 Surveillance Team |
| EPI_ISL_3042368 | Lighthouse Lab in Alderley Park | Wellcome Sanger Institute for the COVID-19 Genomics UK (COG-UK) Consortium | Jacquelyn Wynn, Mairead Hyland, The Lighthouse Lab in Alderley Park and Alex Alderton, Roberto Amato, Jeffrey Barrett, Sonia Goncalves, Ewan Harrison, David K. Jackson, Ian Johnston, Dominic Kwiatkowski, Cordelia Langford, John Sillitoe on behalf of the Wellcome Sanger Institute COVID-19 Surveillance Team |
| EPI_ISL_3042369, EPI_ISL_3042370 | Lighthouse Lab in Milton Keynes | Wellcome Sanger Institute for the COVID-19 Genomics UK (COG-UK) Consortium | The Lighthouse Lab in Milton Keynes and Alex Alderton, Roberto Amato, Jeffrey Barrett, Sonia Goncalves, Ewan Harrison, David K. Jackson, Ian Johnston, Dominic Kwiatkowski, Cordelia Langford, John Sillitoe on behalf of the Wellcome Sanger Institute COVID-19 Surveillance Team |
| EPI_ISL_3042371 | Lighthouse Lab in Alderley Park | Wellcome Sanger Institute for the COVID-19 Genomics UK (COG-UK) Consortium | Jacquelyn Wynn, Mairead Hyland, The Lighthouse Lab in Alderley Park and Alex Alderton, Roberto Amato, Jeffrey Barrett, Sonia Goncalves, Ewan Harrison, David K. Jackson, Ian Johnston, Dominic Kwiatkowski, Cordelia Langford, John Sillitoe on behalf of the Wellcome Sanger Institute COVID-19 Surveillance Team |
| EPI_ISL_3042372 | Lighthouse Laboratory Plymouth | Wellcome Sanger Institute for the COVID-19 Genomics UK (COG-UK) Consortium | Lighthouse Laboratory Plymouth and Alex Alderton, Roberto Amato, Jeffrey Barrett, Sonia Goncalves, Ewan Harrison, David K. Jackson, Ian Johnston, Dominic Kwiatkowski, Cordelia Langford, John Sillitoe on behalf of the Wellcome Sanger Institute COVID-19 Surveillance Team |
| EPI_ISL_3042373 | Berkshire and Surrey Pathology Services Lighthouse Laboratory | Wellcome Sanger Institute for the COVID-19 Genomics UK (COG-UK) Consortium | Berkshire and Surrey Pathology Services Lighthouse Laboratory and Alex Alderton, Roberto Amato, Jeffrey Barrett, Sonia Goncalves, Ewan Harrison, David K. Jackson, Ian Johnston, Dominic Kwiatkowski, Cordelia Langford, John Sillitoe on behalf of the Wellcome Sanger Institute COVID-19 Surveillance Team |
| EPI_ISL_3042374 | Lighthouse Laboratory Plymouth | Wellcome Sanger Institute for the COVID-19 Genomics UK (COG-UK) Consortium | Lighthouse Laboratory Plymouth and Alex Alderton, Roberto Amato, Jeffrey Barrett, Sonia Goncalves, Ewan Harrison, David K. Jackson, Ian Johnston, Dominic Kwiatkowski, Cordelia Langford, John Sillitoe on behalf of the Wellcome Sanger Institute COVID-19 Surveillance Team |
| EPI_ISL_3042375 | Berkshire and Surrey Pathology Services Lighthouse Laboratory | Wellcome Sanger Institute for the COVID-19 Genomics UK (COG-UK) Consortium | Berkshire and Surrey Pathology Services Lighthouse Laboratory and Alex Alderton, Roberto Amato, Jeffrey Barrett, Sonia Goncalves, Ewan Harrison, David K. Jackson, Ian Johnston, Dominic Kwiatkowski, Cordelia Langford, John Sillitoe on behalf of the Wellcome Sanger Institute COVID-19 Surveillance Team |
| EPI_ISL_3042376, EPI_ISL_3042377 | Lighthouse Lab in Milton Keynes | Wellcome Sanger Institute for the COVID-19 Genomics UK (COG-UK) Consortium | The Lighthouse Lab in Milton Keynes and Alex Alderton, Roberto Amato, Jeffrey Barrett, Sonia Goncalves, Ewan Harrison, David K. Jackson, Ian Johnston, Dominic Kwiatkowski, Cordelia Langford, John Sillitoe on behalf of the Wellcome Sanger Institute COVID-19 Surveillance Team |
| EPI_ISL_3042378 | Lighthouse Lab in Alderley Park | Wellcome Sanger Institute for the COVID-19 Genomics UK (COG-UK) Consortium | Jacquelyn Wynn, Mairead Hyland, The Lighthouse Lab in Alderley Park and Alex Alderton, Roberto Amato, Jeffrey Barrett, Sonia Goncalves, Ewan Harrison, David K. Jackson, Ian Johnston, Dominic Kwiatkowski, Cordelia Langford, John Sillitoe on behalf of the Wellcome Sanger Institute COVID-19 Surveillance Team |
| EPI_ISL_3042379 | Lighthouse Laboratory Plymouth | Wellcome Sanger Institute for the COVID-19 Genomics UK (COG-UK) Consortium | Lighthouse Laboratory Plymouth and Alex Alderton, Roberto Amato, Jeffrey Barrett, Sonia Goncalves, Ewan Harrison, David K. Jackson, Ian Johnston, Dominic Kwiatkowski, Cordelia Langford, John Sillitoe on behalf of the Wellcome Sanger Institute COVID-19 Surveillance Team |
| EPI_ISL_3042380 | Lighthouse Lab in Milton Keynes | Wellcome Sanger Institute for the COVID-19 Genomics UK (COG-UK) Consortium | The Lighthouse Lab in Milton Keynes and Alex Alderton, Roberto Amato, Jeffrey Barrett, Sonia Goncalves, Ewan Harrison, David K. Jackson, Ian Johnston, Dominic Kwiatkowski, Cordelia Langford, John Sillitoe on behalf of the Wellcome Sanger Institute COVID-19 Surveillance Team |
| EPI_ISL_3042381 | Lighthouse Lab in Alderley Park | Wellcome Sanger Institute for the COVID-19 Genomics UK (COG-UK) Consortium | Jacquelyn Wynn, Mairead Hyland, The Lighthouse Lab in Alderley Park and Alex Alderton, Roberto Amato, Jeffrey Barrett, Sonia Goncalves, Ewan Harrison, David K. Jackson, Ian Johnston, Dominic Kwiatkowski, Cordelia Langford, John Sillitoe on behalf of the Wellcome Sanger Institute COVID-19 Surveillance Team |
| EPI_ISL_3042382 | Lighthouse Lab in Milton Keynes | Wellcome Sanger Institute for the COVID-19 Genomics UK (COG-UK) Consortium | The Lighthouse Lab in Milton Keynes and Alex Alderton, Roberto Amato, Jeffrey Barrett, Sonia Goncalves, Ewan Harrison, David K. Jackson, Ian Johnston, Dominic Kwiatkowski, Cordelia Langford, John Sillitoe on behalf of the Wellcome Sanger Institute COVID-19 Surveillance Team |
| EPI_ISL_3042383, EPI_ISL_3042384 | Berkshire and Surrey Pathology Services Lighthouse Laboratory | Wellcome Sanger Institute for the COVID-19 Genomics UK (COG-UK) Consortium | Berkshire and Surrey Pathology Services Lighthouse Laboratory and Alex Alderton, Roberto Amato, Jeffrey Barrett, Sonia Goncalves, Ewan Harrison, David K. Jackson, Ian Johnston, Dominic Kwiatkowski, Cordelia Langford, John Sillitoe on behalf of the Wellcome Sanger Institute COVID-19 Surveillance Team |
| EPI_ISL_3042385 | Lighthouse Lab in Alderley Park | Wellcome Sanger Institute for the COVID-19 Genomics UK (COG-UK) Consortium | Jacquelyn Wynn, Mairead Hyland, The Lighthouse Lab in Alderley Park and Alex Alderton, Roberto Amato, Jeffrey Barrett, Sonia Goncalves, Ewan Harrison, David K. Jackson, Ian Johnston, Dominic Kwiatkowski, Cordelia Langford, John Sillitoe on behalf of the Wellcome Sanger Institute COVID-19 Surveillance Team |
| EPI_ISL_3042386, EPI_ISL_3042387 | Berkshire and Surrey Pathology Services Lighthouse Laboratory | Wellcome Sanger Institute for the COVID-19 Genomics UK (COG-UK) Consortium | Berkshire and Surrey Pathology Services Lighthouse Laboratory and Alex Alderton, Roberto Amato, Jeffrey Barrett, Sonia Goncalves, Ewan Harrison, David K. Jackson, Ian Johnston, Dominic Kwiatkowski, Cordelia Langford, John Sillitoe on behalf of the Wellcome Sanger Institute COVID-19 Surveillance Team |
| EPI_ISL_3042388 | Lighthouse Lab in Alderley Park | Wellcome Sanger Institute for the COVID-19 Genomics UK (COG-UK) Consortium | Jacquelyn Wynn, Mairead Hyland, The Lighthouse Lab in Alderley Park and Alex Alderton, Roberto Amato, Jeffrey Barrett, Sonia Goncalves, Ewan Harrison, David K. Jackson, Ian Johnston, Dominic Kwiatkowski, Cordelia Langford, John Sillitoe on behalf of the Wellcome Sanger Institute COVID-19 Surveillance Team |
| EPI_ISL_3042389 | Berkshire and Surrey Pathology Services Lighthouse Laboratory | Wellcome Sanger Institute for the COVID-19 Genomics UK (COG-UK) Consortium | Berkshire and Surrey Pathology Services Lighthouse Laboratory and Alex Alderton, Roberto Amato, Jeffrey Barrett, Sonia Goncalves, Ewan Harrison, David K. Jackson, Ian Johnston, Dominic Kwiatkowski, Cordelia Langford, John Sillitoe on behalf of the Wellcome Sanger Institute COVID-19 Surveillance Team |
| EPI_ISL_3042390, EPI_ISL_3042391 | Lighthouse Lab in Milton Keynes | Wellcome Sanger Institute for the COVID-19 Genomics UK (COG-UK) Consortium | The Lighthouse Lab in Milton Keynes and Alex Alderton, Roberto Amato, Jeffrey Barrett, Sonia Goncalves, Ewan Harrison, David K. Jackson, Ian Johnston, Dominic Kwiatkowski, Cordelia Langford, John Sillitoe on behalf of the Wellcome Sanger Institute COVID-19 Surveillance Team |
| EPI_ISL_3042392, EPI_ISL_3042393, EPI_ISL_3042394, EPI_ISL_3042395 | Lighthouse Laboratory Plymouth | Wellcome Sanger Institute for the COVID-19 Genomics UK (COG-UK) Consortium | Lighthouse Laboratory Plymouth and Alex Alderton, Roberto Amato, Jeffrey Barrett, Sonia Goncalves, Ewan Harrison, David K. Jackson, Ian Johnston, Dominic Kwiatkowski, Cordelia Langford, John Sillitoe on behalf of the Wellcome Sanger Institute COVID-19 Surveillance Team |
| EPI_ISL_3042396 | Lighthouse Lab in Alderley Park | Wellcome Sanger Institute for the COVID-19 Genomics UK (COG-UK) Consortium | Jacquelyn Wynn, Mairead Hyland, The Lighthouse Lab in Alderley Park and Alex Alderton, Roberto Amato, Jeffrey Barrett, Sonia Goncalves, Ewan Harrison, David K. Jackson, Ian Johnston, Dominic Kwiatkowski, Cordelia Langford, John Sillitoe on behalf of the Wellcome Sanger Institute COVID-19 Surveillance Team |
| EPI_ISL_3042397, EPI_ISL_3042398 | Lighthouse Lab in Milton Keynes | Wellcome Sanger Institute for the COVID-19 Genomics UK (COG-UK) Consortium | The Lighthouse Lab in Milton Keynes and Alex Alderton, Roberto Amato, Jeffrey Barrett, Sonia Goncalves, Ewan Harrison, David K. Jackson, Ian Johnston, Dominic Kwiatkowski, Cordelia Langford, John Sillitoe on behalf of the Wellcome Sanger Institute COVID-19 Surveillance Team |
| EPI_ISL_3042399 | Lighthouse Lab in Alderley Park | Wellcome Sanger Institute for the COVID-19 Genomics UK (COG-UK) Consortium | Jacquelyn Wynn, Mairead Hyland, The Lighthouse Lab in Alderley Park and Alex Alderton, Roberto Amato, Jeffrey Barrett, Sonia Goncalves, Ewan Harrison, David K. Jackson, Ian Johnston, Dominic Kwiatkowski, Cordelia Langford, John Sillitoe on behalf of the Wellcome Sanger Institute COVID-19 Surveillance Team |
| EPI_ISL_3042400 | Berkshire and Surrey Pathology Services Lighthouse Laboratory | Wellcome Sanger Institute for the COVID-19 Genomics UK (COG-UK) Consortium | Berkshire and Surrey Pathology Services Lighthouse Laboratory and Alex Alderton, Roberto Amato, Jeffrey Barrett, Sonia Goncalves, Ewan Harrison, David K. Jackson, Ian Johnston, Dominic Kwiatkowski, Cordelia Langford, John Sillitoe on behalf of the Wellcome Sanger Institute COVID-19 Surveillance Team |
| EPI_ISL_3042401 | Lighthouse Laboratory Plymouth | Wellcome Sanger Institute for the COVID-19 Genomics UK (COG-UK) Consortium | Lighthouse Laboratory Plymouth and Alex Alderton, Roberto Amato, Jeffrey Barrett, Sonia Goncalves, Ewan Harrison, David K. Jackson, Ian Johnston, Dominic Kwiatkowski, Cordelia Langford, John Sillitoe on behalf of the Wellcome Sanger Institute COVID-19 Surveillance Team |
| EPI_ISL_3042402 | Lighthouse Lab in Alderley Park | Wellcome Sanger Institute for the COVID-19 Genomics UK (COG-UK) Consortium | Jacquelyn Wynn, Mairead Hyland, The Lighthouse Lab in Alderley Park and Alex Alderton, Roberto Amato, Jeffrey Barrett, Sonia Goncalves, Ewan Harrison, David K. Jackson, Ian Johnston, Dominic Kwiatkowski, Cordelia Langford, John Sillitoe on behalf of the Wellcome Sanger Institute COVID-19 Surveillance Team |
| EPI_ISL_3042403 | Berkshire and Surrey Pathology Services Lighthouse Laboratory | Wellcome Sanger Institute for the COVID-19 Genomics UK (COG-UK) | Berkshire and Surrey Pathology Services Lighthouse Laboratory and Alex Alderton, Roberto Amato, Jeffrey Barrett, Sonia Goncalves, Ewan Harrison, David K. Jackson, Ian Johnston, Dominic Kwiatkowski, Cordelia Langford, John Sillitoe on behalf |

| | | | |
|---|---|---|---|
| | | Consortium | of the Wellcome Sanger Institute COVID-19 Surveillance Team |
| EPI_ISL_3042404 | Lighthouse Laboratory Plymouth | Wellcome Sanger Institute for the COVID-19 Genomics UK (COG-UK) Consortium | Lighthouse Laboratory Plymouth and Alex Alderton, Roberto Amato, Jeffrey Barrett, Sonia Goncalves, Ewan Harrison, David K. Jackson, Ian Johnston, Dominic Kwiatkowski, Cordelia Langford, John Sillitoe on behalf of the Wellcome Sanger Institute COVID-19 Surveillance Team |
| EPI_ISL_3042405 | Berkshire and Surrey Pathology Services Lighthouse Laboratory | Wellcome Sanger Institute for the COVID-19 Genomics UK (COG-UK) Consortium | Berkshire and Surrey Pathology Services Lighthouse Laboratory and Alex Alderton, Roberto Amato, Jeffrey Barrett, Sonia Goncalves, Ewan Harrison, David K. Jackson, Ian Johnston, Dominic Kwiatkowski, Cordelia Langford, John Sillitoe on behalf of the Wellcome Sanger Institute COVID-19 Surveillance Team |
| EPI_ISL_3042406, EPI_ISL_3042407, EPI_ISL_3042408, EPI_ISL_3042409 | Lighthouse Lab in Alderley Park | Wellcome Sanger Institute for the COVID-19 Genomics UK (COG-UK) Consortium | Jacquelyn Wynn, Mairead Hyland, The Lighthouse Lab in Alderley Park and Alex Alderton, Roberto Amato, Jeffrey Barrett, Sonia Goncalves, Ewan Harrison, David K. Jackson, Ian Johnston, Dominic Kwiatkowski, Cordelia Langford, John Sillitoe on behalf of the Wellcome Sanger Institute COVID-19 Surveillance Team |
| EPI_ISL_3042410 | Berkshire and Surrey Pathology Services Lighthouse Laboratory | Wellcome Sanger Institute for the COVID-19 Genomics UK (COG-UK) Consortium | Berkshire and Surrey Pathology Services Lighthouse Laboratory and Alex Alderton, Roberto Amato, Jeffrey Barrett, Sonia Goncalves, Ewan Harrison, David K. Jackson, Ian Johnston, Dominic Kwiatkowski, Cordelia Langford, John Sillitoe on behalf of the Wellcome Sanger Institute COVID-19 Surveillance Team |
| EPI_ISL_3042411, EPI_ISL_3042412 | Lighthouse Laboratory Plymouth | Wellcome Sanger Institute for the COVID-19 Genomics UK (COG-UK) Consortium | Lighthouse Laboratory Plymouth and Alex Alderton, Roberto Amato, Jeffrey Barrett, Sonia Goncalves, Ewan Harrison, David K. Jackson, Ian Johnston, Dominic Kwiatkowski, Cordelia Langford, John Sillitoe on behalf of the Wellcome Sanger Institute COVID-19 Surveillance Team |
| EPI_ISL_3042413 | Lighthouse Lab in Milton Keynes | Wellcome Sanger Institute for the COVID-19 Genomics UK (COG-UK) Consortium | The Lighthouse Lab in Milton Keynes and Alex Alderton, Roberto Amato, Jeffrey Barrett, Sonia Goncalves, Ewan Harrison, David K. Jackson, Ian Johnston, Dominic Kwiatkowski, Cordelia Langford, John Sillitoe on behalf of the Wellcome Sanger Institute COVID-19 Surveillance Team |
| EPI_ISL_3042414 | Lighthouse Laboratory Plymouth | Wellcome Sanger Institute for the COVID-19 Genomics UK (COG-UK) Consortium | Lighthouse Laboratory Plymouth and Alex Alderton, Roberto Amato, Jeffrey Barrett, Sonia Goncalves, Ewan Harrison, David K. Jackson, Ian Johnston, Dominic Kwiatkowski, Cordelia Langford, John Sillitoe on behalf of the Wellcome Sanger Institute COVID-19 Surveillance Team |
| EPI_ISL_3042415 | Berkshire and Surrey Pathology Services Lighthouse Laboratory | Wellcome Sanger Institute for the COVID-19 Genomics UK (COG-UK) Consortium | Berkshire and Surrey Pathology Services Lighthouse Laboratory and Alex Alderton, Roberto Amato, Jeffrey Barrett, Sonia Goncalves, Ewan Harrison, David K. Jackson, Ian Johnston, Dominic Kwiatkowski, Cordelia Langford, John Sillitoe on behalf of the Wellcome Sanger Institute COVID-19 Surveillance Team |
| EPI_ISL_3042416, EPI_ISL_3042417 | Lighthouse Laboratory Plymouth | Wellcome Sanger Institute for the COVID-19 Genomics UK (COG-UK) Consortium | Lighthouse Laboratory Plymouth and Alex Alderton, Roberto Amato, Jeffrey Barrett, Sonia Goncalves, Ewan Harrison, David K. Jackson, Ian Johnston, Dominic Kwiatkowski, Cordelia Langford, John Sillitoe on behalf of the Wellcome Sanger Institute COVID-19 Surveillance Team |
| EPI_ISL_3042418, EPI_ISL_3042419 | Berkshire and Surrey Pathology Services Lighthouse Laboratory | Wellcome Sanger Institute for the COVID-19 Genomics UK (COG-UK) Consortium | Berkshire and Surrey Pathology Services Lighthouse Laboratory and Alex Alderton, Roberto Amato, Jeffrey Barrett, Sonia Goncalves, Ewan Harrison, David K. Jackson, Ian Johnston, Dominic Kwiatkowski, Cordelia Langford, John Sillitoe on behalf of the Wellcome Sanger Institute COVID-19 Surveillance Team |
| EPI_ISL_3042420, EPI_ISL_3042421 | Lighthouse Laboratory Plymouth | Wellcome Sanger Institute for the COVID-19 Genomics UK (COG-UK) Consortium | Lighthouse Laboratory Plymouth and Alex Alderton, Roberto Amato, Jeffrey Barrett, Sonia Goncalves, Ewan Harrison, David K. Jackson, Ian Johnston, Dominic Kwiatkowski, Cordelia Langford, John Sillitoe on behalf of the Wellcome Sanger Institute COVID-19 Surveillance Team |
| EPI_ISL_3042422, EPI_ISL_3042423 | Lighthouse Lab in Alderley Park | Wellcome Sanger Institute for the COVID-19 Genomics UK (COG-UK) Consortium | Jacquelyn Wynn, Mairead Hyland, The Lighthouse Lab in Alderley Park and Alex Alderton, Roberto Amato, Jeffrey Barrett, Sonia Goncalves, Ewan Harrison, David K. Jackson, Ian Johnston, Dominic Kwiatkowski, Cordelia Langford, John Sillitoe on behalf of the Wellcome Sanger Institute COVID-19 Surveillance Team |
| EPI_ISL_3042424 | Berkshire and Surrey Pathology Services Lighthouse Laboratory | Wellcome Sanger Institute for the COVID-19 Genomics UK (COG-UK) Consortium | Berkshire and Surrey Pathology Services Lighthouse Laboratory and Alex Alderton, Roberto Amato, Jeffrey Barrett, Sonia Goncalves, Ewan Harrison, David K. Jackson, Ian Johnston, Dominic Kwiatkowski, Cordelia Langford, John Sillitoe on behalf of the Wellcome Sanger Institute COVID-19 Surveillance Team |
| EPI_ISL_3042425 | Lighthouse Lab in Alderley Park | Wellcome Sanger Institute for the COVID-19 Genomics UK (COG-UK) Consortium | Jacquelyn Wynn, Mairead Hyland, The Lighthouse Lab in Alderley Park and Alex Alderton, Roberto Amato, Jeffrey Barrett, Sonia Goncalves, Ewan Harrison, David K. Jackson, Ian Johnston, Dominic Kwiatkowski, Cordelia Langford, John Sillitoe on behalf of the Wellcome Sanger Institute COVID-19 Surveillance Team |
| EPI_ISL_3042426 | Lighthouse Lab in Milton Keynes | Wellcome Sanger Institute for the COVID-19 Genomics UK (COG-UK) Consortium | The Lighthouse Lab in Milton Keynes and Alex Alderton, Roberto Amato, Jeffrey Barrett, Sonia Goncalves, Ewan Harrison, David K. Jackson, Ian Johnston, Dominic Kwiatkowski, Cordelia Langford, John Sillitoe on behalf of the Wellcome Sanger Institute COVID-19 Surveillance Team |
| EPI_ISL_3042427 | Berkshire and Surrey Pathology Services Lighthouse Laboratory | Wellcome Sanger Institute for the COVID-19 Genomics UK (COG-UK) Consortium | Berkshire and Surrey Pathology Services Lighthouse Laboratory and Alex Alderton, Roberto Amato, Jeffrey Barrett, Sonia Goncalves, Ewan Harrison, David K. Jackson, Ian Johnston, Dominic Kwiatkowski, Cordelia Langford, John Sillitoe on behalf of the Wellcome Sanger Institute COVID-19 Surveillance Team |
| EPI_ISL_3042428, EPI_ISL_3042429 | Lighthouse Lab in Milton Keynes | Wellcome Sanger Institute for the COVID-19 Genomics UK (COG-UK) Consortium | The Lighthouse Lab in Milton Keynes and Alex Alderton, Roberto Amato, Jeffrey Barrett, Sonia Goncalves, Ewan Harrison, David K. Jackson, Ian Johnston, Dominic Kwiatkowski, Cordelia Langford, John Sillitoe on behalf of the Wellcome Sanger Institute COVID-19 Surveillance Team |
| EPI_ISL_3042430 | Lighthouse Lab in Alderley Park | Wellcome Sanger Institute for the COVID-19 Genomics UK (COG-UK) Consortium | Jacquelyn Wynn, Mairead Hyland, The Lighthouse Lab in Alderley Park and Alex Alderton, Roberto Amato, Jeffrey Barrett, Sonia Goncalves, Ewan Harrison, David K. Jackson, Ian Johnston, Dominic Kwiatkowski, Cordelia Langford, John Sillitoe on behalf of the Wellcome Sanger Institute COVID-19 Surveillance Team |
| EPI_ISL_3042431 | Berkshire and Surrey Pathology Services Lighthouse Laboratory | Wellcome Sanger Institute for the COVID-19 Genomics UK (COG-UK) Consortium | Berkshire and Surrey Pathology Services Lighthouse Laboratory and Alex Alderton, Roberto Amato, Jeffrey Barrett, Sonia Goncalves, Ewan Harrison, David K. Jackson, Ian Johnston, Dominic Kwiatkowski, Cordelia Langford, John Sillitoe on behalf of the Wellcome Sanger Institute COVID-19 Surveillance Team |
| EPI_ISL_3042432 | Lighthouse Laboratory Plymouth | Wellcome Sanger Institute for the COVID-19 Genomics UK (COG-UK) Consortium | Lighthouse Laboratory Plymouth and Alex Alderton, Roberto Amato, Jeffrey Barrett, Sonia Goncalves, Ewan Harrison, David K. Jackson, Ian Johnston, Dominic Kwiatkowski, Cordelia Langford, John Sillitoe on behalf of the Wellcome Sanger Institute COVID-19 Surveillance Team |
| EPI_ISL_3042433 | Lighthouse Lab in Alderley Park | Wellcome Sanger Institute for the COVID-19 Genomics UK (COG-UK) Consortium | Jacquelyn Wynn, Mairead Hyland, The Lighthouse Lab in Alderley Park and Alex Alderton, Roberto Amato, Jeffrey Barrett, Sonia Goncalves, Ewan Harrison, David K. Jackson, Ian Johnston, Dominic Kwiatkowski, Cordelia Langford, John Sillitoe on behalf of the Wellcome Sanger Institute COVID-19 Surveillance Team |
| EPI_ISL_3042434 | Lighthouse Lab in Milton Keynes | Wellcome Sanger Institute for the COVID-19 Genomics UK (COG-UK) Consortium | The Lighthouse Lab in Milton Keynes and Alex Alderton, Roberto Amato, Jeffrey Barrett, Sonia Goncalves, Ewan Harrison, David K. Jackson, Ian Johnston, Dominic Kwiatkowski, Cordelia Langford, John Sillitoe on behalf of the Wellcome Sanger Institute COVID-19 Surveillance Team |
| EPI_ISL_3042435 | Lighthouse Lab in Alderley Park | Wellcome Sanger Institute for the COVID-19 Genomics UK (COG-UK) Consortium | Jacquelyn Wynn, Mairead Hyland, The Lighthouse Lab in Alderley Park and Alex Alderton, Roberto Amato, Jeffrey Barrett, Sonia Goncalves, Ewan Harrison, David K. Jackson, Ian Johnston, Dominic Kwiatkowski, Cordelia Langford, John Sillitoe on behalf of the Wellcome Sanger Institute COVID-19 Surveillance Team |
| EPI_ISL_3042436 | Lighthouse Laboratory Plymouth | Wellcome Sanger Institute for the COVID-19 Genomics UK (COG-UK) Consortium | Lighthouse Laboratory Plymouth and Alex Alderton, Roberto Amato, Jeffrey Barrett, Sonia Goncalves, Ewan Harrison, David K. Jackson, Ian Johnston, Dominic Kwiatkowski, Cordelia Langford, John Sillitoe on behalf of the Wellcome Sanger Institute COVID-19 Surveillance Team |
| EPI_ISL_3042437 | Lighthouse Lab in Milton Keynes | Wellcome Sanger Institute for the COVID-19 Genomics UK (COG-UK) Consortium | The Lighthouse Lab in Milton Keynes and Alex Alderton, Roberto Amato, Jeffrey Barrett, Sonia Goncalves, Ewan Harrison, David K. Jackson, Ian Johnston, Dominic Kwiatkowski, Cordelia Langford, John Sillitoe on behalf of the Wellcome Sanger Institute COVID-19 Surveillance Team |
| EPI_ISL_3042438 | Berkshire and Surrey Pathology Services Lighthouse Laboratory | Wellcome Sanger Institute for the COVID-19 Genomics UK (COG-UK) Consortium | Berkshire and Surrey Pathology Services Lighthouse Laboratory and Alex Alderton, Roberto Amato, Jeffrey Barrett, Sonia Goncalves, Ewan Harrison, David K. Jackson, Ian Johnston, Dominic Kwiatkowski, Cordelia Langford, John Sillitoe on behalf of the Wellcome Sanger Institute COVID-19 Surveillance Team |
| EPI_ISL_3042439 | Lighthouse Lab in Milton Keynes | Wellcome Sanger Institute for the COVID-19 Genomics UK (COG-UK) Consortium | The Lighthouse Lab in Milton Keynes and Alex Alderton, Roberto Amato, Jeffrey Barrett, Sonia Goncalves, Ewan Harrison, David K. Jackson, Ian Johnston, Dominic Kwiatkowski, Cordelia Langford, John Sillitoe on behalf of the Wellcome Sanger Institute COVID-19 Surveillance Team |
| EPI_ISL_3042440, EPI_ISL_3042441 | Lighthouse Lab in Alderley Park | Wellcome Sanger Institute for the COVID-19 Genomics UK (COG-UK) Consortium | Jacquelyn Wynn, Mairead Hyland, The Lighthouse Lab in Alderley Park and Alex Alderton, Roberto Amato, Jeffrey Barrett, Sonia Goncalves, Ewan Harrison, David K. Jackson, Ian Johnston, Dominic Kwiatkowski, Cordelia Langford, John Sillitoe on behalf of the Wellcome Sanger Institute COVID-19 Surveillance Team |
| EPI_ISL_3042442 | Lighthouse Lab in Milton Keynes | Wellcome Sanger Institute for the COVID-19 Genomics UK (COG-UK) Consortium | The Lighthouse Lab in Milton Keynes and Alex Alderton, Roberto Amato, Jeffrey Barrett, Sonia Goncalves, Ewan Harrison, David K. Jackson, Ian Johnston, Dominic Kwiatkowski, Cordelia Langford, John Sillitoe on behalf of the Wellcome Sanger Institute COVID-19 Surveillance Team |
| EPI_ISL_3042443 | Lighthouse Laboratory Plymouth | Wellcome Sanger Institute for the COVID-19 Genomics UK (COG-UK) Consortium | Lighthouse Laboratory Plymouth and Alex Alderton, Roberto Amato, Jeffrey Barrett, Sonia Goncalves, Ewan Harrison, David K. Jackson, Ian Johnston, Dominic Kwiatkowski, Cordelia Langford, John Sillitoe on behalf of the Wellcome Sanger Institute COVID-19 Surveillance Team |
| EPI_ISL_3042444, EPI_ISL_3042445, EPI_ISL_3042446 | Lighthouse Lab in Milton Keynes | Wellcome Sanger Institute for the COVID-19 Genomics UK (COG-UK) Consortium | The Lighthouse Lab in Milton Keynes and Alex Alderton, Roberto Amato, Jeffrey Barrett, Sonia Goncalves, Ewan Harrison, David K. Jackson, Ian Johnston, Dominic Kwiatkowski, Cordelia Langford, John Sillitoe on behalf of the Wellcome Sanger Institute COVID-19 Surveillance Team |
| EPI_ISL_3042447 | Berkshire and Surrey Pathology Services Lighthouse Laboratory | Wellcome Sanger Institute for the COVID-19 Genomics UK (COG-UK) Consortium | Berkshire and Surrey Pathology Services Lighthouse Laboratory and Alex Alderton, Roberto Amato, Jeffrey Barrett, Sonia Goncalves, Ewan Harrison, David K. Jackson, Ian Johnston, Dominic Kwiatkowski, Cordelia Langford, John Sillitoe on behalf of the Wellcome Sanger Institute COVID-19 Surveillance Team |
| EPI_ISL_3042448, EPI_ISL_3042449 | Lighthouse Lab in Alderley Park | Wellcome Sanger Institute for the COVID-19 Genomics UK (COG-UK) Consortium | Jacquelyn Wynn, Mairead Hyland, The Lighthouse Lab in Alderley Park and Alex Alderton, Roberto Amato, Jeffrey Barrett, Sonia Goncalves, Ewan Harrison, David K. Jackson, Ian Johnston, Dominic Kwiatkowski, Cordelia Langford, John Sillitoe on behalf of the Wellcome Sanger Institute COVID-19 Surveillance Team |
| EPI_ISL_3042450 | Berkshire and Surrey Pathology Services Lighthouse Laboratory | Wellcome Sanger Institute for the COVID-19 Genomics UK (COG-UK) Consortium | Berkshire and Surrey Pathology Services Lighthouse Laboratory and Alex Alderton, Roberto Amato, Jeffrey Barrett, Sonia Goncalves, Ewan Harrison, David K. Jackson, Ian Johnston, Dominic Kwiatkowski, Cordelia Langford, John Sillitoe on behalf of the Wellcome Sanger Institute COVID-19 Surveillance Team |
| EPI_ISL_3042451, EPI_ISL_3042452 | Lighthouse Laboratory Plymouth | Wellcome Sanger Institute for the COVID-19 Genomics UK (COG-UK) Consortium | Lighthouse Laboratory Plymouth and Alex Alderton, Roberto Amato, Jeffrey Barrett, Sonia Goncalves, Ewan Harrison, David K. Jackson, Ian Johnston, Dominic Kwiatkowski, Cordelia Langford, John Sillitoe on behalf of the Wellcome Sanger Institute COVID-19 Surveillance Team |
| EPI_ISL_3042453 | Lighthouse Lab in Milton Keynes | Wellcome Sanger Institute for the COVID-19 Genomics UK (COG-UK) Consortium | The Lighthouse Lab in Milton Keynes and Alex Alderton, Roberto Amato, Jeffrey Barrett, Sonia Goncalves, Ewan Harrison, David K. Jackson, Ian Johnston, Dominic Kwiatkowski, Cordelia Langford, John Sillitoe on behalf of the Wellcome Sanger Institute COVID-19 Surveillance Team |
| EPI_ISL_3042454, EPI_ISL_3042455 | Berkshire and Surrey Pathology Services Lighthouse Laboratory | Wellcome Sanger Institute for the COVID-19 Genomics UK (COG-UK) Consortium | Berkshire and Surrey Pathology Services Lighthouse Laboratory and Alex Alderton, Roberto Amato, Jeffrey Barrett, Sonia Goncalves, Ewan Harrison, David K. Jackson, Ian Johnston, Dominic Kwiatkowski, Cordelia Langford, John Sillitoe on behalf of the Wellcome Sanger Institute COVID-19 Surveillance Team |
| EPI_ISL_3042456 | Lighthouse Lab in Alderley Park | Wellcome Sanger Institute for the COVID-19 Genomics UK (COG-UK) Consortium | Jacquelyn Wynn, Mairead Hyland, The Lighthouse Lab in Alderley Park and Alex Alderton, Roberto Amato, Jeffrey Barrett, Sonia Goncalves, Ewan Harrison, David K. Jackson, Ian Johnston, Dominic Kwiatkowski, Cordelia Langford, John Sillitoe on behalf of the Wellcome Sanger Institute COVID-19 Surveillance Team |
| EPI_ISL_3042457, EPI_ISL_3042458 | Lighthouse Lab in Milton Keynes | Wellcome Sanger Institute for the COVID-19 Genomics UK (COG-UK) Consortium | The Lighthouse Lab in Milton Keynes and Alex Alderton, Roberto Amato, Jeffrey Barrett, Sonia Goncalves, Ewan Harrison, David K. Jackson, Ian Johnston, Dominic Kwiatkowski, Cordelia Langford, John Sillitoe on behalf of the Wellcome Sanger Institute COVID-19 Surveillance Team |
| EPI_ISL_3042459 | Lighthouse Laboratory Plymouth | Wellcome Sanger Institute for the COVID-19 Genomics UK (COG-UK) Consortium | Lighthouse Laboratory Plymouth and Alex Alderton, Roberto Amato, Jeffrey Barrett, Sonia Goncalves, Ewan Harrison, David K. Jackson, Ian Johnston, Dominic Kwiatkowski, Cordelia Langford, John Sillitoe on behalf of the Wellcome Sanger Institute COVID-19 Surveillance Team |
| EPI_ISL_3042460 | Lighthouse Lab in Milton Keynes | Wellcome Sanger Institute for the COVID-19 Genomics UK (COG-UK) Consortium | The Lighthouse Lab in Milton Keynes and Alex Alderton, Roberto Amato, Jeffrey Barrett, Sonia Goncalves, Ewan Harrison, David K. Jackson, Ian Johnston, Dominic Kwiatkowski, Cordelia Langford, John Sillitoe on behalf of the Wellcome Sanger Institute COVID-19 Surveillance Team |
| EPI_ISL_3042461, EPI_ISL_3042462 | Lighthouse Laboratory Plymouth | Wellcome Sanger Institute for the COVID-19 Genomics UK (COG-UK) Consortium | Lighthouse Laboratory Plymouth and Alex Alderton, Roberto Amato, Jeffrey Barrett, Sonia Goncalves, Ewan Harrison, David K. Jackson, Ian Johnston, Dominic Kwiatkowski, Cordelia Langford, John Sillitoe on behalf of the Wellcome Sanger Institute COVID-19 Surveillance Team |
| EPI_ISL_3042463 | Lighthouse Lab in Milton Keynes | Wellcome Sanger Institute for the COVID-19 Genomics UK (COG-UK) Consortium | The Lighthouse Lab in Milton Keynes and Alex Alderton, Roberto Amato, Jeffrey Barrett, Sonia Goncalves, Ewan Harrison, David K. Jackson, Ian Johnston, Dominic Kwiatkowski, Cordelia Langford, John Sillitoe on behalf of the Wellcome Sanger Institute COVID-19 Surveillance Team |
| EPI_ISL_3042464 | Lighthouse Lab in Alderley Park | Wellcome Sanger Institute for the COVID-19 Genomics UK (COG-UK) | Jacquelyn Wynn, Mairead Hyland, The Lighthouse Lab in Alderley Park and Alex Alderton, Roberto Amato, Jeffrey Barrett, Sonia Goncalves, Ewan Harrison, David K. Jackson, Ian Johnston, Dominic Kwiatkowski, Cordelia Langford, John Sillitoe on |

| | | | |
|---|---|---|---|
| | | Consortium | behalf of the Wellcome Sanger Institute COVID-19 Surveillance Team |
| EPI_ISL_3042465, EPI_ISL_3042466 | Lighthouse Lab in Milton Keynes | Wellcome Sanger Institute for the COVID-19 Genomics UK (COG-UK) Consortium | The Lighthouse Lab in Milton Keynes and Alex Alderton, Roberto Amato, Jeffrey Barrett, Sonia Goncalves, Ewan Harrison, David K. Jackson, Ian Johnston, Dominic Kwiatkowski, Cordelia Langford, John Sillitoe on behalf of the Wellcome Sanger Institute COVID-19 Surveillance Team |
| EPI_ISL_3042467 | Lighthouse Lab in Alderley Park | Wellcome Sanger Institute for the COVID-19 Genomics UK (COG-UK) Consortium | Jacquelyn Wynn, Mairead Hyland, The Lighthouse Lab in Alderley Park and Alex Alderton, Roberto Amato, Jeffrey Barrett, Sonia Goncalves, Ewan Harrison, David K. Jackson, Ian Johnston, Dominic Kwiatkowski, Cordelia Langford, John Sillitoe on behalf of the Wellcome Sanger Institute COVID-19 Surveillance Team |
| EPI_ISL_3042468 | Berkshire and Surrey Pathology Services Lighthouse Laboratory | Wellcome Sanger Institute for the COVID-19 Genomics UK (COG-UK) Consortium | Berkshire and Surrey Pathology Services Lighthouse Laboratory and Alex Alderton, Roberto Amato, Jeffrey Barrett, Sonia Goncalves, Ewan Harrison, David K. Jackson, Ian Johnston, Dominic Kwiatkowski, Cordelia Langford, John Sillitoe on behalf of the Wellcome Sanger Institute COVID-19 Surveillance Team |
| EPI_ISL_3042469 | Lighthouse Lab in Alderley Park | Wellcome Sanger Institute for the COVID-19 Genomics UK (COG-UK) Consortium | Jacquelyn Wynn, Mairead Hyland, The Lighthouse Lab in Alderley Park and Alex Alderton, Roberto Amato, Jeffrey Barrett, Sonia Goncalves, Ewan Harrison, David K. Jackson, Ian Johnston, Dominic Kwiatkowski, Cordelia Langford, John Sillitoe on behalf of the Wellcome Sanger Institute COVID-19 Surveillance Team |
| EPI_ISL_3042470 | Berkshire and Surrey Pathology Services Lighthouse Laboratory | Wellcome Sanger Institute for the COVID-19 Genomics UK (COG-UK) Consortium | Berkshire and Surrey Pathology Services Lighthouse Laboratory and Alex Alderton, Roberto Amato, Jeffrey Barrett, Sonia Goncalves, Ewan Harrison, David K. Jackson, Ian Johnston, Dominic Kwiatkowski, Cordelia Langford, John Sillitoe on behalf of the Wellcome Sanger Institute COVID-19 Surveillance Team |
| EPI_ISL_3042471 | Lighthouse Lab in Milton Keynes | Wellcome Sanger Institute for the COVID-19 Genomics UK (COG-UK) Consortium | The Lighthouse Lab in Milton Keynes and Alex Alderton, Roberto Amato, Jeffrey Barrett, Sonia Goncalves, Ewan Harrison, David K. Jackson, Ian Johnston, Dominic Kwiatkowski, Cordelia Langford, John Sillitoe on behalf of the Wellcome Sanger Institute COVID-19 Surveillance Team |
| EPI_ISL_3042472 | Lighthouse Lab in Alderley Park | Wellcome Sanger Institute for the COVID-19 Genomics UK (COG-UK) Consortium | Jacquelyn Wynn, Mairead Hyland, The Lighthouse Lab in Alderley Park and Alex Alderton, Roberto Amato, Jeffrey Barrett, Sonia Goncalves, Ewan Harrison, David K. Jackson, Ian Johnston, Dominic Kwiatkowski, Cordelia Langford, John Sillitoe on behalf of the Wellcome Sanger Institute COVID-19 Surveillance Team |
| EPI_ISL_3042473 | Lighthouse Laboratory Plymouth | Wellcome Sanger Institute for the COVID-19 Genomics UK (COG-UK) Consortium | Lighthouse Laboratory Plymouth and Alex Alderton, Roberto Amato, Jeffrey Barrett, Sonia Goncalves, Ewan Harrison, David K. Jackson, Ian Johnston, Dominic Kwiatkowski, Cordelia Langford, John Sillitoe on behalf of the Wellcome Sanger Institute COVID-19 Surveillance Team |
| EPI_ISL_3042474, EPI_ISL_3042475 | Lighthouse Lab in Alderley Park | Wellcome Sanger Institute for the COVID-19 Genomics UK (COG-UK) Consortium | Jacquelyn Wynn, Mairead Hyland, The Lighthouse Lab in Alderley Park and Alex Alderton, Roberto Amato, Jeffrey Barrett, Sonia Goncalves, Ewan Harrison, David K. Jackson, Ian Johnston, Dominic Kwiatkowski, Cordelia Langford, John Sillitoe on behalf of the Wellcome Sanger Institute COVID-19 Surveillance Team |
| EPI_ISL_3042476 | Berkshire and Surrey Pathology Services Lighthouse Laboratory | Wellcome Sanger Institute for the COVID-19 Genomics UK (COG-UK) Consortium | Berkshire and Surrey Pathology Services Lighthouse Laboratory and Alex Alderton, Roberto Amato, Jeffrey Barrett, Sonia Goncalves, Ewan Harrison, David K. Jackson, Ian Johnston, Dominic Kwiatkowski, Cordelia Langford, John Sillitoe on behalf of the Wellcome Sanger Institute COVID-19 Surveillance Team |
| EPI_ISL_3042477 | Lighthouse Lab in Alderley Park | Wellcome Sanger Institute for the COVID-19 Genomics UK (COG-UK) Consortium | Jacquelyn Wynn, Mairead Hyland, The Lighthouse Lab in Alderley Park and Alex Alderton, Roberto Amato, Jeffrey Barrett, Sonia Goncalves, Ewan Harrison, David K. Jackson, Ian Johnston, Dominic Kwiatkowski, Cordelia Langford, John Sillitoe on behalf of the Wellcome Sanger Institute COVID-19 Surveillance Team |
| EPI_ISL_3042478 | Lighthouse Laboratory Plymouth | Wellcome Sanger Institute for the COVID-19 Genomics UK (COG-UK) Consortium | Lighthouse Laboratory Plymouth and Alex Alderton, Roberto Amato, Jeffrey Barrett, Sonia Goncalves, Ewan Harrison, David K. Jackson, Ian Johnston, Dominic Kwiatkowski, Cordelia Langford, John Sillitoe on behalf of the Wellcome Sanger Institute COVID-19 Surveillance Team |
| EPI_ISL_3042479 | Lighthouse Lab in Milton Keynes | Wellcome Sanger Institute for the COVID-19 Genomics UK (COG-UK) Consortium | The Lighthouse Lab in Milton Keynes and Alex Alderton, Roberto Amato, Jeffrey Barrett, Sonia Goncalves, Ewan Harrison, David K. Jackson, Ian Johnston, Dominic Kwiatkowski, Cordelia Langford, John Sillitoe on behalf of the Wellcome Sanger Institute COVID-19 Surveillance Team |
| EPI_ISL_3042480 | Lighthouse Laboratory Plymouth | Wellcome Sanger Institute for the COVID-19 Genomics UK (COG-UK) Consortium | Lighthouse Laboratory Plymouth and Alex Alderton, Roberto Amato, Jeffrey Barrett, Sonia Goncalves, Ewan Harrison, David K. Jackson, Ian Johnston, Dominic Kwiatkowski, Cordelia Langford, John Sillitoe on behalf of the Wellcome Sanger Institute COVID-19 Surveillance Team |
| EPI_ISL_3042481 | Lighthouse Lab in Milton Keynes | Wellcome Sanger Institute for the COVID-19 Genomics UK (COG-UK) Consortium | The Lighthouse Lab in Milton Keynes and Alex Alderton, Roberto Amato, Jeffrey Barrett, Sonia Goncalves, Ewan Harrison, David K. Jackson, Ian Johnston, Dominic Kwiatkowski, Cordelia Langford, John Sillitoe on behalf of the Wellcome Sanger Institute COVID-19 Surveillance Team |
| EPI_ISL_3042482, EPI_ISL_3042483 | Berkshire and Surrey Pathology Services Lighthouse Laboratory | Wellcome Sanger Institute for the COVID-19 Genomics UK (COG-UK) Consortium | Berkshire and Surrey Pathology Services Lighthouse Laboratory and Alex Alderton, Roberto Amato, Jeffrey Barrett, Sonia Goncalves, Ewan Harrison, David K. Jackson, Ian Johnston, Dominic Kwiatkowski, Cordelia Langford, John Sillitoe on behalf of the Wellcome Sanger Institute COVID-19 Surveillance Team |
| EPI_ISL_3042484 | Lighthouse Lab in Alderley Park | Wellcome Sanger Institute for the COVID-19 Genomics UK (COG-UK) Consortium | Jacquelyn Wynn, Mairead Hyland, The Lighthouse Lab in Alderley Park and Alex Alderton, Roberto Amato, Jeffrey Barrett, Sonia Goncalves, Ewan Harrison, David K. Jackson, Ian Johnston, Dominic Kwiatkowski, Cordelia Langford, John Sillitoe on behalf of the Wellcome Sanger Institute COVID-19 Surveillance Team |
| EPI_ISL_3042485 | Lighthouse Laboratory Plymouth | Wellcome Sanger Institute for the COVID-19 Genomics UK (COG-UK) Consortium | Lighthouse Laboratory Plymouth and Alex Alderton, Roberto Amato, Jeffrey Barrett, Sonia Goncalves, Ewan Harrison, David K. Jackson, Ian Johnston, Dominic Kwiatkowski, Cordelia Langford, John Sillitoe on behalf of the Wellcome Sanger Institute COVID-19 Surveillance Team |
| EPI_ISL_3042486 | Berkshire and Surrey Pathology Services Lighthouse Laboratory | Wellcome Sanger Institute for the COVID-19 Genomics UK (COG-UK) Consortium | Berkshire and Surrey Pathology Services Lighthouse Laboratory and Alex Alderton, Roberto Amato, Jeffrey Barrett, Sonia Goncalves, Ewan Harrison, David K. Jackson, Ian Johnston, Dominic Kwiatkowski, Cordelia Langford, John Sillitoe on behalf of the Wellcome Sanger Institute COVID-19 Surveillance Team |
| EPI_ISL_3042487 | Lighthouse Lab in Milton Keynes | Wellcome Sanger Institute for the COVID-19 Genomics UK (COG-UK) Consortium | The Lighthouse Lab in Milton Keynes and Alex Alderton, Roberto Amato, Jeffrey Barrett, Sonia Goncalves, Ewan Harrison, David K. Jackson, Ian Johnston, Dominic Kwiatkowski, Cordelia Langford, John Sillitoe on behalf of the Wellcome Sanger Institute COVID-19 Surveillance Team |
| EPI_ISL_3042488, EPI_ISL_3042489 | Lighthouse Laboratory Plymouth | Wellcome Sanger Institute for the COVID-19 Genomics UK (COG-UK) Consortium | Lighthouse Laboratory Plymouth and Alex Alderton, Roberto Amato, Jeffrey Barrett, Sonia Goncalves, Ewan Harrison, David K. Jackson, Ian Johnston, Dominic Kwiatkowski, Cordelia Langford, John Sillitoe on behalf of the Wellcome Sanger Institute COVID-19 Surveillance Team |
| EPI_ISL_3042490 | Lighthouse Lab in Alderley Park | Wellcome Sanger Institute for the COVID-19 Genomics UK (COG-UK) Consortium | Jacquelyn Wynn, Mairead Hyland, The Lighthouse Lab in Alderley Park and Alex Alderton, Roberto Amato, Jeffrey Barrett, Sonia Goncalves, Ewan Harrison, David K. Jackson, Ian Johnston, Dominic Kwiatkowski, Cordelia Langford, John Sillitoe on behalf of the Wellcome Sanger Institute COVID-19 Surveillance Team |
| EPI_ISL_3042491, EPI_ISL_3042492 | Lighthouse Lab in Milton Keynes | Wellcome Sanger Institute for the COVID-19 Genomics UK (COG-UK) Consortium | The Lighthouse Lab in Milton Keynes and Alex Alderton, Roberto Amato, Jeffrey Barrett, Sonia Goncalves, Ewan Harrison, David K. Jackson, Ian Johnston, Dominic Kwiatkowski, Cordelia Langford, John Sillitoe on behalf of the Wellcome Sanger Institute COVID-19 Surveillance Team |
| EPI_ISL_3042493 | Lighthouse Laboratory Plymouth | Wellcome Sanger Institute for the COVID-19 Genomics UK (COG-UK) Consortium | Lighthouse Laboratory Plymouth and Alex Alderton, Roberto Amato, Jeffrey Barrett, Sonia Goncalves, Ewan Harrison, David K. Jackson, Ian Johnston, Dominic Kwiatkowski, Cordelia Langford, John Sillitoe on behalf of the Wellcome Sanger Institute COVID-19 Surveillance Team |
| EPI_ISL_3042494 | Berkshire and Surrey Pathology Services Lighthouse Laboratory | Wellcome Sanger Institute for the COVID-19 Genomics UK (COG-UK) Consortium | Berkshire and Surrey Pathology Services Lighthouse Laboratory and Alex Alderton, Roberto Amato, Jeffrey Barrett, Sonia Goncalves, Ewan Harrison, David K. Jackson, Ian Johnston, Dominic Kwiatkowski, Cordelia Langford, John Sillitoe on behalf of the Wellcome Sanger Institute COVID-19 Surveillance Team |
| EPI_ISL_3042495 | Lighthouse Laboratory Plymouth | Wellcome Sanger Institute for the COVID-19 Genomics UK (COG-UK) Consortium | Lighthouse Laboratory Plymouth and Alex Alderton, Roberto Amato, Jeffrey Barrett, Sonia Goncalves, Ewan Harrison, David K. Jackson, Ian Johnston, Dominic Kwiatkowski, Cordelia Langford, John Sillitoe on behalf of the Wellcome Sanger Institute COVID-19 Surveillance Team |
| EPI_ISL_3042496, EPI_ISL_3042497 | Lighthouse Lab in Milton Keynes | Wellcome Sanger Institute for the COVID-19 Genomics UK (COG-UK) Consortium | The Lighthouse Lab in Milton Keynes and Alex Alderton, Roberto Amato, Jeffrey Barrett, Sonia Goncalves, Ewan Harrison, David K. Jackson, Ian Johnston, Dominic Kwiatkowski, Cordelia Langford, John Sillitoe on behalf of the Wellcome Sanger Institute COVID-19 Surveillance Team |
| EPI_ISL_3042498, EPI_ISL_3042499 | Lighthouse Lab in Alderley Park | Wellcome Sanger Institute for the COVID-19 Genomics UK (COG-UK) Consortium | Jacquelyn Wynn, Mairead Hyland, The Lighthouse Lab in Alderley Park and Alex Alderton, Roberto Amato, Jeffrey Barrett, Sonia Goncalves, Ewan Harrison, David K. Jackson, Ian Johnston, Dominic Kwiatkowski, Cordelia Langford, John Sillitoe on behalf of the Wellcome Sanger Institute COVID-19 Surveillance Team |
| EPI_ISL_3042500 | Berkshire and Surrey Pathology Services Lighthouse Laboratory | Wellcome Sanger Institute for the COVID-19 Genomics UK (COG-UK) Consortium | Berkshire and Surrey Pathology Services Lighthouse Laboratory and Alex Alderton, Roberto Amato, Jeffrey Barrett, Sonia Goncalves, Ewan Harrison, David K. Jackson, Ian Johnston, Dominic Kwiatkowski, Cordelia Langford, John Sillitoe on behalf of the Wellcome Sanger Institute COVID-19 Surveillance Team |
| EPI_ISL_3042501 | Lighthouse Laboratory Plymouth | Wellcome Sanger Institute for the COVID-19 Genomics UK (COG-UK) Consortium | Lighthouse Laboratory Plymouth and Alex Alderton, Roberto Amato, Jeffrey Barrett, Sonia Goncalves, Ewan Harrison, David K. Jackson, Ian Johnston, Dominic Kwiatkowski, Cordelia Langford, John Sillitoe on behalf of the Wellcome Sanger Institute COVID-19 Surveillance Team |
| EPI_ISL_3042502 | Berkshire and Surrey Pathology Services Lighthouse Laboratory | Wellcome Sanger Institute for the COVID-19 Genomics UK (COG-UK) Consortium | Berkshire and Surrey Pathology Services Lighthouse Laboratory and Alex Alderton, Roberto Amato, Jeffrey Barrett, Sonia Goncalves, Ewan Harrison, David K. Jackson, Ian Johnston, Dominic Kwiatkowski, Cordelia Langford, John Sillitoe on behalf of the Wellcome Sanger Institute COVID-19 Surveillance Team |
| EPI_ISL_3042503 | Lighthouse Lab in Milton Keynes | Wellcome Sanger Institute for the COVID-19 Genomics UK (COG-UK) Consortium | The Lighthouse Lab in Milton Keynes and Alex Alderton, Roberto Amato, Jeffrey Barrett, Sonia Goncalves, Ewan Harrison, David K. Jackson, Ian Johnston, Dominic Kwiatkowski, Cordelia Langford, John Sillitoe on behalf of the Wellcome Sanger Institute COVID-19 Surveillance Team |
| EPI_ISL_3042504, EPI_ISL_3042505 | Lighthouse Lab in Alderley Park | Wellcome Sanger Institute for the COVID-19 Genomics UK (COG-UK) Consortium | Jacquelyn Wynn, Mairead Hyland, The Lighthouse Lab in Alderley Park and Alex Alderton, Roberto Amato, Jeffrey Barrett, Sonia Goncalves, Ewan Harrison, David K. Jackson, Ian Johnston, Dominic Kwiatkowski, Cordelia Langford, John Sillitoe on behalf of the Wellcome Sanger Institute COVID-19 Surveillance Team |
| EPI_ISL_3042506 | Lighthouse Laboratory Plymouth | Wellcome Sanger Institute for the COVID-19 Genomics UK (COG-UK) Consortium | Lighthouse Laboratory Plymouth and Alex Alderton, Roberto Amato, Jeffrey Barrett, Sonia Goncalves, Ewan Harrison, David K. Jackson, Ian Johnston, Dominic Kwiatkowski, Cordelia Langford, John Sillitoe on behalf of the Wellcome Sanger Institute COVID-19 Surveillance Team |
| EPI_ISL_3042507 | Lighthouse Lab in Milton Keynes | Wellcome Sanger Institute for the COVID-19 Genomics UK (COG-UK) Consortium | The Lighthouse Lab in Milton Keynes and Alex Alderton, Roberto Amato, Jeffrey Barrett, Sonia Goncalves, Ewan Harrison, David K. Jackson, Ian Johnston, Dominic Kwiatkowski, Cordelia Langford, John Sillitoe on behalf of the Wellcome Sanger Institute COVID-19 Surveillance Team |
| EPI_ISL_3042508 | Berkshire and Surrey Pathology Services Lighthouse Laboratory | Wellcome Sanger Institute for the COVID-19 Genomics UK (COG-UK) Consortium | Berkshire and Surrey Pathology Services Lighthouse Laboratory and Alex Alderton, Roberto Amato, Jeffrey Barrett, Sonia Goncalves, Ewan Harrison, David K. Jackson, Ian Johnston, Dominic Kwiatkowski, Cordelia Langford, John Sillitoe on behalf of the Wellcome Sanger Institute COVID-19 Surveillance Team |
| EPI_ISL_3042509 | Lighthouse Laboratory Plymouth | Wellcome Sanger Institute for the COVID-19 Genomics UK (COG-UK) Consortium | Lighthouse Laboratory Plymouth and Alex Alderton, Roberto Amato, Jeffrey Barrett, Sonia Goncalves, Ewan Harrison, David K. Jackson, Ian Johnston, Dominic Kwiatkowski, Cordelia Langford, John Sillitoe on behalf of the Wellcome Sanger Institute COVID-19 Surveillance Team |
| EPI_ISL_3042510 | Berkshire and Surrey Pathology Services Lighthouse Laboratory | Wellcome Sanger Institute for the COVID-19 Genomics UK (COG-UK) Consortium | Berkshire and Surrey Pathology Services Lighthouse Laboratory and Alex Alderton, Roberto Amato, Jeffrey Barrett, Sonia Goncalves, Ewan Harrison, David K. Jackson, Ian Johnston, Dominic Kwiatkowski, Cordelia Langford, John Sillitoe on behalf of the Wellcome Sanger Institute COVID-19 Surveillance Team |
| EPI_ISL_3042511 | Lighthouse Lab in Alderley Park | Wellcome Sanger Institute for the COVID-19 Genomics UK (COG-UK) Consortium | Jacquelyn Wynn, Mairead Hyland, The Lighthouse Lab in Alderley Park and Alex Alderton, Roberto Amato, Jeffrey Barrett, Sonia Goncalves, Ewan Harrison, David K. Jackson, Ian Johnston, Dominic Kwiatkowski, Cordelia Langford, John Sillitoe on behalf of the Wellcome Sanger Institute COVID-19 Surveillance Team |
| EPI_ISL_3042512 | Lighthouse Lab in Milton Keynes | Wellcome Sanger Institute for the COVID-19 Genomics UK (COG-UK) Consortium | The Lighthouse Lab in Milton Keynes and Alex Alderton, Roberto Amato, Jeffrey Barrett, Sonia Goncalves, Ewan Harrison, David K. Jackson, Ian Johnston, Dominic Kwiatkowski, Cordelia Langford, John Sillitoe on behalf of the Wellcome Sanger Institute COVID-19 Surveillance Team |
| EPI_ISL_3042513 | Lighthouse Laboratory Plymouth | Wellcome Sanger Institute for the COVID-19 Genomics UK (COG-UK) Consortium | Lighthouse Laboratory Plymouth and Alex Alderton, Roberto Amato, Jeffrey Barrett, Sonia Goncalves, Ewan Harrison, David K. Jackson, Ian Johnston, Dominic Kwiatkowski, Cordelia Langford, John Sillitoe on behalf of the Wellcome Sanger Institute COVID-19 Surveillance Team |
| EPI_ISL_3042514 | Lighthouse Lab in Alderley Park | Wellcome Sanger Institute for the COVID-19 Genomics UK (COG-UK) Consortium | Jacquelyn Wynn, Mairead Hyland, The Lighthouse Lab in Alderley Park and Alex Alderton, Roberto Amato, Jeffrey Barrett, Sonia Goncalves, Ewan Harrison, David K. Jackson, Ian Johnston, Dominic Kwiatkowski, Cordelia Langford, John Sillitoe on behalf of the Wellcome Sanger Institute COVID-19 Surveillance Team |
| EPI_ISL_3042515, EPI_ISL_3042516 | Lighthouse Lab in Milton Keynes | Wellcome Sanger Institute for the COVID-19 Genomics UK (COG-UK) Consortium | The Lighthouse Lab in Milton Keynes and Alex Alderton, Roberto Amato, Jeffrey Barrett, Sonia Goncalves, Ewan Harrison, David K. Jackson, Ian Johnston, Dominic Kwiatkowski, Cordelia Langford, John Sillitoe on behalf of the Wellcome Sanger Institute COVID-19 Surveillance Team |
| EPI_ISL_3042517 | Lighthouse Laboratory Plymouth | Wellcome Sanger Institute for the COVID-19 Genomics UK (COG-UK) Consortium | Lighthouse Laboratory Plymouth and Alex Alderton, Roberto Amato, Jeffrey Barrett, Sonia Goncalves, Ewan Harrison, David K. Jackson, Ian Johnston, Dominic Kwiatkowski, Cordelia Langford, John Sillitoe on behalf of the Wellcome Sanger Institute COVID-19 Surveillance Team |
| EPI_ISL_3042518 | Berkshire and Surrey Pathology Services Lighthouse Laboratory | Wellcome Sanger Institute for the COVID-19 Genomics UK (COG-UK) | Berkshire and Surrey Pathology Services Lighthouse Laboratory and Alex Alderton, Roberto Amato, Jeffrey Barrett, Sonia Goncalves, Ewan Harrison, David K. Jackson, Ian Johnston, Dominic Kwiatkowski, Cordelia Langford, John Sillitoe on behalf |

| | | | |
|---|---|---|---|
| | | Consortium | of the Wellcome Sanger Institute COVID-19 Surveillance Team |
| EPI_ISL_3042519 | Lighthouse Lab in Alderley Park | Wellcome Sanger Institute for the COVID-19 Genomics UK (COG-UK) Consortium | Jacquelyn Wynn, Mairead Hyland, The Lighthouse Lab in Alderley Park and Alex Alderton, Roberto Amato, Jeffrey Barrett, Sonia Goncalves, Ewan Harrison, David K. Jackson, Ian Johnston, Dominic Kwiatkowski, Cordelia Langford, John Sillitoe on behalf of the Wellcome Sanger Institute COVID-19 Surveillance Team |
| EPI_ISL_3042520 | Lighthouse Lab in Milton Keynes | Wellcome Sanger Institute for the COVID-19 Genomics UK (COG-UK) Consortium | The Lighthouse Lab in Milton Keynes and Alex Alderton, Roberto Amato, Jeffrey Barrett, Sonia Goncalves, Ewan Harrison, David K. Jackson, Ian Johnston, Dominic Kwiatkowski, Cordelia Langford, John Sillitoe on behalf of the Wellcome Sanger Institute COVID-19 Surveillance Team |
| EPI_ISL_3042521 | Lighthouse Lab in Alderley Park | Wellcome Sanger Institute for the COVID-19 Genomics UK (COG-UK) Consortium | Jacquelyn Wynn, Mairead Hyland, The Lighthouse Lab in Alderley Park and Alex Alderton, Roberto Amato, Jeffrey Barrett, Sonia Goncalves, Ewan Harrison, David K. Jackson, Ian Johnston, Dominic Kwiatkowski, Cordelia Langford, John Sillitoe on behalf of the Wellcome Sanger Institute COVID-19 Surveillance Team |
| EPI_ISL_3042522 | Berkshire and Surrey Pathology Services Lighthouse Laboratory | Wellcome Sanger Institute for the COVID-19 Genomics UK (COG-UK) Consortium | Berkshire and Surrey Pathology Services Lighthouse Laboratory and Alex Alderton, Roberto Amato, Jeffrey Barrett, Sonia Goncalves, Ewan Harrison, David K. Jackson, Ian Johnston, Dominic Kwiatkowski, Cordelia Langford, John Sillitoe on behalf of the Wellcome Sanger Institute COVID-19 Surveillance Team |
| EPI_ISL_3042523 | Lighthouse Lab in Milton Keynes | Wellcome Sanger Institute for the COVID-19 Genomics UK (COG-UK) Consortium | The Lighthouse Lab in Milton Keynes and Alex Alderton, Roberto Amato, Jeffrey Barrett, Sonia Goncalves, Ewan Harrison, David K. Jackson, Ian Johnston, Dominic Kwiatkowski, Cordelia Langford, John Sillitoe on behalf of the Wellcome Sanger Institute COVID-19 Surveillance Team |
| EPI_ISL_3042524 | Lighthouse Laboratory Plymouth | Wellcome Sanger Institute for the COVID-19 Genomics UK (COG-UK) Consortium | Lighthouse Laboratory Plymouth and Alex Alderton, Roberto Amato, Jeffrey Barrett, Sonia Goncalves, Ewan Harrison, David K. Jackson, Ian Johnston, Dominic Kwiatkowski, Cordelia Langford, John Sillitoe on behalf of the Wellcome Sanger Institute COVID-19 Surveillance Team |
| EPI_ISL_3042525 | Berkshire and Surrey Pathology Services Lighthouse Laboratory | Wellcome Sanger Institute for the COVID-19 Genomics UK (COG-UK) Consortium | Berkshire and Surrey Pathology Services Lighthouse Laboratory and Alex Alderton, Roberto Amato, Jeffrey Barrett, Sonia Goncalves, Ewan Harrison, David K. Jackson, Ian Johnston, Dominic Kwiatkowski, Cordelia Langford, John Sillitoe on behalf of the Wellcome Sanger Institute COVID-19 Surveillance Team |
| EPI_ISL_3042526 | Lighthouse Laboratory Plymouth | Wellcome Sanger Institute for the COVID-19 Genomics UK (COG-UK) Consortium | Lighthouse Laboratory Plymouth and Alex Alderton, Roberto Amato, Jeffrey Barrett, Sonia Goncalves, Ewan Harrison, David K. Jackson, Ian Johnston, Dominic Kwiatkowski, Cordelia Langford, John Sillitoe on behalf of the Wellcome Sanger Institute COVID-19 Surveillance Team |
| EPI_ISL_3042527 | Lighthouse Lab in Milton Keynes | Wellcome Sanger Institute for the COVID-19 Genomics UK (COG-UK) Consortium | The Lighthouse Lab in Milton Keynes and Alex Alderton, Roberto Amato, Jeffrey Barrett, Sonia Goncalves, Ewan Harrison, David K. Jackson, Ian Johnston, Dominic Kwiatkowski, Cordelia Langford, John Sillitoe on behalf of the Wellcome Sanger Institute COVID-19 Surveillance Team |
| EPI_ISL_3042528 | Lighthouse Laboratory Plymouth | Wellcome Sanger Institute for the COVID-19 Genomics UK (COG-UK) Consortium | Lighthouse Laboratory Plymouth and Alex Alderton, Roberto Amato, Jeffrey Barrett, Sonia Goncalves, Ewan Harrison, David K. Jackson, Ian Johnston, Dominic Kwiatkowski, Cordelia Langford, John Sillitoe on behalf of the Wellcome Sanger Institute COVID-19 Surveillance Team |
| EPI_ISL_3042529 | Lighthouse Lab in Milton Keynes | Wellcome Sanger Institute for the COVID-19 Genomics UK (COG-UK) Consortium | The Lighthouse Lab in Milton Keynes and Alex Alderton, Roberto Amato, Jeffrey Barrett, Sonia Goncalves, Ewan Harrison, David K. Jackson, Ian Johnston, Dominic Kwiatkowski, Cordelia Langford, John Sillitoe on behalf of the Wellcome Sanger Institute COVID-19 Surveillance Team |
| EPI_ISL_3042530, EPI_ISL_3042531 | Lighthouse Lab in Alderley Park | Wellcome Sanger Institute for the COVID-19 Genomics UK (COG-UK) Consortium | Jacquelyn Wynn, Mairead Hyland, The Lighthouse Lab in Alderley Park and Alex Alderton, Roberto Amato, Jeffrey Barrett, Sonia Goncalves, Ewan Harrison, David K. Jackson, Ian Johnston, Dominic Kwiatkowski, Cordelia Langford, John Sillitoe on behalf of the Wellcome Sanger Institute COVID-19 Surveillance Team |
| EPI_ISL_3042532 | Lighthouse Lab in Milton Keynes | Wellcome Sanger Institute for the COVID-19 Genomics UK (COG-UK) Consortium | The Lighthouse Lab in Milton Keynes and Alex Alderton, Roberto Amato, Jeffrey Barrett, Sonia Goncalves, Ewan Harrison, David K. Jackson, Ian Johnston, Dominic Kwiatkowski, Cordelia Langford, John Sillitoe on behalf of the Wellcome Sanger Institute COVID-19 Surveillance Team |
| EPI_ISL_3042533 | Berkshire and Surrey Pathology Services Lighthouse Laboratory | Wellcome Sanger Institute for the COVID-19 Genomics UK (COG-UK) Consortium | Berkshire and Surrey Pathology Services Lighthouse Laboratory and Alex Alderton, Roberto Amato, Jeffrey Barrett, Sonia Goncalves, Ewan Harrison, David K. Jackson, Ian Johnston, Dominic Kwiatkowski, Cordelia Langford, John Sillitoe on behalf of the Wellcome Sanger Institute COVID-19 Surveillance Team |
| EPI_ISL_3042534 | Lighthouse Lab in Milton Keynes | Wellcome Sanger Institute for the COVID-19 Genomics UK (COG-UK) Consortium | The Lighthouse Lab in Milton Keynes and Alex Alderton, Roberto Amato, Jeffrey Barrett, Sonia Goncalves, Ewan Harrison, David K. Jackson, Ian Johnston, Dominic Kwiatkowski, Cordelia Langford, John Sillitoe on behalf of the Wellcome Sanger Institute COVID-19 Surveillance Team |
| EPI_ISL_3042535 | Lighthouse Laboratory Plymouth | Wellcome Sanger Institute for the COVID-19 Genomics UK (COG-UK) Consortium | Lighthouse Laboratory Plymouth and Alex Alderton, Roberto Amato, Jeffrey Barrett, Sonia Goncalves, Ewan Harrison, David K. Jackson, Ian Johnston, Dominic Kwiatkowski, Cordelia Langford, John Sillitoe on behalf of the Wellcome Sanger Institute COVID-19 Surveillance Team |
| EPI_ISL_3042536 | Lighthouse Lab in Alderley Park | Wellcome Sanger Institute for the COVID-19 Genomics UK (COG-UK) Consortium | Jacquelyn Wynn, Mairead Hyland, The Lighthouse Lab in Alderley Park and Alex Alderton, Roberto Amato, Jeffrey Barrett, Sonia Goncalves, Ewan Harrison, David K. Jackson, Ian Johnston, Dominic Kwiatkowski, Cordelia Langford, John Sillitoe on behalf of the Wellcome Sanger Institute COVID-19 Surveillance Team |
| EPI_ISL_3042537 | Lighthouse Laboratory Plymouth | Wellcome Sanger Institute for the COVID-19 Genomics UK (COG-UK) Consortium | Lighthouse Laboratory Plymouth and Alex Alderton, Roberto Amato, Jeffrey Barrett, Sonia Goncalves, Ewan Harrison, David K. Jackson, Ian Johnston, Dominic Kwiatkowski, Cordelia Langford, John Sillitoe on behalf of the Wellcome Sanger Institute COVID-19 Surveillance Team |
| EPI_ISL_3042538 | Lighthouse Lab in Alderley Park | Wellcome Sanger Institute for the COVID-19 Genomics UK (COG-UK) Consortium | Jacquelyn Wynn, Mairead Hyland, The Lighthouse Lab in Alderley Park and Alex Alderton, Roberto Amato, Jeffrey Barrett, Sonia Goncalves, Ewan Harrison, David K. Jackson, Ian Johnston, Dominic Kwiatkowski, Cordelia Langford, John Sillitoe on behalf of the Wellcome Sanger Institute COVID-19 Surveillance Team |
| EPI_ISL_3042539 | Lighthouse Laboratory Plymouth | Wellcome Sanger Institute for the COVID-19 Genomics UK (COG-UK) Consortium | Lighthouse Laboratory Plymouth and Alex Alderton, Roberto Amato, Jeffrey Barrett, Sonia Goncalves, Ewan Harrison, David K. Jackson, Ian Johnston, Dominic Kwiatkowski, Cordelia Langford, John Sillitoe on behalf of the Wellcome Sanger Institute COVID-19 Surveillance Team |
| EPI_ISL_3042540 | Lighthouse Lab in Alderley Park | Wellcome Sanger Institute for the COVID-19 Genomics UK (COG-UK) Consortium | Jacquelyn Wynn, Mairead Hyland, The Lighthouse Lab in Alderley Park and Alex Alderton, Roberto Amato, Jeffrey Barrett, Sonia Goncalves, Ewan Harrison, David K. Jackson, Ian Johnston, Dominic Kwiatkowski, Cordelia Langford, John Sillitoe on behalf of the Wellcome Sanger Institute COVID-19 Surveillance Team |
| EPI_ISL_3042541, EPI_ISL_3042542 | Lighthouse Laboratory Plymouth | Wellcome Sanger Institute for the COVID-19 Genomics UK (COG-UK) Consortium | Lighthouse Laboratory Plymouth and Alex Alderton, Roberto Amato, Jeffrey Barrett, Sonia Goncalves, Ewan Harrison, David K. Jackson, Ian Johnston, Dominic Kwiatkowski, Cordelia Langford, John Sillitoe on behalf of the Wellcome Sanger Institute COVID-19 Surveillance Team |
| EPI_ISL_3042543 | Berkshire and Surrey Pathology Services Lighthouse Laboratory | Wellcome Sanger Institute for the COVID-19 Genomics UK (COG-UK) Consortium | Berkshire and Surrey Pathology Services Lighthouse Laboratory and Alex Alderton, Roberto Amato, Jeffrey Barrett, Sonia Goncalves, Ewan Harrison, David K. Jackson, Ian Johnston, Dominic Kwiatkowski, Cordelia Langford, John Sillitoe on behalf of the Wellcome Sanger Institute COVID-19 Surveillance Team |
| EPI_ISL_3042544 | Lighthouse Lab in Milton Keynes | Wellcome Sanger Institute for the COVID-19 Genomics UK (COG-UK) Consortium | The Lighthouse Lab in Milton Keynes and Alex Alderton, Roberto Amato, Jeffrey Barrett, Sonia Goncalves, Ewan Harrison, David K. Jackson, Ian Johnston, Dominic Kwiatkowski, Cordelia Langford, John Sillitoe on behalf of the Wellcome Sanger Institute COVID-19 Surveillance Team |
| EPI_ISL_3042545 | Lighthouse Lab in Alderley Park | Wellcome Sanger Institute for the COVID-19 Genomics UK (COG-UK) Consortium | Jacquelyn Wynn, Mairead Hyland, The Lighthouse Lab in Alderley Park and Alex Alderton, Roberto Amato, Jeffrey Barrett, Sonia Goncalves, Ewan Harrison, David K. Jackson, Ian Johnston, Dominic Kwiatkowski, Cordelia Langford, John Sillitoe on behalf of the Wellcome Sanger Institute COVID-19 Surveillance Team |
| EPI_ISL_3042546 | Berkshire and Surrey Pathology Services Lighthouse Laboratory | Wellcome Sanger Institute for the COVID-19 Genomics UK (COG-UK) Consortium | Berkshire and Surrey Pathology Services Lighthouse Laboratory and Alex Alderton, Roberto Amato, Jeffrey Barrett, Sonia Goncalves, Ewan Harrison, David K. Jackson, Ian Johnston, Dominic Kwiatkowski, Cordelia Langford, John Sillitoe on behalf of the Wellcome Sanger Institute COVID-19 Surveillance Team |
| EPI_ISL_3042547, EPI_ISL_3042548 | Lighthouse Lab in Milton Keynes | Wellcome Sanger Institute for the COVID-19 Genomics UK (COG-UK) Consortium | The Lighthouse Lab in Milton Keynes and Alex Alderton, Roberto Amato, Jeffrey Barrett, Sonia Goncalves, Ewan Harrison, David K. Jackson, Ian Johnston, Dominic Kwiatkowski, Cordelia Langford, John Sillitoe on behalf of the Wellcome Sanger Institute COVID-19 Surveillance Team |
| EPI_ISL_3042549 | Lighthouse Lab in Alderley Park | Wellcome Sanger Institute for the COVID-19 Genomics UK (COG-UK) Consortium | Jacquelyn Wynn, Mairead Hyland, The Lighthouse Lab in Alderley Park and Alex Alderton, Roberto Amato, Jeffrey Barrett, Sonia Goncalves, Ewan Harrison, David K. Jackson, Ian Johnston, Dominic Kwiatkowski, Cordelia Langford, John Sillitoe on behalf of the Wellcome Sanger Institute COVID-19 Surveillance Team |
| EPI_ISL_3042550 | Lighthouse Laboratory Plymouth | Wellcome Sanger Institute for the COVID-19 Genomics UK (COG-UK) Consortium | Lighthouse Laboratory Plymouth and Alex Alderton, Roberto Amato, Jeffrey Barrett, Sonia Goncalves, Ewan Harrison, David K. Jackson, Ian Johnston, Dominic Kwiatkowski, Cordelia Langford, John Sillitoe on behalf of the Wellcome Sanger Institute COVID-19 Surveillance Team |
| EPI_ISL_3042551 | Lighthouse Lab in Milton Keynes | Wellcome Sanger Institute for the COVID-19 Genomics UK (COG-UK) Consortium | The Lighthouse Lab in Milton Keynes and Alex Alderton, Roberto Amato, Jeffrey Barrett, Sonia Goncalves, Ewan Harrison, David K. Jackson, Ian Johnston, Dominic Kwiatkowski, Cordelia Langford, John Sillitoe on behalf of the Wellcome Sanger Institute COVID-19 Surveillance Team |
| EPI_ISL_3042552 | Lighthouse Laboratory Plymouth | Wellcome Sanger Institute for the COVID-19 Genomics UK (COG-UK) Consortium | Lighthouse Laboratory Plymouth and Alex Alderton, Roberto Amato, Jeffrey Barrett, Sonia Goncalves, Ewan Harrison, David K. Jackson, Ian Johnston, Dominic Kwiatkowski, Cordelia Langford, John Sillitoe on behalf of the Wellcome Sanger Institute COVID-19 Surveillance Team |
| EPI_ISL_3042553, EPI_ISL_3042554, EPI_ISL_3042555 | Lighthouse Lab in Alderley Park | Wellcome Sanger Institute for the COVID-19 Genomics UK (COG-UK) Consortium | Jacquelyn Wynn, Mairead Hyland, The Lighthouse Lab in Alderley Park and Alex Alderton, Roberto Amato, Jeffrey Barrett, Sonia Goncalves, Ewan Harrison, David K. Jackson, Ian Johnston, Dominic Kwiatkowski, Cordelia Langford, John Sillitoe on behalf of the Wellcome Sanger Institute COVID-19 Surveillance Team |
| EPI_ISL_3042556, EPI_ISL_3042557 | Berkshire and Surrey Pathology Services Lighthouse Laboratory | Wellcome Sanger Institute for the COVID-19 Genomics UK (COG-UK) Consortium | Berkshire and Surrey Pathology Services Lighthouse Laboratory and Alex Alderton, Roberto Amato, Jeffrey Barrett, Sonia Goncalves, Ewan Harrison, David K. Jackson, Ian Johnston, Dominic Kwiatkowski, Cordelia Langford, John Sillitoe on behalf of the Wellcome Sanger Institute COVID-19 Surveillance Team |
| EPI_ISL_3042558 | Lighthouse Lab in Alderley Park | Wellcome Sanger Institute for the COVID-19 Genomics UK (COG-UK) Consortium | Jacquelyn Wynn, Mairead Hyland, The Lighthouse Lab in Alderley Park and Alex Alderton, Roberto Amato, Jeffrey Barrett, Sonia Goncalves, Ewan Harrison, David K. Jackson, Ian Johnston, Dominic Kwiatkowski, Cordelia Langford, John Sillitoe on behalf of the Wellcome Sanger Institute COVID-19 Surveillance Team |
| EPI_ISL_3042559, EPI_ISL_3042560 | Berkshire and Surrey Pathology Services Lighthouse Laboratory | Wellcome Sanger Institute for the COVID-19 Genomics UK (COG-UK) Consortium | Berkshire and Surrey Pathology Services Lighthouse Laboratory and Alex Alderton, Roberto Amato, Jeffrey Barrett, Sonia Goncalves, Ewan Harrison, David K. Jackson, Ian Johnston, Dominic Kwiatkowski, Cordelia Langford, John Sillitoe on behalf of the Wellcome Sanger Institute COVID-19 Surveillance Team |
| EPI_ISL_3042561 | Lighthouse Lab in Milton Keynes | Wellcome Sanger Institute for the COVID-19 Genomics UK (COG-UK) Consortium | The Lighthouse Lab in Milton Keynes and Alex Alderton, Roberto Amato, Jeffrey Barrett, Sonia Goncalves, Ewan Harrison, David K. Jackson, Ian Johnston, Dominic Kwiatkowski, Cordelia Langford, John Sillitoe on behalf of the Wellcome Sanger Institute COVID-19 Surveillance Team |
| EPI_ISL_3042562 | Lighthouse Lab in Alderley Park | Wellcome Sanger Institute for the COVID-19 Genomics UK (COG-UK) Consortium | Jacquelyn Wynn, Mairead Hyland, The Lighthouse Lab in Alderley Park and Alex Alderton, Roberto Amato, Jeffrey Barrett, Sonia Goncalves, Ewan Harrison, David K. Jackson, Ian Johnston, Dominic Kwiatkowski, Cordelia Langford, John Sillitoe on behalf of the Wellcome Sanger Institute COVID-19 Surveillance Team |
| EPI_ISL_3042563 | Lighthouse Laboratory Plymouth | Wellcome Sanger Institute for the COVID-19 Genomics UK (COG-UK) Consortium | Lighthouse Laboratory Plymouth and Alex Alderton, Roberto Amato, Jeffrey Barrett, Sonia Goncalves, Ewan Harrison, David K. Jackson, Ian Johnston, Dominic Kwiatkowski, Cordelia Langford, John Sillitoe on behalf of the Wellcome Sanger Institute COVID-19 Surveillance Team |
| EPI_ISL_3042564 | Berkshire and Surrey Pathology Services Lighthouse Laboratory | Wellcome Sanger Institute for the COVID-19 Genomics UK (COG-UK) Consortium | Berkshire and Surrey Pathology Services Lighthouse Laboratory and Alex Alderton, Roberto Amato, Jeffrey Barrett, Sonia Goncalves, Ewan Harrison, David K. Jackson, Ian Johnston, Dominic Kwiatkowski, Cordelia Langford, John Sillitoe on behalf of the Wellcome Sanger Institute COVID-19 Surveillance Team |
| EPI_ISL_3042565, EPI_ISL_3042566, EPI_ISL_3042567 | Lighthouse Lab in Alderley Park | Wellcome Sanger Institute for the COVID-19 Genomics UK (COG-UK) Consortium | Jacquelyn Wynn, Mairead Hyland, The Lighthouse Lab in Alderley Park and Alex Alderton, Roberto Amato, Jeffrey Barrett, Sonia Goncalves, Ewan Harrison, David K. Jackson, Ian Johnston, Dominic Kwiatkowski, Cordelia Langford, John Sillitoe on behalf of the Wellcome Sanger Institute COVID-19 Surveillance Team |
| EPI_ISL_3042568 | Lighthouse Lab in Milton Keynes | Wellcome Sanger Institute for the COVID-19 Genomics UK (COG-UK) Consortium | The Lighthouse Lab in Milton Keynes and Alex Alderton, Roberto Amato, Jeffrey Barrett, Sonia Goncalves, Ewan Harrison, David K. Jackson, Ian Johnston, Dominic Kwiatkowski, Cordelia Langford, John Sillitoe on behalf of the Wellcome Sanger Institute COVID-19 Surveillance Team |
| EPI_ISL_3042569 | Berkshire and Surrey Pathology Services Lighthouse Laboratory | Wellcome Sanger Institute for the COVID-19 Genomics UK (COG-UK) Consortium | Berkshire and Surrey Pathology Services Lighthouse Laboratory and Alex Alderton, Roberto Amato, Jeffrey Barrett, Sonia Goncalves, Ewan Harrison, David K. Jackson, Ian Johnston, Dominic Kwiatkowski, Cordelia Langford, John Sillitoe on behalf of the Wellcome Sanger Institute COVID-19 Surveillance Team |
| EPI_ISL_3042570, EPI_ISL_3042571 | Lighthouse Laboratory Plymouth | Wellcome Sanger Institute for the COVID-19 Genomics UK (COG-UK) Consortium | Lighthouse Laboratory Plymouth and Alex Alderton, Roberto Amato, Jeffrey Barrett, Sonia Goncalves, Ewan Harrison, David K. Jackson, Ian Johnston, Dominic Kwiatkowski, Cordelia Langford, John Sillitoe on behalf of the Wellcome Sanger Institute COVID-19 Surveillance Team |
| EPI_ISL_3042572 | Lighthouse Lab in Alderley Park | Wellcome Sanger Institute for the COVID-19 Genomics UK (COG-UK) | Jacquelyn Wynn, Mairead Hyland, The Lighthouse Lab in Alderley Park and Alex Alderton, Roberto Amato, Jeffrey Barrett, Sonia Goncalves, Ewan Harrison, David K. Jackson, Ian Johnston, Dominic Kwiatkowski, Cordelia Langford, John Sillitoe on |

| | | | |
|---|---|---|---|
| | | Wellcome Sanger Institute for the COVID-19 Genomics UK (COG-UK) Consortium | behalf of the Wellcome Sanger Institute COVID-19 Surveillance Team |
| EPI_ISL_3042573, EPI_ISL_3042574 | Lighthouse Laboratory Plymouth | Wellcome Sanger Institute for the COVID-19 Genomics UK (COG-UK) Consortium | Lighthouse Laboratory Plymouth and Alex Alderton, Roberto Amato, Jeffrey Barrett, Sonia Goncalves, Ewan Harrison, David K. Jackson, Ian Johnston, Dominic Kwiatkowski, Cordelia Langford, John Sillitoe on behalf of the Wellcome Sanger Institute COVID-19 Surveillance Team |
| EPI_ISL_3042575 | Berkshire and Surrey Pathology Services Lighthouse Laboratory | Wellcome Sanger Institute for the COVID-19 Genomics UK (COG-UK) Consortium | Berkshire and Surrey Pathology Services Lighthouse Laboratory and Alex Alderton, Roberto Amato, Jeffrey Barrett, Sonia Goncalves, Ewan Harrison, David K. Jackson, Ian Johnston, Dominic Kwiatkowski, Cordelia Langford, John Sillitoe on behalf of the Wellcome Sanger Institute COVID-19 Surveillance Team |
| EPI_ISL_3042576 | Lighthouse Lab in Milton Keynes | Wellcome Sanger Institute for the COVID-19 Genomics UK (COG-UK) Consortium | The Lighthouse Lab in Milton Keynes and Alex Alderton, Roberto Amato, Jeffrey Barrett, Sonia Goncalves, Ewan Harrison, David K. Jackson, Ian Johnston, Dominic Kwiatkowski, Cordelia Langford, John Sillitoe on behalf of the Wellcome Sanger Institute COVID-19 Surveillance Team |
| EPI_ISL_3042577, EPI_ISL_3042578 | Lighthouse Lab in Alderley Park | Wellcome Sanger Institute for the COVID-19 Genomics UK (COG-UK) Consortium | Jacquelyn Wynn, Mairead Hyland, The Lighthouse Lab in Alderley Park and Alex Alderton, Roberto Amato, Jeffrey Barrett, Sonia Goncalves, Ewan Harrison, David K. Jackson, Ian Johnston, Dominic Kwiatkowski, Cordelia Langford, John Sillitoe on behalf of the Wellcome Sanger Institute COVID-19 Surveillance Team |
| EPI_ISL_3042579, EPI_ISL_3042580, EPI_ISL_3042581 | Lighthouse Laboratory Plymouth | Wellcome Sanger Institute for the COVID-19 Genomics UK (COG-UK) Consortium | Lighthouse Laboratory Plymouth and Alex Alderton, Roberto Amato, Jeffrey Barrett, Sonia Goncalves, Ewan Harrison, David K. Jackson, Ian Johnston, Dominic Kwiatkowski, Cordelia Langford, John Sillitoe on behalf of the Wellcome Sanger Institute COVID-19 Surveillance Team |
| EPI_ISL_3042582, EPI_ISL_3042583 | Berkshire and Surrey Pathology Services Lighthouse Laboratory | Wellcome Sanger Institute for the COVID-19 Genomics UK (COG-UK) Consortium | Berkshire and Surrey Pathology Services Lighthouse Laboratory and Alex Alderton, Roberto Amato, Jeffrey Barrett, Sonia Goncalves, Ewan Harrison, David K. Jackson, Ian Johnston, Dominic Kwiatkowski, Cordelia Langford, John Sillitoe on behalf of the Wellcome Sanger Institute COVID-19 Surveillance Team |
| EPI_ISL_3042584 | Lighthouse Laboratory Plymouth | Wellcome Sanger Institute for the COVID-19 Genomics UK (COG-UK) Consortium | Lighthouse Laboratory Plymouth and Alex Alderton, Roberto Amato, Jeffrey Barrett, Sonia Goncalves, Ewan Harrison, David K. Jackson, Ian Johnston, Dominic Kwiatkowski, Cordelia Langford, John Sillitoe on behalf of the Wellcome Sanger Institute COVID-19 Surveillance Team |
| EPI_ISL_3042585 | Lighthouse Lab in Milton Keynes | Wellcome Sanger Institute for the COVID-19 Genomics UK (COG-UK) Consortium | The Lighthouse Lab in Milton Keynes and Alex Alderton, Roberto Amato, Jeffrey Barrett, Sonia Goncalves, Ewan Harrison, David K. Jackson, Ian Johnston, Dominic Kwiatkowski, Cordelia Langford, John Sillitoe on behalf of the Wellcome Sanger Institute COVID-19 Surveillance Team |
| EPI_ISL_3042586 | Lighthouse Laboratory Plymouth | Wellcome Sanger Institute for the COVID-19 Genomics UK (COG-UK) Consortium | Lighthouse Laboratory Plymouth and Alex Alderton, Roberto Amato, Jeffrey Barrett, Sonia Goncalves, Ewan Harrison, David K. Jackson, Ian Johnston, Dominic Kwiatkowski, Cordelia Langford, John Sillitoe on behalf of the Wellcome Sanger Institute COVID-19 Surveillance Team |
| EPI_ISL_3042587 | Berkshire and Surrey Pathology Services Lighthouse Laboratory | Wellcome Sanger Institute for the COVID-19 Genomics UK (COG-UK) Consortium | Berkshire and Surrey Pathology Services Lighthouse Laboratory and Alex Alderton, Roberto Amato, Jeffrey Barrett, Sonia Goncalves, Ewan Harrison, David K. Jackson, Ian Johnston, Dominic Kwiatkowski, Cordelia Langford, John Sillitoe on behalf of the Wellcome Sanger Institute COVID-19 Surveillance Team |
| EPI_ISL_3042588 | Lighthouse Lab in Alderley Park | Wellcome Sanger Institute for the COVID-19 Genomics UK (COG-UK) Consortium | Jacquelyn Wynn, Mairead Hyland, The Lighthouse Lab in Alderley Park and Alex Alderton, Roberto Amato, Jeffrey Barrett, Sonia Goncalves, Ewan Harrison, David K. Jackson, Ian Johnston, Dominic Kwiatkowski, Cordelia Langford, John Sillitoe on behalf of the Wellcome Sanger Institute COVID-19 Surveillance Team |
| EPI_ISL_3042589 | Berkshire and Surrey Pathology Services Lighthouse Laboratory | Wellcome Sanger Institute for the COVID-19 Genomics UK (COG-UK) Consortium | Berkshire and Surrey Pathology Services Lighthouse Laboratory and Alex Alderton, Roberto Amato, Jeffrey Barrett, Sonia Goncalves, Ewan Harrison, David K. Jackson, Ian Johnston, Dominic Kwiatkowski, Cordelia Langford, John Sillitoe on behalf of the Wellcome Sanger Institute COVID-19 Surveillance Team |
| EPI_ISL_3042590, EPI_ISL_3042591 | Lighthouse Lab in Alderley Park | Wellcome Sanger Institute for the COVID-19 Genomics UK (COG-UK) Consortium | Jacquelyn Wynn, Mairead Hyland, The Lighthouse Lab in Alderley Park and Alex Alderton, Roberto Amato, Jeffrey Barrett, Sonia Goncalves, Ewan Harrison, David K. Jackson, Ian Johnston, Dominic Kwiatkowski, Cordelia Langford, John Sillitoe on behalf of the Wellcome Sanger Institute COVID-19 Surveillance Team |
| EPI_ISL_3042592 | Lighthouse Laboratory Plymouth | Wellcome Sanger Institute for the COVID-19 Genomics UK (COG-UK) Consortium | Lighthouse Laboratory Plymouth and Alex Alderton, Roberto Amato, Jeffrey Barrett, Sonia Goncalves, Ewan Harrison, David K. Jackson, Ian Johnston, Dominic Kwiatkowski, Cordelia Langford, John Sillitoe on behalf of the Wellcome Sanger Institute COVID-19 Surveillance Team |
| EPI_ISL_3042593 | Lighthouse Lab in Milton Keynes | Wellcome Sanger Institute for the COVID-19 Genomics UK (COG-UK) Consortium | The Lighthouse Lab in Milton Keynes and Alex Alderton, Roberto Amato, Jeffrey Barrett, Sonia Goncalves, Ewan Harrison, David K. Jackson, Ian Johnston, Dominic Kwiatkowski, Cordelia Langford, John Sillitoe on behalf of the Wellcome Sanger Institute COVID-19 Surveillance Team |
| EPI_ISL_3042594 | Lighthouse Laboratory Plymouth | Wellcome Sanger Institute for the COVID-19 Genomics UK (COG-UK) Consortium | Lighthouse Laboratory Plymouth and Alex Alderton, Roberto Amato, Jeffrey Barrett, Sonia Goncalves, Ewan Harrison, David K. Jackson, Ian Johnston, Dominic Kwiatkowski, Cordelia Langford, John Sillitoe on behalf of the Wellcome Sanger Institute COVID-19 Surveillance Team |
| EPI_ISL_3042595 | Berkshire and Surrey Pathology Services Lighthouse Laboratory | Wellcome Sanger Institute for the COVID-19 Genomics UK (COG-UK) Consortium | Berkshire and Surrey Pathology Services Lighthouse Laboratory and Alex Alderton, Roberto Amato, Jeffrey Barrett, Sonia Goncalves, Ewan Harrison, David K. Jackson, Ian Johnston, Dominic Kwiatkowski, Cordelia Langford, John Sillitoe on behalf of the Wellcome Sanger Institute COVID-19 Surveillance Team |
| EPI_ISL_3042596 | Lighthouse Laboratory Plymouth | Wellcome Sanger Institute for the COVID-19 Genomics UK (COG-UK) Consortium | Lighthouse Laboratory Plymouth and Alex Alderton, Roberto Amato, Jeffrey Barrett, Sonia Goncalves, Ewan Harrison, David K. Jackson, Ian Johnston, Dominic Kwiatkowski, Cordelia Langford, John Sillitoe on behalf of the Wellcome Sanger Institute COVID-19 Surveillance Team |
| EPI_ISL_3042597, EPI_ISL_3042598, EPI_ISL_3042599, EPI_ISL_3042600 | Lighthouse Lab in Milton Keynes | Wellcome Sanger Institute for the COVID-19 Genomics UK (COG-UK) Consortium | The Lighthouse Lab in Milton Keynes and Alex Alderton, Roberto Amato, Jeffrey Barrett, Sonia Goncalves, Ewan Harrison, David K. Jackson, Ian Johnston, Dominic Kwiatkowski, Cordelia Langford, John Sillitoe on behalf of the Wellcome Sanger Institute COVID-19 Surveillance Team |
| EPI_ISL_3042601 | Lighthouse Laboratory Plymouth | Wellcome Sanger Institute for the COVID-19 Genomics UK (COG-UK) Consortium | Lighthouse Laboratory Plymouth and Alex Alderton, Roberto Amato, Jeffrey Barrett, Sonia Goncalves, Ewan Harrison, David K. Jackson, Ian Johnston, Dominic Kwiatkowski, Cordelia Langford, John Sillitoe on behalf of the Wellcome Sanger Institute COVID-19 Surveillance Team |
| EPI_ISL_3042602 | Berkshire and Surrey Pathology Services Lighthouse Laboratory | Wellcome Sanger Institute for the COVID-19 Genomics UK (COG-UK) Consortium | Berkshire and Surrey Pathology Services Lighthouse Laboratory and Alex Alderton, Roberto Amato, Jeffrey Barrett, Sonia Goncalves, Ewan Harrison, David K. Jackson, Ian Johnston, Dominic Kwiatkowski, Cordelia Langford, John Sillitoe on behalf of the Wellcome Sanger Institute COVID-19 Surveillance Team |
| EPI_ISL_3042603, EPI_ISL_3042604 | Lighthouse Lab in Milton Keynes | Wellcome Sanger Institute for the COVID-19 Genomics UK (COG-UK) Consortium | The Lighthouse Lab in Milton Keynes and Alex Alderton, Roberto Amato, Jeffrey Barrett, Sonia Goncalves, Ewan Harrison, David K. Jackson, Ian Johnston, Dominic Kwiatkowski, Cordelia Langford, John Sillitoe on behalf of the Wellcome Sanger Institute COVID-19 Surveillance Team |
| EPI_ISL_3042605 | Lighthouse Laboratory Plymouth | Wellcome Sanger Institute for the COVID-19 Genomics UK (COG-UK) Consortium | Lighthouse Laboratory Plymouth and Alex Alderton, Roberto Amato, Jeffrey Barrett, Sonia Goncalves, Ewan Harrison, David K. Jackson, Ian Johnston, Dominic Kwiatkowski, Cordelia Langford, John Sillitoe on behalf of the Wellcome Sanger Institute COVID-19 Surveillance Team |
| EPI_ISL_3042606 | Lighthouse Lab in Alderley Park | Wellcome Sanger Institute for the COVID-19 Genomics UK (COG-UK) Consortium | Jacquelyn Wynn, Mairead Hyland, The Lighthouse Lab in Alderley Park and Alex Alderton, Roberto Amato, Jeffrey Barrett, Sonia Goncalves, Ewan Harrison, David K. Jackson, Ian Johnston, Dominic Kwiatkowski, Cordelia Langford, John Sillitoe on behalf of the Wellcome Sanger Institute COVID-19 Surveillance Team |
| EPI_ISL_3042607 | Lighthouse Lab in Milton Keynes | Wellcome Sanger Institute for the COVID-19 Genomics UK (COG-UK) Consortium | The Lighthouse Lab in Milton Keynes and Alex Alderton, Roberto Amato, Jeffrey Barrett, Sonia Goncalves, Ewan Harrison, David K. Jackson, Ian Johnston, Dominic Kwiatkowski, Cordelia Langford, John Sillitoe on behalf of the Wellcome Sanger Institute COVID-19 Surveillance Team |
| EPI_ISL_3042608, EPI_ISL_3042609 | Berkshire and Surrey Pathology Services Lighthouse Laboratory | Wellcome Sanger Institute for the COVID-19 Genomics UK (COG-UK) Consortium | Berkshire and Surrey Pathology Services Lighthouse Laboratory and Alex Alderton, Roberto Amato, Jeffrey Barrett, Sonia Goncalves, Ewan Harrison, David K. Jackson, Ian Johnston, Dominic Kwiatkowski, Cordelia Langford, John Sillitoe on behalf of the Wellcome Sanger Institute COVID-19 Surveillance Team |
| EPI_ISL_3042610 | Lighthouse Lab in Milton Keynes | Wellcome Sanger Institute for the COVID-19 Genomics UK (COG-UK) Consortium | The Lighthouse Lab in Milton Keynes and Alex Alderton, Roberto Amato, Jeffrey Barrett, Sonia Goncalves, Ewan Harrison, David K. Jackson, Ian Johnston, Dominic Kwiatkowski, Cordelia Langford, John Sillitoe on behalf of the Wellcome Sanger Institute COVID-19 Surveillance Team |
| EPI_ISL_3042611 | Lighthouse Laboratory Plymouth | Wellcome Sanger Institute for the COVID-19 Genomics UK (COG-UK) Consortium | Lighthouse Laboratory Plymouth and Alex Alderton, Roberto Amato, Jeffrey Barrett, Sonia Goncalves, Ewan Harrison, David K. Jackson, Ian Johnston, Dominic Kwiatkowski, Cordelia Langford, John Sillitoe on behalf of the Wellcome Sanger Institute COVID-19 Surveillance Team |
| EPI_ISL_3042612 | Lighthouse Lab in Alderley Park | Wellcome Sanger Institute for the COVID-19 Genomics UK (COG-UK) Consortium | Jacquelyn Wynn, Mairead Hyland, The Lighthouse Lab in Alderley Park and Alex Alderton, Roberto Amato, Jeffrey Barrett, Sonia Goncalves, Ewan Harrison, David K. Jackson, Ian Johnston, Dominic Kwiatkowski, Cordelia Langford, John Sillitoe on behalf of the Wellcome Sanger Institute COVID-19 Surveillance Team |
| EPI_ISL_3042613 | Lighthouse Lab in Milton Keynes | Wellcome Sanger Institute for the COVID-19 Genomics UK (COG-UK) Consortium | The Lighthouse Lab in Milton Keynes and Alex Alderton, Roberto Amato, Jeffrey Barrett, Sonia Goncalves, Ewan Harrison, David K. Jackson, Ian Johnston, Dominic Kwiatkowski, Cordelia Langford, John Sillitoe on behalf of the Wellcome Sanger Institute COVID-19 Surveillance Team |
| EPI_ISL_3042614, EPI_ISL_3042615 | Berkshire and Surrey Pathology Services Lighthouse Laboratory | Wellcome Sanger Institute for the COVID-19 Genomics UK (COG-UK) Consortium | Berkshire and Surrey Pathology Services Lighthouse Laboratory and Alex Alderton, Roberto Amato, Jeffrey Barrett, Sonia Goncalves, Ewan Harrison, David K. Jackson, Ian Johnston, Dominic Kwiatkowski, Cordelia Langford, John Sillitoe on behalf of the Wellcome Sanger Institute COVID-19 Surveillance Team |
| EPI_ISL_3042616 | Lighthouse Laboratory Plymouth | Wellcome Sanger Institute for the COVID-19 Genomics UK (COG-UK) Consortium | Lighthouse Laboratory Plymouth and Alex Alderton, Roberto Amato, Jeffrey Barrett, Sonia Goncalves, Ewan Harrison, David K. Jackson, Ian Johnston, Dominic Kwiatkowski, Cordelia Langford, John Sillitoe on behalf of the Wellcome Sanger Institute COVID-19 Surveillance Team |
| EPI_ISL_3042617 | Lighthouse Lab in Milton Keynes | Wellcome Sanger Institute for the COVID-19 Genomics UK (COG-UK) Consortium | The Lighthouse Lab in Milton Keynes and Alex Alderton, Roberto Amato, Jeffrey Barrett, Sonia Goncalves, Ewan Harrison, David K. Jackson, Ian Johnston, Dominic Kwiatkowski, Cordelia Langford, John Sillitoe on behalf of the Wellcome Sanger Institute COVID-19 Surveillance Team |
| EPI_ISL_3042618 | Berkshire and Surrey Pathology Services Lighthouse Laboratory | Wellcome Sanger Institute for the COVID-19 Genomics UK (COG-UK) Consortium | Berkshire and Surrey Pathology Services Lighthouse Laboratory and Alex Alderton, Roberto Amato, Jeffrey Barrett, Sonia Goncalves, Ewan Harrison, David K. Jackson, Ian Johnston, Dominic Kwiatkowski, Cordelia Langford, John Sillitoe on behalf of the Wellcome Sanger Institute COVID-19 Surveillance Team |
| EPI_ISL_3042619 | Lighthouse Lab in Milton Keynes | Wellcome Sanger Institute for the COVID-19 Genomics UK (COG-UK) Consortium | The Lighthouse Lab in Milton Keynes and Alex Alderton, Roberto Amato, Jeffrey Barrett, Sonia Goncalves, Ewan Harrison, David K. Jackson, Ian Johnston, Dominic Kwiatkowski, Cordelia Langford, John Sillitoe on behalf of the Wellcome Sanger Institute COVID-19 Surveillance Team |
| EPI_ISL_3042620 | Lighthouse Lab in Alderley Park | Wellcome Sanger Institute for the COVID-19 Genomics UK (COG-UK) Consortium | Jacquelyn Wynn, Mairead Hyland, The Lighthouse Lab in Alderley Park and Alex Alderton, Roberto Amato, Jeffrey Barrett, Sonia Goncalves, Ewan Harrison, David K. Jackson, Ian Johnston, Dominic Kwiatkowski, Cordelia Langford, John Sillitoe on behalf of the Wellcome Sanger Institute COVID-19 Surveillance Team |
| EPI_ISL_3042621 | Lighthouse Laboratory Plymouth | Wellcome Sanger Institute for the COVID-19 Genomics UK (COG-UK) Consortium | Lighthouse Laboratory Plymouth and Alex Alderton, Roberto Amato, Jeffrey Barrett, Sonia Goncalves, Ewan Harrison, David K. Jackson, Ian Johnston, Dominic Kwiatkowski, Cordelia Langford, John Sillitoe on behalf of the Wellcome Sanger Institute COVID-19 Surveillance Team |
| EPI_ISL_3042622 | Lighthouse Lab in Milton Keynes | Wellcome Sanger Institute for the COVID-19 Genomics UK (COG-UK) Consortium | The Lighthouse Lab in Milton Keynes and Alex Alderton, Roberto Amato, Jeffrey Barrett, Sonia Goncalves, Ewan Harrison, David K. Jackson, Ian Johnston, Dominic Kwiatkowski, Cordelia Langford, John Sillitoe on behalf of the Wellcome Sanger Institute COVID-19 Surveillance Team |
| EPI_ISL_3042623, EPI_ISL_3042624, EPI_ISL_3042625 | Lighthouse Laboratory Plymouth | Wellcome Sanger Institute for the COVID-19 Genomics UK (COG-UK) Consortium | Lighthouse Laboratory Plymouth and Alex Alderton, Roberto Amato, Jeffrey Barrett, Sonia Goncalves, Ewan Harrison, David K. Jackson, Ian Johnston, Dominic Kwiatkowski, Cordelia Langford, John Sillitoe on behalf of the Wellcome Sanger Institute COVID-19 Surveillance Team |
| EPI_ISL_3042626 | Lighthouse Lab in Milton Keynes | Wellcome Sanger Institute for the COVID-19 Genomics UK (COG-UK) Consortium | The Lighthouse Lab in Milton Keynes and Alex Alderton, Roberto Amato, Jeffrey Barrett, Sonia Goncalves, Ewan Harrison, David K. Jackson, Ian Johnston, Dominic Kwiatkowski, Cordelia Langford, John Sillitoe on behalf of the Wellcome Sanger Institute COVID-19 Surveillance Team |
| EPI_ISL_3042627, EPI_ISL_3042628 | Lighthouse Lab in Alderley Park | Wellcome Sanger Institute for the COVID-19 Genomics UK (COG-UK) Consortium | Jacquelyn Wynn, Mairead Hyland, The Lighthouse Lab in Alderley Park and Alex Alderton, Roberto Amato, Jeffrey Barrett, Sonia Goncalves, Ewan Harrison, David K. Jackson, Ian Johnston, Dominic Kwiatkowski, Cordelia Langford, John Sillitoe on behalf of the Wellcome Sanger Institute COVID-19 Surveillance Team |
| EPI_ISL_3042629 | Lighthouse Lab in Milton Keynes | Wellcome Sanger Institute for the COVID-19 Genomics UK (COG-UK) Consortium | The Lighthouse Lab in Milton Keynes and Alex Alderton, Roberto Amato, Jeffrey Barrett, Sonia Goncalves, Ewan Harrison, David K. Jackson, Ian Johnston, Dominic Kwiatkowski, Cordelia Langford, John Sillitoe on behalf of the Wellcome Sanger Institute COVID-19 Surveillance Team |
| EPI_ISL_3042630 | Lighthouse Laboratory Plymouth | Wellcome Sanger Institute for the COVID-19 Genomics UK (COG-UK) Consortium | Lighthouse Laboratory Plymouth and Alex Alderton, Roberto Amato, Jeffrey Barrett, Sonia Goncalves, Ewan Harrison, David K. Jackson, Ian Johnston, Dominic Kwiatkowski, Cordelia Langford, John Sillitoe on behalf of the Wellcome Sanger Institute COVID-19 Surveillance Team |

| | | | |
|---|---|---|---|
| EPI_ISL_3042631, EPI_ISL_3042632 | Lighthouse Lab in Milton Keynes | Wellcome Sanger Institute for the COVID-19 Genomics UK (COG-UK) Consortium | The Lighthouse Lab in Milton Keynes and Alex Alderton, Roberto Amato, Jeffrey Barrett, Sonia Goncalves, Ewan Harrison, David K. Jackson, Ian Johnston, Dominic Kwiatkowski, Cordelia Langford, John Sillitoe on behalf of the Wellcome Sanger Institute COVID-19 Surveillance Team |
| EPI_ISL_3042633, EPI_ISL_3042634 | Lighthouse Lab in Alderley Park | Wellcome Sanger Institute for the COVID-19 Genomics UK (COG-UK) Consortium | Jacquelyn Wynn, Mairead Hyland, The Lighthouse Lab in Alderley Park and Alex Alderton, Roberto Amato, Jeffrey Barrett, Sonia Goncalves, Ewan Harrison, David K. Jackson, Ian Johnston, Dominic Kwiatkowski, Cordelia Langford, John Sillitoe on behalf of the Wellcome Sanger Institute COVID-19 Surveillance Team |
| EPI_ISL_3042635 | Lighthouse Laboratory Plymouth | Wellcome Sanger Institute for the COVID-19 Genomics UK (COG-UK) Consortium | Lighthouse Laboratory Plymouth and Alex Alderton, Roberto Amato, Jeffrey Barrett, Sonia Goncalves, Ewan Harrison, David K. Jackson, Ian Johnston, Dominic Kwiatkowski, Cordelia Langford, John Sillitoe on behalf of the Wellcome Sanger Institute COVID-19 Surveillance Team |
| EPI_ISL_3042636 | Lighthouse Lab in Milton Keynes | Wellcome Sanger Institute for the COVID-19 Genomics UK (COG-UK) Consortium | The Lighthouse Lab in Milton Keynes and Alex Alderton, Roberto Amato, Jeffrey Barrett, Sonia Goncalves, Ewan Harrison, David K. Jackson, Ian Johnston, Dominic Kwiatkowski, Cordelia Langford, John Sillitoe on behalf of the Wellcome Sanger Institute COVID-19 Surveillance Team |
| EPI_ISL_3042637 | Lighthouse Laboratory Plymouth | Wellcome Sanger Institute for the COVID-19 Genomics UK (COG-UK) Consortium | Lighthouse Laboratory Plymouth and Alex Alderton, Roberto Amato, Jeffrey Barrett, Sonia Goncalves, Ewan Harrison, David K. Jackson, Ian Johnston, Dominic Kwiatkowski, Cordelia Langford, John Sillitoe on behalf of the Wellcome Sanger Institute COVID-19 Surveillance Team |
| EPI_ISL_3042638 | Lighthouse Lab in Milton Keynes | Wellcome Sanger Institute for the COVID-19 Genomics UK (COG-UK) Consortium | The Lighthouse Lab in Milton Keynes and Alex Alderton, Roberto Amato, Jeffrey Barrett, Sonia Goncalves, Ewan Harrison, David K. Jackson, Ian Johnston, Dominic Kwiatkowski, Cordelia Langford, John Sillitoe on behalf of the Wellcome Sanger Institute COVID-19 Surveillance Team |
| EPI_ISL_3042639 | Lighthouse Lab in Alderley Park | Wellcome Sanger Institute for the COVID-19 Genomics UK (COG-UK) Consortium | Jacquelyn Wynn, Mairead Hyland, The Lighthouse Lab in Alderley Park and Alex Alderton, Roberto Amato, Jeffrey Barrett, Sonia Goncalves, Ewan Harrison, David K. Jackson, Ian Johnston, Dominic Kwiatkowski, Cordelia Langford, John Sillitoe on behalf of the Wellcome Sanger Institute COVID-19 Surveillance Team |
| EPI_ISL_3042640 | Berkshire and Surrey Pathology Services Lighthouse Laboratory | Wellcome Sanger Institute for the COVID-19 Genomics UK (COG-UK) Consortium | Berkshire and Surrey Pathology Services Lighthouse Laboratory and Alex Alderton, Roberto Amato, Jeffrey Barrett, Sonia Goncalves, Ewan Harrison, David K. Jackson, Ian Johnston, Dominic Kwiatkowski, Cordelia Langford, John Sillitoe on behalf of the Wellcome Sanger Institute COVID-19 Surveillance Team |
| EPI_ISL_3042641 | Lighthouse Lab in Alderley Park | Wellcome Sanger Institute for the COVID-19 Genomics UK (COG-UK) Consortium | Jacquelyn Wynn, Mairead Hyland, The Lighthouse Lab in Alderley Park and Alex Alderton, Roberto Amato, Jeffrey Barrett, Sonia Goncalves, Ewan Harrison, David K. Jackson, Ian Johnston, Dominic Kwiatkowski, Cordelia Langford, John Sillitoe on behalf of the Wellcome Sanger Institute COVID-19 Surveillance Team |
| EPI_ISL_3042642 | Lighthouse Lab in Glasgow | Wellcome Sanger Institute for the COVID-19 Genomics UK (COG-UK) Consortium | Harper VanSteenhouse, Yumi Kasai, David Gray, Carol Clugston, Anna Dominiczak and Alex Alderton, Roberto Amato, Jeffrey Barrett, Sonia Goncalves, Ewan Harrison, David K. Jackson, Ian Johnston, Dominic Kwiatkowski, Cordelia Langford, John Sillitoe on behalf of the Wellcome Sanger Institute COVID-19 Surveillance Team |
| EPI_ISL_3042643 | Berkshire and Surrey Pathology Services Lighthouse Laboratory | Wellcome Sanger Institute for the COVID-19 Genomics UK (COG-UK) Consortium | Berkshire and Surrey Pathology Services Lighthouse Laboratory and Alex Alderton, Roberto Amato, Jeffrey Barrett, Sonia Goncalves, Ewan Harrison, David K. Jackson, Ian Johnston, Dominic Kwiatkowski, Cordelia Langford, John Sillitoe on behalf of the Wellcome Sanger Institute COVID-19 Surveillance Team |
| EPI_ISL_3042644 | Lighthouse Lab in Alderley Park | Wellcome Sanger Institute for the COVID-19 Genomics UK (COG-UK) Consortium | Jacquelyn Wynn, Mairead Hyland, The Lighthouse Lab in Alderley Park and Alex Alderton, Roberto Amato, Jeffrey Barrett, Sonia Goncalves, Ewan Harrison, David K. Jackson, Ian Johnston, Dominic Kwiatkowski, Cordelia Langford, John Sillitoe on behalf of the Wellcome Sanger Institute COVID-19 Surveillance Team |
| EPI_ISL_3042645 | Lighthouse Lab in Glasgow | Wellcome Sanger Institute for the COVID-19 Genomics UK (COG-UK) Consortium | Harper VanSteenhouse, Yumi Kasai, David Gray, Carol Clugston, Anna Dominiczak and Alex Alderton, Roberto Amato, Jeffrey Barrett, Sonia Goncalves, Ewan Harrison, David K. Jackson, Ian Johnston, Dominic Kwiatkowski, Cordelia Langford, John Sillitoe on behalf of the Wellcome Sanger Institute COVID-19 Surveillance Team |
| EPI_ISL_3042646 | Berkshire and Surrey Pathology Services Lighthouse Laboratory | Wellcome Sanger Institute for the COVID-19 Genomics UK (COG-UK) Consortium | Berkshire and Surrey Pathology Services Lighthouse Laboratory and Alex Alderton, Roberto Amato, Jeffrey Barrett, Sonia Goncalves, Ewan Harrison, David K. Jackson, Ian Johnston, Dominic Kwiatkowski, Cordelia Langford, John Sillitoe on behalf of the Wellcome Sanger Institute COVID-19 Surveillance Team |
| EPI_ISL_3042647 | Lighthouse Lab in Glasgow | Wellcome Sanger Institute for the COVID-19 Genomics UK (COG-UK) Consortium | Harper VanSteenhouse, Yumi Kasai, David Gray, Carol Clugston, Anna Dominiczak and Alex Alderton, Roberto Amato, Jeffrey Barrett, Sonia Goncalves, Ewan Harrison, David K. Jackson, Ian Johnston, Dominic Kwiatkowski, Cordelia Langford, John Sillitoe on behalf of the Wellcome Sanger Institute COVID-19 Surveillance Team |
| EPI_ISL_3042648 | Lighthouse Lab in Alderley Park | Wellcome Sanger Institute for the COVID-19 Genomics UK (COG-UK) Consortium | Jacquelyn Wynn, Mairead Hyland, The Lighthouse Lab in Alderley Park and Alex Alderton, Roberto Amato, Jeffrey Barrett, Sonia Goncalves, Ewan Harrison, David K. Jackson, Ian Johnston, Dominic Kwiatkowski, Cordelia Langford, John Sillitoe on behalf of the Wellcome Sanger Institute COVID-19 Surveillance Team |
| EPI_ISL_3042649 | Lighthouse Lab in Glasgow | Wellcome Sanger Institute for the COVID-19 Genomics UK (COG-UK) Consortium | Harper VanSteenhouse, Yumi Kasai, David Gray, Carol Clugston, Anna Dominiczak and Alex Alderton, Roberto Amato, Jeffrey Barrett, Sonia Goncalves, Ewan Harrison, David K. Jackson, Ian Johnston, Dominic Kwiatkowski, Cordelia Langford, John Sillitoe on behalf of the Wellcome Sanger Institute COVID-19 Surveillance Team |
| EPI_ISL_3042650 | Berkshire and Surrey Pathology Services Lighthouse Laboratory | Wellcome Sanger Institute for the COVID-19 Genomics UK (COG-UK) Consortium | Berkshire and Surrey Pathology Services Lighthouse Laboratory and Alex Alderton, Roberto Amato, Jeffrey Barrett, Sonia Goncalves, Ewan Harrison, David K. Jackson, Ian Johnston, Dominic Kwiatkowski, Cordelia Langford, John Sillitoe on behalf of the Wellcome Sanger Institute COVID-19 Surveillance Team |
| EPI_ISL_3042651 | Lighthouse Lab in Glasgow | Wellcome Sanger Institute for the COVID-19 Genomics UK (COG-UK) Consortium | Harper VanSteenhouse, Yumi Kasai, David Gray, Carol Clugston, Anna Dominiczak and Alex Alderton, Roberto Amato, Jeffrey Barrett, Sonia Goncalves, Ewan Harrison, David K. Jackson, Ian Johnston, Dominic Kwiatkowski, Cordelia Langford, John Sillitoe on behalf of the Wellcome Sanger Institute COVID-19 Surveillance Team |
| EPI_ISL_3042652, EPI_ISL_3042653, EPI_ISL_3042654 | Berkshire and Surrey Pathology Services Lighthouse Laboratory | Wellcome Sanger Institute for the COVID-19 Genomics UK (COG-UK) Consortium | Berkshire and Surrey Pathology Services Lighthouse Laboratory and Alex Alderton, Roberto Amato, Jeffrey Barrett, Sonia Goncalves, Ewan Harrison, David K. Jackson, Ian Johnston, Dominic Kwiatkowski, Cordelia Langford, John Sillitoe on behalf of the Wellcome Sanger Institute COVID-19 Surveillance Team |
| EPI_ISL_3042655 | Lighthouse Lab in Alderley Park | Wellcome Sanger Institute for the COVID-19 Genomics UK (COG-UK) Consortium | Jacquelyn Wynn, Mairead Hyland, The Lighthouse Lab in Alderley Park and Alex Alderton, Roberto Amato, Jeffrey Barrett, Sonia Goncalves, Ewan Harrison, David K. Jackson, Ian Johnston, Dominic Kwiatkowski, Cordelia Langford, John Sillitoe on behalf of the Wellcome Sanger Institute COVID-19 Surveillance Team |
| EPI_ISL_3042656, EPI_ISL_3042657 | Berkshire and Surrey Pathology Services Lighthouse Laboratory | Wellcome Sanger Institute for the COVID-19 Genomics UK (COG-UK) Consortium | Berkshire and Surrey Pathology Services Lighthouse Laboratory and Alex Alderton, Roberto Amato, Jeffrey Barrett, Sonia Goncalves, Ewan Harrison, David K. Jackson, Ian Johnston, Dominic Kwiatkowski, Cordelia Langford, John Sillitoe on behalf of the Wellcome Sanger Institute COVID-19 Surveillance Team |
| EPI_ISL_3042658, EPI_ISL_3042659 | Lighthouse Lab in Glasgow | Wellcome Sanger Institute for the COVID-19 Genomics UK (COG-UK) Consortium | Harper VanSteenhouse, Yumi Kasai, David Gray, Carol Clugston, Anna Dominiczak and Alex Alderton, Roberto Amato, Jeffrey Barrett, Sonia Goncalves, Ewan Harrison, David K. Jackson, Ian Johnston, Dominic Kwiatkowski, Cordelia Langford, John Sillitoe on behalf of the Wellcome Sanger Institute COVID-19 Surveillance Team |
| EPI_ISL_3042660, EPI_ISL_3042661 | Berkshire and Surrey Pathology Services Lighthouse Laboratory | Wellcome Sanger Institute for the COVID-19 Genomics UK (COG-UK) Consortium | Berkshire and Surrey Pathology Services Lighthouse Laboratory and Alex Alderton, Roberto Amato, Jeffrey Barrett, Sonia Goncalves, Ewan Harrison, David K. Jackson, Ian Johnston, Dominic Kwiatkowski, Cordelia Langford, John Sillitoe on behalf of the Wellcome Sanger Institute COVID-19 Surveillance Team |
| EPI_ISL_3042662 | Lighthouse Lab in Alderley Park | Wellcome Sanger Institute for the COVID-19 Genomics UK (COG-UK) Consortium | Jacquelyn Wynn, Mairead Hyland, The Lighthouse Lab in Alderley Park and Alex Alderton, Roberto Amato, Jeffrey Barrett, Sonia Goncalves, Ewan Harrison, David K. Jackson, Ian Johnston, Dominic Kwiatkowski, Cordelia Langford, John Sillitoe on behalf of the Wellcome Sanger Institute COVID-19 Surveillance Team |
| EPI_ISL_3042663 | Lighthouse Lab in Glasgow | Wellcome Sanger Institute for the COVID-19 Genomics UK (COG-UK) Consortium | Harper VanSteenhouse, Yumi Kasai, David Gray, Carol Clugston, Anna Dominiczak and Alex Alderton, Roberto Amato, Jeffrey Barrett, Sonia Goncalves, Ewan Harrison, David K. Jackson, Ian Johnston, Dominic Kwiatkowski, Cordelia Langford, John Sillitoe on behalf of the Wellcome Sanger Institute COVID-19 Surveillance Team |
| EPI_ISL_3042664, EPI_ISL_3042665 | Lighthouse Lab in Alderley Park | Wellcome Sanger Institute for the COVID-19 Genomics UK (COG-UK) Consortium | Jacquelyn Wynn, Mairead Hyland, The Lighthouse Lab in Alderley Park and Alex Alderton, Roberto Amato, Jeffrey Barrett, Sonia Goncalves, Ewan Harrison, David K. Jackson, Ian Johnston, Dominic Kwiatkowski, Cordelia Langford, John Sillitoe on behalf of the Wellcome Sanger Institute COVID-19 Surveillance Team |
| EPI_ISL_3042666 | Berkshire and Surrey Pathology Services Lighthouse Laboratory | Wellcome Sanger Institute for the COVID-19 Genomics UK (COG-UK) Consortium | Berkshire and Surrey Pathology Services Lighthouse Laboratory and Alex Alderton, Roberto Amato, Jeffrey Barrett, Sonia Goncalves, Ewan Harrison, David K. Jackson, Ian Johnston, Dominic Kwiatkowski, Cordelia Langford, John Sillitoe on behalf of the Wellcome Sanger Institute COVID-19 Surveillance Team |
| EPI_ISL_3042667, EPI_ISL_3042668 | Lighthouse Lab in Glasgow | Wellcome Sanger Institute for the COVID-19 Genomics UK (COG-UK) Consortium | Harper VanSteenhouse, Yumi Kasai, David Gray, Carol Clugston, Anna Dominiczak and Alex Alderton, Roberto Amato, Jeffrey Barrett, Sonia Goncalves, Ewan Harrison, David K. Jackson, Ian Johnston, Dominic Kwiatkowski, Cordelia Langford, John Sillitoe on behalf of the Wellcome Sanger Institute COVID-19 Surveillance Team |
| EPI_ISL_3042669 | Lighthouse Lab in Alderley Park | Wellcome Sanger Institute for the COVID-19 Genomics UK (COG-UK) Consortium | Jacquelyn Wynn, Mairead Hyland, The Lighthouse Lab in Alderley Park and Alex Alderton, Roberto Amato, Jeffrey Barrett, Sonia Goncalves, Ewan Harrison, David K. Jackson, Ian Johnston, Dominic Kwiatkowski, Cordelia Langford, John Sillitoe on behalf of the Wellcome Sanger Institute COVID-19 Surveillance Team |
| EPI_ISL_3042670 | Berkshire and Surrey Pathology Services Lighthouse Laboratory | Wellcome Sanger Institute for the COVID-19 Genomics UK (COG-UK) Consortium | Berkshire and Surrey Pathology Services Lighthouse Laboratory and Alex Alderton, Roberto Amato, Jeffrey Barrett, Sonia Goncalves, Ewan Harrison, David K. Jackson, Ian Johnston, Dominic Kwiatkowski, Cordelia Langford, John Sillitoe on behalf of the Wellcome Sanger Institute COVID-19 Surveillance Team |
| EPI_ISL_3042671, EPI_ISL_3042672, EPI_ISL_3042673, EPI_ISL_3042674 | Lighthouse Lab in Alderley Park | Wellcome Sanger Institute for the COVID-19 Genomics UK (COG-UK) Consortium | Jacquelyn Wynn, Mairead Hyland, The Lighthouse Lab in Alderley Park and Alex Alderton, Roberto Amato, Jeffrey Barrett, Sonia Goncalves, Ewan Harrison, David K. Jackson, Ian Johnston, Dominic Kwiatkowski, Cordelia Langford, John Sillitoe on behalf of the Wellcome Sanger Institute COVID-19 Surveillance Team |
| EPI_ISL_3042675, EPI_ISL_3042676 | Berkshire and Surrey Pathology Services Lighthouse Laboratory | Wellcome Sanger Institute for the COVID-19 Genomics UK (COG-UK) Consortium | Berkshire and Surrey Pathology Services Lighthouse Laboratory and Alex Alderton, Roberto Amato, Jeffrey Barrett, Sonia Goncalves, Ewan Harrison, David K. Jackson, Ian Johnston, Dominic Kwiatkowski, Cordelia Langford, John Sillitoe on behalf of the Wellcome Sanger Institute COVID-19 Surveillance Team |
| EPI_ISL_3042677, EPI_ISL_3042678, EPI_ISL_3042679 | Lighthouse Lab in Glasgow | Wellcome Sanger Institute for the COVID-19 Genomics UK (COG-UK) Consortium | Harper VanSteenhouse, Yumi Kasai, David Gray, Carol Clugston, Anna Dominiczak and Alex Alderton, Roberto Amato, Jeffrey Barrett, Sonia Goncalves, Ewan Harrison, David K. Jackson, Ian Johnston, Dominic Kwiatkowski, Cordelia Langford, John Sillitoe on behalf of the Wellcome Sanger Institute COVID-19 Surveillance Team |
| EPI_ISL_3042680 | Berkshire and Surrey Pathology Services Lighthouse Laboratory | Wellcome Sanger Institute for the COVID-19 Genomics UK (COG-UK) Consortium | Berkshire and Surrey Pathology Services Lighthouse Laboratory and Alex Alderton, Roberto Amato, Jeffrey Barrett, Sonia Goncalves, Ewan Harrison, David K. Jackson, Ian Johnston, Dominic Kwiatkowski, Cordelia Langford, John Sillitoe on behalf of the Wellcome Sanger Institute COVID-19 Surveillance Team |
| EPI_ISL_3042681 | Lighthouse Lab in Glasgow | Wellcome Sanger Institute for the COVID-19 Genomics UK (COG-UK) Consortium | Harper VanSteenhouse, Yumi Kasai, David Gray, Carol Clugston, Anna Dominiczak and Alex Alderton, Roberto Amato, Jeffrey Barrett, Sonia Goncalves, Ewan Harrison, David K. Jackson, Ian Johnston, Dominic Kwiatkowski, Cordelia Langford, John Sillitoe on behalf of the Wellcome Sanger Institute COVID-19 Surveillance Team |
| EPI_ISL_3042682 | Lighthouse Lab in Alderley Park | Wellcome Sanger Institute for the COVID-19 Genomics UK (COG-UK) Consortium | Jacquelyn Wynn, Mairead Hyland, The Lighthouse Lab in Alderley Park and Alex Alderton, Roberto Amato, Jeffrey Barrett, Sonia Goncalves, Ewan Harrison, David K. Jackson, Ian Johnston, Dominic Kwiatkowski, Cordelia Langford, John Sillitoe on behalf of the Wellcome Sanger Institute COVID-19 Surveillance Team |
| EPI_ISL_3042683 | Berkshire and Surrey Pathology Services Lighthouse Laboratory | Wellcome Sanger Institute for the COVID-19 Genomics UK (COG-UK) Consortium | Berkshire and Surrey Pathology Services Lighthouse Laboratory and Alex Alderton, Roberto Amato, Jeffrey Barrett, Sonia Goncalves, Ewan Harrison, David K. Jackson, Ian Johnston, Dominic Kwiatkowski, Cordelia Langford, John Sillitoe on behalf of the Wellcome Sanger Institute COVID-19 Surveillance Team |
| EPI_ISL_3042684, EPI_ISL_3042685, EPI_ISL_3042686 | Lighthouse Lab in Glasgow | Wellcome Sanger Institute for the COVID-19 Genomics UK (COG-UK) Consortium | Harper VanSteenhouse, Yumi Kasai, David Gray, Carol Clugston, Anna Dominiczak and Alex Alderton, Roberto Amato, Jeffrey Barrett, Sonia Goncalves, Ewan Harrison, David K. Jackson, Ian Johnston, Dominic Kwiatkowski, Cordelia Langford, John Sillitoe on behalf of the Wellcome Sanger Institute COVID-19 Surveillance Team |
| EPI_ISL_3042687 | Lighthouse Lab in Alderley Park | Wellcome Sanger Institute for the COVID-19 Genomics UK (COG-UK) Consortium | Jacquelyn Wynn, Mairead Hyland, The Lighthouse Lab in Alderley Park and Alex Alderton, Roberto Amato, Jeffrey Barrett, Sonia Goncalves, Ewan Harrison, David K. Jackson, Ian Johnston, Dominic Kwiatkowski, Cordelia Langford, John Sillitoe on behalf of the Wellcome Sanger Institute COVID-19 Surveillance Team |
| EPI_ISL_3042688, EPI_ISL_3042689, EPI_ISL_3042690 | Lighthouse Lab in Glasgow | Wellcome Sanger Institute for the COVID-19 Genomics UK (COG-UK) Consortium | Harper VanSteenhouse, Yumi Kasai, David Gray, Carol Clugston, Anna Dominiczak and Alex Alderton, Roberto Amato, Jeffrey Barrett, Sonia Goncalves, Ewan Harrison, David K. Jackson, Ian Johnston, Dominic Kwiatkowski, Cordelia Langford, John Sillitoe on behalf of the Wellcome Sanger Institute COVID-19 Surveillance Team |
| EPI_ISL_3042691 | Berkshire and Surrey Pathology Services Lighthouse Laboratory | Wellcome Sanger Institute for the COVID-19 Genomics UK (COG-UK) Consortium | Berkshire and Surrey Pathology Services Lighthouse Laboratory and Alex Alderton, Roberto Amato, Jeffrey Barrett, Sonia Goncalves, Ewan Harrison, David K. Jackson, Ian Johnston, Dominic Kwiatkowski, Cordelia Langford, John Sillitoe on behalf of the Wellcome Sanger Institute COVID-19 Surveillance Team |
| EPI_ISL_3042692 | Lighthouse Lab in Glasgow | Wellcome Sanger Institute for the COVID-19 Genomics UK (COG-UK) Consortium | Harper VanSteenhouse, Yumi Kasai, David Gray, Carol Clugston, Anna Dominiczak and Alex Alderton, Roberto Amato, Jeffrey Barrett, Sonia Goncalves, Ewan Harrison, David K. Jackson, Ian Johnston, Dominic Kwiatkowski, Cordelia Langford, John Sillitoe on behalf of the Wellcome Sanger Institute COVID-19 Surveillance Team |

| | | | |
|---|---|---|---|
| EPI_ISL_3042693 | Lighthouse Lab in Alderley Park | Wellcome Sanger Institute for the COVID-19 Genomics UK (COG-UK) Consortium | Jacquelyn Wynn, Mairead Hyland, The Lighthouse Lab in Alderley Park and Alex Alderton, Roberto Amato, Jeffrey Barrett, Sonia Goncalves, Ewan Harrison, David K. Jackson, Ian Johnston, Dominic Kwiatkowski, Cordelia Langford, John Sillitoe on behalf of the Wellcome Sanger Institute COVID-19 Surveillance Team |
| EPI_ISL_3042694, EPI_ISL_3042695 | Berkshire and Surrey Pathology Services Lighthouse Laboratory | Wellcome Sanger Institute for the COVID-19 Genomics UK (COG-UK) Consortium | Berkshire and Surrey Pathology Services Lighthouse Laboratory and Alex Alderton, Roberto Amato, Jeffrey Barrett, Sonia Goncalves, Ewan Harrison, David K. Jackson, Ian Johnston, Dominic Kwiatkowski, Cordelia Langford, John Sillitoe on behalf of the Wellcome Sanger Institute COVID-19 Surveillance Team |
| EPI_ISL_3042696 | Lighthouse Lab in Alderley Park | Wellcome Sanger Institute for the COVID-19 Genomics UK (COG-UK) Consortium | Jacquelyn Wynn, Mairead Hyland, The Lighthouse Lab in Alderley Park and Alex Alderton, Roberto Amato, Jeffrey Barrett, Sonia Goncalves, Ewan Harrison, David K. Jackson, Ian Johnston, Dominic Kwiatkowski, Cordelia Langford, John Sillitoe on behalf of the Wellcome Sanger Institute COVID-19 Surveillance Team |
| EPI_ISL_3042697, EPI_ISL_3042698 | Lighthouse Lab in Glasgow | Wellcome Sanger Institute for the COVID-19 Genomics UK (COG-UK) Consortium | Harper VanSteenhouse, Yumi Kasai, David Gray, Carol Clugston, Anna Dominiczak and Alex Alderton, Roberto Amato, Jeffrey Barrett, Sonia Goncalves, Ewan Harrison, David K. Jackson, Ian Johnston, Dominic Kwiatkowski, Cordelia Langford, John Sillitoe on behalf of the Wellcome Sanger Institute COVID-19 Surveillance Team |
| EPI_ISL_3042699 | Berkshire and Surrey Pathology Services Lighthouse Laboratory | Wellcome Sanger Institute for the COVID-19 Genomics UK (COG-UK) Consortium | Berkshire and Surrey Pathology Services Lighthouse Laboratory and Alex Alderton, Roberto Amato, Jeffrey Barrett, Sonia Goncalves, Ewan Harrison, David K. Jackson, Ian Johnston, Dominic Kwiatkowski, Cordelia Langford, John Sillitoe on behalf of the Wellcome Sanger Institute COVID-19 Surveillance Team |
| EPI_ISL_3042700 | Lighthouse Lab in Glasgow | Wellcome Sanger Institute for the COVID-19 Genomics UK (COG-UK) Consortium | Harper VanSteenhouse, Yumi Kasai, David Gray, Carol Clugston, Anna Dominiczak and Alex Alderton, Roberto Amato, Jeffrey Barrett, Sonia Goncalves, Ewan Harrison, David K. Jackson, Ian Johnston, Dominic Kwiatkowski, Cordelia Langford, John Sillitoe on behalf of the Wellcome Sanger Institute COVID-19 Surveillance Team |
| EPI_ISL_3042701 | Berkshire and Surrey Pathology Services Lighthouse Laboratory | Wellcome Sanger Institute for the COVID-19 Genomics UK (COG-UK) Consortium | Berkshire and Surrey Pathology Services Lighthouse Laboratory and Alex Alderton, Roberto Amato, Jeffrey Barrett, Sonia Goncalves, Ewan Harrison, David K. Jackson, Ian Johnston, Dominic Kwiatkowski, Cordelia Langford, John Sillitoe on behalf of the Wellcome Sanger Institute COVID-19 Surveillance Team |
| EPI_ISL_3042702, EPI_ISL_3042703 | Lighthouse Lab in Alderley Park | Wellcome Sanger Institute for the COVID-19 Genomics UK (COG-UK) Consortium | Jacquelyn Wynn, Mairead Hyland, The Lighthouse Lab in Alderley Park and Alex Alderton, Roberto Amato, Jeffrey Barrett, Sonia Goncalves, Ewan Harrison, David K. Jackson, Ian Johnston, Dominic Kwiatkowski, Cordelia Langford, John Sillitoe on behalf of the Wellcome Sanger Institute COVID-19 Surveillance Team |
| EPI_ISL_3042704 | Berkshire and Surrey Pathology Services Lighthouse Laboratory | Wellcome Sanger Institute for the COVID-19 Genomics UK (COG-UK) Consortium | Berkshire and Surrey Pathology Services Lighthouse Laboratory and Alex Alderton, Roberto Amato, Jeffrey Barrett, Sonia Goncalves, Ewan Harrison, David K. Jackson, Ian Johnston, Dominic Kwiatkowski, Cordelia Langford, John Sillitoe on behalf of the Wellcome Sanger Institute COVID-19 Surveillance Team |
| EPI_ISL_3042705 | Lighthouse Lab in Glasgow | Wellcome Sanger Institute for the COVID-19 Genomics UK (COG-UK) Consortium | Harper VanSteenhouse, Yumi Kasai, David Gray, Carol Clugston, Anna Dominiczak and Alex Alderton, Roberto Amato, Jeffrey Barrett, Sonia Goncalves, Ewan Harrison, David K. Jackson, Ian Johnston, Dominic Kwiatkowski, Cordelia Langford, John Sillitoe on behalf of the Wellcome Sanger Institute COVID-19 Surveillance Team |
| EPI_ISL_3042706, EPI_ISL_3042707 | Lighthouse Lab in Alderley Park | Wellcome Sanger Institute for the COVID-19 Genomics UK (COG-UK) Consortium | Jacquelyn Wynn, Mairead Hyland, The Lighthouse Lab in Alderley Park and Alex Alderton, Roberto Amato, Jeffrey Barrett, Sonia Goncalves, Ewan Harrison, David K. Jackson, Ian Johnston, Dominic Kwiatkowski, Cordelia Langford, John Sillitoe on behalf of the Wellcome Sanger Institute COVID-19 Surveillance Team |
| EPI_ISL_3042708, EPI_ISL_3042709, EPI_ISL_3042710, EPI_ISL_3042711 | Lighthouse Lab in Glasgow | Wellcome Sanger Institute for the COVID-19 Genomics UK (COG-UK) Consortium | Harper VanSteenhouse, Yumi Kasai, David Gray, Carol Clugston, Anna Dominiczak and Alex Alderton, Roberto Amato, Jeffrey Barrett, Sonia Goncalves, Ewan Harrison, David K. Jackson, Ian Johnston, Dominic Kwiatkowski, Cordelia Langford, John Sillitoe on behalf of the Wellcome Sanger Institute COVID-19 Surveillance Team |
| EPI_ISL_3042712 | Lighthouse Lab in Alderley Park | Wellcome Sanger Institute for the COVID-19 Genomics UK (COG-UK) Consortium | Jacquelyn Wynn, Mairead Hyland, The Lighthouse Lab in Alderley Park and Alex Alderton, Roberto Amato, Jeffrey Barrett, Sonia Goncalves, Ewan Harrison, David K. Jackson, Ian Johnston, Dominic Kwiatkowski, Cordelia Langford, John Sillitoe on behalf of the Wellcome Sanger Institute COVID-19 Surveillance Team |
| EPI_ISL_3042713 | Berkshire and Surrey Pathology Services Lighthouse Laboratory | Wellcome Sanger Institute for the COVID-19 Genomics UK (COG-UK) Consortium | Berkshire and Surrey Pathology Services Lighthouse Laboratory and Alex Alderton, Roberto Amato, Jeffrey Barrett, Sonia Goncalves, Ewan Harrison, David K. Jackson, Ian Johnston, Dominic Kwiatkowski, Cordelia Langford, John Sillitoe on behalf of the Wellcome Sanger Institute COVID-19 Surveillance Team |
| EPI_ISL_3042714 | Lighthouse Lab in Alderley Park | Wellcome Sanger Institute for the COVID-19 Genomics UK (COG-UK) Consortium | Jacquelyn Wynn, Mairead Hyland, The Lighthouse Lab in Alderley Park and Alex Alderton, Roberto Amato, Jeffrey Barrett, Sonia Goncalves, Ewan Harrison, David K. Jackson, Ian Johnston, Dominic Kwiatkowski, Cordelia Langford, John Sillitoe on behalf of the Wellcome Sanger Institute COVID-19 Surveillance Team |
| EPI_ISL_3042715 | Lighthouse Lab in Glasgow | Wellcome Sanger Institute for the COVID-19 Genomics UK (COG-UK) Consortium | Harper VanSteenhouse, Yumi Kasai, David Gray, Carol Clugston, Anna Dominiczak and Alex Alderton, Roberto Amato, Jeffrey Barrett, Sonia Goncalves, Ewan Harrison, David K. Jackson, Ian Johnston, Dominic Kwiatkowski, Cordelia Langford, John Sillitoe on behalf of the Wellcome Sanger Institute COVID-19 Surveillance Team |
| EPI_ISL_3042716, EPI_ISL_3042717 | Berkshire and Surrey Pathology Services Lighthouse Laboratory | Wellcome Sanger Institute for the COVID-19 Genomics UK (COG-UK) Consortium | Berkshire and Surrey Pathology Services Lighthouse Laboratory and Alex Alderton, Roberto Amato, Jeffrey Barrett, Sonia Goncalves, Ewan Harrison, David K. Jackson, Ian Johnston, Dominic Kwiatkowski, Cordelia Langford, John Sillitoe on behalf of the Wellcome Sanger Institute COVID-19 Surveillance Team |
| EPI_ISL_3042718, EPI_ISL_3042719 | Lighthouse Lab in Glasgow | Wellcome Sanger Institute for the COVID-19 Genomics UK (COG-UK) Consortium | Harper VanSteenhouse, Yumi Kasai, David Gray, Carol Clugston, Anna Dominiczak and Alex Alderton, Roberto Amato, Jeffrey Barrett, Sonia Goncalves, Ewan Harrison, David K. Jackson, Ian Johnston, Dominic Kwiatkowski, Cordelia Langford, John Sillitoe on behalf of the Wellcome Sanger Institute COVID-19 Surveillance Team |
| EPI_ISL_3042720, EPI_ISL_3042721 | Berkshire and Surrey Pathology Services Lighthouse Laboratory | Wellcome Sanger Institute for the COVID-19 Genomics UK (COG-UK) Consortium | Berkshire and Surrey Pathology Services Lighthouse Laboratory and Alex Alderton, Roberto Amato, Jeffrey Barrett, Sonia Goncalves, Ewan Harrison, David K. Jackson, Ian Johnston, Dominic Kwiatkowski, Cordelia Langford, John Sillitoe on behalf of the Wellcome Sanger Institute COVID-19 Surveillance Team |
| EPI_ISL_3042722 | Lighthouse Lab in Glasgow | Wellcome Sanger Institute for the COVID-19 Genomics UK (COG-UK) Consortium | Harper VanSteenhouse, Yumi Kasai, David Gray, Carol Clugston, Anna Dominiczak and Alex Alderton, Roberto Amato, Jeffrey Barrett, Sonia Goncalves, Ewan Harrison, David K. Jackson, Ian Johnston, Dominic Kwiatkowski, Cordelia Langford, John Sillitoe on behalf of the Wellcome Sanger Institute COVID-19 Surveillance Team |
| EPI_ISL_3042723 | Berkshire and Surrey Pathology Services Lighthouse Laboratory | Wellcome Sanger Institute for the COVID-19 Genomics UK (COG-UK) Consortium | Berkshire and Surrey Pathology Services Lighthouse Laboratory and Alex Alderton, Roberto Amato, Jeffrey Barrett, Sonia Goncalves, Ewan Harrison, David K. Jackson, Ian Johnston, Dominic Kwiatkowski, Cordelia Langford, John Sillitoe on behalf of the Wellcome Sanger Institute COVID-19 Surveillance Team |
| EPI_ISL_3042724 | Lighthouse Lab in Glasgow | Wellcome Sanger Institute for the COVID-19 Genomics UK (COG-UK) Consortium | Harper VanSteenhouse, Yumi Kasai, David Gray, Carol Clugston, Anna Dominiczak and Alex Alderton, Roberto Amato, Jeffrey Barrett, Sonia Goncalves, Ewan Harrison, David K. Jackson, Ian Johnston, Dominic Kwiatkowski, Cordelia Langford, John Sillitoe on behalf of the Wellcome Sanger Institute COVID-19 Surveillance Team |
| EPI_ISL_3042725 | Lighthouse Lab in Alderley Park | Wellcome Sanger Institute for the COVID-19 Genomics UK (COG-UK) Consortium | Jacquelyn Wynn, Mairead Hyland, The Lighthouse Lab in Alderley Park and Alex Alderton, Roberto Amato, Jeffrey Barrett, Sonia Goncalves, Ewan Harrison, David K. Jackson, Ian Johnston, Dominic Kwiatkowski, Cordelia Langford, John Sillitoe on behalf of the Wellcome Sanger Institute COVID-19 Surveillance Team |
| EPI_ISL_3042726, EPI_ISL_3042727, EPI_ISL_3042728 | Berkshire and Surrey Pathology Services Lighthouse Laboratory | Wellcome Sanger Institute for the COVID-19 Genomics UK (COG-UK) Consortium | Berkshire and Surrey Pathology Services Lighthouse Laboratory and Alex Alderton, Roberto Amato, Jeffrey Barrett, Sonia Goncalves, Ewan Harrison, David K. Jackson, Ian Johnston, Dominic Kwiatkowski, Cordelia Langford, John Sillitoe on behalf of the Wellcome Sanger Institute COVID-19 Surveillance Team |
| EPI_ISL_3042729 | Lighthouse Lab in Glasgow | Wellcome Sanger Institute for the COVID-19 Genomics UK (COG-UK) Consortium | Harper VanSteenhouse, Yumi Kasai, David Gray, Carol Clugston, Anna Dominiczak and Alex Alderton, Roberto Amato, Jeffrey Barrett, Sonia Goncalves, Ewan Harrison, David K. Jackson, Ian Johnston, Dominic Kwiatkowski, Cordelia Langford, John Sillitoe on behalf of the Wellcome Sanger Institute COVID-19 Surveillance Team |
| EPI_ISL_3042730 | Berkshire and Surrey Pathology Services Lighthouse Laboratory | Wellcome Sanger Institute for the COVID-19 Genomics UK (COG-UK) Consortium | Berkshire and Surrey Pathology Services Lighthouse Laboratory and Alex Alderton, Roberto Amato, Jeffrey Barrett, Sonia Goncalves, Ewan Harrison, David K. Jackson, Ian Johnston, Dominic Kwiatkowski, Cordelia Langford, John Sillitoe on behalf of the Wellcome Sanger Institute COVID-19 Surveillance Team |
| EPI_ISL_3042731, EPI_ISL_3042732 | Lighthouse Lab in Alderley Park | Wellcome Sanger Institute for the COVID-19 Genomics UK (COG-UK) Consortium | Jacquelyn Wynn, Mairead Hyland, The Lighthouse Lab in Alderley Park and Alex Alderton, Roberto Amato, Jeffrey Barrett, Sonia Goncalves, Ewan Harrison, David K. Jackson, Ian Johnston, Dominic Kwiatkowski, Cordelia Langford, John Sillitoe on behalf of the Wellcome Sanger Institute COVID-19 Surveillance Team |
| EPI_ISL_3042733, EPI_ISL_3042734, EPI_ISL_3042735 | Lighthouse Lab in Glasgow | Wellcome Sanger Institute for the COVID-19 Genomics UK (COG-UK) Consortium | Harper VanSteenhouse, Yumi Kasai, David Gray, Carol Clugston, Anna Dominiczak and Alex Alderton, Roberto Amato, Jeffrey Barrett, Sonia Goncalves, Ewan Harrison, David K. Jackson, Ian Johnston, Dominic Kwiatkowski, Cordelia Langford, John Sillitoe on behalf of the Wellcome Sanger Institute COVID-19 Surveillance Team |
| EPI_ISL_3042736 | Lighthouse Lab in Alderley Park | Wellcome Sanger Institute for the COVID-19 Genomics UK (COG-UK) Consortium | Jacquelyn Wynn, Mairead Hyland, The Lighthouse Lab in Alderley Park and Alex Alderton, Roberto Amato, Jeffrey Barrett, Sonia Goncalves, Ewan Harrison, David K. Jackson, Ian Johnston, Dominic Kwiatkowski, Cordelia Langford, John Sillitoe on behalf of the Wellcome Sanger Institute COVID-19 Surveillance Team |
| EPI_ISL_3042737, EPI_ISL_3042738 | Berkshire and Surrey Pathology Services Lighthouse Laboratory | Wellcome Sanger Institute for the COVID-19 Genomics UK (COG-UK) Consortium | Berkshire and Surrey Pathology Services Lighthouse Laboratory and Alex Alderton, Roberto Amato, Jeffrey Barrett, Sonia Goncalves, Ewan Harrison, David K. Jackson, Ian Johnston, Dominic Kwiatkowski, Cordelia Langford, John Sillitoe on behalf of the Wellcome Sanger Institute COVID-19 Surveillance Team |
| EPI_ISL_3042739, EPI_ISL_3042740, EPI_ISL_3042741 | Lighthouse Lab in Alderley Park | Wellcome Sanger Institute for the COVID-19 Genomics UK (COG-UK) Consortium | Jacquelyn Wynn, Mairead Hyland, The Lighthouse Lab in Alderley Park and Alex Alderton, Roberto Amato, Jeffrey Barrett, Sonia Goncalves, Ewan Harrison, David K. Jackson, Ian Johnston, Dominic Kwiatkowski, Cordelia Langford, John Sillitoe on behalf of the Wellcome Sanger Institute COVID-19 Surveillance Team |
| EPI_ISL_3042742 | Lighthouse Lab in Glasgow | Wellcome Sanger Institute for the COVID-19 Genomics UK (COG-UK) Consortium | Harper VanSteenhouse, Yumi Kasai, David Gray, Carol Clugston, Anna Dominiczak and Alex Alderton, Roberto Amato, Jeffrey Barrett, Sonia Goncalves, Ewan Harrison, David K. Jackson, Ian Johnston, Dominic Kwiatkowski, Cordelia Langford, John Sillitoe on behalf of the Wellcome Sanger Institute COVID-19 Surveillance Team |
| EPI_ISL_3042743 | Lighthouse Lab in Alderley Park | Wellcome Sanger Institute for the COVID-19 Genomics UK (COG-UK) Consortium | Jacquelyn Wynn, Mairead Hyland, The Lighthouse Lab in Alderley Park and Alex Alderton, Roberto Amato, Jeffrey Barrett, Sonia Goncalves, Ewan Harrison, David K. Jackson, Ian Johnston, Dominic Kwiatkowski, Cordelia Langford, John Sillitoe on behalf of the Wellcome Sanger Institute COVID-19 Surveillance Team |
| EPI_ISL_3042744, EPI_ISL_3042745 | Berkshire and Surrey Pathology Services Lighthouse Laboratory | Wellcome Sanger Institute for the COVID-19 Genomics UK (COG-UK) Consortium | Berkshire and Surrey Pathology Services Lighthouse Laboratory and Alex Alderton, Roberto Amato, Jeffrey Barrett, Sonia Goncalves, Ewan Harrison, David K. Jackson, Ian Johnston, Dominic Kwiatkowski, Cordelia Langford, John Sillitoe on behalf of the Wellcome Sanger Institute COVID-19 Surveillance Team |
| EPI_ISL_3042746, EPI_ISL_3042747, EPI_ISL_3042748, EPI_ISL_3042749, EPI_ISL_3042750, EPI_ISL_3042751 | Lighthouse Lab in Glasgow | Wellcome Sanger Institute for the COVID-19 Genomics UK (COG-UK) Consortium | Harper VanSteenhouse, Yumi Kasai, David Gray, Carol Clugston, Anna Dominiczak and Alex Alderton, Roberto Amato, Jeffrey Barrett, Sonia Goncalves, Ewan Harrison, David K. Jackson, Ian Johnston, Dominic Kwiatkowski, Cordelia Langford, John Sillitoe on behalf of the Wellcome Sanger Institute COVID-19 Surveillance Team |
| EPI_ISL_3042752, EPI_ISL_3042753, EPI_ISL_3042754 | Berkshire and Surrey Pathology Services Lighthouse Laboratory | Wellcome Sanger Institute for the COVID-19 Genomics UK (COG-UK) Consortium | Berkshire and Surrey Pathology Services Lighthouse Laboratory and Alex Alderton, Roberto Amato, Jeffrey Barrett, Sonia Goncalves, Ewan Harrison, David K. Jackson, Ian Johnston, Dominic Kwiatkowski, Cordelia Langford, John Sillitoe on behalf of the Wellcome Sanger Institute COVID-19 Surveillance Team |
| EPI_ISL_3042755, EPI_ISL_3042756, EPI_ISL_3042757 | Lighthouse Lab in Glasgow | Wellcome Sanger Institute for the COVID-19 Genomics UK (COG-UK) Consortium | Harper VanSteenhouse, Yumi Kasai, David Gray, Carol Clugston, Anna Dominiczak and Alex Alderton, Roberto Amato, Jeffrey Barrett, Sonia Goncalves, Ewan Harrison, David K. Jackson, Ian Johnston, Dominic Kwiatkowski, Cordelia Langford, John Sillitoe on behalf of the Wellcome Sanger Institute COVID-19 Surveillance Team |
| EPI_ISL_3042758 | Lighthouse Lab in Alderley Park | Wellcome Sanger Institute for the COVID-19 Genomics UK (COG-UK) Consortium | Jacquelyn Wynn, Mairead Hyland, The Lighthouse Lab in Alderley Park and Alex Alderton, Roberto Amato, Jeffrey Barrett, Sonia Goncalves, Ewan Harrison, David K. Jackson, Ian Johnston, Dominic Kwiatkowski, Cordelia Langford, John Sillitoe on behalf of the Wellcome Sanger Institute COVID-19 Surveillance Team |
| EPI_ISL_3042759, EPI_ISL_3042760 | Berkshire and Surrey Pathology Services Lighthouse Laboratory | Wellcome Sanger Institute for the COVID-19 Genomics UK (COG-UK) Consortium | Berkshire and Surrey Pathology Services Lighthouse Laboratory and Alex Alderton, Roberto Amato, Jeffrey Barrett, Sonia Goncalves, Ewan Harrison, David K. Jackson, Ian Johnston, Dominic Kwiatkowski, Cordelia Langford, John Sillitoe on behalf of the Wellcome Sanger Institute COVID-19 Surveillance Team |
| EPI_ISL_3042761, EPI_ISL_3042762 | Lighthouse Lab in Glasgow | Wellcome Sanger Institute for the COVID-19 Genomics UK (COG-UK) Consortium | Harper VanSteenhouse, Yumi Kasai, David Gray, Carol Clugston, Anna Dominiczak and Alex Alderton, Roberto Amato, Jeffrey Barrett, Sonia Goncalves, Ewan Harrison, David K. Jackson, Ian Johnston, Dominic Kwiatkowski, Cordelia Langford, John Sillitoe on behalf of the Wellcome Sanger Institute COVID-19 Surveillance Team |
| EPI_ISL_3042763 | Berkshire and Surrey Pathology Services Lighthouse Laboratory | Wellcome Sanger Institute for the COVID-19 Genomics UK (COG-UK) | Berkshire and Surrey Pathology Services Lighthouse Laboratory and Alex Alderton, Roberto Amato, Jeffrey Barrett, Sonia Goncalves, Ewan Harrison, David K. Jackson, Ian Johnston, Dominic Kwiatkowski, Cordelia Langford, John Sillitoe on behalf |

| | | | |
|---|---|---|---|
| | | Consortium | of the Wellcome Sanger Institute COVID-19 Surveillance Team |
| EPI_ISL_3042764 | Lighthouse Lab in Glasgow | Wellcome Sanger Institute for the COVID-19 Genomics UK (COG-UK) Consortium | Harper VanSteenhouse, Yumi Kasai, David Gray, Carol Clugston, Anna Dominiczak and Alex Alderton, Roberto Amato, Jeffrey Barrett, Sonia Goncalves, Ewan Harrison, David K. Jackson, Ian Johnston, Dominic Kwiatkowski, Cordelia Langford, John Sillitoe on behalf of the Wellcome Sanger Institute COVID-19 Surveillance Team |
| EPI_ISL_3042765 | Berkshire and Surrey Pathology Services Lighthouse Laboratory | Wellcome Sanger Institute for the COVID-19 Genomics UK (COG-UK) Consortium | Berkshire and Surrey Pathology Services Lighthouse Laboratory and Alex Alderton, Roberto Amato, Jeffrey Barrett, Sonia Goncalves, Ewan Harrison, David K. Jackson, Ian Johnston, Dominic Kwiatkowski, Cordelia Langford, John Sillitoe on behalf of the Wellcome Sanger Institute COVID-19 Surveillance Team |
| EPI_ISL_3042766, EPI_ISL_3042767, EPI_ISL_3042768 | Lighthouse Lab in Glasgow | Wellcome Sanger Institute for the COVID-19 Genomics UK (COG-UK) Consortium | Harper VanSteenhouse, Yumi Kasai, David Gray, Carol Clugston, Anna Dominiczak and Alex Alderton, Roberto Amato, Jeffrey Barrett, Sonia Goncalves, Ewan Harrison, David K. Jackson, Ian Johnston, Dominic Kwiatkowski, Cordelia Langford, John Sillitoe on behalf of the Wellcome Sanger Institute COVID-19 Surveillance Team |
| EPI_ISL_3042769, EPI_ISL_3042770 | Berkshire and Surrey Pathology Services Lighthouse Laboratory | Wellcome Sanger Institute for the COVID-19 Genomics UK (COG-UK) Consortium | Berkshire and Surrey Pathology Services Lighthouse Laboratory and Alex Alderton, Roberto Amato, Jeffrey Barrett, Sonia Goncalves, Ewan Harrison, David K. Jackson, Ian Johnston, Dominic Kwiatkowski, Cordelia Langford, John Sillitoe on behalf of the Wellcome Sanger Institute COVID-19 Surveillance Team |
| EPI_ISL_3042771 | Lighthouse Lab in Glasgow | Wellcome Sanger Institute for the COVID-19 Genomics UK (COG-UK) Consortium | Harper VanSteenhouse, Yumi Kasai, David Gray, Carol Clugston, Anna Dominiczak and Alex Alderton, Roberto Amato, Jeffrey Barrett, Sonia Goncalves, Ewan Harrison, David K. Jackson, Ian Johnston, Dominic Kwiatkowski, Cordelia Langford, John Sillitoe on behalf of the Wellcome Sanger Institute COVID-19 Surveillance Team |
| EPI_ISL_3042772 | Lighthouse Lab in Alderley Park | Wellcome Sanger Institute for the COVID-19 Genomics UK (COG-UK) Consortium | Jacquelyn Wynn, Mairead Hyland, The Lighthouse Lab in Alderley Park and Alex Alderton, Roberto Amato, Jeffrey Barrett, Sonia Goncalves, Ewan Harrison, David K. Jackson, Ian Johnston, Dominic Kwiatkowski, Cordelia Langford, John Sillitoe on behalf of the Wellcome Sanger Institute COVID-19 Surveillance Team |
| EPI_ISL_3042773 | Lighthouse Lab in Glasgow | Wellcome Sanger Institute for the COVID-19 Genomics UK (COG-UK) Consortium | Harper VanSteenhouse, Yumi Kasai, David Gray, Carol Clugston, Anna Dominiczak and Alex Alderton, Roberto Amato, Jeffrey Barrett, Sonia Goncalves, Ewan Harrison, David K. Jackson, Ian Johnston, Dominic Kwiatkowski, Cordelia Langford, John Sillitoe on behalf of the Wellcome Sanger Institute COVID-19 Surveillance Team |
| EPI_ISL_3042774 | Lighthouse Lab in Alderley Park | Wellcome Sanger Institute for the COVID-19 Genomics UK (COG-UK) Consortium | Jacquelyn Wynn, Mairead Hyland, The Lighthouse Lab in Alderley Park and Alex Alderton, Roberto Amato, Jeffrey Barrett, Sonia Goncalves, Ewan Harrison, David K. Jackson, Ian Johnston, Dominic Kwiatkowski, Cordelia Langford, John Sillitoe on behalf of the Wellcome Sanger Institute COVID-19 Surveillance Team |
| EPI_ISL_3042775, EPI_ISL_3042776, EPI_ISL_3042777 | Lighthouse Lab in Glasgow | Wellcome Sanger Institute for the COVID-19 Genomics UK (COG-UK) Consortium | Harper VanSteenhouse, Yumi Kasai, David Gray, Carol Clugston, Anna Dominiczak and Alex Alderton, Roberto Amato, Jeffrey Barrett, Sonia Goncalves, Ewan Harrison, David K. Jackson, Ian Johnston, Dominic Kwiatkowski, Cordelia Langford, John Sillitoe on behalf of the Wellcome Sanger Institute COVID-19 Surveillance Team |
| EPI_ISL_3042778 | Lighthouse Lab in Alderley Park | Wellcome Sanger Institute for the COVID-19 Genomics UK (COG-UK) Consortium | Jacquelyn Wynn, Mairead Hyland, The Lighthouse Lab in Alderley Park and Alex Alderton, Roberto Amato, Jeffrey Barrett, Sonia Goncalves, Ewan Harrison, David K. Jackson, Ian Johnston, Dominic Kwiatkowski, Cordelia Langford, John Sillitoe on behalf of the Wellcome Sanger Institute COVID-19 Surveillance Team |
| EPI_ISL_3042779, EPI_ISL_3042780, EPI_ISL_3042781 | Lighthouse Lab in Glasgow | Wellcome Sanger Institute for the COVID-19 Genomics UK (COG-UK) Consortium | Harper VanSteenhouse, Yumi Kasai, David Gray, Carol Clugston, Anna Dominiczak and Alex Alderton, Roberto Amato, Jeffrey Barrett, Sonia Goncalves, Ewan Harrison, David K. Jackson, Ian Johnston, Dominic Kwiatkowski, Cordelia Langford, John Sillitoe on behalf of the Wellcome Sanger Institute COVID-19 Surveillance Team |
| EPI_ISL_3042782, EPI_ISL_3042783, EPI_ISL_3042784 | Lighthouse Lab in Alderley Park | Wellcome Sanger Institute for the COVID-19 Genomics UK (COG-UK) Consortium | Jacquelyn Wynn, Mairead Hyland, The Lighthouse Lab in Alderley Park and Alex Alderton, Roberto Amato, Jeffrey Barrett, Sonia Goncalves, Ewan Harrison, David K. Jackson, Ian Johnston, Dominic Kwiatkowski, Cordelia Langford, John Sillitoe on behalf of the Wellcome Sanger Institute COVID-19 Surveillance Team |
| EPI_ISL_3042785 | Berkshire and Surrey Pathology Services Lighthouse Laboratory | Wellcome Sanger Institute for the COVID-19 Genomics UK (COG-UK) Consortium | Berkshire and Surrey Pathology Services Lighthouse Laboratory and Alex Alderton, Roberto Amato, Jeffrey Barrett, Sonia Goncalves, Ewan Harrison, David K. Jackson, Ian Johnston, Dominic Kwiatkowski, Cordelia Langford, John Sillitoe on behalf of the Wellcome Sanger Institute COVID-19 Surveillance Team |
| EPI_ISL_3042786 | Lighthouse Lab in Alderley Park | Wellcome Sanger Institute for the COVID-19 Genomics UK (COG-UK) Consortium | Jacquelyn Wynn, Mairead Hyland, The Lighthouse Lab in Alderley Park and Alex Alderton, Roberto Amato, Jeffrey Barrett, Sonia Goncalves, Ewan Harrison, David K. Jackson, Ian Johnston, Dominic Kwiatkowski, Cordelia Langford, John Sillitoe on behalf of the Wellcome Sanger Institute COVID-19 Surveillance Team |
| EPI_ISL_3042787 | Lighthouse Lab in Glasgow | Wellcome Sanger Institute for the COVID-19 Genomics UK (COG-UK) Consortium | Harper VanSteenhouse, Yumi Kasai, David Gray, Carol Clugston, Anna Dominiczak and Alex Alderton, Roberto Amato, Jeffrey Barrett, Sonia Goncalves, Ewan Harrison, David K. Jackson, Ian Johnston, Dominic Kwiatkowski, Cordelia Langford, John Sillitoe on behalf of the Wellcome Sanger Institute COVID-19 Surveillance Team |
| EPI_ISL_3042788, EPI_ISL_3042789 | Berkshire and Surrey Pathology Services Lighthouse Laboratory | Wellcome Sanger Institute for the COVID-19 Genomics UK (COG-UK) Consortium | Berkshire and Surrey Pathology Services Lighthouse Laboratory and Alex Alderton, Roberto Amato, Jeffrey Barrett, Sonia Goncalves, Ewan Harrison, David K. Jackson, Ian Johnston, Dominic Kwiatkowski, Cordelia Langford, John Sillitoe on behalf of the Wellcome Sanger Institute COVID-19 Surveillance Team |
| EPI_ISL_3042790 | Lighthouse Lab in Alderley Park | Wellcome Sanger Institute for the COVID-19 Genomics UK (COG-UK) Consortium | Jacquelyn Wynn, Mairead Hyland, The Lighthouse Lab in Alderley Park and Alex Alderton, Roberto Amato, Jeffrey Barrett, Sonia Goncalves, Ewan Harrison, David K. Jackson, Ian Johnston, Dominic Kwiatkowski, Cordelia Langford, John Sillitoe on behalf of the Wellcome Sanger Institute COVID-19 Surveillance Team |
| EPI_ISL_3042791 | Lighthouse Lab in Glasgow | Wellcome Sanger Institute for the COVID-19 Genomics UK (COG-UK) Consortium | Harper VanSteenhouse, Yumi Kasai, David Gray, Carol Clugston, Anna Dominiczak and Alex Alderton, Roberto Amato, Jeffrey Barrett, Sonia Goncalves, Ewan Harrison, David K. Jackson, Ian Johnston, Dominic Kwiatkowski, Cordelia Langford, John Sillitoe on behalf of the Wellcome Sanger Institute COVID-19 Surveillance Team |
| EPI_ISL_3042792 | Lighthouse Lab in Alderley Park | Wellcome Sanger Institute for the COVID-19 Genomics UK (COG-UK) Consortium | Jacquelyn Wynn, Mairead Hyland, The Lighthouse Lab in Alderley Park and Alex Alderton, Roberto Amato, Jeffrey Barrett, Sonia Goncalves, Ewan Harrison, David K. Jackson, Ian Johnston, Dominic Kwiatkowski, Cordelia Langford, John Sillitoe on behalf of the Wellcome Sanger Institute COVID-19 Surveillance Team |
| EPI_ISL_3042793 | Lighthouse Lab in Glasgow | Wellcome Sanger Institute for the COVID-19 Genomics UK (COG-UK) Consortium | Harper VanSteenhouse, Yumi Kasai, David Gray, Carol Clugston, Anna Dominiczak and Alex Alderton, Roberto Amato, Jeffrey Barrett, Sonia Goncalves, Ewan Harrison, David K. Jackson, Ian Johnston, Dominic Kwiatkowski, Cordelia Langford, John Sillitoe on behalf of the Wellcome Sanger Institute COVID-19 Surveillance Team |
| EPI_ISL_3042794 | Lighthouse Lab in Alderley Park | Wellcome Sanger Institute for the COVID-19 Genomics UK (COG-UK) Consortium | Jacquelyn Wynn, Mairead Hyland, The Lighthouse Lab in Alderley Park and Alex Alderton, Roberto Amato, Jeffrey Barrett, Sonia Goncalves, Ewan Harrison, David K. Jackson, Ian Johnston, Dominic Kwiatkowski, Cordelia Langford, John Sillitoe on behalf of the Wellcome Sanger Institute COVID-19 Surveillance Team |
| EPI_ISL_3042795 | Lighthouse Lab in Glasgow | Wellcome Sanger Institute for the COVID-19 Genomics UK (COG-UK) Consortium | Harper VanSteenhouse, Yumi Kasai, David Gray, Carol Clugston, Anna Dominiczak and Alex Alderton, Roberto Amato, Jeffrey Barrett, Sonia Goncalves, Ewan Harrison, David K. Jackson, Ian Johnston, Dominic Kwiatkowski, Cordelia Langford, John Sillitoe on behalf of the Wellcome Sanger Institute COVID-19 Surveillance Team |
| EPI_ISL_3042796 | Lighthouse Lab in Alderley Park | Wellcome Sanger Institute for the COVID-19 Genomics UK (COG-UK) Consortium | Jacquelyn Wynn, Mairead Hyland, The Lighthouse Lab in Alderley Park and Alex Alderton, Roberto Amato, Jeffrey Barrett, Sonia Goncalves, Ewan Harrison, David K. Jackson, Ian Johnston, Dominic Kwiatkowski, Cordelia Langford, John Sillitoe on behalf of the Wellcome Sanger Institute COVID-19 Surveillance Team |
| EPI_ISL_3042797 | Lighthouse Lab in Glasgow | Wellcome Sanger Institute for the COVID-19 Genomics UK (COG-UK) Consortium | Harper VanSteenhouse, Yumi Kasai, David Gray, Carol Clugston, Anna Dominiczak and Alex Alderton, Roberto Amato, Jeffrey Barrett, Sonia Goncalves, Ewan Harrison, David K. Jackson, Ian Johnston, Dominic Kwiatkowski, Cordelia Langford, John Sillitoe on behalf of the Wellcome Sanger Institute COVID-19 Surveillance Team |
| EPI_ISL_3042798, EPI_ISL_3042799 | Lighthouse Lab in Alderley Park | Wellcome Sanger Institute for the COVID-19 Genomics UK (COG-UK) Consortium | Jacquelyn Wynn, Mairead Hyland, The Lighthouse Lab in Alderley Park and Alex Alderton, Roberto Amato, Jeffrey Barrett, Sonia Goncalves, Ewan Harrison, David K. Jackson, Ian Johnston, Dominic Kwiatkowski, Cordelia Langford, John Sillitoe on behalf of the Wellcome Sanger Institute COVID-19 Surveillance Team |
| EPI_ISL_3042800 | Berkshire and Surrey Pathology Services Lighthouse Laboratory | Wellcome Sanger Institute for the COVID-19 Genomics UK (COG-UK) Consortium | Berkshire and Surrey Pathology Services Lighthouse Laboratory and Alex Alderton, Roberto Amato, Jeffrey Barrett, Sonia Goncalves, Ewan Harrison, David K. Jackson, Ian Johnston, Dominic Kwiatkowski, Cordelia Langford, John Sillitoe on behalf of the Wellcome Sanger Institute COVID-19 Surveillance Team |
| EPI_ISL_3042801 | Lighthouse Lab in Glasgow | Wellcome Sanger Institute for the COVID-19 Genomics UK (COG-UK) Consortium | Harper VanSteenhouse, Yumi Kasai, David Gray, Carol Clugston, Anna Dominiczak and Alex Alderton, Roberto Amato, Jeffrey Barrett, Sonia Goncalves, Ewan Harrison, David K. Jackson, Ian Johnston, Dominic Kwiatkowski, Cordelia Langford, John Sillitoe on behalf of the Wellcome Sanger Institute COVID-19 Surveillance Team |
| EPI_ISL_3042802 | Berkshire and Surrey Pathology Services Lighthouse Laboratory | Wellcome Sanger Institute for the COVID-19 Genomics UK (COG-UK) Consortium | Berkshire and Surrey Pathology Services Lighthouse Laboratory and Alex Alderton, Roberto Amato, Jeffrey Barrett, Sonia Goncalves, Ewan Harrison, David K. Jackson, Ian Johnston, Dominic Kwiatkowski, Cordelia Langford, John Sillitoe on behalf of the Wellcome Sanger Institute COVID-19 Surveillance Team |
| EPI_ISL_3042803, EPI_ISL_3042804 | Lighthouse Lab in Glasgow | Wellcome Sanger Institute for the COVID-19 Genomics UK (COG-UK) Consortium | Harper VanSteenhouse, Yumi Kasai, David Gray, Carol Clugston, Anna Dominiczak and Alex Alderton, Roberto Amato, Jeffrey Barrett, Sonia Goncalves, Ewan Harrison, David K. Jackson, Ian Johnston, Dominic Kwiatkowski, Cordelia Langford, John Sillitoe on behalf of the Wellcome Sanger Institute COVID-19 Surveillance Team |
| EPI_ISL_3042805 | Lighthouse Lab in Alderley Park | Wellcome Sanger Institute for the COVID-19 Genomics UK (COG-UK) Consortium | Jacquelyn Wynn, Mairead Hyland, The Lighthouse Lab in Alderley Park and Alex Alderton, Roberto Amato, Jeffrey Barrett, Sonia Goncalves, Ewan Harrison, David K. Jackson, Ian Johnston, Dominic Kwiatkowski, Cordelia Langford, John Sillitoe on behalf of the Wellcome Sanger Institute COVID-19 Surveillance Team |
| EPI_ISL_3042806 | Berkshire and Surrey Pathology Services Lighthouse Laboratory | Wellcome Sanger Institute for the COVID-19 Genomics UK (COG-UK) Consortium | Berkshire and Surrey Pathology Services Lighthouse Laboratory and Alex Alderton, Roberto Amato, Jeffrey Barrett, Sonia Goncalves, Ewan Harrison, David K. Jackson, Ian Johnston, Dominic Kwiatkowski, Cordelia Langford, John Sillitoe on behalf of the Wellcome Sanger Institute COVID-19 Surveillance Team |
| EPI_ISL_3042807 | Lighthouse Lab in Alderley Park | Wellcome Sanger Institute for the COVID-19 Genomics UK (COG-UK) Consortium | Jacquelyn Wynn, Mairead Hyland, The Lighthouse Lab in Alderley Park and Alex Alderton, Roberto Amato, Jeffrey Barrett, Sonia Goncalves, Ewan Harrison, David K. Jackson, Ian Johnston, Dominic Kwiatkowski, Cordelia Langford, John Sillitoe on behalf of the Wellcome Sanger Institute COVID-19 Surveillance Team |
| EPI_ISL_3042808 | Lighthouse Lab in Glasgow | Wellcome Sanger Institute for the COVID-19 Genomics UK (COG-UK) Consortium | Harper VanSteenhouse, Yumi Kasai, David Gray, Carol Clugston, Anna Dominiczak and Alex Alderton, Roberto Amato, Jeffrey Barrett, Sonia Goncalves, Ewan Harrison, David K. Jackson, Ian Johnston, Dominic Kwiatkowski, Cordelia Langford, John Sillitoe on behalf of the Wellcome Sanger Institute COVID-19 Surveillance Team |
| EPI_ISL_3042809 | Lighthouse Lab in Alderley Park | Wellcome Sanger Institute for the COVID-19 Genomics UK (COG-UK) Consortium | Jacquelyn Wynn, Mairead Hyland, The Lighthouse Lab in Alderley Park and Alex Alderton, Roberto Amato, Jeffrey Barrett, Sonia Goncalves, Ewan Harrison, David K. Jackson, Ian Johnston, Dominic Kwiatkowski, Cordelia Langford, John Sillitoe on behalf of the Wellcome Sanger Institute COVID-19 Surveillance Team |
| EPI_ISL_3042810 | Berkshire and Surrey Pathology Services Lighthouse Laboratory | Wellcome Sanger Institute for the COVID-19 Genomics UK (COG-UK) Consortium | Berkshire and Surrey Pathology Services Lighthouse Laboratory and Alex Alderton, Roberto Amato, Jeffrey Barrett, Sonia Goncalves, Ewan Harrison, David K. Jackson, Ian Johnston, Dominic Kwiatkowski, Cordelia Langford, John Sillitoe on behalf of the Wellcome Sanger Institute COVID-19 Surveillance Team |
| EPI_ISL_3042811 | Lighthouse Lab in Glasgow | Wellcome Sanger Institute for the COVID-19 Genomics UK (COG-UK) Consortium | Harper VanSteenhouse, Yumi Kasai, David Gray, Carol Clugston, Anna Dominiczak and Alex Alderton, Roberto Amato, Jeffrey Barrett, Sonia Goncalves, Ewan Harrison, David K. Jackson, Ian Johnston, Dominic Kwiatkowski, Cordelia Langford, John Sillitoe on behalf of the Wellcome Sanger Institute COVID-19 Surveillance Team |
| EPI_ISL_3042812 | Berkshire and Surrey Pathology Services Lighthouse Laboratory | Wellcome Sanger Institute for the COVID-19 Genomics UK (COG-UK) Consortium | Berkshire and Surrey Pathology Services Lighthouse Laboratory and Alex Alderton, Roberto Amato, Jeffrey Barrett, Sonia Goncalves, Ewan Harrison, David K. Jackson, Ian Johnston, Dominic Kwiatkowski, Cordelia Langford, John Sillitoe on behalf of the Wellcome Sanger Institute COVID-19 Surveillance Team |
| EPI_ISL_3042813 | Lighthouse Lab in Alderley Park | Wellcome Sanger Institute for the COVID-19 Genomics UK (COG-UK) Consortium | Jacquelyn Wynn, Mairead Hyland, The Lighthouse Lab in Alderley Park and Alex Alderton, Roberto Amato, Jeffrey Barrett, Sonia Goncalves, Ewan Harrison, David K. Jackson, Ian Johnston, Dominic Kwiatkowski, Cordelia Langford, John Sillitoe on behalf of the Wellcome Sanger Institute COVID-19 Surveillance Team |
| EPI_ISL_3042814, EPI_ISL_3042815, EPI_ISL_3042816 | Lighthouse Lab in Glasgow | Wellcome Sanger Institute for the COVID-19 Genomics UK (COG-UK) Consortium | Harper VanSteenhouse, Yumi Kasai, David Gray, Carol Clugston, Anna Dominiczak and Alex Alderton, Roberto Amato, Jeffrey Barrett, Sonia Goncalves, Ewan Harrison, David K. Jackson, Ian Johnston, Dominic Kwiatkowski, Cordelia Langford, John Sillitoe on behalf of the Wellcome Sanger Institute COVID-19 Surveillance Team |
| EPI_ISL_3042817 | Lighthouse Lab in Alderley Park | Wellcome Sanger Institute for the COVID-19 Genomics UK (COG-UK) Consortium | Jacquelyn Wynn, Mairead Hyland, The Lighthouse Lab in Alderley Park and Alex Alderton, Roberto Amato, Jeffrey Barrett, Sonia Goncalves, Ewan Harrison, David K. Jackson, Ian Johnston, Dominic Kwiatkowski, Cordelia Langford, John Sillitoe on behalf of the Wellcome Sanger Institute COVID-19 Surveillance Team |
| EPI_ISL_3042818, EPI_ISL_3042819, EPI_ISL_3042820 | Lighthouse Lab in Glasgow | Wellcome Sanger Institute for the COVID-19 Genomics UK (COG-UK) Consortium | Harper VanSteenhouse, Yumi Kasai, David Gray, Carol Clugston, Anna Dominiczak and Alex Alderton, Roberto Amato, Jeffrey Barrett, Sonia Goncalves, Ewan Harrison, David K. Jackson, Ian Johnston, Dominic Kwiatkowski, Cordelia Langford, John Sillitoe on behalf of the Wellcome Sanger Institute COVID-19 Surveillance Team |
| EPI_ISL_3042821 | Lighthouse Lab in Alderley Park | Wellcome Sanger Institute for the COVID-19 Genomics UK (COG-UK) Consortium | Jacquelyn Wynn, Mairead Hyland, The Lighthouse Lab in Alderley Park and Alex Alderton, Roberto Amato, Jeffrey Barrett, Sonia Goncalves, Ewan Harrison, David K. Jackson, Ian Johnston, Dominic Kwiatkowski, Cordelia Langford, John Sillitoe on behalf of the Wellcome Sanger Institute COVID-19 Surveillance Team |

| | | | |
|---|---|---|---|
| EPI_ISL_3042822, EPI_ISL_3042823 | Berkshire and Surrey Pathology Services Lighthouse Laboratory | Wellcome Sanger Institute for the COVID-19 Genomics UK (COG-UK) Consortium | Berkshire and Surrey Pathology Services Lighthouse Laboratory and Alex Alderton, Roberto Amato, Jeffrey Barrett, Sonia Goncalves, Ewan Harrison, David K. Jackson, Ian Johnston, Dominic Kwiatkowski, Cordelia Langford, John Sillitoe on behalf of the Wellcome Sanger Institute COVID-19 Surveillance Team |
| EPI_ISL_3042824 | Lighthouse Lab in Glasgow | Wellcome Sanger Institute for the COVID-19 Genomics UK (COG-UK) Consortium | Harper VanSteenhouse, Yumi Kasai, David Gray, Carol Clugston, Anna Dominiczak and Alex Alderton, Roberto Amato, Jeffrey Barrett, Sonia Goncalves, Ewan Harrison, David K. Jackson, Ian Johnston, Dominic Kwiatkowski, Cordelia Langford, John Sillitoe on behalf of the Wellcome Sanger Institute COVID-19 Surveillance Team |
| EPI_ISL_3042825, EPI_ISL_3042826, EPI_ISL_3042827 | Lighthouse Lab in Alderley Park | Wellcome Sanger Institute for the COVID-19 Genomics UK (COG-UK) Consortium | Jacquelyn Wynn, Mairead Hyland, The Lighthouse Lab in Alderley Park and Alex Alderton, Roberto Amato, Jeffrey Barrett, Sonia Goncalves, Ewan Harrison, David K. Jackson, Ian Johnston, Dominic Kwiatkowski, Cordelia Langford, John Sillitoe on behalf of the Wellcome Sanger Institute COVID-19 Surveillance Team |
| EPI_ISL_3042828, EPI_ISL_3042829, EPI_ISL_3042830, EPI_ISL_3042831, EPI_ISL_3042832, EPI_ISL_3042833 | Lighthouse Lab in Glasgow | Wellcome Sanger Institute for the COVID-19 Genomics UK (COG-UK) Consortium | Harper VanSteenhouse, Yumi Kasai, David Gray, Carol Clugston, Anna Dominiczak and Alex Alderton, Roberto Amato, Jeffrey Barrett, Sonia Goncalves, Ewan Harrison, David K. Jackson, Ian Johnston, Dominic Kwiatkowski, Cordelia Langford, John Sillitoe on behalf of the Wellcome Sanger Institute COVID-19 Surveillance Team |
| EPI_ISL_3042834 | Lighthouse Lab in Alderley Park | Wellcome Sanger Institute for the COVID-19 Genomics UK (COG-UK) Consortium | Jacquelyn Wynn, Mairead Hyland, The Lighthouse Lab in Alderley Park and Alex Alderton, Roberto Amato, Jeffrey Barrett, Sonia Goncalves, Ewan Harrison, David K. Jackson, Ian Johnston, Dominic Kwiatkowski, Cordelia Langford, John Sillitoe on behalf of the Wellcome Sanger Institute COVID-19 Surveillance Team |
| EPI_ISL_3042835 | Lighthouse Lab in Glasgow | Wellcome Sanger Institute for the COVID-19 Genomics UK (COG-UK) Consortium | Harper VanSteenhouse, Yumi Kasai, David Gray, Carol Clugston, Anna Dominiczak and Alex Alderton, Roberto Amato, Jeffrey Barrett, Sonia Goncalves, Ewan Harrison, David K. Jackson, Ian Johnston, Dominic Kwiatkowski, Cordelia Langford, John Sillitoe on behalf of the Wellcome Sanger Institute COVID-19 Surveillance Team |
| EPI_ISL_3042836 | Lighthouse Lab in Alderley Park | Wellcome Sanger Institute for the COVID-19 Genomics UK (COG-UK) Consortium | Jacquelyn Wynn, Mairead Hyland, The Lighthouse Lab in Alderley Park and Alex Alderton, Roberto Amato, Jeffrey Barrett, Sonia Goncalves, Ewan Harrison, David K. Jackson, Ian Johnston, Dominic Kwiatkowski, Cordelia Langford, John Sillitoe on behalf of the Wellcome Sanger Institute COVID-19 Surveillance Team |
| EPI_ISL_3042837 | Berkshire and Surrey Pathology Services Lighthouse Laboratory | Wellcome Sanger Institute for the COVID-19 Genomics UK (COG-UK) Consortium | Berkshire and Surrey Pathology Services Lighthouse Laboratory and Alex Alderton, Roberto Amato, Jeffrey Barrett, Sonia Goncalves, Ewan Harrison, David K. Jackson, Ian Johnston, Dominic Kwiatkowski, Cordelia Langford, John Sillitoe on behalf of the Wellcome Sanger Institute COVID-19 Surveillance Team |
| EPI_ISL_3042838 | Lighthouse Lab in Glasgow | Wellcome Sanger Institute for the COVID-19 Genomics UK (COG-UK) Consortium | Harper VanSteenhouse, Yumi Kasai, David Gray, Carol Clugston, Anna Dominiczak and Alex Alderton, Roberto Amato, Jeffrey Barrett, Sonia Goncalves, Ewan Harrison, David K. Jackson, Ian Johnston, Dominic Kwiatkowski, Cordelia Langford, John Sillitoe on behalf of the Wellcome Sanger Institute COVID-19 Surveillance Team |
| EPI_ISL_3042839 | Lighthouse Lab in Alderley Park | Wellcome Sanger Institute for the COVID-19 Genomics UK (COG-UK) Consortium | Jacquelyn Wynn, Mairead Hyland, The Lighthouse Lab in Alderley Park and Alex Alderton, Roberto Amato, Jeffrey Barrett, Sonia Goncalves, Ewan Harrison, David K. Jackson, Ian Johnston, Dominic Kwiatkowski, Cordelia Langford, John Sillitoe on behalf of the Wellcome Sanger Institute COVID-19 Surveillance Team |
| EPI_ISL_3042840 | Lighthouse Lab in Glasgow | Wellcome Sanger Institute for the COVID-19 Genomics UK (COG-UK) Consortium | Harper VanSteenhouse, Yumi Kasai, David Gray, Carol Clugston, Anna Dominiczak and Alex Alderton, Roberto Amato, Jeffrey Barrett, Sonia Goncalves, Ewan Harrison, David K. Jackson, Ian Johnston, Dominic Kwiatkowski, Cordelia Langford, John Sillitoe on behalf of the Wellcome Sanger Institute COVID-19 Surveillance Team |
| EPI_ISL_3042841, EPI_ISL_3042842 | Lighthouse Lab in Alderley Park | Wellcome Sanger Institute for the COVID-19 Genomics UK (COG-UK) Consortium | Jacquelyn Wynn, Mairead Hyland, The Lighthouse Lab in Alderley Park and Alex Alderton, Roberto Amato, Jeffrey Barrett, Sonia Goncalves, Ewan Harrison, David K. Jackson, Ian Johnston, Dominic Kwiatkowski, Cordelia Langford, John Sillitoe on behalf of the Wellcome Sanger Institute COVID-19 Surveillance Team |
| EPI_ISL_3042843 | Lighthouse Lab in Glasgow | Wellcome Sanger Institute for the COVID-19 Genomics UK (COG-UK) Consortium | Harper VanSteenhouse, Yumi Kasai, David Gray, Carol Clugston, Anna Dominiczak and Alex Alderton, Roberto Amato, Jeffrey Barrett, Sonia Goncalves, Ewan Harrison, David K. Jackson, Ian Johnston, Dominic Kwiatkowski, Cordelia Langford, John Sillitoe on behalf of the Wellcome Sanger Institute COVID-19 Surveillance Team |
| EPI_ISL_3042844, EPI_ISL_3042845 | Lighthouse Lab in Alderley Park | Wellcome Sanger Institute for the COVID-19 Genomics UK (COG-UK) Consortium | Jacquelyn Wynn, Mairead Hyland, The Lighthouse Lab in Alderley Park and Alex Alderton, Roberto Amato, Jeffrey Barrett, Sonia Goncalves, Ewan Harrison, David K. Jackson, Ian Johnston, Dominic Kwiatkowski, Cordelia Langford, John Sillitoe on behalf of the Wellcome Sanger Institute COVID-19 Surveillance Team |
| EPI_ISL_3042846, EPI_ISL_3042847 | Lighthouse Lab in Glasgow | Wellcome Sanger Institute for the COVID-19 Genomics UK (COG-UK) Consortium | Harper VanSteenhouse, Yumi Kasai, David Gray, Carol Clugston, Anna Dominiczak and Alex Alderton, Roberto Amato, Jeffrey Barrett, Sonia Goncalves, Ewan Harrison, David K. Jackson, Ian Johnston, Dominic Kwiatkowski, Cordelia Langford, John Sillitoe on behalf of the Wellcome Sanger Institute COVID-19 Surveillance Team |
| EPI_ISL_3042848 | Lighthouse Lab in Alderley Park | Wellcome Sanger Institute for the COVID-19 Genomics UK (COG-UK) Consortium | Jacquelyn Wynn, Mairead Hyland, The Lighthouse Lab in Alderley Park and Alex Alderton, Roberto Amato, Jeffrey Barrett, Sonia Goncalves, Ewan Harrison, David K. Jackson, Ian Johnston, Dominic Kwiatkowski, Cordelia Langford, John Sillitoe on behalf of the Wellcome Sanger Institute COVID-19 Surveillance Team |
| EPI_ISL_3042849, EPI_ISL_3042850 | Lighthouse Lab in Glasgow | Wellcome Sanger Institute for the COVID-19 Genomics UK (COG-UK) Consortium | Harper VanSteenhouse, Yumi Kasai, David Gray, Carol Clugston, Anna Dominiczak and Alex Alderton, Roberto Amato, Jeffrey Barrett, Sonia Goncalves, Ewan Harrison, David K. Jackson, Ian Johnston, Dominic Kwiatkowski, Cordelia Langford, John Sillitoe on behalf of the Wellcome Sanger Institute COVID-19 Surveillance Team |
| EPI_ISL_3042851 | Berkshire and Surrey Pathology Services Lighthouse Laboratory | Wellcome Sanger Institute for the COVID-19 Genomics UK (COG-UK) Consortium | Berkshire and Surrey Pathology Services Lighthouse Laboratory and Alex Alderton, Roberto Amato, Jeffrey Barrett, Sonia Goncalves, Ewan Harrison, David K. Jackson, Ian Johnston, Dominic Kwiatkowski, Cordelia Langford, John Sillitoe on behalf of the Wellcome Sanger Institute COVID-19 Surveillance Team |
| EPI_ISL_3042852, EPI_ISL_3042853, EPI_ISL_3042854 | Lighthouse Lab in Glasgow | Wellcome Sanger Institute for the COVID-19 Genomics UK (COG-UK) Consortium | Harper VanSteenhouse, Yumi Kasai, David Gray, Carol Clugston, Anna Dominiczak and Alex Alderton, Roberto Amato, Jeffrey Barrett, Sonia Goncalves, Ewan Harrison, David K. Jackson, Ian Johnston, Dominic Kwiatkowski, Cordelia Langford, John Sillitoe on behalf of the Wellcome Sanger Institute COVID-19 Surveillance Team |
| EPI_ISL_3042855 | Berkshire and Surrey Pathology Services Lighthouse Laboratory | Wellcome Sanger Institute for the COVID-19 Genomics UK (COG-UK) Consortium | Berkshire and Surrey Pathology Services Lighthouse Laboratory and Alex Alderton, Roberto Amato, Jeffrey Barrett, Sonia Goncalves, Ewan Harrison, David K. Jackson, Ian Johnston, Dominic Kwiatkowski, Cordelia Langford, John Sillitoe on behalf of the Wellcome Sanger Institute COVID-19 Surveillance Team |
| EPI_ISL_3042856 | Lighthouse Lab in Glasgow | Wellcome Sanger Institute for the COVID-19 Genomics UK (COG-UK) Consortium | Harper VanSteenhouse, Yumi Kasai, David Gray, Carol Clugston, Anna Dominiczak and Alex Alderton, Roberto Amato, Jeffrey Barrett, Sonia Goncalves, Ewan Harrison, David K. Jackson, Ian Johnston, Dominic Kwiatkowski, Cordelia Langford, John Sillitoe on behalf of the Wellcome Sanger Institute COVID-19 Surveillance Team |
| EPI_ISL_3042857 | Lighthouse Lab in Alderley Park | Wellcome Sanger Institute for the COVID-19 Genomics UK (COG-UK) Consortium | Jacquelyn Wynn, Mairead Hyland, The Lighthouse Lab in Alderley Park and Alex Alderton, Roberto Amato, Jeffrey Barrett, Sonia Goncalves, Ewan Harrison, David K. Jackson, Ian Johnston, Dominic Kwiatkowski, Cordelia Langford, John Sillitoe on behalf of the Wellcome Sanger Institute COVID-19 Surveillance Team |
| EPI_ISL_3042858 | Lighthouse Lab in Glasgow | Wellcome Sanger Institute for the COVID-19 Genomics UK (COG-UK) Consortium | Harper VanSteenhouse, Yumi Kasai, David Gray, Carol Clugston, Anna Dominiczak and Alex Alderton, Roberto Amato, Jeffrey Barrett, Sonia Goncalves, Ewan Harrison, David K. Jackson, Ian Johnston, Dominic Kwiatkowski, Cordelia Langford, John Sillitoe on behalf of the Wellcome Sanger Institute COVID-19 Surveillance Team |
| EPI_ISL_3042859, EPI_ISL_3042860 | Berkshire and Surrey Pathology Services Lighthouse Laboratory | Wellcome Sanger Institute for the COVID-19 Genomics UK (COG-UK) Consortium | Berkshire and Surrey Pathology Services Lighthouse Laboratory and Alex Alderton, Roberto Amato, Jeffrey Barrett, Sonia Goncalves, Ewan Harrison, David K. Jackson, Ian Johnston, Dominic Kwiatkowski, Cordelia Langford, John Sillitoe on behalf of the Wellcome Sanger Institute COVID-19 Surveillance Team |
| EPI_ISL_3042861, EPI_ISL_3042862 | Lighthouse Lab in Alderley Park | Wellcome Sanger Institute for the COVID-19 Genomics UK (COG-UK) Consortium | Jacquelyn Wynn, Mairead Hyland, The Lighthouse Lab in Alderley Park and Alex Alderton, Roberto Amato, Jeffrey Barrett, Sonia Goncalves, Ewan Harrison, David K. Jackson, Ian Johnston, Dominic Kwiatkowski, Cordelia Langford, John Sillitoe on behalf of the Wellcome Sanger Institute COVID-19 Surveillance Team |
| EPI_ISL_3042863 | Berkshire and Surrey Pathology Services Lighthouse Laboratory | Wellcome Sanger Institute for the COVID-19 Genomics UK (COG-UK) Consortium | Berkshire and Surrey Pathology Services Lighthouse Laboratory and Alex Alderton, Roberto Amato, Jeffrey Barrett, Sonia Goncalves, Ewan Harrison, David K. Jackson, Ian Johnston, Dominic Kwiatkowski, Cordelia Langford, John Sillitoe on behalf of the Wellcome Sanger Institute COVID-19 Surveillance Team |
| EPI_ISL_3042864, EPI_ISL_3042865, EPI_ISL_3042866 | Lighthouse Lab in Glasgow | Wellcome Sanger Institute for the COVID-19 Genomics UK (COG-UK) Consortium | Harper VanSteenhouse, Yumi Kasai, David Gray, Carol Clugston, Anna Dominiczak and Alex Alderton, Roberto Amato, Jeffrey Barrett, Sonia Goncalves, Ewan Harrison, David K. Jackson, Ian Johnston, Dominic Kwiatkowski, Cordelia Langford, John Sillitoe on behalf of the Wellcome Sanger Institute COVID-19 Surveillance Team |
| EPI_ISL_3042867 | Berkshire and Surrey Pathology Services Lighthouse Laboratory | Wellcome Sanger Institute for the COVID-19 Genomics UK (COG-UK) Consortium | Berkshire and Surrey Pathology Services Lighthouse Laboratory and Alex Alderton, Roberto Amato, Jeffrey Barrett, Sonia Goncalves, Ewan Harrison, David K. Jackson, Ian Johnston, Dominic Kwiatkowski, Cordelia Langford, John Sillitoe on behalf of the Wellcome Sanger Institute COVID-19 Surveillance Team |
| EPI_ISL_3042868, EPI_ISL_3042869, EPI_ISL_3042870 | Lighthouse Lab in Glasgow | Wellcome Sanger Institute for the COVID-19 Genomics UK (COG-UK) Consortium | Harper VanSteenhouse, Yumi Kasai, David Gray, Carol Clugston, Anna Dominiczak and Alex Alderton, Roberto Amato, Jeffrey Barrett, Sonia Goncalves, Ewan Harrison, David K. Jackson, Ian Johnston, Dominic Kwiatkowski, Cordelia Langford, John Sillitoe on behalf of the Wellcome Sanger Institute COVID-19 Surveillance Team |
| EPI_ISL_3042871, EPI_ISL_3042872 | Lighthouse Lab in Alderley Park | Wellcome Sanger Institute for the COVID-19 Genomics UK (COG-UK) Consortium | Jacquelyn Wynn, Mairead Hyland, The Lighthouse Lab in Alderley Park and Alex Alderton, Roberto Amato, Jeffrey Barrett, Sonia Goncalves, Ewan Harrison, David K. Jackson, Ian Johnston, Dominic Kwiatkowski, Cordelia Langford, John Sillitoe on behalf of the Wellcome Sanger Institute COVID-19 Surveillance Team |
| EPI_ISL_3042873, EPI_ISL_3042874 | Lighthouse Lab in Glasgow | Wellcome Sanger Institute for the COVID-19 Genomics UK (COG-UK) Consortium | Harper VanSteenhouse, Yumi Kasai, David Gray, Carol Clugston, Anna Dominiczak and Alex Alderton, Roberto Amato, Jeffrey Barrett, Sonia Goncalves, Ewan Harrison, David K. Jackson, Ian Johnston, Dominic Kwiatkowski, Cordelia Langford, John Sillitoe on behalf of the Wellcome Sanger Institute COVID-19 Surveillance Team |
| EPI_ISL_3042875 | Lighthouse Lab in Alderley Park | Wellcome Sanger Institute for the COVID-19 Genomics UK (COG-UK) Consortium | Jacquelyn Wynn, Mairead Hyland, The Lighthouse Lab in Alderley Park and Alex Alderton, Roberto Amato, Jeffrey Barrett, Sonia Goncalves, Ewan Harrison, David K. Jackson, Ian Johnston, Dominic Kwiatkowski, Cordelia Langford, John Sillitoe on behalf of the Wellcome Sanger Institute COVID-19 Surveillance Team |
| EPI_ISL_3042876 | Lighthouse Lab in Glasgow | Wellcome Sanger Institute for the COVID-19 Genomics UK (COG-UK) Consortium | Harper VanSteenhouse, Yumi Kasai, David Gray, Carol Clugston, Anna Dominiczak and Alex Alderton, Roberto Amato, Jeffrey Barrett, Sonia Goncalves, Ewan Harrison, David K. Jackson, Ian Johnston, Dominic Kwiatkowski, Cordelia Langford, John Sillitoe on behalf of the Wellcome Sanger Institute COVID-19 Surveillance Team |
| EPI_ISL_3042877, EPI_ISL_3042878 | Berkshire and Surrey Pathology Services Lighthouse Laboratory | Wellcome Sanger Institute for the COVID-19 Genomics UK (COG-UK) Consortium | Berkshire and Surrey Pathology Services Lighthouse Laboratory and Alex Alderton, Roberto Amato, Jeffrey Barrett, Sonia Goncalves, Ewan Harrison, David K. Jackson, Ian Johnston, Dominic Kwiatkowski, Cordelia Langford, John Sillitoe on behalf of the Wellcome Sanger Institute COVID-19 Surveillance Team |
| EPI_ISL_3042879, EPI_ISL_3042880 | Lighthouse Lab in Glasgow | Wellcome Sanger Institute for the COVID-19 Genomics UK (COG-UK) Consortium | Harper VanSteenhouse, Yumi Kasai, David Gray, Carol Clugston, Anna Dominiczak and Alex Alderton, Roberto Amato, Jeffrey Barrett, Sonia Goncalves, Ewan Harrison, David K. Jackson, Ian Johnston, Dominic Kwiatkowski, Cordelia Langford, John Sillitoe on behalf of the Wellcome Sanger Institute COVID-19 Surveillance Team |
| EPI_ISL_3042881 | Lighthouse Lab in Alderley Park | Wellcome Sanger Institute for the COVID-19 Genomics UK (COG-UK) Consortium | Jacquelyn Wynn, Mairead Hyland, The Lighthouse Lab in Alderley Park and Alex Alderton, Roberto Amato, Jeffrey Barrett, Sonia Goncalves, Ewan Harrison, David K. Jackson, Ian Johnston, Dominic Kwiatkowski, Cordelia Langford, John Sillitoe on behalf of the Wellcome Sanger Institute COVID-19 Surveillance Team |
| EPI_ISL_3042882, EPI_ISL_3042883 | Lighthouse Lab in Glasgow | Wellcome Sanger Institute for the COVID-19 Genomics UK (COG-UK) Consortium | Harper VanSteenhouse, Yumi Kasai, David Gray, Carol Clugston, Anna Dominiczak and Alex Alderton, Roberto Amato, Jeffrey Barrett, Sonia Goncalves, Ewan Harrison, David K. Jackson, Ian Johnston, Dominic Kwiatkowski, Cordelia Langford, John Sillitoe on behalf of the Wellcome Sanger Institute COVID-19 Surveillance Team |
| EPI_ISL_3042884 | Berkshire and Surrey Pathology Services Lighthouse Laboratory | Wellcome Sanger Institute for the COVID-19 Genomics UK (COG-UK) Consortium | Berkshire and Surrey Pathology Services Lighthouse Laboratory and Alex Alderton, Roberto Amato, Jeffrey Barrett, Sonia Goncalves, Ewan Harrison, David K. Jackson, Ian Johnston, Dominic Kwiatkowski, Cordelia Langford, John Sillitoe on behalf of the Wellcome Sanger Institute COVID-19 Surveillance Team |
| EPI_ISL_3042885 | Lighthouse Lab in Alderley Park | Wellcome Sanger Institute for the COVID-19 Genomics UK (COG-UK) Consortium | Jacquelyn Wynn, Mairead Hyland, The Lighthouse Lab in Alderley Park and Alex Alderton, Roberto Amato, Jeffrey Barrett, Sonia Goncalves, Ewan Harrison, David K. Jackson, Ian Johnston, Dominic Kwiatkowski, Cordelia Langford, John Sillitoe on behalf of the Wellcome Sanger Institute COVID-19 Surveillance Team |
| EPI_ISL_3042886, EPI_ISL_3042887 | Berkshire and Surrey Pathology Services Lighthouse Laboratory | Wellcome Sanger Institute for the COVID-19 Genomics UK (COG-UK) Consortium | Berkshire and Surrey Pathology Services Lighthouse Laboratory and Alex Alderton, Roberto Amato, Jeffrey Barrett, Sonia Goncalves, Ewan Harrison, David K. Jackson, Ian Johnston, Dominic Kwiatkowski, Cordelia Langford, John Sillitoe on behalf of the Wellcome Sanger Institute COVID-19 Surveillance Team |
| EPI_ISL_3042888 | Lighthouse Lab in Glasgow | Wellcome Sanger Institute for the COVID-19 Genomics UK (COG-UK) Consortium | Harper VanSteenhouse, Yumi Kasai, David Gray, Carol Clugston, Anna Dominiczak and Alex Alderton, Roberto Amato, Jeffrey Barrett, Sonia Goncalves, Ewan Harrison, David K. Jackson, Ian Johnston, Dominic Kwiatkowski, Cordelia Langford, John Sillitoe on behalf of the Wellcome Sanger Institute COVID-19 Surveillance Team |
| EPI_ISL_3042889 | Lighthouse Lab in Alderley Park | Wellcome Sanger Institute for the COVID-19 Genomics UK (COG-UK) Consortium | Jacquelyn Wynn, Mairead Hyland, The Lighthouse Lab in Alderley Park and Alex Alderton, Roberto Amato, Jeffrey Barrett, Sonia Goncalves, Ewan Harrison, David K. Jackson, Ian Johnston, Dominic Kwiatkowski, Cordelia Langford, John Sillitoe on behalf of the Wellcome Sanger Institute COVID-19 Surveillance Team |

| | | | |
|---|---|---|---|
| EPI_ISL_3042890, EPI_ISL_3042891 | Lighthouse Lab in Glasgow | Wellcome Sanger Institute for the COVID-19 Genomics UK (COG-UK) Consortium | Harper VanSteenhouse, Yumi Kasai, David Gray, Carol Clugston, Anna Dominiczak and Alex Alderton, Roberto Amato, Jeffrey Barrett, Sonia Goncalves, Ewan Harrison, David K. Jackson, Ian Johnston, Dominic Kwiatkowski, Cordelia Langford, John Sillitoe on behalf of the Wellcome Sanger Institute COVID-19 Surveillance Team |
| EPI_ISL_3042892 | Lighthouse Lab in Alderley Park | Wellcome Sanger Institute for the COVID-19 Genomics UK (COG-UK) Consortium | Jacquelyn Wynn, Mairead Hyland, The Lighthouse Lab in Alderley Park and Alex Alderton, Roberto Amato, Jeffrey Barrett, Sonia Goncalves, Ewan Harrison, David K. Jackson, Ian Johnston, Dominic Kwiatkowski, Cordelia Langford, John Sillitoe on behalf of the Wellcome Sanger Institute COVID-19 Surveillance Team |
| EPI_ISL_3042893, EPI_ISL_3042894 | Berkshire and Surrey Pathology Services Lighthouse Laboratory | Wellcome Sanger Institute for the COVID-19 Genomics UK (COG-UK) Consortium | Berkshire and Surrey Pathology Services Lighthouse Laboratory and Alex Alderton, Roberto Amato, Jeffrey Barrett, Sonia Goncalves, Ewan Harrison, David K. Jackson, Ian Johnston, Dominic Kwiatkowski, Cordelia Langford, John Sillitoe on behalf of the Wellcome Sanger Institute COVID-19 Surveillance Team |
| EPI_ISL_3042895 | Lighthouse Lab in Glasgow | Wellcome Sanger Institute for the COVID-19 Genomics UK (COG-UK) Consortium | Harper VanSteenhouse, Yumi Kasai, David Gray, Carol Clugston, Anna Dominiczak and Alex Alderton, Roberto Amato, Jeffrey Barrett, Sonia Goncalves, Ewan Harrison, David K. Jackson, Ian Johnston, Dominic Kwiatkowski, Cordelia Langford, John Sillitoe on behalf of the Wellcome Sanger Institute COVID-19 Surveillance Team |
| EPI_ISL_3042896 | Lighthouse Lab in Alderley Park | Wellcome Sanger Institute for the COVID-19 Genomics UK (COG-UK) Consortium | Jacquelyn Wynn, Mairead Hyland, The Lighthouse Lab in Alderley Park and Alex Alderton, Roberto Amato, Jeffrey Barrett, Sonia Goncalves, Ewan Harrison, David K. Jackson, Ian Johnston, Dominic Kwiatkowski, Cordelia Langford, John Sillitoe on behalf of the Wellcome Sanger Institute COVID-19 Surveillance Team |
| EPI_ISL_3042897, EPI_ISL_3042898 | Lighthouse Lab in Glasgow | Wellcome Sanger Institute for the COVID-19 Genomics UK (COG-UK) Consortium | Harper VanSteenhouse, Yumi Kasai, David Gray, Carol Clugston, Anna Dominiczak and Alex Alderton, Roberto Amato, Jeffrey Barrett, Sonia Goncalves, Ewan Harrison, David K. Jackson, Ian Johnston, Dominic Kwiatkowski, Cordelia Langford, John Sillitoe on behalf of the Wellcome Sanger Institute COVID-19 Surveillance Team |
| EPI_ISL_3042899 | Lighthouse Lab in Alderley Park | Wellcome Sanger Institute for the COVID-19 Genomics UK (COG-UK) Consortium | Jacquelyn Wynn, Mairead Hyland, The Lighthouse Lab in Alderley Park and Alex Alderton, Roberto Amato, Jeffrey Barrett, Sonia Goncalves, Ewan Harrison, David K. Jackson, Ian Johnston, Dominic Kwiatkowski, Cordelia Langford, John Sillitoe on behalf of the Wellcome Sanger Institute COVID-19 Surveillance Team |
| EPI_ISL_3042900 | Berkshire and Surrey Pathology Services Lighthouse Laboratory | Wellcome Sanger Institute for the COVID-19 Genomics UK (COG-UK) Consortium | Berkshire and Surrey Pathology Services Lighthouse Laboratory and Alex Alderton, Roberto Amato, Jeffrey Barrett, Sonia Goncalves, Ewan Harrison, David K. Jackson, Ian Johnston, Dominic Kwiatkowski, Cordelia Langford, John Sillitoe on behalf of the Wellcome Sanger Institute COVID-19 Surveillance Team |
| EPI_ISL_3042901 | Lighthouse Lab in Glasgow | Wellcome Sanger Institute for the COVID-19 Genomics UK (COG-UK) Consortium | Harper VanSteenhouse, Yumi Kasai, David Gray, Carol Clugston, Anna Dominiczak and Alex Alderton, Roberto Amato, Jeffrey Barrett, Sonia Goncalves, Ewan Harrison, David K. Jackson, Ian Johnston, Dominic Kwiatkowski, Cordelia Langford, John Sillitoe on behalf of the Wellcome Sanger Institute COVID-19 Surveillance Team |
| EPI_ISL_3042902, EPI_ISL_3042903 | Lighthouse Lab in Alderley Park | Wellcome Sanger Institute for the COVID-19 Genomics UK (COG-UK) Consortium | Jacquelyn Wynn, Mairead Hyland, The Lighthouse Lab in Alderley Park and Alex Alderton, Roberto Amato, Jeffrey Barrett, Sonia Goncalves, Ewan Harrison, David K. Jackson, Ian Johnston, Dominic Kwiatkowski, Cordelia Langford, John Sillitoe on behalf of the Wellcome Sanger Institute COVID-19 Surveillance Team |
| EPI_ISL_3042904 | Lighthouse Lab in Glasgow | Wellcome Sanger Institute for the COVID-19 Genomics UK (COG-UK) Consortium | Harper VanSteenhouse, Yumi Kasai, David Gray, Carol Clugston, Anna Dominiczak and Alex Alderton, Roberto Amato, Jeffrey Barrett, Sonia Goncalves, Ewan Harrison, David K. Jackson, Ian Johnston, Dominic Kwiatkowski, Cordelia Langford, John Sillitoe on behalf of the Wellcome Sanger Institute COVID-19 Surveillance Team |
| EPI_ISL_3042905 | Lighthouse Lab in Alderley Park | Wellcome Sanger Institute for the COVID-19 Genomics UK (COG-UK) Consortium | Jacquelyn Wynn, Mairead Hyland, The Lighthouse Lab in Alderley Park and Alex Alderton, Roberto Amato, Jeffrey Barrett, Sonia Goncalves, Ewan Harrison, David K. Jackson, Ian Johnston, Dominic Kwiatkowski, Cordelia Langford, John Sillitoe on behalf of the Wellcome Sanger Institute COVID-19 Surveillance Team |
| EPI_ISL_3042906 | Berkshire and Surrey Pathology Services Lighthouse Laboratory | Wellcome Sanger Institute for the COVID-19 Genomics UK (COG-UK) Consortium | Berkshire and Surrey Pathology Services Lighthouse Laboratory and Alex Alderton, Roberto Amato, Jeffrey Barrett, Sonia Goncalves, Ewan Harrison, David K. Jackson, Ian Johnston, Dominic Kwiatkowski, Cordelia Langford, John Sillitoe on behalf of the Wellcome Sanger Institute COVID-19 Surveillance Team |
| EPI_ISL_3042907 | Lighthouse Lab in Alderley Park | Wellcome Sanger Institute for the COVID-19 Genomics UK (COG-UK) Consortium | Jacquelyn Wynn, Mairead Hyland, The Lighthouse Lab in Alderley Park and Alex Alderton, Roberto Amato, Jeffrey Barrett, Sonia Goncalves, Ewan Harrison, David K. Jackson, Ian Johnston, Dominic Kwiatkowski, Cordelia Langford, John Sillitoe on behalf of the Wellcome Sanger Institute COVID-19 Surveillance Team |
| EPI_ISL_3042908 | Lighthouse Lab in Glasgow | Wellcome Sanger Institute for the COVID-19 Genomics UK (COG-UK) Consortium | Harper VanSteenhouse, Yumi Kasai, David Gray, Carol Clugston, Anna Dominiczak and Alex Alderton, Roberto Amato, Jeffrey Barrett, Sonia Goncalves, Ewan Harrison, David K. Jackson, Ian Johnston, Dominic Kwiatkowski, Cordelia Langford, John Sillitoe on behalf of the Wellcome Sanger Institute COVID-19 Surveillance Team |
| EPI_ISL_3042909, EPI_ISL_3042910 | Lighthouse Lab in Alderley Park | Wellcome Sanger Institute for the COVID-19 Genomics UK (COG-UK) Consortium | Jacquelyn Wynn, Mairead Hyland, The Lighthouse Lab in Alderley Park and Alex Alderton, Roberto Amato, Jeffrey Barrett, Sonia Goncalves, Ewan Harrison, David K. Jackson, Ian Johnston, Dominic Kwiatkowski, Cordelia Langford, John Sillitoe on behalf of the Wellcome Sanger Institute COVID-19 Surveillance Team |
| EPI_ISL_3042911 | Lighthouse Lab in Milton Keynes | Wellcome Sanger Institute for the COVID-19 Genomics UK (COG-UK) Consortium | The Lighthouse Lab in Milton Keynes and Alex Alderton, Roberto Amato, Jeffrey Barrett, Sonia Goncalves, Ewan Harrison, David K. Jackson, Ian Johnston, Dominic Kwiatkowski, Cordelia Langford, John Sillitoe on behalf of the Wellcome Sanger Institute COVID-19 Surveillance Team |
| EPI_ISL_3042912 | Randox Laboratories | Wellcome Sanger Institute for the COVID-19 Genomics UK (COG-UK) Consortium | Randox Laboratories and Alex Alderton, Roberto Amato, Jeffrey Barrett, Sonia Goncalves, Ewan Harrison, David K. Jackson, Ian Johnston, Dominic Kwiatkowski, Cordelia Langford, John Sillitoe on behalf of the Wellcome Sanger Institute COVID-19 Surveillance Team |
| EPI_ISL_3042913 | Lighthouse Lab in Milton Keynes | Wellcome Sanger Institute for the COVID-19 Genomics UK (COG-UK) Consortium | The Lighthouse Lab in Milton Keynes and Alex Alderton, Roberto Amato, Jeffrey Barrett, Sonia Goncalves, Ewan Harrison, David K. Jackson, Ian Johnston, Dominic Kwiatkowski, Cordelia Langford, John Sillitoe on behalf of the Wellcome Sanger Institute COVID-19 Surveillance Team |
| EPI_ISL_3042914, EPI_ISL_3042915 | Randox Laboratories | Wellcome Sanger Institute for the COVID-19 Genomics UK (COG-UK) Consortium | Randox Laboratories and Alex Alderton, Roberto Amato, Jeffrey Barrett, Sonia Goncalves, Ewan Harrison, David K. Jackson, Ian Johnston, Dominic Kwiatkowski, Cordelia Langford, John Sillitoe on behalf of the Wellcome Sanger Institute COVID-19 Surveillance Team |
| EPI_ISL_3042916, EPI_ISL_3042917 | Lighthouse Lab in Milton Keynes | Wellcome Sanger Institute for the COVID-19 Genomics UK (COG-UK) Consortium | The Lighthouse Lab in Milton Keynes and Alex Alderton, Roberto Amato, Jeffrey Barrett, Sonia Goncalves, Ewan Harrison, David K. Jackson, Ian Johnston, Dominic Kwiatkowski, Cordelia Langford, John Sillitoe on behalf of the Wellcome Sanger Institute COVID-19 Surveillance Team |
| EPI_ISL_3042918, EPI_ISL_3042919 | Lighthouse Lab in Alderley Park | Wellcome Sanger Institute for the COVID-19 Genomics UK (COG-UK) Consortium | Jacquelyn Wynn, Mairead Hyland, The Lighthouse Lab in Alderley Park and Alex Alderton, Roberto Amato, Jeffrey Barrett, Sonia Goncalves, Ewan Harrison, David K. Jackson, Ian Johnston, Dominic Kwiatkowski, Cordelia Langford, John Sillitoe on behalf of the Wellcome Sanger Institute COVID-19 Surveillance Team |
| EPI_ISL_3042920 | Lighthouse Lab in Glasgow | Wellcome Sanger Institute for the COVID-19 Genomics UK (COG-UK) Consortium | Harper VanSteenhouse, Yumi Kasai, David Gray, Carol Clugston, Anna Dominiczak and Alex Alderton, Roberto Amato, Jeffrey Barrett, Sonia Goncalves, Ewan Harrison, David K. Jackson, Ian Johnston, Dominic Kwiatkowski, Cordelia Langford, John Sillitoe on behalf of the Wellcome Sanger Institute COVID-19 Surveillance Team |
| EPI_ISL_3042921 | Randox Laboratories | Wellcome Sanger Institute for the COVID-19 Genomics UK (COG-UK) Consortium | Randox Laboratories and Alex Alderton, Roberto Amato, Jeffrey Barrett, Sonia Goncalves, Ewan Harrison, David K. Jackson, Ian Johnston, Dominic Kwiatkowski, Cordelia Langford, John Sillitoe on behalf of the Wellcome Sanger Institute COVID-19 Surveillance Team |
| EPI_ISL_3042922 | Lighthouse Lab in Alderley Park | Wellcome Sanger Institute for the COVID-19 Genomics UK (COG-UK) Consortium | Jacquelyn Wynn, Mairead Hyland, The Lighthouse Lab in Alderley Park and Alex Alderton, Roberto Amato, Jeffrey Barrett, Sonia Goncalves, Ewan Harrison, David K. Jackson, Ian Johnston, Dominic Kwiatkowski, Cordelia Langford, John Sillitoe on behalf of the Wellcome Sanger Institute COVID-19 Surveillance Team |
| EPI_ISL_3042923 | Lighthouse Lab in Milton Keynes | Wellcome Sanger Institute for the COVID-19 Genomics UK (COG-UK) Consortium | The Lighthouse Lab in Milton Keynes and Alex Alderton, Roberto Amato, Jeffrey Barrett, Sonia Goncalves, Ewan Harrison, David K. Jackson, Ian Johnston, Dominic Kwiatkowski, Cordelia Langford, John Sillitoe on behalf of the Wellcome Sanger Institute COVID-19 Surveillance Team |
| EPI_ISL_3042924 | Lighthouse Lab in Glasgow | Wellcome Sanger Institute for the COVID-19 Genomics UK (COG-UK) Consortium | Harper VanSteenhouse, Yumi Kasai, David Gray, Carol Clugston, Anna Dominiczak and Alex Alderton, Roberto Amato, Jeffrey Barrett, Sonia Goncalves, Ewan Harrison, David K. Jackson, Ian Johnston, Dominic Kwiatkowski, Cordelia Langford, John Sillitoe on behalf of the Wellcome Sanger Institute COVID-19 Surveillance Team |
| EPI_ISL_3042925 | Randox Laboratories | Wellcome Sanger Institute for the COVID-19 Genomics UK (COG-UK) Consortium | Randox Laboratories and Alex Alderton, Roberto Amato, Jeffrey Barrett, Sonia Goncalves, Ewan Harrison, David K. Jackson, Ian Johnston, Dominic Kwiatkowski, Cordelia Langford, John Sillitoe on behalf of the Wellcome Sanger Institute COVID-19 Surveillance Team |
| EPI_ISL_3042926 | Lighthouse Lab in Alderley Park | Wellcome Sanger Institute for the COVID-19 Genomics UK (COG-UK) Consortium | Jacquelyn Wynn, Mairead Hyland, The Lighthouse Lab in Alderley Park and Alex Alderton, Roberto Amato, Jeffrey Barrett, Sonia Goncalves, Ewan Harrison, David K. Jackson, Ian Johnston, Dominic Kwiatkowski, Cordelia Langford, John Sillitoe on behalf of the Wellcome Sanger Institute COVID-19 Surveillance Team |
| EPI_ISL_3042927, EPI_ISL_3042928 | Lighthouse Lab in Glasgow | Wellcome Sanger Institute for the COVID-19 Genomics UK (COG-UK) Consortium | Harper VanSteenhouse, Yumi Kasai, David Gray, Carol Clugston, Anna Dominiczak and Alex Alderton, Roberto Amato, Jeffrey Barrett, Sonia Goncalves, Ewan Harrison, David K. Jackson, Ian Johnston, Dominic Kwiatkowski, Cordelia Langford, John Sillitoe on behalf of the Wellcome Sanger Institute COVID-19 Surveillance Team |
| EPI_ISL_3042929 | Randox Laboratories | Wellcome Sanger Institute for the COVID-19 Genomics UK (COG-UK) Consortium | Randox Laboratories and Alex Alderton, Roberto Amato, Jeffrey Barrett, Sonia Goncalves, Ewan Harrison, David K. Jackson, Ian Johnston, Dominic Kwiatkowski, Cordelia Langford, John Sillitoe on behalf of the Wellcome Sanger Institute COVID-19 Surveillance Team |
| EPI_ISL_3042930, EPI_ISL_3042931, EPI_ISL_3042932 | Lighthouse Lab in Alderley Park | Wellcome Sanger Institute for the COVID-19 Genomics UK (COG-UK) Consortium | Jacquelyn Wynn, Mairead Hyland, The Lighthouse Lab in Alderley Park and Alex Alderton, Roberto Amato, Jeffrey Barrett, Sonia Goncalves, Ewan Harrison, David K. Jackson, Ian Johnston, Dominic Kwiatkowski, Cordelia Langford, John Sillitoe on behalf of the Wellcome Sanger Institute COVID-19 Surveillance Team |
| EPI_ISL_3042933 | Randox Laboratories | Wellcome Sanger Institute for the COVID-19 Genomics UK (COG-UK) Consortium | Randox Laboratories and Alex Alderton, Roberto Amato, Jeffrey Barrett, Sonia Goncalves, Ewan Harrison, David K. Jackson, Ian Johnston, Dominic Kwiatkowski, Cordelia Langford, John Sillitoe on behalf of the Wellcome Sanger Institute COVID-19 Surveillance Team |
| EPI_ISL_3042934, EPI_ISL_3042935 | Lighthouse Lab in Glasgow | Wellcome Sanger Institute for the COVID-19 Genomics UK (COG-UK) Consortium | Harper VanSteenhouse, Yumi Kasai, David Gray, Carol Clugston, Anna Dominiczak and Alex Alderton, Roberto Amato, Jeffrey Barrett, Sonia Goncalves, Ewan Harrison, David K. Jackson, Ian Johnston, Dominic Kwiatkowski, Cordelia Langford, John Sillitoe on behalf of the Wellcome Sanger Institute COVID-19 Surveillance Team |
| EPI_ISL_3042936, EPI_ISL_3042937 | Lighthouse Lab in Alderley Park | Wellcome Sanger Institute for the COVID-19 Genomics UK (COG-UK) Consortium | Jacquelyn Wynn, Mairead Hyland, The Lighthouse Lab in Alderley Park and Alex Alderton, Roberto Amato, Jeffrey Barrett, Sonia Goncalves, Ewan Harrison, David K. Jackson, Ian Johnston, Dominic Kwiatkowski, Cordelia Langford, John Sillitoe on behalf of the Wellcome Sanger Institute COVID-19 Surveillance Team |
| EPI_ISL_3042938, EPI_ISL_3042939 | Lighthouse Lab in Milton Keynes | Wellcome Sanger Institute for the COVID-19 Genomics UK (COG-UK) Consortium | The Lighthouse Lab in Milton Keynes and Alex Alderton, Roberto Amato, Jeffrey Barrett, Sonia Goncalves, Ewan Harrison, David K. Jackson, Ian Johnston, Dominic Kwiatkowski, Cordelia Langford, John Sillitoe on behalf of the Wellcome Sanger Institute COVID-19 Surveillance Team |
| EPI_ISL_3042940, EPI_ISL_3042941 | Lighthouse Lab in Glasgow | Wellcome Sanger Institute for the COVID-19 Genomics UK (COG-UK) Consortium | Harper VanSteenhouse, Yumi Kasai, David Gray, Carol Clugston, Anna Dominiczak and Alex Alderton, Roberto Amato, Jeffrey Barrett, Sonia Goncalves, Ewan Harrison, David K. Jackson, Ian Johnston, Dominic Kwiatkowski, Cordelia Langford, John Sillitoe on behalf of the Wellcome Sanger Institute COVID-19 Surveillance Team |
| EPI_ISL_3042942 | Lighthouse Lab in Milton Keynes | Wellcome Sanger Institute for the COVID-19 Genomics UK (COG-UK) Consortium | The Lighthouse Lab in Milton Keynes and Alex Alderton, Roberto Amato, Jeffrey Barrett, Sonia Goncalves, Ewan Harrison, David K. Jackson, Ian Johnston, Dominic Kwiatkowski, Cordelia Langford, John Sillitoe on behalf of the Wellcome Sanger Institute COVID-19 Surveillance Team |
| EPI_ISL_3042943, EPI_ISL_3042944 | Lighthouse Lab in Alderley Park | Wellcome Sanger Institute for the COVID-19 Genomics UK (COG-UK) Consortium | Jacquelyn Wynn, Mairead Hyland, The Lighthouse Lab in Alderley Park and Alex Alderton, Roberto Amato, Jeffrey Barrett, Sonia Goncalves, Ewan Harrison, David K. Jackson, Ian Johnston, Dominic Kwiatkowski, Cordelia Langford, John Sillitoe on behalf of the Wellcome Sanger Institute COVID-19 Surveillance Team |
| EPI_ISL_3042945, EPI_ISL_3042946 | Randox Laboratories | Wellcome Sanger Institute for the COVID-19 Genomics UK (COG-UK) Consortium | Randox Laboratories and Alex Alderton, Roberto Amato, Jeffrey Barrett, Sonia Goncalves, Ewan Harrison, David K. Jackson, Ian Johnston, Dominic Kwiatkowski, Cordelia Langford, John Sillitoe on behalf of the Wellcome Sanger Institute COVID-19 Surveillance Team |
| EPI_ISL_3042947, EPI_ISL_3042948 | Lighthouse Lab in Milton Keynes | Wellcome Sanger Institute for the COVID-19 Genomics UK (COG-UK) Consortium | The Lighthouse Lab in Milton Keynes and Alex Alderton, Roberto Amato, Jeffrey Barrett, Sonia Goncalves, Ewan Harrison, David K. Jackson, Ian Johnston, Dominic Kwiatkowski, Cordelia Langford, John Sillitoe on behalf of the Wellcome Sanger Institute COVID-19 Surveillance Team |
| EPI_ISL_3042949, EPI_ISL_3042950 | Lighthouse Lab in Alderley Park | Wellcome Sanger Institute for the COVID-19 Genomics UK (COG-UK) Consortium | Jacquelyn Wynn, Mairead Hyland, The Lighthouse Lab in Alderley Park and Alex Alderton, Roberto Amato, Jeffrey Barrett, Sonia Goncalves, Ewan Harrison, David K. Jackson, Ian Johnston, Dominic Kwiatkowski, Cordelia Langford, John Sillitoe on behalf of the Wellcome Sanger Institute COVID-19 Surveillance Team |
| EPI_ISL_3042951, EPI_ISL_3042952, EPI_ISL_3042953, EPI_ISL_3042954, EPI_ISL_3042955 | Lighthouse Lab in Milton Keynes | Wellcome Sanger Institute for the COVID-19 Genomics UK (COG-UK) Consortium | The Lighthouse Lab in Milton Keynes and Alex Alderton, Roberto Amato, Jeffrey Barrett, Sonia Goncalves, Ewan Harrison, David K. Jackson, Ian Johnston, Dominic Kwiatkowski, Cordelia Langford, John Sillitoe on behalf of the Wellcome Sanger Institute COVID-19 Surveillance Team |

| | | | |
|---|---|---|---|
| EPI_ISL_3042956 | Lighthouse Lab in Alderley Park | Wellcome Sanger Institute for the COVID-19 Genomics UK (COG-UK) Consortium | Jacquelyn Wynn, Mairead Hyland, The Lighthouse Lab in Alderley Park and Alex Alderton, Roberto Amato, Jeffrey Barrett, Sonia Goncalves, Ewan Harrison, David K. Jackson, Ian Johnston, Dominic Kwiatkowski, Cordelia Langford, John Sillitoe on behalf of the Wellcome Sanger Institute COVID-19 Surveillance Team |
| EPI_ISL_3042957 | Lighthouse Lab in Milton Keynes | Wellcome Sanger Institute for the COVID-19 Genomics UK (COG-UK) Consortium | The Lighthouse Lab in Milton Keynes and Alex Alderton, Roberto Amato, Jeffrey Barrett, Sonia Goncalves, Ewan Harrison, David K. Jackson, Ian Johnston, Dominic Kwiatkowski, Cordelia Langford, John Sillitoe on behalf of the Wellcome Sanger Institute COVID-19 Surveillance Team |
| EPI_ISL_3042958, EPI_ISL_3042959, EPI_ISL_3042960 | Lighthouse Lab in Alderley Park | Wellcome Sanger Institute for the COVID-19 Genomics UK (COG-UK) Consortium | Jacquelyn Wynn, Mairead Hyland, The Lighthouse Lab in Alderley Park and Alex Alderton, Roberto Amato, Jeffrey Barrett, Sonia Goncalves, Ewan Harrison, David K. Jackson, Ian Johnston, Dominic Kwiatkowski, Cordelia Langford, John Sillitoe on behalf of the Wellcome Sanger Institute COVID-19 Surveillance Team |
| EPI_ISL_3042961 | Lighthouse Lab in Milton Keynes | Wellcome Sanger Institute for the COVID-19 Genomics UK (COG-UK) Consortium | The Lighthouse Lab in Milton Keynes and Alex Alderton, Roberto Amato, Jeffrey Barrett, Sonia Goncalves, Ewan Harrison, David K. Jackson, Ian Johnston, Dominic Kwiatkowski, Cordelia Langford, John Sillitoe on behalf of the Wellcome Sanger Institute COVID-19 Surveillance Team |
| EPI_ISL_3042962 | Lighthouse Lab in Alderley Park | Wellcome Sanger Institute for the COVID-19 Genomics UK (COG-UK) Consortium | Jacquelyn Wynn, Mairead Hyland, The Lighthouse Lab in Alderley Park and Alex Alderton, Roberto Amato, Jeffrey Barrett, Sonia Goncalves, Ewan Harrison, David K. Jackson, Ian Johnston, Dominic Kwiatkowski, Cordelia Langford, John Sillitoe on behalf of the Wellcome Sanger Institute COVID-19 Surveillance Team |
| EPI_ISL_3042963, EPI_ISL_3042964 | Randox Laboratories | Wellcome Sanger Institute for the COVID-19 Genomics UK (COG-UK) Consortium | Randox Laboratories and Alex Alderton, Roberto Amato, Jeffrey Barrett, Sonia Goncalves, Ewan Harrison, David K. Jackson, Ian Johnston, Dominic Kwiatkowski, Cordelia Langford, John Sillitoe on behalf of the Wellcome Sanger Institute COVID-19 Surveillance Team |
| EPI_ISL_3042965 | Lighthouse Lab in Glasgow | Wellcome Sanger Institute for the COVID-19 Genomics UK (COG-UK) Consortium | Harper VanSteenhouse, Yumi Kasai, David Gray, Carol Clugston, Anna Dominiczak and Alex Alderton, Roberto Amato, Jeffrey Barrett, Sonia Goncalves, Ewan Harrison, David K. Jackson, Ian Johnston, Dominic Kwiatkowski, Cordelia Langford, John Sillitoe on behalf of the Wellcome Sanger Institute COVID-19 Surveillance Team |
| EPI_ISL_3042966 | Lighthouse Lab in Milton Keynes | Wellcome Sanger Institute for the COVID-19 Genomics UK (COG-UK) Consortium | The Lighthouse Lab in Milton Keynes and Alex Alderton, Roberto Amato, Jeffrey Barrett, Sonia Goncalves, Ewan Harrison, David K. Jackson, Ian Johnston, Dominic Kwiatkowski, Cordelia Langford, John Sillitoe on behalf of the Wellcome Sanger Institute COVID-19 Surveillance Team |
| EPI_ISL_3042967 | Randox Laboratories | Wellcome Sanger Institute for the COVID-19 Genomics UK (COG-UK) Consortium | Randox Laboratories and Alex Alderton, Roberto Amato, Jeffrey Barrett, Sonia Goncalves, Ewan Harrison, David K. Jackson, Ian Johnston, Dominic Kwiatkowski, Cordelia Langford, John Sillitoe on behalf of the Wellcome Sanger Institute COVID-19 Surveillance Team |
| EPI_ISL_3042968 | Lighthouse Lab in Milton Keynes | Wellcome Sanger Institute for the COVID-19 Genomics UK (COG-UK) Consortium | The Lighthouse Lab in Milton Keynes and Alex Alderton, Roberto Amato, Jeffrey Barrett, Sonia Goncalves, Ewan Harrison, David K. Jackson, Ian Johnston, Dominic Kwiatkowski, Cordelia Langford, John Sillitoe on behalf of the Wellcome Sanger Institute COVID-19 Surveillance Team |
| EPI_ISL_3042969 | Lighthouse Lab in Alderley Park | Wellcome Sanger Institute for the COVID-19 Genomics UK (COG-UK) Consortium | Jacquelyn Wynn, Mairead Hyland, The Lighthouse Lab in Alderley Park and Alex Alderton, Roberto Amato, Jeffrey Barrett, Sonia Goncalves, Ewan Harrison, David K. Jackson, Ian Johnston, Dominic Kwiatkowski, Cordelia Langford, John Sillitoe on behalf of the Wellcome Sanger Institute COVID-19 Surveillance Team |
| EPI_ISL_3042970 | Lighthouse Lab in Milton Keynes | Wellcome Sanger Institute for the COVID-19 Genomics UK (COG-UK) Consortium | The Lighthouse Lab in Milton Keynes and Alex Alderton, Roberto Amato, Jeffrey Barrett, Sonia Goncalves, Ewan Harrison, David K. Jackson, Ian Johnston, Dominic Kwiatkowski, Cordelia Langford, John Sillitoe on behalf of the Wellcome Sanger Institute COVID-19 Surveillance Team |
| EPI_ISL_3042971, EPI_ISL_3042972 | Lighthouse Lab in Alderley Park | Wellcome Sanger Institute for the COVID-19 Genomics UK (COG-UK) Consortium | Jacquelyn Wynn, Mairead Hyland, The Lighthouse Lab in Alderley Park and Alex Alderton, Roberto Amato, Jeffrey Barrett, Sonia Goncalves, Ewan Harrison, David K. Jackson, Ian Johnston, Dominic Kwiatkowski, Cordelia Langford, John Sillitoe on behalf of the Wellcome Sanger Institute COVID-19 Surveillance Team |
| EPI_ISL_3042973 | Lighthouse Lab in Glasgow | Wellcome Sanger Institute for the COVID-19 Genomics UK (COG-UK) Consortium | Harper VanSteenhouse, Yumi Kasai, David Gray, Carol Clugston, Anna Dominiczak and Alex Alderton, Roberto Amato, Jeffrey Barrett, Sonia Goncalves, Ewan Harrison, David K. Jackson, Ian Johnston, Dominic Kwiatkowski, Cordelia Langford, John Sillitoe on behalf of the Wellcome Sanger Institute COVID-19 Surveillance Team |
| EPI_ISL_3042974, EPI_ISL_3042975 | Randox Laboratories | Wellcome Sanger Institute for the COVID-19 Genomics UK (COG-UK) Consortium | Randox Laboratories and Alex Alderton, Roberto Amato, Jeffrey Barrett, Sonia Goncalves, Ewan Harrison, David K. Jackson, Ian Johnston, Dominic Kwiatkowski, Cordelia Langford, John Sillitoe on behalf of the Wellcome Sanger Institute COVID-19 Surveillance Team |
| EPI_ISL_3042976 | Lighthouse Lab in Alderley Park | Wellcome Sanger Institute for the COVID-19 Genomics UK (COG-UK) Consortium | Jacquelyn Wynn, Mairead Hyland, The Lighthouse Lab in Alderley Park and Alex Alderton, Roberto Amato, Jeffrey Barrett, Sonia Goncalves, Ewan Harrison, David K. Jackson, Ian Johnston, Dominic Kwiatkowski, Cordelia Langford, John Sillitoe on behalf of the Wellcome Sanger Institute COVID-19 Surveillance Team |
| EPI_ISL_3042977 | Randox Laboratories | Wellcome Sanger Institute for the COVID-19 Genomics UK (COG-UK) Consortium | Randox Laboratories and Alex Alderton, Roberto Amato, Jeffrey Barrett, Sonia Goncalves, Ewan Harrison, David K. Jackson, Ian Johnston, Dominic Kwiatkowski, Cordelia Langford, John Sillitoe on behalf of the Wellcome Sanger Institute COVID-19 Surveillance Team |
| EPI_ISL_3042978 | Lighthouse Lab in Alderley Park | Wellcome Sanger Institute for the COVID-19 Genomics UK (COG-UK) Consortium | Jacquelyn Wynn, Mairead Hyland, The Lighthouse Lab in Alderley Park and Alex Alderton, Roberto Amato, Jeffrey Barrett, Sonia Goncalves, Ewan Harrison, David K. Jackson, Ian Johnston, Dominic Kwiatkowski, Cordelia Langford, John Sillitoe on behalf of the Wellcome Sanger Institute COVID-19 Surveillance Team |
| EPI_ISL_3042979 | Lighthouse Lab in Milton Keynes | Wellcome Sanger Institute for the COVID-19 Genomics UK (COG-UK) Consortium | The Lighthouse Lab in Milton Keynes and Alex Alderton, Roberto Amato, Jeffrey Barrett, Sonia Goncalves, Ewan Harrison, David K. Jackson, Ian Johnston, Dominic Kwiatkowski, Cordelia Langford, John Sillitoe on behalf of the Wellcome Sanger Institute COVID-19 Surveillance Team |
| EPI_ISL_3042980 | Lighthouse Lab in Glasgow | Wellcome Sanger Institute for the COVID-19 Genomics UK (COG-UK) Consortium | Harper VanSteenhouse, Yumi Kasai, David Gray, Carol Clugston, Anna Dominiczak and Alex Alderton, Roberto Amato, Jeffrey Barrett, Sonia Goncalves, Ewan Harrison, David K. Jackson, Ian Johnston, Dominic Kwiatkowski, Cordelia Langford, John Sillitoe on behalf of the Wellcome Sanger Institute COVID-19 Surveillance Team |
| EPI_ISL_3042981 | Lighthouse Lab in Milton Keynes | Wellcome Sanger Institute for the COVID-19 Genomics UK (COG-UK) Consortium | The Lighthouse Lab in Milton Keynes and Alex Alderton, Roberto Amato, Jeffrey Barrett, Sonia Goncalves, Ewan Harrison, David K. Jackson, Ian Johnston, Dominic Kwiatkowski, Cordelia Langford, John Sillitoe on behalf of the Wellcome Sanger Institute COVID-19 Surveillance Team |
| EPI_ISL_3042982 | Lighthouse Lab in Alderley Park | Wellcome Sanger Institute for the COVID-19 Genomics UK (COG-UK) Consortium | Jacquelyn Wynn, Mairead Hyland, The Lighthouse Lab in Alderley Park and Alex Alderton, Roberto Amato, Jeffrey Barrett, Sonia Goncalves, Ewan Harrison, David K. Jackson, Ian Johnston, Dominic Kwiatkowski, Cordelia Langford, John Sillitoe on behalf of the Wellcome Sanger Institute COVID-19 Surveillance Team |
| EPI_ISL_3042983, EPI_ISL_3042984 | Lighthouse Lab in Milton Keynes | Wellcome Sanger Institute for the COVID-19 Genomics UK (COG-UK) Consortium | The Lighthouse Lab in Milton Keynes and Alex Alderton, Roberto Amato, Jeffrey Barrett, Sonia Goncalves, Ewan Harrison, David K. Jackson, Ian Johnston, Dominic Kwiatkowski, Cordelia Langford, John Sillitoe on behalf of the Wellcome Sanger Institute COVID-19 Surveillance Team |
| EPI_ISL_3042985, EPI_ISL_3042986 | Lighthouse Lab in Alderley Park | Wellcome Sanger Institute for the COVID-19 Genomics UK (COG-UK) Consortium | Jacquelyn Wynn, Mairead Hyland, The Lighthouse Lab in Alderley Park and Alex Alderton, Roberto Amato, Jeffrey Barrett, Sonia Goncalves, Ewan Harrison, David K. Jackson, Ian Johnston, Dominic Kwiatkowski, Cordelia Langford, John Sillitoe on behalf of the Wellcome Sanger Institute COVID-19 Surveillance Team |
| EPI_ISL_3042987 | Lighthouse Lab in Glasgow | Wellcome Sanger Institute for the COVID-19 Genomics UK (COG-UK) Consortium | Harper VanSteenhouse, Yumi Kasai, David Gray, Carol Clugston, Anna Dominiczak and Alex Alderton, Roberto Amato, Jeffrey Barrett, Sonia Goncalves, Ewan Harrison, David K. Jackson, Ian Johnston, Dominic Kwiatkowski, Cordelia Langford, John Sillitoe on behalf of the Wellcome Sanger Institute COVID-19 Surveillance Team |
| EPI_ISL_3042988 | Randox Laboratories | Wellcome Sanger Institute for the COVID-19 Genomics UK (COG-UK) Consortium | Randox Laboratories and Alex Alderton, Roberto Amato, Jeffrey Barrett, Sonia Goncalves, Ewan Harrison, David K. Jackson, Ian Johnston, Dominic Kwiatkowski, Cordelia Langford, John Sillitoe on behalf of the Wellcome Sanger Institute COVID-19 Surveillance Team |
| EPI_ISL_3042989, EPI_ISL_3042990 | Lighthouse Lab in Alderley Park | Wellcome Sanger Institute for the COVID-19 Genomics UK (COG-UK) Consortium | Jacquelyn Wynn, Mairead Hyland, The Lighthouse Lab in Alderley Park and Alex Alderton, Roberto Amato, Jeffrey Barrett, Sonia Goncalves, Ewan Harrison, David K. Jackson, Ian Johnston, Dominic Kwiatkowski, Cordelia Langford, John Sillitoe on behalf of the Wellcome Sanger Institute COVID-19 Surveillance Team |
| EPI_ISL_3042991 | Randox Laboratories | Wellcome Sanger Institute for the COVID-19 Genomics UK (COG-UK) Consortium | Randox Laboratories and Alex Alderton, Roberto Amato, Jeffrey Barrett, Sonia Goncalves, Ewan Harrison, David K. Jackson, Ian Johnston, Dominic Kwiatkowski, Cordelia Langford, John Sillitoe on behalf of the Wellcome Sanger Institute COVID-19 Surveillance Team |
| EPI_ISL_3042992 | Lighthouse Lab in Alderley Park | Wellcome Sanger Institute for the COVID-19 Genomics UK (COG-UK) Consortium | Jacquelyn Wynn, Mairead Hyland, The Lighthouse Lab in Alderley Park and Alex Alderton, Roberto Amato, Jeffrey Barrett, Sonia Goncalves, Ewan Harrison, David K. Jackson, Ian Johnston, Dominic Kwiatkowski, Cordelia Langford, John Sillitoe on behalf of the Wellcome Sanger Institute COVID-19 Surveillance Team |
| EPI_ISL_3042993 | Lighthouse Lab in Milton Keynes | Wellcome Sanger Institute for the COVID-19 Genomics UK (COG-UK) Consortium | The Lighthouse Lab in Milton Keynes and Alex Alderton, Roberto Amato, Jeffrey Barrett, Sonia Goncalves, Ewan Harrison, David K. Jackson, Ian Johnston, Dominic Kwiatkowski, Cordelia Langford, John Sillitoe on behalf of the Wellcome Sanger Institute COVID-19 Surveillance Team |
| EPI_ISL_3042994 | Lighthouse Lab in Glasgow | Wellcome Sanger Institute for the COVID-19 Genomics UK (COG-UK) Consortium | Harper VanSteenhouse, Yumi Kasai, David Gray, Carol Clugston, Anna Dominiczak and Alex Alderton, Roberto Amato, Jeffrey Barrett, Sonia Goncalves, Ewan Harrison, David K. Jackson, Ian Johnston, Dominic Kwiatkowski, Cordelia Langford, John Sillitoe on behalf of the Wellcome Sanger Institute COVID-19 Surveillance Team |
| EPI_ISL_3042995 | Lighthouse Lab in Alderley Park | Wellcome Sanger Institute for the COVID-19 Genomics UK (COG-UK) Consortium | Jacquelyn Wynn, Mairead Hyland, The Lighthouse Lab in Alderley Park and Alex Alderton, Roberto Amato, Jeffrey Barrett, Sonia Goncalves, Ewan Harrison, David K. Jackson, Ian Johnston, Dominic Kwiatkowski, Cordelia Langford, John Sillitoe on behalf of the Wellcome Sanger Institute COVID-19 Surveillance Team |
| EPI_ISL_3042996 | Lighthouse Lab in Glasgow | Wellcome Sanger Institute for the COVID-19 Genomics UK (COG-UK) Consortium | Harper VanSteenhouse, Yumi Kasai, David Gray, Carol Clugston, Anna Dominiczak and Alex Alderton, Roberto Amato, Jeffrey Barrett, Sonia Goncalves, Ewan Harrison, David K. Jackson, Ian Johnston, Dominic Kwiatkowski, Cordelia Langford, John Sillitoe on behalf of the Wellcome Sanger Institute COVID-19 Surveillance Team |
| EPI_ISL_3042997 | Lighthouse Lab in Alderley Park | Wellcome Sanger Institute for the COVID-19 Genomics UK (COG-UK) Consortium | Jacquelyn Wynn, Mairead Hyland, The Lighthouse Lab in Alderley Park and Alex Alderton, Roberto Amato, Jeffrey Barrett, Sonia Goncalves, Ewan Harrison, David K. Jackson, Ian Johnston, Dominic Kwiatkowski, Cordelia Langford, John Sillitoe on behalf of the Wellcome Sanger Institute COVID-19 Surveillance Team |
| EPI_ISL_3042998 | Lighthouse Lab in Milton Keynes | Wellcome Sanger Institute for the COVID-19 Genomics UK (COG-UK) Consortium | The Lighthouse Lab in Milton Keynes and Alex Alderton, Roberto Amato, Jeffrey Barrett, Sonia Goncalves, Ewan Harrison, David K. Jackson, Ian Johnston, Dominic Kwiatkowski, Cordelia Langford, John Sillitoe on behalf of the Wellcome Sanger Institute COVID-19 Surveillance Team |
| EPI_ISL_3042999 | Lighthouse Lab in Alderley Park | Wellcome Sanger Institute for the COVID-19 Genomics UK (COG-UK) Consortium | Jacquelyn Wynn, Mairead Hyland, The Lighthouse Lab in Alderley Park and Alex Alderton, Roberto Amato, Jeffrey Barrett, Sonia Goncalves, Ewan Harrison, David K. Jackson, Ian Johnston, Dominic Kwiatkowski, Cordelia Langford, John Sillitoe on behalf of the Wellcome Sanger Institute COVID-19 Surveillance Team |
| EPI_ISL_3043000 | Randox Laboratories | Wellcome Sanger Institute for the COVID-19 Genomics UK (COG-UK) Consortium | Randox Laboratories and Alex Alderton, Roberto Amato, Jeffrey Barrett, Sonia Goncalves, Ewan Harrison, David K. Jackson, Ian Johnston, Dominic Kwiatkowski, Cordelia Langford, John Sillitoe on behalf of the Wellcome Sanger Institute COVID-19 Surveillance Team |
| EPI_ISL_3043001 | Lighthouse Lab in Glasgow | Wellcome Sanger Institute for the COVID-19 Genomics UK (COG-UK) Consortium | Harper VanSteenhouse, Yumi Kasai, David Gray, Carol Clugston, Anna Dominiczak and Alex Alderton, Roberto Amato, Jeffrey Barrett, Sonia Goncalves, Ewan Harrison, David K. Jackson, Ian Johnston, Dominic Kwiatkowski, Cordelia Langford, John Sillitoe on behalf of the Wellcome Sanger Institute COVID-19 Surveillance Team |
| EPI_ISL_3043002, EPI_ISL_3043003, EPI_ISL_3043004 | Randox Laboratories | Wellcome Sanger Institute for the COVID-19 Genomics UK (COG-UK) Consortium | Randox Laboratories and Alex Alderton, Roberto Amato, Jeffrey Barrett, Sonia Goncalves, Ewan Harrison, David K. Jackson, Ian Johnston, Dominic Kwiatkowski, Cordelia Langford, John Sillitoe on behalf of the Wellcome Sanger Institute COVID-19 Surveillance Team |
| EPI_ISL_3043005 | Lighthouse Lab in Milton Keynes | Wellcome Sanger Institute for the COVID-19 Genomics UK (COG-UK) Consortium | The Lighthouse Lab in Milton Keynes and Alex Alderton, Roberto Amato, Jeffrey Barrett, Sonia Goncalves, Ewan Harrison, David K. Jackson, Ian Johnston, Dominic Kwiatkowski, Cordelia Langford, John Sillitoe on behalf of the Wellcome Sanger Institute COVID-19 Surveillance Team |
| EPI_ISL_3043006, EPI_ISL_3043007 | Lighthouse Lab in Glasgow | Wellcome Sanger Institute for the COVID-19 Genomics UK (COG-UK) Consortium | Harper VanSteenhouse, Yumi Kasai, David Gray, Carol Clugston, Anna Dominiczak and Alex Alderton, Roberto Amato, Jeffrey Barrett, Sonia Goncalves, Ewan Harrison, David K. Jackson, Ian Johnston, Dominic Kwiatkowski, Cordelia Langford, John Sillitoe on behalf of the Wellcome Sanger Institute COVID-19 Surveillance Team |
| EPI_ISL_3043008 | Randox Laboratories | Wellcome Sanger Institute for the COVID-19 Genomics UK (COG-UK) Consortium | Randox Laboratories and Alex Alderton, Roberto Amato, Jeffrey Barrett, Sonia Goncalves, Ewan Harrison, David K. Jackson, Ian Johnston, Dominic Kwiatkowski, Cordelia Langford, John Sillitoe on behalf of the Wellcome Sanger Institute COVID-19 Surveillance Team |
| EPI_ISL_3043009 | Lighthouse Lab in Milton Keynes | Wellcome Sanger Institute for the COVID-19 Genomics UK (COG-UK) Consortium | The Lighthouse Lab in Milton Keynes and Alex Alderton, Roberto Amato, Jeffrey Barrett, Sonia Goncalves, Ewan Harrison, David K. Jackson, Ian Johnston, Dominic Kwiatkowski, Cordelia Langford, John Sillitoe on behalf of the Wellcome Sanger Institute COVID-19 Surveillance Team |
| EPI_ISL_3043010 | Lighthouse Lab in Glasgow | Wellcome Sanger Institute for the COVID-19 Genomics UK (COG-UK) Consortium | Harper VanSteenhouse, Yumi Kasai, David Gray, Carol Clugston, Anna Dominiczak and Alex Alderton, Roberto Amato, Jeffrey Barrett, Sonia Goncalves, Ewan Harrison, David K. Jackson, Ian Johnston, Dominic Kwiatkowski, Cordelia Langford, John Sillitoe on behalf of the Wellcome Sanger Institute COVID-19 Surveillance Team |

| | | | |
|---|---|---|---|
| EPI_ISL_3043011 | Lighthouse Lab in Milton Keynes | Wellcome Sanger Institute for the COVID-19 Genomics UK (COG-UK) Consortium | The Lighthouse Lab in Milton Keynes and Alex Alderton, Roberto Amato, Jeffrey Barrett, Sonia Goncalves, Ewan Harrison, David K. Jackson, Ian Johnston, Dominic Kwiatkowski, Cordelia Langford, John Sillitoe on behalf of the Wellcome Sanger Institute COVID-19 Surveillance Team |
| EPI_ISL_3043012, EPI_ISL_3043013, EPI_ISL_3043014 | Randox Laboratories | Wellcome Sanger Institute for the COVID-19 Genomics UK (COG-UK) Consortium | Randox Laboratories and Alex Alderton, Roberto Amato, Jeffrey Barrett, Sonia Goncalves, Ewan Harrison, David K. Jackson, Ian Johnston, Dominic Kwiatkowski, Cordelia Langford, John Sillitoe on behalf of the Wellcome Sanger Institute COVID-19 Surveillance Team |
| EPI_ISL_3043015, EPI_ISL_3043016 | Lighthouse Lab in Glasgow | Wellcome Sanger Institute for the COVID-19 Genomics UK (COG-UK) Consortium | Harper VanSteenhouse, Yumi Kasai, David Gray, Carol Clugston, Anna Dominiczak and Alex Alderton, Roberto Amato, Jeffrey Barrett, Sonia Goncalves, Ewan Harrison, David K. Jackson, Ian Johnston, Dominic Kwiatkowski, Cordelia Langford, John Sillitoe on behalf of the Wellcome Sanger Institute COVID-19 Surveillance Team |
| EPI_ISL_3043017 | Lighthouse Lab in Alderley Park | Wellcome Sanger Institute for the COVID-19 Genomics UK (COG-UK) Consortium | Jacquelyn Wynn, Mairead Hyland, The Lighthouse Lab in Alderley Park and Alex Alderton, Roberto Amato, Jeffrey Barrett, Sonia Goncalves, Ewan Harrison, David K. Jackson, Ian Johnston, Dominic Kwiatkowski, Cordelia Langford, John Sillitoe on behalf of the Wellcome Sanger Institute COVID-19 Surveillance Team |
| EPI_ISL_3043018 | Lighthouse Lab in Glasgow | Wellcome Sanger Institute for the COVID-19 Genomics UK (COG-UK) Consortium | Harper VanSteenhouse, Yumi Kasai, David Gray, Carol Clugston, Anna Dominiczak and Alex Alderton, Roberto Amato, Jeffrey Barrett, Sonia Goncalves, Ewan Harrison, David K. Jackson, Ian Johnston, Dominic Kwiatkowski, Cordelia Langford, John Sillitoe on behalf of the Wellcome Sanger Institute COVID-19 Surveillance Team |
| EPI_ISL_3043019, EPI_ISL_3043020 | Lighthouse Lab in Milton Keynes | Wellcome Sanger Institute for the COVID-19 Genomics UK (COG-UK) Consortium | The Lighthouse Lab in Milton Keynes and Alex Alderton, Roberto Amato, Jeffrey Barrett, Sonia Goncalves, Ewan Harrison, David K. Jackson, Ian Johnston, Dominic Kwiatkowski, Cordelia Langford, John Sillitoe on behalf of the Wellcome Sanger Institute COVID-19 Surveillance Team |
| EPI_ISL_3043021 | Lighthouse Lab in Alderley Park | Wellcome Sanger Institute for the COVID-19 Genomics UK (COG-UK) Consortium | Jacquelyn Wynn, Mairead Hyland, The Lighthouse Lab in Alderley Park and Alex Alderton, Roberto Amato, Jeffrey Barrett, Sonia Goncalves, Ewan Harrison, David K. Jackson, Ian Johnston, Dominic Kwiatkowski, Cordelia Langford, John Sillitoe on behalf of the Wellcome Sanger Institute COVID-19 Surveillance Team |
| EPI_ISL_3043022 | Lighthouse Lab in Milton Keynes | Wellcome Sanger Institute for the COVID-19 Genomics UK (COG-UK) Consortium | The Lighthouse Lab in Milton Keynes and Alex Alderton, Roberto Amato, Jeffrey Barrett, Sonia Goncalves, Ewan Harrison, David K. Jackson, Ian Johnston, Dominic Kwiatkowski, Cordelia Langford, John Sillitoe on behalf of the Wellcome Sanger Institute COVID-19 Surveillance Team |
| EPI_ISL_3043023 | Lighthouse Lab in Alderley Park | Wellcome Sanger Institute for the COVID-19 Genomics UK (COG-UK) Consortium | Jacquelyn Wynn, Mairead Hyland, The Lighthouse Lab in Alderley Park and Alex Alderton, Roberto Amato, Jeffrey Barrett, Sonia Goncalves, Ewan Harrison, David K. Jackson, Ian Johnston, Dominic Kwiatkowski, Cordelia Langford, John Sillitoe on behalf of the Wellcome Sanger Institute COVID-19 Surveillance Team |
| EPI_ISL_3043024, EPI_ISL_3043025 | Lighthouse Lab in Glasgow | Wellcome Sanger Institute for the COVID-19 Genomics UK (COG-UK) Consortium | Harper VanSteenhouse, Yumi Kasai, David Gray, Carol Clugston, Anna Dominiczak and Alex Alderton, Roberto Amato, Jeffrey Barrett, Sonia Goncalves, Ewan Harrison, David K. Jackson, Ian Johnston, Dominic Kwiatkowski, Cordelia Langford, John Sillitoe on behalf of the Wellcome Sanger Institute COVID-19 Surveillance Team |
| EPI_ISL_3043026 | Lighthouse Lab in Milton Keynes | Wellcome Sanger Institute for the COVID-19 Genomics UK (COG-UK) Consortium | The Lighthouse Lab in Milton Keynes and Alex Alderton, Roberto Amato, Jeffrey Barrett, Sonia Goncalves, Ewan Harrison, David K. Jackson, Ian Johnston, Dominic Kwiatkowski, Cordelia Langford, John Sillitoe on behalf of the Wellcome Sanger Institute COVID-19 Surveillance Team |
| EPI_ISL_3043027 | Lighthouse Lab in Alderley Park | Wellcome Sanger Institute for the COVID-19 Genomics UK (COG-UK) Consortium | Jacquelyn Wynn, Mairead Hyland, The Lighthouse Lab in Alderley Park and Alex Alderton, Roberto Amato, Jeffrey Barrett, Sonia Goncalves, Ewan Harrison, David K. Jackson, Ian Johnston, Dominic Kwiatkowski, Cordelia Langford, John Sillitoe on behalf of the Wellcome Sanger Institute COVID-19 Surveillance Team |
| EPI_ISL_3043028 | Lighthouse Lab in Milton Keynes | Wellcome Sanger Institute for the COVID-19 Genomics UK (COG-UK) Consortium | The Lighthouse Lab in Milton Keynes and Alex Alderton, Roberto Amato, Jeffrey Barrett, Sonia Goncalves, Ewan Harrison, David K. Jackson, Ian Johnston, Dominic Kwiatkowski, Cordelia Langford, John Sillitoe on behalf of the Wellcome Sanger Institute COVID-19 Surveillance Team |
| EPI_ISL_3043029, EPI_ISL_3043030 | Lighthouse Lab in Glasgow | Wellcome Sanger Institute for the COVID-19 Genomics UK (COG-UK) Consortium | Harper VanSteenhouse, Yumi Kasai, David Gray, Carol Clugston, Anna Dominiczak and Alex Alderton, Roberto Amato, Jeffrey Barrett, Sonia Goncalves, Ewan Harrison, David K. Jackson, Ian Johnston, Dominic Kwiatkowski, Cordelia Langford, John Sillitoe on behalf of the Wellcome Sanger Institute COVID-19 Surveillance Team |
| EPI_ISL_3043031, EPI_ISL_3043032, EPI_ISL_3043033 | Lighthouse Lab in Milton Keynes | Wellcome Sanger Institute for the COVID-19 Genomics UK (COG-UK) Consortium | The Lighthouse Lab in Milton Keynes and Alex Alderton, Roberto Amato, Jeffrey Barrett, Sonia Goncalves, Ewan Harrison, David K. Jackson, Ian Johnston, Dominic Kwiatkowski, Cordelia Langford, John Sillitoe on behalf of the Wellcome Sanger Institute COVID-19 Surveillance Team |
| EPI_ISL_3043034 | Randox Laboratories | Wellcome Sanger Institute for the COVID-19 Genomics UK (COG-UK) Consortium | Randox Laboratories and Alex Alderton, Roberto Amato, Jeffrey Barrett, Sonia Goncalves, Ewan Harrison, David K. Jackson, Ian Johnston, Dominic Kwiatkowski, Cordelia Langford, John Sillitoe on behalf of the Wellcome Sanger Institute COVID-19 Surveillance Team |
| EPI_ISL_3043035 | Lighthouse Lab in Glasgow | Wellcome Sanger Institute for the COVID-19 Genomics UK (COG-UK) Consortium | Harper VanSteenhouse, Yumi Kasai, David Gray, Carol Clugston, Anna Dominiczak and Alex Alderton, Roberto Amato, Jeffrey Barrett, Sonia Goncalves, Ewan Harrison, David K. Jackson, Ian Johnston, Dominic Kwiatkowski, Cordelia Langford, John Sillitoe on behalf of the Wellcome Sanger Institute COVID-19 Surveillance Team |
| EPI_ISL_3043036, EPI_ISL_3043037 | Lighthouse Lab in Milton Keynes | Wellcome Sanger Institute for the COVID-19 Genomics UK (COG-UK) Consortium | The Lighthouse Lab in Milton Keynes and Alex Alderton, Roberto Amato, Jeffrey Barrett, Sonia Goncalves, Ewan Harrison, David K. Jackson, Ian Johnston, Dominic Kwiatkowski, Cordelia Langford, John Sillitoe on behalf of the Wellcome Sanger Institute COVID-19 Surveillance Team |
| EPI_ISL_3043038 | Lighthouse Lab in Glasgow | Wellcome Sanger Institute for the COVID-19 Genomics UK (COG-UK) Consortium | Harper VanSteenhouse, Yumi Kasai, David Gray, Carol Clugston, Anna Dominiczak and Alex Alderton, Roberto Amato, Jeffrey Barrett, Sonia Goncalves, Ewan Harrison, David K. Jackson, Ian Johnston, Dominic Kwiatkowski, Cordelia Langford, John Sillitoe on behalf of the Wellcome Sanger Institute COVID-19 Surveillance Team |
| EPI_ISL_3043039 | Lighthouse Lab in Alderley Park | Wellcome Sanger Institute for the COVID-19 Genomics UK (COG-UK) Consortium | Jacquelyn Wynn, Mairead Hyland, The Lighthouse Lab in Alderley Park and Alex Alderton, Roberto Amato, Jeffrey Barrett, Sonia Goncalves, Ewan Harrison, David K. Jackson, Ian Johnston, Dominic Kwiatkowski, Cordelia Langford, John Sillitoe on behalf of the Wellcome Sanger Institute COVID-19 Surveillance Team |
| EPI_ISL_3043040, EPI_ISL_3043041, EPI_ISL_3043042 | Lighthouse Lab in Glasgow | Wellcome Sanger Institute for the COVID-19 Genomics UK (COG-UK) Consortium | Harper VanSteenhouse, Yumi Kasai, David Gray, Carol Clugston, Anna Dominiczak and Alex Alderton, Roberto Amato, Jeffrey Barrett, Sonia Goncalves, Ewan Harrison, David K. Jackson, Ian Johnston, Dominic Kwiatkowski, Cordelia Langford, John Sillitoe on behalf of the Wellcome Sanger Institute COVID-19 Surveillance Team |
| EPI_ISL_3043043 | Lighthouse Lab in Milton Keynes | Wellcome Sanger Institute for the COVID-19 Genomics UK (COG-UK) Consortium | The Lighthouse Lab in Milton Keynes and Alex Alderton, Roberto Amato, Jeffrey Barrett, Sonia Goncalves, Ewan Harrison, David K. Jackson, Ian Johnston, Dominic Kwiatkowski, Cordelia Langford, John Sillitoe on behalf of the Wellcome Sanger Institute COVID-19 Surveillance Team |
| EPI_ISL_3043044, EPI_ISL_3043045 | Lighthouse Lab in Alderley Park | Wellcome Sanger Institute for the COVID-19 Genomics UK (COG-UK) Consortium | Jacquelyn Wynn, Mairead Hyland, The Lighthouse Lab in Alderley Park and Alex Alderton, Roberto Amato, Jeffrey Barrett, Sonia Goncalves, Ewan Harrison, David K. Jackson, Ian Johnston, Dominic Kwiatkowski, Cordelia Langford, John Sillitoe on behalf of the Wellcome Sanger Institute COVID-19 Surveillance Team |
| EPI_ISL_3043046, EPI_ISL_3043047 | Lighthouse Lab in Milton Keynes | Wellcome Sanger Institute for the COVID-19 Genomics UK (COG-UK) Consortium | The Lighthouse Lab in Milton Keynes and Alex Alderton, Roberto Amato, Jeffrey Barrett, Sonia Goncalves, Ewan Harrison, David K. Jackson, Ian Johnston, Dominic Kwiatkowski, Cordelia Langford, John Sillitoe on behalf of the Wellcome Sanger Institute COVID-19 Surveillance Team |
| EPI_ISL_3043048 | Randox Laboratories | Wellcome Sanger Institute for the COVID-19 Genomics UK (COG-UK) Consortium | Randox Laboratories and Alex Alderton, Roberto Amato, Jeffrey Barrett, Sonia Goncalves, Ewan Harrison, David K. Jackson, Ian Johnston, Dominic Kwiatkowski, Cordelia Langford, John Sillitoe on behalf of the Wellcome Sanger Institute COVID-19 Surveillance Team |
| EPI_ISL_3043049, EPI_ISL_3043050 | Lighthouse Lab in Alderley Park | Wellcome Sanger Institute for the COVID-19 Genomics UK (COG-UK) Consortium | Jacquelyn Wynn, Mairead Hyland, The Lighthouse Lab in Alderley Park and Alex Alderton, Roberto Amato, Jeffrey Barrett, Sonia Goncalves, Ewan Harrison, David K. Jackson, Ian Johnston, Dominic Kwiatkowski, Cordelia Langford, John Sillitoe on behalf of the Wellcome Sanger Institute COVID-19 Surveillance Team |
| EPI_ISL_3043051 | Lighthouse Lab in Milton Keynes | Wellcome Sanger Institute for the COVID-19 Genomics UK (COG-UK) Consortium | The Lighthouse Lab in Milton Keynes and Alex Alderton, Roberto Amato, Jeffrey Barrett, Sonia Goncalves, Ewan Harrison, David K. Jackson, Ian Johnston, Dominic Kwiatkowski, Cordelia Langford, John Sillitoe on behalf of the Wellcome Sanger Institute COVID-19 Surveillance Team |
| EPI_ISL_3043052 | Lighthouse Lab in Glasgow | Wellcome Sanger Institute for the COVID-19 Genomics UK (COG-UK) Consortium | Harper VanSteenhouse, Yumi Kasai, David Gray, Carol Clugston, Anna Dominiczak and Alex Alderton, Roberto Amato, Jeffrey Barrett, Sonia Goncalves, Ewan Harrison, David K. Jackson, Ian Johnston, Dominic Kwiatkowski, Cordelia Langford, John Sillitoe on behalf of the Wellcome Sanger Institute COVID-19 Surveillance Team |
| EPI_ISL_3043053 | Randox Laboratories | Wellcome Sanger Institute for the COVID-19 Genomics UK (COG-UK) Consortium | Randox Laboratories and Alex Alderton, Roberto Amato, Jeffrey Barrett, Sonia Goncalves, Ewan Harrison, David K. Jackson, Ian Johnston, Dominic Kwiatkowski, Cordelia Langford, John Sillitoe on behalf of the Wellcome Sanger Institute COVID-19 Surveillance Team |
| EPI_ISL_3043054 | Lighthouse Lab in Alderley Park | Wellcome Sanger Institute for the COVID-19 Genomics UK (COG-UK) Consortium | Jacquelyn Wynn, Mairead Hyland, The Lighthouse Lab in Alderley Park and Alex Alderton, Roberto Amato, Jeffrey Barrett, Sonia Goncalves, Ewan Harrison, David K. Jackson, Ian Johnston, Dominic Kwiatkowski, Cordelia Langford, John Sillitoe on behalf of the Wellcome Sanger Institute COVID-19 Surveillance Team |
| EPI_ISL_3043055 | Lighthouse Lab in Glasgow | Wellcome Sanger Institute for the COVID-19 Genomics UK (COG-UK) Consortium | Harper VanSteenhouse, Yumi Kasai, David Gray, Carol Clugston, Anna Dominiczak and Alex Alderton, Roberto Amato, Jeffrey Barrett, Sonia Goncalves, Ewan Harrison, David K. Jackson, Ian Johnston, Dominic Kwiatkowski, Cordelia Langford, John Sillitoe on behalf of the Wellcome Sanger Institute COVID-19 Surveillance Team |
| EPI_ISL_3043056 | Lighthouse Lab in Milton Keynes | Wellcome Sanger Institute for the COVID-19 Genomics UK (COG-UK) Consortium | The Lighthouse Lab in Milton Keynes and Alex Alderton, Roberto Amato, Jeffrey Barrett, Sonia Goncalves, Ewan Harrison, David K. Jackson, Ian Johnston, Dominic Kwiatkowski, Cordelia Langford, John Sillitoe on behalf of the Wellcome Sanger Institute COVID-19 Surveillance Team |
| EPI_ISL_3043057 | Lighthouse Lab in Alderley Park | Wellcome Sanger Institute for the COVID-19 Genomics UK (COG-UK) Consortium | Jacquelyn Wynn, Mairead Hyland, The Lighthouse Lab in Alderley Park and Alex Alderton, Roberto Amato, Jeffrey Barrett, Sonia Goncalves, Ewan Harrison, David K. Jackson, Ian Johnston, Dominic Kwiatkowski, Cordelia Langford, John Sillitoe on behalf of the Wellcome Sanger Institute COVID-19 Surveillance Team |
| EPI_ISL_3043058 | Lighthouse Lab in Glasgow | Wellcome Sanger Institute for the COVID-19 Genomics UK (COG-UK) Consortium | Harper VanSteenhouse, Yumi Kasai, David Gray, Carol Clugston, Anna Dominiczak and Alex Alderton, Roberto Amato, Jeffrey Barrett, Sonia Goncalves, Ewan Harrison, David K. Jackson, Ian Johnston, Dominic Kwiatkowski, Cordelia Langford, John Sillitoe on behalf of the Wellcome Sanger Institute COVID-19 Surveillance Team |
| EPI_ISL_3043059 | Lighthouse Lab in Alderley Park | Wellcome Sanger Institute for the COVID-19 Genomics UK (COG-UK) Consortium | Jacquelyn Wynn, Mairead Hyland, The Lighthouse Lab in Alderley Park and Alex Alderton, Roberto Amato, Jeffrey Barrett, Sonia Goncalves, Ewan Harrison, David K. Jackson, Ian Johnston, Dominic Kwiatkowski, Cordelia Langford, John Sillitoe on behalf of the Wellcome Sanger Institute COVID-19 Surveillance Team |
| EPI_ISL_3043060 | Lighthouse Lab in Milton Keynes | Wellcome Sanger Institute for the COVID-19 Genomics UK (COG-UK) Consortium | The Lighthouse Lab in Milton Keynes and Alex Alderton, Roberto Amato, Jeffrey Barrett, Sonia Goncalves, Ewan Harrison, David K. Jackson, Ian Johnston, Dominic Kwiatkowski, Cordelia Langford, John Sillitoe on behalf of the Wellcome Sanger Institute COVID-19 Surveillance Team |
| EPI_ISL_3043061 | Lighthouse Lab in Alderley Park | Wellcome Sanger Institute for the COVID-19 Genomics UK (COG-UK) Consortium | Jacquelyn Wynn, Mairead Hyland, The Lighthouse Lab in Alderley Park and Alex Alderton, Roberto Amato, Jeffrey Barrett, Sonia Goncalves, Ewan Harrison, David K. Jackson, Ian Johnston, Dominic Kwiatkowski, Cordelia Langford, John Sillitoe on behalf of the Wellcome Sanger Institute COVID-19 Surveillance Team |
| EPI_ISL_3043062 | Lighthouse Lab in Glasgow | Wellcome Sanger Institute for the COVID-19 Genomics UK (COG-UK) Consortium | Harper VanSteenhouse, Yumi Kasai, David Gray, Carol Clugston, Anna Dominiczak and Alex Alderton, Roberto Amato, Jeffrey Barrett, Sonia Goncalves, Ewan Harrison, David K. Jackson, Ian Johnston, Dominic Kwiatkowski, Cordelia Langford, John Sillitoe on behalf of the Wellcome Sanger Institute COVID-19 Surveillance Team |
| EPI_ISL_3043063 | Lighthouse Lab in Milton Keynes | Wellcome Sanger Institute for the COVID-19 Genomics UK (COG-UK) Consortium | The Lighthouse Lab in Milton Keynes and Alex Alderton, Roberto Amato, Jeffrey Barrett, Sonia Goncalves, Ewan Harrison, David K. Jackson, Ian Johnston, Dominic Kwiatkowski, Cordelia Langford, John Sillitoe on behalf of the Wellcome Sanger Institute COVID-19 Surveillance Team |
| EPI_ISL_3043064 | Lighthouse Lab in Glasgow | Wellcome Sanger Institute for the COVID-19 Genomics UK (COG-UK) Consortium | Harper VanSteenhouse, Yumi Kasai, David Gray, Carol Clugston, Anna Dominiczak and Alex Alderton, Roberto Amato, Jeffrey Barrett, Sonia Goncalves, Ewan Harrison, David K. Jackson, Ian Johnston, Dominic Kwiatkowski, Cordelia Langford, John Sillitoe on behalf of the Wellcome Sanger Institute COVID-19 Surveillance Team |
| EPI_ISL_3043065 | Lighthouse Lab in Alderley Park | Wellcome Sanger Institute for the COVID-19 Genomics UK (COG-UK) Consortium | Jacquelyn Wynn, Mairead Hyland, The Lighthouse Lab in Alderley Park and Alex Alderton, Roberto Amato, Jeffrey Barrett, Sonia Goncalves, Ewan Harrison, David K. Jackson, Ian Johnston, Dominic Kwiatkowski, Cordelia Langford, John Sillitoe on behalf of the Wellcome Sanger Institute COVID-19 Surveillance Team |
| EPI_ISL_3043066, EPI_ISL_3043067 | Lighthouse Lab in Milton Keynes | Wellcome Sanger Institute for the COVID-19 Genomics UK (COG-UK) Consortium | The Lighthouse Lab in Milton Keynes and Alex Alderton, Roberto Amato, Jeffrey Barrett, Sonia Goncalves, Ewan Harrison, David K. Jackson, Ian Johnston, Dominic Kwiatkowski, Cordelia Langford, John Sillitoe on behalf of the Wellcome Sanger Institute COVID-19 Surveillance Team |
| EPI_ISL_3043068 | Lighthouse Lab in Glasgow | Wellcome Sanger Institute for the COVID-19 Genomics UK (COG-UK) Consortium | Harper VanSteenhouse, Yumi Kasai, David Gray, Carol Clugston, Anna Dominiczak and Alex Alderton, Roberto Amato, Jeffrey Barrett, Sonia Goncalves, Ewan Harrison, David K. Jackson, Ian Johnston, Dominic Kwiatkowski, Cordelia Langford, John Sillitoe on behalf of the Wellcome Sanger Institute COVID-19 Surveillance Team |
| EPI_ISL_3043069 | Lighthouse Lab in Alderley Park | Wellcome Sanger Institute for the COVID-19 Genomics UK (COG-UK) Consortium | Jacquelyn Wynn, Mairead Hyland, The Lighthouse Lab in Alderley Park and Alex Alderton, Roberto Amato, Jeffrey Barrett, Sonia Goncalves, Ewan Harrison, David K. Jackson, Ian Johnston, Dominic Kwiatkowski, Cordelia Langford, John Sillitoe on behalf of the Wellcome Sanger Institute COVID-19 Surveillance Team |

| | | | |
|---|---|---|---|
| EPI_ISL_3043070 | Randox Laboratories | Wellcome Sanger Institute for the COVID-19 Genomics UK (COG-UK) Consortium | Randox Laboratories and Alex Alderton, Roberto Amato, Jeffrey Barrett, Sonia Goncalves, Ewan Harrison, David K. Jackson, Ian Johnston, Dominic Kwiatkowski, Cordelia Langford, John Sillitoe on behalf of the Wellcome Sanger Institute COVID-19 Surveillance Team |
| EPI_ISL_3043071 | Lighthouse Lab in Glasgow | Wellcome Sanger Institute for the COVID-19 Genomics UK (COG-UK) Consortium | Harper VanSteenhouse, Yumi Kasai, David Gray, Carol Clugston, Anna Dominiczak and Alex Alderton, Roberto Amato, Jeffrey Barrett, Sonia Goncalves, Ewan Harrison, David K. Jackson, Ian Johnston, Dominic Kwiatkowski, Cordelia Langford, John Sillitoe on behalf of the Wellcome Sanger Institute COVID-19 Surveillance Team |
| EPI_ISL_3043072, EPI_ISL_3043073 | Randox Laboratories | Wellcome Sanger Institute for the COVID-19 Genomics UK (COG-UK) Consortium | Randox Laboratories and Alex Alderton, Roberto Amato, Jeffrey Barrett, Sonia Goncalves, Ewan Harrison, David K. Jackson, Ian Johnston, Dominic Kwiatkowski, Cordelia Langford, John Sillitoe on behalf of the Wellcome Sanger Institute COVID-19 Surveillance Team |
| EPI_ISL_3043074 | Lighthouse Lab in Alderley Park | Wellcome Sanger Institute for the COVID-19 Genomics UK (COG-UK) Consortium | Jacquelyn Wynn, Mairead Hyland, The Lighthouse Lab in Alderley Park and Alex Alderton, Roberto Amato, Jeffrey Barrett, Sonia Goncalves, Ewan Harrison, David K. Jackson, Ian Johnston, Dominic Kwiatkowski, Cordelia Langford, John Sillitoe on behalf of the Wellcome Sanger Institute COVID-19 Surveillance Team |
| EPI_ISL_3043075 | Randox Laboratories | Wellcome Sanger Institute for the COVID-19 Genomics UK (COG-UK) Consortium | Randox Laboratories and Alex Alderton, Roberto Amato, Jeffrey Barrett, Sonia Goncalves, Ewan Harrison, David K. Jackson, Ian Johnston, Dominic Kwiatkowski, Cordelia Langford, John Sillitoe on behalf of the Wellcome Sanger Institute COVID-19 Surveillance Team |
| EPI_ISL_3043076, EPI_ISL_3043077 | Lighthouse Lab in Alderley Park | Wellcome Sanger Institute for the COVID-19 Genomics UK (COG-UK) Consortium | Jacquelyn Wynn, Mairead Hyland, The Lighthouse Lab in Alderley Park and Alex Alderton, Roberto Amato, Jeffrey Barrett, Sonia Goncalves, Ewan Harrison, David K. Jackson, Ian Johnston, Dominic Kwiatkowski, Cordelia Langford, John Sillitoe on behalf of the Wellcome Sanger Institute COVID-19 Surveillance Team |
| EPI_ISL_3043078 | Lighthouse Lab in Milton Keynes | Wellcome Sanger Institute for the COVID-19 Genomics UK (COG-UK) Consortium | The Lighthouse Lab in Milton Keynes and Alex Alderton, Roberto Amato, Jeffrey Barrett, Sonia Goncalves, Ewan Harrison, David K. Jackson, Ian Johnston, Dominic Kwiatkowski, Cordelia Langford, John Sillitoe on behalf of the Wellcome Sanger Institute COVID-19 Surveillance Team |
| EPI_ISL_3043079 | Randox Laboratories | Wellcome Sanger Institute for the COVID-19 Genomics UK (COG-UK) Consortium | Randox Laboratories and Alex Alderton, Roberto Amato, Jeffrey Barrett, Sonia Goncalves, Ewan Harrison, David K. Jackson, Ian Johnston, Dominic Kwiatkowski, Cordelia Langford, John Sillitoe on behalf of the Wellcome Sanger Institute COVID-19 Surveillance Team |
| EPI_ISL_3043080, EPI_ISL_3043081 | Lighthouse Lab in Alderley Park | Wellcome Sanger Institute for the COVID-19 Genomics UK (COG-UK) Consortium | Jacquelyn Wynn, Mairead Hyland, The Lighthouse Lab in Alderley Park and Alex Alderton, Roberto Amato, Jeffrey Barrett, Sonia Goncalves, Ewan Harrison, David K. Jackson, Ian Johnston, Dominic Kwiatkowski, Cordelia Langford, John Sillitoe on behalf of the Wellcome Sanger Institute COVID-19 Surveillance Team |
| EPI_ISL_3043082 | Lighthouse Lab in Glasgow | Wellcome Sanger Institute for the COVID-19 Genomics UK (COG-UK) Consortium | Harper VanSteenhouse, Yumi Kasai, David Gray, Carol Clugston, Anna Dominiczak and Alex Alderton, Roberto Amato, Jeffrey Barrett, Sonia Goncalves, Ewan Harrison, David K. Jackson, Ian Johnston, Dominic Kwiatkowski, Cordelia Langford, John Sillitoe on behalf of the Wellcome Sanger Institute COVID-19 Surveillance Team |
| EPI_ISL_3043083, EPI_ISL_3043084 | Lighthouse Lab in Alderley Park | Wellcome Sanger Institute for the COVID-19 Genomics UK (COG-UK) Consortium | Jacquelyn Wynn, Mairead Hyland, The Lighthouse Lab in Alderley Park and Alex Alderton, Roberto Amato, Jeffrey Barrett, Sonia Goncalves, Ewan Harrison, David K. Jackson, Ian Johnston, Dominic Kwiatkowski, Cordelia Langford, John Sillitoe on behalf of the Wellcome Sanger Institute COVID-19 Surveillance Team |
| EPI_ISL_3043085, EPI_ISL_3043086 | Lighthouse Lab in Milton Keynes | Wellcome Sanger Institute for the COVID-19 Genomics UK (COG-UK) Consortium | The Lighthouse Lab in Milton Keynes and Alex Alderton, Roberto Amato, Jeffrey Barrett, Sonia Goncalves, Ewan Harrison, David K. Jackson, Ian Johnston, Dominic Kwiatkowski, Cordelia Langford, John Sillitoe on behalf of the Wellcome Sanger Institute COVID-19 Surveillance Team |
| EPI_ISL_3043087 | Lighthouse Lab in Alderley Park | Wellcome Sanger Institute for the COVID-19 Genomics UK (COG-UK) Consortium | Jacquelyn Wynn, Mairead Hyland, The Lighthouse Lab in Alderley Park and Alex Alderton, Roberto Amato, Jeffrey Barrett, Sonia Goncalves, Ewan Harrison, David K. Jackson, Ian Johnston, Dominic Kwiatkowski, Cordelia Langford, John Sillitoe on behalf of the Wellcome Sanger Institute COVID-19 Surveillance Team |
| EPI_ISL_3043088 | Lighthouse Lab in Glasgow | Wellcome Sanger Institute for the COVID-19 Genomics UK (COG-UK) Consortium | Harper VanSteenhouse, Yumi Kasai, David Gray, Carol Clugston, Anna Dominiczak and Alex Alderton, Roberto Amato, Jeffrey Barrett, Sonia Goncalves, Ewan Harrison, David K. Jackson, Ian Johnston, Dominic Kwiatkowski, Cordelia Langford, John Sillitoe on behalf of the Wellcome Sanger Institute COVID-19 Surveillance Team |
| EPI_ISL_3043089 | Randox Laboratories | Wellcome Sanger Institute for the COVID-19 Genomics UK (COG-UK) Consortium | Randox Laboratories and Alex Alderton, Roberto Amato, Jeffrey Barrett, Sonia Goncalves, Ewan Harrison, David K. Jackson, Ian Johnston, Dominic Kwiatkowski, Cordelia Langford, John Sillitoe on behalf of the Wellcome Sanger Institute COVID-19 Surveillance Team |
| EPI_ISL_3043090 | Lighthouse Lab in Milton Keynes | Wellcome Sanger Institute for the COVID-19 Genomics UK (COG-UK) Consortium | The Lighthouse Lab in Milton Keynes and Alex Alderton, Roberto Amato, Jeffrey Barrett, Sonia Goncalves, Ewan Harrison, David K. Jackson, Ian Johnston, Dominic Kwiatkowski, Cordelia Langford, John Sillitoe on behalf of the Wellcome Sanger Institute COVID-19 Surveillance Team |
| EPI_ISL_3043091 | Randox Laboratories | Wellcome Sanger Institute for the COVID-19 Genomics UK (COG-UK) Consortium | Randox Laboratories and Alex Alderton, Roberto Amato, Jeffrey Barrett, Sonia Goncalves, Ewan Harrison, David K. Jackson, Ian Johnston, Dominic Kwiatkowski, Cordelia Langford, John Sillitoe on behalf of the Wellcome Sanger Institute COVID-19 Surveillance Team |
| EPI_ISL_3043092 | Lighthouse Lab in Alderley Park | Wellcome Sanger Institute for the COVID-19 Genomics UK (COG-UK) Consortium | Jacquelyn Wynn, Mairead Hyland, The Lighthouse Lab in Alderley Park and Alex Alderton, Roberto Amato, Jeffrey Barrett, Sonia Goncalves, Ewan Harrison, David K. Jackson, Ian Johnston, Dominic Kwiatkowski, Cordelia Langford, John Sillitoe on behalf of the Wellcome Sanger Institute COVID-19 Surveillance Team |
| EPI_ISL_3043093, EPI_ISL_3043094 | Lighthouse Lab in Glasgow | Wellcome Sanger Institute for the COVID-19 Genomics UK (COG-UK) Consortium | Harper VanSteenhouse, Yumi Kasai, David Gray, Carol Clugston, Anna Dominiczak and Alex Alderton, Roberto Amato, Jeffrey Barrett, Sonia Goncalves, Ewan Harrison, David K. Jackson, Ian Johnston, Dominic Kwiatkowski, Cordelia Langford, John Sillitoe on behalf of the Wellcome Sanger Institute COVID-19 Surveillance Team |
| EPI_ISL_3043095 | Lighthouse Lab in Alderley Park | Wellcome Sanger Institute for the COVID-19 Genomics UK (COG-UK) Consortium | Jacquelyn Wynn, Mairead Hyland, The Lighthouse Lab in Alderley Park and Alex Alderton, Roberto Amato, Jeffrey Barrett, Sonia Goncalves, Ewan Harrison, David K. Jackson, Ian Johnston, Dominic Kwiatkowski, Cordelia Langford, John Sillitoe on behalf of the Wellcome Sanger Institute COVID-19 Surveillance Team |
| EPI_ISL_3043096 | Lighthouse Lab in Milton Keynes | Wellcome Sanger Institute for the COVID-19 Genomics UK (COG-UK) Consortium | The Lighthouse Lab in Milton Keynes and Alex Alderton, Roberto Amato, Jeffrey Barrett, Sonia Goncalves, Ewan Harrison, David K. Jackson, Ian Johnston, Dominic Kwiatkowski, Cordelia Langford, John Sillitoe on behalf of the Wellcome Sanger Institute COVID-19 Surveillance Team |
| EPI_ISL_3043097 | Randox Laboratories | Wellcome Sanger Institute for the COVID-19 Genomics UK (COG-UK) Consortium | Randox Laboratories and Alex Alderton, Roberto Amato, Jeffrey Barrett, Sonia Goncalves, Ewan Harrison, David K. Jackson, Ian Johnston, Dominic Kwiatkowski, Cordelia Langford, John Sillitoe on behalf of the Wellcome Sanger Institute COVID-19 Surveillance Team |
| EPI_ISL_3043098 | Lighthouse Lab in Alderley Park | Wellcome Sanger Institute for the COVID-19 Genomics UK (COG-UK) Consortium | Jacquelyn Wynn, Mairead Hyland, The Lighthouse Lab in Alderley Park and Alex Alderton, Roberto Amato, Jeffrey Barrett, Sonia Goncalves, Ewan Harrison, David K. Jackson, Ian Johnston, Dominic Kwiatkowski, Cordelia Langford, John Sillitoe on behalf of the Wellcome Sanger Institute COVID-19 Surveillance Team |
| EPI_ISL_3043099, EPI_ISL_3043100, EPI_ISL_3043101 | Lighthouse Lab in Milton Keynes | Wellcome Sanger Institute for the COVID-19 Genomics UK (COG-UK) Consortium | The Lighthouse Lab in Milton Keynes and Alex Alderton, Roberto Amato, Jeffrey Barrett, Sonia Goncalves, Ewan Harrison, David K. Jackson, Ian Johnston, Dominic Kwiatkowski, Cordelia Langford, John Sillitoe on behalf of the Wellcome Sanger Institute COVID-19 Surveillance Team |
| EPI_ISL_3043102 | Lighthouse Lab in Glasgow | Wellcome Sanger Institute for the COVID-19 Genomics UK (COG-UK) Consortium | Harper VanSteenhouse, Yumi Kasai, David Gray, Carol Clugston, Anna Dominiczak and Alex Alderton, Roberto Amato, Jeffrey Barrett, Sonia Goncalves, Ewan Harrison, David K. Jackson, Ian Johnston, Dominic Kwiatkowski, Cordelia Langford, John Sillitoe on behalf of the Wellcome Sanger Institute COVID-19 Surveillance Team |
| EPI_ISL_3043103 | Lighthouse Lab in Milton Keynes | Wellcome Sanger Institute for the COVID-19 Genomics UK (COG-UK) Consortium | The Lighthouse Lab in Milton Keynes and Alex Alderton, Roberto Amato, Jeffrey Barrett, Sonia Goncalves, Ewan Harrison, David K. Jackson, Ian Johnston, Dominic Kwiatkowski, Cordelia Langford, John Sillitoe on behalf of the Wellcome Sanger Institute COVID-19 Surveillance Team |
| EPI_ISL_3043104 | Lighthouse Lab in Alderley Park | Wellcome Sanger Institute for the COVID-19 Genomics UK (COG-UK) Consortium | Jacquelyn Wynn, Mairead Hyland, The Lighthouse Lab in Alderley Park and Alex Alderton, Roberto Amato, Jeffrey Barrett, Sonia Goncalves, Ewan Harrison, David K. Jackson, Ian Johnston, Dominic Kwiatkowski, Cordelia Langford, John Sillitoe on behalf of the Wellcome Sanger Institute COVID-19 Surveillance Team |
| EPI_ISL_3043105 | Lighthouse Lab in Milton Keynes | Wellcome Sanger Institute for the COVID-19 Genomics UK (COG-UK) Consortium | The Lighthouse Lab in Milton Keynes and Alex Alderton, Roberto Amato, Jeffrey Barrett, Sonia Goncalves, Ewan Harrison, David K. Jackson, Ian Johnston, Dominic Kwiatkowski, Cordelia Langford, John Sillitoe on behalf of the Wellcome Sanger Institute COVID-19 Surveillance Team |
| EPI_ISL_3043106 | Lighthouse Lab in Alderley Park | Wellcome Sanger Institute for the COVID-19 Genomics UK (COG-UK) Consortium | Jacquelyn Wynn, Mairead Hyland, The Lighthouse Lab in Alderley Park and Alex Alderton, Roberto Amato, Jeffrey Barrett, Sonia Goncalves, Ewan Harrison, David K. Jackson, Ian Johnston, Dominic Kwiatkowski, Cordelia Langford, John Sillitoe on behalf of the Wellcome Sanger Institute COVID-19 Surveillance Team |
| EPI_ISL_3043107 | Lighthouse Lab in Milton Keynes | Wellcome Sanger Institute for the COVID-19 Genomics UK (COG-UK) Consortium | The Lighthouse Lab in Milton Keynes and Alex Alderton, Roberto Amato, Jeffrey Barrett, Sonia Goncalves, Ewan Harrison, David K. Jackson, Ian Johnston, Dominic Kwiatkowski, Cordelia Langford, John Sillitoe on behalf of the Wellcome Sanger Institute COVID-19 Surveillance Team |
| EPI_ISL_3043108 | Randox Laboratories | Wellcome Sanger Institute for the COVID-19 Genomics UK (COG-UK) Consortium | Randox Laboratories and Alex Alderton, Roberto Amato, Jeffrey Barrett, Sonia Goncalves, Ewan Harrison, David K. Jackson, Ian Johnston, Dominic Kwiatkowski, Cordelia Langford, John Sillitoe on behalf of the Wellcome Sanger Institute COVID-19 Surveillance Team |
| EPI_ISL_3043109 | Lighthouse Lab in Milton Keynes | Wellcome Sanger Institute for the COVID-19 Genomics UK (COG-UK) Consortium | The Lighthouse Lab in Milton Keynes and Alex Alderton, Roberto Amato, Jeffrey Barrett, Sonia Goncalves, Ewan Harrison, David K. Jackson, Ian Johnston, Dominic Kwiatkowski, Cordelia Langford, John Sillitoe on behalf of the Wellcome Sanger Institute COVID-19 Surveillance Team |
| EPI_ISL_3043110 | Lighthouse Lab in Alderley Park | Wellcome Sanger Institute for the COVID-19 Genomics UK (COG-UK) Consortium | Jacquelyn Wynn, Mairead Hyland, The Lighthouse Lab in Alderley Park and Alex Alderton, Roberto Amato, Jeffrey Barrett, Sonia Goncalves, Ewan Harrison, David K. Jackson, Ian Johnston, Dominic Kwiatkowski, Cordelia Langford, John Sillitoe on behalf of the Wellcome Sanger Institute COVID-19 Surveillance Team |
| EPI_ISL_3043111 | Lighthouse Lab in Milton Keynes | Wellcome Sanger Institute for the COVID-19 Genomics UK (COG-UK) Consortium | The Lighthouse Lab in Milton Keynes and Alex Alderton, Roberto Amato, Jeffrey Barrett, Sonia Goncalves, Ewan Harrison, David K. Jackson, Ian Johnston, Dominic Kwiatkowski, Cordelia Langford, John Sillitoe on behalf of the Wellcome Sanger Institute COVID-19 Surveillance Team |
| EPI_ISL_3043112 | Lighthouse Lab in Glasgow | Wellcome Sanger Institute for the COVID-19 Genomics UK (COG-UK) Consortium | Harper VanSteenhouse, Yumi Kasai, David Gray, Carol Clugston, Anna Dominiczak and Alex Alderton, Roberto Amato, Jeffrey Barrett, Sonia Goncalves, Ewan Harrison, David K. Jackson, Ian Johnston, Dominic Kwiatkowski, Cordelia Langford, John Sillitoe on behalf of the Wellcome Sanger Institute COVID-19 Surveillance Team |
| EPI_ISL_3043113 | Randox Laboratories | Wellcome Sanger Institute for the COVID-19 Genomics UK (COG-UK) Consortium | Randox Laboratories and Alex Alderton, Roberto Amato, Jeffrey Barrett, Sonia Goncalves, Ewan Harrison, David K. Jackson, Ian Johnston, Dominic Kwiatkowski, Cordelia Langford, John Sillitoe on behalf of the Wellcome Sanger Institute COVID-19 Surveillance Team |
| EPI_ISL_3043114, EPI_ISL_3043115, EPI_ISL_3043116 | Lighthouse Lab in Milton Keynes | Wellcome Sanger Institute for the COVID-19 Genomics UK (COG-UK) Consortium | The Lighthouse Lab in Milton Keynes and Alex Alderton, Roberto Amato, Jeffrey Barrett, Sonia Goncalves, Ewan Harrison, David K. Jackson, Ian Johnston, Dominic Kwiatkowski, Cordelia Langford, John Sillitoe on behalf of the Wellcome Sanger Institute COVID-19 Surveillance Team |
| EPI_ISL_3043117 | Lighthouse Lab in Alderley Park | Wellcome Sanger Institute for the COVID-19 Genomics UK (COG-UK) Consortium | Jacquelyn Wynn, Mairead Hyland, The Lighthouse Lab in Alderley Park and Alex Alderton, Roberto Amato, Jeffrey Barrett, Sonia Goncalves, Ewan Harrison, David K. Jackson, Ian Johnston, Dominic Kwiatkowski, Cordelia Langford, John Sillitoe on behalf of the Wellcome Sanger Institute COVID-19 Surveillance Team |
| EPI_ISL_3043118 | Lighthouse Lab in Glasgow | Wellcome Sanger Institute for the COVID-19 Genomics UK (COG-UK) Consortium | Harper VanSteenhouse, Yumi Kasai, David Gray, Carol Clugston, Anna Dominiczak and Alex Alderton, Roberto Amato, Jeffrey Barrett, Sonia Goncalves, Ewan Harrison, David K. Jackson, Ian Johnston, Dominic Kwiatkowski, Cordelia Langford, John Sillitoe on behalf of the Wellcome Sanger Institute COVID-19 Surveillance Team |
| EPI_ISL_3043119 | Lighthouse Lab in Alderley Park | Wellcome Sanger Institute for the COVID-19 Genomics UK (COG-UK) Consortium | Jacquelyn Wynn, Mairead Hyland, The Lighthouse Lab in Alderley Park and Alex Alderton, Roberto Amato, Jeffrey Barrett, Sonia Goncalves, Ewan Harrison, David K. Jackson, Ian Johnston, Dominic Kwiatkowski, Cordelia Langford, John Sillitoe on behalf of the Wellcome Sanger Institute COVID-19 Surveillance Team |
| EPI_ISL_3043120, EPI_ISL_3043121 | Lighthouse Lab in Glasgow | Wellcome Sanger Institute for the COVID-19 Genomics UK (COG-UK) Consortium | Harper VanSteenhouse, Yumi Kasai, David Gray, Carol Clugston, Anna Dominiczak and Alex Alderton, Roberto Amato, Jeffrey Barrett, Sonia Goncalves, Ewan Harrison, David K. Jackson, Ian Johnston, Dominic Kwiatkowski, Cordelia Langford, John Sillitoe on behalf of the Wellcome Sanger Institute COVID-19 Surveillance Team |
| EPI_ISL_3043122 | Lighthouse Lab in Milton Keynes | Wellcome Sanger Institute for the COVID-19 Genomics UK (COG-UK) Consortium | The Lighthouse Lab in Milton Keynes and Alex Alderton, Roberto Amato, Jeffrey Barrett, Sonia Goncalves, Ewan Harrison, David K. Jackson, Ian Johnston, Dominic Kwiatkowski, Cordelia Langford, John Sillitoe on behalf of the Wellcome Sanger Institute COVID-19 Surveillance Team |
| EPI_ISL_3043123 | Lighthouse Lab in Alderley Park | Wellcome Sanger Institute for the COVID-19 Genomics UK (COG-UK) Consortium | Jacquelyn Wynn, Mairead Hyland, The Lighthouse Lab in Alderley Park and Alex Alderton, Roberto Amato, Jeffrey Barrett, Sonia Goncalves, Ewan Harrison, David K. Jackson, Ian Johnston, Dominic Kwiatkowski, Cordelia Langford, John Sillitoe on behalf of the Wellcome Sanger Institute COVID-19 Surveillance Team |
| EPI_ISL_3043124, EPI_ISL_3043125, EPI_ISL_3043126 | Randox Laboratories | Wellcome Sanger Institute for the COVID-19 Genomics UK (COG-UK) Consortium | Randox Laboratories and Alex Alderton, Roberto Amato, Jeffrey Barrett, Sonia Goncalves, Ewan Harrison, David K. Jackson, Ian Johnston, Dominic Kwiatkowski, Cordelia Langford, John Sillitoe on behalf of the Wellcome Sanger Institute COVID-19 Surveillance Team |

| | | | |
|---|---|---|---|
| EPI_ISL_3043127 | Lighthouse Lab in Glasgow | Wellcome Sanger Institute for the COVID-19 Genomics UK (COG-UK) Consortium | Harper VanSteenhouse, Yumi Kasai, David Gray, Carol Clugston, Anna Dominiczak and Alex Alderton, Roberto Amato, Jeffrey Barrett, Sonia Goncalves, Ewan Harrison, David K. Jackson, Ian Johnston, Dominic Kwiatkowski, Cordelia Langford, John Sillitoe on behalf of the Wellcome Sanger Institute COVID-19 Surveillance Team |
| EPI_ISL_3043128 | Lighthouse Lab in Alderley Park | Wellcome Sanger Institute for the COVID-19 Genomics UK (COG-UK) Consortium | Jacquelyn Wynn, Mairead Hyland, The Lighthouse Lab in Alderley Park and Alex Alderton, Roberto Amato, Jeffrey Barrett, Sonia Goncalves, Ewan Harrison, David K. Jackson, Ian Johnston, Dominic Kwiatkowski, Cordelia Langford, John Sillitoe on behalf of the Wellcome Sanger Institute COVID-19 Surveillance Team |
| EPI_ISL_3043129 | Lighthouse Lab in Milton Keynes | Wellcome Sanger Institute for the COVID-19 Genomics UK (COG-UK) Consortium | The Lighthouse Lab in Milton Keynes and Alex Alderton, Roberto Amato, Jeffrey Barrett, Sonia Goncalves, Ewan Harrison, David K. Jackson, Ian Johnston, Dominic Kwiatkowski, Cordelia Langford, John Sillitoe on behalf of the Wellcome Sanger Institute COVID-19 Surveillance Team |
| EPI_ISL_3043130 | Lighthouse Lab in Glasgow | Wellcome Sanger Institute for the COVID-19 Genomics UK (COG-UK) Consortium | Harper VanSteenhouse, Yumi Kasai, David Gray, Carol Clugston, Anna Dominiczak and Alex Alderton, Roberto Amato, Jeffrey Barrett, Sonia Goncalves, Ewan Harrison, David K. Jackson, Ian Johnston, Dominic Kwiatkowski, Cordelia Langford, John Sillitoe on behalf of the Wellcome Sanger Institute COVID-19 Surveillance Team |
| EPI_ISL_3043131 | Lighthouse Lab in Milton Keynes | Wellcome Sanger Institute for the COVID-19 Genomics UK (COG-UK) Consortium | The Lighthouse Lab in Milton Keynes and Alex Alderton, Roberto Amato, Jeffrey Barrett, Sonia Goncalves, Ewan Harrison, David K. Jackson, Ian Johnston, Dominic Kwiatkowski, Cordelia Langford, John Sillitoe on behalf of the Wellcome Sanger Institute COVID-19 Surveillance Team |
| EPI_ISL_3043132 | Randox Laboratories | Wellcome Sanger Institute for the COVID-19 Genomics UK (COG-UK) Consortium | Randox Laboratories and Alex Alderton, Roberto Amato, Jeffrey Barrett, Sonia Goncalves, Ewan Harrison, David K. Jackson, Ian Johnston, Dominic Kwiatkowski, Cordelia Langford, John Sillitoe on behalf of the Wellcome Sanger Institute COVID-19 Surveillance Team |
| EPI_ISL_3043133, EPI_ISL_3043134, EPI_ISL_3043135 | Lighthouse Lab in Milton Keynes | Wellcome Sanger Institute for the COVID-19 Genomics UK (COG-UK) Consortium | The Lighthouse Lab in Milton Keynes and Alex Alderton, Roberto Amato, Jeffrey Barrett, Sonia Goncalves, Ewan Harrison, David K. Jackson, Ian Johnston, Dominic Kwiatkowski, Cordelia Langford, John Sillitoe on behalf of the Wellcome Sanger Institute COVID-19 Surveillance Team |
| EPI_ISL_3043136, EPI_ISL_3043137, EPI_ISL_3043138 | Lighthouse Lab in Glasgow | Wellcome Sanger Institute for the COVID-19 Genomics UK (COG-UK) Consortium | Harper VanSteenhouse, Yumi Kasai, David Gray, Carol Clugston, Anna Dominiczak and Alex Alderton, Roberto Amato, Jeffrey Barrett, Sonia Goncalves, Ewan Harrison, David K. Jackson, Ian Johnston, Dominic Kwiatkowski, Cordelia Langford, John Sillitoe on behalf of the Wellcome Sanger Institute COVID-19 Surveillance Team |
| EPI_ISL_3043139 | Lighthouse Lab in Milton Keynes | Wellcome Sanger Institute for the COVID-19 Genomics UK (COG-UK) Consortium | The Lighthouse Lab in Milton Keynes and Alex Alderton, Roberto Amato, Jeffrey Barrett, Sonia Goncalves, Ewan Harrison, David K. Jackson, Ian Johnston, Dominic Kwiatkowski, Cordelia Langford, John Sillitoe on behalf of the Wellcome Sanger Institute COVID-19 Surveillance Team |
| EPI_ISL_3043140 | Lighthouse Lab in Glasgow | Wellcome Sanger Institute for the COVID-19 Genomics UK (COG-UK) Consortium | Harper VanSteenhouse, Yumi Kasai, David Gray, Carol Clugston, Anna Dominiczak and Alex Alderton, Roberto Amato, Jeffrey Barrett, Sonia Goncalves, Ewan Harrison, David K. Jackson, Ian Johnston, Dominic Kwiatkowski, Cordelia Langford, John Sillitoe on behalf of the Wellcome Sanger Institute COVID-19 Surveillance Team |
| EPI_ISL_3043141 | Lighthouse Lab in Milton Keynes | Wellcome Sanger Institute for the COVID-19 Genomics UK (COG-UK) Consortium | The Lighthouse Lab in Milton Keynes and Alex Alderton, Roberto Amato, Jeffrey Barrett, Sonia Goncalves, Ewan Harrison, David K. Jackson, Ian Johnston, Dominic Kwiatkowski, Cordelia Langford, John Sillitoe on behalf of the Wellcome Sanger Institute COVID-19 Surveillance Team |
| EPI_ISL_3043142 | Lighthouse Lab in Alderley Park | Wellcome Sanger Institute for the COVID-19 Genomics UK (COG-UK) Consortium | Jacquelyn Wynn, Mairead Hyland, The Lighthouse Lab in Alderley Park and Alex Alderton, Roberto Amato, Jeffrey Barrett, Sonia Goncalves, Ewan Harrison, David K. Jackson, Ian Johnston, Dominic Kwiatkowski, Cordelia Langford, John Sillitoe on behalf of the Wellcome Sanger Institute COVID-19 Surveillance Team |
| EPI_ISL_3043143 | Randox Laboratories | Wellcome Sanger Institute for the COVID-19 Genomics UK (COG-UK) Consortium | Randox Laboratories and Alex Alderton, Roberto Amato, Jeffrey Barrett, Sonia Goncalves, Ewan Harrison, David K. Jackson, Ian Johnston, Dominic Kwiatkowski, Cordelia Langford, John Sillitoe on behalf of the Wellcome Sanger Institute COVID-19 Surveillance Team |
| EPI_ISL_3043144 | Lighthouse Lab in Alderley Park | Wellcome Sanger Institute for the COVID-19 Genomics UK (COG-UK) Consortium | Jacquelyn Wynn, Mairead Hyland, The Lighthouse Lab in Alderley Park and Alex Alderton, Roberto Amato, Jeffrey Barrett, Sonia Goncalves, Ewan Harrison, David K. Jackson, Ian Johnston, Dominic Kwiatkowski, Cordelia Langford, John Sillitoe on behalf of the Wellcome Sanger Institute COVID-19 Surveillance Team |
| EPI_ISL_3043145 | Lighthouse Lab in Glasgow | Wellcome Sanger Institute for the COVID-19 Genomics UK (COG-UK) Consortium | Harper VanSteenhouse, Yumi Kasai, David Gray, Carol Clugston, Anna Dominiczak and Alex Alderton, Roberto Amato, Jeffrey Barrett, Sonia Goncalves, Ewan Harrison, David K. Jackson, Ian Johnston, Dominic Kwiatkowski, Cordelia Langford, John Sillitoe on behalf of the Wellcome Sanger Institute COVID-19 Surveillance Team |
| EPI_ISL_3043146 | Randox Laboratories | Wellcome Sanger Institute for the COVID-19 Genomics UK (COG-UK) Consortium | Randox Laboratories and Alex Alderton, Roberto Amato, Jeffrey Barrett, Sonia Goncalves, Ewan Harrison, David K. Jackson, Ian Johnston, Dominic Kwiatkowski, Cordelia Langford, John Sillitoe on behalf of the Wellcome Sanger Institute COVID-19 Surveillance Team |
| EPI_ISL_3043147, EPI_ISL_3043148 | Lighthouse Lab in Alderley Park | Wellcome Sanger Institute for the COVID-19 Genomics UK (COG-UK) Consortium | Jacquelyn Wynn, Mairead Hyland, The Lighthouse Lab in Alderley Park and Alex Alderton, Roberto Amato, Jeffrey Barrett, Sonia Goncalves, Ewan Harrison, David K. Jackson, Ian Johnston, Dominic Kwiatkowski, Cordelia Langford, John Sillitoe on behalf of the Wellcome Sanger Institute COVID-19 Surveillance Team |
| EPI_ISL_3043149 | Lighthouse Lab in Milton Keynes | Wellcome Sanger Institute for the COVID-19 Genomics UK (COG-UK) Consortium | The Lighthouse Lab in Milton Keynes and Alex Alderton, Roberto Amato, Jeffrey Barrett, Sonia Goncalves, Ewan Harrison, David K. Jackson, Ian Johnston, Dominic Kwiatkowski, Cordelia Langford, John Sillitoe on behalf of the Wellcome Sanger Institute COVID-19 Surveillance Team |
| EPI_ISL_3043150 | Lighthouse Lab in Alderley Park | Wellcome Sanger Institute for the COVID-19 Genomics UK (COG-UK) Consortium | Jacquelyn Wynn, Mairead Hyland, The Lighthouse Lab in Alderley Park and Alex Alderton, Roberto Amato, Jeffrey Barrett, Sonia Goncalves, Ewan Harrison, David K. Jackson, Ian Johnston, Dominic Kwiatkowski, Cordelia Langford, John Sillitoe on behalf of the Wellcome Sanger Institute COVID-19 Surveillance Team |
| EPI_ISL_3043151 | Lighthouse Lab in Glasgow | Wellcome Sanger Institute for the COVID-19 Genomics UK (COG-UK) Consortium | Harper VanSteenhouse, Yumi Kasai, David Gray, Carol Clugston, Anna Dominiczak and Alex Alderton, Roberto Amato, Jeffrey Barrett, Sonia Goncalves, Ewan Harrison, David K. Jackson, Ian Johnston, Dominic Kwiatkowski, Cordelia Langford, John Sillitoe on behalf of the Wellcome Sanger Institute COVID-19 Surveillance Team |
| EPI_ISL_3043152, EPI_ISL_3043153 | Lighthouse Lab in Milton Keynes | Wellcome Sanger Institute for the COVID-19 Genomics UK (COG-UK) Consortium | The Lighthouse Lab in Milton Keynes and Alex Alderton, Roberto Amato, Jeffrey Barrett, Sonia Goncalves, Ewan Harrison, David K. Jackson, Ian Johnston, Dominic Kwiatkowski, Cordelia Langford, John Sillitoe on behalf of the Wellcome Sanger Institute COVID-19 Surveillance Team |
| EPI_ISL_3043154, EPI_ISL_3043155 | Lighthouse Lab in Alderley Park | Wellcome Sanger Institute for the COVID-19 Genomics UK (COG-UK) Consortium | Jacquelyn Wynn, Mairead Hyland, The Lighthouse Lab in Alderley Park and Alex Alderton, Roberto Amato, Jeffrey Barrett, Sonia Goncalves, Ewan Harrison, David K. Jackson, Ian Johnston, Dominic Kwiatkowski, Cordelia Langford, John Sillitoe on behalf of the Wellcome Sanger Institute COVID-19 Surveillance Team |
| EPI_ISL_3043156 | Berkshire and Surrey Pathology Services Lighthouse Laboratory | Wellcome Sanger Institute for the COVID-19 Genomics UK (COG-UK) Consortium | Berkshire and Surrey Pathology Services Lighthouse Laboratory and Alex Alderton, Roberto Amato, Jeffrey Barrett, Sonia Goncalves, Ewan Harrison, David K. Jackson, Ian Johnston, Dominic Kwiatkowski, Cordelia Langford, John Sillitoe on behalf of the Wellcome Sanger Institute COVID-19 Surveillance Team |
| EPI_ISL_3043157 | Lighthouse Lab in Glasgow | Wellcome Sanger Institute for the COVID-19 Genomics UK (COG-UK) Consortium | Harper VanSteenhouse, Yumi Kasai, David Gray, Carol Clugston, Anna Dominiczak and Alex Alderton, Roberto Amato, Jeffrey Barrett, Sonia Goncalves, Ewan Harrison, David K. Jackson, Ian Johnston, Dominic Kwiatkowski, Cordelia Langford, John Sillitoe on behalf of the Wellcome Sanger Institute COVID-19 Surveillance Team |
| EPI_ISL_3043158 | Berkshire and Surrey Pathology Services Lighthouse Laboratory | Wellcome Sanger Institute for the COVID-19 Genomics UK (COG-UK) Consortium | Berkshire and Surrey Pathology Services Lighthouse Laboratory and Alex Alderton, Roberto Amato, Jeffrey Barrett, Sonia Goncalves, Ewan Harrison, David K. Jackson, Ian Johnston, Dominic Kwiatkowski, Cordelia Langford, John Sillitoe on behalf of the Wellcome Sanger Institute COVID-19 Surveillance Team |
| EPI_ISL_3043159, EPI_ISL_3043160 | Lighthouse Lab in Milton Keynes | Wellcome Sanger Institute for the COVID-19 Genomics UK (COG-UK) Consortium | The Lighthouse Lab in Milton Keynes and Alex Alderton, Roberto Amato, Jeffrey Barrett, Sonia Goncalves, Ewan Harrison, David K. Jackson, Ian Johnston, Dominic Kwiatkowski, Cordelia Langford, John Sillitoe on behalf of the Wellcome Sanger Institute COVID-19 Surveillance Team |
| EPI_ISL_3043161, EPI_ISL_3043162, EPI_ISL_3043163, EPI_ISL_3043164, EPI_ISL_3043165, EPI_ISL_3043166 | Lighthouse Lab in Glasgow | Wellcome Sanger Institute for the COVID-19 Genomics UK (COG-UK) Consortium | Harper VanSteenhouse, Yumi Kasai, David Gray, Carol Clugston, Anna Dominiczak and Alex Alderton, Roberto Amato, Jeffrey Barrett, Sonia Goncalves, Ewan Harrison, David K. Jackson, Ian Johnston, Dominic Kwiatkowski, Cordelia Langford, John Sillitoe on behalf of the Wellcome Sanger Institute COVID-19 Surveillance Team |
| EPI_ISL_3043167 | Lighthouse Lab in Milton Keynes | Wellcome Sanger Institute for the COVID-19 Genomics UK (COG-UK) Consortium | The Lighthouse Lab in Milton Keynes and Alex Alderton, Roberto Amato, Jeffrey Barrett, Sonia Goncalves, Ewan Harrison, David K. Jackson, Ian Johnston, Dominic Kwiatkowski, Cordelia Langford, John Sillitoe on behalf of the Wellcome Sanger Institute COVID-19 Surveillance Team |
| EPI_ISL_3043168, EPI_ISL_3043169, EPI_ISL_3043170 | Lighthouse Lab in Glasgow | Wellcome Sanger Institute for the COVID-19 Genomics UK (COG-UK) Consortium | Harper VanSteenhouse, Yumi Kasai, David Gray, Carol Clugston, Anna Dominiczak and Alex Alderton, Roberto Amato, Jeffrey Barrett, Sonia Goncalves, Ewan Harrison, David K. Jackson, Ian Johnston, Dominic Kwiatkowski, Cordelia Langford, John Sillitoe on behalf of the Wellcome Sanger Institute COVID-19 Surveillance Team |
| EPI_ISL_3043171 | Berkshire and Surrey Pathology Services Lighthouse Laboratory | Wellcome Sanger Institute for the COVID-19 Genomics UK (COG-UK) Consortium | Berkshire and Surrey Pathology Services Lighthouse Laboratory and Alex Alderton, Roberto Amato, Jeffrey Barrett, Sonia Goncalves, Ewan Harrison, David K. Jackson, Ian Johnston, Dominic Kwiatkowski, Cordelia Langford, John Sillitoe on behalf of the Wellcome Sanger Institute COVID-19 Surveillance Team |
| EPI_ISL_3043172, EPI_ISL_3043173 | Lighthouse Lab in Milton Keynes | Wellcome Sanger Institute for the COVID-19 Genomics UK (COG-UK) Consortium | The Lighthouse Lab in Milton Keynes and Alex Alderton, Roberto Amato, Jeffrey Barrett, Sonia Goncalves, Ewan Harrison, David K. Jackson, Ian Johnston, Dominic Kwiatkowski, Cordelia Langford, John Sillitoe on behalf of the Wellcome Sanger Institute COVID-19 Surveillance Team |
| EPI_ISL_3043174 | Berkshire and Surrey Pathology Services Lighthouse Laboratory | Wellcome Sanger Institute for the COVID-19 Genomics UK (COG-UK) Consortium | Berkshire and Surrey Pathology Services Lighthouse Laboratory and Alex Alderton, Roberto Amato, Jeffrey Barrett, Sonia Goncalves, Ewan Harrison, David K. Jackson, Ian Johnston, Dominic Kwiatkowski, Cordelia Langford, John Sillitoe on behalf of the Wellcome Sanger Institute COVID-19 Surveillance Team |
| EPI_ISL_3043175, EPI_ISL_3043176, EPI_ISL_3043177 | Lighthouse Lab in Milton Keynes | Wellcome Sanger Institute for the COVID-19 Genomics UK (COG-UK) Consortium | The Lighthouse Lab in Milton Keynes and Alex Alderton, Roberto Amato, Jeffrey Barrett, Sonia Goncalves, Ewan Harrison, David K. Jackson, Ian Johnston, Dominic Kwiatkowski, Cordelia Langford, John Sillitoe on behalf of the Wellcome Sanger Institute COVID-19 Surveillance Team |
| EPI_ISL_3043178 | Lighthouse Lab in Glasgow | Wellcome Sanger Institute for the COVID-19 Genomics UK (COG-UK) Consortium | Harper VanSteenhouse, Yumi Kasai, David Gray, Carol Clugston, Anna Dominiczak and Alex Alderton, Roberto Amato, Jeffrey Barrett, Sonia Goncalves, Ewan Harrison, David K. Jackson, Ian Johnston, Dominic Kwiatkowski, Cordelia Langford, John Sillitoe on behalf of the Wellcome Sanger Institute COVID-19 Surveillance Team |
| EPI_ISL_3043179 | Lighthouse Lab in Milton Keynes | Wellcome Sanger Institute for the COVID-19 Genomics UK (COG-UK) Consortium | The Lighthouse Lab in Milton Keynes and Alex Alderton, Roberto Amato, Jeffrey Barrett, Sonia Goncalves, Ewan Harrison, David K. Jackson, Ian Johnston, Dominic Kwiatkowski, Cordelia Langford, John Sillitoe on behalf of the Wellcome Sanger Institute COVID-19 Surveillance Team |
| EPI_ISL_3043180 | Berkshire and Surrey Pathology Services Lighthouse Laboratory | Wellcome Sanger Institute for the COVID-19 Genomics UK (COG-UK) Consortium | Berkshire and Surrey Pathology Services Lighthouse Laboratory and Alex Alderton, Roberto Amato, Jeffrey Barrett, Sonia Goncalves, Ewan Harrison, David K. Jackson, Ian Johnston, Dominic Kwiatkowski, Cordelia Langford, John Sillitoe on behalf of the Wellcome Sanger Institute COVID-19 Surveillance Team |
| EPI_ISL_3043181, EPI_ISL_3043182 | Lighthouse Lab in Glasgow | Wellcome Sanger Institute for the COVID-19 Genomics UK (COG-UK) Consortium | Harper VanSteenhouse, Yumi Kasai, David Gray, Carol Clugston, Anna Dominiczak and Alex Alderton, Roberto Amato, Jeffrey Barrett, Sonia Goncalves, Ewan Harrison, David K. Jackson, Ian Johnston, Dominic Kwiatkowski, Cordelia Langford, John Sillitoe on behalf of the Wellcome Sanger Institute COVID-19 Surveillance Team |
| EPI_ISL_3043183 | Berkshire and Surrey Pathology Services Lighthouse Laboratory | Wellcome Sanger Institute for the COVID-19 Genomics UK (COG-UK) Consortium | Berkshire and Surrey Pathology Services Lighthouse Laboratory and Alex Alderton, Roberto Amato, Jeffrey Barrett, Sonia Goncalves, Ewan Harrison, David K. Jackson, Ian Johnston, Dominic Kwiatkowski, Cordelia Langford, John Sillitoe on behalf of the Wellcome Sanger Institute COVID-19 Surveillance Team |
| EPI_ISL_3043184 | Lighthouse Lab in Glasgow | Wellcome Sanger Institute for the COVID-19 Genomics UK (COG-UK) Consortium | Harper VanSteenhouse, Yumi Kasai, David Gray, Carol Clugston, Anna Dominiczak and Alex Alderton, Roberto Amato, Jeffrey Barrett, Sonia Goncalves, Ewan Harrison, David K. Jackson, Ian Johnston, Dominic Kwiatkowski, Cordelia Langford, John Sillitoe on behalf of the Wellcome Sanger Institute COVID-19 Surveillance Team |
| EPI_ISL_3043185 | Berkshire and Surrey Pathology Services Lighthouse Laboratory | Wellcome Sanger Institute for the COVID-19 Genomics UK (COG-UK) Consortium | Berkshire and Surrey Pathology Services Lighthouse Laboratory and Alex Alderton, Roberto Amato, Jeffrey Barrett, Sonia Goncalves, Ewan Harrison, David K. Jackson, Ian Johnston, Dominic Kwiatkowski, Cordelia Langford, John Sillitoe on behalf of the Wellcome Sanger Institute COVID-19 Surveillance Team |
| EPI_ISL_3043186 | Lighthouse Lab in Glasgow | Wellcome Sanger Institute for the COVID-19 Genomics UK (COG-UK) Consortium | Harper VanSteenhouse, Yumi Kasai, David Gray, Carol Clugston, Anna Dominiczak and Alex Alderton, Roberto Amato, Jeffrey Barrett, Sonia Goncalves, Ewan Harrison, David K. Jackson, Ian Johnston, Dominic Kwiatkowski, Cordelia Langford, John Sillitoe on behalf of the Wellcome Sanger Institute COVID-19 Surveillance Team |
| EPI_ISL_3043187 | Lighthouse Lab in Milton Keynes | Wellcome Sanger Institute for the COVID-19 Genomics UK (COG-UK) | The Lighthouse Lab in Milton Keynes and Alex Alderton, Roberto Amato, Jeffrey Barrett, Sonia Goncalves, Ewan Harrison, David K. Jackson, Ian Johnston, Dominic Kwiatkowski, Cordelia Langford, John Sillitoe on behalf of the Wellcome Sanger |

| | | | |
|---|---|---|---|
| | | Consortium | Institute COVID-19 Surveillance Team |
| EPI_ISL_3043188, EPI_ISL_3043189 | Lighthouse Lab in Glasgow | Wellcome Sanger Institute for the COVID-19 Genomics UK (COG-UK) Consortium | Harper VanSteenhouse, Yumi Kasai, David Gray, Carol Clugston, Anna Dominiczak and Alex Alderton, Roberto Amato, Jeffrey Barrett, Sonia Goncalves, Ewan Harrison, David K. Jackson, Ian Johnston, Dominic Kwiatkowski, Cordelia Langford, John Sillitoe on behalf of the Wellcome Sanger Institute COVID-19 Surveillance Team |
| EPI_ISL_3043190 | Berkshire and Surrey Pathology Services Lighthouse Laboratory | Wellcome Sanger Institute for the COVID-19 Genomics UK (COG-UK) Consortium | Berkshire and Surrey Pathology Services Lighthouse Laboratory and Alex Alderton, Roberto Amato, Jeffrey Barrett, Sonia Goncalves, Ewan Harrison, David K. Jackson, Ian Johnston, Dominic Kwiatkowski, Cordelia Langford, John Sillitoe on behalf of the Wellcome Sanger Institute COVID-19 Surveillance Team |
| EPI_ISL_3043191 | Lighthouse Lab in Milton Keynes | Wellcome Sanger Institute for the COVID-19 Genomics UK (COG-UK) Consortium | The Lighthouse Lab in Milton Keynes and Alex Alderton, Roberto Amato, Jeffrey Barrett, Sonia Goncalves, Ewan Harrison, David K. Jackson, Ian Johnston, Dominic Kwiatkowski, Cordelia Langford, John Sillitoe on behalf of the Wellcome Sanger Institute COVID-19 Surveillance Team |
| EPI_ISL_3043192 | Berkshire and Surrey Pathology Services Lighthouse Laboratory | Wellcome Sanger Institute for the COVID-19 Genomics UK (COG-UK) Consortium | Berkshire and Surrey Pathology Services Lighthouse Laboratory and Alex Alderton, Roberto Amato, Jeffrey Barrett, Sonia Goncalves, Ewan Harrison, David K. Jackson, Ian Johnston, Dominic Kwiatkowski, Cordelia Langford, John Sillitoe on behalf of the Wellcome Sanger Institute COVID-19 Surveillance Team |
| EPI_ISL_3043193, EPI_ISL_3043194, EPI_ISL_3043195, EPI_ISL_3043196 | Lighthouse Lab in Milton Keynes | Wellcome Sanger Institute for the COVID-19 Genomics UK (COG-UK) Consortium | The Lighthouse Lab in Milton Keynes and Alex Alderton, Roberto Amato, Jeffrey Barrett, Sonia Goncalves, Ewan Harrison, David K. Jackson, Ian Johnston, Dominic Kwiatkowski, Cordelia Langford, John Sillitoe on behalf of the Wellcome Sanger Institute COVID-19 Surveillance Team |
| EPI_ISL_3043197 | Lighthouse Lab in Glasgow | Wellcome Sanger Institute for the COVID-19 Genomics UK (COG-UK) Consortium | Harper VanSteenhouse, Yumi Kasai, David Gray, Carol Clugston, Anna Dominiczak and Alex Alderton, Roberto Amato, Jeffrey Barrett, Sonia Goncalves, Ewan Harrison, David K. Jackson, Ian Johnston, Dominic Kwiatkowski, Cordelia Langford, John Sillitoe on behalf of the Wellcome Sanger Institute COVID-19 Surveillance Team |
| EPI_ISL_3043198 | Berkshire and Surrey Pathology Services Lighthouse Laboratory | Wellcome Sanger Institute for the COVID-19 Genomics UK (COG-UK) Consortium | Berkshire and Surrey Pathology Services Lighthouse Laboratory and Alex Alderton, Roberto Amato, Jeffrey Barrett, Sonia Goncalves, Ewan Harrison, David K. Jackson, Ian Johnston, Dominic Kwiatkowski, Cordelia Langford, John Sillitoe on behalf of the Wellcome Sanger Institute COVID-19 Surveillance Team |
| EPI_ISL_3043199 | Lighthouse Lab in Glasgow | Wellcome Sanger Institute for the COVID-19 Genomics UK (COG-UK) Consortium | Harper VanSteenhouse, Yumi Kasai, David Gray, Carol Clugston, Anna Dominiczak and Alex Alderton, Roberto Amato, Jeffrey Barrett, Sonia Goncalves, Ewan Harrison, David K. Jackson, Ian Johnston, Dominic Kwiatkowski, Cordelia Langford, John Sillitoe on behalf of the Wellcome Sanger Institute COVID-19 Surveillance Team |
| EPI_ISL_3043200 | Berkshire and Surrey Pathology Services Lighthouse Laboratory | Wellcome Sanger Institute for the COVID-19 Genomics UK (COG-UK) Consortium | Berkshire and Surrey Pathology Services Lighthouse Laboratory and Alex Alderton, Roberto Amato, Jeffrey Barrett, Sonia Goncalves, Ewan Harrison, David K. Jackson, Ian Johnston, Dominic Kwiatkowski, Cordelia Langford, John Sillitoe on behalf of the Wellcome Sanger Institute COVID-19 Surveillance Team |
| EPI_ISL_3043201 | Lighthouse Lab in Milton Keynes | Wellcome Sanger Institute for the COVID-19 Genomics UK (COG-UK) Consortium | The Lighthouse Lab in Milton Keynes and Alex Alderton, Roberto Amato, Jeffrey Barrett, Sonia Goncalves, Ewan Harrison, David K. Jackson, Ian Johnston, Dominic Kwiatkowski, Cordelia Langford, John Sillitoe on behalf of the Wellcome Sanger Institute COVID-19 Surveillance Team |
| EPI_ISL_3043202, EPI_ISL_3043203 | Lighthouse Lab in Glasgow | Wellcome Sanger Institute for the COVID-19 Genomics UK (COG-UK) Consortium | Harper VanSteenhouse, Yumi Kasai, David Gray, Carol Clugston, Anna Dominiczak and Alex Alderton, Roberto Amato, Jeffrey Barrett, Sonia Goncalves, Ewan Harrison, David K. Jackson, Ian Johnston, Dominic Kwiatkowski, Cordelia Langford, John Sillitoe on behalf of the Wellcome Sanger Institute COVID-19 Surveillance Team |
| EPI_ISL_3043204 | Lighthouse Lab in Milton Keynes | Wellcome Sanger Institute for the COVID-19 Genomics UK (COG-UK) Consortium | The Lighthouse Lab in Milton Keynes and Alex Alderton, Roberto Amato, Jeffrey Barrett, Sonia Goncalves, Ewan Harrison, David K. Jackson, Ian Johnston, Dominic Kwiatkowski, Cordelia Langford, John Sillitoe on behalf of the Wellcome Sanger Institute COVID-19 Surveillance Team |
| EPI_ISL_3043205 | Lighthouse Lab in Glasgow | Wellcome Sanger Institute for the COVID-19 Genomics UK (COG-UK) Consortium | Harper VanSteenhouse, Yumi Kasai, David Gray, Carol Clugston, Anna Dominiczak and Alex Alderton, Roberto Amato, Jeffrey Barrett, Sonia Goncalves, Ewan Harrison, David K. Jackson, Ian Johnston, Dominic Kwiatkowski, Cordelia Langford, John Sillitoe on behalf of the Wellcome Sanger Institute COVID-19 Surveillance Team |
| EPI_ISL_3043206 | Berkshire and Surrey Pathology Services Lighthouse Laboratory | Wellcome Sanger Institute for the COVID-19 Genomics UK (COG-UK) Consortium | Berkshire and Surrey Pathology Services Lighthouse Laboratory and Alex Alderton, Roberto Amato, Jeffrey Barrett, Sonia Goncalves, Ewan Harrison, David K. Jackson, Ian Johnston, Dominic Kwiatkowski, Cordelia Langford, John Sillitoe on behalf of the Wellcome Sanger Institute COVID-19 Surveillance Team |
| EPI_ISL_3043207, EPI_ISL_3043208 | Lighthouse Lab in Milton Keynes | Wellcome Sanger Institute for the COVID-19 Genomics UK (COG-UK) Consortium | The Lighthouse Lab in Milton Keynes and Alex Alderton, Roberto Amato, Jeffrey Barrett, Sonia Goncalves, Ewan Harrison, David K. Jackson, Ian Johnston, Dominic Kwiatkowski, Cordelia Langford, John Sillitoe on behalf of the Wellcome Sanger Institute COVID-19 Surveillance Team |
| EPI_ISL_3043209 | Lighthouse Lab in Glasgow | Wellcome Sanger Institute for the COVID-19 Genomics UK (COG-UK) Consortium | Harper VanSteenhouse, Yumi Kasai, David Gray, Carol Clugston, Anna Dominiczak and Alex Alderton, Roberto Amato, Jeffrey Barrett, Sonia Goncalves, Ewan Harrison, David K. Jackson, Ian Johnston, Dominic Kwiatkowski, Cordelia Langford, John Sillitoe on behalf of the Wellcome Sanger Institute COVID-19 Surveillance Team |
| EPI_ISL_3043210 | Lighthouse Lab in Milton Keynes | Wellcome Sanger Institute for the COVID-19 Genomics UK (COG-UK) Consortium | The Lighthouse Lab in Milton Keynes and Alex Alderton, Roberto Amato, Jeffrey Barrett, Sonia Goncalves, Ewan Harrison, David K. Jackson, Ian Johnston, Dominic Kwiatkowski, Cordelia Langford, John Sillitoe on behalf of the Wellcome Sanger Institute COVID-19 Surveillance Team |
| EPI_ISL_3043211 | Berkshire and Surrey Pathology Services Lighthouse Laboratory | Wellcome Sanger Institute for the COVID-19 Genomics UK (COG-UK) Consortium | Berkshire and Surrey Pathology Services Lighthouse Laboratory and Alex Alderton, Roberto Amato, Jeffrey Barrett, Sonia Goncalves, Ewan Harrison, David K. Jackson, Ian Johnston, Dominic Kwiatkowski, Cordelia Langford, John Sillitoe on behalf of the Wellcome Sanger Institute COVID-19 Surveillance Team |
| EPI_ISL_3043212 | Lighthouse Lab in Glasgow | Wellcome Sanger Institute for the COVID-19 Genomics UK (COG-UK) Consortium | Harper VanSteenhouse, Yumi Kasai, David Gray, Carol Clugston, Anna Dominiczak and Alex Alderton, Roberto Amato, Jeffrey Barrett, Sonia Goncalves, Ewan Harrison, David K. Jackson, Ian Johnston, Dominic Kwiatkowski, Cordelia Langford, John Sillitoe on behalf of the Wellcome Sanger Institute COVID-19 Surveillance Team |
| EPI_ISL_3043213 | Berkshire and Surrey Pathology Services Lighthouse Laboratory | Wellcome Sanger Institute for the COVID-19 Genomics UK (COG-UK) Consortium | Berkshire and Surrey Pathology Services Lighthouse Laboratory and Alex Alderton, Roberto Amato, Jeffrey Barrett, Sonia Goncalves, Ewan Harrison, David K. Jackson, Ian Johnston, Dominic Kwiatkowski, Cordelia Langford, John Sillitoe on behalf of the Wellcome Sanger Institute COVID-19 Surveillance Team |
| EPI_ISL_3043214, EPI_ISL_3043215, EPI_ISL_3043216, EPI_ISL_3043217 | Lighthouse Lab in Glasgow | Wellcome Sanger Institute for the COVID-19 Genomics UK (COG-UK) Consortium | Harper VanSteenhouse, Yumi Kasai, David Gray, Carol Clugston, Anna Dominiczak and Alex Alderton, Roberto Amato, Jeffrey Barrett, Sonia Goncalves, Ewan Harrison, David K. Jackson, Ian Johnston, Dominic Kwiatkowski, Cordelia Langford, John Sillitoe on behalf of the Wellcome Sanger Institute COVID-19 Surveillance Team |
| EPI_ISL_3043218 | Lighthouse Lab in Milton Keynes | Wellcome Sanger Institute for the COVID-19 Genomics UK (COG-UK) Consortium | The Lighthouse Lab in Milton Keynes and Alex Alderton, Roberto Amato, Jeffrey Barrett, Sonia Goncalves, Ewan Harrison, David K. Jackson, Ian Johnston, Dominic Kwiatkowski, Cordelia Langford, John Sillitoe on behalf of the Wellcome Sanger Institute COVID-19 Surveillance Team |
| EPI_ISL_3043219, EPI_ISL_3043220 | Lighthouse Lab in Glasgow | Wellcome Sanger Institute for the COVID-19 Genomics UK (COG-UK) Consortium | Harper VanSteenhouse, Yumi Kasai, David Gray, Carol Clugston, Anna Dominiczak and Alex Alderton, Roberto Amato, Jeffrey Barrett, Sonia Goncalves, Ewan Harrison, David K. Jackson, Ian Johnston, Dominic Kwiatkowski, Cordelia Langford, John Sillitoe on behalf of the Wellcome Sanger Institute COVID-19 Surveillance Team |
| EPI_ISL_3043221 | Berkshire and Surrey Pathology Services Lighthouse Laboratory | Wellcome Sanger Institute for the COVID-19 Genomics UK (COG-UK) Consortium | Berkshire and Surrey Pathology Services Lighthouse Laboratory and Alex Alderton, Roberto Amato, Jeffrey Barrett, Sonia Goncalves, Ewan Harrison, David K. Jackson, Ian Johnston, Dominic Kwiatkowski, Cordelia Langford, John Sillitoe on behalf of the Wellcome Sanger Institute COVID-19 Surveillance Team |
| EPI_ISL_3043222, EPI_ISL_3043223 | Lighthouse Lab in Glasgow | Wellcome Sanger Institute for the COVID-19 Genomics UK (COG-UK) Consortium | Harper VanSteenhouse, Yumi Kasai, David Gray, Carol Clugston, Anna Dominiczak and Alex Alderton, Roberto Amato, Jeffrey Barrett, Sonia Goncalves, Ewan Harrison, David K. Jackson, Ian Johnston, Dominic Kwiatkowski, Cordelia Langford, John Sillitoe on behalf of the Wellcome Sanger Institute COVID-19 Surveillance Team |
| EPI_ISL_3043224, EPI_ISL_3043225 | Lighthouse Lab in Milton Keynes | Wellcome Sanger Institute for the COVID-19 Genomics UK (COG-UK) Consortium | The Lighthouse Lab in Milton Keynes and Alex Alderton, Roberto Amato, Jeffrey Barrett, Sonia Goncalves, Ewan Harrison, David K. Jackson, Ian Johnston, Dominic Kwiatkowski, Cordelia Langford, John Sillitoe on behalf of the Wellcome Sanger Institute COVID-19 Surveillance Team |
| EPI_ISL_3043226 | Lighthouse Lab in Glasgow | Wellcome Sanger Institute for the COVID-19 Genomics UK (COG-UK) Consortium | Harper VanSteenhouse, Yumi Kasai, David Gray, Carol Clugston, Anna Dominiczak and Alex Alderton, Roberto Amato, Jeffrey Barrett, Sonia Goncalves, Ewan Harrison, David K. Jackson, Ian Johnston, Dominic Kwiatkowski, Cordelia Langford, John Sillitoe on behalf of the Wellcome Sanger Institute COVID-19 Surveillance Team |
| EPI_ISL_3043227 | Berkshire and Surrey Pathology Services Lighthouse Laboratory | Wellcome Sanger Institute for the COVID-19 Genomics UK (COG-UK) Consortium | Berkshire and Surrey Pathology Services Lighthouse Laboratory and Alex Alderton, Roberto Amato, Jeffrey Barrett, Sonia Goncalves, Ewan Harrison, David K. Jackson, Ian Johnston, Dominic Kwiatkowski, Cordelia Langford, John Sillitoe on behalf of the Wellcome Sanger Institute COVID-19 Surveillance Team |
| EPI_ISL_3043228 | Lighthouse Lab in Milton Keynes | Wellcome Sanger Institute for the COVID-19 Genomics UK (COG-UK) Consortium | The Lighthouse Lab in Milton Keynes and Alex Alderton, Roberto Amato, Jeffrey Barrett, Sonia Goncalves, Ewan Harrison, David K. Jackson, Ian Johnston, Dominic Kwiatkowski, Cordelia Langford, John Sillitoe on behalf of the Wellcome Sanger Institute COVID-19 Surveillance Team |
| EPI_ISL_3043229, EPI_ISL_3043230, EPI_ISL_3043231, EPI_ISL_3043232 | Lighthouse Lab in Glasgow | Wellcome Sanger Institute for the COVID-19 Genomics UK (COG-UK) Consortium | Harper VanSteenhouse, Yumi Kasai, David Gray, Carol Clugston, Anna Dominiczak and Alex Alderton, Roberto Amato, Jeffrey Barrett, Sonia Goncalves, Ewan Harrison, David K. Jackson, Ian Johnston, Dominic Kwiatkowski, Cordelia Langford, John Sillitoe on behalf of the Wellcome Sanger Institute COVID-19 Surveillance Team |
| EPI_ISL_3043233 | Berkshire and Surrey Pathology Services Lighthouse Laboratory | Wellcome Sanger Institute for the COVID-19 Genomics UK (COG-UK) Consortium | Berkshire and Surrey Pathology Services Lighthouse Laboratory and Alex Alderton, Roberto Amato, Jeffrey Barrett, Sonia Goncalves, Ewan Harrison, David K. Jackson, Ian Johnston, Dominic Kwiatkowski, Cordelia Langford, John Sillitoe on behalf of the Wellcome Sanger Institute COVID-19 Surveillance Team |
| EPI_ISL_3043234 | Lighthouse Lab in Glasgow | Wellcome Sanger Institute for the COVID-19 Genomics UK (COG-UK) Consortium | Harper VanSteenhouse, Yumi Kasai, David Gray, Carol Clugston, Anna Dominiczak and Alex Alderton, Roberto Amato, Jeffrey Barrett, Sonia Goncalves, Ewan Harrison, David K. Jackson, Ian Johnston, Dominic Kwiatkowski, Cordelia Langford, John Sillitoe on behalf of the Wellcome Sanger Institute COVID-19 Surveillance Team |
| EPI_ISL_3043235, EPI_ISL_3043236 | Lighthouse Lab in Milton Keynes | Wellcome Sanger Institute for the COVID-19 Genomics UK (COG-UK) Consortium | The Lighthouse Lab in Milton Keynes and Alex Alderton, Roberto Amato, Jeffrey Barrett, Sonia Goncalves, Ewan Harrison, David K. Jackson, Ian Johnston, Dominic Kwiatkowski, Cordelia Langford, John Sillitoe on behalf of the Wellcome Sanger Institute COVID-19 Surveillance Team |
| EPI_ISL_3043237 | Lighthouse Lab in Glasgow | Wellcome Sanger Institute for the COVID-19 Genomics UK (COG-UK) Consortium | Harper VanSteenhouse, Yumi Kasai, David Gray, Carol Clugston, Anna Dominiczak and Alex Alderton, Roberto Amato, Jeffrey Barrett, Sonia Goncalves, Ewan Harrison, David K. Jackson, Ian Johnston, Dominic Kwiatkowski, Cordelia Langford, John Sillitoe on behalf of the Wellcome Sanger Institute COVID-19 Surveillance Team |
| EPI_ISL_3043238, EPI_ISL_3043239 | Lighthouse Lab in Milton Keynes | Wellcome Sanger Institute for the COVID-19 Genomics UK (COG-UK) Consortium | The Lighthouse Lab in Milton Keynes and Alex Alderton, Roberto Amato, Jeffrey Barrett, Sonia Goncalves, Ewan Harrison, David K. Jackson, Ian Johnston, Dominic Kwiatkowski, Cordelia Langford, John Sillitoe on behalf of the Wellcome Sanger Institute COVID-19 Surveillance Team |
| EPI_ISL_3043240 | Berkshire and Surrey Pathology Services Lighthouse Laboratory | Wellcome Sanger Institute for the COVID-19 Genomics UK (COG-UK) Consortium | Berkshire and Surrey Pathology Services Lighthouse Laboratory and Alex Alderton, Roberto Amato, Jeffrey Barrett, Sonia Goncalves, Ewan Harrison, David K. Jackson, Ian Johnston, Dominic Kwiatkowski, Cordelia Langford, John Sillitoe on behalf of the Wellcome Sanger Institute COVID-19 Surveillance Team |
| EPI_ISL_3043241 | Lighthouse Lab in Glasgow | Wellcome Sanger Institute for the COVID-19 Genomics UK (COG-UK) Consortium | Harper VanSteenhouse, Yumi Kasai, David Gray, Carol Clugston, Anna Dominiczak and Alex Alderton, Roberto Amato, Jeffrey Barrett, Sonia Goncalves, Ewan Harrison, David K. Jackson, Ian Johnston, Dominic Kwiatkowski, Cordelia Langford, John Sillitoe on behalf of the Wellcome Sanger Institute COVID-19 Surveillance Team |
| EPI_ISL_3043242 | Berkshire and Surrey Pathology Services Lighthouse Laboratory | Wellcome Sanger Institute for the COVID-19 Genomics UK (COG-UK) Consortium | Berkshire and Surrey Pathology Services Lighthouse Laboratory and Alex Alderton, Roberto Amato, Jeffrey Barrett, Sonia Goncalves, Ewan Harrison, David K. Jackson, Ian Johnston, Dominic Kwiatkowski, Cordelia Langford, John Sillitoe on behalf of the Wellcome Sanger Institute COVID-19 Surveillance Team |
| EPI_ISL_3043243 | Lighthouse Lab in Milton Keynes | Wellcome Sanger Institute for the COVID-19 Genomics UK (COG-UK) Consortium | The Lighthouse Lab in Milton Keynes and Alex Alderton, Roberto Amato, Jeffrey Barrett, Sonia Goncalves, Ewan Harrison, David K. Jackson, Ian Johnston, Dominic Kwiatkowski, Cordelia Langford, John Sillitoe on behalf of the Wellcome Sanger Institute COVID-19 Surveillance Team |
| EPI_ISL_3043244 | Berkshire and Surrey Pathology Services Lighthouse Laboratory | Wellcome Sanger Institute for the COVID-19 Genomics UK (COG-UK) Consortium | Berkshire and Surrey Pathology Services Lighthouse Laboratory and Alex Alderton, Roberto Amato, Jeffrey Barrett, Sonia Goncalves, Ewan Harrison, David K. Jackson, Ian Johnston, Dominic Kwiatkowski, Cordelia Langford, John Sillitoe on behalf of the Wellcome Sanger Institute COVID-19 Surveillance Team |
| EPI_ISL_3043245 | Lighthouse Lab in Glasgow | Wellcome Sanger Institute for the COVID-19 Genomics UK (COG-UK) Consortium | Harper VanSteenhouse, Yumi Kasai, David Gray, Carol Clugston, Anna Dominiczak and Alex Alderton, Roberto Amato, Jeffrey Barrett, Sonia Goncalves, Ewan Harrison, David K. Jackson, Ian Johnston, Dominic Kwiatkowski, Cordelia Langford, John Sillitoe on behalf of the Wellcome Sanger Institute COVID-19 Surveillance Team |
| EPI_ISL_3043246 | Lighthouse Lab in Milton Keynes | Wellcome Sanger Institute for the COVID-19 Genomics UK (COG-UK) Consortium | The Lighthouse Lab in Milton Keynes and Alex Alderton, Roberto Amato, Jeffrey Barrett, Sonia Goncalves, Ewan Harrison, David K. Jackson, Ian Johnston, Dominic Kwiatkowski, Cordelia Langford, John Sillitoe on behalf of the Wellcome Sanger Institute COVID-19 Surveillance Team |

| | | | |
|---|---|---|---|
| EPI_ISL_3043247 | Lighthouse Lab in Glasgow | Wellcome Sanger Institute for the COVID-19 Genomics UK (COG-UK) Consortium | Harper VanSteenhouse, Yumi Kasai, David Gray, Carol Clugston, Anna Dominiczak and Alex Alderton, Roberto Amato, Jeffrey Barrett, Sonia Goncalves, Ewan Harrison, David K. Jackson, Ian Johnston, Dominic Kwiatkowski, Cordelia Langford, John Sillitoe on behalf of the Wellcome Sanger Institute COVID-19 Surveillance Team |
| EPI_ISL_3043248 | Lighthouse Lab in Milton Keynes | Wellcome Sanger Institute for the COVID-19 Genomics UK (COG-UK) Consortium | The Lighthouse Lab in Milton Keynes and Alex Alderton, Roberto Amato, Jeffrey Barrett, Sonia Goncalves, Ewan Harrison, David K. Jackson, Ian Johnston, Dominic Kwiatkowski, Cordelia Langford, John Sillitoe on behalf of the Wellcome Sanger Institute COVID-19 Surveillance Team |
| EPI_ISL_3043249 | Lighthouse Lab in Glasgow | Wellcome Sanger Institute for the COVID-19 Genomics UK (COG-UK) Consortium | Harper VanSteenhouse, Yumi Kasai, David Gray, Carol Clugston, Anna Dominiczak and Alex Alderton, Roberto Amato, Jeffrey Barrett, Sonia Goncalves, Ewan Harrison, David K. Jackson, Ian Johnston, Dominic Kwiatkowski, Cordelia Langford, John Sillitoe on behalf of the Wellcome Sanger Institute COVID-19 Surveillance Team |
| EPI_ISL_3043250 | Lighthouse Lab in Milton Keynes | Wellcome Sanger Institute for the COVID-19 Genomics UK (COG-UK) Consortium | The Lighthouse Lab in Milton Keynes and Alex Alderton, Roberto Amato, Jeffrey Barrett, Sonia Goncalves, Ewan Harrison, David K. Jackson, Ian Johnston, Dominic Kwiatkowski, Cordelia Langford, John Sillitoe on behalf of the Wellcome Sanger Institute COVID-19 Surveillance Team |
| EPI_ISL_3043251, EPI_ISL_3043252 | Berkshire and Surrey Pathology Services Lighthouse Laboratory | Wellcome Sanger Institute for the COVID-19 Genomics UK (COG-UK) Consortium | Berkshire and Surrey Pathology Services Lighthouse Laboratory and Alex Alderton, Roberto Amato, Jeffrey Barrett, Sonia Goncalves, Ewan Harrison, David K. Jackson, Ian Johnston, Dominic Kwiatkowski, Cordelia Langford, John Sillitoe on behalf of the Wellcome Sanger Institute COVID-19 Surveillance Team |
| EPI_ISL_3043253 | Lighthouse Lab in Glasgow | Wellcome Sanger Institute for the COVID-19 Genomics UK (COG-UK) Consortium | Harper VanSteenhouse, Yumi Kasai, David Gray, Carol Clugston, Anna Dominiczak and Alex Alderton, Roberto Amato, Jeffrey Barrett, Sonia Goncalves, Ewan Harrison, David K. Jackson, Ian Johnston, Dominic Kwiatkowski, Cordelia Langford, John Sillitoe on behalf of the Wellcome Sanger Institute COVID-19 Surveillance Team |
| EPI_ISL_3043254 | Lighthouse Lab in Milton Keynes | Wellcome Sanger Institute for the COVID-19 Genomics UK (COG-UK) Consortium | The Lighthouse Lab in Milton Keynes and Alex Alderton, Roberto Amato, Jeffrey Barrett, Sonia Goncalves, Ewan Harrison, David K. Jackson, Ian Johnston, Dominic Kwiatkowski, Cordelia Langford, John Sillitoe on behalf of the Wellcome Sanger Institute COVID-19 Surveillance Team |
| EPI_ISL_3043255, EPI_ISL_3043256 | Lighthouse Lab in Glasgow | Wellcome Sanger Institute for the COVID-19 Genomics UK (COG-UK) Consortium | Harper VanSteenhouse, Yumi Kasai, David Gray, Carol Clugston, Anna Dominiczak and Alex Alderton, Roberto Amato, Jeffrey Barrett, Sonia Goncalves, Ewan Harrison, David K. Jackson, Ian Johnston, Dominic Kwiatkowski, Cordelia Langford, John Sillitoe on behalf of the Wellcome Sanger Institute COVID-19 Surveillance Team |
| EPI_ISL_3043257, EPI_ISL_3043258 | Berkshire and Surrey Pathology Services Lighthouse Laboratory | Wellcome Sanger Institute for the COVID-19 Genomics UK (COG-UK) Consortium | Berkshire and Surrey Pathology Services Lighthouse Laboratory and Alex Alderton, Roberto Amato, Jeffrey Barrett, Sonia Goncalves, Ewan Harrison, David K. Jackson, Ian Johnston, Dominic Kwiatkowski, Cordelia Langford, John Sillitoe on behalf of the Wellcome Sanger Institute COVID-19 Surveillance Team |
| EPI_ISL_3043259 | Lighthouse Lab in Glasgow | Wellcome Sanger Institute for the COVID-19 Genomics UK (COG-UK) Consortium | Harper VanSteenhouse, Yumi Kasai, David Gray, Carol Clugston, Anna Dominiczak and Alex Alderton, Roberto Amato, Jeffrey Barrett, Sonia Goncalves, Ewan Harrison, David K. Jackson, Ian Johnston, Dominic Kwiatkowski, Cordelia Langford, John Sillitoe on behalf of the Wellcome Sanger Institute COVID-19 Surveillance Team |
| EPI_ISL_3043260 | Berkshire and Surrey Pathology Services Lighthouse Laboratory | Wellcome Sanger Institute for the COVID-19 Genomics UK (COG-UK) Consortium | Berkshire and Surrey Pathology Services Lighthouse Laboratory and Alex Alderton, Roberto Amato, Jeffrey Barrett, Sonia Goncalves, Ewan Harrison, David K. Jackson, Ian Johnston, Dominic Kwiatkowski, Cordelia Langford, John Sillitoe on behalf of the Wellcome Sanger Institute COVID-19 Surveillance Team |
| EPI_ISL_3043261 | Lighthouse Lab in Milton Keynes | Wellcome Sanger Institute for the COVID-19 Genomics UK (COG-UK) Consortium | The Lighthouse Lab in Milton Keynes and Alex Alderton, Roberto Amato, Jeffrey Barrett, Sonia Goncalves, Ewan Harrison, David K. Jackson, Ian Johnston, Dominic Kwiatkowski, Cordelia Langford, John Sillitoe on behalf of the Wellcome Sanger Institute COVID-19 Surveillance Team |
| EPI_ISL_3043262, EPI_ISL_3043263, EPI_ISL_3043264 | Lighthouse Lab in Glasgow | Wellcome Sanger Institute for the COVID-19 Genomics UK (COG-UK) Consortium | Harper VanSteenhouse, Yumi Kasai, David Gray, Carol Clugston, Anna Dominiczak and Alex Alderton, Roberto Amato, Jeffrey Barrett, Sonia Goncalves, Ewan Harrison, David K. Jackson, Ian Johnston, Dominic Kwiatkowski, Cordelia Langford, John Sillitoe on behalf of the Wellcome Sanger Institute COVID-19 Surveillance Team |
| EPI_ISL_3043265, EPI_ISL_3043266 | Lighthouse Lab in Milton Keynes | Wellcome Sanger Institute for the COVID-19 Genomics UK (COG-UK) Consortium | The Lighthouse Lab in Milton Keynes and Alex Alderton, Roberto Amato, Jeffrey Barrett, Sonia Goncalves, Ewan Harrison, David K. Jackson, Ian Johnston, Dominic Kwiatkowski, Cordelia Langford, John Sillitoe on behalf of the Wellcome Sanger Institute COVID-19 Surveillance Team |
| EPI_ISL_3043267 | Lighthouse Lab in Glasgow | Wellcome Sanger Institute for the COVID-19 Genomics UK (COG-UK) Consortium | Harper VanSteenhouse, Yumi Kasai, David Gray, Carol Clugston, Anna Dominiczak and Alex Alderton, Roberto Amato, Jeffrey Barrett, Sonia Goncalves, Ewan Harrison, David K. Jackson, Ian Johnston, Dominic Kwiatkowski, Cordelia Langford, John Sillitoe on behalf of the Wellcome Sanger Institute COVID-19 Surveillance Team |
| EPI_ISL_3043268 | Lighthouse Lab in Milton Keynes | Wellcome Sanger Institute for the COVID-19 Genomics UK (COG-UK) Consortium | The Lighthouse Lab in Milton Keynes and Alex Alderton, Roberto Amato, Jeffrey Barrett, Sonia Goncalves, Ewan Harrison, David K. Jackson, Ian Johnston, Dominic Kwiatkowski, Cordelia Langford, John Sillitoe on behalf of the Wellcome Sanger Institute COVID-19 Surveillance Team |
| EPI_ISL_3043269, EPI_ISL_3043270, EPI_ISL_3043271, EPI_ISL_3043272 | Lighthouse Lab in Glasgow | Wellcome Sanger Institute for the COVID-19 Genomics UK (COG-UK) Consortium | Harper VanSteenhouse, Yumi Kasai, David Gray, Carol Clugston, Anna Dominiczak and Alex Alderton, Roberto Amato, Jeffrey Barrett, Sonia Goncalves, Ewan Harrison, David K. Jackson, Ian Johnston, Dominic Kwiatkowski, Cordelia Langford, John Sillitoe on behalf of the Wellcome Sanger Institute COVID-19 Surveillance Team |
| EPI_ISL_3043273 | Lighthouse Lab in Milton Keynes | Wellcome Sanger Institute for the COVID-19 Genomics UK (COG-UK) Consortium | The Lighthouse Lab in Milton Keynes and Alex Alderton, Roberto Amato, Jeffrey Barrett, Sonia Goncalves, Ewan Harrison, David K. Jackson, Ian Johnston, Dominic Kwiatkowski, Cordelia Langford, John Sillitoe on behalf of the Wellcome Sanger Institute COVID-19 Surveillance Team |
| EPI_ISL_3043274 | Lighthouse Lab in Glasgow | Wellcome Sanger Institute for the COVID-19 Genomics UK (COG-UK) Consortium | Harper VanSteenhouse, Yumi Kasai, David Gray, Carol Clugston, Anna Dominiczak and Alex Alderton, Roberto Amato, Jeffrey Barrett, Sonia Goncalves, Ewan Harrison, David K. Jackson, Ian Johnston, Dominic Kwiatkowski, Cordelia Langford, John Sillitoe on behalf of the Wellcome Sanger Institute COVID-19 Surveillance Team |
| EPI_ISL_3043275 | Lighthouse Lab in Milton Keynes | Wellcome Sanger Institute for the COVID-19 Genomics UK (COG-UK) Consortium | The Lighthouse Lab in Milton Keynes and Alex Alderton, Roberto Amato, Jeffrey Barrett, Sonia Goncalves, Ewan Harrison, David K. Jackson, Ian Johnston, Dominic Kwiatkowski, Cordelia Langford, John Sillitoe on behalf of the Wellcome Sanger Institute COVID-19 Surveillance Team |
| EPI_ISL_3043276, EPI_ISL_3043277 | Berkshire and Surrey Pathology Services Lighthouse Laboratory | Wellcome Sanger Institute for the COVID-19 Genomics UK (COG-UK) Consortium | Berkshire and Surrey Pathology Services Lighthouse Laboratory and Alex Alderton, Roberto Amato, Jeffrey Barrett, Sonia Goncalves, Ewan Harrison, David K. Jackson, Ian Johnston, Dominic Kwiatkowski, Cordelia Langford, John Sillitoe on behalf of the Wellcome Sanger Institute COVID-19 Surveillance Team |
| EPI_ISL_3043278, EPI_ISL_3043279 | Lighthouse Lab in Glasgow | Wellcome Sanger Institute for the COVID-19 Genomics UK (COG-UK) Consortium | Harper VanSteenhouse, Yumi Kasai, David Gray, Carol Clugston, Anna Dominiczak and Alex Alderton, Roberto Amato, Jeffrey Barrett, Sonia Goncalves, Ewan Harrison, David K. Jackson, Ian Johnston, Dominic Kwiatkowski, Cordelia Langford, John Sillitoe on behalf of the Wellcome Sanger Institute COVID-19 Surveillance Team |
| EPI_ISL_3043280 | Lighthouse Lab in Milton Keynes | Wellcome Sanger Institute for the COVID-19 Genomics UK (COG-UK) Consortium | The Lighthouse Lab in Milton Keynes and Alex Alderton, Roberto Amato, Jeffrey Barrett, Sonia Goncalves, Ewan Harrison, David K. Jackson, Ian Johnston, Dominic Kwiatkowski, Cordelia Langford, John Sillitoe on behalf of the Wellcome Sanger Institute COVID-19 Surveillance Team |
| EPI_ISL_3043281 | Lighthouse Lab in Glasgow | Wellcome Sanger Institute for the COVID-19 Genomics UK (COG-UK) Consortium | Harper VanSteenhouse, Yumi Kasai, David Gray, Carol Clugston, Anna Dominiczak and Alex Alderton, Roberto Amato, Jeffrey Barrett, Sonia Goncalves, Ewan Harrison, David K. Jackson, Ian Johnston, Dominic Kwiatkowski, Cordelia Langford, John Sillitoe on behalf of the Wellcome Sanger Institute COVID-19 Surveillance Team |
| EPI_ISL_3043282, EPI_ISL_3043283 | Lighthouse Lab in Milton Keynes | Wellcome Sanger Institute for the COVID-19 Genomics UK (COG-UK) Consortium | The Lighthouse Lab in Milton Keynes and Alex Alderton, Roberto Amato, Jeffrey Barrett, Sonia Goncalves, Ewan Harrison, David K. Jackson, Ian Johnston, Dominic Kwiatkowski, Cordelia Langford, John Sillitoe on behalf of the Wellcome Sanger Institute COVID-19 Surveillance Team |
| EPI_ISL_3043284 | Berkshire and Surrey Pathology Services Lighthouse Laboratory | Wellcome Sanger Institute for the COVID-19 Genomics UK (COG-UK) Consortium | Berkshire and Surrey Pathology Services Lighthouse Laboratory and Alex Alderton, Roberto Amato, Jeffrey Barrett, Sonia Goncalves, Ewan Harrison, David K. Jackson, Ian Johnston, Dominic Kwiatkowski, Cordelia Langford, John Sillitoe on behalf of the Wellcome Sanger Institute COVID-19 Surveillance Team |
| EPI_ISL_3043285 | Lighthouse Lab in Milton Keynes | Wellcome Sanger Institute for the COVID-19 Genomics UK (COG-UK) Consortium | The Lighthouse Lab in Milton Keynes and Alex Alderton, Roberto Amato, Jeffrey Barrett, Sonia Goncalves, Ewan Harrison, David K. Jackson, Ian Johnston, Dominic Kwiatkowski, Cordelia Langford, John Sillitoe on behalf of the Wellcome Sanger Institute COVID-19 Surveillance Team |
| EPI_ISL_3043286 | Berkshire and Surrey Pathology Services Lighthouse Laboratory | Wellcome Sanger Institute for the COVID-19 Genomics UK (COG-UK) Consortium | Berkshire and Surrey Pathology Services Lighthouse Laboratory and Alex Alderton, Roberto Amato, Jeffrey Barrett, Sonia Goncalves, Ewan Harrison, David K. Jackson, Ian Johnston, Dominic Kwiatkowski, Cordelia Langford, John Sillitoe on behalf of the Wellcome Sanger Institute COVID-19 Surveillance Team |
| EPI_ISL_3043287 | Lighthouse Lab in Glasgow | Wellcome Sanger Institute for the COVID-19 Genomics UK (COG-UK) Consortium | Harper VanSteenhouse, Yumi Kasai, David Gray, Carol Clugston, Anna Dominiczak and Alex Alderton, Roberto Amato, Jeffrey Barrett, Sonia Goncalves, Ewan Harrison, David K. Jackson, Ian Johnston, Dominic Kwiatkowski, Cordelia Langford, John Sillitoe on behalf of the Wellcome Sanger Institute COVID-19 Surveillance Team |
| EPI_ISL_3043288, EPI_ISL_3043289 | Lighthouse Lab in Milton Keynes | Wellcome Sanger Institute for the COVID-19 Genomics UK (COG-UK) Consortium | The Lighthouse Lab in Milton Keynes and Alex Alderton, Roberto Amato, Jeffrey Barrett, Sonia Goncalves, Ewan Harrison, David K. Jackson, Ian Johnston, Dominic Kwiatkowski, Cordelia Langford, John Sillitoe on behalf of the Wellcome Sanger Institute COVID-19 Surveillance Team |
| EPI_ISL_3043290 | Berkshire and Surrey Pathology Services Lighthouse Laboratory | Wellcome Sanger Institute for the COVID-19 Genomics UK (COG-UK) Consortium | Berkshire and Surrey Pathology Services Lighthouse Laboratory and Alex Alderton, Roberto Amato, Jeffrey Barrett, Sonia Goncalves, Ewan Harrison, David K. Jackson, Ian Johnston, Dominic Kwiatkowski, Cordelia Langford, John Sillitoe on behalf of the Wellcome Sanger Institute COVID-19 Surveillance Team |
| EPI_ISL_3043291 | Lighthouse Lab in Milton Keynes | Wellcome Sanger Institute for the COVID-19 Genomics UK (COG-UK) Consortium | The Lighthouse Lab in Milton Keynes and Alex Alderton, Roberto Amato, Jeffrey Barrett, Sonia Goncalves, Ewan Harrison, David K. Jackson, Ian Johnston, Dominic Kwiatkowski, Cordelia Langford, John Sillitoe on behalf of the Wellcome Sanger Institute COVID-19 Surveillance Team |
| EPI_ISL_3043292 | Berkshire and Surrey Pathology Services Lighthouse Laboratory | Wellcome Sanger Institute for the COVID-19 Genomics UK (COG-UK) Consortium | Berkshire and Surrey Pathology Services Lighthouse Laboratory and Alex Alderton, Roberto Amato, Jeffrey Barrett, Sonia Goncalves, Ewan Harrison, David K. Jackson, Ian Johnston, Dominic Kwiatkowski, Cordelia Langford, John Sillitoe on behalf of the Wellcome Sanger Institute COVID-19 Surveillance Team |
| EPI_ISL_3043293, EPI_ISL_3043294 | Lighthouse Lab in Milton Keynes | Wellcome Sanger Institute for the COVID-19 Genomics UK (COG-UK) Consortium | The Lighthouse Lab in Milton Keynes and Alex Alderton, Roberto Amato, Jeffrey Barrett, Sonia Goncalves, Ewan Harrison, David K. Jackson, Ian Johnston, Dominic Kwiatkowski, Cordelia Langford, John Sillitoe on behalf of the Wellcome Sanger Institute COVID-19 Surveillance Team |
| EPI_ISL_3043295 | Lighthouse Lab in Glasgow | Wellcome Sanger Institute for the COVID-19 Genomics UK (COG-UK) Consortium | Harper VanSteenhouse, Yumi Kasai, David Gray, Carol Clugston, Anna Dominiczak and Alex Alderton, Roberto Amato, Jeffrey Barrett, Sonia Goncalves, Ewan Harrison, David K. Jackson, Ian Johnston, Dominic Kwiatkowski, Cordelia Langford, John Sillitoe on behalf of the Wellcome Sanger Institute COVID-19 Surveillance Team |
| EPI_ISL_3043296, EPI_ISL_3043297 | Berkshire and Surrey Pathology Services Lighthouse Laboratory | Wellcome Sanger Institute for the COVID-19 Genomics UK (COG-UK) Consortium | Berkshire and Surrey Pathology Services Lighthouse Laboratory and Alex Alderton, Roberto Amato, Jeffrey Barrett, Sonia Goncalves, Ewan Harrison, David K. Jackson, Ian Johnston, Dominic Kwiatkowski, Cordelia Langford, John Sillitoe on behalf of the Wellcome Sanger Institute COVID-19 Surveillance Team |
| EPI_ISL_3043298 | Lighthouse Lab in Glasgow | Wellcome Sanger Institute for the COVID-19 Genomics UK (COG-UK) Consortium | Harper VanSteenhouse, Yumi Kasai, David Gray, Carol Clugston, Anna Dominiczak and Alex Alderton, Roberto Amato, Jeffrey Barrett, Sonia Goncalves, Ewan Harrison, David K. Jackson, Ian Johnston, Dominic Kwiatkowski, Cordelia Langford, John Sillitoe on behalf of the Wellcome Sanger Institute COVID-19 Surveillance Team |
| EPI_ISL_3043299 | Berkshire and Surrey Pathology Services Lighthouse Laboratory | Wellcome Sanger Institute for the COVID-19 Genomics UK (COG-UK) Consortium | Berkshire and Surrey Pathology Services Lighthouse Laboratory and Alex Alderton, Roberto Amato, Jeffrey Barrett, Sonia Goncalves, Ewan Harrison, David K. Jackson, Ian Johnston, Dominic Kwiatkowski, Cordelia Langford, John Sillitoe on behalf of the Wellcome Sanger Institute COVID-19 Surveillance Team |
| EPI_ISL_3043300, EPI_ISL_3043301, EPI_ISL_3043302 | Lighthouse Lab in Glasgow | Wellcome Sanger Institute for the COVID-19 Genomics UK (COG-UK) Consortium | Harper VanSteenhouse, Yumi Kasai, David Gray, Carol Clugston, Anna Dominiczak and Alex Alderton, Roberto Amato, Jeffrey Barrett, Sonia Goncalves, Ewan Harrison, David K. Jackson, Ian Johnston, Dominic Kwiatkowski, Cordelia Langford, John Sillitoe on behalf of the Wellcome Sanger Institute COVID-19 Surveillance Team |
| EPI_ISL_3043303 | Lighthouse Lab in Milton Keynes | Wellcome Sanger Institute for the COVID-19 Genomics UK (COG-UK) Consortium | The Lighthouse Lab in Milton Keynes and Alex Alderton, Roberto Amato, Jeffrey Barrett, Sonia Goncalves, Ewan Harrison, David K. Jackson, Ian Johnston, Dominic Kwiatkowski, Cordelia Langford, John Sillitoe on behalf of the Wellcome Sanger Institute COVID-19 Surveillance Team |
| EPI_ISL_3043304 | Berkshire and Surrey Pathology Services Lighthouse Laboratory | Wellcome Sanger Institute for the COVID-19 Genomics UK (COG-UK) Consortium | Berkshire and Surrey Pathology Services Lighthouse Laboratory and Alex Alderton, Roberto Amato, Jeffrey Barrett, Sonia Goncalves, Ewan Harrison, David K. Jackson, Ian Johnston, Dominic Kwiatkowski, Cordelia Langford, John Sillitoe on behalf of the Wellcome Sanger Institute COVID-19 Surveillance Team |
| EPI_ISL_3043305 | Lighthouse Lab in Glasgow | Wellcome Sanger Institute for the COVID-19 Genomics UK (COG-UK) Consortium | Harper VanSteenhouse, Yumi Kasai, David Gray, Carol Clugston, Anna Dominiczak and Alex Alderton, Roberto Amato, Jeffrey Barrett, Sonia Goncalves, Ewan Harrison, David K. Jackson, Ian Johnston, Dominic Kwiatkowski, Cordelia Langford, John Sillitoe on behalf of the Wellcome Sanger Institute COVID-19 Surveillance Team |
| EPI_ISL_3043306 | Lighthouse Lab in Milton Keynes | Wellcome Sanger Institute for the COVID-19 Genomics UK (COG-UK) Consortium | The Lighthouse Lab in Milton Keynes and Alex Alderton, Roberto Amato, Jeffrey Barrett, Sonia Goncalves, Ewan Harrison, David K. Jackson, Ian Johnston, Dominic Kwiatkowski, Cordelia Langford, John Sillitoe on behalf of the Wellcome Sanger Institute COVID-19 Surveillance Team |
| EPI_ISL_3043307, | Lighthouse Lab in Glasgow | Wellcome Sanger Institute for the COVID-19 Genomics UK (COG-UK) | Harper VanSteenhouse, Yumi Kasai, David Gray, Carol Clugston, Anna Dominiczak and Alex Alderton, Roberto Amato, Jeffrey Barrett, Sonia Goncalves, Ewan Harrison, David K. Jackson, Ian Johnston, Dominic Kwiatkowski, Cordelia Langford, |

| | | | |
|---|---|---|---|
| EPI_ISL_3043308 | | Consortium | John Sillitoe on behalf of the Wellcome Sanger Institute COVID-19 Surveillance Team |
| EPI_ISL_3043309 | Lighthouse Lab in Milton Keynes | Wellcome Sanger Institute for the COVID-19 Genomics UK (COG-UK) Consortium | The Lighthouse Lab in Milton Keynes and Alex Alderton, Roberto Amato, Jeffrey Barrett, Sonia Goncalves, Ewan Harrison, David K. Jackson, Ian Johnston, Dominic Kwiatkowski, Cordelia Langford, John Sillitoe on behalf of the Wellcome Sanger Institute COVID-19 Surveillance Team |
| EPI_ISL_3043310 | Berkshire and Surrey Pathology Services Lighthouse Laboratory | Wellcome Sanger Institute for the COVID-19 Genomics UK (COG-UK) Consortium | Berkshire and Surrey Pathology Services Lighthouse Laboratory and Alex Alderton, Roberto Amato, Jeffrey Barrett, Sonia Goncalves, Ewan Harrison, David K. Jackson, Ian Johnston, Dominic Kwiatkowski, Cordelia Langford, John Sillitoe on behalf of the Wellcome Sanger Institute COVID-19 Surveillance Team |
| EPI_ISL_3043311 | Lighthouse Lab in Milton Keynes | Wellcome Sanger Institute for the COVID-19 Genomics UK (COG-UK) Consortium | The Lighthouse Lab in Milton Keynes and Alex Alderton, Roberto Amato, Jeffrey Barrett, Sonia Goncalves, Ewan Harrison, David K. Jackson, Ian Johnston, Dominic Kwiatkowski, Cordelia Langford, John Sillitoe on behalf of the Wellcome Sanger Institute COVID-19 Surveillance Team |
| EPI_ISL_3043312, EPI_ISL_3043313, EPI_ISL_3043314 | Berkshire and Surrey Pathology Services Lighthouse Laboratory | Wellcome Sanger Institute for the COVID-19 Genomics UK (COG-UK) Consortium | Berkshire and Surrey Pathology Services Lighthouse Laboratory and Alex Alderton, Roberto Amato, Jeffrey Barrett, Sonia Goncalves, Ewan Harrison, David K. Jackson, Ian Johnston, Dominic Kwiatkowski, Cordelia Langford, John Sillitoe on behalf of the Wellcome Sanger Institute COVID-19 Surveillance Team |
| EPI_ISL_3043315, EPI_ISL_3043316 | Lighthouse Lab in Glasgow | Wellcome Sanger Institute for the COVID-19 Genomics UK (COG-UK) Consortium | Harper VanSteenhouse, Yumi Kasai, David Gray, Carol Clugston, Anna Dominiczak and Alex Alderton, Roberto Amato, Jeffrey Barrett, Sonia Goncalves, Ewan Harrison, David K. Jackson, Ian Johnston, Dominic Kwiatkowski, Cordelia Langford, John Sillitoe on behalf of the Wellcome Sanger Institute COVID-19 Surveillance Team |
| EPI_ISL_3043317 | Lighthouse Lab in Milton Keynes | Wellcome Sanger Institute for the COVID-19 Genomics UK (COG-UK) Consortium | The Lighthouse Lab in Milton Keynes and Alex Alderton, Roberto Amato, Jeffrey Barrett, Sonia Goncalves, Ewan Harrison, David K. Jackson, Ian Johnston, Dominic Kwiatkowski, Cordelia Langford, John Sillitoe on behalf of the Wellcome Sanger Institute COVID-19 Surveillance Team |
| EPI_ISL_3043318 | Berkshire and Surrey Pathology Services Lighthouse Laboratory | Wellcome Sanger Institute for the COVID-19 Genomics UK (COG-UK) Consortium | Berkshire and Surrey Pathology Services Lighthouse Laboratory and Alex Alderton, Roberto Amato, Jeffrey Barrett, Sonia Goncalves, Ewan Harrison, David K. Jackson, Ian Johnston, Dominic Kwiatkowski, Cordelia Langford, John Sillitoe on behalf of the Wellcome Sanger Institute COVID-19 Surveillance Team |
| EPI_ISL_3043319, EPI_ISL_3043320, EPI_ISL_3043321 | Lighthouse Lab in Glasgow | Wellcome Sanger Institute for the COVID-19 Genomics UK (COG-UK) Consortium | Harper VanSteenhouse, Yumi Kasai, David Gray, Carol Clugston, Anna Dominiczak and Alex Alderton, Roberto Amato, Jeffrey Barrett, Sonia Goncalves, Ewan Harrison, David K. Jackson, Ian Johnston, Dominic Kwiatkowski, Cordelia Langford, John Sillitoe on behalf of the Wellcome Sanger Institute COVID-19 Surveillance Team |
| EPI_ISL_3043322, EPI_ISL_3043323 | Berkshire and Surrey Pathology Services Lighthouse Laboratory | Wellcome Sanger Institute for the COVID-19 Genomics UK (COG-UK) Consortium | Berkshire and Surrey Pathology Services Lighthouse Laboratory and Alex Alderton, Roberto Amato, Jeffrey Barrett, Sonia Goncalves, Ewan Harrison, David K. Jackson, Ian Johnston, Dominic Kwiatkowski, Cordelia Langford, John Sillitoe on behalf of the Wellcome Sanger Institute COVID-19 Surveillance Team |
| EPI_ISL_3043324, EPI_ISL_3043325 | Lighthouse Lab in Glasgow | Wellcome Sanger Institute for the COVID-19 Genomics UK (COG-UK) Consortium | Harper VanSteenhouse, Yumi Kasai, David Gray, Carol Clugston, Anna Dominiczak and Alex Alderton, Roberto Amato, Jeffrey Barrett, Sonia Goncalves, Ewan Harrison, David K. Jackson, Ian Johnston, Dominic Kwiatkowski, Cordelia Langford, John Sillitoe on behalf of the Wellcome Sanger Institute COVID-19 Surveillance Team |
| EPI_ISL_3043326 | Lighthouse Lab in Milton Keynes | Wellcome Sanger Institute for the COVID-19 Genomics UK (COG-UK) Consortium | The Lighthouse Lab in Milton Keynes and Alex Alderton, Roberto Amato, Jeffrey Barrett, Sonia Goncalves, Ewan Harrison, David K. Jackson, Ian Johnston, Dominic Kwiatkowski, Cordelia Langford, John Sillitoe on behalf of the Wellcome Sanger Institute COVID-19 Surveillance Team |
| EPI_ISL_3043327 | Berkshire and Surrey Pathology Services Lighthouse Laboratory | Wellcome Sanger Institute for the COVID-19 Genomics UK (COG-UK) Consortium | Berkshire and Surrey Pathology Services Lighthouse Laboratory and Alex Alderton, Roberto Amato, Jeffrey Barrett, Sonia Goncalves, Ewan Harrison, David K. Jackson, Ian Johnston, Dominic Kwiatkowski, Cordelia Langford, John Sillitoe on behalf of the Wellcome Sanger Institute COVID-19 Surveillance Team |
| EPI_ISL_3043328 | Lighthouse Lab in Glasgow | Wellcome Sanger Institute for the COVID-19 Genomics UK (COG-UK) Consortium | Harper VanSteenhouse, Yumi Kasai, David Gray, Carol Clugston, Anna Dominiczak and Alex Alderton, Roberto Amato, Jeffrey Barrett, Sonia Goncalves, Ewan Harrison, David K. Jackson, Ian Johnston, Dominic Kwiatkowski, Cordelia Langford, John Sillitoe on behalf of the Wellcome Sanger Institute COVID-19 Surveillance Team |
| EPI_ISL_3043329 | Berkshire and Surrey Pathology Services Lighthouse Laboratory | Wellcome Sanger Institute for the COVID-19 Genomics UK (COG-UK) Consortium | Berkshire and Surrey Pathology Services Lighthouse Laboratory and Alex Alderton, Roberto Amato, Jeffrey Barrett, Sonia Goncalves, Ewan Harrison, David K. Jackson, Ian Johnston, Dominic Kwiatkowski, Cordelia Langford, John Sillitoe on behalf of the Wellcome Sanger Institute COVID-19 Surveillance Team |
| EPI_ISL_3043330, EPI_ISL_3043331 | Lighthouse Lab in Glasgow | Wellcome Sanger Institute for the COVID-19 Genomics UK (COG-UK) Consortium | Harper VanSteenhouse, Yumi Kasai, David Gray, Carol Clugston, Anna Dominiczak and Alex Alderton, Roberto Amato, Jeffrey Barrett, Sonia Goncalves, Ewan Harrison, David K. Jackson, Ian Johnston, Dominic Kwiatkowski, Cordelia Langford, John Sillitoe on behalf of the Wellcome Sanger Institute COVID-19 Surveillance Team |
| EPI_ISL_3043332, EPI_ISL_3043333 | Berkshire and Surrey Pathology Services Lighthouse Laboratory | Wellcome Sanger Institute for the COVID-19 Genomics UK (COG-UK) Consortium | Berkshire and Surrey Pathology Services Lighthouse Laboratory and Alex Alderton, Roberto Amato, Jeffrey Barrett, Sonia Goncalves, Ewan Harrison, David K. Jackson, Ian Johnston, Dominic Kwiatkowski, Cordelia Langford, John Sillitoe on behalf of the Wellcome Sanger Institute COVID-19 Surveillance Team |
| EPI_ISL_3043334, EPI_ISL_3043335, EPI_ISL_3043336 | Lighthouse Lab in Glasgow | Wellcome Sanger Institute for the COVID-19 Genomics UK (COG-UK) Consortium | Harper VanSteenhouse, Yumi Kasai, David Gray, Carol Clugston, Anna Dominiczak and Alex Alderton, Roberto Amato, Jeffrey Barrett, Sonia Goncalves, Ewan Harrison, David K. Jackson, Ian Johnston, Dominic Kwiatkowski, Cordelia Langford, John Sillitoe on behalf of the Wellcome Sanger Institute COVID-19 Surveillance Team |
| EPI_ISL_3043337 | Lighthouse Lab in Milton Keynes | Wellcome Sanger Institute for the COVID-19 Genomics UK (COG-UK) Consortium | The Lighthouse Lab in Milton Keynes and Alex Alderton, Roberto Amato, Jeffrey Barrett, Sonia Goncalves, Ewan Harrison, David K. Jackson, Ian Johnston, Dominic Kwiatkowski, Cordelia Langford, John Sillitoe on behalf of the Wellcome Sanger Institute COVID-19 Surveillance Team |
| EPI_ISL_3043338, EPI_ISL_3043339, EPI_ISL_3043340, EPI_ISL_3043341, EPI_ISL_3043342, EPI_ISL_3043343 | Lighthouse Lab in Glasgow | Wellcome Sanger Institute for the COVID-19 Genomics UK (COG-UK) Consortium | Harper VanSteenhouse, Yumi Kasai, David Gray, Carol Clugston, Anna Dominiczak and Alex Alderton, Roberto Amato, Jeffrey Barrett, Sonia Goncalves, Ewan Harrison, David K. Jackson, Ian Johnston, Dominic Kwiatkowski, Cordelia Langford, John Sillitoe on behalf of the Wellcome Sanger Institute COVID-19 Surveillance Team |
| EPI_ISL_3043344 | Lighthouse Lab in Milton Keynes | Wellcome Sanger Institute for the COVID-19 Genomics UK (COG-UK) Consortium | The Lighthouse Lab in Milton Keynes and Alex Alderton, Roberto Amato, Jeffrey Barrett, Sonia Goncalves, Ewan Harrison, David K. Jackson, Ian Johnston, Dominic Kwiatkowski, Cordelia Langford, John Sillitoe on behalf of the Wellcome Sanger Institute COVID-19 Surveillance Team |
| EPI_ISL_3043345, EPI_ISL_3043346 | Lighthouse Lab in Glasgow | Wellcome Sanger Institute for the COVID-19 Genomics UK (COG-UK) Consortium | Harper VanSteenhouse, Yumi Kasai, David Gray, Carol Clugston, Anna Dominiczak and Alex Alderton, Roberto Amato, Jeffrey Barrett, Sonia Goncalves, Ewan Harrison, David K. Jackson, Ian Johnston, Dominic Kwiatkowski, Cordelia Langford, John Sillitoe on behalf of the Wellcome Sanger Institute COVID-19 Surveillance Team |
| EPI_ISL_3043347 | Lighthouse Lab in Milton Keynes | Wellcome Sanger Institute for the COVID-19 Genomics UK (COG-UK) Consortium | The Lighthouse Lab in Milton Keynes and Alex Alderton, Roberto Amato, Jeffrey Barrett, Sonia Goncalves, Ewan Harrison, David K. Jackson, Ian Johnston, Dominic Kwiatkowski, Cordelia Langford, John Sillitoe on behalf of the Wellcome Sanger Institute COVID-19 Surveillance Team |
| EPI_ISL_3043348, EPI_ISL_3043349 | Lighthouse Lab in Glasgow | Wellcome Sanger Institute for the COVID-19 Genomics UK (COG-UK) Consortium | Harper VanSteenhouse, Yumi Kasai, David Gray, Carol Clugston, Anna Dominiczak and Alex Alderton, Roberto Amato, Jeffrey Barrett, Sonia Goncalves, Ewan Harrison, David K. Jackson, Ian Johnston, Dominic Kwiatkowski, Cordelia Langford, John Sillitoe on behalf of the Wellcome Sanger Institute COVID-19 Surveillance Team |
| EPI_ISL_3043351 | Berkshire and Surrey Pathology Services Lighthouse Laboratory | Wellcome Sanger Institute for the COVID-19 Genomics UK (COG-UK) Consortium | Berkshire and Surrey Pathology Services Lighthouse Laboratory and Alex Alderton, Roberto Amato, Jeffrey Barrett, Sonia Goncalves, Ewan Harrison, David K. Jackson, Ian Johnston, Dominic Kwiatkowski, Cordelia Langford, John Sillitoe on behalf of the Wellcome Sanger Institute COVID-19 Surveillance Team |
| EPI_ISL_3043352, EPI_ISL_3043353 | Lighthouse Lab in Glasgow | Wellcome Sanger Institute for the COVID-19 Genomics UK (COG-UK) Consortium | Harper VanSteenhouse, Yumi Kasai, David Gray, Carol Clugston, Anna Dominiczak and Alex Alderton, Roberto Amato, Jeffrey Barrett, Sonia Goncalves, Ewan Harrison, David K. Jackson, Ian Johnston, Dominic Kwiatkowski, Cordelia Langford, John Sillitoe on behalf of the Wellcome Sanger Institute COVID-19 Surveillance Team |
| EPI_ISL_3043354 | Berkshire and Surrey Pathology Services Lighthouse Laboratory | Wellcome Sanger Institute for the COVID-19 Genomics UK (COG-UK) Consortium | Berkshire and Surrey Pathology Services Lighthouse Laboratory and Alex Alderton, Roberto Amato, Jeffrey Barrett, Sonia Goncalves, Ewan Harrison, David K. Jackson, Ian Johnston, Dominic Kwiatkowski, Cordelia Langford, John Sillitoe on behalf of the Wellcome Sanger Institute COVID-19 Surveillance Team |
| EPI_ISL_3043355, EPI_ISL_3043356 | Lighthouse Lab in Glasgow | Wellcome Sanger Institute for the COVID-19 Genomics UK (COG-UK) Consortium | Harper VanSteenhouse, Yumi Kasai, David Gray, Carol Clugston, Anna Dominiczak and Alex Alderton, Roberto Amato, Jeffrey Barrett, Sonia Goncalves, Ewan Harrison, David K. Jackson, Ian Johnston, Dominic Kwiatkowski, Cordelia Langford, John Sillitoe on behalf of the Wellcome Sanger Institute COVID-19 Surveillance Team |
| EPI_ISL_3043357 | Berkshire and Surrey Pathology Services Lighthouse Laboratory | Wellcome Sanger Institute for the COVID-19 Genomics UK (COG-UK) Consortium | Berkshire and Surrey Pathology Services Lighthouse Laboratory and Alex Alderton, Roberto Amato, Jeffrey Barrett, Sonia Goncalves, Ewan Harrison, David K. Jackson, Ian Johnston, Dominic Kwiatkowski, Cordelia Langford, John Sillitoe on behalf of the Wellcome Sanger Institute COVID-19 Surveillance Team |
| EPI_ISL_3043358 | Lighthouse Lab in Glasgow | Wellcome Sanger Institute for the COVID-19 Genomics UK (COG-UK) Consortium | Harper VanSteenhouse, Yumi Kasai, David Gray, Carol Clugston, Anna Dominiczak and Alex Alderton, Roberto Amato, Jeffrey Barrett, Sonia Goncalves, Ewan Harrison, David K. Jackson, Ian Johnston, Dominic Kwiatkowski, Cordelia Langford, John Sillitoe on behalf of the Wellcome Sanger Institute COVID-19 Surveillance Team |
| EPI_ISL_3043359 | Berkshire and Surrey Pathology Services Lighthouse Laboratory | Wellcome Sanger Institute for the COVID-19 Genomics UK (COG-UK) Consortium | Berkshire and Surrey Pathology Services Lighthouse Laboratory and Alex Alderton, Roberto Amato, Jeffrey Barrett, Sonia Goncalves, Ewan Harrison, David K. Jackson, Ian Johnston, Dominic Kwiatkowski, Cordelia Langford, John Sillitoe on behalf of the Wellcome Sanger Institute COVID-19 Surveillance Team |
| EPI_ISL_3043360, EPI_ISL_3043361, EPI_ISL_3043362 | Lighthouse Lab in Glasgow | Wellcome Sanger Institute for the COVID-19 Genomics UK (COG-UK) Consortium | Harper VanSteenhouse, Yumi Kasai, David Gray, Carol Clugston, Anna Dominiczak and Alex Alderton, Roberto Amato, Jeffrey Barrett, Sonia Goncalves, Ewan Harrison, David K. Jackson, Ian Johnston, Dominic Kwiatkowski, Cordelia Langford, John Sillitoe on behalf of the Wellcome Sanger Institute COVID-19 Surveillance Team |
| EPI_ISL_3043363 | Lighthouse Lab in Milton Keynes | Wellcome Sanger Institute for the COVID-19 Genomics UK (COG-UK) Consortium | The Lighthouse Lab in Milton Keynes and Alex Alderton, Roberto Amato, Jeffrey Barrett, Sonia Goncalves, Ewan Harrison, David K. Jackson, Ian Johnston, Dominic Kwiatkowski, Cordelia Langford, John Sillitoe on behalf of the Wellcome Sanger Institute COVID-19 Surveillance Team |
| EPI_ISL_3043364, EPI_ISL_3043365 | Lighthouse Lab in Glasgow | Wellcome Sanger Institute for the COVID-19 Genomics UK (COG-UK) Consortium | Harper VanSteenhouse, Yumi Kasai, David Gray, Carol Clugston, Anna Dominiczak and Alex Alderton, Roberto Amato, Jeffrey Barrett, Sonia Goncalves, Ewan Harrison, David K. Jackson, Ian Johnston, Dominic Kwiatkowski, Cordelia Langford, John Sillitoe on behalf of the Wellcome Sanger Institute COVID-19 Surveillance Team |
| EPI_ISL_3043366, EPI_ISL_3043367 | Berkshire and Surrey Pathology Services Lighthouse Laboratory | Wellcome Sanger Institute for the COVID-19 Genomics UK (COG-UK) Consortium | Berkshire and Surrey Pathology Services Lighthouse Laboratory and Alex Alderton, Roberto Amato, Jeffrey Barrett, Sonia Goncalves, Ewan Harrison, David K. Jackson, Ian Johnston, Dominic Kwiatkowski, Cordelia Langford, John Sillitoe on behalf of the Wellcome Sanger Institute COVID-19 Surveillance Team |
| EPI_ISL_3043368, EPI_ISL_3043369, EPI_ISL_3043370, EPI_ISL_3043371 | Lighthouse Lab in Glasgow | Wellcome Sanger Institute for the COVID-19 Genomics UK (COG-UK) Consortium | Harper VanSteenhouse, Yumi Kasai, David Gray, Carol Clugston, Anna Dominiczak and Alex Alderton, Roberto Amato, Jeffrey Barrett, Sonia Goncalves, Ewan Harrison, David K. Jackson, Ian Johnston, Dominic Kwiatkowski, Cordelia Langford, John Sillitoe on behalf of the Wellcome Sanger Institute COVID-19 Surveillance Team |
| EPI_ISL_3043372 | Berkshire and Surrey Pathology Services Lighthouse Laboratory | Wellcome Sanger Institute for the COVID-19 Genomics UK (COG-UK) Consortium | Berkshire and Surrey Pathology Services Lighthouse Laboratory and Alex Alderton, Roberto Amato, Jeffrey Barrett, Sonia Goncalves, Ewan Harrison, David K. Jackson, Ian Johnston, Dominic Kwiatkowski, Cordelia Langford, John Sillitoe on behalf of the Wellcome Sanger Institute COVID-19 Surveillance Team |
| EPI_ISL_3043373 | Lighthouse Lab in Milton Keynes | Wellcome Sanger Institute for the COVID-19 Genomics UK (COG-UK) Consortium | The Lighthouse Lab in Milton Keynes and Alex Alderton, Roberto Amato, Jeffrey Barrett, Sonia Goncalves, Ewan Harrison, David K. Jackson, Ian Johnston, Dominic Kwiatkowski, Cordelia Langford, John Sillitoe on behalf of the Wellcome Sanger Institute COVID-19 Surveillance Team |
| EPI_ISL_3043374 | Berkshire and Surrey Pathology Services Lighthouse Laboratory | Wellcome Sanger Institute for the COVID-19 Genomics UK (COG-UK) Consortium | Berkshire and Surrey Pathology Services Lighthouse Laboratory and Alex Alderton, Roberto Amato, Jeffrey Barrett, Sonia Goncalves, Ewan Harrison, David K. Jackson, Ian Johnston, Dominic Kwiatkowski, Cordelia Langford, John Sillitoe on behalf of the Wellcome Sanger Institute COVID-19 Surveillance Team |
| EPI_ISL_3043375 | Lighthouse Lab in Milton Keynes | Wellcome Sanger Institute for the COVID-19 Genomics UK (COG-UK) Consortium | The Lighthouse Lab in Milton Keynes and Alex Alderton, Roberto Amato, Jeffrey Barrett, Sonia Goncalves, Ewan Harrison, David K. Jackson, Ian Johnston, Dominic Kwiatkowski, Cordelia Langford, John Sillitoe on behalf of the Wellcome Sanger Institute COVID-19 Surveillance Team |
| EPI_ISL_3043376 | Lighthouse Lab in Glasgow | Wellcome Sanger Institute for the COVID-19 Genomics UK (COG-UK) Consortium | Harper VanSteenhouse, Yumi Kasai, David Gray, Carol Clugston, Anna Dominiczak and Alex Alderton, Roberto Amato, Jeffrey Barrett, Sonia Goncalves, Ewan Harrison, David K. Jackson, Ian Johnston, Dominic Kwiatkowski, Cordelia Langford, John Sillitoe on behalf of the Wellcome Sanger Institute COVID-19 Surveillance Team |
| EPI_ISL_3043377 | Berkshire and Surrey Pathology Services Lighthouse Laboratory | Wellcome Sanger Institute for the COVID-19 Genomics UK (COG-UK) | Berkshire and Surrey Pathology Services Lighthouse Laboratory and Alex Alderton, Roberto Amato, Jeffrey Barrett, Sonia Goncalves, Ewan Harrison, David K. Jackson, Ian Johnston, Dominic Kwiatkowski, Cordelia Langford, John Sillitoe on behalf |

| | | | |
|---|---|---|---|
| | | Consortium | of the Wellcome Sanger Institute COVID-19 Surveillance Team |
| EPI_ISL_3043378, EPI_ISL_3043379, EPI_ISL_3043380 | Lighthouse Lab in Milton Keynes | Wellcome Sanger Institute for the COVID-19 Genomics UK (COG-UK) Consortium | The Lighthouse Lab in Milton Keynes and Alex Alderton, Roberto Amato, Jeffrey Barrett, Sonia Goncalves, Ewan Harrison, David K. Jackson, Ian Johnston, Dominic Kwiatkowski, Cordelia Langford, John Sillitoe on behalf of the Wellcome Sanger Institute COVID-19 Surveillance Team |
| EPI_ISL_3043381, EPI_ISL_3043382, EPI_ISL_3043383, EPI_ISL_3043384 | Lighthouse Lab in Glasgow | Wellcome Sanger Institute for the COVID-19 Genomics UK (COG-UK) Consortium | Harper VanSteenhouse, Yumi Kasai, David Gray, Carol Clugston, Anna Dominiczak and Alex Alderton, Roberto Amato, Jeffrey Barrett, Sonia Goncalves, Ewan Harrison, David K. Jackson, Ian Johnston, Dominic Kwiatkowski, Cordelia Langford, John Sillitoe on behalf of the Wellcome Sanger Institute COVID-19 Surveillance Team |
| EPI_ISL_3043385 | Lighthouse Lab in Milton Keynes | Wellcome Sanger Institute for the COVID-19 Genomics UK (COG-UK) Consortium | The Lighthouse Lab in Milton Keynes and Alex Alderton, Roberto Amato, Jeffrey Barrett, Sonia Goncalves, Ewan Harrison, David K. Jackson, Ian Johnston, Dominic Kwiatkowski, Cordelia Langford, John Sillitoe on behalf of the Wellcome Sanger Institute COVID-19 Surveillance Team |
| EPI_ISL_3043386 | Lighthouse Lab in Glasgow | Wellcome Sanger Institute for the COVID-19 Genomics UK (COG-UK) Consortium | Harper VanSteenhouse, Yumi Kasai, David Gray, Carol Clugston, Anna Dominiczak and Alex Alderton, Roberto Amato, Jeffrey Barrett, Sonia Goncalves, Ewan Harrison, David K. Jackson, Ian Johnston, Dominic Kwiatkowski, Cordelia Langford, John Sillitoe on behalf of the Wellcome Sanger Institute COVID-19 Surveillance Team |
| EPI_ISL_3043387 | Berkshire and Surrey Pathology Services Lighthouse Laboratory | Wellcome Sanger Institute for the COVID-19 Genomics UK (COG-UK) Consortium | Berkshire and Surrey Pathology Services Lighthouse Laboratory and Alex Alderton, Roberto Amato, Jeffrey Barrett, Sonia Goncalves, Ewan Harrison, David K. Jackson, Ian Johnston, Dominic Kwiatkowski, Cordelia Langford, John Sillitoe on behalf of the Wellcome Sanger Institute COVID-19 Surveillance Team |
| EPI_ISL_3043388 | Lighthouse Lab in Glasgow | Wellcome Sanger Institute for the COVID-19 Genomics UK (COG-UK) Consortium | Harper VanSteenhouse, Yumi Kasai, David Gray, Carol Clugston, Anna Dominiczak and Alex Alderton, Roberto Amato, Jeffrey Barrett, Sonia Goncalves, Ewan Harrison, David K. Jackson, Ian Johnston, Dominic Kwiatkowski, Cordelia Langford, John Sillitoe on behalf of the Wellcome Sanger Institute COVID-19 Surveillance Team |
| EPI_ISL_3043389, EPI_ISL_3043390, EPI_ISL_3043391 | Lighthouse Lab in Milton Keynes | Wellcome Sanger Institute for the COVID-19 Genomics UK (COG-UK) Consortium | The Lighthouse Lab in Milton Keynes and Alex Alderton, Roberto Amato, Jeffrey Barrett, Sonia Goncalves, Ewan Harrison, David K. Jackson, Ian Johnston, Dominic Kwiatkowski, Cordelia Langford, John Sillitoe on behalf of the Wellcome Sanger Institute COVID-19 Surveillance Team |
| EPI_ISL_3043392 | Lighthouse Lab in Glasgow | Wellcome Sanger Institute for the COVID-19 Genomics UK (COG-UK) Consortium | Harper VanSteenhouse, Yumi Kasai, David Gray, Carol Clugston, Anna Dominiczak and Alex Alderton, Roberto Amato, Jeffrey Barrett, Sonia Goncalves, Ewan Harrison, David K. Jackson, Ian Johnston, Dominic Kwiatkowski, Cordelia Langford, John Sillitoe on behalf of the Wellcome Sanger Institute COVID-19 Surveillance Team |
| EPI_ISL_3043393 | Berkshire and Surrey Pathology Services Lighthouse Laboratory | Wellcome Sanger Institute for the COVID-19 Genomics UK (COG-UK) Consortium | Berkshire and Surrey Pathology Services Lighthouse Laboratory and Alex Alderton, Roberto Amato, Jeffrey Barrett, Sonia Goncalves, Ewan Harrison, David K. Jackson, Ian Johnston, Dominic Kwiatkowski, Cordelia Langford, John Sillitoe on behalf of the Wellcome Sanger Institute COVID-19 Surveillance Team |
| EPI_ISL_3043394 | Lighthouse Lab in Milton Keynes | Wellcome Sanger Institute for the COVID-19 Genomics UK (COG-UK) Consortium | The Lighthouse Lab in Milton Keynes and Alex Alderton, Roberto Amato, Jeffrey Barrett, Sonia Goncalves, Ewan Harrison, David K. Jackson, Ian Johnston, Dominic Kwiatkowski, Cordelia Langford, John Sillitoe on behalf of the Wellcome Sanger Institute COVID-19 Surveillance Team |
| EPI_ISL_3043395 | Berkshire and Surrey Pathology Services Lighthouse Laboratory | Wellcome Sanger Institute for the COVID-19 Genomics UK (COG-UK) Consortium | Berkshire and Surrey Pathology Services Lighthouse Laboratory and Alex Alderton, Roberto Amato, Jeffrey Barrett, Sonia Goncalves, Ewan Harrison, David K. Jackson, Ian Johnston, Dominic Kwiatkowski, Cordelia Langford, John Sillitoe on behalf of the Wellcome Sanger Institute COVID-19 Surveillance Team |
| EPI_ISL_3043396 | Lighthouse Lab in Milton Keynes | Wellcome Sanger Institute for the COVID-19 Genomics UK (COG-UK) Consortium | The Lighthouse Lab in Milton Keynes and Alex Alderton, Roberto Amato, Jeffrey Barrett, Sonia Goncalves, Ewan Harrison, David K. Jackson, Ian Johnston, Dominic Kwiatkowski, Cordelia Langford, John Sillitoe on behalf of the Wellcome Sanger Institute COVID-19 Surveillance Team |
| EPI_ISL_3043397 | Berkshire and Surrey Pathology Services Lighthouse Laboratory | Wellcome Sanger Institute for the COVID-19 Genomics UK (COG-UK) Consortium | Berkshire and Surrey Pathology Services Lighthouse Laboratory and Alex Alderton, Roberto Amato, Jeffrey Barrett, Sonia Goncalves, Ewan Harrison, David K. Jackson, Ian Johnston, Dominic Kwiatkowski, Cordelia Langford, John Sillitoe on behalf of the Wellcome Sanger Institute COVID-19 Surveillance Team |
| EPI_ISL_3043398 | Lighthouse Lab in Glasgow | Wellcome Sanger Institute for the COVID-19 Genomics UK (COG-UK) Consortium | Harper VanSteenhouse, Yumi Kasai, David Gray, Carol Clugston, Anna Dominiczak and Alex Alderton, Roberto Amato, Jeffrey Barrett, Sonia Goncalves, Ewan Harrison, David K. Jackson, Ian Johnston, Dominic Kwiatkowski, Cordelia Langford, John Sillitoe on behalf of the Wellcome Sanger Institute COVID-19 Surveillance Team |
| EPI_ISL_3043399 | Lighthouse Lab in Milton Keynes | Wellcome Sanger Institute for the COVID-19 Genomics UK (COG-UK) Consortium | The Lighthouse Lab in Milton Keynes and Alex Alderton, Roberto Amato, Jeffrey Barrett, Sonia Goncalves, Ewan Harrison, David K. Jackson, Ian Johnston, Dominic Kwiatkowski, Cordelia Langford, John Sillitoe on behalf of the Wellcome Sanger Institute COVID-19 Surveillance Team |
| EPI_ISL_3043400 | Berkshire and Surrey Pathology Services Lighthouse Laboratory | Wellcome Sanger Institute for the COVID-19 Genomics UK (COG-UK) Consortium | Berkshire and Surrey Pathology Services Lighthouse Laboratory and Alex Alderton, Roberto Amato, Jeffrey Barrett, Sonia Goncalves, Ewan Harrison, David K. Jackson, Ian Johnston, Dominic Kwiatkowski, Cordelia Langford, John Sillitoe on behalf of the Wellcome Sanger Institute COVID-19 Surveillance Team |
| EPI_ISL_3043401, EPI_ISL_3043402 | Lighthouse Lab in Glasgow | Wellcome Sanger Institute for the COVID-19 Genomics UK (COG-UK) Consortium | Harper VanSteenhouse, Yumi Kasai, David Gray, Carol Clugston, Anna Dominiczak and Alex Alderton, Roberto Amato, Jeffrey Barrett, Sonia Goncalves, Ewan Harrison, David K. Jackson, Ian Johnston, Dominic Kwiatkowski, Cordelia Langford, John Sillitoe on behalf of the Wellcome Sanger Institute COVID-19 Surveillance Team |
| EPI_ISL_3043403 | Berkshire and Surrey Pathology Services Lighthouse Laboratory | Wellcome Sanger Institute for the COVID-19 Genomics UK (COG-UK) Consortium | Berkshire and Surrey Pathology Services Lighthouse Laboratory and Alex Alderton, Roberto Amato, Jeffrey Barrett, Sonia Goncalves, Ewan Harrison, David K. Jackson, Ian Johnston, Dominic Kwiatkowski, Cordelia Langford, John Sillitoe on behalf of the Wellcome Sanger Institute COVID-19 Surveillance Team |
| EPI_ISL_3043404 | Lighthouse Lab in Glasgow | Wellcome Sanger Institute for the COVID-19 Genomics UK (COG-UK) Consortium | Harper VanSteenhouse, Yumi Kasai, David Gray, Carol Clugston, Anna Dominiczak and Alex Alderton, Roberto Amato, Jeffrey Barrett, Sonia Goncalves, Ewan Harrison, David K. Jackson, Ian Johnston, Dominic Kwiatkowski, Cordelia Langford, John Sillitoe on behalf of the Wellcome Sanger Institute COVID-19 Surveillance Team |
| EPI_ISL_3043405, EPI_ISL_3043406 | Berkshire and Surrey Pathology Services Lighthouse Laboratory | Wellcome Sanger Institute for the COVID-19 Genomics UK (COG-UK) Consortium | Berkshire and Surrey Pathology Services Lighthouse Laboratory and Alex Alderton, Roberto Amato, Jeffrey Barrett, Sonia Goncalves, Ewan Harrison, David K. Jackson, Ian Johnston, Dominic Kwiatkowski, Cordelia Langford, John Sillitoe on behalf of the Wellcome Sanger Institute COVID-19 Surveillance Team |
| EPI_ISL_3043407 | Lighthouse Lab in Glasgow | Wellcome Sanger Institute for the COVID-19 Genomics UK (COG-UK) Consortium | Harper VanSteenhouse, Yumi Kasai, David Gray, Carol Clugston, Anna Dominiczak and Alex Alderton, Roberto Amato, Jeffrey Barrett, Sonia Goncalves, Ewan Harrison, David K. Jackson, Ian Johnston, Dominic Kwiatkowski, Cordelia Langford, John Sillitoe on behalf of the Wellcome Sanger Institute COVID-19 Surveillance Team |
| EPI_ISL_3043408 | Lighthouse Lab in Milton Keynes | Wellcome Sanger Institute for the COVID-19 Genomics UK (COG-UK) Consortium | The Lighthouse Lab in Milton Keynes and Alex Alderton, Roberto Amato, Jeffrey Barrett, Sonia Goncalves, Ewan Harrison, David K. Jackson, Ian Johnston, Dominic Kwiatkowski, Cordelia Langford, John Sillitoe on behalf of the Wellcome Sanger Institute COVID-19 Surveillance Team |
| EPI_ISL_3043409 | Lighthouse Lab in Glasgow | Wellcome Sanger Institute for the COVID-19 Genomics UK (COG-UK) Consortium | Harper VanSteenhouse, Yumi Kasai, David Gray, Carol Clugston, Anna Dominiczak and Alex Alderton, Roberto Amato, Jeffrey Barrett, Sonia Goncalves, Ewan Harrison, David K. Jackson, Ian Johnston, Dominic Kwiatkowski, Cordelia Langford, John Sillitoe on behalf of the Wellcome Sanger Institute COVID-19 Surveillance Team |
| EPI_ISL_3043410 | Berkshire and Surrey Pathology Services Lighthouse Laboratory | Wellcome Sanger Institute for the COVID-19 Genomics UK (COG-UK) Consortium | Berkshire and Surrey Pathology Services Lighthouse Laboratory and Alex Alderton, Roberto Amato, Jeffrey Barrett, Sonia Goncalves, Ewan Harrison, David K. Jackson, Ian Johnston, Dominic Kwiatkowski, Cordelia Langford, John Sillitoe on behalf of the Wellcome Sanger Institute COVID-19 Surveillance Team |
| EPI_ISL_3043411 | Lighthouse Lab in Milton Keynes | Wellcome Sanger Institute for the COVID-19 Genomics UK (COG-UK) Consortium | The Lighthouse Lab in Milton Keynes and Alex Alderton, Roberto Amato, Jeffrey Barrett, Sonia Goncalves, Ewan Harrison, David K. Jackson, Ian Johnston, Dominic Kwiatkowski, Cordelia Langford, John Sillitoe on behalf of the Wellcome Sanger Institute COVID-19 Surveillance Team |
| EPI_ISL_3043412 | Lighthouse Lab in Glasgow | Wellcome Sanger Institute for the COVID-19 Genomics UK (COG-UK) Consortium | Harper VanSteenhouse, Yumi Kasai, David Gray, Carol Clugston, Anna Dominiczak and Alex Alderton, Roberto Amato, Jeffrey Barrett, Sonia Goncalves, Ewan Harrison, David K. Jackson, Ian Johnston, Dominic Kwiatkowski, Cordelia Langford, John Sillitoe on behalf of the Wellcome Sanger Institute COVID-19 Surveillance Team |
| EPI_ISL_3043413 | Berkshire and Surrey Pathology Services Lighthouse Laboratory | Wellcome Sanger Institute for the COVID-19 Genomics UK (COG-UK) Consortium | Berkshire and Surrey Pathology Services Lighthouse Laboratory and Alex Alderton, Roberto Amato, Jeffrey Barrett, Sonia Goncalves, Ewan Harrison, David K. Jackson, Ian Johnston, Dominic Kwiatkowski, Cordelia Langford, John Sillitoe on behalf of the Wellcome Sanger Institute COVID-19 Surveillance Team |
| EPI_ISL_3043414 | Lighthouse Lab in Milton Keynes | Wellcome Sanger Institute for the COVID-19 Genomics UK (COG-UK) Consortium | The Lighthouse Lab in Milton Keynes and Alex Alderton, Roberto Amato, Jeffrey Barrett, Sonia Goncalves, Ewan Harrison, David K. Jackson, Ian Johnston, Dominic Kwiatkowski, Cordelia Langford, John Sillitoe on behalf of the Wellcome Sanger Institute COVID-19 Surveillance Team |
| EPI_ISL_3043415 | Lighthouse Lab in Glasgow | Wellcome Sanger Institute for the COVID-19 Genomics UK (COG-UK) Consortium | Harper VanSteenhouse, Yumi Kasai, David Gray, Carol Clugston, Anna Dominiczak and Alex Alderton, Roberto Amato, Jeffrey Barrett, Sonia Goncalves, Ewan Harrison, David K. Jackson, Ian Johnston, Dominic Kwiatkowski, Cordelia Langford, John Sillitoe on behalf of the Wellcome Sanger Institute COVID-19 Surveillance Team |
| EPI_ISL_3043416 | Berkshire and Surrey Pathology Services Lighthouse Laboratory | Wellcome Sanger Institute for the COVID-19 Genomics UK (COG-UK) Consortium | Berkshire and Surrey Pathology Services Lighthouse Laboratory and Alex Alderton, Roberto Amato, Jeffrey Barrett, Sonia Goncalves, Ewan Harrison, David K. Jackson, Ian Johnston, Dominic Kwiatkowski, Cordelia Langford, John Sillitoe on behalf of the Wellcome Sanger Institute COVID-19 Surveillance Team |
| EPI_ISL_3043417, EPI_ISL_3043418 | Lighthouse Lab in Glasgow | Wellcome Sanger Institute for the COVID-19 Genomics UK (COG-UK) Consortium | Harper VanSteenhouse, Yumi Kasai, David Gray, Carol Clugston, Anna Dominiczak and Alex Alderton, Roberto Amato, Jeffrey Barrett, Sonia Goncalves, Ewan Harrison, David K. Jackson, Ian Johnston, Dominic Kwiatkowski, Cordelia Langford, John Sillitoe on behalf of the Wellcome Sanger Institute COVID-19 Surveillance Team |
| EPI_ISL_3043419 | Lighthouse Lab in Milton Keynes | Wellcome Sanger Institute for the COVID-19 Genomics UK (COG-UK) Consortium | The Lighthouse Lab in Milton Keynes and Alex Alderton, Roberto Amato, Jeffrey Barrett, Sonia Goncalves, Ewan Harrison, David K. Jackson, Ian Johnston, Dominic Kwiatkowski, Cordelia Langford, John Sillitoe on behalf of the Wellcome Sanger Institute COVID-19 Surveillance Team |
| EPI_ISL_3043420, EPI_ISL_3043421 | Lighthouse Lab in Glasgow | Wellcome Sanger Institute for the COVID-19 Genomics UK (COG-UK) Consortium | Harper VanSteenhouse, Yumi Kasai, David Gray, Carol Clugston, Anna Dominiczak and Alex Alderton, Roberto Amato, Jeffrey Barrett, Sonia Goncalves, Ewan Harrison, David K. Jackson, Ian Johnston, Dominic Kwiatkowski, Cordelia Langford, John Sillitoe on behalf of the Wellcome Sanger Institute COVID-19 Surveillance Team |
| EPI_ISL_3043422 | Berkshire and Surrey Pathology Services Lighthouse Laboratory | Wellcome Sanger Institute for the COVID-19 Genomics UK (COG-UK) Consortium | Berkshire and Surrey Pathology Services Lighthouse Laboratory and Alex Alderton, Roberto Amato, Jeffrey Barrett, Sonia Goncalves, Ewan Harrison, David K. Jackson, Ian Johnston, Dominic Kwiatkowski, Cordelia Langford, John Sillitoe on behalf of the Wellcome Sanger Institute COVID-19 Surveillance Team |
| EPI_ISL_3043423 | Lighthouse Lab in Milton Keynes | Wellcome Sanger Institute for the COVID-19 Genomics UK (COG-UK) Consortium | The Lighthouse Lab in Milton Keynes and Alex Alderton, Roberto Amato, Jeffrey Barrett, Sonia Goncalves, Ewan Harrison, David K. Jackson, Ian Johnston, Dominic Kwiatkowski, Cordelia Langford, John Sillitoe on behalf of the Wellcome Sanger Institute COVID-19 Surveillance Team |
| EPI_ISL_3043424 | Lighthouse Lab in Glasgow | Wellcome Sanger Institute for the COVID-19 Genomics UK (COG-UK) Consortium | Harper VanSteenhouse, Yumi Kasai, David Gray, Carol Clugston, Anna Dominiczak and Alex Alderton, Roberto Amato, Jeffrey Barrett, Sonia Goncalves, Ewan Harrison, David K. Jackson, Ian Johnston, Dominic Kwiatkowski, Cordelia Langford, John Sillitoe on behalf of the Wellcome Sanger Institute COVID-19 Surveillance Team |
| EPI_ISL_3043425 | Lighthouse Lab in Milton Keynes | Wellcome Sanger Institute for the COVID-19 Genomics UK (COG-UK) Consortium | The Lighthouse Lab in Milton Keynes and Alex Alderton, Roberto Amato, Jeffrey Barrett, Sonia Goncalves, Ewan Harrison, David K. Jackson, Ian Johnston, Dominic Kwiatkowski, Cordelia Langford, John Sillitoe on behalf of the Wellcome Sanger Institute COVID-19 Surveillance Team |
| EPI_ISL_3043426 | Lighthouse Lab in Glasgow | Wellcome Sanger Institute for the COVID-19 Genomics UK (COG-UK) Consortium | Harper VanSteenhouse, Yumi Kasai, David Gray, Carol Clugston, Anna Dominiczak and Alex Alderton, Roberto Amato, Jeffrey Barrett, Sonia Goncalves, Ewan Harrison, David K. Jackson, Ian Johnston, Dominic Kwiatkowski, Cordelia Langford, John Sillitoe on behalf of the Wellcome Sanger Institute COVID-19 Surveillance Team |
| EPI_ISL_3043427, EPI_ISL_3043428, EPI_ISL_3043429 | Lighthouse Lab in Milton Keynes | Wellcome Sanger Institute for the COVID-19 Genomics UK (COG-UK) Consortium | The Lighthouse Lab in Milton Keynes and Alex Alderton, Roberto Amato, Jeffrey Barrett, Sonia Goncalves, Ewan Harrison, David K. Jackson, Ian Johnston, Dominic Kwiatkowski, Cordelia Langford, John Sillitoe on behalf of the Wellcome Sanger Institute COVID-19 Surveillance Team |
| EPI_ISL_3043430, EPI_ISL_3043431 | Berkshire and Surrey Pathology Services Lighthouse Laboratory | Wellcome Sanger Institute for the COVID-19 Genomics UK (COG-UK) Consortium | Berkshire and Surrey Pathology Services Lighthouse Laboratory and Alex Alderton, Roberto Amato, Jeffrey Barrett, Sonia Goncalves, Ewan Harrison, David K. Jackson, Ian Johnston, Dominic Kwiatkowski, Cordelia Langford, John Sillitoe on behalf of the Wellcome Sanger Institute COVID-19 Surveillance Team |
| EPI_ISL_3043432 | Lighthouse Lab in Milton Keynes | Wellcome Sanger Institute for the COVID-19 Genomics UK (COG-UK) Consortium | The Lighthouse Lab in Milton Keynes and Alex Alderton, Roberto Amato, Jeffrey Barrett, Sonia Goncalves, Ewan Harrison, David K. Jackson, Ian Johnston, Dominic Kwiatkowski, Cordelia Langford, John Sillitoe on behalf of the Wellcome Sanger Institute COVID-19 Surveillance Team |
| EPI_ISL_3043433, EPI_ISL_3043434 | Lighthouse Lab in Glasgow | Wellcome Sanger Institute for the COVID-19 Genomics UK (COG-UK) Consortium | Harper VanSteenhouse, Yumi Kasai, David Gray, Carol Clugston, Anna Dominiczak and Alex Alderton, Roberto Amato, Jeffrey Barrett, Sonia Goncalves, Ewan Harrison, David K. Jackson, Ian Johnston, Dominic Kwiatkowski, Cordelia Langford, John Sillitoe on behalf of the Wellcome Sanger Institute COVID-19 Surveillance Team |
| EPI_ISL_3043435 | Berkshire and Surrey Pathology Services Lighthouse Laboratory | Wellcome Sanger Institute for the COVID-19 Genomics UK (COG-UK) | Berkshire and Surrey Pathology Services Lighthouse Laboratory and Alex Alderton, Roberto Amato, Jeffrey Barrett, Sonia Goncalves, Ewan Harrison, David K. Jackson, Ian Johnston, Dominic Kwiatkowski, Cordelia Langford, John Sillitoe on behalf |

| | | | |
|---|---|---|---|
| | | Consortium | of the Wellcome Sanger Institute COVID-19 Surveillance Team |
| EPI_ISL_3043436, EPI_ISL_3043437 | Lighthouse Lab in Glasgow | Wellcome Sanger Institute for the COVID-19 Genomics UK (COG-UK) Consortium | Harper VanSteenhouse, Yumi Kasai, David Gray, Carol Clugston, Anna Dominiczak and Alex Alderton, Roberto Amato, Jeffrey Barrett, Sonia Goncalves, Ewan Harrison, David K. Jackson, Ian Johnston, Dominic Kwiatkowski, Cordelia Langford, John Sillitoe on behalf of the Wellcome Sanger Institute COVID-19 Surveillance Team |
| EPI_ISL_3043438, EPI_ISL_3043439 | Lighthouse Lab in Milton Keynes | Wellcome Sanger Institute for the COVID-19 Genomics UK (COG-UK) Consortium | The Lighthouse Lab in Milton Keynes and Alex Alderton, Roberto Amato, Jeffrey Barrett, Sonia Goncalves, Ewan Harrison, David K. Jackson, Ian Johnston, Dominic Kwiatkowski, Cordelia Langford, John Sillitoe on behalf of the Wellcome Sanger Institute COVID-19 Surveillance Team |
| EPI_ISL_3043440 | Lighthouse Lab in Glasgow | Wellcome Sanger Institute for the COVID-19 Genomics UK (COG-UK) Consortium | Harper VanSteenhouse, Yumi Kasai, David Gray, Carol Clugston, Anna Dominiczak and Alex Alderton, Roberto Amato, Jeffrey Barrett, Sonia Goncalves, Ewan Harrison, David K. Jackson, Ian Johnston, Dominic Kwiatkowski, Cordelia Langford, John Sillitoe on behalf of the Wellcome Sanger Institute COVID-19 Surveillance Team |
| EPI_ISL_3043441, EPI_ISL_3043442 | Lighthouse Lab in Milton Keynes | Wellcome Sanger Institute for the COVID-19 Genomics UK (COG-UK) Consortium | The Lighthouse Lab in Milton Keynes and Alex Alderton, Roberto Amato, Jeffrey Barrett, Sonia Goncalves, Ewan Harrison, David K. Jackson, Ian Johnston, Dominic Kwiatkowski, Cordelia Langford, John Sillitoe on behalf of the Wellcome Sanger Institute COVID-19 Surveillance Team |
| EPI_ISL_3043443 | Berkshire and Surrey Pathology Services Lighthouse Laboratory | Wellcome Sanger Institute for the COVID-19 Genomics UK (COG-UK) Consortium | Berkshire and Surrey Pathology Services Lighthouse Laboratory and Alex Alderton, Roberto Amato, Jeffrey Barrett, Sonia Goncalves, Ewan Harrison, David K. Jackson, Ian Johnston, Dominic Kwiatkowski, Cordelia Langford, John Sillitoe on behalf of the Wellcome Sanger Institute COVID-19 Surveillance Team |
| EPI_ISL_3043444, EPI_ISL_3043445 | Lighthouse Lab in Milton Keynes | Wellcome Sanger Institute for the COVID-19 Genomics UK (COG-UK) Consortium | The Lighthouse Lab in Milton Keynes and Alex Alderton, Roberto Amato, Jeffrey Barrett, Sonia Goncalves, Ewan Harrison, David K. Jackson, Ian Johnston, Dominic Kwiatkowski, Cordelia Langford, John Sillitoe on behalf of the Wellcome Sanger Institute COVID-19 Surveillance Team |
| EPI_ISL_3043446 | Lighthouse Lab in Alderley Park | Wellcome Sanger Institute for the COVID-19 Genomics UK (COG-UK) Consortium | Jacquelyn Wynn, Mairead Hyland, The Lighthouse Lab in Alderley Park and Alex Alderton, Roberto Amato, Jeffrey Barrett, Sonia Goncalves, Ewan Harrison, David K. Jackson, Ian Johnston, Dominic Kwiatkowski, Cordelia Langford, John Sillitoe on behalf of the Wellcome Sanger Institute COVID-19 Surveillance Team |
| EPI_ISL_3043447, EPI_ISL_3043448, EPI_ISL_3043449, EPI_ISL_3043450, EPI_ISL_3043451, EPI_ISL_3043452 | Lighthouse Lab in Milton Keynes | Wellcome Sanger Institute for the COVID-19 Genomics UK (COG-UK) Consortium | The Lighthouse Lab in Milton Keynes and Alex Alderton, Roberto Amato, Jeffrey Barrett, Sonia Goncalves, Ewan Harrison, David K. Jackson, Ian Johnston, Dominic Kwiatkowski, Cordelia Langford, John Sillitoe on behalf of the Wellcome Sanger Institute COVID-19 Surveillance Team |
| EPI_ISL_3043453 | Lighthouse Lab in Alderley Park | Wellcome Sanger Institute for the COVID-19 Genomics UK (COG-UK) Consortium | Jacquelyn Wynn, Mairead Hyland, The Lighthouse Lab in Alderley Park and Alex Alderton, Roberto Amato, Jeffrey Barrett, Sonia Goncalves, Ewan Harrison, David K. Jackson, Ian Johnston, Dominic Kwiatkowski, Cordelia Langford, John Sillitoe on behalf of the Wellcome Sanger Institute COVID-19 Surveillance Team |
| EPI_ISL_3043454, EPI_ISL_3043455, EPI_ISL_3043456, EPI_ISL_3043457, EPI_ISL_3043458, EPI_ISL_3043459, EPI_ISL_3043460 | Lighthouse Lab in Milton Keynes | Wellcome Sanger Institute for the COVID-19 Genomics UK (COG-UK) Consortium | The Lighthouse Lab in Milton Keynes and Alex Alderton, Roberto Amato, Jeffrey Barrett, Sonia Goncalves, Ewan Harrison, David K. Jackson, Ian Johnston, Dominic Kwiatkowski, Cordelia Langford, John Sillitoe on behalf of the Wellcome Sanger Institute COVID-19 Surveillance Team |
| EPI_ISL_3043461, EPI_ISL_3043462 | Lighthouse Lab in Alderley Park | Wellcome Sanger Institute for the COVID-19 Genomics UK (COG-UK) Consortium | Jacquelyn Wynn, Mairead Hyland, The Lighthouse Lab in Alderley Park and Alex Alderton, Roberto Amato, Jeffrey Barrett, Sonia Goncalves, Ewan Harrison, David K. Jackson, Ian Johnston, Dominic Kwiatkowski, Cordelia Langford, John Sillitoe on behalf of the Wellcome Sanger Institute COVID-19 Surveillance Team |
| EPI_ISL_3043463 | Lighthouse Lab in Milton Keynes | Wellcome Sanger Institute for the COVID-19 Genomics UK (COG-UK) Consortium | The Lighthouse Lab in Milton Keynes and Alex Alderton, Roberto Amato, Jeffrey Barrett, Sonia Goncalves, Ewan Harrison, David K. Jackson, Ian Johnston, Dominic Kwiatkowski, Cordelia Langford, John Sillitoe on behalf of the Wellcome Sanger Institute COVID-19 Surveillance Team |
| EPI_ISL_3043464 | Lighthouse Lab in Alderley Park | Wellcome Sanger Institute for the COVID-19 Genomics UK (COG-UK) Consortium | Jacquelyn Wynn, Mairead Hyland, The Lighthouse Lab in Alderley Park and Alex Alderton, Roberto Amato, Jeffrey Barrett, Sonia Goncalves, Ewan Harrison, David K. Jackson, Ian Johnston, Dominic Kwiatkowski, Cordelia Langford, John Sillitoe on behalf of the Wellcome Sanger Institute COVID-19 Surveillance Team |
| EPI_ISL_3043465, EPI_ISL_3043466, EPI_ISL_3043467, EPI_ISL_3043468, EPI_ISL_3043469, EPI_ISL_3043470 | Lighthouse Lab in Milton Keynes | Wellcome Sanger Institute for the COVID-19 Genomics UK (COG-UK) Consortium | The Lighthouse Lab in Milton Keynes and Alex Alderton, Roberto Amato, Jeffrey Barrett, Sonia Goncalves, Ewan Harrison, David K. Jackson, Ian Johnston, Dominic Kwiatkowski, Cordelia Langford, John Sillitoe on behalf of the Wellcome Sanger Institute COVID-19 Surveillance Team |
| EPI_ISL_3043471 | Lighthouse Lab in Alderley Park | Wellcome Sanger Institute for the COVID-19 Genomics UK (COG-UK) Consortium | Jacquelyn Wynn, Mairead Hyland, The Lighthouse Lab in Alderley Park and Alex Alderton, Roberto Amato, Jeffrey Barrett, Sonia Goncalves, Ewan Harrison, David K. Jackson, Ian Johnston, Dominic Kwiatkowski, Cordelia Langford, John Sillitoe on behalf of the Wellcome Sanger Institute COVID-19 Surveillance Team |
| EPI_ISL_3043472 | Lighthouse Lab in Glasgow | Wellcome Sanger Institute for the COVID-19 Genomics UK (COG-UK) Consortium | Harper VanSteenhouse, Yumi Kasai, David Gray, Carol Clugston, Anna Dominiczak and Alex Alderton, Roberto Amato, Jeffrey Barrett, Sonia Goncalves, Ewan Harrison, David K. Jackson, Ian Johnston, Dominic Kwiatkowski, Cordelia Langford, John Sillitoe on behalf of the Wellcome Sanger Institute COVID-19 Surveillance Team |
| EPI_ISL_3043473, EPI_ISL_3043474 | Lighthouse Lab in Milton Keynes | Wellcome Sanger Institute for the COVID-19 Genomics UK (COG-UK) Consortium | The Lighthouse Lab in Milton Keynes and Alex Alderton, Roberto Amato, Jeffrey Barrett, Sonia Goncalves, Ewan Harrison, David K. Jackson, Ian Johnston, Dominic Kwiatkowski, Cordelia Langford, John Sillitoe on behalf of the Wellcome Sanger Institute COVID-19 Surveillance Team |
| EPI_ISL_3043475 | Lighthouse Lab in Glasgow | Wellcome Sanger Institute for the COVID-19 Genomics UK (COG-UK) Consortium | Harper VanSteenhouse, Yumi Kasai, David Gray, Carol Clugston, Anna Dominiczak and Alex Alderton, Roberto Amato, Jeffrey Barrett, Sonia Goncalves, Ewan Harrison, David K. Jackson, Ian Johnston, Dominic Kwiatkowski, Cordelia Langford, John Sillitoe on behalf of the Wellcome Sanger Institute COVID-19 Surveillance Team |
| EPI_ISL_3043476, EPI_ISL_3043477 | Lighthouse Lab in Milton Keynes | Wellcome Sanger Institute for the COVID-19 Genomics UK (COG-UK) Consortium | The Lighthouse Lab in Milton Keynes and Alex Alderton, Roberto Amato, Jeffrey Barrett, Sonia Goncalves, Ewan Harrison, David K. Jackson, Ian Johnston, Dominic Kwiatkowski, Cordelia Langford, John Sillitoe on behalf of the Wellcome Sanger Institute COVID-19 Surveillance Team |
| EPI_ISL_3043478 | Lighthouse Lab in Alderley Park | Wellcome Sanger Institute for the COVID-19 Genomics UK (COG-UK) Consortium | Jacquelyn Wynn, Mairead Hyland, The Lighthouse Lab in Alderley Park and Alex Alderton, Roberto Amato, Jeffrey Barrett, Sonia Goncalves, Ewan Harrison, David K. Jackson, Ian Johnston, Dominic Kwiatkowski, Cordelia Langford, John Sillitoe on behalf of the Wellcome Sanger Institute COVID-19 Surveillance Team |
| EPI_ISL_3043479, EPI_ISL_3043480 | Lighthouse Lab in Milton Keynes | Wellcome Sanger Institute for the COVID-19 Genomics UK (COG-UK) Consortium | The Lighthouse Lab in Milton Keynes and Alex Alderton, Roberto Amato, Jeffrey Barrett, Sonia Goncalves, Ewan Harrison, David K. Jackson, Ian Johnston, Dominic Kwiatkowski, Cordelia Langford, John Sillitoe on behalf of the Wellcome Sanger Institute COVID-19 Surveillance Team |
| EPI_ISL_3043481 | Lighthouse Lab in Glasgow | Wellcome Sanger Institute for the COVID-19 Genomics UK (COG-UK) Consortium | Harper VanSteenhouse, Yumi Kasai, David Gray, Carol Clugston, Anna Dominiczak and Alex Alderton, Roberto Amato, Jeffrey Barrett, Sonia Goncalves, Ewan Harrison, David K. Jackson, Ian Johnston, Dominic Kwiatkowski, Cordelia Langford, John Sillitoe on behalf of the Wellcome Sanger Institute COVID-19 Surveillance Team |
| EPI_ISL_3043482, EPI_ISL_3043483 | Lighthouse Lab in Milton Keynes | Wellcome Sanger Institute for the COVID-19 Genomics UK (COG-UK) Consortium | The Lighthouse Lab in Milton Keynes and Alex Alderton, Roberto Amato, Jeffrey Barrett, Sonia Goncalves, Ewan Harrison, David K. Jackson, Ian Johnston, Dominic Kwiatkowski, Cordelia Langford, John Sillitoe on behalf of the Wellcome Sanger Institute COVID-19 Surveillance Team |
| EPI_ISL_3043484 | Lighthouse Lab in Glasgow | Wellcome Sanger Institute for the COVID-19 Genomics UK (COG-UK) Consortium | Harper VanSteenhouse, Yumi Kasai, David Gray, Carol Clugston, Anna Dominiczak and Alex Alderton, Roberto Amato, Jeffrey Barrett, Sonia Goncalves, Ewan Harrison, David K. Jackson, Ian Johnston, Dominic Kwiatkowski, Cordelia Langford, John Sillitoe on behalf of the Wellcome Sanger Institute COVID-19 Surveillance Team |
| EPI_ISL_3043485 | Lighthouse Lab in Milton Keynes | Wellcome Sanger Institute for the COVID-19 Genomics UK (COG-UK) Consortium | The Lighthouse Lab in Milton Keynes and Alex Alderton, Roberto Amato, Jeffrey Barrett, Sonia Goncalves, Ewan Harrison, David K. Jackson, Ian Johnston, Dominic Kwiatkowski, Cordelia Langford, John Sillitoe on behalf of the Wellcome Sanger Institute COVID-19 Surveillance Team |
| EPI_ISL_3043486 | Lighthouse Lab in Alderley Park | Wellcome Sanger Institute for the COVID-19 Genomics UK (COG-UK) Consortium | Jacquelyn Wynn, Mairead Hyland, The Lighthouse Lab in Alderley Park and Alex Alderton, Roberto Amato, Jeffrey Barrett, Sonia Goncalves, Ewan Harrison, David K. Jackson, Ian Johnston, Dominic Kwiatkowski, Cordelia Langford, John Sillitoe on behalf of the Wellcome Sanger Institute COVID-19 Surveillance Team |
| EPI_ISL_3043487, EPI_ISL_3043488 | Lighthouse Lab in Milton Keynes | Wellcome Sanger Institute for the COVID-19 Genomics UK (COG-UK) Consortium | The Lighthouse Lab in Milton Keynes and Alex Alderton, Roberto Amato, Jeffrey Barrett, Sonia Goncalves, Ewan Harrison, David K. Jackson, Ian Johnston, Dominic Kwiatkowski, Cordelia Langford, John Sillitoe on behalf of the Wellcome Sanger Institute COVID-19 Surveillance Team |
| EPI_ISL_3043489, EPI_ISL_3043490 | Lighthouse Lab in Glasgow | Wellcome Sanger Institute for the COVID-19 Genomics UK (COG-UK) Consortium | Harper VanSteenhouse, Yumi Kasai, David Gray, Carol Clugston, Anna Dominiczak and Alex Alderton, Roberto Amato, Jeffrey Barrett, Sonia Goncalves, Ewan Harrison, David K. Jackson, Ian Johnston, Dominic Kwiatkowski, Cordelia Langford, John Sillitoe on behalf of the Wellcome Sanger Institute COVID-19 Surveillance Team |
| EPI_ISL_3043491, EPI_ISL_3043492, EPI_ISL_3043493 | Lighthouse Lab in Milton Keynes | Wellcome Sanger Institute for the COVID-19 Genomics UK (COG-UK) Consortium | The Lighthouse Lab in Milton Keynes and Alex Alderton, Roberto Amato, Jeffrey Barrett, Sonia Goncalves, Ewan Harrison, David K. Jackson, Ian Johnston, Dominic Kwiatkowski, Cordelia Langford, John Sillitoe on behalf of the Wellcome Sanger Institute COVID-19 Surveillance Team |
| EPI_ISL_3043494, EPI_ISL_3043495, EPI_ISL_3043496, EPI_ISL_3043497 | Lighthouse Lab in Alderley Park | Wellcome Sanger Institute for the COVID-19 Genomics UK (COG-UK) Consortium | Jacquelyn Wynn, Mairead Hyland, The Lighthouse Lab in Alderley Park and Alex Alderton, Roberto Amato, Jeffrey Barrett, Sonia Goncalves, Ewan Harrison, David K. Jackson, Ian Johnston, Dominic Kwiatkowski, Cordelia Langford, John Sillitoe on behalf of the Wellcome Sanger Institute COVID-19 Surveillance Team |
| EPI_ISL_3043498, EPI_ISL_3043499 | Lighthouse Lab in Milton Keynes | Wellcome Sanger Institute for the COVID-19 Genomics UK (COG-UK) Consortium | The Lighthouse Lab in Milton Keynes and Alex Alderton, Roberto Amato, Jeffrey Barrett, Sonia Goncalves, Ewan Harrison, David K. Jackson, Ian Johnston, Dominic Kwiatkowski, Cordelia Langford, John Sillitoe on behalf of the Wellcome Sanger Institute COVID-19 Surveillance Team |
| EPI_ISL_3043500, EPI_ISL_3043501, EPI_ISL_3043502 | Lighthouse Lab in Alderley Park | Wellcome Sanger Institute for the COVID-19 Genomics UK (COG-UK) Consortium | Jacquelyn Wynn, Mairead Hyland, The Lighthouse Lab in Alderley Park and Alex Alderton, Roberto Amato, Jeffrey Barrett, Sonia Goncalves, Ewan Harrison, David K. Jackson, Ian Johnston, Dominic Kwiatkowski, Cordelia Langford, John Sillitoe on behalf of the Wellcome Sanger Institute COVID-19 Surveillance Team |
| EPI_ISL_3043503, EPI_ISL_3043504, EPI_ISL_3043505, EPI_ISL_3043506, EPI_ISL_3043507 | Lighthouse Lab in Milton Keynes | Wellcome Sanger Institute for the COVID-19 Genomics UK (COG-UK) Consortium | The Lighthouse Lab in Milton Keynes and Alex Alderton, Roberto Amato, Jeffrey Barrett, Sonia Goncalves, Ewan Harrison, David K. Jackson, Ian Johnston, Dominic Kwiatkowski, Cordelia Langford, John Sillitoe on behalf of the Wellcome Sanger Institute COVID-19 Surveillance Team |
| EPI_ISL_3043508 | Lighthouse Lab in Glasgow | Wellcome Sanger Institute for the COVID-19 Genomics UK (COG-UK) Consortium | Harper VanSteenhouse, Yumi Kasai, David Gray, Carol Clugston, Anna Dominiczak and Alex Alderton, Roberto Amato, Jeffrey Barrett, Sonia Goncalves, Ewan Harrison, David K. Jackson, Ian Johnston, Dominic Kwiatkowski, Cordelia Langford, John Sillitoe on behalf of the Wellcome Sanger Institute COVID-19 Surveillance Team |
| EPI_ISL_3043509 | Lighthouse Lab in Milton Keynes | Wellcome Sanger Institute for the COVID-19 Genomics UK (COG-UK) Consortium | The Lighthouse Lab in Milton Keynes and Alex Alderton, Roberto Amato, Jeffrey Barrett, Sonia Goncalves, Ewan Harrison, David K. Jackson, Ian Johnston, Dominic Kwiatkowski, Cordelia Langford, John Sillitoe on behalf of the Wellcome Sanger Institute COVID-19 Surveillance Team |
| EPI_ISL_3043510 | Lighthouse Lab in Alderley Park | Wellcome Sanger Institute for the COVID-19 Genomics UK (COG-UK) Consortium | Jacquelyn Wynn, Mairead Hyland, The Lighthouse Lab in Alderley Park and Alex Alderton, Roberto Amato, Jeffrey Barrett, Sonia Goncalves, Ewan Harrison, David K. Jackson, Ian Johnston, Dominic Kwiatkowski, Cordelia Langford, John Sillitoe on behalf of the Wellcome Sanger Institute COVID-19 Surveillance Team |

| | | | |
|---|---|---|---|
| EPI_ISL_3043511, EPI_ISL_3043512, EPI_ISL_3043513, EPI_ISL_3043514 | Lighthouse Lab in Milton Keynes | Wellcome Sanger Institute for the COVID-19 Genomics UK (COG-UK) Consortium | The Lighthouse Lab in Milton Keynes and Alex Alderton, Roberto Amato, Jeffrey Barrett, Sonia Goncalves, Ewan Harrison, David K. Jackson, Ian Johnston, Dominic Kwiatkowski, Cordelia Langford, John Sillitoe on behalf of the Wellcome Sanger Institute COVID-19 Surveillance Team |
| EPI_ISL_3043515 | Lighthouse Lab in Alderley Park | Wellcome Sanger Institute for the COVID-19 Genomics UK (COG-UK) Consortium | Jacquelyn Wynn, Mairead Hyland, The Lighthouse Lab in Alderley Park and Alex Alderton, Roberto Amato, Jeffrey Barrett, Sonia Goncalves, Ewan Harrison, David K. Jackson, Ian Johnston, Dominic Kwiatkowski, Cordelia Langford, John Sillitoe on behalf of the Wellcome Sanger Institute COVID-19 Surveillance Team |
| EPI_ISL_3043516, EPI_ISL_3043517, EPI_ISL_3043518, EPI_ISL_3043519 | Lighthouse Lab in Milton Keynes | Wellcome Sanger Institute for the COVID-19 Genomics UK (COG-UK) Consortium | The Lighthouse Lab in Milton Keynes and Alex Alderton, Roberto Amato, Jeffrey Barrett, Sonia Goncalves, Ewan Harrison, David K. Jackson, Ian Johnston, Dominic Kwiatkowski, Cordelia Langford, John Sillitoe on behalf of the Wellcome Sanger Institute COVID-19 Surveillance Team |
| EPI_ISL_3043520, EPI_ISL_3043521 | Lighthouse Lab in Glasgow | Wellcome Sanger Institute for the COVID-19 Genomics UK (COG-UK) Consortium | Harper VanSteenhouse, Yumi Kasai, David Gray, Carol Clugston, Anna Dominiczak and Alex Alderton, Roberto Amato, Jeffrey Barrett, Sonia Goncalves, Ewan Harrison, David K. Jackson, Ian Johnston, Dominic Kwiatkowski, Cordelia Langford, John Sillitoe on behalf of the Wellcome Sanger Institute COVID-19 Surveillance Team |
| EPI_ISL_3043522 | Lighthouse Lab in Milton Keynes | Wellcome Sanger Institute for the COVID-19 Genomics UK (COG-UK) Consortium | The Lighthouse Lab in Milton Keynes and Alex Alderton, Roberto Amato, Jeffrey Barrett, Sonia Goncalves, Ewan Harrison, David K. Jackson, Ian Johnston, Dominic Kwiatkowski, Cordelia Langford, John Sillitoe on behalf of the Wellcome Sanger Institute COVID-19 Surveillance Team |
| EPI_ISL_3043523 | Lighthouse Lab in Alderley Park | Wellcome Sanger Institute for the COVID-19 Genomics UK (COG-UK) Consortium | Jacquelyn Wynn, Mairead Hyland, The Lighthouse Lab in Alderley Park and Alex Alderton, Roberto Amato, Jeffrey Barrett, Sonia Goncalves, Ewan Harrison, David K. Jackson, Ian Johnston, Dominic Kwiatkowski, Cordelia Langford, John Sillitoe on behalf of the Wellcome Sanger Institute COVID-19 Surveillance Team |
| EPI_ISL_3043524 | Lighthouse Lab in Milton Keynes | Wellcome Sanger Institute for the COVID-19 Genomics UK (COG-UK) Consortium | The Lighthouse Lab in Milton Keynes and Alex Alderton, Roberto Amato, Jeffrey Barrett, Sonia Goncalves, Ewan Harrison, David K. Jackson, Ian Johnston, Dominic Kwiatkowski, Cordelia Langford, John Sillitoe on behalf of the Wellcome Sanger Institute COVID-19 Surveillance Team |
| EPI_ISL_3043525 | Lighthouse Lab in Alderley Park | Wellcome Sanger Institute for the COVID-19 Genomics UK (COG-UK) Consortium | Jacquelyn Wynn, Mairead Hyland, The Lighthouse Lab in Alderley Park and Alex Alderton, Roberto Amato, Jeffrey Barrett, Sonia Goncalves, Ewan Harrison, David K. Jackson, Ian Johnston, Dominic Kwiatkowski, Cordelia Langford, John Sillitoe on behalf of the Wellcome Sanger Institute COVID-19 Surveillance Team |
| EPI_ISL_3043526 | Lighthouse Lab in Glasgow | Wellcome Sanger Institute for the COVID-19 Genomics UK (COG-UK) Consortium | Harper VanSteenhouse, Yumi Kasai, David Gray, Carol Clugston, Anna Dominiczak and Alex Alderton, Roberto Amato, Jeffrey Barrett, Sonia Goncalves, Ewan Harrison, David K. Jackson, Ian Johnston, Dominic Kwiatkowski, Cordelia Langford, John Sillitoe on behalf of the Wellcome Sanger Institute COVID-19 Surveillance Team |
| EPI_ISL_3043527, EPI_ISL_3043528 | Lighthouse Lab in Alderley Park | Wellcome Sanger Institute for the COVID-19 Genomics UK (COG-UK) Consortium | Jacquelyn Wynn, Mairead Hyland, The Lighthouse Lab in Alderley Park and Alex Alderton, Roberto Amato, Jeffrey Barrett, Sonia Goncalves, Ewan Harrison, David K. Jackson, Ian Johnston, Dominic Kwiatkowski, Cordelia Langford, John Sillitoe on behalf of the Wellcome Sanger Institute COVID-19 Surveillance Team |
| EPI_ISL_3043529 | Lighthouse Lab in Milton Keynes | Wellcome Sanger Institute for the COVID-19 Genomics UK (COG-UK) Consortium | The Lighthouse Lab in Milton Keynes and Alex Alderton, Roberto Amato, Jeffrey Barrett, Sonia Goncalves, Ewan Harrison, David K. Jackson, Ian Johnston, Dominic Kwiatkowski, Cordelia Langford, John Sillitoe on behalf of the Wellcome Sanger Institute COVID-19 Surveillance Team |
| EPI_ISL_3043530 | Lighthouse Lab in Alderley Park | Wellcome Sanger Institute for the COVID-19 Genomics UK (COG-UK) Consortium | Jacquelyn Wynn, Mairead Hyland, The Lighthouse Lab in Alderley Park and Alex Alderton, Roberto Amato, Jeffrey Barrett, Sonia Goncalves, Ewan Harrison, David K. Jackson, Ian Johnston, Dominic Kwiatkowski, Cordelia Langford, John Sillitoe on behalf of the Wellcome Sanger Institute COVID-19 Surveillance Team |
| EPI_ISL_3043531 | Lighthouse Lab in Glasgow | Wellcome Sanger Institute for the COVID-19 Genomics UK (COG-UK) Consortium | Harper VanSteenhouse, Yumi Kasai, David Gray, Carol Clugston, Anna Dominiczak and Alex Alderton, Roberto Amato, Jeffrey Barrett, Sonia Goncalves, Ewan Harrison, David K. Jackson, Ian Johnston, Dominic Kwiatkowski, Cordelia Langford, John Sillitoe on behalf of the Wellcome Sanger Institute COVID-19 Surveillance Team |
| EPI_ISL_3043532, EPI_ISL_3043533 | Lighthouse Lab in Milton Keynes | Wellcome Sanger Institute for the COVID-19 Genomics UK (COG-UK) Consortium | The Lighthouse Lab in Milton Keynes and Alex Alderton, Roberto Amato, Jeffrey Barrett, Sonia Goncalves, Ewan Harrison, David K. Jackson, Ian Johnston, Dominic Kwiatkowski, Cordelia Langford, John Sillitoe on behalf of the Wellcome Sanger Institute COVID-19 Surveillance Team |
| EPI_ISL_3043534 | Lighthouse Lab in Glasgow | Wellcome Sanger Institute for the COVID-19 Genomics UK (COG-UK) Consortium | Harper VanSteenhouse, Yumi Kasai, David Gray, Carol Clugston, Anna Dominiczak and Alex Alderton, Roberto Amato, Jeffrey Barrett, Sonia Goncalves, Ewan Harrison, David K. Jackson, Ian Johnston, Dominic Kwiatkowski, Cordelia Langford, John Sillitoe on behalf of the Wellcome Sanger Institute COVID-19 Surveillance Team |
| EPI_ISL_3043535 | Lighthouse Lab in Milton Keynes | Wellcome Sanger Institute for the COVID-19 Genomics UK (COG-UK) Consortium | The Lighthouse Lab in Milton Keynes and Alex Alderton, Roberto Amato, Jeffrey Barrett, Sonia Goncalves, Ewan Harrison, David K. Jackson, Ian Johnston, Dominic Kwiatkowski, Cordelia Langford, John Sillitoe on behalf of the Wellcome Sanger Institute COVID-19 Surveillance Team |
| EPI_ISL_3043536, EPI_ISL_3043537, EPI_ISL_3043538, EPI_ISL_3043539 | Lighthouse Lab in Alderley Park | Wellcome Sanger Institute for the COVID-19 Genomics UK (COG-UK) Consortium | Jacquelyn Wynn, Mairead Hyland, The Lighthouse Lab in Alderley Park and Alex Alderton, Roberto Amato, Jeffrey Barrett, Sonia Goncalves, Ewan Harrison, David K. Jackson, Ian Johnston, Dominic Kwiatkowski, Cordelia Langford, John Sillitoe on behalf of the Wellcome Sanger Institute COVID-19 Surveillance Team |
| EPI_ISL_3043540, EPI_ISL_3043541, EPI_ISL_3043542, EPI_ISL_3043543 | Lighthouse Lab in Milton Keynes | Wellcome Sanger Institute for the COVID-19 Genomics UK (COG-UK) Consortium | The Lighthouse Lab in Milton Keynes and Alex Alderton, Roberto Amato, Jeffrey Barrett, Sonia Goncalves, Ewan Harrison, David K. Jackson, Ian Johnston, Dominic Kwiatkowski, Cordelia Langford, John Sillitoe on behalf of the Wellcome Sanger Institute COVID-19 Surveillance Team |
| EPI_ISL_3043544 | Lighthouse Lab in Alderley Park | Wellcome Sanger Institute for the COVID-19 Genomics UK (COG-UK) Consortium | Jacquelyn Wynn, Mairead Hyland, The Lighthouse Lab in Alderley Park and Alex Alderton, Roberto Amato, Jeffrey Barrett, Sonia Goncalves, Ewan Harrison, David K. Jackson, Ian Johnston, Dominic Kwiatkowski, Cordelia Langford, John Sillitoe on behalf of the Wellcome Sanger Institute COVID-19 Surveillance Team |
| EPI_ISL_3043545, EPI_ISL_3043546, EPI_ISL_3043547 | Lighthouse Lab in Milton Keynes | Wellcome Sanger Institute for the COVID-19 Genomics UK (COG-UK) Consortium | The Lighthouse Lab in Milton Keynes and Alex Alderton, Roberto Amato, Jeffrey Barrett, Sonia Goncalves, Ewan Harrison, David K. Jackson, Ian Johnston, Dominic Kwiatkowski, Cordelia Langford, John Sillitoe on behalf of the Wellcome Sanger Institute COVID-19 Surveillance Team |
| EPI_ISL_3043548, EPI_ISL_3043549, EPI_ISL_3043550, EPI_ISL_3043551, EPI_ISL_3043552, EPI_ISL_3043553 | Lighthouse Lab in Alderley Park | Wellcome Sanger Institute for the COVID-19 Genomics UK (COG-UK) Consortium | Jacquelyn Wynn, Mairead Hyland, The Lighthouse Lab in Alderley Park and Alex Alderton, Roberto Amato, Jeffrey Barrett, Sonia Goncalves, Ewan Harrison, David K. Jackson, Ian Johnston, Dominic Kwiatkowski, Cordelia Langford, John Sillitoe on behalf of the Wellcome Sanger Institute COVID-19 Surveillance Team |
| EPI_ISL_3043554 | Lighthouse Lab in Milton Keynes | Wellcome Sanger Institute for the COVID-19 Genomics UK (COG-UK) Consortium | The Lighthouse Lab in Milton Keynes and Alex Alderton, Roberto Amato, Jeffrey Barrett, Sonia Goncalves, Ewan Harrison, David K. Jackson, Ian Johnston, Dominic Kwiatkowski, Cordelia Langford, John Sillitoe on behalf of the Wellcome Sanger Institute COVID-19 Surveillance Team |
| EPI_ISL_3043555 | Lighthouse Lab in Alderley Park | Wellcome Sanger Institute for the COVID-19 Genomics UK (COG-UK) Consortium | Jacquelyn Wynn, Mairead Hyland, The Lighthouse Lab in Alderley Park and Alex Alderton, Roberto Amato, Jeffrey Barrett, Sonia Goncalves, Ewan Harrison, David K. Jackson, Ian Johnston, Dominic Kwiatkowski, Cordelia Langford, John Sillitoe on behalf of the Wellcome Sanger Institute COVID-19 Surveillance Team |
| EPI_ISL_3043556, EPI_ISL_3043557, EPI_ISL_3043558, EPI_ISL_3043559 | Lighthouse Lab in Milton Keynes | Wellcome Sanger Institute for the COVID-19 Genomics UK (COG-UK) Consortium | The Lighthouse Lab in Milton Keynes and Alex Alderton, Roberto Amato, Jeffrey Barrett, Sonia Goncalves, Ewan Harrison, David K. Jackson, Ian Johnston, Dominic Kwiatkowski, Cordelia Langford, John Sillitoe on behalf of the Wellcome Sanger Institute COVID-19 Surveillance Team |
| EPI_ISL_3043560 | Lighthouse Lab in Alderley Park | Wellcome Sanger Institute for the COVID-19 Genomics UK (COG-UK) Consortium | Jacquelyn Wynn, Mairead Hyland, The Lighthouse Lab in Alderley Park and Alex Alderton, Roberto Amato, Jeffrey Barrett, Sonia Goncalves, Ewan Harrison, David K. Jackson, Ian Johnston, Dominic Kwiatkowski, Cordelia Langford, John Sillitoe on behalf of the Wellcome Sanger Institute COVID-19 Surveillance Team |
| EPI_ISL_3043561 | Lighthouse Lab in Milton Keynes | Wellcome Sanger Institute for the COVID-19 Genomics UK (COG-UK) Consortium | The Lighthouse Lab in Milton Keynes and Alex Alderton, Roberto Amato, Jeffrey Barrett, Sonia Goncalves, Ewan Harrison, David K. Jackson, Ian Johnston, Dominic Kwiatkowski, Cordelia Langford, John Sillitoe on behalf of the Wellcome Sanger Institute COVID-19 Surveillance Team |
| EPI_ISL_3043562, EPI_ISL_3043563, EPI_ISL_3043564, EPI_ISL_3043565 | Lighthouse Lab in Alderley Park | Wellcome Sanger Institute for the COVID-19 Genomics UK (COG-UK) Consortium | Jacquelyn Wynn, Mairead Hyland, The Lighthouse Lab in Alderley Park and Alex Alderton, Roberto Amato, Jeffrey Barrett, Sonia Goncalves, Ewan Harrison, David K. Jackson, Ian Johnston, Dominic Kwiatkowski, Cordelia Langford, John Sillitoe on behalf of the Wellcome Sanger Institute COVID-19 Surveillance Team |
| EPI_ISL_3043566, EPI_ISL_3043567, EPI_ISL_3043568, EPI_ISL_3043569 | Lighthouse Lab in Milton Keynes | Wellcome Sanger Institute for the COVID-19 Genomics UK (COG-UK) Consortium | The Lighthouse Lab in Milton Keynes and Alex Alderton, Roberto Amato, Jeffrey Barrett, Sonia Goncalves, Ewan Harrison, David K. Jackson, Ian Johnston, Dominic Kwiatkowski, Cordelia Langford, John Sillitoe on behalf of the Wellcome Sanger Institute COVID-19 Surveillance Team |
| EPI_ISL_3043570 | Lighthouse Lab in Glasgow | Wellcome Sanger Institute for the COVID-19 Genomics UK (COG-UK) Consortium | Harper VanSteenhouse, Yumi Kasai, David Gray, Carol Clugston, Anna Dominiczak and Alex Alderton, Roberto Amato, Jeffrey Barrett, Sonia Goncalves, Ewan Harrison, David K. Jackson, Ian Johnston, Dominic Kwiatkowski, Cordelia Langford, John Sillitoe on behalf of the Wellcome Sanger Institute COVID-19 Surveillance Team |
| EPI_ISL_3043571 | Lighthouse Lab in Milton Keynes | Wellcome Sanger Institute for the COVID-19 Genomics UK (COG-UK) Consortium | The Lighthouse Lab in Milton Keynes and Alex Alderton, Roberto Amato, Jeffrey Barrett, Sonia Goncalves, Ewan Harrison, David K. Jackson, Ian Johnston, Dominic Kwiatkowski, Cordelia Langford, John Sillitoe on behalf of the Wellcome Sanger Institute COVID-19 Surveillance Team |
| EPI_ISL_3043572, EPI_ISL_3043573 | Lighthouse Lab in Alderley Park | Wellcome Sanger Institute for the COVID-19 Genomics UK (COG-UK) Consortium | Jacquelyn Wynn, Mairead Hyland, The Lighthouse Lab in Alderley Park and Alex Alderton, Roberto Amato, Jeffrey Barrett, Sonia Goncalves, Ewan Harrison, David K. Jackson, Ian Johnston, Dominic Kwiatkowski, Cordelia Langford, John Sillitoe on behalf of the Wellcome Sanger Institute COVID-19 Surveillance Team |
| EPI_ISL_3043574, EPI_ISL_3043575 | Lighthouse Lab in Glasgow | Wellcome Sanger Institute for the COVID-19 Genomics UK (COG-UK) Consortium | Harper VanSteenhouse, Yumi Kasai, David Gray, Carol Clugston, Anna Dominiczak and Alex Alderton, Roberto Amato, Jeffrey Barrett, Sonia Goncalves, Ewan Harrison, David K. Jackson, Ian Johnston, Dominic Kwiatkowski, Cordelia Langford, John Sillitoe on behalf of the Wellcome Sanger Institute COVID-19 Surveillance Team |
| EPI_ISL_3043576, EPI_ISL_3043577, EPI_ISL_3043578, EPI_ISL_3043579 | Lighthouse Lab in Alderley Park | Wellcome Sanger Institute for the COVID-19 Genomics UK (COG-UK) Consortium | Jacquelyn Wynn, Mairead Hyland, The Lighthouse Lab in Alderley Park and Alex Alderton, Roberto Amato, Jeffrey Barrett, Sonia Goncalves, Ewan Harrison, David K. Jackson, Ian Johnston, Dominic Kwiatkowski, Cordelia Langford, John Sillitoe on behalf of the Wellcome Sanger Institute COVID-19 Surveillance Team |
| EPI_ISL_3043580 | Lighthouse Lab in Milton Keynes | Wellcome Sanger Institute for the COVID-19 Genomics UK (COG-UK) Consortium | The Lighthouse Lab in Milton Keynes and Alex Alderton, Roberto Amato, Jeffrey Barrett, Sonia Goncalves, Ewan Harrison, David K. Jackson, Ian Johnston, Dominic Kwiatkowski, Cordelia Langford, John Sillitoe on behalf of the Wellcome Sanger Institute COVID-19 Surveillance Team |
| EPI_ISL_3043581 | Lighthouse Lab in Alderley Park | Wellcome Sanger Institute for the COVID-19 Genomics UK (COG-UK) Consortium | Jacquelyn Wynn, Mairead Hyland, The Lighthouse Lab in Alderley Park and Alex Alderton, Roberto Amato, Jeffrey Barrett, Sonia Goncalves, Ewan Harrison, David K. Jackson, Ian Johnston, Dominic Kwiatkowski, Cordelia Langford, John Sillitoe on behalf of the Wellcome Sanger Institute COVID-19 Surveillance Team |
| EPI_ISL_3043582 | Lighthouse Lab in Milton Keynes | Wellcome Sanger Institute for the COVID-19 Genomics UK (COG-UK) Consortium | The Lighthouse Lab in Milton Keynes and Alex Alderton, Roberto Amato, Jeffrey Barrett, Sonia Goncalves, Ewan Harrison, David K. Jackson, Ian Johnston, Dominic Kwiatkowski, Cordelia Langford, John Sillitoe on behalf of the Wellcome Sanger Institute COVID-19 Surveillance Team |
| EPI_ISL_3043583 | Lighthouse Lab in Glasgow | Wellcome Sanger Institute for the COVID-19 Genomics UK (COG-UK) | Harper VanSteenhouse, Yumi Kasai, David Gray, Carol Clugston, Anna Dominiczak and Alex Alderton, Roberto Amato, Jeffrey Barrett, Sonia Goncalves, Ewan Harrison, David K. Jackson, Ian Johnston, Dominic Kwiatkowski, Cordelia Langford, |

| | | | |
|---|---|---|---|
| | | Consortium | John Sillitoe on behalf of the Wellcome Sanger Institute COVID-19 Surveillance Team |
| EPI_ISL_3043584, EPI_ISL_3043585 | Lighthouse Lab in Alderley Park | Wellcome Sanger Institute for the COVID-19 Genomics UK (COG-UK) Consortium | Jacquelyn Wynn, Mairead Hyland, The Lighthouse Lab in Alderley Park and Alex Alderton, Roberto Amato, Jeffrey Barrett, Sonia Goncalves, Ewan Harrison, David K. Jackson, Ian Johnston, Dominic Kwiatkowski, Cordelia Langford, John Sillitoe on behalf of the Wellcome Sanger Institute COVID-19 Surveillance Team |
| EPI_ISL_3043586 | Lighthouse Lab in Glasgow | Wellcome Sanger Institute for the COVID-19 Genomics UK (COG-UK) Consortium | Harper VanSteenhouse, Yumi Kasai, David Gray, Carol Clugston, Anna Dominiczak and Alex Alderton, Roberto Amato, Jeffrey Barrett, Sonia Goncalves, Ewan Harrison, David K. Jackson, Ian Johnston, Dominic Kwiatkowski, Cordelia Langford, John Sillitoe on behalf of the Wellcome Sanger Institute COVID-19 Surveillance Team |
| EPI_ISL_3043587 | Lighthouse Lab in Milton Keynes | Wellcome Sanger Institute for the COVID-19 Genomics UK (COG-UK) Consortium | The Lighthouse Lab in Milton Keynes and Alex Alderton, Roberto Amato, Jeffrey Barrett, Sonia Goncalves, Ewan Harrison, David K. Jackson, Ian Johnston, Dominic Kwiatkowski, Cordelia Langford, John Sillitoe on behalf of the Wellcome Sanger Institute COVID-19 Surveillance Team |
| EPI_ISL_3043588, EPI_ISL_3043589 | Lighthouse Lab in Alderley Park | Wellcome Sanger Institute for the COVID-19 Genomics UK (COG-UK) Consortium | Jacquelyn Wynn, Mairead Hyland, The Lighthouse Lab in Alderley Park and Alex Alderton, Roberto Amato, Jeffrey Barrett, Sonia Goncalves, Ewan Harrison, David K. Jackson, Ian Johnston, Dominic Kwiatkowski, Cordelia Langford, John Sillitoe on behalf of the Wellcome Sanger Institute COVID-19 Surveillance Team |
| EPI_ISL_3043590 | Lighthouse Lab in Milton Keynes | Wellcome Sanger Institute for the COVID-19 Genomics UK (COG-UK) Consortium | The Lighthouse Lab in Milton Keynes and Alex Alderton, Roberto Amato, Jeffrey Barrett, Sonia Goncalves, Ewan Harrison, David K. Jackson, Ian Johnston, Dominic Kwiatkowski, Cordelia Langford, John Sillitoe on behalf of the Wellcome Sanger Institute COVID-19 Surveillance Team |
| EPI_ISL_3043591, EPI_ISL_3043592, EPI_ISL_3043593 | Lighthouse Lab in Alderley Park | Wellcome Sanger Institute for the COVID-19 Genomics UK (COG-UK) Consortium | Jacquelyn Wynn, Mairead Hyland, The Lighthouse Lab in Alderley Park and Alex Alderton, Roberto Amato, Jeffrey Barrett, Sonia Goncalves, Ewan Harrison, David K. Jackson, Ian Johnston, Dominic Kwiatkowski, Cordelia Langford, John Sillitoe on behalf of the Wellcome Sanger Institute COVID-19 Surveillance Team |
| EPI_ISL_3043594, EPI_ISL_3043595, EPI_ISL_3043596 | Lighthouse Lab in Milton Keynes | Wellcome Sanger Institute for the COVID-19 Genomics UK (COG-UK) Consortium | The Lighthouse Lab in Milton Keynes and Alex Alderton, Roberto Amato, Jeffrey Barrett, Sonia Goncalves, Ewan Harrison, David K. Jackson, Ian Johnston, Dominic Kwiatkowski, Cordelia Langford, John Sillitoe on behalf of the Wellcome Sanger Institute COVID-19 Surveillance Team |
| EPI_ISL_3043597 | Lighthouse Lab in Alderley Park | Wellcome Sanger Institute for the COVID-19 Genomics UK (COG-UK) Consortium | Jacquelyn Wynn, Mairead Hyland, The Lighthouse Lab in Alderley Park and Alex Alderton, Roberto Amato, Jeffrey Barrett, Sonia Goncalves, Ewan Harrison, David K. Jackson, Ian Johnston, Dominic Kwiatkowski, Cordelia Langford, John Sillitoe on behalf of the Wellcome Sanger Institute COVID-19 Surveillance Team |
| EPI_ISL_3043598 | Lighthouse Lab in Milton Keynes | Wellcome Sanger Institute for the COVID-19 Genomics UK (COG-UK) Consortium | The Lighthouse Lab in Milton Keynes and Alex Alderton, Roberto Amato, Jeffrey Barrett, Sonia Goncalves, Ewan Harrison, David K. Jackson, Ian Johnston, Dominic Kwiatkowski, Cordelia Langford, John Sillitoe on behalf of the Wellcome Sanger Institute COVID-19 Surveillance Team |
| EPI_ISL_3043599 | Lighthouse Lab in Glasgow | Wellcome Sanger Institute for the COVID-19 Genomics UK (COG-UK) Consortium | Harper VanSteenhouse, Yumi Kasai, David Gray, Carol Clugston, Anna Dominiczak and Alex Alderton, Roberto Amato, Jeffrey Barrett, Sonia Goncalves, Ewan Harrison, David K. Jackson, Ian Johnston, Dominic Kwiatkowski, Cordelia Langford, John Sillitoe on behalf of the Wellcome Sanger Institute COVID-19 Surveillance Team |
| EPI_ISL_3043600, EPI_ISL_3043601 | Lighthouse Lab in Milton Keynes | Wellcome Sanger Institute for the COVID-19 Genomics UK (COG-UK) Consortium | The Lighthouse Lab in Milton Keynes and Alex Alderton, Roberto Amato, Jeffrey Barrett, Sonia Goncalves, Ewan Harrison, David K. Jackson, Ian Johnston, Dominic Kwiatkowski, Cordelia Langford, John Sillitoe on behalf of the Wellcome Sanger Institute COVID-19 Surveillance Team |
| EPI_ISL_3043602 | Lighthouse Lab in Alderley Park | Wellcome Sanger Institute for the COVID-19 Genomics UK (COG-UK) Consortium | Jacquelyn Wynn, Mairead Hyland, The Lighthouse Lab in Alderley Park and Alex Alderton, Roberto Amato, Jeffrey Barrett, Sonia Goncalves, Ewan Harrison, David K. Jackson, Ian Johnston, Dominic Kwiatkowski, Cordelia Langford, John Sillitoe on behalf of the Wellcome Sanger Institute COVID-19 Surveillance Team |
| EPI_ISL_3043603, EPI_ISL_3043604 | Lighthouse Lab in Milton Keynes | Wellcome Sanger Institute for the COVID-19 Genomics UK (COG-UK) Consortium | The Lighthouse Lab in Milton Keynes and Alex Alderton, Roberto Amato, Jeffrey Barrett, Sonia Goncalves, Ewan Harrison, David K. Jackson, Ian Johnston, Dominic Kwiatkowski, Cordelia Langford, John Sillitoe on behalf of the Wellcome Sanger Institute COVID-19 Surveillance Team |
| EPI_ISL_3043605 | Lighthouse Lab in Alderley Park | Wellcome Sanger Institute for the COVID-19 Genomics UK (COG-UK) Consortium | Jacquelyn Wynn, Mairead Hyland, The Lighthouse Lab in Alderley Park and Alex Alderton, Roberto Amato, Jeffrey Barrett, Sonia Goncalves, Ewan Harrison, David K. Jackson, Ian Johnston, Dominic Kwiatkowski, Cordelia Langford, John Sillitoe on behalf of the Wellcome Sanger Institute COVID-19 Surveillance Team |
| EPI_ISL_3043606 | Lighthouse Lab in Glasgow | Wellcome Sanger Institute for the COVID-19 Genomics UK (COG-UK) Consortium | Harper VanSteenhouse, Yumi Kasai, David Gray, Carol Clugston, Anna Dominiczak and Alex Alderton, Roberto Amato, Jeffrey Barrett, Sonia Goncalves, Ewan Harrison, David K. Jackson, Ian Johnston, Dominic Kwiatkowski, Cordelia Langford, John Sillitoe on behalf of the Wellcome Sanger Institute COVID-19 Surveillance Team |
| EPI_ISL_3043607, EPI_ISL_3043608 | Lighthouse Lab in Milton Keynes | Wellcome Sanger Institute for the COVID-19 Genomics UK (COG-UK) Consortium | The Lighthouse Lab in Milton Keynes and Alex Alderton, Roberto Amato, Jeffrey Barrett, Sonia Goncalves, Ewan Harrison, David K. Jackson, Ian Johnston, Dominic Kwiatkowski, Cordelia Langford, John Sillitoe on behalf of the Wellcome Sanger Institute COVID-19 Surveillance Team |
| EPI_ISL_3043609, EPI_ISL_3043610, EPI_ISL_3043611, EPI_ISL_3043612 | Lighthouse Lab in Alderley Park | Wellcome Sanger Institute for the COVID-19 Genomics UK (COG-UK) Consortium | Jacquelyn Wynn, Mairead Hyland, The Lighthouse Lab in Alderley Park and Alex Alderton, Roberto Amato, Jeffrey Barrett, Sonia Goncalves, Ewan Harrison, David K. Jackson, Ian Johnston, Dominic Kwiatkowski, Cordelia Langford, John Sillitoe on behalf of the Wellcome Sanger Institute COVID-19 Surveillance Team |
| EPI_ISL_3043613, EPI_ISL_3043614, EPI_ISL_3043615, EPI_ISL_3043616, EPI_ISL_3043617 | Lighthouse Lab in Milton Keynes | Wellcome Sanger Institute for the COVID-19 Genomics UK (COG-UK) Consortium | The Lighthouse Lab in Milton Keynes and Alex Alderton, Roberto Amato, Jeffrey Barrett, Sonia Goncalves, Ewan Harrison, David K. Jackson, Ian Johnston, Dominic Kwiatkowski, Cordelia Langford, John Sillitoe on behalf of the Wellcome Sanger Institute COVID-19 Surveillance Team |
| EPI_ISL_3043618 | Lighthouse Lab in Glasgow | Wellcome Sanger Institute for the COVID-19 Genomics UK (COG-UK) Consortium | Harper VanSteenhouse, Yumi Kasai, David Gray, Carol Clugston, Anna Dominiczak and Alex Alderton, Roberto Amato, Jeffrey Barrett, Sonia Goncalves, Ewan Harrison, David K. Jackson, Ian Johnston, Dominic Kwiatkowski, Cordelia Langford, John Sillitoe on behalf of the Wellcome Sanger Institute COVID-19 Surveillance Team |
| EPI_ISL_3043619 | Lighthouse Lab in Alderley Park | Wellcome Sanger Institute for the COVID-19 Genomics UK (COG-UK) Consortium | Jacquelyn Wynn, Mairead Hyland, The Lighthouse Lab in Alderley Park and Alex Alderton, Roberto Amato, Jeffrey Barrett, Sonia Goncalves, Ewan Harrison, David K. Jackson, Ian Johnston, Dominic Kwiatkowski, Cordelia Langford, John Sillitoe on behalf of the Wellcome Sanger Institute COVID-19 Surveillance Team |
| EPI_ISL_3043620 | Lighthouse Lab in Milton Keynes | Wellcome Sanger Institute for the COVID-19 Genomics UK (COG-UK) Consortium | The Lighthouse Lab in Milton Keynes and Alex Alderton, Roberto Amato, Jeffrey Barrett, Sonia Goncalves, Ewan Harrison, David K. Jackson, Ian Johnston, Dominic Kwiatkowski, Cordelia Langford, John Sillitoe on behalf of the Wellcome Sanger Institute COVID-19 Surveillance Team |
| EPI_ISL_3043621 | Lighthouse Lab in Alderley Park | Wellcome Sanger Institute for the COVID-19 Genomics UK (COG-UK) Consortium | Jacquelyn Wynn, Mairead Hyland, The Lighthouse Lab in Alderley Park and Alex Alderton, Roberto Amato, Jeffrey Barrett, Sonia Goncalves, Ewan Harrison, David K. Jackson, Ian Johnston, Dominic Kwiatkowski, Cordelia Langford, John Sillitoe on behalf of the Wellcome Sanger Institute COVID-19 Surveillance Team |
| EPI_ISL_3043622 | Lighthouse Lab in Milton Keynes | Wellcome Sanger Institute for the COVID-19 Genomics UK (COG-UK) Consortium | The Lighthouse Lab in Milton Keynes and Alex Alderton, Roberto Amato, Jeffrey Barrett, Sonia Goncalves, Ewan Harrison, David K. Jackson, Ian Johnston, Dominic Kwiatkowski, Cordelia Langford, John Sillitoe on behalf of the Wellcome Sanger Institute COVID-19 Surveillance Team |
| EPI_ISL_3043623 | Lighthouse Lab in Alderley Park | Wellcome Sanger Institute for the COVID-19 Genomics UK (COG-UK) Consortium | Jacquelyn Wynn, Mairead Hyland, The Lighthouse Lab in Alderley Park and Alex Alderton, Roberto Amato, Jeffrey Barrett, Sonia Goncalves, Ewan Harrison, David K. Jackson, Ian Johnston, Dominic Kwiatkowski, Cordelia Langford, John Sillitoe on behalf of the Wellcome Sanger Institute COVID-19 Surveillance Team |
| EPI_ISL_3043624 | Lighthouse Lab in Milton Keynes | Wellcome Sanger Institute for the COVID-19 Genomics UK (COG-UK) Consortium | The Lighthouse Lab in Milton Keynes and Alex Alderton, Roberto Amato, Jeffrey Barrett, Sonia Goncalves, Ewan Harrison, David K. Jackson, Ian Johnston, Dominic Kwiatkowski, Cordelia Langford, John Sillitoe on behalf of the Wellcome Sanger Institute COVID-19 Surveillance Team |
| EPI_ISL_3043625, EPI_ISL_3043626 | Lighthouse Lab in Glasgow | Wellcome Sanger Institute for the COVID-19 Genomics UK (COG-UK) Consortium | Harper VanSteenhouse, Yumi Kasai, David Gray, Carol Clugston, Anna Dominiczak and Alex Alderton, Roberto Amato, Jeffrey Barrett, Sonia Goncalves, Ewan Harrison, David K. Jackson, Ian Johnston, Dominic Kwiatkowski, Cordelia Langford, John Sillitoe on behalf of the Wellcome Sanger Institute COVID-19 Surveillance Team |
| EPI_ISL_3043627 | Lighthouse Lab in Milton Keynes | Wellcome Sanger Institute for the COVID-19 Genomics UK (COG-UK) Consortium | The Lighthouse Lab in Milton Keynes and Alex Alderton, Roberto Amato, Jeffrey Barrett, Sonia Goncalves, Ewan Harrison, David K. Jackson, Ian Johnston, Dominic Kwiatkowski, Cordelia Langford, John Sillitoe on behalf of the Wellcome Sanger Institute COVID-19 Surveillance Team |
| EPI_ISL_3043628 | Lighthouse Lab in Alderley Park | Wellcome Sanger Institute for the COVID-19 Genomics UK (COG-UK) Consortium | Jacquelyn Wynn, Mairead Hyland, The Lighthouse Lab in Alderley Park and Alex Alderton, Roberto Amato, Jeffrey Barrett, Sonia Goncalves, Ewan Harrison, David K. Jackson, Ian Johnston, Dominic Kwiatkowski, Cordelia Langford, John Sillitoe on behalf of the Wellcome Sanger Institute COVID-19 Surveillance Team |
| EPI_ISL_3043629, EPI_ISL_3043630 | Lighthouse Lab in Milton Keynes | Wellcome Sanger Institute for the COVID-19 Genomics UK (COG-UK) Consortium | The Lighthouse Lab in Milton Keynes and Alex Alderton, Roberto Amato, Jeffrey Barrett, Sonia Goncalves, Ewan Harrison, David K. Jackson, Ian Johnston, Dominic Kwiatkowski, Cordelia Langford, John Sillitoe on behalf of the Wellcome Sanger Institute COVID-19 Surveillance Team |
| EPI_ISL_3043631 | Lighthouse Lab in Glasgow | Wellcome Sanger Institute for the COVID-19 Genomics UK (COG-UK) Consortium | Harper VanSteenhouse, Yumi Kasai, David Gray, Carol Clugston, Anna Dominiczak and Alex Alderton, Roberto Amato, Jeffrey Barrett, Sonia Goncalves, Ewan Harrison, David K. Jackson, Ian Johnston, Dominic Kwiatkowski, Cordelia Langford, John Sillitoe on behalf of the Wellcome Sanger Institute COVID-19 Surveillance Team |
| EPI_ISL_3043632 | Lighthouse Lab in Alderley Park | Wellcome Sanger Institute for the COVID-19 Genomics UK (COG-UK) Consortium | Jacquelyn Wynn, Mairead Hyland, The Lighthouse Lab in Alderley Park and Alex Alderton, Roberto Amato, Jeffrey Barrett, Sonia Goncalves, Ewan Harrison, David K. Jackson, Ian Johnston, Dominic Kwiatkowski, Cordelia Langford, John Sillitoe on behalf of the Wellcome Sanger Institute COVID-19 Surveillance Team |
| EPI_ISL_3043633 | Lighthouse Lab in Milton Keynes | Wellcome Sanger Institute for the COVID-19 Genomics UK (COG-UK) Consortium | The Lighthouse Lab in Milton Keynes and Alex Alderton, Roberto Amato, Jeffrey Barrett, Sonia Goncalves, Ewan Harrison, David K. Jackson, Ian Johnston, Dominic Kwiatkowski, Cordelia Langford, John Sillitoe on behalf of the Wellcome Sanger Institute COVID-19 Surveillance Team |
| EPI_ISL_3043634 | Lighthouse Lab in Alderley Park | Wellcome Sanger Institute for the COVID-19 Genomics UK (COG-UK) Consortium | Jacquelyn Wynn, Mairead Hyland, The Lighthouse Lab in Alderley Park and Alex Alderton, Roberto Amato, Jeffrey Barrett, Sonia Goncalves, Ewan Harrison, David K. Jackson, Ian Johnston, Dominic Kwiatkowski, Cordelia Langford, John Sillitoe on behalf of the Wellcome Sanger Institute COVID-19 Surveillance Team |
| EPI_ISL_3043635 | Lighthouse Lab in Milton Keynes | Wellcome Sanger Institute for the COVID-19 Genomics UK (COG-UK) Consortium | The Lighthouse Lab in Milton Keynes and Alex Alderton, Roberto Amato, Jeffrey Barrett, Sonia Goncalves, Ewan Harrison, David K. Jackson, Ian Johnston, Dominic Kwiatkowski, Cordelia Langford, John Sillitoe on behalf of the Wellcome Sanger Institute COVID-19 Surveillance Team |
| EPI_ISL_3043636, EPI_ISL_3043637 | Lighthouse Lab in Alderley Park | Wellcome Sanger Institute for the COVID-19 Genomics UK (COG-UK) Consortium | Jacquelyn Wynn, Mairead Hyland, The Lighthouse Lab in Alderley Park and Alex Alderton, Roberto Amato, Jeffrey Barrett, Sonia Goncalves, Ewan Harrison, David K. Jackson, Ian Johnston, Dominic Kwiatkowski, Cordelia Langford, John Sillitoe on behalf of the Wellcome Sanger Institute COVID-19 Surveillance Team |
| EPI_ISL_3043638 | Lighthouse Lab in Milton Keynes | Wellcome Sanger Institute for the COVID-19 Genomics UK (COG-UK) Consortium | The Lighthouse Lab in Milton Keynes and Alex Alderton, Roberto Amato, Jeffrey Barrett, Sonia Goncalves, Ewan Harrison, David K. Jackson, Ian Johnston, Dominic Kwiatkowski, Cordelia Langford, John Sillitoe on behalf of the Wellcome Sanger Institute COVID-19 Surveillance Team |
| EPI_ISL_3043639 | Lighthouse Lab in Alderley Park | Wellcome Sanger Institute for the COVID-19 Genomics UK (COG-UK) Consortium | Jacquelyn Wynn, Mairead Hyland, The Lighthouse Lab in Alderley Park and Alex Alderton, Roberto Amato, Jeffrey Barrett, Sonia Goncalves, Ewan Harrison, David K. Jackson, Ian Johnston, Dominic Kwiatkowski, Cordelia Langford, John Sillitoe on behalf of the Wellcome Sanger Institute COVID-19 Surveillance Team |
| EPI_ISL_3043640 | Lighthouse Lab in Glasgow | Wellcome Sanger Institute for the COVID-19 Genomics UK (COG-UK) Consortium | Harper VanSteenhouse, Yumi Kasai, David Gray, Carol Clugston, Anna Dominiczak and Alex Alderton, Roberto Amato, Jeffrey Barrett, Sonia Goncalves, Ewan Harrison, David K. Jackson, Ian Johnston, Dominic Kwiatkowski, Cordelia Langford, John Sillitoe on behalf of the Wellcome Sanger Institute COVID-19 Surveillance Team |
| EPI_ISL_3043641 | Lighthouse Lab in Milton Keynes | Wellcome Sanger Institute for the COVID-19 Genomics UK (COG-UK) Consortium | The Lighthouse Lab in Milton Keynes and Alex Alderton, Roberto Amato, Jeffrey Barrett, Sonia Goncalves, Ewan Harrison, David K. Jackson, Ian Johnston, Dominic Kwiatkowski, Cordelia Langford, John Sillitoe on behalf of the Wellcome Sanger Institute COVID-19 Surveillance Team |
| EPI_ISL_3043642 | Lighthouse Lab in Glasgow | Wellcome Sanger Institute for the COVID-19 Genomics UK (COG-UK) Consortium | Harper VanSteenhouse, Yumi Kasai, David Gray, Carol Clugston, Anna Dominiczak and Alex Alderton, Roberto Amato, Jeffrey Barrett, Sonia Goncalves, Ewan Harrison, David K. Jackson, Ian Johnston, Dominic Kwiatkowski, Cordelia Langford, John Sillitoe on behalf of the Wellcome Sanger Institute COVID-19 Surveillance Team |
| EPI_ISL_3043643 | Lighthouse Lab in Milton Keynes | Wellcome Sanger Institute for the COVID-19 Genomics UK (COG-UK) Consortium | The Lighthouse Lab in Milton Keynes and Alex Alderton, Roberto Amato, Jeffrey Barrett, Sonia Goncalves, Ewan Harrison, David K. Jackson, Ian Johnston, Dominic Kwiatkowski, Cordelia Langford, John Sillitoe on behalf of the Wellcome Sanger Institute COVID-19 Surveillance Team |
| EPI_ISL_3043644 | Lighthouse Lab in Alderley Park | Wellcome Sanger Institute for the COVID-19 Genomics UK (COG-UK) | Jacquelyn Wynn, Mairead Hyland, The Lighthouse Lab in Alderley Park and Alex Alderton, Roberto Amato, Jeffrey Barrett, Sonia Goncalves, Ewan Harrison, David K. Jackson, Ian Johnston, Dominic Kwiatkowski, Cordelia Langford, John Sillitoe on |

| | | | |
|---|---|---|---|
| | | Consortium | behalf of the Wellcome Sanger Institute COVID-19 Surveillance Team |
| EPI_ISL_3043645, EPI_ISL_3043646, EPI_ISL_3043647 | Lighthouse Lab in Milton Keynes | Wellcome Sanger Institute for the COVID-19 Genomics UK (COG-UK) Consortium | The Lighthouse Lab in Milton Keynes and Alex Alderton, Roberto Amato, Jeffrey Barrett, Sonia Goncalves, Ewan Harrison, David K. Jackson, Ian Johnston, Dominic Kwiatkowski, Cordelia Langford, John Sillitoe on behalf of the Wellcome Sanger Institute COVID-19 Surveillance Team |
| EPI_ISL_3043648 | Lighthouse Lab in Glasgow | Wellcome Sanger Institute for the COVID-19 Genomics UK (COG-UK) Consortium | Harper VanSteenhouse, Yumi Kasai, David Gray, Carol Clugston, Anna Dominiczak and Alex Alderton, Roberto Amato, Jeffrey Barrett, Sonia Goncalves, Ewan Harrison, David K. Jackson, Ian Johnston, Dominic Kwiatkowski, Cordelia Langford, John Sillitoe on behalf of the Wellcome Sanger Institute COVID-19 Surveillance Team |
| EPI_ISL_3043649 | Lighthouse Lab in Alderley Park | Wellcome Sanger Institute for the COVID-19 Genomics UK (COG-UK) Consortium | Jacquelyn Wynn, Mairead Hyland, The Lighthouse Lab in Alderley Park and Alex Alderton, Roberto Amato, Jeffrey Barrett, Sonia Goncalves, Ewan Harrison, David K. Jackson, Ian Johnston, Dominic Kwiatkowski, Cordelia Langford, John Sillitoe on behalf of the Wellcome Sanger Institute COVID-19 Surveillance Team |
| EPI_ISL_3043650, EPI_ISL_3043651, EPI_ISL_3043652, EPI_ISL_3043653 | Lighthouse Lab in Milton Keynes | Wellcome Sanger Institute for the COVID-19 Genomics UK (COG-UK) Consortium | The Lighthouse Lab in Milton Keynes and Alex Alderton, Roberto Amato, Jeffrey Barrett, Sonia Goncalves, Ewan Harrison, David K. Jackson, Ian Johnston, Dominic Kwiatkowski, Cordelia Langford, John Sillitoe on behalf of the Wellcome Sanger Institute COVID-19 Surveillance Team |
| EPI_ISL_3043654, EPI_ISL_3043655, EPI_ISL_3043656 | Lighthouse Lab in Alderley Park | Wellcome Sanger Institute for the COVID-19 Genomics UK (COG-UK) Consortium | Jacquelyn Wynn, Mairead Hyland, The Lighthouse Lab in Alderley Park and Alex Alderton, Roberto Amato, Jeffrey Barrett, Sonia Goncalves, Ewan Harrison, David K. Jackson, Ian Johnston, Dominic Kwiatkowski, Cordelia Langford, John Sillitoe on behalf of the Wellcome Sanger Institute COVID-19 Surveillance Team |
| EPI_ISL_3043657, EPI_ISL_3043658, EPI_ISL_3043659 | Lighthouse Lab in Milton Keynes | Wellcome Sanger Institute for the COVID-19 Genomics UK (COG-UK) Consortium | The Lighthouse Lab in Milton Keynes and Alex Alderton, Roberto Amato, Jeffrey Barrett, Sonia Goncalves, Ewan Harrison, David K. Jackson, Ian Johnston, Dominic Kwiatkowski, Cordelia Langford, John Sillitoe on behalf of the Wellcome Sanger Institute COVID-19 Surveillance Team |
| EPI_ISL_3043660, EPI_ISL_3043661 | Lighthouse Lab in Glasgow | Wellcome Sanger Institute for the COVID-19 Genomics UK (COG-UK) Consortium | Harper VanSteenhouse, Yumi Kasai, David Gray, Carol Clugston, Anna Dominiczak and Alex Alderton, Roberto Amato, Jeffrey Barrett, Sonia Goncalves, Ewan Harrison, David K. Jackson, Ian Johnston, Dominic Kwiatkowski, Cordelia Langford, John Sillitoe on behalf of the Wellcome Sanger Institute COVID-19 Surveillance Team |
| EPI_ISL_3043662, EPI_ISL_3043663, EPI_ISL_3043664 | Lighthouse Lab in Milton Keynes | Wellcome Sanger Institute for the COVID-19 Genomics UK (COG-UK) Consortium | The Lighthouse Lab in Milton Keynes and Alex Alderton, Roberto Amato, Jeffrey Barrett, Sonia Goncalves, Ewan Harrison, David K. Jackson, Ian Johnston, Dominic Kwiatkowski, Cordelia Langford, John Sillitoe on behalf of the Wellcome Sanger Institute COVID-19 Surveillance Team |
| EPI_ISL_3043665 | Lighthouse Lab in Alderley Park | Wellcome Sanger Institute for the COVID-19 Genomics UK (COG-UK) Consortium | Jacquelyn Wynn, Mairead Hyland, The Lighthouse Lab in Alderley Park and Alex Alderton, Roberto Amato, Jeffrey Barrett, Sonia Goncalves, Ewan Harrison, David K. Jackson, Ian Johnston, Dominic Kwiatkowski, Cordelia Langford, John Sillitoe on behalf of the Wellcome Sanger Institute COVID-19 Surveillance Team |
| EPI_ISL_3043666 | Lighthouse Lab in Milton Keynes | Wellcome Sanger Institute for the COVID-19 Genomics UK (COG-UK) Consortium | The Lighthouse Lab in Milton Keynes and Alex Alderton, Roberto Amato, Jeffrey Barrett, Sonia Goncalves, Ewan Harrison, David K. Jackson, Ian Johnston, Dominic Kwiatkowski, Cordelia Langford, John Sillitoe on behalf of the Wellcome Sanger Institute COVID-19 Surveillance Team |
| EPI_ISL_3043667 | Lighthouse Lab in Glasgow | Wellcome Sanger Institute for the COVID-19 Genomics UK (COG-UK) Consortium | Harper VanSteenhouse, Yumi Kasai, David Gray, Carol Clugston, Anna Dominiczak and Alex Alderton, Roberto Amato, Jeffrey Barrett, Sonia Goncalves, Ewan Harrison, David K. Jackson, Ian Johnston, Dominic Kwiatkowski, Cordelia Langford, John Sillitoe on behalf of the Wellcome Sanger Institute COVID-19 Surveillance Team |
| EPI_ISL_3043668, EPI_ISL_3043669, EPI_ISL_3043670 | Lighthouse Lab in Alderley Park | Wellcome Sanger Institute for the COVID-19 Genomics UK (COG-UK) Consortium | Jacquelyn Wynn, Mairead Hyland, The Lighthouse Lab in Alderley Park and Alex Alderton, Roberto Amato, Jeffrey Barrett, Sonia Goncalves, Ewan Harrison, David K. Jackson, Ian Johnston, Dominic Kwiatkowski, Cordelia Langford, John Sillitoe on behalf of the Wellcome Sanger Institute COVID-19 Surveillance Team |
| EPI_ISL_3043671 | Lighthouse Lab in Glasgow | Wellcome Sanger Institute for the COVID-19 Genomics UK (COG-UK) Consortium | Harper VanSteenhouse, Yumi Kasai, David Gray, Carol Clugston, Anna Dominiczak and Alex Alderton, Roberto Amato, Jeffrey Barrett, Sonia Goncalves, Ewan Harrison, David K. Jackson, Ian Johnston, Dominic Kwiatkowski, Cordelia Langford, John Sillitoe on behalf of the Wellcome Sanger Institute COVID-19 Surveillance Team |
| EPI_ISL_3043672, EPI_ISL_3043673, EPI_ISL_3043674 | Lighthouse Lab in Milton Keynes | Wellcome Sanger Institute for the COVID-19 Genomics UK (COG-UK) Consortium | The Lighthouse Lab in Milton Keynes and Alex Alderton, Roberto Amato, Jeffrey Barrett, Sonia Goncalves, Ewan Harrison, David K. Jackson, Ian Johnston, Dominic Kwiatkowski, Cordelia Langford, John Sillitoe on behalf of the Wellcome Sanger Institute COVID-19 Surveillance Team |
| EPI_ISL_3043675 | Lighthouse Lab in Glasgow | Wellcome Sanger Institute for the COVID-19 Genomics UK (COG-UK) Consortium | Harper VanSteenhouse, Yumi Kasai, David Gray, Carol Clugston, Anna Dominiczak and Alex Alderton, Roberto Amato, Jeffrey Barrett, Sonia Goncalves, Ewan Harrison, David K. Jackson, Ian Johnston, Dominic Kwiatkowski, Cordelia Langford, John Sillitoe on behalf of the Wellcome Sanger Institute COVID-19 Surveillance Team |
| EPI_ISL_3043676 | Lighthouse Lab in Milton Keynes | Wellcome Sanger Institute for the COVID-19 Genomics UK (COG-UK) Consortium | The Lighthouse Lab in Milton Keynes and Alex Alderton, Roberto Amato, Jeffrey Barrett, Sonia Goncalves, Ewan Harrison, David K. Jackson, Ian Johnston, Dominic Kwiatkowski, Cordelia Langford, John Sillitoe on behalf of the Wellcome Sanger Institute COVID-19 Surveillance Team |
| EPI_ISL_3043677 | Lighthouse Lab in Alderley Park | Wellcome Sanger Institute for the COVID-19 Genomics UK (COG-UK) Consortium | Jacquelyn Wynn, Mairead Hyland, The Lighthouse Lab in Alderley Park and Alex Alderton, Roberto Amato, Jeffrey Barrett, Sonia Goncalves, Ewan Harrison, David K. Jackson, Ian Johnston, Dominic Kwiatkowski, Cordelia Langford, John Sillitoe on behalf of the Wellcome Sanger Institute COVID-19 Surveillance Team |
| EPI_ISL_3043678 | Lighthouse Lab in Milton Keynes | Wellcome Sanger Institute for the COVID-19 Genomics UK (COG-UK) Consortium | The Lighthouse Lab in Milton Keynes and Alex Alderton, Roberto Amato, Jeffrey Barrett, Sonia Goncalves, Ewan Harrison, David K. Jackson, Ian Johnston, Dominic Kwiatkowski, Cordelia Langford, John Sillitoe on behalf of the Wellcome Sanger Institute COVID-19 Surveillance Team |
| EPI_ISL_3043679, EPI_ISL_3043680, EPI_ISL_3043681, EPI_ISL_3043682, EPI_ISL_3043683, EPI_ISL_3043684, EPI_ISL_3043685, EPI_ISL_3043686, EPI_ISL_3043687, EPI_ISL_3043688, EPI_ISL_3043689 | | | |
| see above | Lighthouse Lab in Alderley Park | Wellcome Sanger Institute for the COVID-19 Genomics UK (COG-UK) Consortium | Jacquelyn Wynn, Mairead Hyland, The Lighthouse Lab in Alderley Park and Alex Alderton, Roberto Amato, Jeffrey Barrett, Sonia Goncalves, Ewan Harrison, David K. Jackson, Ian Johnston, Dominic Kwiatkowski, Cordelia Langford, John Sillitoe on behalf of the Wellcome Sanger Institute COVID-19 Surveillance Team |
| EPI_ISL_3043690 | Lighthouse Lab in Milton Keynes | Wellcome Sanger Institute for the COVID-19 Genomics UK (COG-UK) Consortium | The Lighthouse Lab in Milton Keynes and Alex Alderton, Roberto Amato, Jeffrey Barrett, Sonia Goncalves, Ewan Harrison, David K. Jackson, Ian Johnston, Dominic Kwiatkowski, Cordelia Langford, John Sillitoe on behalf of the Wellcome Sanger Institute COVID-19 Surveillance Team |
| EPI_ISL_3043691, EPI_ISL_3043692, EPI_ISL_3043693, EPI_ISL_3043694, EPI_ISL_3043695, EPI_ISL_3043696, EPI_ISL_3043697, EPI_ISL_3043698, EPI_ISL_3043699, EPI_ISL_3043700 | Lighthouse Lab in Alderley Park | Wellcome Sanger Institute for the COVID-19 Genomics UK (COG-UK) Consortium | Jacquelyn Wynn, Mairead Hyland, The Lighthouse Lab in Alderley Park and Alex Alderton, Roberto Amato, Jeffrey Barrett, Sonia Goncalves, Ewan Harrison, David K. Jackson, Ian Johnston, Dominic Kwiatkowski, Cordelia Langford, John Sillitoe on behalf of the Wellcome Sanger Institute COVID-19 Surveillance Team |
| EPI_ISL_3043701, EPI_ISL_3043702 | Lighthouse Lab in Milton Keynes | Wellcome Sanger Institute for the COVID-19 Genomics UK (COG-UK) Consortium | The Lighthouse Lab in Milton Keynes and Alex Alderton, Roberto Amato, Jeffrey Barrett, Sonia Goncalves, Ewan Harrison, David K. Jackson, Ian Johnston, Dominic Kwiatkowski, Cordelia Langford, John Sillitoe on behalf of the Wellcome Sanger Institute COVID-19 Surveillance Team |
| EPI_ISL_3043703 | Lighthouse Lab in Alderley Park | Wellcome Sanger Institute for the COVID-19 Genomics UK (COG-UK) Consortium | Jacquelyn Wynn, Mairead Hyland, The Lighthouse Lab in Alderley Park and Alex Alderton, Roberto Amato, Jeffrey Barrett, Sonia Goncalves, Ewan Harrison, David K. Jackson, Ian Johnston, Dominic Kwiatkowski, Cordelia Langford, John Sillitoe on behalf of the Wellcome Sanger Institute COVID-19 Surveillance Team |
| EPI_ISL_3043704 | Lighthouse Lab in Milton Keynes | Wellcome Sanger Institute for the COVID-19 Genomics UK (COG-UK) Consortium | The Lighthouse Lab in Milton Keynes and Alex Alderton, Roberto Amato, Jeffrey Barrett, Sonia Goncalves, Ewan Harrison, David K. Jackson, Ian Johnston, Dominic Kwiatkowski, Cordelia Langford, John Sillitoe on behalf of the Wellcome Sanger Institute COVID-19 Surveillance Team |
| EPI_ISL_3043705 | Lighthouse Lab in Glasgow | Wellcome Sanger Institute for the COVID-19 Genomics UK (COG-UK) Consortium | Harper VanSteenhouse, Yumi Kasai, David Gray, Carol Clugston, Anna Dominiczak and Alex Alderton, Roberto Amato, Jeffrey Barrett, Sonia Goncalves, Ewan Harrison, David K. Jackson, Ian Johnston, Dominic Kwiatkowski, Cordelia Langford, John Sillitoe on behalf of the Wellcome Sanger Institute COVID-19 Surveillance Team |
| EPI_ISL_3043706 | Berkshire and Surrey Pathology Services Lighthouse Laboratory | Wellcome Sanger Institute for the COVID-19 Genomics UK (COG-UK) Consortium | Berkshire and Surrey Pathology Services Lighthouse Laboratory and Alex Alderton, Roberto Amato, Jeffrey Barrett, Sonia Goncalves, Ewan Harrison, David K. Jackson, Ian Johnston, Dominic Kwiatkowski, Cordelia Langford, John Sillitoe on behalf of the Wellcome Sanger Institute COVID-19 Surveillance Team |
| EPI_ISL_3043707, EPI_ISL_3043708, EPI_ISL_3043709, EPI_ISL_3043710 | Lighthouse Lab in Alderley Park | Wellcome Sanger Institute for the COVID-19 Genomics UK (COG-UK) Consortium | Jacquelyn Wynn, Mairead Hyland, The Lighthouse Lab in Alderley Park and Alex Alderton, Roberto Amato, Jeffrey Barrett, Sonia Goncalves, Ewan Harrison, David K. Jackson, Ian Johnston, Dominic Kwiatkowski, Cordelia Langford, John Sillitoe on behalf of the Wellcome Sanger Institute COVID-19 Surveillance Team |
| EPI_ISL_3043711 | Berkshire and Surrey Pathology Services Lighthouse Laboratory | Wellcome Sanger Institute for the COVID-19 Genomics UK (COG-UK) Consortium | Berkshire and Surrey Pathology Services Lighthouse Laboratory and Alex Alderton, Roberto Amato, Jeffrey Barrett, Sonia Goncalves, Ewan Harrison, David K. Jackson, Ian Johnston, Dominic Kwiatkowski, Cordelia Langford, John Sillitoe on behalf of the Wellcome Sanger Institute COVID-19 Surveillance Team |
| EPI_ISL_3043712, EPI_ISL_3043713 | Lighthouse Lab in Milton Keynes | Wellcome Sanger Institute for the COVID-19 Genomics UK (COG-UK) Consortium | The Lighthouse Lab in Milton Keynes and Alex Alderton, Roberto Amato, Jeffrey Barrett, Sonia Goncalves, Ewan Harrison, David K. Jackson, Ian Johnston, Dominic Kwiatkowski, Cordelia Langford, John Sillitoe on behalf of the Wellcome Sanger Institute COVID-19 Surveillance Team |
| EPI_ISL_3043714, EPI_ISL_3043715, EPI_ISL_3043716, EPI_ISL_3043717, EPI_ISL_3043718 | Health Services Laboratories | Wellcome Sanger Institute for the COVID-19 Genomics UK (COG-UK) Consortium | Health Services Laboratories and Alex Alderton, Roberto Amato, Jeffrey Barrett, Sonia Goncalves, Ewan Harrison, David K. Jackson, Ian Johnston, Dominic Kwiatkowski, Cordelia Langford, John Sillitoe on behalf of the Wellcome Sanger Institute COVID-19 Surveillance Team |
| EPI_ISL_3043719 | Lighthouse Lab in Milton Keynes | Wellcome Sanger Institute for the COVID-19 Genomics UK (COG-UK) Consortium | The Lighthouse Lab in Milton Keynes and Alex Alderton, Roberto Amato, Jeffrey Barrett, Sonia Goncalves, Ewan Harrison, David K. Jackson, Ian Johnston, Dominic Kwiatkowski, Cordelia Langford, John Sillitoe on behalf of the Wellcome Sanger Institute COVID-19 Surveillance Team |
| EPI_ISL_3043720, EPI_ISL_3043721 | Health Services Laboratories | Wellcome Sanger Institute for the COVID-19 Genomics UK (COG-UK) Consortium | Health Services Laboratories and Alex Alderton, Roberto Amato, Jeffrey Barrett, Sonia Goncalves, Ewan Harrison, David K. Jackson, Ian Johnston, Dominic Kwiatkowski, Cordelia Langford, John Sillitoe on behalf of the Wellcome Sanger Institute COVID-19 Surveillance Team |
| EPI_ISL_3043722 | Lighthouse Lab in Alderley Park | Wellcome Sanger Institute for the COVID-19 Genomics UK (COG-UK) Consortium | Jacquelyn Wynn, Mairead Hyland, The Lighthouse Lab in Alderley Park and Alex Alderton, Roberto Amato, Jeffrey Barrett, Sonia Goncalves, Ewan Harrison, David K. Jackson, Ian Johnston, Dominic Kwiatkowski, Cordelia Langford, John Sillitoe on behalf of the Wellcome Sanger Institute COVID-19 Surveillance Team |
| EPI_ISL_3043723, EPI_ISL_3043724, EPI_ISL_3043725, EPI_ISL_3043726 | Health Services Laboratories | Wellcome Sanger Institute for the COVID-19 Genomics UK (COG-UK) Consortium | Health Services Laboratories and Alex Alderton, Roberto Amato, Jeffrey Barrett, Sonia Goncalves, Ewan Harrison, David K. Jackson, Ian Johnston, Dominic Kwiatkowski, Cordelia Langford, John Sillitoe on behalf of the Wellcome Sanger Institute COVID-19 Surveillance Team |
| EPI_ISL_3043727 | Lighthouse Lab in Glasgow | Wellcome Sanger Institute for the COVID-19 Genomics UK (COG-UK) | Harper VanSteenhouse, Yumi Kasai, David Gray, Carol Clugston, Anna Dominiczak and Alex Alderton, Roberto Amato, Jeffrey Barrett, Sonia Goncalves, Ewan Harrison, David K. Jackson, Ian Johnston, Dominic Kwiatkowski, Cordelia Langford, |

| | | | |
|---|---|---|---|
| | | Consortium | John Sillitoe on behalf of the Wellcome Sanger Institute COVID-19 Surveillance Team |
| EPI_ISL_3043728, EPI_ISL_3043729, EPI_ISL_3043730, EPI_ISL_3043731, EPI_ISL_3043732, EPI_ISL_3043733 | Health Services Laboratories | Wellcome Sanger Institute for the COVID-19 Genomics UK (COG-UK) Consortium | Health Services Laboratories and Alex Alderton, Roberto Amato, Jeffrey Barrett, Sonia Goncalves, Ewan Harrison, David K. Jackson, Ian Johnston, Dominic Kwiatkowski, Cordelia Langford, John Sillitoe on behalf of the Wellcome Sanger Institute COVID-19 Surveillance Team |
| EPI_ISL_3043734 | Lighthouse Lab in Glasgow | Wellcome Sanger Institute for the COVID-19 Genomics UK (COG-UK) Consortium | Harper VanSteenhouse, Yumi Kasai, David Gray, Carol Clugston, Anna Dominiczak and Alex Alderton, Roberto Amato, Jeffrey Barrett, Sonia Goncalves, Ewan Harrison, David K. Jackson, Ian Johnston, Dominic Kwiatkowski, Cordelia Langford, John Sillitoe on behalf of the Wellcome Sanger Institute COVID-19 Surveillance Team |
| EPI_ISL_3043735, EPI_ISL_3043736, EPI_ISL_3043737, EPI_ISL_3043738, EPI_ISL_3043739, EPI_ISL_3043740 | Health Services Laboratories | Wellcome Sanger Institute for the COVID-19 Genomics UK (COG-UK) Consortium | Health Services Laboratories and Alex Alderton, Roberto Amato, Jeffrey Barrett, Sonia Goncalves, Ewan Harrison, David K. Jackson, Ian Johnston, Dominic Kwiatkowski, Cordelia Langford, John Sillitoe on behalf of the Wellcome Sanger Institute COVID-19 Surveillance Team |
| EPI_ISL_3043741 | Lighthouse Lab in Alderley Park | Wellcome Sanger Institute for the COVID-19 Genomics UK (COG-UK) Consortium | Jacquelyn Wynn, Mairead Hyland, The Lighthouse Lab in Alderley Park and Alex Alderton, Roberto Amato, Jeffrey Barrett, Sonia Goncalves, Ewan Harrison, David K. Jackson, Ian Johnston, Dominic Kwiatkowski, Cordelia Langford, John Sillitoe on behalf of the Wellcome Sanger Institute COVID-19 Surveillance Team |
| EPI_ISL_3043742, EPI_ISL_3043743, EPI_ISL_3043744, EPI_ISL_3043745, EPI_ISL_3043746, EPI_ISL_3043747 | Health Services Laboratories | Wellcome Sanger Institute for the COVID-19 Genomics UK (COG-UK) Consortium | Health Services Laboratories and Alex Alderton, Roberto Amato, Jeffrey Barrett, Sonia Goncalves, Ewan Harrison, David K. Jackson, Ian Johnston, Dominic Kwiatkowski, Cordelia Langford, John Sillitoe on behalf of the Wellcome Sanger Institute COVID-19 Surveillance Team |
| EPI_ISL_3043748 | Lighthouse Lab in Glasgow | Wellcome Sanger Institute for the COVID-19 Genomics UK (COG-UK) Consortium | Harper VanSteenhouse, Yumi Kasai, David Gray, Carol Clugston, Anna Dominiczak and Alex Alderton, Roberto Amato, Jeffrey Barrett, Sonia Goncalves, Ewan Harrison, David K. Jackson, Ian Johnston, Dominic Kwiatkowski, Cordelia Langford, John Sillitoe on behalf of the Wellcome Sanger Institute COVID-19 Surveillance Team |
| EPI_ISL_3043749, EPI_ISL_3043750, EPI_ISL_3043751, EPI_ISL_3043752, EPI_ISL_3043753, EPI_ISL_3043754, EPI_ISL_3043755, EPI_ISL_3043756 | Health Services Laboratories | Wellcome Sanger Institute for the COVID-19 Genomics UK (COG-UK) Consortium | Health Services Laboratories and Alex Alderton, Roberto Amato, Jeffrey Barrett, Sonia Goncalves, Ewan Harrison, David K. Jackson, Ian Johnston, Dominic Kwiatkowski, Cordelia Langford, John Sillitoe on behalf of the Wellcome Sanger Institute COVID-19 Surveillance Team |
| EPI_ISL_3043757 | Lighthouse Lab in Milton Keynes | Wellcome Sanger Institute for the COVID-19 Genomics UK (COG-UK) Consortium | The Lighthouse Lab in Milton Keynes and Alex Alderton, Roberto Amato, Jeffrey Barrett, Sonia Goncalves, Ewan Harrison, David K. Jackson, Ian Johnston, Dominic Kwiatkowski, Cordelia Langford, John Sillitoe on behalf of the Wellcome Sanger Institute COVID-19 Surveillance Team |
| EPI_ISL_3043758 | Health Services Laboratories | Wellcome Sanger Institute for the COVID-19 Genomics UK (COG-UK) Consortium | Health Services Laboratories and Alex Alderton, Roberto Amato, Jeffrey Barrett, Sonia Goncalves, Ewan Harrison, David K. Jackson, Ian Johnston, Dominic Kwiatkowski, Cordelia Langford, John Sillitoe on behalf of the Wellcome Sanger Institute COVID-19 Surveillance Team |
| EPI_ISL_3043759 | Lighthouse Lab in Alderley Park | Wellcome Sanger Institute for the COVID-19 Genomics UK (COG-UK) Consortium | Jacquelyn Wynn, Mairead Hyland, The Lighthouse Lab in Alderley Park and Alex Alderton, Roberto Amato, Jeffrey Barrett, Sonia Goncalves, Ewan Harrison, David K. Jackson, Ian Johnston, Dominic Kwiatkowski, Cordelia Langford, John Sillitoe on behalf of the Wellcome Sanger Institute COVID-19 Surveillance Team |
| EPI_ISL_3043760, EPI_ISL_3043761 | Health Services Laboratories | Wellcome Sanger Institute for the COVID-19 Genomics UK (COG-UK) Consortium | Health Services Laboratories and Alex Alderton, Roberto Amato, Jeffrey Barrett, Sonia Goncalves, Ewan Harrison, David K. Jackson, Ian Johnston, Dominic Kwiatkowski, Cordelia Langford, John Sillitoe on behalf of the Wellcome Sanger Institute COVID-19 Surveillance Team |
| EPI_ISL_3043762 | Lighthouse Lab in Glasgow | Wellcome Sanger Institute for the COVID-19 Genomics UK (COG-UK) Consortium | Harper VanSteenhouse, Yumi Kasai, David Gray, Carol Clugston, Anna Dominiczak and Alex Alderton, Roberto Amato, Jeffrey Barrett, Sonia Goncalves, Ewan Harrison, David K. Jackson, Ian Johnston, Dominic Kwiatkowski, Cordelia Langford, John Sillitoe on behalf of the Wellcome Sanger Institute COVID-19 Surveillance Team |
| EPI_ISL_3043763, EPI_ISL_3043764, EPI_ISL_3043765 | Health Services Laboratories | Wellcome Sanger Institute for the COVID-19 Genomics UK (COG-UK) Consortium | Health Services Laboratories and Alex Alderton, Roberto Amato, Jeffrey Barrett, Sonia Goncalves, Ewan Harrison, David K. Jackson, Ian Johnston, Dominic Kwiatkowski, Cordelia Langford, John Sillitoe on behalf of the Wellcome Sanger Institute COVID-19 Surveillance Team |
| EPI_ISL_3043766 | Lighthouse Lab in Glasgow | Wellcome Sanger Institute for the COVID-19 Genomics UK (COG-UK) Consortium | Harper VanSteenhouse, Yumi Kasai, David Gray, Carol Clugston, Anna Dominiczak and Alex Alderton, Roberto Amato, Jeffrey Barrett, Sonia Goncalves, Ewan Harrison, David K. Jackson, Ian Johnston, Dominic Kwiatkowski, Cordelia Langford, John Sillitoe on behalf of the Wellcome Sanger Institute COVID-19 Surveillance Team |
| EPI_ISL_3043767 | Lighthouse Lab in Alderley Park | Wellcome Sanger Institute for the COVID-19 Genomics UK (COG-UK) Consortium | Jacquelyn Wynn, Mairead Hyland, The Lighthouse Lab in Alderley Park and Alex Alderton, Roberto Amato, Jeffrey Barrett, Sonia Goncalves, Ewan Harrison, David K. Jackson, Ian Johnston, Dominic Kwiatkowski, Cordelia Langford, John Sillitoe on behalf of the Wellcome Sanger Institute COVID-19 Surveillance Team |
| EPI_ISL_3043768, EPI_ISL_3043769 | Lighthouse Lab in Glasgow | Wellcome Sanger Institute for the COVID-19 Genomics UK (COG-UK) Consortium | Harper VanSteenhouse, Yumi Kasai, David Gray, Carol Clugston, Anna Dominiczak and Alex Alderton, Roberto Amato, Jeffrey Barrett, Sonia Goncalves, Ewan Harrison, David K. Jackson, Ian Johnston, Dominic Kwiatkowski, Cordelia Langford, John Sillitoe on behalf of the Wellcome Sanger Institute COVID-19 Surveillance Team |
| EPI_ISL_3043770, EPI_ISL_3043771, EPI_ISL_3043772 | Lighthouse Lab in Alderley Park | Wellcome Sanger Institute for the COVID-19 Genomics UK (COG-UK) Consortium | Jacquelyn Wynn, Mairead Hyland, The Lighthouse Lab in Alderley Park and Alex Alderton, Roberto Amato, Jeffrey Barrett, Sonia Goncalves, Ewan Harrison, David K. Jackson, Ian Johnston, Dominic Kwiatkowski, Cordelia Langford, John Sillitoe on behalf of the Wellcome Sanger Institute COVID-19 Surveillance Team |
| EPI_ISL_3043773 | Lighthouse Lab in Glasgow | Wellcome Sanger Institute for the COVID-19 Genomics UK (COG-UK) Consortium | Harper VanSteenhouse, Yumi Kasai, David Gray, Carol Clugston, Anna Dominiczak and Alex Alderton, Roberto Amato, Jeffrey Barrett, Sonia Goncalves, Ewan Harrison, David K. Jackson, Ian Johnston, Dominic Kwiatkowski, Cordelia Langford, John Sillitoe on behalf of the Wellcome Sanger Institute COVID-19 Surveillance Team |
| EPI_ISL_3043774, EPI_ISL_3043775 | Lighthouse Lab in Alderley Park | Wellcome Sanger Institute for the COVID-19 Genomics UK (COG-UK) Consortium | Jacquelyn Wynn, Mairead Hyland, The Lighthouse Lab in Alderley Park and Alex Alderton, Roberto Amato, Jeffrey Barrett, Sonia Goncalves, Ewan Harrison, David K. Jackson, Ian Johnston, Dominic Kwiatkowski, Cordelia Langford, John Sillitoe on behalf of the Wellcome Sanger Institute COVID-19 Surveillance Team |
| EPI_ISL_3043776 | Lighthouse Lab in Glasgow | Wellcome Sanger Institute for the COVID-19 Genomics UK (COG-UK) Consortium | Harper VanSteenhouse, Yumi Kasai, David Gray, Carol Clugston, Anna Dominiczak and Alex Alderton, Roberto Amato, Jeffrey Barrett, Sonia Goncalves, Ewan Harrison, David K. Jackson, Ian Johnston, Dominic Kwiatkowski, Cordelia Langford, John Sillitoe on behalf of the Wellcome Sanger Institute COVID-19 Surveillance Team |
| EPI_ISL_3043777, EPI_ISL_3043778 | Lighthouse Lab in Alderley Park | Wellcome Sanger Institute for the COVID-19 Genomics UK (COG-UK) Consortium | Jacquelyn Wynn, Mairead Hyland, The Lighthouse Lab in Alderley Park and Alex Alderton, Roberto Amato, Jeffrey Barrett, Sonia Goncalves, Ewan Harrison, David K. Jackson, Ian Johnston, Dominic Kwiatkowski, Cordelia Langford, John Sillitoe on behalf of the Wellcome Sanger Institute COVID-19 Surveillance Team |
| EPI_ISL_3043779 | Lighthouse Lab in Glasgow | Wellcome Sanger Institute for the COVID-19 Genomics UK (COG-UK) Consortium | Harper VanSteenhouse, Yumi Kasai, David Gray, Carol Clugston, Anna Dominiczak and Alex Alderton, Roberto Amato, Jeffrey Barrett, Sonia Goncalves, Ewan Harrison, David K. Jackson, Ian Johnston, Dominic Kwiatkowski, Cordelia Langford, John Sillitoe on behalf of the Wellcome Sanger Institute COVID-19 Surveillance Team |
| EPI_ISL_3043780 | Lighthouse Lab in Alderley Park | Wellcome Sanger Institute for the COVID-19 Genomics UK (COG-UK) Consortium | Jacquelyn Wynn, Mairead Hyland, The Lighthouse Lab in Alderley Park and Alex Alderton, Roberto Amato, Jeffrey Barrett, Sonia Goncalves, Ewan Harrison, David K. Jackson, Ian Johnston, Dominic Kwiatkowski, Cordelia Langford, John Sillitoe on behalf of the Wellcome Sanger Institute COVID-19 Surveillance Team |
| EPI_ISL_3043781, EPI_ISL_3043782 | Lighthouse Lab in Glasgow | Wellcome Sanger Institute for the COVID-19 Genomics UK (COG-UK) Consortium | Harper VanSteenhouse, Yumi Kasai, David Gray, Carol Clugston, Anna Dominiczak and Alex Alderton, Roberto Amato, Jeffrey Barrett, Sonia Goncalves, Ewan Harrison, David K. Jackson, Ian Johnston, Dominic Kwiatkowski, Cordelia Langford, John Sillitoe on behalf of the Wellcome Sanger Institute COVID-19 Surveillance Team |
| EPI_ISL_3043783 | Lighthouse Lab in Alderley Park | Wellcome Sanger Institute for the COVID-19 Genomics UK (COG-UK) Consortium | Jacquelyn Wynn, Mairead Hyland, The Lighthouse Lab in Alderley Park and Alex Alderton, Roberto Amato, Jeffrey Barrett, Sonia Goncalves, Ewan Harrison, David K. Jackson, Ian Johnston, Dominic Kwiatkowski, Cordelia Langford, John Sillitoe on behalf of the Wellcome Sanger Institute COVID-19 Surveillance Team |
| EPI_ISL_3043784 | Lighthouse Lab in Glasgow | Wellcome Sanger Institute for the COVID-19 Genomics UK (COG-UK) Consortium | Harper VanSteenhouse, Yumi Kasai, David Gray, Carol Clugston, Anna Dominiczak and Alex Alderton, Roberto Amato, Jeffrey Barrett, Sonia Goncalves, Ewan Harrison, David K. Jackson, Ian Johnston, Dominic Kwiatkowski, Cordelia Langford, John Sillitoe on behalf of the Wellcome Sanger Institute COVID-19 Surveillance Team |
| EPI_ISL_3043785, EPI_ISL_3043786 | Lighthouse Lab in Milton Keynes | Wellcome Sanger Institute for the COVID-19 Genomics UK (COG-UK) Consortium | The Lighthouse Lab in Milton Keynes and Alex Alderton, Roberto Amato, Jeffrey Barrett, Sonia Goncalves, Ewan Harrison, David K. Jackson, Ian Johnston, Dominic Kwiatkowski, Cordelia Langford, John Sillitoe on behalf of the Wellcome Sanger Institute COVID-19 Surveillance Team |
| EPI_ISL_3043787, EPI_ISL_3043788 | Lighthouse Lab in Alderley Park | Wellcome Sanger Institute for the COVID-19 Genomics UK (COG-UK) Consortium | Jacquelyn Wynn, Mairead Hyland, The Lighthouse Lab in Alderley Park and Alex Alderton, Roberto Amato, Jeffrey Barrett, Sonia Goncalves, Ewan Harrison, David K. Jackson, Ian Johnston, Dominic Kwiatkowski, Cordelia Langford, John Sillitoe on behalf of the Wellcome Sanger Institute COVID-19 Surveillance Team |
| EPI_ISL_3043789 | Lighthouse Lab in Glasgow | Wellcome Sanger Institute for the COVID-19 Genomics UK (COG-UK) Consortium | Harper VanSteenhouse, Yumi Kasai, David Gray, Carol Clugston, Anna Dominiczak and Alex Alderton, Roberto Amato, Jeffrey Barrett, Sonia Goncalves, Ewan Harrison, David K. Jackson, Ian Johnston, Dominic Kwiatkowski, Cordelia Langford, John Sillitoe on behalf of the Wellcome Sanger Institute COVID-19 Surveillance Team |
| EPI_ISL_3043790 | Lighthouse Lab in Milton Keynes | Wellcome Sanger Institute for the COVID-19 Genomics UK (COG-UK) Consortium | The Lighthouse Lab in Milton Keynes and Alex Alderton, Roberto Amato, Jeffrey Barrett, Sonia Goncalves, Ewan Harrison, David K. Jackson, Ian Johnston, Dominic Kwiatkowski, Cordelia Langford, John Sillitoe on behalf of the Wellcome Sanger Institute COVID-19 Surveillance Team |
| EPI_ISL_3043791, EPI_ISL_3043792 | Lighthouse Lab in Glasgow | Wellcome Sanger Institute for the COVID-19 Genomics UK (COG-UK) Consortium | Harper VanSteenhouse, Yumi Kasai, David Gray, Carol Clugston, Anna Dominiczak and Alex Alderton, Roberto Amato, Jeffrey Barrett, Sonia Goncalves, Ewan Harrison, David K. Jackson, Ian Johnston, Dominic Kwiatkowski, Cordelia Langford, John Sillitoe on behalf of the Wellcome Sanger Institute COVID-19 Surveillance Team |
| EPI_ISL_3043793, EPI_ISL_3043794 | Lighthouse Lab in Milton Keynes | Wellcome Sanger Institute for the COVID-19 Genomics UK (COG-UK) Consortium | The Lighthouse Lab in Milton Keynes and Alex Alderton, Roberto Amato, Jeffrey Barrett, Sonia Goncalves, Ewan Harrison, David K. Jackson, Ian Johnston, Dominic Kwiatkowski, Cordelia Langford, John Sillitoe on behalf of the Wellcome Sanger Institute COVID-19 Surveillance Team |
| EPI_ISL_3043795 | Lighthouse Lab in Glasgow | Wellcome Sanger Institute for the COVID-19 Genomics UK (COG-UK) Consortium | Harper VanSteenhouse, Yumi Kasai, David Gray, Carol Clugston, Anna Dominiczak and Alex Alderton, Roberto Amato, Jeffrey Barrett, Sonia Goncalves, Ewan Harrison, David K. Jackson, Ian Johnston, Dominic Kwiatkowski, Cordelia Langford, John Sillitoe on behalf of the Wellcome Sanger Institute COVID-19 Surveillance Team |
| EPI_ISL_3043796 | Lighthouse Lab in Alderley Park | Wellcome Sanger Institute for the COVID-19 Genomics UK (COG-UK) Consortium | Jacquelyn Wynn, Mairead Hyland, The Lighthouse Lab in Alderley Park and Alex Alderton, Roberto Amato, Jeffrey Barrett, Sonia Goncalves, Ewan Harrison, David K. Jackson, Ian Johnston, Dominic Kwiatkowski, Cordelia Langford, John Sillitoe on behalf of the Wellcome Sanger Institute COVID-19 Surveillance Team |
| EPI_ISL_3043797 | Lighthouse Lab in Glasgow | Wellcome Sanger Institute for the COVID-19 Genomics UK (COG-UK) Consortium | Harper VanSteenhouse, Yumi Kasai, David Gray, Carol Clugston, Anna Dominiczak and Alex Alderton, Roberto Amato, Jeffrey Barrett, Sonia Goncalves, Ewan Harrison, David K. Jackson, Ian Johnston, Dominic Kwiatkowski, Cordelia Langford, John Sillitoe on behalf of the Wellcome Sanger Institute COVID-19 Surveillance Team |
| EPI_ISL_3043798, EPI_ISL_3043799 | Lighthouse Lab in Alderley Park | Wellcome Sanger Institute for the COVID-19 Genomics UK (COG-UK) Consortium | Jacquelyn Wynn, Mairead Hyland, The Lighthouse Lab in Alderley Park and Alex Alderton, Roberto Amato, Jeffrey Barrett, Sonia Goncalves, Ewan Harrison, David K. Jackson, Ian Johnston, Dominic Kwiatkowski, Cordelia Langford, John Sillitoe on behalf of the Wellcome Sanger Institute COVID-19 Surveillance Team |
| EPI_ISL_3043800, | Lighthouse Lab in Glasgow | Wellcome Sanger Institute for the COVID-19 Genomics UK (COG-UK) | Harper VanSteenhouse, Yumi Kasai, David Gray, Carol Clugston, Anna Dominiczak and Alex Alderton, Roberto Amato, Jeffrey Barrett, Sonia Goncalves, Ewan Harrison, David K. Jackson, Ian Johnston, Dominic Kwiatkowski, Cordelia Langford, |

| | | | |
|---|---|---|---|
| EPI_ISL_3043801 | | Consortium | John Sillitoe on behalf of the Wellcome Sanger Institute COVID-19 Surveillance Team |
| EPI_ISL_3043802 | Health Services Laboratories | Wellcome Sanger Institute for the COVID-19 Genomics UK (COG-UK) Consortium | Health Services Laboratories and Alex Alderton, Roberto Amato, Jeffrey Barrett, Sonia Goncalves, Ewan Harrison, David K. Jackson, Ian Johnston, Dominic Kwiatkowski, Cordelia Langford, John Sillitoe on behalf of the Wellcome Sanger Institute COVID-19 Surveillance Team |
| EPI_ISL_3043803 | Lighthouse Lab in Alderley Park | Wellcome Sanger Institute for the COVID-19 Genomics UK (COG-UK) Consortium | Jacquelyn Wynn, Mairead Hyland, The Lighthouse Lab in Alderley Park and Alex Alderton, Roberto Amato, Jeffrey Barrett, Sonia Goncalves, Ewan Harrison, David K. Jackson, Ian Johnston, Dominic Kwiatkowski, Cordelia Langford, John Sillitoe on behalf of the Wellcome Sanger Institute COVID-19 Surveillance Team |
| EPI_ISL_3043804, EPI_ISL_3043805, EPI_ISL_3043806, EPI_ISL_3043807, EPI_ISL_3043808 | Health Services Laboratories | Wellcome Sanger Institute for the COVID-19 Genomics UK (COG-UK) Consortium | Health Services Laboratories and Alex Alderton, Roberto Amato, Jeffrey Barrett, Sonia Goncalves, Ewan Harrison, David K. Jackson, Ian Johnston, Dominic Kwiatkowski, Cordelia Langford, John Sillitoe on behalf of the Wellcome Sanger Institute COVID-19 Surveillance Team |
| EPI_ISL_3043809 | Lighthouse Lab in Glasgow | Wellcome Sanger Institute for the COVID-19 Genomics UK (COG-UK) Consortium | Harper VanSteenhouse, Yumi Kasai, David Gray, Carol Clugston, Anna Dominiczak and Alex Alderton, Roberto Amato, Jeffrey Barrett, Sonia Goncalves, Ewan Harrison, David K. Jackson, Ian Johnston, Dominic Kwiatkowski, Cordelia Langford, John Sillitoe on behalf of the Wellcome Sanger Institute COVID-19 Surveillance Team |
| EPI_ISL_3043810, EPI_ISL_3043811 | Lighthouse Lab in Alderley Park | Wellcome Sanger Institute for the COVID-19 Genomics UK (COG-UK) Consortium | Jacquelyn Wynn, Mairead Hyland, The Lighthouse Lab in Alderley Park and Alex Alderton, Roberto Amato, Jeffrey Barrett, Sonia Goncalves, Ewan Harrison, David K. Jackson, Ian Johnston, Dominic Kwiatkowski, Cordelia Langford, John Sillitoe on behalf of the Wellcome Sanger Institute COVID-19 Surveillance Team |
| EPI_ISL_3043812, EPI_ISL_3043813, EPI_ISL_3043814, EPI_ISL_3043815, EPI_ISL_3043816, EPI_ISL_3043817, EPI_ISL_3043818, EPI_ISL_3043819, EPI_ISL_3043820, EPI_ISL_3043821, EPI_ISL_3043822 | | | |
| see above | Health Services Laboratories | Wellcome Sanger Institute for the COVID-19 Genomics UK (COG-UK) Consortium | Health Services Laboratories and Alex Alderton, Roberto Amato, Jeffrey Barrett, Sonia Goncalves, Ewan Harrison, David K. Jackson, Ian Johnston, Dominic Kwiatkowski, Cordelia Langford, John Sillitoe on behalf of the Wellcome Sanger Institute COVID-19 Surveillance Team |
| EPI_ISL_3043823 | Lighthouse Lab in Alderley Park | Wellcome Sanger Institute for the COVID-19 Genomics UK (COG-UK) Consortium | Jacquelyn Wynn, Mairead Hyland, The Lighthouse Lab in Alderley Park and Alex Alderton, Roberto Amato, Jeffrey Barrett, Sonia Goncalves, Ewan Harrison, David K. Jackson, Ian Johnston, Dominic Kwiatkowski, Cordelia Langford, John Sillitoe on behalf of the Wellcome Sanger Institute COVID-19 Surveillance Team |
| EPI_ISL_3043824, EPI_ISL_3043825 | Health Services Laboratories | Wellcome Sanger Institute for the COVID-19 Genomics UK (COG-UK) Consortium | Health Services Laboratories and Alex Alderton, Roberto Amato, Jeffrey Barrett, Sonia Goncalves, Ewan Harrison, David K. Jackson, Ian Johnston, Dominic Kwiatkowski, Cordelia Langford, John Sillitoe on behalf of the Wellcome Sanger Institute COVID-19 Surveillance Team |
| EPI_ISL_3043826 | Lighthouse Lab in Alderley Park | Wellcome Sanger Institute for the COVID-19 Genomics UK (COG-UK) Consortium | Jacquelyn Wynn, Mairead Hyland, The Lighthouse Lab in Alderley Park and Alex Alderton, Roberto Amato, Jeffrey Barrett, Sonia Goncalves, Ewan Harrison, David K. Jackson, Ian Johnston, Dominic Kwiatkowski, Cordelia Langford, John Sillitoe on behalf of the Wellcome Sanger Institute COVID-19 Surveillance Team |
| EPI_ISL_3043827, EPI_ISL_3043828 | Health Services Laboratories | Wellcome Sanger Institute for the COVID-19 Genomics UK (COG-UK) Consortium | Health Services Laboratories and Alex Alderton, Roberto Amato, Jeffrey Barrett, Sonia Goncalves, Ewan Harrison, David K. Jackson, Ian Johnston, Dominic Kwiatkowski, Cordelia Langford, John Sillitoe on behalf of the Wellcome Sanger Institute COVID-19 Surveillance Team |
| EPI_ISL_3043829 | Lighthouse Lab in Alderley Park | Wellcome Sanger Institute for the COVID-19 Genomics UK (COG-UK) Consortium | Jacquelyn Wynn, Mairead Hyland, The Lighthouse Lab in Alderley Park and Alex Alderton, Roberto Amato, Jeffrey Barrett, Sonia Goncalves, Ewan Harrison, David K. Jackson, Ian Johnston, Dominic Kwiatkowski, Cordelia Langford, John Sillitoe on behalf of the Wellcome Sanger Institute COVID-19 Surveillance Team |
| EPI_ISL_3043830 | Health Services Laboratories | Wellcome Sanger Institute for the COVID-19 Genomics UK (COG-UK) Consortium | Health Services Laboratories and Alex Alderton, Roberto Amato, Jeffrey Barrett, Sonia Goncalves, Ewan Harrison, David K. Jackson, Ian Johnston, Dominic Kwiatkowski, Cordelia Langford, John Sillitoe on behalf of the Wellcome Sanger Institute COVID-19 Surveillance Team |
| EPI_ISL_3043831 | Lighthouse Lab in Alderley Park | Wellcome Sanger Institute for the COVID-19 Genomics UK (COG-UK) Consortium | Jacquelyn Wynn, Mairead Hyland, The Lighthouse Lab in Alderley Park and Alex Alderton, Roberto Amato, Jeffrey Barrett, Sonia Goncalves, Ewan Harrison, David K. Jackson, Ian Johnston, Dominic Kwiatkowski, Cordelia Langford, John Sillitoe on behalf of the Wellcome Sanger Institute COVID-19 Surveillance Team |
| EPI_ISL_3043832, EPI_ISL_3043833, EPI_ISL_3043834, EPI_ISL_3043835, EPI_ISL_3043836, EPI_ISL_3043837 | Health Services Laboratories | Wellcome Sanger Institute for the COVID-19 Genomics UK (COG-UK) Consortium | Health Services Laboratories and Alex Alderton, Roberto Amato, Jeffrey Barrett, Sonia Goncalves, Ewan Harrison, David K. Jackson, Ian Johnston, Dominic Kwiatkowski, Cordelia Langford, John Sillitoe on behalf of the Wellcome Sanger Institute COVID-19 Surveillance Team |
| EPI_ISL_3043838 | Lighthouse Lab in Alderley Park | Wellcome Sanger Institute for the COVID-19 Genomics UK (COG-UK) Consortium | Jacquelyn Wynn, Mairead Hyland, The Lighthouse Lab in Alderley Park and Alex Alderton, Roberto Amato, Jeffrey Barrett, Sonia Goncalves, Ewan Harrison, David K. Jackson, Ian Johnston, Dominic Kwiatkowski, Cordelia Langford, John Sillitoe on behalf of the Wellcome Sanger Institute COVID-19 Surveillance Team |
| EPI_ISL_3043839 | Berkshire and Surrey Pathology Services Lighthouse Laboratory | Wellcome Sanger Institute for the COVID-19 Genomics UK (COG-UK) Consortium | Berkshire and Surrey Pathology Services Lighthouse Laboratory and Alex Alderton, Roberto Amato, Jeffrey Barrett, Sonia Goncalves, Ewan Harrison, David K. Jackson, Ian Johnston, Dominic Kwiatkowski, Cordelia Langford, John Sillitoe on behalf of the Wellcome Sanger Institute COVID-19 Surveillance Team |
| EPI_ISL_3043840, EPI_ISL_3043841 | Lighthouse Lab in Milton Keynes | Wellcome Sanger Institute for the COVID-19 Genomics UK (COG-UK) Consortium | The Lighthouse Lab in Milton Keynes and Alex Alderton, Roberto Amato, Jeffrey Barrett, Sonia Goncalves, Ewan Harrison, David K. Jackson, Ian Johnston, Dominic Kwiatkowski, Cordelia Langford, John Sillitoe on behalf of the Wellcome Sanger Institute COVID-19 Surveillance Team |
| EPI_ISL_3043842 | Berkshire and Surrey Pathology Services Lighthouse Laboratory | Wellcome Sanger Institute for the COVID-19 Genomics UK (COG-UK) Consortium | Berkshire and Surrey Pathology Services Lighthouse Laboratory and Alex Alderton, Roberto Amato, Jeffrey Barrett, Sonia Goncalves, Ewan Harrison, David K. Jackson, Ian Johnston, Dominic Kwiatkowski, Cordelia Langford, John Sillitoe on behalf of the Wellcome Sanger Institute COVID-19 Surveillance Team |
| EPI_ISL_3043843 | Lighthouse Lab in Alderley Park | Wellcome Sanger Institute for the COVID-19 Genomics UK (COG-UK) Consortium | Jacquelyn Wynn, Mairead Hyland, The Lighthouse Lab in Alderley Park and Alex Alderton, Roberto Amato, Jeffrey Barrett, Sonia Goncalves, Ewan Harrison, David K. Jackson, Ian Johnston, Dominic Kwiatkowski, Cordelia Langford, John Sillitoe on behalf of the Wellcome Sanger Institute COVID-19 Surveillance Team |
| EPI_ISL_3043844 | Lighthouse Lab in Milton Keynes | Wellcome Sanger Institute for the COVID-19 Genomics UK (COG-UK) Consortium | The Lighthouse Lab in Milton Keynes and Alex Alderton, Roberto Amato, Jeffrey Barrett, Sonia Goncalves, Ewan Harrison, David K. Jackson, Ian Johnston, Dominic Kwiatkowski, Cordelia Langford, John Sillitoe on behalf of the Wellcome Sanger Institute COVID-19 Surveillance Team |
| EPI_ISL_3043845 | Berkshire and Surrey Pathology Services Lighthouse Laboratory | Wellcome Sanger Institute for the COVID-19 Genomics UK (COG-UK) Consortium | Berkshire and Surrey Pathology Services Lighthouse Laboratory and Alex Alderton, Roberto Amato, Jeffrey Barrett, Sonia Goncalves, Ewan Harrison, David K. Jackson, Ian Johnston, Dominic Kwiatkowski, Cordelia Langford, John Sillitoe on behalf of the Wellcome Sanger Institute COVID-19 Surveillance Team |
| EPI_ISL_3043846 | Lighthouse Lab in Milton Keynes | Wellcome Sanger Institute for the COVID-19 Genomics UK (COG-UK) Consortium | The Lighthouse Lab in Milton Keynes and Alex Alderton, Roberto Amato, Jeffrey Barrett, Sonia Goncalves, Ewan Harrison, David K. Jackson, Ian Johnston, Dominic Kwiatkowski, Cordelia Langford, John Sillitoe on behalf of the Wellcome Sanger Institute COVID-19 Surveillance Team |
| EPI_ISL_3043847 | Lighthouse Lab in Alderley Park | Wellcome Sanger Institute for the COVID-19 Genomics UK (COG-UK) Consortium | Jacquelyn Wynn, Mairead Hyland, The Lighthouse Lab in Alderley Park and Alex Alderton, Roberto Amato, Jeffrey Barrett, Sonia Goncalves, Ewan Harrison, David K. Jackson, Ian Johnston, Dominic Kwiatkowski, Cordelia Langford, John Sillitoe on behalf of the Wellcome Sanger Institute COVID-19 Surveillance Team |
| EPI_ISL_3043848 | Berkshire and Surrey Pathology Services Lighthouse Laboratory | Wellcome Sanger Institute for the COVID-19 Genomics UK (COG-UK) Consortium | Berkshire and Surrey Pathology Services Lighthouse Laboratory and Alex Alderton, Roberto Amato, Jeffrey Barrett, Sonia Goncalves, Ewan Harrison, David K. Jackson, Ian Johnston, Dominic Kwiatkowski, Cordelia Langford, John Sillitoe on behalf of the Wellcome Sanger Institute COVID-19 Surveillance Team |
| EPI_ISL_3043849 | Lighthouse Lab in Glasgow | Wellcome Sanger Institute for the COVID-19 Genomics UK (COG-UK) Consortium | Harper VanSteenhouse, Yumi Kasai, David Gray, Carol Clugston, Anna Dominiczak and Alex Alderton, Roberto Amato, Jeffrey Barrett, Sonia Goncalves, Ewan Harrison, David K. Jackson, Ian Johnston, Dominic Kwiatkowski, Cordelia Langford, John Sillitoe on behalf of the Wellcome Sanger Institute COVID-19 Surveillance Team |
| EPI_ISL_3043850 | Lighthouse Lab in Alderley Park | Wellcome Sanger Institute for the COVID-19 Genomics UK (COG-UK) Consortium | Jacquelyn Wynn, Mairead Hyland, The Lighthouse Lab in Alderley Park and Alex Alderton, Roberto Amato, Jeffrey Barrett, Sonia Goncalves, Ewan Harrison, David K. Jackson, Ian Johnston, Dominic Kwiatkowski, Cordelia Langford, John Sillitoe on behalf of the Wellcome Sanger Institute COVID-19 Surveillance Team |
| EPI_ISL_3043851, EPI_ISL_3043852 | Lighthouse Lab in Glasgow | Wellcome Sanger Institute for the COVID-19 Genomics UK (COG-UK) Consortium | Harper VanSteenhouse, Yumi Kasai, David Gray, Carol Clugston, Anna Dominiczak and Alex Alderton, Roberto Amato, Jeffrey Barrett, Sonia Goncalves, Ewan Harrison, David K. Jackson, Ian Johnston, Dominic Kwiatkowski, Cordelia Langford, John Sillitoe on behalf of the Wellcome Sanger Institute COVID-19 Surveillance Team |
| EPI_ISL_3043853, EPI_ISL_3043854 | Lighthouse Lab in Milton Keynes | Wellcome Sanger Institute for the COVID-19 Genomics UK (COG-UK) Consortium | The Lighthouse Lab in Milton Keynes and Alex Alderton, Roberto Amato, Jeffrey Barrett, Sonia Goncalves, Ewan Harrison, David K. Jackson, Ian Johnston, Dominic Kwiatkowski, Cordelia Langford, John Sillitoe on behalf of the Wellcome Sanger Institute COVID-19 Surveillance Team |
| EPI_ISL_3043855 | Berkshire and Surrey Pathology Services Lighthouse Laboratory | Wellcome Sanger Institute for the COVID-19 Genomics UK (COG-UK) Consortium | Berkshire and Surrey Pathology Services Lighthouse Laboratory and Alex Alderton, Roberto Amato, Jeffrey Barrett, Sonia Goncalves, Ewan Harrison, David K. Jackson, Ian Johnston, Dominic Kwiatkowski, Cordelia Langford, John Sillitoe on behalf of the Wellcome Sanger Institute COVID-19 Surveillance Team |
| EPI_ISL_3043856 | Lighthouse Lab in Milton Keynes | Wellcome Sanger Institute for the COVID-19 Genomics UK (COG-UK) Consortium | The Lighthouse Lab in Milton Keynes and Alex Alderton, Roberto Amato, Jeffrey Barrett, Sonia Goncalves, Ewan Harrison, David K. Jackson, Ian Johnston, Dominic Kwiatkowski, Cordelia Langford, John Sillitoe on behalf of the Wellcome Sanger Institute COVID-19 Surveillance Team |
| EPI_ISL_3043857 | Lighthouse Lab in Glasgow | Wellcome Sanger Institute for the COVID-19 Genomics UK (COG-UK) Consortium | Harper VanSteenhouse, Yumi Kasai, David Gray, Carol Clugston, Anna Dominiczak and Alex Alderton, Roberto Amato, Jeffrey Barrett, Sonia Goncalves, Ewan Harrison, David K. Jackson, Ian Johnston, Dominic Kwiatkowski, Cordelia Langford, John Sillitoe on behalf of the Wellcome Sanger Institute COVID-19 Surveillance Team |
| EPI_ISL_3043858 | Lighthouse Lab in Milton Keynes | Wellcome Sanger Institute for the COVID-19 Genomics UK (COG-UK) Consortium | The Lighthouse Lab in Milton Keynes and Alex Alderton, Roberto Amato, Jeffrey Barrett, Sonia Goncalves, Ewan Harrison, David K. Jackson, Ian Johnston, Dominic Kwiatkowski, Cordelia Langford, John Sillitoe on behalf of the Wellcome Sanger Institute COVID-19 Surveillance Team |
| EPI_ISL_3043859 | Lighthouse Lab in Glasgow | Wellcome Sanger Institute for the COVID-19 Genomics UK (COG-UK) Consortium | Harper VanSteenhouse, Yumi Kasai, David Gray, Carol Clugston, Anna Dominiczak and Alex Alderton, Roberto Amato, Jeffrey Barrett, Sonia Goncalves, Ewan Harrison, David K. Jackson, Ian Johnston, Dominic Kwiatkowski, Cordelia Langford, John Sillitoe on behalf of the Wellcome Sanger Institute COVID-19 Surveillance Team |
| EPI_ISL_3043860 | Lighthouse Lab in Alderley Park | Wellcome Sanger Institute for the COVID-19 Genomics UK (COG-UK) Consortium | Jacquelyn Wynn, Mairead Hyland, The Lighthouse Lab in Alderley Park and Alex Alderton, Roberto Amato, Jeffrey Barrett, Sonia Goncalves, Ewan Harrison, David K. Jackson, Ian Johnston, Dominic Kwiatkowski, Cordelia Langford, John Sillitoe on behalf of the Wellcome Sanger Institute COVID-19 Surveillance Team |
| EPI_ISL_3043861, EPI_ISL_3043864, EPI_ISL_3043865, EPI_ISL_3043866 | Eurofins MVZ Labor Gelsenkirchen | Robert Koch Institute | unknown |
| EPI_ISL_3043867, EPI_ISL_3043868, EPI_ISL_3043869, EPI_ISL_3043870 | Dianovis GmbH Greiz | Robert Koch Institute | unknown |
| EPI_ISL_3043871, EPI_ISL_3043872, EPI_ISL_3043873, EPI_ISL_3043874, EPI_ISL_3043875, EPI_ISL_3043876, EPI_ISL_3043877, EPI_ISL_3043878, EPI_ISL_3043879, EPI_ISL_3043880, EPI_ISL_3043881, EPI_ISL_3043882, EPI_ISL_3043884, EPI_ISL_3043885, EPI_ISL_3043887, EPI_ISL_3043888, EPI_ISL_3043889, EPI_ISL_3043895, EPI_ISL_3043896, EPI_ISL_3043897, EPI_ISL_3043898, EPI_ISL_3043899, EPI_ISL_3043900, EPI_ISL_3043901, EPI_ISL_3043903, EPI_ISL_3043904, EPI_ISL_3043906, EPI_ISL_3043907, EPI_ISL_3043908, EPI_ISL_3043909, EPI_ISL_3043911, EPI_ISL_3043912, EPI_ISL_3043913, EPI_ISL_3043914, EPI_ISL_3043915, EPI_ISL_3043916, EPI_ISL_3043917, EPI_ISL_3043920, EPI_ISL_3043921, EPI_ISL_3043922, EPI_ISL_3043923, EPI_ISL_3043924, EPI_ISL_3043925, EPI_ISL_3043927, EPI_ISL_3043928, EPI_ISL_3043930, EPI_ISL_3043931, EPI_ISL_3043932, EPI_ISL_3043933, EPI_ISL_3043935, EPI_ISL_3043936, EPI_ISL_3043937, EPI_ISL_3043938, EPI_ISL_3043940, EPI_ISL_3043941 | | | |
| see above | CENTOGENE Frankfurt Laboratory: Niederlassung Industriepark Höchst | Robert Koch Institute | unknown |
| EPI_ISL_3043945, EPI_ISL_3043946, | Klinikum Ernst von Bergmann gemeinnützige GmbH - stationärer Bereich | Robert Koch Institute | unknown |

| | | | |
|---|---|---|---|
| EPI_ISL_3043947 | | | |
| EPI_ISL_3043948 | Institut für Medizinische Mikrobiologie - Universitätsmedizin Göttingen | Robert Koch Institute | unknown |
| EPI_ISL_3043951, EPI_ISL_3043952 | SYNLAB MVZ Weiden | Robert Koch Institute | unknown |
| EPI_ISL_3043953 | SYNLAB Labor München Zentrum LMZ | Robert Koch Institute | unknown |
| EPI_ISL_3043954 | SYNLAB MVZ Weiden | Robert Koch Institute | unknown |
| EPI_ISL_3043955 | SYNLAB Labor München Zentrum LMZ | Robert Koch Institute | unknown |
| EPI_ISL_3043956, EPI_ISL_3043957 | SYNLAB MVZ Weiden | Robert Koch Institute | unknown |
| EPI_ISL_3043958 | SYNLAB MVZ Leverkusen | Robert Koch Institute | unknown |
| EPI_ISL_3043959 | SYNLAB MVZ Weiden | Robert Koch Institute | unknown |
| EPI_ISL_3043961 | SYNLAB MVZ Leinfelden-Echterdingen | Robert Koch Institute | unknown |
| EPI_ISL_3043962, EPI_ISL_3043963 | SYNLAB MVZ Leverkusen | Robert Koch Institute | unknown |
| EPI_ISL_3043964 | SYNLAB MVZ Weiden | Robert Koch Institute | unknown |
| EPI_ISL_3043965 | SYNLAB MVZ Leverkusen | Robert Koch Institute | unknown |
| EPI_ISL_3043966, EPI_ISL_3043967 | Synlab MVZ Augsburg | Robert Koch Institute | unknown |
| EPI_ISL_3043968, EPI_ISL_3043970 | SYNLAB MVZ Weiden | Robert Koch Institute | unknown |
| EPI_ISL_3043971 | SYNLAB MVZ Leinfelden-Echterdingen | Robert Koch Institute | unknown |
| EPI_ISL_3043972 | SYNLAB MVZ Weiden | Robert Koch Institute | unknown |
| EPI_ISL_3043973 | Synlab MVZ Augsburg | Robert Koch Institute | unknown |
| EPI_ISL_3043974 | SYNLAB MVZ Weiden | Robert Koch Institute | unknown |
| EPI_ISL_3043975, EPI_ISL_3043976 | SYNLAB MVZ Leverkusen | Robert Koch Institute | unknown |
| EPI_ISL_3043977 | SYNLAB MVZ Leinfelden-Echterdingen | Robert Koch Institute | unknown |
| EPI_ISL_3043978 | SYNLAB MVZ Weiden | Robert Koch Institute | unknown |
| EPI_ISL_3043980 | Synlab MVZ Augsburg | Robert Koch Institute | unknown |
| EPI_ISL_3043984 | SYNLAB MVZ Leverkusen | Robert Koch Institute | unknown |
| EPI_ISL_3043985 | Synlab MVZ Augsburg | Robert Koch Institute | unknown |
| EPI_ISL_3043986, EPI_ISL_3043987 | SYNLAB MVZ Weiden | Robert Koch Institute | unknown |
| EPI_ISL_3043988, EPI_ISL_3043989 | Synlab MVZ Augsburg | Robert Koch Institute | unknown |
| EPI_ISL_3043990 | SYNLAB MVZ Leverkusen | Robert Koch Institute | unknown |
| EPI_ISL_3043992 | SYNLAB MVZ Weiden | Robert Koch Institute | unknown |
| EPI_ISL_3043993 | SYNLAB MVZ Leverkusen | Robert Koch Institute | unknown |
| EPI_ISL_3043994 | SYNLAB MVZ Weiden | Robert Koch Institute | unknown |
| EPI_ISL_3043995 | SYNLAB Labor München Zentrum LMZ | Robert Koch Institute | unknown |
| EPI_ISL_3043996 | Synlab MVZ Augsburg | Robert Koch Institute | unknown |
| EPI_ISL_3043997, EPI_ISL_3043999 | SYNLAB MVZ Weiden | Robert Koch Institute | unknown |
| EPI_ISL_3044000, EPI_ISL_3044001 | SYNLAB MVZ Leverkusen | Robert Koch Institute | unknown |
| EPI_ISL_3044004, EPI_ISL_3044005 | SYNLAB MVZ Weiden | Robert Koch Institute | unknown |
| EPI_ISL_3044007, EPI_ISL_3044008 | SYNLAB MVZ Leverkusen | Robert Koch Institute | unknown |
| EPI_ISL_3044009 | SYNLAB MVZ Berlin | Robert Koch Institute | unknown |
| EPI_ISL_3044010, EPI_ISL_3044011, EPI_ISL_3044012, EPI_ISL_3044013 | SYNLAB MVZ Leverkusen | Robert Koch Institute | unknown |
| EPI_ISL_3044014, EPI_ISL_3044015 | SYNLAB MVZ Weiden | Robert Koch Institute | unknown |
| EPI_ISL_3044017 | SYNLAB MVZ Leinfelden-Echterdingen | Robert Koch Institute | unknown |
| EPI_ISL_3044018, EPI_ISL_3044020 | Synlab MVZ Augsburg | Robert Koch Institute | unknown |
| EPI_ISL_3044021 | SYNLAB MVZ Weiden | Robert Koch Institute | unknown |
| EPI_ISL_3044023 | SYNLAB MVZ Leverkusen | Robert Koch Institute | unknown |
| EPI_ISL_3044024, EPI_ISL_3044025 | SYNLAB Labor München Zentrum LMZ | Robert Koch Institute | unknown |
| EPI_ISL_3044026 | SYNLAB MVZ Leverkusen | Robert Koch Institute | unknown |
| EPI_ISL_3044027 | SYNLAB MVZ Leinfelden-Echterdingen | Robert Koch Institute | unknown |
| EPI_ISL_3044028 | SYNLAB MVZ Weiden | Robert Koch Institute | unknown |
| EPI_ISL_3044030 | SYNLAB MVZ Leverkusen | Robert Koch Institute | unknown |
| EPI_ISL_3044031 | SYNLAB MVZ Weiden | Robert Koch Institute | unknown |
| EPI_ISL_3044032 | SYNLAB MVZ Trier | Robert Koch Institute | unknown |
| EPI_ISL_3044033, EPI_ISL_3044034 | SYNLAB MVZ Weiden | Robert Koch Institute | unknown |
| EPI_ISL_3044035 | SYNLAB MVZ Leinfelden-Echterdingen | Robert Koch Institute | unknown |
| EPI_ISL_3044036 | SYNLAB MVZ Leverkusen | Robert Koch Institute | unknown |
| EPI_ISL_3044037 | SYNLAB MVZ Weiden | Robert Koch Institute | unknown |
| EPI_ISL_3044038 | SYNLAB MVZ Leverkusen | Robert Koch Institute | unknown |
| EPI_ISL_3044041 | SYNLAB MVZ Trier | Robert Koch Institute | unknown |
| EPI_ISL_3044043 | Synlab MVZ Augsburg | Robert Koch Institute | unknown |
| EPI_ISL_3044045 | SYNLAB MVZ Leinfelden-Echterdingen | Robert Koch Institute | unknown |
| EPI_ISL_3044046 | SYNLAB Jena Oncoscreen | Robert Koch Institute | unknown |
| EPI_ISL_3044047 | SYNLAB MVZ Weiden | Robert Koch Institute | unknown |
| EPI_ISL_3044048 | SYNLAB MVZ Leverkusen | Robert Koch Institute | unknown |

| | | | |
|---|---|---|---|
| EPI_ISL_3044049, EPI_ISL_3044050 | SYNLAB MVZ Weiden | Robert Koch Institute | unknown |
| EPI_ISL_3044051 | SYNLAB MVZ Leverkusen | Robert Koch Institute | unknown |
| EPI_ISL_3044052 | SYNLAB MVZ Leinfelden-Echterdingen | Robert Koch Institute | unknown |
| EPI_ISL_3044053 | SYNLAB MVZ Weiden | Robert Koch Institute | unknown |
| EPI_ISL_3044054 | SYNLAB MVZ Leverkusen | Robert Koch Institute | unknown |
| EPI_ISL_3044055 | SYNLAB Jena Oncoscreen | Robert Koch Institute | unknown |
| EPI_ISL_3044056, EPI_ISL_3044057 | SYNLAB MVZ Leverkusen | Robert Koch Institute | unknown |
| EPI_ISL_3044058 | SYNLAB MVZ Weiden | Robert Koch Institute | unknown |
| EPI_ISL_3044059 | SYNLAB MVZ Dachau | Robert Koch Institute | unknown |
| EPI_ISL_3044060, EPI_ISL_3044061, EPI_ISL_3044062 | SYNLAB MVZ Weiden | Robert Koch Institute | unknown |
| EPI_ISL_3044064 | SYNLAB MVZ Leverkusen | Robert Koch Institute | unknown |
| EPI_ISL_3044065 | SYNLAB MVZ Weiden | Robert Koch Institute | unknown |
| EPI_ISL_3044066 | SYNLAB MVZ Leverkusen | Robert Koch Institute | unknown |
| EPI_ISL_3044067, EPI_ISL_3044068 | SYNLAB MVZ Weiden | Robert Koch Institute | unknown |
| EPI_ISL_3044069 | SYNLAB MVZ Leverkusen | Robert Koch Institute | unknown |
| EPI_ISL_3044072 | SYNLAB MVZ Hamburg | Robert Koch Institute | unknown |
| EPI_ISL_3044073 | SYNLAB MVZ Leverkusen | Robert Koch Institute | unknown |
| EPI_ISL_3044074 | SYNLAB MVZ Weiden | Robert Koch Institute | unknown |
| EPI_ISL_3044075 | SYNLAB MVZ Leverkusen | Robert Koch Institute | unknown |
| EPI_ISL_3044076 | SYNLAB MVZ Trier | Robert Koch Institute | unknown |
| EPI_ISL_3044077 | SYNLAB MVZ Weiden | Robert Koch Institute | unknown |
| EPI_ISL_3044078, EPI_ISL_3044081 | SYNLAB MVZ Leverkusen | Robert Koch Institute | unknown |
| EPI_ISL_3044082 | SYNLAB MVZ Trier | Robert Koch Institute | unknown |
| EPI_ISL_3044083 | SYNLAB MVZ Weiden | Robert Koch Institute | unknown |
| EPI_ISL_3044084, EPI_ISL_3044085 | SYNLAB MVZ Leverkusen | Robert Koch Institute | unknown |
| EPI_ISL_3044086 | SYNLAB MVZ Trier | Robert Koch Institute | unknown |
| EPI_ISL_3044088 | SYNLAB MVZ Weiden | Robert Koch Institute | unknown |
| EPI_ISL_3044089, EPI_ISL_3044090, EPI_ISL_3044091 | SYNLAB MVZ Leverkusen | Robert Koch Institute | unknown |
| EPI_ISL_3044092 | SYNLAB MVZ Weiden | Robert Koch Institute | unknown |
| EPI_ISL_3044094 | Synlab MVZ Augsburg | Robert Koch Institute | unknown |
| EPI_ISL_3044095 | SYNLAB MVZ Leverkusen | Robert Koch Institute | unknown |
| EPI_ISL_3044096 | SYNLAB Labor München Zentrum LMZ | Robert Koch Institute | unknown |
| EPI_ISL_3044097 | SYNLAB MVZ Trier | Robert Koch Institute | unknown |
| EPI_ISL_3044099 | SYNLAB Labor München Zentrum LMZ | Robert Koch Institute | unknown |
| EPI_ISL_3044100 | SYNLAB Jena Oncoscreen | Robert Koch Institute | unknown |
| EPI_ISL_3044101 | SYNLAB MVZ Weiden | Robert Koch Institute | unknown |
| EPI_ISL_3044102, EPI_ISL_3044103 | SYNLAB MVZ Leverkusen | Robert Koch Institute | unknown |
| EPI_ISL_3044104 | Universitätsklinikum Frankfurt - Institut für Medizinische Virologie | Robert Koch Institute | unknown |
| EPI_ISL_3044107, EPI_ISL_3044108 | Bioscientia Labor Wermsdorf | Robert Koch Institute | unknown |
| EPI_ISL_3044109 | Sonic - Dr. Staber & Kollegen GmbH München | Robert Koch Institute | unknown |
| EPI_ISL_3044110, EPI_ISL_3044111, EPI_ISL_3044112, EPI_ISL_3044114 | Bioscientia Labor Wermsdorf | Robert Koch Institute | unknown |
| EPI_ISL_3044115 | LabKom - Labor Hannover MVZ GmbH | Robert Koch Institute | unknown |
| EPI_ISL_3044116 | Bioscientia Labor Wermsdorf | Robert Koch Institute | unknown |
| EPI_ISL_3044118, EPI_ISL_3044119 | Sonic - Labor Dr. von Froreich GmbH | Robert Koch Institute | unknown |
| EPI_ISL_3044120 | Universitätsklinikum Frankfurt - Institut für Medizinische Virologie | Robert Koch Institute | unknown |
| EPI_ISL_3044121, EPI_ISL_3044122, EPI_ISL_3044123, EPI_ISL_3044124 | Bioscientia Labor Wermsdorf | Robert Koch Institute | unknown |
| EPI_ISL_3044127 | Sonic - Bioscientia - MVZ Labor Saar GmbH | Robert Koch Institute | unknown |
| EPI_ISL_3044128 | Bioscientia Labor Wermsdorf | Robert Koch Institute | unknown |
| EPI_ISL_3044129 | Sonic - Labor Dr. von Froreich GmbH | Robert Koch Institute | unknown |
| EPI_ISL_3044130, EPI_ISL_3044132, EPI_ISL_3044133, EPI_ISL_3044134, EPI_ISL_3044135 | Bioscientia Labor Wermsdorf | Robert Koch Institute | unknown |
| EPI_ISL_3044136 | Sonic - Labor Dr. von Froreich GmbH | Robert Koch Institute | unknown |
| EPI_ISL_3044137, EPI_ISL_3044138 | Bioscientia Labor Wermsdorf | Robert Koch Institute | unknown |
| EPI_ISL_3044139 | Sonic - Labor Dr. von Froreich GmbH | Robert Koch Institute | unknown |
| EPI_ISL_3044140, EPI_ISL_3044142, EPI_ISL_3044143 | Bioscientia Labor Wermsdorf | Robert Koch Institute | unknown |
| EPI_ISL_3044144 | Sonic - Labor Staber Nürnberg | Robert Koch Institute | unknown |
| EPI_ISL_3044145 | Bioscientia Labor Wermsdorf | Robert Koch Institute | unknown |
| EPI_ISL_3044146, EPI_ISL_3044147 | alphaomega Labor GbR (Medizinische Kooperationsgemeinschaft Dres. Ackermann/Tennert/Zaiß) | Robert Koch Institute | unknown |

| | | | |
|---|---|---|---|
| EPI_ISL_3044148, EPI_ISL_3044149 | Labor für medizinische Mikrobiologie Bakteriologie Mykologie Virologie & Infektionsserologie - Labor Mölbis | Robert Koch Institute | unknown |
| EPI_ISL_3044150 | MVZ Dr. Eberhard & Partner Dortmund | Robert Koch Institute | unknown |
| EPI_ISL_3044151 | MVZ für Laboratoriumsmedizin und Mikrobiologie Würzburg | Robert Koch Institute | unknown |
| EPI_ISL_3044152, EPI_ISL_3044153 | MVZ Dr. Eberhard & Partner Dortmund | Robert Koch Institute | unknown |
| EPI_ISL_3044154 | Procomcure Biotech Germany GmbH | Robert Koch Institute | unknown |
| EPI_ISL_3044155, EPI_ISL_3044158 | MVZ Dr. Eberhard & Partner Dortmund | Robert Koch Institute | unknown |
| EPI_ISL_3044159, EPI_ISL_3044160, EPI_ISL_3044162 | IMD - Institut für Medizinische Diagnostik Berlin-Potsdam | Robert Koch Institute | unknown |
| EPI_ISL_3044163 | MVZ Labor Dr. Reising-Ackermann und Kollegen | Robert Koch Institute | unknown |
| EPI_ISL_3044164 | Labor Mönchengladbach MVZ Dr. Stein + Kollegen GbR | Robert Koch Institute | unknown |
| EPI_ISL_3044165, EPI_ISL_3044166 | MVZ Labor Dr. Limbach & Kollegen GbR | Robert Koch Institute | unknown |
| EPI_ISL_3044169 | Labor Mönchengladbach MVZ Dr. Stein + Kollegen GbR | Robert Koch Institute | unknown |
| EPI_ISL_3044170 | MVZ Labor Dr. Limbach & Kollegen GbR | Robert Koch Institute | unknown |
| EPI_ISL_3044172 | Limbach - MVZ Labor Eveld & Kollegen Essen | Robert Koch Institute | unknown |
| EPI_ISL_3044173, EPI_ISL_3044174, EPI_ISL_3044175, EPI_ISL_3044176 | MVZ Medizinisches Labor Hannover GmbH | Robert Koch Institute | unknown |
| EPI_ISL_3044177 | MVZ Labor Dr. Limbach & Kollegen GbR | Robert Koch Institute | unknown |
| EPI_ISL_3044179 | MVZ Medizinisches Labor Hannover GmbH | Robert Koch Institute | unknown |
| EPI_ISL_3044180 | MVZ Labor Dr. Limbach & Kollegen GbR | Robert Koch Institute | unknown |
| EPI_ISL_3044181, EPI_ISL_3044182 | MVZ Medizinisches Labor Hannover GmbH | Robert Koch Institute | unknown |
| EPI_ISL_3044183 | MVZ für Laboratoriumsmedizin und Mikrobiologie Koblenz-Mittelrhein (Labor Koblenz) | Robert Koch Institute | unknown |
| EPI_ISL_3044184 | MDI Limbach Berlin GmbH; MVZ Labor Berlin | Robert Koch Institute | unknown |
| EPI_ISL_3044185, EPI_ISL_3044186, EPI_ISL_3044187 | MVZ Medizinisches Labor Hannover GmbH | Robert Koch Institute | unknown |
| EPI_ISL_3044188 | MDI Limbach Berlin GmbH; MVZ Labor Berlin | Robert Koch Institute | unknown |
| EPI_ISL_3044189 | MVZ Medizinisches Labor Hannover GmbH | Robert Koch Institute | unknown |
| EPI_ISL_3044190 | MVZ für Laboratoriumsmedizin und Mikrobiologie Koblenz-Mittelrhein (Labor Koblenz) | Robert Koch Institute | unknown |
| EPI_ISL_3044191 | MVZ Medizinisches Labor Hannover GmbH | Robert Koch Institute | unknown |
| EPI_ISL_3044192, EPI_ISL_3044193 | MDI Limbach Berlin GmbH; MVZ Labor Berlin | Robert Koch Institute | unknown |
| EPI_ISL_3044194, EPI_ISL_3044195 | MVZ für Laboratoriumsmedizin und Mikrobiologie Koblenz-Mittelrhein (Labor Koblenz) | Robert Koch Institute | unknown |
| EPI_ISL_3044196, EPI_ISL_3044197 | MVZ Medizinisches Labor Hannover GmbH | Robert Koch Institute | unknown |
| EPI_ISL_3044198, EPI_ISL_3044199, EPI_ISL_3044200, EPI_ISL_3044201, EPI_ISL_3044202, EPI_ISL_3044203, EPI_ISL_3044204, EPI_ISL_3044205, EPI_ISL_3044206, EPI_ISL_3044207, EPI_ISL_3044208, EPI_ISL_3044209, EPI_ISL_3044210, EPI_ISL_3044211, EPI_ISL_3044212 | | | |
| see above | MVZ für Laboratoriumsmedizin und Mikrobiologie Koblenz-Mittelrhein (Labor Koblenz) | Robert Koch Institute | unknown |
| EPI_ISL_3044213, EPI_ISL_3044214, EPI_ISL_3044215, EPI_ISL_3044216, EPI_ISL_3044217, EPI_ISL_3044218, EPI_ISL_3044219, EPI_ISL_3044220, EPI_ISL_3044221, EPI_ISL_3044222, EPI_ISL_3044223, EPI_ISL_3044224, EPI_ISL_3044225, EPI_ISL_3044226, EPI_ISL_3044227 | | | |
| see above | Labor Dr. Heidrich & Kollegen MVZ GmbH Hamburg | Robert Koch Institute | unknown |
| EPI_ISL_3044228 | MVZ Labor Dr. Limbach & Kollegen GbR | Robert Koch Institute | unknown |
| EPI_ISL_3044229 | MDI Limbach Berlin GmbH; MVZ Labor Berlin | Robert Koch Institute | unknown |
| EPI_ISL_3044230 | MVZ Labor Dr. Limbach & Kollegen GbR | Robert Koch Institute | unknown |
| EPI_ISL_3044231, EPI_ISL_3044232 | MDI Limbach Berlin GmbH; MVZ Labor Berlin | Robert Koch Institute | unknown |
| EPI_ISL_3044233 | MVZ Labor Dr. Limbach & Kollegen GbR | Robert Koch Institute | unknown |
| EPI_ISL_3044234, EPI_ISL_3044235 | MDI Limbach Berlin GmbH; MVZ Labor Berlin | Robert Koch Institute | unknown |
| EPI_ISL_3044236, EPI_ISL_3044237 | MVZ Labor Dr. Limbach & Kollegen GbR | Robert Koch Institute | unknown |
| EPI_ISL_3044238, EPI_ISL_3044239, EPI_ISL_3044240 | MDI Limbach Berlin GmbH; MVZ Labor Berlin | Robert Koch Institute | unknown |
| EPI_ISL_3044241 | MVZ Labor Dr. Limbach & Kollegen GbR | Robert Koch Institute | unknown |
| EPI_ISL_3044242 | MDI Limbach Berlin GmbH; MVZ Labor Berlin | Robert Koch Institute | unknown |
| EPI_ISL_3044243 | MVZ Labor Dr. Limbach & Kollegen GbR | Robert Koch Institute | unknown |
| EPI_ISL_3044244 | MDI Limbach Berlin GmbH; MVZ Labor Berlin | Robert Koch Institute | unknown |
| EPI_ISL_3044246, EPI_ISL_3044249, EPI_ISL_3044250, EPI_ISL_3044251 | Uniklinikum Regensburg; Institut für klinische Mikrobiologie und Hygiene | Robert Koch Institute | unknown |
| EPI_ISL_3044252 | MVZ Medizinisches Labor Hannover GmbH | Robert Koch Institute | unknown |
| EPI_ISL_3044253, EPI_ISL_3044254, EPI_ISL_3044255, EPI_ISL_3044256 | Labormedizin Darmstadt | Robert Koch Institute | unknown |
| EPI_ISL_3044257, EPI_ISL_3044258 | IMD Labor Frankfurt | Robert Koch Institute | unknown |
| EPI_ISL_3044259 | Labormedizin Darmstadt | Robert Koch Institute | unknown |
| EPI_ISL_3044260, EPI_ISL_3044261 | IMD Labor Frankfurt | Robert Koch Institute | unknown |
| EPI_ISL_3044262, EPI_ISL_3044263, EPI_ISL_3044264, EPI_ISL_3044265, EPI_ISL_3044266, EPI_ISL_3044267, EPI_ISL_3044268, EPI_ISL_3044269, EPI_ISL_3044270, EPI_ISL_3044271, EPI_ISL_3044272 | | | |
| see above | Labormedizin Darmstadt | Robert Koch Institute | unknown |
| EPI_ISL_3044273 | IMD Labor Frankfurt | Robert Koch Institute | unknown |
| EPI_ISL_3044274 | Labormedizin Darmstadt | Robert Koch Institute | unknown |
| EPI_ISL_3044275 | IMD Labor Frankfurt | Robert Koch Institute | unknown |

| | | | |
|---|---|---|---|
| EPI_ISL_3044276, EPI_ISL_3044278, EPI_ISL_3044279, EPI_ISL_3044280, EPI_ISL_3044281, EPI_ISL_3044282, EPI_ISL_3044283, EPI_ISL_3044284 | Labormedizin Darmstadt | Robert Koch Institute | unknown |
| EPI_ISL_3044285 | IMD Labor Frankfurt | Robert Koch Institute | unknown |
| EPI_ISL_3044286 | Labormedizin Darmstadt | Robert Koch Institute | unknown |
| EPI_ISL_3044288 | Universitätsklinikum des Saarlandes - Institut für Virologie | Robert Koch Institute | unknown |
| EPI_ISL_3044289 | Labormedizin Darmstadt | Robert Koch Institute | unknown |
| EPI_ISL_3044291 | Carl-Thiem-Klinikum Cottbus - Zentrum für Laboratoriumsmedizin Mikrobiologie und Krankenhaushygiene | Robert Koch Institute | unknown |
| EPI_ISL_3044294, EPI_ISL_3044295, EPI_ISL_3044296, EPI_ISL_3044297, EPI_ISL_3044298, EPI_ISL_3044299, EPI_ISL_3044300, EPI_ISL_3044302, EPI_ISL_3044303, EPI_ISL_3044304, EPI_ISL_3044305, EPI_ISL_3044307, EPI_ISL_3044308, EPI_ISL_3044309, EPI_ISL_3044310, EPI_ISL_3044311, EPI_ISL_3044312, EPI_ISL_3044313, EPI_ISL_3044314, EPI_ISL_3044315, EPI_ISL_3044316, EPI_ISL_3044317, EPI_ISL_3044320, EPI_ISL_3044321, EPI_ISL_3044322, EPI_ISL_3044323, EPI_ISL_3044324, EPI_ISL_3044325, EPI_ISL_3044326, EPI_ISL_3044327, EPI_ISL_3044328, EPI_ISL_3044329 | | | |
| see above | Medizinische Laboratorien Düsseldorf | Robert Koch Institute | unknown |
| EPI_ISL_3044330, EPI_ISL_3044331, EPI_ISL_3044332 | Helios Universitätsklinikum Wuppertal: Universität Witten/Herdecke: Institut für Medizinische Labordiagnostik | Robert Koch Institute | unknown |
| EPI_ISL_3044333, EPI_ISL_3044334 | Philipps-Universität Marburg Fachbereich Medizin Institut für Virologie | Robert Koch Institute | unknown |
| EPI_ISL_3044335, EPI_ISL_3044337, EPI_ISL_3044338, EPI_ISL_3044339, EPI_ISL_3044340, EPI_ISL_3044341, EPI_ISL_3044342, EPI_ISL_3044343, EPI_ISL_3044345, EPI_ISL_3044346, EPI_ISL_3044347, EPI_ISL_3044348, EPI_ISL_3044349, EPI_ISL_3044350 | | | |
| see above | Medizinisch-Diagnostisches Labor Kempten allgäulab | Robert Koch Institute | unknown |
| EPI_ISL_3044351 | Procomcure Biotech Germany GmbH | Robert Koch Institute | unknown |
| EPI_ISL_3044352 | Laborarztpraxis Osnabrück | Robert Koch Institute | unknown |
| EPI_ISL_3044353 | MVZ Dr. Eberhard & Partner Dortmund | Robert Koch Institute | unknown |
| EPI_ISL_3044354 | Labor Lübeck bzw. Laborärztliche Gemeinschaftspraxis Lübeck | Robert Koch Institute | unknown |
| EPI_ISL_3044355 | Laborarztpraxis Osnabrück | Robert Koch Institute | unknown |
| EPI_ISL_3044356 | Labor Lübeck bzw. Laborärztliche Gemeinschaftspraxis Lübeck | Robert Koch Institute | unknown |
| EPI_ISL_3044357, EPI_ISL_3044358, EPI_ISL_3044359 | Laborarztpraxis Osnabrück | Robert Koch Institute | unknown |
| EPI_ISL_3044360 | MVZ Dr. Eberhard & Partner Dortmund | Robert Koch Institute | unknown |
| EPI_ISL_3044361 | Procomcure Biotech Germany GmbH | Robert Koch Institute | unknown |
| EPI_ISL_3044362, EPI_ISL_3044363 | Laborarztpraxis Osnabrück | Robert Koch Institute | unknown |
| EPI_ISL_3044364 | Labor Dr. Wisplinghoff - Köln | Robert Koch Institute | unknown |
| EPI_ISL_3044367, EPI_ISL_3044368, EPI_ISL_3044369, EPI_ISL_3044370, EPI_ISL_3044371 | Niedersächsisches Landesgesundheitsamt (NLGA) | Robert Koch Institute | unknown |
| EPI_ISL_3044373, EPI_ISL_3044374, EPI_ISL_3044375 | Corona-Testzentrum ifp Institut für Produktqualität GmbH | Robert Koch Institute | unknown |
| EPI_ISL_3044376 | MVZ Labor Dr. Quade & Kollegen GmbH | Robert Koch Institute | unknown |
| EPI_ISL_3044378, EPI_ISL_3044379 | Corona-Testzentrum ifp Institut für Produktqualität GmbH | Robert Koch Institute | unknown |
| EPI_ISL_3044380, EPI_ISL_3044381, EPI_ISL_3044382, EPI_ISL_3044383, EPI_ISL_3044384, EPI_ISL_3044385, EPI_ISL_3044386, EPI_ISL_3044387, EPI_ISL_3044389, EPI_ISL_3044390, EPI_ISL_3044391 | | | |
| see above | Eurofins MVZ Labor Gelsenkirchen | Robert Koch Institute | unknown |
| EPI_ISL_3044392 | Labor 28 MVZ GmbH | Robert Koch Institute | unknown |
| EPI_ISL_3044393 | Sonic - MVZ Medizinisches Labor Bremen GmbH | Robert Koch Institute | unknown |
| EPI_ISL_3044397 | Sonic - MVZ Medizinisches Labor Oldenburg | Robert Koch Institute | unknown |
| EPI_ISL_3044398 | Bioscientia MVZ Labor Karlsruhe GmbH | Robert Koch Institute | unknown |
| EPI_ISL_3044399, EPI_ISL_3044400 | Sonic - MVZ Medizinisches Labor Bremen GmbH | Robert Koch Institute | unknown |
| EPI_ISL_3044402 | Sonic - Labor Staber MVZ Kassel | Robert Koch Institute | unknown |
| EPI_ISL_3044403 | Labor 28 MVZ GmbH | Robert Koch Institute | unknown |
| EPI_ISL_3044404 | Bioscientia Labor Wermsdorf | Robert Koch Institute | unknown |
| EPI_ISL_3044405 | Labor 28 MVZ GmbH | Robert Koch Institute | unknown |
| EPI_ISL_3044406 | Bioscientia MVZ Labor Karlsruhe GmbH | Robert Koch Institute | unknown |
| EPI_ISL_3044407, EPI_ISL_3044408 | Bioscientia Labor Wermsdorf | Robert Koch Institute | unknown |
| EPI_ISL_3044409 | Labor 28 MVZ GmbH | Robert Koch Institute | unknown |
| EPI_ISL_3044410 | Sonic - MVZ Medizinisches Labor Bremen GmbH | Robert Koch Institute | unknown |
| EPI_ISL_3044411 | Bioscientia Labor Wermsdorf | Robert Koch Institute | unknown |
| EPI_ISL_3044412 | Labor 28 MVZ GmbH | Robert Koch Institute | unknown |
| EPI_ISL_3044413 | Sonic - MVZ Medizinisches Labor Bremen GmbH | Robert Koch Institute | unknown |
| EPI_ISL_3044416 | Bioscientia Labor Wermsdorf | Robert Koch Institute | unknown |
| EPI_ISL_3044418, EPI_ISL_3044419 | Sonic - MVZ Medizinisches Labor Bremen GmbH | Robert Koch Institute | unknown |
| EPI_ISL_3044420 | Bioscientia Labor Wermsdorf | Robert Koch Institute | unknown |
| EPI_ISL_3044422 | Bioscientia MVZ Labor Karlsruhe GmbH | Robert Koch Institute | unknown |
| EPI_ISL_3044423 | Bioscientia Labor Wermsdorf | Robert Koch Institute | unknown |
| EPI_ISL_3044424 | LabKom - Labor Mainz MVZ GmbH | Robert Koch Institute | unknown |
| EPI_ISL_3044425 | LabKom - Labor an der Salzbrücke MVZ GmbH | Robert Koch Institute | unknown |
| EPI_ISL_3044428, EPI_ISL_3044429, EPI_ISL_3044431 | Bioscientia Labor Wermsdorf | Robert Koch Institute | unknown |
| EPI_ISL_3044432 | Labor 28 MVZ GmbH | Robert Koch Institute | unknown |
| EPI_ISL_3044433, EPI_ISL_3044434 | Bioscientia Labor Wermsdorf | Robert Koch Institute | unknown |
| EPI_ISL_3044435 | Labor 28 MVZ GmbH | Robert Koch Institute | unknown |
| EPI_ISL_3044436, | Bioscientia Labor Wermsdorf | Robert Koch Institute | unknown |

| | | | |
|---|---|---|---|
| EPI_ISL_3044437 | | | |
| EPI_ISL_3044438 | Sonic - MVZ Medizinisches Labor Bremen GmbH | Robert Koch Institute | unknown |
| EPI_ISL_3044439 | Universitätsklinikum Frankfurt - Institut für Medizinische Virologie | Robert Koch Institute | unknown |
| EPI_ISL_3044440, EPI_ISL_3044441 | Sonic - MVZ Medizinisches Labor Bremen GmbH | Robert Koch Institute | unknown |
| EPI_ISL_3044442, EPI_ISL_3044443, EPI_ISL_3044444, EPI_ISL_3044445 | Bioscientia Labor Wermsdorf | Robert Koch Institute | unknown |
| EPI_ISL_3044446 | Sonic - MVZ Medizinisches Labor Bremen GmbH | Robert Koch Institute | unknown |
| EPI_ISL_3044447 | LabKom - MVZ Labor Bochum MLB GmbH | Robert Koch Institute | unknown |
| EPI_ISL_3044448, EPI_ISL_3044449, EPI_ISL_3044450 | Bioscientia Labor Wermsdorf | Robert Koch Institute | unknown |
| EPI_ISL_3044452 | Labor 28 MVZ GmbH | Robert Koch Institute | unknown |
| EPI_ISL_3044453 | Sonic - Bioscientia - MVZ Labor Saar GmbH | Robert Koch Institute | unknown |
| EPI_ISL_3044454 | LabKom - Labor Mainz MVZ GmbH | Robert Koch Institute | unknown |
| EPI_ISL_3044455 | Sonic - MVZ Medizinisches Labor Bremen GmbH | Robert Koch Institute | unknown |
| EPI_ISL_3044456 | Bioscientia Labor Wermsdorf | Robert Koch Institute | unknown |
| EPI_ISL_3044457 | Sonic - MVZ Medizinisches Labor Bremen GmbH | Robert Koch Institute | unknown |
| EPI_ISL_3044458 | Labor 28 MVZ GmbH | Robert Koch Institute | unknown |
| EPI_ISL_3044459, EPI_ISL_3044460 | Bioscientia Labor Wermsdorf | Robert Koch Institute | unknown |
| EPI_ISL_3044461, EPI_ISL_3044464, EPI_ISL_3044465, EPI_ISL_3044466, EPI_ISL_3044467, EPI_ISL_3044469, EPI_ISL_3044470, EPI_ISL_3044472, EPI_ISL_3044473, EPI_ISL_3044474, EPI_ISL_3044476, EPI_ISL_3044477, EPI_ISL_3044478, EPI_ISL_3044479, EPI_ISL_3044480, EPI_ISL_3044481, EPI_ISL_3044482, EPI_ISL_3044483, EPI_ISL_3044484 | | | |
| see above | Chemische und Veterinäruntersuchungsamt Münsterland-Emscher-Lippe (CVUA-MEL) | Robert Koch Institute | unknown |
| EPI_ISL_3044485, EPI_ISL_3044487, EPI_ISL_3044488 | Dianovis GmbH Greiz | Robert Koch Institute | unknown |
| EPI_ISL_3044492, EPI_ISL_3044495, EPI_ISL_3044496, EPI_ISL_3044497, EPI_ISL_3044498, EPI_ISL_3044499, EPI_ISL_3044500, EPI_ISL_3044501, EPI_ISL_3044502, EPI_ISL_3044503, EPI_ISL_3044504, EPI_ISL_3044506, EPI_ISL_3044507, EPI_ISL_3044508, EPI_ISL_3044509, EPI_ISL_3044510, EPI_ISL_3044511, EPI_ISL_3044512, EPI_ISL_3044513, EPI_ISL_3044514, EPI_ISL_3044515, EPI_ISL_3044516, EPI_ISL_3044517, EPI_ISL_3044518, EPI_ISL_3044519, EPI_ISL_3044520, EPI_ISL_3044521, EPI_ISL_3044522 | | | |
| see above | MVZ Labor Krone GbR | Robert Koch Institute | unknown |
| EPI_ISL_3044524 | SYNLAB MVZ Leverkusen | Robert Koch Institute | unknown |
| EPI_ISL_3044525 | SYNLAB Labor München Zentrum LMZ | Robert Koch Institute | unknown |
| EPI_ISL_3044526, EPI_ISL_3044527, EPI_ISL_3044528, EPI_ISL_3044529 | SYNLAB MVZ Leverkusen | Robert Koch Institute | unknown |
| EPI_ISL_3044530 | SYNLAB MVZ Weiden | Robert Koch Institute | unknown |
| EPI_ISL_3044531 | SYNLAB MVZ Leverkusen | Robert Koch Institute | unknown |
| EPI_ISL_3044534 | SYNLAB Labor München Zentrum LMZ | Robert Koch Institute | unknown |
| EPI_ISL_3044536 | SYNLAB MVZ Leverkusen | Robert Koch Institute | unknown |
| EPI_ISL_3044538 | Synlab MVZ Augsburg | Robert Koch Institute | unknown |
| EPI_ISL_3044539 | SYNLAB MVZ Leverkusen | Robert Koch Institute | unknown |
| EPI_ISL_3044540 | SYNLAB MVZ Weiden | Robert Koch Institute | unknown |
| EPI_ISL_3044541 | SYNLAB MVZ Leverkusen | Robert Koch Institute | unknown |
| EPI_ISL_3044542, EPI_ISL_3044543 | SYNLAB MVZ Weiden | Robert Koch Institute | unknown |
| EPI_ISL_3044544 | SYNLAB MVZ Leverkusen | Robert Koch Institute | unknown |
| EPI_ISL_3044545 | SYNLAB MVZ Weiden | Robert Koch Institute | unknown |
| EPI_ISL_3044547 | SYNLAB MVZ Trier | Robert Koch Institute | unknown |
| EPI_ISL_3044548 | SYNLAB MVZ Weiden | Robert Koch Institute | unknown |
| EPI_ISL_3044550 | Synlab MVZ Augsburg | Robert Koch Institute | unknown |
| EPI_ISL_3044551, EPI_ISL_3044553 | SYNLAB MVZ Weiden | Robert Koch Institute | unknown |
| EPI_ISL_3044555 | Synlab MVZ Augsburg | Robert Koch Institute | unknown |
| EPI_ISL_3044556 | SYNLAB MVZ Leverkusen | Robert Koch Institute | unknown |
| EPI_ISL_3044558 | SYNLAB MVZ Weiden | Robert Koch Institute | unknown |
| EPI_ISL_3044559 | SYNLAB MVZ Leverkusen | Robert Koch Institute | unknown |
| EPI_ISL_3044560 | SYNLAB MVZ Weiden | Robert Koch Institute | unknown |
| EPI_ISL_3044561 | SYNLAB Labor München Zentrum LMZ | Robert Koch Institute | unknown |
| EPI_ISL_3044563 | SYNLAB MVZ Dachau | Robert Koch Institute | unknown |
| EPI_ISL_3044564 | SYNLAB MVZ Leverkusen | Robert Koch Institute | unknown |
| EPI_ISL_3044565, EPI_ISL_3044566 | SYNLAB Labor München Zentrum LMZ | Robert Koch Institute | unknown |
| EPI_ISL_3044567, EPI_ISL_3044569 | SYNLAB MVZ Trier | Robert Koch Institute | unknown |
| EPI_ISL_3044570 | SYNLAB MVZ Weiden | Robert Koch Institute | unknown |
| EPI_ISL_3044571 | SYNLAB MVZ Leverkusen | Robert Koch Institute | unknown |
| EPI_ISL_3044572, EPI_ISL_3044573, EPI_ISL_3044574 | SYNLAB Labor München Zentrum LMZ | Robert Koch Institute | unknown |
| EPI_ISL_3044575 | SYNLAB MVZ Leverkusen | Robert Koch Institute | unknown |
| EPI_ISL_3044576 | SYNLAB MVZ Berlin | Robert Koch Institute | unknown |
| EPI_ISL_3044577 | SYNLAB MVZ Trier | Robert Koch Institute | unknown |
| EPI_ISL_3044578 | SYNLAB Labor München Zentrum LMZ | Robert Koch Institute | unknown |
| EPI_ISL_3044579, EPI_ISL_3044581 | SYNLAB MVZ Leverkusen | Robert Koch Institute | unknown |
| EPI_ISL_3044582 | SYNLAB MVZ Berlin | Robert Koch Institute | unknown |
| EPI_ISL_3044583 | Synlab MVZ Augsburg | Robert Koch Institute | unknown |
| EPI_ISL_3044584, EPI_ISL_3044585 | SYNLAB MVZ Leverkusen | Robert Koch Institute | unknown |

| | | | |
|---|---|---|---|
| EPI_ISL_3044586, EPI_ISL_3044587 | SYNLAB MVZ Weiden | Robert Koch Institute | unknown |
| EPI_ISL_3044588 | SYNLAB MVZ Trier | Robert Koch Institute | unknown |
| EPI_ISL_3044589 | SYNLAB MVZ Weiden | Robert Koch Institute | unknown |
| EPI_ISL_3044590 | SYNLAB MVZ Leverkusen | Robert Koch Institute | unknown |
| EPI_ISL_3044591, EPI_ISL_3044593, EPI_ISL_3044594 | SYNLAB MVZ Weiden | Robert Koch Institute | unknown |
| EPI_ISL_3044597 | SYNLAB MVZ Trier | Robert Koch Institute | unknown |
| EPI_ISL_3044598, EPI_ISL_3044599, EPI_ISL_3044600 | SYNLAB MVZ Leverkusen | Robert Koch Institute | unknown |
| EPI_ISL_3044601 | Synlab MVZ Augsburg | Robert Koch Institute | unknown |
| EPI_ISL_3044603 | SYNLAB MVZ Weiden | Robert Koch Institute | unknown |
| EPI_ISL_3044604 | SYNLAB MVZ Leverkusen | Robert Koch Institute | unknown |
| EPI_ISL_3044605 | Synlab MVZ Augsburg | Robert Koch Institute | unknown |
| EPI_ISL_3044606 | SYNLAB MVZ Weiden | Robert Koch Institute | unknown |
| EPI_ISL_3044609 | SYNLAB MVZ Trier | Robert Koch Institute | unknown |
| EPI_ISL_3044610 | SYNLAB MVZ Weiden | Robert Koch Institute | unknown |
| EPI_ISL_3044611, EPI_ISL_3044612, EPI_ISL_3044613 | SYNLAB MVZ Leverkusen | Robert Koch Institute | unknown |
| EPI_ISL_3044614, EPI_ISL_3044615 | Synlab MVZ Augsburg | Robert Koch Institute | unknown |
| EPI_ISL_3044616 | Praxisgemeinschaft für Laboratoriumsmedizin Labor Blumenstraße; Praxis Dr. Sel und Dr. Späte | Robert Koch Institute | unknown |
| EPI_ISL_3044617, EPI_ISL_3044618 | SYNLAB MVZ Weiden | Robert Koch Institute | unknown |
| EPI_ISL_3044619 | SYNLAB MVZ Leverkusen | Robert Koch Institute | unknown |
| EPI_ISL_3044622, EPI_ISL_3044623 | SYNLAB MVZ Weiden | Robert Koch Institute | unknown |
| EPI_ISL_3044625, EPI_ISL_3044626 | SYNLAB MVZ Leverkusen | Robert Koch Institute | unknown |
| EPI_ISL_3044627, EPI_ISL_3044628 | SYNLAB MVZ Weiden | Robert Koch Institute | unknown |
| EPI_ISL_3044629 | Synlab MVZ Augsburg | Robert Koch Institute | unknown |
| EPI_ISL_3044630, EPI_ISL_3044631 | SYNLAB MVZ Weiden | Robert Koch Institute | unknown |
| EPI_ISL_3044632 | SYNLAB MVZ Leverkusen | Robert Koch Institute | unknown |
| EPI_ISL_3044633 | SYNLAB MVZ Trier | Robert Koch Institute | unknown |
| EPI_ISL_3044634, EPI_ISL_3044635, EPI_ISL_3044636 | SYNLAB MVZ Leverkusen | Robert Koch Institute | unknown |
| EPI_ISL_3044637 | SYNLAB MVZ Trier | Robert Koch Institute | unknown |
| EPI_ISL_3044638, EPI_ISL_3044639 | SYNLAB MVZ Leverkusen | Robert Koch Institute | unknown |
| EPI_ISL_3044640 | SYNLAB MVZ Weiden | Robert Koch Institute | unknown |
| EPI_ISL_3044641 | Synlab MVZ Augsburg | Robert Koch Institute | unknown |
| EPI_ISL_3044642 | SYNLAB MVZ Hamburg | Robert Koch Institute | unknown |
| EPI_ISL_3044643, EPI_ISL_3044644 | Synlab MVZ Augsburg | Robert Koch Institute | unknown |
| EPI_ISL_3044645 | SYNLAB MVZ Hamburg | Robert Koch Institute | unknown |
| EPI_ISL_3044646 | SYNLAB MVZ Leverkusen | Robert Koch Institute | unknown |
| EPI_ISL_3044647, EPI_ISL_3044648, EPI_ISL_3044649, EPI_ISL_3044650 | SYNLAB MVZ Weiden | Robert Koch Institute | unknown |
| EPI_ISL_3044651 | SYNLAB MVZ Leverkusen | Robert Koch Institute | unknown |
| EPI_ISL_3044652, EPI_ISL_3044653 | SYNLAB MVZ Weiden | Robert Koch Institute | unknown |
| EPI_ISL_3044654 | SYNLAB MVZ Leverkusen | Robert Koch Institute | unknown |
| EPI_ISL_3044655 | SYNLAB MVZ Weiden | Robert Koch Institute | unknown |
| EPI_ISL_3044656 | SYNLAB MVZ Leverkusen | Robert Koch Institute | unknown |
| EPI_ISL_3044657 | SYNLAB MVZ Trier | Robert Koch Institute | unknown |
| EPI_ISL_3044658, EPI_ISL_3044659, EPI_ISL_3044660 | SYNLAB MVZ Leverkusen | Robert Koch Institute | unknown |
| EPI_ISL_3044661 | Synlab MVZ Augsburg | Robert Koch Institute | unknown |
| EPI_ISL_3044662 | SYNLAB MVZ Weiden | Robert Koch Institute | unknown |
| EPI_ISL_3044663, EPI_ISL_3044666 | SYNLAB MVZ Leverkusen | Robert Koch Institute | unknown |
| EPI_ISL_3044668 | SYNLAB MVZ Hamburg | Robert Koch Institute | unknown |
| EPI_ISL_3044669 | SYNLAB MVZ Leverkusen | Robert Koch Institute | unknown |
| EPI_ISL_3044670 | SYNLAB Labor München Zentrum LMZ | Robert Koch Institute | unknown |
| EPI_ISL_3044673, EPI_ISL_3044675, EPI_ISL_3044676 | SYNLAB MVZ Weiden | Robert Koch Institute | unknown |
| EPI_ISL_3044677 | SYNLAB MVZ Leverkusen | Robert Koch Institute | unknown |
| EPI_ISL_3044678 | SYNLAB MVZ Weiden | Robert Koch Institute | unknown |
| EPI_ISL_3044679 | Synlab MVZ Augsburg | Robert Koch Institute | unknown |
| EPI_ISL_3044680 | SYNLAB MVZ Weiden | Robert Koch Institute | unknown |
| EPI_ISL_3044681 | SYNLAB MVZ Leverkusen | Robert Koch Institute | unknown |
| EPI_ISL_3044682 | SYNLAB Labor München Zentrum LMZ | Robert Koch Institute | unknown |
| EPI_ISL_3044683 | SYNLAB MVZ Weiden | Robert Koch Institute | unknown |

| | | | |
|---|---|---|---|
| EPI_ISL_3044685 | SYNLAB Labor München Zentrum LMZ | Robert Koch Institute | unknown |
| EPI_ISL_3044686 | SYNLAB MVZ Leverkusen | Robert Koch Institute | unknown |
| EPI_ISL_3044687 | Synlab MVZ Augsburg | Robert Koch Institute | unknown |
| EPI_ISL_3044688, EPI_ISL_3044691 | SYNLAB MVZ Leverkusen | Robert Koch Institute | unknown |
| EPI_ISL_3044693 | SYNLAB Jena Oncoscreen | Robert Koch Institute | unknown |
| EPI_ISL_3044694 | SYNLAB MVZ Leverkusen | Robert Koch Institute | unknown |
| EPI_ISL_3044695 | Synlab MVZ Augsburg | Robert Koch Institute | unknown |
| EPI_ISL_3044696 | SYNLAB MVZ Leverkusen | Robert Koch Institute | unknown |
| EPI_ISL_3044697 | SYNLAB MVZ Weiden | Robert Koch Institute | unknown |
| EPI_ISL_3044698 | SYNLAB MVZ Leverkusen | Robert Koch Institute | unknown |
| EPI_ISL_3044701 | SYNLAB MVZ Trier | Robert Koch Institute | unknown |
| EPI_ISL_3044704 | SYNLAB MVZ Leverkusen | Robert Koch Institute | unknown |
| EPI_ISL_3044705, EPI_ISL_3044706 | SYNLAB MVZ Weiden | Robert Koch Institute | unknown |
| EPI_ISL_3044707 | SYNLAB Labor München Zentrum LMZ | Robert Koch Institute | unknown |
| EPI_ISL_3044708 | SYNLAB MVZ Leverkusen | Robert Koch Institute | unknown |
| EPI_ISL_3044709 | SYNLAB MVZ Trier | Robert Koch Institute | unknown |
| EPI_ISL_3044710 | SYNLAB MVZ Hamburg | Robert Koch Institute | unknown |
| EPI_ISL_3044711, EPI_ISL_3044712, EPI_ISL_3044713 | SYNLAB MVZ Weiden | Robert Koch Institute | unknown |
| EPI_ISL_3044714 | SYNLAB MVZ Trier | Robert Koch Institute | unknown |
| EPI_ISL_3044715, EPI_ISL_3044716, EPI_ISL_3044717 | SYNLAB MVZ Leverkusen | Robert Koch Institute | unknown |
| EPI_ISL_3044718 | SYNLAB MVZ Weiden | Robert Koch Institute | unknown |
| EPI_ISL_3044719 | SYNLAB MVZ Hamburg | Robert Koch Institute | unknown |
| EPI_ISL_3044720, EPI_ISL_3044721 | SYNLAB MVZ Leverkusen | Robert Koch Institute | unknown |
| EPI_ISL_3044722 | Synlab MVZ Augsburg | Robert Koch Institute | unknown |
| EPI_ISL_3044724, EPI_ISL_3044725 | SYNLAB MVZ Weiden | Robert Koch Institute | unknown |
| EPI_ISL_3044726, EPI_ISL_3044727 | SYNLAB Labor München Zentrum LMZ | Robert Koch Institute | unknown |
| EPI_ISL_3044728 | Synlab MVZ Augsburg | Robert Koch Institute | unknown |
| EPI_ISL_3044729 | SYNLAB MVZ Trier | Robert Koch Institute | unknown |
| EPI_ISL_3044730 | Synlab MVZ Augsburg | Robert Koch Institute | unknown |
| EPI_ISL_3044731, EPI_ISL_3044733 | SYNLAB MVZ Weiden | Robert Koch Institute | unknown |
| EPI_ISL_3044734 | SYNLAB MVZ Leverkusen | Robert Koch Institute | unknown |
| EPI_ISL_3044735 | SYNLAB MVZ Weiden | Robert Koch Institute | unknown |
| EPI_ISL_3044736 | SYNLAB MVZ Leverkusen | Robert Koch Institute | unknown |
| EPI_ISL_3044737 | Synlab MVZ Augsburg | Robert Koch Institute | unknown |
| EPI_ISL_3044739 | SYNLAB Labor München Zentrum LMZ | Robert Koch Institute | unknown |
| EPI_ISL_3044740 | SYNLAB MVZ Leverkusen | Robert Koch Institute | unknown |
| EPI_ISL_3044741 | SYNLAB Labor München Zentrum LMZ | Robert Koch Institute | unknown |
| EPI_ISL_3044742 | SYNLAB MVZ Leverkusen | Robert Koch Institute | unknown |
| EPI_ISL_3044743 | SYNLAB MVZ Trier | Robert Koch Institute | unknown |
| EPI_ISL_3044744 | SYNLAB MVZ Leverkusen | Robert Koch Institute | unknown |
| EPI_ISL_3044745 | SYNLAB MVZ Trier | Robert Koch Institute | unknown |
| EPI_ISL_3044746 | SYNLAB Labor München Zentrum LMZ | Robert Koch Institute | unknown |
| EPI_ISL_3044747 | SYNLAB MVZ Trier | Robert Koch Institute | unknown |
| EPI_ISL_3044749 | SYNLAB MVZ Leverkusen | Robert Koch Institute | unknown |
| EPI_ISL_3044750 | SYNLAB MVZ Trier | Robert Koch Institute | unknown |
| EPI_ISL_3044751, EPI_ISL_3044752 | SYNLAB MVZ Leverkusen | Robert Koch Institute | unknown |
| EPI_ISL_3044753 | SYNLAB MVZ Weiden | Robert Koch Institute | unknown |
| EPI_ISL_3044754 | SYNLAB MVZ Leverkusen | Robert Koch Institute | unknown |
| EPI_ISL_3044757 | SYNLAB MVZ Weiden | Robert Koch Institute | unknown |
| EPI_ISL_3044758 | SYNLAB Labor München Zentrum LMZ | Robert Koch Institute | unknown |
| EPI_ISL_3044759, EPI_ISL_3044760, EPI_ISL_3044761 | Uniklinikum Jena | Robert Koch Institute | unknown |
| EPI_ISL_3044762 | IMD - MVZ Labor Greifswald | Robert Koch Institute | unknown |
| EPI_ISL_3044763 | IMD - Institut für Medizinische Diagnostik Berlin-Potsdam | Robert Koch Institute | unknown |
| EPI_ISL_3044764, EPI_ISL_3044765 | IMD - Labor Oderland | Robert Koch Institute | unknown |
| EPI_ISL_3044766, EPI_ISL_3044767 | IMD - MVZ Labor Martinsried | Robert Koch Institute | unknown |
| EPI_ISL_3044768 | IMD - Institut für Medizinische Diagnostik Berlin-Potsdam | Robert Koch Institute | unknown |
| EPI_ISL_3044769 | IMD - Labor Oderland | Robert Koch Institute | unknown |
| EPI_ISL_3044770 | IMD - Medizinisches Labor Rostock | Robert Koch Institute | unknown |
| EPI_ISL_3044772, EPI_ISL_3044774, EPI_ISL_3044775, EPI_ISL_3044776, EPI_ISL_3044777, EPI_ISL_3044778, EPI_ISL_3044780, EPI_ISL_3044781, EPI_ISL_3044782, EPI_ISL_3044783 | Labor Becker & Kollegen (Standort München) | Robert Koch Institute | unknown |

| | | | |
|---|---|---|---|
| EPI_ISL_3044784, EPI_ISL_3044785, EPI_ISL_3044786, EPI_ISL_3044787, EPI_ISL_3044788, EPI_ISL_3044789, EPI_ISL_3044790, EPI_ISL_3044791 | Dianovis GmbH Greiz | Robert Koch Institute | unknown |
| EPI_ISL_3044792, EPI_ISL_3044794, EPI_ISL_3044795, EPI_ISL_3044796, EPI_ISL_3044797, EPI_ISL_3044798, EPI_ISL_3044799, EPI_ISL_3044800, EPI_ISL_3044801, EPI_ISL_3044802 | MVZ Labor Dr. Fenner und Kollegen (Standort Hamburg) | Robert Koch Institute | unknown |
| EPI_ISL_3044803, EPI_ISL_3044804, EPI_ISL_3044805, EPI_ISL_3044806, EPI_ISL_3044807, EPI_ISL_3044808, EPI_ISL_3044809, EPI_ISL_3044810, EPI_ISL_3044811, EPI_ISL_3044812, EPI_ISL_3044813, EPI_ISL_3044814, EPI_ISL_3044815, EPI_ISL_3044816, EPI_ISL_3044817, EPI_ISL_3044818, EPI_ISL_3044819, EPI_ISL_3044820, EPI_ISL_3044821, EPI_ISL_3044822, EPI_ISL_3044823, EPI_ISL_3044824, EPI_ISL_3044826, EPI_ISL_3044827, EPI_ISL_3044828, EPI_ISL_3044829, EPI_ISL_3044830, EPI_ISL_3044831, EPI_ISL_3044832, EPI_ISL_3044833, EPI_ISL_3044834, EPI_ISL_3044835, EPI_ISL_3044836, EPI_ISL_3044837, EPI_ISL_3044838, EPI_ISL_3044839, EPI_ISL_3044840, EPI_ISL_3044841, EPI_ISL_3044842, EPI_ISL_3044844, EPI_ISL_3044846, EPI_ISL_3044847, EPI_ISL_3044848, EPI_ISL_3044849, EPI_ISL_3044850, EPI_ISL_3044851, EPI_ISL_3044852, EPI_ISL_3044853, EPI_ISL_3044854, EPI_ISL_3044855, EPI_ISL_3044856, EPI_ISL_3044857, EPI_ISL_3044858, EPI_ISL_3044860, EPI_ISL_3044861, EPI_ISL_3044862, EPI_ISL_3044863, EPI_ISL_3044864, EPI_ISL_3044865, EPI_ISL_3044866, EPI_ISL_3044868, EPI_ISL_3044869, EPI_ISL_3044870, EPI_ISL_3044871, EPI_ISL_3044872, EPI_ISL_3044873, EPI_ISL_3044874, EPI_ISL_3044875, EPI_ISL_3044876, EPI_ISL_3044877, EPI_ISL_3044878 | | | |
| see above | Eurofins LifeCodexx GmbH | Robert Koch Institute | unknown |
| EPI_ISL_3044879, EPI_ISL_3044880, EPI_ISL_3044881, EPI_ISL_3044882, EPI_ISL_3044883, EPI_ISL_3044884, EPI_ISL_3044886, EPI_ISL_3044887, EPI_ISL_3044888, EPI_ISL_3044889, EPI_ISL_3044890, EPI_ISL_3044891, EPI_ISL_3044892, EPI_ISL_3044893, EPI_ISL_3044894, EPI_ISL_3044895, EPI_ISL_3044896, EPI_ISL_3044897, EPI_ISL_3044898, EPI_ISL_3044899, EPI_ISL_3044900, EPI_ISL_3044901, EPI_ISL_3044902, EPI_ISL_3044903, EPI_ISL_3044904 | | | |
| see above | Labor Dr. Wisplinghoff - Köln | Robert Koch Institute | unknown |
| EPI_ISL_3044905, EPI_ISL_3044906 | Landesamt für Verbraucherschutz des Landes Sachsen-Anhalt | Robert Koch Institute | unknown |
| EPI_ISL_3044907, EPI_ISL_3044908, EPI_ISL_3044909, EPI_ISL_3044910, EPI_ISL_3044912, EPI_ISL_3044914, EPI_ISL_3044916, EPI_ISL_3044917, EPI_ISL_3044918 | Bayerisches Landesamt für Gesundheit und Lebensmittelsicherheit (LGL) | Robert Koch Institute | unknown |
| EPI_ISL_3044919 | Universitätsklinikum Essen | Robert Koch Institute | unknown |
| EPI_ISL_3044920, EPI_ISL_3044921, EPI_ISL_3044922, EPI_ISL_3044923, EPI_ISL_3044924, EPI_ISL_3044926, EPI_ISL_3044927, EPI_ISL_3044928, EPI_ISL_3044929, EPI_ISL_3044930, EPI_ISL_3044931, EPI_ISL_3044932, EPI_ISL_3044933, EPI_ISL_3044934, EPI_ISL_3044935, EPI_ISL_3044936 | | | |
| see above | Landesgesundheitsamt Baden-Württemberg | Robert Koch Institute | unknown |
| EPI_ISL_3044937 | San Camillo De Lellis - Rieti | National Institute for Infectious Diseases (INMI) L. Spallanzani I.R.C.C.S | CEM Gruber, F Messina, F Santini, M Rueca, G Bonfiglio, O Butera, B Bartolini, E Giombini, MR Capobianchi, A Di Caro |
| EPI_ISL_3044938, EPI_ISL_3044939, EPI_ISL_3044940, EPI_ISL_3044941, EPI_ISL_3044942, EPI_ISL_3044944, EPI_ISL_3044945 | Landesgesundheitsamt Baden-Württemberg | Robert Koch Institute | unknown |
| EPI_ISL_3044947, EPI_ISL_3044948 | Uniklinikum Carl Gustav Carus an der TU Dresden; Institut für Virologie | Robert Koch Institute | unknown |
| EPI_ISL_3044952 | Landesuntersuchungsanstalt für das Gesundheits- und Veterinärwesen (LUA) Sachsen | Robert Koch Institute | unknown |
| EPI_ISL_3044953 | Uniklinikum Carl Gustav Carus an der TU Dresden; Institut für Virologie | Robert Koch Institute | unknown |
| EPI_ISL_3044954 | Medizinisches Labor Ostsachsen MVZ GbR Nebenbetriebsstätte Dresden | Robert Koch Institute | unknown |
| EPI_ISL_3044956 | Landesuntersuchungsanstalt für das Gesundheits- und Veterinärwesen (LUA) Sachsen | Robert Koch Institute | unknown |
| EPI_ISL_3044958 | EKA Erzgebirgsklinikum Annaberg gGmbH | Robert Koch Institute | unknown |
| EPI_ISL_3044959 | Landesuntersuchungsanstalt für das Gesundheits- und Veterinärwesen (LUA) Sachsen | Robert Koch Institute | unknown |
| EPI_ISL_3044960, EPI_ISL_3044961 | Uniklinikum Carl Gustav Carus an der TU Dresden; Institut für Virologie | Robert Koch Institute | unknown |
| EPI_ISL_3044962 | EKA Erzgebirgsklinikum Annaberg gGmbH | Robert Koch Institute | unknown |
| EPI_ISL_3044964 | Medizinisches Labor Ostsachsen MVZ GbR Nebenbetriebsstätte Dresden | Robert Koch Institute | unknown |
| EPI_ISL_3044966, EPI_ISL_3044967, EPI_ISL_3044970 | Landesuntersuchungsanstalt für das Gesundheits- und Veterinärwesen (LUA) Sachsen | Robert Koch Institute | unknown |
| EPI_ISL_3044973 | Fraunhofer-Institut für Zelltherapie und Immunologie IZI AG Next-Generation Diagnostics | Robert Koch Institute | unknown |
| EPI_ISL_3044975, EPI_ISL_3044976, EPI_ISL_3044977, EPI_ISL_3044980, EPI_ISL_3044981, EPI_ISL_3044982, EPI_ISL_3044983, EPI_ISL_3044984, EPI_ISL_3044985, EPI_ISL_3044987, EPI_ISL_3044988, EPI_ISL_3044990, EPI_ISL_3044991, EPI_ISL_3044993, EPI_ISL_3044994, EPI_ISL_3044995, EPI_ISL_3044996 | | | |
| see above | Labor Mönchengladbach MVZ Dr. Stein + Kollegen GbR | Robert Koch Institute | unknown |
| EPI_ISL_3044997 | MDI Limbach Berlin GmbH; MVZ Labor Berlin | Robert Koch Institute | unknown |
| EPI_ISL_3044998 | Labor Mönchengladbach MVZ Dr. Stein + Kollegen GbR | Robert Koch Institute | unknown |
| EPI_ISL_3045000, EPI_ISL_3045001, EPI_ISL_3045002, EPI_ISL_3045003, EPI_ISL_3045004 | MDI Limbach Berlin GmbH; MVZ Labor Berlin | Robert Koch Institute | unknown |
| EPI_ISL_3045005 | MVZ für Laboratoriumsmedizin und Mikrobiologie Koblenz-Mittelrhein (Labor Koblenz) | Robert Koch Institute | unknown |
| EPI_ISL_3045006 | MDI Limbach Berlin GmbH; MVZ Labor Berlin | Robert Koch Institute | unknown |
| EPI_ISL_3045007 | MVZ Medizinisches Labor Hannover GmbH | Robert Koch Institute | unknown |
| EPI_ISL_3045008, EPI_ISL_3045009 | MDI Limbach Berlin GmbH; MVZ Labor Berlin | Robert Koch Institute | unknown |
| EPI_ISL_3045010 | MVZ Labor Dr. Reising-Ackermann und Kollegen | Robert Koch Institute | unknown |
| EPI_ISL_3045011, EPI_ISL_3045012, EPI_ISL_3045013, EPI_ISL_3045014, EPI_ISL_3045015 | MDI Limbach Berlin GmbH; MVZ Labor Berlin | Robert Koch Institute | unknown |
| EPI_ISL_3045016 | MVZ Medizinisches Labor Hannover GmbH | Robert Koch Institute | unknown |

| | | | |
|---|---|---|---|
| EPI_ISL_3045017 | Limbach - MVZ Gemeinschaftslabor Suhl | Robert Koch Institute | unknown |
| EPI_ISL_3045018, EPI_ISL_3045019, EPI_ISL_3045020 | MVZ Medizinisches Labor Hannover GmbH | Robert Koch Institute | unknown |
| EPI_ISL_3045021 | MDI Limbach Berlin GmbH; MVZ Labor Berlin | Robert Koch Institute | unknown |
| EPI_ISL_3045022 | MVZ Medizinisches Labor Hannover GmbH | Robert Koch Institute | unknown |
| EPI_ISL_3045023 | MDI Limbach Berlin GmbH; MVZ Labor Berlin | Robert Koch Institute | unknown |
| EPI_ISL_3045024 | Limbach - MVZ Gemeinschaftslabor Suhl | Robert Koch Institute | unknown |
| EPI_ISL_3045025, EPI_ISL_3045026 | MDI Limbach Berlin GmbH; MVZ Labor Berlin | Robert Koch Institute | unknown |
| EPI_ISL_3045027 | MVZ Medizinisches Labor Hannover GmbH | Robert Koch Institute | unknown |
| EPI_ISL_3045028 | MDI Limbach Berlin GmbH; MVZ Labor Berlin | Robert Koch Institute | unknown |
| EPI_ISL_3045029, EPI_ISL_3045030 | MVZ Medizinisches Labor Hannover GmbH | Robert Koch Institute | unknown |
| EPI_ISL_3045031 | MDI Limbach Berlin GmbH; MVZ Labor Berlin | Robert Koch Institute | unknown |
| EPI_ISL_3045032 | MVZ Medizinisches Labor Hannover GmbH | Robert Koch Institute | unknown |
| EPI_ISL_3045033 | MDI Limbach Berlin GmbH; MVZ Labor Berlin | Robert Koch Institute | unknown |
| EPI_ISL_3045034 | Limbach - MVZ Gemeinschaftslabor Suhl | Robert Koch Institute | unknown |
| EPI_ISL_3045035, EPI_ISL_3045036, EPI_ISL_3045037 | MVZ Medizinisches Labor Hannover GmbH | Robert Koch Institute | unknown |
| EPI_ISL_3045038, EPI_ISL_3045040 | MDI Limbach Berlin GmbH; MVZ Labor Berlin | Robert Koch Institute | unknown |
| EPI_ISL_3045041, EPI_ISL_3045042, EPI_ISL_3045043, EPI_ISL_3045044 | MVZ Medizinisches Labor Hannover GmbH | Robert Koch Institute | unknown |
| EPI_ISL_3045045 | Labor Prof. Dr. G. Enders MVZ GbR | Robert Koch Institute | unknown |
| EPI_ISL_3045046 | MDI Limbach Berlin GmbH; MVZ Labor Berlin | Robert Koch Institute | unknown |
| EPI_ISL_3045047 | MVZ Medizinisches Labor Hannover GmbH | Robert Koch Institute | unknown |
| EPI_ISL_3045048 | MDI Limbach Berlin GmbH; MVZ Labor Berlin | Robert Koch Institute | unknown |
| EPI_ISL_3045049 | MVZ Medizinisches Labor Hannover GmbH | Robert Koch Institute | unknown |
| EPI_ISL_3045050 | Labor Prof. Dr. G. Enders MVZ GbR | Robert Koch Institute | unknown |
| EPI_ISL_3045052 | MVZ Medizinisches Labor Hannover GmbH | Robert Koch Institute | unknown |
| EPI_ISL_3045053 | Labor Prof. Dr. G. Enders MVZ GbR | Robert Koch Institute | unknown |
| EPI_ISL_3045054 | MDI Limbach Berlin GmbH; MVZ Labor Berlin | Robert Koch Institute | unknown |
| EPI_ISL_3045063, EPI_ISL_3045065, EPI_ISL_3045066, EPI_ISL_3045067, EPI_ISL_3045069, EPI_ISL_3045070, EPI_ISL_3045071, EPI_ISL_3045072, EPI_ISL_3045074, EPI_ISL_3045075, EPI_ISL_3045078, EPI_ISL_3045079, EPI_ISL_3045080, EPI_ISL_3045081, EPI_ISL_3045082, EPI_ISL_3045083, EPI_ISL_3045084, EPI_ISL_3045089, EPI_ISL_3045091, EPI_ISL_3045092, EPI_ISL_3045093, EPI_ISL_3045094, EPI_ISL_3045096, EPI_ISL_3045097, EPI_ISL_3045098, EPI_ISL_3045100, EPI_ISL_3045102, EPI_ISL_3045103 | | | |
| see above | New Mexico Department of Health Scientific Laboratory | New Mexico Department of Health Scientific Laboratory | Ellie Johnson, D'eldra Malone, Jennifer Benoit, Ratheesh Rajan, Linda Salazar, Anastacia Griego-Fisher |
| EPI_ISL_3045104 | Laboratorio de salud publica Puebla | LABOPAT | Luis Mendoza, Cynthia Penaloza, Silvia Montilla |
| EPI_ISL_3045105 | A. Krumbholz, Labor Dr. Krause und Kollegen MVZ GmbH, Kiel | Charité Universitätsmedizin Berlin, Institute of Virology | Victor M Corman, Barbara Mühlemann, Julia Schneider, Jörn Beheim-Schwarzbach, Tobias Bleicker, Julia Tesch, Talitha Veith, Andi Krumbholz, Terry Jones, Christian Drosten |
| EPI_ISL_3045110, EPI_ISL_3045111, EPI_ISL_3045112 | CHU Pontchaillou | CHU Pontchaillou | GROLHIER Claire, DENOUAL Florent, ETCHEVERRY Amandine, SASSI Mohamed, JAGLINE Steven, FEBREAU Christine, PRONIER Charlotte, GALIBERT Marie Dominique, DE TAYRAC Marie, THIBAULT Vincent |
| EPI_ISL_3045136, EPI_ISL_3045211, EPI_ISL_3045246, EPI_ISL_3045272, EPI_ISL_3045273, EPI_ISL_3045274, EPI_ISL_3045275 | Central Public Health Laboratory, National Public Health Organization | Central Public Health Laboratory, National Public Health Organization | Kyriaki Tryfinopoulou, G.Spanakos, A.Katsoulidou et al |
| EPI_ISL_3045290, EPI_ISL_3045291, EPI_ISL_3045292, EPI_ISL_3045293, EPI_ISL_3045294, EPI_ISL_3045295, EPI_ISL_3045296, EPI_ISL_3045297, EPI_ISL_3045298, EPI_ISL_3045299, EPI_ISL_3045300, EPI_ISL_3045301 | | | |
| see above | NCSLPH | NCSLPH | Chase K, Miller MC, Greene S, Glover W |
| EPI_ISL_3045306, EPI_ISL_3045307, EPI_ISL_3045308, EPI_ISL_3045309, EPI_ISL_3045310, EPI_ISL_3045313, EPI_ISL_3045314, EPI_ISL_3045315, EPI_ISL_3045316, EPI_ISL_3045317, EPI_ISL_3045318, EPI_ISL_3045319, EPI_ISL_3045320, EPI_ISL_3045321, EPI_ISL_3045322, EPI_ISL_3045323, EPI_ISL_3045324, EPI_ISL_3045325, EPI_ISL_3045326 | | | |
| see above | Institute for Infectious Diseases | Institute for Infectious Diseases | Stefan Neuenschwander, Christian Baumann, Miguel A Terrazos Miani, Cora SÃ¤gesser, Pascal Bittel, Peter Keller, Franziska Suter-Riniker, Stephen L Leib, Alban Ramette |
| EPI_ISL_3045327 | UniversitätsSpital Zürich | Institute of Medical Virology | Verena Kufner, Gabriela Ziltener, Maryam Zaheri, Stefan Schmutz, Annette Audigé, Maria Grünberg, Kevin Steiner, Jon Huder, Cyril Shah, Riccarda Capaul, Guido Bloemberg, Jürg Böni, Michael Huber, Alexandra Trkola |
| EPI_ISL_3045328 | Spital Männedorf AG | Institute of Medical Virology | Verena Kufner, Gabriela Ziltener, Maryam Zaheri, Stefan Schmutz, Annette Audigé, Maria Grünberg, Kevin Steiner, Jon Huder, Cyril Shah, Riccarda Capaul, Guido Bloemberg, Jürg Böni, Michael Huber, Alexandra Trkola |
| EPI_ISL_3045329, EPI_ISL_3045330 | UniversitätsSpital Zürich | Institute of Medical Virology | Verena Kufner, Gabriela Ziltener, Maryam Zaheri, Stefan Schmutz, Annette Audigé, Maria Grünberg, Kevin Steiner, Jon Huder, Cyril Shah, Riccarda Capaul, Guido Bloemberg, Jürg Böni, Michael Huber, Alexandra Trkola |
| EPI_ISL_3045331 | Kantonsspital Baden AG | Institute of Medical Virology | Verena Kufner, Gabriela Ziltener, Maryam Zaheri, Stefan Schmutz, Annette Audigé, Maria Grünberg, Kevin Steiner, Jon Huder, Cyril Shah, Riccarda Capaul, Guido Bloemberg, Jürg Böni, Michael Huber, Alexandra Trkola |
| EPI_ISL_3045332 | Medica | Institute of Medical Virology | Verena Kufner, Gabriela Ziltener, Maryam Zaheri, Stefan Schmutz, Annette Audigé, Maria Grünberg, Kevin Steiner, Jon Huder, Cyril Shah, Riccarda Capaul, Guido Bloemberg, Jürg Böni, Michael Huber, Alexandra Trkola |
| EPI_ISL_3045333 | Invenimus AG | Institute of Medical Virology | Verena Kufner, Gabriela Ziltener, Maryam Zaheri, Stefan Schmutz, Annette Audigé, Maria Grünberg, Kevin Steiner, Jon Huder, Cyril Shah, Riccarda Capaul, Guido Bloemberg, Jürg Böni, Michael Huber, Alexandra Trkola |
| EPI_ISL_3045334 | Analytica | Institute of Medical Virology | Verena Kufner, Gabriela Ziltener, Maryam Zaheri, Stefan Schmutz, Annette Audigé, Maria Grünberg, Kevin Steiner, Jon Huder, Cyril Shah, Riccarda Capaul, Guido Bloemberg, Jürg Böni, Michael Huber, Alexandra Trkola |
| EPI_ISL_3045335 | Gesundheitsdirektion Zürich | Institute of Medical Virology | Verena Kufner, Gabriela Ziltener, Maryam Zaheri, Stefan Schmutz, Annette Audigé, Maria Grünberg, Kevin Steiner, Jon Huder, Cyril Shah, Riccarda Capaul, Guido Bloemberg, Jürg Böni, Michael Huber, Alexandra Trkola |
| EPI_ISL_3045336 | Zentrallabor Zürich | Institute of Medical Virology | Verena Kufner, Gabriela Ziltener, Maryam Zaheri, Stefan Schmutz, Annette Audigé, Maria Grünberg, Kevin Steiner, Jon Huder, Cyril Shah, Riccarda Capaul, Guido Bloemberg, Jürg Böni, Michael Huber, Alexandra Trkola |
| EPI_ISL_3045337, EPI_ISL_3045338 | Medica | Institute of Medical Virology | Verena Kufner, Gabriela Ziltener, Maryam Zaheri, Stefan Schmutz, Annette Audigé, Maria Grünberg, Kevin Steiner, Jon Huder, Cyril Shah, Riccarda Capaul, Guido Bloemberg, Jürg Böni, Michael Huber, Alexandra Trkola |
| EPI_ISL_3045339 | Unilabs | Institute of Medical Virology | Verena Kufner, Gabriela Ziltener, Maryam Zaheri, Stefan Schmutz, Annette Audigé, Maria Grünberg, Kevin Steiner, Jon Huder, Cyril Shah, Riccarda Capaul, Guido Bloemberg, Jürg Böni, Michael Huber, Alexandra Trkola |
| EPI_ISL_3045340 | GZO Spital Wetzikon | Institute of Medical Virology | Verena Kufner, Gabriela Ziltener, Maryam Zaheri, Stefan Schmutz, Annette Audigé, Maria Grünberg, Kevin Steiner, Jon Huder, Cyril Shah, Riccarda Capaul, Guido Bloemberg, Jürg Böni, Michael Huber, Alexandra Trkola |
| EPI_ISL_3045341 | Analytica | Institute of Medical Virology | Verena Kufner, Gabriela Ziltener, Maryam Zaheri, Stefan Schmutz, Annette Audigé, Maria Grünberg, Kevin Steiner, Jon Huder, Cyril Shah, Riccarda Capaul, Guido Bloemberg, Jürg Böni, Michael Huber, Alexandra Trkola |
| EPI_ISL_3045342 | Kantonsspital Schaffhausen | Institute of Medical Virology | Verena Kufner, Gabriela Ziltener, Maryam Zaheri, Stefan Schmutz, Annette Audigé, Maria Grünberg, Kevin Steiner, Jon Huder, Cyril Shah, Riccarda Capaul, Guido Bloemberg, Jürg Böni, Michael Huber, Alexandra Trkola |
| EPI_ISL_3045343 | Invenimus AG | Institute of Medical Virology | Verena Kufner, Gabriela Ziltener, Maryam Zaheri, Stefan Schmutz, Annette Audigé, Maria Grünberg, Kevin Steiner, Jon Huder, Cyril Shah, Riccarda Capaul, Guido Bloemberg, Jürg Böni, Michael Huber, Alexandra Trkola |
| EPI_ISL_3045344 | Analytica | Institute of Medical Virology | Verena Kufner, Gabriela Ziltener, Maryam Zaheri, Stefan Schmutz, Annette Audigé, Maria Grünberg, Kevin Steiner, Jon Huder, Cyril Shah, Riccarda Capaul, Guido Bloemberg, Jürg Böni, Michael Huber, Alexandra Trkola |
| EPI_ISL_3045345 | Kantonsspital Münsterlingen | Institute of Medical Virology | Verena Kufner, Gabriela Ziltener, Maryam Zaheri, Stefan Schmutz, Annette Audigé, Maria Grünberg, Kevin Steiner, Jon Huder, Cyril Shah, Riccarda Capaul, Guido Bloemberg, Jürg Böni, Michael Huber, Alexandra Trkola |
| EPI_ISL_3045346 | Kantonsspital Baden AG | Institute of Medical Virology | Verena Kufner, Gabriela Ziltener, Maryam Zaheri, Stefan Schmutz, Annette Audigé, Maria Grünberg, Kevin Steiner, Jon Huder, Cyril Shah, Riccarda Capaul, Guido Bloemberg, Jürg Böni, Michael Huber, Alexandra Trkola |
| EPI_ISL_3045348 | Medica | Institute of Medical Virology | Verena Kufner, Gabriela Ziltener, Maryam Zaheri, Stefan Schmutz, Annette Audigé, Maria Grünberg, Kevin Steiner, Jon Huder, Cyril Shah, Riccarda Capaul, Guido Bloemberg, Jürg Böni, Michael Huber, Alexandra Trkola |
| EPI_ISL_3045349 | Unilabs | Institute of Medical Virology | Verena Kufner, Gabriela Ziltener, Maryam Zaheri, Stefan Schmutz, Annette Audigé, Maria Grünberg, Kevin Steiner, Jon Huder, Cyril Shah, Riccarda Capaul, Guido Bloemberg, Jürg Böni, Michael Huber, Alexandra Trkola |
| EPI_ISL_3045350 | USZ Flughafen | Institute of Medical Virology | Verena Kufner, Gabriela Ziltener, Maryam Zaheri, Stefan Schmutz, Annette Audigé, Maria Grünberg, Kevin Steiner, Jon Huder, Cyril Shah, Riccarda Capaul, Guido Bloemberg, Jürg Böni, Michael Huber, Alexandra Trkola |
| EPI_ISL_3045351 | Universitätsspital Zürich | Institute of Medical Virology | Verena Kufner, Gabriela Ziltener, Maryam Zaheri, Stefan Schmutz, Annette Audigé, Maria Grünberg, Kevin Steiner, Jon Huder, Cyril Shah, Riccarda Capaul, Guido Bloemberg, Jürg Böni, Michael Huber, Alexandra Trkola |
| EPI_ISL_3045352 | UniversitätsSpital Zürich | Institute of Medical Virology | Verena Kufner, Gabriela Ziltener, Maryam Zaheri, Stefan Schmutz, Annette Audigé, Maria Grünberg, Kevin Steiner, Jon Huder, Cyril Shah, Riccarda Capaul, Guido Bloemberg, Jürg Böni, Michael Huber, Alexandra Trkola |
| EPI_ISL_3045353 | UniversitätsSpital Zürich 009 | Institute of Medical Virology | Verena Kufner, Gabriela Ziltener, Maryam Zaheri, Stefan Schmutz, Annette Audigé, Maria Grünberg, Kevin Steiner, Jon Huder, Cyril Shah, Riccarda Capaul, Guido Bloemberg, Jürg Böni, Michael Huber, Alexandra Trkola |
| EPI_ISL_3045354, EPI_ISL_3045355, | Spital Männedorf AG | Institute of Medical Virology | Verena Kufner, Gabriela Ziltener, Maryam Zaheri, Stefan Schmutz, Annette Audigé, Maria Grünberg, Kevin Steiner, Jon Huder, Cyril Shah, Riccarda Capaul, Guido Bloemberg, Jürg Böni, Michael Huber, Alexandra Trkola |

| | | | |
|---|---|---|---|
| EPI_ISL_3045356, EPI_ISL_3045357 | | | |
| EPI_ISL_3045358 | Verein Lunge Zürich | Institute of Medical Virology | Verena Kufner, Gabriela Ziltener, Maryam Zaheri, Stefan Schmutz, Annette Audigé, Maria Grünberg, Kevin Steiner, Jon Huder, Cyril Shah, Riccarda Capaul, Guido Bloemberg, Jürg Böni, Michael Huber, Alexandra Trkola |
| EPI_ISL_3045359 | Spital Limmattal | Institute of Medical Virology | Verena Kufner, Gabriela Ziltener, Maryam Zaheri, Stefan Schmutz, Annette Audigé, Maria Grünberg, Kevin Steiner, Jon Huder, Cyril Shah, Riccarda Capaul, Guido Bloemberg, Jürg Böni, Michael Huber, Alexandra Trkola |
| EPI_ISL_3045360, EPI_ISL_3045361, EPI_ISL_3045362, EPI_ISL_3045363, EPI_ISL_3045365, EPI_ISL_3045366 | Stadtspital Triemli | Institute of Medical Virology | Verena Kufner, Gabriela Ziltener, Maryam Zaheri, Stefan Schmutz, Annette Audigé, Maria Grünberg, Kevin Steiner, Jon Huder, Cyril Shah, Riccarda Capaul, Guido Bloemberg, Jürg Böni, Michael Huber, Alexandra Trkola |
| EPI_ISL_3045558, EPI_ISL_3045559, EPI_ISL_3045560, EPI_ISL_3045561 | Banteay Meanchey Regional Laboratory | Virology Unit, Institut Pasteur du Cambodge | Leakhena Pum, Sophoannadedh Rath, Cecile Troupin, Ly Sovann, Kraing Sidonn, Yi Sengdoeurn, Chin Savuth, Chau Darapheak, Veasna Duong, Erik A Karlsson |
| EPI_ISL_3045562, EPI_ISL_3045563, EPI_ISL_3045564 | Battambang Repid Response Team | Virology Unit, Institut Pasteur du Cambodge | Leakhena Pum, Sophoannadedh Rath, Cecile Troupin, Ly Sovann, Kraing Sidonn, Yi Sengdoeurn, Chin Savuth, Chau Darapheak, Veasna Duong, Erik A Karlsson |
| EPI_ISL_3045565 | Oddar Meanchey Rapid Response Team | Virology Unit, Institut Pasteur du Cambodge | Leakhena Pum, Sophoannadedh Rath, Cecile Troupin, Ly Sovann, Kraing Sidonn, Yi Sengdoeurn, Chin Savuth, Chau Darapheak, Veasna Duong, Erik A Karlsson |
| EPI_ISL_3045567 | Battambang Rapid Response Team | Virology Unit, Institut Pasteur du Cambodge | Leakhena Pum, Sophoannadedh Rath, Cecile Troupin, Ly Sovann, Kraing Sidonn, Yi Sengdoeurn, Chin Savuth, Chau Darapheak, Veasna Duong, Erik A Karlsson |
| EPI_ISL_3045575, EPI_ISL_3045593, EPI_ISL_3045594, EPI_ISL_3045595, EPI_ISL_3045596 | Virology Unit, Institut Pasteur du Cambodge | Virology Unit, Institut Pasteur du Cambodge | Leakhena Pum, Sophoannadedh Rath, Cecile Troupin, Ly Sovann, Kraing Sidonn, Yi Sengdoeurn, Chin Savuth, Chau Darapheak, Veasna Duong, Erik A Karlsson |
| EPI_ISL_3045603 | Pailin Rapid Response Team | Virology Unit, Institut Pasteur du Cambodge | Leakhena Pum, Sophoannadedh Rath, Cecile Troupin, Ly Sovann, Kraing Sidonn, Yi Sengdoeurn, Chin Savuth, Chau Darapheak, Veasna Duong, Erik A Karlsson |
| EPI_ISL_3045604, EPI_ISL_3045605, EPI_ISL_3045606, EPI_ISL_3045607, EPI_ISL_3045608, EPI_ISL_3045609, EPI_ISL_3045610, EPI_ISL_3045611, EPI_ISL_3045612, EPI_ISL_3045613, EPI_ISL_3045614, EPI_ISL_3045615, EPI_ISL_3045616, EPI_ISL_3045617, EPI_ISL_3045618, EPI_ISL_3045619, EPI_ISL_3045620, EPI_ISL_3045621, EPI_ISL_3045622, EPI_ISL_3045623, EPI_ISL_3045624, EPI_ISL_3045625, EPI_ISL_3045626, EPI_ISL_3045627, EPI_ISL_3045628, EPI_ISL_3045629, EPI_ISL_3045630, EPI_ISL_3045631, EPI_ISL_3045632, EPI_ISL_3045633, EPI_ISL_3045634, EPI_ISL_3045635, EPI_ISL_3045636, EPI_ISL_3045637, EPI_ISL_3045638, EPI_ISL_3045639, EPI_ISL_3045640, EPI_ISL_3045641, EPI_ISL_3045642, EPI_ISL_3045643, EPI_ISL_3045644, EPI_ISL_3045645, EPI_ISL_3045646, EPI_ISL_3045647, EPI_ISL_3045648, EPI_ISL_3045649, EPI_ISL_3045650, EPI_ISL_3045651, EPI_ISL_3045652, EPI_ISL_3045653, EPI_ISL_3045654, EPI_ISL_3045655, EPI_ISL_3045656, EPI_ISL_3045657, EPI_ISL_3045658, EPI_ISL_3045659, EPI_ISL_3045660, EPI_ISL_3045661, EPI_ISL_3045662, EPI_ISL_3045663, EPI_ISL_3045664, EPI_ISL_3045665, EPI_ISL_3045666, EPI_ISL_3045667, EPI_ISL_3045668, EPI_ISL_3045669, EPI_ISL_3045670, EPI_ISL_3045671, EPI_ISL_3045672, EPI_ISL_3045673, EPI_ISL_3045674, EPI_ISL_3045675, EPI_ISL_3045676, EPI_ISL_3045677, EPI_ISL_3045678, EPI_ISL_3045679, EPI_ISL_3045680, EPI_ISL_3045682, EPI_ISL_3045683, EPI_ISL_3045684, EPI_ISL_3045685, EPI_ISL_3045686, EPI_ISL_3045687, EPI_ISL_3045688, EPI_ISL_3045689, EPI_ISL_3045690, EPI_ISL_3045691, EPI_ISL_3045692, EPI_ISL_3045693, EPI_ISL_3045694, EPI_ISL_3045695, EPI_ISL_3045696, EPI_ISL_3045697, EPI_ISL_3045699, EPI_ISL_3045700, EPI_ISL_3045701, EPI_ISL_3045702, EPI_ISL_3045703, EPI_ISL_3045704, EPI_ISL_3045705, EPI_ISL_3045707, EPI_ISL_3045708, EPI_ISL_3045709, EPI_ISL_3045710, EPI_ISL_3045712, EPI_ISL_3045713, EPI_ISL_3045714, EPI_ISL_3045715, EPI_ISL_3045716, EPI_ISL_3045717, EPI_ISL_3045719, EPI_ISL_3045722, EPI_ISL_3045723, EPI_ISL_3045724, EPI_ISL_3045725, EPI_ISL_3045726, EPI_ISL_3045727, EPI_ISL_3045728, EPI_ISL_3045729, EPI_ISL_3045730, EPI_ISL_3045731, EPI_ISL_3045732, EPI_ISL_3045733, EPI_ISL_3045734, EPI_ISL_3045735, EPI_ISL_3045736, EPI_ISL_3045737, EPI_ISL_3045738, EPI_ISL_3045739, EPI_ISL_3045740, EPI_ISL_3045741, EPI_ISL_3045742, EPI_ISL_3045744, EPI_ISL_3045745, EPI_ISL_3045746, EPI_ISL_3045747, EPI_ISL_3045748, EPI_ISL_3045749, EPI_ISL_3045750, EPI_ISL_3045752, EPI_ISL_3045753, EPI_ISL_3045754, EPI_ISL_3045755, EPI_ISL_3045756, EPI_ISL_3045757, EPI_ISL_3045758, EPI_ISL_3045759, EPI_ISL_3045760, EPI_ISL_3045761, EPI_ISL_3045762, EPI_ISL_3045763, EPI_ISL_3045764, EPI_ISL_3045765, EPI_ISL_3045766, EPI_ISL_3045767, EPI_ISL_3045768, EPI_ISL_3045769, EPI_ISL_3045770, EPI_ISL_3045771 | | | |
| see above | National Virus Reference Laboratory | National Virus Reference Laboratory | Zoe Yandle, Charlene Bennett, Gabriel Gonzalez, Michael Carr, Jonathan Dean, Cillian F De Gascun |
| EPI_ISL_3045840, EPI_ISL_3045843, EPI_ISL_3045844, EPI_ISL_3045845 | Department of Public Health Buzau | National Institute of Infectious Diseases-Prof. Dr. Matei Bals Molecular Diagnostics Laboratory | Corina Casangiu, Leontina Banica, Marius Surleac, Petre Milu, Robert Hohan, Simona Paraschiv, Dan Otelea |
| EPI_ISL_3045850, EPI_ISL_3045851, EPI_ISL_3045852, EPI_ISL_3045853 | Department of Public Health Iasi | National Institute of Infectious Diseases-Prof. Dr. Matei Bals Molecular Diagnostics Laboratory | Corina Casangiu, Leontina Banica, Marius Surleac, Petre Milu, Robert Hohan, Simona Paraschiv, Dan Otelea |
| EPI_ISL_3045857 | National Institute of Infectious Diseases-Prof. Dr. Matei Bals Molecular Diagnostics Laboratory | National Institute of Infectious Diseases-Prof. Dr. Matei Bals Molecular Diagnostics Laboratory | Corina Casangiu, Leontina Banica, Marius Surleac, Petre Milu, Robert Hohan, Simona Paraschiv, Dan Otelea |
| EPI_ISL_3045858 | Department of Public Health Bucharest | National Institute of Infectious Diseases-Prof. Dr. Matei Bals Molecular Diagnostics Laboratory | Corina Casangiu, Leontina Banica, Marius Surleac, Petre Milu, Robert Hohan, Simona Paraschiv, Dan Otelea |
| EPI_ISL_3045859 | National Institute of Infectious Diseases-Prof. Dr. Matei Bals Molecular Diagnostics Laboratory | National Institute of Infectious Diseases-Prof. Dr. Matei Bals Molecular Diagnostics Laboratory | Corina Casangiu, Leontina Banica, Marius Surleac, Petre Milu, Robert Hohan, Simona Paraschiv, Dan Otelea |
| EPI_ISL_3045860 | Department of Public Health Bucharest | National Institute of Infectious Diseases-Prof. Dr. Matei Bals Molecular Diagnostics Laboratory | Corina Casangiu, Leontina Banica, Marius Surleac, Petre Milu, Robert Hohan, Simona Paraschiv, Dan Otelea |
| EPI_ISL_3045862, EPI_ISL_3045863, EPI_ISL_3045864 | National Institute of Infectious Diseases-Prof. Dr. Matei Bals Molecular Diagnostics Laboratory | National Institute of Infectious Diseases-Prof. Dr. Matei Bals Molecular Diagnostics Laboratory | Corina Casangiu, Leontina Banica, Marius Surleac, Petre Milu, Robert Hohan, Simona Paraschiv, Dan Otelea |
| EPI_ISL_3045865, EPI_ISL_3045866, EPI_ISL_3045873, EPI_ISL_3045874, EPI_ISL_3045875, EPI_ISL_3045876, EPI_ISL_3045877, EPI_ISL_3045878 | Department of Public Health Iasi | National Institute of Infectious Diseases-Prof. Dr. Matei Bals Molecular Diagnostics Laboratory | Corina Casangiu, Leontina Banica, Marius Surleac, Petre Milu, Robert Hohan, Simona Paraschiv, Dan Otelea |
| EPI_ISL_3045879, EPI_ISL_3045880, EPI_ISL_3045881, EPI_ISL_3045883, EPI_ISL_3045884, EPI_ISL_3045886, EPI_ISL_3045887, EPI_ISL_3045888, EPI_ISL_3045889, EPI_ISL_3045890, EPI_ISL_3045891, EPI_ISL_3045892, EPI_ISL_3045893, EPI_ISL_3045894, EPI_ISL_3045895, EPI_ISL_3045896, EPI_ISL_3045898, EPI_ISL_3045899, EPI_ISL_3045900, EPI_ISL_3045901, EPI_ISL_3045902, EPI_ISL_3045903 | | | |
| see above | IN State Department of Health Laboratory Services | IN State Department of Health Laboratory Services | Cassandra Campion, Jamie Yeadon, Brian Pope, Lixia Liu, Kyle Brownlee, Melissa Hindenlang, Mark Glazier |
| EPI_ISL_3045914, EPI_ISL_3045915, EPI_ISL_3045916, EPI_ISL_3045917, EPI_ISL_3045920, EPI_ISL_3045921, EPI_ISL_3045922, EPI_ISL_3045924, EPI_ISL_3045925, EPI_ISL_3045926, EPI_ISL_3045927, EPI_ISL_3045928, EPI_ISL_3045929 | | | |
| see above | MD PHL | MD PHL | Maryland Department of Health Laboratories Administration |
| EPI_ISL_3045946, EPI_ISL_3045948 | VIDOH (Virgin Islands Department of Health) | Grubaugh Lab - Yale School of Public Health | Joseph Fauver, Mallery Breban, Isabel Ott, Tara Alpert, Mary Petrone, Anderson Brito, Chantal Vogels, Annie Watkins, Chaney Kalinich, Jessica Rothman, Kendall Billig, Tobias Koch, Jendai Richards, TaLesa Aderohunmu, Esther Ellis, Marlon Lawrence, Brett Ellis, Nathan Grubaugh |
| EPI_ISL_3045965 | PanGenomics International Pvt. Ltd., Ahmedabad | Gujarat Biotechnology Research Centre | Zarna Patel,Nitin Savaliya,Dinesh Kumar,Twinkle Soni,Sonal Sharma ,Zuber Saiyed,Ramesh Pandit,Janvi Raval,Krishna Mohan Singh,Umang Mishra,Nitesh Shah,Chaitanya Joshi,Madhvi Joshi |
| EPI_ISL_3045966 | PanGenomics International Pvt. Ltd., Ahmedabad | Gujarat Biotechnology Research Centre | Nitin Savaliya,Dinesh Kumar,Twinkle Soni,Sonal Sharma ,Zuber Saiyed,Ramesh Pandit,Janvi Raval,Zarna Patel,Krishna Mohan Singh,Umang Mishra,Nitesh Shah,Chaitanya Joshi,Madhvi Joshi |
| EPI_ISL_3045967 | PanGenomics International Pvt. Ltd., Ahmedabad | Gujarat Biotechnology Research Centre | Dinesh Kumar,Twinkle Soni,Sonal Sharma ,Zuber Saiyed,Ramesh Pandit,Janvi Raval,Zarna Patel,Nitin Savaliya,Krishna Mohan Singh,Umang Mishra,Nitesh Shah,Chaitanya Joshi,Madhvi Joshi |
| EPI_ISL_3045968 | PanGenomics International Pvt. Ltd., Ahmedabad | Gujarat Biotechnology Research Centre | Twinkle Soni,Sonal Sharma ,Zuber Saiyed,Ramesh Pandit,Janvi Raval,Zarna Patel,Nitin Savaliya,Dinesh Kumar,Krishna Mohan Singh,Umang Mishra,Nitesh Shah,Chaitanya Joshi,Madhvi Joshi |
| EPI_ISL_3045969 | PanGenomics International Pvt. Ltd., Ahmedabad | Gujarat Biotechnology Research Centre | Sonal Sharma ,Zuber Saiyed,Ramesh Pandit,Janvi Raval,Zarna Patel,Nitin Savaliya,Dinesh Kumar,Twinkle Soni,Krishna Mohan Singh,Umang Mishra,Nitesh Shah,Chaitanya Joshi,Madhvi Joshi |
| EPI_ISL_3045970 | PanGenomics International Pvt. Ltd., Ahmedabad | Gujarat Biotechnology Research Centre | Zuber Saiyed,Ramesh Pandit,Janvi Raval,Zarna Patel,Nitin Savaliya,Dinesh Kumar,Twinkle Soni,Sonal Sharma ,Krishna Mohan Singh,Umang Mishra,Nitesh Shah,Chaitanya Joshi,Madhvi Joshi |
| EPI_ISL_3045971 | PanGenomics International Pvt. Ltd., Ahmedabad | Gujarat Biotechnology Research Centre | Janvi Raval,Zarna Patel,Nitin Savaliya,Dinesh Kumar,Twinkle Soni,Sonal Sharma ,Zuber Saiyed,Ramesh Pandit,Krishna Mohan Singh,Umang Mishra,Nitesh Shah,Chaitanya Joshi,Madhvi Joshi |
| EPI_ISL_3045972 | PanGenomics International Pvt. Ltd., Ahmedabad | Gujarat Biotechnology Research Centre | Nitin Savaliya,Dinesh Kumar,Twinkle Soni,Sonal Sharma ,Zuber Saiyed,Ramesh Pandit,Janvi Raval,Zarna Patel,Krishna Mohan Singh,Umang Mishra,Nitesh Shah,Chaitanya Joshi,Madhvi Joshi |
| EPI_ISL_3045973 | PanGenomics International Pvt. Ltd., Ahmedabad | Gujarat Biotechnology Research Centre | Dinesh Kumar,Twinkle Soni,Sonal Sharma ,Zuber Saiyed,Ramesh Pandit,Janvi Raval,Zarna Patel,Nitin Savaliya,Krishna Mohan Singh,Umang Mishra,Nitesh Shah,Chaitanya Joshi,Madhvi Joshi |
| EPI_ISL_3045974 | PanGenomics International Pvt. Ltd., Ahmedabad | Gujarat Biotechnology Research Centre | Twinkle Soni,Sonal Sharma ,Zuber Saiyed,Ramesh Pandit,Janvi Raval,Zarna Patel,Nitin Savaliya,Dinesh Kumar,Krishna Mohan Singh,Umang Mishra,Nitesh Shah,Chaitanya Joshi,Madhvi Joshi |
| EPI_ISL_3045975 | PanGenomics International Pvt. Ltd., Ahmedabad | Gujarat Biotechnology Research Centre | Sonal Sharma ,Zuber Saiyed,Ramesh Pandit,Janvi Raval,Zarna Patel,Nitin Savaliya,Dinesh Kumar,Twinkle Soni,Krishna Mohan Singh,Umang Mishra,Nitesh Shah,Chaitanya Joshi,Madhvi Joshi |
| EPI_ISL_3045976 | PanGenomics International Pvt. Ltd., Ahmedabad | Gujarat Biotechnology Research Centre | Zuber Saiyed,Ramesh Pandit,Janvi Raval,Zarna Patel,Nitin Savaliya,Dinesh Kumar,Twinkle Soni,Sonal Sharma ,Krishna Mohan Singh,Umang Mishra,Nitesh Shah,Chaitanya Joshi,Madhvi Joshi |
| EPI_ISL_3045977 | PanGenomics International Pvt. Ltd., Ahmedabad | Gujarat Biotechnology Research Centre | Ramesh Pandit,Janvi Raval,Zarna Patel,Nitin Savaliya,Dinesh Kumar,Twinkle Soni,Sonal Sharma ,Zuber Saiyed,Krishna Mohan Singh,Umang Mishra,Nitesh Shah,Chaitanya Joshi,Madhvi Joshi |
| EPI_ISL_3045978 | PanGenomics International Pvt. Ltd., Ahmedabad | Gujarat Biotechnology Research Centre | Janvi Raval,Zarna Patel,Nitin Savaliya,Dinesh Kumar,Twinkle Soni,Sonal Sharma ,Zuber Saiyed,Ramesh Pandit,Krishna Mohan Singh,Umang Mishra,Nitesh Shah,Chaitanya Joshi,Madhvi Joshi |
| EPI_ISL_3046041, EPI_ISL_3046042, EPI_ISL_3046043, EPI_ISL_3046044, EPI_ISL_3046045, EPI_ISL_3046046, EPI_ISL_3046048, EPI_ISL_3046049, EPI_ISL_3046051, EPI_ISL_3046052, EPI_ISL_3046053, EPI_ISL_3046054, EPI_ISL_3046055, EPI_ISL_3046056, EPI_ISL_3046057, EPI_ISL_3046058, EPI_ISL_3046059, EPI_ISL_3046060, EPI_ISL_3046061, EPI_ISL_3046062, EPI_ISL_3046063, EPI_ISL_3046064, EPI_ISL_3046065, EPI_ISL_3046066, EPI_ISL_3046067, EPI_ISL_3046068, EPI_ISL_3046069, EPI_ISL_3046070, EPI_ISL_3046071, EPI_ISL_3046072, EPI_ISL_3046073, EPI_ISL_3046074, EPI_ISL_3046075 | | | |
| see above | Alaska State Virology Laboratory | Alaska State Virology Laboratory | Stephanie DeRonde, Elva House, Jacob Zidek, Lisa Smith, Ph.D., Jack Chen, Ph.D. |
| EPI_ISL_3046077, EPI_ISL_3046078 | LESP Sinaloa | Instituto de Diagnostico y Referencia Epidemiologicos (INDRE) | Claudia Wong-Arambula, Abril Rodriguez-Maldonado, Vanessa Rivero-Arredondo, Ariadna Medina-Benitez, Joaquin Quiroz-Mercado, Sergio Rangel-Guerrero, Maribel Gonzalez-Villa, Natividad Cruz-Ortiz, Tatiana Nunez-Garcia, Gisela Barrera-Badillo, Lucia Hernandez-Rivas, Irma Lopez-Martinez, Ernesto Ramirez-Gonzalez. |
| EPI_ISL_3046079, EPI_ISL_3046080, EPI_ISL_3046081, EPI_ISL_3046082, EPI_ISL_3046083, EPI_ISL_3046084, | LESP Ciudad de Mexico | Instituto de Diagnostico y Referencia Epidemiologicos (INDRE) | Claudia Wong-Arambula, Abril Rodriguez-Maldonado, Vanessa Rivero-Arredondo, Ariadna Medina-Benitez, Joaquin Quiroz-Mercado, Sergio Rangel-Guerrero, Maribel Gonzalez-Villa, Natividad Cruz-Ortiz, Tatiana Nunez-Garcia, Gisela Barrera-Badillo, Lucia Hernandez-Rivas, Irma Lopez-Martinez, Ernesto Ramirez-Gonzalez. |

| | | | |
|---|---|---|---|
| EPI_ISL_3046085, EPI_ISL_3046086, EPI_ISL_3046087 | | | |
| EPI_ISL_3046088, EPI_ISL_3046089, EPI_ISL_3046090, EPI_ISL_3046091, EPI_ISL_3046092, EPI_ISL_3046093, EPI_ISL_3046094, EPI_ISL_3046095, EPI_ISL_3046096 | LESP Queretaro | Instituto de Diagnostico y Referencia Epidemiologicos (INDRE) | Claudia Wong-Arambula, Abril Rodriguez-Maldonado, Vanessa Rivero-Arredondo, Ariadna Medina-Benitez, Joaquin Quiroz-Mercado, Sergio Rangel-Guerrero, Maribel Gonzalez-Villa, Natividad Cruz-Ortiz, Tatiana Nunez-Garcia, Gisela Barrera-Badillo, Lucia Hernandez-Rivas, Irma Lopez-Martinez, Ernesto Ramirez-Gonzalez. |
| EPI_ISL_3046097 | LESP Ciudad de Mexico | Instituto de Diagnostico y Referencia Epidemiologicos (INDRE) | Claudia Wong-Arambula, Abril Rodriguez-Maldonado, Vanessa Rivero-Arredondo, Ariadna Medina-Benitez, Joaquin Quiroz-Mercado, Sergio Rangel-Guerrero, Maribel Gonzalez-Villa, Natividad Cruz-Ortiz, Tatiana Nunez-Garcia, Gisela Barrera-Badillo, Lucia Hernandez-Rivas, Irma Lopez-Martinez, Ernesto Ramirez-Gonzalez. |
| EPI_ISL_3046098, EPI_ISL_3046099 | LESP Queretaro | Instituto de Diagnostico y Referencia Epidemiologicos (INDRE) | Claudia Wong-Arambula, Abril Rodriguez-Maldonado, Vanessa Rivero-Arredondo, Ariadna Medina-Benitez, Joaquin Quiroz-Mercado, Sergio Rangel-Guerrero, Maribel Gonzalez-Villa, Natividad Cruz-Ortiz, Tatiana Nunez-Garcia, Gisela Barrera-Badillo, Lucia Hernandez-Rivas, Irma Lopez-Martinez, Ernesto Ramirez-Gonzalez. |
| EPI_ISL_3046407, EPI_ISL_3046416, EPI_ISL_3046420 | Zavod za javno zdravstvo Međimurske Županije | Hrvatski zavod za javno zdravstvo | Irena Tabain, Ivana Ferenčak |
| EPI_ISL_3046432, EPI_ISL_3046434, EPI_ISL_3046435, EPI_ISL_3046436, EPI_ISL_3046437, EPI_ISL_3046439, EPI_ISL_3046441, EPI_ISL_3046442 | Zavod za javno zdravstvo Šibensko- Kninske Županije | Hrvatski zavod za javno zdravstvo | Irena Tabain, Ivana Ferenčak |
| EPI_ISL_3046521 | CH SAINT-PHILIBERT | CHU Lille - Laboratoire de Virologie | AIT YAHYA Emilie, ALIDJINOU Enagnon Kazali, BOCKET Laurence, CREPIN Michel, DEMAY Christophe, ENGELMANN Ilka, GEFFROY Sandrine, GUIGON Aurélie, LAMBERT Valérie, LAZREK Mouna, NOBILLIAUX Florian, PREVOST Brigitte, THUILLIER Caroline, TINEZ Claire ,TCHANTCHOU NJOSSE YANICK |
| EPI_ISL_3046526, EPI_ISL_3046530, EPI_ISL_3046537 | CHU LILLE | CHU Lille - Laboratoire de Virologie | AIT YAHYA Emilie, ALIDJINOU Enagnon Kazali, BOCKET Laurence, CREPIN Michel, DEMAY Christophe, ENGELMANN Ilka, GEFFROY Sandrine, GUIGON Aurélie, LAMBERT Valérie, LAZREK Mouna, NOBILLIAUX Florian, PREVOST Brigitte, THUILLIER Caroline, TINEZ Claire ,TCHANTCHOU NJOSSE YANICK |
| EPI_ISL_3046540 | CH ROUBAIX | CHU Lille - Laboratoire de Virologie | AIT YAHYA Emilie, ALIDJINOU Enagnon Kazali, BOCKET Laurence, CREPIN Michel, DEMAY Christophe, ENGELMANN Ilka, GEFFROY Sandrine, GUIGON Aurélie, LAMBERT Valérie, LAZREK Mouna, NOBILLIAUX Florian, PREVOST Brigitte, THUILLIER Caroline, TINEZ Claire ,TCHANTCHOU NJOSSE YANICK |
| EPI_ISL_3046541 | CHU LILLE | CHU Lille - Laboratoire de Virologie | AIT YAHYA Emilie, ALIDJINOU Enagnon Kazali, BOCKET Laurence, CREPIN Michel, DEMAY Christophe, ENGELMANN Ilka, GEFFROY Sandrine, GUIGON Aurélie, LAMBERT Valérie, LAZREK Mouna, NOBILLIAUX Florian, PREVOST Brigitte, THUILLIER Caroline, TINEZ Claire ,TCHANTCHOU NJOSSE YANICK |
| EPI_ISL_3046544 | CH DUNKERQUE | CHU Lille - Laboratoire de Virologie | AIT YAHYA Emilie, ALIDJINOU Enagnon Kazali, BOCKET Laurence, CREPIN Michel, DEMAY Christophe, ENGELMANN Ilka, GEFFROY Sandrine, GUIGON Aurélie, LAMBERT Valérie, LAZREK Mouna, NOBILLIAUX Florian, PREVOST Brigitte, THUILLIER Caroline, TINEZ Claire ,TCHANTCHOU NJOSSE YANICK |
| EPI_ISL_3046545 | CH SAINT-PHILIBERT | CHU Lille - Laboratoire de Virologie | AIT YAHYA Emilie, ALIDJINOU Enagnon Kazali, BOCKET Laurence, CREPIN Michel, DEMAY Christophe, ENGELMANN Ilka, GEFFROY Sandrine, GUIGON Aurélie, LAMBERT Valérie, LAZREK Mouna, NOBILLIAUX Florian, PREVOST Brigitte, THUILLIER Caroline, TINEZ Claire ,TCHANTCHOU NJOSSE YANICK |
| EPI_ISL_3046546, EPI_ISL_3046547, EPI_ISL_3046548, EPI_ISL_3046549 | CHU LILLE | CHU Lille - Laboratoire de Virologie | AIT YAHYA Emilie, ALIDJINOU Enagnon Kazali, BOCKET Laurence, CREPIN Michel, DEMAY Christophe, ENGELMANN Ilka, GEFFROY Sandrine, GUIGON Aurélie, LAMBERT Valérie, LAZREK Mouna, NOBILLIAUX Florian, PREVOST Brigitte, THUILLIER Caroline, TINEZ Claire ,TCHANTCHOU NJOSSE YANICK |
| EPI_ISL_3046550 | CH SAINT-PHILIBERT | CHU Lille - Laboratoire de Virologie | AIT YAHYA Emilie, ALIDJINOU Enagnon Kazali, BOCKET Laurence, CREPIN Michel, DEMAY Christophe, ENGELMANN Ilka, GEFFROY Sandrine, GUIGON Aurélie, LAMBERT Valérie, LAZREK Mouna, NOBILLIAUX Florian, PREVOST Brigitte, THUILLIER Caroline, TINEZ Claire ,TCHANTCHOU NJOSSE YANICK |
| EPI_ISL_3046553 | CH TOURCOING | CHU Lille - Laboratoire de Virologie | AIT YAHYA Emilie, ALIDJINOU Enagnon Kazali, BOCKET Laurence, CREPIN Michel, DEMAY Christophe, ENGELMANN Ilka, GEFFROY Sandrine, GUIGON Aurélie, LAMBERT Valérie, LAZREK Mouna, NOBILLIAUX Florian, PREVOST Brigitte, THUILLIER Caroline, TINEZ Claire ,TCHANTCHOU NJOSSE YANICK |
| EPI_ISL_3046555 | CHU LILLE | CHU Lille - Laboratoire de Virologie | AIT YAHYA Emilie, ALIDJINOU Enagnon Kazali, BOCKET Laurence, CREPIN Michel, DEMAY Christophe, ENGELMANN Ilka, GEFFROY Sandrine, GUIGON Aurélie, LAMBERT Valérie, LAZREK Mouna, NOBILLIAUX Florian, PREVOST Brigitte, THUILLIER Caroline, TINEZ Claire ,TCHANTCHOU NJOSSE YANICK |
| EPI_ISL_3046561, EPI_ISL_3046562, EPI_ISL_3046563, EPI_ISL_3046564 | CH VALENCIENNES | CHU Lille - Laboratoire de Virologie | AIT YAHYA Emilie, ALIDJINOU Enagnon Kazali, BOCKET Laurence, CREPIN Michel, DEMAY Christophe, ENGELMANN Ilka, GEFFROY Sandrine, GUIGON Aurélie, LAMBERT Valérie, LAZREK Mouna, NOBILLIAUX Florian, PREVOST Brigitte, THUILLIER Caroline, TINEZ Claire ,TCHANTCHOU NJOSSE YANICK |
| EPI_ISL_3046565, EPI_ISL_3046566, EPI_ISL_3046567, EPI_ISL_3046570 | CHU LILLE | CHU Lille - Laboratoire de Virologie | AIT YAHYA Emilie, ALIDJINOU Enagnon Kazali, BOCKET Laurence, CREPIN Michel, DEMAY Christophe, ENGELMANN Ilka, GEFFROY Sandrine, GUIGON Aurélie, LAMBERT Valérie, LAZREK Mouna, NOBILLIAUX Florian, PREVOST Brigitte, THUILLIER Caroline, TINEZ Claire ,TCHANTCHOU NJOSSE YANICK |
| EPI_ISL_3046575 | CH VALENCIENNES | CHU Lille - Laboratoire de Virologie | AIT YAHYA Emilie, ALIDJINOU Enagnon Kazali, BOCKET Laurence, CREPIN Michel, DEMAY Christophe, ENGELMANN Ilka, GEFFROY Sandrine, GUIGON Aurélie, LAMBERT Valérie, LAZREK Mouna, NOBILLIAUX Florian, PREVOST Brigitte, THUILLIER Caroline, TINEZ Claire ,TCHANTCHOU NJOSSE YANICK |
| EPI_ISL_3046576, EPI_ISL_3046579, EPI_ISL_3046580, EPI_ISL_3046581, EPI_ISL_3046582 | CHU LILLE | CHU Lille - Laboratoire de Virologie | AIT YAHYA Emilie, ALIDJINOU Enagnon Kazali, BOCKET Laurence, CREPIN Michel, DEMAY Christophe, ENGELMANN Ilka, GEFFROY Sandrine, GUIGON Aurélie, LAMBERT Valérie, LAZREK Mouna, NOBILLIAUX Florian, PREVOST Brigitte, THUILLIER Caroline, TINEZ Claire ,TCHANTCHOU NJOSSE YANICK |
| EPI_ISL_3046609, EPI_ISL_3046610, EPI_ISL_3046611, EPI_ISL_3046612, EPI_ISL_3046613, EPI_ISL_3046614, EPI_ISL_3046615, EPI_ISL_3046616, EPI_ISL_3046617, EPI_ISL_3046618, EPI_ISL_3046619, EPI_ISL_3046621, EPI_ISL_3046622, EPI_ISL_3046623, EPI_ISL_3046624, EPI_ISL_3046625, EPI_ISL_3046626, EPI_ISL_3046627, EPI_ISL_3046628, EPI_ISL_3046629, EPI_ISL_3046630, EPI_ISL_3046631, EPI_ISL_3046632, EPI_ISL_3046633, EPI_ISL_3046634, EPI_ISL_3046635, EPI_ISL_3046636, EPI_ISL_3046637 | | | |
| see above | Nebraska Public Health Laboratory | NPHL COVID-19 Response Team | NPHL COVID-19 Response Team |
| EPI_ISL_3046677, EPI_ISL_3046678, EPI_ISL_3046679, EPI_ISL_3046680, EPI_ISL_3046681, EPI_ISL_3046682, EPI_ISL_3046683, EPI_ISL_3046684, EPI_ISL_3046685, EPI_ISL_3046686, EPI_ISL_3046687, EPI_ISL_3046688, EPI_ISL_3046689, EPI_ISL_3046690, EPI_ISL_3046691, EPI_ISL_3046692, EPI_ISL_3046693, EPI_ISL_3046694, EPI_ISL_3046695, EPI_ISL_3046696, EPI_ISL_3046697, EPI_ISL_3046698, EPI_ISL_3046699, EPI_ISL_3046700, EPI_ISL_3046701, EPI_ISL_3046702, EPI_ISL_3046703, EPI_ISL_3046704, EPI_ISL_3046705, EPI_ISL_3046706, EPI_ISL_3046707, EPI_ISL_3046708, EPI_ISL_3046709, EPI_ISL_3046711, EPI_ISL_3046712, EPI_ISL_3046713, EPI_ISL_3046714, EPI_ISL_3046715, EPI_ISL_3046716, EPI_ISL_3046717, EPI_ISL_3046718, EPI_ISL_3046719, EPI_ISL_3046720, EPI_ISL_3046721, EPI_ISL_3046722, EPI_ISL_3046723, EPI_ISL_3046724, EPI_ISL_3046725, EPI_ISL_3046726, EPI_ISL_3046727, EPI_ISL_3046728, EPI_ISL_3046729, EPI_ISL_3046730, EPI_ISL_3046731, EPI_ISL_3046732, EPI_ISL_3046733, EPI_ISL_3046734, EPI_ISL_3046736, EPI_ISL_3046737, EPI_ISL_3046738, EPI_ISL_3046739, EPI_ISL_3046740, EPI_ISL_3046741, EPI_ISL_3046742, EPI_ISL_3046743, EPI_ISL_3046744, EPI_ISL_3046745, EPI_ISL_3046746, EPI_ISL_3046747, EPI_ISL_3046748, EPI_ISL_3046749, EPI_ISL_3046750, EPI_ISL_3046751, EPI_ISL_3046752, EPI_ISL_3046753, EPI_ISL_3046754, EPI_ISL_3046755, EPI_ISL_3046756, EPI_ISL_3046757, EPI_ISL_3046758, EPI_ISL_3046759, EPI_ISL_3046760, EPI_ISL_3046761, EPI_ISL_3046762, EPI_ISL_3046763, EPI_ISL_3046764, EPI_ISL_3046765, EPI_ISL_3046766, EPI_ISL_3046767, EPI_ISL_3046768, EPI_ISL_3046769, EPI_ISL_3046770, EPI_ISL_3046771, EPI_ISL_3046772, EPI_ISL_3046773, EPI_ISL_3046774, EPI_ISL_3046775, EPI_ISL_3046776, EPI_ISL_3046777, EPI_ISL_3046778, EPI_ISL_3046779, EPI_ISL_3046780, EPI_ISL_3046781, EPI_ISL_3046782, EPI_ISL_3046783, EPI_ISL_3046784, EPI_ISL_3046785, EPI_ISL_3046786, EPI_ISL_3046787, EPI_ISL_3046788, EPI_ISL_3046789, EPI_ISL_3046790, EPI_ISL_3046791, EPI_ISL_3046792, EPI_ISL_3046793, EPI_ISL_3046794, EPI_ISL_3046795, EPI_ISL_3046796, EPI_ISL_3046797, EPI_ISL_3046798, EPI_ISL_3046799, EPI_ISL_3046800, EPI_ISL_3046801, EPI_ISL_3046802, EPI_ISL_3046803, EPI_ISL_3046804, EPI_ISL_3046805, EPI_ISL_3046806, EPI_ISL_3046807, EPI_ISL_3046808, EPI_ISL_3046809, EPI_ISL_3046810, EPI_ISL_3046811, EPI_ISL_3046812, EPI_ISL_3046813, EPI_ISL_3046814, EPI_ISL_3046815, EPI_ISL_3046816, EPI_ISL_3046817, EPI_ISL_3046818, EPI_ISL_3046819, EPI_ISL_3046820, EPI_ISL_3046821, EPI_ISL_3046822, EPI_ISL_3046823, EPI_ISL_3046824, EPI_ISL_3046825, EPI_ISL_3046826, EPI_ISL_3046827, EPI_ISL_3046828, EPI_ISL_3046829, EPI_ISL_3046830, EPI_ISL_3046831, EPI_ISL_3046832, EPI_ISL_3046833 | | | |
| see above | Lifebrain Covid Labor GmbH | Dept. of Laboratory Medicine | Harald Esterbauer, Anna Gschaider, Petra Jurkowitsch, Fabian Konig, Sabina Plumer, Robert Strassl, Claudia Weber, Oswald Wagner |
| EPI_ISL_3047110, EPI_ISL_3047111, EPI_ISL_3047112, EPI_ISL_3047113, EPI_ISL_3047114, EPI_ISL_3047115, EPI_ISL_3047116, EPI_ISL_3047117, EPI_ISL_3047118 | Dept. of Laboratory Medicine | Dept. of Laboratory Medicine | Harald Esterbauer, Anna Gschaider, Petra Jurkowitsch, Fabian Konig, Sabina Plumer, Robert Strassl, Claudia Weber, Oswald Wagner |
| EPI_ISL_3047119, EPI_ISL_3047120, EPI_ISL_3047121, EPI_ISL_3047122, EPI_ISL_3047123, EPI_ISL_3047124, EPI_ISL_3047125, EPI_ISL_3047126, EPI_ISL_3047127, EPI_ISL_3047128, EPI_ISL_3047129, EPI_ISL_3047130, EPI_ISL_3047131, EPI_ISL_3047133, EPI_ISL_3047134, EPI_ISL_3047135, EPI_ISL_3047136, EPI_ISL_3047137, EPI_ISL_3047138, EPI_ISL_3047139, EPI_ISL_3047140, EPI_ISL_3047141, EPI_ISL_3047142, EPI_ISL_3047143, EPI_ISL_3047144, EPI_ISL_3047145, EPI_ISL_3047146, EPI_ISL_3047147, EPI_ISL_3047148, EPI_ISL_3047149, EPI_ISL_3047150, EPI_ISL_3047151, EPI_ISL_3047152, EPI_ISL_3047153, EPI_ISL_3047154, EPI_ISL_3047155, EPI_ISL_3047156, EPI_ISL_3047157, EPI_ISL_3047158, EPI_ISL_3047159, EPI_ISL_3047160, EPI_ISL_3047161, EPI_ISL_3047162, EPI_ISL_3047163, EPI_ISL_3047164, EPI_ISL_3047165, EPI_ISL_3047166, EPI_ISL_3047167, EPI_ISL_3047168, EPI_ISL_3047169, EPI_ISL_3047170, EPI_ISL_3047171, EPI_ISL_3047172, EPI_ISL_3047173, EPI_ISL_3047174, EPI_ISL_3047175, EPI_ISL_3047176, EPI_ISL_3047177, EPI_ISL_3047178, EPI_ISL_3047179, EPI_ISL_3047180, EPI_ISL_3047181, EPI_ISL_3047182, EPI_ISL_3047183, EPI_ISL_3047184, EPI_ISL_3047185, EPI_ISL_3047186, EPI_ISL_3047187, EPI_ISL_3047188, EPI_ISL_3047189, EPI_ISL_3047190, EPI_ISL_3047191, EPI_ISL_3047192, EPI_ISL_3047193, EPI_ISL_3047194, EPI_ISL_3047195, EPI_ISL_3047196, EPI_ISL_3047197, EPI_ISL_3047198, EPI_ISL_3047199, EPI_ISL_3047200, EPI_ISL_3047201, EPI_ISL_3047202, EPI_ISL_3047203, EPI_ISL_3047204, EPI_ISL_3047205, EPI_ISL_3047206, EPI_ISL_3047207 | | | |
| see above | Lifebrain Covid Labor GmbH | Dept. of Laboratory Medicine | Harald Esterbauer, Anna Gschaider, Petra Jurkowitsch, Fabian Konig, Sabina Plumer, Robert Strassl, Claudia Weber, Oswald Wagner |
| EPI_ISL_3047211, EPI_ISL_3047212, EPI_ISL_3047213, EPI_ISL_3047214, EPI_ISL_3047215, EPI_ISL_3047216, EPI_ISL_3047217, EPI_ISL_3047218, EPI_ISL_3047219, EPI_ISL_3047220, EPI_ISL_3047221, EPI_ISL_3047222, EPI_ISL_3047223, EPI_ISL_3047224, EPI_ISL_3047225, EPI_ISL_3047226, EPI_ISL_3047227, EPI_ISL_3047228, EPI_ISL_3047229, EPI_ISL_3047230, EPI_ISL_3047231, EPI_ISL_3047232, EPI_ISL_3047233, EPI_ISL_3047234, EPI_ISL_3047235, EPI_ISL_3047236, EPI_ISL_3047237, EPI_ISL_3047238, EPI_ISL_3047239, EPI_ISL_3047240, EPI_ISL_3047241, EPI_ISL_3047243, EPI_ISL_3047244, EPI_ISL_3047245, EPI_ISL_3047246, EPI_ISL_3047247, EPI_ISL_3047248, EPI_ISL_3047249, EPI_ISL_3047250, EPI_ISL_3047251, EPI_ISL_3047252, EPI_ISL_3047253, EPI_ISL_3047254, EPI_ISL_3047255, EPI_ISL_3047256, EPI_ISL_3047257, EPI_ISL_3047258, EPI_ISL_3047259, EPI_ISL_3047260, EPI_ISL_3047261, EPI_ISL_3047262, EPI_ISL_3047263, EPI_ISL_3047264, EPI_ISL_3047265, EPI_ISL_3047266, EPI_ISL_3047267, EPI_ISL_3047268, EPI_ISL_3047269, EPI_ISL_3047270, EPI_ISL_3047271, EPI_ISL_3047272, EPI_ISL_3047273, EPI_ISL_3047274, EPI_ISL_3047275, EPI_ISL_3047276, EPI_ISL_3047277, EPI_ISL_3047278, EPI_ISL_3047279, EPI_ISL_3047280, EPI_ISL_3047281, EPI_ISL_3047282, EPI_ISL_3047283, EPI_ISL_3047284, EPI_ISL_3047285, EPI_ISL_3047286, EPI_ISL_3047287, EPI_ISL_3047288, EPI_ISL_3047289, EPI_ISL_3047290, EPI_ISL_3047291, EPI_ISL_3047292, EPI_ISL_3047293, EPI_ISL_3047294, EPI_ISL_3047295, EPI_ISL_3047296, EPI_ISL_3047297, EPI_ISL_3047298, EPI_ISL_3047299, EPI_ISL_3047300 | | | |
| see above | Hospital Universitari Vall d'Hebron - Vall d'Hebron Institut de | Hospital Universitari Vall d'Hebron - Vall d'Hebron Institut de | Cristina Andrés, Maria Piñana, Alejandra González-Sánchez, Damir Garcia-Cehic, Ariadna Rando, Juliana Esperalba, Maria Gema Codina, Carla Castillo, Karen García, Rodrigo Vásquez, Maria Carmen Martin, Tomàs Pumarola, Josep Quer, Andrés |

| | Recerca | Recerca | Antón |
|---|---|---|---|
| EPI_ISL_3047311 | University of Utah, School of Medicine | Genomics and Discovery, Respiratory Viruses Branch, Division of Viral Diseases, Centers for Disease Control and Prevention | Jing Zhang, Anna Kelleher, Ying Tao, Yan Li, Brian Lynch, Anna Uehara, Peter Cook, Han Jia Justin Ng, Rachel Marine, Clinton R. Paden, Dhwani Batra, Haibin Wang, Tara Coalter, Jasmine Padilla, Morgan Davis, Mili Sheth, Sarah Nobles, Mark Burroughs, Justin Lee, Adam Retchless, Suxiang Tong |
| EPI_ISL_3047314 | Brentwood Biomedical Research | Genomics and Discovery, Respiratory Viruses Branch, Division of Viral Diseases, Centers for Disease Control and Prevention | Jing Zhang, Anna Kelleher, Ying Tao, Yan Li, Brian Lynch, Anna Uehara, Peter Cook, Han Jia Justin Ng, Rachel Marine, Clinton R. Paden, Dhwani Batra, Haibin Wang, Tara Coalter, Jasmine Padilla, Morgan Davis, Mili Sheth, Sarah Nobles, Mark Burroughs, Justin Lee, Adam Retchless, Suxiang Tong |
| EPI_ISL_3047359, EPI_ISL_3047360, EPI_ISL_3047362, EPI_ISL_3047363, EPI_ISL_3047370, EPI_ISL_3047371, EPI_ISL_3047372, EPI_ISL_3047373, EPI_ISL_3047374, EPI_ISL_3047375, EPI_ISL_3047378, EPI_ISL_3047380, EPI_ISL_3047381, EPI_ISL_3047382, EPI_ISL_3047384, EPI_ISL_3047385, EPI_ISL_3047386, EPI_ISL_3047387, EPI_ISL_3047388, EPI_ISL_3047389, EPI_ISL_3047390, EPI_ISL_3047391, EPI_ISL_3047392, EPI_ISL_3047393, EPI_ISL_3047394, EPI_ISL_3047395, EPI_ISL_3047396, EPI_ISL_3047397, EPI_ISL_3047398, EPI_ISL_3047399, EPI_ISL_3047400, EPI_ISL_3047401 | | | |
| see above | Dipartimento di Medicina di Laboratorio, Azienda sanitaria universitaria Friuli Centrale (ASU FC) | Dipartimento di Medicina di Laboratorio, Azienda sanitaria universitaria Friuli Centrale (ASU FC) | Corrado Pipan, Catia Mio, Chiara Dal Secco, Stefania Marzinotto, Francesco Curcio |
| EPI_ISL_3047402, EPI_ISL_3047405, EPI_ISL_3047406, EPI_ISL_3047407, EPI_ISL_3047408, EPI_ISL_3047409, EPI_ISL_3047410, EPI_ISL_3047411, EPI_ISL_3047412, EPI_ISL_3047413, EPI_ISL_3047414, EPI_ISL_3047415, EPI_ISL_3047416, EPI_ISL_3047417, EPI_ISL_3047418, EPI_ISL_3047419, EPI_ISL_3047421, EPI_ISL_3047422, EPI_ISL_3047423, EPI_ISL_3047424, EPI_ISL_3047425, EPI_ISL_3047426, EPI_ISL_3047427, EPI_ISL_3047429, EPI_ISL_3047430, EPI_ISL_3047431, EPI_ISL_3047432, EPI_ISL_3047433, EPI_ISL_3047434, EPI_ISL_3047435, EPI_ISL_3047436, EPI_ISL_3047437, EPI_ISL_3047438, EPI_ISL_3047440, EPI_ISL_3047441, EPI_ISL_3047442, EPI_ISL_3047443, EPI_ISL_3047444, EPI_ISL_3047446, EPI_ISL_3047447, EPI_ISL_3047448, EPI_ISL_3047449, EPI_ISL_3047450, EPI_ISL_3047451, EPI_ISL_3047452, EPI_ISL_3047453, EPI_ISL_3047454, EPI_ISL_3047455, EPI_ISL_3047456, EPI_ISL_3047457, EPI_ISL_3047460, EPI_ISL_3047461, EPI_ISL_3047462, EPI_ISL_3047466, EPI_ISL_3047467, EPI_ISL_3047468, EPI_ISL_3047469, EPI_ISL_3047472, EPI_ISL_3047473, EPI_ISL_3047474, EPI_ISL_3047475, EPI_ISL_3047476, EPI_ISL_3047477, EPI_ISL_3047478, EPI_ISL_3047479, EPI_ISL_3047480, EPI_ISL_3047482, EPI_ISL_3047484, EPI_ISL_3047485, EPI_ISL_3047486, EPI_ISL_3047487, EPI_ISL_3047488, EPI_ISL_3047489, EPI_ISL_3047490, EPI_ISL_3047492, EPI_ISL_3047493, EPI_ISL_3047494, EPI_ISL_3047495, EPI_ISL_3047496, EPI_ISL_3047497, EPI_ISL_3047498, EPI_ISL_3047499, EPI_ISL_3047500, EPI_ISL_3047501, EPI_ISL_3047502, EPI_ISL_3047503, EPI_ISL_3047504, EPI_ISL_3047505, EPI_ISL_3047506, EPI_ISL_3047507, EPI_ISL_3047508, EPI_ISL_3047509, EPI_ISL_3047510, EPI_ISL_3047511, EPI_ISL_3047513, EPI_ISL_3047514, EPI_ISL_3047515, EPI_ISL_3047516, EPI_ISL_3047517, EPI_ISL_3047518, EPI_ISL_3047519, EPI_ISL_3047520, EPI_ISL_3047521, EPI_ISL_3047522, EPI_ISL_3047523, EPI_ISL_3047525, EPI_ISL_3047526, EPI_ISL_3047527, EPI_ISL_3047528, EPI_ISL_3047529, EPI_ISL_3047530, EPI_ISL_3047531, EPI_ISL_3047532, EPI_ISL_3047533 | | | |
| see above | AZDelta | AZ Delta Medical Laboratories in Roeselare, Belgium | Geert Martens, Dieter De Smet, Merijn Vanhee, on behalf of AZ Delta COVID-19 Genomics core (member of Genomic surveillance of SARS-CoV-2 in Belgium network) |
| EPI_ISL_3047542 | Michael E. DeBakey VA Medical Center | Genomics and Discovery, Respiratory Viruses Branch, Division of Viral Diseases, Centers for Disease Control and Prevention | Jing Zhang, Anna Kelleher, Ying Tao, Yan Li, Brian Lynch, Anna Uehara, Peter Cook, Han Jia Justin Ng, Rachel Marine, Clinton R. Paden, Dhwani Batra, Haibin Wang, Tara Coalter, Jasmine Padilla, Morgan Davis, Mili Sheth, Sarah Nobles, Mark Burroughs, Justin Lee, Adam Retchless, Suxiang Tong |
| EPI_ISL_3047545 | Laboratorium Diagnostyczno-Analityczne Ośrodek Profilaktyki i Epidemiologii Nowotworów im. A. Pieńkowskiej | National Institute of Public Health NIH - National Research Institute | Wołkowicz Tomasz, Zacharczuk Katarzyna, Sadkowska-Todys Małgorzata, Gierczyńki Rafał |
| EPI_ISL_3047546 | NZOZ Meditest Diagnostyka Medyczna | National Institute of Public Health NIH - National Research Institute | Wołkowicz Tomasz, Zacharczuk Katarzyna, Sadkowska-Todys Małgorzata, Gierczyńki Rafał |
| EPI_ISL_3047596, EPI_ISL_3047597, EPI_ISL_3047598, EPI_ISL_3047599, EPI_ISL_3047600, EPI_ISL_3047601 | Zavod za javno zdravstvo Međimurske Županije | Hrvatski zavod za javno zdravstvo | Irena Tabain, Ivana Ferenčak |
| EPI_ISL_3047656 | Klinika za infektivne bolesti "Dr. Fran Mihaljević" | Hrvatski zavod za javno zdravstvo | Irena Tabain, Ivana Ferenčak |
| EPI_ISL_3047670, EPI_ISL_3047673 | Nastavni zavod za javno zdravstvo Splitsko- Dalmatinske Županije | Hrvatski zavod za javno zdravstvo | Irena Tabain, Ivana Ferenčak |
| EPI_ISL_3047689, EPI_ISL_3047691, EPI_ISL_3047693 | Zavod za javno zdravstvo Varaždinske županije | Hrvatski zavod za javno zdravstvo | Irena Tabain, Ivana Ferenčak |
| EPI_ISL_3047703, EPI_ISL_3047722 | LATE - Laboratório de Técnicas Especiais - Hospital Israelita Albert Einstein | LATE - Laboratório de Técnicas Especiais - Hospital Israelita Albert Einstein | Deyvid Amgarten, Fernanda de Mello Malta, Raquel Riyuzo, Ana Paula Moreira Salles, Pedro Henrique Sebe Rodrigues, Alexandre Hideaki Takara, Anelisie da Silva Santos, Erick Gustavo Dorlass, Marcio Anunciacao Menezes, João Renato Rebello Pinho |
| EPI_ISL_3047834, EPI_ISL_3047835, EPI_ISL_3047836, EPI_ISL_3047837, EPI_ISL_3047849, EPI_ISL_3047850, EPI_ISL_3047851, EPI_ISL_3047852, EPI_ISL_3047853, EPI_ISL_3047854, EPI_ISL_3047855 | | | |
| see above | Virology Laboratory, Scientific Department, Army Medical Center | Virology Laboratory, Scientific Department, Army Medical Center | Silvia Fillo, Riccardo De Sanctis, Antonella Fortunato, Anella Monte, Rossella Brandi, Giulia Campoli, Marzia Cavalli, Lucia Nicosia, Anna Anselmo, Vanessa Vera Fain, Francesco Giordani, Giandomenico Cerreto, Filippo Molinari, Giancarlo Petralito, Florigio Lista |
| EPI_ISL_3047868, EPI_ISL_3047870, EPI_ISL_3047871, EPI_ISL_3047872, EPI_ISL_3047873, EPI_ISL_3047878, EPI_ISL_3047879, EPI_ISL_3047881, EPI_ISL_3047882, EPI_ISL_3047885, EPI_ISL_3047886, EPI_ISL_3047887, EPI_ISL_3047888, EPI_ISL_3047889, EPI_ISL_3047890, EPI_ISL_3047891, EPI_ISL_3047892, EPI_ISL_3047893, EPI_ISL_3047894, EPI_ISL_3047895, EPI_ISL_3047896, EPI_ISL_3047897, EPI_ISL_3047898, EPI_ISL_3047899, EPI_ISL_3047902, EPI_ISL_3047903, EPI_ISL_3047904, EPI_ISL_3047905, EPI_ISL_3047910, EPI_ISL_3047911, EPI_ISL_3047912, EPI_ISL_3047913, EPI_ISL_3047914, EPI_ISL_3047915, EPI_ISL_3047918, EPI_ISL_3047919, EPI_ISL_3047921, EPI_ISL_3047922, EPI_ISL_3047923, EPI_ISL_3047924, EPI_ISL_3047925, EPI_ISL_3047927, EPI_ISL_3047928, EPI_ISL_3047929, EPI_ISL_3047930, EPI_ISL_3047932, EPI_ISL_3047933, EPI_ISL_3047934, EPI_ISL_3047935, EPI_ISL_3047936, EPI_ISL_3047937, EPI_ISL_3047938, EPI_ISL_3047939, EPI_ISL_3047940 | | | |
| see above | Microvida | Microvida | Suzan D. Pas, Jaco J. Verweij, Joep J. J. M. Stohr |
| EPI_ISL_3047941, EPI_ISL_3047942, EPI_ISL_3047943, EPI_ISL_3047944, EPI_ISL_3047945, EPI_ISL_3047946, EPI_ISL_3047947, EPI_ISL_3047948, EPI_ISL_3047949, EPI_ISL_3047950, EPI_ISL_3047951, EPI_ISL_3047952, EPI_ISL_3047953, EPI_ISL_3047954, EPI_ISL_3047955, EPI_ISL_3047956, EPI_ISL_3047957, EPI_ISL_3047958, EPI_ISL_3047959, EPI_ISL_3047960, EPI_ISL_3047961, EPI_ISL_3047962, EPI_ISL_3047963, EPI_ISL_3047964, EPI_ISL_3047965, EPI_ISL_3047966, EPI_ISL_3047967, EPI_ISL_3047968, EPI_ISL_3047969, EPI_ISL_3047970 | | | |
| see above | Nebraska Public Health Laboratory | NPHL COVID-19 Response Team | NPHL COVID-19 Response Team |
| EPI_ISL_3047971, EPI_ISL_3047972, EPI_ISL_3047974, EPI_ISL_3047975, EPI_ISL_3047977, EPI_ISL_3047978, EPI_ISL_3047979, EPI_ISL_3047980, EPI_ISL_3047982, EPI_ISL_3047983, EPI_ISL_3047984 | | | |
| see above | South Dakota Public Health Laboratory | South Dakota Public Health Laboratory | Chris Carlson |
| EPI_ISL_3048074, EPI_ISL_3048075, EPI_ISL_3048077, EPI_ISL_3048078, EPI_ISL_3048079, EPI_ISL_3048081, EPI_ISL_3048083, EPI_ISL_3048084, EPI_ISL_3048086, EPI_ISL_3048087, EPI_ISL_3048088, EPI_ISL_3048089, EPI_ISL_3048090, EPI_ISL_3048094, EPI_ISL_3048095, EPI_ISL_3048096, EPI_ISL_3048098, EPI_ISL_3048100, EPI_ISL_3048102, EPI_ISL_3048103, EPI_ISL_3048104, EPI_ISL_3048105, EPI_ISL_3048107, EPI_ISL_3048108, EPI_ISL_3048109, EPI_ISL_3048110 | | | |
| see above | Virginia Division of Consolidated Laboratory Services | Virginia Division of Consolidated Laboratory Services | Virginia Division of Consolidated Laboratory Services |
| EPI_ISL_3048113, EPI_ISL_3048114, EPI_ISL_3048115, EPI_ISL_3048116, EPI_ISL_3048117, EPI_ISL_3048119, EPI_ISL_3048121, EPI_ISL_3048122, EPI_ISL_3048123, EPI_ISL_3048124, EPI_ISL_3048125, EPI_ISL_3048126, EPI_ISL_3048127, EPI_ISL_3048128, EPI_ISL_3048131, EPI_ISL_3048134, EPI_ISL_3048137, EPI_ISL_3048138, EPI_ISL_3048139, EPI_ISL_3048140, EPI_ISL_3048141, EPI_ISL_3048142, EPI_ISL_3048143 | | | |
| see above | Oregon State Public Health Laboratory | Oregon State Public Health Laboratory | Rafia Razzaque, Eugene Yeboah, Vanda Makris, Laura Tsaknaridis, John Fontana and Shane Sevey |
| EPI_ISL_3048144, EPI_ISL_3048145, EPI_ISL_3048150, EPI_ISL_3048151, EPI_ISL_3048154, EPI_ISL_3048158, EPI_ISL_3048159, EPI_ISL_3048162, EPI_ISL_3048163 | US Air Force School of Aerospace Medicine | US Air Force School of Aerospace Medicine | Anthony Fries, Jennifer Meyer, William Gruner, Amanda Javorina, Carol Garrett, Sarah Purves, Clarise Starr, Elizabeth Macias |
| EPI_ISL_3048166, EPI_ISL_3048167, EPI_ISL_3048168, EPI_ISL_3048169, EPI_ISL_3048170, EPI_ISL_3048171 | US Air Force School of Aerospace Medicine | US Air Force School of Aerospace Medicine | Anthony Fries, Jennifer Meyer, William Gruner, Amanda Javorina, Carol Garrett, Sarah Purves, Clarise Starr, Elizabeth Macias, Fritz Castillo, Jennifer Black, Cole Anderson |
| EPI_ISL_3048173, EPI_ISL_3048174, EPI_ISL_3048175, EPI_ISL_3048176, EPI_ISL_3048177, EPI_ISL_3048179, EPI_ISL_3048180, EPI_ISL_3048181, EPI_ISL_3048182, EPI_ISL_3048183, EPI_ISL_3048184, EPI_ISL_3048185, EPI_ISL_3048187, EPI_ISL_3048188, EPI_ISL_3048189, EPI_ISL_3048190, EPI_ISL_3048192, EPI_ISL_3048193, EPI_ISL_3048194, EPI_ISL_3048195, EPI_ISL_3048196, EPI_ISL_3048197, EPI_ISL_3048199, EPI_ISL_3048200, EPI_ISL_3048202, EPI_ISL_3048203, EPI_ISL_3048204, EPI_ISL_3048205, EPI_ISL_3048206, EPI_ISL_3048207, EPI_ISL_3048208, EPI_ISL_3048209, EPI_ISL_3048210, EPI_ISL_3048211, EPI_ISL_3048212, EPI_ISL_3048213, EPI_ISL_3048214, EPI_ISL_3048215 | | | |
| see above | Tampa General Hospital Esoteric Lab | Tampa General Hospital Esoteric Research & Development Lab | Jon Faughn, Marci O'Driscoll, Deanna Becker, Dominic Uy, Jorge Mecias-Frias, Elaine Vendrone, Vicki Healer, Amorce Lima, Suzane Silbert |
| EPI_ISL_3048216, EPI_ISL_3048217, EPI_ISL_3048218, EPI_ISL_3048219, EPI_ISL_3048220, EPI_ISL_3048221, EPI_ISL_3048222, EPI_ISL_3048223, EPI_ISL_3048224, EPI_ISL_3048225, EPI_ISL_3048226, EPI_ISL_3048227, EPI_ISL_3048228 | | | |
| see above | Saint Louis County Department of Public Health Environmental Health Laboratory | University of Minnesota Genomics Center | Daryl M. Gohl, John Garbe, Jaquelyn Kuriger-Laber, Corbin Dirkx |
| EPI_ISL_3048232, EPI_ISL_3048234, EPI_ISL_3048235, EPI_ISL_3048236, EPI_ISL_3048237, EPI_ISL_3048238, EPI_ISL_3048239, EPI_ISL_3048241, EPI_ISL_3048243, EPI_ISL_3048247, EPI_ISL_3048250, EPI_ISL_3048253, EPI_ISL_3048254, EPI_ISL_3048255, EPI_ISL_3048256, EPI_ISL_3048257, EPI_ISL_3048259, EPI_ISL_3048260, EPI_ISL_3048261, EPI_ISL_3048263, EPI_ISL_3048267, EPI_ISL_3048272, EPI_ISL_3048273, EPI_ISL_3048274, EPI_ISL_3048294, EPI_ISL_3048295, EPI_ISL_3048296, EPI_ISL_3048297, EPI_ISL_3048298, EPI_ISL_3048310 | | | |
| see above | Colorado Department of Public Health and Environment | Colorado Department of Public Health and Environment | Laura Bankers, Molly C. Hetherington-Rauth, Diana Ir, Alexandria Rossheim, Michael A. Martin, Mandy Waters, Shannon R. Matzinger, Sarah Elizabeth Totten, Emily A. Travanty |
| EPI_ISL_3048312, EPI_ISL_3048313, EPI_ISL_3048314, EPI_ISL_3048315, EPI_ISL_3048316, EPI_ISL_3048317, EPI_ISL_3048318, EPI_ISL_3048319, EPI_ISL_3048320, EPI_ISL_3048321, EPI_ISL_3048322, EPI_ISL_3048323, EPI_ISL_3048324, EPI_ISL_3048325, EPI_ISL_3048326, EPI_ISL_3048327, EPI_ISL_3048328, EPI_ISL_3048329, EPI_ISL_3048330, EPI_ISL_3048331, EPI_ISL_3048334, EPI_ISL_3048335, EPI_ISL_3048336, EPI_ISL_3048337, EPI_ISL_3048338, EPI_ISL_3048339, EPI_ISL_3048340, EPI_ISL_3048341, EPI_ISL_3048342, EPI_ISL_3048343, EPI_ISL_3048344, EPI_ISL_3048345, EPI_ISL_3048346, EPI_ISL_3048347, EPI_ISL_3048348, EPI_ISL_3048349, EPI_ISL_3048354, EPI_ISL_3048355, EPI_ISL_3048356, EPI_ISL_3048357, EPI_ISL_3048358, EPI_ISL_3048359, EPI_ISL_3048360, EPI_ISL_3048361, EPI_ISL_3048362, EPI_ISL_3048363, EPI_ISL_3048365, EPI_ISL_3048366, EPI_ISL_3048367, EPI_ISL_3048369, EPI_ISL_3048370, EPI_ISL_3048371, EPI_ISL_3048372, EPI_ISL_3048373, EPI_ISL_3048374, EPI_ISL_3048375, EPI_ISL_3048376, EPI_ISL_3048377, EPI_ISL_3048378, EPI_ISL_3048379, EPI_ISL_3048380, EPI_ISL_3048381, EPI_ISL_3048382, EPI_ISL_3048383, EPI_ISL_3048384, EPI_ISL_3048385, EPI_ISL_3048386, EPI_ISL_3048387, EPI_ISL_3048388, EPI_ISL_3048389, EPI_ISL_3048390, EPI_ISL_3048391, EPI_ISL_3048392, EPI_ISL_3048393, EPI_ISL_3048394, EPI_ISL_3048395, EPI_ISL_3048396, EPI_ISL_3048397, EPI_ISL_3048398, EPI_ISL_3048399, EPI_ISL_3048400, EPI_ISL_3048401, EPI_ISL_3048402, EPI_ISL_3048403, EPI_ISL_3048404, EPI_ISL_3048405, EPI_ISL_3048406, EPI_ISL_3048407, EPI_ISL_3048408, EPI_ISL_3048409, EPI_ISL_3048410, EPI_ISL_3048411, EPI_ISL_3048412, EPI_ISL_3048413, EPI_ISL_3048414, EPI_ISL_3048415, EPI_ISL_3048416, EPI_ISL_3048417, EPI_ISL_3048418, EPI_ISL_3048419, EPI_ISL_3048420, EPI_ISL_3048421, EPI_ISL_3048422, EPI_ISL_3048423, EPI_ISL_3048424, EPI_ISL_3048425, EPI_ISL_3048426, EPI_ISL_3048427, EPI_ISL_3048428, EPI_ISL_3048429, EPI_ISL_3048430, EPI_ISL_3048431, EPI_ISL_3048432, EPI_ISL_3048433, EPI_ISL_3048435, EPI_ISL_3048436, EPI_ISL_3048437, EPI_ISL_3048438, EPI_ISL_3048439, EPI_ISL_3048440, EPI_ISL_3048441, EPI_ISL_3048442, EPI_ISL_3048443, EPI_ISL_3048444, EPI_ISL_3048445, EPI_ISL_3048446, EPI_ISL_3048447, EPI_ISL_3048448, EPI_ISL_3048449, EPI_ISL_3048450, EPI_ISL_3048451, EPI_ISL_3048452, EPI_ISL_3048453, EPI_ISL_3048454, EPI_ISL_3048455, EPI_ISL_3048456, EPI_ISL_3048457, EPI_ISL_3048458, EPI_ISL_3048459, EPI_ISL_3048460, EPI_ISL_3048461, EPI_ISL_3048462, EPI_ISL_3048463, EPI_ISL_3048464, EPI_ISL_3048465, EPI_ISL_3048466, EPI_ISL_3048467, EPI_ISL_3048468, EPI_ISL_3048469, EPI_ISL_3048470, EPI_ISL_3048471, EPI_ISL_3048472, EPI_ISL_3048473, EPI_ISL_3048474, EPI_ISL_3048475, EPI_ISL_3048476, EPI_ISL_3048478, EPI_ISL_3048481, EPI_ISL_3048482, EPI_ISL_3048484, EPI_ISL_3048485, EPI_ISL_3048486, EPI_ISL_3048487, EPI_ISL_3048488, EPI_ISL_3048489, EPI_ISL_3048490, EPI_ISL_3048491, EPI_ISL_3048492, EPI_ISL_3048494, EPI_ISL_3048495, EPI_ISL_3048496, EPI_ISL_3048497, EPI_ISL_3048498, EPI_ISL_3048499, EPI_ISL_3048500, EPI_ISL_3048501, EPI_ISL_3048502, EPI_ISL_3048503, EPI_ISL_3048504, EPI_ISL_3048505, EPI_ISL_3048506, EPI_ISL_3048507, EPI_ISL_3048508, EPI_ISL_3048509, EPI_ISL_3048510, EPI_ISL_3048511, EPI_ISL_3048512, EPI_ISL_3048513, EPI_ISL_3048514, EPI_ISL_3048515, EPI_ISL_3048516, EPI_ISL_3048517, EPI_ISL_3048518, EPI_ISL_3048519, EPI_ISL_3048520, EPI_ISL_3048521, EPI_ISL_3048522, EPI_ISL_3048523 | | | |
| see above | University Hospitals of Geneva, Laboratory of Virology | HUG, Laboratory of Virology and the Health2030 Genome Center | Samuel Cordey, Ana Rita Goncalves, Laurent Kaiser, Lorenzo Cerutti, Henri Pegeot, Melyssa Elies, Deborah Penet, Keith Harshman, Ioannis Xenarios, Emmanouil Dermitzakis |
| EPI_ISL_3048525, EPI_ISL_3048526, EPI_ISL_3048527, EPI_ISL_3048528, EPI_ISL_3048529, EPI_ISL_3048530, EPI_ISL_3048531, EPI_ISL_3048532, EPI_ISL_3048533, EPI_ISL_3048534, EPI_ISL_3048535, EPI_ISL_3048536, EPI_ISL_3048537, EPI_ISL_3048538, EPI_ISL_3048539, EPI_ISL_3048540, EPI_ISL_3048541, EPI_ISL_3048542, EPI_ISL_3048543, EPI_ISL_3048544, EPI_ISL_3048545, EPI_ISL_3048546, EPI_ISL_3048547, EPI_ISL_3048548, EPI_ISL_3048549, EPI_ISL_3048550, EPI_ISL_3048551, EPI_ISL_3048552, EPI_ISL_3048553, EPI_ISL_3048554, EPI_ISL_3048555, EPI_ISL_3048556, EPI_ISL_3048557, EPI_ISL_3048558, EPI_ISL_3048559, EPI_ISL_3048560, EPI_ISL_3048561, EPI_ISL_3048562, EPI_ISL_3048563, EPI_ISL_3048564, EPI_ISL_3048565, EPI_ISL_3048566, EPI_ISL_3048567, EPI_ISL_3048568, EPI_ISL_3048569, EPI_ISL_3048570, EPI_ISL_3048571, EPI_ISL_3048572, EPI_ISL_3048573, EPI_ISL_3048575, EPI_ISL_3048576, EPI_ISL_3048577, EPI_ISL_3048578, EPI_ISL_3048579, EPI_ISL_3048580, EPI_ISL_3048581, EPI_ISL_3048582, EPI_ISL_3048583, EPI_ISL_3048584, EPI_ISL_3048585, EPI_ISL_3048586, EPI_ISL_3048587, EPI_ISL_3048588, EPI_ISL_3048589, EPI_ISL_3048590, EPI_ISL_3048591, EPI_ISL_3048592, EPI_ISL_3048593, EPI_ISL_3048594, EPI_ISL_3048595, EPI_ISL_3048596, EPI_ISL_3048597, EPI_ISL_3048598, EPI_ISL_3048599, EPI_ISL_3048600, EPI_ISL_3048601, EPI_ISL_3048602, EPI_ISL_3048603, EPI_ISL_3048604, EPI_ISL_3048605, EPI_ISL_3048606, EPI_ISL_3048607, EPI_ISL_3048609, EPI_ISL_3048610, EPI_ISL_3048611, EPI_ISL_3048612, EPI_ISL_3048613, EPI_ISL_3048614, EPI_ISL_3048615, EPI_ISL_3048616, EPI_ISL_3048617, EPI_ISL_3048618, EPI_ISL_3048619, EPI_ISL_3048620, EPI_ISL_3048621, EPI_ISL_3048622, EPI_ISL_3048623, EPI_ISL_3048624, EPI_ISL_3048625, EPI_ISL_3048626, EPI_ISL_3048627, EPI_ISL_3048628, EPI_ISL_3048629, EPI_ISL_3048630, EPI_ISL_3048631, EPI_ISL_3048632, EPI_ISL_3048633, EPI_ISL_3048634, EPI_ISL_3048635, EPI_ISL_3048636, EPI_ISL_3048637, EPI_ISL_3048638, EPI_ISL_3048639, EPI_ISL_3048641, EPI_ISL_3048642, EPI_ISL_3048644, EPI_ISL_3048645, EPI_ISL_3048646, EPI_ISL_3048648, EPI_ISL_3048650, EPI_ISL_3048652, EPI_ISL_3048653, EPI_ISL_3048654, EPI_ISL_3048656, EPI_ISL_3048657, EPI_ISL_3048658, EPI_ISL_3048659, EPI_ISL_3048660, EPI_ISL_3048661, EPI_ISL_3048662, EPI_ISL_3048663, EPI_ISL_3048665, | | | |

| | | | |
|---|---|---|---|
| EPI_ISL_3048667, EPI_ISL_3048668, EPI_ISL_3048669, EPI_ISL_3048670, EPI_ISL_3048671, EPI_ISL_3048672, EPI_ISL_3048673, EPI_ISL_3048674, EPI_ISL_3048676, EPI_ISL_3048677, EPI_ISL_3048678, EPI_ISL_3048679, EPI_ISL_3048680, EPI_ISL_3048681, EPI_ISL_3048682, EPI_ISL_3048683, EPI_ISL_3048684, EPI_ISL_3048685, EPI_ISL_3048686, EPI_ISL_3048687, EPI_ISL_3048689, EPI_ISL_3048690, EPI_ISL_3048691, EPI_ISL_3048692, EPI_ISL_3048693, EPI_ISL_3048694, EPI_ISL_3048695, EPI_ISL_3048696, EPI_ISL_3048697, EPI_ISL_3048698, EPI_ISL_3048699, EPI_ISL_3048700, EPI_ISL_3048701, EPI_ISL_3048703, EPI_ISL_3048704, EPI_ISL_3048705, EPI_ISL_3048706, EPI_ISL_3048707, EPI_ISL_3048708, EPI_ISL_3048709, EPI_ISL_3048710, EPI_ISL_3048711, EPI_ISL_3048712, EPI_ISL_3048713, EPI_ISL_3048714, EPI_ISL_3048716, EPI_ISL_3048717, EPI_ISL_3048718, EPI_ISL_3048719, EPI_ISL_3048720, EPI_ISL_3048721, EPI_ISL_3048722, EPI_ISL_3048723, EPI_ISL_3048724, EPI_ISL_3048725, EPI_ISL_3048726, EPI_ISL_3048727, EPI_ISL_3048728, EPI_ISL_3048729, EPI_ISL_3048730, EPI_ISL_3048732 | | | |
| see above | Arkansas Public Health Laboratory, Arkansas Department of Health | University of Minnesota Genomics Center | Daryl M. Gohl, John Garbe, Jaquelyn Kuriger-Laber, Corbin Dirkx |
| EPI_ISL_3048733, EPI_ISL_3048734, EPI_ISL_3048735, EPI_ISL_3048736, EPI_ISL_3048737, EPI_ISL_3048738, EPI_ISL_3048739, EPI_ISL_3048740, EPI_ISL_3048741, EPI_ISL_3048742, EPI_ISL_3048743, EPI_ISL_3048744, EPI_ISL_3048745, EPI_ISL_3048747, EPI_ISL_3048748, EPI_ISL_3048749, EPI_ISL_3048750, EPI_ISL_3048751 | | | |
| see above | NJDOH, Public Health and Environmental Laboratories | New Jersey Public Health and Environmental Laboratories (NJ PHEL) | Lindsey Bodnar, Shiv K. Verma, Ryan Pachucki, Chelsea San Filippo, Jacquelyn Deverell, Dana Woell, Allison Roder, Byeong Jeong |
| EPI_ISL_3048870 | Santa Clara County Public Health Laboratory | Santa Clara County Public Health Laboratory | Santa Clara County Public Health Department |
| EPI_ISL_3048871, EPI_ISL_3048872, EPI_ISL_3048873, EPI_ISL_3048874, EPI_ISL_3048875, EPI_ISL_3048876, EPI_ISL_3048877, EPI_ISL_3048878, EPI_ISL_3048879, EPI_ISL_3048880, EPI_ISL_3048881, EPI_ISL_3048882, EPI_ISL_3048883, EPI_ISL_3048884, EPI_ISL_3048885, EPI_ISL_3048886, EPI_ISL_3048887, EPI_ISL_3048888, EPI_ISL_3048889, EPI_ISL_3048890, EPI_ISL_3048891, EPI_ISL_3048892, EPI_ISL_3048893, EPI_ISL_3048894, EPI_ISL_3048895, EPI_ISL_3048897, EPI_ISL_3048898, EPI_ISL_3048899, EPI_ISL_3048901, EPI_ISL_3048903, EPI_ISL_3048904, EPI_ISL_3048905, EPI_ISL_3048906, EPI_ISL_3048907 | | | |
| see above | Santa Clara Valley Medical Center | Santa Clara County Public Health Laboratory | Santa Clara County Public Health Department |
| EPI_ISL_3048908 | Santa Clara County Public Health Laboratory | Santa Clara County Public Health Laboratory | Santa Clara County Public Health Department |
| EPI_ISL_3048910, EPI_ISL_3048911, EPI_ISL_3048912, EPI_ISL_3048913, EPI_ISL_3048914, EPI_ISL_3048915, EPI_ISL_3048916, EPI_ISL_3048917, EPI_ISL_3048918, EPI_ISL_3048919, EPI_ISL_3048921, EPI_ISL_3048922, EPI_ISL_3048923, EPI_ISL_3048926, EPI_ISL_3048927, EPI_ISL_3048928, EPI_ISL_3048929, EPI_ISL_3048930, EPI_ISL_3048931, EPI_ISL_3048933, EPI_ISL_3048934, EPI_ISL_3048935, EPI_ISL_3048936, EPI_ISL_3048937, EPI_ISL_3048938, EPI_ISL_3048939, EPI_ISL_3048940, EPI_ISL_3048941, EPI_ISL_3048942, EPI_ISL_3048943, EPI_ISL_3048944, EPI_ISL_3048945, EPI_ISL_3048946 | | | |
| see above | Santa Clara Valley Medical Center | Santa Clara County Public Health Laboratory | Santa Clara County Public Health Department |
| EPI_ISL_3049037, EPI_ISL_3049038 | Utah Public Health Laboratory, Utah Public Health Laboratory Infectious Disease submission group | Utah Public Health Laboratory, Utah Public Health Laboratory Infectious Disease submission group | Linares-Perdomo,O., Oakeson,K., Young,E. |
| EPI_ISL_3049092, EPI_ISL_3049093, EPI_ISL_3049094, EPI_ISL_3049095, EPI_ISL_3049096, EPI_ISL_3049097, EPI_ISL_3049098, EPI_ISL_3049099, EPI_ISL_3049100, EPI_ISL_3049101 | Dianalabs SA | Genesupport | Tanguy ARAUD, Geraldine Jost, Katia Jaton, Nadia Liassine |
| EPI_ISL_3049102 | Dianalabs-Valais SA | Genesupport | Tanguy ARAUD, Geraldine Jost, Katia Jaton, Nadia Liassine |
| EPI_ISL_3049103, EPI_ISL_3049104, EPI_ISL_3049105, EPI_ISL_3049106, EPI_ISL_3049107, EPI_ISL_3049108, EPI_ISL_3049109, EPI_ISL_3049110, EPI_ISL_3049111, EPI_ISL_3049112, EPI_ISL_3049113, EPI_ISL_3049114, EPI_ISL_3049115, EPI_ISL_3049116, EPI_ISL_3049117, EPI_ISL_3049118, EPI_ISL_3049119, EPI_ISL_3049120 | | | |
| see above | Dianalabs SA | Genesupport | Tanguy ARAUD, Geraldine Jost, Katia Jaton, Nadia Liassine |
| EPI_ISL_3049121 | Dianalabs-Valais SA | Genesupport | Tanguy ARAUD, Geraldine Jost, Katia Jaton, Nadia Liassine |
| EPI_ISL_3049122, EPI_ISL_3049123, EPI_ISL_3049126, EPI_ISL_3049127, EPI_ISL_3049128, EPI_ISL_3049129 | Dianalabs SA | Genesupport | Tanguy ARAUD, Geraldine Jost, Katia Jaton, Nadia Liassine |
| EPI_ISL_3049130 | Dianalabs-Valais SA | Genesupport | Tanguy ARAUD, Geraldine Jost, Katia Jaton, Nadia Liassine |
| EPI_ISL_3049131 | Proxilab SA | Genesupport | Tanguy ARAUD, Geraldine Jost, Katia Jaton, Nadia Liassine |
| EPI_ISL_3049132 | Polyanalytic SA | Genesupport | Tanguy ARAUD, Geraldine Jost, Katia Jaton, Nadia Liassine |
| EPI_ISL_3049133, EPI_ISL_3049135 | Dianalabs SA | Genesupport | Tanguy ARAUD, Geraldine Jost, Katia Jaton, Nadia Liassine |
| EPI_ISL_3049136 | Polyanalytic SA | Genesupport | Tanguy ARAUD, Geraldine Jost, Katia Jaton, Nadia Liassine |
| EPI_ISL_3049137, EPI_ISL_3049138 | Dianalabs SA | Genesupport | Tanguy ARAUD, Geraldine Jost, Katia Jaton, Nadia Liassine |
| EPI_ISL_3049139, EPI_ISL_3049140, EPI_ISL_3049141 | Polyanalytic SA | Genesupport | Tanguy ARAUD, Geraldine Jost, Katia Jaton, Nadia Liassine |
| EPI_ISL_3049142 | Dianalabs-Valais SA | Genesupport | Tanguy ARAUD, Geraldine Jost, Katia Jaton, Nadia Liassine |
| EPI_ISL_3049143, EPI_ISL_3049144, EPI_ISL_3049145, EPI_ISL_3049146, EPI_ISL_3049147, EPI_ISL_3049148, EPI_ISL_3049149, EPI_ISL_3049150, EPI_ISL_3049151, EPI_ISL_3049152, EPI_ISL_3049153, EPI_ISL_3049154, EPI_ISL_3049155, EPI_ISL_3049156, EPI_ISL_3049157, EPI_ISL_3049158, EPI_ISL_3049159, EPI_ISL_3049160, EPI_ISL_3049161, EPI_ISL_3049162, EPI_ISL_3049163, EPI_ISL_3049164 | | | |
| see above | Dianalabs SA | Genesupport | Tanguy ARAUD, Geraldine Jost, Katia Jaton, Nadia Liassine |
| EPI_ISL_3049190, EPI_ISL_3049191, EPI_ISL_3049192, EPI_ISL_3049193, EPI_ISL_3049194 | Queensland Health Forensic and Scientific Services | Queensland Health Forensic and Scientific Services | Son Nguyen |
| EPI_ISL_3049299, EPI_ISL_3049303, EPI_ISL_3049321, EPI_ISL_3049322, EPI_ISL_3049323, EPI_ISL_3049324, EPI_ISL_3049325, EPI_ISL_3049333, EPI_ISL_3049344, EPI_ISL_3049346, EPI_ISL_3049347, EPI_ISL_3049358, EPI_ISL_3049362, EPI_ISL_3049363, EPI_ISL_3049364, EPI_ISL_3049366, EPI_ISL_3049367, EPI_ISL_3049368, EPI_ISL_3049369, EPI_ISL_3049370, EPI_ISL_3049373, EPI_ISL_3049374, EPI_ISL_3049375 | | | |
| see above | Tulare County Public Health Laboratory | Tulare County Public Health Laboratory | Denise L Lopez, Joshua Kootstra |
| EPI_ISL_3049428, EPI_ISL_3049430, EPI_ISL_3049435, EPI_ISL_3049438, EPI_ISL_3049439, EPI_ISL_3049455, EPI_ISL_3049457, EPI_ISL_3049458, EPI_ISL_3049463, EPI_ISL_3049464, EPI_ISL_3049465, EPI_ISL_3049466, EPI_ISL_3049467, EPI_ISL_3049468, EPI_ISL_3049470, EPI_ISL_3049471, EPI_ISL_3049474, EPI_ISL_3049475, EPI_ISL_3049476, EPI_ISL_3049477, EPI_ISL_3049479, EPI_ISL_3049480, EPI_ISL_3049481, EPI_ISL_3049482, EPI_ISL_3049483, EPI_ISL_3049484, EPI_ISL_3049494, EPI_ISL_3049497, EPI_ISL_3049498, EPI_ISL_3049500, EPI_ISL_3049501, EPI_ISL_3049502, EPI_ISL_3049503, EPI_ISL_3049504, EPI_ISL_3049505, EPI_ISL_3049506, EPI_ISL_3049507, EPI_ISL_3049508, EPI_ISL_3049509, EPI_ISL_3049511, EPI_ISL_3049512, EPI_ISL_3049513, EPI_ISL_3049524, EPI_ISL_3049526, EPI_ISL_3049527, EPI_ISL_3049528, EPI_ISL_3049529, EPI_ISL_3049530, EPI_ISL_3049531, EPI_ISL_3049532, EPI_ISL_3049533, EPI_ISL_3049534, EPI_ISL_3049535, EPI_ISL_3049536, EPI_ISL_3049538, EPI_ISL_3049539, EPI_ISL_3049540, EPI_ISL_3049541, EPI_ISL_3049542, EPI_ISL_3049543, EPI_ISL_3049545, EPI_ISL_3049546, EPI_ISL_3049547, EPI_ISL_3049548, EPI_ISL_3049549, EPI_ISL_3049550, EPI_ISL_3049551, EPI_ISL_3049552, EPI_ISL_3049554, EPI_ISL_3049555, EPI_ISL_3049556, EPI_ISL_3049557, EPI_ISL_3049558, EPI_ISL_3049559, EPI_ISL_3049560, EPI_ISL_3049561, EPI_ISL_3049562, EPI_ISL_3049563, EPI_ISL_3049565, EPI_ISL_3049566, EPI_ISL_3049567, EPI_ISL_3049569, EPI_ISL_3049570, EPI_ISL_3049576, EPI_ISL_3049578, EPI_ISL_3049579, EPI_ISL_3049581, EPI_ISL_3049582, EPI_ISL_3049583, EPI_ISL_3049586, EPI_ISL_3049591, EPI_ISL_3049592, EPI_ISL_3049596, EPI_ISL_3049597, EPI_ISL_3049598, EPI_ISL_3049600, EPI_ISL_3049602, EPI_ISL_3049603, EPI_ISL_3049604, EPI_ISL_3049605, EPI_ISL_3049606, EPI_ISL_3049607, EPI_ISL_3049608, EPI_ISL_3049609, EPI_ISL_3049610, EPI_ISL_3049612, EPI_ISL_3049613, EPI_ISL_3049614, EPI_ISL_3049616, EPI_ISL_3049617, EPI_ISL_3049618, EPI_ISL_3049619, EPI_ISL_3049621, EPI_ISL_3049622, EPI_ISL_3049626, EPI_ISL_3049629, EPI_ISL_3049632, EPI_ISL_3049633, EPI_ISL_3049635, EPI_ISL_3049636, EPI_ISL_3049640, EPI_ISL_3049641, EPI_ISL_3049645, EPI_ISL_3049646, EPI_ISL_3049650, EPI_ISL_3049656, EPI_ISL_3049658, EPI_ISL_3049659, EPI_ISL_3049660, EPI_ISL_3049661, EPI_ISL_3049662, EPI_ISL_3049663, EPI_ISL_3049664, EPI_ISL_3049665, EPI_ISL_3049666, EPI_ISL_3049667, EPI_ISL_3049668, EPI_ISL_3049669, EPI_ISL_3049670, EPI_ISL_3049671, EPI_ISL_3049672, EPI_ISL_3049673, EPI_ISL_3049674, EPI_ISL_3049675, EPI_ISL_3049676, EPI_ISL_3049677, EPI_ISL_3049678, EPI_ISL_3049679, EPI_ISL_3049680, EPI_ISL_3049681, EPI_ISL_3049682, EPI_ISL_3049683, EPI_ISL_3049684, EPI_ISL_3049685, EPI_ISL_3049688, EPI_ISL_3049689, EPI_ISL_3049690, EPI_ISL_3049691, EPI_ISL_3049692, EPI_ISL_3049693, EPI_ISL_3049695, EPI_ISL_3049696, EPI_ISL_3049697, EPI_ISL_3049698, EPI_ISL_3049699, EPI_ISL_3049700, EPI_ISL_3049701, EPI_ISL_3049702, EPI_ISL_3049703, EPI_ISL_3049704, EPI_ISL_3049705, EPI_ISL_3049706, EPI_ISL_3049707, EPI_ISL_3049710, EPI_ISL_3049711, EPI_ISL_3049712, EPI_ISL_3049713, EPI_ISL_3049718, EPI_ISL_3049732, EPI_ISL_3049735, EPI_ISL_3049736, EPI_ISL_3049737, EPI_ISL_3049739, EPI_ISL_3049740, EPI_ISL_3049741, EPI_ISL_3049742, EPI_ISL_3049743, EPI_ISL_3049744, EPI_ISL_3049745, EPI_ISL_3049746, EPI_ISL_3049747, EPI_ISL_3049748, EPI_ISL_3049749, EPI_ISL_3049750, EPI_ISL_3049751, EPI_ISL_3049752, EPI_ISL_3049756, EPI_ISL_3049757, EPI_ISL_3049758, EPI_ISL_3049759, EPI_ISL_3049760, EPI_ISL_3049761, EPI_ISL_3049762, EPI_ISL_3049763, EPI_ISL_3049764, EPI_ISL_3049765, EPI_ISL_3049766, EPI_ISL_3049767, EPI_ISL_3049768, EPI_ISL_3049769, EPI_ISL_3049770, EPI_ISL_3049771, EPI_ISL_3049772, EPI_ISL_3049773, EPI_ISL_3049774, EPI_ISL_3049775, EPI_ISL_3049777, EPI_ISL_3049778, EPI_ISL_3049779, EPI_ISL_3049780, EPI_ISL_3049781, EPI_ISL_3049782, EPI_ISL_3049783, EPI_ISL_3049784, EPI_ISL_3049785, EPI_ISL_3049786, EPI_ISL_3049787, EPI_ISL_3049788, EPI_ISL_3049789, EPI_ISL_3049790, EPI_ISL_3049791, EPI_ISL_3049792, EPI_ISL_3049793, EPI_ISL_3049794, EPI_ISL_3049795, EPI_ISL_3049796, EPI_ISL_3049797, EPI_ISL_3049798, EPI_ISL_3049800, EPI_ISL_3049801, EPI_ISL_3049802, EPI_ISL_3049803, EPI_ISL_3049804, EPI_ISL_3049805, EPI_ISL_3049806, EPI_ISL_3049807, EPI_ISL_3049808, EPI_ISL_3049809, EPI_ISL_3049810, EPI_ISL_3049811, EPI_ISL_3049812, EPI_ISL_3049813, EPI_ISL_3049814, EPI_ISL_3049815, EPI_ISL_3049816, EPI_ISL_3049817, EPI_ISL_3049819, EPI_ISL_3049820, EPI_ISL_3049821, EPI_ISL_3049822, EPI_ISL_3049823, EPI_ISL_3049824, EPI_ISL_3049825, EPI_ISL_3049827, EPI_ISL_3049828, EPI_ISL_3049829, EPI_ISL_3049830, EPI_ISL_3049831, EPI_ISL_3049832, EPI_ISL_3049833, EPI_ISL_3049834, EPI_ISL_3049835, EPI_ISL_3049836, EPI_ISL_3049837, EPI_ISL_3049838, EPI_ISL_3049839, EPI_ISL_3049840, EPI_ISL_3049841, EPI_ISL_3049842, EPI_ISL_3049843, EPI_ISL_3049844, EPI_ISL_3049845, EPI_ISL_3049846, EPI_ISL_3049847, EPI_ISL_3049849, EPI_ISL_3049850, EPI_ISL_3049851, EPI_ISL_3049852, EPI_ISL_3049853, EPI_ISL_3049854, EPI_ISL_3049855, EPI_ISL_3049856, EPI_ISL_3049857, EPI_ISL_3049858, EPI_ISL_3049859, EPI_ISL_3049860, EPI_ISL_3049861, EPI_ISL_3049862, EPI_ISL_3049863, EPI_ISL_3049864, EPI_ISL_3049865, EPI_ISL_3049866, EPI_ISL_3049867, EPI_ISL_3049868, EPI_ISL_3049869, EPI_ISL_3049871 | | | |
| see above | KEMRI-Wellcome Trust Research Programme,Kilifi | KEMRI-Wellcome Trust Research Programme,Kilifi | Githinji G.,Matoke D.,Mohamed K.S.,de Laurent Z.,Mburu M.W.,Thiongo K. |
| EPI_ISL_3049873, EPI_ISL_3049875, EPI_ISL_3049878, EPI_ISL_3049879, EPI_ISL_3049880, EPI_ISL_3049883, EPI_ISL_3049884, EPI_ISL_3049885, EPI_ISL_3049886, EPI_ISL_3049888, EPI_ISL_3049890, EPI_ISL_3049893, EPI_ISL_3049894, EPI_ISL_3049895, EPI_ISL_3049897, EPI_ISL_3049901, EPI_ISL_3049902, EPI_ISL_3049906, EPI_ISL_3049907, EPI_ISL_3049908, EPI_ISL_3049909, EPI_ISL_3049912, EPI_ISL_3049916, EPI_ISL_3049917, EPI_ISL_3049918, EPI_ISL_3049919, EPI_ISL_3049924, EPI_ISL_3049925, EPI_ISL_3049928, EPI_ISL_3049929, EPI_ISL_3049932, EPI_ISL_3049933, EPI_ISL_3049938, EPI_ISL_3049940, EPI_ISL_3049941, EPI_ISL_3049942, EPI_ISL_3049944, EPI_ISL_3049945, EPI_ISL_3049946, EPI_ISL_3049948, EPI_ISL_3049950, EPI_ISL_3049951, EPI_ISL_3049952, EPI_ISL_3049955, EPI_ISL_3049956, EPI_ISL_3049957, EPI_ISL_3049958, EPI_ISL_3049959, EPI_ISL_3049960, EPI_ISL_3049964, EPI_ISL_3049966, EPI_ISL_3049967, EPI_ISL_3049970, EPI_ISL_3049973, EPI_ISL_3049976, EPI_ISL_3049980, EPI_ISL_3049982, EPI_ISL_3049984, EPI_ISL_3049990, EPI_ISL_3049991, EPI_ISL_3049993, EPI_ISL_3049996, EPI_ISL_3049997, EPI_ISL_3050002, EPI_ISL_3050003, EPI_ISL_3050006, EPI_ISL_3050007, EPI_ISL_3050012, EPI_ISL_3050013, EPI_ISL_3050014, EPI_ISL_3050016, EPI_ISL_3050017, EPI_ISL_3050018, EPI_ISL_3050019, EPI_ISL_3050020, EPI_ISL_3050023, EPI_ISL_3050024, EPI_ISL_3050027, EPI_ISL_3050028, EPI_ISL_3050030, EPI_ISL_3050034, EPI_ISL_3050035, EPI_ISL_3050036, EPI_ISL_3050037, EPI_ISL_3050038, EPI_ISL_3050039, EPI_ISL_3050041, EPI_ISL_3050042, EPI_ISL_3050043, EPI_ISL_3050047, EPI_ISL_3050048, EPI_ISL_3050049, EPI_ISL_3050053, EPI_ISL_3050054, EPI_ISL_3050057, EPI_ISL_3050058, EPI_ISL_3050059, EPI_ISL_3050060, EPI_ISL_3050062, EPI_ISL_3050064, EPI_ISL_3050065, EPI_ISL_3050066, EPI_ISL_3050069, EPI_ISL_3050070, EPI_ISL_3050071, EPI_ISL_3050072, EPI_ISL_3050073, EPI_ISL_3050074, EPI_ISL_3050075, EPI_ISL_3050076, EPI_ISL_3050078, EPI_ISL_3050079, EPI_ISL_3050083, EPI_ISL_3050084, EPI_ISL_3050085, EPI_ISL_3050086, EPI_ISL_3050087, EPI_ISL_3050088, EPI_ISL_3050089, EPI_ISL_3050091, EPI_ISL_3050092, EPI_ISL_3050095, EPI_ISL_3050097, EPI_ISL_3050098, EPI_ISL_3050099, EPI_ISL_3050100, EPI_ISL_3050102, EPI_ISL_3050103, EPI_ISL_3050104, EPI_ISL_3050105, EPI_ISL_3050107, EPI_ISL_3050108, EPI_ISL_3050109, EPI_ISL_3050110, EPI_ISL_3050111, EPI_ISL_3050112, EPI_ISL_3050114, EPI_ISL_3050115, EPI_ISL_3050117, EPI_ISL_3050118, EPI_ISL_3050119, EPI_ISL_3050120, EPI_ISL_3050122, EPI_ISL_3050123, EPI_ISL_3050124, EPI_ISL_3050125, EPI_ISL_3050127, EPI_ISL_3050128, EPI_ISL_3050129, EPI_ISL_3050130, EPI_ISL_3050132, EPI_ISL_3050133, EPI_ISL_3050134, EPI_ISL_3050135 | | | |
| see above | State Testing Facility | Altius Institute for Biomedical Research | Daniel Bates, Rebecca Bruders, Michael Buckley, Mark Frerker, Amanda Gale, Clem Green, Muhammad Halimun, Kneshay Harper, Matt Hartman, Alex Isner, Audra Johnson, Jessica Kunder, Lauren Mitchell, Jemma Nelson, Alex Nguyen, Tobias Ragoczy, Joshua Richards, Jean Robinston, Jacob Rodriguez, John Stamatoyannopoulos, Eric Thorland, Julia Wald |
| EPI_ISL_3050151, EPI_ISL_3050156, EPI_ISL_3050159, EPI_ISL_3050165, EPI_ISL_3050166, EPI_ISL_3050167, EPI_ISL_3050168, EPI_ISL_3050171, EPI_ISL_3050172, EPI_ISL_3050173, EPI_ISL_3050175, EPI_ISL_3050176, EPI_ISL_3050177, EPI_ISL_3050178 | | | |
| see above | Pathology, UAB | Pathology, UAB | Moates,D., Hendrickson,R.C., Hernandez,J., Lefkowitz,E.J., Leal,S.M. |
| EPI_ISL_3050180, EPI_ISL_3050182, EPI_ISL_3050183, EPI_ISL_3050184, EPI_ISL_3050191, EPI_ISL_3050192 | Stanford Health Care | Stanford University School of Medicine, Clinical Virology Laboratory | Mamdouh Sibai, Daniel Solis, Selamawit Bihon, Bernadette Troung, Becky Jiang, Nathan Hammond, James Zehnder, Malaya K. Sahoo, and Benjamin A. Pinsky |
| EPI_ISL_3050193, EPI_ISL_3050195, EPI_ISL_3050196, | Stanford Health Care | Stanford University School of Medicine, Clinical Virology Laboratory | Daniel Solis, Mamdouh Sibai, Selamawit Bihon, Bernadette Troung, Becky Jiang, Nathan Hammond, James Zehnder, Malaya K. Sahoo, and Benjamin A. Pinsky |

| | | | |
|---|---|---|---|
| EPI_ISL_3050199, EPI_ISL_3050200, EPI_ISL_3050201, EPI_ISL_3050202, EPI_ISL_3050203, EPI_ISL_3050204 | | | |
| EPI_ISL_3050208, EPI_ISL_3050211, EPI_ISL_3050216, EPI_ISL_3050217, EPI_ISL_3050218, EPI_ISL_3050219, EPI_ISL_3050221, EPI_ISL_3050222, EPI_ISL_3050223, EPI_ISL_3050224, EPI_ISL_3050225, EPI_ISL_3050226 | | | |
| see above | Stanford Health Care | Stanford University School of Medicine, Clinical Virology Laboratory | Bernadette Troung, Becky Jiang, Daniel Solis, Mamdouh Sibai, Selamawit Bihon, Nathan Hammond, James Zehnder, Malaya K. Sahoo, and Benjamin A. Pinsky |
| EPI_ISL_3050228, EPI_ISL_3050229, EPI_ISL_3050230, EPI_ISL_3050231, EPI_ISL_3050232, EPI_ISL_3050234, EPI_ISL_3050235, EPI_ISL_3050236, EPI_ISL_3050237, EPI_ISL_3050238, EPI_ISL_3050239, EPI_ISL_3050240, EPI_ISL_3050241, EPI_ISL_3050242, EPI_ISL_3050244, EPI_ISL_3050245, EPI_ISL_3050246, EPI_ISL_3050247, EPI_ISL_3050250, EPI_ISL_3050252, EPI_ISL_3050253, EPI_ISL_3050254, EPI_ISL_3050255, EPI_ISL_3050256 | | | |
| see above | Stanford Health Care | Stanford University School of Medicine, Clinical Virology Laboratory | Malaya K. Sahoo, Becky Jiang, Daniel Solis, Mamdouh Sibai, Bernadette Troung, Selamawit Bihon, Nathan Hammond, James Zehnder, and Benjamin A. Pinsky |
| EPI_ISL_3050258 | Shah Alam Hospital | Institute for Medical Research, Infectious Disease Research Centre, National Institutes of Health, Ministry of Health Malaysia | Suppiah J, Kamel K, Mohd Zawawi Z, Azizan MA, Ramly N, Robert F, Anasir MI, Thayan R |
| EPI_ISL_3050770, EPI_ISL_3050771, EPI_ISL_3050772, EPI_ISL_3050773, EPI_ISL_3050774, EPI_ISL_3050775, EPI_ISL_3050776, EPI_ISL_3050778, EPI_ISL_3050779, EPI_ISL_3050780, EPI_ISL_3050781, EPI_ISL_3050782, EPI_ISL_3050783, EPI_ISL_3050784, EPI_ISL_3050785, EPI_ISL_3050786, EPI_ISL_3050787, EPI_ISL_3050788, EPI_ISL_3050789, EPI_ISL_3050790, EPI_ISL_3050791, EPI_ISL_3050792, EPI_ISL_3050793, EPI_ISL_3050794, EPI_ISL_3050795, EPI_ISL_3050796, EPI_ISL_3050798, EPI_ISL_3050799, EPI_ISL_3050800 | | | |
| see above | Microbiological Diagnostic Unit - Public Health Laboratory (MDU-PHL) | MDU-PHL | Seemann T., Sait, M.L., Sherry, N.L. |
| EPI_ISL_3050801 | Victorian Infectious Diseases Reference Laboratory (VIDRL) | VIDRL and MDU-PHL | Caly L., Seemann T., Sait, M.L., Druce J., Sherry, N.L. |
| EPI_ISL_3050802, EPI_ISL_3050803, EPI_ISL_3050804, EPI_ISL_3050805, EPI_ISL_3050806, EPI_ISL_3050808, EPI_ISL_3050809, EPI_ISL_3050810, EPI_ISL_3050811, EPI_ISL_3050812, EPI_ISL_3050813 | | | |
| see above | Microbiological Diagnostic Unit - Public Health Laboratory (MDU-PHL) | MDU-PHL | Seemann T., Sait, M.L., Sherry, N.L. |
| EPI_ISL_3050815 | Lighthouse Lab in Glasgow | Wellcome Sanger Institute for the COVID-19 Genomics UK (COG-UK) Consortium | Harper VanSteenhouse, Yumi Kasai, David Gray, Carol Clugston, Anna Dominiczak and Alex Alderton, Roberto Amato, Jeffrey Barrett, Sonia Goncalves, Ewan Harrison, David K. Jackson, Ian Johnston, Dominic Kwiatkowski, Cordelia Langford, John Sillitoe on behalf of the Wellcome Sanger Institute COVID-19 Surveillance Team |
| EPI_ISL_3050816 | Berkshire and Surrey Pathology Services Lighthouse Laboratory | Wellcome Sanger Institute for the COVID-19 Genomics UK (COG-UK) Consortium | Berkshire and Surrey Pathology Services Lighthouse Laboratory and Alex Alderton, Roberto Amato, Jeffrey Barrett, Sonia Goncalves, Ewan Harrison, David K. Jackson, Ian Johnston, Dominic Kwiatkowski, Cordelia Langford, John Sillitoe on behalf of the Wellcome Sanger Institute COVID-19 Surveillance Team |
| EPI_ISL_3050817 | Randox Laboratories | Wellcome Sanger Institute for the COVID-19 Genomics UK (COG-UK) Consortium | Randox Laboratories and Alex Alderton, Roberto Amato, Jeffrey Barrett, Sonia Goncalves, Ewan Harrison, David K. Jackson, Ian Johnston, Dominic Kwiatkowski, Cordelia Langford, John Sillitoe on behalf of the Wellcome Sanger Institute COVID-19 Surveillance Team |
| EPI_ISL_3050818, EPI_ISL_3050819 | Lighthouse Lab in Glasgow | Wellcome Sanger Institute for the COVID-19 Genomics UK (COG-UK) Consortium | Harper VanSteenhouse, Yumi Kasai, David Gray, Carol Clugston, Anna Dominiczak and Alex Alderton, Roberto Amato, Jeffrey Barrett, Sonia Goncalves, Ewan Harrison, David K. Jackson, Ian Johnston, Dominic Kwiatkowski, Cordelia Langford, John Sillitoe on behalf of the Wellcome Sanger Institute COVID-19 Surveillance Team |
| EPI_ISL_3050820 | Randox Laboratories | Wellcome Sanger Institute for the COVID-19 Genomics UK (COG-UK) Consortium | Randox Laboratories and Alex Alderton, Roberto Amato, Jeffrey Barrett, Sonia Goncalves, Ewan Harrison, David K. Jackson, Ian Johnston, Dominic Kwiatkowski, Cordelia Langford, John Sillitoe on behalf of the Wellcome Sanger Institute COVID-19 Surveillance Team |
| EPI_ISL_3050821 | Lighthouse Lab in Glasgow | Wellcome Sanger Institute for the COVID-19 Genomics UK (COG-UK) Consortium | Harper VanSteenhouse, Yumi Kasai, David Gray, Carol Clugston, Anna Dominiczak and Alex Alderton, Roberto Amato, Jeffrey Barrett, Sonia Goncalves, Ewan Harrison, David K. Jackson, Ian Johnston, Dominic Kwiatkowski, Cordelia Langford, John Sillitoe on behalf of the Wellcome Sanger Institute COVID-19 Surveillance Team |
| EPI_ISL_3050822, EPI_ISL_3050823 | Berkshire and Surrey Pathology Services Lighthouse Laboratory | Wellcome Sanger Institute for the COVID-19 Genomics UK (COG-UK) Consortium | Berkshire and Surrey Pathology Services Lighthouse Laboratory and Alex Alderton, Roberto Amato, Jeffrey Barrett, Sonia Goncalves, Ewan Harrison, David K. Jackson, Ian Johnston, Dominic Kwiatkowski, Cordelia Langford, John Sillitoe on behalf of the Wellcome Sanger Institute COVID-19 Surveillance Team |
| EPI_ISL_3050824, EPI_ISL_3050825, EPI_ISL_3050826, EPI_ISL_3050827 | Lighthouse Lab in Glasgow | Wellcome Sanger Institute for the COVID-19 Genomics UK (COG-UK) Consortium | Harper VanSteenhouse, Yumi Kasai, David Gray, Carol Clugston, Anna Dominiczak and Alex Alderton, Roberto Amato, Jeffrey Barrett, Sonia Goncalves, Ewan Harrison, David K. Jackson, Ian Johnston, Dominic Kwiatkowski, Cordelia Langford, John Sillitoe on behalf of the Wellcome Sanger Institute COVID-19 Surveillance Team |
| EPI_ISL_3050828 | Randox Laboratories | Wellcome Sanger Institute for the COVID-19 Genomics UK (COG-UK) Consortium | Randox Laboratories and Alex Alderton, Roberto Amato, Jeffrey Barrett, Sonia Goncalves, Ewan Harrison, David K. Jackson, Ian Johnston, Dominic Kwiatkowski, Cordelia Langford, John Sillitoe on behalf of the Wellcome Sanger Institute COVID-19 Surveillance Team |
| EPI_ISL_3050829, EPI_ISL_3050830, EPI_ISL_3050831 | Lighthouse Lab in Glasgow | Wellcome Sanger Institute for the COVID-19 Genomics UK (COG-UK) Consortium | Harper VanSteenhouse, Yumi Kasai, David Gray, Carol Clugston, Anna Dominiczak and Alex Alderton, Roberto Amato, Jeffrey Barrett, Sonia Goncalves, Ewan Harrison, David K. Jackson, Ian Johnston, Dominic Kwiatkowski, Cordelia Langford, John Sillitoe on behalf of the Wellcome Sanger Institute COVID-19 Surveillance Team |
| EPI_ISL_3050832 | Berkshire and Surrey Pathology Services Lighthouse Laboratory | Wellcome Sanger Institute for the COVID-19 Genomics UK (COG-UK) Consortium | Berkshire and Surrey Pathology Services Lighthouse Laboratory and Alex Alderton, Roberto Amato, Jeffrey Barrett, Sonia Goncalves, Ewan Harrison, David K. Jackson, Ian Johnston, Dominic Kwiatkowski, Cordelia Langford, John Sillitoe on behalf of the Wellcome Sanger Institute COVID-19 Surveillance Team |
| EPI_ISL_3050833, EPI_ISL_3050834 | Lighthouse Lab in Glasgow | Wellcome Sanger Institute for the COVID-19 Genomics UK (COG-UK) Consortium | Harper VanSteenhouse, Yumi Kasai, David Gray, Carol Clugston, Anna Dominiczak and Alex Alderton, Roberto Amato, Jeffrey Barrett, Sonia Goncalves, Ewan Harrison, David K. Jackson, Ian Johnston, Dominic Kwiatkowski, Cordelia Langford, John Sillitoe on behalf of the Wellcome Sanger Institute COVID-19 Surveillance Team |
| EPI_ISL_3050835 | Berkshire and Surrey Pathology Services Lighthouse Laboratory | Wellcome Sanger Institute for the COVID-19 Genomics UK (COG-UK) Consortium | Berkshire and Surrey Pathology Services Lighthouse Laboratory and Alex Alderton, Roberto Amato, Jeffrey Barrett, Sonia Goncalves, Ewan Harrison, David K. Jackson, Ian Johnston, Dominic Kwiatkowski, Cordelia Langford, John Sillitoe on behalf of the Wellcome Sanger Institute COVID-19 Surveillance Team |
| EPI_ISL_3050836 | Lighthouse Lab in Glasgow | Wellcome Sanger Institute for the COVID-19 Genomics UK (COG-UK) Consortium | Harper VanSteenhouse, Yumi Kasai, David Gray, Carol Clugston, Anna Dominiczak and Alex Alderton, Roberto Amato, Jeffrey Barrett, Sonia Goncalves, Ewan Harrison, David K. Jackson, Ian Johnston, Dominic Kwiatkowski, Cordelia Langford, John Sillitoe on behalf of the Wellcome Sanger Institute COVID-19 Surveillance Team |
| EPI_ISL_3050837, EPI_ISL_3050838, EPI_ISL_3050839 | Randox Laboratories | Wellcome Sanger Institute for the COVID-19 Genomics UK (COG-UK) Consortium | Randox Laboratories and Alex Alderton, Roberto Amato, Jeffrey Barrett, Sonia Goncalves, Ewan Harrison, David K. Jackson, Ian Johnston, Dominic Kwiatkowski, Cordelia Langford, John Sillitoe on behalf of the Wellcome Sanger Institute COVID-19 Surveillance Team |
| EPI_ISL_3050840 | Lighthouse Lab in Glasgow | Wellcome Sanger Institute for the COVID-19 Genomics UK (COG-UK) Consortium | Harper VanSteenhouse, Yumi Kasai, David Gray, Carol Clugston, Anna Dominiczak and Alex Alderton, Roberto Amato, Jeffrey Barrett, Sonia Goncalves, Ewan Harrison, David K. Jackson, Ian Johnston, Dominic Kwiatkowski, Cordelia Langford, John Sillitoe on behalf of the Wellcome Sanger Institute COVID-19 Surveillance Team |
| EPI_ISL_3050841 | Berkshire and Surrey Pathology Services Lighthouse Laboratory | Wellcome Sanger Institute for the COVID-19 Genomics UK (COG-UK) Consortium | Berkshire and Surrey Pathology Services Lighthouse Laboratory and Alex Alderton, Roberto Amato, Jeffrey Barrett, Sonia Goncalves, Ewan Harrison, David K. Jackson, Ian Johnston, Dominic Kwiatkowski, Cordelia Langford, John Sillitoe on behalf of the Wellcome Sanger Institute COVID-19 Surveillance Team |
| EPI_ISL_3050842, EPI_ISL_3050843 | Lighthouse Lab in Glasgow | Wellcome Sanger Institute for the COVID-19 Genomics UK (COG-UK) Consortium | Harper VanSteenhouse, Yumi Kasai, David Gray, Carol Clugston, Anna Dominiczak and Alex Alderton, Roberto Amato, Jeffrey Barrett, Sonia Goncalves, Ewan Harrison, David K. Jackson, Ian Johnston, Dominic Kwiatkowski, Cordelia Langford, John Sillitoe on behalf of the Wellcome Sanger Institute COVID-19 Surveillance Team |
| EPI_ISL_3050844 | Randox Laboratories | Wellcome Sanger Institute for the COVID-19 Genomics UK (COG-UK) Consortium | Randox Laboratories and Alex Alderton, Roberto Amato, Jeffrey Barrett, Sonia Goncalves, Ewan Harrison, David K. Jackson, Ian Johnston, Dominic Kwiatkowski, Cordelia Langford, John Sillitoe on behalf of the Wellcome Sanger Institute COVID-19 Surveillance Team |
| EPI_ISL_3050845 | Berkshire and Surrey Pathology Services Lighthouse Laboratory | Wellcome Sanger Institute for the COVID-19 Genomics UK (COG-UK) Consortium | Berkshire and Surrey Pathology Services Lighthouse Laboratory and Alex Alderton, Roberto Amato, Jeffrey Barrett, Sonia Goncalves, Ewan Harrison, David K. Jackson, Ian Johnston, Dominic Kwiatkowski, Cordelia Langford, John Sillitoe on behalf of the Wellcome Sanger Institute COVID-19 Surveillance Team |
| EPI_ISL_3050846 | Lighthouse Lab in Glasgow | Wellcome Sanger Institute for the COVID-19 Genomics UK (COG-UK) Consortium | Harper VanSteenhouse, Yumi Kasai, David Gray, Carol Clugston, Anna Dominiczak and Alex Alderton, Roberto Amato, Jeffrey Barrett, Sonia Goncalves, Ewan Harrison, David K. Jackson, Ian Johnston, Dominic Kwiatkowski, Cordelia Langford, John Sillitoe on behalf of the Wellcome Sanger Institute COVID-19 Surveillance Team |
| EPI_ISL_3050847 | Randox Laboratories | Wellcome Sanger Institute for the COVID-19 Genomics UK (COG-UK) Consortium | Randox Laboratories and Alex Alderton, Roberto Amato, Jeffrey Barrett, Sonia Goncalves, Ewan Harrison, David K. Jackson, Ian Johnston, Dominic Kwiatkowski, Cordelia Langford, John Sillitoe on behalf of the Wellcome Sanger Institute COVID-19 Surveillance Team |
| EPI_ISL_3050848 | Lighthouse Lab in Glasgow | Wellcome Sanger Institute for the COVID-19 Genomics UK (COG-UK) Consortium | Harper VanSteenhouse, Yumi Kasai, David Gray, Carol Clugston, Anna Dominiczak and Alex Alderton, Roberto Amato, Jeffrey Barrett, Sonia Goncalves, Ewan Harrison, David K. Jackson, Ian Johnston, Dominic Kwiatkowski, Cordelia Langford, John Sillitoe on behalf of the Wellcome Sanger Institute COVID-19 Surveillance Team |
| EPI_ISL_3050849, EPI_ISL_3050850 | Berkshire and Surrey Pathology Services Lighthouse Laboratory | Wellcome Sanger Institute for the COVID-19 Genomics UK (COG-UK) Consortium | Berkshire and Surrey Pathology Services Lighthouse Laboratory and Alex Alderton, Roberto Amato, Jeffrey Barrett, Sonia Goncalves, Ewan Harrison, David K. Jackson, Ian Johnston, Dominic Kwiatkowski, Cordelia Langford, John Sillitoe on behalf of the Wellcome Sanger Institute COVID-19 Surveillance Team |
| EPI_ISL_3050851 | Randox Laboratories | Wellcome Sanger Institute for the COVID-19 Genomics UK (COG-UK) Consortium | Randox Laboratories and Alex Alderton, Roberto Amato, Jeffrey Barrett, Sonia Goncalves, Ewan Harrison, David K. Jackson, Ian Johnston, Dominic Kwiatkowski, Cordelia Langford, John Sillitoe on behalf of the Wellcome Sanger Institute COVID-19 Surveillance Team |
| EPI_ISL_3050852, EPI_ISL_3050853 | Lighthouse Lab in Glasgow | Wellcome Sanger Institute for the COVID-19 Genomics UK (COG-UK) Consortium | Harper VanSteenhouse, Yumi Kasai, David Gray, Carol Clugston, Anna Dominiczak and Alex Alderton, Roberto Amato, Jeffrey Barrett, Sonia Goncalves, Ewan Harrison, David K. Jackson, Ian Johnston, Dominic Kwiatkowski, Cordelia Langford, John Sillitoe on behalf of the Wellcome Sanger Institute COVID-19 Surveillance Team |
| EPI_ISL_3050854 | Randox Laboratories | Wellcome Sanger Institute for the COVID-19 Genomics UK (COG-UK) Consortium | Randox Laboratories and Alex Alderton, Roberto Amato, Jeffrey Barrett, Sonia Goncalves, Ewan Harrison, David K. Jackson, Ian Johnston, Dominic Kwiatkowski, Cordelia Langford, John Sillitoe on behalf of the Wellcome Sanger Institute COVID-19 Surveillance Team |
| EPI_ISL_3050855 | Berkshire and Surrey Pathology Services Lighthouse Laboratory | Wellcome Sanger Institute for the COVID-19 Genomics UK (COG-UK) Consortium | Berkshire and Surrey Pathology Services Lighthouse Laboratory and Alex Alderton, Roberto Amato, Jeffrey Barrett, Sonia Goncalves, Ewan Harrison, David K. Jackson, Ian Johnston, Dominic Kwiatkowski, Cordelia Langford, John Sillitoe on behalf of the Wellcome Sanger Institute COVID-19 Surveillance Team |
| EPI_ISL_3050856, EPI_ISL_3050857 | Randox Laboratories | Wellcome Sanger Institute for the COVID-19 Genomics UK (COG-UK) Consortium | Randox Laboratories and Alex Alderton, Roberto Amato, Jeffrey Barrett, Sonia Goncalves, Ewan Harrison, David K. Jackson, Ian Johnston, Dominic Kwiatkowski, Cordelia Langford, John Sillitoe on behalf of the Wellcome Sanger Institute COVID-19 Surveillance Team |
| EPI_ISL_3050858 | Berkshire and Surrey Pathology Services Lighthouse Laboratory | Wellcome Sanger Institute for the COVID-19 Genomics UK (COG-UK) Consortium | Berkshire and Surrey Pathology Services Lighthouse Laboratory and Alex Alderton, Roberto Amato, Jeffrey Barrett, Sonia Goncalves, Ewan Harrison, David K. Jackson, Ian Johnston, Dominic Kwiatkowski, Cordelia Langford, John Sillitoe on behalf of the Wellcome Sanger Institute COVID-19 Surveillance Team |
| EPI_ISL_3050859, EPI_ISL_3050860 | Randox Laboratories | Wellcome Sanger Institute for the COVID-19 Genomics UK (COG-UK) Consortium | Randox Laboratories and Alex Alderton, Roberto Amato, Jeffrey Barrett, Sonia Goncalves, Ewan Harrison, David K. Jackson, Ian Johnston, Dominic Kwiatkowski, Cordelia Langford, John Sillitoe on behalf of the Wellcome Sanger Institute COVID-19 Surveillance Team |
| EPI_ISL_3050861, EPI_ISL_3050862, EPI_ISL_3050863 | Lighthouse Lab in Glasgow | Wellcome Sanger Institute for the COVID-19 Genomics UK (COG-UK) Consortium | Harper VanSteenhouse, Yumi Kasai, David Gray, Carol Clugston, Anna Dominiczak and Alex Alderton, Roberto Amato, Jeffrey Barrett, Sonia Goncalves, Ewan Harrison, David K. Jackson, Ian Johnston, Dominic Kwiatkowski, Cordelia Langford, John Sillitoe on behalf of the Wellcome Sanger Institute COVID-19 Surveillance Team |
| EPI_ISL_3050864 | Berkshire and Surrey Pathology Services Lighthouse Laboratory | Wellcome Sanger Institute for the COVID-19 Genomics UK (COG-UK) | Berkshire and Surrey Pathology Services Lighthouse Laboratory and Alex Alderton, Roberto Amato, Jeffrey Barrett, Sonia Goncalves, Ewan Harrison, David K. Jackson, Ian Johnston, Dominic Kwiatkowski, Cordelia Langford, John Sillitoe on behalf |

| | | | |
|---|---|---|---|
| | | Consortium | of the Wellcome Sanger Institute COVID-19 Surveillance Team |
| EPI_ISL_3050865 | Randox Laboratories | Wellcome Sanger Institute for the COVID-19 Genomics UK (COG-UK) Consortium | Randox Laboratories and Alex Alderton, Roberto Amato, Jeffrey Barrett, Sonia Goncalves, Ewan Harrison, David K. Jackson, Ian Johnston, Dominic Kwiatkowski, Cordelia Langford, John Sillitoe on behalf of the Wellcome Sanger Institute COVID-19 Surveillance Team |
| EPI_ISL_3050866, EPI_ISL_3050867, EPI_ISL_3050868 | Lighthouse Lab in Glasgow | Wellcome Sanger Institute for the COVID-19 Genomics UK (COG-UK) Consortium | Harper VanSteenhouse, Yumi Kasai, David Gray, Carol Clugston, Anna Dominiczak and Alex Alderton, Roberto Amato, Jeffrey Barrett, Sonia Goncalves, Ewan Harrison, David K. Jackson, Ian Johnston, Dominic Kwiatkowski, Cordelia Langford, John Sillitoe on behalf of the Wellcome Sanger Institute COVID-19 Surveillance Team |
| EPI_ISL_3050869 | Berkshire and Surrey Pathology Services Lighthouse Laboratory | Wellcome Sanger Institute for the COVID-19 Genomics UK (COG-UK) Consortium | Berkshire and Surrey Pathology Services Lighthouse Laboratory and Alex Alderton, Roberto Amato, Jeffrey Barrett, Sonia Goncalves, Ewan Harrison, David K. Jackson, Ian Johnston, Dominic Kwiatkowski, Cordelia Langford, John Sillitoe on behalf of the Wellcome Sanger Institute COVID-19 Surveillance Team |
| EPI_ISL_3050870 | Lighthouse Lab in Glasgow | Wellcome Sanger Institute for the COVID-19 Genomics UK (COG-UK) Consortium | Harper VanSteenhouse, Yumi Kasai, David Gray, Carol Clugston, Anna Dominiczak and Alex Alderton, Roberto Amato, Jeffrey Barrett, Sonia Goncalves, Ewan Harrison, David K. Jackson, Ian Johnston, Dominic Kwiatkowski, Cordelia Langford, John Sillitoe on behalf of the Wellcome Sanger Institute COVID-19 Surveillance Team |
| EPI_ISL_3050871 | Berkshire and Surrey Pathology Services Lighthouse Laboratory | Wellcome Sanger Institute for the COVID-19 Genomics UK (COG-UK) Consortium | Berkshire and Surrey Pathology Services Lighthouse Laboratory and Alex Alderton, Roberto Amato, Jeffrey Barrett, Sonia Goncalves, Ewan Harrison, David K. Jackson, Ian Johnston, Dominic Kwiatkowski, Cordelia Langford, John Sillitoe on behalf of the Wellcome Sanger Institute COVID-19 Surveillance Team |
| EPI_ISL_3050872, EPI_ISL_3050873, EPI_ISL_3050874 | Lighthouse Lab in Glasgow | Wellcome Sanger Institute for the COVID-19 Genomics UK (COG-UK) Consortium | Harper VanSteenhouse, Yumi Kasai, David Gray, Carol Clugston, Anna Dominiczak and Alex Alderton, Roberto Amato, Jeffrey Barrett, Sonia Goncalves, Ewan Harrison, David K. Jackson, Ian Johnston, Dominic Kwiatkowski, Cordelia Langford, John Sillitoe on behalf of the Wellcome Sanger Institute COVID-19 Surveillance Team |
| EPI_ISL_3050875 | Randox Laboratories | Wellcome Sanger Institute for the COVID-19 Genomics UK (COG-UK) Consortium | Randox Laboratories and Alex Alderton, Roberto Amato, Jeffrey Barrett, Sonia Goncalves, Ewan Harrison, David K. Jackson, Ian Johnston, Dominic Kwiatkowski, Cordelia Langford, John Sillitoe on behalf of the Wellcome Sanger Institute COVID-19 Surveillance Team |
| EPI_ISL_3050876, EPI_ISL_3050877 | Berkshire and Surrey Pathology Services Lighthouse Laboratory | Wellcome Sanger Institute for the COVID-19 Genomics UK (COG-UK) Consortium | Berkshire and Surrey Pathology Services Lighthouse Laboratory and Alex Alderton, Roberto Amato, Jeffrey Barrett, Sonia Goncalves, Ewan Harrison, David K. Jackson, Ian Johnston, Dominic Kwiatkowski, Cordelia Langford, John Sillitoe on behalf of the Wellcome Sanger Institute COVID-19 Surveillance Team |
| EPI_ISL_3050878, EPI_ISL_3050879 | Lighthouse Lab in Glasgow | Wellcome Sanger Institute for the COVID-19 Genomics UK (COG-UK) Consortium | Harper VanSteenhouse, Yumi Kasai, David Gray, Carol Clugston, Anna Dominiczak and Alex Alderton, Roberto Amato, Jeffrey Barrett, Sonia Goncalves, Ewan Harrison, David K. Jackson, Ian Johnston, Dominic Kwiatkowski, Cordelia Langford, John Sillitoe on behalf of the Wellcome Sanger Institute COVID-19 Surveillance Team |
| EPI_ISL_3050880 | Randox Laboratories | Wellcome Sanger Institute for the COVID-19 Genomics UK (COG-UK) Consortium | Randox Laboratories and Alex Alderton, Roberto Amato, Jeffrey Barrett, Sonia Goncalves, Ewan Harrison, David K. Jackson, Ian Johnston, Dominic Kwiatkowski, Cordelia Langford, John Sillitoe on behalf of the Wellcome Sanger Institute COVID-19 Surveillance Team |
| EPI_ISL_3050881 | Lighthouse Lab in Glasgow | Wellcome Sanger Institute for the COVID-19 Genomics UK (COG-UK) Consortium | Harper VanSteenhouse, Yumi Kasai, David Gray, Carol Clugston, Anna Dominiczak and Alex Alderton, Roberto Amato, Jeffrey Barrett, Sonia Goncalves, Ewan Harrison, David K. Jackson, Ian Johnston, Dominic Kwiatkowski, Cordelia Langford, John Sillitoe on behalf of the Wellcome Sanger Institute COVID-19 Surveillance Team |
| EPI_ISL_3050882 | Berkshire and Surrey Pathology Services Lighthouse Laboratory | Wellcome Sanger Institute for the COVID-19 Genomics UK (COG-UK) Consortium | Berkshire and Surrey Pathology Services Lighthouse Laboratory and Alex Alderton, Roberto Amato, Jeffrey Barrett, Sonia Goncalves, Ewan Harrison, David K. Jackson, Ian Johnston, Dominic Kwiatkowski, Cordelia Langford, John Sillitoe on behalf of the Wellcome Sanger Institute COVID-19 Surveillance Team |
| EPI_ISL_3050883 | Lighthouse Lab in Glasgow | Wellcome Sanger Institute for the COVID-19 Genomics UK (COG-UK) Consortium | Harper VanSteenhouse, Yumi Kasai, David Gray, Carol Clugston, Anna Dominiczak and Alex Alderton, Roberto Amato, Jeffrey Barrett, Sonia Goncalves, Ewan Harrison, David K. Jackson, Ian Johnston, Dominic Kwiatkowski, Cordelia Langford, John Sillitoe on behalf of the Wellcome Sanger Institute COVID-19 Surveillance Team |
| EPI_ISL_3050884, EPI_ISL_3050885 | Berkshire and Surrey Pathology Services Lighthouse Laboratory | Wellcome Sanger Institute for the COVID-19 Genomics UK (COG-UK) Consortium | Berkshire and Surrey Pathology Services Lighthouse Laboratory and Alex Alderton, Roberto Amato, Jeffrey Barrett, Sonia Goncalves, Ewan Harrison, David K. Jackson, Ian Johnston, Dominic Kwiatkowski, Cordelia Langford, John Sillitoe on behalf of the Wellcome Sanger Institute COVID-19 Surveillance Team |
| EPI_ISL_3050886 | Randox Laboratories | Wellcome Sanger Institute for the COVID-19 Genomics UK (COG-UK) Consortium | Randox Laboratories and Alex Alderton, Roberto Amato, Jeffrey Barrett, Sonia Goncalves, Ewan Harrison, David K. Jackson, Ian Johnston, Dominic Kwiatkowski, Cordelia Langford, John Sillitoe on behalf of the Wellcome Sanger Institute COVID-19 Surveillance Team |
| EPI_ISL_3050887 | Berkshire and Surrey Pathology Services Lighthouse Laboratory | Wellcome Sanger Institute for the COVID-19 Genomics UK (COG-UK) Consortium | Berkshire and Surrey Pathology Services Lighthouse Laboratory and Alex Alderton, Roberto Amato, Jeffrey Barrett, Sonia Goncalves, Ewan Harrison, David K. Jackson, Ian Johnston, Dominic Kwiatkowski, Cordelia Langford, John Sillitoe on behalf of the Wellcome Sanger Institute COVID-19 Surveillance Team |
| EPI_ISL_3050888, EPI_ISL_3050889, EPI_ISL_3050890 | Lighthouse Lab in Glasgow | Wellcome Sanger Institute for the COVID-19 Genomics UK (COG-UK) Consortium | Harper VanSteenhouse, Yumi Kasai, David Gray, Carol Clugston, Anna Dominiczak and Alex Alderton, Roberto Amato, Jeffrey Barrett, Sonia Goncalves, Ewan Harrison, David K. Jackson, Ian Johnston, Dominic Kwiatkowski, Cordelia Langford, John Sillitoe on behalf of the Wellcome Sanger Institute COVID-19 Surveillance Team |
| EPI_ISL_3050891 | Randox Laboratories | Wellcome Sanger Institute for the COVID-19 Genomics UK (COG-UK) Consortium | Randox Laboratories and Alex Alderton, Roberto Amato, Jeffrey Barrett, Sonia Goncalves, Ewan Harrison, David K. Jackson, Ian Johnston, Dominic Kwiatkowski, Cordelia Langford, John Sillitoe on behalf of the Wellcome Sanger Institute COVID-19 Surveillance Team |
| EPI_ISL_3050892, EPI_ISL_3050893, EPI_ISL_3050894 | Lighthouse Lab in Glasgow | Wellcome Sanger Institute for the COVID-19 Genomics UK (COG-UK) Consortium | Harper VanSteenhouse, Yumi Kasai, David Gray, Carol Clugston, Anna Dominiczak and Alex Alderton, Roberto Amato, Jeffrey Barrett, Sonia Goncalves, Ewan Harrison, David K. Jackson, Ian Johnston, Dominic Kwiatkowski, Cordelia Langford, John Sillitoe on behalf of the Wellcome Sanger Institute COVID-19 Surveillance Team |
| EPI_ISL_3050895 | Randox Laboratories | Wellcome Sanger Institute for the COVID-19 Genomics UK (COG-UK) Consortium | Randox Laboratories and Alex Alderton, Roberto Amato, Jeffrey Barrett, Sonia Goncalves, Ewan Harrison, David K. Jackson, Ian Johnston, Dominic Kwiatkowski, Cordelia Langford, John Sillitoe on behalf of the Wellcome Sanger Institute COVID-19 Surveillance Team |
| EPI_ISL_3050896, EPI_ISL_3050897 | Lighthouse Lab in Glasgow | Wellcome Sanger Institute for the COVID-19 Genomics UK (COG-UK) Consortium | Harper VanSteenhouse, Yumi Kasai, David Gray, Carol Clugston, Anna Dominiczak and Alex Alderton, Roberto Amato, Jeffrey Barrett, Sonia Goncalves, Ewan Harrison, David K. Jackson, Ian Johnston, Dominic Kwiatkowski, Cordelia Langford, John Sillitoe on behalf of the Wellcome Sanger Institute COVID-19 Surveillance Team |
| EPI_ISL_3050898, EPI_ISL_3050899, EPI_ISL_3050900 | Berkshire and Surrey Pathology Services Lighthouse Laboratory | Wellcome Sanger Institute for the COVID-19 Genomics UK (COG-UK) Consortium | Berkshire and Surrey Pathology Services Lighthouse Laboratory and Alex Alderton, Roberto Amato, Jeffrey Barrett, Sonia Goncalves, Ewan Harrison, David K. Jackson, Ian Johnston, Dominic Kwiatkowski, Cordelia Langford, John Sillitoe on behalf of the Wellcome Sanger Institute COVID-19 Surveillance Team |
| EPI_ISL_3050901 | Lighthouse Lab in Glasgow | Wellcome Sanger Institute for the COVID-19 Genomics UK (COG-UK) Consortium | Harper VanSteenhouse, Yumi Kasai, David Gray, Carol Clugston, Anna Dominiczak and Alex Alderton, Roberto Amato, Jeffrey Barrett, Sonia Goncalves, Ewan Harrison, David K. Jackson, Ian Johnston, Dominic Kwiatkowski, Cordelia Langford, John Sillitoe on behalf of the Wellcome Sanger Institute COVID-19 Surveillance Team |
| EPI_ISL_3050902 | Berkshire and Surrey Pathology Services Lighthouse Laboratory | Wellcome Sanger Institute for the COVID-19 Genomics UK (COG-UK) Consortium | Berkshire and Surrey Pathology Services Lighthouse Laboratory and Alex Alderton, Roberto Amato, Jeffrey Barrett, Sonia Goncalves, Ewan Harrison, David K. Jackson, Ian Johnston, Dominic Kwiatkowski, Cordelia Langford, John Sillitoe on behalf of the Wellcome Sanger Institute COVID-19 Surveillance Team |
| EPI_ISL_3050903, EPI_ISL_3050904 | Lighthouse Lab in Glasgow | Wellcome Sanger Institute for the COVID-19 Genomics UK (COG-UK) Consortium | Harper VanSteenhouse, Yumi Kasai, David Gray, Carol Clugston, Anna Dominiczak and Alex Alderton, Roberto Amato, Jeffrey Barrett, Sonia Goncalves, Ewan Harrison, David K. Jackson, Ian Johnston, Dominic Kwiatkowski, Cordelia Langford, John Sillitoe on behalf of the Wellcome Sanger Institute COVID-19 Surveillance Team |
| EPI_ISL_3050905 | Berkshire and Surrey Pathology Services Lighthouse Laboratory | Wellcome Sanger Institute for the COVID-19 Genomics UK (COG-UK) Consortium | Berkshire and Surrey Pathology Services Lighthouse Laboratory and Alex Alderton, Roberto Amato, Jeffrey Barrett, Sonia Goncalves, Ewan Harrison, David K. Jackson, Ian Johnston, Dominic Kwiatkowski, Cordelia Langford, John Sillitoe on behalf of the Wellcome Sanger Institute COVID-19 Surveillance Team |
| EPI_ISL_3050906 | Randox Laboratories | Wellcome Sanger Institute for the COVID-19 Genomics UK (COG-UK) Consortium | Randox Laboratories and Alex Alderton, Roberto Amato, Jeffrey Barrett, Sonia Goncalves, Ewan Harrison, David K. Jackson, Ian Johnston, Dominic Kwiatkowski, Cordelia Langford, John Sillitoe on behalf of the Wellcome Sanger Institute COVID-19 Surveillance Team |
| EPI_ISL_3050907, EPI_ISL_3050908 | Lighthouse Lab in Glasgow | Wellcome Sanger Institute for the COVID-19 Genomics UK (COG-UK) Consortium | Harper VanSteenhouse, Yumi Kasai, David Gray, Carol Clugston, Anna Dominiczak and Alex Alderton, Roberto Amato, Jeffrey Barrett, Sonia Goncalves, Ewan Harrison, David K. Jackson, Ian Johnston, Dominic Kwiatkowski, Cordelia Langford, John Sillitoe on behalf of the Wellcome Sanger Institute COVID-19 Surveillance Team |
| EPI_ISL_3050909, EPI_ISL_3050910 | Berkshire and Surrey Pathology Services Lighthouse Laboratory | Wellcome Sanger Institute for the COVID-19 Genomics UK (COG-UK) Consortium | Berkshire and Surrey Pathology Services Lighthouse Laboratory and Alex Alderton, Roberto Amato, Jeffrey Barrett, Sonia Goncalves, Ewan Harrison, David K. Jackson, Ian Johnston, Dominic Kwiatkowski, Cordelia Langford, John Sillitoe on behalf of the Wellcome Sanger Institute COVID-19 Surveillance Team |
| EPI_ISL_3050911, EPI_ISL_3050912 | Lighthouse Lab in Glasgow | Wellcome Sanger Institute for the COVID-19 Genomics UK (COG-UK) Consortium | Harper VanSteenhouse, Yumi Kasai, David Gray, Carol Clugston, Anna Dominiczak and Alex Alderton, Roberto Amato, Jeffrey Barrett, Sonia Goncalves, Ewan Harrison, David K. Jackson, Ian Johnston, Dominic Kwiatkowski, Cordelia Langford, John Sillitoe on behalf of the Wellcome Sanger Institute COVID-19 Surveillance Team |
| EPI_ISL_3050913 | Randox Laboratories | Wellcome Sanger Institute for the COVID-19 Genomics UK (COG-UK) Consortium | Randox Laboratories and Alex Alderton, Roberto Amato, Jeffrey Barrett, Sonia Goncalves, Ewan Harrison, David K. Jackson, Ian Johnston, Dominic Kwiatkowski, Cordelia Langford, John Sillitoe on behalf of the Wellcome Sanger Institute COVID-19 Surveillance Team |
| EPI_ISL_3050914 | Lighthouse Lab in Glasgow | Wellcome Sanger Institute for the COVID-19 Genomics UK (COG-UK) Consortium | Harper VanSteenhouse, Yumi Kasai, David Gray, Carol Clugston, Anna Dominiczak and Alex Alderton, Roberto Amato, Jeffrey Barrett, Sonia Goncalves, Ewan Harrison, David K. Jackson, Ian Johnston, Dominic Kwiatkowski, Cordelia Langford, John Sillitoe on behalf of the Wellcome Sanger Institute COVID-19 Surveillance Team |
| EPI_ISL_3050915 | Randox Laboratories | Wellcome Sanger Institute for the COVID-19 Genomics UK (COG-UK) Consortium | Randox Laboratories and Alex Alderton, Roberto Amato, Jeffrey Barrett, Sonia Goncalves, Ewan Harrison, David K. Jackson, Ian Johnston, Dominic Kwiatkowski, Cordelia Langford, John Sillitoe on behalf of the Wellcome Sanger Institute COVID-19 Surveillance Team |
| EPI_ISL_3050916, EPI_ISL_3050917 | Lighthouse Lab in Glasgow | Wellcome Sanger Institute for the COVID-19 Genomics UK (COG-UK) Consortium | Harper VanSteenhouse, Yumi Kasai, David Gray, Carol Clugston, Anna Dominiczak and Alex Alderton, Roberto Amato, Jeffrey Barrett, Sonia Goncalves, Ewan Harrison, David K. Jackson, Ian Johnston, Dominic Kwiatkowski, Cordelia Langford, John Sillitoe on behalf of the Wellcome Sanger Institute COVID-19 Surveillance Team |
| EPI_ISL_3050918 | Berkshire and Surrey Pathology Services Lighthouse Laboratory | Wellcome Sanger Institute for the COVID-19 Genomics UK (COG-UK) Consortium | Berkshire and Surrey Pathology Services Lighthouse Laboratory and Alex Alderton, Roberto Amato, Jeffrey Barrett, Sonia Goncalves, Ewan Harrison, David K. Jackson, Ian Johnston, Dominic Kwiatkowski, Cordelia Langford, John Sillitoe on behalf of the Wellcome Sanger Institute COVID-19 Surveillance Team |
| EPI_ISL_3050919 | Lighthouse Lab in Glasgow | Wellcome Sanger Institute for the COVID-19 Genomics UK (COG-UK) Consortium | Harper VanSteenhouse, Yumi Kasai, David Gray, Carol Clugston, Anna Dominiczak and Alex Alderton, Roberto Amato, Jeffrey Barrett, Sonia Goncalves, Ewan Harrison, David K. Jackson, Ian Johnston, Dominic Kwiatkowski, Cordelia Langford, John Sillitoe on behalf of the Wellcome Sanger Institute COVID-19 Surveillance Team |
| EPI_ISL_3050920 | Randox Laboratories | Wellcome Sanger Institute for the COVID-19 Genomics UK (COG-UK) Consortium | Randox Laboratories and Alex Alderton, Roberto Amato, Jeffrey Barrett, Sonia Goncalves, Ewan Harrison, David K. Jackson, Ian Johnston, Dominic Kwiatkowski, Cordelia Langford, John Sillitoe on behalf of the Wellcome Sanger Institute COVID-19 Surveillance Team |
| EPI_ISL_3050921, EPI_ISL_3050922 | Lighthouse Lab in Glasgow | Wellcome Sanger Institute for the COVID-19 Genomics UK (COG-UK) Consortium | Harper VanSteenhouse, Yumi Kasai, David Gray, Carol Clugston, Anna Dominiczak and Alex Alderton, Roberto Amato, Jeffrey Barrett, Sonia Goncalves, Ewan Harrison, David K. Jackson, Ian Johnston, Dominic Kwiatkowski, Cordelia Langford, John Sillitoe on behalf of the Wellcome Sanger Institute COVID-19 Surveillance Team |
| EPI_ISL_3050923 | Berkshire and Surrey Pathology Services Lighthouse Laboratory | Wellcome Sanger Institute for the COVID-19 Genomics UK (COG-UK) Consortium | Berkshire and Surrey Pathology Services Lighthouse Laboratory and Alex Alderton, Roberto Amato, Jeffrey Barrett, Sonia Goncalves, Ewan Harrison, David K. Jackson, Ian Johnston, Dominic Kwiatkowski, Cordelia Langford, John Sillitoe on behalf of the Wellcome Sanger Institute COVID-19 Surveillance Team |
| EPI_ISL_3050924 | Randox Laboratories | Wellcome Sanger Institute for the COVID-19 Genomics UK (COG-UK) Consortium | Randox Laboratories and Alex Alderton, Roberto Amato, Jeffrey Barrett, Sonia Goncalves, Ewan Harrison, David K. Jackson, Ian Johnston, Dominic Kwiatkowski, Cordelia Langford, John Sillitoe on behalf of the Wellcome Sanger Institute COVID-19 Surveillance Team |
| EPI_ISL_3050925, EPI_ISL_3050926, EPI_ISL_3050927 | Lighthouse Lab in Glasgow | Wellcome Sanger Institute for the COVID-19 Genomics UK (COG-UK) Consortium | Harper VanSteenhouse, Yumi Kasai, David Gray, Carol Clugston, Anna Dominiczak and Alex Alderton, Roberto Amato, Jeffrey Barrett, Sonia Goncalves, Ewan Harrison, David K. Jackson, Ian Johnston, Dominic Kwiatkowski, Cordelia Langford, John Sillitoe on behalf of the Wellcome Sanger Institute COVID-19 Surveillance Team |
| EPI_ISL_3050928 | Randox Laboratories | Wellcome Sanger Institute for the COVID-19 Genomics UK (COG-UK) Consortium | Randox Laboratories and Alex Alderton, Roberto Amato, Jeffrey Barrett, Sonia Goncalves, Ewan Harrison, David K. Jackson, Ian Johnston, Dominic Kwiatkowski, Cordelia Langford, John Sillitoe on behalf of the Wellcome Sanger Institute COVID-19 Surveillance Team |

| | | | |
|---|---|---|---|
| EPI_ISL_3050929, EPI_ISL_3050930, EPI_ISL_3050931 | Lighthouse Lab in Glasgow | Wellcome Sanger Institute for the COVID-19 Genomics UK (COG-UK) Consortium | Harper VanSteenhouse, Yumi Kasai, David Gray, Carol Clugston, Anna Dominiczak and Alex Alderton, Roberto Amato, Jeffrey Barrett, Sonia Goncalves, Ewan Harrison, David K. Jackson, Ian Johnston, Dominic Kwiatkowski, Cordelia Langford, John Sillitoe on behalf of the Wellcome Sanger Institute COVID-19 Surveillance Team |
| EPI_ISL_3050932 | Randox Laboratories | Wellcome Sanger Institute for the COVID-19 Genomics UK (COG-UK) Consortium | Randox Laboratories and Alex Alderton, Roberto Amato, Jeffrey Barrett, Sonia Goncalves, Ewan Harrison, David K. Jackson, Ian Johnston, Dominic Kwiatkowski, Cordelia Langford, John Sillitoe on behalf of the Wellcome Sanger Institute COVID-19 Surveillance Team |
| EPI_ISL_3050933, EPI_ISL_3050934 | Lighthouse Lab in Glasgow | Wellcome Sanger Institute for the COVID-19 Genomics UK (COG-UK) Consortium | Harper VanSteenhouse, Yumi Kasai, David Gray, Carol Clugston, Anna Dominiczak and Alex Alderton, Roberto Amato, Jeffrey Barrett, Sonia Goncalves, Ewan Harrison, David K. Jackson, Ian Johnston, Dominic Kwiatkowski, Cordelia Langford, John Sillitoe on behalf of the Wellcome Sanger Institute COVID-19 Surveillance Team |
| EPI_ISL_3050935 | Berkshire and Surrey Pathology Services Lighthouse Laboratory | Wellcome Sanger Institute for the COVID-19 Genomics UK (COG-UK) Consortium | Berkshire and Surrey Pathology Services Lighthouse Laboratory and Alex Alderton, Roberto Amato, Jeffrey Barrett, Sonia Goncalves, Ewan Harrison, David K. Jackson, Ian Johnston, Dominic Kwiatkowski, Cordelia Langford, John Sillitoe on behalf of the Wellcome Sanger Institute COVID-19 Surveillance Team |
| EPI_ISL_3050936 | Lighthouse Lab in Glasgow | Wellcome Sanger Institute for the COVID-19 Genomics UK (COG-UK) Consortium | Harper VanSteenhouse, Yumi Kasai, David Gray, Carol Clugston, Anna Dominiczak and Alex Alderton, Roberto Amato, Jeffrey Barrett, Sonia Goncalves, Ewan Harrison, David K. Jackson, Ian Johnston, Dominic Kwiatkowski, Cordelia Langford, John Sillitoe on behalf of the Wellcome Sanger Institute COVID-19 Surveillance Team |
| EPI_ISL_3050937, EPI_ISL_3050938, EPI_ISL_3050939, EPI_ISL_3050940 | Randox Laboratories | Wellcome Sanger Institute for the COVID-19 Genomics UK (COG-UK) Consortium | Randox Laboratories and Alex Alderton, Roberto Amato, Jeffrey Barrett, Sonia Goncalves, Ewan Harrison, David K. Jackson, Ian Johnston, Dominic Kwiatkowski, Cordelia Langford, John Sillitoe on behalf of the Wellcome Sanger Institute COVID-19 Surveillance Team |
| EPI_ISL_3050941, EPI_ISL_3050942, EPI_ISL_3050943, EPI_ISL_3050944 | Lighthouse Lab in Glasgow | Wellcome Sanger Institute for the COVID-19 Genomics UK (COG-UK) Consortium | Harper VanSteenhouse, Yumi Kasai, David Gray, Carol Clugston, Anna Dominiczak and Alex Alderton, Roberto Amato, Jeffrey Barrett, Sonia Goncalves, Ewan Harrison, David K. Jackson, Ian Johnston, Dominic Kwiatkowski, Cordelia Langford, John Sillitoe on behalf of the Wellcome Sanger Institute COVID-19 Surveillance Team |
| EPI_ISL_3050945 | Randox Laboratories | Wellcome Sanger Institute for the COVID-19 Genomics UK (COG-UK) Consortium | Randox Laboratories and Alex Alderton, Roberto Amato, Jeffrey Barrett, Sonia Goncalves, Ewan Harrison, David K. Jackson, Ian Johnston, Dominic Kwiatkowski, Cordelia Langford, John Sillitoe on behalf of the Wellcome Sanger Institute COVID-19 Surveillance Team |
| EPI_ISL_3050946, EPI_ISL_3050947, EPI_ISL_3050948, EPI_ISL_3050949 | Lighthouse Lab in Glasgow | Wellcome Sanger Institute for the COVID-19 Genomics UK (COG-UK) Consortium | Harper VanSteenhouse, Yumi Kasai, David Gray, Carol Clugston, Anna Dominiczak and Alex Alderton, Roberto Amato, Jeffrey Barrett, Sonia Goncalves, Ewan Harrison, David K. Jackson, Ian Johnston, Dominic Kwiatkowski, Cordelia Langford, John Sillitoe on behalf of the Wellcome Sanger Institute COVID-19 Surveillance Team |
| EPI_ISL_3050950 | Randox Laboratories | Wellcome Sanger Institute for the COVID-19 Genomics UK (COG-UK) Consortium | Randox Laboratories and Alex Alderton, Roberto Amato, Jeffrey Barrett, Sonia Goncalves, Ewan Harrison, David K. Jackson, Ian Johnston, Dominic Kwiatkowski, Cordelia Langford, John Sillitoe on behalf of the Wellcome Sanger Institute COVID-19 Surveillance Team |
| EPI_ISL_3050951 | Berkshire and Surrey Pathology Services Lighthouse Laboratory | Wellcome Sanger Institute for the COVID-19 Genomics UK (COG-UK) Consortium | Berkshire and Surrey Pathology Services Lighthouse Laboratory and Alex Alderton, Roberto Amato, Jeffrey Barrett, Sonia Goncalves, Ewan Harrison, David K. Jackson, Ian Johnston, Dominic Kwiatkowski, Cordelia Langford, John Sillitoe on behalf of the Wellcome Sanger Institute COVID-19 Surveillance Team |
| EPI_ISL_3050952 | Randox Laboratories | Wellcome Sanger Institute for the COVID-19 Genomics UK (COG-UK) Consortium | Randox Laboratories and Alex Alderton, Roberto Amato, Jeffrey Barrett, Sonia Goncalves, Ewan Harrison, David K. Jackson, Ian Johnston, Dominic Kwiatkowski, Cordelia Langford, John Sillitoe on behalf of the Wellcome Sanger Institute COVID-19 Surveillance Team |
| EPI_ISL_3050953 | Lighthouse Lab in Glasgow | Wellcome Sanger Institute for the COVID-19 Genomics UK (COG-UK) Consortium | Harper VanSteenhouse, Yumi Kasai, David Gray, Carol Clugston, Anna Dominiczak and Alex Alderton, Roberto Amato, Jeffrey Barrett, Sonia Goncalves, Ewan Harrison, David K. Jackson, Ian Johnston, Dominic Kwiatkowski, Cordelia Langford, John Sillitoe on behalf of the Wellcome Sanger Institute COVID-19 Surveillance Team |
| EPI_ISL_3050954, EPI_ISL_3050955, EPI_ISL_3050956, EPI_ISL_3050957, EPI_ISL_3050958, EPI_ISL_3050959 | Berkshire and Surrey Pathology Services Lighthouse Laboratory | Wellcome Sanger Institute for the COVID-19 Genomics UK (COG-UK) Consortium | Berkshire and Surrey Pathology Services Lighthouse Laboratory and Alex Alderton, Roberto Amato, Jeffrey Barrett, Sonia Goncalves, Ewan Harrison, David K. Jackson, Ian Johnston, Dominic Kwiatkowski, Cordelia Langford, John Sillitoe on behalf of the Wellcome Sanger Institute COVID-19 Surveillance Team |
| EPI_ISL_3050960 | Lighthouse Lab in Glasgow | Wellcome Sanger Institute for the COVID-19 Genomics UK (COG-UK) Consortium | Harper VanSteenhouse, Yumi Kasai, David Gray, Carol Clugston, Anna Dominiczak and Alex Alderton, Roberto Amato, Jeffrey Barrett, Sonia Goncalves, Ewan Harrison, David K. Jackson, Ian Johnston, Dominic Kwiatkowski, Cordelia Langford, John Sillitoe on behalf of the Wellcome Sanger Institute COVID-19 Surveillance Team |
| EPI_ISL_3050961 | Berkshire and Surrey Pathology Services Lighthouse Laboratory | Wellcome Sanger Institute for the COVID-19 Genomics UK (COG-UK) Consortium | Berkshire and Surrey Pathology Services Lighthouse Laboratory and Alex Alderton, Roberto Amato, Jeffrey Barrett, Sonia Goncalves, Ewan Harrison, David K. Jackson, Ian Johnston, Dominic Kwiatkowski, Cordelia Langford, John Sillitoe on behalf of the Wellcome Sanger Institute COVID-19 Surveillance Team |
| EPI_ISL_3050962, EPI_ISL_3050963, EPI_ISL_3050964, EPI_ISL_3050965, EPI_ISL_3050966 | Lighthouse Lab in Glasgow | Wellcome Sanger Institute for the COVID-19 Genomics UK (COG-UK) Consortium | Harper VanSteenhouse, Yumi Kasai, David Gray, Carol Clugston, Anna Dominiczak and Alex Alderton, Roberto Amato, Jeffrey Barrett, Sonia Goncalves, Ewan Harrison, David K. Jackson, Ian Johnston, Dominic Kwiatkowski, Cordelia Langford, John Sillitoe on behalf of the Wellcome Sanger Institute COVID-19 Surveillance Team |
| EPI_ISL_3050967 | Randox Laboratories | Wellcome Sanger Institute for the COVID-19 Genomics UK (COG-UK) Consortium | Randox Laboratories and Alex Alderton, Roberto Amato, Jeffrey Barrett, Sonia Goncalves, Ewan Harrison, David K. Jackson, Ian Johnston, Dominic Kwiatkowski, Cordelia Langford, John Sillitoe on behalf of the Wellcome Sanger Institute COVID-19 Surveillance Team |
| EPI_ISL_3050968 | Berkshire and Surrey Pathology Services Lighthouse Laboratory | Wellcome Sanger Institute for the COVID-19 Genomics UK (COG-UK) Consortium | Berkshire and Surrey Pathology Services Lighthouse Laboratory and Alex Alderton, Roberto Amato, Jeffrey Barrett, Sonia Goncalves, Ewan Harrison, David K. Jackson, Ian Johnston, Dominic Kwiatkowski, Cordelia Langford, John Sillitoe on behalf of the Wellcome Sanger Institute COVID-19 Surveillance Team |
| EPI_ISL_3050969 | Randox Laboratories | Wellcome Sanger Institute for the COVID-19 Genomics UK (COG-UK) Consortium | Randox Laboratories and Alex Alderton, Roberto Amato, Jeffrey Barrett, Sonia Goncalves, Ewan Harrison, David K. Jackson, Ian Johnston, Dominic Kwiatkowski, Cordelia Langford, John Sillitoe on behalf of the Wellcome Sanger Institute COVID-19 Surveillance Team |
| EPI_ISL_3050970, EPI_ISL_3050971 | Berkshire and Surrey Pathology Services Lighthouse Laboratory | Wellcome Sanger Institute for the COVID-19 Genomics UK (COG-UK) Consortium | Berkshire and Surrey Pathology Services Lighthouse Laboratory and Alex Alderton, Roberto Amato, Jeffrey Barrett, Sonia Goncalves, Ewan Harrison, David K. Jackson, Ian Johnston, Dominic Kwiatkowski, Cordelia Langford, John Sillitoe on behalf of the Wellcome Sanger Institute COVID-19 Surveillance Team |
| EPI_ISL_3050972 | Lighthouse Lab in Glasgow | Wellcome Sanger Institute for the COVID-19 Genomics UK (COG-UK) Consortium | Harper VanSteenhouse, Yumi Kasai, David Gray, Carol Clugston, Anna Dominiczak and Alex Alderton, Roberto Amato, Jeffrey Barrett, Sonia Goncalves, Ewan Harrison, David K. Jackson, Ian Johnston, Dominic Kwiatkowski, Cordelia Langford, John Sillitoe on behalf of the Wellcome Sanger Institute COVID-19 Surveillance Team |
| EPI_ISL_3050973 | Randox Laboratories | Wellcome Sanger Institute for the COVID-19 Genomics UK (COG-UK) Consortium | Randox Laboratories and Alex Alderton, Roberto Amato, Jeffrey Barrett, Sonia Goncalves, Ewan Harrison, David K. Jackson, Ian Johnston, Dominic Kwiatkowski, Cordelia Langford, John Sillitoe on behalf of the Wellcome Sanger Institute COVID-19 Surveillance Team |
| EPI_ISL_3050974, EPI_ISL_3050975, EPI_ISL_3050976 | Lighthouse Lab in Glasgow | Wellcome Sanger Institute for the COVID-19 Genomics UK (COG-UK) Consortium | Harper VanSteenhouse, Yumi Kasai, David Gray, Carol Clugston, Anna Dominiczak and Alex Alderton, Roberto Amato, Jeffrey Barrett, Sonia Goncalves, Ewan Harrison, David K. Jackson, Ian Johnston, Dominic Kwiatkowski, Cordelia Langford, John Sillitoe on behalf of the Wellcome Sanger Institute COVID-19 Surveillance Team |
| EPI_ISL_3050977 | Berkshire and Surrey Pathology Services Lighthouse Laboratory | Wellcome Sanger Institute for the COVID-19 Genomics UK (COG-UK) Consortium | Berkshire and Surrey Pathology Services Lighthouse Laboratory and Alex Alderton, Roberto Amato, Jeffrey Barrett, Sonia Goncalves, Ewan Harrison, David K. Jackson, Ian Johnston, Dominic Kwiatkowski, Cordelia Langford, John Sillitoe on behalf of the Wellcome Sanger Institute COVID-19 Surveillance Team |
| EPI_ISL_3050978, EPI_ISL_3050979, EPI_ISL_3050980 | Randox Laboratories | Wellcome Sanger Institute for the COVID-19 Genomics UK (COG-UK) Consortium | Randox Laboratories and Alex Alderton, Roberto Amato, Jeffrey Barrett, Sonia Goncalves, Ewan Harrison, David K. Jackson, Ian Johnston, Dominic Kwiatkowski, Cordelia Langford, John Sillitoe on behalf of the Wellcome Sanger Institute COVID-19 Surveillance Team |
| EPI_ISL_3050981, EPI_ISL_3050982, EPI_ISL_3050983 | Lighthouse Lab in Glasgow | Wellcome Sanger Institute for the COVID-19 Genomics UK (COG-UK) Consortium | Harper VanSteenhouse, Yumi Kasai, David Gray, Carol Clugston, Anna Dominiczak and Alex Alderton, Roberto Amato, Jeffrey Barrett, Sonia Goncalves, Ewan Harrison, David K. Jackson, Ian Johnston, Dominic Kwiatkowski, Cordelia Langford, John Sillitoe on behalf of the Wellcome Sanger Institute COVID-19 Surveillance Team |
| EPI_ISL_3050984 | Berkshire and Surrey Pathology Services Lighthouse Laboratory | Wellcome Sanger Institute for the COVID-19 Genomics UK (COG-UK) Consortium | Berkshire and Surrey Pathology Services Lighthouse Laboratory and Alex Alderton, Roberto Amato, Jeffrey Barrett, Sonia Goncalves, Ewan Harrison, David K. Jackson, Ian Johnston, Dominic Kwiatkowski, Cordelia Langford, John Sillitoe on behalf of the Wellcome Sanger Institute COVID-19 Surveillance Team |
| EPI_ISL_3050985, EPI_ISL_3050986 | Lighthouse Lab in Glasgow | Wellcome Sanger Institute for the COVID-19 Genomics UK (COG-UK) Consortium | Harper VanSteenhouse, Yumi Kasai, David Gray, Carol Clugston, Anna Dominiczak and Alex Alderton, Roberto Amato, Jeffrey Barrett, Sonia Goncalves, Ewan Harrison, David K. Jackson, Ian Johnston, Dominic Kwiatkowski, Cordelia Langford, John Sillitoe on behalf of the Wellcome Sanger Institute COVID-19 Surveillance Team |
| EPI_ISL_3050987, EPI_ISL_3050988 | Randox Laboratories | Wellcome Sanger Institute for the COVID-19 Genomics UK (COG-UK) Consortium | Randox Laboratories and Alex Alderton, Roberto Amato, Jeffrey Barrett, Sonia Goncalves, Ewan Harrison, David K. Jackson, Ian Johnston, Dominic Kwiatkowski, Cordelia Langford, John Sillitoe on behalf of the Wellcome Sanger Institute COVID-19 Surveillance Team |
| EPI_ISL_3050989, EPI_ISL_3050990 | Berkshire and Surrey Pathology Services Lighthouse Laboratory | Wellcome Sanger Institute for the COVID-19 Genomics UK (COG-UK) Consortium | Berkshire and Surrey Pathology Services Lighthouse Laboratory and Alex Alderton, Roberto Amato, Jeffrey Barrett, Sonia Goncalves, Ewan Harrison, David K. Jackson, Ian Johnston, Dominic Kwiatkowski, Cordelia Langford, John Sillitoe on behalf of the Wellcome Sanger Institute COVID-19 Surveillance Team |
| EPI_ISL_3050991 | Lighthouse Lab in Glasgow | Wellcome Sanger Institute for the COVID-19 Genomics UK (COG-UK) Consortium | Harper VanSteenhouse, Yumi Kasai, David Gray, Carol Clugston, Anna Dominiczak and Alex Alderton, Roberto Amato, Jeffrey Barrett, Sonia Goncalves, Ewan Harrison, David K. Jackson, Ian Johnston, Dominic Kwiatkowski, Cordelia Langford, John Sillitoe on behalf of the Wellcome Sanger Institute COVID-19 Surveillance Team |
| EPI_ISL_3050992 | Randox Laboratories | Wellcome Sanger Institute for the COVID-19 Genomics UK (COG-UK) Consortium | Randox Laboratories and Alex Alderton, Roberto Amato, Jeffrey Barrett, Sonia Goncalves, Ewan Harrison, David K. Jackson, Ian Johnston, Dominic Kwiatkowski, Cordelia Langford, John Sillitoe on behalf of the Wellcome Sanger Institute COVID-19 Surveillance Team |
| EPI_ISL_3050993, EPI_ISL_3050994 | Lighthouse Lab in Glasgow | Wellcome Sanger Institute for the COVID-19 Genomics UK (COG-UK) Consortium | Harper VanSteenhouse, Yumi Kasai, David Gray, Carol Clugston, Anna Dominiczak and Alex Alderton, Roberto Amato, Jeffrey Barrett, Sonia Goncalves, Ewan Harrison, David K. Jackson, Ian Johnston, Dominic Kwiatkowski, Cordelia Langford, John Sillitoe on behalf of the Wellcome Sanger Institute COVID-19 Surveillance Team |
| EPI_ISL_3050995 | Berkshire and Surrey Pathology Services Lighthouse Laboratory | Wellcome Sanger Institute for the COVID-19 Genomics UK (COG-UK) Consortium | Berkshire and Surrey Pathology Services Lighthouse Laboratory and Alex Alderton, Roberto Amato, Jeffrey Barrett, Sonia Goncalves, Ewan Harrison, David K. Jackson, Ian Johnston, Dominic Kwiatkowski, Cordelia Langford, John Sillitoe on behalf of the Wellcome Sanger Institute COVID-19 Surveillance Team |
| EPI_ISL_3050996 | Randox Laboratories | Wellcome Sanger Institute for the COVID-19 Genomics UK (COG-UK) Consortium | Randox Laboratories and Alex Alderton, Roberto Amato, Jeffrey Barrett, Sonia Goncalves, Ewan Harrison, David K. Jackson, Ian Johnston, Dominic Kwiatkowski, Cordelia Langford, John Sillitoe on behalf of the Wellcome Sanger Institute COVID-19 Surveillance Team |
| EPI_ISL_3050997, EPI_ISL_3050998, EPI_ISL_3050999 | Berkshire and Surrey Pathology Services Lighthouse Laboratory | Wellcome Sanger Institute for the COVID-19 Genomics UK (COG-UK) Consortium | Berkshire and Surrey Pathology Services Lighthouse Laboratory and Alex Alderton, Roberto Amato, Jeffrey Barrett, Sonia Goncalves, Ewan Harrison, David K. Jackson, Ian Johnston, Dominic Kwiatkowski, Cordelia Langford, John Sillitoe on behalf of the Wellcome Sanger Institute COVID-19 Surveillance Team |
| EPI_ISL_3051000 | Lighthouse Lab in Glasgow | Wellcome Sanger Institute for the COVID-19 Genomics UK (COG-UK) | Harper VanSteenhouse, Yumi Kasai, David Gray, Carol Clugston, Anna Dominiczak and Alex Alderton, Roberto Amato, Jeffrey Barrett, Sonia Goncalves, Ewan Harrison, David K. Jackson, Ian Johnston, Dominic Kwiatkowski, Cordelia Langford, |

| | | | |
|---|---|---|---|
| | | Consortium | John Sillitoe on behalf of the Wellcome Sanger Institute COVID-19 Surveillance Team |
| EPI_ISL_3051001 | Berkshire and Surrey Pathology Services Lighthouse Laboratory | Wellcome Sanger Institute for the COVID-19 Genomics UK (COG-UK) Consortium | Berkshire and Surrey Pathology Services Lighthouse Laboratory and Alex Alderton, Roberto Amato, Jeffrey Barrett, Sonia Goncalves, Ewan Harrison, David K. Jackson, Ian Johnston, Dominic Kwiatkowski, Cordelia Langford, John Sillitoe on behalf of the Wellcome Sanger Institute COVID-19 Surveillance Team |
| EPI_ISL_3051002 | Lighthouse Lab in Glasgow | Wellcome Sanger Institute for the COVID-19 Genomics UK (COG-UK) Consortium | Harper VanSteenhouse, Yumi Kasai, David Gray, Carol Clugston, Anna Dominiczak and Alex Alderton, Roberto Amato, Jeffrey Barrett, Sonia Goncalves, Ewan Harrison, David K. Jackson, Ian Johnston, Dominic Kwiatkowski, Cordelia Langford, John Sillitoe on behalf of the Wellcome Sanger Institute COVID-19 Surveillance Team |
| EPI_ISL_3051003, EPI_ISL_3051004 | Randox Laboratories | Wellcome Sanger Institute for the COVID-19 Genomics UK (COG-UK) Consortium | Randox Laboratories and Alex Alderton, Roberto Amato, Jeffrey Barrett, Sonia Goncalves, Ewan Harrison, David K. Jackson, Ian Johnston, Dominic Kwiatkowski, Cordelia Langford, John Sillitoe on behalf of the Wellcome Sanger Institute COVID-19 Surveillance Team |
| EPI_ISL_3051005 | Lighthouse Lab in Glasgow | Wellcome Sanger Institute for the COVID-19 Genomics UK (COG-UK) Consortium | Harper VanSteenhouse, Yumi Kasai, David Gray, Carol Clugston, Anna Dominiczak and Alex Alderton, Roberto Amato, Jeffrey Barrett, Sonia Goncalves, Ewan Harrison, David K. Jackson, Ian Johnston, Dominic Kwiatkowski, Cordelia Langford, John Sillitoe on behalf of the Wellcome Sanger Institute COVID-19 Surveillance Team |
| EPI_ISL_3051006 | Berkshire and Surrey Pathology Services Lighthouse Laboratory | Wellcome Sanger Institute for the COVID-19 Genomics UK (COG-UK) Consortium | Berkshire and Surrey Pathology Services Lighthouse Laboratory and Alex Alderton, Roberto Amato, Jeffrey Barrett, Sonia Goncalves, Ewan Harrison, David K. Jackson, Ian Johnston, Dominic Kwiatkowski, Cordelia Langford, John Sillitoe on behalf of the Wellcome Sanger Institute COVID-19 Surveillance Team |
| EPI_ISL_3051007 | Randox Laboratories | Wellcome Sanger Institute for the COVID-19 Genomics UK (COG-UK) Consortium | Randox Laboratories and Alex Alderton, Roberto Amato, Jeffrey Barrett, Sonia Goncalves, Ewan Harrison, David K. Jackson, Ian Johnston, Dominic Kwiatkowski, Cordelia Langford, John Sillitoe on behalf of the Wellcome Sanger Institute COVID-19 Surveillance Team |
| EPI_ISL_3051008, EPI_ISL_3051009 | Lighthouse Lab in Glasgow | Wellcome Sanger Institute for the COVID-19 Genomics UK (COG-UK) Consortium | Harper VanSteenhouse, Yumi Kasai, David Gray, Carol Clugston, Anna Dominiczak and Alex Alderton, Roberto Amato, Jeffrey Barrett, Sonia Goncalves, Ewan Harrison, David K. Jackson, Ian Johnston, Dominic Kwiatkowski, Cordelia Langford, John Sillitoe on behalf of the Wellcome Sanger Institute COVID-19 Surveillance Team |
| EPI_ISL_3051010, EPI_ISL_3051011 | Berkshire and Surrey Pathology Services Lighthouse Laboratory | Wellcome Sanger Institute for the COVID-19 Genomics UK (COG-UK) Consortium | Berkshire and Surrey Pathology Services Lighthouse Laboratory and Alex Alderton, Roberto Amato, Jeffrey Barrett, Sonia Goncalves, Ewan Harrison, David K. Jackson, Ian Johnston, Dominic Kwiatkowski, Cordelia Langford, John Sillitoe on behalf of the Wellcome Sanger Institute COVID-19 Surveillance Team |
| EPI_ISL_3051012, EPI_ISL_3051013 | Lighthouse Lab in Glasgow | Wellcome Sanger Institute for the COVID-19 Genomics UK (COG-UK) Consortium | Harper VanSteenhouse, Yumi Kasai, David Gray, Carol Clugston, Anna Dominiczak and Alex Alderton, Roberto Amato, Jeffrey Barrett, Sonia Goncalves, Ewan Harrison, David K. Jackson, Ian Johnston, Dominic Kwiatkowski, Cordelia Langford, John Sillitoe on behalf of the Wellcome Sanger Institute COVID-19 Surveillance Team |
| EPI_ISL_3051014, EPI_ISL_3051015 | Randox Laboratories | Wellcome Sanger Institute for the COVID-19 Genomics UK (COG-UK) Consortium | Randox Laboratories and Alex Alderton, Roberto Amato, Jeffrey Barrett, Sonia Goncalves, Ewan Harrison, David K. Jackson, Ian Johnston, Dominic Kwiatkowski, Cordelia Langford, John Sillitoe on behalf of the Wellcome Sanger Institute COVID-19 Surveillance Team |
| EPI_ISL_3051016 | Berkshire and Surrey Pathology Services Lighthouse Laboratory | Wellcome Sanger Institute for the COVID-19 Genomics UK (COG-UK) Consortium | Berkshire and Surrey Pathology Services Lighthouse Laboratory and Alex Alderton, Roberto Amato, Jeffrey Barrett, Sonia Goncalves, Ewan Harrison, David K. Jackson, Ian Johnston, Dominic Kwiatkowski, Cordelia Langford, John Sillitoe on behalf of the Wellcome Sanger Institute COVID-19 Surveillance Team |
| EPI_ISL_3051017, EPI_ISL_3051018, EPI_ISL_3051019, EPI_ISL_3051020 | Lighthouse Lab in Glasgow | Wellcome Sanger Institute for the COVID-19 Genomics UK (COG-UK) Consortium | Harper VanSteenhouse, Yumi Kasai, David Gray, Carol Clugston, Anna Dominiczak and Alex Alderton, Roberto Amato, Jeffrey Barrett, Sonia Goncalves, Ewan Harrison, David K. Jackson, Ian Johnston, Dominic Kwiatkowski, Cordelia Langford, John Sillitoe on behalf of the Wellcome Sanger Institute COVID-19 Surveillance Team |
| EPI_ISL_3051021 | Berkshire and Surrey Pathology Services Lighthouse Laboratory | Wellcome Sanger Institute for the COVID-19 Genomics UK (COG-UK) Consortium | Berkshire and Surrey Pathology Services Lighthouse Laboratory and Alex Alderton, Roberto Amato, Jeffrey Barrett, Sonia Goncalves, Ewan Harrison, David K. Jackson, Ian Johnston, Dominic Kwiatkowski, Cordelia Langford, John Sillitoe on behalf of the Wellcome Sanger Institute COVID-19 Surveillance Team |
| EPI_ISL_3051022, EPI_ISL_3051023 | Lighthouse Lab in Glasgow | Wellcome Sanger Institute for the COVID-19 Genomics UK (COG-UK) Consortium | Harper VanSteenhouse, Yumi Kasai, David Gray, Carol Clugston, Anna Dominiczak and Alex Alderton, Roberto Amato, Jeffrey Barrett, Sonia Goncalves, Ewan Harrison, David K. Jackson, Ian Johnston, Dominic Kwiatkowski, Cordelia Langford, John Sillitoe on behalf of the Wellcome Sanger Institute COVID-19 Surveillance Team |
| EPI_ISL_3051024, EPI_ISL_3051025 | Berkshire and Surrey Pathology Services Lighthouse Laboratory | Wellcome Sanger Institute for the COVID-19 Genomics UK (COG-UK) Consortium | Berkshire and Surrey Pathology Services Lighthouse Laboratory and Alex Alderton, Roberto Amato, Jeffrey Barrett, Sonia Goncalves, Ewan Harrison, David K. Jackson, Ian Johnston, Dominic Kwiatkowski, Cordelia Langford, John Sillitoe on behalf of the Wellcome Sanger Institute COVID-19 Surveillance Team |
| EPI_ISL_3051026 | Lighthouse Lab in Glasgow | Wellcome Sanger Institute for the COVID-19 Genomics UK (COG-UK) Consortium | Harper VanSteenhouse, Yumi Kasai, David Gray, Carol Clugston, Anna Dominiczak and Alex Alderton, Roberto Amato, Jeffrey Barrett, Sonia Goncalves, Ewan Harrison, David K. Jackson, Ian Johnston, Dominic Kwiatkowski, Cordelia Langford, John Sillitoe on behalf of the Wellcome Sanger Institute COVID-19 Surveillance Team |
| EPI_ISL_3051027 | Berkshire and Surrey Pathology Services Lighthouse Laboratory | Wellcome Sanger Institute for the COVID-19 Genomics UK (COG-UK) Consortium | Berkshire and Surrey Pathology Services Lighthouse Laboratory and Alex Alderton, Roberto Amato, Jeffrey Barrett, Sonia Goncalves, Ewan Harrison, David K. Jackson, Ian Johnston, Dominic Kwiatkowski, Cordelia Langford, John Sillitoe on behalf of the Wellcome Sanger Institute COVID-19 Surveillance Team |
| EPI_ISL_3051028 | Lighthouse Lab in Glasgow | Wellcome Sanger Institute for the COVID-19 Genomics UK (COG-UK) Consortium | Harper VanSteenhouse, Yumi Kasai, David Gray, Carol Clugston, Anna Dominiczak and Alex Alderton, Roberto Amato, Jeffrey Barrett, Sonia Goncalves, Ewan Harrison, David K. Jackson, Ian Johnston, Dominic Kwiatkowski, Cordelia Langford, John Sillitoe on behalf of the Wellcome Sanger Institute COVID-19 Surveillance Team |
| EPI_ISL_3051029 | Berkshire and Surrey Pathology Services Lighthouse Laboratory | Wellcome Sanger Institute for the COVID-19 Genomics UK (COG-UK) Consortium | Berkshire and Surrey Pathology Services Lighthouse Laboratory and Alex Alderton, Roberto Amato, Jeffrey Barrett, Sonia Goncalves, Ewan Harrison, David K. Jackson, Ian Johnston, Dominic Kwiatkowski, Cordelia Langford, John Sillitoe on behalf of the Wellcome Sanger Institute COVID-19 Surveillance Team |
| EPI_ISL_3051030 | Lighthouse Lab in Glasgow | Wellcome Sanger Institute for the COVID-19 Genomics UK (COG-UK) Consortium | Harper VanSteenhouse, Yumi Kasai, David Gray, Carol Clugston, Anna Dominiczak and Alex Alderton, Roberto Amato, Jeffrey Barrett, Sonia Goncalves, Ewan Harrison, David K. Jackson, Ian Johnston, Dominic Kwiatkowski, Cordelia Langford, John Sillitoe on behalf of the Wellcome Sanger Institute COVID-19 Surveillance Team |
| EPI_ISL_3051031 | Berkshire and Surrey Pathology Services Lighthouse Laboratory | Wellcome Sanger Institute for the COVID-19 Genomics UK (COG-UK) Consortium | Berkshire and Surrey Pathology Services Lighthouse Laboratory and Alex Alderton, Roberto Amato, Jeffrey Barrett, Sonia Goncalves, Ewan Harrison, David K. Jackson, Ian Johnston, Dominic Kwiatkowski, Cordelia Langford, John Sillitoe on behalf of the Wellcome Sanger Institute COVID-19 Surveillance Team |
| EPI_ISL_3051032, EPI_ISL_3051033 | Lighthouse Lab in Glasgow | Wellcome Sanger Institute for the COVID-19 Genomics UK (COG-UK) Consortium | Harper VanSteenhouse, Yumi Kasai, David Gray, Carol Clugston, Anna Dominiczak and Alex Alderton, Roberto Amato, Jeffrey Barrett, Sonia Goncalves, Ewan Harrison, David K. Jackson, Ian Johnston, Dominic Kwiatkowski, Cordelia Langford, John Sillitoe on behalf of the Wellcome Sanger Institute COVID-19 Surveillance Team |
| EPI_ISL_3051034 | Randox Laboratories | Wellcome Sanger Institute for the COVID-19 Genomics UK (COG-UK) Consortium | Randox Laboratories and Alex Alderton, Roberto Amato, Jeffrey Barrett, Sonia Goncalves, Ewan Harrison, David K. Jackson, Ian Johnston, Dominic Kwiatkowski, Cordelia Langford, John Sillitoe on behalf of the Wellcome Sanger Institute COVID-19 Surveillance Team |
| EPI_ISL_3051035, EPI_ISL_3051036 | Lighthouse Lab in Glasgow | Wellcome Sanger Institute for the COVID-19 Genomics UK (COG-UK) Consortium | Harper VanSteenhouse, Yumi Kasai, David Gray, Carol Clugston, Anna Dominiczak and Alex Alderton, Roberto Amato, Jeffrey Barrett, Sonia Goncalves, Ewan Harrison, David K. Jackson, Ian Johnston, Dominic Kwiatkowski, Cordelia Langford, John Sillitoe on behalf of the Wellcome Sanger Institute COVID-19 Surveillance Team |
| EPI_ISL_3051037 | Berkshire and Surrey Pathology Services Lighthouse Laboratory | Wellcome Sanger Institute for the COVID-19 Genomics UK (COG-UK) Consortium | Berkshire and Surrey Pathology Services Lighthouse Laboratory and Alex Alderton, Roberto Amato, Jeffrey Barrett, Sonia Goncalves, Ewan Harrison, David K. Jackson, Ian Johnston, Dominic Kwiatkowski, Cordelia Langford, John Sillitoe on behalf of the Wellcome Sanger Institute COVID-19 Surveillance Team |
| EPI_ISL_3051038 | Randox Laboratories | Wellcome Sanger Institute for the COVID-19 Genomics UK (COG-UK) Consortium | Randox Laboratories and Alex Alderton, Roberto Amato, Jeffrey Barrett, Sonia Goncalves, Ewan Harrison, David K. Jackson, Ian Johnston, Dominic Kwiatkowski, Cordelia Langford, John Sillitoe on behalf of the Wellcome Sanger Institute COVID-19 Surveillance Team |
| EPI_ISL_3051039, EPI_ISL_3051040 | Lighthouse Lab in Glasgow | Wellcome Sanger Institute for the COVID-19 Genomics UK (COG-UK) Consortium | Harper VanSteenhouse, Yumi Kasai, David Gray, Carol Clugston, Anna Dominiczak and Alex Alderton, Roberto Amato, Jeffrey Barrett, Sonia Goncalves, Ewan Harrison, David K. Jackson, Ian Johnston, Dominic Kwiatkowski, Cordelia Langford, John Sillitoe on behalf of the Wellcome Sanger Institute COVID-19 Surveillance Team |
| EPI_ISL_3051041 | Berkshire and Surrey Pathology Services Lighthouse Laboratory | Wellcome Sanger Institute for the COVID-19 Genomics UK (COG-UK) Consortium | Berkshire and Surrey Pathology Services Lighthouse Laboratory and Alex Alderton, Roberto Amato, Jeffrey Barrett, Sonia Goncalves, Ewan Harrison, David K. Jackson, Ian Johnston, Dominic Kwiatkowski, Cordelia Langford, John Sillitoe on behalf of the Wellcome Sanger Institute COVID-19 Surveillance Team |
| EPI_ISL_3051042, EPI_ISL_3051043 | Lighthouse Lab in Glasgow | Wellcome Sanger Institute for the COVID-19 Genomics UK (COG-UK) Consortium | Harper VanSteenhouse, Yumi Kasai, David Gray, Carol Clugston, Anna Dominiczak and Alex Alderton, Roberto Amato, Jeffrey Barrett, Sonia Goncalves, Ewan Harrison, David K. Jackson, Ian Johnston, Dominic Kwiatkowski, Cordelia Langford, John Sillitoe on behalf of the Wellcome Sanger Institute COVID-19 Surveillance Team |
| EPI_ISL_3051044 | Berkshire and Surrey Pathology Services Lighthouse Laboratory | Wellcome Sanger Institute for the COVID-19 Genomics UK (COG-UK) Consortium | Berkshire and Surrey Pathology Services Lighthouse Laboratory and Alex Alderton, Roberto Amato, Jeffrey Barrett, Sonia Goncalves, Ewan Harrison, David K. Jackson, Ian Johnston, Dominic Kwiatkowski, Cordelia Langford, John Sillitoe on behalf of the Wellcome Sanger Institute COVID-19 Surveillance Team |
| EPI_ISL_3051045 | Randox Laboratories | Wellcome Sanger Institute for the COVID-19 Genomics UK (COG-UK) Consortium | Randox Laboratories and Alex Alderton, Roberto Amato, Jeffrey Barrett, Sonia Goncalves, Ewan Harrison, David K. Jackson, Ian Johnston, Dominic Kwiatkowski, Cordelia Langford, John Sillitoe on behalf of the Wellcome Sanger Institute COVID-19 Surveillance Team |
| EPI_ISL_3051046, EPI_ISL_3051047, EPI_ISL_3051048, EPI_ISL_3051049 | Lighthouse Lab in Glasgow | Wellcome Sanger Institute for the COVID-19 Genomics UK (COG-UK) Consortium | Harper VanSteenhouse, Yumi Kasai, David Gray, Carol Clugston, Anna Dominiczak and Alex Alderton, Roberto Amato, Jeffrey Barrett, Sonia Goncalves, Ewan Harrison, David K. Jackson, Ian Johnston, Dominic Kwiatkowski, Cordelia Langford, John Sillitoe on behalf of the Wellcome Sanger Institute COVID-19 Surveillance Team |
| EPI_ISL_3051050, EPI_ISL_3051051 | Berkshire and Surrey Pathology Services Lighthouse Laboratory | Wellcome Sanger Institute for the COVID-19 Genomics UK (COG-UK) Consortium | Berkshire and Surrey Pathology Services Lighthouse Laboratory and Alex Alderton, Roberto Amato, Jeffrey Barrett, Sonia Goncalves, Ewan Harrison, David K. Jackson, Ian Johnston, Dominic Kwiatkowski, Cordelia Langford, John Sillitoe on behalf of the Wellcome Sanger Institute COVID-19 Surveillance Team |
| EPI_ISL_3051052, EPI_ISL_3051053 | Randox Laboratories | Wellcome Sanger Institute for the COVID-19 Genomics UK (COG-UK) Consortium | Randox Laboratories and Alex Alderton, Roberto Amato, Jeffrey Barrett, Sonia Goncalves, Ewan Harrison, David K. Jackson, Ian Johnston, Dominic Kwiatkowski, Cordelia Langford, John Sillitoe on behalf of the Wellcome Sanger Institute COVID-19 Surveillance Team |
| EPI_ISL_3051054, EPI_ISL_3051055 | Berkshire and Surrey Pathology Services Lighthouse Laboratory | Wellcome Sanger Institute for the COVID-19 Genomics UK (COG-UK) Consortium | Berkshire and Surrey Pathology Services Lighthouse Laboratory and Alex Alderton, Roberto Amato, Jeffrey Barrett, Sonia Goncalves, Ewan Harrison, David K. Jackson, Ian Johnston, Dominic Kwiatkowski, Cordelia Langford, John Sillitoe on behalf of the Wellcome Sanger Institute COVID-19 Surveillance Team |
| EPI_ISL_3051056 | Randox Laboratories | Wellcome Sanger Institute for the COVID-19 Genomics UK (COG-UK) Consortium | Randox Laboratories and Alex Alderton, Roberto Amato, Jeffrey Barrett, Sonia Goncalves, Ewan Harrison, David K. Jackson, Ian Johnston, Dominic Kwiatkowski, Cordelia Langford, John Sillitoe on behalf of the Wellcome Sanger Institute COVID-19 Surveillance Team |
| EPI_ISL_3051057, EPI_ISL_3051058 | Berkshire and Surrey Pathology Services Lighthouse Laboratory | Wellcome Sanger Institute for the COVID-19 Genomics UK (COG-UK) Consortium | Berkshire and Surrey Pathology Services Lighthouse Laboratory and Alex Alderton, Roberto Amato, Jeffrey Barrett, Sonia Goncalves, Ewan Harrison, David K. Jackson, Ian Johnston, Dominic Kwiatkowski, Cordelia Langford, John Sillitoe on behalf of the Wellcome Sanger Institute COVID-19 Surveillance Team |
| EPI_ISL_3051059 | Lighthouse Lab in Glasgow | Wellcome Sanger Institute for the COVID-19 Genomics UK (COG-UK) Consortium | Harper VanSteenhouse, Yumi Kasai, David Gray, Carol Clugston, Anna Dominiczak and Alex Alderton, Roberto Amato, Jeffrey Barrett, Sonia Goncalves, Ewan Harrison, David K. Jackson, Ian Johnston, Dominic Kwiatkowski, Cordelia Langford, John Sillitoe on behalf of the Wellcome Sanger Institute COVID-19 Surveillance Team |
| EPI_ISL_3051060 | Randox Laboratories | Wellcome Sanger Institute for the COVID-19 Genomics UK (COG-UK) Consortium | Randox Laboratories and Alex Alderton, Roberto Amato, Jeffrey Barrett, Sonia Goncalves, Ewan Harrison, David K. Jackson, Ian Johnston, Dominic Kwiatkowski, Cordelia Langford, John Sillitoe on behalf of the Wellcome Sanger Institute COVID-19 Surveillance Team |
| EPI_ISL_3051061 | Berkshire and Surrey Pathology Services Lighthouse Laboratory | Wellcome Sanger Institute for the COVID-19 Genomics UK (COG-UK) Consortium | Berkshire and Surrey Pathology Services Lighthouse Laboratory and Alex Alderton, Roberto Amato, Jeffrey Barrett, Sonia Goncalves, Ewan Harrison, David K. Jackson, Ian Johnston, Dominic Kwiatkowski, Cordelia Langford, John Sillitoe on behalf of the Wellcome Sanger Institute COVID-19 Surveillance Team |
| EPI_ISL_3051062 | Lighthouse Lab in Glasgow | Wellcome Sanger Institute for the COVID-19 Genomics UK (COG-UK) Consortium | Harper VanSteenhouse, Yumi Kasai, David Gray, Carol Clugston, Anna Dominiczak and Alex Alderton, Roberto Amato, Jeffrey Barrett, Sonia Goncalves, Ewan Harrison, David K. Jackson, Ian Johnston, Dominic Kwiatkowski, Cordelia Langford, John Sillitoe on behalf of the Wellcome Sanger Institute COVID-19 Surveillance Team |
| EPI_ISL_3051063 | Berkshire and Surrey Pathology Services Lighthouse Laboratory | Wellcome Sanger Institute for the COVID-19 Genomics UK (COG-UK) Consortium | Berkshire and Surrey Pathology Services Lighthouse Laboratory and Alex Alderton, Roberto Amato, Jeffrey Barrett, Sonia Goncalves, Ewan Harrison, David K. Jackson, Ian Johnston, Dominic Kwiatkowski, Cordelia Langford, John Sillitoe on behalf of the Wellcome Sanger Institute COVID-19 Surveillance Team |
| EPI_ISL_3051064 | Randox Laboratories | Wellcome Sanger Institute for the COVID-19 Genomics UK (COG-UK) Consortium | Randox Laboratories and Alex Alderton, Roberto Amato, Jeffrey Barrett, Sonia Goncalves, Ewan Harrison, David K. Jackson, Ian Johnston, Dominic Kwiatkowski, Cordelia Langford, John Sillitoe on behalf of the Wellcome Sanger Institute COVID-19 Surveillance Team |

| | | | |
|---|---|---|---|
| EPI_ISL_3051065 | Lighthouse Lab in Glasgow | Wellcome Sanger Institute for the COVID-19 Genomics UK (COG-UK) Consortium | Harper VanSteenhouse, Yumi Kasai, David Gray, Carol Clugston, Anna Dominiczak and Alex Alderton, Roberto Amato, Jeffrey Barrett, Sonia Goncalves, Ewan Harrison, David K. Jackson, Ian Johnston, Dominic Kwiatkowski, Cordelia Langford, John Sillitoe on behalf of the Wellcome Sanger Institute COVID-19 Surveillance Team |
| EPI_ISL_3051066 | Randox Laboratories | Wellcome Sanger Institute for the COVID-19 Genomics UK (COG-UK) Consortium | Randox Laboratories and Alex Alderton, Roberto Amato, Jeffrey Barrett, Sonia Goncalves, Ewan Harrison, David K. Jackson, Ian Johnston, Dominic Kwiatkowski, Cordelia Langford, John Sillitoe on behalf of the Wellcome Sanger Institute COVID-19 Surveillance Team |
| EPI_ISL_3051067 | Lighthouse Lab in Glasgow | Wellcome Sanger Institute for the COVID-19 Genomics UK (COG-UK) Consortium | Harper VanSteenhouse, Yumi Kasai, David Gray, Carol Clugston, Anna Dominiczak and Alex Alderton, Roberto Amato, Jeffrey Barrett, Sonia Goncalves, Ewan Harrison, David K. Jackson, Ian Johnston, Dominic Kwiatkowski, Cordelia Langford, John Sillitoe on behalf of the Wellcome Sanger Institute COVID-19 Surveillance Team |
| EPI_ISL_3051068 | Randox Laboratories | Wellcome Sanger Institute for the COVID-19 Genomics UK (COG-UK) Consortium | Randox Laboratories and Alex Alderton, Roberto Amato, Jeffrey Barrett, Sonia Goncalves, Ewan Harrison, David K. Jackson, Ian Johnston, Dominic Kwiatkowski, Cordelia Langford, John Sillitoe on behalf of the Wellcome Sanger Institute COVID-19 Surveillance Team |
| EPI_ISL_3051069, EPI_ISL_3051070, EPI_ISL_3051071, EPI_ISL_3051072, EPI_ISL_3051073 | Lighthouse Lab in Glasgow | Wellcome Sanger Institute for the COVID-19 Genomics UK (COG-UK) Consortium | Harper VanSteenhouse, Yumi Kasai, David Gray, Carol Clugston, Anna Dominiczak and Alex Alderton, Roberto Amato, Jeffrey Barrett, Sonia Goncalves, Ewan Harrison, David K. Jackson, Ian Johnston, Dominic Kwiatkowski, Cordelia Langford, John Sillitoe on behalf of the Wellcome Sanger Institute COVID-19 Surveillance Team |
| EPI_ISL_3051074 | Randox Laboratories | Wellcome Sanger Institute for the COVID-19 Genomics UK (COG-UK) Consortium | Randox Laboratories and Alex Alderton, Roberto Amato, Jeffrey Barrett, Sonia Goncalves, Ewan Harrison, David K. Jackson, Ian Johnston, Dominic Kwiatkowski, Cordelia Langford, John Sillitoe on behalf of the Wellcome Sanger Institute COVID-19 Surveillance Team |
| EPI_ISL_3051075 | Berkshire and Surrey Pathology Services Lighthouse Laboratory | Wellcome Sanger Institute for the COVID-19 Genomics UK (COG-UK) Consortium | Berkshire and Surrey Pathology Services Lighthouse Laboratory and Alex Alderton, Roberto Amato, Jeffrey Barrett, Sonia Goncalves, Ewan Harrison, David K. Jackson, Ian Johnston, Dominic Kwiatkowski, Cordelia Langford, John Sillitoe on behalf of the Wellcome Sanger Institute COVID-19 Surveillance Team |
| EPI_ISL_3051076 | Randox Laboratories | Wellcome Sanger Institute for the COVID-19 Genomics UK (COG-UK) Consortium | Randox Laboratories and Alex Alderton, Roberto Amato, Jeffrey Barrett, Sonia Goncalves, Ewan Harrison, David K. Jackson, Ian Johnston, Dominic Kwiatkowski, Cordelia Langford, John Sillitoe on behalf of the Wellcome Sanger Institute COVID-19 Surveillance Team |
| EPI_ISL_3051077 | Berkshire and Surrey Pathology Services Lighthouse Laboratory | Wellcome Sanger Institute for the COVID-19 Genomics UK (COG-UK) Consortium | Berkshire and Surrey Pathology Services Lighthouse Laboratory and Alex Alderton, Roberto Amato, Jeffrey Barrett, Sonia Goncalves, Ewan Harrison, David K. Jackson, Ian Johnston, Dominic Kwiatkowski, Cordelia Langford, John Sillitoe on behalf of the Wellcome Sanger Institute COVID-19 Surveillance Team |
| EPI_ISL_3051078, EPI_ISL_3051079, EPI_ISL_3051080 | Lighthouse Lab in Glasgow | Wellcome Sanger Institute for the COVID-19 Genomics UK (COG-UK) Consortium | Harper VanSteenhouse, Yumi Kasai, David Gray, Carol Clugston, Anna Dominiczak and Alex Alderton, Roberto Amato, Jeffrey Barrett, Sonia Goncalves, Ewan Harrison, David K. Jackson, Ian Johnston, Dominic Kwiatkowski, Cordelia Langford, John Sillitoe on behalf of the Wellcome Sanger Institute COVID-19 Surveillance Team |
| EPI_ISL_3051081 | Berkshire and Surrey Pathology Services Lighthouse Laboratory | Wellcome Sanger Institute for the COVID-19 Genomics UK (COG-UK) Consortium | Berkshire and Surrey Pathology Services Lighthouse Laboratory and Alex Alderton, Roberto Amato, Jeffrey Barrett, Sonia Goncalves, Ewan Harrison, David K. Jackson, Ian Johnston, Dominic Kwiatkowski, Cordelia Langford, John Sillitoe on behalf of the Wellcome Sanger Institute COVID-19 Surveillance Team |
| EPI_ISL_3051082 | Lighthouse Lab in Glasgow | Wellcome Sanger Institute for the COVID-19 Genomics UK (COG-UK) Consortium | Harper VanSteenhouse, Yumi Kasai, David Gray, Carol Clugston, Anna Dominiczak and Alex Alderton, Roberto Amato, Jeffrey Barrett, Sonia Goncalves, Ewan Harrison, David K. Jackson, Ian Johnston, Dominic Kwiatkowski, Cordelia Langford, John Sillitoe on behalf of the Wellcome Sanger Institute COVID-19 Surveillance Team |
| EPI_ISL_3051083 | Randox Laboratories | Wellcome Sanger Institute for the COVID-19 Genomics UK (COG-UK) Consortium | Randox Laboratories and Alex Alderton, Roberto Amato, Jeffrey Barrett, Sonia Goncalves, Ewan Harrison, David K. Jackson, Ian Johnston, Dominic Kwiatkowski, Cordelia Langford, John Sillitoe on behalf of the Wellcome Sanger Institute COVID-19 Surveillance Team |
| EPI_ISL_3051084, EPI_ISL_3051085, EPI_ISL_3051086 | Lighthouse Lab in Glasgow | Wellcome Sanger Institute for the COVID-19 Genomics UK (COG-UK) Consortium | Harper VanSteenhouse, Yumi Kasai, David Gray, Carol Clugston, Anna Dominiczak and Alex Alderton, Roberto Amato, Jeffrey Barrett, Sonia Goncalves, Ewan Harrison, David K. Jackson, Ian Johnston, Dominic Kwiatkowski, Cordelia Langford, John Sillitoe on behalf of the Wellcome Sanger Institute COVID-19 Surveillance Team |
| EPI_ISL_3051087 | Randox Laboratories | Wellcome Sanger Institute for the COVID-19 Genomics UK (COG-UK) Consortium | Randox Laboratories and Alex Alderton, Roberto Amato, Jeffrey Barrett, Sonia Goncalves, Ewan Harrison, David K. Jackson, Ian Johnston, Dominic Kwiatkowski, Cordelia Langford, John Sillitoe on behalf of the Wellcome Sanger Institute COVID-19 Surveillance Team |
| EPI_ISL_3051088, EPI_ISL_3051089, EPI_ISL_3051090, EPI_ISL_3051091, EPI_ISL_3051092 | Lighthouse Lab in Glasgow | Wellcome Sanger Institute for the COVID-19 Genomics UK (COG-UK) Consortium | Harper VanSteenhouse, Yumi Kasai, David Gray, Carol Clugston, Anna Dominiczak and Alex Alderton, Roberto Amato, Jeffrey Barrett, Sonia Goncalves, Ewan Harrison, David K. Jackson, Ian Johnston, Dominic Kwiatkowski, Cordelia Langford, John Sillitoe on behalf of the Wellcome Sanger Institute COVID-19 Surveillance Team |
| EPI_ISL_3051093 | Randox Laboratories | Wellcome Sanger Institute for the COVID-19 Genomics UK (COG-UK) Consortium | Randox Laboratories and Alex Alderton, Roberto Amato, Jeffrey Barrett, Sonia Goncalves, Ewan Harrison, David K. Jackson, Ian Johnston, Dominic Kwiatkowski, Cordelia Langford, John Sillitoe on behalf of the Wellcome Sanger Institute COVID-19 Surveillance Team |
| EPI_ISL_3051094, EPI_ISL_3051095 | Lighthouse Lab in Glasgow | Wellcome Sanger Institute for the COVID-19 Genomics UK (COG-UK) Consortium | Harper VanSteenhouse, Yumi Kasai, David Gray, Carol Clugston, Anna Dominiczak and Alex Alderton, Roberto Amato, Jeffrey Barrett, Sonia Goncalves, Ewan Harrison, David K. Jackson, Ian Johnston, Dominic Kwiatkowski, Cordelia Langford, John Sillitoe on behalf of the Wellcome Sanger Institute COVID-19 Surveillance Team |
| EPI_ISL_3051096 | Randox Laboratories | Wellcome Sanger Institute for the COVID-19 Genomics UK (COG-UK) Consortium | Randox Laboratories and Alex Alderton, Roberto Amato, Jeffrey Barrett, Sonia Goncalves, Ewan Harrison, David K. Jackson, Ian Johnston, Dominic Kwiatkowski, Cordelia Langford, John Sillitoe on behalf of the Wellcome Sanger Institute COVID-19 Surveillance Team |
| EPI_ISL_3051097 | Berkshire and Surrey Pathology Services Lighthouse Laboratory | Wellcome Sanger Institute for the COVID-19 Genomics UK (COG-UK) Consortium | Berkshire and Surrey Pathology Services Lighthouse Laboratory and Alex Alderton, Roberto Amato, Jeffrey Barrett, Sonia Goncalves, Ewan Harrison, David K. Jackson, Ian Johnston, Dominic Kwiatkowski, Cordelia Langford, John Sillitoe on behalf of the Wellcome Sanger Institute COVID-19 Surveillance Team |
| EPI_ISL_3051098, EPI_ISL_3051099 | Lighthouse Lab in Glasgow | Wellcome Sanger Institute for the COVID-19 Genomics UK (COG-UK) Consortium | Harper VanSteenhouse, Yumi Kasai, David Gray, Carol Clugston, Anna Dominiczak and Alex Alderton, Roberto Amato, Jeffrey Barrett, Sonia Goncalves, Ewan Harrison, David K. Jackson, Ian Johnston, Dominic Kwiatkowski, Cordelia Langford, John Sillitoe on behalf of the Wellcome Sanger Institute COVID-19 Surveillance Team |
| EPI_ISL_3051100 | Berkshire and Surrey Pathology Services Lighthouse Laboratory | Wellcome Sanger Institute for the COVID-19 Genomics UK (COG-UK) Consortium | Berkshire and Surrey Pathology Services Lighthouse Laboratory and Alex Alderton, Roberto Amato, Jeffrey Barrett, Sonia Goncalves, Ewan Harrison, David K. Jackson, Ian Johnston, Dominic Kwiatkowski, Cordelia Langford, John Sillitoe on behalf of the Wellcome Sanger Institute COVID-19 Surveillance Team |
| EPI_ISL_3051101, EPI_ISL_3051102 | Randox Laboratories | Wellcome Sanger Institute for the COVID-19 Genomics UK (COG-UK) Consortium | Randox Laboratories and Alex Alderton, Roberto Amato, Jeffrey Barrett, Sonia Goncalves, Ewan Harrison, David K. Jackson, Ian Johnston, Dominic Kwiatkowski, Cordelia Langford, John Sillitoe on behalf of the Wellcome Sanger Institute COVID-19 Surveillance Team |
| EPI_ISL_3051103 | Berkshire and Surrey Pathology Services Lighthouse Laboratory | Wellcome Sanger Institute for the COVID-19 Genomics UK (COG-UK) Consortium | Berkshire and Surrey Pathology Services Lighthouse Laboratory and Alex Alderton, Roberto Amato, Jeffrey Barrett, Sonia Goncalves, Ewan Harrison, David K. Jackson, Ian Johnston, Dominic Kwiatkowski, Cordelia Langford, John Sillitoe on behalf of the Wellcome Sanger Institute COVID-19 Surveillance Team |
| EPI_ISL_3051104, EPI_ISL_3051105 | Randox Laboratories | Wellcome Sanger Institute for the COVID-19 Genomics UK (COG-UK) Consortium | Randox Laboratories and Alex Alderton, Roberto Amato, Jeffrey Barrett, Sonia Goncalves, Ewan Harrison, David K. Jackson, Ian Johnston, Dominic Kwiatkowski, Cordelia Langford, John Sillitoe on behalf of the Wellcome Sanger Institute COVID-19 Surveillance Team |
| EPI_ISL_3051106 | Berkshire and Surrey Pathology Services Lighthouse Laboratory | Wellcome Sanger Institute for the COVID-19 Genomics UK (COG-UK) Consortium | Berkshire and Surrey Pathology Services Lighthouse Laboratory and Alex Alderton, Roberto Amato, Jeffrey Barrett, Sonia Goncalves, Ewan Harrison, David K. Jackson, Ian Johnston, Dominic Kwiatkowski, Cordelia Langford, John Sillitoe on behalf of the Wellcome Sanger Institute COVID-19 Surveillance Team |
| EPI_ISL_3051107, EPI_ISL_3051108, EPI_ISL_3051109, EPI_ISL_3051110, EPI_ISL_3051111, EPI_ISL_3051112, EPI_ISL_3051113, EPI_ISL_3051114, EPI_ISL_3051115, EPI_ISL_3051116, EPI_ISL_3051117, EPI_ISL_3051118, EPI_ISL_3051119, EPI_ISL_3051120, EPI_ISL_3051121, EPI_ISL_3051122, EPI_ISL_3051123, EPI_ISL_3051124, EPI_ISL_3051125, EPI_ISL_3051126, EPI_ISL_3051127, EPI_ISL_3051128, EPI_ISL_3051129, EPI_ISL_3051130, EPI_ISL_3051131, EPI_ISL_3051132, EPI_ISL_3051133, EPI_ISL_3051134, EPI_ISL_3051135 | | | |
| see above | Lighthouse Lab in Glasgow | Wellcome Sanger Institute for the COVID-19 Genomics UK (COG-UK) Consortium | Harper VanSteenhouse, Yumi Kasai, David Gray, Carol Clugston, Anna Dominiczak and Alex Alderton, Roberto Amato, Jeffrey Barrett, Sonia Goncalves, Ewan Harrison, David K. Jackson, Ian Johnston, Dominic Kwiatkowski, Cordelia Langford, John Sillitoe on behalf of the Wellcome Sanger Institute COVID-19 Surveillance Team |
| EPI_ISL_3051136 | Lighthouse Lab in Alderley Park | Wellcome Sanger Institute for the COVID-19 Genomics UK (COG-UK) Consortium | Jacquelyn Wynn, Mairead Hyland, The Lighthouse Lab in Alderley Park and Alex Alderton, Roberto Amato, Jeffrey Barrett, Sonia Goncalves, Ewan Harrison, David K. Jackson, Ian Johnston, Dominic Kwiatkowski, Cordelia Langford, John Sillitoe on behalf of the Wellcome Sanger Institute COVID-19 Surveillance Team |
| EPI_ISL_3051137 | Berkshire and Surrey Pathology Services Lighthouse Laboratory | Wellcome Sanger Institute for the COVID-19 Genomics UK (COG-UK) Consortium | Berkshire and Surrey Pathology Services Lighthouse Laboratory and Alex Alderton, Roberto Amato, Jeffrey Barrett, Sonia Goncalves, Ewan Harrison, David K. Jackson, Ian Johnston, Dominic Kwiatkowski, Cordelia Langford, John Sillitoe on behalf of the Wellcome Sanger Institute COVID-19 Surveillance Team |
| EPI_ISL_3051138 | Lighthouse Lab in Alderley Park | Wellcome Sanger Institute for the COVID-19 Genomics UK (COG-UK) Consortium | Jacquelyn Wynn, Mairead Hyland, The Lighthouse Lab in Alderley Park and Alex Alderton, Roberto Amato, Jeffrey Barrett, Sonia Goncalves, Ewan Harrison, David K. Jackson, Ian Johnston, Dominic Kwiatkowski, Cordelia Langford, John Sillitoe on behalf of the Wellcome Sanger Institute COVID-19 Surveillance Team |
| EPI_ISL_3051139 | Randox Laboratories | Wellcome Sanger Institute for the COVID-19 Genomics UK (COG-UK) Consortium | Randox Laboratories and Alex Alderton, Roberto Amato, Jeffrey Barrett, Sonia Goncalves, Ewan Harrison, David K. Jackson, Ian Johnston, Dominic Kwiatkowski, Cordelia Langford, John Sillitoe on behalf of the Wellcome Sanger Institute COVID-19 Surveillance Team |
| EPI_ISL_3051140 | Lighthouse Lab in Milton Keynes | Wellcome Sanger Institute for the COVID-19 Genomics UK (COG-UK) Consortium | The Lighthouse Lab in Milton Keynes and Alex Alderton, Roberto Amato, Jeffrey Barrett, Sonia Goncalves, Ewan Harrison, David K. Jackson, Ian Johnston, Dominic Kwiatkowski, Cordelia Langford, John Sillitoe on behalf of the Wellcome Sanger Institute COVID-19 Surveillance Team |
| EPI_ISL_3051141 | Randox Laboratories | Wellcome Sanger Institute for the COVID-19 Genomics UK (COG-UK) Consortium | Randox Laboratories and Alex Alderton, Roberto Amato, Jeffrey Barrett, Sonia Goncalves, Ewan Harrison, David K. Jackson, Ian Johnston, Dominic Kwiatkowski, Cordelia Langford, John Sillitoe on behalf of the Wellcome Sanger Institute COVID-19 Surveillance Team |
| EPI_ISL_3051142, EPI_ISL_3051143, EPI_ISL_3051144, EPI_ISL_3051145 | Lighthouse Lab in Alderley Park | Wellcome Sanger Institute for the COVID-19 Genomics UK (COG-UK) Consortium | Jacquelyn Wynn, Mairead Hyland, The Lighthouse Lab in Alderley Park and Alex Alderton, Roberto Amato, Jeffrey Barrett, Sonia Goncalves, Ewan Harrison, David K. Jackson, Ian Johnston, Dominic Kwiatkowski, Cordelia Langford, John Sillitoe on behalf of the Wellcome Sanger Institute COVID-19 Surveillance Team |
| EPI_ISL_3051146, EPI_ISL_3051147 | Randox Laboratories | Wellcome Sanger Institute for the COVID-19 Genomics UK (COG-UK) Consortium | Randox Laboratories and Alex Alderton, Roberto Amato, Jeffrey Barrett, Sonia Goncalves, Ewan Harrison, David K. Jackson, Ian Johnston, Dominic Kwiatkowski, Cordelia Langford, John Sillitoe on behalf of the Wellcome Sanger Institute COVID-19 Surveillance Team |
| EPI_ISL_3051148 | Berkshire and Surrey Pathology Services Lighthouse Laboratory | Wellcome Sanger Institute for the COVID-19 Genomics UK (COG-UK) Consortium | Berkshire and Surrey Pathology Services Lighthouse Laboratory and Alex Alderton, Roberto Amato, Jeffrey Barrett, Sonia Goncalves, Ewan Harrison, David K. Jackson, Ian Johnston, Dominic Kwiatkowski, Cordelia Langford, John Sillitoe on behalf of the Wellcome Sanger Institute COVID-19 Surveillance Team |
| EPI_ISL_3051149 | Lighthouse Lab in Glasgow | Wellcome Sanger Institute for the COVID-19 Genomics UK (COG-UK) Consortium | Harper VanSteenhouse, Yumi Kasai, David Gray, Carol Clugston, Anna Dominiczak and Alex Alderton, Roberto Amato, Jeffrey Barrett, Sonia Goncalves, Ewan Harrison, David K. Jackson, Ian Johnston, Dominic Kwiatkowski, Cordelia Langford, John Sillitoe on behalf of the Wellcome Sanger Institute COVID-19 Surveillance Team |
| EPI_ISL_3051150 | Health Services Laboratories | Wellcome Sanger Institute for the COVID-19 Genomics UK (COG-UK) Consortium | Health Services Laboratories and Alex Alderton, Roberto Amato, Jeffrey Barrett, Sonia Goncalves, Ewan Harrison, David K. Jackson, Ian Johnston, Dominic Kwiatkowski, Cordelia Langford, John Sillitoe on behalf of the Wellcome Sanger Institute COVID-19 Surveillance Team |
| EPI_ISL_3051151 | Lighthouse Lab in Alderley Park | Wellcome Sanger Institute for the COVID-19 Genomics UK (COG-UK) Consortium | Jacquelyn Wynn, Mairead Hyland, The Lighthouse Lab in Alderley Park and Alex Alderton, Roberto Amato, Jeffrey Barrett, Sonia Goncalves, Ewan Harrison, David K. Jackson, Ian Johnston, Dominic Kwiatkowski, Cordelia Langford, John Sillitoe on behalf of the Wellcome Sanger Institute COVID-19 Surveillance Team |
| EPI_ISL_3051152, EPI_ISL_3051153, | Health Services Laboratories | Wellcome Sanger Institute for the COVID-19 Genomics UK (COG-UK) Consortium | Health Services Laboratories and Alex Alderton, Roberto Amato, Jeffrey Barrett, Sonia Goncalves, Ewan Harrison, David K. Jackson, Ian Johnston, Dominic Kwiatkowski, Cordelia Langford, John Sillitoe on behalf of the Wellcome Sanger Institute COVID-19 Surveillance Team |

| | | | |
|---|---|---|---|
| EPI_ISL_3051154 | | | |
| EPI_ISL_3051155 | Lighthouse Lab in Glasgow | Wellcome Sanger Institute for the COVID-19 Genomics UK (COG-UK) Consortium | Harper VanSteenhouse, Yumi Kasai, David Gray, Carol Clugston, Anna Dominiczak and Alex Alderton, Roberto Amato, Jeffrey Barrett, Sonia Goncalves, Ewan Harrison, David K. Jackson, Ian Johnston, Dominic Kwiatkowski, Cordelia Langford, John Sillitoe on behalf of the Wellcome Sanger Institute COVID-19 Surveillance Team |
| EPI_ISL_3051156, EPI_ISL_3051157, EPI_ISL_3051158, EPI_ISL_3051159 | Health Services Laboratories | Wellcome Sanger Institute for the COVID-19 Genomics UK (COG-UK) Consortium | Health Services Laboratories and Alex Alderton, Roberto Amato, Jeffrey Barrett, Sonia Goncalves, Ewan Harrison, David K. Jackson, Ian Johnston, Dominic Kwiatkowski, Cordelia Langford, John Sillitoe on behalf of the Wellcome Sanger Institute COVID-19 Surveillance Team |
| EPI_ISL_3051160 | Lighthouse Lab in Glasgow | Wellcome Sanger Institute for the COVID-19 Genomics UK (COG-UK) Consortium | Harper VanSteenhouse, Yumi Kasai, David Gray, Carol Clugston, Anna Dominiczak and Alex Alderton, Roberto Amato, Jeffrey Barrett, Sonia Goncalves, Ewan Harrison, David K. Jackson, Ian Johnston, Dominic Kwiatkowski, Cordelia Langford, John Sillitoe on behalf of the Wellcome Sanger Institute COVID-19 Surveillance Team |
| EPI_ISL_3051161, EPI_ISL_3051162, EPI_ISL_3051163 | Lighthouse Lab in Alderley Park | Wellcome Sanger Institute for the COVID-19 Genomics UK (COG-UK) Consortium | Jacquelyn Wynn, Mairead Hyland, The Lighthouse Lab in Alderley Park and Alex Alderton, Roberto Amato, Jeffrey Barrett, Sonia Goncalves, Ewan Harrison, David K. Jackson, Ian Johnston, Dominic Kwiatkowski, Cordelia Langford, John Sillitoe on behalf of the Wellcome Sanger Institute COVID-19 Surveillance Team |
| EPI_ISL_3051164, EPI_ISL_3051165 | Lighthouse Lab in Glasgow | Wellcome Sanger Institute for the COVID-19 Genomics UK (COG-UK) Consortium | Harper VanSteenhouse, Yumi Kasai, David Gray, Carol Clugston, Anna Dominiczak and Alex Alderton, Roberto Amato, Jeffrey Barrett, Sonia Goncalves, Ewan Harrison, David K. Jackson, Ian Johnston, Dominic Kwiatkowski, Cordelia Langford, John Sillitoe on behalf of the Wellcome Sanger Institute COVID-19 Surveillance Team |
| EPI_ISL_3051166 | Health Services Laboratories | Wellcome Sanger Institute for the COVID-19 Genomics UK (COG-UK) Consortium | Health Services Laboratories and Alex Alderton, Roberto Amato, Jeffrey Barrett, Sonia Goncalves, Ewan Harrison, David K. Jackson, Ian Johnston, Dominic Kwiatkowski, Cordelia Langford, John Sillitoe on behalf of the Wellcome Sanger Institute COVID-19 Surveillance Team |
| EPI_ISL_3051167, EPI_ISL_3051168 | Lighthouse Lab in Glasgow | Wellcome Sanger Institute for the COVID-19 Genomics UK (COG-UK) Consortium | Harper VanSteenhouse, Yumi Kasai, David Gray, Carol Clugston, Anna Dominiczak and Alex Alderton, Roberto Amato, Jeffrey Barrett, Sonia Goncalves, Ewan Harrison, David K. Jackson, Ian Johnston, Dominic Kwiatkowski, Cordelia Langford, John Sillitoe on behalf of the Wellcome Sanger Institute COVID-19 Surveillance Team |
| EPI_ISL_3051169 | Lighthouse Lab in Alderley Park | Wellcome Sanger Institute for the COVID-19 Genomics UK (COG-UK) Consortium | Jacquelyn Wynn, Mairead Hyland, The Lighthouse Lab in Alderley Park and Alex Alderton, Roberto Amato, Jeffrey Barrett, Sonia Goncalves, Ewan Harrison, David K. Jackson, Ian Johnston, Dominic Kwiatkowski, Cordelia Langford, John Sillitoe on behalf of the Wellcome Sanger Institute COVID-19 Surveillance Team |
| EPI_ISL_3051170 | Health Services Laboratories | Wellcome Sanger Institute for the COVID-19 Genomics UK (COG-UK) Consortium | Health Services Laboratories and Alex Alderton, Roberto Amato, Jeffrey Barrett, Sonia Goncalves, Ewan Harrison, David K. Jackson, Ian Johnston, Dominic Kwiatkowski, Cordelia Langford, John Sillitoe on behalf of the Wellcome Sanger Institute COVID-19 Surveillance Team |
| EPI_ISL_3051171 | Lighthouse Lab in Glasgow | Wellcome Sanger Institute for the COVID-19 Genomics UK (COG-UK) Consortium | Harper VanSteenhouse, Yumi Kasai, David Gray, Carol Clugston, Anna Dominiczak and Alex Alderton, Roberto Amato, Jeffrey Barrett, Sonia Goncalves, Ewan Harrison, David K. Jackson, Ian Johnston, Dominic Kwiatkowski, Cordelia Langford, John Sillitoe on behalf of the Wellcome Sanger Institute COVID-19 Surveillance Team |
| EPI_ISL_3051172, EPI_ISL_3051173, EPI_ISL_3051174, EPI_ISL_3051175 | Lighthouse Lab in Alderley Park | Wellcome Sanger Institute for the COVID-19 Genomics UK (COG-UK) Consortium | Jacquelyn Wynn, Mairead Hyland, The Lighthouse Lab in Alderley Park and Alex Alderton, Roberto Amato, Jeffrey Barrett, Sonia Goncalves, Ewan Harrison, David K. Jackson, Ian Johnston, Dominic Kwiatkowski, Cordelia Langford, John Sillitoe on behalf of the Wellcome Sanger Institute COVID-19 Surveillance Team |
| EPI_ISL_3051176, EPI_ISL_3051177 | Lighthouse Lab in Glasgow | Wellcome Sanger Institute for the COVID-19 Genomics UK (COG-UK) Consortium | Harper VanSteenhouse, Yumi Kasai, David Gray, Carol Clugston, Anna Dominiczak and Alex Alderton, Roberto Amato, Jeffrey Barrett, Sonia Goncalves, Ewan Harrison, David K. Jackson, Ian Johnston, Dominic Kwiatkowski, Cordelia Langford, John Sillitoe on behalf of the Wellcome Sanger Institute COVID-19 Surveillance Team |
| EPI_ISL_3051178 | Health Services Laboratories | Wellcome Sanger Institute for the COVID-19 Genomics UK (COG-UK) Consortium | Health Services Laboratories and Alex Alderton, Roberto Amato, Jeffrey Barrett, Sonia Goncalves, Ewan Harrison, David K. Jackson, Ian Johnston, Dominic Kwiatkowski, Cordelia Langford, John Sillitoe on behalf of the Wellcome Sanger Institute COVID-19 Surveillance Team |
| EPI_ISL_3051179 | Lighthouse Lab in Glasgow | Wellcome Sanger Institute for the COVID-19 Genomics UK (COG-UK) Consortium | Harper VanSteenhouse, Yumi Kasai, David Gray, Carol Clugston, Anna Dominiczak and Alex Alderton, Roberto Amato, Jeffrey Barrett, Sonia Goncalves, Ewan Harrison, David K. Jackson, Ian Johnston, Dominic Kwiatkowski, Cordelia Langford, John Sillitoe on behalf of the Wellcome Sanger Institute COVID-19 Surveillance Team |
| EPI_ISL_3051180 | Lighthouse Lab in Alderley Park | Wellcome Sanger Institute for the COVID-19 Genomics UK (COG-UK) Consortium | Jacquelyn Wynn, Mairead Hyland, The Lighthouse Lab in Alderley Park and Alex Alderton, Roberto Amato, Jeffrey Barrett, Sonia Goncalves, Ewan Harrison, David K. Jackson, Ian Johnston, Dominic Kwiatkowski, Cordelia Langford, John Sillitoe on behalf of the Wellcome Sanger Institute COVID-19 Surveillance Team |
| EPI_ISL_3051181 | Lighthouse Lab in Glasgow | Wellcome Sanger Institute for the COVID-19 Genomics UK (COG-UK) Consortium | Harper VanSteenhouse, Yumi Kasai, David Gray, Carol Clugston, Anna Dominiczak and Alex Alderton, Roberto Amato, Jeffrey Barrett, Sonia Goncalves, Ewan Harrison, David K. Jackson, Ian Johnston, Dominic Kwiatkowski, Cordelia Langford, John Sillitoe on behalf of the Wellcome Sanger Institute COVID-19 Surveillance Team |
| EPI_ISL_3051182 | Lighthouse Lab in Alderley Park | Wellcome Sanger Institute for the COVID-19 Genomics UK (COG-UK) Consortium | Jacquelyn Wynn, Mairead Hyland, The Lighthouse Lab in Alderley Park and Alex Alderton, Roberto Amato, Jeffrey Barrett, Sonia Goncalves, Ewan Harrison, David K. Jackson, Ian Johnston, Dominic Kwiatkowski, Cordelia Langford, John Sillitoe on behalf of the Wellcome Sanger Institute COVID-19 Surveillance Team |
| EPI_ISL_3051183 | Health Services Laboratories | Wellcome Sanger Institute for the COVID-19 Genomics UK (COG-UK) Consortium | Health Services Laboratories and Alex Alderton, Roberto Amato, Jeffrey Barrett, Sonia Goncalves, Ewan Harrison, David K. Jackson, Ian Johnston, Dominic Kwiatkowski, Cordelia Langford, John Sillitoe on behalf of the Wellcome Sanger Institute COVID-19 Surveillance Team |
| EPI_ISL_3051184 | Lighthouse Lab in Alderley Park | Wellcome Sanger Institute for the COVID-19 Genomics UK (COG-UK) Consortium | Jacquelyn Wynn, Mairead Hyland, The Lighthouse Lab in Alderley Park and Alex Alderton, Roberto Amato, Jeffrey Barrett, Sonia Goncalves, Ewan Harrison, David K. Jackson, Ian Johnston, Dominic Kwiatkowski, Cordelia Langford, John Sillitoe on behalf of the Wellcome Sanger Institute COVID-19 Surveillance Team |
| EPI_ISL_3051185 | Randox Laboratories | Wellcome Sanger Institute for the COVID-19 Genomics UK (COG-UK) Consortium | Randox Laboratories and Alex Alderton, Roberto Amato, Jeffrey Barrett, Sonia Goncalves, Ewan Harrison, David K. Jackson, Ian Johnston, Dominic Kwiatkowski, Cordelia Langford, John Sillitoe on behalf of the Wellcome Sanger Institute COVID-19 Surveillance Team |
| EPI_ISL_3051186, EPI_ISL_3051187, EPI_ISL_3051188 | Lighthouse Lab in Glasgow | Wellcome Sanger Institute for the COVID-19 Genomics UK (COG-UK) Consortium | Harper VanSteenhouse, Yumi Kasai, David Gray, Carol Clugston, Anna Dominiczak and Alex Alderton, Roberto Amato, Jeffrey Barrett, Sonia Goncalves, Ewan Harrison, David K. Jackson, Ian Johnston, Dominic Kwiatkowski, Cordelia Langford, John Sillitoe on behalf of the Wellcome Sanger Institute COVID-19 Surveillance Team |
| EPI_ISL_3051189 | Lighthouse Lab in Alderley Park | Wellcome Sanger Institute for the COVID-19 Genomics UK (COG-UK) Consortium | Jacquelyn Wynn, Mairead Hyland, The Lighthouse Lab in Alderley Park and Alex Alderton, Roberto Amato, Jeffrey Barrett, Sonia Goncalves, Ewan Harrison, David K. Jackson, Ian Johnston, Dominic Kwiatkowski, Cordelia Langford, John Sillitoe on behalf of the Wellcome Sanger Institute COVID-19 Surveillance Team |
| EPI_ISL_3051190 | Lighthouse Lab in Glasgow | Wellcome Sanger Institute for the COVID-19 Genomics UK (COG-UK) Consortium | Harper VanSteenhouse, Yumi Kasai, David Gray, Carol Clugston, Anna Dominiczak and Alex Alderton, Roberto Amato, Jeffrey Barrett, Sonia Goncalves, Ewan Harrison, David K. Jackson, Ian Johnston, Dominic Kwiatkowski, Cordelia Langford, John Sillitoe on behalf of the Wellcome Sanger Institute COVID-19 Surveillance Team |
| EPI_ISL_3051191 | Lighthouse Lab in Alderley Park | Wellcome Sanger Institute for the COVID-19 Genomics UK (COG-UK) Consortium | Jacquelyn Wynn, Mairead Hyland, The Lighthouse Lab in Alderley Park and Alex Alderton, Roberto Amato, Jeffrey Barrett, Sonia Goncalves, Ewan Harrison, David K. Jackson, Ian Johnston, Dominic Kwiatkowski, Cordelia Langford, John Sillitoe on behalf of the Wellcome Sanger Institute COVID-19 Surveillance Team |
| EPI_ISL_3051192 | Lighthouse Lab in Glasgow | Wellcome Sanger Institute for the COVID-19 Genomics UK (COG-UK) Consortium | Harper VanSteenhouse, Yumi Kasai, David Gray, Carol Clugston, Anna Dominiczak and Alex Alderton, Roberto Amato, Jeffrey Barrett, Sonia Goncalves, Ewan Harrison, David K. Jackson, Ian Johnston, Dominic Kwiatkowski, Cordelia Langford, John Sillitoe on behalf of the Wellcome Sanger Institute COVID-19 Surveillance Team |
| EPI_ISL_3051193, EPI_ISL_3051194 | Health Services Laboratories | Wellcome Sanger Institute for the COVID-19 Genomics UK (COG-UK) Consortium | Health Services Laboratories and Alex Alderton, Roberto Amato, Jeffrey Barrett, Sonia Goncalves, Ewan Harrison, David K. Jackson, Ian Johnston, Dominic Kwiatkowski, Cordelia Langford, John Sillitoe on behalf of the Wellcome Sanger Institute COVID-19 Surveillance Team |
| EPI_ISL_3051195, EPI_ISL_3051196, EPI_ISL_3051197 | Lighthouse Lab in Glasgow | Wellcome Sanger Institute for the COVID-19 Genomics UK (COG-UK) Consortium | Harper VanSteenhouse, Yumi Kasai, David Gray, Carol Clugston, Anna Dominiczak and Alex Alderton, Roberto Amato, Jeffrey Barrett, Sonia Goncalves, Ewan Harrison, David K. Jackson, Ian Johnston, Dominic Kwiatkowski, Cordelia Langford, John Sillitoe on behalf of the Wellcome Sanger Institute COVID-19 Surveillance Team |
| EPI_ISL_3051198, EPI_ISL_3051199, EPI_ISL_3051200 | Health Services Laboratories | Wellcome Sanger Institute for the COVID-19 Genomics UK (COG-UK) Consortium | Health Services Laboratories and Alex Alderton, Roberto Amato, Jeffrey Barrett, Sonia Goncalves, Ewan Harrison, David K. Jackson, Ian Johnston, Dominic Kwiatkowski, Cordelia Langford, John Sillitoe on behalf of the Wellcome Sanger Institute COVID-19 Surveillance Team |
| EPI_ISL_3051201, EPI_ISL_3051202 | Lighthouse Lab in Glasgow | Wellcome Sanger Institute for the COVID-19 Genomics UK (COG-UK) Consortium | Harper VanSteenhouse, Yumi Kasai, David Gray, Carol Clugston, Anna Dominiczak and Alex Alderton, Roberto Amato, Jeffrey Barrett, Sonia Goncalves, Ewan Harrison, David K. Jackson, Ian Johnston, Dominic Kwiatkowski, Cordelia Langford, John Sillitoe on behalf of the Wellcome Sanger Institute COVID-19 Surveillance Team |
| EPI_ISL_3051203 | Lighthouse Lab in Alderley Park | Wellcome Sanger Institute for the COVID-19 Genomics UK (COG-UK) Consortium | Jacquelyn Wynn, Mairead Hyland, The Lighthouse Lab in Alderley Park and Alex Alderton, Roberto Amato, Jeffrey Barrett, Sonia Goncalves, Ewan Harrison, David K. Jackson, Ian Johnston, Dominic Kwiatkowski, Cordelia Langford, John Sillitoe on behalf of the Wellcome Sanger Institute COVID-19 Surveillance Team |
| EPI_ISL_3051204, EPI_ISL_3051205 | Lighthouse Lab in Glasgow | Wellcome Sanger Institute for the COVID-19 Genomics UK (COG-UK) Consortium | Harper VanSteenhouse, Yumi Kasai, David Gray, Carol Clugston, Anna Dominiczak and Alex Alderton, Roberto Amato, Jeffrey Barrett, Sonia Goncalves, Ewan Harrison, David K. Jackson, Ian Johnston, Dominic Kwiatkowski, Cordelia Langford, John Sillitoe on behalf of the Wellcome Sanger Institute COVID-19 Surveillance Team |
| EPI_ISL_3051206 | Lighthouse Lab in Alderley Park | Wellcome Sanger Institute for the COVID-19 Genomics UK (COG-UK) Consortium | Jacquelyn Wynn, Mairead Hyland, The Lighthouse Lab in Alderley Park and Alex Alderton, Roberto Amato, Jeffrey Barrett, Sonia Goncalves, Ewan Harrison, David K. Jackson, Ian Johnston, Dominic Kwiatkowski, Cordelia Langford, John Sillitoe on behalf of the Wellcome Sanger Institute COVID-19 Surveillance Team |
| EPI_ISL_3051207 | Lighthouse Lab in Glasgow | Wellcome Sanger Institute for the COVID-19 Genomics UK (COG-UK) Consortium | Harper VanSteenhouse, Yumi Kasai, David Gray, Carol Clugston, Anna Dominiczak and Alex Alderton, Roberto Amato, Jeffrey Barrett, Sonia Goncalves, Ewan Harrison, David K. Jackson, Ian Johnston, Dominic Kwiatkowski, Cordelia Langford, John Sillitoe on behalf of the Wellcome Sanger Institute COVID-19 Surveillance Team |
| EPI_ISL_3051208, EPI_ISL_3051209, EPI_ISL_3051210, EPI_ISL_3051211 | Lighthouse Lab in Alderley Park | Wellcome Sanger Institute for the COVID-19 Genomics UK (COG-UK) Consortium | Jacquelyn Wynn, Mairead Hyland, The Lighthouse Lab in Alderley Park and Alex Alderton, Roberto Amato, Jeffrey Barrett, Sonia Goncalves, Ewan Harrison, David K. Jackson, Ian Johnston, Dominic Kwiatkowski, Cordelia Langford, John Sillitoe on behalf of the Wellcome Sanger Institute COVID-19 Surveillance Team |
| EPI_ISL_3051212, EPI_ISL_3051213 | Lighthouse Lab in Glasgow | Wellcome Sanger Institute for the COVID-19 Genomics UK (COG-UK) Consortium | Harper VanSteenhouse, Yumi Kasai, David Gray, Carol Clugston, Anna Dominiczak and Alex Alderton, Roberto Amato, Jeffrey Barrett, Sonia Goncalves, Ewan Harrison, David K. Jackson, Ian Johnston, Dominic Kwiatkowski, Cordelia Langford, John Sillitoe on behalf of the Wellcome Sanger Institute COVID-19 Surveillance Team |
| EPI_ISL_3051214, EPI_ISL_3051215 | Lighthouse Lab in Alderley Park | Wellcome Sanger Institute for the COVID-19 Genomics UK (COG-UK) Consortium | Jacquelyn Wynn, Mairead Hyland, The Lighthouse Lab in Alderley Park and Alex Alderton, Roberto Amato, Jeffrey Barrett, Sonia Goncalves, Ewan Harrison, David K. Jackson, Ian Johnston, Dominic Kwiatkowski, Cordelia Langford, John Sillitoe on behalf of the Wellcome Sanger Institute COVID-19 Surveillance Team |
| EPI_ISL_3051216, EPI_ISL_3051217 | Health Services Laboratories | Wellcome Sanger Institute for the COVID-19 Genomics UK (COG-UK) Consortium | Health Services Laboratories and Alex Alderton, Roberto Amato, Jeffrey Barrett, Sonia Goncalves, Ewan Harrison, David K. Jackson, Ian Johnston, Dominic Kwiatkowski, Cordelia Langford, John Sillitoe on behalf of the Wellcome Sanger Institute COVID-19 Surveillance Team |
| EPI_ISL_3051218 | Lighthouse Lab in Glasgow | Wellcome Sanger Institute for the COVID-19 Genomics UK (COG-UK) Consortium | Harper VanSteenhouse, Yumi Kasai, David Gray, Carol Clugston, Anna Dominiczak and Alex Alderton, Roberto Amato, Jeffrey Barrett, Sonia Goncalves, Ewan Harrison, David K. Jackson, Ian Johnston, Dominic Kwiatkowski, Cordelia Langford, John Sillitoe on behalf of the Wellcome Sanger Institute COVID-19 Surveillance Team |
| EPI_ISL_3051219 | Lighthouse Lab in Alderley Park | Wellcome Sanger Institute for the COVID-19 Genomics UK (COG-UK) Consortium | Jacquelyn Wynn, Mairead Hyland, The Lighthouse Lab in Alderley Park and Alex Alderton, Roberto Amato, Jeffrey Barrett, Sonia Goncalves, Ewan Harrison, David K. Jackson, Ian Johnston, Dominic Kwiatkowski, Cordelia Langford, John Sillitoe on behalf of the Wellcome Sanger Institute COVID-19 Surveillance Team |
| EPI_ISL_3051220 | Lighthouse Lab in Glasgow | Wellcome Sanger Institute for the COVID-19 Genomics UK (COG-UK) Consortium | Harper VanSteenhouse, Yumi Kasai, David Gray, Carol Clugston, Anna Dominiczak and Alex Alderton, Roberto Amato, Jeffrey Barrett, Sonia Goncalves, Ewan Harrison, David K. Jackson, Ian Johnston, Dominic Kwiatkowski, Cordelia Langford, John Sillitoe on behalf of the Wellcome Sanger Institute COVID-19 Surveillance Team |
| EPI_ISL_3051221, | Lighthouse Lab in Alderley Park | Wellcome Sanger Institute for the COVID-19 Genomics UK (COG-UK) | Jacquelyn Wynn, Mairead Hyland, The Lighthouse Lab in Alderley Park and Alex Alderton, Roberto Amato, Jeffrey Barrett, Sonia Goncalves, Ewan Harrison, David K. Jackson, Ian Johnston, Dominic Kwiatkowski, Cordelia Langford, John Sillitoe on |

| | | | |
|---|---|---|---|
| EPI_ISL_3051222, EPI_ISL_3051223 | | Consortium | behalf of the Wellcome Sanger Institute COVID-19 Surveillance Team |
| EPI_ISL_3051224 | Lighthouse Lab in Glasgow | Wellcome Sanger Institute for the COVID-19 Genomics UK (COG-UK) Consortium | Harper VanSteenhouse, Yumi Kasai, David Gray, Carol Clugston, Anna Dominiczak and Alex Alderton, Roberto Amato, Jeffrey Barrett, Sonia Goncalves, Ewan Harrison, David K. Jackson, Ian Johnston, Dominic Kwiatkowski, Cordelia Langford, John Sillitoe on behalf of the Wellcome Sanger Institute COVID-19 Surveillance Team |
| EPI_ISL_3051225 | Health Services Laboratories | Wellcome Sanger Institute for the COVID-19 Genomics UK (COG-UK) Consortium | Health Services Laboratories and Alex Alderton, Roberto Amato, Jeffrey Barrett, Sonia Goncalves, Ewan Harrison, David K. Jackson, Ian Johnston, Dominic Kwiatkowski, Cordelia Langford, John Sillitoe on behalf of the Wellcome Sanger Institute COVID-19 Surveillance Team |
| EPI_ISL_3051226 | Lighthouse Lab in Alderley Park | Wellcome Sanger Institute for the COVID-19 Genomics UK (COG-UK) Consortium | Jacquelyn Wynn, Mairead Hyland, The Lighthouse Lab in Alderley Park and Alex Alderton, Roberto Amato, Jeffrey Barrett, Sonia Goncalves, Ewan Harrison, David K. Jackson, Ian Johnston, Dominic Kwiatkowski, Cordelia Langford, John Sillitoe on behalf of the Wellcome Sanger Institute COVID-19 Surveillance Team |
| EPI_ISL_3051227 | Health Services Laboratories | Wellcome Sanger Institute for the COVID-19 Genomics UK (COG-UK) Consortium | Health Services Laboratories and Alex Alderton, Roberto Amato, Jeffrey Barrett, Sonia Goncalves, Ewan Harrison, David K. Jackson, Ian Johnston, Dominic Kwiatkowski, Cordelia Langford, John Sillitoe on behalf of the Wellcome Sanger Institute COVID-19 Surveillance Team |
| EPI_ISL_3051228 | Lighthouse Lab in Glasgow | Wellcome Sanger Institute for the COVID-19 Genomics UK (COG-UK) Consortium | Harper VanSteenhouse, Yumi Kasai, David Gray, Carol Clugston, Anna Dominiczak and Alex Alderton, Roberto Amato, Jeffrey Barrett, Sonia Goncalves, Ewan Harrison, David K. Jackson, Ian Johnston, Dominic Kwiatkowski, Cordelia Langford, John Sillitoe on behalf of the Wellcome Sanger Institute COVID-19 Surveillance Team |
| EPI_ISL_3051229 | Lighthouse Lab in Alderley Park | Wellcome Sanger Institute for the COVID-19 Genomics UK (COG-UK) Consortium | Jacquelyn Wynn, Mairead Hyland, The Lighthouse Lab in Alderley Park and Alex Alderton, Roberto Amato, Jeffrey Barrett, Sonia Goncalves, Ewan Harrison, David K. Jackson, Ian Johnston, Dominic Kwiatkowski, Cordelia Langford, John Sillitoe on behalf of the Wellcome Sanger Institute COVID-19 Surveillance Team |
| EPI_ISL_3051230 | Lighthouse Lab in Glasgow | Wellcome Sanger Institute for the COVID-19 Genomics UK (COG-UK) Consortium | Harper VanSteenhouse, Yumi Kasai, David Gray, Carol Clugston, Anna Dominiczak and Alex Alderton, Roberto Amato, Jeffrey Barrett, Sonia Goncalves, Ewan Harrison, David K. Jackson, Ian Johnston, Dominic Kwiatkowski, Cordelia Langford, John Sillitoe on behalf of the Wellcome Sanger Institute COVID-19 Surveillance Team |
| EPI_ISL_3051231 | Health Services Laboratories | Wellcome Sanger Institute for the COVID-19 Genomics UK (COG-UK) Consortium | Health Services Laboratories and Alex Alderton, Roberto Amato, Jeffrey Barrett, Sonia Goncalves, Ewan Harrison, David K. Jackson, Ian Johnston, Dominic Kwiatkowski, Cordelia Langford, John Sillitoe on behalf of the Wellcome Sanger Institute COVID-19 Surveillance Team |
| EPI_ISL_3051232, EPI_ISL_3051233 | Lighthouse Lab in Glasgow | Wellcome Sanger Institute for the COVID-19 Genomics UK (COG-UK) Consortium | Harper VanSteenhouse, Yumi Kasai, David Gray, Carol Clugston, Anna Dominiczak and Alex Alderton, Roberto Amato, Jeffrey Barrett, Sonia Goncalves, Ewan Harrison, David K. Jackson, Ian Johnston, Dominic Kwiatkowski, Cordelia Langford, John Sillitoe on behalf of the Wellcome Sanger Institute COVID-19 Surveillance Team |
| EPI_ISL_3051234 | Lighthouse Lab in Alderley Park | Wellcome Sanger Institute for the COVID-19 Genomics UK (COG-UK) Consortium | Jacquelyn Wynn, Mairead Hyland, The Lighthouse Lab in Alderley Park and Alex Alderton, Roberto Amato, Jeffrey Barrett, Sonia Goncalves, Ewan Harrison, David K. Jackson, Ian Johnston, Dominic Kwiatkowski, Cordelia Langford, John Sillitoe on behalf of the Wellcome Sanger Institute COVID-19 Surveillance Team |
| EPI_ISL_3051235, EPI_ISL_3051236 | Lighthouse Lab in Glasgow | Wellcome Sanger Institute for the COVID-19 Genomics UK (COG-UK) Consortium | Harper VanSteenhouse, Yumi Kasai, David Gray, Carol Clugston, Anna Dominiczak and Alex Alderton, Roberto Amato, Jeffrey Barrett, Sonia Goncalves, Ewan Harrison, David K. Jackson, Ian Johnston, Dominic Kwiatkowski, Cordelia Langford, John Sillitoe on behalf of the Wellcome Sanger Institute COVID-19 Surveillance Team |
| EPI_ISL_3051237 | Lighthouse Lab in Alderley Park | Wellcome Sanger Institute for the COVID-19 Genomics UK (COG-UK) Consortium | Jacquelyn Wynn, Mairead Hyland, The Lighthouse Lab in Alderley Park and Alex Alderton, Roberto Amato, Jeffrey Barrett, Sonia Goncalves, Ewan Harrison, David K. Jackson, Ian Johnston, Dominic Kwiatkowski, Cordelia Langford, John Sillitoe on behalf of the Wellcome Sanger Institute COVID-19 Surveillance Team |
| EPI_ISL_3051238 | Health Services Laboratories | Wellcome Sanger Institute for the COVID-19 Genomics UK (COG-UK) Consortium | Health Services Laboratories and Alex Alderton, Roberto Amato, Jeffrey Barrett, Sonia Goncalves, Ewan Harrison, David K. Jackson, Ian Johnston, Dominic Kwiatkowski, Cordelia Langford, John Sillitoe on behalf of the Wellcome Sanger Institute COVID-19 Surveillance Team |
| EPI_ISL_3051239, EPI_ISL_3051240 | Lighthouse Lab in Glasgow | Wellcome Sanger Institute for the COVID-19 Genomics UK (COG-UK) Consortium | Harper VanSteenhouse, Yumi Kasai, David Gray, Carol Clugston, Anna Dominiczak and Alex Alderton, Roberto Amato, Jeffrey Barrett, Sonia Goncalves, Ewan Harrison, David K. Jackson, Ian Johnston, Dominic Kwiatkowski, Cordelia Langford, John Sillitoe on behalf of the Wellcome Sanger Institute COVID-19 Surveillance Team |
| EPI_ISL_3051241 | Lighthouse Lab in Alderley Park | Wellcome Sanger Institute for the COVID-19 Genomics UK (COG-UK) Consortium | Jacquelyn Wynn, Mairead Hyland, The Lighthouse Lab in Alderley Park and Alex Alderton, Roberto Amato, Jeffrey Barrett, Sonia Goncalves, Ewan Harrison, David K. Jackson, Ian Johnston, Dominic Kwiatkowski, Cordelia Langford, John Sillitoe on behalf of the Wellcome Sanger Institute COVID-19 Surveillance Team |
| EPI_ISL_3051242 | Lighthouse Lab in Glasgow | Wellcome Sanger Institute for the COVID-19 Genomics UK (COG-UK) Consortium | Harper VanSteenhouse, Yumi Kasai, David Gray, Carol Clugston, Anna Dominiczak and Alex Alderton, Roberto Amato, Jeffrey Barrett, Sonia Goncalves, Ewan Harrison, David K. Jackson, Ian Johnston, Dominic Kwiatkowski, Cordelia Langford, John Sillitoe on behalf of the Wellcome Sanger Institute COVID-19 Surveillance Team |
| EPI_ISL_3051243 | Health Services Laboratories | Wellcome Sanger Institute for the COVID-19 Genomics UK (COG-UK) Consortium | Health Services Laboratories and Alex Alderton, Roberto Amato, Jeffrey Barrett, Sonia Goncalves, Ewan Harrison, David K. Jackson, Ian Johnston, Dominic Kwiatkowski, Cordelia Langford, John Sillitoe on behalf of the Wellcome Sanger Institute COVID-19 Surveillance Team |
| EPI_ISL_3051244 | Lighthouse Lab in Glasgow | Wellcome Sanger Institute for the COVID-19 Genomics UK (COG-UK) Consortium | Harper VanSteenhouse, Yumi Kasai, David Gray, Carol Clugston, Anna Dominiczak and Alex Alderton, Roberto Amato, Jeffrey Barrett, Sonia Goncalves, Ewan Harrison, David K. Jackson, Ian Johnston, Dominic Kwiatkowski, Cordelia Langford, John Sillitoe on behalf of the Wellcome Sanger Institute COVID-19 Surveillance Team |
| EPI_ISL_3051245 | Health Services Laboratories | Wellcome Sanger Institute for the COVID-19 Genomics UK (COG-UK) Consortium | Health Services Laboratories and Alex Alderton, Roberto Amato, Jeffrey Barrett, Sonia Goncalves, Ewan Harrison, David K. Jackson, Ian Johnston, Dominic Kwiatkowski, Cordelia Langford, John Sillitoe on behalf of the Wellcome Sanger Institute COVID-19 Surveillance Team |
| EPI_ISL_3051246, EPI_ISL_3051247 | Lighthouse Lab in Glasgow | Wellcome Sanger Institute for the COVID-19 Genomics UK (COG-UK) Consortium | Harper VanSteenhouse, Yumi Kasai, David Gray, Carol Clugston, Anna Dominiczak and Alex Alderton, Roberto Amato, Jeffrey Barrett, Sonia Goncalves, Ewan Harrison, David K. Jackson, Ian Johnston, Dominic Kwiatkowski, Cordelia Langford, John Sillitoe on behalf of the Wellcome Sanger Institute COVID-19 Surveillance Team |
| EPI_ISL_3051248 | Lighthouse Lab in Alderley Park | Wellcome Sanger Institute for the COVID-19 Genomics UK (COG-UK) Consortium | Jacquelyn Wynn, Mairead Hyland, The Lighthouse Lab in Alderley Park and Alex Alderton, Roberto Amato, Jeffrey Barrett, Sonia Goncalves, Ewan Harrison, David K. Jackson, Ian Johnston, Dominic Kwiatkowski, Cordelia Langford, John Sillitoe on behalf of the Wellcome Sanger Institute COVID-19 Surveillance Team |
| EPI_ISL_3051249 | Health Services Laboratories | Wellcome Sanger Institute for the COVID-19 Genomics UK (COG-UK) Consortium | Health Services Laboratories and Alex Alderton, Roberto Amato, Jeffrey Barrett, Sonia Goncalves, Ewan Harrison, David K. Jackson, Ian Johnston, Dominic Kwiatkowski, Cordelia Langford, John Sillitoe on behalf of the Wellcome Sanger Institute COVID-19 Surveillance Team |
| EPI_ISL_3051250, EPI_ISL_3051251 | Lighthouse Lab in Glasgow | Wellcome Sanger Institute for the COVID-19 Genomics UK (COG-UK) Consortium | Harper VanSteenhouse, Yumi Kasai, David Gray, Carol Clugston, Anna Dominiczak and Alex Alderton, Roberto Amato, Jeffrey Barrett, Sonia Goncalves, Ewan Harrison, David K. Jackson, Ian Johnston, Dominic Kwiatkowski, Cordelia Langford, John Sillitoe on behalf of the Wellcome Sanger Institute COVID-19 Surveillance Team |
| EPI_ISL_3051252 | Health Services Laboratories | Wellcome Sanger Institute for the COVID-19 Genomics UK (COG-UK) Consortium | Health Services Laboratories and Alex Alderton, Roberto Amato, Jeffrey Barrett, Sonia Goncalves, Ewan Harrison, David K. Jackson, Ian Johnston, Dominic Kwiatkowski, Cordelia Langford, John Sillitoe on behalf of the Wellcome Sanger Institute COVID-19 Surveillance Team |
| EPI_ISL_3051253 | Lighthouse Lab in Glasgow | Wellcome Sanger Institute for the COVID-19 Genomics UK (COG-UK) Consortium | Harper VanSteenhouse, Yumi Kasai, David Gray, Carol Clugston, Anna Dominiczak and Alex Alderton, Roberto Amato, Jeffrey Barrett, Sonia Goncalves, Ewan Harrison, David K. Jackson, Ian Johnston, Dominic Kwiatkowski, Cordelia Langford, John Sillitoe on behalf of the Wellcome Sanger Institute COVID-19 Surveillance Team |
| EPI_ISL_3051254 | Health Services Laboratories | Wellcome Sanger Institute for the COVID-19 Genomics UK (COG-UK) Consortium | Health Services Laboratories and Alex Alderton, Roberto Amato, Jeffrey Barrett, Sonia Goncalves, Ewan Harrison, David K. Jackson, Ian Johnston, Dominic Kwiatkowski, Cordelia Langford, John Sillitoe on behalf of the Wellcome Sanger Institute COVID-19 Surveillance Team |
| EPI_ISL_3051255 | Lighthouse Lab in Glasgow | Wellcome Sanger Institute for the COVID-19 Genomics UK (COG-UK) Consortium | Harper VanSteenhouse, Yumi Kasai, David Gray, Carol Clugston, Anna Dominiczak and Alex Alderton, Roberto Amato, Jeffrey Barrett, Sonia Goncalves, Ewan Harrison, David K. Jackson, Ian Johnston, Dominic Kwiatkowski, Cordelia Langford, John Sillitoe on behalf of the Wellcome Sanger Institute COVID-19 Surveillance Team |
| EPI_ISL_3051256 | Lighthouse Lab in Alderley Park | Wellcome Sanger Institute for the COVID-19 Genomics UK (COG-UK) Consortium | Jacquelyn Wynn, Mairead Hyland, The Lighthouse Lab in Alderley Park and Alex Alderton, Roberto Amato, Jeffrey Barrett, Sonia Goncalves, Ewan Harrison, David K. Jackson, Ian Johnston, Dominic Kwiatkowski, Cordelia Langford, John Sillitoe on behalf of the Wellcome Sanger Institute COVID-19 Surveillance Team |
| EPI_ISL_3051257 | Health Services Laboratories | Wellcome Sanger Institute for the COVID-19 Genomics UK (COG-UK) Consortium | Health Services Laboratories and Alex Alderton, Roberto Amato, Jeffrey Barrett, Sonia Goncalves, Ewan Harrison, David K. Jackson, Ian Johnston, Dominic Kwiatkowski, Cordelia Langford, John Sillitoe on behalf of the Wellcome Sanger Institute COVID-19 Surveillance Team |
| EPI_ISL_3051258 | Lighthouse Lab in Alderley Park | Wellcome Sanger Institute for the COVID-19 Genomics UK (COG-UK) Consortium | Jacquelyn Wynn, Mairead Hyland, The Lighthouse Lab in Alderley Park and Alex Alderton, Roberto Amato, Jeffrey Barrett, Sonia Goncalves, Ewan Harrison, David K. Jackson, Ian Johnston, Dominic Kwiatkowski, Cordelia Langford, John Sillitoe on behalf of the Wellcome Sanger Institute COVID-19 Surveillance Team |
| EPI_ISL_3051259 | Health Services Laboratories | Wellcome Sanger Institute for the COVID-19 Genomics UK (COG-UK) Consortium | Health Services Laboratories and Alex Alderton, Roberto Amato, Jeffrey Barrett, Sonia Goncalves, Ewan Harrison, David K. Jackson, Ian Johnston, Dominic Kwiatkowski, Cordelia Langford, John Sillitoe on behalf of the Wellcome Sanger Institute COVID-19 Surveillance Team |
| EPI_ISL_3051260, EPI_ISL_3051261, EPI_ISL_3051262, EPI_ISL_3051263, EPI_ISL_3051264, EPI_ISL_3051265 | Lighthouse Lab in Glasgow | Wellcome Sanger Institute for the COVID-19 Genomics UK (COG-UK) Consortium | Harper VanSteenhouse, Yumi Kasai, David Gray, Carol Clugston, Anna Dominiczak and Alex Alderton, Roberto Amato, Jeffrey Barrett, Sonia Goncalves, Ewan Harrison, David K. Jackson, Ian Johnston, Dominic Kwiatkowski, Cordelia Langford, John Sillitoe on behalf of the Wellcome Sanger Institute COVID-19 Surveillance Team |
| EPI_ISL_3051266, EPI_ISL_3051267 | Health Services Laboratories | Wellcome Sanger Institute for the COVID-19 Genomics UK (COG-UK) Consortium | Health Services Laboratories and Alex Alderton, Roberto Amato, Jeffrey Barrett, Sonia Goncalves, Ewan Harrison, David K. Jackson, Ian Johnston, Dominic Kwiatkowski, Cordelia Langford, John Sillitoe on behalf of the Wellcome Sanger Institute COVID-19 Surveillance Team |
| EPI_ISL_3051268, EPI_ISL_3051269, EPI_ISL_3051270, EPI_ISL_3051271 | Lighthouse Lab in Glasgow | Wellcome Sanger Institute for the COVID-19 Genomics UK (COG-UK) Consortium | Harper VanSteenhouse, Yumi Kasai, David Gray, Carol Clugston, Anna Dominiczak and Alex Alderton, Roberto Amato, Jeffrey Barrett, Sonia Goncalves, Ewan Harrison, David K. Jackson, Ian Johnston, Dominic Kwiatkowski, Cordelia Langford, John Sillitoe on behalf of the Wellcome Sanger Institute COVID-19 Surveillance Team |
| EPI_ISL_3051272, EPI_ISL_3051273 | Lighthouse Lab in Alderley Park | Wellcome Sanger Institute for the COVID-19 Genomics UK (COG-UK) Consortium | Jacquelyn Wynn, Mairead Hyland, The Lighthouse Lab in Alderley Park and Alex Alderton, Roberto Amato, Jeffrey Barrett, Sonia Goncalves, Ewan Harrison, David K. Jackson, Ian Johnston, Dominic Kwiatkowski, Cordelia Langford, John Sillitoe on behalf of the Wellcome Sanger Institute COVID-19 Surveillance Team |
| EPI_ISL_3051274 | Lighthouse Lab in Glasgow | Wellcome Sanger Institute for the COVID-19 Genomics UK (COG-UK) Consortium | Harper VanSteenhouse, Yumi Kasai, David Gray, Carol Clugston, Anna Dominiczak and Alex Alderton, Roberto Amato, Jeffrey Barrett, Sonia Goncalves, Ewan Harrison, David K. Jackson, Ian Johnston, Dominic Kwiatkowski, Cordelia Langford, John Sillitoe on behalf of the Wellcome Sanger Institute COVID-19 Surveillance Team |
| EPI_ISL_3051275, EPI_ISL_3051276 | Lighthouse Lab in Alderley Park | Wellcome Sanger Institute for the COVID-19 Genomics UK (COG-UK) Consortium | Jacquelyn Wynn, Mairead Hyland, The Lighthouse Lab in Alderley Park and Alex Alderton, Roberto Amato, Jeffrey Barrett, Sonia Goncalves, Ewan Harrison, David K. Jackson, Ian Johnston, Dominic Kwiatkowski, Cordelia Langford, John Sillitoe on behalf of the Wellcome Sanger Institute COVID-19 Surveillance Team |
| EPI_ISL_3051277, EPI_ISL_3051278 | Health Services Laboratories | Wellcome Sanger Institute for the COVID-19 Genomics UK (COG-UK) Consortium | Health Services Laboratories and Alex Alderton, Roberto Amato, Jeffrey Barrett, Sonia Goncalves, Ewan Harrison, David K. Jackson, Ian Johnston, Dominic Kwiatkowski, Cordelia Langford, John Sillitoe on behalf of the Wellcome Sanger Institute COVID-19 Surveillance Team |
| EPI_ISL_3051279 | Lighthouse Lab in Glasgow | Wellcome Sanger Institute for the COVID-19 Genomics UK (COG-UK) Consortium | Harper VanSteenhouse, Yumi Kasai, David Gray, Carol Clugston, Anna Dominiczak and Alex Alderton, Roberto Amato, Jeffrey Barrett, Sonia Goncalves, Ewan Harrison, David K. Jackson, Ian Johnston, Dominic Kwiatkowski, Cordelia Langford, John Sillitoe on behalf of the Wellcome Sanger Institute COVID-19 Surveillance Team |
| EPI_ISL_3051280, EPI_ISL_3051281 | Health Services Laboratories | Wellcome Sanger Institute for the COVID-19 Genomics UK (COG-UK) Consortium | Health Services Laboratories and Alex Alderton, Roberto Amato, Jeffrey Barrett, Sonia Goncalves, Ewan Harrison, David K. Jackson, Ian Johnston, Dominic Kwiatkowski, Cordelia Langford, John Sillitoe on behalf of the Wellcome Sanger Institute COVID-19 Surveillance Team |
| EPI_ISL_3051282 | Lighthouse Lab in Alderley Park | Wellcome Sanger Institute for the COVID-19 Genomics UK (COG-UK) Consortium | Jacquelyn Wynn, Mairead Hyland, The Lighthouse Lab in Alderley Park and Alex Alderton, Roberto Amato, Jeffrey Barrett, Sonia Goncalves, Ewan Harrison, David K. Jackson, Ian Johnston, Dominic Kwiatkowski, Cordelia Langford, John Sillitoe on behalf of the Wellcome Sanger Institute COVID-19 Surveillance Team |
| EPI_ISL_3051283 | Lighthouse Lab in Glasgow | Wellcome Sanger Institute for the COVID-19 Genomics UK (COG-UK) | Harper VanSteenhouse, Yumi Kasai, David Gray, Carol Clugston, Anna Dominiczak and Alex Alderton, Roberto Amato, Jeffrey Barrett, Sonia Goncalves, Ewan Harrison, David K. Jackson, Ian Johnston, Dominic Kwiatkowski, Cordelia Langford, |

| | | | |
|---|---|---|---|
| | | Consortium | John Sillitoe on behalf of the Wellcome Sanger Institute COVID-19 Surveillance Team |
| EPI_ISL_3051284 | Lighthouse Lab in Alderley Park | Wellcome Sanger Institute for the COVID-19 Genomics UK (COG-UK) Consortium | Jacquelyn Wynn, Mairead Hyland, The Lighthouse Lab in Alderley Park and Alex Alderton, Roberto Amato, Jeffrey Barrett, Sonia Goncalves, Ewan Harrison, David K. Jackson, Ian Johnston, Dominic Kwiatkowski, Cordelia Langford, John Sillitoe on behalf of the Wellcome Sanger Institute COVID-19 Surveillance Team |
| EPI_ISL_3051285 | Lighthouse Lab in Glasgow | Wellcome Sanger Institute for the COVID-19 Genomics UK (COG-UK) Consortium | Harper VanSteenhouse, Yumi Kasai, David Gray, Carol Clugston, Anna Dominiczak and Alex Alderton, Roberto Amato, Jeffrey Barrett, Sonia Goncalves, Ewan Harrison, David K. Jackson, Ian Johnston, Dominic Kwiatkowski, Cordelia Langford, John Sillitoe on behalf of the Wellcome Sanger Institute COVID-19 Surveillance Team |
| EPI_ISL_3051286, EPI_ISL_3051287 | Health Services Laboratories | Wellcome Sanger Institute for the COVID-19 Genomics UK (COG-UK) Consortium | Health Services Laboratories and Alex Alderton, Roberto Amato, Jeffrey Barrett, Sonia Goncalves, Ewan Harrison, David K. Jackson, Ian Johnston, Dominic Kwiatkowski, Cordelia Langford, John Sillitoe on behalf of the Wellcome Sanger Institute COVID-19 Surveillance Team |
| EPI_ISL_3051288, EPI_ISL_3051289, EPI_ISL_3051290, EPI_ISL_3051291 | Lighthouse Lab in Glasgow | Wellcome Sanger Institute for the COVID-19 Genomics UK (COG-UK) Consortium | Harper VanSteenhouse, Yumi Kasai, David Gray, Carol Clugston, Anna Dominiczak and Alex Alderton, Roberto Amato, Jeffrey Barrett, Sonia Goncalves, Ewan Harrison, David K. Jackson, Ian Johnston, Dominic Kwiatkowski, Cordelia Langford, John Sillitoe on behalf of the Wellcome Sanger Institute COVID-19 Surveillance Team |
| EPI_ISL_3051292 | Health Services Laboratories | Wellcome Sanger Institute for the COVID-19 Genomics UK (COG-UK) Consortium | Health Services Laboratories and Alex Alderton, Roberto Amato, Jeffrey Barrett, Sonia Goncalves, Ewan Harrison, David K. Jackson, Ian Johnston, Dominic Kwiatkowski, Cordelia Langford, John Sillitoe on behalf of the Wellcome Sanger Institute COVID-19 Surveillance Team |
| EPI_ISL_3051293, EPI_ISL_3051294, EPI_ISL_3051295, EPI_ISL_3051296, EPI_ISL_3051297 | Lighthouse Lab in Glasgow | Wellcome Sanger Institute for the COVID-19 Genomics UK (COG-UK) Consortium | Harper VanSteenhouse, Yumi Kasai, David Gray, Carol Clugston, Anna Dominiczak and Alex Alderton, Roberto Amato, Jeffrey Barrett, Sonia Goncalves, Ewan Harrison, David K. Jackson, Ian Johnston, Dominic Kwiatkowski, Cordelia Langford, John Sillitoe on behalf of the Wellcome Sanger Institute COVID-19 Surveillance Team |
| EPI_ISL_3051298, EPI_ISL_3051299, EPI_ISL_3051300 | Health Services Laboratories | Wellcome Sanger Institute for the COVID-19 Genomics UK (COG-UK) Consortium | Health Services Laboratories and Alex Alderton, Roberto Amato, Jeffrey Barrett, Sonia Goncalves, Ewan Harrison, David K. Jackson, Ian Johnston, Dominic Kwiatkowski, Cordelia Langford, John Sillitoe on behalf of the Wellcome Sanger Institute COVID-19 Surveillance Team |
| EPI_ISL_3051301 | Lighthouse Lab in Glasgow | Wellcome Sanger Institute for the COVID-19 Genomics UK (COG-UK) Consortium | Harper VanSteenhouse, Yumi Kasai, David Gray, Carol Clugston, Anna Dominiczak and Alex Alderton, Roberto Amato, Jeffrey Barrett, Sonia Goncalves, Ewan Harrison, David K. Jackson, Ian Johnston, Dominic Kwiatkowski, Cordelia Langford, John Sillitoe on behalf of the Wellcome Sanger Institute COVID-19 Surveillance Team |
| EPI_ISL_3051302 | Lighthouse Lab in Alderley Park | Wellcome Sanger Institute for the COVID-19 Genomics UK (COG-UK) Consortium | Jacquelyn Wynn, Mairead Hyland, The Lighthouse Lab in Alderley Park and Alex Alderton, Roberto Amato, Jeffrey Barrett, Sonia Goncalves, Ewan Harrison, David K. Jackson, Ian Johnston, Dominic Kwiatkowski, Cordelia Langford, John Sillitoe on behalf of the Wellcome Sanger Institute COVID-19 Surveillance Team |
| EPI_ISL_3051303 | Lighthouse Lab in Glasgow | Wellcome Sanger Institute for the COVID-19 Genomics UK (COG-UK) Consortium | Harper VanSteenhouse, Yumi Kasai, David Gray, Carol Clugston, Anna Dominiczak and Alex Alderton, Roberto Amato, Jeffrey Barrett, Sonia Goncalves, Ewan Harrison, David K. Jackson, Ian Johnston, Dominic Kwiatkowski, Cordelia Langford, John Sillitoe on behalf of the Wellcome Sanger Institute COVID-19 Surveillance Team |
| EPI_ISL_3051304 | Health Services Laboratories | Wellcome Sanger Institute for the COVID-19 Genomics UK (COG-UK) Consortium | Health Services Laboratories and Alex Alderton, Roberto Amato, Jeffrey Barrett, Sonia Goncalves, Ewan Harrison, David K. Jackson, Ian Johnston, Dominic Kwiatkowski, Cordelia Langford, John Sillitoe on behalf of the Wellcome Sanger Institute COVID-19 Surveillance Team |
| EPI_ISL_3051305 | Lighthouse Lab in Glasgow | Wellcome Sanger Institute for the COVID-19 Genomics UK (COG-UK) Consortium | Harper VanSteenhouse, Yumi Kasai, David Gray, Carol Clugston, Anna Dominiczak and Alex Alderton, Roberto Amato, Jeffrey Barrett, Sonia Goncalves, Ewan Harrison, David K. Jackson, Ian Johnston, Dominic Kwiatkowski, Cordelia Langford, John Sillitoe on behalf of the Wellcome Sanger Institute COVID-19 Surveillance Team |
| EPI_ISL_3051306, EPI_ISL_3051307 | Health Services Laboratories | Wellcome Sanger Institute for the COVID-19 Genomics UK (COG-UK) Consortium | Health Services Laboratories and Alex Alderton, Roberto Amato, Jeffrey Barrett, Sonia Goncalves, Ewan Harrison, David K. Jackson, Ian Johnston, Dominic Kwiatkowski, Cordelia Langford, John Sillitoe on behalf of the Wellcome Sanger Institute COVID-19 Surveillance Team |
| EPI_ISL_3051308, EPI_ISL_3051309 | Lighthouse Lab in Alderley Park | Wellcome Sanger Institute for the COVID-19 Genomics UK (COG-UK) Consortium | Jacquelyn Wynn, Mairead Hyland, The Lighthouse Lab in Alderley Park and Alex Alderton, Roberto Amato, Jeffrey Barrett, Sonia Goncalves, Ewan Harrison, David K. Jackson, Ian Johnston, Dominic Kwiatkowski, Cordelia Langford, John Sillitoe on behalf of the Wellcome Sanger Institute COVID-19 Surveillance Team |
| EPI_ISL_3051310 | Lighthouse Lab in Glasgow | Wellcome Sanger Institute for the COVID-19 Genomics UK (COG-UK) Consortium | Harper VanSteenhouse, Yumi Kasai, David Gray, Carol Clugston, Anna Dominiczak and Alex Alderton, Roberto Amato, Jeffrey Barrett, Sonia Goncalves, Ewan Harrison, David K. Jackson, Ian Johnston, Dominic Kwiatkowski, Cordelia Langford, John Sillitoe on behalf of the Wellcome Sanger Institute COVID-19 Surveillance Team |
| EPI_ISL_3051311 | Health Services Laboratories | Wellcome Sanger Institute for the COVID-19 Genomics UK (COG-UK) Consortium | Health Services Laboratories and Alex Alderton, Roberto Amato, Jeffrey Barrett, Sonia Goncalves, Ewan Harrison, David K. Jackson, Ian Johnston, Dominic Kwiatkowski, Cordelia Langford, John Sillitoe on behalf of the Wellcome Sanger Institute COVID-19 Surveillance Team |
| EPI_ISL_3051312 | Lighthouse Lab in Alderley Park | Wellcome Sanger Institute for the COVID-19 Genomics UK (COG-UK) Consortium | Jacquelyn Wynn, Mairead Hyland, The Lighthouse Lab in Alderley Park and Alex Alderton, Roberto Amato, Jeffrey Barrett, Sonia Goncalves, Ewan Harrison, David K. Jackson, Ian Johnston, Dominic Kwiatkowski, Cordelia Langford, John Sillitoe on behalf of the Wellcome Sanger Institute COVID-19 Surveillance Team |
| EPI_ISL_3051313 | Lighthouse Lab in Glasgow | Wellcome Sanger Institute for the COVID-19 Genomics UK (COG-UK) Consortium | Harper VanSteenhouse, Yumi Kasai, David Gray, Carol Clugston, Anna Dominiczak and Alex Alderton, Roberto Amato, Jeffrey Barrett, Sonia Goncalves, Ewan Harrison, David K. Jackson, Ian Johnston, Dominic Kwiatkowski, Cordelia Langford, John Sillitoe on behalf of the Wellcome Sanger Institute COVID-19 Surveillance Team |
| EPI_ISL_3051314 | Lighthouse Lab in Alderley Park | Wellcome Sanger Institute for the COVID-19 Genomics UK (COG-UK) Consortium | Jacquelyn Wynn, Mairead Hyland, The Lighthouse Lab in Alderley Park and Alex Alderton, Roberto Amato, Jeffrey Barrett, Sonia Goncalves, Ewan Harrison, David K. Jackson, Ian Johnston, Dominic Kwiatkowski, Cordelia Langford, John Sillitoe on behalf of the Wellcome Sanger Institute COVID-19 Surveillance Team |
| EPI_ISL_3051315 | A.S.L.VERCELLI | Fondazione del Piemonte per l'Oncologia IRCCS | Antonino Sottile, Silvia Brossa, Paola Marino, Giorgio Giardina |
| EPI_ISL_3051316 | A.S.L.NOVARA-BORGOMANERO | Fondazione del Piemonte per l'Oncologia IRCCS | Antonino Sottile, Silvia Brossa, Paola Marino, Giorgio Giardina |
| EPI_ISL_3051317, EPI_ISL_3051318 | A.S.L.TO5 | Fondazione del Piemonte per l'Oncologia IRCCS | Antonino Sottile, Silvia Brossa, Paola Marino, Giorgio Giardina |
| EPI_ISL_3051319, EPI_ISL_3051320 | Lighthouse Lab in Glasgow | Wellcome Sanger Institute for the COVID-19 Genomics UK (COG-UK) Consortium | Harper VanSteenhouse, Yumi Kasai, David Gray, Carol Clugston, Anna Dominiczak and Alex Alderton, Roberto Amato, Jeffrey Barrett, Sonia Goncalves, Ewan Harrison, David K. Jackson, Ian Johnston, Dominic Kwiatkowski, Cordelia Langford, John Sillitoe on behalf of the Wellcome Sanger Institute COVID-19 Surveillance Team |
| EPI_ISL_3051321 | A.S.L.NOVARA-BORGOMANERO | Fondazione del Piemonte per l'Oncologia IRCCS | Antonino Sottile, Silvia Brossa, Paola Marino, Giorgio Giardina |
| EPI_ISL_3051322 | Lighthouse Lab in Glasgow | Wellcome Sanger Institute for the COVID-19 Genomics UK (COG-UK) Consortium | Harper VanSteenhouse, Yumi Kasai, David Gray, Carol Clugston, Anna Dominiczak and Alex Alderton, Roberto Amato, Jeffrey Barrett, Sonia Goncalves, Ewan Harrison, David K. Jackson, Ian Johnston, Dominic Kwiatkowski, Cordelia Langford, John Sillitoe on behalf of the Wellcome Sanger Institute COVID-19 Surveillance Team |
| EPI_ISL_3051323 | Lighthouse Lab in Alderley Park | Wellcome Sanger Institute for the COVID-19 Genomics UK (COG-UK) Consortium | Jacquelyn Wynn, Mairead Hyland, The Lighthouse Lab in Alderley Park and Alex Alderton, Roberto Amato, Jeffrey Barrett, Sonia Goncalves, Ewan Harrison, David K. Jackson, Ian Johnston, Dominic Kwiatkowski, Cordelia Langford, John Sillitoe on behalf of the Wellcome Sanger Institute COVID-19 Surveillance Team |
| EPI_ISL_3051324 | A.S.L.NOVARA-BORGOMANERO | Fondazione del Piemonte per l'Oncologia IRCCS | Antonino Sottile, Silvia Brossa, Paola Marino, Giorgio Giardina |
| EPI_ISL_3051325, EPI_ISL_3051326 | Lighthouse Lab in Glasgow | Wellcome Sanger Institute for the COVID-19 Genomics UK (COG-UK) Consortium | Harper VanSteenhouse, Yumi Kasai, David Gray, Carol Clugston, Anna Dominiczak and Alex Alderton, Roberto Amato, Jeffrey Barrett, Sonia Goncalves, Ewan Harrison, David K. Jackson, Ian Johnston, Dominic Kwiatkowski, Cordelia Langford, John Sillitoe on behalf of the Wellcome Sanger Institute COVID-19 Surveillance Team |
| EPI_ISL_3051327 | Health Services Laboratories | Wellcome Sanger Institute for the COVID-19 Genomics UK (COG-UK) Consortium | Health Services Laboratories and Alex Alderton, Roberto Amato, Jeffrey Barrett, Sonia Goncalves, Ewan Harrison, David K. Jackson, Ian Johnston, Dominic Kwiatkowski, Cordelia Langford, John Sillitoe on behalf of the Wellcome Sanger Institute COVID-19 Surveillance Team |
| EPI_ISL_3051328 | A.S.L.NOVARA-BORGOMANERO | Fondazione del Piemonte per l'Oncologia IRCCS | Antonino Sottile, Silvia Brossa, Paola Marino, Giorgio Giardina |
| EPI_ISL_3051329 | Health Services Laboratories | Wellcome Sanger Institute for the COVID-19 Genomics UK (COG-UK) Consortium | Health Services Laboratories and Alex Alderton, Roberto Amato, Jeffrey Barrett, Sonia Goncalves, Ewan Harrison, David K. Jackson, Ian Johnston, Dominic Kwiatkowski, Cordelia Langford, John Sillitoe on behalf of the Wellcome Sanger Institute COVID-19 Surveillance Team |
| EPI_ISL_3051330 | A.S.L.NOVARA-BORGOMANERO | Fondazione del Piemonte per l'Oncologia IRCCS | Antonino Sottile, Silvia Brossa, Paola Marino, Giorgio Giardina |
| EPI_ISL_3051331 | Health Services Laboratories | Wellcome Sanger Institute for the COVID-19 Genomics UK (COG-UK) Consortium | Health Services Laboratories and Alex Alderton, Roberto Amato, Jeffrey Barrett, Sonia Goncalves, Ewan Harrison, David K. Jackson, Ian Johnston, Dominic Kwiatkowski, Cordelia Langford, John Sillitoe on behalf of the Wellcome Sanger Institute COVID-19 Surveillance Team |
| EPI_ISL_3051332, EPI_ISL_3051333 | Lighthouse Lab in Glasgow | Wellcome Sanger Institute for the COVID-19 Genomics UK (COG-UK) Consortium | Harper VanSteenhouse, Yumi Kasai, David Gray, Carol Clugston, Anna Dominiczak and Alex Alderton, Roberto Amato, Jeffrey Barrett, Sonia Goncalves, Ewan Harrison, David K. Jackson, Ian Johnston, Dominic Kwiatkowski, Cordelia Langford, John Sillitoe on behalf of the Wellcome Sanger Institute COVID-19 Surveillance Team |
| EPI_ISL_3051334 | A.S.L.NOVARA-BORGOMANERO | Fondazione del Piemonte per l'Oncologia IRCCS | Antonino Sottile, Silvia Brossa, Paola Marino, Giorgio Giardina |
| EPI_ISL_3051335, EPI_ISL_3051336 | Lighthouse Lab in Alderley Park | Wellcome Sanger Institute for the COVID-19 Genomics UK (COG-UK) Consortium | Jacquelyn Wynn, Mairead Hyland, The Lighthouse Lab in Alderley Park and Alex Alderton, Roberto Amato, Jeffrey Barrett, Sonia Goncalves, Ewan Harrison, David K. Jackson, Ian Johnston, Dominic Kwiatkowski, Cordelia Langford, John Sillitoe on behalf of the Wellcome Sanger Institute COVID-19 Surveillance Team |
| EPI_ISL_3051337 | A.S.L.NOVARA-BORGOMANERO | Fondazione del Piemonte per l'Oncologia IRCCS | Antonino Sottile, Silvia Brossa, Paola Marino, Giorgio Giardina |
| EPI_ISL_3051338, EPI_ISL_3051339 | Lighthouse Lab in Alderley Park | Wellcome Sanger Institute for the COVID-19 Genomics UK (COG-UK) Consortium | Jacquelyn Wynn, Mairead Hyland, The Lighthouse Lab in Alderley Park and Alex Alderton, Roberto Amato, Jeffrey Barrett, Sonia Goncalves, Ewan Harrison, David K. Jackson, Ian Johnston, Dominic Kwiatkowski, Cordelia Langford, John Sillitoe on behalf of the Wellcome Sanger Institute COVID-19 Surveillance Team |
| EPI_ISL_3051340 | Lighthouse Lab in Glasgow | Wellcome Sanger Institute for the COVID-19 Genomics UK (COG-UK) Consortium | Harper VanSteenhouse, Yumi Kasai, David Gray, Carol Clugston, Anna Dominiczak and Alex Alderton, Roberto Amato, Jeffrey Barrett, Sonia Goncalves, Ewan Harrison, David K. Jackson, Ian Johnston, Dominic Kwiatkowski, Cordelia Langford, John Sillitoe on behalf of the Wellcome Sanger Institute COVID-19 Surveillance Team |
| EPI_ISL_3051341 | A.S.L.NOVARA-BORGOMANERO | Fondazione del Piemonte per l'Oncologia IRCCS | Antonino Sottile, Silvia Brossa, Paola Marino, Giorgio Giardina |
| EPI_ISL_3051342 | Lighthouse Lab in Alderley Park | Wellcome Sanger Institute for the COVID-19 Genomics UK (COG-UK) Consortium | Jacquelyn Wynn, Mairead Hyland, The Lighthouse Lab in Alderley Park and Alex Alderton, Roberto Amato, Jeffrey Barrett, Sonia Goncalves, Ewan Harrison, David K. Jackson, Ian Johnston, Dominic Kwiatkowski, Cordelia Langford, John Sillitoe on behalf of the Wellcome Sanger Institute COVID-19 Surveillance Team |
| EPI_ISL_3051343 | Lighthouse Lab in Glasgow | Wellcome Sanger Institute for the COVID-19 Genomics UK (COG-UK) Consortium | Harper VanSteenhouse, Yumi Kasai, David Gray, Carol Clugston, Anna Dominiczak and Alex Alderton, Roberto Amato, Jeffrey Barrett, Sonia Goncalves, Ewan Harrison, David K. Jackson, Ian Johnston, Dominic Kwiatkowski, Cordelia Langford, John Sillitoe on behalf of the Wellcome Sanger Institute COVID-19 Surveillance Team |
| EPI_ISL_3051344 | Health Services Laboratories | Wellcome Sanger Institute for the COVID-19 Genomics UK (COG-UK) Consortium | Health Services Laboratories and Alex Alderton, Roberto Amato, Jeffrey Barrett, Sonia Goncalves, Ewan Harrison, David K. Jackson, Ian Johnston, Dominic Kwiatkowski, Cordelia Langford, John Sillitoe on behalf of the Wellcome Sanger Institute COVID-19 Surveillance Team |
| EPI_ISL_3051345 | A.S.L.NOVARA-BORGOMANERO | Fondazione del Piemonte per l'Oncologia IRCCS | Antonino Sottile, Silvia Brossa, Paola Marino, Giorgio Giardina |
| EPI_ISL_3051346 | Lighthouse Lab in Glasgow | Wellcome Sanger Institute for the COVID-19 Genomics UK (COG-UK) Consortium | Harper VanSteenhouse, Yumi Kasai, David Gray, Carol Clugston, Anna Dominiczak and Alex Alderton, Roberto Amato, Jeffrey Barrett, Sonia Goncalves, Ewan Harrison, David K. Jackson, Ian Johnston, Dominic Kwiatkowski, Cordelia Langford, John Sillitoe on behalf of the Wellcome Sanger Institute COVID-19 Surveillance Team |
| EPI_ISL_3051347 | Lighthouse Lab in Alderley Park | Wellcome Sanger Institute for the COVID-19 Genomics UK (COG-UK) Consortium | Jacquelyn Wynn, Mairead Hyland, The Lighthouse Lab in Alderley Park and Alex Alderton, Roberto Amato, Jeffrey Barrett, Sonia Goncalves, Ewan Harrison, David K. Jackson, Ian Johnston, Dominic Kwiatkowski, Cordelia Langford, John Sillitoe on behalf of the Wellcome Sanger Institute COVID-19 Surveillance Team |

| | | | |
|---|---|---|---|
| EPI_ISL_3051348 | Lighthouse Lab in Glasgow | Wellcome Sanger Institute for the COVID-19 Genomics UK (COG-UK) Consortium | Harper VanSteenhouse, Yumi Kasai, David Gray, Carol Clugston, Anna Dominiczak and Alex Alderton, Roberto Amato, Jeffrey Barrett, Sonia Goncalves, Ewan Harrison, David K. Jackson, Ian Johnston, Dominic Kwiatkowski, Cordelia Langford, John Sillitoe on behalf of the Wellcome Sanger Institute COVID-19 Surveillance Team |
| EPI_ISL_3051349 | A.S.L.NOVARA-BORGOMANERO | Fondazione del Piemonte per l'Oncologia IRCCS | Antonino Sottile, Silvia Brossa, Paola Marino, Giorgio Giardina |
| EPI_ISL_3051350,<br>EPI_ISL_3051351 | Lighthouse Lab in Glasgow | Wellcome Sanger Institute for the COVID-19 Genomics UK (COG-UK) Consortium | Harper VanSteenhouse, Yumi Kasai, David Gray, Carol Clugston, Anna Dominiczak and Alex Alderton, Roberto Amato, Jeffrey Barrett, Sonia Goncalves, Ewan Harrison, David K. Jackson, Ian Johnston, Dominic Kwiatkowski, Cordelia Langford, John Sillitoe on behalf of the Wellcome Sanger Institute COVID-19 Surveillance Team |
| EPI_ISL_3051352 | A.S.L.NOVARA-BORGOMANERO | Fondazione del Piemonte per l'Oncologia IRCCS | Antonino Sottile, Silvia Brossa, Paola Marino, Giorgio Giardina |
| EPI_ISL_3051353,<br>EPI_ISL_3051354 | Lighthouse Lab in Glasgow | Wellcome Sanger Institute for the COVID-19 Genomics UK (COG-UK) Consortium | Harper VanSteenhouse, Yumi Kasai, David Gray, Carol Clugston, Anna Dominiczak and Alex Alderton, Roberto Amato, Jeffrey Barrett, Sonia Goncalves, Ewan Harrison, David K. Jackson, Ian Johnston, Dominic Kwiatkowski, Cordelia Langford, John Sillitoe on behalf of the Wellcome Sanger Institute COVID-19 Surveillance Team |
| EPI_ISL_3051355 | A.S.L.NOVARA-BORGOMANERO | Fondazione del Piemonte per l'Oncologia IRCCS | Antonino Sottile, Silvia Brossa, Paola Marino, Giorgio Giardina |
| EPI_ISL_3051356,<br>EPI_ISL_3051357 | Lighthouse Lab in Alderley Park | Wellcome Sanger Institute for the COVID-19 Genomics UK (COG-UK) Consortium | Jacquelyn Wynn, Mairead Hyland, The Lighthouse Lab in Alderley Park and Alex Alderton, Roberto Amato, Jeffrey Barrett, Sonia Goncalves, Ewan Harrison, David K. Jackson, Ian Johnston, Dominic Kwiatkowski, Cordelia Langford, John Sillitoe on behalf of the Wellcome Sanger Institute COVID-19 Surveillance Team |
| EPI_ISL_3051358,<br>EPI_ISL_3051360 | Lighthouse Lab in Glasgow | Wellcome Sanger Institute for the COVID-19 Genomics UK (COG-UK) Consortium | Harper VanSteenhouse, Yumi Kasai, David Gray, Carol Clugston, Anna Dominiczak and Alex Alderton, Roberto Amato, Jeffrey Barrett, Sonia Goncalves, Ewan Harrison, David K. Jackson, Ian Johnston, Dominic Kwiatkowski, Cordelia Langford, John Sillitoe on behalf of the Wellcome Sanger Institute COVID-19 Surveillance Team |
| EPI_ISL_3051361 | Health Services Laboratories | Wellcome Sanger Institute for the COVID-19 Genomics UK (COG-UK) Consortium | Health Services Laboratories and Alex Alderton, Roberto Amato, Jeffrey Barrett, Sonia Goncalves, Ewan Harrison, David K. Jackson, Ian Johnston, Dominic Kwiatkowski, Cordelia Langford, John Sillitoe on behalf of the Wellcome Sanger Institute COVID-19 Surveillance Team |
| EPI_ISL_3051362 | Lighthouse Lab in Alderley Park | Wellcome Sanger Institute for the COVID-19 Genomics UK (COG-UK) Consortium | Jacquelyn Wynn, Mairead Hyland, The Lighthouse Lab in Alderley Park and Alex Alderton, Roberto Amato, Jeffrey Barrett, Sonia Goncalves, Ewan Harrison, David K. Jackson, Ian Johnston, Dominic Kwiatkowski, Cordelia Langford, John Sillitoe on behalf of the Wellcome Sanger Institute COVID-19 Surveillance Team |
| EPI_ISL_3051364 | Lighthouse Lab in Glasgow | Wellcome Sanger Institute for the COVID-19 Genomics UK (COG-UK) Consortium | Harper VanSteenhouse, Yumi Kasai, David Gray, Carol Clugston, Anna Dominiczak and Alex Alderton, Roberto Amato, Jeffrey Barrett, Sonia Goncalves, Ewan Harrison, David K. Jackson, Ian Johnston, Dominic Kwiatkowski, Cordelia Langford, John Sillitoe on behalf of the Wellcome Sanger Institute COVID-19 Surveillance Team |
| EPI_ISL_3051365 | Health Services Laboratories | Wellcome Sanger Institute for the COVID-19 Genomics UK (COG-UK) Consortium | Health Services Laboratories and Alex Alderton, Roberto Amato, Jeffrey Barrett, Sonia Goncalves, Ewan Harrison, David K. Jackson, Ian Johnston, Dominic Kwiatkowski, Cordelia Langford, John Sillitoe on behalf of the Wellcome Sanger Institute COVID-19 Surveillance Team |
| EPI_ISL_3051366 | A.S.L.VERCELLI | Fondazione del Piemonte per l'Oncologia IRCCS | Antonino Sottile, Silvia Brossa, Paola Marino, Giorgio Giardina |
| EPI_ISL_3051367 | Lighthouse Lab in Glasgow | Wellcome Sanger Institute for the COVID-19 Genomics UK (COG-UK) Consortium | Harper VanSteenhouse, Yumi Kasai, David Gray, Carol Clugston, Anna Dominiczak and Alex Alderton, Roberto Amato, Jeffrey Barrett, Sonia Goncalves, Ewan Harrison, David K. Jackson, Ian Johnston, Dominic Kwiatkowski, Cordelia Langford, John Sillitoe on behalf of the Wellcome Sanger Institute COVID-19 Surveillance Team |
| EPI_ISL_3051368 | Health Services Laboratories | Wellcome Sanger Institute for the COVID-19 Genomics UK (COG-UK) Consortium | Health Services Laboratories and Alex Alderton, Roberto Amato, Jeffrey Barrett, Sonia Goncalves, Ewan Harrison, David K. Jackson, Ian Johnston, Dominic Kwiatkowski, Cordelia Langford, John Sillitoe on behalf of the Wellcome Sanger Institute COVID-19 Surveillance Team |
| EPI_ISL_3051369 | Lighthouse Lab in Glasgow | Wellcome Sanger Institute for the COVID-19 Genomics UK (COG-UK) Consortium | Harper VanSteenhouse, Yumi Kasai, David Gray, Carol Clugston, Anna Dominiczak and Alex Alderton, Roberto Amato, Jeffrey Barrett, Sonia Goncalves, Ewan Harrison, David K. Jackson, Ian Johnston, Dominic Kwiatkowski, Cordelia Langford, John Sillitoe on behalf of the Wellcome Sanger Institute COVID-19 Surveillance Team |
| EPI_ISL_3051370 | A.S.L.VERCELLI | Fondazione del Piemonte per l'Oncologia IRCCS | Antonino Sottile, Silvia Brossa, Paola Marino, Giorgio Giardina |
| EPI_ISL_3051371 | Lighthouse Lab in Glasgow | Wellcome Sanger Institute for the COVID-19 Genomics UK (COG-UK) Consortium | Harper VanSteenhouse, Yumi Kasai, David Gray, Carol Clugston, Anna Dominiczak and Alex Alderton, Roberto Amato, Jeffrey Barrett, Sonia Goncalves, Ewan Harrison, David K. Jackson, Ian Johnston, Dominic Kwiatkowski, Cordelia Langford, John Sillitoe on behalf of the Wellcome Sanger Institute COVID-19 Surveillance Team |
| EPI_ISL_3051372 | Health Services Laboratories | Wellcome Sanger Institute for the COVID-19 Genomics UK (COG-UK) Consortium | Health Services Laboratories and Alex Alderton, Roberto Amato, Jeffrey Barrett, Sonia Goncalves, Ewan Harrison, David K. Jackson, Ian Johnston, Dominic Kwiatkowski, Cordelia Langford, John Sillitoe on behalf of the Wellcome Sanger Institute COVID-19 Surveillance Team |
| EPI_ISL_3051373 | A.S.L.VERCELLI | Fondazione del Piemonte per l'Oncologia IRCCS | Antonino Sottile, Silvia Brossa, Paola Marino, Giorgio Giardina |
| EPI_ISL_3051374 | Lighthouse Lab in Glasgow | Wellcome Sanger Institute for the COVID-19 Genomics UK (COG-UK) Consortium | Harper VanSteenhouse, Yumi Kasai, David Gray, Carol Clugston, Anna Dominiczak and Alex Alderton, Roberto Amato, Jeffrey Barrett, Sonia Goncalves, Ewan Harrison, David K. Jackson, Ian Johnston, Dominic Kwiatkowski, Cordelia Langford, John Sillitoe on behalf of the Wellcome Sanger Institute COVID-19 Surveillance Team |
| EPI_ISL_3051375 | Lighthouse Lab in Alderley Park | Wellcome Sanger Institute for the COVID-19 Genomics UK (COG-UK) Consortium | Jacquelyn Wynn, Mairead Hyland, The Lighthouse Lab in Alderley Park and Alex Alderton, Roberto Amato, Jeffrey Barrett, Sonia Goncalves, Ewan Harrison, David K. Jackson, Ian Johnston, Dominic Kwiatkowski, Cordelia Langford, John Sillitoe on behalf of the Wellcome Sanger Institute COVID-19 Surveillance Team |
| EPI_ISL_3051376 | Lighthouse Lab in Glasgow | Wellcome Sanger Institute for the COVID-19 Genomics UK (COG-UK) Consortium | Harper VanSteenhouse, Yumi Kasai, David Gray, Carol Clugston, Anna Dominiczak and Alex Alderton, Roberto Amato, Jeffrey Barrett, Sonia Goncalves, Ewan Harrison, David K. Jackson, Ian Johnston, Dominic Kwiatkowski, Cordelia Langford, John Sillitoe on behalf of the Wellcome Sanger Institute COVID-19 Surveillance Team |
| EPI_ISL_3051377 | A.S.L.VERCELLI | Fondazione del Piemonte per l'Oncologia IRCCS | Antonino Sottile, Silvia Brossa, Paola Marino, Giorgio Giardina |
| EPI_ISL_3051378,<br>EPI_ISL_3051379 | Lighthouse Lab in Alderley Park | Wellcome Sanger Institute for the COVID-19 Genomics UK (COG-UK) Consortium | Jacquelyn Wynn, Mairead Hyland, The Lighthouse Lab in Alderley Park and Alex Alderton, Roberto Amato, Jeffrey Barrett, Sonia Goncalves, Ewan Harrison, David K. Jackson, Ian Johnston, Dominic Kwiatkowski, Cordelia Langford, John Sillitoe on behalf of the Wellcome Sanger Institute COVID-19 Surveillance Team |
| EPI_ISL_3051380 | A.S.L.VERCELLI | Fondazione del Piemonte per l'Oncologia IRCCS | Antonino Sottile, Silvia Brossa, Paola Marino, Giorgio Giardina |
| EPI_ISL_3051381 | Lighthouse Lab in Glasgow | Wellcome Sanger Institute for the COVID-19 Genomics UK (COG-UK) Consortium | Harper VanSteenhouse, Yumi Kasai, David Gray, Carol Clugston, Anna Dominiczak and Alex Alderton, Roberto Amato, Jeffrey Barrett, Sonia Goncalves, Ewan Harrison, David K. Jackson, Ian Johnston, Dominic Kwiatkowski, Cordelia Langford, John Sillitoe on behalf of the Wellcome Sanger Institute COVID-19 Surveillance Team |
| EPI_ISL_3051382 | Health Services Laboratories | Wellcome Sanger Institute for the COVID-19 Genomics UK (COG-UK) Consortium | Health Services Laboratories and Alex Alderton, Roberto Amato, Jeffrey Barrett, Sonia Goncalves, Ewan Harrison, David K. Jackson, Ian Johnston, Dominic Kwiatkowski, Cordelia Langford, John Sillitoe on behalf of the Wellcome Sanger Institute COVID-19 Surveillance Team |
| EPI_ISL_3051383 | A.S.L.VERCELLI | Fondazione del Piemonte per l'Oncologia IRCCS | Antonino Sottile, Silvia Brossa, Paola Marino, Giorgio Giardina |
| EPI_ISL_3051384 | Lighthouse Lab in Alderley Park | Wellcome Sanger Institute for the COVID-19 Genomics UK (COG-UK) Consortium | Jacquelyn Wynn, Mairead Hyland, The Lighthouse Lab in Alderley Park and Alex Alderton, Roberto Amato, Jeffrey Barrett, Sonia Goncalves, Ewan Harrison, David K. Jackson, Ian Johnston, Dominic Kwiatkowski, Cordelia Langford, John Sillitoe on behalf of the Wellcome Sanger Institute COVID-19 Surveillance Team |
| EPI_ISL_3051385,<br>EPI_ISL_3051386 | Health Services Laboratories | Wellcome Sanger Institute for the COVID-19 Genomics UK (COG-UK) Consortium | Health Services Laboratories and Alex Alderton, Roberto Amato, Jeffrey Barrett, Sonia Goncalves, Ewan Harrison, David K. Jackson, Ian Johnston, Dominic Kwiatkowski, Cordelia Langford, John Sillitoe on behalf of the Wellcome Sanger Institute COVID-19 Surveillance Team |
| EPI_ISL_3051387,<br>EPI_ISL_3051388 | Lighthouse Lab in Glasgow | Wellcome Sanger Institute for the COVID-19 Genomics UK (COG-UK) Consortium | Harper VanSteenhouse, Yumi Kasai, David Gray, Carol Clugston, Anna Dominiczak and Alex Alderton, Roberto Amato, Jeffrey Barrett, Sonia Goncalves, Ewan Harrison, David K. Jackson, Ian Johnston, Dominic Kwiatkowski, Cordelia Langford, John Sillitoe on behalf of the Wellcome Sanger Institute COVID-19 Surveillance Team |
| EPI_ISL_3051389 | Health Services Laboratories | Wellcome Sanger Institute for the COVID-19 Genomics UK (COG-UK) Consortium | Health Services Laboratories and Alex Alderton, Roberto Amato, Jeffrey Barrett, Sonia Goncalves, Ewan Harrison, David K. Jackson, Ian Johnston, Dominic Kwiatkowski, Cordelia Langford, John Sillitoe on behalf of the Wellcome Sanger Institute COVID-19 Surveillance Team |
| EPI_ISL_3051390,<br>EPI_ISL_3051391,<br>EPI_ISL_3051392,<br>EPI_ISL_3051393,<br>EPI_ISL_3051394,<br>EPI_ISL_3051395,<br>EPI_ISL_3051396 | Lighthouse Lab in Glasgow | Wellcome Sanger Institute for the COVID-19 Genomics UK (COG-UK) Consortium | Harper VanSteenhouse, Yumi Kasai, David Gray, Carol Clugston, Anna Dominiczak and Alex Alderton, Roberto Amato, Jeffrey Barrett, Sonia Goncalves, Ewan Harrison, David K. Jackson, Ian Johnston, Dominic Kwiatkowski, Cordelia Langford, John Sillitoe on behalf of the Wellcome Sanger Institute COVID-19 Surveillance Team |
| EPI_ISL_3051397,<br>EPI_ISL_3051398 | Health Services Laboratories | Wellcome Sanger Institute for the COVID-19 Genomics UK (COG-UK) Consortium | Health Services Laboratories and Alex Alderton, Roberto Amato, Jeffrey Barrett, Sonia Goncalves, Ewan Harrison, David K. Jackson, Ian Johnston, Dominic Kwiatkowski, Cordelia Langford, John Sillitoe on behalf of the Wellcome Sanger Institute COVID-19 Surveillance Team |
| EPI_ISL_3051399 | Randox Laboratories | Wellcome Sanger Institute for the COVID-19 Genomics UK (COG-UK) Consortium | Randox Laboratories and Alex Alderton, Roberto Amato, Jeffrey Barrett, Sonia Goncalves, Ewan Harrison, David K. Jackson, Ian Johnston, Dominic Kwiatkowski, Cordelia Langford, John Sillitoe on behalf of the Wellcome Sanger Institute COVID-19 Surveillance Team |
| EPI_ISL_3051400 | Lighthouse Lab in Glasgow | Wellcome Sanger Institute for the COVID-19 Genomics UK (COG-UK) Consortium | Harper VanSteenhouse, Yumi Kasai, David Gray, Carol Clugston, Anna Dominiczak and Alex Alderton, Roberto Amato, Jeffrey Barrett, Sonia Goncalves, Ewan Harrison, David K. Jackson, Ian Johnston, Dominic Kwiatkowski, Cordelia Langford, John Sillitoe on behalf of the Wellcome Sanger Institute COVID-19 Surveillance Team |
| EPI_ISL_3051401 | Lighthouse Lab in Alderley Park | Wellcome Sanger Institute for the COVID-19 Genomics UK (COG-UK) Consortium | Jacquelyn Wynn, Mairead Hyland, The Lighthouse Lab in Alderley Park and Alex Alderton, Roberto Amato, Jeffrey Barrett, Sonia Goncalves, Ewan Harrison, David K. Jackson, Ian Johnston, Dominic Kwiatkowski, Cordelia Langford, John Sillitoe on behalf of the Wellcome Sanger Institute COVID-19 Surveillance Team |
| EPI_ISL_3051402 | Health Services Laboratories | Wellcome Sanger Institute for the COVID-19 Genomics UK (COG-UK) Consortium | Health Services Laboratories and Alex Alderton, Roberto Amato, Jeffrey Barrett, Sonia Goncalves, Ewan Harrison, David K. Jackson, Ian Johnston, Dominic Kwiatkowski, Cordelia Langford, John Sillitoe on behalf of the Wellcome Sanger Institute COVID-19 Surveillance Team |
| EPI_ISL_3051403 | Lighthouse Lab in Alderley Park | Wellcome Sanger Institute for the COVID-19 Genomics UK (COG-UK) Consortium | Jacquelyn Wynn, Mairead Hyland, The Lighthouse Lab in Alderley Park and Alex Alderton, Roberto Amato, Jeffrey Barrett, Sonia Goncalves, Ewan Harrison, David K. Jackson, Ian Johnston, Dominic Kwiatkowski, Cordelia Langford, John Sillitoe on behalf of the Wellcome Sanger Institute COVID-19 Surveillance Team |
| EPI_ISL_3051404 | Health Services Laboratories | Wellcome Sanger Institute for the COVID-19 Genomics UK (COG-UK) Consortium | Health Services Laboratories and Alex Alderton, Roberto Amato, Jeffrey Barrett, Sonia Goncalves, Ewan Harrison, David K. Jackson, Ian Johnston, Dominic Kwiatkowski, Cordelia Langford, John Sillitoe on behalf of the Wellcome Sanger Institute COVID-19 Surveillance Team |
| EPI_ISL_3051405,<br>EPI_ISL_3051406 | Lighthouse Lab in Alderley Park | Wellcome Sanger Institute for the COVID-19 Genomics UK (COG-UK) Consortium | Jacquelyn Wynn, Mairead Hyland, The Lighthouse Lab in Alderley Park and Alex Alderton, Roberto Amato, Jeffrey Barrett, Sonia Goncalves, Ewan Harrison, David K. Jackson, Ian Johnston, Dominic Kwiatkowski, Cordelia Langford, John Sillitoe on behalf of the Wellcome Sanger Institute COVID-19 Surveillance Team |
| EPI_ISL_3051407,<br>EPI_ISL_3051408 | Lighthouse Lab in Glasgow | Wellcome Sanger Institute for the COVID-19 Genomics UK (COG-UK) Consortium | Harper VanSteenhouse, Yumi Kasai, David Gray, Carol Clugston, Anna Dominiczak and Alex Alderton, Roberto Amato, Jeffrey Barrett, Sonia Goncalves, Ewan Harrison, David K. Jackson, Ian Johnston, Dominic Kwiatkowski, Cordelia Langford, John Sillitoe on behalf of the Wellcome Sanger Institute COVID-19 Surveillance Team |
| EPI_ISL_3051409 | Health Services Laboratories | Wellcome Sanger Institute for the COVID-19 Genomics UK (COG-UK) Consortium | Health Services Laboratories and Alex Alderton, Roberto Amato, Jeffrey Barrett, Sonia Goncalves, Ewan Harrison, David K. Jackson, Ian Johnston, Dominic Kwiatkowski, Cordelia Langford, John Sillitoe on behalf of the Wellcome Sanger Institute COVID-19 Surveillance Team |
| EPI_ISL_3051410 | Lighthouse Lab in Glasgow | Wellcome Sanger Institute for the COVID-19 Genomics UK (COG-UK) Consortium | Harper VanSteenhouse, Yumi Kasai, David Gray, Carol Clugston, Anna Dominiczak and Alex Alderton, Roberto Amato, Jeffrey Barrett, Sonia Goncalves, Ewan Harrison, David K. Jackson, Ian Johnston, Dominic Kwiatkowski, Cordelia Langford, John Sillitoe on behalf of the Wellcome Sanger Institute COVID-19 Surveillance Team |
| EPI_ISL_3051411 | Health Services Laboratories | Wellcome Sanger Institute for the COVID-19 Genomics UK (COG-UK) Consortium | Health Services Laboratories and Alex Alderton, Roberto Amato, Jeffrey Barrett, Sonia Goncalves, Ewan Harrison, David K. Jackson, Ian Johnston, Dominic Kwiatkowski, Cordelia Langford, John Sillitoe on behalf of the Wellcome Sanger Institute COVID-19 Surveillance Team |
| EPI_ISL_3051412 | Lighthouse Lab in Alderley Park | Wellcome Sanger Institute for the COVID-19 Genomics UK (COG-UK) Consortium | Jacquelyn Wynn, Mairead Hyland, The Lighthouse Lab in Alderley Park and Alex Alderton, Roberto Amato, Jeffrey Barrett, Sonia Goncalves, Ewan Harrison, David K. Jackson, Ian Johnston, Dominic Kwiatkowski, Cordelia Langford, John Sillitoe on behalf of the Wellcome Sanger Institute COVID-19 Surveillance Team |

| | | | |
|---|---|---|---|
| EPI_ISL_3051413, EPI_ISL_3051414, EPI_ISL_3051415 | Lighthouse Lab in Glasgow | Wellcome Sanger Institute for the COVID-19 Genomics UK (COG-UK) Consortium | Harper VanSteenhouse, Yumi Kasai, David Gray, Carol Clugston, Anna Dominiczak and Alex Alderton, Roberto Amato, Jeffrey Barrett, Sonia Goncalves, Ewan Harrison, David K. Jackson, Ian Johnston, Dominic Kwiatkowski, Cordelia Langford, John Sillitoe on behalf of the Wellcome Sanger Institute COVID-19 Surveillance Team |
| EPI_ISL_3051416 | Lighthouse Lab in Alderley Park | Wellcome Sanger Institute for the COVID-19 Genomics UK (COG-UK) Consortium | Jacquelyn Wynn, Mairead Hyland, The Lighthouse Lab in Alderley Park and Alex Alderton, Roberto Amato, Jeffrey Barrett, Sonia Goncalves, Ewan Harrison, David K. Jackson, Ian Johnston, Dominic Kwiatkowski, Cordelia Langford, John Sillitoe on behalf of the Wellcome Sanger Institute COVID-19 Surveillance Team |
| EPI_ISL_3051417 | Health Services Laboratories | Wellcome Sanger Institute for the COVID-19 Genomics UK (COG-UK) Consortium | Health Services Laboratories and Alex Alderton, Roberto Amato, Jeffrey Barrett, Sonia Goncalves, Ewan Harrison, David K. Jackson, Ian Johnston, Dominic Kwiatkowski, Cordelia Langford, John Sillitoe on behalf of the Wellcome Sanger Institute COVID-19 Surveillance Team |
| EPI_ISL_3051418 | Lighthouse Lab in Glasgow | Wellcome Sanger Institute for the COVID-19 Genomics UK (COG-UK) Consortium | Harper VanSteenhouse, Yumi Kasai, David Gray, Carol Clugston, Anna Dominiczak and Alex Alderton, Roberto Amato, Jeffrey Barrett, Sonia Goncalves, Ewan Harrison, David K. Jackson, Ian Johnston, Dominic Kwiatkowski, Cordelia Langford, John Sillitoe on behalf of the Wellcome Sanger Institute COVID-19 Surveillance Team |
| EPI_ISL_3051419, EPI_ISL_3051420 | Health Services Laboratories | Wellcome Sanger Institute for the COVID-19 Genomics UK (COG-UK) Consortium | Health Services Laboratories and Alex Alderton, Roberto Amato, Jeffrey Barrett, Sonia Goncalves, Ewan Harrison, David K. Jackson, Ian Johnston, Dominic Kwiatkowski, Cordelia Langford, John Sillitoe on behalf of the Wellcome Sanger Institute COVID-19 Surveillance Team |
| EPI_ISL_3051421, EPI_ISL_3051422, EPI_ISL_3051423 | Lighthouse Lab in Glasgow | Wellcome Sanger Institute for the COVID-19 Genomics UK (COG-UK) Consortium | Harper VanSteenhouse, Yumi Kasai, David Gray, Carol Clugston, Anna Dominiczak and Alex Alderton, Roberto Amato, Jeffrey Barrett, Sonia Goncalves, Ewan Harrison, David K. Jackson, Ian Johnston, Dominic Kwiatkowski, Cordelia Langford, John Sillitoe on behalf of the Wellcome Sanger Institute COVID-19 Surveillance Team |
| EPI_ISL_3051424 | Health Services Laboratories | Wellcome Sanger Institute for the COVID-19 Genomics UK (COG-UK) Consortium | Health Services Laboratories and Alex Alderton, Roberto Amato, Jeffrey Barrett, Sonia Goncalves, Ewan Harrison, David K. Jackson, Ian Johnston, Dominic Kwiatkowski, Cordelia Langford, John Sillitoe on behalf of the Wellcome Sanger Institute COVID-19 Surveillance Team |
| EPI_ISL_3051425 | Lighthouse Lab in Glasgow | Wellcome Sanger Institute for the COVID-19 Genomics UK (COG-UK) Consortium | Harper VanSteenhouse, Yumi Kasai, David Gray, Carol Clugston, Anna Dominiczak and Alex Alderton, Roberto Amato, Jeffrey Barrett, Sonia Goncalves, Ewan Harrison, David K. Jackson, Ian Johnston, Dominic Kwiatkowski, Cordelia Langford, John Sillitoe on behalf of the Wellcome Sanger Institute COVID-19 Surveillance Team |
| EPI_ISL_3051426 | Berkshire and Surrey Pathology Services Lighthouse Laboratory | Wellcome Sanger Institute for the COVID-19 Genomics UK (COG-UK) Consortium | Berkshire and Surrey Pathology Services Lighthouse Laboratory and Alex Alderton, Roberto Amato, Jeffrey Barrett, Sonia Goncalves, Ewan Harrison, David K. Jackson, Ian Johnston, Dominic Kwiatkowski, Cordelia Langford, John Sillitoe on behalf of the Wellcome Sanger Institute COVID-19 Surveillance Team |
| EPI_ISL_3051427 | Lighthouse Lab in Glasgow | Wellcome Sanger Institute for the COVID-19 Genomics UK (COG-UK) Consortium | Harper VanSteenhouse, Yumi Kasai, David Gray, Carol Clugston, Anna Dominiczak and Alex Alderton, Roberto Amato, Jeffrey Barrett, Sonia Goncalves, Ewan Harrison, David K. Jackson, Ian Johnston, Dominic Kwiatkowski, Cordelia Langford, John Sillitoe on behalf of the Wellcome Sanger Institute COVID-19 Surveillance Team |
| EPI_ISL_3051428 | Lighthouse Lab in Milton Keynes | Wellcome Sanger Institute for the COVID-19 Genomics UK (COG-UK) Consortium | The Lighthouse Lab in Milton Keynes and Alex Alderton, Roberto Amato, Jeffrey Barrett, Sonia Goncalves, Ewan Harrison, David K. Jackson, Ian Johnston, Dominic Kwiatkowski, Cordelia Langford, John Sillitoe on behalf of the Wellcome Sanger Institute COVID-19 Surveillance Team |
| EPI_ISL_3051429 | Lighthouse Lab in Glasgow | Wellcome Sanger Institute for the COVID-19 Genomics UK (COG-UK) Consortium | Harper VanSteenhouse, Yumi Kasai, David Gray, Carol Clugston, Anna Dominiczak and Alex Alderton, Roberto Amato, Jeffrey Barrett, Sonia Goncalves, Ewan Harrison, David K. Jackson, Ian Johnston, Dominic Kwiatkowski, Cordelia Langford, John Sillitoe on behalf of the Wellcome Sanger Institute COVID-19 Surveillance Team |
| EPI_ISL_3051430, EPI_ISL_3051431, EPI_ISL_3051432 | Lighthouse Lab in Milton Keynes | Wellcome Sanger Institute for the COVID-19 Genomics UK (COG-UK) Consortium | The Lighthouse Lab in Milton Keynes and Alex Alderton, Roberto Amato, Jeffrey Barrett, Sonia Goncalves, Ewan Harrison, David K. Jackson, Ian Johnston, Dominic Kwiatkowski, Cordelia Langford, John Sillitoe on behalf of the Wellcome Sanger Institute COVID-19 Surveillance Team |
| EPI_ISL_3051433, EPI_ISL_3051434 | Lighthouse Lab in Glasgow | Wellcome Sanger Institute for the COVID-19 Genomics UK (COG-UK) Consortium | Harper VanSteenhouse, Yumi Kasai, David Gray, Carol Clugston, Anna Dominiczak and Alex Alderton, Roberto Amato, Jeffrey Barrett, Sonia Goncalves, Ewan Harrison, David K. Jackson, Ian Johnston, Dominic Kwiatkowski, Cordelia Langford, John Sillitoe on behalf of the Wellcome Sanger Institute COVID-19 Surveillance Team |
| EPI_ISL_3051435, EPI_ISL_3051436, EPI_ISL_3051437 | Lighthouse Lab in Milton Keynes | Wellcome Sanger Institute for the COVID-19 Genomics UK (COG-UK) Consortium | The Lighthouse Lab in Milton Keynes and Alex Alderton, Roberto Amato, Jeffrey Barrett, Sonia Goncalves, Ewan Harrison, David K. Jackson, Ian Johnston, Dominic Kwiatkowski, Cordelia Langford, John Sillitoe on behalf of the Wellcome Sanger Institute COVID-19 Surveillance Team |
| EPI_ISL_3051438 | Lighthouse Lab in Glasgow | Wellcome Sanger Institute for the COVID-19 Genomics UK (COG-UK) Consortium | Harper VanSteenhouse, Yumi Kasai, David Gray, Carol Clugston, Anna Dominiczak and Alex Alderton, Roberto Amato, Jeffrey Barrett, Sonia Goncalves, Ewan Harrison, David K. Jackson, Ian Johnston, Dominic Kwiatkowski, Cordelia Langford, John Sillitoe on behalf of the Wellcome Sanger Institute COVID-19 Surveillance Team |
| EPI_ISL_3051439 | Lighthouse Lab in Alderley Park | Wellcome Sanger Institute for the COVID-19 Genomics UK (COG-UK) Consortium | Jacquelyn Wynn, Mairead Hyland, The Lighthouse Lab in Alderley Park and Alex Alderton, Roberto Amato, Jeffrey Barrett, Sonia Goncalves, Ewan Harrison, David K. Jackson, Ian Johnston, Dominic Kwiatkowski, Cordelia Langford, John Sillitoe on behalf of the Wellcome Sanger Institute COVID-19 Surveillance Team |
| EPI_ISL_3051440 | Lighthouse Lab in Glasgow | Wellcome Sanger Institute for the COVID-19 Genomics UK (COG-UK) Consortium | Harper VanSteenhouse, Yumi Kasai, David Gray, Carol Clugston, Anna Dominiczak and Alex Alderton, Roberto Amato, Jeffrey Barrett, Sonia Goncalves, Ewan Harrison, David K. Jackson, Ian Johnston, Dominic Kwiatkowski, Cordelia Langford, John Sillitoe on behalf of the Wellcome Sanger Institute COVID-19 Surveillance Team |
| EPI_ISL_3051441 | Lighthouse Lab in Alderley Park | Wellcome Sanger Institute for the COVID-19 Genomics UK (COG-UK) Consortium | Jacquelyn Wynn, Mairead Hyland, The Lighthouse Lab in Alderley Park and Alex Alderton, Roberto Amato, Jeffrey Barrett, Sonia Goncalves, Ewan Harrison, David K. Jackson, Ian Johnston, Dominic Kwiatkowski, Cordelia Langford, John Sillitoe on behalf of the Wellcome Sanger Institute COVID-19 Surveillance Team |
| EPI_ISL_3051442, EPI_ISL_3051443 | Berkshire and Surrey Pathology Services Lighthouse Laboratory | Wellcome Sanger Institute for the COVID-19 Genomics UK (COG-UK) Consortium | Berkshire and Surrey Pathology Services Lighthouse Laboratory and Alex Alderton, Roberto Amato, Jeffrey Barrett, Sonia Goncalves, Ewan Harrison, David K. Jackson, Ian Johnston, Dominic Kwiatkowski, Cordelia Langford, John Sillitoe on behalf of the Wellcome Sanger Institute COVID-19 Surveillance Team |
| EPI_ISL_3051444 | Lighthouse Lab in Glasgow | Wellcome Sanger Institute for the COVID-19 Genomics UK (COG-UK) Consortium | Harper VanSteenhouse, Yumi Kasai, David Gray, Carol Clugston, Anna Dominiczak and Alex Alderton, Roberto Amato, Jeffrey Barrett, Sonia Goncalves, Ewan Harrison, David K. Jackson, Ian Johnston, Dominic Kwiatkowski, Cordelia Langford, John Sillitoe on behalf of the Wellcome Sanger Institute COVID-19 Surveillance Team |
| EPI_ISL_3051445 | Lighthouse Lab in Alderley Park | Wellcome Sanger Institute for the COVID-19 Genomics UK (COG-UK) Consortium | Jacquelyn Wynn, Mairead Hyland, The Lighthouse Lab in Alderley Park and Alex Alderton, Roberto Amato, Jeffrey Barrett, Sonia Goncalves, Ewan Harrison, David K. Jackson, Ian Johnston, Dominic Kwiatkowski, Cordelia Langford, John Sillitoe on behalf of the Wellcome Sanger Institute COVID-19 Surveillance Team |
| EPI_ISL_3051446 | Berkshire and Surrey Pathology Services Lighthouse Laboratory | Wellcome Sanger Institute for the COVID-19 Genomics UK (COG-UK) Consortium | Berkshire and Surrey Pathology Services Lighthouse Laboratory and Alex Alderton, Roberto Amato, Jeffrey Barrett, Sonia Goncalves, Ewan Harrison, David K. Jackson, Ian Johnston, Dominic Kwiatkowski, Cordelia Langford, John Sillitoe on behalf of the Wellcome Sanger Institute COVID-19 Surveillance Team |
| EPI_ISL_3051447 | Lighthouse Lab in Glasgow | Wellcome Sanger Institute for the COVID-19 Genomics UK (COG-UK) Consortium | Harper VanSteenhouse, Yumi Kasai, David Gray, Carol Clugston, Anna Dominiczak and Alex Alderton, Roberto Amato, Jeffrey Barrett, Sonia Goncalves, Ewan Harrison, David K. Jackson, Ian Johnston, Dominic Kwiatkowski, Cordelia Langford, John Sillitoe on behalf of the Wellcome Sanger Institute COVID-19 Surveillance Team |
| EPI_ISL_3051448 | Lighthouse Lab in Milton Keynes | Wellcome Sanger Institute for the COVID-19 Genomics UK (COG-UK) Consortium | The Lighthouse Lab in Milton Keynes and Alex Alderton, Roberto Amato, Jeffrey Barrett, Sonia Goncalves, Ewan Harrison, David K. Jackson, Ian Johnston, Dominic Kwiatkowski, Cordelia Langford, John Sillitoe on behalf of the Wellcome Sanger Institute COVID-19 Surveillance Team |
| EPI_ISL_3051449 | Lighthouse Lab in Alderley Park | Wellcome Sanger Institute for the COVID-19 Genomics UK (COG-UK) Consortium | Jacquelyn Wynn, Mairead Hyland, The Lighthouse Lab in Alderley Park and Alex Alderton, Roberto Amato, Jeffrey Barrett, Sonia Goncalves, Ewan Harrison, David K. Jackson, Ian Johnston, Dominic Kwiatkowski, Cordelia Langford, John Sillitoe on behalf of the Wellcome Sanger Institute COVID-19 Surveillance Team |
| EPI_ISL_3051450 | Lighthouse Lab in Milton Keynes | Wellcome Sanger Institute for the COVID-19 Genomics UK (COG-UK) Consortium | The Lighthouse Lab in Milton Keynes and Alex Alderton, Roberto Amato, Jeffrey Barrett, Sonia Goncalves, Ewan Harrison, David K. Jackson, Ian Johnston, Dominic Kwiatkowski, Cordelia Langford, John Sillitoe on behalf of the Wellcome Sanger Institute COVID-19 Surveillance Team |
| EPI_ISL_3051451 | Lighthouse Lab in Glasgow | Wellcome Sanger Institute for the COVID-19 Genomics UK (COG-UK) Consortium | Harper VanSteenhouse, Yumi Kasai, David Gray, Carol Clugston, Anna Dominiczak and Alex Alderton, Roberto Amato, Jeffrey Barrett, Sonia Goncalves, Ewan Harrison, David K. Jackson, Ian Johnston, Dominic Kwiatkowski, Cordelia Langford, John Sillitoe on behalf of the Wellcome Sanger Institute COVID-19 Surveillance Team |
| EPI_ISL_3051452 | Lighthouse Lab in Milton Keynes | Wellcome Sanger Institute for the COVID-19 Genomics UK (COG-UK) Consortium | The Lighthouse Lab in Milton Keynes and Alex Alderton, Roberto Amato, Jeffrey Barrett, Sonia Goncalves, Ewan Harrison, David K. Jackson, Ian Johnston, Dominic Kwiatkowski, Cordelia Langford, John Sillitoe on behalf of the Wellcome Sanger Institute COVID-19 Surveillance Team |
| EPI_ISL_3051453 | Lighthouse Lab in Glasgow | Wellcome Sanger Institute for the COVID-19 Genomics UK (COG-UK) Consortium | Harper VanSteenhouse, Yumi Kasai, David Gray, Carol Clugston, Anna Dominiczak and Alex Alderton, Roberto Amato, Jeffrey Barrett, Sonia Goncalves, Ewan Harrison, David K. Jackson, Ian Johnston, Dominic Kwiatkowski, Cordelia Langford, John Sillitoe on behalf of the Wellcome Sanger Institute COVID-19 Surveillance Team |
| EPI_ISL_3051454 | Lighthouse Lab in Alderley Park | Wellcome Sanger Institute for the COVID-19 Genomics UK (COG-UK) Consortium | Jacquelyn Wynn, Mairead Hyland, The Lighthouse Lab in Alderley Park and Alex Alderton, Roberto Amato, Jeffrey Barrett, Sonia Goncalves, Ewan Harrison, David K. Jackson, Ian Johnston, Dominic Kwiatkowski, Cordelia Langford, John Sillitoe on behalf of the Wellcome Sanger Institute COVID-19 Surveillance Team |
| EPI_ISL_3051455 | Lighthouse Lab in Milton Keynes | Wellcome Sanger Institute for the COVID-19 Genomics UK (COG-UK) Consortium | The Lighthouse Lab in Milton Keynes and Alex Alderton, Roberto Amato, Jeffrey Barrett, Sonia Goncalves, Ewan Harrison, David K. Jackson, Ian Johnston, Dominic Kwiatkowski, Cordelia Langford, John Sillitoe on behalf of the Wellcome Sanger Institute COVID-19 Surveillance Team |
| EPI_ISL_3051456, EPI_ISL_3051457 | Lighthouse Lab in Alderley Park | Wellcome Sanger Institute for the COVID-19 Genomics UK (COG-UK) Consortium | Jacquelyn Wynn, Mairead Hyland, The Lighthouse Lab in Alderley Park and Alex Alderton, Roberto Amato, Jeffrey Barrett, Sonia Goncalves, Ewan Harrison, David K. Jackson, Ian Johnston, Dominic Kwiatkowski, Cordelia Langford, John Sillitoe on behalf of the Wellcome Sanger Institute COVID-19 Surveillance Team |
| EPI_ISL_3051458, EPI_ISL_3051459 | Berkshire and Surrey Pathology Services Lighthouse Laboratory | Wellcome Sanger Institute for the COVID-19 Genomics UK (COG-UK) Consortium | Berkshire and Surrey Pathology Services Lighthouse Laboratory and Alex Alderton, Roberto Amato, Jeffrey Barrett, Sonia Goncalves, Ewan Harrison, David K. Jackson, Ian Johnston, Dominic Kwiatkowski, Cordelia Langford, John Sillitoe on behalf of the Wellcome Sanger Institute COVID-19 Surveillance Team |
| EPI_ISL_3051460 | Lighthouse Lab in Milton Keynes | Wellcome Sanger Institute for the COVID-19 Genomics UK (COG-UK) Consortium | The Lighthouse Lab in Milton Keynes and Alex Alderton, Roberto Amato, Jeffrey Barrett, Sonia Goncalves, Ewan Harrison, David K. Jackson, Ian Johnston, Dominic Kwiatkowski, Cordelia Langford, John Sillitoe on behalf of the Wellcome Sanger Institute COVID-19 Surveillance Team |
| EPI_ISL_3051461, EPI_ISL_3051462 | Lighthouse Lab in Alderley Park | Wellcome Sanger Institute for the COVID-19 Genomics UK (COG-UK) Consortium | Jacquelyn Wynn, Mairead Hyland, The Lighthouse Lab in Alderley Park and Alex Alderton, Roberto Amato, Jeffrey Barrett, Sonia Goncalves, Ewan Harrison, David K. Jackson, Ian Johnston, Dominic Kwiatkowski, Cordelia Langford, John Sillitoe on behalf of the Wellcome Sanger Institute COVID-19 Surveillance Team |
| EPI_ISL_3051463 | Lighthouse Lab in Glasgow | Wellcome Sanger Institute for the COVID-19 Genomics UK (COG-UK) Consortium | Harper VanSteenhouse, Yumi Kasai, David Gray, Carol Clugston, Anna Dominiczak and Alex Alderton, Roberto Amato, Jeffrey Barrett, Sonia Goncalves, Ewan Harrison, David K. Jackson, Ian Johnston, Dominic Kwiatkowski, Cordelia Langford, John Sillitoe on behalf of the Wellcome Sanger Institute COVID-19 Surveillance Team |
| EPI_ISL_3051464 | Lighthouse Lab in Alderley Park | Wellcome Sanger Institute for the COVID-19 Genomics UK (COG-UK) Consortium | Jacquelyn Wynn, Mairead Hyland, The Lighthouse Lab in Alderley Park and Alex Alderton, Roberto Amato, Jeffrey Barrett, Sonia Goncalves, Ewan Harrison, David K. Jackson, Ian Johnston, Dominic Kwiatkowski, Cordelia Langford, John Sillitoe on behalf of the Wellcome Sanger Institute COVID-19 Surveillance Team |
| EPI_ISL_3051465 | Berkshire and Surrey Pathology Services Lighthouse Laboratory | Wellcome Sanger Institute for the COVID-19 Genomics UK (COG-UK) Consortium | Berkshire and Surrey Pathology Services Lighthouse Laboratory and Alex Alderton, Roberto Amato, Jeffrey Barrett, Sonia Goncalves, Ewan Harrison, David K. Jackson, Ian Johnston, Dominic Kwiatkowski, Cordelia Langford, John Sillitoe on behalf of the Wellcome Sanger Institute COVID-19 Surveillance Team |
| EPI_ISL_3051466, EPI_ISL_3051467 | Lighthouse Lab in Alderley Park | Wellcome Sanger Institute for the COVID-19 Genomics UK (COG-UK) Consortium | Jacquelyn Wynn, Mairead Hyland, The Lighthouse Lab in Alderley Park and Alex Alderton, Roberto Amato, Jeffrey Barrett, Sonia Goncalves, Ewan Harrison, David K. Jackson, Ian Johnston, Dominic Kwiatkowski, Cordelia Langford, John Sillitoe on behalf of the Wellcome Sanger Institute COVID-19 Surveillance Team |
| EPI_ISL_3051468 | Berkshire and Surrey Pathology Services Lighthouse Laboratory | Wellcome Sanger Institute for the COVID-19 Genomics UK (COG-UK) Consortium | Berkshire and Surrey Pathology Services Lighthouse Laboratory and Alex Alderton, Roberto Amato, Jeffrey Barrett, Sonia Goncalves, Ewan Harrison, David K. Jackson, Ian Johnston, Dominic Kwiatkowski, Cordelia Langford, John Sillitoe on behalf of the Wellcome Sanger Institute COVID-19 Surveillance Team |
| EPI_ISL_3051469 | Lighthouse Lab in Alderley Park | Wellcome Sanger Institute for the COVID-19 Genomics UK (COG-UK) Consortium | Jacquelyn Wynn, Mairead Hyland, The Lighthouse Lab in Alderley Park and Alex Alderton, Roberto Amato, Jeffrey Barrett, Sonia Goncalves, Ewan Harrison, David K. Jackson, Ian Johnston, Dominic Kwiatkowski, Cordelia Langford, John Sillitoe on behalf of the Wellcome Sanger Institute COVID-19 Surveillance Team |
| EPI_ISL_3051470 | Lighthouse Lab in Glasgow | Wellcome Sanger Institute for the COVID-19 Genomics UK (COG-UK) Consortium | Harper VanSteenhouse, Yumi Kasai, David Gray, Carol Clugston, Anna Dominiczak and Alex Alderton, Roberto Amato, Jeffrey Barrett, Sonia Goncalves, Ewan Harrison, David K. Jackson, Ian Johnston, Dominic Kwiatkowski, Cordelia Langford, John Sillitoe on behalf of the Wellcome Sanger Institute COVID-19 Surveillance Team |

| | | | |
|---|---|---|---|
| EPI_ISL_3051471, EPI_ISL_3051472, EPI_ISL_3051473, EPI_ISL_3051474, EPI_ISL_3051475 | Lighthouse Lab in Alderley Park | Wellcome Sanger Institute for the COVID-19 Genomics UK (COG-UK) Consortium | Jacquelyn Wynn, Mairead Hyland, The Lighthouse Lab in Alderley Park and Alex Alderton, Roberto Amato, Jeffrey Barrett, Sonia Goncalves, Ewan Harrison, David K. Jackson, Ian Johnston, Dominic Kwiatkowski, Cordelia Langford, John Sillitoe on behalf of the Wellcome Sanger Institute COVID-19 Surveillance Team |
| EPI_ISL_3051476 | Lighthouse Lab in Glasgow | Wellcome Sanger Institute for the COVID-19 Genomics UK (COG-UK) Consortium | Harper VanSteenhouse, Yumi Kasai, David Gray, Carol Clugston, Anna Dominiczak and Alex Alderton, Roberto Amato, Jeffrey Barrett, Sonia Goncalves, Ewan Harrison, David K. Jackson, Ian Johnston, Dominic Kwiatkowski, Cordelia Langford, John Sillitoe on behalf of the Wellcome Sanger Institute COVID-19 Surveillance Team |
| EPI_ISL_3051477, EPI_ISL_3051478 | Lighthouse Lab in Alderley Park | Wellcome Sanger Institute for the COVID-19 Genomics UK (COG-UK) Consortium | Jacquelyn Wynn, Mairead Hyland, The Lighthouse Lab in Alderley Park and Alex Alderton, Roberto Amato, Jeffrey Barrett, Sonia Goncalves, Ewan Harrison, David K. Jackson, Ian Johnston, Dominic Kwiatkowski, Cordelia Langford, John Sillitoe on behalf of the Wellcome Sanger Institute COVID-19 Surveillance Team |
| EPI_ISL_3051480 | Lighthouse Lab in Glasgow | Wellcome Sanger Institute for the COVID-19 Genomics UK (COG-UK) Consortium | Harper VanSteenhouse, Yumi Kasai, David Gray, Carol Clugston, Anna Dominiczak and Alex Alderton, Roberto Amato, Jeffrey Barrett, Sonia Goncalves, Ewan Harrison, David K. Jackson, Ian Johnston, Dominic Kwiatkowski, Cordelia Langford, John Sillitoe on behalf of the Wellcome Sanger Institute COVID-19 Surveillance Team |
| EPI_ISL_3051481, EPI_ISL_3051482 | Lighthouse Lab in Alderley Park | Wellcome Sanger Institute for the COVID-19 Genomics UK (COG-UK) Consortium | Jacquelyn Wynn, Mairead Hyland, The Lighthouse Lab in Alderley Park and Alex Alderton, Roberto Amato, Jeffrey Barrett, Sonia Goncalves, Ewan Harrison, David K. Jackson, Ian Johnston, Dominic Kwiatkowski, Cordelia Langford, John Sillitoe on behalf of the Wellcome Sanger Institute COVID-19 Surveillance Team |
| EPI_ISL_3051483, EPI_ISL_3051484 | Lighthouse Lab in Glasgow | Wellcome Sanger Institute for the COVID-19 Genomics UK (COG-UK) Consortium | Harper VanSteenhouse, Yumi Kasai, David Gray, Carol Clugston, Anna Dominiczak and Alex Alderton, Roberto Amato, Jeffrey Barrett, Sonia Goncalves, Ewan Harrison, David K. Jackson, Ian Johnston, Dominic Kwiatkowski, Cordelia Langford, John Sillitoe on behalf of the Wellcome Sanger Institute COVID-19 Surveillance Team |
| EPI_ISL_3051485 | Berkshire and Surrey Pathology Services Lighthouse Laboratory | Wellcome Sanger Institute for the COVID-19 Genomics UK (COG-UK) Consortium | Berkshire and Surrey Pathology Services Lighthouse Laboratory and Alex Alderton, Roberto Amato, Jeffrey Barrett, Sonia Goncalves, Ewan Harrison, David K. Jackson, Ian Johnston, Dominic Kwiatkowski, Cordelia Langford, John Sillitoe on behalf of the Wellcome Sanger Institute COVID-19 Surveillance Team |
| EPI_ISL_3051486 | Lighthouse Lab in Milton Keynes | Wellcome Sanger Institute for the COVID-19 Genomics UK (COG-UK) Consortium | The Lighthouse Lab in Milton Keynes and Alex Alderton, Roberto Amato, Jeffrey Barrett, Sonia Goncalves, Ewan Harrison, David K. Jackson, Ian Johnston, Dominic Kwiatkowski, Cordelia Langford, John Sillitoe on behalf of the Wellcome Sanger Institute COVID-19 Surveillance Team |
| EPI_ISL_3051487 | Berkshire and Surrey Pathology Services Lighthouse Laboratory | Wellcome Sanger Institute for the COVID-19 Genomics UK (COG-UK) Consortium | Berkshire and Surrey Pathology Services Lighthouse Laboratory and Alex Alderton, Roberto Amato, Jeffrey Barrett, Sonia Goncalves, Ewan Harrison, David K. Jackson, Ian Johnston, Dominic Kwiatkowski, Cordelia Langford, John Sillitoe on behalf of the Wellcome Sanger Institute COVID-19 Surveillance Team |
| EPI_ISL_3051488 | Lighthouse Lab in Milton Keynes | Wellcome Sanger Institute for the COVID-19 Genomics UK (COG-UK) Consortium | The Lighthouse Lab in Milton Keynes and Alex Alderton, Roberto Amato, Jeffrey Barrett, Sonia Goncalves, Ewan Harrison, David K. Jackson, Ian Johnston, Dominic Kwiatkowski, Cordelia Langford, John Sillitoe on behalf of the Wellcome Sanger Institute COVID-19 Surveillance Team |
| EPI_ISL_3051489 | Lighthouse Lab in Glasgow | Wellcome Sanger Institute for the COVID-19 Genomics UK (COG-UK) Consortium | Harper VanSteenhouse, Yumi Kasai, David Gray, Carol Clugston, Anna Dominiczak and Alex Alderton, Roberto Amato, Jeffrey Barrett, Sonia Goncalves, Ewan Harrison, David K. Jackson, Ian Johnston, Dominic Kwiatkowski, Cordelia Langford, John Sillitoe on behalf of the Wellcome Sanger Institute COVID-19 Surveillance Team |
| EPI_ISL_3051490 | Lighthouse Lab in Alderley Park | Wellcome Sanger Institute for the COVID-19 Genomics UK (COG-UK) Consortium | Jacquelyn Wynn, Mairead Hyland, The Lighthouse Lab in Alderley Park and Alex Alderton, Roberto Amato, Jeffrey Barrett, Sonia Goncalves, Ewan Harrison, David K. Jackson, Ian Johnston, Dominic Kwiatkowski, Cordelia Langford, John Sillitoe on behalf of the Wellcome Sanger Institute COVID-19 Surveillance Team |
| EPI_ISL_3051491 | Berkshire and Surrey Pathology Services Lighthouse Laboratory | Wellcome Sanger Institute for the COVID-19 Genomics UK (COG-UK) Consortium | Berkshire and Surrey Pathology Services Lighthouse Laboratory and Alex Alderton, Roberto Amato, Jeffrey Barrett, Sonia Goncalves, Ewan Harrison, David K. Jackson, Ian Johnston, Dominic Kwiatkowski, Cordelia Langford, John Sillitoe on behalf of the Wellcome Sanger Institute COVID-19 Surveillance Team |
| EPI_ISL_3051492, EPI_ISL_3051493, EPI_ISL_3051494 | Lighthouse Lab in Glasgow | Wellcome Sanger Institute for the COVID-19 Genomics UK (COG-UK) Consortium | Harper VanSteenhouse, Yumi Kasai, David Gray, Carol Clugston, Anna Dominiczak and Alex Alderton, Roberto Amato, Jeffrey Barrett, Sonia Goncalves, Ewan Harrison, David K. Jackson, Ian Johnston, Dominic Kwiatkowski, Cordelia Langford, John Sillitoe on behalf of the Wellcome Sanger Institute COVID-19 Surveillance Team |
| EPI_ISL_3051495 | Berkshire and Surrey Pathology Services Lighthouse Laboratory | Wellcome Sanger Institute for the COVID-19 Genomics UK (COG-UK) Consortium | Berkshire and Surrey Pathology Services Lighthouse Laboratory and Alex Alderton, Roberto Amato, Jeffrey Barrett, Sonia Goncalves, Ewan Harrison, David K. Jackson, Ian Johnston, Dominic Kwiatkowski, Cordelia Langford, John Sillitoe on behalf of the Wellcome Sanger Institute COVID-19 Surveillance Team |
| EPI_ISL_3051496 | Lighthouse Lab in Glasgow | Wellcome Sanger Institute for the COVID-19 Genomics UK (COG-UK) Consortium | Harper VanSteenhouse, Yumi Kasai, David Gray, Carol Clugston, Anna Dominiczak and Alex Alderton, Roberto Amato, Jeffrey Barrett, Sonia Goncalves, Ewan Harrison, David K. Jackson, Ian Johnston, Dominic Kwiatkowski, Cordelia Langford, John Sillitoe on behalf of the Wellcome Sanger Institute COVID-19 Surveillance Team |
| EPI_ISL_3051497 | Lighthouse Lab in Milton Keynes | Wellcome Sanger Institute for the COVID-19 Genomics UK (COG-UK) Consortium | The Lighthouse Lab in Milton Keynes and Alex Alderton, Roberto Amato, Jeffrey Barrett, Sonia Goncalves, Ewan Harrison, David K. Jackson, Ian Johnston, Dominic Kwiatkowski, Cordelia Langford, John Sillitoe on behalf of the Wellcome Sanger Institute COVID-19 Surveillance Team |
| EPI_ISL_3051498 | Lighthouse Lab in Alderley Park | Wellcome Sanger Institute for the COVID-19 Genomics UK (COG-UK) Consortium | Jacquelyn Wynn, Mairead Hyland, The Lighthouse Lab in Alderley Park and Alex Alderton, Roberto Amato, Jeffrey Barrett, Sonia Goncalves, Ewan Harrison, David K. Jackson, Ian Johnston, Dominic Kwiatkowski, Cordelia Langford, John Sillitoe on behalf of the Wellcome Sanger Institute COVID-19 Surveillance Team |
| EPI_ISL_3051499 | Lighthouse Lab in Glasgow | Wellcome Sanger Institute for the COVID-19 Genomics UK (COG-UK) Consortium | Harper VanSteenhouse, Yumi Kasai, David Gray, Carol Clugston, Anna Dominiczak and Alex Alderton, Roberto Amato, Jeffrey Barrett, Sonia Goncalves, Ewan Harrison, David K. Jackson, Ian Johnston, Dominic Kwiatkowski, Cordelia Langford, John Sillitoe on behalf of the Wellcome Sanger Institute COVID-19 Surveillance Team |
| EPI_ISL_3051500 | Berkshire and Surrey Pathology Services Lighthouse Laboratory | Wellcome Sanger Institute for the COVID-19 Genomics UK (COG-UK) Consortium | Berkshire and Surrey Pathology Services Lighthouse Laboratory and Alex Alderton, Roberto Amato, Jeffrey Barrett, Sonia Goncalves, Ewan Harrison, David K. Jackson, Ian Johnston, Dominic Kwiatkowski, Cordelia Langford, John Sillitoe on behalf of the Wellcome Sanger Institute COVID-19 Surveillance Team |
| EPI_ISL_3051501 | Lighthouse Lab in Glasgow | Wellcome Sanger Institute for the COVID-19 Genomics UK (COG-UK) Consortium | Harper VanSteenhouse, Yumi Kasai, David Gray, Carol Clugston, Anna Dominiczak and Alex Alderton, Roberto Amato, Jeffrey Barrett, Sonia Goncalves, Ewan Harrison, David K. Jackson, Ian Johnston, Dominic Kwiatkowski, Cordelia Langford, John Sillitoe on behalf of the Wellcome Sanger Institute COVID-19 Surveillance Team |
| EPI_ISL_3051502 | Lighthouse Lab in Milton Keynes | Wellcome Sanger Institute for the COVID-19 Genomics UK (COG-UK) Consortium | The Lighthouse Lab in Milton Keynes and Alex Alderton, Roberto Amato, Jeffrey Barrett, Sonia Goncalves, Ewan Harrison, David K. Jackson, Ian Johnston, Dominic Kwiatkowski, Cordelia Langford, John Sillitoe on behalf of the Wellcome Sanger Institute COVID-19 Surveillance Team |
| EPI_ISL_3051503 | Lighthouse Lab in Glasgow | Wellcome Sanger Institute for the COVID-19 Genomics UK (COG-UK) Consortium | Harper VanSteenhouse, Yumi Kasai, David Gray, Carol Clugston, Anna Dominiczak and Alex Alderton, Roberto Amato, Jeffrey Barrett, Sonia Goncalves, Ewan Harrison, David K. Jackson, Ian Johnston, Dominic Kwiatkowski, Cordelia Langford, John Sillitoe on behalf of the Wellcome Sanger Institute COVID-19 Surveillance Team |
| EPI_ISL_3051504, EPI_ISL_3051505, EPI_ISL_3051506 | Lighthouse Lab in Milton Keynes | Wellcome Sanger Institute for the COVID-19 Genomics UK (COG-UK) Consortium | The Lighthouse Lab in Milton Keynes and Alex Alderton, Roberto Amato, Jeffrey Barrett, Sonia Goncalves, Ewan Harrison, David K. Jackson, Ian Johnston, Dominic Kwiatkowski, Cordelia Langford, John Sillitoe on behalf of the Wellcome Sanger Institute COVID-19 Surveillance Team |
| EPI_ISL_3051507 | Lighthouse Lab in Glasgow | Wellcome Sanger Institute for the COVID-19 Genomics UK (COG-UK) Consortium | Harper VanSteenhouse, Yumi Kasai, David Gray, Carol Clugston, Anna Dominiczak and Alex Alderton, Roberto Amato, Jeffrey Barrett, Sonia Goncalves, Ewan Harrison, David K. Jackson, Ian Johnston, Dominic Kwiatkowski, Cordelia Langford, John Sillitoe on behalf of the Wellcome Sanger Institute COVID-19 Surveillance Team |
| EPI_ISL_3051508 | Lighthouse Lab in Milton Keynes | Wellcome Sanger Institute for the COVID-19 Genomics UK (COG-UK) Consortium | The Lighthouse Lab in Milton Keynes and Alex Alderton, Roberto Amato, Jeffrey Barrett, Sonia Goncalves, Ewan Harrison, David K. Jackson, Ian Johnston, Dominic Kwiatkowski, Cordelia Langford, John Sillitoe on behalf of the Wellcome Sanger Institute COVID-19 Surveillance Team |
| EPI_ISL_3051509 | Lighthouse Lab in Alderley Park | Wellcome Sanger Institute for the COVID-19 Genomics UK (COG-UK) Consortium | Jacquelyn Wynn, Mairead Hyland, The Lighthouse Lab in Alderley Park and Alex Alderton, Roberto Amato, Jeffrey Barrett, Sonia Goncalves, Ewan Harrison, David K. Jackson, Ian Johnston, Dominic Kwiatkowski, Cordelia Langford, John Sillitoe on behalf of the Wellcome Sanger Institute COVID-19 Surveillance Team |
| EPI_ISL_3051510, EPI_ISL_3051511 | Lighthouse Lab in Glasgow | Wellcome Sanger Institute for the COVID-19 Genomics UK (COG-UK) Consortium | Harper VanSteenhouse, Yumi Kasai, David Gray, Carol Clugston, Anna Dominiczak and Alex Alderton, Roberto Amato, Jeffrey Barrett, Sonia Goncalves, Ewan Harrison, David K. Jackson, Ian Johnston, Dominic Kwiatkowski, Cordelia Langford, John Sillitoe on behalf of the Wellcome Sanger Institute COVID-19 Surveillance Team |
| EPI_ISL_3051512 | Lighthouse Lab in Alderley Park | Wellcome Sanger Institute for the COVID-19 Genomics UK (COG-UK) Consortium | Jacquelyn Wynn, Mairead Hyland, The Lighthouse Lab in Alderley Park and Alex Alderton, Roberto Amato, Jeffrey Barrett, Sonia Goncalves, Ewan Harrison, David K. Jackson, Ian Johnston, Dominic Kwiatkowski, Cordelia Langford, John Sillitoe on behalf of the Wellcome Sanger Institute COVID-19 Surveillance Team |
| EPI_ISL_3051513, EPI_ISL_3051514 | Berkshire and Surrey Pathology Services Lighthouse Laboratory | Wellcome Sanger Institute for the COVID-19 Genomics UK (COG-UK) Consortium | Berkshire and Surrey Pathology Services Lighthouse Laboratory and Alex Alderton, Roberto Amato, Jeffrey Barrett, Sonia Goncalves, Ewan Harrison, David K. Jackson, Ian Johnston, Dominic Kwiatkowski, Cordelia Langford, John Sillitoe on behalf of the Wellcome Sanger Institute COVID-19 Surveillance Team |
| EPI_ISL_3051515 | Lighthouse Lab in Glasgow | Wellcome Sanger Institute for the COVID-19 Genomics UK (COG-UK) Consortium | Harper VanSteenhouse, Yumi Kasai, David Gray, Carol Clugston, Anna Dominiczak and Alex Alderton, Roberto Amato, Jeffrey Barrett, Sonia Goncalves, Ewan Harrison, David K. Jackson, Ian Johnston, Dominic Kwiatkowski, Cordelia Langford, John Sillitoe on behalf of the Wellcome Sanger Institute COVID-19 Surveillance Team |
| EPI_ISL_3051516 | Lighthouse Lab in Milton Keynes | Wellcome Sanger Institute for the COVID-19 Genomics UK (COG-UK) Consortium | The Lighthouse Lab in Milton Keynes and Alex Alderton, Roberto Amato, Jeffrey Barrett, Sonia Goncalves, Ewan Harrison, David K. Jackson, Ian Johnston, Dominic Kwiatkowski, Cordelia Langford, John Sillitoe on behalf of the Wellcome Sanger Institute COVID-19 Surveillance Team |
| EPI_ISL_3051517 | Berkshire and Surrey Pathology Services Lighthouse Laboratory | Wellcome Sanger Institute for the COVID-19 Genomics UK (COG-UK) Consortium | Berkshire and Surrey Pathology Services Lighthouse Laboratory and Alex Alderton, Roberto Amato, Jeffrey Barrett, Sonia Goncalves, Ewan Harrison, David K. Jackson, Ian Johnston, Dominic Kwiatkowski, Cordelia Langford, John Sillitoe on behalf of the Wellcome Sanger Institute COVID-19 Surveillance Team |
| EPI_ISL_3051518 | Lighthouse Lab in Milton Keynes | Wellcome Sanger Institute for the COVID-19 Genomics UK (COG-UK) Consortium | The Lighthouse Lab in Milton Keynes and Alex Alderton, Roberto Amato, Jeffrey Barrett, Sonia Goncalves, Ewan Harrison, David K. Jackson, Ian Johnston, Dominic Kwiatkowski, Cordelia Langford, John Sillitoe on behalf of the Wellcome Sanger Institute COVID-19 Surveillance Team |
| EPI_ISL_3051519, EPI_ISL_3051520, EPI_ISL_3051521 | Lighthouse Lab in Alderley Park | Wellcome Sanger Institute for the COVID-19 Genomics UK (COG-UK) Consortium | Jacquelyn Wynn, Mairead Hyland, The Lighthouse Lab in Alderley Park and Alex Alderton, Roberto Amato, Jeffrey Barrett, Sonia Goncalves, Ewan Harrison, David K. Jackson, Ian Johnston, Dominic Kwiatkowski, Cordelia Langford, John Sillitoe on behalf of the Wellcome Sanger Institute COVID-19 Surveillance Team |
| EPI_ISL_3051522, EPI_ISL_3051523 | Lighthouse Lab in Milton Keynes | Wellcome Sanger Institute for the COVID-19 Genomics UK (COG-UK) Consortium | The Lighthouse Lab in Milton Keynes and Alex Alderton, Roberto Amato, Jeffrey Barrett, Sonia Goncalves, Ewan Harrison, David K. Jackson, Ian Johnston, Dominic Kwiatkowski, Cordelia Langford, John Sillitoe on behalf of the Wellcome Sanger Institute COVID-19 Surveillance Team |
| EPI_ISL_3051524 | Lighthouse Lab in Alderley Park | Wellcome Sanger Institute for the COVID-19 Genomics UK (COG-UK) Consortium | Jacquelyn Wynn, Mairead Hyland, The Lighthouse Lab in Alderley Park and Alex Alderton, Roberto Amato, Jeffrey Barrett, Sonia Goncalves, Ewan Harrison, David K. Jackson, Ian Johnston, Dominic Kwiatkowski, Cordelia Langford, John Sillitoe on behalf of the Wellcome Sanger Institute COVID-19 Surveillance Team |
| EPI_ISL_3051525 | Lighthouse Lab in Glasgow | Wellcome Sanger Institute for the COVID-19 Genomics UK (COG-UK) Consortium | Harper VanSteenhouse, Yumi Kasai, David Gray, Carol Clugston, Anna Dominiczak and Alex Alderton, Roberto Amato, Jeffrey Barrett, Sonia Goncalves, Ewan Harrison, David K. Jackson, Ian Johnston, Dominic Kwiatkowski, Cordelia Langford, John Sillitoe on behalf of the Wellcome Sanger Institute COVID-19 Surveillance Team |
| EPI_ISL_3051526 | Berkshire and Surrey Pathology Services Lighthouse Laboratory | Wellcome Sanger Institute for the COVID-19 Genomics UK (COG-UK) Consortium | Berkshire and Surrey Pathology Services Lighthouse Laboratory and Alex Alderton, Roberto Amato, Jeffrey Barrett, Sonia Goncalves, Ewan Harrison, David K. Jackson, Ian Johnston, Dominic Kwiatkowski, Cordelia Langford, John Sillitoe on behalf of the Wellcome Sanger Institute COVID-19 Surveillance Team |
| EPI_ISL_3051527 | Lighthouse Lab in Milton Keynes | Wellcome Sanger Institute for the COVID-19 Genomics UK (COG-UK) Consortium | The Lighthouse Lab in Milton Keynes and Alex Alderton, Roberto Amato, Jeffrey Barrett, Sonia Goncalves, Ewan Harrison, David K. Jackson, Ian Johnston, Dominic Kwiatkowski, Cordelia Langford, John Sillitoe on behalf of the Wellcome Sanger Institute COVID-19 Surveillance Team |
| EPI_ISL_3051528 | Lighthouse Lab in Glasgow | Wellcome Sanger Institute for the COVID-19 Genomics UK (COG-UK) Consortium | Harper VanSteenhouse, Yumi Kasai, David Gray, Carol Clugston, Anna Dominiczak and Alex Alderton, Roberto Amato, Jeffrey Barrett, Sonia Goncalves, Ewan Harrison, David K. Jackson, Ian Johnston, Dominic Kwiatkowski, Cordelia Langford, John Sillitoe on behalf of the Wellcome Sanger Institute COVID-19 Surveillance Team |
| EPI_ISL_3051529 | Lighthouse Lab in Milton Keynes | Wellcome Sanger Institute for the COVID-19 Genomics UK (COG-UK) Consortium | The Lighthouse Lab in Milton Keynes and Alex Alderton, Roberto Amato, Jeffrey Barrett, Sonia Goncalves, Ewan Harrison, David K. Jackson, Ian Johnston, Dominic Kwiatkowski, Cordelia Langford, John Sillitoe on behalf of the Wellcome Sanger Institute COVID-19 Surveillance Team |
| EPI_ISL_3051530 | Berkshire and Surrey Pathology Services Lighthouse Laboratory | Wellcome Sanger Institute for the COVID-19 Genomics UK (COG-UK) Consortium | Berkshire and Surrey Pathology Services Lighthouse Laboratory and Alex Alderton, Roberto Amato, Jeffrey Barrett, Sonia Goncalves, Ewan Harrison, David K. Jackson, Ian Johnston, Dominic Kwiatkowski, Cordelia Langford, John Sillitoe on behalf of the Wellcome Sanger Institute COVID-19 Surveillance Team |

| | | | |
|---|---|---|---|
| EPI_ISL_3051531 | Lighthouse Lab in Glasgow | Wellcome Sanger Institute for the COVID-19 Genomics UK (COG-UK) Consortium | Harper VanSteenhouse, Yumi Kasai, David Gray, Carol Clugston, Anna Dominiczak and Alex Alderton, Roberto Amato, Jeffrey Barrett, Sonia Goncalves, Ewan Harrison, David K. Jackson, Ian Johnston, Dominic Kwiatkowski, Cordelia Langford, John Sillitoe on behalf of the Wellcome Sanger Institute COVID-19 Surveillance Team |
| EPI_ISL_3051532, EPI_ISL_3051533 | Lighthouse Lab in Alderley Park | Wellcome Sanger Institute for the COVID-19 Genomics UK (COG-UK) Consortium | Jacquelyn Wynn, Mairead Hyland, The Lighthouse Lab in Alderley Park and Alex Alderton, Roberto Amato, Jeffrey Barrett, Sonia Goncalves, Ewan Harrison, David K. Jackson, Ian Johnston, Dominic Kwiatkowski, Cordelia Langford, John Sillitoe on behalf of the Wellcome Sanger Institute COVID-19 Surveillance Team |
| EPI_ISL_3051534 | Berkshire and Surrey Pathology Services Lighthouse Laboratory | Wellcome Sanger Institute for the COVID-19 Genomics UK (COG-UK) Consortium | Berkshire and Surrey Pathology Services Lighthouse Laboratory and Alex Alderton, Roberto Amato, Jeffrey Barrett, Sonia Goncalves, Ewan Harrison, David K. Jackson, Ian Johnston, Dominic Kwiatkowski, Cordelia Langford, John Sillitoe on behalf of the Wellcome Sanger Institute COVID-19 Surveillance Team |
| EPI_ISL_3051535 | Lighthouse Lab in Alderley Park | Wellcome Sanger Institute for the COVID-19 Genomics UK (COG-UK) Consortium | Jacquelyn Wynn, Mairead Hyland, The Lighthouse Lab in Alderley Park and Alex Alderton, Roberto Amato, Jeffrey Barrett, Sonia Goncalves, Ewan Harrison, David K. Jackson, Ian Johnston, Dominic Kwiatkowski, Cordelia Langford, John Sillitoe on behalf of the Wellcome Sanger Institute COVID-19 Surveillance Team |
| EPI_ISL_3051536 | Berkshire and Surrey Pathology Services Lighthouse Laboratory | Wellcome Sanger Institute for the COVID-19 Genomics UK (COG-UK) Consortium | Berkshire and Surrey Pathology Services Lighthouse Laboratory and Alex Alderton, Roberto Amato, Jeffrey Barrett, Sonia Goncalves, Ewan Harrison, David K. Jackson, Ian Johnston, Dominic Kwiatkowski, Cordelia Langford, John Sillitoe on behalf of the Wellcome Sanger Institute COVID-19 Surveillance Team |
| EPI_ISL_3051537, EPI_ISL_3051538 | Lighthouse Lab in Glasgow | Wellcome Sanger Institute for the COVID-19 Genomics UK (COG-UK) Consortium | Harper VanSteenhouse, Yumi Kasai, David Gray, Carol Clugston, Anna Dominiczak and Alex Alderton, Roberto Amato, Jeffrey Barrett, Sonia Goncalves, Ewan Harrison, David K. Jackson, Ian Johnston, Dominic Kwiatkowski, Cordelia Langford, John Sillitoe on behalf of the Wellcome Sanger Institute COVID-19 Surveillance Team |
| EPI_ISL_3051539 | Berkshire and Surrey Pathology Services Lighthouse Laboratory | Wellcome Sanger Institute for the COVID-19 Genomics UK (COG-UK) Consortium | Berkshire and Surrey Pathology Services Lighthouse Laboratory and Alex Alderton, Roberto Amato, Jeffrey Barrett, Sonia Goncalves, Ewan Harrison, David K. Jackson, Ian Johnston, Dominic Kwiatkowski, Cordelia Langford, John Sillitoe on behalf of the Wellcome Sanger Institute COVID-19 Surveillance Team |
| EPI_ISL_3051540 | Lighthouse Lab in Milton Keynes | Wellcome Sanger Institute for the COVID-19 Genomics UK (COG-UK) Consortium | The Lighthouse Lab in Milton Keynes and Alex Alderton, Roberto Amato, Jeffrey Barrett, Sonia Goncalves, Ewan Harrison, David K. Jackson, Ian Johnston, Dominic Kwiatkowski, Cordelia Langford, John Sillitoe on behalf of the Wellcome Sanger Institute COVID-19 Surveillance Team |
| EPI_ISL_3051541 | Berkshire and Surrey Pathology Services Lighthouse Laboratory | Wellcome Sanger Institute for the COVID-19 Genomics UK (COG-UK) Consortium | Berkshire and Surrey Pathology Services Lighthouse Laboratory and Alex Alderton, Roberto Amato, Jeffrey Barrett, Sonia Goncalves, Ewan Harrison, David K. Jackson, Ian Johnston, Dominic Kwiatkowski, Cordelia Langford, John Sillitoe on behalf of the Wellcome Sanger Institute COVID-19 Surveillance Team |
| EPI_ISL_3051542, EPI_ISL_3051543 | Lighthouse Lab in Glasgow | Wellcome Sanger Institute for the COVID-19 Genomics UK (COG-UK) Consortium | Harper VanSteenhouse, Yumi Kasai, David Gray, Carol Clugston, Anna Dominiczak and Alex Alderton, Roberto Amato, Jeffrey Barrett, Sonia Goncalves, Ewan Harrison, David K. Jackson, Ian Johnston, Dominic Kwiatkowski, Cordelia Langford, John Sillitoe on behalf of the Wellcome Sanger Institute COVID-19 Surveillance Team |
| EPI_ISL_3051544 | Lighthouse Lab in Alderley Park | Wellcome Sanger Institute for the COVID-19 Genomics UK (COG-UK) Consortium | Jacquelyn Wynn, Mairead Hyland, The Lighthouse Lab in Alderley Park and Alex Alderton, Roberto Amato, Jeffrey Barrett, Sonia Goncalves, Ewan Harrison, David K. Jackson, Ian Johnston, Dominic Kwiatkowski, Cordelia Langford, John Sillitoe on behalf of the Wellcome Sanger Institute COVID-19 Surveillance Team |
| EPI_ISL_3051545, EPI_ISL_3051546, EPI_ISL_3051547 | Lighthouse Lab in Glasgow | Wellcome Sanger Institute for the COVID-19 Genomics UK (COG-UK) Consortium | Harper VanSteenhouse, Yumi Kasai, David Gray, Carol Clugston, Anna Dominiczak and Alex Alderton, Roberto Amato, Jeffrey Barrett, Sonia Goncalves, Ewan Harrison, David K. Jackson, Ian Johnston, Dominic Kwiatkowski, Cordelia Langford, John Sillitoe on behalf of the Wellcome Sanger Institute COVID-19 Surveillance Team |
| EPI_ISL_3051548 | Berkshire and Surrey Pathology Services Lighthouse Laboratory | Wellcome Sanger Institute for the COVID-19 Genomics UK (COG-UK) Consortium | Berkshire and Surrey Pathology Services Lighthouse Laboratory and Alex Alderton, Roberto Amato, Jeffrey Barrett, Sonia Goncalves, Ewan Harrison, David K. Jackson, Ian Johnston, Dominic Kwiatkowski, Cordelia Langford, John Sillitoe on behalf of the Wellcome Sanger Institute COVID-19 Surveillance Team |
| EPI_ISL_3051549 | Lighthouse Lab in Glasgow | Wellcome Sanger Institute for the COVID-19 Genomics UK (COG-UK) Consortium | Harper VanSteenhouse, Yumi Kasai, David Gray, Carol Clugston, Anna Dominiczak and Alex Alderton, Roberto Amato, Jeffrey Barrett, Sonia Goncalves, Ewan Harrison, David K. Jackson, Ian Johnston, Dominic Kwiatkowski, Cordelia Langford, John Sillitoe on behalf of the Wellcome Sanger Institute COVID-19 Surveillance Team |
| EPI_ISL_3051550, EPI_ISL_3051551 | Lighthouse Lab in Milton Keynes | Wellcome Sanger Institute for the COVID-19 Genomics UK (COG-UK) Consortium | The Lighthouse Lab in Milton Keynes and Alex Alderton, Roberto Amato, Jeffrey Barrett, Sonia Goncalves, Ewan Harrison, David K. Jackson, Ian Johnston, Dominic Kwiatkowski, Cordelia Langford, John Sillitoe on behalf of the Wellcome Sanger Institute COVID-19 Surveillance Team |
| EPI_ISL_3051552 | Lighthouse Lab in Glasgow | Wellcome Sanger Institute for the COVID-19 Genomics UK (COG-UK) Consortium | Harper VanSteenhouse, Yumi Kasai, David Gray, Carol Clugston, Anna Dominiczak and Alex Alderton, Roberto Amato, Jeffrey Barrett, Sonia Goncalves, Ewan Harrison, David K. Jackson, Ian Johnston, Dominic Kwiatkowski, Cordelia Langford, John Sillitoe on behalf of the Wellcome Sanger Institute COVID-19 Surveillance Team |
| EPI_ISL_3051553 | Lighthouse Lab in Alderley Park | Wellcome Sanger Institute for the COVID-19 Genomics UK (COG-UK) Consortium | Jacquelyn Wynn, Mairead Hyland, The Lighthouse Lab in Alderley Park and Alex Alderton, Roberto Amato, Jeffrey Barrett, Sonia Goncalves, Ewan Harrison, David K. Jackson, Ian Johnston, Dominic Kwiatkowski, Cordelia Langford, John Sillitoe on behalf of the Wellcome Sanger Institute COVID-19 Surveillance Team |
| EPI_ISL_3051554 | Lighthouse Lab in Milton Keynes | Wellcome Sanger Institute for the COVID-19 Genomics UK (COG-UK) Consortium | The Lighthouse Lab in Milton Keynes and Alex Alderton, Roberto Amato, Jeffrey Barrett, Sonia Goncalves, Ewan Harrison, David K. Jackson, Ian Johnston, Dominic Kwiatkowski, Cordelia Langford, John Sillitoe on behalf of the Wellcome Sanger Institute COVID-19 Surveillance Team |
| EPI_ISL_3051555 | Berkshire and Surrey Pathology Services Lighthouse Laboratory | Wellcome Sanger Institute for the COVID-19 Genomics UK (COG-UK) Consortium | Berkshire and Surrey Pathology Services Lighthouse Laboratory and Alex Alderton, Roberto Amato, Jeffrey Barrett, Sonia Goncalves, Ewan Harrison, David K. Jackson, Ian Johnston, Dominic Kwiatkowski, Cordelia Langford, John Sillitoe on behalf of the Wellcome Sanger Institute COVID-19 Surveillance Team |
| EPI_ISL_3051556 | Lighthouse Lab in Milton Keynes | Wellcome Sanger Institute for the COVID-19 Genomics UK (COG-UK) Consortium | The Lighthouse Lab in Milton Keynes and Alex Alderton, Roberto Amato, Jeffrey Barrett, Sonia Goncalves, Ewan Harrison, David K. Jackson, Ian Johnston, Dominic Kwiatkowski, Cordelia Langford, John Sillitoe on behalf of the Wellcome Sanger Institute COVID-19 Surveillance Team |
| EPI_ISL_3051557 | Berkshire and Surrey Pathology Services Lighthouse Laboratory | Wellcome Sanger Institute for the COVID-19 Genomics UK (COG-UK) Consortium | Berkshire and Surrey Pathology Services Lighthouse Laboratory and Alex Alderton, Roberto Amato, Jeffrey Barrett, Sonia Goncalves, Ewan Harrison, David K. Jackson, Ian Johnston, Dominic Kwiatkowski, Cordelia Langford, John Sillitoe on behalf of the Wellcome Sanger Institute COVID-19 Surveillance Team |
| EPI_ISL_3051558, EPI_ISL_3051559 | Lighthouse Lab in Glasgow | Wellcome Sanger Institute for the COVID-19 Genomics UK (COG-UK) Consortium | Harper VanSteenhouse, Yumi Kasai, David Gray, Carol Clugston, Anna Dominiczak and Alex Alderton, Roberto Amato, Jeffrey Barrett, Sonia Goncalves, Ewan Harrison, David K. Jackson, Ian Johnston, Dominic Kwiatkowski, Cordelia Langford, John Sillitoe on behalf of the Wellcome Sanger Institute COVID-19 Surveillance Team |
| EPI_ISL_3051560 | Lighthouse Lab in Alderley Park | Wellcome Sanger Institute for the COVID-19 Genomics UK (COG-UK) Consortium | Jacquelyn Wynn, Mairead Hyland, The Lighthouse Lab in Alderley Park and Alex Alderton, Roberto Amato, Jeffrey Barrett, Sonia Goncalves, Ewan Harrison, David K. Jackson, Ian Johnston, Dominic Kwiatkowski, Cordelia Langford, John Sillitoe on behalf of the Wellcome Sanger Institute COVID-19 Surveillance Team |
| EPI_ISL_3051561 | Lighthouse Lab in Milton Keynes | Wellcome Sanger Institute for the COVID-19 Genomics UK (COG-UK) Consortium | The Lighthouse Lab in Milton Keynes and Alex Alderton, Roberto Amato, Jeffrey Barrett, Sonia Goncalves, Ewan Harrison, David K. Jackson, Ian Johnston, Dominic Kwiatkowski, Cordelia Langford, John Sillitoe on behalf of the Wellcome Sanger Institute COVID-19 Surveillance Team |
| EPI_ISL_3051562 | Lighthouse Lab in Alderley Park | Wellcome Sanger Institute for the COVID-19 Genomics UK (COG-UK) Consortium | Jacquelyn Wynn, Mairead Hyland, The Lighthouse Lab in Alderley Park and Alex Alderton, Roberto Amato, Jeffrey Barrett, Sonia Goncalves, Ewan Harrison, David K. Jackson, Ian Johnston, Dominic Kwiatkowski, Cordelia Langford, John Sillitoe on behalf of the Wellcome Sanger Institute COVID-19 Surveillance Team |
| EPI_ISL_3051563 | Berkshire and Surrey Pathology Services Lighthouse Laboratory | Wellcome Sanger Institute for the COVID-19 Genomics UK (COG-UK) Consortium | Berkshire and Surrey Pathology Services Lighthouse Laboratory and Alex Alderton, Roberto Amato, Jeffrey Barrett, Sonia Goncalves, Ewan Harrison, David K. Jackson, Ian Johnston, Dominic Kwiatkowski, Cordelia Langford, John Sillitoe on behalf of the Wellcome Sanger Institute COVID-19 Surveillance Team |
| EPI_ISL_3051564 | Lighthouse Lab in Milton Keynes | Wellcome Sanger Institute for the COVID-19 Genomics UK (COG-UK) Consortium | The Lighthouse Lab in Milton Keynes and Alex Alderton, Roberto Amato, Jeffrey Barrett, Sonia Goncalves, Ewan Harrison, David K. Jackson, Ian Johnston, Dominic Kwiatkowski, Cordelia Langford, John Sillitoe on behalf of the Wellcome Sanger Institute COVID-19 Surveillance Team |
| EPI_ISL_3051565 | Berkshire and Surrey Pathology Services Lighthouse Laboratory | Wellcome Sanger Institute for the COVID-19 Genomics UK (COG-UK) Consortium | Berkshire and Surrey Pathology Services Lighthouse Laboratory and Alex Alderton, Roberto Amato, Jeffrey Barrett, Sonia Goncalves, Ewan Harrison, David K. Jackson, Ian Johnston, Dominic Kwiatkowski, Cordelia Langford, John Sillitoe on behalf of the Wellcome Sanger Institute COVID-19 Surveillance Team |
| EPI_ISL_3051566 | Lighthouse Lab in Milton Keynes | Wellcome Sanger Institute for the COVID-19 Genomics UK (COG-UK) Consortium | The Lighthouse Lab in Milton Keynes and Alex Alderton, Roberto Amato, Jeffrey Barrett, Sonia Goncalves, Ewan Harrison, David K. Jackson, Ian Johnston, Dominic Kwiatkowski, Cordelia Langford, John Sillitoe on behalf of the Wellcome Sanger Institute COVID-19 Surveillance Team |
| EPI_ISL_3051567, EPI_ISL_3051568 | Lighthouse Lab in Alderley Park | Wellcome Sanger Institute for the COVID-19 Genomics UK (COG-UK) Consortium | Jacquelyn Wynn, Mairead Hyland, The Lighthouse Lab in Alderley Park and Alex Alderton, Roberto Amato, Jeffrey Barrett, Sonia Goncalves, Ewan Harrison, David K. Jackson, Ian Johnston, Dominic Kwiatkowski, Cordelia Langford, John Sillitoe on behalf of the Wellcome Sanger Institute COVID-19 Surveillance Team |
| EPI_ISL_3051569 | Berkshire and Surrey Pathology Services Lighthouse Laboratory | Wellcome Sanger Institute for the COVID-19 Genomics UK (COG-UK) Consortium | Berkshire and Surrey Pathology Services Lighthouse Laboratory and Alex Alderton, Roberto Amato, Jeffrey Barrett, Sonia Goncalves, Ewan Harrison, David K. Jackson, Ian Johnston, Dominic Kwiatkowski, Cordelia Langford, John Sillitoe on behalf of the Wellcome Sanger Institute COVID-19 Surveillance Team |
| EPI_ISL_3051570, EPI_ISL_3051571, EPI_ISL_3051572 | Lighthouse Lab in Glasgow | Wellcome Sanger Institute for the COVID-19 Genomics UK (COG-UK) Consortium | Harper VanSteenhouse, Yumi Kasai, David Gray, Carol Clugston, Anna Dominiczak and Alex Alderton, Roberto Amato, Jeffrey Barrett, Sonia Goncalves, Ewan Harrison, David K. Jackson, Ian Johnston, Dominic Kwiatkowski, Cordelia Langford, John Sillitoe on behalf of the Wellcome Sanger Institute COVID-19 Surveillance Team |
| EPI_ISL_3051573 | Lighthouse Lab in Alderley Park | Wellcome Sanger Institute for the COVID-19 Genomics UK (COG-UK) Consortium | Jacquelyn Wynn, Mairead Hyland, The Lighthouse Lab in Alderley Park and Alex Alderton, Roberto Amato, Jeffrey Barrett, Sonia Goncalves, Ewan Harrison, David K. Jackson, Ian Johnston, Dominic Kwiatkowski, Cordelia Langford, John Sillitoe on behalf of the Wellcome Sanger Institute COVID-19 Surveillance Team |
| EPI_ISL_3051574, EPI_ISL_3051575, EPI_ISL_3051576, EPI_ISL_3051577 | Berkshire and Surrey Pathology Services Lighthouse Laboratory | Wellcome Sanger Institute for the COVID-19 Genomics UK (COG-UK) Consortium | Berkshire and Surrey Pathology Services Lighthouse Laboratory and Alex Alderton, Roberto Amato, Jeffrey Barrett, Sonia Goncalves, Ewan Harrison, David K. Jackson, Ian Johnston, Dominic Kwiatkowski, Cordelia Langford, John Sillitoe on behalf of the Wellcome Sanger Institute COVID-19 Surveillance Team |
| EPI_ISL_3051578 | Lighthouse Lab in Milton Keynes | Wellcome Sanger Institute for the COVID-19 Genomics UK (COG-UK) Consortium | The Lighthouse Lab in Milton Keynes and Alex Alderton, Roberto Amato, Jeffrey Barrett, Sonia Goncalves, Ewan Harrison, David K. Jackson, Ian Johnston, Dominic Kwiatkowski, Cordelia Langford, John Sillitoe on behalf of the Wellcome Sanger Institute COVID-19 Surveillance Team |
| EPI_ISL_3051579 | Lighthouse Lab in Alderley Park | Wellcome Sanger Institute for the COVID-19 Genomics UK (COG-UK) Consortium | Jacquelyn Wynn, Mairead Hyland, The Lighthouse Lab in Alderley Park and Alex Alderton, Roberto Amato, Jeffrey Barrett, Sonia Goncalves, Ewan Harrison, David K. Jackson, Ian Johnston, Dominic Kwiatkowski, Cordelia Langford, John Sillitoe on behalf of the Wellcome Sanger Institute COVID-19 Surveillance Team |
| EPI_ISL_3051580 | Berkshire and Surrey Pathology Services Lighthouse Laboratory | Wellcome Sanger Institute for the COVID-19 Genomics UK (COG-UK) Consortium | Berkshire and Surrey Pathology Services Lighthouse Laboratory and Alex Alderton, Roberto Amato, Jeffrey Barrett, Sonia Goncalves, Ewan Harrison, David K. Jackson, Ian Johnston, Dominic Kwiatkowski, Cordelia Langford, John Sillitoe on behalf of the Wellcome Sanger Institute COVID-19 Surveillance Team |
| EPI_ISL_3051581 | Lighthouse Lab in Glasgow | Wellcome Sanger Institute for the COVID-19 Genomics UK (COG-UK) Consortium | Harper VanSteenhouse, Yumi Kasai, David Gray, Carol Clugston, Anna Dominiczak and Alex Alderton, Roberto Amato, Jeffrey Barrett, Sonia Goncalves, Ewan Harrison, David K. Jackson, Ian Johnston, Dominic Kwiatkowski, Cordelia Langford, John Sillitoe on behalf of the Wellcome Sanger Institute COVID-19 Surveillance Team |
| EPI_ISL_3051582 | Lighthouse Lab in Milton Keynes | Wellcome Sanger Institute for the COVID-19 Genomics UK (COG-UK) Consortium | The Lighthouse Lab in Milton Keynes and Alex Alderton, Roberto Amato, Jeffrey Barrett, Sonia Goncalves, Ewan Harrison, David K. Jackson, Ian Johnston, Dominic Kwiatkowski, Cordelia Langford, John Sillitoe on behalf of the Wellcome Sanger Institute COVID-19 Surveillance Team |
| EPI_ISL_3051583, EPI_ISL_3051584 | Lighthouse Lab in Alderley Park | Wellcome Sanger Institute for the COVID-19 Genomics UK (COG-UK) Consortium | Jacquelyn Wynn, Mairead Hyland, The Lighthouse Lab in Alderley Park and Alex Alderton, Roberto Amato, Jeffrey Barrett, Sonia Goncalves, Ewan Harrison, David K. Jackson, Ian Johnston, Dominic Kwiatkowski, Cordelia Langford, John Sillitoe on behalf of the Wellcome Sanger Institute COVID-19 Surveillance Team |
| EPI_ISL_3051585 | Lighthouse Lab in Milton Keynes | Wellcome Sanger Institute for the COVID-19 Genomics UK (COG-UK) Consortium | The Lighthouse Lab in Milton Keynes and Alex Alderton, Roberto Amato, Jeffrey Barrett, Sonia Goncalves, Ewan Harrison, David K. Jackson, Ian Johnston, Dominic Kwiatkowski, Cordelia Langford, John Sillitoe on behalf of the Wellcome Sanger Institute COVID-19 Surveillance Team |
| EPI_ISL_3051586 | Lighthouse Lab in Glasgow | Wellcome Sanger Institute for the COVID-19 Genomics UK (COG-UK) Consortium | Harper VanSteenhouse, Yumi Kasai, David Gray, Carol Clugston, Anna Dominiczak and Alex Alderton, Roberto Amato, Jeffrey Barrett, Sonia Goncalves, Ewan Harrison, David K. Jackson, Ian Johnston, Dominic Kwiatkowski, Cordelia Langford, John Sillitoe on behalf of the Wellcome Sanger Institute COVID-19 Surveillance Team |
| EPI_ISL_3051587, EPI_ISL_3051588 | Lighthouse Lab in Alderley Park | Wellcome Sanger Institute for the COVID-19 Genomics UK (COG-UK) Consortium | Jacquelyn Wynn, Mairead Hyland, The Lighthouse Lab in Alderley Park and Alex Alderton, Roberto Amato, Jeffrey Barrett, Sonia Goncalves, Ewan Harrison, David K. Jackson, Ian Johnston, Dominic Kwiatkowski, Cordelia Langford, John Sillitoe on behalf of the Wellcome Sanger Institute COVID-19 Surveillance Team |

| | | | |
|---|---|---|---|
| EPI_ISL_3051589 | Lighthouse Lab in Glasgow | Wellcome Sanger Institute for the COVID-19 Genomics UK (COG-UK) Consortium | Harper VanSteenhouse, Yumi Kasai, David Gray, Carol Clugston, Anna Dominiczak and Alex Alderton, Roberto Amato, Jeffrey Barrett, Sonia Goncalves, Ewan Harrison, David K. Jackson, Ian Johnston, Dominic Kwiatkowski, Cordelia Langford, John Sillitoe on behalf of the Wellcome Sanger Institute COVID-19 Surveillance Team |
| EPI_ISL_3051590 | Lighthouse Lab in Alderley Park | Wellcome Sanger Institute for the COVID-19 Genomics UK (COG-UK) Consortium | Jacquelyn Wynn, Mairead Hyland, The Lighthouse Lab in Alderley Park and Alex Alderton, Roberto Amato, Jeffrey Barrett, Sonia Goncalves, Ewan Harrison, David K. Jackson, Ian Johnston, Dominic Kwiatkowski, Cordelia Langford, John Sillitoe on behalf of the Wellcome Sanger Institute COVID-19 Surveillance Team |
| EPI_ISL_3051591 | Lighthouse Lab in Milton Keynes | Wellcome Sanger Institute for the COVID-19 Genomics UK (COG-UK) Consortium | The Lighthouse Lab in Milton Keynes and Alex Alderton, Roberto Amato, Jeffrey Barrett, Sonia Goncalves, Ewan Harrison, David K. Jackson, Ian Johnston, Dominic Kwiatkowski, Cordelia Langford, John Sillitoe on behalf of the Wellcome Sanger Institute COVID-19 Surveillance Team |
| EPI_ISL_3051592 | Lighthouse Lab in Alderley Park | Wellcome Sanger Institute for the COVID-19 Genomics UK (COG-UK) Consortium | Jacquelyn Wynn, Mairead Hyland, The Lighthouse Lab in Alderley Park and Alex Alderton, Roberto Amato, Jeffrey Barrett, Sonia Goncalves, Ewan Harrison, David K. Jackson, Ian Johnston, Dominic Kwiatkowski, Cordelia Langford, John Sillitoe on behalf of the Wellcome Sanger Institute COVID-19 Surveillance Team |
| EPI_ISL_3051593 | Lighthouse Lab in Milton Keynes | Wellcome Sanger Institute for the COVID-19 Genomics UK (COG-UK) Consortium | The Lighthouse Lab in Milton Keynes and Alex Alderton, Roberto Amato, Jeffrey Barrett, Sonia Goncalves, Ewan Harrison, David K. Jackson, Ian Johnston, Dominic Kwiatkowski, Cordelia Langford, John Sillitoe on behalf of the Wellcome Sanger Institute COVID-19 Surveillance Team |
| EPI_ISL_3051594, EPI_ISL_3051595 | Lighthouse Lab in Glasgow | Wellcome Sanger Institute for the COVID-19 Genomics UK (COG-UK) Consortium | Harper VanSteenhouse, Yumi Kasai, David Gray, Carol Clugston, Anna Dominiczak and Alex Alderton, Roberto Amato, Jeffrey Barrett, Sonia Goncalves, Ewan Harrison, David K. Jackson, Ian Johnston, Dominic Kwiatkowski, Cordelia Langford, John Sillitoe on behalf of the Wellcome Sanger Institute COVID-19 Surveillance Team |
| EPI_ISL_3051596 | Berkshire and Surrey Pathology Services Lighthouse Laboratory | Wellcome Sanger Institute for the COVID-19 Genomics UK (COG-UK) Consortium | Berkshire and Surrey Pathology Services Lighthouse Laboratory and Alex Alderton, Roberto Amato, Jeffrey Barrett, Sonia Goncalves, Ewan Harrison, David K. Jackson, Ian Johnston, Dominic Kwiatkowski, Cordelia Langford, John Sillitoe on behalf of the Wellcome Sanger Institute COVID-19 Surveillance Team |
| EPI_ISL_3051597 | Lighthouse Lab in Alderley Park | Wellcome Sanger Institute for the COVID-19 Genomics UK (COG-UK) Consortium | Jacquelyn Wynn, Mairead Hyland, The Lighthouse Lab in Alderley Park and Alex Alderton, Roberto Amato, Jeffrey Barrett, Sonia Goncalves, Ewan Harrison, David K. Jackson, Ian Johnston, Dominic Kwiatkowski, Cordelia Langford, John Sillitoe on behalf of the Wellcome Sanger Institute COVID-19 Surveillance Team |
| EPI_ISL_3051598 | Lighthouse Lab in Milton Keynes | Wellcome Sanger Institute for the COVID-19 Genomics UK (COG-UK) Consortium | The Lighthouse Lab in Milton Keynes and Alex Alderton, Roberto Amato, Jeffrey Barrett, Sonia Goncalves, Ewan Harrison, David K. Jackson, Ian Johnston, Dominic Kwiatkowski, Cordelia Langford, John Sillitoe on behalf of the Wellcome Sanger Institute COVID-19 Surveillance Team |
| EPI_ISL_3051599 | Berkshire and Surrey Pathology Services Lighthouse Laboratory | Wellcome Sanger Institute for the COVID-19 Genomics UK (COG-UK) Consortium | Berkshire and Surrey Pathology Services Lighthouse Laboratory and Alex Alderton, Roberto Amato, Jeffrey Barrett, Sonia Goncalves, Ewan Harrison, David K. Jackson, Ian Johnston, Dominic Kwiatkowski, Cordelia Langford, John Sillitoe on behalf of the Wellcome Sanger Institute COVID-19 Surveillance Team |
| EPI_ISL_3051600 | Lighthouse Lab in Alderley Park | Wellcome Sanger Institute for the COVID-19 Genomics UK (COG-UK) Consortium | Jacquelyn Wynn, Mairead Hyland, The Lighthouse Lab in Alderley Park and Alex Alderton, Roberto Amato, Jeffrey Barrett, Sonia Goncalves, Ewan Harrison, David K. Jackson, Ian Johnston, Dominic Kwiatkowski, Cordelia Langford, John Sillitoe on behalf of the Wellcome Sanger Institute COVID-19 Surveillance Team |
| EPI_ISL_3051601 | Lighthouse Lab in Glasgow | Wellcome Sanger Institute for the COVID-19 Genomics UK (COG-UK) Consortium | Harper VanSteenhouse, Yumi Kasai, David Gray, Carol Clugston, Anna Dominiczak and Alex Alderton, Roberto Amato, Jeffrey Barrett, Sonia Goncalves, Ewan Harrison, David K. Jackson, Ian Johnston, Dominic Kwiatkowski, Cordelia Langford, John Sillitoe on behalf of the Wellcome Sanger Institute COVID-19 Surveillance Team |
| EPI_ISL_3051602 | Lighthouse Lab in Milton Keynes | Wellcome Sanger Institute for the COVID-19 Genomics UK (COG-UK) Consortium | The Lighthouse Lab in Milton Keynes and Alex Alderton, Roberto Amato, Jeffrey Barrett, Sonia Goncalves, Ewan Harrison, David K. Jackson, Ian Johnston, Dominic Kwiatkowski, Cordelia Langford, John Sillitoe on behalf of the Wellcome Sanger Institute COVID-19 Surveillance Team |
| EPI_ISL_3051603, EPI_ISL_3051604, EPI_ISL_3051605, EPI_ISL_3051606 | Lighthouse Lab in Alderley Park | Wellcome Sanger Institute for the COVID-19 Genomics UK (COG-UK) Consortium | Jacquelyn Wynn, Mairead Hyland, The Lighthouse Lab in Alderley Park and Alex Alderton, Roberto Amato, Jeffrey Barrett, Sonia Goncalves, Ewan Harrison, David K. Jackson, Ian Johnston, Dominic Kwiatkowski, Cordelia Langford, John Sillitoe on behalf of the Wellcome Sanger Institute COVID-19 Surveillance Team |
| EPI_ISL_3051607, EPI_ISL_3051608 | Lighthouse Lab in Milton Keynes | Wellcome Sanger Institute for the COVID-19 Genomics UK (COG-UK) Consortium | The Lighthouse Lab in Milton Keynes and Alex Alderton, Roberto Amato, Jeffrey Barrett, Sonia Goncalves, Ewan Harrison, David K. Jackson, Ian Johnston, Dominic Kwiatkowski, Cordelia Langford, John Sillitoe on behalf of the Wellcome Sanger Institute COVID-19 Surveillance Team |
| EPI_ISL_3051609 | Lighthouse Lab in Glasgow | Wellcome Sanger Institute for the COVID-19 Genomics UK (COG-UK) Consortium | Harper VanSteenhouse, Yumi Kasai, David Gray, Carol Clugston, Anna Dominiczak and Alex Alderton, Roberto Amato, Jeffrey Barrett, Sonia Goncalves, Ewan Harrison, David K. Jackson, Ian Johnston, Dominic Kwiatkowski, Cordelia Langford, John Sillitoe on behalf of the Wellcome Sanger Institute COVID-19 Surveillance Team |
| EPI_ISL_3051610 | Lighthouse Lab in Milton Keynes | Wellcome Sanger Institute for the COVID-19 Genomics UK (COG-UK) Consortium | The Lighthouse Lab in Milton Keynes and Alex Alderton, Roberto Amato, Jeffrey Barrett, Sonia Goncalves, Ewan Harrison, David K. Jackson, Ian Johnston, Dominic Kwiatkowski, Cordelia Langford, John Sillitoe on behalf of the Wellcome Sanger Institute COVID-19 Surveillance Team |
| EPI_ISL_3051611 | Lighthouse Lab in Alderley Park | Wellcome Sanger Institute for the COVID-19 Genomics UK (COG-UK) Consortium | Jacquelyn Wynn, Mairead Hyland, The Lighthouse Lab in Alderley Park and Alex Alderton, Roberto Amato, Jeffrey Barrett, Sonia Goncalves, Ewan Harrison, David K. Jackson, Ian Johnston, Dominic Kwiatkowski, Cordelia Langford, John Sillitoe on behalf of the Wellcome Sanger Institute COVID-19 Surveillance Team |
| EPI_ISL_3051612 | Lighthouse Lab in Glasgow | Wellcome Sanger Institute for the COVID-19 Genomics UK (COG-UK) Consortium | Harper VanSteenhouse, Yumi Kasai, David Gray, Carol Clugston, Anna Dominiczak and Alex Alderton, Roberto Amato, Jeffrey Barrett, Sonia Goncalves, Ewan Harrison, David K. Jackson, Ian Johnston, Dominic Kwiatkowski, Cordelia Langford, John Sillitoe on behalf of the Wellcome Sanger Institute COVID-19 Surveillance Team |
| EPI_ISL_3051613, EPI_ISL_3051614 | Lighthouse Lab in Alderley Park | Wellcome Sanger Institute for the COVID-19 Genomics UK (COG-UK) Consortium | Jacquelyn Wynn, Mairead Hyland, The Lighthouse Lab in Alderley Park and Alex Alderton, Roberto Amato, Jeffrey Barrett, Sonia Goncalves, Ewan Harrison, David K. Jackson, Ian Johnston, Dominic Kwiatkowski, Cordelia Langford, John Sillitoe on behalf of the Wellcome Sanger Institute COVID-19 Surveillance Team |
| EPI_ISL_3051615, EPI_ISL_3051616 | Lighthouse Lab in Milton Keynes | Wellcome Sanger Institute for the COVID-19 Genomics UK (COG-UK) Consortium | The Lighthouse Lab in Milton Keynes and Alex Alderton, Roberto Amato, Jeffrey Barrett, Sonia Goncalves, Ewan Harrison, David K. Jackson, Ian Johnston, Dominic Kwiatkowski, Cordelia Langford, John Sillitoe on behalf of the Wellcome Sanger Institute COVID-19 Surveillance Team |
| EPI_ISL_3051617 | Berkshire and Surrey Pathology Services Lighthouse Laboratory | Wellcome Sanger Institute for the COVID-19 Genomics UK (COG-UK) Consortium | Berkshire and Surrey Pathology Services Lighthouse Laboratory and Alex Alderton, Roberto Amato, Jeffrey Barrett, Sonia Goncalves, Ewan Harrison, David K. Jackson, Ian Johnston, Dominic Kwiatkowski, Cordelia Langford, John Sillitoe on behalf of the Wellcome Sanger Institute COVID-19 Surveillance Team |
| EPI_ISL_3051618 | Lighthouse Lab in Alderley Park | Wellcome Sanger Institute for the COVID-19 Genomics UK (COG-UK) Consortium | Jacquelyn Wynn, Mairead Hyland, The Lighthouse Lab in Alderley Park and Alex Alderton, Roberto Amato, Jeffrey Barrett, Sonia Goncalves, Ewan Harrison, David K. Jackson, Ian Johnston, Dominic Kwiatkowski, Cordelia Langford, John Sillitoe on behalf of the Wellcome Sanger Institute COVID-19 Surveillance Team |
| EPI_ISL_3051619 | Lighthouse Lab in Glasgow | Wellcome Sanger Institute for the COVID-19 Genomics UK (COG-UK) Consortium | Harper VanSteenhouse, Yumi Kasai, David Gray, Carol Clugston, Anna Dominiczak and Alex Alderton, Roberto Amato, Jeffrey Barrett, Sonia Goncalves, Ewan Harrison, David K. Jackson, Ian Johnston, Dominic Kwiatkowski, Cordelia Langford, John Sillitoe on behalf of the Wellcome Sanger Institute COVID-19 Surveillance Team |
| EPI_ISL_3051620 | Lighthouse Lab in Alderley Park | Wellcome Sanger Institute for the COVID-19 Genomics UK (COG-UK) Consortium | Jacquelyn Wynn, Mairead Hyland, The Lighthouse Lab in Alderley Park and Alex Alderton, Roberto Amato, Jeffrey Barrett, Sonia Goncalves, Ewan Harrison, David K. Jackson, Ian Johnston, Dominic Kwiatkowski, Cordelia Langford, John Sillitoe on behalf of the Wellcome Sanger Institute COVID-19 Surveillance Team |
| EPI_ISL_3051621 | Lighthouse Lab in Milton Keynes | Wellcome Sanger Institute for the COVID-19 Genomics UK (COG-UK) Consortium | The Lighthouse Lab in Milton Keynes and Alex Alderton, Roberto Amato, Jeffrey Barrett, Sonia Goncalves, Ewan Harrison, David K. Jackson, Ian Johnston, Dominic Kwiatkowski, Cordelia Langford, John Sillitoe on behalf of the Wellcome Sanger Institute COVID-19 Surveillance Team |
| EPI_ISL_3051622 | Lighthouse Lab in Glasgow | Wellcome Sanger Institute for the COVID-19 Genomics UK (COG-UK) Consortium | Harper VanSteenhouse, Yumi Kasai, David Gray, Carol Clugston, Anna Dominiczak and Alex Alderton, Roberto Amato, Jeffrey Barrett, Sonia Goncalves, Ewan Harrison, David K. Jackson, Ian Johnston, Dominic Kwiatkowski, Cordelia Langford, John Sillitoe on behalf of the Wellcome Sanger Institute COVID-19 Surveillance Team |
| EPI_ISL_3051623 | Lighthouse Lab in Milton Keynes | Wellcome Sanger Institute for the COVID-19 Genomics UK (COG-UK) Consortium | The Lighthouse Lab in Milton Keynes and Alex Alderton, Roberto Amato, Jeffrey Barrett, Sonia Goncalves, Ewan Harrison, David K. Jackson, Ian Johnston, Dominic Kwiatkowski, Cordelia Langford, John Sillitoe on behalf of the Wellcome Sanger Institute COVID-19 Surveillance Team |
| EPI_ISL_3051624 | Berkshire and Surrey Pathology Services Lighthouse Laboratory | Wellcome Sanger Institute for the COVID-19 Genomics UK (COG-UK) Consortium | Berkshire and Surrey Pathology Services Lighthouse Laboratory and Alex Alderton, Roberto Amato, Jeffrey Barrett, Sonia Goncalves, Ewan Harrison, David K. Jackson, Ian Johnston, Dominic Kwiatkowski, Cordelia Langford, John Sillitoe on behalf of the Wellcome Sanger Institute COVID-19 Surveillance Team |
| EPI_ISL_3051625 | Lighthouse Lab in Alderley Park | Wellcome Sanger Institute for the COVID-19 Genomics UK (COG-UK) Consortium | Jacquelyn Wynn, Mairead Hyland, The Lighthouse Lab in Alderley Park and Alex Alderton, Roberto Amato, Jeffrey Barrett, Sonia Goncalves, Ewan Harrison, David K. Jackson, Ian Johnston, Dominic Kwiatkowski, Cordelia Langford, John Sillitoe on behalf of the Wellcome Sanger Institute COVID-19 Surveillance Team |
| EPI_ISL_3051626 | Berkshire and Surrey Pathology Services Lighthouse Laboratory | Wellcome Sanger Institute for the COVID-19 Genomics UK (COG-UK) Consortium | Berkshire and Surrey Pathology Services Lighthouse Laboratory and Alex Alderton, Roberto Amato, Jeffrey Barrett, Sonia Goncalves, Ewan Harrison, David K. Jackson, Ian Johnston, Dominic Kwiatkowski, Cordelia Langford, John Sillitoe on behalf of the Wellcome Sanger Institute COVID-19 Surveillance Team |
| EPI_ISL_3051627 | Lighthouse Lab in Alderley Park | Wellcome Sanger Institute for the COVID-19 Genomics UK (COG-UK) Consortium | Jacquelyn Wynn, Mairead Hyland, The Lighthouse Lab in Alderley Park and Alex Alderton, Roberto Amato, Jeffrey Barrett, Sonia Goncalves, Ewan Harrison, David K. Jackson, Ian Johnston, Dominic Kwiatkowski, Cordelia Langford, John Sillitoe on behalf of the Wellcome Sanger Institute COVID-19 Surveillance Team |
| EPI_ISL_3051628 | Berkshire and Surrey Pathology Services Lighthouse Laboratory | Wellcome Sanger Institute for the COVID-19 Genomics UK (COG-UK) Consortium | Berkshire and Surrey Pathology Services Lighthouse Laboratory and Alex Alderton, Roberto Amato, Jeffrey Barrett, Sonia Goncalves, Ewan Harrison, David K. Jackson, Ian Johnston, Dominic Kwiatkowski, Cordelia Langford, John Sillitoe on behalf of the Wellcome Sanger Institute COVID-19 Surveillance Team |
| EPI_ISL_3051629 | Lighthouse Lab in Glasgow | Wellcome Sanger Institute for the COVID-19 Genomics UK (COG-UK) Consortium | Harper VanSteenhouse, Yumi Kasai, David Gray, Carol Clugston, Anna Dominiczak and Alex Alderton, Roberto Amato, Jeffrey Barrett, Sonia Goncalves, Ewan Harrison, David K. Jackson, Ian Johnston, Dominic Kwiatkowski, Cordelia Langford, John Sillitoe on behalf of the Wellcome Sanger Institute COVID-19 Surveillance Team |
| EPI_ISL_3051630 | Lighthouse Lab in Alderley Park | Wellcome Sanger Institute for the COVID-19 Genomics UK (COG-UK) Consortium | Jacquelyn Wynn, Mairead Hyland, The Lighthouse Lab in Alderley Park and Alex Alderton, Roberto Amato, Jeffrey Barrett, Sonia Goncalves, Ewan Harrison, David K. Jackson, Ian Johnston, Dominic Kwiatkowski, Cordelia Langford, John Sillitoe on behalf of the Wellcome Sanger Institute COVID-19 Surveillance Team |
| EPI_ISL_3051631, EPI_ISL_3051632 | Berkshire and Surrey Pathology Services Lighthouse Laboratory | Wellcome Sanger Institute for the COVID-19 Genomics UK (COG-UK) Consortium | Berkshire and Surrey Pathology Services Lighthouse Laboratory and Alex Alderton, Roberto Amato, Jeffrey Barrett, Sonia Goncalves, Ewan Harrison, David K. Jackson, Ian Johnston, Dominic Kwiatkowski, Cordelia Langford, John Sillitoe on behalf of the Wellcome Sanger Institute COVID-19 Surveillance Team |
| EPI_ISL_3051633 | Lighthouse Lab in Alderley Park | Wellcome Sanger Institute for the COVID-19 Genomics UK (COG-UK) Consortium | Jacquelyn Wynn, Mairead Hyland, The Lighthouse Lab in Alderley Park and Alex Alderton, Roberto Amato, Jeffrey Barrett, Sonia Goncalves, Ewan Harrison, David K. Jackson, Ian Johnston, Dominic Kwiatkowski, Cordelia Langford, John Sillitoe on behalf of the Wellcome Sanger Institute COVID-19 Surveillance Team |
| EPI_ISL_3051634, EPI_ISL_3051635 | Berkshire and Surrey Pathology Services Lighthouse Laboratory | Wellcome Sanger Institute for the COVID-19 Genomics UK (COG-UK) Consortium | Berkshire and Surrey Pathology Services Lighthouse Laboratory and Alex Alderton, Roberto Amato, Jeffrey Barrett, Sonia Goncalves, Ewan Harrison, David K. Jackson, Ian Johnston, Dominic Kwiatkowski, Cordelia Langford, John Sillitoe on behalf of the Wellcome Sanger Institute COVID-19 Surveillance Team |
| EPI_ISL_3051636 | Lighthouse Lab in Milton Keynes | Wellcome Sanger Institute for the COVID-19 Genomics UK (COG-UK) Consortium | The Lighthouse Lab in Milton Keynes and Alex Alderton, Roberto Amato, Jeffrey Barrett, Sonia Goncalves, Ewan Harrison, David K. Jackson, Ian Johnston, Dominic Kwiatkowski, Cordelia Langford, John Sillitoe on behalf of the Wellcome Sanger Institute COVID-19 Surveillance Team |
| EPI_ISL_3051637 | Berkshire and Surrey Pathology Services Lighthouse Laboratory | Wellcome Sanger Institute for the COVID-19 Genomics UK (COG-UK) Consortium | Berkshire and Surrey Pathology Services Lighthouse Laboratory and Alex Alderton, Roberto Amato, Jeffrey Barrett, Sonia Goncalves, Ewan Harrison, David K. Jackson, Ian Johnston, Dominic Kwiatkowski, Cordelia Langford, John Sillitoe on behalf of the Wellcome Sanger Institute COVID-19 Surveillance Team |
| EPI_ISL_3051638 | Lighthouse Lab in Alderley Park | Wellcome Sanger Institute for the COVID-19 Genomics UK (COG-UK) Consortium | Jacquelyn Wynn, Mairead Hyland, The Lighthouse Lab in Alderley Park and Alex Alderton, Roberto Amato, Jeffrey Barrett, Sonia Goncalves, Ewan Harrison, David K. Jackson, Ian Johnston, Dominic Kwiatkowski, Cordelia Langford, John Sillitoe on behalf of the Wellcome Sanger Institute COVID-19 Surveillance Team |
| EPI_ISL_3051639 | Berkshire and Surrey Pathology Services Lighthouse Laboratory | Wellcome Sanger Institute for the COVID-19 Genomics UK (COG-UK) Consortium | Berkshire and Surrey Pathology Services Lighthouse Laboratory and Alex Alderton, Roberto Amato, Jeffrey Barrett, Sonia Goncalves, Ewan Harrison, David K. Jackson, Ian Johnston, Dominic Kwiatkowski, Cordelia Langford, John Sillitoe on behalf of the Wellcome Sanger Institute COVID-19 Surveillance Team |
| EPI_ISL_3051640 | Lighthouse Lab in Alderley Park | Wellcome Sanger Institute for the COVID-19 Genomics UK (COG-UK) Consortium | Jacquelyn Wynn, Mairead Hyland, The Lighthouse Lab in Alderley Park and Alex Alderton, Roberto Amato, Jeffrey Barrett, Sonia Goncalves, Ewan Harrison, David K. Jackson, Ian Johnston, Dominic Kwiatkowski, Cordelia Langford, John Sillitoe on behalf of the Wellcome Sanger Institute COVID-19 Surveillance Team |
| EPI_ISL_3051641 | Lighthouse Lab in Glasgow | Wellcome Sanger Institute for the COVID-19 Genomics UK (COG-UK) Consortium | Harper VanSteenhouse, Yumi Kasai, David Gray, Carol Clugston, Anna Dominiczak and Alex Alderton, Roberto Amato, Jeffrey Barrett, Sonia Goncalves, Ewan Harrison, David K. Jackson, Ian Johnston, Dominic Kwiatkowski, Cordelia Langford, John Sillitoe on behalf of the Wellcome Sanger Institute COVID-19 Surveillance Team |
| EPI_ISL_3051642 | Berkshire and Surrey Pathology Services Lighthouse Laboratory | Wellcome Sanger Institute for the COVID-19 Genomics UK (COG-UK) | Berkshire and Surrey Pathology Services Lighthouse Laboratory and Alex Alderton, Roberto Amato, Jeffrey Barrett, Sonia Goncalves, Ewan Harrison, David K. Jackson, Ian Johnston, Dominic Kwiatkowski, Cordelia Langford, John Sillitoe on behalf |

| | | | |
|---|---|---|---|
| | | Consortium | of the Wellcome Sanger Institute COVID-19 Surveillance Team |
| EPI_ISL_3051643, EPI_ISL_3051644 | Lighthouse Lab in Milton Keynes | Wellcome Sanger Institute for the COVID-19 Genomics UK (COG-UK) Consortium | The Lighthouse Lab in Milton Keynes and Alex Alderton, Roberto Amato, Jeffrey Barrett, Sonia Goncalves, Ewan Harrison, David K. Jackson, Ian Johnston, Dominic Kwiatkowski, Cordelia Langford, John Sillitoe on behalf of the Wellcome Sanger Institute COVID-19 Surveillance Team |
| EPI_ISL_3051645 | Berkshire and Surrey Pathology Services Lighthouse Laboratory | Wellcome Sanger Institute for the COVID-19 Genomics UK (COG-UK) Consortium | Berkshire and Surrey Pathology Services Lighthouse Laboratory and Alex Alderton, Roberto Amato, Jeffrey Barrett, Sonia Goncalves, Ewan Harrison, David K. Jackson, Ian Johnston, Dominic Kwiatkowski, Cordelia Langford, John Sillitoe on behalf of the Wellcome Sanger Institute COVID-19 Surveillance Team |
| EPI_ISL_3051646, EPI_ISL_3051647 | Lighthouse Lab in Alderley Park | Wellcome Sanger Institute for the COVID-19 Genomics UK (COG-UK) Consortium | Jacquelyn Wynn, Mairead Hyland, The Lighthouse Lab in Alderley Park and Alex Alderton, Roberto Amato, Jeffrey Barrett, Sonia Goncalves, Ewan Harrison, David K. Jackson, Ian Johnston, Dominic Kwiatkowski, Cordelia Langford, John Sillitoe on behalf of the Wellcome Sanger Institute COVID-19 Surveillance Team |
| EPI_ISL_3051648, EPI_ISL_3051649 | Lighthouse Lab in Glasgow | Wellcome Sanger Institute for the COVID-19 Genomics UK (COG-UK) Consortium | Harper VanSteenhouse, Yumi Kasai, David Gray, Carol Clugston, Anna Dominiczak and Alex Alderton, Roberto Amato, Jeffrey Barrett, Sonia Goncalves, Ewan Harrison, David K. Jackson, Ian Johnston, Dominic Kwiatkowski, Cordelia Langford, John Sillitoe on behalf of the Wellcome Sanger Institute COVID-19 Surveillance Team |
| EPI_ISL_3051650 | Lighthouse Lab in Alderley Park | Wellcome Sanger Institute for the COVID-19 Genomics UK (COG-UK) Consortium | Jacquelyn Wynn, Mairead Hyland, The Lighthouse Lab in Alderley Park and Alex Alderton, Roberto Amato, Jeffrey Barrett, Sonia Goncalves, Ewan Harrison, David K. Jackson, Ian Johnston, Dominic Kwiatkowski, Cordelia Langford, John Sillitoe on behalf of the Wellcome Sanger Institute COVID-19 Surveillance Team |
| EPI_ISL_3051651 | Lighthouse Lab in Glasgow | Wellcome Sanger Institute for the COVID-19 Genomics UK (COG-UK) Consortium | Harper VanSteenhouse, Yumi Kasai, David Gray, Carol Clugston, Anna Dominiczak and Alex Alderton, Roberto Amato, Jeffrey Barrett, Sonia Goncalves, Ewan Harrison, David K. Jackson, Ian Johnston, Dominic Kwiatkowski, Cordelia Langford, John Sillitoe on behalf of the Wellcome Sanger Institute COVID-19 Surveillance Team |
| EPI_ISL_3051652 | Berkshire and Surrey Pathology Services Lighthouse Laboratory | Wellcome Sanger Institute for the COVID-19 Genomics UK (COG-UK) Consortium | Berkshire and Surrey Pathology Services Lighthouse Laboratory and Alex Alderton, Roberto Amato, Jeffrey Barrett, Sonia Goncalves, Ewan Harrison, David K. Jackson, Ian Johnston, Dominic Kwiatkowski, Cordelia Langford, John Sillitoe on behalf of the Wellcome Sanger Institute COVID-19 Surveillance Team |
| EPI_ISL_3051653, EPI_ISL_3051654 | Lighthouse Lab in Glasgow | Wellcome Sanger Institute for the COVID-19 Genomics UK (COG-UK) Consortium | Harper VanSteenhouse, Yumi Kasai, David Gray, Carol Clugston, Anna Dominiczak and Alex Alderton, Roberto Amato, Jeffrey Barrett, Sonia Goncalves, Ewan Harrison, David K. Jackson, Ian Johnston, Dominic Kwiatkowski, Cordelia Langford, John Sillitoe on behalf of the Wellcome Sanger Institute COVID-19 Surveillance Team |
| EPI_ISL_3051655 | Lighthouse Lab in Milton Keynes | Wellcome Sanger Institute for the COVID-19 Genomics UK (COG-UK) Consortium | The Lighthouse Lab in Milton Keynes and Alex Alderton, Roberto Amato, Jeffrey Barrett, Sonia Goncalves, Ewan Harrison, David K. Jackson, Ian Johnston, Dominic Kwiatkowski, Cordelia Langford, John Sillitoe on behalf of the Wellcome Sanger Institute COVID-19 Surveillance Team |
| EPI_ISL_3051656 | Berkshire and Surrey Pathology Services Lighthouse Laboratory | Wellcome Sanger Institute for the COVID-19 Genomics UK (COG-UK) Consortium | Berkshire and Surrey Pathology Services Lighthouse Laboratory and Alex Alderton, Roberto Amato, Jeffrey Barrett, Sonia Goncalves, Ewan Harrison, David K. Jackson, Ian Johnston, Dominic Kwiatkowski, Cordelia Langford, John Sillitoe on behalf of the Wellcome Sanger Institute COVID-19 Surveillance Team |
| EPI_ISL_3051657 | Lighthouse Lab in Milton Keynes | Wellcome Sanger Institute for the COVID-19 Genomics UK (COG-UK) Consortium | The Lighthouse Lab in Milton Keynes and Alex Alderton, Roberto Amato, Jeffrey Barrett, Sonia Goncalves, Ewan Harrison, David K. Jackson, Ian Johnston, Dominic Kwiatkowski, Cordelia Langford, John Sillitoe on behalf of the Wellcome Sanger Institute COVID-19 Surveillance Team |
| EPI_ISL_3051658, EPI_ISL_3051659 | Berkshire and Surrey Pathology Services Lighthouse Laboratory | Wellcome Sanger Institute for the COVID-19 Genomics UK (COG-UK) Consortium | Berkshire and Surrey Pathology Services Lighthouse Laboratory and Alex Alderton, Roberto Amato, Jeffrey Barrett, Sonia Goncalves, Ewan Harrison, David K. Jackson, Ian Johnston, Dominic Kwiatkowski, Cordelia Langford, John Sillitoe on behalf of the Wellcome Sanger Institute COVID-19 Surveillance Team |
| EPI_ISL_3051660 | Lighthouse Lab in Alderley Park | Wellcome Sanger Institute for the COVID-19 Genomics UK (COG-UK) Consortium | Jacquelyn Wynn, Mairead Hyland, The Lighthouse Lab in Alderley Park and Alex Alderton, Roberto Amato, Jeffrey Barrett, Sonia Goncalves, Ewan Harrison, David K. Jackson, Ian Johnston, Dominic Kwiatkowski, Cordelia Langford, John Sillitoe on behalf of the Wellcome Sanger Institute COVID-19 Surveillance Team |
| EPI_ISL_3051661, EPI_ISL_3051662 | Berkshire and Surrey Pathology Services Lighthouse Laboratory | Wellcome Sanger Institute for the COVID-19 Genomics UK (COG-UK) Consortium | Berkshire and Surrey Pathology Services Lighthouse Laboratory and Alex Alderton, Roberto Amato, Jeffrey Barrett, Sonia Goncalves, Ewan Harrison, David K. Jackson, Ian Johnston, Dominic Kwiatkowski, Cordelia Langford, John Sillitoe on behalf of the Wellcome Sanger Institute COVID-19 Surveillance Team |
| EPI_ISL_3051663 | Lighthouse Lab in Milton Keynes | Wellcome Sanger Institute for the COVID-19 Genomics UK (COG-UK) Consortium | The Lighthouse Lab in Milton Keynes and Alex Alderton, Roberto Amato, Jeffrey Barrett, Sonia Goncalves, Ewan Harrison, David K. Jackson, Ian Johnston, Dominic Kwiatkowski, Cordelia Langford, John Sillitoe on behalf of the Wellcome Sanger Institute COVID-19 Surveillance Team |
| EPI_ISL_3051664 | Lighthouse Lab in Glasgow | Wellcome Sanger Institute for the COVID-19 Genomics UK (COG-UK) Consortium | Harper VanSteenhouse, Yumi Kasai, David Gray, Carol Clugston, Anna Dominiczak and Alex Alderton, Roberto Amato, Jeffrey Barrett, Sonia Goncalves, Ewan Harrison, David K. Jackson, Ian Johnston, Dominic Kwiatkowski, Cordelia Langford, John Sillitoe on behalf of the Wellcome Sanger Institute COVID-19 Surveillance Team |
| EPI_ISL_3051665, EPI_ISL_3051666 | Lighthouse Lab in Alderley Park | Wellcome Sanger Institute for the COVID-19 Genomics UK (COG-UK) Consortium | Jacquelyn Wynn, Mairead Hyland, The Lighthouse Lab in Alderley Park and Alex Alderton, Roberto Amato, Jeffrey Barrett, Sonia Goncalves, Ewan Harrison, David K. Jackson, Ian Johnston, Dominic Kwiatkowski, Cordelia Langford, John Sillitoe on behalf of the Wellcome Sanger Institute COVID-19 Surveillance Team |
| EPI_ISL_3051667 | Berkshire and Surrey Pathology Services Lighthouse Laboratory | Wellcome Sanger Institute for the COVID-19 Genomics UK (COG-UK) Consortium | Berkshire and Surrey Pathology Services Lighthouse Laboratory and Alex Alderton, Roberto Amato, Jeffrey Barrett, Sonia Goncalves, Ewan Harrison, David K. Jackson, Ian Johnston, Dominic Kwiatkowski, Cordelia Langford, John Sillitoe on behalf of the Wellcome Sanger Institute COVID-19 Surveillance Team |
| EPI_ISL_3051668 | Lighthouse Lab in Alderley Park | Wellcome Sanger Institute for the COVID-19 Genomics UK (COG-UK) Consortium | Jacquelyn Wynn, Mairead Hyland, The Lighthouse Lab in Alderley Park and Alex Alderton, Roberto Amato, Jeffrey Barrett, Sonia Goncalves, Ewan Harrison, David K. Jackson, Ian Johnston, Dominic Kwiatkowski, Cordelia Langford, John Sillitoe on behalf of the Wellcome Sanger Institute COVID-19 Surveillance Team |
| EPI_ISL_3051669, EPI_ISL_3051670, EPI_ISL_3051671 | Lighthouse Lab in Milton Keynes | Wellcome Sanger Institute for the COVID-19 Genomics UK (COG-UK) Consortium | The Lighthouse Lab in Milton Keynes and Alex Alderton, Roberto Amato, Jeffrey Barrett, Sonia Goncalves, Ewan Harrison, David K. Jackson, Ian Johnston, Dominic Kwiatkowski, Cordelia Langford, John Sillitoe on behalf of the Wellcome Sanger Institute COVID-19 Surveillance Team |
| EPI_ISL_3051672, EPI_ISL_3051673 | Lighthouse Lab in Glasgow | Wellcome Sanger Institute for the COVID-19 Genomics UK (COG-UK) Consortium | Harper VanSteenhouse, Yumi Kasai, David Gray, Carol Clugston, Anna Dominiczak and Alex Alderton, Roberto Amato, Jeffrey Barrett, Sonia Goncalves, Ewan Harrison, David K. Jackson, Ian Johnston, Dominic Kwiatkowski, Cordelia Langford, John Sillitoe on behalf of the Wellcome Sanger Institute COVID-19 Surveillance Team |
| EPI_ISL_3051674, EPI_ISL_3051675 | Lighthouse Lab in Alderley Park | Wellcome Sanger Institute for the COVID-19 Genomics UK (COG-UK) Consortium | Jacquelyn Wynn, Mairead Hyland, The Lighthouse Lab in Alderley Park and Alex Alderton, Roberto Amato, Jeffrey Barrett, Sonia Goncalves, Ewan Harrison, David K. Jackson, Ian Johnston, Dominic Kwiatkowski, Cordelia Langford, John Sillitoe on behalf of the Wellcome Sanger Institute COVID-19 Surveillance Team |
| EPI_ISL_3051676 | Lighthouse Lab in Milton Keynes | Wellcome Sanger Institute for the COVID-19 Genomics UK (COG-UK) Consortium | The Lighthouse Lab in Milton Keynes and Alex Alderton, Roberto Amato, Jeffrey Barrett, Sonia Goncalves, Ewan Harrison, David K. Jackson, Ian Johnston, Dominic Kwiatkowski, Cordelia Langford, John Sillitoe on behalf of the Wellcome Sanger Institute COVID-19 Surveillance Team |
| EPI_ISL_3051677 | Lighthouse Lab in Alderley Park | Wellcome Sanger Institute for the COVID-19 Genomics UK (COG-UK) Consortium | Jacquelyn Wynn, Mairead Hyland, The Lighthouse Lab in Alderley Park and Alex Alderton, Roberto Amato, Jeffrey Barrett, Sonia Goncalves, Ewan Harrison, David K. Jackson, Ian Johnston, Dominic Kwiatkowski, Cordelia Langford, John Sillitoe on behalf of the Wellcome Sanger Institute COVID-19 Surveillance Team |
| EPI_ISL_3051678 | Lighthouse Lab in Glasgow | Wellcome Sanger Institute for the COVID-19 Genomics UK (COG-UK) Consortium | Harper VanSteenhouse, Yumi Kasai, David Gray, Carol Clugston, Anna Dominiczak and Alex Alderton, Roberto Amato, Jeffrey Barrett, Sonia Goncalves, Ewan Harrison, David K. Jackson, Ian Johnston, Dominic Kwiatkowski, Cordelia Langford, John Sillitoe on behalf of the Wellcome Sanger Institute COVID-19 Surveillance Team |
| EPI_ISL_3051679 | Berkshire and Surrey Pathology Services Lighthouse Laboratory | Wellcome Sanger Institute for the COVID-19 Genomics UK (COG-UK) Consortium | Berkshire and Surrey Pathology Services Lighthouse Laboratory and Alex Alderton, Roberto Amato, Jeffrey Barrett, Sonia Goncalves, Ewan Harrison, David K. Jackson, Ian Johnston, Dominic Kwiatkowski, Cordelia Langford, John Sillitoe on behalf of the Wellcome Sanger Institute COVID-19 Surveillance Team |
| EPI_ISL_3051680, EPI_ISL_3051681 | Lighthouse Lab in Milton Keynes | Wellcome Sanger Institute for the COVID-19 Genomics UK (COG-UK) Consortium | The Lighthouse Lab in Milton Keynes and Alex Alderton, Roberto Amato, Jeffrey Barrett, Sonia Goncalves, Ewan Harrison, David K. Jackson, Ian Johnston, Dominic Kwiatkowski, Cordelia Langford, John Sillitoe on behalf of the Wellcome Sanger Institute COVID-19 Surveillance Team |
| EPI_ISL_3051682, EPI_ISL_3051683 | Lighthouse Lab in Glasgow | Wellcome Sanger Institute for the COVID-19 Genomics UK (COG-UK) Consortium | Harper VanSteenhouse, Yumi Kasai, David Gray, Carol Clugston, Anna Dominiczak and Alex Alderton, Roberto Amato, Jeffrey Barrett, Sonia Goncalves, Ewan Harrison, David K. Jackson, Ian Johnston, Dominic Kwiatkowski, Cordelia Langford, John Sillitoe on behalf of the Wellcome Sanger Institute COVID-19 Surveillance Team |
| EPI_ISL_3051684, EPI_ISL_3051685 | Lighthouse Lab in Milton Keynes | Wellcome Sanger Institute for the COVID-19 Genomics UK (COG-UK) Consortium | The Lighthouse Lab in Milton Keynes and Alex Alderton, Roberto Amato, Jeffrey Barrett, Sonia Goncalves, Ewan Harrison, David K. Jackson, Ian Johnston, Dominic Kwiatkowski, Cordelia Langford, John Sillitoe on behalf of the Wellcome Sanger Institute COVID-19 Surveillance Team |
| EPI_ISL_3051686 | Lighthouse Lab in Alderley Park | Wellcome Sanger Institute for the COVID-19 Genomics UK (COG-UK) Consortium | Jacquelyn Wynn, Mairead Hyland, The Lighthouse Lab in Alderley Park and Alex Alderton, Roberto Amato, Jeffrey Barrett, Sonia Goncalves, Ewan Harrison, David K. Jackson, Ian Johnston, Dominic Kwiatkowski, Cordelia Langford, John Sillitoe on behalf of the Wellcome Sanger Institute COVID-19 Surveillance Team |
| EPI_ISL_3051687 | Berkshire and Surrey Pathology Services Lighthouse Laboratory | Wellcome Sanger Institute for the COVID-19 Genomics UK (COG-UK) Consortium | Berkshire and Surrey Pathology Services Lighthouse Laboratory and Alex Alderton, Roberto Amato, Jeffrey Barrett, Sonia Goncalves, Ewan Harrison, David K. Jackson, Ian Johnston, Dominic Kwiatkowski, Cordelia Langford, John Sillitoe on behalf of the Wellcome Sanger Institute COVID-19 Surveillance Team |
| EPI_ISL_3051688, EPI_ISL_3051689 | Lighthouse Lab in Alderley Park | Wellcome Sanger Institute for the COVID-19 Genomics UK (COG-UK) Consortium | Jacquelyn Wynn, Mairead Hyland, The Lighthouse Lab in Alderley Park and Alex Alderton, Roberto Amato, Jeffrey Barrett, Sonia Goncalves, Ewan Harrison, David K. Jackson, Ian Johnston, Dominic Kwiatkowski, Cordelia Langford, John Sillitoe on behalf of the Wellcome Sanger Institute COVID-19 Surveillance Team |
| EPI_ISL_3051690 | Lighthouse Lab in Glasgow | Wellcome Sanger Institute for the COVID-19 Genomics UK (COG-UK) Consortium | Harper VanSteenhouse, Yumi Kasai, David Gray, Carol Clugston, Anna Dominiczak and Alex Alderton, Roberto Amato, Jeffrey Barrett, Sonia Goncalves, Ewan Harrison, David K. Jackson, Ian Johnston, Dominic Kwiatkowski, Cordelia Langford, John Sillitoe on behalf of the Wellcome Sanger Institute COVID-19 Surveillance Team |
| EPI_ISL_3051691 | Lighthouse Lab in Milton Keynes | Wellcome Sanger Institute for the COVID-19 Genomics UK (COG-UK) Consortium | The Lighthouse Lab in Milton Keynes and Alex Alderton, Roberto Amato, Jeffrey Barrett, Sonia Goncalves, Ewan Harrison, David K. Jackson, Ian Johnston, Dominic Kwiatkowski, Cordelia Langford, John Sillitoe on behalf of the Wellcome Sanger Institute COVID-19 Surveillance Team |
| EPI_ISL_3051692, EPI_ISL_3051693 | Berkshire and Surrey Pathology Services Lighthouse Laboratory | Wellcome Sanger Institute for the COVID-19 Genomics UK (COG-UK) Consortium | Berkshire and Surrey Pathology Services Lighthouse Laboratory and Alex Alderton, Roberto Amato, Jeffrey Barrett, Sonia Goncalves, Ewan Harrison, David K. Jackson, Ian Johnston, Dominic Kwiatkowski, Cordelia Langford, John Sillitoe on behalf of the Wellcome Sanger Institute COVID-19 Surveillance Team |
| EPI_ISL_3051694 | Lighthouse Lab in Milton Keynes | Wellcome Sanger Institute for the COVID-19 Genomics UK (COG-UK) Consortium | The Lighthouse Lab in Milton Keynes and Alex Alderton, Roberto Amato, Jeffrey Barrett, Sonia Goncalves, Ewan Harrison, David K. Jackson, Ian Johnston, Dominic Kwiatkowski, Cordelia Langford, John Sillitoe on behalf of the Wellcome Sanger Institute COVID-19 Surveillance Team |
| EPI_ISL_3051695 | Lighthouse Lab in Alderley Park | Wellcome Sanger Institute for the COVID-19 Genomics UK (COG-UK) Consortium | Jacquelyn Wynn, Mairead Hyland, The Lighthouse Lab in Alderley Park and Alex Alderton, Roberto Amato, Jeffrey Barrett, Sonia Goncalves, Ewan Harrison, David K. Jackson, Ian Johnston, Dominic Kwiatkowski, Cordelia Langford, John Sillitoe on behalf of the Wellcome Sanger Institute COVID-19 Surveillance Team |
| EPI_ISL_3051696, EPI_ISL_3051697 | Berkshire and Surrey Pathology Services Lighthouse Laboratory | Wellcome Sanger Institute for the COVID-19 Genomics UK (COG-UK) Consortium | Berkshire and Surrey Pathology Services Lighthouse Laboratory and Alex Alderton, Roberto Amato, Jeffrey Barrett, Sonia Goncalves, Ewan Harrison, David K. Jackson, Ian Johnston, Dominic Kwiatkowski, Cordelia Langford, John Sillitoe on behalf of the Wellcome Sanger Institute COVID-19 Surveillance Team |
| EPI_ISL_3051698 | Lighthouse Lab in Milton Keynes | Wellcome Sanger Institute for the COVID-19 Genomics UK (COG-UK) Consortium | The Lighthouse Lab in Milton Keynes and Alex Alderton, Roberto Amato, Jeffrey Barrett, Sonia Goncalves, Ewan Harrison, David K. Jackson, Ian Johnston, Dominic Kwiatkowski, Cordelia Langford, John Sillitoe on behalf of the Wellcome Sanger Institute COVID-19 Surveillance Team |
| EPI_ISL_3051699 | Lighthouse Lab in Glasgow | Wellcome Sanger Institute for the COVID-19 Genomics UK (COG-UK) Consortium | Harper VanSteenhouse, Yumi Kasai, David Gray, Carol Clugston, Anna Dominiczak and Alex Alderton, Roberto Amato, Jeffrey Barrett, Sonia Goncalves, Ewan Harrison, David K. Jackson, Ian Johnston, Dominic Kwiatkowski, Cordelia Langford, John Sillitoe on behalf of the Wellcome Sanger Institute COVID-19 Surveillance Team |
| EPI_ISL_3051700 | Berkshire and Surrey Pathology Services Lighthouse Laboratory | Wellcome Sanger Institute for the COVID-19 Genomics UK (COG-UK) Consortium | Berkshire and Surrey Pathology Services Lighthouse Laboratory and Alex Alderton, Roberto Amato, Jeffrey Barrett, Sonia Goncalves, Ewan Harrison, David K. Jackson, Ian Johnston, Dominic Kwiatkowski, Cordelia Langford, John Sillitoe on behalf of the Wellcome Sanger Institute COVID-19 Surveillance Team |
| EPI_ISL_3051701 | Lighthouse Lab in Milton Keynes | Wellcome Sanger Institute for the COVID-19 Genomics UK (COG-UK) Consortium | The Lighthouse Lab in Milton Keynes and Alex Alderton, Roberto Amato, Jeffrey Barrett, Sonia Goncalves, Ewan Harrison, David K. Jackson, Ian Johnston, Dominic Kwiatkowski, Cordelia Langford, John Sillitoe on behalf of the Wellcome Sanger Institute COVID-19 Surveillance Team |
| EPI_ISL_3051702 | Lighthouse Lab in Alderley Park | Wellcome Sanger Institute for the COVID-19 Genomics UK (COG-UK) Consortium | Jacquelyn Wynn, Mairead Hyland, The Lighthouse Lab in Alderley Park and Alex Alderton, Roberto Amato, Jeffrey Barrett, Sonia Goncalves, Ewan Harrison, David K. Jackson, Ian Johnston, Dominic Kwiatkowski, Cordelia Langford, John Sillitoe on behalf of the Wellcome Sanger Institute COVID-19 Surveillance Team |
| EPI_ISL_3051703 | Berkshire and Surrey Pathology Services Lighthouse Laboratory | Wellcome Sanger Institute for the COVID-19 Genomics UK (COG-UK) Consortium | Berkshire and Surrey Pathology Services Lighthouse Laboratory and Alex Alderton, Roberto Amato, Jeffrey Barrett, Sonia Goncalves, Ewan Harrison, David K. Jackson, Ian Johnston, Dominic Kwiatkowski, Cordelia Langford, John Sillitoe on behalf of the Wellcome Sanger Institute COVID-19 Surveillance Team |

| | | | |
|---|---|---|---|
| EPI_ISL_3051704 | Lighthouse Lab in Alderley Park | Wellcome Sanger Institute for the COVID-19 Genomics UK (COG-UK) Consortium | Jacquelyn Wynn, Mairead Hyland, The Lighthouse Lab in Alderley Park and Alex Alderton, Roberto Amato, Jeffrey Barrett, Sonia Goncalves, Ewan Harrison, David K. Jackson, Ian Johnston, Dominic Kwiatkowski, Cordelia Langford, John Sillitoe on behalf of the Wellcome Sanger Institute COVID-19 Surveillance Team |
| EPI_ISL_3051705 | Lighthouse Lab in Glasgow | Wellcome Sanger Institute for the COVID-19 Genomics UK (COG-UK) Consortium | Harper VanSteenhouse, Yumi Kasai, David Gray, Carol Clugston, Anna Dominiczak and Alex Alderton, Roberto Amato, Jeffrey Barrett, Sonia Goncalves, Ewan Harrison, David K. Jackson, Ian Johnston, Dominic Kwiatkowski, Cordelia Langford, John Sillitoe on behalf of the Wellcome Sanger Institute COVID-19 Surveillance Team |
| EPI_ISL_3051706 | Lighthouse Lab in Alderley Park | Wellcome Sanger Institute for the COVID-19 Genomics UK (COG-UK) Consortium | Jacquelyn Wynn, Mairead Hyland, The Lighthouse Lab in Alderley Park and Alex Alderton, Roberto Amato, Jeffrey Barrett, Sonia Goncalves, Ewan Harrison, David K. Jackson, Ian Johnston, Dominic Kwiatkowski, Cordelia Langford, John Sillitoe on behalf of the Wellcome Sanger Institute COVID-19 Surveillance Team |
| EPI_ISL_3051707 | Berkshire and Surrey Pathology Services Lighthouse Laboratory | Wellcome Sanger Institute for the COVID-19 Genomics UK (COG-UK) Consortium | Berkshire and Surrey Pathology Services Lighthouse Laboratory and Alex Alderton, Roberto Amato, Jeffrey Barrett, Sonia Goncalves, Ewan Harrison, David K. Jackson, Ian Johnston, Dominic Kwiatkowski, Cordelia Langford, John Sillitoe on behalf of the Wellcome Sanger Institute COVID-19 Surveillance Team |
| EPI_ISL_3051708 | Lighthouse Lab in Milton Keynes | Wellcome Sanger Institute for the COVID-19 Genomics UK (COG-UK) Consortium | The Lighthouse Lab in Milton Keynes and Alex Alderton, Roberto Amato, Jeffrey Barrett, Sonia Goncalves, Ewan Harrison, David K. Jackson, Ian Johnston, Dominic Kwiatkowski, Cordelia Langford, John Sillitoe on behalf of the Wellcome Sanger Institute COVID-19 Surveillance Team |
| EPI_ISL_3051709 | Berkshire and Surrey Pathology Services Lighthouse Laboratory | Wellcome Sanger Institute for the COVID-19 Genomics UK (COG-UK) Consortium | Berkshire and Surrey Pathology Services Lighthouse Laboratory and Alex Alderton, Roberto Amato, Jeffrey Barrett, Sonia Goncalves, Ewan Harrison, David K. Jackson, Ian Johnston, Dominic Kwiatkowski, Cordelia Langford, John Sillitoe on behalf of the Wellcome Sanger Institute COVID-19 Surveillance Team |
| EPI_ISL_3051710 | Lighthouse Lab in Glasgow | Wellcome Sanger Institute for the COVID-19 Genomics UK (COG-UK) Consortium | Harper VanSteenhouse, Yumi Kasai, David Gray, Carol Clugston, Anna Dominiczak and Alex Alderton, Roberto Amato, Jeffrey Barrett, Sonia Goncalves, Ewan Harrison, David K. Jackson, Ian Johnston, Dominic Kwiatkowski, Cordelia Langford, John Sillitoe on behalf of the Wellcome Sanger Institute COVID-19 Surveillance Team |
| EPI_ISL_3051711, EPI_ISL_3051712 | Lighthouse Lab in Alderley Park | Wellcome Sanger Institute for the COVID-19 Genomics UK (COG-UK) Consortium | Jacquelyn Wynn, Mairead Hyland, The Lighthouse Lab in Alderley Park and Alex Alderton, Roberto Amato, Jeffrey Barrett, Sonia Goncalves, Ewan Harrison, David K. Jackson, Ian Johnston, Dominic Kwiatkowski, Cordelia Langford, John Sillitoe on behalf of the Wellcome Sanger Institute COVID-19 Surveillance Team |
| EPI_ISL_3051713 | Lighthouse Lab in Glasgow | Wellcome Sanger Institute for the COVID-19 Genomics UK (COG-UK) Consortium | Harper VanSteenhouse, Yumi Kasai, David Gray, Carol Clugston, Anna Dominiczak and Alex Alderton, Roberto Amato, Jeffrey Barrett, Sonia Goncalves, Ewan Harrison, David K. Jackson, Ian Johnston, Dominic Kwiatkowski, Cordelia Langford, John Sillitoe on behalf of the Wellcome Sanger Institute COVID-19 Surveillance Team |
| EPI_ISL_3051714 | Berkshire and Surrey Pathology Services Lighthouse Laboratory | Wellcome Sanger Institute for the COVID-19 Genomics UK (COG-UK) Consortium | Berkshire and Surrey Pathology Services Lighthouse Laboratory and Alex Alderton, Roberto Amato, Jeffrey Barrett, Sonia Goncalves, Ewan Harrison, David K. Jackson, Ian Johnston, Dominic Kwiatkowski, Cordelia Langford, John Sillitoe on behalf of the Wellcome Sanger Institute COVID-19 Surveillance Team |
| EPI_ISL_3051715 | Lighthouse Lab in Alderley Park | Wellcome Sanger Institute for the COVID-19 Genomics UK (COG-UK) Consortium | Jacquelyn Wynn, Mairead Hyland, The Lighthouse Lab in Alderley Park and Alex Alderton, Roberto Amato, Jeffrey Barrett, Sonia Goncalves, Ewan Harrison, David K. Jackson, Ian Johnston, Dominic Kwiatkowski, Cordelia Langford, John Sillitoe on behalf of the Wellcome Sanger Institute COVID-19 Surveillance Team |
| EPI_ISL_3051716 | Berkshire and Surrey Pathology Services Lighthouse Laboratory | Wellcome Sanger Institute for the COVID-19 Genomics UK (COG-UK) Consortium | Berkshire and Surrey Pathology Services Lighthouse Laboratory and Alex Alderton, Roberto Amato, Jeffrey Barrett, Sonia Goncalves, Ewan Harrison, David K. Jackson, Ian Johnston, Dominic Kwiatkowski, Cordelia Langford, John Sillitoe on behalf of the Wellcome Sanger Institute COVID-19 Surveillance Team |
| EPI_ISL_3051717, EPI_ISL_3051718 | Lighthouse Lab in Alderley Park | Wellcome Sanger Institute for the COVID-19 Genomics UK (COG-UK) Consortium | Jacquelyn Wynn, Mairead Hyland, The Lighthouse Lab in Alderley Park and Alex Alderton, Roberto Amato, Jeffrey Barrett, Sonia Goncalves, Ewan Harrison, David K. Jackson, Ian Johnston, Dominic Kwiatkowski, Cordelia Langford, John Sillitoe on behalf of the Wellcome Sanger Institute COVID-19 Surveillance Team |
| EPI_ISL_3051719 | Lighthouse Lab in Glasgow | Wellcome Sanger Institute for the COVID-19 Genomics UK (COG-UK) Consortium | Harper VanSteenhouse, Yumi Kasai, David Gray, Carol Clugston, Anna Dominiczak and Alex Alderton, Roberto Amato, Jeffrey Barrett, Sonia Goncalves, Ewan Harrison, David K. Jackson, Ian Johnston, Dominic Kwiatkowski, Cordelia Langford, John Sillitoe on behalf of the Wellcome Sanger Institute COVID-19 Surveillance Team |
| EPI_ISL_3051720 | Lighthouse Lab in Alderley Park | Wellcome Sanger Institute for the COVID-19 Genomics UK (COG-UK) Consortium | Jacquelyn Wynn, Mairead Hyland, The Lighthouse Lab in Alderley Park and Alex Alderton, Roberto Amato, Jeffrey Barrett, Sonia Goncalves, Ewan Harrison, David K. Jackson, Ian Johnston, Dominic Kwiatkowski, Cordelia Langford, John Sillitoe on behalf of the Wellcome Sanger Institute COVID-19 Surveillance Team |
| EPI_ISL_3051721, EPI_ISL_3051722, EPI_ISL_3051723, EPI_ISL_3051724 | Lighthouse Lab in Milton Keynes | Wellcome Sanger Institute for the COVID-19 Genomics UK (COG-UK) Consortium | The Lighthouse Lab in Milton Keynes and Alex Alderton, Roberto Amato, Jeffrey Barrett, Sonia Goncalves, Ewan Harrison, David K. Jackson, Ian Johnston, Dominic Kwiatkowski, Cordelia Langford, John Sillitoe on behalf of the Wellcome Sanger Institute COVID-19 Surveillance Team |
| EPI_ISL_3051725 | Lighthouse Lab in Glasgow | Wellcome Sanger Institute for the COVID-19 Genomics UK (COG-UK) Consortium | Harper VanSteenhouse, Yumi Kasai, David Gray, Carol Clugston, Anna Dominiczak and Alex Alderton, Roberto Amato, Jeffrey Barrett, Sonia Goncalves, Ewan Harrison, David K. Jackson, Ian Johnston, Dominic Kwiatkowski, Cordelia Langford, John Sillitoe on behalf of the Wellcome Sanger Institute COVID-19 Surveillance Team |
| EPI_ISL_3051726, EPI_ISL_3051727 | Berkshire and Surrey Pathology Services Lighthouse Laboratory | Wellcome Sanger Institute for the COVID-19 Genomics UK (COG-UK) Consortium | Berkshire and Surrey Pathology Services Lighthouse Laboratory and Alex Alderton, Roberto Amato, Jeffrey Barrett, Sonia Goncalves, Ewan Harrison, David K. Jackson, Ian Johnston, Dominic Kwiatkowski, Cordelia Langford, John Sillitoe on behalf of the Wellcome Sanger Institute COVID-19 Surveillance Team |
| EPI_ISL_3051728 | Lighthouse Lab in Glasgow | Wellcome Sanger Institute for the COVID-19 Genomics UK (COG-UK) Consortium | Harper VanSteenhouse, Yumi Kasai, David Gray, Carol Clugston, Anna Dominiczak and Alex Alderton, Roberto Amato, Jeffrey Barrett, Sonia Goncalves, Ewan Harrison, David K. Jackson, Ian Johnston, Dominic Kwiatkowski, Cordelia Langford, John Sillitoe on behalf of the Wellcome Sanger Institute COVID-19 Surveillance Team |
| EPI_ISL_3051729 | Lighthouse Lab in Alderley Park | Wellcome Sanger Institute for the COVID-19 Genomics UK (COG-UK) Consortium | Jacquelyn Wynn, Mairead Hyland, The Lighthouse Lab in Alderley Park and Alex Alderton, Roberto Amato, Jeffrey Barrett, Sonia Goncalves, Ewan Harrison, David K. Jackson, Ian Johnston, Dominic Kwiatkowski, Cordelia Langford, John Sillitoe on behalf of the Wellcome Sanger Institute COVID-19 Surveillance Team |
| EPI_ISL_3051730 | Lighthouse Lab in Glasgow | Wellcome Sanger Institute for the COVID-19 Genomics UK (COG-UK) Consortium | Harper VanSteenhouse, Yumi Kasai, David Gray, Carol Clugston, Anna Dominiczak and Alex Alderton, Roberto Amato, Jeffrey Barrett, Sonia Goncalves, Ewan Harrison, David K. Jackson, Ian Johnston, Dominic Kwiatkowski, Cordelia Langford, John Sillitoe on behalf of the Wellcome Sanger Institute COVID-19 Surveillance Team |
| EPI_ISL_3051731 | Lighthouse Lab in Milton Keynes | Wellcome Sanger Institute for the COVID-19 Genomics UK (COG-UK) Consortium | The Lighthouse Lab in Milton Keynes and Alex Alderton, Roberto Amato, Jeffrey Barrett, Sonia Goncalves, Ewan Harrison, David K. Jackson, Ian Johnston, Dominic Kwiatkowski, Cordelia Langford, John Sillitoe on behalf of the Wellcome Sanger Institute COVID-19 Surveillance Team |
| EPI_ISL_3051732 | Berkshire and Surrey Pathology Services Lighthouse Laboratory | Wellcome Sanger Institute for the COVID-19 Genomics UK (COG-UK) Consortium | Berkshire and Surrey Pathology Services Lighthouse Laboratory and Alex Alderton, Roberto Amato, Jeffrey Barrett, Sonia Goncalves, Ewan Harrison, David K. Jackson, Ian Johnston, Dominic Kwiatkowski, Cordelia Langford, John Sillitoe on behalf of the Wellcome Sanger Institute COVID-19 Surveillance Team |
| EPI_ISL_3051733 | Lighthouse Lab in Milton Keynes | Wellcome Sanger Institute for the COVID-19 Genomics UK (COG-UK) Consortium | The Lighthouse Lab in Milton Keynes and Alex Alderton, Roberto Amato, Jeffrey Barrett, Sonia Goncalves, Ewan Harrison, David K. Jackson, Ian Johnston, Dominic Kwiatkowski, Cordelia Langford, John Sillitoe on behalf of the Wellcome Sanger Institute COVID-19 Surveillance Team |
| EPI_ISL_3051734 | Berkshire and Surrey Pathology Services Lighthouse Laboratory | Wellcome Sanger Institute for the COVID-19 Genomics UK (COG-UK) Consortium | Berkshire and Surrey Pathology Services Lighthouse Laboratory and Alex Alderton, Roberto Amato, Jeffrey Barrett, Sonia Goncalves, Ewan Harrison, David K. Jackson, Ian Johnston, Dominic Kwiatkowski, Cordelia Langford, John Sillitoe on behalf of the Wellcome Sanger Institute COVID-19 Surveillance Team |
| EPI_ISL_3051735 | Lighthouse Lab in Milton Keynes | Wellcome Sanger Institute for the COVID-19 Genomics UK (COG-UK) Consortium | The Lighthouse Lab in Milton Keynes and Alex Alderton, Roberto Amato, Jeffrey Barrett, Sonia Goncalves, Ewan Harrison, David K. Jackson, Ian Johnston, Dominic Kwiatkowski, Cordelia Langford, John Sillitoe on behalf of the Wellcome Sanger Institute COVID-19 Surveillance Team |
| EPI_ISL_3051736 | Lighthouse Lab in Glasgow | Wellcome Sanger Institute for the COVID-19 Genomics UK (COG-UK) Consortium | Harper VanSteenhouse, Yumi Kasai, David Gray, Carol Clugston, Anna Dominiczak and Alex Alderton, Roberto Amato, Jeffrey Barrett, Sonia Goncalves, Ewan Harrison, David K. Jackson, Ian Johnston, Dominic Kwiatkowski, Cordelia Langford, John Sillitoe on behalf of the Wellcome Sanger Institute COVID-19 Surveillance Team |
| EPI_ISL_3051737, EPI_ISL_3051738 | Berkshire and Surrey Pathology Services Lighthouse Laboratory | Wellcome Sanger Institute for the COVID-19 Genomics UK (COG-UK) Consortium | Berkshire and Surrey Pathology Services Lighthouse Laboratory and Alex Alderton, Roberto Amato, Jeffrey Barrett, Sonia Goncalves, Ewan Harrison, David K. Jackson, Ian Johnston, Dominic Kwiatkowski, Cordelia Langford, John Sillitoe on behalf of the Wellcome Sanger Institute COVID-19 Surveillance Team |
| EPI_ISL_3051739, EPI_ISL_3051740, EPI_ISL_3051741 | Lighthouse Lab in Alderley Park | Wellcome Sanger Institute for the COVID-19 Genomics UK (COG-UK) Consortium | Jacquelyn Wynn, Mairead Hyland, The Lighthouse Lab in Alderley Park and Alex Alderton, Roberto Amato, Jeffrey Barrett, Sonia Goncalves, Ewan Harrison, David K. Jackson, Ian Johnston, Dominic Kwiatkowski, Cordelia Langford, John Sillitoe on behalf of the Wellcome Sanger Institute COVID-19 Surveillance Team |
| EPI_ISL_3051742 | Lighthouse Laboratory Plymouth | Wellcome Sanger Institute for the COVID-19 Genomics UK (COG-UK) Consortium | Lighthouse Laboratory Plymouth and Alex Alderton, Roberto Amato, Jeffrey Barrett, Sonia Goncalves, Ewan Harrison, David K. Jackson, Ian Johnston, Dominic Kwiatkowski, Cordelia Langford, John Sillitoe on behalf of the Wellcome Sanger Institute COVID-19 Surveillance Team |
| EPI_ISL_3051743 | Berkshire and Surrey Pathology Services Lighthouse Laboratory | Wellcome Sanger Institute for the COVID-19 Genomics UK (COG-UK) Consortium | Berkshire and Surrey Pathology Services Lighthouse Laboratory and Alex Alderton, Roberto Amato, Jeffrey Barrett, Sonia Goncalves, Ewan Harrison, David K. Jackson, Ian Johnston, Dominic Kwiatkowski, Cordelia Langford, John Sillitoe on behalf of the Wellcome Sanger Institute COVID-19 Surveillance Team |
| EPI_ISL_3051744, EPI_ISL_3051745 | Lighthouse Lab in Alderley Park | Wellcome Sanger Institute for the COVID-19 Genomics UK (COG-UK) Consortium | Jacquelyn Wynn, Mairead Hyland, The Lighthouse Lab in Alderley Park and Alex Alderton, Roberto Amato, Jeffrey Barrett, Sonia Goncalves, Ewan Harrison, David K. Jackson, Ian Johnston, Dominic Kwiatkowski, Cordelia Langford, John Sillitoe on behalf of the Wellcome Sanger Institute COVID-19 Surveillance Team |
| EPI_ISL_3051746 | Lighthouse Lab in Milton Keynes | Wellcome Sanger Institute for the COVID-19 Genomics UK (COG-UK) Consortium | The Lighthouse Lab in Milton Keynes and Alex Alderton, Roberto Amato, Jeffrey Barrett, Sonia Goncalves, Ewan Harrison, David K. Jackson, Ian Johnston, Dominic Kwiatkowski, Cordelia Langford, John Sillitoe on behalf of the Wellcome Sanger Institute COVID-19 Surveillance Team |
| EPI_ISL_3051747 | Berkshire and Surrey Pathology Services Lighthouse Laboratory | Wellcome Sanger Institute for the COVID-19 Genomics UK (COG-UK) Consortium | Berkshire and Surrey Pathology Services Lighthouse Laboratory and Alex Alderton, Roberto Amato, Jeffrey Barrett, Sonia Goncalves, Ewan Harrison, David K. Jackson, Ian Johnston, Dominic Kwiatkowski, Cordelia Langford, John Sillitoe on behalf of the Wellcome Sanger Institute COVID-19 Surveillance Team |
| EPI_ISL_3051748 | Lighthouse Lab in Alderley Park | Wellcome Sanger Institute for the COVID-19 Genomics UK (COG-UK) Consortium | Jacquelyn Wynn, Mairead Hyland, The Lighthouse Lab in Alderley Park and Alex Alderton, Roberto Amato, Jeffrey Barrett, Sonia Goncalves, Ewan Harrison, David K. Jackson, Ian Johnston, Dominic Kwiatkowski, Cordelia Langford, John Sillitoe on behalf of the Wellcome Sanger Institute COVID-19 Surveillance Team |
| EPI_ISL_3051749, EPI_ISL_3051750 | Berkshire and Surrey Pathology Services Lighthouse Laboratory | Wellcome Sanger Institute for the COVID-19 Genomics UK (COG-UK) Consortium | Berkshire and Surrey Pathology Services Lighthouse Laboratory and Alex Alderton, Roberto Amato, Jeffrey Barrett, Sonia Goncalves, Ewan Harrison, David K. Jackson, Ian Johnston, Dominic Kwiatkowski, Cordelia Langford, John Sillitoe on behalf of the Wellcome Sanger Institute COVID-19 Surveillance Team |
| EPI_ISL_3051751, EPI_ISL_3051752 | Lighthouse Lab in Alderley Park | Wellcome Sanger Institute for the COVID-19 Genomics UK (COG-UK) Consortium | Jacquelyn Wynn, Mairead Hyland, The Lighthouse Lab in Alderley Park and Alex Alderton, Roberto Amato, Jeffrey Barrett, Sonia Goncalves, Ewan Harrison, David K. Jackson, Ian Johnston, Dominic Kwiatkowski, Cordelia Langford, John Sillitoe on behalf of the Wellcome Sanger Institute COVID-19 Surveillance Team |
| EPI_ISL_3051753 | Lighthouse Lab in Milton Keynes | Wellcome Sanger Institute for the COVID-19 Genomics UK (COG-UK) Consortium | The Lighthouse Lab in Milton Keynes and Alex Alderton, Roberto Amato, Jeffrey Barrett, Sonia Goncalves, Ewan Harrison, David K. Jackson, Ian Johnston, Dominic Kwiatkowski, Cordelia Langford, John Sillitoe on behalf of the Wellcome Sanger Institute COVID-19 Surveillance Team |
| EPI_ISL_3051754, EPI_ISL_3051755 | Lighthouse Lab in Alderley Park | Wellcome Sanger Institute for the COVID-19 Genomics UK (COG-UK) Consortium | Jacquelyn Wynn, Mairead Hyland, The Lighthouse Lab in Alderley Park and Alex Alderton, Roberto Amato, Jeffrey Barrett, Sonia Goncalves, Ewan Harrison, David K. Jackson, Ian Johnston, Dominic Kwiatkowski, Cordelia Langford, John Sillitoe on behalf of the Wellcome Sanger Institute COVID-19 Surveillance Team |
| EPI_ISL_3051756 | Berkshire and Surrey Pathology Services Lighthouse Laboratory | Wellcome Sanger Institute for the COVID-19 Genomics UK (COG-UK) Consortium | Berkshire and Surrey Pathology Services Lighthouse Laboratory and Alex Alderton, Roberto Amato, Jeffrey Barrett, Sonia Goncalves, Ewan Harrison, David K. Jackson, Ian Johnston, Dominic Kwiatkowski, Cordelia Langford, John Sillitoe on behalf of the Wellcome Sanger Institute COVID-19 Surveillance Team |
| EPI_ISL_3051757 | Lighthouse Laboratory Plymouth | Wellcome Sanger Institute for the COVID-19 Genomics UK (COG-UK) Consortium | Lighthouse Laboratory Plymouth and Alex Alderton, Roberto Amato, Jeffrey Barrett, Sonia Goncalves, Ewan Harrison, David K. Jackson, Ian Johnston, Dominic Kwiatkowski, Cordelia Langford, John Sillitoe on behalf of the Wellcome Sanger Institute COVID-19 Surveillance Team |
| EPI_ISL_3051758 | Lighthouse Lab in Alderley Park | Wellcome Sanger Institute for the COVID-19 Genomics UK (COG-UK) Consortium | Jacquelyn Wynn, Mairead Hyland, The Lighthouse Lab in Alderley Park and Alex Alderton, Roberto Amato, Jeffrey Barrett, Sonia Goncalves, Ewan Harrison, David K. Jackson, Ian Johnston, Dominic Kwiatkowski, Cordelia Langford, John Sillitoe on behalf of the Wellcome Sanger Institute COVID-19 Surveillance Team |
| EPI_ISL_3051759 | Lighthouse Lab in Milton Keynes | Wellcome Sanger Institute for the COVID-19 Genomics UK (COG-UK) Consortium | The Lighthouse Lab in Milton Keynes and Alex Alderton, Roberto Amato, Jeffrey Barrett, Sonia Goncalves, Ewan Harrison, David K. Jackson, Ian Johnston, Dominic Kwiatkowski, Cordelia Langford, John Sillitoe on behalf of the Wellcome Sanger Institute COVID-19 Surveillance Team |
| EPI_ISL_3051760, EPI_ISL_3051761 | Berkshire and Surrey Pathology Services Lighthouse Laboratory | Wellcome Sanger Institute for the COVID-19 Genomics UK (COG-UK) Consortium | Berkshire and Surrey Pathology Services Lighthouse Laboratory and Alex Alderton, Roberto Amato, Jeffrey Barrett, Sonia Goncalves, Ewan Harrison, David K. Jackson, Ian Johnston, Dominic Kwiatkowski, Cordelia Langford, John Sillitoe on behalf of the Wellcome Sanger Institute COVID-19 Surveillance Team |

| | | | |
|---|---|---|---|
| EPI_ISL_3051762 | Lighthouse Lab in Alderley Park | Wellcome Sanger Institute for the COVID-19 Genomics UK (COG-UK) Consortium | Jacquelyn Wynn, Mairead Hyland, The Lighthouse Lab in Alderley Park and Alex Alderton, Roberto Amato, Jeffrey Barrett, Sonia Goncalves, Ewan Harrison, David K. Jackson, Ian Johnston, Dominic Kwiatkowski, Cordelia Langford, John Sillitoe on behalf of the Wellcome Sanger Institute COVID-19 Surveillance Team |
| EPI_ISL_3051763, EPI_ISL_3051764 | Lighthouse Lab in Milton Keynes | Wellcome Sanger Institute for the COVID-19 Genomics UK (COG-UK) Consortium | The Lighthouse Lab in Milton Keynes and Alex Alderton, Roberto Amato, Jeffrey Barrett, Sonia Goncalves, Ewan Harrison, David K. Jackson, Ian Johnston, Dominic Kwiatkowski, Cordelia Langford, John Sillitoe on behalf of the Wellcome Sanger Institute COVID-19 Surveillance Team |
| EPI_ISL_3051765 | Lighthouse Lab in Alderley Park | Wellcome Sanger Institute for the COVID-19 Genomics UK (COG-UK) Consortium | Jacquelyn Wynn, Mairead Hyland, The Lighthouse Lab in Alderley Park and Alex Alderton, Roberto Amato, Jeffrey Barrett, Sonia Goncalves, Ewan Harrison, David K. Jackson, Ian Johnston, Dominic Kwiatkowski, Cordelia Langford, John Sillitoe on behalf of the Wellcome Sanger Institute COVID-19 Surveillance Team |
| EPI_ISL_3051766, EPI_ISL_3051767, EPI_ISL_3051768, EPI_ISL_3051769 | Lighthouse Lab in Glasgow | Wellcome Sanger Institute for the COVID-19 Genomics UK (COG-UK) Consortium | Harper VanSteenhouse, Yumi Kasai, David Gray, Carol Clugston, Anna Dominiczak and Alex Alderton, Roberto Amato, Jeffrey Barrett, Sonia Goncalves, Ewan Harrison, David K. Jackson, Ian Johnston, Dominic Kwiatkowski, Cordelia Langford, John Sillitoe on behalf of the Wellcome Sanger Institute COVID-19 Surveillance Team |
| EPI_ISL_3051770 | Lighthouse Lab in Alderley Park | Wellcome Sanger Institute for the COVID-19 Genomics UK (COG-UK) Consortium | Jacquelyn Wynn, Mairead Hyland, The Lighthouse Lab in Alderley Park and Alex Alderton, Roberto Amato, Jeffrey Barrett, Sonia Goncalves, Ewan Harrison, David K. Jackson, Ian Johnston, Dominic Kwiatkowski, Cordelia Langford, John Sillitoe on behalf of the Wellcome Sanger Institute COVID-19 Surveillance Team |
| EPI_ISL_3051771 | Lighthouse Lab in Glasgow | Wellcome Sanger Institute for the COVID-19 Genomics UK (COG-UK) Consortium | Harper VanSteenhouse, Yumi Kasai, David Gray, Carol Clugston, Anna Dominiczak and Alex Alderton, Roberto Amato, Jeffrey Barrett, Sonia Goncalves, Ewan Harrison, David K. Jackson, Ian Johnston, Dominic Kwiatkowski, Cordelia Langford, John Sillitoe on behalf of the Wellcome Sanger Institute COVID-19 Surveillance Team |
| EPI_ISL_3051772, EPI_ISL_3051773 | Lighthouse Lab in Alderley Park | Wellcome Sanger Institute for the COVID-19 Genomics UK (COG-UK) Consortium | Jacquelyn Wynn, Mairead Hyland, The Lighthouse Lab in Alderley Park and Alex Alderton, Roberto Amato, Jeffrey Barrett, Sonia Goncalves, Ewan Harrison, David K. Jackson, Ian Johnston, Dominic Kwiatkowski, Cordelia Langford, John Sillitoe on behalf of the Wellcome Sanger Institute COVID-19 Surveillance Team |
| EPI_ISL_3051774 | Lighthouse Lab in Glasgow | Wellcome Sanger Institute for the COVID-19 Genomics UK (COG-UK) Consortium | Harper VanSteenhouse, Yumi Kasai, David Gray, Carol Clugston, Anna Dominiczak and Alex Alderton, Roberto Amato, Jeffrey Barrett, Sonia Goncalves, Ewan Harrison, David K. Jackson, Ian Johnston, Dominic Kwiatkowski, Cordelia Langford, John Sillitoe on behalf of the Wellcome Sanger Institute COVID-19 Surveillance Team |
| EPI_ISL_3051775 | Lighthouse Lab in Alderley Park | Wellcome Sanger Institute for the COVID-19 Genomics UK (COG-UK) Consortium | Jacquelyn Wynn, Mairead Hyland, The Lighthouse Lab in Alderley Park and Alex Alderton, Roberto Amato, Jeffrey Barrett, Sonia Goncalves, Ewan Harrison, David K. Jackson, Ian Johnston, Dominic Kwiatkowski, Cordelia Langford, John Sillitoe on behalf of the Wellcome Sanger Institute COVID-19 Surveillance Team |
| EPI_ISL_3051776, EPI_ISL_3051777, EPI_ISL_3051778 | Lighthouse Lab in Glasgow | Wellcome Sanger Institute for the COVID-19 Genomics UK (COG-UK) Consortium | Harper VanSteenhouse, Yumi Kasai, David Gray, Carol Clugston, Anna Dominiczak and Alex Alderton, Roberto Amato, Jeffrey Barrett, Sonia Goncalves, Ewan Harrison, David K. Jackson, Ian Johnston, Dominic Kwiatkowski, Cordelia Langford, John Sillitoe on behalf of the Wellcome Sanger Institute COVID-19 Surveillance Team |
| EPI_ISL_3051779, EPI_ISL_3051780 | Lighthouse Lab in Alderley Park | Wellcome Sanger Institute for the COVID-19 Genomics UK (COG-UK) Consortium | Jacquelyn Wynn, Mairead Hyland, The Lighthouse Lab in Alderley Park and Alex Alderton, Roberto Amato, Jeffrey Barrett, Sonia Goncalves, Ewan Harrison, David K. Jackson, Ian Johnston, Dominic Kwiatkowski, Cordelia Langford, John Sillitoe on behalf of the Wellcome Sanger Institute COVID-19 Surveillance Team |
| EPI_ISL_3051781, EPI_ISL_3051782 | Lighthouse Lab in Glasgow | Wellcome Sanger Institute for the COVID-19 Genomics UK (COG-UK) Consortium | Harper VanSteenhouse, Yumi Kasai, David Gray, Carol Clugston, Anna Dominiczak and Alex Alderton, Roberto Amato, Jeffrey Barrett, Sonia Goncalves, Ewan Harrison, David K. Jackson, Ian Johnston, Dominic Kwiatkowski, Cordelia Langford, John Sillitoe on behalf of the Wellcome Sanger Institute COVID-19 Surveillance Team |
| EPI_ISL_3051783, EPI_ISL_3051784, EPI_ISL_3051785, EPI_ISL_3051786, EPI_ISL_3051787 | Lighthouse Lab in Alderley Park | Wellcome Sanger Institute for the COVID-19 Genomics UK (COG-UK) Consortium | Jacquelyn Wynn, Mairead Hyland, The Lighthouse Lab in Alderley Park and Alex Alderton, Roberto Amato, Jeffrey Barrett, Sonia Goncalves, Ewan Harrison, David K. Jackson, Ian Johnston, Dominic Kwiatkowski, Cordelia Langford, John Sillitoe on behalf of the Wellcome Sanger Institute COVID-19 Surveillance Team |
| EPI_ISL_3051788, EPI_ISL_3051789 | Lighthouse Lab in Glasgow | Wellcome Sanger Institute for the COVID-19 Genomics UK (COG-UK) Consortium | Harper VanSteenhouse, Yumi Kasai, David Gray, Carol Clugston, Anna Dominiczak and Alex Alderton, Roberto Amato, Jeffrey Barrett, Sonia Goncalves, Ewan Harrison, David K. Jackson, Ian Johnston, Dominic Kwiatkowski, Cordelia Langford, John Sillitoe on behalf of the Wellcome Sanger Institute COVID-19 Surveillance Team |
| EPI_ISL_3051790, EPI_ISL_3051791 | Lighthouse Lab in Alderley Park | Wellcome Sanger Institute for the COVID-19 Genomics UK (COG-UK) Consortium | Jacquelyn Wynn, Mairead Hyland, The Lighthouse Lab in Alderley Park and Alex Alderton, Roberto Amato, Jeffrey Barrett, Sonia Goncalves, Ewan Harrison, David K. Jackson, Ian Johnston, Dominic Kwiatkowski, Cordelia Langford, John Sillitoe on behalf of the Wellcome Sanger Institute COVID-19 Surveillance Team |
| EPI_ISL_3051792, EPI_ISL_3051793, EPI_ISL_3051794 | Lighthouse Lab in Glasgow | Wellcome Sanger Institute for the COVID-19 Genomics UK (COG-UK) Consortium | Harper VanSteenhouse, Yumi Kasai, David Gray, Carol Clugston, Anna Dominiczak and Alex Alderton, Roberto Amato, Jeffrey Barrett, Sonia Goncalves, Ewan Harrison, David K. Jackson, Ian Johnston, Dominic Kwiatkowski, Cordelia Langford, John Sillitoe on behalf of the Wellcome Sanger Institute COVID-19 Surveillance Team |
| EPI_ISL_3051795, EPI_ISL_3051796 | Lighthouse Lab in Alderley Park | Wellcome Sanger Institute for the COVID-19 Genomics UK (COG-UK) Consortium | Jacquelyn Wynn, Mairead Hyland, The Lighthouse Lab in Alderley Park and Alex Alderton, Roberto Amato, Jeffrey Barrett, Sonia Goncalves, Ewan Harrison, David K. Jackson, Ian Johnston, Dominic Kwiatkowski, Cordelia Langford, John Sillitoe on behalf of the Wellcome Sanger Institute COVID-19 Surveillance Team |
| EPI_ISL_3051797 | Lighthouse Lab in Glasgow | Wellcome Sanger Institute for the COVID-19 Genomics UK (COG-UK) Consortium | Harper VanSteenhouse, Yumi Kasai, David Gray, Carol Clugston, Anna Dominiczak and Alex Alderton, Roberto Amato, Jeffrey Barrett, Sonia Goncalves, Ewan Harrison, David K. Jackson, Ian Johnston, Dominic Kwiatkowski, Cordelia Langford, John Sillitoe on behalf of the Wellcome Sanger Institute COVID-19 Surveillance Team |
| EPI_ISL_3051798 | Lighthouse Lab in Alderley Park | Wellcome Sanger Institute for the COVID-19 Genomics UK (COG-UK) Consortium | Jacquelyn Wynn, Mairead Hyland, The Lighthouse Lab in Alderley Park and Alex Alderton, Roberto Amato, Jeffrey Barrett, Sonia Goncalves, Ewan Harrison, David K. Jackson, Ian Johnston, Dominic Kwiatkowski, Cordelia Langford, John Sillitoe on behalf of the Wellcome Sanger Institute COVID-19 Surveillance Team |
| EPI_ISL_3051799, EPI_ISL_3051800, EPI_ISL_3051801 | Lighthouse Lab in Glasgow | Wellcome Sanger Institute for the COVID-19 Genomics UK (COG-UK) Consortium | Harper VanSteenhouse, Yumi Kasai, David Gray, Carol Clugston, Anna Dominiczak and Alex Alderton, Roberto Amato, Jeffrey Barrett, Sonia Goncalves, Ewan Harrison, David K. Jackson, Ian Johnston, Dominic Kwiatkowski, Cordelia Langford, John Sillitoe on behalf of the Wellcome Sanger Institute COVID-19 Surveillance Team |
| EPI_ISL_3051802, EPI_ISL_3051803 | Lighthouse Lab in Alderley Park | Wellcome Sanger Institute for the COVID-19 Genomics UK (COG-UK) Consortium | Jacquelyn Wynn, Mairead Hyland, The Lighthouse Lab in Alderley Park and Alex Alderton, Roberto Amato, Jeffrey Barrett, Sonia Goncalves, Ewan Harrison, David K. Jackson, Ian Johnston, Dominic Kwiatkowski, Cordelia Langford, John Sillitoe on behalf of the Wellcome Sanger Institute COVID-19 Surveillance Team |
| EPI_ISL_3051804, EPI_ISL_3051805, EPI_ISL_3051806, EPI_ISL_3051807, EPI_ISL_3051808 | Lighthouse Lab in Glasgow | Wellcome Sanger Institute for the COVID-19 Genomics UK (COG-UK) Consortium | Harper VanSteenhouse, Yumi Kasai, David Gray, Carol Clugston, Anna Dominiczak and Alex Alderton, Roberto Amato, Jeffrey Barrett, Sonia Goncalves, Ewan Harrison, David K. Jackson, Ian Johnston, Dominic Kwiatkowski, Cordelia Langford, John Sillitoe on behalf of the Wellcome Sanger Institute COVID-19 Surveillance Team |
| EPI_ISL_3051809 | Lighthouse Lab in Alderley Park | Wellcome Sanger Institute for the COVID-19 Genomics UK (COG-UK) Consortium | Jacquelyn Wynn, Mairead Hyland, The Lighthouse Lab in Alderley Park and Alex Alderton, Roberto Amato, Jeffrey Barrett, Sonia Goncalves, Ewan Harrison, David K. Jackson, Ian Johnston, Dominic Kwiatkowski, Cordelia Langford, John Sillitoe on behalf of the Wellcome Sanger Institute COVID-19 Surveillance Team |
| EPI_ISL_3051810, EPI_ISL_3051811 | Lighthouse Lab in Glasgow | Wellcome Sanger Institute for the COVID-19 Genomics UK (COG-UK) Consortium | Harper VanSteenhouse, Yumi Kasai, David Gray, Carol Clugston, Anna Dominiczak and Alex Alderton, Roberto Amato, Jeffrey Barrett, Sonia Goncalves, Ewan Harrison, David K. Jackson, Ian Johnston, Dominic Kwiatkowski, Cordelia Langford, John Sillitoe on behalf of the Wellcome Sanger Institute COVID-19 Surveillance Team |
| EPI_ISL_3051812 | Lighthouse Lab in Alderley Park | Wellcome Sanger Institute for the COVID-19 Genomics UK (COG-UK) Consortium | Jacquelyn Wynn, Mairead Hyland, The Lighthouse Lab in Alderley Park and Alex Alderton, Roberto Amato, Jeffrey Barrett, Sonia Goncalves, Ewan Harrison, David K. Jackson, Ian Johnston, Dominic Kwiatkowski, Cordelia Langford, John Sillitoe on behalf of the Wellcome Sanger Institute COVID-19 Surveillance Team |
| EPI_ISL_3051813 | Lighthouse Lab in Glasgow | Wellcome Sanger Institute for the COVID-19 Genomics UK (COG-UK) Consortium | Harper VanSteenhouse, Yumi Kasai, David Gray, Carol Clugston, Anna Dominiczak and Alex Alderton, Roberto Amato, Jeffrey Barrett, Sonia Goncalves, Ewan Harrison, David K. Jackson, Ian Johnston, Dominic Kwiatkowski, Cordelia Langford, John Sillitoe on behalf of the Wellcome Sanger Institute COVID-19 Surveillance Team |
| EPI_ISL_3051814, EPI_ISL_3051815 | Lighthouse Lab in Alderley Park | Wellcome Sanger Institute for the COVID-19 Genomics UK (COG-UK) Consortium | Jacquelyn Wynn, Mairead Hyland, The Lighthouse Lab in Alderley Park and Alex Alderton, Roberto Amato, Jeffrey Barrett, Sonia Goncalves, Ewan Harrison, David K. Jackson, Ian Johnston, Dominic Kwiatkowski, Cordelia Langford, John Sillitoe on behalf of the Wellcome Sanger Institute COVID-19 Surveillance Team |
| EPI_ISL_3051816 | Lighthouse Lab in Glasgow | Wellcome Sanger Institute for the COVID-19 Genomics UK (COG-UK) Consortium | Harper VanSteenhouse, Yumi Kasai, David Gray, Carol Clugston, Anna Dominiczak and Alex Alderton, Roberto Amato, Jeffrey Barrett, Sonia Goncalves, Ewan Harrison, David K. Jackson, Ian Johnston, Dominic Kwiatkowski, Cordelia Langford, John Sillitoe on behalf of the Wellcome Sanger Institute COVID-19 Surveillance Team |
| EPI_ISL_3051817, EPI_ISL_3051818 | Lighthouse Lab in Alderley Park | Wellcome Sanger Institute for the COVID-19 Genomics UK (COG-UK) Consortium | Jacquelyn Wynn, Mairead Hyland, The Lighthouse Lab in Alderley Park and Alex Alderton, Roberto Amato, Jeffrey Barrett, Sonia Goncalves, Ewan Harrison, David K. Jackson, Ian Johnston, Dominic Kwiatkowski, Cordelia Langford, John Sillitoe on behalf of the Wellcome Sanger Institute COVID-19 Surveillance Team |
| EPI_ISL_3051819 | Lighthouse Lab in Glasgow | Wellcome Sanger Institute for the COVID-19 Genomics UK (COG-UK) Consortium | Harper VanSteenhouse, Yumi Kasai, David Gray, Carol Clugston, Anna Dominiczak and Alex Alderton, Roberto Amato, Jeffrey Barrett, Sonia Goncalves, Ewan Harrison, David K. Jackson, Ian Johnston, Dominic Kwiatkowski, Cordelia Langford, John Sillitoe on behalf of the Wellcome Sanger Institute COVID-19 Surveillance Team |
| EPI_ISL_3051820 | Lighthouse Lab in Alderley Park | Wellcome Sanger Institute for the COVID-19 Genomics UK (COG-UK) Consortium | Jacquelyn Wynn, Mairead Hyland, The Lighthouse Lab in Alderley Park and Alex Alderton, Roberto Amato, Jeffrey Barrett, Sonia Goncalves, Ewan Harrison, David K. Jackson, Ian Johnston, Dominic Kwiatkowski, Cordelia Langford, John Sillitoe on behalf of the Wellcome Sanger Institute COVID-19 Surveillance Team |
| EPI_ISL_3051821, EPI_ISL_3051822, EPI_ISL_3051823, EPI_ISL_3051824, EPI_ISL_3051825, EPI_ISL_3051826 | Lighthouse Lab in Glasgow | Wellcome Sanger Institute for the COVID-19 Genomics UK (COG-UK) Consortium | Harper VanSteenhouse, Yumi Kasai, David Gray, Carol Clugston, Anna Dominiczak and Alex Alderton, Roberto Amato, Jeffrey Barrett, Sonia Goncalves, Ewan Harrison, David K. Jackson, Ian Johnston, Dominic Kwiatkowski, Cordelia Langford, John Sillitoe on behalf of the Wellcome Sanger Institute COVID-19 Surveillance Team |
| EPI_ISL_3051827, EPI_ISL_3051828 | Lighthouse Lab in Alderley Park | Wellcome Sanger Institute for the COVID-19 Genomics UK (COG-UK) Consortium | Jacquelyn Wynn, Mairead Hyland, The Lighthouse Lab in Alderley Park and Alex Alderton, Roberto Amato, Jeffrey Barrett, Sonia Goncalves, Ewan Harrison, David K. Jackson, Ian Johnston, Dominic Kwiatkowski, Cordelia Langford, John Sillitoe on behalf of the Wellcome Sanger Institute COVID-19 Surveillance Team |
| EPI_ISL_3051829, EPI_ISL_3051830, EPI_ISL_3051831, EPI_ISL_3051832 | Lighthouse Lab in Glasgow | Wellcome Sanger Institute for the COVID-19 Genomics UK (COG-UK) Consortium | Harper VanSteenhouse, Yumi Kasai, David Gray, Carol Clugston, Anna Dominiczak and Alex Alderton, Roberto Amato, Jeffrey Barrett, Sonia Goncalves, Ewan Harrison, David K. Jackson, Ian Johnston, Dominic Kwiatkowski, Cordelia Langford, John Sillitoe on behalf of the Wellcome Sanger Institute COVID-19 Surveillance Team |
| EPI_ISL_3051833 | Lighthouse Lab in Alderley Park | Wellcome Sanger Institute for the COVID-19 Genomics UK (COG-UK) Consortium | Jacquelyn Wynn, Mairead Hyland, The Lighthouse Lab in Alderley Park and Alex Alderton, Roberto Amato, Jeffrey Barrett, Sonia Goncalves, Ewan Harrison, David K. Jackson, Ian Johnston, Dominic Kwiatkowski, Cordelia Langford, John Sillitoe on behalf of the Wellcome Sanger Institute COVID-19 Surveillance Team |
| EPI_ISL_3051834, EPI_ISL_3051835, EPI_ISL_3051836 | Lighthouse Lab in Glasgow | Wellcome Sanger Institute for the COVID-19 Genomics UK (COG-UK) Consortium | Harper VanSteenhouse, Yumi Kasai, David Gray, Carol Clugston, Anna Dominiczak and Alex Alderton, Roberto Amato, Jeffrey Barrett, Sonia Goncalves, Ewan Harrison, David K. Jackson, Ian Johnston, Dominic Kwiatkowski, Cordelia Langford, John Sillitoe on behalf of the Wellcome Sanger Institute COVID-19 Surveillance Team |
| EPI_ISL_3051837, EPI_ISL_3051838, EPI_ISL_3051839, | Lighthouse Lab in Alderley Park | Wellcome Sanger Institute for the COVID-19 Genomics UK (COG-UK) Consortium | Jacquelyn Wynn, Mairead Hyland, The Lighthouse Lab in Alderley Park and Alex Alderton, Roberto Amato, Jeffrey Barrett, Sonia Goncalves, Ewan Harrison, David K. Jackson, Ian Johnston, Dominic Kwiatkowski, Cordelia Langford, John Sillitoe on behalf of the Wellcome Sanger Institute COVID-19 Surveillance Team |

| | | | |
|---|---|---|---|
| EPI_ISL_3051840, EPI_ISL_3051841, EPI_ISL_3051842, EPI_ISL_3051843, EPI_ISL_3051844 | | | |
| EPI_ISL_3051845 | Lighthouse Lab in Glasgow | Wellcome Sanger Institute for the COVID-19 Genomics UK (COG-UK) Consortium | Harper VanSteenhouse, Yumi Kasai, David Gray, Carol Clugston, Anna Dominiczak and Alex Alderton, Roberto Amato, Jeffrey Barrett, Sonia Goncalves, Ewan Harrison, David K. Jackson, Ian Johnston, Dominic Kwiatkowski, Cordelia Langford, John Sillitoe on behalf of the Wellcome Sanger Institute COVID-19 Surveillance Team |
| EPI_ISL_3051846 | Lighthouse Lab in Alderley Park | Wellcome Sanger Institute for the COVID-19 Genomics UK (COG-UK) Consortium | Jacquelyn Wynn, Mairead Hyland, The Lighthouse Lab in Alderley Park and Alex Alderton, Roberto Amato, Jeffrey Barrett, Sonia Goncalves, Ewan Harrison, David K. Jackson, Ian Johnston, Dominic Kwiatkowski, Cordelia Langford, John Sillitoe on behalf of the Wellcome Sanger Institute COVID-19 Surveillance Team |
| EPI_ISL_3051847, EPI_ISL_3051848 | Lighthouse Lab in Glasgow | Wellcome Sanger Institute for the COVID-19 Genomics UK (COG-UK) Consortium | Harper VanSteenhouse, Yumi Kasai, David Gray, Carol Clugston, Anna Dominiczak and Alex Alderton, Roberto Amato, Jeffrey Barrett, Sonia Goncalves, Ewan Harrison, David K. Jackson, Ian Johnston, Dominic Kwiatkowski, Cordelia Langford, John Sillitoe on behalf of the Wellcome Sanger Institute COVID-19 Surveillance Team |
| EPI_ISL_3051849 | Lighthouse Lab in Alderley Park | Wellcome Sanger Institute for the COVID-19 Genomics UK (COG-UK) Consortium | Jacquelyn Wynn, Mairead Hyland, The Lighthouse Lab in Alderley Park and Alex Alderton, Roberto Amato, Jeffrey Barrett, Sonia Goncalves, Ewan Harrison, David K. Jackson, Ian Johnston, Dominic Kwiatkowski, Cordelia Langford, John Sillitoe on behalf of the Wellcome Sanger Institute COVID-19 Surveillance Team |
| EPI_ISL_3051850, EPI_ISL_3051851, EPI_ISL_3051852 | Lighthouse Lab in Glasgow | Wellcome Sanger Institute for the COVID-19 Genomics UK (COG-UK) Consortium | Harper VanSteenhouse, Yumi Kasai, David Gray, Carol Clugston, Anna Dominiczak and Alex Alderton, Roberto Amato, Jeffrey Barrett, Sonia Goncalves, Ewan Harrison, David K. Jackson, Ian Johnston, Dominic Kwiatkowski, Cordelia Langford, John Sillitoe on behalf of the Wellcome Sanger Institute COVID-19 Surveillance Team |
| EPI_ISL_3051853 | Lighthouse Lab in Alderley Park | Wellcome Sanger Institute for the COVID-19 Genomics UK (COG-UK) Consortium | Jacquelyn Wynn, Mairead Hyland, The Lighthouse Lab in Alderley Park and Alex Alderton, Roberto Amato, Jeffrey Barrett, Sonia Goncalves, Ewan Harrison, David K. Jackson, Ian Johnston, Dominic Kwiatkowski, Cordelia Langford, John Sillitoe on behalf of the Wellcome Sanger Institute COVID-19 Surveillance Team |
| EPI_ISL_3051854 | Lighthouse Lab in Glasgow | Wellcome Sanger Institute for the COVID-19 Genomics UK (COG-UK) Consortium | Harper VanSteenhouse, Yumi Kasai, David Gray, Carol Clugston, Anna Dominiczak and Alex Alderton, Roberto Amato, Jeffrey Barrett, Sonia Goncalves, Ewan Harrison, David K. Jackson, Ian Johnston, Dominic Kwiatkowski, Cordelia Langford, John Sillitoe on behalf of the Wellcome Sanger Institute COVID-19 Surveillance Team |
| EPI_ISL_3051855, EPI_ISL_3051856, EPI_ISL_3051857, EPI_ISL_3051858 | Lighthouse Lab in Alderley Park | Wellcome Sanger Institute for the COVID-19 Genomics UK (COG-UK) Consortium | Jacquelyn Wynn, Mairead Hyland, The Lighthouse Lab in Alderley Park and Alex Alderton, Roberto Amato, Jeffrey Barrett, Sonia Goncalves, Ewan Harrison, David K. Jackson, Ian Johnston, Dominic Kwiatkowski, Cordelia Langford, John Sillitoe on behalf of the Wellcome Sanger Institute COVID-19 Surveillance Team |
| EPI_ISL_3051859 | Lighthouse Lab in Glasgow | Wellcome Sanger Institute for the COVID-19 Genomics UK (COG-UK) Consortium | Harper VanSteenhouse, Yumi Kasai, David Gray, Carol Clugston, Anna Dominiczak and Alex Alderton, Roberto Amato, Jeffrey Barrett, Sonia Goncalves, Ewan Harrison, David K. Jackson, Ian Johnston, Dominic Kwiatkowski, Cordelia Langford, John Sillitoe on behalf of the Wellcome Sanger Institute COVID-19 Surveillance Team |
| EPI_ISL_3051860 | Lighthouse Lab in Alderley Park | Wellcome Sanger Institute for the COVID-19 Genomics UK (COG-UK) Consortium | Jacquelyn Wynn, Mairead Hyland, The Lighthouse Lab in Alderley Park and Alex Alderton, Roberto Amato, Jeffrey Barrett, Sonia Goncalves, Ewan Harrison, David K. Jackson, Ian Johnston, Dominic Kwiatkowski, Cordelia Langford, John Sillitoe on behalf of the Wellcome Sanger Institute COVID-19 Surveillance Team |
| EPI_ISL_3051861, EPI_ISL_3051862 | Lighthouse Lab in Glasgow | Wellcome Sanger Institute for the COVID-19 Genomics UK (COG-UK) Consortium | Harper VanSteenhouse, Yumi Kasai, David Gray, Carol Clugston, Anna Dominiczak and Alex Alderton, Roberto Amato, Jeffrey Barrett, Sonia Goncalves, Ewan Harrison, David K. Jackson, Ian Johnston, Dominic Kwiatkowski, Cordelia Langford, John Sillitoe on behalf of the Wellcome Sanger Institute COVID-19 Surveillance Team |
| EPI_ISL_3051863 | Lighthouse Lab in Alderley Park | Wellcome Sanger Institute for the COVID-19 Genomics UK (COG-UK) Consortium | Jacquelyn Wynn, Mairead Hyland, The Lighthouse Lab in Alderley Park and Alex Alderton, Roberto Amato, Jeffrey Barrett, Sonia Goncalves, Ewan Harrison, David K. Jackson, Ian Johnston, Dominic Kwiatkowski, Cordelia Langford, John Sillitoe on behalf of the Wellcome Sanger Institute COVID-19 Surveillance Team |
| EPI_ISL_3051864 | Lighthouse Lab in Glasgow | Wellcome Sanger Institute for the COVID-19 Genomics UK (COG-UK) Consortium | Harper VanSteenhouse, Yumi Kasai, David Gray, Carol Clugston, Anna Dominiczak and Alex Alderton, Roberto Amato, Jeffrey Barrett, Sonia Goncalves, Ewan Harrison, David K. Jackson, Ian Johnston, Dominic Kwiatkowski, Cordelia Langford, John Sillitoe on behalf of the Wellcome Sanger Institute COVID-19 Surveillance Team |
| EPI_ISL_3051865, EPI_ISL_3051866 | Lighthouse Lab in Alderley Park | Wellcome Sanger Institute for the COVID-19 Genomics UK (COG-UK) Consortium | Jacquelyn Wynn, Mairead Hyland, The Lighthouse Lab in Alderley Park and Alex Alderton, Roberto Amato, Jeffrey Barrett, Sonia Goncalves, Ewan Harrison, David K. Jackson, Ian Johnston, Dominic Kwiatkowski, Cordelia Langford, John Sillitoe on behalf of the Wellcome Sanger Institute COVID-19 Surveillance Team |
| EPI_ISL_3051867 | Lighthouse Lab in Glasgow | Wellcome Sanger Institute for the COVID-19 Genomics UK (COG-UK) Consortium | Harper VanSteenhouse, Yumi Kasai, David Gray, Carol Clugston, Anna Dominiczak and Alex Alderton, Roberto Amato, Jeffrey Barrett, Sonia Goncalves, Ewan Harrison, David K. Jackson, Ian Johnston, Dominic Kwiatkowski, Cordelia Langford, John Sillitoe on behalf of the Wellcome Sanger Institute COVID-19 Surveillance Team |
| EPI_ISL_3051868, EPI_ISL_3051869, EPI_ISL_3051870, EPI_ISL_3051871, EPI_ISL_3051872, EPI_ISL_3051873, EPI_ISL_3051874 | Lighthouse Lab in Alderley Park | Wellcome Sanger Institute for the COVID-19 Genomics UK (COG-UK) Consortium | Jacquelyn Wynn, Mairead Hyland, The Lighthouse Lab in Alderley Park and Alex Alderton, Roberto Amato, Jeffrey Barrett, Sonia Goncalves, Ewan Harrison, David K. Jackson, Ian Johnston, Dominic Kwiatkowski, Cordelia Langford, John Sillitoe on behalf of the Wellcome Sanger Institute COVID-19 Surveillance Team |
| EPI_ISL_3051875, EPI_ISL_3051876 | Lighthouse Lab in Glasgow | Wellcome Sanger Institute for the COVID-19 Genomics UK (COG-UK) Consortium | Harper VanSteenhouse, Yumi Kasai, David Gray, Carol Clugston, Anna Dominiczak and Alex Alderton, Roberto Amato, Jeffrey Barrett, Sonia Goncalves, Ewan Harrison, David K. Jackson, Ian Johnston, Dominic Kwiatkowski, Cordelia Langford, John Sillitoe on behalf of the Wellcome Sanger Institute COVID-19 Surveillance Team |
| EPI_ISL_3051877 | Lighthouse Lab in Alderley Park | Wellcome Sanger Institute for the COVID-19 Genomics UK (COG-UK) Consortium | Jacquelyn Wynn, Mairead Hyland, The Lighthouse Lab in Alderley Park and Alex Alderton, Roberto Amato, Jeffrey Barrett, Sonia Goncalves, Ewan Harrison, David K. Jackson, Ian Johnston, Dominic Kwiatkowski, Cordelia Langford, John Sillitoe on behalf of the Wellcome Sanger Institute COVID-19 Surveillance Team |
| EPI_ISL_3051878 | Lighthouse Lab in Glasgow | Wellcome Sanger Institute for the COVID-19 Genomics UK (COG-UK) Consortium | Harper VanSteenhouse, Yumi Kasai, David Gray, Carol Clugston, Anna Dominiczak and Alex Alderton, Roberto Amato, Jeffrey Barrett, Sonia Goncalves, Ewan Harrison, David K. Jackson, Ian Johnston, Dominic Kwiatkowski, Cordelia Langford, John Sillitoe on behalf of the Wellcome Sanger Institute COVID-19 Surveillance Team |
| EPI_ISL_3051879, EPI_ISL_3051880, EPI_ISL_3051881 | Lighthouse Lab in Alderley Park | Wellcome Sanger Institute for the COVID-19 Genomics UK (COG-UK) Consortium | Jacquelyn Wynn, Mairead Hyland, The Lighthouse Lab in Alderley Park and Alex Alderton, Roberto Amato, Jeffrey Barrett, Sonia Goncalves, Ewan Harrison, David K. Jackson, Ian Johnston, Dominic Kwiatkowski, Cordelia Langford, John Sillitoe on behalf of the Wellcome Sanger Institute COVID-19 Surveillance Team |
| EPI_ISL_3051882 | Lighthouse Lab in Glasgow | Wellcome Sanger Institute for the COVID-19 Genomics UK (COG-UK) Consortium | Harper VanSteenhouse, Yumi Kasai, David Gray, Carol Clugston, Anna Dominiczak and Alex Alderton, Roberto Amato, Jeffrey Barrett, Sonia Goncalves, Ewan Harrison, David K. Jackson, Ian Johnston, Dominic Kwiatkowski, Cordelia Langford, John Sillitoe on behalf of the Wellcome Sanger Institute COVID-19 Surveillance Team |
| EPI_ISL_3051883, EPI_ISL_3051884 | Lighthouse Lab in Alderley Park | Wellcome Sanger Institute for the COVID-19 Genomics UK (COG-UK) Consortium | Jacquelyn Wynn, Mairead Hyland, The Lighthouse Lab in Alderley Park and Alex Alderton, Roberto Amato, Jeffrey Barrett, Sonia Goncalves, Ewan Harrison, David K. Jackson, Ian Johnston, Dominic Kwiatkowski, Cordelia Langford, John Sillitoe on behalf of the Wellcome Sanger Institute COVID-19 Surveillance Team |
| EPI_ISL_3051885, EPI_ISL_3051886 | Lighthouse Lab in Glasgow | Wellcome Sanger Institute for the COVID-19 Genomics UK (COG-UK) Consortium | Harper VanSteenhouse, Yumi Kasai, David Gray, Carol Clugston, Anna Dominiczak and Alex Alderton, Roberto Amato, Jeffrey Barrett, Sonia Goncalves, Ewan Harrison, David K. Jackson, Ian Johnston, Dominic Kwiatkowski, Cordelia Langford, John Sillitoe on behalf of the Wellcome Sanger Institute COVID-19 Surveillance Team |
| EPI_ISL_3051887, EPI_ISL_3051888, EPI_ISL_3051889 | Lighthouse Lab in Alderley Park | Wellcome Sanger Institute for the COVID-19 Genomics UK (COG-UK) Consortium | Jacquelyn Wynn, Mairead Hyland, The Lighthouse Lab in Alderley Park and Alex Alderton, Roberto Amato, Jeffrey Barrett, Sonia Goncalves, Ewan Harrison, David K. Jackson, Ian Johnston, Dominic Kwiatkowski, Cordelia Langford, John Sillitoe on behalf of the Wellcome Sanger Institute COVID-19 Surveillance Team |
| EPI_ISL_3051890 | Lighthouse Lab in Glasgow | Wellcome Sanger Institute for the COVID-19 Genomics UK (COG-UK) Consortium | Harper VanSteenhouse, Yumi Kasai, David Gray, Carol Clugston, Anna Dominiczak and Alex Alderton, Roberto Amato, Jeffrey Barrett, Sonia Goncalves, Ewan Harrison, David K. Jackson, Ian Johnston, Dominic Kwiatkowski, Cordelia Langford, John Sillitoe on behalf of the Wellcome Sanger Institute COVID-19 Surveillance Team |
| EPI_ISL_3051891, EPI_ISL_3051892, EPI_ISL_3051893 | Lighthouse Lab in Alderley Park | Wellcome Sanger Institute for the COVID-19 Genomics UK (COG-UK) Consortium | Jacquelyn Wynn, Mairead Hyland, The Lighthouse Lab in Alderley Park and Alex Alderton, Roberto Amato, Jeffrey Barrett, Sonia Goncalves, Ewan Harrison, David K. Jackson, Ian Johnston, Dominic Kwiatkowski, Cordelia Langford, John Sillitoe on behalf of the Wellcome Sanger Institute COVID-19 Surveillance Team |
| EPI_ISL_3051894 | Lighthouse Lab in Glasgow | Wellcome Sanger Institute for the COVID-19 Genomics UK (COG-UK) Consortium | Harper VanSteenhouse, Yumi Kasai, David Gray, Carol Clugston, Anna Dominiczak and Alex Alderton, Roberto Amato, Jeffrey Barrett, Sonia Goncalves, Ewan Harrison, David K. Jackson, Ian Johnston, Dominic Kwiatkowski, Cordelia Langford, John Sillitoe on behalf of the Wellcome Sanger Institute COVID-19 Surveillance Team |
| EPI_ISL_3051895, EPI_ISL_3051896, EPI_ISL_3051897, EPI_ISL_3051898, EPI_ISL_3051899, EPI_ISL_3051900 | Lighthouse Lab in Alderley Park | Wellcome Sanger Institute for the COVID-19 Genomics UK (COG-UK) Consortium | Jacquelyn Wynn, Mairead Hyland, The Lighthouse Lab in Alderley Park and Alex Alderton, Roberto Amato, Jeffrey Barrett, Sonia Goncalves, Ewan Harrison, David K. Jackson, Ian Johnston, Dominic Kwiatkowski, Cordelia Langford, John Sillitoe on behalf of the Wellcome Sanger Institute COVID-19 Surveillance Team |
| EPI_ISL_3051901, EPI_ISL_3051902, EPI_ISL_3051903, EPI_ISL_3051904, EPI_ISL_3051905, EPI_ISL_3051906, EPI_ISL_3051907 | Lighthouse Lab in Glasgow | Wellcome Sanger Institute for the COVID-19 Genomics UK (COG-UK) Consortium | Harper VanSteenhouse, Yumi Kasai, David Gray, Carol Clugston, Anna Dominiczak and Alex Alderton, Roberto Amato, Jeffrey Barrett, Sonia Goncalves, Ewan Harrison, David K. Jackson, Ian Johnston, Dominic Kwiatkowski, Cordelia Langford, John Sillitoe on behalf of the Wellcome Sanger Institute COVID-19 Surveillance Team |
| EPI_ISL_3051908 | Lighthouse Lab in Alderley Park | Wellcome Sanger Institute for the COVID-19 Genomics UK (COG-UK) Consortium | Jacquelyn Wynn, Mairead Hyland, The Lighthouse Lab in Alderley Park and Alex Alderton, Roberto Amato, Jeffrey Barrett, Sonia Goncalves, Ewan Harrison, David K. Jackson, Ian Johnston, Dominic Kwiatkowski, Cordelia Langford, John Sillitoe on behalf of the Wellcome Sanger Institute COVID-19 Surveillance Team |
| EPI_ISL_3051909 | Lighthouse Lab in Glasgow | Wellcome Sanger Institute for the COVID-19 Genomics UK (COG-UK) Consortium | Harper VanSteenhouse, Yumi Kasai, David Gray, Carol Clugston, Anna Dominiczak and Alex Alderton, Roberto Amato, Jeffrey Barrett, Sonia Goncalves, Ewan Harrison, David K. Jackson, Ian Johnston, Dominic Kwiatkowski, Cordelia Langford, John Sillitoe on behalf of the Wellcome Sanger Institute COVID-19 Surveillance Team |
| EPI_ISL_3051910, EPI_ISL_3051911, EPI_ISL_3051912 | Lighthouse Lab in Alderley Park | Wellcome Sanger Institute for the COVID-19 Genomics UK (COG-UK) Consortium | Jacquelyn Wynn, Mairead Hyland, The Lighthouse Lab in Alderley Park and Alex Alderton, Roberto Amato, Jeffrey Barrett, Sonia Goncalves, Ewan Harrison, David K. Jackson, Ian Johnston, Dominic Kwiatkowski, Cordelia Langford, John Sillitoe on behalf of the Wellcome Sanger Institute COVID-19 Surveillance Team |
| EPI_ISL_3051913 | Lighthouse Lab in Glasgow | Wellcome Sanger Institute for the COVID-19 Genomics UK (COG-UK) Consortium | Harper VanSteenhouse, Yumi Kasai, David Gray, Carol Clugston, Anna Dominiczak and Alex Alderton, Roberto Amato, Jeffrey Barrett, Sonia Goncalves, Ewan Harrison, David K. Jackson, Ian Johnston, Dominic Kwiatkowski, Cordelia Langford, John Sillitoe on behalf of the Wellcome Sanger Institute COVID-19 Surveillance Team |
| EPI_ISL_3051914, EPI_ISL_3051915 | Lighthouse Lab in Alderley Park | Wellcome Sanger Institute for the COVID-19 Genomics UK (COG-UK) Consortium | Jacquelyn Wynn, Mairead Hyland, The Lighthouse Lab in Alderley Park and Alex Alderton, Roberto Amato, Jeffrey Barrett, Sonia Goncalves, Ewan Harrison, David K. Jackson, Ian Johnston, Dominic Kwiatkowski, Cordelia Langford, John Sillitoe on behalf of the Wellcome Sanger Institute COVID-19 Surveillance Team |
| EPI_ISL_3051916, | Lighthouse Lab in Glasgow | Wellcome Sanger Institute for the COVID-19 Genomics UK (COG-UK) | Harper VanSteenhouse, Yumi Kasai, David Gray, Carol Clugston, Anna Dominiczak and Alex Alderton, Roberto Amato, Jeffrey Barrett, Sonia Goncalves, Ewan Harrison, David K. Jackson, Ian Johnston, Dominic Kwiatkowski, Cordelia Langford, |

| | | | |
|---|---|---|---|
| EPI_ISL_3051917, EPI_ISL_3051918, EPI_ISL_3051919 | | Consortium | John Sillitoe on behalf of the Wellcome Sanger Institute COVID-19 Surveillance Team |
| EPI_ISL_3051920 | Lighthouse Lab in Alderley Park | Wellcome Sanger Institute for the COVID-19 Genomics UK (COG-UK) Consortium | Jacquelyn Wynn, Mairead Hyland, The Lighthouse Lab in Alderley Park and Alex Alderton, Roberto Amato, Jeffrey Barrett, Sonia Goncalves, Ewan Harrison, David K. Jackson, Ian Johnston, Dominic Kwiatkowski, Cordelia Langford, John Sillitoe on behalf of the Wellcome Sanger Institute COVID-19 Surveillance Team |
| EPI_ISL_3051921, EPI_ISL_3051922, EPI_ISL_3051923 | Lighthouse Lab in Glasgow | Wellcome Sanger Institute for the COVID-19 Genomics UK (COG-UK) Consortium | Harper VanSteenhouse, Yumi Kasai, David Gray, Carol Clugston, Anna Dominiczak and Alex Alderton, Roberto Amato, Jeffrey Barrett, Sonia Goncalves, Ewan Harrison, David K. Jackson, Ian Johnston, Dominic Kwiatkowski, Cordelia Langford, John Sillitoe on behalf of the Wellcome Sanger Institute COVID-19 Surveillance Team |
| EPI_ISL_3051924 | Lighthouse Lab in Alderley Park | Wellcome Sanger Institute for the COVID-19 Genomics UK (COG-UK) Consortium | Jacquelyn Wynn, Mairead Hyland, The Lighthouse Lab in Alderley Park and Alex Alderton, Roberto Amato, Jeffrey Barrett, Sonia Goncalves, Ewan Harrison, David K. Jackson, Ian Johnston, Dominic Kwiatkowski, Cordelia Langford, John Sillitoe on behalf of the Wellcome Sanger Institute COVID-19 Surveillance Team |
| EPI_ISL_3051925, EPI_ISL_3051926 | Lighthouse Lab in Glasgow | Wellcome Sanger Institute for the COVID-19 Genomics UK (COG-UK) Consortium | Harper VanSteenhouse, Yumi Kasai, David Gray, Carol Clugston, Anna Dominiczak and Alex Alderton, Roberto Amato, Jeffrey Barrett, Sonia Goncalves, Ewan Harrison, David K. Jackson, Ian Johnston, Dominic Kwiatkowski, Cordelia Langford, John Sillitoe on behalf of the Wellcome Sanger Institute COVID-19 Surveillance Team |
| EPI_ISL_3051927 | Lighthouse Lab in Alderley Park | Wellcome Sanger Institute for the COVID-19 Genomics UK (COG-UK) Consortium | Jacquelyn Wynn, Mairead Hyland, The Lighthouse Lab in Alderley Park and Alex Alderton, Roberto Amato, Jeffrey Barrett, Sonia Goncalves, Ewan Harrison, David K. Jackson, Ian Johnston, Dominic Kwiatkowski, Cordelia Langford, John Sillitoe on behalf of the Wellcome Sanger Institute COVID-19 Surveillance Team |
| EPI_ISL_3051928 | Lighthouse Lab in Glasgow | Wellcome Sanger Institute for the COVID-19 Genomics UK (COG-UK) Consortium | Harper VanSteenhouse, Yumi Kasai, David Gray, Carol Clugston, Anna Dominiczak and Alex Alderton, Roberto Amato, Jeffrey Barrett, Sonia Goncalves, Ewan Harrison, David K. Jackson, Ian Johnston, Dominic Kwiatkowski, Cordelia Langford, John Sillitoe on behalf of the Wellcome Sanger Institute COVID-19 Surveillance Team |
| EPI_ISL_3051929 | Lighthouse Lab in Alderley Park | Wellcome Sanger Institute for the COVID-19 Genomics UK (COG-UK) Consortium | Jacquelyn Wynn, Mairead Hyland, The Lighthouse Lab in Alderley Park and Alex Alderton, Roberto Amato, Jeffrey Barrett, Sonia Goncalves, Ewan Harrison, David K. Jackson, Ian Johnston, Dominic Kwiatkowski, Cordelia Langford, John Sillitoe on behalf of the Wellcome Sanger Institute COVID-19 Surveillance Team |
| EPI_ISL_3051930 | Lighthouse Lab in Glasgow | Wellcome Sanger Institute for the COVID-19 Genomics UK (COG-UK) Consortium | Harper VanSteenhouse, Yumi Kasai, David Gray, Carol Clugston, Anna Dominiczak and Alex Alderton, Roberto Amato, Jeffrey Barrett, Sonia Goncalves, Ewan Harrison, David K. Jackson, Ian Johnston, Dominic Kwiatkowski, Cordelia Langford, John Sillitoe on behalf of the Wellcome Sanger Institute COVID-19 Surveillance Team |
| EPI_ISL_3051931, EPI_ISL_3051932 | Lighthouse Lab in Alderley Park | Wellcome Sanger Institute for the COVID-19 Genomics UK (COG-UK) Consortium | Jacquelyn Wynn, Mairead Hyland, The Lighthouse Lab in Alderley Park and Alex Alderton, Roberto Amato, Jeffrey Barrett, Sonia Goncalves, Ewan Harrison, David K. Jackson, Ian Johnston, Dominic Kwiatkowski, Cordelia Langford, John Sillitoe on behalf of the Wellcome Sanger Institute COVID-19 Surveillance Team |
| EPI_ISL_3051933, EPI_ISL_3051934 | Lighthouse Lab in Glasgow | Wellcome Sanger Institute for the COVID-19 Genomics UK (COG-UK) Consortium | Harper VanSteenhouse, Yumi Kasai, David Gray, Carol Clugston, Anna Dominiczak and Alex Alderton, Roberto Amato, Jeffrey Barrett, Sonia Goncalves, Ewan Harrison, David K. Jackson, Ian Johnston, Dominic Kwiatkowski, Cordelia Langford, John Sillitoe on behalf of the Wellcome Sanger Institute COVID-19 Surveillance Team |
| EPI_ISL_3051935 | Lighthouse Lab in Alderley Park | Wellcome Sanger Institute for the COVID-19 Genomics UK (COG-UK) Consortium | Jacquelyn Wynn, Mairead Hyland, The Lighthouse Lab in Alderley Park and Alex Alderton, Roberto Amato, Jeffrey Barrett, Sonia Goncalves, Ewan Harrison, David K. Jackson, Ian Johnston, Dominic Kwiatkowski, Cordelia Langford, John Sillitoe on behalf of the Wellcome Sanger Institute COVID-19 Surveillance Team |
| EPI_ISL_3051936 | Lighthouse Lab in Glasgow | Wellcome Sanger Institute for the COVID-19 Genomics UK (COG-UK) Consortium | Harper VanSteenhouse, Yumi Kasai, David Gray, Carol Clugston, Anna Dominiczak and Alex Alderton, Roberto Amato, Jeffrey Barrett, Sonia Goncalves, Ewan Harrison, David K. Jackson, Ian Johnston, Dominic Kwiatkowski, Cordelia Langford, John Sillitoe on behalf of the Wellcome Sanger Institute COVID-19 Surveillance Team |
| EPI_ISL_3051937 | Lighthouse Lab in Alderley Park | Wellcome Sanger Institute for the COVID-19 Genomics UK (COG-UK) Consortium | Jacquelyn Wynn, Mairead Hyland, The Lighthouse Lab in Alderley Park and Alex Alderton, Roberto Amato, Jeffrey Barrett, Sonia Goncalves, Ewan Harrison, David K. Jackson, Ian Johnston, Dominic Kwiatkowski, Cordelia Langford, John Sillitoe on behalf of the Wellcome Sanger Institute COVID-19 Surveillance Team |
| EPI_ISL_3051938, EPI_ISL_3051939, EPI_ISL_3051940, EPI_ISL_3051941 | Lighthouse Lab in Glasgow | Wellcome Sanger Institute for the COVID-19 Genomics UK (COG-UK) Consortium | Harper VanSteenhouse, Yumi Kasai, David Gray, Carol Clugston, Anna Dominiczak and Alex Alderton, Roberto Amato, Jeffrey Barrett, Sonia Goncalves, Ewan Harrison, David K. Jackson, Ian Johnston, Dominic Kwiatkowski, Cordelia Langford, John Sillitoe on behalf of the Wellcome Sanger Institute COVID-19 Surveillance Team |
| EPI_ISL_3051942, EPI_ISL_3051943, EPI_ISL_3051944 | Lighthouse Lab in Alderley Park | Wellcome Sanger Institute for the COVID-19 Genomics UK (COG-UK) Consortium | Jacquelyn Wynn, Mairead Hyland, The Lighthouse Lab in Alderley Park and Alex Alderton, Roberto Amato, Jeffrey Barrett, Sonia Goncalves, Ewan Harrison, David K. Jackson, Ian Johnston, Dominic Kwiatkowski, Cordelia Langford, John Sillitoe on behalf of the Wellcome Sanger Institute COVID-19 Surveillance Team |
| EPI_ISL_3051945 | Lighthouse Lab in Glasgow | Wellcome Sanger Institute for the COVID-19 Genomics UK (COG-UK) Consortium | Harper VanSteenhouse, Yumi Kasai, David Gray, Carol Clugston, Anna Dominiczak and Alex Alderton, Roberto Amato, Jeffrey Barrett, Sonia Goncalves, Ewan Harrison, David K. Jackson, Ian Johnston, Dominic Kwiatkowski, Cordelia Langford, John Sillitoe on behalf of the Wellcome Sanger Institute COVID-19 Surveillance Team |
| EPI_ISL_3051946, EPI_ISL_3051947 | Lighthouse Lab in Alderley Park | Wellcome Sanger Institute for the COVID-19 Genomics UK (COG-UK) Consortium | Jacquelyn Wynn, Mairead Hyland, The Lighthouse Lab in Alderley Park and Alex Alderton, Roberto Amato, Jeffrey Barrett, Sonia Goncalves, Ewan Harrison, David K. Jackson, Ian Johnston, Dominic Kwiatkowski, Cordelia Langford, John Sillitoe on behalf of the Wellcome Sanger Institute COVID-19 Surveillance Team |
| EPI_ISL_3051948, EPI_ISL_3051949 | Lighthouse Lab in Glasgow | Wellcome Sanger Institute for the COVID-19 Genomics UK (COG-UK) Consortium | Harper VanSteenhouse, Yumi Kasai, David Gray, Carol Clugston, Anna Dominiczak and Alex Alderton, Roberto Amato, Jeffrey Barrett, Sonia Goncalves, Ewan Harrison, David K. Jackson, Ian Johnston, Dominic Kwiatkowski, Cordelia Langford, John Sillitoe on behalf of the Wellcome Sanger Institute COVID-19 Surveillance Team |
| EPI_ISL_3051950, EPI_ISL_3051951 | Lighthouse Lab in Alderley Park | Wellcome Sanger Institute for the COVID-19 Genomics UK (COG-UK) Consortium | Jacquelyn Wynn, Mairead Hyland, The Lighthouse Lab in Alderley Park and Alex Alderton, Roberto Amato, Jeffrey Barrett, Sonia Goncalves, Ewan Harrison, David K. Jackson, Ian Johnston, Dominic Kwiatkowski, Cordelia Langford, John Sillitoe on behalf of the Wellcome Sanger Institute COVID-19 Surveillance Team |
| EPI_ISL_3051952 | Lighthouse Lab in Glasgow | Wellcome Sanger Institute for the COVID-19 Genomics UK (COG-UK) Consortium | Harper VanSteenhouse, Yumi Kasai, David Gray, Carol Clugston, Anna Dominiczak and Alex Alderton, Roberto Amato, Jeffrey Barrett, Sonia Goncalves, Ewan Harrison, David K. Jackson, Ian Johnston, Dominic Kwiatkowski, Cordelia Langford, John Sillitoe on behalf of the Wellcome Sanger Institute COVID-19 Surveillance Team |
| EPI_ISL_3051953, EPI_ISL_3051954, EPI_ISL_3051955, EPI_ISL_3051956 | Lighthouse Lab in Alderley Park | Wellcome Sanger Institute for the COVID-19 Genomics UK (COG-UK) Consortium | Jacquelyn Wynn, Mairead Hyland, The Lighthouse Lab in Alderley Park and Alex Alderton, Roberto Amato, Jeffrey Barrett, Sonia Goncalves, Ewan Harrison, David K. Jackson, Ian Johnston, Dominic Kwiatkowski, Cordelia Langford, John Sillitoe on behalf of the Wellcome Sanger Institute COVID-19 Surveillance Team |
| EPI_ISL_3051957, EPI_ISL_3051958, EPI_ISL_3051959 | Lighthouse Lab in Glasgow | Wellcome Sanger Institute for the COVID-19 Genomics UK (COG-UK) Consortium | Harper VanSteenhouse, Yumi Kasai, David Gray, Carol Clugston, Anna Dominiczak and Alex Alderton, Roberto Amato, Jeffrey Barrett, Sonia Goncalves, Ewan Harrison, David K. Jackson, Ian Johnston, Dominic Kwiatkowski, Cordelia Langford, John Sillitoe on behalf of the Wellcome Sanger Institute COVID-19 Surveillance Team |
| EPI_ISL_3051960, EPI_ISL_3051961, EPI_ISL_3051962, EPI_ISL_3051963, EPI_ISL_3051964, EPI_ISL_3051965 | Lighthouse Lab in Alderley Park | Wellcome Sanger Institute for the COVID-19 Genomics UK (COG-UK) Consortium | Jacquelyn Wynn, Mairead Hyland, The Lighthouse Lab in Alderley Park and Alex Alderton, Roberto Amato, Jeffrey Barrett, Sonia Goncalves, Ewan Harrison, David K. Jackson, Ian Johnston, Dominic Kwiatkowski, Cordelia Langford, John Sillitoe on behalf of the Wellcome Sanger Institute COVID-19 Surveillance Team |
| EPI_ISL_3051966 | Lighthouse Lab in Glasgow | Wellcome Sanger Institute for the COVID-19 Genomics UK (COG-UK) Consortium | Harper VanSteenhouse, Yumi Kasai, David Gray, Carol Clugston, Anna Dominiczak and Alex Alderton, Roberto Amato, Jeffrey Barrett, Sonia Goncalves, Ewan Harrison, David K. Jackson, Ian Johnston, Dominic Kwiatkowski, Cordelia Langford, John Sillitoe on behalf of the Wellcome Sanger Institute COVID-19 Surveillance Team |
| EPI_ISL_3051967 | Lighthouse Lab in Alderley Park | Wellcome Sanger Institute for the COVID-19 Genomics UK (COG-UK) Consortium | Jacquelyn Wynn, Mairead Hyland, The Lighthouse Lab in Alderley Park and Alex Alderton, Roberto Amato, Jeffrey Barrett, Sonia Goncalves, Ewan Harrison, David K. Jackson, Ian Johnston, Dominic Kwiatkowski, Cordelia Langford, John Sillitoe on behalf of the Wellcome Sanger Institute COVID-19 Surveillance Team |
| EPI_ISL_3051968, EPI_ISL_3051969 | Lighthouse Lab in Glasgow | Wellcome Sanger Institute for the COVID-19 Genomics UK (COG-UK) Consortium | Harper VanSteenhouse, Yumi Kasai, David Gray, Carol Clugston, Anna Dominiczak and Alex Alderton, Roberto Amato, Jeffrey Barrett, Sonia Goncalves, Ewan Harrison, David K. Jackson, Ian Johnston, Dominic Kwiatkowski, Cordelia Langford, John Sillitoe on behalf of the Wellcome Sanger Institute COVID-19 Surveillance Team |
| EPI_ISL_3051970 | Lighthouse Lab in Alderley Park | Wellcome Sanger Institute for the COVID-19 Genomics UK (COG-UK) Consortium | Jacquelyn Wynn, Mairead Hyland, The Lighthouse Lab in Alderley Park and Alex Alderton, Roberto Amato, Jeffrey Barrett, Sonia Goncalves, Ewan Harrison, David K. Jackson, Ian Johnston, Dominic Kwiatkowski, Cordelia Langford, John Sillitoe on behalf of the Wellcome Sanger Institute COVID-19 Surveillance Team |
| EPI_ISL_3051971 | Lighthouse Lab in Glasgow | Wellcome Sanger Institute for the COVID-19 Genomics UK (COG-UK) Consortium | Harper VanSteenhouse, Yumi Kasai, David Gray, Carol Clugston, Anna Dominiczak and Alex Alderton, Roberto Amato, Jeffrey Barrett, Sonia Goncalves, Ewan Harrison, David K. Jackson, Ian Johnston, Dominic Kwiatkowski, Cordelia Langford, John Sillitoe on behalf of the Wellcome Sanger Institute COVID-19 Surveillance Team |
| EPI_ISL_3051972, EPI_ISL_3051973, EPI_ISL_3051974, EPI_ISL_3051975, EPI_ISL_3051976 | Lighthouse Lab in Alderley Park | Wellcome Sanger Institute for the COVID-19 Genomics UK (COG-UK) Consortium | Jacquelyn Wynn, Mairead Hyland, The Lighthouse Lab in Alderley Park and Alex Alderton, Roberto Amato, Jeffrey Barrett, Sonia Goncalves, Ewan Harrison, David K. Jackson, Ian Johnston, Dominic Kwiatkowski, Cordelia Langford, John Sillitoe on behalf of the Wellcome Sanger Institute COVID-19 Surveillance Team |
| EPI_ISL_3051977, EPI_ISL_3051978, EPI_ISL_3051979 | Lighthouse Lab in Glasgow | Wellcome Sanger Institute for the COVID-19 Genomics UK (COG-UK) Consortium | Harper VanSteenhouse, Yumi Kasai, David Gray, Carol Clugston, Anna Dominiczak and Alex Alderton, Roberto Amato, Jeffrey Barrett, Sonia Goncalves, Ewan Harrison, David K. Jackson, Ian Johnston, Dominic Kwiatkowski, Cordelia Langford, John Sillitoe on behalf of the Wellcome Sanger Institute COVID-19 Surveillance Team |
| EPI_ISL_3051980 | Lighthouse Lab in Alderley Park | Wellcome Sanger Institute for the COVID-19 Genomics UK (COG-UK) Consortium | Jacquelyn Wynn, Mairead Hyland, The Lighthouse Lab in Alderley Park and Alex Alderton, Roberto Amato, Jeffrey Barrett, Sonia Goncalves, Ewan Harrison, David K. Jackson, Ian Johnston, Dominic Kwiatkowski, Cordelia Langford, John Sillitoe on behalf of the Wellcome Sanger Institute COVID-19 Surveillance Team |
| EPI_ISL_3051981, EPI_ISL_3051982 | Lighthouse Lab in Glasgow | Wellcome Sanger Institute for the COVID-19 Genomics UK (COG-UK) Consortium | Harper VanSteenhouse, Yumi Kasai, David Gray, Carol Clugston, Anna Dominiczak and Alex Alderton, Roberto Amato, Jeffrey Barrett, Sonia Goncalves, Ewan Harrison, David K. Jackson, Ian Johnston, Dominic Kwiatkowski, Cordelia Langford, John Sillitoe on behalf of the Wellcome Sanger Institute COVID-19 Surveillance Team |
| EPI_ISL_3051983, EPI_ISL_3051984, EPI_ISL_3051985, EPI_ISL_3051986 | Lighthouse Lab in Alderley Park | Wellcome Sanger Institute for the COVID-19 Genomics UK (COG-UK) Consortium | Jacquelyn Wynn, Mairead Hyland, The Lighthouse Lab in Alderley Park and Alex Alderton, Roberto Amato, Jeffrey Barrett, Sonia Goncalves, Ewan Harrison, David K. Jackson, Ian Johnston, Dominic Kwiatkowski, Cordelia Langford, John Sillitoe on behalf of the Wellcome Sanger Institute COVID-19 Surveillance Team |
| EPI_ISL_3051987 | Lighthouse Lab in Glasgow | Wellcome Sanger Institute for the COVID-19 Genomics UK (COG-UK) Consortium | Harper VanSteenhouse, Yumi Kasai, David Gray, Carol Clugston, Anna Dominiczak and Alex Alderton, Roberto Amato, Jeffrey Barrett, Sonia Goncalves, Ewan Harrison, David K. Jackson, Ian Johnston, Dominic Kwiatkowski, Cordelia Langford, John Sillitoe on behalf of the Wellcome Sanger Institute COVID-19 Surveillance Team |
| EPI_ISL_3051988, EPI_ISL_3051989, EPI_ISL_3051990, EPI_ISL_3051991, EPI_ISL_3051992, EPI_ISL_3051993, EPI_ISL_3051994, | Lighthouse Lab in Alderley Park | Wellcome Sanger Institute for the COVID-19 Genomics UK (COG-UK) Consortium | Jacquelyn Wynn, Mairead Hyland, The Lighthouse Lab in Alderley Park and Alex Alderton, Roberto Amato, Jeffrey Barrett, Sonia Goncalves, Ewan Harrison, David K. Jackson, Ian Johnston, Dominic Kwiatkowski, Cordelia Langford, John Sillitoe on behalf of the Wellcome Sanger Institute COVID-19 Surveillance Team |

| | | | |
|---|---|---|---|
| EPI_ISL_3051995 | | | |
| EPI_ISL_3051996 | Lighthouse Lab in Glasgow | Wellcome Sanger Institute for the COVID-19 Genomics UK (COG-UK) Consortium | Harper VanSteenhouse, Yumi Kasai, David Gray, Carol Clugston, Anna Dominiczak and Alex Alderton, Roberto Amato, Jeffrey Barrett, Sonia Goncalves, Ewan Harrison, David K. Jackson, Ian Johnston, Dominic Kwiatkowski, Cordelia Langford, John Sillitoe on behalf of the Wellcome Sanger Institute COVID-19 Surveillance Team |
| EPI_ISL_3051997, EPI_ISL_3051998 | Lighthouse Lab in Alderley Park | Wellcome Sanger Institute for the COVID-19 Genomics UK (COG-UK) Consortium | Jacquelyn Wynn, Mairead Hyland, The Lighthouse Lab in Alderley Park and Alex Alderton, Roberto Amato, Jeffrey Barrett, Sonia Goncalves, Ewan Harrison, David K. Jackson, Ian Johnston, Dominic Kwiatkowski, Cordelia Langford, John Sillitoe on behalf of the Wellcome Sanger Institute COVID-19 Surveillance Team |
| EPI_ISL_3051999, EPI_ISL_3052000 | Lighthouse Lab in Glasgow | Wellcome Sanger Institute for the COVID-19 Genomics UK (COG-UK) Consortium | Harper VanSteenhouse, Yumi Kasai, David Gray, Carol Clugston, Anna Dominiczak and Alex Alderton, Roberto Amato, Jeffrey Barrett, Sonia Goncalves, Ewan Harrison, David K. Jackson, Ian Johnston, Dominic Kwiatkowski, Cordelia Langford, John Sillitoe on behalf of the Wellcome Sanger Institute COVID-19 Surveillance Team |
| EPI_ISL_3052001, EPI_ISL_3052002 | Lighthouse Lab in Alderley Park | Wellcome Sanger Institute for the COVID-19 Genomics UK (COG-UK) Consortium | Jacquelyn Wynn, Mairead Hyland, The Lighthouse Lab in Alderley Park and Alex Alderton, Roberto Amato, Jeffrey Barrett, Sonia Goncalves, Ewan Harrison, David K. Jackson, Ian Johnston, Dominic Kwiatkowski, Cordelia Langford, John Sillitoe on behalf of the Wellcome Sanger Institute COVID-19 Surveillance Team |
| EPI_ISL_3052003 | Lighthouse Lab in Glasgow | Wellcome Sanger Institute for the COVID-19 Genomics UK (COG-UK) Consortium | Harper VanSteenhouse, Yumi Kasai, David Gray, Carol Clugston, Anna Dominiczak and Alex Alderton, Roberto Amato, Jeffrey Barrett, Sonia Goncalves, Ewan Harrison, David K. Jackson, Ian Johnston, Dominic Kwiatkowski, Cordelia Langford, John Sillitoe on behalf of the Wellcome Sanger Institute COVID-19 Surveillance Team |
| EPI_ISL_3052004, EPI_ISL_3052005, EPI_ISL_3052006 | Lighthouse Lab in Alderley Park | Wellcome Sanger Institute for the COVID-19 Genomics UK (COG-UK) Consortium | Jacquelyn Wynn, Mairead Hyland, The Lighthouse Lab in Alderley Park and Alex Alderton, Roberto Amato, Jeffrey Barrett, Sonia Goncalves, Ewan Harrison, David K. Jackson, Ian Johnston, Dominic Kwiatkowski, Cordelia Langford, John Sillitoe on behalf of the Wellcome Sanger Institute COVID-19 Surveillance Team |
| EPI_ISL_3052007 | Lighthouse Lab in Glasgow | Wellcome Sanger Institute for the COVID-19 Genomics UK (COG-UK) Consortium | Harper VanSteenhouse, Yumi Kasai, David Gray, Carol Clugston, Anna Dominiczak and Alex Alderton, Roberto Amato, Jeffrey Barrett, Sonia Goncalves, Ewan Harrison, David K. Jackson, Ian Johnston, Dominic Kwiatkowski, Cordelia Langford, John Sillitoe on behalf of the Wellcome Sanger Institute COVID-19 Surveillance Team |
| EPI_ISL_3052008, EPI_ISL_3052009, EPI_ISL_3052010, EPI_ISL_3052011 | Lighthouse Lab in Alderley Park | Wellcome Sanger Institute for the COVID-19 Genomics UK (COG-UK) Consortium | Jacquelyn Wynn, Mairead Hyland, The Lighthouse Lab in Alderley Park and Alex Alderton, Roberto Amato, Jeffrey Barrett, Sonia Goncalves, Ewan Harrison, David K. Jackson, Ian Johnston, Dominic Kwiatkowski, Cordelia Langford, John Sillitoe on behalf of the Wellcome Sanger Institute COVID-19 Surveillance Team |
| EPI_ISL_3052012 | Lighthouse Lab in Glasgow | Wellcome Sanger Institute for the COVID-19 Genomics UK (COG-UK) Consortium | Harper VanSteenhouse, Yumi Kasai, David Gray, Carol Clugston, Anna Dominiczak and Alex Alderton, Roberto Amato, Jeffrey Barrett, Sonia Goncalves, Ewan Harrison, David K. Jackson, Ian Johnston, Dominic Kwiatkowski, Cordelia Langford, John Sillitoe on behalf of the Wellcome Sanger Institute COVID-19 Surveillance Team |
| EPI_ISL_3052013, EPI_ISL_3052014 | Lighthouse Lab in Alderley Park | Wellcome Sanger Institute for the COVID-19 Genomics UK (COG-UK) Consortium | Jacquelyn Wynn, Mairead Hyland, The Lighthouse Lab in Alderley Park and Alex Alderton, Roberto Amato, Jeffrey Barrett, Sonia Goncalves, Ewan Harrison, David K. Jackson, Ian Johnston, Dominic Kwiatkowski, Cordelia Langford, John Sillitoe on behalf of the Wellcome Sanger Institute COVID-19 Surveillance Team |
| EPI_ISL_3052015, EPI_ISL_3052016, EPI_ISL_3052017, EPI_ISL_3052018 | Lighthouse Lab in Glasgow | Wellcome Sanger Institute for the COVID-19 Genomics UK (COG-UK) Consortium | Harper VanSteenhouse, Yumi Kasai, David Gray, Carol Clugston, Anna Dominiczak and Alex Alderton, Roberto Amato, Jeffrey Barrett, Sonia Goncalves, Ewan Harrison, David K. Jackson, Ian Johnston, Dominic Kwiatkowski, Cordelia Langford, John Sillitoe on behalf of the Wellcome Sanger Institute COVID-19 Surveillance Team |
| EPI_ISL_3052019 | Lighthouse Lab in Alderley Park | Wellcome Sanger Institute for the COVID-19 Genomics UK (COG-UK) Consortium | Jacquelyn Wynn, Mairead Hyland, The Lighthouse Lab in Alderley Park and Alex Alderton, Roberto Amato, Jeffrey Barrett, Sonia Goncalves, Ewan Harrison, David K. Jackson, Ian Johnston, Dominic Kwiatkowski, Cordelia Langford, John Sillitoe on behalf of the Wellcome Sanger Institute COVID-19 Surveillance Team |
| EPI_ISL_3052020 | Lighthouse Lab in Glasgow | Wellcome Sanger Institute for the COVID-19 Genomics UK (COG-UK) Consortium | Harper VanSteenhouse, Yumi Kasai, David Gray, Carol Clugston, Anna Dominiczak and Alex Alderton, Roberto Amato, Jeffrey Barrett, Sonia Goncalves, Ewan Harrison, David K. Jackson, Ian Johnston, Dominic Kwiatkowski, Cordelia Langford, John Sillitoe on behalf of the Wellcome Sanger Institute COVID-19 Surveillance Team |
| EPI_ISL_3052021 | Lighthouse Lab in Alderley Park | Wellcome Sanger Institute for the COVID-19 Genomics UK (COG-UK) Consortium | Jacquelyn Wynn, Mairead Hyland, The Lighthouse Lab in Alderley Park and Alex Alderton, Roberto Amato, Jeffrey Barrett, Sonia Goncalves, Ewan Harrison, David K. Jackson, Ian Johnston, Dominic Kwiatkowski, Cordelia Langford, John Sillitoe on behalf of the Wellcome Sanger Institute COVID-19 Surveillance Team |
| EPI_ISL_3052022 | Lighthouse Lab in Glasgow | Wellcome Sanger Institute for the COVID-19 Genomics UK (COG-UK) Consortium | Harper VanSteenhouse, Yumi Kasai, David Gray, Carol Clugston, Anna Dominiczak and Alex Alderton, Roberto Amato, Jeffrey Barrett, Sonia Goncalves, Ewan Harrison, David K. Jackson, Ian Johnston, Dominic Kwiatkowski, Cordelia Langford, John Sillitoe on behalf of the Wellcome Sanger Institute COVID-19 Surveillance Team |
| EPI_ISL_3052023, EPI_ISL_3052024, EPI_ISL_3052025, EPI_ISL_3052026, EPI_ISL_3052027 | Lighthouse Lab in Alderley Park | Wellcome Sanger Institute for the COVID-19 Genomics UK (COG-UK) Consortium | Jacquelyn Wynn, Mairead Hyland, The Lighthouse Lab in Alderley Park and Alex Alderton, Roberto Amato, Jeffrey Barrett, Sonia Goncalves, Ewan Harrison, David K. Jackson, Ian Johnston, Dominic Kwiatkowski, Cordelia Langford, John Sillitoe on behalf of the Wellcome Sanger Institute COVID-19 Surveillance Team |
| EPI_ISL_3052028 | Lighthouse Lab in Glasgow | Wellcome Sanger Institute for the COVID-19 Genomics UK (COG-UK) Consortium | Harper VanSteenhouse, Yumi Kasai, David Gray, Carol Clugston, Anna Dominiczak and Alex Alderton, Roberto Amato, Jeffrey Barrett, Sonia Goncalves, Ewan Harrison, David K. Jackson, Ian Johnston, Dominic Kwiatkowski, Cordelia Langford, John Sillitoe on behalf of the Wellcome Sanger Institute COVID-19 Surveillance Team |
| EPI_ISL_3052029, EPI_ISL_3052030, EPI_ISL_3052031 | Lighthouse Lab in Alderley Park | Wellcome Sanger Institute for the COVID-19 Genomics UK (COG-UK) Consortium | Jacquelyn Wynn, Mairead Hyland, The Lighthouse Lab in Alderley Park and Alex Alderton, Roberto Amato, Jeffrey Barrett, Sonia Goncalves, Ewan Harrison, David K. Jackson, Ian Johnston, Dominic Kwiatkowski, Cordelia Langford, John Sillitoe on behalf of the Wellcome Sanger Institute COVID-19 Surveillance Team |
| EPI_ISL_3052032, EPI_ISL_3052033 | Lighthouse Lab in Glasgow | Wellcome Sanger Institute for the COVID-19 Genomics UK (COG-UK) Consortium | Harper VanSteenhouse, Yumi Kasai, David Gray, Carol Clugston, Anna Dominiczak and Alex Alderton, Roberto Amato, Jeffrey Barrett, Sonia Goncalves, Ewan Harrison, David K. Jackson, Ian Johnston, Dominic Kwiatkowski, Cordelia Langford, John Sillitoe on behalf of the Wellcome Sanger Institute COVID-19 Surveillance Team |
| EPI_ISL_3052034, EPI_ISL_3052035, EPI_ISL_3052036 | Lighthouse Lab in Alderley Park | Wellcome Sanger Institute for the COVID-19 Genomics UK (COG-UK) Consortium | Jacquelyn Wynn, Mairead Hyland, The Lighthouse Lab in Alderley Park and Alex Alderton, Roberto Amato, Jeffrey Barrett, Sonia Goncalves, Ewan Harrison, David K. Jackson, Ian Johnston, Dominic Kwiatkowski, Cordelia Langford, John Sillitoe on behalf of the Wellcome Sanger Institute COVID-19 Surveillance Team |
| EPI_ISL_3052037, EPI_ISL_3052038, EPI_ISL_3052039 | Lighthouse Lab in Glasgow | Wellcome Sanger Institute for the COVID-19 Genomics UK (COG-UK) Consortium | Harper VanSteenhouse, Yumi Kasai, David Gray, Carol Clugston, Anna Dominiczak and Alex Alderton, Roberto Amato, Jeffrey Barrett, Sonia Goncalves, Ewan Harrison, David K. Jackson, Ian Johnston, Dominic Kwiatkowski, Cordelia Langford, John Sillitoe on behalf of the Wellcome Sanger Institute COVID-19 Surveillance Team |
| EPI_ISL_3052040, EPI_ISL_3052041 | Lighthouse Lab in Alderley Park | Wellcome Sanger Institute for the COVID-19 Genomics UK (COG-UK) Consortium | Jacquelyn Wynn, Mairead Hyland, The Lighthouse Lab in Alderley Park and Alex Alderton, Roberto Amato, Jeffrey Barrett, Sonia Goncalves, Ewan Harrison, David K. Jackson, Ian Johnston, Dominic Kwiatkowski, Cordelia Langford, John Sillitoe on behalf of the Wellcome Sanger Institute COVID-19 Surveillance Team |
| EPI_ISL_3052042 | Lighthouse Lab in Glasgow | Wellcome Sanger Institute for the COVID-19 Genomics UK (COG-UK) Consortium | Harper VanSteenhouse, Yumi Kasai, David Gray, Carol Clugston, Anna Dominiczak and Alex Alderton, Roberto Amato, Jeffrey Barrett, Sonia Goncalves, Ewan Harrison, David K. Jackson, Ian Johnston, Dominic Kwiatkowski, Cordelia Langford, John Sillitoe on behalf of the Wellcome Sanger Institute COVID-19 Surveillance Team |
| EPI_ISL_3052043, EPI_ISL_3052044, EPI_ISL_3052045, EPI_ISL_3052046 | Lighthouse Lab in Alderley Park | Wellcome Sanger Institute for the COVID-19 Genomics UK (COG-UK) Consortium | Jacquelyn Wynn, Mairead Hyland, The Lighthouse Lab in Alderley Park and Alex Alderton, Roberto Amato, Jeffrey Barrett, Sonia Goncalves, Ewan Harrison, David K. Jackson, Ian Johnston, Dominic Kwiatkowski, Cordelia Langford, John Sillitoe on behalf of the Wellcome Sanger Institute COVID-19 Surveillance Team |
| EPI_ISL_3052047, EPI_ISL_3052048, EPI_ISL_3052049 | Lighthouse Lab in Glasgow | Wellcome Sanger Institute for the COVID-19 Genomics UK (COG-UK) Consortium | Harper VanSteenhouse, Yumi Kasai, David Gray, Carol Clugston, Anna Dominiczak and Alex Alderton, Roberto Amato, Jeffrey Barrett, Sonia Goncalves, Ewan Harrison, David K. Jackson, Ian Johnston, Dominic Kwiatkowski, Cordelia Langford, John Sillitoe on behalf of the Wellcome Sanger Institute COVID-19 Surveillance Team |
| EPI_ISL_3052050 | Lighthouse Lab in Alderley Park | Wellcome Sanger Institute for the COVID-19 Genomics UK (COG-UK) Consortium | Jacquelyn Wynn, Mairead Hyland, The Lighthouse Lab in Alderley Park and Alex Alderton, Roberto Amato, Jeffrey Barrett, Sonia Goncalves, Ewan Harrison, David K. Jackson, Ian Johnston, Dominic Kwiatkowski, Cordelia Langford, John Sillitoe on behalf of the Wellcome Sanger Institute COVID-19 Surveillance Team |
| EPI_ISL_3052051 | Lighthouse Lab in Glasgow | Wellcome Sanger Institute for the COVID-19 Genomics UK (COG-UK) Consortium | Harper VanSteenhouse, Yumi Kasai, David Gray, Carol Clugston, Anna Dominiczak and Alex Alderton, Roberto Amato, Jeffrey Barrett, Sonia Goncalves, Ewan Harrison, David K. Jackson, Ian Johnston, Dominic Kwiatkowski, Cordelia Langford, John Sillitoe on behalf of the Wellcome Sanger Institute COVID-19 Surveillance Team |
| EPI_ISL_3052052, EPI_ISL_3052053 | Lighthouse Lab in Alderley Park | Wellcome Sanger Institute for the COVID-19 Genomics UK (COG-UK) Consortium | Jacquelyn Wynn, Mairead Hyland, The Lighthouse Lab in Alderley Park and Alex Alderton, Roberto Amato, Jeffrey Barrett, Sonia Goncalves, Ewan Harrison, David K. Jackson, Ian Johnston, Dominic Kwiatkowski, Cordelia Langford, John Sillitoe on behalf of the Wellcome Sanger Institute COVID-19 Surveillance Team |
| EPI_ISL_3052054 | Lighthouse Lab in Glasgow | Wellcome Sanger Institute for the COVID-19 Genomics UK (COG-UK) Consortium | Harper VanSteenhouse, Yumi Kasai, David Gray, Carol Clugston, Anna Dominiczak and Alex Alderton, Roberto Amato, Jeffrey Barrett, Sonia Goncalves, Ewan Harrison, David K. Jackson, Ian Johnston, Dominic Kwiatkowski, Cordelia Langford, John Sillitoe on behalf of the Wellcome Sanger Institute COVID-19 Surveillance Team |
| EPI_ISL_3052055 | Lighthouse Lab in Alderley Park | Wellcome Sanger Institute for the COVID-19 Genomics UK (COG-UK) Consortium | Jacquelyn Wynn, Mairead Hyland, The Lighthouse Lab in Alderley Park and Alex Alderton, Roberto Amato, Jeffrey Barrett, Sonia Goncalves, Ewan Harrison, David K. Jackson, Ian Johnston, Dominic Kwiatkowski, Cordelia Langford, John Sillitoe on behalf of the Wellcome Sanger Institute COVID-19 Surveillance Team |
| EPI_ISL_3052056, EPI_ISL_3052057, EPI_ISL_3052058, EPI_ISL_3052059 | Lighthouse Lab in Glasgow | Wellcome Sanger Institute for the COVID-19 Genomics UK (COG-UK) Consortium | Harper VanSteenhouse, Yumi Kasai, David Gray, Carol Clugston, Anna Dominiczak and Alex Alderton, Roberto Amato, Jeffrey Barrett, Sonia Goncalves, Ewan Harrison, David K. Jackson, Ian Johnston, Dominic Kwiatkowski, Cordelia Langford, John Sillitoe on behalf of the Wellcome Sanger Institute COVID-19 Surveillance Team |
| EPI_ISL_3052060 | Lighthouse Lab in Alderley Park | Wellcome Sanger Institute for the COVID-19 Genomics UK (COG-UK) Consortium | Jacquelyn Wynn, Mairead Hyland, The Lighthouse Lab in Alderley Park and Alex Alderton, Roberto Amato, Jeffrey Barrett, Sonia Goncalves, Ewan Harrison, David K. Jackson, Ian Johnston, Dominic Kwiatkowski, Cordelia Langford, John Sillitoe on behalf of the Wellcome Sanger Institute COVID-19 Surveillance Team |
| EPI_ISL_3052061, EPI_ISL_3052062, EPI_ISL_3052063, EPI_ISL_3052064 | Lighthouse Lab in Glasgow | Wellcome Sanger Institute for the COVID-19 Genomics UK (COG-UK) Consortium | Harper VanSteenhouse, Yumi Kasai, David Gray, Carol Clugston, Anna Dominiczak and Alex Alderton, Roberto Amato, Jeffrey Barrett, Sonia Goncalves, Ewan Harrison, David K. Jackson, Ian Johnston, Dominic Kwiatkowski, Cordelia Langford, John Sillitoe on behalf of the Wellcome Sanger Institute COVID-19 Surveillance Team |
| EPI_ISL_3052065, EPI_ISL_3052066, EPI_ISL_3052067, EPI_ISL_3052068 | Lighthouse Lab in Alderley Park | Wellcome Sanger Institute for the COVID-19 Genomics UK (COG-UK) Consortium | Jacquelyn Wynn, Mairead Hyland, The Lighthouse Lab in Alderley Park and Alex Alderton, Roberto Amato, Jeffrey Barrett, Sonia Goncalves, Ewan Harrison, David K. Jackson, Ian Johnston, Dominic Kwiatkowski, Cordelia Langford, John Sillitoe on behalf of the Wellcome Sanger Institute COVID-19 Surveillance Team |
| EPI_ISL_3052069 | Lighthouse Lab in Glasgow | Wellcome Sanger Institute for the COVID-19 Genomics UK (COG-UK) Consortium | Harper VanSteenhouse, Yumi Kasai, David Gray, Carol Clugston, Anna Dominiczak and Alex Alderton, Roberto Amato, Jeffrey Barrett, Sonia Goncalves, Ewan Harrison, David K. Jackson, Ian Johnston, Dominic Kwiatkowski, Cordelia Langford, John Sillitoe on behalf of the Wellcome Sanger Institute COVID-19 Surveillance Team |
| EPI_ISL_3052070, EPI_ISL_3052071 | Lighthouse Lab in Alderley Park | Wellcome Sanger Institute for the COVID-19 Genomics UK (COG-UK) Consortium | Jacquelyn Wynn, Mairead Hyland, The Lighthouse Lab in Alderley Park and Alex Alderton, Roberto Amato, Jeffrey Barrett, Sonia Goncalves, Ewan Harrison, David K. Jackson, Ian Johnston, Dominic Kwiatkowski, Cordelia Langford, John Sillitoe on behalf of the Wellcome Sanger Institute COVID-19 Surveillance Team |
| EPI_ISL_3052072 | Lighthouse Lab in Glasgow | Wellcome Sanger Institute for the COVID-19 Genomics UK (COG-UK) | Harper VanSteenhouse, Yumi Kasai, David Gray, Carol Clugston, Anna Dominiczak and Alex Alderton, Roberto Amato, Jeffrey Barrett, Sonia Goncalves, Ewan Harrison, David K. Jackson, Ian Johnston, Dominic Kwiatkowski, Cordelia Langford, |

| | | | |
|---|---|---|---|
| | | Consortium | John Sillitoe on behalf of the Wellcome Sanger Institute COVID-19 Surveillance Team |
| EPI_ISL_3052073 | Lighthouse Lab in Alderley Park | Wellcome Sanger Institute for the COVID-19 Genomics UK (COG-UK) Consortium | Jacquelyn Wynn, Mairead Hyland, The Lighthouse Lab in Alderley Park and Alex Alderton, Roberto Amato, Jeffrey Barrett, Sonia Goncalves, Ewan Harrison, David K. Jackson, Ian Johnston, Dominic Kwiatkowski, Cordelia Langford, John Sillitoe on behalf of the Wellcome Sanger Institute COVID-19 Surveillance Team |
| EPI_ISL_3052074, EPI_ISL_3052075 | Lighthouse Lab in Glasgow | Wellcome Sanger Institute for the COVID-19 Genomics UK (COG-UK) Consortium | Harper VanSteenhouse, Yumi Kasai, David Gray, Carol Clugston, Anna Dominiczak and Alex Alderton, Roberto Amato, Jeffrey Barrett, Sonia Goncalves, Ewan Harrison, David K. Jackson, Ian Johnston, Dominic Kwiatkowski, Cordelia Langford, John Sillitoe on behalf of the Wellcome Sanger Institute COVID-19 Surveillance Team |
| EPI_ISL_3052076, EPI_ISL_3052077 | Lighthouse Lab in Alderley Park | Wellcome Sanger Institute for the COVID-19 Genomics UK (COG-UK) Consortium | Jacquelyn Wynn, Mairead Hyland, The Lighthouse Lab in Alderley Park and Alex Alderton, Roberto Amato, Jeffrey Barrett, Sonia Goncalves, Ewan Harrison, David K. Jackson, Ian Johnston, Dominic Kwiatkowski, Cordelia Langford, John Sillitoe on behalf of the Wellcome Sanger Institute COVID-19 Surveillance Team |
| EPI_ISL_3052078 | Lighthouse Lab in Glasgow | Wellcome Sanger Institute for the COVID-19 Genomics UK (COG-UK) Consortium | Harper VanSteenhouse, Yumi Kasai, David Gray, Carol Clugston, Anna Dominiczak and Alex Alderton, Roberto Amato, Jeffrey Barrett, Sonia Goncalves, Ewan Harrison, David K. Jackson, Ian Johnston, Dominic Kwiatkowski, Cordelia Langford, John Sillitoe on behalf of the Wellcome Sanger Institute COVID-19 Surveillance Team |
| EPI_ISL_3052079 | Lighthouse Lab in Alderley Park | Wellcome Sanger Institute for the COVID-19 Genomics UK (COG-UK) Consortium | Jacquelyn Wynn, Mairead Hyland, The Lighthouse Lab in Alderley Park and Alex Alderton, Roberto Amato, Jeffrey Barrett, Sonia Goncalves, Ewan Harrison, David K. Jackson, Ian Johnston, Dominic Kwiatkowski, Cordelia Langford, John Sillitoe on behalf of the Wellcome Sanger Institute COVID-19 Surveillance Team |
| EPI_ISL_3052080 | Lighthouse Lab in Glasgow | Wellcome Sanger Institute for the COVID-19 Genomics UK (COG-UK) Consortium | Harper VanSteenhouse, Yumi Kasai, David Gray, Carol Clugston, Anna Dominiczak and Alex Alderton, Roberto Amato, Jeffrey Barrett, Sonia Goncalves, Ewan Harrison, David K. Jackson, Ian Johnston, Dominic Kwiatkowski, Cordelia Langford, John Sillitoe on behalf of the Wellcome Sanger Institute COVID-19 Surveillance Team |
| EPI_ISL_3052081, EPI_ISL_3052082, EPI_ISL_3052083, EPI_ISL_3052084, EPI_ISL_3052085 | Lighthouse Lab in Alderley Park | Wellcome Sanger Institute for the COVID-19 Genomics UK (COG-UK) Consortium | Jacquelyn Wynn, Mairead Hyland, The Lighthouse Lab in Alderley Park and Alex Alderton, Roberto Amato, Jeffrey Barrett, Sonia Goncalves, Ewan Harrison, David K. Jackson, Ian Johnston, Dominic Kwiatkowski, Cordelia Langford, John Sillitoe on behalf of the Wellcome Sanger Institute COVID-19 Surveillance Team |
| EPI_ISL_3052086 | Lighthouse Lab in Glasgow | Wellcome Sanger Institute for the COVID-19 Genomics UK (COG-UK) Consortium | Harper VanSteenhouse, Yumi Kasai, David Gray, Carol Clugston, Anna Dominiczak and Alex Alderton, Roberto Amato, Jeffrey Barrett, Sonia Goncalves, Ewan Harrison, David K. Jackson, Ian Johnston, Dominic Kwiatkowski, Cordelia Langford, John Sillitoe on behalf of the Wellcome Sanger Institute COVID-19 Surveillance Team |
| EPI_ISL_3052087 | Lighthouse Lab in Alderley Park | Wellcome Sanger Institute for the COVID-19 Genomics UK (COG-UK) Consortium | Jacquelyn Wynn, Mairead Hyland, The Lighthouse Lab in Alderley Park and Alex Alderton, Roberto Amato, Jeffrey Barrett, Sonia Goncalves, Ewan Harrison, David K. Jackson, Ian Johnston, Dominic Kwiatkowski, Cordelia Langford, John Sillitoe on behalf of the Wellcome Sanger Institute COVID-19 Surveillance Team |
| EPI_ISL_3052088, EPI_ISL_3052089, EPI_ISL_3052090, EPI_ISL_3052091 | Lighthouse Lab in Glasgow | Wellcome Sanger Institute for the COVID-19 Genomics UK (COG-UK) Consortium | Harper VanSteenhouse, Yumi Kasai, David Gray, Carol Clugston, Anna Dominiczak and Alex Alderton, Roberto Amato, Jeffrey Barrett, Sonia Goncalves, Ewan Harrison, David K. Jackson, Ian Johnston, Dominic Kwiatkowski, Cordelia Langford, John Sillitoe on behalf of the Wellcome Sanger Institute COVID-19 Surveillance Team |
| EPI_ISL_3052092, EPI_ISL_3052093, EPI_ISL_3052094, EPI_ISL_3052095, EPI_ISL_3052096, EPI_ISL_3052097, EPI_ISL_3052098, EPI_ISL_3052099, EPI_ISL_3052100, EPI_ISL_3052101 | Lighthouse Lab in Alderley Park | Wellcome Sanger Institute for the COVID-19 Genomics UK (COG-UK) Consortium | Jacquelyn Wynn, Mairead Hyland, The Lighthouse Lab in Alderley Park and Alex Alderton, Roberto Amato, Jeffrey Barrett, Sonia Goncalves, Ewan Harrison, David K. Jackson, Ian Johnston, Dominic Kwiatkowski, Cordelia Langford, John Sillitoe on behalf of the Wellcome Sanger Institute COVID-19 Surveillance Team |
| EPI_ISL_3052102 | Lighthouse Lab in Glasgow | Wellcome Sanger Institute for the COVID-19 Genomics UK (COG-UK) Consortium | Harper VanSteenhouse, Yumi Kasai, David Gray, Carol Clugston, Anna Dominiczak and Alex Alderton, Roberto Amato, Jeffrey Barrett, Sonia Goncalves, Ewan Harrison, David K. Jackson, Ian Johnston, Dominic Kwiatkowski, Cordelia Langford, John Sillitoe on behalf of the Wellcome Sanger Institute COVID-19 Surveillance Team |
| EPI_ISL_3052103, EPI_ISL_3052104, EPI_ISL_3052105 | Lighthouse Lab in Alderley Park | Wellcome Sanger Institute for the COVID-19 Genomics UK (COG-UK) Consortium | Jacquelyn Wynn, Mairead Hyland, The Lighthouse Lab in Alderley Park and Alex Alderton, Roberto Amato, Jeffrey Barrett, Sonia Goncalves, Ewan Harrison, David K. Jackson, Ian Johnston, Dominic Kwiatkowski, Cordelia Langford, John Sillitoe on behalf of the Wellcome Sanger Institute COVID-19 Surveillance Team |
| EPI_ISL_3052106 | Lighthouse Lab in Glasgow | Wellcome Sanger Institute for the COVID-19 Genomics UK (COG-UK) Consortium | Harper VanSteenhouse, Yumi Kasai, David Gray, Carol Clugston, Anna Dominiczak and Alex Alderton, Roberto Amato, Jeffrey Barrett, Sonia Goncalves, Ewan Harrison, David K. Jackson, Ian Johnston, Dominic Kwiatkowski, Cordelia Langford, John Sillitoe on behalf of the Wellcome Sanger Institute COVID-19 Surveillance Team |
| EPI_ISL_3052107 | Lighthouse Lab in Alderley Park | Wellcome Sanger Institute for the COVID-19 Genomics UK (COG-UK) Consortium | Jacquelyn Wynn, Mairead Hyland, The Lighthouse Lab in Alderley Park and Alex Alderton, Roberto Amato, Jeffrey Barrett, Sonia Goncalves, Ewan Harrison, David K. Jackson, Ian Johnston, Dominic Kwiatkowski, Cordelia Langford, John Sillitoe on behalf of the Wellcome Sanger Institute COVID-19 Surveillance Team |
| EPI_ISL_3052108 | Lighthouse Lab in Glasgow | Wellcome Sanger Institute for the COVID-19 Genomics UK (COG-UK) Consortium | Harper VanSteenhouse, Yumi Kasai, David Gray, Carol Clugston, Anna Dominiczak and Alex Alderton, Roberto Amato, Jeffrey Barrett, Sonia Goncalves, Ewan Harrison, David K. Jackson, Ian Johnston, Dominic Kwiatkowski, Cordelia Langford, John Sillitoe on behalf of the Wellcome Sanger Institute COVID-19 Surveillance Team |
| EPI_ISL_3052109, EPI_ISL_3052110, EPI_ISL_3052111 | Lighthouse Lab in Alderley Park | Wellcome Sanger Institute for the COVID-19 Genomics UK (COG-UK) Consortium | Jacquelyn Wynn, Mairead Hyland, The Lighthouse Lab in Alderley Park and Alex Alderton, Roberto Amato, Jeffrey Barrett, Sonia Goncalves, Ewan Harrison, David K. Jackson, Ian Johnston, Dominic Kwiatkowski, Cordelia Langford, John Sillitoe on behalf of the Wellcome Sanger Institute COVID-19 Surveillance Team |
| EPI_ISL_3052112 | Lighthouse Lab in Glasgow | Wellcome Sanger Institute for the COVID-19 Genomics UK (COG-UK) Consortium | Harper VanSteenhouse, Yumi Kasai, David Gray, Carol Clugston, Anna Dominiczak and Alex Alderton, Roberto Amato, Jeffrey Barrett, Sonia Goncalves, Ewan Harrison, David K. Jackson, Ian Johnston, Dominic Kwiatkowski, Cordelia Langford, John Sillitoe on behalf of the Wellcome Sanger Institute COVID-19 Surveillance Team |
| EPI_ISL_3052113 | Lighthouse Lab in Alderley Park | Wellcome Sanger Institute for the COVID-19 Genomics UK (COG-UK) Consortium | Jacquelyn Wynn, Mairead Hyland, The Lighthouse Lab in Alderley Park and Alex Alderton, Roberto Amato, Jeffrey Barrett, Sonia Goncalves, Ewan Harrison, David K. Jackson, Ian Johnston, Dominic Kwiatkowski, Cordelia Langford, John Sillitoe on behalf of the Wellcome Sanger Institute COVID-19 Surveillance Team |
| EPI_ISL_3052114 | Lighthouse Lab in Glasgow | Wellcome Sanger Institute for the COVID-19 Genomics UK (COG-UK) Consortium | Harper VanSteenhouse, Yumi Kasai, David Gray, Carol Clugston, Anna Dominiczak and Alex Alderton, Roberto Amato, Jeffrey Barrett, Sonia Goncalves, Ewan Harrison, David K. Jackson, Ian Johnston, Dominic Kwiatkowski, Cordelia Langford, John Sillitoe on behalf of the Wellcome Sanger Institute COVID-19 Surveillance Team |
| EPI_ISL_3052115 | Lighthouse Lab in Alderley Park | Wellcome Sanger Institute for the COVID-19 Genomics UK (COG-UK) Consortium | Jacquelyn Wynn, Mairead Hyland, The Lighthouse Lab in Alderley Park and Alex Alderton, Roberto Amato, Jeffrey Barrett, Sonia Goncalves, Ewan Harrison, David K. Jackson, Ian Johnston, Dominic Kwiatkowski, Cordelia Langford, John Sillitoe on behalf of the Wellcome Sanger Institute COVID-19 Surveillance Team |
| EPI_ISL_3052116, EPI_ISL_3052117, EPI_ISL_3052118 | Lighthouse Lab in Glasgow | Wellcome Sanger Institute for the COVID-19 Genomics UK (COG-UK) Consortium | Harper VanSteenhouse, Yumi Kasai, David Gray, Carol Clugston, Anna Dominiczak and Alex Alderton, Roberto Amato, Jeffrey Barrett, Sonia Goncalves, Ewan Harrison, David K. Jackson, Ian Johnston, Dominic Kwiatkowski, Cordelia Langford, John Sillitoe on behalf of the Wellcome Sanger Institute COVID-19 Surveillance Team |
| EPI_ISL_3052119 | Lighthouse Lab in Alderley Park | Wellcome Sanger Institute for the COVID-19 Genomics UK (COG-UK) Consortium | Jacquelyn Wynn, Mairead Hyland, The Lighthouse Lab in Alderley Park and Alex Alderton, Roberto Amato, Jeffrey Barrett, Sonia Goncalves, Ewan Harrison, David K. Jackson, Ian Johnston, Dominic Kwiatkowski, Cordelia Langford, John Sillitoe on behalf of the Wellcome Sanger Institute COVID-19 Surveillance Team |
| EPI_ISL_3052120, EPI_ISL_3052121 | Lighthouse Lab in Glasgow | Wellcome Sanger Institute for the COVID-19 Genomics UK (COG-UK) Consortium | Harper VanSteenhouse, Yumi Kasai, David Gray, Carol Clugston, Anna Dominiczak and Alex Alderton, Roberto Amato, Jeffrey Barrett, Sonia Goncalves, Ewan Harrison, David K. Jackson, Ian Johnston, Dominic Kwiatkowski, Cordelia Langford, John Sillitoe on behalf of the Wellcome Sanger Institute COVID-19 Surveillance Team |
| EPI_ISL_3052122 | Lighthouse Lab in Alderley Park | Wellcome Sanger Institute for the COVID-19 Genomics UK (COG-UK) Consortium | Jacquelyn Wynn, Mairead Hyland, The Lighthouse Lab in Alderley Park and Alex Alderton, Roberto Amato, Jeffrey Barrett, Sonia Goncalves, Ewan Harrison, David K. Jackson, Ian Johnston, Dominic Kwiatkowski, Cordelia Langford, John Sillitoe on behalf of the Wellcome Sanger Institute COVID-19 Surveillance Team |
| EPI_ISL_3052123, EPI_ISL_3052124 | Lighthouse Lab in Glasgow | Wellcome Sanger Institute for the COVID-19 Genomics UK (COG-UK) Consortium | Harper VanSteenhouse, Yumi Kasai, David Gray, Carol Clugston, Anna Dominiczak and Alex Alderton, Roberto Amato, Jeffrey Barrett, Sonia Goncalves, Ewan Harrison, David K. Jackson, Ian Johnston, Dominic Kwiatkowski, Cordelia Langford, John Sillitoe on behalf of the Wellcome Sanger Institute COVID-19 Surveillance Team |
| EPI_ISL_3052125, EPI_ISL_3052126 | Lighthouse Lab in Alderley Park | Wellcome Sanger Institute for the COVID-19 Genomics UK (COG-UK) Consortium | Jacquelyn Wynn, Mairead Hyland, The Lighthouse Lab in Alderley Park and Alex Alderton, Roberto Amato, Jeffrey Barrett, Sonia Goncalves, Ewan Harrison, David K. Jackson, Ian Johnston, Dominic Kwiatkowski, Cordelia Langford, John Sillitoe on behalf of the Wellcome Sanger Institute COVID-19 Surveillance Team |
| EPI_ISL_3052127 | Lighthouse Lab in Glasgow | Wellcome Sanger Institute for the COVID-19 Genomics UK (COG-UK) Consortium | Harper VanSteenhouse, Yumi Kasai, David Gray, Carol Clugston, Anna Dominiczak and Alex Alderton, Roberto Amato, Jeffrey Barrett, Sonia Goncalves, Ewan Harrison, David K. Jackson, Ian Johnston, Dominic Kwiatkowski, Cordelia Langford, John Sillitoe on behalf of the Wellcome Sanger Institute COVID-19 Surveillance Team |
| EPI_ISL_3052128, EPI_ISL_3052129, EPI_ISL_3052130 | Lighthouse Lab in Alderley Park | Wellcome Sanger Institute for the COVID-19 Genomics UK (COG-UK) Consortium | Jacquelyn Wynn, Mairead Hyland, The Lighthouse Lab in Alderley Park and Alex Alderton, Roberto Amato, Jeffrey Barrett, Sonia Goncalves, Ewan Harrison, David K. Jackson, Ian Johnston, Dominic Kwiatkowski, Cordelia Langford, John Sillitoe on behalf of the Wellcome Sanger Institute COVID-19 Surveillance Team |
| EPI_ISL_3052131, EPI_ISL_3052132 | Lighthouse Lab in Glasgow | Wellcome Sanger Institute for the COVID-19 Genomics UK (COG-UK) Consortium | Harper VanSteenhouse, Yumi Kasai, David Gray, Carol Clugston, Anna Dominiczak and Alex Alderton, Roberto Amato, Jeffrey Barrett, Sonia Goncalves, Ewan Harrison, David K. Jackson, Ian Johnston, Dominic Kwiatkowski, Cordelia Langford, John Sillitoe on behalf of the Wellcome Sanger Institute COVID-19 Surveillance Team |
| EPI_ISL_3052133, EPI_ISL_3052134, EPI_ISL_3052135 | Lighthouse Lab in Alderley Park | Wellcome Sanger Institute for the COVID-19 Genomics UK (COG-UK) Consortium | Jacquelyn Wynn, Mairead Hyland, The Lighthouse Lab in Alderley Park and Alex Alderton, Roberto Amato, Jeffrey Barrett, Sonia Goncalves, Ewan Harrison, David K. Jackson, Ian Johnston, Dominic Kwiatkowski, Cordelia Langford, John Sillitoe on behalf of the Wellcome Sanger Institute COVID-19 Surveillance Team |
| EPI_ISL_3052136 | Lighthouse Lab in Glasgow | Wellcome Sanger Institute for the COVID-19 Genomics UK (COG-UK) Consortium | Harper VanSteenhouse, Yumi Kasai, David Gray, Carol Clugston, Anna Dominiczak and Alex Alderton, Roberto Amato, Jeffrey Barrett, Sonia Goncalves, Ewan Harrison, David K. Jackson, Ian Johnston, Dominic Kwiatkowski, Cordelia Langford, John Sillitoe on behalf of the Wellcome Sanger Institute COVID-19 Surveillance Team |
| EPI_ISL_3052137, EPI_ISL_3052138 | Lighthouse Lab in Alderley Park | Wellcome Sanger Institute for the COVID-19 Genomics UK (COG-UK) Consortium | Jacquelyn Wynn, Mairead Hyland, The Lighthouse Lab in Alderley Park and Alex Alderton, Roberto Amato, Jeffrey Barrett, Sonia Goncalves, Ewan Harrison, David K. Jackson, Ian Johnston, Dominic Kwiatkowski, Cordelia Langford, John Sillitoe on behalf of the Wellcome Sanger Institute COVID-19 Surveillance Team |
| EPI_ISL_3052139, EPI_ISL_3052140 | Lighthouse Lab in Glasgow | Wellcome Sanger Institute for the COVID-19 Genomics UK (COG-UK) Consortium | Harper VanSteenhouse, Yumi Kasai, David Gray, Carol Clugston, Anna Dominiczak and Alex Alderton, Roberto Amato, Jeffrey Barrett, Sonia Goncalves, Ewan Harrison, David K. Jackson, Ian Johnston, Dominic Kwiatkowski, Cordelia Langford, John Sillitoe on behalf of the Wellcome Sanger Institute COVID-19 Surveillance Team |
| EPI_ISL_3052141, EPI_ISL_3052142, EPI_ISL_3052143, EPI_ISL_3052144 | Lighthouse Lab in Alderley Park | Wellcome Sanger Institute for the COVID-19 Genomics UK (COG-UK) Consortium | Jacquelyn Wynn, Mairead Hyland, The Lighthouse Lab in Alderley Park and Alex Alderton, Roberto Amato, Jeffrey Barrett, Sonia Goncalves, Ewan Harrison, David K. Jackson, Ian Johnston, Dominic Kwiatkowski, Cordelia Langford, John Sillitoe on behalf of the Wellcome Sanger Institute COVID-19 Surveillance Team |
| EPI_ISL_3052145, EPI_ISL_3052146, EPI_ISL_3052147 | Lighthouse Lab in Glasgow | Wellcome Sanger Institute for the COVID-19 Genomics UK (COG-UK) Consortium | Harper VanSteenhouse, Yumi Kasai, David Gray, Carol Clugston, Anna Dominiczak and Alex Alderton, Roberto Amato, Jeffrey Barrett, Sonia Goncalves, Ewan Harrison, David K. Jackson, Ian Johnston, Dominic Kwiatkowski, Cordelia Langford, John Sillitoe on behalf of the Wellcome Sanger Institute COVID-19 Surveillance Team |

| | | | |
|---|---|---|---|
| EPI_ISL_3052148, EPI_ISL_3052149 | Lighthouse Lab in Alderley Park | Wellcome Sanger Institute for the COVID-19 Genomics UK (COG-UK) Consortium | Jacquelyn Wynn, Mairead Hyland, The Lighthouse Lab in Alderley Park and Alex Alderton, Roberto Amato, Jeffrey Barrett, Sonia Goncalves, Ewan Harrison, David K. Jackson, Ian Johnston, Dominic Kwiatkowski, Cordelia Langford, John Sillitoe on behalf of the Wellcome Sanger Institute COVID-19 Surveillance Team |
| EPI_ISL_3052150, EPI_ISL_3052151, EPI_ISL_3052152 | Lighthouse Lab in Glasgow | Wellcome Sanger Institute for the COVID-19 Genomics UK (COG-UK) Consortium | Harper VanSteenhouse, Yumi Kasai, David Gray, Carol Clugston, Anna Dominiczak and Alex Alderton, Roberto Amato, Jeffrey Barrett, Sonia Goncalves, Ewan Harrison, David K. Jackson, Ian Johnston, Dominic Kwiatkowski, Cordelia Langford, John Sillitoe on behalf of the Wellcome Sanger Institute COVID-19 Surveillance Team |
| EPI_ISL_3052153, EPI_ISL_3052154 | Lighthouse Lab in Alderley Park | Wellcome Sanger Institute for the COVID-19 Genomics UK (COG-UK) Consortium | Jacquelyn Wynn, Mairead Hyland, The Lighthouse Lab in Alderley Park and Alex Alderton, Roberto Amato, Jeffrey Barrett, Sonia Goncalves, Ewan Harrison, David K. Jackson, Ian Johnston, Dominic Kwiatkowski, Cordelia Langford, John Sillitoe on behalf of the Wellcome Sanger Institute COVID-19 Surveillance Team |
| EPI_ISL_3052155 | Lighthouse Lab in Glasgow | Wellcome Sanger Institute for the COVID-19 Genomics UK (COG-UK) Consortium | Harper VanSteenhouse, Yumi Kasai, David Gray, Carol Clugston, Anna Dominiczak and Alex Alderton, Roberto Amato, Jeffrey Barrett, Sonia Goncalves, Ewan Harrison, David K. Jackson, Ian Johnston, Dominic Kwiatkowski, Cordelia Langford, John Sillitoe on behalf of the Wellcome Sanger Institute COVID-19 Surveillance Team |
| EPI_ISL_3052156, EPI_ISL_3052157, EPI_ISL_3052158, EPI_ISL_3052159 | Lighthouse Lab in Alderley Park | Wellcome Sanger Institute for the COVID-19 Genomics UK (COG-UK) Consortium | Jacquelyn Wynn, Mairead Hyland, The Lighthouse Lab in Alderley Park and Alex Alderton, Roberto Amato, Jeffrey Barrett, Sonia Goncalves, Ewan Harrison, David K. Jackson, Ian Johnston, Dominic Kwiatkowski, Cordelia Langford, John Sillitoe on behalf of the Wellcome Sanger Institute COVID-19 Surveillance Team |
| EPI_ISL_3052160 | Lighthouse Lab in Glasgow | Wellcome Sanger Institute for the COVID-19 Genomics UK (COG-UK) Consortium | Harper VanSteenhouse, Yumi Kasai, David Gray, Carol Clugston, Anna Dominiczak and Alex Alderton, Roberto Amato, Jeffrey Barrett, Sonia Goncalves, Ewan Harrison, David K. Jackson, Ian Johnston, Dominic Kwiatkowski, Cordelia Langford, John Sillitoe on behalf of the Wellcome Sanger Institute COVID-19 Surveillance Team |
| EPI_ISL_3052161 | Lighthouse Lab in Alderley Park | Wellcome Sanger Institute for the COVID-19 Genomics UK (COG-UK) Consortium | Jacquelyn Wynn, Mairead Hyland, The Lighthouse Lab in Alderley Park and Alex Alderton, Roberto Amato, Jeffrey Barrett, Sonia Goncalves, Ewan Harrison, David K. Jackson, Ian Johnston, Dominic Kwiatkowski, Cordelia Langford, John Sillitoe on behalf of the Wellcome Sanger Institute COVID-19 Surveillance Team |
| EPI_ISL_3052162 | Lighthouse Lab in Glasgow | Wellcome Sanger Institute for the COVID-19 Genomics UK (COG-UK) Consortium | Harper VanSteenhouse, Yumi Kasai, David Gray, Carol Clugston, Anna Dominiczak and Alex Alderton, Roberto Amato, Jeffrey Barrett, Sonia Goncalves, Ewan Harrison, David K. Jackson, Ian Johnston, Dominic Kwiatkowski, Cordelia Langford, John Sillitoe on behalf of the Wellcome Sanger Institute COVID-19 Surveillance Team |
| EPI_ISL_3052163, EPI_ISL_3052164, EPI_ISL_3052165, EPI_ISL_3052166, EPI_ISL_3052167 | Lighthouse Lab in Alderley Park | Wellcome Sanger Institute for the COVID-19 Genomics UK (COG-UK) Consortium | Jacquelyn Wynn, Mairead Hyland, The Lighthouse Lab in Alderley Park and Alex Alderton, Roberto Amato, Jeffrey Barrett, Sonia Goncalves, Ewan Harrison, David K. Jackson, Ian Johnston, Dominic Kwiatkowski, Cordelia Langford, John Sillitoe on behalf of the Wellcome Sanger Institute COVID-19 Surveillance Team |
| EPI_ISL_3052168 | Lighthouse Lab in Glasgow | Wellcome Sanger Institute for the COVID-19 Genomics UK (COG-UK) Consortium | Harper VanSteenhouse, Yumi Kasai, David Gray, Carol Clugston, Anna Dominiczak and Alex Alderton, Roberto Amato, Jeffrey Barrett, Sonia Goncalves, Ewan Harrison, David K. Jackson, Ian Johnston, Dominic Kwiatkowski, Cordelia Langford, John Sillitoe on behalf of the Wellcome Sanger Institute COVID-19 Surveillance Team |
| EPI_ISL_3052169, EPI_ISL_3052170 | Lighthouse Lab in Alderley Park | Wellcome Sanger Institute for the COVID-19 Genomics UK (COG-UK) Consortium | Jacquelyn Wynn, Mairead Hyland, The Lighthouse Lab in Alderley Park and Alex Alderton, Roberto Amato, Jeffrey Barrett, Sonia Goncalves, Ewan Harrison, David K. Jackson, Ian Johnston, Dominic Kwiatkowski, Cordelia Langford, John Sillitoe on behalf of the Wellcome Sanger Institute COVID-19 Surveillance Team |
| EPI_ISL_3052171, EPI_ISL_3052172 | Lighthouse Lab in Glasgow | Wellcome Sanger Institute for the COVID-19 Genomics UK (COG-UK) Consortium | Harper VanSteenhouse, Yumi Kasai, David Gray, Carol Clugston, Anna Dominiczak and Alex Alderton, Roberto Amato, Jeffrey Barrett, Sonia Goncalves, Ewan Harrison, David K. Jackson, Ian Johnston, Dominic Kwiatkowski, Cordelia Langford, John Sillitoe on behalf of the Wellcome Sanger Institute COVID-19 Surveillance Team |
| EPI_ISL_3052173, EPI_ISL_3052174, EPI_ISL_3052175 | Lighthouse Lab in Alderley Park | Wellcome Sanger Institute for the COVID-19 Genomics UK (COG-UK) Consortium | Jacquelyn Wynn, Mairead Hyland, The Lighthouse Lab in Alderley Park and Alex Alderton, Roberto Amato, Jeffrey Barrett, Sonia Goncalves, Ewan Harrison, David K. Jackson, Ian Johnston, Dominic Kwiatkowski, Cordelia Langford, John Sillitoe on behalf of the Wellcome Sanger Institute COVID-19 Surveillance Team |
| EPI_ISL_3052176, EPI_ISL_3052177 | Lighthouse Lab in Glasgow | Wellcome Sanger Institute for the COVID-19 Genomics UK (COG-UK) Consortium | Harper VanSteenhouse, Yumi Kasai, David Gray, Carol Clugston, Anna Dominiczak and Alex Alderton, Roberto Amato, Jeffrey Barrett, Sonia Goncalves, Ewan Harrison, David K. Jackson, Ian Johnston, Dominic Kwiatkowski, Cordelia Langford, John Sillitoe on behalf of the Wellcome Sanger Institute COVID-19 Surveillance Team |
| EPI_ISL_3052178 | Lighthouse Lab in Alderley Park | Wellcome Sanger Institute for the COVID-19 Genomics UK (COG-UK) Consortium | Jacquelyn Wynn, Mairead Hyland, The Lighthouse Lab in Alderley Park and Alex Alderton, Roberto Amato, Jeffrey Barrett, Sonia Goncalves, Ewan Harrison, David K. Jackson, Ian Johnston, Dominic Kwiatkowski, Cordelia Langford, John Sillitoe on behalf of the Wellcome Sanger Institute COVID-19 Surveillance Team |
| EPI_ISL_3052179 | Lighthouse Lab in Glasgow | Wellcome Sanger Institute for the COVID-19 Genomics UK (COG-UK) Consortium | Harper VanSteenhouse, Yumi Kasai, David Gray, Carol Clugston, Anna Dominiczak and Alex Alderton, Roberto Amato, Jeffrey Barrett, Sonia Goncalves, Ewan Harrison, David K. Jackson, Ian Johnston, Dominic Kwiatkowski, Cordelia Langford, John Sillitoe on behalf of the Wellcome Sanger Institute COVID-19 Surveillance Team |
| EPI_ISL_3052180 | Lighthouse Lab in Alderley Park | Wellcome Sanger Institute for the COVID-19 Genomics UK (COG-UK) Consortium | Jacquelyn Wynn, Mairead Hyland, The Lighthouse Lab in Alderley Park and Alex Alderton, Roberto Amato, Jeffrey Barrett, Sonia Goncalves, Ewan Harrison, David K. Jackson, Ian Johnston, Dominic Kwiatkowski, Cordelia Langford, John Sillitoe on behalf of the Wellcome Sanger Institute COVID-19 Surveillance Team |
| EPI_ISL_3052181, EPI_ISL_3052182, EPI_ISL_3052183 | Lighthouse Lab in Glasgow | Wellcome Sanger Institute for the COVID-19 Genomics UK (COG-UK) Consortium | Harper VanSteenhouse, Yumi Kasai, David Gray, Carol Clugston, Anna Dominiczak and Alex Alderton, Roberto Amato, Jeffrey Barrett, Sonia Goncalves, Ewan Harrison, David K. Jackson, Ian Johnston, Dominic Kwiatkowski, Cordelia Langford, John Sillitoe on behalf of the Wellcome Sanger Institute COVID-19 Surveillance Team |
| EPI_ISL_3052184 | Lighthouse Lab in Alderley Park | Wellcome Sanger Institute for the COVID-19 Genomics UK (COG-UK) Consortium | Jacquelyn Wynn, Mairead Hyland, The Lighthouse Lab in Alderley Park and Alex Alderton, Roberto Amato, Jeffrey Barrett, Sonia Goncalves, Ewan Harrison, David K. Jackson, Ian Johnston, Dominic Kwiatkowski, Cordelia Langford, John Sillitoe on behalf of the Wellcome Sanger Institute COVID-19 Surveillance Team |
| EPI_ISL_3052185, EPI_ISL_3052186 | Lighthouse Lab in Glasgow | Wellcome Sanger Institute for the COVID-19 Genomics UK (COG-UK) Consortium | Harper VanSteenhouse, Yumi Kasai, David Gray, Carol Clugston, Anna Dominiczak and Alex Alderton, Roberto Amato, Jeffrey Barrett, Sonia Goncalves, Ewan Harrison, David K. Jackson, Ian Johnston, Dominic Kwiatkowski, Cordelia Langford, John Sillitoe on behalf of the Wellcome Sanger Institute COVID-19 Surveillance Team |
| EPI_ISL_3052187 | Lighthouse Lab in Alderley Park | Wellcome Sanger Institute for the COVID-19 Genomics UK (COG-UK) Consortium | Jacquelyn Wynn, Mairead Hyland, The Lighthouse Lab in Alderley Park and Alex Alderton, Roberto Amato, Jeffrey Barrett, Sonia Goncalves, Ewan Harrison, David K. Jackson, Ian Johnston, Dominic Kwiatkowski, Cordelia Langford, John Sillitoe on behalf of the Wellcome Sanger Institute COVID-19 Surveillance Team |
| EPI_ISL_3052188, EPI_ISL_3052189 | Lighthouse Lab in Glasgow | Wellcome Sanger Institute for the COVID-19 Genomics UK (COG-UK) Consortium | Harper VanSteenhouse, Yumi Kasai, David Gray, Carol Clugston, Anna Dominiczak and Alex Alderton, Roberto Amato, Jeffrey Barrett, Sonia Goncalves, Ewan Harrison, David K. Jackson, Ian Johnston, Dominic Kwiatkowski, Cordelia Langford, John Sillitoe on behalf of the Wellcome Sanger Institute COVID-19 Surveillance Team |
| EPI_ISL_3052190 | Lighthouse Lab in Alderley Park | Wellcome Sanger Institute for the COVID-19 Genomics UK (COG-UK) Consortium | Jacquelyn Wynn, Mairead Hyland, The Lighthouse Lab in Alderley Park and Alex Alderton, Roberto Amato, Jeffrey Barrett, Sonia Goncalves, Ewan Harrison, David K. Jackson, Ian Johnston, Dominic Kwiatkowski, Cordelia Langford, John Sillitoe on behalf of the Wellcome Sanger Institute COVID-19 Surveillance Team |
| EPI_ISL_3052191 | Lighthouse Lab in Glasgow | Wellcome Sanger Institute for the COVID-19 Genomics UK (COG-UK) Consortium | Harper VanSteenhouse, Yumi Kasai, David Gray, Carol Clugston, Anna Dominiczak and Alex Alderton, Roberto Amato, Jeffrey Barrett, Sonia Goncalves, Ewan Harrison, David K. Jackson, Ian Johnston, Dominic Kwiatkowski, Cordelia Langford, John Sillitoe on behalf of the Wellcome Sanger Institute COVID-19 Surveillance Team |
| EPI_ISL_3052192 | Lighthouse Lab in Alderley Park | Wellcome Sanger Institute for the COVID-19 Genomics UK (COG-UK) Consortium | Jacquelyn Wynn, Mairead Hyland, The Lighthouse Lab in Alderley Park and Alex Alderton, Roberto Amato, Jeffrey Barrett, Sonia Goncalves, Ewan Harrison, David K. Jackson, Ian Johnston, Dominic Kwiatkowski, Cordelia Langford, John Sillitoe on behalf of the Wellcome Sanger Institute COVID-19 Surveillance Team |
| EPI_ISL_3052193, EPI_ISL_3052194, EPI_ISL_3052195 | Lighthouse Lab in Glasgow | Wellcome Sanger Institute for the COVID-19 Genomics UK (COG-UK) Consortium | Harper VanSteenhouse, Yumi Kasai, David Gray, Carol Clugston, Anna Dominiczak and Alex Alderton, Roberto Amato, Jeffrey Barrett, Sonia Goncalves, Ewan Harrison, David K. Jackson, Ian Johnston, Dominic Kwiatkowski, Cordelia Langford, John Sillitoe on behalf of the Wellcome Sanger Institute COVID-19 Surveillance Team |
| EPI_ISL_3052196, EPI_ISL_3052197 | Lighthouse Lab in Alderley Park | Wellcome Sanger Institute for the COVID-19 Genomics UK (COG-UK) Consortium | Jacquelyn Wynn, Mairead Hyland, The Lighthouse Lab in Alderley Park and Alex Alderton, Roberto Amato, Jeffrey Barrett, Sonia Goncalves, Ewan Harrison, David K. Jackson, Ian Johnston, Dominic Kwiatkowski, Cordelia Langford, John Sillitoe on behalf of the Wellcome Sanger Institute COVID-19 Surveillance Team |
| EPI_ISL_3052198 | Lighthouse Lab in Glasgow | Wellcome Sanger Institute for the COVID-19 Genomics UK (COG-UK) Consortium | Harper VanSteenhouse, Yumi Kasai, David Gray, Carol Clugston, Anna Dominiczak and Alex Alderton, Roberto Amato, Jeffrey Barrett, Sonia Goncalves, Ewan Harrison, David K. Jackson, Ian Johnston, Dominic Kwiatkowski, Cordelia Langford, John Sillitoe on behalf of the Wellcome Sanger Institute COVID-19 Surveillance Team |
| EPI_ISL_3052199 | Lighthouse Lab in Alderley Park | Wellcome Sanger Institute for the COVID-19 Genomics UK (COG-UK) Consortium | Jacquelyn Wynn, Mairead Hyland, The Lighthouse Lab in Alderley Park and Alex Alderton, Roberto Amato, Jeffrey Barrett, Sonia Goncalves, Ewan Harrison, David K. Jackson, Ian Johnston, Dominic Kwiatkowski, Cordelia Langford, John Sillitoe on behalf of the Wellcome Sanger Institute COVID-19 Surveillance Team |
| EPI_ISL_3052200, EPI_ISL_3052201 | Lighthouse Lab in Glasgow | Wellcome Sanger Institute for the COVID-19 Genomics UK (COG-UK) Consortium | Harper VanSteenhouse, Yumi Kasai, David Gray, Carol Clugston, Anna Dominiczak and Alex Alderton, Roberto Amato, Jeffrey Barrett, Sonia Goncalves, Ewan Harrison, David K. Jackson, Ian Johnston, Dominic Kwiatkowski, Cordelia Langford, John Sillitoe on behalf of the Wellcome Sanger Institute COVID-19 Surveillance Team |
| EPI_ISL_3052202 | Lighthouse Lab in Alderley Park | Wellcome Sanger Institute for the COVID-19 Genomics UK (COG-UK) Consortium | Jacquelyn Wynn, Mairead Hyland, The Lighthouse Lab in Alderley Park and Alex Alderton, Roberto Amato, Jeffrey Barrett, Sonia Goncalves, Ewan Harrison, David K. Jackson, Ian Johnston, Dominic Kwiatkowski, Cordelia Langford, John Sillitoe on behalf of the Wellcome Sanger Institute COVID-19 Surveillance Team |
| EPI_ISL_3052203, EPI_ISL_3052204 | Lighthouse Lab in Glasgow | Wellcome Sanger Institute for the COVID-19 Genomics UK (COG-UK) Consortium | Harper VanSteenhouse, Yumi Kasai, David Gray, Carol Clugston, Anna Dominiczak and Alex Alderton, Roberto Amato, Jeffrey Barrett, Sonia Goncalves, Ewan Harrison, David K. Jackson, Ian Johnston, Dominic Kwiatkowski, Cordelia Langford, John Sillitoe on behalf of the Wellcome Sanger Institute COVID-19 Surveillance Team |
| EPI_ISL_3052205 | Lighthouse Lab in Alderley Park | Wellcome Sanger Institute for the COVID-19 Genomics UK (COG-UK) Consortium | Jacquelyn Wynn, Mairead Hyland, The Lighthouse Lab in Alderley Park and Alex Alderton, Roberto Amato, Jeffrey Barrett, Sonia Goncalves, Ewan Harrison, David K. Jackson, Ian Johnston, Dominic Kwiatkowski, Cordelia Langford, John Sillitoe on behalf of the Wellcome Sanger Institute COVID-19 Surveillance Team |
| EPI_ISL_3052206, EPI_ISL_3052207 | Lighthouse Lab in Glasgow | Wellcome Sanger Institute for the COVID-19 Genomics UK (COG-UK) Consortium | Harper VanSteenhouse, Yumi Kasai, David Gray, Carol Clugston, Anna Dominiczak and Alex Alderton, Roberto Amato, Jeffrey Barrett, Sonia Goncalves, Ewan Harrison, David K. Jackson, Ian Johnston, Dominic Kwiatkowski, Cordelia Langford, John Sillitoe on behalf of the Wellcome Sanger Institute COVID-19 Surveillance Team |
| EPI_ISL_3052208 | Lighthouse Lab in Alderley Park | Wellcome Sanger Institute for the COVID-19 Genomics UK (COG-UK) Consortium | Jacquelyn Wynn, Mairead Hyland, The Lighthouse Lab in Alderley Park and Alex Alderton, Roberto Amato, Jeffrey Barrett, Sonia Goncalves, Ewan Harrison, David K. Jackson, Ian Johnston, Dominic Kwiatkowski, Cordelia Langford, John Sillitoe on behalf of the Wellcome Sanger Institute COVID-19 Surveillance Team |
| EPI_ISL_3052209, EPI_ISL_3052210 | Lighthouse Lab in Glasgow | Wellcome Sanger Institute for the COVID-19 Genomics UK (COG-UK) Consortium | Harper VanSteenhouse, Yumi Kasai, David Gray, Carol Clugston, Anna Dominiczak and Alex Alderton, Roberto Amato, Jeffrey Barrett, Sonia Goncalves, Ewan Harrison, David K. Jackson, Ian Johnston, Dominic Kwiatkowski, Cordelia Langford, John Sillitoe on behalf of the Wellcome Sanger Institute COVID-19 Surveillance Team |
| EPI_ISL_3052211, EPI_ISL_3052212, EPI_ISL_3052213, EPI_ISL_3052214 | Lighthouse Lab in Alderley Park | Wellcome Sanger Institute for the COVID-19 Genomics UK (COG-UK) Consortium | Jacquelyn Wynn, Mairead Hyland, The Lighthouse Lab in Alderley Park and Alex Alderton, Roberto Amato, Jeffrey Barrett, Sonia Goncalves, Ewan Harrison, David K. Jackson, Ian Johnston, Dominic Kwiatkowski, Cordelia Langford, John Sillitoe on behalf of the Wellcome Sanger Institute COVID-19 Surveillance Team |
| EPI_ISL_3052215 | Lighthouse Lab in Glasgow | Wellcome Sanger Institute for the COVID-19 Genomics UK (COG-UK) Consortium | Harper VanSteenhouse, Yumi Kasai, David Gray, Carol Clugston, Anna Dominiczak and Alex Alderton, Roberto Amato, Jeffrey Barrett, Sonia Goncalves, Ewan Harrison, David K. Jackson, Ian Johnston, Dominic Kwiatkowski, Cordelia Langford, John Sillitoe on behalf of the Wellcome Sanger Institute COVID-19 Surveillance Team |
| EPI_ISL_3052216 | Lighthouse Lab in Alderley Park | Wellcome Sanger Institute for the COVID-19 Genomics UK (COG-UK) Consortium | Jacquelyn Wynn, Mairead Hyland, The Lighthouse Lab in Alderley Park and Alex Alderton, Roberto Amato, Jeffrey Barrett, Sonia Goncalves, Ewan Harrison, David K. Jackson, Ian Johnston, Dominic Kwiatkowski, Cordelia Langford, John Sillitoe on behalf of the Wellcome Sanger Institute COVID-19 Surveillance Team |
| EPI_ISL_3052217, EPI_ISL_3052218 | Lighthouse Lab in Glasgow | Wellcome Sanger Institute for the COVID-19 Genomics UK (COG-UK) Consortium | Harper VanSteenhouse, Yumi Kasai, David Gray, Carol Clugston, Anna Dominiczak and Alex Alderton, Roberto Amato, Jeffrey Barrett, Sonia Goncalves, Ewan Harrison, David K. Jackson, Ian Johnston, Dominic Kwiatkowski, Cordelia Langford, John Sillitoe on behalf of the Wellcome Sanger Institute COVID-19 Surveillance Team |
| EPI_ISL_3052219, | Lighthouse Lab in Alderley Park | Wellcome Sanger Institute for the COVID-19 Genomics UK (COG-UK) | Jacquelyn Wynn, Mairead Hyland, The Lighthouse Lab in Alderley Park and Alex Alderton, Roberto Amato, Jeffrey Barrett, Sonia Goncalves, Ewan Harrison, David K. Jackson, Ian Johnston, Dominic Kwiatkowski, Cordelia Langford, John Sillitoe on |

| | | | |
|---|---|---|---|
| EPI_ISL_3052220, EPI_ISL_3052221, EPI_ISL_3052222 | | Consortium | behalf of the Wellcome Sanger Institute COVID-19 Surveillance Team |
| EPI_ISL_3052223 | Lighthouse Lab in Glasgow | Wellcome Sanger Institute for the COVID-19 Genomics UK (COG-UK) Consortium | Harper VanSteenhouse, Yumi Kasai, David Gray, Carol Clugston, Anna Dominiczak and Alex Alderton, Roberto Amato, Jeffrey Barrett, Sonia Goncalves, Ewan Harrison, David K. Jackson, Ian Johnston, Dominic Kwiatkowski, Cordelia Langford, John Sillitoe on behalf of the Wellcome Sanger Institute COVID-19 Surveillance Team |
| EPI_ISL_3052224 | Lighthouse Lab in Alderley Park | Wellcome Sanger Institute for the COVID-19 Genomics UK (COG-UK) Consortium | Jacquelyn Wynn, Mairead Hyland, The Lighthouse Lab in Alderley Park and Alex Alderton, Roberto Amato, Jeffrey Barrett, Sonia Goncalves, Ewan Harrison, David K. Jackson, Ian Johnston, Dominic Kwiatkowski, Cordelia Langford, John Sillitoe on behalf of the Wellcome Sanger Institute COVID-19 Surveillance Team |
| EPI_ISL_3052225, EPI_ISL_3052226, EPI_ISL_3052227 | Lighthouse Lab in Glasgow | Wellcome Sanger Institute for the COVID-19 Genomics UK (COG-UK) Consortium | Harper VanSteenhouse, Yumi Kasai, David Gray, Carol Clugston, Anna Dominiczak and Alex Alderton, Roberto Amato, Jeffrey Barrett, Sonia Goncalves, Ewan Harrison, David K. Jackson, Ian Johnston, Dominic Kwiatkowski, Cordelia Langford, John Sillitoe on behalf of the Wellcome Sanger Institute COVID-19 Surveillance Team |
| EPI_ISL_3052228, EPI_ISL_3052229, EPI_ISL_3052230 | Lighthouse Lab in Alderley Park | Wellcome Sanger Institute for the COVID-19 Genomics UK (COG-UK) Consortium | Jacquelyn Wynn, Mairead Hyland, The Lighthouse Lab in Alderley Park and Alex Alderton, Roberto Amato, Jeffrey Barrett, Sonia Goncalves, Ewan Harrison, David K. Jackson, Ian Johnston, Dominic Kwiatkowski, Cordelia Langford, John Sillitoe on behalf of the Wellcome Sanger Institute COVID-19 Surveillance Team |
| EPI_ISL_3052231 | Lighthouse Lab in Glasgow | Wellcome Sanger Institute for the COVID-19 Genomics UK (COG-UK) Consortium | Harper VanSteenhouse, Yumi Kasai, David Gray, Carol Clugston, Anna Dominiczak and Alex Alderton, Roberto Amato, Jeffrey Barrett, Sonia Goncalves, Ewan Harrison, David K. Jackson, Ian Johnston, Dominic Kwiatkowski, Cordelia Langford, John Sillitoe on behalf of the Wellcome Sanger Institute COVID-19 Surveillance Team |
| EPI_ISL_3052232 | Lighthouse Lab in Alderley Park | Wellcome Sanger Institute for the COVID-19 Genomics UK (COG-UK) Consortium | Jacquelyn Wynn, Mairead Hyland, The Lighthouse Lab in Alderley Park and Alex Alderton, Roberto Amato, Jeffrey Barrett, Sonia Goncalves, Ewan Harrison, David K. Jackson, Ian Johnston, Dominic Kwiatkowski, Cordelia Langford, John Sillitoe on behalf of the Wellcome Sanger Institute COVID-19 Surveillance Team |
| EPI_ISL_3052233 | Lighthouse Lab in Glasgow | Wellcome Sanger Institute for the COVID-19 Genomics UK (COG-UK) Consortium | Harper VanSteenhouse, Yumi Kasai, David Gray, Carol Clugston, Anna Dominiczak and Alex Alderton, Roberto Amato, Jeffrey Barrett, Sonia Goncalves, Ewan Harrison, David K. Jackson, Ian Johnston, Dominic Kwiatkowski, Cordelia Langford, John Sillitoe on behalf of the Wellcome Sanger Institute COVID-19 Surveillance Team |
| EPI_ISL_3052234, EPI_ISL_3052235 | Lighthouse Lab in Alderley Park | Wellcome Sanger Institute for the COVID-19 Genomics UK (COG-UK) Consortium | Jacquelyn Wynn, Mairead Hyland, The Lighthouse Lab in Alderley Park and Alex Alderton, Roberto Amato, Jeffrey Barrett, Sonia Goncalves, Ewan Harrison, David K. Jackson, Ian Johnston, Dominic Kwiatkowski, Cordelia Langford, John Sillitoe on behalf of the Wellcome Sanger Institute COVID-19 Surveillance Team |
| EPI_ISL_3052236, EPI_ISL_3052237 | Lighthouse Lab in Glasgow | Wellcome Sanger Institute for the COVID-19 Genomics UK (COG-UK) Consortium | Harper VanSteenhouse, Yumi Kasai, David Gray, Carol Clugston, Anna Dominiczak and Alex Alderton, Roberto Amato, Jeffrey Barrett, Sonia Goncalves, Ewan Harrison, David K. Jackson, Ian Johnston, Dominic Kwiatkowski, Cordelia Langford, John Sillitoe on behalf of the Wellcome Sanger Institute COVID-19 Surveillance Team |
| EPI_ISL_3052238 | Lighthouse Lab in Alderley Park | Wellcome Sanger Institute for the COVID-19 Genomics UK (COG-UK) Consortium | Jacquelyn Wynn, Mairead Hyland, The Lighthouse Lab in Alderley Park and Alex Alderton, Roberto Amato, Jeffrey Barrett, Sonia Goncalves, Ewan Harrison, David K. Jackson, Ian Johnston, Dominic Kwiatkowski, Cordelia Langford, John Sillitoe on behalf of the Wellcome Sanger Institute COVID-19 Surveillance Team |
| EPI_ISL_3052239 | Lighthouse Lab in Glasgow | Wellcome Sanger Institute for the COVID-19 Genomics UK (COG-UK) Consortium | Harper VanSteenhouse, Yumi Kasai, David Gray, Carol Clugston, Anna Dominiczak and Alex Alderton, Roberto Amato, Jeffrey Barrett, Sonia Goncalves, Ewan Harrison, David K. Jackson, Ian Johnston, Dominic Kwiatkowski, Cordelia Langford, John Sillitoe on behalf of the Wellcome Sanger Institute COVID-19 Surveillance Team |
| EPI_ISL_3052240 | Lighthouse Lab in Alderley Park | Wellcome Sanger Institute for the COVID-19 Genomics UK (COG-UK) Consortium | Jacquelyn Wynn, Mairead Hyland, The Lighthouse Lab in Alderley Park and Alex Alderton, Roberto Amato, Jeffrey Barrett, Sonia Goncalves, Ewan Harrison, David K. Jackson, Ian Johnston, Dominic Kwiatkowski, Cordelia Langford, John Sillitoe on behalf of the Wellcome Sanger Institute COVID-19 Surveillance Team |
| EPI_ISL_3052241, EPI_ISL_3052242 | Lighthouse Lab in Glasgow | Wellcome Sanger Institute for the COVID-19 Genomics UK (COG-UK) Consortium | Harper VanSteenhouse, Yumi Kasai, David Gray, Carol Clugston, Anna Dominiczak and Alex Alderton, Roberto Amato, Jeffrey Barrett, Sonia Goncalves, Ewan Harrison, David K. Jackson, Ian Johnston, Dominic Kwiatkowski, Cordelia Langford, John Sillitoe on behalf of the Wellcome Sanger Institute COVID-19 Surveillance Team |
| EPI_ISL_3052243, EPI_ISL_3052244 | Lighthouse Lab in Alderley Park | Wellcome Sanger Institute for the COVID-19 Genomics UK (COG-UK) Consortium | Jacquelyn Wynn, Mairead Hyland, The Lighthouse Lab in Alderley Park and Alex Alderton, Roberto Amato, Jeffrey Barrett, Sonia Goncalves, Ewan Harrison, David K. Jackson, Ian Johnston, Dominic Kwiatkowski, Cordelia Langford, John Sillitoe on behalf of the Wellcome Sanger Institute COVID-19 Surveillance Team |
| EPI_ISL_3052245 | Lighthouse Lab in Glasgow | Wellcome Sanger Institute for the COVID-19 Genomics UK (COG-UK) Consortium | Harper VanSteenhouse, Yumi Kasai, David Gray, Carol Clugston, Anna Dominiczak and Alex Alderton, Roberto Amato, Jeffrey Barrett, Sonia Goncalves, Ewan Harrison, David K. Jackson, Ian Johnston, Dominic Kwiatkowski, Cordelia Langford, John Sillitoe on behalf of the Wellcome Sanger Institute COVID-19 Surveillance Team |
| EPI_ISL_3052246 | Lighthouse Lab in Alderley Park | Wellcome Sanger Institute for the COVID-19 Genomics UK (COG-UK) Consortium | Jacquelyn Wynn, Mairead Hyland, The Lighthouse Lab in Alderley Park and Alex Alderton, Roberto Amato, Jeffrey Barrett, Sonia Goncalves, Ewan Harrison, David K. Jackson, Ian Johnston, Dominic Kwiatkowski, Cordelia Langford, John Sillitoe on behalf of the Wellcome Sanger Institute COVID-19 Surveillance Team |
| EPI_ISL_3052247, EPI_ISL_3052248, EPI_ISL_3052249, EPI_ISL_3052250 | Lighthouse Lab in Glasgow | Wellcome Sanger Institute for the COVID-19 Genomics UK (COG-UK) Consortium | Harper VanSteenhouse, Yumi Kasai, David Gray, Carol Clugston, Anna Dominiczak and Alex Alderton, Roberto Amato, Jeffrey Barrett, Sonia Goncalves, Ewan Harrison, David K. Jackson, Ian Johnston, Dominic Kwiatkowski, Cordelia Langford, John Sillitoe on behalf of the Wellcome Sanger Institute COVID-19 Surveillance Team |
| EPI_ISL_3052251 | Lighthouse Lab in Alderley Park | Wellcome Sanger Institute for the COVID-19 Genomics UK (COG-UK) Consortium | Jacquelyn Wynn, Mairead Hyland, The Lighthouse Lab in Alderley Park and Alex Alderton, Roberto Amato, Jeffrey Barrett, Sonia Goncalves, Ewan Harrison, David K. Jackson, Ian Johnston, Dominic Kwiatkowski, Cordelia Langford, John Sillitoe on behalf of the Wellcome Sanger Institute COVID-19 Surveillance Team |
| EPI_ISL_3052252 | Lighthouse Lab in Glasgow | Wellcome Sanger Institute for the COVID-19 Genomics UK (COG-UK) Consortium | Harper VanSteenhouse, Yumi Kasai, David Gray, Carol Clugston, Anna Dominiczak and Alex Alderton, Roberto Amato, Jeffrey Barrett, Sonia Goncalves, Ewan Harrison, David K. Jackson, Ian Johnston, Dominic Kwiatkowski, Cordelia Langford, John Sillitoe on behalf of the Wellcome Sanger Institute COVID-19 Surveillance Team |
| EPI_ISL_3052253 | Lighthouse Lab in Alderley Park | Wellcome Sanger Institute for the COVID-19 Genomics UK (COG-UK) Consortium | Jacquelyn Wynn, Mairead Hyland, The Lighthouse Lab in Alderley Park and Alex Alderton, Roberto Amato, Jeffrey Barrett, Sonia Goncalves, Ewan Harrison, David K. Jackson, Ian Johnston, Dominic Kwiatkowski, Cordelia Langford, John Sillitoe on behalf of the Wellcome Sanger Institute COVID-19 Surveillance Team |
| EPI_ISL_3052254 | Lighthouse Lab in Glasgow | Wellcome Sanger Institute for the COVID-19 Genomics UK (COG-UK) Consortium | Harper VanSteenhouse, Yumi Kasai, David Gray, Carol Clugston, Anna Dominiczak and Alex Alderton, Roberto Amato, Jeffrey Barrett, Sonia Goncalves, Ewan Harrison, David K. Jackson, Ian Johnston, Dominic Kwiatkowski, Cordelia Langford, John Sillitoe on behalf of the Wellcome Sanger Institute COVID-19 Surveillance Team |
| EPI_ISL_3052255, EPI_ISL_3052256, EPI_ISL_3052257, EPI_ISL_3052258, EPI_ISL_3052259, EPI_ISL_3052260 | Lighthouse Lab in Alderley Park | Wellcome Sanger Institute for the COVID-19 Genomics UK (COG-UK) Consortium | Jacquelyn Wynn, Mairead Hyland, The Lighthouse Lab in Alderley Park and Alex Alderton, Roberto Amato, Jeffrey Barrett, Sonia Goncalves, Ewan Harrison, David K. Jackson, Ian Johnston, Dominic Kwiatkowski, Cordelia Langford, John Sillitoe on behalf of the Wellcome Sanger Institute COVID-19 Surveillance Team |
| EPI_ISL_3052261 | Lighthouse Lab in Glasgow | Wellcome Sanger Institute for the COVID-19 Genomics UK (COG-UK) Consortium | Harper VanSteenhouse, Yumi Kasai, David Gray, Carol Clugston, Anna Dominiczak and Alex Alderton, Roberto Amato, Jeffrey Barrett, Sonia Goncalves, Ewan Harrison, David K. Jackson, Ian Johnston, Dominic Kwiatkowski, Cordelia Langford, John Sillitoe on behalf of the Wellcome Sanger Institute COVID-19 Surveillance Team |
| EPI_ISL_3052262, EPI_ISL_3052263 | Lighthouse Lab in Alderley Park | Wellcome Sanger Institute for the COVID-19 Genomics UK (COG-UK) Consortium | Jacquelyn Wynn, Mairead Hyland, The Lighthouse Lab in Alderley Park and Alex Alderton, Roberto Amato, Jeffrey Barrett, Sonia Goncalves, Ewan Harrison, David K. Jackson, Ian Johnston, Dominic Kwiatkowski, Cordelia Langford, John Sillitoe on behalf of the Wellcome Sanger Institute COVID-19 Surveillance Team |
| EPI_ISL_3052264 | Lighthouse Lab in Glasgow | Wellcome Sanger Institute for the COVID-19 Genomics UK (COG-UK) Consortium | Harper VanSteenhouse, Yumi Kasai, David Gray, Carol Clugston, Anna Dominiczak and Alex Alderton, Roberto Amato, Jeffrey Barrett, Sonia Goncalves, Ewan Harrison, David K. Jackson, Ian Johnston, Dominic Kwiatkowski, Cordelia Langford, John Sillitoe on behalf of the Wellcome Sanger Institute COVID-19 Surveillance Team |
| EPI_ISL_3052265 | Lighthouse Lab in Alderley Park | Wellcome Sanger Institute for the COVID-19 Genomics UK (COG-UK) Consortium | Jacquelyn Wynn, Mairead Hyland, The Lighthouse Lab in Alderley Park and Alex Alderton, Roberto Amato, Jeffrey Barrett, Sonia Goncalves, Ewan Harrison, David K. Jackson, Ian Johnston, Dominic Kwiatkowski, Cordelia Langford, John Sillitoe on behalf of the Wellcome Sanger Institute COVID-19 Surveillance Team |
| EPI_ISL_3052266, EPI_ISL_3052267, EPI_ISL_3052268, EPI_ISL_3052269 | Lighthouse Lab in Glasgow | Wellcome Sanger Institute for the COVID-19 Genomics UK (COG-UK) Consortium | Harper VanSteenhouse, Yumi Kasai, David Gray, Carol Clugston, Anna Dominiczak and Alex Alderton, Roberto Amato, Jeffrey Barrett, Sonia Goncalves, Ewan Harrison, David K. Jackson, Ian Johnston, Dominic Kwiatkowski, Cordelia Langford, John Sillitoe on behalf of the Wellcome Sanger Institute COVID-19 Surveillance Team |
| EPI_ISL_3052270 | Lighthouse Lab in Alderley Park | Wellcome Sanger Institute for the COVID-19 Genomics UK (COG-UK) Consortium | Jacquelyn Wynn, Mairead Hyland, The Lighthouse Lab in Alderley Park and Alex Alderton, Roberto Amato, Jeffrey Barrett, Sonia Goncalves, Ewan Harrison, David K. Jackson, Ian Johnston, Dominic Kwiatkowski, Cordelia Langford, John Sillitoe on behalf of the Wellcome Sanger Institute COVID-19 Surveillance Team |
| EPI_ISL_3052271, EPI_ISL_3052272, EPI_ISL_3052273 | Lighthouse Lab in Glasgow | Wellcome Sanger Institute for the COVID-19 Genomics UK (COG-UK) Consortium | Harper VanSteenhouse, Yumi Kasai, David Gray, Carol Clugston, Anna Dominiczak and Alex Alderton, Roberto Amato, Jeffrey Barrett, Sonia Goncalves, Ewan Harrison, David K. Jackson, Ian Johnston, Dominic Kwiatkowski, Cordelia Langford, John Sillitoe on behalf of the Wellcome Sanger Institute COVID-19 Surveillance Team |
| EPI_ISL_3052274, EPI_ISL_3052275 | Lighthouse Lab in Alderley Park | Wellcome Sanger Institute for the COVID-19 Genomics UK (COG-UK) Consortium | Jacquelyn Wynn, Mairead Hyland, The Lighthouse Lab in Alderley Park and Alex Alderton, Roberto Amato, Jeffrey Barrett, Sonia Goncalves, Ewan Harrison, David K. Jackson, Ian Johnston, Dominic Kwiatkowski, Cordelia Langford, John Sillitoe on behalf of the Wellcome Sanger Institute COVID-19 Surveillance Team |
| EPI_ISL_3052276, EPI_ISL_3052277 | Lighthouse Lab in Glasgow | Wellcome Sanger Institute for the COVID-19 Genomics UK (COG-UK) Consortium | Harper VanSteenhouse, Yumi Kasai, David Gray, Carol Clugston, Anna Dominiczak and Alex Alderton, Roberto Amato, Jeffrey Barrett, Sonia Goncalves, Ewan Harrison, David K. Jackson, Ian Johnston, Dominic Kwiatkowski, Cordelia Langford, John Sillitoe on behalf of the Wellcome Sanger Institute COVID-19 Surveillance Team |
| EPI_ISL_3052278, EPI_ISL_3052279, EPI_ISL_3052280 | Lighthouse Lab in Alderley Park | Wellcome Sanger Institute for the COVID-19 Genomics UK (COG-UK) Consortium | Jacquelyn Wynn, Mairead Hyland, The Lighthouse Lab in Alderley Park and Alex Alderton, Roberto Amato, Jeffrey Barrett, Sonia Goncalves, Ewan Harrison, David K. Jackson, Ian Johnston, Dominic Kwiatkowski, Cordelia Langford, John Sillitoe on behalf of the Wellcome Sanger Institute COVID-19 Surveillance Team |
| EPI_ISL_3052281 | Lighthouse Lab in Glasgow | Wellcome Sanger Institute for the COVID-19 Genomics UK (COG-UK) Consortium | Harper VanSteenhouse, Yumi Kasai, David Gray, Carol Clugston, Anna Dominiczak and Alex Alderton, Roberto Amato, Jeffrey Barrett, Sonia Goncalves, Ewan Harrison, David K. Jackson, Ian Johnston, Dominic Kwiatkowski, Cordelia Langford, John Sillitoe on behalf of the Wellcome Sanger Institute COVID-19 Surveillance Team |
| EPI_ISL_3052282, EPI_ISL_3052283 | Lighthouse Lab in Alderley Park | Wellcome Sanger Institute for the COVID-19 Genomics UK (COG-UK) Consortium | Jacquelyn Wynn, Mairead Hyland, The Lighthouse Lab in Alderley Park and Alex Alderton, Roberto Amato, Jeffrey Barrett, Sonia Goncalves, Ewan Harrison, David K. Jackson, Ian Johnston, Dominic Kwiatkowski, Cordelia Langford, John Sillitoe on behalf of the Wellcome Sanger Institute COVID-19 Surveillance Team |
| EPI_ISL_3052284, EPI_ISL_3052285 | Lighthouse Lab in Glasgow | Wellcome Sanger Institute for the COVID-19 Genomics UK (COG-UK) Consortium | Harper VanSteenhouse, Yumi Kasai, David Gray, Carol Clugston, Anna Dominiczak and Alex Alderton, Roberto Amato, Jeffrey Barrett, Sonia Goncalves, Ewan Harrison, David K. Jackson, Ian Johnston, Dominic Kwiatkowski, Cordelia Langford, John Sillitoe on behalf of the Wellcome Sanger Institute COVID-19 Surveillance Team |
| EPI_ISL_3052286, EPI_ISL_3052287, EPI_ISL_3052288 | Lighthouse Lab in Alderley Park | Wellcome Sanger Institute for the COVID-19 Genomics UK (COG-UK) Consortium | Jacquelyn Wynn, Mairead Hyland, The Lighthouse Lab in Alderley Park and Alex Alderton, Roberto Amato, Jeffrey Barrett, Sonia Goncalves, Ewan Harrison, David K. Jackson, Ian Johnston, Dominic Kwiatkowski, Cordelia Langford, John Sillitoe on behalf of the Wellcome Sanger Institute COVID-19 Surveillance Team |
| EPI_ISL_3052289 | Lighthouse Lab in Glasgow | Wellcome Sanger Institute for the COVID-19 Genomics UK (COG-UK) Consortium | Harper VanSteenhouse, Yumi Kasai, David Gray, Carol Clugston, Anna Dominiczak and Alex Alderton, Roberto Amato, Jeffrey Barrett, Sonia Goncalves, Ewan Harrison, David K. Jackson, Ian Johnston, Dominic Kwiatkowski, Cordelia Langford, John Sillitoe on behalf of the Wellcome Sanger Institute COVID-19 Surveillance Team |
| EPI_ISL_3052290 | Lighthouse Lab in Alderley Park | Wellcome Sanger Institute for the COVID-19 Genomics UK (COG-UK) Consortium | Jacquelyn Wynn, Mairead Hyland, The Lighthouse Lab in Alderley Park and Alex Alderton, Roberto Amato, Jeffrey Barrett, Sonia Goncalves, Ewan Harrison, David K. Jackson, Ian Johnston, Dominic Kwiatkowski, Cordelia Langford, John Sillitoe on behalf of the Wellcome Sanger Institute COVID-19 Surveillance Team |
| EPI_ISL_3052291, | Lighthouse Lab in Glasgow | Wellcome Sanger Institute for the COVID-19 Genomics UK (COG-UK) | Harper VanSteenhouse, Yumi Kasai, David Gray, Carol Clugston, Anna Dominiczak and Alex Alderton, Roberto Amato, Jeffrey Barrett, Sonia Goncalves, Ewan Harrison, David K. Jackson, Ian Johnston, Dominic Kwiatkowski, Cordelia Langford, |

| | | | |
|---|---|---|---|
| EPI_ISL_3052292 | | Consortium | John Sillitoe on behalf of the Wellcome Sanger Institute COVID-19 Surveillance Team |
| EPI_ISL_3052293, EPI_ISL_3052294, EPI_ISL_3052295, EPI_ISL_3052296 | Lighthouse Lab in Alderley Park | Wellcome Sanger Institute for the COVID-19 Genomics UK (COG-UK) Consortium | Jacquelyn Wynn, Mairead Hyland, The Lighthouse Lab in Alderley Park and Alex Alderton, Roberto Amato, Jeffrey Barrett, Sonia Goncalves, Ewan Harrison, David K. Jackson, Ian Johnston, Dominic Kwiatkowski, Cordelia Langford, John Sillitoe on behalf of the Wellcome Sanger Institute COVID-19 Surveillance Team |
| EPI_ISL_3052297, EPI_ISL_3052298, EPI_ISL_3052299 | Lighthouse Lab in Glasgow | Wellcome Sanger Institute for the COVID-19 Genomics UK (COG-UK) Consortium | Harper VanSteenhouse, Yumi Kasai, David Gray, Carol Clugston, Anna Dominiczak and Alex Alderton, Roberto Amato, Jeffrey Barrett, Sonia Goncalves, Ewan Harrison, David K. Jackson, Ian Johnston, Dominic Kwiatkowski, Cordelia Langford, John Sillitoe on behalf of the Wellcome Sanger Institute COVID-19 Surveillance Team |
| EPI_ISL_3052300, EPI_ISL_3052301, EPI_ISL_3052302 | Lighthouse Lab in Alderley Park | Wellcome Sanger Institute for the COVID-19 Genomics UK (COG-UK) Consortium | Jacquelyn Wynn, Mairead Hyland, The Lighthouse Lab in Alderley Park and Alex Alderton, Roberto Amato, Jeffrey Barrett, Sonia Goncalves, Ewan Harrison, David K. Jackson, Ian Johnston, Dominic Kwiatkowski, Cordelia Langford, John Sillitoe on behalf of the Wellcome Sanger Institute COVID-19 Surveillance Team |
| EPI_ISL_3052303, EPI_ISL_3052304, EPI_ISL_3052305 | Lighthouse Lab in Glasgow | Wellcome Sanger Institute for the COVID-19 Genomics UK (COG-UK) Consortium | Harper VanSteenhouse, Yumi Kasai, David Gray, Carol Clugston, Anna Dominiczak and Alex Alderton, Roberto Amato, Jeffrey Barrett, Sonia Goncalves, Ewan Harrison, David K. Jackson, Ian Johnston, Dominic Kwiatkowski, Cordelia Langford, John Sillitoe on behalf of the Wellcome Sanger Institute COVID-19 Surveillance Team |
| EPI_ISL_3052306 | Lighthouse Lab in Alderley Park | Wellcome Sanger Institute for the COVID-19 Genomics UK (COG-UK) Consortium | Jacquelyn Wynn, Mairead Hyland, The Lighthouse Lab in Alderley Park and Alex Alderton, Roberto Amato, Jeffrey Barrett, Sonia Goncalves, Ewan Harrison, David K. Jackson, Ian Johnston, Dominic Kwiatkowski, Cordelia Langford, John Sillitoe on behalf of the Wellcome Sanger Institute COVID-19 Surveillance Team |
| EPI_ISL_3052307, EPI_ISL_3052308, EPI_ISL_3052309 | Lighthouse Lab in Glasgow | Wellcome Sanger Institute for the COVID-19 Genomics UK (COG-UK) Consortium | Harper VanSteenhouse, Yumi Kasai, David Gray, Carol Clugston, Anna Dominiczak and Alex Alderton, Roberto Amato, Jeffrey Barrett, Sonia Goncalves, Ewan Harrison, David K. Jackson, Ian Johnston, Dominic Kwiatkowski, Cordelia Langford, John Sillitoe on behalf of the Wellcome Sanger Institute COVID-19 Surveillance Team |
| EPI_ISL_3052310, EPI_ISL_3052311, EPI_ISL_3052312, EPI_ISL_3052313 | Lighthouse Lab in Alderley Park | Wellcome Sanger Institute for the COVID-19 Genomics UK (COG-UK) Consortium | Jacquelyn Wynn, Mairead Hyland, The Lighthouse Lab in Alderley Park and Alex Alderton, Roberto Amato, Jeffrey Barrett, Sonia Goncalves, Ewan Harrison, David K. Jackson, Ian Johnston, Dominic Kwiatkowski, Cordelia Langford, John Sillitoe on behalf of the Wellcome Sanger Institute COVID-19 Surveillance Team |
| EPI_ISL_3052314, EPI_ISL_3052315, EPI_ISL_3052316 | Lighthouse Lab in Glasgow | Wellcome Sanger Institute for the COVID-19 Genomics UK (COG-UK) Consortium | Harper VanSteenhouse, Yumi Kasai, David Gray, Carol Clugston, Anna Dominiczak and Alex Alderton, Roberto Amato, Jeffrey Barrett, Sonia Goncalves, Ewan Harrison, David K. Jackson, Ian Johnston, Dominic Kwiatkowski, Cordelia Langford, John Sillitoe on behalf of the Wellcome Sanger Institute COVID-19 Surveillance Team |
| EPI_ISL_3052317, EPI_ISL_3052318 | Lighthouse Lab in Alderley Park | Wellcome Sanger Institute for the COVID-19 Genomics UK (COG-UK) Consortium | Jacquelyn Wynn, Mairead Hyland, The Lighthouse Lab in Alderley Park and Alex Alderton, Roberto Amato, Jeffrey Barrett, Sonia Goncalves, Ewan Harrison, David K. Jackson, Ian Johnston, Dominic Kwiatkowski, Cordelia Langford, John Sillitoe on behalf of the Wellcome Sanger Institute COVID-19 Surveillance Team |
| EPI_ISL_3052319 | Lighthouse Lab in Glasgow | Wellcome Sanger Institute for the COVID-19 Genomics UK (COG-UK) Consortium | Harper VanSteenhouse, Yumi Kasai, David Gray, Carol Clugston, Anna Dominiczak and Alex Alderton, Roberto Amato, Jeffrey Barrett, Sonia Goncalves, Ewan Harrison, David K. Jackson, Ian Johnston, Dominic Kwiatkowski, Cordelia Langford, John Sillitoe on behalf of the Wellcome Sanger Institute COVID-19 Surveillance Team |
| EPI_ISL_3052320, EPI_ISL_3052321, EPI_ISL_3052322, EPI_ISL_3052323 | Lighthouse Lab in Alderley Park | Wellcome Sanger Institute for the COVID-19 Genomics UK (COG-UK) Consortium | Jacquelyn Wynn, Mairead Hyland, The Lighthouse Lab in Alderley Park and Alex Alderton, Roberto Amato, Jeffrey Barrett, Sonia Goncalves, Ewan Harrison, David K. Jackson, Ian Johnston, Dominic Kwiatkowski, Cordelia Langford, John Sillitoe on behalf of the Wellcome Sanger Institute COVID-19 Surveillance Team |
| EPI_ISL_3052324, EPI_ISL_3052325, EPI_ISL_3052326, EPI_ISL_3052327 | Lighthouse Lab in Glasgow | Wellcome Sanger Institute for the COVID-19 Genomics UK (COG-UK) Consortium | Harper VanSteenhouse, Yumi Kasai, David Gray, Carol Clugston, Anna Dominiczak and Alex Alderton, Roberto Amato, Jeffrey Barrett, Sonia Goncalves, Ewan Harrison, David K. Jackson, Ian Johnston, Dominic Kwiatkowski, Cordelia Langford, John Sillitoe on behalf of the Wellcome Sanger Institute COVID-19 Surveillance Team |
| EPI_ISL_3052328 | Lighthouse Lab in Alderley Park | Wellcome Sanger Institute for the COVID-19 Genomics UK (COG-UK) Consortium | Jacquelyn Wynn, Mairead Hyland, The Lighthouse Lab in Alderley Park and Alex Alderton, Roberto Amato, Jeffrey Barrett, Sonia Goncalves, Ewan Harrison, David K. Jackson, Ian Johnston, Dominic Kwiatkowski, Cordelia Langford, John Sillitoe on behalf of the Wellcome Sanger Institute COVID-19 Surveillance Team |
| EPI_ISL_3052329, EPI_ISL_3052330, EPI_ISL_3052331 | Lighthouse Lab in Glasgow | Wellcome Sanger Institute for the COVID-19 Genomics UK (COG-UK) Consortium | Harper VanSteenhouse, Yumi Kasai, David Gray, Carol Clugston, Anna Dominiczak and Alex Alderton, Roberto Amato, Jeffrey Barrett, Sonia Goncalves, Ewan Harrison, David K. Jackson, Ian Johnston, Dominic Kwiatkowski, Cordelia Langford, John Sillitoe on behalf of the Wellcome Sanger Institute COVID-19 Surveillance Team |
| EPI_ISL_3052332, EPI_ISL_3052333, EPI_ISL_3052334 | Lighthouse Lab in Alderley Park | Wellcome Sanger Institute for the COVID-19 Genomics UK (COG-UK) Consortium | Jacquelyn Wynn, Mairead Hyland, The Lighthouse Lab in Alderley Park and Alex Alderton, Roberto Amato, Jeffrey Barrett, Sonia Goncalves, Ewan Harrison, David K. Jackson, Ian Johnston, Dominic Kwiatkowski, Cordelia Langford, John Sillitoe on behalf of the Wellcome Sanger Institute COVID-19 Surveillance Team |
| EPI_ISL_3052335, EPI_ISL_3052336, EPI_ISL_3052337, EPI_ISL_3052338, EPI_ISL_3052339, EPI_ISL_3052340, EPI_ISL_3052341, EPI_ISL_3052342, EPI_ISL_3052343 | Lighthouse Lab in Glasgow | Wellcome Sanger Institute for the COVID-19 Genomics UK (COG-UK) Consortium | Harper VanSteenhouse, Yumi Kasai, David Gray, Carol Clugston, Anna Dominiczak and Alex Alderton, Roberto Amato, Jeffrey Barrett, Sonia Goncalves, Ewan Harrison, David K. Jackson, Ian Johnston, Dominic Kwiatkowski, Cordelia Langford, John Sillitoe on behalf of the Wellcome Sanger Institute COVID-19 Surveillance Team |
| EPI_ISL_3052344 | Lighthouse Lab in Alderley Park | Wellcome Sanger Institute for the COVID-19 Genomics UK (COG-UK) Consortium | Jacquelyn Wynn, Mairead Hyland, The Lighthouse Lab in Alderley Park and Alex Alderton, Roberto Amato, Jeffrey Barrett, Sonia Goncalves, Ewan Harrison, David K. Jackson, Ian Johnston, Dominic Kwiatkowski, Cordelia Langford, John Sillitoe on behalf of the Wellcome Sanger Institute COVID-19 Surveillance Team |
| EPI_ISL_3052345 | Lighthouse Lab in Glasgow | Wellcome Sanger Institute for the COVID-19 Genomics UK (COG-UK) Consortium | Harper VanSteenhouse, Yumi Kasai, David Gray, Carol Clugston, Anna Dominiczak and Alex Alderton, Roberto Amato, Jeffrey Barrett, Sonia Goncalves, Ewan Harrison, David K. Jackson, Ian Johnston, Dominic Kwiatkowski, Cordelia Langford, John Sillitoe on behalf of the Wellcome Sanger Institute COVID-19 Surveillance Team |
| EPI_ISL_3052346, EPI_ISL_3052347 | Lighthouse Lab in Alderley Park | Wellcome Sanger Institute for the COVID-19 Genomics UK (COG-UK) Consortium | Jacquelyn Wynn, Mairead Hyland, The Lighthouse Lab in Alderley Park and Alex Alderton, Roberto Amato, Jeffrey Barrett, Sonia Goncalves, Ewan Harrison, David K. Jackson, Ian Johnston, Dominic Kwiatkowski, Cordelia Langford, John Sillitoe on behalf of the Wellcome Sanger Institute COVID-19 Surveillance Team |
| EPI_ISL_3052348, EPI_ISL_3052349 | Lighthouse Lab in Glasgow | Wellcome Sanger Institute for the COVID-19 Genomics UK (COG-UK) Consortium | Harper VanSteenhouse, Yumi Kasai, David Gray, Carol Clugston, Anna Dominiczak and Alex Alderton, Roberto Amato, Jeffrey Barrett, Sonia Goncalves, Ewan Harrison, David K. Jackson, Ian Johnston, Dominic Kwiatkowski, Cordelia Langford, John Sillitoe on behalf of the Wellcome Sanger Institute COVID-19 Surveillance Team |
| EPI_ISL_3052350, EPI_ISL_3052351 | Lighthouse Lab in Alderley Park | Wellcome Sanger Institute for the COVID-19 Genomics UK (COG-UK) Consortium | Jacquelyn Wynn, Mairead Hyland, The Lighthouse Lab in Alderley Park and Alex Alderton, Roberto Amato, Jeffrey Barrett, Sonia Goncalves, Ewan Harrison, David K. Jackson, Ian Johnston, Dominic Kwiatkowski, Cordelia Langford, John Sillitoe on behalf of the Wellcome Sanger Institute COVID-19 Surveillance Team |
| EPI_ISL_3052352 | Lighthouse Lab in Glasgow | Wellcome Sanger Institute for the COVID-19 Genomics UK (COG-UK) Consortium | Harper VanSteenhouse, Yumi Kasai, David Gray, Carol Clugston, Anna Dominiczak and Alex Alderton, Roberto Amato, Jeffrey Barrett, Sonia Goncalves, Ewan Harrison, David K. Jackson, Ian Johnston, Dominic Kwiatkowski, Cordelia Langford, John Sillitoe on behalf of the Wellcome Sanger Institute COVID-19 Surveillance Team |
| EPI_ISL_3052353, EPI_ISL_3052354, EPI_ISL_3052355, EPI_ISL_3052356, EPI_ISL_3052357, EPI_ISL_3052358, EPI_ISL_3052359 | Lighthouse Lab in Alderley Park | Wellcome Sanger Institute for the COVID-19 Genomics UK (COG-UK) Consortium | Jacquelyn Wynn, Mairead Hyland, The Lighthouse Lab in Alderley Park and Alex Alderton, Roberto Amato, Jeffrey Barrett, Sonia Goncalves, Ewan Harrison, David K. Jackson, Ian Johnston, Dominic Kwiatkowski, Cordelia Langford, John Sillitoe on behalf of the Wellcome Sanger Institute COVID-19 Surveillance Team |
| EPI_ISL_3052360, EPI_ISL_3052361, EPI_ISL_3052362, EPI_ISL_3052363, EPI_ISL_3052364 | Lighthouse Lab in Glasgow | Wellcome Sanger Institute for the COVID-19 Genomics UK (COG-UK) Consortium | Harper VanSteenhouse, Yumi Kasai, David Gray, Carol Clugston, Anna Dominiczak and Alex Alderton, Roberto Amato, Jeffrey Barrett, Sonia Goncalves, Ewan Harrison, David K. Jackson, Ian Johnston, Dominic Kwiatkowski, Cordelia Langford, John Sillitoe on behalf of the Wellcome Sanger Institute COVID-19 Surveillance Team |
| EPI_ISL_3052365, EPI_ISL_3052366, EPI_ISL_3052367 | Lighthouse Lab in Alderley Park | Wellcome Sanger Institute for the COVID-19 Genomics UK (COG-UK) Consortium | Jacquelyn Wynn, Mairead Hyland, The Lighthouse Lab in Alderley Park and Alex Alderton, Roberto Amato, Jeffrey Barrett, Sonia Goncalves, Ewan Harrison, David K. Jackson, Ian Johnston, Dominic Kwiatkowski, Cordelia Langford, John Sillitoe on behalf of the Wellcome Sanger Institute COVID-19 Surveillance Team |
| EPI_ISL_3052368, EPI_ISL_3052369 | Lighthouse Lab in Glasgow | Wellcome Sanger Institute for the COVID-19 Genomics UK (COG-UK) Consortium | Harper VanSteenhouse, Yumi Kasai, David Gray, Carol Clugston, Anna Dominiczak and Alex Alderton, Roberto Amato, Jeffrey Barrett, Sonia Goncalves, Ewan Harrison, David K. Jackson, Ian Johnston, Dominic Kwiatkowski, Cordelia Langford, John Sillitoe on behalf of the Wellcome Sanger Institute COVID-19 Surveillance Team |
| EPI_ISL_3052370 | Lighthouse Lab in Alderley Park | Wellcome Sanger Institute for the COVID-19 Genomics UK (COG-UK) Consortium | Jacquelyn Wynn, Mairead Hyland, The Lighthouse Lab in Alderley Park and Alex Alderton, Roberto Amato, Jeffrey Barrett, Sonia Goncalves, Ewan Harrison, David K. Jackson, Ian Johnston, Dominic Kwiatkowski, Cordelia Langford, John Sillitoe on behalf of the Wellcome Sanger Institute COVID-19 Surveillance Team |
| EPI_ISL_3052371 | Lighthouse Lab in Glasgow | Wellcome Sanger Institute for the COVID-19 Genomics UK (COG-UK) Consortium | Harper VanSteenhouse, Yumi Kasai, David Gray, Carol Clugston, Anna Dominiczak and Alex Alderton, Roberto Amato, Jeffrey Barrett, Sonia Goncalves, Ewan Harrison, David K. Jackson, Ian Johnston, Dominic Kwiatkowski, Cordelia Langford, John Sillitoe on behalf of the Wellcome Sanger Institute COVID-19 Surveillance Team |
| EPI_ISL_3052372 | Lighthouse Lab in Alderley Park | Wellcome Sanger Institute for the COVID-19 Genomics UK (COG-UK) Consortium | Jacquelyn Wynn, Mairead Hyland, The Lighthouse Lab in Alderley Park and Alex Alderton, Roberto Amato, Jeffrey Barrett, Sonia Goncalves, Ewan Harrison, David K. Jackson, Ian Johnston, Dominic Kwiatkowski, Cordelia Langford, John Sillitoe on behalf of the Wellcome Sanger Institute COVID-19 Surveillance Team |
| EPI_ISL_3052373 | Lighthouse Lab in Glasgow | Wellcome Sanger Institute for the COVID-19 Genomics UK (COG-UK) Consortium | Harper VanSteenhouse, Yumi Kasai, David Gray, Carol Clugston, Anna Dominiczak and Alex Alderton, Roberto Amato, Jeffrey Barrett, Sonia Goncalves, Ewan Harrison, David K. Jackson, Ian Johnston, Dominic Kwiatkowski, Cordelia Langford, John Sillitoe on behalf of the Wellcome Sanger Institute COVID-19 Surveillance Team |
| EPI_ISL_3052374, EPI_ISL_3052375 | Lighthouse Lab in Alderley Park | Wellcome Sanger Institute for the COVID-19 Genomics UK (COG-UK) Consortium | Jacquelyn Wynn, Mairead Hyland, The Lighthouse Lab in Alderley Park and Alex Alderton, Roberto Amato, Jeffrey Barrett, Sonia Goncalves, Ewan Harrison, David K. Jackson, Ian Johnston, Dominic Kwiatkowski, Cordelia Langford, John Sillitoe on behalf of the Wellcome Sanger Institute COVID-19 Surveillance Team |
| EPI_ISL_3052376 | Berkshire and Surrey Pathology Services Lighthouse Laboratory | Wellcome Sanger Institute for the COVID-19 Genomics UK (COG-UK) | Berkshire and Surrey Pathology Services Lighthouse Laboratory and Alex Alderton, Roberto Amato, Jeffrey Barrett, Sonia Goncalves, Ewan Harrison, David K. Jackson, Ian Johnston, Dominic Kwiatkowski, Cordelia Langford, John Sillitoe on behalf |

| | | | |
|---|---|---|---|
| | | Consortium | of the Wellcome Sanger Institute COVID-19 Surveillance Team |

EPI_ISL_3052377, EPI_ISL_3052378, EPI_ISL_3052379, EPI_ISL_3052380, EPI_ISL_3052381, EPI_ISL_3052382, EPI_ISL_3052383, EPI_ISL_3052384, EPI_ISL_3052385, EPI_ISL_3052386, EPI_ISL_3052387, EPI_ISL_3052388, EPI_ISL_3052389, EPI_ISL_3052390, EPI_ISL_3052391, EPI_ISL_3052392, EPI_ISL_3052393, EPI_ISL_3052394, EPI_ISL_3052395, EPI_ISL_3052396, EPI_ISL_3052397, EPI_ISL_3052398, EPI_ISL_3052399, EPI_ISL_3052400, EPI_ISL_3052401, EPI_ISL_3052402, EPI_ISL_3052403, EPI_ISL_3052404, EPI_ISL_3052405, EPI_ISL_3052406, EPI_ISL_3052407, EPI_ISL_3052408, EPI_ISL_3052409, EPI_ISL_3052410, EPI_ISL_3052411, EPI_ISL_3052412

| | | | |
|---|---|---|---|
| see above | Lighthouse Lab in Glasgow | Wellcome Sanger Institute for the COVID-19 Genomics UK (COG-UK) Consortium | Harper VanSteenhouse, Yumi Kasai, David Gray, Carol Clugston, Anna Dominiczak and Alex Alderton, Roberto Amato, Jeffrey Barrett, Sonia Goncalves, Ewan Harrison, David K. Jackson, Ian Johnston, Dominic Kwiatkowski, Cordelia Langford, John Sillitoe on behalf of the Wellcome Sanger Institute COVID-19 Surveillance Team |
| EPI_ISL_3052413 | Lighthouse Lab in Milton Keynes | Wellcome Sanger Institute for the COVID-19 Genomics UK (COG-UK) Consortium | The Lighthouse Lab in Milton Keynes and Alex Alderton, Roberto Amato, Jeffrey Barrett, Sonia Goncalves, Ewan Harrison, David K. Jackson, Ian Johnston, Dominic Kwiatkowski, Cordelia Langford, John Sillitoe on behalf of the Wellcome Sanger Institute COVID-19 Surveillance Team |
| EPI_ISL_3052414 | Randox Laboratories | Wellcome Sanger Institute for the COVID-19 Genomics UK (COG-UK) Consortium | Randox Laboratories and Alex Alderton, Roberto Amato, Jeffrey Barrett, Sonia Goncalves, Ewan Harrison, David K. Jackson, Ian Johnston, Dominic Kwiatkowski, Cordelia Langford, John Sillitoe on behalf of the Wellcome Sanger Institute COVID-19 Surveillance Team |
| EPI_ISL_3052415 | Lighthouse Lab in Glasgow | Wellcome Sanger Institute for the COVID-19 Genomics UK (COG-UK) Consortium | Harper VanSteenhouse, Yumi Kasai, David Gray, Carol Clugston, Anna Dominiczak and Alex Alderton, Roberto Amato, Jeffrey Barrett, Sonia Goncalves, Ewan Harrison, David K. Jackson, Ian Johnston, Dominic Kwiatkowski, Cordelia Langford, John Sillitoe on behalf of the Wellcome Sanger Institute COVID-19 Surveillance Team |
| EPI_ISL_3052416 | Lighthouse Lab in Alderley Park | Wellcome Sanger Institute for the COVID-19 Genomics UK (COG-UK) Consortium | Jacquelyn Wynn, Mairead Hyland, The Lighthouse Lab in Alderley Park and Alex Alderton, Roberto Amato, Jeffrey Barrett, Sonia Goncalves, Ewan Harrison, David K. Jackson, Ian Johnston, Dominic Kwiatkowski, Cordelia Langford, John Sillitoe on behalf of the Wellcome Sanger Institute COVID-19 Surveillance Team |
| EPI_ISL_3052417 | Lighthouse Lab in Glasgow | Wellcome Sanger Institute for the COVID-19 Genomics UK (COG-UK) Consortium | Harper VanSteenhouse, Yumi Kasai, David Gray, Carol Clugston, Anna Dominiczak and Alex Alderton, Roberto Amato, Jeffrey Barrett, Sonia Goncalves, Ewan Harrison, David K. Jackson, Ian Johnston, Dominic Kwiatkowski, Cordelia Langford, John Sillitoe on behalf of the Wellcome Sanger Institute COVID-19 Surveillance Team |
| EPI_ISL_3052418 | Lighthouse Lab in Milton Keynes | Wellcome Sanger Institute for the COVID-19 Genomics UK (COG-UK) Consortium | The Lighthouse Lab in Milton Keynes and Alex Alderton, Roberto Amato, Jeffrey Barrett, Sonia Goncalves, Ewan Harrison, David K. Jackson, Ian Johnston, Dominic Kwiatkowski, Cordelia Langford, John Sillitoe on behalf of the Wellcome Sanger Institute COVID-19 Surveillance Team |
| EPI_ISL_3052419, EPI_ISL_3052420 | Lighthouse Lab in Glasgow | Wellcome Sanger Institute for the COVID-19 Genomics UK (COG-UK) Consortium | Harper VanSteenhouse, Yumi Kasai, David Gray, Carol Clugston, Anna Dominiczak and Alex Alderton, Roberto Amato, Jeffrey Barrett, Sonia Goncalves, Ewan Harrison, David K. Jackson, Ian Johnston, Dominic Kwiatkowski, Cordelia Langford, John Sillitoe on behalf of the Wellcome Sanger Institute COVID-19 Surveillance Team |
| EPI_ISL_3052421 | Lighthouse Lab in Milton Keynes | Wellcome Sanger Institute for the COVID-19 Genomics UK (COG-UK) Consortium | The Lighthouse Lab in Milton Keynes and Alex Alderton, Roberto Amato, Jeffrey Barrett, Sonia Goncalves, Ewan Harrison, David K. Jackson, Ian Johnston, Dominic Kwiatkowski, Cordelia Langford, John Sillitoe on behalf of the Wellcome Sanger Institute COVID-19 Surveillance Team |
| EPI_ISL_3052422, EPI_ISL_3052423 | Lighthouse Lab in Glasgow | Wellcome Sanger Institute for the COVID-19 Genomics UK (COG-UK) Consortium | Harper VanSteenhouse, Yumi Kasai, David Gray, Carol Clugston, Anna Dominiczak and Alex Alderton, Roberto Amato, Jeffrey Barrett, Sonia Goncalves, Ewan Harrison, David K. Jackson, Ian Johnston, Dominic Kwiatkowski, Cordelia Langford, John Sillitoe on behalf of the Wellcome Sanger Institute COVID-19 Surveillance Team |
| EPI_ISL_3052424 | Randox Laboratories | Wellcome Sanger Institute for the COVID-19 Genomics UK (COG-UK) Consortium | Randox Laboratories and Alex Alderton, Roberto Amato, Jeffrey Barrett, Sonia Goncalves, Ewan Harrison, David K. Jackson, Ian Johnston, Dominic Kwiatkowski, Cordelia Langford, John Sillitoe on behalf of the Wellcome Sanger Institute COVID-19 Surveillance Team |
| EPI_ISL_3052425 | Lighthouse Lab in Glasgow | Wellcome Sanger Institute for the COVID-19 Genomics UK (COG-UK) Consortium | Harper VanSteenhouse, Yumi Kasai, David Gray, Carol Clugston, Anna Dominiczak and Alex Alderton, Roberto Amato, Jeffrey Barrett, Sonia Goncalves, Ewan Harrison, David K. Jackson, Ian Johnston, Dominic Kwiatkowski, Cordelia Langford, John Sillitoe on behalf of the Wellcome Sanger Institute COVID-19 Surveillance Team |
| EPI_ISL_3052426 | Lighthouse Lab in Alderley Park | Wellcome Sanger Institute for the COVID-19 Genomics UK (COG-UK) Consortium | Jacquelyn Wynn, Mairead Hyland, The Lighthouse Lab in Alderley Park and Alex Alderton, Roberto Amato, Jeffrey Barrett, Sonia Goncalves, Ewan Harrison, David K. Jackson, Ian Johnston, Dominic Kwiatkowski, Cordelia Langford, John Sillitoe on behalf of the Wellcome Sanger Institute COVID-19 Surveillance Team |
| EPI_ISL_3052427, EPI_ISL_3052428, EPI_ISL_3052429, EPI_ISL_3052430 | Lighthouse Lab in Glasgow | Wellcome Sanger Institute for the COVID-19 Genomics UK (COG-UK) Consortium | Harper VanSteenhouse, Yumi Kasai, David Gray, Carol Clugston, Anna Dominiczak and Alex Alderton, Roberto Amato, Jeffrey Barrett, Sonia Goncalves, Ewan Harrison, David K. Jackson, Ian Johnston, Dominic Kwiatkowski, Cordelia Langford, John Sillitoe on behalf of the Wellcome Sanger Institute COVID-19 Surveillance Team |
| EPI_ISL_3052431 | Lighthouse Lab in Alderley Park | Wellcome Sanger Institute for the COVID-19 Genomics UK (COG-UK) Consortium | Jacquelyn Wynn, Mairead Hyland, The Lighthouse Lab in Alderley Park and Alex Alderton, Roberto Amato, Jeffrey Barrett, Sonia Goncalves, Ewan Harrison, David K. Jackson, Ian Johnston, Dominic Kwiatkowski, Cordelia Langford, John Sillitoe on behalf of the Wellcome Sanger Institute COVID-19 Surveillance Team |
| EPI_ISL_3052432, EPI_ISL_3052433 | Lighthouse Lab in Glasgow | Wellcome Sanger Institute for the COVID-19 Genomics UK (COG-UK) Consortium | Harper VanSteenhouse, Yumi Kasai, David Gray, Carol Clugston, Anna Dominiczak and Alex Alderton, Roberto Amato, Jeffrey Barrett, Sonia Goncalves, Ewan Harrison, David K. Jackson, Ian Johnston, Dominic Kwiatkowski, Cordelia Langford, John Sillitoe on behalf of the Wellcome Sanger Institute COVID-19 Surveillance Team |
| EPI_ISL_3052434 | Lighthouse Lab in Milton Keynes | Wellcome Sanger Institute for the COVID-19 Genomics UK (COG-UK) Consortium | The Lighthouse Lab in Milton Keynes and Alex Alderton, Roberto Amato, Jeffrey Barrett, Sonia Goncalves, Ewan Harrison, David K. Jackson, Ian Johnston, Dominic Kwiatkowski, Cordelia Langford, John Sillitoe on behalf of the Wellcome Sanger Institute COVID-19 Surveillance Team |
| EPI_ISL_3052435 | Lighthouse Lab in Alderley Park | Wellcome Sanger Institute for the COVID-19 Genomics UK (COG-UK) Consortium | Jacquelyn Wynn, Mairead Hyland, The Lighthouse Lab in Alderley Park and Alex Alderton, Roberto Amato, Jeffrey Barrett, Sonia Goncalves, Ewan Harrison, David K. Jackson, Ian Johnston, Dominic Kwiatkowski, Cordelia Langford, John Sillitoe on behalf of the Wellcome Sanger Institute COVID-19 Surveillance Team |
| EPI_ISL_3052436 | Lighthouse Lab in Glasgow | Wellcome Sanger Institute for the COVID-19 Genomics UK (COG-UK) Consortium | Harper VanSteenhouse, Yumi Kasai, David Gray, Carol Clugston, Anna Dominiczak and Alex Alderton, Roberto Amato, Jeffrey Barrett, Sonia Goncalves, Ewan Harrison, David K. Jackson, Ian Johnston, Dominic Kwiatkowski, Cordelia Langford, John Sillitoe on behalf of the Wellcome Sanger Institute COVID-19 Surveillance Team |
| EPI_ISL_3052437 | Lighthouse Lab in Milton Keynes | Wellcome Sanger Institute for the COVID-19 Genomics UK (COG-UK) Consortium | The Lighthouse Lab in Milton Keynes and Alex Alderton, Roberto Amato, Jeffrey Barrett, Sonia Goncalves, Ewan Harrison, David K. Jackson, Ian Johnston, Dominic Kwiatkowski, Cordelia Langford, John Sillitoe on behalf of the Wellcome Sanger Institute COVID-19 Surveillance Team |
| EPI_ISL_3052438 | Lighthouse Lab in Alderley Park | Wellcome Sanger Institute for the COVID-19 Genomics UK (COG-UK) Consortium | Jacquelyn Wynn, Mairead Hyland, The Lighthouse Lab in Alderley Park and Alex Alderton, Roberto Amato, Jeffrey Barrett, Sonia Goncalves, Ewan Harrison, David K. Jackson, Ian Johnston, Dominic Kwiatkowski, Cordelia Langford, John Sillitoe on behalf of the Wellcome Sanger Institute COVID-19 Surveillance Team |
| EPI_ISL_3052439, EPI_ISL_3052440, EPI_ISL_3052441, EPI_ISL_3052442, EPI_ISL_3052443, EPI_ISL_3052444, EPI_ISL_3052445 | Lighthouse Lab in Glasgow | Wellcome Sanger Institute for the COVID-19 Genomics UK (COG-UK) Consortium | Harper VanSteenhouse, Yumi Kasai, David Gray, Carol Clugston, Anna Dominiczak and Alex Alderton, Roberto Amato, Jeffrey Barrett, Sonia Goncalves, Ewan Harrison, David K. Jackson, Ian Johnston, Dominic Kwiatkowski, Cordelia Langford, John Sillitoe on behalf of the Wellcome Sanger Institute COVID-19 Surveillance Team |
| EPI_ISL_3052446 | Berkshire and Surrey Pathology Services Lighthouse Laboratory | Wellcome Sanger Institute for the COVID-19 Genomics UK (COG-UK) Consortium | Berkshire and Surrey Pathology Services Lighthouse Laboratory and Alex Alderton, Roberto Amato, Jeffrey Barrett, Sonia Goncalves, Ewan Harrison, David K. Jackson, Ian Johnston, Dominic Kwiatkowski, Cordelia Langford, John Sillitoe on behalf of the Wellcome Sanger Institute COVID-19 Surveillance Team |
| EPI_ISL_3052447 | Lighthouse Lab in Glasgow | Wellcome Sanger Institute for the COVID-19 Genomics UK (COG-UK) Consortium | Harper VanSteenhouse, Yumi Kasai, David Gray, Carol Clugston, Anna Dominiczak and Alex Alderton, Roberto Amato, Jeffrey Barrett, Sonia Goncalves, Ewan Harrison, David K. Jackson, Ian Johnston, Dominic Kwiatkowski, Cordelia Langford, John Sillitoe on behalf of the Wellcome Sanger Institute COVID-19 Surveillance Team |
| EPI_ISL_3052448 | Berkshire and Surrey Pathology Services Lighthouse Laboratory | Wellcome Sanger Institute for the COVID-19 Genomics UK (COG-UK) Consortium | Berkshire and Surrey Pathology Services Lighthouse Laboratory and Alex Alderton, Roberto Amato, Jeffrey Barrett, Sonia Goncalves, Ewan Harrison, David K. Jackson, Ian Johnston, Dominic Kwiatkowski, Cordelia Langford, John Sillitoe on behalf of the Wellcome Sanger Institute COVID-19 Surveillance Team |

EPI_ISL_3052449, EPI_ISL_3052450, EPI_ISL_3052451, EPI_ISL_3052452, EPI_ISL_3052453, EPI_ISL_3052454, EPI_ISL_3052455, EPI_ISL_3052456, EPI_ISL_3052457, EPI_ISL_3052458, EPI_ISL_3052459, EPI_ISL_3052460, EPI_ISL_3052461, EPI_ISL_3052462, EPI_ISL_3052463, EPI_ISL_3052464, EPI_ISL_3052465, EPI_ISL_3052466, EPI_ISL_3052467, EPI_ISL_3052468, EPI_ISL_3052469, EPI_ISL_3052470, EPI_ISL_3052471, EPI_ISL_3052472, EPI_ISL_3052473, EPI_ISL_3052474, EPI_ISL_3052475, EPI_ISL_3052476, EPI_ISL_3052477, EPI_ISL_3052478, EPI_ISL_3052479, EPI_ISL_3052480, EPI_ISL_3052481, EPI_ISL_3052482, EPI_ISL_3052483, EPI_ISL_3052484

| | | | |
|---|---|---|---|
| see above | Lighthouse Lab in Glasgow | Wellcome Sanger Institute for the COVID-19 Genomics UK (COG-UK) Consortium | Harper VanSteenhouse, Yumi Kasai, David Gray, Carol Clugston, Anna Dominiczak and Alex Alderton, Roberto Amato, Jeffrey Barrett, Sonia Goncalves, Ewan Harrison, David K. Jackson, Ian Johnston, Dominic Kwiatkowski, Cordelia Langford, John Sillitoe on behalf of the Wellcome Sanger Institute COVID-19 Surveillance Team |
| EPI_ISL_3052485 | Lighthouse Lab in Milton Keynes | Wellcome Sanger Institute for the COVID-19 Genomics UK (COG-UK) Consortium | The Lighthouse Lab in Milton Keynes and Alex Alderton, Roberto Amato, Jeffrey Barrett, Sonia Goncalves, Ewan Harrison, David K. Jackson, Ian Johnston, Dominic Kwiatkowski, Cordelia Langford, John Sillitoe on behalf of the Wellcome Sanger Institute COVID-19 Surveillance Team |
| EPI_ISL_3052486, EPI_ISL_3052487, EPI_ISL_3052488 | Lighthouse Lab in Glasgow | Wellcome Sanger Institute for the COVID-19 Genomics UK (COG-UK) Consortium | Harper VanSteenhouse, Yumi Kasai, David Gray, Carol Clugston, Anna Dominiczak and Alex Alderton, Roberto Amato, Jeffrey Barrett, Sonia Goncalves, Ewan Harrison, David K. Jackson, Ian Johnston, Dominic Kwiatkowski, Cordelia Langford, John Sillitoe on behalf of the Wellcome Sanger Institute COVID-19 Surveillance Team |
| EPI_ISL_3052489 | Lighthouse Lab in Milton Keynes | Wellcome Sanger Institute for the COVID-19 Genomics UK (COG-UK) Consortium | The Lighthouse Lab in Milton Keynes and Alex Alderton, Roberto Amato, Jeffrey Barrett, Sonia Goncalves, Ewan Harrison, David K. Jackson, Ian Johnston, Dominic Kwiatkowski, Cordelia Langford, John Sillitoe on behalf of the Wellcome Sanger Institute COVID-19 Surveillance Team |
| EPI_ISL_3052490, EPI_ISL_3052491, EPI_ISL_3052492, EPI_ISL_3052493, EPI_ISL_3052494 | Lighthouse Lab in Glasgow | Wellcome Sanger Institute for the COVID-19 Genomics UK (COG-UK) Consortium | Harper VanSteenhouse, Yumi Kasai, David Gray, Carol Clugston, Anna Dominiczak and Alex Alderton, Roberto Amato, Jeffrey Barrett, Sonia Goncalves, Ewan Harrison, David K. Jackson, Ian Johnston, Dominic Kwiatkowski, Cordelia Langford, John Sillitoe on behalf of the Wellcome Sanger Institute COVID-19 Surveillance Team |
| EPI_ISL_3052495 | Lighthouse Lab in Milton Keynes | Wellcome Sanger Institute for the COVID-19 Genomics UK (COG-UK) Consortium | The Lighthouse Lab in Milton Keynes and Alex Alderton, Roberto Amato, Jeffrey Barrett, Sonia Goncalves, Ewan Harrison, David K. Jackson, Ian Johnston, Dominic Kwiatkowski, Cordelia Langford, John Sillitoe on behalf of the Wellcome Sanger Institute COVID-19 Surveillance Team |
| EPI_ISL_3052496, EPI_ISL_3052497, EPI_ISL_3052498, EPI_ISL_3052499, EPI_ISL_3052500, EPI_ISL_3052501 | Lighthouse Lab in Glasgow | Wellcome Sanger Institute for the COVID-19 Genomics UK (COG-UK) Consortium | Harper VanSteenhouse, Yumi Kasai, David Gray, Carol Clugston, Anna Dominiczak and Alex Alderton, Roberto Amato, Jeffrey Barrett, Sonia Goncalves, Ewan Harrison, David K. Jackson, Ian Johnston, Dominic Kwiatkowski, Cordelia Langford, John Sillitoe on behalf of the Wellcome Sanger Institute COVID-19 Surveillance Team |
| EPI_ISL_3052502 | Lighthouse Lab in Milton Keynes | Wellcome Sanger Institute for the COVID-19 Genomics UK (COG-UK) Consortium | The Lighthouse Lab in Milton Keynes and Alex Alderton, Roberto Amato, Jeffrey Barrett, Sonia Goncalves, Ewan Harrison, David K. Jackson, Ian Johnston, Dominic Kwiatkowski, Cordelia Langford, John Sillitoe on behalf of the Wellcome Sanger Institute COVID-19 Surveillance Team |
| EPI_ISL_3052503, EPI_ISL_3052504, EPI_ISL_3052505, EPI_ISL_3052506, EPI_ISL_3052507, EPI_ISL_3052508, EPI_ISL_3052509 | Lighthouse Lab in Glasgow | Wellcome Sanger Institute for the COVID-19 Genomics UK (COG-UK) Consortium | Harper VanSteenhouse, Yumi Kasai, David Gray, Carol Clugston, Anna Dominiczak and Alex Alderton, Roberto Amato, Jeffrey Barrett, Sonia Goncalves, Ewan Harrison, David K. Jackson, Ian Johnston, Dominic Kwiatkowski, Cordelia Langford, John Sillitoe on behalf of the Wellcome Sanger Institute COVID-19 Surveillance Team |

| | | | |
|---|---|---|---|
| EPI_ISL_3052510 | Lighthouse Lab in Milton Keynes | Wellcome Sanger Institute for the COVID-19 Genomics UK (COG-UK) Consortium | The Lighthouse Lab in Milton Keynes and Alex Alderton, Roberto Amato, Jeffrey Barrett, Sonia Goncalves, Ewan Harrison, David K. Jackson, Ian Johnston, Dominic Kwiatkowski, Cordelia Langford, John Sillitoe on behalf of the Wellcome Sanger Institute COVID-19 Surveillance Team |
| EPI_ISL_3052511, EPI_ISL_3052512 | Lighthouse Lab in Glasgow | Wellcome Sanger Institute for the COVID-19 Genomics UK (COG-UK) Consortium | Harper VanSteenhouse, Yumi Kasai, David Gray, Carol Clugston, Anna Dominiczak and Alex Alderton, Roberto Amato, Jeffrey Barrett, Sonia Goncalves, Ewan Harrison, David K. Jackson, Ian Johnston, Dominic Kwiatkowski, Cordelia Langford, John Sillitoe on behalf of the Wellcome Sanger Institute COVID-19 Surveillance Team |
| EPI_ISL_3052513, EPI_ISL_3052514 | Lighthouse Lab in Milton Keynes | Wellcome Sanger Institute for the COVID-19 Genomics UK (COG-UK) Consortium | The Lighthouse Lab in Milton Keynes and Alex Alderton, Roberto Amato, Jeffrey Barrett, Sonia Goncalves, Ewan Harrison, David K. Jackson, Ian Johnston, Dominic Kwiatkowski, Cordelia Langford, John Sillitoe on behalf of the Wellcome Sanger Institute COVID-19 Surveillance Team |
| EPI_ISL_3052515 | Lighthouse Lab in Glasgow | Wellcome Sanger Institute for the COVID-19 Genomics UK (COG-UK) Consortium | Harper VanSteenhouse, Yumi Kasai, David Gray, Carol Clugston, Anna Dominiczak and Alex Alderton, Roberto Amato, Jeffrey Barrett, Sonia Goncalves, Ewan Harrison, David K. Jackson, Ian Johnston, Dominic Kwiatkowski, Cordelia Langford, John Sillitoe on behalf of the Wellcome Sanger Institute COVID-19 Surveillance Team |
| EPI_ISL_3052516 | Lighthouse Lab in Milton Keynes | Wellcome Sanger Institute for the COVID-19 Genomics UK (COG-UK) Consortium | The Lighthouse Lab in Milton Keynes and Alex Alderton, Roberto Amato, Jeffrey Barrett, Sonia Goncalves, Ewan Harrison, David K. Jackson, Ian Johnston, Dominic Kwiatkowski, Cordelia Langford, John Sillitoe on behalf of the Wellcome Sanger Institute COVID-19 Surveillance Team |
| EPI_ISL_3052517, EPI_ISL_3052518, EPI_ISL_3052519, EPI_ISL_3052520 | Lighthouse Lab in Glasgow | Wellcome Sanger Institute for the COVID-19 Genomics UK (COG-UK) Consortium | Harper VanSteenhouse, Yumi Kasai, David Gray, Carol Clugston, Anna Dominiczak and Alex Alderton, Roberto Amato, Jeffrey Barrett, Sonia Goncalves, Ewan Harrison, David K. Jackson, Ian Johnston, Dominic Kwiatkowski, Cordelia Langford, John Sillitoe on behalf of the Wellcome Sanger Institute COVID-19 Surveillance Team |
| EPI_ISL_3052521 | Lighthouse Lab in Alderley Park | Wellcome Sanger Institute for the COVID-19 Genomics UK (COG-UK) Consortium | Jacquelyn Wynn, Mairead Hyland, The Lighthouse Lab in Alderley Park and Alex Alderton, Roberto Amato, Jeffrey Barrett, Sonia Goncalves, Ewan Harrison, David K. Jackson, Ian Johnston, Dominic Kwiatkowski, Cordelia Langford, John Sillitoe on behalf of the Wellcome Sanger Institute COVID-19 Surveillance Team |
| EPI_ISL_3052522, EPI_ISL_3052523, EPI_ISL_3052524, EPI_ISL_3052525 | Lighthouse Lab in Milton Keynes | Wellcome Sanger Institute for the COVID-19 Genomics UK (COG-UK) Consortium | The Lighthouse Lab in Milton Keynes and Alex Alderton, Roberto Amato, Jeffrey Barrett, Sonia Goncalves, Ewan Harrison, David K. Jackson, Ian Johnston, Dominic Kwiatkowski, Cordelia Langford, John Sillitoe on behalf of the Wellcome Sanger Institute COVID-19 Surveillance Team |
| EPI_ISL_3052526 | Lighthouse Lab in Glasgow | Wellcome Sanger Institute for the COVID-19 Genomics UK (COG-UK) Consortium | Harper VanSteenhouse, Yumi Kasai, David Gray, Carol Clugston, Anna Dominiczak and Alex Alderton, Roberto Amato, Jeffrey Barrett, Sonia Goncalves, Ewan Harrison, David K. Jackson, Ian Johnston, Dominic Kwiatkowski, Cordelia Langford, John Sillitoe on behalf of the Wellcome Sanger Institute COVID-19 Surveillance Team |
| EPI_ISL_3052527 | Lighthouse Lab in Alderley Park | Wellcome Sanger Institute for the COVID-19 Genomics UK (COG-UK) Consortium | Jacquelyn Wynn, Mairead Hyland, The Lighthouse Lab in Alderley Park and Alex Alderton, Roberto Amato, Jeffrey Barrett, Sonia Goncalves, Ewan Harrison, David K. Jackson, Ian Johnston, Dominic Kwiatkowski, Cordelia Langford, John Sillitoe on behalf of the Wellcome Sanger Institute COVID-19 Surveillance Team |
| EPI_ISL_3052528, EPI_ISL_3052529 | Lighthouse Lab in Milton Keynes | Wellcome Sanger Institute for the COVID-19 Genomics UK (COG-UK) Consortium | The Lighthouse Lab in Milton Keynes and Alex Alderton, Roberto Amato, Jeffrey Barrett, Sonia Goncalves, Ewan Harrison, David K. Jackson, Ian Johnston, Dominic Kwiatkowski, Cordelia Langford, John Sillitoe on behalf of the Wellcome Sanger Institute COVID-19 Surveillance Team |
| EPI_ISL_3052530, EPI_ISL_3052531, EPI_ISL_3052532 | Lighthouse Lab in Glasgow | Wellcome Sanger Institute for the COVID-19 Genomics UK (COG-UK) Consortium | Harper VanSteenhouse, Yumi Kasai, David Gray, Carol Clugston, Anna Dominiczak and Alex Alderton, Roberto Amato, Jeffrey Barrett, Sonia Goncalves, Ewan Harrison, David K. Jackson, Ian Johnston, Dominic Kwiatkowski, Cordelia Langford, John Sillitoe on behalf of the Wellcome Sanger Institute COVID-19 Surveillance Team |
| EPI_ISL_3052533 | Lighthouse Lab in Milton Keynes | Wellcome Sanger Institute for the COVID-19 Genomics UK (COG-UK) Consortium | The Lighthouse Lab in Milton Keynes and Alex Alderton, Roberto Amato, Jeffrey Barrett, Sonia Goncalves, Ewan Harrison, David K. Jackson, Ian Johnston, Dominic Kwiatkowski, Cordelia Langford, John Sillitoe on behalf of the Wellcome Sanger Institute COVID-19 Surveillance Team |
| EPI_ISL_3052534, EPI_ISL_3052535, EPI_ISL_3052536, EPI_ISL_3052537, EPI_ISL_3052538 | Lighthouse Lab in Glasgow | Wellcome Sanger Institute for the COVID-19 Genomics UK (COG-UK) Consortium | Harper VanSteenhouse, Yumi Kasai, David Gray, Carol Clugston, Anna Dominiczak and Alex Alderton, Roberto Amato, Jeffrey Barrett, Sonia Goncalves, Ewan Harrison, David K. Jackson, Ian Johnston, Dominic Kwiatkowski, Cordelia Langford, John Sillitoe on behalf of the Wellcome Sanger Institute COVID-19 Surveillance Team |
| EPI_ISL_3052539, EPI_ISL_3052540 | Lighthouse Lab in Milton Keynes | Wellcome Sanger Institute for the COVID-19 Genomics UK (COG-UK) Consortium | The Lighthouse Lab in Milton Keynes and Alex Alderton, Roberto Amato, Jeffrey Barrett, Sonia Goncalves, Ewan Harrison, David K. Jackson, Ian Johnston, Dominic Kwiatkowski, Cordelia Langford, John Sillitoe on behalf of the Wellcome Sanger Institute COVID-19 Surveillance Team |
| EPI_ISL_3052541, EPI_ISL_3052542, EPI_ISL_3052543, EPI_ISL_3052544 | Lighthouse Lab in Glasgow | Wellcome Sanger Institute for the COVID-19 Genomics UK (COG-UK) Consortium | Harper VanSteenhouse, Yumi Kasai, David Gray, Carol Clugston, Anna Dominiczak and Alex Alderton, Roberto Amato, Jeffrey Barrett, Sonia Goncalves, Ewan Harrison, David K. Jackson, Ian Johnston, Dominic Kwiatkowski, Cordelia Langford, John Sillitoe on behalf of the Wellcome Sanger Institute COVID-19 Surveillance Team |
| EPI_ISL_3052545, EPI_ISL_3052546 | Lighthouse Lab in Milton Keynes | Wellcome Sanger Institute for the COVID-19 Genomics UK (COG-UK) Consortium | The Lighthouse Lab in Milton Keynes and Alex Alderton, Roberto Amato, Jeffrey Barrett, Sonia Goncalves, Ewan Harrison, David K. Jackson, Ian Johnston, Dominic Kwiatkowski, Cordelia Langford, John Sillitoe on behalf of the Wellcome Sanger Institute COVID-19 Surveillance Team |
| EPI_ISL_3052547, EPI_ISL_3052548, EPI_ISL_3052549, EPI_ISL_3052550, EPI_ISL_3052551, EPI_ISL_3052552, EPI_ISL_3052553, EPI_ISL_3052554, EPI_ISL_3052555, EPI_ISL_3052556, EPI_ISL_3052557, EPI_ISL_3052558, EPI_ISL_3052559, EPI_ISL_3052560, EPI_ISL_3052561, EPI_ISL_3052562, EPI_ISL_3052563, EPI_ISL_3052564, EPI_ISL_3052565, EPI_ISL_3052566, EPI_ISL_3052567, EPI_ISL_3052568, EPI_ISL_3052569, EPI_ISL_3052570, EPI_ISL_3052571, EPI_ISL_3052572, EPI_ISL_3052573, EPI_ISL_3052574, EPI_ISL_3052575, EPI_ISL_3052576, EPI_ISL_3052577, EPI_ISL_3052578, EPI_ISL_3052579, EPI_ISL_3052580, EPI_ISL_3052582, EPI_ISL_3052583, EPI_ISL_3052584, EPI_ISL_3052585, EPI_ISL_3052586, EPI_ISL_3052587, EPI_ISL_3052588, EPI_ISL_3052589, EPI_ISL_3052590, EPI_ISL_3052591, EPI_ISL_3052592, EPI_ISL_3052593, EPI_ISL_3052594, EPI_ISL_3052595, EPI_ISL_3052596, EPI_ISL_3052597, EPI_ISL_3052598, EPI_ISL_3052599, EPI_ISL_3052600, EPI_ISL_3052601, EPI_ISL_3052602, EPI_ISL_3052603, EPI_ISL_3052604, EPI_ISL_3052605, EPI_ISL_3052606, EPI_ISL_3052607, EPI_ISL_3052608, EPI_ISL_3052609, EPI_ISL_3052610, EPI_ISL_3052611, EPI_ISL_3052612, EPI_ISL_3052613, EPI_ISL_3052614, EPI_ISL_3052615, EPI_ISL_3052616, EPI_ISL_3052617, EPI_ISL_3052618, EPI_ISL_3052619, EPI_ISL_3052620, EPI_ISL_3052621, EPI_ISL_3052622, EPI_ISL_3052623, EPI_ISL_3052624, EPI_ISL_3052625, EPI_ISL_3052626, EPI_ISL_3052627, EPI_ISL_3052628, EPI_ISL_3052629, EPI_ISL_3052630, EPI_ISL_3052631 | | | |
| see above | Lighthouse Lab in Glasgow | Wellcome Sanger Institute for the COVID-19 Genomics UK (COG-UK) Consortium | Harper VanSteenhouse, Yumi Kasai, David Gray, Carol Clugston, Anna Dominiczak and Alex Alderton, Roberto Amato, Jeffrey Barrett, Sonia Goncalves, Ewan Harrison, David K. Jackson, Ian Johnston, Dominic Kwiatkowski, Cordelia Langford, John Sillitoe on behalf of the Wellcome Sanger Institute COVID-19 Surveillance Team |
| EPI_ISL_3052638 | Genomik Solidaritas Indonesia Laboratorium | Genomik Solidaritas Indonesia Laboratorium | Vania Gavrila Wikasa, Carissa Sintca Wijaya, Anna Christina Brazia, Annisa Muthiah Sukirman, Gracia Felias Enos Korompis, Anuraj Shankar, Louisa Markus, Meutia Ayuputeri Kumaheri, Ariel Pradipta |
| EPI_ISL_3052639, EPI_ISL_3052640, EPI_ISL_3052641, EPI_ISL_3052642, EPI_ISL_3052643, EPI_ISL_3052644, EPI_ISL_3052646, EPI_ISL_3052647, EPI_ISL_3052649, EPI_ISL_3052650, EPI_ISL_3052651, EPI_ISL_3052652, EPI_ISL_3052653, EPI_ISL_3052654 | | | |
| see above | ASL BARI | University of Bari Biomedical Sciences and Human Oncology | Maria Chironna |
| EPI_ISL_3052666, EPI_ISL_3052667, EPI_ISL_3052668, EPI_ISL_3052669, EPI_ISL_3052670, EPI_ISL_3052671, EPI_ISL_3052673, EPI_ISL_3052680, EPI_ISL_3052681, EPI_ISL_3052682, EPI_ISL_3052683 | | | |
| see above | Molecular Diagnostics Pathology Department Mater Dei Hospital Malta | Molecular Diagnostics Pathology Department Mater Dei Hospital Malta | G Zahra, R Borg, C Cilia, L Grech, M Briffa |
| EPI_ISL_3052685, EPI_ISL_3052686, EPI_ISL_3052687, EPI_ISL_3052688, EPI_ISL_3052689, EPI_ISL_3052690, EPI_ISL_3052691, EPI_ISL_3052692, EPI_ISL_3052693, EPI_ISL_3052694, EPI_ISL_3052695, EPI_ISL_3052696, EPI_ISL_3052697, EPI_ISL_3052698, EPI_ISL_3052699, EPI_ISL_3052700, EPI_ISL_3052701, EPI_ISL_3052702, EPI_ISL_3052704, EPI_ISL_3052705, EPI_ISL_3052706, EPI_ISL_3052707, EPI_ISL_3052708, EPI_ISL_3052709, EPI_ISL_3052710, EPI_ISL_3052711, EPI_ISL_3052712, EPI_ISL_3052713, EPI_ISL_3052714, EPI_ISL_3052715, EPI_ISL_3052716, EPI_ISL_3052717, EPI_ISL_3052718, EPI_ISL_3052719, EPI_ISL_3052720, EPI_ISL_3052721, EPI_ISL_3052722, EPI_ISL_3052723, EPI_ISL_3052724, EPI_ISL_3052725, EPI_ISL_3052726, EPI_ISL_3052727, EPI_ISL_3052728, EPI_ISL_3052729, EPI_ISL_3052730, EPI_ISL_3052731, EPI_ISL_3052732, EPI_ISL_3052733, EPI_ISL_3052734, EPI_ISL_3052735, EPI_ISL_3052736, EPI_ISL_3052737, EPI_ISL_3052738, EPI_ISL_3052739, EPI_ISL_3052740, EPI_ISL_3052741, EPI_ISL_3052742, EPI_ISL_3052743, EPI_ISL_3052744, EPI_ISL_3052745, EPI_ISL_3052746, EPI_ISL_3052747, EPI_ISL_3052748, EPI_ISL_3052749, EPI_ISL_3052750, EPI_ISL_3052751, EPI_ISL_3052752, EPI_ISL_3052753, EPI_ISL_3052754, EPI_ISL_3052755, EPI_ISL_3052756, EPI_ISL_3052757, EPI_ISL_3052758, EPI_ISL_3052759, EPI_ISL_3052760, EPI_ISL_3052761, EPI_ISL_3052762, EPI_ISL_3052763, EPI_ISL_3052764, EPI_ISL_3052765, EPI_ISL_3052766, EPI_ISL_3052767, EPI_ISL_3052768, EPI_ISL_3052769, EPI_ISL_3052770, EPI_ISL_3052771, EPI_ISL_3052772, EPI_ISL_3052773, EPI_ISL_3052774, EPI_ISL_3052775, EPI_ISL_3052776, EPI_ISL_3052777, EPI_ISL_3052778, EPI_ISL_3052779, EPI_ISL_3052780, EPI_ISL_3052781, EPI_ISL_3052782, EPI_ISL_3052783, EPI_ISL_3052784, EPI_ISL_3052785, EPI_ISL_3052786, EPI_ISL_3052787, EPI_ISL_3052788, EPI_ISL_3052789, EPI_ISL_3052790, EPI_ISL_3052791, EPI_ISL_3052792, EPI_ISL_3052793, EPI_ISL_3052794, EPI_ISL_3052795, EPI_ISL_3052796, EPI_ISL_3052797, EPI_ISL_3052798, EPI_ISL_3052799, EPI_ISL_3052800, EPI_ISL_3052801, EPI_ISL_3052802, EPI_ISL_3052803, EPI_ISL_3052804, EPI_ISL_3052805, EPI_ISL_3052806, EPI_ISL_3052807, EPI_ISL_3052808, EPI_ISL_3052809, EPI_ISL_3052810, EPI_ISL_3052811, EPI_ISL_3052812, EPI_ISL_3052813, EPI_ISL_3052814, EPI_ISL_3052815, EPI_ISL_3052816, EPI_ISL_3052817, EPI_ISL_3052818, EPI_ISL_3052819, EPI_ISL_3052820, EPI_ISL_3052821, EPI_ISL_3052822, EPI_ISL_3052823, EPI_ISL_3052824, EPI_ISL_3052825, EPI_ISL_3052826, EPI_ISL_3052827, EPI_ISL_3052828, EPI_ISL_3052829, EPI_ISL_3052830, EPI_ISL_3052831, EPI_ISL_3052832, EPI_ISL_3052833, EPI_ISL_3052834, EPI_ISL_3052835, EPI_ISL_3052836, EPI_ISL_3052837, EPI_ISL_3052838, EPI_ISL_3052839, EPI_ISL_3052840, EPI_ISL_3052841, EPI_ISL_3052842, EPI_ISL_3052843, EPI_ISL_3052844, EPI_ISL_3052845, EPI_ISL_3052846, EPI_ISL_3052847, EPI_ISL_3052848, EPI_ISL_3052849, EPI_ISL_3052850, EPI_ISL_3052851, EPI_ISL_3052852, EPI_ISL_3052853, EPI_ISL_3052854, EPI_ISL_3052855, EPI_ISL_3052856, EPI_ISL_3052857, EPI_ISL_3052858, EPI_ISL_3052859, EPI_ISL_3052860, EPI_ISL_3052861, EPI_ISL_3052862, EPI_ISL_3052863, EPI_ISL_3052864, EPI_ISL_3052865, EPI_ISL_3052866, EPI_ISL_3052867, EPI_ISL_3052868, EPI_ISL_3052869, EPI_ISL_3052870, EPI_ISL_3052871, EPI_ISL_3052872, EPI_ISL_3052873, EPI_ISL_3052874, EPI_ISL_3052875, EPI_ISL_3052876, EPI_ISL_3052877, EPI_ISL_3052878, EPI_ISL_3052879, EPI_ISL_3052880, EPI_ISL_3052881, EPI_ISL_3052882, EPI_ISL_3052883, EPI_ISL_3052884, EPI_ISL_3052885, EPI_ISL_3052886, EPI_ISL_3052887, EPI_ISL_3052888, EPI_ISL_3052889, EPI_ISL_3052890, EPI_ISL_3052891, EPI_ISL_3052892, EPI_ISL_3052893, EPI_ISL_3052894, EPI_ISL_3052895, EPI_ISL_3052896, EPI_ISL_3052897, EPI_ISL_3052898, EPI_ISL_3052899, EPI_ISL_3052900, EPI_ISL_3052901, EPI_ISL_3052902, EPI_ISL_3052903, EPI_ISL_3052904 | | | |
| see above | National Public Health Laboratory, National Centre for Infectious Diseases | National Public Health Laboratory, National Centre for Infectious Diseases | Tze Minn Mak, Katherine Ching, Zhenyang Zhou, Royce Ang, Lin Cui, Raymond Tzer Pin Lin |
| EPI_ISL_3052906, EPI_ISL_3052908, EPI_ISL_3052910, EPI_ISL_3052911, EPI_ISL_3052912, EPI_ISL_3052913, EPI_ISL_3052914, EPI_ISL_3052915, EPI_ISL_3052916, EPI_ISL_3052917, EPI_ISL_3052918, EPI_ISL_3052922, EPI_ISL_3052924 | | | |
| see above | Molecular Diagnostics Pathology Department Mater Dei Hospital Malta | Molecular Diagnostics Pathology Department Mater Dei Hospital Malta | G Zahra, R Borg, C Cilia, L Grech, M Briffa |
| EPI_ISL_3052925, EPI_ISL_3052927, EPI_ISL_3052928, EPI_ISL_3052929, EPI_ISL_3052930, EPI_ISL_3052931, EPI_ISL_3052932, EPI_ISL_3052933, EPI_ISL_3052934, EPI_ISL_3052935, EPI_ISL_3052936, EPI_ISL_3052937, EPI_ISL_3052938, EPI_ISL_3052939, EPI_ISL_3052940, EPI_ISL_3052941, EPI_ISL_3052942, EPI_ISL_3052943, EPI_ISL_3052944, EPI_ISL_3052945, EPI_ISL_3052946, EPI_ISL_3052947, EPI_ISL_3052948, EPI_ISL_3052949, EPI_ISL_3052950, EPI_ISL_3052951, EPI_ISL_3052952, EPI_ISL_3052953, EPI_ISL_3052954, EPI_ISL_3052955, EPI_ISL_3052956, EPI_ISL_3052957, EPI_ISL_3052958, EPI_ISL_3052959, EPI_ISL_3052960, EPI_ISL_3052961, EPI_ISL_3052962, EPI_ISL_3052963, EPI_ISL_3052964, EPI_ISL_3052965, EPI_ISL_3052966, EPI_ISL_3052967, EPI_ISL_3052968, EPI_ISL_3052969, EPI_ISL_3052970, EPI_ISL_3052971, EPI_ISL_3052972, EPI_ISL_3052974, EPI_ISL_3052975, EPI_ISL_3052976, EPI_ISL_3052978, EPI_ISL_3052980, EPI_ISL_3052981, EPI_ISL_3052982, EPI_ISL_3052983, EPI_ISL_3052984, EPI_ISL_3052985, EPI_ISL_3052986 | | | |
| see above | Department of Bacteria, Parasites and Fungi, Statens Serum Institut, Copenhagen, Denmark | Statens Serum Institut Bioinformatics and Microbial Genomics | Danish Covid-19 Genome Consortium |
| EPI_ISL_3052987 | Molecular Diagnostics Pathology Department Mater Dei Hospital Malta | Molecular Diagnostics Pathology Department Mater Dei Hospital Malta | G Zahra, R Borg, C Cilia, L Grech, M Briffa |
| EPI_ISL_3052988, EPI_ISL_3052989, EPI_ISL_3052990, EPI_ISL_3052991, EPI_ISL_3052992, EPI_ISL_3052993, EPI_ISL_3052994, EPI_ISL_3052995, EPI_ISL_3052996, EPI_ISL_3052997, EPI_ISL_3052998, EPI_ISL_3052999, EPI_ISL_3053000, EPI_ISL_3053001, EPI_ISL_3053002, EPI_ISL_3053003, EPI_ISL_3053004, EPI_ISL_3053005, EPI_ISL_3053006, EPI_ISL_3053008, EPI_ISL_3053009, EPI_ISL_3053011, EPI_ISL_3053012, EPI_ISL_3053013, EPI_ISL_3053014, EPI_ISL_3053015, EPI_ISL_3053016, EPI_ISL_3053017, EPI_ISL_3053018, EPI_ISL_3053019, EPI_ISL_3053020, EPI_ISL_3053021, EPI_ISL_3053022, EPI_ISL_3053023, EPI_ISL_3053026, EPI_ISL_3053027, EPI_ISL_3053028, EPI_ISL_3053029, EPI_ISL_3053030, EPI_ISL_3053031, EPI_ISL_3053032, EPI_ISL_3053033, EPI_ISL_3053034, EPI_ISL_3053035, EPI_ISL_3053036, EPI_ISL_3053037, EPI_ISL_3053038, EPI_ISL_3053039, EPI_ISL_3053040, EPI_ISL_3053041, EPI_ISL_3053042, EPI_ISL_3053043, EPI_ISL_3053044, EPI_ISL_3053045, EPI_ISL_3053046, EPI_ISL_3053047, EPI_ISL_3053048, EPI_ISL_3053049, EPI_ISL_3053050, EPI_ISL_3053051, EPI_ISL_3053052, EPI_ISL_3053054, EPI_ISL_3053055, EPI_ISL_3053056, EPI_ISL_3053057, EPI_ISL_3053058, EPI_ISL_3053059, EPI_ISL_3053060, EPI_ISL_3053061, EPI_ISL_3053062, EPI_ISL_3053063, EPI_ISL_3053064, EPI_ISL_3053065, EPI_ISL_3053066, EPI_ISL_3053067, EPI_ISL_3053068, EPI_ISL_3053069, EPI_ISL_3053070, EPI_ISL_3053071, EPI_ISL_3053072, EPI_ISL_3053073, EPI_ISL_3053074, EPI_ISL_3053075, EPI_ISL_3053076, EPI_ISL_3053077, EPI_ISL_3053078, EPI_ISL_3053079, EPI_ISL_3053080, EPI_ISL_3053081, EPI_ISL_3053082, EPI_ISL_3053083, EPI_ISL_3053084, EPI_ISL_3053085, EPI_ISL_3053086, EPI_ISL_3053087, EPI_ISL_3053088, EPI_ISL_3053089, EPI_ISL_3053090, EPI_ISL_3053091, EPI_ISL_3053092, EPI_ISL_3053093, EPI_ISL_3053094, EPI_ISL_3053095, EPI_ISL_3053096, EPI_ISL_3053097, EPI_ISL_3053098, EPI_ISL_3053099, EPI_ISL_3053100, EPI_ISL_3053101, EPI_ISL_3053102, EPI_ISL_3053103, EPI_ISL_3053104, EPI_ISL_3053105, EPI_ISL_3053106, EPI_ISL_3053107, EPI_ISL_3053108, EPI_ISL_3053109, EPI_ISL_3053110, EPI_ISL_3053111, EPI_ISL_3053112, EPI_ISL_3053113, EPI_ISL_3053114, EPI_ISL_3053115, EPI_ISL_3053116, EPI_ISL_3053117, EPI_ISL_3053118, EPI_ISL_3053119, EPI_ISL_3053120, EPI_ISL_3053121, EPI_ISL_3053122, EPI_ISL_3053123, EPI_ISL_3053124, EPI_ISL_3053125, EPI_ISL_3053126, EPI_ISL_3053127, EPI_ISL_3053128, EPI_ISL_3053129, EPI_ISL_3053130, EPI_ISL_3053131, EPI_ISL_3053132, EPI_ISL_3053134, EPI_ISL_3053135, EPI_ISL_3053136, EPI_ISL_3053137, EPI_ISL_3053138, EPI_ISL_3053139, EPI_ISL_3053140, EPI_ISL_3053141, EPI_ISL_3053142, EPI_ISL_3053143, EPI_ISL_3053144, EPI_ISL_3053145, EPI_ISL_3053146, EPI_ISL_3053147, EPI_ISL_3053148, EPI_ISL_3053149, EPI_ISL_3053150, EPI_ISL_3053151, EPI_ISL_3053153, EPI_ISL_3053155, EPI_ISL_3053156, EPI_ISL_3053157, EPI_ISL_3053158, EPI_ISL_3053159, EPI_ISL_3053160, EPI_ISL_3053162, EPI_ISL_3053163, EPI_ISL_3053164, EPI_ISL_3053165, EPI_ISL_3053166, EPI_ISL_3053167, EPI_ISL_3053168, EPI_ISL_3053169, EPI_ISL_3053171, EPI_ISL_3053172, EPI_ISL_3053173, EPI_ISL_3053174, EPI_ISL_3053175, EPI_ISL_3053176, EPI_ISL_3053177, EPI_ISL_3053178, EPI_ISL_3053179, EPI_ISL_3053180, EPI_ISL_3053181, EPI_ISL_3053182, EPI_ISL_3053183, EPI_ISL_3053184 | | | |
| see above | Department of Bacteria, Parasites and Fungi, Statens Serum Institut, Copenhagen, Denmark | Statens Serum Institut Bioinformatics and Microbial Genomics | Danish Covid-19 Genome Consortium |
| EPI_ISL_3053185 | Department of Clinical Microbiology, Odense University Hospital, Odense, Denmark | Statens Serum Institut Bioinformatics and Microbial Genomics | Danish Covid-19 Genome Consortium |
| EPI_ISL_3053186, EPI_ISL_3053187, EPI_ISL_3053188, EPI_ISL_3053189, EPI_ISL_3053190, EPI_ISL_3053191, EPI_ISL_3053192, EPI_ISL_3053193, EPI_ISL_3053194, EPI_ISL_3053195, EPI_ISL_3053196, EPI_ISL_3053197, EPI_ISL_3053198, EPI_ISL_3053199, EPI_ISL_3053200, EPI_ISL_3053201, EPI_ISL_3053202, EPI_ISL_3053203, EPI_ISL_3053204, EPI_ISL_3053205 | | | |
| see above | Department of Bacteria, Parasites and Fungi, Statens Serum Institut, Copenhagen, Denmark | Statens Serum Institut Bioinformatics and Microbial Genomics | Danish Covid-19 Genome Consortium |

EPI_ISL_3053206 Department of Clinical Microbiology, Odense University Hospital, Odense, Denmark Statens Serum Institut Bioinformatics and Microbial Genomics Danish Covid-19 Genome Consortium

EPI_ISL_3053207, EPI_ISL_3053208, EPI_ISL_3053209, EPI_ISL_3053210, EPI_ISL_3053211, EPI_ISL_3053212, EPI_ISL_3053213, EPI_ISL_3053214, EPI_ISL_3053215, EPI_ISL_3053216, EPI_ISL_3053217, EPI_ISL_3053218, EPI_ISL_3053219, EPI_ISL_3053220, EPI_ISL_3053221, EPI_ISL_3053222, EPI_ISL_3053223, EPI_ISL_3053224, EPI_ISL_3053225, EPI_ISL_3053226, EPI_ISL_3053227, EPI_ISL_3053228,
EPI_ISL_3053229, EPI_ISL_3053230, EPI_ISL_3053231, EPI_ISL_3053232, EPI_ISL_3053233, EPI_ISL_3053234, EPI_ISL_3053235, EPI_ISL_3053236, EPI_ISL_3053237, EPI_ISL_3053238, EPI_ISL_3053240, EPI_ISL_3053241, EPI_ISL_3053242, EPI_ISL_3053243, EPI_ISL_3053244, EPI_ISL_3053245, EPI_ISL_3053246, EPI_ISL_3053247, EPI_ISL_3053248, EPI_ISL_3053249, EPI_ISL_3053250, EPI_ISL_3053251,
EPI_ISL_3053252, EPI_ISL_3053253, EPI_ISL_3053254, EPI_ISL_3053255, EPI_ISL_3053256, EPI_ISL_3053258, EPI_ISL_3053259, EPI_ISL_3053260, EPI_ISL_3053261, EPI_ISL_3053262, EPI_ISL_3053263, EPI_ISL_3053264, EPI_ISL_3053265, EPI_ISL_3053266, EPI_ISL_3053267, EPI_ISL_3053268, EPI_ISL_3053270, EPI_ISL_3053271, EPI_ISL_3053272, EPI_ISL_3053273, EPI_ISL_3053274, EPI_ISL_3053275,
EPI_ISL_3053276, EPI_ISL_3053277, EPI_ISL_3053278, EPI_ISL_3053279, EPI_ISL_3053280, EPI_ISL_3053281, EPI_ISL_3053282, EPI_ISL_3053283, EPI_ISL_3053284, EPI_ISL_3053285, EPI_ISL_3053286, EPI_ISL_3053287, EPI_ISL_3053288, EPI_ISL_3053289, EPI_ISL_3053290, EPI_ISL_3053291, EPI_ISL_3053292, EPI_ISL_3053293, EPI_ISL_3053294, EPI_ISL_3053295, EPI_ISL_3053296, EPI_ISL_3053297,
EPI_ISL_3053298, EPI_ISL_3053299, EPI_ISL_3053300, EPI_ISL_3053301, EPI_ISL_3053302, EPI_ISL_3053303, EPI_ISL_3053304, EPI_ISL_3053305, EPI_ISL_3053306, EPI_ISL_3053307, EPI_ISL_3053308, EPI_ISL_3053309, EPI_ISL_3053310, EPI_ISL_3053311, EPI_ISL_3053312, EPI_ISL_3053313, EPI_ISL_3053314, EPI_ISL_3053315, EPI_ISL_3053316, EPI_ISL_3053318, EPI_ISL_3053319, EPI_ISL_3053320,
EPI_ISL_3053321, EPI_ISL_3053322, EPI_ISL_3053323, EPI_ISL_3053324, EPI_ISL_3053325, EPI_ISL_3053327, EPI_ISL_3053328, EPI_ISL_3053329, EPI_ISL_3053330, EPI_ISL_3053331, EPI_ISL_3053332, EPI_ISL_3053333, EPI_ISL_3053335, EPI_ISL_3053336, EPI_ISL_3053337, EPI_ISL_3053338, EPI_ISL_3053339, EPI_ISL_3053340, EPI_ISL_3053341, EPI_ISL_3053342, EPI_ISL_3053343, EPI_ISL_3053344,
EPI_ISL_3053345, EPI_ISL_3053346, EPI_ISL_3053347, EPI_ISL_3053348, EPI_ISL_3053349, EPI_ISL_3053350, EPI_ISL_3053351, EPI_ISL_3053352, EPI_ISL_3053353, EPI_ISL_3053354, EPI_ISL_3053355, EPI_ISL_3053356, EPI_ISL_3053357, EPI_ISL_3053358, EPI_ISL_3053359, EPI_ISL_3053360, EPI_ISL_3053361, EPI_ISL_3053362, EPI_ISL_3053363, EPI_ISL_3053364, EPI_ISL_3053365, EPI_ISL_3053366,
EPI_ISL_3053367, EPI_ISL_3053368, EPI_ISL_3053369, EPI_ISL_3053370, EPI_ISL_3053371, EPI_ISL_3053372, EPI_ISL_3053374, EPI_ISL_3053375, EPI_ISL_3053376, EPI_ISL_3053377, EPI_ISL_3053378, EPI_ISL_3053379, EPI_ISL_3053380, EPI_ISL_3053381, EPI_ISL_3053382, EPI_ISL_3053383, EPI_ISL_3053384, EPI_ISL_3053385, EPI_ISL_3053386, EPI_ISL_3053387, EPI_ISL_3053388, EPI_ISL_3053389,
EPI_ISL_3053390, EPI_ISL_3053391, EPI_ISL_3053392, EPI_ISL_3053393, EPI_ISL_3053394, EPI_ISL_3053395, EPI_ISL_3053396, EPI_ISL_3053397, EPI_ISL_3053398, EPI_ISL_3053399, EPI_ISL_3053400, EPI_ISL_3053401, EPI_ISL_3053402, EPI_ISL_3053403, EPI_ISL_3053404, EPI_ISL_3053405, EPI_ISL_3053406, EPI_ISL_3053407, EPI_ISL_3053408, EPI_ISL_3053409, EPI_ISL_3053410, EPI_ISL_3053411,
EPI_ISL_3053412, EPI_ISL_3053413, EPI_ISL_3053414, EPI_ISL_3053415, EPI_ISL_3053416, EPI_ISL_3053417, EPI_ISL_3053418, EPI_ISL_3053419, EPI_ISL_3053420, EPI_ISL_3053421, EPI_ISL_3053422, EPI_ISL_3053423, EPI_ISL_3053424, EPI_ISL_3053425, EPI_ISL_3053426, EPI_ISL_3053427, EPI_ISL_3053428, EPI_ISL_3053429, EPI_ISL_3053430, EPI_ISL_3053431, EPI_ISL_3053432, EPI_ISL_3053433,
EPI_ISL_3053434, EPI_ISL_3053435, EPI_ISL_3053436, EPI_ISL_3053437, EPI_ISL_3053438, EPI_ISL_3053439, EPI_ISL_3053440, EPI_ISL_3053441, EPI_ISL_3053442, EPI_ISL_3053443, EPI_ISL_3053444, EPI_ISL_3053445, EPI_ISL_3053447, EPI_ISL_3053448, EPI_ISL_3053449, EPI_ISL_3053450, EPI_ISL_3053451, EPI_ISL_3053452, EPI_ISL_3053453, EPI_ISL_3053454, EPI_ISL_3053456, EPI_ISL_3053457,
EPI_ISL_3053458, EPI_ISL_3053459, EPI_ISL_3053460, EPI_ISL_3053461, EPI_ISL_3053462, EPI_ISL_3053464, EPI_ISL_3053465, EPI_ISL_3053466, EPI_ISL_3053467, EPI_ISL_3053468, EPI_ISL_3053469, EPI_ISL_3053470, EPI_ISL_3053471, EPI_ISL_3053472, EPI_ISL_3053473, EPI_ISL_3053474, EPI_ISL_3053475, EPI_ISL_3053476, EPI_ISL_3053477, EPI_ISL_3053478, EPI_ISL_3053479, EPI_ISL_3053480,
EPI_ISL_3053481, EPI_ISL_3053482, EPI_ISL_3053483, EPI_ISL_3053484, EPI_ISL_3053485, EPI_ISL_3053486, EPI_ISL_3053487, EPI_ISL_3053488, EPI_ISL_3053489, EPI_ISL_3053490, EPI_ISL_3053491, EPI_ISL_3053492, EPI_ISL_3053493, EPI_ISL_3053494, EPI_ISL_3053495, EPI_ISL_3053496, EPI_ISL_3053497, EPI_ISL_3053498, EPI_ISL_3053499, EPI_ISL_3053500, EPI_ISL_3053501, EPI_ISL_3053502,
EPI_ISL_3053503, EPI_ISL_3053504, EPI_ISL_3053505, EPI_ISL_3053506, EPI_ISL_3053507, EPI_ISL_3053508, EPI_ISL_3053509, EPI_ISL_3053510, EPI_ISL_3053511, EPI_ISL_3053512, EPI_ISL_3053513, EPI_ISL_3053514, EPI_ISL_3053515, EPI_ISL_3053516, EPI_ISL_3053517, EPI_ISL_3053518, EPI_ISL_3053519, EPI_ISL_3053520, EPI_ISL_3053521, EPI_ISL_3053522, EPI_ISL_3053523, EPI_ISL_3053524,
EPI_ISL_3053525, EPI_ISL_3053526, EPI_ISL_3053527, EPI_ISL_3053528, EPI_ISL_3053529, EPI_ISL_3053530, EPI_ISL_3053531, EPI_ISL_3053532, EPI_ISL_3053533, EPI_ISL_3053534, EPI_ISL_3053535, EPI_ISL_3053536, EPI_ISL_3053537, EPI_ISL_3053538, EPI_ISL_3053539, EPI_ISL_3053540, EPI_ISL_3053541, EPI_ISL_3053542, EPI_ISL_3053543, EPI_ISL_3053544, EPI_ISL_3053545, EPI_ISL_3053546,
EPI_ISL_3053547, EPI_ISL_3053548, EPI_ISL_3053549, EPI_ISL_3053550, EPI_ISL_3053551, EPI_ISL_3053553, EPI_ISL_3053554, EPI_ISL_3053555, EPI_ISL_3053556, EPI_ISL_3053557, EPI_ISL_3053558, EPI_ISL_3053559, EPI_ISL_3053561, EPI_ISL_3053562, EPI_ISL_3053563, EPI_ISL_3053564, EPI_ISL_3053565, EPI_ISL_3053566, EPI_ISL_3053567, EPI_ISL_3053568, EPI_ISL_3053569, EPI_ISL_3053570,
EPI_ISL_3053571, EPI_ISL_3053572, EPI_ISL_3053574, EPI_ISL_3053576, EPI_ISL_3053577, EPI_ISL_3053578, EPI_ISL_3053579, EPI_ISL_3053580, EPI_ISL_3053581, EPI_ISL_3053582, EPI_ISL_3053583, EPI_ISL_3053584, EPI_ISL_3053585, EPI_ISL_3053586, EPI_ISL_3053587, EPI_ISL_3053588, EPI_ISL_3053589, EPI_ISL_3053591, EPI_ISL_3053592, EPI_ISL_3053593, EPI_ISL_3053595, EPI_ISL_3053596,
EPI_ISL_3053597, EPI_ISL_3053598, EPI_ISL_3053599, EPI_ISL_3053600, EPI_ISL_3053601, EPI_ISL_3053602, EPI_ISL_3053603, EPI_ISL_3053604, EPI_ISL_3053605, EPI_ISL_3053606, EPI_ISL_3053607, EPI_ISL_3053608, EPI_ISL_3053609, EPI_ISL_3053610, EPI_ISL_3053611, EPI_ISL_3053612, EPI_ISL_3053613, EPI_ISL_3053614, EPI_ISL_3053615, EPI_ISL_3053616, EPI_ISL_3053617, EPI_ISL_3053618,
EPI_ISL_3053619, EPI_ISL_3053620, EPI_ISL_3053621, EPI_ISL_3053622, EPI_ISL_3053624, EPI_ISL_3053625, EPI_ISL_3053626, EPI_ISL_3053627, EPI_ISL_3053628, EPI_ISL_3053629, EPI_ISL_3053630, EPI_ISL_3053631, EPI_ISL_3053632, EPI_ISL_3053633, EPI_ISL_3053634, EPI_ISL_3053635, EPI_ISL_3053636, EPI_ISL_3053637, EPI_ISL_3053638, EPI_ISL_3053639, EPI_ISL_3053640, EPI_ISL_3053641,
EPI_ISL_3053642, EPI_ISL_3053643, EPI_ISL_3053644, EPI_ISL_3053645, EPI_ISL_3053646, EPI_ISL_3053647, EPI_ISL_3053648, EPI_ISL_3053649, EPI_ISL_3053650, EPI_ISL_3053651, EPI_ISL_3053652, EPI_ISL_3053653, EPI_ISL_3053654, EPI_ISL_3053655, EPI_ISL_3053656, EPI_ISL_3053657, EPI_ISL_3053658, EPI_ISL_3053659, EPI_ISL_3053660, EPI_ISL_3053661, EPI_ISL_3053662, EPI_ISL_3053663,
EPI_ISL_3053664, EPI_ISL_3053665, EPI_ISL_3053666, EPI_ISL_3053667, EPI_ISL_3053668, EPI_ISL_3053669, EPI_ISL_3053670, EPI_ISL_3053671, EPI_ISL_3053672, EPI_ISL_3053673, EPI_ISL_3053674, EPI_ISL_3053675, EPI_ISL_3053676, EPI_ISL_3053677, EPI_ISL_3053679, EPI_ISL_3053680, EPI_ISL_3053681, EPI_ISL_3053682, EPI_ISL_3053683, EPI_ISL_3053684, EPI_ISL_3053685, EPI_ISL_3053686,
EPI_ISL_3053688, EPI_ISL_3053689, EPI_ISL_3053690, EPI_ISL_3053691, EPI_ISL_3053692, EPI_ISL_3053693, EPI_ISL_3053694, EPI_ISL_3053695, EPI_ISL_3053696, EPI_ISL_3053697, EPI_ISL_3053698, EPI_ISL_3053699, EPI_ISL_3053700, EPI_ISL_3053701, EPI_ISL_3053702, EPI_ISL_3053703, EPI_ISL_3053704, EPI_ISL_3053705, EPI_ISL_3053706, EPI_ISL_3053707, EPI_ISL_3053708, EPI_ISL_3053709,
EPI_ISL_3053710, EPI_ISL_3053711, EPI_ISL_3053712, EPI_ISL_3053713, EPI_ISL_3053714, EPI_ISL_3053715, EPI_ISL_3053717, EPI_ISL_3053718, EPI_ISL_3053719, EPI_ISL_3053720, EPI_ISL_3053721, EPI_ISL_3053722, EPI_ISL_3053723, EPI_ISL_3053724, EPI_ISL_3053725, EPI_ISL_3053726, EPI_ISL_3053727, EPI_ISL_3053728, EPI_ISL_3053729, EPI_ISL_3053730, EPI_ISL_3053731, EPI_ISL_3053732,
EPI_ISL_3053733, EPI_ISL_3053734, EPI_ISL_3053735, EPI_ISL_3053736, EPI_ISL_3053737, EPI_ISL_3053738, EPI_ISL_3053739, EPI_ISL_3053740, EPI_ISL_3053741, EPI_ISL_3053742, EPI_ISL_3053743, EPI_ISL_3053744, EPI_ISL_3053745, EPI_ISL_3053746, EPI_ISL_3053747, EPI_ISL_3053748, EPI_ISL_3053749, EPI_ISL_3053750, EPI_ISL_3053751, EPI_ISL_3053752, EPI_ISL_3053753, EPI_ISL_3053754,
EPI_ISL_3053755, EPI_ISL_3053756, EPI_ISL_3053757, EPI_ISL_3053758, EPI_ISL_3053760, EPI_ISL_3053761, EPI_ISL_3053762, EPI_ISL_3053763, EPI_ISL_3053764, EPI_ISL_3053765, EPI_ISL_3053766, EPI_ISL_3053767, EPI_ISL_3053768, EPI_ISL_3053769, EPI_ISL_3053770, EPI_ISL_3053771, EPI_ISL_3053772, EPI_ISL_3053773, EPI_ISL_3053774, EPI_ISL_3053775, EPI_ISL_3053776, EPI_ISL_3053777,
EPI_ISL_3053778, EPI_ISL_3053779, EPI_ISL_3053780, EPI_ISL_3053781, EPI_ISL_3053782, EPI_ISL_3053783, EPI_ISL_3053784, EPI_ISL_3053785, EPI_ISL_3053786, EPI_ISL_3053787, EPI_ISL_3053788, EPI_ISL_3053789, EPI_ISL_3053790

see above Department of Bacteria, Parasites and Fungi, Statens Serum Institut, Copenhagen, Denmark Statens Serum Institut Bioinformatics and Microbial Genomics Danish Covid-19 Genome Consortium

EPI_ISL_3053791 Department of Clinical Microbiology, Odense University Hospital, Odense, Denmark Statens Serum Institut Bioinformatics and Microbial Genomics Danish Covid-19 Genome Consortium

EPI_ISL_3053792, EPI_ISL_3053793, EPI_ISL_3053794, EPI_ISL_3053795, EPI_ISL_3053796, EPI_ISL_3053797, EPI_ISL_3053798, EPI_ISL_3053800, EPI_ISL_3053801, EPI_ISL_3053802, EPI_ISL_3053804, EPI_ISL_3053805, EPI_ISL_3053806, EPI_ISL_3053807, EPI_ISL_3053808, EPI_ISL_3053809, EPI_ISL_3053810, EPI_ISL_3053811, EPI_ISL_3053812, EPI_ISL_3053813, EPI_ISL_3053814, EPI_ISL_3053815,
EPI_ISL_3053816, EPI_ISL_3053817, EPI_ISL_3053818, EPI_ISL_3053819, EPI_ISL_3053820

see above Department of Bacteria, Parasites and Fungi, Statens Serum Institut, Copenhagen, Denmark Statens Serum Institut Bioinformatics and Microbial Genomics Danish Covid-19 Genome Consortium

EPI_ISL_3053833, EPI_ISL_3053849, EPI_ISL_3053851, EPI_ISL_3053853, EPI_ISL_3053857, EPI_ISL_3053867, EPI_ISL_3053872, EPI_ISL_3053877, EPI_ISL_3053884, EPI_ISL_3053886, EPI_ISL_3053889, EPI_ISL_3053890, EPI_ISL_3053893, EPI_ISL_3053896, EPI_ISL_3053897, EPI_ISL_3053907, EPI_ISL_3053916, EPI_ISL_3053924, EPI_ISL_3053927, EPI_ISL_3053929, EPI_ISL_3053930, EPI_ISL_3053937,
EPI_ISL_3053942, EPI_ISL_3053943, EPI_ISL_3053946, EPI_ISL_3053948, EPI_ISL_3053949, EPI_ISL_3053950, EPI_ISL_3053951, EPI_ISL_3053952, EPI_ISL_3053955, EPI_ISL_3053957, EPI_ISL_3053958, EPI_ISL_3053960, EPI_ISL_3053962, EPI_ISL_3053963, EPI_ISL_3053966, EPI_ISL_3053969, EPI_ISL_3053977, EPI_ISL_3053978, EPI_ISL_3053986, EPI_ISL_3053990, EPI_ISL_3053997, EPI_ISL_3054007,
EPI_ISL_3054009, EPI_ISL_3054010, EPI_ISL_3054011, EPI_ISL_3054013, EPI_ISL_3054016, EPI_ISL_3054022, EPI_ISL_3054025, EPI_ISL_3054026, EPI_ISL_3054029, EPI_ISL_3054030, EPI_ISL_3054032, EPI_ISL_3054033, EPI_ISL_3054038, EPI_ISL_3054042, EPI_ISL_3054048, EPI_ISL_3054049, EPI_ISL_3054050, EPI_ISL_3054054, EPI_ISL_3054055, EPI_ISL_3054059, EPI_ISL_3054061, EPI_ISL_3054062,
EPI_ISL_3054068, EPI_ISL_3054076, EPI_ISL_3054078, EPI_ISL_3054083, EPI_ISL_3054085, EPI_ISL_3054091, EPI_ISL_3054095, EPI_ISL_3054096, EPI_ISL_3054105, EPI_ISL_3054108, EPI_ISL_3054114, EPI_ISL_3054117, EPI_ISL_3054118, EPI_ISL_3054119, EPI_ISL_3054120, EPI_ISL_3054121, EPI_ISL_3054123, EPI_ISL_3054124, EPI_ISL_3054128, EPI_ISL_3054129, EPI_ISL_3054130, EPI_ISL_3054131,
EPI_ISL_3054132, EPI_ISL_3054134, EPI_ISL_3054135, EPI_ISL_3054137, EPI_ISL_3054144, EPI_ISL_3054146, EPI_ISL_3054149, EPI_ISL_3054151, EPI_ISL_3054152, EPI_ISL_3054153, EPI_ISL_3054154, EPI_ISL_3054159, EPI_ISL_3054160, EPI_ISL_3054179, EPI_ISL_3054186, EPI_ISL_3054206, EPI_ISL_3054220, EPI_ISL_3054234, EPI_ISL_3054236, EPI_ISL_3054238, EPI_ISL_3054241, EPI_ISL_3054249,
EPI_ISL_3054268, EPI_ISL_3054275, EPI_ISL_3054377, EPI_ISL_3054388, EPI_ISL_3054409, EPI_ISL_3054412, EPI_ISL_3054434, EPI_ISL_3054436, EPI_ISL_3054437, EPI_ISL_3054442, EPI_ISL_3054457, EPI_ISL_3054459, EPI_ISL_3054473, EPI_ISL_3054475, EPI_ISL_3054483, EPI_ISL_3054670, EPI_ISL_3054698, EPI_ISL_3054727, EPI_ISL_3054735, EPI_ISL_3054736, EPI_ISL_3054737, EPI_ISL_3054738,
EPI_ISL_3054739, EPI_ISL_3054740, EPI_ISL_3054741, EPI_ISL_3054743, EPI_ISL_3054744, EPI_ISL_3054759, EPI_ISL_3054760, EPI_ISL_3054761, EPI_ISL_3054762, EPI_ISL_3054763, EPI_ISL_3054764, EPI_ISL_3054765, EPI_ISL_3054766, EPI_ISL_3054767, EPI_ISL_3054768, EPI_ISL_3054769, EPI_ISL_3054770, EPI_ISL_3054771, EPI_ISL_3054772, EPI_ISL_3054773, EPI_ISL_3054774, EPI_ISL_3054775,
EPI_ISL_3054776, EPI_ISL_3054777, EPI_ISL_3054778, EPI_ISL_3054779, EPI_ISL_3054780, EPI_ISL_3054781, EPI_ISL_3054782, EPI_ISL_3054783, EPI_ISL_3054784, EPI_ISL_3054785, EPI_ISL_3054786, EPI_ISL_3054787, EPI_ISL_3054788, EPI_ISL_3054789, EPI_ISL_3054790, EPI_ISL_3054791, EPI_ISL_3054792, EPI_ISL_3054793, EPI_ISL_3054794, EPI_ISL_3054795, EPI_ISL_3054796, EPI_ISL_3054797,
EPI_ISL_3054798, EPI_ISL_3054799, EPI_ISL_3054800, EPI_ISL_3054801, EPI_ISL_3054802, EPI_ISL_3054803, EPI_ISL_3054804, EPI_ISL_3054805, EPI_ISL_3054806, EPI_ISL_3054807, EPI_ISL_3054808, EPI_ISL_3054809, EPI_ISL_3054810, EPI_ISL_3054811, EPI_ISL_3054812, EPI_ISL_3054813, EPI_ISL_3054814, EPI_ISL_3054815, EPI_ISL_3054816, EPI_ISL_3054817, EPI_ISL_3054818, EPI_ISL_3054819,
EPI_ISL_3054820, EPI_ISL_3054821, EPI_ISL_3054822, EPI_ISL_3054823, EPI_ISL_3054824, EPI_ISL_3054825, EPI_ISL_3054826, EPI_ISL_3054827, EPI_ISL_3054828, EPI_ISL_3054829, EPI_ISL_3054830, EPI_ISL_3054831, EPI_ISL_3054832, EPI_ISL_3054833, EPI_ISL_3054834, EPI_ISL_3054835, EPI_ISL_3054836, EPI_ISL_3054837, EPI_ISL_3054838, EPI_ISL_3054839, EPI_ISL_3054840, EPI_ISL_3054841,
EPI_ISL_3054842, EPI_ISL_3054843, EPI_ISL_3054844, EPI_ISL_3054845, EPI_ISL_3054846, EPI_ISL_3054847, EPI_ISL_3054848, EPI_ISL_3054849, EPI_ISL_3054850, EPI_ISL_3054851, EPI_ISL_3054852, EPI_ISL_3054853, EPI_ISL_3054854, EPI_ISL_3054855, EPI_ISL_3054856, EPI_ISL_3054857, EPI_ISL_3054858, EPI_ISL_3054859, EPI_ISL_3054860, EPI_ISL_3054861, EPI_ISL_3054862, EPI_ISL_3054863,
EPI_ISL_3054864, EPI_ISL_3054865, EPI_ISL_3054866, EPI_ISL_3054867, EPI_ISL_3054868, EPI_ISL_3054869, EPI_ISL_3054870, EPI_ISL_3054871, EPI_ISL_3054872, EPI_ISL_3054873, EPI_ISL_3054874, EPI_ISL_3054875, EPI_ISL_3054876, EPI_ISL_3054877, EPI_ISL_3054878, EPI_ISL_3054879, EPI_ISL_3054882, EPI_ISL_3055392

see above MEPHI, Aix Marseille University MEPHI, Aix Marseille University Anthony LEVASSEUR

EPI_ISL_3055536 Dipartimento di Medicina di Laboratorio, Azienda sanitaria universitaria Friuli Centrale (ASU FC) Dipartimento di Medicina di Laboratorio, Azienda sanitaria universitaria Friuli Centrale (ASU FC) Corrado Pipan, Catia Mio, Chiara Dal Secco, Stefania Marzinotto, Francesco Curcio

EPI_ISL_3055541 National Institute of Infectious Diseases-Prof. Dr. Matei Bals Molecular Diagnostics Laboratory National Institute of Infectious Diseases-Prof. Dr. Matei Bals Molecular Diagnostics Laboratory Corina Casangiu, Leontina Banica, Marius Surleac, Petre Milu, Robert Hohan, Simona Paraschiv, Dan Otelea

EPI_ISL_3055551, EPI_ISL_3055555, EPI_ISL_3055562, EPI_ISL_3055563 Laboratorio de salud publica Puebla LABOPAT Luis Mendoza, Cynthia Penaloza, Silvia Montilla

EPI_ISL_3055584, EPI_ISL_3055589, EPI_ISL_3055593, EPI_ISL_3055594, EPI_ISL_3055596, EPI_ISL_3055598, EPI_ISL_3055599, EPI_ISL_3055605, EPI_ISL_3055606, EPI_ISL_3055607, EPI_ISL_3055608, EPI_ISL_3055609, EPI_ISL_3055610, EPI_ISL_3055611, EPI_ISL_3055612, EPI_ISL_3055613, EPI_ISL_3055614, EPI_ISL_3055616, EPI_ISL_3055617, EPI_ISL_3055620, EPI_ISL_3055621, EPI_ISL_3055624,
EPI_ISL_3055625, EPI_ISL_3055628, EPI_ISL_3055631, EPI_ISL_3055642, EPI_ISL_3055644, EPI_ISL_3055652, EPI_ISL_3055655, EPI_ISL_3055660, EPI_ISL_3055665, EPI_ISL_3055676, EPI_ISL_3055682, EPI_ISL_3055684, EPI_ISL_3055696, EPI_ISL_3055700, EPI_ISL_3055704, EPI_ISL_3055706, EPI_ISL_3055900, EPI_ISL_3056106, EPI_ISL_3056107, EPI_ISL_3056108, EPI_ISL_3056109, EPI_ISL_3056110,
EPI_ISL_3056111, EPI_ISL_3056112, EPI_ISL_3056113, EPI_ISL_3056116, EPI_ISL_3056119, EPI_ISL_3056125, EPI_ISL_3056127, EPI_ISL_3056128, EPI_ISL_3056129, EPI_ISL_3056130, EPI_ISL_3056208, EPI_ISL_3056209, EPI_ISL_3056218, EPI_ISL_3056242, EPI_ISL_3056255, EPI_ISL_3056502, EPI_ISL_3056532, EPI_ISL_3056539, EPI_ISL_3056540, EPI_ISL_3056543, EPI_ISL_3056544, EPI_ISL_3056552,
EPI_ISL_3056553, EPI_ISL_3056554, EPI_ISL_3056555, EPI_ISL_3056556, EPI_ISL_3056557, EPI_ISL_3056558, EPI_ISL_3056559, EPI_ISL_3056560, EPI_ISL_3056561, EPI_ISL_3056562, EPI_ISL_3056563, EPI_ISL_3056564, EPI_ISL_3056565, EPI_ISL_3056566, EPI_ISL_3056567, EPI_ISL_3056568, EPI_ISL_3056569, EPI_ISL_3056570, EPI_ISL_3056571, EPI_ISL_3056572, EPI_ISL_3056573, EPI_ISL_3056574,
EPI_ISL_3056575, EPI_ISL_3056576, EPI_ISL_3056577, EPI_ISL_3056578, EPI_ISL_3056579, EPI_ISL_3056580, EPI_ISL_3056581, EPI_ISL_3056582, EPI_ISL_3056583, EPI_ISL_3056584, EPI_ISL_3056585, EPI_ISL_3056586, EPI_ISL_3056587, EPI_ISL_3056588, EPI_ISL_3056589, EPI_ISL_3056590, EPI_ISL_3056591, EPI_ISL_3056592, EPI_ISL_3056593, EPI_ISL_3056594, EPI_ISL_3056595, EPI_ISL_3056596,
EPI_ISL_3056597, EPI_ISL_3056598, EPI_ISL_3056599, EPI_ISL_3056600, EPI_ISL_3056601, EPI_ISL_3056602, EPI_ISL_3056603, EPI_ISL_3056604, EPI_ISL_3056605, EPI_ISL_3056606, EPI_ISL_3056607, EPI_ISL_3056608, EPI_ISL_3056609, EPI_ISL_3056610, EPI_ISL_3056611, EPI_ISL_3056612, EPI_ISL_3056613, EPI_ISL_3056614, EPI_ISL_3056615, EPI_ISL_3056616, EPI_ISL_3056617, EPI_ISL_3056618,
EPI_ISL_3056619, EPI_ISL_3056620, EPI_ISL_3056621, EPI_ISL_3056622, EPI_ISL_3056623, EPI_ISL_3056624, EPI_ISL_3056625, EPI_ISL_3056626, EPI_ISL_3056627, EPI_ISL_3056628, EPI_ISL_3056629, EPI_ISL_3056630, EPI_ISL_3056631, EPI_ISL_3056632, EPI_ISL_3056633, EPI_ISL_3056634, EPI_ISL_3056635, EPI_ISL_3056636, EPI_ISL_3056637, EPI_ISL_3056638, EPI_ISL_3056639, EPI_ISL_3056640,
EPI_ISL_3056641, EPI_ISL_3056642, EPI_ISL_3056643, EPI_ISL_3056644, EPI_ISL_3056645, EPI_ISL_3056646, EPI_ISL_3056647, EPI_ISL_3056648, EPI_ISL_3056649, EPI_ISL_3056650, EPI_ISL_3056651, EPI_ISL_3056652, EPI_ISL_3056653, EPI_ISL_3056667, EPI_ISL_3056668, EPI_ISL_3056669, EPI_ISL_3056670, EPI_ISL_3056671, EPI_ISL_3056672, EPI_ISL_3056673, EPI_ISL_3056674, EPI_ISL_3056675,
EPI_ISL_3056676, EPI_ISL_3056677, EPI_ISL_3056678, EPI_ISL_3056679, EPI_ISL_3056680, EPI_ISL_3056681, EPI_ISL_3056682, EPI_ISL_3056683, EPI_ISL_3056684, EPI_ISL_3056685, EPI_ISL_3056686, EPI_ISL_3056687, EPI_ISL_3056688, EPI_ISL_3056689, EPI_ISL_3056690, EPI_ISL_3056691, EPI_ISL_3056692, EPI_ISL_3056693, EPI_ISL_3056694, EPI_ISL_3056695, EPI_ISL_3056696, EPI_ISL_3056697,
EPI_ISL_3056698, EPI_ISL_3056699

see above SARS-CoV-2 Sequencing Castilla y Leon-Spain Consortium SARS-CoV-2 Sequencing Castilla y Leon-Spain Consortium Antonio Orduña-Domingo, Marta Hernandez, David Abad, Marta Dominguez-Gil, Silvia Rojo, Gabriel March Rosello, Sonsoles Garcinuño Pérez, Carmen Aldea-Mansilla, Mª Fe Brezmes-Valdivieso, Gregoria Megías Lobón, María Antonia García Castro, Carmen Gimeno Crespo, Noelia Arenal Andrés, Carlos Fuster Foz, M. Isabel Fernandez-Natal, Jose María Eiros Bouza

EPI_ISL_3056721, EPI_ISL_3056722, EPI_ISL_3056723, EPI_ISL_3056725, EPI_ISL_3056726, EPI_ISL_3056727, EPI_ISL_3056728, EPI_ISL_3056729, EPI_ISL_3056730, EPI_ISL_3056732, EPI_ISL_3056733, EPI_ISL_3056734, EPI_ISL_3056735, EPI_ISL_3056736, EPI_ISL_3056737, EPI_ISL_3056738, EPI_ISL_3056739, EPI_ISL_3056741, EPI_ISL_3056742, EPI_ISL_3056743, EPI_ISL_3056745, EPI_ISL_3056746,
EPI_ISL_3056747, EPI_ISL_3056748, EPI_ISL_3056751, EPI_ISL_3056752, EPI_ISL_3056753, EPI_ISL_3056754, EPI_ISL_3056755, EPI_ISL_3056756, EPI_ISL_3056757, EPI_ISL_3056759, EPI_ISL_3056760, EPI_ISL_3056761

see above Dutch COVID-19 response team National Institute for Public Health and the Environment (RIVM) Adam Meijer, Harry Vennema, Dirk Eggink, Jeroen Cremer, Sharon van den Brink, Bas van der Veer, AnneMarie van den Brandt, Lisa Wijsman, Kim Freriks, Ryanne Jaarsma, Euníce Then, Lynn Aarts, Sanne Bos, Melissa van Tuil, Florian Zwagemaker, Dennis Schmitz, Annelies Kroneman, Karim Hajji, Chantal Reusken, on behalf of the national COVID-19 response team

EPI_ISL_3056762 CT Department of Public Health CT Department of Public Health Claire_Pearson,Tu_N_Nguyen

EPI_ISL_3056763, EPI_ISL_3056764, EPI_ISL_3056765, EPI_ISL_3056766, EPI_ISL_3056767, EPI_ISL_3056768, EPI_ISL_3056769, EPI_ISL_3056770, EPI_ISL_3056771, EPI_ISL_3056772, EPI_ISL_3056774, EPI_ISL_3056775, EPI_ISL_3056776, EPI_ISL_3056777, EPI_ISL_3056778, EPI_ISL_3056780, EPI_ISL_3056781, EPI_ISL_3056782, EPI_ISL_3056784, EPI_ISL_3056785, EPI_ISL_3056787, EPI_ISL_3056788,
EPI_ISL_3056790, EPI_ISL_3056792, EPI_ISL_3056793, EPI_ISL_3056794, EPI_ISL_3056795, EPI_ISL_3056797, EPI_ISL_3056798, EPI_ISL_3056799, EPI_ISL_3056800, EPI_ISL_3056802, EPI_ISL_3056803, EPI_ISL_3056804, EPI_ISL_3056805, EPI_ISL_3056806, EPI_ISL_3056807, EPI_ISL_3056808, EPI_ISL_3056809, EPI_ISL_3056810, EPI_ISL_3056811, EPI_ISL_3056812, EPI_ISL_3056813, EPI_ISL_3056814,
EPI_ISL_3056815, EPI_ISL_3056816, EPI_ISL_3056817, EPI_ISL_3056818, EPI_ISL_3056819, EPI_ISL_3056820, EPI_ISL_3056822, EPI_ISL_3056823, EPI_ISL_3056824, EPI_ISL_3056825, EPI_ISL_3056826, EPI_ISL_3056827, EPI_ISL_3056828, EPI_ISL_3056829, EPI_ISL_3056830, EPI_ISL_3056831, EPI_ISL_3056832, EPI_ISL_3056833, EPI_ISL_3056834, EPI_ISL_3056835, EPI_ISL_3056836, EPI_ISL_3056837,
EPI_ISL_3056838, EPI_ISL_3056839, EPI_ISL_3056840, EPI_ISL_3056841, EPI_ISL_3056842, EPI_ISL_3056843, EPI_ISL_3056844, EPI_ISL_3056845, EPI_ISL_3056846, EPI_ISL_3056847, EPI_ISL_3056848, EPI_ISL_3056850, EPI_ISL_3056851, EPI_ISL_3056852, EPI_ISL_3056853, EPI_ISL_3056854, EPI_ISL_3056857, EPI_ISL_3056858, EPI_ISL_3056859, EPI_ISL_3056860, EPI_ISL_3056861, EPI_ISL_3056862,
EPI_ISL_3056863, EPI_ISL_3056865, EPI_ISL_3056866, EPI_ISL_3056867, EPI_ISL_3056869, EPI_ISL_3056870, EPI_ISL_3056871, EPI_ISL_3056872, EPI_ISL_3056873, EPI_ISL_3056874, EPI_ISL_3056875, EPI_ISL_3056876, EPI_ISL_3056877, EPI_ISL_3056878, EPI_ISL_3056879, EPI_ISL_3056880, EPI_ISL_3056881, EPI_ISL_3056882, EPI_ISL_3056883, EPI_ISL_3056884, EPI_ISL_3056885, EPI_ISL_3056886,
EPI_ISL_3056887, EPI_ISL_3056888, EPI_ISL_3056889, EPI_ISL_3056892, EPI_ISL_3056893, EPI_ISL_3056894, EPI_ISL_3056895, EPI_ISL_3056896, EPI_ISL_3056897, EPI_ISL_3056898, EPI_ISL_3056899, EPI_ISL_3056900, EPI_ISL_3056901, EPI_ISL_3056902, EPI_ISL_3056903, EPI_ISL_3056904, EPI_ISL_3056906, EPI_ISL_3056907, EPI_ISL_3056908, EPI_ISL_3056909, EPI_ISL_3056910, EPI_ISL_3056911,
EPI_ISL_3056912, EPI_ISL_3056913, EPI_ISL_3056914, EPI_ISL_3056915, EPI_ISL_3056916, EPI_ISL_3056917, EPI_ISL_3056918, EPI_ISL_3056919, EPI_ISL_3056920, EPI_ISL_3056921, EPI_ISL_3056922, EPI_ISL_3056923, EPI_ISL_3056924, EPI_ISL_3056925, EPI_ISL_3056926, EPI_ISL_3056927, EPI_ISL_3056928, EPI_ISL_3056929, EPI_ISL_3056930, EPI_ISL_3056931, EPI_ISL_3056932, EPI_ISL_3056933,
EPI_ISL_3056934, EPI_ISL_3056935, EPI_ISL_3056936, EPI_ISL_3056937, EPI_ISL_3056938, EPI_ISL_3056939, EPI_ISL_3056940, EPI_ISL_3056941, EPI_ISL_3056942, EPI_ISL_3056944, EPI_ISL_3056945, EPI_ISL_3056946, EPI_ISL_3056947, EPI_ISL_3056948, EPI_ISL_3056949, EPI_ISL_3056950, EPI_ISL_3056951, EPI_ISL_3056952, EPI_ISL_3056953, EPI_ISL_3056954, EPI_ISL_3056956, EPI_ISL_3056957,
EPI_ISL_3056958, EPI_ISL_3056959, EPI_ISL_3056960, EPI_ISL_3056961, EPI_ISL_3056962, EPI_ISL_3056963, EPI_ISL_3056964, EPI_ISL_3056965, EPI_ISL_3056966, EPI_ISL_3056967, EPI_ISL_3056968, EPI_ISL_3056969, EPI_ISL_3056970, EPI_ISL_3056971, EPI_ISL_3056972, EPI_ISL_3056973, EPI_ISL_3056974, EPI_ISL_3056975, EPI_ISL_3056976, EPI_ISL_3056977, EPI_ISL_3056978, EPI_ISL_3056979,
EPI_ISL_3056980, EPI_ISL_3056981, EPI_ISL_3056982, EPI_ISL_3056984, EPI_ISL_3056985, EPI_ISL_3056986, EPI_ISL_3056987, EPI_ISL_3056988, EPI_ISL_3056990, EPI_ISL_3056991, EPI_ISL_3056992, EPI_ISL_3056993, EPI_ISL_3056994, EPI_ISL_3056995, EPI_ISL_3056996, EPI_ISL_3056997, EPI_ISL_3056998, EPI_ISL_3056999, EPI_ISL_3057000, EPI_ISL_3057002, EPI_ISL_3057003, EPI_ISL_3057004,
EPI_ISL_3057005, EPI_ISL_3057006, EPI_ISL_3057007, EPI_ISL_3057008, EPI_ISL_3057009, EPI_ISL_3057010, EPI_ISL_3057011, EPI_ISL_3057012, EPI_ISL_3057013, EPI_ISL_3057014, EPI_ISL_3057015, EPI_ISL_3057016, EPI_ISL_3057017, EPI_ISL_3057018, EPI_ISL_3057019, EPI_ISL_3057020, EPI_ISL_3057021, EPI_ISL_3057022, EPI_ISL_3057023, EPI_ISL_3057024, EPI_ISL_3057025, EPI_ISL_3057026,
EPI_ISL_3057028, EPI_ISL_3057029, EPI_ISL_3057030, EPI_ISL_3057031, EPI_ISL_3057033, EPI_ISL_3057034, EPI_ISL_3057035, EPI_ISL_3057036, EPI_ISL_3057037, EPI_ISL_3057038, EPI_ISL_3057039, EPI_ISL_3057040, EPI_ISL_3057041, EPI_ISL_3057042, EPI_ISL_3057043, EPI_ISL_3057044, EPI_ISL_3057046, EPI_ISL_3057047, EPI_ISL_3057048, EPI_ISL_3057049, EPI_ISL_3057050, EPI_ISL_3057051,
EPI_ISL_3057052, EPI_ISL_3057053, EPI_ISL_3057054, EPI_ISL_3057055, EPI_ISL_3057056, EPI_ISL_3057057, EPI_ISL_3057058, EPI_ISL_3057059, EPI_ISL_3057060, EPI_ISL_3057061, EPI_ISL_3057062, EPI_ISL_3057063, EPI_ISL_3057064, EPI_ISL_3057065, EPI_ISL_3057066, EPI_ISL_3057067, EPI_ISL_3057068, EPI_ISL_3057069, EPI_ISL_3057070, EPI_ISL_3057071, EPI_ISL_3057072, EPI_ISL_3057073,
EPI_ISL_3057074, EPI_ISL_3057075, EPI_ISL_3057076, EPI_ISL_3057077, EPI_ISL_3057078, EPI_ISL_3057079, EPI_ISL_3057080, EPI_ISL_3057081, EPI_ISL_3057082, EPI_ISL_3057083, EPI_ISL_3057084, EPI_ISL_3057085, EPI_ISL_3057086, EPI_ISL_3057087, EPI_ISL_3057088, EPI_ISL_3057089, EPI_ISL_3057090, EPI_ISL_3057091, EPI_ISL_3057092, EPI_ISL_3057093, EPI_ISL_3057094, EPI_ISL_3057095,
EPI_ISL_3057096, EPI_ISL_3057097, EPI_ISL_3057098, EPI_ISL_3057099, EPI_ISL_3057100, EPI_ISL_3057101, EPI_ISL_3057102, EPI_ISL_3057103, EPI_ISL_3057104, EPI_ISL_3057105, EPI_ISL_3057106, EPI_ISL_3057108, EPI_ISL_3057109, EPI_ISL_3057110, EPI_ISL_3057111, EPI_ISL_3057112, EPI_ISL_3057113, EPI_ISL_3057114, EPI_ISL_3057115, EPI_ISL_3057116, EPI_ISL_3057117, EPI_ISL_3057118,
EPI_ISL_3057119, EPI_ISL_3057120, EPI_ISL_3057121, EPI_ISL_3057122, EPI_ISL_3057123, EPI_ISL_3057124, EPI_ISL_3057125, EPI_ISL_3057126, EPI_ISL_3057127, EPI_ISL_3057128, EPI_ISL_3057129, EPI_ISL_3057130, EPI_ISL_3057131, EPI_ISL_3057132, EPI_ISL_3057133, EPI_ISL_3057134, EPI_ISL_3057135, EPI_ISL_3057136, EPI_ISL_3057137, EPI_ISL_3057138, EPI_ISL_3057140, EPI_ISL_3057142,
EPI_ISL_3057143, EPI_ISL_3057144, EPI_ISL_3057145, EPI_ISL_3057146, EPI_ISL_3057147, EPI_ISL_3057149, EPI_ISL_3057150, EPI_ISL_3057151, EPI_ISL_3057152, EPI_ISL_3057153, EPI_ISL_3057154, EPI_ISL_3057155, EPI_ISL_3057156, EPI_ISL_3057158, EPI_ISL_3057159, EPI_ISL_3057160, EPI_ISL_3057161, EPI_ISL_3057162, EPI_ISL_3057163, EPI_ISL_3057164, EPI_ISL_3057165, EPI_ISL_3057166,
EPI_ISL_3057168, EPI_ISL_3057169, EPI_ISL_3057170, EPI_ISL_3057171, EPI_ISL_3057172, EPI_ISL_3057174, EPI_ISL_3057175, EPI_ISL_3057176, EPI_ISL_3057178, EPI_ISL_3057179, EPI_ISL_3057180, EPI_ISL_3057181, EPI_ISL_3057182, EPI_ISL_3057183, EPI_ISL_3057184, EPI_ISL_3057185, EPI_ISL_3057186, EPI_ISL_3057187, EPI_ISL_3057190, EPI_ISL_3057191, EPI_ISL_3057192, EPI_ISL_3057193,
EPI_ISL_3057194, EPI_ISL_3057198, EPI_ISL_3057199, EPI_ISL_3057200, EPI_ISL_3057201, EPI_ISL_3057202, EPI_ISL_3057203, EPI_ISL_3057204, EPI_ISL_3057205, EPI_ISL_3057206, EPI_ISL_3057212, EPI_ISL_3057213, EPI_ISL_3057214, EPI_ISL_3057215, EPI_ISL_3057216, EPI_ISL_3057217, EPI_ISL_3057218, EPI_ISL_3057220, EPI_ISL_3057221, EPI_ISL_3057222, EPI_ISL_3057223, EPI_ISL_3057224,
EPI_ISL_3057225, EPI_ISL_3057226, EPI_ISL_3057227, EPI_ISL_3057228, EPI_ISL_3057229, EPI_ISL_3057230, EPI_ISL_3057231, EPI_ISL_3057232, EPI_ISL_3057233, EPI_ISL_3057234, EPI_ISL_3057235, EPI_ISL_3057236, EPI_ISL_3057237, EPI_ISL_3057238, EPI_ISL_3057239, EPI_ISL_3057240, EPI_ISL_3057241, EPI_ISL_3057243, EPI_ISL_3057245, EPI_ISL_3057246, EPI_ISL_3057247, EPI_ISL_3057248,
EPI_ISL_3057251, EPI_ISL_3057252, EPI_ISL_3057255, EPI_ISL_3057256, EPI_ISL_3057257, EPI_ISL_3057258, EPI_ISL_3057259, EPI_ISL_3057260, EPI_ISL_3057261, EPI_ISL_3057262, EPI_ISL_3057264, EPI_ISL_3057265, EPI_ISL_3057266, EPI_ISL_3057267, EPI_ISL_3057269, EPI_ISL_3057273, EPI_ISL_3057274, EPI_ISL_3057275, EPI_ISL_3057276, EPI_ISL_3057277, EPI_ISL_3057278, EPI_ISL_3057280,
EPI_ISL_3057282, EPI_ISL_3057283, EPI_ISL_3057284, EPI_ISL_3057285, EPI_ISL_3057286, EPI_ISL_3057287, EPI_ISL_3057288, EPI_ISL_3057289, EPI_ISL_3057291, EPI_ISL_3057292, EPI_ISL_3057294, EPI_ISL_3057295, EPI_ISL_3057296, EPI_ISL_3057297, EPI_ISL_3057300, EPI_ISL_3057303, EPI_ISL_3057304, EPI_ISL_3057305, EPI_ISL_3057306, EPI_ISL_3057307, EPI_ISL_3057310, EPI_ISL_3057312,
EPI_ISL_3057313, EPI_ISL_3057315, EPI_ISL_3057319, EPI_ISL_3057320, EPI_ISL_3057321, EPI_ISL_3057322, EPI_ISL_3057323, EPI_ISL_3057326, EPI_ISL_3057327, EPI_ISL_3057328, EPI_ISL_3057329, EPI_ISL_3057331, EPI_ISL_3057332, EPI_ISL_3057335, EPI_ISL_3057336, EPI_ISL_3057337, EPI_ISL_3057338, EPI_ISL_3057339, EPI_ISL_3057340, EPI_ISL_3057341, EPI_ISL_3057342, EPI_ISL_3057343,
EPI_ISL_3057344, EPI_ISL_3057345, EPI_ISL_3057347, EPI_ISL_3057348, EPI_ISL_3057350, EPI_ISL_3057351, EPI_ISL_3057352, EPI_ISL_3057353, EPI_ISL_3057354, EPI_ISL_3057355, EPI_ISL_3057356, EPI_ISL_3057358, EPI_ISL_3057359, EPI_ISL_3057360, EPI_ISL_3057361, EPI_ISL_3057363, EPI_ISL_3057364, EPI_ISL_3057365, EPI_ISL_3057366, EPI_ISL_3057367, EPI_ISL_3057369, EPI_ISL_3057371,
EPI_ISL_3057372, EPI_ISL_3057373, EPI_ISL_3057374, EPI_ISL_3057375, EPI_ISL_3057376, EPI_ISL_3057377, EPI_ISL_3057378, EPI_ISL_3057379, EPI_ISL_3057380, EPI_ISL_3057381, EPI_ISL_3057382, EPI_ISL_3057383, EPI_ISL_3057384, EPI_ISL_3057385, EPI_ISL_3057388, EPI_ISL_3057389, EPI_ISL_3057390, EPI_ISL_3057391, EPI_ISL_3057392, EPI_ISL_3057393, EPI_ISL_3057394, EPI_ISL_3057398,
EPI_ISL_3057399, EPI_ISL_3057400, EPI_ISL_3057402, EPI_ISL_3057403, EPI_ISL_3057405, EPI_ISL_3057406, EPI_ISL_3057409, EPI_ISL_3057410, EPI_ISL_3057411, EPI_ISL_3057413, EPI_ISL_3057414, EPI_ISL_3057415, EPI_ISL_3057416, EPI_ISL_3057417, EPI_ISL_3057418, EPI_ISL_3057419, EPI_ISL_3057420, EPI_ISL_3057421, EPI_ISL_3057422, EPI_ISL_3057424, EPI_ISL_3057425, EPI_ISL_3057427,
EPI_ISL_3057428, EPI_ISL_3057430, EPI_ISL_3057431, EPI_ISL_3057434, EPI_ISL_3057436, EPI_ISL_3057438, EPI_ISL_3057439, EPI_ISL_3057442, EPI_ISL_3057443, EPI_ISL_3057444, EPI_ISL_3057445, EPI_ISL_3057448, EPI_ISL_3057449, EPI_ISL_3057450, EPI_ISL_3057451, EPI_ISL_3057452, EPI_ISL_3057453, EPI_ISL_3057454, EPI_ISL_3057456, EPI_ISL_3057458, EPI_ISL_3057459, EPI_ISL_3057460,
EPI_ISL_3057461, EPI_ISL_3057462, EPI_ISL_3057464, EPI_ISL_3057465, EPI_ISL_3057466, EPI_ISL_3057475, EPI_ISL_3057477, EPI_ISL_3057485, EPI_ISL_3057486, EPI_ISL_3057488, EPI_ISL_3057489, EPI_ISL_3057490, EPI_ISL_3057491, EPI_ISL_3057493, EPI_ISL_3057494, EPI_ISL_3057496, EPI_ISL_3057497, EPI_ISL_3057498, EPI_ISL_3057500, EPI_ISL_3057503, EPI_ISL_3057504, EPI_ISL_3057505,
EPI_ISL_3057506, EPI_ISL_3057507, EPI_ISL_3057508, EPI_ISL_3057509, EPI_ISL_3057510, EPI_ISL_3057514, EPI_ISL_3057518, EPI_ISL_3057523, EPI_ISL_3057524, EPI_ISL_3057525, EPI_ISL_3057526, EPI_ISL_3057528, EPI_ISL_3057529, EPI_ISL_3057530, EPI_ISL_3057533, EPI_ISL_3057534, EPI_ISL_3057535, EPI_ISL_3057536, EPI_ISL_3057537, EPI_ISL_3057538, EPI_ISL_3057539, EPI_ISL_3057540,

| | | | |
|---|---|---|---|
| EPI_ISL_3057543, EPI_ISL_3057544, EPI_ISL_3057545, EPI_ISL_3057547, EPI_ISL_3057548, EPI_ISL_3057549, EPI_ISL_3057551, EPI_ISL_3057552, EPI_ISL_3057555, EPI_ISL_3057556, EPI_ISL_3057559, EPI_ISL_3057560, EPI_ISL_3057563, EPI_ISL_3057564, EPI_ISL_3057565, EPI_ISL_3057567, EPI_ISL_3057568, EPI_ISL_3057570, EPI_ISL_3057571, EPI_ISL_3057572, EPI_ISL_3057573, EPI_ISL_3057574, EPI_ISL_3057576, EPI_ISL_3057578, EPI_ISL_3057579, EPI_ISL_3057580, EPI_ISL_3057583, EPI_ISL_3057584, EPI_ISL_3057585, EPI_ISL_3057586, EPI_ISL_3057587, EPI_ISL_3057588, EPI_ISL_3057589, EPI_ISL_3057590, EPI_ISL_3057592, EPI_ISL_3057593, EPI_ISL_3057594, EPI_ISL_3057595, EPI_ISL_3057596, EPI_ISL_3057597, EPI_ISL_3057598, EPI_ISL_3057599, EPI_ISL_3057600, EPI_ISL_3057601, EPI_ISL_3057602, EPI_ISL_3057603, EPI_ISL_3057605, EPI_ISL_3057606, EPI_ISL_3057607, EPI_ISL_3057608, EPI_ISL_3057609, EPI_ISL_3057610, EPI_ISL_3057611, EPI_ISL_3057613, EPI_ISL_3057614, EPI_ISL_3057615, EPI_ISL_3057616, EPI_ISL_3057617, EPI_ISL_3057618, EPI_ISL_3057619, EPI_ISL_3057620, EPI_ISL_3057621, EPI_ISL_3057622, EPI_ISL_3057623, EPI_ISL_3057624, EPI_ISL_3057625, EPI_ISL_3057626, EPI_ISL_3057627, EPI_ISL_3057628, EPI_ISL_3057629, EPI_ISL_3057631, EPI_ISL_3057632, EPI_ISL_3057633, EPI_ISL_3057634, EPI_ISL_3057636, EPI_ISL_3057637, EPI_ISL_3057638, EPI_ISL_3057641, EPI_ISL_3057644, EPI_ISL_3057645, EPI_ISL_3057646, EPI_ISL_3057647, EPI_ISL_3057649, EPI_ISL_3057650, EPI_ISL_3057651, EPI_ISL_3057652, EPI_ISL_3057654, EPI_ISL_3057655, EPI_ISL_3057656, EPI_ISL_3057657, EPI_ISL_3057658, EPI_ISL_3057659, EPI_ISL_3057660, EPI_ISL_3057661, EPI_ISL_3057662, EPI_ISL_3057663, EPI_ISL_3057664, EPI_ISL_3057665, EPI_ISL_3057667, EPI_ISL_3057669, EPI_ISL_3057670, EPI_ISL_3057671, EPI_ISL_3057672, EPI_ISL_3057673, EPI_ISL_3057674, EPI_ISL_3057675, EPI_ISL_3057677, EPI_ISL_3057678, EPI_ISL_3057679, EPI_ISL_3057680, EPI_ISL_3057682, EPI_ISL_3057683, EPI_ISL_3057684, EPI_ISL_3057685, EPI_ISL_3057686, EPI_ISL_3057687, EPI_ISL_3057688, EPI_ISL_3057689, EPI_ISL_3057690, EPI_ISL_3057692, EPI_ISL_3057693, EPI_ISL_3057696, EPI_ISL_3057699, EPI_ISL_3057700, EPI_ISL_3057701, EPI_ISL_3057702, EPI_ISL_3057703, EPI_ISL_3057704, EPI_ISL_3057705, EPI_ISL_3057706, EPI_ISL_3057708, EPI_ISL_3057709, EPI_ISL_3057710, EPI_ISL_3057711, EPI_ISL_3057712, EPI_ISL_3057713, EPI_ISL_3057714, EPI_ISL_3057715, EPI_ISL_3057716, EPI_ISL_3057720, EPI_ISL_3057721, EPI_ISL_3057722, EPI_ISL_3057726, EPI_ISL_3057727, EPI_ISL_3057728, EPI_ISL_3057729, EPI_ISL_3057731, EPI_ISL_3057732, EPI_ISL_3057733, EPI_ISL_3057734, EPI_ISL_3057736, EPI_ISL_3057737, EPI_ISL_3057739, EPI_ISL_3057740, EPI_ISL_3057741, EPI_ISL_3057742, EPI_ISL_3057743, EPI_ISL_3057745, EPI_ISL_3057746, EPI_ISL_3057747, EPI_ISL_3057748, EPI_ISL_3057749, EPI_ISL_3057750, EPI_ISL_3057751, EPI_ISL_3057752, EPI_ISL_3057755, EPI_ISL_3057756, EPI_ISL_3057757, EPI_ISL_3057760, EPI_ISL_3057761, EPI_ISL_3057762, EPI_ISL_3057764, EPI_ISL_3057768, EPI_ISL_3057770, EPI_ISL_3057771, EPI_ISL_3057772, EPI_ISL_3057774, EPI_ISL_3057775, EPI_ISL_3057776, EPI_ISL_3057777, EPI_ISL_3057778, EPI_ISL_3057779, EPI_ISL_3057780, EPI_ISL_3057781, EPI_ISL_3057782, EPI_ISL_3057783, EPI_ISL_3057784, EPI_ISL_3057785, EPI_ISL_3057786, EPI_ISL_3057787, EPI_ISL_3057788, EPI_ISL_3057792, EPI_ISL_3057793, EPI_ISL_3057794, EPI_ISL_3057795, EPI_ISL_3057797, EPI_ISL_3057798, EPI_ISL_3057799, EPI_ISL_3057800, EPI_ISL_3057801, EPI_ISL_3057802, EPI_ISL_3057803, EPI_ISL_3057806, EPI_ISL_3057807, EPI_ISL_3057808, EPI_ISL_3057809, EPI_ISL_3057810, EPI_ISL_3057811, EPI_ISL_3057812, EPI_ISL_3057813, EPI_ISL_3057815, EPI_ISL_3057816, EPI_ISL_3057817, EPI_ISL_3057818, EPI_ISL_3057819, EPI_ISL_3057820, EPI_ISL_3057821, EPI_ISL_3057822, EPI_ISL_3057823, EPI_ISL_3057824, EPI_ISL_3057825, EPI_ISL_3057826, EPI_ISL_3057827, EPI_ISL_3057828, EPI_ISL_3057833, EPI_ISL_3057835, EPI_ISL_3057836, EPI_ISL_3057837, EPI_ISL_3057838, EPI_ISL_3057839, EPI_ISL_3057840, EPI_ISL_3057841, EPI_ISL_3057844, EPI_ISL_3057845, EPI_ISL_3057846, EPI_ISL_3057847, EPI_ISL_3057848, EPI_ISL_3057850, EPI_ISL_3057851, EPI_ISL_3057852, EPI_ISL_3057853, EPI_ISL_3057854, EPI_ISL_3057858, EPI_ISL_3057860, EPI_ISL_3057861, EPI_ISL_3057863, EPI_ISL_3057866, EPI_ISL_3057867, EPI_ISL_3057868, EPI_ISL_3057870, EPI_ISL_3057874, EPI_ISL_3057875, EPI_ISL_3057879, EPI_ISL_3057880, EPI_ISL_3057881, EPI_ISL_3057882, EPI_ISL_3057888, EPI_ISL_3057889, EPI_ISL_3057890, EPI_ISL_3057891, EPI_ISL_3057892, EPI_ISL_3057895, EPI_ISL_3057900, EPI_ISL_3057901, EPI_ISL_3057902, EPI_ISL_3057903, EPI_ISL_3057909, EPI_ISL_3057910, EPI_ISL_3057912, EPI_ISL_3057913, EPI_ISL_3057915, EPI_ISL_3057917, EPI_ISL_3057919, EPI_ISL_3057920, EPI_ISL_3057922, EPI_ISL_3057924, EPI_ISL_3057926, EPI_ISL_3057929, EPI_ISL_3057931, EPI_ISL_3057932, EPI_ISL_3057933, EPI_ISL_3057935, EPI_ISL_3057939, EPI_ISL_3057940, EPI_ISL_3057943, EPI_ISL_3057944, EPI_ISL_3057945, EPI_ISL_3057946, EPI_ISL_3057947, EPI_ISL_3057949, EPI_ISL_3057950, EPI_ISL_3057951, EPI_ISL_3057952, EPI_ISL_3057953, EPI_ISL_3057954, EPI_ISL_3057955, EPI_ISL_3057956, EPI_ISL_3057957, EPI_ISL_3057958, EPI_ISL_3057959, EPI_ISL_3057961, EPI_ISL_3057962, EPI_ISL_3057964, EPI_ISL_3057965, EPI_ISL_3057966, EPI_ISL_3057967, EPI_ISL_3057968, EPI_ISL_3057970, EPI_ISL_3057971, EPI_ISL_3057973, EPI_ISL_3057974, EPI_ISL_3057975, EPI_ISL_3057976, EPI_ISL_3057977, EPI_ISL_3057978, EPI_ISL_3057979, EPI_ISL_3057981, EPI_ISL_3057982, EPI_ISL_3057983, EPI_ISL_3057984, EPI_ISL_3057985, EPI_ISL_3057986, EPI_ISL_3057987, EPI_ISL_3057989, EPI_ISL_3057990, EPI_ISL_3057991, EPI_ISL_3057992, EPI_ISL_3057995, EPI_ISL_3057996, EPI_ISL_3057997, EPI_ISL_3057998, EPI_ISL_3057999, EPI_ISL_3058000, EPI_ISL_3058001, EPI_ISL_3058002, EPI_ISL_3058003, EPI_ISL_3058005, EPI_ISL_3058006, EPI_ISL_3058007, EPI_ISL_3058008, EPI_ISL_3058009, EPI_ISL_3058010, EPI_ISL_3058011, EPI_ISL_3058012, EPI_ISL_3058013, EPI_ISL_3058014, EPI_ISL_3058015, EPI_ISL_3058017, EPI_ISL_3058018, EPI_ISL_3058020, EPI_ISL_3058021, EPI_ISL_3058022, EPI_ISL_3058023, EPI_ISL_3058024, EPI_ISL_3058025, EPI_ISL_3058026, EPI_ISL_3058027, EPI_ISL_3058028, EPI_ISL_3058029, EPI_ISL_3058031, EPI_ISL_3058032, EPI_ISL_3058033, EPI_ISL_3058036, EPI_ISL_3058037, EPI_ISL_3058038, EPI_ISL_3058039, EPI_ISL_3058040, EPI_ISL_3058041, EPI_ISL_3058042, EPI_ISL_3058046, EPI_ISL_3058049, EPI_ISL_3058052, EPI_ISL_3058053, EPI_ISL_3058054, EPI_ISL_3058055, EPI_ISL_3058058, EPI_ISL_3058059, EPI_ISL_3058060, EPI_ISL_3058061, EPI_ISL_3058064, EPI_ISL_3058065, EPI_ISL_3058066, EPI_ISL_3058067, EPI_ISL_3058069, EPI_ISL_3058073, EPI_ISL_3058074, EPI_ISL_3058077, EPI_ISL_3058078, EPI_ISL_3058079, EPI_ISL_3058080, EPI_ISL_3058081, EPI_ISL_3058085, EPI_ISL_3058086, EPI_ISL_3058088, EPI_ISL_3058089, EPI_ISL_3058091, EPI_ISL_3058092, EPI_ISL_3058094, EPI_ISL_3058095, EPI_ISL_3058096, EPI_ISL_3058097, EPI_ISL_3058098, EPI_ISL_3058101, EPI_ISL_3058102, EPI_ISL_3058103, EPI_ISL_3058104, EPI_ISL_3058106 | | | |
| see above | Dutch COVID-19 response team | National Institute for Public Health and the Environment (RIVM) | Adam Meijer, Harry Vennema, Dirk Eggink, Jeroen Cremer, Sharon van den Brink, Bas van der Veer, AnneMarie van den Brandt, Lisa Wijsman, Kim Freriks, Ryanne Jaarsma, Euníce Then, Lynn Aarts, Sanne Bos, Melissa van Tuil, Florian Zwagemaker, Dennis Schmitz, Annelies Kroneman, Karim Hajji, Chantal Reusken, on behalf of the national COVID-19 response team |
| EPI_ISL_3058107, EPI_ISL_3058109, EPI_ISL_3058110, EPI_ISL_3058113, EPI_ISL_3058115, EPI_ISL_3058122, EPI_ISL_3058123, EPI_ISL_3058125, EPI_ISL_3058126, EPI_ISL_3058129, EPI_ISL_3058130, EPI_ISL_3058132, EPI_ISL_3058133 | | | |
| see above | Nastavni zavod za javno zdravstvo Splitsko- Dalmatinske Županije | Hrvatski zavod za javno zdravstvo | Irena Tabain, Ivana Ferenčak |
| EPI_ISL_3058139, EPI_ISL_3058141, EPI_ISL_3058142, EPI_ISL_3058143 | Zavod za javno zdravstvo Zadarske Županije | Hrvatski zavod za javno zdravstvo | Irena Tabain, Ivana Ferenčak |
| EPI_ISL_3058144, EPI_ISL_3058145, EPI_ISL_3058146 | Nastavni zavod za javno zdravstvo Splitsko- Dalmatinske Županije | Hrvatski zavod za javno zdravstvo | Irena Tabain, Ivana Ferenčak |
| EPI_ISL_3058152, EPI_ISL_3058153, EPI_ISL_3058154, EPI_ISL_3058156, EPI_ISL_3058157, EPI_ISL_3058158 | Zavod za javno zdravstvo Međimurske Županije | Hrvatski zavod za javno zdravstvo | Irena Tabain, Ivana Ferenčak |
| EPI_ISL_3058167, EPI_ISL_3058168 | Hrvatski zavod za javno zdravstvo | Hrvatski zavod za javno zdravstvo | Irena Tabain, Ivana Ferenčak |
| EPI_ISL_3058175, EPI_ISL_3058176, EPI_ISL_3058177, EPI_ISL_3058178, EPI_ISL_3058179, EPI_ISL_3058180, EPI_ISL_3058181, EPI_ISL_3058182, EPI_ISL_3058183, EPI_ISL_3058184, EPI_ISL_3058185, EPI_ISL_3058186, EPI_ISL_3058187 | | | |
| see above | Zavod za javno zdravstvo Šibensko- Kninske Županije | Hrvatski zavod za javno zdravstvo | Irena Tabain, Ivana Ferenčak |
| EPI_ISL_3058188 | Klinički bolnički centar Split | Hrvatski zavod za javno zdravstvo | Irena Tabain, Ivana Ferenčak |
| EPI_ISL_3058192, EPI_ISL_3058193, EPI_ISL_3058194, EPI_ISL_3058195, EPI_ISL_3058196, EPI_ISL_3058200 | Nastavni zavod za javno zdravstvo Splitsko- Dalmatinske Županije | Hrvatski zavod za javno zdravstvo | Irena Tabain, Ivana Ferenčak |
| EPI_ISL_3058202 | Hospital | National Reference Center for Viruses of Respiratory Infections, Institut Pasteur, Paris | Marion Barbet, Sylvie Behillil, Méline Bizard, Angela Brisebarre, Camille Capel, Vincent Enouf, Louise Lefrançois, Frédéric Lemoine, Christophe Malabat, Corinne Maufrais, Victoire Baillet, Etienne Simon-Lorière, Maud Vanpeene, Sylvie Van der Werf ,Anne Holstein |
| EPI_ISL_3058203 | Labo Analyses Med | National Reference Center for Viruses of Respiratory Infections, Institut Pasteur, Paris | Marion Barbet, Sylvie Behillil, Méline Bizard, Angela Brisebarre, Camille Capel, Vincent Enouf, Louise Lefrançois, Frédéric Lemoine, Christophe Malabat, Corinne Maufrais, Victoire Baillet, Etienne Simon-Lorière, Maud Vanpeene, Sylvie Van der Werf ,VéRonique Bourgois |
| EPI_ISL_3058204 | Labo Analyses Med | National Reference Center for Viruses of Respiratory Infections, Institut Pasteur, Paris | Marion Barbet, Sylvie Behillil, Méline Bizard, Angela Brisebarre, Camille Capel, Vincent Enouf, Louise Lefrançois, Frédéric Lemoine, Christophe Malabat, Corinne Maufrais, Hub Bioinformatique Biostatistiques, Etienne Simon-Lorière, Maud Vanpeene, Sylvie Van der Werf ,VéRonique Bourgois |
| EPI_ISL_3058206, EPI_ISL_3058207, EPI_ISL_3058208 | Labo Analyses Med | National Reference Center for Viruses of Respiratory Infections, Institut Pasteur, Paris | Marion Barbet, Sylvie Behillil, Méline Bizard, Angela Brisebarre, Camille Capel, Vincent Enouf, Louise Lefrançois, Frédéric Lemoine, Christophe Malabat, Corinne Maufrais, Hub Bioinformatique Biostatistiques, Etienne Simon-Lorière, Maud Vanpeene, Sylvie Van der Werf ,BéNéDicte Roquebert |
| EPI_ISL_3058209 | Labo Analyses Med | National Reference Center for Viruses of Respiratory Infections, Institut Pasteur, Paris | Marion Barbet, Sylvie Behillil, Méline Bizard, Angela Brisebarre, Camille Capel, Vincent Enouf, Louise Lefrançois, Frédéric Lemoine, Christophe Malabat, Corinne Maufrais, Victoire Baillet, Etienne Simon-Lorière, Maud Vanpeene, Sylvie Van der Werf ,BéNéDicte Roquebert |
| EPI_ISL_3058211 | Labo Analyses Med | National Reference Center for Viruses of Respiratory Infections, Institut Pasteur, Paris | Marion Barbet, Sylvie Behillil, Méline Bizard, Angela Brisebarre, Camille Capel, Vincent Enouf, Louise Lefrançois, Frédéric Lemoine, Christophe Malabat, Corinne Maufrais, Hub Bioinformatique Biostatistiques, Etienne Simon-Lorière, Maud Vanpeene, Sylvie Van der Werf ,BéNéDicte Roquebert |
| EPI_ISL_3058212 | Labo Analyses Med | National Reference Center for Viruses of Respiratory Infections, Institut Pasteur, Paris | Marion Barbet, Sylvie Behillil, Méline Bizard, Angela Brisebarre, Camille Capel, Vincent Enouf, Louise Lefrançois, Frédéric Lemoine, Christophe Malabat, Corinne Maufrais, Hub Bioinformatique Biostatistiques, Etienne Simon-Lorière, Maud Vanpeene, Sylvie Van der Werf ,Jean-Baptiste Foulquier |
| EPI_ISL_3058213, EPI_ISL_3058214, EPI_ISL_3058215 | Labo Analyses Med | National Reference Center for Viruses of Respiratory Infections, Institut Pasteur, Paris | Marion Barbet, Sylvie Behillil, Méline Bizard, Angela Brisebarre, Camille Capel, Vincent Enouf, Louise Lefrançois, Frédéric Lemoine, Christophe Malabat, Corinne Maufrais, Victoire Baillet, Etienne Simon-Lorière, Maud Vanpeene, Sylvie Van der Werf ,Jean-Baptiste Foulquier |
| EPI_ISL_3058218 | Hospital | National Reference Center for Viruses of Respiratory Infections, Institut Pasteur, Paris | Marion Barbet, Sylvie Behillil, Méline Bizard, Angela Brisebarre, Camille Capel, Vincent Enouf, Louise Lefrançois, Frédéric Lemoine, Christophe Malabat, Corinne Maufrais, Hub Bioinformatique Biostatistiques, Etienne Simon-Lorière, Maud Vanpeene, Sylvie Van der Werf ,Eric Farfour |
| EPI_ISL_3058221, EPI_ISL_3058223, EPI_ISL_3058224, EPI_ISL_3058225, EPI_ISL_3058226, EPI_ISL_3058227, EPI_ISL_3058228, EPI_ISL_3058229, EPI_ISL_3058230, EPI_ISL_3058232, EPI_ISL_3058233, EPI_ISL_3058234, EPI_ISL_3058235 | | | |
| see above | Hospital | National Reference Center for Viruses of Respiratory Infections, Institut Pasteur, Paris | Marion Barbet, Sylvie Behillil, Méline Bizard, Angela Brisebarre, Camille Capel, Vincent Enouf, Louise Lefrançois, Frédéric Lemoine, Christophe Malabat, Corinne Maufrais, Hub Bioinformatique Biostatistiques, Etienne Simon-Lorière, Maud Vanpeene, Sylvie Van der Werf ,Nabil Gastli |
| EPI_ISL_3058240 | Hospital | National Reference Center for Viruses of Respiratory Infections, Institut Pasteur, Paris | Marion Barbet, Sylvie Behillil, Méline Bizard, Angela Brisebarre, Camille Capel, Vincent Enouf, Louise Lefrançois, Frédéric Lemoine, Christophe Malabat, Corinne Maufrais, Hub Bioinformatique Biostatistiques, Etienne Simon-Lorière, Maud Vanpeene, Sylvie Van der Werf ,Marine Desroches |
| EPI_ISL_3058241, EPI_ISL_3058243, EPI_ISL_3058244 | Labo Analyses Med | National Reference Center for Viruses of Respiratory Infections, Institut Pasteur, Paris | Marion Barbet, Sylvie Behillil, Méline Bizard, Angela Brisebarre, Camille Capel, Vincent Enouf, Louise Lefrançois, Frédéric Lemoine, Christophe Malabat, Corinne Maufrais, Victoire Baillet, Etienne Simon-Lorière, Maud Vanpeene, Sylvie Van der Werf ,Arnaud Serpentini |
| EPI_ISL_3058245 | Hospital | National Reference Center for Viruses of Respiratory Infections, Institut Pasteur, Paris | Marion Barbet, Sylvie Behillil, Méline Bizard, Angela Brisebarre, Camille Capel, Vincent Enouf, Louise Lefrançois, Frédéric Lemoine, Christophe Malabat, Corinne Maufrais, Hub Bioinformatique Biostatistiques, Etienne Simon-Lorière, Maud Vanpeene, Sylvie Van der Werf ,Mj Dufour |
| EPI_ISL_3058246 | Hospital | National Reference Center for Viruses of Respiratory Infections, Institut Pasteur, Paris | Marion Barbet, Sylvie Behillil, Méline Bizard, Angela Brisebarre, Camille Capel, Vincent Enouf, Louise Lefrançois, Frédéric Lemoine, Christophe Malabat, Corinne Maufrais, Victoire Baillet, Etienne Simon-Lorière, Maud Vanpeene, Sylvie Van der Werf ,Mj Dufour |
| EPI_ISL_3058247 | Hospital | National Reference Center for Viruses of Respiratory Infections, Institut Pasteur, Paris | Marion Barbet, Sylvie Behillil, Méline Bizard, Angela Brisebarre, Camille Capel, Vincent Enouf, Louise Lefrançois, Frédéric Lemoine, Christophe Malabat, Corinne Maufrais, Hub Bioinformatique Biostatistiques, Etienne Simon-Lorière, Maud Vanpeene, Sylvie Van der Werf ,CéCile Hermann |
| EPI_ISL_3058267, EPI_ISL_3058268, EPI_ISL_3058269, EPI_ISL_3058270, EPI_ISL_3058271, EPI_ISL_3058272, EPI_ISL_3058273, EPI_ISL_3058274, EPI_ISL_3058275, EPI_ISL_3058276, EPI_ISL_3058277, EPI_ISL_3058278, EPI_ISL_3058279, EPI_ISL_3058280, EPI_ISL_3058281, EPI_ISL_3058282, EPI_ISL_3058283, EPI_ISL_3058284, EPI_ISL_3058285, EPI_ISL_3058286, EPI_ISL_3058287, EPI_ISL_3058288, EPI_ISL_3058289, EPI_ISL_3058290, EPI_ISL_3058291, EPI_ISL_3058292, EPI_ISL_3058293, EPI_ISL_3058294, EPI_ISL_3058295, EPI_ISL_3058296, EPI_ISL_3058297, EPI_ISL_3058298, EPI_ISL_3058299, EPI_ISL_3058301, EPI_ISL_3058302, EPI_ISL_3058303, EPI_ISL_3058304, EPI_ISL_3058305, EPI_ISL_3058306, EPI_ISL_3058307, EPI_ISL_3058308, EPI_ISL_3058309, EPI_ISL_3058310, EPI_ISL_3058311, EPI_ISL_3058312, EPI_ISL_3058313, EPI_ISL_3058314, EPI_ISL_3058315, EPI_ISL_3058316, EPI_ISL_3058317, EPI_ISL_3058318, EPI_ISL_3058319, EPI_ISL_3058320, EPI_ISL_3058321, EPI_ISL_3058322, EPI_ISL_3058323, EPI_ISL_3058324, EPI_ISL_3058325, EPI_ISL_3058326, EPI_ISL_3058327, EPI_ISL_3058328, EPI_ISL_3058329, EPI_ISL_3058330, EPI_ISL_3058331, EPI_ISL_3058332, EPI_ISL_3058333, EPI_ISL_3058334, EPI_ISL_3058335, EPI_ISL_3058336, EPI_ISL_3058337, EPI_ISL_3058338, EPI_ISL_3058339, EPI_ISL_3058340, EPI_ISL_3058341, EPI_ISL_3058342, EPI_ISL_3058343, EPI_ISL_3058344, EPI_ISL_3058345, EPI_ISL_3058346, EPI_ISL_3058347, EPI_ISL_3058348, EPI_ISL_3058349, EPI_ISL_3058350, EPI_ISL_3058351, EPI_ISL_3058352, EPI_ISL_3058353, EPI_ISL_3058354, EPI_ISL_3058355, EPI_ISL_3058356, EPI_ISL_3058357, EPI_ISL_3058358, EPI_ISL_3058359, EPI_ISL_3058360, EPI_ISL_3058361, EPI_ISL_3058362, EPI_ISL_3058363, EPI_ISL_3058364, EPI_ISL_3058365, EPI_ISL_3058366, EPI_ISL_3058367, EPI_ISL_3058368, EPI_ISL_3058369, EPI_ISL_3058371, EPI_ISL_3058372, EPI_ISL_3058373, EPI_ISL_3058374, EPI_ISL_3058375, EPI_ISL_3058376, EPI_ISL_3058377, EPI_ISL_3058378, EPI_ISL_3058379, EPI_ISL_3058380, EPI_ISL_3058381, EPI_ISL_3058382, EPI_ISL_3058383, EPI_ISL_3058384, EPI_ISL_3058385, EPI_ISL_3058386, EPI_ISL_3058387, EPI_ISL_3058388, EPI_ISL_3058389, EPI_ISL_3058390, EPI_ISL_3058391, EPI_ISL_3058392, EPI_ISL_3058393, EPI_ISL_3058394, EPI_ISL_3058395, EPI_ISL_3058396, EPI_ISL_3058397, EPI_ISL_3058398, EPI_ISL_3058399, EPI_ISL_3058400, EPI_ISL_3058401, EPI_ISL_3058402, EPI_ISL_3058403, EPI_ISL_3058404, EPI_ISL_3058405, EPI_ISL_3058406, EPI_ISL_3058407, EPI_ISL_3058408, EPI_ISL_3058409, EPI_ISL_3058410, EPI_ISL_3058411, EPI_ISL_3058412, EPI_ISL_3058413, EPI_ISL_3058414, EPI_ISL_3058415, EPI_ISL_3058416, EPI_ISL_3058417, EPI_ISL_3058418, EPI_ISL_3058419, EPI_ISL_3058420, EPI_ISL_3058421, EPI_ISL_3058422, EPI_ISL_3058423, EPI_ISL_3058424, EPI_ISL_3058425, EPI_ISL_3058426, EPI_ISL_3058427, EPI_ISL_3058428, EPI_ISL_3058429, EPI_ISL_3058430, EPI_ISL_3058431, EPI_ISL_3058432, EPI_ISL_3058433, EPI_ISL_3058434, EPI_ISL_3058435, EPI_ISL_3058436, EPI_ISL_3058437, EPI_ISL_3058438, EPI_ISL_3058439, EPI_ISL_3058440, EPI_ISL_3058441, EPI_ISL_3058442, EPI_ISL_3058443, EPI_ISL_3058444, EPI_ISL_3058445, EPI_ISL_3058446, EPI_ISL_3058447, EPI_ISL_3058448, EPI_ISL_3058449, EPI_ISL_3058450, EPI_ISL_3058451, EPI_ISL_3058452, EPI_ISL_3058453, EPI_ISL_3058454, EPI_ISL_3058455, EPI_ISL_3058456, EPI_ISL_3058457, EPI_ISL_3058458, EPI_ISL_3058459, EPI_ISL_3058460, EPI_ISL_3058461, EPI_ISL_3058462, EPI_ISL_3058463, EPI_ISL_3058464, EPI_ISL_3058465, EPI_ISL_3058466, EPI_ISL_3058467, EPI_ISL_3058468, EPI_ISL_3058469, EPI_ISL_3058470, EPI_ISL_3058471, EPI_ISL_3058472, EPI_ISL_3058473, EPI_ISL_3058474, EPI_ISL_3058475, EPI_ISL_3058476, EPI_ISL_3058477, EPI_ISL_3058478, EPI_ISL_3058479, EPI_ISL_3058480, EPI_ISL_3058481, EPI_ISL_3058482, EPI_ISL_3058483, EPI_ISL_3058484, EPI_ISL_3058485, EPI_ISL_3058486, EPI_ISL_3058487, EPI_ISL_3058488, EPI_ISL_3058489, EPI_ISL_3058490, EPI_ISL_3058491, EPI_ISL_3058492, EPI_ISL_3058493, EPI_ISL_3058494, EPI_ISL_3058495 | | | |
| see above | National Virus Reference Laboratory | National Virus Reference Laboratory | Zoe Yandle, Charlene Bennett, Gabriel Gonzalez, Michael Carr, Jonathan Dean, Cillian F De Gascun |
| EPI_ISL_3060002, EPI_ISL_3060003, | UAB "Rezus.lt" | Institute of Biotechnology, Life Sciences Center, Vilnius University | Emilija Vasiliunaite; Milda Norkiene; Danguole Ziogiene; Albertas Timinskas; Alma Gedvilaite |

| | | | |
|---|---|---|---|
| EPI_ISL_3060004 | | | |
| EPI_ISL_3060005, EPI_ISL_3060006, EPI_ISL_3060007 | Viešoji įstaiga Klaipėdos universitetinė ligoninė | Institute of Biotechnology, Life Sciences Center, Vilnius University | Emilija Vasiliunaite; Milda Norkiene; Danguole Ziogiene; Albertas Timinskas; Alma Gedvilaite |
| EPI_ISL_3060008, EPI_ISL_3060009 | UAB "Rezus.lt" | Institute of Biotechnology, Life Sciences Center, Vilnius University | Emilija Vasiliunaite; Milda Norkiene; Danguole Ziogiene; Albertas Timinskas; Alma Gedvilaite |
| EPI_ISL_3060010, EPI_ISL_3060011 | Viešoji įstaiga Klaipėdos universitetinė ligoninė | Institute of Biotechnology, Life Sciences Center, Vilnius University | Emilija Vasiliunaite; Milda Norkiene; Danguole Ziogiene; Albertas Timinskas; Alma Gedvilaite |
| EPI_ISL_3060012 | Viešoji įstaiga Vilniaus universiteto ligoninė Santaros klinikos | Institute of Biotechnology, Life Sciences Center, Vilnius University | Emilija Vasiliunaite; Milda Norkiene; Danguole Ziogiene; Albertas Timinskas; Alma Gedvilaite |
| EPI_ISL_3060015, EPI_ISL_3060016, EPI_ISL_3060018, EPI_ISL_3060019 | UAB "Rezus.lt" | Institute of Biotechnology, Life Sciences Center, Vilnius University | Emilija Vasiliunaite; Milda Norkiene; Danguole Ziogiene; Albertas Timinskas; Alma Gedvilaite |
| EPI_ISL_3060020 | UAB Diagnostikos laboratorija | Institute of Biotechnology, Life Sciences Center, Vilnius University | Emilija Vasiliunaite; Milda Norkiene; Danguole Ziogiene; Albertas Timinskas; Alma Gedvilaite |
| EPI_ISL_3060021, EPI_ISL_3060022 | Viešoji įstaiga Klaipėdos universitetinė ligoninė | Institute of Biotechnology, Life Sciences Center, Vilnius University | Emilija Vasiliunaite; Milda Norkiene; Danguole Ziogiene; Albertas Timinskas; Alma Gedvilaite |
| EPI_ISL_3060023 | Viešoji įstaiga Vilniaus universiteto ligoninė Santaros klinikos | Institute of Biotechnology, Life Sciences Center, Vilnius University | Emilija Vasiliunaite; Milda Norkiene; Danguole Ziogiene; Albertas Timinskas; Alma Gedvilaite |
| EPI_ISL_3060024, EPI_ISL_3060028, EPI_ISL_3060029 | UAB "Rezus.lt" | Institute of Biotechnology, Life Sciences Center, Vilnius University | Emilija Vasiliunaite; Milda Norkiene; Danguole Ziogiene; Albertas Timinskas; Alma Gedvilaite |
| EPI_ISL_3060030, EPI_ISL_3060031 | Viešoji įstaiga Klaipėdos universitetinė ligoninė | Institute of Biotechnology, Life Sciences Center, Vilnius University | Emilija Vasiliunaite; Milda Norkiene; Danguole Ziogiene; Albertas Timinskas; Alma Gedvilaite |
| EPI_ISL_3060032 | Viešoji įstaiga Vilniaus universiteto ligoninė Santaros klinikos | Institute of Biotechnology, Life Sciences Center, Vilnius University | Emilija Vasiliunaite; Milda Norkiene; Danguole Ziogiene; Albertas Timinskas; Alma Gedvilaite |
| EPI_ISL_3060033 | Viešoji įstaiga Klaipėdos universitetinė ligoninė | Institute of Biotechnology, Life Sciences Center, Vilnius University | Emilija Vasiliunaite; Milda Norkiene; Danguole Ziogiene; Albertas Timinskas; Alma Gedvilaite |
| EPI_ISL_3060034 | UAB "Rezus.lt" | Institute of Biotechnology, Life Sciences Center, Vilnius University | Emilija Vasiliunaite; Milda Norkiene; Danguole Ziogiene; Albertas Timinskas; Alma Gedvilaite |
| EPI_ISL_3060035, EPI_ISL_3060037, EPI_ISL_3060038, EPI_ISL_3060039, EPI_ISL_3060040 | Viešoji įstaiga Klaipėdos universitetinė ligoninė | Institute of Biotechnology, Life Sciences Center, Vilnius University | Emilija Vasiliunaite; Milda Norkiene; Danguole Ziogiene; Albertas Timinskas; Alma Gedvilaite |
| EPI_ISL_3060041 | UAB "Rezus.lt" | Institute of Biotechnology, Life Sciences Center, Vilnius University | Emilija Vasiliunaite; Milda Norkiene; Danguole Ziogiene; Albertas Timinskas; Alma Gedvilaite |
| EPI_ISL_3060042 | Viešoji įstaiga Klaipėdos universitetinė ligoninė | Institute of Biotechnology, Life Sciences Center, Vilnius University | Emilija Vasiliunaite; Milda Norkiene; Danguole Ziogiene; Albertas Timinskas; Alma Gedvilaite |
| EPI_ISL_3060043 | UAB Medicina practica laboratorija | unknown | Rasa Ugenskiene, Darius Cereskevicius, Inga Nasvytiene, Zivile Zemeckiene, Mantas Sarauskas, Marius Sukys, Rima Vainoriene, Astra Vitkauskiene, Renaldas Jurkevicius |
| EPI_ISL_3060044 | Viešoji įstaiga Vilniaus universiteto ligoninė Santaros klinikos | unknown | Rasa Ugenskiene, Darius Cereskevicius, Inga Nasvytiene, Zivile Zemeckiene, Mantas Sarauskas, Marius Sukys, Rima Vainoriene, Astra Vitkauskiene, Renaldas Jurkevicius |
| EPI_ISL_3060045 | UAB Diagnostikos laboratorija | unknown | Rasa Ugenskiene, Darius Cereskevicius, Inga Nasvytiene, Zivile Zemeckiene, Mantas Sarauskas, Marius Sukys, Rima Vainoriene, Astra Vitkauskiene, Renaldas Jurkevicius |
| EPI_ISL_3060046, EPI_ISL_3060047 | UAB Medicina practica laboratorija | unknown | Rasa Ugenskiene, Darius Cereskevicius, Inga Nasvytiene, Zivile Zemeckiene, Mantas Sarauskas, Marius Sukys, Rima Vainoriene, Astra Vitkauskiene, Renaldas Jurkevicius |
| EPI_ISL_3060048, EPI_ISL_3060049, EPI_ISL_3060050, EPI_ISL_3060051 | UAB InMedica | unknown | Rasa Ugenskiene, Darius Cereskevicius, Inga Nasvytiene, Zivile Zemeckiene, Mantas Sarauskas, Marius Sukys, Rima Vainoriene, Astra Vitkauskiene, Renaldas Jurkevicius |
| EPI_ISL_3060052 | UAB "Baltic Medics" | unknown | Rasa Ugenskiene, Darius Cereskevicius, Inga Nasvytiene, Zivile Zemeckiene, Mantas Sarauskas, Marius Sukys, Rima Vainoriene, Astra Vitkauskiene, Renaldas Jurkevicius |
| EPI_ISL_3060053, EPI_ISL_3060054, EPI_ISL_3060055, EPI_ISL_3060056 | UAB Medicina practica laboratorija | unknown | Rasa Ugenskiene, Darius Cereskevicius, Inga Nasvytiene, Zivile Zemeckiene, Mantas Sarauskas, Marius Sukys, Rima Vainoriene, Astra Vitkauskiene, Renaldas Jurkevicius |
| EPI_ISL_3060057, EPI_ISL_3060058, EPI_ISL_3060059 | UAB Diagnostikos laboratorija | unknown | Rasa Ugenskiene, Darius Cereskevicius, Inga Nasvytiene, Zivile Zemeckiene, Mantas Sarauskas, Marius Sukys, Rima Vainoriene, Astra Vitkauskiene, Renaldas Jurkevicius |
| EPI_ISL_3060060, EPI_ISL_3060061 | UAB InMedica | unknown | Rasa Ugenskiene, Darius Cereskevicius, Inga Nasvytiene, Zivile Zemeckiene, Mantas Sarauskas, Marius Sukys, Rima Vainoriene, Astra Vitkauskiene, Renaldas Jurkevicius |
| EPI_ISL_3060062 | UAB "Rezus.lt" | unknown | Rasa Ugenskiene, Darius Cereskevicius, Inga Nasvytiene, Zivile Zemeckiene, Mantas Sarauskas, Marius Sukys, Rima Vainoriene, Astra Vitkauskiene, Renaldas Jurkevicius |
| EPI_ISL_3060063, EPI_ISL_3060064, EPI_ISL_3060065 | UAB Medicina practica laboratorija | unknown | Rasa Ugenskiene, Darius Cereskevicius, Inga Nasvytiene, Zivile Zemeckiene, Mantas Sarauskas, Marius Sukys, Rima Vainoriene, Astra Vitkauskiene, Renaldas Jurkevicius |
| EPI_ISL_3060069 | UAB "Rezus.lt" | Vilnius University Hospital Santaros Klinikos, Center of Laboratory Medicine | Dovile Juozapaite, Ingrida Olendraite, Gytis Dudas, Daniel Naumovas, Ligita Raugaite, Monika Katenaite, Rimvydas Norvilas |
| EPI_ISL_3060071 | Viešoji įstaiga Respublikinė Šiaulių ligoninė | Vilnius University Hospital Santaros Klinikos, Center of Laboratory Medicine | Dovile Juozapaite, Ingrida Olendraite, Gytis Dudas, Daniel Naumovas, Ligita Raugaite, Monika Katenaite, Rimvydas Norvilas |
| EPI_ISL_3060073 | Viešoji įstaiga Klaipėdos universitetinė ligoninė | Vilnius University Hospital Santaros Klinikos, Center of Laboratory Medicine | Dovile Juozapaite, Ingrida Olendraite, Gytis Dudas, Daniel Naumovas, Ligita Raugaite, Monika Katenaite, Rimvydas Norvilas |
| EPI_ISL_3060079 | UAB "Baltic Medics" | Vilnius University Hospital Santaros Klinikos, Center of Laboratory Medicine | Dovile Juozapaite, Ingrida Olendraite, Gytis Dudas, Daniel Naumovas, Ligita Raugaite, Monika Katenaite, Rimvydas Norvilas |
| EPI_ISL_3060081 | UAB "Rezus.lt" | Vilnius University Hospital Santaros Klinikos, Center of Laboratory Medicine | Dovile Juozapaite, Ingrida Olendraite, Gytis Dudas, Daniel Naumovas, Ligita Raugaite, Monika Katenaite, Rimvydas Norvilas |
| EPI_ISL_3060082 | Nacionalinis maisto ir veterinarijos rizikos vertinimo institutas | Vilnius University Hospital Santaros Klinikos, Center of Laboratory Medicine | Dovile Juozapaite, Ingrida Olendraite, Gytis Dudas, Daniel Naumovas, Ligita Raugaite, Monika Katenaite, Rimvydas Norvilas |
| EPI_ISL_3060086 | Viešoji įstaiga Klaipėdos universitetinė ligoninė | Vilnius University Hospital Santaros Klinikos, Center of Laboratory Medicine | Dovile Juozapaite, Ingrida Olendraite, Gytis Dudas, Daniel Naumovas, Ligita Raugaite, Monika Katenaite, Rimvydas Norvilas |
| EPI_ISL_3060089 | UAB "Rezus.lt" | Vilnius University Hospital Santaros Klinikos, Center of Laboratory Medicine | Dovile Juozapaite, Ingrida Olendraite, Gytis Dudas, Daniel Naumovas, Ligita Raugaite, Monika Katenaite, Rimvydas Norvilas |
| EPI_ISL_3060091, EPI_ISL_3060092 | Nacionalinis maisto ir veterinarijos rizikos vertinimo institutas | Vilnius University Hospital Santaros Klinikos, Center of Laboratory Medicine | Dovile Juozapaite, Ingrida Olendraite, Gytis Dudas, Daniel Naumovas, Ligita Raugaite, Monika Katenaite, Rimvydas Norvilas |
| EPI_ISL_3060093 | SYNLAB Lietuva UAB | Vilnius University Hospital Santaros Klinikos, Center of Laboratory Medicine | Dovile Juozapaite, Ingrida Olendraite, Gytis Dudas, Daniel Naumovas, Ligita Raugaite, Monika Katenaite, Rimvydas Norvilas |
| EPI_ISL_3060094 | UAB "Rezus.lt" | Vilnius University Hospital Santaros Klinikos, Center of Laboratory Medicine | Dovile Juozapaite, Ingrida Olendraite, Gytis Dudas, Daniel Naumovas, Ligita Raugaite, Monika Katenaite, Rimvydas Norvilas |
| EPI_ISL_3060096 | Nacionalinis maisto ir veterinarijos rizikos vertinimo institutas | Vilnius University Hospital Santaros Klinikos, Center of Laboratory Medicine | Dovile Juozapaite, Ingrida Olendraite, Gytis Dudas, Daniel Naumovas, Ligita Raugaite, Monika Katenaite, Rimvydas Norvilas |
| EPI_ISL_3060100 | UAB "Baltic Medics" | Vilnius University Hospital Santaros Klinikos, Center of Laboratory Medicine | Dovile Juozapaite, Ingrida Olendraite, Gytis Dudas, Daniel Naumovas, Ligita Raugaite, Monika Katenaite, Rimvydas Norvilas |
| EPI_ISL_3060102, EPI_ISL_3060107, EPI_ISL_3060108, EPI_ISL_3060109 | Nacionalinis maisto ir veterinarijos rizikos vertinimo institutas | Vilnius University Hospital Santaros Klinikos, Center of Laboratory Medicine | Dovile Juozapaite, Ingrida Olendraite, Gytis Dudas, Daniel Naumovas, Ligita Raugaite, Monika Katenaite, Rimvydas Norvilas |
| EPI_ISL_3060112 | Viešoji įstaiga Klaipėdos universitetinė ligoninė | Vilnius University Hospital Santaros Klinikos, Center of Laboratory Medicine | Dovile Juozapaite, Ingrida Olendraite, Gytis Dudas, Daniel Naumovas, Ligita Raugaite, Monika Katenaite, Rimvydas Norvilas |
| EPI_ISL_3060114 | Nacionalinis maisto ir veterinarijos rizikos vertinimo institutas | Vilnius University Hospital Santaros Klinikos, Center of Laboratory Medicine | Dovile Juozapaite, Ingrida Olendraite, Gytis Dudas, Daniel Naumovas, Ligita Raugaite, Monika Katenaite, Rimvydas Norvilas |
| EPI_ISL_3060118 | UAB "Rezus.lt" | Vilnius University Hospital Santaros Klinikos, Center of Laboratory | Dovile Juozapaite, Ingrida Olendraite, Gytis Dudas, Daniel Naumovas, Ligita Raugaite, Monika Katenaite, Rimvydas Norvilas |

| | | | |
|---|---|---|---|
| | | Medicine | |
| EPI_ISL_3060119 | Viešoji įstaiga Vilniaus universiteto ligoninė Santaros klinikos | Vilnius University Hospital Santaros Klinikos, Center of Laboratory Medicine | Dovile Juozapaite, Ingrida Olendraite, Gytis Dudas, Daniel Naumovas, Ligita Raugaite, Monika Katenaite, Rimvydas Norvilas |
| EPI_ISL_3060121 | Nacionalinis maisto ir veterinarijos rizikos vertinimo institutas | Vilnius University Hospital Santaros Klinikos, Center of Laboratory Medicine | Dovile Juozapaite, Ingrida Olendraite, Gytis Dudas, Daniel Naumovas, Ligita Raugaite, Monika Katenaite, Rimvydas Norvilas |
| EPI_ISL_3060123 | UAB "Rezus.lt" | Vilnius University Hospital Santaros Klinikos, Center of Laboratory Medicine | Dovile Juozapaite, Ingrida Olendraite, Gytis Dudas, Daniel Naumovas, Ligita Raugaite, Monika Katenaite, Rimvydas Norvilas |
| EPI_ISL_3060130 | Nacionalinis maisto ir veterinarijos rizikos vertinimo institutas | Vilnius University Hospital Santaros Klinikos, Center of Laboratory Medicine | Rimvydas Norvilas__ Dovile Juozapaite__ Ingrida Olendraite__ Gytis Dudas__ Daniel Naumovas__ Ligita Raugaite__ Monika Katenaite |
| EPI_ISL_3060131 | Viešoji įstaiga Vilniaus universiteto ligoninė Santaros klinikos | Vilnius University Hospital Santaros Klinikos, Center of Laboratory Medicine | Rimvydas Norvilas__ Dovile Juozapaite__ Ingrida Olendraite__ Gytis Dudas__ Daniel Naumovas__ Ligita Raugaite__ Monika Katenaite |
| EPI_ISL_3060138 | Nacionalinis maisto ir veterinarijos rizikos vertinimo institutas | Vilnius University Hospital Santaros Klinikos, Center of Laboratory Medicine | Rimvydas Norvilas__ Dovile Juozapaite__ Ingrida Olendraite__ Gytis Dudas__ Daniel Naumovas__ Ligita Raugaite__ Monika Katenaite |
| EPI_ISL_3060143 | Viešoji įstaiga Vilniaus universiteto ligoninė Santaros klinikos | Vilnius University Hospital Santaros Klinikos, Center of Laboratory Medicine | Rimvydas Norvilas__ Dovile Juozapaite__ Ingrida Olendraite__ Gytis Dudas__ Daniel Naumovas__ Ligita Raugaite__ Monika Katenaite |
| EPI_ISL_3060148, EPI_ISL_3060162 | Nacionalinis maisto ir veterinarijos rizikos vertinimo institutas | Vilnius University Hospital Santaros Klinikos, Center of Laboratory Medicine | Rimvydas Norvilas__ Dovile Juozapaite__ Ingrida Olendraite__ Gytis Dudas__ Daniel Naumovas__ Ligita Raugaite__ Monika Katenaite |
| EPI_ISL_3060174 | UAB Diagnostikos laboratorija | Vilnius University Hospital Santaros Klinikos, Center of Laboratory Medicine | Rimvydas Norvilas__ Dovile Juozapaite__ Ingrida Olendraite__ Gytis Dudas__ Daniel Naumovas__ Ligita Raugaite__ Monika Katenaite |
| EPI_ISL_3060179 | Viešoji įstaiga Vilniaus universiteto ligoninė Santaros klinikos | Vilnius University Hospital Santaros Klinikos, Center of Laboratory Medicine | Rimvydas Norvilas__ Dovile Juozapaite__ Ingrida Olendraite__ Gytis Dudas__ Daniel Naumovas__ Ligita Raugaite__ Monika Katenaite |
| EPI_ISL_3060180 | Nacionalinis maisto ir veterinarijos rizikos vertinimo institutas | Vilnius University Hospital Santaros Klinikos, Center of Laboratory Medicine | Rimvydas Norvilas__ Dovile Juozapaite__ Ingrida Olendraite__ Gytis Dudas__ Daniel Naumovas__ Ligita Raugaite__ Monika Katenaite |
| EPI_ISL_3060183 | UAB Diagnostikos laboratorija | Vilnius University Hospital Santaros Klinikos, Center of Laboratory Medicine | Rimvydas Norvilas__ Dovile Juozapaite__ Ingrida Olendraite__ Gytis Dudas__ Daniel Naumovas__ Ligita Raugaite__ Monika Katenaite |
| EPI_ISL_3060201 | Nacionalinis maisto ir veterinarijos rizikos vertinimo institutas | Vilnius University Hospital Santaros Klinikos, Center of Laboratory Medicine | Rimvydas Norvilas__ Dovile Juozapaite__ Ingrida Olendraite__ Gytis Dudas__ Daniel Naumovas__ Ligita Raugaite__ Monika Katenaite |
| EPI_ISL_3060202, EPI_ISL_3060203, EPI_ISL_3060209, EPI_ISL_3060212, EPI_ISL_3060215, EPI_ISL_3060216, EPI_ISL_3060221, EPI_ISL_3060222 | UAB Diagnostikos laboratorija | unknown | Rasa Ugenskiene, Darius Cereskevicius, Inga Nasvytiene, Zivile Zemeckiene, Mantas Sarauskas, Marius Sukys, Rima Vainoriene, Astra Vitkauskiene, Renaldas Jurkevicius |
| EPI_ISL_3060275 | MKK Mühlenkreiskliniken, Johannes Wesling Klinikum Minden | Hannover Medical School, Institute of Virology | Lars Steinbrück |
| EPI_ISL_3060291 | Molecular Biology Laboratory, Westchester Medical Center | Genomics Core Laboratory, New York Medical College | Weihua Huang, Utsav Pandey, Jian Zhuge, Changhong Yin |
| EPI_ISL_3060306, EPI_ISL_3060307, EPI_ISL_3060308, EPI_ISL_3060309, EPI_ISL_3060310, EPI_ISL_3060311, EPI_ISL_3060312, EPI_ISL_3060313, EPI_ISL_3060314, EPI_ISL_3060315, EPI_ISL_3060316, EPI_ISL_3060317, EPI_ISL_3060318, EPI_ISL_3060319, EPI_ISL_3060320, EPI_ISL_3060321, EPI_ISL_3060322, EPI_ISL_3060324, EPI_ISL_3060325, EPI_ISL_3060326, EPI_ISL_3060327, EPI_ISL_3060328, EPI_ISL_3060329, EPI_ISL_3060330, EPI_ISL_3060331, EPI_ISL_3060332, EPI_ISL_3060333, EPI_ISL_3060334, EPI_ISL_3060335, EPI_ISL_3060336, EPI_ISL_3060337, EPI_ISL_3060339, EPI_ISL_3060340, EPI_ISL_3060341, EPI_ISL_3060342, EPI_ISL_3060343, EPI_ISL_3060344, EPI_ISL_3060345, EPI_ISL_3060346, EPI_ISL_3060347, EPI_ISL_3060348, EPI_ISL_3060349, EPI_ISL_3060350, EPI_ISL_3060351, EPI_ISL_3060352, EPI_ISL_3060353, EPI_ISL_3060354, EPI_ISL_3060355, EPI_ISL_3060356, EPI_ISL_3060357, EPI_ISL_3060358, EPI_ISL_3060359, EPI_ISL_3060360, EPI_ISL_3060361, EPI_ISL_3060362, EPI_ISL_3060363, EPI_ISL_3060364, EPI_ISL_3060365, EPI_ISL_3060367, EPI_ISL_3060368, EPI_ISL_3060369, EPI_ISL_3060370, EPI_ISL_3060371, EPI_ISL_3060372, EPI_ISL_3060373, EPI_ISL_3060374, EPI_ISL_3060375, EPI_ISL_3060376 | | | |
| see above | Hospital Universitari Vall d'Hebron - Vall d'Hebron Institut de Recerca | Hospital Universitari Vall d'Hebron - Vall d'Hebron Institut de Recerca | Cristina Andrés, Maria Piñana, Alejandra González-Sánchez, Damir Garcia-Cehic, Ariadna Rando, Juliana Esperalba, Maria Gema Codina, Carla Castillo, Karen García, Rodrigo Vásquez, Maria Carmen Martin, Tomàs Pumarola, Josep Quer, Andrés Antón |
| EPI_ISL_3060377, EPI_ISL_3060378, EPI_ISL_3060379, EPI_ISL_3060380, EPI_ISL_3060381, EPI_ISL_3060382, EPI_ISL_3060383, EPI_ISL_3060384, EPI_ISL_3060386, EPI_ISL_3060387, EPI_ISL_3060388, EPI_ISL_3060389, EPI_ISL_3060390, EPI_ISL_3060391, EPI_ISL_3060392, EPI_ISL_3060393, EPI_ISL_3060394 | | | |
| see above | Hospital Universitari Arnau de Vilanova | Hospital Universitari Vall d'Hebron - Vall d'Hebron Institut de Recerca | Cristina Andrés, Maria Piñana, Alejandra González-Sánchez, Damir Garcia-Cehic, Ariadna Rando, Juliana Esperalba, Maria Gema Codina, Carla Castillo, Karen García, Rodrigo Vásquez, Maria Carmen Martin, Tomàs Pumarola, Josep Quer, Andrés Antón |
| EPI_ISL_3060395, EPI_ISL_3060396, EPI_ISL_3060397, EPI_ISL_3060398, EPI_ISL_3060399, EPI_ISL_3060400, EPI_ISL_3060401, EPI_ISL_3060402 | AP SSO | CSIR-Centre for Cellular and Molecular Biology-INSACOG | Shreekant Verma, Amareshwar Vodapalli ,Ara Sreenivas,B Himasri,Valli Nagalakshmi Undamatla,Payel Mukherjee, Lamuk Zaveri, Onkar Kulkarni,Tulasi Nagabandi, Sofia Banu, Jandhyala Sai Krishna,Sumedha Avadhanula, Priya Nurkurthy, Vidhyadhari Methuku, Archana Bharadwaj Siva, Karthik Bharadwaj Tallapaka,Rakesh K Mishra,Divya Tej Sowpati |
| EPI_ISL_3060416, EPI_ISL_3060418, EPI_ISL_3060419, EPI_ISL_3060420, EPI_ISL_3060421, EPI_ISL_3060422, EPI_ISL_3060423, EPI_ISL_3060424, EPI_ISL_3060425, EPI_ISL_3060426, EPI_ISL_3060427, EPI_ISL_3060428, EPI_ISL_3060429, EPI_ISL_3060430, EPI_ISL_3060431, EPI_ISL_3060432, EPI_ISL_3060433 | | | |
| see above | AP SSO | CSIR-Centre for Cellular and Molecular Biology-INSACOG | Amareshwar Vodapalli,Ara Sreenivas,Shreekant Verma, B Himasri,Valli Nagalakshmi Undamatla,Payel Mukherjee, Lamuk Zaveri, Onkar Kulkarni,Tulasi Nagabandi, Sofia Banu, Jandhyala Sai Krishna,Priya Nurkurthy, Vidhyadhari Methuku, Sumedha Avadhanula, Archana Bharadwaj Siva, Karthik Bharadwaj Tallapaka,Rakesh K Mishra,Divya Tej Sowpati |
| EPI_ISL_3060434 | CSIR-Centre for Cellular and Molecular Biology | CSIR-Centre for Cellular and Molecular Biology-INSACOG | Amareshwar Vodapalli,Ara Sreenivas,Shreekant Verma, B Himasri,Valli Nagalakshmi Undamatla,Payel Mukherjee, Lamuk Zaveri, Onkar Kulkarni,Tulasi Nagabandi, Sofia Banu, Jandhyala Sai Krishna,Priya Nurkurthy, Vidhyadhari Methuku, Sumedha Avadhanula, Archana Bharadwaj Siva, Karthik Bharadwaj Tallapaka,Rakesh K Mishra,Divya Tej Sowpati |
| EPI_ISL_3060435 | CSIR-Centre for Cellular and Molecular Biology | CSIR-Centre for Cellular and Molecular Biology-INSACOG | B Himasri,Amareshwar Vodapalli,Ara Sreenivas,Shreekant Verma,Valli Nagalakshmi Undamatla,Payel Mukherjee, Lamuk Zaveri, Onkar Kulkarni,Tulasi Nagabandi, Sofia Banu, Vidhyadhari Methuku,Priya Nurkurthy, Jandhyala Sai Krishna,Sumedha Avadhanula, Archana Bharadwaj Siva, Karthik Bharadwaj Tallapaka,Rakesh K Mishra,Divya Tej Sowpati |
| EPI_ISL_3060444, EPI_ISL_3060445, EPI_ISL_3060446, EPI_ISL_3060447, EPI_ISL_3060448, EPI_ISL_3060449, EPI_ISL_3060450 | AP SSO | CSIR-Centre for Cellular and Molecular Biology-INSACOG | B Himasri,Amareshwar Vodapalli,Ara Sreenivas,Shreekant Verma,Valli Nagalakshmi Undamatla,Payel Mukherjee, Lamuk Zaveri, Onkar Kulkarni,Tulasi Nagabandi, Sofia Banu, Vidhyadhari Methuku,Priya Nurkurthy, Jandhyala Sai Krishna,Sumedha Avadhanula, Archana Bharadwaj Siva, Karthik Bharadwaj Tallapaka,Rakesh K Mishra,Divya Tej Sowpati |
| EPI_ISL_3060452, EPI_ISL_3060453, EPI_ISL_3060454, EPI_ISL_3060456, EPI_ISL_3060457, EPI_ISL_3060458, EPI_ISL_3060459, EPI_ISL_3060460, EPI_ISL_3060461, EPI_ISL_3060462, EPI_ISL_3060463 | | | |
| see above | AP SSO | CSIR-Centre for Cellular and Molecular Biology-INSACOG | Sofia Banu,Payel Mukherjee, Lamuk Zaveri, Onkar Kulkarni,Tulasi Nagabandi, Jandhyala Sai Krishna, Ara Sreenivas,Shreekant Verma, Amareshwar Vodapalli ,B Himasri,Valli Nagalakshmi Undamatla, Sumedha Avadhanula, Vidhyadhari Methuku, Priya Nurkurthy, Archana Bharadwaj Siva, Karthik Bharadwaj Tallapaka,Rakesh K Mishra,Divya Tej Sowpati |
| EPI_ISL_3060464, EPI_ISL_3060465, EPI_ISL_3060466, EPI_ISL_3060467, EPI_ISL_3060468, EPI_ISL_3060469, EPI_ISL_3060471, EPI_ISL_3060472, EPI_ISL_3060473, EPI_ISL_3060474 | AP SSO | CSIR-Centre for Cellular and Molecular Biology-INSACOG | Sumedha Avadhanula, Lamuk Zaveri, Tulasi Nagabandi, Ara Sreenivas,Shreekant Verma, Amareshwar Vodapalli ,B Himasri,Valli Nagalakshmi Undamatla, Vidhyadhari Methuku, Priya Nurkurthy, Payel Mukherjee,Jandhyala Sai Krishna,Onkar Kulkarni, Sofia Banu, Archana Bharadwaj Siva, Karthik Bharadwaj Tallapaka,Rakesh K Mishra,Divya Tej Sowpati |
| EPI_ISL_3060475, EPI_ISL_3060476, EPI_ISL_3060477, EPI_ISL_3060478, EPI_ISL_3060479, EPI_ISL_3060480, EPI_ISL_3060481, EPI_ISL_3060482, EPI_ISL_3060483, EPI_ISL_3060484, EPI_ISL_3060485 | | | |
| see above | AP SSO | CSIR-Centre for Cellular and Molecular Biology-INSACOG | Priya Nurkurthy,Ara Sreenivas,Shreekant Verma, Amareshwar Vodapalli ,Lamuk Zaveri,Tulasi Nagabandi,B Himasri,Valli Nagalakshmi Undamatla, Sumedha Avadhanula, Vidhyadhari Methuku,Payel Mukherjee,Onkar Kulkarni, Jandhyala Sai Krishna,Sofia Banu, Archana Bharadwaj Siva, Karthik Bharadwaj Tallapaka,Rakesh K Mishra,Divya Tej Sowpati |
| EPI_ISL_3060486, EPI_ISL_3060487, EPI_ISL_3060488, EPI_ISL_3060489, EPI_ISL_3060490, EPI_ISL_3060491, EPI_ISL_3060492, EPI_ISL_3060493, EPI_ISL_3060494, EPI_ISL_3060495, EPI_ISL_3060496, EPI_ISL_3060497, EPI_ISL_3060498, EPI_ISL_3060499, EPI_ISL_3060500, EPI_ISL_3060501, EPI_ISL_3060502, EPI_ISL_3060503, EPI_ISL_3060504 | | | |
| see above | AP SSO | CSIR-Centre for Cellular and Molecular Biology-INSACOG | Vidhyadhari Methuku,Tulasi Nagabandi,Lamuk Zaveri, Ara Sreenivas,Shreekant Verma, Amareshwar Vodapalli ,B Himasri,Valli Nagalakshmi Undamatla, Priya Nurkurthy,Sumedha Avadhanula, Archana Bharadwaj Siva,Payel Mukherjee,Onkar Kulkarni, Jandhyala Sai Krishna,Sofia Banu, Karthik Bharadwaj Tallapaka,Rakesh K Mishra,Divya Tej Sowpati |
| EPI_ISL_3060505, EPI_ISL_3060506, EPI_ISL_3060507, EPI_ISL_3060508, EPI_ISL_3060509, EPI_ISL_3060510, EPI_ISL_3060511, EPI_ISL_3060512, EPI_ISL_3060513, EPI_ISL_3060514, EPI_ISL_3060515, EPI_ISL_3060516, EPI_ISL_3060517, EPI_ISL_3060518, EPI_ISL_3060519, EPI_ISL_3060520, EPI_ISL_3060521, EPI_ISL_3060522 | | | |
| see above | AP SSO | CSIR-Centre for Cellular and Molecular Biology-INSACOG | Payel Mukherjee, Lamuk Zaveri, Onkar Kulkarni,Jandhyala Sai Krishna,Tulasi Nagabandi, Sofia Banu, Ara Sreenivas,Shreekant Verma, Amareshwar Vodapalli ,B Himasri,Valli Nagalakshmi Undamatla, Sumedha Avadhanula, Vidhyadhari Methuku, Priya Nurkurthy, Archana Bharadwaj Siva, Karthik Bharadwaj Tallapaka,Rakesh K Mishra,Divya Tej Sowpati |
| EPI_ISL_3060523, EPI_ISL_3060524, EPI_ISL_3060525, | AP SSO | CSIR-Centre for Cellular and Molecular Biology-INSACOG | Lamuk Zaveri,Tulasi Nagabandi,Payel Mukherjee,Onkar Kulkarni, Sofia Banu, Jandhyala Sai Krishna,Ara Sreenivas,Shreekant Verma, Amareshwar Vodapalli ,B Himasri,Valli Nagalakshmi Undamatla, Sumedha Avadhanula, Priya Nurkurthy, Vidhyadhari Methuku, Archana Bharadwaj Siva, Karthik Bharadwaj Tallapaka,Rakesh K Mishra,Divya Tej Sowpati |

| | | | |
|---|---|---|---|
| EPI_ISL_3060526, EPI_ISL_3060527, EPI_ISL_3060528, EPI_ISL_3060529, EPI_ISL_3060530 | | | |
| EPI_ISL_3060531, EPI_ISL_3060532, EPI_ISL_3060533, EPI_ISL_3060534, EPI_ISL_3060535, EPI_ISL_3060536, EPI_ISL_3060537, EPI_ISL_3060538, EPI_ISL_3060539, EPI_ISL_3060540, EPI_ISL_3060541 | | | |
| see above | CSIR-Centre for Cellular and Molecular Biology | CSIR-Centre for Cellular and Molecular Biology-INSACOG | Lamuk Zaveri,Tulasi Nagabandi,Payel Mukherjee,Onkar Kulkarni, Sofia Banu, Jandhyala Sai Krishna,Ara Sreenivas,Shreekant Verma, Amareshwar Vodapalli ,B Himasri,Valli Nagalakshmi Undamatla, Sumedha Avadhanula, Priya Nurkurthy, Vidhyadhari Methuku, Archana Bharadwaj Siva, Karthik Bharadwaj Tallapaka,Rakesh K Mishra,Divya Tej Sowpati |
| EPI_ISL_3060542 | Mapmygenome | CSIR-Centre for Cellular and Molecular Biology-INSACOG | Lamuk Zaveri,Tulasi Nagabandi,Payel Mukherjee,Onkar Kulkarni, Sofia Banu, Jandhyala Sai Krishna,Ara Sreenivas,Shreekant Verma, Amareshwar Vodapalli ,B Himasri,Valli Nagalakshmi Undamatla, Sumedha Avadhanula, Priya Nurkurthy, Vidhyadhari Methuku, Archana Bharadwaj Siva, Karthik Bharadwaj Tallapaka,Rakesh K Mishra,Divya Tej Sowpati |
| EPI_ISL_3060543, EPI_ISL_3060544, EPI_ISL_3060545, EPI_ISL_3060546, EPI_ISL_3060547, EPI_ISL_3060548, EPI_ISL_3060549, EPI_ISL_3060551, EPI_ISL_3060552, EPI_ISL_3060553, EPI_ISL_3060554 | | | |
| see above | Mapmygenome | CSIR-Centre for Cellular and Molecular Biology-INSACOG | Tulasi Nagabandi,Lamuk Zaveri,Payel Mukherjee,Onkar Kulkarni,Jandhyala Sai Krishna, Sofia Banu, Ara Sreenivas,Shreekant Verma, Amareshwar Vodapalli ,B Himasri,Valli Nagalakshmi Undamatla, Priya Nurkurthy,Sumedha Avadhanula, Vidhyadhari Methuku, Archana Bharadwaj Siva, Karthik Bharadwaj Tallapaka,Rakesh K Mishra,Divya Tej Sowpati |
| EPI_ISL_3060555, EPI_ISL_3060556, EPI_ISL_3060557, EPI_ISL_3060558, EPI_ISL_3060559, EPI_ISL_3060560, EPI_ISL_3060561 | CSIR-National Environmental Engineering Research Institute | CSIR-Centre for Cellular and Molecular Biology-INSACOG | Payel Mukherjee, Lamuk Zaveri, Onkar Kulkarni,Tulasi Nagabandi, Sofia Banu, Ara Sreenivas,Shreekant Verma, Amareshwar Vodapalli ,B Himasri,Valli Nagalakshmi Undamatla, Sumedha Avadhanula, Vidhyadhari Methuku, Priya Nurkurthy, Archana Bharadwaj Siva, Karthik Bharadwaj Tallapaka,Krishna Khairnar,Rakesh K Mishra,Divya Tej Sowpati |
| EPI_ISL_3060562, EPI_ISL_3060563, EPI_ISL_3060564, EPI_ISL_3060565, EPI_ISL_3060568, EPI_ISL_3060569, EPI_ISL_3060570 | AP SSO | CSIR-Centre for Cellular and Molecular Biology-INSACOG | B Himasri,Amareshwar Vodapalli,Ara Sreenivas,Shreekant Verma,Valli Nagalakshmi Undamatla,Payel Mukherjee, Lamuk Zaveri, Onkar Kulkarni,Tulasi Nagabandi, Sofia Banu, Vidhyadhari Methuku,Priya Nurkurthy, Jandhyala Sai Krishna,Sumedha Avadhanula, Archana Bharadwaj Siva, Karthik Bharadwaj Tallapaka,Rakesh K Mishra,Divya Tej Sowpati |
| EPI_ISL_3060571, EPI_ISL_3060572, EPI_ISL_3060573, EPI_ISL_3060574, EPI_ISL_3060575, EPI_ISL_3060576, EPI_ISL_3060577, EPI_ISL_3060578, EPI_ISL_3060581, EPI_ISL_3060582, EPI_ISL_3060583, EPI_ISL_3060584, EPI_ISL_3060589, EPI_ISL_3060590, EPI_ISL_3060592, EPI_ISL_3060593, EPI_ISL_3060594 | | | |
| see above | AP SSO | CSIR-Centre for Cellular and Molecular Biology-INSACOG | Sofia Banu,Payel Mukherjee, Lamuk Zaveri, Onkar Kulkarni,Tulasi Nagabandi, Jandhyala Sai Krishna, Ara Sreenivas,Shreekant Verma, Amareshwar Vodapalli ,B Himasri,Valli Nagalakshmi Undamatla, Sumedha Avadhanula, Vidhyadhari Methuku, Priya Nurkurthy, Archana Bharadwaj Siva, Karthik Bharadwaj Tallapaka,Rakesh K Mishra,Divya Tej Sowpati |
| EPI_ISL_3060595, EPI_ISL_3060597, EPI_ISL_3060598, EPI_ISL_3060599, EPI_ISL_3060600, EPI_ISL_3060601 | AP SSO | CSIR-Centre for Cellular and Molecular Biology-INSACOG | Sumedha Avadhanula, Lamuk Zaveri, Tulasi Nagabandi, Ara Sreenivas,Shreekant Verma, Amareshwar Vodapalli ,B Himasri,Valli Nagalakshmi Undamatla, Vidhyadhari Methuku, Priya Nurkurthy, Payel Mukherjee,Jandhyala Sai Krishna,Onkar Kulkarni, Sofia Banu, Archana Bharadwaj Siva, Karthik Bharadwaj Tallapaka,Rakesh K Mishra,Divya Tej Sowpati |
| EPI_ISL_3060603 | Mapmygenome | CSIR-Centre for Cellular and Molecular Biology-INSACOG | Ara Sreenivas,Shreekant Verma, Amareshwar Vodapalli ,B Himasri,Valli Nagalakshmi Undamatla,Payel Mukherjee, Lamuk Zaveri, Onkar Kulkarni,Jandhyala Sai Krishna,Tulasi Nagabandi, Sofia Banu, ,Priya Nurkurthy, Vidhyadhari Methuku, Sumedha Avadhanula, Archana Bharadwaj Siva, Karthik Bharadwaj Tallapaka,Rakesh K Mishra,Divya Tej Sowpati |
| EPI_ISL_3060607, EPI_ISL_3060608, EPI_ISL_3060609, EPI_ISL_3060610, EPI_ISL_3060611, EPI_ISL_3060612, EPI_ISL_3060615, EPI_ISL_3060616 | AP SSO | CSIR-Centre for Cellular and Molecular Biology-INSACOG | Tulasi Nagabandi,Lamuk Zaveri,Payel Mukherjee,Onkar Kulkarni,Jandhyala Sai Krishna, Sofia Banu, Ara Sreenivas,Shreekant Verma, Amareshwar Vodapalli ,B Himasri,Valli Nagalakshmi Undamatla, Priya Nurkurthy,Sumedha Avadhanula, Vidhyadhari Methuku, Archana Bharadwaj Siva, Karthik Bharadwaj Tallapaka,Rakesh K Mishra,Divya Tej Sowpati |
| EPI_ISL_3060617, EPI_ISL_3060619 | AP SSO | CSIR-Centre for Cellular and Molecular Biology-INSACOG | Ara Sreenivas,Shreekant Verma, Amareshwar Vodapalli ,B Himasri,Valli Nagalakshmi Undamatla,Payel Mukherjee, Lamuk Zaveri, Onkar Kulkarni,Jandhyala Sai Krishna,Tulasi Nagabandi, Sofia Banu, ,Priya Nurkurthy, Vidhyadhari Methuku, Sumedha Avadhanula, Archana Bharadwaj Siva, Karthik Bharadwaj Tallapaka,Rakesh K Mishra,Divya Tej Sowpati |
| EPI_ISL_3060621, EPI_ISL_3060622 | Mapmygenome | CSIR-Centre for Cellular and Molecular Biology-INSACOG | Shreekant Verma, Amareshwar Vodapalli ,Ara Sreenivas,B Himasri,Valli Nagalakshmi Undamatla,Payel Mukherjee, Lamuk Zaveri, Onkar Kulkarni,Tulasi Nagabandi, Sofia Banu, Jandhyala Sai Krishna,Sumedha Avadhanula, Priya Nurkurthy, Vidhyadhari Methuku, Archana Bharadwaj Siva, Karthik Bharadwaj Tallapaka,Rakesh K Mishra,Divya Tej Sowpati |
| EPI_ISL_3060626, EPI_ISL_3060627, EPI_ISL_3060630, EPI_ISL_3060631, EPI_ISL_3060632, EPI_ISL_3060633, EPI_ISL_3060634, EPI_ISL_3060635, EPI_ISL_3060636, EPI_ISL_3060637, EPI_ISL_3060638, EPI_ISL_3060639 | | | |
| see above | Mapmygenome | CSIR-Centre for Cellular and Molecular Biology-INSACOG | Amareshwar Vodapalli,Ara Sreenivas,Shreekant Verma, B Himasri,Valli Nagalakshmi Undamatla,Payel Mukherjee, Lamuk Zaveri, Onkar Kulkarni,Tulasi Nagabandi, Sofia Banu, Jandhyala Sai Krishna,Priya Nurkurthy, Vidhyadhari Methuku, Sumedha Avadhanula, Archana Bharadwaj Siva, Karthik Bharadwaj Tallapaka,Rakesh K Mishra,Divya Tej Sowpati |
| EPI_ISL_3060641, EPI_ISL_3060642, EPI_ISL_3060644, EPI_ISL_3060645, EPI_ISL_3060646, EPI_ISL_3060650, EPI_ISL_3060651, EPI_ISL_3060652 | Mapmygenome | CSIR-Centre for Cellular and Molecular Biology-INSACOG | B Himasri,Amareshwar Vodapalli,Ara Sreenivas,Shreekant Verma,Valli Nagalakshmi Undamatla,Payel Mukherjee, Lamuk Zaveri, Onkar Kulkarni,Tulasi Nagabandi, Sofia Banu, Vidhyadhari Methuku,Priya Nurkurthy, Jandhyala Sai Krishna,Sumedha Avadhanula, Archana Bharadwaj Siva, Karthik Bharadwaj Tallapaka,Rakesh K Mishra,Divya Tej Sowpati |
| EPI_ISL_3060655, EPI_ISL_3060656, EPI_ISL_3060657, EPI_ISL_3060658, EPI_ISL_3060659, EPI_ISL_3060660, EPI_ISL_3060661, EPI_ISL_3060662, EPI_ISL_3060663, EPI_ISL_3060664, EPI_ISL_3060665, EPI_ISL_3060666, EPI_ISL_3060667, EPI_ISL_3060668, EPI_ISL_3060669, EPI_ISL_3060670 | | | |
| see above | AP SSO | CSIR-Centre for Cellular and Molecular Biology-INSACOG | Sumedha Avadhanula, Lamuk Zaveri, Tulasi Nagabandi, Ara Sreenivas,Shreekant Verma, Amareshwar Vodapalli ,B Himasri,Valli Nagalakshmi Undamatla, Vidhyadhari Methuku, Priya Nurkurthy, Payel Mukherjee,Jandhyala Sai Krishna,Onkar Kulkarni, Sofia Banu, Archana Bharadwaj Siva, Karthik Bharadwaj Tallapaka,Rakesh K Mishra,Divya Tej Sowpati |
| EPI_ISL_3060671, EPI_ISL_3060672 | AP SSO | CSIR-Centre for Cellular and Molecular Biology-INSACOG | Priya Nurkurthy,Ara Sreenivas,Shreekant Verma, Amareshwar Vodapalli ,Lamuk Zaveri,Tulasi Nagabandi,B Himasri,Valli Nagalakshmi Undamatla, Sumedha Avadhanula, Vidhyadhari Methuku,Payel Mukherjee,Onkar Kulkarni, Jandhyala Sai Krishna,Sofia Banu, Archana Bharadwaj Siva, Karthik Bharadwaj Tallapaka,Rakesh K Mishra,Divya Tej Sowpati |
| EPI_ISL_3060673, EPI_ISL_3060676, EPI_ISL_3060677, EPI_ISL_3060678, EPI_ISL_3060679, EPI_ISL_3060680, EPI_ISL_3060681 | CSIR-National Environmental Engineering Research Institute | CSIR-Centre for Cellular and Molecular Biology-INSACOG | Payel Mukherjee, Lamuk Zaveri, Onkar Kulkarni,Tulasi Nagabandi, Sofia Banu, Ara Sreenivas,Shreekant Verma, Amareshwar Vodapalli ,B Himasri,Valli Nagalakshmi Undamatla, Sumedha Avadhanula, Vidhyadhari Methuku, Priya Nurkurthy, Archana Bharadwaj Siva, Karthik Bharadwaj Tallapaka,Krishna Khairnar,Rakesh K Mishra,Divya Tej Sowpati |
| EPI_ISL_3060683, EPI_ISL_3060684 | CSIR-Centre for Cellular and Molecular Biology | CSIR-Centre for Cellular and Molecular Biology-INSACOG | Payel Mukherjee, Lamuk Zaveri, Onkar Kulkarni,Jandhyala Sai Krishna,Tulasi Nagabandi, Sofia Banu, Ara Sreenivas,Shreekant Verma, Amareshwar Vodapalli ,B Himasri,Valli Nagalakshmi Undamatla, Sumedha Avadhanula, Vidhyadhari Methuku, Priya Nurkurthy, Archana Bharadwaj Siva, Karthik Bharadwaj Tallapaka,Rakesh K Mishra,Divya Tej Sowpati |
| EPI_ISL_3060685, EPI_ISL_3060686 | AP SSO | CSIR-Centre for Cellular and Molecular Biology-INSACOG | Payel Mukherjee, Lamuk Zaveri, Onkar Kulkarni,Jandhyala Sai Krishna,Tulasi Nagabandi, Sofia Banu, Ara Sreenivas,Shreekant Verma, Amareshwar Vodapalli ,B Himasri,Valli Nagalakshmi Undamatla, Sumedha Avadhanula, Vidhyadhari Methuku, Priya Nurkurthy, Archana Bharadwaj Siva, Karthik Bharadwaj Tallapaka,Rakesh K Mishra,Divya Tej Sowpati |
| EPI_ISL_3060687, EPI_ISL_3060688, EPI_ISL_3060689, EPI_ISL_3060690 | AP SSO | CSIR-Centre for Cellular and Molecular Biology-INSACOG | Lamuk Zaveri,Tulasi Nagabandi,Payel Mukherjee,Onkar Kulkarni, Sofia Banu, Jandhyala Sai Krishna,Ara Sreenivas,Shreekant Verma, Amareshwar Vodapalli ,B Himasri,Valli Nagalakshmi Undamatla, Sumedha Avadhanula, Priya Nurkurthy, Vidhyadhari Methuku, Archana Bharadwaj Siva, Karthik Bharadwaj Tallapaka,Rakesh K Mishra,Divya Tej Sowpati |
| EPI_ISL_3060691, EPI_ISL_3060692, EPI_ISL_3060693, EPI_ISL_3060694, EPI_ISL_3060695 | AP SSO | CSIR-Centre for Cellular and Molecular Biology-INSACOG | Tulasi Nagabandi,Lamuk Zaveri,Payel Mukherjee,Onkar Kulkarni,Jandhyala Sai Krishna, Sofia Banu, Ara Sreenivas,Shreekant Verma, Amareshwar Vodapalli ,B Himasri,Valli Nagalakshmi Undamatla, Priya Nurkurthy,Sumedha Avadhanula, Vidhyadhari Methuku, Archana Bharadwaj Siva, Karthik Bharadwaj Tallapaka,Rakesh K Mishra,Divya Tej Sowpati |
| EPI_ISL_3060696, EPI_ISL_3060697, EPI_ISL_3060698, EPI_ISL_3060699 | AP SSO | CSIR-Centre for Cellular and Molecular Biology-INSACOG | Ara Sreenivas,Shreekant Verma, Amareshwar Vodapalli ,B Himasri,Valli Nagalakshmi Undamatla,Payel Mukherjee, Lamuk Zaveri, Onkar Kulkarni,Jandhyala Sai Krishna,Tulasi Nagabandi, Sofia Banu, ,Priya Nurkurthy, Vidhyadhari Methuku, Sumedha Avadhanula, Archana Bharadwaj Siva, Karthik Bharadwaj Tallapaka,Rakesh K Mishra,Divya Tej Sowpati |
| EPI_ISL_3060700, EPI_ISL_3060701, EPI_ISL_3060703, EPI_ISL_3060704 | AP SSO | CSIR-Centre for Cellular and Molecular Biology-INSACOG | Shreekant Verma, Amareshwar Vodapalli ,Ara Sreenivas,B Himasri,Valli Nagalakshmi Undamatla,Payel Mukherjee, Lamuk Zaveri, Onkar Kulkarni,Tulasi Nagabandi, Sofia Banu, Jandhyala Sai Krishna,Sumedha Avadhanula, Priya Nurkurthy, Vidhyadhari Methuku, Archana Bharadwaj Siva, Karthik Bharadwaj Tallapaka,Rakesh K Mishra,Divya Tej Sowpati |
| EPI_ISL_3060705, EPI_ISL_3060706, EPI_ISL_3060707, | AP SSO | CSIR-Centre for Cellular and Molecular Biology-INSACOG | Amareshwar Vodapalli,Ara Sreenivas,Shreekant Verma, B Himasri,Valli Nagalakshmi Undamatla,Payel Mukherjee, Lamuk Zaveri, Onkar Kulkarni,Tulasi Nagabandi, Sofia Banu, Jandhyala Sai Krishna,Priya Nurkurthy, Vidhyadhari Methuku, Sumedha Avadhanula, Archana Bharadwaj Siva, Karthik Bharadwaj Tallapaka,Rakesh K Mishra,Divya Tej Sowpati |

| | | | |
|---|---|---|---|
| EPI_ISL_3060708 | | | |
| EPI_ISL_3060709, EPI_ISL_3060710, EPI_ISL_3060711, EPI_ISL_3060712 | AP SSO | CSIR-Centre for Cellular and Molecular Biology-INSACOG | B Himasri,Amareshwar Vodapalli,Ara Sreenivas,Shreekant Verma,Valli Nagalakshmi Undamatla,Payel Mukherjee, Lamuk Zaveri, Onkar Kulkarni,Tulasi Nagabandi, Sofia Banu, Vidhyadhari Methuku,Priya Nurkurthy, Jandhyala Sai Krishna,Sumedha Avadhanula, Archana Bharadwaj Siva, Karthik Bharadwaj Tallapaka,Rakesh K Mishra,Divya Tej Sowpati |
| EPI_ISL_3060713, EPI_ISL_3060714, EPI_ISL_3060715, EPI_ISL_3060716 | AP SSO | CSIR-Centre for Cellular and Molecular Biology-INSACOG | Sofia Banu,Payel Mukherjee, Lamuk Zaveri, Onkar Kulkarni,Tulasi Nagabandi, Jandhyala Sai Krishna, Ara Sreenivas,Shreekant Verma, Amareshwar Vodapalli ,B Himasri,Valli Nagalakshmi Undamatla, Sumedha Avadhanula, Vidhyadhari Methuku, Priya Nurkurthy, Archana Bharadwaj Siva, Karthik Bharadwaj Tallapaka,Rakesh K Mishra,Divya Tej Sowpati |
| EPI_ISL_3060717, EPI_ISL_3060718, EPI_ISL_3060720, EPI_ISL_3060721 | AP SSO | CSIR-Centre for Cellular and Molecular Biology-INSACOG | Sumedha Avadhanula, Lamuk Zaveri, Tulasi Nagabandi, Ara Sreenivas,Shreekant Verma, Amareshwar Vodapalli ,B Himasri,Valli Nagalakshmi Undamatla, Vidhyadhari Methuku, Priya Nurkurthy, Payel Mukherjee,Jandhyala Sai Krishna,Onkar Kulkarni, Sofia Banu, Archana Bharadwaj Siva, Karthik Bharadwaj Tallapaka,Rakesh K Mishra,Divya Tej Sowpati |
| EPI_ISL_3060722, EPI_ISL_3060724, EPI_ISL_3060725, EPI_ISL_3060726 | AP SSO | CSIR-Centre for Cellular and Molecular Biology-INSACOG | Priya Nurkurthy,Ara Sreenivas,Shreekant Verma, Amareshwar Vodapalli ,Lamuk Zaveri,Tulasi Nagabandi,B Himasri,Valli Nagalakshmi Undamatla, Sumedha Avadhanula, Vidhyadhari Methuku,Payel Mukherjee,Onkar Kulkarni, Jandhyala Sai Krishna,Sofia Banu, Archana Bharadwaj Siva, Karthik Bharadwaj Tallapaka,Rakesh K Mishra,Divya Tej Sowpati |
| EPI_ISL_3060727, EPI_ISL_3060728 | AP SSO | CSIR-Centre for Cellular and Molecular Biology-INSACOG | Vidhyadhari Methuku,Tulasi Nagabandi,Lamuk Zaveri, Ara Sreenivas,Shreekant Verma, Amareshwar Vodapalli ,B Himasri,Valli Nagalakshmi Undamatla, Priya Nurkurthy,Sumedha Avadhanula, Archana Bharadwaj Siva,Payel Mukherjee,Onkar Kulkarni, Jandhyala Sai Krishna,Sofia Banu, Karthik Bharadwaj Tallapaka,Rakesh K Mishra,Divya Tej Sowpati |
| EPI_ISL_3060732 | AP SSO | CSIR-Centre for Cellular and Molecular Biology-INSACOG | Payel Mukherjee, Lamuk Zaveri, Onkar Kulkarni,Jandhyala Sai Krishna,Tulasi Nagabandi, Sofia Banu, Ara Sreenivas,Shreekant Verma, Amareshwar Vodapalli ,B Himasri,Valli Nagalakshmi Undamatla, Sumedha Avadhanula, Vidhyadhari Methuku, Priya Nurkurthy, Archana Bharadwaj Siva, Karthik Bharadwaj Tallapaka,Rakesh K Mishra,Divya Tej Sowpati |
| EPI_ISL_3060738, EPI_ISL_3060739 | AP SSO | CSIR-Centre for Cellular and Molecular Biology-INSACOG | Lamuk Zaveri,Tulasi Nagabandi,Payel Mukherjee,Onkar Kulkarni, Sofia Banu, Jandhyala Sai Krishna,Ara Sreenivas,Shreekant Verma, Amareshwar Vodapalli ,B Himasri,Valli Nagalakshmi Undamatla, Sumedha Avadhanula, Priya Nurkurthy, Vidhyadhari Methuku, Archana Bharadwaj Siva, Karthik Bharadwaj Tallapaka,Rakesh K Mishra,Divya Tej Sowpati |
| EPI_ISL_3060740, EPI_ISL_3060741, EPI_ISL_3060742, EPI_ISL_3060743, EPI_ISL_3060744 | AP SSO | CSIR-Centre for Cellular and Molecular Biology-INSACOG | Tulasi Nagabandi,Lamuk Zaveri,Payel Mukherjee,Onkar Kulkarni,Jandhyala Sai Krishna, Sofia Banu, Ara Sreenivas,Shreekant Verma, Amareshwar Vodapalli ,B Himasri,Valli Nagalakshmi Undamatla, Priya Nurkurthy,Sumedha Avadhanula, Vidhyadhari Methuku, Archana Bharadwaj Siva, Karthik Bharadwaj Tallapaka,Rakesh K Mishra,Divya Tej Sowpati |
| EPI_ISL_3060745, EPI_ISL_3060746, EPI_ISL_3060747, EPI_ISL_3060748 | AP SSO | CSIR-Centre for Cellular and Molecular Biology-INSACOG | Ara Sreenivas,Shreekant Verma, Amareshwar Vodapalli ,B Himasri,Valli Nagalakshmi Undamatla,Payel Mukherjee, Lamuk Zaveri, Onkar Kulkarni,Jandhyala Sai Krishna,Tulasi Nagabandi, Sofia Banu, ,Priya Nurkurthy, Vidhyadhari Methuku, Sumedha Avadhanula, Archana Bharadwaj Siva, Karthik Bharadwaj Tallapaka,Rakesh K Mishra,Divya Tej Sowpati |
| EPI_ISL_3060750, EPI_ISL_3060751, EPI_ISL_3060752 | AP SSO | CSIR-Centre for Cellular and Molecular Biology-INSACOG | Shreekant Verma, Amareshwar Vodapalli ,Ara Sreenivas,B Himasri,Valli Nagalakshmi Undamatla,Payel Mukherjee, Lamuk Zaveri, Onkar Kulkarni,Tulasi Nagabandi, Sofia Banu, Jandhyala Sai Krishna,Sumedha Avadhanula, Priya Nurkurthy, Vidhyadhari Methuku, Archana Bharadwaj Siva, Karthik Bharadwaj Tallapaka,Rakesh K Mishra,Divya Tej Sowpati |
| EPI_ISL_3060753, EPI_ISL_3060754, EPI_ISL_3060756 | AP SSO | CSIR-Centre for Cellular and Molecular Biology-INSACOG | Amareshwar Vodapalli,Ara Sreenivas,Shreekant Verma, B Himasri,Valli Nagalakshmi Undamatla,Payel Mukherjee, Lamuk Zaveri, Onkar Kulkarni,Tulasi Nagabandi, Sofia Banu, Jandhyala Sai Krishna,Priya Nurkurthy, Vidhyadhari Methuku, Sumedha Avadhanula, Archana Bharadwaj Siva, Karthik Bharadwaj Tallapaka,Rakesh K Mishra,Divya Tej Sowpati |
| EPI_ISL_3060757, EPI_ISL_3060758, EPI_ISL_3060759 | AP SSO | CSIR-Centre for Cellular and Molecular Biology-INSACOG | B Himasri,Amareshwar Vodapalli,Ara Sreenivas,Shreekant Verma,Valli Nagalakshmi Undamatla,Payel Mukherjee, Lamuk Zaveri, Onkar Kulkarni,Tulasi Nagabandi, Sofia Banu, Vidhyadhari Methuku,Priya Nurkurthy, Jandhyala Sai Krishna,Sumedha Avadhanula, Archana Bharadwaj Siva, Karthik Bharadwaj Tallapaka,Rakesh K Mishra,Divya Tej Sowpati |
| EPI_ISL_3060761, EPI_ISL_3060763, EPI_ISL_3060764 | AP SSO | CSIR-Centre for Cellular and Molecular Biology-INSACOG | Sofia Banu,Payel Mukherjee, Lamuk Zaveri, Onkar Kulkarni,Tulasi Nagabandi, Jandhyala Sai Krishna, Ara Sreenivas,Shreekant Verma, Amareshwar Vodapalli ,B Himasri,Valli Nagalakshmi Undamatla, Sumedha Avadhanula, Vidhyadhari Methuku, Priya Nurkurthy, Archana Bharadwaj Siva, Karthik Bharadwaj Tallapaka,Rakesh K Mishra,Divya Tej Sowpati |
| EPI_ISL_3060765, EPI_ISL_3060767, EPI_ISL_3060768 | AP SSO | CSIR-Centre for Cellular and Molecular Biology-INSACOG | Sumedha Avadhanula, Lamuk Zaveri, Tulasi Nagabandi, Ara Sreenivas,Shreekant Verma, Amareshwar Vodapalli ,B Himasri,Valli Nagalakshmi Undamatla, Vidhyadhari Methuku, Priya Nurkurthy, Payel Mukherjee,Jandhyala Sai Krishna,Onkar Kulkarni, Sofia Banu, Archana Bharadwaj Siva, Karthik Bharadwaj Tallapaka,Rakesh K Mishra,Divya Tej Sowpati |
| EPI_ISL_3060769 | Mapmygenome | CSIR-Centre for Cellular and Molecular Biology-INSACOG | Sumedha Avadhanula, Lamuk Zaveri, Tulasi Nagabandi, Ara Sreenivas,Shreekant Verma, Amareshwar Vodapalli ,B Himasri,Valli Nagalakshmi Undamatla, Vidhyadhari Methuku, Priya Nurkurthy, Payel Mukherjee,Jandhyala Sai Krishna,Onkar Kulkarni, Sofia Banu, Archana Bharadwaj Siva, Karthik Bharadwaj Tallapaka,Rakesh K Mishra,Divya Tej Sowpati |
| EPI_ISL_3060770, EPI_ISL_3060771, EPI_ISL_3060772, EPI_ISL_3060773, EPI_ISL_3060774 | Mapmygenome | CSIR-Centre for Cellular and Molecular Biology-INSACOG | Priya Nurkurthy,Ara Sreenivas,Shreekant Verma, Amareshwar Vodapalli ,Lamuk Zaveri,Tulasi Nagabandi,B Himasri,Valli Nagalakshmi Undamatla, Sumedha Avadhanula, Vidhyadhari Methuku,Payel Mukherjee,Onkar Kulkarni, Jandhyala Sai Krishna,Sofia Banu, Archana Bharadwaj Siva, Karthik Bharadwaj Tallapaka,Rakesh K Mishra,Divya Tej Sowpati |
| EPI_ISL_3060775, EPI_ISL_3060776, EPI_ISL_3060777 | Mapmygenome | CSIR-Centre for Cellular and Molecular Biology-INSACOG | Vidhyadhari Methuku,Tulasi Nagabandi,Lamuk Zaveri, Ara Sreenivas,Shreekant Verma, Amareshwar Vodapalli ,B Himasri,Valli Nagalakshmi Undamatla, Priya Nurkurthy,Sumedha Avadhanula, Archana Bharadwaj Siva,Payel Mukherjee,Onkar Kulkarni, Jandhyala Sai Krishna,Sofia Banu, Karthik Bharadwaj Tallapaka,Rakesh K Mishra,Divya Tej Sowpati |
| EPI_ISL_3060778, EPI_ISL_3060779, EPI_ISL_3060780, EPI_ISL_3060781, EPI_ISL_3060782, EPI_ISL_3060783, EPI_ISL_3060784, EPI_ISL_3060785, EPI_ISL_3060786, EPI_ISL_3060787, EPI_ISL_3060788, EPI_ISL_3060789, EPI_ISL_3060790, EPI_ISL_3060791, EPI_ISL_3060792, EPI_ISL_3060793, EPI_ISL_3060794, EPI_ISL_3060795 | | | |
| see above | AP SSO | CSIR-Centre for Cellular and Molecular Biology-INSACOG | Payel Mukherjee, Lamuk Zaveri, Onkar Kulkarni,Jandhyala Sai Krishna,Tulasi Nagabandi, Sofia Banu, Ara Sreenivas,Shreekant Verma, Amareshwar Vodapalli ,B Himasri,Valli Nagalakshmi Undamatla, Sumedha Avadhanula, Vidhyadhari Methuku, Priya Nurkurthy, Archana Bharadwaj Siva, Karthik Bharadwaj Tallapaka,Rakesh K Mishra,Divya Tej Sowpati |
| EPI_ISL_3060796, EPI_ISL_3060797, EPI_ISL_3060798, EPI_ISL_3060799, EPI_ISL_3060800, EPI_ISL_3060801, EPI_ISL_3060802, EPI_ISL_3060803, EPI_ISL_3060804, EPI_ISL_3060805, EPI_ISL_3060806, EPI_ISL_3060807, EPI_ISL_3060808, EPI_ISL_3060809, EPI_ISL_3060810 | | | |
| see above | AP SSO | CSIR-Centre for Cellular and Molecular Biology-INSACOG | Lamuk Zaveri,Tulasi Nagabandi,Payel Mukherjee,Onkar Kulkarni, Sofia Banu, Jandhyala Sai Krishna,Ara Sreenivas,Shreekant Verma, Amareshwar Vodapalli ,B Himasri,Valli Nagalakshmi Undamatla, Sumedha Avadhanula, Priya Nurkurthy, Vidhyadhari Methuku, Archana Bharadwaj Siva, Karthik Bharadwaj Tallapaka,Rakesh K Mishra,Divya Tej Sowpati |
| EPI_ISL_3060812, EPI_ISL_3060813, EPI_ISL_3060814, EPI_ISL_3060815, EPI_ISL_3060816, EPI_ISL_3060821, EPI_ISL_3060824, EPI_ISL_3060825, EPI_ISL_3060827 | AP SSO | CSIR-Centre for Cellular and Molecular Biology-INSACOG | Ara Sreenivas,Shreekant Verma, Amareshwar Vodapalli ,B Himasri,Valli Nagalakshmi Undamatla,Payel Mukherjee, Lamuk Zaveri, Onkar Kulkarni,Jandhyala Sai Krishna,Tulasi Nagabandi, Sofia Banu, ,Priya Nurkurthy, Vidhyadhari Methuku, Sumedha Avadhanula, Archana Bharadwaj Siva, Karthik Bharadwaj Tallapaka,Rakesh K Mishra,Divya Tej Sowpati |
| EPI_ISL_3060830 | AP SSO | CSIR-Centre for Cellular and Molecular Biology-INSACOG | Shreekant Verma, Amareshwar Vodapalli ,Ara Sreenivas,B Himasri,Valli Nagalakshmi Undamatla,Payel Mukherjee, Lamuk Zaveri, Onkar Kulkarni,Tulasi Nagabandi, Sofia Banu, Jandhyala Sai Krishna,Sumedha Avadhanula, Priya Nurkurthy, Vidhyadhari Methuku, Archana Bharadwaj Siva, Karthik Bharadwaj Tallapaka,Rakesh K Mishra,Divya Tej Sowpati |
| EPI_ISL_3060831, EPI_ISL_3060832, EPI_ISL_3060833, EPI_ISL_3060834, EPI_ISL_3060835, EPI_ISL_3060836 | CSIR-Centre for Cellular and Molecular Biology | CSIR-Centre for Cellular and Molecular Biology-INSACOG | Shreekant Verma, Amareshwar Vodapalli ,Ara Sreenivas,B Himasri,Valli Nagalakshmi Undamatla,Payel Mukherjee, Lamuk Zaveri, Onkar Kulkarni,Tulasi Nagabandi, Sofia Banu, Jandhyala Sai Krishna,Sumedha Avadhanula, Priya Nurkurthy, Vidhyadhari Methuku, Archana Bharadwaj Siva, Karthik Bharadwaj Tallapaka,Rakesh K Mishra,Divya Tej Sowpati |
| EPI_ISL_3060837, EPI_ISL_3060838, EPI_ISL_3060840, EPI_ISL_3060841, EPI_ISL_3060842 | AP SSO | CSIR-Centre for Cellular and Molecular Biology-INSACOG | Shreekant Verma, Amareshwar Vodapalli ,Ara Sreenivas,B Himasri,Valli Nagalakshmi Undamatla,Payel Mukherjee, Lamuk Zaveri, Onkar Kulkarni,Tulasi Nagabandi, Sofia Banu, Jandhyala Sai Krishna,Sumedha Avadhanula, Priya Nurkurthy, Vidhyadhari Methuku, Archana Bharadwaj Siva, Karthik Bharadwaj Tallapaka,Rakesh K Mishra,Divya Tej Sowpati |
| EPI_ISL_3060848, EPI_ISL_3060849, EPI_ISL_3060850 | AP SSO | CSIR-Centre for Cellular and Molecular Biology-INSACOG | Amareshwar Vodapalli,Ara Sreenivas,Shreekant Verma, B Himasri,Valli Nagalakshmi Undamatla,Payel Mukherjee, Lamuk Zaveri, Onkar Kulkarni,Tulasi Nagabandi, Sofia Banu, Jandhyala Sai Krishna,Priya Nurkurthy, Vidhyadhari Methuku, Sumedha Avadhanula, Archana Bharadwaj Siva, Karthik Bharadwaj Tallapaka,Rakesh K Mishra,Divya Tej Sowpati |
| EPI_ISL_3060852, EPI_ISL_3060853, EPI_ISL_3060854, EPI_ISL_3060855, EPI_ISL_3060856, EPI_ISL_3060857, EPI_ISL_3060858, EPI_ISL_3060859, EPI_ISL_3060860, EPI_ISL_3060861, EPI_ISL_3060862, EPI_ISL_3060863, EPI_ISL_3060864, EPI_ISL_3060865, EPI_ISL_3060866, EPI_ISL_3060867 | | | |
| see above | AP SSO | CSIR-Centre for Cellular and Molecular Biology-INSACOG | Priya Nurkurthy,Ara Sreenivas,Shreekant Verma, Amareshwar Vodapalli ,Lamuk Zaveri,Tulasi Nagabandi,B Himasri,Valli Nagalakshmi Undamatla, Sumedha Avadhanula, Vidhyadhari Methuku,Payel Mukherjee,Onkar Kulkarni, Jandhyala Sai Krishna,Sofia Banu, Archana Bharadwaj Siva, Karthik Bharadwaj Tallapaka,Rakesh K Mishra,Divya Tej Sowpati |
| EPI_ISL_3060868, EPI_ISL_3060869, EPI_ISL_3060870, EPI_ISL_3060871, EPI_ISL_3060872, EPI_ISL_3060873, | AP SSO | CSIR-Centre for Cellular and Molecular Biology-INSACOG | Vidhyadhari Methuku,Tulasi Nagabandi,Lamuk Zaveri, Ara Sreenivas,Shreekant Verma, Amareshwar Vodapalli ,B Himasri,Valli Nagalakshmi Undamatla, Priya Nurkurthy,Sumedha Avadhanula, Archana Bharadwaj Siva,Payel Mukherjee,Onkar Kulkarni, Jandhyala Sai Krishna,Sofia Banu, Karthik Bharadwaj Tallapaka,Rakesh K Mishra,Divya Tej Sowpati |

| | | | |
|---|---|---|---|
| EPI_ISL_3060874, EPI_ISL_3060875, EPI_ISL_3060876, EPI_ISL_3060877 | | | |
| EPI_ISL_3060879, EPI_ISL_3060880, EPI_ISL_3060882, EPI_ISL_3060883, EPI_ISL_3060885, EPI_ISL_3060886, EPI_ISL_3060887, EPI_ISL_3060888 | CSIR-National Environmental Engineering Research Institute | CSIR-Centre for Cellular and Molecular Biology-INSACOG | Payel Mukherjee, Lamuk Zaveri, Onkar Kulkarni,Tulasi Nagabandi, Sofia Banu, Ara Sreenivas,Shreekant Verma, Amareshwar Vodapalli ,B Himasri,Valli Nagalakshmi Undamatla, Sumedha Avadhanula, Vidhyadhari Methuku, Priya Nurkurthy, Archana Bharadwaj Siva, Karthik Bharadwaj Tallapaka,Krishna Khairnar,Rakesh K Mishra,Divya Tej Sowpati |
| EPI_ISL_3060890, EPI_ISL_3060898 | CSIR-Centre for Cellular and Molecular Biology | CSIR-Centre for Cellular and Molecular Biology-INSACOG | Vidhyadhari Methuku,Tulasi Nagabandi,Lamuk Zaveri, Ara Sreenivas,Shreekant Verma, Amareshwar Vodapalli ,B Himasri,Valli Nagalakshmi Undamatla, Priya Nurkurthy,Sumedha Avadhanula, Archana Bharadwaj Siva,Payel Mukherjee,Onkar Kulkarni, Jandhyala Sai Krishna,Sofia Banu, Karthik Bharadwaj Tallapaka,Rakesh K Mishra,Divya Tej Sowpati |
| EPI_ISL_3060902, EPI_ISL_3060903, EPI_ISL_3060904, EPI_ISL_3060906, EPI_ISL_3060907, EPI_ISL_3060908, EPI_ISL_3060910, EPI_ISL_3060911, EPI_ISL_3060912, EPI_ISL_3060913 | CSIR-National Environmental Engineering Research Institute | CSIR-Centre for Cellular and Molecular Biology - INSACOG | Jandhyala Sai Krishna,Lamuk Zaveri,Onkar Kulkarni,Tulasi Nagabandi, Payel Mukherjee, Sofia Banu, Ara Sreenivas,Shreekant Verma, Amareshwar Vodapalli ,B Himasri,Valli Nagalakshmi Undamatla, Sumedha Avadhanula, Vidhyadhari Methuku, Priya Nurkurthy, Archana Bharadwaj Siva, Karthik Bharadwaj Tallapaka,Krishna Khairnar,Rakesh K Mishra,Divya Tej Sowpati |
| EPI_ISL_3060914, EPI_ISL_3060916 | LKO | Jessa | Berden et al. on behalf of the Jessa_cmdLab |
| EPI_ISL_3060917 | Jessa | Jessa | Berden et al. on behalf of the Jessa_cmdLab |
| EPI_ISL_3060918, EPI_ISL_3060919, EPI_ISL_3060920, EPI_ISL_3060922, EPI_ISL_3060923, EPI_ISL_3060924, EPI_ISL_3060925, EPI_ISL_3060926, EPI_ISL_3060927, EPI_ISL_3060928, EPI_ISL_3060929, EPI_ISL_3060930, EPI_ISL_3060931, EPI_ISL_3060933, EPI_ISL_3060934, EPI_ISL_3060935, EPI_ISL_3060936, EPI_ISL_3060937, EPI_ISL_3060938, EPI_ISL_3060939, EPI_ISL_3060941, EPI_ISL_3060942, EPI_ISL_3060943, EPI_ISL_3060945, EPI_ISL_3060946, EPI_ISL_3060947, EPI_ISL_3060948, EPI_ISL_3060949, EPI_ISL_3060950, EPI_ISL_3060951, EPI_ISL_3060952, EPI_ISL_3060953, EPI_ISL_3060954, EPI_ISL_3060955, EPI_ISL_3060956 | | | |
| see above | ZOL | Jessa | Berden et al. on behalf of the Jessa_cmdLab |
| EPI_ISL_3060957, EPI_ISL_3060958, EPI_ISL_3060959, EPI_ISL_3060960, EPI_ISL_3060961, EPI_ISL_3060962, EPI_ISL_3060963, EPI_ISL_3060964, EPI_ISL_3060966, EPI_ISL_3060967, EPI_ISL_3060968, EPI_ISL_3060969, EPI_ISL_3060970, EPI_ISL_3060971, EPI_ISL_3060974, EPI_ISL_3060975, EPI_ISL_3060976, EPI_ISL_3060978 | | | |
| see above | Jessa | Jessa | Berden et al. on behalf of the Jessa_cmdLab |
| EPI_ISL_3060979, EPI_ISL_3060980, EPI_ISL_3060981, EPI_ISL_3060983, EPI_ISL_3060985, EPI_ISL_3060986, EPI_ISL_3060988, EPI_ISL_3060989, EPI_ISL_3060991, EPI_ISL_3060992, EPI_ISL_3060993, EPI_ISL_3060994, EPI_ISL_3060995, EPI_ISL_3060996, EPI_ISL_3060997, EPI_ISL_3060998, EPI_ISL_3061000, EPI_ISL_3061001, EPI_ISL_3061002, EPI_ISL_3061003, EPI_ISL_3061004, EPI_ISL_3061006, EPI_ISL_3061007, EPI_ISL_3061008, EPI_ISL_3061009, EPI_ISL_3061011, EPI_ISL_3061012, EPI_ISL_3061013, EPI_ISL_3061014, EPI_ISL_3061015, EPI_ISL_3061016, EPI_ISL_3061017, EPI_ISL_3061018, EPI_ISL_3061019, EPI_ISL_3061020, EPI_ISL_3061021, EPI_ISL_3061022, EPI_ISL_3061023, EPI_ISL_3061024, EPI_ISL_3061025, EPI_ISL_3061026, EPI_ISL_3061027, EPI_ISL_3061028, EPI_ISL_3061029, EPI_ISL_3061030, EPI_ISL_3061031, EPI_ISL_3061032, EPI_ISL_3061033, EPI_ISL_3061034, EPI_ISL_3061035, EPI_ISL_3061036, EPI_ISL_3061037, EPI_ISL_3061038, EPI_ISL_3061039, EPI_ISL_3061040, EPI_ISL_3061041, EPI_ISL_3061042, EPI_ISL_3061043, EPI_ISL_3061044, EPI_ISL_3061045, EPI_ISL_3061046, EPI_ISL_3061047, EPI_ISL_3061048, EPI_ISL_3061049, EPI_ISL_3061050, EPI_ISL_3061051, EPI_ISL_3061053, EPI_ISL_3061054, EPI_ISL_3061055, EPI_ISL_3061056, EPI_ISL_3061057, EPI_ISL_3061058 | | | |
| see above | Servicio de Microbiología. Consorcio Hospital General Universitario de Valencia | SeqCOVID-SPAIN consortium/IBV(CSIC) | María Dolores Ocete, Begoña Fuster Escrivá, Carme Salvador García, Rafael Medina González, Concepción Gimeno Cardona and SeqCOVID-SPAIN consortium |
| EPI_ISL_3061059 | VIDYMED LAUSANNE | Laboratory of genomics and metagenomics | Trestan Pillonel, Damien Jacot, Sébastien Aeby, Gilbert Greub, Claire Bertelli |
| EPI_ISL_3061063 | CHUV | Laboratory of genomics and metagenomics | Trestan Pillonel, Damien Jacot, Sébastien Aeby, Gilbert Greub, Claire Bertelli |
| EPI_ISL_3061064 | VIDYMED LAUSANNE | Laboratory of genomics and metagenomics | Trestan Pillonel, Damien Jacot, Sébastien Aeby, Gilbert Greub, Claire Bertelli |
| EPI_ISL_3061065, EPI_ISL_3061066 | CHUV | Laboratory of genomics and metagenomics | Trestan Pillonel, Damien Jacot, Sébastien Aeby, Gilbert Greub, Claire Bertelli |
| EPI_ISL_3061067 | VIDYMED LAUSANNE | Laboratory of genomics and metagenomics | Trestan Pillonel, Damien Jacot, Sébastien Aeby, Gilbert Greub, Claire Bertelli |
| EPI_ISL_3061068 | CHUV | Laboratory of genomics and metagenomics | Trestan Pillonel, Damien Jacot, Sébastien Aeby, Gilbert Greub, Claire Bertelli |
| EPI_ISL_3061070 | VIDYMED LAUSANNE | Laboratory of genomics and metagenomics | Trestan Pillonel, Damien Jacot, Sébastien Aeby, Gilbert Greub, Claire Bertelli |
| EPI_ISL_3061071, EPI_ISL_3061072 | CHUV | Laboratory of genomics and metagenomics | Trestan Pillonel, Damien Jacot, Sébastien Aeby, Gilbert Greub, Claire Bertelli |
| EPI_ISL_3061073 | LA SOURCE | Laboratory of genomics and metagenomics | Trestan Pillonel, Damien Jacot, Sébastien Aeby, Gilbert Greub, Claire Bertelli |
| EPI_ISL_3061075 | VIDYMED LAUSANNE | Laboratory of genomics and metagenomics | Trestan Pillonel, Damien Jacot, Sébastien Aeby, Gilbert Greub, Claire Bertelli |
| EPI_ISL_3061076, EPI_ISL_3061077, EPI_ISL_3061078, EPI_ISL_3061079, EPI_ISL_3061080, EPI_ISL_3061081 | EHNV | Laboratory of genomics and metagenomics | Trestan Pillonel, Damien Jacot, Sébastien Aeby, Gilbert Greub, Claire Bertelli |
| EPI_ISL_3061082, EPI_ISL_3061083, EPI_ISL_3061084, EPI_ISL_3061085 | CHUV | Laboratory of genomics and metagenomics | Trestan Pillonel, Damien Jacot, Sébastien Aeby, Gilbert Greub, Claire Bertelli |
| EPI_ISL_3061086 | EHC MORGES | Laboratory of genomics and metagenomics | Trestan Pillonel, Damien Jacot, Sébastien Aeby, Gilbert Greub, Claire Bertelli |
| EPI_ISL_3061087, EPI_ISL_3061088 | VIDYMED LAUSANNE | Laboratory of genomics and metagenomics | Trestan Pillonel, Damien Jacot, Sébastien Aeby, Gilbert Greub, Claire Bertelli |
| EPI_ISL_3061089 | CHUV | Laboratory of genomics and metagenomics | Trestan Pillonel, Damien Jacot, Sébastien Aeby, Gilbert Greub, Claire Bertelli |
| EPI_ISL_3061090, EPI_ISL_3061091 | VIDYMED LAUSANNE | Laboratory of genomics and metagenomics | Trestan Pillonel, Damien Jacot, Sébastien Aeby, Gilbert Greub, Claire Bertelli |
| EPI_ISL_3061092 | EHC MORGES | Laboratory of genomics and metagenomics | Trestan Pillonel, Damien Jacot, Sébastien Aeby, Gilbert Greub, Claire Bertelli |
| EPI_ISL_3061093 | CHUV | Laboratory of genomics and metagenomics | Trestan Pillonel, Damien Jacot, Sébastien Aeby, Gilbert Greub, Claire Bertelli |
| EPI_ISL_3061094, EPI_ISL_3061095, EPI_ISL_3061096, EPI_ISL_3061097, EPI_ISL_3061098 | VIDYMED LAUSANNE | Laboratory of genomics and metagenomics | Trestan Pillonel, Damien Jacot, Sébastien Aeby, Gilbert Greub, Claire Bertelli |
| EPI_ISL_3061099 | CHUV | Laboratory of genomics and metagenomics | Trestan Pillonel, Damien Jacot, Sébastien Aeby, Gilbert Greub, Claire Bertelli |
| EPI_ISL_3061100, EPI_ISL_3061101 | VIDYMED LAUSANNE | Laboratory of genomics and metagenomics | Trestan Pillonel, Damien Jacot, Sébastien Aeby, Gilbert Greub, Claire Bertelli |
| EPI_ISL_3061102 | CHUV | Laboratory of genomics and metagenomics | Trestan Pillonel, Damien Jacot, Sébastien Aeby, Gilbert Greub, Claire Bertelli |
| EPI_ISL_3061103 | VIDYMED LAUSANNE | Laboratory of genomics and metagenomics | Trestan Pillonel, Damien Jacot, Sébastien Aeby, Gilbert Greub, Claire Bertelli |
| EPI_ISL_3061104, EPI_ISL_3061105, EPI_ISL_3061106, EPI_ISL_3061107, EPI_ISL_3061108, EPI_ISL_3061109, EPI_ISL_3061110, EPI_ISL_3061111, EPI_ISL_3061112, EPI_ISL_3061113, EPI_ISL_3061114, EPI_ISL_3061115, EPI_ISL_3061116 | | | |
| see above | EHNV | Laboratory of genomics and metagenomics | Trestan Pillonel, Damien Jacot, Sébastien Aeby, Gilbert Greub, Claire Bertelli |
| EPI_ISL_3061117 | CHUV | Laboratory of genomics and metagenomics | Trestan Pillonel, Damien Jacot, Sébastien Aeby, Gilbert Greub, Claire Bertelli |
| EPI_ISL_3061118 | VIDYMED LAUSANNE | Laboratory of genomics and metagenomics | Trestan Pillonel, Damien Jacot, Sébastien Aeby, Gilbert Greub, Claire Bertelli |
| EPI_ISL_3061119 | EHNV | Laboratory of genomics and metagenomics | Trestan Pillonel, Damien Jacot, Sébastien Aeby, Gilbert Greub, Claire Bertelli |
| EPI_ISL_3061120, EPI_ISL_3061121 | VIDYMED LAUSANNE | Laboratory of genomics and metagenomics | Trestan Pillonel, Damien Jacot, Sébastien Aeby, Gilbert Greub, Claire Bertelli |
| EPI_ISL_3061122, EPI_ISL_3061123 | CHUV | Laboratory of genomics and metagenomics | Trestan Pillonel, Damien Jacot, Sébastien Aeby, Gilbert Greub, Claire Bertelli |
| EPI_ISL_3061124, EPI_ISL_3061125 | VIDYMED LAUSANNE | Laboratory of genomics and metagenomics | Trestan Pillonel, Damien Jacot, Sébastien Aeby, Gilbert Greub, Claire Bertelli |
| EPI_ISL_3061126 | Kantonsspital Winterthur | Institute of Medical Virology | Daniel Ehrsam, Isabel Stürmer, Catharine Aquino, Joel Wirz, Weihong Qi, Hubert Rehrauer, Verena Kufner, Gabriela Ziltener, Maryam Zaheri, Stefan Schmutz, Annette Audigé, Maria Grünberg, Kevin Steiner, Jon Huder, Cyril Shah, Riccarda Capaul, Guido Bloemberg, Jürg Böni, Michael Huber, Alexandra Trkola |

| | | | |
|---|---|---|---|
| EPI_ISL_3061127 | UniversitätsSpital Zürich | Institute of Medical Virology | Daniel Ehrsam, Isabel Stürmer, Catharine Aquino, Joel Wirz, Weihong Qi, Hubert Rehrauer, Verena Kufner, Gabriela Ziltener, Maryam Zaheri, Stefan Schmutz, Annette Audigé, Maria Grünberg, Kevin Steiner, Jon Huder, Cyril Shah, Riccarda Capaul, Guido Bloemberg, Jürg Böni, Michael Huber, Alexandra Trkola |
| EPI_ISL_3061128 | Kantonsspital Aarau | Institute of Medical Virology | Daniel Ehrsam, Isabel Stürmer, Catharine Aquino, Joel Wirz, Weihong Qi, Hubert Rehrauer, Verena Kufner, Gabriela Ziltener, Maryam Zaheri, Stefan Schmutz, Annette Audigé, Maria Grünberg, Kevin Steiner, Jon Huder, Cyril Shah, Riccarda Capaul, Guido Bloemberg, Jürg Böni, Michael Huber, Alexandra Trkola |
| EPI_ISL_3061130, EPI_ISL_3061131, EPI_ISL_3061132, EPI_ISL_3061133, EPI_ISL_3061134 | UniversitätsSpital Zürich | Institute of Medical Virology | Daniel Ehrsam, Isabel Stürmer, Catharine Aquino, Joel Wirz, Weihong Qi, Hubert Rehrauer, Verena Kufner, Gabriela Ziltener, Maryam Zaheri, Stefan Schmutz, Annette Audigé, Maria Grünberg, Kevin Steiner, Jon Huder, Cyril Shah, Riccarda Capaul, Guido Bloemberg, Jürg Böni, Michael Huber, Alexandra Trkola |
| EPI_ISL_3061135, EPI_ISL_3061136, EPI_ISL_3061137, EPI_ISL_3061139, EPI_ISL_3061140, EPI_ISL_3061141, EPI_ISL_3061142, EPI_ISL_3061143, EPI_ISL_3061144, EPI_ISL_3061145, EPI_ISL_3061146, EPI_ISL_3061147 | | | |
| see above | Kantonsspital Winterthur | Institute of Medical Virology | Daniel Ehrsam, Isabel Stürmer, Catharine Aquino, Joel Wirz, Weihong Qi, Hubert Rehrauer, Verena Kufner, Gabriela Ziltener, Maryam Zaheri, Stefan Schmutz, Annette Audigé, Maria Grünberg, Kevin Steiner, Jon Huder, Cyril Shah, Riccarda Capaul, Guido Bloemberg, Jürg Böni, Michael Huber, Alexandra Trkola |
| EPI_ISL_3061148, EPI_ISL_3061149, EPI_ISL_3061150, EPI_ISL_3061152, EPI_ISL_3061153, EPI_ISL_3061154, EPI_ISL_3061156, EPI_ISL_3061157, EPI_ISL_3061158, EPI_ISL_3061159, EPI_ISL_3061160, EPI_ISL_3061161 | | | |
| see above | Stadtspital Triemli | Institute of Medical Virology | Daniel Ehrsam, Isabel Stürmer, Catharine Aquino, Joel Wirz, Weihong Qi, Hubert Rehrauer, Verena Kufner, Gabriela Ziltener, Maryam Zaheri, Stefan Schmutz, Annette Audigé, Maria Grünberg, Kevin Steiner, Jon Huder, Cyril Shah, Riccarda Capaul, Guido Bloemberg, Jürg Böni, Michael Huber, Alexandra Trkola |
| EPI_ISL_3061162, EPI_ISL_3061163 | UniversitätsSpital Zürich | Institute of Medical Virology | Daniel Ehrsam, Isabel Stürmer, Catharine Aquino, Joel Wirz, Weihong Qi, Hubert Rehrauer, Verena Kufner, Gabriela Ziltener, Maryam Zaheri, Stefan Schmutz, Annette Audigé, Maria Grünberg, Kevin Steiner, Jon Huder, Cyril Shah, Riccarda Capaul, Guido Bloemberg, Jürg Böni, Michael Huber, Alexandra Trkola |
| EPI_ISL_3061164, EPI_ISL_3061165, EPI_ISL_3061166 | Spital Limmattal | Institute of Medical Virology | Daniel Ehrsam, Isabel Stürmer, Catharine Aquino, Joel Wirz, Weihong Qi, Hubert Rehrauer, Verena Kufner, Gabriela Ziltener, Maryam Zaheri, Stefan Schmutz, Annette Audigé, Maria Grünberg, Kevin Steiner, Jon Huder, Cyril Shah, Riccarda Capaul, Guido Bloemberg, Jürg Böni, Michael Huber, Alexandra Trkola |
| EPI_ISL_3061167 | Spital Männedorf AG | Institute of Medical Virology | Daniel Ehrsam, Isabel Stürmer, Catharine Aquino, Joel Wirz, Weihong Qi, Hubert Rehrauer, Verena Kufner, Gabriela Ziltener, Maryam Zaheri, Stefan Schmutz, Annette Audigé, Maria Grünberg, Kevin Steiner, Jon Huder, Cyril Shah, Riccarda Capaul, Guido Bloemberg, Jürg Böni, Michael Huber, Alexandra Trkola |
| EPI_ISL_3061168, EPI_ISL_3061169, EPI_ISL_3061170 | Spital Limmattal | Institute of Medical Virology | Daniel Ehrsam, Isabel Stürmer, Catharine Aquino, Joel Wirz, Weihong Qi, Hubert Rehrauer, Verena Kufner, Gabriela Ziltener, Maryam Zaheri, Stefan Schmutz, Annette Audigé, Maria Grünberg, Kevin Steiner, Jon Huder, Cyril Shah, Riccarda Capaul, Guido Bloemberg, Jürg Böni, Michael Huber, Alexandra Trkola |
| EPI_ISL_3061171 | Verein Lunge Zürich | Institute of Medical Virology | Daniel Ehrsam, Isabel Stürmer, Catharine Aquino, Joel Wirz, Weihong Qi, Hubert Rehrauer, Verena Kufner, Gabriela Ziltener, Maryam Zaheri, Stefan Schmutz, Annette Audigé, Maria Grünberg, Kevin Steiner, Jon Huder, Cyril Shah, Riccarda Capaul, Guido Bloemberg, Jürg Böni, Michael Huber, Alexandra Trkola |
| EPI_ISL_3061172, EPI_ISL_3061173 | UniversitätsSpital Zürich | Institute of Medical Virology | Daniel Ehrsam, Isabel Stürmer, Catharine Aquino, Joel Wirz, Weihong Qi, Hubert Rehrauer, Verena Kufner, Gabriela Ziltener, Maryam Zaheri, Stefan Schmutz, Annette Audigé, Maria Grünberg, Kevin Steiner, Jon Huder, Cyril Shah, Riccarda Capaul, Guido Bloemberg, Jürg Böni, Michael Huber, Alexandra Trkola |
| EPI_ISL_3061174 | Kinderspital Zürich | Institute of Medical Virology | Daniel Ehrsam, Isabel Stürmer, Catharine Aquino, Joel Wirz, Weihong Qi, Hubert Rehrauer, Verena Kufner, Gabriela Ziltener, Maryam Zaheri, Stefan Schmutz, Annette Audigé, Maria Grünberg, Kevin Steiner, Jon Huder, Cyril Shah, Riccarda Capaul, Guido Bloemberg, Jürg Böni, Michael Huber, Alexandra Trkola |
| EPI_ISL_3061175, EPI_ISL_3061176, EPI_ISL_3061177 | UniversitätsSpital Zürich | Institute of Medical Virology | Daniel Ehrsam, Isabel Stürmer, Catharine Aquino, Joel Wirz, Weihong Qi, Hubert Rehrauer, Verena Kufner, Gabriela Ziltener, Maryam Zaheri, Stefan Schmutz, Annette Audigé, Maria Grünberg, Kevin Steiner, Jon Huder, Cyril Shah, Riccarda Capaul, Guido Bloemberg, Jürg Böni, Michael Huber, Alexandra Trkola |
| EPI_ISL_3061178, EPI_ISL_3061179, EPI_ISL_3061180, EPI_ISL_3061181 | Stadtspital Triemli | Institute of Medical Virology | Daniel Ehrsam, Isabel Stürmer, Catharine Aquino, Joel Wirz, Weihong Qi, Hubert Rehrauer, Verena Kufner, Gabriela Ziltener, Maryam Zaheri, Stefan Schmutz, Annette Audigé, Maria Grünberg, Kevin Steiner, Jon Huder, Cyril Shah, Riccarda Capaul, Guido Bloemberg, Jürg Böni, Michael Huber, Alexandra Trkola |
| EPI_ISL_3061182, EPI_ISL_3061183 | UniversitätsSpital Zürich | Institute of Medical Virology | Daniel Ehrsam, Isabel Stürmer, Catharine Aquino, Joel Wirz, Weihong Qi, Hubert Rehrauer, Verena Kufner, Gabriela Ziltener, Maryam Zaheri, Stefan Schmutz, Annette Audigé, Maria Grünberg, Kevin Steiner, Jon Huder, Cyril Shah, Riccarda Capaul, Guido Bloemberg, Jürg Böni, Michael Huber, Alexandra Trkola |
| EPI_ISL_3061190, EPI_ISL_3061196, EPI_ISL_3061197 | ASST FBF Sacco | Laboratory of Clinical Microbiology, Virology and Bioemergencies, ASST Fatebenefratelli Sacco - Sacco Hospital | Valeria Micheli |
| EPI_ISL_3061208, EPI_ISL_3061209, EPI_ISL_3061211, EPI_ISL_3061212, EPI_ISL_3061214, EPI_ISL_3061215, EPI_ISL_3061217, EPI_ISL_3061218, EPI_ISL_3061219, EPI_ISL_3061221, EPI_ISL_3061222, EPI_ISL_3061223, EPI_ISL_3061224, EPI_ISL_3061225, EPI_ISL_3061226, EPI_ISL_3061227, EPI_ISL_3061228, EPI_ISL_3061229, EPI_ISL_3061230, EPI_ISL_3061231, EPI_ISL_3061232, EPI_ISL_3061233, EPI_ISL_3061234, EPI_ISL_3061235, EPI_ISL_3061236, EPI_ISL_3061237, EPI_ISL_3061238, EPI_ISL_3061239, EPI_ISL_3061240, EPI_ISL_3061241, EPI_ISL_3061242 | | | |
| see above | Dept. of Microbiology and Infection Control, Akershus University Hospital HF | Dept. of Microbiology and Infection Control, Akershus University Hospital HF | Hege Vangstein Aamot, Alexander Hesselberg Løvestad |
| EPI_ISL_3061244, EPI_ISL_3061245, EPI_ISL_3061249, EPI_ISL_3061250, EPI_ISL_3061251, EPI_ISL_3061252, EPI_ISL_3061253, EPI_ISL_3061254, EPI_ISL_3061255, EPI_ISL_3061256, EPI_ISL_3061258, EPI_ISL_3061259, EPI_ISL_3061260, EPI_ISL_3061261 | | | |
| see above | IN State Department of Health Laboratory Services | IN State Department of Health Laboratory Services | Cassandra Campion, Jamie Yeadon, Brian Pope, Lixia Liu, Kyle Brownlee, Melissa Hindenlang, Mark Glazier |
| EPI_ISL_3061262 | Nastavni zavod za javno zdravstvo Splitsko- Dalmatinske Županije | Hrvatski zavod za javno zdravstvo | Irena Tabain, Ivana Ferenčak |
| EPI_ISL_3061264, EPI_ISL_3061265, EPI_ISL_3061266, EPI_ISL_3061267, EPI_ISL_3061268 | Zavod za javno zdravstvo Zadarske Županije | Hrvatski zavod za javno zdravstvo | Irena Tabain, Ivana Ferenčak |
| EPI_ISL_3061270, EPI_ISL_3061277 | Klinika za infektivne bolesti "Dr. Fran Mihaljević" | Hrvatski zavod za javno zdravstvo | Irena Tabain, Ivana Ferenčak |
| EPI_ISL_3061278, EPI_ISL_3061280, EPI_ISL_3061281, EPI_ISL_3061284, EPI_ISL_3061285, EPI_ISL_3061286, EPI_ISL_3061287, EPI_ISL_3061288, EPI_ISL_3061289, EPI_ISL_3061290, EPI_ISL_3061292, EPI_ISL_3061293, EPI_ISL_3061294 | | | |
| see above | Zavod za javno zdravstvo Međimurske Županije | Hrvatski zavod za javno zdravstvo | Irena Tabain, Ivana Ferenčak |
| EPI_ISL_3061296, EPI_ISL_3061297, EPI_ISL_3061298, EPI_ISL_3061299, EPI_ISL_3061300, EPI_ISL_3061301, EPI_ISL_3061303, EPI_ISL_3061304, EPI_ISL_3061305, EPI_ISL_3061306, EPI_ISL_3061307, EPI_ISL_3061308, EPI_ISL_3061309, EPI_ISL_3061310, EPI_ISL_3061315, EPI_ISL_3061317 | | | |
| see above | Nastavni zavod za javno zdravstvo Primorsko- Goranske Županije | Hrvatski zavod za javno zdravstvo | Irena Tabain, Ivana Ferenčak |
| EPI_ISL_3061327, EPI_ISL_3061328, EPI_ISL_3061329, EPI_ISL_3061333, EPI_ISL_3061334 | Hrvatski zavod za javno zdravstvo | Hrvatski zavod za javno zdravstvo | Irena Tabain, Ivana Ferenčak |
| EPI_ISL_3061348, EPI_ISL_3061350, EPI_ISL_3061352, EPI_ISL_3061353, EPI_ISL_3061354, EPI_ISL_3061356, EPI_ISL_3061358, EPI_ISL_3061359, EPI_ISL_3061360, EPI_ISL_3061361, EPI_ISL_3061362, EPI_ISL_3061363, EPI_ISL_3061364, EPI_ISL_3061365, EPI_ISL_3061366, EPI_ISL_3061368, EPI_ISL_3061369, EPI_ISL_3061370, EPI_ISL_3061371 | | | |
| see above | National Laboratory for Health, Environment and Food, OMM, Maribor | NLZOH (National Laboratory for Health, Environment and Food) / CISLD (Clinical Institute of Special Laboratory Diagnostics), University Children's Hospital, University Medical Center Ljubljana | Sandra Janezic, Aleksander Mahnic, Maja Rupnik, Tjaša Žohar Čretnik, Alenka Štorman, Nika Gobec, Aleksander Kocuvan, Kaja Tominc, Maša Jarčič, Mojca Cimerman, Nika Volmajer, Andrej Golle / Jernej Kovač, Barbara Jenko Bizjan, Tine Tesovnik, Robert Šket, Katarina Kozmos, Ana Grom, Maruša Debeljak, Marko Pokorn, Tadej Battelino |
| EPI_ISL_3061372, EPI_ISL_3061373, EPI_ISL_3061375, EPI_ISL_3061376, EPI_ISL_3061377, EPI_ISL_3061378, EPI_ISL_3061379, EPI_ISL_3061380, EPI_ISL_3061381, EPI_ISL_3061382, EPI_ISL_3061383, EPI_ISL_3061384, EPI_ISL_3061385, EPI_ISL_3061386, EPI_ISL_3061387, EPI_ISL_3061388, EPI_ISL_3061389, EPI_ISL_3061390, EPI_ISL_3061391, EPI_ISL_3061392 | | | |
| see above | National Laboratory for Health, Environment and Food, OMM, Kranj | NLZOH (National Laboratory for Health, Environment and Food) / CISLD (Clinical Institute of Special Laboratory Diagnostics), University Children's Hospital, University Medical Center Ljubljana | Sandra Janezic, Aleksander Mahnic, Maja Rupnik, Tjaša Žohar Čretnik, Alenka Štorman, Nika Gobec, Aleksander Kocuvan, Kaja Tominc, Maša Jarčič, Mateja Ravnik, Monika Korošec, Marjana Petrevčič / Jernej Kovač, Barbara Jenko Bizjan, Tine Tesovnik, Robert Šket, Katarina Kozmos, Ana Grom, Maruša Debeljak, Marko Pokorn, Tadej Battelino |
| EPI_ISL_3061394, EPI_ISL_3061395, EPI_ISL_3061396 | National Laboratory for Health, Environment and Food, OMM, Celje | NLZOH (National Laboratory for Health, Environment and Food) / CISLD (Clinical Institute of Special Laboratory Diagnostics), University Children's Hospital, University Medical Center Ljubljana | Sandra Janezic, Aleksander Mahnic, Maja Rupnik, Tjaša Žohar Čretnik, Alenka Štorman, Nika Gobec, Aleksander Kocuvan, Kaja Tominc, Maša Jarčič, Mateja Borinc, Daša Kavka / Jernej Kovač, Barbara Jenko Bizjan, Tine Tesovnik, Robert Šket, Katarina Kozmos, Ana Grom, Maruša Debeljak, Marko Pokorn, Tadej Battelino |
| EPI_ISL_3061400, EPI_ISL_3061404, EPI_ISL_3061407 | National Laboratory for Health, Environment and Food, OMM, Koper | NLZOH (National Laboratory for Health, Environment and Food) / CISLD (Clinical Institute of Special Laboratory Diagnostics), University Children's Hospital, University Medical Center Ljubljana | Sandra Janezic, Aleksander Mahnic, Maja Rupnik, Tjaša Žohar Čretnik, Alenka Štorman, Nika Gobec, Aleksander Kocuvan, Kaja Tominc, Maša Jarčič, Gašper Strugar, Tina Cvetković, Mitja Rak / Jernej Kovač, Barbara Jenko Bizjan, Tine Tesovnik, Robert Šket, Katarina Kozmos, Ana Grom, Maruša Debeljak, Marko Pokorn, Tadej Battelino |
| EPI_ISL_3061412, EPI_ISL_3061413, EPI_ISL_3061414, EPI_ISL_3061415, EPI_ISL_3061416, EPI_ISL_3061417, EPI_ISL_3061419, EPI_ISL_3061422, EPI_ISL_3061423, EPI_ISL_3061428, EPI_ISL_3061429, EPI_ISL_3061431, EPI_ISL_3061433, EPI_ISL_3061436, EPI_ISL_3061438, EPI_ISL_3061439, EPI_ISL_3061442, EPI_ISL_3061443, EPI_ISL_3061444, EPI_ISL_3061447, EPI_ISL_3061448, EPI_ISL_3061450, EPI_ISL_3061452 | | | |
| see above | National Laboratory for Health, Environment and Food, OMM, Novo mesto | NLZOH (National Laboratory for Health, Environment and Food) / CISLD (Clinical Institute of Special Laboratory Diagnostics), University Children's Hospital, University Medical Center Ljubljana | Sandra Janezic, Aleksander Mahnic, Maja Rupnik, Tjaša Žohar Čretnik, Alenka Štorman, Nika Gobec, Aleksander Kocuvan, Kaja Tominc, Maša Jarčič, David Cvetko, Tatjana Harlander, Matjaž Retelj / Jernej Kovač, Barbara Jenko Bizjan, Tine Tesovnik, Robert Šket, Katarina Kozmos, Ana Grom, Maruša Debeljak, Marko Pokorn, Tadej Battelino |
| EPI_ISL_3061453 | A.S.L.TO3 | Fondazione del Piemonte per l'Oncologia IRCCS | Antonino Sottile, Silvia Brossa, Paola Marino, Giorgio Giardina |
| EPI_ISL_3061454 | National Laboratory for Health, Environment and Food, OMM, Novo mesto | NLZOH (National Laboratory for Health, Environment and Food) / CISLD (Clinical Institute of Special Laboratory Diagnostics), University Children's Hospital, University Medical Center Ljubljana | Sandra Janezic, Aleksander Mahnic, Maja Rupnik, Tjaša Žohar Čretnik, Alenka Štorman, Nika Gobec, Aleksander Kocuvan, Kaja Tominc, Maša Jarčič, David Cvetko, Tatjana Harlander, Matjaž Retelj / Jernej Kovač, Barbara Jenko Bizjan, Tine Tesovnik, Robert Šket, Katarina Kozmos, Ana Grom, Maruša Debeljak, Marko Pokorn, Tadej Battelino |
| EPI_ISL_3061455 | A.S.L.IVREA | Fondazione del Piemonte per l'Oncologia IRCCS | Antonino Sottile, Silvia Brossa, Paola Marino, Giorgio Giardina |

| | | | |
|---|---|---|---|
| EPI_ISL_3061456, EPI_ISL_3061457, EPI_ISL_3061459 | National Laboratory for Health, Environment and Food, OMM, Novo mesto | NLZOH (National Laboratory for Health, Environment and Food) / CISLD (Clinical Institute of Special Laboratory Diagnostics), University Children's Hospital, University Medical Center Ljubljana | Sandra Janezic, Aleksander Mahnic, Maja Rupnik, Tjaša Žohar Čretnik, Alenka Štorman, Nika Gobec, Aleksander Kocuvan, Kaja Tominc, Maša Jarčič, David Cvetko, Tatjana Harlander, Matjaž Retelj / Jernej Kovač, Barbara Jenko Bizjan, Tine Tesovnik, Robert Šket, Katarina Kozmos, Ana Grom, Maruša Debeljak, Marko Pokorn, Tadej Battelino |
| EPI_ISL_3061460 | A.S.L.IVREA | Fondazione del Piemonte per l'Oncologia IRCCS | Antonino Sottile, Silvia Brossa, Paola Marino, Giorgio Giardina |
| EPI_ISL_3061461 | National Laboratory for Health, Environment and Food, OMM, Novo mesto | NLZOH (National Laboratory for Health, Environment and Food) / CISLD (Clinical Institute of Special Laboratory Diagnostics), University Children's Hospital, University Medical Center Ljubljana | Sandra Janezic, Aleksander Mahnic, Maja Rupnik, Tjaša Žohar Čretnik, Alenka Štorman, Nika Gobec, Aleksander Kocuvan, Kaja Tominc, Maša Jarčič, David Cvetko, Tatjana Harlander, Matjaž Retelj / Jernej Kovač, Barbara Jenko Bizjan, Tine Tesovnik, Robert Šket, Katarina Kozmos, Ana Grom, Maruša Debeljak, Marko Pokorn, Tadej Battelino |
| EPI_ISL_3061462, EPI_ISL_3061466 | A.S.L.IVREA | Fondazione del Piemonte per l'Oncologia IRCCS | Antonino Sottile, Silvia Brossa, Paola Marino, Giorgio Giardina |
| EPI_ISL_3061468 | A.S.L.TO3 | Fondazione del Piemonte per l'Oncologia IRCCS | Antonino Sottile, Silvia Brossa, Paola Marino, Giorgio Giardina |
| EPI_ISL_3061469 | National Laboratory for Health, Environment and Food, OMM, Novo mesto | NLZOH (National Laboratory for Health, Environment and Food) / CISLD (Clinical Institute of Special Laboratory Diagnostics), University Children's Hospital, University Medical Center Ljubljana | Sandra Janezic, Aleksander Mahnic, Maja Rupnik, Tjaša Žohar Čretnik, Alenka Štorman, Nika Gobec, Aleksander Kocuvan, Kaja Tominc, Maša Jarčič, David Cvetko, Tatjana Harlander, Matjaž Retelj / Jernej Kovač, Barbara Jenko Bizjan, Tine Tesovnik, Robert Šket, Katarina Kozmos, Ana Grom, Maruša Debeljak, Marko Pokorn, Tadej Battelino |
| EPI_ISL_3061470, EPI_ISL_3061472 | A.S.L.TO3 | Fondazione del Piemonte per l'Oncologia IRCCS | Antonino Sottile, Silvia Brossa, Paola Marino, Giorgio Giardina |
| EPI_ISL_3061473, EPI_ISL_3061474 | National Laboratory for Health, Environment and Food, OMM, Novo mesto | NLZOH (National Laboratory for Health, Environment and Food) / CISLD (Clinical Institute of Special Laboratory Diagnostics), University Children's Hospital, University Medical Center Ljubljana | Sandra Janezic, Aleksander Mahnic, Maja Rupnik, Tjaša Žohar Čretnik, Alenka Štorman, Nika Gobec, Aleksander Kocuvan, Kaja Tominc, Maša Jarčič, David Cvetko, Tatjana Harlander, Matjaž Retelj / Jernej Kovač, Barbara Jenko Bizjan, Tine Tesovnik, Robert Šket, Katarina Kozmos, Ana Grom, Maruša Debeljak, Marko Pokorn, Tadej Battelino |
| EPI_ISL_3061475 | A.S.L.TO3 | Fondazione del Piemonte per l'Oncologia IRCCS | Antonino Sottile, Silvia Brossa, Paola Marino, Giorgio Giardina |
| EPI_ISL_3061476 | National Laboratory for Health, Environment and Food, OMM, Novo mesto | NLZOH (National Laboratory for Health, Environment and Food) / CISLD (Clinical Institute of Special Laboratory Diagnostics), University Children's Hospital, University Medical Center Ljubljana | Sandra Janezic, Aleksander Mahnic, Maja Rupnik, Tjaša Žohar Čretnik, Alenka Štorman, Nika Gobec, Aleksander Kocuvan, Kaja Tominc, Maša Jarčič, David Cvetko, Tatjana Harlander, Matjaž Retelj / Jernej Kovač, Barbara Jenko Bizjan, Tine Tesovnik, Robert Šket, Katarina Kozmos, Ana Grom, Maruša Debeljak, Marko Pokorn, Tadej Battelino |
| EPI_ISL_3061477, EPI_ISL_3061479 | A.S.L.TO3 | Fondazione del Piemonte per l'Oncologia IRCCS | Antonino Sottile, Silvia Brossa, Paola Marino, Giorgio Giardina |
| EPI_ISL_3061480 | National Laboratory for Health, Environment and Food, OMM, Novo mesto | NLZOH (National Laboratory for Health, Environment and Food) / CISLD (Clinical Institute of Special Laboratory Diagnostics), University Children's Hospital, University Medical Center Ljubljana | Sandra Janezic, Aleksander Mahnic, Maja Rupnik, Tjaša Žohar Čretnik, Alenka Štorman, Nika Gobec, Aleksander Kocuvan, Kaja Tominc, Maša Jarčič, David Cvetko, Tatjana Harlander, Matjaž Retelj / Jernej Kovač, Barbara Jenko Bizjan, Tine Tesovnik, Robert Šket, Katarina Kozmos, Ana Grom, Maruša Debeljak, Marko Pokorn, Tadej Battelino |
| EPI_ISL_3061481 | A.S.L.TO3 | Fondazione del Piemonte per l'Oncologia IRCCS | Antonino Sottile, Silvia Brossa, Paola Marino, Giorgio Giardina |
| EPI_ISL_3061482 | National Laboratory for Health, Environment and Food, OMM, Novo mesto | NLZOH (National Laboratory for Health, Environment and Food) / CISLD (Clinical Institute of Special Laboratory Diagnostics), University Children's Hospital, University Medical Center Ljubljana | Sandra Janezic, Aleksander Mahnic, Maja Rupnik, Tjaša Žohar Čretnik, Alenka Štorman, Nika Gobec, Aleksander Kocuvan, Kaja Tominc, Maša Jarčič, David Cvetko, Tatjana Harlander, Matjaž Retelj / Jernej Kovač, Barbara Jenko Bizjan, Tine Tesovnik, Robert Šket, Katarina Kozmos, Ana Grom, Maruša Debeljak, Marko Pokorn, Tadej Battelino |
| EPI_ISL_3061483 | A.S.L.TO3 | Fondazione del Piemonte per l'Oncologia IRCCS | Antonino Sottile, Silvia Brossa, Paola Marino, Giorgio Giardina |
| EPI_ISL_3061484 | National Laboratory for Health, Environment and Food, OMM, Novo mesto | NLZOH (National Laboratory for Health, Environment and Food) / CISLD (Clinical Institute of Special Laboratory Diagnostics), University Children's Hospital, University Medical Center Ljubljana | Sandra Janezic, Aleksander Mahnic, Maja Rupnik, Tjaša Žohar Čretnik, Alenka Štorman, Nika Gobec, Aleksander Kocuvan, Kaja Tominc, Maša Jarčič, David Cvetko, Tatjana Harlander, Matjaž Retelj / Jernej Kovač, Barbara Jenko Bizjan, Tine Tesovnik, Robert Šket, Katarina Kozmos, Ana Grom, Maruša Debeljak, Marko Pokorn, Tadej Battelino |
| EPI_ISL_3061486, EPI_ISL_3061488, EPI_ISL_3061489 | A.S.L.TO3 | Fondazione del Piemonte per l'Oncologia IRCCS | Antonino Sottile, Silvia Brossa, Paola Marino, Giorgio Giardina |
| EPI_ISL_3061493, EPI_ISL_3061496, EPI_ISL_3061497, EPI_ISL_3061498, EPI_ISL_3061499, EPI_ISL_3061501, EPI_ISL_3061502, EPI_ISL_3061505, EPI_ISL_3061506, EPI_ISL_3061507, EPI_ISL_3061508, EPI_ISL_3061510, EPI_ISL_3061512, EPI_ISL_3061513 | | | |
| see above | National Laboratory for Health, Environment and Food, OMM, Novo mesto | NLZOH (National Laboratory for Health, Environment and Food) / CISLD (Clinical Institute of Special Laboratory Diagnostics), University Children's Hospital, University Medical Center Ljubljana | Sandra Janezic, Aleksander Mahnic, Maja Rupnik, Tjaša Žohar Čretnik, Alenka Štorman, Nika Gobec, Aleksander Kocuvan, Kaja Tominc, Maša Jarčič, David Cvetko, Tatjana Harlander, Matjaž Retelj / Jernej Kovač, Barbara Jenko Bizjan, Tine Tesovnik, Robert Šket, Katarina Kozmos, Ana Grom, Maruša Debeljak, Marko Pokorn, Tadej Battelino |
| EPI_ISL_3061562, EPI_ISL_3061563 | UNC-CH COVID Surveillance Lab | Jeremy Wang | Jeremy Wang, Melissa Miller, Corbin Jones, Amy James Loftis, Amir Barzin, Susan Fiscus |
| EPI_ISL_3061570, EPI_ISL_3061580, EPI_ISL_3061583, EPI_ISL_3061584, EPI_ISL_3061586, EPI_ISL_3061587, EPI_ISL_3061589, EPI_ISL_3061590, EPI_ISL_3061591, EPI_ISL_3061593, EPI_ISL_3061595, EPI_ISL_3061596, EPI_ISL_3061597, EPI_ISL_3061598, EPI_ISL_3061599, EPI_ISL_3061600, EPI_ISL_3061601, EPI_ISL_3061602, EPI_ISL_3061603, EPI_ISL_3061605, EPI_ISL_3061606, EPI_ISL_3061607, EPI_ISL_3061608, EPI_ISL_3061609, EPI_ISL_3061610, EPI_ISL_3061611, EPI_ISL_3061612, EPI_ISL_3061613, EPI_ISL_3061614, EPI_ISL_3061615, EPI_ISL_3061617, EPI_ISL_3061618, EPI_ISL_3061619, EPI_ISL_3061620, EPI_ISL_3061621, EPI_ISL_3061622, EPI_ISL_3061624, EPI_ISL_3061625, EPI_ISL_3061626, EPI_ISL_3061627, EPI_ISL_3061628, EPI_ISL_3061629 | | | |
| see above | Clinical Molecular Microbiology Laboratory, UNC Hospitals | Jeremy Wang | Jeremy Wang, Shawn Hawken, Corbin Jones, Melissa Miller |
| EPI_ISL_3061631, EPI_ISL_3061633, EPI_ISL_3061634 | UNC-CH COVID Surveillance Lab | Jeremy Wang | Jeremy Wang, Melissa Miller, Corbin Jones, Amy James Loftis, Amir Barzin, Susan Fiscus |
| EPI_ISL_3061683, EPI_ISL_3061684, EPI_ISL_3061687, EPI_ISL_3061688, EPI_ISL_3061689, EPI_ISL_3061690, EPI_ISL_3061691, EPI_ISL_3061692, EPI_ISL_3061693, EPI_ISL_3061694, EPI_ISL_3061695, EPI_ISL_3061696, EPI_ISL_3061697, EPI_ISL_3061698, EPI_ISL_3061699, EPI_ISL_3061700, EPI_ISL_3061701 | | | |
| see above | Zavod za javno zdravstvo Varaždinske županije | Hrvatski zavod za javno zdravstvo | Irena Tabain, Ivana Ferenčak |
| EPI_ISL_3061702, EPI_ISL_3061703, EPI_ISL_3061704, EPI_ISL_3061706, EPI_ISL_3061707, EPI_ISL_3061708, EPI_ISL_3061709, EPI_ISL_3061710, EPI_ISL_3061711, EPI_ISL_3061712, EPI_ISL_3061713 | | | |
| see above | Klinika za infektivne bolesti "Dr. Fran Mihaljević" | Hrvatski zavod za javno zdravstvo | Irena Tabain, Ivana Ferenčak |
| EPI_ISL_3061714, EPI_ISL_3061715, EPI_ISL_3061717, EPI_ISL_3061720, EPI_ISL_3061722 | Zavod za javno zdravstvo Varaždinske županije | Hrvatski zavod za javno zdravstvo | Irena Tabain, Ivana Ferenčak |
| EPI_ISL_3061723, EPI_ISL_3061725, EPI_ISL_3061726, EPI_ISL_3061727, EPI_ISL_3061728, EPI_ISL_3061729 | Nastavni zavod za javno zdravstvo Splitsko- Dalmatinske Županije | Hrvatski zavod za javno zdravstvo | Irena Tabain, Ivana Ferenčak |
| EPI_ISL_3061730, EPI_ISL_3061731, EPI_ISL_3061732, EPI_ISL_3061733, EPI_ISL_3061734, EPI_ISL_3061735, EPI_ISL_3061736, EPI_ISL_3061737, EPI_ISL_3061738 | Zavod za javno zdravstvo Međimurske Županije | Hrvatski zavod za javno zdravstvo | Irena Tabain, Ivana Ferenčak |
| EPI_ISL_3061739, EPI_ISL_3061740, EPI_ISL_3061741, EPI_ISL_3061742, EPI_ISL_3061743, EPI_ISL_3061744 | Zavod za javno zdravstvo Zadarske Županije | Hrvatski zavod za javno zdravstvo | Irena Tabain, Ivana Ferenčak |
| EPI_ISL_3061748 | Zavod za javno zdravstvo Istarske Županije | Hrvatski zavod za javno zdravstvo | Irena Tabain, Ivana Ferenčak |
| EPI_ISL_3061749 | Zavod za javno zdravstvo Ličko-senjske županije | Hrvatski zavod za javno zdravstvo | Irena Tabain, Ivana Ferenčak |
| EPI_ISL_3061751 | Zavod za javno zdravstvo Karlovačke Županije | Hrvatski zavod za javno zdravstvo | Irena Tabain, Ivana Ferenčak |
| EPI_ISL_3061770, EPI_ISL_3061771, EPI_ISL_3061772, EPI_ISL_3061773, EPI_ISL_3061774, EPI_ISL_3061775, EPI_ISL_3061776, EPI_ISL_3061777, EPI_ISL_3061778, EPI_ISL_3061779, EPI_ISL_3061780, EPI_ISL_3061781, EPI_ISL_3061782, EPI_ISL_3061783, EPI_ISL_3061784, EPI_ISL_3061785 | | | |
| see above | Zavod za javno zdravstvo Šibensko- Kninske Županije | Hrvatski zavod za javno zdravstvo | Irena Tabain, Ivana Ferenčak |
| EPI_ISL_3061794, EPI_ISL_3061795, EPI_ISL_3061796 | Zavod za javno zdravstvo Zagrebačke Županije | Hrvatski zavod za javno zdravstvo | Irena Tabain, Ivana Ferenčak |
| EPI_ISL_3061797 | Hrvatski zavod za javno zdravstvo | Hrvatski zavod za javno zdravstvo | Irena Tabain, Ivana Ferenčak |
| EPI_ISL_3061798, EPI_ISL_3061799, EPI_ISL_3061800, EPI_ISL_3061801, EPI_ISL_3061802, EPI_ISL_3061803, EPI_ISL_3061804, EPI_ISL_3061806, EPI_ISL_3061807, EPI_ISL_3061808, EPI_ISL_3061809, EPI_ISL_3061810, EPI_ISL_3061811, EPI_ISL_3061812, EPI_ISL_3061813, EPI_ISL_3061814, EPI_ISL_3061815, EPI_ISL_3061816, EPI_ISL_3061817, EPI_ISL_3061818, EPI_ISL_3061819, EPI_ISL_3061820, EPI_ISL_3061821, EPI_ISL_3061822, EPI_ISL_3061823, EPI_ISL_3061824, EPI_ISL_3061825, EPI_ISL_3061826, EPI_ISL_3061828, EPI_ISL_3061829, EPI_ISL_3061830 | | | |
| see above | Nastavni zavod za javno zdravstvo Primorsko- Goranske | Hrvatski zavod za javno zdravstvo | Irena Tabain, Ivana Ferenčak |

| | | | |
|---|---|---|---|
| | Županije | | |
| EPI_ISL_3061831, EPI_ISL_3061832, EPI_ISL_3061833, EPI_ISL_3061834, EPI_ISL_3061835, EPI_ISL_3061836, EPI_ISL_3061837, EPI_ISL_3061838, EPI_ISL_3061839, EPI_ISL_3061840, EPI_ISL_3061841, EPI_ISL_3061842, EPI_ISL_3061843, EPI_ISL_3061844, EPI_ISL_3061845, EPI_ISL_3061846, EPI_ISL_3061847, EPI_ISL_3061848, EPI_ISL_3061849, EPI_ISL_3061850, EPI_ISL_3061851, EPI_ISL_3061852, EPI_ISL_3061853 | | | |
| see above | National Platform bis UMONS/Jolimont | National Platform bis UMONS/Jolimont | François Dufrasne, Guillaume Bayon-Vicente, Florian Juszczak, Laetitia Gheysen, Eric Tarantino, Gautier Detry, Ruddy Wattiez |
| EPI_ISL_3061857 | Labo Analyses Med | National Reference Center for Viruses of Respiratory Infections, Institut Pasteur, Paris | Marion Barbet, Sylvie Behillil, Méline Bizard, Angela Brisebarre, Camille Capel, Vincent Enouf, Louise Lefrançois, Frédéric Lemoine, Christophe Malabat, Corinne Maufrais, Hub de Bioinformatique et Biostatistique, Etienne Simon-Lorière, Maud Vanpeene, Sylvie Van der Werf ,Alexandra Ducancelle |
| EPI_ISL_3061865 | CH VALENCIENNES | CHU Lille - Laboratoire de Virologie | AIT YAHYA Emilie, ALIDJINOU Enagnon Kazali, BOCKET Laurence, CREPIN Michel, DEMAY Christophe, ENGELMANN Ilka, GEFFROY Sandrine, GUIGON Aurélie, LAMBERT Valérie, LAZREK Mouna, NOBILLIAUX Florian, PREVOST Brigitte, THUILLIER Caroline, TINEZ Claire ,TCHANTCHOU NJOSSE YANICK |
| EPI_ISL_3061866 | CHU LILLE | CHU Lille - Laboratoire de Virologie | AIT YAHYA Emilie, ALIDJINOU Enagnon Kazali, BOCKET Laurence, CREPIN Michel, DEMAY Christophe, ENGELMANN Ilka, GEFFROY Sandrine, GUIGON Aurélie, LAMBERT Valérie, LAZREK Mouna, NOBILLIAUX Florian, PREVOST Brigitte, THUILLIER Caroline, TINEZ Claire ,TCHANTCHOU NJOSSE YANICK |
| EPI_ISL_3061867, EPI_ISL_3061868, EPI_ISL_3061870, EPI_ISL_3061871 | CT Department of Public Health | CT Department of Public Health | Claire_Pearson,Tu_N_Nguyen |
| EPI_ISL_3061872, EPI_ISL_3061873, EPI_ISL_3061874, EPI_ISL_3061875, EPI_ISL_3061876 | Laboratorio Biologia Molecolare Sars Cov2 – UOC Laboratorio Analisi – Servizio Medicina di Laboratorio, Ospedale "San Francesco" - ATS-ASSL Nuoro | Laboratorio specialistico UOC Ematologia - Ospedale "San Francesco" - ATS-ASSL Nuoro | Rosanna Asproni, Antonio Manai,Giovanna Piras,Maura Fiamma, Maria Cristina Garau,Giuseppe Mameli |
| EPI_ISL_3061904, EPI_ISL_3061905, EPI_ISL_3061906, EPI_ISL_3061907, EPI_ISL_3061908 | UW Virology Lab | UW Virology Lab | Pavitra Roychoudhury, Hong Xie, Lasata Shrestha, Shah Mohamed Bakhash, Tien V. Nguyen, Noah R. Baker, Sean Ellis, Meei-Li Huang, Keith R Jerome, Alexander Greninger |
| EPI_ISL_3061912, EPI_ISL_3061913, EPI_ISL_3061914, EPI_ISL_3061915, EPI_ISL_3061916, EPI_ISL_3061917, EPI_ISL_3061918, EPI_ISL_3061919, EPI_ISL_3061920, EPI_ISL_3061921, EPI_ISL_3061922, EPI_ISL_3061923, EPI_ISL_3061924, EPI_ISL_3061925 | | | |
| see above | Kansas Health and Environmental Lab | Kansas Health and Environmental Lab | Katherine Wiggins, Mike Grose, Jonathan Barnell, Ben Olsen, and Phil Adam |
| EPI_ISL_3061926, EPI_ISL_3061927 | Wichita State University - Molecular Diagnostics Lab | Kansas Health and Environmental Lab | Katherine Wiggins, Mike Grose, Jonathan Barnell, Ben Olsen, and Phil Adam |
| EPI_ISL_3061930 | Oregon State Public Health Laboratory | Oregon State Public Health Laboratory | Rafia Razzaque, Eugene Yeboah, Vanda Makris, Laura Tsaknaridis, John Fontana and Shane Sevey |
| EPI_ISL_3061932, EPI_ISL_3061933, EPI_ISL_3061934 | UCSC Genomics Institute | UCSC Genomics Institute | Mark Akeson, Ikenna Anigbogu, Eric Beraut, Isabel Bjork, Molly Cassatt-Johnstone, Terren Chang, Russell Corbett-Detig, Namrita Dhillon, Maximilian Haeussler, David Haussler, Angie Hinrichs, Miten Jain, Joshua Kapp, A. Marm Kilpatrick, Jakob McBroome, Hugh Olsen, Jeremy Sanford, Beth Shapiro, Michael Stone, Bryan Thornlow, Yatish Turakhia, Ciara Wanket |
| EPI_ISL_3061938, EPI_ISL_3061939, EPI_ISL_3061940, EPI_ISL_3061941, EPI_ISL_3061942, EPI_ISL_3061944, EPI_ISL_3061945, EPI_ISL_3061946, EPI_ISL_3061948, EPI_ISL_3061949, EPI_ISL_3061950, EPI_ISL_3061952, EPI_ISL_3061953 | | | |
| see above | Plateforme de testing Namuroise | Plateforme de testing Namuroise | Lesly Nyinkeu Kemamen; Demars Aurore ; Maschietto Céline ; Nobis Chloé ; Otto Gaetan ; Denis Olivier ; Degosserie Jonathan ; Mullier François |
| EPI_ISL_3061955, EPI_ISL_3061956 | Plateforme de testing Namuroise | Arlon | Lesly Nyinkeu Kemamen; Demars Aurore ; Maschietto Céline ; Nobis Chloé ; Otto Gaetan ; Denis Olivier ; Degosserie Jonathan ; Mullier François |
| EPI_ISL_3061959, EPI_ISL_3061960, EPI_ISL_3061961, EPI_ISL_3061964, EPI_ISL_3061970, EPI_ISL_3061973, EPI_ISL_3061974, EPI_ISL_3061975, EPI_ISL_3061977, EPI_ISL_3061978, EPI_ISL_3061980, EPI_ISL_3061981, EPI_ISL_3061982, EPI_ISL_3061985, EPI_ISL_3061986, EPI_ISL_3061987, EPI_ISL_3061988 | | | |
| see above | Plateforme de testing Namuroise | Plateforme de testing Namuroise | Lesly Nyinkeu Kemamen; Demars Aurore ; Maschietto Céline ; Nobis Chloé ; Otto Gaetan ; Denis Olivier ; Degosserie Jonathan ; Mullier François |
| EPI_ISL_3061990, EPI_ISL_3061991, EPI_ISL_3061992, EPI_ISL_3061993, EPI_ISL_3061994, EPI_ISL_3061995, EPI_ISL_3061996, EPI_ISL_3061997, EPI_ISL_3061998, EPI_ISL_3061999, EPI_ISL_3062000, EPI_ISL_3062001, EPI_ISL_3062002, EPI_ISL_3062003, EPI_ISL_3062004, EPI_ISL_3062005, EPI_ISL_3062006, EPI_ISL_3062007, EPI_ISL_3062008, EPI_ISL_3062009, EPI_ISL_3062010, EPI_ISL_3062011, EPI_ISL_3062012, EPI_ISL_3062013, EPI_ISL_3062014, EPI_ISL_3062015, EPI_ISL_3062016, EPI_ISL_3062017, EPI_ISL_3062018, EPI_ISL_3062019, EPI_ISL_3062020 | | | |
| see above | Nebraska Public Health Laboratory | NPHL COVID-19 Response Team | NPHL COVID-19 Response Team |
| EPI_ISL_3062024, EPI_ISL_3062025, EPI_ISL_3062026, EPI_ISL_3062027, EPI_ISL_3062029, EPI_ISL_3062030, EPI_ISL_3062031, EPI_ISL_3062032, EPI_ISL_3062034, EPI_ISL_3062035, EPI_ISL_3062036, EPI_ISL_3062037, EPI_ISL_3062038, EPI_ISL_3062040, EPI_ISL_3062041, EPI_ISL_3062042, EPI_ISL_3062043, EPI_ISL_3062044, EPI_ISL_3062045, EPI_ISL_3062046, EPI_ISL_3062047, EPI_ISL_3062048, EPI_ISL_3062049, EPI_ISL_3062050, EPI_ISL_3062051, EPI_ISL_3062053, EPI_ISL_3062054, EPI_ISL_3062055, EPI_ISL_3062056, EPI_ISL_3062057, EPI_ISL_3062059, EPI_ISL_3062060, EPI_ISL_3062061, EPI_ISL_3062062, EPI_ISL_3062063, EPI_ISL_3062064, EPI_ISL_3062065, EPI_ISL_3062066, EPI_ISL_3062067, EPI_ISL_3062068, EPI_ISL_3062069, EPI_ISL_3062071, EPI_ISL_3062072, EPI_ISL_3062073, EPI_ISL_3062075, EPI_ISL_3062076, EPI_ISL_3062077, EPI_ISL_3062078, EPI_ISL_3062079, EPI_ISL_3062080, EPI_ISL_3062081, EPI_ISL_3062082, EPI_ISL_3062083, EPI_ISL_3062084, EPI_ISL_3062085 | | | |
| see above | ASL Napoli 1 Centro | AMES Centro Polidiagnostico Strumentale S.r.l. | Giovanni Savarese, Raffaella Ruggiero, Eloisa Evangelista, Antonella Di Carlo, Luisa Circelli, Luigi D'Amore, Roberto Sirica,Maurizio D'Amora Antonio Fico |
| EPI_ISL_3062106, EPI_ISL_3062107, EPI_ISL_3062109, EPI_ISL_3062110, EPI_ISL_3062111, EPI_ISL_3062112, EPI_ISL_3062113, EPI_ISL_3062114, EPI_ISL_3062115, EPI_ISL_3062116, EPI_ISL_3062117, EPI_ISL_3062118, EPI_ISL_3062119, EPI_ISL_3062120, EPI_ISL_3062121, EPI_ISL_3062122, EPI_ISL_3062124, EPI_ISL_3062125, EPI_ISL_3062126, EPI_ISL_3062127, EPI_ISL_3062128, EPI_ISL_3062129, EPI_ISL_3062130, EPI_ISL_3062131, EPI_ISL_3062132, EPI_ISL_3062133, EPI_ISL_3062134, EPI_ISL_3062135, EPI_ISL_3062136 | | | |
| see above | Wichita State University - Molecular Diagnostics Lab | Kansas Health and Environmental Lab | Katherine Wiggins, Mike Grose, Jonathan Barnell, Ben Olsen, and Phil Adam |
| EPI_ISL_3062146, EPI_ISL_3062148, EPI_ISL_3062149, EPI_ISL_3062150, EPI_ISL_3062151, EPI_ISL_3062152, EPI_ISL_3062153, EPI_ISL_3062154, EPI_ISL_3062155, EPI_ISL_3062318, EPI_ISL_3062319, EPI_ISL_3062320, EPI_ISL_3062321, EPI_ISL_3062322 | | | |
| see above | North Dakota Department of Health, Public Health Laboratory | North Dakota Department of Health, Public Health Laboratory | Lisa Wingerter |
| EPI_ISL_3062324, EPI_ISL_3062325 | St James's Hospital, Dublin 8, Ireland. | ELDA biotech | Deirdre M. Broderick, Peter R. Flanagan, Brendan Crowley |
| EPI_ISL_3062326, EPI_ISL_3062328 | CAP Vic Nord | Banc de Sang i Teixits | Francisco Vidal, Irene Corrales, Nina Borràs, Carlos Hobeich, Natália Comes, Lorena Ramírez, Noemí Gonzalez, Silvia Sauleda, Maria Glòria Soria |
| EPI_ISL_3062329 | Fundacio Althaia-Manresa | Banc de Sang i Teixits | Francisco Vidal, Irene Corrales, Nina Borràs, Carlos Hobeich, Natália Comes, Lorena Ramírez, Noemí Gonzalez, Silvia Sauleda, Maria Glòria Soria |
| EPI_ISL_3062331 | CAP Folgueroles | Banc de Sang i Teixits | Francisco Vidal, Irene Corrales, Nina Borràs, Carlos Hobeich, Natália Comes, Lorena Ramírez, Noemí Gonzalez, Silvia Sauleda, Maria Glòria Soria |
| EPI_ISL_3062332 | CAP Manlleu | Banc de Sang i Teixits | Francisco Vidal, Irene Corrales, Nina Borràs, Carlos Hobeich, Natália Comes, Lorena Ramírez, Noemí Gonzalez, Silvia Sauleda, Maria Glòria Soria |
| EPI_ISL_3062333 | Fundacio Althaia-Manresa | Banc de Sang i Teixits | Francisco Vidal, Irene Corrales, Nina Borràs, Carlos Hobeich, Natália Comes, Lorena Ramírez, Noemí Gonzalez, Silvia Sauleda, Maria Glòria Soria |
| EPI_ISL_3062334 | CAP Roda de Ter | Banc de Sang i Teixits | Francisco Vidal, Irene Corrales, Nina Borràs, Carlos Hobeich, Natália Comes, Lorena Ramírez, Noemí Gonzalez, Silvia Sauleda, Maria Glòria Soria |
| EPI_ISL_3062336 | CAP Tona | Banc de Sang i Teixits | Francisco Vidal, Irene Corrales, Nina Borràs, Carlos Hobeich, Natália Comes, Lorena Ramírez, Noemí Gonzalez, Silvia Sauleda, Maria Glòria Soria |
| EPI_ISL_3062337 | CAP El Remei | Banc de Sang i Teixits | Francisco Vidal, Irene Corrales, Nina Borràs, Carlos Hobeich, Natália Comes, Lorena Ramírez, Noemí Gonzalez, Silvia Sauleda, Maria Glòria Soria |
| EPI_ISL_3062340 | CAP Roda de Ter | Banc de Sang i Teixits | Francisco Vidal, Irene Corrales, Nina Borràs, Carlos Hobeich, Natália Comes, Lorena Ramírez, Noemí Gonzalez, Silvia Sauleda, Maria Glòria Soria |
| EPI_ISL_3062342 | CAP Vic Nord | Banc de Sang i Teixits | Francisco Vidal, Irene Corrales, Nina Borràs, Carlos Hobeich, Natália Comes, Lorena Ramírez, Noemí Gonzalez, Silvia Sauleda, Maria Glòria Soria |
| EPI_ISL_3062343, EPI_ISL_3062344, EPI_ISL_3062345, EPI_ISL_3062346, EPI_ISL_3062347, EPI_ISL_3062348 | Laboratory of Molecular Biology and Cancer Immunology, Faculty of Sciences, Lebanese University | Microbial Pathogenomics Lab - LAU | Sima tokajian, Georgi Merhi, Jad Koweyes, Bassam Badran, Fadi Abdel Sater, Rawan Makki, Tatiana Tohmeh, Nada Ghosn, Hamad Hassan |
| EPI_ISL_3062349, EPI_ISL_3062350, EPI_ISL_3062351 | Laboratory of Molecular Biology and Cancer Immunology, Faculty of Sciences, Lebanese University | Microbial Pathogenomics Lab - LAU | Sima tokajian, Georgi Merhi, Jad Koweyes, Bassam Badran, Fadi Abdel Sater, Rawan Makki, Hamad Hassan |
| EPI_ISL_3062352, EPI_ISL_3062353 | Wichita State University - Molecular Diagnostics Lab | Kansas Health and Environmental Lab | Mike Grose, Jonathan Barnell, Ben Olsen, and Phil Adam |
| EPI_ISL_3062354, EPI_ISL_3062355, EPI_ISL_3062356, EPI_ISL_3062357, EPI_ISL_3062358, EPI_ISL_3062359, EPI_ISL_3062360, EPI_ISL_3062361, EPI_ISL_3062362, EPI_ISL_3062363, EPI_ISL_3062364, EPI_ISL_3062365, EPI_ISL_3062366, EPI_ISL_3062367, EPI_ISL_3062368, EPI_ISL_3062369, EPI_ISL_3062370, EPI_ISL_3062371, EPI_ISL_3062372 | | | |
| see above | Clinical Reference Laboratory | Kansas Health and Environmental Lab | Mike Grose, Jonathan Barnell, Ben Olsen, and Phil Adam |
| EPI_ISL_3062373 | Department of Virology, Istituto Zooprofilattico Sperimentale del Lazio e della Toscana (IZSLT) | Department of General Diagnostics; Department of Virology; Istituto Zooprofilattico Sperimentale del Lazio e della Toscana (IZSLT) | Patricia Alba, Giuseppe Manna, Elena L. Diaconu, Fabiola Feltrin, Raffaela Conti, Teresa Scicluna, Virginia Carfora, Antonella Cersini, Alessia Franco, Antonio Battisti. |
| EPI_ISL_3062377 | A.S.L. ROMA 1 - SISP | Department of General Diagnostics; Department of Virology; Istituto Zooprofilattico Sperimentale del Lazio e della Toscana (IZSLT) | Patricia Alba, Giuseppe Manna, Elena L. Diaconu, Fabiola Feltrin, Raffaela Conti, Teresa Scicluna, Virginia Carfora, Antonella Cersini, Alessia Franco, Antonio Battisti. |
| EPI_ISL_3062379 | Sant'Eugenio/CTO ASL Roma 2 | Department of General Diagnostics; Department of Virology; Istituto Zooprofilattico Sperimentale del Lazio e della Toscana (IZSLT) | Patricia Alba, Giuseppe Manna, Elena L. Diaconu, Fabiola Feltrin, Raffaela Conti, Teresa Scicluna, Virginia Carfora, Antonella Cersini, Alessia Franco, Antonio Battisti. |
| EPI_ISL_3062380 | A.S.L. ROMA 2 - SISP | Department of General Diagnostics; Department of Virology; Istituto Zooprofilattico Sperimentale del Lazio e della Toscana (IZSLT) | Patricia Alba, Giuseppe Manna, Elena L. Diaconu, Fabiola Feltrin, Raffaela Conti, Teresa Scicluna, Virginia Carfora, Antonella Cersini, Alessia Franco, Antonio Battisti. |
| EPI_ISL_3062382 | Sant'Eugenio/CTO ASL Roma 2 | Department of General Diagnostics; Department of Virology; Istituto Zooprofilattico Sperimentale del Lazio e della Toscana (IZSLT) | Patricia Alba, Giuseppe Manna, Elena L. Diaconu, Fabiola Feltrin, Raffaela Conti, Teresa Scicluna, Virginia Carfora, Antonella Cersini, Alessia Franco, Antonio Battisti. |
| EPI_ISL_3062383, EPI_ISL_3062384, EPI_ISL_3062385, | Azienda Ospedaliera San Camillo Forlanini | Department of General Diagnostics; Department of Virology; Istituto Zooprofilattico Sperimentale del Lazio e della Toscana (IZSLT) | Patricia Alba, Giuseppe Manna, Elena L. Diaconu, Fabiola Feltrin, Raffaela Conti, Teresa Scicluna, Virginia Carfora, Antonella Cersini, Alessia Franco, Antonio Battisti. |

| | | | |
|---|---|---|---|
| EPI_ISL_3062386, EPI_ISL_3062388 | | | |
| EPI_ISL_3062389 | Ospedale di Genzano - ASL RM 6 | Department of General Diagnostics; Department of Virology; Istituto Zooprofilattico Sperimentale del Lazio e della Toscana (IZSLT) | Patricia Alba, Giuseppe Manna, Elena L. Diaconu, Fabiola Feltrin, Raffaela Conti, Teresa Scicluna, Virginia Carfora, Antonella Cersini, Alessia Franco, Antonio Battisti. |
| EPI_ISL_3062390 | University of Rome Tor Vergata: Departm Experim Medicine Chair of Virology | Department of General Diagnostics; Department of Virology; Istituto Zooprofilattico Sperimentale del Lazio e della Toscana (IZSLT) | Patricia Alba, Giuseppe Manna, Elena L. Diaconu, Fabiola Feltrin, Raffaela Conti, Teresa Scicluna, Virginia Carfora, Antonella Cersini, Alessia Franco, Antonio Battisti. |
| EPI_ISL_3062394 | AOU Policlinico Umberto I; Sapienza Università di Roma | Department of General Diagnostics; Department of Virology; Istituto Zooprofilattico Sperimentale del Lazio e della Toscana (IZSLT) | Patricia Alba, Giuseppe Manna, Elena L. Diaconu, Fabiola Feltrin, Raffaela Conti, Teresa Scicluna, Virginia Carfora, Antonella Cersini, Alessia Franco, Antonio Battisti. |
| EPI_ISL_3062395, EPI_ISL_3062399, EPI_ISL_3062401, EPI_ISL_3062402, EPI_ISL_3062403 | Ospedale Sandro Pertini ASL Roma2 | Department of General Diagnostics; Department of Virology; Istituto Zooprofilattico Sperimentale del Lazio e della Toscana (IZSLT) | Patricia Alba, Giuseppe Manna, Elena L. Diaconu, Fabiola Feltrin, Raffaela Conti, Teresa Scicluna, Virginia Carfora, Antonella Cersini, Alessia Franco, Antonio Battisti. |
| EPI_ISL_3062404, EPI_ISL_3062405, EPI_ISL_3062406, EPI_ISL_3062414, EPI_ISL_3062415, EPI_ISL_3062416, EPI_ISL_3062417 | Ospedale San Filippo Neri | Department of General Diagnostics; Department of Virology; Istituto Zooprofilattico Sperimentale del Lazio e della Toscana (IZSLT) | Patricia Alba, Giuseppe Manna, Elena L. Diaconu, Fabiola Feltrin, Raffaela Conti, Teresa Scicluna, Virginia Carfora, Antonella Cersini, Alessia Franco, Antonio Battisti. |
| EPI_ISL_3062422 | Ospedale S.M. Goretti di Latina - ASL Latina presidio ospedaliero Nord | Department of General Diagnostics; Department of Virology; Istituto Zooprofilattico Sperimentale del Lazio e della Toscana (IZSLT) | Patricia Alba, Giuseppe Manna, Elena L. Diaconu, Fabiola Feltrin, Raffaela Conti, Teresa Scicluna, Virginia Carfora, Antonella Cersini, Alessia Franco, Antonio Battisti. |
| EPI_ISL_3062426 | Policlinico Universitario Campus Bio-Medico | Department of General Diagnostics; Department of Virology; Istituto Zooprofilattico Sperimentale del Lazio e della Toscana (IZSLT) | Patricia Alba, Giuseppe Manna, Elena L. Diaconu, Fabiola Feltrin, Raffaela Conti, Teresa Scicluna, Virginia Carfora, Antonella Cersini, Alessia Franco, Antonio Battisti. |
| EPI_ISL_3062427 | Ospedale San Filippo Neri | Department of General Diagnostics; Department of Virology; Istituto Zooprofilattico Sperimentale del Lazio e della Toscana (IZSLT) | Patricia Alba, Giuseppe Manna, Elena L. Diaconu, Fabiola Feltrin, Raffaela Conti, Teresa Scicluna, Virginia Carfora, Antonella Cersini, Alessia Franco, Antonio Battisti. |
| EPI_ISL_3062428 | DOHMH PHL | New York City Public Health Laboratory | Jade Wang, et al. |
| EPI_ISL_3062429 | DOHMH Crown Heights | New York City Public Health Laboratory | Jade Wang, et al. |
| EPI_ISL_3062430 | DOHMH Corona | New York City Public Health Laboratory | Jade Wang, et al. |
| EPI_ISL_3062432 | DOHMH Riverside | New York City Public Health Laboratory | Jade Wang, et al. |
| EPI_ISL_3062433 | DOHMH Corona | New York City Public Health Laboratory | Jade Wang, et al. |
| EPI_ISL_3062434, EPI_ISL_3062435 | DOHMH Chelsea | New York City Public Health Laboratory | Jade Wang, et al. |
| EPI_ISL_3062436, EPI_ISL_3062437, EPI_ISL_3062438, EPI_ISL_3062439 | DOHMH PHL | New York City Public Health Laboratory | Jade Wang, et al. |
| EPI_ISL_3062440, EPI_ISL_3062441 | DOHMH Crown Heights | New York City Public Health Laboratory | Jade Wang, et al. |
| EPI_ISL_3062442 | DOHMH Riverside | New York City Public Health Laboratory | Jade Wang, et al. |
| EPI_ISL_3062443 | DOHMH PHL | New York City Public Health Laboratory | Jade Wang, et al. |
| EPI_ISL_3062445 | DOHMH Jamaica | New York City Public Health Laboratory | Jade Wang, et al. |
| EPI_ISL_3062447 | DOHMH Crown Heights | New York City Public Health Laboratory | Jade Wang, et al. |
| EPI_ISL_3062448 | DOHMH Corona | New York City Public Health Laboratory | Jade Wang, et al. |
| EPI_ISL_3062449 | DOHMH Chelsea | New York City Public Health Laboratory | Jade Wang, et al. |
| EPI_ISL_3062450 | DOHMH Crown Heights | New York City Public Health Laboratory | Jade Wang, et al. |
| EPI_ISL_3062451, EPI_ISL_3062452 | DOHMH Morrisania | New York City Public Health Laboratory | Jade Wang, et al. |
| EPI_ISL_3062453 | DOHMH Fort Greene | New York City Public Health Laboratory | Jade Wang, et al. |
| EPI_ISL_3062455, EPI_ISL_3062456, EPI_ISL_3062457 | Quest Diagnostics | New York City Public Health Laboratory | Jade Wang, et al. |
| EPI_ISL_3062459 | DOHMH Chelsea | New York City Public Health Laboratory | Jade Wang, et al. |
| EPI_ISL_3062460 | DOHMH Crown Heights | New York City Public Health Laboratory | Jade Wang, et al. |
| EPI_ISL_3062461 | DOHMH Chelsea | New York City Public Health Laboratory | Jade Wang, et al. |
| EPI_ISL_3062462 | DOHMH Corona | New York City Public Health Laboratory | Jade Wang, et al. |
| EPI_ISL_3062463 | Department of Homeless Services | New York City Public Health Laboratory | Jade Wang, et al. |
| EPI_ISL_3062466, EPI_ISL_3062467, EPI_ISL_3062468, EPI_ISL_3062469, EPI_ISL_3062470, EPI_ISL_3062471, EPI_ISL_3062472, EPI_ISL_3062473, EPI_ISL_3062474, EPI_ISL_3062476, EPI_ISL_3062477, EPI_ISL_3062478, EPI_ISL_3062479, EPI_ISL_3062480, EPI_ISL_3062481, EPI_ISL_3062482, EPI_ISL_3062483, EPI_ISL_3062484, EPI_ISL_3062485 | | | |
| see above | Wyoming Public Health Laboratory | Wyoming Public Health Laboratory | Jim Mildenberger, Taylor Fearing, Channing Weber, Ashley Norberg, Chayse Rowley, Marley Goetz, Brian Dominguez, Elliot Thomasson, Sam Britz, Cari Sloma, Robert Petit, and Rob Christensen |
| EPI_ISL_3062567, EPI_ISL_3062570, EPI_ISL_3062571, EPI_ISL_3062572, EPI_ISL_3062573, EPI_ISL_3062574, EPI_ISL_3062575 | University of Michigan Clinical Microbiology Laboratory | Lauring Lab, University of Michigan, Department of Microbiology and Immunology | Valesano |
| EPI_ISL_3062576, EPI_ISL_3062577, EPI_ISL_3062578, EPI_ISL_3062579, EPI_ISL_3062580, EPI_ISL_3062581, EPI_ISL_3062582, EPI_ISL_3062583, EPI_ISL_3062584, EPI_ISL_3062585, EPI_ISL_3062586, EPI_ISL_3062587, EPI_ISL_3062588, EPI_ISL_3062591, EPI_ISL_3062592, EPI_ISL_3062593, EPI_ISL_3062594, EPI_ISL_3062595, EPI_ISL_3062596, EPI_ISL_3062597, EPI_ISL_3062598, EPI_ISL_3062599, EPI_ISL_3062600, EPI_ISL_3062601, EPI_ISL_3062602, EPI_ISL_3062603, EPI_ISL_3062604, EPI_ISL_3062605, EPI_ISL_3062606, EPI_ISL_3062607, EPI_ISL_3062608, EPI_ISL_3062609, EPI_ISL_3062610, EPI_ISL_3062611, EPI_ISL_3062612, EPI_ISL_3062613, EPI_ISL_3062614, EPI_ISL_3062615, EPI_ISL_3062616, EPI_ISL_3062617, EPI_ISL_3062618, EPI_ISL_3062619, EPI_ISL_3062620, EPI_ISL_3062621, EPI_ISL_3062623, EPI_ISL_3062624, EPI_ISL_3062625, EPI_ISL_3062626, EPI_ISL_3062627, EPI_ISL_3062628, EPI_ISL_3062629, EPI_ISL_3062630, EPI_ISL_3062631, EPI_ISL_3062632, EPI_ISL_3062633, EPI_ISL_3062634, EPI_ISL_3062635, EPI_ISL_3062636, EPI_ISL_3062637, EPI_ISL_3062638, EPI_ISL_3062639, EPI_ISL_3062640, EPI_ISL_3062641, EPI_ISL_3062642, EPI_ISL_3062643, EPI_ISL_3062644, EPI_ISL_3062645, EPI_ISL_3062646, EPI_ISL_3062650, EPI_ISL_3062651, EPI_ISL_3062652, EPI_ISL_3062653, EPI_ISL_3062654, EPI_ISL_3062655, EPI_ISL_3062656, EPI_ISL_3062657, EPI_ISL_3062658, EPI_ISL_3062659, EPI_ISL_3062660, EPI_ISL_3062661, EPI_ISL_3062662, EPI_ISL_3062663 | | | |
| see above | UMC Groningen, Clinical Virology, Department of Medical Microbiology and Infection Prevention | UMC Groningen, Clinical Virology, Department of Medical Microbiology and Infection Prevention | Hubert Niesters, Alexander Friedrich, Erley Lizarazo-Forero, Monika Fliss, Lilli Gard, Sigrid Rosema, Coretta Van Leer-Buter, Xuewei Zhou, Marjolein Knoester |
| EPI_ISL_3062664, EPI_ISL_3062665, EPI_ISL_3062666, EPI_ISL_3062667, EPI_ISL_3062668, EPI_ISL_3062669, EPI_ISL_3062670, EPI_ISL_3062671, EPI_ISL_3062672 | Singapore General Hospital | Department of Microbiology | Nurdyana Abdul Rahman, Kenneth Xin Long Chan, Chayaporn Suphavilai, Kun Lee Lim, Sui Sin Goh, Kian Sing Chan, Lynette Oon, Kern Rei Chng, James Sim Heng Chiak, Niranjan Nagarajan, Karrie Ko |
| EPI_ISL_3062673, EPI_ISL_3062674, EPI_ISL_3062675, EPI_ISL_3062676, EPI_ISL_3062677, EPI_ISL_3062678, EPI_ISL_3062679, EPI_ISL_3062680, EPI_ISL_3062681, EPI_ISL_3062682 | Tampa General Hospital Esoteric Lab | Tampa General Hospital Esoteric Research & Development Lab | Jon Faughn, Marci O'Driscoll, Deanna Becker, Dominic Uy, Jorge Mecias-Frias, Elaine Vendrone, Vicki Healer, Amorce Lima, Suzane Silbert |
| EPI_ISL_3062683, EPI_ISL_3062684, EPI_ISL_3062685, EPI_ISL_3062686, EPI_ISL_3062687, EPI_ISL_3062688, EPI_ISL_3062689, EPI_ISL_3062690, EPI_ISL_3062691, EPI_ISL_3062692, EPI_ISL_3062693, EPI_ISL_3062694, EPI_ISL_3062695, EPI_ISL_3062696, EPI_ISL_3062697, EPI_ISL_3062698, EPI_ISL_3062699, EPI_ISL_3062700, EPI_ISL_3062701, EPI_ISL_3062702, EPI_ISL_3062703, EPI_ISL_3062704, EPI_ISL_3062705, EPI_ISL_3062706, EPI_ISL_3062707, EPI_ISL_3062708, EPI_ISL_3062709, EPI_ISL_3062710, EPI_ISL_3062711, EPI_ISL_3062712, EPI_ISL_3062713, EPI_ISL_3062714, EPI_ISL_3062715, EPI_ISL_3062716, EPI_ISL_3062717, EPI_ISL_3062718, EPI_ISL_3062719, EPI_ISL_3062720, EPI_ISL_3062721, EPI_ISL_3062722, EPI_ISL_3062723, EPI_ISL_3062724, EPI_ISL_3062725, EPI_ISL_3062726, | | | |

| | | | |
|---|---|---|---|
| EPI_ISL_3062727, EPI_ISL_3062728, EPI_ISL_3062729, EPI_ISL_3062730, EPI_ISL_3062731, EPI_ISL_3062734, EPI_ISL_3062735, EPI_ISL_3062736, EPI_ISL_3062737, EPI_ISL_3062738, EPI_ISL_3062739, EPI_ISL_3062740, EPI_ISL_3062741, EPI_ISL_3062742, EPI_ISL_3062743, EPI_ISL_3062744, EPI_ISL_3062745, EPI_ISL_3062746, EPI_ISL_3062747, EPI_ISL_3062748, EPI_ISL_3062749, EPI_ISL_3062750, EPI_ISL_3062751, EPI_ISL_3062752, EPI_ISL_3062753, EPI_ISL_3062754, EPI_ISL_3062755, EPI_ISL_3062756, EPI_ISL_3062757, EPI_ISL_3062758, EPI_ISL_3062759, EPI_ISL_3062760, EPI_ISL_3062761, EPI_ISL_3062762, EPI_ISL_3062763, EPI_ISL_3062764, EPI_ISL_3062765, EPI_ISL_3062766, EPI_ISL_3062767, EPI_ISL_3062768, EPI_ISL_3062769, EPI_ISL_3062770, EPI_ISL_3062771, EPI_ISL_3062772, EPI_ISL_3062773, EPI_ISL_3062774, EPI_ISL_3062775, EPI_ISL_3062776, EPI_ISL_3062777, EPI_ISL_3062778, EPI_ISL_3062779, EPI_ISL_3062780, EPI_ISL_3062781, EPI_ISL_3062782, EPI_ISL_3062783, EPI_ISL_3062784, EPI_ISL_3062785, EPI_ISL_3062786, EPI_ISL_3062787, EPI_ISL_3062788, EPI_ISL_3062789, EPI_ISL_3062790, EPI_ISL_3062791, EPI_ISL_3062792, EPI_ISL_3062793, EPI_ISL_3062794, EPI_ISL_3062795, EPI_ISL_3062796, EPI_ISL_3062797, EPI_ISL_3062798, EPI_ISL_3062799, EPI_ISL_3062801, EPI_ISL_3062802, EPI_ISL_3062803, EPI_ISL_3062804, EPI_ISL_3062805, EPI_ISL_3062806, EPI_ISL_3062807, EPI_ISL_3062808, EPI_ISL_3062809, EPI_ISL_3062810, EPI_ISL_3062811, EPI_ISL_3062812, EPI_ISL_3062813, EPI_ISL_3062814, EPI_ISL_3062815, EPI_ISL_3062816, EPI_ISL_3062817, EPI_ISL_3062818, EPI_ISL_3062819, EPI_ISL_3062820, EPI_ISL_3062821, EPI_ISL_3062822, EPI_ISL_3062823, EPI_ISL_3062824, EPI_ISL_3062825, EPI_ISL_3062826, EPI_ISL_3062827, EPI_ISL_3062828, EPI_ISL_3062829, EPI_ISL_3062830, EPI_ISL_3062831, EPI_ISL_3062832, EPI_ISL_3062833, EPI_ISL_3062834, EPI_ISL_3062835 | | | |
| see above | University of Mississippi Medical Center, Department of Pathology | University of Mississippi Medical Center, Molecular and Genomics Core Facility | Ashley C. Johnson, Wenjie Wu, Ithiel J. Frame, Krishna K. Ayyalasomayajula, Michael R. Garrett, D. Ashley Robinson |
| EPI_ISL_3062839 | Salud Digna | Instituto Nacional de Medicina Genomica | Cedro-Tanda A, Hidalgo-Miranda A, Cisneros-Villanueva M, Blancas S, Mendoza-Vargas A, Reyes-Grajeda JP, Abraham Campos-Romero, Moreno-Camacho José Luis, Rodriguez-Gallegos Jorge, Luna-Ruiz Marco, Gonzalez-Barrera D, Rangel-DeLeon D, Munguia-Garza P, Ramirez-Vega O, Escobar-Arrazola, M, Herrera-Montalvo LA. |
| EPI_ISL_3062848 | Instituto Nacional de Medicina Genomica | Instituto Nacional de Medicina Genomica | Cedro-Tanda A, Hidalgo-Miranda A, Cisneros-Villanueva M, Blancas S, Mendoza-Vargas A, Reyes-Grajeda JP, Cisneros- Villanueva M, Gonzalez-Barrera D, Rangel-DeLeon D, Munguia-Garza P, Ramirez-Vega O, Escobar-Arrazola, M, Herrera-Montalvo LA. |
| EPI_ISL_3062860 | Salud Digna | Instituto Nacional de Medicina Genomica | Cedro-Tanda A, Hidalgo-Miranda A, Cisneros-Villanueva M, Blancas S, Mendoza-Vargas A, Reyes-Grajeda JP, Abraham Campos-Romero, Moreno-Camacho José Luis, Rodriguez-Gallegos Jorge, Luna-Ruiz Marco, Gonzalez-Barrera D, Rangel-DeLeon D, Munguia-Garza P, Ramirez-Vega O, Escobar-Arrazola, M, Herrera-Montalvo LA. |
| EPI_ISL_3062869, EPI_ISL_3062870, EPI_ISL_3062871, EPI_ISL_3062872, EPI_ISL_3062873, EPI_ISL_3062876, EPI_ISL_3062877, EPI_ISL_3062878, EPI_ISL_3062879, EPI_ISL_3062881, EPI_ISL_3062882, EPI_ISL_3062883, EPI_ISL_3062884 | | | |
| see above | ASL Napoli 1 Centro | AMES Centro Polidiagnostico Strumentale S.r.l. | Giovanni Savarese, Raffaella Ruggiero, Eloisa Evangelista, Antonella Di Carlo, Luisa Circelli, Luigi D'Amore, Roberto Sirica,Maurizio D'Amora Antonio Fico |
| EPI_ISL_3062885, EPI_ISL_3062886, EPI_ISL_3062887, EPI_ISL_3062888, EPI_ISL_3062889, EPI_ISL_3062890, EPI_ISL_3062891, EPI_ISL_3062892, EPI_ISL_3062893, EPI_ISL_3062896, EPI_ISL_3062897, EPI_ISL_3062898, EPI_ISL_3062899, EPI_ISL_3062900, EPI_ISL_3062901, EPI_ISL_3062902, EPI_ISL_3062903, EPI_ISL_3062904, EPI_ISL_3062905, EPI_ISL_3062906, EPI_ISL_3062908, EPI_ISL_3062909, EPI_ISL_3062911, EPI_ISL_3062913, EPI_ISL_3062914, EPI_ISL_3062915, EPI_ISL_3062916, EPI_ISL_3062919, EPI_ISL_3062921, EPI_ISL_3062922, EPI_ISL_3062923, EPI_ISL_3062924, EPI_ISL_3062925, EPI_ISL_3062926, EPI_ISL_3062927, EPI_ISL_3062928, EPI_ISL_3062929, EPI_ISL_3062930, EPI_ISL_3062931, EPI_ISL_3062932, EPI_ISL_3062933, EPI_ISL_3062935, EPI_ISL_3062936, EPI_ISL_3062937, EPI_ISL_3062938, EPI_ISL_3062940, EPI_ISL_3062941, EPI_ISL_3062943, EPI_ISL_3062944, EPI_ISL_3062946, EPI_ISL_3062948, EPI_ISL_3062949, EPI_ISL_3062955, EPI_ISL_3062956, EPI_ISL_3062958, EPI_ISL_3062961, EPI_ISL_3062962 | | | |
| see above | ASL Napoli 1 Centro | AMES Centro Polidiagnostico Strumentale S.r.l. | "Giovanni Savarese, Raffaella Ruggiero, Eloisa Evangelista, Antonella Di Carlo, Luisa Circelli, Luigi D'Amore, Roberto Sirica,Maurizio D'Amora Antonio Fico" |
| EPI_ISL_3062963, EPI_ISL_3062964, EPI_ISL_3062965, EPI_ISL_3062966, EPI_ISL_3062967, EPI_ISL_3062968, EPI_ISL_3062969, EPI_ISL_3062970, EPI_ISL_3062971, EPI_ISL_3062972, EPI_ISL_3062973, EPI_ISL_3062974, EPI_ISL_3062975, EPI_ISL_3062976, EPI_ISL_3062977, EPI_ISL_3062978, EPI_ISL_3062980, EPI_ISL_3062981, EPI_ISL_3062982, EPI_ISL_3062983, EPI_ISL_3062985, EPI_ISL_3062986, EPI_ISL_3062989, EPI_ISL_3062990, EPI_ISL_3062991, EPI_ISL_3062992, EPI_ISL_3062993, EPI_ISL_3062994, EPI_ISL_3062995, EPI_ISL_3062996, EPI_ISL_3062997, EPI_ISL_3062998, EPI_ISL_3062999, EPI_ISL_3063000, EPI_ISL_3063002, EPI_ISL_3063003, EPI_ISL_3063005, EPI_ISL_3063006, EPI_ISL_3063007, EPI_ISL_3063008, EPI_ISL_3063009, EPI_ISL_3063010, EPI_ISL_3063011, EPI_ISL_3063012, EPI_ISL_3063013, EPI_ISL_3063014, EPI_ISL_3063015, EPI_ISL_3063017, EPI_ISL_3063018, EPI_ISL_3063019, EPI_ISL_3063023, EPI_ISL_3063024, EPI_ISL_3063025, EPI_ISL_3063026, EPI_ISL_3063027, EPI_ISL_3063030, EPI_ISL_3063034, EPI_ISL_3063036, EPI_ISL_3063039, EPI_ISL_3063041 | | | |
| see above | ASL Napoli 1 Centro | AMES Centro Polidiagnostico Strumentale S.r.l. | Giovanni Savarese, Raffaella Ruggiero, Eloisa Evangelista, Antonella Di Carlo, Luisa Circelli, Luigi D'Amore, Roberto Sirica,Maurizio D'Amora Antonio Fico |
| EPI_ISL_3063042, EPI_ISL_3063043, EPI_ISL_3063044, EPI_ISL_3063045, EPI_ISL_3063047, EPI_ISL_3063048, EPI_ISL_3063049, EPI_ISL_3063050, EPI_ISL_3063051, EPI_ISL_3063052, EPI_ISL_3063053, EPI_ISL_3063054, EPI_ISL_3063055, EPI_ISL_3063057, EPI_ISL_3063060, EPI_ISL_3063061, EPI_ISL_3063062, EPI_ISL_3063063, EPI_ISL_3063064, EPI_ISL_3063065, EPI_ISL_3063066, EPI_ISL_3063067, EPI_ISL_3063068, EPI_ISL_3063069, EPI_ISL_3063070, EPI_ISL_3063071, EPI_ISL_3063072, EPI_ISL_3063073, EPI_ISL_3063074, EPI_ISL_3063075, EPI_ISL_3063076, EPI_ISL_3063078, EPI_ISL_3063079, EPI_ISL_3063081, EPI_ISL_3063082, EPI_ISL_3063083, EPI_ISL_3063084, EPI_ISL_3063085, EPI_ISL_3063086, EPI_ISL_3063089, EPI_ISL_3063090, EPI_ISL_3063091, EPI_ISL_3063092, EPI_ISL_3063093, EPI_ISL_3063094, EPI_ISL_3063095, EPI_ISL_3063096, EPI_ISL_3063097, EPI_ISL_3063098, EPI_ISL_3063099, EPI_ISL_3063100, EPI_ISL_3063101, EPI_ISL_3063102, EPI_ISL_3063103, EPI_ISL_3063104 | | | |
| see above | Orange County Public Health Laboratory | Orange County Public Health Laboratory | Angelica Torres, Julia Wolfe |
| EPI_ISL_3063108, EPI_ISL_3063114, EPI_ISL_3063118, EPI_ISL_3063119, EPI_ISL_3063121, EPI_ISL_3063122, EPI_ISL_3063123, EPI_ISL_3063126, EPI_ISL_3063128, EPI_ISL_3063132, EPI_ISL_3063133, EPI_ISL_3063139, EPI_ISL_3063141, EPI_ISL_3063142, EPI_ISL_3063144, EPI_ISL_3063146, EPI_ISL_3063147, EPI_ISL_3063148, EPI_ISL_3063149, EPI_ISL_3063150, EPI_ISL_3063151, EPI_ISL_3063152, EPI_ISL_3063153, EPI_ISL_3063154, EPI_ISL_3063155, EPI_ISL_3063157, EPI_ISL_3063158, EPI_ISL_3063161, EPI_ISL_3063162, EPI_ISL_3063163, EPI_ISL_3063164, EPI_ISL_3063165, EPI_ISL_3063166, EPI_ISL_3063167, EPI_ISL_3063168, EPI_ISL_3063169, EPI_ISL_3063170, EPI_ISL_3063171, EPI_ISL_3063186, EPI_ISL_3063187, EPI_ISL_3063191, EPI_ISL_3063192, EPI_ISL_3063193, EPI_ISL_3063194, EPI_ISL_3063195, EPI_ISL_3063196, EPI_ISL_3063198, EPI_ISL_3063199, EPI_ISL_3063201, EPI_ISL_3063202, EPI_ISL_3063203, EPI_ISL_3063204, EPI_ISL_3063208, EPI_ISL_3063209, EPI_ISL_3063210, EPI_ISL_3063211, EPI_ISL_3063212, EPI_ISL_3063213, EPI_ISL_3063214, EPI_ISL_3063215, EPI_ISL_3063216, EPI_ISL_3063218, EPI_ISL_3063219, EPI_ISL_3063220, EPI_ISL_3063222, EPI_ISL_3063223, EPI_ISL_3063224, EPI_ISL_3063225, EPI_ISL_3063226, EPI_ISL_3063227, EPI_ISL_3063231, EPI_ISL_3063232, EPI_ISL_3063233, EPI_ISL_3063234, EPI_ISL_3063235, EPI_ISL_3063236, EPI_ISL_3063237, EPI_ISL_3063242, EPI_ISL_3063243, EPI_ISL_3063244, EPI_ISL_3063245, EPI_ISL_3063246, EPI_ISL_3063247, EPI_ISL_3063248, EPI_ISL_3063249, EPI_ISL_3063250, EPI_ISL_3063251, EPI_ISL_3063252, EPI_ISL_3063253, EPI_ISL_3063254, EPI_ISL_3063255, EPI_ISL_3063256, EPI_ISL_3063257, EPI_ISL_3063258, EPI_ISL_3063259, EPI_ISL_3063260, EPI_ISL_3063261, EPI_ISL_3063262, EPI_ISL_3063263, EPI_ISL_3063264, EPI_ISL_3063265, EPI_ISL_3063266 | | | |
| see above | National Institute of Public Health | National Institute of Public Health | Helena Jirincova, Jaromira Vecerova, Timotej Suri, Dusan Trnka, Alexander Nagy |
| EPI_ISL_3063274, EPI_ISL_3063275, EPI_ISL_3063276, EPI_ISL_3063277, EPI_ISL_3063278, EPI_ISL_3063279 | Ospedale "San Francesco", ASSL Nuoro | ATS SARDEGNA - ASSL Nuoro, Ospedale San Francesco, Laboratorio COVID e Laboratorio Specialistico UOC Ematologia | Aspróni Rosanna, Piras Giovanna, Malune Paolo, Pira Giovanna, Fancello Patrizia, Rosu Valentina, Carta Franco, Fiamma Maura, Mameli Giuseppe, Garau Maria Cristina , Palmas Angelo Domenico, Caterina Serra, Elena Rimini, Salvatore Rubino |
| EPI_ISL_3063280 | National Platform bis UMONS/Jolimont | National Platform bis UMONS/Jolimont | François Dufrasne, Guillaume Bayon-Vicente, Florian Juszczak, Laetitia Gheysen, Eric Tarantino, Gautier Detry, Ruddy Wattiez |
| EPI_ISL_3063283, EPI_ISL_3063284 | Ospedale Giovanni Paolo II Ragusa RG | AOU Polclinico "G Rodolico-S.Marco" PO Rodolico and BIOMETEC UNICT CATANIA | STEFANI AND SCALIA |
| EPI_ISL_3063286, EPI_ISL_3063289 | AOU Cannizzaro Catania | AOU Polclinico "G Rodolico-S.Marco" PO Rodolico and BIOMETEC UNICT CATANIA | STEFANI AND SCALIA |
| EPI_ISL_3063291 | Ospedale Giovanni Paolo II Ragusa RG | AOU Polclinico "G Rodolico-S.Marco" PO Rodolico and BIOMETEC UNICT CATANIA | STEFANI AND SCALIA |
| EPI_ISL_3063293, EPI_ISL_3063296, EPI_ISL_3063343, EPI_ISL_3063344, EPI_ISL_3063346, EPI_ISL_3063347, EPI_ISL_3063352, EPI_ISL_3063354, EPI_ISL_3063356, EPI_ISL_3063357, EPI_ISL_3063359, EPI_ISL_3063360, EPI_ISL_3063362, EPI_ISL_3063363, EPI_ISL_3063364, EPI_ISL_3063367, EPI_ISL_3063370, EPI_ISL_3063371, EPI_ISL_3063372, EPI_ISL_3063373, EPI_ISL_3063374, EPI_ISL_3063375, EPI_ISL_3063376, EPI_ISL_3063377, EPI_ISL_3063379, EPI_ISL_3063380, EPI_ISL_3063383, EPI_ISL_3063384, EPI_ISL_3063385, EPI_ISL_3063387, EPI_ISL_3063388, EPI_ISL_3063389, EPI_ISL_3063390, EPI_ISL_3063391, EPI_ISL_3063392, EPI_ISL_3063393, EPI_ISL_3063394, EPI_ISL_3063395, EPI_ISL_3063396, EPI_ISL_3063397 | | | |
| see above | Ministry of Health Turkey | Ministry of Health Turkey | Fatma Bayrakdar, Yasemin Cosgun, Suleyman Yalcin, Gulay Korukluoglu |
| EPI_ISL_3063487, EPI_ISL_3063488, EPI_ISL_3063489, EPI_ISL_3063490, EPI_ISL_3063491, EPI_ISL_3063492, EPI_ISL_3063493, EPI_ISL_3063494, EPI_ISL_3063496, EPI_ISL_3063497, EPI_ISL_3063499, EPI_ISL_3063500, EPI_ISL_3063501, EPI_ISL_3063502, EPI_ISL_3063504, EPI_ISL_3063505, EPI_ISL_3063506, EPI_ISL_3063507, EPI_ISL_3063508, EPI_ISL_3063509, EPI_ISL_3063510, EPI_ISL_3063511, EPI_ISL_3063512, EPI_ISL_3063513, EPI_ISL_3063514, EPI_ISL_3063515, EPI_ISL_3063516, EPI_ISL_3063517, EPI_ISL_3063519, EPI_ISL_3063521, EPI_ISL_3063522, EPI_ISL_3063523, EPI_ISL_3063525, EPI_ISL_3063526, EPI_ISL_3063527, EPI_ISL_3063528, EPI_ISL_3063529, EPI_ISL_3063530, EPI_ISL_3063531, EPI_ISL_3063532, EPI_ISL_3063533, EPI_ISL_3063535, EPI_ISL_3063537, EPI_ISL_3063538, EPI_ISL_3063539, EPI_ISL_3063540, EPI_ISL_3063541, EPI_ISL_3063542, EPI_ISL_3063543, EPI_ISL_3063544, EPI_ISL_3063546, EPI_ISL_3063547, EPI_ISL_3063548, EPI_ISL_3063549, EPI_ISL_3063550, EPI_ISL_3063551, EPI_ISL_3063552, EPI_ISL_3063553, EPI_ISL_3063554, EPI_ISL_3063555, EPI_ISL_3063556, EPI_ISL_3063557, EPI_ISL_3063558, EPI_ISL_3063559, EPI_ISL_3063561, EPI_ISL_3063562, EPI_ISL_3063564, EPI_ISL_3063565, EPI_ISL_3063566, EPI_ISL_3063567, EPI_ISL_3063568, EPI_ISL_3063569, EPI_ISL_3063570, EPI_ISL_3063571, EPI_ISL_3063572, EPI_ISL_3063573, EPI_ISL_3063574, EPI_ISL_3063575, EPI_ISL_3063578, EPI_ISL_3063579, EPI_ISL_3063580, EPI_ISL_3063582, EPI_ISL_3063583, EPI_ISL_3063584, EPI_ISL_3063585, EPI_ISL_3063586, EPI_ISL_3063587, EPI_ISL_3063589, EPI_ISL_3063590, EPI_ISL_3063591, EPI_ISL_3063592, EPI_ISL_3063593, EPI_ISL_3063594, EPI_ISL_3063595, EPI_ISL_3063596, EPI_ISL_3063597, EPI_ISL_3063598, EPI_ISL_3063599, EPI_ISL_3063600, EPI_ISL_3063601, EPI_ISL_3063602, EPI_ISL_3063604, EPI_ISL_3063605, EPI_ISL_3063606, EPI_ISL_3063608, EPI_ISL_3063610, EPI_ISL_3063611, EPI_ISL_3063612, EPI_ISL_3063613, EPI_ISL_3063614, EPI_ISL_3063615, EPI_ISL_3063616, EPI_ISL_3063617, EPI_ISL_3063618, EPI_ISL_3063619, EPI_ISL_3063620, EPI_ISL_3063621, EPI_ISL_3063622, EPI_ISL_3063623, EPI_ISL_3063624, EPI_ISL_3063625, EPI_ISL_3063626, EPI_ISL_3063627, EPI_ISL_3063628, EPI_ISL_3063631, EPI_ISL_3063632, EPI_ISL_3063633, EPI_ISL_3063635, EPI_ISL_3063636, EPI_ISL_3063637, EPI_ISL_3063638, EPI_ISL_3063639, EPI_ISL_3063640, EPI_ISL_3063641, EPI_ISL_3063642, EPI_ISL_3063643, EPI_ISL_3063645, EPI_ISL_3063646, EPI_ISL_3063647, EPI_ISL_3063648, EPI_ISL_3063649, EPI_ISL_3063650, EPI_ISL_3063651, EPI_ISL_3063652, EPI_ISL_3063653, EPI_ISL_3063654, EPI_ISL_3063655, EPI_ISL_3063658, EPI_ISL_3063659, EPI_ISL_3063660, EPI_ISL_3063661, EPI_ISL_3063662, EPI_ISL_3063663, EPI_ISL_3063664, EPI_ISL_3063666, EPI_ISL_3063667, EPI_ISL_3063668, EPI_ISL_3063670, EPI_ISL_3063671, EPI_ISL_3063673, EPI_ISL_3063674, EPI_ISL_3063675, EPI_ISL_3063676, EPI_ISL_3063677, EPI_ISL_3063678, EPI_ISL_3063679, EPI_ISL_3063681, EPI_ISL_3063682, EPI_ISL_3063683, EPI_ISL_3063686, EPI_ISL_3063687, EPI_ISL_3063688, EPI_ISL_3063689, EPI_ISL_3063690, EPI_ISL_3063691, EPI_ISL_3063692, EPI_ISL_3063693, EPI_ISL_3063694, EPI_ISL_3063695, EPI_ISL_3063696, EPI_ISL_3063697, EPI_ISL_3063698, EPI_ISL_3063699, EPI_ISL_3063700, EPI_ISL_3063701, EPI_ISL_3063703, EPI_ISL_3063704, EPI_ISL_3063705, EPI_ISL_3063706, EPI_ISL_3063707, EPI_ISL_3063708, EPI_ISL_3063709, EPI_ISL_3063710, EPI_ISL_3063712, EPI_ISL_3063713, EPI_ISL_3063715, EPI_ISL_3063716, EPI_ISL_3063717, EPI_ISL_3063718, EPI_ISL_3063719, EPI_ISL_3063720, EPI_ISL_3063721, EPI_ISL_3063722, EPI_ISL_3063723, EPI_ISL_3063724, EPI_ISL_3063725, EPI_ISL_3063727, EPI_ISL_3063728, EPI_ISL_3063729, EPI_ISL_3063730, EPI_ISL_3063731, EPI_ISL_3063732, EPI_ISL_3063733, EPI_ISL_3063734, EPI_ISL_3063735, EPI_ISL_3063736, EPI_ISL_3063737, EPI_ISL_3063739, EPI_ISL_3063740 | | | |
| see above | UW Virology Lab | UW Virology Lab | Pavitra Roychoudhury, Shah Mohamed Bakhash, Ricardo Perez, Hong Xie, Tien V. Nguyen, Maria Lukes, Lasata Shrestha, Sean Ellis, Nathan Breit, Robert J. Livingston, Meei-Li Huang, Keith R Jerome, Patrick Mathias, Alexander Greninger |
| EPI_ISL_3063741, EPI_ISL_3063743, EPI_ISL_3063744, EPI_ISL_3063745, EPI_ISL_3063746, EPI_ISL_3063747, EPI_ISL_3063748, EPI_ISL_3063749, EPI_ISL_3063750, EPI_ISL_3063752 | Servicio de Microbiología. Hospital Clínico Universitario de Valencia | SeqCOVID-SPAIN consortium/IBV(CSIC) | David Navarro Ortega, Eliseo Albert Vicent, Ignacio Torres and SeqCOVID-SPAIN consortium |
| EPI_ISL_3063753, EPI_ISL_3063754, EPI_ISL_3063755, EPI_ISL_3063756, EPI_ISL_3063759, EPI_ISL_3063760, EPI_ISL_3063761, EPI_ISL_3063762, EPI_ISL_3063765, EPI_ISL_3063766, EPI_ISL_3063769, EPI_ISL_3063771, EPI_ISL_3063772, EPI_ISL_3063774, EPI_ISL_3063775, EPI_ISL_3063776, EPI_ISL_3063778, EPI_ISL_3063779, EPI_ISL_3063780, EPI_ISL_3063781, EPI_ISL_3063782, EPI_ISL_3063784, EPI_ISL_3063785, EPI_ISL_3063789, EPI_ISL_3063790, EPI_ISL_3063791, EPI_ISL_3063792, EPI_ISL_3063793, EPI_ISL_3063794, EPI_ISL_3063795, EPI_ISL_3063796, EPI_ISL_3063797, EPI_ISL_3063798, EPI_ISL_3063800, EPI_ISL_3063801 | | | |
| see above | Servicio de Microbiología. Hospital Universitario Doctor Peset | SeqCOVID-SPAIN consortium/IBV(CSIC) | Juan Alberola Enguídanos, Juan José Camarena Miñana, Rosa González Pellicer, José Miguel Nogueira Coito and SeqCOVID-SPAIN consortium |
| EPI_ISL_3063802 | Hospital Universitario de La Ribera (Alzira, València) | SeqCOVID-SPAIN consortium/IBV(CSIC) | Olalla Martínez Macias, Julia González Cantó and SeqCOVID-SPAIN consortium |
| EPI_ISL_3063803, EPI_ISL_3063804, EPI_ISL_3063805, EPI_ISL_3063806, EPI_ISL_3063807, EPI_ISL_3063808 | Hospital Comarcal de Vinaròs | SeqCOVID-SPAIN consortium/IBV(CSIC) | Mercedes Roig Cardells and SeqCOVID-SPAIN consortium |
| EPI_ISL_3063809, EPI_ISL_3063810 | Laboratorio de Biología Molecular. Servicio de Análisis Clínicos y Microbiología. Hospital de Sagunto | SeqCOVID-SPAIN consortium/IBV(CSIC) | Dra. Victoria Simón García, Gonzalo Llop Furquet, Agustín Irazo Tatay, Sandra Moreno Marro, Noelia Lozano Rodríguez and SeqCOVID-SPAIN consortium |
| EPI_ISL_3063811, EPI_ISL_3063812, EPI_ISL_3063813, EPI_ISL_3063814, EPI_ISL_3063815, EPI_ISL_3063816, EPI_ISL_3063817, EPI_ISL_3063818, EPI_ISL_3063819, EPI_ISL_3063820 | Hospital Universitari i Politècnic La Fe de València | SeqCOVID-SPAIN consortium/IBV(CSIC) | María Dolores Gómez Ruiz, Eva González Barberá, Ana Gil Brusola, Salvador Giner Almaraz, José Luis López Hontangas and SeqCOVID-SPAIN consortium |
| EPI_ISL_3064129, EPI_ISL_3064130, EPI_ISL_3064132, EPI_ISL_3064133, EPI_ISL_3064135, EPI_ISL_3064136, EPI_ISL_3064137, EPI_ISL_3064138, EPI_ISL_3064139, EPI_ISL_3064140, EPI_ISL_3064141, EPI_ISL_3064142, EPI_ISL_3064143, EPI_ISL_3064144, EPI_ISL_3064145, EPI_ISL_3064146, EPI_ISL_3064147 | | | |
| see above | CHU Pontchaillou | CHU Pontchaillou | GROLHIER Claire, DENOUAL Florent, ETCHEVERRY Amandine, SASSI Mohamed, JAGLINE Steven, FEBREAU Christine, PRONIER Charlotte, GALIBERT Marie Dominique, DE TAYRAC Marie, THIBAULT Vincent |
| EPI_ISL_3064242, EPI_ISL_3064243, EPI_ISL_3064244, EPI_ISL_3064245, EPI_ISL_3064246, EPI_ISL_3064247, EPI_ISL_3064248, EPI_ISL_3064249, EPI_ISL_3064250, EPI_ISL_3064251, EPI_ISL_3064252, EPI_ISL_3064253, EPI_ISL_3064254, EPI_ISL_3064255, EPI_ISL_3064256, EPI_ISL_3064257, EPI_ISL_3064258, EPI_ISL_3064259, EPI_ISL_3064260, EPI_ISL_3064261, EPI_ISL_3064262, EPI_ISL_3064263, EPI_ISL_3064264, EPI_ISL_3064265, EPI_ISL_3064266, EPI_ISL_3064267, EPI_ISL_3064268, EPI_ISL_3064269, EPI_ISL_3064270, EPI_ISL_3064271, EPI_ISL_3064272, EPI_ISL_3064273, EPI_ISL_3064274, EPI_ISL_3064275, EPI_ISL_3064276, EPI_ISL_3064277, EPI_ISL_3064278, EPI_ISL_3064279, EPI_ISL_3064280, EPI_ISL_3064281, EPI_ISL_3064282, EPI_ISL_3064283, EPI_ISL_3064284, EPI_ISL_3064285, | | | |

EPI_ISL_3064286, EPI_ISL_3064287, EPI_ISL_3064288, EPI_ISL_3064289, EPI_ISL_3064290, EPI_ISL_3064291, EPI_ISL_3064292, EPI_ISL_3064293, EPI_ISL_3064294, EPI_ISL_3064295, EPI_ISL_3064296, EPI_ISL_3064297, EPI_ISL_3064298, EPI_ISL_3064299, EPI_ISL_3064300, EPI_ISL_3064301

| | | | |
|---|---|---|---|
| see above | Pathology and Laboratory Medicine Institute, Cleveland Clinic, Ohio, USA | Pathology and Laboratory Medicine Institute, Cleveland Clinic, Ohio, USA | Jessica Spildener, Concetta Peck, Jennifer Starbuck, Joy Nakitandwe, Kristen McDonnell, David Plunkett, Zheng Jin Tu, Jay Brock, Yu-Wei Cheng, Gary Procop, Daniel Rhoads, Daniel H. Farkas, David Bosler |

EPI_ISL_3064316, EPI_ISL_3064317, EPI_ISL_3064319, EPI_ISL_3064320, EPI_ISL_3064321, EPI_ISL_3064322, EPI_ISL_3064325, EPI_ISL_3064326, EPI_ISL_3064327, EPI_ISL_3064328, EPI_ISL_3064329, EPI_ISL_3064330, EPI_ISL_3064333, EPI_ISL_3064334, EPI_ISL_3064335, EPI_ISL_3064336, EPI_ISL_3064337, EPI_ISL_3064338, EPI_ISL_3064339, EPI_ISL_3064340, EPI_ISL_3064342, EPI_ISL_3064343, EPI_ISL_3064344, EPI_ISL_3064345, EPI_ISL_3064347, EPI_ISL_3064348, EPI_ISL_3064349, EPI_ISL_3064350, EPI_ISL_3064351, EPI_ISL_3064352, EPI_ISL_3064353, EPI_ISL_3064354, EPI_ISL_3064356, EPI_ISL_3064358, EPI_ISL_3064359, EPI_ISL_3064361, EPI_ISL_3064362, EPI_ISL_3064363, EPI_ISL_3064364, EPI_ISL_3064366, EPI_ISL_3064367, EPI_ISL_3064368, EPI_ISL_3064369, EPI_ISL_3064370, EPI_ISL_3064371, EPI_ISL_3064373, EPI_ISL_3064374, EPI_ISL_3064375, EPI_ISL_3064376

| | | | |
|---|---|---|---|
| see above | Arizona State University | Arizona State University | Ajeet Bains, Peter T. Skidmore, LaRinda A. Holland, Matthew F. Smith, Rabia Maqsood, Nicholas J. Mellor, Joy M. Blain, Valerie Harris, Joshua LaBaer, Vel Murugan, Efrem S. Lim |

EPI_ISL_3064386, EPI_ISL_3064406, EPI_ISL_3064409, EPI_ISL_3064413, EPI_ISL_3064416, EPI_ISL_3064417, EPI_ISL_3064418, EPI_ISL_3064419, EPI_ISL_3064420, EPI_ISL_3064422, EPI_ISL_3064423, EPI_ISL_3064424, EPI_ISL_3064425, EPI_ISL_3064427, EPI_ISL_3064428, EPI_ISL_3064429, EPI_ISL_3064430, EPI_ISL_3064431, EPI_ISL_3064432, EPI_ISL_3064433, EPI_ISL_3064434, EPI_ISL_3064435, EPI_ISL_3064436, EPI_ISL_3064437, EPI_ISL_3064438, EPI_ISL_3064440, EPI_ISL_3064441, EPI_ISL_3064442, EPI_ISL_3064443, EPI_ISL_3064444, EPI_ISL_3064446, EPI_ISL_3064447, EPI_ISL_3064448, EPI_ISL_3064449, EPI_ISL_3064451, EPI_ISL_3064452, EPI_ISL_3064453, EPI_ISL_3064454, EPI_ISL_3064456, EPI_ISL_3064457, EPI_ISL_3064458, EPI_ISL_3064459, EPI_ISL_3064460, EPI_ISL_3064461, EPI_ISL_3064462, EPI_ISL_3064463, EPI_ISL_3064467, EPI_ISL_3064468, EPI_ISL_3064469, EPI_ISL_3064470, EPI_ISL_3064473, EPI_ISL_3064477, EPI_ISL_3064479, EPI_ISL_3064480, EPI_ISL_3064481, EPI_ISL_3064482, EPI_ISL_3064484, EPI_ISL_3064486, EPI_ISL_3064487, EPI_ISL_3064488, EPI_ISL_3064490, EPI_ISL_3064492, EPI_ISL_3064493, EPI_ISL_3064494, EPI_ISL_3064496, EPI_ISL_3064497, EPI_ISL_3064498, EPI_ISL_3064500, EPI_ISL_3064505, EPI_ISL_3064506, EPI_ISL_3064507, EPI_ISL_3064508, EPI_ISL_3064509, EPI_ISL_3064510, EPI_ISL_3064512, EPI_ISL_3064513, EPI_ISL_3064514, EPI_ISL_3064519, EPI_ISL_3064520, EPI_ISL_3064521, EPI_ISL_3064522, EPI_ISL_3064527, EPI_ISL_3064528, EPI_ISL_3064529, EPI_ISL_3064533, EPI_ISL_3064534, EPI_ISL_3064536, EPI_ISL_3064538, EPI_ISL_3064543, EPI_ISL_3064544, EPI_ISL_3064545, EPI_ISL_3064546, EPI_ISL_3064547, EPI_ISL_3064548, EPI_ISL_3064549, EPI_ISL_3064550, EPI_ISL_3064551, EPI_ISL_3064552, EPI_ISL_3064554, EPI_ISL_3064555, EPI_ISL_3064556, EPI_ISL_3064557, EPI_ISL_3064558, EPI_ISL_3064559, EPI_ISL_3064561, EPI_ISL_3064562, EPI_ISL_3064563, EPI_ISL_3064564, EPI_ISL_3064565, EPI_ISL_3064566, EPI_ISL_3064567, EPI_ISL_3064568, EPI_ISL_3064569, EPI_ISL_3064570, EPI_ISL_3064571, EPI_ISL_3064572, EPI_ISL_3064573, EPI_ISL_3064575, EPI_ISL_3064576, EPI_ISL_3064577, EPI_ISL_3064578, EPI_ISL_3064579, EPI_ISL_3064580, EPI_ISL_3064581, EPI_ISL_3064583, EPI_ISL_3064584, EPI_ISL_3064585, EPI_ISL_3064586, EPI_ISL_3064588, EPI_ISL_3064589, EPI_ISL_3064591, EPI_ISL_3064592, EPI_ISL_3064593, EPI_ISL_3064594, EPI_ISL_3064595, EPI_ISL_3064628, EPI_ISL_3064634, EPI_ISL_3064638, EPI_ISL_3064646, EPI_ISL_3064651, EPI_ISL_3064656, EPI_ISL_3064658, EPI_ISL_3064659, EPI_ISL_3064664, EPI_ISL_3064665, EPI_ISL_3064666, EPI_ISL_3064669, EPI_ISL_3064671, EPI_ISL_3064673, EPI_ISL_3064674, EPI_ISL_3064675, EPI_ISL_3064676, EPI_ISL_3064677, EPI_ISL_3064678, EPI_ISL_3064680, EPI_ISL_3064682, EPI_ISL_3064683, EPI_ISL_3064685, EPI_ISL_3064686, EPI_ISL_3064687, EPI_ISL_3064688, EPI_ISL_3064689, EPI_ISL_3064691, EPI_ISL_3064692, EPI_ISL_3064693, EPI_ISL_3064695, EPI_ISL_3064696, EPI_ISL_3064697, EPI_ISL_3064698, EPI_ISL_3064699, EPI_ISL_3064700

| | | | |
|---|---|---|---|
| see above | Curative Labs | Curative Labs | Mikhail Hanewich-Hollatz, J.Cesar Ignacio-Espinoza, Elias L. Salfati, Nabjot Sandhu, George Way, Zhiyi Xie, Eugenia Khorosheva, Sophia Quasem, Janet Chen, Vladimir Slepnev |

EPI_ISL_3064701, EPI_ISL_3064702, EPI_ISL_3064705, EPI_ISL_3064706, EPI_ISL_3064707, EPI_ISL_3064708, EPI_ISL_3064709, EPI_ISL_3064710, EPI_ISL_3064711, EPI_ISL_3064712, EPI_ISL_3064713

| | | | |
|---|---|---|---|
| see above | National Agency for Public Health, Republic of Moldov | Charité Universitätsmedizin Berlin, Institut für Virologie | Victor M Corman, Barbara Mühlemann, Jörn Beheim-Schwarzbach, Talitha Veith, Julia Schneider, Maria Apostol, Ala Halacu, Terry Jones, Christian Drosten |
| EPI_ISL_3064714, EPI_ISL_3064715, EPI_ISL_3064716, EPI_ISL_3064717, EPI_ISL_3064718 | "OK Public Health Laboratory, Oklahoma State DOH" | Centers for Disease Control and Prevention Division of Viral Diseases, Pathogen Discovery | Mili Sheth, Sarah Nobles, Jasmine Padilla, Alex Burgin, Meghan Bentz, Mark Burroughs, Peter Cook, Clinton R. Paden, Dhwani Batra, Krista Queen, Kristen Knipe, Dakota Howard, Yvette Unoarumhi, Matthew Schmerer, Ben L. Rambo-Martin, Kristine Lacek, Sam Shepard, Dave Wentworth, Vivien Dugan, Suxiang Tong, Justin Lee |
| EPI_ISL_3064719 | NJ Public Health and Environmental Laboratories | Centers for Disease Control and Prevention Division of Viral Diseases, Pathogen Discovery | Mili Sheth, Sarah Nobles, Jasmine Padilla, Alex Burgin, Meghan Bentz, Mark Burroughs, Peter Cook, Clinton R. Paden, Dhwani Batra, Krista Queen, Kristen Knipe, Dakota Howard, Yvette Unoarumhi, Matthew Schmerer, Ben L. Rambo-Martin, Kristine Lacek, Sam Shepard, Dave Wentworth, Vivien Dugan, Suxiang Tong, Justin Lee |

EPI_ISL_3064720, EPI_ISL_3064722, EPI_ISL_3064725, EPI_ISL_3064726, EPI_ISL_3064727, EPI_ISL_3064729, EPI_ISL_3064731, EPI_ISL_3064734, EPI_ISL_3064735, EPI_ISL_3064737, EPI_ISL_3064738, EPI_ISL_3064739, EPI_ISL_3064740

| | | | |
|---|---|---|---|
| see above | "AR Dept. of Health-PHL, Molecular Diagnostics" | Centers for Disease Control and Prevention Division of Viral Diseases, Pathogen Discovery | Mili Sheth, Sarah Nobles, Jasmine Padilla, Alex Burgin, Meghan Bentz, Mark Burroughs, Peter Cook, Clinton R. Paden, Dhwani Batra, Krista Queen, Kristen Knipe, Dakota Howard, Yvette Unoarumhi, Matthew Schmerer, Ben L. Rambo-Martin, Kristine Lacek, Sam Shepard, Dave Wentworth, Vivien Dugan, Suxiang Tong, Justin Lee |
| EPI_ISL_3064743 | LA Office of Public Health Laboratories | Centers for Disease Control and Prevention Division of Viral Diseases, Pathogen Discovery | Mili Sheth, Sarah Nobles, Jasmine Padilla, Alex Burgin, Meghan Bentz, Mark Burroughs, Peter Cook, Clinton R. Paden, Dhwani Batra, Krista Queen, Kristen Knipe, Dakota Howard, Yvette Unoarumhi, Matthew Schmerer, Ben L. Rambo-Martin, Kristine Lacek, Sam Shepard, Dave Wentworth, Vivien Dugan, Suxiang Tong, Justin Lee |
| EPI_ISL_3064744, EPI_ISL_3064746 | "NM Dept. Health, Scientific Laboratory Division " | Centers for Disease Control and Prevention Division of Viral Diseases, Pathogen Discovery | Mili Sheth, Sarah Nobles, Jasmine Padilla, Alex Burgin, Meghan Bentz, Mark Burroughs, Peter Cook, Clinton R. Paden, Dhwani Batra, Krista Queen, Kristen Knipe, Dakota Howard, Yvette Unoarumhi, Matthew Schmerer, Ben L. Rambo-Martin, Kristine Lacek, Sam Shepard, Dave Wentworth, Vivien Dugan, Suxiang Tong, Justin Lee |
| EPI_ISL_3064749 | TN Division of Laboratory Services | Centers for Disease Control and Prevention Division of Viral Diseases, Pathogen Discovery | Mili Sheth, Sarah Nobles, Jasmine Padilla, Alex Burgin, Meghan Bentz, Mark Burroughs, Peter Cook, Clinton R. Paden, Dhwani Batra, Krista Queen, Kristen Knipe, Dakota Howard, Yvette Unoarumhi, Matthew Schmerer, Ben L. Rambo-Martin, Kristine Lacek, Sam Shepard, Dave Wentworth, Vivien Dugan, Suxiang Tong, Justin Lee |

EPI_ISL_3064754, EPI_ISL_3064755, EPI_ISL_3064756, EPI_ISL_3064757, EPI_ISL_3064761, EPI_ISL_3064762, EPI_ISL_3064763, EPI_ISL_3064765, EPI_ISL_3064767, EPI_ISL_3064768, EPI_ISL_3064769

| | | | |
|---|---|---|---|
| see above | "OK Public Health Laboratory, Oklahoma State DOH" | Centers for Disease Control and Prevention Division of Viral Diseases, Pathogen Discovery | Mili Sheth, Sarah Nobles, Jasmine Padilla, Alex Burgin, Meghan Bentz, Mark Burroughs, Peter Cook, Clinton R. Paden, Dhwani Batra, Krista Queen, Kristen Knipe, Dakota Howard, Yvette Unoarumhi, Matthew Schmerer, Ben L. Rambo-Martin, Kristine Lacek, Sam Shepard, Dave Wentworth, Vivien Dugan, Suxiang Tong, Justin Lee |
| EPI_ISL_3064770, EPI_ISL_3064771 | SD Public Health Laboratory | Centers for Disease Control and Prevention Division of Viral Diseases, Pathogen Discovery | Mili Sheth, Sarah Nobles, Jasmine Padilla, Alex Burgin, Meghan Bentz, Mark Burroughs, Peter Cook, Clinton R. Paden, Dhwani Batra, Krista Queen, Kristen Knipe, Dakota Howard, Yvette Unoarumhi, Matthew Schmerer, Ben L. Rambo-Martin, Kristine Lacek, Sam Shepard, Dave Wentworth, Vivien Dugan, Suxiang Tong, Justin Lee |
| EPI_ISL_3064774, EPI_ISL_3064775 | NC State Laboratory of Public Health | Centers for Disease Control and Prevention Division of Viral Diseases, Pathogen Discovery | Mili Sheth, Sarah Nobles, Jasmine Padilla, Alex Burgin, Meghan Bentz, Mark Burroughs, Peter Cook, Clinton R. Paden, Dhwani Batra, Krista Queen, Kristen Knipe, Dakota Howard, Yvette Unoarumhi, Matthew Schmerer, Ben L. Rambo-Martin, Kristine Lacek, Sam Shepard, Dave Wentworth, Vivien Dugan, Suxiang Tong, Justin Lee |

EPI_ISL_3064778, EPI_ISL_3064779, EPI_ISL_3064780, EPI_ISL_3064781, EPI_ISL_3064784, EPI_ISL_3064785, EPI_ISL_3064787, EPI_ISL_3064788, EPI_ISL_3064789, EPI_ISL_3064790, EPI_ISL_3064791, EPI_ISL_3064792, EPI_ISL_3064793, EPI_ISL_3064794, EPI_ISL_3064795, EPI_ISL_3064797, EPI_ISL_3064798, EPI_ISL_3064799, EPI_ISL_3064800, EPI_ISL_3064801, EPI_ISL_3064802, EPI_ISL_3064803, EPI_ISL_3064804, EPI_ISL_3064805, EPI_ISL_3064806, EPI_ISL_3064807, EPI_ISL_3064808, EPI_ISL_3064809, EPI_ISL_3064810, EPI_ISL_3064811, EPI_ISL_3064814, EPI_ISL_3064819, EPI_ISL_3064820, EPI_ISL_3064821, EPI_ISL_3064822, EPI_ISL_3064823, EPI_ISL_3064824, EPI_ISL_3064825, EPI_ISL_3064826, EPI_ISL_3064827, EPI_ISL_3064828, EPI_ISL_3064829, EPI_ISL_3064830, EPI_ISL_3064831, EPI_ISL_3064832, EPI_ISL_3064833, EPI_ISL_3064834, EPI_ISL_3064835, EPI_ISL_3064836, EPI_ISL_3064837, EPI_ISL_3064838, EPI_ISL_3064839, EPI_ISL_3064840, EPI_ISL_3064841, EPI_ISL_3064842, EPI_ISL_3064843, EPI_ISL_3064844, EPI_ISL_3064847, EPI_ISL_3064849, EPI_ISL_3064850, EPI_ISL_3064851, EPI_ISL_3064852, EPI_ISL_3064853, EPI_ISL_3064854, EPI_ISL_3064855, EPI_ISL_3064856, EPI_ISL_3064857, EPI_ISL_3064858, EPI_ISL_3064859, EPI_ISL_3064860, EPI_ISL_3064862, EPI_ISL_3064863, EPI_ISL_3064864, EPI_ISL_3064865, EPI_ISL_3064869, EPI_ISL_3064871, EPI_ISL_3064872, EPI_ISL_3064873, EPI_ISL_3064874, EPI_ISL_3064876, EPI_ISL_3064877, EPI_ISL_3064878, EPI_ISL_3064879, EPI_ISL_3064880, EPI_ISL_3064881, EPI_ISL_3064882, EPI_ISL_3064883, EPI_ISL_3064884, EPI_ISL_3064885, EPI_ISL_3064886, EPI_ISL_3064887, EPI_ISL_3064888, EPI_ISL_3064889, EPI_ISL_3064890, EPI_ISL_3064891, EPI_ISL_3064892, EPI_ISL_3064894, EPI_ISL_3064895, EPI_ISL_3064896, EPI_ISL_3064898, EPI_ISL_3064899, EPI_ISL_3064900, EPI_ISL_3064901, EPI_ISL_3064902, EPI_ISL_3064903, EPI_ISL_3064904, EPI_ISL_3064905, EPI_ISL_3064906, EPI_ISL_3064907, EPI_ISL_3064908, EPI_ISL_3064909, EPI_ISL_3064910, EPI_ISL_3064911, EPI_ISL_3064912, EPI_ISL_3064913, EPI_ISL_3064914, EPI_ISL_3064915, EPI_ISL_3064916, EPI_ISL_3064917, EPI_ISL_3064918, EPI_ISL_3064919, EPI_ISL_3064920, EPI_ISL_3064921, EPI_ISL_3064922, EPI_ISL_3064923, EPI_ISL_3064924, EPI_ISL_3064925, EPI_ISL_3064926, EPI_ISL_3064927, EPI_ISL_3064928, EPI_ISL_3064929, EPI_ISL_3064930, EPI_ISL_3064931, EPI_ISL_3064932, EPI_ISL_3064933, EPI_ISL_3064934, EPI_ISL_3064935, EPI_ISL_3064936, EPI_ISL_3064937, EPI_ISL_3064939, EPI_ISL_3064940, EPI_ISL_3064941, EPI_ISL_3064942, EPI_ISL_3064943, EPI_ISL_3064944, EPI_ISL_3064945, EPI_ISL_3064946, EPI_ISL_3064947, EPI_ISL_3064948, EPI_ISL_3064949, EPI_ISL_3064950, EPI_ISL_3064951, EPI_ISL_3064952, EPI_ISL_3064953, EPI_ISL_3064954, EPI_ISL_3064955, EPI_ISL_3064957, EPI_ISL_3064958, EPI_ISL_3064959, EPI_ISL_3064960, EPI_ISL_3064961, EPI_ISL_3064962, EPI_ISL_3064964, EPI_ISL_3064965, EPI_ISL_3064966, EPI_ISL_3064967, EPI_ISL_3064968, EPI_ISL_3064969, EPI_ISL_3064971, EPI_ISL_3064972, EPI_ISL_3064973, EPI_ISL_3064974, EPI_ISL_3064975, EPI_ISL_3064976, EPI_ISL_3064977, EPI_ISL_3064978, EPI_ISL_3064979, EPI_ISL_3064980, EPI_ISL_3064981, EPI_ISL_3064983, EPI_ISL_3064984, EPI_ISL_3064985, EPI_ISL_3064986, EPI_ISL_3064987, EPI_ISL_3064988, EPI_ISL_3064989, EPI_ISL_3064990, EPI_ISL_3064992, EPI_ISL_3064993, EPI_ISL_3064994, EPI_ISL_3064995, EPI_ISL_3064996, EPI_ISL_3064997, EPI_ISL_3064998, EPI_ISL_3065000, EPI_ISL_3065001, EPI_ISL_3065002, EPI_ISL_3065003, EPI_ISL_3065004, EPI_ISL_3065005, EPI_ISL_3065006, EPI_ISL_3065008, EPI_ISL_3065009, EPI_ISL_3065010, EPI_ISL_3065011, EPI_ISL_3065012, EPI_ISL_3065013, EPI_ISL_3065014, EPI_ISL_3065015, EPI_ISL_3065016, EPI_ISL_3065017, EPI_ISL_3065019, EPI_ISL_3065020, EPI_ISL_3065021, EPI_ISL_3065022, EPI_ISL_3065023, EPI_ISL_3065024, EPI_ISL_3065025, EPI_ISL_3065026, EPI_ISL_3065027, EPI_ISL_3065028, EPI_ISL_3065031, EPI_ISL_3065032, EPI_ISL_3065033, EPI_ISL_3065034, EPI_ISL_3065035, EPI_ISL_3065036, EPI_ISL_3065037, EPI_ISL_3065038

| | | | |
|---|---|---|---|
| see above | Fulgent Genetics | Centers for Disease Control and Prevention Division of Viral Diseases, Pathogen Discovery | Dakota Howard, Dhwani Batra, Peter W. Cook, Kara Moser, Adrian Paskey, Jason Caravas, Benjamin Rambo-Martin, Shatavia Morrison, Christopher Gulvick, Scott Sammons, Yvette Unoarumhi, Darlene Wagner, Matthew Schmerer, Harry Gao, Mickey Li, John Gao, Joseph Fierro, Benafsh Sapra, Becky Tsai, Yan Meng, Doreen Ng, James Xie, Clinton R. Paden, Duncan MacCannell |

EPI_ISL_3065039, EPI_ISL_3065040, EPI_ISL_3065041, EPI_ISL_3065042, EPI_ISL_3065043, EPI_ISL_3065044, EPI_ISL_3065045, EPI_ISL_3065046, EPI_ISL_3065047, EPI_ISL_3065048, EPI_ISL_3065049, EPI_ISL_3065050, EPI_ISL_3065051, EPI_ISL_3065052, EPI_ISL_3065053, EPI_ISL_3065054, EPI_ISL_3065055, EPI_ISL_3065056, EPI_ISL_3065057, EPI_ISL_3065059, EPI_ISL_3065061, EPI_ISL_3065063, EPI_ISL_3065065, EPI_ISL_3065066, EPI_ISL_3065067, EPI_ISL_3065068, EPI_ISL_3065069, EPI_ISL_3065070, EPI_ISL_3065071, EPI_ISL_3065072, EPI_ISL_3065073, EPI_ISL_3065074, EPI_ISL_3065075, EPI_ISL_3065076, EPI_ISL_3065077, EPI_ISL_3065078, EPI_ISL_3065080, EPI_ISL_3065081, EPI_ISL_3065082, EPI_ISL_3065083, EPI_ISL_3065084, EPI_ISL_3065085, EPI_ISL_3065086, EPI_ISL_3065087, EPI_ISL_3065088, EPI_ISL_3065089, EPI_ISL_3065090, EPI_ISL_3065092, EPI_ISL_3065093, EPI_ISL_3065094, EPI_ISL_3065095, EPI_ISL_3065096, EPI_ISL_3065097, EPI_ISL_3065098, EPI_ISL_3065099, EPI_ISL_3065100, EPI_ISL_3065101, EPI_ISL_3065102, EPI_ISL_3065103, EPI_ISL_3065104, EPI_ISL_3065105, EPI_ISL_3065106, EPI_ISL_3065107, EPI_ISL_3065108, EPI_ISL_3065109, EPI_ISL_3065110, EPI_ISL_3065111, EPI_ISL_3065112, EPI_ISL_3065113, EPI_ISL_3065114, EPI_ISL_3065115, EPI_ISL_3065117, EPI_ISL_3065118, EPI_ISL_3065119, EPI_ISL_3065120, EPI_ISL_3065121, EPI_ISL_3065122, EPI_ISL_3065123, EPI_ISL_3065125, EPI_ISL_3065126, EPI_ISL_3065127, EPI_ISL_3065128, EPI_ISL_3065129, EPI_ISL_3065130, EPI_ISL_3065131, EPI_ISL_3065132, EPI_ISL_3065133, EPI_ISL_3065134, EPI_ISL_3065135, EPI_ISL_3065136, EPI_ISL_3065137, EPI_ISL_3065138, EPI_ISL_3065139, EPI_ISL_3065140, EPI_ISL_3065141, EPI_ISL_3065142, EPI_ISL_3065143, EPI_ISL_3065145, EPI_ISL_3065146, EPI_ISL_3065147, EPI_ISL_3065149, EPI_ISL_3065151, EPI_ISL_3065153, EPI_ISL_3065154, EPI_ISL_3065157, EPI_ISL_3065158, EPI_ISL_3065160, EPI_ISL_3065161, EPI_ISL_3065164, EPI_ISL_3065165, EPI_ISL_3065166, EPI_ISL_3065167, EPI_ISL_3065168, EPI_ISL_3065169, EPI_ISL_3065170, EPI_ISL_3065171, EPI_ISL_3065174, EPI_ISL_3065177, EPI_ISL_3065179, EPI_ISL_3065182, EPI_ISL_3065183, EPI_ISL_3065185, EPI_ISL_3065186, EPI_ISL_3065187, EPI_ISL_3065189, EPI_ISL_3065190, EPI_ISL_3065191, EPI_ISL_3065192, EPI_ISL_3065193, EPI_ISL_3065194, EPI_ISL_3065195, EPI_ISL_3065196, EPI_ISL_3065197, EPI_ISL_3065198, EPI_ISL_3065199, EPI_ISL_3065200, EPI_ISL_3065201, EPI_ISL_3065202, EPI_ISL_3065204, EPI_ISL_3065207, EPI_ISL_3065209, EPI_ISL_3065211, EPI_ISL_3065212, EPI_ISL_3065213, EPI_ISL_3065215, EPI_ISL_3065217, EPI_ISL_3065218, EPI_ISL_3065219, EPI_ISL_3065220, EPI_ISL_3065221, EPI_ISL_3065222, EPI_ISL_3065223, EPI_ISL_3065225, EPI_ISL_3065226, EPI_ISL_3065227, EPI_ISL_3065228, EPI_ISL_3065229, EPI_ISL_3065231, EPI_ISL_3065232, EPI_ISL_3065233, EPI_ISL_3065234, EPI_ISL_3065235, EPI_ISL_3065236, EPI_ISL_3065237, EPI_ISL_3065238, EPI_ISL_3065240, EPI_ISL_3065241, EPI_ISL_3065242, EPI_ISL_3065243, EPI_ISL_3065244, EPI_ISL_3065245, EPI_ISL_3065246, EPI_ISL_3065247, EPI_ISL_3065249, EPI_ISL_3065250, EPI_ISL_3065251, EPI_ISL_3065252, EPI_ISL_3065253, EPI_ISL_3065254, EPI_ISL_3065255, EPI_ISL_3065256, EPI_ISL_3065257, EPI_ISL_3065258, EPI_ISL_3065259, EPI_ISL_3065260, EPI_ISL_3065261, EPI_ISL_3065262, EPI_ISL_3065263, EPI_ISL_3065265, EPI_ISL_3065266, EPI_ISL_3065267, EPI_ISL_3065268, EPI_ISL_3065269, EPI_ISL_3065270, EPI_ISL_3065272, EPI_ISL_3065274, EPI_ISL_3065275, EPI_ISL_3065276, EPI_ISL_3065277, EPI_ISL_3065279, EPI_ISL_3065280, EPI_ISL_3065286, EPI_ISL_3065287, EPI_ISL_3065288, EPI_ISL_3065289, EPI_ISL_3065290, EPI_ISL_3065292, EPI_ISL_3065293, EPI_ISL_3065294, EPI_ISL_3065295, EPI_ISL_3065297, EPI_ISL_3065298, EPI_ISL_3065299, EPI_ISL_3065301, EPI_ISL_3065302, EPI_ISL_3065303, EPI_ISL_3065304, EPI_ISL_3065305, EPI_ISL_3065307, EPI_ISL_3065308, EPI_ISL_3065309, EPI_ISL_3065310, EPI_ISL_3065311, EPI_ISL_3065312, EPI_ISL_3065313, EPI_ISL_3065314, EPI_ISL_3065316, EPI_ISL_3065317, EPI_ISL_3065318, EPI_ISL_3065319, EPI_ISL_3065320, EPI_ISL_3065321, EPI_ISL_3065322, EPI_ISL_3065325, EPI_ISL_3065326, EPI_ISL_3065329, EPI_ISL_3065331, EPI_ISL_3065332, EPI_ISL_3065333, EPI_ISL_3065334, EPI_ISL_3065335, EPI_ISL_3065336, EPI_ISL_3065337, EPI_ISL_3065338, EPI_ISL_3065339, EPI_ISL_3065340, EPI_ISL_3065341, EPI_ISL_3065342, EPI_ISL_3065343, EPI_ISL_3065345, EPI_ISL_3065346, EPI_ISL_3065347, EPI_ISL_3065348, EPI_ISL_3065349, EPI_ISL_3065350, EPI_ISL_3065352, EPI_ISL_3065353, EPI_ISL_3065354, EPI_ISL_3065355, EPI_ISL_3065356, EPI_ISL_3065357, EPI_ISL_3065358, EPI_ISL_3065359, EPI_ISL_3065360, EPI_ISL_3065363, EPI_ISL_3065364, EPI_ISL_3065365, EPI_ISL_3065366, EPI_ISL_3065368, EPI_ISL_3065369, EPI_ISL_3065371, EPI_ISL_3065372, EPI_ISL_3065373, EPI_ISL_3065374, EPI_ISL_3065375, EPI_ISL_3065376, EPI_ISL_3065377, EPI_ISL_3065379, EPI_ISL_3065381, EPI_ISL_3065382, EPI_ISL_3065383, EPI_ISL_3065384, EPI_ISL_3065385, EPI_ISL_3065386, EPI_ISL_3065387, EPI_ISL_3065388, EPI_ISL_3065390, EPI_ISL_3065391, EPI_ISL_3065392, EPI_ISL_3065393, EPI_ISL_3065394, EPI_ISL_3065395, EPI_ISL_3065396, EPI_ISL_3065397, EPI_ISL_3065398, EPI_ISL_3065399, EPI_ISL_3065400, EPI_ISL_3065401, EPI_ISL_3065402, EPI_ISL_3065403, EPI_ISL_3065404, EPI_ISL_3065406, EPI_ISL_3065407, EPI_ISL_3065408, EPI_ISL_3065409, EPI_ISL_3065410, EPI_ISL_3065411, EPI_ISL_3065412, EPI_ISL_3065414, EPI_ISL_3065415, EPI_ISL_3065416, EPI_ISL_3065417, EPI_ISL_3065418, EPI_ISL_3065419, EPI_ISL_3065420, EPI_ISL_3065421, EPI_ISL_3065422, EPI_ISL_3065423, EPI_ISL_3065424, EPI_ISL_3065425, EPI_ISL_3065426, EPI_ISL_3065427, EPI_ISL_3065428, EPI_ISL_3065429, EPI_ISL_3065430, EPI_ISL_3065431, EPI_ISL_3065432, EPI_ISL_3065433, EPI_ISL_3065434, EPI_ISL_3065435, EPI_ISL_3065436, EPI_ISL_3065437, EPI_ISL_3065438, EPI_ISL_3065439, EPI_ISL_3065440, EPI_ISL_3065441, EPI_ISL_3065442, EPI_ISL_3065443, EPI_ISL_3065444, EPI_ISL_3065445, EPI_ISL_3065446, EPI_ISL_3065447, EPI_ISL_3065448, EPI_ISL_3065450, EPI_ISL_3065451, EPI_ISL_3065452, EPI_ISL_3065454, EPI_ISL_3065455, EPI_ISL_3065456, EPI_ISL_3065457, EPI_ISL_3065458, EPI_ISL_3065459, EPI_ISL_3065460, EPI_ISL_3065461, EPI_ISL_3065462, EPI_ISL_3065463, EPI_ISL_3065465, EPI_ISL_3065466, EPI_ISL_3065467, EPI_ISL_3065468, EPI_ISL_3065469, EPI_ISL_3065470, EPI_ISL_3065471, EPI_ISL_3065472, EPI_ISL_3065473, EPI_ISL_3065475, EPI_ISL_3065476, EPI_ISL_3065477, EPI_ISL_3065478, EPI_ISL_3065480, EPI_ISL_3065482, EPI_ISL_3065483, EPI_ISL_3065485, EPI_ISL_3065486, EPI_ISL_3065488, EPI_ISL_3065489, EPI_ISL_3065490, EPI_ISL_3065491, EPI_ISL_3065492, EPI_ISL_3065494, EPI_ISL_3065495, EPI_ISL_3065496, EPI_ISL_3065498, EPI_ISL_3065500, EPI_ISL_3065501, EPI_ISL_3065502, EPI_ISL_3065503, EPI_ISL_3065504

| | | | |
|---|---|---|---|
| see above | Mako Medical | Centers for Disease Control and Prevention Division of Viral Diseases, Pathogen Discovery | Dakota Howard, Dhwani Batra, Peter W. Cook, Kara Moser, Adrian Paskey, Jason Caravas, Benjamin Rambo-Martin, Shatavia Morrison, Christopher Gulvick, Scott Sammons, Yvette Unoarumhi, Darlene Wagner, Matthew Schmerer,Matthew Tugwell, Lauren Moon, Clinton R. Paden, Duncan MacCannell |
| EPI_ISL_3065505 | FUNDACIÓN VALLE DEL LILI | Laboratorio de Biotecnología, Universidad Icesi | María I. Gutiérrez López, Diana López-Álvarez, Ludwig Luis Albornoz-Tovar, Marcela Mercado-Reyes, Andrés M. Castillo-Giraldo, Beatriz Parra, María F. Villegas-Torres, Paola A. Caicedo |

EPI_ISL_3065507, EPI_ISL_3065508, EPI_ISL_3065509, EPI_ISL_3065510, EPI_ISL_3065511, EPI_ISL_3065512, EPI_ISL_3065513, EPI_ISL_3065514, EPI_ISL_3065515, EPI_ISL_3065516, EPI_ISL_3065517, EPI_ISL_3065518, EPI_ISL_3065519, EPI_ISL_3065520

| | | | |
|---|---|---|---|
| see above | Monterey County Public Health Laboratory | Monterey County Public Health Laboratory | Monterey County Public Health Laboratory |

EPI_ISL_3065521, EPI_ISL_3065522, EPI_ISL_3065523, EPI_ISL_3065524, EPI_ISL_3065525, EPI_ISL_3065526, EPI_ISL_3065527, EPI_ISL_3065528, EPI_ISL_3065529, EPI_ISL_3065530, EPI_ISL_3065531, EPI_ISL_3065532, EPI_ISL_3065533, EPI_ISL_3065534, EPI_ISL_3065535, EPI_ISL_3065536, EPI_ISL_3065537, EPI_ISL_3065538, EPI_ISL_3065539, EPI_ISL_3065540, EPI_ISL_3065541, EPI_ISL_3065542, EPI_ISL_3065543, EPI_ISL_3065544, EPI_ISL_3065545, EPI_ISL_3065546, EPI_ISL_3065547, EPI_ISL_3065548, EPI_ISL_3065549, EPI_ISL_3065550, EPI_ISL_3065551, EPI_ISL_3065552, EPI_ISL_3065553, EPI_ISL_3065554, EPI_ISL_3065555, EPI_ISL_3065556, EPI_ISL_3065557, EPI_ISL_3065558, EPI_ISL_3065561, EPI_ISL_3065562, EPI_ISL_3065563, EPI_ISL_3065564, EPI_ISL_3065565, EPI_ISL_3065566, EPI_ISL_3065567, EPI_ISL_3065568, EPI_ISL_3065569, EPI_ISL_3065570, EPI_ISL_3065571, EPI_ISL_3065572, EPI_ISL_3065573, EPI_ISL_3065574, EPI_ISL_3065575, EPI_ISL_3065576, EPI_ISL_3065577, EPI_ISL_3065578, EPI_ISL_3065579, EPI_ISL_3065580, EPI_ISL_3065581, EPI_ISL_3065582, EPI_ISL_3065583, EPI_ISL_3065584, EPI_ISL_3065585, EPI_ISL_3065586, EPI_ISL_3065587, EPI_ISL_3065588, EPI_ISL_3065589, EPI_ISL_3065590, EPI_ISL_3065591, EPI_ISL_3065592, EPI_ISL_3065593, EPI_ISL_3065594, EPI_ISL_3065595, EPI_ISL_3065596, EPI_ISL_3065597, EPI_ISL_3065598

| | | | |
|---|---|---|---|
| see above | Missouri State Public Health Laboratory | Missouri State Public Health Laboratory | Matthew Sinn, Joshua Barry, Ashley New |
| EPI_ISL_3065600, EPI_ISL_3065617 | OHSU-MM Lab | OHSU-MM Lab | Rebecca O'Gara |

EPI_ISL_3065859, EPI_ISL_3065860, EPI_ISL_3065861, EPI_ISL_3065862, EPI_ISL_3065863, EPI_ISL_3065864, EPI_ISL_3065865, EPI_ISL_3065866, EPI_ISL_3065867, EPI_ISL_3065868, EPI_ISL_3065869, EPI_ISL_3065870, EPI_ISL_3065871, EPI_ISL_3065872, EPI_ISL_3065873, EPI_ISL_3065874, EPI_ISL_3065875

| | | | |
|---|---|---|---|
| see above | Platform BIS UZA/UAntwerpen | Labo Klinische Biologie, UZA | Jasmine Coppens, Marie Le Mercier, Kathleen Holemans, Ines Verbesselt, Basil Britto Xavier, Christine Lammens, Veerle Matheeussen, Herman Goossens |
| EPI_ISL_3065876, EPI_ISL_3065877 | Centrum voor Medische Analyse | Labo Klinische Biologie, UZA | Jasmine Coppens, Marie Le Mercier, Kathleen Holemans, Ines Verbesselt, Basil Britto Xavier, Christine Lammens, Veerle Matheeussen, Herman Goossens |
| EPI_ISL_3065878 | Algemeen Klinisch Labo | Labo Klinische Biologie, UZA | Jasmine Coppens, Marie Le Mercier, Kathleen Holemans, Ines Verbesselt, Basil Britto Xavier, Christine Lammens, Veerle Matheeussen, Herman Goossens |
| EPI_ISL_3065879, EPI_ISL_3065880 | Centrum voor Medische Analyse | Labo Klinische Biologie, UZA | Jasmine Coppens, Marie Le Mercier, Kathleen Holemans, Ines Verbesselt, Basil Britto Xavier, Christine Lammens, Veerle Matheeussen, Herman Goossens |
| EPI_ISL_3065881 | Algemeen Klinisch Labo | Labo Klinische Biologie, UZA | Jasmine Coppens, Marie Le Mercier, Kathleen Holemans, Ines Verbesselt, Basil Britto Xavier, Christine Lammens, Veerle Matheeussen, Herman Goossens |
| EPI_ISL_3065882, EPI_ISL_3065883, EPI_ISL_3065884, | Centrum voor Medische Analyse | Labo Klinische Biologie, UZA | Jasmine Coppens, Marie Le Mercier, Kathleen Holemans, Ines Verbesselt, Basil Britto Xavier, Christine Lammens, Veerle Matheeussen, Herman Goossens |

| | | | |
|---|---|---|---|
| EPI_ISL_3065885 | | | |
| EPI_ISL_3065887, EPI_ISL_3065888, EPI_ISL_3065889, EPI_ISL_3065890, EPI_ISL_3065891, EPI_ISL_3065892, EPI_ISL_3065893, EPI_ISL_3065894, EPI_ISL_3065895, EPI_ISL_3065896, EPI_ISL_3065897, EPI_ISL_3065898, EPI_ISL_3065899, EPI_ISL_3065900, EPI_ISL_3065901, EPI_ISL_3065902, EPI_ISL_3065903, EPI_ISL_3065904, EPI_ISL_3065905, EPI_ISL_3065906, EPI_ISL_3065907, EPI_ISL_3065908 | | | |
| see above | Platform BIS UZA/UAntwerpen | Labo Klinische Biologie, UZA | Jasmine Coppens, Marie Le Mercier, Kathleen Holemans, Ines Verbesselt, Basil Britto Xavier, Christine Lammens, Veerle Matheeussen, Herman Goossens |
| EPI_ISL_3065909, EPI_ISL_3065910, EPI_ISL_3065911, EPI_ISL_3065912, EPI_ISL_3065913, EPI_ISL_3065914, EPI_ISL_3065915 | Algemeen Medisch Labo | Labo Klinische Biologie, UZA | Jasmine Coppens, Marie Le Mercier, Kathleen Holemans, Ines Verbesselt, Basil Britto Xavier, Christine Lammens, Veerle Matheeussen, Herman Goossens |
| EPI_ISL_3065916, EPI_ISL_3065917, EPI_ISL_3065918, EPI_ISL_3065919, EPI_ISL_3065920, EPI_ISL_3065921, EPI_ISL_3065922, EPI_ISL_3065923, EPI_ISL_3065924, EPI_ISL_3065925, EPI_ISL_3065926, EPI_ISL_3065927, EPI_ISL_3065928 | | | |
| see above | Platform BIS UZA/UAntwerpen | Labo Klinische Biologie, UZA | Jasmine Coppens, Marie Le Mercier, Kathleen Holemans, Ines Verbesselt, Basil Britto Xavier, Christine Lammens, Veerle Matheeussen, Herman Goossens |
| EPI_ISL_3065929, EPI_ISL_3065930, EPI_ISL_3065931, EPI_ISL_3065932, EPI_ISL_3065933, EPI_ISL_3065934, EPI_ISL_3065935, EPI_ISL_3065936, EPI_ISL_3065937, EPI_ISL_3065938, EPI_ISL_3065940, EPI_ISL_3065941, EPI_ISL_3065942, EPI_ISL_3065943, EPI_ISL_3065944, EPI_ISL_3065945 | | | |
| see above | Algemeen Medisch Labo | Labo Klinische Biologie, UZA | Jasmine Coppens, Marie Le Mercier, Kathleen Holemans, Ines Verbesselt, Basil Britto Xavier, Christine Lammens, Veerle Matheeussen, Herman Goossens |
| EPI_ISL_3065946 | PanGenomics International Pvt. Ltd., Ahmedabad | Gujarat Biotechnology Research Centre | Ramesh Pandit,Janvi Raval,Zarna Patel,Nitin Savaliya,Dinesh Kumar,Twinkle Soni,Sonal Sharma ,Zuber Saiyed,Krishna Mohan Singh,Umang Mishra,Nitesh Shah,Chaitanya Joshi,Madhvi Joshi |
| EPI_ISL_3065947, EPI_ISL_3066395 | PanGenomics International Pvt. Ltd., Ahmedabad | Gujarat Biotechnology Research Centre | Zarna Patel,Nitin Savaliya,Dinesh Kumar,Twinkle Soni,Sonal Sharma ,Zuber Saiyed,Ramesh Pandit,Janvi Raval,Krishna Mohan Singh,Umang Mishra,Nitesh Shah,Chaitanya Joshi,Madhvi Joshi |
| EPI_ISL_3066396 | PanGenomics International Pvt. Ltd., Ahmedabad | Gujarat Biotechnology Research Centre | Nitin Savaliya,Dinesh Kumar,Twinkle Soni,Sonal Sharma ,Zuber Saiyed,Ramesh Pandit,Janvi Raval,Zarna Patel,Krishna Mohan Singh,Umang Mishra,Nitesh Shah,Chaitanya Joshi,Madhvi Joshi |
| EPI_ISL_3066397 | PanGenomics International Pvt. Ltd., Ahmedabad | Gujarat Biotechnology Research Centre | Dinesh Kumar,Twinkle Soni,Sonal Sharma ,Zuber Saiyed,Ramesh Pandit,Janvi Raval,Zarna Patel,Nitin Savaliya,Krishna Mohan Singh,Umang Mishra,Nitesh Shah,Chaitanya Joshi,Madhvi Joshi |
| EPI_ISL_3066398 | PanGenomics International Pvt. Ltd., Ahmedabad | Gujarat Biotechnology Research Centre | Twinkle Soni, Sonal Sharma, Zuber Saiyed, Ramesh Pandit, Janvi Raval, Zarna Patel, Nitin Savaliya, Dinesh Kumar, Krishna Mohan Singh, Umang Mishra, Nitesh Shah, Chaitanya Joshi, Madhvi Joshi |
| EPI_ISL_3066399 | PanGenomics International Pvt. Ltd., Ahmedabad | Gujarat Biotechnology Research Centre | Sonal Sharma ,Zuber Saiyed,Ramesh Pandit,Janvi Raval,Zarna Patel,Nitin Savaliya,Dinesh Kumar,Twinkle Soni,Krishna Mohan Singh,Umang Mishra,Nitesh Shah,Chaitanya Joshi,Madhvi Joshi |
| EPI_ISL_3066400 | PanGenomics International Pvt. Ltd., Ahmedabad | Gujarat Biotechnology Research Centre | Zuber Saiyed,Ramesh Pandit,Janvi Raval,Zarna Patel,Nitin Savaliya,Dinesh Kumar,Twinkle Soni,Sonal Sharma ,Krishna Mohan Singh,Umang Mishra,Nitesh Shah,Chaitanya Joshi,Madhvi Joshi |
| EPI_ISL_3066401 | PanGenomics International Pvt. Ltd., Ahmedabad | Gujarat Biotechnology Research Centre | Ramesh Pandit,Janvi Raval,Zarna Patel,Nitin Savaliya,Dinesh Kumar,Twinkle Soni,Sonal Sharma ,Zuber Saiyed,Krishna Mohan Singh,Umang Mishra,Nitesh Shah,Chaitanya Joshi,Madhvi Joshi |
| EPI_ISL_3066402 | Supratech Micropath Laboratory Research Institute Pvt Ltd, Ahmedabad | Gujarat Biotechnology Research Centre | Janvi Raval,Zarna Patel,Nitin Savaliya,Dinesh Kumar,Twinkle Soni,Sonal Sharma ,Zuber Saiyed,Ramesh Pandit,Mahendra Parikh,Umang Mishra,Nitesh Shah,Chaitanya Joshi,Madhvi Joshi |
| EPI_ISL_3066403 | NHL Medical College, Ahmedabad | Gujarat Biotechnology Research Centre | Zarna Patel,Nitin Savaliya,Dinesh Kumar,Twinkle Soni,Sonal Sharma ,Zuber Saiyed,Ramesh Pandit,Janvi Raval,Jayshri Pethani,Umang Mishra,Nitesh Shah,Chaitanya Joshi,Madhvi Joshi |
| EPI_ISL_3066404 | NHL Medical College, Ahmedabad | Gujarat Biotechnology Research Centre | Nitin Savaliya,Dinesh Kumar,Twinkle Soni,Sonal Sharma ,Zuber Saiyed,Ramesh Pandit,Janvi Raval,Zarna Patel,Jayshri Pethani,Umang Mishra,Nitesh Shah,Chaitanya Joshi,Madhvi Joshi |
| EPI_ISL_3066405 | NHL Medical College, Ahmedabad | Gujarat Biotechnology Research Centre | Dinesh Kumar,Twinkle Soni,Sonal Sharma ,Zuber Saiyed,Ramesh Pandit,Janvi Raval,Zarna Patel,Nitin Savaliya,Jayshri Pethani,Umang Mishra,Nitesh Shah,Chaitanya Joshi,Madhvi Joshi |
| EPI_ISL_3066406 | NHL Medical College, Ahmedabad | Gujarat Biotechnology Research Centre | Twinkle Soni,Sonal Sharma ,Zuber Saiyed,Ramesh Pandit,Janvi Raval,Zarna Patel,Nitin Savaliya,Dinesh Kumar,Jayshri Pethani,Umang Mishra,Nitesh Shah,Chaitanya Joshi,Madhvi Joshi |
| EPI_ISL_3066407 | NHL Medical College, Ahmedabad | Gujarat Biotechnology Research Centre | Sonal Sharma ,Zuber Saiyed,Ramesh Pandit,Janvi Raval,Zarna Patel,Nitin Savaliya,Dinesh Kumar,Twinkle Soni,Jayshri Pethani,Umang Mishra,Nitesh Shah,Chaitanya Joshi,Madhvi Joshi |
| EPI_ISL_3066408 | NHL Medical College, Ahmedabad | Gujarat Biotechnology Research Centre | Zuber Saiyed,Ramesh Pandit,Janvi Raval,Zarna Patel,Nitin Savaliya,Dinesh Kumar,Twinkle Soni,Sonal Sharma ,Jayshri Pethani,Umang Mishra,Nitesh Shah,Chaitanya Joshi,Madhvi Joshi |
| EPI_ISL_3066409 | BJ Medical College Ahmedabad | Gujarat Biotechnology Research Centre | Ramesh Pandit,Janvi Raval,Zarna Patel,Nitin Savaliya,Dinesh Kumar,Twinkle Soni,Sonal Sharma ,Zuber Saiyed,Pranay Shah,Umang Mishra,Nitesh Shah,Chaitanya Joshi,Madhvi Joshi |
| EPI_ISL_3066410 | BJ Medical College Ahmedabad | Gujarat Biotechnology Research Centre | Janvi Raval,Zarna Patel,Nitin Savaliya,Dinesh Kumar,Twinkle Soni,Sonal Sharma ,Zuber Saiyed,Ramesh Pandit,Pranay Shah,Umang Mishra,Nitesh Shah,Chaitanya Joshi,Madhvi Joshi |
| EPI_ISL_3066411 | BJ Medical College Ahmedabad | Gujarat Biotechnology Research Centre | Zarna Patel,Nitin Savaliya,Dinesh Kumar,Twinkle Soni,Sonal Sharma ,Zuber Saiyed,Ramesh Pandit,Janvi Raval,Pranay Shah,Umang Mishra,Nitesh Shah,Chaitanya Joshi,Madhvi Joshi |
| EPI_ISL_3066412 | GMERS, Government Medical College, Himmatnagar | Gujarat Biotechnology Research Centre | Dinesh Kumar,Twinkle Soni,Sonal Sharma ,Zuber Saiyed,Ramesh Pandit,Janvi Raval,Zarna Patel,Nitin Savaliya,Himanshu Khatri,Umang Mishra,Nitesh Shah,Chaitanya Joshi,Madhvi Joshi |
| EPI_ISL_3066413 | GMERS, Government Medical College, Himmatnagar | Gujarat Biotechnology Research Centre | Twinkle Soni,Sonal Sharma ,Zuber Saiyed,Ramesh Pandit,Janvi Raval,Zarna Patel,Nitin Savaliya,Dinesh Kumar,Himanshu Khatri,Umang Mishra,Nitesh Shah,Chaitanya Joshi,Madhvi Joshi |
| EPI_ISL_3066415 | GMERS, Government Medical College, Himmatnagar | Gujarat Biotechnology Research Centre | Zuber Saiyed,Ramesh Pandit,Janvi Raval,Zarna Patel,Nitin Savaliya,Dinesh Kumar,Twinkle Soni,Sonal Sharma ,Himanshu Khatri,Umang Mishra,Nitesh Shah,Chaitanya Joshi,Madhvi Joshi |
| EPI_ISL_3066416 | GMERS, Government Medical College, Himmatnagar | Gujarat Biotechnology Research Centre | Ramesh Pandit,Janvi Raval,Zarna Patel,Nitin Savaliya,Dinesh Kumar,Twinkle Soni,Sonal Sharma ,Zuber Saiyed,Himanshu Khatri,Umang Mishra,Nitesh Shah,Chaitanya Joshi,Madhvi Joshi |
| EPI_ISL_3066417 | GMERS, Government Medical College, Himmatnagar | Gujarat Biotechnology Research Centre | Janvi Raval,Zarna Patel,Nitin Savaliya,Dinesh Kumar,Twinkle Soni,Sonal Sharma ,Zuber Saiyed,Ramesh Pandit,Himanshu Khatri,Umang Mishra,Nitesh Shah,Chaitanya Joshi,Madhvi Joshi |
| EPI_ISL_3066418 | GMERS, Government Medical College, Himmatnagar | Gujarat Biotechnology Research Centre | Nitin Savaliya,Dinesh Kumar,Twinkle Soni,Sonal Sharma ,Zuber Saiyed,Ramesh Pandit,Janvi Raval,Zarna Patel,Himanshu Khatri,Umang Mishra,Nitesh Shah,Chaitanya Joshi,Madhvi Joshi |
| EPI_ISL_3066419 | GMERS, Government Medical College, Himmatnagar | Gujarat Biotechnology Research Centre | Dinesh Kumar,Twinkle Soni,Sonal Sharma ,Zuber Saiyed,Ramesh Pandit,Janvi Raval,Zarna Patel,Nitin Savaliya,Himanshu Khatri,Umang Mishra,Nitesh Shah,Chaitanya Joshi,Madhvi Joshi |
| EPI_ISL_3066420 | GMERS, Government Medical College, Himmatnagar | Gujarat Biotechnology Research Centre | Twinkle Soni,Sonal Sharma ,Zuber Saiyed,Ramesh Pandit,Janvi Raval,Zarna Patel,Nitin Savaliya,Dinesh Kumar,Himanshu Khatri,Umang Mishra,Nitesh Shah,Chaitanya Joshi,Madhvi Joshi |
| EPI_ISL_3066421 | GMERS, Government Medical College, Himmatnagar | Gujarat Biotechnology Research Centre | Sonal Sharma ,Zuber Saiyed,Ramesh Pandit,Janvi Raval,Zarna Patel,Nitin Savaliya,Dinesh Kumar,Twinkle Soni,Himanshu Khatri,Umang Mishra,Nitesh Shah,Chaitanya Joshi,Madhvi Joshi |
| EPI_ISL_3066422 | GMERS, Government Medical College, Himmatnagar | Gujarat Biotechnology Research Centre | Zuber Saiyed,Ramesh Pandit,Janvi Raval,Zarna Patel,Nitin Savaliya,Dinesh Kumar,Twinkle Soni,Sonal Sharma ,Himanshu Khatri,Umang Mishra,Nitesh Shah,Chaitanya Joshi,Madhvi Joshi |
| EPI_ISL_3066423 | GMERS, Government Medical College, Himmatnagar | Gujarat Biotechnology Research Centre | Ramesh Pandit,Janvi Raval,Zarna Patel,Nitin Savaliya,Dinesh Kumar,Twinkle Soni,Sonal Sharma ,Zuber Saiyed,Himanshu Khatri,Umang Mishra,Nitesh Shah,Chaitanya Joshi,Madhvi Joshi |
| EPI_ISL_3066424 | GMERS, Government Medical College, Himmatnagar | Gujarat Biotechnology Research Centre | Janvi Raval,Zarna Patel,Nitin Savaliya,Dinesh Kumar,Twinkle Soni,Sonal Sharma ,Zuber Saiyed,Ramesh Pandit,Himanshu Khatri,Umang Mishra,Nitesh Shah,Chaitanya Joshi,Madhvi Joshi |
| EPI_ISL_3066425 | GMERS, Government Medical College, Himmatnagar | Gujarat Biotechnology Research Centre | Zarna Patel,Nitin Savaliya,Dinesh Kumar,Twinkle Soni,Sonal Sharma ,Zuber Saiyed,Ramesh Pandit,Janvi Raval,Himanshu Khatri,Umang Mishra,Nitesh Shah,Chaitanya Joshi,Madhvi Joshi |
| EPI_ISL_3066426 | GMERS, Government Medical College, Himmatnagar | Gujarat Biotechnology Research Centre | Nitin Savaliya,Dinesh Kumar,Twinkle Soni,Sonal Sharma ,Zuber Saiyed,Ramesh Pandit,Janvi Raval,Zarna Patel,Himanshu Khatri,Umang Mishra,Nitesh Shah,Chaitanya Joshi,Madhvi Joshi |
| EPI_ISL_3066428 | GMERS, Government Medical College, Himmatnagar | Gujarat Biotechnology Research Centre | Twinkle Soni,Sonal Sharma ,Zuber Saiyed,Ramesh Pandit,Janvi Raval,Zarna Patel,Nitin Savaliya,Dinesh Kumar,Himanshu Khatri,Umang Mishra,Nitesh Shah,Chaitanya Joshi,Madhvi Joshi |
| EPI_ISL_3066429 | GMERS, Government Medical College, Himmatnagar | Gujarat Biotechnology Research Centre | Zuber Saiyed,Ramesh Pandit,Janvi Raval,Zarna Patel,Nitin Savaliya,Dinesh Kumar,Twinkle Soni,Sonal Sharma ,Himanshu Khatri,Umang Mishra,Nitesh Shah,Chaitanya Joshi,Madhvi Joshi |
| EPI_ISL_3066430 | GMERS, Government Medical College, Himmatnagar | Gujarat Biotechnology Research Centre | Ramesh Pandit,Janvi Raval,Zarna Patel,Nitin Savaliya,Dinesh Kumar,Twinkle Soni,Sonal Sharma ,Zuber Saiyed,Himanshu Khatri,Umang Mishra,Nitesh Shah,Chaitanya Joshi,Madhvi Joshi |
| EPI_ISL_3066431 | GMERS, Government Medical College, Himmatnagar | Gujarat Biotechnology Research Centre | Janvi Raval,Zarna Patel,Nitin Savaliya,Dinesh Kumar,Twinkle Soni,Sonal Sharma ,Zuber Saiyed,Ramesh Pandit,Himanshu Khatri,Umang Mishra,Nitesh Shah,Chaitanya Joshi,Madhvi Joshi |
| EPI_ISL_3066432 | GMERS, Government Medical College, Himmatnagar | Gujarat Biotechnology Research Centre | Zarna Patel,Nitin Savaliya,Dinesh Kumar,Twinkle Soni,Sonal Sharma ,Zuber Saiyed,Ramesh Pandit,Janvi Raval,Himanshu Khatri,Umang Mishra,Nitesh Shah,Chaitanya Joshi,Madhvi Joshi |
| EPI_ISL_3066433 | GMERS, Government Medical College, Himmatnagar | Gujarat Biotechnology Research Centre | Nitin Savaliya,Dinesh Kumar,Twinkle Soni,Sonal Sharma ,Zuber Saiyed,Ramesh Pandit,Janvi Raval,Zarna Patel,Himanshu Khatri,Umang Mishra,Nitesh Shah,Chaitanya Joshi,Madhvi Joshi |
| EPI_ISL_3066434 | GMERS, Government Medical College, Himmatnagar | Gujarat Biotechnology Research Centre | Dinesh Kumar,Twinkle Soni,Sonal Sharma ,Zuber Saiyed,Ramesh Pandit,Janvi Raval,Zarna Patel,Nitin Savaliya,Himanshu Khatri,Umang Mishra,Nitesh Shah,Chaitanya Joshi,Madhvi Joshi |
| EPI_ISL_3066435 | GMERS, Government Medical College, Himmatnagar | Gujarat Biotechnology Research Centre | Twinkle Soni,Sonal Sharma ,Zuber Saiyed,Ramesh Pandit,Janvi Raval,Zarna Patel,Nitin Savaliya,Dinesh Kumar,Himanshu Khatri,Umang Mishra,Nitesh Shah,Chaitanya Joshi,Madhvi Joshi |
| EPI_ISL_3066437 | GMERS, Government Medical College, Himmatnagar | Gujarat Biotechnology Research Centre | Ramesh Pandit,Janvi Raval,Zarna Patel,Nitin Savaliya,Dinesh Kumar,Twinkle Soni,Sonal Sharma ,Zuber Saiyed,Himanshu Khatri,Umang Mishra,Nitesh Shah,Chaitanya Joshi,Madhvi Joshi |
| EPI_ISL_3066438 | GMERS, Government Medical College, Himmatnagar | Gujarat Biotechnology Research Centre | Janvi Raval,Zarna Patel,Nitin Savaliya,Dinesh Kumar,Twinkle Soni,Sonal Sharma ,Zuber Saiyed,Ramesh Pandit,Himanshu Khatri,Umang Mishra,Nitesh Shah,Chaitanya Joshi,Madhvi Joshi |
| EPI_ISL_3066439 | GMERS, Government Medical College, Himmatnagar | Gujarat Biotechnology Research Centre | Zarna Patel,Nitin Savaliya,Dinesh Kumar,Twinkle Soni,Sonal Sharma ,Zuber Saiyed,Ramesh Pandit,Janvi Raval,Himanshu Khatri,Umang Mishra,Nitesh Shah,Chaitanya Joshi,Madhvi Joshi |
| EPI_ISL_3066440 | GMERS, Government Medical College, Himmatnagar | Gujarat Biotechnology Research Centre | Twinkle Soni,Sonal Sharma ,Zuber Saiyed,Ramesh Pandit,Janvi Raval,Zarna Patel,Nitin Savaliya,Dinesh Kumar,Himanshu Khatri,Umang Mishra,Nitesh Shah,Chaitanya Joshi,Madhvi Joshi |
| EPI_ISL_3066441 | GMERS, Government Medical College, Himmatnagar | Gujarat Biotechnology Research Centre | Sonal Sharma ,Zuber Saiyed,Ramesh Pandit,Janvi Raval,Zarna Patel,Nitin Savaliya,Dinesh Kumar,Twinkle Soni,Himanshu Khatri,Umang Mishra,Nitesh Shah,Chaitanya Joshi,Madhvi Joshi |
| EPI_ISL_3066442 | GMERS, Government Medical College, Himmatnagar | Gujarat Biotechnology Research Centre | Ramesh Pandit,Janvi Raval,Zarna Patel,Nitin Savaliya,Dinesh Kumar,Twinkle Soni,Sonal Sharma ,Zuber Saiyed,Himanshu Khatri,Umang Mishra,Nitesh Shah,Chaitanya Joshi,Madhvi Joshi |
| EPI_ISL_3066443 | GMERS, Government Medical College, Himmatnagar | Gujarat Biotechnology Research Centre | Janvi Raval,Zarna Patel,Nitin Savaliya,Dinesh Kumar,Twinkle Soni,Sonal Sharma ,Zuber Saiyed,Ramesh Pandit,Himanshu Khatri,Umang Mishra,Nitesh Shah,Chaitanya Joshi,Madhvi Joshi |
| EPI_ISL_3066444 | Neuberg Supratech Reference Laboratory, Surat | Gujarat Biotechnology Research Centre | Zarna Patel,Nitin Savaliya,Dinesh Kumar,Twinkle Soni,Sonal Sharma ,Zuber Saiyed,Ramesh Pandit,Janvi Raval,Athmanathan Ganeshan,Umang Mishra,Nitesh Shah,Chaitanya Joshi,Madhvi Joshi |
| EPI_ISL_3066445 | Neuberg Supratech Reference Laboratory, Surat | Gujarat Biotechnology Research Centre | Nitin Savaliya,Dinesh Kumar,Twinkle Soni,Sonal Sharma ,Zuber Saiyed,Ramesh Pandit,Janvi Raval,Zarna Patel,Athmanathan Ganeshan,Umang Mishra,Nitesh Shah,Chaitanya Joshi,Madhvi Joshi |
| EPI_ISL_3066446 | Neuberg Supratech Reference Laboratory, Surat | Gujarat Biotechnology Research Centre | Dinesh Kumar,Twinkle Soni,Sonal Sharma ,Zuber Saiyed,Ramesh Pandit,Janvi Raval,Zarna Patel,Nitin Savaliya,Athmanathan Ganeshan,Umang Mishra,Nitesh Shah,Chaitanya Joshi,Madhvi Joshi |
| EPI_ISL_3066447 | Neuberg Supratech Reference Laboratory, Surat | Gujarat Biotechnology Research Centre | Twinkle Soni,Sonal Sharma ,Zuber Saiyed,Ramesh Pandit,Janvi Raval,Zarna Patel,Nitin Savaliya,Dinesh Kumar,Athmanathan Ganeshan,Umang Mishra,Nitesh Shah,Chaitanya Joshi,Madhvi Joshi |
| EPI_ISL_3066448 | Neuberg Supratech Reference Laboratory, Surat | Gujarat Biotechnology Research Centre | Sonal Sharma ,Zuber Saiyed,Ramesh Pandit,Janvi Raval,Zarna Patel,Nitin Savaliya,Dinesh Kumar,Twinkle Soni,Athmanathan Ganeshan,Umang Mishra,Nitesh Shah,Chaitanya Joshi,Madhvi Joshi |
| EPI_ISL_3066449 | GMERS, Medical College, Gotri | Gujarat Biotechnology Research Centre | Zuber Saiyed,Ramesh Pandit,Janvi Raval,Zarna Patel,Nitin Savaliya,Dinesh Kumar,Twinkle Soni,Sonal Sharma ,Bithika Duttaroy,Umang Mishra,Nitesh Shah,Chaitanya Joshi,Madhvi Joshi |
| EPI_ISL_3066532, EPI_ISL_3066540, EPI_ISL_3066544, EPI_ISL_3066670, EPI_ISL_3066671, EPI_ISL_3066731, EPI_ISL_3066732, EPI_ISL_3066750, EPI_ISL_3066752, EPI_ISL_3066753 | Ministry of Health Turkey | Ministry of Health Turkey | Fatma Bayrakdar, Yasemin Cosgun, Suleyman Yalcin, Gulay Korukluoglu |
| EPI_ISL_3066772, EPI_ISL_3066775, EPI_ISL_3066776, EPI_ISL_3066777, EPI_ISL_3066778, EPI_ISL_3066779, EPI_ISL_3066780, EPI_ISL_3066781, EPI_ISL_3066782, EPI_ISL_3066783, EPI_ISL_3066784, EPI_ISL_3066785, EPI_ISL_3066786, EPI_ISL_3066787, EPI_ISL_3066788, EPI_ISL_3066789, EPI_ISL_3066792, EPI_ISL_3066793, EPI_ISL_3066794, EPI_ISL_3066796, EPI_ISL_3066797, EPI_ISL_3066798, EPI_ISL_3066799, EPI_ISL_3066800, EPI_ISL_3066802, EPI_ISL_3066807, EPI_ISL_3066815, EPI_ISL_3066816, EPI_ISL_3066817 | | | |
| see above | Diagnostyka. Laboratoria Medyczne. | 1. ViroGenetics - BSL3 Laboratory of Virology, Małopolska Centre of | Rabalski L., Gromowski,T., Rogalska-Kupiec M., Swadzba J., Wydmanski,W., Mazur-Panasiuk,N., Kowalski,M., Maciej Kosinski, Natalia Derewonko, Szulc,P., Sylwia Januszczak, Labaj,P.P., Pyrc,K. |

| | | | |
|---|---|---|---|
| | | Biotechnology, Jagiellonian University; 2. Diagtron Laboratoria Łukasz Rąbalski | |
| EPI_ISL_3066819, EPI_ISL_3066820, EPI_ISL_3066823, EPI_ISL_3066825, EPI_ISL_3066826, EPI_ISL_3066827 | Indraprastha Apollo Hospital, New Delhi, India | NCDC Delhi, Biotechnology Division INSACOG | Meena Datta, Priyanka Singh, Uma Sharma, Hemlata Lall, Manoj K Singh, Radhakrishnan V. S, Robin Marwal, Mahesh S Dhar, Kalaiarasan Ponnusamy, Hema Gogia, Preeti Madan, Sandhya Kabra, Sujeet K Singh, Partha Rakshit |
| EPI_ISL_3066829, EPI_ISL_3066830, EPI_ISL_3066832, EPI_ISL_3066833, EPI_ISL_3066834, EPI_ISL_3066835, EPI_ISL_3066837 | Indraprastha Apollo Hospital, New Delhi, India | NCDC Delhi, Biotechnology Division INSACOG | Priyanka Singh, Uma Sharma, Hemlata Lall, Manoj K Singh, Radhakrishnan V. S, Robin Marwal, Mahesh S Dhar, Kalaiarasan Ponnusamy, Meena Datta, Hema Gogia, Preeti Madan, Sandhya Kabra, Sujeet K Singh, Partha Rakshit |
| EPI_ISL_3066838 | National Centre for Disease Control (NCDC) Biotechnology Division, Delhi | NCDC Delhi, Biotechnology Division INSACOG | Priyanka Singh, Uma Sharma, Hemlata Lall, Manoj K Singh, Radhakrishnan V. S, Robin Marwal, Mahesh S Dhar, Kalaiarasan Ponnusamy, Meena Datta, Hema Gogia, Preeti Madan, Sandhya Kabra, Sujeet K Singh, Partha Rakshit |
| EPI_ISL_3066843, EPI_ISL_3066848, EPI_ISL_3066849, EPI_ISL_3066850, EPI_ISL_3066851, EPI_ISL_3066852, EPI_ISL_3066853 | Northern Railway Central Hospital, New Delhi, India | NCDC Delhi, Biotechnology Division INSACOG | Priyanka Singh, Uma Sharma, Hemlata Lall, Manoj K Singh, Radhakrishnan V. S, Robin Marwal, Mahesh S Dhar, Kalaiarasan Ponnusamy, Meena Datta, Hema Gogia, Preeti Madan, Sandhya Kabra, Sujeet K Singh, Partha Rakshit |
| EPI_ISL_3066855 | NYU Langone Health | Departments of Pathology and Medicine, New York University School of Medicine | Adriana Heguy, Dacia Dimartino, Emily Guzman, Christian Marier, Peter Meyn, Sitharam Ramaswami, Gael Westby, Paul Zappile, Yutong Zhang, Paolo Cotzia, Guiqing Wang |
| EPI_ISL_3066877, EPI_ISL_3066879, EPI_ISL_3066880, EPI_ISL_3066881, EPI_ISL_3066882 | The Jackson Laboratory | The Jackson Laboratory | Long J, Renzette N, Adams M, Omerza G, Kelly K, Li L |
| EPI_ISL_3066883, EPI_ISL_3066884, EPI_ISL_3066885, EPI_ISL_3066887, EPI_ISL_3066888, EPI_ISL_3066889, EPI_ISL_3066890, EPI_ISL_3066891, EPI_ISL_3066892, EPI_ISL_3066893, EPI_ISL_3066894, EPI_ISL_3066895, EPI_ISL_3066896, EPI_ISL_3066897, EPI_ISL_3066899, EPI_ISL_3066901, EPI_ISL_3066903 | | | |
| see above | Wyoming Public Health Laboratory | Wyoming Public Health Laboratory | Jim Mildenberger, Taylor Fearing, Channing Weber, Ashley Norberg, Chayse Rowley, Marley Goetz, Brian Dominguez, Elliot Thomasson, Sam Britz, Cari Sloma, Robert Petit, and Rob Christensen |
| EPI_ISL_3066981, EPI_ISL_3066982, EPI_ISL_3066983, EPI_ISL_3066984, EPI_ISL_3066985, EPI_ISL_3066986 | Idaho Bureau of Laboratories | Center for Global Health, University of New Mexico Health Sciences Center | Daryl Domman, Kurt Schwalm, Valerie Morley, Matthew Burns, Robert Voermans, Christopher Ball, Darrell Dinwiddie |
| EPI_ISL_3066988 | Pathology and Laboratory Medicine Institute, Cleveland Clinic, Ohio, USA | Pathology and Laboratory Medicine Institute, Cleveland Clinic, Ohio, USA | Concetta Peck, Jennifer Starbuck, Joy Nakitandwe, Kristen McDonnell, David Plunkett, Zheng Jin Tu, Jay Brock, Yu-Wei Cheng, Gary Procop, Daniel Rhoads, Daniel H. Farkas, David Bosler |
| EPI_ISL_3067362, EPI_ISL_3067363, EPI_ISL_3067364, EPI_ISL_3067365, EPI_ISL_3067366, EPI_ISL_3067367, EPI_ISL_3067368, EPI_ISL_3067369, EPI_ISL_3067370, EPI_ISL_3067371, EPI_ISL_3067372, EPI_ISL_3067373, EPI_ISL_3067374, EPI_ISL_3067375, EPI_ISL_3067376, EPI_ISL_3067377, EPI_ISL_3067378, EPI_ISL_3067379, EPI_ISL_3067380, EPI_ISL_3067381, EPI_ISL_3067382, EPI_ISL_3067383, EPI_ISL_3067384, EPI_ISL_3067385, EPI_ISL_3067386, EPI_ISL_3067394 | | | |
| see above | Oregon State Public Health Laboratory | Oregon State Public Health Laboratory | Rafia Razzaque, Eugene Yeboah, Vanda Makris, Laura Tsaknaridis, John Fontana and Shane Sevey |
| EPI_ISL_3067398, EPI_ISL_3067399, EPI_ISL_3067400, EPI_ISL_3067401, EPI_ISL_3067402, EPI_ISL_3067403, EPI_ISL_3067405, EPI_ISL_3067407, EPI_ISL_3067408, EPI_ISL_3067409, EPI_ISL_3067410, EPI_ISL_3067411, EPI_ISL_3067412, EPI_ISL_3067413, EPI_ISL_3067414, EPI_ISL_3067415 | | | |
| see above | TXDSHS | TXDSHS | Rashmi Tuladhar, Bonnie Oh, Jenny Zhang, Maliha Rahman, Mayela Pedrueza, Anita Pokharel, Karen Bobier, Lorraine Rodriguez, Myong Koag, Chun Wang, Rachel Lee, Grace Kubin |
| EPI_ISL_3067421 | Southern Nevada Public Health Laboratory | Southern Nevada Public Health Laboratory | Michael Picker |
| EPI_ISL_3067496, EPI_ISL_3067497, EPI_ISL_3067498, EPI_ISL_3067499, EPI_ISL_3067500, EPI_ISL_3067501, EPI_ISL_3067502, EPI_ISL_3067503, EPI_ISL_3067504, EPI_ISL_3067505, EPI_ISL_3067507, EPI_ISL_3067508, EPI_ISL_3067510, EPI_ISL_3067511, EPI_ISL_3067512, EPI_ISL_3067513, EPI_ISL_3067514, EPI_ISL_3067515, EPI_ISL_3067516, EPI_ISL_3067517, EPI_ISL_3067518, EPI_ISL_3067519, EPI_ISL_3067520, EPI_ISL_3067522, EPI_ISL_3067523, EPI_ISL_3067524, EPI_ISL_3067525, EPI_ISL_3067526, EPI_ISL_3067527, EPI_ISL_3067528, EPI_ISL_3067529, EPI_ISL_3067530, EPI_ISL_3067531, EPI_ISL_3067532, EPI_ISL_3067533, EPI_ISL_3067534, EPI_ISL_3067535, EPI_ISL_3067536, EPI_ISL_3067537, EPI_ISL_3067538, EPI_ISL_3067539, EPI_ISL_3067540, EPI_ISL_3067541, EPI_ISL_3067542, EPI_ISL_3067543, EPI_ISL_3067544, EPI_ISL_3067545, EPI_ISL_3067546, EPI_ISL_3067547, EPI_ISL_3067548, EPI_ISL_3067549, EPI_ISL_3067550, EPI_ISL_3067553, EPI_ISL_3067554, EPI_ISL_3067555, EPI_ISL_3067556, EPI_ISL_3067557, EPI_ISL_3067558, EPI_ISL_3067559, EPI_ISL_3067560, EPI_ISL_3067562, EPI_ISL_3067563, EPI_ISL_3067564, EPI_ISL_3067565, EPI_ISL_3067566, EPI_ISL_3067567, EPI_ISL_3067568, EPI_ISL_3067569, EPI_ISL_3067570, EPI_ISL_3067571 | | | |
| see above | Alaska State Virology Laboratory | Alaska State Virology Laboratory | Stephanie DeRonde, Elva House, Jacob Zidek, Lisa Smith, Ph.D., Jack Chen, Ph.D. |
| EPI_ISL_3067574, EPI_ISL_3067575, EPI_ISL_3067576, EPI_ISL_3067577, EPI_ISL_3067578, EPI_ISL_3067579, EPI_ISL_3067580, EPI_ISL_3067581, EPI_ISL_3067582, EPI_ISL_3067583, EPI_ISL_3067585, EPI_ISL_3067586, EPI_ISL_3067587, EPI_ISL_3067588, EPI_ISL_3067589, EPI_ISL_3067590, EPI_ISL_3067591, EPI_ISL_3067592, EPI_ISL_3067593, EPI_ISL_3067594, EPI_ISL_3067595, EPI_ISL_3067596, EPI_ISL_3067597, EPI_ISL_3067598, EPI_ISL_3067603, EPI_ISL_3067604, EPI_ISL_3067606, EPI_ISL_3067607, EPI_ISL_3067608, EPI_ISL_3067609, EPI_ISL_3067610, EPI_ISL_3067611, EPI_ISL_3067612, EPI_ISL_3067614, EPI_ISL_3067615, EPI_ISL_3067616, EPI_ISL_3067617, EPI_ISL_3067618, EPI_ISL_3067619, EPI_ISL_3067620, EPI_ISL_3067621, EPI_ISL_3067622, EPI_ISL_3067623, EPI_ISL_3067624, EPI_ISL_3067625, EPI_ISL_3067626, EPI_ISL_3067627, EPI_ISL_3067628, EPI_ISL_3067629, EPI_ISL_3067630, EPI_ISL_3067631, EPI_ISL_3067632, EPI_ISL_3067633, EPI_ISL_3067634, EPI_ISL_3067635, EPI_ISL_3067636, EPI_ISL_3067637, EPI_ISL_3067638, EPI_ISL_3067639, EPI_ISL_3067640, EPI_ISL_3067641, EPI_ISL_3067642, EPI_ISL_3067643, EPI_ISL_3067644, EPI_ISL_3067645, EPI_ISL_3067646, EPI_ISL_3067647, EPI_ISL_3067648, EPI_ISL_3067649, EPI_ISL_3067650, EPI_ISL_3067651 | | | |
| see above | Santa Clara Valley Medical Center | Santa Clara County Public Health Laboratory | Santa Clara County Public Health Department |
| EPI_ISL_3067652, EPI_ISL_3067653 | Instituto Nacional de Medicina Genomica | Instituto Nacional de Medicina Genomica | Cedro-Tanda A, Hidalgo-Miranda A, Mendoza-Vargas A, Reyes-Grajeda JP, Abraham Campos-Romero, Moreno-Camacho José Luis, Rodriguez-Gallegos Jorge, Luna-Ruiz Marco, Gonzalez-Barrera D, Rangel-DeLeon D, Munguia-Garza P, Ramirez-Vega O, Escobar-Arrazola, M, Herrera-Montalvo LA. |
| EPI_ISL_3067654 | Centro Medico ABC | Instituto Nacional de Medicina Genomica | Cedro-Tanda A, Hidalgo-Miranda A, Mendoza-Vargas A, Reyes-Grajeda JP, Roxana Trejo-González, Cesar Lara, Jacomine Reyes-Carrasco, Daniel Aguirre, Gonzalez-Barrera D, Rangel-DeLeon D, Munguia-Garza P, Ramirez-Vega O, Escobar-Arrazola, M, Herrera-Montalvo LA. |
| EPI_ISL_3067655 | Instituto Nacional de Medicina Genomica | Instituto Nacional de Medicina Genomica | Cedro-Tanda A, Hidalgo-Miranda A, Mendoza-Vargas A, Reyes-Grajeda JP, Abraham Campos-Romero, Moreno-Camacho José Luis, Rodriguez-Gallegos Jorge, Luna-Ruiz Marco, Gonzalez-Barrera D, Rangel-DeLeon D, Munguia-Garza P, Ramirez-Vega O, Escobar-Arrazola, M, Herrera-Montalvo LA. |
| EPI_ISL_3067657 | INCMNSZ | Instituto Nacional de Medicina Genomica | Cedro-Tanda A, Hidalgo-Miranda A, Mendoza-Vargas A, Reyes-Grajeda JP, Abraham Campos-Romero, Moreno-Camacho José Luis, Rodriguez-Gallegos Jorge, Luna-Ruiz Marco, Gonzalez-Barrera D, Rangel-DeLeon D, Munguia-Garza P, Ramirez-Vega O, Escobar-Arrazola, M, Herrera-Montalvo LA. |
| EPI_ISL_3067658 | Salud Digna | Instituto Nacional de Medicina Genomica | Cedro-Tanda A, Hidalgo-Miranda A, Mendoza-Vargas A, Reyes-Grajeda JP, Abraham Campos-Romero, Moreno-Camacho José Luis, Rodriguez-Gallegos Jorge, Luna-Ruiz Marco, Gonzalez-Barrera D, Rangel-DeLeon D, Munguia-Garza P, Ramirez-Vega O, Escobar-Arrazola, M, Herrera-Montalvo LA. |
| EPI_ISL_3067660 | Centro Medico ABC | Instituto Nacional de Medicina Genomica | Cedro-Tanda A, Hidalgo-Miranda A, Mendoza-Vargas A, Reyes-Grajeda JP, Roxana Trejo-González, Cesar Lara, Jacomine Reyes-Carrasco, Daniel Aguirre, Gonzalez-Barrera D, Rangel-DeLeon D, Munguia-Garza P, Ramirez-Vega O, Escobar-Arrazola, M, Herrera-Montalvo LA. |
| EPI_ISL_3067662, EPI_ISL_3067664 | Instituto Nacional de Medicina Genomica | Instituto Nacional de Medicina Genomica | Cedro-Tanda A, Hidalgo-Miranda A, Mendoza-Vargas A, Reyes-Grajeda JP, Abraham Campos-Romero, Moreno-Camacho José Luis, Rodriguez-Gallegos Jorge, Luna-Ruiz Marco, Gonzalez-Barrera D, Rangel-DeLeon D, Munguia-Garza P, Ramirez-Vega O, Escobar-Arrazola, M, Herrera-Montalvo LA. |
| EPI_ISL_3067665 | INCMNSZ | Instituto Nacional de Medicina Genomica | Cedro-Tanda A, Hidalgo-Miranda A, Mendoza-Vargas A, Reyes-Grajeda JP, Abraham Campos-Romero, Moreno-Camacho José Luis, Rodriguez-Gallegos Jorge, Luna-Ruiz Marco, Gonzalez-Barrera D, Rangel-DeLeon D, Munguia-Garza P, Ramirez-Vega O, Escobar-Arrazola, M, Herrera-Montalvo LA. |
| EPI_ISL_3067666, EPI_ISL_3067667, EPI_ISL_3067668, EPI_ISL_3067669, EPI_ISL_3067671 | Instituto Nacional de Medicina Genomica | Instituto Nacional de Medicina Genomica | Cedro-Tanda A, Hidalgo-Miranda A, Mendoza-Vargas A, Reyes-Grajeda JP, Abraham Campos-Romero, Moreno-Camacho José Luis, Rodriguez-Gallegos Jorge, Luna-Ruiz Marco, Gonzalez-Barrera D, Rangel-DeLeon D, Munguia-Garza P, Ramirez-Vega O, Escobar-Arrazola, M, Herrera-Montalvo LA. |
| EPI_ISL_3067672 | Salud Digna | Instituto Nacional de Medicina Genomica | Cedro-Tanda A, Hidalgo-Miranda A, Mendoza-Vargas A, Reyes-Grajeda JP, Abraham Campos-Romero, Moreno-Camacho José Luis, Rodriguez-Gallegos Jorge, Luna-Ruiz Marco, Gonzalez-Barrera D, Rangel-DeLeon D, Munguia-Garza P, Ramirez-Vega O, Escobar-Arrazola, M, Herrera-Montalvo LA. |
| EPI_ISL_3067673 | INCMNSZ | Instituto Nacional de Medicina Genomica | Cedro-Tanda A, Hidalgo-Miranda A, Mendoza-Vargas A, Reyes-Grajeda JP, Abraham Campos-Romero, Moreno-Camacho José Luis, Rodriguez-Gallegos Jorge, Luna-Ruiz Marco, Gonzalez-Barrera D, Rangel-DeLeon D, Munguia-Garza P, Ramirez-Vega O, Escobar-Arrazola, M, Herrera-Montalvo LA. |
| EPI_ISL_3067674 | Instituto Nacional de Medicina Genomica | Instituto Nacional de Medicina Genomica | Cedro-Tanda A, Hidalgo-Miranda A, Mendoza-Vargas A, Reyes-Grajeda JP, Abraham Campos-Romero, Moreno-Camacho José Luis, Rodriguez-Gallegos Jorge, Luna-Ruiz Marco, Gonzalez-Barrera D, Rangel-DeLeon D, Munguia-Garza P, Ramirez-Vega O, Escobar-Arrazola, M, Herrera-Montalvo LA. |
| EPI_ISL_3067675 | Salud Digna | Instituto Nacional de Medicina Genomica | Cedro-Tanda A, Hidalgo-Miranda A, Mendoza-Vargas A, Reyes-Grajeda JP, Abraham Campos-Romero, Moreno-Camacho José Luis, Rodriguez-Gallegos Jorge, Luna-Ruiz Marco, Gonzalez-Barrera D, Rangel-DeLeon D, Munguia-Garza P, Ramirez-Vega O, Escobar-Arrazola, M, Herrera-Montalvo LA. |
| EPI_ISL_3067676 | Instituto Nacional de Medicina Genomica | Instituto Nacional de Medicina Genomica | Cedro-Tanda A, Hidalgo-Miranda A, Mendoza-Vargas A, Reyes-Grajeda JP, Abraham Campos-Romero, Moreno-Camacho José Luis, Rodriguez-Gallegos Jorge, Luna-Ruiz Marco, Gonzalez-Barrera D, Rangel-DeLeon D, Munguia-Garza P, Ramirez-Vega O, Escobar-Arrazola, M, Herrera-Montalvo LA. |
| EPI_ISL_3067679 | Salud Digna | Instituto Nacional de Medicina Genomica | Cedro-Tanda A, Hidalgo-Miranda A, Mendoza-Vargas A, Reyes-Grajeda JP, Abraham Campos-Romero, Moreno-Camacho José Luis, Rodriguez-Gallegos Jorge, Luna-Ruiz Marco, Gonzalez-Barrera D, Rangel-DeLeon D, Munguia-Garza P, Ramirez-Vega O, Escobar-Arrazola, M, Herrera-Montalvo LA. |
| EPI_ISL_3067680 | Instituto Nacional de Medicina Genomica | Instituto Nacional de Medicina Genomica | Cedro-Tanda A, Hidalgo-Miranda A, Mendoza-Vargas A, Reyes-Grajeda JP, Abraham Campos-Romero, Moreno-Camacho José Luis, Rodriguez-Gallegos Jorge, Luna-Ruiz Marco, Gonzalez-Barrera D, Rangel-DeLeon D, Munguia-Garza P, Ramirez-Vega O, Escobar-Arrazola, M, Herrera-Montalvo LA. |
| EPI_ISL_3067681 | Salud Digna | Instituto Nacional de Medicina Genomica | Cedro-Tanda A, Hidalgo-Miranda A, Mendoza-Vargas A, Reyes-Grajeda JP, Abraham Campos-Romero, Moreno-Camacho José Luis, Rodriguez-Gallegos Jorge, Luna-Ruiz Marco, Gonzalez-Barrera D, Rangel-DeLeon D, Munguia-Garza P, Ramirez-Vega O, Escobar-Arrazola, M, Herrera-Montalvo LA. |

| | | | |
|---|---|---|---|
| EPI_ISL_3067682, EPI_ISL_3067683 | Instituto Nacional de Medicina Genomica | Instituto Nacional de Medicina Genomica | Cedro-Tanda A, Hidalgo-Miranda A, Mendoza-Vargas A, Reyes-Grajeda JP, Abraham Campos-Romero, Moreno-Camacho José Luis, Rodriguez-Gallegos Jorge, Luna-Ruiz Marco, Gonzalez-Barrera D, Rangel-DeLeon D, Munguia-Garza P, Ramirez-Vega O, Escobar-Arrazola, M, Herrera-Montalvo LA. |
| EPI_ISL_3067684, EPI_ISL_3067686, EPI_ISL_3067687 | Salud Digna | Instituto Nacional de Medicina Genomica | Cedro-Tanda A, Hidalgo-Miranda A, Mendoza-Vargas A, Reyes-Grajeda JP, Abraham Campos-Romero, Moreno-Camacho José Luis, Rodriguez-Gallegos Jorge, Luna-Ruiz Marco, Gonzalez-Barrera D, Rangel-DeLeon D, Munguia-Garza P, Ramirez-Vega O, Escobar-Arrazola, M, Herrera-Montalvo LA. |
| EPI_ISL_3067688 | Instituto Nacional de Medicina Genomica | Instituto Nacional de Medicina Genomica | Cedro-Tanda A, Hidalgo-Miranda A, Mendoza-Vargas A, Reyes-Grajeda JP, Abraham Campos-Romero, Moreno-Camacho José Luis, Rodriguez-Gallegos Jorge, Luna-Ruiz Marco, Gonzalez-Barrera D, Rangel-DeLeon D, Munguia-Garza P, Ramirez-Vega O, Escobar-Arrazola, M, Herrera-Montalvo LA. |
| EPI_ISL_3067689 | Salud Digna | Instituto Nacional de Medicina Genomica | Cedro-Tanda A, Hidalgo-Miranda A, Mendoza-Vargas A, Reyes-Grajeda JP, Abraham Campos-Romero, Moreno-Camacho José Luis, Rodriguez-Gallegos Jorge, Luna-Ruiz Marco, Gonzalez-Barrera D, Rangel-DeLeon D, Munguia-Garza P, Ramirez-Vega O, Escobar-Arrazola, M, Herrera-Montalvo LA. |
| EPI_ISL_3067690 | INCMNSZ | Instituto Nacional de Medicina Genomica | Cedro-Tanda A, Hidalgo-Miranda A, Mendoza-Vargas A, Reyes-Grajeda JP, Abraham Campos-Romero, Moreno-Camacho José Luis, Rodriguez-Gallegos Jorge, Luna-Ruiz Marco, Gonzalez-Barrera D, Rangel-DeLeon D, Munguia-Garza P, Ramirez-Vega O, Escobar-Arrazola, M, Herrera-Montalvo LA. |
| EPI_ISL_3067691 | Salud Digna | Instituto Nacional de Medicina Genomica | Cedro-Tanda A, Hidalgo-Miranda A, Mendoza-Vargas A, Reyes-Grajeda JP, Abraham Campos-Romero, Moreno-Camacho José Luis, Rodriguez-Gallegos Jorge, Luna-Ruiz Marco, Gonzalez-Barrera D, Rangel-DeLeon D, Munguia-Garza P, Ramirez-Vega O, Escobar-Arrazola, M, Herrera-Montalvo LA. |
| EPI_ISL_3067692 | Instituto Nacional de Medicina Genomica | Instituto Nacional de Medicina Genomica | Cedro-Tanda A, Hidalgo-Miranda A, Mendoza-Vargas A, Reyes-Grajeda JP, Abraham Campos-Romero, Moreno-Camacho José Luis, Rodriguez-Gallegos Jorge, Luna-Ruiz Marco, Gonzalez-Barrera D, Rangel-DeLeon D, Munguia-Garza P, Ramirez-Vega O, Escobar-Arrazola, M, Herrera-Montalvo LA. |
| EPI_ISL_3067693 | INCMNSZ | Instituto Nacional de Medicina Genomica | Cedro-Tanda A, Hidalgo-Miranda A, Mendoza-Vargas A, Reyes-Grajeda JP, Abraham Campos-Romero, Moreno-Camacho José Luis, Rodriguez-Gallegos Jorge, Luna-Ruiz Marco, Gonzalez-Barrera D, Rangel-DeLeon D, Munguia-Garza P, Ramirez-Vega O, Escobar-Arrazola, M, Herrera-Montalvo LA. |
| EPI_ISL_3067694 | Instituto Nacional de Medicina Genomica | Instituto Nacional de Medicina Genomica | Cedro-Tanda A, Hidalgo-Miranda A, Mendoza-Vargas A, Reyes-Grajeda JP, Abraham Campos-Romero, Moreno-Camacho José Luis, Rodriguez-Gallegos Jorge, Luna-Ruiz Marco, Gonzalez-Barrera D, Rangel-DeLeon D, Munguia-Garza P, Ramirez-Vega O, Escobar-Arrazola, M, Herrera-Montalvo LA. |
| EPI_ISL_3067695 | Salud Digna | Instituto Nacional de Medicina Genomica | Cedro-Tanda A, Hidalgo-Miranda A, Mendoza-Vargas A, Reyes-Grajeda JP, Abraham Campos-Romero, Moreno-Camacho José Luis, Rodriguez-Gallegos Jorge, Luna-Ruiz Marco, Gonzalez-Barrera D, Rangel-DeLeon D, Munguia-Garza P, Ramirez-Vega O, Escobar-Arrazola, M, Herrera-Montalvo LA. |
| EPI_ISL_3067696 | Instituto Nacional de Medicina Genomica | Instituto Nacional de Medicina Genomica | Cedro-Tanda A, Hidalgo-Miranda A, Mendoza-Vargas A, Reyes-Grajeda JP, Abraham Campos-Romero, Moreno-Camacho José Luis, Rodriguez-Gallegos Jorge, Luna-Ruiz Marco, Gonzalez-Barrera D, Rangel-DeLeon D, Munguia-Garza P, Ramirez-Vega O, Escobar-Arrazola, M, Herrera-Montalvo LA. |
| EPI_ISL_3067697, EPI_ISL_3067698 | Salud Digna | Instituto Nacional de Medicina Genomica | Cedro-Tanda A, Hidalgo-Miranda A, Mendoza-Vargas A, Reyes-Grajeda JP, Abraham Campos-Romero, Moreno-Camacho José Luis, Rodriguez-Gallegos Jorge, Luna-Ruiz Marco, Gonzalez-Barrera D, Rangel-DeLeon D, Munguia-Garza P, Ramirez-Vega O, Escobar-Arrazola, M, Herrera-Montalvo LA. |
| EPI_ISL_3067699, EPI_ISL_3067700, EPI_ISL_3067701 | Instituto Nacional de Medicina Genomica | Instituto Nacional de Medicina Genomica | Cedro-Tanda A, Hidalgo-Miranda A, Mendoza-Vargas A, Reyes-Grajeda JP, Abraham Campos-Romero, Moreno-Camacho José Luis, Rodriguez-Gallegos Jorge, Luna-Ruiz Marco, Gonzalez-Barrera D, Rangel-DeLeon D, Munguia-Garza P, Ramirez-Vega O, Escobar-Arrazola, M, Herrera-Montalvo LA. |
| EPI_ISL_3067702, EPI_ISL_3067703 | Salud Digna | Instituto Nacional de Medicina Genomica | Cedro-Tanda A, Hidalgo-Miranda A, Mendoza-Vargas A, Reyes-Grajeda JP, Abraham Campos-Romero, Moreno-Camacho José Luis, Rodriguez-Gallegos Jorge, Luna-Ruiz Marco, Gonzalez-Barrera D, Rangel-DeLeon D, Munguia-Garza P, Ramirez-Vega O, Escobar-Arrazola, M, Herrera-Montalvo LA. |
| EPI_ISL_3067704, EPI_ISL_3067705, EPI_ISL_3067706 | INCMNSZ | Instituto Nacional de Medicina Genomica | Cedro-Tanda A, Hidalgo-Miranda A, Mendoza-Vargas A, Reyes-Grajeda JP, Abraham Campos-Romero, Moreno-Camacho José Luis, Rodriguez-Gallegos Jorge, Luna-Ruiz Marco, Gonzalez-Barrera D, Rangel-DeLeon D, Munguia-Garza P, Ramirez-Vega O, Escobar-Arrazola, M, Herrera-Montalvo LA. |
| EPI_ISL_3067707 | Salud Digna | Instituto Nacional de Medicina Genomica | Cedro-Tanda A, Hidalgo-Miranda A, Mendoza-Vargas A, Reyes-Grajeda JP, Abraham Campos-Romero, Moreno-Camacho José Luis, Rodriguez-Gallegos Jorge, Luna-Ruiz Marco, Gonzalez-Barrera D, Rangel-DeLeon D, Munguia-Garza P, Ramirez-Vega O, Escobar-Arrazola, M, Herrera-Montalvo LA. |
| EPI_ISL_3067708, EPI_ISL_3067709 | INCMNSZ | Instituto Nacional de Medicina Genomica | Cedro-Tanda A, Hidalgo-Miranda A, Mendoza-Vargas A, Reyes-Grajeda JP, Abraham Campos-Romero, Moreno-Camacho José Luis, Rodriguez-Gallegos Jorge, Luna-Ruiz Marco, Gonzalez-Barrera D, Rangel-DeLeon D, Munguia-Garza P, Ramirez-Vega O, Escobar-Arrazola, M, Herrera-Montalvo LA. |
| EPI_ISL_3067711 | Instituto Nacional de Medicina Genomica | Instituto Nacional de Medicina Genomica | Cedro-Tanda A, Hidalgo-Miranda A, Mendoza-Vargas A, Reyes-Grajeda JP, Abraham Campos-Romero, Moreno-Camacho José Luis, Rodriguez-Gallegos Jorge, Luna-Ruiz Marco, Gonzalez-Barrera D, Rangel-DeLeon D, Munguia-Garza P, Ramirez-Vega O, Escobar-Arrazola, M, Herrera-Montalvo LA. |
| EPI_ISL_3067713, EPI_ISL_3067714 | INCMNSZ | Instituto Nacional de Medicina Genomica | Cedro-Tanda A, Hidalgo-Miranda A, Mendoza-Vargas A, Reyes-Grajeda JP, Abraham Campos-Romero, Moreno-Camacho José Luis, Rodriguez-Gallegos Jorge, Luna-Ruiz Marco, Gonzalez-Barrera D, Rangel-DeLeon D, Munguia-Garza P, Ramirez-Vega O, Escobar-Arrazola, M, Herrera-Montalvo LA. |
| EPI_ISL_3067715 | Instituto Nacional de Medicina Genomica | Instituto Nacional de Medicina Genomica | Cedro-Tanda A, Hidalgo-Miranda A, Mendoza-Vargas A, Reyes-Grajeda JP, Abraham Campos-Romero, Moreno-Camacho José Luis, Rodriguez-Gallegos Jorge, Luna-Ruiz Marco, Gonzalez-Barrera D, Rangel-DeLeon D, Munguia-Garza P, Ramirez-Vega O, Escobar-Arrazola, M, Herrera-Montalvo LA. |
| EPI_ISL_3067716 | Salud Digna | Instituto Nacional de Medicina Genomica | Cedro-Tanda A, Hidalgo-Miranda A, Mendoza-Vargas A, Reyes-Grajeda JP, Abraham Campos-Romero, Moreno-Camacho José Luis, Rodriguez-Gallegos Jorge, Luna-Ruiz Marco, Gonzalez-Barrera D, Rangel-DeLeon D, Munguia-Garza P, Ramirez-Vega O, Escobar-Arrazola, M, Herrera-Montalvo LA. |
| EPI_ISL_3067717 | INCMNSZ | Instituto Nacional de Medicina Genomica | Cedro-Tanda A, Hidalgo-Miranda A, Mendoza-Vargas A, Reyes-Grajeda JP, Abraham Campos-Romero, Moreno-Camacho José Luis, Rodriguez-Gallegos Jorge, Luna-Ruiz Marco, Gonzalez-Barrera D, Rangel-DeLeon D, Munguia-Garza P, Ramirez-Vega O, Escobar-Arrazola, M, Herrera-Montalvo LA. |
| EPI_ISL_3067718 | Instituto Nacional de Medicina Genomica | Instituto Nacional de Medicina Genomica | Cedro-Tanda A, Hidalgo-Miranda A, Mendoza-Vargas A, Reyes-Grajeda JP, Abraham Campos-Romero, Moreno-Camacho José Luis, Rodriguez-Gallegos Jorge, Luna-Ruiz Marco, Gonzalez-Barrera D, Rangel-DeLeon D, Munguia-Garza P, Ramirez-Vega O, Escobar-Arrazola, M, Herrera-Montalvo LA. |
| EPI_ISL_3067719, EPI_ISL_3067720 | Salud Digna | Instituto Nacional de Medicina Genomica | Cedro-Tanda A, Hidalgo-Miranda A, Mendoza-Vargas A, Reyes-Grajeda JP, Abraham Campos-Romero, Moreno-Camacho José Luis, Rodriguez-Gallegos Jorge, Luna-Ruiz Marco, Gonzalez-Barrera D, Rangel-DeLeon D, Munguia-Garza P, Ramirez-Vega O, Escobar-Arrazola, M, Herrera-Montalvo LA. |
| EPI_ISL_3067721, EPI_ISL_3067722, EPI_ISL_3067724 | Instituto Nacional de Medicina Genomica | Instituto Nacional de Medicina Genomica | Cedro-Tanda A, Hidalgo-Miranda A, Mendoza-Vargas A, Reyes-Grajeda JP, Abraham Campos-Romero, Moreno-Camacho José Luis, Rodriguez-Gallegos Jorge, Luna-Ruiz Marco, Gonzalez-Barrera D, Rangel-DeLeon D, Munguia-Garza P, Ramirez-Vega O, Escobar-Arrazola, M, Herrera-Montalvo LA. |
| EPI_ISL_3067725 | Salud Digna | Instituto Nacional de Medicina Genomica | Cedro-Tanda A, Hidalgo-Miranda A, Mendoza-Vargas A, Reyes-Grajeda JP, Abraham Campos-Romero, Moreno-Camacho José Luis, Rodriguez-Gallegos Jorge, Luna-Ruiz Marco, Gonzalez-Barrera D, Rangel-DeLeon D, Munguia-Garza P, Ramirez-Vega O, Escobar-Arrazola, M, Herrera-Montalvo LA. |
| EPI_ISL_3067726 | INCMNSZ | Instituto Nacional de Medicina Genomica | Cedro-Tanda A, Hidalgo-Miranda A, Mendoza-Vargas A, Reyes-Grajeda JP, Abraham Campos-Romero, Moreno-Camacho José Luis, Rodriguez-Gallegos Jorge, Luna-Ruiz Marco, Gonzalez-Barrera D, Rangel-DeLeon D, Munguia-Garza P, Ramirez-Vega O, Escobar-Arrazola, M, Herrera-Montalvo LA. |
| EPI_ISL_3067727, EPI_ISL_3067728 | Instituto Nacional de Medicina Genomica | Instituto Nacional de Medicina Genomica | Cedro-Tanda A, Hidalgo-Miranda A, Mendoza-Vargas A, Reyes-Grajeda JP, Abraham Campos-Romero, Moreno-Camacho José Luis, Rodriguez-Gallegos Jorge, Luna-Ruiz Marco, Gonzalez-Barrera D, Rangel-DeLeon D, Munguia-Garza P, Ramirez-Vega O, Escobar-Arrazola, M, Herrera-Montalvo LA. |
| EPI_ISL_3067729 | INCMNSZ | Instituto Nacional de Medicina Genomica | Cedro-Tanda A, Hidalgo-Miranda A, Mendoza-Vargas A, Reyes-Grajeda JP, Abraham Campos-Romero, Moreno-Camacho José Luis, Rodriguez-Gallegos Jorge, Luna-Ruiz Marco, Gonzalez-Barrera D, Rangel-DeLeon D, Munguia-Garza P, Ramirez-Vega O, Escobar-Arrazola, M, Herrera-Montalvo LA. |
| EPI_ISL_3067731, EPI_ISL_3067732 | Instituto Nacional de Medicina Genomica | Instituto Nacional de Medicina Genomica | Cedro-Tanda A, Hidalgo-Miranda A, Mendoza-Vargas A, Reyes-Grajeda JP, Abraham Campos-Romero, Moreno-Camacho José Luis, Rodriguez-Gallegos Jorge, Luna-Ruiz Marco, Gonzalez-Barrera D, Rangel-DeLeon D, Munguia-Garza P, Ramirez-Vega O, Escobar-Arrazola, M, Herrera-Montalvo LA. |
| EPI_ISL_3067733, EPI_ISL_3067734, EPI_ISL_3067735 | Salud Digna | Instituto Nacional de Medicina Genomica | Cedro-Tanda A, Hidalgo-Miranda A, Mendoza-Vargas A, Reyes-Grajeda JP, Abraham Campos-Romero, Moreno-Camacho José Luis, Rodriguez-Gallegos Jorge, Luna-Ruiz Marco, Gonzalez-Barrera D, Rangel-DeLeon D, Munguia-Garza P, Ramirez-Vega O, Escobar-Arrazola, M, Herrera-Montalvo LA. |
| EPI_ISL_3067736, EPI_ISL_3067737, EPI_ISL_3067738 | Instituto Nacional de Medicina Genomica | Instituto Nacional de Medicina Genomica | Cedro-Tanda A, Hidalgo-Miranda A, Mendoza-Vargas A, Reyes-Grajeda JP, Abraham Campos-Romero, Moreno-Camacho José Luis, Rodriguez-Gallegos Jorge, Luna-Ruiz Marco, Gonzalez-Barrera D, Rangel-DeLeon D, Munguia-Garza P, Ramirez-Vega O, Escobar-Arrazola, M, Herrera-Montalvo LA. |
| EPI_ISL_3067739, EPI_ISL_3067740 | INCMNSZ | Instituto Nacional de Medicina Genomica | Cedro-Tanda A, Hidalgo-Miranda A, Mendoza-Vargas A, Reyes-Grajeda JP, Abraham Campos-Romero, Moreno-Camacho José Luis, Rodriguez-Gallegos Jorge, Luna-Ruiz Marco, Gonzalez-Barrera D, Rangel-DeLeon D, Munguia-Garza P, Ramirez-Vega O, Escobar-Arrazola, M, Herrera-Montalvo LA. |
| EPI_ISL_3067741 | Instituto Nacional de Medicina Genomica | Instituto Nacional de Medicina Genomica | Cedro-Tanda A, Hidalgo-Miranda A, Mendoza-Vargas A, Reyes-Grajeda JP, Abraham Campos-Romero, Moreno-Camacho José Luis, Rodriguez-Gallegos Jorge, Luna-Ruiz Marco, Gonzalez-Barrera D, Rangel-DeLeon D, Munguia-Garza P, Ramirez-Vega O, Escobar-Arrazola, M, Herrera-Montalvo LA. |
| EPI_ISL_3067742 | Salud Digna | Instituto Nacional de Medicina Genomica | Cedro-Tanda A, Hidalgo-Miranda A, Mendoza-Vargas A, Reyes-Grajeda JP, Abraham Campos-Romero, Moreno-Camacho José Luis, Rodriguez-Gallegos Jorge, Luna-Ruiz Marco, Gonzalez-Barrera D, Rangel-DeLeon D, Munguia-Garza P, Ramirez-Vega O, Escobar-Arrazola, M, Herrera-Montalvo LA. |
| EPI_ISL_3067743 | Instituto Nacional de Medicina Genomica | Instituto Nacional de Medicina Genomica | Cedro-Tanda A, Hidalgo-Miranda A, Mendoza-Vargas A, Reyes-Grajeda JP, Abraham Campos-Romero, Moreno-Camacho José Luis, Rodriguez-Gallegos Jorge, Luna-Ruiz Marco, Gonzalez-Barrera D, Rangel-DeLeon D, Munguia-Garza P, Ramirez-Vega O, Escobar-Arrazola, M, Herrera-Montalvo LA. |
| EPI_ISL_3067744, EPI_ISL_3067745 | Salud Digna | Instituto Nacional de Medicina Genomica | Cedro-Tanda A, Hidalgo-Miranda A, Mendoza-Vargas A, Reyes-Grajeda JP, Abraham Campos-Romero, Moreno-Camacho José Luis, Rodriguez-Gallegos Jorge, Luna-Ruiz Marco, Gonzalez-Barrera D, Rangel-DeLeon D, Munguia-Garza P, Ramirez-Vega O, Escobar-Arrazola, M, Herrera-Montalvo LA. |
| EPI_ISL_3067746, EPI_ISL_3067748 | INCMNSZ | Instituto Nacional de Medicina Genomica | Cedro-Tanda A, Hidalgo-Miranda A, Mendoza-Vargas A, Reyes-Grajeda JP, Abraham Campos-Romero, Moreno-Camacho José Luis, Rodriguez-Gallegos Jorge, Luna-Ruiz Marco, Gonzalez-Barrera D, Rangel-DeLeon D, Munguia-Garza P, Ramirez-Vega O, Escobar-Arrazola, M, Herrera-Montalvo LA. |
| EPI_ISL_3067750, EPI_ISL_3067751, EPI_ISL_3067752 | Instituto Nacional de Medicina Genomica | Instituto Nacional de Medicina Genomica | Cedro-Tanda A, Hidalgo-Miranda A, Mendoza-Vargas A, Reyes-Grajeda JP, Abraham Campos-Romero, Moreno-Camacho José Luis, Rodriguez-Gallegos Jorge, Luna-Ruiz Marco, Gonzalez-Barrera D, Rangel-DeLeon D, Munguia-Garza P, Ramirez-Vega O, Escobar-Arrazola, M, Herrera-Montalvo LA. |
| EPI_ISL_3067753 | Salud Digna | Instituto Nacional de Medicina Genomica | Cedro-Tanda A, Hidalgo-Miranda A, Mendoza-Vargas A, Reyes-Grajeda JP, Abraham Campos-Romero, Moreno-Camacho José Luis, Rodriguez-Gallegos Jorge, Luna-Ruiz Marco, Gonzalez-Barrera D, Rangel-DeLeon D, Munguia-Garza P, Ramirez-Vega O, Escobar-Arrazola, M, Herrera-Montalvo LA. |
| EPI_ISL_3067754 | INCMNSZ | Instituto Nacional de Medicina Genomica | Cedro-Tanda A, Hidalgo-Miranda A, Mendoza-Vargas A, Reyes-Grajeda JP, Abraham Campos-Romero, Moreno-Camacho José Luis, Rodriguez-Gallegos Jorge, Luna-Ruiz Marco, Gonzalez-Barrera D, Rangel-DeLeon D, Munguia-Garza P, Ramirez-Vega O, Escobar-Arrazola, M, Herrera-Montalvo LA. |
| EPI_ISL_3067755 | Instituto Nacional de Medicina Genomica | Instituto Nacional de Medicina Genomica | Cedro-Tanda A, Hidalgo-Miranda A, Mendoza-Vargas A, Reyes-Grajeda JP, Abraham Campos-Romero, Moreno-Camacho José Luis, Rodriguez-Gallegos Jorge, Luna-Ruiz Marco, Gonzalez-Barrera D, Rangel-DeLeon D, Munguia-Garza P, Ramirez-Vega O, Escobar-Arrazola, M, Herrera-Montalvo LA. |

| | | | |
|---|---|---|---|
| EPI_ISL_3067756, EPI_ISL_3067757, EPI_ISL_3067758 | Salud Digna | Instituto Nacional de Medicina Genomica | Cedro-Tanda A, Hidalgo-Miranda A, Mendoza-Vargas A, Reyes-Grajeda JP, Abraham Campos-Romero, Moreno-Camacho José Luis, Rodriguez-Gallegos Jorge, Luna-Ruiz Marco, Gonzalez-Barrera D, Rangel-DeLeon D, Munguia-Garza P, Ramirez-Vega O, Escobar-Arrazola, M, Herrera-Montalvo LA. |
| EPI_ISL_3067759, EPI_ISL_3067760 | INCMNSZ | Instituto Nacional de Medicina Genomica | Cedro-Tanda A, Hidalgo-Miranda A, Mendoza-Vargas A, Reyes-Grajeda JP, Abraham Campos-Romero, Moreno-Camacho José Luis, Rodriguez-Gallegos Jorge, Luna-Ruiz Marco, Gonzalez-Barrera D, Rangel-DeLeon D, Munguia-Garza P, Ramirez-Vega O, Escobar-Arrazola, M, Herrera-Montalvo LA. |
| EPI_ISL_3067761, EPI_ISL_3067762 | Salud Digna | Instituto Nacional de Medicina Genomica | Cedro-Tanda A, Hidalgo-Miranda A, Mendoza-Vargas A, Reyes-Grajeda JP, Abraham Campos-Romero, Moreno-Camacho José Luis, Rodriguez-Gallegos Jorge, Luna-Ruiz Marco, Gonzalez-Barrera D, Rangel-DeLeon D, Munguia-Garza P, Ramirez-Vega O, Escobar-Arrazola, M, Herrera-Montalvo LA. |
| EPI_ISL_3067764, EPI_ISL_3067765 | INCMNSZ | Instituto Nacional de Medicina Genomica | Cedro-Tanda A, Hidalgo-Miranda A, Mendoza-Vargas A, Reyes-Grajeda JP, Abraham Campos-Romero, Moreno-Camacho José Luis, Rodriguez-Gallegos Jorge, Luna-Ruiz Marco, Gonzalez-Barrera D, Rangel-DeLeon D, Munguia-Garza P, Ramirez-Vega O, Escobar-Arrazola, M, Herrera-Montalvo LA. |
| EPI_ISL_3067766, EPI_ISL_3067767, EPI_ISL_3067768 | Salud Digna | Instituto Nacional de Medicina Genomica | Cedro-Tanda A, Hidalgo-Miranda A, Mendoza-Vargas A, Reyes-Grajeda JP, Abraham Campos-Romero, Moreno-Camacho José Luis, Rodriguez-Gallegos Jorge, Luna-Ruiz Marco, Gonzalez-Barrera D, Rangel-DeLeon D, Munguia-Garza P, Ramirez-Vega O, Escobar-Arrazola, M, Herrera-Montalvo LA. |
| EPI_ISL_3067770 | INCMNSZ | Instituto Nacional de Medicina Genomica | Cedro-Tanda A, Hidalgo-Miranda A, Mendoza-Vargas A, Reyes-Grajeda JP, Abraham Campos-Romero, Moreno-Camacho José Luis, Rodriguez-Gallegos Jorge, Luna-Ruiz Marco, Gonzalez-Barrera D, Rangel-DeLeon D, Munguia-Garza P, Ramirez-Vega O, Escobar-Arrazola, M, Herrera-Montalvo LA. |
| EPI_ISL_3067771, EPI_ISL_3067772, EPI_ISL_3067773 | Salud Digna | Instituto Nacional de Medicina Genomica | Cedro-Tanda A, Hidalgo-Miranda A, Mendoza-Vargas A, Reyes-Grajeda JP, Abraham Campos-Romero, Moreno-Camacho José Luis, Rodriguez-Gallegos Jorge, Luna-Ruiz Marco, Gonzalez-Barrera D, Rangel-DeLeon D, Munguia-Garza P, Ramirez-Vega O, Escobar-Arrazola, M, Herrera-Montalvo LA. |
| EPI_ISL_3067774 | INCMNSZ | Instituto Nacional de Medicina Genomica | Cedro-Tanda A, Hidalgo-Miranda A, Mendoza-Vargas A, Reyes-Grajeda JP, Abraham Campos-Romero, Moreno-Camacho José Luis, Rodriguez-Gallegos Jorge, Luna-Ruiz Marco, Gonzalez-Barrera D, Rangel-DeLeon D, Munguia-Garza P, Ramirez-Vega O, Escobar-Arrazola, M, Herrera-Montalvo LA. |
| EPI_ISL_3067775, EPI_ISL_3067777, EPI_ISL_3067778, EPI_ISL_3067779, EPI_ISL_3067780 | Salud Digna | Instituto Nacional de Medicina Genomica | Cedro-Tanda A, Hidalgo-Miranda A, Mendoza-Vargas A, Reyes-Grajeda JP, Abraham Campos-Romero, Moreno-Camacho José Luis, Rodriguez-Gallegos Jorge, Luna-Ruiz Marco, Gonzalez-Barrera D, Rangel-DeLeon D, Munguia-Garza P, Ramirez-Vega O, Escobar-Arrazola, M, Herrera-Montalvo LA. |
| EPI_ISL_3067781 | Instituto Nacional de Cancerología | Instituto Nacional de Medicina Genomica | Cedro-Tanda A, Hidalgo-Miranda A, Mendoza-Vargas A, Reyes-Grajeda JP, Abraham Campos-Romero, Moreno-Camacho José Luis, Rodriguez-Gallegos Jorge, Luna-Ruiz Marco, Gonzalez-Barrera D, Rangel-DeLeon D, Munguia-Garza P, Ramirez-Vega O, Escobar-Arrazola, M, Herrera-Montalvo LA. |
| EPI_ISL_3067783 | INCMNSZ | Instituto Nacional de Medicina Genomica | Cedro-Tanda A, Hidalgo-Miranda A, Mendoza-Vargas A, Reyes-Grajeda JP, Abraham Campos-Romero, Moreno-Camacho José Luis, Rodriguez-Gallegos Jorge, Luna-Ruiz Marco, Gonzalez-Barrera D, Rangel-DeLeon D, Munguia-Garza P, Ramirez-Vega O, Escobar-Arrazola, M, Herrera-Montalvo LA. |
| EPI_ISL_3067784 | Instituto Nacional de Medicina Genomica | Instituto Nacional de Medicina Genomica | Cedro-Tanda A, Hidalgo-Miranda A, Mendoza-Vargas A, Reyes-Grajeda JP, Abraham Campos-Romero, Moreno-Camacho José Luis, Rodriguez-Gallegos Jorge, Luna-Ruiz Marco, Gonzalez-Barrera D, Rangel-DeLeon D, Munguia-Garza P, Ramirez-Vega O, Escobar-Arrazola, M, Herrera-Montalvo LA. |
| EPI_ISL_3067785, EPI_ISL_3067786 | Salud Digna | Instituto Nacional de Medicina Genomica | Cedro-Tanda A, Hidalgo-Miranda A, Mendoza-Vargas A, Reyes-Grajeda JP, Abraham Campos-Romero, Moreno-Camacho José Luis, Rodriguez-Gallegos Jorge, Luna-Ruiz Marco, Gonzalez-Barrera D, Rangel-DeLeon D, Munguia-Garza P, Ramirez-Vega O, Escobar-Arrazola, M, Herrera-Montalvo LA. |
| EPI_ISL_3067787 | Instituto Nacional de Medicina Genomica | Instituto Nacional de Medicina Genomica | Cedro-Tanda A, Hidalgo-Miranda A, Mendoza-Vargas A, Reyes-Grajeda JP, Abraham Campos-Romero, Moreno-Camacho José Luis, Rodriguez-Gallegos Jorge, Luna-Ruiz Marco, Gonzalez-Barrera D, Rangel-DeLeon D, Munguia-Garza P, Ramirez-Vega O, Escobar-Arrazola, M, Herrera-Montalvo LA. |
| EPI_ISL_3067788 | Salud Digna | Instituto Nacional de Medicina Genomica | Cedro-Tanda A, Hidalgo-Miranda A, Mendoza-Vargas A, Reyes-Grajeda JP, Abraham Campos-Romero, Moreno-Camacho José Luis, Rodriguez-Gallegos Jorge, Luna-Ruiz Marco, Gonzalez-Barrera D, Rangel-DeLeon D, Munguia-Garza P, Ramirez-Vega O, Escobar-Arrazola, M, Herrera-Montalvo LA. |
| EPI_ISL_3067789, EPI_ISL_3067790 | Instituto Nacional de Medicina Genomica | Instituto Nacional de Medicina Genomica | Cedro-Tanda A, Hidalgo-Miranda A, Mendoza-Vargas A, Reyes-Grajeda JP, Abraham Campos-Romero, Moreno-Camacho José Luis, Rodriguez-Gallegos Jorge, Luna-Ruiz Marco, Gonzalez-Barrera D, Rangel-DeLeon D, Munguia-Garza P, Ramirez-Vega O, Escobar-Arrazola, M, Herrera-Montalvo LA. |
| EPI_ISL_3067791, EPI_ISL_3067792 | Salud Digna | Instituto Nacional de Medicina Genomica | Cedro-Tanda A, Hidalgo-Miranda A, Mendoza-Vargas A, Reyes-Grajeda JP, Abraham Campos-Romero, Moreno-Camacho José Luis, Rodriguez-Gallegos Jorge, Luna-Ruiz Marco, Gonzalez-Barrera D, Rangel-DeLeon D, Munguia-Garza P, Ramirez-Vega O, Escobar-Arrazola, M, Herrera-Montalvo LA. |
| EPI_ISL_3067793 | Instituto Nacional de Medicina Genomica | Instituto Nacional de Medicina Genomica | Cedro-Tanda A, Hidalgo-Miranda A, Mendoza-Vargas A, Reyes-Grajeda JP, Abraham Campos-Romero, Moreno-Camacho José Luis, Rodriguez-Gallegos Jorge, Luna-Ruiz Marco, Gonzalez-Barrera D, Rangel-DeLeon D, Munguia-Garza P, Ramirez-Vega O, Escobar-Arrazola, M, Herrera-Montalvo LA. |
| EPI_ISL_3067794, EPI_ISL_3067795, EPI_ISL_3067796 | Salud Digna | Instituto Nacional de Medicina Genomica | Cedro-Tanda A, Hidalgo-Miranda A, Mendoza-Vargas A, Reyes-Grajeda JP, Abraham Campos-Romero, Moreno-Camacho José Luis, Rodriguez-Gallegos Jorge, Luna-Ruiz Marco, Gonzalez-Barrera D, Rangel-DeLeon D, Munguia-Garza P, Ramirez-Vega O, Escobar-Arrazola, M, Herrera-Montalvo LA. |
| EPI_ISL_3067797 | Instituto Nacional de Medicina Genomica | Instituto Nacional de Medicina Genomica | Cedro-Tanda A, Hidalgo-Miranda A, Mendoza-Vargas A, Reyes-Grajeda JP, Abraham Campos-Romero, Moreno-Camacho José Luis, Rodriguez-Gallegos Jorge, Luna-Ruiz Marco, Gonzalez-Barrera D, Rangel-DeLeon D, Munguia-Garza P, Ramirez-Vega O, Escobar-Arrazola, M, Herrera-Montalvo LA. |
| EPI_ISL_3067798 | INCMNSZ | Instituto Nacional de Medicina Genomica | Cedro-Tanda A, Hidalgo-Miranda A, Mendoza-Vargas A, Reyes-Grajeda JP, Abraham Campos-Romero, Moreno-Camacho José Luis, Rodriguez-Gallegos Jorge, Luna-Ruiz Marco, Gonzalez-Barrera D, Rangel-DeLeon D, Munguia-Garza P, Ramirez-Vega O, Escobar-Arrazola, M, Herrera-Montalvo LA. |
| EPI_ISL_3067799, EPI_ISL_3067800, EPI_ISL_3067801, EPI_ISL_3067802, EPI_ISL_3067803, EPI_ISL_3067804, EPI_ISL_3067805, EPI_ISL_3067806 | Salud Digna | Instituto Nacional de Medicina Genomica | Cedro-Tanda A, Hidalgo-Miranda A, Mendoza-Vargas A, Reyes-Grajeda JP, Abraham Campos-Romero, Moreno-Camacho José Luis, Rodriguez-Gallegos Jorge, Luna-Ruiz Marco, Gonzalez-Barrera D, Rangel-DeLeon D, Munguia-Garza P, Ramirez-Vega O, Escobar-Arrazola, M, Herrera-Montalvo LA. |
| EPI_ISL_3067807 | INCMNSZ | Instituto Nacional de Medicina Genomica | Cedro-Tanda A, Hidalgo-Miranda A, Mendoza-Vargas A, Reyes-Grajeda JP, Abraham Campos-Romero, Moreno-Camacho José Luis, Rodriguez-Gallegos Jorge, Luna-Ruiz Marco, Gonzalez-Barrera D, Rangel-DeLeon D, Munguia-Garza P, Ramirez-Vega O, Escobar-Arrazola, M, Herrera-Montalvo LA. |
| EPI_ISL_3067808 | Instituto Nacional de Medicina Genomica | Instituto Nacional de Medicina Genomica | Cedro-Tanda A, Hidalgo-Miranda A, Mendoza-Vargas A, Reyes-Grajeda JP, Abraham Campos-Romero, Moreno-Camacho José Luis, Rodriguez-Gallegos Jorge, Luna-Ruiz Marco, Gonzalez-Barrera D, Rangel-DeLeon D, Munguia-Garza P, Ramirez-Vega O, Escobar-Arrazola, M, Herrera-Montalvo LA. |
| EPI_ISL_3067809, EPI_ISL_3067810, EPI_ISL_3067811, EPI_ISL_3067812, EPI_ISL_3067813 | Salud Digna | Instituto Nacional de Medicina Genomica | Cedro-Tanda A, Hidalgo-Miranda A, Mendoza-Vargas A, Reyes-Grajeda JP, Abraham Campos-Romero, Moreno-Camacho José Luis, Rodriguez-Gallegos Jorge, Luna-Ruiz Marco, Gonzalez-Barrera D, Rangel-DeLeon D, Munguia-Garza P, Ramirez-Vega O, Escobar-Arrazola, M, Herrera-Montalvo LA. |
| EPI_ISL_3067814, EPI_ISL_3067815 | Instituto Nacional de Medicina Genomica | Instituto Nacional de Medicina Genomica | Cedro-Tanda A, Hidalgo-Miranda A, Mendoza-Vargas A, Reyes-Grajeda JP, Abraham Campos-Romero, Moreno-Camacho José Luis, Rodriguez-Gallegos Jorge, Luna-Ruiz Marco, Gonzalez-Barrera D, Rangel-DeLeon D, Munguia-Garza P, Ramirez-Vega O, Escobar-Arrazola, M, Herrera-Montalvo LA. |
| EPI_ISL_3067816, EPI_ISL_3067817 | INCMNSZ | Instituto Nacional de Medicina Genomica | Cedro-Tanda A, Hidalgo-Miranda A, Mendoza-Vargas A, Reyes-Grajeda JP, Abraham Campos-Romero, Moreno-Camacho José Luis, Rodriguez-Gallegos Jorge, Luna-Ruiz Marco, Gonzalez-Barrera D, Rangel-DeLeon D, Munguia-Garza P, Ramirez-Vega O, Escobar-Arrazola, M, Herrera-Montalvo LA. |
| EPI_ISL_3067818, EPI_ISL_3067819, EPI_ISL_3067820 | Instituto Nacional de Medicina Genomica | Instituto Nacional de Medicina Genomica | Cedro-Tanda A, Hidalgo-Miranda A, Mendoza-Vargas A, Reyes-Grajeda JP, Abraham Campos-Romero, Moreno-Camacho José Luis, Rodriguez-Gallegos Jorge, Luna-Ruiz Marco, Gonzalez-Barrera D, Rangel-DeLeon D, Munguia-Garza P, Ramirez-Vega O, Escobar-Arrazola, M, Herrera-Montalvo LA. |
| EPI_ISL_3067821 | INCMNSZ | Instituto Nacional de Medicina Genomica | Cedro-Tanda A, Hidalgo-Miranda A, Mendoza-Vargas A, Reyes-Grajeda JP, Abraham Campos-Romero, Moreno-Camacho José Luis, Rodriguez-Gallegos Jorge, Luna-Ruiz Marco, Gonzalez-Barrera D, Rangel-DeLeon D, Munguia-Garza P, Ramirez-Vega O, Escobar-Arrazola, M, Herrera-Montalvo LA. |
| EPI_ISL_3067822 | Instituto Nacional de Medicina Genomica | Instituto Nacional de Medicina Genomica | Cedro-Tanda A, Hidalgo-Miranda A, Mendoza-Vargas A, Reyes-Grajeda JP, Abraham Campos-Romero, Moreno-Camacho José Luis, Rodriguez-Gallegos Jorge, Luna-Ruiz Marco, Gonzalez-Barrera D, Rangel-DeLeon D, Munguia-Garza P, Ramirez-Vega O, Escobar-Arrazola, M, Herrera-Montalvo LA. |
| EPI_ISL_3067823 | Salud Digna | Instituto Nacional de Medicina Genomica | Cedro-Tanda A, Hidalgo-Miranda A, Mendoza-Vargas A, Reyes-Grajeda JP, Abraham Campos-Romero, Moreno-Camacho José Luis, Rodriguez-Gallegos Jorge, Luna-Ruiz Marco, Gonzalez-Barrera D, Rangel-DeLeon D, Munguia-Garza P, Ramirez-Vega O, Escobar-Arrazola, M, Herrera-Montalvo LA. |
| EPI_ISL_3067824 | Instituto Nacional de Medicina Genomica | Instituto Nacional de Medicina Genomica | Cedro-Tanda A, Hidalgo-Miranda A, Mendoza-Vargas A, Reyes-Grajeda JP, Abraham Campos-Romero, Moreno-Camacho José Luis, Rodriguez-Gallegos Jorge, Luna-Ruiz Marco, Gonzalez-Barrera D, Rangel-DeLeon D, Munguia-Garza P, Ramirez-Vega O, Escobar-Arrazola, M, Herrera-Montalvo LA. |
| EPI_ISL_3067825 | INCMNSZ | Instituto Nacional de Medicina Genomica | Cedro-Tanda A, Hidalgo-Miranda A, Mendoza-Vargas A, Reyes-Grajeda JP, Abraham Campos-Romero, Moreno-Camacho José Luis, Rodriguez-Gallegos Jorge, Luna-Ruiz Marco, Gonzalez-Barrera D, Rangel-DeLeon D, Munguia-Garza P, Ramirez-Vega O, Escobar-Arrazola, M, Herrera-Montalvo LA. |
| EPI_ISL_3067826, EPI_ISL_3067827 | Instituto Nacional de Medicina Genomica | Instituto Nacional de Medicina Genomica | Cedro-Tanda A, Hidalgo-Miranda A, Mendoza-Vargas A, Reyes-Grajeda JP, Abraham Campos-Romero, Moreno-Camacho José Luis, Rodriguez-Gallegos Jorge, Luna-Ruiz Marco, Gonzalez-Barrera D, Rangel-DeLeon D, Munguia-Garza P, Ramirez-Vega O, Escobar-Arrazola, M, Herrera-Montalvo LA. |
| EPI_ISL_3067828, EPI_ISL_3067829, EPI_ISL_3067830, EPI_ISL_3067831 | Salud Digna | Instituto Nacional de Medicina Genomica | Cedro-Tanda A, Hidalgo-Miranda A, Mendoza-Vargas A, Reyes-Grajeda JP, Abraham Campos-Romero, Moreno-Camacho José Luis, Rodriguez-Gallegos Jorge, Luna-Ruiz Marco, Gonzalez-Barrera D, Rangel-DeLeon D, Munguia-Garza P, Ramirez-Vega O, Escobar-Arrazola, M, Herrera-Montalvo LA. |
| EPI_ISL_3067832 | Instituto Nacional de Cancerología | Instituto Nacional de Medicina Genomica | Cedro-Tanda A, Hidalgo-Miranda A, Mendoza-Vargas A, Reyes-Grajeda JP, Abraham Campos-Romero, Moreno-Camacho José Luis, Rodriguez-Gallegos Jorge, Luna-Ruiz Marco, Gonzalez-Barrera D, Rangel-DeLeon D, Munguia-Garza P, Ramirez-Vega O, Escobar-Arrazola, M, Herrera-Montalvo LA. |
| EPI_ISL_3067833, EPI_ISL_3067834 | Instituto Nacional de Medicina Genomica | Instituto Nacional de Medicina Genomica | Cedro-Tanda A, Hidalgo-Miranda A, Mendoza-Vargas A, Reyes-Grajeda JP, Abraham Campos-Romero, Moreno-Camacho José Luis, Rodriguez-Gallegos Jorge, Luna-Ruiz Marco, Gonzalez-Barrera D, Rangel-DeLeon D, Munguia-Garza P, Ramirez-Vega O, Escobar-Arrazola, M, Herrera-Montalvo LA. |
| EPI_ISL_3067835, | Salud Digna | Instituto Nacional de Medicina Genomica | Cedro-Tanda A, Hidalgo-Miranda A, Mendoza-Vargas A, Reyes-Grajeda JP, Abraham Campos-Romero, Moreno-Camacho José Luis, Rodriguez-Gallegos Jorge, Luna-Ruiz Marco, Gonzalez-Barrera D, Rangel-DeLeon D, Munguia-Garza P, Ramirez- |

| | | | |
|---|---|---|---|
| EPI_ISL_3067836, EPI_ISL_3067837 | | | Vega O, Escobar-Arrazola, M, Herrera-Montalvo LA. |
| EPI_ISL_3067839 | INCMNSZ | Instituto Nacional de Medicina Genomica | Cedro-Tanda A, Hidalgo-Miranda A, Mendoza-Vargas A, Reyes-Grajeda JP, Abraham Campos-Romero, Moreno-Camacho José Luis, Rodriguez-Gallegos Jorge, Luna-Ruiz Marco, Gonzalez-Barrera D, Rangel-DeLeon D, Munguia-Garza P, Ramirez-Vega O, Escobar-Arrazola, M, Herrera-Montalvo LA. |
| EPI_ISL_3067840, EPI_ISL_3067841, EPI_ISL_3067842, EPI_ISL_3067843 | Salud Digna | Instituto Nacional de Medicina Genomica | Cedro-Tanda A, Hidalgo-Miranda A, Mendoza-Vargas A, Reyes-Grajeda JP, Abraham Campos-Romero, Moreno-Camacho José Luis, Rodriguez-Gallegos Jorge, Luna-Ruiz Marco, Gonzalez-Barrera D, Rangel-DeLeon D, Munguia-Garza P, Ramirez-Vega O, Escobar-Arrazola, M, Herrera-Montalvo LA. |
| EPI_ISL_3067844 | Instituto Nacional de Medicina Genomica | Instituto Nacional de Medicina Genomica | Cedro-Tanda A, Hidalgo-Miranda A, Mendoza-Vargas A, Reyes-Grajeda JP, Abraham Campos-Romero, Moreno-Camacho José Luis, Rodriguez-Gallegos Jorge, Luna-Ruiz Marco, Gonzalez-Barrera D, Rangel-DeLeon D, Munguia-Garza P, Ramirez-Vega O, Escobar-Arrazola, M, Herrera-Montalvo LA. |
| EPI_ISL_3067845 | Salud Digna | Instituto Nacional de Medicina Genomica | Cedro-Tanda A, Hidalgo-Miranda A, Mendoza-Vargas A, Reyes-Grajeda JP, Abraham Campos-Romero, Moreno-Camacho José Luis, Rodriguez-Gallegos Jorge, Luna-Ruiz Marco, Gonzalez-Barrera D, Rangel-DeLeon D, Munguia-Garza P, Ramirez-Vega O, Escobar-Arrazola, M, Herrera-Montalvo LA. |
| EPI_ISL_3067846 | Instituto Nacional de Medicina Genomica | Instituto Nacional de Medicina Genomica | Cedro-Tanda A, Hidalgo-Miranda A, Mendoza-Vargas A, Reyes-Grajeda JP, Abraham Campos-Romero, Moreno-Camacho José Luis, Rodriguez-Gallegos Jorge, Luna-Ruiz Marco, Gonzalez-Barrera D, Rangel-DeLeon D, Munguia-Garza P, Ramirez-Vega O, Escobar-Arrazola, M, Herrera-Montalvo LA. |
| EPI_ISL_3067847 | Salud Digna | Instituto Nacional de Medicina Genomica | Cedro-Tanda A, Hidalgo-Miranda A, Mendoza-Vargas A, Reyes-Grajeda JP, Abraham Campos-Romero, Moreno-Camacho José Luis, Rodriguez-Gallegos Jorge, Luna-Ruiz Marco, Gonzalez-Barrera D, Rangel-DeLeon D, Munguia-Garza P, Ramirez-Vega O, Escobar-Arrazola, M, Herrera-Montalvo LA. |
| EPI_ISL_3067848, EPI_ISL_3067850 | Instituto Nacional de Medicina Genomica | Instituto Nacional de Medicina Genomica | Cedro-Tanda A, Hidalgo-Miranda A, Mendoza-Vargas A, Reyes-Grajeda JP, Abraham Campos-Romero, Moreno-Camacho José Luis, Rodriguez-Gallegos Jorge, Luna-Ruiz Marco, Gonzalez-Barrera D, Rangel-DeLeon D, Munguia-Garza P, Ramirez-Vega O, Escobar-Arrazola, M, Herrera-Montalvo LA. |
| EPI_ISL_3067851 | Salud Digna | Instituto Nacional de Medicina Genomica | Cedro-Tanda A, Hidalgo-Miranda A, Mendoza-Vargas A, Reyes-Grajeda JP, Abraham Campos-Romero, Moreno-Camacho José Luis, Rodriguez-Gallegos Jorge, Luna-Ruiz Marco, Gonzalez-Barrera D, Rangel-DeLeon D, Munguia-Garza P, Ramirez-Vega O, Escobar-Arrazola, M, Herrera-Montalvo LA. |
| EPI_ISL_3067852 | INCMNSZ | Instituto Nacional de Medicina Genomica | Cedro-Tanda A, Hidalgo-Miranda A, Mendoza-Vargas A, Reyes-Grajeda JP, Abraham Campos-Romero, Moreno-Camacho José Luis, Rodriguez-Gallegos Jorge, Luna-Ruiz Marco, Gonzalez-Barrera D, Rangel-DeLeon D, Munguia-Garza P, Ramirez-Vega O, Escobar-Arrazola, M, Herrera-Montalvo LA. |
| EPI_ISL_3067853 | Instituto Nacional de Medicina Genomica | Instituto Nacional de Medicina Genomica | Cedro-Tanda A, Hidalgo-Miranda A, Mendoza-Vargas A, Reyes-Grajeda JP, Abraham Campos-Romero, Moreno-Camacho José Luis, Rodriguez-Gallegos Jorge, Luna-Ruiz Marco, Gonzalez-Barrera D, Rangel-DeLeon D, Munguia-Garza P, Ramirez-Vega O, Escobar-Arrazola, M, Herrera-Montalvo LA. |
| EPI_ISL_3067854, EPI_ISL_3067855 | Salud Digna | Instituto Nacional de Medicina Genomica | Cedro-Tanda A, Hidalgo-Miranda A, Mendoza-Vargas A, Reyes-Grajeda JP, Abraham Campos-Romero, Moreno-Camacho José Luis, Rodriguez-Gallegos Jorge, Luna-Ruiz Marco, Gonzalez-Barrera D, Rangel-DeLeon D, Munguia-Garza P, Ramirez-Vega O, Escobar-Arrazola, M, Herrera-Montalvo LA. |
| EPI_ISL_3067856 | INCMNSZ | Instituto Nacional de Medicina Genomica | Cedro-Tanda A, Hidalgo-Miranda A, Mendoza-Vargas A, Reyes-Grajeda JP, Abraham Campos-Romero, Moreno-Camacho José Luis, Rodriguez-Gallegos Jorge, Luna-Ruiz Marco, Gonzalez-Barrera D, Rangel-DeLeon D, Munguia-Garza P, Ramirez-Vega O, Escobar-Arrazola, M, Herrera-Montalvo LA. |
| EPI_ISL_3067857 | Salud Digna | Instituto Nacional de Medicina Genomica | Cedro-Tanda A, Hidalgo-Miranda A, Mendoza-Vargas A, Reyes-Grajeda JP, Abraham Campos-Romero, Moreno-Camacho José Luis, Rodriguez-Gallegos Jorge, Luna-Ruiz Marco, Gonzalez-Barrera D, Rangel-DeLeon D, Munguia-Garza P, Ramirez-Vega O, Escobar-Arrazola, M, Herrera-Montalvo LA. |
| EPI_ISL_3067858, EPI_ISL_3067859, EPI_ISL_3067860 | Instituto Nacional de Medicina Genomica | Instituto Nacional de Medicina Genomica | Cedro-Tanda A, Hidalgo-Miranda A, Mendoza-Vargas A, Reyes-Grajeda JP, Abraham Campos-Romero, Moreno-Camacho José Luis, Rodriguez-Gallegos Jorge, Luna-Ruiz Marco, Gonzalez-Barrera D, Rangel-DeLeon D, Munguia-Garza P, Ramirez-Vega O, Escobar-Arrazola, M, Herrera-Montalvo LA. |
| EPI_ISL_3067861, EPI_ISL_3067862 | Salud Digna | Instituto Nacional de Medicina Genomica | Cedro-Tanda A, Hidalgo-Miranda A, Mendoza-Vargas A, Reyes-Grajeda JP, Abraham Campos-Romero, Moreno-Camacho José Luis, Rodriguez-Gallegos Jorge, Luna-Ruiz Marco, Gonzalez-Barrera D, Rangel-DeLeon D, Munguia-Garza P, Ramirez-Vega O, Escobar-Arrazola, M, Herrera-Montalvo LA. |
| EPI_ISL_3067863 | INCMNSZ | Instituto Nacional de Medicina Genomica | Cedro-Tanda A, Hidalgo-Miranda A, Mendoza-Vargas A, Reyes-Grajeda JP, Abraham Campos-Romero, Moreno-Camacho José Luis, Rodriguez-Gallegos Jorge, Luna-Ruiz Marco, Gonzalez-Barrera D, Rangel-DeLeon D, Munguia-Garza P, Ramirez-Vega O, Escobar-Arrazola, M, Herrera-Montalvo LA. |
| EPI_ISL_3067864, EPI_ISL_3067865 | Salud Digna | Instituto Nacional de Medicina Genomica | Cedro-Tanda A, Hidalgo-Miranda A, Mendoza-Vargas A, Reyes-Grajeda JP, Abraham Campos-Romero, Moreno-Camacho José Luis, Rodriguez-Gallegos Jorge, Luna-Ruiz Marco, Gonzalez-Barrera D, Rangel-DeLeon D, Munguia-Garza P, Ramirez-Vega O, Escobar-Arrazola, M, Herrera-Montalvo LA. |
| EPI_ISL_3067866 | Instituto Nacional de Medicina Genomica | Instituto Nacional de Medicina Genomica | Cedro-Tanda A, Hidalgo-Miranda A, Mendoza-Vargas A, Reyes-Grajeda JP, Abraham Campos-Romero, Moreno-Camacho José Luis, Rodriguez-Gallegos Jorge, Luna-Ruiz Marco, Gonzalez-Barrera D, Rangel-DeLeon D, Munguia-Garza P, Ramirez-Vega O, Escobar-Arrazola, M, Herrera-Montalvo LA. |
| EPI_ISL_3067867 | Centro Medico ABC | Instituto Nacional de Medicina Genomica | Cedro-Tanda A, Hidalgo-Miranda A, Mendoza-Vargas A, Reyes-Grajeda JP, Roxana Trejo-González, Cesar Lara, Jacomine Reyes-Carrasco, Daniel Aguirre, Gonzalez-Barrera D, Rangel-DeLeon D, Munguia-Garza P, Ramirez-Vega O, Escobar-Arrazola, M, Herrera-Montalvo LA. |
| EPI_ISL_3067868 | Salud Digna | Instituto Nacional de Medicina Genomica | Cedro-Tanda A, Hidalgo-Miranda A, Mendoza-Vargas A, Reyes-Grajeda JP, Abraham Campos-Romero, Moreno-Camacho José Luis, Rodriguez-Gallegos Jorge, Luna-Ruiz Marco, Gonzalez-Barrera D, Rangel-DeLeon D, Munguia-Garza P, Ramirez-Vega O, Escobar-Arrazola, M, Herrera-Montalvo LA. |
| EPI_ISL_3067869 | Instituto Nacional de Medicina Genomica | Instituto Nacional de Medicina Genomica | Cedro-Tanda A, Hidalgo-Miranda A, Mendoza-Vargas A, Reyes-Grajeda JP, Abraham Campos-Romero, Moreno-Camacho José Luis, Rodriguez-Gallegos Jorge, Luna-Ruiz Marco, Gonzalez-Barrera D, Rangel-DeLeon D, Munguia-Garza P, Ramirez-Vega O, Escobar-Arrazola, M, Herrera-Montalvo LA. |
| EPI_ISL_3067870 | Salud Digna | Instituto Nacional de Medicina Genomica | Cedro-Tanda A, Hidalgo-Miranda A, Mendoza-Vargas A, Reyes-Grajeda JP, Abraham Campos-Romero, Moreno-Camacho José Luis, Rodriguez-Gallegos Jorge, Luna-Ruiz Marco, Gonzalez-Barrera D, Rangel-DeLeon D, Munguia-Garza P, Ramirez-Vega O, Escobar-Arrazola, M, Herrera-Montalvo LA. |
| EPI_ISL_3067871 | Instituto Nacional de Medicina Genomica | Instituto Nacional de Medicina Genomica | Cedro-Tanda A, Hidalgo-Miranda A, Mendoza-Vargas A, Reyes-Grajeda JP, Abraham Campos-Romero, Moreno-Camacho José Luis, Rodriguez-Gallegos Jorge, Luna-Ruiz Marco, Gonzalez-Barrera D, Rangel-DeLeon D, Munguia-Garza P, Ramirez-Vega O, Escobar-Arrazola, M, Herrera-Montalvo LA. |
| EPI_ISL_3067872 | INCMNSZ | Instituto Nacional de Medicina Genomica | Cedro-Tanda A, Hidalgo-Miranda A, Mendoza-Vargas A, Reyes-Grajeda JP, Abraham Campos-Romero, Moreno-Camacho José Luis, Rodriguez-Gallegos Jorge, Luna-Ruiz Marco, Gonzalez-Barrera D, Rangel-DeLeon D, Munguia-Garza P, Ramirez-Vega O, Escobar-Arrazola, M, Herrera-Montalvo LA. |
| EPI_ISL_3067874, EPI_ISL_3067875, EPI_ISL_3067877 | Salud Digna | Instituto Nacional de Medicina Genomica | Cedro-Tanda A, Hidalgo-Miranda A, Mendoza-Vargas A, Reyes-Grajeda JP, Abraham Campos-Romero, Moreno-Camacho José Luis, Rodriguez-Gallegos Jorge, Luna-Ruiz Marco, Gonzalez-Barrera D, Rangel-DeLeon D, Munguia-Garza P, Ramirez-Vega O, Escobar-Arrazola, M, Herrera-Montalvo LA. |
| EPI_ISL_3067878 | Centro Medico ABC | Instituto Nacional de Medicina Genomica | Cedro-Tanda A, Hidalgo-Miranda A, Mendoza-Vargas A, Reyes-Grajeda JP, Roxana Trejo-González, Cesar Lara, Jacomine Reyes-Carrasco, Daniel Aguirre, Gonzalez-Barrera D, Rangel-DeLeon D, Munguia-Garza P, Ramirez-Vega O, Escobar-Arrazola, M, Herrera-Montalvo LA. |
| EPI_ISL_3067879, EPI_ISL_3067880 | Salud Digna | Instituto Nacional de Medicina Genomica | Cedro-Tanda A, Hidalgo-Miranda A, Mendoza-Vargas A, Reyes-Grajeda JP, Abraham Campos-Romero, Moreno-Camacho José Luis, Rodriguez-Gallegos Jorge, Luna-Ruiz Marco, Gonzalez-Barrera D, Rangel-DeLeon D, Munguia-Garza P, Ramirez-Vega O, Escobar-Arrazola, M, Herrera-Montalvo LA. |
| EPI_ISL_3067881 | Instituto Nacional de Medicina Genomica | Instituto Nacional de Medicina Genomica | Cedro-Tanda A, Hidalgo-Miranda A, Mendoza-Vargas A, Reyes-Grajeda JP, Abraham Campos-Romero, Moreno-Camacho José Luis, Rodriguez-Gallegos Jorge, Luna-Ruiz Marco, Gonzalez-Barrera D, Rangel-DeLeon D, Munguia-Garza P, Ramirez-Vega O, Escobar-Arrazola, M, Herrera-Montalvo LA. |
| EPI_ISL_3067882, EPI_ISL_3067883 | Salud Digna | Instituto Nacional de Medicina Genomica | Cedro-Tanda A, Hidalgo-Miranda A, Mendoza-Vargas A, Reyes-Grajeda JP, Abraham Campos-Romero, Moreno-Camacho José Luis, Rodriguez-Gallegos Jorge, Luna-Ruiz Marco, Gonzalez-Barrera D, Rangel-DeLeon D, Munguia-Garza P, Ramirez-Vega O, Escobar-Arrazola, M, Herrera-Montalvo LA. |
| EPI_ISL_3067884 | Instituto Nacional de Medicina Genomica | Instituto Nacional de Medicina Genomica | Cedro-Tanda A, Hidalgo-Miranda A, Mendoza-Vargas A, Reyes-Grajeda JP, Abraham Campos-Romero, Moreno-Camacho José Luis, Rodriguez-Gallegos Jorge, Luna-Ruiz Marco, Gonzalez-Barrera D, Rangel-DeLeon D, Munguia-Garza P, Ramirez-Vega O, Escobar-Arrazola, M, Herrera-Montalvo LA. |
| EPI_ISL_3067885 | Salud Digna | Instituto Nacional de Medicina Genomica | Cedro-Tanda A, Hidalgo-Miranda A, Mendoza-Vargas A, Reyes-Grajeda JP, Abraham Campos-Romero, Moreno-Camacho José Luis, Rodriguez-Gallegos Jorge, Luna-Ruiz Marco, Gonzalez-Barrera D, Rangel-DeLeon D, Munguia-Garza P, Ramirez-Vega O, Escobar-Arrazola, M, Herrera-Montalvo LA. |
| EPI_ISL_3067886 | Instituto Nacional de Medicina Genomica | Instituto Nacional de Medicina Genomica | Cedro-Tanda A, Hidalgo-Miranda A, Mendoza-Vargas A, Reyes-Grajeda JP, Abraham Campos-Romero, Moreno-Camacho José Luis, Rodriguez-Gallegos Jorge, Luna-Ruiz Marco, Gonzalez-Barrera D, Rangel-DeLeon D, Munguia-Garza P, Ramirez-Vega O, Escobar-Arrazola, M, Herrera-Montalvo LA. |
| EPI_ISL_3067887 | INCMNSZ | Instituto Nacional de Medicina Genomica | Cedro-Tanda A, Hidalgo-Miranda A, Mendoza-Vargas A, Reyes-Grajeda JP, Abraham Campos-Romero, Moreno-Camacho José Luis, Rodriguez-Gallegos Jorge, Luna-Ruiz Marco, Gonzalez-Barrera D, Rangel-DeLeon D, Munguia-Garza P, Ramirez-Vega O, Escobar-Arrazola, M, Herrera-Montalvo LA. |
| EPI_ISL_3067888, EPI_ISL_3067889, EPI_ISL_3067890, EPI_ISL_3067891, EPI_ISL_3067892, EPI_ISL_3067893, EPI_ISL_3067894 | Salud Digna | Instituto Nacional de Medicina Genomica | Cedro-Tanda A, Hidalgo-Miranda A, Mendoza-Vargas A, Reyes-Grajeda JP, Abraham Campos-Romero, Moreno-Camacho José Luis, Rodriguez-Gallegos Jorge, Luna-Ruiz Marco, Gonzalez-Barrera D, Rangel-DeLeon D, Munguia-Garza P, Ramirez-Vega O, Escobar-Arrazola, M, Herrera-Montalvo LA. |
| EPI_ISL_3067895 | INCMNSZ | Instituto Nacional de Medicina Genomica | Cedro-Tanda A, Hidalgo-Miranda A, Mendoza-Vargas A, Reyes-Grajeda JP, Abraham Campos-Romero, Moreno-Camacho José Luis, Rodriguez-Gallegos Jorge, Luna-Ruiz Marco, Gonzalez-Barrera D, Rangel-DeLeon D, Munguia-Garza P, Ramirez-Vega O, Escobar-Arrazola, M, Herrera-Montalvo LA. |
| EPI_ISL_3067897 | Salud Digna | Instituto Nacional de Medicina Genomica | Cedro-Tanda A, Hidalgo-Miranda A, Mendoza-Vargas A, Reyes-Grajeda JP, Abraham Campos-Romero, Moreno-Camacho José Luis, Rodriguez-Gallegos Jorge, Luna-Ruiz Marco, Gonzalez-Barrera D, Rangel-DeLeon D, Munguia-Garza P, Ramirez-Vega O, Escobar-Arrazola, M, Herrera-Montalvo LA. |
| EPI_ISL_3067898 | Instituto Nacional de Medicina Genomica | Instituto Nacional de Medicina Genomica | Cedro-Tanda A, Hidalgo-Miranda A, Mendoza-Vargas A, Reyes-Grajeda JP, Abraham Campos-Romero, Moreno-Camacho José Luis, Rodriguez-Gallegos Jorge, Luna-Ruiz Marco, Gonzalez-Barrera D, Rangel-DeLeon D, Munguia-Garza P, Ramirez-Vega O, Escobar-Arrazola, M, Herrera-Montalvo LA. |
| EPI_ISL_3067900 | Salud Digna | Instituto Nacional de Medicina Genomica | Cedro-Tanda A, Hidalgo-Miranda A, Mendoza-Vargas A, Reyes-Grajeda JP, Abraham Campos-Romero, Moreno-Camacho José Luis, Rodriguez-Gallegos Jorge, Luna-Ruiz Marco, Gonzalez-Barrera D, Rangel-DeLeon D, Munguia-Garza P, Ramirez-Vega O, Escobar-Arrazola, M, Herrera-Montalvo LA. |
| EPI_ISL_3067901 | Instituto Nacional de Medicina Genomica | Instituto Nacional de Medicina Genomica | Cedro-Tanda A, Hidalgo-Miranda A, Mendoza-Vargas A, Reyes-Grajeda JP, Abraham Campos-Romero, Moreno-Camacho José Luis, Rodriguez-Gallegos Jorge, Luna-Ruiz Marco, Gonzalez-Barrera D, Rangel-DeLeon D, Munguia-Garza P, Ramirez-Vega O, Escobar-Arrazola, M, Herrera-Montalvo LA. |
| EPI_ISL_3067903, EPI_ISL_3067904, EPI_ISL_3067905 | Salud Digna | Instituto Nacional de Medicina Genomica | Cedro-Tanda A, Hidalgo-Miranda A, Mendoza-Vargas A, Reyes-Grajeda JP, Abraham Campos-Romero, Moreno-Camacho José Luis, Rodriguez-Gallegos Jorge, Luna-Ruiz Marco, Gonzalez-Barrera D, Rangel-DeLeon D, Munguia-Garza P, Ramirez-Vega O, Escobar-Arrazola, M, Herrera-Montalvo LA. |

| | | | |
|---|---|---|---|
| EPI_ISL_3067906 | INCMNSZ | Instituto Nacional de Medicina Genomica | Cedro-Tanda A, Hidalgo-Miranda A, Mendoza-Vargas A, Reyes-Grajeda JP, Abraham Campos-Romero, Moreno-Camacho José Luis, Rodriguez-Gallegos Jorge, Luna-Ruiz Marco, Gonzalez-Barrera D, Rangel-DeLeon D, Munguia-Garza P, Ramirez-Vega O, Escobar-Arrazola, M, Herrera-Montalvo LA. |
| EPI_ISL_3067918 | Salud Digna | Instituto Nacional de Medicina Genomica | Cedro-Tanda A, Hidalgo-Miranda A, Mendoza-Vargas A, Reyes-Grajeda JP, Abraham Campos-Romero, Moreno-Camacho José Luis, Rodriguez-Gallegos Jorge, Luna-Ruiz Marco, Gonzalez-Barrera D, Rangel-DeLeon D, Munguia-Garza P, Ramirez-Vega O, Escobar-Arrazola, M, Herrera-Montalvo LA. |
| EPI_ISL_3067919 | Centro Medico ABC | Instituto Nacional de Medicina Genomica | Cedro-Tanda A, Hidalgo-Miranda A, Mendoza-Vargas A, Reyes-Grajeda JP, Roxana Trejo-González, Cesar Lara, Jacomine Reyes-Carrasco, Daniel Aguirre, Gonzalez-Barrera D, Rangel-DeLeon D, Munguia-Garza P, Ramirez-Vega O, Escobar-Arrazola, M, Herrera-Montalvo LA. |
| EPI_ISL_3067920, EPI_ISL_3067921 | Instituto Nacional de Medicina Genomica | Instituto Nacional de Medicina Genomica | Cedro-Tanda A, Hidalgo-Miranda A, Mendoza-Vargas A, Reyes-Grajeda JP, Abraham Campos-Romero, Moreno-Camacho José Luis, Rodriguez-Gallegos Jorge, Luna-Ruiz Marco, Gonzalez-Barrera D, Rangel-DeLeon D, Munguia-Garza P, Ramirez-Vega O, Escobar-Arrazola, M, Herrera-Montalvo LA. |
| EPI_ISL_3067922 | Instituto Nacional de Cancerología | Instituto Nacional de Medicina Genomica | Cedro-Tanda A, Hidalgo-Miranda A, Mendoza-Vargas A, Reyes-Grajeda JP, Abraham Campos-Romero, Moreno-Camacho José Luis, Rodriguez-Gallegos Jorge, Luna-Ruiz Marco, Gonzalez-Barrera D, Rangel-DeLeon D, Munguia-Garza P, Ramirez-Vega O, Escobar-Arrazola, M, Herrera-Montalvo LA. |
| EPI_ISL_3067923, EPI_ISL_3067924 | Instituto Nacional de Medicina Genomica | Instituto Nacional de Medicina Genomica | Cedro-Tanda A, Hidalgo-Miranda A, Mendoza-Vargas A, Reyes-Grajeda JP, Abraham Campos-Romero, Moreno-Camacho José Luis, Rodriguez-Gallegos Jorge, Luna-Ruiz Marco, Gonzalez-Barrera D, Rangel-DeLeon D, Munguia-Garza P, Ramirez-Vega O, Escobar-Arrazola, M, Herrera-Montalvo LA. |
| EPI_ISL_3067925 | Salud Digna | Instituto Nacional de Medicina Genomica | Cedro-Tanda A, Hidalgo-Miranda A, Mendoza-Vargas A, Reyes-Grajeda JP, Abraham Campos-Romero, Moreno-Camacho José Luis, Rodriguez-Gallegos Jorge, Luna-Ruiz Marco, Gonzalez-Barrera D, Rangel-DeLeon D, Munguia-Garza P, Ramirez-Vega O, Escobar-Arrazola, M, Herrera-Montalvo LA. |
| EPI_ISL_3067939 | Instituto Nacional de Medicina Genomica | Instituto Nacional de Medicina Genomica | Cedro-Tanda A, Hidalgo-Miranda A, Mendoza-Vargas A, Reyes-Grajeda JP, Abraham Campos-Romero, Moreno-Camacho José Luis, Rodriguez-Gallegos Jorge, Luna-Ruiz Marco, Gonzalez-Barrera D, Rangel-DeLeon D, Munguia-Garza P, Ramirez-Vega O, Escobar-Arrazola, M, Herrera-Montalvo LA. |
| EPI_ISL_3067942 | Salud Digna | Instituto Nacional de Medicina Genomica | Cedro-Tanda A, Hidalgo-Miranda A, Mendoza-Vargas A, Reyes-Grajeda JP, Abraham Campos-Romero, Moreno-Camacho José Luis, Rodriguez-Gallegos Jorge, Luna-Ruiz Marco, Gonzalez-Barrera D, Rangel-DeLeon D, Munguia-Garza P, Ramirez-Vega O, Escobar-Arrazola, M, Herrera-Montalvo LA. |
| EPI_ISL_3067943 | INCMNSZ | Instituto Nacional de Medicina Genomica | Cedro-Tanda A, Hidalgo-Miranda A, Mendoza-Vargas A, Reyes-Grajeda JP, Abraham Campos-Romero, Moreno-Camacho José Luis, Rodriguez-Gallegos Jorge, Luna-Ruiz Marco, Gonzalez-Barrera D, Rangel-DeLeon D, Munguia-Garza P, Ramirez-Vega O, Escobar-Arrazola, M, Herrera-Montalvo LA. |
| EPI_ISL_3067944, EPI_ISL_3067945, EPI_ISL_3067946, EPI_ISL_3067947, EPI_ISL_3067948, EPI_ISL_3067949, EPI_ISL_3067950, EPI_ISL_3067951, EPI_ISL_3067952 | Salud Digna | Instituto Nacional de Medicina Genomica | Cedro-Tanda A, Hidalgo-Miranda A, Mendoza-Vargas A, Reyes-Grajeda JP, Abraham Campos-Romero, Moreno-Camacho José Luis, Rodriguez-Gallegos Jorge, Luna-Ruiz Marco, Gonzalez-Barrera D, Rangel-DeLeon D, Munguia-Garza P, Ramirez-Vega O, Escobar-Arrazola, M, Herrera-Montalvo LA. |
| EPI_ISL_3067953, EPI_ISL_3067954, EPI_ISL_3067955, EPI_ISL_3067956 | INCMNSZ | Instituto Nacional de Medicina Genomica | Cedro-Tanda A, Hidalgo-Miranda A, Mendoza-Vargas A, Reyes-Grajeda JP, Abraham Campos-Romero, Moreno-Camacho José Luis, Rodriguez-Gallegos Jorge, Luna-Ruiz Marco, Gonzalez-Barrera D, Rangel-DeLeon D, Munguia-Garza P, Ramirez-Vega O, Escobar-Arrazola, M, Herrera-Montalvo LA. |
| EPI_ISL_3067957, EPI_ISL_3067958, EPI_ISL_3067959, EPI_ISL_3067960 | Instituto Nacional de Medicina Genomica | Instituto Nacional de Medicina Genomica | Cedro-Tanda A, Hidalgo-Miranda A, Mendoza-Vargas A, Reyes-Grajeda JP, Abraham Campos-Romero, Moreno-Camacho José Luis, Rodriguez-Gallegos Jorge, Luna-Ruiz Marco, Gonzalez-Barrera D, Rangel-DeLeon D, Munguia-Garza P, Ramirez-Vega O, Escobar-Arrazola, M, Herrera-Montalvo LA. |
| EPI_ISL_3067961 | INCMNSZ | Instituto Nacional de Medicina Genomica | Cedro-Tanda A, Hidalgo-Miranda A, Mendoza-Vargas A, Reyes-Grajeda JP, Abraham Campos-Romero, Moreno-Camacho José Luis, Rodriguez-Gallegos Jorge, Luna-Ruiz Marco, Gonzalez-Barrera D, Rangel-DeLeon D, Munguia-Garza P, Ramirez-Vega O, Escobar-Arrazola, M, Herrera-Montalvo LA. |
| EPI_ISL_3067962, EPI_ISL_3067963 | Instituto Nacional de Medicina Genomica | Instituto Nacional de Medicina Genomica | Cedro-Tanda A, Hidalgo-Miranda A, Mendoza-Vargas A, Reyes-Grajeda JP, Abraham Campos-Romero, Moreno-Camacho José Luis, Rodriguez-Gallegos Jorge, Luna-Ruiz Marco, Gonzalez-Barrera D, Rangel-DeLeon D, Munguia-Garza P, Ramirez-Vega O, Escobar-Arrazola, M, Herrera-Montalvo LA. |
| EPI_ISL_3067964, EPI_ISL_3067965, EPI_ISL_3067966, EPI_ISL_3067967 | INCMNSZ | Instituto Nacional de Medicina Genomica | Cedro-Tanda A, Hidalgo-Miranda A, Mendoza-Vargas A, Reyes-Grajeda JP, Abraham Campos-Romero, Moreno-Camacho José Luis, Rodriguez-Gallegos Jorge, Luna-Ruiz Marco, Gonzalez-Barrera D, Rangel-DeLeon D, Munguia-Garza P, Ramirez-Vega O, Escobar-Arrazola, M, Herrera-Montalvo LA. |
| EPI_ISL_3067968, EPI_ISL_3067969 | Centro Medico ABC | Instituto Nacional de Medicina Genomica | Cedro-Tanda A, Hidalgo-Miranda A, Mendoza-Vargas A, Reyes-Grajeda JP, Roxana Trejo-González, Cesar Lara, Jacomine Reyes-Carrasco, Daniel Aguirre, Gonzalez-Barrera D, Rangel-DeLeon D, Munguia-Garza P, Ramirez-Vega O, Escobar-Arrazola, M, Herrera-Montalvo LA. |
| EPI_ISL_3067970 | Instituto Nacional de Medicina Genomica | Instituto Nacional de Medicina Genomica | Cedro-Tanda A, Hidalgo-Miranda A, Mendoza-Vargas A, Reyes-Grajeda JP, Abraham Campos-Romero, Moreno-Camacho José Luis, Rodriguez-Gallegos Jorge, Luna-Ruiz Marco, Gonzalez-Barrera D, Rangel-DeLeon D, Munguia-Garza P, Ramirez-Vega O, Escobar-Arrazola, M, Herrera-Montalvo LA. |
| EPI_ISL_3067971, EPI_ISL_3067972, EPI_ISL_3067973 | INCMNSZ | Instituto Nacional de Medicina Genomica | Cedro-Tanda A, Hidalgo-Miranda A, Mendoza-Vargas A, Reyes-Grajeda JP, Abraham Campos-Romero, Moreno-Camacho José Luis, Rodriguez-Gallegos Jorge, Luna-Ruiz Marco, Gonzalez-Barrera D, Rangel-DeLeon D, Munguia-Garza P, Ramirez-Vega O, Escobar-Arrazola, M, Herrera-Montalvo LA. |
| EPI_ISL_3067974, EPI_ISL_3067975, EPI_ISL_3067976, EPI_ISL_3067977, EPI_ISL_3067978, EPI_ISL_3067979 | Instituto Nacional de Medicina Genomica | Instituto Nacional de Medicina Genomica | Cedro-Tanda A, Hidalgo-Miranda A, Mendoza-Vargas A, Reyes-Grajeda JP, Abraham Campos-Romero, Moreno-Camacho José Luis, Rodriguez-Gallegos Jorge, Luna-Ruiz Marco, Gonzalez-Barrera D, Rangel-DeLeon D, Munguia-Garza P, Ramirez-Vega O, Escobar-Arrazola, M, Herrera-Montalvo LA. |
| EPI_ISL_3068244 | IRCCS San Gallicano Dermatological Institute | IRCCS Regina Elena National Cancer Institute | Sabrina Strano, Sara Donzelli, Matteo Pallocca, Grazia Prignano, Fabrizio Ensoli, Fulvia Pimpinelli, Maurizio Fanciulli, Giovanni Blandino, Aldo Morrone, Gennaro Ciliberto |
| EPI_ISL_3068245 | IRCCS San Gallicano Dermatological Institute | IRCCS Regina Elena National Cancer Institute | Francesca De Nicola, Giulia Orlandi, Eleonora Sperandio, Elisabetta Trento, Fabrizio Ensoli, Fulvia Pimpinelli, Maurizio Fanciulli, Giovanni Blandino, Aldo Morrone, Gennaro Ciliberto |
| EPI_ISL_3068246 | IRCCS San Gallicano Dermatological Institute | IRCCS Regina Elena National Cancer Institute | Ludovica Ciuffreda, Sara Donzelli, Alice Massacci, Arianna Mastrofrancesco, Fabrizio Ensoli, Fulvia Pimpinelli, Maurizio Fanciulli, Giovanni Blandino, Aldo Morrone, Gennaro Ciliberto |
| EPI_ISL_3068247 | IRCCS San Gallicano Dermatological Institute | IRCCS Regina Elena National Cancer Institute | Frauke Goeman, Giulia Orlandi, Matteo Pallocca, Giovanna D’agosto, Fabrizio Ensoli, Fulvia Pimpinelli, Maurizio Fanciulli, Giovanni Blandino, Aldo Morrone, Gennaro Ciliberto |
| EPI_ISL_3068248 | IRCCS San Gallicano Dermatological Institute | IRCCS Regina Elena National Cancer Institute | Sabrina Strano, Sara Donzelli, Eleonora Sperandio, Carla Mottini, Fabrizio Ensoli, Fulvia Pimpinelli, Maurizio Fanciulli, Giovanni Blandino, Aldo Morrone, Gennaro Ciliberto |
| EPI_ISL_3068249 | IRCCS San Gallicano Dermatological Institute | IRCCS Regina Elena National Cancer Institute | Francesca De Nicola, Giulia Orlandi, Alice Massacci, Martina Pontone, Fabrizio Ensoli, Fulvia Pimpinelli, Maurizio Fanciulli, Giovanni Blandino, Aldo Morrone, Gennaro Ciliberto |
| EPI_ISL_3068251 | IRCCS San Gallicano Dermatological Institute | IRCCS Regina Elena National Cancer Institute | Frauke Goeman, Giulia Orlandi, Eleonora Sperandio, Ilaria Cavallo, Fabrizio Ensoli, Fulvia Pimpinelli, Maurizio Fanciulli, Giovanni Blandino, Aldo Morrone, Gennaro Ciliberto |
| EPI_ISL_3068252 | IRCCS San Gallicano Dermatological Institute | IRCCS Regina Elena National Cancer Institute | Sabrina Strano, Sara Donzelli, Alice Massacci, Francesca Sivori, Fabrizio Ensoli, Fulvia Pimpinelli, Maurizio Fanciulli, Giovanni Blandino, Aldo Morrone, Gennaro Ciliberto |
| EPI_ISL_3068253 | IRCCS San Gallicano Dermatological Institute | IRCCS Regina Elena National Cancer Institute | Francesca De Nicola, Giulia Orlandi, Matteo Pallocca, Andrea Cazzani, Fabrizio Ensoli, Fulvia Pimpinelli, Maurizio Fanciulli, Giovanni Blandino, Aldo Morrone, Gennaro Ciliberto |
| EPI_ISL_3068255 | IRCCS San Gallicano Dermatological Institute | IRCCS Regina Elena National Cancer Institute | Frauke Goeman, Giulia Orlandi, Alice Massacci, Martina Diano, Fabrizio Ensoli, Fulvia Pimpinelli, Maurizio Fanciulli, Giovanni Blandino, Aldo Morrone, Gennaro Ciliberto |
| EPI_ISL_3068256 | IRCCS San Gallicano Dermatological Institute | IRCCS Regina Elena National Cancer Institute | Sabrina Strano, Sara Donzelli, Matteo Pallocca, Antonio Federico, Fabrizio Ensoli, Fulvia Pimpinelli, Maurizio Fanciulli, Giovanni Blandino, Aldo Morrone, Gennaro Ciliberto |
| EPI_ISL_3068257 | IRCCS San Gallicano Dermatological Institute | IRCCS Regina Elena National Cancer Institute | Francesca De Nicola, Giulia Orlandi, Eleonora Sperandio, Fulvia Fraticelli, Fabrizio Ensoli, Fulvia Pimpinelli, Maurizio Fanciulli, Giovanni Blandino, Aldo Morrone, Gennaro Ciliberto |
| EPI_ISL_3068258 | IRCCS San Gallicano Dermatological Institute | IRCCS Regina Elena National Cancer Institute | Ludovica Ciuffreda, Sara Donzelli, Alice Massacci, Lorenzo Furzi, Fabrizio Ensoli, Fulvia Pimpinelli, Maurizio Fanciulli, Giovanni Blandino, Aldo Morrone, Gennaro Ciliberto |
| EPI_ISL_3068259 | IRCCS San Gallicano Dermatological Institute | IRCCS Regina Elena National Cancer Institute | Frauke Goeman, Giulia Orlandi, Matteo Pallocca, Alessia Lauretti, Fabrizio Ensoli, Fulvia Pimpinelli, Maurizio Fanciulli, Giovanni Blandino, Aldo Morrone, Gennaro Ciliberto |
| EPI_ISL_3068261 | IRCCS San Gallicano Dermatological Institute | IRCCS Regina Elena National Cancer Institute | Francesca De Nicola, Giulia Orlandi, Alice Massacci, Francisco Obregon, Fabrizio Ensoli, Fulvia Pimpinelli, Maurizio Fanciulli, Giovanni Blandino, Aldo Morrone, Gennaro Ciliberto |
| EPI_ISL_3068262 | IRCCS San Gallicano Dermatological Institute | IRCCS Regina Elena National Cancer Institute | Ludovica Ciuffreda, Sara Donzelli, Matteo Pallocca, Silvia Paluzzi, Fabrizio Ensoli, Fulvia Pimpinelli, Maurizio Fanciulli, Giovanni Blandino, Aldo Morrone, Gennaro Ciliberto |
| EPI_ISL_3068263 | IRCCS San Gallicano Dermatological Institute | IRCCS Regina Elena National Cancer Institute | Frauke Goeman, Giulia Orlandi, Eleonora Sperandio, Sara Petrolo, Fabrizio Ensoli, Fulvia Pimpinelli, Maurizio Fanciulli, Giovanni Blandino, Aldo Morrone, Gennaro Ciliberto |
| EPI_ISL_3068264 | IRCCS San Gallicano Dermatological Institute | IRCCS Regina Elena National Cancer Institute | Sabrina Strano, Sara Donzelli, Alice Massacci, Valentina Ricca, Fabrizio Ensoli, Fulvia Pimpinelli, Maurizio Fanciulli, Giovanni Blandino, Aldo Morrone, Gennaro Ciliberto |
| EPI_ISL_3068265 | IRCCS San Gallicano Dermatological Institute | IRCCS Regina Elena National Cancer Institute | Francesca De Nicola, Giulia Orlandi, Matteo Pallocca, Serena Salvo, Fabrizio Ensoli, Fulvia Pimpinelli, Maurizio Fanciulli, Giovanni Blandino, Aldo Morrone, Gennaro Ciliberto |
| EPI_ISL_3068266 | IRCCS San Gallicano Dermatological Institute | IRCCS Regina Elena National Cancer Institute | Ludovica Ciuffreda, Sara Donzelli, Eleonora Sperandio, Grazia Prignano, Fabrizio Ensoli, Fulvia Pimpinelli, Maurizio Fanciulli, Giovanni Blandino, Aldo Morrone, Gennaro Ciliberto |
| EPI_ISL_3068267 | IRCCS San Gallicano Dermatological Institute | IRCCS Regina Elena National Cancer Institute | Frauke Goeman, Giulia Orlandi, Alice Massacci, Elisabetta Trento, Fabrizio Ensoli, Fulvia Pimpinelli, Maurizio Fanciulli, Giovanni Blandino, Aldo Morrone, Gennaro Ciliberto |
| EPI_ISL_3068268 | IRCCS San Gallicano Dermatological Institute | IRCCS Regina Elena National Cancer Institute | Sabrina Strano, Sara Donzelli, Matteo Pallocca, Arianna Mastrofrancesco, Fabrizio Ensoli, Fulvia Pimpinelli, Maurizio Fanciulli, Giovanni Blandino, Aldo Morrone, Gennaro Ciliberto |
| EPI_ISL_3068269 | IRCCS San Gallicano Dermatological Institute | IRCCS Regina Elena National Cancer Institute | Francesca De Nicola, Giulia Orlandi, Eleonora Sperandio, Giovanna D’agosto, Fabrizio Ensoli, Fulvia Pimpinelli, Maurizio Fanciulli, Giovanni Blandino, Aldo Morrone, Gennaro Ciliberto |
| EPI_ISL_3068270 | IRCCS San Gallicano Dermatological Institute | IRCCS Regina Elena National Cancer Institute | Ludovica Ciuffreda, Sara Donzelli, Alice Massacci, Carla Mottini, Fabrizio Ensoli, Fulvia Pimpinelli, Maurizio Fanciulli, Giovanni Blandino, Aldo Morrone, Gennaro Ciliberto |
| EPI_ISL_3068271 | IRCCS San Gallicano Dermatological Institute | IRCCS Regina Elena National Cancer Institute | Frauke Goeman, Giulia Orlandi, Matteo Pallocca, Martina Pontone, Fabrizio Ensoli, Fulvia Pimpinelli, Maurizio Fanciulli, Giovanni Blandino, Aldo Morrone, Gennaro Ciliberto |
| EPI_ISL_3068272 | IRCCS San Gallicano Dermatological Institute | IRCCS Regina Elena National Cancer Institute | Sabrina Strano, Sara Donzelli, Eleonora Sperandio, di Domenico Enea Gino, Fabrizio Ensoli, Fulvia Pimpinelli, Maurizio Fanciulli, Giovanni Blandino, Aldo Morrone, Gennaro Ciliberto |
| EPI_ISL_3068273 | IRCCS San Gallicano Dermatological Institute | IRCCS Regina Elena National Cancer Institute | Francesca De Nicola, Giulia Orlandi, Alice Massacci, Ilaria Cavallo, Fabrizio Ensoli, Fulvia Pimpinelli, Maurizio Fanciulli, Giovanni Blandino, Aldo Morrone, Gennaro Ciliberto |
| EPI_ISL_3068274 | IRCCS San Gallicano Dermatological Institute | IRCCS Regina Elena National Cancer Institute | Ludovica Ciuffreda, Sara Donzelli, Matteo Pallocca, Francesca Sivori, Fabrizio Ensoli, Fulvia Pimpinelli, Maurizio Fanciulli, Giovanni Blandino, Aldo Morrone, Gennaro Ciliberto |
| EPI_ISL_3068275 | IRCCS San Gallicano Dermatological Institute | IRCCS Regina Elena National Cancer Institute | Frauke Goeman, Giulia Orlandi, Eleonora Sperandio, Andrea Cazzani, Fabrizio Ensoli, Fulvia Pimpinelli, Maurizio Fanciulli, Giovanni Blandino, Aldo Morrone, Gennaro Ciliberto |

| | | | |
|---|---|---|---|
| EPI_ISL_3068276 | IRCCS San Gallicano Dermatological Institute | IRCCS Regina Elena National Cancer Institute | Sabrina Strano, Sara Donzelli, Alice Massacci, Ilaria Celesti, Fabrizio Ensoli, Fulvia Pimpinelli, Maurizio Fanciulli, Giovanni Blandino, Aldo Morrone, Gennaro Ciliberto |
| EPI_ISL_3068277 | IRCCS San Gallicano Dermatological Institute | IRCCS Regina Elena National Cancer Institute | Francesca De Nicola, Giulia Orlandi, Matteo Pallocca, Martina Diano, Fabrizio Ensoli, Fulvia Pimpinelli, Maurizio Fanciulli, Giovanni Blandino, Aldo Morrone, Gennaro Ciliberto |
| EPI_ISL_3068278 | IRCCS San Gallicano Dermatological Institute | IRCCS Regina Elena National Cancer Institute | Ludovica Ciuffreda, Sara Donzelli, Eleonora Sperandio, Antonio Federico, Fabrizio Ensoli, Fulvia Pimpinelli, Maurizio Fanciulli, Giovanni Blandino, Aldo Morrone, Gennaro Ciliberto |
| EPI_ISL_3068279 | IRCCS San Gallicano Dermatological Institute | IRCCS Regina Elena National Cancer Institute | Frauke Goeman, Giulia Orlandi, Alice Massacci, Fulvia Fraticelli, Fabrizio Ensoli, Fulvia Pimpinelli, Maurizio Fanciulli, Giovanni Blandino, Aldo Morrone, Gennaro Ciliberto |
| EPI_ISL_3068280 | IRCCS San Gallicano Dermatological Institute | IRCCS Regina Elena National Cancer Institute | Sabrina Strano, Sara Donzelli, Matteo Pallocca, Lorenzo Furzi, Fabrizio Ensoli, Fulvia Pimpinelli, Maurizio Fanciulli, Giovanni Blandino, Aldo Morrone, Gennaro Ciliberto |
| EPI_ISL_3068281 | IRCCS San Gallicano Dermatological Institute | IRCCS Regina Elena National Cancer Institute | Francesca De Nicola, Giulia Orlandi, Eleonora Sperandio, Alessia Lauretti, Fabrizio Ensoli, Fulvia Pimpinelli, Maurizio Fanciulli, Giovanni Blandino, Aldo Morrone, Gennaro Ciliberto |
| EPI_ISL_3068283 | IRCCS San Gallicano Dermatological Institute | IRCCS Regina Elena National Cancer Institute | Frauke Goeman, Giulia Orlandi, Matteo Pallocca, Francisco Obregon, Fabrizio Ensoli, Fulvia Pimpinelli, Maurizio Fanciulli, Giovanni Blandino, Aldo Morrone, Gennaro Ciliberto |
| EPI_ISL_3068284 | IRCCS San Gallicano Dermatological Institute | IRCCS Regina Elena National Cancer Institute | Sabrina Strano, Sara Donzelli, Eleonora Sperandio, Silvia Paluzzi, Fabrizio Ensoli, Fulvia Pimpinelli, Maurizio Fanciulli, Giovanni Blandino, Aldo Morrone, Gennaro Ciliberto |
| EPI_ISL_3068286 | IRCCS San Gallicano Dermatological Institute | IRCCS Regina Elena National Cancer Institute | Ludovica Ciuffreda, Sara Donzelli, Matteo Pallocca, Valentina Ricca, Fabrizio Ensoli, Fulvia Pimpinelli, Maurizio Fanciulli, Giovanni Blandino, Aldo Morrone, Gennaro Ciliberto |
| EPI_ISL_3068287 | IRCCS San Gallicano Dermatological Institute | IRCCS Regina Elena National Cancer Institute | Frauke Goeman, Giulia Orlandi, Eleonora Sperandio, Serena Salvo, Fabrizio Ensoli, Fulvia Pimpinelli, Maurizio Fanciulli, Giovanni Blandino, Aldo Morrone, Gennaro Ciliberto |
| EPI_ISL_3068288 | IRCCS San Gallicano Dermatological Institute | IRCCS Regina Elena National Cancer Institute | Sabrina Strano, Sara Donzelli, Alice Massacci, Grazia Prignano, Fabrizio Ensoli, Fulvia Pimpinelli, Maurizio Fanciulli, Giovanni Blandino, Aldo Morrone, Gennaro Ciliberto |
| EPI_ISL_3068289 | IRCCS San Gallicano Dermatological Institute | IRCCS Regina Elena National Cancer Institute | Francesca De Nicola, Giulia Orlandi, Matteo Pallocca, Elisabetta Trento, Fabrizio Ensoli, Fulvia Pimpinelli, Maurizio Fanciulli, Giovanni Blandino, Aldo Morrone, Gennaro Ciliberto |
| EPI_ISL_3068290 | IRCCS San Gallicano Dermatological Institute | IRCCS Regina Elena National Cancer Institute | Ludovica Ciuffreda, Sara Donzelli, Eleonora Sperandio, Arianna Mastrofrancesco, Fabrizio Ensoli, Fulvia Pimpinelli, Maurizio Fanciulli, Giovanni Blandino, Aldo Morrone, Gennaro Ciliberto |
| EPI_ISL_3068291 | IRCCS San Gallicano Dermatological Institute | IRCCS Regina Elena National Cancer Institute | Frauke Goeman, Giulia Orlandi, Alice Massacci, Giovanna D'agosto, Fabrizio Ensoli, Fulvia Pimpinelli, Maurizio Fanciulli, Giovanni Blandino, Aldo Morrone, Gennaro Ciliberto |
| EPI_ISL_3068292 | IRCCS San Gallicano Dermatological Institute | IRCCS Regina Elena National Cancer Institute | Sabrina Strano, Sara Donzelli, Matteo Pallocca, Carla Mottini, Fabrizio Ensoli, Fulvia Pimpinelli, Maurizio Fanciulli, Giovanni Blandino, Aldo Morrone, Gennaro Ciliberto |
| EPI_ISL_3068293 | IRCCS San Gallicano Dermatological Institute | IRCCS Regina Elena National Cancer Institute | Francesca De Nicola, Giulia Orlandi, Eleonora Sperandio, Martina Pontone, Fabrizio Ensoli, Fulvia Pimpinelli, Maurizio Fanciulli, Giovanni Blandino, Aldo Morrone, Gennaro Ciliberto |
| EPI_ISL_3068294 | IRCCS San Gallicano Dermatological Institute | IRCCS Regina Elena National Cancer Institute | Ludovica Ciuffreda, Sara Donzelli, Alice Massacci, di Domenico Enea Gino, Fabrizio Ensoli, Fulvia Pimpinelli, Maurizio Fanciulli, Giovanni Blandino, Aldo Morrone, Gennaro Ciliberto |
| EPI_ISL_3068296 | IRCCS San Gallicano Dermatological Institute | IRCCS Regina Elena National Cancer Institute | Sabrina Strano, Sara Donzelli, Eleonora Sperandio, Francesca Sivori, Fabrizio Ensoli, Fulvia Pimpinelli, Maurizio Fanciulli, Giovanni Blandino, Aldo Morrone, Gennaro Ciliberto |
| EPI_ISL_3068297 | IRCCS San Gallicano Dermatological Institute | IRCCS Regina Elena National Cancer Institute | Francesca De Nicola, Giulia Orlandi, Alice Massacci, Andrea Cazzani, Fabrizio Ensoli, Fulvia Pimpinelli, Maurizio Fanciulli, Giovanni Blandino, Aldo Morrone, Gennaro Ciliberto |
| EPI_ISL_3068298 | IRCCS San Gallicano Dermatological Institute | IRCCS Regina Elena National Cancer Institute | Ludovica Ciuffreda, Sara Donzelli, Matteo Pallocca, Ilaria Celesti, Fabrizio Ensoli, Fulvia Pimpinelli, Maurizio Fanciulli, Giovanni Blandino, Aldo Morrone, Gennaro Ciliberto |
| EPI_ISL_3068299 | IRCCS San Gallicano Dermatological Institute | IRCCS Regina Elena National Cancer Institute | Frauke Goeman, Giulia Orlandi, Eleonora Sperandio, Martina Diano, Fabrizio Ensoli, Fulvia Pimpinelli, Maurizio Fanciulli, Giovanni Blandino, Aldo Morrone, Gennaro Ciliberto |
| EPI_ISL_3068300 | IRCCS San Gallicano Dermatological Institute | IRCCS Regina Elena National Cancer Institute | Sabrina Strano, Sara Donzelli, Alice Massacci, Antonio Federico, Fabrizio Ensoli, Fulvia Pimpinelli, Maurizio Fanciulli, Giovanni Blandino, Aldo Morrone, Gennaro Ciliberto |
| EPI_ISL_3068303 | IRCCS San Gallicano Dermatological Institute | IRCCS Regina Elena National Cancer Institute | Frauke Goeman, Giulia Orlandi, Alice Massacci, Alessia Lauretti, Fabrizio Ensoli, Fulvia Pimpinelli, Maurizio Fanciulli, Giovanni Blandino, Aldo Morrone, Gennaro Ciliberto |
| EPI_ISL_3068304 | IRCCS San Gallicano Dermatological Institute | IRCCS Regina Elena National Cancer Institute | Sabrina Strano, Sara Donzelli, Matteo Pallocca, Francesca Maione, Fabrizio Ensoli, Fulvia Pimpinelli, Maurizio Fanciulli, Giovanni Blandino, Aldo Morrone, Gennaro Ciliberto |
| EPI_ISL_3068305 | IRCCS San Gallicano Dermatological Institute | IRCCS Regina Elena National Cancer Institute | Francesca De Nicola, Giulia Orlandi, Eleonora Sperandio, Francisco Obregon, Fabrizio Ensoli, Fulvia Pimpinelli, Maurizio Fanciulli, Giovanni Blandino, Aldo Morrone, Gennaro Ciliberto |
| EPI_ISL_3068306 | IRCCS San Gallicano Dermatological Institute | IRCCS Regina Elena National Cancer Institute | Ludovica Ciuffreda, Sara Donzelli, Alice Massacci, Silvia Paluzzi, Fabrizio Ensoli, Fulvia Pimpinelli, Maurizio Fanciulli, Giovanni Blandino, Aldo Morrone, Gennaro Ciliberto |
| EPI_ISL_3068307 | IRCCS San Gallicano Dermatological Institute | IRCCS Regina Elena National Cancer Institute | Frauke Goeman, Giulia Orlandi, Matteo Pallocca, Sara Petrolo, Fabrizio Ensoli, Fulvia Pimpinelli, Maurizio Fanciulli, Giovanni Blandino, Aldo Morrone, Gennaro Ciliberto |
| EPI_ISL_3068308 | IRCCS San Gallicano Dermatological Institute | IRCCS Regina Elena National Cancer Institute | Sabrina Strano, Sara Donzelli, Eleonora Sperandio, Valentina Ricca, Fabrizio Ensoli, Fulvia Pimpinelli, Maurizio Fanciulli, Giovanni Blandino, Aldo Morrone, Gennaro Ciliberto |
| EPI_ISL_3068310 | IRCCS San Gallicano Dermatological Institute | IRCCS Regina Elena National Cancer Institute | Ludovica Ciuffreda, Sara Donzelli, Matteo Pallocca, Grazia Prignano, Fabrizio Ensoli, Fulvia Pimpinelli, Maurizio Fanciulli, Giovanni Blandino, Aldo Morrone, Gennaro Ciliberto |
| EPI_ISL_3068311 | IRCCS San Gallicano Dermatological Institute | IRCCS Regina Elena National Cancer Institute | Frauke Goeman, Giulia Orlandi, Eleonora Sperandio, Elisabetta Trento, Fabrizio Ensoli, Fulvia Pimpinelli, Maurizio Fanciulli, Giovanni Blandino, Aldo Morrone, Gennaro Ciliberto |
| EPI_ISL_3068312 | IRCCS San Gallicano Dermatological Institute | IRCCS Regina Elena National Cancer Institute | Sabrina Strano, Sara Donzelli, Alice Massacci, Arianna Mastrofrancesco, Fabrizio Ensoli, Fulvia Pimpinelli, Maurizio Fanciulli, Giovanni Blandino, Aldo Morrone, Gennaro Ciliberto |
| EPI_ISL_3068313 | IRCCS San Gallicano Dermatological Institute | IRCCS Regina Elena National Cancer Institute | Francesca De Nicola, Giulia Orlandi, Matteo Pallocca, Giovanna D'agosto, Fabrizio Ensoli, Fulvia Pimpinelli, Maurizio Fanciulli, Giovanni Blandino, Aldo Morrone, Gennaro Ciliberto |
| EPI_ISL_3068314 | IRCCS San Gallicano Dermatological Institute | IRCCS Regina Elena National Cancer Institute | Ludovica Ciuffreda, Sara Donzelli, Eleonora Sperandio, Carla Mottini, Fabrizio Ensoli, Fulvia Pimpinelli, Maurizio Fanciulli, Giovanni Blandino, Aldo Morrone, Gennaro Ciliberto |
| EPI_ISL_3068315 | IRCCS San Gallicano Dermatological Institute | IRCCS Regina Elena National Cancer Institute | Frauke Goeman, Giulia Orlandi, Alice Massacci, Martina Pontone, Fabrizio Ensoli, Fulvia Pimpinelli, Maurizio Fanciulli, Giovanni Blandino, Aldo Morrone, Gennaro Ciliberto |
| EPI_ISL_3068316 | IRCCS San Gallicano Dermatological Institute | IRCCS Regina Elena National Cancer Institute | Sabrina Strano, Sara Donzelli, Matteo Pallocca, di Domenico Enea Gino, Fabrizio Ensoli, Fulvia Pimpinelli, Maurizio Fanciulli, Giovanni Blandino, Aldo Morrone, Gennaro Ciliberto |
| EPI_ISL_3068317 | IRCCS San Gallicano Dermatological Institute | IRCCS Regina Elena National Cancer Institute | Francesca De Nicola, Giulia Orlandi, Eleonora Sperandio, Ilaria Cavallo, Fabrizio Ensoli, Fulvia Pimpinelli, Maurizio Fanciulli, Giovanni Blandino, Aldo Morrone, Gennaro Ciliberto |
| EPI_ISL_3068318 | IRCCS San Gallicano Dermatological Institute | IRCCS Regina Elena National Cancer Institute | Ludovica Ciuffreda, Sara Donzelli, Alice Massacci, Francesca Sivori, Fabrizio Ensoli, Fulvia Pimpinelli, Maurizio Fanciulli, Giovanni Blandino, Aldo Morrone, Gennaro Ciliberto |
| EPI_ISL_3068319 | IRCCS San Gallicano Dermatological Institute | IRCCS Regina Elena National Cancer Institute | Frauke Goeman, Giulia Orlandi, Matteo Pallocca, Andrea Cazzani, Fabrizio Ensoli, Fulvia Pimpinelli, Maurizio Fanciulli, Giovanni Blandino, Aldo Morrone, Gennaro Ciliberto |
| EPI_ISL_3068320 | IRCCS San Gallicano Dermatological Institute | IRCCS Regina Elena National Cancer Institute | Sabrina Strano, Sara Donzelli, Eleonora Sperandio, Ilaria Celesti, Fabrizio Ensoli, Fulvia Pimpinelli, Maurizio Fanciulli, Giovanni Blandino, Aldo Morrone, Gennaro Ciliberto |
| EPI_ISL_3068321 | IRCCS San Gallicano Dermatological Institute | IRCCS Regina Elena National Cancer Institute | Francesca De Nicola, Giulia Orlandi, Alice Massacci, Martina Diano, Fabrizio Ensoli, Fulvia Pimpinelli, Maurizio Fanciulli, Giovanni Blandino, Aldo Morrone, Gennaro Ciliberto |
| EPI_ISL_3068322 | IRCCS San Gallicano Dermatological Institute | IRCCS Regina Elena National Cancer Institute | Ludovica Ciuffreda, Sara Donzelli, Matteo Pallocca, Antonio Federico, Fabrizio Ensoli, Fulvia Pimpinelli, Maurizio Fanciulli, Giovanni Blandino, Aldo Morrone, Gennaro Ciliberto |
| EPI_ISL_3068323 | IRCCS San Gallicano Dermatological Institute | IRCCS Regina Elena National Cancer Institute | Frauke Goeman, Giulia Orlandi, Eleonora Sperandio, Fulvia Fraticelli, Fabrizio Ensoli, Fulvia Pimpinelli, Maurizio Fanciulli, Giovanni Blandino, Aldo Morrone, Gennaro Ciliberto |
| EPI_ISL_3068324 | IRCCS San Gallicano Dermatological Institute | IRCCS Regina Elena National Cancer Institute | Sabrina Strano, Sara Donzelli, Alice Massacci, Lorenzo Furzi, Fabrizio Ensoli, Fulvia Pimpinelli, Maurizio Fanciulli, Giovanni Blandino, Aldo Morrone, Gennaro Ciliberto |
| EPI_ISL_3068325 | IRCCS San Gallicano Dermatological Institute | IRCCS Regina Elena National Cancer Institute | Francesca De Nicola, Giulia Orlandi, Matteo Pallocca, Alessia Lauretti, Fabrizio Ensoli, Fulvia Pimpinelli, Maurizio Fanciulli, Giovanni Blandino, Aldo Morrone, Gennaro Ciliberto |
| EPI_ISL_3068327 | IRCCS San Gallicano Dermatological Institute | IRCCS Regina Elena National Cancer Institute | Frauke Goeman, Giulia Orlandi, Alice Massacci, Francisco Obregon, Fabrizio Ensoli, Fulvia Pimpinelli, Maurizio Fanciulli, Giovanni Blandino, Aldo Morrone, Gennaro Ciliberto |
| EPI_ISL_3068328 | IRCCS San Gallicano Dermatological Institute | IRCCS Regina Elena National Cancer Institute | Sabrina Strano, Sara Donzelli, Matteo Pallocca, Silvia Paluzzi, Fabrizio Ensoli, Fulvia Pimpinelli, Maurizio Fanciulli, Giovanni Blandino, Aldo Morrone, Gennaro Ciliberto |
| EPI_ISL_3068330 | IRCCS San Gallicano Dermatological Institute | IRCCS Regina Elena National Cancer Institute | Ludovica Ciuffreda, Sara Donzelli, Alice Massacci, Valentina Ricca, Fabrizio Ensoli, Fulvia Pimpinelli, Maurizio Fanciulli, Giovanni Blandino, Aldo Morrone, Gennaro Ciliberto |
| EPI_ISL_3068331 | IRCCS San Gallicano Dermatological Institute | IRCCS Regina Elena National Cancer Institute | Frauke Goeman, Giulia Orlandi, Matteo Pallocca, Serena Salvo, Fabrizio Ensoli, Fulvia Pimpinelli, Maurizio Fanciulli, Giovanni Blandino, Aldo Morrone, Gennaro Ciliberto |
| EPI_ISL_3068332 | IRCCS San Gallicano Dermatological Institute | IRCCS Regina Elena National Cancer Institute | Sabrina Strano, Sara Donzelli, Eleonora Sperandio, Grazia Prignano, Fabrizio Ensoli, Fulvia Pimpinelli, Maurizio Fanciulli, Giovanni Blandino, Aldo Morrone, Gennaro Ciliberto |
| EPI_ISL_3068333 | IRCCS San Gallicano Dermatological Institute | IRCCS Regina Elena National Cancer Institute | Francesca De Nicola, Giulia Orlandi, Alice Massacci, Elisabetta Trento, Fabrizio Ensoli, Fulvia Pimpinelli, Maurizio Fanciulli, Giovanni Blandino, Aldo Morrone, Gennaro Ciliberto |
| EPI_ISL_3068334 | IRCCS San Gallicano Dermatological Institute | IRCCS Regina Elena National Cancer Institute | Ludovica Ciuffreda, Sara Donzelli, Matteo Pallocca, Arianna Mastrofrancesco, Fabrizio Ensoli, Fulvia Pimpinelli, Maurizio Fanciulli, Giovanni Blandino, Aldo Morrone, Gennaro Ciliberto |
| EPI_ISL_3068335 | IRCCS San Gallicano Dermatological Institute | IRCCS Regina Elena National Cancer Institute | Frauke Goeman, Giulia Orlandi, Eleonora Sperandio, Giovanna D'agosto, Fabrizio Ensoli, Fulvia Pimpinelli, Maurizio Fanciulli, Giovanni Blandino, Aldo Morrone, Gennaro Ciliberto |
| EPI_ISL_3068336 | IRCCS San Gallicano Dermatological Institute | IRCCS Regina Elena National Cancer Institute | Sabrina Strano, Sara Donzelli, Alice Massacci, Carla Mottini, Fabrizio Ensoli, Fulvia Pimpinelli, Maurizio Fanciulli, Giovanni Blandino, Aldo Morrone, Gennaro Ciliberto |
| EPI_ISL_3068337 | IRCCS San Gallicano Dermatological Institute | IRCCS Regina Elena National Cancer Institute | Francesca De Nicola, Giulia Orlandi, Matteo Pallocca, Martina Pontone, Fabrizio Ensoli, Fulvia Pimpinelli, Maurizio Fanciulli, Giovanni Blandino, Aldo Morrone, Gennaro Ciliberto |
| EPI_ISL_3068338 | IRCCS San Gallicano Dermatological Institute | IRCCS Regina Elena National Cancer Institute | Ludovica Ciuffreda, Sara Donzelli, Eleonora Sperandio, di Domenico Enea Gino, Fabrizio Ensoli, Fulvia Pimpinelli, Maurizio Fanciulli, Giovanni Blandino, Aldo Morrone, Gennaro Ciliberto |
| EPI_ISL_3068339 | IRCCS San Gallicano Dermatological Institute | IRCCS Regina Elena National Cancer Institute | Frauke Goeman, Giulia Orlandi, Alice Massacci, Ilaria Cavallo, Fabrizio Ensoli, Fulvia Pimpinelli, Maurizio Fanciulli, Giovanni Blandino, Aldo Morrone, Gennaro Ciliberto |
| EPI_ISL_3068340 | IRCCS San Gallicano Dermatological Institute | IRCCS Regina Elena National Cancer Institute | Sabrina Strano, Sara Donzelli, Matteo Pallocca, Francesca Sivori, Fabrizio Ensoli, Fulvia Pimpinelli, Maurizio Fanciulli, Giovanni Blandino, Aldo Morrone, Gennaro Ciliberto |
| EPI_ISL_3068341 | IRCCS San Gallicano Dermatological Institute | IRCCS Regina Elena National Cancer Institute | Francesca De Nicola, Giulia Orlandi, Eleonora Sperandio, Andrea Cazzani, Fabrizio Ensoli, Fulvia Pimpinelli, Maurizio Fanciulli, Giovanni Blandino, Aldo Morrone, Gennaro Ciliberto |
| EPI_ISL_3068342 | IRCCS San Gallicano Dermatological Institute | IRCCS Regina Elena National Cancer Institute | Ludovica Ciuffreda, Sara Donzelli, Alice Massacci, Ilaria Celesti, Fabrizio Ensoli, Fulvia Pimpinelli, Maurizio Fanciulli, Giovanni Blandino, Aldo Morrone, Gennaro Ciliberto |
| EPI_ISL_3068343 | IRCCS San Gallicano Dermatological Institute | IRCCS Regina Elena National Cancer Institute | Frauke Goeman, Giulia Orlandi, Matteo Pallocca, Martina Diano, Fabrizio Ensoli, Fulvia Pimpinelli, Maurizio Fanciulli, Giovanni Blandino, Aldo Morrone, Gennaro Ciliberto |
| EPI_ISL_3068344 | IRCCS San Gallicano Dermatological Institute | IRCCS Regina Elena National Cancer Institute | Sabrina Strano, Sara Donzelli, Eleonora Sperandio, Antonio Federico, Fabrizio Ensoli, Fulvia Pimpinelli, Maurizio Fanciulli, Giovanni Blandino, Aldo Morrone, Gennaro Ciliberto |
| EPI_ISL_3068345 | IRCCS San Gallicano Dermatological Institute | IRCCS Regina Elena National Cancer Institute | Francesca De Nicola, Giulia Orlandi, Alice Massacci, Fulvia Fraticelli, Fabrizio Ensoli, Fulvia Pimpinelli, Maurizio Fanciulli, Giovanni Blandino, Aldo Morrone, Gennaro Ciliberto |
| EPI_ISL_3068346 | IRCCS San Gallicano Dermatological Institute | IRCCS Regina Elena National Cancer Institute | Ludovica Ciuffreda, Sara Donzelli, Matteo Pallocca, Lorenzo Furzi, Fabrizio Ensoli, Fulvia Pimpinelli, Maurizio Fanciulli, Giovanni Blandino, Aldo Morrone, Gennaro Ciliberto |
| EPI_ISL_3068347 | IRCCS San Gallicano Dermatological Institute | IRCCS Regina Elena National Cancer Institute | Frauke Goeman, Giulia Orlandi, Eleonora Sperandio, Alessia Lauretti, Fabrizio Ensoli, Fulvia Pimpinelli, Maurizio Fanciulli, Giovanni Blandino, Aldo Morrone, Gennaro Ciliberto |
| EPI_ISL_3068348 | IRCCS San Gallicano Dermatological Institute | IRCCS Regina Elena National Cancer Institute | Sabrina Strano, Sara Donzelli, Alice Massacci, Francesca Maione, Fabrizio Ensoli, Fulvia Pimpinelli, Maurizio Fanciulli, Giovanni Blandino, Aldo Morrone, Gennaro Ciliberto |
| EPI_ISL_3068349 | IRCCS San Gallicano Dermatological Institute | IRCCS Regina Elena National Cancer Institute | Francesca De Nicola, Giulia Orlandi, Matteo Pallocca, Francisco Obregon, Fabrizio Ensoli, Fulvia Pimpinelli, Maurizio Fanciulli, Giovanni Blandino, Aldo Morrone, Gennaro Ciliberto |
| EPI_ISL_3068350 | IRCCS San Gallicano Dermatological Institute | IRCCS Regina Elena National Cancer Institute | Ludovica Ciuffreda, Sara Donzelli, Eleonora Sperandio, Silvia Paluzzi, Fabrizio Ensoli, Fulvia Pimpinelli, Maurizio Fanciulli, Giovanni Blandino, Aldo Morrone, Gennaro Ciliberto |
| EPI_ISL_3068351 | IRCCS San Gallicano Dermatological Institute | IRCCS Regina Elena National Cancer Institute | Frauke Goeman, Giulia Orlandi, Alice Massacci, Sara Petrolo, Fabrizio Ensoli, Fulvia Pimpinelli, Maurizio Fanciulli, Giovanni Blandino, Aldo Morrone, Gennaro Ciliberto |
| EPI_ISL_3068352 | IRCCS San Gallicano Dermatological Institute | IRCCS Regina Elena National Cancer Institute | Sabrina Strano, Sara Donzelli, Matteo Pallocca, Valentina Ricca, Fabrizio Ensoli, Fulvia Pimpinelli, Maurizio Fanciulli, Giovanni Blandino, Aldo Morrone, Gennaro Ciliberto |
| EPI_ISL_3068353 | IRCCS San Gallicano Dermatological Institute | IRCCS Regina Elena National Cancer Institute | Francesca De Nicola, Giulia Orlandi, Eleonora Sperandio, Serena Salvo, Fabrizio Ensoli, Fulvia Pimpinelli, Maurizio Fanciulli, Giovanni Blandino, Aldo Morrone, Gennaro Ciliberto |
| EPI_ISL_3068355 | IRCCS San Gallicano Dermatological Institute | IRCCS Regina Elena National Cancer Institute | Frauke Goeman, Giulia Orlandi, Matteo Pallocca, Elisabetta Trento, Fabrizio Ensoli, Fulvia Pimpinelli, Maurizio Fanciulli, Giovanni Blandino, Aldo Morrone, Gennaro Ciliberto |
| EPI_ISL_3068356 | IRCCS San Gallicano Dermatological Institute | IRCCS Regina Elena National Cancer Institute | Sabrina Strano, Sara Donzelli, Eleonora Sperandio, Arianna Mastrofrancesco, Fabrizio Ensoli, Fulvia Pimpinelli, Maurizio Fanciulli, Giovanni Blandino, Aldo Morrone, Gennaro Ciliberto |
| EPI_ISL_3068357 | IRCCS San Gallicano Dermatological Institute | IRCCS Regina Elena National Cancer Institute | Francesca De Nicola, Giulia Orlandi, Alice Massacci, Giovanna D'agosto, Fabrizio Ensoli, Fulvia Pimpinelli, Maurizio Fanciulli, Giovanni Blandino, Aldo Morrone, Gennaro Ciliberto |
| EPI_ISL_3068358 | IRCCS San Gallicano Dermatological Institute | IRCCS Regina Elena National Cancer Institute | Ludovica Ciuffreda, Sara Donzelli, Matteo Pallocca, Carla Mottini, Fabrizio Ensoli, Fulvia Pimpinelli, Maurizio Fanciulli, Giovanni Blandino, Aldo Morrone, Gennaro Ciliberto |
| EPI_ISL_3068359 | IRCCS San Gallicano Dermatological Institute | IRCCS Regina Elena National Cancer Institute | Frauke Goeman, Giulia Orlandi, Eleonora Sperandio, Martina Pontone, Fabrizio Ensoli, Fulvia Pimpinelli, Maurizio Fanciulli, Giovanni Blandino, Aldo Morrone, Gennaro Ciliberto |
| EPI_ISL_3068360 | IRCCS San Gallicano Dermatological Institute | IRCCS Regina Elena National Cancer Institute | Sabrina Strano, Sara Donzelli, Alice Massacci, di Domenico Enea Gino, Fabrizio Ensoli, Fulvia Pimpinelli, Maurizio Fanciulli, Giovanni Blandino, Aldo Morrone, Gennaro Ciliberto |
| EPI_ISL_3068361 | IRCCS San Gallicano Dermatological Institute | IRCCS Regina Elena National Cancer Institute | Francesca De Nicola, Giulia Orlandi, Matteo Pallocca, Ilaria Cavallo, Fabrizio Ensoli, Fulvia Pimpinelli, Maurizio Fanciulli, Giovanni Blandino, Aldo Morrone, Gennaro Ciliberto |

| | | | |
|---|---|---|---|
| EPI_ISL_3068362 | IRCCS San Gallicano Dermatological Institute | IRCCS Regina Elena National Cancer Institute | Ludovica Ciuffreda, Sara Donzelli, Eleonora Sperandio, Francesca Sivori, Fabrizio Ensoli, Fulvia Pimpinelli, Maurizio Fanciulli, Giovanni Blandino, Aldo Morrone, Gennaro Ciliberto |
| EPI_ISL_3068363 | IRCCS San Gallicano Dermatological Institute | IRCCS Regina Elena National Cancer Institute | Frauke Goeman, Giulia Orlandi, Alice Massacci, Andrea Cazzani, Fabrizio Ensoli, Fulvia Pimpinelli, Maurizio Fanciulli, Giovanni Blandino, Aldo Morrone, Gennaro Ciliberto |
| EPI_ISL_3068364 | IRCCS San Gallicano Dermatological Institute | IRCCS Regina Elena National Cancer Institute | Sabrina Strano, Sara Donzelli, Matteo Pallocca, Ilaria Celesti, Fabrizio Ensoli, Fulvia Pimpinelli, Maurizio Fanciulli, Giovanni Blandino, Aldo Morrone, Gennaro Ciliberto |
| EPI_ISL_3068365 | IRCCS San Gallicano Dermatological Institute | IRCCS Regina Elena National Cancer Institute | Francesca De Nicola, Giulia Orlandi, Eleonora Sperandio, Martina Diano, Fabrizio Ensoli, Fulvia Pimpinelli, Maurizio Fanciulli, Giovanni Blandino, Aldo Morrone, Gennaro Ciliberto |
| EPI_ISL_3068366 | IRCCS San Gallicano Dermatological Institute | IRCCS Regina Elena National Cancer Institute | Ludovica Ciuffreda, Sara Donzelli, Alice Massacci, Antonio Federico, Fabrizio Ensoli, Fulvia Pimpinelli, Maurizio Fanciulli, Giovanni Blandino, Aldo Morrone, Gennaro Ciliberto |
| EPI_ISL_3068367 | IRCCS San Gallicano Dermatological Institute | IRCCS Regina Elena National Cancer Institute | Frauke Goeman, Giulia Orlandi, Matteo Pallocca, Fulvia Fraticelli, Fabrizio Ensoli, Fulvia Pimpinelli, Maurizio Fanciulli, Giovanni Blandino, Aldo Morrone, Gennaro Ciliberto |
| EPI_ISL_3068368 | IRCCS San Gallicano Dermatological Institute | IRCCS Regina Elena National Cancer Institute | Sabrina Strano, Sara Donzelli, Eleonora Sperandio, Lorenzo Furzi, Fabrizio Ensoli, Fulvia Pimpinelli, Maurizio Fanciulli, Giovanni Blandino, Aldo Morrone, Gennaro Ciliberto |
| EPI_ISL_3068369 | IRCCS San Gallicano Dermatological Institute | IRCCS Regina Elena National Cancer Institute | Francesca De Nicola, Giulia Orlandi, Alice Massacci, Alessia Lauretti, Fabrizio Ensoli, Fulvia Pimpinelli, Maurizio Fanciulli, Giovanni Blandino, Aldo Morrone, Gennaro Ciliberto |
| EPI_ISL_3068370 | IRCCS San Gallicano Dermatological Institute | IRCCS Regina Elena National Cancer Institute | Ludovica Ciuffreda, Sara Donzelli, Matteo Pallocca, Francesca Maione, Fabrizio Ensoli, Fulvia Pimpinelli, Maurizio Fanciulli, Giovanni Blandino, Aldo Morrone, Gennaro Ciliberto |
| EPI_ISL_3068372 | IRCCS San Gallicano Dermatological Institute | IRCCS Regina Elena National Cancer Institute | Sabrina Strano, Sara Donzelli, Alice Massacci, Silvia Paluzzi, Fabrizio Ensoli, Fulvia Pimpinelli, Maurizio Fanciulli, Giovanni Blandino, Aldo Morrone, Gennaro Ciliberto |
| EPI_ISL_3068373 | IRCCS San Gallicano Dermatological Institute | IRCCS Regina Elena National Cancer Institute | Francesca De Nicola, Giulia Orlandi, Matteo Pallocca, Sara Petrolo, Fabrizio Ensoli, Fulvia Pimpinelli, Maurizio Fanciulli, Giovanni Blandino, Aldo Morrone, Gennaro Ciliberto |
| EPI_ISL_3068374 | IRCCS San Gallicano Dermatological Institute | IRCCS Regina Elena National Cancer Institute | Ludovica Ciuffreda, Sara Donzelli, Eleonora Sperandio, Valentina Ricca, Fabrizio Ensoli, Fulvia Pimpinelli, Maurizio Fanciulli, Giovanni Blandino, Aldo Morrone, Gennaro Ciliberto |
| EPI_ISL_3068375 | IRCCS San Gallicano Dermatological Institute | IRCCS Regina Elena National Cancer Institute | Frauke Goeman, Giulia Orlandi, Alice Massacci, Serena Salvo, Fabrizio Ensoli, Fulvia Pimpinelli, Maurizio Fanciulli, Giovanni Blandino, Aldo Morrone, Gennaro Ciliberto |
| EPI_ISL_3068376 | IRCCS San Gallicano Dermatological Institute | IRCCS Regina Elena National Cancer Institute | Sabrina Strano, Sara Donzelli, Matteo Pallocca, Grazia Prignano, Fabrizio Ensoli, Fulvia Pimpinelli, Maurizio Fanciulli, Giovanni Blandino, Aldo Morrone, Gennaro Ciliberto |
| EPI_ISL_3068377 | IRCCS San Gallicano Dermatological Institute | IRCCS Regina Elena National Cancer Institute | Francesca De Nicola, Giulia Orlandi, Eleonora Sperandio, Elisabetta Trento, Fabrizio Ensoli, Fulvia Pimpinelli, Maurizio Fanciulli, Giovanni Blandino, Aldo Morrone, Gennaro Ciliberto |
| EPI_ISL_3068378 | IRCCS San Gallicano Dermatological Institute | IRCCS Regina Elena National Cancer Institute | Ludovica Ciuffreda, Sara Donzelli, Alice Massacci, Arianna Mastrofrancesco, Fabrizio Ensoli, Fulvia Pimpinelli, Maurizio Fanciulli, Giovanni Blandino, Aldo Morrone, Gennaro Ciliberto |
| EPI_ISL_3068379 | IRCCS San Gallicano Dermatological Institute | IRCCS Regina Elena National Cancer Institute | Frauke Goeman, Giulia Orlandi, Matteo Pallocca, Giovanna D'agosto, Fabrizio Ensoli, Fulvia Pimpinelli, Maurizio Fanciulli, Giovanni Blandino, Aldo Morrone, Gennaro Ciliberto |
| EPI_ISL_3068380 | IRCCS San Gallicano Dermatological Institute | IRCCS Regina Elena National Cancer Institute | Sabrina Strano, Sara Donzelli, Eleonora Sperandio, Carla Mottini, Fabrizio Ensoli, Fulvia Pimpinelli, Maurizio Fanciulli, Giovanni Blandino, Aldo Morrone, Gennaro Ciliberto |
| EPI_ISL_3068381 | IRCCS San Gallicano Dermatological Institute | IRCCS Regina Elena National Cancer Institute | Francesca De Nicola, Giulia Orlandi, Alice Massacci, Martina Pontone, Fabrizio Ensoli, Fulvia Pimpinelli, Maurizio Fanciulli, Giovanni Blandino, Aldo Morrone, Gennaro Ciliberto |
| EPI_ISL_3068382 | IRCCS San Gallicano Dermatological Institute | IRCCS Regina Elena National Cancer Institute | Ludovica Ciuffreda, Sara Donzelli, Matteo Pallocca, di Domenico Enea Gino, Fabrizio Ensoli, Fulvia Pimpinelli, Maurizio Fanciulli, Giovanni Blandino, Aldo Morrone, Gennaro Ciliberto |
| EPI_ISL_3068383 | IRCCS San Gallicano Dermatological Institute | IRCCS Regina Elena National Cancer Institute | Frauke Goeman, Giulia Orlandi, Eleonora Sperandio, Ilaria Cavallo, Fabrizio Ensoli, Fulvia Pimpinelli, Maurizio Fanciulli, Giovanni Blandino, Aldo Morrone, Gennaro Ciliberto |
| EPI_ISL_3068384 | IRCCS San Gallicano Dermatological Institute | IRCCS Regina Elena National Cancer Institute | Sabrina Strano, Sara Donzelli, Alice Massacci, Francesca Sivori, Fabrizio Ensoli, Fulvia Pimpinelli, Maurizio Fanciulli, Giovanni Blandino, Aldo Morrone, Gennaro Ciliberto |
| EPI_ISL_3068385 | IRCCS San Gallicano Dermatological Institute | IRCCS Regina Elena National Cancer Institute | Francesca De Nicola, Giulia Orlandi, Matteo Pallocca, Andrea Cazzani, Fabrizio Ensoli, Fulvia Pimpinelli, Maurizio Fanciulli, Giovanni Blandino, Aldo Morrone, Gennaro Ciliberto |
| EPI_ISL_3068386 | IRCCS San Gallicano Dermatological Institute | IRCCS Regina Elena National Cancer Institute | Ludovica Ciuffreda, Sara Donzelli, Eleonora Sperandio, Ilaria Celesti, Fabrizio Ensoli, Fulvia Pimpinelli, Maurizio Fanciulli, Giovanni Blandino, Aldo Morrone, Gennaro Ciliberto |
| EPI_ISL_3068387 | IRCCS San Gallicano Dermatological Institute | IRCCS Regina Elena National Cancer Institute | Frauke Goeman, Giulia Orlandi, Alice Massacci, Martina Diano, Fabrizio Ensoli, Fulvia Pimpinelli, Maurizio Fanciulli, Giovanni Blandino, Aldo Morrone, Gennaro Ciliberto |
| EPI_ISL_3068388 | IRCCS San Gallicano Dermatological Institute | IRCCS Regina Elena National Cancer Institute | Sabrina Strano, Sara Donzelli, Matteo Pallocca, Antonio Federico, Fabrizio Ensoli, Fulvia Pimpinelli, Maurizio Fanciulli, Giovanni Blandino, Aldo Morrone, Gennaro Ciliberto |
| EPI_ISL_3068389 | IRCCS San Gallicano Dermatological Institute | IRCCS Regina Elena National Cancer Institute | Francesca De Nicola, Giulia Orlandi, Eleonora Sperandio, Fulvia Fraticelli, Fabrizio Ensoli, Fulvia Pimpinelli, Maurizio Fanciulli, Giovanni Blandino, Aldo Morrone, Gennaro Ciliberto |
| EPI_ISL_3068390 | IRCCS San Gallicano Dermatological Institute | IRCCS Regina Elena National Cancer Institute | Ludovica Ciuffreda, Sara Donzelli, Alice Massacci, Lorenzo Furzi, Fabrizio Ensoli, Fulvia Pimpinelli, Maurizio Fanciulli, Giovanni Blandino, Aldo Morrone, Gennaro Ciliberto |
| EPI_ISL_3068391 | IRCCS San Gallicano Dermatological Institute | IRCCS Regina Elena National Cancer Institute | Frauke Goeman, Giulia Orlandi, Matteo Pallocca, Alessia Lauretti, Fabrizio Ensoli, Fulvia Pimpinelli, Maurizio Fanciulli, Giovanni Blandino, Aldo Morrone, Gennaro Ciliberto |
| EPI_ISL_3068392 | IRCCS San Gallicano Dermatological Institute | IRCCS Regina Elena National Cancer Institute | Sabrina Strano, Sara Donzelli, Eleonora Sperandio, Francesca Maione, Fabrizio Ensoli, Fulvia Pimpinelli, Maurizio Fanciulli, Giovanni Blandino, Aldo Morrone, Gennaro Ciliberto |
| EPI_ISL_3068393 | IRCCS San Gallicano Dermatological Institute | IRCCS Regina Elena National Cancer Institute | Francesca De Nicola, Giulia Orlandi, Alice Massacci, Francisco Obregon, Fabrizio Ensoli, Fulvia Pimpinelli, Maurizio Fanciulli, Giovanni Blandino, Aldo Morrone, Gennaro Ciliberto |
| EPI_ISL_3068394 | IRCCS San Gallicano Dermatological Institute | IRCCS Regina Elena National Cancer Institute | Ludovica Ciuffreda, Sara Donzelli, Matteo Pallocca, Silvia Paluzzi, Fabrizio Ensoli, Fulvia Pimpinelli, Maurizio Fanciulli, Giovanni Blandino, Aldo Morrone, Gennaro Ciliberto |
| EPI_ISL_3068395 | IRCCS San Gallicano Dermatological Institute | IRCCS Regina Elena National Cancer Institute | Frauke Goeman, Giulia Orlandi, Eleonora Sperandio, Sara Petrolo, Fabrizio Ensoli, Fulvia Pimpinelli, Maurizio Fanciulli, Giovanni Blandino, Aldo Morrone, Gennaro Ciliberto |
| EPI_ISL_3068396 | IRCCS San Gallicano Dermatological Institute | IRCCS Regina Elena National Cancer Institute | Sabrina Strano, Sara Donzelli, Alice Massacci, Valentina Ricca, Fabrizio Ensoli, Fulvia Pimpinelli, Maurizio Fanciulli, Giovanni Blandino, Aldo Morrone, Gennaro Ciliberto |
| EPI_ISL_3068397 | IRCCS San Gallicano Dermatological Institute | IRCCS Regina Elena National Cancer Institute | Francesca De Nicola, Giulia Orlandi, Matteo Pallocca, Serena Salvo, Fabrizio Ensoli, Fulvia Pimpinelli, Maurizio Fanciulli, Giovanni Blandino, Aldo Morrone, Gennaro Ciliberto |
| EPI_ISL_3068398 | IRCCS San Gallicano Dermatological Institute | IRCCS Regina Elena National Cancer Institute | Ludovica Ciuffreda, Sara Donzelli, Eleonora Sperandio, Grazia Prignano, Fabrizio Ensoli, Fulvia Pimpinelli, Maurizio Fanciulli, Giovanni Blandino, Aldo Morrone, Gennaro Ciliberto |
| EPI_ISL_3068399 | IRCCS San Gallicano Dermatological Institute | IRCCS Regina Elena National Cancer Institute | Frauke Goeman, Giulia Orlandi, Alice Massacci, Elisabetta Trento, Fabrizio Ensoli, Fulvia Pimpinelli, Maurizio Fanciulli, Giovanni Blandino, Aldo Morrone, Gennaro Ciliberto |
| EPI_ISL_3068400 | IRCCS San Gallicano Dermatological Institute | IRCCS Regina Elena National Cancer Institute | Francesca De Nicola, Giulia Orlandi, Eleonora Sperandio, Giovanna D'agosto, Fabrizio Ensoli, Fulvia Pimpinelli, Maurizio Fanciulli, Giovanni Blandino, Aldo Morrone, Gennaro Ciliberto |
| EPI_ISL_3068401 | IRCCS San Gallicano Dermatological Institute | IRCCS Regina Elena National Cancer Institute | Ludovica Ciuffreda, Sara Donzelli, Alice Massacci, Carla Mottini, Fabrizio Ensoli, Fulvia Pimpinelli, Maurizio Fanciulli, Giovanni Blandino, Aldo Morrone, Gennaro Ciliberto |
| EPI_ISL_3068402 | IRCCS San Gallicano Dermatological Institute | IRCCS Regina Elena National Cancer Institute | Frauke Goeman, Giulia Orlandi, Matteo Pallocca, Martina Pontone, Fabrizio Ensoli, Fulvia Pimpinelli, Maurizio Fanciulli, Giovanni Blandino, Aldo Morrone, Gennaro Ciliberto |
| EPI_ISL_3068403 | IRCCS San Gallicano Dermatological Institute | IRCCS Regina Elena National Cancer Institute | Sabrina Strano, Sara Donzelli, Eleonora Sperandio, di Domenico Enea Gino, Fabrizio Ensoli, Fulvia Pimpinelli, Maurizio Fanciulli, Giovanni Blandino, Aldo Morrone, Gennaro Ciliberto |
| EPI_ISL_3068404 | IRCCS San Gallicano Dermatological Institute | IRCCS Regina Elena National Cancer Institute | Francesca De Nicola, Giulia Orlandi, Alice Massacci, Ilaria Cavallo, Fabrizio Ensoli, Fulvia Pimpinelli, Maurizio Fanciulli, Giovanni Blandino, Aldo Morrone, Gennaro Ciliberto |
| EPI_ISL_3068405 | IRCCS San Gallicano Dermatological Institute | IRCCS Regina Elena National Cancer Institute | Ludovica Ciuffreda, Sara Donzelli, Matteo Pallocca, Francesca Sivori, Fabrizio Ensoli, Fulvia Pimpinelli, Maurizio Fanciulli, Giovanni Blandino, Aldo Morrone, Gennaro Ciliberto |
| EPI_ISL_3068406 | IRCCS San Gallicano Dermatological Institute | IRCCS Regina Elena National Cancer Institute | Frauke Goeman, Giulia Orlandi, Eleonora Sperandio, Andrea Cazzani, Fabrizio Ensoli, Fulvia Pimpinelli, Maurizio Fanciulli, Giovanni Blandino, Aldo Morrone, Gennaro Ciliberto |
| EPI_ISL_3068407 | IRCCS San Gallicano Dermatological Institute | IRCCS Regina Elena National Cancer Institute | Sabrina Strano, Sara Donzelli, Alice Massacci, Ilaria Celesti, Fabrizio Ensoli, Fulvia Pimpinelli, Maurizio Fanciulli, Giovanni Blandino, Aldo Morrone, Gennaro Ciliberto |
| EPI_ISL_3068408 | IRCCS San Gallicano Dermatological Institute | IRCCS Regina Elena National Cancer Institute | Francesca De Nicola, Giulia Orlandi, Matteo Pallocca, Martina Diano, Fabrizio Ensoli, Fulvia Pimpinelli, Maurizio Fanciulli, Giovanni Blandino, Aldo Morrone, Gennaro Ciliberto |
| EPI_ISL_3068409 | IRCCS San Gallicano Dermatological Institute | IRCCS Regina Elena National Cancer Institute | Ludovica Ciuffreda, Sara Donzelli, Eleonora Sperandio, Antonio Federico, Fabrizio Ensoli, Fulvia Pimpinelli, Maurizio Fanciulli, Giovanni Blandino, Aldo Morrone, Gennaro Ciliberto |
| EPI_ISL_3068410 | IRCCS San Gallicano Dermatological Institute | IRCCS Regina Elena National Cancer Institute | Frauke Goeman, Giulia Orlandi, Alice Massacci, Fulvia Fraticelli, Fabrizio Ensoli, Fulvia Pimpinelli, Maurizio Fanciulli, Giovanni Blandino, Aldo Morrone, Gennaro Ciliberto |
| EPI_ISL_3068411 | IRCCS San Gallicano Dermatological Institute | IRCCS Regina Elena National Cancer Institute | Sabrina Strano, Sara Donzelli, Matteo Pallocca, Lorenzo Furzi, Fabrizio Ensoli, Fulvia Pimpinelli, Maurizio Fanciulli, Giovanni Blandino, Aldo Morrone, Gennaro Ciliberto |
| EPI_ISL_3068413 | IRCCS San Gallicano Dermatological Institute | IRCCS Regina Elena National Cancer Institute | Ludovica Ciuffreda, Sara Donzelli, Alice Massacci, Francesca Maione, Fabrizio Ensoli, Fulvia Pimpinelli, Maurizio Fanciulli, Giovanni Blandino, Aldo Morrone, Gennaro Ciliberto |
| EPI_ISL_3068414 | IRCCS San Gallicano Dermatological Institute | IRCCS Regina Elena National Cancer Institute | Frauke Goeman, Giulia Orlandi, Matteo Pallocca, Francisco Obregon, Fabrizio Ensoli, Fulvia Pimpinelli, Maurizio Fanciulli, Giovanni Blandino, Aldo Morrone, Gennaro Ciliberto |
| EPI_ISL_3068415 | IRCCS San Gallicano Dermatological Institute | IRCCS Regina Elena National Cancer Institute | Sabrina Strano, Sara Donzelli, Eleonora Sperandio, Silvia Paluzzi, Fabrizio Ensoli, Fulvia Pimpinelli, Maurizio Fanciulli, Giovanni Blandino, Aldo Morrone, Gennaro Ciliberto |
| EPI_ISL_3068416 | IRCCS San Gallicano Dermatological Institute | IRCCS Regina Elena National Cancer Institute | Francesca De Nicola, Giulia Orlandi, Alice Massacci, Sara Petrolo, Fabrizio Ensoli, Fulvia Pimpinelli, Maurizio Fanciulli, Giovanni Blandino, Aldo Morrone, Gennaro Ciliberto |
| EPI_ISL_3068417 | IRCCS San Gallicano Dermatological Institute | IRCCS Regina Elena National Cancer Institute | Ludovica Ciuffreda, Sara Donzelli, Matteo Pallocca, Valentina Ricca, Fabrizio Ensoli, Fulvia Pimpinelli, Maurizio Fanciulli, Giovanni Blandino, Aldo Morrone, Gennaro Ciliberto |
| EPI_ISL_3068418 | IRCCS San Gallicano Dermatological Institute | IRCCS Regina Elena National Cancer Institute | Frauke Goeman, Giulia Orlandi, Eleonora Sperandio, Serena Salvo, Fabrizio Ensoli, Fulvia Pimpinelli, Maurizio Fanciulli, Giovanni Blandino, Aldo Morrone, Gennaro Ciliberto |
| EPI_ISL_3068419 | IRCCS San Gallicano Dermatological Institute | IRCCS Regina Elena National Cancer Institute | Sabrina Strano, Sara Donzelli, Alice Massacci, Grazia Prignano, Fabrizio Ensoli, Fulvia Pimpinelli, Maurizio Fanciulli, Giovanni Blandino, Aldo Morrone, Gennaro Ciliberto |
| EPI_ISL_3068420 | IRCCS San Gallicano Dermatological Institute | IRCCS Regina Elena National Cancer Institute | Francesca De Nicola, Giulia Orlandi, Matteo Pallocca, Elisabetta Trento, Fabrizio Ensoli, Fulvia Pimpinelli, Maurizio Fanciulli, Giovanni Blandino, Aldo Morrone, Gennaro Ciliberto |
| EPI_ISL_3068421 | IRCCS San Gallicano Dermatological Institute | IRCCS Regina Elena National Cancer Institute | Ludovica Ciuffreda, Sara Donzelli, Eleonora Sperandio, Arianna Mastrofrancesco, Fabrizio Ensoli, Fulvia Pimpinelli, Maurizio Fanciulli, Giovanni Blandino, Aldo Morrone, Gennaro Ciliberto |
| EPI_ISL_3068422 | IRCCS San Gallicano Dermatological Institute | IRCCS Regina Elena National Cancer Institute | Frauke Goeman, Giulia Orlandi, Alice Massacci, Giovanna D'agosto, Fabrizio Ensoli, Fulvia Pimpinelli, Maurizio Fanciulli, Giovanni Blandino, Aldo Morrone, Gennaro Ciliberto |
| EPI_ISL_3068423 | IRCCS San Gallicano Dermatological Institute | IRCCS Regina Elena National Cancer Institute | Sabrina Strano, Sara Donzelli, Matteo Pallocca, Carla Mottini, Fabrizio Ensoli, Fulvia Pimpinelli, Maurizio Fanciulli, Giovanni Blandino, Aldo Morrone, Gennaro Ciliberto |
| EPI_ISL_3068424 | IRCCS San Gallicano Dermatological Institute | IRCCS Regina Elena National Cancer Institute | Francesca De Nicola, Giulia Orlandi, Eleonora Sperandio, Martina Pontone, Fabrizio Ensoli, Fulvia Pimpinelli, Maurizio Fanciulli, Giovanni Blandino, Aldo Morrone, Gennaro Ciliberto |
| EPI_ISL_3068425 | IRCCS San Gallicano Dermatological Institute | IRCCS Regina Elena National Cancer Institute | Ludovica Ciuffreda, Sara Donzelli, Alice Massacci, di Domenico Enea Gino, Fabrizio Ensoli, Fulvia Pimpinelli, Maurizio Fanciulli, Giovanni Blandino, Aldo Morrone, Gennaro Ciliberto |
| EPI_ISL_3068426 | IRCCS San Gallicano Dermatological Institute | IRCCS Regina Elena National Cancer Institute | Frauke Goeman, Giulia Orlandi, Matteo Pallocca, Ilaria Cavallo, Fabrizio Ensoli, Fulvia Pimpinelli, Maurizio Fanciulli, Giovanni Blandino, Aldo Morrone, Gennaro Ciliberto |
| EPI_ISL_3068427 | IRCCS San Gallicano Dermatological Institute | IRCCS Regina Elena National Cancer Institute | Sabrina Strano, Sara Donzelli, Eleonora Sperandio, Francesca Sivori, Fabrizio Ensoli, Fulvia Pimpinelli, Maurizio Fanciulli, Giovanni Blandino, Aldo Morrone, Gennaro Ciliberto |
| EPI_ISL_3068428 | IRCCS San Gallicano Dermatological Institute | IRCCS Regina Elena National Cancer Institute | Francesca De Nicola, Giulia Orlandi, Alice Massacci, Andrea Cazzani, Fabrizio Ensoli, Fulvia Pimpinelli, Maurizio Fanciulli, Giovanni Blandino, Aldo Morrone, Gennaro Ciliberto |
| EPI_ISL_3068429 | IRCCS San Gallicano Dermatological Institute | IRCCS Regina Elena National Cancer Institute | Ludovica Ciuffreda, Sara Donzelli, Matteo Pallocca, Ilaria Celesti, Fabrizio Ensoli, Fulvia Pimpinelli, Maurizio Fanciulli, Giovanni Blandino, Aldo Morrone, Gennaro Ciliberto |
| EPI_ISL_3068430 | IRCCS San Gallicano Dermatological Institute | IRCCS Regina Elena National Cancer Institute | Frauke Goeman, Giulia Orlandi, Eleonora Sperandio, Martina Diano, Fabrizio Ensoli, Fulvia Pimpinelli, Maurizio Fanciulli, Giovanni Blandino, Aldo Morrone, Gennaro Ciliberto |
| EPI_ISL_3068431 | IRCCS San Gallicano Dermatological Institute | IRCCS Regina Elena National Cancer Institute | Sabrina Strano, Sara Donzelli, Alice Massacci, Antonio Federico, Fabrizio Ensoli, Fulvia Pimpinelli, Maurizio Fanciulli, Giovanni Blandino, Aldo Morrone, Gennaro Ciliberto |
| EPI_ISL_3068432 | IRCCS San Gallicano Dermatological Institute | IRCCS Regina Elena National Cancer Institute | Francesca De Nicola, Giulia Orlandi, Matteo Pallocca, Fulvia Fraticelli, Fabrizio Ensoli, Fulvia Pimpinelli, Maurizio Fanciulli, Giovanni Blandino, Aldo Morrone, Gennaro Ciliberto |
| EPI_ISL_3068433 | IRCCS San Gallicano Dermatological Institute | IRCCS Regina Elena National Cancer Institute | Ludovica Ciuffreda, Sara Donzelli, Eleonora Sperandio, Lorenzo Furzi, Fabrizio Ensoli, Fulvia Pimpinelli, Maurizio Fanciulli, Giovanni Blandino, Aldo Morrone, Gennaro Ciliberto |
| EPI_ISL_3068434 | IRCCS San Gallicano Dermatological Institute | IRCCS Regina Elena National Cancer Institute | Frauke Goeman, Giulia Orlandi, Alice Massacci, Alessia Lauretti, Fabrizio Ensoli, Fulvia Pimpinelli, Maurizio Fanciulli, Giovanni Blandino, Aldo Morrone, Gennaro Ciliberto |
| EPI_ISL_3068435 | IRCCS San Gallicano Dermatological Institute | IRCCS Regina Elena National Cancer Institute | Sabrina Strano, Sara Donzelli, Matteo Pallocca, Francesca Maione, Fabrizio Ensoli, Fulvia Pimpinelli, Maurizio Fanciulli, Giovanni Blandino, Aldo Morrone, Gennaro Ciliberto |
| EPI_ISL_3068436 | IRCCS San Gallicano Dermatological Institute | IRCCS Regina Elena National Cancer Institute | Francesca De Nicola, Giulia Orlandi, Eleonora Sperandio, Francisco Obregon, Fabrizio Ensoli, Fulvia Pimpinelli, Maurizio Fanciulli, Giovanni Blandino, Aldo Morrone, Gennaro Ciliberto |
| EPI_ISL_3068437 | IRCCS San Gallicano Dermatological Institute | IRCCS Regina Elena National Cancer Institute | Ludovica Ciuffreda, Sara Donzelli, Alice Massacci, Silvia Paluzzi, Fabrizio Ensoli, Fulvia Pimpinelli, Maurizio Fanciulli, Giovanni Blandino, Aldo Morrone, Gennaro Ciliberto |
| EPI_ISL_3068438 | IRCCS San Gallicano Dermatological Institute | IRCCS Regina Elena National Cancer Institute | Frauke Goeman, Giulia Orlandi, Matteo Pallocca, Sara Petrolo, Fabrizio Ensoli, Fulvia Pimpinelli, Maurizio Fanciulli, Giovanni Blandino, Aldo Morrone, Gennaro Ciliberto |
| EPI_ISL_3068439 | IRCCS San Gallicano Dermatological Institute | IRCCS Regina Elena National Cancer Institute | Sabrina Strano, Sara Donzelli, Eleonora Sperandio, Valentina Ricca, Fabrizio Ensoli, Fulvia Pimpinelli, Maurizio Fanciulli, Giovanni Blandino, Aldo Morrone, Gennaro Ciliberto |

| | | | |
|---|---|---|---|
| EPI_ISL_3068440 | IRCCS San Gallicano Dermatological Institute | IRCCS Regina Elena National Cancer Institute | Francesca De Nicola, Giulia Orlandi, Alice Massacci, Serena Salvo, Fabrizio Ensoli, Fulvia Pimpinelli, Maurizio Fanciulli, Giovanni Blandino, Aldo Morrone, Gennaro Ciliberto |
| EPI_ISL_3068442 | IRCCS San Gallicano Dermatological Institute | IRCCS Regina Elena National Cancer Institute | Frauke Goeman, Giulia Orlandi, Eleonora Sperandio, Elisabetta Trento, Fabrizio Ensoli, Fulvia Pimpinelli, Maurizio Fanciulli, Giovanni Blandino, Aldo Morrone, Gennaro Ciliberto |
| EPI_ISL_3068444 | IRCCS San Gallicano Dermatological Institute | IRCCS Regina Elena National Cancer Institute | Francesca De Nicola, Giulia Orlandi, Matteo Pallocca, Giovanna D'agosto, Fabrizio Ensoli, Fulvia Pimpinelli, Maurizio Fanciulli, Giovanni Blandino, Aldo Morrone, Gennaro Ciliberto |
| EPI_ISL_3068445 | IRCCS San Gallicano Dermatological Institute | IRCCS Regina Elena National Cancer Institute | Ludovica Ciuffreda, Sara Donzelli, Eleonora Sperandio, Carla Mottini, Fabrizio Ensoli, Fulvia Pimpinelli, Maurizio Fanciulli, Giovanni Blandino, Aldo Morrone, Gennaro Ciliberto |
| EPI_ISL_3068446 | IRCCS San Gallicano Dermatological Institute | IRCCS Regina Elena National Cancer Institute | Frauke Goeman, Giulia Orlandi, Alice Massacci, Martina Pontone, Fabrizio Ensoli, Fulvia Pimpinelli, Maurizio Fanciulli, Giovanni Blandino, Aldo Morrone, Gennaro Ciliberto |
| EPI_ISL_3068447 | IRCCS San Gallicano Dermatological Institute | IRCCS Regina Elena National Cancer Institute | Sabrina Strano, Sara Donzelli, Matteo Pallocca, di Domenico Enea Gino, Fabrizio Ensoli, Fulvia Pimpinelli, Maurizio Fanciulli, Giovanni Blandino, Aldo Morrone, Gennaro Ciliberto |
| EPI_ISL_3068448 | IRCCS San Gallicano Dermatological Institute | IRCCS Regina Elena National Cancer Institute | Francesca De Nicola, Giulia Orlandi, Eleonora Sperandio, Ilaria Cavallo, Fabrizio Ensoli, Fulvia Pimpinelli, Maurizio Fanciulli, Giovanni Blandino, Aldo Morrone, Gennaro Ciliberto |
| EPI_ISL_3068449 | IRCCS San Gallicano Dermatological Institute | IRCCS Regina Elena National Cancer Institute | Ludovica Ciuffreda, Sara Donzelli, Alice Massacci, Francesca Sivori, Fabrizio Ensoli, Fulvia Pimpinelli, Maurizio Fanciulli, Giovanni Blandino, Aldo Morrone, Gennaro Ciliberto |
| EPI_ISL_3068450 | IRCCS San Gallicano Dermatological Institute | IRCCS Regina Elena National Cancer Institute | Frauke Goeman, Giulia Orlandi, Matteo Pallocca, Andrea Cazzani, Fabrizio Ensoli, Fulvia Pimpinelli, Maurizio Fanciulli, Giovanni Blandino, Aldo Morrone, Gennaro Ciliberto |
| EPI_ISL_3068451 | IRCCS San Gallicano Dermatological Institute | IRCCS Regina Elena National Cancer Institute | Sabrina Strano, Sara Donzelli, Eleonora Sperandio, Ilaria Celesti, Fabrizio Ensoli, Fulvia Pimpinelli, Maurizio Fanciulli, Giovanni Blandino, Aldo Morrone, Gennaro Ciliberto |
| EPI_ISL_3068452 | IRCCS San Gallicano Dermatological Institute | IRCCS Regina Elena National Cancer Institute | Francesca De Nicola, Giulia Orlandi, Alice Massacci, Martina Diano, Fabrizio Ensoli, Fulvia Pimpinelli, Maurizio Fanciulli, Giovanni Blandino, Aldo Morrone, Gennaro Ciliberto |
| EPI_ISL_3068453 | IRCCS San Gallicano Dermatological Institute | IRCCS Regina Elena National Cancer Institute | Ludovica Ciuffreda, Sara Donzelli, Matteo Pallocca, Antonio Federico, Fabrizio Ensoli, Fulvia Pimpinelli, Maurizio Fanciulli, Giovanni Blandino, Aldo Morrone, Gennaro Ciliberto |
| EPI_ISL_3068454 | IRCCS San Gallicano Dermatological Institute | IRCCS Regina Elena National Cancer Institute | Frauke Goeman, Giulia Orlandi, Eleonora Sperandio, Fulvia Fraticelli, Fabrizio Ensoli, Fulvia Pimpinelli, Maurizio Fanciulli, Giovanni Blandino, Aldo Morrone, Gennaro Ciliberto |
| EPI_ISL_3068455 | IRCCS San Gallicano Dermatological Institute | IRCCS Regina Elena National Cancer Institute | Sabrina Strano, Sara Donzelli, Alice Massacci, Lorenzo Furzi, Fabrizio Ensoli, Fulvia Pimpinelli, Maurizio Fanciulli, Giovanni Blandino, Aldo Morrone, Gennaro Ciliberto |
| EPI_ISL_3068456 | IRCCS San Gallicano Dermatological Institute | IRCCS Regina Elena National Cancer Institute | Francesca De Nicola, Giulia Orlandi, Matteo Pallocca, Alessia Lauretti, Fabrizio Ensoli, Fulvia Pimpinelli, Maurizio Fanciulli, Giovanni Blandino, Aldo Morrone, Gennaro Ciliberto |
| EPI_ISL_3068457 | IRCCS San Gallicano Dermatological Institute | IRCCS Regina Elena National Cancer Institute | Ludovica Ciuffreda, Sara Donzelli, Eleonora Sperandio, Francesca Maione, Fabrizio Ensoli, Fulvia Pimpinelli, Maurizio Fanciulli, Giovanni Blandino, Aldo Morrone, Gennaro Ciliberto |
| EPI_ISL_3068458 | IRCCS San Gallicano Dermatological Institute | IRCCS Regina Elena National Cancer Institute | Frauke Goeman, Giulia Orlandi, Alice Massacci, Francisco Obregon, Fabrizio Ensoli, Fulvia Pimpinelli, Maurizio Fanciulli, Giovanni Blandino, Aldo Morrone, Gennaro Ciliberto |
| EPI_ISL_3068459 | IRCCS San Gallicano Dermatological Institute | IRCCS Regina Elena National Cancer Institute | Sabrina Strano, Sara Donzelli, Matteo Pallocca, Silvia Paluzzi, Fabrizio Ensoli, Fulvia Pimpinelli, Maurizio Fanciulli, Giovanni Blandino, Aldo Morrone, Gennaro Ciliberto |
| EPI_ISL_3068460 | IRCCS San Gallicano Dermatological Institute | IRCCS Regina Elena National Cancer Institute | Francesca De Nicola, Giulia Orlandi, Eleonora Sperandio, Sara Petrolo, Fabrizio Ensoli, Fulvia Pimpinelli, Maurizio Fanciulli, Giovanni Blandino, Aldo Morrone, Gennaro Ciliberto |
| EPI_ISL_3068461 | IRCCS San Gallicano Dermatological Institute | IRCCS Regina Elena National Cancer Institute | Ludovica Ciuffreda, Sara Donzelli, Alice Massacci, Valentina Ricca, Fabrizio Ensoli, Fulvia Pimpinelli, Maurizio Fanciulli, Giovanni Blandino, Aldo Morrone, Gennaro Ciliberto |
| EPI_ISL_3068462 | IRCCS San Gallicano Dermatological Institute | IRCCS Regina Elena National Cancer Institute | Frauke Goeman, Giulia Orlandi, Matteo Pallocca, Serena Salvo, Fabrizio Ensoli, Fulvia Pimpinelli, Maurizio Fanciulli, Giovanni Blandino, Aldo Morrone, Gennaro Ciliberto |
| EPI_ISL_3068463 | IRCCS San Gallicano Dermatological Institute | IRCCS Regina Elena National Cancer Institute | Sabrina Strano, Sara Donzelli, Eleonora Sperandio, Grazia Prignano, Fabrizio Ensoli, Fulvia Pimpinelli, Maurizio Fanciulli, Giovanni Blandino, Aldo Morrone, Gennaro Ciliberto |
| EPI_ISL_3068464 | IRCCS San Gallicano Dermatological Institute | IRCCS Regina Elena National Cancer Institute | Francesca De Nicola, Giulia Orlandi, Alice Massacci, Elisabetta Trento, Fabrizio Ensoli, Fulvia Pimpinelli, Maurizio Fanciulli, Giovanni Blandino, Aldo Morrone, Gennaro Ciliberto |
| EPI_ISL_3068465 | IRCCS San Gallicano Dermatological Institute | IRCCS Regina Elena National Cancer Institute | Ludovica Ciuffreda, Sara Donzelli, Matteo Pallocca, Arianna Mastrofrancesco, Fabrizio Ensoli, Fulvia Pimpinelli, Maurizio Fanciulli, Giovanni Blandino, Aldo Morrone, Gennaro Ciliberto |
| EPI_ISL_3068466 | IRCCS San Gallicano Dermatological Institute | IRCCS Regina Elena National Cancer Institute | Frauke Goeman, Giulia Orlandi, Eleonora Sperandio, Giovanna D'agosto, Fabrizio Ensoli, Fulvia Pimpinelli, Maurizio Fanciulli, Giovanni Blandino, Aldo Morrone, Gennaro Ciliberto |
| EPI_ISL_3068467 | IRCCS San Gallicano Dermatological Institute | IRCCS Regina Elena National Cancer Institute | Sabrina Strano, Sara Donzelli, Alice Massacci, Carla Mottini, Fabrizio Ensoli, Fulvia Pimpinelli, Maurizio Fanciulli, Giovanni Blandino, Aldo Morrone, Gennaro Ciliberto |
| EPI_ISL_3068468 | IRCCS San Gallicano Dermatological Institute | IRCCS Regina Elena National Cancer Institute | Francesca De Nicola, Giulia Orlandi, Matteo Pallocca, Martina Pontone, Fabrizio Ensoli, Fulvia Pimpinelli, Maurizio Fanciulli, Giovanni Blandino, Aldo Morrone, Gennaro Ciliberto |
| EPI_ISL_3068469 | IRCCS San Gallicano Dermatological Institute | IRCCS Regina Elena National Cancer Institute | Ludovica Ciuffreda, Sara Donzelli, Eleonora Sperandio, di Domenico Enea Gino, Fabrizio Ensoli, Fulvia Pimpinelli, Maurizio Fanciulli, Giovanni Blandino, Aldo Morrone, Gennaro Ciliberto |
| EPI_ISL_3068470 | IRCCS San Gallicano Dermatological Institute | IRCCS Regina Elena National Cancer Institute | Frauke Goeman, Giulia Orlandi, Alice Massacci, Ilaria Cavallo, Fabrizio Ensoli, Fulvia Pimpinelli, Maurizio Fanciulli, Giovanni Blandino, Aldo Morrone, Gennaro Ciliberto |
| EPI_ISL_3068471 | IRCCS San Gallicano Dermatological Institute | IRCCS Regina Elena National Cancer Institute | Sabrina Strano, Sara Donzelli, Matteo Pallocca, Francesca Sivori, Fabrizio Ensoli, Fulvia Pimpinelli, Maurizio Fanciulli, Giovanni Blandino, Aldo Morrone, Gennaro Ciliberto |
| EPI_ISL_3068472 | IRCCS San Gallicano Dermatological Institute | IRCCS Regina Elena National Cancer Institute | Francesca De Nicola, Giulia Orlandi, Eleonora Sperandio, Andrea Cazzani, Fabrizio Ensoli, Fulvia Pimpinelli, Maurizio Fanciulli, Giovanni Blandino, Aldo Morrone, Gennaro Ciliberto |
| EPI_ISL_3068473 | IRCCS San Gallicano Dermatological Institute | IRCCS Regina Elena National Cancer Institute | Ludovica Ciuffreda, Sara Donzelli, Alice Massacci, Ilaria Celesti, Fabrizio Ensoli, Fulvia Pimpinelli, Maurizio Fanciulli, Giovanni Blandino, Aldo Morrone, Gennaro Ciliberto |
| EPI_ISL_3068474 | IRCCS San Gallicano Dermatological Institute | IRCCS Regina Elena National Cancer Institute | Frauke Goeman, Giulia Orlandi, Matteo Pallocca, Martina Diano, Fabrizio Ensoli, Fulvia Pimpinelli, Maurizio Fanciulli, Giovanni Blandino, Aldo Morrone, Gennaro Ciliberto |
| EPI_ISL_3068475 | IRCCS San Gallicano Dermatological Institute | IRCCS Regina Elena National Cancer Institute | Sabrina Strano, Sara Donzelli, Eleonora Sperandio, Antonio Federico, Fabrizio Ensoli, Fulvia Pimpinelli, Maurizio Fanciulli, Giovanni Blandino, Aldo Morrone, Gennaro Ciliberto |
| EPI_ISL_3068476 | IRCCS San Gallicano Dermatological Institute | IRCCS Regina Elena National Cancer Institute | Francesca De Nicola, Giulia Orlandi, Alice Massacci, Fulvia Fraticelli, Fabrizio Ensoli, Fulvia Pimpinelli, Maurizio Fanciulli, Giovanni Blandino, Aldo Morrone, Gennaro Ciliberto |
| EPI_ISL_3068478 | IRCCS San Gallicano Dermatological Institute | IRCCS Regina Elena National Cancer Institute | Frauke Goeman, Giulia Orlandi, Eleonora Sperandio, Alessia Lauretti, Fabrizio Ensoli, Fulvia Pimpinelli, Maurizio Fanciulli, Giovanni Blandino, Aldo Morrone, Gennaro Ciliberto |
| EPI_ISL_3068479 | IRCCS San Gallicano Dermatological Institute | IRCCS Regina Elena National Cancer Institute | Sabrina Strano, Sara Donzelli, Alice Massacci, Francesca Maione, Fabrizio Ensoli, Fulvia Pimpinelli, Maurizio Fanciulli, Giovanni Blandino, Aldo Morrone, Gennaro Ciliberto |
| EPI_ISL_3068480 | IRCCS San Gallicano Dermatological Institute | IRCCS Regina Elena National Cancer Institute | Francesca De Nicola, Giulia Orlandi, Matteo Pallocca, Francisco Obregon, Fabrizio Ensoli, Fulvia Pimpinelli, Maurizio Fanciulli, Giovanni Blandino, Aldo Morrone, Gennaro Ciliberto |
| EPI_ISL_3068481 | IRCCS San Gallicano Dermatological Institute | IRCCS Regina Elena National Cancer Institute | Ludovica Ciuffreda, Sara Donzelli, Eleonora Sperandio, Silvia Paluzzi, Fabrizio Ensoli, Fulvia Pimpinelli, Maurizio Fanciulli, Giovanni Blandino, Aldo Morrone, Gennaro Ciliberto |
| EPI_ISL_3068482 | IRCCS San Gallicano Dermatological Institute | IRCCS Regina Elena National Cancer Institute | Frauke Goeman, Giulia Orlandi, Alice Massacci, Sara Petrolo, Fabrizio Ensoli, Fulvia Pimpinelli, Maurizio Fanciulli, Giovanni Blandino, Aldo Morrone, Gennaro Ciliberto |
| EPI_ISL_3068483 | IRCCS San Gallicano Dermatological Institute | IRCCS Regina Elena National Cancer Institute | Sabrina Strano, Sara Donzelli, Matteo Pallocca, Valentina Ricca, Fabrizio Ensoli, Fulvia Pimpinelli, Maurizio Fanciulli, Giovanni Blandino, Aldo Morrone, Gennaro Ciliberto |
| EPI_ISL_3068484 | IRCCS San Gallicano Dermatological Institute | IRCCS Regina Elena National Cancer Institute | Francesca De Nicola, Giulia Orlandi, Eleonora Sperandio, Serena Salvo, Fabrizio Ensoli, Fulvia Pimpinelli, Maurizio Fanciulli, Giovanni Blandino, Aldo Morrone, Gennaro Ciliberto |
| EPI_ISL_3068485 | IRCCS San Gallicano Dermatological Institute | IRCCS Regina Elena National Cancer Institute | Ludovica Ciuffreda, Sara Donzelli, Alice Massacci, Grazia Prignano, Fabrizio Ensoli, Fulvia Pimpinelli, Maurizio Fanciulli, Giovanni Blandino, Aldo Morrone, Gennaro Ciliberto |
| EPI_ISL_3068486 | IRCCS San Gallicano Dermatological Institute | IRCCS Regina Elena National Cancer Institute | Frauke Goeman, Giulia Orlandi, Matteo Pallocca, Elisabetta Trento, Fabrizio Ensoli, Fulvia Pimpinelli, Maurizio Fanciulli, Giovanni Blandino, Aldo Morrone, Gennaro Ciliberto |
| EPI_ISL_3068487 | IRCCS San Gallicano Dermatological Institute | IRCCS Regina Elena National Cancer Institute | Sabrina Strano, Sara Donzelli, Eleonora Sperandio, Arianna Mastrofrancesco, Fabrizio Ensoli, Fulvia Pimpinelli, Maurizio Fanciulli, Giovanni Blandino, Aldo Morrone, Gennaro Ciliberto |
| EPI_ISL_3068488 | IRCCS San Gallicano Dermatological Institute | IRCCS Regina Elena National Cancer Institute | Francesca De Nicola, Giulia Orlandi, Alice Massacci, Giovanna D'agosto, Fabrizio Ensoli, Fulvia Pimpinelli, Maurizio Fanciulli, Giovanni Blandino, Aldo Morrone, Gennaro Ciliberto |
| EPI_ISL_3068489 | IRCCS San Gallicano Dermatological Institute | IRCCS Regina Elena National Cancer Institute | Ludovica Ciuffreda, Sara Donzelli, Matteo Pallocca, Carla Mottini, Fabrizio Ensoli, Fulvia Pimpinelli, Maurizio Fanciulli, Giovanni Blandino, Aldo Morrone, Gennaro Ciliberto |
| EPI_ISL_3068490 | IRCCS San Gallicano Dermatological Institute | IRCCS Regina Elena National Cancer Institute | Frauke Goeman, Giulia Orlandi, Eleonora Sperandio, Martina Pontone, Fabrizio Ensoli, Fulvia Pimpinelli, Maurizio Fanciulli, Giovanni Blandino, Aldo Morrone, Gennaro Ciliberto |
| EPI_ISL_3068491 | IRCCS San Gallicano Dermatological Institute | IRCCS Regina Elena National Cancer Institute | Sabrina Strano, Sara Donzelli, Alice Massacci, di Domenico Enea Gino, Fabrizio Ensoli, Fulvia Pimpinelli, Maurizio Fanciulli, Giovanni Blandino, Aldo Morrone, Gennaro Ciliberto |
| EPI_ISL_3068493 | IRCCS San Gallicano Dermatological Institute | IRCCS Regina Elena National Cancer Institute | Ludovica Ciuffreda, Sara Donzelli, Eleonora Sperandio, Francesca Sivori, Fabrizio Ensoli, Fulvia Pimpinelli, Maurizio Fanciulli, Giovanni Blandino, Aldo Morrone, Gennaro Ciliberto |
| EPI_ISL_3068494 | IRCCS San Gallicano Dermatological Institute | IRCCS Regina Elena National Cancer Institute | Frauke Goeman, Giulia Orlandi, Alice Massacci, Andrea Cazzani, Fabrizio Ensoli, Fulvia Pimpinelli, Maurizio Fanciulli, Giovanni Blandino, Aldo Morrone, Gennaro Ciliberto |
| EPI_ISL_3068495 | IRCCS San Gallicano Dermatological Institute | IRCCS Regina Elena National Cancer Institute | Sabrina Strano, Sara Donzelli, Matteo Pallocca, Ilaria Celesti, Fabrizio Ensoli, Fulvia Pimpinelli, Maurizio Fanciulli, Giovanni Blandino, Aldo Morrone, Gennaro Ciliberto |
| EPI_ISL_3068496 | IRCCS San Gallicano Dermatological Institute | IRCCS Regina Elena National Cancer Institute | Ludovica Ciuffreda, Sara Donzelli, Alice Massacci, Antonio Federico, Fabrizio Ensoli, Fulvia Pimpinelli, Maurizio Fanciulli, Giovanni Blandino, Aldo Morrone, Gennaro Ciliberto |
| EPI_ISL_3068497 | IRCCS San Gallicano Dermatological Institute | IRCCS Regina Elena National Cancer Institute | Frauke Goeman, Giulia Orlandi, Matteo Pallocca, Fulvia Fraticelli, Fabrizio Ensoli, Fulvia Pimpinelli, Maurizio Fanciulli, Giovanni Blandino, Aldo Morrone, Gennaro Ciliberto |
| EPI_ISL_3068498 | IRCCS San Gallicano Dermatological Institute | IRCCS Regina Elena National Cancer Institute | Sabrina Strano, Sara Donzelli, Eleonora Sperandio, Lorenzo Furzi, Fabrizio Ensoli, Fulvia Pimpinelli, Maurizio Fanciulli, Giovanni Blandino, Aldo Morrone, Gennaro Ciliberto |
| EPI_ISL_3068499 | IRCCS San Gallicano Dermatological Institute | IRCCS Regina Elena National Cancer Institute | Francesca De Nicola, Giulia Orlandi, Alice Massacci, Alessia Lauretti, Fabrizio Ensoli, Fulvia Pimpinelli, Maurizio Fanciulli, Giovanni Blandino, Aldo Morrone, Gennaro Ciliberto |
| EPI_ISL_3068500 | IRCCS San Gallicano Dermatological Institute | IRCCS Regina Elena National Cancer Institute | Ludovica Ciuffreda, Sara Donzelli, Matteo Pallocca, Francesca Maione, Fabrizio Ensoli, Fulvia Pimpinelli, Maurizio Fanciulli, Giovanni Blandino, Aldo Morrone, Gennaro Ciliberto |
| EPI_ISL_3068501 | IRCCS San Gallicano Dermatological Institute | IRCCS Regina Elena National Cancer Institute | Frauke Goeman, Giulia Orlandi, Eleonora Sperandio, Francisco Obregon, Fabrizio Ensoli, Fulvia Pimpinelli, Maurizio Fanciulli, Giovanni Blandino, Aldo Morrone, Gennaro Ciliberto |
| EPI_ISL_3068502 | IRCCS San Gallicano Dermatological Institute | IRCCS Regina Elena National Cancer Institute | Sabrina Strano, Sara Donzelli, Alice Massacci, Silvia Paluzzi, Fabrizio Ensoli, Fulvia Pimpinelli, Maurizio Fanciulli, Giovanni Blandino, Aldo Morrone, Gennaro Ciliberto |
| EPI_ISL_3068503 | IRCCS San Gallicano Dermatological Institute | IRCCS Regina Elena National Cancer Institute | Francesca De Nicola, Giulia Orlandi, Matteo Pallocca, Sara Petrolo, Fabrizio Ensoli, Fulvia Pimpinelli, Maurizio Fanciulli, Giovanni Blandino, Aldo Morrone, Gennaro Ciliberto |
| EPI_ISL_3068504 | IRCCS San Gallicano Dermatological Institute | IRCCS Regina Elena National Cancer Institute | Ludovica Ciuffreda, Sara Donzelli, Eleonora Sperandio, Valentina Ricca, Fabrizio Ensoli, Fulvia Pimpinelli, Maurizio Fanciulli, Giovanni Blandino, Aldo Morrone, Gennaro Ciliberto |
| EPI_ISL_3068505 | IRCCS San Gallicano Dermatological Institute | IRCCS Regina Elena National Cancer Institute | Frauke Goeman, Giulia Orlandi, Alice Massacci, Serena Salvo, Fabrizio Ensoli, Fulvia Pimpinelli, Maurizio Fanciulli, Giovanni Blandino, Aldo Morrone, Gennaro Ciliberto |
| EPI_ISL_3068506 | IRCCS San Gallicano Dermatological Institute | IRCCS Regina Elena National Cancer Institute | Sabrina Strano, Sara Donzelli, Matteo Pallocca, Grazia Prignano, Fabrizio Ensoli, Fulvia Pimpinelli, Maurizio Fanciulli, Giovanni Blandino, Aldo Morrone, Gennaro Ciliberto |
| EPI_ISL_3068507 | IRCCS San Gallicano Dermatological Institute | IRCCS Regina Elena National Cancer Institute | Francesca De Nicola, Giulia Orlandi, Eleonora Sperandio, Elisabetta Trento, Fabrizio Ensoli, Fulvia Pimpinelli, Maurizio Fanciulli, Giovanni Blandino, Aldo Morrone, Gennaro Ciliberto |
| EPI_ISL_3068508 | IRCCS San Gallicano Dermatological Institute | IRCCS Regina Elena National Cancer Institute | Ludovica Ciuffreda, Sara Donzelli, Alice Massacci, Arianna Mastrofrancesco, Fabrizio Ensoli, Fulvia Pimpinelli, Maurizio Fanciulli, Giovanni Blandino, Aldo Morrone, Gennaro Ciliberto |
| EPI_ISL_3068509 | IRCCS San Gallicano Dermatological Institute | IRCCS Regina Elena National Cancer Institute | Frauke Goeman, Giulia Orlandi, Matteo Pallocca, Giovanna D'agosto, Fabrizio Ensoli, Fulvia Pimpinelli, Maurizio Fanciulli, Giovanni Blandino, Aldo Morrone, Gennaro Ciliberto |
| EPI_ISL_3068510 | IRCCS San Gallicano Dermatological Institute | IRCCS Regina Elena National Cancer Institute | Sabrina Strano, Sara Donzelli, Eleonora Sperandio, Carla Mottini, Fabrizio Ensoli, Fulvia Pimpinelli, Maurizio Fanciulli, Giovanni Blandino, Aldo Morrone, Gennaro Ciliberto |
| EPI_ISL_3068511 | IRCCS San Gallicano Dermatological Institute | IRCCS Regina Elena National Cancer Institute | Francesca De Nicola, Giulia Orlandi, Alice Massacci, Martina Pontone, Fabrizio Ensoli, Fulvia Pimpinelli, Maurizio Fanciulli, Giovanni Blandino, Aldo Morrone, Gennaro Ciliberto |
| EPI_ISL_3068512 | IRCCS San Gallicano Dermatological Institute | IRCCS Regina Elena National Cancer Institute | Ludovica Ciuffreda, Sara Donzelli, Matteo Pallocca, di Domenico Enea Gino, Fabrizio Ensoli, Fulvia Pimpinelli, Maurizio Fanciulli, Giovanni Blandino, Aldo Morrone, Gennaro Ciliberto |
| EPI_ISL_3068513 | IRCCS San Gallicano Dermatological Institute | IRCCS Regina Elena National Cancer Institute | Frauke Goeman, Giulia Orlandi, Eleonora Sperandio, Ilaria Cavallo, Fabrizio Ensoli, Fulvia Pimpinelli, Maurizio Fanciulli, Giovanni Blandino, Aldo Morrone, Gennaro Ciliberto |
| EPI_ISL_3068514 | IRCCS San Gallicano Dermatological Institute | IRCCS Regina Elena National Cancer Institute | Sabrina Strano, Sara Donzelli, Alice Massacci, Francesca Sivori, Fabrizio Ensoli, Fulvia Pimpinelli, Maurizio Fanciulli, Giovanni Blandino, Aldo Morrone, Gennaro Ciliberto |
| EPI_ISL_3068515 | IRCCS San Gallicano Dermatological Institute | IRCCS Regina Elena National Cancer Institute | Francesca De Nicola, Giulia Orlandi, Matteo Pallocca, Andrea Cazzani, Fabrizio Ensoli, Fulvia Pimpinelli, Maurizio Fanciulli, Giovanni Blandino, Aldo Morrone, Gennaro Ciliberto |
| EPI_ISL_3068516 | IRCCS San Gallicano Dermatological Institute | IRCCS Regina Elena National Cancer Institute | Ludovica Ciuffreda, Sara Donzelli, Eleonora Sperandio, Ilaria Celesti, Fabrizio Ensoli, Fulvia Pimpinelli, Maurizio Fanciulli, Giovanni Blandino, Aldo Morrone, Gennaro Ciliberto |
| EPI_ISL_3068517 | IRCCS San Gallicano Dermatological Institute | IRCCS Regina Elena National Cancer Institute | Frauke Goeman, Giulia Orlandi, Alice Massacci, Martina Diano, Fabrizio Ensoli, Fulvia Pimpinelli, Maurizio Fanciulli, Giovanni Blandino, Aldo Morrone, Gennaro Ciliberto |
| EPI_ISL_3068518 | IRCCS San Gallicano Dermatological Institute | IRCCS Regina Elena National Cancer Institute | Sabrina Strano, Sara Donzelli, Matteo Pallocca, Antonio Federico, Fabrizio Ensoli, Fulvia Pimpinelli, Maurizio Fanciulli, Giovanni Blandino, Aldo Morrone, Gennaro Ciliberto |
| EPI_ISL_3068519 | IRCCS San Gallicano Dermatological Institute | IRCCS Regina Elena National Cancer Institute | Francesca De Nicola, Giulia Orlandi, Eleonora Sperandio, Fulvia Fraticelli, Fabrizio Ensoli, Fulvia Pimpinelli, Maurizio Fanciulli, Giovanni Blandino, Aldo Morrone, Gennaro Ciliberto |
| EPI_ISL_3068520 | IRCCS San Gallicano Dermatological Institute | IRCCS Regina Elena National Cancer Institute | Ludovica Ciuffreda, Sara Donzelli, Alice Massacci, Lorenzo Furzi, Fabrizio Ensoli, Fulvia Pimpinelli, Maurizio Fanciulli, Giovanni Blandino, Aldo Morrone, Gennaro Ciliberto |

| | | | |
|---|---|---|---|
| EPI_ISL_3068521 | IRCCS San Gallicano Dermatological Institute | IRCCS Regina Elena National Cancer Institute | Frauke Goeman, Giulia Orlandi, Matteo Pallocca, Alessia Lauretti, Fabrizio Ensoli, Fulvia Pimpinelli, Maurizio Fanciulli, Giovanni Blandino, Aldo Morrone, Gennaro Ciliberto |
| EPI_ISL_3068522 | IRCCS San Gallicano Dermatological Institute | IRCCS Regina Elena National Cancer Institute | Sabrina Strano, Sara Donzelli, Eleonora Sperandio, Francesca Maione, Fabrizio Ensoli, Fulvia Pimpinelli, Maurizio Fanciulli, Giovanni Blandino, Aldo Morrone, Gennaro Ciliberto |
| EPI_ISL_3068523 | IRCCS San Gallicano Dermatological Institute | IRCCS Regina Elena National Cancer Institute | Francesca De Nicola, Giulia Orlandi, Alice Massacci, Francisco Obregon, Fabrizio Ensoli, Fulvia Pimpinelli, Maurizio Fanciulli, Giovanni Blandino, Aldo Morrone, Gennaro Ciliberto |
| EPI_ISL_3068524 | IRCCS San Gallicano Dermatological Institute | IRCCS Regina Elena National Cancer Institute | Ludovica Ciuffreda, Sara Donzelli, Matteo Pallocca, Silvia Paluzzi, Fabrizio Ensoli, Fulvia Pimpinelli, Maurizio Fanciulli, Giovanni Blandino, Aldo Morrone, Gennaro Ciliberto |
| EPI_ISL_3068525 | IRCCS San Gallicano Dermatological Institute | IRCCS Regina Elena National Cancer Institute | Frauke Goeman, Giulia Orlandi, Eleonora Sperandio, Sara Petrolo, Fabrizio Ensoli, Fulvia Pimpinelli, Maurizio Fanciulli, Giovanni Blandino, Aldo Morrone, Gennaro Ciliberto |
| EPI_ISL_3068526 | IRCCS San Gallicano Dermatological Institute | IRCCS Regina Elena National Cancer Institute | Francesca De Nicola, Giulia Orlandi, Matteo Pallocca, Serena Salvo, Fabrizio Ensoli, Fulvia Pimpinelli, Maurizio Fanciulli, Giovanni Blandino, Aldo Morrone, Gennaro Ciliberto |
| EPI_ISL_3068527 | IRCCS San Gallicano Dermatological Institute | IRCCS Regina Elena National Cancer Institute | Ludovica Ciuffreda, Sara Donzelli, Eleonora Sperandio, Grazia Prignano, Fabrizio Ensoli, Fulvia Pimpinelli, Maurizio Fanciulli, Giovanni Blandino, Aldo Morrone, Gennaro Ciliberto |
| EPI_ISL_3068528 | IRCCS San Gallicano Dermatological Institute | IRCCS Regina Elena National Cancer Institute | Frauke Goeman, Giulia Orlandi, Alice Massacci, Elisabetta Trento, Fabrizio Ensoli, Fulvia Pimpinelli, Maurizio Fanciulli, Giovanni Blandino, Aldo Morrone, Gennaro Ciliberto |
| EPI_ISL_3068529 | IRCCS San Gallicano Dermatological Institute | IRCCS Regina Elena National Cancer Institute | Sabrina Strano, Sara Donzelli, Matteo Pallocca, Arianna Mastrofrancesco, Fabrizio Ensoli, Fulvia Pimpinelli, Maurizio Fanciulli, Giovanni Blandino, Aldo Morrone, Gennaro Ciliberto |
| EPI_ISL_3068531 | IRCCS San Gallicano Dermatological Institute | IRCCS Regina Elena National Cancer Institute | Ludovica Ciuffreda, Sara Donzelli, Alice Massacci, Carla Mottini, Fabrizio Ensoli, Fulvia Pimpinelli, Maurizio Fanciulli, Giovanni Blandino, Aldo Morrone, Gennaro Ciliberto |
| EPI_ISL_3068532 | IRCCS San Gallicano Dermatological Institute | IRCCS Regina Elena National Cancer Institute | Frauke Goeman, Giulia Orlandi, Matteo Pallocca, Martina Pontone, Fabrizio Ensoli, Fulvia Pimpinelli, Maurizio Fanciulli, Giovanni Blandino, Aldo Morrone, Gennaro Ciliberto |
| EPI_ISL_3068533 | IRCCS San Gallicano Dermatological Institute | IRCCS Regina Elena National Cancer Institute | Sabrina Strano, Sara Donzelli, Eleonora Sperandio, di Domenico Enea Gino, Fabrizio Ensoli, Fulvia Pimpinelli, Maurizio Fanciulli, Giovanni Blandino, Aldo Morrone, Gennaro Ciliberto |
| EPI_ISL_3068534 | IRCCS San Gallicano Dermatological Institute | IRCCS Regina Elena National Cancer Institute | Francesca De Nicola, Giulia Orlandi, Alice Massacci, Ilaria Cavallo, Fabrizio Ensoli, Fulvia Pimpinelli, Maurizio Fanciulli, Giovanni Blandino, Aldo Morrone, Gennaro Ciliberto |
| EPI_ISL_3068535 | IRCCS San Gallicano Dermatological Institute | IRCCS Regina Elena National Cancer Institute | Ludovica Ciuffreda, Sara Donzelli, Matteo Pallocca, Francesca Sivori, Fabrizio Ensoli, Fulvia Pimpinelli, Maurizio Fanciulli, Giovanni Blandino, Aldo Morrone, Gennaro Ciliberto |
| EPI_ISL_3068536 | IRCCS San Gallicano Dermatological Institute | IRCCS Regina Elena National Cancer Institute | Frauke Goeman, Giulia Orlandi, Eleonora Sperandio, Andrea Cazzani, Fabrizio Ensoli, Fulvia Pimpinelli, Maurizio Fanciulli, Giovanni Blandino, Aldo Morrone, Gennaro Ciliberto |
| EPI_ISL_3068537 | IRCCS San Gallicano Dermatological Institute | IRCCS Regina Elena National Cancer Institute | Sabrina Strano, Sara Donzelli, Alice Massacci, Ilaria Celesti, Fabrizio Ensoli, Fulvia Pimpinelli, Maurizio Fanciulli, Giovanni Blandino, Aldo Morrone, Gennaro Ciliberto |
| EPI_ISL_3068538 | IRCCS San Gallicano Dermatological Institute | IRCCS Regina Elena National Cancer Institute | Francesca De Nicola, Giulia Orlandi, Matteo Pallocca, Martina Diano, Fabrizio Ensoli, Fulvia Pimpinelli, Maurizio Fanciulli, Giovanni Blandino, Aldo Morrone, Gennaro Ciliberto |
| EPI_ISL_3068539 | IRCCS San Gallicano Dermatological Institute | IRCCS Regina Elena National Cancer Institute | Ludovica Ciuffreda, Sara Donzelli, Eleonora Sperandio, Antonio Federico, Fabrizio Ensoli, Fulvia Pimpinelli, Maurizio Fanciulli, Giovanni Blandino, Aldo Morrone, Gennaro Ciliberto |
| EPI_ISL_3068540 | IRCCS San Gallicano Dermatological Institute | IRCCS Regina Elena National Cancer Institute | Frauke Goeman, Giulia Orlandi, Alice Massacci, Fulvia Fraticelli, Fabrizio Ensoli, Fulvia Pimpinelli, Maurizio Fanciulli, Giovanni Blandino, Aldo Morrone, Gennaro Ciliberto |
| EPI_ISL_3068541 | IRCCS San Gallicano Dermatological Institute | IRCCS Regina Elena National Cancer Institute | Sabrina Strano, Sara Donzelli, Matteo Pallocca, Lorenzo Furzi, Fabrizio Ensoli, Fulvia Pimpinelli, Maurizio Fanciulli, Giovanni Blandino, Aldo Morrone, Gennaro Ciliberto |
| EPI_ISL_3068542 | IRCCS San Gallicano Dermatological Institute | IRCCS Regina Elena National Cancer Institute | Francesca De Nicola, Giulia Orlandi, Eleonora Sperandio, Alessia Lauretti, Fabrizio Ensoli, Fulvia Pimpinelli, Maurizio Fanciulli, Giovanni Blandino, Aldo Morrone, Gennaro Ciliberto |
| EPI_ISL_3068543 | IRCCS San Gallicano Dermatological Institute | IRCCS Regina Elena National Cancer Institute | Ludovica Ciuffreda, Sara Donzelli, Alice Massacci, Francesca Maione, Fabrizio Ensoli, Fulvia Pimpinelli, Maurizio Fanciulli, Giovanni Blandino, Aldo Morrone, Gennaro Ciliberto |
| EPI_ISL_3068544 | IRCCS San Gallicano Dermatological Institute | IRCCS Regina Elena National Cancer Institute | Frauke Goeman, Giulia Orlandi, Matteo Pallocca, Francisco Obregon, Fabrizio Ensoli, Fulvia Pimpinelli, Maurizio Fanciulli, Giovanni Blandino, Aldo Morrone, Gennaro Ciliberto |
| EPI_ISL_3068545 | IRCCS San Gallicano Dermatological Institute | IRCCS Regina Elena National Cancer Institute | Sabrina Strano, Sara Donzelli, Eleonora Sperandio, Silvia Paluzzi, Fabrizio Ensoli, Fulvia Pimpinelli, Maurizio Fanciulli, Giovanni Blandino, Aldo Morrone, Gennaro Ciliberto |
| EPI_ISL_3068546 | IRCCS San Gallicano Dermatological Institute | IRCCS Regina Elena National Cancer Institute | Francesca De Nicola, Giulia Orlandi, Alice Massacci, Sara Petrolo, Fabrizio Ensoli, Fulvia Pimpinelli, Maurizio Fanciulli, Giovanni Blandino, Aldo Morrone, Gennaro Ciliberto |
| EPI_ISL_3068547 | IRCCS San Gallicano Dermatological Institute | IRCCS Regina Elena National Cancer Institute | Ludovica Ciuffreda, Sara Donzelli, Matteo Pallocca, Valentina Ricca, Fabrizio Ensoli, Fulvia Pimpinelli, Maurizio Fanciulli, Giovanni Blandino, Aldo Morrone, Gennaro Ciliberto |
| EPI_ISL_3068548 | IRCCS San Gallicano Dermatological Institute | IRCCS Regina Elena National Cancer Institute | Frauke Goeman, Giulia Orlandi, Eleonora Sperandio, Serena Salvo, Fabrizio Ensoli, Fulvia Pimpinelli, Maurizio Fanciulli, Giovanni Blandino, Aldo Morrone, Gennaro Ciliberto |
| EPI_ISL_3068549 | IRCCS San Gallicano Dermatological Institute | IRCCS Regina Elena National Cancer Institute | Sabrina Strano, Sara Donzelli, Alice Massacci, Grazia Prignano, Fabrizio Ensoli, Fulvia Pimpinelli, Maurizio Fanciulli, Giovanni Blandino, Aldo Morrone, Gennaro Ciliberto |
| EPI_ISL_3068550 | IRCCS San Gallicano Dermatological Institute | IRCCS Regina Elena National Cancer Institute | Ludovica Ciuffreda, Sara Donzelli, Eleonora Sperandio, Arianna Mastrofrancesco, Fabrizio Ensoli, Fulvia Pimpinelli, Maurizio Fanciulli, Giovanni Blandino, Aldo Morrone, Gennaro Ciliberto |
| EPI_ISL_3068551 | IRCCS San Gallicano Dermatological Institute | IRCCS Regina Elena National Cancer Institute | Frauke Goeman, Giulia Orlandi, Alice Massacci, Giovanna D’agosto, Fabrizio Ensoli, Fulvia Pimpinelli, Maurizio Fanciulli, Giovanni Blandino, Aldo Morrone, Gennaro Ciliberto |
| EPI_ISL_3068552 | IRCCS San Gallicano Dermatological Institute | IRCCS Regina Elena National Cancer Institute | Sabrina Strano, Sara Donzelli, Matteo Pallocca, Carla Mottini, Fabrizio Ensoli, Fulvia Pimpinelli, Maurizio Fanciulli, Giovanni Blandino, Aldo Morrone, Gennaro Ciliberto |
| EPI_ISL_3068553 | IRCCS San Gallicano Dermatological Institute | IRCCS Regina Elena National Cancer Institute | Francesca De Nicola, Giulia Orlandi, Eleonora Sperandio, Martina Pontone, Fabrizio Ensoli, Fulvia Pimpinelli, Maurizio Fanciulli, Giovanni Blandino, Aldo Morrone, Gennaro Ciliberto |
| EPI_ISL_3068554 | IRCCS San Gallicano Dermatological Institute | IRCCS Regina Elena National Cancer Institute | Ludovica Ciuffreda, Sara Donzelli, Alice Massacci, di Domenico Enea Gino, Fabrizio Ensoli, Fulvia Pimpinelli, Maurizio Fanciulli, Giovanni Blandino, Aldo Morrone, Gennaro Ciliberto |
| EPI_ISL_3068555 | IRCCS San Gallicano Dermatological Institute | IRCCS Regina Elena National Cancer Institute | Frauke Goeman, Giulia Orlandi, Matteo Pallocca, Ilaria Cavallo, Fabrizio Ensoli, Fulvia Pimpinelli, Maurizio Fanciulli, Giovanni Blandino, Aldo Morrone, Gennaro Ciliberto |
| EPI_ISL_3068556 | IRCCS San Gallicano Dermatological Institute | IRCCS Regina Elena National Cancer Institute | Sabrina Strano, Sara Donzelli, Eleonora Sperandio, Francesca Sivori, Fabrizio Ensoli, Fulvia Pimpinelli, Maurizio Fanciulli, Giovanni Blandino, Aldo Morrone, Gennaro Ciliberto |
| EPI_ISL_3068557 | IRCCS San Gallicano Dermatological Institute | IRCCS Regina Elena National Cancer Institute | Francesca De Nicola, Giulia Orlandi, Alice Massacci, Andrea Cazzani, Fabrizio Ensoli, Fulvia Pimpinelli, Maurizio Fanciulli, Giovanni Blandino, Aldo Morrone, Gennaro Ciliberto |
| EPI_ISL_3068558 | IRCCS San Gallicano Dermatological Institute | IRCCS Regina Elena National Cancer Institute | Ludovica Ciuffreda, Sara Donzelli, Matteo Pallocca, Ilaria Celesti, Fabrizio Ensoli, Fulvia Pimpinelli, Maurizio Fanciulli, Giovanni Blandino, Aldo Morrone, Gennaro Ciliberto |
| EPI_ISL_3068559 | IRCCS San Gallicano Dermatological Institute | IRCCS Regina Elena National Cancer Institute | Frauke Goeman, Giulia Orlandi, Eleonora Sperandio, Martina Diano, Fabrizio Ensoli, Fulvia Pimpinelli, Maurizio Fanciulli, Giovanni Blandino, Aldo Morrone, Gennaro Ciliberto |
| EPI_ISL_3068560 | IRCCS San Gallicano Dermatological Institute | IRCCS Regina Elena National Cancer Institute | Sabrina Strano, Sara Donzelli, Alice Massacci, Antonio Federico, Fabrizio Ensoli, Fulvia Pimpinelli, Maurizio Fanciulli, Giovanni Blandino, Aldo Morrone, Gennaro Ciliberto |
| EPI_ISL_3068561 | IRCCS San Gallicano Dermatological Institute | IRCCS Regina Elena National Cancer Institute | Francesca De Nicola, Giulia Orlandi, Matteo Pallocca, Fulvia Fraticelli, Fabrizio Ensoli, Fulvia Pimpinelli, Maurizio Fanciulli, Giovanni Blandino, Aldo Morrone, Gennaro Ciliberto |
| EPI_ISL_3068563 | IRCCS San Gallicano Dermatological Institute | IRCCS Regina Elena National Cancer Institute | Frauke Goeman, Giulia Orlandi, Alice Massacci, Alessia Lauretti, Fabrizio Ensoli, Fulvia Pimpinelli, Maurizio Fanciulli, Giovanni Blandino, Aldo Morrone, Gennaro Ciliberto |
| EPI_ISL_3068564 | IRCCS San Gallicano Dermatological Institute | IRCCS Regina Elena National Cancer Institute | Sabrina Strano, Sara Donzelli, Matteo Pallocca, Francesca Maione, Fabrizio Ensoli, Fulvia Pimpinelli, Maurizio Fanciulli, Giovanni Blandino, Aldo Morrone, Gennaro Ciliberto |
| EPI_ISL_3068565 | IRCCS San Gallicano Dermatological Institute | IRCCS Regina Elena National Cancer Institute | Francesca De Nicola, Giulia Orlandi, Eleonora Sperandio, Francisco Obregon, Fabrizio Ensoli, Fulvia Pimpinelli, Maurizio Fanciulli, Giovanni Blandino, Aldo Morrone, Gennaro Ciliberto |
| EPI_ISL_3068566 | IRCCS San Gallicano Dermatological Institute | IRCCS Regina Elena National Cancer Institute | Ludovica Ciuffreda, Sara Donzelli, Alice Massacci, Silvia Paluzzi, Fabrizio Ensoli, Fulvia Pimpinelli, Maurizio Fanciulli, Giovanni Blandino, Aldo Morrone, Gennaro Ciliberto |
| EPI_ISL_3068567 | IRCCS San Gallicano Dermatological Institute | IRCCS Regina Elena National Cancer Institute | Frauke Goeman, Giulia Orlandi, Matteo Pallocca, Sara Petrolo, Fabrizio Ensoli, Fulvia Pimpinelli, Maurizio Fanciulli, Giovanni Blandino, Aldo Morrone, Gennaro Ciliberto |
| EPI_ISL_3068568 | IRCCS San Gallicano Dermatological Institute | IRCCS Regina Elena National Cancer Institute | Sabrina Strano, Sara Donzelli, Eleonora Sperandio, Valentina Ricca, Fabrizio Ensoli, Fulvia Pimpinelli, Maurizio Fanciulli, Giovanni Blandino, Aldo Morrone, Gennaro Ciliberto |
| EPI_ISL_3068569 | IRCCS San Gallicano Dermatological Institute | IRCCS Regina Elena National Cancer Institute | Francesca De Nicola, Giulia Orlandi, Alice Massacci, Serena Salvo, Fabrizio Ensoli, Fulvia Pimpinelli, Maurizio Fanciulli, Giovanni Blandino, Aldo Morrone, Gennaro Ciliberto |
| EPI_ISL_3068570 | IRCCS San Gallicano Dermatological Institute | IRCCS Regina Elena National Cancer Institute | Ludovica Ciuffreda, Sara Donzelli, Matteo Pallocca, Grazia Prignano, Fabrizio Ensoli, Fulvia Pimpinelli, Maurizio Fanciulli, Giovanni Blandino, Aldo Morrone, Gennaro Ciliberto |
| EPI_ISL_3068571 | IRCCS San Gallicano Dermatological Institute | IRCCS Regina Elena National Cancer Institute | Frauke Goeman, Giulia Orlandi, Eleonora Sperandio, Elisabetta Trento, Fabrizio Ensoli, Fulvia Pimpinelli, Maurizio Fanciulli, Giovanni Blandino, Aldo Morrone, Gennaro Ciliberto |
| EPI_ISL_3068572 | IRCCS San Gallicano Dermatological Institute | IRCCS Regina Elena National Cancer Institute | Sabrina Strano, Sara Donzelli, Alice Massacci, Arianna Mastrofrancesco, Fabrizio Ensoli, Fulvia Pimpinelli, Maurizio Fanciulli, Giovanni Blandino, Aldo Morrone, Gennaro Ciliberto |
| EPI_ISL_3068573 | IRCCS San Gallicano Dermatological Institute | IRCCS Regina Elena National Cancer Institute | Francesca De Nicola, Giulia Orlandi, Matteo Pallocca, Giovanna D’agosto, Fabrizio Ensoli, Fulvia Pimpinelli, Maurizio Fanciulli, Giovanni Blandino, Aldo Morrone, Gennaro Ciliberto |
| EPI_ISL_3068574 | IRCCS San Gallicano Dermatological Institute | IRCCS Regina Elena National Cancer Institute | Ludovica Ciuffreda, Sara Donzelli, Eleonora Sperandio, Carla Mottini, Fabrizio Ensoli, Fulvia Pimpinelli, Maurizio Fanciulli, Giovanni Blandino, Aldo Morrone, Gennaro Ciliberto |
| EPI_ISL_3068575 | IRCCS San Gallicano Dermatological Institute | IRCCS Regina Elena National Cancer Institute | Frauke Goeman, Giulia Orlandi, Alice Massacci, Martina Pontone, Fabrizio Ensoli, Fulvia Pimpinelli, Maurizio Fanciulli, Giovanni Blandino, Aldo Morrone, Gennaro Ciliberto |
| EPI_ISL_3068576 | IRCCS San Gallicano Dermatological Institute | IRCCS Regina Elena National Cancer Institute | Sabrina Strano, Sara Donzelli, Matteo Pallocca, di Domenico Enea Gino, Fabrizio Ensoli, Fulvia Pimpinelli, Maurizio Fanciulli, Giovanni Blandino, Aldo Morrone, Gennaro Ciliberto |
| EPI_ISL_3068577 | IRCCS San Gallicano Dermatological Institute | IRCCS Regina Elena National Cancer Institute | Francesca De Nicola, Giulia Orlandi, Eleonora Sperandio, Ilaria Cavallo, Fabrizio Ensoli, Fulvia Pimpinelli, Maurizio Fanciulli, Giovanni Blandino, Aldo Morrone, Gennaro Ciliberto |
| EPI_ISL_3068578 | IRCCS San Gallicano Dermatological Institute | IRCCS Regina Elena National Cancer Institute | Ludovica Ciuffreda, Sara Donzelli, Alice Massacci, Francesca Sivori, Fabrizio Ensoli, Fulvia Pimpinelli, Maurizio Fanciulli, Giovanni Blandino, Aldo Morrone, Gennaro Ciliberto |
| EPI_ISL_3068579 | IRCCS San Gallicano Dermatological Institute | IRCCS Regina Elena National Cancer Institute | Frauke Goeman, Giulia Orlandi, Matteo Pallocca, Andrea Cazzani, Fabrizio Ensoli, Fulvia Pimpinelli, Maurizio Fanciulli, Giovanni Blandino, Aldo Morrone, Gennaro Ciliberto |
| EPI_ISL_3068580 | IRCCS San Gallicano Dermatological Institute | IRCCS Regina Elena National Cancer Institute | Sabrina Strano, Sara Donzelli, Eleonora Sperandio, Ilaria Celesti, Fabrizio Ensoli, Fulvia Pimpinelli, Maurizio Fanciulli, Giovanni Blandino, Aldo Morrone, Gennaro Ciliberto |
| EPI_ISL_3068581 | IRCCS San Gallicano Dermatological Institute | IRCCS Regina Elena National Cancer Institute | Francesca De Nicola, Giulia Orlandi, Alice Massacci, Martina Diano, Fabrizio Ensoli, Fulvia Pimpinelli, Maurizio Fanciulli, Giovanni Blandino, Aldo Morrone, Gennaro Ciliberto |
| EPI_ISL_3068582 | IRCCS San Gallicano Dermatological Institute | IRCCS Regina Elena National Cancer Institute | Ludovica Ciuffreda, Sara Donzelli, Matteo Pallocca, Antonio Federico, Fabrizio Ensoli, Fulvia Pimpinelli, Maurizio Fanciulli, Giovanni Blandino, Aldo Morrone, Gennaro Ciliberto |
| EPI_ISL_3068583 | IRCCS San Gallicano Dermatological Institute | IRCCS Regina Elena National Cancer Institute | Frauke Goeman, Giulia Orlandi, Eleonora Sperandio, Fulvia Fraticelli, Fabrizio Ensoli, Fulvia Pimpinelli, Maurizio Fanciulli, Giovanni Blandino, Aldo Morrone, Gennaro Ciliberto |
| EPI_ISL_3068584 | IRCCS San Gallicano Dermatological Institute | IRCCS Regina Elena National Cancer Institute | Sabrina Strano, Sara Donzelli, Alice Massacci, Lorenzo Furzi, Fabrizio Ensoli, Fulvia Pimpinelli, Maurizio Fanciulli, Giovanni Blandino, Aldo Morrone, Gennaro Ciliberto |
| EPI_ISL_3068585 | IRCCS San Gallicano Dermatological Institute | IRCCS Regina Elena National Cancer Institute | Francesca De Nicola, Giulia Orlandi, Matteo Pallocca, Alessia Lauretti, Fabrizio Ensoli, Fulvia Pimpinelli, Maurizio Fanciulli, Giovanni Blandino, Aldo Morrone, Gennaro Ciliberto |
| EPI_ISL_3068587 | IRCCS San Gallicano Dermatological Institute | IRCCS Regina Elena National Cancer Institute | Frauke Goeman, Giulia Orlandi, Alice Massacci, Francisco Obregon, Fabrizio Ensoli, Fulvia Pimpinelli, Maurizio Fanciulli, Giovanni Blandino, Aldo Morrone, Gennaro Ciliberto |
| EPI_ISL_3068591, EPI_ISL_3068592 | Lighthouse Lab in Alderley Park | Wellcome Sanger Institute for the COVID-19 Genomics UK (COG-UK) Consortium | Jacquelyn Wynn, Mairead Hyland, The Lighthouse Lab in Alderley Park and Alex Alderton, Roberto Amato, Jeffrey Barrett, Sonia Goncalves, Ewan Harrison, David K. Jackson, Ian Johnston, Dominic Kwiatkowski, Cordelia Langford, John Sillitoe on behalf of the Wellcome Sanger Institute COVID-19 Surveillance Team |
| EPI_ISL_3068593, EPI_ISL_3068594 | Lighthouse Laboratory Plymouth | Wellcome Sanger Institute for the COVID-19 Genomics UK (COG-UK) Consortium | Lighthouse Laboratory Plymouth and Alex Alderton, Roberto Amato, Jeffrey Barrett, Sonia Goncalves, Ewan Harrison, David K. Jackson, Ian Johnston, Dominic Kwiatkowski, Cordelia Langford, John Sillitoe on behalf of the Wellcome Sanger Institute COVID-19 Surveillance Team |
| EPI_ISL_3068595 | Health Services Laboratories | Wellcome Sanger Institute for the COVID-19 Genomics UK (COG-UK) Consortium | Health Services Laboratories and Alex Alderton, Roberto Amato, Jeffrey Barrett, Sonia Goncalves, Ewan Harrison, David K. Jackson, Ian Johnston, Dominic Kwiatkowski, Cordelia Langford, John Sillitoe on behalf of the Wellcome Sanger Institute COVID-19 Surveillance Team |
| EPI_ISL_3068596 | Lighthouse Lab in Alderley Park | Wellcome Sanger Institute for the COVID-19 Genomics UK (COG-UK) Consortium | Jacquelyn Wynn, Mairead Hyland, The Lighthouse Lab in Alderley Park and Alex Alderton, Roberto Amato, Jeffrey Barrett, Sonia Goncalves, Ewan Harrison, David K. Jackson, Ian Johnston, Dominic Kwiatkowski, Cordelia Langford, John Sillitoe on behalf of the Wellcome Sanger Institute COVID-19 Surveillance Team |
| EPI_ISL_3068597, EPI_ISL_3068598 | Lighthouse Laboratory Plymouth | Wellcome Sanger Institute for the COVID-19 Genomics UK (COG-UK) Consortium | Lighthouse Laboratory Plymouth and Alex Alderton, Roberto Amato, Jeffrey Barrett, Sonia Goncalves, Ewan Harrison, David K. Jackson, Ian Johnston, Dominic Kwiatkowski, Cordelia Langford, John Sillitoe on behalf of the Wellcome Sanger Institute COVID-19 Surveillance Team |
| EPI_ISL_3068599 | Lighthouse Lab in Alderley Park | Wellcome Sanger Institute for the COVID-19 Genomics UK (COG-UK) Consortium | Jacquelyn Wynn, Mairead Hyland, The Lighthouse Lab in Alderley Park and Alex Alderton, Roberto Amato, Jeffrey Barrett, Sonia Goncalves, Ewan Harrison, David K. Jackson, Ian Johnston, Dominic Kwiatkowski, Cordelia Langford, John Sillitoe on behalf of the Wellcome Sanger Institute COVID-19 Surveillance Team |
| EPI_ISL_3068600 | Lighthouse Laboratory Plymouth | Wellcome Sanger Institute for the COVID-19 Genomics UK (COG-UK) Consortium | Lighthouse Laboratory Plymouth and Alex Alderton, Roberto Amato, Jeffrey Barrett, Sonia Goncalves, Ewan Harrison, David K. Jackson, Ian Johnston, Dominic Kwiatkowski, Cordelia Langford, John Sillitoe on behalf of the Wellcome Sanger Institute COVID-19 Surveillance Team |

| | | | |
|---|---|---|---|
| EPI_ISL_3068601, EPI_ISL_3068602 | Lighthouse Lab in Alderley Park | Wellcome Sanger Institute for the COVID-19 Genomics UK (COG-UK) Consortium | Jacquelyn Wynn, Mairead Hyland, The Lighthouse Lab in Alderley Park and Alex Alderton, Roberto Amato, Jeffrey Barrett, Sonia Goncalves, Ewan Harrison, David K. Jackson, Ian Johnston, Dominic Kwiatkowski, Cordelia Langford, John Sillitoe on behalf of the Wellcome Sanger Institute COVID-19 Surveillance Team |
| EPI_ISL_3068603 | Health Services Laboratories | Wellcome Sanger Institute for the COVID-19 Genomics UK (COG-UK) Consortium | Health Services Laboratories and Alex Alderton, Roberto Amato, Jeffrey Barrett, Sonia Goncalves, Ewan Harrison, David K. Jackson, Ian Johnston, Dominic Kwiatkowski, Cordelia Langford, John Sillitoe on behalf of the Wellcome Sanger Institute COVID-19 Surveillance Team |
| EPI_ISL_3068604 | Berkshire and Surrey Pathology Services Lighthouse Laboratory | Wellcome Sanger Institute for the COVID-19 Genomics UK (COG-UK) Consortium | Berkshire and Surrey Pathology Services Lighthouse Laboratory and Alex Alderton, Roberto Amato, Jeffrey Barrett, Sonia Goncalves, Ewan Harrison, David K. Jackson, Ian Johnston, Dominic Kwiatkowski, Cordelia Langford, John Sillitoe on behalf of the Wellcome Sanger Institute COVID-19 Surveillance Team |
| EPI_ISL_3068605, EPI_ISL_3068606 | Lighthouse Laboratory Plymouth | Wellcome Sanger Institute for the COVID-19 Genomics UK (COG-UK) Consortium | Lighthouse Laboratory Plymouth and Alex Alderton, Roberto Amato, Jeffrey Barrett, Sonia Goncalves, Ewan Harrison, David K. Jackson, Ian Johnston, Dominic Kwiatkowski, Cordelia Langford, John Sillitoe on behalf of the Wellcome Sanger Institute COVID-19 Surveillance Team |
| EPI_ISL_3068607, EPI_ISL_3068608, EPI_ISL_3068609 | Lighthouse Lab in Alderley Park | Wellcome Sanger Institute for the COVID-19 Genomics UK (COG-UK) Consortium | Jacquelyn Wynn, Mairead Hyland, The Lighthouse Lab in Alderley Park and Alex Alderton, Roberto Amato, Jeffrey Barrett, Sonia Goncalves, Ewan Harrison, David K. Jackson, Ian Johnston, Dominic Kwiatkowski, Cordelia Langford, John Sillitoe on behalf of the Wellcome Sanger Institute COVID-19 Surveillance Team |
| EPI_ISL_3068610, EPI_ISL_3068611 | Lighthouse Laboratory Plymouth | Wellcome Sanger Institute for the COVID-19 Genomics UK (COG-UK) Consortium | Lighthouse Laboratory Plymouth and Alex Alderton, Roberto Amato, Jeffrey Barrett, Sonia Goncalves, Ewan Harrison, David K. Jackson, Ian Johnston, Dominic Kwiatkowski, Cordelia Langford, John Sillitoe on behalf of the Wellcome Sanger Institute COVID-19 Surveillance Team |
| EPI_ISL_3068612 | Lighthouse Lab in Alderley Park | Wellcome Sanger Institute for the COVID-19 Genomics UK (COG-UK) Consortium | Jacquelyn Wynn, Mairead Hyland, The Lighthouse Lab in Alderley Park and Alex Alderton, Roberto Amato, Jeffrey Barrett, Sonia Goncalves, Ewan Harrison, David K. Jackson, Ian Johnston, Dominic Kwiatkowski, Cordelia Langford, John Sillitoe on behalf of the Wellcome Sanger Institute COVID-19 Surveillance Team |
| EPI_ISL_3068613 | Lighthouse Laboratory Plymouth | Wellcome Sanger Institute for the COVID-19 Genomics UK (COG-UK) Consortium | Lighthouse Laboratory Plymouth and Alex Alderton, Roberto Amato, Jeffrey Barrett, Sonia Goncalves, Ewan Harrison, David K. Jackson, Ian Johnston, Dominic Kwiatkowski, Cordelia Langford, John Sillitoe on behalf of the Wellcome Sanger Institute COVID-19 Surveillance Team |
| EPI_ISL_3068614 | Lighthouse Lab in Alderley Park | Wellcome Sanger Institute for the COVID-19 Genomics UK (COG-UK) Consortium | Jacquelyn Wynn, Mairead Hyland, The Lighthouse Lab in Alderley Park and Alex Alderton, Roberto Amato, Jeffrey Barrett, Sonia Goncalves, Ewan Harrison, David K. Jackson, Ian Johnston, Dominic Kwiatkowski, Cordelia Langford, John Sillitoe on behalf of the Wellcome Sanger Institute COVID-19 Surveillance Team |
| EPI_ISL_3068615 | Health Services Laboratories | Wellcome Sanger Institute for the COVID-19 Genomics UK (COG-UK) Consortium | Health Services Laboratories and Alex Alderton, Roberto Amato, Jeffrey Barrett, Sonia Goncalves, Ewan Harrison, David K. Jackson, Ian Johnston, Dominic Kwiatkowski, Cordelia Langford, John Sillitoe on behalf of the Wellcome Sanger Institute COVID-19 Surveillance Team |
| EPI_ISL_3068616, EPI_ISL_3068617, EPI_ISL_3068618, EPI_ISL_3068619, EPI_ISL_3068620 | Lighthouse Laboratory Plymouth | Wellcome Sanger Institute for the COVID-19 Genomics UK (COG-UK) Consortium | Lighthouse Laboratory Plymouth and Alex Alderton, Roberto Amato, Jeffrey Barrett, Sonia Goncalves, Ewan Harrison, David K. Jackson, Ian Johnston, Dominic Kwiatkowski, Cordelia Langford, John Sillitoe on behalf of the Wellcome Sanger Institute COVID-19 Surveillance Team |
| EPI_ISL_3068621, EPI_ISL_3068622 | Lighthouse Lab in Alderley Park | Wellcome Sanger Institute for the COVID-19 Genomics UK (COG-UK) Consortium | Jacquelyn Wynn, Mairead Hyland, The Lighthouse Lab in Alderley Park and Alex Alderton, Roberto Amato, Jeffrey Barrett, Sonia Goncalves, Ewan Harrison, David K. Jackson, Ian Johnston, Dominic Kwiatkowski, Cordelia Langford, John Sillitoe on behalf of the Wellcome Sanger Institute COVID-19 Surveillance Team |
| EPI_ISL_3068623 | Lighthouse Laboratory Plymouth | Wellcome Sanger Institute for the COVID-19 Genomics UK (COG-UK) Consortium | Lighthouse Laboratory Plymouth and Alex Alderton, Roberto Amato, Jeffrey Barrett, Sonia Goncalves, Ewan Harrison, David K. Jackson, Ian Johnston, Dominic Kwiatkowski, Cordelia Langford, John Sillitoe on behalf of the Wellcome Sanger Institute COVID-19 Surveillance Team |
| EPI_ISL_3068624 | Lighthouse Lab in Alderley Park | Wellcome Sanger Institute for the COVID-19 Genomics UK (COG-UK) Consortium | Jacquelyn Wynn, Mairead Hyland, The Lighthouse Lab in Alderley Park and Alex Alderton, Roberto Amato, Jeffrey Barrett, Sonia Goncalves, Ewan Harrison, David K. Jackson, Ian Johnston, Dominic Kwiatkowski, Cordelia Langford, John Sillitoe on behalf of the Wellcome Sanger Institute COVID-19 Surveillance Team |
| EPI_ISL_3068625 | Health Services Laboratories | Wellcome Sanger Institute for the COVID-19 Genomics UK (COG-UK) Consortium | Health Services Laboratories and Alex Alderton, Roberto Amato, Jeffrey Barrett, Sonia Goncalves, Ewan Harrison, David K. Jackson, Ian Johnston, Dominic Kwiatkowski, Cordelia Langford, John Sillitoe on behalf of the Wellcome Sanger Institute COVID-19 Surveillance Team |
| EPI_ISL_3068626 | Lighthouse Laboratory Plymouth | Wellcome Sanger Institute for the COVID-19 Genomics UK (COG-UK) Consortium | Lighthouse Laboratory Plymouth and Alex Alderton, Roberto Amato, Jeffrey Barrett, Sonia Goncalves, Ewan Harrison, David K. Jackson, Ian Johnston, Dominic Kwiatkowski, Cordelia Langford, John Sillitoe on behalf of the Wellcome Sanger Institute COVID-19 Surveillance Team |
| EPI_ISL_3068627 | Health Services Laboratories | Wellcome Sanger Institute for the COVID-19 Genomics UK (COG-UK) Consortium | Health Services Laboratories and Alex Alderton, Roberto Amato, Jeffrey Barrett, Sonia Goncalves, Ewan Harrison, David K. Jackson, Ian Johnston, Dominic Kwiatkowski, Cordelia Langford, John Sillitoe on behalf of the Wellcome Sanger Institute COVID-19 Surveillance Team |
| EPI_ISL_3068628 | Lighthouse Lab in Alderley Park | Wellcome Sanger Institute for the COVID-19 Genomics UK (COG-UK) Consortium | Jacquelyn Wynn, Mairead Hyland, The Lighthouse Lab in Alderley Park and Alex Alderton, Roberto Amato, Jeffrey Barrett, Sonia Goncalves, Ewan Harrison, David K. Jackson, Ian Johnston, Dominic Kwiatkowski, Cordelia Langford, John Sillitoe on behalf of the Wellcome Sanger Institute COVID-19 Surveillance Team |
| EPI_ISL_3068629 | Lighthouse Laboratory Plymouth | Wellcome Sanger Institute for the COVID-19 Genomics UK (COG-UK) Consortium | Lighthouse Laboratory Plymouth and Alex Alderton, Roberto Amato, Jeffrey Barrett, Sonia Goncalves, Ewan Harrison, David K. Jackson, Ian Johnston, Dominic Kwiatkowski, Cordelia Langford, John Sillitoe on behalf of the Wellcome Sanger Institute COVID-19 Surveillance Team |
| EPI_ISL_3068630 | Health Services Laboratories | Wellcome Sanger Institute for the COVID-19 Genomics UK (COG-UK) Consortium | Health Services Laboratories and Alex Alderton, Roberto Amato, Jeffrey Barrett, Sonia Goncalves, Ewan Harrison, David K. Jackson, Ian Johnston, Dominic Kwiatkowski, Cordelia Langford, John Sillitoe on behalf of the Wellcome Sanger Institute COVID-19 Surveillance Team |
| EPI_ISL_3068631, EPI_ISL_3068632, EPI_ISL_3068633 | Lighthouse Lab in Alderley Park | Wellcome Sanger Institute for the COVID-19 Genomics UK (COG-UK) Consortium | Jacquelyn Wynn, Mairead Hyland, The Lighthouse Lab in Alderley Park and Alex Alderton, Roberto Amato, Jeffrey Barrett, Sonia Goncalves, Ewan Harrison, David K. Jackson, Ian Johnston, Dominic Kwiatkowski, Cordelia Langford, John Sillitoe on behalf of the Wellcome Sanger Institute COVID-19 Surveillance Team |
| EPI_ISL_3068634, EPI_ISL_3068635, EPI_ISL_3068636, EPI_ISL_3068637 | Lighthouse Laboratory Plymouth | Wellcome Sanger Institute for the COVID-19 Genomics UK (COG-UK) Consortium | Lighthouse Laboratory Plymouth and Alex Alderton, Roberto Amato, Jeffrey Barrett, Sonia Goncalves, Ewan Harrison, David K. Jackson, Ian Johnston, Dominic Kwiatkowski, Cordelia Langford, John Sillitoe on behalf of the Wellcome Sanger Institute COVID-19 Surveillance Team |
| EPI_ISL_3068638 | Health Services Laboratories | Wellcome Sanger Institute for the COVID-19 Genomics UK (COG-UK) Consortium | Health Services Laboratories and Alex Alderton, Roberto Amato, Jeffrey Barrett, Sonia Goncalves, Ewan Harrison, David K. Jackson, Ian Johnston, Dominic Kwiatkowski, Cordelia Langford, John Sillitoe on behalf of the Wellcome Sanger Institute COVID-19 Surveillance Team |
| EPI_ISL_3068639 | Lighthouse Laboratory Plymouth | Wellcome Sanger Institute for the COVID-19 Genomics UK (COG-UK) Consortium | Lighthouse Laboratory Plymouth and Alex Alderton, Roberto Amato, Jeffrey Barrett, Sonia Goncalves, Ewan Harrison, David K. Jackson, Ian Johnston, Dominic Kwiatkowski, Cordelia Langford, John Sillitoe on behalf of the Wellcome Sanger Institute COVID-19 Surveillance Team |
| EPI_ISL_3068640 | Lighthouse Lab in Alderley Park | Wellcome Sanger Institute for the COVID-19 Genomics UK (COG-UK) Consortium | Jacquelyn Wynn, Mairead Hyland, The Lighthouse Lab in Alderley Park and Alex Alderton, Roberto Amato, Jeffrey Barrett, Sonia Goncalves, Ewan Harrison, David K. Jackson, Ian Johnston, Dominic Kwiatkowski, Cordelia Langford, John Sillitoe on behalf of the Wellcome Sanger Institute COVID-19 Surveillance Team |
| EPI_ISL_3068641, EPI_ISL_3068642, EPI_ISL_3068643 | Lighthouse Laboratory Plymouth | Wellcome Sanger Institute for the COVID-19 Genomics UK (COG-UK) Consortium | Lighthouse Laboratory Plymouth and Alex Alderton, Roberto Amato, Jeffrey Barrett, Sonia Goncalves, Ewan Harrison, David K. Jackson, Ian Johnston, Dominic Kwiatkowski, Cordelia Langford, John Sillitoe on behalf of the Wellcome Sanger Institute COVID-19 Surveillance Team |
| EPI_ISL_3068644 | Lighthouse Lab in Alderley Park | Wellcome Sanger Institute for the COVID-19 Genomics UK (COG-UK) Consortium | Jacquelyn Wynn, Mairead Hyland, The Lighthouse Lab in Alderley Park and Alex Alderton, Roberto Amato, Jeffrey Barrett, Sonia Goncalves, Ewan Harrison, David K. Jackson, Ian Johnston, Dominic Kwiatkowski, Cordelia Langford, John Sillitoe on behalf of the Wellcome Sanger Institute COVID-19 Surveillance Team |
| EPI_ISL_3068645 | Lighthouse Laboratory Plymouth | Wellcome Sanger Institute for the COVID-19 Genomics UK (COG-UK) Consortium | Lighthouse Laboratory Plymouth and Alex Alderton, Roberto Amato, Jeffrey Barrett, Sonia Goncalves, Ewan Harrison, David K. Jackson, Ian Johnston, Dominic Kwiatkowski, Cordelia Langford, John Sillitoe on behalf of the Wellcome Sanger Institute COVID-19 Surveillance Team |
| EPI_ISL_3068646 | Lighthouse Lab in Alderley Park | Wellcome Sanger Institute for the COVID-19 Genomics UK (COG-UK) Consortium | Jacquelyn Wynn, Mairead Hyland, The Lighthouse Lab in Alderley Park and Alex Alderton, Roberto Amato, Jeffrey Barrett, Sonia Goncalves, Ewan Harrison, David K. Jackson, Ian Johnston, Dominic Kwiatkowski, Cordelia Langford, John Sillitoe on behalf of the Wellcome Sanger Institute COVID-19 Surveillance Team |
| EPI_ISL_3068647 | Health Services Laboratories | Wellcome Sanger Institute for the COVID-19 Genomics UK (COG-UK) Consortium | Health Services Laboratories and Alex Alderton, Roberto Amato, Jeffrey Barrett, Sonia Goncalves, Ewan Harrison, David K. Jackson, Ian Johnston, Dominic Kwiatkowski, Cordelia Langford, John Sillitoe on behalf of the Wellcome Sanger Institute COVID-19 Surveillance Team |
| EPI_ISL_3068648 | Lighthouse Laboratory Plymouth | Wellcome Sanger Institute for the COVID-19 Genomics UK (COG-UK) Consortium | Lighthouse Laboratory Plymouth and Alex Alderton, Roberto Amato, Jeffrey Barrett, Sonia Goncalves, Ewan Harrison, David K. Jackson, Ian Johnston, Dominic Kwiatkowski, Cordelia Langford, John Sillitoe on behalf of the Wellcome Sanger Institute COVID-19 Surveillance Team |
| EPI_ISL_3068649 | Health Services Laboratories | Wellcome Sanger Institute for the COVID-19 Genomics UK (COG-UK) Consortium | Health Services Laboratories and Alex Alderton, Roberto Amato, Jeffrey Barrett, Sonia Goncalves, Ewan Harrison, David K. Jackson, Ian Johnston, Dominic Kwiatkowski, Cordelia Langford, John Sillitoe on behalf of the Wellcome Sanger Institute COVID-19 Surveillance Team |
| EPI_ISL_3068650 | Lighthouse Laboratory Plymouth | Wellcome Sanger Institute for the COVID-19 Genomics UK (COG-UK) Consortium | Lighthouse Laboratory Plymouth and Alex Alderton, Roberto Amato, Jeffrey Barrett, Sonia Goncalves, Ewan Harrison, David K. Jackson, Ian Johnston, Dominic Kwiatkowski, Cordelia Langford, John Sillitoe on behalf of the Wellcome Sanger Institute COVID-19 Surveillance Team |
| EPI_ISL_3068651 | Health Services Laboratories | Wellcome Sanger Institute for the COVID-19 Genomics UK (COG-UK) Consortium | Health Services Laboratories and Alex Alderton, Roberto Amato, Jeffrey Barrett, Sonia Goncalves, Ewan Harrison, David K. Jackson, Ian Johnston, Dominic Kwiatkowski, Cordelia Langford, John Sillitoe on behalf of the Wellcome Sanger Institute COVID-19 Surveillance Team |
| EPI_ISL_3068652 | Lighthouse Lab in Alderley Park | Wellcome Sanger Institute for the COVID-19 Genomics UK (COG-UK) Consortium | Jacquelyn Wynn, Mairead Hyland, The Lighthouse Lab in Alderley Park and Alex Alderton, Roberto Amato, Jeffrey Barrett, Sonia Goncalves, Ewan Harrison, David K. Jackson, Ian Johnston, Dominic Kwiatkowski, Cordelia Langford, John Sillitoe on behalf of the Wellcome Sanger Institute COVID-19 Surveillance Team |
| EPI_ISL_3068653 | Health Services Laboratories | Wellcome Sanger Institute for the COVID-19 Genomics UK (COG-UK) Consortium | Health Services Laboratories and Alex Alderton, Roberto Amato, Jeffrey Barrett, Sonia Goncalves, Ewan Harrison, David K. Jackson, Ian Johnston, Dominic Kwiatkowski, Cordelia Langford, John Sillitoe on behalf of the Wellcome Sanger Institute COVID-19 Surveillance Team |
| EPI_ISL_3068654, EPI_ISL_3068655 | Lighthouse Laboratory Plymouth | Wellcome Sanger Institute for the COVID-19 Genomics UK (COG-UK) Consortium | Lighthouse Laboratory Plymouth and Alex Alderton, Roberto Amato, Jeffrey Barrett, Sonia Goncalves, Ewan Harrison, David K. Jackson, Ian Johnston, Dominic Kwiatkowski, Cordelia Langford, John Sillitoe on behalf of the Wellcome Sanger Institute COVID-19 Surveillance Team |
| EPI_ISL_3068656, EPI_ISL_3068657 | Lighthouse Lab in Alderley Park | Wellcome Sanger Institute for the COVID-19 Genomics UK (COG-UK) Consortium | Jacquelyn Wynn, Mairead Hyland, The Lighthouse Lab in Alderley Park and Alex Alderton, Roberto Amato, Jeffrey Barrett, Sonia Goncalves, Ewan Harrison, David K. Jackson, Ian Johnston, Dominic Kwiatkowski, Cordelia Langford, John Sillitoe on behalf of the Wellcome Sanger Institute COVID-19 Surveillance Team |
| EPI_ISL_3068658 | Health Services Laboratories | Wellcome Sanger Institute for the COVID-19 Genomics UK (COG-UK) Consortium | Health Services Laboratories and Alex Alderton, Roberto Amato, Jeffrey Barrett, Sonia Goncalves, Ewan Harrison, David K. Jackson, Ian Johnston, Dominic Kwiatkowski, Cordelia Langford, John Sillitoe on behalf of the Wellcome Sanger Institute COVID-19 Surveillance Team |
| EPI_ISL_3068659 | Lighthouse Laboratory Plymouth | Wellcome Sanger Institute for the COVID-19 Genomics UK (COG-UK) Consortium | Lighthouse Laboratory Plymouth and Alex Alderton, Roberto Amato, Jeffrey Barrett, Sonia Goncalves, Ewan Harrison, David K. Jackson, Ian Johnston, Dominic Kwiatkowski, Cordelia Langford, John Sillitoe on behalf of the Wellcome Sanger Institute COVID-19 Surveillance Team |
| EPI_ISL_3068660 | Lighthouse Lab in Alderley Park | Wellcome Sanger Institute for the COVID-19 Genomics UK (COG-UK) Consortium | Jacquelyn Wynn, Mairead Hyland, The Lighthouse Lab in Alderley Park and Alex Alderton, Roberto Amato, Jeffrey Barrett, Sonia Goncalves, Ewan Harrison, David K. Jackson, Ian Johnston, Dominic Kwiatkowski, Cordelia Langford, John Sillitoe on behalf of the Wellcome Sanger Institute COVID-19 Surveillance Team |
| EPI_ISL_3068661, EPI_ISL_3068662, | Lighthouse Laboratory Plymouth | Wellcome Sanger Institute for the COVID-19 Genomics UK (COG-UK) Consortium | Lighthouse Laboratory Plymouth and Alex Alderton, Roberto Amato, Jeffrey Barrett, Sonia Goncalves, Ewan Harrison, David K. Jackson, Ian Johnston, Dominic Kwiatkowski, Cordelia Langford, John Sillitoe on behalf of the Wellcome Sanger Institute COVID-19 Surveillance Team |

| | | | |
|---|---|---|---|
| EPI_ISL_3068663 | | | |
| EPI_ISL_3068664 | Lighthouse Lab in Alderley Park | Wellcome Sanger Institute for the COVID-19 Genomics UK (COG-UK) Consortium | Jacquelyn Wynn, Mairead Hyland, The Lighthouse Lab in Alderley Park and Alex Alderton, Roberto Amato, Jeffrey Barrett, Sonia Goncalves, Ewan Harrison, David K. Jackson, Ian Johnston, Dominic Kwiatkowski, Cordelia Langford, John Sillitoe on behalf of the Wellcome Sanger Institute COVID-19 Surveillance Team |
| EPI_ISL_3068665 | Lighthouse Laboratory Plymouth | Wellcome Sanger Institute for the COVID-19 Genomics UK (COG-UK) Consortium | Lighthouse Laboratory Plymouth and Alex Alderton, Roberto Amato, Jeffrey Barrett, Sonia Goncalves, Ewan Harrison, David K. Jackson, Ian Johnston, Dominic Kwiatkowski, Cordelia Langford, John Sillitoe on behalf of the Wellcome Sanger Institute COVID-19 Surveillance Team |
| EPI_ISL_3068666 | Berkshire and Surrey Pathology Services Lighthouse Laboratory | Wellcome Sanger Institute for the COVID-19 Genomics UK (COG-UK) Consortium | Berkshire and Surrey Pathology Services Lighthouse Laboratory and Alex Alderton, Roberto Amato, Jeffrey Barrett, Sonia Goncalves, Ewan Harrison, David K. Jackson, Ian Johnston, Dominic Kwiatkowski, Cordelia Langford, John Sillitoe on behalf of the Wellcome Sanger Institute COVID-19 Surveillance Team |
| EPI_ISL_3068667, EPI_ISL_3068668 | Lighthouse Laboratory Plymouth | Wellcome Sanger Institute for the COVID-19 Genomics UK (COG-UK) Consortium | Lighthouse Laboratory Plymouth and Alex Alderton, Roberto Amato, Jeffrey Barrett, Sonia Goncalves, Ewan Harrison, David K. Jackson, Ian Johnston, Dominic Kwiatkowski, Cordelia Langford, John Sillitoe on behalf of the Wellcome Sanger Institute COVID-19 Surveillance Team |
| EPI_ISL_3068669 | Lighthouse Lab in Alderley Park | Wellcome Sanger Institute for the COVID-19 Genomics UK (COG-UK) Consortium | Jacquelyn Wynn, Mairead Hyland, The Lighthouse Lab in Alderley Park and Alex Alderton, Roberto Amato, Jeffrey Barrett, Sonia Goncalves, Ewan Harrison, David K. Jackson, Ian Johnston, Dominic Kwiatkowski, Cordelia Langford, John Sillitoe on behalf of the Wellcome Sanger Institute COVID-19 Surveillance Team |
| EPI_ISL_3068670, EPI_ISL_3068671 | Lighthouse Laboratory Plymouth | Wellcome Sanger Institute for the COVID-19 Genomics UK (COG-UK) Consortium | Lighthouse Laboratory Plymouth and Alex Alderton, Roberto Amato, Jeffrey Barrett, Sonia Goncalves, Ewan Harrison, David K. Jackson, Ian Johnston, Dominic Kwiatkowski, Cordelia Langford, John Sillitoe on behalf of the Wellcome Sanger Institute COVID-19 Surveillance Team |
| EPI_ISL_3068672 | Berkshire and Surrey Pathology Services Lighthouse Laboratory | Wellcome Sanger Institute for the COVID-19 Genomics UK (COG-UK) Consortium | Berkshire and Surrey Pathology Services Lighthouse Laboratory and Alex Alderton, Roberto Amato, Jeffrey Barrett, Sonia Goncalves, Ewan Harrison, David K. Jackson, Ian Johnston, Dominic Kwiatkowski, Cordelia Langford, John Sillitoe on behalf of the Wellcome Sanger Institute COVID-19 Surveillance Team |
| EPI_ISL_3068673 | Lighthouse Laboratory Plymouth | Wellcome Sanger Institute for the COVID-19 Genomics UK (COG-UK) Consortium | Lighthouse Laboratory Plymouth and Alex Alderton, Roberto Amato, Jeffrey Barrett, Sonia Goncalves, Ewan Harrison, David K. Jackson, Ian Johnston, Dominic Kwiatkowski, Cordelia Langford, John Sillitoe on behalf of the Wellcome Sanger Institute COVID-19 Surveillance Team |
| EPI_ISL_3068674 | Health Services Laboratories | Wellcome Sanger Institute for the COVID-19 Genomics UK (COG-UK) Consortium | Health Services Laboratories and Alex Alderton, Roberto Amato, Jeffrey Barrett, Sonia Goncalves, Ewan Harrison, David K. Jackson, Ian Johnston, Dominic Kwiatkowski, Cordelia Langford, John Sillitoe on behalf of the Wellcome Sanger Institute COVID-19 Surveillance Team |
| EPI_ISL_3068675, EPI_ISL_3068676 | Lighthouse Lab in Alderley Park | Wellcome Sanger Institute for the COVID-19 Genomics UK (COG-UK) Consortium | Jacquelyn Wynn, Mairead Hyland, The Lighthouse Lab in Alderley Park and Alex Alderton, Roberto Amato, Jeffrey Barrett, Sonia Goncalves, Ewan Harrison, David K. Jackson, Ian Johnston, Dominic Kwiatkowski, Cordelia Langford, John Sillitoe on behalf of the Wellcome Sanger Institute COVID-19 Surveillance Team |
| EPI_ISL_3068677, EPI_ISL_3068678 | Lighthouse Laboratory Plymouth | Wellcome Sanger Institute for the COVID-19 Genomics UK (COG-UK) Consortium | Lighthouse Laboratory Plymouth and Alex Alderton, Roberto Amato, Jeffrey Barrett, Sonia Goncalves, Ewan Harrison, David K. Jackson, Ian Johnston, Dominic Kwiatkowski, Cordelia Langford, John Sillitoe on behalf of the Wellcome Sanger Institute COVID-19 Surveillance Team |
| EPI_ISL_3068679 | Lighthouse Lab in Alderley Park | Wellcome Sanger Institute for the COVID-19 Genomics UK (COG-UK) Consortium | Jacquelyn Wynn, Mairead Hyland, The Lighthouse Lab in Alderley Park and Alex Alderton, Roberto Amato, Jeffrey Barrett, Sonia Goncalves, Ewan Harrison, David K. Jackson, Ian Johnston, Dominic Kwiatkowski, Cordelia Langford, John Sillitoe on behalf of the Wellcome Sanger Institute COVID-19 Surveillance Team |
| EPI_ISL_3068680, EPI_ISL_3068681 | Lighthouse Laboratory Plymouth | Wellcome Sanger Institute for the COVID-19 Genomics UK (COG-UK) Consortium | Lighthouse Laboratory Plymouth and Alex Alderton, Roberto Amato, Jeffrey Barrett, Sonia Goncalves, Ewan Harrison, David K. Jackson, Ian Johnston, Dominic Kwiatkowski, Cordelia Langford, John Sillitoe on behalf of the Wellcome Sanger Institute COVID-19 Surveillance Team |
| EPI_ISL_3068682 | Lighthouse Lab in Alderley Park | Wellcome Sanger Institute for the COVID-19 Genomics UK (COG-UK) Consortium | Jacquelyn Wynn, Mairead Hyland, The Lighthouse Lab in Alderley Park and Alex Alderton, Roberto Amato, Jeffrey Barrett, Sonia Goncalves, Ewan Harrison, David K. Jackson, Ian Johnston, Dominic Kwiatkowski, Cordelia Langford, John Sillitoe on behalf of the Wellcome Sanger Institute COVID-19 Surveillance Team |
| EPI_ISL_3068683 | Lighthouse Laboratory Plymouth | Wellcome Sanger Institute for the COVID-19 Genomics UK (COG-UK) Consortium | Lighthouse Laboratory Plymouth and Alex Alderton, Roberto Amato, Jeffrey Barrett, Sonia Goncalves, Ewan Harrison, David K. Jackson, Ian Johnston, Dominic Kwiatkowski, Cordelia Langford, John Sillitoe on behalf of the Wellcome Sanger Institute COVID-19 Surveillance Team |
| EPI_ISL_3068684 | Lighthouse Lab in Alderley Park | Wellcome Sanger Institute for the COVID-19 Genomics UK (COG-UK) Consortium | Jacquelyn Wynn, Mairead Hyland, The Lighthouse Lab in Alderley Park and Alex Alderton, Roberto Amato, Jeffrey Barrett, Sonia Goncalves, Ewan Harrison, David K. Jackson, Ian Johnston, Dominic Kwiatkowski, Cordelia Langford, John Sillitoe on behalf of the Wellcome Sanger Institute COVID-19 Surveillance Team |
| EPI_ISL_3068685 | Lighthouse Laboratory Plymouth | Wellcome Sanger Institute for the COVID-19 Genomics UK (COG-UK) Consortium | Lighthouse Laboratory Plymouth and Alex Alderton, Roberto Amato, Jeffrey Barrett, Sonia Goncalves, Ewan Harrison, David K. Jackson, Ian Johnston, Dominic Kwiatkowski, Cordelia Langford, John Sillitoe on behalf of the Wellcome Sanger Institute COVID-19 Surveillance Team |
| EPI_ISL_3068686, EPI_ISL_3068687 | Health Services Laboratories | Wellcome Sanger Institute for the COVID-19 Genomics UK (COG-UK) Consortium | Health Services Laboratories and Alex Alderton, Roberto Amato, Jeffrey Barrett, Sonia Goncalves, Ewan Harrison, David K. Jackson, Ian Johnston, Dominic Kwiatkowski, Cordelia Langford, John Sillitoe on behalf of the Wellcome Sanger Institute COVID-19 Surveillance Team |
| EPI_ISL_3068688, EPI_ISL_3068689, EPI_ISL_3068690 | Lighthouse Laboratory Plymouth | Wellcome Sanger Institute for the COVID-19 Genomics UK (COG-UK) Consortium | Lighthouse Laboratory Plymouth and Alex Alderton, Roberto Amato, Jeffrey Barrett, Sonia Goncalves, Ewan Harrison, David K. Jackson, Ian Johnston, Dominic Kwiatkowski, Cordelia Langford, John Sillitoe on behalf of the Wellcome Sanger Institute COVID-19 Surveillance Team |
| EPI_ISL_3068691, EPI_ISL_3068692 | Lighthouse Lab in Alderley Park | Wellcome Sanger Institute for the COVID-19 Genomics UK (COG-UK) Consortium | Jacquelyn Wynn, Mairead Hyland, The Lighthouse Lab in Alderley Park and Alex Alderton, Roberto Amato, Jeffrey Barrett, Sonia Goncalves, Ewan Harrison, David K. Jackson, Ian Johnston, Dominic Kwiatkowski, Cordelia Langford, John Sillitoe on behalf of the Wellcome Sanger Institute COVID-19 Surveillance Team |
| EPI_ISL_3068693, EPI_ISL_3068694, EPI_ISL_3068695 | Lighthouse Laboratory Plymouth | Wellcome Sanger Institute for the COVID-19 Genomics UK (COG-UK) Consortium | Lighthouse Laboratory Plymouth and Alex Alderton, Roberto Amato, Jeffrey Barrett, Sonia Goncalves, Ewan Harrison, David K. Jackson, Ian Johnston, Dominic Kwiatkowski, Cordelia Langford, John Sillitoe on behalf of the Wellcome Sanger Institute COVID-19 Surveillance Team |
| EPI_ISL_3068696 | Lighthouse Lab in Alderley Park | Wellcome Sanger Institute for the COVID-19 Genomics UK (COG-UK) Consortium | Jacquelyn Wynn, Mairead Hyland, The Lighthouse Lab in Alderley Park and Alex Alderton, Roberto Amato, Jeffrey Barrett, Sonia Goncalves, Ewan Harrison, David K. Jackson, Ian Johnston, Dominic Kwiatkowski, Cordelia Langford, John Sillitoe on behalf of the Wellcome Sanger Institute COVID-19 Surveillance Team |
| EPI_ISL_3068697, EPI_ISL_3068698, EPI_ISL_3068699, EPI_ISL_3068700, EPI_ISL_3068701 | Lighthouse Laboratory Plymouth | Wellcome Sanger Institute for the COVID-19 Genomics UK (COG-UK) Consortium | Lighthouse Laboratory Plymouth and Alex Alderton, Roberto Amato, Jeffrey Barrett, Sonia Goncalves, Ewan Harrison, David K. Jackson, Ian Johnston, Dominic Kwiatkowski, Cordelia Langford, John Sillitoe on behalf of the Wellcome Sanger Institute COVID-19 Surveillance Team |
| EPI_ISL_3068702 | Health Services Laboratories | Wellcome Sanger Institute for the COVID-19 Genomics UK (COG-UK) Consortium | Health Services Laboratories and Alex Alderton, Roberto Amato, Jeffrey Barrett, Sonia Goncalves, Ewan Harrison, David K. Jackson, Ian Johnston, Dominic Kwiatkowski, Cordelia Langford, John Sillitoe on behalf of the Wellcome Sanger Institute COVID-19 Surveillance Team |
| EPI_ISL_3068703, EPI_ISL_3068704, EPI_ISL_3068705, EPI_ISL_3068706 | Lighthouse Laboratory Plymouth | Wellcome Sanger Institute for the COVID-19 Genomics UK (COG-UK) Consortium | Lighthouse Laboratory Plymouth and Alex Alderton, Roberto Amato, Jeffrey Barrett, Sonia Goncalves, Ewan Harrison, David K. Jackson, Ian Johnston, Dominic Kwiatkowski, Cordelia Langford, John Sillitoe on behalf of the Wellcome Sanger Institute COVID-19 Surveillance Team |
| EPI_ISL_3068707 | Health Services Laboratories | Wellcome Sanger Institute for the COVID-19 Genomics UK (COG-UK) Consortium | Health Services Laboratories and Alex Alderton, Roberto Amato, Jeffrey Barrett, Sonia Goncalves, Ewan Harrison, David K. Jackson, Ian Johnston, Dominic Kwiatkowski, Cordelia Langford, John Sillitoe on behalf of the Wellcome Sanger Institute COVID-19 Surveillance Team |
| EPI_ISL_3068708, EPI_ISL_3068709 | Lighthouse Laboratory Plymouth | Wellcome Sanger Institute for the COVID-19 Genomics UK (COG-UK) Consortium | Lighthouse Laboratory Plymouth and Alex Alderton, Roberto Amato, Jeffrey Barrett, Sonia Goncalves, Ewan Harrison, David K. Jackson, Ian Johnston, Dominic Kwiatkowski, Cordelia Langford, John Sillitoe on behalf of the Wellcome Sanger Institute COVID-19 Surveillance Team |
| EPI_ISL_3068710 | Lighthouse Lab in Alderley Park | Wellcome Sanger Institute for the COVID-19 Genomics UK (COG-UK) Consortium | Jacquelyn Wynn, Mairead Hyland, The Lighthouse Lab in Alderley Park and Alex Alderton, Roberto Amato, Jeffrey Barrett, Sonia Goncalves, Ewan Harrison, David K. Jackson, Ian Johnston, Dominic Kwiatkowski, Cordelia Langford, John Sillitoe on behalf of the Wellcome Sanger Institute COVID-19 Surveillance Team |
| EPI_ISL_3068711 | Lighthouse Laboratory Plymouth | Wellcome Sanger Institute for the COVID-19 Genomics UK (COG-UK) Consortium | Lighthouse Laboratory Plymouth and Alex Alderton, Roberto Amato, Jeffrey Barrett, Sonia Goncalves, Ewan Harrison, David K. Jackson, Ian Johnston, Dominic Kwiatkowski, Cordelia Langford, John Sillitoe on behalf of the Wellcome Sanger Institute COVID-19 Surveillance Team |
| EPI_ISL_3068712, EPI_ISL_3068713 | Health Services Laboratories | Wellcome Sanger Institute for the COVID-19 Genomics UK (COG-UK) Consortium | Health Services Laboratories and Alex Alderton, Roberto Amato, Jeffrey Barrett, Sonia Goncalves, Ewan Harrison, David K. Jackson, Ian Johnston, Dominic Kwiatkowski, Cordelia Langford, John Sillitoe on behalf of the Wellcome Sanger Institute COVID-19 Surveillance Team |
| EPI_ISL_3068714, EPI_ISL_3068715 | Lighthouse Lab in Alderley Park | Wellcome Sanger Institute for the COVID-19 Genomics UK (COG-UK) Consortium | Jacquelyn Wynn, Mairead Hyland, The Lighthouse Lab in Alderley Park and Alex Alderton, Roberto Amato, Jeffrey Barrett, Sonia Goncalves, Ewan Harrison, David K. Jackson, Ian Johnston, Dominic Kwiatkowski, Cordelia Langford, John Sillitoe on behalf of the Wellcome Sanger Institute COVID-19 Surveillance Team |
| EPI_ISL_3068716 | Berkshire and Surrey Pathology Services Lighthouse Laboratory | Wellcome Sanger Institute for the COVID-19 Genomics UK (COG-UK) Consortium | Berkshire and Surrey Pathology Services Lighthouse Laboratory and Alex Alderton, Roberto Amato, Jeffrey Barrett, Sonia Goncalves, Ewan Harrison, David K. Jackson, Ian Johnston, Dominic Kwiatkowski, Cordelia Langford, John Sillitoe on behalf of the Wellcome Sanger Institute COVID-19 Surveillance Team |
| EPI_ISL_3068717 | Health Services Laboratories | Wellcome Sanger Institute for the COVID-19 Genomics UK (COG-UK) Consortium | Health Services Laboratories and Alex Alderton, Roberto Amato, Jeffrey Barrett, Sonia Goncalves, Ewan Harrison, David K. Jackson, Ian Johnston, Dominic Kwiatkowski, Cordelia Langford, John Sillitoe on behalf of the Wellcome Sanger Institute COVID-19 Surveillance Team |
| EPI_ISL_3068718 | Lighthouse Laboratory Plymouth | Wellcome Sanger Institute for the COVID-19 Genomics UK (COG-UK) Consortium | Lighthouse Laboratory Plymouth and Alex Alderton, Roberto Amato, Jeffrey Barrett, Sonia Goncalves, Ewan Harrison, David K. Jackson, Ian Johnston, Dominic Kwiatkowski, Cordelia Langford, John Sillitoe on behalf of the Wellcome Sanger Institute COVID-19 Surveillance Team |
| EPI_ISL_3068719 | Lighthouse Lab in Alderley Park | Wellcome Sanger Institute for the COVID-19 Genomics UK (COG-UK) Consortium | Jacquelyn Wynn, Mairead Hyland, The Lighthouse Lab in Alderley Park and Alex Alderton, Roberto Amato, Jeffrey Barrett, Sonia Goncalves, Ewan Harrison, David K. Jackson, Ian Johnston, Dominic Kwiatkowski, Cordelia Langford, John Sillitoe on behalf of the Wellcome Sanger Institute COVID-19 Surveillance Team |
| EPI_ISL_3068720 | Health Services Laboratories | Wellcome Sanger Institute for the COVID-19 Genomics UK (COG-UK) Consortium | Health Services Laboratories and Alex Alderton, Roberto Amato, Jeffrey Barrett, Sonia Goncalves, Ewan Harrison, David K. Jackson, Ian Johnston, Dominic Kwiatkowski, Cordelia Langford, John Sillitoe on behalf of the Wellcome Sanger Institute COVID-19 Surveillance Team |
| EPI_ISL_3068721 | Lighthouse Laboratory Plymouth | Wellcome Sanger Institute for the COVID-19 Genomics UK (COG-UK) Consortium | Lighthouse Laboratory Plymouth and Alex Alderton, Roberto Amato, Jeffrey Barrett, Sonia Goncalves, Ewan Harrison, David K. Jackson, Ian Johnston, Dominic Kwiatkowski, Cordelia Langford, John Sillitoe on behalf of the Wellcome Sanger Institute COVID-19 Surveillance Team |
| EPI_ISL_3068722 | Health Services Laboratories | Wellcome Sanger Institute for the COVID-19 Genomics UK (COG-UK) Consortium | Health Services Laboratories and Alex Alderton, Roberto Amato, Jeffrey Barrett, Sonia Goncalves, Ewan Harrison, David K. Jackson, Ian Johnston, Dominic Kwiatkowski, Cordelia Langford, John Sillitoe on behalf of the Wellcome Sanger Institute COVID-19 Surveillance Team |
| EPI_ISL_3068723, EPI_ISL_3068724, EPI_ISL_3068725, EPI_ISL_3068726, EPI_ISL_3068727 | Lighthouse Laboratory Plymouth | Wellcome Sanger Institute for the COVID-19 Genomics UK (COG-UK) Consortium | Lighthouse Laboratory Plymouth and Alex Alderton, Roberto Amato, Jeffrey Barrett, Sonia Goncalves, Ewan Harrison, David K. Jackson, Ian Johnston, Dominic Kwiatkowski, Cordelia Langford, John Sillitoe on behalf of the Wellcome Sanger Institute COVID-19 Surveillance Team |
| EPI_ISL_3068728 | Lighthouse Lab in Alderley Park | Wellcome Sanger Institute for the COVID-19 Genomics UK (COG-UK) Consortium | Jacquelyn Wynn, Mairead Hyland, The Lighthouse Lab in Alderley Park and Alex Alderton, Roberto Amato, Jeffrey Barrett, Sonia Goncalves, Ewan Harrison, David K. Jackson, Ian Johnston, Dominic Kwiatkowski, Cordelia Langford, John Sillitoe on behalf of the Wellcome Sanger Institute COVID-19 Surveillance Team |
| EPI_ISL_3068729, | Lighthouse Laboratory Plymouth | Wellcome Sanger Institute for the COVID-19 Genomics UK (COG-UK) | Lighthouse Laboratory Plymouth and Alex Alderton, Roberto Amato, Jeffrey Barrett, Sonia Goncalves, Ewan Harrison, David K. Jackson, Ian Johnston, Dominic Kwiatkowski, Cordelia Langford, John Sillitoe on behalf of the Wellcome Sanger |

| | | | |
|---|---|---|---|
| EPI_ISL_3068730 | | Consortium | Institute COVID-19 Surveillance Team |
| EPI_ISL_3068731 | Health Services Laboratories | Wellcome Sanger Institute for the COVID-19 Genomics UK (COG-UK) Consortium | Health Services Laboratories and Alex Alderton, Roberto Amato, Jeffrey Barrett, Sonia Goncalves, Ewan Harrison, David K. Jackson, Ian Johnston, Dominic Kwiatkowski, Cordelia Langford, John Sillitoe on behalf of the Wellcome Sanger Institute COVID-19 Surveillance Team |
| EPI_ISL_3068732, EPI_ISL_3068733, EPI_ISL_3068734 | Lighthouse Laboratory Plymouth | Wellcome Sanger Institute for the COVID-19 Genomics UK (COG-UK) Consortium | Lighthouse Laboratory Plymouth and Alex Alderton, Roberto Amato, Jeffrey Barrett, Sonia Goncalves, Ewan Harrison, David K. Jackson, Ian Johnston, Dominic Kwiatkowski, Cordelia Langford, John Sillitoe on behalf of the Wellcome Sanger Institute COVID-19 Surveillance Team |
| EPI_ISL_3068735, EPI_ISL_3068736 | Lighthouse Lab in Alderley Park | Wellcome Sanger Institute for the COVID-19 Genomics UK (COG-UK) Consortium | Jacquelyn Wynn, Mairead Hyland, The Lighthouse Lab in Alderley Park and Alex Alderton, Roberto Amato, Jeffrey Barrett, Sonia Goncalves, Ewan Harrison, David K. Jackson, Ian Johnston, Dominic Kwiatkowski, Cordelia Langford, John Sillitoe on behalf of the Wellcome Sanger Institute COVID-19 Surveillance Team |
| EPI_ISL_3068737 | Health Services Laboratories | Wellcome Sanger Institute for the COVID-19 Genomics UK (COG-UK) Consortium | Health Services Laboratories and Alex Alderton, Roberto Amato, Jeffrey Barrett, Sonia Goncalves, Ewan Harrison, David K. Jackson, Ian Johnston, Dominic Kwiatkowski, Cordelia Langford, John Sillitoe on behalf of the Wellcome Sanger Institute COVID-19 Surveillance Team |
| EPI_ISL_3068738, EPI_ISL_3068739, EPI_ISL_3068740, EPI_ISL_3068741, EPI_ISL_3068742, EPI_ISL_3068743 | Lighthouse Laboratory Plymouth | Wellcome Sanger Institute for the COVID-19 Genomics UK (COG-UK) Consortium | Lighthouse Laboratory Plymouth and Alex Alderton, Roberto Amato, Jeffrey Barrett, Sonia Goncalves, Ewan Harrison, David K. Jackson, Ian Johnston, Dominic Kwiatkowski, Cordelia Langford, John Sillitoe on behalf of the Wellcome Sanger Institute COVID-19 Surveillance Team |
| EPI_ISL_3068744 | Health Services Laboratories | Wellcome Sanger Institute for the COVID-19 Genomics UK (COG-UK) Consortium | Health Services Laboratories and Alex Alderton, Roberto Amato, Jeffrey Barrett, Sonia Goncalves, Ewan Harrison, David K. Jackson, Ian Johnston, Dominic Kwiatkowski, Cordelia Langford, John Sillitoe on behalf of the Wellcome Sanger Institute COVID-19 Surveillance Team |
| EPI_ISL_3068745 | Lighthouse Lab in Alderley Park | Wellcome Sanger Institute for the COVID-19 Genomics UK (COG-UK) Consortium | Jacquelyn Wynn, Mairead Hyland, The Lighthouse Lab in Alderley Park and Alex Alderton, Roberto Amato, Jeffrey Barrett, Sonia Goncalves, Ewan Harrison, David K. Jackson, Ian Johnston, Dominic Kwiatkowski, Cordelia Langford, John Sillitoe on behalf of the Wellcome Sanger Institute COVID-19 Surveillance Team |
| EPI_ISL_3068746 | Lighthouse Laboratory Plymouth | Wellcome Sanger Institute for the COVID-19 Genomics UK (COG-UK) Consortium | Lighthouse Laboratory Plymouth and Alex Alderton, Roberto Amato, Jeffrey Barrett, Sonia Goncalves, Ewan Harrison, David K. Jackson, Ian Johnston, Dominic Kwiatkowski, Cordelia Langford, John Sillitoe on behalf of the Wellcome Sanger Institute COVID-19 Surveillance Team |
| EPI_ISL_3068747 | Lighthouse Lab in Alderley Park | Wellcome Sanger Institute for the COVID-19 Genomics UK (COG-UK) Consortium | Jacquelyn Wynn, Mairead Hyland, The Lighthouse Lab in Alderley Park and Alex Alderton, Roberto Amato, Jeffrey Barrett, Sonia Goncalves, Ewan Harrison, David K. Jackson, Ian Johnston, Dominic Kwiatkowski, Cordelia Langford, John Sillitoe on behalf of the Wellcome Sanger Institute COVID-19 Surveillance Team |
| EPI_ISL_3068748 | Lighthouse Laboratory Plymouth | Wellcome Sanger Institute for the COVID-19 Genomics UK (COG-UK) Consortium | Lighthouse Laboratory Plymouth and Alex Alderton, Roberto Amato, Jeffrey Barrett, Sonia Goncalves, Ewan Harrison, David K. Jackson, Ian Johnston, Dominic Kwiatkowski, Cordelia Langford, John Sillitoe on behalf of the Wellcome Sanger Institute COVID-19 Surveillance Team |
| EPI_ISL_3068749, EPI_ISL_3068750 | Lighthouse Lab in Alderley Park | Wellcome Sanger Institute for the COVID-19 Genomics UK (COG-UK) Consortium | Jacquelyn Wynn, Mairead Hyland, The Lighthouse Lab in Alderley Park and Alex Alderton, Roberto Amato, Jeffrey Barrett, Sonia Goncalves, Ewan Harrison, David K. Jackson, Ian Johnston, Dominic Kwiatkowski, Cordelia Langford, John Sillitoe on behalf of the Wellcome Sanger Institute COVID-19 Surveillance Team |
| EPI_ISL_3068751 | Health Services Laboratories | Wellcome Sanger Institute for the COVID-19 Genomics UK (COG-UK) Consortium | Health Services Laboratories and Alex Alderton, Roberto Amato, Jeffrey Barrett, Sonia Goncalves, Ewan Harrison, David K. Jackson, Ian Johnston, Dominic Kwiatkowski, Cordelia Langford, John Sillitoe on behalf of the Wellcome Sanger Institute COVID-19 Surveillance Team |
| EPI_ISL_3068752 | Berkshire and Surrey Pathology Services Lighthouse Laboratory | Wellcome Sanger Institute for the COVID-19 Genomics UK (COG-UK) Consortium | Berkshire and Surrey Pathology Services Lighthouse Laboratory and Alex Alderton, Roberto Amato, Jeffrey Barrett, Sonia Goncalves, Ewan Harrison, David K. Jackson, Ian Johnston, Dominic Kwiatkowski, Cordelia Langford, John Sillitoe on behalf of the Wellcome Sanger Institute COVID-19 Surveillance Team |
| EPI_ISL_3068753, EPI_ISL_3068754, EPI_ISL_3068755 | Lighthouse Laboratory Plymouth | Wellcome Sanger Institute for the COVID-19 Genomics UK (COG-UK) Consortium | Lighthouse Laboratory Plymouth and Alex Alderton, Roberto Amato, Jeffrey Barrett, Sonia Goncalves, Ewan Harrison, David K. Jackson, Ian Johnston, Dominic Kwiatkowski, Cordelia Langford, John Sillitoe on behalf of the Wellcome Sanger Institute COVID-19 Surveillance Team |
| EPI_ISL_3068756 | Lighthouse Lab in Alderley Park | Wellcome Sanger Institute for the COVID-19 Genomics UK (COG-UK) Consortium | Jacquelyn Wynn, Mairead Hyland, The Lighthouse Lab in Alderley Park and Alex Alderton, Roberto Amato, Jeffrey Barrett, Sonia Goncalves, Ewan Harrison, David K. Jackson, Ian Johnston, Dominic Kwiatkowski, Cordelia Langford, John Sillitoe on behalf of the Wellcome Sanger Institute COVID-19 Surveillance Team |
| EPI_ISL_3068757, EPI_ISL_3068758, EPI_ISL_3068759, EPI_ISL_3068760 | Lighthouse Laboratory Plymouth | Wellcome Sanger Institute for the COVID-19 Genomics UK (COG-UK) Consortium | Lighthouse Laboratory Plymouth and Alex Alderton, Roberto Amato, Jeffrey Barrett, Sonia Goncalves, Ewan Harrison, David K. Jackson, Ian Johnston, Dominic Kwiatkowski, Cordelia Langford, John Sillitoe on behalf of the Wellcome Sanger Institute COVID-19 Surveillance Team |
| EPI_ISL_3068761 | Lighthouse Lab in Alderley Park | Wellcome Sanger Institute for the COVID-19 Genomics UK (COG-UK) Consortium | Jacquelyn Wynn, Mairead Hyland, The Lighthouse Lab in Alderley Park and Alex Alderton, Roberto Amato, Jeffrey Barrett, Sonia Goncalves, Ewan Harrison, David K. Jackson, Ian Johnston, Dominic Kwiatkowski, Cordelia Langford, John Sillitoe on behalf of the Wellcome Sanger Institute COVID-19 Surveillance Team |
| EPI_ISL_3068762 | Lighthouse Laboratory Plymouth | Wellcome Sanger Institute for the COVID-19 Genomics UK (COG-UK) Consortium | Lighthouse Laboratory Plymouth and Alex Alderton, Roberto Amato, Jeffrey Barrett, Sonia Goncalves, Ewan Harrison, David K. Jackson, Ian Johnston, Dominic Kwiatkowski, Cordelia Langford, John Sillitoe on behalf of the Wellcome Sanger Institute COVID-19 Surveillance Team |
| EPI_ISL_3068763 | Lighthouse Lab in Alderley Park | Wellcome Sanger Institute for the COVID-19 Genomics UK (COG-UK) Consortium | Jacquelyn Wynn, Mairead Hyland, The Lighthouse Lab in Alderley Park and Alex Alderton, Roberto Amato, Jeffrey Barrett, Sonia Goncalves, Ewan Harrison, David K. Jackson, Ian Johnston, Dominic Kwiatkowski, Cordelia Langford, John Sillitoe on behalf of the Wellcome Sanger Institute COVID-19 Surveillance Team |
| EPI_ISL_3068764 | Lighthouse Laboratory Plymouth | Wellcome Sanger Institute for the COVID-19 Genomics UK (COG-UK) Consortium | Lighthouse Laboratory Plymouth and Alex Alderton, Roberto Amato, Jeffrey Barrett, Sonia Goncalves, Ewan Harrison, David K. Jackson, Ian Johnston, Dominic Kwiatkowski, Cordelia Langford, John Sillitoe on behalf of the Wellcome Sanger Institute COVID-19 Surveillance Team |
| EPI_ISL_3068765, EPI_ISL_3068766 | Lighthouse Lab in Alderley Park | Wellcome Sanger Institute for the COVID-19 Genomics UK (COG-UK) Consortium | Jacquelyn Wynn, Mairead Hyland, The Lighthouse Lab in Alderley Park and Alex Alderton, Roberto Amato, Jeffrey Barrett, Sonia Goncalves, Ewan Harrison, David K. Jackson, Ian Johnston, Dominic Kwiatkowski, Cordelia Langford, John Sillitoe on behalf of the Wellcome Sanger Institute COVID-19 Surveillance Team |
| EPI_ISL_3068767, EPI_ISL_3068768, EPI_ISL_3068769 | Lighthouse Laboratory Plymouth | Wellcome Sanger Institute for the COVID-19 Genomics UK (COG-UK) Consortium | Lighthouse Laboratory Plymouth and Alex Alderton, Roberto Amato, Jeffrey Barrett, Sonia Goncalves, Ewan Harrison, David K. Jackson, Ian Johnston, Dominic Kwiatkowski, Cordelia Langford, John Sillitoe on behalf of the Wellcome Sanger Institute COVID-19 Surveillance Team |
| EPI_ISL_3068770 | Lighthouse Lab in Alderley Park | Wellcome Sanger Institute for the COVID-19 Genomics UK (COG-UK) Consortium | Jacquelyn Wynn, Mairead Hyland, The Lighthouse Lab in Alderley Park and Alex Alderton, Roberto Amato, Jeffrey Barrett, Sonia Goncalves, Ewan Harrison, David K. Jackson, Ian Johnston, Dominic Kwiatkowski, Cordelia Langford, John Sillitoe on behalf of the Wellcome Sanger Institute COVID-19 Surveillance Team |
| EPI_ISL_3068771, EPI_ISL_3068772 | Lighthouse Laboratory Plymouth | Wellcome Sanger Institute for the COVID-19 Genomics UK (COG-UK) Consortium | Lighthouse Laboratory Plymouth and Alex Alderton, Roberto Amato, Jeffrey Barrett, Sonia Goncalves, Ewan Harrison, David K. Jackson, Ian Johnston, Dominic Kwiatkowski, Cordelia Langford, John Sillitoe on behalf of the Wellcome Sanger Institute COVID-19 Surveillance Team |
| EPI_ISL_3068773 | Health Services Laboratories | Wellcome Sanger Institute for the COVID-19 Genomics UK (COG-UK) Consortium | Health Services Laboratories and Alex Alderton, Roberto Amato, Jeffrey Barrett, Sonia Goncalves, Ewan Harrison, David K. Jackson, Ian Johnston, Dominic Kwiatkowski, Cordelia Langford, John Sillitoe on behalf of the Wellcome Sanger Institute COVID-19 Surveillance Team |
| EPI_ISL_3068774 | Lighthouse Lab in Alderley Park | Wellcome Sanger Institute for the COVID-19 Genomics UK (COG-UK) Consortium | Jacquelyn Wynn, Mairead Hyland, The Lighthouse Lab in Alderley Park and Alex Alderton, Roberto Amato, Jeffrey Barrett, Sonia Goncalves, Ewan Harrison, David K. Jackson, Ian Johnston, Dominic Kwiatkowski, Cordelia Langford, John Sillitoe on behalf of the Wellcome Sanger Institute COVID-19 Surveillance Team |
| EPI_ISL_3068775, EPI_ISL_3068776, EPI_ISL_3068777 | Lighthouse Laboratory Plymouth | Wellcome Sanger Institute for the COVID-19 Genomics UK (COG-UK) Consortium | Lighthouse Laboratory Plymouth and Alex Alderton, Roberto Amato, Jeffrey Barrett, Sonia Goncalves, Ewan Harrison, David K. Jackson, Ian Johnston, Dominic Kwiatkowski, Cordelia Langford, John Sillitoe on behalf of the Wellcome Sanger Institute COVID-19 Surveillance Team |
| EPI_ISL_3068778 | Lighthouse Lab in Alderley Park | Wellcome Sanger Institute for the COVID-19 Genomics UK (COG-UK) Consortium | Jacquelyn Wynn, Mairead Hyland, The Lighthouse Lab in Alderley Park and Alex Alderton, Roberto Amato, Jeffrey Barrett, Sonia Goncalves, Ewan Harrison, David K. Jackson, Ian Johnston, Dominic Kwiatkowski, Cordelia Langford, John Sillitoe on behalf of the Wellcome Sanger Institute COVID-19 Surveillance Team |
| EPI_ISL_3068779, EPI_ISL_3068780, EPI_ISL_3068781 | Lighthouse Laboratory Plymouth | Wellcome Sanger Institute for the COVID-19 Genomics UK (COG-UK) Consortium | Lighthouse Laboratory Plymouth and Alex Alderton, Roberto Amato, Jeffrey Barrett, Sonia Goncalves, Ewan Harrison, David K. Jackson, Ian Johnston, Dominic Kwiatkowski, Cordelia Langford, John Sillitoe on behalf of the Wellcome Sanger Institute COVID-19 Surveillance Team |
| EPI_ISL_3068782 | Health Services Laboratories | Wellcome Sanger Institute for the COVID-19 Genomics UK (COG-UK) Consortium | Health Services Laboratories and Alex Alderton, Roberto Amato, Jeffrey Barrett, Sonia Goncalves, Ewan Harrison, David K. Jackson, Ian Johnston, Dominic Kwiatkowski, Cordelia Langford, John Sillitoe on behalf of the Wellcome Sanger Institute COVID-19 Surveillance Team |
| EPI_ISL_3068783, EPI_ISL_3068784 | Lighthouse Laboratory Plymouth | Wellcome Sanger Institute for the COVID-19 Genomics UK (COG-UK) Consortium | Lighthouse Laboratory Plymouth and Alex Alderton, Roberto Amato, Jeffrey Barrett, Sonia Goncalves, Ewan Harrison, David K. Jackson, Ian Johnston, Dominic Kwiatkowski, Cordelia Langford, John Sillitoe on behalf of the Wellcome Sanger Institute COVID-19 Surveillance Team |
| EPI_ISL_3068785, EPI_ISL_3068786 | Lighthouse Lab in Alderley Park | Wellcome Sanger Institute for the COVID-19 Genomics UK (COG-UK) Consortium | Jacquelyn Wynn, Mairead Hyland, The Lighthouse Lab in Alderley Park and Alex Alderton, Roberto Amato, Jeffrey Barrett, Sonia Goncalves, Ewan Harrison, David K. Jackson, Ian Johnston, Dominic Kwiatkowski, Cordelia Langford, John Sillitoe on behalf of the Wellcome Sanger Institute COVID-19 Surveillance Team |
| EPI_ISL_3068787 | Health Services Laboratories | Wellcome Sanger Institute for the COVID-19 Genomics UK (COG-UK) Consortium | Health Services Laboratories and Alex Alderton, Roberto Amato, Jeffrey Barrett, Sonia Goncalves, Ewan Harrison, David K. Jackson, Ian Johnston, Dominic Kwiatkowski, Cordelia Langford, John Sillitoe on behalf of the Wellcome Sanger Institute COVID-19 Surveillance Team |
| EPI_ISL_3068788 | Lighthouse Lab in Alderley Park | Wellcome Sanger Institute for the COVID-19 Genomics UK (COG-UK) Consortium | Jacquelyn Wynn, Mairead Hyland, The Lighthouse Lab in Alderley Park and Alex Alderton, Roberto Amato, Jeffrey Barrett, Sonia Goncalves, Ewan Harrison, David K. Jackson, Ian Johnston, Dominic Kwiatkowski, Cordelia Langford, John Sillitoe on behalf of the Wellcome Sanger Institute COVID-19 Surveillance Team |
| EPI_ISL_3068789 | Lighthouse Laboratory Plymouth | Wellcome Sanger Institute for the COVID-19 Genomics UK (COG-UK) Consortium | Lighthouse Laboratory Plymouth and Alex Alderton, Roberto Amato, Jeffrey Barrett, Sonia Goncalves, Ewan Harrison, David K. Jackson, Ian Johnston, Dominic Kwiatkowski, Cordelia Langford, John Sillitoe on behalf of the Wellcome Sanger Institute COVID-19 Surveillance Team |
| EPI_ISL_3068790 | Lighthouse Lab in Alderley Park | Wellcome Sanger Institute for the COVID-19 Genomics UK (COG-UK) Consortium | Jacquelyn Wynn, Mairead Hyland, The Lighthouse Lab in Alderley Park and Alex Alderton, Roberto Amato, Jeffrey Barrett, Sonia Goncalves, Ewan Harrison, David K. Jackson, Ian Johnston, Dominic Kwiatkowski, Cordelia Langford, John Sillitoe on behalf of the Wellcome Sanger Institute COVID-19 Surveillance Team |
| EPI_ISL_3068791, EPI_ISL_3068792, EPI_ISL_3068793, EPI_ISL_3068794 | Lighthouse Laboratory Plymouth | Wellcome Sanger Institute for the COVID-19 Genomics UK (COG-UK) Consortium | Lighthouse Laboratory Plymouth and Alex Alderton, Roberto Amato, Jeffrey Barrett, Sonia Goncalves, Ewan Harrison, David K. Jackson, Ian Johnston, Dominic Kwiatkowski, Cordelia Langford, John Sillitoe on behalf of the Wellcome Sanger Institute COVID-19 Surveillance Team |
| EPI_ISL_3068795 | Lighthouse Lab in Alderley Park | Wellcome Sanger Institute for the COVID-19 Genomics UK (COG-UK) Consortium | Jacquelyn Wynn, Mairead Hyland, The Lighthouse Lab in Alderley Park and Alex Alderton, Roberto Amato, Jeffrey Barrett, Sonia Goncalves, Ewan Harrison, David K. Jackson, Ian Johnston, Dominic Kwiatkowski, Cordelia Langford, John Sillitoe on behalf of the Wellcome Sanger Institute COVID-19 Surveillance Team |
| EPI_ISL_3068796 | Lighthouse Laboratory Plymouth | Wellcome Sanger Institute for the COVID-19 Genomics UK (COG-UK) Consortium | Lighthouse Laboratory Plymouth and Alex Alderton, Roberto Amato, Jeffrey Barrett, Sonia Goncalves, Ewan Harrison, David K. Jackson, Ian Johnston, Dominic Kwiatkowski, Cordelia Langford, John Sillitoe on behalf of the Wellcome Sanger Institute COVID-19 Surveillance Team |

| | | | |
|---|---|---|---|
| EPI_ISL_3068797 | Lighthouse Lab in Alderley Park | Wellcome Sanger Institute for the COVID-19 Genomics UK (COG-UK) Consortium | Jacquelyn Wynn, Mairead Hyland, The Lighthouse Lab in Alderley Park and Alex Alderton, Roberto Amato, Jeffrey Barrett, Sonia Goncalves, Ewan Harrison, David K. Jackson, Ian Johnston, Dominic Kwiatkowski, Cordelia Langford, John Sillitoe on behalf of the Wellcome Sanger Institute COVID-19 Surveillance Team |
| EPI_ISL_3068798 | Lighthouse Laboratory Plymouth | Wellcome Sanger Institute for the COVID-19 Genomics UK (COG-UK) Consortium | Lighthouse Laboratory Plymouth and Alex Alderton, Roberto Amato, Jeffrey Barrett, Sonia Goncalves, Ewan Harrison, David K. Jackson, Ian Johnston, Dominic Kwiatkowski, Cordelia Langford, John Sillitoe on behalf of the Wellcome Sanger Institute COVID-19 Surveillance Team |
| EPI_ISL_3068799 | Lighthouse Lab in Alderley Park | Wellcome Sanger Institute for the COVID-19 Genomics UK (COG-UK) Consortium | Jacquelyn Wynn, Mairead Hyland, The Lighthouse Lab in Alderley Park and Alex Alderton, Roberto Amato, Jeffrey Barrett, Sonia Goncalves, Ewan Harrison, David K. Jackson, Ian Johnston, Dominic Kwiatkowski, Cordelia Langford, John Sillitoe on behalf of the Wellcome Sanger Institute COVID-19 Surveillance Team |
| EPI_ISL_3068800, EPI_ISL_3068801, EPI_ISL_3068802 | Lighthouse Laboratory Plymouth | Wellcome Sanger Institute for the COVID-19 Genomics UK (COG-UK) Consortium | Lighthouse Laboratory Plymouth and Alex Alderton, Roberto Amato, Jeffrey Barrett, Sonia Goncalves, Ewan Harrison, David K. Jackson, Ian Johnston, Dominic Kwiatkowski, Cordelia Langford, John Sillitoe on behalf of the Wellcome Sanger Institute COVID-19 Surveillance Team |
| EPI_ISL_3068803, EPI_ISL_3068804 | Lighthouse Lab in Alderley Park | Wellcome Sanger Institute for the COVID-19 Genomics UK (COG-UK) Consortium | Jacquelyn Wynn, Mairead Hyland, The Lighthouse Lab in Alderley Park and Alex Alderton, Roberto Amato, Jeffrey Barrett, Sonia Goncalves, Ewan Harrison, David K. Jackson, Ian Johnston, Dominic Kwiatkowski, Cordelia Langford, John Sillitoe on behalf of the Wellcome Sanger Institute COVID-19 Surveillance Team |
| EPI_ISL_3068805 | Lighthouse Laboratory Plymouth | Wellcome Sanger Institute for the COVID-19 Genomics UK (COG-UK) Consortium | Lighthouse Laboratory Plymouth and Alex Alderton, Roberto Amato, Jeffrey Barrett, Sonia Goncalves, Ewan Harrison, David K. Jackson, Ian Johnston, Dominic Kwiatkowski, Cordelia Langford, John Sillitoe on behalf of the Wellcome Sanger Institute COVID-19 Surveillance Team |
| EPI_ISL_3068806 | Lighthouse Lab in Alderley Park | Wellcome Sanger Institute for the COVID-19 Genomics UK (COG-UK) Consortium | Jacquelyn Wynn, Mairead Hyland, The Lighthouse Lab in Alderley Park and Alex Alderton, Roberto Amato, Jeffrey Barrett, Sonia Goncalves, Ewan Harrison, David K. Jackson, Ian Johnston, Dominic Kwiatkowski, Cordelia Langford, John Sillitoe on behalf of the Wellcome Sanger Institute COVID-19 Surveillance Team |
| EPI_ISL_3068807, EPI_ISL_3068808, EPI_ISL_3068809, EPI_ISL_3068810, EPI_ISL_3068811 | Lighthouse Laboratory Plymouth | Wellcome Sanger Institute for the COVID-19 Genomics UK (COG-UK) Consortium | Lighthouse Laboratory Plymouth and Alex Alderton, Roberto Amato, Jeffrey Barrett, Sonia Goncalves, Ewan Harrison, David K. Jackson, Ian Johnston, Dominic Kwiatkowski, Cordelia Langford, John Sillitoe on behalf of the Wellcome Sanger Institute COVID-19 Surveillance Team |
| EPI_ISL_3068812 | Health Services Laboratories | Wellcome Sanger Institute for the COVID-19 Genomics UK (COG-UK) Consortium | Health Services Laboratories and Alex Alderton, Roberto Amato, Jeffrey Barrett, Sonia Goncalves, Ewan Harrison, David K. Jackson, Ian Johnston, Dominic Kwiatkowski, Cordelia Langford, John Sillitoe on behalf of the Wellcome Sanger Institute COVID-19 Surveillance Team |
| EPI_ISL_3068813 | Lighthouse Lab in Alderley Park | Wellcome Sanger Institute for the COVID-19 Genomics UK (COG-UK) Consortium | Jacquelyn Wynn, Mairead Hyland, The Lighthouse Lab in Alderley Park and Alex Alderton, Roberto Amato, Jeffrey Barrett, Sonia Goncalves, Ewan Harrison, David K. Jackson, Ian Johnston, Dominic Kwiatkowski, Cordelia Langford, John Sillitoe on behalf of the Wellcome Sanger Institute COVID-19 Surveillance Team |
| EPI_ISL_3068814 | Health Services Laboratories | Wellcome Sanger Institute for the COVID-19 Genomics UK (COG-UK) Consortium | Health Services Laboratories and Alex Alderton, Roberto Amato, Jeffrey Barrett, Sonia Goncalves, Ewan Harrison, David K. Jackson, Ian Johnston, Dominic Kwiatkowski, Cordelia Langford, John Sillitoe on behalf of the Wellcome Sanger Institute COVID-19 Surveillance Team |
| EPI_ISL_3068815 | Lighthouse Lab in Alderley Park | Wellcome Sanger Institute for the COVID-19 Genomics UK (COG-UK) Consortium | Jacquelyn Wynn, Mairead Hyland, The Lighthouse Lab in Alderley Park and Alex Alderton, Roberto Amato, Jeffrey Barrett, Sonia Goncalves, Ewan Harrison, David K. Jackson, Ian Johnston, Dominic Kwiatkowski, Cordelia Langford, John Sillitoe on behalf of the Wellcome Sanger Institute COVID-19 Surveillance Team |
| EPI_ISL_3068816, EPI_ISL_3068817, EPI_ISL_3068818 | Lighthouse Laboratory Plymouth | Wellcome Sanger Institute for the COVID-19 Genomics UK (COG-UK) Consortium | Lighthouse Laboratory Plymouth and Alex Alderton, Roberto Amato, Jeffrey Barrett, Sonia Goncalves, Ewan Harrison, David K. Jackson, Ian Johnston, Dominic Kwiatkowski, Cordelia Langford, John Sillitoe on behalf of the Wellcome Sanger Institute COVID-19 Surveillance Team |
| EPI_ISL_3068819 | Lighthouse Lab in Alderley Park | Wellcome Sanger Institute for the COVID-19 Genomics UK (COG-UK) Consortium | Jacquelyn Wynn, Mairead Hyland, The Lighthouse Lab in Alderley Park and Alex Alderton, Roberto Amato, Jeffrey Barrett, Sonia Goncalves, Ewan Harrison, David K. Jackson, Ian Johnston, Dominic Kwiatkowski, Cordelia Langford, John Sillitoe on behalf of the Wellcome Sanger Institute COVID-19 Surveillance Team |
| EPI_ISL_3068820, EPI_ISL_3068821, EPI_ISL_3068822 | Lighthouse Laboratory Plymouth | Wellcome Sanger Institute for the COVID-19 Genomics UK (COG-UK) Consortium | Lighthouse Laboratory Plymouth and Alex Alderton, Roberto Amato, Jeffrey Barrett, Sonia Goncalves, Ewan Harrison, David K. Jackson, Ian Johnston, Dominic Kwiatkowski, Cordelia Langford, John Sillitoe on behalf of the Wellcome Sanger Institute COVID-19 Surveillance Team |
| EPI_ISL_3068823 | Lighthouse Lab in Alderley Park | Wellcome Sanger Institute for the COVID-19 Genomics UK (COG-UK) Consortium | Jacquelyn Wynn, Mairead Hyland, The Lighthouse Lab in Alderley Park and Alex Alderton, Roberto Amato, Jeffrey Barrett, Sonia Goncalves, Ewan Harrison, David K. Jackson, Ian Johnston, Dominic Kwiatkowski, Cordelia Langford, John Sillitoe on behalf of the Wellcome Sanger Institute COVID-19 Surveillance Team |
| EPI_ISL_3068824 | Lighthouse Laboratory Plymouth | Wellcome Sanger Institute for the COVID-19 Genomics UK (COG-UK) Consortium | Lighthouse Laboratory Plymouth and Alex Alderton, Roberto Amato, Jeffrey Barrett, Sonia Goncalves, Ewan Harrison, David K. Jackson, Ian Johnston, Dominic Kwiatkowski, Cordelia Langford, John Sillitoe on behalf of the Wellcome Sanger Institute COVID-19 Surveillance Team |
| EPI_ISL_3068825 | Health Services Laboratories | Wellcome Sanger Institute for the COVID-19 Genomics UK (COG-UK) Consortium | Health Services Laboratories and Alex Alderton, Roberto Amato, Jeffrey Barrett, Sonia Goncalves, Ewan Harrison, David K. Jackson, Ian Johnston, Dominic Kwiatkowski, Cordelia Langford, John Sillitoe on behalf of the Wellcome Sanger Institute COVID-19 Surveillance Team |
| EPI_ISL_3068826, EPI_ISL_3068827 | Lighthouse Lab in Alderley Park | Wellcome Sanger Institute for the COVID-19 Genomics UK (COG-UK) Consortium | Jacquelyn Wynn, Mairead Hyland, The Lighthouse Lab in Alderley Park and Alex Alderton, Roberto Amato, Jeffrey Barrett, Sonia Goncalves, Ewan Harrison, David K. Jackson, Ian Johnston, Dominic Kwiatkowski, Cordelia Langford, John Sillitoe on behalf of the Wellcome Sanger Institute COVID-19 Surveillance Team |
| EPI_ISL_3068828 | Lighthouse Laboratory Plymouth | Wellcome Sanger Institute for the COVID-19 Genomics UK (COG-UK) Consortium | Lighthouse Laboratory Plymouth and Alex Alderton, Roberto Amato, Jeffrey Barrett, Sonia Goncalves, Ewan Harrison, David K. Jackson, Ian Johnston, Dominic Kwiatkowski, Cordelia Langford, John Sillitoe on behalf of the Wellcome Sanger Institute COVID-19 Surveillance Team |
| EPI_ISL_3068829 | Lighthouse Lab in Alderley Park | Wellcome Sanger Institute for the COVID-19 Genomics UK (COG-UK) Consortium | Jacquelyn Wynn, Mairead Hyland, The Lighthouse Lab in Alderley Park and Alex Alderton, Roberto Amato, Jeffrey Barrett, Sonia Goncalves, Ewan Harrison, David K. Jackson, Ian Johnston, Dominic Kwiatkowski, Cordelia Langford, John Sillitoe on behalf of the Wellcome Sanger Institute COVID-19 Surveillance Team |
| EPI_ISL_3068830 | Health Services Laboratories | Wellcome Sanger Institute for the COVID-19 Genomics UK (COG-UK) Consortium | Health Services Laboratories and Alex Alderton, Roberto Amato, Jeffrey Barrett, Sonia Goncalves, Ewan Harrison, David K. Jackson, Ian Johnston, Dominic Kwiatkowski, Cordelia Langford, John Sillitoe on behalf of the Wellcome Sanger Institute COVID-19 Surveillance Team |
| EPI_ISL_3068831, EPI_ISL_3068832, EPI_ISL_3068833, EPI_ISL_3068834, EPI_ISL_3068835 | Lighthouse Laboratory Plymouth | Wellcome Sanger Institute for the COVID-19 Genomics UK (COG-UK) Consortium | Lighthouse Laboratory Plymouth and Alex Alderton, Roberto Amato, Jeffrey Barrett, Sonia Goncalves, Ewan Harrison, David K. Jackson, Ian Johnston, Dominic Kwiatkowski, Cordelia Langford, John Sillitoe on behalf of the Wellcome Sanger Institute COVID-19 Surveillance Team |
| EPI_ISL_3068836 | Lighthouse Lab in Alderley Park | Wellcome Sanger Institute for the COVID-19 Genomics UK (COG-UK) Consortium | Jacquelyn Wynn, Mairead Hyland, The Lighthouse Lab in Alderley Park and Alex Alderton, Roberto Amato, Jeffrey Barrett, Sonia Goncalves, Ewan Harrison, David K. Jackson, Ian Johnston, Dominic Kwiatkowski, Cordelia Langford, John Sillitoe on behalf of the Wellcome Sanger Institute COVID-19 Surveillance Team |
| EPI_ISL_3068837 | Lighthouse Laboratory Plymouth | Wellcome Sanger Institute for the COVID-19 Genomics UK (COG-UK) Consortium | Lighthouse Laboratory Plymouth and Alex Alderton, Roberto Amato, Jeffrey Barrett, Sonia Goncalves, Ewan Harrison, David K. Jackson, Ian Johnston, Dominic Kwiatkowski, Cordelia Langford, John Sillitoe on behalf of the Wellcome Sanger Institute COVID-19 Surveillance Team |
| EPI_ISL_3068838 | Health Services Laboratories | Wellcome Sanger Institute for the COVID-19 Genomics UK (COG-UK) Consortium | Health Services Laboratories and Alex Alderton, Roberto Amato, Jeffrey Barrett, Sonia Goncalves, Ewan Harrison, David K. Jackson, Ian Johnston, Dominic Kwiatkowski, Cordelia Langford, John Sillitoe on behalf of the Wellcome Sanger Institute COVID-19 Surveillance Team |
| EPI_ISL_3068839, EPI_ISL_3068840, EPI_ISL_3068841 | Lighthouse Laboratory Plymouth | Wellcome Sanger Institute for the COVID-19 Genomics UK (COG-UK) Consortium | Lighthouse Laboratory Plymouth and Alex Alderton, Roberto Amato, Jeffrey Barrett, Sonia Goncalves, Ewan Harrison, David K. Jackson, Ian Johnston, Dominic Kwiatkowski, Cordelia Langford, John Sillitoe on behalf of the Wellcome Sanger Institute COVID-19 Surveillance Team |
| EPI_ISL_3068842 | Health Services Laboratories | Wellcome Sanger Institute for the COVID-19 Genomics UK (COG-UK) Consortium | Health Services Laboratories and Alex Alderton, Roberto Amato, Jeffrey Barrett, Sonia Goncalves, Ewan Harrison, David K. Jackson, Ian Johnston, Dominic Kwiatkowski, Cordelia Langford, John Sillitoe on behalf of the Wellcome Sanger Institute COVID-19 Surveillance Team |
| EPI_ISL_3068843 | Lighthouse Lab in Alderley Park | Wellcome Sanger Institute for the COVID-19 Genomics UK (COG-UK) Consortium | Jacquelyn Wynn, Mairead Hyland, The Lighthouse Lab in Alderley Park and Alex Alderton, Roberto Amato, Jeffrey Barrett, Sonia Goncalves, Ewan Harrison, David K. Jackson, Ian Johnston, Dominic Kwiatkowski, Cordelia Langford, John Sillitoe on behalf of the Wellcome Sanger Institute COVID-19 Surveillance Team |
| EPI_ISL_3068844, EPI_ISL_3068845, EPI_ISL_3068846 | Lighthouse Laboratory Plymouth | Wellcome Sanger Institute for the COVID-19 Genomics UK (COG-UK) Consortium | Lighthouse Laboratory Plymouth and Alex Alderton, Roberto Amato, Jeffrey Barrett, Sonia Goncalves, Ewan Harrison, David K. Jackson, Ian Johnston, Dominic Kwiatkowski, Cordelia Langford, John Sillitoe on behalf of the Wellcome Sanger Institute COVID-19 Surveillance Team |
| EPI_ISL_3068847 | Berkshire and Surrey Pathology Services Lighthouse Laboratory | Wellcome Sanger Institute for the COVID-19 Genomics UK (COG-UK) Consortium | Berkshire and Surrey Pathology Services Lighthouse Laboratory and Alex Alderton, Roberto Amato, Jeffrey Barrett, Sonia Goncalves, Ewan Harrison, David K. Jackson, Ian Johnston, Dominic Kwiatkowski, Cordelia Langford, John Sillitoe on behalf of the Wellcome Sanger Institute COVID-19 Surveillance Team |
| EPI_ISL_3068848 | Lighthouse Laboratory Plymouth | Wellcome Sanger Institute for the COVID-19 Genomics UK (COG-UK) Consortium | Lighthouse Laboratory Plymouth and Alex Alderton, Roberto Amato, Jeffrey Barrett, Sonia Goncalves, Ewan Harrison, David K. Jackson, Ian Johnston, Dominic Kwiatkowski, Cordelia Langford, John Sillitoe on behalf of the Wellcome Sanger Institute COVID-19 Surveillance Team |
| EPI_ISL_3068849 | Lighthouse Lab in Alderley Park | Wellcome Sanger Institute for the COVID-19 Genomics UK (COG-UK) Consortium | Jacquelyn Wynn, Mairead Hyland, The Lighthouse Lab in Alderley Park and Alex Alderton, Roberto Amato, Jeffrey Barrett, Sonia Goncalves, Ewan Harrison, David K. Jackson, Ian Johnston, Dominic Kwiatkowski, Cordelia Langford, John Sillitoe on behalf of the Wellcome Sanger Institute COVID-19 Surveillance Team |
| EPI_ISL_3068850, EPI_ISL_3068851 | Lighthouse Laboratory Plymouth | Wellcome Sanger Institute for the COVID-19 Genomics UK (COG-UK) Consortium | Lighthouse Laboratory Plymouth and Alex Alderton, Roberto Amato, Jeffrey Barrett, Sonia Goncalves, Ewan Harrison, David K. Jackson, Ian Johnston, Dominic Kwiatkowski, Cordelia Langford, John Sillitoe on behalf of the Wellcome Sanger Institute COVID-19 Surveillance Team |
| EPI_ISL_3068852 | Lighthouse Lab in Alderley Park | Wellcome Sanger Institute for the COVID-19 Genomics UK (COG-UK) Consortium | Jacquelyn Wynn, Mairead Hyland, The Lighthouse Lab in Alderley Park and Alex Alderton, Roberto Amato, Jeffrey Barrett, Sonia Goncalves, Ewan Harrison, David K. Jackson, Ian Johnston, Dominic Kwiatkowski, Cordelia Langford, John Sillitoe on behalf of the Wellcome Sanger Institute COVID-19 Surveillance Team |
| EPI_ISL_3068853 | Lighthouse Laboratory Plymouth | Wellcome Sanger Institute for the COVID-19 Genomics UK (COG-UK) Consortium | Lighthouse Laboratory Plymouth and Alex Alderton, Roberto Amato, Jeffrey Barrett, Sonia Goncalves, Ewan Harrison, David K. Jackson, Ian Johnston, Dominic Kwiatkowski, Cordelia Langford, John Sillitoe on behalf of the Wellcome Sanger Institute COVID-19 Surveillance Team |
| EPI_ISL_3068854, EPI_ISL_3068855 | Lighthouse Lab in Alderley Park | Wellcome Sanger Institute for the COVID-19 Genomics UK (COG-UK) Consortium | Jacquelyn Wynn, Mairead Hyland, The Lighthouse Lab in Alderley Park and Alex Alderton, Roberto Amato, Jeffrey Barrett, Sonia Goncalves, Ewan Harrison, David K. Jackson, Ian Johnston, Dominic Kwiatkowski, Cordelia Langford, John Sillitoe on behalf of the Wellcome Sanger Institute COVID-19 Surveillance Team |
| EPI_ISL_3068856, EPI_ISL_3068857 | Lighthouse Laboratory Plymouth | Wellcome Sanger Institute for the COVID-19 Genomics UK (COG-UK) Consortium | Lighthouse Laboratory Plymouth and Alex Alderton, Roberto Amato, Jeffrey Barrett, Sonia Goncalves, Ewan Harrison, David K. Jackson, Ian Johnston, Dominic Kwiatkowski, Cordelia Langford, John Sillitoe on behalf of the Wellcome Sanger Institute COVID-19 Surveillance Team |
| EPI_ISL_3068858 | Lighthouse Lab in Alderley Park | Wellcome Sanger Institute for the COVID-19 Genomics UK (COG-UK) Consortium | Jacquelyn Wynn, Mairead Hyland, The Lighthouse Lab in Alderley Park and Alex Alderton, Roberto Amato, Jeffrey Barrett, Sonia Goncalves, Ewan Harrison, David K. Jackson, Ian Johnston, Dominic Kwiatkowski, Cordelia Langford, John Sillitoe on behalf of the Wellcome Sanger Institute COVID-19 Surveillance Team |
| EPI_ISL_3068859, EPI_ISL_3068860 | Lighthouse Laboratory Plymouth | Wellcome Sanger Institute for the COVID-19 Genomics UK (COG-UK) Consortium | Lighthouse Laboratory Plymouth and Alex Alderton, Roberto Amato, Jeffrey Barrett, Sonia Goncalves, Ewan Harrison, David K. Jackson, Ian Johnston, Dominic Kwiatkowski, Cordelia Langford, John Sillitoe on behalf of the Wellcome Sanger Institute COVID-19 Surveillance Team |
| EPI_ISL_3068861 | Health Services Laboratories | Wellcome Sanger Institute for the COVID-19 Genomics UK (COG-UK) Consortium | Health Services Laboratories and Alex Alderton, Roberto Amato, Jeffrey Barrett, Sonia Goncalves, Ewan Harrison, David K. Jackson, Ian Johnston, Dominic Kwiatkowski, Cordelia Langford, John Sillitoe on behalf of the Wellcome Sanger Institute COVID-19 Surveillance Team |

| | | | |
|---|---|---|---|
| EPI_ISL_3068862 | Lighthouse Laboratory Plymouth | Wellcome Sanger Institute for the COVID-19 Genomics UK (COG-UK) Consortium | Lighthouse Laboratory Plymouth and Alex Alderton, Roberto Amato, Jeffrey Barrett, Sonia Goncalves, Ewan Harrison, David K. Jackson, Ian Johnston, Dominic Kwiatkowski, Cordelia Langford, John Sillitoe on behalf of the Wellcome Sanger Institute COVID-19 Surveillance Team |
| EPI_ISL_3068863 | Health Services Laboratories | Wellcome Sanger Institute for the COVID-19 Genomics UK (COG-UK) Consortium | Health Services Laboratories and Alex Alderton, Roberto Amato, Jeffrey Barrett, Sonia Goncalves, Ewan Harrison, David K. Jackson, Ian Johnston, Dominic Kwiatkowski, Cordelia Langford, John Sillitoe on behalf of the Wellcome Sanger Institute COVID-19 Surveillance Team |
| EPI_ISL_3068864 | Lighthouse Laboratory Plymouth | Wellcome Sanger Institute for the COVID-19 Genomics UK (COG-UK) Consortium | Lighthouse Laboratory Plymouth and Alex Alderton, Roberto Amato, Jeffrey Barrett, Sonia Goncalves, Ewan Harrison, David K. Jackson, Ian Johnston, Dominic Kwiatkowski, Cordelia Langford, John Sillitoe on behalf of the Wellcome Sanger Institute COVID-19 Surveillance Team |
| EPI_ISL_3068865 | Health Services Laboratories | Wellcome Sanger Institute for the COVID-19 Genomics UK (COG-UK) Consortium | Health Services Laboratories and Alex Alderton, Roberto Amato, Jeffrey Barrett, Sonia Goncalves, Ewan Harrison, David K. Jackson, Ian Johnston, Dominic Kwiatkowski, Cordelia Langford, John Sillitoe on behalf of the Wellcome Sanger Institute COVID-19 Surveillance Team |
| EPI_ISL_3068866 | Lighthouse Lab in Alderley Park | Wellcome Sanger Institute for the COVID-19 Genomics UK (COG-UK) Consortium | Jacquelyn Wynn, Mairead Hyland, The Lighthouse Lab in Alderley Park and Alex Alderton, Roberto Amato, Jeffrey Barrett, Sonia Goncalves, Ewan Harrison, David K. Jackson, Ian Johnston, Dominic Kwiatkowski, Cordelia Langford, John Sillitoe on behalf of the Wellcome Sanger Institute COVID-19 Surveillance Team |
| EPI_ISL_3068867, EPI_ISL_3068868, EPI_ISL_3068869, EPI_ISL_3068870 | Lighthouse Laboratory Plymouth | Wellcome Sanger Institute for the COVID-19 Genomics UK (COG-UK) Consortium | Lighthouse Laboratory Plymouth and Alex Alderton, Roberto Amato, Jeffrey Barrett, Sonia Goncalves, Ewan Harrison, David K. Jackson, Ian Johnston, Dominic Kwiatkowski, Cordelia Langford, John Sillitoe on behalf of the Wellcome Sanger Institute COVID-19 Surveillance Team |
| EPI_ISL_3068871, EPI_ISL_3068872 | Health Services Laboratories | Wellcome Sanger Institute for the COVID-19 Genomics UK (COG-UK) Consortium | Health Services Laboratories and Alex Alderton, Roberto Amato, Jeffrey Barrett, Sonia Goncalves, Ewan Harrison, David K. Jackson, Ian Johnston, Dominic Kwiatkowski, Cordelia Langford, John Sillitoe on behalf of the Wellcome Sanger Institute COVID-19 Surveillance Team |
| EPI_ISL_3068873, EPI_ISL_3068874 | Lighthouse Laboratory Plymouth | Wellcome Sanger Institute for the COVID-19 Genomics UK (COG-UK) Consortium | Lighthouse Laboratory Plymouth and Alex Alderton, Roberto Amato, Jeffrey Barrett, Sonia Goncalves, Ewan Harrison, David K. Jackson, Ian Johnston, Dominic Kwiatkowski, Cordelia Langford, John Sillitoe on behalf of the Wellcome Sanger Institute COVID-19 Surveillance Team |
| EPI_ISL_3068875, EPI_ISL_3068876 | Health Services Laboratories | Wellcome Sanger Institute for the COVID-19 Genomics UK (COG-UK) Consortium | Health Services Laboratories and Alex Alderton, Roberto Amato, Jeffrey Barrett, Sonia Goncalves, Ewan Harrison, David K. Jackson, Ian Johnston, Dominic Kwiatkowski, Cordelia Langford, John Sillitoe on behalf of the Wellcome Sanger Institute COVID-19 Surveillance Team |
| EPI_ISL_3068877, EPI_ISL_3068878, EPI_ISL_3068879 | Lighthouse Laboratory Plymouth | Wellcome Sanger Institute for the COVID-19 Genomics UK (COG-UK) Consortium | Lighthouse Laboratory Plymouth and Alex Alderton, Roberto Amato, Jeffrey Barrett, Sonia Goncalves, Ewan Harrison, David K. Jackson, Ian Johnston, Dominic Kwiatkowski, Cordelia Langford, John Sillitoe on behalf of the Wellcome Sanger Institute COVID-19 Surveillance Team |
| EPI_ISL_3068880 | Lighthouse Lab in Alderley Park | Wellcome Sanger Institute for the COVID-19 Genomics UK (COG-UK) Consortium | Jacquelyn Wynn, Mairead Hyland, The Lighthouse Lab in Alderley Park and Alex Alderton, Roberto Amato, Jeffrey Barrett, Sonia Goncalves, Ewan Harrison, David K. Jackson, Ian Johnston, Dominic Kwiatkowski, Cordelia Langford, John Sillitoe on behalf of the Wellcome Sanger Institute COVID-19 Surveillance Team |
| EPI_ISL_3068881 | Lighthouse Laboratory Plymouth | Wellcome Sanger Institute for the COVID-19 Genomics UK (COG-UK) Consortium | Lighthouse Laboratory Plymouth and Alex Alderton, Roberto Amato, Jeffrey Barrett, Sonia Goncalves, Ewan Harrison, David K. Jackson, Ian Johnston, Dominic Kwiatkowski, Cordelia Langford, John Sillitoe on behalf of the Wellcome Sanger Institute COVID-19 Surveillance Team |
| EPI_ISL_3068882 | Health Services Laboratories | Wellcome Sanger Institute for the COVID-19 Genomics UK (COG-UK) Consortium | Health Services Laboratories and Alex Alderton, Roberto Amato, Jeffrey Barrett, Sonia Goncalves, Ewan Harrison, David K. Jackson, Ian Johnston, Dominic Kwiatkowski, Cordelia Langford, John Sillitoe on behalf of the Wellcome Sanger Institute COVID-19 Surveillance Team |
| EPI_ISL_3068883 | Lighthouse Laboratory Plymouth | Wellcome Sanger Institute for the COVID-19 Genomics UK (COG-UK) Consortium | Lighthouse Laboratory Plymouth and Alex Alderton, Roberto Amato, Jeffrey Barrett, Sonia Goncalves, Ewan Harrison, David K. Jackson, Ian Johnston, Dominic Kwiatkowski, Cordelia Langford, John Sillitoe on behalf of the Wellcome Sanger Institute COVID-19 Surveillance Team |
| EPI_ISL_3068884, EPI_ISL_3068885 | Lighthouse Lab in Alderley Park | Wellcome Sanger Institute for the COVID-19 Genomics UK (COG-UK) Consortium | Jacquelyn Wynn, Mairead Hyland, The Lighthouse Lab in Alderley Park and Alex Alderton, Roberto Amato, Jeffrey Barrett, Sonia Goncalves, Ewan Harrison, David K. Jackson, Ian Johnston, Dominic Kwiatkowski, Cordelia Langford, John Sillitoe on behalf of the Wellcome Sanger Institute COVID-19 Surveillance Team |
| EPI_ISL_3068886, EPI_ISL_3068887, EPI_ISL_3068888 | Lighthouse Laboratory Plymouth | Wellcome Sanger Institute for the COVID-19 Genomics UK (COG-UK) Consortium | Lighthouse Laboratory Plymouth and Alex Alderton, Roberto Amato, Jeffrey Barrett, Sonia Goncalves, Ewan Harrison, David K. Jackson, Ian Johnston, Dominic Kwiatkowski, Cordelia Langford, John Sillitoe on behalf of the Wellcome Sanger Institute COVID-19 Surveillance Team |
| EPI_ISL_3068889 | Lighthouse Lab in Alderley Park | Wellcome Sanger Institute for the COVID-19 Genomics UK (COG-UK) Consortium | Jacquelyn Wynn, Mairead Hyland, The Lighthouse Lab in Alderley Park and Alex Alderton, Roberto Amato, Jeffrey Barrett, Sonia Goncalves, Ewan Harrison, David K. Jackson, Ian Johnston, Dominic Kwiatkowski, Cordelia Langford, John Sillitoe on behalf of the Wellcome Sanger Institute COVID-19 Surveillance Team |
| EPI_ISL_3068890 | Lighthouse Laboratory Plymouth | Wellcome Sanger Institute for the COVID-19 Genomics UK (COG-UK) Consortium | Lighthouse Laboratory Plymouth and Alex Alderton, Roberto Amato, Jeffrey Barrett, Sonia Goncalves, Ewan Harrison, David K. Jackson, Ian Johnston, Dominic Kwiatkowski, Cordelia Langford, John Sillitoe on behalf of the Wellcome Sanger Institute COVID-19 Surveillance Team |
| EPI_ISL_3068891, EPI_ISL_3068892, EPI_ISL_3068893 | Lighthouse Lab in Alderley Park | Wellcome Sanger Institute for the COVID-19 Genomics UK (COG-UK) Consortium | Jacquelyn Wynn, Mairead Hyland, The Lighthouse Lab in Alderley Park and Alex Alderton, Roberto Amato, Jeffrey Barrett, Sonia Goncalves, Ewan Harrison, David K. Jackson, Ian Johnston, Dominic Kwiatkowski, Cordelia Langford, John Sillitoe on behalf of the Wellcome Sanger Institute COVID-19 Surveillance Team |
| EPI_ISL_3068894 | Lighthouse Laboratory Plymouth | Wellcome Sanger Institute for the COVID-19 Genomics UK (COG-UK) Consortium | Lighthouse Laboratory Plymouth and Alex Alderton, Roberto Amato, Jeffrey Barrett, Sonia Goncalves, Ewan Harrison, David K. Jackson, Ian Johnston, Dominic Kwiatkowski, Cordelia Langford, John Sillitoe on behalf of the Wellcome Sanger Institute COVID-19 Surveillance Team |
| EPI_ISL_3068895 | Health Services Laboratories | Wellcome Sanger Institute for the COVID-19 Genomics UK (COG-UK) Consortium | Health Services Laboratories and Alex Alderton, Roberto Amato, Jeffrey Barrett, Sonia Goncalves, Ewan Harrison, David K. Jackson, Ian Johnston, Dominic Kwiatkowski, Cordelia Langford, John Sillitoe on behalf of the Wellcome Sanger Institute COVID-19 Surveillance Team |
| EPI_ISL_3068896 | Lighthouse Lab in Alderley Park | Wellcome Sanger Institute for the COVID-19 Genomics UK (COG-UK) Consortium | Jacquelyn Wynn, Mairead Hyland, The Lighthouse Lab in Alderley Park and Alex Alderton, Roberto Amato, Jeffrey Barrett, Sonia Goncalves, Ewan Harrison, David K. Jackson, Ian Johnston, Dominic Kwiatkowski, Cordelia Langford, John Sillitoe on behalf of the Wellcome Sanger Institute COVID-19 Surveillance Team |
| EPI_ISL_3068897 | Lighthouse Laboratory Plymouth | Wellcome Sanger Institute for the COVID-19 Genomics UK (COG-UK) Consortium | Lighthouse Laboratory Plymouth and Alex Alderton, Roberto Amato, Jeffrey Barrett, Sonia Goncalves, Ewan Harrison, David K. Jackson, Ian Johnston, Dominic Kwiatkowski, Cordelia Langford, John Sillitoe on behalf of the Wellcome Sanger Institute COVID-19 Surveillance Team |
| EPI_ISL_3068898 | Berkshire and Surrey Pathology Services Lighthouse Laboratory | Wellcome Sanger Institute for the COVID-19 Genomics UK (COG-UK) Consortium | Berkshire and Surrey Pathology Services Lighthouse Laboratory and Alex Alderton, Roberto Amato, Jeffrey Barrett, Sonia Goncalves, Ewan Harrison, David K. Jackson, Ian Johnston, Dominic Kwiatkowski, Cordelia Langford, John Sillitoe on behalf of the Wellcome Sanger Institute COVID-19 Surveillance Team |
| EPI_ISL_3068899, EPI_ISL_3068900 | Lighthouse Lab in Glasgow | Wellcome Sanger Institute for the COVID-19 Genomics UK (COG-UK) Consortium | Harper VanSteenhouse, Yumi Kasai, David Gray, Carol Clugston, Anna Dominiczak and Alex Alderton, Roberto Amato, Jeffrey Barrett, Sonia Goncalves, Ewan Harrison, David K. Jackson, Ian Johnston, Dominic Kwiatkowski, Cordelia Langford, John Sillitoe on behalf of the Wellcome Sanger Institute COVID-19 Surveillance Team |
| EPI_ISL_3068901 | Lighthouse Lab in Alderley Park | Wellcome Sanger Institute for the COVID-19 Genomics UK (COG-UK) Consortium | Jacquelyn Wynn, Mairead Hyland, The Lighthouse Lab in Alderley Park and Alex Alderton, Roberto Amato, Jeffrey Barrett, Sonia Goncalves, Ewan Harrison, David K. Jackson, Ian Johnston, Dominic Kwiatkowski, Cordelia Langford, John Sillitoe on behalf of the Wellcome Sanger Institute COVID-19 Surveillance Team |
| EPI_ISL_3068902 | Lighthouse Lab in Glasgow | Wellcome Sanger Institute for the COVID-19 Genomics UK (COG-UK) Consortium | Harper VanSteenhouse, Yumi Kasai, David Gray, Carol Clugston, Anna Dominiczak and Alex Alderton, Roberto Amato, Jeffrey Barrett, Sonia Goncalves, Ewan Harrison, David K. Jackson, Ian Johnston, Dominic Kwiatkowski, Cordelia Langford, John Sillitoe on behalf of the Wellcome Sanger Institute COVID-19 Surveillance Team |
| EPI_ISL_3068903 | Berkshire and Surrey Pathology Services Lighthouse Laboratory | Wellcome Sanger Institute for the COVID-19 Genomics UK (COG-UK) Consortium | Berkshire and Surrey Pathology Services Lighthouse Laboratory and Alex Alderton, Roberto Amato, Jeffrey Barrett, Sonia Goncalves, Ewan Harrison, David K. Jackson, Ian Johnston, Dominic Kwiatkowski, Cordelia Langford, John Sillitoe on behalf of the Wellcome Sanger Institute COVID-19 Surveillance Team |
| EPI_ISL_3068904, EPI_ISL_3068905 | Lighthouse Lab in Glasgow | Wellcome Sanger Institute for the COVID-19 Genomics UK (COG-UK) Consortium | Harper VanSteenhouse, Yumi Kasai, David Gray, Carol Clugston, Anna Dominiczak and Alex Alderton, Roberto Amato, Jeffrey Barrett, Sonia Goncalves, Ewan Harrison, David K. Jackson, Ian Johnston, Dominic Kwiatkowski, Cordelia Langford, John Sillitoe on behalf of the Wellcome Sanger Institute COVID-19 Surveillance Team |
| EPI_ISL_3068906 | Berkshire and Surrey Pathology Services Lighthouse Laboratory | Wellcome Sanger Institute for the COVID-19 Genomics UK (COG-UK) Consortium | Berkshire and Surrey Pathology Services Lighthouse Laboratory and Alex Alderton, Roberto Amato, Jeffrey Barrett, Sonia Goncalves, Ewan Harrison, David K. Jackson, Ian Johnston, Dominic Kwiatkowski, Cordelia Langford, John Sillitoe on behalf of the Wellcome Sanger Institute COVID-19 Surveillance Team |
| EPI_ISL_3068907 | Lighthouse Lab in Alderley Park | Wellcome Sanger Institute for the COVID-19 Genomics UK (COG-UK) Consortium | Jacquelyn Wynn, Mairead Hyland, The Lighthouse Lab in Alderley Park and Alex Alderton, Roberto Amato, Jeffrey Barrett, Sonia Goncalves, Ewan Harrison, David K. Jackson, Ian Johnston, Dominic Kwiatkowski, Cordelia Langford, John Sillitoe on behalf of the Wellcome Sanger Institute COVID-19 Surveillance Team |
| EPI_ISL_3068908, EPI_ISL_3068909 | Lighthouse Lab in Glasgow | Wellcome Sanger Institute for the COVID-19 Genomics UK (COG-UK) Consortium | Harper VanSteenhouse, Yumi Kasai, David Gray, Carol Clugston, Anna Dominiczak and Alex Alderton, Roberto Amato, Jeffrey Barrett, Sonia Goncalves, Ewan Harrison, David K. Jackson, Ian Johnston, Dominic Kwiatkowski, Cordelia Langford, John Sillitoe on behalf of the Wellcome Sanger Institute COVID-19 Surveillance Team |
| EPI_ISL_3068910, EPI_ISL_3068911 | Berkshire and Surrey Pathology Services Lighthouse Laboratory | Wellcome Sanger Institute for the COVID-19 Genomics UK (COG-UK) Consortium | Berkshire and Surrey Pathology Services Lighthouse Laboratory and Alex Alderton, Roberto Amato, Jeffrey Barrett, Sonia Goncalves, Ewan Harrison, David K. Jackson, Ian Johnston, Dominic Kwiatkowski, Cordelia Langford, John Sillitoe on behalf of the Wellcome Sanger Institute COVID-19 Surveillance Team |
| EPI_ISL_3068912 | Lighthouse Lab in Alderley Park | Wellcome Sanger Institute for the COVID-19 Genomics UK (COG-UK) Consortium | Jacquelyn Wynn, Mairead Hyland, The Lighthouse Lab in Alderley Park and Alex Alderton, Roberto Amato, Jeffrey Barrett, Sonia Goncalves, Ewan Harrison, David K. Jackson, Ian Johnston, Dominic Kwiatkowski, Cordelia Langford, John Sillitoe on behalf of the Wellcome Sanger Institute COVID-19 Surveillance Team |
| EPI_ISL_3068913, EPI_ISL_3068914 | Lighthouse Lab in Glasgow | Wellcome Sanger Institute for the COVID-19 Genomics UK (COG-UK) Consortium | Harper VanSteenhouse, Yumi Kasai, David Gray, Carol Clugston, Anna Dominiczak and Alex Alderton, Roberto Amato, Jeffrey Barrett, Sonia Goncalves, Ewan Harrison, David K. Jackson, Ian Johnston, Dominic Kwiatkowski, Cordelia Langford, John Sillitoe on behalf of the Wellcome Sanger Institute COVID-19 Surveillance Team |
| EPI_ISL_3068915, EPI_ISL_3068916 | Lighthouse Lab in Alderley Park | Wellcome Sanger Institute for the COVID-19 Genomics UK (COG-UK) Consortium | Jacquelyn Wynn, Mairead Hyland, The Lighthouse Lab in Alderley Park and Alex Alderton, Roberto Amato, Jeffrey Barrett, Sonia Goncalves, Ewan Harrison, David K. Jackson, Ian Johnston, Dominic Kwiatkowski, Cordelia Langford, John Sillitoe on behalf of the Wellcome Sanger Institute COVID-19 Surveillance Team |
| EPI_ISL_3068917 | Berkshire and Surrey Pathology Services Lighthouse Laboratory | Wellcome Sanger Institute for the COVID-19 Genomics UK (COG-UK) Consortium | Berkshire and Surrey Pathology Services Lighthouse Laboratory and Alex Alderton, Roberto Amato, Jeffrey Barrett, Sonia Goncalves, Ewan Harrison, David K. Jackson, Ian Johnston, Dominic Kwiatkowski, Cordelia Langford, John Sillitoe on behalf of the Wellcome Sanger Institute COVID-19 Surveillance Team |
| EPI_ISL_3068918 | Lighthouse Lab in Glasgow | Wellcome Sanger Institute for the COVID-19 Genomics UK (COG-UK) Consortium | Harper VanSteenhouse, Yumi Kasai, David Gray, Carol Clugston, Anna Dominiczak and Alex Alderton, Roberto Amato, Jeffrey Barrett, Sonia Goncalves, Ewan Harrison, David K. Jackson, Ian Johnston, Dominic Kwiatkowski, Cordelia Langford, John Sillitoe on behalf of the Wellcome Sanger Institute COVID-19 Surveillance Team |
| EPI_ISL_3068919 | Berkshire and Surrey Pathology Services Lighthouse Laboratory | Wellcome Sanger Institute for the COVID-19 Genomics UK (COG-UK) Consortium | Berkshire and Surrey Pathology Services Lighthouse Laboratory and Alex Alderton, Roberto Amato, Jeffrey Barrett, Sonia Goncalves, Ewan Harrison, David K. Jackson, Ian Johnston, Dominic Kwiatkowski, Cordelia Langford, John Sillitoe on behalf of the Wellcome Sanger Institute COVID-19 Surveillance Team |
| EPI_ISL_3068920, EPI_ISL_3068921, EPI_ISL_3068922 | Lighthouse Lab in Glasgow | Wellcome Sanger Institute for the COVID-19 Genomics UK (COG-UK) Consortium | Harper VanSteenhouse, Yumi Kasai, David Gray, Carol Clugston, Anna Dominiczak and Alex Alderton, Roberto Amato, Jeffrey Barrett, Sonia Goncalves, Ewan Harrison, David K. Jackson, Ian Johnston, Dominic Kwiatkowski, Cordelia Langford, John Sillitoe on behalf of the Wellcome Sanger Institute COVID-19 Surveillance Team |
| EPI_ISL_3068923 | Lighthouse Lab in Alderley Park | Wellcome Sanger Institute for the COVID-19 Genomics UK (COG-UK) Consortium | Jacquelyn Wynn, Mairead Hyland, The Lighthouse Lab in Alderley Park and Alex Alderton, Roberto Amato, Jeffrey Barrett, Sonia Goncalves, Ewan Harrison, David K. Jackson, Ian Johnston, Dominic Kwiatkowski, Cordelia Langford, John Sillitoe on behalf of the Wellcome Sanger Institute COVID-19 Surveillance Team |
| EPI_ISL_3068924 | Berkshire and Surrey Pathology Services Lighthouse Laboratory | Wellcome Sanger Institute for the COVID-19 Genomics UK (COG-UK) Consortium | Berkshire and Surrey Pathology Services Lighthouse Laboratory and Alex Alderton, Roberto Amato, Jeffrey Barrett, Sonia Goncalves, Ewan Harrison, David K. Jackson, Ian Johnston, Dominic Kwiatkowski, Cordelia Langford, John Sillitoe on behalf of the Wellcome Sanger Institute COVID-19 Surveillance Team |
| EPI_ISL_3068925 | Lighthouse Lab in Glasgow | Wellcome Sanger Institute for the COVID-19 Genomics UK (COG-UK) | Harper VanSteenhouse, Yumi Kasai, David Gray, Carol Clugston, Anna Dominiczak and Alex Alderton, Roberto Amato, Jeffrey Barrett, Sonia Goncalves, Ewan Harrison, David K. Jackson, Ian Johnston, Dominic Kwiatkowski, Cordelia Langford, |

| | | | |
|---|---|---|---|
| | | Consortium | John Sillitoe on behalf of the Wellcome Sanger Institute COVID-19 Surveillance Team |
| EPI_ISL_3068926 | Lighthouse Lab in Alderley Park | Wellcome Sanger Institute for the COVID-19 Genomics UK (COG-UK) Consortium | Jacquelyn Wynn, Mairead Hyland, The Lighthouse Lab in Alderley Park and Alex Alderton, Roberto Amato, Jeffrey Barrett, Sonia Goncalves, Ewan Harrison, David K. Jackson, Ian Johnston, Dominic Kwiatkowski, Cordelia Langford, John Sillitoe on behalf of the Wellcome Sanger Institute COVID-19 Surveillance Team |
| EPI_ISL_3068927 | Lighthouse Lab in Glasgow | Wellcome Sanger Institute for the COVID-19 Genomics UK (COG-UK) Consortium | Harper VanSteenhouse, Yumi Kasai, David Gray, Carol Clugston, Anna Dominiczak and Alex Alderton, Roberto Amato, Jeffrey Barrett, Sonia Goncalves, Ewan Harrison, David K. Jackson, Ian Johnston, Dominic Kwiatkowski, Cordelia Langford, John Sillitoe on behalf of the Wellcome Sanger Institute COVID-19 Surveillance Team |
| EPI_ISL_3068928, EPI_ISL_3068929, EPI_ISL_3068930 | Berkshire and Surrey Pathology Services Lighthouse Laboratory | Wellcome Sanger Institute for the COVID-19 Genomics UK (COG-UK) Consortium | Berkshire and Surrey Pathology Services Lighthouse Laboratory and Alex Alderton, Roberto Amato, Jeffrey Barrett, Sonia Goncalves, Ewan Harrison, David K. Jackson, Ian Johnston, Dominic Kwiatkowski, Cordelia Langford, John Sillitoe on behalf of the Wellcome Sanger Institute COVID-19 Surveillance Team |
| EPI_ISL_3068931, EPI_ISL_3068932, EPI_ISL_3068933 | Lighthouse Lab in Glasgow | Wellcome Sanger Institute for the COVID-19 Genomics UK (COG-UK) Consortium | Harper VanSteenhouse, Yumi Kasai, David Gray, Carol Clugston, Anna Dominiczak and Alex Alderton, Roberto Amato, Jeffrey Barrett, Sonia Goncalves, Ewan Harrison, David K. Jackson, Ian Johnston, Dominic Kwiatkowski, Cordelia Langford, John Sillitoe on behalf of the Wellcome Sanger Institute COVID-19 Surveillance Team |
| EPI_ISL_3068934 | Berkshire and Surrey Pathology Services Lighthouse Laboratory | Wellcome Sanger Institute for the COVID-19 Genomics UK (COG-UK) Consortium | Berkshire and Surrey Pathology Services Lighthouse Laboratory and Alex Alderton, Roberto Amato, Jeffrey Barrett, Sonia Goncalves, Ewan Harrison, David K. Jackson, Ian Johnston, Dominic Kwiatkowski, Cordelia Langford, John Sillitoe on behalf of the Wellcome Sanger Institute COVID-19 Surveillance Team |
| EPI_ISL_3068935 | Lighthouse Lab in Alderley Park | Wellcome Sanger Institute for the COVID-19 Genomics UK (COG-UK) Consortium | Jacquelyn Wynn, Mairead Hyland, The Lighthouse Lab in Alderley Park and Alex Alderton, Roberto Amato, Jeffrey Barrett, Sonia Goncalves, Ewan Harrison, David K. Jackson, Ian Johnston, Dominic Kwiatkowski, Cordelia Langford, John Sillitoe on behalf of the Wellcome Sanger Institute COVID-19 Surveillance Team |
| EPI_ISL_3068936 | Berkshire and Surrey Pathology Services Lighthouse Laboratory | Wellcome Sanger Institute for the COVID-19 Genomics UK (COG-UK) Consortium | Berkshire and Surrey Pathology Services Lighthouse Laboratory and Alex Alderton, Roberto Amato, Jeffrey Barrett, Sonia Goncalves, Ewan Harrison, David K. Jackson, Ian Johnston, Dominic Kwiatkowski, Cordelia Langford, John Sillitoe on behalf of the Wellcome Sanger Institute COVID-19 Surveillance Team |
| EPI_ISL_3068937, EPI_ISL_3068938, EPI_ISL_3068939 | Lighthouse Lab in Alderley Park | Wellcome Sanger Institute for the COVID-19 Genomics UK (COG-UK) Consortium | Jacquelyn Wynn, Mairead Hyland, The Lighthouse Lab in Alderley Park and Alex Alderton, Roberto Amato, Jeffrey Barrett, Sonia Goncalves, Ewan Harrison, David K. Jackson, Ian Johnston, Dominic Kwiatkowski, Cordelia Langford, John Sillitoe on behalf of the Wellcome Sanger Institute COVID-19 Surveillance Team |
| EPI_ISL_3068940, EPI_ISL_3068941 | Lighthouse Lab in Glasgow | Wellcome Sanger Institute for the COVID-19 Genomics UK (COG-UK) Consortium | Harper VanSteenhouse, Yumi Kasai, David Gray, Carol Clugston, Anna Dominiczak and Alex Alderton, Roberto Amato, Jeffrey Barrett, Sonia Goncalves, Ewan Harrison, David K. Jackson, Ian Johnston, Dominic Kwiatkowski, Cordelia Langford, John Sillitoe on behalf of the Wellcome Sanger Institute COVID-19 Surveillance Team |
| EPI_ISL_3068942 | Berkshire and Surrey Pathology Services Lighthouse Laboratory | Wellcome Sanger Institute for the COVID-19 Genomics UK (COG-UK) Consortium | Berkshire and Surrey Pathology Services Lighthouse Laboratory and Alex Alderton, Roberto Amato, Jeffrey Barrett, Sonia Goncalves, Ewan Harrison, David K. Jackson, Ian Johnston, Dominic Kwiatkowski, Cordelia Langford, John Sillitoe on behalf of the Wellcome Sanger Institute COVID-19 Surveillance Team |
| EPI_ISL_3068943 | Lighthouse Lab in Glasgow | Wellcome Sanger Institute for the COVID-19 Genomics UK (COG-UK) Consortium | Harper VanSteenhouse, Yumi Kasai, David Gray, Carol Clugston, Anna Dominiczak and Alex Alderton, Roberto Amato, Jeffrey Barrett, Sonia Goncalves, Ewan Harrison, David K. Jackson, Ian Johnston, Dominic Kwiatkowski, Cordelia Langford, John Sillitoe on behalf of the Wellcome Sanger Institute COVID-19 Surveillance Team |
| EPI_ISL_3068944 | Lighthouse Lab in Alderley Park | Wellcome Sanger Institute for the COVID-19 Genomics UK (COG-UK) Consortium | Jacquelyn Wynn, Mairead Hyland, The Lighthouse Lab in Alderley Park and Alex Alderton, Roberto Amato, Jeffrey Barrett, Sonia Goncalves, Ewan Harrison, David K. Jackson, Ian Johnston, Dominic Kwiatkowski, Cordelia Langford, John Sillitoe on behalf of the Wellcome Sanger Institute COVID-19 Surveillance Team |
| EPI_ISL_3068945 | Berkshire and Surrey Pathology Services Lighthouse Laboratory | Wellcome Sanger Institute for the COVID-19 Genomics UK (COG-UK) Consortium | Berkshire and Surrey Pathology Services Lighthouse Laboratory and Alex Alderton, Roberto Amato, Jeffrey Barrett, Sonia Goncalves, Ewan Harrison, David K. Jackson, Ian Johnston, Dominic Kwiatkowski, Cordelia Langford, John Sillitoe on behalf of the Wellcome Sanger Institute COVID-19 Surveillance Team |
| EPI_ISL_3068946 | Lighthouse Lab in Glasgow | Wellcome Sanger Institute for the COVID-19 Genomics UK (COG-UK) Consortium | Harper VanSteenhouse, Yumi Kasai, David Gray, Carol Clugston, Anna Dominiczak and Alex Alderton, Roberto Amato, Jeffrey Barrett, Sonia Goncalves, Ewan Harrison, David K. Jackson, Ian Johnston, Dominic Kwiatkowski, Cordelia Langford, John Sillitoe on behalf of the Wellcome Sanger Institute COVID-19 Surveillance Team |
| EPI_ISL_3068947 | Berkshire and Surrey Pathology Services Lighthouse Laboratory | Wellcome Sanger Institute for the COVID-19 Genomics UK (COG-UK) Consortium | Berkshire and Surrey Pathology Services Lighthouse Laboratory and Alex Alderton, Roberto Amato, Jeffrey Barrett, Sonia Goncalves, Ewan Harrison, David K. Jackson, Ian Johnston, Dominic Kwiatkowski, Cordelia Langford, John Sillitoe on behalf of the Wellcome Sanger Institute COVID-19 Surveillance Team |
| EPI_ISL_3068948 | Lighthouse Lab in Glasgow | Wellcome Sanger Institute for the COVID-19 Genomics UK (COG-UK) Consortium | Harper VanSteenhouse, Yumi Kasai, David Gray, Carol Clugston, Anna Dominiczak and Alex Alderton, Roberto Amato, Jeffrey Barrett, Sonia Goncalves, Ewan Harrison, David K. Jackson, Ian Johnston, Dominic Kwiatkowski, Cordelia Langford, John Sillitoe on behalf of the Wellcome Sanger Institute COVID-19 Surveillance Team |
| EPI_ISL_3068949 | Berkshire and Surrey Pathology Services Lighthouse Laboratory | Wellcome Sanger Institute for the COVID-19 Genomics UK (COG-UK) Consortium | Berkshire and Surrey Pathology Services Lighthouse Laboratory and Alex Alderton, Roberto Amato, Jeffrey Barrett, Sonia Goncalves, Ewan Harrison, David K. Jackson, Ian Johnston, Dominic Kwiatkowski, Cordelia Langford, John Sillitoe on behalf of the Wellcome Sanger Institute COVID-19 Surveillance Team |
| EPI_ISL_3068950 | Lighthouse Lab in Glasgow | Wellcome Sanger Institute for the COVID-19 Genomics UK (COG-UK) Consortium | Harper VanSteenhouse, Yumi Kasai, David Gray, Carol Clugston, Anna Dominiczak and Alex Alderton, Roberto Amato, Jeffrey Barrett, Sonia Goncalves, Ewan Harrison, David K. Jackson, Ian Johnston, Dominic Kwiatkowski, Cordelia Langford, John Sillitoe on behalf of the Wellcome Sanger Institute COVID-19 Surveillance Team |
| EPI_ISL_3068951 | Lighthouse Lab in Alderley Park | Wellcome Sanger Institute for the COVID-19 Genomics UK (COG-UK) Consortium | Jacquelyn Wynn, Mairead Hyland, The Lighthouse Lab in Alderley Park and Alex Alderton, Roberto Amato, Jeffrey Barrett, Sonia Goncalves, Ewan Harrison, David K. Jackson, Ian Johnston, Dominic Kwiatkowski, Cordelia Langford, John Sillitoe on behalf of the Wellcome Sanger Institute COVID-19 Surveillance Team |
| EPI_ISL_3068952, EPI_ISL_3068953, EPI_ISL_3068954, EPI_ISL_3068955, EPI_ISL_3068956, EPI_ISL_3068957, EPI_ISL_3068958, EPI_ISL_3068959, EPI_ISL_3068960, EPI_ISL_3068961 | Lighthouse Lab in Glasgow | Wellcome Sanger Institute for the COVID-19 Genomics UK (COG-UK) Consortium | Harper VanSteenhouse, Yumi Kasai, David Gray, Carol Clugston, Anna Dominiczak and Alex Alderton, Roberto Amato, Jeffrey Barrett, Sonia Goncalves, Ewan Harrison, David K. Jackson, Ian Johnston, Dominic Kwiatkowski, Cordelia Langford, John Sillitoe on behalf of the Wellcome Sanger Institute COVID-19 Surveillance Team |
| EPI_ISL_3068962 | Berkshire and Surrey Pathology Services Lighthouse Laboratory | Wellcome Sanger Institute for the COVID-19 Genomics UK (COG-UK) Consortium | Berkshire and Surrey Pathology Services Lighthouse Laboratory and Alex Alderton, Roberto Amato, Jeffrey Barrett, Sonia Goncalves, Ewan Harrison, David K. Jackson, Ian Johnston, Dominic Kwiatkowski, Cordelia Langford, John Sillitoe on behalf of the Wellcome Sanger Institute COVID-19 Surveillance Team |
| EPI_ISL_3068963 | Lighthouse Lab in Alderley Park | Wellcome Sanger Institute for the COVID-19 Genomics UK (COG-UK) Consortium | Jacquelyn Wynn, Mairead Hyland, The Lighthouse Lab in Alderley Park and Alex Alderton, Roberto Amato, Jeffrey Barrett, Sonia Goncalves, Ewan Harrison, David K. Jackson, Ian Johnston, Dominic Kwiatkowski, Cordelia Langford, John Sillitoe on behalf of the Wellcome Sanger Institute COVID-19 Surveillance Team |
| EPI_ISL_3068964, EPI_ISL_3068965 | Lighthouse Lab in Glasgow | Wellcome Sanger Institute for the COVID-19 Genomics UK (COG-UK) Consortium | Harper VanSteenhouse, Yumi Kasai, David Gray, Carol Clugston, Anna Dominiczak and Alex Alderton, Roberto Amato, Jeffrey Barrett, Sonia Goncalves, Ewan Harrison, David K. Jackson, Ian Johnston, Dominic Kwiatkowski, Cordelia Langford, John Sillitoe on behalf of the Wellcome Sanger Institute COVID-19 Surveillance Team |
| EPI_ISL_3068966 | Berkshire and Surrey Pathology Services Lighthouse Laboratory | Wellcome Sanger Institute for the COVID-19 Genomics UK (COG-UK) Consortium | Berkshire and Surrey Pathology Services Lighthouse Laboratory and Alex Alderton, Roberto Amato, Jeffrey Barrett, Sonia Goncalves, Ewan Harrison, David K. Jackson, Ian Johnston, Dominic Kwiatkowski, Cordelia Langford, John Sillitoe on behalf of the Wellcome Sanger Institute COVID-19 Surveillance Team |
| EPI_ISL_3068967 | Lighthouse Lab in Glasgow | Wellcome Sanger Institute for the COVID-19 Genomics UK (COG-UK) Consortium | Harper VanSteenhouse, Yumi Kasai, David Gray, Carol Clugston, Anna Dominiczak and Alex Alderton, Roberto Amato, Jeffrey Barrett, Sonia Goncalves, Ewan Harrison, David K. Jackson, Ian Johnston, Dominic Kwiatkowski, Cordelia Langford, John Sillitoe on behalf of the Wellcome Sanger Institute COVID-19 Surveillance Team |
| EPI_ISL_3068968 | Lighthouse Lab in Alderley Park | Wellcome Sanger Institute for the COVID-19 Genomics UK (COG-UK) Consortium | Jacquelyn Wynn, Mairead Hyland, The Lighthouse Lab in Alderley Park and Alex Alderton, Roberto Amato, Jeffrey Barrett, Sonia Goncalves, Ewan Harrison, David K. Jackson, Ian Johnston, Dominic Kwiatkowski, Cordelia Langford, John Sillitoe on behalf of the Wellcome Sanger Institute COVID-19 Surveillance Team |
| EPI_ISL_3068969, EPI_ISL_3068970, EPI_ISL_3068971 | Lighthouse Lab in Glasgow | Wellcome Sanger Institute for the COVID-19 Genomics UK (COG-UK) Consortium | Harper VanSteenhouse, Yumi Kasai, David Gray, Carol Clugston, Anna Dominiczak and Alex Alderton, Roberto Amato, Jeffrey Barrett, Sonia Goncalves, Ewan Harrison, David K. Jackson, Ian Johnston, Dominic Kwiatkowski, Cordelia Langford, John Sillitoe on behalf of the Wellcome Sanger Institute COVID-19 Surveillance Team |
| EPI_ISL_3068972 | Lighthouse Lab in Alderley Park | Wellcome Sanger Institute for the COVID-19 Genomics UK (COG-UK) Consortium | Jacquelyn Wynn, Mairead Hyland, The Lighthouse Lab in Alderley Park and Alex Alderton, Roberto Amato, Jeffrey Barrett, Sonia Goncalves, Ewan Harrison, David K. Jackson, Ian Johnston, Dominic Kwiatkowski, Cordelia Langford, John Sillitoe on behalf of the Wellcome Sanger Institute COVID-19 Surveillance Team |
| EPI_ISL_3068973 | Berkshire and Surrey Pathology Services Lighthouse Laboratory | Wellcome Sanger Institute for the COVID-19 Genomics UK (COG-UK) Consortium | Berkshire and Surrey Pathology Services Lighthouse Laboratory and Alex Alderton, Roberto Amato, Jeffrey Barrett, Sonia Goncalves, Ewan Harrison, David K. Jackson, Ian Johnston, Dominic Kwiatkowski, Cordelia Langford, John Sillitoe on behalf of the Wellcome Sanger Institute COVID-19 Surveillance Team |
| EPI_ISL_3068974, EPI_ISL_3068975, EPI_ISL_3068976 | Lighthouse Lab in Glasgow | Wellcome Sanger Institute for the COVID-19 Genomics UK (COG-UK) Consortium | Harper VanSteenhouse, Yumi Kasai, David Gray, Carol Clugston, Anna Dominiczak and Alex Alderton, Roberto Amato, Jeffrey Barrett, Sonia Goncalves, Ewan Harrison, David K. Jackson, Ian Johnston, Dominic Kwiatkowski, Cordelia Langford, John Sillitoe on behalf of the Wellcome Sanger Institute COVID-19 Surveillance Team |
| EPI_ISL_3068977 | Lighthouse Lab in Alderley Park | Wellcome Sanger Institute for the COVID-19 Genomics UK (COG-UK) Consortium | Jacquelyn Wynn, Mairead Hyland, The Lighthouse Lab in Alderley Park and Alex Alderton, Roberto Amato, Jeffrey Barrett, Sonia Goncalves, Ewan Harrison, David K. Jackson, Ian Johnston, Dominic Kwiatkowski, Cordelia Langford, John Sillitoe on behalf of the Wellcome Sanger Institute COVID-19 Surveillance Team |
| EPI_ISL_3068978 | Berkshire and Surrey Pathology Services Lighthouse Laboratory | Wellcome Sanger Institute for the COVID-19 Genomics UK (COG-UK) Consortium | Berkshire and Surrey Pathology Services Lighthouse Laboratory and Alex Alderton, Roberto Amato, Jeffrey Barrett, Sonia Goncalves, Ewan Harrison, David K. Jackson, Ian Johnston, Dominic Kwiatkowski, Cordelia Langford, John Sillitoe on behalf of the Wellcome Sanger Institute COVID-19 Surveillance Team |
| EPI_ISL_3068979 | Lighthouse Lab in Alderley Park | Wellcome Sanger Institute for the COVID-19 Genomics UK (COG-UK) Consortium | Jacquelyn Wynn, Mairead Hyland, The Lighthouse Lab in Alderley Park and Alex Alderton, Roberto Amato, Jeffrey Barrett, Sonia Goncalves, Ewan Harrison, David K. Jackson, Ian Johnston, Dominic Kwiatkowski, Cordelia Langford, John Sillitoe on behalf of the Wellcome Sanger Institute COVID-19 Surveillance Team |
| EPI_ISL_3068980 | Lighthouse Lab in Glasgow | Wellcome Sanger Institute for the COVID-19 Genomics UK (COG-UK) Consortium | Harper VanSteenhouse, Yumi Kasai, David Gray, Carol Clugston, Anna Dominiczak and Alex Alderton, Roberto Amato, Jeffrey Barrett, Sonia Goncalves, Ewan Harrison, David K. Jackson, Ian Johnston, Dominic Kwiatkowski, Cordelia Langford, John Sillitoe on behalf of the Wellcome Sanger Institute COVID-19 Surveillance Team |
| EPI_ISL_3068981, EPI_ISL_3068982, EPI_ISL_3068983 | Lighthouse Lab in Alderley Park | Wellcome Sanger Institute for the COVID-19 Genomics UK (COG-UK) Consortium | Jacquelyn Wynn, Mairead Hyland, The Lighthouse Lab in Alderley Park and Alex Alderton, Roberto Amato, Jeffrey Barrett, Sonia Goncalves, Ewan Harrison, David K. Jackson, Ian Johnston, Dominic Kwiatkowski, Cordelia Langford, John Sillitoe on behalf of the Wellcome Sanger Institute COVID-19 Surveillance Team |
| EPI_ISL_3068984 | Lighthouse Lab in Glasgow | Wellcome Sanger Institute for the COVID-19 Genomics UK (COG-UK) Consortium | Harper VanSteenhouse, Yumi Kasai, David Gray, Carol Clugston, Anna Dominiczak and Alex Alderton, Roberto Amato, Jeffrey Barrett, Sonia Goncalves, Ewan Harrison, David K. Jackson, Ian Johnston, Dominic Kwiatkowski, Cordelia Langford, John Sillitoe on behalf of the Wellcome Sanger Institute COVID-19 Surveillance Team |
| EPI_ISL_3068985 | Lighthouse Lab in Alderley Park | Wellcome Sanger Institute for the COVID-19 Genomics UK (COG-UK) Consortium | Jacquelyn Wynn, Mairead Hyland, The Lighthouse Lab in Alderley Park and Alex Alderton, Roberto Amato, Jeffrey Barrett, Sonia Goncalves, Ewan Harrison, David K. Jackson, Ian Johnston, Dominic Kwiatkowski, Cordelia Langford, John Sillitoe on behalf of the Wellcome Sanger Institute COVID-19 Surveillance Team |
| EPI_ISL_3068986 | Berkshire and Surrey Pathology Services Lighthouse Laboratory | Wellcome Sanger Institute for the COVID-19 Genomics UK (COG-UK) Consortium | Berkshire and Surrey Pathology Services Lighthouse Laboratory and Alex Alderton, Roberto Amato, Jeffrey Barrett, Sonia Goncalves, Ewan Harrison, David K. Jackson, Ian Johnston, Dominic Kwiatkowski, Cordelia Langford, John Sillitoe on behalf of the Wellcome Sanger Institute COVID-19 Surveillance Team |
| EPI_ISL_3068987 | Lighthouse Lab in Glasgow | Wellcome Sanger Institute for the COVID-19 Genomics UK (COG-UK) | Harper VanSteenhouse, Yumi Kasai, David Gray, Carol Clugston, Anna Dominiczak and Alex Alderton, Roberto Amato, Jeffrey Barrett, Sonia Goncalves, Ewan Harrison, David K. Jackson, Ian Johnston, Dominic Kwiatkowski, Cordelia Langford, |

| | | | |
|---|---|---|---|
| | | Consortium | John Sillitoe on behalf of the Wellcome Sanger Institute COVID-19 Surveillance Team |
| EPI_ISL_3068988, EPI_ISL_3068989 | Berkshire and Surrey Pathology Services Lighthouse Laboratory | Wellcome Sanger Institute for the COVID-19 Genomics UK (COG-UK) Consortium | Berkshire and Surrey Pathology Services Lighthouse Laboratory and Alex Alderton, Roberto Amato, Jeffrey Barrett, Sonia Goncalves, Ewan Harrison, David K. Jackson, Ian Johnston, Dominic Kwiatkowski, Cordelia Langford, John Sillitoe on behalf of the Wellcome Sanger Institute COVID-19 Surveillance Team |
| EPI_ISL_3068990 | Lighthouse Lab in Glasgow | Wellcome Sanger Institute for the COVID-19 Genomics UK (COG-UK) Consortium | Harper VanSteenhouse, Yumi Kasai, David Gray, Carol Clugston, Anna Dominiczak and Alex Alderton, Roberto Amato, Jeffrey Barrett, Sonia Goncalves, Ewan Harrison, David K. Jackson, Ian Johnston, Dominic Kwiatkowski, Cordelia Langford, John Sillitoe on behalf of the Wellcome Sanger Institute COVID-19 Surveillance Team |
| EPI_ISL_3068991, EPI_ISL_3068992 | Berkshire and Surrey Pathology Services Lighthouse Laboratory | Wellcome Sanger Institute for the COVID-19 Genomics UK (COG-UK) Consortium | Berkshire and Surrey Pathology Services Lighthouse Laboratory and Alex Alderton, Roberto Amato, Jeffrey Barrett, Sonia Goncalves, Ewan Harrison, David K. Jackson, Ian Johnston, Dominic Kwiatkowski, Cordelia Langford, John Sillitoe on behalf of the Wellcome Sanger Institute COVID-19 Surveillance Team |
| EPI_ISL_3068993, EPI_ISL_3068994 | Lighthouse Lab in Glasgow | Wellcome Sanger Institute for the COVID-19 Genomics UK (COG-UK) Consortium | Harper VanSteenhouse, Yumi Kasai, David Gray, Carol Clugston, Anna Dominiczak and Alex Alderton, Roberto Amato, Jeffrey Barrett, Sonia Goncalves, Ewan Harrison, David K. Jackson, Ian Johnston, Dominic Kwiatkowski, Cordelia Langford, John Sillitoe on behalf of the Wellcome Sanger Institute COVID-19 Surveillance Team |
| EPI_ISL_3068995, EPI_ISL_3068996 | Lighthouse Lab in Alderley Park | Wellcome Sanger Institute for the COVID-19 Genomics UK (COG-UK) Consortium | Jacquelyn Wynn, Mairead Hyland, The Lighthouse Lab in Alderley Park and Alex Alderton, Roberto Amato, Jeffrey Barrett, Sonia Goncalves, Ewan Harrison, David K. Jackson, Ian Johnston, Dominic Kwiatkowski, Cordelia Langford, John Sillitoe on behalf of the Wellcome Sanger Institute COVID-19 Surveillance Team |
| EPI_ISL_3068997, EPI_ISL_3068998 | Berkshire and Surrey Pathology Services Lighthouse Laboratory | Wellcome Sanger Institute for the COVID-19 Genomics UK (COG-UK) Consortium | Berkshire and Surrey Pathology Services Lighthouse Laboratory and Alex Alderton, Roberto Amato, Jeffrey Barrett, Sonia Goncalves, Ewan Harrison, David K. Jackson, Ian Johnston, Dominic Kwiatkowski, Cordelia Langford, John Sillitoe on behalf of the Wellcome Sanger Institute COVID-19 Surveillance Team |
| EPI_ISL_3068999 | Lighthouse Lab in Glasgow | Wellcome Sanger Institute for the COVID-19 Genomics UK (COG-UK) Consortium | Harper VanSteenhouse, Yumi Kasai, David Gray, Carol Clugston, Anna Dominiczak and Alex Alderton, Roberto Amato, Jeffrey Barrett, Sonia Goncalves, Ewan Harrison, David K. Jackson, Ian Johnston, Dominic Kwiatkowski, Cordelia Langford, John Sillitoe on behalf of the Wellcome Sanger Institute COVID-19 Surveillance Team |
| EPI_ISL_3069000, EPI_ISL_3069001 | Berkshire and Surrey Pathology Services Lighthouse Laboratory | Wellcome Sanger Institute for the COVID-19 Genomics UK (COG-UK) Consortium | Berkshire and Surrey Pathology Services Lighthouse Laboratory and Alex Alderton, Roberto Amato, Jeffrey Barrett, Sonia Goncalves, Ewan Harrison, David K. Jackson, Ian Johnston, Dominic Kwiatkowski, Cordelia Langford, John Sillitoe on behalf of the Wellcome Sanger Institute COVID-19 Surveillance Team |
| EPI_ISL_3069002 | Lighthouse Lab in Glasgow | Wellcome Sanger Institute for the COVID-19 Genomics UK (COG-UK) Consortium | Harper VanSteenhouse, Yumi Kasai, David Gray, Carol Clugston, Anna Dominiczak and Alex Alderton, Roberto Amato, Jeffrey Barrett, Sonia Goncalves, Ewan Harrison, David K. Jackson, Ian Johnston, Dominic Kwiatkowski, Cordelia Langford, John Sillitoe on behalf of the Wellcome Sanger Institute COVID-19 Surveillance Team |
| EPI_ISL_3069003 | Lighthouse Lab in Alderley Park | Wellcome Sanger Institute for the COVID-19 Genomics UK (COG-UK) Consortium | Jacquelyn Wynn, Mairead Hyland, The Lighthouse Lab in Alderley Park and Alex Alderton, Roberto Amato, Jeffrey Barrett, Sonia Goncalves, Ewan Harrison, David K. Jackson, Ian Johnston, Dominic Kwiatkowski, Cordelia Langford, John Sillitoe on behalf of the Wellcome Sanger Institute COVID-19 Surveillance Team |
| EPI_ISL_3069004 | Berkshire and Surrey Pathology Services Lighthouse Laboratory | Wellcome Sanger Institute for the COVID-19 Genomics UK (COG-UK) Consortium | Berkshire and Surrey Pathology Services Lighthouse Laboratory and Alex Alderton, Roberto Amato, Jeffrey Barrett, Sonia Goncalves, Ewan Harrison, David K. Jackson, Ian Johnston, Dominic Kwiatkowski, Cordelia Langford, John Sillitoe on behalf of the Wellcome Sanger Institute COVID-19 Surveillance Team |
| EPI_ISL_3069005 | Lighthouse Lab in Glasgow | Wellcome Sanger Institute for the COVID-19 Genomics UK (COG-UK) Consortium | Harper VanSteenhouse, Yumi Kasai, David Gray, Carol Clugston, Anna Dominiczak and Alex Alderton, Roberto Amato, Jeffrey Barrett, Sonia Goncalves, Ewan Harrison, David K. Jackson, Ian Johnston, Dominic Kwiatkowski, Cordelia Langford, John Sillitoe on behalf of the Wellcome Sanger Institute COVID-19 Surveillance Team |
| EPI_ISL_3069006 | Berkshire and Surrey Pathology Services Lighthouse Laboratory | Wellcome Sanger Institute for the COVID-19 Genomics UK (COG-UK) Consortium | Berkshire and Surrey Pathology Services Lighthouse Laboratory and Alex Alderton, Roberto Amato, Jeffrey Barrett, Sonia Goncalves, Ewan Harrison, David K. Jackson, Ian Johnston, Dominic Kwiatkowski, Cordelia Langford, John Sillitoe on behalf of the Wellcome Sanger Institute COVID-19 Surveillance Team |
| EPI_ISL_3069007 | Lighthouse Lab in Alderley Park | Wellcome Sanger Institute for the COVID-19 Genomics UK (COG-UK) Consortium | Jacquelyn Wynn, Mairead Hyland, The Lighthouse Lab in Alderley Park and Alex Alderton, Roberto Amato, Jeffrey Barrett, Sonia Goncalves, Ewan Harrison, David K. Jackson, Ian Johnston, Dominic Kwiatkowski, Cordelia Langford, John Sillitoe on behalf of the Wellcome Sanger Institute COVID-19 Surveillance Team |
| EPI_ISL_3069008, EPI_ISL_3069009, EPI_ISL_3069010 | Lighthouse Lab in Glasgow | Wellcome Sanger Institute for the COVID-19 Genomics UK (COG-UK) Consortium | Harper VanSteenhouse, Yumi Kasai, David Gray, Carol Clugston, Anna Dominiczak and Alex Alderton, Roberto Amato, Jeffrey Barrett, Sonia Goncalves, Ewan Harrison, David K. Jackson, Ian Johnston, Dominic Kwiatkowski, Cordelia Langford, John Sillitoe on behalf of the Wellcome Sanger Institute COVID-19 Surveillance Team |
| EPI_ISL_3069011 | Berkshire and Surrey Pathology Services Lighthouse Laboratory | Wellcome Sanger Institute for the COVID-19 Genomics UK (COG-UK) Consortium | Berkshire and Surrey Pathology Services Lighthouse Laboratory and Alex Alderton, Roberto Amato, Jeffrey Barrett, Sonia Goncalves, Ewan Harrison, David K. Jackson, Ian Johnston, Dominic Kwiatkowski, Cordelia Langford, John Sillitoe on behalf of the Wellcome Sanger Institute COVID-19 Surveillance Team |
| EPI_ISL_3069012, EPI_ISL_3069013, EPI_ISL_3069014 | Lighthouse Lab in Glasgow | Wellcome Sanger Institute for the COVID-19 Genomics UK (COG-UK) Consortium | Harper VanSteenhouse, Yumi Kasai, David Gray, Carol Clugston, Anna Dominiczak and Alex Alderton, Roberto Amato, Jeffrey Barrett, Sonia Goncalves, Ewan Harrison, David K. Jackson, Ian Johnston, Dominic Kwiatkowski, Cordelia Langford, John Sillitoe on behalf of the Wellcome Sanger Institute COVID-19 Surveillance Team |
| EPI_ISL_3069015 | Lighthouse Lab in Alderley Park | Wellcome Sanger Institute for the COVID-19 Genomics UK (COG-UK) Consortium | Jacquelyn Wynn, Mairead Hyland, The Lighthouse Lab in Alderley Park and Alex Alderton, Roberto Amato, Jeffrey Barrett, Sonia Goncalves, Ewan Harrison, David K. Jackson, Ian Johnston, Dominic Kwiatkowski, Cordelia Langford, John Sillitoe on behalf of the Wellcome Sanger Institute COVID-19 Surveillance Team |
| EPI_ISL_3069016 | Lighthouse Lab in Glasgow | Wellcome Sanger Institute for the COVID-19 Genomics UK (COG-UK) Consortium | Harper VanSteenhouse, Yumi Kasai, David Gray, Carol Clugston, Anna Dominiczak and Alex Alderton, Roberto Amato, Jeffrey Barrett, Sonia Goncalves, Ewan Harrison, David K. Jackson, Ian Johnston, Dominic Kwiatkowski, Cordelia Langford, John Sillitoe on behalf of the Wellcome Sanger Institute COVID-19 Surveillance Team |
| EPI_ISL_3069017 | Berkshire and Surrey Pathology Services Lighthouse Laboratory | Wellcome Sanger Institute for the COVID-19 Genomics UK (COG-UK) Consortium | Berkshire and Surrey Pathology Services Lighthouse Laboratory and Alex Alderton, Roberto Amato, Jeffrey Barrett, Sonia Goncalves, Ewan Harrison, David K. Jackson, Ian Johnston, Dominic Kwiatkowski, Cordelia Langford, John Sillitoe on behalf of the Wellcome Sanger Institute COVID-19 Surveillance Team |
| EPI_ISL_3069018 | Lighthouse Lab in Glasgow | Wellcome Sanger Institute for the COVID-19 Genomics UK (COG-UK) Consortium | Harper VanSteenhouse, Yumi Kasai, David Gray, Carol Clugston, Anna Dominiczak and Alex Alderton, Roberto Amato, Jeffrey Barrett, Sonia Goncalves, Ewan Harrison, David K. Jackson, Ian Johnston, Dominic Kwiatkowski, Cordelia Langford, John Sillitoe on behalf of the Wellcome Sanger Institute COVID-19 Surveillance Team |
| EPI_ISL_3069019, EPI_ISL_3069020 | Berkshire and Surrey Pathology Services Lighthouse Laboratory | Wellcome Sanger Institute for the COVID-19 Genomics UK (COG-UK) Consortium | Berkshire and Surrey Pathology Services Lighthouse Laboratory and Alex Alderton, Roberto Amato, Jeffrey Barrett, Sonia Goncalves, Ewan Harrison, David K. Jackson, Ian Johnston, Dominic Kwiatkowski, Cordelia Langford, John Sillitoe on behalf of the Wellcome Sanger Institute COVID-19 Surveillance Team |
| EPI_ISL_3069021 | Lighthouse Lab in Alderley Park | Wellcome Sanger Institute for the COVID-19 Genomics UK (COG-UK) Consortium | Jacquelyn Wynn, Mairead Hyland, The Lighthouse Lab in Alderley Park and Alex Alderton, Roberto Amato, Jeffrey Barrett, Sonia Goncalves, Ewan Harrison, David K. Jackson, Ian Johnston, Dominic Kwiatkowski, Cordelia Langford, John Sillitoe on behalf of the Wellcome Sanger Institute COVID-19 Surveillance Team |
| EPI_ISL_3069022, EPI_ISL_3069023 | Berkshire and Surrey Pathology Services Lighthouse Laboratory | Wellcome Sanger Institute for the COVID-19 Genomics UK (COG-UK) Consortium | Berkshire and Surrey Pathology Services Lighthouse Laboratory and Alex Alderton, Roberto Amato, Jeffrey Barrett, Sonia Goncalves, Ewan Harrison, David K. Jackson, Ian Johnston, Dominic Kwiatkowski, Cordelia Langford, John Sillitoe on behalf of the Wellcome Sanger Institute COVID-19 Surveillance Team |
| EPI_ISL_3069024 | Lighthouse Lab in Alderley Park | Wellcome Sanger Institute for the COVID-19 Genomics UK (COG-UK) Consortium | Jacquelyn Wynn, Mairead Hyland, The Lighthouse Lab in Alderley Park and Alex Alderton, Roberto Amato, Jeffrey Barrett, Sonia Goncalves, Ewan Harrison, David K. Jackson, Ian Johnston, Dominic Kwiatkowski, Cordelia Langford, John Sillitoe on behalf of the Wellcome Sanger Institute COVID-19 Surveillance Team |
| EPI_ISL_3069025 | Berkshire and Surrey Pathology Services Lighthouse Laboratory | Wellcome Sanger Institute for the COVID-19 Genomics UK (COG-UK) Consortium | Berkshire and Surrey Pathology Services Lighthouse Laboratory and Alex Alderton, Roberto Amato, Jeffrey Barrett, Sonia Goncalves, Ewan Harrison, David K. Jackson, Ian Johnston, Dominic Kwiatkowski, Cordelia Langford, John Sillitoe on behalf of the Wellcome Sanger Institute COVID-19 Surveillance Team |
| EPI_ISL_3069026 | Lighthouse Lab in Alderley Park | Wellcome Sanger Institute for the COVID-19 Genomics UK (COG-UK) Consortium | Jacquelyn Wynn, Mairead Hyland, The Lighthouse Lab in Alderley Park and Alex Alderton, Roberto Amato, Jeffrey Barrett, Sonia Goncalves, Ewan Harrison, David K. Jackson, Ian Johnston, Dominic Kwiatkowski, Cordelia Langford, John Sillitoe on behalf of the Wellcome Sanger Institute COVID-19 Surveillance Team |
| EPI_ISL_3069027 | Berkshire and Surrey Pathology Services Lighthouse Laboratory | Wellcome Sanger Institute for the COVID-19 Genomics UK (COG-UK) Consortium | Berkshire and Surrey Pathology Services Lighthouse Laboratory and Alex Alderton, Roberto Amato, Jeffrey Barrett, Sonia Goncalves, Ewan Harrison, David K. Jackson, Ian Johnston, Dominic Kwiatkowski, Cordelia Langford, John Sillitoe on behalf of the Wellcome Sanger Institute COVID-19 Surveillance Team |
| EPI_ISL_3069028 | Lighthouse Lab in Glasgow | Wellcome Sanger Institute for the COVID-19 Genomics UK (COG-UK) Consortium | Harper VanSteenhouse, Yumi Kasai, David Gray, Carol Clugston, Anna Dominiczak and Alex Alderton, Roberto Amato, Jeffrey Barrett, Sonia Goncalves, Ewan Harrison, David K. Jackson, Ian Johnston, Dominic Kwiatkowski, Cordelia Langford, John Sillitoe on behalf of the Wellcome Sanger Institute COVID-19 Surveillance Team |
| EPI_ISL_3069029 | Lighthouse Lab in Alderley Park | Wellcome Sanger Institute for the COVID-19 Genomics UK (COG-UK) Consortium | Jacquelyn Wynn, Mairead Hyland, The Lighthouse Lab in Alderley Park and Alex Alderton, Roberto Amato, Jeffrey Barrett, Sonia Goncalves, Ewan Harrison, David K. Jackson, Ian Johnston, Dominic Kwiatkowski, Cordelia Langford, John Sillitoe on behalf of the Wellcome Sanger Institute COVID-19 Surveillance Team |
| EPI_ISL_3069030 | Berkshire and Surrey Pathology Services Lighthouse Laboratory | Wellcome Sanger Institute for the COVID-19 Genomics UK (COG-UK) Consortium | Berkshire and Surrey Pathology Services Lighthouse Laboratory and Alex Alderton, Roberto Amato, Jeffrey Barrett, Sonia Goncalves, Ewan Harrison, David K. Jackson, Ian Johnston, Dominic Kwiatkowski, Cordelia Langford, John Sillitoe on behalf of the Wellcome Sanger Institute COVID-19 Surveillance Team |
| EPI_ISL_3069031 | Lighthouse Lab in Glasgow | Wellcome Sanger Institute for the COVID-19 Genomics UK (COG-UK) Consortium | Harper VanSteenhouse, Yumi Kasai, David Gray, Carol Clugston, Anna Dominiczak and Alex Alderton, Roberto Amato, Jeffrey Barrett, Sonia Goncalves, Ewan Harrison, David K. Jackson, Ian Johnston, Dominic Kwiatkowski, Cordelia Langford, John Sillitoe on behalf of the Wellcome Sanger Institute COVID-19 Surveillance Team |
| EPI_ISL_3069032, EPI_ISL_3069033, EPI_ISL_3069034, EPI_ISL_3069035 | Lighthouse Lab in Alderley Park | Wellcome Sanger Institute for the COVID-19 Genomics UK (COG-UK) Consortium | Jacquelyn Wynn, Mairead Hyland, The Lighthouse Lab in Alderley Park and Alex Alderton, Roberto Amato, Jeffrey Barrett, Sonia Goncalves, Ewan Harrison, David K. Jackson, Ian Johnston, Dominic Kwiatkowski, Cordelia Langford, John Sillitoe on behalf of the Wellcome Sanger Institute COVID-19 Surveillance Team |
| EPI_ISL_3069036 | Lighthouse Lab in Glasgow | Wellcome Sanger Institute for the COVID-19 Genomics UK (COG-UK) Consortium | Harper VanSteenhouse, Yumi Kasai, David Gray, Carol Clugston, Anna Dominiczak and Alex Alderton, Roberto Amato, Jeffrey Barrett, Sonia Goncalves, Ewan Harrison, David K. Jackson, Ian Johnston, Dominic Kwiatkowski, Cordelia Langford, John Sillitoe on behalf of the Wellcome Sanger Institute COVID-19 Surveillance Team |
| EPI_ISL_3069037 | Lighthouse Lab in Alderley Park | Wellcome Sanger Institute for the COVID-19 Genomics UK (COG-UK) Consortium | Jacquelyn Wynn, Mairead Hyland, The Lighthouse Lab in Alderley Park and Alex Alderton, Roberto Amato, Jeffrey Barrett, Sonia Goncalves, Ewan Harrison, David K. Jackson, Ian Johnston, Dominic Kwiatkowski, Cordelia Langford, John Sillitoe on behalf of the Wellcome Sanger Institute COVID-19 Surveillance Team |
| EPI_ISL_3069038 | Berkshire and Surrey Pathology Services Lighthouse Laboratory | Wellcome Sanger Institute for the COVID-19 Genomics UK (COG-UK) Consortium | Berkshire and Surrey Pathology Services Lighthouse Laboratory and Alex Alderton, Roberto Amato, Jeffrey Barrett, Sonia Goncalves, Ewan Harrison, David K. Jackson, Ian Johnston, Dominic Kwiatkowski, Cordelia Langford, John Sillitoe on behalf of the Wellcome Sanger Institute COVID-19 Surveillance Team |
| EPI_ISL_3069039 | Lighthouse Lab in Alderley Park | Wellcome Sanger Institute for the COVID-19 Genomics UK (COG-UK) Consortium | Jacquelyn Wynn, Mairead Hyland, The Lighthouse Lab in Alderley Park and Alex Alderton, Roberto Amato, Jeffrey Barrett, Sonia Goncalves, Ewan Harrison, David K. Jackson, Ian Johnston, Dominic Kwiatkowski, Cordelia Langford, John Sillitoe on behalf of the Wellcome Sanger Institute COVID-19 Surveillance Team |
| EPI_ISL_3069040 | Lighthouse Lab in Glasgow | Wellcome Sanger Institute for the COVID-19 Genomics UK (COG-UK) Consortium | Harper VanSteenhouse, Yumi Kasai, David Gray, Carol Clugston, Anna Dominiczak and Alex Alderton, Roberto Amato, Jeffrey Barrett, Sonia Goncalves, Ewan Harrison, David K. Jackson, Ian Johnston, Dominic Kwiatkowski, Cordelia Langford, John Sillitoe on behalf of the Wellcome Sanger Institute COVID-19 Surveillance Team |
| EPI_ISL_3069041 | Berkshire and Surrey Pathology Services Lighthouse Laboratory | Wellcome Sanger Institute for the COVID-19 Genomics UK (COG-UK) Consortium | Berkshire and Surrey Pathology Services Lighthouse Laboratory and Alex Alderton, Roberto Amato, Jeffrey Barrett, Sonia Goncalves, Ewan Harrison, David K. Jackson, Ian Johnston, Dominic Kwiatkowski, Cordelia Langford, John Sillitoe on behalf of the Wellcome Sanger Institute COVID-19 Surveillance Team |
| EPI_ISL_3069042 | Lighthouse Lab in Alderley Park | Wellcome Sanger Institute for the COVID-19 Genomics UK (COG-UK) Consortium | Jacquelyn Wynn, Mairead Hyland, The Lighthouse Lab in Alderley Park and Alex Alderton, Roberto Amato, Jeffrey Barrett, Sonia Goncalves, Ewan Harrison, David K. Jackson, Ian Johnston, Dominic Kwiatkowski, Cordelia Langford, John Sillitoe on behalf of the Wellcome Sanger Institute COVID-19 Surveillance Team |
| EPI_ISL_3069043 | Berkshire and Surrey Pathology Services Lighthouse Laboratory | Wellcome Sanger Institute for the COVID-19 Genomics UK (COG-UK) Consortium | Berkshire and Surrey Pathology Services Lighthouse Laboratory and Alex Alderton, Roberto Amato, Jeffrey Barrett, Sonia Goncalves, Ewan Harrison, David K. Jackson, Ian Johnston, Dominic Kwiatkowski, Cordelia Langford, John Sillitoe on behalf of the Wellcome Sanger Institute COVID-19 Surveillance Team |
| EPI_ISL_3069044 | Lighthouse Lab in Alderley Park | Wellcome Sanger Institute for the COVID-19 Genomics UK (COG-UK) Consortium | Jacquelyn Wynn, Mairead Hyland, The Lighthouse Lab in Alderley Park and Alex Alderton, Roberto Amato, Jeffrey Barrett, Sonia Goncalves, Ewan Harrison, David K. Jackson, Ian Johnston, Dominic Kwiatkowski, Cordelia Langford, John Sillitoe on behalf of the Wellcome Sanger Institute COVID-19 Surveillance Team |
| EPI_ISL_3069045 | Lighthouse Lab in Glasgow | Wellcome Sanger Institute for the COVID-19 Genomics UK (COG-UK) Consortium | Harper VanSteenhouse, Yumi Kasai, David Gray, Carol Clugston, Anna Dominiczak and Alex Alderton, Roberto Amato, Jeffrey Barrett, Sonia Goncalves, Ewan Harrison, David K. Jackson, Ian Johnston, Dominic Kwiatkowski, Cordelia Langford, John Sillitoe on behalf of the Wellcome Sanger Institute COVID-19 Surveillance Team |

| | | | |
|---|---|---|---|
| EPI_ISL_3069046 | Berkshire and Surrey Pathology Services Lighthouse Laboratory | Wellcome Sanger Institute for the COVID-19 Genomics UK (COG-UK) Consortium | Berkshire and Surrey Pathology Services Lighthouse Laboratory and Alex Alderton, Roberto Amato, Jeffrey Barrett, Sonia Goncalves, Ewan Harrison, David K. Jackson, Ian Johnston, Dominic Kwiatkowski, Cordelia Langford, John Sillitoe on behalf of the Wellcome Sanger Institute COVID-19 Surveillance Team |
| EPI_ISL_3069047, EPI_ISL_3069048 | Lighthouse Lab in Glasgow | Wellcome Sanger Institute for the COVID-19 Genomics UK (COG-UK) Consortium | Harper VanSteenhouse, Yumi Kasai, David Gray, Carol Clugston, Anna Dominiczak and Alex Alderton, Roberto Amato, Jeffrey Barrett, Sonia Goncalves, Ewan Harrison, David K. Jackson, Ian Johnston, Dominic Kwiatkowski, Cordelia Langford, John Sillitoe on behalf of the Wellcome Sanger Institute COVID-19 Surveillance Team |
| EPI_ISL_3069049 | Berkshire and Surrey Pathology Services Lighthouse Laboratory | Wellcome Sanger Institute for the COVID-19 Genomics UK (COG-UK) Consortium | Berkshire and Surrey Pathology Services Lighthouse Laboratory and Alex Alderton, Roberto Amato, Jeffrey Barrett, Sonia Goncalves, Ewan Harrison, David K. Jackson, Ian Johnston, Dominic Kwiatkowski, Cordelia Langford, John Sillitoe on behalf of the Wellcome Sanger Institute COVID-19 Surveillance Team |
| EPI_ISL_3069050 | Lighthouse Lab in Glasgow | Wellcome Sanger Institute for the COVID-19 Genomics UK (COG-UK) Consortium | Harper VanSteenhouse, Yumi Kasai, David Gray, Carol Clugston, Anna Dominiczak and Alex Alderton, Roberto Amato, Jeffrey Barrett, Sonia Goncalves, Ewan Harrison, David K. Jackson, Ian Johnston, Dominic Kwiatkowski, Cordelia Langford, John Sillitoe on behalf of the Wellcome Sanger Institute COVID-19 Surveillance Team |
| EPI_ISL_3069051, EPI_ISL_3069052 | Lighthouse Lab in Alderley Park | Wellcome Sanger Institute for the COVID-19 Genomics UK (COG-UK) Consortium | Jacquelyn Wynn, Mairead Hyland, The Lighthouse Lab in Alderley Park and Alex Alderton, Roberto Amato, Jeffrey Barrett, Sonia Goncalves, Ewan Harrison, David K. Jackson, Ian Johnston, Dominic Kwiatkowski, Cordelia Langford, John Sillitoe on behalf of the Wellcome Sanger Institute COVID-19 Surveillance Team |
| EPI_ISL_3069053, EPI_ISL_3069054 | Lighthouse Lab in Glasgow | Wellcome Sanger Institute for the COVID-19 Genomics UK (COG-UK) Consortium | Harper VanSteenhouse, Yumi Kasai, David Gray, Carol Clugston, Anna Dominiczak and Alex Alderton, Roberto Amato, Jeffrey Barrett, Sonia Goncalves, Ewan Harrison, David K. Jackson, Ian Johnston, Dominic Kwiatkowski, Cordelia Langford, John Sillitoe on behalf of the Wellcome Sanger Institute COVID-19 Surveillance Team |
| EPI_ISL_3069055 | Berkshire and Surrey Pathology Services Lighthouse Laboratory | Wellcome Sanger Institute for the COVID-19 Genomics UK (COG-UK) Consortium | Berkshire and Surrey Pathology Services Lighthouse Laboratory and Alex Alderton, Roberto Amato, Jeffrey Barrett, Sonia Goncalves, Ewan Harrison, David K. Jackson, Ian Johnston, Dominic Kwiatkowski, Cordelia Langford, John Sillitoe on behalf of the Wellcome Sanger Institute COVID-19 Surveillance Team |
| EPI_ISL_3069056, EPI_ISL_3069057 | Lighthouse Lab in Glasgow | Wellcome Sanger Institute for the COVID-19 Genomics UK (COG-UK) Consortium | Harper VanSteenhouse, Yumi Kasai, David Gray, Carol Clugston, Anna Dominiczak and Alex Alderton, Roberto Amato, Jeffrey Barrett, Sonia Goncalves, Ewan Harrison, David K. Jackson, Ian Johnston, Dominic Kwiatkowski, Cordelia Langford, John Sillitoe on behalf of the Wellcome Sanger Institute COVID-19 Surveillance Team |
| EPI_ISL_3069058 | Lighthouse Lab in Alderley Park | Wellcome Sanger Institute for the COVID-19 Genomics UK (COG-UK) Consortium | Jacquelyn Wynn, Mairead Hyland, The Lighthouse Lab in Alderley Park and Alex Alderton, Roberto Amato, Jeffrey Barrett, Sonia Goncalves, Ewan Harrison, David K. Jackson, Ian Johnston, Dominic Kwiatkowski, Cordelia Langford, John Sillitoe on behalf of the Wellcome Sanger Institute COVID-19 Surveillance Team |
| EPI_ISL_3069059 | Lighthouse Lab in Glasgow | Wellcome Sanger Institute for the COVID-19 Genomics UK (COG-UK) Consortium | Harper VanSteenhouse, Yumi Kasai, David Gray, Carol Clugston, Anna Dominiczak and Alex Alderton, Roberto Amato, Jeffrey Barrett, Sonia Goncalves, Ewan Harrison, David K. Jackson, Ian Johnston, Dominic Kwiatkowski, Cordelia Langford, John Sillitoe on behalf of the Wellcome Sanger Institute COVID-19 Surveillance Team |
| EPI_ISL_3069060 | Lighthouse Lab in Alderley Park | Wellcome Sanger Institute for the COVID-19 Genomics UK (COG-UK) Consortium | Jacquelyn Wynn, Mairead Hyland, The Lighthouse Lab in Alderley Park and Alex Alderton, Roberto Amato, Jeffrey Barrett, Sonia Goncalves, Ewan Harrison, David K. Jackson, Ian Johnston, Dominic Kwiatkowski, Cordelia Langford, John Sillitoe on behalf of the Wellcome Sanger Institute COVID-19 Surveillance Team |
| EPI_ISL_3069061 | Berkshire and Surrey Pathology Services Lighthouse Laboratory | Wellcome Sanger Institute for the COVID-19 Genomics UK (COG-UK) Consortium | Berkshire and Surrey Pathology Services Lighthouse Laboratory and Alex Alderton, Roberto Amato, Jeffrey Barrett, Sonia Goncalves, Ewan Harrison, David K. Jackson, Ian Johnston, Dominic Kwiatkowski, Cordelia Langford, John Sillitoe on behalf of the Wellcome Sanger Institute COVID-19 Surveillance Team |
| EPI_ISL_3069062 | Lighthouse Lab in Alderley Park | Wellcome Sanger Institute for the COVID-19 Genomics UK (COG-UK) Consortium | Jacquelyn Wynn, Mairead Hyland, The Lighthouse Lab in Alderley Park and Alex Alderton, Roberto Amato, Jeffrey Barrett, Sonia Goncalves, Ewan Harrison, David K. Jackson, Ian Johnston, Dominic Kwiatkowski, Cordelia Langford, John Sillitoe on behalf of the Wellcome Sanger Institute COVID-19 Surveillance Team |
| EPI_ISL_3069063, EPI_ISL_3069064, EPI_ISL_3069065, EPI_ISL_3069066 | Lighthouse Lab in Glasgow | Wellcome Sanger Institute for the COVID-19 Genomics UK (COG-UK) Consortium | Harper VanSteenhouse, Yumi Kasai, David Gray, Carol Clugston, Anna Dominiczak and Alex Alderton, Roberto Amato, Jeffrey Barrett, Sonia Goncalves, Ewan Harrison, David K. Jackson, Ian Johnston, Dominic Kwiatkowski, Cordelia Langford, John Sillitoe on behalf of the Wellcome Sanger Institute COVID-19 Surveillance Team |
| EPI_ISL_3069067 | Berkshire and Surrey Pathology Services Lighthouse Laboratory | Wellcome Sanger Institute for the COVID-19 Genomics UK (COG-UK) Consortium | Berkshire and Surrey Pathology Services Lighthouse Laboratory and Alex Alderton, Roberto Amato, Jeffrey Barrett, Sonia Goncalves, Ewan Harrison, David K. Jackson, Ian Johnston, Dominic Kwiatkowski, Cordelia Langford, John Sillitoe on behalf of the Wellcome Sanger Institute COVID-19 Surveillance Team |
| EPI_ISL_3069068 | Lighthouse Lab in Glasgow | Wellcome Sanger Institute for the COVID-19 Genomics UK (COG-UK) Consortium | Harper VanSteenhouse, Yumi Kasai, David Gray, Carol Clugston, Anna Dominiczak and Alex Alderton, Roberto Amato, Jeffrey Barrett, Sonia Goncalves, Ewan Harrison, David K. Jackson, Ian Johnston, Dominic Kwiatkowski, Cordelia Langford, John Sillitoe on behalf of the Wellcome Sanger Institute COVID-19 Surveillance Team |
| EPI_ISL_3069069 | Lighthouse Lab in Alderley Park | Wellcome Sanger Institute for the COVID-19 Genomics UK (COG-UK) Consortium | Jacquelyn Wynn, Mairead Hyland, The Lighthouse Lab in Alderley Park and Alex Alderton, Roberto Amato, Jeffrey Barrett, Sonia Goncalves, Ewan Harrison, David K. Jackson, Ian Johnston, Dominic Kwiatkowski, Cordelia Langford, John Sillitoe on behalf of the Wellcome Sanger Institute COVID-19 Surveillance Team |
| EPI_ISL_3069070 | Berkshire and Surrey Pathology Services Lighthouse Laboratory | Wellcome Sanger Institute for the COVID-19 Genomics UK (COG-UK) Consortium | Berkshire and Surrey Pathology Services Lighthouse Laboratory and Alex Alderton, Roberto Amato, Jeffrey Barrett, Sonia Goncalves, Ewan Harrison, David K. Jackson, Ian Johnston, Dominic Kwiatkowski, Cordelia Langford, John Sillitoe on behalf of the Wellcome Sanger Institute COVID-19 Surveillance Team |
| EPI_ISL_3069071 | Lighthouse Lab in Alderley Park | Wellcome Sanger Institute for the COVID-19 Genomics UK (COG-UK) Consortium | Jacquelyn Wynn, Mairead Hyland, The Lighthouse Lab in Alderley Park and Alex Alderton, Roberto Amato, Jeffrey Barrett, Sonia Goncalves, Ewan Harrison, David K. Jackson, Ian Johnston, Dominic Kwiatkowski, Cordelia Langford, John Sillitoe on behalf of the Wellcome Sanger Institute COVID-19 Surveillance Team |
| EPI_ISL_3069072 | Berkshire and Surrey Pathology Services Lighthouse Laboratory | Wellcome Sanger Institute for the COVID-19 Genomics UK (COG-UK) Consortium | Berkshire and Surrey Pathology Services Lighthouse Laboratory and Alex Alderton, Roberto Amato, Jeffrey Barrett, Sonia Goncalves, Ewan Harrison, David K. Jackson, Ian Johnston, Dominic Kwiatkowski, Cordelia Langford, John Sillitoe on behalf of the Wellcome Sanger Institute COVID-19 Surveillance Team |
| EPI_ISL_3069073, EPI_ISL_3069074, EPI_ISL_3069075, EPI_ISL_3069076 | Lighthouse Lab in Glasgow | Wellcome Sanger Institute for the COVID-19 Genomics UK (COG-UK) Consortium | Harper VanSteenhouse, Yumi Kasai, David Gray, Carol Clugston, Anna Dominiczak and Alex Alderton, Roberto Amato, Jeffrey Barrett, Sonia Goncalves, Ewan Harrison, David K. Jackson, Ian Johnston, Dominic Kwiatkowski, Cordelia Langford, John Sillitoe on behalf of the Wellcome Sanger Institute COVID-19 Surveillance Team |
| EPI_ISL_3069077, EPI_ISL_3069078 | Lighthouse Lab in Alderley Park | Wellcome Sanger Institute for the COVID-19 Genomics UK (COG-UK) Consortium | Jacquelyn Wynn, Mairead Hyland, The Lighthouse Lab in Alderley Park and Alex Alderton, Roberto Amato, Jeffrey Barrett, Sonia Goncalves, Ewan Harrison, David K. Jackson, Ian Johnston, Dominic Kwiatkowski, Cordelia Langford, John Sillitoe on behalf of the Wellcome Sanger Institute COVID-19 Surveillance Team |
| EPI_ISL_3069079, EPI_ISL_3069080, EPI_ISL_3069081 | Berkshire and Surrey Pathology Services Lighthouse Laboratory | Wellcome Sanger Institute for the COVID-19 Genomics UK (COG-UK) Consortium | Berkshire and Surrey Pathology Services Lighthouse Laboratory and Alex Alderton, Roberto Amato, Jeffrey Barrett, Sonia Goncalves, Ewan Harrison, David K. Jackson, Ian Johnston, Dominic Kwiatkowski, Cordelia Langford, John Sillitoe on behalf of the Wellcome Sanger Institute COVID-19 Surveillance Team |
| EPI_ISL_3069082, EPI_ISL_3069083 | Lighthouse Lab in Glasgow | Wellcome Sanger Institute for the COVID-19 Genomics UK (COG-UK) Consortium | Harper VanSteenhouse, Yumi Kasai, David Gray, Carol Clugston, Anna Dominiczak and Alex Alderton, Roberto Amato, Jeffrey Barrett, Sonia Goncalves, Ewan Harrison, David K. Jackson, Ian Johnston, Dominic Kwiatkowski, Cordelia Langford, John Sillitoe on behalf of the Wellcome Sanger Institute COVID-19 Surveillance Team |
| EPI_ISL_3069084 | Lighthouse Lab in Alderley Park | Wellcome Sanger Institute for the COVID-19 Genomics UK (COG-UK) Consortium | Jacquelyn Wynn, Mairead Hyland, The Lighthouse Lab in Alderley Park and Alex Alderton, Roberto Amato, Jeffrey Barrett, Sonia Goncalves, Ewan Harrison, David K. Jackson, Ian Johnston, Dominic Kwiatkowski, Cordelia Langford, John Sillitoe on behalf of the Wellcome Sanger Institute COVID-19 Surveillance Team |
| EPI_ISL_3069085, EPI_ISL_3069086, EPI_ISL_3069087 | Lighthouse Lab in Glasgow | Wellcome Sanger Institute for the COVID-19 Genomics UK (COG-UK) Consortium | Harper VanSteenhouse, Yumi Kasai, David Gray, Carol Clugston, Anna Dominiczak and Alex Alderton, Roberto Amato, Jeffrey Barrett, Sonia Goncalves, Ewan Harrison, David K. Jackson, Ian Johnston, Dominic Kwiatkowski, Cordelia Langford, John Sillitoe on behalf of the Wellcome Sanger Institute COVID-19 Surveillance Team |
| EPI_ISL_3069088, EPI_ISL_3069089 | Lighthouse Lab in Alderley Park | Wellcome Sanger Institute for the COVID-19 Genomics UK (COG-UK) Consortium | Jacquelyn Wynn, Mairead Hyland, The Lighthouse Lab in Alderley Park and Alex Alderton, Roberto Amato, Jeffrey Barrett, Sonia Goncalves, Ewan Harrison, David K. Jackson, Ian Johnston, Dominic Kwiatkowski, Cordelia Langford, John Sillitoe on behalf of the Wellcome Sanger Institute COVID-19 Surveillance Team |
| EPI_ISL_3069090, EPI_ISL_3069091 | Lighthouse Lab in Glasgow | Wellcome Sanger Institute for the COVID-19 Genomics UK (COG-UK) Consortium | Harper VanSteenhouse, Yumi Kasai, David Gray, Carol Clugston, Anna Dominiczak and Alex Alderton, Roberto Amato, Jeffrey Barrett, Sonia Goncalves, Ewan Harrison, David K. Jackson, Ian Johnston, Dominic Kwiatkowski, Cordelia Langford, John Sillitoe on behalf of the Wellcome Sanger Institute COVID-19 Surveillance Team |
| EPI_ISL_3069092 | Lighthouse Lab in Alderley Park | Wellcome Sanger Institute for the COVID-19 Genomics UK (COG-UK) Consortium | Jacquelyn Wynn, Mairead Hyland, The Lighthouse Lab in Alderley Park and Alex Alderton, Roberto Amato, Jeffrey Barrett, Sonia Goncalves, Ewan Harrison, David K. Jackson, Ian Johnston, Dominic Kwiatkowski, Cordelia Langford, John Sillitoe on behalf of the Wellcome Sanger Institute COVID-19 Surveillance Team |
| EPI_ISL_3069093, EPI_ISL_3069094, EPI_ISL_3069095 | Lighthouse Lab in Glasgow | Wellcome Sanger Institute for the COVID-19 Genomics UK (COG-UK) Consortium | Harper VanSteenhouse, Yumi Kasai, David Gray, Carol Clugston, Anna Dominiczak and Alex Alderton, Roberto Amato, Jeffrey Barrett, Sonia Goncalves, Ewan Harrison, David K. Jackson, Ian Johnston, Dominic Kwiatkowski, Cordelia Langford, John Sillitoe on behalf of the Wellcome Sanger Institute COVID-19 Surveillance Team |
| EPI_ISL_3069096 | Lighthouse Lab in Alderley Park | Wellcome Sanger Institute for the COVID-19 Genomics UK (COG-UK) Consortium | Jacquelyn Wynn, Mairead Hyland, The Lighthouse Lab in Alderley Park and Alex Alderton, Roberto Amato, Jeffrey Barrett, Sonia Goncalves, Ewan Harrison, David K. Jackson, Ian Johnston, Dominic Kwiatkowski, Cordelia Langford, John Sillitoe on behalf of the Wellcome Sanger Institute COVID-19 Surveillance Team |
| EPI_ISL_3069097 | Lighthouse Lab in Glasgow | Wellcome Sanger Institute for the COVID-19 Genomics UK (COG-UK) Consortium | Harper VanSteenhouse, Yumi Kasai, David Gray, Carol Clugston, Anna Dominiczak and Alex Alderton, Roberto Amato, Jeffrey Barrett, Sonia Goncalves, Ewan Harrison, David K. Jackson, Ian Johnston, Dominic Kwiatkowski, Cordelia Langford, John Sillitoe on behalf of the Wellcome Sanger Institute COVID-19 Surveillance Team |
| EPI_ISL_3069098 | Berkshire and Surrey Pathology Services Lighthouse Laboratory | Wellcome Sanger Institute for the COVID-19 Genomics UK (COG-UK) Consortium | Berkshire and Surrey Pathology Services Lighthouse Laboratory and Alex Alderton, Roberto Amato, Jeffrey Barrett, Sonia Goncalves, Ewan Harrison, David K. Jackson, Ian Johnston, Dominic Kwiatkowski, Cordelia Langford, John Sillitoe on behalf of the Wellcome Sanger Institute COVID-19 Surveillance Team |
| EPI_ISL_3069099 | Lighthouse Lab in Glasgow | Wellcome Sanger Institute for the COVID-19 Genomics UK (COG-UK) Consortium | Harper VanSteenhouse, Yumi Kasai, David Gray, Carol Clugston, Anna Dominiczak and Alex Alderton, Roberto Amato, Jeffrey Barrett, Sonia Goncalves, Ewan Harrison, David K. Jackson, Ian Johnston, Dominic Kwiatkowski, Cordelia Langford, John Sillitoe on behalf of the Wellcome Sanger Institute COVID-19 Surveillance Team |
| EPI_ISL_3069100 | Lighthouse Lab in Alderley Park | Wellcome Sanger Institute for the COVID-19 Genomics UK (COG-UK) Consortium | Jacquelyn Wynn, Mairead Hyland, The Lighthouse Lab in Alderley Park and Alex Alderton, Roberto Amato, Jeffrey Barrett, Sonia Goncalves, Ewan Harrison, David K. Jackson, Ian Johnston, Dominic Kwiatkowski, Cordelia Langford, John Sillitoe on behalf of the Wellcome Sanger Institute COVID-19 Surveillance Team |
| EPI_ISL_3069101 | Berkshire and Surrey Pathology Services Lighthouse Laboratory | Wellcome Sanger Institute for the COVID-19 Genomics UK (COG-UK) Consortium | Berkshire and Surrey Pathology Services Lighthouse Laboratory and Alex Alderton, Roberto Amato, Jeffrey Barrett, Sonia Goncalves, Ewan Harrison, David K. Jackson, Ian Johnston, Dominic Kwiatkowski, Cordelia Langford, John Sillitoe on behalf of the Wellcome Sanger Institute COVID-19 Surveillance Team |
| EPI_ISL_3069102 | Lighthouse Lab in Glasgow | Wellcome Sanger Institute for the COVID-19 Genomics UK (COG-UK) Consortium | Harper VanSteenhouse, Yumi Kasai, David Gray, Carol Clugston, Anna Dominiczak and Alex Alderton, Roberto Amato, Jeffrey Barrett, Sonia Goncalves, Ewan Harrison, David K. Jackson, Ian Johnston, Dominic Kwiatkowski, Cordelia Langford, John Sillitoe on behalf of the Wellcome Sanger Institute COVID-19 Surveillance Team |
| EPI_ISL_3069103 | Lighthouse Lab in Alderley Park | Wellcome Sanger Institute for the COVID-19 Genomics UK (COG-UK) Consortium | Jacquelyn Wynn, Mairead Hyland, The Lighthouse Lab in Alderley Park and Alex Alderton, Roberto Amato, Jeffrey Barrett, Sonia Goncalves, Ewan Harrison, David K. Jackson, Ian Johnston, Dominic Kwiatkowski, Cordelia Langford, John Sillitoe on behalf of the Wellcome Sanger Institute COVID-19 Surveillance Team |
| EPI_ISL_3069104 | Berkshire and Surrey Pathology Services Lighthouse Laboratory | Wellcome Sanger Institute for the COVID-19 Genomics UK (COG-UK) Consortium | Berkshire and Surrey Pathology Services Lighthouse Laboratory and Alex Alderton, Roberto Amato, Jeffrey Barrett, Sonia Goncalves, Ewan Harrison, David K. Jackson, Ian Johnston, Dominic Kwiatkowski, Cordelia Langford, John Sillitoe on behalf of the Wellcome Sanger Institute COVID-19 Surveillance Team |
| EPI_ISL_3069105, EPI_ISL_3069106 | Lighthouse Lab in Alderley Park | Wellcome Sanger Institute for the COVID-19 Genomics UK (COG-UK) Consortium | Jacquelyn Wynn, Mairead Hyland, The Lighthouse Lab in Alderley Park and Alex Alderton, Roberto Amato, Jeffrey Barrett, Sonia Goncalves, Ewan Harrison, David K. Jackson, Ian Johnston, Dominic Kwiatkowski, Cordelia Langford, John Sillitoe on behalf of the Wellcome Sanger Institute COVID-19 Surveillance Team |
| EPI_ISL_3069107 | Berkshire and Surrey Pathology Services Lighthouse Laboratory | Wellcome Sanger Institute for the COVID-19 Genomics UK (COG-UK) Consortium | Berkshire and Surrey Pathology Services Lighthouse Laboratory and Alex Alderton, Roberto Amato, Jeffrey Barrett, Sonia Goncalves, Ewan Harrison, David K. Jackson, Ian Johnston, Dominic Kwiatkowski, Cordelia Langford, John Sillitoe on behalf of the Wellcome Sanger Institute COVID-19 Surveillance Team |
| EPI_ISL_3069108 | Lighthouse Lab in Glasgow | Wellcome Sanger Institute for the COVID-19 Genomics UK (COG-UK) Consortium | Harper VanSteenhouse, Yumi Kasai, David Gray, Carol Clugston, Anna Dominiczak and Alex Alderton, Roberto Amato, Jeffrey Barrett, Sonia Goncalves, Ewan Harrison, David K. Jackson, Ian Johnston, Dominic Kwiatkowski, Cordelia Langford, John Sillitoe on behalf of the Wellcome Sanger Institute COVID-19 Surveillance Team |
| EPI_ISL_3069109 | Berkshire and Surrey Pathology Services Lighthouse Laboratory | Wellcome Sanger Institute for the COVID-19 Genomics UK (COG-UK) | Berkshire and Surrey Pathology Services Lighthouse Laboratory and Alex Alderton, Roberto Amato, Jeffrey Barrett, Sonia Goncalves, Ewan Harrison, David K. Jackson, Ian Johnston, Dominic Kwiatkowski, Cordelia Langford, John Sillitoe on behalf |

| | | | |
|---|---|---|---|
| | | Consortium | of the Wellcome Sanger Institute COVID-19 Surveillance Team |
| EPI_ISL_3069110 | Lighthouse Lab in Glasgow | Wellcome Sanger Institute for the COVID-19 Genomics UK (COG-UK) Consortium | Harper VanSteenhouse, Yumi Kasai, David Gray, Carol Clugston, Anna Dominiczak and Alex Alderton, Roberto Amato, Jeffrey Barrett, Sonia Goncalves, Ewan Harrison, David K. Jackson, Ian Johnston, Dominic Kwiatkowski, Cordelia Langford, John Sillitoe on behalf of the Wellcome Sanger Institute COVID-19 Surveillance Team |
| EPI_ISL_3069111 | Berkshire and Surrey Pathology Services Lighthouse Laboratory | Wellcome Sanger Institute for the COVID-19 Genomics UK (COG-UK) Consortium | Berkshire and Surrey Pathology Services Lighthouse Laboratory and Alex Alderton, Roberto Amato, Jeffrey Barrett, Sonia Goncalves, Ewan Harrison, David K. Jackson, Ian Johnston, Dominic Kwiatkowski, Cordelia Langford, John Sillitoe on behalf of the Wellcome Sanger Institute COVID-19 Surveillance Team |
| EPI_ISL_3069112 | Lighthouse Lab in Glasgow | Wellcome Sanger Institute for the COVID-19 Genomics UK (COG-UK) Consortium | Harper VanSteenhouse, Yumi Kasai, David Gray, Carol Clugston, Anna Dominiczak and Alex Alderton, Roberto Amato, Jeffrey Barrett, Sonia Goncalves, Ewan Harrison, David K. Jackson, Ian Johnston, Dominic Kwiatkowski, Cordelia Langford, John Sillitoe on behalf of the Wellcome Sanger Institute COVID-19 Surveillance Team |
| EPI_ISL_3069113, EPI_ISL_3069114 | Lighthouse Lab in Alderley Park | Wellcome Sanger Institute for the COVID-19 Genomics UK (COG-UK) Consortium | Jacquelyn Wynn, Mairead Hyland, The Lighthouse Lab in Alderley Park and Alex Alderton, Roberto Amato, Jeffrey Barrett, Sonia Goncalves, Ewan Harrison, David K. Jackson, Ian Johnston, Dominic Kwiatkowski, Cordelia Langford, John Sillitoe on behalf of the Wellcome Sanger Institute COVID-19 Surveillance Team |
| EPI_ISL_3069115 | Lighthouse Lab in Glasgow | Wellcome Sanger Institute for the COVID-19 Genomics UK (COG-UK) Consortium | Harper VanSteenhouse, Yumi Kasai, David Gray, Carol Clugston, Anna Dominiczak and Alex Alderton, Roberto Amato, Jeffrey Barrett, Sonia Goncalves, Ewan Harrison, David K. Jackson, Ian Johnston, Dominic Kwiatkowski, Cordelia Langford, John Sillitoe on behalf of the Wellcome Sanger Institute COVID-19 Surveillance Team |
| EPI_ISL_3069116, EPI_ISL_3069117 | Lighthouse Lab in Alderley Park | Wellcome Sanger Institute for the COVID-19 Genomics UK (COG-UK) Consortium | Jacquelyn Wynn, Mairead Hyland, The Lighthouse Lab in Alderley Park and Alex Alderton, Roberto Amato, Jeffrey Barrett, Sonia Goncalves, Ewan Harrison, David K. Jackson, Ian Johnston, Dominic Kwiatkowski, Cordelia Langford, John Sillitoe on behalf of the Wellcome Sanger Institute COVID-19 Surveillance Team |
| EPI_ISL_3069118 | Lighthouse Lab in Glasgow | Wellcome Sanger Institute for the COVID-19 Genomics UK (COG-UK) Consortium | Harper VanSteenhouse, Yumi Kasai, David Gray, Carol Clugston, Anna Dominiczak and Alex Alderton, Roberto Amato, Jeffrey Barrett, Sonia Goncalves, Ewan Harrison, David K. Jackson, Ian Johnston, Dominic Kwiatkowski, Cordelia Langford, John Sillitoe on behalf of the Wellcome Sanger Institute COVID-19 Surveillance Team |
| EPI_ISL_3069119 | Lighthouse Lab in Alderley Park | Wellcome Sanger Institute for the COVID-19 Genomics UK (COG-UK) Consortium | Jacquelyn Wynn, Mairead Hyland, The Lighthouse Lab in Alderley Park and Alex Alderton, Roberto Amato, Jeffrey Barrett, Sonia Goncalves, Ewan Harrison, David K. Jackson, Ian Johnston, Dominic Kwiatkowski, Cordelia Langford, John Sillitoe on behalf of the Wellcome Sanger Institute COVID-19 Surveillance Team |
| EPI_ISL_3069120 | Lighthouse Lab in Glasgow | Wellcome Sanger Institute for the COVID-19 Genomics UK (COG-UK) Consortium | Harper VanSteenhouse, Yumi Kasai, David Gray, Carol Clugston, Anna Dominiczak and Alex Alderton, Roberto Amato, Jeffrey Barrett, Sonia Goncalves, Ewan Harrison, David K. Jackson, Ian Johnston, Dominic Kwiatkowski, Cordelia Langford, John Sillitoe on behalf of the Wellcome Sanger Institute COVID-19 Surveillance Team |
| EPI_ISL_3069121 | Berkshire and Surrey Pathology Services Lighthouse Laboratory | Wellcome Sanger Institute for the COVID-19 Genomics UK (COG-UK) Consortium | Berkshire and Surrey Pathology Services Lighthouse Laboratory and Alex Alderton, Roberto Amato, Jeffrey Barrett, Sonia Goncalves, Ewan Harrison, David K. Jackson, Ian Johnston, Dominic Kwiatkowski, Cordelia Langford, John Sillitoe on behalf of the Wellcome Sanger Institute COVID-19 Surveillance Team |
| EPI_ISL_3069122 | Lighthouse Lab in Glasgow | Wellcome Sanger Institute for the COVID-19 Genomics UK (COG-UK) Consortium | Harper VanSteenhouse, Yumi Kasai, David Gray, Carol Clugston, Anna Dominiczak and Alex Alderton, Roberto Amato, Jeffrey Barrett, Sonia Goncalves, Ewan Harrison, David K. Jackson, Ian Johnston, Dominic Kwiatkowski, Cordelia Langford, John Sillitoe on behalf of the Wellcome Sanger Institute COVID-19 Surveillance Team |
| EPI_ISL_3069123 | Berkshire and Surrey Pathology Services Lighthouse Laboratory | Wellcome Sanger Institute for the COVID-19 Genomics UK (COG-UK) Consortium | Berkshire and Surrey Pathology Services Lighthouse Laboratory and Alex Alderton, Roberto Amato, Jeffrey Barrett, Sonia Goncalves, Ewan Harrison, David K. Jackson, Ian Johnston, Dominic Kwiatkowski, Cordelia Langford, John Sillitoe on behalf of the Wellcome Sanger Institute COVID-19 Surveillance Team |
| EPI_ISL_3069124 | Lighthouse Lab in Alderley Park | Wellcome Sanger Institute for the COVID-19 Genomics UK (COG-UK) Consortium | Jacquelyn Wynn, Mairead Hyland, The Lighthouse Lab in Alderley Park and Alex Alderton, Roberto Amato, Jeffrey Barrett, Sonia Goncalves, Ewan Harrison, David K. Jackson, Ian Johnston, Dominic Kwiatkowski, Cordelia Langford, John Sillitoe on behalf of the Wellcome Sanger Institute COVID-19 Surveillance Team |
| EPI_ISL_3069125, EPI_ISL_3069126, EPI_ISL_3069127, EPI_ISL_3069129, EPI_ISL_3069130, EPI_ISL_3069131, EPI_ISL_3069132, EPI_ISL_3069133, EPI_ISL_3069134 | Lighthouse Lab in Glasgow | Wellcome Sanger Institute for the COVID-19 Genomics UK (COG-UK) Consortium | Harper VanSteenhouse, Yumi Kasai, David Gray, Carol Clugston, Anna Dominiczak and Alex Alderton, Roberto Amato, Jeffrey Barrett, Sonia Goncalves, Ewan Harrison, David K. Jackson, Ian Johnston, Dominic Kwiatkowski, Cordelia Langford, John Sillitoe on behalf of the Wellcome Sanger Institute COVID-19 Surveillance Team |
| EPI_ISL_3069135, EPI_ISL_3069136 | Lighthouse Lab in Alderley Park | Wellcome Sanger Institute for the COVID-19 Genomics UK (COG-UK) Consortium | Jacquelyn Wynn, Mairead Hyland, The Lighthouse Lab in Alderley Park and Alex Alderton, Roberto Amato, Jeffrey Barrett, Sonia Goncalves, Ewan Harrison, David K. Jackson, Ian Johnston, Dominic Kwiatkowski, Cordelia Langford, John Sillitoe on behalf of the Wellcome Sanger Institute COVID-19 Surveillance Team |
| EPI_ISL_3069137 | Lighthouse Lab in Glasgow | Wellcome Sanger Institute for the COVID-19 Genomics UK (COG-UK) Consortium | Harper VanSteenhouse, Yumi Kasai, David Gray, Carol Clugston, Anna Dominiczak and Alex Alderton, Roberto Amato, Jeffrey Barrett, Sonia Goncalves, Ewan Harrison, David K. Jackson, Ian Johnston, Dominic Kwiatkowski, Cordelia Langford, John Sillitoe on behalf of the Wellcome Sanger Institute COVID-19 Surveillance Team |
| EPI_ISL_3069138 | Berkshire and Surrey Pathology Services Lighthouse Laboratory | Wellcome Sanger Institute for the COVID-19 Genomics UK (COG-UK) Consortium | Berkshire and Surrey Pathology Services Lighthouse Laboratory and Alex Alderton, Roberto Amato, Jeffrey Barrett, Sonia Goncalves, Ewan Harrison, David K. Jackson, Ian Johnston, Dominic Kwiatkowski, Cordelia Langford, John Sillitoe on behalf of the Wellcome Sanger Institute COVID-19 Surveillance Team |
| EPI_ISL_3069139 | Lighthouse Lab in Glasgow | Wellcome Sanger Institute for the COVID-19 Genomics UK (COG-UK) Consortium | Harper VanSteenhouse, Yumi Kasai, David Gray, Carol Clugston, Anna Dominiczak and Alex Alderton, Roberto Amato, Jeffrey Barrett, Sonia Goncalves, Ewan Harrison, David K. Jackson, Ian Johnston, Dominic Kwiatkowski, Cordelia Langford, John Sillitoe on behalf of the Wellcome Sanger Institute COVID-19 Surveillance Team |
| EPI_ISL_3069140 | Berkshire and Surrey Pathology Services Lighthouse Laboratory | Wellcome Sanger Institute for the COVID-19 Genomics UK (COG-UK) Consortium | Berkshire and Surrey Pathology Services Lighthouse Laboratory and Alex Alderton, Roberto Amato, Jeffrey Barrett, Sonia Goncalves, Ewan Harrison, David K. Jackson, Ian Johnston, Dominic Kwiatkowski, Cordelia Langford, John Sillitoe on behalf of the Wellcome Sanger Institute COVID-19 Surveillance Team |
| EPI_ISL_3069141 | Lighthouse Lab in Glasgow | Wellcome Sanger Institute for the COVID-19 Genomics UK (COG-UK) Consortium | Harper VanSteenhouse, Yumi Kasai, David Gray, Carol Clugston, Anna Dominiczak and Alex Alderton, Roberto Amato, Jeffrey Barrett, Sonia Goncalves, Ewan Harrison, David K. Jackson, Ian Johnston, Dominic Kwiatkowski, Cordelia Langford, John Sillitoe on behalf of the Wellcome Sanger Institute COVID-19 Surveillance Team |
| EPI_ISL_3069142 | Berkshire and Surrey Pathology Services Lighthouse Laboratory | Wellcome Sanger Institute for the COVID-19 Genomics UK (COG-UK) Consortium | Berkshire and Surrey Pathology Services Lighthouse Laboratory and Alex Alderton, Roberto Amato, Jeffrey Barrett, Sonia Goncalves, Ewan Harrison, David K. Jackson, Ian Johnston, Dominic Kwiatkowski, Cordelia Langford, John Sillitoe on behalf of the Wellcome Sanger Institute COVID-19 Surveillance Team |
| EPI_ISL_3069143 | Lighthouse Lab in Glasgow | Wellcome Sanger Institute for the COVID-19 Genomics UK (COG-UK) Consortium | Harper VanSteenhouse, Yumi Kasai, David Gray, Carol Clugston, Anna Dominiczak and Alex Alderton, Roberto Amato, Jeffrey Barrett, Sonia Goncalves, Ewan Harrison, David K. Jackson, Ian Johnston, Dominic Kwiatkowski, Cordelia Langford, John Sillitoe on behalf of the Wellcome Sanger Institute COVID-19 Surveillance Team |
| EPI_ISL_3069144 | Lighthouse Lab in Alderley Park | Wellcome Sanger Institute for the COVID-19 Genomics UK (COG-UK) Consortium | Jacquelyn Wynn, Mairead Hyland, The Lighthouse Lab in Alderley Park and Alex Alderton, Roberto Amato, Jeffrey Barrett, Sonia Goncalves, Ewan Harrison, David K. Jackson, Ian Johnston, Dominic Kwiatkowski, Cordelia Langford, John Sillitoe on behalf of the Wellcome Sanger Institute COVID-19 Surveillance Team |
| EPI_ISL_3069145, EPI_ISL_3069146, EPI_ISL_3069147, EPI_ISL_3069148, EPI_ISL_3069149, EPI_ISL_3069150 | Lighthouse Lab in Glasgow | Wellcome Sanger Institute for the COVID-19 Genomics UK (COG-UK) Consortium | Harper VanSteenhouse, Yumi Kasai, David Gray, Carol Clugston, Anna Dominiczak and Alex Alderton, Roberto Amato, Jeffrey Barrett, Sonia Goncalves, Ewan Harrison, David K. Jackson, Ian Johnston, Dominic Kwiatkowski, Cordelia Langford, John Sillitoe on behalf of the Wellcome Sanger Institute COVID-19 Surveillance Team |
| EPI_ISL_3069151 | Lighthouse Lab in Alderley Park | Wellcome Sanger Institute for the COVID-19 Genomics UK (COG-UK) Consortium | Jacquelyn Wynn, Mairead Hyland, The Lighthouse Lab in Alderley Park and Alex Alderton, Roberto Amato, Jeffrey Barrett, Sonia Goncalves, Ewan Harrison, David K. Jackson, Ian Johnston, Dominic Kwiatkowski, Cordelia Langford, John Sillitoe on behalf of the Wellcome Sanger Institute COVID-19 Surveillance Team |
| EPI_ISL_3069152 | Berkshire and Surrey Pathology Services Lighthouse Laboratory | Wellcome Sanger Institute for the COVID-19 Genomics UK (COG-UK) Consortium | Berkshire and Surrey Pathology Services Lighthouse Laboratory and Alex Alderton, Roberto Amato, Jeffrey Barrett, Sonia Goncalves, Ewan Harrison, David K. Jackson, Ian Johnston, Dominic Kwiatkowski, Cordelia Langford, John Sillitoe on behalf of the Wellcome Sanger Institute COVID-19 Surveillance Team |
| EPI_ISL_3069153 | Lighthouse Lab in Alderley Park | Wellcome Sanger Institute for the COVID-19 Genomics UK (COG-UK) Consortium | Jacquelyn Wynn, Mairead Hyland, The Lighthouse Lab in Alderley Park and Alex Alderton, Roberto Amato, Jeffrey Barrett, Sonia Goncalves, Ewan Harrison, David K. Jackson, Ian Johnston, Dominic Kwiatkowski, Cordelia Langford, John Sillitoe on behalf of the Wellcome Sanger Institute COVID-19 Surveillance Team |
| EPI_ISL_3069154 | Lighthouse Lab in Glasgow | Wellcome Sanger Institute for the COVID-19 Genomics UK (COG-UK) Consortium | Harper VanSteenhouse, Yumi Kasai, David Gray, Carol Clugston, Anna Dominiczak and Alex Alderton, Roberto Amato, Jeffrey Barrett, Sonia Goncalves, Ewan Harrison, David K. Jackson, Ian Johnston, Dominic Kwiatkowski, Cordelia Langford, John Sillitoe on behalf of the Wellcome Sanger Institute COVID-19 Surveillance Team |
| EPI_ISL_3069155 | Berkshire and Surrey Pathology Services Lighthouse Laboratory | Wellcome Sanger Institute for the COVID-19 Genomics UK (COG-UK) Consortium | Berkshire and Surrey Pathology Services Lighthouse Laboratory and Alex Alderton, Roberto Amato, Jeffrey Barrett, Sonia Goncalves, Ewan Harrison, David K. Jackson, Ian Johnston, Dominic Kwiatkowski, Cordelia Langford, John Sillitoe on behalf of the Wellcome Sanger Institute COVID-19 Surveillance Team |
| EPI_ISL_3069156, EPI_ISL_3069157 | Lighthouse Lab in Alderley Park | Wellcome Sanger Institute for the COVID-19 Genomics UK (COG-UK) Consortium | Jacquelyn Wynn, Mairead Hyland, The Lighthouse Lab in Alderley Park and Alex Alderton, Roberto Amato, Jeffrey Barrett, Sonia Goncalves, Ewan Harrison, David K. Jackson, Ian Johnston, Dominic Kwiatkowski, Cordelia Langford, John Sillitoe on behalf of the Wellcome Sanger Institute COVID-19 Surveillance Team |
| EPI_ISL_3069158 | Lighthouse Lab in Glasgow | Wellcome Sanger Institute for the COVID-19 Genomics UK (COG-UK) Consortium | Harper VanSteenhouse, Yumi Kasai, David Gray, Carol Clugston, Anna Dominiczak and Alex Alderton, Roberto Amato, Jeffrey Barrett, Sonia Goncalves, Ewan Harrison, David K. Jackson, Ian Johnston, Dominic Kwiatkowski, Cordelia Langford, John Sillitoe on behalf of the Wellcome Sanger Institute COVID-19 Surveillance Team |
| EPI_ISL_3069159 | Berkshire and Surrey Pathology Services Lighthouse Laboratory | Wellcome Sanger Institute for the COVID-19 Genomics UK (COG-UK) Consortium | Berkshire and Surrey Pathology Services Lighthouse Laboratory and Alex Alderton, Roberto Amato, Jeffrey Barrett, Sonia Goncalves, Ewan Harrison, David K. Jackson, Ian Johnston, Dominic Kwiatkowski, Cordelia Langford, John Sillitoe on behalf of the Wellcome Sanger Institute COVID-19 Surveillance Team |
| EPI_ISL_3069160, EPI_ISL_3069161 | Lighthouse Lab in Glasgow | Wellcome Sanger Institute for the COVID-19 Genomics UK (COG-UK) Consortium | Harper VanSteenhouse, Yumi Kasai, David Gray, Carol Clugston, Anna Dominiczak and Alex Alderton, Roberto Amato, Jeffrey Barrett, Sonia Goncalves, Ewan Harrison, David K. Jackson, Ian Johnston, Dominic Kwiatkowski, Cordelia Langford, John Sillitoe on behalf of the Wellcome Sanger Institute COVID-19 Surveillance Team |
| EPI_ISL_3069162 | Berkshire and Surrey Pathology Services Lighthouse Laboratory | Wellcome Sanger Institute for the COVID-19 Genomics UK (COG-UK) Consortium | Berkshire and Surrey Pathology Services Lighthouse Laboratory and Alex Alderton, Roberto Amato, Jeffrey Barrett, Sonia Goncalves, Ewan Harrison, David K. Jackson, Ian Johnston, Dominic Kwiatkowski, Cordelia Langford, John Sillitoe on behalf of the Wellcome Sanger Institute COVID-19 Surveillance Team |
| EPI_ISL_3069163, EPI_ISL_3069164, EPI_ISL_3069165 | Lighthouse Lab in Alderley Park | Wellcome Sanger Institute for the COVID-19 Genomics UK (COG-UK) Consortium | Jacquelyn Wynn, Mairead Hyland, The Lighthouse Lab in Alderley Park and Alex Alderton, Roberto Amato, Jeffrey Barrett, Sonia Goncalves, Ewan Harrison, David K. Jackson, Ian Johnston, Dominic Kwiatkowski, Cordelia Langford, John Sillitoe on behalf of the Wellcome Sanger Institute COVID-19 Surveillance Team |
| EPI_ISL_3069166 | Berkshire and Surrey Pathology Services Lighthouse Laboratory | Wellcome Sanger Institute for the COVID-19 Genomics UK (COG-UK) Consortium | Berkshire and Surrey Pathology Services Lighthouse Laboratory and Alex Alderton, Roberto Amato, Jeffrey Barrett, Sonia Goncalves, Ewan Harrison, David K. Jackson, Ian Johnston, Dominic Kwiatkowski, Cordelia Langford, John Sillitoe on behalf of the Wellcome Sanger Institute COVID-19 Surveillance Team |
| EPI_ISL_3069167 | Lighthouse Lab in Alderley Park | Wellcome Sanger Institute for the COVID-19 Genomics UK (COG-UK) Consortium | Jacquelyn Wynn, Mairead Hyland, The Lighthouse Lab in Alderley Park and Alex Alderton, Roberto Amato, Jeffrey Barrett, Sonia Goncalves, Ewan Harrison, David K. Jackson, Ian Johnston, Dominic Kwiatkowski, Cordelia Langford, John Sillitoe on behalf of the Wellcome Sanger Institute COVID-19 Surveillance Team |
| EPI_ISL_3069168 | Lighthouse Lab in Glasgow | Wellcome Sanger Institute for the COVID-19 Genomics UK (COG-UK) Consortium | Harper VanSteenhouse, Yumi Kasai, David Gray, Carol Clugston, Anna Dominiczak and Alex Alderton, Roberto Amato, Jeffrey Barrett, Sonia Goncalves, Ewan Harrison, David K. Jackson, Ian Johnston, Dominic Kwiatkowski, Cordelia Langford, John Sillitoe on behalf of the Wellcome Sanger Institute COVID-19 Surveillance Team |
| EPI_ISL_3069169 | Berkshire and Surrey Pathology Services Lighthouse Laboratory | Wellcome Sanger Institute for the COVID-19 Genomics UK (COG-UK) Consortium | Berkshire and Surrey Pathology Services Lighthouse Laboratory and Alex Alderton, Roberto Amato, Jeffrey Barrett, Sonia Goncalves, Ewan Harrison, David K. Jackson, Ian Johnston, Dominic Kwiatkowski, Cordelia Langford, John Sillitoe on behalf of the Wellcome Sanger Institute COVID-19 Surveillance Team |
| EPI_ISL_3069170 | Lighthouse Lab in Alderley Park | Wellcome Sanger Institute for the COVID-19 Genomics UK (COG-UK) Consortium | Jacquelyn Wynn, Mairead Hyland, The Lighthouse Lab in Alderley Park and Alex Alderton, Roberto Amato, Jeffrey Barrett, Sonia Goncalves, Ewan Harrison, David K. Jackson, Ian Johnston, Dominic Kwiatkowski, Cordelia Langford, John Sillitoe on behalf of the Wellcome Sanger Institute COVID-19 Surveillance Team |

| | | | |
|---|---|---|---|
| EPI_ISL_3069171, EPI_ISL_3069172, EPI_ISL_3069173, EPI_ISL_3069174 | Lighthouse Lab in Glasgow | Wellcome Sanger Institute for the COVID-19 Genomics UK (COG-UK) Consortium | Harper VanSteenhouse, Yumi Kasai, David Gray, Carol Clugston, Anna Dominiczak and Alex Alderton, Roberto Amato, Jeffrey Barrett, Sonia Goncalves, Ewan Harrison, David K. Jackson, Ian Johnston, Dominic Kwiatkowski, Cordelia Langford, John Sillitoe on behalf of the Wellcome Sanger Institute COVID-19 Surveillance Team |
| EPI_ISL_3069175 | Berkshire and Surrey Pathology Services Lighthouse Laboratory | Wellcome Sanger Institute for the COVID-19 Genomics UK (COG-UK) Consortium | Berkshire and Surrey Pathology Services Lighthouse Laboratory and Alex Alderton, Roberto Amato, Jeffrey Barrett, Sonia Goncalves, Ewan Harrison, David K. Jackson, Ian Johnston, Dominic Kwiatkowski, Cordelia Langford, John Sillitoe on behalf of the Wellcome Sanger Institute COVID-19 Surveillance Team |
| EPI_ISL_3069176 | Lighthouse Lab in Glasgow | Wellcome Sanger Institute for the COVID-19 Genomics UK (COG-UK) Consortium | Harper VanSteenhouse, Yumi Kasai, David Gray, Carol Clugston, Anna Dominiczak and Alex Alderton, Roberto Amato, Jeffrey Barrett, Sonia Goncalves, Ewan Harrison, David K. Jackson, Ian Johnston, Dominic Kwiatkowski, Cordelia Langford, John Sillitoe on behalf of the Wellcome Sanger Institute COVID-19 Surveillance Team |
| EPI_ISL_3069177 | Lighthouse Lab in Alderley Park | Wellcome Sanger Institute for the COVID-19 Genomics UK (COG-UK) Consortium | Jacquelyn Wynn, Mairead Hyland, The Lighthouse Lab in Alderley Park and Alex Alderton, Roberto Amato, Jeffrey Barrett, Sonia Goncalves, Ewan Harrison, David K. Jackson, Ian Johnston, Dominic Kwiatkowski, Cordelia Langford, John Sillitoe on behalf of the Wellcome Sanger Institute COVID-19 Surveillance Team |
| EPI_ISL_3069178, EPI_ISL_3069179 | Berkshire and Surrey Pathology Services Lighthouse Laboratory | Wellcome Sanger Institute for the COVID-19 Genomics UK (COG-UK) Consortium | Berkshire and Surrey Pathology Services Lighthouse Laboratory and Alex Alderton, Roberto Amato, Jeffrey Barrett, Sonia Goncalves, Ewan Harrison, David K. Jackson, Ian Johnston, Dominic Kwiatkowski, Cordelia Langford, John Sillitoe on behalf of the Wellcome Sanger Institute COVID-19 Surveillance Team |
| EPI_ISL_3069180, EPI_ISL_3069181, EPI_ISL_3069182, EPI_ISL_3069183 | Lighthouse Lab in Glasgow | Wellcome Sanger Institute for the COVID-19 Genomics UK (COG-UK) Consortium | Harper VanSteenhouse, Yumi Kasai, David Gray, Carol Clugston, Anna Dominiczak and Alex Alderton, Roberto Amato, Jeffrey Barrett, Sonia Goncalves, Ewan Harrison, David K. Jackson, Ian Johnston, Dominic Kwiatkowski, Cordelia Langford, John Sillitoe on behalf of the Wellcome Sanger Institute COVID-19 Surveillance Team |
| EPI_ISL_3069184, EPI_ISL_3069185 | Lighthouse Lab in Alderley Park | Wellcome Sanger Institute for the COVID-19 Genomics UK (COG-UK) Consortium | Jacquelyn Wynn, Mairead Hyland, The Lighthouse Lab in Alderley Park and Alex Alderton, Roberto Amato, Jeffrey Barrett, Sonia Goncalves, Ewan Harrison, David K. Jackson, Ian Johnston, Dominic Kwiatkowski, Cordelia Langford, John Sillitoe on behalf of the Wellcome Sanger Institute COVID-19 Surveillance Team |
| EPI_ISL_3069186 | Berkshire and Surrey Pathology Services Lighthouse Laboratory | Wellcome Sanger Institute for the COVID-19 Genomics UK (COG-UK) Consortium | Berkshire and Surrey Pathology Services Lighthouse Laboratory and Alex Alderton, Roberto Amato, Jeffrey Barrett, Sonia Goncalves, Ewan Harrison, David K. Jackson, Ian Johnston, Dominic Kwiatkowski, Cordelia Langford, John Sillitoe on behalf of the Wellcome Sanger Institute COVID-19 Surveillance Team |
| EPI_ISL_3069187 | Lighthouse Lab in Alderley Park | Wellcome Sanger Institute for the COVID-19 Genomics UK (COG-UK) Consortium | Jacquelyn Wynn, Mairead Hyland, The Lighthouse Lab in Alderley Park and Alex Alderton, Roberto Amato, Jeffrey Barrett, Sonia Goncalves, Ewan Harrison, David K. Jackson, Ian Johnston, Dominic Kwiatkowski, Cordelia Langford, John Sillitoe on behalf of the Wellcome Sanger Institute COVID-19 Surveillance Team |
| EPI_ISL_3069188 | Lighthouse Lab in Glasgow | Wellcome Sanger Institute for the COVID-19 Genomics UK (COG-UK) Consortium | Harper VanSteenhouse, Yumi Kasai, David Gray, Carol Clugston, Anna Dominiczak and Alex Alderton, Roberto Amato, Jeffrey Barrett, Sonia Goncalves, Ewan Harrison, David K. Jackson, Ian Johnston, Dominic Kwiatkowski, Cordelia Langford, John Sillitoe on behalf of the Wellcome Sanger Institute COVID-19 Surveillance Team |
| EPI_ISL_3069189 | Berkshire and Surrey Pathology Services Lighthouse Laboratory | Wellcome Sanger Institute for the COVID-19 Genomics UK (COG-UK) Consortium | Berkshire and Surrey Pathology Services Lighthouse Laboratory and Alex Alderton, Roberto Amato, Jeffrey Barrett, Sonia Goncalves, Ewan Harrison, David K. Jackson, Ian Johnston, Dominic Kwiatkowski, Cordelia Langford, John Sillitoe on behalf of the Wellcome Sanger Institute COVID-19 Surveillance Team |
| EPI_ISL_3069190 | Lighthouse Lab in Glasgow | Wellcome Sanger Institute for the COVID-19 Genomics UK (COG-UK) Consortium | Harper VanSteenhouse, Yumi Kasai, David Gray, Carol Clugston, Anna Dominiczak and Alex Alderton, Roberto Amato, Jeffrey Barrett, Sonia Goncalves, Ewan Harrison, David K. Jackson, Ian Johnston, Dominic Kwiatkowski, Cordelia Langford, John Sillitoe on behalf of the Wellcome Sanger Institute COVID-19 Surveillance Team |
| EPI_ISL_3069191, EPI_ISL_3069192 | Lighthouse Lab in Alderley Park | Wellcome Sanger Institute for the COVID-19 Genomics UK (COG-UK) Consortium | Jacquelyn Wynn, Mairead Hyland, The Lighthouse Lab in Alderley Park and Alex Alderton, Roberto Amato, Jeffrey Barrett, Sonia Goncalves, Ewan Harrison, David K. Jackson, Ian Johnston, Dominic Kwiatkowski, Cordelia Langford, John Sillitoe on behalf of the Wellcome Sanger Institute COVID-19 Surveillance Team |
| EPI_ISL_3069193 | Lighthouse Lab in Glasgow | Wellcome Sanger Institute for the COVID-19 Genomics UK (COG-UK) Consortium | Harper VanSteenhouse, Yumi Kasai, David Gray, Carol Clugston, Anna Dominiczak and Alex Alderton, Roberto Amato, Jeffrey Barrett, Sonia Goncalves, Ewan Harrison, David K. Jackson, Ian Johnston, Dominic Kwiatkowski, Cordelia Langford, John Sillitoe on behalf of the Wellcome Sanger Institute COVID-19 Surveillance Team |
| EPI_ISL_3069194 | Berkshire and Surrey Pathology Services Lighthouse Laboratory | Wellcome Sanger Institute for the COVID-19 Genomics UK (COG-UK) Consortium | Berkshire and Surrey Pathology Services Lighthouse Laboratory and Alex Alderton, Roberto Amato, Jeffrey Barrett, Sonia Goncalves, Ewan Harrison, David K. Jackson, Ian Johnston, Dominic Kwiatkowski, Cordelia Langford, John Sillitoe on behalf of the Wellcome Sanger Institute COVID-19 Surveillance Team |
| EPI_ISL_3069195 | Lighthouse Lab in Glasgow | Wellcome Sanger Institute for the COVID-19 Genomics UK (COG-UK) Consortium | Harper VanSteenhouse, Yumi Kasai, David Gray, Carol Clugston, Anna Dominiczak and Alex Alderton, Roberto Amato, Jeffrey Barrett, Sonia Goncalves, Ewan Harrison, David K. Jackson, Ian Johnston, Dominic Kwiatkowski, Cordelia Langford, John Sillitoe on behalf of the Wellcome Sanger Institute COVID-19 Surveillance Team |
| EPI_ISL_3069196, EPI_ISL_3069197, EPI_ISL_3069198 | Lighthouse Lab in Alderley Park | Wellcome Sanger Institute for the COVID-19 Genomics UK (COG-UK) Consortium | Jacquelyn Wynn, Mairead Hyland, The Lighthouse Lab in Alderley Park and Alex Alderton, Roberto Amato, Jeffrey Barrett, Sonia Goncalves, Ewan Harrison, David K. Jackson, Ian Johnston, Dominic Kwiatkowski, Cordelia Langford, John Sillitoe on behalf of the Wellcome Sanger Institute COVID-19 Surveillance Team |
| EPI_ISL_3069199 | Berkshire and Surrey Pathology Services Lighthouse Laboratory | Wellcome Sanger Institute for the COVID-19 Genomics UK (COG-UK) Consortium | Berkshire and Surrey Pathology Services Lighthouse Laboratory and Alex Alderton, Roberto Amato, Jeffrey Barrett, Sonia Goncalves, Ewan Harrison, David K. Jackson, Ian Johnston, Dominic Kwiatkowski, Cordelia Langford, John Sillitoe on behalf of the Wellcome Sanger Institute COVID-19 Surveillance Team |
| EPI_ISL_3069200, EPI_ISL_3069201 | Lighthouse Lab in Glasgow | Wellcome Sanger Institute for the COVID-19 Genomics UK (COG-UK) Consortium | Harper VanSteenhouse, Yumi Kasai, David Gray, Carol Clugston, Anna Dominiczak and Alex Alderton, Roberto Amato, Jeffrey Barrett, Sonia Goncalves, Ewan Harrison, David K. Jackson, Ian Johnston, Dominic Kwiatkowski, Cordelia Langford, John Sillitoe on behalf of the Wellcome Sanger Institute COVID-19 Surveillance Team |
| EPI_ISL_3069202 | Lighthouse Lab in Alderley Park | Wellcome Sanger Institute for the COVID-19 Genomics UK (COG-UK) Consortium | Jacquelyn Wynn, Mairead Hyland, The Lighthouse Lab in Alderley Park and Alex Alderton, Roberto Amato, Jeffrey Barrett, Sonia Goncalves, Ewan Harrison, David K. Jackson, Ian Johnston, Dominic Kwiatkowski, Cordelia Langford, John Sillitoe on behalf of the Wellcome Sanger Institute COVID-19 Surveillance Team |
| EPI_ISL_3069203 | Lighthouse Lab in Glasgow | Wellcome Sanger Institute for the COVID-19 Genomics UK (COG-UK) Consortium | Harper VanSteenhouse, Yumi Kasai, David Gray, Carol Clugston, Anna Dominiczak and Alex Alderton, Roberto Amato, Jeffrey Barrett, Sonia Goncalves, Ewan Harrison, David K. Jackson, Ian Johnston, Dominic Kwiatkowski, Cordelia Langford, John Sillitoe on behalf of the Wellcome Sanger Institute COVID-19 Surveillance Team |
| EPI_ISL_3069204 | Berkshire and Surrey Pathology Services Lighthouse Laboratory | Wellcome Sanger Institute for the COVID-19 Genomics UK (COG-UK) Consortium | Berkshire and Surrey Pathology Services Lighthouse Laboratory and Alex Alderton, Roberto Amato, Jeffrey Barrett, Sonia Goncalves, Ewan Harrison, David K. Jackson, Ian Johnston, Dominic Kwiatkowski, Cordelia Langford, John Sillitoe on behalf of the Wellcome Sanger Institute COVID-19 Surveillance Team |
| EPI_ISL_3069205 | Lighthouse Lab in Glasgow | Wellcome Sanger Institute for the COVID-19 Genomics UK (COG-UK) Consortium | Harper VanSteenhouse, Yumi Kasai, David Gray, Carol Clugston, Anna Dominiczak and Alex Alderton, Roberto Amato, Jeffrey Barrett, Sonia Goncalves, Ewan Harrison, David K. Jackson, Ian Johnston, Dominic Kwiatkowski, Cordelia Langford, John Sillitoe on behalf of the Wellcome Sanger Institute COVID-19 Surveillance Team |
| EPI_ISL_3069206 | Lighthouse Lab in Alderley Park | Wellcome Sanger Institute for the COVID-19 Genomics UK (COG-UK) Consortium | Jacquelyn Wynn, Mairead Hyland, The Lighthouse Lab in Alderley Park and Alex Alderton, Roberto Amato, Jeffrey Barrett, Sonia Goncalves, Ewan Harrison, David K. Jackson, Ian Johnston, Dominic Kwiatkowski, Cordelia Langford, John Sillitoe on behalf of the Wellcome Sanger Institute COVID-19 Surveillance Team |
| EPI_ISL_3069207 | Berkshire and Surrey Pathology Services Lighthouse Laboratory | Wellcome Sanger Institute for the COVID-19 Genomics UK (COG-UK) Consortium | Berkshire and Surrey Pathology Services Lighthouse Laboratory and Alex Alderton, Roberto Amato, Jeffrey Barrett, Sonia Goncalves, Ewan Harrison, David K. Jackson, Ian Johnston, Dominic Kwiatkowski, Cordelia Langford, John Sillitoe on behalf of the Wellcome Sanger Institute COVID-19 Surveillance Team |
| EPI_ISL_3069208 | Lighthouse Lab in Alderley Park | Wellcome Sanger Institute for the COVID-19 Genomics UK (COG-UK) Consortium | Jacquelyn Wynn, Mairead Hyland, The Lighthouse Lab in Alderley Park and Alex Alderton, Roberto Amato, Jeffrey Barrett, Sonia Goncalves, Ewan Harrison, David K. Jackson, Ian Johnston, Dominic Kwiatkowski, Cordelia Langford, John Sillitoe on behalf of the Wellcome Sanger Institute COVID-19 Surveillance Team |
| EPI_ISL_3069209, EPI_ISL_3069210 | Berkshire and Surrey Pathology Services Lighthouse Laboratory | Wellcome Sanger Institute for the COVID-19 Genomics UK (COG-UK) Consortium | Berkshire and Surrey Pathology Services Lighthouse Laboratory and Alex Alderton, Roberto Amato, Jeffrey Barrett, Sonia Goncalves, Ewan Harrison, David K. Jackson, Ian Johnston, Dominic Kwiatkowski, Cordelia Langford, John Sillitoe on behalf of the Wellcome Sanger Institute COVID-19 Surveillance Team |
| EPI_ISL_3069211, EPI_ISL_3069212 | Lighthouse Lab in Glasgow | Wellcome Sanger Institute for the COVID-19 Genomics UK (COG-UK) Consortium | Harper VanSteenhouse, Yumi Kasai, David Gray, Carol Clugston, Anna Dominiczak and Alex Alderton, Roberto Amato, Jeffrey Barrett, Sonia Goncalves, Ewan Harrison, David K. Jackson, Ian Johnston, Dominic Kwiatkowski, Cordelia Langford, John Sillitoe on behalf of the Wellcome Sanger Institute COVID-19 Surveillance Team |
| EPI_ISL_3069213 | Berkshire and Surrey Pathology Services Lighthouse Laboratory | Wellcome Sanger Institute for the COVID-19 Genomics UK (COG-UK) Consortium | Berkshire and Surrey Pathology Services Lighthouse Laboratory and Alex Alderton, Roberto Amato, Jeffrey Barrett, Sonia Goncalves, Ewan Harrison, David K. Jackson, Ian Johnston, Dominic Kwiatkowski, Cordelia Langford, John Sillitoe on behalf of the Wellcome Sanger Institute COVID-19 Surveillance Team |
| EPI_ISL_3069214 | Lighthouse Lab in Alderley Park | Wellcome Sanger Institute for the COVID-19 Genomics UK (COG-UK) Consortium | Jacquelyn Wynn, Mairead Hyland, The Lighthouse Lab in Alderley Park and Alex Alderton, Roberto Amato, Jeffrey Barrett, Sonia Goncalves, Ewan Harrison, David K. Jackson, Ian Johnston, Dominic Kwiatkowski, Cordelia Langford, John Sillitoe on behalf of the Wellcome Sanger Institute COVID-19 Surveillance Team |
| EPI_ISL_3069215 | Lighthouse Lab in Glasgow | Wellcome Sanger Institute for the COVID-19 Genomics UK (COG-UK) Consortium | Harper VanSteenhouse, Yumi Kasai, David Gray, Carol Clugston, Anna Dominiczak and Alex Alderton, Roberto Amato, Jeffrey Barrett, Sonia Goncalves, Ewan Harrison, David K. Jackson, Ian Johnston, Dominic Kwiatkowski, Cordelia Langford, John Sillitoe on behalf of the Wellcome Sanger Institute COVID-19 Surveillance Team |
| EPI_ISL_3069216 | Lighthouse Lab in Alderley Park | Wellcome Sanger Institute for the COVID-19 Genomics UK (COG-UK) Consortium | Jacquelyn Wynn, Mairead Hyland, The Lighthouse Lab in Alderley Park and Alex Alderton, Roberto Amato, Jeffrey Barrett, Sonia Goncalves, Ewan Harrison, David K. Jackson, Ian Johnston, Dominic Kwiatkowski, Cordelia Langford, John Sillitoe on behalf of the Wellcome Sanger Institute COVID-19 Surveillance Team |
| EPI_ISL_3069217, EPI_ISL_3069218 | Lighthouse Lab in Glasgow | Wellcome Sanger Institute for the COVID-19 Genomics UK (COG-UK) Consortium | Harper VanSteenhouse, Yumi Kasai, David Gray, Carol Clugston, Anna Dominiczak and Alex Alderton, Roberto Amato, Jeffrey Barrett, Sonia Goncalves, Ewan Harrison, David K. Jackson, Ian Johnston, Dominic Kwiatkowski, Cordelia Langford, John Sillitoe on behalf of the Wellcome Sanger Institute COVID-19 Surveillance Team |
| EPI_ISL_3069219 | Berkshire and Surrey Pathology Services Lighthouse Laboratory | Wellcome Sanger Institute for the COVID-19 Genomics UK (COG-UK) Consortium | Berkshire and Surrey Pathology Services Lighthouse Laboratory and Alex Alderton, Roberto Amato, Jeffrey Barrett, Sonia Goncalves, Ewan Harrison, David K. Jackson, Ian Johnston, Dominic Kwiatkowski, Cordelia Langford, John Sillitoe on behalf of the Wellcome Sanger Institute COVID-19 Surveillance Team |
| EPI_ISL_3069220 | Lighthouse Lab in Glasgow | Wellcome Sanger Institute for the COVID-19 Genomics UK (COG-UK) Consortium | Harper VanSteenhouse, Yumi Kasai, David Gray, Carol Clugston, Anna Dominiczak and Alex Alderton, Roberto Amato, Jeffrey Barrett, Sonia Goncalves, Ewan Harrison, David K. Jackson, Ian Johnston, Dominic Kwiatkowski, Cordelia Langford, John Sillitoe on behalf of the Wellcome Sanger Institute COVID-19 Surveillance Team |
| EPI_ISL_3069221 | Lighthouse Lab in Alderley Park | Wellcome Sanger Institute for the COVID-19 Genomics UK (COG-UK) Consortium | Jacquelyn Wynn, Mairead Hyland, The Lighthouse Lab in Alderley Park and Alex Alderton, Roberto Amato, Jeffrey Barrett, Sonia Goncalves, Ewan Harrison, David K. Jackson, Ian Johnston, Dominic Kwiatkowski, Cordelia Langford, John Sillitoe on behalf of the Wellcome Sanger Institute COVID-19 Surveillance Team |
| EPI_ISL_3069222 | Lighthouse Lab in Glasgow | Wellcome Sanger Institute for the COVID-19 Genomics UK (COG-UK) Consortium | Harper VanSteenhouse, Yumi Kasai, David Gray, Carol Clugston, Anna Dominiczak and Alex Alderton, Roberto Amato, Jeffrey Barrett, Sonia Goncalves, Ewan Harrison, David K. Jackson, Ian Johnston, Dominic Kwiatkowski, Cordelia Langford, John Sillitoe on behalf of the Wellcome Sanger Institute COVID-19 Surveillance Team |
| EPI_ISL_3069223, EPI_ISL_3069224 | Lighthouse Lab in Alderley Park | Wellcome Sanger Institute for the COVID-19 Genomics UK (COG-UK) Consortium | Jacquelyn Wynn, Mairead Hyland, The Lighthouse Lab in Alderley Park and Alex Alderton, Roberto Amato, Jeffrey Barrett, Sonia Goncalves, Ewan Harrison, David K. Jackson, Ian Johnston, Dominic Kwiatkowski, Cordelia Langford, John Sillitoe on behalf of the Wellcome Sanger Institute COVID-19 Surveillance Team |
| EPI_ISL_3069225, EPI_ISL_3069226, EPI_ISL_3069227 | Berkshire and Surrey Pathology Services Lighthouse Laboratory | Wellcome Sanger Institute for the COVID-19 Genomics UK (COG-UK) Consortium | Berkshire and Surrey Pathology Services Lighthouse Laboratory and Alex Alderton, Roberto Amato, Jeffrey Barrett, Sonia Goncalves, Ewan Harrison, David K. Jackson, Ian Johnston, Dominic Kwiatkowski, Cordelia Langford, John Sillitoe on behalf of the Wellcome Sanger Institute COVID-19 Surveillance Team |
| EPI_ISL_3069228 | Lighthouse Lab in Glasgow | Wellcome Sanger Institute for the COVID-19 Genomics UK (COG-UK) Consortium | Harper VanSteenhouse, Yumi Kasai, David Gray, Carol Clugston, Anna Dominiczak and Alex Alderton, Roberto Amato, Jeffrey Barrett, Sonia Goncalves, Ewan Harrison, David K. Jackson, Ian Johnston, Dominic Kwiatkowski, Cordelia Langford, John Sillitoe on behalf of the Wellcome Sanger Institute COVID-19 Surveillance Team |
| EPI_ISL_3069229 | Lighthouse Lab in Alderley Park | Wellcome Sanger Institute for the COVID-19 Genomics UK (COG-UK) Consortium | Jacquelyn Wynn, Mairead Hyland, The Lighthouse Lab in Alderley Park and Alex Alderton, Roberto Amato, Jeffrey Barrett, Sonia Goncalves, Ewan Harrison, David K. Jackson, Ian Johnston, Dominic Kwiatkowski, Cordelia Langford, John Sillitoe on behalf of the Wellcome Sanger Institute COVID-19 Surveillance Team |
| EPI_ISL_3069230 | Berkshire and Surrey Pathology Services Lighthouse Laboratory | Wellcome Sanger Institute for the COVID-19 Genomics UK (COG-UK) Consortium | Berkshire and Surrey Pathology Services Lighthouse Laboratory and Alex Alderton, Roberto Amato, Jeffrey Barrett, Sonia Goncalves, Ewan Harrison, David K. Jackson, Ian Johnston, Dominic Kwiatkowski, Cordelia Langford, John Sillitoe on behalf of the Wellcome Sanger Institute COVID-19 Surveillance Team |
| EPI_ISL_3069231, | Lighthouse Lab in Alderley Park | Wellcome Sanger Institute for the COVID-19 Genomics UK (COG-UK) | Jacquelyn Wynn, Mairead Hyland, The Lighthouse Lab in Alderley Park and Alex Alderton, Roberto Amato, Jeffrey Barrett, Sonia Goncalves, Ewan Harrison, David K. Jackson, Ian Johnston, Dominic Kwiatkowski, Cordelia Langford, John Sillitoe on |

| | | | |
|---|---|---|---|
| EPI_ISL_3069232 | | Consortium | behalf of the Wellcome Sanger Institute COVID-19 Surveillance Team |
| EPI_ISL_3069233 | Lighthouse Lab in Glasgow | Wellcome Sanger Institute for the COVID-19 Genomics UK (COG-UK) Consortium | Harper VanSteenhouse, Yumi Kasai, David Gray, Carol Clugston, Anna Dominiczak and Alex Alderton, Roberto Amato, Jeffrey Barrett, Sonia Goncalves, Ewan Harrison, David K. Jackson, Ian Johnston, Dominic Kwiatkowski, Cordelia Langford, John Sillitoe on behalf of the Wellcome Sanger Institute COVID-19 Surveillance Team |
| EPI_ISL_3069234 | Lighthouse Lab in Alderley Park | Wellcome Sanger Institute for the COVID-19 Genomics UK (COG-UK) Consortium | Jacquelyn Wynn, Mairead Hyland, The Lighthouse Lab in Alderley Park and Alex Alderton, Roberto Amato, Jeffrey Barrett, Sonia Goncalves, Ewan Harrison, David K. Jackson, Ian Johnston, Dominic Kwiatkowski, Cordelia Langford, John Sillitoe on behalf of the Wellcome Sanger Institute COVID-19 Surveillance Team |
| EPI_ISL_3069235, EPI_ISL_3069236 | Berkshire and Surrey Pathology Services Lighthouse Laboratory | Wellcome Sanger Institute for the COVID-19 Genomics UK (COG-UK) Consortium | Berkshire and Surrey Pathology Services Lighthouse Laboratory and Alex Alderton, Roberto Amato, Jeffrey Barrett, Sonia Goncalves, Ewan Harrison, David K. Jackson, Ian Johnston, Dominic Kwiatkowski, Cordelia Langford, John Sillitoe on behalf of the Wellcome Sanger Institute COVID-19 Surveillance Team |
| EPI_ISL_3069237 | Lighthouse Lab in Glasgow | Wellcome Sanger Institute for the COVID-19 Genomics UK (COG-UK) Consortium | Harper VanSteenhouse, Yumi Kasai, David Gray, Carol Clugston, Anna Dominiczak and Alex Alderton, Roberto Amato, Jeffrey Barrett, Sonia Goncalves, Ewan Harrison, David K. Jackson, Ian Johnston, Dominic Kwiatkowski, Cordelia Langford, John Sillitoe on behalf of the Wellcome Sanger Institute COVID-19 Surveillance Team |
| EPI_ISL_3069238 | Berkshire and Surrey Pathology Services Lighthouse Laboratory | Wellcome Sanger Institute for the COVID-19 Genomics UK (COG-UK) Consortium | Berkshire and Surrey Pathology Services Lighthouse Laboratory and Alex Alderton, Roberto Amato, Jeffrey Barrett, Sonia Goncalves, Ewan Harrison, David K. Jackson, Ian Johnston, Dominic Kwiatkowski, Cordelia Langford, John Sillitoe on behalf of the Wellcome Sanger Institute COVID-19 Surveillance Team |
| EPI_ISL_3069239 | Lighthouse Lab in Alderley Park | Wellcome Sanger Institute for the COVID-19 Genomics UK (COG-UK) Consortium | Jacquelyn Wynn, Mairead Hyland, The Lighthouse Lab in Alderley Park and Alex Alderton, Roberto Amato, Jeffrey Barrett, Sonia Goncalves, Ewan Harrison, David K. Jackson, Ian Johnston, Dominic Kwiatkowski, Cordelia Langford, John Sillitoe on behalf of the Wellcome Sanger Institute COVID-19 Surveillance Team |
| EPI_ISL_3069240, EPI_ISL_3069241, EPI_ISL_3069242, EPI_ISL_3069243 | Berkshire and Surrey Pathology Services Lighthouse Laboratory | Wellcome Sanger Institute for the COVID-19 Genomics UK (COG-UK) Consortium | Berkshire and Surrey Pathology Services Lighthouse Laboratory and Alex Alderton, Roberto Amato, Jeffrey Barrett, Sonia Goncalves, Ewan Harrison, David K. Jackson, Ian Johnston, Dominic Kwiatkowski, Cordelia Langford, John Sillitoe on behalf of the Wellcome Sanger Institute COVID-19 Surveillance Team |
| EPI_ISL_3069244 | Lighthouse Lab in Glasgow | Wellcome Sanger Institute for the COVID-19 Genomics UK (COG-UK) Consortium | Harper VanSteenhouse, Yumi Kasai, David Gray, Carol Clugston, Anna Dominiczak and Alex Alderton, Roberto Amato, Jeffrey Barrett, Sonia Goncalves, Ewan Harrison, David K. Jackson, Ian Johnston, Dominic Kwiatkowski, Cordelia Langford, John Sillitoe on behalf of the Wellcome Sanger Institute COVID-19 Surveillance Team |
| EPI_ISL_3069245 | Berkshire and Surrey Pathology Services Lighthouse Laboratory | Wellcome Sanger Institute for the COVID-19 Genomics UK (COG-UK) Consortium | Berkshire and Surrey Pathology Services Lighthouse Laboratory and Alex Alderton, Roberto Amato, Jeffrey Barrett, Sonia Goncalves, Ewan Harrison, David K. Jackson, Ian Johnston, Dominic Kwiatkowski, Cordelia Langford, John Sillitoe on behalf of the Wellcome Sanger Institute COVID-19 Surveillance Team |
| EPI_ISL_3069246 | Lighthouse Lab in Glasgow | Wellcome Sanger Institute for the COVID-19 Genomics UK (COG-UK) Consortium | Harper VanSteenhouse, Yumi Kasai, David Gray, Carol Clugston, Anna Dominiczak and Alex Alderton, Roberto Amato, Jeffrey Barrett, Sonia Goncalves, Ewan Harrison, David K. Jackson, Ian Johnston, Dominic Kwiatkowski, Cordelia Langford, John Sillitoe on behalf of the Wellcome Sanger Institute COVID-19 Surveillance Team |
| EPI_ISL_3069247 | Berkshire and Surrey Pathology Services Lighthouse Laboratory | Wellcome Sanger Institute for the COVID-19 Genomics UK (COG-UK) Consortium | Berkshire and Surrey Pathology Services Lighthouse Laboratory and Alex Alderton, Roberto Amato, Jeffrey Barrett, Sonia Goncalves, Ewan Harrison, David K. Jackson, Ian Johnston, Dominic Kwiatkowski, Cordelia Langford, John Sillitoe on behalf of the Wellcome Sanger Institute COVID-19 Surveillance Team |
| EPI_ISL_3069248 | Lighthouse Lab in Glasgow | Wellcome Sanger Institute for the COVID-19 Genomics UK (COG-UK) Consortium | Harper VanSteenhouse, Yumi Kasai, David Gray, Carol Clugston, Anna Dominiczak and Alex Alderton, Roberto Amato, Jeffrey Barrett, Sonia Goncalves, Ewan Harrison, David K. Jackson, Ian Johnston, Dominic Kwiatkowski, Cordelia Langford, John Sillitoe on behalf of the Wellcome Sanger Institute COVID-19 Surveillance Team |
| EPI_ISL_3069249 | Lighthouse Lab in Alderley Park | Wellcome Sanger Institute for the COVID-19 Genomics UK (COG-UK) Consortium | Jacquelyn Wynn, Mairead Hyland, The Lighthouse Lab in Alderley Park and Alex Alderton, Roberto Amato, Jeffrey Barrett, Sonia Goncalves, Ewan Harrison, David K. Jackson, Ian Johnston, Dominic Kwiatkowski, Cordelia Langford, John Sillitoe on behalf of the Wellcome Sanger Institute COVID-19 Surveillance Team |
| EPI_ISL_3069250 | Berkshire and Surrey Pathology Services Lighthouse Laboratory | Wellcome Sanger Institute for the COVID-19 Genomics UK (COG-UK) Consortium | Berkshire and Surrey Pathology Services Lighthouse Laboratory and Alex Alderton, Roberto Amato, Jeffrey Barrett, Sonia Goncalves, Ewan Harrison, David K. Jackson, Ian Johnston, Dominic Kwiatkowski, Cordelia Langford, John Sillitoe on behalf of the Wellcome Sanger Institute COVID-19 Surveillance Team |
| EPI_ISL_3069251, EPI_ISL_3069252 | Lighthouse Lab in Alderley Park | Wellcome Sanger Institute for the COVID-19 Genomics UK (COG-UK) Consortium | Jacquelyn Wynn, Mairead Hyland, The Lighthouse Lab in Alderley Park and Alex Alderton, Roberto Amato, Jeffrey Barrett, Sonia Goncalves, Ewan Harrison, David K. Jackson, Ian Johnston, Dominic Kwiatkowski, Cordelia Langford, John Sillitoe on behalf of the Wellcome Sanger Institute COVID-19 Surveillance Team |
| EPI_ISL_3069253 | Berkshire and Surrey Pathology Services Lighthouse Laboratory | Wellcome Sanger Institute for the COVID-19 Genomics UK (COG-UK) Consortium | Berkshire and Surrey Pathology Services Lighthouse Laboratory and Alex Alderton, Roberto Amato, Jeffrey Barrett, Sonia Goncalves, Ewan Harrison, David K. Jackson, Ian Johnston, Dominic Kwiatkowski, Cordelia Langford, John Sillitoe on behalf of the Wellcome Sanger Institute COVID-19 Surveillance Team |
| EPI_ISL_3069254, EPI_ISL_3069255, EPI_ISL_3069256 | Lighthouse Lab in Glasgow | Wellcome Sanger Institute for the COVID-19 Genomics UK (COG-UK) Consortium | Harper VanSteenhouse, Yumi Kasai, David Gray, Carol Clugston, Anna Dominiczak and Alex Alderton, Roberto Amato, Jeffrey Barrett, Sonia Goncalves, Ewan Harrison, David K. Jackson, Ian Johnston, Dominic Kwiatkowski, Cordelia Langford, John Sillitoe on behalf of the Wellcome Sanger Institute COVID-19 Surveillance Team |
| EPI_ISL_3069257 | Lighthouse Lab in Alderley Park | Wellcome Sanger Institute for the COVID-19 Genomics UK (COG-UK) Consortium | Jacquelyn Wynn, Mairead Hyland, The Lighthouse Lab in Alderley Park and Alex Alderton, Roberto Amato, Jeffrey Barrett, Sonia Goncalves, Ewan Harrison, David K. Jackson, Ian Johnston, Dominic Kwiatkowski, Cordelia Langford, John Sillitoe on behalf of the Wellcome Sanger Institute COVID-19 Surveillance Team |
| EPI_ISL_3069258, EPI_ISL_3069259 | Berkshire and Surrey Pathology Services Lighthouse Laboratory | Wellcome Sanger Institute for the COVID-19 Genomics UK (COG-UK) Consortium | Berkshire and Surrey Pathology Services Lighthouse Laboratory and Alex Alderton, Roberto Amato, Jeffrey Barrett, Sonia Goncalves, Ewan Harrison, David K. Jackson, Ian Johnston, Dominic Kwiatkowski, Cordelia Langford, John Sillitoe on behalf of the Wellcome Sanger Institute COVID-19 Surveillance Team |
| EPI_ISL_3069260 | Lighthouse Lab in Alderley Park | Wellcome Sanger Institute for the COVID-19 Genomics UK (COG-UK) Consortium | Jacquelyn Wynn, Mairead Hyland, The Lighthouse Lab in Alderley Park and Alex Alderton, Roberto Amato, Jeffrey Barrett, Sonia Goncalves, Ewan Harrison, David K. Jackson, Ian Johnston, Dominic Kwiatkowski, Cordelia Langford, John Sillitoe on behalf of the Wellcome Sanger Institute COVID-19 Surveillance Team |
| EPI_ISL_3069261, EPI_ISL_3069262 | Berkshire and Surrey Pathology Services Lighthouse Laboratory | Wellcome Sanger Institute for the COVID-19 Genomics UK (COG-UK) Consortium | Berkshire and Surrey Pathology Services Lighthouse Laboratory and Alex Alderton, Roberto Amato, Jeffrey Barrett, Sonia Goncalves, Ewan Harrison, David K. Jackson, Ian Johnston, Dominic Kwiatkowski, Cordelia Langford, John Sillitoe on behalf of the Wellcome Sanger Institute COVID-19 Surveillance Team |
| EPI_ISL_3069263 | Lighthouse Lab in Alderley Park | Wellcome Sanger Institute for the COVID-19 Genomics UK (COG-UK) Consortium | Jacquelyn Wynn, Mairead Hyland, The Lighthouse Lab in Alderley Park and Alex Alderton, Roberto Amato, Jeffrey Barrett, Sonia Goncalves, Ewan Harrison, David K. Jackson, Ian Johnston, Dominic Kwiatkowski, Cordelia Langford, John Sillitoe on behalf of the Wellcome Sanger Institute COVID-19 Surveillance Team |
| EPI_ISL_3069264 | Lighthouse Lab in Glasgow | Wellcome Sanger Institute for the COVID-19 Genomics UK (COG-UK) Consortium | Harper VanSteenhouse, Yumi Kasai, David Gray, Carol Clugston, Anna Dominiczak and Alex Alderton, Roberto Amato, Jeffrey Barrett, Sonia Goncalves, Ewan Harrison, David K. Jackson, Ian Johnston, Dominic Kwiatkowski, Cordelia Langford, John Sillitoe on behalf of the Wellcome Sanger Institute COVID-19 Surveillance Team |
| EPI_ISL_3069265 | Lighthouse Lab in Alderley Park | Wellcome Sanger Institute for the COVID-19 Genomics UK (COG-UK) Consortium | Jacquelyn Wynn, Mairead Hyland, The Lighthouse Lab in Alderley Park and Alex Alderton, Roberto Amato, Jeffrey Barrett, Sonia Goncalves, Ewan Harrison, David K. Jackson, Ian Johnston, Dominic Kwiatkowski, Cordelia Langford, John Sillitoe on behalf of the Wellcome Sanger Institute COVID-19 Surveillance Team |
| EPI_ISL_3069266 | Lighthouse Lab in Glasgow | Wellcome Sanger Institute for the COVID-19 Genomics UK (COG-UK) Consortium | Harper VanSteenhouse, Yumi Kasai, David Gray, Carol Clugston, Anna Dominiczak and Alex Alderton, Roberto Amato, Jeffrey Barrett, Sonia Goncalves, Ewan Harrison, David K. Jackson, Ian Johnston, Dominic Kwiatkowski, Cordelia Langford, John Sillitoe on behalf of the Wellcome Sanger Institute COVID-19 Surveillance Team |
| EPI_ISL_3069267, EPI_ISL_3069268 | Berkshire and Surrey Pathology Services Lighthouse Laboratory | Wellcome Sanger Institute for the COVID-19 Genomics UK (COG-UK) Consortium | Berkshire and Surrey Pathology Services Lighthouse Laboratory and Alex Alderton, Roberto Amato, Jeffrey Barrett, Sonia Goncalves, Ewan Harrison, David K. Jackson, Ian Johnston, Dominic Kwiatkowski, Cordelia Langford, John Sillitoe on behalf of the Wellcome Sanger Institute COVID-19 Surveillance Team |
| EPI_ISL_3069269, EPI_ISL_3069270 | Lighthouse Lab in Glasgow | Wellcome Sanger Institute for the COVID-19 Genomics UK (COG-UK) Consortium | Harper VanSteenhouse, Yumi Kasai, David Gray, Carol Clugston, Anna Dominiczak and Alex Alderton, Roberto Amato, Jeffrey Barrett, Sonia Goncalves, Ewan Harrison, David K. Jackson, Ian Johnston, Dominic Kwiatkowski, Cordelia Langford, John Sillitoe on behalf of the Wellcome Sanger Institute COVID-19 Surveillance Team |
| EPI_ISL_3069271, EPI_ISL_3069272 | Berkshire and Surrey Pathology Services Lighthouse Laboratory | Wellcome Sanger Institute for the COVID-19 Genomics UK (COG-UK) Consortium | Berkshire and Surrey Pathology Services Lighthouse Laboratory and Alex Alderton, Roberto Amato, Jeffrey Barrett, Sonia Goncalves, Ewan Harrison, David K. Jackson, Ian Johnston, Dominic Kwiatkowski, Cordelia Langford, John Sillitoe on behalf of the Wellcome Sanger Institute COVID-19 Surveillance Team |
| EPI_ISL_3069273, EPI_ISL_3069274 | Lighthouse Lab in Alderley Park | Wellcome Sanger Institute for the COVID-19 Genomics UK (COG-UK) Consortium | Jacquelyn Wynn, Mairead Hyland, The Lighthouse Lab in Alderley Park and Alex Alderton, Roberto Amato, Jeffrey Barrett, Sonia Goncalves, Ewan Harrison, David K. Jackson, Ian Johnston, Dominic Kwiatkowski, Cordelia Langford, John Sillitoe on behalf of the Wellcome Sanger Institute COVID-19 Surveillance Team |
| EPI_ISL_3069275, EPI_ISL_3069276, EPI_ISL_3069277 | Berkshire and Surrey Pathology Services Lighthouse Laboratory | Wellcome Sanger Institute for the COVID-19 Genomics UK (COG-UK) Consortium | Berkshire and Surrey Pathology Services Lighthouse Laboratory and Alex Alderton, Roberto Amato, Jeffrey Barrett, Sonia Goncalves, Ewan Harrison, David K. Jackson, Ian Johnston, Dominic Kwiatkowski, Cordelia Langford, John Sillitoe on behalf of the Wellcome Sanger Institute COVID-19 Surveillance Team |
| EPI_ISL_3069278 | Lighthouse Lab in Alderley Park | Wellcome Sanger Institute for the COVID-19 Genomics UK (COG-UK) Consortium | Jacquelyn Wynn, Mairead Hyland, The Lighthouse Lab in Alderley Park and Alex Alderton, Roberto Amato, Jeffrey Barrett, Sonia Goncalves, Ewan Harrison, David K. Jackson, Ian Johnston, Dominic Kwiatkowski, Cordelia Langford, John Sillitoe on behalf of the Wellcome Sanger Institute COVID-19 Surveillance Team |
| EPI_ISL_3069279, EPI_ISL_3069280, EPI_ISL_3069281, EPI_ISL_3069282 | Berkshire and Surrey Pathology Services Lighthouse Laboratory | Wellcome Sanger Institute for the COVID-19 Genomics UK (COG-UK) Consortium | Berkshire and Surrey Pathology Services Lighthouse Laboratory and Alex Alderton, Roberto Amato, Jeffrey Barrett, Sonia Goncalves, Ewan Harrison, David K. Jackson, Ian Johnston, Dominic Kwiatkowski, Cordelia Langford, John Sillitoe on behalf of the Wellcome Sanger Institute COVID-19 Surveillance Team |
| EPI_ISL_3069283 | Lighthouse Lab in Glasgow | Wellcome Sanger Institute for the COVID-19 Genomics UK (COG-UK) Consortium | Harper VanSteenhouse, Yumi Kasai, David Gray, Carol Clugston, Anna Dominiczak and Alex Alderton, Roberto Amato, Jeffrey Barrett, Sonia Goncalves, Ewan Harrison, David K. Jackson, Ian Johnston, Dominic Kwiatkowski, Cordelia Langford, John Sillitoe on behalf of the Wellcome Sanger Institute COVID-19 Surveillance Team |
| EPI_ISL_3069284 | Berkshire and Surrey Pathology Services Lighthouse Laboratory | Wellcome Sanger Institute for the COVID-19 Genomics UK (COG-UK) Consortium | Berkshire and Surrey Pathology Services Lighthouse Laboratory and Alex Alderton, Roberto Amato, Jeffrey Barrett, Sonia Goncalves, Ewan Harrison, David K. Jackson, Ian Johnston, Dominic Kwiatkowski, Cordelia Langford, John Sillitoe on behalf of the Wellcome Sanger Institute COVID-19 Surveillance Team |
| EPI_ISL_3069285 | Lighthouse Lab in Alderley Park | Wellcome Sanger Institute for the COVID-19 Genomics UK (COG-UK) Consortium | Jacquelyn Wynn, Mairead Hyland, The Lighthouse Lab in Alderley Park and Alex Alderton, Roberto Amato, Jeffrey Barrett, Sonia Goncalves, Ewan Harrison, David K. Jackson, Ian Johnston, Dominic Kwiatkowski, Cordelia Langford, John Sillitoe on behalf of the Wellcome Sanger Institute COVID-19 Surveillance Team |
| EPI_ISL_3069286, EPI_ISL_3069287, EPI_ISL_3069288 | Berkshire and Surrey Pathology Services Lighthouse Laboratory | Wellcome Sanger Institute for the COVID-19 Genomics UK (COG-UK) Consortium | Berkshire and Surrey Pathology Services Lighthouse Laboratory and Alex Alderton, Roberto Amato, Jeffrey Barrett, Sonia Goncalves, Ewan Harrison, David K. Jackson, Ian Johnston, Dominic Kwiatkowski, Cordelia Langford, John Sillitoe on behalf of the Wellcome Sanger Institute COVID-19 Surveillance Team |
| EPI_ISL_3069289 | Lighthouse Lab in Alderley Park | Wellcome Sanger Institute for the COVID-19 Genomics UK (COG-UK) Consortium | Jacquelyn Wynn, Mairead Hyland, The Lighthouse Lab in Alderley Park and Alex Alderton, Roberto Amato, Jeffrey Barrett, Sonia Goncalves, Ewan Harrison, David K. Jackson, Ian Johnston, Dominic Kwiatkowski, Cordelia Langford, John Sillitoe on behalf of the Wellcome Sanger Institute COVID-19 Surveillance Team |
| EPI_ISL_3069290, EPI_ISL_3069291, EPI_ISL_3069292 | Berkshire and Surrey Pathology Services Lighthouse Laboratory | Wellcome Sanger Institute for the COVID-19 Genomics UK (COG-UK) Consortium | Berkshire and Surrey Pathology Services Lighthouse Laboratory and Alex Alderton, Roberto Amato, Jeffrey Barrett, Sonia Goncalves, Ewan Harrison, David K. Jackson, Ian Johnston, Dominic Kwiatkowski, Cordelia Langford, John Sillitoe on behalf of the Wellcome Sanger Institute COVID-19 Surveillance Team |
| EPI_ISL_3069293 | Lighthouse Lab in Glasgow | Wellcome Sanger Institute for the COVID-19 Genomics UK (COG-UK) Consortium | Harper VanSteenhouse, Yumi Kasai, David Gray, Carol Clugston, Anna Dominiczak and Alex Alderton, Roberto Amato, Jeffrey Barrett, Sonia Goncalves, Ewan Harrison, David K. Jackson, Ian Johnston, Dominic Kwiatkowski, Cordelia Langford, John Sillitoe on behalf of the Wellcome Sanger Institute COVID-19 Surveillance Team |
| EPI_ISL_3069294 | Berkshire and Surrey Pathology Services Lighthouse Laboratory | Wellcome Sanger Institute for the COVID-19 Genomics UK (COG-UK) Consortium | Berkshire and Surrey Pathology Services Lighthouse Laboratory and Alex Alderton, Roberto Amato, Jeffrey Barrett, Sonia Goncalves, Ewan Harrison, David K. Jackson, Ian Johnston, Dominic Kwiatkowski, Cordelia Langford, John Sillitoe on behalf of the Wellcome Sanger Institute COVID-19 Surveillance Team |
| EPI_ISL_3069295, EPI_ISL_3069296, EPI_ISL_3069297 | Lighthouse Lab in Glasgow | Wellcome Sanger Institute for the COVID-19 Genomics UK (COG-UK) Consortium | Harper VanSteenhouse, Yumi Kasai, David Gray, Carol Clugston, Anna Dominiczak and Alex Alderton, Roberto Amato, Jeffrey Barrett, Sonia Goncalves, Ewan Harrison, David K. Jackson, Ian Johnston, Dominic Kwiatkowski, Cordelia Langford, John Sillitoe on behalf of the Wellcome Sanger Institute COVID-19 Surveillance Team |

| | | | |
|---|---|---|---|
| EPI_ISL_3069298, EPI_ISL_3069299, EPI_ISL_3069300 | Berkshire and Surrey Pathology Services Lighthouse Laboratory | Wellcome Sanger Institute for the COVID-19 Genomics UK (COG-UK) Consortium | Berkshire and Surrey Pathology Services Lighthouse Laboratory and Alex Alderton, Roberto Amato, Jeffrey Barrett, Sonia Goncalves, Ewan Harrison, David K. Jackson, Ian Johnston, Dominic Kwiatkowski, Cordelia Langford, John Sillitoe on behalf of the Wellcome Sanger Institute COVID-19 Surveillance Team |
| EPI_ISL_3069301, EPI_ISL_3069302 | Lighthouse Lab in Glasgow | Wellcome Sanger Institute for the COVID-19 Genomics UK (COG-UK) Consortium | Harper VanSteenhouse, Yumi Kasai, David Gray, Carol Clugston, Anna Dominiczak and Alex Alderton, Roberto Amato, Jeffrey Barrett, Sonia Goncalves, Ewan Harrison, David K. Jackson, Ian Johnston, Dominic Kwiatkowski, Cordelia Langford, John Sillitoe on behalf of the Wellcome Sanger Institute COVID-19 Surveillance Team |
| EPI_ISL_3069303 | Lighthouse Lab in Alderley Park | Wellcome Sanger Institute for the COVID-19 Genomics UK (COG-UK) Consortium | Jacquelyn Wynn, Mairead Hyland, The Lighthouse Lab in Alderley Park and Alex Alderton, Roberto Amato, Jeffrey Barrett, Sonia Goncalves, Ewan Harrison, David K. Jackson, Ian Johnston, Dominic Kwiatkowski, Cordelia Langford, John Sillitoe on behalf of the Wellcome Sanger Institute COVID-19 Surveillance Team |
| EPI_ISL_3069304 | Berkshire and Surrey Pathology Services Lighthouse Laboratory | Wellcome Sanger Institute for the COVID-19 Genomics UK (COG-UK) Consortium | Berkshire and Surrey Pathology Services Lighthouse Laboratory and Alex Alderton, Roberto Amato, Jeffrey Barrett, Sonia Goncalves, Ewan Harrison, David K. Jackson, Ian Johnston, Dominic Kwiatkowski, Cordelia Langford, John Sillitoe on behalf of the Wellcome Sanger Institute COVID-19 Surveillance Team |
| EPI_ISL_3069305, EPI_ISL_3069306 | Lighthouse Lab in Alderley Park | Wellcome Sanger Institute for the COVID-19 Genomics UK (COG-UK) Consortium | Jacquelyn Wynn, Mairead Hyland, The Lighthouse Lab in Alderley Park and Alex Alderton, Roberto Amato, Jeffrey Barrett, Sonia Goncalves, Ewan Harrison, David K. Jackson, Ian Johnston, Dominic Kwiatkowski, Cordelia Langford, John Sillitoe on behalf of the Wellcome Sanger Institute COVID-19 Surveillance Team |
| EPI_ISL_3069307 | Lighthouse Lab in Glasgow | Wellcome Sanger Institute for the COVID-19 Genomics UK (COG-UK) Consortium | Harper VanSteenhouse, Yumi Kasai, David Gray, Carol Clugston, Anna Dominiczak and Alex Alderton, Roberto Amato, Jeffrey Barrett, Sonia Goncalves, Ewan Harrison, David K. Jackson, Ian Johnston, Dominic Kwiatkowski, Cordelia Langford, John Sillitoe on behalf of the Wellcome Sanger Institute COVID-19 Surveillance Team |
| EPI_ISL_3069308 | Berkshire and Surrey Pathology Services Lighthouse Laboratory | Wellcome Sanger Institute for the COVID-19 Genomics UK (COG-UK) Consortium | Berkshire and Surrey Pathology Services Lighthouse Laboratory and Alex Alderton, Roberto Amato, Jeffrey Barrett, Sonia Goncalves, Ewan Harrison, David K. Jackson, Ian Johnston, Dominic Kwiatkowski, Cordelia Langford, John Sillitoe on behalf of the Wellcome Sanger Institute COVID-19 Surveillance Team |
| EPI_ISL_3069309 | Lighthouse Lab in Alderley Park | Wellcome Sanger Institute for the COVID-19 Genomics UK (COG-UK) Consortium | Jacquelyn Wynn, Mairead Hyland, The Lighthouse Lab in Alderley Park and Alex Alderton, Roberto Amato, Jeffrey Barrett, Sonia Goncalves, Ewan Harrison, David K. Jackson, Ian Johnston, Dominic Kwiatkowski, Cordelia Langford, John Sillitoe on behalf of the Wellcome Sanger Institute COVID-19 Surveillance Team |
| EPI_ISL_3069310 | Berkshire and Surrey Pathology Services Lighthouse Laboratory | Wellcome Sanger Institute for the COVID-19 Genomics UK (COG-UK) Consortium | Berkshire and Surrey Pathology Services Lighthouse Laboratory and Alex Alderton, Roberto Amato, Jeffrey Barrett, Sonia Goncalves, Ewan Harrison, David K. Jackson, Ian Johnston, Dominic Kwiatkowski, Cordelia Langford, John Sillitoe on behalf of the Wellcome Sanger Institute COVID-19 Surveillance Team |
| EPI_ISL_3069311, EPI_ISL_3069312 | Lighthouse Lab in Glasgow | Wellcome Sanger Institute for the COVID-19 Genomics UK (COG-UK) Consortium | Harper VanSteenhouse, Yumi Kasai, David Gray, Carol Clugston, Anna Dominiczak and Alex Alderton, Roberto Amato, Jeffrey Barrett, Sonia Goncalves, Ewan Harrison, David K. Jackson, Ian Johnston, Dominic Kwiatkowski, Cordelia Langford, John Sillitoe on behalf of the Wellcome Sanger Institute COVID-19 Surveillance Team |
| EPI_ISL_3069313, EPI_ISL_3069314 | Lighthouse Lab in Alderley Park | Wellcome Sanger Institute for the COVID-19 Genomics UK (COG-UK) Consortium | Jacquelyn Wynn, Mairead Hyland, The Lighthouse Lab in Alderley Park and Alex Alderton, Roberto Amato, Jeffrey Barrett, Sonia Goncalves, Ewan Harrison, David K. Jackson, Ian Johnston, Dominic Kwiatkowski, Cordelia Langford, John Sillitoe on behalf of the Wellcome Sanger Institute COVID-19 Surveillance Team |
| EPI_ISL_3069315, EPI_ISL_3069316 | Berkshire and Surrey Pathology Services Lighthouse Laboratory | Wellcome Sanger Institute for the COVID-19 Genomics UK (COG-UK) Consortium | Berkshire and Surrey Pathology Services Lighthouse Laboratory and Alex Alderton, Roberto Amato, Jeffrey Barrett, Sonia Goncalves, Ewan Harrison, David K. Jackson, Ian Johnston, Dominic Kwiatkowski, Cordelia Langford, John Sillitoe on behalf of the Wellcome Sanger Institute COVID-19 Surveillance Team |
| EPI_ISL_3069317, EPI_ISL_3069318, EPI_ISL_3069319 | Lighthouse Lab in Alderley Park | Wellcome Sanger Institute for the COVID-19 Genomics UK (COG-UK) Consortium | Jacquelyn Wynn, Mairead Hyland, The Lighthouse Lab in Alderley Park and Alex Alderton, Roberto Amato, Jeffrey Barrett, Sonia Goncalves, Ewan Harrison, David K. Jackson, Ian Johnston, Dominic Kwiatkowski, Cordelia Langford, John Sillitoe on behalf of the Wellcome Sanger Institute COVID-19 Surveillance Team |
| EPI_ISL_3069320 | Lighthouse Lab in Glasgow | Wellcome Sanger Institute for the COVID-19 Genomics UK (COG-UK) Consortium | Harper VanSteenhouse, Yumi Kasai, David Gray, Carol Clugston, Anna Dominiczak and Alex Alderton, Roberto Amato, Jeffrey Barrett, Sonia Goncalves, Ewan Harrison, David K. Jackson, Ian Johnston, Dominic Kwiatkowski, Cordelia Langford, John Sillitoe on behalf of the Wellcome Sanger Institute COVID-19 Surveillance Team |
| EPI_ISL_3069321, EPI_ISL_3069322 | Berkshire and Surrey Pathology Services Lighthouse Laboratory | Wellcome Sanger Institute for the COVID-19 Genomics UK (COG-UK) Consortium | Berkshire and Surrey Pathology Services Lighthouse Laboratory and Alex Alderton, Roberto Amato, Jeffrey Barrett, Sonia Goncalves, Ewan Harrison, David K. Jackson, Ian Johnston, Dominic Kwiatkowski, Cordelia Langford, John Sillitoe on behalf of the Wellcome Sanger Institute COVID-19 Surveillance Team |
| EPI_ISL_3069323 | Lighthouse Lab in Glasgow | Wellcome Sanger Institute for the COVID-19 Genomics UK (COG-UK) Consortium | Harper VanSteenhouse, Yumi Kasai, David Gray, Carol Clugston, Anna Dominiczak and Alex Alderton, Roberto Amato, Jeffrey Barrett, Sonia Goncalves, Ewan Harrison, David K. Jackson, Ian Johnston, Dominic Kwiatkowski, Cordelia Langford, John Sillitoe on behalf of the Wellcome Sanger Institute COVID-19 Surveillance Team |
| EPI_ISL_3069324, EPI_ISL_3069325 | Berkshire and Surrey Pathology Services Lighthouse Laboratory | Wellcome Sanger Institute for the COVID-19 Genomics UK (COG-UK) Consortium | Berkshire and Surrey Pathology Services Lighthouse Laboratory and Alex Alderton, Roberto Amato, Jeffrey Barrett, Sonia Goncalves, Ewan Harrison, David K. Jackson, Ian Johnston, Dominic Kwiatkowski, Cordelia Langford, John Sillitoe on behalf of the Wellcome Sanger Institute COVID-19 Surveillance Team |
| EPI_ISL_3069326, EPI_ISL_3069327 | Lighthouse Lab in Alderley Park | Wellcome Sanger Institute for the COVID-19 Genomics UK (COG-UK) Consortium | Jacquelyn Wynn, Mairead Hyland, The Lighthouse Lab in Alderley Park and Alex Alderton, Roberto Amato, Jeffrey Barrett, Sonia Goncalves, Ewan Harrison, David K. Jackson, Ian Johnston, Dominic Kwiatkowski, Cordelia Langford, John Sillitoe on behalf of the Wellcome Sanger Institute COVID-19 Surveillance Team |
| EPI_ISL_3069328 | Lighthouse Lab in Glasgow | Wellcome Sanger Institute for the COVID-19 Genomics UK (COG-UK) Consortium | Harper VanSteenhouse, Yumi Kasai, David Gray, Carol Clugston, Anna Dominiczak and Alex Alderton, Roberto Amato, Jeffrey Barrett, Sonia Goncalves, Ewan Harrison, David K. Jackson, Ian Johnston, Dominic Kwiatkowski, Cordelia Langford, John Sillitoe on behalf of the Wellcome Sanger Institute COVID-19 Surveillance Team |
| EPI_ISL_3069329 | Berkshire and Surrey Pathology Services Lighthouse Laboratory | Wellcome Sanger Institute for the COVID-19 Genomics UK (COG-UK) Consortium | Berkshire and Surrey Pathology Services Lighthouse Laboratory and Alex Alderton, Roberto Amato, Jeffrey Barrett, Sonia Goncalves, Ewan Harrison, David K. Jackson, Ian Johnston, Dominic Kwiatkowski, Cordelia Langford, John Sillitoe on behalf of the Wellcome Sanger Institute COVID-19 Surveillance Team |
| EPI_ISL_3069330, EPI_ISL_3069331, EPI_ISL_3069332 | Lighthouse Lab in Alderley Park | Wellcome Sanger Institute for the COVID-19 Genomics UK (COG-UK) Consortium | Jacquelyn Wynn, Mairead Hyland, The Lighthouse Lab in Alderley Park and Alex Alderton, Roberto Amato, Jeffrey Barrett, Sonia Goncalves, Ewan Harrison, David K. Jackson, Ian Johnston, Dominic Kwiatkowski, Cordelia Langford, John Sillitoe on behalf of the Wellcome Sanger Institute COVID-19 Surveillance Team |
| EPI_ISL_3069333, EPI_ISL_3069334 | Berkshire and Surrey Pathology Services Lighthouse Laboratory | Wellcome Sanger Institute for the COVID-19 Genomics UK (COG-UK) Consortium | Berkshire and Surrey Pathology Services Lighthouse Laboratory and Alex Alderton, Roberto Amato, Jeffrey Barrett, Sonia Goncalves, Ewan Harrison, David K. Jackson, Ian Johnston, Dominic Kwiatkowski, Cordelia Langford, John Sillitoe on behalf of the Wellcome Sanger Institute COVID-19 Surveillance Team |
| EPI_ISL_3069335, EPI_ISL_3069336 | Lighthouse Lab in Glasgow | Wellcome Sanger Institute for the COVID-19 Genomics UK (COG-UK) Consortium | Harper VanSteenhouse, Yumi Kasai, David Gray, Carol Clugston, Anna Dominiczak and Alex Alderton, Roberto Amato, Jeffrey Barrett, Sonia Goncalves, Ewan Harrison, David K. Jackson, Ian Johnston, Dominic Kwiatkowski, Cordelia Langford, John Sillitoe on behalf of the Wellcome Sanger Institute COVID-19 Surveillance Team |
| EPI_ISL_3069337 | Berkshire and Surrey Pathology Services Lighthouse Laboratory | Wellcome Sanger Institute for the COVID-19 Genomics UK (COG-UK) Consortium | Berkshire and Surrey Pathology Services Lighthouse Laboratory and Alex Alderton, Roberto Amato, Jeffrey Barrett, Sonia Goncalves, Ewan Harrison, David K. Jackson, Ian Johnston, Dominic Kwiatkowski, Cordelia Langford, John Sillitoe on behalf of the Wellcome Sanger Institute COVID-19 Surveillance Team |
| EPI_ISL_3069338, EPI_ISL_3069339 | Lighthouse Lab in Alderley Park | Wellcome Sanger Institute for the COVID-19 Genomics UK (COG-UK) Consortium | Jacquelyn Wynn, Mairead Hyland, The Lighthouse Lab in Alderley Park and Alex Alderton, Roberto Amato, Jeffrey Barrett, Sonia Goncalves, Ewan Harrison, David K. Jackson, Ian Johnston, Dominic Kwiatkowski, Cordelia Langford, John Sillitoe on behalf of the Wellcome Sanger Institute COVID-19 Surveillance Team |
| EPI_ISL_3069340, EPI_ISL_3069341 | Berkshire and Surrey Pathology Services Lighthouse Laboratory | Wellcome Sanger Institute for the COVID-19 Genomics UK (COG-UK) Consortium | Berkshire and Surrey Pathology Services Lighthouse Laboratory and Alex Alderton, Roberto Amato, Jeffrey Barrett, Sonia Goncalves, Ewan Harrison, David K. Jackson, Ian Johnston, Dominic Kwiatkowski, Cordelia Langford, John Sillitoe on behalf of the Wellcome Sanger Institute COVID-19 Surveillance Team |
| EPI_ISL_3069342 | Lighthouse Lab in Alderley Park | Wellcome Sanger Institute for the COVID-19 Genomics UK (COG-UK) Consortium | Jacquelyn Wynn, Mairead Hyland, The Lighthouse Lab in Alderley Park and Alex Alderton, Roberto Amato, Jeffrey Barrett, Sonia Goncalves, Ewan Harrison, David K. Jackson, Ian Johnston, Dominic Kwiatkowski, Cordelia Langford, John Sillitoe on behalf of the Wellcome Sanger Institute COVID-19 Surveillance Team |
| EPI_ISL_3069343, EPI_ISL_3069344, EPI_ISL_3069345 | Berkshire and Surrey Pathology Services Lighthouse Laboratory | Wellcome Sanger Institute for the COVID-19 Genomics UK (COG-UK) Consortium | Berkshire and Surrey Pathology Services Lighthouse Laboratory and Alex Alderton, Roberto Amato, Jeffrey Barrett, Sonia Goncalves, Ewan Harrison, David K. Jackson, Ian Johnston, Dominic Kwiatkowski, Cordelia Langford, John Sillitoe on behalf of the Wellcome Sanger Institute COVID-19 Surveillance Team |
| EPI_ISL_3069346 | Lighthouse Lab in Alderley Park | Wellcome Sanger Institute for the COVID-19 Genomics UK (COG-UK) Consortium | Jacquelyn Wynn, Mairead Hyland, The Lighthouse Lab in Alderley Park and Alex Alderton, Roberto Amato, Jeffrey Barrett, Sonia Goncalves, Ewan Harrison, David K. Jackson, Ian Johnston, Dominic Kwiatkowski, Cordelia Langford, John Sillitoe on behalf of the Wellcome Sanger Institute COVID-19 Surveillance Team |
| EPI_ISL_3069347 | Lighthouse Lab in Glasgow | Wellcome Sanger Institute for the COVID-19 Genomics UK (COG-UK) Consortium | Harper VanSteenhouse, Yumi Kasai, David Gray, Carol Clugston, Anna Dominiczak and Alex Alderton, Roberto Amato, Jeffrey Barrett, Sonia Goncalves, Ewan Harrison, David K. Jackson, Ian Johnston, Dominic Kwiatkowski, Cordelia Langford, John Sillitoe on behalf of the Wellcome Sanger Institute COVID-19 Surveillance Team |
| EPI_ISL_3069348 | Lighthouse Lab in Alderley Park | Wellcome Sanger Institute for the COVID-19 Genomics UK (COG-UK) Consortium | Jacquelyn Wynn, Mairead Hyland, The Lighthouse Lab in Alderley Park and Alex Alderton, Roberto Amato, Jeffrey Barrett, Sonia Goncalves, Ewan Harrison, David K. Jackson, Ian Johnston, Dominic Kwiatkowski, Cordelia Langford, John Sillitoe on behalf of the Wellcome Sanger Institute COVID-19 Surveillance Team |
| EPI_ISL_3069349 | Lighthouse Lab in Glasgow | Wellcome Sanger Institute for the COVID-19 Genomics UK (COG-UK) Consortium | Harper VanSteenhouse, Yumi Kasai, David Gray, Carol Clugston, Anna Dominiczak and Alex Alderton, Roberto Amato, Jeffrey Barrett, Sonia Goncalves, Ewan Harrison, David K. Jackson, Ian Johnston, Dominic Kwiatkowski, Cordelia Langford, John Sillitoe on behalf of the Wellcome Sanger Institute COVID-19 Surveillance Team |
| EPI_ISL_3069350, EPI_ISL_3069351 | Lighthouse Lab in Alderley Park | Wellcome Sanger Institute for the COVID-19 Genomics UK (COG-UK) Consortium | Jacquelyn Wynn, Mairead Hyland, The Lighthouse Lab in Alderley Park and Alex Alderton, Roberto Amato, Jeffrey Barrett, Sonia Goncalves, Ewan Harrison, David K. Jackson, Ian Johnston, Dominic Kwiatkowski, Cordelia Langford, John Sillitoe on behalf of the Wellcome Sanger Institute COVID-19 Surveillance Team |
| EPI_ISL_3069352 | Berkshire and Surrey Pathology Services Lighthouse Laboratory | Wellcome Sanger Institute for the COVID-19 Genomics UK (COG-UK) Consortium | Berkshire and Surrey Pathology Services Lighthouse Laboratory and Alex Alderton, Roberto Amato, Jeffrey Barrett, Sonia Goncalves, Ewan Harrison, David K. Jackson, Ian Johnston, Dominic Kwiatkowski, Cordelia Langford, John Sillitoe on behalf of the Wellcome Sanger Institute COVID-19 Surveillance Team |
| EPI_ISL_3069353 | Lighthouse Lab in Glasgow | Wellcome Sanger Institute for the COVID-19 Genomics UK (COG-UK) Consortium | Harper VanSteenhouse, Yumi Kasai, David Gray, Carol Clugston, Anna Dominiczak and Alex Alderton, Roberto Amato, Jeffrey Barrett, Sonia Goncalves, Ewan Harrison, David K. Jackson, Ian Johnston, Dominic Kwiatkowski, Cordelia Langford, John Sillitoe on behalf of the Wellcome Sanger Institute COVID-19 Surveillance Team |
| EPI_ISL_3069354 | Lighthouse Lab in Alderley Park | Wellcome Sanger Institute for the COVID-19 Genomics UK (COG-UK) Consortium | Jacquelyn Wynn, Mairead Hyland, The Lighthouse Lab in Alderley Park and Alex Alderton, Roberto Amato, Jeffrey Barrett, Sonia Goncalves, Ewan Harrison, David K. Jackson, Ian Johnston, Dominic Kwiatkowski, Cordelia Langford, John Sillitoe on behalf of the Wellcome Sanger Institute COVID-19 Surveillance Team |
| EPI_ISL_3069355, EPI_ISL_3069356, EPI_ISL_3069357, EPI_ISL_3069358, EPI_ISL_3069359 | Berkshire and Surrey Pathology Services Lighthouse Laboratory | Wellcome Sanger Institute for the COVID-19 Genomics UK (COG-UK) Consortium | Berkshire and Surrey Pathology Services Lighthouse Laboratory and Alex Alderton, Roberto Amato, Jeffrey Barrett, Sonia Goncalves, Ewan Harrison, David K. Jackson, Ian Johnston, Dominic Kwiatkowski, Cordelia Langford, John Sillitoe on behalf of the Wellcome Sanger Institute COVID-19 Surveillance Team |
| EPI_ISL_3069360 | Lighthouse Lab in Glasgow | Wellcome Sanger Institute for the COVID-19 Genomics UK (COG-UK) Consortium | Harper VanSteenhouse, Yumi Kasai, David Gray, Carol Clugston, Anna Dominiczak and Alex Alderton, Roberto Amato, Jeffrey Barrett, Sonia Goncalves, Ewan Harrison, David K. Jackson, Ian Johnston, Dominic Kwiatkowski, Cordelia Langford, John Sillitoe on behalf of the Wellcome Sanger Institute COVID-19 Surveillance Team |
| EPI_ISL_3069361 | Berkshire and Surrey Pathology Services Lighthouse Laboratory | Wellcome Sanger Institute for the COVID-19 Genomics UK (COG-UK) Consortium | Berkshire and Surrey Pathology Services Lighthouse Laboratory and Alex Alderton, Roberto Amato, Jeffrey Barrett, Sonia Goncalves, Ewan Harrison, David K. Jackson, Ian Johnston, Dominic Kwiatkowski, Cordelia Langford, John Sillitoe on behalf of the Wellcome Sanger Institute COVID-19 Surveillance Team |
| EPI_ISL_3069362, EPI_ISL_3069363 | Lighthouse Lab in Glasgow | Wellcome Sanger Institute for the COVID-19 Genomics UK (COG-UK) Consortium | Harper VanSteenhouse, Yumi Kasai, David Gray, Carol Clugston, Anna Dominiczak and Alex Alderton, Roberto Amato, Jeffrey Barrett, Sonia Goncalves, Ewan Harrison, David K. Jackson, Ian Johnston, Dominic Kwiatkowski, Cordelia Langford, John Sillitoe on behalf of the Wellcome Sanger Institute COVID-19 Surveillance Team |
| EPI_ISL_3069364, EPI_ISL_3069365 | Berkshire and Surrey Pathology Services Lighthouse Laboratory | Wellcome Sanger Institute for the COVID-19 Genomics UK (COG-UK) Consortium | Berkshire and Surrey Pathology Services Lighthouse Laboratory and Alex Alderton, Roberto Amato, Jeffrey Barrett, Sonia Goncalves, Ewan Harrison, David K. Jackson, Ian Johnston, Dominic Kwiatkowski, Cordelia Langford, John Sillitoe on behalf of the Wellcome Sanger Institute COVID-19 Surveillance Team |
| EPI_ISL_3069366 | Lighthouse Lab in Glasgow | Wellcome Sanger Institute for the COVID-19 Genomics UK (COG-UK) Consortium | Harper VanSteenhouse, Yumi Kasai, David Gray, Carol Clugston, Anna Dominiczak and Alex Alderton, Roberto Amato, Jeffrey Barrett, Sonia Goncalves, Ewan Harrison, David K. Jackson, Ian Johnston, Dominic Kwiatkowski, Cordelia Langford, John Sillitoe on behalf of the Wellcome Sanger Institute COVID-19 Surveillance Team |
| EPI_ISL_3069367, | Berkshire and Surrey Pathology Services Lighthouse Laboratory | Wellcome Sanger Institute for the COVID-19 Genomics UK (COG-UK) | Berkshire and Surrey Pathology Services Lighthouse Laboratory and Alex Alderton, Roberto Amato, Jeffrey Barrett, Sonia Goncalves, Ewan Harrison, David K. Jackson, Ian Johnston, Dominic Kwiatkowski, Cordelia Langford, John Sillitoe on behalf |

| | | | |
|---|---|---|---|
| EPI_ISL_3069368 | | Consortium | of the Wellcome Sanger Institute COVID-19 Surveillance Team |
| EPI_ISL_3069369, EPI_ISL_3069370 | Lighthouse Lab in Alderley Park | Wellcome Sanger Institute for the COVID-19 Genomics UK (COG-UK) Consortium | Jacquelyn Wynn, Mairead Hyland, The Lighthouse Lab in Alderley Park and Alex Alderton, Roberto Amato, Jeffrey Barrett, Sonia Goncalves, Ewan Harrison, David K. Jackson, Ian Johnston, Dominic Kwiatkowski, Cordelia Langford, John Sillitoe on behalf of the Wellcome Sanger Institute COVID-19 Surveillance Team |
| EPI_ISL_3069371, EPI_ISL_3069372 | Berkshire and Surrey Pathology Services Lighthouse Laboratory | Wellcome Sanger Institute for the COVID-19 Genomics UK (COG-UK) Consortium | Berkshire and Surrey Pathology Services Lighthouse Laboratory and Alex Alderton, Roberto Amato, Jeffrey Barrett, Sonia Goncalves, Ewan Harrison, David K. Jackson, Ian Johnston, Dominic Kwiatkowski, Cordelia Langford, John Sillitoe on behalf of the Wellcome Sanger Institute COVID-19 Surveillance Team |
| EPI_ISL_3069373 | Lighthouse Lab in Glasgow | Wellcome Sanger Institute for the COVID-19 Genomics UK (COG-UK) Consortium | Harper VanSteenhouse, Yumi Kasai, David Gray, Carol Clugston, Anna Dominiczak and Alex Alderton, Roberto Amato, Jeffrey Barrett, Sonia Goncalves, Ewan Harrison, David K. Jackson, Ian Johnston, Dominic Kwiatkowski, Cordelia Langford, John Sillitoe on behalf of the Wellcome Sanger Institute COVID-19 Surveillance Team |
| EPI_ISL_3069374, EPI_ISL_3069375, EPI_ISL_3069376 | Berkshire and Surrey Pathology Services Lighthouse Laboratory | Wellcome Sanger Institute for the COVID-19 Genomics UK (COG-UK) Consortium | Berkshire and Surrey Pathology Services Lighthouse Laboratory and Alex Alderton, Roberto Amato, Jeffrey Barrett, Sonia Goncalves, Ewan Harrison, David K. Jackson, Ian Johnston, Dominic Kwiatkowski, Cordelia Langford, John Sillitoe on behalf of the Wellcome Sanger Institute COVID-19 Surveillance Team |
| EPI_ISL_3069377, EPI_ISL_3069378 | Lighthouse Lab in Glasgow | Wellcome Sanger Institute for the COVID-19 Genomics UK (COG-UK) Consortium | Harper VanSteenhouse, Yumi Kasai, David Gray, Carol Clugston, Anna Dominiczak and Alex Alderton, Roberto Amato, Jeffrey Barrett, Sonia Goncalves, Ewan Harrison, David K. Jackson, Ian Johnston, Dominic Kwiatkowski, Cordelia Langford, John Sillitoe on behalf of the Wellcome Sanger Institute COVID-19 Surveillance Team |
| EPI_ISL_3069379 | Berkshire and Surrey Pathology Services Lighthouse Laboratory | Wellcome Sanger Institute for the COVID-19 Genomics UK (COG-UK) Consortium | Berkshire and Surrey Pathology Services Lighthouse Laboratory and Alex Alderton, Roberto Amato, Jeffrey Barrett, Sonia Goncalves, Ewan Harrison, David K. Jackson, Ian Johnston, Dominic Kwiatkowski, Cordelia Langford, John Sillitoe on behalf of the Wellcome Sanger Institute COVID-19 Surveillance Team |
| EPI_ISL_3069380 | Lighthouse Lab in Alderley Park | Wellcome Sanger Institute for the COVID-19 Genomics UK (COG-UK) Consortium | Jacquelyn Wynn, Mairead Hyland, The Lighthouse Lab in Alderley Park and Alex Alderton, Roberto Amato, Jeffrey Barrett, Sonia Goncalves, Ewan Harrison, David K. Jackson, Ian Johnston, Dominic Kwiatkowski, Cordelia Langford, John Sillitoe on behalf of the Wellcome Sanger Institute COVID-19 Surveillance Team |
| EPI_ISL_3069381, EPI_ISL_3069382, EPI_ISL_3069383, EPI_ISL_3069384 | Lighthouse Lab in Glasgow | Wellcome Sanger Institute for the COVID-19 Genomics UK (COG-UK) Consortium | Harper VanSteenhouse, Yumi Kasai, David Gray, Carol Clugston, Anna Dominiczak and Alex Alderton, Roberto Amato, Jeffrey Barrett, Sonia Goncalves, Ewan Harrison, David K. Jackson, Ian Johnston, Dominic Kwiatkowski, Cordelia Langford, John Sillitoe on behalf of the Wellcome Sanger Institute COVID-19 Surveillance Team |
| EPI_ISL_3069385, EPI_ISL_3069386 | Lighthouse Lab in Alderley Park | Wellcome Sanger Institute for the COVID-19 Genomics UK (COG-UK) Consortium | Jacquelyn Wynn, Mairead Hyland, The Lighthouse Lab in Alderley Park and Alex Alderton, Roberto Amato, Jeffrey Barrett, Sonia Goncalves, Ewan Harrison, David K. Jackson, Ian Johnston, Dominic Kwiatkowski, Cordelia Langford, John Sillitoe on behalf of the Wellcome Sanger Institute COVID-19 Surveillance Team |
| EPI_ISL_3069387, EPI_ISL_3069388, EPI_ISL_3069389 | Lighthouse Lab in Glasgow | Wellcome Sanger Institute for the COVID-19 Genomics UK (COG-UK) Consortium | Harper VanSteenhouse, Yumi Kasai, David Gray, Carol Clugston, Anna Dominiczak and Alex Alderton, Roberto Amato, Jeffrey Barrett, Sonia Goncalves, Ewan Harrison, David K. Jackson, Ian Johnston, Dominic Kwiatkowski, Cordelia Langford, John Sillitoe on behalf of the Wellcome Sanger Institute COVID-19 Surveillance Team |
| EPI_ISL_3069390 | Lighthouse Lab in Alderley Park | Wellcome Sanger Institute for the COVID-19 Genomics UK (COG-UK) Consortium | Jacquelyn Wynn, Mairead Hyland, The Lighthouse Lab in Alderley Park and Alex Alderton, Roberto Amato, Jeffrey Barrett, Sonia Goncalves, Ewan Harrison, David K. Jackson, Ian Johnston, Dominic Kwiatkowski, Cordelia Langford, John Sillitoe on behalf of the Wellcome Sanger Institute COVID-19 Surveillance Team |
| EPI_ISL_3069391 | Lighthouse Lab in Glasgow | Wellcome Sanger Institute for the COVID-19 Genomics UK (COG-UK) Consortium | Harper VanSteenhouse, Yumi Kasai, David Gray, Carol Clugston, Anna Dominiczak and Alex Alderton, Roberto Amato, Jeffrey Barrett, Sonia Goncalves, Ewan Harrison, David K. Jackson, Ian Johnston, Dominic Kwiatkowski, Cordelia Langford, John Sillitoe on behalf of the Wellcome Sanger Institute COVID-19 Surveillance Team |
| EPI_ISL_3069392 | Lighthouse Lab in Alderley Park | Wellcome Sanger Institute for the COVID-19 Genomics UK (COG-UK) Consortium | Jacquelyn Wynn, Mairead Hyland, The Lighthouse Lab in Alderley Park and Alex Alderton, Roberto Amato, Jeffrey Barrett, Sonia Goncalves, Ewan Harrison, David K. Jackson, Ian Johnston, Dominic Kwiatkowski, Cordelia Langford, John Sillitoe on behalf of the Wellcome Sanger Institute COVID-19 Surveillance Team |
| EPI_ISL_3069393 | Berkshire and Surrey Pathology Services Lighthouse Laboratory | Wellcome Sanger Institute for the COVID-19 Genomics UK (COG-UK) Consortium | Berkshire and Surrey Pathology Services Lighthouse Laboratory and Alex Alderton, Roberto Amato, Jeffrey Barrett, Sonia Goncalves, Ewan Harrison, David K. Jackson, Ian Johnston, Dominic Kwiatkowski, Cordelia Langford, John Sillitoe on behalf of the Wellcome Sanger Institute COVID-19 Surveillance Team |
| EPI_ISL_3069394, EPI_ISL_3069395, EPI_ISL_3069396 | Lighthouse Lab in Alderley Park | Wellcome Sanger Institute for the COVID-19 Genomics UK (COG-UK) Consortium | Jacquelyn Wynn, Mairead Hyland, The Lighthouse Lab in Alderley Park and Alex Alderton, Roberto Amato, Jeffrey Barrett, Sonia Goncalves, Ewan Harrison, David K. Jackson, Ian Johnston, Dominic Kwiatkowski, Cordelia Langford, John Sillitoe on behalf of the Wellcome Sanger Institute COVID-19 Surveillance Team |
| EPI_ISL_3069397 | Berkshire and Surrey Pathology Services Lighthouse Laboratory | Wellcome Sanger Institute for the COVID-19 Genomics UK (COG-UK) Consortium | Berkshire and Surrey Pathology Services Lighthouse Laboratory and Alex Alderton, Roberto Amato, Jeffrey Barrett, Sonia Goncalves, Ewan Harrison, David K. Jackson, Ian Johnston, Dominic Kwiatkowski, Cordelia Langford, John Sillitoe on behalf of the Wellcome Sanger Institute COVID-19 Surveillance Team |
| EPI_ISL_3069398, EPI_ISL_3069399 | Lighthouse Lab in Alderley Park | Wellcome Sanger Institute for the COVID-19 Genomics UK (COG-UK) Consortium | Jacquelyn Wynn, Mairead Hyland, The Lighthouse Lab in Alderley Park and Alex Alderton, Roberto Amato, Jeffrey Barrett, Sonia Goncalves, Ewan Harrison, David K. Jackson, Ian Johnston, Dominic Kwiatkowski, Cordelia Langford, John Sillitoe on behalf of the Wellcome Sanger Institute COVID-19 Surveillance Team |
| EPI_ISL_3069400 | Berkshire and Surrey Pathology Services Lighthouse Laboratory | Wellcome Sanger Institute for the COVID-19 Genomics UK (COG-UK) Consortium | Berkshire and Surrey Pathology Services Lighthouse Laboratory and Alex Alderton, Roberto Amato, Jeffrey Barrett, Sonia Goncalves, Ewan Harrison, David K. Jackson, Ian Johnston, Dominic Kwiatkowski, Cordelia Langford, John Sillitoe on behalf of the Wellcome Sanger Institute COVID-19 Surveillance Team |
| EPI_ISL_3069401, EPI_ISL_3069402, EPI_ISL_3069403 | Lighthouse Lab in Alderley Park | Wellcome Sanger Institute for the COVID-19 Genomics UK (COG-UK) Consortium | Jacquelyn Wynn, Mairead Hyland, The Lighthouse Lab in Alderley Park and Alex Alderton, Roberto Amato, Jeffrey Barrett, Sonia Goncalves, Ewan Harrison, David K. Jackson, Ian Johnston, Dominic Kwiatkowski, Cordelia Langford, John Sillitoe on behalf of the Wellcome Sanger Institute COVID-19 Surveillance Team |
| EPI_ISL_3069404 | Lighthouse Lab in Glasgow | Wellcome Sanger Institute for the COVID-19 Genomics UK (COG-UK) Consortium | Harper VanSteenhouse, Yumi Kasai, David Gray, Carol Clugston, Anna Dominiczak and Alex Alderton, Roberto Amato, Jeffrey Barrett, Sonia Goncalves, Ewan Harrison, David K. Jackson, Ian Johnston, Dominic Kwiatkowski, Cordelia Langford, John Sillitoe on behalf of the Wellcome Sanger Institute COVID-19 Surveillance Team |
| EPI_ISL_3069405 | Berkshire and Surrey Pathology Services Lighthouse Laboratory | Wellcome Sanger Institute for the COVID-19 Genomics UK (COG-UK) Consortium | Berkshire and Surrey Pathology Services Lighthouse Laboratory and Alex Alderton, Roberto Amato, Jeffrey Barrett, Sonia Goncalves, Ewan Harrison, David K. Jackson, Ian Johnston, Dominic Kwiatkowski, Cordelia Langford, John Sillitoe on behalf of the Wellcome Sanger Institute COVID-19 Surveillance Team |
| EPI_ISL_3069406, EPI_ISL_3069407, EPI_ISL_3069408 | Lighthouse Lab in Alderley Park | Wellcome Sanger Institute for the COVID-19 Genomics UK (COG-UK) Consortium | Jacquelyn Wynn, Mairead Hyland, The Lighthouse Lab in Alderley Park and Alex Alderton, Roberto Amato, Jeffrey Barrett, Sonia Goncalves, Ewan Harrison, David K. Jackson, Ian Johnston, Dominic Kwiatkowski, Cordelia Langford, John Sillitoe on behalf of the Wellcome Sanger Institute COVID-19 Surveillance Team |
| EPI_ISL_3069409 | Berkshire and Surrey Pathology Services Lighthouse Laboratory | Wellcome Sanger Institute for the COVID-19 Genomics UK (COG-UK) Consortium | Berkshire and Surrey Pathology Services Lighthouse Laboratory and Alex Alderton, Roberto Amato, Jeffrey Barrett, Sonia Goncalves, Ewan Harrison, David K. Jackson, Ian Johnston, Dominic Kwiatkowski, Cordelia Langford, John Sillitoe on behalf of the Wellcome Sanger Institute COVID-19 Surveillance Team |
| EPI_ISL_3069410, EPI_ISL_3069411 | Lighthouse Lab in Alderley Park | Wellcome Sanger Institute for the COVID-19 Genomics UK (COG-UK) Consortium | Jacquelyn Wynn, Mairead Hyland, The Lighthouse Lab in Alderley Park and Alex Alderton, Roberto Amato, Jeffrey Barrett, Sonia Goncalves, Ewan Harrison, David K. Jackson, Ian Johnston, Dominic Kwiatkowski, Cordelia Langford, John Sillitoe on behalf of the Wellcome Sanger Institute COVID-19 Surveillance Team |
| EPI_ISL_3069412, EPI_ISL_3069413, EPI_ISL_3069414 | Lighthouse Lab in Glasgow | Wellcome Sanger Institute for the COVID-19 Genomics UK (COG-UK) Consortium | Harper VanSteenhouse, Yumi Kasai, David Gray, Carol Clugston, Anna Dominiczak and Alex Alderton, Roberto Amato, Jeffrey Barrett, Sonia Goncalves, Ewan Harrison, David K. Jackson, Ian Johnston, Dominic Kwiatkowski, Cordelia Langford, John Sillitoe on behalf of the Wellcome Sanger Institute COVID-19 Surveillance Team |
| EPI_ISL_3069415, EPI_ISL_3069416, EPI_ISL_3069417 | Berkshire and Surrey Pathology Services Lighthouse Laboratory | Wellcome Sanger Institute for the COVID-19 Genomics UK (COG-UK) Consortium | Berkshire and Surrey Pathology Services Lighthouse Laboratory and Alex Alderton, Roberto Amato, Jeffrey Barrett, Sonia Goncalves, Ewan Harrison, David K. Jackson, Ian Johnston, Dominic Kwiatkowski, Cordelia Langford, John Sillitoe on behalf of the Wellcome Sanger Institute COVID-19 Surveillance Team |
| EPI_ISL_3069418 | Lighthouse Lab in Alderley Park | Wellcome Sanger Institute for the COVID-19 Genomics UK (COG-UK) Consortium | Jacquelyn Wynn, Mairead Hyland, The Lighthouse Lab in Alderley Park and Alex Alderton, Roberto Amato, Jeffrey Barrett, Sonia Goncalves, Ewan Harrison, David K. Jackson, Ian Johnston, Dominic Kwiatkowski, Cordelia Langford, John Sillitoe on behalf of the Wellcome Sanger Institute COVID-19 Surveillance Team |
| EPI_ISL_3069419, EPI_ISL_3069420 | Lighthouse Lab in Glasgow | Wellcome Sanger Institute for the COVID-19 Genomics UK (COG-UK) Consortium | Harper VanSteenhouse, Yumi Kasai, David Gray, Carol Clugston, Anna Dominiczak and Alex Alderton, Roberto Amato, Jeffrey Barrett, Sonia Goncalves, Ewan Harrison, David K. Jackson, Ian Johnston, Dominic Kwiatkowski, Cordelia Langford, John Sillitoe on behalf of the Wellcome Sanger Institute COVID-19 Surveillance Team |
| EPI_ISL_3069421 | Lighthouse Lab in Alderley Park | Wellcome Sanger Institute for the COVID-19 Genomics UK (COG-UK) Consortium | Jacquelyn Wynn, Mairead Hyland, The Lighthouse Lab in Alderley Park and Alex Alderton, Roberto Amato, Jeffrey Barrett, Sonia Goncalves, Ewan Harrison, David K. Jackson, Ian Johnston, Dominic Kwiatkowski, Cordelia Langford, John Sillitoe on behalf of the Wellcome Sanger Institute COVID-19 Surveillance Team |
| EPI_ISL_3069422 | Berkshire and Surrey Pathology Services Lighthouse Laboratory | Wellcome Sanger Institute for the COVID-19 Genomics UK (COG-UK) Consortium | Berkshire and Surrey Pathology Services Lighthouse Laboratory and Alex Alderton, Roberto Amato, Jeffrey Barrett, Sonia Goncalves, Ewan Harrison, David K. Jackson, Ian Johnston, Dominic Kwiatkowski, Cordelia Langford, John Sillitoe on behalf of the Wellcome Sanger Institute COVID-19 Surveillance Team |
| EPI_ISL_3069423, EPI_ISL_3069424 | Lighthouse Lab in Alderley Park | Wellcome Sanger Institute for the COVID-19 Genomics UK (COG-UK) Consortium | Jacquelyn Wynn, Mairead Hyland, The Lighthouse Lab in Alderley Park and Alex Alderton, Roberto Amato, Jeffrey Barrett, Sonia Goncalves, Ewan Harrison, David K. Jackson, Ian Johnston, Dominic Kwiatkowski, Cordelia Langford, John Sillitoe on behalf of the Wellcome Sanger Institute COVID-19 Surveillance Team |
| EPI_ISL_3069425, EPI_ISL_3069426, EPI_ISL_3069427, EPI_ISL_3069428, EPI_ISL_3069429, EPI_ISL_3069430, EPI_ISL_3069431, EPI_ISL_3069432 | Berkshire and Surrey Pathology Services Lighthouse Laboratory | Wellcome Sanger Institute for the COVID-19 Genomics UK (COG-UK) Consortium | Berkshire and Surrey Pathology Services Lighthouse Laboratory and Alex Alderton, Roberto Amato, Jeffrey Barrett, Sonia Goncalves, Ewan Harrison, David K. Jackson, Ian Johnston, Dominic Kwiatkowski, Cordelia Langford, John Sillitoe on behalf of the Wellcome Sanger Institute COVID-19 Surveillance Team |
| EPI_ISL_3069433 | Lighthouse Lab in Glasgow | Wellcome Sanger Institute for the COVID-19 Genomics UK (COG-UK) Consortium | Harper VanSteenhouse, Yumi Kasai, David Gray, Carol Clugston, Anna Dominiczak and Alex Alderton, Roberto Amato, Jeffrey Barrett, Sonia Goncalves, Ewan Harrison, David K. Jackson, Ian Johnston, Dominic Kwiatkowski, Cordelia Langford, John Sillitoe on behalf of the Wellcome Sanger Institute COVID-19 Surveillance Team |
| EPI_ISL_3069434, EPI_ISL_3069435, EPI_ISL_3069436 | Berkshire and Surrey Pathology Services Lighthouse Laboratory | Wellcome Sanger Institute for the COVID-19 Genomics UK (COG-UK) Consortium | Berkshire and Surrey Pathology Services Lighthouse Laboratory and Alex Alderton, Roberto Amato, Jeffrey Barrett, Sonia Goncalves, Ewan Harrison, David K. Jackson, Ian Johnston, Dominic Kwiatkowski, Cordelia Langford, John Sillitoe on behalf of the Wellcome Sanger Institute COVID-19 Surveillance Team |
| EPI_ISL_3069437 | Lighthouse Lab in Alderley Park | Wellcome Sanger Institute for the COVID-19 Genomics UK (COG-UK) Consortium | Jacquelyn Wynn, Mairead Hyland, The Lighthouse Lab in Alderley Park and Alex Alderton, Roberto Amato, Jeffrey Barrett, Sonia Goncalves, Ewan Harrison, David K. Jackson, Ian Johnston, Dominic Kwiatkowski, Cordelia Langford, John Sillitoe on behalf of the Wellcome Sanger Institute COVID-19 Surveillance Team |
| EPI_ISL_3069438 | Lighthouse Lab in Glasgow | Wellcome Sanger Institute for the COVID-19 Genomics UK (COG-UK) Consortium | Harper VanSteenhouse, Yumi Kasai, David Gray, Carol Clugston, Anna Dominiczak and Alex Alderton, Roberto Amato, Jeffrey Barrett, Sonia Goncalves, Ewan Harrison, David K. Jackson, Ian Johnston, Dominic Kwiatkowski, Cordelia Langford, John Sillitoe on behalf of the Wellcome Sanger Institute COVID-19 Surveillance Team |
| EPI_ISL_3069439 | Berkshire and Surrey Pathology Services Lighthouse Laboratory | Wellcome Sanger Institute for the COVID-19 Genomics UK (COG-UK) Consortium | Berkshire and Surrey Pathology Services Lighthouse Laboratory and Alex Alderton, Roberto Amato, Jeffrey Barrett, Sonia Goncalves, Ewan Harrison, David K. Jackson, Ian Johnston, Dominic Kwiatkowski, Cordelia Langford, John Sillitoe on behalf of the Wellcome Sanger Institute COVID-19 Surveillance Team |
| EPI_ISL_3069440 | Lighthouse Lab in Glasgow | Wellcome Sanger Institute for the COVID-19 Genomics UK (COG-UK) Consortium | Harper VanSteenhouse, Yumi Kasai, David Gray, Carol Clugston, Anna Dominiczak and Alex Alderton, Roberto Amato, Jeffrey Barrett, Sonia Goncalves, Ewan Harrison, David K. Jackson, Ian Johnston, Dominic Kwiatkowski, Cordelia Langford, John Sillitoe on behalf of the Wellcome Sanger Institute COVID-19 Surveillance Team |

| | | | |
|---|---|---|---|
| EPI_ISL_3069441, EPI_ISL_3069442 | Lighthouse Lab in Alderley Park | Wellcome Sanger Institute for the COVID-19 Genomics UK (COG-UK) Consortium | Jacquelyn Wynn, Mairead Hyland, The Lighthouse Lab in Alderley Park and Alex Alderton, Roberto Amato, Jeffrey Barrett, Sonia Goncalves, Ewan Harrison, David K. Jackson, Ian Johnston, Dominic Kwiatkowski, Cordelia Langford, John Sillitoe on behalf of the Wellcome Sanger Institute COVID-19 Surveillance Team |
| EPI_ISL_3069443 | Berkshire and Surrey Pathology Services Lighthouse Laboratory | Wellcome Sanger Institute for the COVID-19 Genomics UK (COG-UK) Consortium | Berkshire and Surrey Pathology Services Lighthouse Laboratory and Alex Alderton, Roberto Amato, Jeffrey Barrett, Sonia Goncalves, Ewan Harrison, David K. Jackson, Ian Johnston, Dominic Kwiatkowski, Cordelia Langford, John Sillitoe on behalf of the Wellcome Sanger Institute COVID-19 Surveillance Team |
| EPI_ISL_3069444, EPI_ISL_3069445 | Lighthouse Lab in Glasgow | Wellcome Sanger Institute for the COVID-19 Genomics UK (COG-UK) Consortium | Harper VanSteenhouse, Yumi Kasai, David Gray, Carol Clugston, Anna Dominiczak and Alex Alderton, Roberto Amato, Jeffrey Barrett, Sonia Goncalves, Ewan Harrison, David K. Jackson, Ian Johnston, Dominic Kwiatkowski, Cordelia Langford, John Sillitoe on behalf of the Wellcome Sanger Institute COVID-19 Surveillance Team |
| EPI_ISL_3069446 | Lighthouse Lab in Alderley Park | Wellcome Sanger Institute for the COVID-19 Genomics UK (COG-UK) Consortium | Jacquelyn Wynn, Mairead Hyland, The Lighthouse Lab in Alderley Park and Alex Alderton, Roberto Amato, Jeffrey Barrett, Sonia Goncalves, Ewan Harrison, David K. Jackson, Ian Johnston, Dominic Kwiatkowski, Cordelia Langford, John Sillitoe on behalf of the Wellcome Sanger Institute COVID-19 Surveillance Team |
| EPI_ISL_3069447, EPI_ISL_3069448 | Berkshire and Surrey Pathology Services Lighthouse Laboratory | Wellcome Sanger Institute for the COVID-19 Genomics UK (COG-UK) Consortium | Berkshire and Surrey Pathology Services Lighthouse Laboratory and Alex Alderton, Roberto Amato, Jeffrey Barrett, Sonia Goncalves, Ewan Harrison, David K. Jackson, Ian Johnston, Dominic Kwiatkowski, Cordelia Langford, John Sillitoe on behalf of the Wellcome Sanger Institute COVID-19 Surveillance Team |
| EPI_ISL_3069449 | Lighthouse Lab in Alderley Park | Wellcome Sanger Institute for the COVID-19 Genomics UK (COG-UK) Consortium | Jacquelyn Wynn, Mairead Hyland, The Lighthouse Lab in Alderley Park and Alex Alderton, Roberto Amato, Jeffrey Barrett, Sonia Goncalves, Ewan Harrison, David K. Jackson, Ian Johnston, Dominic Kwiatkowski, Cordelia Langford, John Sillitoe on behalf of the Wellcome Sanger Institute COVID-19 Surveillance Team |
| EPI_ISL_3069450, EPI_ISL_3069451, EPI_ISL_3069452 | Berkshire and Surrey Pathology Services Lighthouse Laboratory | Wellcome Sanger Institute for the COVID-19 Genomics UK (COG-UK) Consortium | Berkshire and Surrey Pathology Services Lighthouse Laboratory and Alex Alderton, Roberto Amato, Jeffrey Barrett, Sonia Goncalves, Ewan Harrison, David K. Jackson, Ian Johnston, Dominic Kwiatkowski, Cordelia Langford, John Sillitoe on behalf of the Wellcome Sanger Institute COVID-19 Surveillance Team |
| EPI_ISL_3069453 | Lighthouse Lab in Alderley Park | Wellcome Sanger Institute for the COVID-19 Genomics UK (COG-UK) Consortium | Jacquelyn Wynn, Mairead Hyland, The Lighthouse Lab in Alderley Park and Alex Alderton, Roberto Amato, Jeffrey Barrett, Sonia Goncalves, Ewan Harrison, David K. Jackson, Ian Johnston, Dominic Kwiatkowski, Cordelia Langford, John Sillitoe on behalf of the Wellcome Sanger Institute COVID-19 Surveillance Team |
| EPI_ISL_3069454 | Lighthouse Lab in Glasgow | Wellcome Sanger Institute for the COVID-19 Genomics UK (COG-UK) Consortium | Harper VanSteenhouse, Yumi Kasai, David Gray, Carol Clugston, Anna Dominiczak and Alex Alderton, Roberto Amato, Jeffrey Barrett, Sonia Goncalves, Ewan Harrison, David K. Jackson, Ian Johnston, Dominic Kwiatkowski, Cordelia Langford, John Sillitoe on behalf of the Wellcome Sanger Institute COVID-19 Surveillance Team |
| EPI_ISL_3069455 | Lighthouse Lab in Alderley Park | Wellcome Sanger Institute for the COVID-19 Genomics UK (COG-UK) Consortium | Jacquelyn Wynn, Mairead Hyland, The Lighthouse Lab in Alderley Park and Alex Alderton, Roberto Amato, Jeffrey Barrett, Sonia Goncalves, Ewan Harrison, David K. Jackson, Ian Johnston, Dominic Kwiatkowski, Cordelia Langford, John Sillitoe on behalf of the Wellcome Sanger Institute COVID-19 Surveillance Team |
| EPI_ISL_3069456 | Berkshire and Surrey Pathology Services Lighthouse Laboratory | Wellcome Sanger Institute for the COVID-19 Genomics UK (COG-UK) Consortium | Berkshire and Surrey Pathology Services Lighthouse Laboratory and Alex Alderton, Roberto Amato, Jeffrey Barrett, Sonia Goncalves, Ewan Harrison, David K. Jackson, Ian Johnston, Dominic Kwiatkowski, Cordelia Langford, John Sillitoe on behalf of the Wellcome Sanger Institute COVID-19 Surveillance Team |
| EPI_ISL_3069457 | Lighthouse Lab in Glasgow | Wellcome Sanger Institute for the COVID-19 Genomics UK (COG-UK) Consortium | Harper VanSteenhouse, Yumi Kasai, David Gray, Carol Clugston, Anna Dominiczak and Alex Alderton, Roberto Amato, Jeffrey Barrett, Sonia Goncalves, Ewan Harrison, David K. Jackson, Ian Johnston, Dominic Kwiatkowski, Cordelia Langford, John Sillitoe on behalf of the Wellcome Sanger Institute COVID-19 Surveillance Team |
| EPI_ISL_3069458 | Lighthouse Lab in Alderley Park | Wellcome Sanger Institute for the COVID-19 Genomics UK (COG-UK) Consortium | Jacquelyn Wynn, Mairead Hyland, The Lighthouse Lab in Alderley Park and Alex Alderton, Roberto Amato, Jeffrey Barrett, Sonia Goncalves, Ewan Harrison, David K. Jackson, Ian Johnston, Dominic Kwiatkowski, Cordelia Langford, John Sillitoe on behalf of the Wellcome Sanger Institute COVID-19 Surveillance Team |
| EPI_ISL_3069459 | Lighthouse Lab in Glasgow | Wellcome Sanger Institute for the COVID-19 Genomics UK (COG-UK) Consortium | Harper VanSteenhouse, Yumi Kasai, David Gray, Carol Clugston, Anna Dominiczak and Alex Alderton, Roberto Amato, Jeffrey Barrett, Sonia Goncalves, Ewan Harrison, David K. Jackson, Ian Johnston, Dominic Kwiatkowski, Cordelia Langford, John Sillitoe on behalf of the Wellcome Sanger Institute COVID-19 Surveillance Team |
| EPI_ISL_3069460 | Lighthouse Lab in Alderley Park | Wellcome Sanger Institute for the COVID-19 Genomics UK (COG-UK) Consortium | Jacquelyn Wynn, Mairead Hyland, The Lighthouse Lab in Alderley Park and Alex Alderton, Roberto Amato, Jeffrey Barrett, Sonia Goncalves, Ewan Harrison, David K. Jackson, Ian Johnston, Dominic Kwiatkowski, Cordelia Langford, John Sillitoe on behalf of the Wellcome Sanger Institute COVID-19 Surveillance Team |
| EPI_ISL_3069461 | Berkshire and Surrey Pathology Services Lighthouse Laboratory | Wellcome Sanger Institute for the COVID-19 Genomics UK (COG-UK) Consortium | Berkshire and Surrey Pathology Services Lighthouse Laboratory and Alex Alderton, Roberto Amato, Jeffrey Barrett, Sonia Goncalves, Ewan Harrison, David K. Jackson, Ian Johnston, Dominic Kwiatkowski, Cordelia Langford, John Sillitoe on behalf of the Wellcome Sanger Institute COVID-19 Surveillance Team |
| EPI_ISL_3069462, EPI_ISL_3069463 | Lighthouse Lab in Glasgow | Wellcome Sanger Institute for the COVID-19 Genomics UK (COG-UK) Consortium | Harper VanSteenhouse, Yumi Kasai, David Gray, Carol Clugston, Anna Dominiczak and Alex Alderton, Roberto Amato, Jeffrey Barrett, Sonia Goncalves, Ewan Harrison, David K. Jackson, Ian Johnston, Dominic Kwiatkowski, Cordelia Langford, John Sillitoe on behalf of the Wellcome Sanger Institute COVID-19 Surveillance Team |
| EPI_ISL_3069464 | Berkshire and Surrey Pathology Services Lighthouse Laboratory | Wellcome Sanger Institute for the COVID-19 Genomics UK (COG-UK) Consortium | Berkshire and Surrey Pathology Services Lighthouse Laboratory and Alex Alderton, Roberto Amato, Jeffrey Barrett, Sonia Goncalves, Ewan Harrison, David K. Jackson, Ian Johnston, Dominic Kwiatkowski, Cordelia Langford, John Sillitoe on behalf of the Wellcome Sanger Institute COVID-19 Surveillance Team |
| EPI_ISL_3069465 | Lighthouse Lab in Alderley Park | Wellcome Sanger Institute for the COVID-19 Genomics UK (COG-UK) Consortium | Jacquelyn Wynn, Mairead Hyland, The Lighthouse Lab in Alderley Park and Alex Alderton, Roberto Amato, Jeffrey Barrett, Sonia Goncalves, Ewan Harrison, David K. Jackson, Ian Johnston, Dominic Kwiatkowski, Cordelia Langford, John Sillitoe on behalf of the Wellcome Sanger Institute COVID-19 Surveillance Team |
| EPI_ISL_3069466, EPI_ISL_3069467, EPI_ISL_3069468 | Berkshire and Surrey Pathology Services Lighthouse Laboratory | Wellcome Sanger Institute for the COVID-19 Genomics UK (COG-UK) Consortium | Berkshire and Surrey Pathology Services Lighthouse Laboratory and Alex Alderton, Roberto Amato, Jeffrey Barrett, Sonia Goncalves, Ewan Harrison, David K. Jackson, Ian Johnston, Dominic Kwiatkowski, Cordelia Langford, John Sillitoe on behalf of the Wellcome Sanger Institute COVID-19 Surveillance Team |
| EPI_ISL_3069469 | Lighthouse Lab in Alderley Park | Wellcome Sanger Institute for the COVID-19 Genomics UK (COG-UK) Consortium | Jacquelyn Wynn, Mairead Hyland, The Lighthouse Lab in Alderley Park and Alex Alderton, Roberto Amato, Jeffrey Barrett, Sonia Goncalves, Ewan Harrison, David K. Jackson, Ian Johnston, Dominic Kwiatkowski, Cordelia Langford, John Sillitoe on behalf of the Wellcome Sanger Institute COVID-19 Surveillance Team |
| EPI_ISL_3069470 | Berkshire and Surrey Pathology Services Lighthouse Laboratory | Wellcome Sanger Institute for the COVID-19 Genomics UK (COG-UK) Consortium | Berkshire and Surrey Pathology Services Lighthouse Laboratory and Alex Alderton, Roberto Amato, Jeffrey Barrett, Sonia Goncalves, Ewan Harrison, David K. Jackson, Ian Johnston, Dominic Kwiatkowski, Cordelia Langford, John Sillitoe on behalf of the Wellcome Sanger Institute COVID-19 Surveillance Team |
| EPI_ISL_3069471 | Lighthouse Lab in Glasgow | Wellcome Sanger Institute for the COVID-19 Genomics UK (COG-UK) Consortium | Harper VanSteenhouse, Yumi Kasai, David Gray, Carol Clugston, Anna Dominiczak and Alex Alderton, Roberto Amato, Jeffrey Barrett, Sonia Goncalves, Ewan Harrison, David K. Jackson, Ian Johnston, Dominic Kwiatkowski, Cordelia Langford, John Sillitoe on behalf of the Wellcome Sanger Institute COVID-19 Surveillance Team |
| EPI_ISL_3069472, EPI_ISL_3069473, EPI_ISL_3069474, EPI_ISL_3069475, EPI_ISL_3069476 | Berkshire and Surrey Pathology Services Lighthouse Laboratory | Wellcome Sanger Institute for the COVID-19 Genomics UK (COG-UK) Consortium | Berkshire and Surrey Pathology Services Lighthouse Laboratory and Alex Alderton, Roberto Amato, Jeffrey Barrett, Sonia Goncalves, Ewan Harrison, David K. Jackson, Ian Johnston, Dominic Kwiatkowski, Cordelia Langford, John Sillitoe on behalf of the Wellcome Sanger Institute COVID-19 Surveillance Team |
| EPI_ISL_3069477 | Lighthouse Lab in Alderley Park | Wellcome Sanger Institute for the COVID-19 Genomics UK (COG-UK) Consortium | Jacquelyn Wynn, Mairead Hyland, The Lighthouse Lab in Alderley Park and Alex Alderton, Roberto Amato, Jeffrey Barrett, Sonia Goncalves, Ewan Harrison, David K. Jackson, Ian Johnston, Dominic Kwiatkowski, Cordelia Langford, John Sillitoe on behalf of the Wellcome Sanger Institute COVID-19 Surveillance Team |
| EPI_ISL_3069478, EPI_ISL_3069479, EPI_ISL_3069480, EPI_ISL_3069481, EPI_ISL_3069482, EPI_ISL_3069483, EPI_ISL_3069484, EPI_ISL_3069485, EPI_ISL_3069486, EPI_ISL_3069487, EPI_ISL_3069488, EPI_ISL_3069489, EPI_ISL_3069490, EPI_ISL_3069491, EPI_ISL_3069492, EPI_ISL_3069493, EPI_ISL_3069494, EPI_ISL_3069495, EPI_ISL_3069496, EPI_ISL_3069497, EPI_ISL_3069498, EPI_ISL_3069499, EPI_ISL_3069500, EPI_ISL_3069501, EPI_ISL_3069502, EPI_ISL_3069503, EPI_ISL_3069504, EPI_ISL_3069505, EPI_ISL_3069506, EPI_ISL_3069507, EPI_ISL_3069508, EPI_ISL_3069509, EPI_ISL_3069510, EPI_ISL_3069511, EPI_ISL_3069512, EPI_ISL_3069513, EPI_ISL_3069514, EPI_ISL_3069515, EPI_ISL_3069516, EPI_ISL_3069517, EPI_ISL_3069518, EPI_ISL_3069519, EPI_ISL_3069520, EPI_ISL_3069521, EPI_ISL_3069522, EPI_ISL_3069523, EPI_ISL_3069524, EPI_ISL_3069525, EPI_ISL_3069526, EPI_ISL_3069527, EPI_ISL_3069528, EPI_ISL_3069529, EPI_ISL_3069530, EPI_ISL_3069531, EPI_ISL_3069532, EPI_ISL_3069533, EPI_ISL_3069534, EPI_ISL_3069535, EPI_ISL_3069536, EPI_ISL_3069537, EPI_ISL_3069538, EPI_ISL_3069539, EPI_ISL_3069540, EPI_ISL_3069541, EPI_ISL_3069542, EPI_ISL_3069543, EPI_ISL_3069544, EPI_ISL_3069545, EPI_ISL_3069546, EPI_ISL_3069547, EPI_ISL_3069548, EPI_ISL_3069549, EPI_ISL_3069550, EPI_ISL_3069551, EPI_ISL_3069552, EPI_ISL_3069553, EPI_ISL_3069554, EPI_ISL_3069555, EPI_ISL_3069556, EPI_ISL_3069557, EPI_ISL_3069558, EPI_ISL_3069559, EPI_ISL_3069560, EPI_ISL_3069561, EPI_ISL_3069562, EPI_ISL_3069563, EPI_ISL_3069564, EPI_ISL_3069565, EPI_ISL_3069566, EPI_ISL_3069567, EPI_ISL_3069568, EPI_ISL_3069569, EPI_ISL_3069570, EPI_ISL_3069571, EPI_ISL_3069572, EPI_ISL_3069573, EPI_ISL_3069574, EPI_ISL_3069575, EPI_ISL_3069576, EPI_ISL_3069577, EPI_ISL_3069578, EPI_ISL_3069579, EPI_ISL_3069580, EPI_ISL_3069581, EPI_ISL_3069582, EPI_ISL_3069583, EPI_ISL_3069584, EPI_ISL_3069585, EPI_ISL_3069586, EPI_ISL_3069587, EPI_ISL_3069588, EPI_ISL_3069589, EPI_ISL_3069590, EPI_ISL_3069591, EPI_ISL_3069592, EPI_ISL_3069593, EPI_ISL_3069594, EPI_ISL_3069595, EPI_ISL_3069596, EPI_ISL_3069597, EPI_ISL_3069598, EPI_ISL_3069599, EPI_ISL_3069600, EPI_ISL_3069601, EPI_ISL_3069602, EPI_ISL_3069603, EPI_ISL_3069604, EPI_ISL_3069605, EPI_ISL_3069606, EPI_ISL_3069607, EPI_ISL_3069608, EPI_ISL_3069609, EPI_ISL_3069610, EPI_ISL_3069611, EPI_ISL_3069612, EPI_ISL_3069613, EPI_ISL_3069614, EPI_ISL_3069615, EPI_ISL_3069616, EPI_ISL_3069617, EPI_ISL_3069618, EPI_ISL_3069619, EPI_ISL_3069620, EPI_ISL_3069621, EPI_ISL_3069622, EPI_ISL_3069623, EPI_ISL_3069624, EPI_ISL_3069625, EPI_ISL_3069626, EPI_ISL_3069627, EPI_ISL_3069628, EPI_ISL_3069629, EPI_ISL_3069630, EPI_ISL_3069631, EPI_ISL_3069632, EPI_ISL_3069633, EPI_ISL_3069634, EPI_ISL_3069635, EPI_ISL_3069636, EPI_ISL_3069637, EPI_ISL_3069638, EPI_ISL_3069639, EPI_ISL_3069640, EPI_ISL_3069641, EPI_ISL_3069642, EPI_ISL_3069643, EPI_ISL_3069644, EPI_ISL_3069645, EPI_ISL_3069646, EPI_ISL_3069647, EPI_ISL_3069648, EPI_ISL_3069649, EPI_ISL_3069650, EPI_ISL_3069651, EPI_ISL_3069652, EPI_ISL_3069653, EPI_ISL_3069654, EPI_ISL_3069655, EPI_ISL_3069656, EPI_ISL_3069657, EPI_ISL_3069658, EPI_ISL_3069659, EPI_ISL_3069660, EPI_ISL_3069661, EPI_ISL_3069662, EPI_ISL_3069663, EPI_ISL_3069664, EPI_ISL_3069665, EPI_ISL_3069666, EPI_ISL_3069667, EPI_ISL_3069668, EPI_ISL_3069669, EPI_ISL_3069670, EPI_ISL_3069671, EPI_ISL_3069672, EPI_ISL_3069673, EPI_ISL_3069674, EPI_ISL_3069675, EPI_ISL_3069676, EPI_ISL_3069677, EPI_ISL_3069678, EPI_ISL_3069679, EPI_ISL_3069680, EPI_ISL_3069681, EPI_ISL_3069682, EPI_ISL_3069683, EPI_ISL_3069684, EPI_ISL_3069685, EPI_ISL_3069686, EPI_ISL_3069687, EPI_ISL_3069688, EPI_ISL_3069689, EPI_ISL_3069690, EPI_ISL_3069691, EPI_ISL_3069692, EPI_ISL_3069693, EPI_ISL_3069694, EPI_ISL_3069695, EPI_ISL_3069696, EPI_ISL_3069697, EPI_ISL_3069698, EPI_ISL_3069700, EPI_ISL_3069701, EPI_ISL_3069702, EPI_ISL_3069703, EPI_ISL_3069704, EPI_ISL_3069705, EPI_ISL_3069706, EPI_ISL_3069707, EPI_ISL_3069708, EPI_ISL_3069709, EPI_ISL_3069710, EPI_ISL_3069711, EPI_ISL_3069712, EPI_ISL_3069713, EPI_ISL_3069714, EPI_ISL_3069715, EPI_ISL_3069716, EPI_ISL_3069717, EPI_ISL_3069718, EPI_ISL_3069719, EPI_ISL_3069720, EPI_ISL_3069721, EPI_ISL_3069722, EPI_ISL_3069723, EPI_ISL_3069724, EPI_ISL_3069725, EPI_ISL_3069726, EPI_ISL_3069727, EPI_ISL_3069728, EPI_ISL_3069729, EPI_ISL_3069730, EPI_ISL_3069731, EPI_ISL_3069732, EPI_ISL_3069733, EPI_ISL_3069734, EPI_ISL_3069735, EPI_ISL_3069736, EPI_ISL_3069737, EPI_ISL_3069738, EPI_ISL_3069739, EPI_ISL_3069740, EPI_ISL_3069741, EPI_ISL_3069742, EPI_ISL_3069743, EPI_ISL_3069744, EPI_ISL_3069745, EPI_ISL_3069746, EPI_ISL_3069747, EPI_ISL_3069748, EPI_ISL_3069749, EPI_ISL_3069750, EPI_ISL_3069751, EPI_ISL_3069752, EPI_ISL_3069753, EPI_ISL_3069754, EPI_ISL_3069755, EPI_ISL_3069756 | | | |
| see above | Berkshire and Surrey Pathology Services Lighthouse Laboratory | Wellcome Sanger Institute for the COVID-19 Genomics UK (COG-UK) Consortium | Berkshire and Surrey Pathology Services Lighthouse Laboratory and Alex Alderton, Roberto Amato, Jeffrey Barrett, Sonia Goncalves, Ewan Harrison, David K. Jackson, Ian Johnston, Dominic Kwiatkowski, Cordelia Langford, John Sillitoe on behalf of the Wellcome Sanger Institute COVID-19 Surveillance Team |
| EPI_ISL_3069757 | Lighthouse Lab in Alderley Park | Wellcome Sanger Institute for the COVID-19 Genomics UK (COG-UK) Consortium | Jacquelyn Wynn, Mairead Hyland, The Lighthouse Lab in Alderley Park and Alex Alderton, Roberto Amato, Jeffrey Barrett, Sonia Goncalves, Ewan Harrison, David K. Jackson, Ian Johnston, Dominic Kwiatkowski, Cordelia Langford, John Sillitoe on behalf of the Wellcome Sanger Institute COVID-19 Surveillance Team |
| EPI_ISL_3069758 | Lighthouse Lab in Glasgow | Wellcome Sanger Institute for the COVID-19 Genomics UK (COG-UK) Consortium | Harper VanSteenhouse, Yumi Kasai, David Gray, Carol Clugston, Anna Dominiczak and Alex Alderton, Roberto Amato, Jeffrey Barrett, Sonia Goncalves, Ewan Harrison, David K. Jackson, Ian Johnston, Dominic Kwiatkowski, Cordelia Langford, John Sillitoe on behalf of the Wellcome Sanger Institute COVID-19 Surveillance Team |
| EPI_ISL_3069759, EPI_ISL_3069760 | Berkshire and Surrey Pathology Services Lighthouse Laboratory | Wellcome Sanger Institute for the COVID-19 Genomics UK (COG-UK) Consortium | Berkshire and Surrey Pathology Services Lighthouse Laboratory and Alex Alderton, Roberto Amato, Jeffrey Barrett, Sonia Goncalves, Ewan Harrison, David K. Jackson, Ian Johnston, Dominic Kwiatkowski, Cordelia Langford, John Sillitoe on behalf of the Wellcome Sanger Institute COVID-19 Surveillance Team |
| EPI_ISL_3069761, EPI_ISL_3069762 | Lighthouse Lab in Glasgow | Wellcome Sanger Institute for the COVID-19 Genomics UK (COG-UK) Consortium | Harper VanSteenhouse, Yumi Kasai, David Gray, Carol Clugston, Anna Dominiczak and Alex Alderton, Roberto Amato, Jeffrey Barrett, Sonia Goncalves, Ewan Harrison, David K. Jackson, Ian Johnston, Dominic Kwiatkowski, Cordelia Langford, John Sillitoe on behalf of the Wellcome Sanger Institute COVID-19 Surveillance Team |
| EPI_ISL_3069763 | Lighthouse Lab in Alderley Park | Wellcome Sanger Institute for the COVID-19 Genomics UK (COG-UK) Consortium | Jacquelyn Wynn, Mairead Hyland, The Lighthouse Lab in Alderley Park and Alex Alderton, Roberto Amato, Jeffrey Barrett, Sonia Goncalves, Ewan Harrison, David K. Jackson, Ian Johnston, Dominic Kwiatkowski, Cordelia Langford, John Sillitoe on behalf of the Wellcome Sanger Institute COVID-19 Surveillance Team |
| EPI_ISL_3069764, EPI_ISL_3069765 | Lighthouse Lab in Glasgow | Wellcome Sanger Institute for the COVID-19 Genomics UK (COG-UK) Consortium | Harper VanSteenhouse, Yumi Kasai, David Gray, Carol Clugston, Anna Dominiczak and Alex Alderton, Roberto Amato, Jeffrey Barrett, Sonia Goncalves, Ewan Harrison, David K. Jackson, Ian Johnston, Dominic Kwiatkowski, Cordelia Langford, John Sillitoe on behalf of the Wellcome Sanger Institute COVID-19 Surveillance Team |
| EPI_ISL_3069766, EPI_ISL_3069767 | Berkshire and Surrey Pathology Services Lighthouse Laboratory | Wellcome Sanger Institute for the COVID-19 Genomics UK (COG-UK) Consortium | Berkshire and Surrey Pathology Services Lighthouse Laboratory and Alex Alderton, Roberto Amato, Jeffrey Barrett, Sonia Goncalves, Ewan Harrison, David K. Jackson, Ian Johnston, Dominic Kwiatkowski, Cordelia Langford, John Sillitoe on behalf of the Wellcome Sanger Institute COVID-19 Surveillance Team |
| EPI_ISL_3069768 | Lighthouse Lab in Glasgow | Wellcome Sanger Institute for the COVID-19 Genomics UK (COG-UK) Consortium | Harper VanSteenhouse, Yumi Kasai, David Gray, Carol Clugston, Anna Dominiczak and Alex Alderton, Roberto Amato, Jeffrey Barrett, Sonia Goncalves, Ewan Harrison, David K. Jackson, Ian Johnston, Dominic Kwiatkowski, Cordelia Langford, John Sillitoe on behalf of the Wellcome Sanger Institute COVID-19 Surveillance Team |
| EPI_ISL_3069769 | Lighthouse Lab in Alderley Park | Wellcome Sanger Institute for the COVID-19 Genomics UK (COG-UK) Consortium | Jacquelyn Wynn, Mairead Hyland, The Lighthouse Lab in Alderley Park and Alex Alderton, Roberto Amato, Jeffrey Barrett, Sonia Goncalves, Ewan Harrison, David K. Jackson, Ian Johnston, Dominic Kwiatkowski, Cordelia Langford, John Sillitoe on behalf of the Wellcome Sanger Institute COVID-19 Surveillance Team |
| EPI_ISL_3069770 | Lighthouse Lab in Glasgow | Wellcome Sanger Institute for the COVID-19 Genomics UK (COG-UK) Consortium | Harper VanSteenhouse, Yumi Kasai, David Gray, Carol Clugston, Anna Dominiczak and Alex Alderton, Roberto Amato, Jeffrey Barrett, Sonia Goncalves, Ewan Harrison, David K. Jackson, Ian Johnston, Dominic Kwiatkowski, Cordelia Langford, John Sillitoe on behalf of the Wellcome Sanger Institute COVID-19 Surveillance Team |
| EPI_ISL_3069771, EPI_ISL_3069772, EPI_ISL_3069773 | Lighthouse Lab in Alderley Park | Wellcome Sanger Institute for the COVID-19 Genomics UK (COG-UK) Consortium | Jacquelyn Wynn, Mairead Hyland, The Lighthouse Lab in Alderley Park and Alex Alderton, Roberto Amato, Jeffrey Barrett, Sonia Goncalves, Ewan Harrison, David K. Jackson, Ian Johnston, Dominic Kwiatkowski, Cordelia Langford, John Sillitoe on behalf of the Wellcome Sanger Institute COVID-19 Surveillance Team |

| | | | |
|---|---|---|---|
| EPI_ISL_3069774, EPI_ISL_3069775, EPI_ISL_3069776, EPI_ISL_3069777 | Lighthouse Lab in Glasgow | Wellcome Sanger Institute for the COVID-19 Genomics UK (COG-UK) Consortium | Harper VanSteenhouse, Yumi Kasai, David Gray, Carol Clugston, Anna Dominiczak and Alex Alderton, Roberto Amato, Jeffrey Barrett, Sonia Goncalves, Ewan Harrison, David K. Jackson, Ian Johnston, Dominic Kwiatkowski, Cordelia Langford, John Sillitoe on behalf of the Wellcome Sanger Institute COVID-19 Surveillance Team |
| EPI_ISL_3069778, EPI_ISL_3069779, EPI_ISL_3069780 | Lighthouse Lab in Alderley Park | Wellcome Sanger Institute for the COVID-19 Genomics UK (COG-UK) Consortium | Jacquelyn Wynn, Mairead Hyland, The Lighthouse Lab in Alderley Park and Alex Alderton, Roberto Amato, Jeffrey Barrett, Sonia Goncalves, Ewan Harrison, David K. Jackson, Ian Johnston, Dominic Kwiatkowski, Cordelia Langford, John Sillitoe on behalf of the Wellcome Sanger Institute COVID-19 Surveillance Team |
| EPI_ISL_3069781, EPI_ISL_3069782, EPI_ISL_3069783, EPI_ISL_3069784 | Lighthouse Lab in Glasgow | Wellcome Sanger Institute for the COVID-19 Genomics UK (COG-UK) Consortium | Harper VanSteenhouse, Yumi Kasai, David Gray, Carol Clugston, Anna Dominiczak and Alex Alderton, Roberto Amato, Jeffrey Barrett, Sonia Goncalves, Ewan Harrison, David K. Jackson, Ian Johnston, Dominic Kwiatkowski, Cordelia Langford, John Sillitoe on behalf of the Wellcome Sanger Institute COVID-19 Surveillance Team |
| EPI_ISL_3069785 | Lighthouse Lab in Alderley Park | Wellcome Sanger Institute for the COVID-19 Genomics UK (COG-UK) Consortium | Jacquelyn Wynn, Mairead Hyland, The Lighthouse Lab in Alderley Park and Alex Alderton, Roberto Amato, Jeffrey Barrett, Sonia Goncalves, Ewan Harrison, David K. Jackson, Ian Johnston, Dominic Kwiatkowski, Cordelia Langford, John Sillitoe on behalf of the Wellcome Sanger Institute COVID-19 Surveillance Team |
| EPI_ISL_3069786 | Berkshire and Surrey Pathology Services Lighthouse Laboratory | Wellcome Sanger Institute for the COVID-19 Genomics UK (COG-UK) Consortium | Berkshire and Surrey Pathology Services Lighthouse Laboratory and Alex Alderton, Roberto Amato, Jeffrey Barrett, Sonia Goncalves, Ewan Harrison, David K. Jackson, Ian Johnston, Dominic Kwiatkowski, Cordelia Langford, John Sillitoe on behalf of the Wellcome Sanger Institute COVID-19 Surveillance Team |
| EPI_ISL_3069787 | Lighthouse Lab in Alderley Park | Wellcome Sanger Institute for the COVID-19 Genomics UK (COG-UK) Consortium | Jacquelyn Wynn, Mairead Hyland, The Lighthouse Lab in Alderley Park and Alex Alderton, Roberto Amato, Jeffrey Barrett, Sonia Goncalves, Ewan Harrison, David K. Jackson, Ian Johnston, Dominic Kwiatkowski, Cordelia Langford, John Sillitoe on behalf of the Wellcome Sanger Institute COVID-19 Surveillance Team |
| EPI_ISL_3069788 | Lighthouse Lab in Glasgow | Wellcome Sanger Institute for the COVID-19 Genomics UK (COG-UK) Consortium | Harper VanSteenhouse, Yumi Kasai, David Gray, Carol Clugston, Anna Dominiczak and Alex Alderton, Roberto Amato, Jeffrey Barrett, Sonia Goncalves, Ewan Harrison, David K. Jackson, Ian Johnston, Dominic Kwiatkowski, Cordelia Langford, John Sillitoe on behalf of the Wellcome Sanger Institute COVID-19 Surveillance Team |
| EPI_ISL_3069789 | Lighthouse Lab in Alderley Park | Wellcome Sanger Institute for the COVID-19 Genomics UK (COG-UK) Consortium | Jacquelyn Wynn, Mairead Hyland, The Lighthouse Lab in Alderley Park and Alex Alderton, Roberto Amato, Jeffrey Barrett, Sonia Goncalves, Ewan Harrison, David K. Jackson, Ian Johnston, Dominic Kwiatkowski, Cordelia Langford, John Sillitoe on behalf of the Wellcome Sanger Institute COVID-19 Surveillance Team |
| EPI_ISL_3069790, EPI_ISL_3069791, EPI_ISL_3069792, EPI_ISL_3069793, EPI_ISL_3069794, EPI_ISL_3069795 | Lighthouse Lab in Glasgow | Wellcome Sanger Institute for the COVID-19 Genomics UK (COG-UK) Consortium | Harper VanSteenhouse, Yumi Kasai, David Gray, Carol Clugston, Anna Dominiczak and Alex Alderton, Roberto Amato, Jeffrey Barrett, Sonia Goncalves, Ewan Harrison, David K. Jackson, Ian Johnston, Dominic Kwiatkowski, Cordelia Langford, John Sillitoe on behalf of the Wellcome Sanger Institute COVID-19 Surveillance Team |
| EPI_ISL_3069796 | Lighthouse Lab in Alderley Park | Wellcome Sanger Institute for the COVID-19 Genomics UK (COG-UK) Consortium | Jacquelyn Wynn, Mairead Hyland, The Lighthouse Lab in Alderley Park and Alex Alderton, Roberto Amato, Jeffrey Barrett, Sonia Goncalves, Ewan Harrison, David K. Jackson, Ian Johnston, Dominic Kwiatkowski, Cordelia Langford, John Sillitoe on behalf of the Wellcome Sanger Institute COVID-19 Surveillance Team |
| EPI_ISL_3069797 | Berkshire and Surrey Pathology Services Lighthouse Laboratory | Wellcome Sanger Institute for the COVID-19 Genomics UK (COG-UK) Consortium | Berkshire and Surrey Pathology Services Lighthouse Laboratory and Alex Alderton, Roberto Amato, Jeffrey Barrett, Sonia Goncalves, Ewan Harrison, David K. Jackson, Ian Johnston, Dominic Kwiatkowski, Cordelia Langford, John Sillitoe on behalf of the Wellcome Sanger Institute COVID-19 Surveillance Team |
| EPI_ISL_3069798, EPI_ISL_3069799 | Lighthouse Lab in Glasgow | Wellcome Sanger Institute for the COVID-19 Genomics UK (COG-UK) Consortium | Harper VanSteenhouse, Yumi Kasai, David Gray, Carol Clugston, Anna Dominiczak and Alex Alderton, Roberto Amato, Jeffrey Barrett, Sonia Goncalves, Ewan Harrison, David K. Jackson, Ian Johnston, Dominic Kwiatkowski, Cordelia Langford, John Sillitoe on behalf of the Wellcome Sanger Institute COVID-19 Surveillance Team |
| EPI_ISL_3069800 | Berkshire and Surrey Pathology Services Lighthouse Laboratory | Wellcome Sanger Institute for the COVID-19 Genomics UK (COG-UK) Consortium | Berkshire and Surrey Pathology Services Lighthouse Laboratory and Alex Alderton, Roberto Amato, Jeffrey Barrett, Sonia Goncalves, Ewan Harrison, David K. Jackson, Ian Johnston, Dominic Kwiatkowski, Cordelia Langford, John Sillitoe on behalf of the Wellcome Sanger Institute COVID-19 Surveillance Team |
| EPI_ISL_3069801 | Lighthouse Lab in Alderley Park | Wellcome Sanger Institute for the COVID-19 Genomics UK (COG-UK) Consortium | Jacquelyn Wynn, Mairead Hyland, The Lighthouse Lab in Alderley Park and Alex Alderton, Roberto Amato, Jeffrey Barrett, Sonia Goncalves, Ewan Harrison, David K. Jackson, Ian Johnston, Dominic Kwiatkowski, Cordelia Langford, John Sillitoe on behalf of the Wellcome Sanger Institute COVID-19 Surveillance Team |
| EPI_ISL_3069802, EPI_ISL_3069803 | Lighthouse Lab in Glasgow | Wellcome Sanger Institute for the COVID-19 Genomics UK (COG-UK) Consortium | Harper VanSteenhouse, Yumi Kasai, David Gray, Carol Clugston, Anna Dominiczak and Alex Alderton, Roberto Amato, Jeffrey Barrett, Sonia Goncalves, Ewan Harrison, David K. Jackson, Ian Johnston, Dominic Kwiatkowski, Cordelia Langford, John Sillitoe on behalf of the Wellcome Sanger Institute COVID-19 Surveillance Team |
| EPI_ISL_3069804, EPI_ISL_3069805, EPI_ISL_3069806, EPI_ISL_3069807 | Lighthouse Lab in Alderley Park | Wellcome Sanger Institute for the COVID-19 Genomics UK (COG-UK) Consortium | Jacquelyn Wynn, Mairead Hyland, The Lighthouse Lab in Alderley Park and Alex Alderton, Roberto Amato, Jeffrey Barrett, Sonia Goncalves, Ewan Harrison, David K. Jackson, Ian Johnston, Dominic Kwiatkowski, Cordelia Langford, John Sillitoe on behalf of the Wellcome Sanger Institute COVID-19 Surveillance Team |
| EPI_ISL_3069808, EPI_ISL_3069809 | Lighthouse Lab in Glasgow | Wellcome Sanger Institute for the COVID-19 Genomics UK (COG-UK) Consortium | Harper VanSteenhouse, Yumi Kasai, David Gray, Carol Clugston, Anna Dominiczak and Alex Alderton, Roberto Amato, Jeffrey Barrett, Sonia Goncalves, Ewan Harrison, David K. Jackson, Ian Johnston, Dominic Kwiatkowski, Cordelia Langford, John Sillitoe on behalf of the Wellcome Sanger Institute COVID-19 Surveillance Team |
| EPI_ISL_3069810, EPI_ISL_3069811 | Berkshire and Surrey Pathology Services Lighthouse Laboratory | Wellcome Sanger Institute for the COVID-19 Genomics UK (COG-UK) Consortium | Berkshire and Surrey Pathology Services Lighthouse Laboratory and Alex Alderton, Roberto Amato, Jeffrey Barrett, Sonia Goncalves, Ewan Harrison, David K. Jackson, Ian Johnston, Dominic Kwiatkowski, Cordelia Langford, John Sillitoe on behalf of the Wellcome Sanger Institute COVID-19 Surveillance Team |
| EPI_ISL_3069812 | Lighthouse Lab in Glasgow | Wellcome Sanger Institute for the COVID-19 Genomics UK (COG-UK) Consortium | Harper VanSteenhouse, Yumi Kasai, David Gray, Carol Clugston, Anna Dominiczak and Alex Alderton, Roberto Amato, Jeffrey Barrett, Sonia Goncalves, Ewan Harrison, David K. Jackson, Ian Johnston, Dominic Kwiatkowski, Cordelia Langford, John Sillitoe on behalf of the Wellcome Sanger Institute COVID-19 Surveillance Team |
| EPI_ISL_3069813 | Lighthouse Lab in Alderley Park | Wellcome Sanger Institute for the COVID-19 Genomics UK (COG-UK) Consortium | Jacquelyn Wynn, Mairead Hyland, The Lighthouse Lab in Alderley Park and Alex Alderton, Roberto Amato, Jeffrey Barrett, Sonia Goncalves, Ewan Harrison, David K. Jackson, Ian Johnston, Dominic Kwiatkowski, Cordelia Langford, John Sillitoe on behalf of the Wellcome Sanger Institute COVID-19 Surveillance Team |
| EPI_ISL_3069814 | Lighthouse Lab in Glasgow | Wellcome Sanger Institute for the COVID-19 Genomics UK (COG-UK) Consortium | Harper VanSteenhouse, Yumi Kasai, David Gray, Carol Clugston, Anna Dominiczak and Alex Alderton, Roberto Amato, Jeffrey Barrett, Sonia Goncalves, Ewan Harrison, David K. Jackson, Ian Johnston, Dominic Kwiatkowski, Cordelia Langford, John Sillitoe on behalf of the Wellcome Sanger Institute COVID-19 Surveillance Team |
| EPI_ISL_3069815 | Lighthouse Lab in Alderley Park | Wellcome Sanger Institute for the COVID-19 Genomics UK (COG-UK) Consortium | Jacquelyn Wynn, Mairead Hyland, The Lighthouse Lab in Alderley Park and Alex Alderton, Roberto Amato, Jeffrey Barrett, Sonia Goncalves, Ewan Harrison, David K. Jackson, Ian Johnston, Dominic Kwiatkowski, Cordelia Langford, John Sillitoe on behalf of the Wellcome Sanger Institute COVID-19 Surveillance Team |
| EPI_ISL_3069816, EPI_ISL_3069817 | Berkshire and Surrey Pathology Services Lighthouse Laboratory | Wellcome Sanger Institute for the COVID-19 Genomics UK (COG-UK) Consortium | Berkshire and Surrey Pathology Services Lighthouse Laboratory and Alex Alderton, Roberto Amato, Jeffrey Barrett, Sonia Goncalves, Ewan Harrison, David K. Jackson, Ian Johnston, Dominic Kwiatkowski, Cordelia Langford, John Sillitoe on behalf of the Wellcome Sanger Institute COVID-19 Surveillance Team |
| EPI_ISL_3069818 | Lighthouse Lab in Alderley Park | Wellcome Sanger Institute for the COVID-19 Genomics UK (COG-UK) Consortium | Jacquelyn Wynn, Mairead Hyland, The Lighthouse Lab in Alderley Park and Alex Alderton, Roberto Amato, Jeffrey Barrett, Sonia Goncalves, Ewan Harrison, David K. Jackson, Ian Johnston, Dominic Kwiatkowski, Cordelia Langford, John Sillitoe on behalf of the Wellcome Sanger Institute COVID-19 Surveillance Team |
| EPI_ISL_3069819 | Berkshire and Surrey Pathology Services Lighthouse Laboratory | Wellcome Sanger Institute for the COVID-19 Genomics UK (COG-UK) Consortium | Berkshire and Surrey Pathology Services Lighthouse Laboratory and Alex Alderton, Roberto Amato, Jeffrey Barrett, Sonia Goncalves, Ewan Harrison, David K. Jackson, Ian Johnston, Dominic Kwiatkowski, Cordelia Langford, John Sillitoe on behalf of the Wellcome Sanger Institute COVID-19 Surveillance Team |
| EPI_ISL_3069820 | Lighthouse Lab in Alderley Park | Wellcome Sanger Institute for the COVID-19 Genomics UK (COG-UK) Consortium | Jacquelyn Wynn, Mairead Hyland, The Lighthouse Lab in Alderley Park and Alex Alderton, Roberto Amato, Jeffrey Barrett, Sonia Goncalves, Ewan Harrison, David K. Jackson, Ian Johnston, Dominic Kwiatkowski, Cordelia Langford, John Sillitoe on behalf of the Wellcome Sanger Institute COVID-19 Surveillance Team |
| EPI_ISL_3069821, EPI_ISL_3069822 | Lighthouse Lab in Glasgow | Wellcome Sanger Institute for the COVID-19 Genomics UK (COG-UK) Consortium | Harper VanSteenhouse, Yumi Kasai, David Gray, Carol Clugston, Anna Dominiczak and Alex Alderton, Roberto Amato, Jeffrey Barrett, Sonia Goncalves, Ewan Harrison, David K. Jackson, Ian Johnston, Dominic Kwiatkowski, Cordelia Langford, John Sillitoe on behalf of the Wellcome Sanger Institute COVID-19 Surveillance Team |
| EPI_ISL_3069823 | Lighthouse Lab in Alderley Park | Wellcome Sanger Institute for the COVID-19 Genomics UK (COG-UK) Consortium | Jacquelyn Wynn, Mairead Hyland, The Lighthouse Lab in Alderley Park and Alex Alderton, Roberto Amato, Jeffrey Barrett, Sonia Goncalves, Ewan Harrison, David K. Jackson, Ian Johnston, Dominic Kwiatkowski, Cordelia Langford, John Sillitoe on behalf of the Wellcome Sanger Institute COVID-19 Surveillance Team |
| EPI_ISL_3069824 | Berkshire and Surrey Pathology Services Lighthouse Laboratory | Wellcome Sanger Institute for the COVID-19 Genomics UK (COG-UK) Consortium | Berkshire and Surrey Pathology Services Lighthouse Laboratory and Alex Alderton, Roberto Amato, Jeffrey Barrett, Sonia Goncalves, Ewan Harrison, David K. Jackson, Ian Johnston, Dominic Kwiatkowski, Cordelia Langford, John Sillitoe on behalf of the Wellcome Sanger Institute COVID-19 Surveillance Team |
| EPI_ISL_3069825, EPI_ISL_3069826 | Lighthouse Lab in Glasgow | Wellcome Sanger Institute for the COVID-19 Genomics UK (COG-UK) Consortium | Harper VanSteenhouse, Yumi Kasai, David Gray, Carol Clugston, Anna Dominiczak and Alex Alderton, Roberto Amato, Jeffrey Barrett, Sonia Goncalves, Ewan Harrison, David K. Jackson, Ian Johnston, Dominic Kwiatkowski, Cordelia Langford, John Sillitoe on behalf of the Wellcome Sanger Institute COVID-19 Surveillance Team |
| EPI_ISL_3069827, EPI_ISL_3069828 | Lighthouse Lab in Alderley Park | Wellcome Sanger Institute for the COVID-19 Genomics UK (COG-UK) Consortium | Jacquelyn Wynn, Mairead Hyland, The Lighthouse Lab in Alderley Park and Alex Alderton, Roberto Amato, Jeffrey Barrett, Sonia Goncalves, Ewan Harrison, David K. Jackson, Ian Johnston, Dominic Kwiatkowski, Cordelia Langford, John Sillitoe on behalf of the Wellcome Sanger Institute COVID-19 Surveillance Team |
| EPI_ISL_3069829 | Berkshire and Surrey Pathology Services Lighthouse Laboratory | Wellcome Sanger Institute for the COVID-19 Genomics UK (COG-UK) Consortium | Berkshire and Surrey Pathology Services Lighthouse Laboratory and Alex Alderton, Roberto Amato, Jeffrey Barrett, Sonia Goncalves, Ewan Harrison, David K. Jackson, Ian Johnston, Dominic Kwiatkowski, Cordelia Langford, John Sillitoe on behalf of the Wellcome Sanger Institute COVID-19 Surveillance Team |
| EPI_ISL_3069830 | Lighthouse Lab in Glasgow | Wellcome Sanger Institute for the COVID-19 Genomics UK (COG-UK) Consortium | Harper VanSteenhouse, Yumi Kasai, David Gray, Carol Clugston, Anna Dominiczak and Alex Alderton, Roberto Amato, Jeffrey Barrett, Sonia Goncalves, Ewan Harrison, David K. Jackson, Ian Johnston, Dominic Kwiatkowski, Cordelia Langford, John Sillitoe on behalf of the Wellcome Sanger Institute COVID-19 Surveillance Team |
| EPI_ISL_3069831 | Berkshire and Surrey Pathology Services Lighthouse Laboratory | Wellcome Sanger Institute for the COVID-19 Genomics UK (COG-UK) Consortium | Berkshire and Surrey Pathology Services Lighthouse Laboratory and Alex Alderton, Roberto Amato, Jeffrey Barrett, Sonia Goncalves, Ewan Harrison, David K. Jackson, Ian Johnston, Dominic Kwiatkowski, Cordelia Langford, John Sillitoe on behalf of the Wellcome Sanger Institute COVID-19 Surveillance Team |
| EPI_ISL_3069832 | Lighthouse Lab in Glasgow | Wellcome Sanger Institute for the COVID-19 Genomics UK (COG-UK) Consortium | Harper VanSteenhouse, Yumi Kasai, David Gray, Carol Clugston, Anna Dominiczak and Alex Alderton, Roberto Amato, Jeffrey Barrett, Sonia Goncalves, Ewan Harrison, David K. Jackson, Ian Johnston, Dominic Kwiatkowski, Cordelia Langford, John Sillitoe on behalf of the Wellcome Sanger Institute COVID-19 Surveillance Team |
| EPI_ISL_3069833, EPI_ISL_3069834, EPI_ISL_3069835, EPI_ISL_3069836 | Berkshire and Surrey Pathology Services Lighthouse Laboratory | Wellcome Sanger Institute for the COVID-19 Genomics UK (COG-UK) Consortium | Berkshire and Surrey Pathology Services Lighthouse Laboratory and Alex Alderton, Roberto Amato, Jeffrey Barrett, Sonia Goncalves, Ewan Harrison, David K. Jackson, Ian Johnston, Dominic Kwiatkowski, Cordelia Langford, John Sillitoe on behalf of the Wellcome Sanger Institute COVID-19 Surveillance Team |
| EPI_ISL_3069837 | Lighthouse Lab in Glasgow | Wellcome Sanger Institute for the COVID-19 Genomics UK (COG-UK) Consortium | Harper VanSteenhouse, Yumi Kasai, David Gray, Carol Clugston, Anna Dominiczak and Alex Alderton, Roberto Amato, Jeffrey Barrett, Sonia Goncalves, Ewan Harrison, David K. Jackson, Ian Johnston, Dominic Kwiatkowski, Cordelia Langford, John Sillitoe on behalf of the Wellcome Sanger Institute COVID-19 Surveillance Team |
| EPI_ISL_3069838 | Berkshire and Surrey Pathology Services Lighthouse Laboratory | Wellcome Sanger Institute for the COVID-19 Genomics UK (COG-UK) Consortium | Berkshire and Surrey Pathology Services Lighthouse Laboratory and Alex Alderton, Roberto Amato, Jeffrey Barrett, Sonia Goncalves, Ewan Harrison, David K. Jackson, Ian Johnston, Dominic Kwiatkowski, Cordelia Langford, John Sillitoe on behalf of the Wellcome Sanger Institute COVID-19 Surveillance Team |
| EPI_ISL_3069839 | Lighthouse Lab in Glasgow | Wellcome Sanger Institute for the COVID-19 Genomics UK (COG-UK) Consortium | Harper VanSteenhouse, Yumi Kasai, David Gray, Carol Clugston, Anna Dominiczak and Alex Alderton, Roberto Amato, Jeffrey Barrett, Sonia Goncalves, Ewan Harrison, David K. Jackson, Ian Johnston, Dominic Kwiatkowski, Cordelia Langford, John Sillitoe on behalf of the Wellcome Sanger Institute COVID-19 Surveillance Team |
| EPI_ISL_3069840 | Berkshire and Surrey Pathology Services Lighthouse Laboratory | Wellcome Sanger Institute for the COVID-19 Genomics UK (COG-UK) | Berkshire and Surrey Pathology Services Lighthouse Laboratory and Alex Alderton, Roberto Amato, Jeffrey Barrett, Sonia Goncalves, Ewan Harrison, David K. Jackson, Ian Johnston, Dominic Kwiatkowski, Cordelia Langford, John Sillitoe on behalf |

| | | | |
|---|---|---|---|
| | | Consortium | of the Wellcome Sanger Institute COVID-19 Surveillance Team |
| EPI_ISL_3069841, EPI_ISL_3069842 | Lighthouse Lab in Glasgow | Wellcome Sanger Institute for the COVID-19 Genomics UK (COG-UK) Consortium | Harper VanSteenhouse, Yumi Kasai, David Gray, Carol Clugston, Anna Dominiczak and Alex Alderton, Roberto Amato, Jeffrey Barrett, Sonia Goncalves, Ewan Harrison, David K. Jackson, Ian Johnston, Dominic Kwiatkowski, Cordelia Langford, John Sillitoe on behalf of the Wellcome Sanger Institute COVID-19 Surveillance Team |
| EPI_ISL_3069843 | Berkshire and Surrey Pathology Services Lighthouse Laboratory | Wellcome Sanger Institute for the COVID-19 Genomics UK (COG-UK) Consortium | Berkshire and Surrey Pathology Services Lighthouse Laboratory and Alex Alderton, Roberto Amato, Jeffrey Barrett, Sonia Goncalves, Ewan Harrison, David K. Jackson, Ian Johnston, Dominic Kwiatkowski, Cordelia Langford, John Sillitoe on behalf of the Wellcome Sanger Institute COVID-19 Surveillance Team |
| EPI_ISL_3069844 | Lighthouse Lab in Glasgow | Wellcome Sanger Institute for the COVID-19 Genomics UK (COG-UK) Consortium | Harper VanSteenhouse, Yumi Kasai, David Gray, Carol Clugston, Anna Dominiczak and Alex Alderton, Roberto Amato, Jeffrey Barrett, Sonia Goncalves, Ewan Harrison, David K. Jackson, Ian Johnston, Dominic Kwiatkowski, Cordelia Langford, John Sillitoe on behalf of the Wellcome Sanger Institute COVID-19 Surveillance Team |
| EPI_ISL_3069845 | Berkshire and Surrey Pathology Services Lighthouse Laboratory | Wellcome Sanger Institute for the COVID-19 Genomics UK (COG-UK) Consortium | Berkshire and Surrey Pathology Services Lighthouse Laboratory and Alex Alderton, Roberto Amato, Jeffrey Barrett, Sonia Goncalves, Ewan Harrison, David K. Jackson, Ian Johnston, Dominic Kwiatkowski, Cordelia Langford, John Sillitoe on behalf of the Wellcome Sanger Institute COVID-19 Surveillance Team |
| EPI_ISL_3069846, EPI_ISL_3069847, EPI_ISL_3069848 | Lighthouse Lab in Alderley Park | Wellcome Sanger Institute for the COVID-19 Genomics UK (COG-UK) Consortium | Jacquelyn Wynn, Mairead Hyland, The Lighthouse Lab in Alderley Park and Alex Alderton, Roberto Amato, Jeffrey Barrett, Sonia Goncalves, Ewan Harrison, David K. Jackson, Ian Johnston, Dominic Kwiatkowski, Cordelia Langford, John Sillitoe on behalf of the Wellcome Sanger Institute COVID-19 Surveillance Team |
| EPI_ISL_3069849 | Berkshire and Surrey Pathology Services Lighthouse Laboratory | Wellcome Sanger Institute for the COVID-19 Genomics UK (COG-UK) Consortium | Berkshire and Surrey Pathology Services Lighthouse Laboratory and Alex Alderton, Roberto Amato, Jeffrey Barrett, Sonia Goncalves, Ewan Harrison, David K. Jackson, Ian Johnston, Dominic Kwiatkowski, Cordelia Langford, John Sillitoe on behalf of the Wellcome Sanger Institute COVID-19 Surveillance Team |
| EPI_ISL_3069850, EPI_ISL_3069851 | Lighthouse Lab in Glasgow | Wellcome Sanger Institute for the COVID-19 Genomics UK (COG-UK) Consortium | Harper VanSteenhouse, Yumi Kasai, David Gray, Carol Clugston, Anna Dominiczak and Alex Alderton, Roberto Amato, Jeffrey Barrett, Sonia Goncalves, Ewan Harrison, David K. Jackson, Ian Johnston, Dominic Kwiatkowski, Cordelia Langford, John Sillitoe on behalf of the Wellcome Sanger Institute COVID-19 Surveillance Team |
| EPI_ISL_3069852 | Berkshire and Surrey Pathology Services Lighthouse Laboratory | Wellcome Sanger Institute for the COVID-19 Genomics UK (COG-UK) Consortium | Berkshire and Surrey Pathology Services Lighthouse Laboratory and Alex Alderton, Roberto Amato, Jeffrey Barrett, Sonia Goncalves, Ewan Harrison, David K. Jackson, Ian Johnston, Dominic Kwiatkowski, Cordelia Langford, John Sillitoe on behalf of the Wellcome Sanger Institute COVID-19 Surveillance Team |
| EPI_ISL_3069853, EPI_ISL_3069854 | Lighthouse Lab in Glasgow | Wellcome Sanger Institute for the COVID-19 Genomics UK (COG-UK) Consortium | Harper VanSteenhouse, Yumi Kasai, David Gray, Carol Clugston, Anna Dominiczak and Alex Alderton, Roberto Amato, Jeffrey Barrett, Sonia Goncalves, Ewan Harrison, David K. Jackson, Ian Johnston, Dominic Kwiatkowski, Cordelia Langford, John Sillitoe on behalf of the Wellcome Sanger Institute COVID-19 Surveillance Team |
| EPI_ISL_3069855 | Lighthouse Lab in Alderley Park | Wellcome Sanger Institute for the COVID-19 Genomics UK (COG-UK) Consortium | Jacquelyn Wynn, Mairead Hyland, The Lighthouse Lab in Alderley Park and Alex Alderton, Roberto Amato, Jeffrey Barrett, Sonia Goncalves, Ewan Harrison, David K. Jackson, Ian Johnston, Dominic Kwiatkowski, Cordelia Langford, John Sillitoe on behalf of the Wellcome Sanger Institute COVID-19 Surveillance Team |
| EPI_ISL_3069856 | Berkshire and Surrey Pathology Services Lighthouse Laboratory | Wellcome Sanger Institute for the COVID-19 Genomics UK (COG-UK) Consortium | Berkshire and Surrey Pathology Services Lighthouse Laboratory and Alex Alderton, Roberto Amato, Jeffrey Barrett, Sonia Goncalves, Ewan Harrison, David K. Jackson, Ian Johnston, Dominic Kwiatkowski, Cordelia Langford, John Sillitoe on behalf of the Wellcome Sanger Institute COVID-19 Surveillance Team |
| EPI_ISL_3069857 | Lighthouse Lab in Alderley Park | Wellcome Sanger Institute for the COVID-19 Genomics UK (COG-UK) Consortium | Jacquelyn Wynn, Mairead Hyland, The Lighthouse Lab in Alderley Park and Alex Alderton, Roberto Amato, Jeffrey Barrett, Sonia Goncalves, Ewan Harrison, David K. Jackson, Ian Johnston, Dominic Kwiatkowski, Cordelia Langford, John Sillitoe on behalf of the Wellcome Sanger Institute COVID-19 Surveillance Team |
| EPI_ISL_3069858 | Lighthouse Lab in Glasgow | Wellcome Sanger Institute for the COVID-19 Genomics UK (COG-UK) Consortium | Harper VanSteenhouse, Yumi Kasai, David Gray, Carol Clugston, Anna Dominiczak and Alex Alderton, Roberto Amato, Jeffrey Barrett, Sonia Goncalves, Ewan Harrison, David K. Jackson, Ian Johnston, Dominic Kwiatkowski, Cordelia Langford, John Sillitoe on behalf of the Wellcome Sanger Institute COVID-19 Surveillance Team |
| EPI_ISL_3069859 | Berkshire and Surrey Pathology Services Lighthouse Laboratory | Wellcome Sanger Institute for the COVID-19 Genomics UK (COG-UK) Consortium | Berkshire and Surrey Pathology Services Lighthouse Laboratory and Alex Alderton, Roberto Amato, Jeffrey Barrett, Sonia Goncalves, Ewan Harrison, David K. Jackson, Ian Johnston, Dominic Kwiatkowski, Cordelia Langford, John Sillitoe on behalf of the Wellcome Sanger Institute COVID-19 Surveillance Team |
| EPI_ISL_3069860 | Lighthouse Lab in Alderley Park | Wellcome Sanger Institute for the COVID-19 Genomics UK (COG-UK) Consortium | Jacquelyn Wynn, Mairead Hyland, The Lighthouse Lab in Alderley Park and Alex Alderton, Roberto Amato, Jeffrey Barrett, Sonia Goncalves, Ewan Harrison, David K. Jackson, Ian Johnston, Dominic Kwiatkowski, Cordelia Langford, John Sillitoe on behalf of the Wellcome Sanger Institute COVID-19 Surveillance Team |
| EPI_ISL_3069861 | Berkshire and Surrey Pathology Services Lighthouse Laboratory | Wellcome Sanger Institute for the COVID-19 Genomics UK (COG-UK) Consortium | Berkshire and Surrey Pathology Services Lighthouse Laboratory and Alex Alderton, Roberto Amato, Jeffrey Barrett, Sonia Goncalves, Ewan Harrison, David K. Jackson, Ian Johnston, Dominic Kwiatkowski, Cordelia Langford, John Sillitoe on behalf of the Wellcome Sanger Institute COVID-19 Surveillance Team |
| EPI_ISL_3069862, EPI_ISL_3069863, EPI_ISL_3069864 | Lighthouse Lab in Alderley Park | Wellcome Sanger Institute for the COVID-19 Genomics UK (COG-UK) Consortium | Jacquelyn Wynn, Mairead Hyland, The Lighthouse Lab in Alderley Park and Alex Alderton, Roberto Amato, Jeffrey Barrett, Sonia Goncalves, Ewan Harrison, David K. Jackson, Ian Johnston, Dominic Kwiatkowski, Cordelia Langford, John Sillitoe on behalf of the Wellcome Sanger Institute COVID-19 Surveillance Team |
| EPI_ISL_3069865, EPI_ISL_3069866 | Lighthouse Lab in Glasgow | Wellcome Sanger Institute for the COVID-19 Genomics UK (COG-UK) Consortium | Harper VanSteenhouse, Yumi Kasai, David Gray, Carol Clugston, Anna Dominiczak and Alex Alderton, Roberto Amato, Jeffrey Barrett, Sonia Goncalves, Ewan Harrison, David K. Jackson, Ian Johnston, Dominic Kwiatkowski, Cordelia Langford, John Sillitoe on behalf of the Wellcome Sanger Institute COVID-19 Surveillance Team |
| EPI_ISL_3069867 | Lighthouse Lab in Alderley Park | Wellcome Sanger Institute for the COVID-19 Genomics UK (COG-UK) Consortium | Jacquelyn Wynn, Mairead Hyland, The Lighthouse Lab in Alderley Park and Alex Alderton, Roberto Amato, Jeffrey Barrett, Sonia Goncalves, Ewan Harrison, David K. Jackson, Ian Johnston, Dominic Kwiatkowski, Cordelia Langford, John Sillitoe on behalf of the Wellcome Sanger Institute COVID-19 Surveillance Team |
| EPI_ISL_3069868, EPI_ISL_3069869, EPI_ISL_3069870, EPI_ISL_3069871 | Lighthouse Lab in Glasgow | Wellcome Sanger Institute for the COVID-19 Genomics UK (COG-UK) Consortium | Harper VanSteenhouse, Yumi Kasai, David Gray, Carol Clugston, Anna Dominiczak and Alex Alderton, Roberto Amato, Jeffrey Barrett, Sonia Goncalves, Ewan Harrison, David K. Jackson, Ian Johnston, Dominic Kwiatkowski, Cordelia Langford, John Sillitoe on behalf of the Wellcome Sanger Institute COVID-19 Surveillance Team |
| EPI_ISL_3069872, EPI_ISL_3069873 | Berkshire and Surrey Pathology Services Lighthouse Laboratory | Wellcome Sanger Institute for the COVID-19 Genomics UK (COG-UK) Consortium | Berkshire and Surrey Pathology Services Lighthouse Laboratory and Alex Alderton, Roberto Amato, Jeffrey Barrett, Sonia Goncalves, Ewan Harrison, David K. Jackson, Ian Johnston, Dominic Kwiatkowski, Cordelia Langford, John Sillitoe on behalf of the Wellcome Sanger Institute COVID-19 Surveillance Team |
| EPI_ISL_3069874, EPI_ISL_3069875 | Lighthouse Lab in Glasgow | Wellcome Sanger Institute for the COVID-19 Genomics UK (COG-UK) Consortium | Harper VanSteenhouse, Yumi Kasai, David Gray, Carol Clugston, Anna Dominiczak and Alex Alderton, Roberto Amato, Jeffrey Barrett, Sonia Goncalves, Ewan Harrison, David K. Jackson, Ian Johnston, Dominic Kwiatkowski, Cordelia Langford, John Sillitoe on behalf of the Wellcome Sanger Institute COVID-19 Surveillance Team |
| EPI_ISL_3069876, EPI_ISL_3069877, EPI_ISL_3069878 | Berkshire and Surrey Pathology Services Lighthouse Laboratory | Wellcome Sanger Institute for the COVID-19 Genomics UK (COG-UK) Consortium | Berkshire and Surrey Pathology Services Lighthouse Laboratory and Alex Alderton, Roberto Amato, Jeffrey Barrett, Sonia Goncalves, Ewan Harrison, David K. Jackson, Ian Johnston, Dominic Kwiatkowski, Cordelia Langford, John Sillitoe on behalf of the Wellcome Sanger Institute COVID-19 Surveillance Team |
| EPI_ISL_3069879 | Lighthouse Lab in Alderley Park | Wellcome Sanger Institute for the COVID-19 Genomics UK (COG-UK) Consortium | Jacquelyn Wynn, Mairead Hyland, The Lighthouse Lab in Alderley Park and Alex Alderton, Roberto Amato, Jeffrey Barrett, Sonia Goncalves, Ewan Harrison, David K. Jackson, Ian Johnston, Dominic Kwiatkowski, Cordelia Langford, John Sillitoe on behalf of the Wellcome Sanger Institute COVID-19 Surveillance Team |
| EPI_ISL_3069880, EPI_ISL_3069881, EPI_ISL_3069882, EPI_ISL_3069883 | Lighthouse Lab in Glasgow | Wellcome Sanger Institute for the COVID-19 Genomics UK (COG-UK) Consortium | Harper VanSteenhouse, Yumi Kasai, David Gray, Carol Clugston, Anna Dominiczak and Alex Alderton, Roberto Amato, Jeffrey Barrett, Sonia Goncalves, Ewan Harrison, David K. Jackson, Ian Johnston, Dominic Kwiatkowski, Cordelia Langford, John Sillitoe on behalf of the Wellcome Sanger Institute COVID-19 Surveillance Team |
| EPI_ISL_3069884 | Lighthouse Lab in Alderley Park | Wellcome Sanger Institute for the COVID-19 Genomics UK (COG-UK) Consortium | Jacquelyn Wynn, Mairead Hyland, The Lighthouse Lab in Alderley Park and Alex Alderton, Roberto Amato, Jeffrey Barrett, Sonia Goncalves, Ewan Harrison, David K. Jackson, Ian Johnston, Dominic Kwiatkowski, Cordelia Langford, John Sillitoe on behalf of the Wellcome Sanger Institute COVID-19 Surveillance Team |
| EPI_ISL_3069885, EPI_ISL_3069886 | Lighthouse Lab in Glasgow | Wellcome Sanger Institute for the COVID-19 Genomics UK (COG-UK) Consortium | Harper VanSteenhouse, Yumi Kasai, David Gray, Carol Clugston, Anna Dominiczak and Alex Alderton, Roberto Amato, Jeffrey Barrett, Sonia Goncalves, Ewan Harrison, David K. Jackson, Ian Johnston, Dominic Kwiatkowski, Cordelia Langford, John Sillitoe on behalf of the Wellcome Sanger Institute COVID-19 Surveillance Team |
| EPI_ISL_3069887 | Lighthouse Lab in Alderley Park | Wellcome Sanger Institute for the COVID-19 Genomics UK (COG-UK) Consortium | Jacquelyn Wynn, Mairead Hyland, The Lighthouse Lab in Alderley Park and Alex Alderton, Roberto Amato, Jeffrey Barrett, Sonia Goncalves, Ewan Harrison, David K. Jackson, Ian Johnston, Dominic Kwiatkowski, Cordelia Langford, John Sillitoe on behalf of the Wellcome Sanger Institute COVID-19 Surveillance Team |
| EPI_ISL_3069888 | Lighthouse Lab in Glasgow | Wellcome Sanger Institute for the COVID-19 Genomics UK (COG-UK) Consortium | Harper VanSteenhouse, Yumi Kasai, David Gray, Carol Clugston, Anna Dominiczak and Alex Alderton, Roberto Amato, Jeffrey Barrett, Sonia Goncalves, Ewan Harrison, David K. Jackson, Ian Johnston, Dominic Kwiatkowski, Cordelia Langford, John Sillitoe on behalf of the Wellcome Sanger Institute COVID-19 Surveillance Team |
| EPI_ISL_3069889 | Lighthouse Lab in Alderley Park | Wellcome Sanger Institute for the COVID-19 Genomics UK (COG-UK) Consortium | Jacquelyn Wynn, Mairead Hyland, The Lighthouse Lab in Alderley Park and Alex Alderton, Roberto Amato, Jeffrey Barrett, Sonia Goncalves, Ewan Harrison, David K. Jackson, Ian Johnston, Dominic Kwiatkowski, Cordelia Langford, John Sillitoe on behalf of the Wellcome Sanger Institute COVID-19 Surveillance Team |
| EPI_ISL_3069890 | Lighthouse Lab in Glasgow | Wellcome Sanger Institute for the COVID-19 Genomics UK (COG-UK) Consortium | Harper VanSteenhouse, Yumi Kasai, David Gray, Carol Clugston, Anna Dominiczak and Alex Alderton, Roberto Amato, Jeffrey Barrett, Sonia Goncalves, Ewan Harrison, David K. Jackson, Ian Johnston, Dominic Kwiatkowski, Cordelia Langford, John Sillitoe on behalf of the Wellcome Sanger Institute COVID-19 Surveillance Team |
| EPI_ISL_3069891, EPI_ISL_3069892, EPI_ISL_3069893 | Lighthouse Lab in Alderley Park | Wellcome Sanger Institute for the COVID-19 Genomics UK (COG-UK) Consortium | Jacquelyn Wynn, Mairead Hyland, The Lighthouse Lab in Alderley Park and Alex Alderton, Roberto Amato, Jeffrey Barrett, Sonia Goncalves, Ewan Harrison, David K. Jackson, Ian Johnston, Dominic Kwiatkowski, Cordelia Langford, John Sillitoe on behalf of the Wellcome Sanger Institute COVID-19 Surveillance Team |
| EPI_ISL_3069894, EPI_ISL_3069895 | Berkshire and Surrey Pathology Services Lighthouse Laboratory | Wellcome Sanger Institute for the COVID-19 Genomics UK (COG-UK) Consortium | Berkshire and Surrey Pathology Services Lighthouse Laboratory and Alex Alderton, Roberto Amato, Jeffrey Barrett, Sonia Goncalves, Ewan Harrison, David K. Jackson, Ian Johnston, Dominic Kwiatkowski, Cordelia Langford, John Sillitoe on behalf of the Wellcome Sanger Institute COVID-19 Surveillance Team |
| EPI_ISL_3069896 | Lighthouse Lab in Glasgow | Wellcome Sanger Institute for the COVID-19 Genomics UK (COG-UK) Consortium | Harper VanSteenhouse, Yumi Kasai, David Gray, Carol Clugston, Anna Dominiczak and Alex Alderton, Roberto Amato, Jeffrey Barrett, Sonia Goncalves, Ewan Harrison, David K. Jackson, Ian Johnston, Dominic Kwiatkowski, Cordelia Langford, John Sillitoe on behalf of the Wellcome Sanger Institute COVID-19 Surveillance Team |
| EPI_ISL_3069897 | Berkshire and Surrey Pathology Services Lighthouse Laboratory | Wellcome Sanger Institute for the COVID-19 Genomics UK (COG-UK) Consortium | Berkshire and Surrey Pathology Services Lighthouse Laboratory and Alex Alderton, Roberto Amato, Jeffrey Barrett, Sonia Goncalves, Ewan Harrison, David K. Jackson, Ian Johnston, Dominic Kwiatkowski, Cordelia Langford, John Sillitoe on behalf of the Wellcome Sanger Institute COVID-19 Surveillance Team |
| EPI_ISL_3069898, EPI_ISL_3069899 | Lighthouse Lab in Alderley Park | Wellcome Sanger Institute for the COVID-19 Genomics UK (COG-UK) Consortium | Jacquelyn Wynn, Mairead Hyland, The Lighthouse Lab in Alderley Park and Alex Alderton, Roberto Amato, Jeffrey Barrett, Sonia Goncalves, Ewan Harrison, David K. Jackson, Ian Johnston, Dominic Kwiatkowski, Cordelia Langford, John Sillitoe on behalf of the Wellcome Sanger Institute COVID-19 Surveillance Team |
| EPI_ISL_3069900 | Lighthouse Lab in Glasgow | Wellcome Sanger Institute for the COVID-19 Genomics UK (COG-UK) Consortium | Harper VanSteenhouse, Yumi Kasai, David Gray, Carol Clugston, Anna Dominiczak and Alex Alderton, Roberto Amato, Jeffrey Barrett, Sonia Goncalves, Ewan Harrison, David K. Jackson, Ian Johnston, Dominic Kwiatkowski, Cordelia Langford, John Sillitoe on behalf of the Wellcome Sanger Institute COVID-19 Surveillance Team |
| EPI_ISL_3069901 | Berkshire and Surrey Pathology Services Lighthouse Laboratory | Wellcome Sanger Institute for the COVID-19 Genomics UK (COG-UK) Consortium | Berkshire and Surrey Pathology Services Lighthouse Laboratory and Alex Alderton, Roberto Amato, Jeffrey Barrett, Sonia Goncalves, Ewan Harrison, David K. Jackson, Ian Johnston, Dominic Kwiatkowski, Cordelia Langford, John Sillitoe on behalf of the Wellcome Sanger Institute COVID-19 Surveillance Team |
| EPI_ISL_3069902, EPI_ISL_3069903 | Lighthouse Lab in Glasgow | Wellcome Sanger Institute for the COVID-19 Genomics UK (COG-UK) Consortium | Harper VanSteenhouse, Yumi Kasai, David Gray, Carol Clugston, Anna Dominiczak and Alex Alderton, Roberto Amato, Jeffrey Barrett, Sonia Goncalves, Ewan Harrison, David K. Jackson, Ian Johnston, Dominic Kwiatkowski, Cordelia Langford, John Sillitoe on behalf of the Wellcome Sanger Institute COVID-19 Surveillance Team |
| EPI_ISL_3069904 | Berkshire and Surrey Pathology Services Lighthouse Laboratory | Wellcome Sanger Institute for the COVID-19 Genomics UK (COG-UK) Consortium | Berkshire and Surrey Pathology Services Lighthouse Laboratory and Alex Alderton, Roberto Amato, Jeffrey Barrett, Sonia Goncalves, Ewan Harrison, David K. Jackson, Ian Johnston, Dominic Kwiatkowski, Cordelia Langford, John Sillitoe on behalf of the Wellcome Sanger Institute COVID-19 Surveillance Team |
| EPI_ISL_3069905, EPI_ISL_3069906, EPI_ISL_3069907 | Lighthouse Lab in Glasgow | Wellcome Sanger Institute for the COVID-19 Genomics UK (COG-UK) Consortium | Harper VanSteenhouse, Yumi Kasai, David Gray, Carol Clugston, Anna Dominiczak and Alex Alderton, Roberto Amato, Jeffrey Barrett, Sonia Goncalves, Ewan Harrison, David K. Jackson, Ian Johnston, Dominic Kwiatkowski, Cordelia Langford, John Sillitoe on behalf of the Wellcome Sanger Institute COVID-19 Surveillance Team |

| | | | |
|---|---|---|---|
| EPI_ISL_3069908 | Lighthouse Lab in Alderley Park | Wellcome Sanger Institute for the COVID-19 Genomics UK (COG-UK) Consortium | Jacquelyn Wynn, Mairead Hyland, The Lighthouse Lab in Alderley Park and Alex Alderton, Roberto Amato, Jeffrey Barrett, Sonia Goncalves, Ewan Harrison, David K. Jackson, Ian Johnston, Dominic Kwiatkowski, Cordelia Langford, John Sillitoe on behalf of the Wellcome Sanger Institute COVID-19 Surveillance Team |
| EPI_ISL_3069909 | Berkshire and Surrey Pathology Services Lighthouse Laboratory | Wellcome Sanger Institute for the COVID-19 Genomics UK (COG-UK) Consortium | Berkshire and Surrey Pathology Services Lighthouse Laboratory and Alex Alderton, Roberto Amato, Jeffrey Barrett, Sonia Goncalves, Ewan Harrison, David K. Jackson, Ian Johnston, Dominic Kwiatkowski, Cordelia Langford, John Sillitoe on behalf of the Wellcome Sanger Institute COVID-19 Surveillance Team |
| EPI_ISL_3069910 | Lighthouse Lab in Glasgow | Wellcome Sanger Institute for the COVID-19 Genomics UK (COG-UK) Consortium | Harper VanSteenhouse, Yumi Kasai, David Gray, Carol Clugston, Anna Dominiczak and Alex Alderton, Roberto Amato, Jeffrey Barrett, Sonia Goncalves, Ewan Harrison, David K. Jackson, Ian Johnston, Dominic Kwiatkowski, Cordelia Langford, John Sillitoe on behalf of the Wellcome Sanger Institute COVID-19 Surveillance Team |
| EPI_ISL_3069911 | Lighthouse Lab in Alderley Park | Wellcome Sanger Institute for the COVID-19 Genomics UK (COG-UK) Consortium | Jacquelyn Wynn, Mairead Hyland, The Lighthouse Lab in Alderley Park and Alex Alderton, Roberto Amato, Jeffrey Barrett, Sonia Goncalves, Ewan Harrison, David K. Jackson, Ian Johnston, Dominic Kwiatkowski, Cordelia Langford, John Sillitoe on behalf of the Wellcome Sanger Institute COVID-19 Surveillance Team |
| EPI_ISL_3069912, EPI_ISL_3069913, EPI_ISL_3069914 | Lighthouse Lab in Glasgow | Wellcome Sanger Institute for the COVID-19 Genomics UK (COG-UK) Consortium | Harper VanSteenhouse, Yumi Kasai, David Gray, Carol Clugston, Anna Dominiczak and Alex Alderton, Roberto Amato, Jeffrey Barrett, Sonia Goncalves, Ewan Harrison, David K. Jackson, Ian Johnston, Dominic Kwiatkowski, Cordelia Langford, John Sillitoe on behalf of the Wellcome Sanger Institute COVID-19 Surveillance Team |
| EPI_ISL_3069915, EPI_ISL_3069916, EPI_ISL_3069917 | Lighthouse Lab in Alderley Park | Wellcome Sanger Institute for the COVID-19 Genomics UK (COG-UK) Consortium | Jacquelyn Wynn, Mairead Hyland, The Lighthouse Lab in Alderley Park and Alex Alderton, Roberto Amato, Jeffrey Barrett, Sonia Goncalves, Ewan Harrison, David K. Jackson, Ian Johnston, Dominic Kwiatkowski, Cordelia Langford, John Sillitoe on behalf of the Wellcome Sanger Institute COVID-19 Surveillance Team |
| EPI_ISL_3069918 | Berkshire and Surrey Pathology Services Lighthouse Laboratory | Wellcome Sanger Institute for the COVID-19 Genomics UK (COG-UK) Consortium | Berkshire and Surrey Pathology Services Lighthouse Laboratory and Alex Alderton, Roberto Amato, Jeffrey Barrett, Sonia Goncalves, Ewan Harrison, David K. Jackson, Ian Johnston, Dominic Kwiatkowski, Cordelia Langford, John Sillitoe on behalf of the Wellcome Sanger Institute COVID-19 Surveillance Team |
| EPI_ISL_3069919 | Lighthouse Lab in Glasgow | Wellcome Sanger Institute for the COVID-19 Genomics UK (COG-UK) Consortium | Harper VanSteenhouse, Yumi Kasai, David Gray, Carol Clugston, Anna Dominiczak and Alex Alderton, Roberto Amato, Jeffrey Barrett, Sonia Goncalves, Ewan Harrison, David K. Jackson, Ian Johnston, Dominic Kwiatkowski, Cordelia Langford, John Sillitoe on behalf of the Wellcome Sanger Institute COVID-19 Surveillance Team |
| EPI_ISL_3069920 | Berkshire and Surrey Pathology Services Lighthouse Laboratory | Wellcome Sanger Institute for the COVID-19 Genomics UK (COG-UK) Consortium | Berkshire and Surrey Pathology Services Lighthouse Laboratory and Alex Alderton, Roberto Amato, Jeffrey Barrett, Sonia Goncalves, Ewan Harrison, David K. Jackson, Ian Johnston, Dominic Kwiatkowski, Cordelia Langford, John Sillitoe on behalf of the Wellcome Sanger Institute COVID-19 Surveillance Team |
| EPI_ISL_3069921 | Lighthouse Lab in Alderley Park | Wellcome Sanger Institute for the COVID-19 Genomics UK (COG-UK) Consortium | Jacquelyn Wynn, Mairead Hyland, The Lighthouse Lab in Alderley Park and Alex Alderton, Roberto Amato, Jeffrey Barrett, Sonia Goncalves, Ewan Harrison, David K. Jackson, Ian Johnston, Dominic Kwiatkowski, Cordelia Langford, John Sillitoe on behalf of the Wellcome Sanger Institute COVID-19 Surveillance Team |
| EPI_ISL_3069922, EPI_ISL_3069923, EPI_ISL_3069924 | Lighthouse Lab in Glasgow | Wellcome Sanger Institute for the COVID-19 Genomics UK (COG-UK) Consortium | Harper VanSteenhouse, Yumi Kasai, David Gray, Carol Clugston, Anna Dominiczak and Alex Alderton, Roberto Amato, Jeffrey Barrett, Sonia Goncalves, Ewan Harrison, David K. Jackson, Ian Johnston, Dominic Kwiatkowski, Cordelia Langford, John Sillitoe on behalf of the Wellcome Sanger Institute COVID-19 Surveillance Team |
| EPI_ISL_3069925 | Berkshire and Surrey Pathology Services Lighthouse Laboratory | Wellcome Sanger Institute for the COVID-19 Genomics UK (COG-UK) Consortium | Berkshire and Surrey Pathology Services Lighthouse Laboratory and Alex Alderton, Roberto Amato, Jeffrey Barrett, Sonia Goncalves, Ewan Harrison, David K. Jackson, Ian Johnston, Dominic Kwiatkowski, Cordelia Langford, John Sillitoe on behalf of the Wellcome Sanger Institute COVID-19 Surveillance Team |
| EPI_ISL_3069926 | Lighthouse Lab in Alderley Park | Wellcome Sanger Institute for the COVID-19 Genomics UK (COG-UK) Consortium | Jacquelyn Wynn, Mairead Hyland, The Lighthouse Lab in Alderley Park and Alex Alderton, Roberto Amato, Jeffrey Barrett, Sonia Goncalves, Ewan Harrison, David K. Jackson, Ian Johnston, Dominic Kwiatkowski, Cordelia Langford, John Sillitoe on behalf of the Wellcome Sanger Institute COVID-19 Surveillance Team |
| EPI_ISL_3069927 | Berkshire and Surrey Pathology Services Lighthouse Laboratory | Wellcome Sanger Institute for the COVID-19 Genomics UK (COG-UK) Consortium | Berkshire and Surrey Pathology Services Lighthouse Laboratory and Alex Alderton, Roberto Amato, Jeffrey Barrett, Sonia Goncalves, Ewan Harrison, David K. Jackson, Ian Johnston, Dominic Kwiatkowski, Cordelia Langford, John Sillitoe on behalf of the Wellcome Sanger Institute COVID-19 Surveillance Team |
| EPI_ISL_3069928, EPI_ISL_3069929, EPI_ISL_3069930, EPI_ISL_3069931 | Lighthouse Lab in Glasgow | Wellcome Sanger Institute for the COVID-19 Genomics UK (COG-UK) Consortium | Harper VanSteenhouse, Yumi Kasai, David Gray, Carol Clugston, Anna Dominiczak and Alex Alderton, Roberto Amato, Jeffrey Barrett, Sonia Goncalves, Ewan Harrison, David K. Jackson, Ian Johnston, Dominic Kwiatkowski, Cordelia Langford, John Sillitoe on behalf of the Wellcome Sanger Institute COVID-19 Surveillance Team |
| EPI_ISL_3069932 | Lighthouse Lab in Alderley Park | Wellcome Sanger Institute for the COVID-19 Genomics UK (COG-UK) Consortium | Jacquelyn Wynn, Mairead Hyland, The Lighthouse Lab in Alderley Park and Alex Alderton, Roberto Amato, Jeffrey Barrett, Sonia Goncalves, Ewan Harrison, David K. Jackson, Ian Johnston, Dominic Kwiatkowski, Cordelia Langford, John Sillitoe on behalf of the Wellcome Sanger Institute COVID-19 Surveillance Team |
| EPI_ISL_3069933, EPI_ISL_3069934 | Lighthouse Lab in Glasgow | Wellcome Sanger Institute for the COVID-19 Genomics UK (COG-UK) Consortium | Harper VanSteenhouse, Yumi Kasai, David Gray, Carol Clugston, Anna Dominiczak and Alex Alderton, Roberto Amato, Jeffrey Barrett, Sonia Goncalves, Ewan Harrison, David K. Jackson, Ian Johnston, Dominic Kwiatkowski, Cordelia Langford, John Sillitoe on behalf of the Wellcome Sanger Institute COVID-19 Surveillance Team |
| EPI_ISL_3069935 | Lighthouse Lab in Alderley Park | Wellcome Sanger Institute for the COVID-19 Genomics UK (COG-UK) Consortium | Jacquelyn Wynn, Mairead Hyland, The Lighthouse Lab in Alderley Park and Alex Alderton, Roberto Amato, Jeffrey Barrett, Sonia Goncalves, Ewan Harrison, David K. Jackson, Ian Johnston, Dominic Kwiatkowski, Cordelia Langford, John Sillitoe on behalf of the Wellcome Sanger Institute COVID-19 Surveillance Team |
| EPI_ISL_3069936 | Berkshire and Surrey Pathology Services Lighthouse Laboratory | Wellcome Sanger Institute for the COVID-19 Genomics UK (COG-UK) Consortium | Berkshire and Surrey Pathology Services Lighthouse Laboratory and Alex Alderton, Roberto Amato, Jeffrey Barrett, Sonia Goncalves, Ewan Harrison, David K. Jackson, Ian Johnston, Dominic Kwiatkowski, Cordelia Langford, John Sillitoe on behalf of the Wellcome Sanger Institute COVID-19 Surveillance Team |
| EPI_ISL_3069937 | Lighthouse Lab in Glasgow | Wellcome Sanger Institute for the COVID-19 Genomics UK (COG-UK) Consortium | Harper VanSteenhouse, Yumi Kasai, David Gray, Carol Clugston, Anna Dominiczak and Alex Alderton, Roberto Amato, Jeffrey Barrett, Sonia Goncalves, Ewan Harrison, David K. Jackson, Ian Johnston, Dominic Kwiatkowski, Cordelia Langford, John Sillitoe on behalf of the Wellcome Sanger Institute COVID-19 Surveillance Team |
| EPI_ISL_3069938 | Lighthouse Lab in Alderley Park | Wellcome Sanger Institute for the COVID-19 Genomics UK (COG-UK) Consortium | Jacquelyn Wynn, Mairead Hyland, The Lighthouse Lab in Alderley Park and Alex Alderton, Roberto Amato, Jeffrey Barrett, Sonia Goncalves, Ewan Harrison, David K. Jackson, Ian Johnston, Dominic Kwiatkowski, Cordelia Langford, John Sillitoe on behalf of the Wellcome Sanger Institute COVID-19 Surveillance Team |
| EPI_ISL_3069939 | Lighthouse Lab in Glasgow | Wellcome Sanger Institute for the COVID-19 Genomics UK (COG-UK) Consortium | Harper VanSteenhouse, Yumi Kasai, David Gray, Carol Clugston, Anna Dominiczak and Alex Alderton, Roberto Amato, Jeffrey Barrett, Sonia Goncalves, Ewan Harrison, David K. Jackson, Ian Johnston, Dominic Kwiatkowski, Cordelia Langford, John Sillitoe on behalf of the Wellcome Sanger Institute COVID-19 Surveillance Team |
| EPI_ISL_3069940, EPI_ISL_3069941, EPI_ISL_3069942 | Berkshire and Surrey Pathology Services Lighthouse Laboratory | Wellcome Sanger Institute for the COVID-19 Genomics UK (COG-UK) Consortium | Berkshire and Surrey Pathology Services Lighthouse Laboratory and Alex Alderton, Roberto Amato, Jeffrey Barrett, Sonia Goncalves, Ewan Harrison, David K. Jackson, Ian Johnston, Dominic Kwiatkowski, Cordelia Langford, John Sillitoe on behalf of the Wellcome Sanger Institute COVID-19 Surveillance Team |
| EPI_ISL_3069943, EPI_ISL_3069944 | Lighthouse Lab in Glasgow | Wellcome Sanger Institute for the COVID-19 Genomics UK (COG-UK) Consortium | Harper VanSteenhouse, Yumi Kasai, David Gray, Carol Clugston, Anna Dominiczak and Alex Alderton, Roberto Amato, Jeffrey Barrett, Sonia Goncalves, Ewan Harrison, David K. Jackson, Ian Johnston, Dominic Kwiatkowski, Cordelia Langford, John Sillitoe on behalf of the Wellcome Sanger Institute COVID-19 Surveillance Team |
| EPI_ISL_3069945, EPI_ISL_3069946 | Berkshire and Surrey Pathology Services Lighthouse Laboratory | Wellcome Sanger Institute for the COVID-19 Genomics UK (COG-UK) Consortium | Berkshire and Surrey Pathology Services Lighthouse Laboratory and Alex Alderton, Roberto Amato, Jeffrey Barrett, Sonia Goncalves, Ewan Harrison, David K. Jackson, Ian Johnston, Dominic Kwiatkowski, Cordelia Langford, John Sillitoe on behalf of the Wellcome Sanger Institute COVID-19 Surveillance Team |
| EPI_ISL_3069947 | Lighthouse Lab in Glasgow | Wellcome Sanger Institute for the COVID-19 Genomics UK (COG-UK) Consortium | Harper VanSteenhouse, Yumi Kasai, David Gray, Carol Clugston, Anna Dominiczak and Alex Alderton, Roberto Amato, Jeffrey Barrett, Sonia Goncalves, Ewan Harrison, David K. Jackson, Ian Johnston, Dominic Kwiatkowski, Cordelia Langford, John Sillitoe on behalf of the Wellcome Sanger Institute COVID-19 Surveillance Team |
| EPI_ISL_3069948 | Lighthouse Lab in Alderley Park | Wellcome Sanger Institute for the COVID-19 Genomics UK (COG-UK) Consortium | Jacquelyn Wynn, Mairead Hyland, The Lighthouse Lab in Alderley Park and Alex Alderton, Roberto Amato, Jeffrey Barrett, Sonia Goncalves, Ewan Harrison, David K. Jackson, Ian Johnston, Dominic Kwiatkowski, Cordelia Langford, John Sillitoe on behalf of the Wellcome Sanger Institute COVID-19 Surveillance Team |
| EPI_ISL_3069949, EPI_ISL_3069950, EPI_ISL_3069951 | Lighthouse Lab in Glasgow | Wellcome Sanger Institute for the COVID-19 Genomics UK (COG-UK) Consortium | Harper VanSteenhouse, Yumi Kasai, David Gray, Carol Clugston, Anna Dominiczak and Alex Alderton, Roberto Amato, Jeffrey Barrett, Sonia Goncalves, Ewan Harrison, David K. Jackson, Ian Johnston, Dominic Kwiatkowski, Cordelia Langford, John Sillitoe on behalf of the Wellcome Sanger Institute COVID-19 Surveillance Team |
| EPI_ISL_3069952 | Berkshire and Surrey Pathology Services Lighthouse Laboratory | Wellcome Sanger Institute for the COVID-19 Genomics UK (COG-UK) Consortium | Berkshire and Surrey Pathology Services Lighthouse Laboratory and Alex Alderton, Roberto Amato, Jeffrey Barrett, Sonia Goncalves, Ewan Harrison, David K. Jackson, Ian Johnston, Dominic Kwiatkowski, Cordelia Langford, John Sillitoe on behalf of the Wellcome Sanger Institute COVID-19 Surveillance Team |
| EPI_ISL_3069953 | Lighthouse Lab in Alderley Park | Wellcome Sanger Institute for the COVID-19 Genomics UK (COG-UK) Consortium | Jacquelyn Wynn, Mairead Hyland, The Lighthouse Lab in Alderley Park and Alex Alderton, Roberto Amato, Jeffrey Barrett, Sonia Goncalves, Ewan Harrison, David K. Jackson, Ian Johnston, Dominic Kwiatkowski, Cordelia Langford, John Sillitoe on behalf of the Wellcome Sanger Institute COVID-19 Surveillance Team |
| EPI_ISL_3069954, EPI_ISL_3069955 | Berkshire and Surrey Pathology Services Lighthouse Laboratory | Wellcome Sanger Institute for the COVID-19 Genomics UK (COG-UK) Consortium | Berkshire and Surrey Pathology Services Lighthouse Laboratory and Alex Alderton, Roberto Amato, Jeffrey Barrett, Sonia Goncalves, Ewan Harrison, David K. Jackson, Ian Johnston, Dominic Kwiatkowski, Cordelia Langford, John Sillitoe on behalf of the Wellcome Sanger Institute COVID-19 Surveillance Team |
| EPI_ISL_3069956 | Lighthouse Lab in Glasgow | Wellcome Sanger Institute for the COVID-19 Genomics UK (COG-UK) Consortium | Harper VanSteenhouse, Yumi Kasai, David Gray, Carol Clugston, Anna Dominiczak and Alex Alderton, Roberto Amato, Jeffrey Barrett, Sonia Goncalves, Ewan Harrison, David K. Jackson, Ian Johnston, Dominic Kwiatkowski, Cordelia Langford, John Sillitoe on behalf of the Wellcome Sanger Institute COVID-19 Surveillance Team |
| EPI_ISL_3069957 | Lighthouse Lab in Alderley Park | Wellcome Sanger Institute for the COVID-19 Genomics UK (COG-UK) Consortium | Jacquelyn Wynn, Mairead Hyland, The Lighthouse Lab in Alderley Park and Alex Alderton, Roberto Amato, Jeffrey Barrett, Sonia Goncalves, Ewan Harrison, David K. Jackson, Ian Johnston, Dominic Kwiatkowski, Cordelia Langford, John Sillitoe on behalf of the Wellcome Sanger Institute COVID-19 Surveillance Team |
| EPI_ISL_3069958 | Lighthouse Lab in Glasgow | Wellcome Sanger Institute for the COVID-19 Genomics UK (COG-UK) Consortium | Harper VanSteenhouse, Yumi Kasai, David Gray, Carol Clugston, Anna Dominiczak and Alex Alderton, Roberto Amato, Jeffrey Barrett, Sonia Goncalves, Ewan Harrison, David K. Jackson, Ian Johnston, Dominic Kwiatkowski, Cordelia Langford, John Sillitoe on behalf of the Wellcome Sanger Institute COVID-19 Surveillance Team |
| EPI_ISL_3069959 | Lighthouse Lab in Alderley Park | Wellcome Sanger Institute for the COVID-19 Genomics UK (COG-UK) Consortium | Jacquelyn Wynn, Mairead Hyland, The Lighthouse Lab in Alderley Park and Alex Alderton, Roberto Amato, Jeffrey Barrett, Sonia Goncalves, Ewan Harrison, David K. Jackson, Ian Johnston, Dominic Kwiatkowski, Cordelia Langford, John Sillitoe on behalf of the Wellcome Sanger Institute COVID-19 Surveillance Team |
| EPI_ISL_3069960 | Berkshire and Surrey Pathology Services Lighthouse Laboratory | Wellcome Sanger Institute for the COVID-19 Genomics UK (COG-UK) Consortium | Berkshire and Surrey Pathology Services Lighthouse Laboratory and Alex Alderton, Roberto Amato, Jeffrey Barrett, Sonia Goncalves, Ewan Harrison, David K. Jackson, Ian Johnston, Dominic Kwiatkowski, Cordelia Langford, John Sillitoe on behalf of the Wellcome Sanger Institute COVID-19 Surveillance Team |
| EPI_ISL_3069961 | Lighthouse Lab in Glasgow | Wellcome Sanger Institute for the COVID-19 Genomics UK (COG-UK) Consortium | Harper VanSteenhouse, Yumi Kasai, David Gray, Carol Clugston, Anna Dominiczak and Alex Alderton, Roberto Amato, Jeffrey Barrett, Sonia Goncalves, Ewan Harrison, David K. Jackson, Ian Johnston, Dominic Kwiatkowski, Cordelia Langford, John Sillitoe on behalf of the Wellcome Sanger Institute COVID-19 Surveillance Team |
| EPI_ISL_3069962 | Lighthouse Lab in Alderley Park | Wellcome Sanger Institute for the COVID-19 Genomics UK (COG-UK) Consortium | Jacquelyn Wynn, Mairead Hyland, The Lighthouse Lab in Alderley Park and Alex Alderton, Roberto Amato, Jeffrey Barrett, Sonia Goncalves, Ewan Harrison, David K. Jackson, Ian Johnston, Dominic Kwiatkowski, Cordelia Langford, John Sillitoe on behalf of the Wellcome Sanger Institute COVID-19 Surveillance Team |
| EPI_ISL_3069963 | Lighthouse Lab in Glasgow | Wellcome Sanger Institute for the COVID-19 Genomics UK (COG-UK) Consortium | Harper VanSteenhouse, Yumi Kasai, David Gray, Carol Clugston, Anna Dominiczak and Alex Alderton, Roberto Amato, Jeffrey Barrett, Sonia Goncalves, Ewan Harrison, David K. Jackson, Ian Johnston, Dominic Kwiatkowski, Cordelia Langford, John Sillitoe on behalf of the Wellcome Sanger Institute COVID-19 Surveillance Team |
| EPI_ISL_3069964 | Lighthouse Lab in Alderley Park | Wellcome Sanger Institute for the COVID-19 Genomics UK (COG-UK) Consortium | Jacquelyn Wynn, Mairead Hyland, The Lighthouse Lab in Alderley Park and Alex Alderton, Roberto Amato, Jeffrey Barrett, Sonia Goncalves, Ewan Harrison, David K. Jackson, Ian Johnston, Dominic Kwiatkowski, Cordelia Langford, John Sillitoe on behalf of the Wellcome Sanger Institute COVID-19 Surveillance Team |
| EPI_ISL_3069965, EPI_ISL_3069966 | Lighthouse Lab in Glasgow | Wellcome Sanger Institute for the COVID-19 Genomics UK (COG-UK) Consortium | Harper VanSteenhouse, Yumi Kasai, David Gray, Carol Clugston, Anna Dominiczak and Alex Alderton, Roberto Amato, Jeffrey Barrett, Sonia Goncalves, Ewan Harrison, David K. Jackson, Ian Johnston, Dominic Kwiatkowski, Cordelia Langford, John Sillitoe on behalf of the Wellcome Sanger Institute COVID-19 Surveillance Team |
| EPI_ISL_3069967, | Lighthouse Lab in Alderley Park | Wellcome Sanger Institute for the COVID-19 Genomics UK (COG-UK) | Jacquelyn Wynn, Mairead Hyland, The Lighthouse Lab in Alderley Park and Alex Alderton, Roberto Amato, Jeffrey Barrett, Sonia Goncalves, Ewan Harrison, David K. Jackson, Ian Johnston, Dominic Kwiatkowski, Cordelia Langford, John Sillitoe on |

| | | | |
|---|---|---|---|
| EPI_ISL_3069968 | | Consortium | behalf of the Wellcome Sanger Institute COVID-19 Surveillance Team |
| EPI_ISL_3069969, EPI_ISL_3069970 | Lighthouse Lab in Glasgow | Wellcome Sanger Institute for the COVID-19 Genomics UK (COG-UK) Consortium | Harper VanSteenhouse, Yumi Kasai, David Gray, Carol Clugston, Anna Dominiczak and Alex Alderton, Roberto Amato, Jeffrey Barrett, Sonia Goncalves, Ewan Harrison, David K. Jackson, Ian Johnston, Dominic Kwiatkowski, Cordelia Langford, John Sillitoe on behalf of the Wellcome Sanger Institute COVID-19 Surveillance Team |
| EPI_ISL_3069971 | Lighthouse Lab in Alderley Park | Wellcome Sanger Institute for the COVID-19 Genomics UK (COG-UK) Consortium | Jacquelyn Wynn, Mairead Hyland, The Lighthouse Lab in Alderley Park and Alex Alderton, Roberto Amato, Jeffrey Barrett, Sonia Goncalves, Ewan Harrison, David K. Jackson, Ian Johnston, Dominic Kwiatkowski, Cordelia Langford, John Sillitoe on behalf of the Wellcome Sanger Institute COVID-19 Surveillance Team |
| EPI_ISL_3069972 | Lighthouse Lab in Glasgow | Wellcome Sanger Institute for the COVID-19 Genomics UK (COG-UK) Consortium | Harper VanSteenhouse, Yumi Kasai, David Gray, Carol Clugston, Anna Dominiczak and Alex Alderton, Roberto Amato, Jeffrey Barrett, Sonia Goncalves, Ewan Harrison, David K. Jackson, Ian Johnston, Dominic Kwiatkowski, Cordelia Langford, John Sillitoe on behalf of the Wellcome Sanger Institute COVID-19 Surveillance Team |
| EPI_ISL_3069973, EPI_ISL_3069974 | Lighthouse Lab in Alderley Park | Wellcome Sanger Institute for the COVID-19 Genomics UK (COG-UK) Consortium | Jacquelyn Wynn, Mairead Hyland, The Lighthouse Lab in Alderley Park and Alex Alderton, Roberto Amato, Jeffrey Barrett, Sonia Goncalves, Ewan Harrison, David K. Jackson, Ian Johnston, Dominic Kwiatkowski, Cordelia Langford, John Sillitoe on behalf of the Wellcome Sanger Institute COVID-19 Surveillance Team |
| EPI_ISL_3069975, EPI_ISL_3069976, EPI_ISL_3069977 | Lighthouse Lab in Glasgow | Wellcome Sanger Institute for the COVID-19 Genomics UK (COG-UK) Consortium | Harper VanSteenhouse, Yumi Kasai, David Gray, Carol Clugston, Anna Dominiczak and Alex Alderton, Roberto Amato, Jeffrey Barrett, Sonia Goncalves, Ewan Harrison, David K. Jackson, Ian Johnston, Dominic Kwiatkowski, Cordelia Langford, John Sillitoe on behalf of the Wellcome Sanger Institute COVID-19 Surveillance Team |
| EPI_ISL_3069978, EPI_ISL_3069979, EPI_ISL_3069980 | Berkshire and Surrey Pathology Services Lighthouse Laboratory | Wellcome Sanger Institute for the COVID-19 Genomics UK (COG-UK) Consortium | Berkshire and Surrey Pathology Services Lighthouse Laboratory and Alex Alderton, Roberto Amato, Jeffrey Barrett, Sonia Goncalves, Ewan Harrison, David K. Jackson, Ian Johnston, Dominic Kwiatkowski, Cordelia Langford, John Sillitoe on behalf of the Wellcome Sanger Institute COVID-19 Surveillance Team |
| EPI_ISL_3069981 | Lighthouse Lab in Glasgow | Wellcome Sanger Institute for the COVID-19 Genomics UK (COG-UK) Consortium | Harper VanSteenhouse, Yumi Kasai, David Gray, Carol Clugston, Anna Dominiczak and Alex Alderton, Roberto Amato, Jeffrey Barrett, Sonia Goncalves, Ewan Harrison, David K. Jackson, Ian Johnston, Dominic Kwiatkowski, Cordelia Langford, John Sillitoe on behalf of the Wellcome Sanger Institute COVID-19 Surveillance Team |
| EPI_ISL_3069982 | Lighthouse Lab in Alderley Park | Wellcome Sanger Institute for the COVID-19 Genomics UK (COG-UK) Consortium | Jacquelyn Wynn, Mairead Hyland, The Lighthouse Lab in Alderley Park and Alex Alderton, Roberto Amato, Jeffrey Barrett, Sonia Goncalves, Ewan Harrison, David K. Jackson, Ian Johnston, Dominic Kwiatkowski, Cordelia Langford, John Sillitoe on behalf of the Wellcome Sanger Institute COVID-19 Surveillance Team |
| EPI_ISL_3069983 | Berkshire and Surrey Pathology Services Lighthouse Laboratory | Wellcome Sanger Institute for the COVID-19 Genomics UK (COG-UK) Consortium | Berkshire and Surrey Pathology Services Lighthouse Laboratory and Alex Alderton, Roberto Amato, Jeffrey Barrett, Sonia Goncalves, Ewan Harrison, David K. Jackson, Ian Johnston, Dominic Kwiatkowski, Cordelia Langford, John Sillitoe on behalf of the Wellcome Sanger Institute COVID-19 Surveillance Team |
| EPI_ISL_3069984 | Lighthouse Lab in Alderley Park | Wellcome Sanger Institute for the COVID-19 Genomics UK (COG-UK) Consortium | Jacquelyn Wynn, Mairead Hyland, The Lighthouse Lab in Alderley Park and Alex Alderton, Roberto Amato, Jeffrey Barrett, Sonia Goncalves, Ewan Harrison, David K. Jackson, Ian Johnston, Dominic Kwiatkowski, Cordelia Langford, John Sillitoe on behalf of the Wellcome Sanger Institute COVID-19 Surveillance Team |
| EPI_ISL_3069985 | Berkshire and Surrey Pathology Services Lighthouse Laboratory | Wellcome Sanger Institute for the COVID-19 Genomics UK (COG-UK) Consortium | Berkshire and Surrey Pathology Services Lighthouse Laboratory and Alex Alderton, Roberto Amato, Jeffrey Barrett, Sonia Goncalves, Ewan Harrison, David K. Jackson, Ian Johnston, Dominic Kwiatkowski, Cordelia Langford, John Sillitoe on behalf of the Wellcome Sanger Institute COVID-19 Surveillance Team |
| EPI_ISL_3069986, EPI_ISL_3069987 | Lighthouse Lab in Alderley Park | Wellcome Sanger Institute for the COVID-19 Genomics UK (COG-UK) Consortium | Jacquelyn Wynn, Mairead Hyland, The Lighthouse Lab in Alderley Park and Alex Alderton, Roberto Amato, Jeffrey Barrett, Sonia Goncalves, Ewan Harrison, David K. Jackson, Ian Johnston, Dominic Kwiatkowski, Cordelia Langford, John Sillitoe on behalf of the Wellcome Sanger Institute COVID-19 Surveillance Team |
| EPI_ISL_3069988, EPI_ISL_3069989, EPI_ISL_3069990 | Lighthouse Lab in Glasgow | Wellcome Sanger Institute for the COVID-19 Genomics UK (COG-UK) Consortium | Harper VanSteenhouse, Yumi Kasai, David Gray, Carol Clugston, Anna Dominiczak and Alex Alderton, Roberto Amato, Jeffrey Barrett, Sonia Goncalves, Ewan Harrison, David K. Jackson, Ian Johnston, Dominic Kwiatkowski, Cordelia Langford, John Sillitoe on behalf of the Wellcome Sanger Institute COVID-19 Surveillance Team |
| EPI_ISL_3069991 | Berkshire and Surrey Pathology Services Lighthouse Laboratory | Wellcome Sanger Institute for the COVID-19 Genomics UK (COG-UK) Consortium | Berkshire and Surrey Pathology Services Lighthouse Laboratory and Alex Alderton, Roberto Amato, Jeffrey Barrett, Sonia Goncalves, Ewan Harrison, David K. Jackson, Ian Johnston, Dominic Kwiatkowski, Cordelia Langford, John Sillitoe on behalf of the Wellcome Sanger Institute COVID-19 Surveillance Team |
| EPI_ISL_3069992 | Lighthouse Lab in Glasgow | Wellcome Sanger Institute for the COVID-19 Genomics UK (COG-UK) Consortium | Harper VanSteenhouse, Yumi Kasai, David Gray, Carol Clugston, Anna Dominiczak and Alex Alderton, Roberto Amato, Jeffrey Barrett, Sonia Goncalves, Ewan Harrison, David K. Jackson, Ian Johnston, Dominic Kwiatkowski, Cordelia Langford, John Sillitoe on behalf of the Wellcome Sanger Institute COVID-19 Surveillance Team |
| EPI_ISL_3069993, EPI_ISL_3069994 | Lighthouse Lab in Alderley Park | Wellcome Sanger Institute for the COVID-19 Genomics UK (COG-UK) Consortium | Jacquelyn Wynn, Mairead Hyland, The Lighthouse Lab in Alderley Park and Alex Alderton, Roberto Amato, Jeffrey Barrett, Sonia Goncalves, Ewan Harrison, David K. Jackson, Ian Johnston, Dominic Kwiatkowski, Cordelia Langford, John Sillitoe on behalf of the Wellcome Sanger Institute COVID-19 Surveillance Team |
| EPI_ISL_3069995 | Berkshire and Surrey Pathology Services Lighthouse Laboratory | Wellcome Sanger Institute for the COVID-19 Genomics UK (COG-UK) Consortium | Berkshire and Surrey Pathology Services Lighthouse Laboratory and Alex Alderton, Roberto Amato, Jeffrey Barrett, Sonia Goncalves, Ewan Harrison, David K. Jackson, Ian Johnston, Dominic Kwiatkowski, Cordelia Langford, John Sillitoe on behalf of the Wellcome Sanger Institute COVID-19 Surveillance Team |
| EPI_ISL_3069996 | Lighthouse Lab in Alderley Park | Wellcome Sanger Institute for the COVID-19 Genomics UK (COG-UK) Consortium | Jacquelyn Wynn, Mairead Hyland, The Lighthouse Lab in Alderley Park and Alex Alderton, Roberto Amato, Jeffrey Barrett, Sonia Goncalves, Ewan Harrison, David K. Jackson, Ian Johnston, Dominic Kwiatkowski, Cordelia Langford, John Sillitoe on behalf of the Wellcome Sanger Institute COVID-19 Surveillance Team |
| EPI_ISL_3069997 | Berkshire and Surrey Pathology Services Lighthouse Laboratory | Wellcome Sanger Institute for the COVID-19 Genomics UK (COG-UK) Consortium | Berkshire and Surrey Pathology Services Lighthouse Laboratory and Alex Alderton, Roberto Amato, Jeffrey Barrett, Sonia Goncalves, Ewan Harrison, David K. Jackson, Ian Johnston, Dominic Kwiatkowski, Cordelia Langford, John Sillitoe on behalf of the Wellcome Sanger Institute COVID-19 Surveillance Team |
| EPI_ISL_3069998, EPI_ISL_3069999 | Lighthouse Lab in Glasgow | Wellcome Sanger Institute for the COVID-19 Genomics UK (COG-UK) Consortium | Harper VanSteenhouse, Yumi Kasai, David Gray, Carol Clugston, Anna Dominiczak and Alex Alderton, Roberto Amato, Jeffrey Barrett, Sonia Goncalves, Ewan Harrison, David K. Jackson, Ian Johnston, Dominic Kwiatkowski, Cordelia Langford, John Sillitoe on behalf of the Wellcome Sanger Institute COVID-19 Surveillance Team |
| EPI_ISL_3070000, EPI_ISL_3070001 | Berkshire and Surrey Pathology Services Lighthouse Laboratory | Wellcome Sanger Institute for the COVID-19 Genomics UK (COG-UK) Consortium | Berkshire and Surrey Pathology Services Lighthouse Laboratory and Alex Alderton, Roberto Amato, Jeffrey Barrett, Sonia Goncalves, Ewan Harrison, David K. Jackson, Ian Johnston, Dominic Kwiatkowski, Cordelia Langford, John Sillitoe on behalf of the Wellcome Sanger Institute COVID-19 Surveillance Team |
| EPI_ISL_3070002 | Lighthouse Lab in Glasgow | Wellcome Sanger Institute for the COVID-19 Genomics UK (COG-UK) Consortium | Harper VanSteenhouse, Yumi Kasai, David Gray, Carol Clugston, Anna Dominiczak and Alex Alderton, Roberto Amato, Jeffrey Barrett, Sonia Goncalves, Ewan Harrison, David K. Jackson, Ian Johnston, Dominic Kwiatkowski, Cordelia Langford, John Sillitoe on behalf of the Wellcome Sanger Institute COVID-19 Surveillance Team |
| EPI_ISL_3070003 | Berkshire and Surrey Pathology Services Lighthouse Laboratory | Wellcome Sanger Institute for the COVID-19 Genomics UK (COG-UK) Consortium | Berkshire and Surrey Pathology Services Lighthouse Laboratory and Alex Alderton, Roberto Amato, Jeffrey Barrett, Sonia Goncalves, Ewan Harrison, David K. Jackson, Ian Johnston, Dominic Kwiatkowski, Cordelia Langford, John Sillitoe on behalf of the Wellcome Sanger Institute COVID-19 Surveillance Team |
| EPI_ISL_3070004, EPI_ISL_3070005 | Lighthouse Lab in Alderley Park | Wellcome Sanger Institute for the COVID-19 Genomics UK (COG-UK) Consortium | Jacquelyn Wynn, Mairead Hyland, The Lighthouse Lab in Alderley Park and Alex Alderton, Roberto Amato, Jeffrey Barrett, Sonia Goncalves, Ewan Harrison, David K. Jackson, Ian Johnston, Dominic Kwiatkowski, Cordelia Langford, John Sillitoe on behalf of the Wellcome Sanger Institute COVID-19 Surveillance Team |
| EPI_ISL_3070006, EPI_ISL_3070007 | Lighthouse Lab in Glasgow | Wellcome Sanger Institute for the COVID-19 Genomics UK (COG-UK) Consortium | Harper VanSteenhouse, Yumi Kasai, David Gray, Carol Clugston, Anna Dominiczak and Alex Alderton, Roberto Amato, Jeffrey Barrett, Sonia Goncalves, Ewan Harrison, David K. Jackson, Ian Johnston, Dominic Kwiatkowski, Cordelia Langford, John Sillitoe on behalf of the Wellcome Sanger Institute COVID-19 Surveillance Team |
| EPI_ISL_3070008 | Lighthouse Lab in Alderley Park | Wellcome Sanger Institute for the COVID-19 Genomics UK (COG-UK) Consortium | Jacquelyn Wynn, Mairead Hyland, The Lighthouse Lab in Alderley Park and Alex Alderton, Roberto Amato, Jeffrey Barrett, Sonia Goncalves, Ewan Harrison, David K. Jackson, Ian Johnston, Dominic Kwiatkowski, Cordelia Langford, John Sillitoe on behalf of the Wellcome Sanger Institute COVID-19 Surveillance Team |
| EPI_ISL_3070009 | Lighthouse Lab in Glasgow | Wellcome Sanger Institute for the COVID-19 Genomics UK (COG-UK) Consortium | Harper VanSteenhouse, Yumi Kasai, David Gray, Carol Clugston, Anna Dominiczak and Alex Alderton, Roberto Amato, Jeffrey Barrett, Sonia Goncalves, Ewan Harrison, David K. Jackson, Ian Johnston, Dominic Kwiatkowski, Cordelia Langford, John Sillitoe on behalf of the Wellcome Sanger Institute COVID-19 Surveillance Team |
| EPI_ISL_3070010, EPI_ISL_3070011, EPI_ISL_3070012 | Berkshire and Surrey Pathology Services Lighthouse Laboratory | Wellcome Sanger Institute for the COVID-19 Genomics UK (COG-UK) Consortium | Berkshire and Surrey Pathology Services Lighthouse Laboratory and Alex Alderton, Roberto Amato, Jeffrey Barrett, Sonia Goncalves, Ewan Harrison, David K. Jackson, Ian Johnston, Dominic Kwiatkowski, Cordelia Langford, John Sillitoe on behalf of the Wellcome Sanger Institute COVID-19 Surveillance Team |
| EPI_ISL_3070013, EPI_ISL_3070014 | Lighthouse Lab in Glasgow | Wellcome Sanger Institute for the COVID-19 Genomics UK (COG-UK) Consortium | Harper VanSteenhouse, Yumi Kasai, David Gray, Carol Clugston, Anna Dominiczak and Alex Alderton, Roberto Amato, Jeffrey Barrett, Sonia Goncalves, Ewan Harrison, David K. Jackson, Ian Johnston, Dominic Kwiatkowski, Cordelia Langford, John Sillitoe on behalf of the Wellcome Sanger Institute COVID-19 Surveillance Team |
| EPI_ISL_3070015 | Lighthouse Lab in Alderley Park | Wellcome Sanger Institute for the COVID-19 Genomics UK (COG-UK) Consortium | Jacquelyn Wynn, Mairead Hyland, The Lighthouse Lab in Alderley Park and Alex Alderton, Roberto Amato, Jeffrey Barrett, Sonia Goncalves, Ewan Harrison, David K. Jackson, Ian Johnston, Dominic Kwiatkowski, Cordelia Langford, John Sillitoe on behalf of the Wellcome Sanger Institute COVID-19 Surveillance Team |
| EPI_ISL_3070016 | Berkshire and Surrey Pathology Services Lighthouse Laboratory | Wellcome Sanger Institute for the COVID-19 Genomics UK (COG-UK) Consortium | Berkshire and Surrey Pathology Services Lighthouse Laboratory and Alex Alderton, Roberto Amato, Jeffrey Barrett, Sonia Goncalves, Ewan Harrison, David K. Jackson, Ian Johnston, Dominic Kwiatkowski, Cordelia Langford, John Sillitoe on behalf of the Wellcome Sanger Institute COVID-19 Surveillance Team |
| EPI_ISL_3070017, EPI_ISL_3070018 | Lighthouse Lab in Glasgow | Wellcome Sanger Institute for the COVID-19 Genomics UK (COG-UK) Consortium | Harper VanSteenhouse, Yumi Kasai, David Gray, Carol Clugston, Anna Dominiczak and Alex Alderton, Roberto Amato, Jeffrey Barrett, Sonia Goncalves, Ewan Harrison, David K. Jackson, Ian Johnston, Dominic Kwiatkowski, Cordelia Langford, John Sillitoe on behalf of the Wellcome Sanger Institute COVID-19 Surveillance Team |
| EPI_ISL_3070019, EPI_ISL_3070020 | Lighthouse Lab in Alderley Park | Wellcome Sanger Institute for the COVID-19 Genomics UK (COG-UK) Consortium | Jacquelyn Wynn, Mairead Hyland, The Lighthouse Lab in Alderley Park and Alex Alderton, Roberto Amato, Jeffrey Barrett, Sonia Goncalves, Ewan Harrison, David K. Jackson, Ian Johnston, Dominic Kwiatkowski, Cordelia Langford, John Sillitoe on behalf of the Wellcome Sanger Institute COVID-19 Surveillance Team |
| EPI_ISL_3070021 | Berkshire and Surrey Pathology Services Lighthouse Laboratory | Wellcome Sanger Institute for the COVID-19 Genomics UK (COG-UK) Consortium | Berkshire and Surrey Pathology Services Lighthouse Laboratory and Alex Alderton, Roberto Amato, Jeffrey Barrett, Sonia Goncalves, Ewan Harrison, David K. Jackson, Ian Johnston, Dominic Kwiatkowski, Cordelia Langford, John Sillitoe on behalf of the Wellcome Sanger Institute COVID-19 Surveillance Team |
| EPI_ISL_3070022, EPI_ISL_3070023, EPI_ISL_3070024 | Lighthouse Lab in Alderley Park | Wellcome Sanger Institute for the COVID-19 Genomics UK (COG-UK) Consortium | Jacquelyn Wynn, Mairead Hyland, The Lighthouse Lab in Alderley Park and Alex Alderton, Roberto Amato, Jeffrey Barrett, Sonia Goncalves, Ewan Harrison, David K. Jackson, Ian Johnston, Dominic Kwiatkowski, Cordelia Langford, John Sillitoe on behalf of the Wellcome Sanger Institute COVID-19 Surveillance Team |
| EPI_ISL_3070025 | Berkshire and Surrey Pathology Services Lighthouse Laboratory | Wellcome Sanger Institute for the COVID-19 Genomics UK (COG-UK) Consortium | Berkshire and Surrey Pathology Services Lighthouse Laboratory and Alex Alderton, Roberto Amato, Jeffrey Barrett, Sonia Goncalves, Ewan Harrison, David K. Jackson, Ian Johnston, Dominic Kwiatkowski, Cordelia Langford, John Sillitoe on behalf of the Wellcome Sanger Institute COVID-19 Surveillance Team |
| EPI_ISL_3070026 | Lighthouse Lab in Glasgow | Wellcome Sanger Institute for the COVID-19 Genomics UK (COG-UK) Consortium | Harper VanSteenhouse, Yumi Kasai, David Gray, Carol Clugston, Anna Dominiczak and Alex Alderton, Roberto Amato, Jeffrey Barrett, Sonia Goncalves, Ewan Harrison, David K. Jackson, Ian Johnston, Dominic Kwiatkowski, Cordelia Langford, John Sillitoe on behalf of the Wellcome Sanger Institute COVID-19 Surveillance Team |
| EPI_ISL_3070027 | Lighthouse Lab in Alderley Park | Wellcome Sanger Institute for the COVID-19 Genomics UK (COG-UK) Consortium | Jacquelyn Wynn, Mairead Hyland, The Lighthouse Lab in Alderley Park and Alex Alderton, Roberto Amato, Jeffrey Barrett, Sonia Goncalves, Ewan Harrison, David K. Jackson, Ian Johnston, Dominic Kwiatkowski, Cordelia Langford, John Sillitoe on behalf of the Wellcome Sanger Institute COVID-19 Surveillance Team |
| EPI_ISL_3070028 | Lighthouse Lab in Glasgow | Wellcome Sanger Institute for the COVID-19 Genomics UK (COG-UK) Consortium | Harper VanSteenhouse, Yumi Kasai, David Gray, Carol Clugston, Anna Dominiczak and Alex Alderton, Roberto Amato, Jeffrey Barrett, Sonia Goncalves, Ewan Harrison, David K. Jackson, Ian Johnston, Dominic Kwiatkowski, Cordelia Langford, John Sillitoe on behalf of the Wellcome Sanger Institute COVID-19 Surveillance Team |
| EPI_ISL_3070029 | Berkshire and Surrey Pathology Services Lighthouse Laboratory | Wellcome Sanger Institute for the COVID-19 Genomics UK (COG-UK) Consortium | Berkshire and Surrey Pathology Services Lighthouse Laboratory and Alex Alderton, Roberto Amato, Jeffrey Barrett, Sonia Goncalves, Ewan Harrison, David K. Jackson, Ian Johnston, Dominic Kwiatkowski, Cordelia Langford, John Sillitoe on behalf of the Wellcome Sanger Institute COVID-19 Surveillance Team |
| EPI_ISL_3070030 | Lighthouse Lab in Alderley Park | Wellcome Sanger Institute for the COVID-19 Genomics UK (COG-UK) Consortium | Jacquelyn Wynn, Mairead Hyland, The Lighthouse Lab in Alderley Park and Alex Alderton, Roberto Amato, Jeffrey Barrett, Sonia Goncalves, Ewan Harrison, David K. Jackson, Ian Johnston, Dominic Kwiatkowski, Cordelia Langford, John Sillitoe on behalf of the Wellcome Sanger Institute COVID-19 Surveillance Team |
| EPI_ISL_3070031 | Berkshire and Surrey Pathology Services Lighthouse Laboratory | Wellcome Sanger Institute for the COVID-19 Genomics UK (COG-UK) Consortium | Berkshire and Surrey Pathology Services Lighthouse Laboratory and Alex Alderton, Roberto Amato, Jeffrey Barrett, Sonia Goncalves, Ewan Harrison, David K. Jackson, Ian Johnston, Dominic Kwiatkowski, Cordelia Langford, John Sillitoe on behalf of the Wellcome Sanger Institute COVID-19 Surveillance Team |
| EPI_ISL_3070032 | Lighthouse Lab in Alderley Park | Wellcome Sanger Institute for the COVID-19 Genomics UK (COG-UK) | Jacquelyn Wynn, Mairead Hyland, The Lighthouse Lab in Alderley Park and Alex Alderton, Roberto Amato, Jeffrey Barrett, Sonia Goncalves, Ewan Harrison, David K. Jackson, Ian Johnston, Dominic Kwiatkowski, Cordelia Langford, John Sillitoe on |

| | | | |
|---|---|---|---|
| | | Consortium | behalf of the Wellcome Sanger Institute COVID-19 Surveillance Team |
| EPI_ISL_3070033 | Berkshire and Surrey Pathology Services Lighthouse Laboratory | Wellcome Sanger Institute for the COVID-19 Genomics UK (COG-UK) Consortium | Berkshire and Surrey Pathology Services Lighthouse Laboratory and Alex Alderton, Roberto Amato, Jeffrey Barrett, Sonia Goncalves, Ewan Harrison, David K. Jackson, Ian Johnston, Dominic Kwiatkowski, Cordelia Langford, John Sillitoe on behalf of the Wellcome Sanger Institute COVID-19 Surveillance Team |
| EPI_ISL_3070034, EPI_ISL_3070035, EPI_ISL_3070036, EPI_ISL_3070037 | Lighthouse Lab in Glasgow | Wellcome Sanger Institute for the COVID-19 Genomics UK (COG-UK) Consortium | Harper VanSteenhouse, Yumi Kasai, David Gray, Carol Clugston, Anna Dominiczak and Alex Alderton, Roberto Amato, Jeffrey Barrett, Sonia Goncalves, Ewan Harrison, David K. Jackson, Ian Johnston, Dominic Kwiatkowski, Cordelia Langford, John Sillitoe on behalf of the Wellcome Sanger Institute COVID-19 Surveillance Team |
| EPI_ISL_3070038, EPI_ISL_3070039, EPI_ISL_3070040 | Berkshire and Surrey Pathology Services Lighthouse Laboratory | Wellcome Sanger Institute for the COVID-19 Genomics UK (COG-UK) Consortium | Berkshire and Surrey Pathology Services Lighthouse Laboratory and Alex Alderton, Roberto Amato, Jeffrey Barrett, Sonia Goncalves, Ewan Harrison, David K. Jackson, Ian Johnston, Dominic Kwiatkowski, Cordelia Langford, John Sillitoe on behalf of the Wellcome Sanger Institute COVID-19 Surveillance Team |
| EPI_ISL_3070041 | Lighthouse Lab in Glasgow | Wellcome Sanger Institute for the COVID-19 Genomics UK (COG-UK) Consortium | Harper VanSteenhouse, Yumi Kasai, David Gray, Carol Clugston, Anna Dominiczak and Alex Alderton, Roberto Amato, Jeffrey Barrett, Sonia Goncalves, Ewan Harrison, David K. Jackson, Ian Johnston, Dominic Kwiatkowski, Cordelia Langford, John Sillitoe on behalf of the Wellcome Sanger Institute COVID-19 Surveillance Team |
| EPI_ISL_3070042 | Lighthouse Lab in Alderley Park | Wellcome Sanger Institute for the COVID-19 Genomics UK (COG-UK) Consortium | Jacquelyn Wynn, Mairead Hyland, The Lighthouse Lab in Alderley Park and Alex Alderton, Roberto Amato, Jeffrey Barrett, Sonia Goncalves, Ewan Harrison, David K. Jackson, Ian Johnston, Dominic Kwiatkowski, Cordelia Langford, John Sillitoe on behalf of the Wellcome Sanger Institute COVID-19 Surveillance Team |
| EPI_ISL_3070043, EPI_ISL_3070044 | Lighthouse Lab in Glasgow | Wellcome Sanger Institute for the COVID-19 Genomics UK (COG-UK) Consortium | Harper VanSteenhouse, Yumi Kasai, David Gray, Carol Clugston, Anna Dominiczak and Alex Alderton, Roberto Amato, Jeffrey Barrett, Sonia Goncalves, Ewan Harrison, David K. Jackson, Ian Johnston, Dominic Kwiatkowski, Cordelia Langford, John Sillitoe on behalf of the Wellcome Sanger Institute COVID-19 Surveillance Team |
| EPI_ISL_3070045, EPI_ISL_3070046 | Lighthouse Lab in Alderley Park | Wellcome Sanger Institute for the COVID-19 Genomics UK (COG-UK) Consortium | Jacquelyn Wynn, Mairead Hyland, The Lighthouse Lab in Alderley Park and Alex Alderton, Roberto Amato, Jeffrey Barrett, Sonia Goncalves, Ewan Harrison, David K. Jackson, Ian Johnston, Dominic Kwiatkowski, Cordelia Langford, John Sillitoe on behalf of the Wellcome Sanger Institute COVID-19 Surveillance Team |
| EPI_ISL_3070047, EPI_ISL_3070048 | Lighthouse Lab in Glasgow | Wellcome Sanger Institute for the COVID-19 Genomics UK (COG-UK) Consortium | Harper VanSteenhouse, Yumi Kasai, David Gray, Carol Clugston, Anna Dominiczak and Alex Alderton, Roberto Amato, Jeffrey Barrett, Sonia Goncalves, Ewan Harrison, David K. Jackson, Ian Johnston, Dominic Kwiatkowski, Cordelia Langford, John Sillitoe on behalf of the Wellcome Sanger Institute COVID-19 Surveillance Team |
| EPI_ISL_3070049 | Berkshire and Surrey Pathology Services Lighthouse Laboratory | Wellcome Sanger Institute for the COVID-19 Genomics UK (COG-UK) Consortium | Berkshire and Surrey Pathology Services Lighthouse Laboratory and Alex Alderton, Roberto Amato, Jeffrey Barrett, Sonia Goncalves, Ewan Harrison, David K. Jackson, Ian Johnston, Dominic Kwiatkowski, Cordelia Langford, John Sillitoe on behalf of the Wellcome Sanger Institute COVID-19 Surveillance Team |
| EPI_ISL_3070050, EPI_ISL_3070051, EPI_ISL_3070052, EPI_ISL_3070053, EPI_ISL_3070054, EPI_ISL_3070055 | Lighthouse Lab in Glasgow | Wellcome Sanger Institute for the COVID-19 Genomics UK (COG-UK) Consortium | Harper VanSteenhouse, Yumi Kasai, David Gray, Carol Clugston, Anna Dominiczak and Alex Alderton, Roberto Amato, Jeffrey Barrett, Sonia Goncalves, Ewan Harrison, David K. Jackson, Ian Johnston, Dominic Kwiatkowski, Cordelia Langford, John Sillitoe on behalf of the Wellcome Sanger Institute COVID-19 Surveillance Team |
| EPI_ISL_3070056 | Lighthouse Lab in Alderley Park | Wellcome Sanger Institute for the COVID-19 Genomics UK (COG-UK) Consortium | Jacquelyn Wynn, Mairead Hyland, The Lighthouse Lab in Alderley Park and Alex Alderton, Roberto Amato, Jeffrey Barrett, Sonia Goncalves, Ewan Harrison, David K. Jackson, Ian Johnston, Dominic Kwiatkowski, Cordelia Langford, John Sillitoe on behalf of the Wellcome Sanger Institute COVID-19 Surveillance Team |
| EPI_ISL_3070057 | Lighthouse Lab in Glasgow | Wellcome Sanger Institute for the COVID-19 Genomics UK (COG-UK) Consortium | Harper VanSteenhouse, Yumi Kasai, David Gray, Carol Clugston, Anna Dominiczak and Alex Alderton, Roberto Amato, Jeffrey Barrett, Sonia Goncalves, Ewan Harrison, David K. Jackson, Ian Johnston, Dominic Kwiatkowski, Cordelia Langford, John Sillitoe on behalf of the Wellcome Sanger Institute COVID-19 Surveillance Team |
| EPI_ISL_3070058 | Berkshire and Surrey Pathology Services Lighthouse Laboratory | Wellcome Sanger Institute for the COVID-19 Genomics UK (COG-UK) Consortium | Berkshire and Surrey Pathology Services Lighthouse Laboratory and Alex Alderton, Roberto Amato, Jeffrey Barrett, Sonia Goncalves, Ewan Harrison, David K. Jackson, Ian Johnston, Dominic Kwiatkowski, Cordelia Langford, John Sillitoe on behalf of the Wellcome Sanger Institute COVID-19 Surveillance Team |
| EPI_ISL_3070059 | Lighthouse Lab in Alderley Park | Wellcome Sanger Institute for the COVID-19 Genomics UK (COG-UK) Consortium | Jacquelyn Wynn, Mairead Hyland, The Lighthouse Lab in Alderley Park and Alex Alderton, Roberto Amato, Jeffrey Barrett, Sonia Goncalves, Ewan Harrison, David K. Jackson, Ian Johnston, Dominic Kwiatkowski, Cordelia Langford, John Sillitoe on behalf of the Wellcome Sanger Institute COVID-19 Surveillance Team |
| EPI_ISL_3070060 | Berkshire and Surrey Pathology Services Lighthouse Laboratory | Wellcome Sanger Institute for the COVID-19 Genomics UK (COG-UK) Consortium | Berkshire and Surrey Pathology Services Lighthouse Laboratory and Alex Alderton, Roberto Amato, Jeffrey Barrett, Sonia Goncalves, Ewan Harrison, David K. Jackson, Ian Johnston, Dominic Kwiatkowski, Cordelia Langford, John Sillitoe on behalf of the Wellcome Sanger Institute COVID-19 Surveillance Team |
| EPI_ISL_3070061 | Lighthouse Lab in Alderley Park | Wellcome Sanger Institute for the COVID-19 Genomics UK (COG-UK) Consortium | Jacquelyn Wynn, Mairead Hyland, The Lighthouse Lab in Alderley Park and Alex Alderton, Roberto Amato, Jeffrey Barrett, Sonia Goncalves, Ewan Harrison, David K. Jackson, Ian Johnston, Dominic Kwiatkowski, Cordelia Langford, John Sillitoe on behalf of the Wellcome Sanger Institute COVID-19 Surveillance Team |
| EPI_ISL_3070062 | Lighthouse Lab in Glasgow | Wellcome Sanger Institute for the COVID-19 Genomics UK (COG-UK) Consortium | Harper VanSteenhouse, Yumi Kasai, David Gray, Carol Clugston, Anna Dominiczak and Alex Alderton, Roberto Amato, Jeffrey Barrett, Sonia Goncalves, Ewan Harrison, David K. Jackson, Ian Johnston, Dominic Kwiatkowski, Cordelia Langford, John Sillitoe on behalf of the Wellcome Sanger Institute COVID-19 Surveillance Team |
| EPI_ISL_3070063, EPI_ISL_3070064 | Lighthouse Lab in Alderley Park | Wellcome Sanger Institute for the COVID-19 Genomics UK (COG-UK) Consortium | Jacquelyn Wynn, Mairead Hyland, The Lighthouse Lab in Alderley Park and Alex Alderton, Roberto Amato, Jeffrey Barrett, Sonia Goncalves, Ewan Harrison, David K. Jackson, Ian Johnston, Dominic Kwiatkowski, Cordelia Langford, John Sillitoe on behalf of the Wellcome Sanger Institute COVID-19 Surveillance Team |
| EPI_ISL_3070065 | Berkshire and Surrey Pathology Services Lighthouse Laboratory | Wellcome Sanger Institute for the COVID-19 Genomics UK (COG-UK) Consortium | Berkshire and Surrey Pathology Services Lighthouse Laboratory and Alex Alderton, Roberto Amato, Jeffrey Barrett, Sonia Goncalves, Ewan Harrison, David K. Jackson, Ian Johnston, Dominic Kwiatkowski, Cordelia Langford, John Sillitoe on behalf of the Wellcome Sanger Institute COVID-19 Surveillance Team |
| EPI_ISL_3070066, EPI_ISL_3070067 | Lighthouse Lab in Glasgow | Wellcome Sanger Institute for the COVID-19 Genomics UK (COG-UK) Consortium | Harper VanSteenhouse, Yumi Kasai, David Gray, Carol Clugston, Anna Dominiczak and Alex Alderton, Roberto Amato, Jeffrey Barrett, Sonia Goncalves, Ewan Harrison, David K. Jackson, Ian Johnston, Dominic Kwiatkowski, Cordelia Langford, John Sillitoe on behalf of the Wellcome Sanger Institute COVID-19 Surveillance Team |
| EPI_ISL_3070068, EPI_ISL_3070069 | Lighthouse Lab in Alderley Park | Wellcome Sanger Institute for the COVID-19 Genomics UK (COG-UK) Consortium | Jacquelyn Wynn, Mairead Hyland, The Lighthouse Lab in Alderley Park and Alex Alderton, Roberto Amato, Jeffrey Barrett, Sonia Goncalves, Ewan Harrison, David K. Jackson, Ian Johnston, Dominic Kwiatkowski, Cordelia Langford, John Sillitoe on behalf of the Wellcome Sanger Institute COVID-19 Surveillance Team |
| EPI_ISL_3070070 | Berkshire and Surrey Pathology Services Lighthouse Laboratory | Wellcome Sanger Institute for the COVID-19 Genomics UK (COG-UK) Consortium | Berkshire and Surrey Pathology Services Lighthouse Laboratory and Alex Alderton, Roberto Amato, Jeffrey Barrett, Sonia Goncalves, Ewan Harrison, David K. Jackson, Ian Johnston, Dominic Kwiatkowski, Cordelia Langford, John Sillitoe on behalf of the Wellcome Sanger Institute COVID-19 Surveillance Team |
| EPI_ISL_3070071 | Lighthouse Laboratory Plymouth | Wellcome Sanger Institute for the COVID-19 Genomics UK (COG-UK) Consortium | Lighthouse Laboratory Plymouth and Alex Alderton, Roberto Amato, Jeffrey Barrett, Sonia Goncalves, Ewan Harrison, David K. Jackson, Ian Johnston, Dominic Kwiatkowski, Cordelia Langford, John Sillitoe on behalf of the Wellcome Sanger Institute COVID-19 Surveillance Team |
| EPI_ISL_3070072 | Berkshire and Surrey Pathology Services Lighthouse Laboratory | Wellcome Sanger Institute for the COVID-19 Genomics UK (COG-UK) Consortium | Berkshire and Surrey Pathology Services Lighthouse Laboratory and Alex Alderton, Roberto Amato, Jeffrey Barrett, Sonia Goncalves, Ewan Harrison, David K. Jackson, Ian Johnston, Dominic Kwiatkowski, Cordelia Langford, John Sillitoe on behalf of the Wellcome Sanger Institute COVID-19 Surveillance Team |
| EPI_ISL_3070073 | Lighthouse Lab in Alderley Park | Wellcome Sanger Institute for the COVID-19 Genomics UK (COG-UK) Consortium | Jacquelyn Wynn, Mairead Hyland, The Lighthouse Lab in Alderley Park and Alex Alderton, Roberto Amato, Jeffrey Barrett, Sonia Goncalves, Ewan Harrison, David K. Jackson, Ian Johnston, Dominic Kwiatkowski, Cordelia Langford, John Sillitoe on behalf of the Wellcome Sanger Institute COVID-19 Surveillance Team |
| EPI_ISL_3070074, EPI_ISL_3070075 | Lighthouse Laboratory Plymouth | Wellcome Sanger Institute for the COVID-19 Genomics UK (COG-UK) Consortium | Lighthouse Laboratory Plymouth and Alex Alderton, Roberto Amato, Jeffrey Barrett, Sonia Goncalves, Ewan Harrison, David K. Jackson, Ian Johnston, Dominic Kwiatkowski, Cordelia Langford, John Sillitoe on behalf of the Wellcome Sanger Institute COVID-19 Surveillance Team |
| EPI_ISL_3070076, EPI_ISL_3070077 | Berkshire and Surrey Pathology Services Lighthouse Laboratory | Wellcome Sanger Institute for the COVID-19 Genomics UK (COG-UK) Consortium | Berkshire and Surrey Pathology Services Lighthouse Laboratory and Alex Alderton, Roberto Amato, Jeffrey Barrett, Sonia Goncalves, Ewan Harrison, David K. Jackson, Ian Johnston, Dominic Kwiatkowski, Cordelia Langford, John Sillitoe on behalf of the Wellcome Sanger Institute COVID-19 Surveillance Team |
| EPI_ISL_3070078 | Lighthouse Lab in Alderley Park | Wellcome Sanger Institute for the COVID-19 Genomics UK (COG-UK) Consortium | Jacquelyn Wynn, Mairead Hyland, The Lighthouse Lab in Alderley Park and Alex Alderton, Roberto Amato, Jeffrey Barrett, Sonia Goncalves, Ewan Harrison, David K. Jackson, Ian Johnston, Dominic Kwiatkowski, Cordelia Langford, John Sillitoe on behalf of the Wellcome Sanger Institute COVID-19 Surveillance Team |
| EPI_ISL_3070079 | Berkshire and Surrey Pathology Services Lighthouse Laboratory | Wellcome Sanger Institute for the COVID-19 Genomics UK (COG-UK) Consortium | Berkshire and Surrey Pathology Services Lighthouse Laboratory and Alex Alderton, Roberto Amato, Jeffrey Barrett, Sonia Goncalves, Ewan Harrison, David K. Jackson, Ian Johnston, Dominic Kwiatkowski, Cordelia Langford, John Sillitoe on behalf of the Wellcome Sanger Institute COVID-19 Surveillance Team |
| EPI_ISL_3070080, EPI_ISL_3070081 | Lighthouse Lab in Alderley Park | Wellcome Sanger Institute for the COVID-19 Genomics UK (COG-UK) Consortium | Jacquelyn Wynn, Mairead Hyland, The Lighthouse Lab in Alderley Park and Alex Alderton, Roberto Amato, Jeffrey Barrett, Sonia Goncalves, Ewan Harrison, David K. Jackson, Ian Johnston, Dominic Kwiatkowski, Cordelia Langford, John Sillitoe on behalf of the Wellcome Sanger Institute COVID-19 Surveillance Team |
| EPI_ISL_3070082 | Berkshire and Surrey Pathology Services Lighthouse Laboratory | Wellcome Sanger Institute for the COVID-19 Genomics UK (COG-UK) Consortium | Berkshire and Surrey Pathology Services Lighthouse Laboratory and Alex Alderton, Roberto Amato, Jeffrey Barrett, Sonia Goncalves, Ewan Harrison, David K. Jackson, Ian Johnston, Dominic Kwiatkowski, Cordelia Langford, John Sillitoe on behalf of the Wellcome Sanger Institute COVID-19 Surveillance Team |
| EPI_ISL_3070083, EPI_ISL_3070084 | Lighthouse Lab in Alderley Park | Wellcome Sanger Institute for the COVID-19 Genomics UK (COG-UK) Consortium | Jacquelyn Wynn, Mairead Hyland, The Lighthouse Lab in Alderley Park and Alex Alderton, Roberto Amato, Jeffrey Barrett, Sonia Goncalves, Ewan Harrison, David K. Jackson, Ian Johnston, Dominic Kwiatkowski, Cordelia Langford, John Sillitoe on behalf of the Wellcome Sanger Institute COVID-19 Surveillance Team |
| EPI_ISL_3070085, EPI_ISL_3070086 | Berkshire and Surrey Pathology Services Lighthouse Laboratory | Wellcome Sanger Institute for the COVID-19 Genomics UK (COG-UK) Consortium | Berkshire and Surrey Pathology Services Lighthouse Laboratory and Alex Alderton, Roberto Amato, Jeffrey Barrett, Sonia Goncalves, Ewan Harrison, David K. Jackson, Ian Johnston, Dominic Kwiatkowski, Cordelia Langford, John Sillitoe on behalf of the Wellcome Sanger Institute COVID-19 Surveillance Team |
| EPI_ISL_3070088 | Lighthouse Laboratory Plymouth | Wellcome Sanger Institute for the COVID-19 Genomics UK (COG-UK) Consortium | Lighthouse Laboratory Plymouth and Alex Alderton, Roberto Amato, Jeffrey Barrett, Sonia Goncalves, Ewan Harrison, David K. Jackson, Ian Johnston, Dominic Kwiatkowski, Cordelia Langford, John Sillitoe on behalf of the Wellcome Sanger Institute COVID-19 Surveillance Team |
| EPI_ISL_3070089 | Berkshire and Surrey Pathology Services Lighthouse Laboratory | Wellcome Sanger Institute for the COVID-19 Genomics UK (COG-UK) Consortium | Berkshire and Surrey Pathology Services Lighthouse Laboratory and Alex Alderton, Roberto Amato, Jeffrey Barrett, Sonia Goncalves, Ewan Harrison, David K. Jackson, Ian Johnston, Dominic Kwiatkowski, Cordelia Langford, John Sillitoe on behalf of the Wellcome Sanger Institute COVID-19 Surveillance Team |
| EPI_ISL_3070090 | Lighthouse Laboratory Plymouth | Wellcome Sanger Institute for the COVID-19 Genomics UK (COG-UK) Consortium | Lighthouse Laboratory Plymouth and Alex Alderton, Roberto Amato, Jeffrey Barrett, Sonia Goncalves, Ewan Harrison, David K. Jackson, Ian Johnston, Dominic Kwiatkowski, Cordelia Langford, John Sillitoe on behalf of the Wellcome Sanger Institute COVID-19 Surveillance Team |
| EPI_ISL_3070091 | Berkshire and Surrey Pathology Services Lighthouse Laboratory | Wellcome Sanger Institute for the COVID-19 Genomics UK (COG-UK) Consortium | Berkshire and Surrey Pathology Services Lighthouse Laboratory and Alex Alderton, Roberto Amato, Jeffrey Barrett, Sonia Goncalves, Ewan Harrison, David K. Jackson, Ian Johnston, Dominic Kwiatkowski, Cordelia Langford, John Sillitoe on behalf of the Wellcome Sanger Institute COVID-19 Surveillance Team |
| EPI_ISL_3070092 | Lighthouse Lab in Alderley Park | Wellcome Sanger Institute for the COVID-19 Genomics UK (COG-UK) Consortium | Jacquelyn Wynn, Mairead Hyland, The Lighthouse Lab in Alderley Park and Alex Alderton, Roberto Amato, Jeffrey Barrett, Sonia Goncalves, Ewan Harrison, David K. Jackson, Ian Johnston, Dominic Kwiatkowski, Cordelia Langford, John Sillitoe on behalf of the Wellcome Sanger Institute COVID-19 Surveillance Team |
| EPI_ISL_3070093 | Berkshire and Surrey Pathology Services Lighthouse Laboratory | Wellcome Sanger Institute for the COVID-19 Genomics UK (COG-UK) Consortium | Berkshire and Surrey Pathology Services Lighthouse Laboratory and Alex Alderton, Roberto Amato, Jeffrey Barrett, Sonia Goncalves, Ewan Harrison, David K. Jackson, Ian Johnston, Dominic Kwiatkowski, Cordelia Langford, John Sillitoe on behalf of the Wellcome Sanger Institute COVID-19 Surveillance Team |
| EPI_ISL_3070094 | Lighthouse Laboratory Plymouth | Wellcome Sanger Institute for the COVID-19 Genomics UK (COG-UK) Consortium | Lighthouse Laboratory Plymouth and Alex Alderton, Roberto Amato, Jeffrey Barrett, Sonia Goncalves, Ewan Harrison, David K. Jackson, Ian Johnston, Dominic Kwiatkowski, Cordelia Langford, John Sillitoe on behalf of the Wellcome Sanger Institute COVID-19 Surveillance Team |
| EPI_ISL_3070095 | Berkshire and Surrey Pathology Services Lighthouse Laboratory | Wellcome Sanger Institute for the COVID-19 Genomics UK (COG-UK) Consortium | Berkshire and Surrey Pathology Services Lighthouse Laboratory and Alex Alderton, Roberto Amato, Jeffrey Barrett, Sonia Goncalves, Ewan Harrison, David K. Jackson, Ian Johnston, Dominic Kwiatkowski, Cordelia Langford, John Sillitoe on behalf of the Wellcome Sanger Institute COVID-19 Surveillance Team |
| EPI_ISL_3070096 | Lighthouse Lab in Alderley Park | Wellcome Sanger Institute for the COVID-19 Genomics UK (COG-UK) Consortium | Jacquelyn Wynn, Mairead Hyland, The Lighthouse Lab in Alderley Park and Alex Alderton, Roberto Amato, Jeffrey Barrett, Sonia Goncalves, Ewan Harrison, David K. Jackson, Ian Johnston, Dominic Kwiatkowski, Cordelia Langford, John Sillitoe on behalf of the Wellcome Sanger Institute COVID-19 Surveillance Team |

| | | | |
|---|---|---|---|
| EPI_ISL_3070097 | Lighthouse Laboratory Plymouth | Wellcome Sanger Institute for the COVID-19 Genomics UK (COG-UK) Consortium | Lighthouse Laboratory Plymouth and Alex Alderton, Roberto Amato, Jeffrey Barrett, Sonia Goncalves, Ewan Harrison, David K. Jackson, Ian Johnston, Dominic Kwiatkowski, Cordelia Langford, John Sillitoe on behalf of the Wellcome Sanger Institute COVID-19 Surveillance Team |
| EPI_ISL_3070098, EPI_ISL_3070099 | Lighthouse Lab in Alderley Park | Wellcome Sanger Institute for the COVID-19 Genomics UK (COG-UK) Consortium | Jacquelyn Wynn, Mairead Hyland, The Lighthouse Lab in Alderley Park and Alex Alderton, Roberto Amato, Jeffrey Barrett, Sonia Goncalves, Ewan Harrison, David K. Jackson, Ian Johnston, Dominic Kwiatkowski, Cordelia Langford, John Sillitoe on behalf of the Wellcome Sanger Institute COVID-19 Surveillance Team |
| EPI_ISL_3070100, EPI_ISL_3070101 | Berkshire and Surrey Pathology Services Lighthouse Laboratory | Wellcome Sanger Institute for the COVID-19 Genomics UK (COG-UK) Consortium | Berkshire and Surrey Pathology Services Lighthouse Laboratory and Alex Alderton, Roberto Amato, Jeffrey Barrett, Sonia Goncalves, Ewan Harrison, David K. Jackson, Ian Johnston, Dominic Kwiatkowski, Cordelia Langford, John Sillitoe on behalf of the Wellcome Sanger Institute COVID-19 Surveillance Team |
| EPI_ISL_3070102 | Lighthouse Laboratory Plymouth | Wellcome Sanger Institute for the COVID-19 Genomics UK (COG-UK) Consortium | Lighthouse Laboratory Plymouth and Alex Alderton, Roberto Amato, Jeffrey Barrett, Sonia Goncalves, Ewan Harrison, David K. Jackson, Ian Johnston, Dominic Kwiatkowski, Cordelia Langford, John Sillitoe on behalf of the Wellcome Sanger Institute COVID-19 Surveillance Team |
| EPI_ISL_3070103 | Berkshire and Surrey Pathology Services Lighthouse Laboratory | Wellcome Sanger Institute for the COVID-19 Genomics UK (COG-UK) Consortium | Berkshire and Surrey Pathology Services Lighthouse Laboratory and Alex Alderton, Roberto Amato, Jeffrey Barrett, Sonia Goncalves, Ewan Harrison, David K. Jackson, Ian Johnston, Dominic Kwiatkowski, Cordelia Langford, John Sillitoe on behalf of the Wellcome Sanger Institute COVID-19 Surveillance Team |
| EPI_ISL_3070104 | Lighthouse Laboratory Plymouth | Wellcome Sanger Institute for the COVID-19 Genomics UK (COG-UK) Consortium | Lighthouse Laboratory Plymouth and Alex Alderton, Roberto Amato, Jeffrey Barrett, Sonia Goncalves, Ewan Harrison, David K. Jackson, Ian Johnston, Dominic Kwiatkowski, Cordelia Langford, John Sillitoe on behalf of the Wellcome Sanger Institute COVID-19 Surveillance Team |
| EPI_ISL_3070105 | Berkshire and Surrey Pathology Services Lighthouse Laboratory | Wellcome Sanger Institute for the COVID-19 Genomics UK (COG-UK) Consortium | Berkshire and Surrey Pathology Services Lighthouse Laboratory and Alex Alderton, Roberto Amato, Jeffrey Barrett, Sonia Goncalves, Ewan Harrison, David K. Jackson, Ian Johnston, Dominic Kwiatkowski, Cordelia Langford, John Sillitoe on behalf of the Wellcome Sanger Institute COVID-19 Surveillance Team |
| EPI_ISL_3070106 | Lighthouse Laboratory Plymouth | Wellcome Sanger Institute for the COVID-19 Genomics UK (COG-UK) Consortium | Lighthouse Laboratory Plymouth and Alex Alderton, Roberto Amato, Jeffrey Barrett, Sonia Goncalves, Ewan Harrison, David K. Jackson, Ian Johnston, Dominic Kwiatkowski, Cordelia Langford, John Sillitoe on behalf of the Wellcome Sanger Institute COVID-19 Surveillance Team |
| EPI_ISL_3070107 | Lighthouse Lab in Alderley Park | Wellcome Sanger Institute for the COVID-19 Genomics UK (COG-UK) Consortium | Jacquelyn Wynn, Mairead Hyland, The Lighthouse Lab in Alderley Park and Alex Alderton, Roberto Amato, Jeffrey Barrett, Sonia Goncalves, Ewan Harrison, David K. Jackson, Ian Johnston, Dominic Kwiatkowski, Cordelia Langford, John Sillitoe on behalf of the Wellcome Sanger Institute COVID-19 Surveillance Team |
| EPI_ISL_3070108, EPI_ISL_3070109 | Lighthouse Laboratory Plymouth | Wellcome Sanger Institute for the COVID-19 Genomics UK (COG-UK) Consortium | Lighthouse Laboratory Plymouth and Alex Alderton, Roberto Amato, Jeffrey Barrett, Sonia Goncalves, Ewan Harrison, David K. Jackson, Ian Johnston, Dominic Kwiatkowski, Cordelia Langford, John Sillitoe on behalf of the Wellcome Sanger Institute COVID-19 Surveillance Team |
| EPI_ISL_3070110, EPI_ISL_3070111 | Berkshire and Surrey Pathology Services Lighthouse Laboratory | Wellcome Sanger Institute for the COVID-19 Genomics UK (COG-UK) Consortium | Berkshire and Surrey Pathology Services Lighthouse Laboratory and Alex Alderton, Roberto Amato, Jeffrey Barrett, Sonia Goncalves, Ewan Harrison, David K. Jackson, Ian Johnston, Dominic Kwiatkowski, Cordelia Langford, John Sillitoe on behalf of the Wellcome Sanger Institute COVID-19 Surveillance Team |
| EPI_ISL_3070112, EPI_ISL_3070113 | Lighthouse Lab in Alderley Park | Wellcome Sanger Institute for the COVID-19 Genomics UK (COG-UK) Consortium | Jacquelyn Wynn, Mairead Hyland, The Lighthouse Lab in Alderley Park and Alex Alderton, Roberto Amato, Jeffrey Barrett, Sonia Goncalves, Ewan Harrison, David K. Jackson, Ian Johnston, Dominic Kwiatkowski, Cordelia Langford, John Sillitoe on behalf of the Wellcome Sanger Institute COVID-19 Surveillance Team |
| EPI_ISL_3070114 | Lighthouse Laboratory Plymouth | Wellcome Sanger Institute for the COVID-19 Genomics UK (COG-UK) Consortium | Lighthouse Laboratory Plymouth and Alex Alderton, Roberto Amato, Jeffrey Barrett, Sonia Goncalves, Ewan Harrison, David K. Jackson, Ian Johnston, Dominic Kwiatkowski, Cordelia Langford, John Sillitoe on behalf of the Wellcome Sanger Institute COVID-19 Surveillance Team |
| EPI_ISL_3070115 | Lighthouse Lab in Alderley Park | Wellcome Sanger Institute for the COVID-19 Genomics UK (COG-UK) Consortium | Jacquelyn Wynn, Mairead Hyland, The Lighthouse Lab in Alderley Park and Alex Alderton, Roberto Amato, Jeffrey Barrett, Sonia Goncalves, Ewan Harrison, David K. Jackson, Ian Johnston, Dominic Kwiatkowski, Cordelia Langford, John Sillitoe on behalf of the Wellcome Sanger Institute COVID-19 Surveillance Team |
| EPI_ISL_3070116, EPI_ISL_3070117, EPI_ISL_3070118 | Lighthouse Laboratory Plymouth | Wellcome Sanger Institute for the COVID-19 Genomics UK (COG-UK) Consortium | Lighthouse Laboratory Plymouth and Alex Alderton, Roberto Amato, Jeffrey Barrett, Sonia Goncalves, Ewan Harrison, David K. Jackson, Ian Johnston, Dominic Kwiatkowski, Cordelia Langford, John Sillitoe on behalf of the Wellcome Sanger Institute COVID-19 Surveillance Team |
| EPI_ISL_3070119 | Lighthouse Lab in Alderley Park | Wellcome Sanger Institute for the COVID-19 Genomics UK (COG-UK) Consortium | Jacquelyn Wynn, Mairead Hyland, The Lighthouse Lab in Alderley Park and Alex Alderton, Roberto Amato, Jeffrey Barrett, Sonia Goncalves, Ewan Harrison, David K. Jackson, Ian Johnston, Dominic Kwiatkowski, Cordelia Langford, John Sillitoe on behalf of the Wellcome Sanger Institute COVID-19 Surveillance Team |
| EPI_ISL_3070120 | Lighthouse Laboratory Plymouth | Wellcome Sanger Institute for the COVID-19 Genomics UK (COG-UK) Consortium | Lighthouse Laboratory Plymouth and Alex Alderton, Roberto Amato, Jeffrey Barrett, Sonia Goncalves, Ewan Harrison, David K. Jackson, Ian Johnston, Dominic Kwiatkowski, Cordelia Langford, John Sillitoe on behalf of the Wellcome Sanger Institute COVID-19 Surveillance Team |
| EPI_ISL_3070121 | Lighthouse Lab in Alderley Park | Wellcome Sanger Institute for the COVID-19 Genomics UK (COG-UK) Consortium | Jacquelyn Wynn, Mairead Hyland, The Lighthouse Lab in Alderley Park and Alex Alderton, Roberto Amato, Jeffrey Barrett, Sonia Goncalves, Ewan Harrison, David K. Jackson, Ian Johnston, Dominic Kwiatkowski, Cordelia Langford, John Sillitoe on behalf of the Wellcome Sanger Institute COVID-19 Surveillance Team |
| EPI_ISL_3070123, EPI_ISL_3070124, EPI_ISL_3070125 | Berkshire and Surrey Pathology Services Lighthouse Laboratory | Wellcome Sanger Institute for the COVID-19 Genomics UK (COG-UK) Consortium | Berkshire and Surrey Pathology Services Lighthouse Laboratory and Alex Alderton, Roberto Amato, Jeffrey Barrett, Sonia Goncalves, Ewan Harrison, David K. Jackson, Ian Johnston, Dominic Kwiatkowski, Cordelia Langford, John Sillitoe on behalf of the Wellcome Sanger Institute COVID-19 Surveillance Team |
| EPI_ISL_3070126 | Lighthouse Laboratory Plymouth | Wellcome Sanger Institute for the COVID-19 Genomics UK (COG-UK) Consortium | Lighthouse Laboratory Plymouth and Alex Alderton, Roberto Amato, Jeffrey Barrett, Sonia Goncalves, Ewan Harrison, David K. Jackson, Ian Johnston, Dominic Kwiatkowski, Cordelia Langford, John Sillitoe on behalf of the Wellcome Sanger Institute COVID-19 Surveillance Team |
| EPI_ISL_3070127 | Lighthouse Lab in Alderley Park | Wellcome Sanger Institute for the COVID-19 Genomics UK (COG-UK) Consortium | Jacquelyn Wynn, Mairead Hyland, The Lighthouse Lab in Alderley Park and Alex Alderton, Roberto Amato, Jeffrey Barrett, Sonia Goncalves, Ewan Harrison, David K. Jackson, Ian Johnston, Dominic Kwiatkowski, Cordelia Langford, John Sillitoe on behalf of the Wellcome Sanger Institute COVID-19 Surveillance Team |
| EPI_ISL_3070128 | Lighthouse Laboratory Plymouth | Wellcome Sanger Institute for the COVID-19 Genomics UK (COG-UK) Consortium | Lighthouse Laboratory Plymouth and Alex Alderton, Roberto Amato, Jeffrey Barrett, Sonia Goncalves, Ewan Harrison, David K. Jackson, Ian Johnston, Dominic Kwiatkowski, Cordelia Langford, John Sillitoe on behalf of the Wellcome Sanger Institute COVID-19 Surveillance Team |
| EPI_ISL_3070129, EPI_ISL_3070130 | Berkshire and Surrey Pathology Services Lighthouse Laboratory | Wellcome Sanger Institute for the COVID-19 Genomics UK (COG-UK) Consortium | Berkshire and Surrey Pathology Services Lighthouse Laboratory and Alex Alderton, Roberto Amato, Jeffrey Barrett, Sonia Goncalves, Ewan Harrison, David K. Jackson, Ian Johnston, Dominic Kwiatkowski, Cordelia Langford, John Sillitoe on behalf of the Wellcome Sanger Institute COVID-19 Surveillance Team |
| EPI_ISL_3070131, EPI_ISL_3070132 | Lighthouse Laboratory Plymouth | Wellcome Sanger Institute for the COVID-19 Genomics UK (COG-UK) Consortium | Lighthouse Laboratory Plymouth and Alex Alderton, Roberto Amato, Jeffrey Barrett, Sonia Goncalves, Ewan Harrison, David K. Jackson, Ian Johnston, Dominic Kwiatkowski, Cordelia Langford, John Sillitoe on behalf of the Wellcome Sanger Institute COVID-19 Surveillance Team |
| EPI_ISL_3070133 | Lighthouse Lab in Alderley Park | Wellcome Sanger Institute for the COVID-19 Genomics UK (COG-UK) Consortium | Jacquelyn Wynn, Mairead Hyland, The Lighthouse Lab in Alderley Park and Alex Alderton, Roberto Amato, Jeffrey Barrett, Sonia Goncalves, Ewan Harrison, David K. Jackson, Ian Johnston, Dominic Kwiatkowski, Cordelia Langford, John Sillitoe on behalf of the Wellcome Sanger Institute COVID-19 Surveillance Team |
| EPI_ISL_3070134 | Berkshire and Surrey Pathology Services Lighthouse Laboratory | Wellcome Sanger Institute for the COVID-19 Genomics UK (COG-UK) Consortium | Berkshire and Surrey Pathology Services Lighthouse Laboratory and Alex Alderton, Roberto Amato, Jeffrey Barrett, Sonia Goncalves, Ewan Harrison, David K. Jackson, Ian Johnston, Dominic Kwiatkowski, Cordelia Langford, John Sillitoe on behalf of the Wellcome Sanger Institute COVID-19 Surveillance Team |
| EPI_ISL_3070135 | Lighthouse Laboratory Plymouth | Wellcome Sanger Institute for the COVID-19 Genomics UK (COG-UK) Consortium | Lighthouse Laboratory Plymouth and Alex Alderton, Roberto Amato, Jeffrey Barrett, Sonia Goncalves, Ewan Harrison, David K. Jackson, Ian Johnston, Dominic Kwiatkowski, Cordelia Langford, John Sillitoe on behalf of the Wellcome Sanger Institute COVID-19 Surveillance Team |
| EPI_ISL_3070136 | Lighthouse Lab in Alderley Park | Wellcome Sanger Institute for the COVID-19 Genomics UK (COG-UK) Consortium | Jacquelyn Wynn, Mairead Hyland, The Lighthouse Lab in Alderley Park and Alex Alderton, Roberto Amato, Jeffrey Barrett, Sonia Goncalves, Ewan Harrison, David K. Jackson, Ian Johnston, Dominic Kwiatkowski, Cordelia Langford, John Sillitoe on behalf of the Wellcome Sanger Institute COVID-19 Surveillance Team |
| EPI_ISL_3070137 | Berkshire and Surrey Pathology Services Lighthouse Laboratory | Wellcome Sanger Institute for the COVID-19 Genomics UK (COG-UK) Consortium | Berkshire and Surrey Pathology Services Lighthouse Laboratory and Alex Alderton, Roberto Amato, Jeffrey Barrett, Sonia Goncalves, Ewan Harrison, David K. Jackson, Ian Johnston, Dominic Kwiatkowski, Cordelia Langford, John Sillitoe on behalf of the Wellcome Sanger Institute COVID-19 Surveillance Team |
| EPI_ISL_3070138 | Lighthouse Laboratory Plymouth | Wellcome Sanger Institute for the COVID-19 Genomics UK (COG-UK) Consortium | Lighthouse Laboratory Plymouth and Alex Alderton, Roberto Amato, Jeffrey Barrett, Sonia Goncalves, Ewan Harrison, David K. Jackson, Ian Johnston, Dominic Kwiatkowski, Cordelia Langford, John Sillitoe on behalf of the Wellcome Sanger Institute COVID-19 Surveillance Team |
| EPI_ISL_3070139 | Lighthouse Lab in Alderley Park | Wellcome Sanger Institute for the COVID-19 Genomics UK (COG-UK) Consortium | Jacquelyn Wynn, Mairead Hyland, The Lighthouse Lab in Alderley Park and Alex Alderton, Roberto Amato, Jeffrey Barrett, Sonia Goncalves, Ewan Harrison, David K. Jackson, Ian Johnston, Dominic Kwiatkowski, Cordelia Langford, John Sillitoe on behalf of the Wellcome Sanger Institute COVID-19 Surveillance Team |
| EPI_ISL_3070140, EPI_ISL_3070141, EPI_ISL_3070142, EPI_ISL_3070143, EPI_ISL_3070144, EPI_ISL_3070145, EPI_ISL_3070146 | Berkshire and Surrey Pathology Services Lighthouse Laboratory | Wellcome Sanger Institute for the COVID-19 Genomics UK (COG-UK) Consortium | Berkshire and Surrey Pathology Services Lighthouse Laboratory and Alex Alderton, Roberto Amato, Jeffrey Barrett, Sonia Goncalves, Ewan Harrison, David K. Jackson, Ian Johnston, Dominic Kwiatkowski, Cordelia Langford, John Sillitoe on behalf of the Wellcome Sanger Institute COVID-19 Surveillance Team |
| EPI_ISL_3070147, EPI_ISL_3070148 | Lighthouse Lab in Alderley Park | Wellcome Sanger Institute for the COVID-19 Genomics UK (COG-UK) Consortium | Jacquelyn Wynn, Mairead Hyland, The Lighthouse Lab in Alderley Park and Alex Alderton, Roberto Amato, Jeffrey Barrett, Sonia Goncalves, Ewan Harrison, David K. Jackson, Ian Johnston, Dominic Kwiatkowski, Cordelia Langford, John Sillitoe on behalf of the Wellcome Sanger Institute COVID-19 Surveillance Team |
| EPI_ISL_3070149 | Berkshire and Surrey Pathology Services Lighthouse Laboratory | Wellcome Sanger Institute for the COVID-19 Genomics UK (COG-UK) Consortium | Berkshire and Surrey Pathology Services Lighthouse Laboratory and Alex Alderton, Roberto Amato, Jeffrey Barrett, Sonia Goncalves, Ewan Harrison, David K. Jackson, Ian Johnston, Dominic Kwiatkowski, Cordelia Langford, John Sillitoe on behalf of the Wellcome Sanger Institute COVID-19 Surveillance Team |
| EPI_ISL_3070150 | Lighthouse Laboratory Plymouth | Wellcome Sanger Institute for the COVID-19 Genomics UK (COG-UK) Consortium | Lighthouse Laboratory Plymouth and Alex Alderton, Roberto Amato, Jeffrey Barrett, Sonia Goncalves, Ewan Harrison, David K. Jackson, Ian Johnston, Dominic Kwiatkowski, Cordelia Langford, John Sillitoe on behalf of the Wellcome Sanger Institute COVID-19 Surveillance Team |
| EPI_ISL_3070151 | Lighthouse Lab in Alderley Park | Wellcome Sanger Institute for the COVID-19 Genomics UK (COG-UK) Consortium | Jacquelyn Wynn, Mairead Hyland, The Lighthouse Lab in Alderley Park and Alex Alderton, Roberto Amato, Jeffrey Barrett, Sonia Goncalves, Ewan Harrison, David K. Jackson, Ian Johnston, Dominic Kwiatkowski, Cordelia Langford, John Sillitoe on behalf of the Wellcome Sanger Institute COVID-19 Surveillance Team |
| EPI_ISL_3070152, EPI_ISL_3070153, EPI_ISL_3070154, EPI_ISL_3070155 | Berkshire and Surrey Pathology Services Lighthouse Laboratory | Wellcome Sanger Institute for the COVID-19 Genomics UK (COG-UK) Consortium | Berkshire and Surrey Pathology Services Lighthouse Laboratory and Alex Alderton, Roberto Amato, Jeffrey Barrett, Sonia Goncalves, Ewan Harrison, David K. Jackson, Ian Johnston, Dominic Kwiatkowski, Cordelia Langford, John Sillitoe on behalf of the Wellcome Sanger Institute COVID-19 Surveillance Team |
| EPI_ISL_3070156 | Lighthouse Laboratory Plymouth | Wellcome Sanger Institute for the COVID-19 Genomics UK (COG-UK) Consortium | Lighthouse Laboratory Plymouth and Alex Alderton, Roberto Amato, Jeffrey Barrett, Sonia Goncalves, Ewan Harrison, David K. Jackson, Ian Johnston, Dominic Kwiatkowski, Cordelia Langford, John Sillitoe on behalf of the Wellcome Sanger Institute COVID-19 Surveillance Team |
| EPI_ISL_3070157 | Lighthouse Lab in Alderley Park | Wellcome Sanger Institute for the COVID-19 Genomics UK (COG-UK) Consortium | Jacquelyn Wynn, Mairead Hyland, The Lighthouse Lab in Alderley Park and Alex Alderton, Roberto Amato, Jeffrey Barrett, Sonia Goncalves, Ewan Harrison, David K. Jackson, Ian Johnston, Dominic Kwiatkowski, Cordelia Langford, John Sillitoe on behalf of the Wellcome Sanger Institute COVID-19 Surveillance Team |
| EPI_ISL_3070158, EPI_ISL_3070159 | Lighthouse Laboratory Plymouth | Wellcome Sanger Institute for the COVID-19 Genomics UK (COG-UK) Consortium | Lighthouse Laboratory Plymouth and Alex Alderton, Roberto Amato, Jeffrey Barrett, Sonia Goncalves, Ewan Harrison, David K. Jackson, Ian Johnston, Dominic Kwiatkowski, Cordelia Langford, John Sillitoe on behalf of the Wellcome Sanger Institute COVID-19 Surveillance Team |
| EPI_ISL_3070160, EPI_ISL_3070161 | Berkshire and Surrey Pathology Services Lighthouse Laboratory | Wellcome Sanger Institute for the COVID-19 Genomics UK (COG-UK) Consortium | Berkshire and Surrey Pathology Services Lighthouse Laboratory and Alex Alderton, Roberto Amato, Jeffrey Barrett, Sonia Goncalves, Ewan Harrison, David K. Jackson, Ian Johnston, Dominic Kwiatkowski, Cordelia Langford, John Sillitoe on behalf of the Wellcome Sanger Institute COVID-19 Surveillance Team |

| | | | |
|---|---|---|---|
| EPI_ISL_3070162, EPI_ISL_3070163 | Lighthouse Laboratory Plymouth | Wellcome Sanger Institute for the COVID-19 Genomics UK (COG-UK) Consortium | Lighthouse Laboratory Plymouth and Alex Alderton, Roberto Amato, Jeffrey Barrett, Sonia Goncalves, Ewan Harrison, David K. Jackson, Ian Johnston, Dominic Kwiatkowski, Cordelia Langford, John Sillitoe on behalf of the Wellcome Sanger Institute COVID-19 Surveillance Team |
| EPI_ISL_3070164, EPI_ISL_3070165, EPI_ISL_3070166 | Lighthouse Lab in Alderley Park | Wellcome Sanger Institute for the COVID-19 Genomics UK (COG-UK) Consortium | Jacquelyn Wynn, Mairead Hyland, The Lighthouse Lab in Alderley Park and Alex Alderton, Roberto Amato, Jeffrey Barrett, Sonia Goncalves, Ewan Harrison, David K. Jackson, Ian Johnston, Dominic Kwiatkowski, Cordelia Langford, John Sillitoe on behalf of the Wellcome Sanger Institute COVID-19 Surveillance Team |
| EPI_ISL_3070167 | Berkshire and Surrey Pathology Services Lighthouse Laboratory | Wellcome Sanger Institute for the COVID-19 Genomics UK (COG-UK) Consortium | Berkshire and Surrey Pathology Services Lighthouse Laboratory and Alex Alderton, Roberto Amato, Jeffrey Barrett, Sonia Goncalves, Ewan Harrison, David K. Jackson, Ian Johnston, Dominic Kwiatkowski, Cordelia Langford, John Sillitoe on behalf of the Wellcome Sanger Institute COVID-19 Surveillance Team |
| EPI_ISL_3070168 | Lighthouse Laboratory Plymouth | Wellcome Sanger Institute for the COVID-19 Genomics UK (COG-UK) Consortium | Lighthouse Laboratory Plymouth and Alex Alderton, Roberto Amato, Jeffrey Barrett, Sonia Goncalves, Ewan Harrison, David K. Jackson, Ian Johnston, Dominic Kwiatkowski, Cordelia Langford, John Sillitoe on behalf of the Wellcome Sanger Institute COVID-19 Surveillance Team |
| EPI_ISL_3070169 | Lighthouse Lab in Alderley Park | Wellcome Sanger Institute for the COVID-19 Genomics UK (COG-UK) Consortium | Jacquelyn Wynn, Mairead Hyland, The Lighthouse Lab in Alderley Park and Alex Alderton, Roberto Amato, Jeffrey Barrett, Sonia Goncalves, Ewan Harrison, David K. Jackson, Ian Johnston, Dominic Kwiatkowski, Cordelia Langford, John Sillitoe on behalf of the Wellcome Sanger Institute COVID-19 Surveillance Team |
| EPI_ISL_3070170, EPI_ISL_3070171, EPI_ISL_3070172 | Berkshire and Surrey Pathology Services Lighthouse Laboratory | Wellcome Sanger Institute for the COVID-19 Genomics UK (COG-UK) Consortium | Berkshire and Surrey Pathology Services Lighthouse Laboratory and Alex Alderton, Roberto Amato, Jeffrey Barrett, Sonia Goncalves, Ewan Harrison, David K. Jackson, Ian Johnston, Dominic Kwiatkowski, Cordelia Langford, John Sillitoe on behalf of the Wellcome Sanger Institute COVID-19 Surveillance Team |
| EPI_ISL_3070173 | Lighthouse Laboratory Plymouth | Wellcome Sanger Institute for the COVID-19 Genomics UK (COG-UK) Consortium | Lighthouse Laboratory Plymouth and Alex Alderton, Roberto Amato, Jeffrey Barrett, Sonia Goncalves, Ewan Harrison, David K. Jackson, Ian Johnston, Dominic Kwiatkowski, Cordelia Langford, John Sillitoe on behalf of the Wellcome Sanger Institute COVID-19 Surveillance Team |
| EPI_ISL_3070174, EPI_ISL_3070175 | Berkshire and Surrey Pathology Services Lighthouse Laboratory | Wellcome Sanger Institute for the COVID-19 Genomics UK (COG-UK) Consortium | Berkshire and Surrey Pathology Services Lighthouse Laboratory and Alex Alderton, Roberto Amato, Jeffrey Barrett, Sonia Goncalves, Ewan Harrison, David K. Jackson, Ian Johnston, Dominic Kwiatkowski, Cordelia Langford, John Sillitoe on behalf of the Wellcome Sanger Institute COVID-19 Surveillance Team |
| EPI_ISL_3070176 | Lighthouse Lab in Alderley Park | Wellcome Sanger Institute for the COVID-19 Genomics UK (COG-UK) Consortium | Jacquelyn Wynn, Mairead Hyland, The Lighthouse Lab in Alderley Park and Alex Alderton, Roberto Amato, Jeffrey Barrett, Sonia Goncalves, Ewan Harrison, David K. Jackson, Ian Johnston, Dominic Kwiatkowski, Cordelia Langford, John Sillitoe on behalf of the Wellcome Sanger Institute COVID-19 Surveillance Team |
| EPI_ISL_3070177, EPI_ISL_3070178 | Berkshire and Surrey Pathology Services Lighthouse Laboratory | Wellcome Sanger Institute for the COVID-19 Genomics UK (COG-UK) Consortium | Berkshire and Surrey Pathology Services Lighthouse Laboratory and Alex Alderton, Roberto Amato, Jeffrey Barrett, Sonia Goncalves, Ewan Harrison, David K. Jackson, Ian Johnston, Dominic Kwiatkowski, Cordelia Langford, John Sillitoe on behalf of the Wellcome Sanger Institute COVID-19 Surveillance Team |
| EPI_ISL_3070179, EPI_ISL_3070180, EPI_ISL_3070181 | Lighthouse Lab in Alderley Park | Wellcome Sanger Institute for the COVID-19 Genomics UK (COG-UK) Consortium | Jacquelyn Wynn, Mairead Hyland, The Lighthouse Lab in Alderley Park and Alex Alderton, Roberto Amato, Jeffrey Barrett, Sonia Goncalves, Ewan Harrison, David K. Jackson, Ian Johnston, Dominic Kwiatkowski, Cordelia Langford, John Sillitoe on behalf of the Wellcome Sanger Institute COVID-19 Surveillance Team |
| EPI_ISL_3070182 | Lighthouse Laboratory Plymouth | Wellcome Sanger Institute for the COVID-19 Genomics UK (COG-UK) Consortium | Lighthouse Laboratory Plymouth and Alex Alderton, Roberto Amato, Jeffrey Barrett, Sonia Goncalves, Ewan Harrison, David K. Jackson, Ian Johnston, Dominic Kwiatkowski, Cordelia Langford, John Sillitoe on behalf of the Wellcome Sanger Institute COVID-19 Surveillance Team |
| EPI_ISL_3070183, EPI_ISL_3070184, EPI_ISL_3070185 | Berkshire and Surrey Pathology Services Lighthouse Laboratory | Wellcome Sanger Institute for the COVID-19 Genomics UK (COG-UK) Consortium | Berkshire and Surrey Pathology Services Lighthouse Laboratory and Alex Alderton, Roberto Amato, Jeffrey Barrett, Sonia Goncalves, Ewan Harrison, David K. Jackson, Ian Johnston, Dominic Kwiatkowski, Cordelia Langford, John Sillitoe on behalf of the Wellcome Sanger Institute COVID-19 Surveillance Team |
| EPI_ISL_3070186 | Lighthouse Lab in Alderley Park | Wellcome Sanger Institute for the COVID-19 Genomics UK (COG-UK) Consortium | Jacquelyn Wynn, Mairead Hyland, The Lighthouse Lab in Alderley Park and Alex Alderton, Roberto Amato, Jeffrey Barrett, Sonia Goncalves, Ewan Harrison, David K. Jackson, Ian Johnston, Dominic Kwiatkowski, Cordelia Langford, John Sillitoe on behalf of the Wellcome Sanger Institute COVID-19 Surveillance Team |
| EPI_ISL_3070187, EPI_ISL_3070188, EPI_ISL_3070189 | Berkshire and Surrey Pathology Services Lighthouse Laboratory | Wellcome Sanger Institute for the COVID-19 Genomics UK (COG-UK) Consortium | Berkshire and Surrey Pathology Services Lighthouse Laboratory and Alex Alderton, Roberto Amato, Jeffrey Barrett, Sonia Goncalves, Ewan Harrison, David K. Jackson, Ian Johnston, Dominic Kwiatkowski, Cordelia Langford, John Sillitoe on behalf of the Wellcome Sanger Institute COVID-19 Surveillance Team |
| EPI_ISL_3070190 | Lighthouse Laboratory Plymouth | Wellcome Sanger Institute for the COVID-19 Genomics UK (COG-UK) Consortium | Lighthouse Laboratory Plymouth and Alex Alderton, Roberto Amato, Jeffrey Barrett, Sonia Goncalves, Ewan Harrison, David K. Jackson, Ian Johnston, Dominic Kwiatkowski, Cordelia Langford, John Sillitoe on behalf of the Wellcome Sanger Institute COVID-19 Surveillance Team |
| EPI_ISL_3070191, EPI_ISL_3070192, EPI_ISL_3070193 | Lighthouse Lab in Alderley Park | Wellcome Sanger Institute for the COVID-19 Genomics UK (COG-UK) Consortium | Jacquelyn Wynn, Mairead Hyland, The Lighthouse Lab in Alderley Park and Alex Alderton, Roberto Amato, Jeffrey Barrett, Sonia Goncalves, Ewan Harrison, David K. Jackson, Ian Johnston, Dominic Kwiatkowski, Cordelia Langford, John Sillitoe on behalf of the Wellcome Sanger Institute COVID-19 Surveillance Team |
| EPI_ISL_3070194 | Berkshire and Surrey Pathology Services Lighthouse Laboratory | Wellcome Sanger Institute for the COVID-19 Genomics UK (COG-UK) Consortium | Berkshire and Surrey Pathology Services Lighthouse Laboratory and Alex Alderton, Roberto Amato, Jeffrey Barrett, Sonia Goncalves, Ewan Harrison, David K. Jackson, Ian Johnston, Dominic Kwiatkowski, Cordelia Langford, John Sillitoe on behalf of the Wellcome Sanger Institute COVID-19 Surveillance Team |
| EPI_ISL_3070195 | Lighthouse Laboratory Plymouth | Wellcome Sanger Institute for the COVID-19 Genomics UK (COG-UK) Consortium | Lighthouse Laboratory Plymouth and Alex Alderton, Roberto Amato, Jeffrey Barrett, Sonia Goncalves, Ewan Harrison, David K. Jackson, Ian Johnston, Dominic Kwiatkowski, Cordelia Langford, John Sillitoe on behalf of the Wellcome Sanger Institute COVID-19 Surveillance Team |
| EPI_ISL_3070196 | Berkshire and Surrey Pathology Services Lighthouse Laboratory | Wellcome Sanger Institute for the COVID-19 Genomics UK (COG-UK) Consortium | Berkshire and Surrey Pathology Services Lighthouse Laboratory and Alex Alderton, Roberto Amato, Jeffrey Barrett, Sonia Goncalves, Ewan Harrison, David K. Jackson, Ian Johnston, Dominic Kwiatkowski, Cordelia Langford, John Sillitoe on behalf of the Wellcome Sanger Institute COVID-19 Surveillance Team |
| EPI_ISL_3070197 | Lighthouse Laboratory Plymouth | Wellcome Sanger Institute for the COVID-19 Genomics UK (COG-UK) Consortium | Lighthouse Laboratory Plymouth and Alex Alderton, Roberto Amato, Jeffrey Barrett, Sonia Goncalves, Ewan Harrison, David K. Jackson, Ian Johnston, Dominic Kwiatkowski, Cordelia Langford, John Sillitoe on behalf of the Wellcome Sanger Institute COVID-19 Surveillance Team |
| EPI_ISL_3070198 | Berkshire and Surrey Pathology Services Lighthouse Laboratory | Wellcome Sanger Institute for the COVID-19 Genomics UK (COG-UK) Consortium | Berkshire and Surrey Pathology Services Lighthouse Laboratory and Alex Alderton, Roberto Amato, Jeffrey Barrett, Sonia Goncalves, Ewan Harrison, David K. Jackson, Ian Johnston, Dominic Kwiatkowski, Cordelia Langford, John Sillitoe on behalf of the Wellcome Sanger Institute COVID-19 Surveillance Team |
| EPI_ISL_3070199 | Lighthouse Laboratory Plymouth | Wellcome Sanger Institute for the COVID-19 Genomics UK (COG-UK) Consortium | Lighthouse Laboratory Plymouth and Alex Alderton, Roberto Amato, Jeffrey Barrett, Sonia Goncalves, Ewan Harrison, David K. Jackson, Ian Johnston, Dominic Kwiatkowski, Cordelia Langford, John Sillitoe on behalf of the Wellcome Sanger Institute COVID-19 Surveillance Team |
| EPI_ISL_3070200 | Lighthouse Lab in Alderley Park | Wellcome Sanger Institute for the COVID-19 Genomics UK (COG-UK) Consortium | Jacquelyn Wynn, Mairead Hyland, The Lighthouse Lab in Alderley Park and Alex Alderton, Roberto Amato, Jeffrey Barrett, Sonia Goncalves, Ewan Harrison, David K. Jackson, Ian Johnston, Dominic Kwiatkowski, Cordelia Langford, John Sillitoe on behalf of the Wellcome Sanger Institute COVID-19 Surveillance Team |
| EPI_ISL_3070201 | Berkshire and Surrey Pathology Services Lighthouse Laboratory | Wellcome Sanger Institute for the COVID-19 Genomics UK (COG-UK) Consortium | Berkshire and Surrey Pathology Services Lighthouse Laboratory and Alex Alderton, Roberto Amato, Jeffrey Barrett, Sonia Goncalves, Ewan Harrison, David K. Jackson, Ian Johnston, Dominic Kwiatkowski, Cordelia Langford, John Sillitoe on behalf of the Wellcome Sanger Institute COVID-19 Surveillance Team |
| EPI_ISL_3070202 | Lighthouse Laboratory Plymouth | Wellcome Sanger Institute for the COVID-19 Genomics UK (COG-UK) Consortium | Lighthouse Laboratory Plymouth and Alex Alderton, Roberto Amato, Jeffrey Barrett, Sonia Goncalves, Ewan Harrison, David K. Jackson, Ian Johnston, Dominic Kwiatkowski, Cordelia Langford, John Sillitoe on behalf of the Wellcome Sanger Institute COVID-19 Surveillance Team |
| EPI_ISL_3070203 | Lighthouse Lab in Alderley Park | Wellcome Sanger Institute for the COVID-19 Genomics UK (COG-UK) Consortium | Jacquelyn Wynn, Mairead Hyland, The Lighthouse Lab in Alderley Park and Alex Alderton, Roberto Amato, Jeffrey Barrett, Sonia Goncalves, Ewan Harrison, David K. Jackson, Ian Johnston, Dominic Kwiatkowski, Cordelia Langford, John Sillitoe on behalf of the Wellcome Sanger Institute COVID-19 Surveillance Team |
| EPI_ISL_3070204, EPI_ISL_3070205 | Berkshire and Surrey Pathology Services Lighthouse Laboratory | Wellcome Sanger Institute for the COVID-19 Genomics UK (COG-UK) Consortium | Berkshire and Surrey Pathology Services Lighthouse Laboratory and Alex Alderton, Roberto Amato, Jeffrey Barrett, Sonia Goncalves, Ewan Harrison, David K. Jackson, Ian Johnston, Dominic Kwiatkowski, Cordelia Langford, John Sillitoe on behalf of the Wellcome Sanger Institute COVID-19 Surveillance Team |
| EPI_ISL_3070206, EPI_ISL_3070207 | Lighthouse Lab in Alderley Park | Wellcome Sanger Institute for the COVID-19 Genomics UK (COG-UK) Consortium | Jacquelyn Wynn, Mairead Hyland, The Lighthouse Lab in Alderley Park and Alex Alderton, Roberto Amato, Jeffrey Barrett, Sonia Goncalves, Ewan Harrison, David K. Jackson, Ian Johnston, Dominic Kwiatkowski, Cordelia Langford, John Sillitoe on behalf of the Wellcome Sanger Institute COVID-19 Surveillance Team |
| EPI_ISL_3070208, EPI_ISL_3070209 | Lighthouse Laboratory Plymouth | Wellcome Sanger Institute for the COVID-19 Genomics UK (COG-UK) Consortium | Lighthouse Laboratory Plymouth and Alex Alderton, Roberto Amato, Jeffrey Barrett, Sonia Goncalves, Ewan Harrison, David K. Jackson, Ian Johnston, Dominic Kwiatkowski, Cordelia Langford, John Sillitoe on behalf of the Wellcome Sanger Institute COVID-19 Surveillance Team |
| EPI_ISL_3070210, EPI_ISL_3070211 | Lighthouse Lab in Alderley Park | Wellcome Sanger Institute for the COVID-19 Genomics UK (COG-UK) Consortium | Jacquelyn Wynn, Mairead Hyland, The Lighthouse Lab in Alderley Park and Alex Alderton, Roberto Amato, Jeffrey Barrett, Sonia Goncalves, Ewan Harrison, David K. Jackson, Ian Johnston, Dominic Kwiatkowski, Cordelia Langford, John Sillitoe on behalf of the Wellcome Sanger Institute COVID-19 Surveillance Team |
| EPI_ISL_3070212, EPI_ISL_3070213 | Berkshire and Surrey Pathology Services Lighthouse Laboratory | Wellcome Sanger Institute for the COVID-19 Genomics UK (COG-UK) Consortium | Berkshire and Surrey Pathology Services Lighthouse Laboratory and Alex Alderton, Roberto Amato, Jeffrey Barrett, Sonia Goncalves, Ewan Harrison, David K. Jackson, Ian Johnston, Dominic Kwiatkowski, Cordelia Langford, John Sillitoe on behalf of the Wellcome Sanger Institute COVID-19 Surveillance Team |
| EPI_ISL_3070214 | Lighthouse Laboratory Plymouth | Wellcome Sanger Institute for the COVID-19 Genomics UK (COG-UK) Consortium | Lighthouse Laboratory Plymouth and Alex Alderton, Roberto Amato, Jeffrey Barrett, Sonia Goncalves, Ewan Harrison, David K. Jackson, Ian Johnston, Dominic Kwiatkowski, Cordelia Langford, John Sillitoe on behalf of the Wellcome Sanger Institute COVID-19 Surveillance Team |
| EPI_ISL_3070215 | Berkshire and Surrey Pathology Services Lighthouse Laboratory | Wellcome Sanger Institute for the COVID-19 Genomics UK (COG-UK) Consortium | Berkshire and Surrey Pathology Services Lighthouse Laboratory and Alex Alderton, Roberto Amato, Jeffrey Barrett, Sonia Goncalves, Ewan Harrison, David K. Jackson, Ian Johnston, Dominic Kwiatkowski, Cordelia Langford, John Sillitoe on behalf of the Wellcome Sanger Institute COVID-19 Surveillance Team |
| EPI_ISL_3070216 | Lighthouse Lab in Alderley Park | Wellcome Sanger Institute for the COVID-19 Genomics UK (COG-UK) Consortium | Jacquelyn Wynn, Mairead Hyland, The Lighthouse Lab in Alderley Park and Alex Alderton, Roberto Amato, Jeffrey Barrett, Sonia Goncalves, Ewan Harrison, David K. Jackson, Ian Johnston, Dominic Kwiatkowski, Cordelia Langford, John Sillitoe on behalf of the Wellcome Sanger Institute COVID-19 Surveillance Team |
| EPI_ISL_3070217 | Berkshire and Surrey Pathology Services Lighthouse Laboratory | Wellcome Sanger Institute for the COVID-19 Genomics UK (COG-UK) Consortium | Berkshire and Surrey Pathology Services Lighthouse Laboratory and Alex Alderton, Roberto Amato, Jeffrey Barrett, Sonia Goncalves, Ewan Harrison, David K. Jackson, Ian Johnston, Dominic Kwiatkowski, Cordelia Langford, John Sillitoe on behalf of the Wellcome Sanger Institute COVID-19 Surveillance Team |
| EPI_ISL_3070218, EPI_ISL_3070219 | Lighthouse Laboratory Plymouth | Wellcome Sanger Institute for the COVID-19 Genomics UK (COG-UK) Consortium | Lighthouse Laboratory Plymouth and Alex Alderton, Roberto Amato, Jeffrey Barrett, Sonia Goncalves, Ewan Harrison, David K. Jackson, Ian Johnston, Dominic Kwiatkowski, Cordelia Langford, John Sillitoe on behalf of the Wellcome Sanger Institute COVID-19 Surveillance Team |
| EPI_ISL_3070220 | Lighthouse Lab in Alderley Park | Wellcome Sanger Institute for the COVID-19 Genomics UK (COG-UK) Consortium | Jacquelyn Wynn, Mairead Hyland, The Lighthouse Lab in Alderley Park and Alex Alderton, Roberto Amato, Jeffrey Barrett, Sonia Goncalves, Ewan Harrison, David K. Jackson, Ian Johnston, Dominic Kwiatkowski, Cordelia Langford, John Sillitoe on behalf of the Wellcome Sanger Institute COVID-19 Surveillance Team |
| EPI_ISL_3070221, EPI_ISL_3070222 | Berkshire and Surrey Pathology Services Lighthouse Laboratory | Wellcome Sanger Institute for the COVID-19 Genomics UK (COG-UK) Consortium | Berkshire and Surrey Pathology Services Lighthouse Laboratory and Alex Alderton, Roberto Amato, Jeffrey Barrett, Sonia Goncalves, Ewan Harrison, David K. Jackson, Ian Johnston, Dominic Kwiatkowski, Cordelia Langford, John Sillitoe on behalf of the Wellcome Sanger Institute COVID-19 Surveillance Team |
| EPI_ISL_3070223 | Lighthouse Lab in Alderley Park | Wellcome Sanger Institute for the COVID-19 Genomics UK (COG-UK) Consortium | Jacquelyn Wynn, Mairead Hyland, The Lighthouse Lab in Alderley Park and Alex Alderton, Roberto Amato, Jeffrey Barrett, Sonia Goncalves, Ewan Harrison, David K. Jackson, Ian Johnston, Dominic Kwiatkowski, Cordelia Langford, John Sillitoe on behalf of the Wellcome Sanger Institute COVID-19 Surveillance Team |
| EPI_ISL_3070224 | Lighthouse Laboratory Plymouth | Wellcome Sanger Institute for the COVID-19 Genomics UK (COG-UK) Consortium | Lighthouse Laboratory Plymouth and Alex Alderton, Roberto Amato, Jeffrey Barrett, Sonia Goncalves, Ewan Harrison, David K. Jackson, Ian Johnston, Dominic Kwiatkowski, Cordelia Langford, John Sillitoe on behalf of the Wellcome Sanger Institute COVID-19 Surveillance Team |
| EPI_ISL_3070225 | Berkshire and Surrey Pathology Services Lighthouse Laboratory | Wellcome Sanger Institute for the COVID-19 Genomics UK (COG-UK) Consortium | Berkshire and Surrey Pathology Services Lighthouse Laboratory and Alex Alderton, Roberto Amato, Jeffrey Barrett, Sonia Goncalves, Ewan Harrison, David K. Jackson, Ian Johnston, Dominic Kwiatkowski, Cordelia Langford, John Sillitoe on behalf of the Wellcome Sanger Institute COVID-19 Surveillance Team |
| EPI_ISL_3070226 | Lighthouse Lab in Alderley Park | Wellcome Sanger Institute for the COVID-19 Genomics UK (COG-UK) | Jacquelyn Wynn, Mairead Hyland, The Lighthouse Lab in Alderley Park and Alex Alderton, Roberto Amato, Jeffrey Barrett, Sonia Goncalves, Ewan Harrison, David K. Jackson, Ian Johnston, Dominic Kwiatkowski, Cordelia Langford, John Sillitoe on |

| | | | |
|---|---|---|---|
| | | Consortium | behalf of the Wellcome Sanger Institute COVID-19 Surveillance Team |
| EPI_ISL_3070227, EPI_ISL_3070228 | Berkshire and Surrey Pathology Services Lighthouse Laboratory | Wellcome Sanger Institute for the COVID-19 Genomics UK (COG-UK) Consortium | Berkshire and Surrey Pathology Services Lighthouse Laboratory and Alex Alderton, Roberto Amato, Jeffrey Barrett, Sonia Goncalves, Ewan Harrison, David K. Jackson, Ian Johnston, Dominic Kwiatkowski, Cordelia Langford, John Sillitoe on behalf of the Wellcome Sanger Institute COVID-19 Surveillance Team |
| EPI_ISL_3070229 | Lighthouse Lab in Alderley Park | Wellcome Sanger Institute for the COVID-19 Genomics UK (COG-UK) Consortium | Jacquelyn Wynn, Mairead Hyland, The Lighthouse Lab in Alderley Park and Alex Alderton, Roberto Amato, Jeffrey Barrett, Sonia Goncalves, Ewan Harrison, David K. Jackson, Ian Johnston, Dominic Kwiatkowski, Cordelia Langford, John Sillitoe on behalf of the Wellcome Sanger Institute COVID-19 Surveillance Team |
| EPI_ISL_3070230 | Berkshire and Surrey Pathology Services Lighthouse Laboratory | Wellcome Sanger Institute for the COVID-19 Genomics UK (COG-UK) Consortium | Berkshire and Surrey Pathology Services Lighthouse Laboratory and Alex Alderton, Roberto Amato, Jeffrey Barrett, Sonia Goncalves, Ewan Harrison, David K. Jackson, Ian Johnston, Dominic Kwiatkowski, Cordelia Langford, John Sillitoe on behalf of the Wellcome Sanger Institute COVID-19 Surveillance Team |
| EPI_ISL_3070231 | Lighthouse Lab in Alderley Park | Wellcome Sanger Institute for the COVID-19 Genomics UK (COG-UK) Consortium | Jacquelyn Wynn, Mairead Hyland, The Lighthouse Lab in Alderley Park and Alex Alderton, Roberto Amato, Jeffrey Barrett, Sonia Goncalves, Ewan Harrison, David K. Jackson, Ian Johnston, Dominic Kwiatkowski, Cordelia Langford, John Sillitoe on behalf of the Wellcome Sanger Institute COVID-19 Surveillance Team |
| EPI_ISL_3070232 | Berkshire and Surrey Pathology Services Lighthouse Laboratory | Wellcome Sanger Institute for the COVID-19 Genomics UK (COG-UK) Consortium | Berkshire and Surrey Pathology Services Lighthouse Laboratory and Alex Alderton, Roberto Amato, Jeffrey Barrett, Sonia Goncalves, Ewan Harrison, David K. Jackson, Ian Johnston, Dominic Kwiatkowski, Cordelia Langford, John Sillitoe on behalf of the Wellcome Sanger Institute COVID-19 Surveillance Team |
| EPI_ISL_3070233 | Lighthouse Laboratory Plymouth | Wellcome Sanger Institute for the COVID-19 Genomics UK (COG-UK) Consortium | Lighthouse Laboratory Plymouth and Alex Alderton, Roberto Amato, Jeffrey Barrett, Sonia Goncalves, Ewan Harrison, David K. Jackson, Ian Johnston, Dominic Kwiatkowski, Cordelia Langford, John Sillitoe on behalf of the Wellcome Sanger Institute COVID-19 Surveillance Team |
| EPI_ISL_3070234, EPI_ISL_3070235 | Berkshire and Surrey Pathology Services Lighthouse Laboratory | Wellcome Sanger Institute for the COVID-19 Genomics UK (COG-UK) Consortium | Berkshire and Surrey Pathology Services Lighthouse Laboratory and Alex Alderton, Roberto Amato, Jeffrey Barrett, Sonia Goncalves, Ewan Harrison, David K. Jackson, Ian Johnston, Dominic Kwiatkowski, Cordelia Langford, John Sillitoe on behalf of the Wellcome Sanger Institute COVID-19 Surveillance Team |
| EPI_ISL_3070236 | Lighthouse Lab in Alderley Park | Wellcome Sanger Institute for the COVID-19 Genomics UK (COG-UK) Consortium | Jacquelyn Wynn, Mairead Hyland, The Lighthouse Lab in Alderley Park and Alex Alderton, Roberto Amato, Jeffrey Barrett, Sonia Goncalves, Ewan Harrison, David K. Jackson, Ian Johnston, Dominic Kwiatkowski, Cordelia Langford, John Sillitoe on behalf of the Wellcome Sanger Institute COVID-19 Surveillance Team |
| EPI_ISL_3070237, EPI_ISL_3070238 | Berkshire and Surrey Pathology Services Lighthouse Laboratory | Wellcome Sanger Institute for the COVID-19 Genomics UK (COG-UK) Consortium | Berkshire and Surrey Pathology Services Lighthouse Laboratory and Alex Alderton, Roberto Amato, Jeffrey Barrett, Sonia Goncalves, Ewan Harrison, David K. Jackson, Ian Johnston, Dominic Kwiatkowski, Cordelia Langford, John Sillitoe on behalf of the Wellcome Sanger Institute COVID-19 Surveillance Team |
| EPI_ISL_3070239 | Lighthouse Laboratory Plymouth | Wellcome Sanger Institute for the COVID-19 Genomics UK (COG-UK) Consortium | Lighthouse Laboratory Plymouth and Alex Alderton, Roberto Amato, Jeffrey Barrett, Sonia Goncalves, Ewan Harrison, David K. Jackson, Ian Johnston, Dominic Kwiatkowski, Cordelia Langford, John Sillitoe on behalf of the Wellcome Sanger Institute COVID-19 Surveillance Team |
| EPI_ISL_3070240, EPI_ISL_3070241 | Berkshire and Surrey Pathology Services Lighthouse Laboratory | Wellcome Sanger Institute for the COVID-19 Genomics UK (COG-UK) Consortium | Berkshire and Surrey Pathology Services Lighthouse Laboratory and Alex Alderton, Roberto Amato, Jeffrey Barrett, Sonia Goncalves, Ewan Harrison, David K. Jackson, Ian Johnston, Dominic Kwiatkowski, Cordelia Langford, John Sillitoe on behalf of the Wellcome Sanger Institute COVID-19 Surveillance Team |
| EPI_ISL_3070242 | Lighthouse Lab in Alderley Park | Wellcome Sanger Institute for the COVID-19 Genomics UK (COG-UK) Consortium | Jacquelyn Wynn, Mairead Hyland, The Lighthouse Lab in Alderley Park and Alex Alderton, Roberto Amato, Jeffrey Barrett, Sonia Goncalves, Ewan Harrison, David K. Jackson, Ian Johnston, Dominic Kwiatkowski, Cordelia Langford, John Sillitoe on behalf of the Wellcome Sanger Institute COVID-19 Surveillance Team |
| EPI_ISL_3070243, EPI_ISL_3070244, EPI_ISL_3070245, EPI_ISL_3070246, EPI_ISL_3070247, EPI_ISL_3070248 | Berkshire and Surrey Pathology Services Lighthouse Laboratory | Wellcome Sanger Institute for the COVID-19 Genomics UK (COG-UK) Consortium | Berkshire and Surrey Pathology Services Lighthouse Laboratory and Alex Alderton, Roberto Amato, Jeffrey Barrett, Sonia Goncalves, Ewan Harrison, David K. Jackson, Ian Johnston, Dominic Kwiatkowski, Cordelia Langford, John Sillitoe on behalf of the Wellcome Sanger Institute COVID-19 Surveillance Team |
| EPI_ISL_3070249 | Lighthouse Laboratory Plymouth | Wellcome Sanger Institute for the COVID-19 Genomics UK (COG-UK) Consortium | Lighthouse Laboratory Plymouth and Alex Alderton, Roberto Amato, Jeffrey Barrett, Sonia Goncalves, Ewan Harrison, David K. Jackson, Ian Johnston, Dominic Kwiatkowski, Cordelia Langford, John Sillitoe on behalf of the Wellcome Sanger Institute COVID-19 Surveillance Team |
| EPI_ISL_3070250 | Lighthouse Lab in Alderley Park | Wellcome Sanger Institute for the COVID-19 Genomics UK (COG-UK) Consortium | Jacquelyn Wynn, Mairead Hyland, The Lighthouse Lab in Alderley Park and Alex Alderton, Roberto Amato, Jeffrey Barrett, Sonia Goncalves, Ewan Harrison, David K. Jackson, Ian Johnston, Dominic Kwiatkowski, Cordelia Langford, John Sillitoe on behalf of the Wellcome Sanger Institute COVID-19 Surveillance Team |
| EPI_ISL_3070251 | Berkshire and Surrey Pathology Services Lighthouse Laboratory | Wellcome Sanger Institute for the COVID-19 Genomics UK (COG-UK) Consortium | Berkshire and Surrey Pathology Services Lighthouse Laboratory and Alex Alderton, Roberto Amato, Jeffrey Barrett, Sonia Goncalves, Ewan Harrison, David K. Jackson, Ian Johnston, Dominic Kwiatkowski, Cordelia Langford, John Sillitoe on behalf of the Wellcome Sanger Institute COVID-19 Surveillance Team |
| EPI_ISL_3070252 | Lighthouse Lab in Alderley Park | Wellcome Sanger Institute for the COVID-19 Genomics UK (COG-UK) Consortium | Jacquelyn Wynn, Mairead Hyland, The Lighthouse Lab in Alderley Park and Alex Alderton, Roberto Amato, Jeffrey Barrett, Sonia Goncalves, Ewan Harrison, David K. Jackson, Ian Johnston, Dominic Kwiatkowski, Cordelia Langford, John Sillitoe on behalf of the Wellcome Sanger Institute COVID-19 Surveillance Team |
| EPI_ISL_3070253 | Berkshire and Surrey Pathology Services Lighthouse Laboratory | Wellcome Sanger Institute for the COVID-19 Genomics UK (COG-UK) Consortium | Berkshire and Surrey Pathology Services Lighthouse Laboratory and Alex Alderton, Roberto Amato, Jeffrey Barrett, Sonia Goncalves, Ewan Harrison, David K. Jackson, Ian Johnston, Dominic Kwiatkowski, Cordelia Langford, John Sillitoe on behalf of the Wellcome Sanger Institute COVID-19 Surveillance Team |
| EPI_ISL_3070254 | Lighthouse Lab in Alderley Park | Wellcome Sanger Institute for the COVID-19 Genomics UK (COG-UK) Consortium | Jacquelyn Wynn, Mairead Hyland, The Lighthouse Lab in Alderley Park and Alex Alderton, Roberto Amato, Jeffrey Barrett, Sonia Goncalves, Ewan Harrison, David K. Jackson, Ian Johnston, Dominic Kwiatkowski, Cordelia Langford, John Sillitoe on behalf of the Wellcome Sanger Institute COVID-19 Surveillance Team |
| EPI_ISL_3070255 | Berkshire and Surrey Pathology Services Lighthouse Laboratory | Wellcome Sanger Institute for the COVID-19 Genomics UK (COG-UK) Consortium | Berkshire and Surrey Pathology Services Lighthouse Laboratory and Alex Alderton, Roberto Amato, Jeffrey Barrett, Sonia Goncalves, Ewan Harrison, David K. Jackson, Ian Johnston, Dominic Kwiatkowski, Cordelia Langford, John Sillitoe on behalf of the Wellcome Sanger Institute COVID-19 Surveillance Team |
| EPI_ISL_3070256, EPI_ISL_3070257, EPI_ISL_3070258 | Lighthouse Laboratory Plymouth | Wellcome Sanger Institute for the COVID-19 Genomics UK (COG-UK) Consortium | Lighthouse Laboratory Plymouth and Alex Alderton, Roberto Amato, Jeffrey Barrett, Sonia Goncalves, Ewan Harrison, David K. Jackson, Ian Johnston, Dominic Kwiatkowski, Cordelia Langford, John Sillitoe on behalf of the Wellcome Sanger Institute COVID-19 Surveillance Team |
| EPI_ISL_3070259 | Berkshire and Surrey Pathology Services Lighthouse Laboratory | Wellcome Sanger Institute for the COVID-19 Genomics UK (COG-UK) Consortium | Berkshire and Surrey Pathology Services Lighthouse Laboratory and Alex Alderton, Roberto Amato, Jeffrey Barrett, Sonia Goncalves, Ewan Harrison, David K. Jackson, Ian Johnston, Dominic Kwiatkowski, Cordelia Langford, John Sillitoe on behalf of the Wellcome Sanger Institute COVID-19 Surveillance Team |
| EPI_ISL_3070260, EPI_ISL_3070261, EPI_ISL_3070262 | Lighthouse Laboratory Plymouth | Wellcome Sanger Institute for the COVID-19 Genomics UK (COG-UK) Consortium | Lighthouse Laboratory Plymouth and Alex Alderton, Roberto Amato, Jeffrey Barrett, Sonia Goncalves, Ewan Harrison, David K. Jackson, Ian Johnston, Dominic Kwiatkowski, Cordelia Langford, John Sillitoe on behalf of the Wellcome Sanger Institute COVID-19 Surveillance Team |
| EPI_ISL_3070263 | Lighthouse Lab in Alderley Park | Wellcome Sanger Institute for the COVID-19 Genomics UK (COG-UK) Consortium | Jacquelyn Wynn, Mairead Hyland, The Lighthouse Lab in Alderley Park and Alex Alderton, Roberto Amato, Jeffrey Barrett, Sonia Goncalves, Ewan Harrison, David K. Jackson, Ian Johnston, Dominic Kwiatkowski, Cordelia Langford, John Sillitoe on behalf of the Wellcome Sanger Institute COVID-19 Surveillance Team |
| EPI_ISL_3070264, EPI_ISL_3070265 | Berkshire and Surrey Pathology Services Lighthouse Laboratory | Wellcome Sanger Institute for the COVID-19 Genomics UK (COG-UK) Consortium | Berkshire and Surrey Pathology Services Lighthouse Laboratory and Alex Alderton, Roberto Amato, Jeffrey Barrett, Sonia Goncalves, Ewan Harrison, David K. Jackson, Ian Johnston, Dominic Kwiatkowski, Cordelia Langford, John Sillitoe on behalf of the Wellcome Sanger Institute COVID-19 Surveillance Team |
| EPI_ISL_3070266, EPI_ISL_3070267 | Lighthouse Laboratory Plymouth | Wellcome Sanger Institute for the COVID-19 Genomics UK (COG-UK) Consortium | Lighthouse Laboratory Plymouth and Alex Alderton, Roberto Amato, Jeffrey Barrett, Sonia Goncalves, Ewan Harrison, David K. Jackson, Ian Johnston, Dominic Kwiatkowski, Cordelia Langford, John Sillitoe on behalf of the Wellcome Sanger Institute COVID-19 Surveillance Team |
| EPI_ISL_3070268 | Berkshire and Surrey Pathology Services Lighthouse Laboratory | Wellcome Sanger Institute for the COVID-19 Genomics UK (COG-UK) Consortium | Berkshire and Surrey Pathology Services Lighthouse Laboratory and Alex Alderton, Roberto Amato, Jeffrey Barrett, Sonia Goncalves, Ewan Harrison, David K. Jackson, Ian Johnston, Dominic Kwiatkowski, Cordelia Langford, John Sillitoe on behalf of the Wellcome Sanger Institute COVID-19 Surveillance Team |
| EPI_ISL_3070269 | Lighthouse Laboratory Plymouth | Wellcome Sanger Institute for the COVID-19 Genomics UK (COG-UK) Consortium | Lighthouse Laboratory Plymouth and Alex Alderton, Roberto Amato, Jeffrey Barrett, Sonia Goncalves, Ewan Harrison, David K. Jackson, Ian Johnston, Dominic Kwiatkowski, Cordelia Langford, John Sillitoe on behalf of the Wellcome Sanger Institute COVID-19 Surveillance Team |
| EPI_ISL_3070270 | Berkshire and Surrey Pathology Services Lighthouse Laboratory | Wellcome Sanger Institute for the COVID-19 Genomics UK (COG-UK) Consortium | Berkshire and Surrey Pathology Services Lighthouse Laboratory and Alex Alderton, Roberto Amato, Jeffrey Barrett, Sonia Goncalves, Ewan Harrison, David K. Jackson, Ian Johnston, Dominic Kwiatkowski, Cordelia Langford, John Sillitoe on behalf of the Wellcome Sanger Institute COVID-19 Surveillance Team |
| EPI_ISL_3070271 | Lighthouse Lab in Alderley Park | Wellcome Sanger Institute for the COVID-19 Genomics UK (COG-UK) Consortium | Jacquelyn Wynn, Mairead Hyland, The Lighthouse Lab in Alderley Park and Alex Alderton, Roberto Amato, Jeffrey Barrett, Sonia Goncalves, Ewan Harrison, David K. Jackson, Ian Johnston, Dominic Kwiatkowski, Cordelia Langford, John Sillitoe on behalf of the Wellcome Sanger Institute COVID-19 Surveillance Team |
| EPI_ISL_3070272, EPI_ISL_3070273 | Berkshire and Surrey Pathology Services Lighthouse Laboratory | Wellcome Sanger Institute for the COVID-19 Genomics UK (COG-UK) Consortium | Berkshire and Surrey Pathology Services Lighthouse Laboratory and Alex Alderton, Roberto Amato, Jeffrey Barrett, Sonia Goncalves, Ewan Harrison, David K. Jackson, Ian Johnston, Dominic Kwiatkowski, Cordelia Langford, John Sillitoe on behalf of the Wellcome Sanger Institute COVID-19 Surveillance Team |
| EPI_ISL_3070274 | Lighthouse Lab in Alderley Park | Wellcome Sanger Institute for the COVID-19 Genomics UK (COG-UK) Consortium | Jacquelyn Wynn, Mairead Hyland, The Lighthouse Lab in Alderley Park and Alex Alderton, Roberto Amato, Jeffrey Barrett, Sonia Goncalves, Ewan Harrison, David K. Jackson, Ian Johnston, Dominic Kwiatkowski, Cordelia Langford, John Sillitoe on behalf of the Wellcome Sanger Institute COVID-19 Surveillance Team |
| EPI_ISL_3070275, EPI_ISL_3070276 | Lighthouse Laboratory Plymouth | Wellcome Sanger Institute for the COVID-19 Genomics UK (COG-UK) Consortium | Lighthouse Laboratory Plymouth and Alex Alderton, Roberto Amato, Jeffrey Barrett, Sonia Goncalves, Ewan Harrison, David K. Jackson, Ian Johnston, Dominic Kwiatkowski, Cordelia Langford, John Sillitoe on behalf of the Wellcome Sanger Institute COVID-19 Surveillance Team |
| EPI_ISL_3070277, EPI_ISL_3070278 | Berkshire and Surrey Pathology Services Lighthouse Laboratory | Wellcome Sanger Institute for the COVID-19 Genomics UK (COG-UK) Consortium | Berkshire and Surrey Pathology Services Lighthouse Laboratory and Alex Alderton, Roberto Amato, Jeffrey Barrett, Sonia Goncalves, Ewan Harrison, David K. Jackson, Ian Johnston, Dominic Kwiatkowski, Cordelia Langford, John Sillitoe on behalf of the Wellcome Sanger Institute COVID-19 Surveillance Team |
| EPI_ISL_3070279 | Lighthouse Laboratory Plymouth | Wellcome Sanger Institute for the COVID-19 Genomics UK (COG-UK) Consortium | Lighthouse Laboratory Plymouth and Alex Alderton, Roberto Amato, Jeffrey Barrett, Sonia Goncalves, Ewan Harrison, David K. Jackson, Ian Johnston, Dominic Kwiatkowski, Cordelia Langford, John Sillitoe on behalf of the Wellcome Sanger Institute COVID-19 Surveillance Team |
| EPI_ISL_3070280, EPI_ISL_3070281, EPI_ISL_3070282, EPI_ISL_3070283, EPI_ISL_3070284, EPI_ISL_3070285, EPI_ISL_3070286, EPI_ISL_3070287 | Berkshire and Surrey Pathology Services Lighthouse Laboratory | Wellcome Sanger Institute for the COVID-19 Genomics UK (COG-UK) Consortium | Berkshire and Surrey Pathology Services Lighthouse Laboratory and Alex Alderton, Roberto Amato, Jeffrey Barrett, Sonia Goncalves, Ewan Harrison, David K. Jackson, Ian Johnston, Dominic Kwiatkowski, Cordelia Langford, John Sillitoe on behalf of the Wellcome Sanger Institute COVID-19 Surveillance Team |
| EPI_ISL_3070288, EPI_ISL_3070289 | Lighthouse Lab in Alderley Park | Wellcome Sanger Institute for the COVID-19 Genomics UK (COG-UK) Consortium | Jacquelyn Wynn, Mairead Hyland, The Lighthouse Lab in Alderley Park and Alex Alderton, Roberto Amato, Jeffrey Barrett, Sonia Goncalves, Ewan Harrison, David K. Jackson, Ian Johnston, Dominic Kwiatkowski, Cordelia Langford, John Sillitoe on behalf of the Wellcome Sanger Institute COVID-19 Surveillance Team |
| EPI_ISL_3070290 | Lighthouse Laboratory Plymouth | Wellcome Sanger Institute for the COVID-19 Genomics UK (COG-UK) Consortium | Lighthouse Laboratory Plymouth and Alex Alderton, Roberto Amato, Jeffrey Barrett, Sonia Goncalves, Ewan Harrison, David K. Jackson, Ian Johnston, Dominic Kwiatkowski, Cordelia Langford, John Sillitoe on behalf of the Wellcome Sanger Institute COVID-19 Surveillance Team |
| EPI_ISL_3070291 | Lighthouse Lab in Alderley Park | Wellcome Sanger Institute for the COVID-19 Genomics UK (COG-UK) Consortium | Jacquelyn Wynn, Mairead Hyland, The Lighthouse Lab in Alderley Park and Alex Alderton, Roberto Amato, Jeffrey Barrett, Sonia Goncalves, Ewan Harrison, David K. Jackson, Ian Johnston, Dominic Kwiatkowski, Cordelia Langford, John Sillitoe on behalf of the Wellcome Sanger Institute COVID-19 Surveillance Team |

| | | | |
|---|---|---|---|
| EPI_ISL_3070292 | Lighthouse Laboratory Plymouth | Wellcome Sanger Institute for the COVID-19 Genomics UK (COG-UK) Consortium | Lighthouse Laboratory Plymouth and Alex Alderton, Roberto Amato, Jeffrey Barrett, Sonia Goncalves, Ewan Harrison, David K. Jackson, Ian Johnston, Dominic Kwiatkowski, Cordelia Langford, John Sillitoe on behalf of the Wellcome Sanger Institute COVID-19 Surveillance Team |
| EPI_ISL_3070293 | Berkshire and Surrey Pathology Services Lighthouse Laboratory | Wellcome Sanger Institute for the COVID-19 Genomics UK (COG-UK) Consortium | Berkshire and Surrey Pathology Services Lighthouse Laboratory and Alex Alderton, Roberto Amato, Jeffrey Barrett, Sonia Goncalves, Ewan Harrison, David K. Jackson, Ian Johnston, Dominic Kwiatkowski, Cordelia Langford, John Sillitoe on behalf of the Wellcome Sanger Institute COVID-19 Surveillance Team |
| EPI_ISL_3070294 | Lighthouse Laboratory Plymouth | Wellcome Sanger Institute for the COVID-19 Genomics UK (COG-UK) Consortium | Lighthouse Laboratory Plymouth and Alex Alderton, Roberto Amato, Jeffrey Barrett, Sonia Goncalves, Ewan Harrison, David K. Jackson, Ian Johnston, Dominic Kwiatkowski, Cordelia Langford, John Sillitoe on behalf of the Wellcome Sanger Institute COVID-19 Surveillance Team |
| EPI_ISL_3070295, EPI_ISL_3070296 | Lighthouse Lab in Alderley Park | Wellcome Sanger Institute for the COVID-19 Genomics UK (COG-UK) Consortium | Jacquelyn Wynn, Mairead Hyland, The Lighthouse Lab in Alderley Park and Alex Alderton, Roberto Amato, Jeffrey Barrett, Sonia Goncalves, Ewan Harrison, David K. Jackson, Ian Johnston, Dominic Kwiatkowski, Cordelia Langford, John Sillitoe on behalf of the Wellcome Sanger Institute COVID-19 Surveillance Team |
| EPI_ISL_3070297, EPI_ISL_3070298 | Berkshire and Surrey Pathology Services Lighthouse Laboratory | Wellcome Sanger Institute for the COVID-19 Genomics UK (COG-UK) Consortium | Berkshire and Surrey Pathology Services Lighthouse Laboratory and Alex Alderton, Roberto Amato, Jeffrey Barrett, Sonia Goncalves, Ewan Harrison, David K. Jackson, Ian Johnston, Dominic Kwiatkowski, Cordelia Langford, John Sillitoe on behalf of the Wellcome Sanger Institute COVID-19 Surveillance Team |
| EPI_ISL_3070299 | Lighthouse Laboratory Plymouth | Wellcome Sanger Institute for the COVID-19 Genomics UK (COG-UK) Consortium | Lighthouse Laboratory Plymouth and Alex Alderton, Roberto Amato, Jeffrey Barrett, Sonia Goncalves, Ewan Harrison, David K. Jackson, Ian Johnston, Dominic Kwiatkowski, Cordelia Langford, John Sillitoe on behalf of the Wellcome Sanger Institute COVID-19 Surveillance Team |
| EPI_ISL_3070300 | Berkshire and Surrey Pathology Services Lighthouse Laboratory | Wellcome Sanger Institute for the COVID-19 Genomics UK (COG-UK) Consortium | Berkshire and Surrey Pathology Services Lighthouse Laboratory and Alex Alderton, Roberto Amato, Jeffrey Barrett, Sonia Goncalves, Ewan Harrison, David K. Jackson, Ian Johnston, Dominic Kwiatkowski, Cordelia Langford, John Sillitoe on behalf of the Wellcome Sanger Institute COVID-19 Surveillance Team |
| EPI_ISL_3070301 | Lighthouse Laboratory Plymouth | Wellcome Sanger Institute for the COVID-19 Genomics UK (COG-UK) Consortium | Lighthouse Laboratory Plymouth and Alex Alderton, Roberto Amato, Jeffrey Barrett, Sonia Goncalves, Ewan Harrison, David K. Jackson, Ian Johnston, Dominic Kwiatkowski, Cordelia Langford, John Sillitoe on behalf of the Wellcome Sanger Institute COVID-19 Surveillance Team |
| EPI_ISL_3070302, EPI_ISL_3070303, EPI_ISL_3070304, EPI_ISL_3070305 | Berkshire and Surrey Pathology Services Lighthouse Laboratory | Wellcome Sanger Institute for the COVID-19 Genomics UK (COG-UK) Consortium | Berkshire and Surrey Pathology Services Lighthouse Laboratory and Alex Alderton, Roberto Amato, Jeffrey Barrett, Sonia Goncalves, Ewan Harrison, David K. Jackson, Ian Johnston, Dominic Kwiatkowski, Cordelia Langford, John Sillitoe on behalf of the Wellcome Sanger Institute COVID-19 Surveillance Team |
| EPI_ISL_3070306, EPI_ISL_3070307, EPI_ISL_3070308 | Lighthouse Lab in Alderley Park | Wellcome Sanger Institute for the COVID-19 Genomics UK (COG-UK) Consortium | Jacquelyn Wynn, Mairead Hyland, The Lighthouse Lab in Alderley Park and Alex Alderton, Roberto Amato, Jeffrey Barrett, Sonia Goncalves, Ewan Harrison, David K. Jackson, Ian Johnston, Dominic Kwiatkowski, Cordelia Langford, John Sillitoe on behalf of the Wellcome Sanger Institute COVID-19 Surveillance Team |
| EPI_ISL_3070309, EPI_ISL_3070310 | Berkshire and Surrey Pathology Services Lighthouse Laboratory | Wellcome Sanger Institute for the COVID-19 Genomics UK (COG-UK) Consortium | Berkshire and Surrey Pathology Services Lighthouse Laboratory and Alex Alderton, Roberto Amato, Jeffrey Barrett, Sonia Goncalves, Ewan Harrison, David K. Jackson, Ian Johnston, Dominic Kwiatkowski, Cordelia Langford, John Sillitoe on behalf of the Wellcome Sanger Institute COVID-19 Surveillance Team |
| EPI_ISL_3070311 | Lighthouse Lab in Alderley Park | Wellcome Sanger Institute for the COVID-19 Genomics UK (COG-UK) Consortium | Jacquelyn Wynn, Mairead Hyland, The Lighthouse Lab in Alderley Park and Alex Alderton, Roberto Amato, Jeffrey Barrett, Sonia Goncalves, Ewan Harrison, David K. Jackson, Ian Johnston, Dominic Kwiatkowski, Cordelia Langford, John Sillitoe on behalf of the Wellcome Sanger Institute COVID-19 Surveillance Team |
| EPI_ISL_3070312 | Berkshire and Surrey Pathology Services Lighthouse Laboratory | Wellcome Sanger Institute for the COVID-19 Genomics UK (COG-UK) Consortium | Berkshire and Surrey Pathology Services Lighthouse Laboratory and Alex Alderton, Roberto Amato, Jeffrey Barrett, Sonia Goncalves, Ewan Harrison, David K. Jackson, Ian Johnston, Dominic Kwiatkowski, Cordelia Langford, John Sillitoe on behalf of the Wellcome Sanger Institute COVID-19 Surveillance Team |
| EPI_ISL_3070313 | Lighthouse Lab in Alderley Park | Wellcome Sanger Institute for the COVID-19 Genomics UK (COG-UK) Consortium | Jacquelyn Wynn, Mairead Hyland, The Lighthouse Lab in Alderley Park and Alex Alderton, Roberto Amato, Jeffrey Barrett, Sonia Goncalves, Ewan Harrison, David K. Jackson, Ian Johnston, Dominic Kwiatkowski, Cordelia Langford, John Sillitoe on behalf of the Wellcome Sanger Institute COVID-19 Surveillance Team |
| EPI_ISL_3070314 | Lighthouse Laboratory Plymouth | Wellcome Sanger Institute for the COVID-19 Genomics UK (COG-UK) Consortium | Lighthouse Laboratory Plymouth and Alex Alderton, Roberto Amato, Jeffrey Barrett, Sonia Goncalves, Ewan Harrison, David K. Jackson, Ian Johnston, Dominic Kwiatkowski, Cordelia Langford, John Sillitoe on behalf of the Wellcome Sanger Institute COVID-19 Surveillance Team |
| EPI_ISL_3070315, EPI_ISL_3070316 | Lighthouse Lab in Alderley Park | Wellcome Sanger Institute for the COVID-19 Genomics UK (COG-UK) Consortium | Jacquelyn Wynn, Mairead Hyland, The Lighthouse Lab in Alderley Park and Alex Alderton, Roberto Amato, Jeffrey Barrett, Sonia Goncalves, Ewan Harrison, David K. Jackson, Ian Johnston, Dominic Kwiatkowski, Cordelia Langford, John Sillitoe on behalf of the Wellcome Sanger Institute COVID-19 Surveillance Team |
| EPI_ISL_3070317 | Berkshire and Surrey Pathology Services Lighthouse Laboratory | Wellcome Sanger Institute for the COVID-19 Genomics UK (COG-UK) Consortium | Berkshire and Surrey Pathology Services Lighthouse Laboratory and Alex Alderton, Roberto Amato, Jeffrey Barrett, Sonia Goncalves, Ewan Harrison, David K. Jackson, Ian Johnston, Dominic Kwiatkowski, Cordelia Langford, John Sillitoe on behalf of the Wellcome Sanger Institute COVID-19 Surveillance Team |
| EPI_ISL_3070318, EPI_ISL_3070319 | Lighthouse Laboratory Plymouth | Wellcome Sanger Institute for the COVID-19 Genomics UK (COG-UK) Consortium | Lighthouse Laboratory Plymouth and Alex Alderton, Roberto Amato, Jeffrey Barrett, Sonia Goncalves, Ewan Harrison, David K. Jackson, Ian Johnston, Dominic Kwiatkowski, Cordelia Langford, John Sillitoe on behalf of the Wellcome Sanger Institute COVID-19 Surveillance Team |
| EPI_ISL_3070320 | Lighthouse Lab in Alderley Park | Wellcome Sanger Institute for the COVID-19 Genomics UK (COG-UK) Consortium | Jacquelyn Wynn, Mairead Hyland, The Lighthouse Lab in Alderley Park and Alex Alderton, Roberto Amato, Jeffrey Barrett, Sonia Goncalves, Ewan Harrison, David K. Jackson, Ian Johnston, Dominic Kwiatkowski, Cordelia Langford, John Sillitoe on behalf of the Wellcome Sanger Institute COVID-19 Surveillance Team |
| EPI_ISL_3070321 | Berkshire and Surrey Pathology Services Lighthouse Laboratory | Wellcome Sanger Institute for the COVID-19 Genomics UK (COG-UK) Consortium | Berkshire and Surrey Pathology Services Lighthouse Laboratory and Alex Alderton, Roberto Amato, Jeffrey Barrett, Sonia Goncalves, Ewan Harrison, David K. Jackson, Ian Johnston, Dominic Kwiatkowski, Cordelia Langford, John Sillitoe on behalf of the Wellcome Sanger Institute COVID-19 Surveillance Team |
| EPI_ISL_3070322 | Lighthouse Laboratory Plymouth | Wellcome Sanger Institute for the COVID-19 Genomics UK (COG-UK) Consortium | Lighthouse Laboratory Plymouth and Alex Alderton, Roberto Amato, Jeffrey Barrett, Sonia Goncalves, Ewan Harrison, David K. Jackson, Ian Johnston, Dominic Kwiatkowski, Cordelia Langford, John Sillitoe on behalf of the Wellcome Sanger Institute COVID-19 Surveillance Team |
| EPI_ISL_3070323, EPI_ISL_3070324, EPI_ISL_3070325, EPI_ISL_3070326 | Berkshire and Surrey Pathology Services Lighthouse Laboratory | Wellcome Sanger Institute for the COVID-19 Genomics UK (COG-UK) Consortium | Berkshire and Surrey Pathology Services Lighthouse Laboratory and Alex Alderton, Roberto Amato, Jeffrey Barrett, Sonia Goncalves, Ewan Harrison, David K. Jackson, Ian Johnston, Dominic Kwiatkowski, Cordelia Langford, John Sillitoe on behalf of the Wellcome Sanger Institute COVID-19 Surveillance Team |
| EPI_ISL_3070327 | Lighthouse Laboratory Plymouth | Wellcome Sanger Institute for the COVID-19 Genomics UK (COG-UK) Consortium | Lighthouse Laboratory Plymouth and Alex Alderton, Roberto Amato, Jeffrey Barrett, Sonia Goncalves, Ewan Harrison, David K. Jackson, Ian Johnston, Dominic Kwiatkowski, Cordelia Langford, John Sillitoe on behalf of the Wellcome Sanger Institute COVID-19 Surveillance Team |
| EPI_ISL_3070328 | Lighthouse Lab in Alderley Park | Wellcome Sanger Institute for the COVID-19 Genomics UK (COG-UK) Consortium | Jacquelyn Wynn, Mairead Hyland, The Lighthouse Lab in Alderley Park and Alex Alderton, Roberto Amato, Jeffrey Barrett, Sonia Goncalves, Ewan Harrison, David K. Jackson, Ian Johnston, Dominic Kwiatkowski, Cordelia Langford, John Sillitoe on behalf of the Wellcome Sanger Institute COVID-19 Surveillance Team |
| EPI_ISL_3070329 | Berkshire and Surrey Pathology Services Lighthouse Laboratory | Wellcome Sanger Institute for the COVID-19 Genomics UK (COG-UK) Consortium | Berkshire and Surrey Pathology Services Lighthouse Laboratory and Alex Alderton, Roberto Amato, Jeffrey Barrett, Sonia Goncalves, Ewan Harrison, David K. Jackson, Ian Johnston, Dominic Kwiatkowski, Cordelia Langford, John Sillitoe on behalf of the Wellcome Sanger Institute COVID-19 Surveillance Team |
| EPI_ISL_3070330 | Lighthouse Laboratory Plymouth | Wellcome Sanger Institute for the COVID-19 Genomics UK (COG-UK) Consortium | Lighthouse Laboratory Plymouth and Alex Alderton, Roberto Amato, Jeffrey Barrett, Sonia Goncalves, Ewan Harrison, David K. Jackson, Ian Johnston, Dominic Kwiatkowski, Cordelia Langford, John Sillitoe on behalf of the Wellcome Sanger Institute COVID-19 Surveillance Team |
| EPI_ISL_3070331, EPI_ISL_3070332 | Berkshire and Surrey Pathology Services Lighthouse Laboratory | Wellcome Sanger Institute for the COVID-19 Genomics UK (COG-UK) Consortium | Berkshire and Surrey Pathology Services Lighthouse Laboratory and Alex Alderton, Roberto Amato, Jeffrey Barrett, Sonia Goncalves, Ewan Harrison, David K. Jackson, Ian Johnston, Dominic Kwiatkowski, Cordelia Langford, John Sillitoe on behalf of the Wellcome Sanger Institute COVID-19 Surveillance Team |
| EPI_ISL_3070333 | Lighthouse Laboratory Plymouth | Wellcome Sanger Institute for the COVID-19 Genomics UK (COG-UK) Consortium | Lighthouse Laboratory Plymouth and Alex Alderton, Roberto Amato, Jeffrey Barrett, Sonia Goncalves, Ewan Harrison, David K. Jackson, Ian Johnston, Dominic Kwiatkowski, Cordelia Langford, John Sillitoe on behalf of the Wellcome Sanger Institute COVID-19 Surveillance Team |
| EPI_ISL_3070334 | Lighthouse Lab in Alderley Park | Wellcome Sanger Institute for the COVID-19 Genomics UK (COG-UK) Consortium | Jacquelyn Wynn, Mairead Hyland, The Lighthouse Lab in Alderley Park and Alex Alderton, Roberto Amato, Jeffrey Barrett, Sonia Goncalves, Ewan Harrison, David K. Jackson, Ian Johnston, Dominic Kwiatkowski, Cordelia Langford, John Sillitoe on behalf of the Wellcome Sanger Institute COVID-19 Surveillance Team |
| EPI_ISL_3070335, EPI_ISL_3070336, EPI_ISL_3070337 | Berkshire and Surrey Pathology Services Lighthouse Laboratory | Wellcome Sanger Institute for the COVID-19 Genomics UK (COG-UK) Consortium | Berkshire and Surrey Pathology Services Lighthouse Laboratory and Alex Alderton, Roberto Amato, Jeffrey Barrett, Sonia Goncalves, Ewan Harrison, David K. Jackson, Ian Johnston, Dominic Kwiatkowski, Cordelia Langford, John Sillitoe on behalf of the Wellcome Sanger Institute COVID-19 Surveillance Team |
| EPI_ISL_3070338 | Lighthouse Laboratory Plymouth | Wellcome Sanger Institute for the COVID-19 Genomics UK (COG-UK) Consortium | Lighthouse Laboratory Plymouth and Alex Alderton, Roberto Amato, Jeffrey Barrett, Sonia Goncalves, Ewan Harrison, David K. Jackson, Ian Johnston, Dominic Kwiatkowski, Cordelia Langford, John Sillitoe on behalf of the Wellcome Sanger Institute COVID-19 Surveillance Team |
| EPI_ISL_3070339 | Berkshire and Surrey Pathology Services Lighthouse Laboratory | Wellcome Sanger Institute for the COVID-19 Genomics UK (COG-UK) Consortium | Berkshire and Surrey Pathology Services Lighthouse Laboratory and Alex Alderton, Roberto Amato, Jeffrey Barrett, Sonia Goncalves, Ewan Harrison, David K. Jackson, Ian Johnston, Dominic Kwiatkowski, Cordelia Langford, John Sillitoe on behalf of the Wellcome Sanger Institute COVID-19 Surveillance Team |
| EPI_ISL_3070340 | Lighthouse Laboratory Plymouth | Wellcome Sanger Institute for the COVID-19 Genomics UK (COG-UK) Consortium | Lighthouse Laboratory Plymouth and Alex Alderton, Roberto Amato, Jeffrey Barrett, Sonia Goncalves, Ewan Harrison, David K. Jackson, Ian Johnston, Dominic Kwiatkowski, Cordelia Langford, John Sillitoe on behalf of the Wellcome Sanger Institute COVID-19 Surveillance Team |
| EPI_ISL_3070341, EPI_ISL_3070342, EPI_ISL_3070343, EPI_ISL_3070344 | Berkshire and Surrey Pathology Services Lighthouse Laboratory | Wellcome Sanger Institute for the COVID-19 Genomics UK (COG-UK) Consortium | Berkshire and Surrey Pathology Services Lighthouse Laboratory and Alex Alderton, Roberto Amato, Jeffrey Barrett, Sonia Goncalves, Ewan Harrison, David K. Jackson, Ian Johnston, Dominic Kwiatkowski, Cordelia Langford, John Sillitoe on behalf of the Wellcome Sanger Institute COVID-19 Surveillance Team |
| EPI_ISL_3070345, EPI_ISL_3070346 | Lighthouse Laboratory Plymouth | Wellcome Sanger Institute for the COVID-19 Genomics UK (COG-UK) Consortium | Lighthouse Laboratory Plymouth and Alex Alderton, Roberto Amato, Jeffrey Barrett, Sonia Goncalves, Ewan Harrison, David K. Jackson, Ian Johnston, Dominic Kwiatkowski, Cordelia Langford, John Sillitoe on behalf of the Wellcome Sanger Institute COVID-19 Surveillance Team |
| EPI_ISL_3070347 | Berkshire and Surrey Pathology Services Lighthouse Laboratory | Wellcome Sanger Institute for the COVID-19 Genomics UK (COG-UK) Consortium | Berkshire and Surrey Pathology Services Lighthouse Laboratory and Alex Alderton, Roberto Amato, Jeffrey Barrett, Sonia Goncalves, Ewan Harrison, David K. Jackson, Ian Johnston, Dominic Kwiatkowski, Cordelia Langford, John Sillitoe on behalf of the Wellcome Sanger Institute COVID-19 Surveillance Team |
| EPI_ISL_3070348 | Lighthouse Lab in Alderley Park | Wellcome Sanger Institute for the COVID-19 Genomics UK (COG-UK) Consortium | Jacquelyn Wynn, Mairead Hyland, The Lighthouse Lab in Alderley Park and Alex Alderton, Roberto Amato, Jeffrey Barrett, Sonia Goncalves, Ewan Harrison, David K. Jackson, Ian Johnston, Dominic Kwiatkowski, Cordelia Langford, John Sillitoe on behalf of the Wellcome Sanger Institute COVID-19 Surveillance Team |
| EPI_ISL_3070349, EPI_ISL_3070350 | Berkshire and Surrey Pathology Services Lighthouse Laboratory | Wellcome Sanger Institute for the COVID-19 Genomics UK (COG-UK) Consortium | Berkshire and Surrey Pathology Services Lighthouse Laboratory and Alex Alderton, Roberto Amato, Jeffrey Barrett, Sonia Goncalves, Ewan Harrison, David K. Jackson, Ian Johnston, Dominic Kwiatkowski, Cordelia Langford, John Sillitoe on behalf of the Wellcome Sanger Institute COVID-19 Surveillance Team |
| EPI_ISL_3070351 | Lighthouse Laboratory Plymouth | Wellcome Sanger Institute for the COVID-19 Genomics UK (COG-UK) Consortium | Lighthouse Laboratory Plymouth and Alex Alderton, Roberto Amato, Jeffrey Barrett, Sonia Goncalves, Ewan Harrison, David K. Jackson, Ian Johnston, Dominic Kwiatkowski, Cordelia Langford, John Sillitoe on behalf of the Wellcome Sanger Institute COVID-19 Surveillance Team |
| EPI_ISL_3070352 | Lighthouse Lab in Alderley Park | Wellcome Sanger Institute for the COVID-19 Genomics UK (COG-UK) Consortium | Jacquelyn Wynn, Mairead Hyland, The Lighthouse Lab in Alderley Park and Alex Alderton, Roberto Amato, Jeffrey Barrett, Sonia Goncalves, Ewan Harrison, David K. Jackson, Ian Johnston, Dominic Kwiatkowski, Cordelia Langford, John Sillitoe on behalf of the Wellcome Sanger Institute COVID-19 Surveillance Team |
| EPI_ISL_3070353 | Berkshire and Surrey Pathology Services Lighthouse Laboratory | Wellcome Sanger Institute for the COVID-19 Genomics UK (COG-UK) Consortium | Berkshire and Surrey Pathology Services Lighthouse Laboratory and Alex Alderton, Roberto Amato, Jeffrey Barrett, Sonia Goncalves, Ewan Harrison, David K. Jackson, Ian Johnston, Dominic Kwiatkowski, Cordelia Langford, John Sillitoe on behalf of the Wellcome Sanger Institute COVID-19 Surveillance Team |
| EPI_ISL_3070354, | Lighthouse Laboratory Plymouth | Wellcome Sanger Institute for the COVID-19 Genomics UK (COG-UK) | Lighthouse Laboratory Plymouth and Alex Alderton, Roberto Amato, Jeffrey Barrett, Sonia Goncalves, Ewan Harrison, David K. Jackson, Ian Johnston, Dominic Kwiatkowski, Cordelia Langford, John Sillitoe on behalf of the Wellcome Sanger |

| | | | |
|---|---|---|---|
| EPI_ISL_3070355 | | Consortium | Institute COVID-19 Surveillance Team |
| EPI_ISL_3070356 | Berkshire and Surrey Pathology Services Lighthouse Laboratory | Wellcome Sanger Institute for the COVID-19 Genomics UK (COG-UK) Consortium | Berkshire and Surrey Pathology Services Lighthouse Laboratory and Alex Alderton, Roberto Amato, Jeffrey Barrett, Sonia Goncalves, Ewan Harrison, David K. Jackson, Ian Johnston, Dominic Kwiatkowski, Cordelia Langford, John Sillitoe on behalf of the Wellcome Sanger Institute COVID-19 Surveillance Team |
| EPI_ISL_3070357 | Lighthouse Lab in Alderley Park | Wellcome Sanger Institute for the COVID-19 Genomics UK (COG-UK) Consortium | Jacquelyn Wynn, Mairead Hyland, The Lighthouse Lab in Alderley Park and Alex Alderton, Roberto Amato, Jeffrey Barrett, Sonia Goncalves, Ewan Harrison, David K. Jackson, Ian Johnston, Dominic Kwiatkowski, Cordelia Langford, John Sillitoe on behalf of the Wellcome Sanger Institute COVID-19 Surveillance Team |
| EPI_ISL_3070358, EPI_ISL_3070359, EPI_ISL_3070360, EPI_ISL_3070361, EPI_ISL_3070362, EPI_ISL_3070363, EPI_ISL_3070364, EPI_ISL_3070365 | Berkshire and Surrey Pathology Services Lighthouse Laboratory | Wellcome Sanger Institute for the COVID-19 Genomics UK (COG-UK) Consortium | Berkshire and Surrey Pathology Services Lighthouse Laboratory and Alex Alderton, Roberto Amato, Jeffrey Barrett, Sonia Goncalves, Ewan Harrison, David K. Jackson, Ian Johnston, Dominic Kwiatkowski, Cordelia Langford, John Sillitoe on behalf of the Wellcome Sanger Institute COVID-19 Surveillance Team |
| EPI_ISL_3070366 | Lighthouse Lab in Alderley Park | Wellcome Sanger Institute for the COVID-19 Genomics UK (COG-UK) Consortium | Jacquelyn Wynn, Mairead Hyland, The Lighthouse Lab in Alderley Park and Alex Alderton, Roberto Amato, Jeffrey Barrett, Sonia Goncalves, Ewan Harrison, David K. Jackson, Ian Johnston, Dominic Kwiatkowski, Cordelia Langford, John Sillitoe on behalf of the Wellcome Sanger Institute COVID-19 Surveillance Team |
| EPI_ISL_3070367, EPI_ISL_3070368 | Berkshire and Surrey Pathology Services Lighthouse Laboratory | Wellcome Sanger Institute for the COVID-19 Genomics UK (COG-UK) Consortium | Berkshire and Surrey Pathology Services Lighthouse Laboratory and Alex Alderton, Roberto Amato, Jeffrey Barrett, Sonia Goncalves, Ewan Harrison, David K. Jackson, Ian Johnston, Dominic Kwiatkowski, Cordelia Langford, John Sillitoe on behalf of the Wellcome Sanger Institute COVID-19 Surveillance Team |
| EPI_ISL_3070369 | Lighthouse Lab in Milton Keynes | Wellcome Sanger Institute for the COVID-19 Genomics UK (COG-UK) Consortium | The Lighthouse Lab in Milton Keynes and Alex Alderton, Roberto Amato, Jeffrey Barrett, Sonia Goncalves, Ewan Harrison, David K. Jackson, Ian Johnston, Dominic Kwiatkowski, Cordelia Langford, John Sillitoe on behalf of the Wellcome Sanger Institute COVID-19 Surveillance Team |
| EPI_ISL_3070370 | Lighthouse Lab in Alderley Park | Wellcome Sanger Institute for the COVID-19 Genomics UK (COG-UK) Consortium | Jacquelyn Wynn, Mairead Hyland, The Lighthouse Lab in Alderley Park and Alex Alderton, Roberto Amato, Jeffrey Barrett, Sonia Goncalves, Ewan Harrison, David K. Jackson, Ian Johnston, Dominic Kwiatkowski, Cordelia Langford, John Sillitoe on behalf of the Wellcome Sanger Institute COVID-19 Surveillance Team |
| EPI_ISL_3070371, EPI_ISL_3070372 | Lighthouse Lab in Milton Keynes | Wellcome Sanger Institute for the COVID-19 Genomics UK (COG-UK) Consortium | The Lighthouse Lab in Milton Keynes and Alex Alderton, Roberto Amato, Jeffrey Barrett, Sonia Goncalves, Ewan Harrison, David K. Jackson, Ian Johnston, Dominic Kwiatkowski, Cordelia Langford, John Sillitoe on behalf of the Wellcome Sanger Institute COVID-19 Surveillance Team |
| EPI_ISL_3070373 | Berkshire and Surrey Pathology Services Lighthouse Laboratory | Wellcome Sanger Institute for the COVID-19 Genomics UK (COG-UK) Consortium | Berkshire and Surrey Pathology Services Lighthouse Laboratory and Alex Alderton, Roberto Amato, Jeffrey Barrett, Sonia Goncalves, Ewan Harrison, David K. Jackson, Ian Johnston, Dominic Kwiatkowski, Cordelia Langford, John Sillitoe on behalf of the Wellcome Sanger Institute COVID-19 Surveillance Team |
| EPI_ISL_3070374 | Lighthouse Lab in Milton Keynes | Wellcome Sanger Institute for the COVID-19 Genomics UK (COG-UK) Consortium | The Lighthouse Lab in Milton Keynes and Alex Alderton, Roberto Amato, Jeffrey Barrett, Sonia Goncalves, Ewan Harrison, David K. Jackson, Ian Johnston, Dominic Kwiatkowski, Cordelia Langford, John Sillitoe on behalf of the Wellcome Sanger Institute COVID-19 Surveillance Team |
| EPI_ISL_3070375, EPI_ISL_3070376, EPI_ISL_3070377 | Health Services Laboratories | Wellcome Sanger Institute for the COVID-19 Genomics UK (COG-UK) Consortium | Health Services Laboratories and Alex Alderton, Roberto Amato, Jeffrey Barrett, Sonia Goncalves, Ewan Harrison, David K. Jackson, Ian Johnston, Dominic Kwiatkowski, Cordelia Langford, John Sillitoe on behalf of the Wellcome Sanger Institute COVID-19 Surveillance Team |
| EPI_ISL_3070378 | Lighthouse Lab in Glasgow | Wellcome Sanger Institute for the COVID-19 Genomics UK (COG-UK) Consortium | Harper VanSteenhouse, Yumi Kasai, David Gray, Carol Clugston, Anna Dominiczak and Alex Alderton, Roberto Amato, Jeffrey Barrett, Sonia Goncalves, Ewan Harrison, David K. Jackson, Ian Johnston, Dominic Kwiatkowski, Cordelia Langford, John Sillitoe on behalf of the Wellcome Sanger Institute COVID-19 Surveillance Team |
| EPI_ISL_3070379 | Health Services Laboratories | Wellcome Sanger Institute for the COVID-19 Genomics UK (COG-UK) Consortium | Health Services Laboratories and Alex Alderton, Roberto Amato, Jeffrey Barrett, Sonia Goncalves, Ewan Harrison, David K. Jackson, Ian Johnston, Dominic Kwiatkowski, Cordelia Langford, John Sillitoe on behalf of the Wellcome Sanger Institute COVID-19 Surveillance Team |
| EPI_ISL_3070380, EPI_ISL_3070381, EPI_ISL_3070382, EPI_ISL_3070383, EPI_ISL_3070384, EPI_ISL_3070385 | Lighthouse Lab in Glasgow | Wellcome Sanger Institute for the COVID-19 Genomics UK (COG-UK) Consortium | Harper VanSteenhouse, Yumi Kasai, David Gray, Carol Clugston, Anna Dominiczak and Alex Alderton, Roberto Amato, Jeffrey Barrett, Sonia Goncalves, Ewan Harrison, David K. Jackson, Ian Johnston, Dominic Kwiatkowski, Cordelia Langford, John Sillitoe on behalf of the Wellcome Sanger Institute COVID-19 Surveillance Team |
| EPI_ISL_3070386 | Health Services Laboratories | Wellcome Sanger Institute for the COVID-19 Genomics UK (COG-UK) Consortium | Health Services Laboratories and Alex Alderton, Roberto Amato, Jeffrey Barrett, Sonia Goncalves, Ewan Harrison, David K. Jackson, Ian Johnston, Dominic Kwiatkowski, Cordelia Langford, John Sillitoe on behalf of the Wellcome Sanger Institute COVID-19 Surveillance Team |
| EPI_ISL_3070387, EPI_ISL_3070388, EPI_ISL_3070389, EPI_ISL_3070390 | Lighthouse Lab in Glasgow | Wellcome Sanger Institute for the COVID-19 Genomics UK (COG-UK) Consortium | Harper VanSteenhouse, Yumi Kasai, David Gray, Carol Clugston, Anna Dominiczak and Alex Alderton, Roberto Amato, Jeffrey Barrett, Sonia Goncalves, Ewan Harrison, David K. Jackson, Ian Johnston, Dominic Kwiatkowski, Cordelia Langford, John Sillitoe on behalf of the Wellcome Sanger Institute COVID-19 Surveillance Team |
| EPI_ISL_3070391 | Health Services Laboratories | Wellcome Sanger Institute for the COVID-19 Genomics UK (COG-UK) Consortium | Health Services Laboratories and Alex Alderton, Roberto Amato, Jeffrey Barrett, Sonia Goncalves, Ewan Harrison, David K. Jackson, Ian Johnston, Dominic Kwiatkowski, Cordelia Langford, John Sillitoe on behalf of the Wellcome Sanger Institute COVID-19 Surveillance Team |
| EPI_ISL_3070392 | Lighthouse Lab in Alderley Park | Wellcome Sanger Institute for the COVID-19 Genomics UK (COG-UK) Consortium | Jacquelyn Wynn, Mairead Hyland, The Lighthouse Lab in Alderley Park and Alex Alderton, Roberto Amato, Jeffrey Barrett, Sonia Goncalves, Ewan Harrison, David K. Jackson, Ian Johnston, Dominic Kwiatkowski, Cordelia Langford, John Sillitoe on behalf of the Wellcome Sanger Institute COVID-19 Surveillance Team |
| EPI_ISL_3070393 | Health Services Laboratories | Wellcome Sanger Institute for the COVID-19 Genomics UK (COG-UK) Consortium | Health Services Laboratories and Alex Alderton, Roberto Amato, Jeffrey Barrett, Sonia Goncalves, Ewan Harrison, David K. Jackson, Ian Johnston, Dominic Kwiatkowski, Cordelia Langford, John Sillitoe on behalf of the Wellcome Sanger Institute COVID-19 Surveillance Team |
| EPI_ISL_3070394 | Lighthouse Lab in Glasgow | Wellcome Sanger Institute for the COVID-19 Genomics UK (COG-UK) Consortium | Harper VanSteenhouse, Yumi Kasai, David Gray, Carol Clugston, Anna Dominiczak and Alex Alderton, Roberto Amato, Jeffrey Barrett, Sonia Goncalves, Ewan Harrison, David K. Jackson, Ian Johnston, Dominic Kwiatkowski, Cordelia Langford, John Sillitoe on behalf of the Wellcome Sanger Institute COVID-19 Surveillance Team |
| EPI_ISL_3070395, EPI_ISL_3070396 | Health Services Laboratories | Wellcome Sanger Institute for the COVID-19 Genomics UK (COG-UK) Consortium | Health Services Laboratories and Alex Alderton, Roberto Amato, Jeffrey Barrett, Sonia Goncalves, Ewan Harrison, David K. Jackson, Ian Johnston, Dominic Kwiatkowski, Cordelia Langford, John Sillitoe on behalf of the Wellcome Sanger Institute COVID-19 Surveillance Team |
| EPI_ISL_3070397, EPI_ISL_3070398 | Lighthouse Lab in Glasgow | Wellcome Sanger Institute for the COVID-19 Genomics UK (COG-UK) Consortium | Harper VanSteenhouse, Yumi Kasai, David Gray, Carol Clugston, Anna Dominiczak and Alex Alderton, Roberto Amato, Jeffrey Barrett, Sonia Goncalves, Ewan Harrison, David K. Jackson, Ian Johnston, Dominic Kwiatkowski, Cordelia Langford, John Sillitoe on behalf of the Wellcome Sanger Institute COVID-19 Surveillance Team |
| EPI_ISL_3070399, EPI_ISL_3070400, EPI_ISL_3070401, EPI_ISL_3070402 | Health Services Laboratories | Wellcome Sanger Institute for the COVID-19 Genomics UK (COG-UK) Consortium | Health Services Laboratories and Alex Alderton, Roberto Amato, Jeffrey Barrett, Sonia Goncalves, Ewan Harrison, David K. Jackson, Ian Johnston, Dominic Kwiatkowski, Cordelia Langford, John Sillitoe on behalf of the Wellcome Sanger Institute COVID-19 Surveillance Team |
| EPI_ISL_3070403, EPI_ISL_3070404 | Lighthouse Lab in Glasgow | Wellcome Sanger Institute for the COVID-19 Genomics UK (COG-UK) Consortium | Harper VanSteenhouse, Yumi Kasai, David Gray, Carol Clugston, Anna Dominiczak and Alex Alderton, Roberto Amato, Jeffrey Barrett, Sonia Goncalves, Ewan Harrison, David K. Jackson, Ian Johnston, Dominic Kwiatkowski, Cordelia Langford, John Sillitoe on behalf of the Wellcome Sanger Institute COVID-19 Surveillance Team |
| EPI_ISL_3070405 | Health Services Laboratories | Wellcome Sanger Institute for the COVID-19 Genomics UK (COG-UK) Consortium | Health Services Laboratories and Alex Alderton, Roberto Amato, Jeffrey Barrett, Sonia Goncalves, Ewan Harrison, David K. Jackson, Ian Johnston, Dominic Kwiatkowski, Cordelia Langford, John Sillitoe on behalf of the Wellcome Sanger Institute COVID-19 Surveillance Team |
| EPI_ISL_3070406, EPI_ISL_3070407 | Lighthouse Lab in Glasgow | Wellcome Sanger Institute for the COVID-19 Genomics UK (COG-UK) Consortium | Harper VanSteenhouse, Yumi Kasai, David Gray, Carol Clugston, Anna Dominiczak and Alex Alderton, Roberto Amato, Jeffrey Barrett, Sonia Goncalves, Ewan Harrison, David K. Jackson, Ian Johnston, Dominic Kwiatkowski, Cordelia Langford, John Sillitoe on behalf of the Wellcome Sanger Institute COVID-19 Surveillance Team |
| EPI_ISL_3070408, EPI_ISL_3070409 | Health Services Laboratories | Wellcome Sanger Institute for the COVID-19 Genomics UK (COG-UK) Consortium | Health Services Laboratories and Alex Alderton, Roberto Amato, Jeffrey Barrett, Sonia Goncalves, Ewan Harrison, David K. Jackson, Ian Johnston, Dominic Kwiatkowski, Cordelia Langford, John Sillitoe on behalf of the Wellcome Sanger Institute COVID-19 Surveillance Team |
| EPI_ISL_3070410, EPI_ISL_3070411, EPI_ISL_3070412, EPI_ISL_3070413, EPI_ISL_3070414, EPI_ISL_3070415 | Berkshire and Surrey Pathology Services Lighthouse Laboratory | Wellcome Sanger Institute for the COVID-19 Genomics UK (COG-UK) Consortium | Berkshire and Surrey Pathology Services Lighthouse Laboratory and Alex Alderton, Roberto Amato, Jeffrey Barrett, Sonia Goncalves, Ewan Harrison, David K. Jackson, Ian Johnston, Dominic Kwiatkowski, Cordelia Langford, John Sillitoe on behalf of the Wellcome Sanger Institute COVID-19 Surveillance Team |
| EPI_ISL_3070416, EPI_ISL_3070417 | Lighthouse Lab in Milton Keynes | Wellcome Sanger Institute for the COVID-19 Genomics UK (COG-UK) Consortium | The Lighthouse Lab in Milton Keynes and Alex Alderton, Roberto Amato, Jeffrey Barrett, Sonia Goncalves, Ewan Harrison, David K. Jackson, Ian Johnston, Dominic Kwiatkowski, Cordelia Langford, John Sillitoe on behalf of the Wellcome Sanger Institute COVID-19 Surveillance Team |
| EPI_ISL_3070418, EPI_ISL_3070419 | Berkshire and Surrey Pathology Services Lighthouse Laboratory | Wellcome Sanger Institute for the COVID-19 Genomics UK (COG-UK) Consortium | Berkshire and Surrey Pathology Services Lighthouse Laboratory and Alex Alderton, Roberto Amato, Jeffrey Barrett, Sonia Goncalves, Ewan Harrison, David K. Jackson, Ian Johnston, Dominic Kwiatkowski, Cordelia Langford, John Sillitoe on behalf of the Wellcome Sanger Institute COVID-19 Surveillance Team |
| EPI_ISL_3070420 | Lighthouse Lab in Milton Keynes | Wellcome Sanger Institute for the COVID-19 Genomics UK (COG-UK) Consortium | The Lighthouse Lab in Milton Keynes and Alex Alderton, Roberto Amato, Jeffrey Barrett, Sonia Goncalves, Ewan Harrison, David K. Jackson, Ian Johnston, Dominic Kwiatkowski, Cordelia Langford, John Sillitoe on behalf of the Wellcome Sanger Institute COVID-19 Surveillance Team |
| EPI_ISL_3070421, EPI_ISL_3070422, EPI_ISL_3070423, EPI_ISL_3070424, EPI_ISL_3070425 | Berkshire and Surrey Pathology Services Lighthouse Laboratory | Wellcome Sanger Institute for the COVID-19 Genomics UK (COG-UK) Consortium | Berkshire and Surrey Pathology Services Lighthouse Laboratory and Alex Alderton, Roberto Amato, Jeffrey Barrett, Sonia Goncalves, Ewan Harrison, David K. Jackson, Ian Johnston, Dominic Kwiatkowski, Cordelia Langford, John Sillitoe on behalf of the Wellcome Sanger Institute COVID-19 Surveillance Team |
| EPI_ISL_3070426 | Lighthouse Lab in Milton Keynes | Wellcome Sanger Institute for the COVID-19 Genomics UK (COG-UK) Consortium | The Lighthouse Lab in Milton Keynes and Alex Alderton, Roberto Amato, Jeffrey Barrett, Sonia Goncalves, Ewan Harrison, David K. Jackson, Ian Johnston, Dominic Kwiatkowski, Cordelia Langford, John Sillitoe on behalf of the Wellcome Sanger Institute COVID-19 Surveillance Team |
| EPI_ISL_3070427, EPI_ISL_3070428, EPI_ISL_3070429, EPI_ISL_3070430, EPI_ISL_3070431, EPI_ISL_3070432, | Berkshire and Surrey Pathology Services Lighthouse Laboratory | Wellcome Sanger Institute for the COVID-19 Genomics UK (COG-UK) Consortium | Berkshire and Surrey Pathology Services Lighthouse Laboratory and Alex Alderton, Roberto Amato, Jeffrey Barrett, Sonia Goncalves, Ewan Harrison, David K. Jackson, Ian Johnston, Dominic Kwiatkowski, Cordelia Langford, John Sillitoe on behalf of the Wellcome Sanger Institute COVID-19 Surveillance Team |

| | | | |
|---|---|---|---|
| EPI_ISL_3070433, EPI_ISL_3070434, EPI_ISL_3070435, EPI_ISL_3070436 | | | |
| EPI_ISL_3070437 | Lighthouse Lab in Milton Keynes | Wellcome Sanger Institute for the COVID-19 Genomics UK (COG-UK) Consortium | The Lighthouse Lab in Milton Keynes and Alex Alderton, Roberto Amato, Jeffrey Barrett, Sonia Goncalves, Ewan Harrison, David K. Jackson, Ian Johnston, Dominic Kwiatkowski, Cordelia Langford, John Sillitoe on behalf of the Wellcome Sanger Institute COVID-19 Surveillance Team |
| EPI_ISL_3070438, EPI_ISL_3070439, EPI_ISL_3070440 | Berkshire and Surrey Pathology Services Lighthouse Laboratory | Wellcome Sanger Institute for the COVID-19 Genomics UK (COG-UK) Consortium | Berkshire and Surrey Pathology Services Lighthouse Laboratory and Alex Alderton, Roberto Amato, Jeffrey Barrett, Sonia Goncalves, Ewan Harrison, David K. Jackson, Ian Johnston, Dominic Kwiatkowski, Cordelia Langford, John Sillitoe on behalf of the Wellcome Sanger Institute COVID-19 Surveillance Team |
| EPI_ISL_3070441, EPI_ISL_3070442 | Lighthouse Lab in Milton Keynes | Wellcome Sanger Institute for the COVID-19 Genomics UK (COG-UK) Consortium | The Lighthouse Lab in Milton Keynes and Alex Alderton, Roberto Amato, Jeffrey Barrett, Sonia Goncalves, Ewan Harrison, David K. Jackson, Ian Johnston, Dominic Kwiatkowski, Cordelia Langford, John Sillitoe on behalf of the Wellcome Sanger Institute COVID-19 Surveillance Team |
| EPI_ISL_3070443, EPI_ISL_3070444, EPI_ISL_3070445, EPI_ISL_3070446, EPI_ISL_3070447, EPI_ISL_3070448, EPI_ISL_3070449, EPI_ISL_3070450, EPI_ISL_3070451, EPI_ISL_3070452, EPI_ISL_3070453, EPI_ISL_3070454 | | | |
| see above | Berkshire and Surrey Pathology Services Lighthouse Laboratory | Wellcome Sanger Institute for the COVID-19 Genomics UK (COG-UK) Consortium | Berkshire and Surrey Pathology Services Lighthouse Laboratory and Alex Alderton, Roberto Amato, Jeffrey Barrett, Sonia Goncalves, Ewan Harrison, David K. Jackson, Ian Johnston, Dominic Kwiatkowski, Cordelia Langford, John Sillitoe on behalf of the Wellcome Sanger Institute COVID-19 Surveillance Team |
| EPI_ISL_3070455 | Lighthouse Lab in Milton Keynes | Wellcome Sanger Institute for the COVID-19 Genomics UK (COG-UK) Consortium | The Lighthouse Lab in Milton Keynes and Alex Alderton, Roberto Amato, Jeffrey Barrett, Sonia Goncalves, Ewan Harrison, David K. Jackson, Ian Johnston, Dominic Kwiatkowski, Cordelia Langford, John Sillitoe on behalf of the Wellcome Sanger Institute COVID-19 Surveillance Team |
| EPI_ISL_3070456, EPI_ISL_3070457, EPI_ISL_3070458, EPI_ISL_3070459, EPI_ISL_3070460, EPI_ISL_3070461, EPI_ISL_3070462, EPI_ISL_3070463, EPI_ISL_3070464, EPI_ISL_3070465, EPI_ISL_3070466, EPI_ISL_3070467 | | | |
| see above | Berkshire and Surrey Pathology Services Lighthouse Laboratory | Wellcome Sanger Institute for the COVID-19 Genomics UK (COG-UK) Consortium | Berkshire and Surrey Pathology Services Lighthouse Laboratory and Alex Alderton, Roberto Amato, Jeffrey Barrett, Sonia Goncalves, Ewan Harrison, David K. Jackson, Ian Johnston, Dominic Kwiatkowski, Cordelia Langford, John Sillitoe on behalf of the Wellcome Sanger Institute COVID-19 Surveillance Team |
| EPI_ISL_3070468 | Lighthouse Lab in Milton Keynes | Wellcome Sanger Institute for the COVID-19 Genomics UK (COG-UK) Consortium | The Lighthouse Lab in Milton Keynes and Alex Alderton, Roberto Amato, Jeffrey Barrett, Sonia Goncalves, Ewan Harrison, David K. Jackson, Ian Johnston, Dominic Kwiatkowski, Cordelia Langford, John Sillitoe on behalf of the Wellcome Sanger Institute COVID-19 Surveillance Team |
| EPI_ISL_3070469, EPI_ISL_3070470 | Berkshire and Surrey Pathology Services Lighthouse Laboratory | Wellcome Sanger Institute for the COVID-19 Genomics UK (COG-UK) Consortium | Berkshire and Surrey Pathology Services Lighthouse Laboratory and Alex Alderton, Roberto Amato, Jeffrey Barrett, Sonia Goncalves, Ewan Harrison, David K. Jackson, Ian Johnston, Dominic Kwiatkowski, Cordelia Langford, John Sillitoe on behalf of the Wellcome Sanger Institute COVID-19 Surveillance Team |
| EPI_ISL_3070471 | Lighthouse Lab in Milton Keynes | Wellcome Sanger Institute for the COVID-19 Genomics UK (COG-UK) Consortium | The Lighthouse Lab in Milton Keynes and Alex Alderton, Roberto Amato, Jeffrey Barrett, Sonia Goncalves, Ewan Harrison, David K. Jackson, Ian Johnston, Dominic Kwiatkowski, Cordelia Langford, John Sillitoe on behalf of the Wellcome Sanger Institute COVID-19 Surveillance Team |
| EPI_ISL_3070472, EPI_ISL_3070473, EPI_ISL_3070474 | Berkshire and Surrey Pathology Services Lighthouse Laboratory | Wellcome Sanger Institute for the COVID-19 Genomics UK (COG-UK) Consortium | Berkshire and Surrey Pathology Services Lighthouse Laboratory and Alex Alderton, Roberto Amato, Jeffrey Barrett, Sonia Goncalves, Ewan Harrison, David K. Jackson, Ian Johnston, Dominic Kwiatkowski, Cordelia Langford, John Sillitoe on behalf of the Wellcome Sanger Institute COVID-19 Surveillance Team |
| EPI_ISL_3070475 | Lighthouse Lab in Milton Keynes | Wellcome Sanger Institute for the COVID-19 Genomics UK (COG-UK) Consortium | The Lighthouse Lab in Milton Keynes and Alex Alderton, Roberto Amato, Jeffrey Barrett, Sonia Goncalves, Ewan Harrison, David K. Jackson, Ian Johnston, Dominic Kwiatkowski, Cordelia Langford, John Sillitoe on behalf of the Wellcome Sanger Institute COVID-19 Surveillance Team |
| EPI_ISL_3070476, EPI_ISL_3070477, EPI_ISL_3070478, EPI_ISL_3070479, EPI_ISL_3070480, EPI_ISL_3070481, EPI_ISL_3070482, EPI_ISL_3070483 | Berkshire and Surrey Pathology Services Lighthouse Laboratory | Wellcome Sanger Institute for the COVID-19 Genomics UK (COG-UK) Consortium | Berkshire and Surrey Pathology Services Lighthouse Laboratory and Alex Alderton, Roberto Amato, Jeffrey Barrett, Sonia Goncalves, Ewan Harrison, David K. Jackson, Ian Johnston, Dominic Kwiatkowski, Cordelia Langford, John Sillitoe on behalf of the Wellcome Sanger Institute COVID-19 Surveillance Team |
| EPI_ISL_3070484 | Lighthouse Lab in Milton Keynes | Wellcome Sanger Institute for the COVID-19 Genomics UK (COG-UK) Consortium | The Lighthouse Lab in Milton Keynes and Alex Alderton, Roberto Amato, Jeffrey Barrett, Sonia Goncalves, Ewan Harrison, David K. Jackson, Ian Johnston, Dominic Kwiatkowski, Cordelia Langford, John Sillitoe on behalf of the Wellcome Sanger Institute COVID-19 Surveillance Team |
| EPI_ISL_3070485, EPI_ISL_3070486 | Berkshire and Surrey Pathology Services Lighthouse Laboratory | Wellcome Sanger Institute for the COVID-19 Genomics UK (COG-UK) Consortium | Berkshire and Surrey Pathology Services Lighthouse Laboratory and Alex Alderton, Roberto Amato, Jeffrey Barrett, Sonia Goncalves, Ewan Harrison, David K. Jackson, Ian Johnston, Dominic Kwiatkowski, Cordelia Langford, John Sillitoe on behalf of the Wellcome Sanger Institute COVID-19 Surveillance Team |
| EPI_ISL_3070487, EPI_ISL_3070488 | Lighthouse Lab in Milton Keynes | Wellcome Sanger Institute for the COVID-19 Genomics UK (COG-UK) Consortium | The Lighthouse Lab in Milton Keynes and Alex Alderton, Roberto Amato, Jeffrey Barrett, Sonia Goncalves, Ewan Harrison, David K. Jackson, Ian Johnston, Dominic Kwiatkowski, Cordelia Langford, John Sillitoe on behalf of the Wellcome Sanger Institute COVID-19 Surveillance Team |
| EPI_ISL_3070489, EPI_ISL_3070490, EPI_ISL_3070491, EPI_ISL_3070492, EPI_ISL_3070493 | Berkshire and Surrey Pathology Services Lighthouse Laboratory | Wellcome Sanger Institute for the COVID-19 Genomics UK (COG-UK) Consortium | Berkshire and Surrey Pathology Services Lighthouse Laboratory and Alex Alderton, Roberto Amato, Jeffrey Barrett, Sonia Goncalves, Ewan Harrison, David K. Jackson, Ian Johnston, Dominic Kwiatkowski, Cordelia Langford, John Sillitoe on behalf of the Wellcome Sanger Institute COVID-19 Surveillance Team |
| EPI_ISL_3070494 | Lighthouse Lab in Milton Keynes | Wellcome Sanger Institute for the COVID-19 Genomics UK (COG-UK) Consortium | The Lighthouse Lab in Milton Keynes and Alex Alderton, Roberto Amato, Jeffrey Barrett, Sonia Goncalves, Ewan Harrison, David K. Jackson, Ian Johnston, Dominic Kwiatkowski, Cordelia Langford, John Sillitoe on behalf of the Wellcome Sanger Institute COVID-19 Surveillance Team |
| EPI_ISL_3070495, EPI_ISL_3070496, EPI_ISL_3070497, EPI_ISL_3070498 | Berkshire and Surrey Pathology Services Lighthouse Laboratory | Wellcome Sanger Institute for the COVID-19 Genomics UK (COG-UK) Consortium | Berkshire and Surrey Pathology Services Lighthouse Laboratory and Alex Alderton, Roberto Amato, Jeffrey Barrett, Sonia Goncalves, Ewan Harrison, David K. Jackson, Ian Johnston, Dominic Kwiatkowski, Cordelia Langford, John Sillitoe on behalf of the Wellcome Sanger Institute COVID-19 Surveillance Team |
| EPI_ISL_3070499, EPI_ISL_3070500 | Lighthouse Lab in Milton Keynes | Wellcome Sanger Institute for the COVID-19 Genomics UK (COG-UK) Consortium | The Lighthouse Lab in Milton Keynes and Alex Alderton, Roberto Amato, Jeffrey Barrett, Sonia Goncalves, Ewan Harrison, David K. Jackson, Ian Johnston, Dominic Kwiatkowski, Cordelia Langford, John Sillitoe on behalf of the Wellcome Sanger Institute COVID-19 Surveillance Team |
| EPI_ISL_3070501, EPI_ISL_3070502, EPI_ISL_3070503, EPI_ISL_3070504, EPI_ISL_3070505, EPI_ISL_3070506, EPI_ISL_3070507 | Berkshire and Surrey Pathology Services Lighthouse Laboratory | Wellcome Sanger Institute for the COVID-19 Genomics UK (COG-UK) Consortium | Berkshire and Surrey Pathology Services Lighthouse Laboratory and Alex Alderton, Roberto Amato, Jeffrey Barrett, Sonia Goncalves, Ewan Harrison, David K. Jackson, Ian Johnston, Dominic Kwiatkowski, Cordelia Langford, John Sillitoe on behalf of the Wellcome Sanger Institute COVID-19 Surveillance Team |
| EPI_ISL_3070508, EPI_ISL_3070509, EPI_ISL_3070510 | Lighthouse Lab in Milton Keynes | Wellcome Sanger Institute for the COVID-19 Genomics UK (COG-UK) Consortium | The Lighthouse Lab in Milton Keynes and Alex Alderton, Roberto Amato, Jeffrey Barrett, Sonia Goncalves, Ewan Harrison, David K. Jackson, Ian Johnston, Dominic Kwiatkowski, Cordelia Langford, John Sillitoe on behalf of the Wellcome Sanger Institute COVID-19 Surveillance Team |
| EPI_ISL_3070511, EPI_ISL_3070512, EPI_ISL_3070513 | Berkshire and Surrey Pathology Services Lighthouse Laboratory | Wellcome Sanger Institute for the COVID-19 Genomics UK (COG-UK) Consortium | Berkshire and Surrey Pathology Services Lighthouse Laboratory and Alex Alderton, Roberto Amato, Jeffrey Barrett, Sonia Goncalves, Ewan Harrison, David K. Jackson, Ian Johnston, Dominic Kwiatkowski, Cordelia Langford, John Sillitoe on behalf of the Wellcome Sanger Institute COVID-19 Surveillance Team |
| EPI_ISL_3070514, EPI_ISL_3070515 | Lighthouse Lab in Milton Keynes | Wellcome Sanger Institute for the COVID-19 Genomics UK (COG-UK) Consortium | The Lighthouse Lab in Milton Keynes and Alex Alderton, Roberto Amato, Jeffrey Barrett, Sonia Goncalves, Ewan Harrison, David K. Jackson, Ian Johnston, Dominic Kwiatkowski, Cordelia Langford, John Sillitoe on behalf of the Wellcome Sanger Institute COVID-19 Surveillance Team |
| EPI_ISL_3070516, EPI_ISL_3070517, EPI_ISL_3070518 | Berkshire and Surrey Pathology Services Lighthouse Laboratory | Wellcome Sanger Institute for the COVID-19 Genomics UK (COG-UK) Consortium | Berkshire and Surrey Pathology Services Lighthouse Laboratory and Alex Alderton, Roberto Amato, Jeffrey Barrett, Sonia Goncalves, Ewan Harrison, David K. Jackson, Ian Johnston, Dominic Kwiatkowski, Cordelia Langford, John Sillitoe on behalf of the Wellcome Sanger Institute COVID-19 Surveillance Team |
| EPI_ISL_3070519 | Lighthouse Lab in Milton Keynes | Wellcome Sanger Institute for the COVID-19 Genomics UK (COG-UK) Consortium | The Lighthouse Lab in Milton Keynes and Alex Alderton, Roberto Amato, Jeffrey Barrett, Sonia Goncalves, Ewan Harrison, David K. Jackson, Ian Johnston, Dominic Kwiatkowski, Cordelia Langford, John Sillitoe on behalf of the Wellcome Sanger Institute COVID-19 Surveillance Team |
| EPI_ISL_3070520, EPI_ISL_3070521, EPI_ISL_3070522, EPI_ISL_3070523, EPI_ISL_3070524 | Berkshire and Surrey Pathology Services Lighthouse Laboratory | Wellcome Sanger Institute for the COVID-19 Genomics UK (COG-UK) Consortium | Berkshire and Surrey Pathology Services Lighthouse Laboratory and Alex Alderton, Roberto Amato, Jeffrey Barrett, Sonia Goncalves, Ewan Harrison, David K. Jackson, Ian Johnston, Dominic Kwiatkowski, Cordelia Langford, John Sillitoe on behalf of the Wellcome Sanger Institute COVID-19 Surveillance Team |
| EPI_ISL_3070525 | Lighthouse Lab in Milton Keynes | Wellcome Sanger Institute for the COVID-19 Genomics UK (COG-UK) Consortium | The Lighthouse Lab in Milton Keynes and Alex Alderton, Roberto Amato, Jeffrey Barrett, Sonia Goncalves, Ewan Harrison, David K. Jackson, Ian Johnston, Dominic Kwiatkowski, Cordelia Langford, John Sillitoe on behalf of the Wellcome Sanger Institute COVID-19 Surveillance Team |
| EPI_ISL_3070526 | Berkshire and Surrey Pathology Services Lighthouse Laboratory | Wellcome Sanger Institute for the COVID-19 Genomics UK (COG-UK) Consortium | Berkshire and Surrey Pathology Services Lighthouse Laboratory and Alex Alderton, Roberto Amato, Jeffrey Barrett, Sonia Goncalves, Ewan Harrison, David K. Jackson, Ian Johnston, Dominic Kwiatkowski, Cordelia Langford, John Sillitoe on behalf of the Wellcome Sanger Institute COVID-19 Surveillance Team |
| EPI_ISL_3070527, EPI_ISL_3070528, EPI_ISL_3070529 | Lighthouse Lab in Milton Keynes | Wellcome Sanger Institute for the COVID-19 Genomics UK (COG-UK) Consortium | The Lighthouse Lab in Milton Keynes and Alex Alderton, Roberto Amato, Jeffrey Barrett, Sonia Goncalves, Ewan Harrison, David K. Jackson, Ian Johnston, Dominic Kwiatkowski, Cordelia Langford, John Sillitoe on behalf of the Wellcome Sanger Institute COVID-19 Surveillance Team |
| EPI_ISL_3070530 | Berkshire and Surrey Pathology Services Lighthouse Laboratory | Wellcome Sanger Institute for the COVID-19 Genomics UK (COG-UK) Consortium | Berkshire and Surrey Pathology Services Lighthouse Laboratory and Alex Alderton, Roberto Amato, Jeffrey Barrett, Sonia Goncalves, Ewan Harrison, David K. Jackson, Ian Johnston, Dominic Kwiatkowski, Cordelia Langford, John Sillitoe on behalf of the Wellcome Sanger Institute COVID-19 Surveillance Team |
| EPI_ISL_3070531 | Lighthouse Lab in Milton Keynes | Wellcome Sanger Institute for the COVID-19 Genomics UK (COG-UK) Consortium | The Lighthouse Lab in Milton Keynes and Alex Alderton, Roberto Amato, Jeffrey Barrett, Sonia Goncalves, Ewan Harrison, David K. Jackson, Ian Johnston, Dominic Kwiatkowski, Cordelia Langford, John Sillitoe on behalf of the Wellcome Sanger Institute COVID-19 Surveillance Team |
| EPI_ISL_3070532 | Berkshire and Surrey Pathology Services Lighthouse Laboratory | Wellcome Sanger Institute for the COVID-19 Genomics UK (COG-UK) Consortium | Berkshire and Surrey Pathology Services Lighthouse Laboratory and Alex Alderton, Roberto Amato, Jeffrey Barrett, Sonia Goncalves, Ewan Harrison, David K. Jackson, Ian Johnston, Dominic Kwiatkowski, Cordelia Langford, John Sillitoe on behalf of the Wellcome Sanger Institute COVID-19 Surveillance Team |

| | | | |
|---|---|---|---|
| EPI_ISL_3070533 | Lighthouse Lab in Milton Keynes | Wellcome Sanger Institute for the COVID-19 Genomics UK (COG-UK) Consortium | The Lighthouse Lab in Milton Keynes and Alex Alderton, Roberto Amato, Jeffrey Barrett, Sonia Goncalves, Ewan Harrison, David K. Jackson, Ian Johnston, Dominic Kwiatkowski, Cordelia Langford, John Sillitoe on behalf of the Wellcome Sanger Institute COVID-19 Surveillance Team |
| EPI_ISL_3070534 | Berkshire and Surrey Pathology Services Lighthouse Laboratory | Wellcome Sanger Institute for the COVID-19 Genomics UK (COG-UK) Consortium | Berkshire and Surrey Pathology Services Lighthouse Laboratory and Alex Alderton, Roberto Amato, Jeffrey Barrett, Sonia Goncalves, Ewan Harrison, David K. Jackson, Ian Johnston, Dominic Kwiatkowski, Cordelia Langford, John Sillitoe on behalf of the Wellcome Sanger Institute COVID-19 Surveillance Team |
| EPI_ISL_3070535 | Lighthouse Lab in Milton Keynes | Wellcome Sanger Institute for the COVID-19 Genomics UK (COG-UK) Consortium | The Lighthouse Lab in Milton Keynes and Alex Alderton, Roberto Amato, Jeffrey Barrett, Sonia Goncalves, Ewan Harrison, David K. Jackson, Ian Johnston, Dominic Kwiatkowski, Cordelia Langford, John Sillitoe on behalf of the Wellcome Sanger Institute COVID-19 Surveillance Team |
| EPI_ISL_3070536, EPI_ISL_3070537 | Berkshire and Surrey Pathology Services Lighthouse Laboratory | Wellcome Sanger Institute for the COVID-19 Genomics UK (COG-UK) Consortium | Berkshire and Surrey Pathology Services Lighthouse Laboratory and Alex Alderton, Roberto Amato, Jeffrey Barrett, Sonia Goncalves, Ewan Harrison, David K. Jackson, Ian Johnston, Dominic Kwiatkowski, Cordelia Langford, John Sillitoe on behalf of the Wellcome Sanger Institute COVID-19 Surveillance Team |
| EPI_ISL_3070538, EPI_ISL_3070539 | Lighthouse Lab in Milton Keynes | Wellcome Sanger Institute for the COVID-19 Genomics UK (COG-UK) Consortium | The Lighthouse Lab in Milton Keynes and Alex Alderton, Roberto Amato, Jeffrey Barrett, Sonia Goncalves, Ewan Harrison, David K. Jackson, Ian Johnston, Dominic Kwiatkowski, Cordelia Langford, John Sillitoe on behalf of the Wellcome Sanger Institute COVID-19 Surveillance Team |
| EPI_ISL_3070540, EPI_ISL_3070541, EPI_ISL_3070542 | Berkshire and Surrey Pathology Services Lighthouse Laboratory | Wellcome Sanger Institute for the COVID-19 Genomics UK (COG-UK) Consortium | Berkshire and Surrey Pathology Services Lighthouse Laboratory and Alex Alderton, Roberto Amato, Jeffrey Barrett, Sonia Goncalves, Ewan Harrison, David K. Jackson, Ian Johnston, Dominic Kwiatkowski, Cordelia Langford, John Sillitoe on behalf of the Wellcome Sanger Institute COVID-19 Surveillance Team |
| EPI_ISL_3070543 | Lighthouse Lab in Milton Keynes | Wellcome Sanger Institute for the COVID-19 Genomics UK (COG-UK) Consortium | The Lighthouse Lab in Milton Keynes and Alex Alderton, Roberto Amato, Jeffrey Barrett, Sonia Goncalves, Ewan Harrison, David K. Jackson, Ian Johnston, Dominic Kwiatkowski, Cordelia Langford, John Sillitoe on behalf of the Wellcome Sanger Institute COVID-19 Surveillance Team |
| EPI_ISL_3070544, EPI_ISL_3070545, EPI_ISL_3070546, EPI_ISL_3070547, EPI_ISL_3070548, EPI_ISL_3070549, EPI_ISL_3070550, EPI_ISL_3070551, EPI_ISL_3070552 | Berkshire and Surrey Pathology Services Lighthouse Laboratory | Wellcome Sanger Institute for the COVID-19 Genomics UK (COG-UK) Consortium | Berkshire and Surrey Pathology Services Lighthouse Laboratory and Alex Alderton, Roberto Amato, Jeffrey Barrett, Sonia Goncalves, Ewan Harrison, David K. Jackson, Ian Johnston, Dominic Kwiatkowski, Cordelia Langford, John Sillitoe on behalf of the Wellcome Sanger Institute COVID-19 Surveillance Team |
| EPI_ISL_3070553 | Lighthouse Lab in Milton Keynes | Wellcome Sanger Institute for the COVID-19 Genomics UK (COG-UK) Consortium | The Lighthouse Lab in Milton Keynes and Alex Alderton, Roberto Amato, Jeffrey Barrett, Sonia Goncalves, Ewan Harrison, David K. Jackson, Ian Johnston, Dominic Kwiatkowski, Cordelia Langford, John Sillitoe on behalf of the Wellcome Sanger Institute COVID-19 Surveillance Team |
| EPI_ISL_3070554, EPI_ISL_3070555, EPI_ISL_3070556, EPI_ISL_3070557, EPI_ISL_3070558 | Berkshire and Surrey Pathology Services Lighthouse Laboratory | Wellcome Sanger Institute for the COVID-19 Genomics UK (COG-UK) Consortium | Berkshire and Surrey Pathology Services Lighthouse Laboratory and Alex Alderton, Roberto Amato, Jeffrey Barrett, Sonia Goncalves, Ewan Harrison, David K. Jackson, Ian Johnston, Dominic Kwiatkowski, Cordelia Langford, John Sillitoe on behalf of the Wellcome Sanger Institute COVID-19 Surveillance Team |
| EPI_ISL_3070559 | Lighthouse Lab in Milton Keynes | Wellcome Sanger Institute for the COVID-19 Genomics UK (COG-UK) Consortium | The Lighthouse Lab in Milton Keynes and Alex Alderton, Roberto Amato, Jeffrey Barrett, Sonia Goncalves, Ewan Harrison, David K. Jackson, Ian Johnston, Dominic Kwiatkowski, Cordelia Langford, John Sillitoe on behalf of the Wellcome Sanger Institute COVID-19 Surveillance Team |
| EPI_ISL_3070560, EPI_ISL_3070561, EPI_ISL_3070562 | Berkshire and Surrey Pathology Services Lighthouse Laboratory | Wellcome Sanger Institute for the COVID-19 Genomics UK (COG-UK) Consortium | Berkshire and Surrey Pathology Services Lighthouse Laboratory and Alex Alderton, Roberto Amato, Jeffrey Barrett, Sonia Goncalves, Ewan Harrison, David K. Jackson, Ian Johnston, Dominic Kwiatkowski, Cordelia Langford, John Sillitoe on behalf of the Wellcome Sanger Institute COVID-19 Surveillance Team |
| EPI_ISL_3070563 | Lighthouse Lab in Milton Keynes | Wellcome Sanger Institute for the COVID-19 Genomics UK (COG-UK) Consortium | The Lighthouse Lab in Milton Keynes and Alex Alderton, Roberto Amato, Jeffrey Barrett, Sonia Goncalves, Ewan Harrison, David K. Jackson, Ian Johnston, Dominic Kwiatkowski, Cordelia Langford, John Sillitoe on behalf of the Wellcome Sanger Institute COVID-19 Surveillance Team |
| EPI_ISL_3070564, EPI_ISL_3070565, EPI_ISL_3070566 | Berkshire and Surrey Pathology Services Lighthouse Laboratory | Wellcome Sanger Institute for the COVID-19 Genomics UK (COG-UK) Consortium | Berkshire and Surrey Pathology Services Lighthouse Laboratory and Alex Alderton, Roberto Amato, Jeffrey Barrett, Sonia Goncalves, Ewan Harrison, David K. Jackson, Ian Johnston, Dominic Kwiatkowski, Cordelia Langford, John Sillitoe on behalf of the Wellcome Sanger Institute COVID-19 Surveillance Team |
| EPI_ISL_3070567 | Lighthouse Lab in Milton Keynes | Wellcome Sanger Institute for the COVID-19 Genomics UK (COG-UK) Consortium | The Lighthouse Lab in Milton Keynes and Alex Alderton, Roberto Amato, Jeffrey Barrett, Sonia Goncalves, Ewan Harrison, David K. Jackson, Ian Johnston, Dominic Kwiatkowski, Cordelia Langford, John Sillitoe on behalf of the Wellcome Sanger Institute COVID-19 Surveillance Team |
| EPI_ISL_3070568 | Berkshire and Surrey Pathology Services Lighthouse Laboratory | Wellcome Sanger Institute for the COVID-19 Genomics UK (COG-UK) Consortium | Berkshire and Surrey Pathology Services Lighthouse Laboratory and Alex Alderton, Roberto Amato, Jeffrey Barrett, Sonia Goncalves, Ewan Harrison, David K. Jackson, Ian Johnston, Dominic Kwiatkowski, Cordelia Langford, John Sillitoe on behalf of the Wellcome Sanger Institute COVID-19 Surveillance Team |
| EPI_ISL_3070569 | Lighthouse Lab in Milton Keynes | Wellcome Sanger Institute for the COVID-19 Genomics UK (COG-UK) Consortium | The Lighthouse Lab in Milton Keynes and Alex Alderton, Roberto Amato, Jeffrey Barrett, Sonia Goncalves, Ewan Harrison, David K. Jackson, Ian Johnston, Dominic Kwiatkowski, Cordelia Langford, John Sillitoe on behalf of the Wellcome Sanger Institute COVID-19 Surveillance Team |
| EPI_ISL_3070570, EPI_ISL_3070571 | Berkshire and Surrey Pathology Services Lighthouse Laboratory | Wellcome Sanger Institute for the COVID-19 Genomics UK (COG-UK) Consortium | Berkshire and Surrey Pathology Services Lighthouse Laboratory and Alex Alderton, Roberto Amato, Jeffrey Barrett, Sonia Goncalves, Ewan Harrison, David K. Jackson, Ian Johnston, Dominic Kwiatkowski, Cordelia Langford, John Sillitoe on behalf of the Wellcome Sanger Institute COVID-19 Surveillance Team |
| EPI_ISL_3070572 | Lighthouse Lab in Milton Keynes | Wellcome Sanger Institute for the COVID-19 Genomics UK (COG-UK) Consortium | The Lighthouse Lab in Milton Keynes and Alex Alderton, Roberto Amato, Jeffrey Barrett, Sonia Goncalves, Ewan Harrison, David K. Jackson, Ian Johnston, Dominic Kwiatkowski, Cordelia Langford, John Sillitoe on behalf of the Wellcome Sanger Institute COVID-19 Surveillance Team |
| EPI_ISL_3070573 | Berkshire and Surrey Pathology Services Lighthouse Laboratory | Wellcome Sanger Institute for the COVID-19 Genomics UK (COG-UK) Consortium | Berkshire and Surrey Pathology Services Lighthouse Laboratory and Alex Alderton, Roberto Amato, Jeffrey Barrett, Sonia Goncalves, Ewan Harrison, David K. Jackson, Ian Johnston, Dominic Kwiatkowski, Cordelia Langford, John Sillitoe on behalf of the Wellcome Sanger Institute COVID-19 Surveillance Team |
| EPI_ISL_3070574 | Lighthouse Lab in Milton Keynes | Wellcome Sanger Institute for the COVID-19 Genomics UK (COG-UK) Consortium | The Lighthouse Lab in Milton Keynes and Alex Alderton, Roberto Amato, Jeffrey Barrett, Sonia Goncalves, Ewan Harrison, David K. Jackson, Ian Johnston, Dominic Kwiatkowski, Cordelia Langford, John Sillitoe on behalf of the Wellcome Sanger Institute COVID-19 Surveillance Team |
| EPI_ISL_3070575 | Berkshire and Surrey Pathology Services Lighthouse Laboratory | Wellcome Sanger Institute for the COVID-19 Genomics UK (COG-UK) Consortium | Berkshire and Surrey Pathology Services Lighthouse Laboratory and Alex Alderton, Roberto Amato, Jeffrey Barrett, Sonia Goncalves, Ewan Harrison, David K. Jackson, Ian Johnston, Dominic Kwiatkowski, Cordelia Langford, John Sillitoe on behalf of the Wellcome Sanger Institute COVID-19 Surveillance Team |
| EPI_ISL_3070576 | Lighthouse Lab in Milton Keynes | Wellcome Sanger Institute for the COVID-19 Genomics UK (COG-UK) Consortium | The Lighthouse Lab in Milton Keynes and Alex Alderton, Roberto Amato, Jeffrey Barrett, Sonia Goncalves, Ewan Harrison, David K. Jackson, Ian Johnston, Dominic Kwiatkowski, Cordelia Langford, John Sillitoe on behalf of the Wellcome Sanger Institute COVID-19 Surveillance Team |
| EPI_ISL_3070577, EPI_ISL_3070578, EPI_ISL_3070579 | Berkshire and Surrey Pathology Services Lighthouse Laboratory | Wellcome Sanger Institute for the COVID-19 Genomics UK (COG-UK) Consortium | Berkshire and Surrey Pathology Services Lighthouse Laboratory and Alex Alderton, Roberto Amato, Jeffrey Barrett, Sonia Goncalves, Ewan Harrison, David K. Jackson, Ian Johnston, Dominic Kwiatkowski, Cordelia Langford, John Sillitoe on behalf of the Wellcome Sanger Institute COVID-19 Surveillance Team |
| EPI_ISL_3070580, EPI_ISL_3070581 | Lighthouse Lab in Milton Keynes | Wellcome Sanger Institute for the COVID-19 Genomics UK (COG-UK) Consortium | The Lighthouse Lab in Milton Keynes and Alex Alderton, Roberto Amato, Jeffrey Barrett, Sonia Goncalves, Ewan Harrison, David K. Jackson, Ian Johnston, Dominic Kwiatkowski, Cordelia Langford, John Sillitoe on behalf of the Wellcome Sanger Institute COVID-19 Surveillance Team |
| EPI_ISL_3070582, EPI_ISL_3070583, EPI_ISL_3070584, EPI_ISL_3070585, EPI_ISL_3070586 | Berkshire and Surrey Pathology Services Lighthouse Laboratory | Wellcome Sanger Institute for the COVID-19 Genomics UK (COG-UK) Consortium | Berkshire and Surrey Pathology Services Lighthouse Laboratory and Alex Alderton, Roberto Amato, Jeffrey Barrett, Sonia Goncalves, Ewan Harrison, David K. Jackson, Ian Johnston, Dominic Kwiatkowski, Cordelia Langford, John Sillitoe on behalf of the Wellcome Sanger Institute COVID-19 Surveillance Team |
| EPI_ISL_3070587 | Lighthouse Lab in Milton Keynes | Wellcome Sanger Institute for the COVID-19 Genomics UK (COG-UK) Consortium | The Lighthouse Lab in Milton Keynes and Alex Alderton, Roberto Amato, Jeffrey Barrett, Sonia Goncalves, Ewan Harrison, David K. Jackson, Ian Johnston, Dominic Kwiatkowski, Cordelia Langford, John Sillitoe on behalf of the Wellcome Sanger Institute COVID-19 Surveillance Team |
| EPI_ISL_3070588 | Berkshire and Surrey Pathology Services Lighthouse Laboratory | Wellcome Sanger Institute for the COVID-19 Genomics UK (COG-UK) Consortium | Berkshire and Surrey Pathology Services Lighthouse Laboratory and Alex Alderton, Roberto Amato, Jeffrey Barrett, Sonia Goncalves, Ewan Harrison, David K. Jackson, Ian Johnston, Dominic Kwiatkowski, Cordelia Langford, John Sillitoe on behalf of the Wellcome Sanger Institute COVID-19 Surveillance Team |
| EPI_ISL_3070589 | Lighthouse Lab in Milton Keynes | Wellcome Sanger Institute for the COVID-19 Genomics UK (COG-UK) Consortium | The Lighthouse Lab in Milton Keynes and Alex Alderton, Roberto Amato, Jeffrey Barrett, Sonia Goncalves, Ewan Harrison, David K. Jackson, Ian Johnston, Dominic Kwiatkowski, Cordelia Langford, John Sillitoe on behalf of the Wellcome Sanger Institute COVID-19 Surveillance Team |
| EPI_ISL_3070590, EPI_ISL_3070591, EPI_ISL_3070592, EPI_ISL_3070593, EPI_ISL_3070594 | Berkshire and Surrey Pathology Services Lighthouse Laboratory | Wellcome Sanger Institute for the COVID-19 Genomics UK (COG-UK) Consortium | Berkshire and Surrey Pathology Services Lighthouse Laboratory and Alex Alderton, Roberto Amato, Jeffrey Barrett, Sonia Goncalves, Ewan Harrison, David K. Jackson, Ian Johnston, Dominic Kwiatkowski, Cordelia Langford, John Sillitoe on behalf of the Wellcome Sanger Institute COVID-19 Surveillance Team |
| EPI_ISL_3070595 | Lighthouse Lab in Milton Keynes | Wellcome Sanger Institute for the COVID-19 Genomics UK (COG-UK) Consortium | The Lighthouse Lab in Milton Keynes and Alex Alderton, Roberto Amato, Jeffrey Barrett, Sonia Goncalves, Ewan Harrison, David K. Jackson, Ian Johnston, Dominic Kwiatkowski, Cordelia Langford, John Sillitoe on behalf of the Wellcome Sanger Institute COVID-19 Surveillance Team |
| EPI_ISL_3070596, EPI_ISL_3070597 | Berkshire and Surrey Pathology Services Lighthouse Laboratory | Wellcome Sanger Institute for the COVID-19 Genomics UK (COG-UK) Consortium | Berkshire and Surrey Pathology Services Lighthouse Laboratory and Alex Alderton, Roberto Amato, Jeffrey Barrett, Sonia Goncalves, Ewan Harrison, David K. Jackson, Ian Johnston, Dominic Kwiatkowski, Cordelia Langford, John Sillitoe on behalf of the Wellcome Sanger Institute COVID-19 Surveillance Team |
| EPI_ISL_3070598 | Lighthouse Lab in Milton Keynes | Wellcome Sanger Institute for the COVID-19 Genomics UK (COG-UK) Consortium | The Lighthouse Lab in Milton Keynes and Alex Alderton, Roberto Amato, Jeffrey Barrett, Sonia Goncalves, Ewan Harrison, David K. Jackson, Ian Johnston, Dominic Kwiatkowski, Cordelia Langford, John Sillitoe on behalf of the Wellcome Sanger Institute COVID-19 Surveillance Team |
| EPI_ISL_3070599 | Berkshire and Surrey Pathology Services Lighthouse Laboratory | Wellcome Sanger Institute for the COVID-19 Genomics UK (COG-UK) Consortium | Berkshire and Surrey Pathology Services Lighthouse Laboratory and Alex Alderton, Roberto Amato, Jeffrey Barrett, Sonia Goncalves, Ewan Harrison, David K. Jackson, Ian Johnston, Dominic Kwiatkowski, Cordelia Langford, John Sillitoe on behalf of the Wellcome Sanger Institute COVID-19 Surveillance Team |
| EPI_ISL_3070600 | Lighthouse Lab in Milton Keynes | Wellcome Sanger Institute for the COVID-19 Genomics UK (COG-UK) Consortium | The Lighthouse Lab in Milton Keynes and Alex Alderton, Roberto Amato, Jeffrey Barrett, Sonia Goncalves, Ewan Harrison, David K. Jackson, Ian Johnston, Dominic Kwiatkowski, Cordelia Langford, John Sillitoe on behalf of the Wellcome Sanger Institute COVID-19 Surveillance Team |
| EPI_ISL_3070601, EPI_ISL_3070602, EPI_ISL_3070603, EPI_ISL_3070604, | Berkshire and Surrey Pathology Services Lighthouse Laboratory | Wellcome Sanger Institute for the COVID-19 Genomics UK (COG-UK) Consortium | Berkshire and Surrey Pathology Services Lighthouse Laboratory and Alex Alderton, Roberto Amato, Jeffrey Barrett, Sonia Goncalves, Ewan Harrison, David K. Jackson, Ian Johnston, Dominic Kwiatkowski, Cordelia Langford, John Sillitoe on behalf of the Wellcome Sanger Institute COVID-19 Surveillance Team |

| | | | |
|---|---|---|---|
| EPI_ISL_3070605, EPI_ISL_3070606 | | | |
| EPI_ISL_3070607 | Lighthouse Lab in Milton Keynes | Wellcome Sanger Institute for the COVID-19 Genomics UK (COG-UK) Consortium | The Lighthouse Lab in Milton Keynes and Alex Alderton, Roberto Amato, Jeffrey Barrett, Sonia Goncalves, Ewan Harrison, David K. Jackson, Ian Johnston, Dominic Kwiatkowski, Cordelia Langford, John Sillitoe on behalf of the Wellcome Sanger Institute COVID-19 Surveillance Team |
| EPI_ISL_3070608 | Berkshire and Surrey Pathology Services Lighthouse Laboratory | Wellcome Sanger Institute for the COVID-19 Genomics UK (COG-UK) Consortium | Berkshire and Surrey Pathology Services Lighthouse Laboratory and Alex Alderton, Roberto Amato, Jeffrey Barrett, Sonia Goncalves, Ewan Harrison, David K. Jackson, Ian Johnston, Dominic Kwiatkowski, Cordelia Langford, John Sillitoe on behalf of the Wellcome Sanger Institute COVID-19 Surveillance Team |
| EPI_ISL_3070609 | Lighthouse Lab in Milton Keynes | Wellcome Sanger Institute for the COVID-19 Genomics UK (COG-UK) Consortium | The Lighthouse Lab in Milton Keynes and Alex Alderton, Roberto Amato, Jeffrey Barrett, Sonia Goncalves, Ewan Harrison, David K. Jackson, Ian Johnston, Dominic Kwiatkowski, Cordelia Langford, John Sillitoe on behalf of the Wellcome Sanger Institute COVID-19 Surveillance Team |
| EPI_ISL_3070610 | Berkshire and Surrey Pathology Services Lighthouse Laboratory | Wellcome Sanger Institute for the COVID-19 Genomics UK (COG-UK) Consortium | Berkshire and Surrey Pathology Services Lighthouse Laboratory and Alex Alderton, Roberto Amato, Jeffrey Barrett, Sonia Goncalves, Ewan Harrison, David K. Jackson, Ian Johnston, Dominic Kwiatkowski, Cordelia Langford, John Sillitoe on behalf of the Wellcome Sanger Institute COVID-19 Surveillance Team |
| EPI_ISL_3070611, EPI_ISL_3070612, EPI_ISL_3070613 | Lighthouse Lab in Milton Keynes | Wellcome Sanger Institute for the COVID-19 Genomics UK (COG-UK) Consortium | The Lighthouse Lab in Milton Keynes and Alex Alderton, Roberto Amato, Jeffrey Barrett, Sonia Goncalves, Ewan Harrison, David K. Jackson, Ian Johnston, Dominic Kwiatkowski, Cordelia Langford, John Sillitoe on behalf of the Wellcome Sanger Institute COVID-19 Surveillance Team |
| EPI_ISL_3070614, EPI_ISL_3070615, EPI_ISL_3070616, EPI_ISL_3070617, EPI_ISL_3070618, EPI_ISL_3070619 | Berkshire and Surrey Pathology Services Lighthouse Laboratory | Wellcome Sanger Institute for the COVID-19 Genomics UK (COG-UK) Consortium | Berkshire and Surrey Pathology Services Lighthouse Laboratory and Alex Alderton, Roberto Amato, Jeffrey Barrett, Sonia Goncalves, Ewan Harrison, David K. Jackson, Ian Johnston, Dominic Kwiatkowski, Cordelia Langford, John Sillitoe on behalf of the Wellcome Sanger Institute COVID-19 Surveillance Team |
| EPI_ISL_3070620, EPI_ISL_3070621 | Lighthouse Lab in Milton Keynes | Wellcome Sanger Institute for the COVID-19 Genomics UK (COG-UK) Consortium | The Lighthouse Lab in Milton Keynes and Alex Alderton, Roberto Amato, Jeffrey Barrett, Sonia Goncalves, Ewan Harrison, David K. Jackson, Ian Johnston, Dominic Kwiatkowski, Cordelia Langford, John Sillitoe on behalf of the Wellcome Sanger Institute COVID-19 Surveillance Team |
| EPI_ISL_3070622 | Berkshire and Surrey Pathology Services Lighthouse Laboratory | Wellcome Sanger Institute for the COVID-19 Genomics UK (COG-UK) Consortium | Berkshire and Surrey Pathology Services Lighthouse Laboratory and Alex Alderton, Roberto Amato, Jeffrey Barrett, Sonia Goncalves, Ewan Harrison, David K. Jackson, Ian Johnston, Dominic Kwiatkowski, Cordelia Langford, John Sillitoe on behalf of the Wellcome Sanger Institute COVID-19 Surveillance Team |
| EPI_ISL_3070623 | Lighthouse Lab in Milton Keynes | Wellcome Sanger Institute for the COVID-19 Genomics UK (COG-UK) Consortium | The Lighthouse Lab in Milton Keynes and Alex Alderton, Roberto Amato, Jeffrey Barrett, Sonia Goncalves, Ewan Harrison, David K. Jackson, Ian Johnston, Dominic Kwiatkowski, Cordelia Langford, John Sillitoe on behalf of the Wellcome Sanger Institute COVID-19 Surveillance Team |
| EPI_ISL_3070624, EPI_ISL_3070625, EPI_ISL_3070626 | Berkshire and Surrey Pathology Services Lighthouse Laboratory | Wellcome Sanger Institute for the COVID-19 Genomics UK (COG-UK) Consortium | Berkshire and Surrey Pathology Services Lighthouse Laboratory and Alex Alderton, Roberto Amato, Jeffrey Barrett, Sonia Goncalves, Ewan Harrison, David K. Jackson, Ian Johnston, Dominic Kwiatkowski, Cordelia Langford, John Sillitoe on behalf of the Wellcome Sanger Institute COVID-19 Surveillance Team |
| EPI_ISL_3070627 | Lighthouse Lab in Milton Keynes | Wellcome Sanger Institute for the COVID-19 Genomics UK (COG-UK) Consortium | The Lighthouse Lab in Milton Keynes and Alex Alderton, Roberto Amato, Jeffrey Barrett, Sonia Goncalves, Ewan Harrison, David K. Jackson, Ian Johnston, Dominic Kwiatkowski, Cordelia Langford, John Sillitoe on behalf of the Wellcome Sanger Institute COVID-19 Surveillance Team |
| EPI_ISL_3070628, EPI_ISL_3070629, EPI_ISL_3070630, EPI_ISL_3070631, EPI_ISL_3070632 | Berkshire and Surrey Pathology Services Lighthouse Laboratory | Wellcome Sanger Institute for the COVID-19 Genomics UK (COG-UK) Consortium | Berkshire and Surrey Pathology Services Lighthouse Laboratory and Alex Alderton, Roberto Amato, Jeffrey Barrett, Sonia Goncalves, Ewan Harrison, David K. Jackson, Ian Johnston, Dominic Kwiatkowski, Cordelia Langford, John Sillitoe on behalf of the Wellcome Sanger Institute COVID-19 Surveillance Team |
| EPI_ISL_3070633 | Lighthouse Lab in Milton Keynes | Wellcome Sanger Institute for the COVID-19 Genomics UK (COG-UK) Consortium | The Lighthouse Lab in Milton Keynes and Alex Alderton, Roberto Amato, Jeffrey Barrett, Sonia Goncalves, Ewan Harrison, David K. Jackson, Ian Johnston, Dominic Kwiatkowski, Cordelia Langford, John Sillitoe on behalf of the Wellcome Sanger Institute COVID-19 Surveillance Team |
| EPI_ISL_3070634, EPI_ISL_3070635, EPI_ISL_3070636, EPI_ISL_3070637 | Berkshire and Surrey Pathology Services Lighthouse Laboratory | Wellcome Sanger Institute for the COVID-19 Genomics UK (COG-UK) Consortium | Berkshire and Surrey Pathology Services Lighthouse Laboratory and Alex Alderton, Roberto Amato, Jeffrey Barrett, Sonia Goncalves, Ewan Harrison, David K. Jackson, Ian Johnston, Dominic Kwiatkowski, Cordelia Langford, John Sillitoe on behalf of the Wellcome Sanger Institute COVID-19 Surveillance Team |
| EPI_ISL_3070638 | Lighthouse Lab in Milton Keynes | Wellcome Sanger Institute for the COVID-19 Genomics UK (COG-UK) Consortium | The Lighthouse Lab in Milton Keynes and Alex Alderton, Roberto Amato, Jeffrey Barrett, Sonia Goncalves, Ewan Harrison, David K. Jackson, Ian Johnston, Dominic Kwiatkowski, Cordelia Langford, John Sillitoe on behalf of the Wellcome Sanger Institute COVID-19 Surveillance Team |
| EPI_ISL_3070639 | Berkshire and Surrey Pathology Services Lighthouse Laboratory | Wellcome Sanger Institute for the COVID-19 Genomics UK (COG-UK) Consortium | Berkshire and Surrey Pathology Services Lighthouse Laboratory and Alex Alderton, Roberto Amato, Jeffrey Barrett, Sonia Goncalves, Ewan Harrison, David K. Jackson, Ian Johnston, Dominic Kwiatkowski, Cordelia Langford, John Sillitoe on behalf of the Wellcome Sanger Institute COVID-19 Surveillance Team |
| EPI_ISL_3070640 | Lighthouse Lab in Milton Keynes | Wellcome Sanger Institute for the COVID-19 Genomics UK (COG-UK) Consortium | The Lighthouse Lab in Milton Keynes and Alex Alderton, Roberto Amato, Jeffrey Barrett, Sonia Goncalves, Ewan Harrison, David K. Jackson, Ian Johnston, Dominic Kwiatkowski, Cordelia Langford, John Sillitoe on behalf of the Wellcome Sanger Institute COVID-19 Surveillance Team |
| EPI_ISL_3070641 | Berkshire and Surrey Pathology Services Lighthouse Laboratory | Wellcome Sanger Institute for the COVID-19 Genomics UK (COG-UK) Consortium | Berkshire and Surrey Pathology Services Lighthouse Laboratory and Alex Alderton, Roberto Amato, Jeffrey Barrett, Sonia Goncalves, Ewan Harrison, David K. Jackson, Ian Johnston, Dominic Kwiatkowski, Cordelia Langford, John Sillitoe on behalf of the Wellcome Sanger Institute COVID-19 Surveillance Team |
| EPI_ISL_3070642 | Lighthouse Lab in Milton Keynes | Wellcome Sanger Institute for the COVID-19 Genomics UK (COG-UK) Consortium | The Lighthouse Lab in Milton Keynes and Alex Alderton, Roberto Amato, Jeffrey Barrett, Sonia Goncalves, Ewan Harrison, David K. Jackson, Ian Johnston, Dominic Kwiatkowski, Cordelia Langford, John Sillitoe on behalf of the Wellcome Sanger Institute COVID-19 Surveillance Team |
| EPI_ISL_3070643, EPI_ISL_3070644, EPI_ISL_3070645, EPI_ISL_3070646, EPI_ISL_3070647, EPI_ISL_3070648, EPI_ISL_3070649 | Berkshire and Surrey Pathology Services Lighthouse Laboratory | Wellcome Sanger Institute for the COVID-19 Genomics UK (COG-UK) Consortium | Berkshire and Surrey Pathology Services Lighthouse Laboratory and Alex Alderton, Roberto Amato, Jeffrey Barrett, Sonia Goncalves, Ewan Harrison, David K. Jackson, Ian Johnston, Dominic Kwiatkowski, Cordelia Langford, John Sillitoe on behalf of the Wellcome Sanger Institute COVID-19 Surveillance Team |
| EPI_ISL_3070650, EPI_ISL_3070651 | Lighthouse Lab in Milton Keynes | Wellcome Sanger Institute for the COVID-19 Genomics UK (COG-UK) Consortium | The Lighthouse Lab in Milton Keynes and Alex Alderton, Roberto Amato, Jeffrey Barrett, Sonia Goncalves, Ewan Harrison, David K. Jackson, Ian Johnston, Dominic Kwiatkowski, Cordelia Langford, John Sillitoe on behalf of the Wellcome Sanger Institute COVID-19 Surveillance Team |
| EPI_ISL_3070652, EPI_ISL_3070653, EPI_ISL_3070654, EPI_ISL_3070655 | Berkshire and Surrey Pathology Services Lighthouse Laboratory | Wellcome Sanger Institute for the COVID-19 Genomics UK (COG-UK) Consortium | Berkshire and Surrey Pathology Services Lighthouse Laboratory and Alex Alderton, Roberto Amato, Jeffrey Barrett, Sonia Goncalves, Ewan Harrison, David K. Jackson, Ian Johnston, Dominic Kwiatkowski, Cordelia Langford, John Sillitoe on behalf of the Wellcome Sanger Institute COVID-19 Surveillance Team |
| EPI_ISL_3070656, EPI_ISL_3070657 | Lighthouse Lab in Milton Keynes | Wellcome Sanger Institute for the COVID-19 Genomics UK (COG-UK) Consortium | The Lighthouse Lab in Milton Keynes and Alex Alderton, Roberto Amato, Jeffrey Barrett, Sonia Goncalves, Ewan Harrison, David K. Jackson, Ian Johnston, Dominic Kwiatkowski, Cordelia Langford, John Sillitoe on behalf of the Wellcome Sanger Institute COVID-19 Surveillance Team |
| EPI_ISL_3070658 | Berkshire and Surrey Pathology Services Lighthouse Laboratory | Wellcome Sanger Institute for the COVID-19 Genomics UK (COG-UK) Consortium | Berkshire and Surrey Pathology Services Lighthouse Laboratory and Alex Alderton, Roberto Amato, Jeffrey Barrett, Sonia Goncalves, Ewan Harrison, David K. Jackson, Ian Johnston, Dominic Kwiatkowski, Cordelia Langford, John Sillitoe on behalf of the Wellcome Sanger Institute COVID-19 Surveillance Team |
| EPI_ISL_3070659, EPI_ISL_3070660, EPI_ISL_3070661 | Lighthouse Lab in Milton Keynes | Wellcome Sanger Institute for the COVID-19 Genomics UK (COG-UK) Consortium | The Lighthouse Lab in Milton Keynes and Alex Alderton, Roberto Amato, Jeffrey Barrett, Sonia Goncalves, Ewan Harrison, David K. Jackson, Ian Johnston, Dominic Kwiatkowski, Cordelia Langford, John Sillitoe on behalf of the Wellcome Sanger Institute COVID-19 Surveillance Team |
| EPI_ISL_3070662, EPI_ISL_3070663, EPI_ISL_3070664, EPI_ISL_3070665 | Berkshire and Surrey Pathology Services Lighthouse Laboratory | Wellcome Sanger Institute for the COVID-19 Genomics UK (COG-UK) Consortium | Berkshire and Surrey Pathology Services Lighthouse Laboratory and Alex Alderton, Roberto Amato, Jeffrey Barrett, Sonia Goncalves, Ewan Harrison, David K. Jackson, Ian Johnston, Dominic Kwiatkowski, Cordelia Langford, John Sillitoe on behalf of the Wellcome Sanger Institute COVID-19 Surveillance Team |
| EPI_ISL_3070666 | Lighthouse Lab in Milton Keynes | Wellcome Sanger Institute for the COVID-19 Genomics UK (COG-UK) Consortium | The Lighthouse Lab in Milton Keynes and Alex Alderton, Roberto Amato, Jeffrey Barrett, Sonia Goncalves, Ewan Harrison, David K. Jackson, Ian Johnston, Dominic Kwiatkowski, Cordelia Langford, John Sillitoe on behalf of the Wellcome Sanger Institute COVID-19 Surveillance Team |
| EPI_ISL_3070667, EPI_ISL_3070668, EPI_ISL_3070669, EPI_ISL_3070670 | Berkshire and Surrey Pathology Services Lighthouse Laboratory | Wellcome Sanger Institute for the COVID-19 Genomics UK (COG-UK) Consortium | Berkshire and Surrey Pathology Services Lighthouse Laboratory and Alex Alderton, Roberto Amato, Jeffrey Barrett, Sonia Goncalves, Ewan Harrison, David K. Jackson, Ian Johnston, Dominic Kwiatkowski, Cordelia Langford, John Sillitoe on behalf of the Wellcome Sanger Institute COVID-19 Surveillance Team |
| EPI_ISL_3070671 | Lighthouse Lab in Milton Keynes | Wellcome Sanger Institute for the COVID-19 Genomics UK (COG-UK) Consortium | The Lighthouse Lab in Milton Keynes and Alex Alderton, Roberto Amato, Jeffrey Barrett, Sonia Goncalves, Ewan Harrison, David K. Jackson, Ian Johnston, Dominic Kwiatkowski, Cordelia Langford, John Sillitoe on behalf of the Wellcome Sanger Institute COVID-19 Surveillance Team |
| EPI_ISL_3070672, EPI_ISL_3070673, EPI_ISL_3070674, EPI_ISL_3070675, EPI_ISL_3070676, EPI_ISL_3070677, EPI_ISL_3070678, EPI_ISL_3070679, EPI_ISL_3070680, EPI_ISL_3070681, EPI_ISL_3070682, EPI_ISL_3070683 | | | |
| see above | Berkshire and Surrey Pathology Services Lighthouse Laboratory | Wellcome Sanger Institute for the COVID-19 Genomics UK (COG-UK) Consortium | Berkshire and Surrey Pathology Services Lighthouse Laboratory and Alex Alderton, Roberto Amato, Jeffrey Barrett, Sonia Goncalves, Ewan Harrison, David K. Jackson, Ian Johnston, Dominic Kwiatkowski, Cordelia Langford, John Sillitoe on behalf of the Wellcome Sanger Institute COVID-19 Surveillance Team |
| EPI_ISL_3070684 | Lighthouse Lab in Milton Keynes | Wellcome Sanger Institute for the COVID-19 Genomics UK (COG-UK) Consortium | The Lighthouse Lab in Milton Keynes and Alex Alderton, Roberto Amato, Jeffrey Barrett, Sonia Goncalves, Ewan Harrison, David K. Jackson, Ian Johnston, Dominic Kwiatkowski, Cordelia Langford, John Sillitoe on behalf of the Wellcome Sanger Institute COVID-19 Surveillance Team |
| EPI_ISL_3070685 | Berkshire and Surrey Pathology Services Lighthouse Laboratory | Wellcome Sanger Institute for the COVID-19 Genomics UK (COG-UK) Consortium | Berkshire and Surrey Pathology Services Lighthouse Laboratory and Alex Alderton, Roberto Amato, Jeffrey Barrett, Sonia Goncalves, Ewan Harrison, David K. Jackson, Ian Johnston, Dominic Kwiatkowski, Cordelia Langford, John Sillitoe on behalf of the Wellcome Sanger Institute COVID-19 Surveillance Team |
| EPI_ISL_3070686 | Lighthouse Lab in Milton Keynes | Wellcome Sanger Institute for the COVID-19 Genomics UK (COG-UK) Consortium | The Lighthouse Lab in Milton Keynes and Alex Alderton, Roberto Amato, Jeffrey Barrett, Sonia Goncalves, Ewan Harrison, David K. Jackson, Ian Johnston, Dominic Kwiatkowski, Cordelia Langford, John Sillitoe on behalf of the Wellcome Sanger Institute COVID-19 Surveillance Team |
| EPI_ISL_3070687, EPI_ISL_3070688, | Berkshire and Surrey Pathology Services Lighthouse Laboratory | Wellcome Sanger Institute for the COVID-19 Genomics UK (COG-UK) Consortium | Berkshire and Surrey Pathology Services Lighthouse Laboratory and Alex Alderton, Roberto Amato, Jeffrey Barrett, Sonia Goncalves, Ewan Harrison, David K. Jackson, Ian Johnston, Dominic Kwiatkowski, Cordelia Langford, John Sillitoe on behalf of the Wellcome Sanger Institute COVID-19 Surveillance Team |

| | | | |
|---|---|---|---|
| EPI_ISL_3070689 | | | |
| EPI_ISL_3070690, EPI_ISL_3070691 | Lighthouse Lab in Milton Keynes | Wellcome Sanger Institute for the COVID-19 Genomics UK (COG-UK) Consortium | The Lighthouse Lab in Milton Keynes and Alex Alderton, Roberto Amato, Jeffrey Barrett, Sonia Goncalves, Ewan Harrison, David K. Jackson, Ian Johnston, Dominic Kwiatkowski, Cordelia Langford, John Sillitoe on behalf of the Wellcome Sanger Institute COVID-19 Surveillance Team |
| EPI_ISL_3070692, EPI_ISL_3070693 | Berkshire and Surrey Pathology Services Lighthouse Laboratory | Wellcome Sanger Institute for the COVID-19 Genomics UK (COG-UK) Consortium | Berkshire and Surrey Pathology Services Lighthouse Laboratory and Alex Alderton, Roberto Amato, Jeffrey Barrett, Sonia Goncalves, Ewan Harrison, David K. Jackson, Ian Johnston, Dominic Kwiatkowski, Cordelia Langford, John Sillitoe on behalf of the Wellcome Sanger Institute COVID-19 Surveillance Team |
| EPI_ISL_3070694, EPI_ISL_3070695, EPI_ISL_3070696 | Lighthouse Lab in Milton Keynes | Wellcome Sanger Institute for the COVID-19 Genomics UK (COG-UK) Consortium | The Lighthouse Lab in Milton Keynes and Alex Alderton, Roberto Amato, Jeffrey Barrett, Sonia Goncalves, Ewan Harrison, David K. Jackson, Ian Johnston, Dominic Kwiatkowski, Cordelia Langford, John Sillitoe on behalf of the Wellcome Sanger Institute COVID-19 Surveillance Team |
| EPI_ISL_3070697, EPI_ISL_3070698, EPI_ISL_3070699, EPI_ISL_3070700, EPI_ISL_3070701, EPI_ISL_3070702, EPI_ISL_3070703, EPI_ISL_3070704 | Berkshire and Surrey Pathology Services Lighthouse Laboratory | Wellcome Sanger Institute for the COVID-19 Genomics UK (COG-UK) Consortium | Berkshire and Surrey Pathology Services Lighthouse Laboratory and Alex Alderton, Roberto Amato, Jeffrey Barrett, Sonia Goncalves, Ewan Harrison, David K. Jackson, Ian Johnston, Dominic Kwiatkowski, Cordelia Langford, John Sillitoe on behalf of the Wellcome Sanger Institute COVID-19 Surveillance Team |
| EPI_ISL_3070705 | Lighthouse Lab in Milton Keynes | Wellcome Sanger Institute for the COVID-19 Genomics UK (COG-UK) Consortium | The Lighthouse Lab in Milton Keynes and Alex Alderton, Roberto Amato, Jeffrey Barrett, Sonia Goncalves, Ewan Harrison, David K. Jackson, Ian Johnston, Dominic Kwiatkowski, Cordelia Langford, John Sillitoe on behalf of the Wellcome Sanger Institute COVID-19 Surveillance Team |
| EPI_ISL_3070706 | Berkshire and Surrey Pathology Services Lighthouse Laboratory | Wellcome Sanger Institute for the COVID-19 Genomics UK (COG-UK) Consortium | Berkshire and Surrey Pathology Services Lighthouse Laboratory and Alex Alderton, Roberto Amato, Jeffrey Barrett, Sonia Goncalves, Ewan Harrison, David K. Jackson, Ian Johnston, Dominic Kwiatkowski, Cordelia Langford, John Sillitoe on behalf of the Wellcome Sanger Institute COVID-19 Surveillance Team |
| EPI_ISL_3070707, EPI_ISL_3070708, EPI_ISL_3070709, EPI_ISL_3070710 | Lighthouse Lab in Milton Keynes | Wellcome Sanger Institute for the COVID-19 Genomics UK (COG-UK) Consortium | The Lighthouse Lab in Milton Keynes and Alex Alderton, Roberto Amato, Jeffrey Barrett, Sonia Goncalves, Ewan Harrison, David K. Jackson, Ian Johnston, Dominic Kwiatkowski, Cordelia Langford, John Sillitoe on behalf of the Wellcome Sanger Institute COVID-19 Surveillance Team |
| EPI_ISL_3070711, EPI_ISL_3070712 | Berkshire and Surrey Pathology Services Lighthouse Laboratory | Wellcome Sanger Institute for the COVID-19 Genomics UK (COG-UK) Consortium | Berkshire and Surrey Pathology Services Lighthouse Laboratory and Alex Alderton, Roberto Amato, Jeffrey Barrett, Sonia Goncalves, Ewan Harrison, David K. Jackson, Ian Johnston, Dominic Kwiatkowski, Cordelia Langford, John Sillitoe on behalf of the Wellcome Sanger Institute COVID-19 Surveillance Team |
| EPI_ISL_3070713 | Lighthouse Lab in Milton Keynes | Wellcome Sanger Institute for the COVID-19 Genomics UK (COG-UK) Consortium | The Lighthouse Lab in Milton Keynes and Alex Alderton, Roberto Amato, Jeffrey Barrett, Sonia Goncalves, Ewan Harrison, David K. Jackson, Ian Johnston, Dominic Kwiatkowski, Cordelia Langford, John Sillitoe on behalf of the Wellcome Sanger Institute COVID-19 Surveillance Team |
| EPI_ISL_3070714 | Berkshire and Surrey Pathology Services Lighthouse Laboratory | Wellcome Sanger Institute for the COVID-19 Genomics UK (COG-UK) Consortium | Berkshire and Surrey Pathology Services Lighthouse Laboratory and Alex Alderton, Roberto Amato, Jeffrey Barrett, Sonia Goncalves, Ewan Harrison, David K. Jackson, Ian Johnston, Dominic Kwiatkowski, Cordelia Langford, John Sillitoe on behalf of the Wellcome Sanger Institute COVID-19 Surveillance Team |
| EPI_ISL_3070715 | Lighthouse Lab in Milton Keynes | Wellcome Sanger Institute for the COVID-19 Genomics UK (COG-UK) Consortium | The Lighthouse Lab in Milton Keynes and Alex Alderton, Roberto Amato, Jeffrey Barrett, Sonia Goncalves, Ewan Harrison, David K. Jackson, Ian Johnston, Dominic Kwiatkowski, Cordelia Langford, John Sillitoe on behalf of the Wellcome Sanger Institute COVID-19 Surveillance Team |
| EPI_ISL_3070716, EPI_ISL_3070717 | Berkshire and Surrey Pathology Services Lighthouse Laboratory | Wellcome Sanger Institute for the COVID-19 Genomics UK (COG-UK) Consortium | Berkshire and Surrey Pathology Services Lighthouse Laboratory and Alex Alderton, Roberto Amato, Jeffrey Barrett, Sonia Goncalves, Ewan Harrison, David K. Jackson, Ian Johnston, Dominic Kwiatkowski, Cordelia Langford, John Sillitoe on behalf of the Wellcome Sanger Institute COVID-19 Surveillance Team |
| EPI_ISL_3070718 | Lighthouse Lab in Alderley Park | Wellcome Sanger Institute for the COVID-19 Genomics UK (COG-UK) Consortium | Jacquelyn Wynn, Mairead Hyland, The Lighthouse Lab in Alderley Park and Alex Alderton, Roberto Amato, Jeffrey Barrett, Sonia Goncalves, Ewan Harrison, David K. Jackson, Ian Johnston, Dominic Kwiatkowski, Cordelia Langford, John Sillitoe on behalf of the Wellcome Sanger Institute COVID-19 Surveillance Team |
| EPI_ISL_3070719 | Lighthouse Lab in Glasgow | Wellcome Sanger Institute for the COVID-19 Genomics UK (COG-UK) Consortium | Harper VanSteenhouse, Yumi Kasai, David Gray, Carol Clugston, Anna Dominiczak and Alex Alderton, Roberto Amato, Jeffrey Barrett, Sonia Goncalves, Ewan Harrison, David K. Jackson, Ian Johnston, Dominic Kwiatkowski, Cordelia Langford, John Sillitoe on behalf of the Wellcome Sanger Institute COVID-19 Surveillance Team |
| EPI_ISL_3070720 | Lighthouse Lab in Alderley Park | Wellcome Sanger Institute for the COVID-19 Genomics UK (COG-UK) Consortium | Jacquelyn Wynn, Mairead Hyland, The Lighthouse Lab in Alderley Park and Alex Alderton, Roberto Amato, Jeffrey Barrett, Sonia Goncalves, Ewan Harrison, David K. Jackson, Ian Johnston, Dominic Kwiatkowski, Cordelia Langford, John Sillitoe on behalf of the Wellcome Sanger Institute COVID-19 Surveillance Team |
| EPI_ISL_3070721 | Lighthouse Lab in Glasgow | Wellcome Sanger Institute for the COVID-19 Genomics UK (COG-UK) Consortium | Harper VanSteenhouse, Yumi Kasai, David Gray, Carol Clugston, Anna Dominiczak and Alex Alderton, Roberto Amato, Jeffrey Barrett, Sonia Goncalves, Ewan Harrison, David K. Jackson, Ian Johnston, Dominic Kwiatkowski, Cordelia Langford, John Sillitoe on behalf of the Wellcome Sanger Institute COVID-19 Surveillance Team |
| EPI_ISL_3070722, EPI_ISL_3070723, EPI_ISL_3070724, EPI_ISL_3070725, EPI_ISL_3070726, EPI_ISL_3070727, EPI_ISL_3070728 | Lighthouse Lab in Alderley Park | Wellcome Sanger Institute for the COVID-19 Genomics UK (COG-UK) Consortium | Jacquelyn Wynn, Mairead Hyland, The Lighthouse Lab in Alderley Park and Alex Alderton, Roberto Amato, Jeffrey Barrett, Sonia Goncalves, Ewan Harrison, David K. Jackson, Ian Johnston, Dominic Kwiatkowski, Cordelia Langford, John Sillitoe on behalf of the Wellcome Sanger Institute COVID-19 Surveillance Team |
| EPI_ISL_3070729 | Lighthouse Lab in Glasgow | Wellcome Sanger Institute for the COVID-19 Genomics UK (COG-UK) Consortium | Harper VanSteenhouse, Yumi Kasai, David Gray, Carol Clugston, Anna Dominiczak and Alex Alderton, Roberto Amato, Jeffrey Barrett, Sonia Goncalves, Ewan Harrison, David K. Jackson, Ian Johnston, Dominic Kwiatkowski, Cordelia Langford, John Sillitoe on behalf of the Wellcome Sanger Institute COVID-19 Surveillance Team |
| EPI_ISL_3070730, EPI_ISL_3070731, EPI_ISL_3070732, EPI_ISL_3070733, EPI_ISL_3070734, EPI_ISL_3070735 | Lighthouse Lab in Alderley Park | Wellcome Sanger Institute for the COVID-19 Genomics UK (COG-UK) Consortium | Jacquelyn Wynn, Mairead Hyland, The Lighthouse Lab in Alderley Park and Alex Alderton, Roberto Amato, Jeffrey Barrett, Sonia Goncalves, Ewan Harrison, David K. Jackson, Ian Johnston, Dominic Kwiatkowski, Cordelia Langford, John Sillitoe on behalf of the Wellcome Sanger Institute COVID-19 Surveillance Team |
| EPI_ISL_3070736, EPI_ISL_3070737, EPI_ISL_3070738, EPI_ISL_3070739 | Lighthouse Lab in Glasgow | Wellcome Sanger Institute for the COVID-19 Genomics UK (COG-UK) Consortium | Harper VanSteenhouse, Yumi Kasai, David Gray, Carol Clugston, Anna Dominiczak and Alex Alderton, Roberto Amato, Jeffrey Barrett, Sonia Goncalves, Ewan Harrison, David K. Jackson, Ian Johnston, Dominic Kwiatkowski, Cordelia Langford, John Sillitoe on behalf of the Wellcome Sanger Institute COVID-19 Surveillance Team |
| EPI_ISL_3070740, EPI_ISL_3070741 | Lighthouse Lab in Alderley Park | Wellcome Sanger Institute for the COVID-19 Genomics UK (COG-UK) Consortium | Jacquelyn Wynn, Mairead Hyland, The Lighthouse Lab in Alderley Park and Alex Alderton, Roberto Amato, Jeffrey Barrett, Sonia Goncalves, Ewan Harrison, David K. Jackson, Ian Johnston, Dominic Kwiatkowski, Cordelia Langford, John Sillitoe on behalf of the Wellcome Sanger Institute COVID-19 Surveillance Team |
| EPI_ISL_3070742 | Berkshire and Surrey Pathology Services Lighthouse Laboratory | Wellcome Sanger Institute for the COVID-19 Genomics UK (COG-UK) Consortium | Berkshire and Surrey Pathology Services Lighthouse Laboratory and Alex Alderton, Roberto Amato, Jeffrey Barrett, Sonia Goncalves, Ewan Harrison, David K. Jackson, Ian Johnston, Dominic Kwiatkowski, Cordelia Langford, John Sillitoe on behalf of the Wellcome Sanger Institute COVID-19 Surveillance Team |
| EPI_ISL_3070743, EPI_ISL_3070744, EPI_ISL_3070745, EPI_ISL_3070746, EPI_ISL_3070747 | Lighthouse Lab in Glasgow | Wellcome Sanger Institute for the COVID-19 Genomics UK (COG-UK) Consortium | Harper VanSteenhouse, Yumi Kasai, David Gray, Carol Clugston, Anna Dominiczak and Alex Alderton, Roberto Amato, Jeffrey Barrett, Sonia Goncalves, Ewan Harrison, David K. Jackson, Ian Johnston, Dominic Kwiatkowski, Cordelia Langford, John Sillitoe on behalf of the Wellcome Sanger Institute COVID-19 Surveillance Team |
| EPI_ISL_3070748 | Berkshire and Surrey Pathology Services Lighthouse Laboratory | Wellcome Sanger Institute for the COVID-19 Genomics UK (COG-UK) Consortium | Berkshire and Surrey Pathology Services Lighthouse Laboratory and Alex Alderton, Roberto Amato, Jeffrey Barrett, Sonia Goncalves, Ewan Harrison, David K. Jackson, Ian Johnston, Dominic Kwiatkowski, Cordelia Langford, John Sillitoe on behalf of the Wellcome Sanger Institute COVID-19 Surveillance Team |
| EPI_ISL_3070749 | Randox Laboratories | Wellcome Sanger Institute for the COVID-19 Genomics UK (COG-UK) Consortium | Randox Laboratories and Alex Alderton, Roberto Amato, Jeffrey Barrett, Sonia Goncalves, Ewan Harrison, David K. Jackson, Ian Johnston, Dominic Kwiatkowski, Cordelia Langford, John Sillitoe on behalf of the Wellcome Sanger Institute COVID-19 Surveillance Team |
| EPI_ISL_3070750, EPI_ISL_3070751, EPI_ISL_3070752, EPI_ISL_3070753, EPI_ISL_3070754, EPI_ISL_3070755, EPI_ISL_3070756, EPI_ISL_3070757, EPI_ISL_3070758, EPI_ISL_3070759 | Lighthouse Lab in Glasgow | Wellcome Sanger Institute for the COVID-19 Genomics UK (COG-UK) Consortium | Harper VanSteenhouse, Yumi Kasai, David Gray, Carol Clugston, Anna Dominiczak and Alex Alderton, Roberto Amato, Jeffrey Barrett, Sonia Goncalves, Ewan Harrison, David K. Jackson, Ian Johnston, Dominic Kwiatkowski, Cordelia Langford, John Sillitoe on behalf of the Wellcome Sanger Institute COVID-19 Surveillance Team |
| EPI_ISL_3070760 | Lighthouse Lab in Alderley Park | Wellcome Sanger Institute for the COVID-19 Genomics UK (COG-UK) Consortium | Jacquelyn Wynn, Mairead Hyland, The Lighthouse Lab in Alderley Park and Alex Alderton, Roberto Amato, Jeffrey Barrett, Sonia Goncalves, Ewan Harrison, David K. Jackson, Ian Johnston, Dominic Kwiatkowski, Cordelia Langford, John Sillitoe on behalf of the Wellcome Sanger Institute COVID-19 Surveillance Team |
| EPI_ISL_3070761, EPI_ISL_3070762 | Randox Laboratories | Wellcome Sanger Institute for the COVID-19 Genomics UK (COG-UK) Consortium | Randox Laboratories and Alex Alderton, Roberto Amato, Jeffrey Barrett, Sonia Goncalves, Ewan Harrison, David K. Jackson, Ian Johnston, Dominic Kwiatkowski, Cordelia Langford, John Sillitoe on behalf of the Wellcome Sanger Institute COVID-19 Surveillance Team |
| EPI_ISL_3070763 | Lighthouse Lab in Alderley Park | Wellcome Sanger Institute for the COVID-19 Genomics UK (COG-UK) Consortium | Jacquelyn Wynn, Mairead Hyland, The Lighthouse Lab in Alderley Park and Alex Alderton, Roberto Amato, Jeffrey Barrett, Sonia Goncalves, Ewan Harrison, David K. Jackson, Ian Johnston, Dominic Kwiatkowski, Cordelia Langford, John Sillitoe on behalf of the Wellcome Sanger Institute COVID-19 Surveillance Team |
| EPI_ISL_3070775, EPI_ISL_3070776, EPI_ISL_3070777, EPI_ISL_3070778, EPI_ISL_3070779, EPI_ISL_3070780, EPI_ISL_3070781, EPI_ISL_3070782, EPI_ISL_3070783, EPI_ISL_3070786, EPI_ISL_3070787, EPI_ISL_3070788, EPI_ISL_3070789, EPI_ISL_3070790, EPI_ISL_3070791, EPI_ISL_3070792, EPI_ISL_3070845, EPI_ISL_3070846, EPI_ISL_3070847, EPI_ISL_3070848, EPI_ISL_3070849, EPI_ISL_3070850, EPI_ISL_3070851, EPI_ISL_3070852, EPI_ISL_3070853, EPI_ISL_3070854, EPI_ISL_3070855 | | | |
| see above | Allergy, Immunology and Cell Biology Unit (AICBU) | Allergy, Immunology and Cell Biology Unit (AICBU) | Chandima Jeewandara, Deshni Jayathilaka, Dumni Gunasinghe, Tibutius Thanesh, Dinuka Ariyarathne, Ayesha Wijesinghe, Diyanath Ranasinghe, Gathsaurie Neelika Malavige |
| EPI_ISL_3070856, EPI_ISL_3070857 | RSUP Dr. Kariadi | Eijkman Institute for Molecular Biology, National Research and Innovation Agency | Iskandar Adnan, Sukma Oktavianthi, Lidwina Priliani, Edison Johar, Frilasita A Yudhaputri, Muhammad Rezki Rasyak, Willy Agustine, Hidayat Trimarsanto, Lydia V. Panggalo, Safarina G Malik, Khin Saw Myint, Amin Soebandrio |
| EPI_ISL_3070858 | RS Bogor Senior | Eijkman Institute for Molecular Biology, National Research and | Muhammad Rezki Rasyak, Willy Agustine, Hidayat Trimarsanto, Lydia V. Panggalo, Iskandar Adnan, Sukma Oktavianthi, Lidwina Priliani, Edison Johar, Frilasita A Yudhaputri, Safarina G Malik, Khin Saw Myint, Amin Soebandrio |

| | | | |
|---|---|---|---|
| | | Innovation Agency | |
| EPI_ISL_3070859 | RS Mitra Keluarga Gading Serpong | Eijkman Institute for Molecular Biology, National Research and Innovation Agency | Lidwina Priliani, Edison Johar, Frilasita A Yudhaputri, Muhammad Rezki Rasyak, Willy Agustine, Hidayat Trimarsanto, Lydia V. Panggalo, Iskandar Adnan, Sukma Oktavianthi, Safarina G Malik, Khin Saw Myint, Amin Soebandrio |
| EPI_ISL_3070860 | Dinas Kesehatan Provinsi Nusa Tenggara Timur | Eijkman Institute for Molecular Biology, National Research and Innovation Agency | Iskandar Adnan, Sukma Oktavianthi, Lidwina Priliani, Edison Johar, Frilasita A Yudhaputri, Muhammad Rezki Rasyak, Willy Agustine, Hidayat Trimarsanto, Lydia V. Panggalo, Safarina G Malik, Khin Saw Myint, Amin Soebandrio |
| EPI_ISL_3070862 | RS Rawa Lumbu | Eijkman Institute for Molecular Biology, National Research and Innovation Agency | Edison Johar, Frilasita A Yudhaputri, Muhammad Rezki Rasyak, Willy Agustine, Hidayat Trimarsanto, Lydia V. Panggalo, Iskandar Adnan, Sukma Oktavianthi, Lidwina Priliani, Safarina G Malik, Khin Saw Myint, Amin Soebandrio |
| EPI_ISL_3070863 | RSU Dinda Tangerang | Eijkman Institute for Molecular Biology, National Research and Innovation Agency | Frilasita A Yudhaputri, Muhammad Rezki Rasyak, Willy Agustine, Hidayat Trimarsanto, Lydia V. Panggalo, Iskandar Adnan, Sukma Oktavianthi, Lidwina Priliani, Edison Johar, Safarina G Malik, Khin Saw Myint, Amin Soebandrio |
| EPI_ISL_3070864 | RSU Dinda Tangerang | Eijkman Institute for Molecular Biology, National Research and Innovation Agency | Muhammad Rezki Rasyak, Willy Agustine, Hidayat Trimarsanto, Lydia V. Panggalo, Iskandar Adnan, Sukma Oktavianthi, Lidwina Priliani, Edison Johar, Frilasita A Yudhaputri, Safarina G Malik, Khin Saw Myint, Amin Soebandrio |
| EPI_ISL_3070865 | RSU Dinda Tangerang | Eijkman Institute for Molecular Biology, National Research and Innovation Agency | Willy Agustine, Hidayat Trimarsanto, Lydia V. Panggalo, Iskandar Adnan, Sukma Oktavianthi, Lidwina Priliani, Edison Johar, Frilasita A Yudhaputri, Muhammad Rezki Rasyak, Safarina G Malik, Khin Saw Myint, Amin Soebandrio |
| EPI_ISL_3070866 | RSU Dinda Tangerang | Eijkman Institute for Molecular Biology, National Research and Innovation Agency | Hidayat Trimarsanto, Lydia V. Panggalo, Iskandar Adnan, Sukma Oktavianthi, Lidwina Priliani, Edison Johar, Frilasita A Yudhaputri, Muhammad Rezki Rasyak, Willy Agustine, Safarina G Malik, Khin Saw Myint, Amin Soebandrio |
| EPI_ISL_3070867 | RSUP dr. Mohammad Hoesin/Fakultas Kedokteran Universitas Sriwijaya | Eijkman Institute for Molecular Biology, National Research and Innovation Agency | Lydia V. Panggalo, Iskandar Adnan, Sukma Oktavianthi, Lidwina Priliani, Edison Johar, Frilasita A Yudhaputri, Muhammad Rezki Rasyak, Willy Agustine, Hidayat Trimarsanto, Chairil Anwar, Eny Rahmawati, Phey Liana, Nelda Aprilia Salim, Yulia Iriani, Iche Andriani Liberty, Ahmad Ghiffari, Safarina G Malik, Khin Saw Myint, Amin Soebandrio |
| EPI_ISL_3070871 | RSUP dr. Mohammad Hoesin/Fakultas Kedokteran Universitas Sriwijaya | Eijkman Institute for Molecular Biology, National Research and Innovation Agency | Willy Agustine, Hidayat Trimarsanto, Lydia V. Panggalo, Iskandar Adnan, Sukma Oktavianthi, Lidwina Priliani, Edison Johar, Frilasita A Yudhaputri, Muhammad Rezki Rasyak, Chairil Anwar, Eny Rahmawati, Phey Liana, Nelda Aprilia Salim, Yulia Iriani, Iche Andriani Liberty, Ahmad Ghiffari, Safarina G Malik, Khin Saw Myint, Amin Soebandrio |
| EPI_ISL_3070940 | BTKLPP Kelas I Batam | Eijkman Institute for Molecular Biology, National Research and Innovation Agency; National Institute of Health Research and Development | Frilasita A Yudhaputri, Muhammad Rezki Rasyak, Willy Agustine, Hidayat Trimarsanto, Lydia V. Panggalo, Iskandar Adnan, Sukma Oktavianthi, Lidwina Priliani, Edison Johar, Vivi Setiawaty, Hana Apsari Pawestri, Subangkit, Safarina G Malik, Khin Saw Myint, Amin Soebandrio |
| EPI_ISL_3070963 | RS Mitra Kasih | Eijkman Institute for Molecular Biology, National Research and Innovation Agency; West Java Health Laboratory | Muhammad Rezki Rasyak, Willy Agustine, Hidayat Trimarsanto, Lydia V. Panggalo, Iskandar Adnan, Sukma Oktavianthi, Lidwina Priliani, Edison Johar, Frilasita A Yudhaputri, Azzania Fibriani, Ema Rahmawati, Ryan Bayusantika Ristandi, Rifky Waluyajati Rachman, Cut Nur Cinthia Alamanda, Safarina G Malik, Khin Saw Myint, Amin Soebandrio |
| EPI_ISL_3071075, EPI_ISL_3071076, EPI_ISL_3071077, EPI_ISL_3071079, EPI_ISL_3071080, EPI_ISL_3071081, EPI_ISL_3071088, EPI_ISL_3071089, EPI_ISL_3071090, EPI_ISL_3071091, EPI_ISL_3071092, EPI_ISL_3071093, EPI_ISL_3071094, EPI_ISL_3071096, EPI_ISL_3071097, EPI_ISL_3071098, EPI_ISL_3071099, EPI_ISL_3071100, EPI_ISL_3071101, EPI_ISL_3071102, EPI_ISL_3071103, EPI_ISL_3071104, EPI_ISL_3071105, EPI_ISL_3071106, EPI_ISL_3071108, EPI_ISL_3071109, EPI_ISL_3071110, EPI_ISL_3071113, EPI_ISL_3071114, EPI_ISL_3071115, EPI_ISL_3071116, EPI_ISL_3071117, EPI_ISL_3071118, EPI_ISL_3071119, EPI_ISL_3071120, EPI_ISL_3071121, EPI_ISL_3071123, EPI_ISL_3071124, EPI_ISL_3071125, EPI_ISL_3071126, EPI_ISL_3071127, EPI_ISL_3071128, EPI_ISL_3071129, EPI_ISL_3071130 | | | |
| see above | Child Health Research Foundation | Child Health Research Foundation | CHRF Bangladesh Genomics Team |
| EPI_ISL_3071133 | Shimantik Pathology and Diagnostic Center | Child Health Research Foundation | CHRF Bangladesh Genomics Team, Md. Parvej Alam, Md. Mobarok Karim |
| EPI_ISL_3071139, EPI_ISL_3071140, EPI_ISL_3071141, EPI_ISL_3071142, EPI_ISL_3071143, EPI_ISL_3071144, EPI_ISL_3071145, EPI_ISL_3071146, EPI_ISL_3071147, EPI_ISL_3071148, EPI_ISL_3071149, EPI_ISL_3071150, EPI_ISL_3071151, EPI_ISL_3071152, EPI_ISL_3071153, EPI_ISL_3071154, EPI_ISL_3071155, EPI_ISL_3071156, EPI_ISL_3071157, EPI_ISL_3071158, EPI_ISL_3071159, EPI_ISL_3071160, EPI_ISL_3071161, EPI_ISL_3071162, EPI_ISL_3071163, EPI_ISL_3071164, EPI_ISL_3071165, EPI_ISL_3071166, EPI_ISL_3071167, EPI_ISL_3071168, EPI_ISL_3071169, EPI_ISL_3071170, EPI_ISL_3071171, EPI_ISL_3071172, EPI_ISL_3071173, EPI_ISL_3071174, EPI_ISL_3071175, EPI_ISL_3071176, EPI_ISL_3071177, EPI_ISL_3071178, EPI_ISL_3071179, EPI_ISL_3071180, EPI_ISL_3071181, EPI_ISL_3071182, EPI_ISL_3071183, EPI_ISL_3071184, EPI_ISL_3071185, EPI_ISL_3071186, EPI_ISL_3071187, EPI_ISL_3071188, EPI_ISL_3071189, EPI_ISL_3071190, EPI_ISL_3071191, EPI_ISL_3071192, EPI_ISL_3071193, EPI_ISL_3071194, EPI_ISL_3071195, EPI_ISL_3071196, EPI_ISL_3071197, EPI_ISL_3071198, EPI_ISL_3071199, EPI_ISL_3071200, EPI_ISL_3071201, EPI_ISL_3071202, EPI_ISL_3071203, EPI_ISL_3071204, EPI_ISL_3071205, EPI_ISL_3071206, EPI_ISL_3071207, EPI_ISL_3071208, EPI_ISL_3071209, EPI_ISL_3071210, EPI_ISL_3071211, EPI_ISL_3071212, EPI_ISL_3071213, EPI_ISL_3071214, EPI_ISL_3071215, EPI_ISL_3071216, EPI_ISL_3071217, EPI_ISL_3071218, EPI_ISL_3071219, EPI_ISL_3071220, EPI_ISL_3071221, EPI_ISL_3071222, EPI_ISL_3071223, EPI_ISL_3071224, EPI_ISL_3071225, EPI_ISL_3071226, EPI_ISL_3071227, EPI_ISL_3071228, EPI_ISL_3071229, EPI_ISL_3071230, EPI_ISL_3071231, EPI_ISL_3071232 | | | |
| see above | UNILABS | Instituto Nacional de Saude (INSA) and Instituto Gulbenkian de Ciencia (IGC) | Borges et al |
| EPI_ISL_3071233, EPI_ISL_3071234, EPI_ISL_3071235, EPI_ISL_3071236, EPI_ISL_3071237, EPI_ISL_3071238, EPI_ISL_3071239, EPI_ISL_3071240, EPI_ISL_3071241, EPI_ISL_3071242, EPI_ISL_3071243 | | | |
| see above | H Evora | Instituto Nacional de Saude (INSA) and Institute of Biomedicine (iBiMed), Universidade de Aveiro | Borges et al |
| EPI_ISL_3071244, EPI_ISL_3071245, EPI_ISL_3071246, EPI_ISL_3071247, EPI_ISL_3071248, EPI_ISL_3071249, EPI_ISL_3071250, EPI_ISL_3071251 | H Santarem | Instituto Nacional de Saude (INSA) and Institute of Biomedicine (iBiMed), Universidade de Aveiro | Borges et al |
| EPI_ISL_3071252, EPI_ISL_3071253, EPI_ISL_3071254, EPI_ISL_3071255, EPI_ISL_3071256, EPI_ISL_3071257, EPI_ISL_3071258, EPI_ISL_3071259, EPI_ISL_3071260, EPI_ISL_3071261, EPI_ISL_3071262, EPI_ISL_3071263, EPI_ISL_3071264 | | | |
| see above | CHLC | Instituto Nacional de Saude (INSA) and Institute of Biomedicine (iBiMed), Universidade de Aveiro | Borges et al |
| EPI_ISL_3071265, EPI_ISL_3071266, EPI_ISL_3071267, EPI_ISL_3071268, EPI_ISL_3071269, EPI_ISL_3071270, EPI_ISL_3071271, EPI_ISL_3071272, EPI_ISL_3071273, EPI_ISL_3071274 | LAC UCoimbra | Instituto Nacional de Saude (INSA) and Institute of Biomedicine (iBiMed), Universidade de Aveiro | Borges et al |
| EPI_ISL_3071275, EPI_ISL_3071276, EPI_ISL_3071277, EPI_ISL_3071278, EPI_ISL_3071279, EPI_ISL_3071280, EPI_ISL_3071281, EPI_ISL_3071282, EPI_ISL_3071283, EPI_ISL_3071284 | ULS Norte Alentejano | Instituto Nacional de Saude (INSA) and Institute of Biomedicine (iBiMed), Universidade de Aveiro | Borges et al |
| EPI_ISL_3071285 | ARS Algarve - Laboratorio Laura Ayres | Instituto Nacional de Saude (INSA) | Borges et al |
| EPI_ISL_3071286, EPI_ISL_3071287, EPI_ISL_3071288, EPI_ISL_3071289, EPI_ISL_3071290, EPI_ISL_3071292, EPI_ISL_3071293, EPI_ISL_3071294, EPI_ISL_3071295, EPI_ISL_3071296, EPI_ISL_3071297, EPI_ISL_3071299, EPI_ISL_3071300, EPI_ISL_3071301 | | | |
| see above | Synlab | Instituto Nacional de Saude (INSA) | Borges et al |
| EPI_ISL_3071302, EPI_ISL_3071303, EPI_ISL_3071304, EPI_ISL_3071305, EPI_ISL_3071306, EPI_ISL_3071307, EPI_ISL_3071308, EPI_ISL_3071309, EPI_ISL_3071310, EPI_ISL_3071311, EPI_ISL_3071312, EPI_ISL_3071313, EPI_ISL_3071314, EPI_ISL_3071315 | | | |
| see above | ULS Litoral Alentejano | Instituto Nacional de Saude (INSA) | Borges et al |
| EPI_ISL_3071316, EPI_ISL_3071317, EPI_ISL_3071318, EPI_ISL_3071319, EPI_ISL_3071320, EPI_ISL_3071321, EPI_ISL_3071322, EPI_ISL_3071323, EPI_ISL_3071324, EPI_ISL_3071325, EPI_ISL_3071326, EPI_ISL_3071327, EPI_ISL_3071328, EPI_ISL_3071329, EPI_ISL_3071330 | | | |
| see above | H Vila Franca Xira | Instituto Nacional de Saude (INSA) | Borges et al |
| EPI_ISL_3071331, EPI_ISL_3071332, EPI_ISL_3071333, EPI_ISL_3071334, EPI_ISL_3071335, EPI_ISL_3071336, EPI_ISL_3071337, EPI_ISL_3071338, EPI_ISL_3071339, EPI_ISL_3071340, EPI_ISL_3071341, EPI_ISL_3071342, EPI_ISL_3071343, EPI_ISL_3071344 | | | |
| see above | H Fernando Fonseca | Instituto Nacional de Saude (INSA) | Borges et al |
| EPI_ISL_3071345, EPI_ISL_3071346, EPI_ISL_3071347, EPI_ISL_3071348, EPI_ISL_3071349, EPI_ISL_3071350, EPI_ISL_3071351, EPI_ISL_3071352, EPI_ISL_3071353, EPI_ISL_3071354, EPI_ISL_3071355, EPI_ISL_3071356, EPI_ISL_3071357, EPI_ISL_3071358, EPI_ISL_3071359, EPI_ISL_3071360, EPI_ISL_3071361, EPI_ISL_3071362, EPI_ISL_3071363, EPI_ISL_3071365, EPI_ISL_3071366, EPI_ISL_3071367, EPI_ISL_3071368, EPI_ISL_3071369, EPI_ISL_3071370, EPI_ISL_3071371, EPI_ISL_3071372, EPI_ISL_3071374 | | | |
| see above | ABC Algarve | Instituto Nacional de Saude (INSA) | Borges et al |
| EPI_ISL_3071375, EPI_ISL_3071376, EPI_ISL_3071378, EPI_ISL_3071379, EPI_ISL_3071380, EPI_ISL_3071381, EPI_ISL_3071382, EPI_ISL_3071383, EPI_ISL_3071384, EPI_ISL_3071385, EPI_ISL_3071386, EPI_ISL_3071387, EPI_ISL_3071388, EPI_ISL_3071389, EPI_ISL_3071390, EPI_ISL_3071391, EPI_ISL_3071392 | | | |
| see above | ULS Castelo Branco | Instituto Nacional de Saude (INSA) | Borges et al |
| EPI_ISL_3071393, EPI_ISL_3071394, EPI_ISL_3071395, EPI_ISL_3071396, EPI_ISL_3071397, EPI_ISL_3071398, EPI_ISL_3071399, EPI_ISL_3071400, EPI_ISL_3071401, EPI_ISL_3071402, EPI_ISL_3071403, EPI_ISL_3071405, EPI_ISL_3071406 | | | |
| see above | CHLO | Instituto Nacional de Saude (INSA) | Borges et al |
| EPI_ISL_3071407, EPI_ISL_3071408, EPI_ISL_3071409, EPI_ISL_3071410, EPI_ISL_3071412, EPI_ISL_3071413, EPI_ISL_3071414, EPI_ISL_3071416, EPI_ISL_3071417, EPI_ISL_3071418, EPI_ISL_3071419, EPI_ISL_3071420, EPI_ISL_3071421 | | | |
| see above | H Braga | Instituto Nacional de Saude (INSA) | Borges et al |
| EPI_ISL_3071422, EPI_ISL_3071423, EPI_ISL_3071424, EPI_ISL_3071425, EPI_ISL_3071426, EPI_ISL_3071427, EPI_ISL_3071428, EPI_ISL_3071429, EPI_ISL_3071430, EPI_ISL_3071431, EPI_ISL_3071432, EPI_ISL_3071433, EPI_ISL_3071434, EPI_ISL_3071435 | | | |
| see above | Labeto - CAB - Leiria | Instituto Nacional de Saude (INSA) | Borges et al |
| EPI_ISL_3071436, EPI_ISL_3071437, | CH Setubal | Instituto Nacional de Saude (INSA) | Borges et al |

| | | | |
|---|---|---|---|
| EPI_ISL_3071438, EPI_ISL_3071439 | | | |
| EPI_ISL_3071440, EPI_ISL_3071441, EPI_ISL_3071442, EPI_ISL_3071443, EPI_ISL_3071444, EPI_ISL_3071445, EPI_ISL_3071446, EPI_ISL_3071447, EPI_ISL_3071448, EPI_ISL_3071449, EPI_ISL_3071450, EPI_ISL_3071451, EPI_ISL_3071452, EPI_ISL_3071453, EPI_ISL_3071454, EPI_ISL_3071455, EPI_ISL_3071456, EPI_ISL_3071457, EPI_ISL_3071458, EPI_ISL_3071459, EPI_ISL_3071460, EPI_ISL_3071461, EPI_ISL_3071462, EPI_ISL_3071463, EPI_ISL_3071464, EPI_ISL_3071465, EPI_ISL_3071466, EPI_ISL_3071467, EPI_ISL_3071468, EPI_ISL_3071469, EPI_ISL_3071470, EPI_ISL_3071471, EPI_ISL_3071472, EPI_ISL_3071473, EPI_ISL_3071474, EPI_ISL_3071475, EPI_ISL_3071476, EPI_ISL_3071477, EPI_ISL_3071478, EPI_ISL_3071479, EPI_ISL_3071480, EPI_ISL_3071481, EPI_ISL_3071482, EPI_ISL_3071483, EPI_ISL_3071484, EPI_ISL_3071485, EPI_ISL_3071486, EPI_ISL_3071487, EPI_ISL_3071488, EPI_ISL_3071489, EPI_ISL_3071490, EPI_ISL_3071491, EPI_ISL_3071492, EPI_ISL_3071493, EPI_ISL_3071494, EPI_ISL_3071495, EPI_ISL_3071496, EPI_ISL_3071497, EPI_ISL_3071498 | | | |
| see above | ARS Algarve - Laboratorio Laura Ayres | Instituto Nacional de Saude (INSA) | Borges et al |
| EPI_ISL_3071499, EPI_ISL_3071500, EPI_ISL_3071501, EPI_ISL_3071502, EPI_ISL_3071503, EPI_ISL_3071504, EPI_ISL_3071505, EPI_ISL_3071506, EPI_ISL_3071507, EPI_ISL_3071508, EPI_ISL_3071509, EPI_ISL_3071510, EPI_ISL_3071511, EPI_ISL_3071512, EPI_ISL_3071513 | | | |
| see above | CHU Sao Joao, Porto | Instituto Nacional de Saude (INSA) | Borges et al |
| EPI_ISL_3071514, EPI_ISL_3071515, EPI_ISL_3071516, EPI_ISL_3071517, EPI_ISL_3071518, EPI_ISL_3071519, EPI_ISL_3071520, EPI_ISL_3071521, EPI_ISL_3071522, EPI_ISL_3071523, EPI_ISL_3071524, EPI_ISL_3071525, EPI_ISL_3071526, EPI_ISL_3071527, EPI_ISL_3071528 | | | |
| see above | CHTMAD | Instituto Nacional de Saude (INSA) | Borges et al |
| EPI_ISL_3071529, EPI_ISL_3071530, EPI_ISL_3071531, EPI_ISL_3071532, EPI_ISL_3071533, EPI_ISL_3071534, EPI_ISL_3071535, EPI_ISL_3071536, EPI_ISL_3071537, EPI_ISL_3071538, EPI_ISL_3071539, EPI_ISL_3071540 | | | |
| see above | CH Tondela Viseu | Instituto Nacional de Saude (INSA) | Borges et al |
| EPI_ISL_3071541, EPI_ISL_3071542, EPI_ISL_3071543, EPI_ISL_3071544, EPI_ISL_3071545, EPI_ISL_3071546, EPI_ISL_3071547, EPI_ISL_3071548, EPI_ISL_3071549, EPI_ISL_3071550, EPI_ISL_3071551, EPI_ISL_3071552, EPI_ISL_3071553, EPI_ISL_3071554, EPI_ISL_3071555 | | | |
| see above | CHEDV | Instituto Nacional de Saude (INSA) | Borges et al |
| EPI_ISL_3071556, EPI_ISL_3071557, EPI_ISL_3071558, EPI_ISL_3071559, EPI_ISL_3071560, EPI_ISL_3071561, EPI_ISL_3071562, EPI_ISL_3071563, EPI_ISL_3071564, EPI_ISL_3071565, EPI_ISL_3071566, EPI_ISL_3071567, EPI_ISL_3071568 | | | |
| see above | ULS Guarda | Instituto Nacional de Saude (INSA) | Borges et al |
| EPI_ISL_3071569, EPI_ISL_3071570, EPI_ISL_3071571, EPI_ISL_3071572, EPI_ISL_3071573, EPI_ISL_3071574, EPI_ISL_3071575, EPI_ISL_3071576, EPI_ISL_3071577, EPI_ISL_3071578, EPI_ISL_3071579, EPI_ISL_3071580, EPI_ISL_3071581, EPI_ISL_3071582, EPI_ISL_3071583, EPI_ISL_3071584, EPI_ISL_3071585, EPI_ISL_3071586, EPI_ISL_3071587, EPI_ISL_3071588, EPI_ISL_3071589, EPI_ISL_3071590, EPI_ISL_3071591, EPI_ISL_3071592, EPI_ISL_3071593 | | | |
| see above | H Garcia de Orta | Instituto Nacional de Saude (INSA) | Borges et al |
| EPI_ISL_3071594, EPI_ISL_3071595, EPI_ISL_3071596, EPI_ISL_3071597, EPI_ISL_3071598, EPI_ISL_3071599, EPI_ISL_3071600, EPI_ISL_3071601, EPI_ISL_3071602, EPI_ISL_3071603, EPI_ISL_3071604, EPI_ISL_3071605, EPI_ISL_3071606, EPI_ISL_3071607, EPI_ISL_3071608 | | | |
| see above | CH Porto - H Sto Antonio | Instituto Nacional de Saude (INSA) | Borges et al |
| EPI_ISL_3071609, EPI_ISL_3071610, EPI_ISL_3071611, EPI_ISL_3071612, EPI_ISL_3071613, EPI_ISL_3071614, EPI_ISL_3071616, EPI_ISL_3071617, EPI_ISL_3071618, EPI_ISL_3071619, EPI_ISL_3071620, EPI_ISL_3071621, EPI_ISL_3071622, EPI_ISL_3071623, EPI_ISL_3071624, EPI_ISL_3071625, EPI_ISL_3071626, EPI_ISL_3071627, EPI_ISL_3071628, EPI_ISL_3071629, EPI_ISL_3071630, EPI_ISL_3071631, EPI_ISL_3071632, EPI_ISL_3071633, EPI_ISL_3071634, EPI_ISL_3071635, EPI_ISL_3071636, EPI_ISL_3071637 | | | |
| see above | H Divino Espirito Santo - Ponta Delgada | Instituto Nacional de Saude (INSA) | Borges et al |
| EPI_ISL_3071638, EPI_ISL_3071639, EPI_ISL_3071640, EPI_ISL_3071641, EPI_ISL_3071642, EPI_ISL_3071643, EPI_ISL_3071644, EPI_ISL_3071645, EPI_ISL_3071648, EPI_ISL_3071649, EPI_ISL_3071650, EPI_ISL_3071651, EPI_ISL_3071652, EPI_ISL_3071653, EPI_ISL_3071654, EPI_ISL_3071655, EPI_ISL_3071657, EPI_ISL_3071658, EPI_ISL_3071659, EPI_ISL_3071660, EPI_ISL_3071661, EPI_ISL_3071662, EPI_ISL_3071663, EPI_ISL_3071665, EPI_ISL_3071666 | | | |
| see above | SEEBMO | Instituto Nacional de Saude (INSA) | Borges et al |
| EPI_ISL_3071667, EPI_ISL_3071668, EPI_ISL_3071669, EPI_ISL_3071670, EPI_ISL_3071671, EPI_ISL_3071672, EPI_ISL_3071673, EPI_ISL_3071674, EPI_ISL_3071675, EPI_ISL_3071676, EPI_ISL_3071677, EPI_ISL_3071678, EPI_ISL_3071679, EPI_ISL_3071680, EPI_ISL_3071681 | | | |
| see above | CHULN - H Santa Maria | Instituto Nacional de Saude (INSA) | Borges et al |
| EPI_ISL_3071682, EPI_ISL_3071683, EPI_ISL_3071684, EPI_ISL_3071685, EPI_ISL_3071686, EPI_ISL_3071687, EPI_ISL_3071688, EPI_ISL_3071689, EPI_ISL_3071690, EPI_ISL_3071691, EPI_ISL_3071692, EPI_ISL_3071693, EPI_ISL_3071694, EPI_ISL_3071695, EPI_ISL_3071696 | | | |
| see above | Avelab | Instituto Nacional de Saude (INSA) | Borges et al |
| EPI_ISL_3071697, EPI_ISL_3071698, EPI_ISL_3071699, EPI_ISL_3071700, EPI_ISL_3071701, EPI_ISL_3071702, EPI_ISL_3071703, EPI_ISL_3071704, EPI_ISL_3071705, EPI_ISL_3071706, EPI_ISL_3071707, EPI_ISL_3071708, EPI_ISL_3071709, EPI_ISL_3071710, EPI_ISL_3071711, EPI_ISL_3071712, EPI_ISL_3071714, EPI_ISL_3071715, EPI_ISL_3071716, EPI_ISL_3071717, EPI_ISL_3071718, EPI_ISL_3071719, EPI_ISL_3071720, EPI_ISL_3071721, EPI_ISL_3071722, EPI_ISL_3071723, EPI_ISL_3071724, EPI_ISL_3071726, EPI_ISL_3071727, EPI_ISL_3071728, EPI_ISL_3071729, EPI_ISL_3071730, EPI_ISL_3071731, EPI_ISL_3071732, EPI_ISL_3071733, EPI_ISL_3071734 | | | |
| see above | SESARAM | Instituto Nacional de Saude (INSA) | Borges et al |
| EPI_ISL_3071735 | Infogene | Instituto Nacional de Saude (INSA) | Borges et al |
| EPI_ISL_3071736 | Labovedras | Instituto Nacional de Saude (INSA) | Borges et al |
| EPI_ISL_3071737, EPI_ISL_3071738, EPI_ISL_3071739, EPI_ISL_3071740 | CHMT | Instituto Nacional de Saude (INSA) | Borges et al |
| EPI_ISL_3071741, EPI_ISL_3071742, EPI_ISL_3071743, EPI_ISL_3071744, EPI_ISL_3071745, EPI_ISL_3071746, EPI_ISL_3071747, EPI_ISL_3071748, EPI_ISL_3071749, EPI_ISL_3071750, EPI_ISL_3071751, EPI_ISL_3071752, EPI_ISL_3071753, EPI_ISL_3071754, EPI_ISL_3071755, EPI_ISL_3071756, EPI_ISL_3071757, EPI_ISL_3071758, EPI_ISL_3071759, EPI_ISL_3071760, EPI_ISL_3071761, EPI_ISL_3071762, EPI_ISL_3071763, EPI_ISL_3071764, EPI_ISL_3071765, EPI_ISL_3071766, EPI_ISL_3071767, EPI_ISL_3071768, EPI_ISL_3071769, EPI_ISL_3071770, EPI_ISL_3071771, EPI_ISL_3071772, EPI_ISL_3071773, EPI_ISL_3071774, EPI_ISL_3071775, EPI_ISL_3071776, EPI_ISL_3071777, EPI_ISL_3071778, EPI_ISL_3071779, EPI_ISL_3071780, EPI_ISL_3071781, EPI_ISL_3071782, EPI_ISL_3071783, EPI_ISL_3071784, EPI_ISL_3071785, EPI_ISL_3071786, EPI_ISL_3071787, EPI_ISL_3071788, EPI_ISL_3071789, EPI_ISL_3071790, EPI_ISL_3071791, EPI_ISL_3071792, EPI_ISL_3071793, EPI_ISL_3071794, EPI_ISL_3071795, EPI_ISL_3071796, EPI_ISL_3071797, EPI_ISL_3071798, EPI_ISL_3071799, EPI_ISL_3071800, EPI_ISL_3071801, EPI_ISL_3071802, EPI_ISL_3071803, EPI_ISL_3071804, EPI_ISL_3071805, EPI_ISL_3071806, EPI_ISL_3071807, EPI_ISL_3071808, EPI_ISL_3071809, EPI_ISL_3071810, EPI_ISL_3071811, EPI_ISL_3071812, EPI_ISL_3071813, EPI_ISL_3071814, EPI_ISL_3071815, EPI_ISL_3071816, EPI_ISL_3071817, EPI_ISL_3071818, EPI_ISL_3071819, EPI_ISL_3071820, EPI_ISL_3071821, EPI_ISL_3071822, EPI_ISL_3071823, EPI_ISL_3071824, EPI_ISL_3071825, EPI_ISL_3071826, EPI_ISL_3071827, EPI_ISL_3071828, EPI_ISL_3071829, EPI_ISL_3071830, EPI_ISL_3071831, EPI_ISL_3071832, EPI_ISL_3071833, EPI_ISL_3071834 | | | |
| see above | UNILABS | Instituto Nacional de Saude (INSA) | Borges et al |
| EPI_ISL_3071835, EPI_ISL_3071836 | WSSE w Olsztynie | Wojewodzka Stacja Sanitarno-Epidemiologiczna w Olsztynie, Laboratorium Badan Epidemiologiczno-Klinicznych | Sylwia Krzetowska, Monika Czerminska, Ewa Liszewska, Tomasz Jakubczak, Marta Lukian, Patryk Bielecki, Aleksandra Kobiatko, Emilia Tarabasz, Paulina Rozycka, Barbara Dolinska |
| EPI_ISL_3071838, EPI_ISL_3071839 | WSSE w Bydgoszczy | Wojewodzka Stacja Sanitarno-Epidemiologiczna w Olsztynie, Laboratorium Badan Epidemiologiczno-Klinicznych | Sylwia Krzetowska, Monika Czerminska, Ewa Liszewska, Tomasz Jakubczak, Marta Lukian, Patryk Bielecki, Aleksandra Kobiatko, Emilia Tarabasz, Paulina Rozycka, Barbara Dolinska |
| EPI_ISL_3071840, EPI_ISL_3071841 | WSSE w Bialymstoku | Wojewodzka Stacja Sanitarno-Epidemiologiczna w Olsztynie, Laboratorium Badan Epidemiologiczno-Klinicznych | Sylwia Krzetowska, Monika Czerminska, Ewa Liszewska, Tomasz Jakubczak, Marta Lukian, Patryk Bielecki, Aleksandra Kobiatko, Emilia Tarabasz, Paulina Rozycka, Barbara Dolinska |
| EPI_ISL_3071842, EPI_ISL_3071843 | WSSE w Olsztynie | Wojewodzka Stacja Sanitarno-Epidemiologiczna w Olsztynie, Laboratorium Badan Epidemiologiczno-Klinicznych | Sylwia Krzetowska, Monika Czerminska, Ewa Liszewska, Tomasz Jakubczak, Marta Lukian, Patryk Bielecki, Aleksandra Kobiatko, Emilia Tarabasz, Paulina Rozycka, Barbara Dolinska |
| EPI_ISL_3071844, EPI_ISL_3071845, EPI_ISL_3071846, EPI_ISL_3071847, EPI_ISL_3071848, EPI_ISL_3071849, EPI_ISL_3071850, EPI_ISL_3071851, EPI_ISL_3071852, EPI_ISL_3071853, EPI_ISL_3071854 | | | |
| see above | Renegade.bio | DeRisi Lab, University of California, San Francisco | Sabrina Mann, Anthea Mitchell, Matthias Hapte-Selassie, Jamin Liu, Sara Sunshine, Matthew Laurie, Genay Pilarowski, Amy Kistler, Manu Vanaerschot, Patrick Ayscue, Lucy Li, Eric Chow, IDseq Team, James Peng, Carina Marquez, Luis Rubio, Gabriel Chamie, Diane Jones, Jon Jacobo, Susana Rojas, Susy Rojas, Valerie Tulier-Laiwa, Douglas Black, Jackie Martinez, Maya Petersen, Salu Ribeiro, Diane Havlir, and Joseph DeRisi |
| EPI_ISL_3071855, EPI_ISL_3071856 | Melaka Hospital | Institute for Medical Research, Infectious Disease Research Centre, National Institutes of Health, Ministry of Health Malaysia | Suppiah J, Kamel K, Mohd Zawawi Z, Ramly N, Robert F, Anasir MI, Thayan R |
| EPI_ISL_3071857, EPI_ISL_3071858 | Tengku Ampuan Rahimah Hospital | Institute for Medical Research, Infectious Disease Research Centre, National Institutes of Health, Ministry of Health Malaysia | Suppiah J, Kamel K, Mohd Zawawi Z, Ramly N, Robert F, Anasir MI, Thayan R |
| EPI_ISL_3071861, EPI_ISL_3071862, EPI_ISL_3071863, EPI_ISL_3071864, EPI_ISL_3071866 | SA Pathology | SA Pathology | Caitlin Selway, Lex Leong, Chuan Kok Lim, Mark Turra, Ivan Bastian, Geoff Higgins |
| EPI_ISL_3071867, EPI_ISL_3071868, EPI_ISL_3071869, EPI_ISL_3071870, EPI_ISL_3071871, EPI_ISL_3071872, EPI_ISL_3071873, EPI_ISL_3071874, EPI_ISL_3071875, EPI_ISL_3071876, EPI_ISL_3071877, EPI_ISL_3071878, EPI_ISL_3071879, EPI_ISL_3071880, EPI_ISL_3071883, EPI_ISL_3071884, EPI_ISL_3071885, EPI_ISL_3071886, EPI_ISL_3071887, EPI_ISL_3071888, EPI_ISL_3071889, EPI_ISL_3071890, EPI_ISL_3071891, EPI_ISL_3071892, EPI_ISL_3071894, EPI_ISL_3071895, EPI_ISL_3071896, EPI_ISL_3071897, EPI_ISL_3071898, EPI_ISL_3071899, EPI_ISL_3071900, EPI_ISL_3071901, EPI_ISL_3071902, EPI_ISL_3071903, EPI_ISL_3071904, EPI_ISL_3071905, EPI_ISL_3071907, EPI_ISL_3071908, EPI_ISL_3071909, EPI_ISL_3071910, EPI_ISL_3071911, EPI_ISL_3071912, EPI_ISL_3071913, EPI_ISL_3071915, EPI_ISL_3071916, EPI_ISL_3071917, EPI_ISL_3071918, EPI_ISL_3071919, EPI_ISL_3071920, EPI_ISL_3071921, EPI_ISL_3071922, EPI_ISL_3071923, EPI_ISL_3071924, EPI_ISL_3071925, EPI_ISL_3071926, EPI_ISL_3071927, EPI_ISL_3071928, EPI_ISL_3071929, EPI_ISL_3071930, EPI_ISL_3071931, EPI_ISL_3071932, EPI_ISL_3071934, EPI_ISL_3071936, EPI_ISL_3071937, EPI_ISL_3071938, EPI_ISL_3071939, EPI_ISL_3071940, EPI_ISL_3071941, EPI_ISL_3071942, EPI_ISL_3071944, EPI_ISL_3071945, EPI_ISL_3071946, EPI_ISL_3071947, EPI_ISL_3071948, EPI_ISL_3071949, EPI_ISL_3071950, EPI_ISL_3071951, EPI_ISL_3071952, EPI_ISL_3071953, EPI_ISL_3071954, EPI_ISL_3071955, EPI_ISL_3071956, EPI_ISL_3071957, EPI_ISL_3071958, EPI_ISL_3071959, EPI_ISL_3071961, EPI_ISL_3071962, EPI_ISL_3071963, EPI_ISL_3071964, EPI_ISL_3071966, EPI_ISL_3071967, EPI_ISL_3071968, EPI_ISL_3071969, EPI_ISL_3071970, EPI_ISL_3071971, EPI_ISL_3071972, EPI_ISL_3071973, EPI_ISL_3071974, EPI_ISL_3071975, EPI_ISL_3071976, EPI_ISL_3071977, EPI_ISL_3071978, EPI_ISL_3071979, EPI_ISL_3071980, EPI_ISL_3071981, EPI_ISL_3071982, EPI_ISL_3071983, EPI_ISL_3071984, EPI_ISL_3071985 | | | |
| see above | Pandemic Response Lab - NYC | Pandemic Response Lab, R&D | Henry Lee, Michael Hammerling, Melissa Hopkins, Cybill del Castillo, Shinyoung Clair Kang, William Ward, Pradeep Bugga, Sol Rey, Dylan Law, Isabel Fernandez Escapa, Haiping Hao, Jon Laurent |
| EPI_ISL_3071986 | Ipoh Public Health Laboratory | Institute for Medical Research, Infectious Disease Research Centre, National Institutes of Health, Ministry of Health Malaysia | Suppiah J, Kamel K, Mohd Zawawi Z, Ramly N, Robert F, Anasir MI, Thayan R |
| EPI_ISL_3071988 | Kuala Pilah District Health Office | Institute for Medical Research, Infectious Disease Research Centre, National Institutes of Health, Ministry of Health Malaysia | Suppiah J, Kamel K, Mohd Zawawi Z, Ramly N, Robert F, Anasir MI, Thayan R |
| EPI_ISL_3071992, EPI_ISL_3071993, EPI_ISL_3071994, EPI_ISL_3071995, EPI_ISL_3071996, EPI_ISL_3071997, EPI_ISL_3071998, EPI_ISL_3071999, EPI_ISL_3072000, EPI_ISL_3072001, EPI_ISL_3072002, EPI_ISL_3072003, EPI_ISL_3072005, EPI_ISL_3072006, EPI_ISL_3072007, EPI_ISL_3072008, EPI_ISL_3072009, EPI_ISL_3072010, EPI_ISL_3072011, EPI_ISL_3072012, EPI_ISL_3072013, EPI_ISL_3072014, EPI_ISL_3072015, EPI_ISL_3072016, EPI_ISL_3072017, EPI_ISL_3072018, EPI_ISL_3072019, EPI_ISL_3072020, EPI_ISL_3072021, EPI_ISL_3072022, EPI_ISL_3072023, EPI_ISL_3072024, EPI_ISL_3072025, EPI_ISL_3072026, EPI_ISL_3072027, EPI_ISL_3072028, EPI_ISL_3072029, EPI_ISL_3072030, EPI_ISL_3072031, EPI_ISL_3072032, EPI_ISL_3072033, EPI_ISL_3072034, EPI_ISL_3072036, EPI_ISL_3072037, EPI_ISL_3072038, EPI_ISL_3072039, EPI_ISL_3072042, EPI_ISL_3072043, EPI_ISL_3072044, EPI_ISL_3072045, EPI_ISL_3072046, EPI_ISL_3072047, EPI_ISL_3072048, EPI_ISL_3072049, EPI_ISL_3072050, EPI_ISL_3072051, EPI_ISL_3072052, EPI_ISL_3072053, EPI_ISL_3072054, EPI_ISL_3072056, EPI_ISL_3072057, EPI_ISL_3072058, EPI_ISL_3072059, EPI_ISL_3072060, EPI_ISL_3072062, EPI_ISL_3072063, EPI_ISL_3072064, EPI_ISL_3072065, EPI_ISL_3072066, EPI_ISL_3072067, EPI_ISL_3072068, EPI_ISL_3072069, EPI_ISL_3072070, EPI_ISL_3072071, EPI_ISL_3072073, EPI_ISL_3072074, EPI_ISL_3072075, EPI_ISL_3072076, EPI_ISL_3072077, EPI_ISL_3072078, EPI_ISL_3072079, EPI_ISL_3072080, EPI_ISL_3072081, EPI_ISL_3072082, EPI_ISL_3072083, EPI_ISL_3072084, EPI_ISL_3072085, EPI_ISL_3072086, EPI_ISL_3072088, EPI_ISL_3072089, EPI_ISL_3072090, EPI_ISL_3072091, EPI_ISL_3072092, EPI_ISL_3072093, EPI_ISL_3072095, EPI_ISL_3072096, EPI_ISL_3072097, EPI_ISL_3072099, EPI_ISL_3072100, EPI_ISL_3072102, EPI_ISL_3072103, EPI_ISL_3072106, EPI_ISL_3072107, EPI_ISL_3072108, EPI_ISL_3072110, EPI_ISL_3072112, EPI_ISL_3072113, EPI_ISL_3072115, EPI_ISL_3072116, EPI_ISL_3072117, EPI_ISL_3072118, EPI_ISL_3072119, EPI_ISL_3072120, EPI_ISL_3072121, EPI_ISL_3072123, EPI_ISL_3072124, EPI_ISL_3072125, EPI_ISL_3072126, EPI_ISL_3072127, EPI_ISL_3072128, EPI_ISL_3072129, EPI_ISL_3072130, EPI_ISL_3072131, EPI_ISL_3072132, EPI_ISL_3072133, EPI_ISL_3072134, EPI_ISL_3072135, EPI_ISL_3072136, EPI_ISL_3072137, EPI_ISL_3072138, EPI_ISL_3072140, EPI_ISL_3072141, EPI_ISL_3072142, EPI_ISL_3072145, EPI_ISL_3072146, EPI_ISL_3072148 | | | |
| see above | New South Wales Health Pathology Royal Prince Alfred Hospital | Microbiology RPAH | Foster, C.; Au, J.; Ruiz Silva, M.; Deveson, I.; Bull, R.; Van Hal, S.; Rawlinson, W. |
| EPI_ISL_3072328 | Molekylær Medicinsk Afdeling, Aarhus University Hospital, Aarhus, Denmark | Statens Serum Institut Bioinformatics and Microbial Genomics | Danish Covid-19 Genome Consortium |
| EPI_ISL_3072330, EPI_ISL_3072332, EPI_ISL_3072334, EPI_ISL_3072336, EPI_ISL_3072338, EPI_ISL_3072340, EPI_ISL_3072342, EPI_ISL_3072344, EPI_ISL_3072346, EPI_ISL_3072348, EPI_ISL_3072350, EPI_ISL_3072352, EPI_ISL_3072354, EPI_ISL_3072358, EPI_ISL_3072360, EPI_ISL_3072361, EPI_ISL_3072362, EPI_ISL_3072363, EPI_ISL_3072364, EPI_ISL_3072365, EPI_ISL_3072367, EPI_ISL_3072368, EPI_ISL_3072369, EPI_ISL_3072371, EPI_ISL_3072372, EPI_ISL_3072373, EPI_ISL_3072375, EPI_ISL_3072376, EPI_ISL_3072377, EPI_ISL_3072379, EPI_ISL_3072380, EPI_ISL_3072382, EPI_ISL_3072383, EPI_ISL_3072384, EPI_ISL_3072385, EPI_ISL_3072386, EPI_ISL_3072387, EPI_ISL_3072388, EPI_ISL_3072389, EPI_ISL_3072390, EPI_ISL_3072391, EPI_ISL_3072392, EPI_ISL_3072393, EPI_ISL_3072395, EPI_ISL_3072397, EPI_ISL_3072398, EPI_ISL_3072400, EPI_ISL_3072402, EPI_ISL_3072404, EPI_ISL_3072406, EPI_ISL_3072408, EPI_ISL_3072410, EPI_ISL_3072412, EPI_ISL_3072414, EPI_ISL_3072416, EPI_ISL_3072418, EPI_ISL_3072420, EPI_ISL_3072422, EPI_ISL_3072424, EPI_ISL_3072426, EPI_ISL_3072428, EPI_ISL_3072429, EPI_ISL_3072433, EPI_ISL_3072435, EPI_ISL_3072437, EPI_ISL_3072439, | | | |

| | | | |
|---|---|---|---|
| EPI_ISL_3072441, EPI_ISL_3072443, EPI_ISL_3072445, EPI_ISL_3072447, EPI_ISL_3072449, EPI_ISL_3072453, EPI_ISL_3072455, EPI_ISL_3072457, EPI_ISL_3072459, EPI_ISL_3072461, EPI_ISL_3072466, EPI_ISL_3072468, EPI_ISL_3072470, EPI_ISL_3072472, EPI_ISL_3072474, EPI_ISL_3072478, EPI_ISL_3072480, EPI_ISL_3072482, EPI_ISL_3072486, EPI_ISL_3072488, EPI_ISL_3072490, EPI_ISL_3072492, EPI_ISL_3072493, EPI_ISL_3072496, EPI_ISL_3072498, EPI_ISL_3072500, EPI_ISL_3072502, EPI_ISL_3072504, EPI_ISL_3072508, EPI_ISL_3072510, EPI_ISL_3072512, EPI_ISL_3072516, EPI_ISL_3072518, EPI_ISL_3072520, EPI_ISL_3072522, EPI_ISL_3072524, EPI_ISL_3072526, EPI_ISL_3072528, EPI_ISL_3072530, EPI_ISL_3072532, EPI_ISL_3072534, EPI_ISL_3072537, EPI_ISL_3072538, EPI_ISL_3072540, EPI_ISL_3072544, EPI_ISL_3072546, EPI_ISL_3072548, EPI_ISL_3072550, EPI_ISL_3072552, EPI_ISL_3072554, EPI_ISL_3072556, EPI_ISL_3072558, EPI_ISL_3072560, EPI_ISL_3072562, EPI_ISL_3072564, EPI_ISL_3072566, EPI_ISL_3072568, EPI_ISL_3072570, EPI_ISL_3072572, EPI_ISL_3072574, EPI_ISL_3072576, EPI_ISL_3072578, EPI_ISL_3072580, EPI_ISL_3072582 | | | |
| see above | Department of Bacteria, Parasites and Fungi, Statens Serum Institut, Copenhagen, Denmark | Statens Serum Institut Bioinformatics and Microbial Genomics | Danish Covid-19 Genome Consortium |
| EPI_ISL_3072583 | Department of Clinical Microbiology, Odense University Hospital, Odense, Denmark | Statens Serum Institut Bioinformatics and Microbial Genomics | Danish Covid-19 Genome Consortium |
| EPI_ISL_3072585, EPI_ISL_3072587, EPI_ISL_3072589, EPI_ISL_3072591, EPI_ISL_3072593, EPI_ISL_3072594, EPI_ISL_3072596, EPI_ISL_3072599, EPI_ISL_3072601, EPI_ISL_3072602, EPI_ISL_3072605, EPI_ISL_3072607, EPI_ISL_3072609 | | | |
| see above | Department of Bacteria, Parasites and Fungi, Statens Serum Institut, Copenhagen, Denmark | Statens Serum Institut Bioinformatics and Microbial Genomics | Danish Covid-19 Genome Consortium |
| EPI_ISL_3072611 | Molekylær Medicinsk Afdeling, Aarhus University Hospital, Aarhus, Denmark | Statens Serum Institut Bioinformatics and Microbial Genomics | Danish Covid-19 Genome Consortium |
| EPI_ISL_3072613, EPI_ISL_3072615, EPI_ISL_3072617, EPI_ISL_3072619, EPI_ISL_3072621, EPI_ISL_3072623, EPI_ISL_3072625, EPI_ISL_3072627, EPI_ISL_3072628, EPI_ISL_3072629, EPI_ISL_3072630, EPI_ISL_3072631, EPI_ISL_3072632, EPI_ISL_3072633, EPI_ISL_3072634, EPI_ISL_3072635, EPI_ISL_3072636, EPI_ISL_3072637, EPI_ISL_3072638, EPI_ISL_3072639, EPI_ISL_3072640, EPI_ISL_3072641, EPI_ISL_3072642, EPI_ISL_3072643, EPI_ISL_3072644, EPI_ISL_3072645, EPI_ISL_3072646, EPI_ISL_3072647, EPI_ISL_3072648, EPI_ISL_3072649, EPI_ISL_3072650, EPI_ISL_3072651, EPI_ISL_3072652, EPI_ISL_3072654, EPI_ISL_3072655, EPI_ISL_3072656, EPI_ISL_3072657, EPI_ISL_3072658, EPI_ISL_3072659, EPI_ISL_3072660, EPI_ISL_3072661, EPI_ISL_3072662, EPI_ISL_3072663, EPI_ISL_3072664, EPI_ISL_3072665, EPI_ISL_3072666, EPI_ISL_3072667, EPI_ISL_3072668, EPI_ISL_3072669, EPI_ISL_3072670, EPI_ISL_3072671, EPI_ISL_3072672, EPI_ISL_3072673, EPI_ISL_3072674, EPI_ISL_3072675, EPI_ISL_3072676, EPI_ISL_3072677, EPI_ISL_3072678, EPI_ISL_3072679, EPI_ISL_3072680, EPI_ISL_3072681, EPI_ISL_3072682, EPI_ISL_3072683, EPI_ISL_3072684, EPI_ISL_3072685, EPI_ISL_3072686, EPI_ISL_3072687, EPI_ISL_3072688, EPI_ISL_3072689 | | | |
| see above | Department of Bacteria, Parasites and Fungi, Statens Serum Institut, Copenhagen, Denmark | Statens Serum Institut Bioinformatics and Microbial Genomics | Danish Covid-19 Genome Consortium |
| EPI_ISL_3072690 | Molekylær Medicinsk Afdeling, Aarhus University Hospital, Aarhus, Denmark | Statens Serum Institut Bioinformatics and Microbial Genomics | Danish Covid-19 Genome Consortium |
| EPI_ISL_3072691, EPI_ISL_3072692, EPI_ISL_3072694, EPI_ISL_3072695, EPI_ISL_3072696, EPI_ISL_3072697, EPI_ISL_3072698, EPI_ISL_3072699, EPI_ISL_3072700, EPI_ISL_3072701, EPI_ISL_3072702, EPI_ISL_3072703, EPI_ISL_3072705, EPI_ISL_3072706, EPI_ISL_3072707, EPI_ISL_3072708, EPI_ISL_3072710, EPI_ISL_3072711, EPI_ISL_3072712, EPI_ISL_3072713, EPI_ISL_3072714, EPI_ISL_3072715, EPI_ISL_3072716, EPI_ISL_3072717, EPI_ISL_3072718, EPI_ISL_3072719, EPI_ISL_3072720, EPI_ISL_3072721, EPI_ISL_3072722, EPI_ISL_3072723, EPI_ISL_3072724, EPI_ISL_3072725, EPI_ISL_3072726, EPI_ISL_3072727, EPI_ISL_3072728, EPI_ISL_3072729, EPI_ISL_3072730, EPI_ISL_3072731, EPI_ISL_3072732, EPI_ISL_3072733, EPI_ISL_3072734, EPI_ISL_3072735, EPI_ISL_3072736 | | | |
| see above | Department of Bacteria, Parasites and Fungi, Statens Serum Institut, Copenhagen, Denmark | Statens Serum Institut Bioinformatics and Microbial Genomics | Danish Covid-19 Genome Consortium |
| EPI_ISL_3072737 | Department of Clinical Microbiology, Odense University Hospital, Odense, Denmark | Statens Serum Institut Bioinformatics and Microbial Genomics | Danish Covid-19 Genome Consortium |
| EPI_ISL_3072738, EPI_ISL_3072739, EPI_ISL_3072740, EPI_ISL_3072741, EPI_ISL_3072742, EPI_ISL_3072743, EPI_ISL_3072745, EPI_ISL_3072746, EPI_ISL_3072747, EPI_ISL_3072748, EPI_ISL_3072749, EPI_ISL_3072750, EPI_ISL_3072751, EPI_ISL_3072752, EPI_ISL_3072753, EPI_ISL_3072754, EPI_ISL_3072755, EPI_ISL_3072756, EPI_ISL_3072757, EPI_ISL_3072758, EPI_ISL_3072759, EPI_ISL_3072760, EPI_ISL_3072761, EPI_ISL_3072762, EPI_ISL_3072763, EPI_ISL_3072764, EPI_ISL_3072765, EPI_ISL_3072766, EPI_ISL_3072767, EPI_ISL_3072768, EPI_ISL_3072769, EPI_ISL_3072770, EPI_ISL_3072771, EPI_ISL_3072772, EPI_ISL_3072773, EPI_ISL_3072774, EPI_ISL_3072775, EPI_ISL_3072776, EPI_ISL_3072777, EPI_ISL_3072778, EPI_ISL_3072779, EPI_ISL_3072780, EPI_ISL_3072781, EPI_ISL_3072782 | | | |
| see above | Department of Bacteria, Parasites and Fungi, Statens Serum Institut, Copenhagen, Denmark | Statens Serum Institut Bioinformatics and Microbial Genomics | Danish Covid-19 Genome Consortium |
| EPI_ISL_3072783 | Department of Clinical Microbiology, Odense University Hospital, Odense, Denmark | Statens Serum Institut Bioinformatics and Microbial Genomics | Danish Covid-19 Genome Consortium |
| EPI_ISL_3072784 | Department of Bacteria, Parasites and Fungi, Statens Serum Institut, Copenhagen, Denmark | Statens Serum Institut Bioinformatics and Microbial Genomics | Danish Covid-19 Genome Consortium |
| EPI_ISL_3072785 | Molekylær Medicinsk Afdeling, Aarhus University Hospital, Aarhus, Denmark | Statens Serum Institut Bioinformatics and Microbial Genomics | Danish Covid-19 Genome Consortium |
| EPI_ISL_3072786, EPI_ISL_3072787, EPI_ISL_3072788, EPI_ISL_3072789, EPI_ISL_3072790, EPI_ISL_3072791, EPI_ISL_3072792, EPI_ISL_3072793, EPI_ISL_3072794, EPI_ISL_3072795, EPI_ISL_3072796, EPI_ISL_3072797, EPI_ISL_3072798 | | | |
| see above | Department of Bacteria, Parasites and Fungi, Statens Serum Institut, Copenhagen, Denmark | Statens Serum Institut Bioinformatics and Microbial Genomics | Danish Covid-19 Genome Consortium |
| EPI_ISL_3072799 | Molekylær Medicinsk Afdeling, Aarhus University Hospital, Aarhus, Denmark | Statens Serum Institut Bioinformatics and Microbial Genomics | Danish Covid-19 Genome Consortium |
| EPI_ISL_3072800, EPI_ISL_3072801, EPI_ISL_3072802, EPI_ISL_3072803, EPI_ISL_3072804, EPI_ISL_3072805, EPI_ISL_3072806, EPI_ISL_3072807, EPI_ISL_3072808, EPI_ISL_3072809 | Department of Bacteria, Parasites and Fungi, Statens Serum Institut, Copenhagen, Denmark | Statens Serum Institut Bioinformatics and Microbial Genomics | Danish Covid-19 Genome Consortium |
| EPI_ISL_3072810 | Molekylær Medicinsk Afdeling, Aarhus University Hospital, Aarhus, Denmark | Statens Serum Institut Bioinformatics and Microbial Genomics | Danish Covid-19 Genome Consortium |
| EPI_ISL_3072811, EPI_ISL_3072812, EPI_ISL_3072813 | Department of Bacteria, Parasites and Fungi, Statens Serum Institut, Copenhagen, Denmark | Statens Serum Institut Bioinformatics and Microbial Genomics | Danish Covid-19 Genome Consortium |
| EPI_ISL_3072814 | Department of Clinical Microbiology, Odense University Hospital, Odense, Denmark | Statens Serum Institut Bioinformatics and Microbial Genomics | Danish Covid-19 Genome Consortium |
| EPI_ISL_3072815, EPI_ISL_3072816, EPI_ISL_3072817 | Department of Bacteria, Parasites and Fungi, Statens Serum Institut, Copenhagen, Denmark | Statens Serum Institut Bioinformatics and Microbial Genomics | Danish Covid-19 Genome Consortium |
| EPI_ISL_3072818 | Department of Clinical Microbiology and Center for Genomic Medicine, Rigshospitalet, Copenhagen, Denmark | Statens Serum Institut Bioinformatics and Microbial Genomics | Danish Covid-19 Genome Consortium |
| EPI_ISL_3072819, EPI_ISL_3072820, EPI_ISL_3072821, EPI_ISL_3072822, EPI_ISL_3072823, EPI_ISL_3072824, EPI_ISL_3072825 | Department of Bacteria, Parasites and Fungi, Statens Serum Institut, Copenhagen, Denmark | Statens Serum Institut Bioinformatics and Microbial Genomics | Danish Covid-19 Genome Consortium |
| EPI_ISL_3072826 | Department of Clinical Microbiology and Center for Genomic Medicine, Rigshospitalet, Copenhagen, Denmark | Statens Serum Institut Bioinformatics and Microbial Genomics | Danish Covid-19 Genome Consortium |
| EPI_ISL_3072827, EPI_ISL_3072828, EPI_ISL_3072829, EPI_ISL_3072830, EPI_ISL_3072831, EPI_ISL_3072832, EPI_ISL_3072833, EPI_ISL_3072834, EPI_ISL_3072835 | Department of Bacteria, Parasites and Fungi, Statens Serum Institut, Copenhagen, Denmark | Statens Serum Institut Bioinformatics and Microbial Genomics | Danish Covid-19 Genome Consortium |
| EPI_ISL_3072836 | Molekylær Medicinsk Afdeling, Aarhus University Hospital, Aarhus, Denmark | Statens Serum Institut Bioinformatics and Microbial Genomics | Danish Covid-19 Genome Consortium |
| EPI_ISL_3072837, EPI_ISL_3072838, EPI_ISL_3072839, EPI_ISL_3072840, EPI_ISL_3072841, EPI_ISL_3072842, EPI_ISL_3072843, EPI_ISL_3072844, EPI_ISL_3072845, EPI_ISL_3072846, EPI_ISL_3072847, EPI_ISL_3072848, EPI_ISL_3072849, EPI_ISL_3072850, EPI_ISL_3072851, EPI_ISL_3072852, EPI_ISL_3072853, EPI_ISL_3072854, EPI_ISL_3072855, EPI_ISL_3072856, EPI_ISL_3072857, EPI_ISL_3072858, EPI_ISL_3072859, EPI_ISL_3072860, EPI_ISL_3072861, EPI_ISL_3072862, EPI_ISL_3072863, EPI_ISL_3072864, EPI_ISL_3072865, EPI_ISL_3072866, EPI_ISL_3072867, EPI_ISL_3072868, EPI_ISL_3072869, EPI_ISL_3072870, EPI_ISL_3072871, EPI_ISL_3072872, EPI_ISL_3072873, EPI_ISL_3072874, EPI_ISL_3072875, EPI_ISL_3072876, EPI_ISL_3072877, EPI_ISL_3072878, EPI_ISL_3072879, EPI_ISL_3072880, EPI_ISL_3072881 | | | |
| see above | Department of Bacteria, Parasites and Fungi, Statens Serum Institut, Copenhagen, Denmark | Statens Serum Institut Bioinformatics and Microbial Genomics | Danish Covid-19 Genome Consortium |
| EPI_ISL_3072882 | Molekylær Medicinsk Afdeling, Aarhus University Hospital, Aarhus, Denmark | Statens Serum Institut Bioinformatics and Microbial Genomics | Danish Covid-19 Genome Consortium |
| EPI_ISL_3072883, EPI_ISL_3072885, EPI_ISL_3072886, EPI_ISL_3072887, EPI_ISL_3072888, EPI_ISL_3072889, | Department of Bacteria, Parasites and Fungi, Statens Serum Institut, Copenhagen, Denmark | Statens Serum Institut Bioinformatics and Microbial Genomics | Danish Covid-19 Genome Consortium |

| | | | |
|---|---|---|---|
| EPI_ISL_3072890, EPI_ISL_3072891, EPI_ISL_3072892, EPI_ISL_3072893 | | | |
| EPI_ISL_3072894 | Department of Clinical Microbiology and Center for Genomic Medicine, Rigshospitalet, Copenhagen, Denmark | Statens Serum Institut Bioinformatics and Microbial Genomics | Danish Covid-19 Genome Consortium |
| EPI_ISL_3072895, EPI_ISL_3072896, EPI_ISL_3072897, EPI_ISL_3072898, EPI_ISL_3072899, EPI_ISL_3072900, EPI_ISL_3072902, EPI_ISL_3072903, EPI_ISL_3072904, EPI_ISL_3072905, EPI_ISL_3072906, EPI_ISL_3072907, EPI_ISL_3072909, EPI_ISL_3072910, EPI_ISL_3072911, EPI_ISL_3072912, EPI_ISL_3072913, EPI_ISL_3072914, EPI_ISL_3072915, EPI_ISL_3072916, EPI_ISL_3072917, EPI_ISL_3072918, EPI_ISL_3072919, EPI_ISL_3072920, EPI_ISL_3072921, EPI_ISL_3072922 | | | |
| see above | Department of Bacteria, Parasites and Fungi, Statens Serum Institut, Copenhagen, Denmark | Statens Serum Institut Bioinformatics and Microbial Genomics | Danish Covid-19 Genome Consortium |
| EPI_ISL_3072923 | Department of Clinical Microbiology, Odense University Hospital, Odense, Denmark | Statens Serum Institut Bioinformatics and Microbial Genomics | Danish Covid-19 Genome Consortium |
| EPI_ISL_3072925, EPI_ISL_3072926, EPI_ISL_3072927, EPI_ISL_3072928, EPI_ISL_3072929 | Department of Bacteria, Parasites and Fungi, Statens Serum Institut, Copenhagen, Denmark | Statens Serum Institut Bioinformatics and Microbial Genomics | Danish Covid-19 Genome Consortium |
| EPI_ISL_3072930 | Molekylær Medicinsk Afdeling, Aarhus University Hospital, Aarhus, Denmark | Statens Serum Institut Bioinformatics and Microbial Genomics | Danish Covid-19 Genome Consortium |
| EPI_ISL_3072931, EPI_ISL_3072932, EPI_ISL_3072933, EPI_ISL_3072934, EPI_ISL_3072935, EPI_ISL_3072936, EPI_ISL_3072937, EPI_ISL_3072938, EPI_ISL_3072939, EPI_ISL_3072940, EPI_ISL_3072941, EPI_ISL_3072942, EPI_ISL_3072943, EPI_ISL_3072944, EPI_ISL_3072945, EPI_ISL_3072946, EPI_ISL_3072947, EPI_ISL_3072948, EPI_ISL_3072949, EPI_ISL_3072950, EPI_ISL_3072951, EPI_ISL_3072952, EPI_ISL_3072953, EPI_ISL_3072954, EPI_ISL_3072955, EPI_ISL_3072956, EPI_ISL_3072957, EPI_ISL_3072958, EPI_ISL_3072959, EPI_ISL_3072960, EPI_ISL_3072961, EPI_ISL_3072962, EPI_ISL_3072963, EPI_ISL_3072964, EPI_ISL_3072965, EPI_ISL_3072966, EPI_ISL_3072967, EPI_ISL_3072968, EPI_ISL_3072969, EPI_ISL_3072970, EPI_ISL_3072971, EPI_ISL_3072972, EPI_ISL_3072973, EPI_ISL_3072974, EPI_ISL_3072975 | | | |
| see above | Department of Bacteria, Parasites and Fungi, Statens Serum Institut, Copenhagen, Denmark | Statens Serum Institut Bioinformatics and Microbial Genomics | Danish Covid-19 Genome Consortium |
| EPI_ISL_3072976 | Department of Clinical Microbiology and Center for Genomic Medicine, Rigshospitalet, Copenhagen, Denmark | Statens Serum Institut Bioinformatics and Microbial Genomics | Danish Covid-19 Genome Consortium |
| EPI_ISL_3072977, EPI_ISL_3072978, EPI_ISL_3072979, EPI_ISL_3072980, EPI_ISL_3072982, EPI_ISL_3072983, EPI_ISL_3072984, EPI_ISL_3072985, EPI_ISL_3072986 | Department of Bacteria, Parasites and Fungi, Statens Serum Institut, Copenhagen, Denmark | Statens Serum Institut Bioinformatics and Microbial Genomics | Danish Covid-19 Genome Consortium |
| EPI_ISL_3072987 | Molekylær Medicinsk Afdeling, Aarhus University Hospital, Aarhus, Denmark | Statens Serum Institut Bioinformatics and Microbial Genomics | Danish Covid-19 Genome Consortium |
| EPI_ISL_3072988, EPI_ISL_3072989, EPI_ISL_3072991, EPI_ISL_3072992, EPI_ISL_3072993, EPI_ISL_3072994, EPI_ISL_3072995, EPI_ISL_3072996, EPI_ISL_3072998, EPI_ISL_3072999, EPI_ISL_3073000, EPI_ISL_3073001, EPI_ISL_3073002, EPI_ISL_3073003, EPI_ISL_3073004, EPI_ISL_3073005, EPI_ISL_3073006, EPI_ISL_3073007, EPI_ISL_3073008, EPI_ISL_3073009, EPI_ISL_3073010, EPI_ISL_3073011, EPI_ISL_3073012, EPI_ISL_3073013, EPI_ISL_3073014, EPI_ISL_3073015, EPI_ISL_3073016, EPI_ISL_3073017, EPI_ISL_3073018, EPI_ISL_3073020, EPI_ISL_3073021, EPI_ISL_3073022, EPI_ISL_3073023, EPI_ISL_3073024, EPI_ISL_3073025, EPI_ISL_3073026, EPI_ISL_3073027, EPI_ISL_3073028, EPI_ISL_3073029, EPI_ISL_3073030, EPI_ISL_3073031, EPI_ISL_3073032, EPI_ISL_3073033, EPI_ISL_3073034, EPI_ISL_3073035, EPI_ISL_3073036, EPI_ISL_3073037, EPI_ISL_3073038, EPI_ISL_3073039, EPI_ISL_3073040, EPI_ISL_3073041, EPI_ISL_3073042, EPI_ISL_3073043, EPI_ISL_3073044, EPI_ISL_3073045, EPI_ISL_3073046, EPI_ISL_3073047, EPI_ISL_3073048, EPI_ISL_3073049, EPI_ISL_3073050, EPI_ISL_3073051, EPI_ISL_3073052, EPI_ISL_3073053, EPI_ISL_3073054, EPI_ISL_3073055, EPI_ISL_3073056, EPI_ISL_3073057, EPI_ISL_3073058, EPI_ISL_3073059, EPI_ISL_3073060, EPI_ISL_3073061, EPI_ISL_3073062, EPI_ISL_3073063, EPI_ISL_3073064, EPI_ISL_3073065, EPI_ISL_3073066, EPI_ISL_3073067, EPI_ISL_3073068 | | | |
| see above | Department of Bacteria, Parasites and Fungi, Statens Serum Institut, Copenhagen, Denmark | Statens Serum Institut Bioinformatics and Microbial Genomics | Danish Covid-19 Genome Consortium |
| EPI_ISL_3073069 | Molekylær Medicinsk Afdeling, Aarhus University Hospital, Aarhus, Denmark | Statens Serum Institut Bioinformatics and Microbial Genomics | Danish Covid-19 Genome Consortium |
| EPI_ISL_3073070, EPI_ISL_3073071, EPI_ISL_3073072, EPI_ISL_3073073, EPI_ISL_3073074, EPI_ISL_3073075, EPI_ISL_3073076, EPI_ISL_3073077, EPI_ISL_3073079, EPI_ISL_3073080, EPI_ISL_3073081, EPI_ISL_3073082, EPI_ISL_3073083, EPI_ISL_3073084, EPI_ISL_3073085, EPI_ISL_3073086, EPI_ISL_3073087, EPI_ISL_3073088, EPI_ISL_3073089, EPI_ISL_3073090, EPI_ISL_3073091, EPI_ISL_3073092, EPI_ISL_3073093, EPI_ISL_3073094, EPI_ISL_3073095, EPI_ISL_3073096, EPI_ISL_3073097, EPI_ISL_3073098, EPI_ISL_3073099, EPI_ISL_3073100, EPI_ISL_3073101, EPI_ISL_3073102, EPI_ISL_3073103, EPI_ISL_3073104, EPI_ISL_3073105, EPI_ISL_3073106, EPI_ISL_3073107, EPI_ISL_3073108, EPI_ISL_3073110, EPI_ISL_3073111, EPI_ISL_3073112, EPI_ISL_3073113, EPI_ISL_3073114, EPI_ISL_3073115, EPI_ISL_3073116, EPI_ISL_3073117, EPI_ISL_3073118, EPI_ISL_3073119, EPI_ISL_3073120, EPI_ISL_3073121, EPI_ISL_3073122, EPI_ISL_3073123, EPI_ISL_3073124, EPI_ISL_3073125, EPI_ISL_3073126, EPI_ISL_3073127, EPI_ISL_3073128, EPI_ISL_3073129, EPI_ISL_3073130, EPI_ISL_3073131, EPI_ISL_3073132 | | | |
| see above | Department of Bacteria, Parasites and Fungi, Statens Serum Institut, Copenhagen, Denmark | Statens Serum Institut Bioinformatics and Microbial Genomics | Danish Covid-19 Genome Consortium |
| EPI_ISL_3073133 | Molekylær Medicinsk Afdeling, Aarhus University Hospital, Aarhus, Denmark | Statens Serum Institut Bioinformatics and Microbial Genomics | Danish Covid-19 Genome Consortium |
| EPI_ISL_3073134, EPI_ISL_3073135, EPI_ISL_3073136, EPI_ISL_3073137, EPI_ISL_3073138, EPI_ISL_3073139, EPI_ISL_3073140, EPI_ISL_3073141, EPI_ISL_3073142, EPI_ISL_3073143, EPI_ISL_3073144, EPI_ISL_3073145, EPI_ISL_3073146, EPI_ISL_3073147, EPI_ISL_3073148, EPI_ISL_3073149, EPI_ISL_3073150, EPI_ISL_3073151, EPI_ISL_3073152, EPI_ISL_3073153, EPI_ISL_3073155, EPI_ISL_3073156, EPI_ISL_3073157, EPI_ISL_3073158, EPI_ISL_3073159, EPI_ISL_3073160, EPI_ISL_3073161, EPI_ISL_3073162, EPI_ISL_3073163, EPI_ISL_3073164, EPI_ISL_3073165, EPI_ISL_3073166, EPI_ISL_3073167, EPI_ISL_3073168, EPI_ISL_3073169, EPI_ISL_3073170, EPI_ISL_3073171, EPI_ISL_3073172, EPI_ISL_3073173, EPI_ISL_3073174, EPI_ISL_3073175, EPI_ISL_3073176, EPI_ISL_3073177, EPI_ISL_3073178, EPI_ISL_3073179, EPI_ISL_3073180, EPI_ISL_3073181, EPI_ISL_3073182, EPI_ISL_3073183, EPI_ISL_3073184, EPI_ISL_3073185, EPI_ISL_3073186, EPI_ISL_3073187, EPI_ISL_3073188, EPI_ISL_3073190, EPI_ISL_3073191, EPI_ISL_3073192, EPI_ISL_3073193, EPI_ISL_3073194, EPI_ISL_3073195, EPI_ISL_3073196, EPI_ISL_3073197, EPI_ISL_3073198, EPI_ISL_3073199, EPI_ISL_3073200, EPI_ISL_3073201, EPI_ISL_3073202, EPI_ISL_3073203, EPI_ISL_3073204, EPI_ISL_3073205, EPI_ISL_3073206, EPI_ISL_3073207, EPI_ISL_3073208, EPI_ISL_3073209, EPI_ISL_3073210, EPI_ISL_3073212, EPI_ISL_3073213, EPI_ISL_3073214, EPI_ISL_3073215, EPI_ISL_3073216, EPI_ISL_3073217, EPI_ISL_3073218, EPI_ISL_3073219, EPI_ISL_3073220, EPI_ISL_3073221, EPI_ISL_3073222, EPI_ISL_3073223, EPI_ISL_3073224 | | | |
| see above | Department of Bacteria, Parasites and Fungi, Statens Serum Institut, Copenhagen, Denmark | Statens Serum Institut Bioinformatics and Microbial Genomics | Danish Covid-19 Genome Consortium |
| EPI_ISL_3073225 | Department of Clinical Microbiology, Odense University Hospital, Odense, Denmark | Statens Serum Institut Bioinformatics and Microbial Genomics | Danish Covid-19 Genome Consortium |
| EPI_ISL_3073226, EPI_ISL_3073227, EPI_ISL_3073228, EPI_ISL_3073229, EPI_ISL_3073230, EPI_ISL_3073231, EPI_ISL_3073232, EPI_ISL_3073233, EPI_ISL_3073234, EPI_ISL_3073235, EPI_ISL_3073236, EPI_ISL_3073237, EPI_ISL_3073238, EPI_ISL_3073239, EPI_ISL_3073240, EPI_ISL_3073241, EPI_ISL_3073242, EPI_ISL_3073243, EPI_ISL_3073244, EPI_ISL_3073245, EPI_ISL_3073246, EPI_ISL_3073247, EPI_ISL_3073248, EPI_ISL_3073249, EPI_ISL_3073250, EPI_ISL_3073251, EPI_ISL_3073252, EPI_ISL_3073253, EPI_ISL_3073254, EPI_ISL_3073255, EPI_ISL_3073256, EPI_ISL_3073257, EPI_ISL_3073258, EPI_ISL_3073259, EPI_ISL_3073260, EPI_ISL_3073261, EPI_ISL_3073262, EPI_ISL_3073263, EPI_ISL_3073264, EPI_ISL_3073265, EPI_ISL_3073266 | | | |
| see above | Department of Bacteria, Parasites and Fungi, Statens Serum Institut, Copenhagen, Denmark | Statens Serum Institut Bioinformatics and Microbial Genomics | Danish Covid-19 Genome Consortium |
| EPI_ISL_3073267 | Department of Clinical Microbiology and Center for Genomic Medicine, Rigshospitalet, Copenhagen, Denmark | Statens Serum Institut Bioinformatics and Microbial Genomics | Danish Covid-19 Genome Consortium |
| EPI_ISL_3073268, EPI_ISL_3073269, EPI_ISL_3073270, EPI_ISL_3073271, EPI_ISL_3073272, EPI_ISL_3073273, EPI_ISL_3073274, EPI_ISL_3073275, EPI_ISL_3073276, EPI_ISL_3073277, EPI_ISL_3073278, EPI_ISL_3073279, EPI_ISL_3073280, EPI_ISL_3073281, EPI_ISL_3073282, EPI_ISL_3073283, EPI_ISL_3073284, EPI_ISL_3073285, EPI_ISL_3073286, EPI_ISL_3073287, EPI_ISL_3073288, EPI_ISL_3073289, EPI_ISL_3073290, EPI_ISL_3073291, EPI_ISL_3073292, EPI_ISL_3073293, EPI_ISL_3073294, EPI_ISL_3073295, EPI_ISL_3073296, EPI_ISL_3073297, EPI_ISL_3073298, EPI_ISL_3073299, EPI_ISL_3073300, EPI_ISL_3073301, EPI_ISL_3073302, EPI_ISL_3073303, EPI_ISL_3073304, EPI_ISL_3073305, EPI_ISL_3073306, EPI_ISL_3073307, EPI_ISL_3073308, EPI_ISL_3073310, EPI_ISL_3073311, EPI_ISL_3073312, EPI_ISL_3073313, EPI_ISL_3073314, EPI_ISL_3073315, EPI_ISL_3073316, EPI_ISL_3073317, EPI_ISL_3073318, EPI_ISL_3073319, EPI_ISL_3073320, EPI_ISL_3073321, EPI_ISL_3073323, EPI_ISL_3073324, EPI_ISL_3073325 | | | |
| see above | Department of Bacteria, Parasites and Fungi, Statens Serum Institut, Copenhagen, Denmark | Statens Serum Institut Bioinformatics and Microbial Genomics | Danish Covid-19 Genome Consortium |
| EPI_ISL_3073326 | Department of Clinical Microbiology and Center for Genomic Medicine, Rigshospitalet, Copenhagen, Denmark | Statens Serum Institut Bioinformatics and Microbial Genomics | Danish Covid-19 Genome Consortium |
| EPI_ISL_3073327, EPI_ISL_3073328, EPI_ISL_3073329, EPI_ISL_3073330, EPI_ISL_3073331, EPI_ISL_3073332, EPI_ISL_3073333, EPI_ISL_3073334 | Department of Bacteria, Parasites and Fungi, Statens Serum Institut, Copenhagen, Denmark | Statens Serum Institut Bioinformatics and Microbial Genomics | Danish Covid-19 Genome Consortium |
| EPI_ISL_3073335 | Department of Clinical Microbiology and Center for Genomic Medicine, Rigshospitalet, Copenhagen, Denmark | Statens Serum Institut Bioinformatics and Microbial Genomics | Danish Covid-19 Genome Consortium |
| EPI_ISL_3073336, EPI_ISL_3073337, EPI_ISL_3073338, EPI_ISL_3073339, EPI_ISL_3073340, EPI_ISL_3073341, EPI_ISL_3073342, EPI_ISL_3073343, EPI_ISL_3073344, EPI_ISL_3073345, EPI_ISL_3073346, EPI_ISL_3073347, EPI_ISL_3073348, EPI_ISL_3073349, EPI_ISL_3073350, EPI_ISL_3073351, EPI_ISL_3073352, EPI_ISL_3073353, EPI_ISL_3073354, EPI_ISL_3073355, EPI_ISL_3073356, EPI_ISL_3073357, EPI_ISL_3073358, EPI_ISL_3073359, EPI_ISL_3073360, EPI_ISL_3073361, EPI_ISL_3073362, EPI_ISL_3073363, EPI_ISL_3073364, EPI_ISL_3073365, EPI_ISL_3073366, EPI_ISL_3073367, EPI_ISL_3073368, EPI_ISL_3073369, EPI_ISL_3073370, EPI_ISL_3073371, EPI_ISL_3073372, EPI_ISL_3073373, EPI_ISL_3073374, EPI_ISL_3073375, EPI_ISL_3073376, EPI_ISL_3073377, EPI_ISL_3073378, EPI_ISL_3073379, EPI_ISL_3073380, EPI_ISL_3073381, EPI_ISL_3073382, EPI_ISL_3073383, EPI_ISL_3073384, EPI_ISL_3073385, EPI_ISL_3073386, EPI_ISL_3073387, EPI_ISL_3073388, EPI_ISL_3073389, EPI_ISL_3073390, EPI_ISL_3073391, EPI_ISL_3073392, EPI_ISL_3073393, EPI_ISL_3073394, EPI_ISL_3073395, EPI_ISL_3073396, EPI_ISL_3073397, EPI_ISL_3073398, EPI_ISL_3073399, EPI_ISL_3073400, EPI_ISL_3073401, EPI_ISL_3073402, EPI_ISL_3073403, EPI_ISL_3073404, EPI_ISL_3073405, EPI_ISL_3073406, EPI_ISL_3073407, EPI_ISL_3073408, EPI_ISL_3073409, EPI_ISL_3073410, EPI_ISL_3073411, EPI_ISL_3073412, EPI_ISL_3073413, EPI_ISL_3073415, EPI_ISL_3073416, EPI_ISL_3073417 | | | |
| see above | Department of Bacteria, Parasites and Fungi, Statens Serum Institut, Copenhagen, Denmark | Statens Serum Institut Bioinformatics and Microbial Genomics | Danish Covid-19 Genome Consortium |
| EPI_ISL_3073418 | Department of Clinical Microbiology and Center for Genomic Medicine, Rigshospitalet, Copenhagen, Denmark | Statens Serum Institut Bioinformatics and Microbial Genomics | Danish Covid-19 Genome Consortium |
| EPI_ISL_3073419, EPI_ISL_3073420, EPI_ISL_3073421, EPI_ISL_3073422, EPI_ISL_3073423, EPI_ISL_3073424, EPI_ISL_3073425, EPI_ISL_3073426, EPI_ISL_3073427, EPI_ISL_3073428, EPI_ISL_3073429, EPI_ISL_3073430, EPI_ISL_3073431, EPI_ISL_3073432, EPI_ISL_3073433, EPI_ISL_3073434, EPI_ISL_3073435, EPI_ISL_3073436, EPI_ISL_3073437, EPI_ISL_3073438, EPI_ISL_3073439, EPI_ISL_3073440, EPI_ISL_3073441, EPI_ISL_3073442, EPI_ISL_3073443, EPI_ISL_3073444, EPI_ISL_3073445, EPI_ISL_3073446, EPI_ISL_3073447, EPI_ISL_3073448, EPI_ISL_3073449, EPI_ISL_3073450, EPI_ISL_3073451, EPI_ISL_3073452, EPI_ISL_3073453, EPI_ISL_3073454, EPI_ISL_3073455, EPI_ISL_3073456, EPI_ISL_3073457, EPI_ISL_3073458, EPI_ISL_3073459, EPI_ISL_3073460, EPI_ISL_3073461, EPI_ISL_3073462, EPI_ISL_3073463, EPI_ISL_3073464, EPI_ISL_3073465, EPI_ISL_3073466, EPI_ISL_3073467, EPI_ISL_3073468, EPI_ISL_3073470, EPI_ISL_3073471, EPI_ISL_3073472, EPI_ISL_3073473, EPI_ISL_3073474, EPI_ISL_3073475, EPI_ISL_3073476, EPI_ISL_3073477, EPI_ISL_3073478, EPI_ISL_3073479, EPI_ISL_3073480, EPI_ISL_3073481, EPI_ISL_3073482, EPI_ISL_3073483, EPI_ISL_3073484, EPI_ISL_3073485, EPI_ISL_3073486, EPI_ISL_3073487, EPI_ISL_3073488, EPI_ISL_3073489, EPI_ISL_3073490, EPI_ISL_3073491, EPI_ISL_3073492, EPI_ISL_3073493, EPI_ISL_3073494, EPI_ISL_3073495, EPI_ISL_3073496, EPI_ISL_3073497, EPI_ISL_3073498, EPI_ISL_3073499, EPI_ISL_3073500, EPI_ISL_3073501, EPI_ISL_3073502, EPI_ISL_3073503, EPI_ISL_3073504, EPI_ISL_3073505, EPI_ISL_3073506, EPI_ISL_3073507, EPI_ISL_3073508, EPI_ISL_3073509, EPI_ISL_3073510, EPI_ISL_3073511, EPI_ISL_3073512, EPI_ISL_3073513, EPI_ISL_3073514, EPI_ISL_3073515, EPI_ISL_3073516, EPI_ISL_3073517, EPI_ISL_3073518, EPI_ISL_3073519, EPI_ISL_3073520, EPI_ISL_3073521, EPI_ISL_3073522, EPI_ISL_3073523, EPI_ISL_3073524, EPI_ISL_3073525, EPI_ISL_3073526, EPI_ISL_3073527 | | | |

| | | | |
|---|---|---|---|
| see above | Department of Bacteria, Parasites and Fungi, Statens Serum Institut, Copenhagen, Denmark | Statens Serum Institut Bioinformatics and Microbial Genomics | Danish Covid-19 Genome Consortium |
| EPI_ISL_3073528 | Department of Clinical Microbiology, Odense University Hospital, Odense, Denmark | Statens Serum Institut Bioinformatics and Microbial Genomics | Danish Covid-19 Genome Consortium |
| EPI_ISL_3073529, EPI_ISL_3073530, EPI_ISL_3073531, EPI_ISL_3073532, EPI_ISL_3073533, EPI_ISL_3073534, EPI_ISL_3073535, EPI_ISL_3073536, EPI_ISL_3073537, EPI_ISL_3073538, EPI_ISL_3073539, EPI_ISL_3073540, EPI_ISL_3073541, EPI_ISL_3073542, EPI_ISL_3073543, EPI_ISL_3073544, EPI_ISL_3073545, EPI_ISL_3073546, EPI_ISL_3073547, EPI_ISL_3073548, EPI_ISL_3073549, EPI_ISL_3073551, EPI_ISL_3073552, EPI_ISL_3073553, EPI_ISL_3073554, EPI_ISL_3073555, EPI_ISL_3073556, EPI_ISL_3073557, EPI_ISL_3073558, EPI_ISL_3073559, EPI_ISL_3073560, EPI_ISL_3073561, EPI_ISL_3073562, EPI_ISL_3073563, EPI_ISL_3073564, EPI_ISL_3073565, EPI_ISL_3073566, EPI_ISL_3073567, EPI_ISL_3073568, EPI_ISL_3073569, EPI_ISL_3073570, EPI_ISL_3073571, EPI_ISL_3073572, EPI_ISL_3073573, EPI_ISL_3073574, EPI_ISL_3073575, EPI_ISL_3073576, EPI_ISL_3073577, EPI_ISL_3073578, EPI_ISL_3073579, EPI_ISL_3073580, EPI_ISL_3073581, EPI_ISL_3073582, EPI_ISL_3073583, EPI_ISL_3073584, EPI_ISL_3073585, EPI_ISL_3073586, EPI_ISL_3073587, EPI_ISL_3073588, EPI_ISL_3073589, EPI_ISL_3073590, EPI_ISL_3073591, EPI_ISL_3073592, EPI_ISL_3073593, EPI_ISL_3073594, EPI_ISL_3073595, EPI_ISL_3073597, EPI_ISL_3073598, EPI_ISL_3073599, EPI_ISL_3073600, EPI_ISL_3073601, EPI_ISL_3073602, EPI_ISL_3073603, EPI_ISL_3073604, EPI_ISL_3073605, EPI_ISL_3073606, EPI_ISL_3073607, EPI_ISL_3073608, EPI_ISL_3073609, EPI_ISL_3073610, EPI_ISL_3073611, EPI_ISL_3073612, EPI_ISL_3073613, EPI_ISL_3073614, EPI_ISL_3073615, EPI_ISL_3073616, EPI_ISL_3073617, EPI_ISL_3073618, EPI_ISL_3073619, EPI_ISL_3073620, EPI_ISL_3073621, EPI_ISL_3073622, EPI_ISL_3073623, EPI_ISL_3073624, EPI_ISL_3073625, EPI_ISL_3073626, EPI_ISL_3073627, EPI_ISL_3073628, EPI_ISL_3073629, EPI_ISL_3073630, EPI_ISL_3073631, EPI_ISL_3073632, EPI_ISL_3073633, EPI_ISL_3073634, EPI_ISL_3073635, EPI_ISL_3073636, EPI_ISL_3073637, EPI_ISL_3073638, EPI_ISL_3073639, EPI_ISL_3073640, EPI_ISL_3073641, EPI_ISL_3073642, EPI_ISL_3073643, EPI_ISL_3073644, EPI_ISL_3073645, EPI_ISL_3073646, EPI_ISL_3073647, EPI_ISL_3073648, EPI_ISL_3073649, EPI_ISL_3073650, EPI_ISL_3073651, EPI_ISL_3073652 | | | |
| see above | Department of Bacteria, Parasites and Fungi, Statens Serum Institut, Copenhagen, Denmark | Statens Serum Institut Bioinformatics and Microbial Genomics | Danish Covid-19 Genome Consortium |
| EPI_ISL_3073653 | Molekylær Medicinsk Afdeling, Aarhus University Hospital, Aarhus, Denmark | Statens Serum Institut Bioinformatics and Microbial Genomics | Danish Covid-19 Genome Consortium |
| EPI_ISL_3073654, EPI_ISL_3073655, EPI_ISL_3073657, EPI_ISL_3073658, EPI_ISL_3073659, EPI_ISL_3073660, EPI_ISL_3073661, EPI_ISL_3073662, EPI_ISL_3073663, EPI_ISL_3073664, EPI_ISL_3073665, EPI_ISL_3073666, EPI_ISL_3073667, EPI_ISL_3073668, EPI_ISL_3073669, EPI_ISL_3073670, EPI_ISL_3073671, EPI_ISL_3073672, EPI_ISL_3073673, EPI_ISL_3073674, EPI_ISL_3073675, EPI_ISL_3073676, EPI_ISL_3073677, EPI_ISL_3073678, EPI_ISL_3073679, EPI_ISL_3073680, EPI_ISL_3073681, EPI_ISL_3073682, EPI_ISL_3073683, EPI_ISL_3073684, EPI_ISL_3073685, EPI_ISL_3073686, EPI_ISL_3073687, EPI_ISL_3073688, EPI_ISL_3073689, EPI_ISL_3073690, EPI_ISL_3073691, EPI_ISL_3073692, EPI_ISL_3073693, EPI_ISL_3073694, EPI_ISL_3073695, EPI_ISL_3073696, EPI_ISL_3073697, EPI_ISL_3073698, EPI_ISL_3073699, EPI_ISL_3073700, EPI_ISL_3073701, EPI_ISL_3073702 | | | |
| see above | Department of Bacteria, Parasites and Fungi, Statens Serum Institut, Copenhagen, Denmark | Statens Serum Institut Bioinformatics and Microbial Genomics | Danish Covid-19 Genome Consortium |
| EPI_ISL_3073703 | Molekylær Medicinsk Afdeling, Aarhus University Hospital, Aarhus, Denmark | Statens Serum Institut Bioinformatics and Microbial Genomics | Danish Covid-19 Genome Consortium |
| EPI_ISL_3073704, EPI_ISL_3073705, EPI_ISL_3073706, EPI_ISL_3073707, EPI_ISL_3073708, EPI_ISL_3073709, EPI_ISL_3073710, EPI_ISL_3073711, EPI_ISL_3073712, EPI_ISL_3073713, EPI_ISL_3073714, EPI_ISL_3073715, EPI_ISL_3073716, EPI_ISL_3073717, EPI_ISL_3073718, EPI_ISL_3073719, EPI_ISL_3073720, EPI_ISL_3073721, EPI_ISL_3073722, EPI_ISL_3073723, EPI_ISL_3073724, EPI_ISL_3073725, EPI_ISL_3073726, EPI_ISL_3073727, EPI_ISL_3073728, EPI_ISL_3073729, EPI_ISL_3073730, EPI_ISL_3073731, EPI_ISL_3073732, EPI_ISL_3073733, EPI_ISL_3073734, EPI_ISL_3073735, EPI_ISL_3073736, EPI_ISL_3073737, EPI_ISL_3073738, EPI_ISL_3073739, EPI_ISL_3073740, EPI_ISL_3073741, EPI_ISL_3073742, EPI_ISL_3073743, EPI_ISL_3073744, EPI_ISL_3073745, EPI_ISL_3073746, EPI_ISL_3073747, EPI_ISL_3073748, EPI_ISL_3073749, EPI_ISL_3073750, EPI_ISL_3073751, EPI_ISL_3073752, EPI_ISL_3073753, EPI_ISL_3073754, EPI_ISL_3073755, EPI_ISL_3073756, EPI_ISL_3073757, EPI_ISL_3073760, EPI_ISL_3073761, EPI_ISL_3073762, EPI_ISL_3073763, EPI_ISL_3073764, EPI_ISL_3073765, EPI_ISL_3073766, EPI_ISL_3073767, EPI_ISL_3073768, EPI_ISL_3073769, EPI_ISL_3073770, EPI_ISL_3073771, EPI_ISL_3073773, EPI_ISL_3073774, EPI_ISL_3073775, EPI_ISL_3073776, EPI_ISL_3073777, EPI_ISL_3073778, EPI_ISL_3073779, EPI_ISL_3073780, EPI_ISL_3073781, EPI_ISL_3073782, EPI_ISL_3073783, EPI_ISL_3073784, EPI_ISL_3073785, EPI_ISL_3073786, EPI_ISL_3073787, EPI_ISL_3073788, EPI_ISL_3073789, EPI_ISL_3073790, EPI_ISL_3073791 | | | |
| see above | Department of Bacteria, Parasites and Fungi, Statens Serum Institut, Copenhagen, Denmark | Statens Serum Institut Bioinformatics and Microbial Genomics | Danish Covid-19 Genome Consortium |
| EPI_ISL_3073792 | Molekylær Medicinsk Afdeling, Aarhus University Hospital, Aarhus, Denmark | Statens Serum Institut Bioinformatics and Microbial Genomics | Danish Covid-19 Genome Consortium |
| EPI_ISL_3073793, EPI_ISL_3073794, EPI_ISL_3073795, EPI_ISL_3073797, EPI_ISL_3073798, EPI_ISL_3073799, EPI_ISL_3073800 | Department of Bacteria, Parasites and Fungi, Statens Serum Institut, Copenhagen, Denmark | Statens Serum Institut Bioinformatics and Microbial Genomics | Danish Covid-19 Genome Consortium |
| EPI_ISL_3073801 | Molekylær Medicinsk Afdeling, Aarhus University Hospital, Aarhus, Denmark | Statens Serum Institut Bioinformatics and Microbial Genomics | Danish Covid-19 Genome Consortium |
| EPI_ISL_3073802, EPI_ISL_3073803, EPI_ISL_3073804, EPI_ISL_3073805, EPI_ISL_3073806, EPI_ISL_3073807, EPI_ISL_3073808, EPI_ISL_3073809, EPI_ISL_3073810, EPI_ISL_3073811, EPI_ISL_3073812, EPI_ISL_3073813, EPI_ISL_3073814, EPI_ISL_3073815, EPI_ISL_3073816, EPI_ISL_3073817, EPI_ISL_3073818, EPI_ISL_3073819, EPI_ISL_3073820, EPI_ISL_3073821, EPI_ISL_3073822, EPI_ISL_3073823, EPI_ISL_3073824, EPI_ISL_3073825, EPI_ISL_3073826, EPI_ISL_3073827, EPI_ISL_3073828, EPI_ISL_3073829, EPI_ISL_3073830, EPI_ISL_3073831, EPI_ISL_3073832, EPI_ISL_3073833, EPI_ISL_3073834, EPI_ISL_3073835, EPI_ISL_3073836, EPI_ISL_3073837, EPI_ISL_3073838, EPI_ISL_3073839, EPI_ISL_3073840, EPI_ISL_3073841, EPI_ISL_3073842, EPI_ISL_3073843, EPI_ISL_3073844, EPI_ISL_3073845, EPI_ISL_3073846, EPI_ISL_3073847, EPI_ISL_3073848, EPI_ISL_3073849, EPI_ISL_3073850, EPI_ISL_3073851, EPI_ISL_3073852, EPI_ISL_3073853, EPI_ISL_3073854, EPI_ISL_3073855, EPI_ISL_3073856, EPI_ISL_3073857 | | | |
| see above | Department of Bacteria, Parasites and Fungi, Statens Serum Institut, Copenhagen, Denmark | Statens Serum Institut Bioinformatics and Microbial Genomics | Danish Covid-19 Genome Consortium |
| EPI_ISL_3073861 | Department of Microbiology, Yamanashi Institute for Public Health and Environment | Genome Analysis Center, Yamanashi Central Hospital | Yosuke Hirotsu |
| EPI_ISL_3073863, EPI_ISL_3073869, EPI_ISL_3073871, EPI_ISL_3073872, EPI_ISL_3073873 | Genome Analysis Center, Yamanashi Central Hospital | Genome Analysis Center, Yamanashi Central Hospital | Yosuke Hirotsu |
| EPI_ISL_3073874, EPI_ISL_3073877, EPI_ISL_3073878, EPI_ISL_3073879 | Zuckerberg San Francisco General Hospital | DeRisi Lab, University of California, San Francisco | Jeffery Whitman, Sabrina Mann, Anthea Mitchell, Matthias Hapte-Selassie, Jamin Liu, Sara Sunshine, Matthew Laurie, Genay Pilarowski, Amy Kistler, Manu Vanaerschot, Patrick Ayscue, Lucy Li, Eric Chow, IDseq Team, James Peng, Carina Marquez, Luis Rubio, Gabriel Chamie, Diane Jones, Jon Jacobo, Susana Rojas, Susy Rojas, Valerie Tulier-Laiwa, Douglas Black, Jackie Martinez, Maya Petersen, Diane Havlir, and Joseph DeRisi |
| EPI_ISL_3073886 | Clínica Rotger Laboratorio de Análisis Clínicos Dpto. Biología Molecular | Clínica Rotger Laboratorio de Análisis Clínicos Dpto. Biología Molecular | Francisco Aliaga-Lozano, Chaymaá Gammani-Chairi, Laura Gómez, Juana Gonzalez-Calderón, Marina Barceló-Vives, Catalina Vidal-Gómez, Marina Kleber-Pol, Esperança Frau-Fuster, Carmen Morales-González, Esther Nieto-Molina |
| EPI_ISL_3073888, EPI_ISL_3073889, EPI_ISL_3073890, EPI_ISL_3073891, EPI_ISL_3073892, EPI_ISL_3073893, EPI_ISL_3073894, EPI_ISL_3073895, EPI_ISL_3073896, EPI_ISL_3073897, EPI_ISL_3073898, EPI_ISL_3073899, EPI_ISL_3073900, EPI_ISL_3073901, EPI_ISL_3073903, EPI_ISL_3073906, EPI_ISL_3073907, EPI_ISL_3073909, EPI_ISL_3073910, EPI_ISL_3073911 | | | |
| see above | Lancet Laboratory | National Institute for Communicable Diseases of the National Health Laboratory Service | Amoako DG, Everatt J, Scheepers C, Mohale T, Ntuli N, Mahlangu B, Mnguni A, Ismail A, Bhiman JN |
| EPI_ISL_3073912, EPI_ISL_3073913, EPI_ISL_3073914, EPI_ISL_3073915, EPI_ISL_3073916, EPI_ISL_3073917, EPI_ISL_3073918, EPI_ISL_3073919, EPI_ISL_3073921, EPI_ISL_3073922, EPI_ISL_3073923, EPI_ISL_3073924, EPI_ISL_3073926, EPI_ISL_3073927, EPI_ISL_3073929, EPI_ISL_3073930, EPI_ISL_3073931, EPI_ISL_3073932, EPI_ISL_3073933, EPI_ISL_3073934, EPI_ISL_3073935, EPI_ISL_3073936, EPI_ISL_3073937, EPI_ISL_3073939, EPI_ISL_3073940, EPI_ISL_3073941, EPI_ISL_3073942, EPI_ISL_3073944, EPI_ISL_3073945, EPI_ISL_3073946, EPI_ISL_3073947, EPI_ISL_3073948, EPI_ISL_3073949, EPI_ISL_3073950, EPI_ISL_3073951, EPI_ISL_3073952, EPI_ISL_3073953, EPI_ISL_3073954, EPI_ISL_3073955, EPI_ISL_3073957, EPI_ISL_3073958, EPI_ISL_3073959, EPI_ISL_3073960, EPI_ISL_3073961, EPI_ISL_3073962, EPI_ISL_3073963, EPI_ISL_3073964, EPI_ISL_3073965, EPI_ISL_3073966 | | | |
| see above | MANKWENG LABORATORY | National Institute for Communicable Diseases of the National Health Laboratory Service | Amoako DG, Everatt J, Scheepers C, Mohale T, Ntuli N, Mahlangu B, Mnguni A, Ismail A, Bhiman JN |
| EPI_ISL_3073967, EPI_ISL_3073968, EPI_ISL_3073970, EPI_ISL_3073971, EPI_ISL_3073972, EPI_ISL_3073973, EPI_ISL_3073974, EPI_ISL_3073977, EPI_ISL_3073980, EPI_ISL_3073981, EPI_ISL_3073983, EPI_ISL_3073984, EPI_ISL_3073985, EPI_ISL_3073986, EPI_ISL_3073987 | | | |
| see above | TSHEPONG LABORATORY | National Institute for Communicable Diseases of the National Health Laboratory Service | Amoako DG, Everatt J, Scheepers C, Mohale T, Ntuli N, Mahlangu B, Mnguni A, Ismail A, Bhiman JN |
| EPI_ISL_3073990, EPI_ISL_3073991, EPI_ISL_3073992, EPI_ISL_3073993, EPI_ISL_3073994, EPI_ISL_3073995, EPI_ISL_3073996, EPI_ISL_3073997, EPI_ISL_3073998, EPI_ISL_3073999, EPI_ISL_3074000, EPI_ISL_3074001, EPI_ISL_3074002, EPI_ISL_3074003, EPI_ISL_3074004 | | | |
| see above | MANKWENG LABORATORY | National Institute for Communicable Diseases of the National Health Laboratory Service | Amoako DG, Everatt J, Scheepers C, Mohale T, Ntuli N, Mahlangu B, Mnguni A, Ismail A, Bhiman JN |
| EPI_ISL_3074005, EPI_ISL_3074006, EPI_ISL_3074007, EPI_ISL_3074008, EPI_ISL_3074009, EPI_ISL_3074010, EPI_ISL_3074011, EPI_ISL_3074012, EPI_ISL_3074013 | LEBOWAKGOMO LABORATORY | National Institute for Communicable Diseases of the National Health Laboratory Service | Amoako DG, Everatt J, Scheepers C, Mohale T, Ntuli N, Mahlangu B, Mnguni A, Ismail A, Bhiman JN |
| EPI_ISL_3074014 | LEBOWAKGOMO LABORATORY | National Institute for Communicable Diseases of the National Health Laboratory Service | Sisonke Team, Amoako DG, Everatt J, Scheepers C, Mohale T, Ntuli N, Mahlangu B, Mnguni A, Ismail A, Bhiman JN |
| EPI_ISL_3074015, EPI_ISL_3074016, EPI_ISL_3074017, EPI_ISL_3074018, EPI_ISL_3074019, EPI_ISL_3074022, EPI_ISL_3074023, EPI_ISL_3074024, EPI_ISL_3074025, EPI_ISL_3074026, EPI_ISL_3074027, EPI_ISL_3074028, EPI_ISL_3074029 | | | |
| see above | LEBOWAKGOMO LABORATORY | National Institute for Communicable Diseases of the National Health Laboratory Service | Amoako DG, Everatt J, Scheepers C, Mohale T, Ntuli N, Mahlangu B, Mnguni A, Ismail A, Bhiman JN |
| EPI_ISL_3074030, EPI_ISL_3074032, EPI_ISL_3074034, EPI_ISL_3074035, EPI_ISL_3074036, EPI_ISL_3074037, EPI_ISL_3074038, EPI_ISL_3074039, EPI_ISL_3074040, EPI_ISL_3074041 | TAMBO MEMORIAL LABORATORY | National Institute for Communicable Diseases of the National Health Laboratory Service | Amoako DG, Everatt J, Scheepers C, Mohale T, Ntuli N, Mahlangu B, Mnguni A, Ismail A, Bhiman JN |
| EPI_ISL_3074049, EPI_ISL_3074050, | Lancet Laboratory | National Institute for Communicable Diseases of the National Health Laboratory Service | Amoako DG, Everatt J, Scheepers C, Mohale T, Ntuli N, Mahlangu B, Mnguni A, Ismail A, Bhiman JN |

| | | | |
|---|---|---|---|
| EPI_ISL_3074055 | | | |
| EPI_ISL_3074107, EPI_ISL_3074108 | Department of Virology I, National Institute of Infectious Diseases | Department of Veterinary Science, National Institute of Infectious Diseases | Yudai Kuroda, Tsukasa Yamamoto, Keita Ishijima, Tsuguto Fujimoto, Souichi Yamada, Shuetsu Fukushi, Ken Maeda |
| EPI_ISL_3075899, EPI_ISL_3075903, EPI_ISL_3076006, EPI_ISL_3076055, EPI_ISL_3076102, EPI_ISL_3076202, EPI_ISL_3076327, EPI_ISL_3076490, EPI_ISL_3076500, EPI_ISL_3076568 | Department of Bacteria, Parasites and Fungi, Statens Serum Institut, Copenhagen, Denmark | Statens Serum Institut Bioinformatics and Microbial Genomics | Danish Covid-19 Genome Consortium |
| EPI_ISL_3076600 | SARS-CoV-2 testing team, National Institute of Infectious Diseases | Pathogen Genomics Center, National Institute of Infectious Diseases | Tsuyoshi Sekizuka, Kentaro Itokawa, Rina Tanaka, Masanori Hashino, Nozomu Hanaoka, Masumichi Saito, Naomi Nojiri, Hazuka Y Furihata, Hiromizu Takahashi, Tsuguto Fujimoto, Makoto Kuroda |
| EPI_ISL_3076603 | Department of Bacteria, Parasites and Fungi, Statens Serum Institut, Copenhagen, Denmark | Statens Serum Institut Bioinformatics and Microbial Genomics | Danish Covid-19 Genome Consortium |
| EPI_ISL_3076606, EPI_ISL_3076608, EPI_ISL_3076610, EPI_ISL_3076614, EPI_ISL_3076618, EPI_ISL_3076620, EPI_ISL_3076622, EPI_ISL_3076624, EPI_ISL_3076626, EPI_ISL_3076628, EPI_ISL_3076629, EPI_ISL_3076631, EPI_ISL_3076632, EPI_ISL_3076633, EPI_ISL_3076634, EPI_ISL_3076636, EPI_ISL_3076637, EPI_ISL_3076638, EPI_ISL_3076639, EPI_ISL_3076640, EPI_ISL_3076641, EPI_ISL_3076642, EPI_ISL_3076643, EPI_ISL_3076644, EPI_ISL_3076645, EPI_ISL_3076646, EPI_ISL_3076648, EPI_ISL_3076649, EPI_ISL_3076650, EPI_ISL_3076651, EPI_ISL_3076652, EPI_ISL_3076653, EPI_ISL_3076654, EPI_ISL_3076656, EPI_ISL_3076657, EPI_ISL_3076658, EPI_ISL_3076659, EPI_ISL_3076660, EPI_ISL_3076661, EPI_ISL_3076662, EPI_ISL_3076663, EPI_ISL_3076664, EPI_ISL_3076665, EPI_ISL_3076666, EPI_ISL_3076667, EPI_ISL_3076668, EPI_ISL_3076669, EPI_ISL_3076670, EPI_ISL_3076671, EPI_ISL_3076672, EPI_ISL_3076673, EPI_ISL_3076674, EPI_ISL_3076675, EPI_ISL_3076676, EPI_ISL_3076677, EPI_ISL_3076678, EPI_ISL_3076679, EPI_ISL_3076680, EPI_ISL_3076681, EPI_ISL_3076682, EPI_ISL_3076684, EPI_ISL_3076685, EPI_ISL_3076686, EPI_ISL_3076687, EPI_ISL_3076688, EPI_ISL_3076689, EPI_ISL_3076690, EPI_ISL_3076691, EPI_ISL_3076692, EPI_ISL_3076693, EPI_ISL_3076695, EPI_ISL_3076696, EPI_ISL_3076697, EPI_ISL_3076698, EPI_ISL_3076699, EPI_ISL_3076700, EPI_ISL_3076701, EPI_ISL_3076702, EPI_ISL_3076703, EPI_ISL_3076704, EPI_ISL_3076705, EPI_ISL_3076706, EPI_ISL_3076707, EPI_ISL_3076708, EPI_ISL_3076709, EPI_ISL_3076710, EPI_ISL_3076711, EPI_ISL_3076714, EPI_ISL_3076716, EPI_ISL_3076717, EPI_ISL_3076718, EPI_ISL_3076719, EPI_ISL_3076720, EPI_ISL_3076722, EPI_ISL_3076723 | | | |
| see above | SARS-CoV-2 testing team, National Institute of Infectious Diseases | Pathogen Genomics Center, National Institute of Infectious Diseases | Tsuyoshi Sekizuka, Kentaro Itokawa, Rina Tanaka, Masanori Hashino, Nozomu Hanaoka, Masumichi Saito, Naomi Nojiri, Hazuka Y Furihata, Hiromizu Takahashi, Tsuguto Fujimoto, Makoto Kuroda |
| EPI_ISL_3076724, EPI_ISL_3076725 | AP SSO | CSIR-Centre for Cellular and Molecular Biology-INSACOG | Tulasi Nagabandi,Lamuk Zaveri,Payel Mukherjee,Onkar Kulkarni,Jandhyala Sai Krishna, Sofia Banu, Ara Sreenivas,Shreekant Verma, Amareshwar Vodapalli ,B Himasri,Valli Nagalakshmi Undamatla, Priya Nurkurthy,Sumedha Avadhanula, Vidhyadhari Methuku, Archana Bharadwaj Siva, Karthik Bharadwaj Tallapaka,Rakesh K Mishra,Divya Tej Sowpati |
| EPI_ISL_3076727, EPI_ISL_3076728, EPI_ISL_3076729 | AP SSO | CSIR-Centre for Cellular and Molecular Biology-INSACOG | Priya Nurkurthy,Ara Sreenivas,Shreekant Verma, Amareshwar Vodapalli ,Lamuk Zaveri,Tulasi Nagabandi,B Himasri,Valli Nagalakshmi Undamatla, Sumedha Avadhanula, Vidhyadhari Methuku,Payel Mukherjee,Onkar Kulkarni, Jandhyala Sai Krishna,Sofia Banu, Archana Bharadwaj Siva, Karthik Bharadwaj Tallapaka,Rakesh K Mishra,Divya Tej Sowpati |
| EPI_ISL_3076731 | Hospital Universitari Arnau de Vilanova | Hospital Universitari Vall d'Hebron - Vall d'Hebron Institut de Recerca | Cristina Andrés, Maria Piñana, Alejandra González-Sánchez, Damir Garcia-Cehic, Ariadna Rando, Juliana Esperalba, Maria Gema Codina, Carla Castillo, Karen García, Rodrigo Vásquez, Maria Carmen Martin, Tomàs Pumarola, Josep Quer, Andrés Antón |
| EPI_ISL_3076736, EPI_ISL_3076737, EPI_ISL_3076739, EPI_ISL_3076740, EPI_ISL_3076741, EPI_ISL_3076742, EPI_ISL_3076743, EPI_ISL_3076745, EPI_ISL_3076746, EPI_ISL_3076747, EPI_ISL_3076750, EPI_ISL_3076752, EPI_ISL_3076753, EPI_ISL_3076754, EPI_ISL_3076756, EPI_ISL_3076757, EPI_ISL_3076758, EPI_ISL_3076759, EPI_ISL_3076761, EPI_ISL_3076762, EPI_ISL_3076763, EPI_ISL_3076764, EPI_ISL_3076765, EPI_ISL_3076766, EPI_ISL_3076767, EPI_ISL_3076768, EPI_ISL_3076770, EPI_ISL_3076775, EPI_ISL_3076777, EPI_ISL_3076778, EPI_ISL_3076779, EPI_ISL_3076780, EPI_ISL_3076782, EPI_ISL_3076783, EPI_ISL_3076784, EPI_ISL_3076785, EPI_ISL_3076786, EPI_ISL_3076787, EPI_ISL_3076789, EPI_ISL_3076790, EPI_ISL_3076793, EPI_ISL_3076796, EPI_ISL_3076799, EPI_ISL_3076801, EPI_ISL_3076802, EPI_ISL_3076804, EPI_ISL_3076805, EPI_ISL_3076809, EPI_ISL_3076813, EPI_ISL_3076814, EPI_ISL_3076816 | | | |
| see above | National Center of Infectious and Parasitic Diseases | National Center of Infectious and Parasitic Diseases | Alexiev et al |
| EPI_ISL_3076820, EPI_ISL_3076821, EPI_ISL_3076822, EPI_ISL_3076823, EPI_ISL_3076824, EPI_ISL_3076825, EPI_ISL_3076826, EPI_ISL_3076827, EPI_ISL_3076829, EPI_ISL_3076830, EPI_ISL_3076831, EPI_ISL_3076832, EPI_ISL_3076833, EPI_ISL_3076834, EPI_ISL_3076835, EPI_ISL_3076838, EPI_ISL_3076839, EPI_ISL_3076840, EPI_ISL_3076841, EPI_ISL_3076842, EPI_ISL_3076844, EPI_ISL_3076845, EPI_ISL_3076846, EPI_ISL_3076847, EPI_ISL_3076848, EPI_ISL_3076849, EPI_ISL_3076850, EPI_ISL_3076851, EPI_ISL_3076852, EPI_ISL_3076853, EPI_ISL_3076854, EPI_ISL_3076855, EPI_ISL_3076856, EPI_ISL_3076857, EPI_ISL_3076858, EPI_ISL_3076859, EPI_ISL_3076860, EPI_ISL_3076861, EPI_ISL_3076862, EPI_ISL_3076863, EPI_ISL_3076864, EPI_ISL_3076865, EPI_ISL_3076866, EPI_ISL_3076868, EPI_ISL_3076869, EPI_ISL_3076870, EPI_ISL_3076871, EPI_ISL_3076872, EPI_ISL_3076873, EPI_ISL_3076874, EPI_ISL_3076875, EPI_ISL_3076876, EPI_ISL_3076877, EPI_ISL_3076878, EPI_ISL_3076879, EPI_ISL_3076880, EPI_ISL_3076881, EPI_ISL_3076882, EPI_ISL_3076883, EPI_ISL_3076884, EPI_ISL_3076885, EPI_ISL_3076886, EPI_ISL_3076887, EPI_ISL_3076888, EPI_ISL_3076889, EPI_ISL_3076890, EPI_ISL_3076891, EPI_ISL_3076892, EPI_ISL_3076894, EPI_ISL_3076895, EPI_ISL_3076896, EPI_ISL_3076897 | | | |
| see above | Clinical Microbiology, Infection Prevention and Control | Section for Molecular Diagnostics | Björn Hallström, Jonas Björkman |
| EPI_ISL_3076899 | PZU “Eurofarm - Centar Poliklinika” | University of Sarajevo, Veterinary Faculty, Laboratory for Molecular Diagnostic and Research Laboratory | Goletic S, Goletic T, Prguda-Mujic J, Huseinbegovic E, Softic A, Sabic E., Jazic A., Nicevic M., Hodzic A., Terzic I. |
| EPI_ISL_3076900 | University Clinical Hospital of Mostar, Department of Microbiology and Molecular Diagnostics | University of Sarajevo, Veterinary Faculty, Laboratory for Molecular Diagnostic and Research Laboratory | Goletic S, Goletic T, Ostojic M, Softic A, Sabic E., Jazic A., Nicevic M., Hodzic A., Terzic I. |
| EPI_ISL_3076901 | University Clinical Hospital of Mostar, Department of Microbiology and Molecular Diagnostics | University of Sarajevo, Veterinary Faculty, Laboratory for Molecular Diagnostic and Research Laboratory | Goletic S, Goletic T, Ostojic M, Softic A, Terzic I., Sabic E., Hodzic A., Jazic A., Nicevic M. |
| EPI_ISL_3076902 | University Clinical Hospital of Mostar, Department of Microbiology and Molecular Diagnostics | University of Sarajevo, Veterinary Faculty, Laboratory for Molecular Diagnostic and Research Laboratory | Goletic S, Goletic T, Ostojic M, Softic A, Hodzic A., Jazic A., Terzic I., Nicevic M., Sabic E. |
| EPI_ISL_3076905 | Hospital del Oriente de Asturias | Laboratorio de Virología HUCA | Sandoval M, Castelló C, Gómez de Oña J, Boga JA, Rojo S, Alvarez-Arguelles ME, Abreu F, Costales I, Perez-Martínez Z, Martín-Rodríguez G, Coto E, Melón S |
| EPI_ISL_3076907 | Hospital de Oriente de Asturias | Laboratorio de Virología HUCA | Sandoval M, Castelló C, Gómez de Oña J, Boga JA, Rojo S, Alvarez-Arguelles ME, Abreu F, Costales I, Perez-Martínez Z, Martín-Rodríguez G, Coto E, Melón S |
| EPI_ISL_3076909, EPI_ISL_3076912 | Hospital del Oriente de Asturias | Laboratorio de Virología HUCA | Sandoval M, Castelló C, Gómez de Oña J, Boga JA, Rojo S, Alvarez-Arguelles ME, Abreu F, Costales I, Perez-Martínez Z, Martín-Rodríguez G, Coto E, Melón S |
| EPI_ISL_3076914 | Clínica Rotger Laboratorio de Análisis Clínicos Dpto. Biología Molecular | Clínica Rotger Laboratorio de Análisis Clínicos Dpto. Biología Molecular | Francisco Aliaga-Lozano, Chaymaá Gammani-Chairi, Laura Gómez, Juana Gonzalez-Calderón, Marina Barceló-Vives, Catalina Vidal-Gómez, Marina Kleber-Pol, Esperança Frau-Fuster, Carmen Morales-González, Esther Nieto-Molina |
| EPI_ISL_3076915 | Department for Virology, Molecular Biology and Genome Research, R. G. Lugar Center for Public Health Research, National Center for Disease Control and Public Health (NCDC) of Georgia. | Department for Virology, Molecular Biology and Genome Research, R. G. Lugar Center for Public Health Research, National Center for Disease Control and Public Health (NCDC) of Georgia. | Giorgi Gogoladze, Giorgi Tomashvili, Meri Pantsulaia, Gvantsa Brachveli, Nino Berishvili, Tata Imnadze, Ana Papkiauri, Gvantsa Chanturia, Ann Machablishvili, Nato Kotaria, Marine Murtskhvaladze, Lela Sabadze, Mari Gavashelidze, Tamar Jashiashvili, Tea Tevdoradze, Ketevan Sidamonidze, Ekaterine Khmaladze, Ekaterine Zhghenti, Roena Sukhiashvili, Mariam Zakalashvili, Lela Urushadze, Magda Dgebuadze, Davit Tsaguria, Ekaterine Zangaladze, Adam Kotorashvili, Maia Alkhazashvili, Irma Burjanadze, Anna Kasradze, Khatuna Zakhashvili, Paata Imnadze, Amiran Gamkrelidze. |
| EPI_ISL_3076916 | Hospital del Oriente de Asturias | Laboratorio de Virología HUCA | Sandoval M, Castelló C, Gómez de Oña J, Boga JA, Rojo S, Alvarez-Arguelles ME, Abreu F, Costales I, Perez-Martínez Z, Martín-Rodríguez G, Coto E, Melón S |
| EPI_ISL_3076917 | Department for Virology, Molecular Biology and Genome Research, R. G. Lugar Center for Public Health Research, National Center for Disease Control and Public Health (NCDC) of Georgia. | Department for Virology, Molecular Biology and Genome Research, R. G. Lugar Center for Public Health Research, National Center for Disease Control and Public Health (NCDC) of Georgia. | Giorgi Tomashvili, Meri Pantsulaia, Gvantsa Brachveli, Giorgi Gogoladze, Nino Berishvili, Tata Imnadze, Ana Papkiauri, Gvantsa Chanturia, Ann Machablishvili, Nato Kotaria, Marine Murtskhvaladze, Lela Sabadze, Mari Gavashelidze, Tamar Jashiashvili, Tea Tevdoradze, Ketevan Sidamonidze, Ekaterine Khmaladze, Ekaterine Zhghenti, Roena Sukhiashvili, Mariam Zakalashvili, Lela Urushadze, Magda Dgebuadze, Davit Tsaguria, Ekaterine Zangaladze, Adam Kotorashvili, Maia Alkhazashvili, Irma Burjanadze, Anna Kasradze, Khatuna Zakhashvili, Paata Imnadze, Amiran Gamkrelidze. |
| EPI_ISL_3076918, EPI_ISL_3076919 | Hospital del Oriente de Asturias | Laboratorio de Virología HUCA | Sandoval M, Castelló C, Gómez de Oña J, Boga JA, Rojo S, Alvarez-Arguelles ME, Abreu F, Costales I, Perez-Martínez Z, Martín-Rodríguez G, Coto E, Melón S |
| EPI_ISL_3076921 | Department for Virology, Molecular Biology and Genome Research, R. G. Lugar Center for Public Health Research, National Center for Disease Control and Public Health (NCDC) of Georgia. | Department for Virology, Molecular Biology and Genome Research, R. G. Lugar Center for Public Health Research, National Center for Disease Control and Public Health (NCDC) of Georgia. | Giorgi Tomashvili, Meri Pantsulaia, Gvantsa Brachveli, Giorgi Gogoladze, Nino Berishvili, Tata Imnadze, Ana Papkiauri, Gvantsa Chanturia, Ann Machablishvili, Nato Kotaria, Marine Murtskhvaladze, Lela Sabadze, Mari Gavashelidze, Tamar Jashiashvili, Tea Tevdoradze, Ketevan Sidamonidze, Ekaterine Khmaladze, Ekaterine Zhghenti, Roena Sukhiashvili, Mariam Zakalashvili, Lela Urushadze, Magda Dgebuadze, Davit Tsaguria, Ekaterine Zangaladze, Adam Kotorashvili, Maia Alkhazashvili, Irma Burjanadze, Anna Kasradze, Khatuna Zakhashvili, Paata Imnadze, Amiran Gamkrelidze. |
| EPI_ISL_3076922 | Clínica Rotger Laboratorio de Análisis Clínicos Dpto. Biología Molecular | Clínica Rotger Laboratorio de Análisis Clínicos Dpto. Biología Molecular | Francisco Aliaga-Lozano, Chaymaá Gammani-Chairi, Laura Gómez, Juana Gonzalez-Calderón, Marina Barceló-Vives, Catalina Vidal-Gómez, Marina Kleber-Pol, Esperança Frau-Fuster, Carmen Morales-González, Esther Nieto-Molina |
| EPI_ISL_3076923 | Hospital del Oriente de Asturias | Laboratorio de Virología HUCA | Sandoval M, Castelló C, Gómez de Oña J, Boga JA, Rojo S, Alvarez-Arguelles ME, Abreu F, Costales I, Perez-Martínez Z, Martín-Rodríguez G, Coto E, Melón S |
| EPI_ISL_3076924 | Department for Virology, Molecular Biology and Genome Research, R. G. Lugar Center for Public Health Research, National Center for Disease Control and Public Health (NCDC) of Georgia. | Department for Virology, Molecular Biology and Genome Research, R. G. Lugar Center for Public Health Research, National Center for Disease Control and Public Health (NCDC) of Georgia. | Gvantsa Brachveli, Giorgi Tomashvili, Meri Pantsulaia, Giorgi Gogoladze, Nino Berishvili, Tata Imnadze, Ana Papkiauri, Gvantsa Chanturia, Ann Machablishvili, Nato Kotaria, Marine Murtskhvaladze, Lela Sabadze, Mari Gavashelidze, Tamar Jashiashvili, Tea Tevdoradze, Ketevan Sidamonidze, Ekaterine Khmaladze, Ekaterine Zhghenti, Roena Sukhiashvili, Mariam Zakalashvili, Lela Urushadze, Magda Dgebuadze, Davit Tsaguria, Ekaterine Zangaladze, Adam Kotorashvili, Maia Alkhazashvili, Irma Burjanadze, Anna Kasradze, Khatuna Zakhashvili, Paata Imnadze, Amiran Gamkrelidze. |
| EPI_ISL_3076925 | Department for Virology, Molecular Biology and Genome Research, R. G. Lugar Center for Public Health Research, National Center for Disease Control and Public Health (NCDC) of Georgia. | Department for Virology, Molecular Biology and Genome Research, R. G. Lugar Center for Public Health Research, National Center for Disease Control and Public Health (NCDC) of Georgia. | Ana Papkiauri, Giorgi Tomashvili, Meri Pantsulaia, Gvantsa Brachveli, Giorgi Gogoladze, Nino Berishvili, Tata Imnadze, Gvantsa Chanturia, Ann Machablishvili, Nato Kotaria, Marine Murtskhvaladze, Lela Sabadze, Mari Gavashelidze, Tamar Jashiashvili, Tea Tevdoradze, Ketevan Sidamonidze, Ekaterine Khmaladze, Ekaterine Zhghenti, Roena Sukhiashvili, Mariam Zakalashvili, Lela Urushadze, Magda Dgebuadze, Davit Tsaguria, Ekaterine Zangaladze, Adam Kotorashvili, Maia Alkhazashvili, Irma Burjanadze, Anna Kasradze, Khatuna Zakhashvili, Paata Imnadze, Amiran Gamkrelidze. |
| EPI_ISL_3076927, EPI_ISL_3076929 | Department for Virology, Molecular Biology and Genome Research, R. G. Lugar Center for Public Health Research, National Center for Disease Control and Public Health (NCDC) of Georgia. | Department for Virology, Molecular Biology and Genome Research, R. G. Lugar Center for Public Health Research, National Center for Disease Control and Public Health (NCDC) of Georgia. | Giorgi Tomashvili, Meri Pantsulaia, Gvantsa Brachveli, Giorgi Gogoladze, Nino Berishvili, Tata Imnadze, Ana Papkiauri, Gvantsa Chanturia, Ann Machablishvili, Nato Kotaria, Marine Murtskhvaladze, Lela Sabadze, Mari Gavashelidze, Tamar Jashiashvili, Tea Tevdoradze, Ketevan Sidamonidze, Ekaterine Khmaladze, Ekaterine Zhghenti, Roena Sukhiashvili, Mariam Zakalashvili, Lela Urushadze, Magda Dgebuadze, Davit Tsaguria, Ekaterine Zangaladze, Adam Kotorashvili, Maia Alkhazashvili, Irma Burjanadze, Anna Kasradze, Khatuna Zakhashvili, Paata Imnadze, Amiran Gamkrelidze. |
| EPI_ISL_3076937 | University Clinical Hospital of Mostar, Department of Microbiology and Molecular Diagnostics | University of Sarajevo, Veterinary Faculty, Laboratory for Molecular Diagnostic and Research Laboratory | Goletic S, Goletic T, Ostojic M, Softic A, Nicevic M., Terzic I., Sabic E., Hodzic A., Jazic A. |
| EPI_ISL_3076941 | University Clinical Hospital of Mostar, Department of Microbiology and Molecular Diagnostics | University of Sarajevo, Veterinary Faculty, Laboratory for Molecular Diagnostic and Research Laboratory | Goletic S, Goletic T, Ostojic M, Softic A, Terzic I., Sabic E., Hodzic A., Jazic A., Nicevic M. |
| EPI_ISL_3076942 | Hospital del Oriente de Asturias | Laboratorio de Virología HUCA | Sandoval M, Castelló C, Gómez de Oña J, Boga JA, Rojo S, Alvarez-Arguelles ME, Abreu F, Costales I, Perez-Martínez Z, Martín-Rodríguez G, Coto E, Melón S |
| EPI_ISL_3076943 | Tuzla University Clinical Center, Department of Microbiology | University of Sarajevo, Veterinary Faculty, Laboratory for Molecular Diagnostic and Research Laboratory | Goletic S, Goletic T, Ostojic M, Softic A, Sabic E., Jazic A., Nicevic M., Hodzic A., Terzic I. |
| EPI_ISL_3076946, EPI_ISL_3076947, EPI_ISL_3076948, EPI_ISL_3076949, EPI_ISL_3076950, EPI_ISL_3076951, EPI_ISL_3076952, EPI_ISL_3076953, EPI_ISL_3076954, EPI_ISL_3076955, EPI_ISL_3076956, EPI_ISL_3076957 | | | |
| see above | Quadram Institute Bioscience | COVID-19 Genomics UK (COG-UK) Consortium | Dave J. Baker, Gemma L. Kay, Alp Aydin, Thanh Le-Viet, Steven Rudder, Ana P. Tedim, Anastasia Kolyva, Maria Diaz, Leonardo de Oliveira Martins, Nabil-Fareed Alikhan, Lizzie Meadows, Rachael Stanley, Rose K Davidson, Ngozi Elumogo, |

| | | | |
|---|---|---|---|
| | | | Muhammed Yasir, Sophie J. Prosolek, Ebenezer Foster-Nyarko, Martin Lott, Christopher Jeanes, Lewis G. Spurgin, Nicholas M. Thomson, Alexander J Trotter, Rachel Gilroy, Luke Bedford, Samuel Bloomfield, Claire Stuart, Andrew Bell, Reenesh Prakash, Lindsay Coupland, Samir Dervisevic, Alison E. Mather, John Wain, Mark Webber, Andrew J. Page, Justin O'Grady |
| EPI_ISL_3077001, EPI_ISL_3077002, EPI_ISL_3077003, EPI_ISL_3077004, EPI_ISL_3077005, EPI_ISL_3077006, EPI_ISL_3077007, EPI_ISL_3077008, EPI_ISL_3077009, EPI_ISL_3077010, EPI_ISL_3077011, EPI_ISL_3077012, EPI_ISL_3077013, EPI_ISL_3077014, EPI_ISL_3077015, EPI_ISL_3077016, EPI_ISL_3077017, EPI_ISL_3077018, EPI_ISL_3077019, EPI_ISL_3077020, EPI_ISL_3077021, EPI_ISL_3077022, EPI_ISL_3077023, EPI_ISL_3077024, EPI_ISL_3077025, EPI_ISL_3077026, EPI_ISL_3077027, EPI_ISL_3077028, EPI_ISL_3077029, EPI_ISL_3077030, EPI_ISL_3077031, EPI_ISL_3077032, EPI_ISL_3077033, EPI_ISL_3077034, EPI_ISL_3077035, EPI_ISL_3077036, EPI_ISL_3077037, EPI_ISL_3077038, EPI_ISL_3077039, EPI_ISL_3077040, EPI_ISL_3077041, EPI_ISL_3077042, EPI_ISL_3077043, EPI_ISL_3077044, EPI_ISL_3077045, EPI_ISL_3077046, EPI_ISL_3077047, EPI_ISL_3077048, EPI_ISL_3077049, EPI_ISL_3077050, EPI_ISL_3077051, EPI_ISL_3077052, EPI_ISL_3077053, EPI_ISL_3077054, EPI_ISL_3077055, EPI_ISL_3077056, EPI_ISL_3077057, EPI_ISL_3077058, EPI_ISL_3077059, EPI_ISL_3077060, EPI_ISL_3077061, EPI_ISL_3077062, EPI_ISL_3077063, EPI_ISL_3077064, EPI_ISL_3077065, EPI_ISL_3077066, EPI_ISL_3077067, EPI_ISL_3077068, EPI_ISL_3077069, EPI_ISL_3077070, EPI_ISL_3077071, EPI_ISL_3077072, EPI_ISL_3077073, EPI_ISL_3077074, EPI_ISL_3077075, EPI_ISL_3077076, EPI_ISL_3077077, EPI_ISL_3077078, EPI_ISL_3077079, EPI_ISL_3077080, EPI_ISL_3077081, EPI_ISL_3077082, EPI_ISL_3077083, EPI_ISL_3077084, EPI_ISL_3077085, EPI_ISL_3077086, EPI_ISL_3077087, EPI_ISL_3077088, EPI_ISL_3077089, EPI_ISL_3077090, EPI_ISL_3077091, EPI_ISL_3077092, EPI_ISL_3077093, EPI_ISL_3077094, EPI_ISL_3077095, EPI_ISL_3077096, EPI_ISL_3077097, EPI_ISL_3077098, EPI_ISL_3077099, EPI_ISL_3077100, EPI_ISL_3077101, EPI_ISL_3077102, EPI_ISL_3077103, EPI_ISL_3077104, EPI_ISL_3077105, EPI_ISL_3077106, EPI_ISL_3077107, EPI_ISL_3077108, EPI_ISL_3077109, EPI_ISL_3077110, EPI_ISL_3077111, EPI_ISL_3077112, EPI_ISL_3077113, EPI_ISL_3077114, EPI_ISL_3077115, EPI_ISL_3077116, EPI_ISL_3077117, EPI_ISL_3077118, EPI_ISL_3077119, EPI_ISL_3077120, EPI_ISL_3077121, EPI_ISL_3077122, EPI_ISL_3077123, EPI_ISL_3077124, EPI_ISL_3077125, EPI_ISL_3077126, EPI_ISL_3077127, EPI_ISL_3077128, EPI_ISL_3077129, EPI_ISL_3077130, EPI_ISL_3077131, EPI_ISL_3077132, EPI_ISL_3077133, EPI_ISL_3077134, EPI_ISL_3077135, EPI_ISL_3077136, EPI_ISL_3077137, EPI_ISL_3077138, EPI_ISL_3077139, EPI_ISL_3077140, EPI_ISL_3077141, EPI_ISL_3077142, EPI_ISL_3077143, EPI_ISL_3077144, EPI_ISL_3077145, EPI_ISL_3077146, EPI_ISL_3077147, EPI_ISL_3077148, EPI_ISL_3077149, EPI_ISL_3077150 | | | |
| see above | Department of Pathology, University of Cambridge | COVID-19 Genomics UK (COG-UK) Consortium | Aminu S. Jahun, Yasmin Chaudhry, Iliana Georgana, Myra Hosmillo, Rhys Izuagbe, William L. Hamilton, Martin D. Curran, Surendra Parmar, Ian Goodfellow |
| EPI_ISL_3077151, EPI_ISL_3077152, EPI_ISL_3077153, EPI_ISL_3077154, EPI_ISL_3077155, EPI_ISL_3077156, EPI_ISL_3077157, EPI_ISL_3077158, EPI_ISL_3077159, EPI_ISL_3077160, EPI_ISL_3077161, EPI_ISL_3077162, EPI_ISL_3077163, EPI_ISL_3077164, EPI_ISL_3077165, EPI_ISL_3077166, EPI_ISL_3077167, EPI_ISL_3077168, EPI_ISL_3077169, EPI_ISL_3077170, EPI_ISL_3077171, EPI_ISL_3077172, EPI_ISL_3077173, EPI_ISL_3077174, EPI_ISL_3077175, EPI_ISL_3077176, EPI_ISL_3077177, EPI_ISL_3077178, EPI_ISL_3077179, EPI_ISL_3077180, EPI_ISL_3077181, EPI_ISL_3077182, EPI_ISL_3077183, EPI_ISL_3077184, EPI_ISL_3077185, EPI_ISL_3077186, EPI_ISL_3077187, EPI_ISL_3077188, EPI_ISL_3077189, EPI_ISL_3077190, EPI_ISL_3077191, EPI_ISL_3077192, EPI_ISL_3077193, EPI_ISL_3077194, EPI_ISL_3077195, EPI_ISL_3077196, EPI_ISL_3077197, EPI_ISL_3077198, EPI_ISL_3077199, EPI_ISL_3077200, EPI_ISL_3077201, EPI_ISL_3077202, EPI_ISL_3077203, EPI_ISL_3077204, EPI_ISL_3077205, EPI_ISL_3077206, EPI_ISL_3077207, EPI_ISL_3077208, EPI_ISL_3077209, EPI_ISL_3077210, EPI_ISL_3077211, EPI_ISL_3077212, EPI_ISL_3077213, EPI_ISL_3077214, EPI_ISL_3077215, EPI_ISL_3077216, EPI_ISL_3077217, EPI_ISL_3077218, EPI_ISL_3077219, EPI_ISL_3077220, EPI_ISL_3077221, EPI_ISL_3077222, EPI_ISL_3077223, EPI_ISL_3077224, EPI_ISL_3077225, EPI_ISL_3077226, EPI_ISL_3077227, EPI_ISL_3077228, EPI_ISL_3077229, EPI_ISL_3077230, EPI_ISL_3077231, EPI_ISL_3077232, EPI_ISL_3077233, EPI_ISL_3077234, EPI_ISL_3077235, EPI_ISL_3077236, EPI_ISL_3077237, EPI_ISL_3077238, EPI_ISL_3077239, EPI_ISL_3077240, EPI_ISL_3077241, EPI_ISL_3077242, EPI_ISL_3077243, EPI_ISL_3077244, EPI_ISL_3077245, EPI_ISL_3077246, EPI_ISL_3077247, EPI_ISL_3077248, EPI_ISL_3077249, EPI_ISL_3077250, EPI_ISL_3077251, EPI_ISL_3077252, EPI_ISL_3077253, EPI_ISL_3077254, EPI_ISL_3077255, EPI_ISL_3077256, EPI_ISL_3077257, EPI_ISL_3077258, EPI_ISL_3077259, EPI_ISL_3077260, EPI_ISL_3077261, EPI_ISL_3077262, EPI_ISL_3077263, EPI_ISL_3077264, EPI_ISL_3077265, EPI_ISL_3077266, EPI_ISL_3077267, EPI_ISL_3077268, EPI_ISL_3077269, EPI_ISL_3077270, EPI_ISL_3077271, EPI_ISL_3077272, EPI_ISL_3077273, EPI_ISL_3077274, EPI_ISL_3077275, EPI_ISL_3077276, EPI_ISL_3077277, EPI_ISL_3077278, EPI_ISL_3077279, EPI_ISL_3077280, EPI_ISL_3077281, EPI_ISL_3077282, EPI_ISL_3077283, EPI_ISL_3077284, EPI_ISL_3077285, EPI_ISL_3077286, EPI_ISL_3077287, EPI_ISL_3077288, EPI_ISL_3077289, EPI_ISL_3077290, EPI_ISL_3077291, EPI_ISL_3077292, EPI_ISL_3077293, EPI_ISL_3077294, EPI_ISL_3077295, EPI_ISL_3077296, EPI_ISL_3077297, EPI_ISL_3077298, EPI_ISL_3077299, EPI_ISL_3077300, EPI_ISL_3077301, EPI_ISL_3077302, EPI_ISL_3077303, EPI_ISL_3077304, EPI_ISL_3077305, EPI_ISL_3077306, EPI_ISL_3077307 | | | |
| see above | West of Scotland Specialist Virology Centre, NHSGGC / MRC-University of Glasgow Centre for Virus Research | COVID-19 Genomics UK (COG-UK) Consortium | Ana da Silva Filipe, Natasha Johnson, Kathy Smollett, Daniel Mair, Stephen Carmichael, Alice Broos, Lily Tong, Jenna Nichols, Kyriaki Nomikou; Sarah McDonald; Ioulia Tsatsani, Richard Orton, Joseph Hughes, Sreenu Vattipally, David L Robertson; Alasdair MacLean, Rory Gunson; Sharif Shaaban, Matthew Holden; Rachel Blacow, Guy Mollett, Kathy Li, James Shepherd, Antonia Ho, Emma Thomson |
| EPI_ISL_3077308, EPI_ISL_3077309, EPI_ISL_3077310, EPI_ISL_3077311, EPI_ISL_3077312, EPI_ISL_3077313, EPI_ISL_3077314, EPI_ISL_3077315, EPI_ISL_3077316, EPI_ISL_3077317, EPI_ISL_3077318, EPI_ISL_3077319, EPI_ISL_3077320, EPI_ISL_3077321, EPI_ISL_3077322, EPI_ISL_3077323, EPI_ISL_3077324, EPI_ISL_3077325, EPI_ISL_3077326, EPI_ISL_3077327, EPI_ISL_3077328, EPI_ISL_3077329, EPI_ISL_3077330, EPI_ISL_3077331, EPI_ISL_3077332, EPI_ISL_3077333, EPI_ISL_3077334, EPI_ISL_3077335, EPI_ISL_3077336, EPI_ISL_3077337, EPI_ISL_3077338, EPI_ISL_3077339, EPI_ISL_3077340, EPI_ISL_3077341, EPI_ISL_3077342, EPI_ISL_3077343, EPI_ISL_3077344, EPI_ISL_3077345, EPI_ISL_3077346, EPI_ISL_3077347, EPI_ISL_3077348, EPI_ISL_3077349, EPI_ISL_3077350 | | | |
| see above | The Roslin Institute and R(D)SVS, University of Edinburgh / Virology Department, Royal Infirmary of Edinburgh, NHS Lothian / School of Biological Sciences, University of Edinburgh | COVID-19 Genomics UK (COG-UK) Consortium | Warr A, Newman, C, Tait-Burkhard C, McHugh M, Dewar R, Cotton S, Gallagher A, Maloney D, Fernandez G, O'Toole Á, Scher E, Hill V, McCrone JT, Colquhoun R, Yu X, Jackson B, Rambaut A, Templeton K |
| EPI_ISL_3077351, EPI_ISL_3077352 | Virology Department, Royal Infirmary of Edinburgh, NHS Lothian / School of Biological Sciences, University of Edinburgh | COVID-19 Genomics UK (COG-UK) Consortium | McHugh M, Dewar R, Cotton S, Gallagher A, Maloney D, Fernandez G, O'Toole Á, Scher E, Hill V, McCrone JT, Colquhoun R, Yu X, Jackson B, Rambaut A, Templeton K |
| EPI_ISL_3077353, EPI_ISL_3077354, EPI_ISL_3077355, EPI_ISL_3077356, EPI_ISL_3077357, EPI_ISL_3077358, EPI_ISL_3077359, EPI_ISL_3077360, EPI_ISL_3077361, EPI_ISL_3077362 | The Roslin Institute and R(D)SVS, University of Edinburgh / Virology Department, Royal Infirmary of Edinburgh, NHS Lothian / School of Biological Sciences, University of Edinburgh | COVID-19 Genomics UK (COG-UK) Consortium | Warr A, Newman, C, Tait-Burkhard C, McHugh M, Dewar R, Cotton S, Gallagher A, Maloney D, Fernandez G, O'Toole Á, Scher E, Hill V, McCrone JT, Colquhoun R, Yu X, Jackson B, Rambaut A, Templeton K |
| EPI_ISL_3077363, EPI_ISL_3077364, EPI_ISL_3077365, EPI_ISL_3077366, EPI_ISL_3077367, EPI_ISL_3077368, EPI_ISL_3077369, EPI_ISL_3077370, EPI_ISL_3077371, EPI_ISL_3077372, EPI_ISL_3077373, EPI_ISL_3077374, EPI_ISL_3077375, EPI_ISL_3077376, EPI_ISL_3077377, EPI_ISL_3077378, EPI_ISL_3077379, EPI_ISL_3077380, EPI_ISL_3077381, EPI_ISL_3077383, EPI_ISL_3077384, EPI_ISL_3077385, EPI_ISL_3077386, EPI_ISL_3077387, EPI_ISL_3077388, EPI_ISL_3077389, EPI_ISL_3077390, EPI_ISL_3077391, EPI_ISL_3077392, EPI_ISL_3077393, EPI_ISL_3077394, EPI_ISL_3077395, EPI_ISL_3077396, EPI_ISL_3077397, EPI_ISL_3077398, EPI_ISL_3077399, EPI_ISL_3077400, EPI_ISL_3077401, EPI_ISL_3077402, EPI_ISL_3077403, EPI_ISL_3077404, EPI_ISL_3077405, EPI_ISL_3077406, EPI_ISL_3077407, EPI_ISL_3077408, EPI_ISL_3077409, EPI_ISL_3077410, EPI_ISL_3077411, EPI_ISL_3077412, EPI_ISL_3077413, EPI_ISL_3077414, EPI_ISL_3077415, EPI_ISL_3077416, EPI_ISL_3077417, EPI_ISL_3077418, EPI_ISL_3077419, EPI_ISL_3077420, EPI_ISL_3077421, EPI_ISL_3077422, EPI_ISL_3077423, EPI_ISL_3077424, EPI_ISL_3077425, EPI_ISL_3077426, EPI_ISL_3077427, EPI_ISL_3077428, EPI_ISL_3077429, EPI_ISL_3077430, EPI_ISL_3077431, EPI_ISL_3077432, EPI_ISL_3077433, EPI_ISL_3077434, EPI_ISL_3077435, EPI_ISL_3077436, EPI_ISL_3077437, EPI_ISL_3077438, EPI_ISL_3077439, EPI_ISL_3077440, EPI_ISL_3077441, EPI_ISL_3077442, EPI_ISL_3077443, EPI_ISL_3077444, EPI_ISL_3077445, EPI_ISL_3077446, EPI_ISL_3077447, EPI_ISL_3077448, EPI_ISL_3077449, EPI_ISL_3077451, EPI_ISL_3077452, EPI_ISL_3077453, EPI_ISL_3077454, EPI_ISL_3077455, EPI_ISL_3077456, EPI_ISL_3077457, EPI_ISL_3077458, EPI_ISL_3077459, EPI_ISL_3077460, EPI_ISL_3077461, EPI_ISL_3077462, EPI_ISL_3077463, EPI_ISL_3077464, EPI_ISL_3077465, EPI_ISL_3077466, EPI_ISL_3077467, EPI_ISL_3077468, EPI_ISL_3077469, EPI_ISL_3077470, EPI_ISL_3077471, EPI_ISL_3077472, EPI_ISL_3077473, EPI_ISL_3077474, EPI_ISL_3077475, EPI_ISL_3077476, EPI_ISL_3077477, EPI_ISL_3077478, EPI_ISL_3077479 | | | |
| see above | Virology Department, Royal Infirmary of Edinburgh, NHS Lothian / School of Biological Sciences, University of Edinburgh | COVID-19 Genomics UK (COG-UK) Consortium | McHugh M, Dewar R, Cotton S, Gallagher A, Maloney D, Fernandez G, O'Toole Á, Scher E, Hill V, McCrone JT, Colquhoun R, Yu X, Jackson B, Rambaut A, Templeton K |
| EPI_ISL_3077480, EPI_ISL_3077481, EPI_ISL_3077482, EPI_ISL_3077483, EPI_ISL_3077484, EPI_ISL_3077485, EPI_ISL_3077486, EPI_ISL_3077487, EPI_ISL_3077488, EPI_ISL_3077489, EPI_ISL_3077490, EPI_ISL_3077491, EPI_ISL_3077492, EPI_ISL_3077493, EPI_ISL_3077494, EPI_ISL_3077495, EPI_ISL_3077496, EPI_ISL_3077497, EPI_ISL_3077498, EPI_ISL_3077499, EPI_ISL_3077500, EPI_ISL_3077501, EPI_ISL_3077502, EPI_ISL_3077503, EPI_ISL_3077504, EPI_ISL_3077505, EPI_ISL_3077506, EPI_ISL_3077507, EPI_ISL_3077508, EPI_ISL_3077509, EPI_ISL_3077510, EPI_ISL_3077511, EPI_ISL_3077512, EPI_ISL_3077513, EPI_ISL_3077514, EPI_ISL_3077515, EPI_ISL_3077516, EPI_ISL_3077517, EPI_ISL_3077518, EPI_ISL_3077519, EPI_ISL_3077520, EPI_ISL_3077521, EPI_ISL_3077522, EPI_ISL_3077523, EPI_ISL_3077524, EPI_ISL_3077525, EPI_ISL_3077526, EPI_ISL_3077527, EPI_ISL_3077528, EPI_ISL_3077529, EPI_ISL_3077530, EPI_ISL_3077531, EPI_ISL_3077532, EPI_ISL_3077533, EPI_ISL_3077534, EPI_ISL_3077535, EPI_ISL_3077536, EPI_ISL_3077537, EPI_ISL_3077538, EPI_ISL_3077539, EPI_ISL_3077540, EPI_ISL_3077541, EPI_ISL_3077542, EPI_ISL_3077543, EPI_ISL_3077544, EPI_ISL_3077545, EPI_ISL_3077546, EPI_ISL_3077547, EPI_ISL_3077548, EPI_ISL_3077549, EPI_ISL_3077550, EPI_ISL_3077551, EPI_ISL_3077552, EPI_ISL_3077553, EPI_ISL_3077554, EPI_ISL_3077555, EPI_ISL_3077556, EPI_ISL_3077557, EPI_ISL_3077558, EPI_ISL_3077559, EPI_ISL_3077560, EPI_ISL_3077561, EPI_ISL_3077562, EPI_ISL_3077563, EPI_ISL_3077564, EPI_ISL_3077565, EPI_ISL_3077566, EPI_ISL_3077567, EPI_ISL_3077568, EPI_ISL_3077569, EPI_ISL_3077570, EPI_ISL_3077571, EPI_ISL_3077572, EPI_ISL_3077573, EPI_ISL_3077574, EPI_ISL_3077575, EPI_ISL_3077576, EPI_ISL_3077577, EPI_ISL_3077578, EPI_ISL_3077579, EPI_ISL_3077580, EPI_ISL_3077581, EPI_ISL_3077582, EPI_ISL_3077584, EPI_ISL_3077585, EPI_ISL_3077586, EPI_ISL_3077587, EPI_ISL_3077589, EPI_ISL_3077590, EPI_ISL_3077591, EPI_ISL_3077592, EPI_ISL_3077593, EPI_ISL_3077594, EPI_ISL_3077595, EPI_ISL_3077596, EPI_ISL_3077597, EPI_ISL_3077598, EPI_ISL_3077599, EPI_ISL_3077600, EPI_ISL_3077601, EPI_ISL_3077602, EPI_ISL_3077603, EPI_ISL_3077604, EPI_ISL_3077605, EPI_ISL_3077606, EPI_ISL_3077607, EPI_ISL_3077608, EPI_ISL_3077609, EPI_ISL_3077610, EPI_ISL_3077611, EPI_ISL_3077612, EPI_ISL_3077613, EPI_ISL_3077614, EPI_ISL_3077615, EPI_ISL_3077616, EPI_ISL_3077617, EPI_ISL_3077618, EPI_ISL_3077619, EPI_ISL_3077620, EPI_ISL_3077621, EPI_ISL_3077622, EPI_ISL_3077623, EPI_ISL_3077624, EPI_ISL_3077625, EPI_ISL_3077626, EPI_ISL_3077627, EPI_ISL_3077628, EPI_ISL_3077629, EPI_ISL_3077630, EPI_ISL_3077631, EPI_ISL_3077632, EPI_ISL_3077633, EPI_ISL_3077634, EPI_ISL_3077635, EPI_ISL_3077636, EPI_ISL_3077637, EPI_ISL_3077638, EPI_ISL_3077639, EPI_ISL_3077640, EPI_ISL_3077641, EPI_ISL_3077642, EPI_ISL_3077643, EPI_ISL_3077644, EPI_ISL_3077645, EPI_ISL_3077646, EPI_ISL_3077647, EPI_ISL_3077648, EPI_ISL_3077649, EPI_ISL_3077650, EPI_ISL_3077651, EPI_ISL_3077652, EPI_ISL_3077653, EPI_ISL_3077654, EPI_ISL_3077655, EPI_ISL_3077656, EPI_ISL_3077657, EPI_ISL_3077658, EPI_ISL_3077659, EPI_ISL_3077660, EPI_ISL_3077661, EPI_ISL_3077662, EPI_ISL_3077663, EPI_ISL_3077664, EPI_ISL_3077665, EPI_ISL_3077666, EPI_ISL_3077667, EPI_ISL_3077668, EPI_ISL_3077669, EPI_ISL_3077670, EPI_ISL_3077671, EPI_ISL_3077672, EPI_ISL_3077673, EPI_ISL_3077674, EPI_ISL_3077675, EPI_ISL_3077676, EPI_ISL_3077677, EPI_ISL_3077678, EPI_ISL_3077679, EPI_ISL_3077680, EPI_ISL_3077681, EPI_ISL_3077682, EPI_ISL_3077683, EPI_ISL_3077684, EPI_ISL_3077685, EPI_ISL_3077686, EPI_ISL_3077687, EPI_ISL_3077688, EPI_ISL_3077689, EPI_ISL_3077690, EPI_ISL_3077691, EPI_ISL_3077692, EPI_ISL_3077693, EPI_ISL_3077694, EPI_ISL_3077695, EPI_ISL_3077696, EPI_ISL_3077697, EPI_ISL_3077699, EPI_ISL_3077700, EPI_ISL_3077701, EPI_ISL_3077702, EPI_ISL_3077703, EPI_ISL_3077704, EPI_ISL_3077705, EPI_ISL_3077706, EPI_ISL_3077707, EPI_ISL_3077708, EPI_ISL_3077709, EPI_ISL_3077710, EPI_ISL_3077711, EPI_ISL_3077712, EPI_ISL_3077713, EPI_ISL_3077714, EPI_ISL_3077715, EPI_ISL_3077716, EPI_ISL_3077717, EPI_ISL_3077718, EPI_ISL_3077719, EPI_ISL_3077720, EPI_ISL_3077721, EPI_ISL_3077722, EPI_ISL_3077723, EPI_ISL_3077724, EPI_ISL_3077725, EPI_ISL_3077726, EPI_ISL_3077727, EPI_ISL_3077728, EPI_ISL_3077729, EPI_ISL_3077730, EPI_ISL_3077731, EPI_ISL_3077732, EPI_ISL_3077733, EPI_ISL_3077734, EPI_ISL_3077735, EPI_ISL_3077736, EPI_ISL_3077737, EPI_ISL_3077739, EPI_ISL_3077740, EPI_ISL_3077741, EPI_ISL_3077742, EPI_ISL_3077743, EPI_ISL_3077744, EPI_ISL_3077745, EPI_ISL_3077746, EPI_ISL_3077747, EPI_ISL_3077748, EPI_ISL_3077749, EPI_ISL_3077750, EPI_ISL_3077751, EPI_ISL_3077752, EPI_ISL_3077753, EPI_ISL_3077754, EPI_ISL_3077755, EPI_ISL_3077756, EPI_ISL_3077757, EPI_ISL_3077758, EPI_ISL_3077759, EPI_ISL_3077761, EPI_ISL_3077762, EPI_ISL_3077763, EPI_ISL_3077764, EPI_ISL_3077765, EPI_ISL_3077766, EPI_ISL_3077767, EPI_ISL_3077768, EPI_ISL_3077769, EPI_ISL_3077770, EPI_ISL_3077771, EPI_ISL_3077772, EPI_ISL_3077773, EPI_ISL_3077774, EPI_ISL_3077775, EPI_ISL_3077776, EPI_ISL_3077778, EPI_ISL_3077779, EPI_ISL_3077780, EPI_ISL_3077781, EPI_ISL_3077782, EPI_ISL_3077783, EPI_ISL_3077784, EPI_ISL_3077785, EPI_ISL_3077786, EPI_ISL_3077787, EPI_ISL_3077788, EPI_ISL_3077789, EPI_ISL_3077790, EPI_ISL_3077791, EPI_ISL_3077792, EPI_ISL_3077793, EPI_ISL_3077794, EPI_ISL_3077795, EPI_ISL_3077796, EPI_ISL_3077797, EPI_ISL_3077798, EPI_ISL_3077799, EPI_ISL_3077800, EPI_ISL_3077801, EPI_ISL_3077802, EPI_ISL_3077803, EPI_ISL_3077804, EPI_ISL_3077805, EPI_ISL_3077806, EPI_ISL_3077807, EPI_ISL_3077808, EPI_ISL_3077809, EPI_ISL_3077810, EPI_ISL_3077811, EPI_ISL_3077812, EPI_ISL_3077813, EPI_ISL_3077814, EPI_ISL_3077815, EPI_ISL_3077816, EPI_ISL_3077817, EPI_ISL_3077818, EPI_ISL_3077819, EPI_ISL_3077820, EPI_ISL_3077821, EPI_ISL_3077822, EPI_ISL_3077823, EPI_ISL_3077824, EPI_ISL_3077825, EPI_ISL_3077826, EPI_ISL_3077827, EPI_ISL_3077828, EPI_ISL_3077829, EPI_ISL_3077830, EPI_ISL_3077831, EPI_ISL_3077832, EPI_ISL_3077833, EPI_ISL_3077834, EPI_ISL_3077835, EPI_ISL_3077836, EPI_ISL_3077837, EPI_ISL_3077838, EPI_ISL_3077839, EPI_ISL_3077840, EPI_ISL_3077841, EPI_ISL_3077842, EPI_ISL_3077843, EPI_ISL_3077844, EPI_ISL_3077845, EPI_ISL_3077846, EPI_ISL_3077847, EPI_ISL_3077848, EPI_ISL_3077849, EPI_ISL_3077850, EPI_ISL_3077851, EPI_ISL_3077852, EPI_ISL_3077853, EPI_ISL_3077855, EPI_ISL_3077856, EPI_ISL_3077857, EPI_ISL_3077858, EPI_ISL_3077859, EPI_ISL_3077860, EPI_ISL_3077861, EPI_ISL_3077862, EPI_ISL_3077863, EPI_ISL_3077864, EPI_ISL_3077865, EPI_ISL_3077866, EPI_ISL_3077867, EPI_ISL_3077868, EPI_ISL_3077869, EPI_ISL_3077870, EPI_ISL_3077871, EPI_ISL_3077872, EPI_ISL_3077873, EPI_ISL_3077874, EPI_ISL_3077875, EPI_ISL_3077876, EPI_ISL_3077877, EPI_ISL_3077878, EPI_ISL_3077879, EPI_ISL_3077880, EPI_ISL_3077881, EPI_ISL_3077882, EPI_ISL_3077883, EPI_ISL_3077884, EPI_ISL_3077885, EPI_ISL_3077886, EPI_ISL_3077887, EPI_ISL_3077888, EPI_ISL_3077889, EPI_ISL_3077890, EPI_ISL_3077891, EPI_ISL_3077892, EPI_ISL_3077893, EPI_ISL_3077894, EPI_ISL_3077895, EPI_ISL_3077896, EPI_ISL_3077897, EPI_ISL_3077898, EPI_ISL_3077899, EPI_ISL_3077901, EPI_ISL_3077902, EPI_ISL_3077903, EPI_ISL_3077904, EPI_ISL_3077905, EPI_ISL_3077906, EPI_ISL_3077907, EPI_ISL_3077908, EPI_ISL_3077909, EPI_ISL_3077910, EPI_ISL_3077911, EPI_ISL_3077912, EPI_ISL_3077913, EPI_ISL_3077914, EPI_ISL_3077915, EPI_ISL_3077916, EPI_ISL_3077917, EPI_ISL_3077918, EPI_ISL_3077919, EPI_ISL_3077920, EPI_ISL_3077921, EPI_ISL_3077922, EPI_ISL_3077923, EPI_ISL_3077924, EPI_ISL_3077925, EPI_ISL_3077926, EPI_ISL_3077927, EPI_ISL_3077928, EPI_ISL_3077929, EPI_ISL_3077930, EPI_ISL_3077931, EPI_ISL_3077932, EPI_ISL_3077933, EPI_ISL_3077934, EPI_ISL_3077935, EPI_ISL_3077936, EPI_ISL_3077937, EPI_ISL_3077938, EPI_ISL_3077939, EPI_ISL_3077940, EPI_ISL_3077941, EPI_ISL_3077942, EPI_ISL_3077943, EPI_ISL_3077944, EPI_ISL_3077945, EPI_ISL_3077946, EPI_ISL_3077947, EPI_ISL_3077948, EPI_ISL_3077949, EPI_ISL_3077950, EPI_ISL_3077951, EPI_ISL_3077952, EPI_ISL_3077953, EPI_ISL_3077954, EPI_ISL_3077955, EPI_ISL_3077956, EPI_ISL_3077957, EPI_ISL_3077958, EPI_ISL_3077959, EPI_ISL_3077960, EPI_ISL_3077961, EPI_ISL_3077962, EPI_ISL_3077963, EPI_ISL_3077964, EPI_ISL_3077965, EPI_ISL_3077966, EPI_ISL_3077967, EPI_ISL_3077968, EPI_ISL_3077969, EPI_ISL_3077970, EPI_ISL_3077971, EPI_ISL_3077972, EPI_ISL_3077973, EPI_ISL_3077974, EPI_ISL_3077975, EPI_ISL_3077976, EPI_ISL_3077977, EPI_ISL_3077978 | | | |
| see above | Liverpool Clinical Laboratories | COVID-19 Genomics UK (COG-UK) Consortium | Sam Haldenby, Alistair Darby, Steve Paterson, Anita Lucaci, Julian Hiscox, M Almsaud, A Alrezaihi, Muhannad Alruwaili, Stuart D Armstrong, Jones Benjamin, Eleanor G Bentley, Anu Chawla, Jordan J Clark, Angela Cowell, Richard Eccles, Isabel García-Dorival, Matthew Gemmell, Alessandro Gerada, PKF Gilmore, Richard Gregory, Ximeng Han, Catherine Hartley, Margaret Hughes, Miren Iturriza-Gomara, James Johnson, L Luu, Jenifer Manson, Charlotte Nelson, Elaine O'Toole, Cassie Olateju, Rebekah Penrice-Randal , Lucille Rainbow, N.P Randle, Trevor Ian Robinson, Parul Sharma, Ghada T Shawli, James P Stewart, Neil Swainston, Ecaterina Vamos, Joanne Watts, Mark Whitehead, Hermione Webster |
| EPI_ISL_3077979, EPI_ISL_3077980, EPI_ISL_3077981, EPI_ISL_3077982, EPI_ISL_3077983, EPI_ISL_3077984, EPI_ISL_3077985, EPI_ISL_3077986, EPI_ISL_3077987, EPI_ISL_3077988, EPI_ISL_3077989, EPI_ISL_3077990, EPI_ISL_3077991, EPI_ISL_3077992, EPI_ISL_3077993, EPI_ISL_3077994, EPI_ISL_3077995, EPI_ISL_3077996, EPI_ISL_3077997, EPI_ISL_3077998, EPI_ISL_3077999, EPI_ISL_3078000, EPI_ISL_3078001, EPI_ISL_3078002, EPI_ISL_3078003, EPI_ISL_3078004, EPI_ISL_3078005, EPI_ISL_3078006, EPI_ISL_3078007, EPI_ISL_3078008, EPI_ISL_3078009, EPI_ISL_3078010, EPI_ISL_3078011, EPI_ISL_3078012, EPI_ISL_3078013, EPI_ISL_3078014, EPI_ISL_3078015, EPI_ISL_3078016, EPI_ISL_3078017, EPI_ISL_3078018, EPI_ISL_3078019, EPI_ISL_3078020, EPI_ISL_3078021, EPI_ISL_3078022, EPI_ISL_3078023, EPI_ISL_3078024, EPI_ISL_3078025, EPI_ISL_3078026, EPI_ISL_3078027, EPI_ISL_3078028, EPI_ISL_3078029, EPI_ISL_3078030, EPI_ISL_3078031, EPI_ISL_3078032, EPI_ISL_3078033, EPI_ISL_3078034, EPI_ISL_3078035, EPI_ISL_3078036, EPI_ISL_3078037, EPI_ISL_3078038, EPI_ISL_3078039, EPI_ISL_3078040, EPI_ISL_3078041, EPI_ISL_3078042, EPI_ISL_3078043, EPI_ISL_3078044, EPI_ISL_3078045, EPI_ISL_3078046, EPI_ISL_3078047, EPI_ISL_3078048, EPI_ISL_3078049, EPI_ISL_3078050, EPI_ISL_3078051, EPI_ISL_3078052, EPI_ISL_3078053, EPI_ISL_3078054, EPI_ISL_3078055, EPI_ISL_3078056, EPI_ISL_3078057, EPI_ISL_3078058, EPI_ISL_3078059, EPI_ISL_3078060, EPI_ISL_3078061, EPI_ISL_3078062, EPI_ISL_3078063, EPI_ISL_3078064, EPI_ISL_3078065, EPI_ISL_3078066, EPI_ISL_3078067, EPI_ISL_3078068, EPI_ISL_3078069, EPI_ISL_3078070, EPI_ISL_3078072, EPI_ISL_3078073, EPI_ISL_3078074, EPI_ISL_3078075, EPI_ISL_3078076, EPI_ISL_3078077, EPI_ISL_3078078, EPI_ISL_3078079, EPI_ISL_3078080, EPI_ISL_3078081, EPI_ISL_3078082, EPI_ISL_3078083, EPI_ISL_3078084, EPI_ISL_3078085, EPI_ISL_3078086, EPI_ISL_3078087, EPI_ISL_3078088, EPI_ISL_3078089, EPI_ISL_3078090, EPI_ISL_3078091, EPI_ISL_3078092, EPI_ISL_3078093, EPI_ISL_3078094, EPI_ISL_3078095, EPI_ISL_3078096, EPI_ISL_3078097, EPI_ISL_3078098, EPI_ISL_3078099, EPI_ISL_3078100, EPI_ISL_3078101, EPI_ISL_3078102, EPI_ISL_3078103, EPI_ISL_3078104, EPI_ISL_3078105, EPI_ISL_3078106, EPI_ISL_3078107, EPI_ISL_3078108, EPI_ISL_3078109, EPI_ISL_3078110, EPI_ISL_3078111, EPI_ISL_3078112, EPI_ISL_3078113, EPI_ISL_3078114, EPI_ISL_3078115, EPI_ISL_3078116, EPI_ISL_3078117, EPI_ISL_3078118, EPI_ISL_3078119 | | | |
| see above | University College London, Great Ormond Street Hospital for Children NHS Foundation Trust, Imperial College Healthcare NHS Trust | COVID-19 Genomics UK (COG-UK) Consortium | Sergi Castellano, Rachel Williams, Mark Kristiansen, Sunando Roy, Charlotte Williams, Nadua Bayzid, Marius Cotic, Helena Tutill, Patricia Dyal, Judith Breuer |
| EPI_ISL_3078120, EPI_ISL_3078121, EPI_ISL_3078122, EPI_ISL_3078123, EPI_ISL_3078124, EPI_ISL_3078125, EPI_ISL_3078126, EPI_ISL_3078127, EPI_ISL_3078128, EPI_ISL_3078129, EPI_ISL_3078130, EPI_ISL_3078131, EPI_ISL_3078132, EPI_ISL_3078133, EPI_ISL_3078134, EPI_ISL_3078135, EPI_ISL_3078136, EPI_ISL_3078137, EPI_ISL_3078138, EPI_ISL_3078139, EPI_ISL_3078140, EPI_ISL_3078141, EPI_ISL_3078142, EPI_ISL_3078143, EPI_ISL_3078144, EPI_ISL_3078145, EPI_ISL_3078146, EPI_ISL_3078147, EPI_ISL_3078148, EPI_ISL_3078149, EPI_ISL_3078150, EPI_ISL_3078151, EPI_ISL_3078152, EPI_ISL_3078153, EPI_ISL_3078154, EPI_ISL_3078155, EPI_ISL_3078156, EPI_ISL_3078157, EPI_ISL_3078158, EPI_ISL_3078160 | | | |
| see above | Northumbria University / South Tees Hospitals NHS Foundation Trust / North Cumbria Integrated Care NHS Foundation Trust / North Tees and Hartlepool NHS Foundation Trust / Newcastle Hospitals NHS Foundation Trust | COVID-19 Genomics UK (COG-UK) Consortium | Darren L Smith,Andrew Nelson,Matthew Bashton,Greg R Young,Joshua Loh,John Allan,Mohammad A Tariq,Giles S Holt,Gary Black,Wen C Yew,Lynn Dover,Paul Baker,Steve Liggett,Sarah Essex,Jane Greenaway,Debra Padgett,Clive Graham,Garren Scott,Edward Barton,Emma Swindells,Brendan Payne,Jennifer Collins,Yusri Taha,Gary Eltringham |
| EPI_ISL_3078161 | Presidio Ospedaliero "Madonna delle Grazie" di Matera | ASM - P.O. Madonna delle Grazie - Matera | Massimo Dell'Edera |
| EPI_ISL_3078162 | Northumbria University / South Tees Hospitals NHS Foundation Trust / North Cumbria Integrated Care NHS Foundation Trust / North Tees and Hartlepool NHS Foundation Trust / Newcastle Hospitals NHS Foundation Trust | COVID-19 Genomics UK (COG-UK) Consortium | Darren L Smith,Andrew Nelson,Matthew Bashton,Greg R Young,Joshua Loh,John Allan,Mohammad A Tariq,Giles S Holt,Gary Black,Wen C Yew,Lynn Dover,Paul Baker,Steve Liggett,Sarah Essex,Jane Greenaway,Debra Padgett,Clive Graham,Garren Scott,Edward Barton,Emma Swindells,Brendan Payne,Jennifer Collins,Yusri Taha,Gary Eltringham |
| EPI_ISL_3078163 | Presidio Ospedaliero "Madonna delle Grazie" di Matera | ASM - P.O. Madonna delle Grazie - Matera | Massimo Dell'Edera |
| EPI_ISL_3078164 | Northumbria University / South Tees Hospitals NHS Foundation Trust / North Cumbria Integrated Care NHS Foundation Trust / North Tees and Hartlepool NHS Foundation Trust / Newcastle Hospitals NHS Foundation Trust | COVID-19 Genomics UK (COG-UK) Consortium | Darren L Smith,Andrew Nelson,Matthew Bashton,Greg R Young,Joshua Loh,John Allan,Mohammad A Tariq,Giles S Holt,Gary Black,Wen C Yew,Lynn Dover,Paul Baker,Steve Liggett,Sarah Essex,Jane Greenaway,Debra Padgett,Clive Graham,Garren Scott,Edward Barton,Emma Swindells,Brendan Payne,Jennifer Collins,Yusri Taha,Gary Eltringham |
| EPI_ISL_3078165 | Presidio Ospedaliero "Madonna delle Grazie" di Matera | ASM - P.O. Madonna delle Grazie - Matera | Massimo Dell'Edera |
| EPI_ISL_3078166, EPI_ISL_3078168 | Northumbria University / South Tees Hospitals NHS Foundation Trust / North Cumbria Integrated Care NHS Foundation Trust / | COVID-19 Genomics UK (COG-UK) Consortium | Darren L Smith,Andrew Nelson,Matthew Bashton,Greg R Young,Joshua Loh,John Allan,Mohammad A Tariq,Giles S Holt,Gary Black,Wen C Yew,Lynn Dover,Paul Baker,Steve Liggett,Sarah Essex,Jane Greenaway,Debra Padgett,Clive Graham,Garren Scott,Edward Barton,Emma Swindells,Brendan Payne,Jennifer Collins,Yusri Taha,Gary Eltringham |

| | | | |
|---|---|---|---|
| | North Tees and Hartlepool NHS Foundation Trust / Newcastle Hospitals NHS Foundation Trust | | |
| EPI_ISL_3078169 | Presidio Ospedaliero "Madonna delle Grazie" di Matera | ASM - P.O. Madonna delle Grazie - Matera | Massimo Dell'Edera |

EPI_ISL_3078170, EPI_ISL_3078171, EPI_ISL_3078174, EPI_ISL_3078175, EPI_ISL_3078176, EPI_ISL_3078177, EPI_ISL_3078178, EPI_ISL_3078179, EPI_ISL_3078180, EPI_ISL_3078181, EPI_ISL_3078182, EPI_ISL_3078183, EPI_ISL_3078184, EPI_ISL_3078185, EPI_ISL_3078186, EPI_ISL_3078187, EPI_ISL_3078188, EPI_ISL_3078189, EPI_ISL_3078190, EPI_ISL_3078191, EPI_ISL_3078192, EPI_ISL_3078193, EPI_ISL_3078194, EPI_ISL_3078195, EPI_ISL_3078196, EPI_ISL_3078197, EPI_ISL_3078198, EPI_ISL_3078199, EPI_ISL_3078200, EPI_ISL_3078201, EPI_ISL_3078202, EPI_ISL_3078203, EPI_ISL_3078204, EPI_ISL_3078205, EPI_ISL_3078206, EPI_ISL_3078207, EPI_ISL_3078208, EPI_ISL_3078209, EPI_ISL_3078210, EPI_ISL_3078211, EPI_ISL_3078212, EPI_ISL_3078213, EPI_ISL_3078214, EPI_ISL_3078215, EPI_ISL_3078216, EPI_ISL_3078217, EPI_ISL_3078218, EPI_ISL_3078219, EPI_ISL_3078220, EPI_ISL_3078221, EPI_ISL_3078222, EPI_ISL_3078223, EPI_ISL_3078224, EPI_ISL_3078225, EPI_ISL_3078226, EPI_ISL_3078227, EPI_ISL_3078228, EPI_ISL_3078229, EPI_ISL_3078230, EPI_ISL_3078231, EPI_ISL_3078232, EPI_ISL_3078233, EPI_ISL_3078234, EPI_ISL_3078235, EPI_ISL_3078236, EPI_ISL_3078237, EPI_ISL_3078238, EPI_ISL_3078239, EPI_ISL_3078240, EPI_ISL_3078241, EPI_ISL_3078242, EPI_ISL_3078243, EPI_ISL_3078244, EPI_ISL_3078245, EPI_ISL_3078246, EPI_ISL_3078247, EPI_ISL_3078248, EPI_ISL_3078249, EPI_ISL_3078250, EPI_ISL_3078251, EPI_ISL_3078252, EPI_ISL_3078253, EPI_ISL_3078254, EPI_ISL_3078255, EPI_ISL_3078256, EPI_ISL_3078257, EPI_ISL_3078258, EPI_ISL_3078259, EPI_ISL_3078260, EPI_ISL_3078261, EPI_ISL_3078262, EPI_ISL_3078263, EPI_ISL_3078264, EPI_ISL_3078265, EPI_ISL_3078266, EPI_ISL_3078267, EPI_ISL_3078268, EPI_ISL_3078269, EPI_ISL_3078270, EPI_ISL_3078271, EPI_ISL_3078272, EPI_ISL_3078273, EPI_ISL_3078274, EPI_ISL_3078275, EPI_ISL_3078276, EPI_ISL_3078277, EPI_ISL_3078278, EPI_ISL_3078279, EPI_ISL_3078280, EPI_ISL_3078281, EPI_ISL_3078282, EPI_ISL_3078283, EPI_ISL_3078284, EPI_ISL_3078285, EPI_ISL_3078286, EPI_ISL_3078287, EPI_ISL_3078288, EPI_ISL_3078289, EPI_ISL_3078290, EPI_ISL_3078291, EPI_ISL_3078292, EPI_ISL_3078293, EPI_ISL_3078294, EPI_ISL_3078295, EPI_ISL_3078296, EPI_ISL_3078297, EPI_ISL_3078298, EPI_ISL_3078299, EPI_ISL_3078300, EPI_ISL_3078301, EPI_ISL_3078302, EPI_ISL_3078303, EPI_ISL_3078304, EPI_ISL_3078305, EPI_ISL_3078306, EPI_ISL_3078307, EPI_ISL_3078308, EPI_ISL_3078309, EPI_ISL_3078310, EPI_ISL_3078311, EPI_ISL_3078312, EPI_ISL_3078313, EPI_ISL_3078314, EPI_ISL_3078315, EPI_ISL_3078316, EPI_ISL_3078317, EPI_ISL_3078318, EPI_ISL_3078319, EPI_ISL_3078320, EPI_ISL_3078321, EPI_ISL_3078322, EPI_ISL_3078323, EPI_ISL_3078324, EPI_ISL_3078325, EPI_ISL_3078326, EPI_ISL_3078327, EPI_ISL_3078328, EPI_ISL_3078329, EPI_ISL_3078330, EPI_ISL_3078331, EPI_ISL_3078332, EPI_ISL_3078333, EPI_ISL_3078334, EPI_ISL_3078335, EPI_ISL_3078336, EPI_ISL_3078337, EPI_ISL_3078338, EPI_ISL_3078339, EPI_ISL_3078340, EPI_ISL_3078341, EPI_ISL_3078342, EPI_ISL_3078343, EPI_ISL_3078344, EPI_ISL_3078345, EPI_ISL_3078346, EPI_ISL_3078347, EPI_ISL_3078348, EPI_ISL_3078349, EPI_ISL_3078350, EPI_ISL_3078351, EPI_ISL_3078352, EPI_ISL_3078353, EPI_ISL_3078354, EPI_ISL_3078355, EPI_ISL_3078356, EPI_ISL_3078357, EPI_ISL_3078358, EPI_ISL_3078359, EPI_ISL_3078360, EPI_ISL_3078361, EPI_ISL_3078362, EPI_ISL_3078363, EPI_ISL_3078364, EPI_ISL_3078365, EPI_ISL_3078366, EPI_ISL_3078367, EPI_ISL_3078368, EPI_ISL_3078369, EPI_ISL_3078370, EPI_ISL_3078371, EPI_ISL_3078372, EPI_ISL_3078373, EPI_ISL_3078374, EPI_ISL_3078375, EPI_ISL_3078376, EPI_ISL_3078377, EPI_ISL_3078378, EPI_ISL_3078379, EPI_ISL_3078380, EPI_ISL_3078381, EPI_ISL_3078382, EPI_ISL_3078383, EPI_ISL_3078384, EPI_ISL_3078385, EPI_ISL_3078386, EPI_ISL_3078387, EPI_ISL_3078388, EPI_ISL_3078389, EPI_ISL_3078390, EPI_ISL_3078391, EPI_ISL_3078392, EPI_ISL_3078393, EPI_ISL_3078394, EPI_ISL_3078395, EPI_ISL_3078396, EPI_ISL_3078397, EPI_ISL_3078398, EPI_ISL_3078399, EPI_ISL_3078400, EPI_ISL_3078401, EPI_ISL_3078402, EPI_ISL_3078403, EPI_ISL_3078404, EPI_ISL_3078405, EPI_ISL_3078406, EPI_ISL_3078407, EPI_ISL_3078408, EPI_ISL_3078409, EPI_ISL_3078410, EPI_ISL_3078411, EPI_ISL_3078412, EPI_ISL_3078413, EPI_ISL_3078414, EPI_ISL_3078415, EPI_ISL_3078416, EPI_ISL_3078417, EPI_ISL_3078418, EPI_ISL_3078419, EPI_ISL_3078420, EPI_ISL_3078421, EPI_ISL_3078422, EPI_ISL_3078423, EPI_ISL_3078424, EPI_ISL_3078425, EPI_ISL_3078426, EPI_ISL_3078427, EPI_ISL_3078428, EPI_ISL_3078429, EPI_ISL_3078430, EPI_ISL_3078431, EPI_ISL_3078432, EPI_ISL_3078433, EPI_ISL_3078434, EPI_ISL_3078435, EPI_ISL_3078436, EPI_ISL_3078437, EPI_ISL_3078438, EPI_ISL_3078439, EPI_ISL_3078440, EPI_ISL_3078441, EPI_ISL_3078442, EPI_ISL_3078443, EPI_ISL_3078444, EPI_ISL_3078445, EPI_ISL_3078446, EPI_ISL_3078447, EPI_ISL_3078448, EPI_ISL_3078449, EPI_ISL_3078450, EPI_ISL_3078451, EPI_ISL_3078452, EPI_ISL_3078453, EPI_ISL_3078454, EPI_ISL_3078455, EPI_ISL_3078456, EPI_ISL_3078457, EPI_ISL_3078458, EPI_ISL_3078459, EPI_ISL_3078460, EPI_ISL_3078461, EPI_ISL_3078462, EPI_ISL_3078463, EPI_ISL_3078464, EPI_ISL_3078465, EPI_ISL_3078466, EPI_ISL_3078467, EPI_ISL_3078468, EPI_ISL_3078469, EPI_ISL_3078470, EPI_ISL_3078471, EPI_ISL_3078472, EPI_ISL_3078473, EPI_ISL_3078474, EPI_ISL_3078475, EPI_ISL_3078476, EPI_ISL_3078477, EPI_ISL_3078478, EPI_ISL_3078479, EPI_ISL_3078480, EPI_ISL_3078481, EPI_ISL_3078482, EPI_ISL_3078483, EPI_ISL_3078484, EPI_ISL_3078485, EPI_ISL_3078486, EPI_ISL_3078487, EPI_ISL_3078488, EPI_ISL_3078489, EPI_ISL_3078490, EPI_ISL_3078491, EPI_ISL_3078492, EPI_ISL_3078493, EPI_ISL_3078494, EPI_ISL_3078495, EPI_ISL_3078496, EPI_ISL_3078497, EPI_ISL_3078498, EPI_ISL_3078499, EPI_ISL_3078500, EPI_ISL_3078501, EPI_ISL_3078502

| | | | |
|---|---|---|---|
| see above | Northumbria University / South Tees Hospitals NHS Foundation Trust / North Cumbria Integrated Care NHS Foundation Trust / North Tees and Hartlepool NHS Foundation Trust / Newcastle Hospitals NHS Foundation Trust | COVID-19 Genomics UK (COG-UK) Consortium | Darren L Smith,Andrew Nelson,Matthew Bashton,Greg R Young,Joshua Loh,John Allan,Mohammad A Tariq,Giles S Holt,Gary Black,Wen C Yew,Lynn Dover,Paul Baker,Steve Liggett,Sarah Essex,Jane Greenaway,Debra Padgett,Clive Graham,Garren Scott,Edward Barton,Emma Swindells,Brendan Payne,Jennifer Collins,Yusri Taha,Gary Eltringham |

EPI_ISL_3078503, EPI_ISL_3078504, EPI_ISL_3078505, EPI_ISL_3078506, EPI_ISL_3078507, EPI_ISL_3078508, EPI_ISL_3078509, EPI_ISL_3078510, EPI_ISL_3078511, EPI_ISL_3078512, EPI_ISL_3078513, EPI_ISL_3078514, EPI_ISL_3078515, EPI_ISL_3078516, EPI_ISL_3078517, EPI_ISL_3078518, EPI_ISL_3078519, EPI_ISL_3078520, EPI_ISL_3078521, EPI_ISL_3078522, EPI_ISL_3078523, EPI_ISL_3078524, EPI_ISL_3078525, EPI_ISL_3078526, EPI_ISL_3078527, EPI_ISL_3078528, EPI_ISL_3078529, EPI_ISL_3078530, EPI_ISL_3078531, EPI_ISL_3078532, EPI_ISL_3078533, EPI_ISL_3078534, EPI_ISL_3078535, EPI_ISL_3078536, EPI_ISL_3078537, EPI_ISL_3078538, EPI_ISL_3078539, EPI_ISL_3078540, EPI_ISL_3078541, EPI_ISL_3078542, EPI_ISL_3078543, EPI_ISL_3078544, EPI_ISL_3078545, EPI_ISL_3078546, EPI_ISL_3078547, EPI_ISL_3078548, EPI_ISL_3078549, EPI_ISL_3078550, EPI_ISL_3078551, EPI_ISL_3078552, EPI_ISL_3078553, EPI_ISL_3078554, EPI_ISL_3078555, EPI_ISL_3078556, EPI_ISL_3078557, EPI_ISL_3078558, EPI_ISL_3078559, EPI_ISL_3078560, EPI_ISL_3078561, EPI_ISL_3078562, EPI_ISL_3078563, EPI_ISL_3078564, EPI_ISL_3078565, EPI_ISL_3078566, EPI_ISL_3078567, EPI_ISL_3078568, EPI_ISL_3078569, EPI_ISL_3078570, EPI_ISL_3078571, EPI_ISL_3078572, EPI_ISL_3078573, EPI_ISL_3078574, EPI_ISL_3078575, EPI_ISL_3078576, EPI_ISL_3078577, EPI_ISL_3078578, EPI_ISL_3078579, EPI_ISL_3078580, EPI_ISL_3078581, EPI_ISL_3078582, EPI_ISL_3078583, EPI_ISL_3078584, EPI_ISL_3078585, EPI_ISL_3078586, EPI_ISL_3078587, EPI_ISL_3078588, EPI_ISL_3078589, EPI_ISL_3078590, EPI_ISL_3078591, EPI_ISL_3078592, EPI_ISL_3078593, EPI_ISL_3078594, EPI_ISL_3078595, EPI_ISL_3078596, EPI_ISL_3078597, EPI_ISL_3078598, EPI_ISL_3078599, EPI_ISL_3078600, EPI_ISL_3078601, EPI_ISL_3078602, EPI_ISL_3078603, EPI_ISL_3078604, EPI_ISL_3078605, EPI_ISL_3078606, EPI_ISL_3078607, EPI_ISL_3078608, EPI_ISL_3078609, EPI_ISL_3078610, EPI_ISL_3078611, EPI_ISL_3078612, EPI_ISL_3078613, EPI_ISL_3078614, EPI_ISL_3078615, EPI_ISL_3078616, EPI_ISL_3078617, EPI_ISL_3078618, EPI_ISL_3078619, EPI_ISL_3078620, EPI_ISL_3078621, EPI_ISL_3078622, EPI_ISL_3078623, EPI_ISL_3078624, EPI_ISL_3078625, EPI_ISL_3078626, EPI_ISL_3078627, EPI_ISL_3078628, EPI_ISL_3078629, EPI_ISL_3078630, EPI_ISL_3078631, EPI_ISL_3078632, EPI_ISL_3078633, EPI_ISL_3078634, EPI_ISL_3078635, EPI_ISL_3078636, EPI_ISL_3078637, EPI_ISL_3078638, EPI_ISL_3078639, EPI_ISL_3078640, EPI_ISL_3078641

| | | | |
|---|---|---|---|
| see above | Quadram Institute Bioscience | COVID-19 Genomics UK (COG-UK) Consortium | Dave J. Baker, Gemma L. Kay, Alp Aydin, Thanh Le-Viet, Steven Rudder, Ana P. Tedim, Anastasia Kolyva, Maria Diaz, Leonardo de Oliveira Martins, Nabil-Fareed Alikhan, Lizzie Meadows, Rachael Stanley, Rose K Davidson, Ngozi Elumogo, Muhammed Yasir, Sophie J. Prosolek, Ebenezer Foster-Nyarko, Martin Lott, Christopher Jeanes, Lewis G. Spurgin, Nicholas M. Thomson, Alexander J Trotter, Rachel Gilroy, Luke Bedford, Samuel Bloomfield, Claire Stuart, Andrew Bell, Reenesh Prakash, Lindsay Coupland, Samir Dervisevic, Alison E. Mather, John Wain, Mark Webber, Andrew J. Page, Justin O'Grady |

EPI_ISL_3078642, EPI_ISL_3078643, EPI_ISL_3078644, EPI_ISL_3078645, EPI_ISL_3078646, EPI_ISL_3078647, EPI_ISL_3078648, EPI_ISL_3078649, EPI_ISL_3078650, EPI_ISL_3078651, EPI_ISL_3078652, EPI_ISL_3078653, EPI_ISL_3078654, EPI_ISL_3078655, EPI_ISL_3078656, EPI_ISL_3078657, EPI_ISL_3078658, EPI_ISL_3078659, EPI_ISL_3078660, EPI_ISL_3078661, EPI_ISL_3078662, EPI_ISL_3078663, EPI_ISL_3078664, EPI_ISL_3078665, EPI_ISL_3078666, EPI_ISL_3078667, EPI_ISL_3078668, EPI_ISL_3078669, EPI_ISL_3078670, EPI_ISL_3078671, EPI_ISL_3078672, EPI_ISL_3078673, EPI_ISL_3078674, EPI_ISL_3078675, EPI_ISL_3078676, EPI_ISL_3078677, EPI_ISL_3078678, EPI_ISL_3078679, EPI_ISL_3078680, EPI_ISL_3078681, EPI_ISL_3078682, EPI_ISL_3078683, EPI_ISL_3078684, EPI_ISL_3078685, EPI_ISL_3078686, EPI_ISL_3078687, EPI_ISL_3078688, EPI_ISL_3078689, EPI_ISL_3078690, EPI_ISL_3078691, EPI_ISL_3078692, EPI_ISL_3078693, EPI_ISL_3078694, EPI_ISL_3078695, EPI_ISL_3078696, EPI_ISL_3078697, EPI_ISL_3078698, EPI_ISL_3078699, EPI_ISL_3078700, EPI_ISL_3078701, EPI_ISL_3078702, EPI_ISL_3078703, EPI_ISL_3078704, EPI_ISL_3078705, EPI_ISL_3078706, EPI_ISL_3078707, EPI_ISL_3078708, EPI_ISL_3078709, EPI_ISL_3078710, EPI_ISL_3078711, EPI_ISL_3078712, EPI_ISL_3078713, EPI_ISL_3078714, EPI_ISL_3078715, EPI_ISL_3078716, EPI_ISL_3078717, EPI_ISL_3078718, EPI_ISL_3078719, EPI_ISL_3078720, EPI_ISL_3078721, EPI_ISL_3078722, EPI_ISL_3078723, EPI_ISL_3078724, EPI_ISL_3078725, EPI_ISL_3078726, EPI_ISL_3078727, EPI_ISL_3078728, EPI_ISL_3078729, EPI_ISL_3078730, EPI_ISL_3078731, EPI_ISL_3078732, EPI_ISL_3078733, EPI_ISL_3078734, EPI_ISL_3078735, EPI_ISL_3078736, EPI_ISL_3078737, EPI_ISL_3078738, EPI_ISL_3078739, EPI_ISL_3078740, EPI_ISL_3078741, EPI_ISL_3078742, EPI_ISL_3078743, EPI_ISL_3078744, EPI_ISL_3078745, EPI_ISL_3078746, EPI_ISL_3078747, EPI_ISL_3078748, EPI_ISL_3078749, EPI_ISL_3078750, EPI_ISL_3078751, EPI_ISL_3078752, EPI_ISL_3078753, EPI_ISL_3078754, EPI_ISL_3078755, EPI_ISL_3078756, EPI_ISL_3078757, EPI_ISL_3078758, EPI_ISL_3078759, EPI_ISL_3078760, EPI_ISL_3078761, EPI_ISL_3078762, EPI_ISL_3078763, EPI_ISL_3078764, EPI_ISL_3078765, EPI_ISL_3078766, EPI_ISL_3078767, EPI_ISL_3078768, EPI_ISL_3078769, EPI_ISL_3078770, EPI_ISL_3078771, EPI_ISL_3078772, EPI_ISL_3078773, EPI_ISL_3078774, EPI_ISL_3078775, EPI_ISL_3078776, EPI_ISL_3078777, EPI_ISL_3078778, EPI_ISL_3078779, EPI_ISL_3078780, EPI_ISL_3078781, EPI_ISL_3078782, EPI_ISL_3078783, EPI_ISL_3078784, EPI_ISL_3078785, EPI_ISL_3078786, EPI_ISL_3078787, EPI_ISL_3078788, EPI_ISL_3078789, EPI_ISL_3078790, EPI_ISL_3078791, EPI_ISL_3078792, EPI_ISL_3078793, EPI_ISL_3078794, EPI_ISL_3078795, EPI_ISL_3078796, EPI_ISL_3078797, EPI_ISL_3078798, EPI_ISL_3078799, EPI_ISL_3078800, EPI_ISL_3078801, EPI_ISL_3078802, EPI_ISL_3078803, EPI_ISL_3078804, EPI_ISL_3078805, EPI_ISL_3078806, EPI_ISL_3078807, EPI_ISL_3078808, EPI_ISL_3078809, EPI_ISL_3078810, EPI_ISL_3078811, EPI_ISL_3078812, EPI_ISL_3078813, EPI_ISL_3078814, EPI_ISL_3078815

| | | | |
|---|---|---|---|
| see above | Queens Medical Centre, Clinical Microbiology Department / DeepSeq Nottingham | COVID-19 Genomics UK (COG-UK) Consortium | Gemma Clark, Wendy Smith, Manjinder Khakh, Vicki M Fleming, Michelle M Lister, Hannah Howson-Wells, Jonathan Ball, Timothy Byaruhanga, Jayasree Dey, Emily Park, Jack Hill, Patrick McClure, Joseph Chappell, Theocharis Tsoleridis, Nadine Holmes, Matthew Carlisle, Christopher Moore, Fei Sang, Johnny Debebe, Victoria Wright, Matthew Loose |

EPI_ISL_3078816, EPI_ISL_3078817, EPI_ISL_3078818, EPI_ISL_3078820, EPI_ISL_3078821, EPI_ISL_3078822, EPI_ISL_3078823, EPI_ISL_3078825, EPI_ISL_3078827, EPI_ISL_3078828, EPI_ISL_3078829, EPI_ISL_3078830, EPI_ISL_3078831, EPI_ISL_3078833, EPI_ISL_3078835, EPI_ISL_3078836, EPI_ISL_3078837, EPI_ISL_3078838, EPI_ISL_3078839, EPI_ISL_3078840, EPI_ISL_3078841, EPI_ISL_3078843, EPI_ISL_3078845, EPI_ISL_3078846, EPI_ISL_3078847, EPI_ISL_3078848, EPI_ISL_3078849, EPI_ISL_3078850, EPI_ISL_3078851, EPI_ISL_3078852, EPI_ISL_3078854, EPI_ISL_3078855, EPI_ISL_3078856, EPI_ISL_3078858, EPI_ISL_3078859, EPI_ISL_3078861, EPI_ISL_3078862, EPI_ISL_3078863, EPI_ISL_3078865, EPI_ISL_3078867, EPI_ISL_3078869, EPI_ISL_3078870, EPI_ISL_3078874, EPI_ISL_3078876, EPI_ISL_3078877, EPI_ISL_3078878, EPI_ISL_3078881, EPI_ISL_3078882, EPI_ISL_3078883, EPI_ISL_3078888, EPI_ISL_3078891, EPI_ISL_3078892, EPI_ISL_3078893, EPI_ISL_3078895, EPI_ISL_3078899, EPI_ISL_3078902, EPI_ISL_3078903, EPI_ISL_3078906, EPI_ISL_3078907, EPI_ISL_3078908, EPI_ISL_3078910, EPI_ISL_3078911, EPI_ISL_3078914, EPI_ISL_3078915, EPI_ISL_3078916, EPI_ISL_3078918, EPI_ISL_3078919, EPI_ISL_3078992, EPI_ISL_3079008, EPI_ISL_3079010, EPI_ISL_3079011, EPI_ISL_3079012, EPI_ISL_3079013, EPI_ISL_3079014, EPI_ISL_3079015, EPI_ISL_3079016, EPI_ISL_3079017, EPI_ISL_3079019, EPI_ISL_3079020, EPI_ISL_3079022, EPI_ISL_3079025, EPI_ISL_3079026, EPI_ISL_3079027, EPI_ISL_3079028, EPI_ISL_3079029, EPI_ISL_3079031, EPI_ISL_3079033, EPI_ISL_3079034, EPI_ISL_3079035, EPI_ISL_3079036, EPI_ISL_3079038, EPI_ISL_3079040, EPI_ISL_3079041, EPI_ISL_3079042, EPI_ISL_3079044, EPI_ISL_3079045, EPI_ISL_3079046, EPI_ISL_3079047, EPI_ISL_3079048, EPI_ISL_3079050, EPI_ISL_3079051, EPI_ISL_3079053, EPI_ISL_3079054, EPI_ISL_3079055, EPI_ISL_3079056, EPI_ISL_3079058, EPI_ISL_3079059, EPI_ISL_3079060, EPI_ISL_3079061, EPI_ISL_3079063, EPI_ISL_3079066, EPI_ISL_3079067, EPI_ISL_3079068, EPI_ISL_3079069, EPI_ISL_3079070, EPI_ISL_3079071, EPI_ISL_3079072, EPI_ISL_3079073, EPI_ISL_3079074, EPI_ISL_3079075, EPI_ISL_3079076, EPI_ISL_3079078, EPI_ISL_3079079, EPI_ISL_3079080, EPI_ISL_3079083, EPI_ISL_3079085, EPI_ISL_3079086, EPI_ISL_3079087, EPI_ISL_3079089, EPI_ISL_3079090, EPI_ISL_3079091, EPI_ISL_3079092, EPI_ISL_3079093, EPI_ISL_3079095, EPI_ISL_3079096, EPI_ISL_3079097, EPI_ISL_3079099, EPI_ISL_3079100, EPI_ISL_3079101, EPI_ISL_3079103, EPI_ISL_3079104, EPI_ISL_3079105, EPI_ISL_3079106, EPI_ISL_3079108, EPI_ISL_3079109, EPI_ISL_3079110, EPI_ISL_3079111, EPI_ISL_3079112, EPI_ISL_3079113, EPI_ISL_3079114, EPI_ISL_3079116, EPI_ISL_3079117, EPI_ISL_3079118, EPI_ISL_3079119, EPI_ISL_3079121, EPI_ISL_3079122, EPI_ISL_3079123, EPI_ISL_3079124, EPI_ISL_3079125, EPI_ISL_3079126, EPI_ISL_3079127, EPI_ISL_3079128, EPI_ISL_3079129, EPI_ISL_3079131, EPI_ISL_3079132, EPI_ISL_3079133, EPI_ISL_3079134, EPI_ISL_3079135, EPI_ISL_3079136, EPI_ISL_3079138, EPI_ISL_3079139, EPI_ISL_3079140, EPI_ISL_3079141, EPI_ISL_3079142, EPI_ISL_3079143, EPI_ISL_3079144, EPI_ISL_3079145, EPI_ISL_3079146, EPI_ISL_3079147, EPI_ISL_3079148, EPI_ISL_3079149, EPI_ISL_3079150, EPI_ISL_3079151, EPI_ISL_3079152, EPI_ISL_3079153, EPI_ISL_3079154, EPI_ISL_3079155, EPI_ISL_3079156, EPI_ISL_3079157, EPI_ISL_3079158, EPI_ISL_3079159, EPI_ISL_3079160, EPI_ISL_3079161, EPI_ISL_3079162, EPI_ISL_3079163, EPI_ISL_3079164, EPI_ISL_3079165, EPI_ISL_3079166, EPI_ISL_3079167, EPI_ISL_3079169, EPI_ISL_3079170, EPI_ISL_3079171, EPI_ISL_3079172, EPI_ISL_3079173, EPI_ISL_3079175, EPI_ISL_3079176, EPI_ISL_3079177, EPI_ISL_3079182, EPI_ISL_3079183, EPI_ISL_3079184, EPI_ISL_3079187, EPI_ISL_3079188, EPI_ISL_3079189, EPI_ISL_3079191, EPI_ISL_3079192, EPI_ISL_3079193, EPI_ISL_3079194, EPI_ISL_3079195, EPI_ISL_3079196, EPI_ISL_3079197, EPI_ISL_3079199, EPI_ISL_3079200, EPI_ISL_3079201, EPI_ISL_3079202, EPI_ISL_3079203, EPI_ISL_3079205, EPI_ISL_3079206, EPI_ISL_3079207, EPI_ISL_3079208, EPI_ISL_3079209, EPI_ISL_3079210, EPI_ISL_3079211, EPI_ISL_3079213, EPI_ISL_3079214, EPI_ISL_3079215, EPI_ISL_3079217, EPI_ISL_3079218, EPI_ISL_3079219, EPI_ISL_3079220, EPI_ISL_3079221, EPI_ISL_3079223, EPI_ISL_3079224, EPI_ISL_3079225, EPI_ISL_3079227, EPI_ISL_3079229, EPI_ISL_3079230, EPI_ISL_3079231, EPI_ISL_3079232, EPI_ISL_3079234, EPI_ISL_3079235, EPI_ISL_3079236, EPI_ISL_3079237, EPI_ISL_3079242, EPI_ISL_3079243, EPI_ISL_3079244, EPI_ISL_3079245, EPI_ISL_3079246, EPI_ISL_3079248, EPI_ISL_3079249, EPI_ISL_3079250, EPI_ISL_3079251, EPI_ISL_3079252, EPI_ISL_3079253, EPI_ISL_3079254, EPI_ISL_3079255, EPI_ISL_3079258, EPI_ISL_3079259, EPI_ISL_3079260, EPI_ISL_3079262, EPI_ISL_3079264, EPI_ISL_3079265, EPI_ISL_3079266

| | | | |
|---|---|---|---|
| see above | Oxford Viromics, NDM, University of Oxford; Oxford University Hospitals; Basingstoke and North Hampshire Hospital | COVID-19 Genomics UK (COG-UK) Consortium | Tanya Golubchik, David Bonsall, George Macintyre, Amy Trebes, Mariateresa de Cesare, Catrin Moore, Alex Mobbs, Anita Justice, Robert Shaw, Monique Andersson, Timothy Peto, Emma Wise, Nathan Moore, Jessica Lynch, Nick Cortes, Matilde Mori, Stephen Kidd, David Buck, John Todd, Christophe Fraser |

EPI_ISL_3079267, EPI_ISL_3079268, EPI_ISL_3079269, EPI_ISL_3079270, EPI_ISL_3079272, EPI_ISL_3079273, EPI_ISL_3079274, EPI_ISL_3079275, EPI_ISL_3079276, EPI_ISL_3079277, EPI_ISL_3079278, EPI_ISL_3079279, EPI_ISL_3079280, EPI_ISL_3079281, EPI_ISL_3079282, EPI_ISL_3079283, EPI_ISL_3079284, EPI_ISL_3079285, EPI_ISL_3079286, EPI_ISL_3079287, EPI_ISL_3079288, EPI_ISL_3079289, EPI_ISL_3079290, EPI_ISL_3079291, EPI_ISL_3079292, EPI_ISL_3079293, EPI_ISL_3079294, EPI_ISL_3079295, EPI_ISL_3079296, EPI_ISL_3079297, EPI_ISL_3079298, EPI_ISL_3079299, EPI_ISL_3079300, EPI_ISL_3079301, EPI_ISL_3079302, EPI_ISL_3079303, EPI_ISL_3079304, EPI_ISL_3079305, EPI_ISL_3079306, EPI_ISL_3079307, EPI_ISL_3079308, EPI_ISL_3079309, EPI_ISL_3079310, EPI_ISL_3079311, EPI_ISL_3079312, EPI_ISL_3079313, EPI_ISL_3079314, EPI_ISL_3079315, EPI_ISL_3079316, EPI_ISL_3079317, EPI_ISL_3079318, EPI_ISL_3079319, EPI_ISL_3079320, EPI_ISL_3079321, EPI_ISL_3079322, EPI_ISL_3079323, EPI_ISL_3079324, EPI_ISL_3079325, EPI_ISL_3079326, EPI_ISL_3079327, EPI_ISL_3079328, EPI_ISL_3079329, EPI_ISL_3079330, EPI_ISL_3079331, EPI_ISL_3079332, EPI_ISL_3079333, EPI_ISL_3079334, EPI_ISL_3079335, EPI_ISL_3079336, EPI_ISL_3079337, EPI_ISL_3079338, EPI_ISL_3079339, EPI_ISL_3079340, EPI_ISL_3079341, EPI_ISL_3079342, EPI_ISL_3079343, EPI_ISL_3079344, EPI_ISL_3079345, EPI_ISL_3079346, EPI_ISL_3079347, EPI_ISL_3079348, EPI_ISL_3079349, EPI_ISL_3079350, EPI_ISL_3079351, EPI_ISL_3079352, EPI_ISL_3079353, EPI_ISL_3079354, EPI_ISL_3079355, EPI_ISL_3079356, EPI_ISL_3079357, EPI_ISL_3079358, EPI_ISL_3079359, EPI_ISL_3079360, EPI_ISL_3079361, EPI_ISL_3079362, EPI_ISL_3079363, EPI_ISL_3079364, EPI_ISL_3079365, EPI_ISL_3079366, EPI_ISL_3079367, EPI_ISL_3079368, EPI_ISL_3079369, EPI_ISL_3079370, EPI_ISL_3079371, EPI_ISL_3079372, EPI_ISL_3079373, EPI_ISL_3079374, EPI_ISL_3079375, EPI_ISL_3079376, EPI_ISL_3079377, EPI_ISL_3079378, EPI_ISL_3079379, EPI_ISL_3079380, EPI_ISL_3079382, EPI_ISL_3079383, EPI_ISL_3079384, EPI_ISL_3079385, EPI_ISL_3079386, EPI_ISL_3079387, EPI_ISL_3079388, EPI_ISL_3079389, EPI_ISL_3079390, EPI_ISL_3079391, EPI_ISL_3079392, EPI_ISL_3079393, EPI_ISL_3079394, EPI_ISL_3079395, EPI_ISL_3079396, EPI_ISL_3079397, EPI_ISL_3079398, EPI_ISL_3079399, EPI_ISL_3079400, EPI_ISL_3079401, EPI_ISL_3079402, EPI_ISL_3079403, EPI_ISL_3079404, EPI_ISL_3079405, EPI_ISL_3079406, EPI_ISL_3079407, EPI_ISL_3079408, EPI_ISL_3079409, EPI_ISL_3079410, EPI_ISL_3079411, EPI_ISL_3079412, EPI_ISL_3079413, EPI_ISL_3079414, EPI_ISL_3079415, EPI_ISL_3079416, EPI_ISL_3079417, EPI_ISL_3079418, EPI_ISL_3079419, EPI_ISL_3079420, EPI_ISL_3079421, EPI_ISL_3079422, EPI_ISL_3079423, EPI_ISL_3079424, EPI_ISL_3079425, EPI_ISL_3079426, EPI_ISL_3079427, EPI_ISL_3079428, EPI_ISL_3079429, EPI_ISL_3079430, EPI_ISL_3079431, EPI_ISL_3079432, EPI_ISL_3079433, EPI_ISL_3079434, EPI_ISL_3079435, EPI_ISL_3079436, EPI_ISL_3079437, EPI_ISL_3079438, EPI_ISL_3079439, EPI_ISL_3079440, EPI_ISL_3079441, EPI_ISL_3079442, EPI_ISL_3079443, EPI_ISL_3079444, EPI_ISL_3079445, EPI_ISL_3079446, EPI_ISL_3079447, EPI_ISL_3079448, EPI_ISL_3079449, EPI_ISL_3079450, EPI_ISL_3079451, EPI_ISL_3079452, EPI_ISL_3079453, EPI_ISL_3079454, EPI_ISL_3079455, EPI_ISL_3079456, EPI_ISL_3079457, EPI_ISL_3079458, EPI_ISL_3079459, EPI_ISL_3079460, EPI_ISL_3079461, EPI_ISL_3079462, EPI_ISL_3079463, EPI_ISL_3079464, EPI_ISL_3079465, EPI_ISL_3079466, EPI_ISL_3079467, EPI_ISL_3079468, EPI_ISL_3079469, EPI_ISL_3079470, EPI_ISL_3079471, EPI_ISL_3079472, EPI_ISL_3079473, EPI_ISL_3079474, EPI_ISL_3079475, EPI_ISL_3079476, EPI_ISL_3079477, EPI_ISL_3079478, EPI_ISL_3079479, EPI_ISL_3079480, EPI_ISL_3079481, EPI_ISL_3079482, EPI_ISL_3079483, EPI_ISL_3079484, EPI_ISL_3079485, EPI_ISL_3079486, EPI_ISL_3079487, EPI_ISL_3079488, EPI_ISL_3079489, EPI_ISL_3079490, EPI_ISL_3079491, EPI_ISL_3079492, EPI_ISL_3079493, EPI_ISL_3079494, EPI_ISL_3079495, EPI_ISL_3079496, EPI_ISL_3079497, EPI_ISL_3079498, EPI_ISL_3079499, EPI_ISL_3079500, EPI_ISL_3079501, EPI_ISL_3079502, EPI_ISL_3079503, EPI_ISL_3079504, EPI_ISL_3079505, EPI_ISL_3079506, EPI_ISL_3079507, EPI_ISL_3079508, EPI_ISL_3079509, EPI_ISL_3079510, EPI_ISL_3079511, EPI_ISL_3079512, EPI_ISL_3079513, EPI_ISL_3079514, EPI_ISL_3079515, EPI_ISL_3079516, EPI_ISL_3079517, EPI_ISL_3079518, EPI_ISL_3079519, EPI_ISL_3079520, EPI_ISL_3079521, EPI_ISL_3079522, EPI_ISL_3079523, EPI_ISL_3079524, EPI_ISL_3079525, EPI_ISL_3079526, EPI_ISL_3079527, EPI_ISL_3079528, EPI_ISL_3079529, EPI_ISL_3079530, EPI_ISL_3079531, EPI_ISL_3079532, EPI_ISL_3079533, EPI_ISL_3079534, EPI_ISL_3079535, EPI_ISL_3079536, EPI_ISL_3079537, EPI_ISL_3079538, EPI_ISL_3079539, EPI_ISL_3079540, EPI_ISL_3079541, EPI_ISL_3079542, EPI_ISL_3079543, EPI_ISL_3079544, EPI_ISL_3079545, EPI_ISL_3079546, EPI_ISL_3079547, EPI_ISL_3079548, EPI_ISL_3079549, EPI_ISL_3079550, EPI_ISL_3079551, EPI_ISL_3079552, EPI_ISL_3079553, EPI_ISL_3079554, EPI_ISL_3079555, EPI_ISL_3079556, EPI_ISL_3079557, EPI_ISL_3079558, EPI_ISL_3079559, EPI_ISL_3079560, EPI_ISL_3079561, EPI_ISL_3079562, EPI_ISL_3079563, EPI_ISL_3079564, EPI_ISL_3079565, EPI_ISL_3079566, EPI_ISL_3079567, EPI_ISL_3079568, EPI_ISL_3079569, EPI_ISL_3079570, EPI_ISL_3079571, EPI_ISL_3079572, EPI_ISL_3079573, EPI_ISL_3079574, EPI_ISL_3079575, EPI_ISL_3079576, EPI_ISL_3079577, EPI_ISL_3079578, EPI_ISL_3079579, EPI_ISL_3079580, EPI_ISL_3079581, EPI_ISL_3079582, EPI_ISL_3079583, EPI_ISL_3079584, EPI_ISL_3079585, EPI_ISL_3079586, EPI_ISL_3079587, EPI_ISL_3079588, EPI_ISL_3079589, EPI_ISL_3079590, EPI_ISL_3079591, EPI_ISL_3079592, EPI_ISL_3079593, EPI_ISL_3079594, EPI_ISL_3079595, EPI_ISL_3079596, EPI_ISL_3079597, EPI_ISL_3079598, EPI_ISL_3079599, EPI_ISL_3079600, EPI_ISL_3079601, EPI_ISL_3079602, EPI_ISL_3079603, EPI_ISL_3079604, EPI_ISL_3079605, EPI_ISL_3079606, EPI_ISL_3079607, EPI_ISL_3079608, EPI_ISL_3079609, EPI_ISL_3079610, EPI_ISL_3079611, EPI_ISL_3079612, EPI_ISL_3079613, EPI_ISL_3079614, EPI_ISL_3079615, EPI_ISL_3079616, EPI_ISL_3079617, EPI_ISL_3079619, EPI_ISL_3079620, EPI_ISL_3079621, EPI_ISL_3079622, EPI_ISL_3079623, EPI_ISL_3079624, EPI_ISL_3079625, EPI_ISL_3079626, EPI_ISL_3079627, EPI_ISL_3079628, EPI_ISL_3079629, EPI_ISL_3079630, EPI_ISL_3079631, EPI_ISL_3079632, EPI_ISL_3079633, EPI_ISL_3079634, EPI_ISL_3079635, EPI_ISL_3079636, EPI_ISL_3079637, EPI_ISL_3079638, EPI_ISL_3079639, EPI_ISL_3079640, EPI_ISL_3079641, EPI_ISL_3079642, EPI_ISL_3079643, EPI_ISL_3079644, EPI_ISL_3079645, EPI_ISL_3079646, EPI_ISL_3079647, EPI_ISL_3079648, EPI_ISL_3079649, EPI_ISL_3079650, EPI_ISL_3079651, EPI_ISL_3079652, EPI_ISL_3079653, EPI_ISL_3079654, EPI_ISL_3079655, EPI_ISL_3079656, EPI_ISL_3079657, EPI_ISL_3079658, EPI_ISL_3079659, EPI_ISL_3079661, EPI_ISL_3079662, EPI_ISL_3079663, EPI_ISL_3079665, EPI_ISL_3079666, EPI_ISL_3079667, EPI_ISL_3079668, EPI_ISL_3079669, EPI_ISL_3079670, EPI_ISL_3079671, EPI_ISL_3079672, EPI_ISL_3079673, EPI_ISL_3079674, EPI_ISL_3079675, EPI_ISL_3079676, EPI_ISL_3079677, EPI_ISL_3079678, EPI_ISL_3079679, EPI_ISL_3079682, EPI_ISL_3079683, EPI_ISL_3079684, EPI_ISL_3079686, EPI_ISL_3079687, EPI_ISL_3079688, EPI_ISL_3079689, EPI_ISL_3079690, EPI_ISL_3079691, EPI_ISL_3079692, EPI_ISL_3079693, EPI_ISL_3079694, EPI_ISL_3079695, EPI_ISL_3079696, EPI_ISL_3079697, EPI_ISL_3079698, EPI_ISL_3079699, EPI_ISL_3079700, EPI_ISL_3079701, EPI_ISL_3079702, EPI_ISL_3079703, EPI_ISL_3079704, EPI_ISL_3079705, EPI_ISL_3079706, EPI_ISL_3079707, EPI_ISL_3079709, EPI_ISL_3079710, EPI_ISL_3079711, EPI_ISL_3079712, EPI_ISL_3079713, EPI_ISL_3079714, EPI_ISL_3079715, EPI_ISL_3079716, EPI_ISL_3079717, EPI_ISL_3079718, EPI_ISL_3079719, EPI_ISL_3079720, EPI_ISL_3079721, EPI_ISL_3079722, EPI_ISL_3079723, EPI_ISL_3079724, EPI_ISL_3079725, EPI_ISL_3079726, EPI_ISL_3079727, EPI_ISL_3079728, EPI_ISL_3079729, EPI_ISL_3079730, EPI_ISL_3079731, EPI_ISL_3079732, EPI_ISL_3079733, EPI_ISL_3079734, EPI_ISL_3079735, EPI_ISL_3079736, EPI_ISL_3079737, EPI_ISL_3079738, EPI_ISL_3079739, EPI_ISL_3079740, EPI_ISL_3079741, EPI_ISL_3079742, EPI_ISL_3079743, EPI_ISL_3079744, EPI_ISL_3079745, EPI_ISL_3079746, EPI_ISL_3079747, EPI_ISL_3079748, EPI_ISL_3079749, EPI_ISL_3079750, EPI_ISL_3079751, EPI_ISL_3079752, EPI_ISL_3079753, EPI_ISL_3079754, EPI_ISL_3079755, EPI_ISL_3079756, EPI_ISL_3079757, EPI_ISL_3079758, EPI_ISL_3079759, EPI_ISL_3079760, EPI_ISL_3079761, EPI_ISL_3079762, EPI_ISL_3079763, EPI_ISL_3079764, EPI_ISL_3079765, EPI_ISL_3079766, EPI_ISL_3079767, EPI_ISL_3079768, EPI_ISL_3079769, EPI_ISL_3079770, EPI_ISL_3079771, EPI_ISL_3079772, EPI_ISL_3079773, EPI_ISL_3079774, EPI_ISL_3079775, EPI_ISL_3079776, EPI_ISL_3079777, EPI_ISL_3079778, EPI_ISL_3079779, EPI_ISL_3079780, EPI_ISL_3079781, EPI_ISL_3079782, EPI_ISL_3079783, EPI_ISL_3079784, EPI_ISL_3079785, EPI_ISL_3079786, EPI_ISL_3079787, EPI_ISL_3079788, EPI_ISL_3079789, EPI_ISL_3079790, EPI_ISL_3079791, EPI_ISL_3079792, EPI_ISL_3079793, EPI_ISL_3079794, EPI_ISL_3079795, EPI_ISL_3079796, EPI_ISL_3079797, EPI_ISL_3079798, EPI_ISL_3079799, EPI_ISL_3079800, EPI_ISL_3079801, EPI_ISL_3079802, EPI_ISL_3079803, EPI_ISL_3079804, EPI_ISL_3079805, EPI_ISL_3079806, EPI_ISL_3079807, EPI_ISL_3079808, EPI_ISL_3079809, EPI_ISL_3079810, EPI_ISL_3079811, EPI_ISL_3079812, EPI_ISL_3079813, EPI_ISL_3079814, EPI_ISL_3079815, EPI_ISL_3079816, EPI_ISL_3079817, EPI_ISL_3079818, EPI_ISL_3079819, EPI_ISL_3079820, EPI_ISL_3079821, EPI_ISL_3079822, EPI_ISL_3079823, EPI_ISL_3079824, EPI_ISL_3079825, EPI_ISL_3079826, EPI_ISL_3079827, EPI_ISL_3079828, EPI_ISL_3079829, EPI_ISL_3079830, EPI_ISL_3079831, EPI_ISL_3079832, EPI_ISL_3079833, EPI_ISL_3079834, EPI_ISL_3079835, EPI_ISL_3079836, EPI_ISL_3079837, EPI_ISL_3079838, EPI_ISL_3079839, EPI_ISL_3079840, EPI_ISL_3079841, EPI_ISL_3079842, EPI_ISL_3079843, EPI_ISL_3079844, EPI_ISL_3079845, EPI_ISL_3079846, EPI_ISL_3079847, EPI_ISL_3079848, EPI_ISL_3079849, EPI_ISL_3079851, EPI_ISL_3079852, EPI_ISL_3079853, EPI_ISL_3079854, EPI_ISL_3079855, EPI_ISL_3079856, EPI_ISL_3079857, EPI_ISL_3079858, EPI_ISL_3079859, EPI_ISL_3079860, EPI_ISL_3079861, EPI_ISL_3079862, EPI_ISL_3079863, EPI_ISL_3079864, EPI_ISL_3079865, EPI_ISL_3079866, EPI_ISL_3079867, EPI_ISL_3079868, EPI_ISL_3079869, EPI_ISL_3079870, EPI_ISL_3079871, EPI_ISL_3079872, EPI_ISL_3079873, EPI_ISL_3079874, EPI_ISL_3079875, EPI_ISL_3079876, EPI_ISL_3079877, EPI_ISL_3079878, EPI_ISL_3079879, EPI_ISL_3079880, EPI_ISL_3079881, EPI_ISL_3079882, EPI_ISL_3079883, EPI_ISL_3079884, EPI_ISL_3079885, EPI_ISL_3079886, EPI_ISL_3079887, EPI_ISL_3079888, EPI_ISL_3079889, EPI_ISL_3079890, EPI_ISL_3079891, EPI_ISL_3079892, EPI_ISL_3079893, EPI_ISL_3079894, EPI_ISL_3079895, EPI_ISL_3079896, EPI_ISL_3079897, EPI_ISL_3079898, EPI_ISL_3079899, EPI_ISL_3079900, EPI_ISL_3079901, EPI_ISL_3079902, EPI_ISL_3079903, EPI_ISL_3079904, EPI_ISL_3079905, EPI_ISL_3079906, EPI_ISL_3079908, EPI_ISL_3079909, EPI_ISL_3079910, EPI_ISL_3079911, EPI_ISL_3079912, EPI_ISL_3079913, EPI_ISL_3079914, EPI_ISL_3079915, EPI_ISL_3079916, EPI_ISL_3079917, EPI_ISL_3079918, EPI_ISL_3079919, EPI_ISL_3079920, EPI_ISL_3079921, EPI_ISL_3079922, EPI_ISL_3079923, EPI_ISL_3079924, EPI_ISL_3079925, EPI_ISL_3079926, EPI_ISL_3079927, EPI_ISL_3079928, EPI_ISL_3079929, EPI_ISL_3079930, EPI_ISL_3079931, EPI_ISL_3079932, EPI_ISL_3079933, EPI_ISL_3079934, EPI_ISL_3079935, EPI_ISL_3079936, EPI_ISL_3079937, EPI_ISL_3079938, EPI_ISL_3079939, EPI_ISL_3079940, EPI_ISL_3079941, EPI_ISL_3079942, EPI_ISL_3079943, EPI_ISL_3079944, EPI_ISL_3079945, EPI_ISL_3079946, EPI_ISL_3079947, EPI_ISL_3079948, EPI_ISL_3079949, EPI_ISL_3079950, EPI_ISL_3079951, EPI_ISL_3079952, EPI_ISL_3079953, EPI_ISL_3079954, EPI_ISL_3079955, EPI_ISL_3079956, EPI_ISL_3079957, EPI_ISL_3079958, EPI_ISL_3079959, EPI_ISL_3079960, EPI_ISL_3079961, EPI_ISL_3079962, EPI_ISL_3079963, EPI_ISL_3079964, EPI_ISL_3079965, EPI_ISL_3079966, EPI_ISL_3079967, EPI_ISL_3079968, EPI_ISL_3079969, EPI_ISL_3079970, EPI_ISL_3079971, EPI_ISL_3079972, EPI_ISL_3079973, EPI_ISL_3079974, EPI_ISL_3079975, EPI_ISL_3079976, EPI_ISL_3079977, EPI_ISL_3079978, EPI_ISL_3079979, EPI_ISL_3079980, EPI_ISL_3079981, EPI_ISL_3079982, EPI_ISL_3079983, EPI_ISL_3079984, EPI_ISL_3079985, EPI_ISL_3079986, EPI_ISL_3079987, EPI_ISL_3079988, EPI_ISL_3079989, EPI_ISL_3079990, EPI_ISL_3079991, EPI_ISL_3079992, EPI_ISL_3079993, EPI_ISL_3079994, EPI_ISL_3079996, EPI_ISL_3079997, EPI_ISL_3079998, EPI_ISL_3079999, EPI_ISL_3080000, EPI_ISL_3080001, EPI_ISL_3080002, EPI_ISL_3080003, EPI_ISL_3080004, EPI_ISL_3080005, EPI_ISL_3080006, EPI_ISL_3080007, EPI_ISL_3080008, EPI_ISL_3080009, EPI_ISL_3080010, EPI_ISL_3080011, EPI_ISL_3080012, EPI_ISL_3080013, EPI_ISL_3080014, EPI_ISL_3080015, EPI_ISL_3080016, EPI_ISL_3080017, EPI_ISL_3080018, EPI_ISL_3080019, EPI_ISL_3080020, EPI_ISL_3080021, EPI_ISL_3080022, EPI_ISL_3080023, EPI_ISL_3080024, EPI_ISL_3080025, EPI_ISL_3080026, EPI_ISL_3080027, EPI_ISL_3080028, EPI_ISL_3080029, EPI_ISL_3080030, EPI_ISL_3080031, EPI_ISL_3080032, EPI_ISL_3080033, EPI_ISL_3080034, EPI_ISL_3080035, EPI_ISL_3080036, EPI_ISL_3080037, EPI_ISL_3080038, EPI_ISL_3080039, EPI_ISL_3080040, EPI_ISL_3080041, EPI_ISL_3080042, EPI_ISL_3080043, EPI_ISL_3080044, EPI_ISL_3080045, EPI_ISL_3080046, EPI_ISL_3080047, EPI_ISL_3080048, EPI_ISL_3080049, EPI_ISL_3080050, EPI_ISL_3080051, EPI_ISL_3080052, EPI_ISL_3080053, EPI_ISL_3080054, EPI_ISL_3080055, EPI_ISL_3080056, EPI_ISL_3080057, EPI_ISL_3080058, EPI_ISL_3080059, EPI_ISL_3080060, EPI_ISL_3080061, EPI_ISL_3080062, EPI_ISL_3080063, EPI_ISL_3080065, EPI_ISL_3080066, EPI_ISL_3080067, EPI_ISL_3080068, EPI_ISL_3080069, EPI_ISL_3080070, EPI_ISL_3080071, EPI_ISL_3080072, EPI_ISL_3080073, EPI_ISL_3080074, EPI_ISL_3080075, EPI_ISL_3080076, EPI_ISL_3080077, EPI_ISL_3080078, EPI_ISL_3080079, EPI_ISL_3080080, EPI_ISL_3080081, EPI_ISL_3080082, EPI_ISL_3080083, EPI_ISL_3080084, EPI_ISL_3080085, EPI_ISL_3080086, EPI_ISL_3080087, EPI_ISL_3080088, EPI_ISL_3080089, EPI_ISL_3080090, EPI_ISL_3080091, EPI_ISL_3080092, EPI_ISL_3080093, EPI_ISL_3080094, EPI_ISL_3080095, EPI_ISL_3080096, EPI_ISL_3080097, EPI_ISL_3080098, EPI_ISL_3080099, EPI_ISL_3080100, EPI_ISL_3080101, EPI_ISL_3080102, EPI_ISL_3080103, EPI_ISL_3080104, EPI_ISL_3080105, EPI_ISL_3080106, EPI_ISL_3080107, EPI_ISL_3080108, EPI_ISL_3080109, EPI_ISL_3080110, EPI_ISL_3080111, EPI_ISL_3080112, EPI_ISL_3080113, EPI_ISL_3080114, EPI_ISL_3080115, EPI_ISL_3080116, EPI_ISL_3080117, EPI_ISL_3080118, EPI_ISL_3080119, EPI_ISL_3080120, EPI_ISL_3080121, EPI_ISL_3080122, EPI_ISL_3080123, EPI_ISL_3080124, EPI_ISL_3080125, EPI_ISL_3080126, EPI_ISL_3080127, EPI_ISL_3080128, EPI_ISL_3080129, EPI_ISL_3080130, EPI_ISL_3080131, EPI_ISL_3080132, EPI_ISL_3080133, EPI_ISL_3080134, EPI_ISL_3080135, EPI_ISL_3080136, EPI_ISL_3080137, EPI_ISL_3080138, EPI_ISL_3080139, EPI_ISL_3080140, EPI_ISL_3080141, EPI_ISL_3080142, EPI_ISL_3080143, EPI_ISL_3080144, EPI_ISL_3080145, EPI_ISL_3080146, EPI_ISL_3080147, EPI_ISL_3080148, EPI_ISL_3080149, EPI_ISL_3080150, EPI_ISL_3080151, EPI_ISL_3080152, EPI_ISL_3080153, EPI_ISL_3080154, EPI_ISL_3080155, EPI_ISL_3080156, EPI_ISL_3080157, EPI_ISL_3080158, EPI_ISL_3080159, EPI_ISL_3080160, EPI_ISL_3080161, EPI_ISL_3080162, EPI_ISL_3080163, EPI_ISL_3080164, EPI_ISL_3080165, EPI_ISL_3080166, EPI_ISL_3080167, EPI_ISL_3080168, EPI_ISL_3080169, EPI_ISL_3080170, EPI_ISL_3080171, EPI_ISL_3080172, EPI_ISL_3080173, EPI_ISL_3080174, EPI_ISL_3080175, EPI_ISL_3080176, EPI_ISL_3080177, EPI_ISL_3080178, EPI_ISL_3080179, EPI_ISL_3080180, EPI_ISL_3080181, EPI_ISL_3080182, EPI_ISL_3080184, EPI_ISL_3080185, EPI_ISL_3080187, EPI_ISL_3080188, EPI_ISL_3080189, EPI_ISL_3080191, EPI_ISL_3080192, EPI_ISL_3080193, EPI_ISL_3080194, EPI_ISL_3080195, EPI_ISL_3080196, EPI_ISL_3080198, EPI_ISL_3080199, EPI_ISL_3080201, EPI_ISL_3080202, EPI_ISL_3080203, EPI_ISL_3080204, EPI_ISL_3080205, EPI_ISL_3080206, EPI_ISL_3080209, EPI_ISL_3080210, EPI_ISL_3080211, EPI_ISL_3080212, EPI_ISL_3080213, EPI_ISL_3080214, EPI_ISL_3080215, EPI_ISL_3080216, EPI_ISL_3080217, EPI_ISL_3080218, EPI_ISL_3080219, EPI_ISL_3080220, EPI_ISL_3080221, EPI_ISL_3080222, EPI_ISL_3080223, EPI_ISL_3080224, EPI_ISL_3080225, EPI_ISL_3080226, EPI_ISL_3080227, EPI_ISL_3080228, EPI_ISL_3080229, EPI_ISL_3080230, EPI_ISL_3080231, EPI_ISL_3080232, EPI_ISL_3080234, EPI_ISL_3080235, EPI_ISL_3080236, EPI_ISL_3080237, EPI_ISL_3080238, EPI_ISL_3080239, EPI_ISL_3080241, EPI_ISL_3080242, EPI_ISL_3080243, EPI_ISL_3080244, EPI_ISL_3080246, EPI_ISL_3080248, EPI_ISL_3080249, EPI_ISL_3080250, EPI_ISL_3080251, EPI_ISL_3080252, EPI_ISL_3080253, EPI_ISL_3080254, EPI_ISL_3080255, EPI_ISL_3080257, EPI_ISL_3080258, EPI_ISL_3080259, EPI_ISL_3080260, EPI_ISL_3080261, EPI_ISL_3080262, EPI_ISL_3080263, EPI_ISL_3080264, EPI_ISL_3080265, EPI_ISL_3080266, EPI_ISL_3080267, EPI_ISL_3080268, EPI_ISL_3080269, EPI_ISL_3080270, EPI_ISL_3080271, EPI_ISL_3080272, EPI_ISL_3080273, EPI_ISL_3080274, EPI_ISL_3080275, EPI_ISL_3080276, EPI_ISL_3080277, EPI_ISL_3080278, EPI_ISL_3080279, EPI_ISL_3080280, EPI_ISL_3080281, EPI_ISL_3080282, EPI_ISL_3080283, EPI_ISL_3080284, EPI_ISL_3080285, EPI_ISL_3080286, EPI_ISL_3080287, EPI_ISL_3080288, EPI_ISL_3080289, EPI_ISL_3080290, EPI_ISL_3080291, EPI_ISL_3080292, EPI_ISL_3080293, EPI_ISL_3080294, EPI_ISL_3080295, EPI_ISL_3080296, EPI_ISL_3080297, EPI_ISL_3080298, EPI_ISL_3080299, EPI_ISL_3080301, EPI_ISL_3080302, EPI_ISL_3080303, EPI_ISL_3080304, EPI_ISL_3080305, EPI_ISL_3080306, EPI_ISL_3080307, EPI_ISL_3080308, EPI_ISL_3080309, EPI_ISL_3080310, EPI_ISL_3080311, EPI_ISL_3080312, EPI_ISL_3080313, EPI_ISL_3080314, EPI_ISL_3080315, EPI_ISL_3080316, EPI_ISL_3080317, EPI_ISL_3080318, EPI_ISL_3080320, EPI_ISL_3080321, EPI_ISL_3080322, EPI_ISL_3080323, EPI_ISL_3080325, EPI_ISL_3080326, EPI_ISL_3080327, EPI_ISL_3080328, EPI_ISL_3080329, EPI_ISL_3080330, EPI_ISL_3080331, EPI_ISL_3080332, EPI_ISL_3080333, EPI_ISL_3080334, EPI_ISL_3080335, EPI_ISL_3080336, EPI_ISL_3080337, EPI_ISL_3080338, EPI_ISL_3080340, EPI_ISL_3080341, EPI_ISL_3080342, EPI_ISL_3080343, EPI_ISL_3080344, EPI_ISL_3080345, EPI_ISL_3080346, EPI_ISL_3080347, EPI_ISL_3080348, EPI_ISL_3080349, EPI_ISL_3080351, EPI_ISL_3080352,

| | | | |
|---|---|---|---|
| EPI_ISL_3080353, EPI_ISL_3080354, EPI_ISL_3080355, EPI_ISL_3080357, EPI_ISL_3080358, EPI_ISL_3080359, EPI_ISL_3080360, EPI_ISL_3080361, EPI_ISL_3080362, EPI_ISL_3080363, EPI_ISL_3080364, EPI_ISL_3080365, EPI_ISL_3080367, EPI_ISL_3080368, EPI_ISL_3080369, EPI_ISL_3080370, EPI_ISL_3080371, EPI_ISL_3080372, EPI_ISL_3080373, EPI_ISL_3080374, EPI_ISL_3080375, EPI_ISL_3080376, EPI_ISL_3080378, EPI_ISL_3080379, EPI_ISL_3080380, EPI_ISL_3080381, EPI_ISL_3080382, EPI_ISL_3080383, EPI_ISL_3080384, EPI_ISL_3080385, EPI_ISL_3080386, EPI_ISL_3080387, EPI_ISL_3080388, EPI_ISL_3080390, EPI_ISL_3080391, EPI_ISL_3080392, EPI_ISL_3080393, EPI_ISL_3080394, EPI_ISL_3080406, EPI_ISL_3080395, EPI_ISL_3080396, EPI_ISL_3080398, EPI_ISL_3080399, EPI_ISL_3080400, EPI_ISL_3080401, EPI_ISL_3080402, EPI_ISL_3080403, EPI_ISL_3080404, EPI_ISL_3080405, EPI_ISL_3080406, EPI_ISL_3080407, EPI_ISL_3080408, EPI_ISL_3080409, EPI_ISL_3080410, EPI_ISL_3080411, EPI_ISL_3080413, EPI_ISL_3080414, EPI_ISL_3080415, EPI_ISL_3080416, EPI_ISL_3080417, EPI_ISL_3080418, EPI_ISL_3080419, EPI_ISL_3080420, EPI_ISL_3080422, EPI_ISL_3080423, EPI_ISL_3080424, EPI_ISL_3080425, EPI_ISL_3080426, EPI_ISL_3080427, EPI_ISL_3080428, EPI_ISL_3080429, EPI_ISL_3080430, EPI_ISL_3080431, EPI_ISL_3080432, EPI_ISL_3080433, EPI_ISL_3080435, EPI_ISL_3080437, EPI_ISL_3080438, EPI_ISL_3080439, EPI_ISL_3080441, EPI_ISL_3080442, EPI_ISL_3080443, EPI_ISL_3080444, EPI_ISL_3080445, EPI_ISL_3080446, EPI_ISL_3080447, EPI_ISL_3080449, EPI_ISL_3080450, EPI_ISL_3080452, EPI_ISL_3080453, EPI_ISL_3080454, EPI_ISL_3080455, EPI_ISL_3080456, EPI_ISL_3080457, EPI_ISL_3080458, EPI_ISL_3080459, EPI_ISL_3080460, EPI_ISL_3080461, EPI_ISL_3080462, EPI_ISL_3080463, EPI_ISL_3080464, EPI_ISL_3080465, EPI_ISL_3080466, EPI_ISL_3080467, EPI_ISL_3080468, EPI_ISL_3080469, EPI_ISL_3080470, EPI_ISL_3080471, EPI_ISL_3080472, EPI_ISL_3080473, EPI_ISL_3080474, EPI_ISL_3080475, EPI_ISL_3080476, EPI_ISL_3080477, EPI_ISL_3080478, EPI_ISL_3080479, EPI_ISL_3080480, EPI_ISL_3080481, EPI_ISL_3080482, EPI_ISL_3080483, EPI_ISL_3080484, EPI_ISL_3080485, EPI_ISL_3080486, EPI_ISL_3080487, EPI_ISL_3080488, EPI_ISL_3080489, EPI_ISL_3080490, EPI_ISL_3080493, EPI_ISL_3080494, EPI_ISL_3080495, EPI_ISL_3080496, EPI_ISL_3080498, EPI_ISL_3080499, EPI_ISL_3080500, EPI_ISL_3080501, EPI_ISL_3080502, EPI_ISL_3080503, EPI_ISL_3080504, EPI_ISL_3080505, EPI_ISL_3080506, EPI_ISL_3080507, EPI_ISL_3080508, EPI_ISL_3080509, EPI_ISL_3080510, EPI_ISL_3080511, EPI_ISL_3080512, EPI_ISL_3080513, EPI_ISL_3080514, EPI_ISL_3080515, EPI_ISL_3080516, EPI_ISL_3080517, EPI_ISL_3080518, EPI_ISL_3080519, EPI_ISL_3080520, EPI_ISL_3080521, EPI_ISL_3080522, EPI_ISL_3080523, EPI_ISL_3080524, EPI_ISL_3080525, EPI_ISL_3080526, EPI_ISL_3080527, EPI_ISL_3080528, EPI_ISL_3080529, EPI_ISL_3080530, EPI_ISL_3080531, EPI_ISL_3080532, EPI_ISL_3080533, EPI_ISL_3080534, EPI_ISL_3080535, EPI_ISL_3080536, EPI_ISL_3080537, EPI_ISL_3080538, EPI_ISL_3080539, EPI_ISL_3080540, EPI_ISL_3080541, EPI_ISL_3080542, EPI_ISL_3080543, EPI_ISL_3080544, EPI_ISL_3080546, EPI_ISL_3080547, EPI_ISL_3080548, EPI_ISL_3080549, EPI_ISL_3080550, EPI_ISL_3080551, EPI_ISL_3080552, EPI_ISL_3080553, EPI_ISL_3080554, EPI_ISL_3080555, EPI_ISL_3080556, EPI_ISL_3080557, EPI_ISL_3080558, EPI_ISL_3080559, EPI_ISL_3080560, EPI_ISL_3080561, EPI_ISL_3080562, EPI_ISL_3080563, EPI_ISL_3080564, EPI_ISL_3080566, EPI_ISL_3080567, EPI_ISL_3080568, EPI_ISL_3080569, EPI_ISL_3080570, EPI_ISL_3080571, EPI_ISL_3080572, EPI_ISL_3080573, EPI_ISL_3080574, EPI_ISL_3080575, EPI_ISL_3080576, EPI_ISL_3080577, EPI_ISL_3080578, EPI_ISL_3080579, EPI_ISL_3080580, EPI_ISL_3080581, EPI_ISL_3080582, EPI_ISL_3080583, EPI_ISL_3080584, EPI_ISL_3080585, EPI_ISL_3080586, EPI_ISL_3080587, EPI_ISL_3080589, EPI_ISL_3080590, EPI_ISL_3080591, EPI_ISL_3080592, EPI_ISL_3080593, EPI_ISL_3080594, EPI_ISL_3080595, EPI_ISL_3080596, EPI_ISL_3080597, EPI_ISL_3080598, EPI_ISL_3080599, EPI_ISL_3080600, EPI_ISL_3080601, EPI_ISL_3080602, EPI_ISL_3080603, EPI_ISL_3080604, EPI_ISL_3080605, EPI_ISL_3080606, EPI_ISL_3080607, EPI_ISL_3080608, EPI_ISL_3080609, EPI_ISL_3080610, EPI_ISL_3080611, EPI_ISL_3080612, EPI_ISL_3080613, EPI_ISL_3080614, EPI_ISL_3080615, EPI_ISL_3080616, EPI_ISL_3080617, EPI_ISL_3080618, EPI_ISL_3080619, EPI_ISL_3080620, EPI_ISL_3080621, EPI_ISL_3080622, EPI_ISL_3080623, EPI_ISL_3080624, EPI_ISL_3080625, EPI_ISL_3080626, EPI_ISL_3080627, EPI_ISL_3080628, EPI_ISL_3080629, EPI_ISL_3080630, EPI_ISL_3080631, EPI_ISL_3080632, EPI_ISL_3080633, EPI_ISL_3080634, EPI_ISL_3080635, EPI_ISL_3080636, EPI_ISL_3080637, EPI_ISL_3080638, EPI_ISL_3080639, EPI_ISL_3080640, EPI_ISL_3080641, EPI_ISL_3080642, EPI_ISL_3080643, EPI_ISL_3080644, EPI_ISL_3080645, EPI_ISL_3080646, EPI_ISL_3080647, EPI_ISL_3080648, EPI_ISL_3080649, EPI_ISL_3080650, EPI_ISL_3080651, EPI_ISL_3080653, EPI_ISL_3080654, EPI_ISL_3080655, EPI_ISL_3080656, EPI_ISL_3080657, EPI_ISL_3080658, EPI_ISL_3080659, EPI_ISL_3080660, EPI_ISL_3080661, EPI_ISL_3080663, EPI_ISL_3080664, EPI_ISL_3080665, EPI_ISL_3080666, EPI_ISL_3080667, EPI_ISL_3080668, EPI_ISL_3080669, EPI_ISL_3080670, EPI_ISL_3080671, EPI_ISL_3080672, EPI_ISL_3080673, EPI_ISL_3080674, EPI_ISL_3080675, EPI_ISL_3080676, EPI_ISL_3080677, EPI_ISL_3080678, EPI_ISL_3080679, EPI_ISL_3080680, EPI_ISL_3080681, EPI_ISL_3080682, EPI_ISL_3080683, EPI_ISL_3080684, EPI_ISL_3080685, EPI_ISL_3080686, EPI_ISL_3080687, EPI_ISL_3080688, EPI_ISL_3080689, EPI_ISL_3080690, EPI_ISL_3080691, EPI_ISL_3080692, EPI_ISL_3080693, EPI_ISL_3080694, EPI_ISL_3080695, EPI_ISL_3080696, EPI_ISL_3080697, EPI_ISL_3080698, EPI_ISL_3080699, EPI_ISL_3080700, EPI_ISL_3080701, EPI_ISL_3080702, EPI_ISL_3080703, EPI_ISL_3080704, EPI_ISL_3080706, EPI_ISL_3080707, EPI_ISL_3080708, EPI_ISL_3080709, EPI_ISL_3080710, EPI_ISL_3080711, EPI_ISL_3080712, EPI_ISL_3080713, EPI_ISL_3080714, EPI_ISL_3080715, EPI_ISL_3080716, EPI_ISL_3080717, EPI_ISL_3080718, EPI_ISL_3080719, EPI_ISL_3080720, EPI_ISL_3080721, EPI_ISL_3080722, EPI_ISL_3080723, EPI_ISL_3080724, EPI_ISL_3080725, EPI_ISL_3080726, EPI_ISL_3080727, EPI_ISL_3080728, EPI_ISL_3080729, EPI_ISL_3080730, EPI_ISL_3080731, EPI_ISL_3080732, EPI_ISL_3080733, EPI_ISL_3080734, EPI_ISL_3080735, EPI_ISL_3080736, EPI_ISL_3080737, EPI_ISL_3080738, EPI_ISL_3080740, EPI_ISL_3080741, EPI_ISL_3080742, EPI_ISL_3080744, EPI_ISL_3080746, EPI_ISL_3080747, EPI_ISL_3080748, EPI_ISL_3080750, EPI_ISL_3080751, EPI_ISL_3080752, EPI_ISL_3080753, EPI_ISL_3080754, EPI_ISL_3080755, EPI_ISL_3080756, EPI_ISL_3080757, EPI_ISL_3080758, EPI_ISL_3080759, EPI_ISL_3080760, EPI_ISL_3080761, EPI_ISL_3080762, EPI_ISL_3080763, EPI_ISL_3080764, EPI_ISL_3080765, EPI_ISL_3080766, EPI_ISL_3080767, EPI_ISL_3080768, EPI_ISL_3080769, EPI_ISL_3080770, EPI_ISL_3080771, EPI_ISL_3080772, EPI_ISL_3080773, EPI_ISL_3080774, EPI_ISL_3080775, EPI_ISL_3080776, EPI_ISL_3080777, EPI_ISL_3080778, EPI_ISL_3080779, EPI_ISL_3080781, EPI_ISL_3080782, EPI_ISL_3080783, EPI_ISL_3080784, EPI_ISL_3080785, EPI_ISL_3080786, EPI_ISL_3080787, EPI_ISL_3080788, EPI_ISL_3080789, EPI_ISL_3080791, EPI_ISL_3080792, EPI_ISL_3080793, EPI_ISL_3080794, EPI_ISL_3080795, EPI_ISL_3080796, EPI_ISL_3080797, EPI_ISL_3080798, EPI_ISL_3080799, EPI_ISL_3080800, EPI_ISL_3080802, EPI_ISL_3080803, EPI_ISL_3080804, EPI_ISL_3080805, EPI_ISL_3080806, EPI_ISL_3080807, EPI_ISL_3080808, EPI_ISL_3080809, EPI_ISL_3080810, EPI_ISL_3080811, EPI_ISL_3080812, EPI_ISL_3080813, EPI_ISL_3080814, EPI_ISL_3080815, EPI_ISL_3080816, EPI_ISL_3080817, EPI_ISL_3080818, EPI_ISL_3080819, EPI_ISL_3080820, EPI_ISL_3080821, EPI_ISL_3080822, EPI_ISL_3080823, EPI_ISL_3080824, EPI_ISL_3080825, EPI_ISL_3080826, EPI_ISL_3080827, EPI_ISL_3080828, EPI_ISL_3080829, EPI_ISL_3080830, EPI_ISL_3080831, EPI_ISL_3080832, EPI_ISL_3080833, EPI_ISL_3080834, EPI_ISL_3080835, EPI_ISL_3080836, EPI_ISL_3080837, EPI_ISL_3080838, EPI_ISL_3080839, EPI_ISL_3080840, EPI_ISL_3080841, EPI_ISL_3080842, EPI_ISL_3080843, EPI_ISL_3080844, EPI_ISL_3080847, EPI_ISL_3080848, EPI_ISL_3080849, EPI_ISL_3080850, EPI_ISL_3080851, EPI_ISL_3080852, EPI_ISL_3080853, EPI_ISL_3080854, EPI_ISL_3080855, EPI_ISL_3080856, EPI_ISL_3080857, EPI_ISL_3080858, EPI_ISL_3080859, EPI_ISL_3080860, EPI_ISL_3080861, EPI_ISL_3080862, EPI_ISL_3080863, EPI_ISL_3080864, EPI_ISL_3080865, EPI_ISL_3080866, EPI_ISL_3080867, EPI_ISL_3080868, EPI_ISL_3080869, EPI_ISL_3080871, EPI_ISL_3080873, EPI_ISL_3080874, EPI_ISL_3080875, EPI_ISL_3080876, EPI_ISL_3080877, EPI_ISL_3080878, EPI_ISL_3080879, EPI_ISL_3080880, EPI_ISL_3080881, EPI_ISL_3080882, EPI_ISL_3080883, EPI_ISL_3080884, EPI_ISL_3080885, EPI_ISL_3080887, EPI_ISL_3080889, EPI_ISL_3080890, EPI_ISL_3080891, EPI_ISL_3080892, EPI_ISL_3080893, EPI_ISL_3080894, EPI_ISL_3080895, EPI_ISL_3080897, EPI_ISL_3080898, EPI_ISL_3080899, EPI_ISL_3080900, EPI_ISL_3080901, EPI_ISL_3080902, EPI_ISL_3080903, EPI_ISL_3080904, EPI_ISL_3080905, EPI_ISL_3080906, EPI_ISL_3080907, EPI_ISL_3080908, EPI_ISL_3080909, EPI_ISL_3080910, EPI_ISL_3080911, EPI_ISL_3080912, EPI_ISL_3080913, EPI_ISL_3080914, EPI_ISL_3080915, EPI_ISL_3080916, EPI_ISL_3080917, EPI_ISL_3080918, EPI_ISL_3080919, EPI_ISL_3080920, EPI_ISL_3080921, EPI_ISL_3080922, EPI_ISL_3080923, EPI_ISL_3080924, EPI_ISL_3080925, EPI_ISL_3080926, EPI_ISL_3080927, EPI_ISL_3080928, EPI_ISL_3080929, EPI_ISL_3080930, EPI_ISL_3080931, EPI_ISL_3080932, EPI_ISL_3080933, EPI_ISL_3080935, EPI_ISL_3080936, EPI_ISL_3080937, EPI_ISL_3080938, EPI_ISL_3080939, EPI_ISL_3080940, EPI_ISL_3080941, EPI_ISL_3080942, EPI_ISL_3080943, EPI_ISL_3080944, EPI_ISL_3080945, EPI_ISL_3080946, EPI_ISL_3080947, EPI_ISL_3080948, EPI_ISL_3080949, EPI_ISL_3080950, EPI_ISL_3080951, EPI_ISL_3080952, EPI_ISL_3080953, EPI_ISL_3080954, EPI_ISL_3080955, EPI_ISL_3080956, EPI_ISL_3080957, EPI_ISL_3080958, EPI_ISL_3080959, EPI_ISL_3080960, EPI_ISL_3080961, EPI_ISL_3080962, EPI_ISL_3080964, EPI_ISL_3080966, EPI_ISL_3080967, EPI_ISL_3080968, EPI_ISL_3080969, EPI_ISL_3080970, EPI_ISL_3080971, EPI_ISL_3080972, EPI_ISL_3080973, EPI_ISL_3080975, EPI_ISL_3080976, EPI_ISL_3080977, EPI_ISL_3080978, EPI_ISL_3080979, EPI_ISL_3080981, EPI_ISL_3080982, EPI_ISL_3080983, EPI_ISL_3080984, EPI_ISL_3080985, EPI_ISL_3080986, EPI_ISL_3080987, EPI_ISL_3080988, EPI_ISL_3080989, EPI_ISL_3080990, EPI_ISL_3080991, EPI_ISL_3080992, EPI_ISL_3080993, EPI_ISL_3080994, EPI_ISL_3080995, EPI_ISL_3080996, EPI_ISL_3080997, EPI_ISL_3080998, EPI_ISL_3080999, EPI_ISL_3081000, EPI_ISL_3081001, EPI_ISL_3081002, EPI_ISL_3081004, EPI_ISL_3081005, EPI_ISL_3081006, EPI_ISL_3081007, EPI_ISL_3081008, EPI_ISL_3081009, EPI_ISL_3081010, EPI_ISL_3081011, EPI_ISL_3081012, EPI_ISL_3081013, EPI_ISL_3081014, EPI_ISL_3081015, EPI_ISL_3081016, EPI_ISL_3081017, EPI_ISL_3081018, EPI_ISL_3081019, EPI_ISL_3081020, EPI_ISL_3081021, EPI_ISL_3081022, EPI_ISL_3081023, EPI_ISL_3081024, EPI_ISL_3081025, EPI_ISL_3081026, EPI_ISL_3081027, EPI_ISL_3081028, EPI_ISL_3081029, EPI_ISL_3081030, EPI_ISL_3081031, EPI_ISL_3081032, EPI_ISL_3081033, EPI_ISL_3081034, EPI_ISL_3081035, EPI_ISL_3081036, EPI_ISL_3081037, EPI_ISL_3081038, EPI_ISL_3081039, EPI_ISL_3081040, EPI_ISL_3081041, EPI_ISL_3081042, EPI_ISL_3081043, EPI_ISL_3081044, EPI_ISL_3081045, EPI_ISL_3081046, EPI_ISL_3081047, EPI_ISL_3081048, EPI_ISL_3081049, EPI_ISL_3081050, EPI_ISL_3081051, EPI_ISL_3081052, EPI_ISL_3081053, EPI_ISL_3081054, EPI_ISL_3081055, EPI_ISL_3081056, EPI_ISL_3081057, EPI_ISL_3081058, EPI_ISL_3081059, EPI_ISL_3081060, EPI_ISL_3081061, EPI_ISL_3081062, EPI_ISL_3081064, EPI_ISL_3081066, EPI_ISL_3081067, EPI_ISL_3081068, EPI_ISL_3081069, EPI_ISL_3081070, EPI_ISL_3081071, EPI_ISL_3081072, EPI_ISL_3081073, EPI_ISL_3081074, EPI_ISL_3081075, EPI_ISL_3081076, EPI_ISL_3081077, EPI_ISL_3081078, EPI_ISL_3081079, EPI_ISL_3081080, EPI_ISL_3081081, EPI_ISL_3081082, EPI_ISL_3081083, EPI_ISL_3081084, EPI_ISL_3081085, EPI_ISL_3081086, EPI_ISL_3081087, EPI_ISL_3081088, EPI_ISL_3081089, EPI_ISL_3081090, EPI_ISL_3081091, EPI_ISL_3081092, EPI_ISL_3081093, EPI_ISL_3081094, EPI_ISL_3081095, EPI_ISL_3081096, EPI_ISL_3081097, EPI_ISL_3081098, EPI_ISL_3081099, EPI_ISL_3081100, EPI_ISL_3081101, EPI_ISL_3081102, EPI_ISL_3081103, EPI_ISL_3081104, EPI_ISL_3081105, EPI_ISL_3081106, EPI_ISL_3081107, EPI_ISL_3081108, EPI_ISL_3081109, EPI_ISL_3081110, EPI_ISL_3081111, EPI_ISL_3081112, EPI_ISL_3081113, EPI_ISL_3081114, EPI_ISL_3081115, EPI_ISL_3081116, EPI_ISL_3081117, EPI_ISL_3081118, EPI_ISL_3081119, EPI_ISL_3081120, EPI_ISL_3081121, EPI_ISL_3081123, EPI_ISL_3081124, EPI_ISL_3081125, EPI_ISL_3081126, EPI_ISL_3081127, EPI_ISL_3081129, EPI_ISL_3081130, EPI_ISL_3081132, EPI_ISL_3081133, EPI_ISL_3081134, EPI_ISL_3081135, EPI_ISL_3081136, EPI_ISL_3081137, EPI_ISL_3081138, EPI_ISL_3081139, EPI_ISL_3081140, EPI_ISL_3081141, EPI_ISL_3081142, EPI_ISL_3081143, EPI_ISL_3081144, EPI_ISL_3081145, EPI_ISL_3081146, EPI_ISL_3081147, EPI_ISL_3081148, EPI_ISL_3081149, EPI_ISL_3081150, EPI_ISL_3081151, EPI_ISL_3081152, EPI_ISL_3081153, EPI_ISL_3081154, EPI_ISL_3081155, EPI_ISL_3081157, EPI_ISL_3081158, EPI_ISL_3081159, EPI_ISL_3081160, EPI_ISL_3081161, EPI_ISL_3081162, EPI_ISL_3081163, EPI_ISL_3081164, EPI_ISL_3081165, EPI_ISL_3081166, EPI_ISL_3081167, EPI_ISL_3081168, EPI_ISL_3081169, EPI_ISL_3081170, EPI_ISL_3081171, EPI_ISL_3081172, EPI_ISL_3081173, EPI_ISL_3081174, EPI_ISL_3081175, EPI_ISL_3081176, EPI_ISL_3081177, EPI_ISL_3081178, EPI_ISL_3081179, EPI_ISL_3081180, EPI_ISL_3081181, EPI_ISL_3081182, EPI_ISL_3081183, EPI_ISL_3081184, EPI_ISL_3081185, EPI_ISL_3081186, EPI_ISL_3081187, EPI_ISL_3081188, EPI_ISL_3081189, EPI_ISL_3081190, EPI_ISL_3081191, EPI_ISL_3081192, EPI_ISL_3081193, EPI_ISL_3081194, EPI_ISL_3081195, EPI_ISL_3081196, EPI_ISL_3081197, EPI_ISL_3081198, EPI_ISL_3081199, EPI_ISL_3081200, EPI_ISL_3081201, EPI_ISL_3081202, EPI_ISL_3081203, EPI_ISL_3081204, EPI_ISL_3081205, EPI_ISL_3081206, EPI_ISL_3081207, EPI_ISL_3081208, EPI_ISL_3081209, EPI_ISL_3081210, EPI_ISL_3081212, EPI_ISL_3081213, EPI_ISL_3081215, EPI_ISL_3081216, EPI_ISL_3081217, EPI_ISL_3081218, EPI_ISL_3081219, EPI_ISL_3081220, EPI_ISL_3081221, EPI_ISL_3081222, EPI_ISL_3081223, EPI_ISL_3081224, EPI_ISL_3081225, EPI_ISL_3081226, EPI_ISL_3081227, EPI_ISL_3081228, EPI_ISL_3081229, EPI_ISL_3081230, EPI_ISL_3081231, EPI_ISL_3081232, EPI_ISL_3081233, EPI_ISL_3081234, EPI_ISL_3081235, EPI_ISL_3081236, EPI_ISL_3081237, EPI_ISL_3081238, EPI_ISL_3081239, EPI_ISL_3081240, EPI_ISL_3081241, EPI_ISL_3081242, EPI_ISL_3081243, EPI_ISL_3081244, EPI_ISL_3081245, EPI_ISL_3081246, EPI_ISL_3081247, EPI_ISL_3081248, EPI_ISL_3081249, EPI_ISL_3081250, EPI_ISL_3081251, EPI_ISL_3081252, EPI_ISL_3081253, EPI_ISL_3081254, EPI_ISL_3081255, EPI_ISL_3081256, EPI_ISL_3081257, EPI_ISL_3081258, EPI_ISL_3081259, EPI_ISL_3081260, EPI_ISL_3081261, EPI_ISL_3081262, EPI_ISL_3081263, EPI_ISL_3081264, EPI_ISL_3081265, EPI_ISL_3081266, EPI_ISL_3081267, EPI_ISL_3081268, EPI_ISL_3081269, EPI_ISL_3081270, EPI_ISL_3081271, EPI_ISL_3081272, EPI_ISL_3081273, EPI_ISL_3081274, EPI_ISL_3081275, EPI_ISL_3081276, EPI_ISL_3081277, EPI_ISL_3081278, EPI_ISL_3081279, EPI_ISL_3081280, EPI_ISL_3081281, EPI_ISL_3081282, EPI_ISL_3081283, EPI_ISL_3081284, EPI_ISL_3081285, EPI_ISL_3081286, EPI_ISL_3081287, EPI_ISL_3081288, EPI_ISL_3081289, EPI_ISL_3081290, EPI_ISL_3081291, EPI_ISL_3081292, EPI_ISL_3081293, EPI_ISL_3081294, EPI_ISL_3081295, EPI_ISL_3081296, EPI_ISL_3081297, EPI_ISL_3081298, EPI_ISL_3081299, EPI_ISL_3081300, EPI_ISL_3081301, EPI_ISL_3081302, EPI_ISL_3081303, EPI_ISL_3081304, EPI_ISL_3081305, EPI_ISL_3081306, EPI_ISL_3081307, EPI_ISL_3081308, EPI_ISL_3081309, EPI_ISL_3081310, EPI_ISL_3081311, EPI_ISL_3081312, EPI_ISL_3081313, EPI_ISL_3081314, EPI_ISL_3081315, EPI_ISL_3081316, EPI_ISL_3081317, EPI_ISL_3081318, EPI_ISL_3081319, EPI_ISL_3081320, EPI_ISL_3081321, EPI_ISL_3081322, EPI_ISL_3081323, EPI_ISL_3081324, EPI_ISL_3081325, EPI_ISL_3081326, EPI_ISL_3081327, EPI_ISL_3081328, EPI_ISL_3081329, EPI_ISL_3081330, EPI_ISL_3081331, EPI_ISL_3081332, EPI_ISL_3081333, EPI_ISL_3081334, EPI_ISL_3081335, EPI_ISL_3081336, EPI_ISL_3081337, EPI_ISL_3081338, EPI_ISL_3081339, EPI_ISL_3081340, EPI_ISL_3081341, EPI_ISL_3081342, EPI_ISL_3081343, EPI_ISL_3081344, EPI_ISL_3081345, EPI_ISL_3081346, EPI_ISL_3081347, EPI_ISL_3081348, EPI_ISL_3081349, EPI_ISL_3081350, EPI_ISL_3081351, EPI_ISL_3081352, EPI_ISL_3081353, EPI_ISL_3081354, EPI_ISL_3081355, EPI_ISL_3081356, EPI_ISL_3081357, EPI_ISL_3081358, EPI_ISL_3081359, EPI_ISL_3081360, EPI_ISL_3081361, EPI_ISL_3081362, EPI_ISL_3081363, EPI_ISL_3081364, EPI_ISL_3081365, EPI_ISL_3081366, EPI_ISL_3081367, EPI_ISL_3081368, EPI_ISL_3081369, EPI_ISL_3081370, EPI_ISL_3081371, EPI_ISL_3081372, EPI_ISL_3081373, EPI_ISL_3081374, EPI_ISL_3081375, EPI_ISL_3081376, EPI_ISL_3081377, EPI_ISL_3081378, EPI_ISL_3081379, EPI_ISL_3081380, EPI_ISL_3081381, EPI_ISL_3081382, EPI_ISL_3081383, EPI_ISL_3081384, EPI_ISL_3081385, EPI_ISL_3081386, EPI_ISL_3081387, EPI_ISL_3081388, EPI_ISL_3081389, EPI_ISL_3081390, EPI_ISL_3081391, EPI_ISL_3081392, EPI_ISL_3081393, EPI_ISL_3081394 | | | |
| see above | Respiratory Virus Unit, Microbiology Services Colindale, Public Health England | COVID-19 Genomics UK (COG-UK) Consortium | PHE Covid Sequencing Team |
| EPI_ISL_3081395, EPI_ISL_3081396, EPI_ISL_3081397, EPI_ISL_3081398, EPI_ISL_3081399, EPI_ISL_3081400, EPI_ISL_3081401, EPI_ISL_3081402, EPI_ISL_3081403, EPI_ISL_3081404, EPI_ISL_3081405, EPI_ISL_3081406, EPI_ISL_3081407, EPI_ISL_3081408, EPI_ISL_3081409, EPI_ISL_3081410, EPI_ISL_3081411, EPI_ISL_3081412 | | | |
| see above | Originating lab: Wales Specialist Virology Centre Sequencing lab: Pathogen Genomics Unit | Public Health Wales Microbiology Cardiff Wales Specialist Virology Centre | Catherine Moore, Johnathan Evans, Laura Gifford, Malorie Perry, Simon Cottrell, Angela Marchbank, Alec Birchley, Alexander Adams, Amy Gaskin, Bree Gatica-Wilcox, Jason Coombes, Joel Southgate, Lauren Gilbert, Lee Graham, Nicole Pacchiarini, Sara Kumziene-Summerhayes, Sarah Taylor, Sophie Jones, Sara Rey, Matthew Bull, Joanne Watkins, Sally Corden, Tom Connor |
| EPI_ISL_3081413 | Ospedale Regionale Umberto Parini | U.O. Genomica - Istituto Zooprofilattico Sperimentale del Piemonte, Liguria e Valle d'Aosta | Simone Peletto |
| EPI_ISL_3081414 | Originating lab: Wales Specialist Virology Centre Sequencing lab: Pathogen Genomics Unit | Public Health Wales Microbiology Cardiff Wales Specialist Virology Centre | Catherine Moore, Johnathan Evans, Laura Gifford, Malorie Perry, Simon Cottrell, Angela Marchbank, Alec Birchley, Alexander Adams, Amy Gaskin, Bree Gatica-Wilcox, Jason Coombes, Joel Southgate, Lauren Gilbert, Lee Graham, Nicole Pacchiarini, Sara Kumziene-Summerhayes, Sarah Taylor, Sophie Jones, Sara Rey, Matthew Bull, Joanne Watkins, Sally Corden, Tom Connor |
| EPI_ISL_3081415 | Ospedale Regionale Umberto Parini | U.O. Genomica - Istituto Zooprofilattico Sperimentale del Piemonte, Liguria e Valle d'Aosta | Simone Peletto |
| EPI_ISL_3081416, EPI_ISL_3081418, EPI_ISL_3081419, EPI_ISL_3081420, EPI_ISL_3081421, EPI_ISL_3081422, EPI_ISL_3081423, EPI_ISL_3081424, EPI_ISL_3081425, EPI_ISL_3081426, EPI_ISL_3081427, EPI_ISL_3081428, EPI_ISL_3081429, EPI_ISL_3081430, EPI_ISL_3081431, EPI_ISL_3081432, EPI_ISL_3081433, EPI_ISL_3081434, EPI_ISL_3081435, EPI_ISL_3081436, EPI_ISL_3081437, EPI_ISL_3081438, EPI_ISL_3081439, EPI_ISL_3081440, EPI_ISL_3081441, EPI_ISL_3081442, EPI_ISL_3081443, EPI_ISL_3081444, EPI_ISL_3081445, EPI_ISL_3081446, EPI_ISL_3081447, EPI_ISL_3081449, EPI_ISL_3081450, EPI_ISL_3081451, EPI_ISL_3081452, EPI_ISL_3081453, EPI_ISL_3081454, EPI_ISL_3081455, EPI_ISL_3081456, EPI_ISL_3081457, EPI_ISL_3081458, EPI_ISL_3081459, EPI_ISL_3081460, EPI_ISL_3081461, EPI_ISL_3081462, EPI_ISL_3081463, EPI_ISL_3081464, EPI_ISL_3081465, EPI_ISL_3081466, EPI_ISL_3081467, EPI_ISL_3081468, EPI_ISL_3081469, EPI_ISL_3081470, EPI_ISL_3081471, EPI_ISL_3081472, EPI_ISL_3081473, EPI_ISL_3081474, EPI_ISL_3081475, EPI_ISL_3081476, EPI_ISL_3081477, EPI_ISL_3081478, EPI_ISL_3081479, EPI_ISL_3081480, EPI_ISL_3081481, EPI_ISL_3081482, EPI_ISL_3081483, EPI_ISL_3081484, EPI_ISL_3081485, EPI_ISL_3081486, EPI_ISL_3081487, EPI_ISL_3081488, EPI_ISL_3081489, EPI_ISL_3081490, EPI_ISL_3081491, EPI_ISL_3081492, EPI_ISL_3081493, EPI_ISL_3081494, EPI_ISL_3081495, EPI_ISL_3081496, EPI_ISL_3081497, EPI_ISL_3081498, EPI_ISL_3081499, EPI_ISL_3081500, EPI_ISL_3081501, EPI_ISL_3081502, EPI_ISL_3081503, EPI_ISL_3081504, EPI_ISL_3081505, EPI_ISL_3081506, EPI_ISL_3081507, EPI_ISL_3081508, EPI_ISL_3081509, EPI_ISL_3081510, EPI_ISL_3081511, EPI_ISL_3081512, EPI_ISL_3081513, EPI_ISL_3081514, EPI_ISL_3081515, EPI_ISL_3081516, EPI_ISL_3081517, EPI_ISL_3081518, EPI_ISL_3081519, EPI_ISL_3081520, EPI_ISL_3081521, EPI_ISL_3081522, EPI_ISL_3081523, EPI_ISL_3081524, EPI_ISL_3081525, EPI_ISL_3081526, EPI_ISL_3081527, EPI_ISL_3081528, EPI_ISL_3081529, EPI_ISL_3081530, EPI_ISL_3081531, EPI_ISL_3081532, EPI_ISL_3081533, EPI_ISL_3081534, EPI_ISL_3081535, EPI_ISL_3081536, EPI_ISL_3081537, EPI_ISL_3081538, EPI_ISL_3081539, EPI_ISL_3081540, EPI_ISL_3081541, EPI_ISL_3081542, EPI_ISL_3081543, EPI_ISL_3081544, EPI_ISL_3081545, EPI_ISL_3081546, EPI_ISL_3081547, EPI_ISL_3081548, EPI_ISL_3081549, EPI_ISL_3081550, EPI_ISL_3081551, EPI_ISL_3081552, EPI_ISL_3081553, EPI_ISL_3081554, EPI_ISL_3081555, EPI_ISL_3081556, EPI_ISL_3081557, EPI_ISL_3081558, EPI_ISL_3081559, EPI_ISL_3081560, EPI_ISL_3081561, EPI_ISL_3081562, EPI_ISL_3081563, EPI_ISL_3081564, EPI_ISL_3081565, EPI_ISL_3081566, EPI_ISL_3081567, EPI_ISL_3081568, EPI_ISL_3081569, EPI_ISL_3081570, EPI_ISL_3081571, EPI_ISL_3081572, EPI_ISL_3081573, EPI_ISL_3081574, EPI_ISL_3081575, EPI_ISL_3081576, EPI_ISL_3081577, EPI_ISL_3081578, EPI_ISL_3081579, EPI_ISL_3081580, EPI_ISL_3081581, EPI_ISL_3081582, EPI_ISL_3081583, EPI_ISL_3081585, EPI_ISL_3081586, EPI_ISL_3081587, EPI_ISL_3081588, EPI_ISL_3081589, EPI_ISL_3081590, EPI_ISL_3081591, EPI_ISL_3081592, EPI_ISL_3081593, EPI_ISL_3081594, EPI_ISL_3081595, EPI_ISL_3081596, EPI_ISL_3081597, EPI_ISL_3081598, EPI_ISL_3081599, EPI_ISL_3081600, EPI_ISL_3081601, EPI_ISL_3081602, EPI_ISL_3081603, EPI_ISL_3081604, EPI_ISL_3081605, EPI_ISL_3081606, EPI_ISL_3081607, EPI_ISL_3081608, EPI_ISL_3081609, EPI_ISL_3081610, EPI_ISL_3081611, EPI_ISL_3081612, EPI_ISL_3081613, EPI_ISL_3081614, EPI_ISL_3081615, EPI_ISL_3081616, EPI_ISL_3081617, EPI_ISL_3081618, EPI_ISL_3081619, EPI_ISL_3081620, EPI_ISL_3081621, EPI_ISL_3081622, EPI_ISL_3081623, EPI_ISL_3081624, EPI_ISL_3081625, EPI_ISL_3081626, EPI_ISL_3081627, EPI_ISL_3081628, EPI_ISL_3081629, EPI_ISL_3081630, EPI_ISL_3081631, EPI_ISL_3081632, EPI_ISL_3081633, EPI_ISL_3081634, EPI_ISL_3081635, EPI_ISL_3081636, EPI_ISL_3081637, EPI_ISL_3081638, EPI_ISL_3081639, EPI_ISL_3081640, EPI_ISL_3081641, EPI_ISL_3081642, EPI_ISL_3081643, EPI_ISL_3081644, EPI_ISL_3081645, EPI_ISL_3081646, EPI_ISL_3081647, EPI_ISL_3081648, EPI_ISL_3081649, EPI_ISL_3081650, EPI_ISL_3081651, EPI_ISL_3081652, EPI_ISL_3081653, EPI_ISL_3081654, EPI_ISL_3081655, EPI_ISL_3081656, EPI_ISL_3081657, EPI_ISL_3081658, EPI_ISL_3081659, EPI_ISL_3081660, EPI_ISL_3081661, EPI_ISL_3081662, EPI_ISL_3081663, EPI_ISL_3081664, EPI_ISL_3081665, EPI_ISL_3081666, EPI_ISL_3081667, EPI_ISL_3081668, EPI_ISL_3081669, EPI_ISL_3081670, EPI_ISL_3081671, EPI_ISL_3081672, EPI_ISL_3081673, EPI_ISL_3081674, EPI_ISL_3081675, EPI_ISL_3081676, EPI_ISL_3081677, EPI_ISL_3081678, EPI_ISL_3081679, EPI_ISL_3081680, EPI_ISL_3081681, EPI_ISL_3081682, EPI_ISL_3081683, EPI_ISL_3081684, EPI_ISL_3081685, EPI_ISL_3081686, EPI_ISL_3081687, EPI_ISL_3081688, EPI_ISL_3081689, EPI_ISL_3081690, EPI_ISL_3081691, EPI_ISL_3081692, EPI_ISL_3081693, EPI_ISL_3081694, EPI_ISL_3081695, EPI_ISL_3081696, EPI_ISL_3081697, EPI_ISL_3081698, EPI_ISL_3081699, EPI_ISL_3081700, EPI_ISL_3081701, EPI_ISL_3081702, EPI_ISL_3081703, EPI_ISL_3081704, EPI_ISL_3081705, EPI_ISL_3081706, EPI_ISL_3081707, EPI_ISL_3081708, EPI_ISL_3081709, EPI_ISL_3081710, EPI_ISL_3081711, EPI_ISL_3081712, EPI_ISL_3081713, EPI_ISL_3081714, EPI_ISL_3081715 | | | |
| see above | Originating lab: Wales Specialist Virology Centre Sequencing lab: Pathogen Genomics Unit | Public Health Wales Microbiology Cardiff Wales Specialist Virology Centre | Catherine Moore, Johnathan Evans, Laura Gifford, Malorie Perry, Simon Cottrell, Angela Marchbank, Alec Birchley, Alexander Adams, Amy Gaskin, Bree Gatica-Wilcox, Jason Coombes, Joel Southgate, Lauren Gilbert, Lee Graham, Nicole Pacchiarini, Sara Kumziene-Summerhayes, Sarah Taylor, Sophie Jones, Sara Rey, Matthew Bull, Joanne Watkins, Sally Corden, Tom Connor |
| EPI_ISL_3081716 | Tuzla University Clinical Center, Department of Microbiology | University of Sarajevo, Veterinary Faculty, Laboratory for Molecular Diagnostic and Research Laboratory | Goletic S, Goletic T, Tihic N, Softic A, Terzic I., Sabic E., Hodzic A., Jazic A., Nicevic M. |
| EPI_ISL_3081717, EPI_ISL_3081718, EPI_ISL_3081719, EPI_ISL_3081720, EPI_ISL_3081721, EPI_ISL_3081722, EPI_ISL_3081723, EPI_ISL_3081724, EPI_ISL_3081725, EPI_ISL_3081726, EPI_ISL_3081727, EPI_ISL_3081728, EPI_ISL_3081729, EPI_ISL_3081730, EPI_ISL_3081731, EPI_ISL_3081732, EPI_ISL_3081733, EPI_ISL_3081734, EPI_ISL_3081735, EPI_ISL_3081736, EPI_ISL_3081737, EPI_ISL_3081738, EPI_ISL_3081739, EPI_ISL_3081740, EPI_ISL_3081741, EPI_ISL_3081742, EPI_ISL_3081743, EPI_ISL_3081744, EPI_ISL_3081745, EPI_ISL_3081746, EPI_ISL_3081747, EPI_ISL_3081748, EPI_ISL_3081749, EPI_ISL_3081750, EPI_ISL_3081751, EPI_ISL_3081752, EPI_ISL_3081753, EPI_ISL_3081754, EPI_ISL_3081755, EPI_ISL_3081756, EPI_ISL_3081757, EPI_ISL_3081758, EPI_ISL_3081759, EPI_ISL_3081760, EPI_ISL_3081761, EPI_ISL_3081762, EPI_ISL_3081763, EPI_ISL_3081764, EPI_ISL_3081765, EPI_ISL_3081766, EPI_ISL_3081767, EPI_ISL_3081768, EPI_ISL_3081769, EPI_ISL_3081770, EPI_ISL_3081771, EPI_ISL_3081772, EPI_ISL_3081773, EPI_ISL_3081774, EPI_ISL_3081775, EPI_ISL_3081776, EPI_ISL_3081777, EPI_ISL_3081778, EPI_ISL_3081779, EPI_ISL_3081780, EPI_ISL_3081781, EPI_ISL_3081782, EPI_ISL_3081783, EPI_ISL_3081784, EPI_ISL_3081785, EPI_ISL_3081786, EPI_ISL_3081787, EPI_ISL_3081788, EPI_ISL_3081789, EPI_ISL_3081790, EPI_ISL_3081791, EPI_ISL_3081792, EPI_ISL_3081793, EPI_ISL_3081794, EPI_ISL_3081795, EPI_ISL_3081796, EPI_ISL_3081797, EPI_ISL_3081798, EPI_ISL_3081799, EPI_ISL_3081800, EPI_ISL_3081801, EPI_ISL_3081802, EPI_ISL_3081803, EPI_ISL_3081804, EPI_ISL_3081805, EPI_ISL_3081806, EPI_ISL_3081807, EPI_ISL_3081808, EPI_ISL_3081809, EPI_ISL_3081810, EPI_ISL_3081811, EPI_ISL_3081812, EPI_ISL_3081813, EPI_ISL_3081814, EPI_ISL_3081815, EPI_ISL_3081816, EPI_ISL_3081817, EPI_ISL_3081818, EPI_ISL_3081819, EPI_ISL_3081820, EPI_ISL_3081821, EPI_ISL_3081822, EPI_ISL_3081823, EPI_ISL_3081824, EPI_ISL_3081825, EPI_ISL_3081826, EPI_ISL_3081827, EPI_ISL_3081828, EPI_ISL_3081829, EPI_ISL_3081830, EPI_ISL_3081831, EPI_ISL_3081832, EPI_ISL_3081833, EPI_ISL_3081834, EPI_ISL_3081835, EPI_ISL_3081836, EPI_ISL_3081837, EPI_ISL_3081838, EPI_ISL_3081839, EPI_ISL_3081840, EPI_ISL_3081841, EPI_ISL_3081842, EPI_ISL_3081843, EPI_ISL_3081844, EPI_ISL_3081845, EPI_ISL_3081846, EPI_ISL_3081847, EPI_ISL_3081848, EPI_ISL_3081849, EPI_ISL_3081850, EPI_ISL_3081851, EPI_ISL_3081852, EPI_ISL_3081853, EPI_ISL_3081854, EPI_ISL_3081855, EPI_ISL_3081856, EPI_ISL_3081857, EPI_ISL_3081858, EPI_ISL_3081859, EPI_ISL_3081860, EPI_ISL_3081861, EPI_ISL_3081862, EPI_ISL_3081863, EPI_ISL_3081864, EPI_ISL_3081865, EPI_ISL_3081866, EPI_ISL_3081867, EPI_ISL_3081868, EPI_ISL_3081869, EPI_ISL_3081870, EPI_ISL_3081871, EPI_ISL_3081872, EPI_ISL_3081873, EPI_ISL_3081874, EPI_ISL_3081875, EPI_ISL_3081876, EPI_ISL_3081877, EPI_ISL_3081878, EPI_ISL_3081879, EPI_ISL_3081880, EPI_ISL_3081881, EPI_ISL_3081882, EPI_ISL_3081883, EPI_ISL_3081884, EPI_ISL_3081885, EPI_ISL_3081886, EPI_ISL_3081887, EPI_ISL_3081888, EPI_ISL_3081889, EPI_ISL_3081890, EPI_ISL_3081891, EPI_ISL_3081892, EPI_ISL_3081893, EPI_ISL_3081894, EPI_ISL_3081895, EPI_ISL_3081896, EPI_ISL_3081897, EPI_ISL_3081898, EPI_ISL_3081899, EPI_ISL_3081900, EPI_ISL_3081901, EPI_ISL_3081902, EPI_ISL_3081903, EPI_ISL_3081904, EPI_ISL_3081905, EPI_ISL_3081906, EPI_ISL_3081907, EPI_ISL_3081908, EPI_ISL_3081909, EPI_ISL_3081910, EPI_ISL_3081911, EPI_ISL_3081912, EPI_ISL_3081913, EPI_ISL_3081914, EPI_ISL_3081915, EPI_ISL_3081916, EPI_ISL_3081917, EPI_ISL_3081918, EPI_ISL_3081919, EPI_ISL_3081920, EPI_ISL_3081921, EPI_ISL_3081922, EPI_ISL_3081923, EPI_ISL_3081924, EPI_ISL_3081925, EPI_ISL_3081926, EPI_ISL_3081927, EPI_ISL_3081928, EPI_ISL_3081929, EPI_ISL_3081930, EPI_ISL_3081931, EPI_ISL_3081932, EPI_ISL_3081933, EPI_ISL_3081934, EPI_ISL_3081935, EPI_ISL_3081936, EPI_ISL_3081937, EPI_ISL_3081938, EPI_ISL_3081939, EPI_ISL_3081940, EPI_ISL_3081941, EPI_ISL_3081942, EPI_ISL_3081943, EPI_ISL_3081944, EPI_ISL_3081945, EPI_ISL_3081946, EPI_ISL_3081947, EPI_ISL_3081948, EPI_ISL_3081949, EPI_ISL_3081950, EPI_ISL_3081951, EPI_ISL_3081952, EPI_ISL_3081953, EPI_ISL_3081954, EPI_ISL_3081955, EPI_ISL_3081956, EPI_ISL_3081957, EPI_ISL_3081958, EPI_ISL_3081959, EPI_ISL_3081960, EPI_ISL_3081961, EPI_ISL_3081962, EPI_ISL_3081963, EPI_ISL_3081964, EPI_ISL_3081965, EPI_ISL_3081966, EPI_ISL_3081967, EPI_ISL_3081968, EPI_ISL_3081969, EPI_ISL_3081970, EPI_ISL_3081971, EPI_ISL_3081972, EPI_ISL_3081973, EPI_ISL_3081974, EPI_ISL_3081975, EPI_ISL_3081976, EPI_ISL_3081977, EPI_ISL_3081978, EPI_ISL_3081979, EPI_ISL_3081980, EPI_ISL_3081981, EPI_ISL_3081982, EPI_ISL_3081983, EPI_ISL_3081984, EPI_ISL_3081985, EPI_ISL_3081986, EPI_ISL_3081987, EPI_ISL_3081988, EPI_ISL_3081989, EPI_ISL_3081990, EPI_ISL_3081991, EPI_ISL_3081992, EPI_ISL_3081993, EPI_ISL_3081994, EPI_ISL_3081995, EPI_ISL_3081996, EPI_ISL_3081997, EPI_ISL_3081998, EPI_ISL_3081999, EPI_ISL_3082000, EPI_ISL_3082001, EPI_ISL_3082002, EPI_ISL_3082003, EPI_ISL_3082004, EPI_ISL_3082005, EPI_ISL_3082006, EPI_ISL_3082007, EPI_ISL_3082008, EPI_ISL_3082009, EPI_ISL_3082010, EPI_ISL_3082011, EPI_ISL_3082012, EPI_ISL_3082013, EPI_ISL_3082014, EPI_ISL_3082015, EPI_ISL_3082016, EPI_ISL_3082017, EPI_ISL_3082018, EPI_ISL_3082019, EPI_ISL_3082020, EPI_ISL_3082021, EPI_ISL_3082022, EPI_ISL_3082023, EPI_ISL_3082024, EPI_ISL_3082025, EPI_ISL_3082026, EPI_ISL_3082027, EPI_ISL_3082028, EPI_ISL_3082029, EPI_ISL_3082030, EPI_ISL_3082031, EPI_ISL_3082032, EPI_ISL_3082033, EPI_ISL_3082034, EPI_ISL_3082035, EPI_ISL_3082036, EPI_ISL_3082037, EPI_ISL_3082038, EPI_ISL_3082039, EPI_ISL_3082040, EPI_ISL_3082041, EPI_ISL_3082042, EPI_ISL_3082043, EPI_ISL_3082044, EPI_ISL_3082045, EPI_ISL_3082046, EPI_ISL_3082047, EPI_ISL_3082048, EPI_ISL_3082049, EPI_ISL_3082050, EPI_ISL_3082051, EPI_ISL_3082052, EPI_ISL_3082053, EPI_ISL_3082054, EPI_ISL_3082055, EPI_ISL_3082056, EPI_ISL_3082057, EPI_ISL_3082058, EPI_ISL_3082059, EPI_ISL_3082060, EPI_ISL_3082061, EPI_ISL_3082062, EPI_ISL_3082063, EPI_ISL_3082064, EPI_ISL_3082065, EPI_ISL_3082066, EPI_ISL_3082067, EPI_ISL_3082068, EPI_ISL_3082069, EPI_ISL_3082070, EPI_ISL_3082071, EPI_ISL_3082072, EPI_ISL_3082073, EPI_ISL_3082074, EPI_ISL_3082075, EPI_ISL_3082076, EPI_ISL_3082077, EPI_ISL_3082078, EPI_ISL_3082079, EPI_ISL_3082080, EPI_ISL_3082081, EPI_ISL_3082082, EPI_ISL_3082083, EPI_ISL_3082084, EPI_ISL_3082085, EPI_ISL_3082086, EPI_ISL_3082087, EPI_ISL_3082088, EPI_ISL_3082089, EPI_ISL_3082090, EPI_ISL_3082091, EPI_ISL_3082092, EPI_ISL_3082093, EPI_ISL_3082094, EPI_ISL_3082095, EPI_ISL_3082096, EPI_ISL_3082097, EPI_ISL_3082098, EPI_ISL_3082099, EPI_ISL_3082100, EPI_ISL_3082101, EPI_ISL_3082102, EPI_ISL_3082103, EPI_ISL_3082104, EPI_ISL_3082105, EPI_ISL_3082106, EPI_ISL_3082107, EPI_ISL_3082108, EPI_ISL_3082109, EPI_ISL_3082110, EPI_ISL_3082111, EPI_ISL_3082112, EPI_ISL_3082113, EPI_ISL_3082114, EPI_ISL_3082115, EPI_ISL_3082116, EPI_ISL_3082117, EPI_ISL_3082118, EPI_ISL_3082119, EPI_ISL_3082120, EPI_ISL_3082121, EPI_ISL_3082122, EPI_ISL_3082123, EPI_ISL_3082124, EPI_ISL_3082125, EPI_ISL_3082126, EPI_ISL_3082127, EPI_ISL_3082128, EPI_ISL_3082129, EPI_ISL_3082130, EPI_ISL_3082131, EPI_ISL_3082132, EPI_ISL_3082133, EPI_ISL_3082134, EPI_ISL_3082135, EPI_ISL_3082136, EPI_ISL_3082137, EPI_ISL_3082138, EPI_ISL_3082139, EPI_ISL_3082140, EPI_ISL_3082141, EPI_ISL_3082142, EPI_ISL_3082143, EPI_ISL_3082144, EPI_ISL_3082145, EPI_ISL_3082146, EPI_ISL_3082147, EPI_ISL_3082148, EPI_ISL_3082149, EPI_ISL_3082150, EPI_ISL_3082151, EPI_ISL_3082152, EPI_ISL_3082153, EPI_ISL_3082154, EPI_ISL_3082155, EPI_ISL_3082156, EPI_ISL_3082157, EPI_ISL_3082158, EPI_ISL_3082159, EPI_ISL_3082160, EPI_ISL_3082161, EPI_ISL_3082162, EPI_ISL_3082163, EPI_ISL_3082164, EPI_ISL_3082165, EPI_ISL_3082166, EPI_ISL_3082167, EPI_ISL_3082168, EPI_ISL_3082169, EPI_ISL_3082170, EPI_ISL_3082171, EPI_ISL_3082172, EPI_ISL_3082173, EPI_ISL_3082174, EPI_ISL_3082175, EPI_ISL_3082176, EPI_ISL_3082177, EPI_ISL_3082178, EPI_ISL_3082179, EPI_ISL_3082180, EPI_ISL_3082181, EPI_ISL_3082182, EPI_ISL_3082183, EPI_ISL_3082184, EPI_ISL_3082185, EPI_ISL_3082186, EPI_ISL_3082187, EPI_ISL_3082188, EPI_ISL_3082189, EPI_ISL_3082190, EPI_ISL_3082191, EPI_ISL_3082192, EPI_ISL_3082193, EPI_ISL_3082194, EPI_ISL_3082195, EPI_ISL_3082196, EPI_ISL_3082197, EPI_ISL_3082198, EPI_ISL_3082199, EPI_ISL_3082200, EPI_ISL_3082201, EPI_ISL_3082202, EPI_ISL_3082203, EPI_ISL_3082204, EPI_ISL_3082205, EPI_ISL_3082206, EPI_ISL_3082207, EPI_ISL_3082208, EPI_ISL_3082209, EPI_ISL_3082210, EPI_ISL_3082211, EPI_ISL_3082212, EPI_ISL_3082213 | | | |
| see above | Originating lab: Wales Specialist Virology Centre Sequencing lab: Pathogen Genomics Unit | Public Health Wales Microbiology Cardiff Wales Specialist Virology Centre | Catherine Moore, Johnathan Evans, Laura Gifford, Malorie Perry, Simon Cottrell, Angela Marchbank, Alec Birchley, Alexander Adams, Amy Gaskin, Bree Gatica-Wilcox, Jason Coombes, Joel Southgate, Lauren Gilbert, Lee Graham, Nicole Pacchiarini, Sara Kumziene-Summerhayes, Sarah Taylor, Sophie Jones, Sara Rey, Matthew Bull, Joanne Watkins, Sally Corden, Tom Connor |
| EPI_ISL_3082214 | Tuzla University Clinical Center, Department of Microbiology | University of Sarajevo, Veterinary Faculty, Laboratory for Molecular Diagnostic and Research Laboratory | Goletic S, Goletic T, Tihic N, Softic A, Hodzic A., Jazic A., Terzic I., Nicevic M., Sabic E. |
| EPI_ISL_3082215, EPI_ISL_3082216, EPI_ISL_3082217, EPI_ISL_3082218, EPI_ISL_3082219, EPI_ISL_3082220, EPI_ISL_3082221, EPI_ISL_3082222, EPI_ISL_3082223, EPI_ISL_3082224, EPI_ISL_3082225, EPI_ISL_3082226, EPI_ISL_3082227, EPI_ISL_3082228, EPI_ISL_3082229, EPI_ISL_3082230, EPI_ISL_3082231, EPI_ISL_3082232, EPI_ISL_3082233, EPI_ISL_3082234, EPI_ISL_3082235, EPI_ISL_3082236, EPI_ISL_3082237, EPI_ISL_3082238, EPI_ISL_3082239, EPI_ISL_3082240, EPI_ISL_3082241, EPI_ISL_3082242, EPI_ISL_3082243, EPI_ISL_3082244, EPI_ISL_3082245, EPI_ISL_3082246, EPI_ISL_3082247, EPI_ISL_3082248, EPI_ISL_3082249, EPI_ISL_3082250, EPI_ISL_3082251, EPI_ISL_3082252, EPI_ISL_3082253, EPI_ISL_3082254, EPI_ISL_3082255, EPI_ISL_3082256, EPI_ISL_3082257, EPI_ISL_3082258, | | | |

EPI_ISL_3082259, EPI_ISL_3082260, EPI_ISL_3082261, EPI_ISL_3082262, EPI_ISL_3082263, EPI_ISL_3082264, EPI_ISL_3082265, EPI_ISL_3082266, EPI_ISL_3082267, EPI_ISL_3082268, EPI_ISL_3082269, EPI_ISL_3082270, EPI_ISL_3082271, EPI_ISL_3082272, EPI_ISL_3082273, EPI_ISL_3082274, EPI_ISL_3082275, EPI_ISL_3082276, EPI_ISL_3082277, EPI_ISL_3082278, EPI_ISL_3082279, EPI_ISL_3082280, EPI_ISL_3082281, EPI_ISL_3082282, EPI_ISL_3082283, EPI_ISL_3082284, EPI_ISL_3082285, EPI_ISL_3082286, EPI_ISL_3082287, EPI_ISL_3082288, EPI_ISL_3082289, EPI_ISL_3082290, EPI_ISL_3082291, EPI_ISL_3082292, EPI_ISL_3082293, EPI_ISL_3082294, EPI_ISL_3082295, EPI_ISL_3082296, EPI_ISL_3082297, EPI_ISL_3082298, EPI_ISL_3082299, EPI_ISL_3082300, EPI_ISL_3082301, EPI_ISL_3082302, EPI_ISL_3082303, EPI_ISL_3082304, EPI_ISL_3082305, EPI_ISL_3082306, EPI_ISL_3082307, EPI_ISL_3082308, EPI_ISL_3082309, EPI_ISL_3082310, EPI_ISL_3082311, EPI_ISL_3082312, EPI_ISL_3082313, EPI_ISL_3082314, EPI_ISL_3082315, EPI_ISL_3082316, EPI_ISL_3082317, EPI_ISL_3082318, EPI_ISL_3082319, EPI_ISL_3082320, EPI_ISL_3082321, EPI_ISL_3082322, EPI_ISL_3082323, EPI_ISL_3082324, EPI_ISL_3082325, EPI_ISL_3082326, EPI_ISL_3082327, EPI_ISL_3082328, EPI_ISL_3082329, EPI_ISL_3082330, EPI_ISL_3082331, EPI_ISL_3082332, EPI_ISL_3082333, EPI_ISL_3082334, EPI_ISL_3082335, EPI_ISL_3082336, EPI_ISL_3082337, EPI_ISL_3082338, EPI_ISL_3082339, EPI_ISL_3082340, EPI_ISL_3082341, EPI_ISL_3082342, EPI_ISL_3082343, EPI_ISL_3082344, EPI_ISL_3082345, EPI_ISL_3082346, EPI_ISL_3082347, EPI_ISL_3082348, EPI_ISL_3082349, EPI_ISL_3082350, EPI_ISL_3082351, EPI_ISL_3082352, EPI_ISL_3082353, EPI_ISL_3082354, EPI_ISL_3082355, EPI_ISL_3082356, EPI_ISL_3082357, EPI_ISL_3082358, EPI_ISL_3082359, EPI_ISL_3082360, EPI_ISL_3082361, EPI_ISL_3082362, EPI_ISL_3082363, EPI_ISL_3082364, EPI_ISL_3082365, EPI_ISL_3082366, EPI_ISL_3082367, EPI_ISL_3082368, EPI_ISL_3082369, EPI_ISL_3082370, EPI_ISL_3082371, EPI_ISL_3082372, EPI_ISL_3082373, EPI_ISL_3082374, EPI_ISL_3082375, EPI_ISL_3082376, EPI_ISL_3082377, EPI_ISL_3082378, EPI_ISL_3082379, EPI_ISL_3082380, EPI_ISL_3082381, EPI_ISL_3082382, EPI_ISL_3082383, EPI_ISL_3082384, EPI_ISL_3082385, EPI_ISL_3082386, EPI_ISL_3082387, EPI_ISL_3082388, EPI_ISL_3082389, EPI_ISL_3082390, EPI_ISL_3082391, EPI_ISL_3082392, EPI_ISL_3082393, EPI_ISL_3082394, EPI_ISL_3082395, EPI_ISL_3082396, EPI_ISL_3082397, EPI_ISL_3082398, EPI_ISL_3082399, EPI_ISL_3082400, EPI_ISL_3082401, EPI_ISL_3082402, EPI_ISL_3082403, EPI_ISL_3082404, EPI_ISL_3082405, EPI_ISL_3082406, EPI_ISL_3082407, EPI_ISL_3082408, EPI_ISL_3082409, EPI_ISL_3082410, EPI_ISL_3082411, EPI_ISL_3082412, EPI_ISL_3082413, EPI_ISL_3082414, EPI_ISL_3082415, EPI_ISL_3082416, EPI_ISL_3082417, EPI_ISL_3082418, EPI_ISL_3082419, EPI_ISL_3082420, EPI_ISL_3082421, EPI_ISL_3082422, EPI_ISL_3082423, EPI_ISL_3082424, EPI_ISL_3082425, EPI_ISL_3082426, EPI_ISL_3082427, EPI_ISL_3082428, EPI_ISL_3082429, EPI_ISL_3082430, EPI_ISL_3082431, EPI_ISL_3082432, EPI_ISL_3082433, EPI_ISL_3082434, EPI_ISL_3082435, EPI_ISL_3082436, EPI_ISL_3082437, EPI_ISL_3082438, EPI_ISL_3082439, EPI_ISL_3082440, EPI_ISL_3082441, EPI_ISL_3082442, EPI_ISL_3082443, EPI_ISL_3082444, EPI_ISL_3082445, EPI_ISL_3082446, EPI_ISL_3082447, EPI_ISL_3082448, EPI_ISL_3082449, EPI_ISL_3082450, EPI_ISL_3082451, EPI_ISL_3082452, EPI_ISL_3082453, EPI_ISL_3082454, EPI_ISL_3082455, EPI_ISL_3082456, EPI_ISL_3082457, EPI_ISL_3082458, EPI_ISL_3082459, EPI_ISL_3082460, EPI_ISL_3082461, EPI_ISL_3082462, EPI_ISL_3082463, EPI_ISL_3082464, EPI_ISL_3082465, EPI_ISL_3082466, EPI_ISL_3082467, EPI_ISL_3082468, EPI_ISL_3082469, EPI_ISL_3082470, EPI_ISL_3082471, EPI_ISL_3082472, EPI_ISL_3082473, EPI_ISL_3082474, EPI_ISL_3082475, EPI_ISL_3082476, EPI_ISL_3082477, EPI_ISL_3082478, EPI_ISL_3082479, EPI_ISL_3082480, EPI_ISL_3082481, EPI_ISL_3082482, EPI_ISL_3082483, EPI_ISL_3082484, EPI_ISL_3082485, EPI_ISL_3082486, EPI_ISL_3082487, EPI_ISL_3082488, EPI_ISL_3082489, EPI_ISL_3082490, EPI_ISL_3082491, EPI_ISL_3082492, EPI_ISL_3082493, EPI_ISL_3082494, EPI_ISL_3082495, EPI_ISL_3082496, EPI_ISL_3082497, EPI_ISL_3082498, EPI_ISL_3082499, EPI_ISL_3082500, EPI_ISL_3082501, EPI_ISL_3082502, EPI_ISL_3082503, EPI_ISL_3082504, EPI_ISL_3082505, EPI_ISL_3082506, EPI_ISL_3082507, EPI_ISL_3082508, EPI_ISL_3082509, EPI_ISL_3082510, EPI_ISL_3082511, EPI_ISL_3082512, EPI_ISL_3082513, EPI_ISL_3082514, EPI_ISL_3082515, EPI_ISL_3082516, EPI_ISL_3082517, EPI_ISL_3082518, EPI_ISL_3082519, EPI_ISL_3082520, EPI_ISL_3082521, EPI_ISL_3082522, EPI_ISL_3082523, EPI_ISL_3082524, EPI_ISL_3082525, EPI_ISL_3082526, EPI_ISL_3082527, EPI_ISL_3082528, EPI_ISL_3082529, EPI_ISL_3082530, EPI_ISL_3082531, EPI_ISL_3082532, EPI_ISL_3082533, EPI_ISL_3082534, EPI_ISL_3082535, EPI_ISL_3082536, EPI_ISL_3082537, EPI_ISL_3082538, EPI_ISL_3082539, EPI_ISL_3082540, EPI_ISL_3082541, EPI_ISL_3082542, EPI_ISL_3082543, EPI_ISL_3082544, EPI_ISL_3082545, EPI_ISL_3082546, EPI_ISL_3082547, EPI_ISL_3082548, EPI_ISL_3082549, EPI_ISL_3082550, EPI_ISL_3082551, EPI_ISL_3082552, EPI_ISL_3082553, EPI_ISL_3082554, EPI_ISL_3082555, EPI_ISL_3082556, EPI_ISL_3082557, EPI_ISL_3082558, EPI_ISL_3082559, EPI_ISL_3082560, EPI_ISL_3082561, EPI_ISL_3082562, EPI_ISL_3082563, EPI_ISL_3082564, EPI_ISL_3082565, EPI_ISL_3082566, EPI_ISL_3082567, EPI_ISL_3082568, EPI_ISL_3082569, EPI_ISL_3082570, EPI_ISL_3082571, EPI_ISL_3082572, EPI_ISL_3082573, EPI_ISL_3082574, EPI_ISL_3082575, EPI_ISL_3082576, EPI_ISL_3082577, EPI_ISL_3082578, EPI_ISL_3082579, EPI_ISL_3082580, EPI_ISL_3082581, EPI_ISL_3082582, EPI_ISL_3082583, EPI_ISL_3082584, EPI_ISL_3082585, EPI_ISL_3082586, EPI_ISL_3082587, EPI_ISL_3082588, EPI_ISL_3082589, EPI_ISL_3082590, EPI_ISL_3082591, EPI_ISL_3082592, EPI_ISL_3082593, EPI_ISL_3082594, EPI_ISL_3082595, EPI_ISL_3082596, EPI_ISL_3082597, EPI_ISL_3082598, EPI_ISL_3082599, EPI_ISL_3082600, EPI_ISL_3082601, EPI_ISL_3082602, EPI_ISL_3082603, EPI_ISL_3082604, EPI_ISL_3082605, EPI_ISL_3082606, EPI_ISL_3082607, EPI_ISL_3082608, EPI_ISL_3082609, EPI_ISL_3082610, EPI_ISL_3082611, EPI_ISL_3082612, EPI_ISL_3082613, EPI_ISL_3082614, EPI_ISL_3082615, EPI_ISL_3082616, EPI_ISL_3082617, EPI_ISL_3082618, EPI_ISL_3082619, EPI_ISL_3082620, EPI_ISL_3082621, EPI_ISL_3082622, EPI_ISL_3082623, EPI_ISL_3082624, EPI_ISL_3082625, EPI_ISL_3082626, EPI_ISL_3082627, EPI_ISL_3082628, EPI_ISL_3082629, EPI_ISL_3082630, EPI_ISL_3082631, EPI_ISL_3082632, EPI_ISL_3082633, EPI_ISL_3082634, EPI_ISL_3082635, EPI_ISL_3082636, EPI_ISL_3082637, EPI_ISL_3082638, EPI_ISL_3082639, EPI_ISL_3082640

| | | | |
|---|---|---|---|
| see above | Originating lab: Wales Specialist Virology Centre Sequencing lab: Pathogen Genomics Unit | Public Health Wales Microbiology Cardiff Wales Specialist Virology Centre | Catherine Moore, Johnathan Evans, Laura Gifford, Malorie Perry, Simon Cottrell, Angela Marchbank, Alec Birchley, Alexander Adams, Amy Gaskin, Bree Gatica-Wilcox, Jason Coombes, Joel Southgate, Lauren Gilbert, Lee Graham, Nicole Pacchiarini, Sara Kumziene-Summerhayes, Sarah Taylor, Sophie Jones, Sara Rey, Matthew Bull, Joanne Watkins, Sally Corden, Tom Connor |
| EPI_ISL_3082641 | Tuzla University Clinical Center, Department of Microbiology | University of Sarajevo, Veterinary Faculty, Laboratory for Molecular Diagnostic and Research Laboratory | Goletic S, Goletic T, Tihic N, Softic A, Nicevic M., Terzic I., Sabic E., Hodzic A., Jazic A. |

EPI_ISL_3082642, EPI_ISL_3082643, EPI_ISL_3082644, EPI_ISL_3082645, EPI_ISL_3082646, EPI_ISL_3082647, EPI_ISL_3082648, EPI_ISL_3082649, EPI_ISL_3082650, EPI_ISL_3082651, EPI_ISL_3082652, EPI_ISL_3082653, EPI_ISL_3082654, EPI_ISL_3082655, EPI_ISL_3082656, EPI_ISL_3082657, EPI_ISL_3082658, EPI_ISL_3082659, EPI_ISL_3082660, EPI_ISL_3082661, EPI_ISL_3082662, EPI_ISL_3082663, EPI_ISL_3082664, EPI_ISL_3082665, EPI_ISL_3082666, EPI_ISL_3082667, EPI_ISL_3082668, EPI_ISL_3082669, EPI_ISL_3082670, EPI_ISL_3082671, EPI_ISL_3082672, EPI_ISL_3082673, EPI_ISL_3082674, EPI_ISL_3082675, EPI_ISL_3082676, EPI_ISL_3082677, EPI_ISL_3082678, EPI_ISL_3082679, EPI_ISL_3082680, EPI_ISL_3082681, EPI_ISL_3082682, EPI_ISL_3082683, EPI_ISL_3082684, EPI_ISL_3082685, EPI_ISL_3082686, EPI_ISL_3082687, EPI_ISL_3082688, EPI_ISL_3082689, EPI_ISL_3082690, EPI_ISL_3082691, EPI_ISL_3082692, EPI_ISL_3082693, EPI_ISL_3082694, EPI_ISL_3082695, EPI_ISL_3082696, EPI_ISL_3082697, EPI_ISL_3082698, EPI_ISL_3082699, EPI_ISL_3082700, EPI_ISL_3082701, EPI_ISL_3082702, EPI_ISL_3082703, EPI_ISL_3082704, EPI_ISL_3082705, EPI_ISL_3082706, EPI_ISL_3082707, EPI_ISL_3082708, EPI_ISL_3082709, EPI_ISL_3082710, EPI_ISL_3082711, EPI_ISL_3082712, EPI_ISL_3082713, EPI_ISL_3082714, EPI_ISL_3082715, EPI_ISL_3082716, EPI_ISL_3082717, EPI_ISL_3082718, EPI_ISL_3082719, EPI_ISL_3082720, EPI_ISL_3082721, EPI_ISL_3082722, EPI_ISL_3082723, EPI_ISL_3082724, EPI_ISL_3082725, EPI_ISL_3082726, EPI_ISL_3082727, EPI_ISL_3082728, EPI_ISL_3082729, EPI_ISL_3082730, EPI_ISL_3082731, EPI_ISL_3082732, EPI_ISL_3082733, EPI_ISL_3082734, EPI_ISL_3082735, EPI_ISL_3082736, EPI_ISL_3082737, EPI_ISL_3082738, EPI_ISL_3082739, EPI_ISL_3082740, EPI_ISL_3082741, EPI_ISL_3082742, EPI_ISL_3082743, EPI_ISL_3082744, EPI_ISL_3082745, EPI_ISL_3082746, EPI_ISL_3082747, EPI_ISL_3082748, EPI_ISL_3082749, EPI_ISL_3082750, EPI_ISL_3082751, EPI_ISL_3082752, EPI_ISL_3082753, EPI_ISL_3082754, EPI_ISL_3082755, EPI_ISL_3082756, EPI_ISL_3082757, EPI_ISL_3082758, EPI_ISL_3082759, EPI_ISL_3082760, EPI_ISL_3082761, EPI_ISL_3082762, EPI_ISL_3082763, EPI_ISL_3082764, EPI_ISL_3082765, EPI_ISL_3082766, EPI_ISL_3082767, EPI_ISL_3082768, EPI_ISL_3082769, EPI_ISL_3082770, EPI_ISL_3082771, EPI_ISL_3082772, EPI_ISL_3082773, EPI_ISL_3082774, EPI_ISL_3082775, EPI_ISL_3082776, EPI_ISL_3082777, EPI_ISL_3082778, EPI_ISL_3082779, EPI_ISL_3082780, EPI_ISL_3082781, EPI_ISL_3082782, EPI_ISL_3082783, EPI_ISL_3082784, EPI_ISL_3082785, EPI_ISL_3082786, EPI_ISL_3082787, EPI_ISL_3082788, EPI_ISL_3082789, EPI_ISL_3082790, EPI_ISL_3082791, EPI_ISL_3082792, EPI_ISL_3082793, EPI_ISL_3082794, EPI_ISL_3082795, EPI_ISL_3082796, EPI_ISL_3082797, EPI_ISL_3082798, EPI_ISL_3082799, EPI_ISL_3082800, EPI_ISL_3082801, EPI_ISL_3082802, EPI_ISL_3082803, EPI_ISL_3082804, EPI_ISL_3082805, EPI_ISL_3082806, EPI_ISL_3082807, EPI_ISL_3082808, EPI_ISL_3082809, EPI_ISL_3082810, EPI_ISL_3082811, EPI_ISL_3082812, EPI_ISL_3082813, EPI_ISL_3082814, EPI_ISL_3082815, EPI_ISL_3082816, EPI_ISL_3082817, EPI_ISL_3082818, EPI_ISL_3082819, EPI_ISL_3082820, EPI_ISL_3082821, EPI_ISL_3082822, EPI_ISL_3082823, EPI_ISL_3082824, EPI_ISL_3082825, EPI_ISL_3082826, EPI_ISL_3082827, EPI_ISL_3082828, EPI_ISL_3082829, EPI_ISL_3082830, EPI_ISL_3082831, EPI_ISL_3082832, EPI_ISL_3082833, EPI_ISL_3082834, EPI_ISL_3082835, EPI_ISL_3082836, EPI_ISL_3082837, EPI_ISL_3082838, EPI_ISL_3082839, EPI_ISL_3082840, EPI_ISL_3082841, EPI_ISL_3082842, EPI_ISL_3082843, EPI_ISL_3082844, EPI_ISL_3082845, EPI_ISL_3082846, EPI_ISL_3082847, EPI_ISL_3082848, EPI_ISL_3082849, EPI_ISL_3082850, EPI_ISL_3082851, EPI_ISL_3082852, EPI_ISL_3082853, EPI_ISL_3082854, EPI_ISL_3082855, EPI_ISL_3082856, EPI_ISL_3082857, EPI_ISL_3082858, EPI_ISL_3082859, EPI_ISL_3082860, EPI_ISL_3082861, EPI_ISL_3082862, EPI_ISL_3082863, EPI_ISL_3082864, EPI_ISL_3082865, EPI_ISL_3082866, EPI_ISL_3082867, EPI_ISL_3082868, EPI_ISL_3082869, EPI_ISL_3082870, EPI_ISL_3082871, EPI_ISL_3082872, EPI_ISL_3082873, EPI_ISL_3082874, EPI_ISL_3082875, EPI_ISL_3082876, EPI_ISL_3082877, EPI_ISL_3082878, EPI_ISL_3082879, EPI_ISL_3082880, EPI_ISL_3082881, EPI_ISL_3082882, EPI_ISL_3082883, EPI_ISL_3082884, EPI_ISL_3082885, EPI_ISL_3082886, EPI_ISL_3082887, EPI_ISL_3082888, EPI_ISL_3082889, EPI_ISL_3082890, EPI_ISL_3082891, EPI_ISL_3082892, EPI_ISL_3082893, EPI_ISL_3082894, EPI_ISL_3082895, EPI_ISL_3082896, EPI_ISL_3082897, EPI_ISL_3082898, EPI_ISL_3082899, EPI_ISL_3082900, EPI_ISL_3082901, EPI_ISL_3082902, EPI_ISL_3082903, EPI_ISL_3082904, EPI_ISL_3082905, EPI_ISL_3082906, EPI_ISL_3082907, EPI_ISL_3082908, EPI_ISL_3082909, EPI_ISL_3082910, EPI_ISL_3082911, EPI_ISL_3082912, EPI_ISL_3082913, EPI_ISL_3082914, EPI_ISL_3082915, EPI_ISL_3082916, EPI_ISL_3082917, EPI_ISL_3082918, EPI_ISL_3082919, EPI_ISL_3082920, EPI_ISL_3082921, EPI_ISL_3082922, EPI_ISL_3082923, EPI_ISL_3082924, EPI_ISL_3082925, EPI_ISL_3082926, EPI_ISL_3082927, EPI_ISL_3082928, EPI_ISL_3082929, EPI_ISL_3082930, EPI_ISL_3082931, EPI_ISL_3082932, EPI_ISL_3082933, EPI_ISL_3082934, EPI_ISL_3082935, EPI_ISL_3082936, EPI_ISL_3082937, EPI_ISL_3082938, EPI_ISL_3082939, EPI_ISL_3082940, EPI_ISL_3082941, EPI_ISL_3082942, EPI_ISL_3082943, EPI_ISL_3082944, EPI_ISL_3082945, EPI_ISL_3082946, EPI_ISL_3082947, EPI_ISL_3082948, EPI_ISL_3082949, EPI_ISL_3082950, EPI_ISL_3082951, EPI_ISL_3082952, EPI_ISL_3082953, EPI_ISL_3082954, EPI_ISL_3082955, EPI_ISL_3082956, EPI_ISL_3082957, EPI_ISL_3082958, EPI_ISL_3082959, EPI_ISL_3082960, EPI_ISL_3082961, EPI_ISL_3082962, EPI_ISL_3082963, EPI_ISL_3082964, EPI_ISL_3082965, EPI_ISL_3082966, EPI_ISL_3082967, EPI_ISL_3082968, EPI_ISL_3082969, EPI_ISL_3082970, EPI_ISL_3082971, EPI_ISL_3082972, EPI_ISL_3082973, EPI_ISL_3082974, EPI_ISL_3082975, EPI_ISL_3082976, EPI_ISL_3082977, EPI_ISL_3082978, EPI_ISL_3082979, EPI_ISL_3082980, EPI_ISL_3082981, EPI_ISL_3082982, EPI_ISL_3082983, EPI_ISL_3082984, EPI_ISL_3082985, EPI_ISL_3082986, EPI_ISL_3082987, EPI_ISL_3082988, EPI_ISL_3082989, EPI_ISL_3082990, EPI_ISL_3082991, EPI_ISL_3082992, EPI_ISL_3082993, EPI_ISL_3082994, EPI_ISL_3082995, EPI_ISL_3082996, EPI_ISL_3082997, EPI_ISL_3082998, EPI_ISL_3082999, EPI_ISL_3083000, EPI_ISL_3083001, EPI_ISL_3083002, EPI_ISL_3083003, EPI_ISL_3083004, EPI_ISL_3083005, EPI_ISL_3083006, EPI_ISL_3083007, EPI_ISL_3083008, EPI_ISL_3083009, EPI_ISL_3083010, EPI_ISL_3083011, EPI_ISL_3083012, EPI_ISL_3083013, EPI_ISL_3083014, EPI_ISL_3083015, EPI_ISL_3083016, EPI_ISL_3083017, EPI_ISL_3083018, EPI_ISL_3083019, EPI_ISL_3083020, EPI_ISL_3083021, EPI_ISL_3083022, EPI_ISL_3083023, EPI_ISL_3083024, EPI_ISL_3083025, EPI_ISL_3083026, EPI_ISL_3083027, EPI_ISL_3083028, EPI_ISL_3083029, EPI_ISL_3083030, EPI_ISL_3083031, EPI_ISL_3083032, EPI_ISL_3083033, EPI_ISL_3083034, EPI_ISL_3083035, EPI_ISL_3083036, EPI_ISL_3083037, EPI_ISL_3083038, EPI_ISL_3083039, EPI_ISL_3083040, EPI_ISL_3083041, EPI_ISL_3083042, EPI_ISL_3083043, EPI_ISL_3083044, EPI_ISL_3083045, EPI_ISL_3083046, EPI_ISL_3083047, EPI_ISL_3083048, EPI_ISL_3083049, EPI_ISL_3083050, EPI_ISL_3083051, EPI_ISL_3083052, EPI_ISL_3083053, EPI_ISL_3083054, EPI_ISL_3083055, EPI_ISL_3083056, EPI_ISL_3083057, EPI_ISL_3083058, EPI_ISL_3083059, EPI_ISL_3083060, EPI_ISL_3083061, EPI_ISL_3083062, EPI_ISL_3083063, EPI_ISL_3083064, EPI_ISL_3083065, EPI_ISL_3083066, EPI_ISL_3083067, EPI_ISL_3083068, EPI_ISL_3083069, EPI_ISL_3083070, EPI_ISL_3083071, EPI_ISL_3083072, EPI_ISL_3083073, EPI_ISL_3083074, EPI_ISL_3083075, EPI_ISL_3083076, EPI_ISL_3083077, EPI_ISL_3083078, EPI_ISL_3083079, EPI_ISL_3083080, EPI_ISL_3083081, EPI_ISL_3083082, EPI_ISL_3083083, EPI_ISL_3083084, EPI_ISL_3083085, EPI_ISL_3083086, EPI_ISL_3083087, EPI_ISL_3083088, EPI_ISL_3083089, EPI_ISL_3083090, EPI_ISL_3083091, EPI_ISL_3083092, EPI_ISL_3083093, EPI_ISL_3083094, EPI_ISL_3083095, EPI_ISL_3083096, EPI_ISL_3083097, EPI_ISL_3083098, EPI_ISL_3083099, EPI_ISL_3083100, EPI_ISL_3083101, EPI_ISL_3083102, EPI_ISL_3083103, EPI_ISL_3083104, EPI_ISL_3083105, EPI_ISL_3083106, EPI_ISL_3083107, EPI_ISL_3083108, EPI_ISL_3083109, EPI_ISL_3083110, EPI_ISL_3083111, EPI_ISL_3083112, EPI_ISL_3083113, EPI_ISL_3083114, EPI_ISL_3083115, EPI_ISL_3083116, EPI_ISL_3083117, EPI_ISL_3083118, EPI_ISL_3083119, EPI_ISL_3083120, EPI_ISL_3083121, EPI_ISL_3083122, EPI_ISL_3083123, EPI_ISL_3083124, EPI_ISL_3083125, EPI_ISL_3083126, EPI_ISL_3083127, EPI_ISL_3083128, EPI_ISL_3083129, EPI_ISL_3083130, EPI_ISL_3083131, EPI_ISL_3083132, EPI_ISL_3083133, EPI_ISL_3083134, EPI_ISL_3083135, EPI_ISL_3083136, EPI_ISL_3083137, EPI_ISL_3083138, EPI_ISL_3083139, EPI_ISL_3083140, EPI_ISL_3083141, EPI_ISL_3083142, EPI_ISL_3083143, EPI_ISL_3083144, EPI_ISL_3083145, EPI_ISL_3083146, EPI_ISL_3083147, EPI_ISL_3083148, EPI_ISL_3083149, EPI_ISL_3083150, EPI_ISL_3083151, EPI_ISL_3083152, EPI_ISL_3083153, EPI_ISL_3083154, EPI_ISL_3083155, EPI_ISL_3083156, EPI_ISL_3083157, EPI_ISL_3083158, EPI_ISL_3083159, EPI_ISL_3083160, EPI_ISL_3083161, EPI_ISL_3083162, EPI_ISL_3083163, EPI_ISL_3083164, EPI_ISL_3083165, EPI_ISL_3083166, EPI_ISL_3083167, EPI_ISL_3083168, EPI_ISL_3083169, EPI_ISL_3083170, EPI_ISL_3083171, EPI_ISL_3083172, EPI_ISL_3083173, EPI_ISL_3083174, EPI_ISL_3083175, EPI_ISL_3083176, EPI_ISL_3083177, EPI_ISL_3083178, EPI_ISL_3083179, EPI_ISL_3083180, EPI_ISL_3083181, EPI_ISL_3083182, EPI_ISL_3083183, EPI_ISL_3083184, EPI_ISL_3083185, EPI_ISL_3083186, EPI_ISL_3083187, EPI_ISL_3083188, EPI_ISL_3083189, EPI_ISL_3083190, EPI_ISL_3083191, EPI_ISL_3083192, EPI_ISL_3083193, EPI_ISL_3083194, EPI_ISL_3083195, EPI_ISL_3083196, EPI_ISL_3083197, EPI_ISL_3083198, EPI_ISL_3083199, EPI_ISL_3083200, EPI_ISL_3083201, EPI_ISL_3083202, EPI_ISL_3083203, EPI_ISL_3083204, EPI_ISL_3083205, EPI_ISL_3083206, EPI_ISL_3083207, EPI_ISL_3083208, EPI_ISL_3083209, EPI_ISL_3083210, EPI_ISL_3083211, EPI_ISL_3083212, EPI_ISL_3083213, EPI_ISL_3083214, EPI_ISL_3083215, EPI_ISL_3083216, EPI_ISL_3083217, EPI_ISL_3083218, EPI_ISL_3083219, EPI_ISL_3083220, EPI_ISL_3083221, EPI_ISL_3083222, EPI_ISL_3083223, EPI_ISL_3083224, EPI_ISL_3083225, EPI_ISL_3083226, EPI_ISL_3083227, EPI_ISL_3083228, EPI_ISL_3083229, EPI_ISL_3083230, EPI_ISL_3083231, EPI_ISL_3083232, EPI_ISL_3083233, EPI_ISL_3083234, EPI_ISL_3083235, EPI_ISL_3083236, EPI_ISL_3083237, EPI_ISL_3083238, EPI_ISL_3083239, EPI_ISL_3083240, EPI_ISL_3083241, EPI_ISL_3083242, EPI_ISL_3083243, EPI_ISL_3083244, EPI_ISL_3083245, EPI_ISL_3083246, EPI_ISL_3083247, EPI_ISL_3083248, EPI_ISL_3083249, EPI_ISL_3083250, EPI_ISL_3083251, EPI_ISL_3083252, EPI_ISL_3083253, EPI_ISL_3083254, EPI_ISL_3083255, EPI_ISL_3083256, EPI_ISL_3083257, EPI_ISL_3083258, EPI_ISL_3083259, EPI_ISL_3083260, EPI_ISL_3083261, EPI_ISL_3083262, EPI_ISL_3083263, EPI_ISL_3083264, EPI_ISL_3083265, EPI_ISL_3083266, EPI_ISL_3083267, EPI_ISL_3083268, EPI_ISL_3083269, EPI_ISL_3083270, EPI_ISL_3083271, EPI_ISL_3083272, EPI_ISL_3083273, EPI_ISL_3083274, EPI_ISL_3083275, EPI_ISL_3083276, EPI_ISL_3083277, EPI_ISL_3083278, EPI_ISL_3083279, EPI_ISL_3083280, EPI_ISL_3083281, EPI_ISL_3083282, EPI_ISL_3083283, EPI_ISL_3083284, EPI_ISL_3083285, EPI_ISL_3083286, EPI_ISL_3083287, EPI_ISL_3083288, EPI_ISL_3083289, EPI_ISL_3083290, EPI_ISL_3083291, EPI_ISL_3083292, EPI_ISL_3083293, EPI_ISL_3083294, EPI_ISL_3083295, EPI_ISL_3083296, EPI_ISL_3083297, EPI_ISL_3083298, EPI_ISL_3083299, EPI_ISL_3083300, EPI_ISL_3083301, EPI_ISL_3083302, EPI_ISL_3083303, EPI_ISL_3083304, EPI_ISL_3083305, EPI_ISL_3083306, EPI_ISL_3083307, EPI_ISL_3083308, EPI_ISL_3083309, EPI_ISL_3083310, EPI_ISL_3083311, EPI_ISL_3083312, EPI_ISL_3083313, EPI_ISL_3083314, EPI_ISL_3083315, EPI_ISL_3083316, EPI_ISL_3083317, EPI_ISL_3083318, EPI_ISL_3083319, EPI_ISL_3083320, EPI_ISL_3083321, EPI_ISL_3083322, EPI_ISL_3083323, EPI_ISL_3083324, EPI_ISL_3083325, EPI_ISL_3083326, EPI_ISL_3083327, EPI_ISL_3083328, EPI_ISL_3083329, EPI_ISL_3083330, EPI_ISL_3083331, EPI_ISL_3083332, EPI_ISL_3083333, EPI_ISL_3083334, EPI_ISL_3083335, EPI_ISL_3083336, EPI_ISL_3083337, EPI_ISL_3083338, EPI_ISL_3083339, EPI_ISL_3083340, EPI_ISL_3083341, EPI_ISL_3083342, EPI_ISL_3083343, EPI_ISL_3083344, EPI_ISL_3083345, EPI_ISL_3083346, EPI_ISL_3083347, EPI_ISL_3083348, EPI_ISL_3083349, EPI_ISL_3083350, EPI_ISL_3083351, EPI_ISL_3083352, EPI_ISL_3083353, EPI_ISL_3083354, EPI_ISL_3083355, EPI_ISL_3083356, EPI_ISL_3083357, EPI_ISL_3083358, EPI_ISL_3083359, EPI_ISL_3083360, EPI_ISL_3083361, EPI_ISL_3083362, EPI_ISL_3083363, EPI_ISL_3083364, EPI_ISL_3083365, EPI_ISL_3083366, EPI_ISL_3083367, EPI_ISL_3083368, EPI_ISL_3083369, EPI_ISL_3083370, EPI_ISL_3083371, EPI_ISL_3083372, EPI_ISL_3083373, EPI_ISL_3083374, EPI_ISL_3083375, EPI_ISL_3083376, EPI_ISL_3083377, EPI_ISL_3083378, EPI_ISL_3083379, EPI_ISL_3083380, EPI_ISL_3083381, EPI_ISL_3083382, EPI_ISL_3083383, EPI_ISL_3083384, EPI_ISL_3083385, EPI_ISL_3083386, EPI_ISL_3083387, EPI_ISL_3083388, EPI_ISL_3083389, EPI_ISL_3083390, EPI_ISL_3083391, EPI_ISL_3083392, EPI_ISL_3083393, EPI_ISL_3083394, EPI_ISL_3083395, EPI_ISL_3083396, EPI_ISL_3083397, EPI_ISL_3083398, EPI_ISL_3083399, EPI_ISL_3083400, EPI_ISL_3083401, EPI_ISL_3083402, EPI_ISL_3083403, EPI_ISL_3083404, EPI_ISL_3083405, EPI_ISL_3083406, EPI_ISL_3083407, EPI_ISL_3083408, EPI_ISL_3083409, EPI_ISL_3083410, EPI_ISL_3083411, EPI_ISL_3083412, EPI_ISL_3083413, EPI_ISL_3083414, EPI_ISL_3083415, EPI_ISL_3083416, EPI_ISL_3083417, EPI_ISL_3083418, EPI_ISL_3083419, EPI_ISL_3083420, EPI_ISL_3083421, EPI_ISL_3083422, EPI_ISL_3083423, EPI_ISL_3083424, EPI_ISL_3083425, EPI_ISL_3083426, EPI_ISL_3083427, EPI_ISL_3083428, EPI_ISL_3083429, EPI_ISL_3083430, EPI_ISL_3083431, EPI_ISL_3083432, EPI_ISL_3083433, EPI_ISL_3083434, EPI_ISL_3083435, EPI_ISL_3083436, EPI_ISL_3083437, EPI_ISL_3083438, EPI_ISL_3083439, EPI_ISL_3083440, EPI_ISL_3083441, EPI_ISL_3083442, EPI_ISL_3083443, EPI_ISL_3083444, EPI_ISL_3083445, EPI_ISL_3083446, EPI_ISL_3083447, EPI_ISL_3083448, EPI_ISL_3083449, EPI_ISL_3083450, EPI_ISL_3083451, EPI_ISL_3083452, EPI_ISL_3083453, EPI_ISL_3083454, EPI_ISL_3083455, EPI_ISL_3083456, EPI_ISL_3083457, EPI_ISL_3083458, EPI_ISL_3083459, EPI_ISL_3083460, EPI_ISL_3083461, EPI_ISL_3083462, EPI_ISL_3083463, EPI_ISL_3083464, EPI_ISL_3083465, EPI_ISL_3083466, EPI_ISL_3083467, EPI_ISL_3083468, EPI_ISL_3083469, EPI_ISL_3083470, EPI_ISL_3083471, EPI_ISL_3083472, EPI_ISL_3083473, EPI_ISL_3083474, EPI_ISL_3083475, EPI_ISL_3083476, EPI_ISL_3083477, EPI_ISL_3083478, EPI_ISL_3083479, EPI_ISL_3083480, EPI_ISL_3083481, EPI_ISL_3083482, EPI_ISL_3083483, EPI_ISL_3083484, EPI_ISL_3083485, EPI_ISL_3083486, EPI_ISL_3083487, EPI_ISL_3083488, EPI_ISL_3083489, EPI_ISL_3083490, EPI_ISL_3083491, EPI_ISL_3083492, EPI_ISL_3083493, EPI_ISL_3083494, EPI_ISL_3083495, EPI_ISL_3083496, EPI_ISL_3083497, EPI_ISL_3083498, EPI_ISL_3083499, EPI_ISL_3083500, EPI_ISL_3083501, EPI_ISL_3083502, EPI_ISL_3083503, EPI_ISL_3083504, EPI_ISL_3083505, EPI_ISL_3083506, EPI_ISL_3083507, EPI_ISL_3083508, EPI_ISL_3083509, EPI_ISL_3083510, EPI_ISL_3083511, EPI_ISL_3083512, EPI_ISL_3083513, EPI_ISL_3083514, EPI_ISL_3083515, EPI_ISL_3083516, EPI_ISL_3083517, EPI_ISL_3083518, EPI_ISL_3083519, EPI_ISL_3083520, EPI_ISL_3083521, EPI_ISL_3083522, EPI_ISL_3083523, EPI_ISL_3083524, EPI_ISL_3083525, EPI_ISL_3083526, EPI_ISL_3083527, EPI_ISL_3083528, EPI_ISL_3083529, EPI_ISL_3083530, EPI_ISL_3083531, EPI_ISL_3083532, EPI_ISL_3083533, EPI_ISL_3083534, EPI_ISL_3083535, EPI_ISL_3083536, EPI_ISL_3083537, EPI_ISL_3083538, EPI_ISL_3083539, EPI_ISL_3083540, EPI_ISL_3083541, EPI_ISL_3083542, EPI_ISL_3083543, EPI_ISL_3083544, EPI_ISL_3083545, EPI_ISL_3083546, EPI_ISL_3083547, EPI_ISL_3083548, EPI_ISL_3083549, EPI_ISL_3083550, EPI_ISL_3083551, EPI_ISL_3083552, EPI_ISL_3083553, EPI_ISL_3083554, EPI_ISL_3083555, EPI_ISL_3083556, EPI_ISL_3083557, EPI_ISL_3083558, EPI_ISL_3083559, EPI_ISL_3083560, EPI_ISL_3083561, EPI_ISL_3083562, EPI_ISL_3083563, EPI_ISL_3083564, EPI_ISL_3083565, EPI_ISL_3083566, EPI_ISL_3083567, EPI_ISL_3083568, EPI_ISL_3083569, EPI_ISL_3083570, EPI_ISL_3083571, EPI_ISL_3083572, EPI_ISL_3083573, EPI_ISL_3083574, EPI_ISL_3083575, EPI_ISL_3083576, EPI_ISL_3083577, EPI_ISL_3083578, EPI_ISL_3083579, EPI_ISL_3083580, EPI_ISL_3083581, EPI_ISL_3083582, EPI_ISL_3083583, EPI_ISL_3083584, EPI_ISL_3083585, EPI_ISL_3083586, EPI_ISL_3083587, EPI_ISL_3083588, EPI_ISL_3083589, EPI_ISL_3083590, EPI_ISL_3083591, EPI_ISL_3083592, EPI_ISL_3083593, EPI_ISL_3083594, EPI_ISL_3083595, EPI_ISL_3083596, EPI_ISL_3083597, EPI_ISL_3083598, EPI_ISL_3083599, EPI_ISL_3083600, EPI_ISL_3083601, EPI_ISL_3083602, EPI_ISL_3083603, EPI_ISL_3083604, EPI_ISL_3083605, EPI_ISL_3083606, EPI_ISL_3083607, EPI_ISL_3083608, EPI_ISL_3083609, EPI_ISL_3083610, EPI_ISL_3083611, EPI_ISL_3083612, EPI_ISL_3083613, EPI_ISL_3083614, EPI_ISL_3083615, EPI_ISL_3083616, EPI_ISL_3083617, EPI_ISL_3083618, EPI_ISL_3083619, EPI_ISL_3083620, EPI_ISL_3083621, EPI_ISL_3083622, EPI_ISL_3083623, EPI_ISL_3083624, EPI_ISL_3083625, EPI_ISL_3083626, EPI_ISL_3083627, EPI_ISL_3083628, EPI_ISL_3083629, EPI_ISL_3083630, EPI_ISL_3083631, EPI_ISL_3083632, EPI_ISL_3083633, EPI_ISL_3083634, EPI_ISL_3083635, EPI_ISL_3083636, EPI_ISL_3083637, EPI_ISL_3083638, EPI_ISL_3083639, EPI_ISL_3083640, EPI_ISL_3083641, EPI_ISL_3083642, EPI_ISL_3083643, EPI_ISL_3083644, EPI_ISL_3083645, EPI_ISL_3083646, EPI_ISL_3083647, EPI_ISL_3083648, EPI_ISL_3083649, EPI_ISL_3083650, EPI_ISL_3083651, EPI_ISL_3083652, EPI_ISL_3083653, EPI_ISL_3083654, EPI_ISL_3083655, EPI_ISL_3083656, EPI_ISL_3083657, EPI_ISL_3083658, EPI_ISL_3083659, EPI_ISL_3083660, EPI_ISL_3083661, EPI_ISL_3083662, EPI_ISL_3083663, EPI_ISL_3083664, EPI_ISL_3083665, EPI_ISL_3083666, EPI_ISL_3083667, EPI_ISL_3083668, EPI_ISL_3083669, EPI_ISL_3083670, EPI_ISL_3083671, EPI_ISL_3083672, EPI_ISL_3083673, EPI_ISL_3083674, EPI_ISL_3083675, EPI_ISL_3083676, EPI_ISL_3083677, EPI_ISL_3083678, EPI_ISL_3083679, EPI_ISL_3083680, EPI_ISL_3083681, EPI_ISL_3083682, EPI_ISL_3083683, EPI_ISL_3083684, EPI_ISL_3083685, EPI_ISL_3083686, EPI_ISL_3083687, EPI_ISL_3083688, EPI_ISL_3083689, EPI_ISL_3083690, EPI_ISL_3083691, EPI_ISL_3083692, EPI_ISL_3083693, EPI_ISL_3083694, EPI_ISL_3083695, EPI_ISL_3083696, EPI_ISL_3083697, EPI_ISL_3083698, EPI_ISL_3083699, EPI_ISL_3083700, EPI_ISL_3083701, EPI_ISL_3083702, EPI_ISL_3083703, EPI_ISL_3083704, EPI_ISL_3083705, EPI_ISL_3083706, EPI_ISL_3083707, EPI_ISL_3083708, EPI_ISL_3083709, EPI_ISL_3083710, EPI_ISL_3083711, EPI_ISL_3083712, EPI_ISL_3083713, EPI_ISL_3083714, EPI_ISL_3083715, EPI_ISL_3083716, EPI_ISL_3083717, EPI_ISL_3083718, EPI_ISL_3083719, EPI_ISL_3083720, EPI_ISL_3083721, EPI_ISL_3083722, EPI_ISL_3083723, EPI_ISL_3083724, EPI_ISL_3083725, EPI_ISL_3083726, EPI_ISL_3083727, EPI_ISL_3083728, EPI_ISL_3083729, EPI_ISL_3083730, EPI_ISL_3083731, EPI_ISL_3083732, EPI_ISL_3083733, EPI_ISL_3083734, EPI_ISL_3083735, EPI_ISL_3083736, EPI_ISL_3083737, EPI_ISL_3083738, EPI_ISL_3083739, EPI_ISL_3083740, EPI_ISL_3083741, EPI_ISL_3083742, EPI_ISL_3083743, EPI_ISL_3083744, EPI_ISL_3083745, EPI_ISL_3083746, EPI_ISL_3083747, EPI_ISL_3083748, EPI_ISL_3083749, EPI_ISL_3083750, EPI_ISL_3083751, EPI_ISL_3083752, EPI_ISL_3083753, EPI_ISL_3083754, EPI_ISL_3083755, EPI_ISL_3083756, EPI_ISL_3083757, EPI_ISL_3083758, EPI_ISL_3083759, EPI_ISL_3083760, EPI_ISL_3083761, EPI_ISL_3083762, EPI_ISL_3083763, EPI_ISL_3083764, EPI_ISL_3083765, EPI_ISL_3083766, EPI_ISL_3083767, EPI_ISL_3083768, EPI_ISL_3083769, EPI_ISL_3083770, EPI_ISL_3083771, EPI_ISL_3083772, EPI_ISL_3083773, EPI_ISL_3083774, EPI_ISL_3083775, EPI_ISL_3083776, EPI_ISL_3083777, EPI_ISL_3083778, EPI_ISL_3083779, EPI_ISL_3083780, EPI_ISL_3083781, EPI_ISL_3083782, EPI_ISL_3083783, EPI_ISL_3083784, EPI_ISL_3083785, EPI_ISL_3083786, EPI_ISL_3083787, EPI_ISL_3083788, EPI_ISL_3083789, EPI_ISL_3083790, EPI_ISL_3083791, EPI_ISL_3083792, EPI_ISL_3083793, EPI_ISL_3083794, EPI_ISL_3083795, EPI_ISL_3083796, EPI_ISL_3083797, EPI_ISL_3083798, EPI_ISL_3083799, EPI_ISL_3083800, EPI_ISL_3083801, EPI_ISL_3083802, EPI_ISL_3083803, EPI_ISL_3083804, EPI_ISL_3083805, EPI_ISL_3083806, EPI_ISL_3083807, EPI_ISL_3083808, EPI_ISL_3083809, EPI_ISL_3083810, EPI_ISL_3083811, EPI_ISL_3083812, EPI_ISL_3083813, EPI_ISL_3083814, EPI_ISL_3083815, EPI_ISL_3083816, EPI_ISL_3083817, EPI_ISL_3083818, EPI_ISL_3083819, EPI_ISL_3083820, EPI_ISL_3083821, EPI_ISL_3083822, EPI_ISL_3083823, EPI_ISL_3083824, EPI_ISL_3083825, EPI_ISL_3083826, EPI_ISL_3083827, EPI_ISL_3083828, EPI_ISL_3083829, EPI_ISL_3083830, EPI_ISL_3083831, EPI_ISL_3083832, EPI_ISL_3083833, EPI_ISL_3083834, EPI_ISL_3083835, EPI_ISL_3083836, EPI_ISL_3083837, EPI_ISL_3083838, EPI_ISL_3083839, EPI_ISL_3083840, EPI_ISL_3083841, EPI_ISL_3083842, EPI_ISL_3083843, EPI_ISL_3083844, EPI_ISL_3083845, EPI_ISL_3083846, EPI_ISL_3083847, EPI_ISL_3083848, EPI_ISL_3083849, EPI_ISL_3083850, EPI_ISL_3083851, EPI_ISL_3083852, EPI_ISL_3083853, EPI_ISL_3083854, EPI_ISL_3083855, EPI_ISL_3083856, EPI_ISL_3083857, EPI_ISL_3083858, EPI_ISL_3083859, EPI_ISL_3083860, EPI_ISL_3083861, EPI_ISL_3083862, EPI_ISL_3083863, EPI_ISL_3083864, EPI_ISL_3083865, EPI_ISL_3083866, EPI_ISL_3083867, EPI_ISL_3083868, EPI_ISL_3083869, EPI_ISL_3083870, EPI_ISL_3083871, EPI_ISL_3083872, EPI_ISL_3083873, EPI_ISL_3083874, EPI_ISL_3083875, EPI_ISL_3083876, EPI_ISL_3083877, EPI_ISL_3083878, EPI_ISL_3083879, EPI_ISL_3083880, EPI_ISL_3083881, EPI_ISL_3083882, EPI_ISL_3083883, EPI_ISL_3083884, EPI_ISL_3083885, EPI_ISL_3083886, EPI_ISL_3083887, EPI_ISL_3083888, EPI_ISL_3083889, EPI_ISL_3083890, EPI_ISL_3083891, EPI_ISL_3083892, EPI_ISL_3083893, EPI_ISL_3083894, EPI_ISL_3083895, EPI_ISL_3083896, EPI_ISL_3083897, EPI_ISL_3083898, EPI_ISL_3083899, EPI_ISL_3083900, EPI_ISL_3083901, EPI_ISL_3083902, EPI_ISL_3083903, EPI_ISL_3083904, EPI_ISL_3083905, EPI_ISL_3083906, EPI_ISL_3083907, EPI_ISL_3083908, EPI_ISL_3083909, EPI_ISL_3083910, EPI_ISL_3083911, EPI_ISL_3083912, EPI_ISL_3083913, EPI_ISL_3083914, EPI_ISL_3083915, EPI_ISL_3083916, EPI_ISL_3083917, EPI_ISL_3083918, EPI_ISL_3083919, EPI_ISL_3083920, EPI_ISL_3083921, EPI_ISL_3083922, EPI_ISL_3083923, EPI_ISL_3083924, EPI_ISL_3083925, EPI_ISL_3083926, EPI_ISL_3083927, EPI_ISL_3083928, EPI_ISL_3083929, EPI_ISL_3083930, EPI_ISL_3083931, EPI_ISL_3083932, EPI_ISL_3083933, EPI_ISL_3083934, EPI_ISL_3083935, EPI_ISL_3083936, EPI_ISL_3083937, EPI_ISL_3083938, EPI_ISL_3083939, EPI_ISL_3083940, EPI_ISL_3083941, EPI_ISL_3083942, EPI_ISL_3083943, EPI_ISL_3083944, EPI_ISL_3083945, EPI_ISL_3083946, EPI_ISL_3083947, EPI_ISL_3083948, EPI_ISL_3083949, EPI_ISL_3083950, EPI_ISL_3083951, EPI_ISL_3083952, EPI_ISL_3083953, EPI_ISL_3083954, EPI_ISL_3083955, EPI_ISL_3083956, EPI_ISL_3083957, EPI_ISL_3083958, EPI_ISL_3083959, EPI_ISL_3083960, EPI_ISL_3083961, EPI_ISL_3083962, EPI_ISL_3083963, EPI_ISL_3083964, EPI_ISL_3083965, EPI_ISL_3083966, EPI_ISL_3083967, EPI_ISL_3083968, EPI_ISL_3083969, EPI_ISL_3083970, EPI_ISL_3083971, EPI_ISL_3083972, EPI_ISL_3083973, EPI_ISL_3083974, EPI_ISL_3083975, EPI_ISL_3083976, EPI_ISL_3083977, EPI_ISL_3083978, EPI_ISL_3083979, EPI_ISL_3083980, EPI_ISL_3083981, EPI_ISL_3083982, EPI_ISL_3083983, EPI_ISL_3083984, EPI_ISL_3083985, EPI_ISL_3083986, EPI_ISL_3083987, EPI_ISL_3083988, EPI_ISL_3083989, EPI_ISL_3083990, EPI_ISL_3083991, EPI_ISL_3083992, EPI_ISL_3083993, EPI_ISL_3083994, EPI_ISL_3083995, EPI_ISL_3083996, EPI_ISL_3083997, EPI_ISL_3083998, EPI_ISL_3083999, EPI_ISL_3084000, EPI_ISL_3084001, EPI_ISL_3084002, EPI_ISL_3084003, EPI_ISL_3084004, EPI_ISL_3084005, EPI_ISL_3084006, EPI_ISL_3084007, EPI_ISL_3084008, EPI_ISL_3084009, EPI_ISL_3084010, EPI_ISL_3084011, EPI_ISL_3084012, EPI_ISL_3084013, EPI_ISL_3084014, EPI_ISL_3084015, EPI_ISL_3084016, EPI_ISL_3084017, EPI_ISL_3084018, EPI_ISL_3084019, EPI_ISL_3084020, EPI_ISL_3084021, EPI_ISL_3084022, EPI_ISL_3084023, EPI_ISL_3084024, EPI_ISL_3084025, EPI_ISL_3084026, EPI_ISL_3084027, EPI_ISL_3084028, EPI_ISL_3084029, EPI_ISL_3084030, EPI_ISL_3084031, EPI_ISL_3084032, EPI_ISL_3084033, EPI_ISL_3084034, EPI_ISL_3084035, EPI_ISL_3084036, EPI_ISL_3084037, EPI_ISL_3084038, EPI_ISL_3084039, EPI_ISL_3084040, EPI_ISL_3084041, EPI_ISL_3084042, EPI_ISL_3084043, EPI_ISL_3084044, EPI_ISL_3084045, EPI_ISL_3084046, EPI_ISL_3084047, EPI_ISL_3084048, EPI_ISL_3084049, EPI_ISL_3084050, EPI_ISL_3084051, EPI_ISL_3084052, EPI_ISL_3084053, EPI_ISL_3084054, EPI_ISL_3084055, EPI_ISL_3084056, EPI_ISL_3084057, EPI_ISL_3084058, EPI_ISL_3084059, EPI_ISL_3084060, EPI_ISL_3084061, EPI_ISL_3084062, EPI_ISL_3084063, EPI_ISL_3084064, EPI_ISL_3084065, EPI_ISL_3084066, EPI_ISL_3084067, EPI_ISL_3084068, EPI_ISL_3084069, EPI_ISL_3084070, EPI_ISL_3084071, EPI_ISL_3084072, EPI_ISL_3084073, EPI_ISL_3084074, EPI_ISL_3084075, EPI_ISL_3084076, EPI_ISL_3084077, EPI_ISL_3084078, EPI_ISL_3084079, EPI_ISL_3084080, EPI_ISL_3084081, EPI_ISL_3084082, EPI_ISL_3084083, EPI_ISL_3084084, EPI_ISL_3084085, EPI_ISL_3084086, EPI_ISL_3084087, EPI_ISL_3084088, EPI_ISL_3084089, EPI_ISL_3084090, EPI_ISL_3084091, EPI_ISL_3084092, EPI_ISL_3084093, EPI_ISL_3084094, EPI_ISL_3084095, EPI_ISL_3084096, EPI_ISL_3084097, EPI_ISL_3084098, EPI_ISL_3084099, EPI_ISL_3084100, EPI_ISL_3084101, EPI_ISL_3084102, EPI_ISL_3084103, EPI_ISL_3084104, EPI_ISL_3084105, EPI_ISL_3084106, EPI_ISL_3084107, EPI_ISL_3084108, EPI_ISL_3084109, EPI_ISL_3084110, EPI_ISL_3084111, EPI_ISL_3084112, EPI_ISL_3084113, EPI_ISL_3084114, EPI_ISL_3084115, EPI_ISL_3084116, EPI_ISL_3084117, EPI_ISL_3084118, EPI_ISL_3084119, EPI_ISL_3084120, EPI_ISL_3084121, EPI_ISL_3084122, EPI_ISL_3084123, EPI_ISL_3084124, EPI_ISL_3084125, EPI_ISL_3084126, EPI_ISL_3084127, EPI_ISL_3084128, EPI_ISL_3084129, EPI_ISL_3084130, EPI_ISL_3084131, EPI_ISL_3084132, EPI_ISL_3084133, EPI_ISL_3084134, EPI_ISL_3084135, EPI_ISL_3084136, EPI_ISL_3084137, EPI_ISL_3084138, EPI_ISL_3084139, EPI_ISL_3084140, EPI_ISL_3084141, EPI_ISL_3084142, EPI_ISL_3084143, EPI_ISL_3084144, EPI_ISL_3084145, EPI_ISL_3084146, EPI_ISL_3084147, EPI_ISL_3084148, EPI_ISL_3084149, EPI_ISL_3084150, EPI_ISL_3084151, EPI_ISL_3084152, EPI_ISL_3084153, EPI_ISL_3084154, EPI_ISL_3084155, EPI_ISL_3084156, EPI_ISL_3084157, EPI_ISL_3084158, EPI_ISL_3084159, EPI_ISL_3084160

| | | | |
|---|---|---|---|
| see above | Originating lab: Wales Specialist Virology Centre Sequencing lab: Pathogen Genomics Unit | Public Health Wales Microbiology Cardiff Wales Specialist Virology Centre | Catherine Moore, Johnathan Evans, Laura Gifford, Malorie Perry, Simon Cottrell, Angela Marchbank, Alec Birchley, Alexander Adams, Amy Gaskin, Bree Gatica-Wilcox, Jason Coombes, Joel Southgate, Lauren Gilbert, Lee Graham, Nicole Pacchiarini, Sara Kumziene-Summerhayes, Sarah Taylor, Sophie Jones, Sara Rey, Matthew Bull, Joanne Watkins, Sally Corden, Tom Connor |

EPI_ISL_3084161, EPI_ISL_3084162, EPI_ISL_3084163, EPI_ISL_3084164, EPI_ISL_3084165, EPI_ISL_3084166, EPI_ISL_3084167, EPI_ISL_3084168, EPI_ISL_3084169, EPI_ISL_3084170, EPI_ISL_3084171, EPI_ISL_3084172, EPI_ISL_3084173, EPI_ISL_3084174, EPI_ISL_3084175, EPI_ISL_3084176, EPI_ISL_3084177, EPI_ISL_3084178, EPI_ISL_3084179, EPI_ISL_3084180, EPI_ISL_3084181, EPI_ISL_3084182, EPI_ISL_3084183, EPI_ISL_3084184, EPI_ISL_3084185, EPI_ISL_3084186, EPI_ISL_3084187, EPI_ISL_3084188, EPI_ISL_3084189, EPI_ISL_3084190, EPI_ISL_3084191, EPI_ISL_3084192, EPI_ISL_3084193, EPI_ISL_3084194, EPI_ISL_3084195, EPI_ISL_3084196, EPI_ISL_3084197, EPI_ISL_3084198, EPI_ISL_3084199, EPI_ISL_3084200, EPI_ISL_3084201, EPI_ISL_3084202, EPI_ISL_3084203, EPI_ISL_3084204, EPI_ISL_3084205, EPI_ISL_3084206, EPI_ISL_3084207, EPI_ISL_3084208, EPI_ISL_3084209, EPI_ISL_3084210, EPI_ISL_3084211, EPI_ISL_3084212, EPI_ISL_3084213, EPI_ISL_3084214, EPI_ISL_3084215, EPI_ISL_3084216, EPI_ISL_3084217, EPI_ISL_3084218, EPI_ISL_3084219, EPI_ISL_3084220, EPI_ISL_3084221, EPI_ISL_3084222, EPI_ISL_3084223, EPI_ISL_3084224, EPI_ISL_3084225, EPI_ISL_3084226, EPI_ISL_3084227, EPI_ISL_3084228, EPI_ISL_3084229, EPI_ISL_3084230, EPI_ISL_3084231, EPI_ISL_3084232, EPI_ISL_3084233, EPI_ISL_3084234, EPI_ISL_3084235, EPI_ISL_3084236, EPI_ISL_3084237, EPI_ISL_3084238, EPI_ISL_3084239, EPI_ISL_3084240, EPI_ISL_3084241, EPI_ISL_3084242, EPI_ISL_3084243, EPI_ISL_3084244, EPI_ISL_3084245, EPI_ISL_3084246, EPI_ISL_3084247, EPI_ISL_3084248, EPI_ISL_3084249, EPI_ISL_3084250, EPI_ISL_3084251, EPI_ISL_3084252, EPI_ISL_3084253, EPI_ISL_3084254, EPI_ISL_3084255, EPI_ISL_3084256, EPI_ISL_3084257, EPI_ISL_3084258, EPI_ISL_3084259, EPI_ISL_3084260, EPI_ISL_3084261, EPI_ISL_3084262, EPI_ISL_3084263, EPI_ISL_3084264, EPI_ISL_3084265, EPI_ISL_3084266, EPI_ISL_3084267, EPI_ISL_3084268, EPI_ISL_3084269, EPI_ISL_3084270, EPI_ISL_3084271, EPI_ISL_3084272, EPI_ISL_3084273, EPI_ISL_3084274, EPI_ISL_3084275, EPI_ISL_3084276, EPI_ISL_3084277, EPI_ISL_3084278, EPI_ISL_3084279, EPI_ISL_3084280, EPI_ISL_3084281, EPI_ISL_3084282, EPI_ISL_3084283, EPI_ISL_3084284, EPI_ISL_3084285, EPI_ISL_3084286, EPI_ISL_3084287, EPI_ISL_3084288

| | | | |
|---|---|---|---|
| see above | Centre for Enzyme Innovation, University of Portsmouth / Translational Research Laboratory, Portsmouth Hospitals NHS Trust / Hampshire Hospitals NHS Foundation Trust / Bournemouth University / University Hospitals Dorset NHS Foundation Trust / Brighton and Sussex University Hospitals NHS Trust / Dartford and Gravesham NHS Trust / Isle of Wight NHS Trust / Maidstone and Tunbridge Wells NHS Trust / University | COVID-19 Genomics UK (COG-UK) Consortium | Samuel Robson, Angela Beckett, Salman Goudarzi, Christopher Fearn, Kate Cook, Katie Loveson, Sharon Glaysher, Scott Elliott, Kelly Bicknell, Sarah Wyllie, Allyson Lloyd, Robert Impey, Anoop Chauhan, Stephen Kidd, Nathan Moore, Nick Cortes, Claire Thomas, Anna Mantzaouratou, Sarah Buchan, Magdalena Barrow, Andrew Butt, Liz Sheridan, Jonnie Seymour, Dorian Crudgington, Ben Macklin, Mohammed Hassan-Ibrahim, Cassandra Malone, Benjamin Cogger, Kevin Tucker, Samirakhon Raupova, Rachael Jeremiah, Anibolina Castigador, Emily Macnaughton, Karen Withell, Kordo Saeed, Jacqui Prieto, Adhyana Mahanama, Buddhini Samaraweera, Siona Silvieri, Emanuela Pelosi, Eleri Wilson-Davies, Sarah Jeremiah, Helen Wheeler, Matthew Harvey, Thea Sass, Helen Umpleby, Stephen Aplin |

| | | | |
|---|---|---|---|
| | Hospital Southampton NHS Foundation Trust | | |
| EPI_ISL_3084289, EPI_ISL_3084290, EPI_ISL_3084291, EPI_ISL_3084292, EPI_ISL_3084293, EPI_ISL_3084294, EPI_ISL_3084295, EPI_ISL_3084296, EPI_ISL_3084297, EPI_ISL_3084298, EPI_ISL_3084299, EPI_ISL_3084300, EPI_ISL_3084301, EPI_ISL_3084302, EPI_ISL_3084303, EPI_ISL_3084304, EPI_ISL_3084305, EPI_ISL_3084306, EPI_ISL_3084307, EPI_ISL_3084308, EPI_ISL_3084309, EPI_ISL_3084310, EPI_ISL_3084311, EPI_ISL_3084312, EPI_ISL_3084313, EPI_ISL_3084314, EPI_ISL_3084315, EPI_ISL_3084316, EPI_ISL_3084317, EPI_ISL_3084318, EPI_ISL_3084319, EPI_ISL_3084320, EPI_ISL_3084321, EPI_ISL_3084322, EPI_ISL_3084323, EPI_ISL_3084324, EPI_ISL_3084325, EPI_ISL_3084326, EPI_ISL_3084327, EPI_ISL_3084328, EPI_ISL_3084329, EPI_ISL_3084330, EPI_ISL_3084331, EPI_ISL_3084332, EPI_ISL_3084333, EPI_ISL_3084334, EPI_ISL_3084335, EPI_ISL_3084336, EPI_ISL_3084337, EPI_ISL_3084338, EPI_ISL_3084339, EPI_ISL_3084340, EPI_ISL_3084341, EPI_ISL_3084342, EPI_ISL_3084343, EPI_ISL_3084344, EPI_ISL_3084345, EPI_ISL_3084346, EPI_ISL_3084347, EPI_ISL_3084348, EPI_ISL_3084349, EPI_ISL_3084350, EPI_ISL_3084351, EPI_ISL_3084352, EPI_ISL_3084353, EPI_ISL_3084354, EPI_ISL_3084355, EPI_ISL_3084356, EPI_ISL_3084357, EPI_ISL_3084358, EPI_ISL_3084359, EPI_ISL_3084360, EPI_ISL_3084361, EPI_ISL_3084362, EPI_ISL_3084363, EPI_ISL_3084364, EPI_ISL_3084365, EPI_ISL_3084366, EPI_ISL_3084367, EPI_ISL_3084368, EPI_ISL_3084369, EPI_ISL_3084370, EPI_ISL_3084371, EPI_ISL_3084372, EPI_ISL_3084373, EPI_ISL_3084374, EPI_ISL_3084375, EPI_ISL_3084376, EPI_ISL_3084377, EPI_ISL_3084378, EPI_ISL_3084379, EPI_ISL_3084380, EPI_ISL_3084381, EPI_ISL_3084382, EPI_ISL_3084383, EPI_ISL_3084384, EPI_ISL_3084385, EPI_ISL_3084386, EPI_ISL_3084387, EPI_ISL_3084388, EPI_ISL_3084389, EPI_ISL_3084390, EPI_ISL_3084391, EPI_ISL_3084392, EPI_ISL_3084393, EPI_ISL_3084394, EPI_ISL_3084395, EPI_ISL_3084396, EPI_ISL_3084397, EPI_ISL_3084398, EPI_ISL_3084399, EPI_ISL_3084400, EPI_ISL_3084401, EPI_ISL_3084402, EPI_ISL_3084403, EPI_ISL_3084404, EPI_ISL_3084405, EPI_ISL_3084406, EPI_ISL_3084407, EPI_ISL_3084408, EPI_ISL_3084409, EPI_ISL_3084410, EPI_ISL_3084411, EPI_ISL_3084412, EPI_ISL_3084413, EPI_ISL_3084414, EPI_ISL_3084415, EPI_ISL_3084416, EPI_ISL_3084417, EPI_ISL_3084418, EPI_ISL_3084419, EPI_ISL_3084420, EPI_ISL_3084421, EPI_ISL_3084422, EPI_ISL_3084423, EPI_ISL_3084424, EPI_ISL_3084425, EPI_ISL_3084426, EPI_ISL_3084427, EPI_ISL_3084428, EPI_ISL_3084429, EPI_ISL_3084430, EPI_ISL_3084431, EPI_ISL_3084432, EPI_ISL_3084433, EPI_ISL_3084434, EPI_ISL_3084435, EPI_ISL_3084436, EPI_ISL_3084437, EPI_ISL_3084438, EPI_ISL_3084439, EPI_ISL_3084440, EPI_ISL_3084441, EPI_ISL_3084442, EPI_ISL_3084443, EPI_ISL_3084444, EPI_ISL_3084445, EPI_ISL_3084446, EPI_ISL_3084447, EPI_ISL_3084448, EPI_ISL_3084449, EPI_ISL_3084450, EPI_ISL_3084451, EPI_ISL_3084452, EPI_ISL_3084453, EPI_ISL_3084454, EPI_ISL_3084455, EPI_ISL_3084456, EPI_ISL_3084457, EPI_ISL_3084458, EPI_ISL_3084459, EPI_ISL_3084460, EPI_ISL_3084461, EPI_ISL_3084462, EPI_ISL_3084463, EPI_ISL_3084464, EPI_ISL_3084465, EPI_ISL_3084466, EPI_ISL_3084467, EPI_ISL_3084468, EPI_ISL_3084469, EPI_ISL_3084470, EPI_ISL_3084471, EPI_ISL_3084472, EPI_ISL_3084473, EPI_ISL_3084474, EPI_ISL_3084475, EPI_ISL_3084476, EPI_ISL_3084477, EPI_ISL_3084478, EPI_ISL_3084479, EPI_ISL_3084480, EPI_ISL_3084481, EPI_ISL_3084482, EPI_ISL_3084483, EPI_ISL_3084484, EPI_ISL_3084485, EPI_ISL_3084486, EPI_ISL_3084487, EPI_ISL_3084488, EPI_ISL_3084489, EPI_ISL_3084490, EPI_ISL_3084491, EPI_ISL_3084492, EPI_ISL_3084493, EPI_ISL_3084494, EPI_ISL_3084495, EPI_ISL_3084496, EPI_ISL_3084497, EPI_ISL_3084498, EPI_ISL_3084499, EPI_ISL_3084500, EPI_ISL_3084501, EPI_ISL_3084502, EPI_ISL_3084503, EPI_ISL_3084504, EPI_ISL_3084505, EPI_ISL_3084506, EPI_ISL_3084507, EPI_ISL_3084508, EPI_ISL_3084509, EPI_ISL_3084510, EPI_ISL_3084511, EPI_ISL_3084512, EPI_ISL_3084513, EPI_ISL_3084514, EPI_ISL_3084515, EPI_ISL_3084516, EPI_ISL_3084517, EPI_ISL_3084518, EPI_ISL_3084519, EPI_ISL_3084520, EPI_ISL_3084521, EPI_ISL_3084522, EPI_ISL_3084523, EPI_ISL_3084524, EPI_ISL_3084525, EPI_ISL_3084526, EPI_ISL_3084527, EPI_ISL_3084528, EPI_ISL_3084529, EPI_ISL_3084530, EPI_ISL_3084531, EPI_ISL_3084532, EPI_ISL_3084533, EPI_ISL_3084534, EPI_ISL_3084535, EPI_ISL_3084536, EPI_ISL_3084537, EPI_ISL_3084538, EPI_ISL_3084539, EPI_ISL_3084540, EPI_ISL_3084541, EPI_ISL_3084542, EPI_ISL_3084543, EPI_ISL_3084544, EPI_ISL_3084545, EPI_ISL_3084546, EPI_ISL_3084547, EPI_ISL_3084548, EPI_ISL_3084549, EPI_ISL_3084550 | | | |
| see above | Virology Department, Sheffield Teaching Hospitals NHS Foundation Trust/Department of Infection, Immunity and Cardiovascular Disease, The Medical School, University of Sheffield | COVID-19 Genomics UK (COG-UK) Consortium | Thushan de Silva, Matthew Parker, Nikki Smith, Adri Angyal, Rebecca Brown, Luke Green, Rachel Tucker, Paul Parsons, Danielle Groves, Katie Johnson, Laura Carrilero, Alex Keeley, Dave Partridge, Matthew Wyles, Benjamin Lindsey, Mehmet Yavuz, Mohammad Raza, Cariad Evans |
| EPI_ISL_3084573 | Tivoli | University Campus Bio-Medico of Rome (UCBM) | Angeletti Silvia, Riva Elisabetta |
| EPI_ISL_3084575, EPI_ISL_3084576 | Tivoli | University Campus Bio-Medico of Rome | Angeletti Silvia, Riva Elisabetta |
| EPI_ISL_3084578 | Tivoli | University Campus Bio-Medico of Rome | Angeletti S, Riva E |
| EPI_ISL_3084579 | Ospedale Sandro Pertini | University Campus Bio-Medico of Rome | Angeletti Silvia, Riva Elisabetta |
| EPI_ISL_3084580 | Ospedale San Pietro FatebeneFratelli of Rome | University Campus Bio-Medico of Rome | Angeletti Silvia, Riva Elisabetta |
| EPI_ISL_3084582 | ASL LATINA | University Campus Bio-Medico of Rome | Angeletti Silvia, Riva Elisabetta |
| EPI_ISL_3084584 | ASL Latina | University Campus Bio-Medico of Rome | Angeletti Silvia, Riva Elisabetta |
| EPI_ISL_3084585 | ASL LATINA | University Campus Bio-Medico of Rome | Angeletti Silvia, Riva Elisabetta |
| EPI_ISL_3084586, EPI_ISL_3084587, EPI_ISL_3084588, EPI_ISL_3084589, EPI_ISL_3084590, EPI_ISL_3084591, EPI_ISL_3084592, EPI_ISL_3084593, EPI_ISL_3084594, EPI_ISL_3084595, EPI_ISL_3084596, EPI_ISL_3084597, EPI_ISL_3084598, EPI_ISL_3084599, EPI_ISL_3084600, EPI_ISL_3084601, EPI_ISL_3084602, EPI_ISL_3084603, EPI_ISL_3084604, EPI_ISL_3084605, EPI_ISL_3084606, EPI_ISL_3084607, EPI_ISL_3084608, EPI_ISL_3084609, EPI_ISL_3084610, EPI_ISL_3084611, EPI_ISL_3084612, EPI_ISL_3084613, EPI_ISL_3084614, EPI_ISL_3084615, EPI_ISL_3084616, EPI_ISL_3084617, EPI_ISL_3084618, EPI_ISL_3084619, EPI_ISL_3084620, EPI_ISL_3084621, EPI_ISL_3084622, EPI_ISL_3084624, EPI_ISL_3084625, EPI_ISL_3084626, EPI_ISL_3084627, EPI_ISL_3084628, EPI_ISL_3084629, EPI_ISL_3084630, EPI_ISL_3084631, EPI_ISL_3084632, EPI_ISL_3084633, EPI_ISL_3084634, EPI_ISL_3084635, EPI_ISL_3084637, EPI_ISL_3084638, EPI_ISL_3084639, EPI_ISL_3084640, EPI_ISL_3084641, EPI_ISL_3084642, EPI_ISL_3084643, EPI_ISL_3084644, EPI_ISL_3084645, EPI_ISL_3084646, EPI_ISL_3084647, EPI_ISL_3084648, EPI_ISL_3084649, EPI_ISL_3084650, EPI_ISL_3084651, EPI_ISL_3084652, EPI_ISL_3084653, EPI_ISL_3084654, EPI_ISL_3084655, EPI_ISL_3084656, EPI_ISL_3084657, EPI_ISL_3084658, EPI_ISL_3084659, EPI_ISL_3084660, EPI_ISL_3084661, EPI_ISL_3084662, EPI_ISL_3084664, EPI_ISL_3084665, EPI_ISL_3084666, EPI_ISL_3084667, EPI_ISL_3084668, EPI_ISL_3084669, EPI_ISL_3084670, EPI_ISL_3084671, EPI_ISL_3084672, EPI_ISL_3084673, EPI_ISL_3084674, EPI_ISL_3084676, EPI_ISL_3084677, EPI_ISL_3084678, EPI_ISL_3084679, EPI_ISL_3084680, EPI_ISL_3084681, EPI_ISL_3084682, EPI_ISL_3084683, EPI_ISL_3084684, EPI_ISL_3084685, EPI_ISL_3084686, EPI_ISL_3084687, EPI_ISL_3084688, EPI_ISL_3084689, EPI_ISL_3084690, EPI_ISL_3084691, EPI_ISL_3084692, EPI_ISL_3084693, EPI_ISL_3084694, EPI_ISL_3084695, EPI_ISL_3084696, EPI_ISL_3084697, EPI_ISL_3084698, EPI_ISL_3084699, EPI_ISL_3084700, EPI_ISL_3084701, EPI_ISL_3084702, EPI_ISL_3084703, EPI_ISL_3084704, EPI_ISL_3084705, EPI_ISL_3084706, EPI_ISL_3084707, EPI_ISL_3084708, EPI_ISL_3084709, EPI_ISL_3084710, EPI_ISL_3084711, EPI_ISL_3084712, EPI_ISL_3084713, EPI_ISL_3084714, EPI_ISL_3084715, EPI_ISL_3084716, EPI_ISL_3084717, EPI_ISL_3084718, EPI_ISL_3084719, EPI_ISL_3084720, EPI_ISL_3084721, EPI_ISL_3084722, EPI_ISL_3084723, EPI_ISL_3084724, EPI_ISL_3084725, EPI_ISL_3084726, EPI_ISL_3084727, EPI_ISL_3084728, EPI_ISL_3084729, EPI_ISL_3084731, EPI_ISL_3084732, EPI_ISL_3084733, EPI_ISL_3084734, EPI_ISL_3084735, EPI_ISL_3084736, EPI_ISL_3084737, EPI_ISL_3084738, EPI_ISL_3084739, EPI_ISL_3084740, EPI_ISL_3084741, EPI_ISL_3084742, EPI_ISL_3084743, EPI_ISL_3084744, EPI_ISL_3084745, EPI_ISL_3084746, EPI_ISL_3084747, EPI_ISL_3084748, EPI_ISL_3084749, EPI_ISL_3084751, EPI_ISL_3084752, EPI_ISL_3084753, EPI_ISL_3084754, EPI_ISL_3084755, EPI_ISL_3084756, EPI_ISL_3084757, EPI_ISL_3084758, EPI_ISL_3084759, EPI_ISL_3084760, EPI_ISL_3084761, EPI_ISL_3084762, EPI_ISL_3084763, EPI_ISL_3084764, EPI_ISL_3084765, EPI_ISL_3084766, EPI_ISL_3084767, EPI_ISL_3084768, EPI_ISL_3084769, EPI_ISL_3084770, EPI_ISL_3084772, EPI_ISL_3084773, EPI_ISL_3084774, EPI_ISL_3084775, EPI_ISL_3084776, EPI_ISL_3084777, EPI_ISL_3084778, EPI_ISL_3084779, EPI_ISL_3084780, EPI_ISL_3084781, EPI_ISL_3084782, EPI_ISL_3084783, EPI_ISL_3084784, EPI_ISL_3084785, EPI_ISL_3084786, EPI_ISL_3084787, EPI_ISL_3084788, EPI_ISL_3084789, EPI_ISL_3084790, EPI_ISL_3084791, EPI_ISL_3084792, EPI_ISL_3084793, EPI_ISL_3084794, EPI_ISL_3084795, EPI_ISL_3084796, EPI_ISL_3084797, EPI_ISL_3084798, EPI_ISL_3084799, EPI_ISL_3084800, EPI_ISL_3084801, EPI_ISL_3084802, EPI_ISL_3084803, EPI_ISL_3084804, EPI_ISL_3084805, EPI_ISL_3084806, EPI_ISL_3084807, EPI_ISL_3084808, EPI_ISL_3084809, EPI_ISL_3084810, EPI_ISL_3084811, EPI_ISL_3084812, EPI_ISL_3084813, EPI_ISL_3084814, EPI_ISL_3084816, EPI_ISL_3084817, EPI_ISL_3084818, EPI_ISL_3084819, EPI_ISL_3084820, EPI_ISL_3084821, EPI_ISL_3084822, EPI_ISL_3084823, EPI_ISL_3084824, EPI_ISL_3084825, EPI_ISL_3084826, EPI_ISL_3084827, EPI_ISL_3084828, EPI_ISL_3084829, EPI_ISL_3084830, EPI_ISL_3084831, EPI_ISL_3084832, EPI_ISL_3084833, EPI_ISL_3084834, EPI_ISL_3084835, EPI_ISL_3084836, EPI_ISL_3084837, EPI_ISL_3084838, EPI_ISL_3084839, EPI_ISL_3084840, EPI_ISL_3084841, EPI_ISL_3084842, EPI_ISL_3084843, EPI_ISL_3084844, EPI_ISL_3084845, EPI_ISL_3084846, EPI_ISL_3084847, EPI_ISL_3084848, EPI_ISL_3084849, EPI_ISL_3084850, EPI_ISL_3084852, EPI_ISL_3084853, EPI_ISL_3084855, EPI_ISL_3084856, EPI_ISL_3084857, EPI_ISL_3084858, EPI_ISL_3084859, EPI_ISL_3084860, EPI_ISL_3084861, EPI_ISL_3084862, EPI_ISL_3084863, EPI_ISL_3084864, EPI_ISL_3084865, EPI_ISL_3084866, EPI_ISL_3084867, EPI_ISL_3084868, EPI_ISL_3084869, EPI_ISL_3084870, EPI_ISL_3084871, EPI_ISL_3084872, EPI_ISL_3084873, EPI_ISL_3084875, EPI_ISL_3084876, EPI_ISL_3084878, EPI_ISL_3084879, EPI_ISL_3084880, EPI_ISL_3084883, EPI_ISL_3084884, EPI_ISL_3084885, EPI_ISL_3084886, EPI_ISL_3084887, EPI_ISL_3084888, EPI_ISL_3084889, EPI_ISL_3084890, EPI_ISL_3084891, EPI_ISL_3084892, EPI_ISL_3084893, EPI_ISL_3084894, EPI_ISL_3084895, EPI_ISL_3084896, EPI_ISL_3084897, EPI_ISL_3084898, EPI_ISL_3084899, EPI_ISL_3084900, EPI_ISL_3084902, EPI_ISL_3084904, EPI_ISL_3084905, EPI_ISL_3084907, EPI_ISL_3084908, EPI_ISL_3084909, EPI_ISL_3084910, EPI_ISL_3084911, EPI_ISL_3084912, EPI_ISL_3084913, EPI_ISL_3084914, EPI_ISL_3084915, EPI_ISL_3084916, EPI_ISL_3084917, EPI_ISL_3084918, EPI_ISL_3084919, EPI_ISL_3084920, EPI_ISL_3084921, EPI_ISL_3084922, EPI_ISL_3084923, EPI_ISL_3084924, EPI_ISL_3084925, EPI_ISL_3084926, EPI_ISL_3084927, EPI_ISL_3084928, EPI_ISL_3084929, EPI_ISL_3084930, EPI_ISL_3084931, EPI_ISL_3084932, EPI_ISL_3084933, EPI_ISL_3084934, EPI_ISL_3084935, EPI_ISL_3084936, EPI_ISL_3084937, EPI_ISL_3084938, EPI_ISL_3084939, EPI_ISL_3084940, EPI_ISL_3084941, EPI_ISL_3084942, EPI_ISL_3084943, EPI_ISL_3084944, EPI_ISL_3084946, EPI_ISL_3084947, EPI_ISL_3084948, EPI_ISL_3084949, EPI_ISL_3084950, EPI_ISL_3084951, EPI_ISL_3084952, EPI_ISL_3084953, EPI_ISL_3084954, EPI_ISL_3084955, EPI_ISL_3084956, EPI_ISL_3084957, EPI_ISL_3084958, EPI_ISL_3084959, EPI_ISL_3084960, EPI_ISL_3084961, EPI_ISL_3084962, EPI_ISL_3084963, EPI_ISL_3084964, EPI_ISL_3084965, EPI_ISL_3084966, EPI_ISL_3084967, EPI_ISL_3084968, EPI_ISL_3084969, EPI_ISL_3084970, EPI_ISL_3084972, EPI_ISL_3084973, EPI_ISL_3084974, EPI_ISL_3084975, EPI_ISL_3084976, EPI_ISL_3084977, EPI_ISL_3084978, EPI_ISL_3084979, EPI_ISL_3084980, EPI_ISL_3084981, EPI_ISL_3084982, EPI_ISL_3084983, EPI_ISL_3084984, EPI_ISL_3084985, EPI_ISL_3084986, EPI_ISL_3084987, EPI_ISL_3084988, EPI_ISL_3084989, EPI_ISL_3084990, EPI_ISL_3084991, EPI_ISL_3084993, EPI_ISL_3084994, EPI_ISL_3084995, EPI_ISL_3084996, EPI_ISL_3084997, EPI_ISL_3084998, EPI_ISL_3084999, EPI_ISL_3085000, EPI_ISL_3085001, EPI_ISL_3085002, EPI_ISL_3085003, EPI_ISL_3085004, EPI_ISL_3085005, EPI_ISL_3085006, EPI_ISL_3085007, EPI_ISL_3085008, EPI_ISL_3085009, EPI_ISL_3085010, EPI_ISL_3085011, EPI_ISL_3085012, EPI_ISL_3085013, EPI_ISL_3085014, EPI_ISL_3085015, EPI_ISL_3085016, EPI_ISL_3085017, EPI_ISL_3085018, EPI_ISL_3085019, EPI_ISL_3085020, EPI_ISL_3085021, EPI_ISL_3085023, EPI_ISL_3085024, EPI_ISL_3085025, EPI_ISL_3085026, EPI_ISL_3085027, EPI_ISL_3085028, EPI_ISL_3085029, EPI_ISL_3085030, EPI_ISL_3085032, EPI_ISL_3085033, EPI_ISL_3085034, EPI_ISL_3085035, EPI_ISL_3085036, EPI_ISL_3085037, EPI_ISL_3085038, EPI_ISL_3085039, EPI_ISL_3085040, EPI_ISL_3085042, EPI_ISL_3085043, EPI_ISL_3085044, EPI_ISL_3085045, EPI_ISL_3085046, EPI_ISL_3085048, EPI_ISL_3085049, EPI_ISL_3085050, EPI_ISL_3085051, EPI_ISL_3085052, EPI_ISL_3085053, EPI_ISL_3085054, EPI_ISL_3085055, EPI_ISL_3085057, EPI_ISL_3085058, EPI_ISL_3085059, EPI_ISL_3085060, EPI_ISL_3085062, EPI_ISL_3085063, EPI_ISL_3085064, EPI_ISL_3085065, EPI_ISL_3085066, EPI_ISL_3085067, EPI_ISL_3085068, EPI_ISL_3085069, EPI_ISL_3085070, EPI_ISL_3085071, EPI_ISL_3085072, EPI_ISL_3085073, EPI_ISL_3085074, EPI_ISL_3085075, EPI_ISL_3085076, EPI_ISL_3085077, EPI_ISL_3085078, EPI_ISL_3085079, EPI_ISL_3085081, EPI_ISL_3085082, EPI_ISL_3085083, EPI_ISL_3085084, EPI_ISL_3085085, EPI_ISL_3085086, EPI_ISL_3085087, EPI_ISL_3085088, EPI_ISL_3085089, EPI_ISL_3085090, EPI_ISL_3085091, EPI_ISL_3085092, EPI_ISL_3085093, EPI_ISL_3085094, EPI_ISL_3085095, EPI_ISL_3085096, EPI_ISL_3085097, EPI_ISL_3085098, EPI_ISL_3085099, EPI_ISL_3085100, EPI_ISL_3085101, EPI_ISL_3085102, EPI_ISL_3085103, EPI_ISL_3085104, EPI_ISL_3085105, EPI_ISL_3085106, EPI_ISL_3085107, EPI_ISL_3085108, EPI_ISL_3085109, EPI_ISL_3085110, EPI_ISL_3085111, EPI_ISL_3085112, EPI_ISL_3085113, EPI_ISL_3085114, EPI_ISL_3085115, EPI_ISL_3085116, EPI_ISL_3085117, EPI_ISL_3085118, EPI_ISL_3085119, EPI_ISL_3085120, EPI_ISL_3085121 | | | |
| see above | Broad Institute Clinical Research Sequencing Platform | Infectious Disease Program, Broad Institute of Harvard and MIT | Siddle,K.J., Adams,G., Pearlman,L., Gladden-Young,A., Vicente,G., Blumenstiel,B., DeFelice,M., Lee,M., McGovern,S., Lagerborg,K., Rudy,M., DeRuff,K., Carter,A., Normandin,E., Bauer,M., Reilly,S., Tomkins-Tinch,C., Loreth,C., Chaluvadi,S., Meldrim,J., Granger,B., Lemieux,J.E., Birren,B.W., Sabeti,P.C., Larkin,K., Dodge,S., Lennon,N., Madoff,L., Brown,C., Gallagher,G., Smole,S., Park,D.J., Gabriel,S., and MacInnis,B.L. |
| EPI_ISL_3085361 | Karolinska University Hospital Solna | Karolinska University Hospital | Jan Albert ,Tobias Allander ,Annelie Bjerkner ,Sandra Broddesson ,Robert Dyrdak ,Martin Ekman ,Lynda Eneh ,Lina Guerra Blomqvist ,Karolina Ininbergs ,Tanja Normark ,Isak Sylvin ,Zhibing Yun ,Martina Wahlund ,Valtteri Wirta |
| EPI_ISL_3085362 | Unilabs | Karolinska University Hospital | Jan Albert ,Tobias Allander ,Annelie Bjerkner ,Sandra Broddesson ,Robert Dyrdak ,Martin Ekman ,Lynda Eneh ,Lina Guerra Blomqvist ,Karolina Ininbergs ,Tanja Normark ,Isak Sylvin ,Zhibing Yun ,Martina Wahlund ,Valtteri Wirta |
| EPI_ISL_3085367 | Karolinska University Hospital Solna | Karolinska University Hospital | Jan Albert ,Tobias Allander ,Annelie Bjerkner ,Sandra Broddesson ,Robert Dyrdak ,Martin Ekman ,Lynda Eneh ,Lina Guerra Blomqvist ,Karolina Ininbergs ,Tanja Normark ,Isak Sylvin ,Zhibing Yun ,Martina Wahlund ,Valtteri Wirta |
| EPI_ISL_3085370 | Unilabs | Karolinska University Hospital | Jan Albert ,Tobias Allander ,Annelie Bjerkner ,Sandra Broddesson ,Robert Dyrdak ,Martin Ekman ,Lynda Eneh ,Lina Guerra Blomqvist ,Karolina Ininbergs ,Tanja Normark ,Isak Sylvin ,Zhibing Yun ,Martina Wahlund ,Valtteri Wirta |
| EPI_ISL_3085371, EPI_ISL_3085372 | Karolinska University Hospital Solna | Karolinska University Hospital | Jan Albert ,Tobias Allander ,Annelie Bjerkner ,Sandra Broddesson ,Robert Dyrdak ,Martin Ekman ,Lynda Eneh ,Lina Guerra Blomqvist ,Karolina Ininbergs ,Tanja Normark ,Isak Sylvin ,Zhibing Yun ,Martina Wahlund ,Valtteri Wirta |
| EPI_ISL_3085375 | Unilabs | Karolinska University Hospital | Jan Albert ,Tobias Allander ,Annelie Bjerkner ,Sandra Broddesson ,Robert Dyrdak ,Martin Ekman ,Lynda Eneh ,Lina Guerra Blomqvist ,Karolina Ininbergs ,Tanja Normark ,Isak Sylvin ,Zhibing Yun ,Martina Wahlund ,Valtteri Wirta |
| EPI_ISL_3085376 | Karolinska University Hospital Solna | Karolinska University Hospital | Jan Albert ,Tobias Allander ,Annelie Bjerkner ,Sandra Broddesson ,Robert Dyrdak ,Martin Ekman ,Lynda Eneh ,Lina Guerra Blomqvist ,Karolina Ininbergs ,Tanja Normark ,Isak Sylvin ,Zhibing Yun ,Martina Wahlund ,Valtteri Wirta |
| EPI_ISL_3085378, EPI_ISL_3085380 | Unilabs | Karolinska University Hospital | Jan Albert ,Tobias Allander ,Annelie Bjerkner ,Sandra Broddesson ,Robert Dyrdak ,Martin Ekman ,Lynda Eneh ,Lina Guerra Blomqvist ,Karolina Ininbergs ,Tanja Normark ,Isak Sylvin ,Zhibing Yun ,Martina Wahlund ,Valtteri Wirta |
| EPI_ISL_3085384 | Karolinska University Hospital Solna | Karolinska University Hospital | Jan Albert ,Tobias Allander ,Annelie Bjerkner ,Sandra Broddesson ,Robert Dyrdak ,Martin Ekman ,Lynda Eneh ,Lina Guerra Blomqvist ,Karolina Ininbergs ,Tanja Normark ,Isak Sylvin ,Zhibing Yun ,Martina Wahlund ,Valtteri Wirta |
| EPI_ISL_3085386, EPI_ISL_3085387, EPI_ISL_3085389, EPI_ISL_3085392 | Unilabs | Karolinska University Hospital | Jan Albert ,Tobias Allander ,Annelie Bjerkner ,Sandra Broddesson ,Robert Dyrdak ,Martin Ekman ,Lynda Eneh ,Lina Guerra Blomqvist ,Karolina Ininbergs ,Tanja Normark ,Isak Sylvin ,Zhibing Yun ,Martina Wahlund ,Valtteri Wirta |
| EPI_ISL_3085394 | Karolinska University Hospital Solna | Karolinska University Hospital | Jan Albert ,Tobias Allander ,Annelie Bjerkner ,Sandra Broddesson ,Robert Dyrdak ,Martin Ekman ,Lynda Eneh ,Lina Guerra Blomqvist ,Karolina Ininbergs ,Tanja Normark ,Isak Sylvin ,Zhibing Yun ,Martina Wahlund ,Valtteri Wirta |
| EPI_ISL_3085395, EPI_ISL_3085397, EPI_ISL_3085399, EPI_ISL_3085400 | Unilabs | Karolinska University Hospital | Jan Albert ,Tobias Allander ,Annelie Bjerkner ,Sandra Broddesson ,Robert Dyrdak ,Martin Ekman ,Lynda Eneh ,Lina Guerra Blomqvist ,Karolina Ininbergs ,Tanja Normark ,Isak Sylvin ,Zhibing Yun ,Martina Wahlund ,Valtteri Wirta |
| EPI_ISL_3085401 | Karolinska University Hospital Solna | Karolinska University Hospital | Jan Albert ,Tobias Allander ,Annelie Bjerkner ,Sandra Broddesson ,Robert Dyrdak ,Martin Ekman ,Lynda Eneh ,Lina Guerra Blomqvist ,Karolina Ininbergs ,Tanja Normark ,Isak Sylvin ,Zhibing Yun ,Martina Wahlund ,Valtteri Wirta |
| EPI_ISL_3085402, EPI_ISL_3085404 | Unilabs | Karolinska University Hospital | Jan Albert ,Tobias Allander ,Annelie Bjerkner ,Sandra Broddesson ,Robert Dyrdak ,Martin Ekman ,Lynda Eneh ,Lina Guerra Blomqvist ,Karolina Ininbergs ,Tanja Normark ,Isak Sylvin ,Zhibing Yun ,Martina Wahlund ,Valtteri Wirta |
| EPI_ISL_3085407 | Karolinska University Hospital Solna | Karolinska University Hospital | Jan Albert ,Tobias Allander ,Annelie Bjerkner ,Sandra Broddesson ,Robert Dyrdak ,Martin Ekman ,Lynda Eneh ,Lina Guerra Blomqvist ,Karolina Ininbergs ,Tanja Normark ,Isak Sylvin ,Zhibing Yun ,Martina Wahlund ,Valtteri Wirta |
| EPI_ISL_3085408 | Unilabs | Karolinska University Hospital | Jan Albert ,Tobias Allander ,Annelie Bjerkner ,Sandra Broddesson ,Robert Dyrdak ,Martin Ekman ,Lynda Eneh ,Lina Guerra Blomqvist ,Karolina Ininbergs ,Tanja Normark ,Isak Sylvin ,Zhibing Yun ,Martina Wahlund ,Valtteri Wirta |
| EPI_ISL_3085409 | Karolinska University Hospital Solna | Karolinska University Hospital | Jan Albert ,Tobias Allander ,Annelie Bjerkner ,Sandra Broddesson ,Robert Dyrdak ,Martin Ekman ,Lynda Eneh ,Lina Guerra Blomqvist ,Karolina Ininbergs ,Tanja Normark ,Isak Sylvin ,Zhibing Yun ,Martina Wahlund ,Valtteri Wirta |
| EPI_ISL_3085411 | Synlab Medilab | Karolinska University Hospital | Jan Albert ,Tobias Allander ,Annelie Bjerkner ,Sandra Broddesson ,Robert Dyrdak ,Martin Ekman ,Lynda Eneh ,Lina Guerra Blomqvist ,Karolina Ininbergs ,Tanja Normark ,Isak Sylvin ,Zhibing Yun ,Martina Wahlund ,Valtteri Wirta |
| EPI_ISL_3085412 | Unilabs | Karolinska University Hospital | Jan Albert ,Tobias Allander ,Annelie Bjerkner ,Sandra Broddesson ,Robert Dyrdak ,Martin Ekman ,Lynda Eneh ,Lina Guerra Blomqvist ,Karolina Ininbergs ,Tanja Normark ,Isak Sylvin ,Zhibing Yun ,Martina Wahlund ,Valtteri Wirta |
| EPI_ISL_3085413 | Karolinska University Hospital Solna | Karolinska University Hospital | Jan Albert ,Tobias Allander ,Annelie Bjerkner ,Sandra Broddesson ,Robert Dyrdak ,Martin Ekman ,Lynda Eneh ,Lina Guerra Blomqvist ,Karolina Ininbergs ,Tanja Normark ,Isak Sylvin ,Zhibing Yun ,Martina Wahlund ,Valtteri Wirta |
| EPI_ISL_3085414, EPI_ISL_3085415, EPI_ISL_3085420, EPI_ISL_3085421, EPI_ISL_3085429 | Unilabs | Karolinska University Hospital | Jan Albert ,Tobias Allander ,Annelie Bjerkner ,Sandra Broddesson ,Robert Dyrdak ,Martin Ekman ,Lynda Eneh ,Lina Guerra Blomqvist ,Karolina Ininbergs ,Tanja Normark ,Isak Sylvin ,Zhibing Yun ,Martina Wahlund ,Valtteri Wirta |
| EPI_ISL_3085431 | Karolinska University Hospital Solna | Karolinska University Hospital | Jan Albert ,Tobias Allander ,Annelie Bjerkner ,Sandra Broddesson ,Robert Dyrdak ,Martin Ekman ,Lynda Eneh ,Lina Guerra Blomqvist ,Karolina Ininbergs ,Tanja Normark ,Isak Sylvin ,Zhibing Yun ,Martina Wahlund ,Valtteri Wirta |
| EPI_ISL_3085432 | Unilabs | Karolinska University Hospital | Jan Albert ,Tobias Allander ,Annelie Bjerkner ,Sandra Broddesson ,Robert Dyrdak ,Martin Ekman ,Lynda Eneh ,Lina Guerra Blomqvist ,Karolina Ininbergs ,Tanja Normark ,Isak Sylvin ,Zhibing Yun ,Martina Wahlund ,Valtteri Wirta |
| EPI_ISL_3085433, EPI_ISL_3085434 | Karolinska University Hospital Solna | Karolinska University Hospital | Jan Albert ,Tobias Allander ,Annelie Bjerkner ,Sandra Broddesson ,Robert Dyrdak ,Martin Ekman ,Lynda Eneh ,Lina Guerra Blomqvist ,Karolina Ininbergs ,Tanja Normark ,Isak Sylvin ,Zhibing Yun ,Martina Wahlund ,Valtteri Wirta |
| EPI_ISL_3085435 | Unilabs | Karolinska University Hospital | Jan Albert ,Tobias Allander ,Annelie Bjerkner ,Sandra Broddesson ,Robert Dyrdak ,Martin Ekman ,Lynda Eneh ,Lina Guerra Blomqvist ,Karolina Ininbergs ,Tanja Normark ,Isak Sylvin ,Zhibing Yun ,Martina Wahlund ,Valtteri Wirta |
| EPI_ISL_3085438, | Karolinska University Hospital Solna | Karolinska University Hospital | Jan Albert ,Tobias Allander ,Annelie Bjerkner ,Sandra Broddesson ,Robert Dyrdak ,Martin Ekman ,Lynda Eneh ,Lina Guerra Blomqvist ,Karolina Ininbergs ,Tanja Normark ,Isak Sylvin ,Zhibing Yun ,Martina Wahlund ,Valtteri Wirta |

| | | | |
|---|---|---|---|
| EPI_ISL_3085439 | | | |
| EPI_ISL_3085440 | Unilabs | Karolinska University Hospital | Jan Albert ,Tobias Allander ,Annelie Bjerkner ,Sandra Broddesson ,Robert Dyrdak ,Martin Ekman ,Lynda Eneh ,Lina Guerra Blomqvist ,Karolina Ininbergs ,Tanja Normark ,Isak Sylvin ,Zhibing Yun ,Martina Wahlund ,Valtteri Wirta |
| EPI_ISL_3085441, EPI_ISL_3085442 | Karolinska University Hospital Solna | Karolinska University Hospital | Jan Albert ,Tobias Allander ,Annelie Bjerkner ,Sandra Broddesson ,Robert Dyrdak ,Martin Ekman ,Lynda Eneh ,Lina Guerra Blomqvist ,Karolina Ininbergs ,Tanja Normark ,Isak Sylvin ,Zhibing Yun ,Martina Wahlund ,Valtteri Wirta |
| EPI_ISL_3085443 | Unilabs | Karolinska University Hospital | Jan Albert ,Tobias Allander ,Annelie Bjerkner ,Sandra Broddesson ,Robert Dyrdak ,Martin Ekman ,Lynda Eneh ,Lina Guerra Blomqvist ,Karolina Ininbergs ,Tanja Normark ,Isak Sylvin ,Zhibing Yun ,Martina Wahlund ,Valtteri Wirta |
| EPI_ISL_3085444, EPI_ISL_3085445 | Karolinska University Hospital Solna | Karolinska University Hospital | Jan Albert ,Tobias Allander ,Annelie Bjerkner ,Sandra Broddesson ,Robert Dyrdak ,Martin Ekman ,Lynda Eneh ,Lina Guerra Blomqvist ,Karolina Ininbergs ,Tanja Normark ,Isak Sylvin ,Zhibing Yun ,Martina Wahlund ,Valtteri Wirta |
| EPI_ISL_3085446, EPI_ISL_3085447, EPI_ISL_3085448, EPI_ISL_3085449 | Unilabs | Karolinska University Hospital | Jan Albert ,Tobias Allander ,Annelie Bjerkner ,Sandra Broddesson ,Robert Dyrdak ,Martin Ekman ,Lynda Eneh ,Lina Guerra Blomqvist ,Karolina Ininbergs ,Tanja Normark ,Isak Sylvin ,Zhibing Yun ,Martina Wahlund ,Valtteri Wirta |
| EPI_ISL_3085450 | Karolinska University Hospital Solna | Karolinska University Hospital | Jan Albert ,Tobias Allander ,Annelie Bjerkner ,Sandra Broddesson ,Robert Dyrdak ,Martin Ekman ,Lynda Eneh ,Lina Guerra Blomqvist ,Karolina Ininbergs ,Tanja Normark ,Isak Sylvin ,Zhibing Yun ,Martina Wahlund ,Valtteri Wirta |
| EPI_ISL_3085451, EPI_ISL_3085452, EPI_ISL_3085453 | Unilabs | Karolinska University Hospital | Jan Albert ,Tobias Allander ,Annelie Bjerkner ,Sandra Broddesson ,Robert Dyrdak ,Martin Ekman ,Lynda Eneh ,Lina Guerra Blomqvist ,Karolina Ininbergs ,Tanja Normark ,Isak Sylvin ,Zhibing Yun ,Martina Wahlund ,Valtteri Wirta |
| EPI_ISL_3085454 | Karolinska University Hospital Solna | Karolinska University Hospital | Jan Albert ,Tobias Allander ,Annelie Bjerkner ,Sandra Broddesson ,Robert Dyrdak ,Martin Ekman ,Lynda Eneh ,Lina Guerra Blomqvist ,Karolina Ininbergs ,Tanja Normark ,Isak Sylvin ,Zhibing Yun ,Martina Wahlund ,Valtteri Wirta |
| EPI_ISL_3085455 | Unilabs | Karolinska University Hospital | Jan Albert ,Tobias Allander ,Annelie Bjerkner ,Sandra Broddesson ,Robert Dyrdak ,Martin Ekman ,Lynda Eneh ,Lina Guerra Blomqvist ,Karolina Ininbergs ,Tanja Normark ,Isak Sylvin ,Zhibing Yun ,Martina Wahlund ,Valtteri Wirta |
| EPI_ISL_3085456, EPI_ISL_3085457, EPI_ISL_3085458 | Karolinska University Hospital Solna | Karolinska University Hospital | Jan Albert ,Tobias Allander ,Annelie Bjerkner ,Sandra Broddesson ,Robert Dyrdak ,Martin Ekman ,Lynda Eneh ,Lina Guerra Blomqvist ,Karolina Ininbergs ,Tanja Normark ,Isak Sylvin ,Zhibing Yun ,Martina Wahlund ,Valtteri Wirta |
| EPI_ISL_3085459 | Unilabs | Karolinska University Hospital | Jan Albert ,Tobias Allander ,Annelie Bjerkner ,Sandra Broddesson ,Robert Dyrdak ,Martin Ekman ,Lynda Eneh ,Lina Guerra Blomqvist ,Karolina Ininbergs ,Tanja Normark ,Isak Sylvin ,Zhibing Yun ,Martina Wahlund ,Valtteri Wirta |
| EPI_ISL_3085460 | Karolinska University Hospital Solna | Karolinska University Hospital | Jan Albert ,Tobias Allander ,Annelie Bjerkner ,Sandra Broddesson ,Robert Dyrdak ,Martin Ekman ,Lynda Eneh ,Lina Guerra Blomqvist ,Karolina Ininbergs ,Tanja Normark ,Isak Sylvin ,Zhibing Yun ,Martina Wahlund ,Valtteri Wirta |
| EPI_ISL_3085461 | Unilabs | Karolinska University Hospital | Jan Albert ,Tobias Allander ,Annelie Bjerkner ,Sandra Broddesson ,Robert Dyrdak ,Martin Ekman ,Lynda Eneh ,Lina Guerra Blomqvist ,Karolina Ininbergs ,Tanja Normark ,Isak Sylvin ,Zhibing Yun ,Martina Wahlund ,Valtteri Wirta |
| EPI_ISL_3085462, EPI_ISL_3085463, EPI_ISL_3085464, EPI_ISL_3085465, EPI_ISL_3085467, EPI_ISL_3085468, EPI_ISL_3085469, EPI_ISL_3085470, EPI_ISL_3085471 | Karolinska University Hospital Solna | Karolinska University Hospital | Jan Albert ,Tobias Allander ,Annelie Bjerkner ,Sandra Broddesson ,Robert Dyrdak ,Martin Ekman ,Lynda Eneh ,Lina Guerra Blomqvist ,Karolina Ininbergs ,Tanja Normark ,Isak Sylvin ,Zhibing Yun ,Martina Wahlund ,Valtteri Wirta |
| EPI_ISL_3085472 | Unilabs | Karolinska University Hospital | Jan Albert ,Tobias Allander ,Annelie Bjerkner ,Sandra Broddesson ,Robert Dyrdak ,Martin Ekman ,Lynda Eneh ,Lina Guerra Blomqvist ,Karolina Ininbergs ,Tanja Normark ,Isak Sylvin ,Zhibing Yun ,Martina Wahlund ,Valtteri Wirta |
| EPI_ISL_3085473, EPI_ISL_3085474, EPI_ISL_3085475, EPI_ISL_3085476, EPI_ISL_3085477 | Karolinska University Hospital Solna | Karolinska University Hospital | Jan Albert ,Tobias Allander ,Annelie Bjerkner ,Sandra Broddesson ,Robert Dyrdak ,Martin Ekman ,Lynda Eneh ,Lina Guerra Blomqvist ,Karolina Ininbergs ,Tanja Normark ,Isak Sylvin ,Zhibing Yun ,Martina Wahlund ,Valtteri Wirta |
| EPI_ISL_3085478 | Unilabs | Karolinska University Hospital | Jan Albert ,Tobias Allander ,Annelie Bjerkner ,Sandra Broddesson ,Robert Dyrdak ,Martin Ekman ,Lynda Eneh ,Lina Guerra Blomqvist ,Karolina Ininbergs ,Tanja Normark ,Isak Sylvin ,Zhibing Yun ,Martina Wahlund ,Valtteri Wirta |
| EPI_ISL_3085479 | Karolinska University Hospital Solna | Karolinska University Hospital | Jan Albert ,Tobias Allander ,Annelie Bjerkner ,Sandra Broddesson ,Robert Dyrdak ,Martin Ekman ,Lynda Eneh ,Lina Guerra Blomqvist ,Karolina Ininbergs ,Tanja Normark ,Isak Sylvin ,Zhibing Yun ,Martina Wahlund ,Valtteri Wirta |
| EPI_ISL_3085480 | Unilabs | Karolinska University Hospital | Jan Albert ,Tobias Allander ,Annelie Bjerkner ,Sandra Broddesson ,Robert Dyrdak ,Martin Ekman ,Lynda Eneh ,Lina Guerra Blomqvist ,Karolina Ininbergs ,Tanja Normark ,Isak Sylvin ,Zhibing Yun ,Martina Wahlund ,Valtteri Wirta |
| EPI_ISL_3085481 | Karolinska University Hospital Solna | Karolinska University Hospital | Jan Albert ,Tobias Allander ,Annelie Bjerkner ,Sandra Broddesson ,Robert Dyrdak ,Martin Ekman ,Lynda Eneh ,Lina Guerra Blomqvist ,Karolina Ininbergs ,Tanja Normark ,Isak Sylvin ,Zhibing Yun ,Martina Wahlund ,Valtteri Wirta |
| EPI_ISL_3085482 | Unilabs | Karolinska University Hospital | Jan Albert ,Tobias Allander ,Annelie Bjerkner ,Sandra Broddesson ,Robert Dyrdak ,Martin Ekman ,Lynda Eneh ,Lina Guerra Blomqvist ,Karolina Ininbergs ,Tanja Normark ,Isak Sylvin ,Zhibing Yun ,Martina Wahlund ,Valtteri Wirta |
| EPI_ISL_3085483 | Karolinska University Hospital Solna | Karolinska University Hospital | Jan Albert ,Tobias Allander ,Annelie Bjerkner ,Sandra Broddesson ,Robert Dyrdak ,Martin Ekman ,Lynda Eneh ,Lina Guerra Blomqvist ,Karolina Ininbergs ,Tanja Normark ,Isak Sylvin ,Zhibing Yun ,Martina Wahlund ,Valtteri Wirta |
| EPI_ISL_3085485, EPI_ISL_3085486 | Unilabs | Karolinska University Hospital | Jan Albert ,Tobias Allander ,Annelie Bjerkner ,Sandra Broddesson ,Robert Dyrdak ,Martin Ekman ,Lynda Eneh ,Lina Guerra Blomqvist ,Karolina Ininbergs ,Tanja Normark ,Isak Sylvin ,Zhibing Yun ,Martina Wahlund ,Valtteri Wirta |
| EPI_ISL_3085487 | Karolinska University Hospital Solna | Karolinska University Hospital | Jan Albert ,Tobias Allander ,Annelie Bjerkner ,Sandra Broddesson ,Robert Dyrdak ,Martin Ekman ,Lynda Eneh ,Lina Guerra Blomqvist ,Karolina Ininbergs ,Tanja Normark ,Isak Sylvin ,Zhibing Yun ,Martina Wahlund ,Valtteri Wirta |
| EPI_ISL_3085488 | Unilabs | Karolinska University Hospital | Jan Albert ,Tobias Allander ,Annelie Bjerkner ,Sandra Broddesson ,Robert Dyrdak ,Martin Ekman ,Lynda Eneh ,Lina Guerra Blomqvist ,Karolina Ininbergs ,Tanja Normark ,Isak Sylvin ,Zhibing Yun ,Martina Wahlund ,Valtteri Wirta |
| EPI_ISL_3085489, EPI_ISL_3085490, EPI_ISL_3085491, EPI_ISL_3085492, EPI_ISL_3085493 | Karolinska University Hospital Solna | Karolinska University Hospital | Jan Albert ,Tobias Allander ,Annelie Bjerkner ,Sandra Broddesson ,Robert Dyrdak ,Martin Ekman ,Lynda Eneh ,Lina Guerra Blomqvist ,Karolina Ininbergs ,Tanja Normark ,Isak Sylvin ,Zhibing Yun ,Martina Wahlund ,Valtteri Wirta |
| EPI_ISL_3085494, EPI_ISL_3085495, EPI_ISL_3085496, EPI_ISL_3085497, EPI_ISL_3085498, EPI_ISL_3085499, EPI_ISL_3085500, EPI_ISL_3085501, EPI_ISL_3085502, EPI_ISL_3085503, EPI_ISL_3085504, EPI_ISL_3085505, EPI_ISL_3085506, EPI_ISL_3085507, EPI_ISL_3085508, EPI_ISL_3085509, EPI_ISL_3085510 | | | |
| see above | Karolinska University Hospital Huddinge | Karolinska University Hospital | Jan Albert ,Tobias Allander ,Annelie Bjerkner ,Sandra Broddesson ,Robert Dyrdak ,Martin Ekman ,Lynda Eneh ,Lina Guerra Blomqvist ,Karolina Ininbergs ,Tanja Normark ,Isak Sylvin ,Zhibing Yun ,Martina Wahlund ,Valtteri Wirta |
| EPI_ISL_3085511 | Unilabs | Karolinska University Hospital | Jan Albert ,Tobias Allander ,Annelie Bjerkner ,Sandra Broddesson ,Robert Dyrdak ,Martin Ekman ,Lynda Eneh ,Lina Guerra Blomqvist ,Karolina Ininbergs ,Tanja Normark ,Isak Sylvin ,Zhibing Yun ,Martina Wahlund ,Valtteri Wirta |
| EPI_ISL_3085512 | Karolinska University Hospital Solna | Karolinska University Hospital | Jan Albert ,Tobias Allander ,Annelie Bjerkner ,Sandra Broddesson ,Robert Dyrdak ,Martin Ekman ,Lynda Eneh ,Lina Guerra Blomqvist ,Karolina Ininbergs ,Tanja Normark ,Isak Sylvin ,Zhibing Yun ,Martina Wahlund ,Valtteri Wirta |
| EPI_ISL_3085513, EPI_ISL_3085514 | Unilabs | Karolinska University Hospital | Jan Albert ,Tobias Allander ,Annelie Bjerkner ,Sandra Broddesson ,Robert Dyrdak ,Martin Ekman ,Lynda Eneh ,Lina Guerra Blomqvist ,Karolina Ininbergs ,Tanja Normark ,Isak Sylvin ,Zhibing Yun ,Martina Wahlund ,Valtteri Wirta |
| EPI_ISL_3085515 | Karolinska University Hospital Solna | Karolinska University Hospital | Jan Albert ,Tobias Allander ,Annelie Bjerkner ,Sandra Broddesson ,Robert Dyrdak ,Martin Ekman ,Lynda Eneh ,Lina Guerra Blomqvist ,Karolina Ininbergs ,Tanja Normark ,Isak Sylvin ,Zhibing Yun ,Martina Wahlund ,Valtteri Wirta |
| EPI_ISL_3085516, EPI_ISL_3085517 | Synlab Medilab | Karolinska University Hospital | Jan Albert ,Tobias Allander ,Annelie Bjerkner ,Sandra Broddesson ,Robert Dyrdak ,Martin Ekman ,Lynda Eneh ,Lina Guerra Blomqvist ,Karolina Ininbergs ,Tanja Normark ,Isak Sylvin ,Zhibing Yun ,Martina Wahlund ,Valtteri Wirta |
| EPI_ISL_3085518 | Karolinska University Hospital Solna | Karolinska University Hospital | Jan Albert ,Tobias Allander ,Annelie Bjerkner ,Sandra Broddesson ,Robert Dyrdak ,Martin Ekman ,Lynda Eneh ,Lina Guerra Blomqvist ,Karolina Ininbergs ,Tanja Normark ,Isak Sylvin ,Zhibing Yun ,Martina Wahlund ,Valtteri Wirta |
| EPI_ISL_3085519, EPI_ISL_3085520, EPI_ISL_3085522 | Synlab Medilab | Karolinska University Hospital | Jan Albert ,Tobias Allander ,Annelie Bjerkner ,Sandra Broddesson ,Robert Dyrdak ,Martin Ekman ,Lynda Eneh ,Lina Guerra Blomqvist ,Karolina Ininbergs ,Tanja Normark ,Isak Sylvin ,Zhibing Yun ,Martina Wahlund ,Valtteri Wirta |
| EPI_ISL_3085523 | Unilabs | Karolinska University Hospital | Jan Albert ,Tobias Allander ,Annelie Bjerkner ,Sandra Broddesson ,Robert Dyrdak ,Martin Ekman ,Lynda Eneh ,Lina Guerra Blomqvist ,Karolina Ininbergs ,Tanja Normark ,Isak Sylvin ,Zhibing Yun ,Martina Wahlund ,Valtteri Wirta |
| EPI_ISL_3085524, EPI_ISL_3085525, EPI_ISL_3085526 | Karolinska University Hospital Solna | Karolinska University Hospital | Jan Albert ,Tobias Allander ,Annelie Bjerkner ,Sandra Broddesson ,Robert Dyrdak ,Martin Ekman ,Lynda Eneh ,Lina Guerra Blomqvist ,Karolina Ininbergs ,Tanja Normark ,Isak Sylvin ,Zhibing Yun ,Martina Wahlund ,Valtteri Wirta |
| EPI_ISL_3085527 | Unilabs | Karolinska University Hospital | Jan Albert ,Tobias Allander ,Annelie Bjerkner ,Sandra Broddesson ,Robert Dyrdak ,Martin Ekman ,Lynda Eneh ,Lina Guerra Blomqvist ,Karolina Ininbergs ,Tanja Normark ,Isak Sylvin ,Zhibing Yun ,Martina Wahlund ,Valtteri Wirta |
| EPI_ISL_3085528 | Synlab Medilab | Karolinska University Hospital | Jan Albert ,Tobias Allander ,Annelie Bjerkner ,Sandra Broddesson ,Robert Dyrdak ,Martin Ekman ,Lynda Eneh ,Lina Guerra Blomqvist ,Karolina Ininbergs ,Tanja Normark ,Isak Sylvin ,Zhibing Yun ,Martina Wahlund ,Valtteri Wirta |
| EPI_ISL_3085529, EPI_ISL_3085530 | Unilabs | Karolinska University Hospital | Jan Albert ,Tobias Allander ,Annelie Bjerkner ,Sandra Broddesson ,Robert Dyrdak ,Martin Ekman ,Lynda Eneh ,Lina Guerra Blomqvist ,Karolina Ininbergs ,Tanja Normark ,Isak Sylvin ,Zhibing Yun ,Martina Wahlund ,Valtteri Wirta |
| EPI_ISL_3085531 | Karolinska University Hospital Solna | Karolinska University Hospital | Jan Albert ,Tobias Allander ,Annelie Bjerkner ,Sandra Broddesson ,Robert Dyrdak ,Martin Ekman ,Lynda Eneh ,Lina Guerra Blomqvist ,Karolina Ininbergs ,Tanja Normark ,Isak Sylvin ,Zhibing Yun ,Martina Wahlund ,Valtteri Wirta |
| EPI_ISL_3085532, EPI_ISL_3085533, EPI_ISL_3085534 | Unilabs | Karolinska University Hospital | Jan Albert ,Tobias Allander ,Annelie Bjerkner ,Sandra Broddesson ,Robert Dyrdak ,Martin Ekman ,Lynda Eneh ,Lina Guerra Blomqvist ,Karolina Ininbergs ,Tanja Normark ,Isak Sylvin ,Zhibing Yun ,Martina Wahlund ,Valtteri Wirta |
| EPI_ISL_3085535 | Karolinska University Hospital Solna | Karolinska University Hospital | Jan Albert ,Tobias Allander ,Annelie Bjerkner ,Sandra Broddesson ,Robert Dyrdak ,Martin Ekman ,Lynda Eneh ,Lina Guerra Blomqvist ,Karolina Ininbergs ,Tanja Normark ,Isak Sylvin ,Zhibing Yun ,Martina Wahlund ,Valtteri Wirta |
| EPI_ISL_3085536 | Synlab Medilab | Karolinska University Hospital | Jan Albert ,Tobias Allander ,Annelie Bjerkner ,Sandra Broddesson ,Robert Dyrdak ,Martin Ekman ,Lynda Eneh ,Lina Guerra Blomqvist ,Karolina Ininbergs ,Tanja Normark ,Isak Sylvin ,Zhibing Yun ,Martina Wahlund ,Valtteri Wirta |
| EPI_ISL_3085537, EPI_ISL_3085538, EPI_ISL_3085539, EPI_ISL_3085540, EPI_ISL_3085541 | Karolinska University Hospital Solna | Karolinska University Hospital | Jan Albert ,Tobias Allander ,Annelie Bjerkner ,Sandra Broddesson ,Robert Dyrdak ,Martin Ekman ,Lynda Eneh ,Lina Guerra Blomqvist ,Karolina Ininbergs ,Tanja Normark ,Isak Sylvin ,Zhibing Yun ,Martina Wahlund ,Valtteri Wirta |
| EPI_ISL_3085543, EPI_ISL_3085544 | Unilabs | Karolinska University Hospital | Jan Albert ,Tobias Allander ,Annelie Bjerkner ,Sandra Broddesson ,Robert Dyrdak ,Martin Ekman ,Lynda Eneh ,Lina Guerra Blomqvist ,Karolina Ininbergs ,Tanja Normark ,Isak Sylvin ,Zhibing Yun ,Martina Wahlund ,Valtteri Wirta |
| EPI_ISL_3085546, EPI_ISL_3085547 | Karolinska University Hospital Solna | Karolinska University Hospital | Jan Albert ,Tobias Allander ,Annelie Bjerkner ,Sandra Broddesson ,Robert Dyrdak ,Martin Ekman ,Lynda Eneh ,Lina Guerra Blomqvist ,Karolina Ininbergs ,Tanja Normark ,Isak Sylvin ,Zhibing Yun ,Martina Wahlund ,Valtteri Wirta |

| | | | |
|---|---|---|---|
| EPI_ISL_3085548, EPI_ISL_3085549 | Unilabs | Karolinska University Hospital | Jan Albert ,Tobias Allander ,Annelie Bjerkner ,Sandra Broddesson ,Robert Dyrdak ,Martin Ekman ,Lynda Eneh ,Lina Guerra Blomqvist ,Karolina Ininbergs ,Tanja Normark ,Isak Sylvin ,Zhibing Yun ,Martina Wahlund ,Valtteri Wirta |
| EPI_ISL_3085550 | Karolinska University Hospital Solna | Karolinska University Hospital | Jan Albert ,Tobias Allander ,Annelie Bjerkner ,Sandra Broddesson ,Robert Dyrdak ,Martin Ekman ,Lynda Eneh ,Lina Guerra Blomqvist ,Karolina Ininbergs ,Tanja Normark ,Isak Sylvin ,Zhibing Yun ,Martina Wahlund ,Valtteri Wirta |
| EPI_ISL_3085551 | Synlab Medilab | Karolinska University Hospital | Jan Albert ,Tobias Allander ,Annelie Bjerkner ,Sandra Broddesson ,Robert Dyrdak ,Martin Ekman ,Lynda Eneh ,Lina Guerra Blomqvist ,Karolina Ininbergs ,Tanja Normark ,Isak Sylvin ,Zhibing Yun ,Martina Wahlund ,Valtteri Wirta |
| EPI_ISL_3085552 | Unilabs | Karolinska University Hospital | Jan Albert ,Tobias Allander ,Annelie Bjerkner ,Sandra Broddesson ,Robert Dyrdak ,Martin Ekman ,Lynda Eneh ,Lina Guerra Blomqvist ,Karolina Ininbergs ,Tanja Normark ,Isak Sylvin ,Zhibing Yun ,Martina Wahlund ,Valtteri Wirta |
| EPI_ISL_3085553 | Synlab Medilab | Karolinska University Hospital | Jan Albert ,Tobias Allander ,Annelie Bjerkner ,Sandra Broddesson ,Robert Dyrdak ,Martin Ekman ,Lynda Eneh ,Lina Guerra Blomqvist ,Karolina Ininbergs ,Tanja Normark ,Isak Sylvin ,Zhibing Yun ,Martina Wahlund ,Valtteri Wirta |
| EPI_ISL_3085554 | Unilabs | Karolinska University Hospital | Jan Albert ,Tobias Allander ,Annelie Bjerkner ,Sandra Broddesson ,Robert Dyrdak ,Martin Ekman ,Lynda Eneh ,Lina Guerra Blomqvist ,Karolina Ininbergs ,Tanja Normark ,Isak Sylvin ,Zhibing Yun ,Martina Wahlund ,Valtteri Wirta |
| EPI_ISL_3085555 | Karolinska University Hospital Solna | Karolinska University Hospital | Jan Albert ,Tobias Allander ,Annelie Bjerkner ,Sandra Broddesson ,Robert Dyrdak ,Martin Ekman ,Lynda Eneh ,Lina Guerra Blomqvist ,Karolina Ininbergs ,Tanja Normark ,Isak Sylvin ,Zhibing Yun ,Martina Wahlund ,Valtteri Wirta |
| EPI_ISL_3085557 | Unilabs | Karolinska University Hospital | Jan Albert ,Tobias Allander ,Annelie Bjerkner ,Sandra Broddesson ,Robert Dyrdak ,Martin Ekman ,Lynda Eneh ,Lina Guerra Blomqvist ,Karolina Ininbergs ,Tanja Normark ,Isak Sylvin ,Zhibing Yun ,Martina Wahlund ,Valtteri Wirta |
| EPI_ISL_3085558 | Karolinska University Hospital Solna | Karolinska University Hospital | Jan Albert ,Tobias Allander ,Annelie Bjerkner ,Sandra Broddesson ,Robert Dyrdak ,Martin Ekman ,Lynda Eneh ,Lina Guerra Blomqvist ,Karolina Ininbergs ,Tanja Normark ,Isak Sylvin ,Zhibing Yun ,Martina Wahlund ,Valtteri Wirta |
| EPI_ISL_3085559 | Synlab Medilab | Karolinska University Hospital | Jan Albert ,Tobias Allander ,Annelie Bjerkner ,Sandra Broddesson ,Robert Dyrdak ,Martin Ekman ,Lynda Eneh ,Lina Guerra Blomqvist ,Karolina Ininbergs ,Tanja Normark ,Isak Sylvin ,Zhibing Yun ,Martina Wahlund ,Valtteri Wirta |
| EPI_ISL_3085560 | Karolinska University Hospital Solna | Karolinska University Hospital | Jan Albert ,Tobias Allander ,Annelie Bjerkner ,Sandra Broddesson ,Robert Dyrdak ,Martin Ekman ,Lynda Eneh ,Lina Guerra Blomqvist ,Karolina Ininbergs ,Tanja Normark ,Isak Sylvin ,Zhibing Yun ,Martina Wahlund ,Valtteri Wirta |
| EPI_ISL_3085561, EPI_ISL_3085563 | Synlab Medilab | Karolinska University Hospital | Jan Albert ,Tobias Allander ,Annelie Bjerkner ,Sandra Broddesson ,Robert Dyrdak ,Martin Ekman ,Lynda Eneh ,Lina Guerra Blomqvist ,Karolina Ininbergs ,Tanja Normark ,Isak Sylvin ,Zhibing Yun ,Martina Wahlund ,Valtteri Wirta |
| EPI_ISL_3085564, EPI_ISL_3085565 | Unilabs | Karolinska University Hospital | Jan Albert ,Tobias Allander ,Annelie Bjerkner ,Sandra Broddesson ,Robert Dyrdak ,Martin Ekman ,Lynda Eneh ,Lina Guerra Blomqvist ,Karolina Ininbergs ,Tanja Normark ,Isak Sylvin ,Zhibing Yun ,Martina Wahlund ,Valtteri Wirta |
| EPI_ISL_3085566, EPI_ISL_3085567 | Synlab Medilab | Karolinska University Hospital | Jan Albert ,Tobias Allander ,Annelie Bjerkner ,Sandra Broddesson ,Robert Dyrdak ,Martin Ekman ,Lynda Eneh ,Lina Guerra Blomqvist ,Karolina Ininbergs ,Tanja Normark ,Isak Sylvin ,Zhibing Yun ,Martina Wahlund ,Valtteri Wirta |
| EPI_ISL_3085568 | Unilabs | Karolinska University Hospital | Jan Albert ,Tobias Allander ,Annelie Bjerkner ,Sandra Broddesson ,Robert Dyrdak ,Martin Ekman ,Lynda Eneh ,Lina Guerra Blomqvist ,Karolina Ininbergs ,Tanja Normark ,Isak Sylvin ,Zhibing Yun ,Martina Wahlund ,Valtteri Wirta |
| EPI_ISL_3085569 | Karolinska University Hospital Solna | Karolinska University Hospital | Jan Albert ,Tobias Allander ,Annelie Bjerkner ,Sandra Broddesson ,Robert Dyrdak ,Martin Ekman ,Lynda Eneh ,Lina Guerra Blomqvist ,Karolina Ininbergs ,Tanja Normark ,Isak Sylvin ,Zhibing Yun ,Martina Wahlund ,Valtteri Wirta |
| EPI_ISL_3085570, EPI_ISL_3085571, EPI_ISL_3085572, EPI_ISL_3085573 | Unilabs | Karolinska University Hospital | Jan Albert ,Tobias Allander ,Annelie Bjerkner ,Sandra Broddesson ,Robert Dyrdak ,Martin Ekman ,Lynda Eneh ,Lina Guerra Blomqvist ,Karolina Ininbergs ,Tanja Normark ,Isak Sylvin ,Zhibing Yun ,Martina Wahlund ,Valtteri Wirta |
| EPI_ISL_3085574, EPI_ISL_3085575 | Synlab Medilab | Karolinska University Hospital | Jan Albert ,Tobias Allander ,Annelie Bjerkner ,Sandra Broddesson ,Robert Dyrdak ,Martin Ekman ,Lynda Eneh ,Lina Guerra Blomqvist ,Karolina Ininbergs ,Tanja Normark ,Isak Sylvin ,Zhibing Yun ,Martina Wahlund ,Valtteri Wirta |
| EPI_ISL_3085576, EPI_ISL_3085577 | Unilabs | Karolinska University Hospital | Jan Albert ,Tobias Allander ,Annelie Bjerkner ,Sandra Broddesson ,Robert Dyrdak ,Martin Ekman ,Lynda Eneh ,Lina Guerra Blomqvist ,Karolina Ininbergs ,Tanja Normark ,Isak Sylvin ,Zhibing Yun ,Martina Wahlund ,Valtteri Wirta |
| EPI_ISL_3085578 | Synlab Medilab | Karolinska University Hospital | Jan Albert ,Tobias Allander ,Annelie Bjerkner ,Sandra Broddesson ,Robert Dyrdak ,Martin Ekman ,Lynda Eneh ,Lina Guerra Blomqvist ,Karolina Ininbergs ,Tanja Normark ,Isak Sylvin ,Zhibing Yun ,Martina Wahlund ,Valtteri Wirta |
| EPI_ISL_3085579 | Karolinska University Hospital Solna | Karolinska University Hospital | Jan Albert ,Tobias Allander ,Annelie Bjerkner ,Sandra Broddesson ,Robert Dyrdak ,Martin Ekman ,Lynda Eneh ,Lina Guerra Blomqvist ,Karolina Ininbergs ,Tanja Normark ,Isak Sylvin ,Zhibing Yun ,Martina Wahlund ,Valtteri Wirta |
| EPI_ISL_3085580 | Synlab Medilab | Karolinska University Hospital | Jan Albert ,Tobias Allander ,Annelie Bjerkner ,Sandra Broddesson ,Robert Dyrdak ,Martin Ekman ,Lynda Eneh ,Lina Guerra Blomqvist ,Karolina Ininbergs ,Tanja Normark ,Isak Sylvin ,Zhibing Yun ,Martina Wahlund ,Valtteri Wirta |
| EPI_ISL_3085581 | Unilabs | Karolinska University Hospital | Jan Albert ,Tobias Allander ,Annelie Bjerkner ,Sandra Broddesson ,Robert Dyrdak ,Martin Ekman ,Lynda Eneh ,Lina Guerra Blomqvist ,Karolina Ininbergs ,Tanja Normark ,Isak Sylvin ,Zhibing Yun ,Martina Wahlund ,Valtteri Wirta |
| EPI_ISL_3085582 | Synlab Medilab | Karolinska University Hospital | Jan Albert ,Tobias Allander ,Annelie Bjerkner ,Sandra Broddesson ,Robert Dyrdak ,Martin Ekman ,Lynda Eneh ,Lina Guerra Blomqvist ,Karolina Ininbergs ,Tanja Normark ,Isak Sylvin ,Zhibing Yun ,Martina Wahlund ,Valtteri Wirta |
| EPI_ISL_3085583, EPI_ISL_3085584, EPI_ISL_3085585, EPI_ISL_3085586, EPI_ISL_3085587, EPI_ISL_3085588, EPI_ISL_3085655, EPI_ISL_3085657, EPI_ISL_3085658, EPI_ISL_3085659, EPI_ISL_3085660, EPI_ISL_3085661 | | | |
| see above | Unilabs | Karolinska University Hospital | Jan Albert ,Tobias Allander ,Annelie Bjerkner ,Sandra Broddesson ,Robert Dyrdak ,Martin Ekman ,Lynda Eneh ,Lina Guerra Blomqvist ,Karolina Ininbergs ,Tanja Normark ,Isak Sylvin ,Zhibing Yun ,Martina Wahlund ,Valtteri Wirta |
| EPI_ISL_3085662 | Karolinska University Hospital Solna | Karolinska University Hospital | Jan Albert ,Tobias Allander ,Annelie Bjerkner ,Sandra Broddesson ,Robert Dyrdak ,Martin Ekman ,Lynda Eneh ,Lina Guerra Blomqvist ,Karolina Ininbergs ,Tanja Normark ,Isak Sylvin ,Zhibing Yun ,Martina Wahlund ,Valtteri Wirta |
| EPI_ISL_3085663 | Unilabs | Karolinska University Hospital | Jan Albert ,Tobias Allander ,Annelie Bjerkner ,Sandra Broddesson ,Robert Dyrdak ,Martin Ekman ,Lynda Eneh ,Lina Guerra Blomqvist ,Karolina Ininbergs ,Tanja Normark ,Isak Sylvin ,Zhibing Yun ,Martina Wahlund ,Valtteri Wirta |
| EPI_ISL_3085664 | Karolinska University Hospital Solna | Karolinska University Hospital | Jan Albert ,Tobias Allander ,Annelie Bjerkner ,Sandra Broddesson ,Robert Dyrdak ,Martin Ekman ,Lynda Eneh ,Lina Guerra Blomqvist ,Karolina Ininbergs ,Tanja Normark ,Isak Sylvin ,Zhibing Yun ,Martina Wahlund ,Valtteri Wirta |
| EPI_ISL_3085665, EPI_ISL_3085666 | Unilabs | Karolinska University Hospital | Jan Albert ,Tobias Allander ,Annelie Bjerkner ,Sandra Broddesson ,Robert Dyrdak ,Martin Ekman ,Lynda Eneh ,Lina Guerra Blomqvist ,Karolina Ininbergs ,Tanja Normark ,Isak Sylvin ,Zhibing Yun ,Martina Wahlund ,Valtteri Wirta |
| EPI_ISL_3085667 | Karolinska University Hospital Solna | Karolinska University Hospital | Jan Albert ,Tobias Allander ,Annelie Bjerkner ,Sandra Broddesson ,Robert Dyrdak ,Martin Ekman ,Lynda Eneh ,Lina Guerra Blomqvist ,Karolina Ininbergs ,Tanja Normark ,Isak Sylvin ,Zhibing Yun ,Martina Wahlund ,Valtteri Wirta |
| EPI_ISL_3085669, EPI_ISL_3085670, EPI_ISL_3085671, EPI_ISL_3085672, EPI_ISL_3085673, EPI_ISL_3085674, EPI_ISL_3085675, EPI_ISL_3085676, EPI_ISL_3085677 | Unilabs | Karolinska University Hospital | Jan Albert ,Tobias Allander ,Annelie Bjerkner ,Sandra Broddesson ,Robert Dyrdak ,Martin Ekman ,Lynda Eneh ,Lina Guerra Blomqvist ,Karolina Ininbergs ,Tanja Normark ,Isak Sylvin ,Zhibing Yun ,Martina Wahlund ,Valtteri Wirta |
| EPI_ISL_3085678 | Karolinska University Hospital Solna | Karolinska University Hospital | Jan Albert ,Tobias Allander ,Annelie Bjerkner ,Sandra Broddesson ,Robert Dyrdak ,Martin Ekman ,Lynda Eneh ,Lina Guerra Blomqvist ,Karolina Ininbergs ,Tanja Normark ,Isak Sylvin ,Zhibing Yun ,Martina Wahlund ,Valtteri Wirta |
| EPI_ISL_3085679, EPI_ISL_3085681, EPI_ISL_3085683 | Unilabs | Karolinska University Hospital | Jan Albert ,Tobias Allander ,Annelie Bjerkner ,Sandra Broddesson ,Robert Dyrdak ,Martin Ekman ,Lynda Eneh ,Lina Guerra Blomqvist ,Karolina Ininbergs ,Tanja Normark ,Isak Sylvin ,Zhibing Yun ,Martina Wahlund ,Valtteri Wirta |
| EPI_ISL_3085684, EPI_ISL_3085685, EPI_ISL_3085686 | Karolinska University Hospital Solna | Karolinska University Hospital | Jan Albert ,Tobias Allander ,Annelie Bjerkner ,Sandra Broddesson ,Robert Dyrdak ,Martin Ekman ,Lynda Eneh ,Lina Guerra Blomqvist ,Karolina Ininbergs ,Tanja Normark ,Isak Sylvin ,Zhibing Yun ,Martina Wahlund ,Valtteri Wirta |
| EPI_ISL_3085688, EPI_ISL_3085689, EPI_ISL_3085690, EPI_ISL_3085691, EPI_ISL_3085692, EPI_ISL_3085693 | Unilabs | Karolinska University Hospital | Jan Albert ,Tobias Allander ,Annelie Bjerkner ,Sandra Broddesson ,Robert Dyrdak ,Martin Ekman ,Lynda Eneh ,Lina Guerra Blomqvist ,Karolina Ininbergs ,Tanja Normark ,Isak Sylvin ,Zhibing Yun ,Martina Wahlund ,Valtteri Wirta |
| EPI_ISL_3085694 | Karolinska University Hospital Solna | Karolinska University Hospital | Jan Albert ,Tobias Allander ,Annelie Bjerkner ,Sandra Broddesson ,Robert Dyrdak ,Martin Ekman ,Lynda Eneh ,Lina Guerra Blomqvist ,Karolina Ininbergs ,Tanja Normark ,Isak Sylvin ,Zhibing Yun ,Martina Wahlund ,Valtteri Wirta |
| EPI_ISL_3085695, EPI_ISL_3085697, EPI_ISL_3085698, EPI_ISL_3085699, EPI_ISL_3085700 | Unilabs | Karolinska University Hospital | Jan Albert ,Tobias Allander ,Annelie Bjerkner ,Sandra Broddesson ,Robert Dyrdak ,Martin Ekman ,Lynda Eneh ,Lina Guerra Blomqvist ,Karolina Ininbergs ,Tanja Normark ,Isak Sylvin ,Zhibing Yun ,Martina Wahlund ,Valtteri Wirta |
| EPI_ISL_3085702 | Karolinska University Hospital Solna | Karolinska University Hospital | Jan Albert ,Tobias Allander ,Annelie Bjerkner ,Sandra Broddesson ,Robert Dyrdak ,Martin Ekman ,Lynda Eneh ,Lina Guerra Blomqvist ,Karolina Ininbergs ,Tanja Normark ,Isak Sylvin ,Zhibing Yun ,Martina Wahlund ,Valtteri Wirta |
| EPI_ISL_3085703, EPI_ISL_3085704, EPI_ISL_3085705, EPI_ISL_3085706, EPI_ISL_3085707 | Unilabs | Karolinska University Hospital | Jan Albert ,Tobias Allander ,Annelie Bjerkner ,Sandra Broddesson ,Robert Dyrdak ,Martin Ekman ,Lynda Eneh ,Lina Guerra Blomqvist ,Karolina Ininbergs ,Tanja Normark ,Isak Sylvin ,Zhibing Yun ,Martina Wahlund ,Valtteri Wirta |
| EPI_ISL_3085708 | Karolinska University Hospital Solna | Karolinska University Hospital | Jan Albert ,Tobias Allander ,Annelie Bjerkner ,Sandra Broddesson ,Robert Dyrdak ,Martin Ekman ,Lynda Eneh ,Lina Guerra Blomqvist ,Karolina Ininbergs ,Tanja Normark ,Isak Sylvin ,Zhibing Yun ,Martina Wahlund ,Valtteri Wirta |
| EPI_ISL_3085709, EPI_ISL_3085710 | Unilabs | Karolinska University Hospital | Jan Albert ,Tobias Allander ,Annelie Bjerkner ,Sandra Broddesson ,Robert Dyrdak ,Martin Ekman ,Lynda Eneh ,Lina Guerra Blomqvist ,Karolina Ininbergs ,Tanja Normark ,Isak Sylvin ,Zhibing Yun ,Martina Wahlund ,Valtteri Wirta |
| EPI_ISL_3085711 | Karolinska University Hospital Solna | Karolinska University Hospital | Jan Albert ,Tobias Allander ,Annelie Bjerkner ,Sandra Broddesson ,Robert Dyrdak ,Martin Ekman ,Lynda Eneh ,Lina Guerra Blomqvist ,Karolina Ininbergs ,Tanja Normark ,Isak Sylvin ,Zhibing Yun ,Martina Wahlund ,Valtteri Wirta |
| EPI_ISL_3085712, EPI_ISL_3085713, EPI_ISL_3085715, EPI_ISL_3085716, EPI_ISL_3085717, EPI_ISL_3085718, EPI_ISL_3085720, EPI_ISL_3085721, EPI_ISL_3085722, EPI_ISL_3085723, EPI_ISL_3085724, EPI_ISL_3085725, EPI_ISL_3085726, EPI_ISL_3085727, EPI_ISL_3085728, EPI_ISL_3085729, EPI_ISL_3085730, EPI_ISL_3085731, EPI_ISL_3085732, EPI_ISL_3085733, EPI_ISL_3085734 | | | |
| see above | Unilabs | Karolinska University Hospital | Jan Albert ,Tobias Allander ,Annelie Bjerkner ,Sandra Broddesson ,Robert Dyrdak ,Martin Ekman ,Lynda Eneh ,Lina Guerra Blomqvist ,Karolina Ininbergs ,Tanja Normark ,Isak Sylvin ,Zhibing Yun ,Martina Wahlund ,Valtteri Wirta |
| EPI_ISL_3085735, EPI_ISL_3085736, EPI_ISL_3085737, EPI_ISL_3085738, EPI_ISL_3085739, EPI_ISL_3085740, EPI_ISL_3085741, EPI_ISL_3085742, EPI_ISL_3085743, EPI_ISL_3085744, EPI_ISL_3085746, EPI_ISL_3085747, EPI_ISL_3085748 | | | |
| see above | Karolinska University Hospital Huddinge | Karolinska University Hospital | Jan Albert ,Tobias Allander ,Annelie Bjerkner ,Sandra Broddesson ,Robert Dyrdak ,Martin Ekman ,Lynda Eneh ,Lina Guerra Blomqvist ,Karolina Ininbergs ,Tanja Normark ,Isak Sylvin ,Zhibing Yun ,Martina Wahlund ,Valtteri Wirta |
| EPI_ISL_3085749, EPI_ISL_3085750, EPI_ISL_3085751, EPI_ISL_3085753 | Karolinska University Hospital Solna | Karolinska University Hospital | Jan Albert ,Tobias Allander ,Annelie Bjerkner ,Sandra Broddesson ,Robert Dyrdak ,Martin Ekman ,Lynda Eneh ,Lina Guerra Blomqvist ,Karolina Ininbergs ,Tanja Normark ,Isak Sylvin ,Zhibing Yun ,Martina Wahlund ,Valtteri Wirta |
| EPI_ISL_3085754 | Synlab Medilab | Karolinska University Hospital | Jan Albert ,Tobias Allander ,Annelie Bjerkner ,Sandra Broddesson ,Robert Dyrdak ,Martin Ekman ,Lynda Eneh ,Lina Guerra Blomqvist ,Karolina Ininbergs ,Tanja Normark ,Isak Sylvin ,Zhibing Yun ,Martina Wahlund ,Valtteri Wirta |

| | | | |
|---|---|---|---|
| EPI_ISL_3085755, EPI_ISL_3085756, EPI_ISL_3085757 | Karolinska University Hospital Solna | Karolinska University Hospital | Jan Albert ,Tobias Allander ,Annelie Bjerkner ,Sandra Broddesson ,Robert Dyrdak ,Martin Ekman ,Lynda Eneh ,Lina Guerra Blomqvist ,Karolina Ininbergs ,Tanja Normark ,Isak Sylvin ,Zhibing Yun ,Martina Wahlund ,Valtteri Wirta |
| EPI_ISL_3085758 | Synlab Medilab | Karolinska University Hospital | Jan Albert ,Tobias Allander ,Annelie Bjerkner ,Sandra Broddesson ,Robert Dyrdak ,Martin Ekman ,Lynda Eneh ,Lina Guerra Blomqvist ,Karolina Ininbergs ,Tanja Normark ,Isak Sylvin ,Zhibing Yun ,Martina Wahlund ,Valtteri Wirta |
| EPI_ISL_3085759, EPI_ISL_3085760, EPI_ISL_3085761, EPI_ISL_3085762 | Karolinska University Hospital Solna | Karolinska University Hospital | Jan Albert ,Tobias Allander ,Annelie Bjerkner ,Sandra Broddesson ,Robert Dyrdak ,Martin Ekman ,Lynda Eneh ,Lina Guerra Blomqvist ,Karolina Ininbergs ,Tanja Normark ,Isak Sylvin ,Zhibing Yun ,Martina Wahlund ,Valtteri Wirta |
| EPI_ISL_3085763 | Unilabs | Karolinska University Hospital | Jan Albert ,Tobias Allander ,Annelie Bjerkner ,Sandra Broddesson ,Robert Dyrdak ,Martin Ekman ,Lynda Eneh ,Lina Guerra Blomqvist ,Karolina Ininbergs ,Tanja Normark ,Isak Sylvin ,Zhibing Yun ,Martina Wahlund ,Valtteri Wirta |
| EPI_ISL_3085764 | Synlab Medilab | Karolinska University Hospital | Jan Albert ,Tobias Allander ,Annelie Bjerkner ,Sandra Broddesson ,Robert Dyrdak ,Martin Ekman ,Lynda Eneh ,Lina Guerra Blomqvist ,Karolina Ininbergs ,Tanja Normark ,Isak Sylvin ,Zhibing Yun ,Martina Wahlund ,Valtteri Wirta |
| EPI_ISL_3085765, EPI_ISL_3085766, EPI_ISL_3085767 | Karolinska University Hospital Solna | Karolinska University Hospital | Jan Albert ,Tobias Allander ,Annelie Bjerkner ,Sandra Broddesson ,Robert Dyrdak ,Martin Ekman ,Lynda Eneh ,Lina Guerra Blomqvist ,Karolina Ininbergs ,Tanja Normark ,Isak Sylvin ,Zhibing Yun ,Martina Wahlund ,Valtteri Wirta |
| EPI_ISL_3085768 | Synlab Medilab | Karolinska University Hospital | Jan Albert ,Tobias Allander ,Annelie Bjerkner ,Sandra Broddesson ,Robert Dyrdak ,Martin Ekman ,Lynda Eneh ,Lina Guerra Blomqvist ,Karolina Ininbergs ,Tanja Normark ,Isak Sylvin ,Zhibing Yun ,Martina Wahlund ,Valtteri Wirta |
| EPI_ISL_3085769 | Unilabs | Karolinska University Hospital | Jan Albert ,Tobias Allander ,Annelie Bjerkner ,Sandra Broddesson ,Robert Dyrdak ,Martin Ekman ,Lynda Eneh ,Lina Guerra Blomqvist ,Karolina Ininbergs ,Tanja Normark ,Isak Sylvin ,Zhibing Yun ,Martina Wahlund ,Valtteri Wirta |
| EPI_ISL_3085770, EPI_ISL_3085771, EPI_ISL_3085772, EPI_ISL_3085773, EPI_ISL_3085774 | Karolinska University Hospital Solna | Karolinska University Hospital | Jan Albert ,Tobias Allander ,Annelie Bjerkner ,Sandra Broddesson ,Robert Dyrdak ,Martin Ekman ,Lynda Eneh ,Lina Guerra Blomqvist ,Karolina Ininbergs ,Tanja Normark ,Isak Sylvin ,Zhibing Yun ,Martina Wahlund ,Valtteri Wirta |
| EPI_ISL_3085776 | Unilabs | Karolinska University Hospital | Jan Albert ,Tobias Allander ,Annelie Bjerkner ,Sandra Broddesson ,Robert Dyrdak ,Martin Ekman ,Lynda Eneh ,Lina Guerra Blomqvist ,Karolina Ininbergs ,Tanja Normark ,Isak Sylvin ,Zhibing Yun ,Martina Wahlund ,Valtteri Wirta |
| EPI_ISL_3085777 | Karolinska University Hospital Solna | Karolinska University Hospital | Jan Albert ,Tobias Allander ,Annelie Bjerkner ,Sandra Broddesson ,Robert Dyrdak ,Martin Ekman ,Lynda Eneh ,Lina Guerra Blomqvist ,Karolina Ininbergs ,Tanja Normark ,Isak Sylvin ,Zhibing Yun ,Martina Wahlund ,Valtteri Wirta |
| EPI_ISL_3085778, EPI_ISL_3085779 | Unilabs | Karolinska University Hospital | Jan Albert ,Tobias Allander ,Annelie Bjerkner ,Sandra Broddesson ,Robert Dyrdak ,Martin Ekman ,Lynda Eneh ,Lina Guerra Blomqvist ,Karolina Ininbergs ,Tanja Normark ,Isak Sylvin ,Zhibing Yun ,Martina Wahlund ,Valtteri Wirta |
| EPI_ISL_3085780, EPI_ISL_3085781 | Karolinska University Hospital Solna | Karolinska University Hospital | Jan Albert ,Tobias Allander ,Annelie Bjerkner ,Sandra Broddesson ,Robert Dyrdak ,Martin Ekman ,Lynda Eneh ,Lina Guerra Blomqvist ,Karolina Ininbergs ,Tanja Normark ,Isak Sylvin ,Zhibing Yun ,Martina Wahlund ,Valtteri Wirta |
| EPI_ISL_3085782 | Unilabs | Karolinska University Hospital | Jan Albert ,Tobias Allander ,Annelie Bjerkner ,Sandra Broddesson ,Robert Dyrdak ,Martin Ekman ,Lynda Eneh ,Lina Guerra Blomqvist ,Karolina Ininbergs ,Tanja Normark ,Isak Sylvin ,Zhibing Yun ,Martina Wahlund ,Valtteri Wirta |
| EPI_ISL_3085783, EPI_ISL_3085784, EPI_ISL_3085786, EPI_ISL_3085787, EPI_ISL_3085789, EPI_ISL_3085790, EPI_ISL_3085791, EPI_ISL_3085792 | Karolinska University Hospital Solna | Karolinska University Hospital | Jan Albert ,Tobias Allander ,Annelie Bjerkner ,Sandra Broddesson ,Robert Dyrdak ,Martin Ekman ,Lynda Eneh ,Lina Guerra Blomqvist ,Karolina Ininbergs ,Tanja Normark ,Isak Sylvin ,Zhibing Yun ,Martina Wahlund ,Valtteri Wirta |
| EPI_ISL_3085793 | Unilabs | Karolinska University Hospital | Jan Albert ,Tobias Allander ,Annelie Bjerkner ,Sandra Broddesson ,Robert Dyrdak ,Martin Ekman ,Lynda Eneh ,Lina Guerra Blomqvist ,Karolina Ininbergs ,Tanja Normark ,Isak Sylvin ,Zhibing Yun ,Martina Wahlund ,Valtteri Wirta |
| EPI_ISL_3085794, EPI_ISL_3085795, EPI_ISL_3085796, EPI_ISL_3085797, EPI_ISL_3085798, EPI_ISL_3085799, EPI_ISL_3085800, EPI_ISL_3085801, EPI_ISL_3085802 | Karolinska University Hospital Solna | Karolinska University Hospital | Jan Albert ,Tobias Allander ,Annelie Bjerkner ,Sandra Broddesson ,Robert Dyrdak ,Martin Ekman ,Lynda Eneh ,Lina Guerra Blomqvist ,Karolina Ininbergs ,Tanja Normark ,Isak Sylvin ,Zhibing Yun ,Martina Wahlund ,Valtteri Wirta |
| EPI_ISL_3085803 | Synlab Medilab | Karolinska University Hospital | Jan Albert ,Tobias Allander ,Annelie Bjerkner ,Sandra Broddesson ,Robert Dyrdak ,Martin Ekman ,Lynda Eneh ,Lina Guerra Blomqvist ,Karolina Ininbergs ,Tanja Normark ,Isak Sylvin ,Zhibing Yun ,Martina Wahlund ,Valtteri Wirta |
| EPI_ISL_3085804 | Karolinska University Hospital Solna | Karolinska University Hospital | Jan Albert ,Tobias Allander ,Annelie Bjerkner ,Sandra Broddesson ,Robert Dyrdak ,Martin Ekman ,Lynda Eneh ,Lina Guerra Blomqvist ,Karolina Ininbergs ,Tanja Normark ,Isak Sylvin ,Zhibing Yun ,Martina Wahlund ,Valtteri Wirta |
| EPI_ISL_3085805 | Unilabs | Karolinska University Hospital | Jan Albert ,Tobias Allander ,Annelie Bjerkner ,Sandra Broddesson ,Robert Dyrdak ,Martin Ekman ,Lynda Eneh ,Lina Guerra Blomqvist ,Karolina Ininbergs ,Tanja Normark ,Isak Sylvin ,Zhibing Yun ,Martina Wahlund ,Valtteri Wirta |
| EPI_ISL_3085807, EPI_ISL_3085808, EPI_ISL_3085809, EPI_ISL_3085811, EPI_ISL_3085813, EPI_ISL_3085814 | Karolinska University Hospital Solna | Karolinska University Hospital | Jan Albert ,Tobias Allander ,Annelie Bjerkner ,Sandra Broddesson ,Robert Dyrdak ,Martin Ekman ,Lynda Eneh ,Lina Guerra Blomqvist ,Karolina Ininbergs ,Tanja Normark ,Isak Sylvin ,Zhibing Yun ,Martina Wahlund ,Valtteri Wirta |
| EPI_ISL_3085815 | Synlab Medilab | Karolinska University Hospital | Jan Albert ,Tobias Allander ,Annelie Bjerkner ,Sandra Broddesson ,Robert Dyrdak ,Martin Ekman ,Lynda Eneh ,Lina Guerra Blomqvist ,Karolina Ininbergs ,Tanja Normark ,Isak Sylvin ,Zhibing Yun ,Martina Wahlund ,Valtteri Wirta |
| EPI_ISL_3085816, EPI_ISL_3085817 | Karolinska University Hospital Solna | Karolinska University Hospital | Jan Albert ,Tobias Allander ,Annelie Bjerkner ,Sandra Broddesson ,Robert Dyrdak ,Martin Ekman ,Lynda Eneh ,Lina Guerra Blomqvist ,Karolina Ininbergs ,Tanja Normark ,Isak Sylvin ,Zhibing Yun ,Martina Wahlund ,Valtteri Wirta |
| EPI_ISL_3085819 | Unilabs | Karolinska University Hospital | Jan Albert ,Tobias Allander ,Annelie Bjerkner ,Sandra Broddesson ,Robert Dyrdak ,Martin Ekman ,Lynda Eneh ,Lina Guerra Blomqvist ,Karolina Ininbergs ,Tanja Normark ,Isak Sylvin ,Zhibing Yun ,Martina Wahlund ,Valtteri Wirta |
| EPI_ISL_3085827 | Laboratory Corporation of America | Centers for Disease Control and Prevention Division of Viral Diseases, Pathogen Discovery | Dakota Howard, Dhwani Batra, Peter W. Cook, Kara Moser, Adrian Paskey, Jason Caravas, Benjamin Rambo-Martin, Shatavia Morrison, Christopher Gulvick, Scott Sammons, Yvette Unoarumhi, Darlene Wagner, Matthew Schmerer, Minoo Agarwal, Eyad Almasri, Debbie Boles, Ayla Burns, Nuthawin Charoensri, Oren Cohen, Susan Countryman, Mary Ann Cristobal, Bobbi Croy, Suzanne Dale, Hrushikesh Deshmukh, Amanda Douglas, Vincent Drouillon, Marcia Eisenberg, Howard Engler, Rama Ghatti, Prashant Gupta, Susan Hicks, Jake Humphrey, Lax Iyer, Lisa Pfefferle, Manoj Jain, Matthew Robinson, Mohan Kolli, Brian Krueger, Tim Kuphal, Stanley Letovsky, Michael Levandoski, Craig Lukasik, Jonathan Meltzer, Brian Norvell, Mindy Nye, Scott Parker, Christos Petropoulos, John Pruitt, Steven Ragan, Scott Ryan, Mike Sapeta, Jana Schroth, Suresh Babu Selvaraju, Goran Stevovic, Amanda Suchanek, Andrea Throop, Lyndon Tilson, Thomas Urban, Joe Voshell, Kimberly Wagner, Jonathan Williams, Mary Williamson, Qian Zeng, Tricia Zwiefelhofer, Clinton R. Paden, Duncan MacCannell |
| EPI_ISL_3085868, EPI_ISL_3085870, EPI_ISL_3085873, EPI_ISL_3085874, EPI_ISL_3085875, EPI_ISL_3085876 | Infinity Biologix | Centers for Disease Control and Prevention Division of Viral Diseases, Pathogen Discovery | Dakota Howard, Dhwani Batra, Peter W. Cook, Kara Moser, Adrian Paskey, Jason Caravas, Benjamin Rambo-Martin, Shatavia Morrison, Christopher Gulvick, Scott Sammons, Yvette Unoarumhi, Darlene Wagner, Matthew Schmerer, Christian Bixby, Yihe Wang, Jonathan Schultz, Chirayu Goswami, Russ Hager, Robin Grimwood, Clinton R. Paden, Duncan MacCannell |
| EPI_ISL_3085882, EPI_ISL_3086027, EPI_ISL_3086109, EPI_ISL_3086111, EPI_ISL_3086120, EPI_ISL_3086121, EPI_ISL_3086124, EPI_ISL_3086125, EPI_ISL_3086126, EPI_ISL_3086127, EPI_ISL_3086130, EPI_ISL_3086133, EPI_ISL_3086134, EPI_ISL_3086135, EPI_ISL_3086142 | | | |
| see above | Maine Health and Environmental Testing Laboratory | Tewhey Lab, The Jackson Laboratory | Matluk,N., Dewey,H., Iosue,F., Barter,M., Lynch,R., Munger,H. and Tewhey,R. |
| EPI_ISL_3086143, EPI_ISL_3086147, EPI_ISL_3086149, EPI_ISL_3086153, EPI_ISL_3086154 | Curative Labs | New Mexico Department of Health Scientific Laboratory | Ellie Johnson, D'eldra Malone, Jennifer Benoit, Ratheesh Rajan, Linda Salazar, Anastacia Griego-Fisher |
| EPI_ISL_3086157, EPI_ISL_3086159, EPI_ISL_3086160, EPI_ISL_3086161, EPI_ISL_3086162, EPI_ISL_3086166, EPI_ISL_3086168, EPI_ISL_3086169, EPI_ISL_3086170, EPI_ISL_3086171, EPI_ISL_3086172, EPI_ISL_3086173, EPI_ISL_3086174, EPI_ISL_3086175, EPI_ISL_3086176, EPI_ISL_3086177 | | | |
| see above | New Mexico Department of Health Scientific Laboratory | New Mexico Department of Health Scientific Laboratory | Ellie Johnson, D'eldra Malone, Jennifer Benoit, Ratheesh Rajan, Linda Salazar, Anastacia Griego-Fisher |
| EPI_ISL_3086178, EPI_ISL_3086179, EPI_ISL_3086182, EPI_ISL_3086210 | Curative Labs | New Mexico Department of Health Scientific Laboratory | Ellie Johnson, D'eldra Malone, Jennifer Benoit, Ratheesh Rajan, Linda Salazar, Anastacia Griego-Fisher |
| EPI_ISL_3086245, EPI_ISL_3086246, EPI_ISL_3086248, EPI_ISL_3086249, EPI_ISL_3086250, EPI_ISL_3086251, EPI_ISL_3086252, EPI_ISL_3086253, EPI_ISL_3086254, EPI_ISL_3086255, EPI_ISL_3086256, EPI_ISL_3086257, EPI_ISL_3086258, EPI_ISL_3086261, EPI_ISL_3086264 | | | |
| see above | IN State Department of Health Laboratory Services | IN State Department of Health Laboratory Services | Cassandra Campion, Jamie Yeadon, Brian Pope, Lixia Liu, Kyle Brownlee, Melissa Hindenlang, Mark Glazier |
| EPI_ISL_3086265 | Clinical Virology | Clinical Bacteriology | Tim Roloff, Fanny Wegner, Helena MB Seth-Smith, Alfredo Mari, Karoline Leuzinger, Julia Bielicki, Manuel Battegay, Hans Hirsch, Adrian Egli |
| EPI_ISL_3086266 | IN State Department of Health Laboratory Services | IN State Department of Health Laboratory Services | Cassandra Campion, Jamie Yeadon, Brian Pope, Lixia Liu, Kyle Brownlee, Melissa Hindenlang, Mark Glazier |
| EPI_ISL_3086268 | Clinical Virology | Clinical Bacteriology | Tim Roloff, Fanny Wegner, Helena MB Seth-Smith, Alfredo Mari, Karoline Leuzinger, Julia Bielicki, Manuel Battegay, Hans Hirsch, Adrian Egli |
| EPI_ISL_3086270 | IN State Department of Health Laboratory Services | IN State Department of Health Laboratory Services | Cassandra Campion, Jamie Yeadon, Brian Pope, Lixia Liu, Kyle Brownlee, Melissa Hindenlang, Mark Glazier |
| EPI_ISL_3086271 | Clinical Virology | Clinical Bacteriology | Tim Roloff, Fanny Wegner, Helena MB Seth-Smith, Alfredo Mari, Karoline Leuzinger, Julia Bielicki, Manuel Battegay, Hans Hirsch, Adrian Egli |
| EPI_ISL_3086272 | IN State Department of Health Laboratory Services | IN State Department of Health Laboratory Services | Cassandra Campion, Jamie Yeadon, Brian Pope, Lixia Liu, Kyle Brownlee, Melissa Hindenlang, Mark Glazier |
| EPI_ISL_3086273, EPI_ISL_3086274 | Clinical Virology | Clinical Bacteriology | Tim Roloff, Fanny Wegner, Helena MB Seth-Smith, Alfredo Mari, Karoline Leuzinger, Julia Bielicki, Manuel Battegay, Hans Hirsch, Adrian Egli |

| | | | |
|---|---|---|---|
| EPI_ISL_3086275 | IN State Department of Health Laboratory Services | IN State Department of Health Laboratory Services | Cassandra Campion, Jamie Yeadon, Brian Pope, Lixia Liu, Kyle Brownlee, Melissa Hindenlang, Mark Glazier |
| EPI_ISL_3086276, EPI_ISL_3086277, EPI_ISL_3086279, EPI_ISL_3086280, EPI_ISL_3086282, EPI_ISL_3086284, EPI_ISL_3086285 | Clinical Virology | Clinical Bacteriology | Tim Roloff, Fanny Wegner, Helena MB Seth-Smith, Alfredo Mari, Karoline Leuzinger, Julia Bielicki, Manuel Battegay, Hans Hirsch, Adrian Egli |
| EPI_ISL_3086286 | IN State Department of Health Laboratory Services | IN State Department of Health Laboratory Services | Cassandra Campion, Jamie Yeadon, Brian Pope, Lixia Liu, Kyle Brownlee, Melissa Hindenlang, Mark Glazier |
| EPI_ISL_3086287 | Clinical Virology | Clinical Bacteriology | Tim Roloff, Fanny Wegner, Helena MB Seth-Smith, Alfredo Mari, Karoline Leuzinger, Julia Bielicki, Manuel Battegay, Hans Hirsch, Adrian Egli |
| EPI_ISL_3086288 | IN State Department of Health Laboratory Services | IN State Department of Health Laboratory Services | Cassandra Campion, Jamie Yeadon, Brian Pope, Lixia Liu, Kyle Brownlee, Melissa Hindenlang, Mark Glazier |
| EPI_ISL_3086289, EPI_ISL_3086290 | Clinical Virology | Clinical Bacteriology | Tim Roloff, Fanny Wegner, Helena MB Seth-Smith, Alfredo Mari, Karoline Leuzinger, Julia Bielicki, Manuel Battegay, Hans Hirsch, Adrian Egli |
| EPI_ISL_3086291 | IN State Department of Health Laboratory Services | IN State Department of Health Laboratory Services | Cassandra Campion, Jamie Yeadon, Brian Pope, Lixia Liu, Kyle Brownlee, Melissa Hindenlang, Mark Glazier |
| EPI_ISL_3086292 | Clinical Virology | Clinical Bacteriology | Tim Roloff, Fanny Wegner, Helena MB Seth-Smith, Alfredo Mari, Karoline Leuzinger, Julia Bielicki, Manuel Battegay, Hans Hirsch, Adrian Egli |
| EPI_ISL_3086293 | IN State Department of Health Laboratory Services | IN State Department of Health Laboratory Services | Cassandra Campion, Jamie Yeadon, Brian Pope, Lixia Liu, Kyle Brownlee, Melissa Hindenlang, Mark Glazier |
| EPI_ISL_3086294, EPI_ISL_3086295 | Clinical Virology | Clinical Bacteriology | Tim Roloff, Fanny Wegner, Helena MB Seth-Smith, Alfredo Mari, Karoline Leuzinger, Julia Bielicki, Manuel Battegay, Hans Hirsch, Adrian Egli |
| EPI_ISL_3086296 | IN State Department of Health Laboratory Services | IN State Department of Health Laboratory Services | Cassandra Campion, Jamie Yeadon, Brian Pope, Lixia Liu, Kyle Brownlee, Melissa Hindenlang, Mark Glazier |
| EPI_ISL_3086297 | Clinical Virology | Clinical Bacteriology | Tim Roloff, Fanny Wegner, Helena MB Seth-Smith, Alfredo Mari, Karoline Leuzinger, Julia Bielicki, Manuel Battegay, Hans Hirsch, Adrian Egli |
| EPI_ISL_3086298 | IN State Department of Health Laboratory Services | IN State Department of Health Laboratory Services | Cassandra Campion, Jamie Yeadon, Brian Pope, Lixia Liu, Kyle Brownlee, Melissa Hindenlang, Mark Glazier |
| EPI_ISL_3086299, EPI_ISL_3086300 | Clinical Virology | Clinical Bacteriology | Tim Roloff, Fanny Wegner, Helena MB Seth-Smith, Alfredo Mari, Karoline Leuzinger, Julia Bielicki, Manuel Battegay, Hans Hirsch, Adrian Egli |
| EPI_ISL_3086301 | IN State Department of Health Laboratory Services | IN State Department of Health Laboratory Services | Cassandra Campion, Jamie Yeadon, Brian Pope, Lixia Liu, Kyle Brownlee, Melissa Hindenlang, Mark Glazier |
| EPI_ISL_3086302, EPI_ISL_3086303, EPI_ISL_3086304, EPI_ISL_3086305, EPI_ISL_3086306, EPI_ISL_3086308, EPI_ISL_3086309 | Clinical Virology | Clinical Bacteriology | Tim Roloff, Fanny Wegner, Helena MB Seth-Smith, Alfredo Mari, Karoline Leuzinger, Julia Bielicki, Manuel Battegay, Hans Hirsch, Adrian Egli |
| EPI_ISL_3086310, EPI_ISL_3086311, EPI_ISL_3086312, EPI_ISL_3086314, EPI_ISL_3086315 | Synlab Suisse SA | Clinical Bacteriology | Tim Roloff, Fanny Wegner, Helena MB Seth-Smith, Alfredo Mari, Karoline Leuzinger, Julia Bielicki, Manuel Battegay, Hans Hirsch, Adrian Egli |
| EPI_ISL_3086316, EPI_ISL_3086317 | Rothen Medizinische Laboratorien AG | Clinical Bacteriology | Tim Roloff, Fanny Wegner, Helena MB Seth-Smith, Alfredo Mari, Karoline Leuzinger, Julia Bielicki, Ingrid Steffen, Manuel Battegay, Hans Hirsch, Adrian Egli |
| EPI_ISL_3086318, EPI_ISL_3086319 | Viollier AG | Clinical Bacteriology | Tim Roloff, Fanny Wegner, Helena MB Seth-Smith, Alfredo Mari, Karoline Leuzinger, Julia Bielicki, Christiane Beckmann, Manuel Battegay, Hans Hirsch, Adrian Egli |
| EPI_ISL_3086320, EPI_ISL_3086321, EPI_ISL_3086322 | Hopital du Jura: Delemont | Clinical Bacteriology | Tim Roloff, Fanny Wegner, Helena MB Seth-Smith, Alfredo Mari, Karoline Leuzinger, Julia Bielicki, Manuel Battegay, Hans Hirsch, Adrian Egli |
| EPI_ISL_3086331, EPI_ISL_3086332, EPI_ISL_3086334, EPI_ISL_3086335, EPI_ISL_3086337, EPI_ISL_3086338, EPI_ISL_3086341, EPI_ISL_3086342, EPI_ISL_3086344, EPI_ISL_3086345, EPI_ISL_3086347, EPI_ISL_3086348, EPI_ISL_3086349, EPI_ISL_3086351, EPI_ISL_3086352, EPI_ISL_3086355, EPI_ISL_3086357, EPI_ISL_3086358, EPI_ISL_3086360, EPI_ISL_3086361, EPI_ISL_3086364, EPI_ISL_3086366, EPI_ISL_3086367, EPI_ISL_3086369, EPI_ISL_3086370, EPI_ISL_3086371, EPI_ISL_3086373, EPI_ISL_3086374, EPI_ISL_3086376, EPI_ISL_3086377, EPI_ISL_3086379, EPI_ISL_3086380, EPI_ISL_3086382, EPI_ISL_3086383, EPI_ISL_3086385, EPI_ISL_3086386, EPI_ISL_3086387, EPI_ISL_3086389, EPI_ISL_3086390, EPI_ISL_3086392, EPI_ISL_3086393, EPI_ISL_3086395, EPI_ISL_3086396, EPI_ISL_3086398, EPI_ISL_3086399, EPI_ISL_3086401, EPI_ISL_3086402 | | | |
| see above | Laboratorio di Microbiologia | Laboratorio di Microbiologia | Martinetti Lucchini Gladys, Valeria Spina |
| EPI_ISL_3086445, EPI_ISL_3086446, EPI_ISL_3086447, EPI_ISL_3086448 | Louisiana Office of Public Heath Laboratory | LSUHS Emerging Viral Threat Laboratory | Gregory L. Ware, Alexander Mijalis, Jeremy P. Kamil, Jennifer L. Carroll, Maarten Van Diest, Rona S. Scott, Andrew D. Yurochko, Christopher G. Kevil, John A. Vanchiere |
| EPI_ISL_3086449 | LSUHS Emerging Viral Threat Laboratory | LSUHS Emerging Viral Threat Laboratory | Gregory L. Ware, Alexander Mijalis, Jeremy P. Kamil, Jennifer L. Carroll, Maarten Van Diest, Rona S. Scott, Andrew D. Yurochko, Christopher G. Kevil, John A. Vanchiere |
| EPI_ISL_3086450, EPI_ISL_3086451, EPI_ISL_3086452, EPI_ISL_3086453, EPI_ISL_3086454, EPI_ISL_3086455, EPI_ISL_3086456, EPI_ISL_3086457 | Louisiana Office of Public Heath Laboratory | LSUHS Emerging Viral Threat Laboratory | Gregory L. Ware, Alexander Mijalis, Jeremy P. Kamil, Jennifer L. Carroll, Maarten Van Diest, Rona S. Scott, Andrew D. Yurochko, Christopher G. Kevil, John A. Vanchiere |
| EPI_ISL_3086458 | LSUHS Emerging Viral Threat Laboratory | LSUHS Emerging Viral Threat Laboratory | Gregory L. Ware, Alexander Mijalis, Jeremy P. Kamil, Jennifer L. Carroll, Maarten Van Diest, Rona S. Scott, Andrew D. Yurochko, Christopher G. Kevil, John A. Vanchiere |
| EPI_ISL_3086459 | Louisiana Office of Public Heath Laboratory | LSUHS Emerging Viral Threat Laboratory | Gregory L. Ware, Alexander Mijalis, Jeremy P. Kamil, Jennifer L. Carroll, Maarten Van Diest, Rona S. Scott, Andrew D. Yurochko, Christopher G. Kevil, John A. Vanchiere |
| EPI_ISL_3086460 | LSUHS Emerging Viral Threat Laboratory | LSUHS Emerging Viral Threat Laboratory | Gregory L. Ware, Alexander Mijalis, Jeremy P. Kamil, Jennifer L. Carroll, Maarten Van Diest, Rona S. Scott, Andrew D. Yurochko, Christopher G. Kevil, John A. Vanchiere |
| EPI_ISL_3086461, EPI_ISL_3086462 | Louisiana Office of Public Heath Laboratory | LSUHS Emerging Viral Threat Laboratory | Gregory L. Ware, Alexander Mijalis, Jeremy P. Kamil, Jennifer L. Carroll, Maarten Van Diest, Rona S. Scott, Andrew D. Yurochko, Christopher G. Kevil, John A. Vanchiere |
| EPI_ISL_3086463 | LSUHS Emerging Viral Threat Laboratory | LSUHS Emerging Viral Threat Laboratory | Gregory L. Ware, Alexander Mijalis, Jeremy P. Kamil, Jennifer L. Carroll, Maarten Van Diest, Rona S. Scott, Andrew D. Yurochko, Christopher G. Kevil, John A. Vanchiere |
| EPI_ISL_3086464 | Louisiana Office of Public Heath Laboratory | LSUHS Emerging Viral Threat Laboratory | Gregory L. Ware, Alexander Mijalis, Jeremy P. Kamil, Jennifer L. Carroll, Maarten Van Diest, Rona S. Scott, Andrew D. Yurochko, Christopher G. Kevil, John A. Vanchiere |
| EPI_ISL_3086465, EPI_ISL_3086466 | LSUHS Emerging Viral Threat Laboratory | LSUHS Emerging Viral Threat Laboratory | Gregory L. Ware, Alexander Mijalis, Jeremy P. Kamil, Jennifer L. Carroll, Maarten Van Diest, Rona S. Scott, Andrew D. Yurochko, Christopher G. Kevil, John A. Vanchiere |
| EPI_ISL_3086467, EPI_ISL_3086468 | Louisiana Office of Public Heath Laboratory | LSUHS Emerging Viral Threat Laboratory | Gregory L. Ware, Alexander Mijalis, Jeremy P. Kamil, Jennifer L. Carroll, Maarten Van Diest, Rona S. Scott, Andrew D. Yurochko, Christopher G. Kevil, John A. Vanchiere |
| EPI_ISL_3086469 | LSUHS Emerging Viral Threat Laboratory | LSUHS Emerging Viral Threat Laboratory | Gregory L. Ware, Alexander Mijalis, Jeremy P. Kamil, Jennifer L. Carroll, Maarten Van Diest, Rona S. Scott, Andrew D. Yurochko, Christopher G. Kevil, John A. Vanchiere |
| EPI_ISL_3086470 | Louisiana Office of Public Heath Laboratory | LSUHS Emerging Viral Threat Laboratory | Gregory L. Ware, Alexander Mijalis, Jeremy P. Kamil, Jennifer L. Carroll, Maarten Van Diest, Rona S. Scott, Andrew D. Yurochko, Christopher G. Kevil, John A. Vanchiere |
| EPI_ISL_3086471 | LSUHS Emerging Viral Threat Laboratory | LSUHS Emerging Viral Threat Laboratory | Gregory L. Ware, Alexander Mijalis, Jeremy P. Kamil, Jennifer L. Carroll, Maarten Van Diest, Rona S. Scott, Andrew D. Yurochko, Christopher G. Kevil, John A. Vanchiere |
| EPI_ISL_3086472, EPI_ISL_3086473, EPI_ISL_3086474, EPI_ISL_3086475, EPI_ISL_3086476, EPI_ISL_3086477 | Willis-Knighton Medical Center Hospital Laboratory | LSUHS Emerging Viral Threat Laboratory | Gregory L. Ware, Alexander Mijalis, Jeremy P. Kamil, Jennifer L. Carroll, Maarten Van Diest, Rona S. Scott, Andrew D. Yurochko, Christopher G. Kevil, John A. Vanchiere |
| EPI_ISL_3086478 | Louisiana Office of Public Heath Laboratory | LSUHS Emerging Viral Threat Laboratory | Gregory L. Ware, Alexander Mijalis, Jeremy P. Kamil, Jennifer L. Carroll, Maarten Van Diest, Rona S. Scott, Andrew D. Yurochko, Christopher G. Kevil, John A. Vanchiere |
| EPI_ISL_3086479 | LSUHS Emerging Viral Threat Laboratory | LSUHS Emerging Viral Threat Laboratory | Gregory L. Ware, Alexander Mijalis, Jeremy P. Kamil, Jennifer L. Carroll, Maarten Van Diest, Rona S. Scott, Andrew D. Yurochko, Christopher G. Kevil, John A. Vanchiere |
| EPI_ISL_3086480, EPI_ISL_3086481, EPI_ISL_3086482, EPI_ISL_3086483, EPI_ISL_3086484, EPI_ISL_3086485, EPI_ISL_3086486 | Louisiana Office of Public Heath Laboratory | LSUHS Emerging Viral Threat Laboratory | Gregory L. Ware, Alexander Mijalis, Jeremy P. Kamil, Jennifer L. Carroll, Maarten Van Diest, Rona S. Scott, Andrew D. Yurochko, Christopher G. Kevil, John A. Vanchiere |
| EPI_ISL_3086487 | LSUHS Emerging Viral Threat Laboratory | LSUHS Emerging Viral Threat Laboratory | Gregory L. Ware, Alexander Mijalis, Jeremy P. Kamil, Jennifer L. Carroll, Maarten Van Diest, Rona S. Scott, Andrew D. Yurochko, Christopher G. Kevil, John A. Vanchiere |
| EPI_ISL_3086488, EPI_ISL_3086489 | Louisiana Office of Public Heath Laboratory | LSUHS Emerging Viral Threat Laboratory | Gregory L. Ware, Alexander Mijalis, Jeremy P. Kamil, Jennifer L. Carroll, Maarten Van Diest, Rona S. Scott, Andrew D. Yurochko, Christopher G. Kevil, John A. Vanchiere |

| | | | |
|---|---|---|---|
| EPI_ISL_3086490 | LSUHS Emerging Viral Threat Laboratory | LSUHS Emerging Viral Threat Laboratory | Gregory L. Ware, Alexander Mijalis, Jeremy P. Kamil, Jennifer L. Carroll, Maarten Van Diest, Rona S. Scott, Andrew D. Yurochko, Christopher G. Kevil, John A. Vanchiere |
| EPI_ISL_3086491, EPI_ISL_3086492, EPI_ISL_3086493, EPI_ISL_3086494, EPI_ISL_3086495 | Louisiana Office of Public Heath Laboratory | LSUHS Emerging Viral Threat Laboratory | Gregory L. Ware, Alexander Mijalis, Jeremy P. Kamil, Jennifer L. Carroll, Maarten Van Diest, Rona S. Scott, Andrew D. Yurochko, Christopher G. Kevil, John A. Vanchiere |
| EPI_ISL_3086496 | LSUHS Emerging Viral Threat Laboratory | LSUHS Emerging Viral Threat Laboratory | Gregory L. Ware, Alexander Mijalis, Jeremy P. Kamil, Jennifer L. Carroll, Maarten Van Diest, Rona S. Scott, Andrew D. Yurochko, Christopher G. Kevil, John A. Vanchiere |
| EPI_ISL_3086497, EPI_ISL_3086498, EPI_ISL_3086499, EPI_ISL_3086500, EPI_ISL_3086501, EPI_ISL_3086502 | Louisiana Office of Public Heath Laboratory | LSUHS Emerging Viral Threat Laboratory | Gregory L. Ware, Alexander Mijalis, Jeremy P. Kamil, Jennifer L. Carroll, Maarten Van Diest, Rona S. Scott, Andrew D. Yurochko, Christopher G. Kevil, John A. Vanchiere |
| EPI_ISL_3086503 | LSUHS Emerging Viral Threat Laboratory | LSUHS Emerging Viral Threat Laboratory | Gregory L. Ware, Alexander Mijalis, Jeremy P. Kamil, Jennifer L. Carroll, Maarten Van Diest, Rona S. Scott, Andrew D. Yurochko, Christopher G. Kevil, John A. Vanchiere |
| EPI_ISL_3086504 | Louisiana Office of Public Heath Laboratory | LSUHS Emerging Viral Threat Laboratory | Gregory L. Ware, Alexander Mijalis, Jeremy P. Kamil, Jennifer L. Carroll, Maarten Van Diest, Rona S. Scott, Andrew D. Yurochko, Christopher G. Kevil, John A. Vanchiere |
| EPI_ISL_3086505 | LSUHS Emerging Viral Threat Laboratory | LSUHS Emerging Viral Threat Laboratory | Gregory L. Ware, Alexander Mijalis, Jeremy P. Kamil, Jennifer L. Carroll, Maarten Van Diest, Rona S. Scott, Andrew D. Yurochko, Christopher G. Kevil, John A. Vanchiere |
| EPI_ISL_3086506, EPI_ISL_3086507, EPI_ISL_3086508, EPI_ISL_3086509, EPI_ISL_3086510, EPI_ISL_3086511, EPI_ISL_3086512, EPI_ISL_3086513, EPI_ISL_3086514, EPI_ISL_3086515, EPI_ISL_3086516, EPI_ISL_3086517, EPI_ISL_3086518, EPI_ISL_3086519, EPI_ISL_3086520, EPI_ISL_3086521, EPI_ISL_3086522, EPI_ISL_3086523, EPI_ISL_3086524, EPI_ISL_3086525 | | | |
| see above | Willis-Knighton Medical Center Hospital Laboratory | LSUHS Emerging Viral Threat Laboratory | Gregory L. Ware, Alexander Mijalis, Jeremy P. Kamil, Jennifer L. Carroll, Maarten Van Diest, Rona S. Scott, Andrew D. Yurochko, Christopher G. Kevil, John A. Vanchiere |
| EPI_ISL_3086526 | Louisiana Office of Public Heath Laboratory | LSUHS Emerging Viral Threat Laboratory | Gregory L. Ware, Alexander Mijalis, Jeremy P. Kamil, Jennifer L. Carroll, Maarten Van Diest, Rona S. Scott, Andrew D. Yurochko, Christopher G. Kevil, John A. Vanchiere |
| EPI_ISL_3086533, EPI_ISL_3086534, EPI_ISL_3086535, EPI_ISL_3086536, EPI_ISL_3086537, EPI_ISL_3086538, EPI_ISL_3086539, EPI_ISL_3086540, EPI_ISL_3086541, EPI_ISL_3086542, EPI_ISL_3086543, EPI_ISL_3086544, EPI_ISL_3086545, EPI_ISL_3086546, EPI_ISL_3086547, EPI_ISL_3086548, EPI_ISL_3086549, EPI_ISL_3086550, EPI_ISL_3086551, EPI_ISL_3086552, EPI_ISL_3086553, EPI_ISL_3086554, EPI_ISL_3086555, EPI_ISL_3086556, EPI_ISL_3086557, EPI_ISL_3086558, EPI_ISL_3086559, EPI_ISL_3086560, EPI_ISL_3086561, EPI_ISL_3086562, EPI_ISL_3086563, EPI_ISL_3086564, EPI_ISL_3086565, EPI_ISL_3086566, EPI_ISL_3086567, EPI_ISL_3086568, EPI_ISL_3086569, EPI_ISL_3086570, EPI_ISL_3086571, EPI_ISL_3086572, EPI_ISL_3086573, EPI_ISL_3086574, EPI_ISL_3086575, EPI_ISL_3086576, EPI_ISL_3086577, EPI_ISL_3086578, EPI_ISL_3086579, EPI_ISL_3086580, EPI_ISL_3086581, EPI_ISL_3086582, EPI_ISL_3086583, EPI_ISL_3086584, EPI_ISL_3086585, EPI_ISL_3086586, EPI_ISL_3086587, EPI_ISL_3086588, EPI_ISL_3086589, EPI_ISL_3086590, EPI_ISL_3086591, EPI_ISL_3086592, EPI_ISL_3086593, EPI_ISL_3086594, EPI_ISL_3086595, EPI_ISL_3086596, EPI_ISL_3086597, EPI_ISL_3086598, EPI_ISL_3086599, EPI_ISL_3086600, EPI_ISL_3086601, EPI_ISL_3086602, EPI_ISL_3086603, EPI_ISL_3086604, EPI_ISL_3086605, EPI_ISL_3086606, EPI_ISL_3086607, EPI_ISL_3086608, EPI_ISL_3086609, EPI_ISL_3086610, EPI_ISL_3086611, EPI_ISL_3086612, EPI_ISL_3086613, EPI_ISL_3086614, EPI_ISL_3086615, EPI_ISL_3086616, EPI_ISL_3086617, EPI_ISL_3086618, EPI_ISL_3086619, EPI_ISL_3086620, EPI_ISL_3086621, EPI_ISL_3086622, EPI_ISL_3086623, EPI_ISL_3086624, EPI_ISL_3086625, EPI_ISL_3086626, EPI_ISL_3086627, EPI_ISL_3086628, EPI_ISL_3086629, EPI_ISL_3086630, EPI_ISL_3086631, EPI_ISL_3086632, EPI_ISL_3086633, EPI_ISL_3086634, EPI_ISL_3086635, EPI_ISL_3086636, EPI_ISL_3086637, EPI_ISL_3086638, EPI_ISL_3086639, EPI_ISL_3086640, EPI_ISL_3086641, EPI_ISL_3086642, EPI_ISL_3086643, EPI_ISL_3086644, EPI_ISL_3086645, EPI_ISL_3086646, EPI_ISL_3086647, EPI_ISL_3086648, EPI_ISL_3086649, EPI_ISL_3086650, EPI_ISL_3086651, EPI_ISL_3086652, EPI_ISL_3086653, EPI_ISL_3086654, EPI_ISL_3086655, EPI_ISL_3086657, EPI_ISL_3086658, EPI_ISL_3086659, EPI_ISL_3086660, EPI_ISL_3086661, EPI_ISL_3086662, EPI_ISL_3086663, EPI_ISL_3086664, EPI_ISL_3086665, EPI_ISL_3086666, EPI_ISL_3086667, EPI_ISL_3086668, EPI_ISL_3086669, EPI_ISL_3086670, EPI_ISL_3086671, EPI_ISL_3086672, EPI_ISL_3086673, EPI_ISL_3086674, EPI_ISL_3086675, EPI_ISL_3086676, EPI_ISL_3086677, EPI_ISL_3086678, EPI_ISL_3086679, EPI_ISL_3086680, EPI_ISL_3086681, EPI_ISL_3086682, EPI_ISL_3086683, EPI_ISL_3086684, EPI_ISL_3086685, EPI_ISL_3086686, EPI_ISL_3086687, EPI_ISL_3086688, EPI_ISL_3086689, EPI_ISL_3086690 | | | |
| see above | N.F. Gamaleya Research Center for Epidemiology and Microbiology | WHO National Influenza Centre Russian Federation | Andrey Komissarov, Artem Fadeev, Kseniya Komissarova, Oula Mansour, Kirill Varchenko, Mikhail Bakaev, Tamila Musaeva, Veronika Eder, Maria Pisareva, Nikita Yolshin, Daria Danilenko, Elizaveta Divisenko, Alina Odintsova, Elena Shidlovskaya, Ludmila Vasilchenko, Nadezhda Kuznetsova, Maria Nikiforova, Alexey Shchetinin, Artem Tkachuk, Andrei Pochtovyi, Andrei Siniavin, Denis Kleymenov, Evgeniia Bykonia, Liubov Popova, Elena Mazunina, Inna Dolzhikova, Anna Kovyrshina, Andrei Botikov, Dmitry Shcheblyakov, Vladimir Gushchin, Denis Logunov, Alexander Gintsburg, Ksenia Safina, Elena Nabieva, Georgii Bazykin, Dmitry Lioznov |
| EPI_ISL_3086691 | Hospital Universitari Vall d'Hebron - Vall d'Hebron Institut de Recerca | Hospital Universitari Vall d'Hebron - Vall d'Hebron Institut de Recerca | Cristina Andrés, Maria Piñana, Alejandra González-Sánchez, Damir Garcia-Cehic, Ariadna Rando, Juliana Esperalba, Maria Gema Codina, Carla Castillo, Karen García, Rodrigo Vásquez, Maria Carmen Martin, Tomàs Pumarola, Josep Quer, Andrés Antón |
| EPI_ISL_3086692, EPI_ISL_3086693 | N.F. Gamaleya Research Center for Epidemiology and Microbiology | WHO National Influenza Centre Russian Federation | Andrey Komissarov, Artem Fadeev, Kseniya Komissarova, Oula Mansour, Kirill Varchenko, Mikhail Bakaev, Tamila Musaeva, Veronika Eder, Maria Pisareva, Nikita Yolshin, Daria Danilenko, Elizaveta Divisenko, Alina Odintsova, Elena Shidlovskaya, Ludmila Vasilchenko, Nadezhda Kuznetsova, Maria Nikiforova, Alexey Shchetinin, Artem Tkachuk, Andrei Pochtovyi, Andrei Siniavin, Denis Kleymenov, Evgeniia Bykonia, Liubov Popova, Elena Mazunina, Inna Dolzhikova, Anna Kovyrshina, Andrei Botikov, Dmitry Shcheblyakov, Vladimir Gushchin, Denis Logunov, Alexander Gintsburg, Ksenia Safina, Elena Nabieva, Georgii Bazykin, Dmitry Lioznov |
| EPI_ISL_3086694 | Hospital Universitari Vall d'Hebron - Vall d'Hebron Institut de Recerca | Hospital Universitari Vall d'Hebron - Vall d'Hebron Institut de Recerca | Cristina Andrés, Maria Piñana, Alejandra González-Sánchez, Damir Garcia-Cehic, Ariadna Rando, Juliana Esperalba, Maria Gema Codina, Carla Castillo, Karen García, Rodrigo Vásquez, Maria Carmen Martin, Tomàs Pumarola, Josep Quer, Andrés Antón |
| EPI_ISL_3086695 | N.F. Gamaleya Research Center for Epidemiology and Microbiology | WHO National Influenza Centre Russian Federation | Andrey Komissarov, Artem Fadeev, Kseniya Komissarova, Oula Mansour, Kirill Varchenko, Mikhail Bakaev, Tamila Musaeva, Veronika Eder, Maria Pisareva, Nikita Yolshin, Daria Danilenko, Elizaveta Divisenko, Alina Odintsova, Elena Shidlovskaya, Ludmila Vasilchenko, Nadezhda Kuznetsova, Maria Nikiforova, Alexey Shchetinin, Artem Tkachuk, Andrei Pochtovyi, Andrei Siniavin, Denis Kleymenov, Evgeniia Bykonia, Liubov Popova, Elena Mazunina, Inna Dolzhikova, Anna Kovyrshina, Andrei Botikov, Dmitry Shcheblyakov, Vladimir Gushchin, Denis Logunov, Alexander Gintsburg, Ksenia Safina, Elena Nabieva, Georgii Bazykin, Dmitry Lioznov |
| EPI_ISL_3086696 | Hospital Universitari Vall d'Hebron - Vall d'Hebron Institut de Recerca | Hospital Universitari Vall d'Hebron - Vall d'Hebron Institut de Recerca | Cristina Andrés, Maria Piñana, Alejandra González-Sánchez, Damir Garcia-Cehic, Ariadna Rando, Juliana Esperalba, Maria Gema Codina, Carla Castillo, Karen García, Rodrigo Vásquez, Maria Carmen Martin, Tomàs Pumarola, Josep Quer, Andrés Antón |
| EPI_ISL_3086697 | N.F. Gamaleya Research Center for Epidemiology and Microbiology | WHO National Influenza Centre Russian Federation | Andrey Komissarov, Artem Fadeev, Kseniya Komissarova, Oula Mansour, Kirill Varchenko, Mikhail Bakaev, Tamila Musaeva, Veronika Eder, Maria Pisareva, Nikita Yolshin, Daria Danilenko, Elizaveta Divisenko, Alina Odintsova, Elena Shidlovskaya, Ludmila Vasilchenko, Nadezhda Kuznetsova, Maria Nikiforova, Alexey Shchetinin, Artem Tkachuk, Andrei Pochtovyi, Andrei Siniavin, Denis Kleymenov, Evgeniia Bykonia, Liubov Popova, Elena Mazunina, Inna Dolzhikova, Anna Kovyrshina, Andrei Botikov, Dmitry Shcheblyakov, Vladimir Gushchin, Denis Logunov, Alexander Gintsburg, Ksenia Safina, Elena Nabieva, Georgii Bazykin, Dmitry Lioznov |
| EPI_ISL_3086698, EPI_ISL_3086699 | Hospital Universitari Vall d'Hebron - Vall d'Hebron Institut de Recerca | Hospital Universitari Vall d'Hebron - Vall d'Hebron Institut de Recerca | Cristina Andrés, Maria Piñana, Alejandra González-Sánchez, Damir Garcia-Cehic, Ariadna Rando, Juliana Esperalba, Maria Gema Codina, Carla Castillo, Karen García, Rodrigo Vásquez, Maria Carmen Martin, Tomàs Pumarola, Josep Quer, Andrés Antón |
| EPI_ISL_3086700 | N.F. Gamaleya Research Center for Epidemiology and Microbiology | WHO National Influenza Centre Russian Federation | Andrey Komissarov, Artem Fadeev, Kseniya Komissarova, Oula Mansour, Kirill Varchenko, Mikhail Bakaev, Tamila Musaeva, Veronika Eder, Maria Pisareva, Nikita Yolshin, Daria Danilenko, Elizaveta Divisenko, Alina Odintsova, Elena Shidlovskaya, Ludmila Vasilchenko, Nadezhda Kuznetsova, Maria Nikiforova, Alexey Shchetinin, Artem Tkachuk, Andrei Pochtovyi, Andrei Siniavin, Denis Kleymenov, Evgeniia Bykonia, Liubov Popova, Elena Mazunina, Inna Dolzhikova, Anna Kovyrshina, Andrei Botikov, Dmitry Shcheblyakov, Vladimir Gushchin, Denis Logunov, Alexander Gintsburg, Ksenia Safina, Elena Nabieva, Georgii Bazykin, Dmitry Lioznov |
| EPI_ISL_3086701 | Hospital Universitari Vall d'Hebron - Vall d'Hebron Institut de Recerca | Hospital Universitari Vall d'Hebron - Vall d'Hebron Institut de Recerca | Cristina Andrés, Maria Piñana, Alejandra González-Sánchez, Damir Garcia-Cehic, Ariadna Rando, Juliana Esperalba, Maria Gema Codina, Carla Castillo, Karen García, Rodrigo Vásquez, Maria Carmen Martin, Tomàs Pumarola, Josep Quer, Andrés Antón |
| EPI_ISL_3086702 | N.F. Gamaleya Research Center for Epidemiology and Microbiology | WHO National Influenza Centre Russian Federation | Andrey Komissarov, Artem Fadeev, Kseniya Komissarova, Oula Mansour, Kirill Varchenko, Mikhail Bakaev, Tamila Musaeva, Veronika Eder, Maria Pisareva, Nikita Yolshin, Daria Danilenko, Elizaveta Divisenko, Alina Odintsova, Elena Shidlovskaya, Ludmila Vasilchenko, Nadezhda Kuznetsova, Maria Nikiforova, Alexey Shchetinin, Artem Tkachuk, Andrei Pochtovyi, Andrei Siniavin, Denis Kleymenov, Evgeniia Bykonia, Liubov Popova, Elena Mazunina, Inna Dolzhikova, Anna Kovyrshina, Andrei Botikov, Dmitry Shcheblyakov, Vladimir Gushchin, Denis Logunov, Alexander Gintsburg, Ksenia Safina, Elena Nabieva, Georgii Bazykin, Dmitry Lioznov |
| EPI_ISL_3086703 | Hospital Universitari Vall d'Hebron - Vall d'Hebron Institut de Recerca | Hospital Universitari Vall d'Hebron - Vall d'Hebron Institut de Recerca | Cristina Andrés, Maria Piñana, Alejandra González-Sánchez, Damir Garcia-Cehic, Ariadna Rando, Juliana Esperalba, Maria Gema Codina, Carla Castillo, Karen García, Rodrigo Vásquez, Maria Carmen Martin, Tomàs Pumarola, Josep Quer, Andrés Antón |
| EPI_ISL_3086704, EPI_ISL_3086706 | N.F. Gamaleya Research Center for Epidemiology and Microbiology | WHO National Influenza Centre Russian Federation | Andrey Komissarov, Artem Fadeev, Kseniya Komissarova, Oula Mansour, Kirill Varchenko, Mikhail Bakaev, Tamila Musaeva, Veronika Eder, Maria Pisareva, Nikita Yolshin, Daria Danilenko, Elizaveta Divisenko, Alina Odintsova, Elena Shidlovskaya, Ludmila Vasilchenko, Nadezhda Kuznetsova, Maria Nikiforova, Alexey Shchetinin, Artem Tkachuk, Andrei Pochtovyi, Andrei Siniavin, Denis Kleymenov, Evgeniia Bykonia, Liubov Popova, Elena Mazunina, Inna Dolzhikova, Anna Kovyrshina, Andrei Botikov, Dmitry Shcheblyakov, Vladimir Gushchin, Denis Logunov, Alexander Gintsburg, Ksenia Safina, Elena Nabieva, Georgii Bazykin, Dmitry Lioznov |
| EPI_ISL_3086707, EPI_ISL_3086708 | Hospital Universitari Vall d'Hebron - Vall d'Hebron Institut de Recerca | Hospital Universitari Vall d'Hebron - Vall d'Hebron Institut de Recerca | Cristina Andrés, Maria Piñana, Alejandra González-Sánchez, Damir Garcia-Cehic, Ariadna Rando, Juliana Esperalba, Maria Gema Codina, Carla Castillo, Karen García, Rodrigo Vásquez, Maria Carmen Martin, Tomàs Pumarola, Josep Quer, Andrés Antón |
| EPI_ISL_3086709 | N.F. Gamaleya Research Center for Epidemiology and Microbiology | WHO National Influenza Centre Russian Federation | Andrey Komissarov, Artem Fadeev, Kseniya Komissarova, Oula Mansour, Kirill Varchenko, Mikhail Bakaev, Tamila Musaeva, Veronika Eder, Maria Pisareva, Nikita Yolshin, Daria Danilenko, Elizaveta Divisenko, Alina Odintsova, Elena Shidlovskaya, Ludmila Vasilchenko, Nadezhda Kuznetsova, Maria Nikiforova, Alexey Shchetinin, Artem Tkachuk, Andrei Pochtovyi, Andrei Siniavin, Denis Kleymenov, Evgeniia Bykonia, Liubov Popova, Elena Mazunina, Inna Dolzhikova, Anna Kovyrshina, Andrei Botikov, Dmitry Shcheblyakov, Vladimir Gushchin, Denis Logunov, Alexander Gintsburg, Ksenia Safina, Elena Nabieva, Georgii Bazykin, Dmitry Lioznov |
| EPI_ISL_3086710 | Hospital Universitari Vall d'Hebron - Vall d'Hebron Institut de Recerca | Hospital Universitari Vall d'Hebron - Vall d'Hebron Institut de Recerca | Cristina Andrés, Maria Piñana, Alejandra González-Sánchez, Damir Garcia-Cehic, Ariadna Rando, Juliana Esperalba, Maria Gema Codina, Carla Castillo, Karen García, Rodrigo Vásquez, Maria Carmen Martin, Tomàs Pumarola, Josep Quer, Andrés Antón |
| EPI_ISL_3086711 | N.F. Gamaleya Research Center for Epidemiology and Microbiology | WHO National Influenza Centre Russian Federation | Andrey Komissarov, Artem Fadeev, Kseniya Komissarova, Oula Mansour, Kirill Varchenko, Mikhail Bakaev, Tamila Musaeva, Veronika Eder, Maria Pisareva, Nikita Yolshin, Daria Danilenko, Elizaveta Divisenko, Alina Odintsova, Elena Shidlovskaya, Ludmila Vasilchenko, Nadezhda Kuznetsova, Maria Nikiforova, Alexey Shchetinin, Artem Tkachuk, Andrei Pochtovyi, Andrei Siniavin, Denis Kleymenov, Evgeniia Bykonia, Liubov Popova, Elena Mazunina, Inna Dolzhikova, Anna Kovyrshina, Andrei Botikov, Dmitry Shcheblyakov, Vladimir Gushchin, Denis Logunov, Alexander Gintsburg, Ksenia Safina, Elena Nabieva, Georgii Bazykin, Dmitry Lioznov |
| EPI_ISL_3086712, EPI_ISL_3086713, EPI_ISL_3086714, EPI_ISL_3086715, EPI_ISL_3086716, EPI_ISL_3086717, EPI_ISL_3086718, EPI_ISL_3086719, EPI_ISL_3086720, EPI_ISL_3086721, EPI_ISL_3086722, EPI_ISL_3086723, EPI_ISL_3086724, EPI_ISL_3086725, EPI_ISL_3086726, EPI_ISL_3086727, EPI_ISL_3086728, EPI_ISL_3086729, EPI_ISL_3086730, EPI_ISL_3086731, EPI_ISL_3086732, EPI_ISL_3086733, EPI_ISL_3086734, EPI_ISL_3086735, EPI_ISL_3086736, EPI_ISL_3086737, EPI_ISL_3086738, EPI_ISL_3086739, EPI_ISL_3086740, EPI_ISL_3086741, EPI_ISL_3086742, EPI_ISL_3086743, EPI_ISL_3086744, EPI_ISL_3086745, EPI_ISL_3086746, EPI_ISL_3086747, EPI_ISL_3086748, EPI_ISL_3086749, EPI_ISL_3086750, EPI_ISL_3086751, EPI_ISL_3086752, EPI_ISL_3086753, EPI_ISL_3086754 | | | |
| see above | Hospital Universitari Vall d'Hebron - Vall d'Hebron Institut de Recerca | Hospital Universitari Vall d'Hebron - Vall d'Hebron Institut de Recerca | Cristina Andrés, Maria Piñana, Alejandra González-Sánchez, Damir Garcia-Cehic, Ariadna Rando, Juliana Esperalba, Maria Gema Codina, Carla Castillo, Karen García, Rodrigo Vásquez, Maria Carmen Martin, Tomàs Pumarola, Josep Quer, Andrés Antón |
| EPI_ISL_3086755, EPI_ISL_3086756, EPI_ISL_3086757, EPI_ISL_3086758, EPI_ISL_3086759, EPI_ISL_3086760, EPI_ISL_3086761, EPI_ISL_3086762, EPI_ISL_3086763, EPI_ISL_3086764, EPI_ISL_3086765, EPI_ISL_3086766, EPI_ISL_3086770, EPI_ISL_3086771, EPI_ISL_3086772, EPI_ISL_3086773, EPI_ISL_3086774, EPI_ISL_3086775, EPI_ISL_3086776, EPI_ISL_3086777, EPI_ISL_3086778, EPI_ISL_3086779, EPI_ISL_3086780, EPI_ISL_3086781 | | | |
| see above | Hospital Universitari Arnau de Vilanova | Hospital Universitari Vall d'Hebron - Vall d'Hebron Institut de Recerca | Cristina Andrés, Maria Piñana, Alejandra González-Sánchez, Damir Garcia-Cehic, Ariadna Rando, Juliana Esperalba, Maria Gema Codina, Carla Castillo, Karen García, Rodrigo Vásquez, Maria Carmen Martin, Tomàs Pumarola, Josep Quer, Andrés Antón |
| EPI_ISL_3086782 | Neuberg Supratech Reference Laboratory, Surat | Gujarat Biotechnology Research Centre | Ramesh Pandit,Janvi Raval,Zarna Patel,Nitin Savaliya,Dinesh Kumar,Twinkle Soni,Sonal Sharma ,Zuber Saiyed,Manthan Shah ,Umang Mishra,Nitesh Shah,Chaitanya Joshi,Madhvi Joshi |
| EPI_ISL_3086783 | Neuberg Supratech Reference Laboratory, Surat | Gujarat Biotechnology Research Centre | Janvi Raval,Zarna Patel,Nitin Savaliya,Dinesh Kumar,Twinkle Soni,Sonal Sharma ,Zuber Saiyed,Ramesh Pandit,Manthan Shah ,Umang Mishra,Nitesh Shah,Chaitanya Joshi,Madhvi Joshi |
| EPI_ISL_3086784 | BJ Medical College, Ahmedabad | Gujarat Biotechnology Research Centre | Zarna Patel,Nitin Savaliya,Dinesh Kumar,Twinkle Soni,Sonal Sharma ,Zuber Saiyed,Ramesh Pandit,Janvi Raval,Pranay Shah,Umang Mishra,Nitesh Shah,Chaitanya Joshi,Madhvi Joshi |
| EPI_ISL_3086785 | BJ Medical College, Ahmedabad | Gujarat Biotechnology Research Centre | Nitin Savaliya,Dinesh Kumar,Twinkle Soni,Sonal Sharma ,Zuber Saiyed,Ramesh Pandit,Janvi Raval,Zarna Patel,Pranay Shah,Umang Mishra,Nitesh Shah,Chaitanya Joshi,Madhvi Joshi |
| EPI_ISL_3086786 | College of Veterinary Science and A.H Dantiwada | Gujarat Biotechnology Research Centre | Dinesh Kumar,Twinkle Soni,Sonal Sharma ,Zuber Saiyed,Ramesh Pandit,Janvi Raval,Zarna Patel,Nitin Savaliya,Hemlatta Sunsara,Umang Mishra,Nitesh Shah,Chaitanya Joshi,Madhvi Joshi |
| EPI_ISL_3086787 | College of Veterinary Science and A.H Dantiwada | Gujarat Biotechnology Research Centre | Twinkle Soni,Sonal Sharma ,Zuber Saiyed,Ramesh Pandit,Janvi Raval,Zarna Patel,Nitin Savaliya,Dinesh Kumar,Hemlatta Sunsara,Umang Mishra,Nitesh Shah,Chaitanya Joshi,Madhvi Joshi |
| EPI_ISL_3086788 | Neuberg Supratech Reference Laboratory, Surat | Gujarat Biotechnology Research Centre | Sonal Sharma ,Zuber Saiyed,Ramesh Pandit,Janvi Raval,Zarna Patel,Nitin Savaliya,Dinesh Kumar,Twinkle Soni,Manthan Shah ,Umang Mishra,Nitesh Shah,Chaitanya Joshi,Madhvi Joshi |
| EPI_ISL_3086789 | Supratech Micropath Laboratory Research Institute Pvt Ltd, Ahmedabad | Gujarat Biotechnology Research Centre | Zuber Saiyed,Ramesh Pandit,Janvi Raval,Zarna Patel,Nitin Savaliya,Dinesh Kumar,Twinkle Soni,Sonal Sharma ,Mahendra Parikh,Umang Mishra,Nitesh Shah,Chaitanya Joshi,Madhvi Joshi |
| EPI_ISL_3086790 | GMERS Medical College, Valsad | Gujarat Biotechnology Research Centre | Ramesh Pandit,Janvi Raval,Zarna Patel,Nitin Savaliya,Dinesh Kumar,Twinkle Soni,Sonal Sharma ,Zuber Saiyed,Vicky Gandhi,Umang Mishra,Nitesh Shah,Chaitanya Joshi,Madhvi Joshi |
| EPI_ISL_3086791 | GMERS Medical College, Valsad | Gujarat Biotechnology Research Centre | Janvi Raval,Zarna Patel,Nitin Savaliya,Dinesh Kumar,Twinkle Soni,Sonal Sharma ,Zuber Saiyed,Ramesh Pandit,Vicky Gandhi,Umang Mishra,Nitesh Shah,Chaitanya Joshi,Madhvi Joshi |
| EPI_ISL_3086792 | GMERS Medical College, Valsad | Gujarat Biotechnology Research Centre | Zarna Patel,Nitin Savaliya,Dinesh Kumar,Twinkle Soni,Sonal Sharma ,Zuber Saiyed,Ramesh Pandit,Janvi Raval,Vicky Gandhi,Umang Mishra,Nitesh Shah,Chaitanya Joshi,Madhvi Joshi |
| EPI_ISL_3086793 | GMERS Medical College, Valsad | Gujarat Biotechnology Research Centre | Nitin Savaliya,Dinesh Kumar,Twinkle Soni,Sonal Sharma ,Zuber Saiyed,Ramesh Pandit,Janvi Raval,Zarna Patel,Vicky Gandhi,Umang Mishra,Nitesh Shah,Chaitanya Joshi,Madhvi Joshi |

| | | | |
|---|---|---|---|
| EPI_ISL_3086794 | Banas Medical College & Research Institute (BMCRI), Palanpur | Gujarat Biotechnology Research Centre | Dinesh Kumar,Twinkle Soni,Sonal Sharma ,Zuber Saiyed,Ramesh Pandit,Janvi Raval,Zarna Patel,Nitin Savaliya,Anil Choudhary ,Amar Sajjan,Umang Mishra,Nitesh Shah,Chaitanya Joshi,Madhvi Joshi |
| EPI_ISL_3086795 | Banas Medical College & Research Institute (BMCRI), Palanpur | Gujarat Biotechnology Research Centre | Twinkle Soni,Sonal Sharma ,Zuber Saiyed,Ramesh Pandit,Janvi Raval,Zarna Patel,Nitin Savaliya,Dinesh Kumar,Anil Choudhary ,Amar Sajjan,Umang Mishra,Nitesh Shah,Chaitanya Joshi,Madhvi Joshi |
| EPI_ISL_3086796 | Banas Medical College & Research Institute (BMCRI), Palanpur | Gujarat Biotechnology Research Centre | Sonal Sharma ,Zuber Saiyed,Ramesh Pandit,Janvi Raval,Zarna Patel,Nitin Savaliya,Dinesh Kumar,Twinkle Soni,Anil Choudhary ,Amar Sajjan,Umang Mishra,Nitesh Shah,Chaitanya Joshi,Madhvi Joshi |
| EPI_ISL_3086797 | Banas Medical College & Research Institute (BMCRI), Palanpur | Gujarat Biotechnology Research Centre | Zuber Saiyed,Ramesh Pandit,Janvi Raval,Zarna Patel,Nitin Savaliya,Dinesh Kumar,Twinkle Soni,Sonal Sharma ,Anil Choudhary ,Amar Sajjan,Umang Mishra,Nitesh Shah,Chaitanya Joshi,Madhvi Joshi |
| EPI_ISL_3086798 | College of Veterinary Science and A.H Dantiwada | Gujarat Biotechnology Research Centre | Ramesh Pandit,Janvi Raval,Zarna Patel,Nitin Savaliya,Dinesh Kumar,Twinkle Soni,Sonal Sharma ,Zuber Saiyed,Hemlatta Sunsara,Umang Mishra,Nitesh Shah,Chaitanya Joshi,Madhvi Joshi |
| EPI_ISL_3086799 | College of Veterinary Science and A.H Dantiwada | Gujarat Biotechnology Research Centre | Janvi Raval,Zarna Patel,Nitin Savaliya,Dinesh Kumar,Twinkle Soni,Sonal Sharma ,Zuber Saiyed,Ramesh Pandit,Hemlatta Sunsara,Umang Mishra,Nitesh Shah,Chaitanya Joshi,Madhvi Joshi |
| EPI_ISL_3086800 | College of Veterinary Science and A.H Dantiwada | Gujarat Biotechnology Research Centre | Zarna Patel,Nitin Savaliya,Dinesh Kumar,Twinkle Soni,Sonal Sharma ,Zuber Saiyed,Ramesh Pandit,Janvi Raval,Hemlatta Sunsara,Umang Mishra,Nitesh Shah,Chaitanya Joshi,Madhvi Joshi |
| EPI_ISL_3086801 | College of Veterinary Science and A.H Dantiwada | Gujarat Biotechnology Research Centre | Nitin Savaliya,Dinesh Kumar,Twinkle Soni,Sonal Sharma ,Zuber Saiyed,Ramesh Pandit,Janvi Raval,Zarna Patel,Hemlatta Sunsara,Umang Mishra,Nitesh Shah,Chaitanya Joshi,Madhvi Joshi |
| EPI_ISL_3086802 | College of Veterinary Science and A.H Dantiwada | Gujarat Biotechnology Research Centre | Dinesh Kumar,Twinkle Soni,Sonal Sharma ,Zuber Saiyed,Ramesh Pandit,Janvi Raval,Zarna Patel,Nitin Savaliya,Hemlatta Sunsara,Umang Mishra,Nitesh Shah,Chaitanya Joshi,Madhvi Joshi |
| EPI_ISL_3086851, EPI_ISL_3086852, EPI_ISL_3086853, EPI_ISL_3086854, EPI_ISL_3086855, EPI_ISL_3086856, EPI_ISL_3086857, EPI_ISL_3086858, EPI_ISL_3086859, EPI_ISL_3086860, EPI_ISL_3086861, EPI_ISL_3086862, EPI_ISL_3086863, EPI_ISL_3086864 | | | |
| see above | Plateforme de testing Namuroise | Plateforme de testing Namuroise | Lesly Nyinkeu Kemamen; Demars Aurore ; Maschietto Céline ; Nobis Chloé ; Otto Gaetan ; Denis Olivier ; Degosserie Jonathan ; Mullier François |
| EPI_ISL_3086865 | UPTD Puskesmas Limo | Indonesian Institute of Sciences (LIPI) and Laboratorium Kesehatan Daerah (Labkesda) Kota Depok | Masrukhin, Syam B. Iryanto, Idris, Syaiful Rizal, Erwin F. Hasrianda, Yuliawati, Listiana Oktavia, Ruby Setiawan, Herjuno A. Nugroho, Elin Herliana, Kania R. Astuti, Kurniyawati, Harto W. Tamara, Sugeng Suhadi, Anggia Prasetyoputri, Ratih A. Ningrum |
| EPI_ISL_3086866 | UPTD Puskesmas Mampang | Indonesian Institute of Sciences (LIPI) and Laboratorium Kesehatan Daerah (Labkesda) Kota Depok | Masrukhin, Syam B. Iryanto, Idris, Syaiful Rizal, Erwin F. Hasrianda, Yuliawati, Listiana Oktavia, Ruby Setiawan, Herjuno A. Nugroho, Elin Herliana, Kania R. Astuti, Kurniyawati, Harto W. Tamara, Sugeng Suhadi, Anggia Prasetyoputri, Ratih A. Ningrum |
| EPI_ISL_3086867 | UPTD Puskesmas Limo | Indonesian Institute of Sciences (LIPI) and Laboratorium Kesehatan Daerah (Labkesda) Kota Depok | Masrukhin, Syam B. Iryanto, Idris, Syaiful Rizal, Erwin F. Hasrianda, Yuliawati, Listiana Oktavia, Ruby Setiawan, Herjuno A. Nugroho, Elin Herliana, Kania R. Astuti, Kurniyawati, Harto W. Tamara, Sugeng Suhadi, Anggia Prasetyoputri, Ratih A. Ningrum |
| EPI_ISL_3086868 | UPTD Puskesmas Pondok Sukmajaya | Indonesian Institute of Sciences (LIPI) and Laboratorium Kesehatan Daerah (Labkesda) Kota Depok | Yuliawati, Listiana Oktavia, Herjuno A. Nugroho, Ruby Setiawan, Masrukhin, Syam B. Iryanto, Idris, Syaiful Rizal, Erwin F. Hasrianda, Elin Herliana, Kania R. Astuti, Kurniyawati, Harto W. Tamara, Sugeng Suhadi, Andri Wardiana, Ratih A. Ningrum |
| EPI_ISL_3086869 | UPTD Puskesmas Cimanggis | Indonesian Institute of Sciences (LIPI) and Laboratorium Kesehatan Daerah (Labkesda) Kota Depok | Yuliawati, Listiana Oktavia, Herjuno A. Nugroho, Ruby Setiawan, Masrukhin, Syam B. Iryanto, Idris, Syaiful Rizal, Erwin F. Hasrianda, Elin Herliana, Kania R. Astuti, Kurniyawati, Harto W. Tamara, Sugeng Suhadi, Andri Wardiana, Ratih A. Ningrum |
| EPI_ISL_3086870 | UPTD Puskesmas Sawangan | Indonesian Institute of Sciences (LIPI) and Laboratorium Kesehatan Daerah (Labkesda) Kota Depok | Yuliawati, Listiana Oktavia, Herjuno A. Nugroho, Ruby Setiawan, Masrukhin, Syam B. Iryanto, Idris, Syaiful Rizal, Erwin F. Hasrianda, Elin Herliana, Kania R. Astuti, Kurniyawati, Harto W. Tamara, Sugeng Suhadi, Andri Wardiana, Ratih A. Ningrum |
| EPI_ISL_3086871 | UPTD Puskesmas Mampang | Indonesian Institute of Sciences (LIPI) and Laboratorium Kesehatan Daerah (Labkesda) Kota Depok | Listiana Oktavia, Herjuno A. Nugroho, Ruby Setiawan, Yuliawati, Masrukhin, Syam B. Iryanto, Idris, Syaiful Rizal, Erwin F. Hasrianda, Elin Herliana, Kania R. Astuti, Kurniyawati, Harto W. Tamara, Sugeng Suhadi, Sugiyono Saputra, Ratih A. Ningrum |
| EPI_ISL_3086873 | UPTD Puskesmas Limo | Indonesian Institute of Sciences (LIPI) and Laboratorium Kesehatan Daerah (Labkesda) Kota Depok | Ruby Setiawan, Listiana Oktavia, Yuliawati, Masrukhin, Syam B. Iryanto, Herjuno A. Nugroho, Idris, Syaiful Rizal, Erwin F. Hasrianda, Elin Herliana, Kania R. Astuti, Kurniyawati, Harto W. Tamara, Sugeng Suhadi, Ratih A. Ningrum, Puspita Lisdiyanti |
| EPI_ISL_3086874 | UPTD Puskesmas Mampang | Indonesian Institute of Sciences (LIPI) and Laboratorium Kesehatan Daerah (Labkesda) Kota Depok | Ruby Setiawan, Listiana Oktavia, Yuliawati, Masrukhin, Syam B. Iryanto, Herjuno A. Nugroho, Idris, Syaiful Rizal, Erwin F. Hasrianda, Elin Herliana, Kania R. Astuti, Kurniyawati, Harto W. Tamara, Sugeng Suhadi, Ratih A. Ningrum, Puspita Lisdiyanti |
| EPI_ISL_3086875 | UPTD Puskesmas Tanah Baru | Indonesian Institute of Sciences (LIPI) and Laboratorium Kesehatan Daerah (Labkesda) Kota Depok | Herjuno A. Nugroho,Ruby Setiawan, Listiana Oktavia, Yuliawati, Masrukhin, Syam B. Iryanto, Idris, Syaiful Rizal, Erwin F. Hasrianda, Elin Herliana, Kania R. Astuti, Kurniyawati, Harto W. Tamara, Sugeng Suhadi, Ratih A. Ningrum, Sugiyono Saputra |
| EPI_ISL_3086876, EPI_ISL_3086877 | UPTD Puskesmas Cimanggis | Indonesian Institute of Sciences (LIPI) and Laboratorium Kesehatan Daerah (Labkesda) Kota Depok | Herjuno A. Nugroho,Ruby Setiawan, Listiana Oktavia, Yuliawati, Masrukhin, Syam B. Iryanto, Idris, Syaiful Rizal, Erwin F. Hasrianda, Elin Herliana, Kania R. Astuti, Kurniyawati, Harto W. Tamara, Sugeng Suhadi, Ratih A. Ningrum, Sugiyono Saputra |
| EPI_ISL_3086878 | UPTD Labkesda Depok | Indonesian Institute of Sciences (LIPI) and Laboratorium Kesehatan Daerah (Labkesda) Kota Depok | Syam B. Iryanto, Herjuno A. Nugroho,Ruby Setiawan, Listiana Oktavia, Yuliawati, Masrukhin, Idris, Syaiful Rizal, Erwin F. Hasrianda, Elin Herliana, Kania R. Astuti, Kurniyawati, Harto W. Tamara, Sugeng Suhadi, Ratih A. Ningrum, Anik B. Dharmayanthi |
| EPI_ISL_3086879 | UPTD Puskesmas Limo | Indonesian Institute of Sciences (LIPI) and Laboratorium Kesehatan Daerah (Labkesda) Kota Depok | Syam B. Iryanto, Herjuno A. Nugroho,Ruby Setiawan, Listiana Oktavia, Yuliawati, Masrukhin, Idris, Syaiful Rizal, Erwin F. Hasrianda, Elin Herliana, Kania R. Astuti, Kurniyawati, Harto W. Tamara, Sugeng Suhadi, Ratih A. Ningrum, Anik B. Dharmayanthi |
| EPI_ISL_3086880 | RS Citra Medika | Indonesian Institute of Sciences (LIPI) | Indriawati, Isa Nuryana, Syaiful Rizal, Yuhoniz A. Maryam, Ocha P. Meitha, Willery Yeo, Annisa D. Septiani, Lusiana, Maya Komalasari, Michael Saut Parulian, Asep M. Ridwanulloh, Sugiyono Saputra |
| EPI_ISL_3086881, EPI_ISL_3086882 | RS Umum Trimitra | Indonesian Institute of Sciences (LIPI) | Isa Nuryana, Syaiful Rizal, Indriawati, Yuhoniz A. Maryam, Ocha P. Meitha, Willery Yeo, Annisa D. Septiani, Lusiana, Maya Komalasari, Michael Saut Parulian, Syam B. Iryanto, Anggia Prasetyoputri |
| EPI_ISL_3086883 | RS Umum Trimitra | Indonesian Institute of Sciences (LIPI) | Syaiful Rizal, Indriawati, Isa Nuryana, Yuhoniz A. Maryam, Ocha P. Meitha, Willery Yeo, Annisa D. Septiani, Lusiana, Maya Komalasari, Michael Saut Parulian, Anik B. Dharmayanthi, Sugiyono Saputra |
| EPI_ISL_3086884 | RS Umum Trimitra | Indonesian Institute of Sciences (LIPI) | Syaiful Rizal, Indriawati, Isa Nuryana, Yuhoniz A. Maryam, Ocha P. Meitha, Willery Yeo, Annisa D. Septiani, Lusiana, Maya Komalasari, Michael Saut Parulian, Anik B. Dharmayanthi, Andri Wardiana |
| EPI_ISL_3086885 | RS Umum Trimitra | Indonesian Institute of Sciences (LIPI) | Syam B. Iryanto, Syaiful Rizal, Indriawati, Isa Nuryana, Yuhoniz A. Maryam, Ocha P. Meitha, Willery Yeo, Annisa D. Septiani, Lusiana, Maya Komalasari, Michael Saut Parulian, Anik B. Dharmayanthi, Ratih A. Ningrum |
| EPI_ISL_3086886 | Laboratorium Riset Unsoed | Indonesian Institute of Sciences (LIPI) and Laboratorium Riset Universitas Jenderal Soedirman | Syaiful Rizal, Idris, Masrukhin, Erwin F. Hasrianda, Herjuno A. Nugroho, Yuliawati, Listiana Oktavia, Ruby Setiawan, Endang S. Kusmintarsih, M. Z. Syamsul Hidayat, M. M. Rudi Prihatno, Dody Novrial, Syam B. Iryanto, Ratih A. Ningrum, Sugiyono Saputra |
| EPI_ISL_3086887 | Laboratorium Riset Unsoed | Indonesian Institute of Sciences (LIPI) and Laboratorium Riset Universitas Jenderal Soedirman | Idris, Herjuno A. Nugroho, Syaiful Rizal, Masrukhin, Erwin F. Hasrianda, Yuliawati, Listiana Oktavia, Ruby Setiawan, Endang S. Kusmintarsih, M. Z. Syamsul Hidayat, M. M. Rudi Prihatno, Dody Novrial, Syam B. Iryanto, Anggia Prasetyoputri, Sugiyono Saputra |
| EPI_ISL_3086888 | Laboratorium Riset Unsoed | Indonesian Institute of Sciences (LIPI) and Laboratorium Riset Universitas Jenderal Soedirman | Isa Nuryana, Syaiful Rizal, Indriawati, Idris, Endang S. Kusmintarsih, Wahyu Siswandari, Norman A. Prayogi, Dhadhang W. Kurniawan, Suprayogi, Sugiyono Saputra, Syam B. Iryanto |
| EPI_ISL_3086889 | Laboratorium Riset Unsoed | Indonesian Institute of Sciences (LIPI) and Laboratorium Riset Universitas Jenderal Soedirman | Isa Nuryana, Syaiful Rizal, Indriawati, Idris, Endang S. Kusmintarsih, Wahyu Siswandari, Norman A. Prayogi, Dhadhang W. Kurniawan, Suprayogi, Anik B. Dharmayanthi, Sugiyono Saputra |
| EPI_ISL_3086890 | Laboratorium Riset Unsoed | Indonesian Institute of Sciences (LIPI) and Laboratorium Riset Universitas Jenderal Soedirman | Syaiful Rizal, Isa Nuryana, Indriawati, Idris, Endang S. Kusmintarsih, Wahyu Siswandari, Norman A. Prayogi, Dhadhang W. Kurniawan, Suprayogi, Sugiyono Saputra, Syam B. Iryanto, Anggia Prasetyoputri |
| EPI_ISL_3086891, EPI_ISL_3086892 | Laboratorium Riset Unsoed | Indonesian Institute of Sciences (LIPI) and Laboratorium Riset Universitas Jenderal Soedirman | Indriawati, Isa Nuryana, Syaiful Rizal, Idris, Endang S. Kusmintarsih, Wahyu Siswandari, Norman A. Prayogi, Dhadhang W. Kurniawan, Suprayogi, Sugiyono Saputra, Ratih A. Ningrum |
| EPI_ISL_3086893 | Laboratorium Riset Unsoed | Indonesian Institute of Sciences (LIPI) and Laboratorium Riset Universitas Jenderal Soedirman | Syam B. Iryanto, Syaiful Rizal, Indriawati, Isa Nuryana, Idris, Endang S. Kusmintarsih, M. Z. Syamsul Hidayat, M. M. Rudi Prihatno, Dody Novrial, Puspita Lisdiyanti, Sugiyono Saputra |
| EPI_ISL_3086897, EPI_ISL_3086898, EPI_ISL_3086899, EPI_ISL_3086901, EPI_ISL_3086902, EPI_ISL_3086903, EPI_ISL_3086904, EPI_ISL_3086905, EPI_ISL_3086906, EPI_ISL_3086907, EPI_ISL_3086908, EPI_ISL_3086909, EPI_ISL_3086910, EPI_ISL_3086911, EPI_ISL_3086912, EPI_ISL_3086913, EPI_ISL_3086914, EPI_ISL_3086916, EPI_ISL_3086919, EPI_ISL_3086920, EPI_ISL_3086921, EPI_ISL_3086922, EPI_ISL_3086923, EPI_ISL_3086924, EPI_ISL_3086925, EPI_ISL_3086926, EPI_ISL_3086927, EPI_ISL_3086928, EPI_ISL_3086929, EPI_ISL_3086930, EPI_ISL_3086932, EPI_ISL_3086933, EPI_ISL_3086934, EPI_ISL_3086935, EPI_ISL_3086936, EPI_ISL_3086937, EPI_ISL_3086938, EPI_ISL_3086939, EPI_ISL_3086940, EPI_ISL_3086941, EPI_ISL_3086942, EPI_ISL_3086943, EPI_ISL_3086944, EPI_ISL_3086945, EPI_ISL_3086946, EPI_ISL_3086947 | | | |
| see above | Viral Respiratory Lab, National Institute for Biomedical Research (INRB) | Pathogen Sequencing Lab, National Institute for Biomedical Research (INRB) | Placide Mbala-Kingebeni, Edith Nkwembe, Eddy Kinganda-Lusamaki, Amuri Aziza, Francisca Muyembe Mawete, Emmanuel Lokilo Lofiko, Jean Claude Makangara, Raphaël Lumembe, Gabriel Kabamba, Catherine Pratt, Matthias Pauthner, Josh Quick, Allison Black, James Hadfield, Trevor Bedford, Ian Goodfellow, Andrew Rambaut, Nick Loman, Kristian Andersen, Michael Wiley, Steve Ahuka-Mundeke, Jean-Jacques Muyembe Tamfum |
| EPI_ISL_3086948, EPI_ISL_3086949, EPI_ISL_3086950, EPI_ISL_3086951, EPI_ISL_3086952, EPI_ISL_3086953, EPI_ISL_3086954, EPI_ISL_3086955 | Hospital Universitari Joan XXIII | Hospital Universitari Joan XXIII | Gemma Recio, Cristina Gutiérrez, Clara Benavent, Carla Martín, Margarida Terrón, Ester Picó, Natalia Bastón |
| EPI_ISL_3086956, EPI_ISL_3086959, EPI_ISL_3086960, EPI_ISL_3086962, EPI_ISL_3086963, EPI_ISL_3086964, EPI_ISL_3086968, EPI_ISL_3086969, EPI_ISL_3086970, EPI_ISL_3086971, EPI_ISL_3086972, EPI_ISL_3086973, EPI_ISL_3086975, EPI_ISL_3086976, EPI_ISL_3086977, EPI_ISL_3086978, EPI_ISL_3086979, EPI_ISL_3086981, EPI_ISL_3086982, EPI_ISL_3086983, EPI_ISL_3086984, EPI_ISL_3086985, EPI_ISL_3086986, EPI_ISL_3086987, EPI_ISL_3086988, EPI_ISL_3086989, EPI_ISL_3086990, EPI_ISL_3086991, EPI_ISL_3086992, EPI_ISL_3086993, EPI_ISL_3086994 | | | |
| see above | ADPH | ADPH | Adrianna Walker |
| EPI_ISL_3086995 | BJ Medical College Ahmedabad | Gujarat Biotechnology Research Centre | Nitin Savaliya,Dinesh Kumar,Twinkle Soni,Sonal Sharma ,Zuber Saiyed,Ramesh Pandit,Janvi Raval,Zarna Patel,Pranay Shah,Umang Mishra,Nitesh Shah,Chaitanya Joshi,Madhvi Joshi |
| EPI_ISL_3086996 | GMERS, Government Medical College, Himmatnagar | Gujarat Biotechnology Research Centre | Zarna Patel,Nitin Savaliya,Dinesh Kumar,Twinkle Soni,Sonal Sharma ,Zuber Saiyed,Ramesh Pandit,Janvi Raval,Himanshu Khatri,Umang Mishra,Nitesh Shah,Chaitanya Joshi,Madhvi Joshi |
| EPI_ISL_3086997 | GMERS, Government Medical College, Himmatnagar | Gujarat Biotechnology Research Centre | Sonal Sharma ,Zuber Saiyed,Ramesh Pandit,Janvi Raval,Zarna Patel,Nitin Savaliya,Dinesh Kumar,Twinkle Soni,Himanshu Khatri,Umang Mishra,Nitesh Shah,Chaitanya Joshi,Madhvi Joshi |
| EPI_ISL_3086998 | GMERS, Government Medical College, Himmatnagar | Gujarat Biotechnology Research Centre | Zuber Saiyed,Ramesh Pandit,Janvi Raval,Zarna Patel,Nitin Savaliya,Dinesh Kumar,Twinkle Soni,Sonal Sharma ,Himanshu Khatri,Umang Mishra,Nitesh Shah,Chaitanya Joshi,Madhvi Joshi |
| EPI_ISL_3086999 | GMERS, Government Medical College, Himmatnagar | Gujarat Biotechnology Research Centre | Nitin Savaliya,Dinesh Kumar,Twinkle Soni,Sonal Sharma ,Zuber Saiyed,Ramesh Pandit,Janvi Raval,Zarna Patel,Himanshu Khatri,Umang Mishra,Nitesh Shah,Chaitanya Joshi,Madhvi Joshi |
| EPI_ISL_3087000 | GMERS, Government Medical College, Himmatnagar | Gujarat Biotechnology Research Centre | Dinesh Kumar,Twinkle Soni,Sonal Sharma ,Zuber Saiyed,Ramesh Pandit,Janvi Raval,Zarna Patel,Nitin Savaliya,Himanshu Khatri,Umang Mishra,Nitesh Shah,Chaitanya Joshi,Madhvi Joshi |
| EPI_ISL_3087001 | GMERS, Government Medical College, Himmatnagar | Gujarat Biotechnology Research Centre | Zuber Saiyed,Ramesh Pandit,Janvi Raval,Zarna Patel,Nitin Savaliya,Dinesh Kumar,Twinkle Soni,Sonal Sharma ,Himanshu Khatri,Umang Mishra,Nitesh Shah,Chaitanya Joshi,Madhvi Joshi |
| EPI_ISL_3087002, EPI_ISL_3087003, EPI_ISL_3087004, EPI_ISL_3087005, EPI_ISL_3087006, EPI_ISL_3087008, EPI_ISL_3087009 | Orebro University Hospital, Dept Laboratory Medicine, Clinical Microbiology | Orebro University Hospital | Sundqvist M et al |
| EPI_ISL_3087011, EPI_ISL_3087012, EPI_ISL_3087013, EPI_ISL_3087014, EPI_ISL_3087015, EPI_ISL_3087016, EPI_ISL_3087017, EPI_ISL_3087018, EPI_ISL_3087019, EPI_ISL_3087020, EPI_ISL_3087021, EPI_ISL_3087023, EPI_ISL_3087024, EPI_ISL_3087025, EPI_ISL_3087026, EPI_ISL_3087027, EPI_ISL_3087028, EPI_ISL_3087029, EPI_ISL_3087030, EPI_ISL_3087031, EPI_ISL_3087032, EPI_ISL_3087033, EPI_ISL_3087034, EPI_ISL_3087035, EPI_ISL_3087036, EPI_ISL_3087037, EPI_ISL_3087038, EPI_ISL_3087040, EPI_ISL_3087041, EPI_ISL_3087042, EPI_ISL_3087043, EPI_ISL_3087044, EPI_ISL_3087045, EPI_ISL_3087046, EPI_ISL_3087047, EPI_ISL_3087048, EPI_ISL_3087049, EPI_ISL_3087050, EPI_ISL_3087051, EPI_ISL_3087052, EPI_ISL_3087053, EPI_ISL_3087054, EPI_ISL_3087055, EPI_ISL_3087056, | | | |

EPI_ISL_3087057, EPI_ISL_3087058, EPI_ISL_3087059, EPI_ISL_3087060, EPI_ISL_3087061, EPI_ISL_3087062, EPI_ISL_3087063, EPI_ISL_3087064, EPI_ISL_3087065, EPI_ISL_3087066, EPI_ISL_3087067, EPI_ISL_3087068, EPI_ISL_3087069, EPI_ISL_3087070, EPI_ISL_3087071, EPI_ISL_3087072, EPI_ISL_3087073, EPI_ISL_3087074, EPI_ISL_3087075, EPI_ISL_3087076, EPI_ISL_3087077, EPI_ISL_3087078, EPI_ISL_3087079, EPI_ISL_3087080, EPI_ISL_3087081, EPI_ISL_3087082, EPI_ISL_3087083, EPI_ISL_3087084, EPI_ISL_3087085, EPI_ISL_3087086, EPI_ISL_3087087, EPI_ISL_3087089

| | | | |
|---|---|---|---|
| see above | Baylor Scott & White-Temple | Baylor Scott & White-Temple | Ari Rao, Linden Morales, Kimberly Walker, Marcus Volz, Shelby Johnson |
| EPI_ISL_3087092 | HOSPITAL UNIVERSITARIO SON ESPASES | HOSPITAL UNIVERSITARIO SON ESPASES | Servicio de Microbiología, Hospital Universitario Son Espases |
| EPI_ISL_3087093 | HOSPITAL UNIVERSITARIO SON LLATZER | HOSPITAL UNIVERSITARIO SON ESPASES | Servicio de Microbiología, Hospital Universitario Son Espases |
| EPI_ISL_3087094, EPI_ISL_3087095 | HOSPITAL MANACOR | HOSPITAL UNIVERSITARIO SON ESPASES | Servicio de Microbiología, Hospital Universitario Son Espases |
| EPI_ISL_3087096 | HOSPITAL CAN MISSES | HOSPITAL UNIVERSITARIO SON ESPASES | Servicio de Microbiología, Hospital Universitario Son Espases |
| EPI_ISL_3087097, EPI_ISL_3087098 | HOSPITAL UNIVERSITARIO SON ESPASES | HOSPITAL UNIVERSITARIO SON ESPASES | Servicio de Microbiología, Hospital Universitario Son Espases |
| EPI_ISL_3087099, EPI_ISL_3087100 | HOSPITAL CAN MISSES | HOSPITAL UNIVERSITARIO SON ESPASES | Servicio de Microbiología, Hospital Universitario Son Espases |
| EPI_ISL_3087101, EPI_ISL_3087102, EPI_ISL_3087103 | HOSPITAL UNIVERSITARIO SON ESPASES | HOSPITAL UNIVERSITARIO SON ESPASES | Servicio de Microbiología, Hospital Universitario Son Espases |
| EPI_ISL_3087104 | HOSPITAL MANACOR | HOSPITAL UNIVERSITARIO SON ESPASES | Servicio de Microbiología, Hospital Universitario Son Espases |
| EPI_ISL_3087105 | HOSPITAL UNIVERSITARIO SON ESPASES | HOSPITAL UNIVERSITARIO SON ESPASES | Servicio de Microbiología, Hospital Universitario Son Espases |
| EPI_ISL_3087106 | HOSPITAL CAN MISSES | HOSPITAL UNIVERSITARIO SON ESPASES | Servicio de Microbiología, Hospital Universitario Son Espases |
| EPI_ISL_3087107, EPI_ISL_3087108 | HOSPITAL UNIVERSITARIO SON ESPASES | HOSPITAL UNIVERSITARIO SON ESPASES | Servicio de Microbiología, Hospital Universitario Son Espases |
| EPI_ISL_3087109, EPI_ISL_3087110 | HOSPITAL CAN MISSES | HOSPITAL UNIVERSITARIO SON ESPASES | Servicio de Microbiología, Hospital Universitario Son Espases |
| EPI_ISL_3087111, EPI_ISL_3087112 | HOSPITAL UNIVERSITARIO SON ESPASES | HOSPITAL UNIVERSITARIO SON ESPASES | Servicio de Microbiología, Hospital Universitario Son Espases |
| EPI_ISL_3087113, EPI_ISL_3087114 | HOSPITAL MANACOR | HOSPITAL UNIVERSITARIO SON ESPASES | Servicio de Microbiología, Hospital Universitario Son Espases |
| EPI_ISL_3087115 | HOSPITAL CAN MISSES | HOSPITAL UNIVERSITARIO SON ESPASES | Servicio de Microbiología, Hospital Universitario Son Espases |
| EPI_ISL_3087116, EPI_ISL_3087117 | HOSPITAL UNIVERSITARIO SON ESPASES | HOSPITAL UNIVERSITARIO SON ESPASES | Servicio de Microbiología, Hospital Universitario Son Espases |
| EPI_ISL_3087118, EPI_ISL_3087119 | HOSPITAL CAN MISSES | HOSPITAL UNIVERSITARIO SON ESPASES | Servicio de Microbiología, Hospital Universitario Son Espases |
| EPI_ISL_3087121, EPI_ISL_3087122, EPI_ISL_3087123 | HOSPITAL UNIVERSITARIO SON ESPASES | HOSPITAL UNIVERSITARIO SON ESPASES | Servicio de Microbiología, Hospital Universitario Son Espases |
| EPI_ISL_3087124 | HOSPITAL MANACOR | HOSPITAL UNIVERSITARIO SON ESPASES | Servicio de Microbiología, Hospital Universitario Son Espases |
| EPI_ISL_3087125 | HOSPITAL UNIVERSITARIO SON ESPASES | HOSPITAL UNIVERSITARIO SON ESPASES | Servicio de Microbiología, Hospital Universitario Son Espases |
| EPI_ISL_3087126 | HOSPITAL CAN MISSES | HOSPITAL UNIVERSITARIO SON ESPASES | Servicio de Microbiología, Hospital Universitario Son Espases |
| EPI_ISL_3087127 | HOSPITAL UNIVERSITARIO SON ESPASES | HOSPITAL UNIVERSITARIO SON ESPASES | Servicio de Microbiología, Hospital Universitario Son Espases |
| EPI_ISL_3087128 | HOSPITAL CAN MISSES | HOSPITAL UNIVERSITARIO SON ESPASES | Servicio de Microbiología, Hospital Universitario Son Espases |
| EPI_ISL_3087130 | HOSPITAL UNIVERSITARIO SON ESPASES | HOSPITAL UNIVERSITARIO SON ESPASES | Servicio de Microbiología, Hospital Universitario Son Espases |
| EPI_ISL_3087131 | HOSPITAL UNIVERSITARIO SON LLATZER | HOSPITAL UNIVERSITARIO SON ESPASES | Servicio de Microbiología, Hospital Universitario Son Espases |
| EPI_ISL_3087132 | HOSPITAL MANACOR | HOSPITAL UNIVERSITARIO SON ESPASES | Servicio de Microbiología, Hospital Universitario Son Espases |
| EPI_ISL_3087133 | HOSPITAL UNIVERSITARIO SON LLATZER | HOSPITAL UNIVERSITARIO SON ESPASES | Servicio de Microbiología, Hospital Universitario Son Espases |
| EPI_ISL_3087134 | HOSPITAL MANACOR | HOSPITAL UNIVERSITARIO SON ESPASES | Servicio de Microbiología, Hospital Universitario Son Espases |
| EPI_ISL_3087135 | HOSPITAL UNIVERSITARIO SON ESPASES | HOSPITAL UNIVERSITARIO SON ESPASES | Servicio de Microbiología, Hospital Universitario Son Espases |
| EPI_ISL_3087136 | HOSPITAL CAN MISSES | HOSPITAL UNIVERSITARIO SON ESPASES | Servicio de Microbiología, Hospital Universitario Son Espases |
| EPI_ISL_3087137 | HOSPITAL UNIVERSITARIO SON ESPASES | HOSPITAL UNIVERSITARIO SON ESPASES | Servicio de Microbiología, Hospital Universitario Son Espases |
| EPI_ISL_3087139 | HOSPITAL CAN MISSES | HOSPITAL UNIVERSITARIO SON ESPASES | Servicio de Microbiología, Hospital Universitario Son Espases |
| EPI_ISL_3087140, EPI_ISL_3087141, EPI_ISL_3087143 | HOSPITAL UNIVERSITARIO SON ESPASES | HOSPITAL UNIVERSITARIO SON ESPASES | Servicio de Microbiología, Hospital Universitario Son Espases |
| EPI_ISL_3087144, EPI_ISL_3087145 | HOSPITAL MANACOR | HOSPITAL UNIVERSITARIO SON ESPASES | Servicio de Microbiología, Hospital Universitario Son Espases |
| EPI_ISL_3087146 | HOSPITAL UNIVERSITARIO SON ESPASES | HOSPITAL UNIVERSITARIO SON ESPASES | Servicio de Microbiología, Hospital Universitario Son Espases |
| EPI_ISL_3087147 | HOSPITAL UNIVERSITARIO SON LLATZER | HOSPITAL UNIVERSITARIO SON ESPASES | Servicio de Microbiología, Hospital Universitario Son Espases |
| EPI_ISL_3087148 | HOSPITAL CAN MISSES | HOSPITAL UNIVERSITARIO SON ESPASES | Servicio de Microbiología, Hospital Universitario Son Espases |
| EPI_ISL_3087149 | HOSPITAL UNIVERSITARIO SON ESPASES | HOSPITAL UNIVERSITARIO SON ESPASES | Servicio de Microbiología, Hospital Universitario Son Espases |
| EPI_ISL_3087150 | HOSPITAL CAN MISSES | HOSPITAL UNIVERSITARIO SON ESPASES | Servicio de Microbiología, Hospital Universitario Son Espases |
| EPI_ISL_3087151, EPI_ISL_3087152, EPI_ISL_3087153 | HOSPITAL UNIVERSITARIO SON ESPASES | HOSPITAL UNIVERSITARIO SON ESPASES | Servicio de Microbiología, Hospital Universitario Son Espases |
| EPI_ISL_3087154 | HOSPITAL MANACOR | HOSPITAL UNIVERSITARIO SON ESPASES | Servicio de Microbiología, Hospital Universitario Son Espases |
| EPI_ISL_3087156 | HOSPITAL UNIVERSITARIO SON ESPASES | HOSPITAL UNIVERSITARIO SON ESPASES | Servicio de Microbiología, Hospital Universitario Son Espases |
| EPI_ISL_3087157 | HOSPITAL CAN MISSES | HOSPITAL UNIVERSITARIO SON ESPASES | Servicio de Microbiología, Hospital Universitario Son Espases |
| EPI_ISL_3087158 | HOSPITAL UNIVERSITARIO SON ESPASES | HOSPITAL UNIVERSITARIO SON ESPASES | Servicio de Microbiología, Hospital Universitario Son Espases |
| EPI_ISL_3087159, EPI_ISL_3087160 | HOSPITAL CAN MISSES | HOSPITAL UNIVERSITARIO SON ESPASES | Servicio de Microbiología, Hospital Universitario Son Espases |
| EPI_ISL_3087161, EPI_ISL_3087163 | HOSPITAL UNIVERSITARIO SON ESPASES | HOSPITAL UNIVERSITARIO SON ESPASES | Servicio de Microbiología, Hospital Universitario Son Espases |
| EPI_ISL_3087164, EPI_ISL_3087165 | HOSPITAL MANACOR | HOSPITAL UNIVERSITARIO SON ESPASES | Servicio de Microbiología, Hospital Universitario Son Espases |
| EPI_ISL_3087166 | HOSPITAL UNIVERSITARIO SON ESPASES | HOSPITAL UNIVERSITARIO SON ESPASES | Servicio de Microbiología, Hospital Universitario Son Espases |
| EPI_ISL_3087167 | HOSPITAL CAN MISSES | HOSPITAL UNIVERSITARIO SON ESPASES | Servicio de Microbiología, Hospital Universitario Son Espases |
| EPI_ISL_3087168, EPI_ISL_3087169 | HOSPITAL UNIVERSITARIO SON ESPASES | HOSPITAL UNIVERSITARIO SON ESPASES | Servicio de Microbiología, Hospital Universitario Son Espases |
| EPI_ISL_3087170 | HOSPITAL CAN MISSES | HOSPITAL UNIVERSITARIO SON ESPASES | Servicio de Microbiología, Hospital Universitario Son Espases |
| EPI_ISL_3087174, EPI_ISL_3087175, EPI_ISL_3087176, EPI_ISL_3087177, EPI_ISL_3087178, EPI_ISL_3087179, EPI_ISL_3087182, EPI_ISL_3087185, EPI_ISL_3087186, EPI_ISL_3087197 | URMC LABS | URMC LABS | Ghinwa Dumyati, Dwight Hardy, Andrew Cameron, Stepan Voljehn, Kelly DeLary |

| | | | |
|---|---|---|---|
| EPI_ISL_3087199, EPI_ISL_3087201, EPI_ISL_3087202, EPI_ISL_3087203, EPI_ISL_3087204, EPI_ISL_3087205, EPI_ISL_3087206, EPI_ISL_3087207, EPI_ISL_3087240, EPI_ISL_3087241, EPI_ISL_3087242, EPI_ISL_3087243, EPI_ISL_3087244, EPI_ISL_3087245, EPI_ISL_3087246, EPI_ISL_3087247, EPI_ISL_3087248, EPI_ISL_3087249 | | | |
| see above | University of Wisconsin-Madison AIDS Vaccine Research Laboratories | University of Wisconsin-Madison AIDS Vaccine Research Laboratories | Gage Moreno, Katarina Braun, et al. AIDS Vaccine Research Laboratories |
| EPI_ISL_3087250, EPI_ISL_3087251, EPI_ISL_3087253 | Department of Clinical Microbiology | GIGA Medical Genomics | Keith Durkin, Maria Artesi, Bouchra Boujemla, Nathalie Renotte, Nadine Cambisano, Sébastien Bontems, Cécile Meex, Claire Gourzonès, Olivier Ek, Laurent Gillet, Marie-Pierre Hayette, Vincent Bours |
| EPI_ISL_3087254 | C.H.Bois de l'Abbaye | GIGA Medical Genomics | Keith Durkin, Maria Artesi, Bouchra Boujemla, Nathalie Renotte, Nadine Cambisano, Sébastien Bontems, Cécile Meex, Claire Gourzonès, Olivier Ek, Laurent Gillet, Marie-Pierre Hayette, Vincent Bours |
| EPI_ISL_3087255, EPI_ISL_3087256, EPI_ISL_3087257, EPI_ISL_3087259, EPI_ISL_3087260, EPI_ISL_3087261, EPI_ISL_3087262, EPI_ISL_3087265, EPI_ISL_3087266, EPI_ISL_3087267, EPI_ISL_3087268, EPI_ISL_3087269, EPI_ISL_3087270, EPI_ISL_3087272, EPI_ISL_3087273, EPI_ISL_3087274, EPI_ISL_3087276, EPI_ISL_3087277, EPI_ISL_3087280, EPI_ISL_3087283, EPI_ISL_3087285 | | | |
| see above | SYNLAB | GIGA Medical Genomics | Keith Durkin, Maria Artesi, Bouchra Boujemla, Nathalie Renotte, Nadine Cambisano, Sébastien Bontems, Cécile Meex, Claire Gourzonès, Olivier Ek, Laurent Gillet, Marie-Pierre Hayette, Vincent Bours |
| EPI_ISL_3087286, EPI_ISL_3087287, EPI_ISL_3087288, EPI_ISL_3087290, EPI_ISL_3087291 | Department of Clinical Microbiology | GIGA Medical Genomics | Keith Durkin, Maria Artesi, Bouchra Boujemla, Nathalie Renotte, Nadine Cambisano, Sébastien Bontems, Cécile Meex, Claire Gourzonès, Olivier Ek, Laurent Gillet, Marie-Pierre Hayette, Vincent Bours |
| EPI_ISL_3087294, EPI_ISL_3087295, EPI_ISL_3087296, EPI_ISL_3087297, EPI_ISL_3087299, EPI_ISL_3087301, EPI_ISL_3087303 | SYNLAB | GIGA Medical Genomics | Keith Durkin, Maria Artesi, Bouchra Boujemla, Nathalie Renotte, Nadine Cambisano, Sébastien Bontems, Cécile Meex, Claire Gourzonès, Olivier Ek, Laurent Gillet, Marie-Pierre Hayette, Vincent Bours |
| EPI_ISL_3087305 | Department of Clinical Microbiology | GIGA Medical Genomics | Keith Durkin, Maria Artesi, Bouchra Boujemla, Nathalie Renotte, Nadine Cambisano, Sébastien Bontems, Cécile Meex, Claire Gourzonès, Olivier Ek, Laurent Gillet, Marie-Pierre Hayette, Vincent Bours |
| EPI_ISL_3087306, EPI_ISL_3087307 | SYNLAB | GIGA Medical Genomics | Keith Durkin, Maria Artesi, Bouchra Boujemla, Nathalie Renotte, Nadine Cambisano, Sébastien Bontems, Cécile Meex, Claire Gourzonès, Olivier Ek, Laurent Gillet, Marie-Pierre Hayette, Vincent Bours |
| EPI_ISL_3087308 | C.H.C MontLégia | GIGA Medical Genomics | Keith Durkin, Maria Artesi, Bouchra Boujemla, Nathalie Renotte, Nadine Cambisano, Sébastien Bontems, Cécile Meex, Claire Gourzonès, Olivier Ek, Laurent Gillet, Marie-Pierre Hayette, Vincent Bours |
| EPI_ISL_3087309, EPI_ISL_3087310, EPI_ISL_3087311, EPI_ISL_3087312, EPI_ISL_3087313, EPI_ISL_3087314, EPI_ISL_3087315, EPI_ISL_3087317, EPI_ISL_3087318, EPI_ISL_3087320 | SYNLAB | GIGA Medical Genomics | Keith Durkin, Maria Artesi, Bouchra Boujemla, Nathalie Renotte, Nadine Cambisano, Sébastien Bontems, Cécile Meex, Claire Gourzonès, Olivier Ek, Laurent Gillet, Marie-Pierre Hayette, Vincent Bours |
| EPI_ISL_3087322, EPI_ISL_3087323, EPI_ISL_3087325, EPI_ISL_3087326, EPI_ISL_3087327, EPI_ISL_3087329, EPI_ISL_3087330, EPI_ISL_3087333, EPI_ISL_3087334 | University of Liège COVID-19 testing center | GIGA Medical Genomics | Keith Durkin, Maria Artesi, Bouchra Boujemla, Nathalie Renotte, Nadine Cambisano, Sébastien Bontems, Cécile Meex, Claire Gourzonès, Olivier Ek, Laurent Gillet, Marie-Pierre Hayette, Vincent Bours |
| EPI_ISL_3087335, EPI_ISL_3087336, EPI_ISL_3087337, EPI_ISL_3087338, EPI_ISL_3087339, EPI_ISL_3087340, EPI_ISL_3087341, EPI_ISL_3087342, EPI_ISL_3087343, EPI_ISL_3087344, EPI_ISL_3087345, EPI_ISL_3087346, EPI_ISL_3087347, EPI_ISL_3087348, EPI_ISL_3087349, EPI_ISL_3087350, EPI_ISL_3087351, EPI_ISL_3087352, EPI_ISL_3087353, EPI_ISL_3087354, EPI_ISL_3087355, EPI_ISL_3087356, EPI_ISL_3087357, EPI_ISL_3087358, EPI_ISL_3087359, EPI_ISL_3087360, EPI_ISL_3087361, EPI_ISL_3087362, EPI_ISL_3087363, EPI_ISL_3087364, EPI_ISL_3087365, EPI_ISL_3087366, EPI_ISL_3087367, EPI_ISL_3087368, EPI_ISL_3087369, EPI_ISL_3087370, EPI_ISL_3087371, EPI_ISL_3087372, EPI_ISL_3087373, EPI_ISL_3087374, EPI_ISL_3087375, EPI_ISL_3087376, EPI_ISL_3087377, EPI_ISL_3087378, EPI_ISL_3087379, EPI_ISL_3087380, EPI_ISL_3087381, EPI_ISL_3087382, EPI_ISL_3087383, EPI_ISL_3087384, EPI_ISL_3087385, EPI_ISL_3087386, EPI_ISL_3087387, EPI_ISL_3087388, EPI_ISL_3087389, EPI_ISL_3087390, EPI_ISL_3087391, EPI_ISL_3087392, EPI_ISL_3087393, EPI_ISL_3087394, EPI_ISL_3087395, EPI_ISL_3087396, EPI_ISL_3087397, EPI_ISL_3087398, EPI_ISL_3087399, EPI_ISL_3087400, EPI_ISL_3087401, EPI_ISL_3087402, EPI_ISL_3087403, EPI_ISL_3087404, EPI_ISL_3087405, EPI_ISL_3087406, EPI_ISL_3087407, EPI_ISL_3087408, EPI_ISL_3087409, EPI_ISL_3087410, EPI_ISL_3087411, EPI_ISL_3087412, EPI_ISL_3087413, EPI_ISL_3087414, EPI_ISL_3087415, EPI_ISL_3087416, EPI_ISL_3087417, EPI_ISL_3087418, EPI_ISL_3087419, EPI_ISL_3087420, EPI_ISL_3087421, EPI_ISL_3087422, EPI_ISL_3087423, EPI_ISL_3087424, EPI_ISL_3087425, EPI_ISL_3087426, EPI_ISL_3087427, EPI_ISL_3087428, EPI_ISL_3087429, EPI_ISL_3087430, EPI_ISL_3087431, EPI_ISL_3087432, EPI_ISL_3087433, EPI_ISL_3087434, EPI_ISL_3087435, EPI_ISL_3087436, EPI_ISL_3087437, EPI_ISL_3087438, EPI_ISL_3087439, EPI_ISL_3087440, EPI_ISL_3087441, EPI_ISL_3087442, EPI_ISL_3087443, EPI_ISL_3087444, EPI_ISL_3087445, EPI_ISL_3087446, EPI_ISL_3087447, EPI_ISL_3087448, EPI_ISL_3087449, EPI_ISL_3087450, EPI_ISL_3087451, EPI_ISL_3087452, EPI_ISL_3087453, EPI_ISL_3087454, EPI_ISL_3087455, EPI_ISL_3087456, EPI_ISL_3087457, EPI_ISL_3087458, EPI_ISL_3087459, EPI_ISL_3087460, EPI_ISL_3087461, EPI_ISL_3087462, EPI_ISL_3087463, EPI_ISL_3087464, EPI_ISL_3087465, EPI_ISL_3087466, EPI_ISL_3087467, EPI_ISL_3087468, EPI_ISL_3087469, EPI_ISL_3087470, EPI_ISL_3087471, EPI_ISL_3087472, EPI_ISL_3087473, EPI_ISL_3087474, EPI_ISL_3087475, EPI_ISL_3087476, EPI_ISL_3087477, EPI_ISL_3087478, EPI_ISL_3087479, EPI_ISL_3087480, EPI_ISL_3087481, EPI_ISL_3087482, EPI_ISL_3087483, EPI_ISL_3087484, EPI_ISL_3087485, EPI_ISL_3087486, EPI_ISL_3087487, EPI_ISL_3087488, EPI_ISL_3087489, EPI_ISL_3087490, EPI_ISL_3087491, EPI_ISL_3087492, EPI_ISL_3087493, EPI_ISL_3087494, EPI_ISL_3087495, EPI_ISL_3087496, EPI_ISL_3087497, EPI_ISL_3087498, EPI_ISL_3087499, EPI_ISL_3087500, EPI_ISL_3087501, EPI_ISL_3087502, EPI_ISL_3087503, EPI_ISL_3087504, EPI_ISL_3087505, EPI_ISL_3087506, EPI_ISL_3087507, EPI_ISL_3087508, EPI_ISL_3087509, EPI_ISL_3087510, EPI_ISL_3087511, EPI_ISL_3087512, EPI_ISL_3087513, EPI_ISL_3087514, EPI_ISL_3087515, EPI_ISL_3087516, EPI_ISL_3087517, EPI_ISL_3087518, EPI_ISL_3087519, EPI_ISL_3087520, EPI_ISL_3087521, EPI_ISL_3087522, EPI_ISL_3087523, EPI_ISL_3087524, EPI_ISL_3087525, EPI_ISL_3087526, EPI_ISL_3087527, EPI_ISL_3087528, EPI_ISL_3087529, EPI_ISL_3087530, EPI_ISL_3087531, EPI_ISL_3087532, EPI_ISL_3087533, EPI_ISL_3087534, EPI_ISL_3087535, EPI_ISL_3087536, EPI_ISL_3087537, EPI_ISL_3087538, EPI_ISL_3087539, EPI_ISL_3087540, EPI_ISL_3087541, EPI_ISL_3087542, EPI_ISL_3087543, EPI_ISL_3087544, EPI_ISL_3087545, EPI_ISL_3087546, EPI_ISL_3087547, EPI_ISL_3087548, EPI_ISL_3087549, EPI_ISL_3087550, EPI_ISL_3087551, EPI_ISL_3087552, EPI_ISL_3087553, EPI_ISL_3087554, EPI_ISL_3087555, EPI_ISL_3087556, EPI_ISL_3087557, EPI_ISL_3087558, EPI_ISL_3087559, EPI_ISL_3087560, EPI_ISL_3087561 | | | |
| see above | National Public Health Laboratory, National Centre for Infectious Diseases | National Public Health Laboratory, National Centre for Infectious Diseases | Tze Minn Mak, Katherine Ching, Zhenyang Zhou, Royce Ang, Lin Cui, Raymond Tzer Pin Lin |
| EPI_ISL_3087562, EPI_ISL_3087563, EPI_ISL_3087564, EPI_ISL_3087565, EPI_ISL_3087566, EPI_ISL_3087567, EPI_ISL_3087568, EPI_ISL_3087569, EPI_ISL_3087571, EPI_ISL_3087572 | University of Wisconsin-Madison AIDS Vaccine Research Laboratories | University of Wisconsin-Madison AIDS Vaccine Research Laboratories | Gage Moreno, Katarina Braun, et al. AIDS Vaccine Research Laboratories |
| EPI_ISL_3087573, EPI_ISL_3087574, EPI_ISL_3087575, EPI_ISL_3087576 | Jaber Al Ahmad Hospital | Virology Lab, Jaber Al Ahmad Hospital | Dr. Zahrah Buhamad, Dr. Ebaa Al-Awadhi, Haroon Masih, Khubaid-ur-Rehman, Adil Afridi |
| EPI_ISL_3087577, EPI_ISL_3087578, EPI_ISL_3087579, EPI_ISL_3087580, EPI_ISL_3087581, EPI_ISL_3087582, EPI_ISL_3087583, EPI_ISL_3087584, EPI_ISL_3087585, EPI_ISL_3087586, EPI_ISL_3087587, EPI_ISL_3087588, EPI_ISL_3087589 | | | |
| see above | Jaber Quarantine Station | Virology Lab, Jaber Al Ahmad Hospital | Dr. Zahrah Buhamad, Dr. Ebaa Al-Awadhi, Haroon Masih, Khubaid-ur-Rehman, Adil Afridi |
| EPI_ISL_3087590 | Jaber Al Ahmad Hospital | Virology Lab, Jaber Al Ahmad Hospital | Dr. Zahrah Buhamad, Dr. Ebaa Al-Awadhi, Haroon Masih, Khubaid-ur-Rehman, Adil Afridi |
| EPI_ISL_3087591, EPI_ISL_3087592, EPI_ISL_3087593, EPI_ISL_3087594, EPI_ISL_3087596, EPI_ISL_3087597, EPI_ISL_3087598, EPI_ISL_3087599, EPI_ISL_3087600, EPI_ISL_3087601 | Jaber Quarantine Station | Virology Lab, Jaber Al Ahmad Hospital | Dr. Zahrah Buhamad, Dr. Ebaa Al-Awadhi, Haroon Masih, Khubaid-ur-Rehman, Adil Afridi |
| EPI_ISL_3087602 | Jaber Al Ahmad Hospital | Virology Lab, Jaber Al Ahmad Hospital | Dr. Zahrah Buhamad, Dr. Ebaa Al-Awadhi, Haroon Masih, Khubaid-ur-Rehman, Adil Afridi |
| EPI_ISL_3087682, EPI_ISL_3087683, EPI_ISL_3087685 | Clínica Rotger Laboratorio de Análisis Clínicos Dpto. Biología Molecular | Clínica Rotger Laboratorio de Análisis Clínicos Dpto. Biología Molecular | Francisco Aliaga-Lozano, Chaymaá Gammani-Chairi, Laura Gómez, Juana González-Calderón, Marina Barceló-Vives, Catalina Vidal-Gómez, Marina Kleber-Pol, Esperança Frau-Fuster, Carmen Morales-González, Esther Nieto-Molina |
| EPI_ISL_3087778 | Landesamt für Verbraucherschutz Sachsen Anhalt, Magdeburg | Institute of Medical Microbiology and Hospital Hygiene | Prof. Dr. Achim Kaasch, Aljoscha Tersteegen |
| EPI_ISL_3087819 | Institute of Medical Microbiology and Hospital Hygiene | Institute of Medical Microbiology and Hospital Hygiene | Prof. Dr. Achim Kaasch, Aljoscha Tersteegen |
| EPI_ISL_3087865, EPI_ISL_3087866, EPI_ISL_3087867, EPI_ISL_3087868, EPI_ISL_3087869, EPI_ISL_3087870, EPI_ISL_3087871, EPI_ISL_3087872, EPI_ISL_3087873, EPI_ISL_3087874, EPI_ISL_3087875, EPI_ISL_3087876, EPI_ISL_3087877, EPI_ISL_3087878, EPI_ISL_3087879, EPI_ISL_3087880, EPI_ISL_3087881, EPI_ISL_3087882, EPI_ISL_3087883, EPI_ISL_3087884, EPI_ISL_3087885, EPI_ISL_3087886, EPI_ISL_3087887, EPI_ISL_3087888, EPI_ISL_3087890, EPI_ISL_3087891, EPI_ISL_3087892, EPI_ISL_3087893, EPI_ISL_3087894, EPI_ISL_3087895, EPI_ISL_3087896 | | | |
| see above | Kansas Health and Environmental Lab | Kansas Health and Environmental Lab | Katherine Wiggins, Mike Grose, Jonathan Barnell, Ben Olsen, and Phil Adam |
| EPI_ISL_3087899, EPI_ISL_3087900, EPI_ISL_3087901, EPI_ISL_3087902, EPI_ISL_3087903, EPI_ISL_3087904, EPI_ISL_3087905, EPI_ISL_3087906, EPI_ISL_3087907, EPI_ISL_3087908, EPI_ISL_3087909, EPI_ISL_3087910, EPI_ISL_3087911, EPI_ISL_3087912, EPI_ISL_3087913, EPI_ISL_3087914, EPI_ISL_3087915, EPI_ISL_3087916, EPI_ISL_3087918, EPI_ISL_3087919, EPI_ISL_3087920, EPI_ISL_3087921, EPI_ISL_3087922, EPI_ISL_3087923, EPI_ISL_3087925, EPI_ISL_3087926, EPI_ISL_3087927, EPI_ISL_3087928, EPI_ISL_3087930, EPI_ISL_3087931, EPI_ISL_3087932, EPI_ISL_3087933, EPI_ISL_3087934, EPI_ISL_3087935, EPI_ISL_3087936, EPI_ISL_3087937, EPI_ISL_3087938, EPI_ISL_3087939, EPI_ISL_3087941, EPI_ISL_3087942, EPI_ISL_3087944, EPI_ISL_3087945, EPI_ISL_3087948, EPI_ISL_3087949, EPI_ISL_3087950, EPI_ISL_3087951, EPI_ISL_3087952, EPI_ISL_3087953, EPI_ISL_3087954, EPI_ISL_3087955, EPI_ISL_3087956, EPI_ISL_3087957, EPI_ISL_3087958, EPI_ISL_3087959, EPI_ISL_3087960, EPI_ISL_3087961, EPI_ISL_3087962, EPI_ISL_3087963, EPI_ISL_3087964, EPI_ISL_3087965, EPI_ISL_3087966, EPI_ISL_3087967, EPI_ISL_3087968, EPI_ISL_3087969, EPI_ISL_3087970, EPI_ISL_3087971, EPI_ISL_3087972, EPI_ISL_3087973, EPI_ISL_3087974, EPI_ISL_3087975, EPI_ISL_3087976, EPI_ISL_3087977, EPI_ISL_3087978, EPI_ISL_3087979 | | | |
| see above | Hospital Universitari i Politècnic La Fe de València | SeqCOVID-SPAIN consortium/IBV(CSIC) | María Dolores Gómez Ruiz, Eva González Barberá, Ana Gil Brusola, Salvador Giner Almaraz, José Luis López Hontangas and SeqCOVID-SPAIN consortium |

| | | | |
|---|---|---|---|
| EPI_ISL_3087980 | Hospital Universitario de La Ribera (Alzira, València) | SeqCOVID-SPAIN consortium/IBV(CSIC) | Olalla Martínez Macias, Julia González Cantó and SeqCOVID-SPAIN consortium |
| EPI_ISL_3087981, EPI_ISL_3087982, EPI_ISL_3087983, EPI_ISL_3087984, EPI_ISL_3087985, EPI_ISL_3087986, EPI_ISL_3087988 | Servicio de Microbiología. Hospital General Universitario de Castellón | SeqCOVID-SPAIN consortium/IBV(CSIC) | Rosario Moreno Muñoz, María Dolores Tirado Balaguer and SeqCOVID-SPAIN consortium |
| EPI_ISL_3087989, EPI_ISL_3087990, EPI_ISL_3087991, EPI_ISL_3087992, EPI_ISL_3087993, EPI_ISL_3087994, EPI_ISL_3087995, EPI_ISL_3087996, EPI_ISL_3087997, EPI_ISL_3087998, EPI_ISL_3087999, EPI_ISL_3088000, EPI_ISL_3088001, EPI_ISL_3088002, EPI_ISL_3088003, EPI_ISL_3088004, EPI_ISL_3088005, EPI_ISL_3088006, EPI_ISL_3088007, EPI_ISL_3088008, EPI_ISL_3088009, EPI_ISL_3088010, EPI_ISL_3088011, EPI_ISL_3088012, EPI_ISL_3088013, EPI_ISL_3088014, EPI_ISL_3088015, EPI_ISL_3088016, EPI_ISL_3088017, EPI_ISL_3088018, EPI_ISL_3088019, EPI_ISL_3088020, EPI_ISL_3088021, EPI_ISL_3088022, EPI_ISL_3088023, EPI_ISL_3088024, EPI_ISL_3088025, EPI_ISL_3088026, EPI_ISL_3088027, EPI_ISL_3088028, EPI_ISL_3088029, EPI_ISL_3088030, EPI_ISL_3088031, EPI_ISL_3088032, EPI_ISL_3088033, EPI_ISL_3088034, EPI_ISL_3088035, EPI_ISL_3088036, EPI_ISL_3088037, EPI_ISL_3088038, EPI_ISL_3088039, EPI_ISL_3088040, EPI_ISL_3088041, EPI_ISL_3088042, EPI_ISL_3088043, EPI_ISL_3088044, EPI_ISL_3088045, EPI_ISL_3088046, EPI_ISL_3088047, EPI_ISL_3088048, EPI_ISL_3088049, EPI_ISL_3088050, EPI_ISL_3088051, EPI_ISL_3088052, EPI_ISL_3088053, EPI_ISL_3088054, EPI_ISL_3088055, EPI_ISL_3088056, EPI_ISL_3088057, EPI_ISL_3088058, EPI_ISL_3088059, EPI_ISL_3088060, EPI_ISL_3088061, EPI_ISL_3088062, EPI_ISL_3088063, EPI_ISL_3088064, EPI_ISL_3088065, EPI_ISL_3088066, EPI_ISL_3088067, EPI_ISL_3088068, EPI_ISL_3088069, EPI_ISL_3088070, EPI_ISL_3088071, EPI_ISL_3088072, EPI_ISL_3088073, EPI_ISL_3088074, EPI_ISL_3088075, EPI_ISL_3088076, EPI_ISL_3088077, EPI_ISL_3088078, EPI_ISL_3088079, EPI_ISL_3088080, EPI_ISL_3088081, EPI_ISL_3088082, EPI_ISL_3088083, EPI_ISL_3088084, EPI_ISL_3088085, EPI_ISL_3088086, EPI_ISL_3088087, EPI_ISL_3088088, EPI_ISL_3088089, EPI_ISL_3088090, EPI_ISL_3088091, EPI_ISL_3088092, EPI_ISL_3088093, EPI_ISL_3088094, EPI_ISL_3088095, EPI_ISL_3088096, EPI_ISL_3088097, EPI_ISL_3088098, EPI_ISL_3088099, EPI_ISL_3088100, EPI_ISL_3088101, EPI_ISL_3088102, EPI_ISL_3088103, EPI_ISL_3088104, EPI_ISL_3088105, EPI_ISL_3088106, EPI_ISL_3088107, EPI_ISL_3088108, EPI_ISL_3088109, EPI_ISL_3088110, EPI_ISL_3088111, EPI_ISL_3088112, EPI_ISL_3088113, EPI_ISL_3088114, EPI_ISL_3088115, EPI_ISL_3088116, EPI_ISL_3088117, EPI_ISL_3088118, EPI_ISL_3088119, EPI_ISL_3088120, EPI_ISL_3088121, EPI_ISL_3088122, EPI_ISL_3088123, EPI_ISL_3088124, EPI_ISL_3088125, EPI_ISL_3088126, EPI_ISL_3088127, EPI_ISL_3088128, EPI_ISL_3088129, EPI_ISL_3088130, EPI_ISL_3088131, EPI_ISL_3088132, EPI_ISL_3088133, EPI_ISL_3088134, EPI_ISL_3088135, EPI_ISL_3088136, EPI_ISL_3088137, EPI_ISL_3088138, EPI_ISL_3088139, EPI_ISL_3088140, EPI_ISL_3088141, EPI_ISL_3088142, EPI_ISL_3088143, EPI_ISL_3088144, EPI_ISL_3088145, EPI_ISL_3088146, EPI_ISL_3088147, EPI_ISL_3088148, EPI_ISL_3088149, EPI_ISL_3088150, EPI_ISL_3088151, EPI_ISL_3088152, EPI_ISL_3088153, EPI_ISL_3088154, EPI_ISL_3088155, EPI_ISL_3088156, EPI_ISL_3088157, EPI_ISL_3088158, EPI_ISL_3088159, EPI_ISL_3088160, EPI_ISL_3088161, EPI_ISL_3088162, EPI_ISL_3088163, EPI_ISL_3088164, EPI_ISL_3088165, EPI_ISL_3088166, EPI_ISL_3088167, EPI_ISL_3088168, EPI_ISL_3088169, EPI_ISL_3088170, EPI_ISL_3088171, EPI_ISL_3088172, EPI_ISL_3088173, EPI_ISL_3088174, EPI_ISL_3088175, EPI_ISL_3088176, EPI_ISL_3088177, EPI_ISL_3088178, EPI_ISL_3088179, EPI_ISL_3088180, EPI_ISL_3088181, EPI_ISL_3088182, EPI_ISL_3088183, EPI_ISL_3088184, EPI_ISL_3088185, EPI_ISL_3088186, EPI_ISL_3088187, EPI_ISL_3088188, EPI_ISL_3088189, EPI_ISL_3088190, EPI_ISL_3088191, EPI_ISL_3088192, EPI_ISL_3088193, EPI_ISL_3088194, EPI_ISL_3088195, EPI_ISL_3088196, EPI_ISL_3088197, EPI_ISL_3088198, EPI_ISL_3088199, EPI_ISL_3088200, EPI_ISL_3088201, EPI_ISL_3088202, EPI_ISL_3088203, EPI_ISL_3088204, EPI_ISL_3088205, EPI_ISL_3088206, EPI_ISL_3088207, EPI_ISL_3088208, EPI_ISL_3088209, EPI_ISL_3088210, EPI_ISL_3088211, EPI_ISL_3088212, EPI_ISL_3088213, EPI_ISL_3088214, EPI_ISL_3088215, EPI_ISL_3088216, EPI_ISL_3088217, EPI_ISL_3088218, EPI_ISL_3088219, EPI_ISL_3088220, EPI_ISL_3088221, EPI_ISL_3088222, EPI_ISL_3088223, EPI_ISL_3088224, EPI_ISL_3088225, EPI_ISL_3088226, EPI_ISL_3088227, EPI_ISL_3088228, EPI_ISL_3088229, EPI_ISL_3088230, EPI_ISL_3088231, EPI_ISL_3088232, EPI_ISL_3088233, EPI_ISL_3088234, EPI_ISL_3088235, EPI_ISL_3088236, EPI_ISL_3088237, EPI_ISL_3088238, EPI_ISL_3088239, EPI_ISL_3088240, EPI_ISL_3088241, EPI_ISL_3088242, EPI_ISL_3088243, EPI_ISL_3088244, EPI_ISL_3088245, EPI_ISL_3088246, EPI_ISL_3088247, EPI_ISL_3088248, EPI_ISL_3088249, EPI_ISL_3088250, EPI_ISL_3088251, EPI_ISL_3088252, EPI_ISL_3088253, EPI_ISL_3088254, EPI_ISL_3088255, EPI_ISL_3088256, EPI_ISL_3088257, EPI_ISL_3088258, EPI_ISL_3088259, EPI_ISL_3088260, EPI_ISL_3088261, EPI_ISL_3088262, EPI_ISL_3088263, EPI_ISL_3088264, EPI_ISL_3088265, EPI_ISL_3088266, EPI_ISL_3088267, EPI_ISL_3088268, EPI_ISL_3088269, EPI_ISL_3088270, EPI_ISL_3088271, EPI_ISL_3088272, EPI_ISL_3088273, EPI_ISL_3088274, EPI_ISL_3088275, EPI_ISL_3088276, EPI_ISL_3088277, EPI_ISL_3088278, EPI_ISL_3088279, EPI_ISL_3088280, EPI_ISL_3088281, EPI_ISL_3088282, EPI_ISL_3088283, EPI_ISL_3088284, EPI_ISL_3088285, EPI_ISL_3088286, EPI_ISL_3088287, EPI_ISL_3088288, EPI_ISL_3088289, EPI_ISL_3088290, EPI_ISL_3088291, EPI_ISL_3088292, EPI_ISL_3088293, EPI_ISL_3088294, EPI_ISL_3088295, EPI_ISL_3088296, EPI_ISL_3088297, EPI_ISL_3088298, EPI_ISL_3088299, EPI_ISL_3088300 | | | |
| see above | National Virus Reference Laboratory | National Virus Reference Laboratory | Zoe Yandle, Charlene Bennett, Gabriel Gonzalez, Michael Carr, Jonathan Dean, Cillian F De Gascun |
| EPI_ISL_3088307, EPI_ISL_3088310 | Omics Sciences Laboratory | Omics Sciences Laboratory | Derly Andrade Molina, Gabriel Morey León, Darlyn Amaya, Rubén Armas González, Juan Carlos Fernández Cadena |
| EPI_ISL_3088324, EPI_ISL_3088329, EPI_ISL_3088330, EPI_ISL_3088332 | INTERLAB | Omics Sciences Laboratory | Derly Andrade Molina, Gabriel Morey León, Darlyn Amaya, Rubén Armas González, Juan Carlos Fernández Cadena |
| EPI_ISL_3088374, EPI_ISL_3088375, EPI_ISL_3088376, EPI_ISL_3088377, EPI_ISL_3088378, EPI_ISL_3088379, EPI_ISL_3088380, EPI_ISL_3088381, EPI_ISL_3088382, EPI_ISL_3088383, EPI_ISL_3088384, EPI_ISL_3088385, EPI_ISL_3088386 | | | |
| see above | Kansas Health and Environmental Lab | Kansas Health and Environmental Lab | Katherine Wiggins, Mike Grose, Jonathan Barnell, Ben Olsen, and Phil Adam |
| EPI_ISL_3088387, EPI_ISL_3088388, EPI_ISL_3088389, EPI_ISL_3088390, EPI_ISL_3088391, EPI_ISL_3088392, EPI_ISL_3088393, EPI_ISL_3088394, EPI_ISL_3088395, EPI_ISL_3088396, EPI_ISL_3088397, EPI_ISL_3088398, EPI_ISL_3088399, EPI_ISL_3088400, EPI_ISL_3088401, EPI_ISL_3088402, EPI_ISL_3088403, EPI_ISL_3088404, EPI_ISL_3088405 | | | |
| see above | Wichita State University - Molecular Diagnostics Lab | Kansas Health and Environmental Lab | Katherine Wiggins, Mike Grose, Jonathan Barnell, Ben Olsen, and Phil Adam |
| EPI_ISL_3088406, EPI_ISL_3088407, EPI_ISL_3088408, EPI_ISL_3088409, EPI_ISL_3088410, EPI_ISL_3088411, EPI_ISL_3088412, EPI_ISL_3088413, EPI_ISL_3088414, EPI_ISL_3088415, EPI_ISL_3088416, EPI_ISL_3088417, EPI_ISL_3088418, EPI_ISL_3088419, EPI_ISL_3088420, EPI_ISL_3088421, EPI_ISL_3088422, EPI_ISL_3088423, EPI_ISL_3088424, EPI_ISL_3088425, EPI_ISL_3088426, EPI_ISL_3088427, EPI_ISL_3088428, EPI_ISL_3088429, EPI_ISL_3088430, EPI_ISL_3088431, EPI_ISL_3088432, EPI_ISL_3088433, EPI_ISL_3088434, EPI_ISL_3088435, EPI_ISL_3088436, EPI_ISL_3088437, EPI_ISL_3088438, EPI_ISL_3088439, EPI_ISL_3088440, EPI_ISL_3088441, EPI_ISL_3088442, EPI_ISL_3088443, EPI_ISL_3088444, EPI_ISL_3088445, EPI_ISL_3088446, EPI_ISL_3088447, EPI_ISL_3088448, EPI_ISL_3088450, EPI_ISL_3088451, EPI_ISL_3088452, EPI_ISL_3088453, EPI_ISL_3088456, EPI_ISL_3088457, EPI_ISL_3088458, EPI_ISL_3088459, EPI_ISL_3088462, EPI_ISL_3088463, EPI_ISL_3088464, EPI_ISL_3088465, EPI_ISL_3088466, EPI_ISL_3088467, EPI_ISL_3088468, EPI_ISL_3088469, EPI_ISL_3088470, EPI_ISL_3088471, EPI_ISL_3088472, EPI_ISL_3088473, EPI_ISL_3088474, EPI_ISL_3088475, EPI_ISL_3088476, EPI_ISL_3088477, EPI_ISL_3088478, EPI_ISL_3088479, EPI_ISL_3088480, EPI_ISL_3088481, EPI_ISL_3088482, EPI_ISL_3088483, EPI_ISL_3088484, EPI_ISL_3088485, EPI_ISL_3088487, EPI_ISL_3088488, EPI_ISL_3088489, EPI_ISL_3088490, EPI_ISL_3088491, EPI_ISL_3088492, EPI_ISL_3088493, EPI_ISL_3088494, EPI_ISL_3088495 | | | |
| see above | Illinois Department of Public Health - Springfield Lab | Illinois Department of Public Health - Springfield Lab | Bryan Sim, Marshall Smock |
| EPI_ISL_3088497, EPI_ISL_3088498, EPI_ISL_3088499, EPI_ISL_3088500, EPI_ISL_3088501, EPI_ISL_3088503, EPI_ISL_3088504, EPI_ISL_3088505, EPI_ISL_3088506 | Wichita State University - Molecular Diagnostics Lab | Kansas Health and Environmental Lab | Katherine Wiggins, Mike Grose, Jonathan Barnell, Ben Olsen, and Phil Adam |
| EPI_ISL_3088507, EPI_ISL_3088508, EPI_ISL_3088509, EPI_ISL_3088510, EPI_ISL_3088511, EPI_ISL_3088512, EPI_ISL_3088513, EPI_ISL_3088514, EPI_ISL_3088515, EPI_ISL_3088516, EPI_ISL_3088517, EPI_ISL_3088518, EPI_ISL_3088519, EPI_ISL_3088520, EPI_ISL_3088521, EPI_ISL_3088522, EPI_ISL_3088523, EPI_ISL_3088524, EPI_ISL_3088525, EPI_ISL_3088526, EPI_ISL_3088527, EPI_ISL_3088528, EPI_ISL_3088529, EPI_ISL_3088530, EPI_ISL_3088531, EPI_ISL_3088532, EPI_ISL_3088533, EPI_ISL_3088534, EPI_ISL_3088535, EPI_ISL_3088536, EPI_ISL_3088537, EPI_ISL_3088538, EPI_ISL_3088539, EPI_ISL_3088540, EPI_ISL_3088541, EPI_ISL_3088542, EPI_ISL_3088543, EPI_ISL_3088544, EPI_ISL_3088545, EPI_ISL_3088546, EPI_ISL_3088547, EPI_ISL_3088548, EPI_ISL_3088549, EPI_ISL_3088550, EPI_ISL_3088551, EPI_ISL_3088552 | | | |
| see above | Kansas Health and Environmental Lab | Kansas Health and Environmental Lab | Katherine Wiggins, Mike Grose, Jonathan Barnell, Ben Olsen, and Phil Adam |
| EPI_ISL_3088556, EPI_ISL_3088557, EPI_ISL_3088558, EPI_ISL_3088561, EPI_ISL_3088569, EPI_ISL_3088578, EPI_ISL_3088581, EPI_ISL_3088589, EPI_ISL_3088594, EPI_ISL_3088604, EPI_ISL_3088616, EPI_ISL_3088621, EPI_ISL_3088629, EPI_ISL_3088636, EPI_ISL_3088638, EPI_ISL_3088643, EPI_ISL_3088646, EPI_ISL_3088649, EPI_ISL_3088652, EPI_ISL_3088653, EPI_ISL_3088656, EPI_ISL_3088661, EPI_ISL_3088664, EPI_ISL_3088670, EPI_ISL_3088673, EPI_ISL_3088689, EPI_ISL_3088690, EPI_ISL_3088691, EPI_ISL_3088697, EPI_ISL_3088707, EPI_ISL_3088709, EPI_ISL_3088713, EPI_ISL_3088715, EPI_ISL_3088718, EPI_ISL_3088719, EPI_ISL_3088721, EPI_ISL_3088749, EPI_ISL_3088754, EPI_ISL_3088760, EPI_ISL_3088761, EPI_ISL_3088762, EPI_ISL_3088769, EPI_ISL_3088770, EPI_ISL_3088772, EPI_ISL_3088775, EPI_ISL_3088777, EPI_ISL_3088779, EPI_ISL_3088782, EPI_ISL_3088784, EPI_ISL_3088785, EPI_ISL_3088789, EPI_ISL_3088807, EPI_ISL_3088809, EPI_ISL_3088820 | | | |
| see above | TGen North | TGen North | "Jolene Bowers, Heather Centner, Brett Van Tassel, Chris French, Hayley Yaglom, Darrin Lemmer, Dave Engelthaler" |
| EPI_ISL_3088821, EPI_ISL_3088822 | Kansas Health and Environmental Lab | Kansas Health and Environmental Lab | Katherine Wiggins, Mike Grose, Jonathan Barnell, Ben Olsen, and Phil Adam |
| EPI_ISL_3088823, EPI_ISL_3088824, EPI_ISL_3088825, EPI_ISL_3088826, EPI_ISL_3088827, EPI_ISL_3088828, EPI_ISL_3088830, EPI_ISL_3088831, EPI_ISL_3088832, EPI_ISL_3088833, EPI_ISL_3088834, EPI_ISL_3088835, EPI_ISL_3088836, EPI_ISL_3088837, EPI_ISL_3088838, EPI_ISL_3088839, EPI_ISL_3088840, EPI_ISL_3088843, EPI_ISL_3088844, EPI_ISL_3088845, EPI_ISL_3088846 | | | |
| see above | Wichita State University - Molecular Diagnostics Lab | Kansas Health and Environmental Lab | Katherine Wiggins, Mike Grose, Jonathan Barnell, Ben Olsen, and Phil Adam |
| EPI_ISL_3088847, EPI_ISL_3088849, EPI_ISL_3088850, EPI_ISL_3088851, EPI_ISL_3088852, EPI_ISL_3088853, EPI_ISL_3088854, EPI_ISL_3088855, EPI_ISL_3088856, EPI_ISL_3088857, EPI_ISL_3088858, EPI_ISL_3088859, EPI_ISL_3088860, EPI_ISL_3088861, EPI_ISL_3088862, EPI_ISL_3088863, EPI_ISL_3088864, EPI_ISL_3088865, EPI_ISL_3088866, EPI_ISL_3088867, EPI_ISL_3088869, EPI_ISL_3088870, EPI_ISL_3088871, EPI_ISL_3088872, EPI_ISL_3088873, EPI_ISL_3088874, EPI_ISL_3088875, EPI_ISL_3088876, EPI_ISL_3088878, EPI_ISL_3088879, EPI_ISL_3088880, EPI_ISL_3088881, EPI_ISL_3088882, EPI_ISL_3088883, EPI_ISL_3088884, EPI_ISL_3088885, EPI_ISL_3088888, EPI_ISL_3088889, EPI_ISL_3088890, EPI_ISL_3088891, EPI_ISL_3088892, EPI_ISL_3088893, EPI_ISL_3088894, EPI_ISL_3088895, EPI_ISL_3088896, EPI_ISL_3088897, EPI_ISL_3088898, EPI_ISL_3088899, EPI_ISL_3088900, EPI_ISL_3088901, EPI_ISL_3088902, EPI_ISL_3088903, EPI_ISL_3088904, EPI_ISL_3088905, EPI_ISL_3088906, EPI_ISL_3088907, EPI_ISL_3088908, EPI_ISL_3088909, EPI_ISL_3088910, EPI_ISL_3088911, EPI_ISL_3088912, EPI_ISL_3088913, EPI_ISL_3088914, EPI_ISL_3088915, EPI_ISL_3088916, EPI_ISL_3088917, EPI_ISL_3088918, EPI_ISL_3088919, EPI_ISL_3088920, EPI_ISL_3088921, EPI_ISL_3088922, EPI_ISL_3088923, EPI_ISL_3088924, EPI_ISL_3088926, EPI_ISL_3088927, EPI_ISL_3088928, EPI_ISL_3088929, EPI_ISL_3088930, EPI_ISL_3088931, EPI_ISL_3088932, EPI_ISL_3088933, EPI_ISL_3088934, EPI_ISL_3088935, EPI_ISL_3088936, EPI_ISL_3088937, EPI_ISL_3088938, EPI_ISL_3088939, EPI_ISL_3088940, EPI_ISL_3088942, EPI_ISL_3088943, EPI_ISL_3088944, EPI_ISL_3088945, EPI_ISL_3088946, EPI_ISL_3088947, EPI_ISL_3088948, EPI_ISL_3088949, EPI_ISL_3088950, EPI_ISL_3088951, EPI_ISL_3088952, EPI_ISL_3088953, EPI_ISL_3088954, EPI_ISL_3088955, EPI_ISL_3088956, EPI_ISL_3088957, EPI_ISL_3088958, EPI_ISL_3088959, EPI_ISL_3088960, EPI_ISL_3088961, EPI_ISL_3088962, EPI_ISL_3088963, EPI_ISL_3088964, EPI_ISL_3088965, EPI_ISL_3088966, EPI_ISL_3088967, EPI_ISL_3088968, EPI_ISL_3088969, EPI_ISL_3088970, EPI_ISL_3088971, EPI_ISL_3088972, EPI_ISL_3088973, EPI_ISL_3088974, EPI_ISL_3088975, EPI_ISL_3088976, EPI_ISL_3088977, EPI_ISL_3088978, EPI_ISL_3088979, EPI_ISL_3088980, EPI_ISL_3088981, EPI_ISL_3088983, EPI_ISL_3088985, EPI_ISL_3088986, EPI_ISL_3088987, EPI_ISL_3088988, EPI_ISL_3088990 | | | |
| see above | Ochsner Health | Bioinfoexperts, LLC | Judy Crabtree, Amy Feehan, Rebecca Rose, David J. Nolan, Tessa LaFleur, Julia-Garcia-Diaz, Lucio Miele, Susanna L. Lamers |
| EPI_ISL_3088992, EPI_ISL_3088993, EPI_ISL_3088994, EPI_ISL_3088995, EPI_ISL_3088996, EPI_ISL_3088997, EPI_ISL_3088998, EPI_ISL_3088999, EPI_ISL_3089000, EPI_ISL_3089001, EPI_ISL_3089002, EPI_ISL_3089003, EPI_ISL_3089004 | | | |
| see above | Kansas Health and Environmental Lab | Kansas Health and Environmental Lab | Katherine Wiggins, Mike Grose, Jonathan Barnell, Ben Olsen, and Phil Adam |
| EPI_ISL_3089005, EPI_ISL_3089006, EPI_ISL_3089007, EPI_ISL_3089008, EPI_ISL_3089009, EPI_ISL_3089010, EPI_ISL_3089011, EPI_ISL_3089012, EPI_ISL_3089013, EPI_ISL_3089014, EPI_ISL_3089015 | | | |
| see above | Wichita State University - Molecular Diagnostics Lab | Kansas Health and Environmental Lab | Katherine Wiggins, Mike Grose, Jonathan Barnell, Ben Olsen, and Phil Adam |
| EPI_ISL_3089016, EPI_ISL_3089017, EPI_ISL_3089018, EPI_ISL_3089020, EPI_ISL_3089021, EPI_ISL_3089022, EPI_ISL_3089023, EPI_ISL_3089024, EPI_ISL_3089025, EPI_ISL_3089026, EPI_ISL_3089027, EPI_ISL_3089028, EPI_ISL_3089030, EPI_ISL_3089031, EPI_ISL_3089032, EPI_ISL_3089033, EPI_ISL_3089034, EPI_ISL_3089035, EPI_ISL_3089036, EPI_ISL_3089037, EPI_ISL_3089038, EPI_ISL_3089039, EPI_ISL_3089040, EPI_ISL_3089041, EPI_ISL_3089042, EPI_ISL_3089043, EPI_ISL_3089044, EPI_ISL_3089045, EPI_ISL_3089046, EPI_ISL_3089047 | | | |
| see above | Clinique Saint-Pierre Ottignies | UCLouvain/IREC/MBLG | Jean Ruelle, Bertrand Bearzatto, Benoit Kabamba Mukadi |
| EPI_ISL_3089049, EPI_ISL_3089050, EPI_ISL_3089051, EPI_ISL_3089052, EPI_ISL_3089053, EPI_ISL_3089054, EPI_ISL_3089055, EPI_ISL_3089056, EPI_ISL_3089057, EPI_ISL_3089058, EPI_ISL_3089059, EPI_ISL_3089060, EPI_ISL_3089061, EPI_ISL_3089062, EPI_ISL_3089063, EPI_ISL_3089064, EPI_ISL_3089065, EPI_ISL_3089066, EPI_ISL_3089067, EPI_ISL_3089068, EPI_ISL_3089069, EPI_ISL_3089070, EPI_ISL_3089071, EPI_ISL_3089072, EPI_ISL_3089073, EPI_ISL_3089074, EPI_ISL_3089075, EPI_ISL_3089076, EPI_ISL_3089078, EPI_ISL_3089079, EPI_ISL_3089080, EPI_ISL_3089081, EPI_ISL_3089082, EPI_ISL_3089084, EPI_ISL_3089085, EPI_ISL_3089086, EPI_ISL_3089087, EPI_ISL_3089088, EPI_ISL_3089089, EPI_ISL_3089091, EPI_ISL_3089092, EPI_ISL_3089093, EPI_ISL_3089094, EPI_ISL_3089095, EPI_ISL_3089096, EPI_ISL_3089097, EPI_ISL_3089098, EPI_ISL_3089099, EPI_ISL_3089100, EPI_ISL_3089101, EPI_ISL_3089102, EPI_ISL_3089103, EPI_ISL_3089104, EPI_ISL_3089105, EPI_ISL_3089106, EPI_ISL_3089107, EPI_ISL_3089108, EPI_ISL_3089109 | | | |
| see above | Cliniques universitaires Saint-Luc | UCLouvain/IREC/MBLG | Jean Ruelle, Bertrand Bearzatto, Benoit Kabamba Mukadi |
| EPI_ISL_3089110, EPI_ISL_3089111, EPI_ISL_3089112, EPI_ISL_3089113, EPI_ISL_3089114, EPI_ISL_3089115, EPI_ISL_3089116, EPI_ISL_3089117, EPI_ISL_3089118, EPI_ISL_3089119, EPI_ISL_3089120, EPI_ISL_3089121, EPI_ISL_3089122, EPI_ISL_3089123, EPI_ISL_3089124, EPI_ISL_3089125, EPI_ISL_3089126, EPI_ISL_3089127, EPI_ISL_3089128, EPI_ISL_3089129, EPI_ISL_3089130, EPI_ISL_3089131, EPI_ISL_3089132 | | | |
| see above | University of Wisconsin-Madison AIDS Vaccine Research Laboratories | University of Wisconsin-Madison AIDS Vaccine Research Laboratories | Gage Moreno, Katarina Braun, et al. AIDS Vaccine Research Laboratories |
| EPI_ISL_3089133, EPI_ISL_3089134, EPI_ISL_3089135, EPI_ISL_3089136, EPI_ISL_3089137, EPI_ISL_3089138, EPI_ISL_3089139, EPI_ISL_3089140, EPI_ISL_3089141, EPI_ISL_3089142, EPI_ISL_3089143, EPI_ISL_3089144, EPI_ISL_3089145, EPI_ISL_3089146, EPI_ISL_3089147, EPI_ISL_3089148, EPI_ISL_3089149, EPI_ISL_3089150, EPI_ISL_3089151, EPI_ISL_3089152, EPI_ISL_3089153, EPI_ISL_3089154, EPI_ISL_3089155, EPI_ISL_3089156, EPI_ISL_3089157, EPI_ISL_3089158, EPI_ISL_3089159, EPI_ISL_3089160, EPI_ISL_3089161, EPI_ISL_3089162, EPI_ISL_3089163 | | | |
| see above | Nebraska Public Health Laboratory | NPHL COVID-19 Response Team | NPHL COVID-19 Response Team |
| EPI_ISL_3089170, EPI_ISL_3089171, EPI_ISL_3089172, EPI_ISL_3089174, EPI_ISL_3089175, EPI_ISL_3089176, EPI_ISL_3089177, EPI_ISL_3089178, EPI_ISL_3089179, EPI_ISL_3089180, EPI_ISL_3089181, EPI_ISL_3089182, EPI_ISL_3089184, EPI_ISL_3089185, EPI_ISL_3089186, EPI_ISL_3089187, EPI_ISL_3089188, EPI_ISL_3089189, EPI_ISL_3089190, EPI_ISL_3089192, EPI_ISL_3089193, EPI_ISL_3089194, EPI_ISL_3089195, EPI_ISL_3089197, EPI_ISL_3089198, EPI_ISL_3089207, EPI_ISL_3089208, EPI_ISL_3089209, EPI_ISL_3089211, EPI_ISL_3089212, EPI_ISL_3089213, EPI_ISL_3089214 | | | |
| see above | Arizona State Public Health Laboratory | Arizona State Public Health Laboratory | Trung Huynh, Jessica Escobar, Katherine Fullerton, Nobuko Fukushima, Matthew Contursi, Stacy White, Linda Getsinger, Victor Waddell |
| EPI_ISL_3089215, EPI_ISL_3089216, EPI_ISL_3089217, EPI_ISL_3089219, | Rhode Island State Health Laboratory | Rhode Island State Health Laboratory | Ewa King, Richard C. Huard, Kristin Carpenter-Azevedo |

| | | | |
|---|---|---|---|
| EPI_ISL_3089220,<br>EPI_ISL_3089221,<br>EPI_ISL_3089222,<br>EPI_ISL_3089223,<br>EPI_ISL_3089250 | | | |
| EPI_ISL_3089272, EPI_ISL_3089273, EPI_ISL_3089279, EPI_ISL_3089285, EPI_ISL_3089287, EPI_ISL_3089288, EPI_ISL_3089289, EPI_ISL_3089290, EPI_ISL_3089291, EPI_ISL_3089294, EPI_ISL_3089295, EPI_ISL_3089296, EPI_ISL_3089299, EPI_ISL_3089305, EPI_ISL_3089307, EPI_ISL_3089314, EPI_ISL_3089316, EPI_ISL_3089320, EPI_ISL_3089323, EPI_ISL_3089327, EPI_ISL_3089328, EPI_ISL_3089334, EPI_ISL_3089346, EPI_ISL_3089347, EPI_ISL_3089351, EPI_ISL_3089352, EPI_ISL_3089353, EPI_ISL_3089354, EPI_ISL_3089355, EPI_ISL_3089356, EPI_ISL_3089357, EPI_ISL_3089358, EPI_ISL_3089359, EPI_ISL_3089360, EPI_ISL_3089363, EPI_ISL_3089371, EPI_ISL_3089374, EPI_ISL_3089375, EPI_ISL_3089376, EPI_ISL_3089377, EPI_ISL_3089379, EPI_ISL_3089380, EPI_ISL_3089382, EPI_ISL_3089383, EPI_ISL_3089384, EPI_ISL_3089385, EPI_ISL_3089386, EPI_ISL_3089387, EPI_ISL_3089388, EPI_ISL_3089389, EPI_ISL_3089390, EPI_ISL_3089391, EPI_ISL_3089392, EPI_ISL_3089393, EPI_ISL_3089394, EPI_ISL_3089395, EPI_ISL_3089396, EPI_ISL_3089397, EPI_ISL_3089398, EPI_ISL_3089399, EPI_ISL_3089400, EPI_ISL_3089401, EPI_ISL_3089402, EPI_ISL_3089403, EPI_ISL_3089404, EPI_ISL_3089405, EPI_ISL_3089406, EPI_ISL_3089407, EPI_ISL_3089408, EPI_ISL_3089409, EPI_ISL_3089410, EPI_ISL_3089411, EPI_ISL_3089412, EPI_ISL_3089413, EPI_ISL_3089414, EPI_ISL_3089415, EPI_ISL_3089416, EPI_ISL_3089417, EPI_ISL_3089418, EPI_ISL_3089419, EPI_ISL_3089420, EPI_ISL_3089421, EPI_ISL_3089424, EPI_ISL_3089425, EPI_ISL_3089426, EPI_ISL_3089427, EPI_ISL_3089428, EPI_ISL_3089430, EPI_ISL_3089431, EPI_ISL_3089432, EPI_ISL_3089434, EPI_ISL_3089435, EPI_ISL_3089436, EPI_ISL_3089438, EPI_ISL_3089439, EPI_ISL_3089440, EPI_ISL_3089442, EPI_ISL_3089443, EPI_ISL_3089444, EPI_ISL_3089445, EPI_ISL_3089450, EPI_ISL_3089451, EPI_ISL_3089452, EPI_ISL_3089453, EPI_ISL_3089454, EPI_ISL_3089455, EPI_ISL_3089456, EPI_ISL_3089457, EPI_ISL_3089458, EPI_ISL_3089459, EPI_ISL_3089460, EPI_ISL_3089461, EPI_ISL_3089462, EPI_ISL_3089463, EPI_ISL_3089464, EPI_ISL_3089465, EPI_ISL_3089466, EPI_ISL_3089467, EPI_ISL_3089468, EPI_ISL_3089469, EPI_ISL_3089470, EPI_ISL_3089471, EPI_ISL_3089472, EPI_ISL_3089473, EPI_ISL_3089474, EPI_ISL_3089475, EPI_ISL_3089476, EPI_ISL_3089477, EPI_ISL_3089478, EPI_ISL_3089479, EPI_ISL_3089482, EPI_ISL_3089483, EPI_ISL_3089484, EPI_ISL_3089485, EPI_ISL_3089486, EPI_ISL_3089487, EPI_ISL_3089488, EPI_ISL_3089491, EPI_ISL_3089492, EPI_ISL_3089493, EPI_ISL_3089494, EPI_ISL_3089495, EPI_ISL_3089496, EPI_ISL_3089497, EPI_ISL_3089498, EPI_ISL_3089499, EPI_ISL_3089500, EPI_ISL_3089501, EPI_ISL_3089502, EPI_ISL_3089503, EPI_ISL_3089504, EPI_ISL_3089505, EPI_ISL_3089506, EPI_ISL_3089507, EPI_ISL_3089508, EPI_ISL_3089509, EPI_ISL_3089510, EPI_ISL_3089511, EPI_ISL_3089512, EPI_ISL_3089513, EPI_ISL_3089514, EPI_ISL_3089515, EPI_ISL_3089516, EPI_ISL_3089518, EPI_ISL_3089521, EPI_ISL_3089522, EPI_ISL_3089523, EPI_ISL_3089524, EPI_ISL_3089525, EPI_ISL_3089526, EPI_ISL_3089527, EPI_ISL_3089528, EPI_ISL_3089529, EPI_ISL_3089530, EPI_ISL_3089535, EPI_ISL_3089536, EPI_ISL_3089537, EPI_ISL_3089538, EPI_ISL_3089539, EPI_ISL_3089540, EPI_ISL_3089541, EPI_ISL_3089544, EPI_ISL_3089545, EPI_ISL_3089546, EPI_ISL_3089547, EPI_ISL_3089548, EPI_ISL_3089549, EPI_ISL_3089550, EPI_ISL_3089551, EPI_ISL_3089552, EPI_ISL_3089553, EPI_ISL_3089554, EPI_ISL_3089555, EPI_ISL_3089556, EPI_ISL_3089558, EPI_ISL_3089559, EPI_ISL_3089561, EPI_ISL_3089566, EPI_ISL_3089569, EPI_ISL_3089570, EPI_ISL_3089571, EPI_ISL_3089572, EPI_ISL_3089573, EPI_ISL_3089575, EPI_ISL_3089576, EPI_ISL_3089577, EPI_ISL_3089579, EPI_ISL_3089580, EPI_ISL_3089581, EPI_ISL_3089582, EPI_ISL_3089583, EPI_ISL_3089584, EPI_ISL_3089585, EPI_ISL_3089587, EPI_ISL_3089588, EPI_ISL_3089589, EPI_ISL_3089590, EPI_ISL_3089591, EPI_ISL_3089592, EPI_ISL_3089593, EPI_ISL_3089594, EPI_ISL_3089595, EPI_ISL_3089596, EPI_ISL_3089597, EPI_ISL_3089598, EPI_ISL_3089599, EPI_ISL_3089600, EPI_ISL_3089601, EPI_ISL_3089602, EPI_ISL_3089603, EPI_ISL_3089604, EPI_ISL_3089605, EPI_ISL_3089606, EPI_ISL_3089607, EPI_ISL_3089608, EPI_ISL_3089609, EPI_ISL_3089610, EPI_ISL_3089611, EPI_ISL_3089612, EPI_ISL_3089613, EPI_ISL_3089616, EPI_ISL_3089617, EPI_ISL_3089618, EPI_ISL_3089619, EPI_ISL_3089620, EPI_ISL_3089623, EPI_ISL_3089624, EPI_ISL_3089625, EPI_ISL_3089627, EPI_ISL_3089628, EPI_ISL_3089629, EPI_ISL_3089632, EPI_ISL_3089633, EPI_ISL_3089637, EPI_ISL_3089638, EPI_ISL_3089640, EPI_ISL_3089642, EPI_ISL_3089644, EPI_ISL_3089645, EPI_ISL_3089647, EPI_ISL_3089648, EPI_ISL_3089649, EPI_ISL_3089650, EPI_ISL_3089651, EPI_ISL_3089652, EPI_ISL_3089654, EPI_ISL_3089655, EPI_ISL_3089656, EPI_ISL_3089657, EPI_ISL_3089658, EPI_ISL_3089662, EPI_ISL_3089663, EPI_ISL_3089664, EPI_ISL_3089665, EPI_ISL_3089666, EPI_ISL_3089667, EPI_ISL_3089668, EPI_ISL_3089669, EPI_ISL_3089670, EPI_ISL_3089671, EPI_ISL_3089672, EPI_ISL_3089673, EPI_ISL_3089674, EPI_ISL_3089676, EPI_ISL_3089677, EPI_ISL_3089678, EPI_ISL_3089679, EPI_ISL_3089680, EPI_ISL_3089681, EPI_ISL_3089682, EPI_ISL_3089683, EPI_ISL_3089684, EPI_ISL_3089685, EPI_ISL_3089686, EPI_ISL_3089687, EPI_ISL_3089690, EPI_ISL_3089691, EPI_ISL_3089692, EPI_ISL_3089693, EPI_ISL_3089694, EPI_ISL_3089695, EPI_ISL_3089696, EPI_ISL_3089697, EPI_ISL_3089698, EPI_ISL_3089700, EPI_ISL_3089701, EPI_ISL_3089703, EPI_ISL_3089704, EPI_ISL_3089706, EPI_ISL_3089707, EPI_ISL_3089709, EPI_ISL_3089710, EPI_ISL_3089711, EPI_ISL_3089712, EPI_ISL_3089713, EPI_ISL_3089714, EPI_ISL_3089715, EPI_ISL_3089716, EPI_ISL_3089717, EPI_ISL_3089718, EPI_ISL_3089719, EPI_ISL_3089720, EPI_ISL_3089721, EPI_ISL_3089722, EPI_ISL_3089723, EPI_ISL_3089724, EPI_ISL_3089725, EPI_ISL_3089726, EPI_ISL_3089727, EPI_ISL_3089728, EPI_ISL_3089729, EPI_ISL_3089730, EPI_ISL_3089731, EPI_ISL_3089732, EPI_ISL_3089733, EPI_ISL_3089734, EPI_ISL_3089735, EPI_ISL_3089736, EPI_ISL_3089737, EPI_ISL_3089740, EPI_ISL_3089743, EPI_ISL_3089744, EPI_ISL_3089745, EPI_ISL_3089746, EPI_ISL_3089748, EPI_ISL_3089749, EPI_ISL_3089750, EPI_ISL_3089751, EPI_ISL_3089752, EPI_ISL_3089753, EPI_ISL_3089754, EPI_ISL_3089755, EPI_ISL_3089756, EPI_ISL_3089757, EPI_ISL_3089758, EPI_ISL_3089759, EPI_ISL_3089762, EPI_ISL_3089763, EPI_ISL_3089764, EPI_ISL_3089765, EPI_ISL_3089766, EPI_ISL_3089767, EPI_ISL_3089768, EPI_ISL_3089769, EPI_ISL_3089770, EPI_ISL_3089772, EPI_ISL_3089773, EPI_ISL_3089774, EPI_ISL_3089777, EPI_ISL_3089778, EPI_ISL_3089779, EPI_ISL_3089780, EPI_ISL_3089781, EPI_ISL_3089782, EPI_ISL_3089783, EPI_ISL_3089784, EPI_ISL_3089785, EPI_ISL_3089786, EPI_ISL_3089787, EPI_ISL_3089788, EPI_ISL_3089789, EPI_ISL_3089790, EPI_ISL_3089791, EPI_ISL_3089793, EPI_ISL_3089794, EPI_ISL_3089795, EPI_ISL_3089796, EPI_ISL_3089797, EPI_ISL_3089798, EPI_ISL_3089799, EPI_ISL_3089800, EPI_ISL_3089801, EPI_ISL_3089802, EPI_ISL_3089803, EPI_ISL_3089804, EPI_ISL_3089805, EPI_ISL_3089806, EPI_ISL_3089807, EPI_ISL_3089808, EPI_ISL_3089809, EPI_ISL_3089810, EPI_ISL_3089811, EPI_ISL_3089812, EPI_ISL_3089816, EPI_ISL_3089818, EPI_ISL_3089819, EPI_ISL_3089822, EPI_ISL_3089823, EPI_ISL_3089824, EPI_ISL_3089825, EPI_ISL_3089826, EPI_ISL_3089829, EPI_ISL_3089830, EPI_ISL_3089831, EPI_ISL_3089832, EPI_ISL_3089833, EPI_ISL_3089834, EPI_ISL_3089835, EPI_ISL_3089836, EPI_ISL_3089838, EPI_ISL_3089839, EPI_ISL_3089843, EPI_ISL_3089847, EPI_ISL_3089848, EPI_ISL_3089849, EPI_ISL_3089850, EPI_ISL_3089855, EPI_ISL_3089856, EPI_ISL_3089857, EPI_ISL_3089858, EPI_ISL_3089859, EPI_ISL_3089860, EPI_ISL_3089861, EPI_ISL_3089862, EPI_ISL_3089864, EPI_ISL_3089866, EPI_ISL_3089867, EPI_ISL_3089868, EPI_ISL_3089869, EPI_ISL_3089870, EPI_ISL_3089871, EPI_ISL_3089872, EPI_ISL_3089873, EPI_ISL_3089874, EPI_ISL_3089875, EPI_ISL_3089876, EPI_ISL_3089877, EPI_ISL_3089878, EPI_ISL_3089879, EPI_ISL_3089881, EPI_ISL_3089882, EPI_ISL_3089883, EPI_ISL_3089884, EPI_ISL_3089885, EPI_ISL_3089886, EPI_ISL_3089887, EPI_ISL_3089888, EPI_ISL_3089889, EPI_ISL_3089890, EPI_ISL_3089891, EPI_ISL_3089892, EPI_ISL_3089893, EPI_ISL_3089894, EPI_ISL_3089895, EPI_ISL_3089896, EPI_ISL_3089897, EPI_ISL_3089898, EPI_ISL_3089899, EPI_ISL_3089900, EPI_ISL_3089901, EPI_ISL_3089903, EPI_ISL_3089904, EPI_ISL_3089905, EPI_ISL_3089906, EPI_ISL_3089907, EPI_ISL_3089908, EPI_ISL_3089909, EPI_ISL_3089910, EPI_ISL_3089912, EPI_ISL_3089913, EPI_ISL_3089914, EPI_ISL_3089915, EPI_ISL_3089916, EPI_ISL_3089917, EPI_ISL_3089918, EPI_ISL_3089919, EPI_ISL_3089920, EPI_ISL_3089921, EPI_ISL_3089922, EPI_ISL_3089923, EPI_ISL_3089924, EPI_ISL_3089925, EPI_ISL_3089926, EPI_ISL_3089927, EPI_ISL_3089928, EPI_ISL_3089929, EPI_ISL_3089930, EPI_ISL_3089931, EPI_ISL_3089932, EPI_ISL_3089933, EPI_ISL_3089934, EPI_ISL_3089935, EPI_ISL_3089937, EPI_ISL_3089938, EPI_ISL_3089939, EPI_ISL_3089940, EPI_ISL_3089941, EPI_ISL_3089942, EPI_ISL_3089944, EPI_ISL_3089946, EPI_ISL_3089947, EPI_ISL_3089948, EPI_ISL_3089949, EPI_ISL_3089951, EPI_ISL_3089952, EPI_ISL_3089953, EPI_ISL_3089955, EPI_ISL_3089956, EPI_ISL_3089957, EPI_ISL_3089958, EPI_ISL_3089959, EPI_ISL_3089960, EPI_ISL_3089961, EPI_ISL_3089963, EPI_ISL_3089964, EPI_ISL_3089965, EPI_ISL_3089966, EPI_ISL_3089967, EPI_ISL_3089968, EPI_ISL_3089969, EPI_ISL_3089970, EPI_ISL_3089971, EPI_ISL_3089972, EPI_ISL_3089974 | | | |
| see above | Public Health Ontario Laboratory | Public Health Ontario Laboratory | Vanessa G Allen, Philip Banh, Yao Chen, Richard de Borja, Alireza Eshaghi, Nahuel Fittipaldi, Christine Frantz, Jonathan B Gubbay, Jennifer L Guthrie, Lawrence Heisler, Esha Joshi, Michael Laszloffy, Aimin Li, Michael CY Li, Dean Maxwell, Sandeep Nagra, Samir N Patel, Jared Simpson, Karthikeyan Sivaraman, Ashleigh Sullivan, Yogi Sundaravadanam, Sarah Teatero, Andre Villegas, Matthew Watson, Sandra Zittermann |
| EPI_ISL_3090041,<br>EPI_ISL_3090042 | County of San Luis Obispo Public Health Laboratory | County of San Luis Obispo Public Health Laboratory | Katrina Erwin, Frances Sidhu |
| EPI_ISL_3090043 | Marian Medical Center Lab | County of San Luis Obispo Public Health Laboratory | Katrina Erwin, Frances Sidhu |
| EPI_ISL_3090044,<br>EPI_ISL_3090045,<br>EPI_ISL_3090046,<br>EPI_ISL_3090047,<br>EPI_ISL_3090048 | County of San Luis Obispo Public Health Laboratory | County of San Luis Obispo Public Health Laboratory | Katrina Erwin, Frances Sidhu |
| EPI_ISL_3090049,<br>EPI_ISL_3090050,<br>EPI_ISL_3090051,<br>EPI_ISL_3090052,<br>EPI_ISL_3090053,<br>EPI_ISL_3090054,<br>EPI_ISL_3090056,<br>EPI_ISL_3090057,<br>EPI_ISL_3090058,<br>EPI_ISL_3090059 | UW Virology Lab | UW Virology Lab | Pavitra Roychoudhury, Hong Xie, Lasata Shrestha, Shah Mohamed Bakhash, Tien V. Nguyen, Noah R. Baker, Sean Ellis, Meei-Li Huang, Keith R Jerome, Alexander Greninger |
| EPI_ISL_3090060, EPI_ISL_3090061, EPI_ISL_3090062, EPI_ISL_3090063, EPI_ISL_3090064, EPI_ISL_3090067, EPI_ISL_3090068, EPI_ISL_3090070, EPI_ISL_3090071, EPI_ISL_3090072, EPI_ISL_3090074, EPI_ISL_3090075, EPI_ISL_3090076, EPI_ISL_3090077, EPI_ISL_3090078, EPI_ISL_3090079, EPI_ISL_3090080, EPI_ISL_3090081, EPI_ISL_3090082, EPI_ISL_3090083, EPI_ISL_3090084, EPI_ISL_3090085, EPI_ISL_3090087, EPI_ISL_3090088, EPI_ISL_3090089, EPI_ISL_3090090, EPI_ISL_3090091, EPI_ISL_3090092, EPI_ISL_3090093, EPI_ISL_3090096, EPI_ISL_3090098 | | | |
| see above | Arizona State Public Health Laboratory | Arizona State Public Health Laboratory | Trung Huynh, Jessica Escobar, Katherine Fullerton, Nobuko Fukushima, Matthew Contursi, Stacy White, Linda Getsinger, Victor Waddell |
| EPI_ISL_3090728, EPI_ISL_3090729, EPI_ISL_3090730, EPI_ISL_3090731, EPI_ISL_3090732, EPI_ISL_3090733, EPI_ISL_3090734, EPI_ISL_3090735, EPI_ISL_3090736, EPI_ISL_3090737, EPI_ISL_3090738, EPI_ISL_3090739, EPI_ISL_3090740, EPI_ISL_3090741, EPI_ISL_3090742, EPI_ISL_3090743, EPI_ISL_3090745, EPI_ISL_3090746, EPI_ISL_3090747, EPI_ISL_3090748, EPI_ISL_3090749, EPI_ISL_3090750 | | | |
| see above | Oregon State Public Health Laboratory | Oregon State Public Health Laboratory | Rafia Razzaque, Eugene Yeboah, Vanda Makris, Laura Tsaknaridis, John Fontana and Shane Sevey |
| EPI_ISL_3090751,<br>EPI_ISL_3090752,<br>EPI_ISL_3090753,<br>EPI_ISL_3090754,<br>EPI_ISL_3090755,<br>EPI_ISL_3090756 | Queensland Health Forensic and Scientific Services | Queensland Health Forensic and Scientific Services | Chenwei-Wang |
| EPI_ISL_3090842,<br>EPI_ISL_3090843 | Idaho Bureau of Laboratories | Boise VA Medical Center, PALMS | Matthew Burns, Aimee Ceniseros, Christian Loera, Dr. Christopher Ball, Ying Pei, Cheri Lamb McFarlane, James Razor, Emily Bartlette |
| EPI_ISL_3090849 | Centracare Laboratory Services | Minnesota Department of Health, Public Health Laboratory | Alexandra Lorentz, Jacob Garfin, Matt Plumb, and Xiong Wang |
| EPI_ISL_3090850,<br>EPI_ISL_3090851,<br>EPI_ISL_3090852 | Allina Health Laboratory | Minnesota Department of Health, Public Health Laboratory | Alexandra Lorentz, Jacob Garfin, Matt Plumb, and Xiong Wang |
| EPI_ISL_3090853 | Minnesota Department of Health, Public Health Laboratory | Minnesota Department of Health, Public Health Laboratory | Alexandra Lorentz, Jacob Garfin, Matt Plumb, and Xiong Wang |
| EPI_ISL_3090854,<br>EPI_ISL_3090855,<br>EPI_ISL_3090856,<br>EPI_ISL_3090857,<br>EPI_ISL_3090858,<br>EPI_ISL_3090859 | Allina Health Laboratory | Minnesota Department of Health, Public Health Laboratory | Alexandra Lorentz, Jacob Garfin, Matt Plumb, and Xiong Wang |
| EPI_ISL_3090861,<br>EPI_ISL_3090862,<br>EPI_ISL_3090863,<br>EPI_ISL_3090864,<br>EPI_ISL_3090865,<br>EPI_ISL_3090867,<br>EPI_ISL_3090868,<br>EPI_ISL_3090869 | HealthPartners Central Lab | Minnesota Department of Health, Public Health Laboratory | Alexandra Lorentz, Jacob Garfin, Matt Plumb, and Xiong Wang |
| EPI_ISL_3090870, EPI_ISL_3090871, EPI_ISL_3090872, EPI_ISL_3090873, EPI_ISL_3090874, EPI_ISL_3090875, EPI_ISL_3090876, EPI_ISL_3090877, EPI_ISL_3090878, EPI_ISL_3090879, EPI_ISL_3090880, EPI_ISL_3090882, EPI_ISL_3090883, EPI_ISL_3090884, EPI_ISL_3090885, EPI_ISL_3090886, EPI_ISL_3090888, EPI_ISL_3090889, EPI_ISL_3090890, EPI_ISL_3090891, EPI_ISL_3090892, EPI_ISL_3090893, EPI_ISL_3090894, EPI_ISL_3090895, EPI_ISL_3090896, EPI_ISL_3090897, EPI_ISL_3090898, EPI_ISL_3090899, EPI_ISL_3090900, EPI_ISL_3090901, EPI_ISL_3090902, EPI_ISL_3090903, EPI_ISL_3090904, EPI_ISL_3090905, EPI_ISL_3090906, EPI_ISL_3090907, EPI_ISL_3090908, EPI_ISL_3090909, EPI_ISL_3090910, EPI_ISL_3090911, EPI_ISL_3090912, EPI_ISL_3090913, EPI_ISL_3090914, EPI_ISL_3090915, EPI_ISL_3090916, EPI_ISL_3090917, EPI_ISL_3090918, EPI_ISL_3090919, EPI_ISL_3090920, EPI_ISL_3090921, EPI_ISL_3090922, EPI_ISL_3090923, EPI_ISL_3090924, EPI_ISL_3090925, EPI_ISL_3090926, EPI_ISL_3090927, EPI_ISL_3090928, EPI_ISL_3090929, EPI_ISL_3090930, EPI_ISL_3090931, EPI_ISL_3090932, EPI_ISL_3090933, EPI_ISL_3090934, EPI_ISL_3090936, EPI_ISL_3090937, EPI_ISL_3090938, EPI_ISL_3090939, EPI_ISL_3090940, EPI_ISL_3090942, EPI_ISL_3090943, EPI_ISL_3090944, EPI_ISL_3090945, EPI_ISL_3090946, EPI_ISL_3090947, EPI_ISL_3090948, EPI_ISL_3090950, EPI_ISL_3090951, EPI_ISL_3090952, EPI_ISL_3090954, EPI_ISL_3090955, EPI_ISL_3090956, EPI_ISL_3090957, EPI_ISL_3090958, EPI_ISL_3090959, EPI_ISL_3090960, EPI_ISL_3090962, EPI_ISL_3090963, EPI_ISL_3090964, EPI_ISL_3090965, EPI_ISL_3090966, EPI_ISL_3090967, EPI_ISL_3090968, EPI_ISL_3090969, EPI_ISL_3090970, EPI_ISL_3090972, EPI_ISL_3090973, EPI_ISL_3090974, EPI_ISL_3090975, EPI_ISL_3090976, EPI_ISL_3090977, EPI_ISL_3090978, EPI_ISL_3090979, EPI_ISL_3090980, EPI_ISL_3090981, EPI_ISL_3090982, EPI_ISL_3090984, EPI_ISL_3090985, EPI_ISL_3090986, EPI_ISL_3090987, EPI_ISL_3090988, EPI_ISL_3090989 | | | |
| see above | Ochsner Health | Bioinfoexperts, LLC | Judy Crabtree, Amy Feehan, Rebecca Rose, David J. Nolan, Tessa LaFleur, Julia-Garcia-Diaz, Lucio Miele, Susanna L. Lamers |
| EPI_ISL_3090993,<br>EPI_ISL_3090996,<br>EPI_ISL_3090997,<br>EPI_ISL_3090998,<br>EPI_ISL_3090999,<br>EPI_ISL_3091000,<br>EPI_ISL_3091001,<br>EPI_ISL_3091002,<br>EPI_ISL_3091003 | UW Virology Lab | UW Virology Lab | Pavitra Roychoudhury, Hong Xie, Lasata Shrestha, Shah Mohamed Bakhash, Tien V. Nguyen, Noah R. Baker, Sean Ellis, Meei-Li Huang, Keith R Jerome, Alexander Greninger |
| EPI_ISL_3091055,<br>EPI_ISL_3091057 | UCSC Genomics Institute | UCSC Genomics Institute | Mark Akeson, Ikenna Anigbogu, Eric Beraut, Isabel Bjork, Molly Cassatt-Johnstone, Terren Chang, Russell Corbett-Detig, Namrita Dhillon, Maximilian Haeussler, David Haussler, Angie Hinrichs, Miten Jain, Joshua Kapp, A. Marm Kilpatrick, Jakob McBroome, Hugh Olsen, Jeremy Sanford, Beth Shapiro, Michael Stone, Bryan Thornlow, Yatish Turakhia, Ciara Wanket |

| | | | |
|---|---|---|---|
| EPI_ISL_3091171, EPI_ISL_3091172, EPI_ISL_3091173, EPI_ISL_3091174, EPI_ISL_3091175, EPI_ISL_3091176, EPI_ISL_3091178, EPI_ISL_3091179, EPI_ISL_3091180, EPI_ISL_3091181, EPI_ISL_3091182, EPI_ISL_3091183, EPI_ISL_3091184, EPI_ISL_3091185, EPI_ISL_3091186, EPI_ISL_3091187, EPI_ISL_3091188, EPI_ISL_3091189, EPI_ISL_3091190, EPI_ISL_3091191, EPI_ISL_3091192, EPI_ISL_3091193, EPI_ISL_3091194, EPI_ISL_3091195, EPI_ISL_3091196, EPI_ISL_3091197, EPI_ISL_3091200, EPI_ISL_3091201, EPI_ISL_3091202, EPI_ISL_3091203, EPI_ISL_3091204, EPI_ISL_3091206, EPI_ISL_3091207, EPI_ISL_3091208, EPI_ISL_3091209, EPI_ISL_3091210, EPI_ISL_3091211, EPI_ISL_3091212, EPI_ISL_3091213, EPI_ISL_3091214, EPI_ISL_3091215, EPI_ISL_3091216, EPI_ISL_3091217, EPI_ISL_3091218, EPI_ISL_3091221, EPI_ISL_3091222, EPI_ISL_3091223, EPI_ISL_3091224, EPI_ISL_3091225 | | | |
| see above | Houston Health Dept. | Houston Health Dept. | Ryker Penn, Pamela Brown, Adolpho Lara, Yanlai Lai |
| EPI_ISL_3091517, EPI_ISL_3091518, EPI_ISL_3091519, EPI_ISL_3091520, EPI_ISL_3091522, EPI_ISL_3091523, EPI_ISL_3091524, EPI_ISL_3091525, EPI_ISL_3091526, EPI_ISL_3091527, EPI_ISL_3091528, EPI_ISL_3091529, EPI_ISL_3091530, EPI_ISL_3091531, EPI_ISL_3091532, EPI_ISL_3091536, EPI_ISL_3091537, EPI_ISL_3091538, EPI_ISL_3091539, EPI_ISL_3091540, EPI_ISL_3091541, EPI_ISL_3091542 | | | |
| see above | Wexner Medical Center | The Ohio State University College of Medicine | Koenig,S.,Aidja,Z. |
| EPI_ISL_3091566, EPI_ISL_3091570, EPI_ISL_3091571, EPI_ISL_3091572, EPI_ISL_3091573, EPI_ISL_3091574, EPI_ISL_3091575, EPI_ISL_3091576, EPI_ISL_3091577, EPI_ISL_3091580, EPI_ISL_3091581, EPI_ISL_3091582, EPI_ISL_3091592, EPI_ISL_3091595, EPI_ISL_3091596, EPI_ISL_3091597, EPI_ISL_3091601, EPI_ISL_3091602, EPI_ISL_3091603, EPI_ISL_3091604, EPI_ISL_3091606, EPI_ISL_3091609, EPI_ISL_3091612, EPI_ISL_3091613, EPI_ISL_3091615, EPI_ISL_3091617, EPI_ISL_3091621, EPI_ISL_3091623, EPI_ISL_3091624, EPI_ISL_3091626, EPI_ISL_3091627, EPI_ISL_3091631, EPI_ISL_3091633, EPI_ISL_3091634, EPI_ISL_3091635, EPI_ISL_3091636, EPI_ISL_3091637, EPI_ISL_3091638, EPI_ISL_3091640, EPI_ISL_3091646, EPI_ISL_3091648, EPI_ISL_3091651, EPI_ISL_3091653, EPI_ISL_3091654, EPI_ISL_3091656, EPI_ISL_3091658, EPI_ISL_3091659, EPI_ISL_3091660, EPI_ISL_3091661, EPI_ISL_3091662, EPI_ISL_3091663, EPI_ISL_3091664, EPI_ISL_3091666, EPI_ISL_3091668, EPI_ISL_3091669, EPI_ISL_3091670, EPI_ISL_3091671, EPI_ISL_3091672, EPI_ISL_3091673, EPI_ISL_3091674, EPI_ISL_3091676, EPI_ISL_3091683, EPI_ISL_3091686, EPI_ISL_3091689, EPI_ISL_3091693, EPI_ISL_3091695, EPI_ISL_3091696, EPI_ISL_3091698, EPI_ISL_3091703, EPI_ISL_3091704, EPI_ISL_3091706, EPI_ISL_3091707, EPI_ISL_3091708, EPI_ISL_3091711, EPI_ISL_3091713, EPI_ISL_3091716, EPI_ISL_3091720, EPI_ISL_3091721, EPI_ISL_3091722, EPI_ISL_3091723, EPI_ISL_3091725, EPI_ISL_3091727, EPI_ISL_3091729, EPI_ISL_3091734, EPI_ISL_3091736, EPI_ISL_3091738, EPI_ISL_3091739, EPI_ISL_3091742, EPI_ISL_3091743, EPI_ISL_3091744, EPI_ISL_3091745, EPI_ISL_3091747, EPI_ISL_3091749, EPI_ISL_3091750, EPI_ISL_3091752, EPI_ISL_3091755, EPI_ISL_3091756, EPI_ISL_3091758, EPI_ISL_3091760, EPI_ISL_3091766, EPI_ISL_3091768, EPI_ISL_3091769, EPI_ISL_3091770, EPI_ISL_3091775, EPI_ISL_3091782, EPI_ISL_3091783, EPI_ISL_3091784, EPI_ISL_3091785, EPI_ISL_3091786, EPI_ISL_3091787, EPI_ISL_3091788, EPI_ISL_3091791, EPI_ISL_3091792, EPI_ISL_3091793, EPI_ISL_3091794, EPI_ISL_3091798, EPI_ISL_3091799, EPI_ISL_3091800, EPI_ISL_3091803, EPI_ISL_3091805, EPI_ISL_3091808, EPI_ISL_3091810, EPI_ISL_3091812, EPI_ISL_3091814, EPI_ISL_3091824, EPI_ISL_3091826, EPI_ISL_3091830, EPI_ISL_3091834, EPI_ISL_3091836, EPI_ISL_3091840, EPI_ISL_3091843, EPI_ISL_3091844, EPI_ISL_3091845, EPI_ISL_3091852, EPI_ISL_3091853, EPI_ISL_3091854, EPI_ISL_3091858, EPI_ISL_3091859, EPI_ISL_3091861, EPI_ISL_3091864, EPI_ISL_3091865, EPI_ISL_3091866, EPI_ISL_3091867, EPI_ISL_3091868, EPI_ISL_3091870, EPI_ISL_3091875, EPI_ISL_3091876, EPI_ISL_3091877, EPI_ISL_3091881, EPI_ISL_3091883, EPI_ISL_3091884, EPI_ISL_3091885, EPI_ISL_3091888, EPI_ISL_3091893, EPI_ISL_3091894, EPI_ISL_3091898, EPI_ISL_3091899, EPI_ISL_3091900, EPI_ISL_3091903, EPI_ISL_3091915, EPI_ISL_3091918, EPI_ISL_3091921, EPI_ISL_3091923, EPI_ISL_3091926, EPI_ISL_3091929, EPI_ISL_3091930, EPI_ISL_3091931, EPI_ISL_3091932, EPI_ISL_3091936, EPI_ISL_3091939, EPI_ISL_3091942, EPI_ISL_3091946, EPI_ISL_3091947, EPI_ISL_3091948, EPI_ISL_3091949, EPI_ISL_3091951, EPI_ISL_3091953, EPI_ISL_3091955, EPI_ISL_3091958, EPI_ISL_3091959, EPI_ISL_3091961, EPI_ISL_3091963, EPI_ISL_3091964, EPI_ISL_3091966, EPI_ISL_3091968, EPI_ISL_3091972, EPI_ISL_3091973, EPI_ISL_3091979, EPI_ISL_3091980, EPI_ISL_3091981, EPI_ISL_3091982, EPI_ISL_3091984, EPI_ISL_3091987, EPI_ISL_3091993, EPI_ISL_3091995, EPI_ISL_3091997, EPI_ISL_3091998, EPI_ISL_3092000, EPI_ISL_3092003, EPI_ISL_3092004, EPI_ISL_3092008, EPI_ISL_3092009, EPI_ISL_3092014, EPI_ISL_3092018, EPI_ISL_3092023, EPI_ISL_3092024, EPI_ISL_3092025, EPI_ISL_3092028, EPI_ISL_3092030, EPI_ISL_3092031, EPI_ISL_3092033, EPI_ISL_3092034, EPI_ISL_3092035, EPI_ISL_3092037, EPI_ISL_3092038, EPI_ISL_3092039, EPI_ISL_3092040, EPI_ISL_3092053 | | | |
| see above | Respiratory Viruses Branch, Centers for Disease Control and Prevention | Respiratory Viruses Branch, Centers for Disease Control and Prevention | Howard,D., Batra,D., Cook,P.W., Moser,K., Paskey,A., Caravas,J., Rambo-Martin,B., Morrison,S., Gulvick,C., Sammons,S., Unoarumhi,Y., Wagner,D., Schmerer,M., Rosenthal,S.H., Gerasimova,A., Kagan,R.M., Anderson,B., Hua,M., Liu,Y., Bernstein,L., Livingston,K., Perez,A., Shlyakhter,I.A., Rolando,R.V., Owen,R., Tanpaiboon,P., Lacbawan,F., Paden,C.R., MacCannell,D. |
| EPI_ISL_3092056, EPI_ISL_3092057 | Hospital del Oriente de Asturias | Laboratorio de Virología HUCA | Sandoval M, Castelló C, Gómez de Oña J, Boga JA, Rojo S, Alvarez-Arguelles ME, Abreu F, Costales I, Perez-Martínez Z, Martín-Rodríguez G, Coto E, Melón S |
| EPI_ISL_3092058, EPI_ISL_3092059 | Hospital de Cabueñes | Laboratorio de Virología HUCA | Sandoval M, Castelló C, Gómez de Oña J, Boga JA, Rojo S, Alvarez-Arguelles ME, Abreu F, Costales I, Perez-Martínez Z, Martín-Rodríguez G, Coto E, Melón S |
| EPI_ISL_3092060, EPI_ISL_3092061, EPI_ISL_3092062, EPI_ISL_3092063, EPI_ISL_3092064, EPI_ISL_3092065, EPI_ISL_3092066, EPI_ISL_3092067, EPI_ISL_3092068, EPI_ISL_3092069, EPI_ISL_3092070, EPI_ISL_3092071, EPI_ISL_3092072, EPI_ISL_3092073, EPI_ISL_3092074, EPI_ISL_3092075, EPI_ISL_3092076, EPI_ISL_3092077, EPI_ISL_3092080, EPI_ISL_3092081, EPI_ISL_3092083, EPI_ISL_3092084, EPI_ISL_3092085, EPI_ISL_3092086, EPI_ISL_3092087, EPI_ISL_3092088, EPI_ISL_3092089 | | | |
| see above | Wyoming Public Health Laboratory | Wyoming Public Health Laboratory | Jim Mildenberger, Taylor Fearing, Channing Weber, Ashley Norberg, Chayse Rowley, Marley Goetz, Brian Dominguez, Elliot Thomasson, Sam Britz, Cari Sloma, Robert Petit, and Rob Christensen |
| EPI_ISL_3092090 | Hospital de Cabueñes | Laboratorio de Virología HUCA | Sandoval M, Castelló C, Gómez de Oña J, Boga JA, Rojo S, Alvarez-Arguelles ME, Abreu F, Costales I, Perez-Martínez Z, Martín-Rodríguez G, Coto E, Melón S |
| EPI_ISL_3092091, EPI_ISL_3092092, EPI_ISL_3092093, EPI_ISL_3092094, EPI_ISL_3092095, EPI_ISL_3092097, EPI_ISL_3092098, EPI_ISL_3092099, EPI_ISL_3092100, EPI_ISL_3092101, EPI_ISL_3092102, EPI_ISL_3092103, EPI_ISL_3092104, EPI_ISL_3092105, EPI_ISL_3092106, EPI_ISL_3092107, EPI_ISL_3092108, EPI_ISL_3092109, EPI_ISL_3092110, EPI_ISL_3092111, EPI_ISL_3092112, EPI_ISL_3092113, EPI_ISL_3092114, EPI_ISL_3092115, EPI_ISL_3092116, EPI_ISL_3092117, EPI_ISL_3092118, EPI_ISL_3092119, EPI_ISL_3092120, EPI_ISL_3092121, EPI_ISL_3092122, EPI_ISL_3092123, EPI_ISL_3092124, EPI_ISL_3092125, EPI_ISL_3092126, EPI_ISL_3092127, EPI_ISL_3092128, EPI_ISL_3092130, EPI_ISL_3092132, EPI_ISL_3092135, EPI_ISL_3092136, EPI_ISL_3092137, EPI_ISL_3092138, EPI_ISL_3092139, EPI_ISL_3092140, EPI_ISL_3092141, EPI_ISL_3092142, EPI_ISL_3092143, EPI_ISL_3092144, EPI_ISL_3092145, EPI_ISL_3092146, EPI_ISL_3092147, EPI_ISL_3092148, EPI_ISL_3092149, EPI_ISL_3092150, EPI_ISL_3092151, EPI_ISL_3092152, EPI_ISL_3092153, EPI_ISL_3092154, EPI_ISL_3092155, EPI_ISL_3092156, EPI_ISL_3092157, EPI_ISL_3092158, EPI_ISL_3092159, EPI_ISL_3092160, EPI_ISL_3092161, EPI_ISL_3092162, EPI_ISL_3092164, EPI_ISL_3092168, EPI_ISL_3092169, EPI_ISL_3092170, EPI_ISL_3092171, EPI_ISL_3092172, EPI_ISL_3092173, EPI_ISL_3092174, EPI_ISL_3092175, EPI_ISL_3092176, EPI_ISL_3092177, EPI_ISL_3092178, EPI_ISL_3092179, EPI_ISL_3092180, EPI_ISL_3092181, EPI_ISL_3092182, EPI_ISL_3092183, EPI_ISL_3092184, EPI_ISL_3092185, EPI_ISL_3092186, EPI_ISL_3092187, EPI_ISL_3092188, EPI_ISL_3092189, EPI_ISL_3092190, EPI_ISL_3092191, EPI_ISL_3092192, EPI_ISL_3092193, EPI_ISL_3092194, EPI_ISL_3092195, EPI_ISL_3092196, EPI_ISL_3092197, EPI_ISL_3092198, EPI_ISL_3092199, EPI_ISL_3092200, EPI_ISL_3092201, EPI_ISL_3092202, EPI_ISL_3092203, EPI_ISL_3092204, EPI_ISL_3092205, EPI_ISL_3092206, EPI_ISL_3092207, EPI_ISL_3092208, EPI_ISL_3092209, EPI_ISL_3092212, EPI_ISL_3092213, EPI_ISL_3092214, EPI_ISL_3092215, EPI_ISL_3092216, EPI_ISL_3092217, EPI_ISL_3092218, EPI_ISL_3092219, EPI_ISL_3092220, EPI_ISL_3092221, EPI_ISL_3092222, EPI_ISL_3092223, EPI_ISL_3092224, EPI_ISL_3092225, EPI_ISL_3092227, EPI_ISL_3092228, EPI_ISL_3092229, EPI_ISL_3092230, EPI_ISL_3092231, EPI_ISL_3092232, EPI_ISL_3092233, EPI_ISL_3092234, EPI_ISL_3092235, EPI_ISL_3092236, EPI_ISL_3092237, EPI_ISL_3092238, EPI_ISL_3092239, EPI_ISL_3092240, EPI_ISL_3092241, EPI_ISL_3092242, EPI_ISL_3092243, EPI_ISL_3092244, EPI_ISL_3092245, EPI_ISL_3092246, EPI_ISL_3092247, EPI_ISL_3092248, EPI_ISL_3092249, EPI_ISL_3092250, EPI_ISL_3092251, EPI_ISL_3092252, EPI_ISL_3092253, EPI_ISL_3092254, EPI_ISL_3092255, EPI_ISL_3092256, EPI_ISL_3092257, EPI_ISL_3092258, EPI_ISL_3092259, EPI_ISL_3092260, EPI_ISL_3092261, EPI_ISL_3092262, EPI_ISL_3092263, EPI_ISL_3092264, EPI_ISL_3092265, EPI_ISL_3092266, EPI_ISL_3092267, EPI_ISL_3092268, EPI_ISL_3092269, EPI_ISL_3092271, EPI_ISL_3092272, EPI_ISL_3092273, EPI_ISL_3092274, EPI_ISL_3092275, EPI_ISL_3092276, EPI_ISL_3092277, EPI_ISL_3092278, EPI_ISL_3092279, EPI_ISL_3092280, EPI_ISL_3092282, EPI_ISL_3092283, EPI_ISL_3092284, EPI_ISL_3092285, EPI_ISL_3092286, EPI_ISL_3092287, EPI_ISL_3092288, EPI_ISL_3092289, EPI_ISL_3092290, EPI_ISL_3092291, EPI_ISL_3092292, EPI_ISL_3092293, EPI_ISL_3092294, EPI_ISL_3092295, EPI_ISL_3092296, EPI_ISL_3092297, EPI_ISL_3092298, EPI_ISL_3092299, EPI_ISL_3092300, EPI_ISL_3092301, EPI_ISL_3092302, EPI_ISL_3092303, EPI_ISL_3092304, EPI_ISL_3092305, EPI_ISL_3092306, EPI_ISL_3092307, EPI_ISL_3092308, EPI_ISL_3092311, EPI_ISL_3092312, EPI_ISL_3092313, EPI_ISL_3092316, EPI_ISL_3092317, EPI_ISL_3092318, EPI_ISL_3092319, EPI_ISL_3092320, EPI_ISL_3092322, EPI_ISL_3092323, EPI_ISL_3092324, EPI_ISL_3092325, EPI_ISL_3092326, EPI_ISL_3092327, EPI_ISL_3092328, EPI_ISL_3092329, EPI_ISL_3092330, EPI_ISL_3092331, EPI_ISL_3092332, EPI_ISL_3092333, EPI_ISL_3092334, EPI_ISL_3092335, EPI_ISL_3092337, EPI_ISL_3092338, EPI_ISL_3092339, EPI_ISL_3092340, EPI_ISL_3092341, EPI_ISL_3092342, EPI_ISL_3092343, EPI_ISL_3092344, EPI_ISL_3092345, EPI_ISL_3092346, EPI_ISL_3092347, EPI_ISL_3092348, EPI_ISL_3092349, EPI_ISL_3092350, EPI_ISL_3092351, EPI_ISL_3092353, EPI_ISL_3092354, EPI_ISL_3092355, EPI_ISL_3092356, EPI_ISL_3092357, EPI_ISL_3092358, EPI_ISL_3092359, EPI_ISL_3092360, EPI_ISL_3092361, EPI_ISL_3092362, EPI_ISL_3092364, EPI_ISL_3092365, EPI_ISL_3092366, EPI_ISL_3092367, EPI_ISL_3092368, EPI_ISL_3092369, EPI_ISL_3092370, EPI_ISL_3092371, EPI_ISL_3092372, EPI_ISL_3092373, EPI_ISL_3092374, EPI_ISL_3092375, EPI_ISL_3092376, EPI_ISL_3092378, EPI_ISL_3092379, EPI_ISL_3092380, EPI_ISL_3092381, EPI_ISL_3092382, EPI_ISL_3092383, EPI_ISL_3092384, EPI_ISL_3092385, EPI_ISL_3092386, EPI_ISL_3092387, EPI_ISL_3092388, EPI_ISL_3092389, EPI_ISL_3092390, EPI_ISL_3092391, EPI_ISL_3092392, EPI_ISL_3092393, EPI_ISL_3092394, EPI_ISL_3092395, EPI_ISL_3092396, EPI_ISL_3092398, EPI_ISL_3092399, EPI_ISL_3092400, EPI_ISL_3092401, EPI_ISL_3092402, EPI_ISL_3092403, EPI_ISL_3092404, EPI_ISL_3092405, EPI_ISL_3092407, EPI_ISL_3092408, EPI_ISL_3092409, EPI_ISL_3092410, EPI_ISL_3092411, EPI_ISL_3092412, EPI_ISL_3092413, EPI_ISL_3092414, EPI_ISL_3092415, EPI_ISL_3092416, EPI_ISL_3092417, EPI_ISL_3092418 | | | |
| see above | Cotugno | TIGEM | Antonio Grimaldi Patrizia Annunziata Francesco Panariello Biancamaria Pierri Claudia Tiberio Teresa Giuliano Valentina Bouche Chiara Colantuono Maria Concetta Cuomo Denise Di Concilio Lucio Di Filippo Anna Manfredi Marcello Salvi Antonio Limone Luigi Atripaldi Pellegrino Cerino Andrea Ballabio Davide Cacchiarelli |
| EPI_ISL_3092453 | Hospital del Oriente de Asturias | Laboratorio de Virología HUCA | Sandoval M, Castelló C, Gómez de Oña J, Boga JA, Rojo S, Alvarez-Arguelles ME, Abreu F, Costales I, Perez-Martínez Z, Martín-Rodríguez G, Coto E, Melón S |
| EPI_ISL_3092454, EPI_ISL_3092455, EPI_ISL_3092458 | Laboratorio de Virología HUCA | Laboratorio de Virología HUCA | Sandoval M, Castelló C, Gómez de Oña J, Boga JA, Rojo S, Alvarez-Arguelles ME, Abreu F, Costales I, Perez-Martínez Z, Martín-Rodríguez G, Coto E, Melón S |
| EPI_ISL_3092464 | DINAS KESEHATAN DKI JAKARTA | Genomik Solidaritas Indonesia Laboratorium | Normastuti Adhini Tantyo, Dhahlia Agustina Cahyono, Alfin Mohammad Abdillah, Vania Gavrila Wikasa, Carissa Sintca Wijaya, Mohammad Anang Syarifudin, Annisa Muthiah Sukirman, Gracia Felias Enos Korompis, Widyastuti, Dwi Oktavia, Tiranti Vindha, Ngabila Salama, Anuraj Shankar, Meutia Ayuputeri Kumaheri, Ariel Pradipta |
| EPI_ISL_3092580 | Laboratorio de Virología HUCA | Laboratorio de Virología HUCA | Sandoval M, Castelló C, Gómez de Oña J, Boga JA, Rojo S, Alvarez-Arguelles ME, Abreu F, Costales I, Perez-Martínez Z, Martín-Rodríguez G, Coto E, Melón S |
| EPI_ISL_3092582, EPI_ISL_3092583, EPI_ISL_3092584, EPI_ISL_3092585, EPI_ISL_3092586, EPI_ISL_3092587, EPI_ISL_3092588, EPI_ISL_3092589, EPI_ISL_3092590, EPI_ISL_3092591, EPI_ISL_3092592, EPI_ISL_3092593, EPI_ISL_3092594, EPI_ISL_3092595, EPI_ISL_3092596, EPI_ISL_3092597, EPI_ISL_3092598, EPI_ISL_3092599, EPI_ISL_3092600, EPI_ISL_3092601, EPI_ISL_3092602, EPI_ISL_3092603, EPI_ISL_3092604, EPI_ISL_3092605, EPI_ISL_3092606, EPI_ISL_3092607, EPI_ISL_3092608, EPI_ISL_3092609, EPI_ISL_3092610, EPI_ISL_3092611, EPI_ISL_3092612, EPI_ISL_3092613 | | | |
| see above | Lighthouse Lab in Milton Keynes | Wellcome Sanger Institute for the COVID-19 Genomics UK (COG-UK) Consortium | The Lighthouse Lab in Milton Keynes and Alex Alderton, Roberto Amato, Jeffrey Barrett, Sonia Goncalves, Ewan Harrison, David K. Jackson, Ian Johnston, Dominic Kwiatkowski, Cordelia Langford, John Sillitoe on behalf of the Wellcome Sanger Institute COVID-19 Surveillance Team |
| EPI_ISL_3092614 | Hospital del Oriente de Asturias | Laboratorio de Virología HUCA | Sandoval M, Castelló C, Gómez de Oña J, Boga JA, Rojo S, Alvarez-Arguelles ME, Abreu F, Costales I, Perez-Martínez Z, Martín-Rodríguez G, Coto E, Melón S |
| EPI_ISL_3092615, EPI_ISL_3092616, EPI_ISL_3092617, EPI_ISL_3092618, EPI_ISL_3092619, EPI_ISL_3092620, EPI_ISL_3092621, EPI_ISL_3092622, EPI_ISL_3092623, EPI_ISL_3092624, EPI_ISL_3092625, EPI_ISL_3092626, EPI_ISL_3092627, EPI_ISL_3092628, EPI_ISL_3092629, EPI_ISL_3092630, EPI_ISL_3092631, EPI_ISL_3092632, EPI_ISL_3092633, EPI_ISL_3092634, EPI_ISL_3092635, EPI_ISL_3092636, EPI_ISL_3092637, EPI_ISL_3092638, EPI_ISL_3092639, EPI_ISL_3092640, EPI_ISL_3092641, EPI_ISL_3092642, EPI_ISL_3092643, EPI_ISL_3092644, EPI_ISL_3092645, EPI_ISL_3092646, EPI_ISL_3092647, EPI_ISL_3092648, EPI_ISL_3092649, EPI_ISL_3092650, EPI_ISL_3092651, EPI_ISL_3092652, EPI_ISL_3092653, EPI_ISL_3092654, EPI_ISL_3092655, EPI_ISL_3092656, EPI_ISL_3092657, EPI_ISL_3092658, EPI_ISL_3092659, EPI_ISL_3092660, EPI_ISL_3092661, EPI_ISL_3092662, EPI_ISL_3092663, EPI_ISL_3092664, EPI_ISL_3092665, EPI_ISL_3092666, EPI_ISL_3092667, EPI_ISL_3092668, EPI_ISL_3092669, EPI_ISL_3092670, EPI_ISL_3092671, EPI_ISL_3092672, EPI_ISL_3092673, EPI_ISL_3092674, EPI_ISL_3092675, EPI_ISL_3092676, EPI_ISL_3092677, EPI_ISL_3092678, EPI_ISL_3092679, EPI_ISL_3092680, EPI_ISL_3092681, EPI_ISL_3092682, EPI_ISL_3092683, EPI_ISL_3092684, EPI_ISL_3092685, EPI_ISL_3092686, EPI_ISL_3092687, EPI_ISL_3092688, EPI_ISL_3092689, EPI_ISL_3092690, EPI_ISL_3092691, EPI_ISL_3092692, EPI_ISL_3092693, EPI_ISL_3092694, EPI_ISL_3092695, EPI_ISL_3092696, EPI_ISL_3092697, EPI_ISL_3092698, EPI_ISL_3092699, EPI_ISL_3092700, EPI_ISL_3092701, EPI_ISL_3092702, EPI_ISL_3092703, EPI_ISL_3092704, EPI_ISL_3092705, EPI_ISL_3092706, EPI_ISL_3092707, EPI_ISL_3092708, EPI_ISL_3092709, EPI_ISL_3092710, EPI_ISL_3092711, EPI_ISL_3092712, EPI_ISL_3092713, EPI_ISL_3092714, EPI_ISL_3092715, EPI_ISL_3092716, EPI_ISL_3092717, EPI_ISL_3092718, EPI_ISL_3092719, EPI_ISL_3092720, EPI_ISL_3092721, EPI_ISL_3092722, EPI_ISL_3092723, EPI_ISL_3092724, EPI_ISL_3092725, EPI_ISL_3092726, EPI_ISL_3092727, EPI_ISL_3092728, EPI_ISL_3092729, EPI_ISL_3092730, EPI_ISL_3092731, EPI_ISL_3092732, EPI_ISL_3092733, EPI_ISL_3092734, EPI_ISL_3092735, EPI_ISL_3092736, EPI_ISL_3092737, EPI_ISL_3092738, EPI_ISL_3092740, EPI_ISL_3092741, EPI_ISL_3092742, EPI_ISL_3092743, EPI_ISL_3092744, EPI_ISL_3092745, EPI_ISL_3092746, EPI_ISL_3092747, EPI_ISL_3092748, EPI_ISL_3092749, EPI_ISL_3092750, EPI_ISL_3092751, EPI_ISL_3092752, EPI_ISL_3092753, EPI_ISL_3092754, EPI_ISL_3092755, EPI_ISL_3092756, EPI_ISL_3092757, EPI_ISL_3092758, EPI_ISL_3092759, EPI_ISL_3092760, EPI_ISL_3092761, EPI_ISL_3092762, EPI_ISL_3092763, EPI_ISL_3092764, EPI_ISL_3092765, EPI_ISL_3092766, EPI_ISL_3092767, EPI_ISL_3092768, EPI_ISL_3092769, EPI_ISL_3092770, EPI_ISL_3092771, EPI_ISL_3092772, EPI_ISL_3092773, EPI_ISL_3092774, EPI_ISL_3092775, EPI_ISL_3092776, EPI_ISL_3092777, EPI_ISL_3092778, EPI_ISL_3092779, EPI_ISL_3092780, EPI_ISL_3092781, EPI_ISL_3092782, EPI_ISL_3092783, EPI_ISL_3092784, EPI_ISL_3092785, EPI_ISL_3092786, EPI_ISL_3092787, EPI_ISL_3092788, EPI_ISL_3092789, EPI_ISL_3092790, EPI_ISL_3092791, EPI_ISL_3092792, EPI_ISL_3092793, EPI_ISL_3092794, EPI_ISL_3092795, EPI_ISL_3092796, EPI_ISL_3092797, EPI_ISL_3092798, EPI_ISL_3092799, EPI_ISL_3092800, EPI_ISL_3092801, EPI_ISL_3092802, EPI_ISL_3092803, EPI_ISL_3092804, EPI_ISL_3092805, EPI_ISL_3092806, EPI_ISL_3092807, EPI_ISL_3092808, EPI_ISL_3092809, EPI_ISL_3092810, EPI_ISL_3092811, EPI_ISL_3092812, EPI_ISL_3092813, EPI_ISL_3092814, EPI_ISL_3092815, EPI_ISL_3092816, EPI_ISL_3092817, EPI_ISL_3092818, EPI_ISL_3092819, EPI_ISL_3092820, EPI_ISL_3092821, EPI_ISL_3092822, EPI_ISL_3092823, EPI_ISL_3092824, EPI_ISL_3092825, EPI_ISL_3092826, EPI_ISL_3092827, EPI_ISL_3092828, EPI_ISL_3092829, EPI_ISL_3092830, EPI_ISL_3092831, EPI_ISL_3092832, EPI_ISL_3092833, EPI_ISL_3092834, EPI_ISL_3092835, EPI_ISL_3092836, EPI_ISL_3092837, EPI_ISL_3092838, EPI_ISL_3092839, EPI_ISL_3092840, EPI_ISL_3092841, EPI_ISL_3092842, EPI_ISL_3092843, EPI_ISL_3092844, EPI_ISL_3092845, EPI_ISL_3092846, EPI_ISL_3092847, EPI_ISL_3092848, EPI_ISL_3092849, EPI_ISL_3092850, EPI_ISL_3092851, EPI_ISL_3092852, EPI_ISL_3092853, EPI_ISL_3092854, EPI_ISL_3092855, EPI_ISL_3092856, EPI_ISL_3092857, EPI_ISL_3092858, EPI_ISL_3092859, EPI_ISL_3092860, EPI_ISL_3092861, EPI_ISL_3092862, EPI_ISL_3092863, EPI_ISL_3092864, EPI_ISL_3092865, EPI_ISL_3092866, EPI_ISL_3092867, EPI_ISL_3092868, EPI_ISL_3092869, EPI_ISL_3092870, EPI_ISL_3092871, EPI_ISL_3092872, EPI_ISL_3092873, EPI_ISL_3092874, EPI_ISL_3092875, EPI_ISL_3092876 | | | |
| see above | Lighthouse Lab in Milton Keynes | Wellcome Sanger Institute for the COVID-19 Genomics UK (COG-UK) Consortium | The Lighthouse Lab in Milton Keynes and Alex Alderton, Roberto Amato, Jeffrey Barrett, Sonia Goncalves, Ewan Harrison, David K. Jackson, Ian Johnston, Dominic Kwiatkowski, Cordelia Langford, John Sillitoe on behalf of the Wellcome Sanger Institute COVID-19 Surveillance Team |
| EPI_ISL_3092877 | Berkshire and Surrey Pathology Services Lighthouse Laboratory | Wellcome Sanger Institute for the COVID-19 Genomics UK (COG-UK) Consortium | Berkshire and Surrey Pathology Services Lighthouse Laboratory and Alex Alderton, Roberto Amato, Jeffrey Barrett, Sonia Goncalves, Ewan Harrison, David K. Jackson, Ian Johnston, Dominic Kwiatkowski, Cordelia Langford, John Sillitoe on behalf of the Wellcome Sanger Institute COVID-19 Surveillance Team |
| EPI_ISL_3092878 | Lighthouse Lab in Alderley Park | Wellcome Sanger Institute for the COVID-19 Genomics UK (COG-UK) Consortium | Jacquelyn Wynn, Mairead Hyland, The Lighthouse Lab in Alderley Park and Alex Alderton, Roberto Amato, Jeffrey Barrett, Sonia Goncalves, Ewan Harrison, David K. Jackson, Ian Johnston, Dominic Kwiatkowski, Cordelia Langford, John Sillitoe on behalf of the Wellcome Sanger Institute COVID-19 Surveillance Team |
| EPI_ISL_3092879 | Berkshire and Surrey Pathology Services Lighthouse Laboratory | Wellcome Sanger Institute for the COVID-19 Genomics UK (COG-UK) Consortium | Berkshire and Surrey Pathology Services Lighthouse Laboratory and Alex Alderton, Roberto Amato, Jeffrey Barrett, Sonia Goncalves, Ewan Harrison, David K. Jackson, Ian Johnston, Dominic Kwiatkowski, Cordelia Langford, John Sillitoe on behalf of the Wellcome Sanger Institute COVID-19 Surveillance Team |
| EPI_ISL_3092880, EPI_ISL_3092881 | Lighthouse Lab in Alderley Park | Wellcome Sanger Institute for the COVID-19 Genomics UK (COG-UK) Consortium | Jacquelyn Wynn, Mairead Hyland, The Lighthouse Lab in Alderley Park and Alex Alderton, Roberto Amato, Jeffrey Barrett, Sonia Goncalves, Ewan Harrison, David K. Jackson, Ian Johnston, Dominic Kwiatkowski, Cordelia Langford, John Sillitoe on behalf of the Wellcome Sanger Institute COVID-19 Surveillance Team |
| EPI_ISL_3092882, EPI_ISL_3092883 | Lighthouse Lab in Milton Keynes | Wellcome Sanger Institute for the COVID-19 Genomics UK (COG-UK) Consortium | The Lighthouse Lab in Milton Keynes and Alex Alderton, Roberto Amato, Jeffrey Barrett, Sonia Goncalves, Ewan Harrison, David K. Jackson, Ian Johnston, Dominic Kwiatkowski, Cordelia Langford, John Sillitoe on behalf of the Wellcome Sanger Institute COVID-19 Surveillance Team |
| EPI_ISL_3092884 | Lighthouse Lab in Glasgow | Wellcome Sanger Institute for the COVID-19 Genomics UK (COG-UK) Consortium | Harper VanSteenhouse, Yumi Kasai, David Gray, Carol Clugston, Anna Dominiczak and Alex Alderton, Roberto Amato, Jeffrey Barrett, Sonia Goncalves, Ewan Harrison, David K. Jackson, Ian Johnston, Dominic Kwiatkowski, Cordelia Langford, John Sillitoe on behalf of the Wellcome Sanger Institute COVID-19 Surveillance Team |
| EPI_ISL_3092885 | Lighthouse Lab in Alderley Park | Wellcome Sanger Institute for the COVID-19 Genomics UK (COG-UK) Consortium | Jacquelyn Wynn, Mairead Hyland, The Lighthouse Lab in Alderley Park and Alex Alderton, Roberto Amato, Jeffrey Barrett, Sonia Goncalves, Ewan Harrison, David K. Jackson, Ian Johnston, Dominic Kwiatkowski, Cordelia Langford, John Sillitoe on behalf of the Wellcome Sanger Institute COVID-19 Surveillance Team |
| EPI_ISL_3092886 | Berkshire and Surrey Pathology Services Lighthouse Laboratory | Wellcome Sanger Institute for the COVID-19 Genomics UK (COG-UK) Consortium | Berkshire and Surrey Pathology Services Lighthouse Laboratory and Alex Alderton, Roberto Amato, Jeffrey Barrett, Sonia Goncalves, Ewan Harrison, David K. Jackson, Ian Johnston, Dominic Kwiatkowski, Cordelia Langford, John Sillitoe on behalf of the Wellcome Sanger Institute COVID-19 Surveillance Team |
| EPI_ISL_3092887 | Lighthouse Lab in Milton Keynes | Wellcome Sanger Institute for the COVID-19 Genomics UK (COG-UK) Consortium | The Lighthouse Lab in Milton Keynes and Alex Alderton, Roberto Amato, Jeffrey Barrett, Sonia Goncalves, Ewan Harrison, David K. Jackson, Ian Johnston, Dominic Kwiatkowski, Cordelia Langford, John Sillitoe on behalf of the Wellcome Sanger Institute COVID-19 Surveillance Team |
| EPI_ISL_3092888 | Lighthouse Lab in Glasgow | Wellcome Sanger Institute for the COVID-19 Genomics UK (COG-UK) Consortium | Harper VanSteenhouse, Yumi Kasai, David Gray, Carol Clugston, Anna Dominiczak and Alex Alderton, Roberto Amato, Jeffrey Barrett, Sonia Goncalves, Ewan Harrison, David K. Jackson, Ian Johnston, Dominic Kwiatkowski, Cordelia Langford, John Sillitoe on behalf of the Wellcome Sanger Institute COVID-19 Surveillance Team |
| EPI_ISL_3092889 | Lighthouse Lab in Alderley Park | Wellcome Sanger Institute for the COVID-19 Genomics UK (COG-UK) Consortium | Jacquelyn Wynn, Mairead Hyland, The Lighthouse Lab in Alderley Park and Alex Alderton, Roberto Amato, Jeffrey Barrett, Sonia Goncalves, Ewan Harrison, David K. Jackson, Ian Johnston, Dominic Kwiatkowski, Cordelia Langford, John Sillitoe on behalf of the Wellcome Sanger Institute COVID-19 Surveillance Team |
| EPI_ISL_3092890 | Berkshire and Surrey Pathology Services Lighthouse Laboratory | Wellcome Sanger Institute for the COVID-19 Genomics UK (COG-UK) Consortium | Berkshire and Surrey Pathology Services Lighthouse Laboratory and Alex Alderton, Roberto Amato, Jeffrey Barrett, Sonia Goncalves, Ewan Harrison, David K. Jackson, Ian Johnston, Dominic Kwiatkowski, Cordelia Langford, John Sillitoe on behalf of the Wellcome Sanger Institute COVID-19 Surveillance Team |
| EPI_ISL_3092891, EPI_ISL_3092892 | Lighthouse Lab in Milton Keynes | Wellcome Sanger Institute for the COVID-19 Genomics UK (COG-UK) Consortium | The Lighthouse Lab in Milton Keynes and Alex Alderton, Roberto Amato, Jeffrey Barrett, Sonia Goncalves, Ewan Harrison, David K. Jackson, Ian Johnston, Dominic Kwiatkowski, Cordelia Langford, John Sillitoe on behalf of the Wellcome Sanger Institute COVID-19 Surveillance Team |
| EPI_ISL_3092893 | Berkshire and Surrey Pathology Services Lighthouse Laboratory | Wellcome Sanger Institute for the COVID-19 Genomics UK (COG-UK) | Berkshire and Surrey Pathology Services Lighthouse Laboratory and Alex Alderton, Roberto Amato, Jeffrey Barrett, Sonia Goncalves, Ewan Harrison, David K. Jackson, Ian Johnston, Dominic Kwiatkowski, Cordelia Langford, John Sillitoe on behalf |

| | | | |
|---|---|---|---|
| | | Consortium | of the Wellcome Sanger Institute COVID-19 Surveillance Team |
| EPI_ISL_3092894, EPI_ISL_3092895, EPI_ISL_3092896, EPI_ISL_3092897, EPI_ISL_3092898 | Lighthouse Lab in Milton Keynes | Wellcome Sanger Institute for the COVID-19 Genomics UK (COG-UK) Consortium | The Lighthouse Lab in Milton Keynes and Alex Alderton, Roberto Amato, Jeffrey Barrett, Sonia Goncalves, Ewan Harrison, David K. Jackson, Ian Johnston, Dominic Kwiatkowski, Cordelia Langford, John Sillitoe on behalf of the Wellcome Sanger Institute COVID-19 Surveillance Team |
| EPI_ISL_3092899 | Lighthouse Lab in Alderley Park | Wellcome Sanger Institute for the COVID-19 Genomics UK (COG-UK) Consortium | Jacquelyn Wynn, Mairead Hyland, The Lighthouse Lab in Alderley Park and Alex Alderton, Roberto Amato, Jeffrey Barrett, Sonia Goncalves, Ewan Harrison, David K. Jackson, Ian Johnston, Dominic Kwiatkowski, Cordelia Langford, John Sillitoe on behalf of the Wellcome Sanger Institute COVID-19 Surveillance Team |
| EPI_ISL_3092900, EPI_ISL_3092901 | Lighthouse Lab in Milton Keynes | Wellcome Sanger Institute for the COVID-19 Genomics UK (COG-UK) Consortium | The Lighthouse Lab in Milton Keynes and Alex Alderton, Roberto Amato, Jeffrey Barrett, Sonia Goncalves, Ewan Harrison, David K. Jackson, Ian Johnston, Dominic Kwiatkowski, Cordelia Langford, John Sillitoe on behalf of the Wellcome Sanger Institute COVID-19 Surveillance Team |
| EPI_ISL_3092902 | Lighthouse Lab in Alderley Park | Wellcome Sanger Institute for the COVID-19 Genomics UK (COG-UK) Consortium | Jacquelyn Wynn, Mairead Hyland, The Lighthouse Lab in Alderley Park and Alex Alderton, Roberto Amato, Jeffrey Barrett, Sonia Goncalves, Ewan Harrison, David K. Jackson, Ian Johnston, Dominic Kwiatkowski, Cordelia Langford, John Sillitoe on behalf of the Wellcome Sanger Institute COVID-19 Surveillance Team |
| EPI_ISL_3092903 | Lighthouse Lab in Glasgow | Wellcome Sanger Institute for the COVID-19 Genomics UK (COG-UK) Consortium | Harper VanSteenhouse, Yumi Kasai, David Gray, Carol Clugston, Anna Dominiczak and Alex Alderton, Roberto Amato, Jeffrey Barrett, Sonia Goncalves, Ewan Harrison, David K. Jackson, Ian Johnston, Dominic Kwiatkowski, Cordelia Langford, John Sillitoe on behalf of the Wellcome Sanger Institute COVID-19 Surveillance Team |
| EPI_ISL_3092904, EPI_ISL_3092905 | Lighthouse Lab in Alderley Park | Wellcome Sanger Institute for the COVID-19 Genomics UK (COG-UK) Consortium | Jacquelyn Wynn, Mairead Hyland, The Lighthouse Lab in Alderley Park and Alex Alderton, Roberto Amato, Jeffrey Barrett, Sonia Goncalves, Ewan Harrison, David K. Jackson, Ian Johnston, Dominic Kwiatkowski, Cordelia Langford, John Sillitoe on behalf of the Wellcome Sanger Institute COVID-19 Surveillance Team |
| EPI_ISL_3092906 | Lighthouse Lab in Glasgow | Wellcome Sanger Institute for the COVID-19 Genomics UK (COG-UK) Consortium | Harper VanSteenhouse, Yumi Kasai, David Gray, Carol Clugston, Anna Dominiczak and Alex Alderton, Roberto Amato, Jeffrey Barrett, Sonia Goncalves, Ewan Harrison, David K. Jackson, Ian Johnston, Dominic Kwiatkowski, Cordelia Langford, John Sillitoe on behalf of the Wellcome Sanger Institute COVID-19 Surveillance Team |
| EPI_ISL_3092907 | Lighthouse Lab in Alderley Park | Wellcome Sanger Institute for the COVID-19 Genomics UK (COG-UK) Consortium | Jacquelyn Wynn, Mairead Hyland, The Lighthouse Lab in Alderley Park and Alex Alderton, Roberto Amato, Jeffrey Barrett, Sonia Goncalves, Ewan Harrison, David K. Jackson, Ian Johnston, Dominic Kwiatkowski, Cordelia Langford, John Sillitoe on behalf of the Wellcome Sanger Institute COVID-19 Surveillance Team |
| EPI_ISL_3092908 | Lighthouse Lab in Glasgow | Wellcome Sanger Institute for the COVID-19 Genomics UK (COG-UK) Consortium | Harper VanSteenhouse, Yumi Kasai, David Gray, Carol Clugston, Anna Dominiczak and Alex Alderton, Roberto Amato, Jeffrey Barrett, Sonia Goncalves, Ewan Harrison, David K. Jackson, Ian Johnston, Dominic Kwiatkowski, Cordelia Langford, John Sillitoe on behalf of the Wellcome Sanger Institute COVID-19 Surveillance Team |
| EPI_ISL_3092909, EPI_ISL_3092910 | Berkshire and Surrey Pathology Services Lighthouse Laboratory | Wellcome Sanger Institute for the COVID-19 Genomics UK (COG-UK) Consortium | Berkshire and Surrey Pathology Services Lighthouse Laboratory and Alex Alderton, Roberto Amato, Jeffrey Barrett, Sonia Goncalves, Ewan Harrison, David K. Jackson, Ian Johnston, Dominic Kwiatkowski, Cordelia Langford, John Sillitoe on behalf of the Wellcome Sanger Institute COVID-19 Surveillance Team |
| EPI_ISL_3092911 | Lighthouse Lab in Glasgow | Wellcome Sanger Institute for the COVID-19 Genomics UK (COG-UK) Consortium | Harper VanSteenhouse, Yumi Kasai, David Gray, Carol Clugston, Anna Dominiczak and Alex Alderton, Roberto Amato, Jeffrey Barrett, Sonia Goncalves, Ewan Harrison, David K. Jackson, Ian Johnston, Dominic Kwiatkowski, Cordelia Langford, John Sillitoe on behalf of the Wellcome Sanger Institute COVID-19 Surveillance Team |
| EPI_ISL_3092912 | Lighthouse Lab in Milton Keynes | Wellcome Sanger Institute for the COVID-19 Genomics UK (COG-UK) Consortium | The Lighthouse Lab in Milton Keynes and Alex Alderton, Roberto Amato, Jeffrey Barrett, Sonia Goncalves, Ewan Harrison, David K. Jackson, Ian Johnston, Dominic Kwiatkowski, Cordelia Langford, John Sillitoe on behalf of the Wellcome Sanger Institute COVID-19 Surveillance Team |
| EPI_ISL_3092913 | Lighthouse Lab in Glasgow | Wellcome Sanger Institute for the COVID-19 Genomics UK (COG-UK) Consortium | Harper VanSteenhouse, Yumi Kasai, David Gray, Carol Clugston, Anna Dominiczak and Alex Alderton, Roberto Amato, Jeffrey Barrett, Sonia Goncalves, Ewan Harrison, David K. Jackson, Ian Johnston, Dominic Kwiatkowski, Cordelia Langford, John Sillitoe on behalf of the Wellcome Sanger Institute COVID-19 Surveillance Team |
| EPI_ISL_3092914, EPI_ISL_3092915, EPI_ISL_3092916 | Lighthouse Lab in Alderley Park | Wellcome Sanger Institute for the COVID-19 Genomics UK (COG-UK) Consortium | Jacquelyn Wynn, Mairead Hyland, The Lighthouse Lab in Alderley Park and Alex Alderton, Roberto Amato, Jeffrey Barrett, Sonia Goncalves, Ewan Harrison, David K. Jackson, Ian Johnston, Dominic Kwiatkowski, Cordelia Langford, John Sillitoe on behalf of the Wellcome Sanger Institute COVID-19 Surveillance Team |
| EPI_ISL_3092917 | Lighthouse Lab in Glasgow | Wellcome Sanger Institute for the COVID-19 Genomics UK (COG-UK) Consortium | Harper VanSteenhouse, Yumi Kasai, David Gray, Carol Clugston, Anna Dominiczak and Alex Alderton, Roberto Amato, Jeffrey Barrett, Sonia Goncalves, Ewan Harrison, David K. Jackson, Ian Johnston, Dominic Kwiatkowski, Cordelia Langford, John Sillitoe on behalf of the Wellcome Sanger Institute COVID-19 Surveillance Team |
| EPI_ISL_3092918, EPI_ISL_3092919 | Berkshire and Surrey Pathology Services Lighthouse Laboratory | Wellcome Sanger Institute for the COVID-19 Genomics UK (COG-UK) Consortium | Berkshire and Surrey Pathology Services Lighthouse Laboratory and Alex Alderton, Roberto Amato, Jeffrey Barrett, Sonia Goncalves, Ewan Harrison, David K. Jackson, Ian Johnston, Dominic Kwiatkowski, Cordelia Langford, John Sillitoe on behalf of the Wellcome Sanger Institute COVID-19 Surveillance Team |
| EPI_ISL_3092920, EPI_ISL_3092921 | Lighthouse Lab in Glasgow | Wellcome Sanger Institute for the COVID-19 Genomics UK (COG-UK) Consortium | Harper VanSteenhouse, Yumi Kasai, David Gray, Carol Clugston, Anna Dominiczak and Alex Alderton, Roberto Amato, Jeffrey Barrett, Sonia Goncalves, Ewan Harrison, David K. Jackson, Ian Johnston, Dominic Kwiatkowski, Cordelia Langford, John Sillitoe on behalf of the Wellcome Sanger Institute COVID-19 Surveillance Team |
| EPI_ISL_3092922, EPI_ISL_3092923, EPI_ISL_3092924 | Lighthouse Lab in Alderley Park | Wellcome Sanger Institute for the COVID-19 Genomics UK (COG-UK) Consortium | Jacquelyn Wynn, Mairead Hyland, The Lighthouse Lab in Alderley Park and Alex Alderton, Roberto Amato, Jeffrey Barrett, Sonia Goncalves, Ewan Harrison, David K. Jackson, Ian Johnston, Dominic Kwiatkowski, Cordelia Langford, John Sillitoe on behalf of the Wellcome Sanger Institute COVID-19 Surveillance Team |
| EPI_ISL_3092925 | Lighthouse Lab in Glasgow | Wellcome Sanger Institute for the COVID-19 Genomics UK (COG-UK) Consortium | Harper VanSteenhouse, Yumi Kasai, David Gray, Carol Clugston, Anna Dominiczak and Alex Alderton, Roberto Amato, Jeffrey Barrett, Sonia Goncalves, Ewan Harrison, David K. Jackson, Ian Johnston, Dominic Kwiatkowski, Cordelia Langford, John Sillitoe on behalf of the Wellcome Sanger Institute COVID-19 Surveillance Team |
| EPI_ISL_3092926 | Berkshire and Surrey Pathology Services Lighthouse Laboratory | Wellcome Sanger Institute for the COVID-19 Genomics UK (COG-UK) Consortium | Berkshire and Surrey Pathology Services Lighthouse Laboratory and Alex Alderton, Roberto Amato, Jeffrey Barrett, Sonia Goncalves, Ewan Harrison, David K. Jackson, Ian Johnston, Dominic Kwiatkowski, Cordelia Langford, John Sillitoe on behalf of the Wellcome Sanger Institute COVID-19 Surveillance Team |
| EPI_ISL_3092927 | Lighthouse Lab in Milton Keynes | Wellcome Sanger Institute for the COVID-19 Genomics UK (COG-UK) Consortium | The Lighthouse Lab in Milton Keynes and Alex Alderton, Roberto Amato, Jeffrey Barrett, Sonia Goncalves, Ewan Harrison, David K. Jackson, Ian Johnston, Dominic Kwiatkowski, Cordelia Langford, John Sillitoe on behalf of the Wellcome Sanger Institute COVID-19 Surveillance Team |
| EPI_ISL_3092928, EPI_ISL_3092929 | Lighthouse Lab in Glasgow | Wellcome Sanger Institute for the COVID-19 Genomics UK (COG-UK) Consortium | Harper VanSteenhouse, Yumi Kasai, David Gray, Carol Clugston, Anna Dominiczak and Alex Alderton, Roberto Amato, Jeffrey Barrett, Sonia Goncalves, Ewan Harrison, David K. Jackson, Ian Johnston, Dominic Kwiatkowski, Cordelia Langford, John Sillitoe on behalf of the Wellcome Sanger Institute COVID-19 Surveillance Team |
| EPI_ISL_3092930 | Lighthouse Lab in Alderley Park | Wellcome Sanger Institute for the COVID-19 Genomics UK (COG-UK) Consortium | Jacquelyn Wynn, Mairead Hyland, The Lighthouse Lab in Alderley Park and Alex Alderton, Roberto Amato, Jeffrey Barrett, Sonia Goncalves, Ewan Harrison, David K. Jackson, Ian Johnston, Dominic Kwiatkowski, Cordelia Langford, John Sillitoe on behalf of the Wellcome Sanger Institute COVID-19 Surveillance Team |
| EPI_ISL_3092931, EPI_ISL_3092932 | Lighthouse Lab in Glasgow | Wellcome Sanger Institute for the COVID-19 Genomics UK (COG-UK) Consortium | Harper VanSteenhouse, Yumi Kasai, David Gray, Carol Clugston, Anna Dominiczak and Alex Alderton, Roberto Amato, Jeffrey Barrett, Sonia Goncalves, Ewan Harrison, David K. Jackson, Ian Johnston, Dominic Kwiatkowski, Cordelia Langford, John Sillitoe on behalf of the Wellcome Sanger Institute COVID-19 Surveillance Team |
| EPI_ISL_3092933 | Lighthouse Lab in Alderley Park | Wellcome Sanger Institute for the COVID-19 Genomics UK (COG-UK) Consortium | Jacquelyn Wynn, Mairead Hyland, The Lighthouse Lab in Alderley Park and Alex Alderton, Roberto Amato, Jeffrey Barrett, Sonia Goncalves, Ewan Harrison, David K. Jackson, Ian Johnston, Dominic Kwiatkowski, Cordelia Langford, John Sillitoe on behalf of the Wellcome Sanger Institute COVID-19 Surveillance Team |
| EPI_ISL_3092934 | Lighthouse Lab in Glasgow | Wellcome Sanger Institute for the COVID-19 Genomics UK (COG-UK) Consortium | Harper VanSteenhouse, Yumi Kasai, David Gray, Carol Clugston, Anna Dominiczak and Alex Alderton, Roberto Amato, Jeffrey Barrett, Sonia Goncalves, Ewan Harrison, David K. Jackson, Ian Johnston, Dominic Kwiatkowski, Cordelia Langford, John Sillitoe on behalf of the Wellcome Sanger Institute COVID-19 Surveillance Team |
| EPI_ISL_3092935, EPI_ISL_3092936, EPI_ISL_3092937 | Berkshire and Surrey Pathology Services Lighthouse Laboratory | Wellcome Sanger Institute for the COVID-19 Genomics UK (COG-UK) Consortium | Berkshire and Surrey Pathology Services Lighthouse Laboratory and Alex Alderton, Roberto Amato, Jeffrey Barrett, Sonia Goncalves, Ewan Harrison, David K. Jackson, Ian Johnston, Dominic Kwiatkowski, Cordelia Langford, John Sillitoe on behalf of the Wellcome Sanger Institute COVID-19 Surveillance Team |
| EPI_ISL_3092938, EPI_ISL_3092939 | Lighthouse Lab in Glasgow | Wellcome Sanger Institute for the COVID-19 Genomics UK (COG-UK) Consortium | Harper VanSteenhouse, Yumi Kasai, David Gray, Carol Clugston, Anna Dominiczak and Alex Alderton, Roberto Amato, Jeffrey Barrett, Sonia Goncalves, Ewan Harrison, David K. Jackson, Ian Johnston, Dominic Kwiatkowski, Cordelia Langford, John Sillitoe on behalf of the Wellcome Sanger Institute COVID-19 Surveillance Team |
| EPI_ISL_3092940 | Lighthouse Lab in Milton Keynes | Wellcome Sanger Institute for the COVID-19 Genomics UK (COG-UK) Consortium | The Lighthouse Lab in Milton Keynes and Alex Alderton, Roberto Amato, Jeffrey Barrett, Sonia Goncalves, Ewan Harrison, David K. Jackson, Ian Johnston, Dominic Kwiatkowski, Cordelia Langford, John Sillitoe on behalf of the Wellcome Sanger Institute COVID-19 Surveillance Team |
| EPI_ISL_3092941, EPI_ISL_3092942 | Lighthouse Lab in Glasgow | Wellcome Sanger Institute for the COVID-19 Genomics UK (COG-UK) Consortium | Harper VanSteenhouse, Yumi Kasai, David Gray, Carol Clugston, Anna Dominiczak and Alex Alderton, Roberto Amato, Jeffrey Barrett, Sonia Goncalves, Ewan Harrison, David K. Jackson, Ian Johnston, Dominic Kwiatkowski, Cordelia Langford, John Sillitoe on behalf of the Wellcome Sanger Institute COVID-19 Surveillance Team |
| EPI_ISL_3092943 | Lighthouse Lab in Alderley Park | Wellcome Sanger Institute for the COVID-19 Genomics UK (COG-UK) Consortium | Jacquelyn Wynn, Mairead Hyland, The Lighthouse Lab in Alderley Park and Alex Alderton, Roberto Amato, Jeffrey Barrett, Sonia Goncalves, Ewan Harrison, David K. Jackson, Ian Johnston, Dominic Kwiatkowski, Cordelia Langford, John Sillitoe on behalf of the Wellcome Sanger Institute COVID-19 Surveillance Team |
| EPI_ISL_3092944 | Berkshire and Surrey Pathology Services Lighthouse Laboratory | Wellcome Sanger Institute for the COVID-19 Genomics UK (COG-UK) Consortium | Berkshire and Surrey Pathology Services Lighthouse Laboratory and Alex Alderton, Roberto Amato, Jeffrey Barrett, Sonia Goncalves, Ewan Harrison, David K. Jackson, Ian Johnston, Dominic Kwiatkowski, Cordelia Langford, John Sillitoe on behalf of the Wellcome Sanger Institute COVID-19 Surveillance Team |
| EPI_ISL_3092945, EPI_ISL_3092946 | Lighthouse Lab in Alderley Park | Wellcome Sanger Institute for the COVID-19 Genomics UK (COG-UK) Consortium | Jacquelyn Wynn, Mairead Hyland, The Lighthouse Lab in Alderley Park and Alex Alderton, Roberto Amato, Jeffrey Barrett, Sonia Goncalves, Ewan Harrison, David K. Jackson, Ian Johnston, Dominic Kwiatkowski, Cordelia Langford, John Sillitoe on behalf of the Wellcome Sanger Institute COVID-19 Surveillance Team |
| EPI_ISL_3092947 | Lighthouse Lab in Glasgow | Wellcome Sanger Institute for the COVID-19 Genomics UK (COG-UK) Consortium | Harper VanSteenhouse, Yumi Kasai, David Gray, Carol Clugston, Anna Dominiczak and Alex Alderton, Roberto Amato, Jeffrey Barrett, Sonia Goncalves, Ewan Harrison, David K. Jackson, Ian Johnston, Dominic Kwiatkowski, Cordelia Langford, John Sillitoe on behalf of the Wellcome Sanger Institute COVID-19 Surveillance Team |
| EPI_ISL_3092948 | Lighthouse Lab in Milton Keynes | Wellcome Sanger Institute for the COVID-19 Genomics UK (COG-UK) Consortium | The Lighthouse Lab in Milton Keynes and Alex Alderton, Roberto Amato, Jeffrey Barrett, Sonia Goncalves, Ewan Harrison, David K. Jackson, Ian Johnston, Dominic Kwiatkowski, Cordelia Langford, John Sillitoe on behalf of the Wellcome Sanger Institute COVID-19 Surveillance Team |
| EPI_ISL_3092949 | Berkshire and Surrey Pathology Services Lighthouse Laboratory | Wellcome Sanger Institute for the COVID-19 Genomics UK (COG-UK) Consortium | Berkshire and Surrey Pathology Services Lighthouse Laboratory and Alex Alderton, Roberto Amato, Jeffrey Barrett, Sonia Goncalves, Ewan Harrison, David K. Jackson, Ian Johnston, Dominic Kwiatkowski, Cordelia Langford, John Sillitoe on behalf of the Wellcome Sanger Institute COVID-19 Surveillance Team |
| EPI_ISL_3092950, EPI_ISL_3092951 | Lighthouse Lab in Alderley Park | Wellcome Sanger Institute for the COVID-19 Genomics UK (COG-UK) Consortium | Jacquelyn Wynn, Mairead Hyland, The Lighthouse Lab in Alderley Park and Alex Alderton, Roberto Amato, Jeffrey Barrett, Sonia Goncalves, Ewan Harrison, David K. Jackson, Ian Johnston, Dominic Kwiatkowski, Cordelia Langford, John Sillitoe on behalf of the Wellcome Sanger Institute COVID-19 Surveillance Team |
| EPI_ISL_3092952 | Berkshire and Surrey Pathology Services Lighthouse Laboratory | Wellcome Sanger Institute for the COVID-19 Genomics UK (COG-UK) Consortium | Berkshire and Surrey Pathology Services Lighthouse Laboratory and Alex Alderton, Roberto Amato, Jeffrey Barrett, Sonia Goncalves, Ewan Harrison, David K. Jackson, Ian Johnston, Dominic Kwiatkowski, Cordelia Langford, John Sillitoe on behalf of the Wellcome Sanger Institute COVID-19 Surveillance Team |
| EPI_ISL_3092953 | Lighthouse Lab in Glasgow | Wellcome Sanger Institute for the COVID-19 Genomics UK (COG-UK) Consortium | Harper VanSteenhouse, Yumi Kasai, David Gray, Carol Clugston, Anna Dominiczak and Alex Alderton, Roberto Amato, Jeffrey Barrett, Sonia Goncalves, Ewan Harrison, David K. Jackson, Ian Johnston, Dominic Kwiatkowski, Cordelia Langford, John Sillitoe on behalf of the Wellcome Sanger Institute COVID-19 Surveillance Team |
| EPI_ISL_3092954 | Lighthouse Lab in Milton Keynes | Wellcome Sanger Institute for the COVID-19 Genomics UK (COG-UK) Consortium | The Lighthouse Lab in Milton Keynes and Alex Alderton, Roberto Amato, Jeffrey Barrett, Sonia Goncalves, Ewan Harrison, David K. Jackson, Ian Johnston, Dominic Kwiatkowski, Cordelia Langford, John Sillitoe on behalf of the Wellcome Sanger Institute COVID-19 Surveillance Team |
| EPI_ISL_3092955 | Lighthouse Lab in Alderley Park | Wellcome Sanger Institute for the COVID-19 Genomics UK (COG-UK) Consortium | Jacquelyn Wynn, Mairead Hyland, The Lighthouse Lab in Alderley Park and Alex Alderton, Roberto Amato, Jeffrey Barrett, Sonia Goncalves, Ewan Harrison, David K. Jackson, Ian Johnston, Dominic Kwiatkowski, Cordelia Langford, John Sillitoe on behalf of the Wellcome Sanger Institute COVID-19 Surveillance Team |
| EPI_ISL_3092956 | Lighthouse Lab in Milton Keynes | Wellcome Sanger Institute for the COVID-19 Genomics UK (COG-UK) Consortium | The Lighthouse Lab in Milton Keynes and Alex Alderton, Roberto Amato, Jeffrey Barrett, Sonia Goncalves, Ewan Harrison, David K. Jackson, Ian Johnston, Dominic Kwiatkowski, Cordelia Langford, John Sillitoe on behalf of the Wellcome Sanger Institute COVID-19 Surveillance Team |

| | | | |
|---|---|---|---|
| EPI_ISL_3092957 | Lighthouse Lab in Glasgow | Wellcome Sanger Institute for the COVID-19 Genomics UK (COG-UK) Consortium | Harper VanSteenhouse, Yumi Kasai, David Gray, Carol Clugston, Anna Dominiczak and Alex Alderton, Roberto Amato, Jeffrey Barrett, Sonia Goncalves, Ewan Harrison, David K. Jackson, Ian Johnston, Dominic Kwiatkowski, Cordelia Langford, John Sillitoe on behalf of the Wellcome Sanger Institute COVID-19 Surveillance Team |
| EPI_ISL_3092958 | Lighthouse Lab in Milton Keynes | Wellcome Sanger Institute for the COVID-19 Genomics UK (COG-UK) Consortium | The Lighthouse Lab in Milton Keynes and Alex Alderton, Roberto Amato, Jeffrey Barrett, Sonia Goncalves, Ewan Harrison, David K. Jackson, Ian Johnston, Dominic Kwiatkowski, Cordelia Langford, John Sillitoe on behalf of the Wellcome Sanger Institute COVID-19 Surveillance Team |
| EPI_ISL_3092959 | Lighthouse Lab in Glasgow | Wellcome Sanger Institute for the COVID-19 Genomics UK (COG-UK) Consortium | Harper VanSteenhouse, Yumi Kasai, David Gray, Carol Clugston, Anna Dominiczak and Alex Alderton, Roberto Amato, Jeffrey Barrett, Sonia Goncalves, Ewan Harrison, David K. Jackson, Ian Johnston, Dominic Kwiatkowski, Cordelia Langford, John Sillitoe on behalf of the Wellcome Sanger Institute COVID-19 Surveillance Team |
| EPI_ISL_3092960 | Lighthouse Lab in Milton Keynes | Wellcome Sanger Institute for the COVID-19 Genomics UK (COG-UK) Consortium | The Lighthouse Lab in Milton Keynes and Alex Alderton, Roberto Amato, Jeffrey Barrett, Sonia Goncalves, Ewan Harrison, David K. Jackson, Ian Johnston, Dominic Kwiatkowski, Cordelia Langford, John Sillitoe on behalf of the Wellcome Sanger Institute COVID-19 Surveillance Team |
| EPI_ISL_3092961 | Lighthouse Lab in Glasgow | Wellcome Sanger Institute for the COVID-19 Genomics UK (COG-UK) Consortium | Harper VanSteenhouse, Yumi Kasai, David Gray, Carol Clugston, Anna Dominiczak and Alex Alderton, Roberto Amato, Jeffrey Barrett, Sonia Goncalves, Ewan Harrison, David K. Jackson, Ian Johnston, Dominic Kwiatkowski, Cordelia Langford, John Sillitoe on behalf of the Wellcome Sanger Institute COVID-19 Surveillance Team |
| EPI_ISL_3092962, EPI_ISL_3092963 | Lighthouse Lab in Milton Keynes | Wellcome Sanger Institute for the COVID-19 Genomics UK (COG-UK) Consortium | The Lighthouse Lab in Milton Keynes and Alex Alderton, Roberto Amato, Jeffrey Barrett, Sonia Goncalves, Ewan Harrison, David K. Jackson, Ian Johnston, Dominic Kwiatkowski, Cordelia Langford, John Sillitoe on behalf of the Wellcome Sanger Institute COVID-19 Surveillance Team |
| EPI_ISL_3092964, EPI_ISL_3092965 | Lighthouse Lab in Glasgow | Wellcome Sanger Institute for the COVID-19 Genomics UK (COG-UK) Consortium | Harper VanSteenhouse, Yumi Kasai, David Gray, Carol Clugston, Anna Dominiczak and Alex Alderton, Roberto Amato, Jeffrey Barrett, Sonia Goncalves, Ewan Harrison, David K. Jackson, Ian Johnston, Dominic Kwiatkowski, Cordelia Langford, John Sillitoe on behalf of the Wellcome Sanger Institute COVID-19 Surveillance Team |
| EPI_ISL_3092966 | Berkshire and Surrey Pathology Services Lighthouse Laboratory | Wellcome Sanger Institute for the COVID-19 Genomics UK (COG-UK) Consortium | Berkshire and Surrey Pathology Services Lighthouse Laboratory and Alex Alderton, Roberto Amato, Jeffrey Barrett, Sonia Goncalves, Ewan Harrison, David K. Jackson, Ian Johnston, Dominic Kwiatkowski, Cordelia Langford, John Sillitoe on behalf of the Wellcome Sanger Institute COVID-19 Surveillance Team |
| EPI_ISL_3092967 | Lighthouse Lab in Glasgow | Wellcome Sanger Institute for the COVID-19 Genomics UK (COG-UK) Consortium | Harper VanSteenhouse, Yumi Kasai, David Gray, Carol Clugston, Anna Dominiczak and Alex Alderton, Roberto Amato, Jeffrey Barrett, Sonia Goncalves, Ewan Harrison, David K. Jackson, Ian Johnston, Dominic Kwiatkowski, Cordelia Langford, John Sillitoe on behalf of the Wellcome Sanger Institute COVID-19 Surveillance Team |
| EPI_ISL_3092968 | Lighthouse Lab in Milton Keynes | Wellcome Sanger Institute for the COVID-19 Genomics UK (COG-UK) Consortium | The Lighthouse Lab in Milton Keynes and Alex Alderton, Roberto Amato, Jeffrey Barrett, Sonia Goncalves, Ewan Harrison, David K. Jackson, Ian Johnston, Dominic Kwiatkowski, Cordelia Langford, John Sillitoe on behalf of the Wellcome Sanger Institute COVID-19 Surveillance Team |
| EPI_ISL_3092969 | Lighthouse Lab in Alderley Park | Wellcome Sanger Institute for the COVID-19 Genomics UK (COG-UK) Consortium | Jacquelyn Wynn, Mairead Hyland, The Lighthouse Lab in Alderley Park and Alex Alderton, Roberto Amato, Jeffrey Barrett, Sonia Goncalves, Ewan Harrison, David K. Jackson, Ian Johnston, Dominic Kwiatkowski, Cordelia Langford, John Sillitoe on behalf of the Wellcome Sanger Institute COVID-19 Surveillance Team |
| EPI_ISL_3092970 | Lighthouse Lab in Milton Keynes | Wellcome Sanger Institute for the COVID-19 Genomics UK (COG-UK) Consortium | The Lighthouse Lab in Milton Keynes and Alex Alderton, Roberto Amato, Jeffrey Barrett, Sonia Goncalves, Ewan Harrison, David K. Jackson, Ian Johnston, Dominic Kwiatkowski, Cordelia Langford, John Sillitoe on behalf of the Wellcome Sanger Institute COVID-19 Surveillance Team |
| EPI_ISL_3092971 | Lighthouse Lab in Alderley Park | Wellcome Sanger Institute for the COVID-19 Genomics UK (COG-UK) Consortium | Jacquelyn Wynn, Mairead Hyland, The Lighthouse Lab in Alderley Park and Alex Alderton, Roberto Amato, Jeffrey Barrett, Sonia Goncalves, Ewan Harrison, David K. Jackson, Ian Johnston, Dominic Kwiatkowski, Cordelia Langford, John Sillitoe on behalf of the Wellcome Sanger Institute COVID-19 Surveillance Team |
| EPI_ISL_3092972, EPI_ISL_3092973 | Lighthouse Lab in Milton Keynes | Wellcome Sanger Institute for the COVID-19 Genomics UK (COG-UK) Consortium | The Lighthouse Lab in Milton Keynes and Alex Alderton, Roberto Amato, Jeffrey Barrett, Sonia Goncalves, Ewan Harrison, David K. Jackson, Ian Johnston, Dominic Kwiatkowski, Cordelia Langford, John Sillitoe on behalf of the Wellcome Sanger Institute COVID-19 Surveillance Team |
| EPI_ISL_3092974 | Berkshire and Surrey Pathology Services Lighthouse Laboratory | Wellcome Sanger Institute for the COVID-19 Genomics UK (COG-UK) Consortium | Berkshire and Surrey Pathology Services Lighthouse Laboratory and Alex Alderton, Roberto Amato, Jeffrey Barrett, Sonia Goncalves, Ewan Harrison, David K. Jackson, Ian Johnston, Dominic Kwiatkowski, Cordelia Langford, John Sillitoe on behalf of the Wellcome Sanger Institute COVID-19 Surveillance Team |
| EPI_ISL_3092975 | Lighthouse Lab in Milton Keynes | Wellcome Sanger Institute for the COVID-19 Genomics UK (COG-UK) Consortium | The Lighthouse Lab in Milton Keynes and Alex Alderton, Roberto Amato, Jeffrey Barrett, Sonia Goncalves, Ewan Harrison, David K. Jackson, Ian Johnston, Dominic Kwiatkowski, Cordelia Langford, John Sillitoe on behalf of the Wellcome Sanger Institute COVID-19 Surveillance Team |
| EPI_ISL_3092976 | Berkshire and Surrey Pathology Services Lighthouse Laboratory | Wellcome Sanger Institute for the COVID-19 Genomics UK (COG-UK) Consortium | Berkshire and Surrey Pathology Services Lighthouse Laboratory and Alex Alderton, Roberto Amato, Jeffrey Barrett, Sonia Goncalves, Ewan Harrison, David K. Jackson, Ian Johnston, Dominic Kwiatkowski, Cordelia Langford, John Sillitoe on behalf of the Wellcome Sanger Institute COVID-19 Surveillance Team |
| EPI_ISL_3092977 | Lighthouse Lab in Milton Keynes | Wellcome Sanger Institute for the COVID-19 Genomics UK (COG-UK) Consortium | The Lighthouse Lab in Milton Keynes and Alex Alderton, Roberto Amato, Jeffrey Barrett, Sonia Goncalves, Ewan Harrison, David K. Jackson, Ian Johnston, Dominic Kwiatkowski, Cordelia Langford, John Sillitoe on behalf of the Wellcome Sanger Institute COVID-19 Surveillance Team |
| EPI_ISL_3092978 | Lighthouse Lab in Glasgow | Wellcome Sanger Institute for the COVID-19 Genomics UK (COG-UK) Consortium | Harper VanSteenhouse, Yumi Kasai, David Gray, Carol Clugston, Anna Dominiczak and Alex Alderton, Roberto Amato, Jeffrey Barrett, Sonia Goncalves, Ewan Harrison, David K. Jackson, Ian Johnston, Dominic Kwiatkowski, Cordelia Langford, John Sillitoe on behalf of the Wellcome Sanger Institute COVID-19 Surveillance Team |
| EPI_ISL_3092979 | Lighthouse Lab in Milton Keynes | Wellcome Sanger Institute for the COVID-19 Genomics UK (COG-UK) Consortium | The Lighthouse Lab in Milton Keynes and Alex Alderton, Roberto Amato, Jeffrey Barrett, Sonia Goncalves, Ewan Harrison, David K. Jackson, Ian Johnston, Dominic Kwiatkowski, Cordelia Langford, John Sillitoe on behalf of the Wellcome Sanger Institute COVID-19 Surveillance Team |
| EPI_ISL_3092980 | Lighthouse Lab in Alderley Park | Wellcome Sanger Institute for the COVID-19 Genomics UK (COG-UK) Consortium | Jacquelyn Wynn, Mairead Hyland, The Lighthouse Lab in Alderley Park and Alex Alderton, Roberto Amato, Jeffrey Barrett, Sonia Goncalves, Ewan Harrison, David K. Jackson, Ian Johnston, Dominic Kwiatkowski, Cordelia Langford, John Sillitoe on behalf of the Wellcome Sanger Institute COVID-19 Surveillance Team |
| EPI_ISL_3092981 | Lighthouse Lab in Milton Keynes | Wellcome Sanger Institute for the COVID-19 Genomics UK (COG-UK) Consortium | The Lighthouse Lab in Milton Keynes and Alex Alderton, Roberto Amato, Jeffrey Barrett, Sonia Goncalves, Ewan Harrison, David K. Jackson, Ian Johnston, Dominic Kwiatkowski, Cordelia Langford, John Sillitoe on behalf of the Wellcome Sanger Institute COVID-19 Surveillance Team |
| EPI_ISL_3092982 | Berkshire and Surrey Pathology Services Lighthouse Laboratory | Wellcome Sanger Institute for the COVID-19 Genomics UK (COG-UK) Consortium | Berkshire and Surrey Pathology Services Lighthouse Laboratory and Alex Alderton, Roberto Amato, Jeffrey Barrett, Sonia Goncalves, Ewan Harrison, David K. Jackson, Ian Johnston, Dominic Kwiatkowski, Cordelia Langford, John Sillitoe on behalf of the Wellcome Sanger Institute COVID-19 Surveillance Team |
| EPI_ISL_3092983, EPI_ISL_3092984 | Lighthouse Lab in Milton Keynes | Wellcome Sanger Institute for the COVID-19 Genomics UK (COG-UK) Consortium | The Lighthouse Lab in Milton Keynes and Alex Alderton, Roberto Amato, Jeffrey Barrett, Sonia Goncalves, Ewan Harrison, David K. Jackson, Ian Johnston, Dominic Kwiatkowski, Cordelia Langford, John Sillitoe on behalf of the Wellcome Sanger Institute COVID-19 Surveillance Team |
| EPI_ISL_3092985 | Lighthouse Lab in Alderley Park | Wellcome Sanger Institute for the COVID-19 Genomics UK (COG-UK) Consortium | Jacquelyn Wynn, Mairead Hyland, The Lighthouse Lab in Alderley Park and Alex Alderton, Roberto Amato, Jeffrey Barrett, Sonia Goncalves, Ewan Harrison, David K. Jackson, Ian Johnston, Dominic Kwiatkowski, Cordelia Langford, John Sillitoe on behalf of the Wellcome Sanger Institute COVID-19 Surveillance Team |
| EPI_ISL_3092986 | Berkshire and Surrey Pathology Services Lighthouse Laboratory | Wellcome Sanger Institute for the COVID-19 Genomics UK (COG-UK) Consortium | Berkshire and Surrey Pathology Services Lighthouse Laboratory and Alex Alderton, Roberto Amato, Jeffrey Barrett, Sonia Goncalves, Ewan Harrison, David K. Jackson, Ian Johnston, Dominic Kwiatkowski, Cordelia Langford, John Sillitoe on behalf of the Wellcome Sanger Institute COVID-19 Surveillance Team |
| EPI_ISL_3092987, EPI_ISL_3092988 | Lighthouse Lab in Glasgow | Wellcome Sanger Institute for the COVID-19 Genomics UK (COG-UK) Consortium | Harper VanSteenhouse, Yumi Kasai, David Gray, Carol Clugston, Anna Dominiczak and Alex Alderton, Roberto Amato, Jeffrey Barrett, Sonia Goncalves, Ewan Harrison, David K. Jackson, Ian Johnston, Dominic Kwiatkowski, Cordelia Langford, John Sillitoe on behalf of the Wellcome Sanger Institute COVID-19 Surveillance Team |
| EPI_ISL_3092989, EPI_ISL_3092990, EPI_ISL_3092991 | Berkshire and Surrey Pathology Services Lighthouse Laboratory | Wellcome Sanger Institute for the COVID-19 Genomics UK (COG-UK) Consortium | Berkshire and Surrey Pathology Services Lighthouse Laboratory and Alex Alderton, Roberto Amato, Jeffrey Barrett, Sonia Goncalves, Ewan Harrison, David K. Jackson, Ian Johnston, Dominic Kwiatkowski, Cordelia Langford, John Sillitoe on behalf of the Wellcome Sanger Institute COVID-19 Surveillance Team |
| EPI_ISL_3092992 | Lighthouse Lab in Milton Keynes | Wellcome Sanger Institute for the COVID-19 Genomics UK (COG-UK) Consortium | The Lighthouse Lab in Milton Keynes and Alex Alderton, Roberto Amato, Jeffrey Barrett, Sonia Goncalves, Ewan Harrison, David K. Jackson, Ian Johnston, Dominic Kwiatkowski, Cordelia Langford, John Sillitoe on behalf of the Wellcome Sanger Institute COVID-19 Surveillance Team |
| EPI_ISL_3092993, EPI_ISL_3092994 | Lighthouse Lab in Glasgow | Wellcome Sanger Institute for the COVID-19 Genomics UK (COG-UK) Consortium | Harper VanSteenhouse, Yumi Kasai, David Gray, Carol Clugston, Anna Dominiczak and Alex Alderton, Roberto Amato, Jeffrey Barrett, Sonia Goncalves, Ewan Harrison, David K. Jackson, Ian Johnston, Dominic Kwiatkowski, Cordelia Langford, John Sillitoe on behalf of the Wellcome Sanger Institute COVID-19 Surveillance Team |
| EPI_ISL_3092995 | Lighthouse Lab in Alderley Park | Wellcome Sanger Institute for the COVID-19 Genomics UK (COG-UK) Consortium | Jacquelyn Wynn, Mairead Hyland, The Lighthouse Lab in Alderley Park and Alex Alderton, Roberto Amato, Jeffrey Barrett, Sonia Goncalves, Ewan Harrison, David K. Jackson, Ian Johnston, Dominic Kwiatkowski, Cordelia Langford, John Sillitoe on behalf of the Wellcome Sanger Institute COVID-19 Surveillance Team |
| EPI_ISL_3092996, EPI_ISL_3092997, EPI_ISL_3092998 | Berkshire and Surrey Pathology Services Lighthouse Laboratory | Wellcome Sanger Institute for the COVID-19 Genomics UK (COG-UK) Consortium | Berkshire and Surrey Pathology Services Lighthouse Laboratory and Alex Alderton, Roberto Amato, Jeffrey Barrett, Sonia Goncalves, Ewan Harrison, David K. Jackson, Ian Johnston, Dominic Kwiatkowski, Cordelia Langford, John Sillitoe on behalf of the Wellcome Sanger Institute COVID-19 Surveillance Team |
| EPI_ISL_3092999 | Lighthouse Lab in Alderley Park | Wellcome Sanger Institute for the COVID-19 Genomics UK (COG-UK) Consortium | Jacquelyn Wynn, Mairead Hyland, The Lighthouse Lab in Alderley Park and Alex Alderton, Roberto Amato, Jeffrey Barrett, Sonia Goncalves, Ewan Harrison, David K. Jackson, Ian Johnston, Dominic Kwiatkowski, Cordelia Langford, John Sillitoe on behalf of the Wellcome Sanger Institute COVID-19 Surveillance Team |
| EPI_ISL_3093000, EPI_ISL_3093001 | Lighthouse Lab in Glasgow | Wellcome Sanger Institute for the COVID-19 Genomics UK (COG-UK) Consortium | Harper VanSteenhouse, Yumi Kasai, David Gray, Carol Clugston, Anna Dominiczak and Alex Alderton, Roberto Amato, Jeffrey Barrett, Sonia Goncalves, Ewan Harrison, David K. Jackson, Ian Johnston, Dominic Kwiatkowski, Cordelia Langford, John Sillitoe on behalf of the Wellcome Sanger Institute COVID-19 Surveillance Team |
| EPI_ISL_3093002 | Lighthouse Lab in Alderley Park | Wellcome Sanger Institute for the COVID-19 Genomics UK (COG-UK) Consortium | Jacquelyn Wynn, Mairead Hyland, The Lighthouse Lab in Alderley Park and Alex Alderton, Roberto Amato, Jeffrey Barrett, Sonia Goncalves, Ewan Harrison, David K. Jackson, Ian Johnston, Dominic Kwiatkowski, Cordelia Langford, John Sillitoe on behalf of the Wellcome Sanger Institute COVID-19 Surveillance Team |
| EPI_ISL_3093003, EPI_ISL_3093004 | Berkshire and Surrey Pathology Services Lighthouse Laboratory | Wellcome Sanger Institute for the COVID-19 Genomics UK (COG-UK) Consortium | Berkshire and Surrey Pathology Services Lighthouse Laboratory and Alex Alderton, Roberto Amato, Jeffrey Barrett, Sonia Goncalves, Ewan Harrison, David K. Jackson, Ian Johnston, Dominic Kwiatkowski, Cordelia Langford, John Sillitoe on behalf of the Wellcome Sanger Institute COVID-19 Surveillance Team |
| EPI_ISL_3093005, EPI_ISL_3093006 | Lighthouse Lab in Milton Keynes | Wellcome Sanger Institute for the COVID-19 Genomics UK (COG-UK) Consortium | The Lighthouse Lab in Milton Keynes and Alex Alderton, Roberto Amato, Jeffrey Barrett, Sonia Goncalves, Ewan Harrison, David K. Jackson, Ian Johnston, Dominic Kwiatkowski, Cordelia Langford, John Sillitoe on behalf of the Wellcome Sanger Institute COVID-19 Surveillance Team |
| EPI_ISL_3093007 | Lighthouse Lab in Alderley Park | Wellcome Sanger Institute for the COVID-19 Genomics UK (COG-UK) Consortium | Jacquelyn Wynn, Mairead Hyland, The Lighthouse Lab in Alderley Park and Alex Alderton, Roberto Amato, Jeffrey Barrett, Sonia Goncalves, Ewan Harrison, David K. Jackson, Ian Johnston, Dominic Kwiatkowski, Cordelia Langford, John Sillitoe on behalf of the Wellcome Sanger Institute COVID-19 Surveillance Team |
| EPI_ISL_3093008 | Berkshire and Surrey Pathology Services Lighthouse Laboratory | Wellcome Sanger Institute for the COVID-19 Genomics UK (COG-UK) Consortium | Berkshire and Surrey Pathology Services Lighthouse Laboratory and Alex Alderton, Roberto Amato, Jeffrey Barrett, Sonia Goncalves, Ewan Harrison, David K. Jackson, Ian Johnston, Dominic Kwiatkowski, Cordelia Langford, John Sillitoe on behalf of the Wellcome Sanger Institute COVID-19 Surveillance Team |
| EPI_ISL_3093009, EPI_ISL_3093010 | Lighthouse Lab in Alderley Park | Wellcome Sanger Institute for the COVID-19 Genomics UK (COG-UK) Consortium | Jacquelyn Wynn, Mairead Hyland, The Lighthouse Lab in Alderley Park and Alex Alderton, Roberto Amato, Jeffrey Barrett, Sonia Goncalves, Ewan Harrison, David K. Jackson, Ian Johnston, Dominic Kwiatkowski, Cordelia Langford, John Sillitoe on behalf of the Wellcome Sanger Institute COVID-19 Surveillance Team |
| EPI_ISL_3093011, EPI_ISL_3093012 | Lighthouse Lab in Milton Keynes | Wellcome Sanger Institute for the COVID-19 Genomics UK (COG-UK) Consortium | The Lighthouse Lab in Milton Keynes and Alex Alderton, Roberto Amato, Jeffrey Barrett, Sonia Goncalves, Ewan Harrison, David K. Jackson, Ian Johnston, Dominic Kwiatkowski, Cordelia Langford, John Sillitoe on behalf of the Wellcome Sanger Institute COVID-19 Surveillance Team |
| EPI_ISL_3093013 | Lighthouse Lab in Glasgow | Wellcome Sanger Institute for the COVID-19 Genomics UK (COG-UK) Consortium | Harper VanSteenhouse, Yumi Kasai, David Gray, Carol Clugston, Anna Dominiczak and Alex Alderton, Roberto Amato, Jeffrey Barrett, Sonia Goncalves, Ewan Harrison, David K. Jackson, Ian Johnston, Dominic Kwiatkowski, Cordelia Langford, John Sillitoe on behalf of the Wellcome Sanger Institute COVID-19 Surveillance Team |
| EPI_ISL_3093014 | Lighthouse Lab in Milton Keynes | Wellcome Sanger Institute for the COVID-19 Genomics UK (COG-UK) Consortium | The Lighthouse Lab in Milton Keynes and Alex Alderton, Roberto Amato, Jeffrey Barrett, Sonia Goncalves, Ewan Harrison, David K. Jackson, Ian Johnston, Dominic Kwiatkowski, Cordelia Langford, John Sillitoe on behalf of the Wellcome Sanger Institute COVID-19 Surveillance Team |
| EPI_ISL_3093015 | Berkshire and Surrey Pathology Services Lighthouse Laboratory | Wellcome Sanger Institute for the COVID-19 Genomics UK (COG-UK) Consortium | Berkshire and Surrey Pathology Services Lighthouse Laboratory and Alex Alderton, Roberto Amato, Jeffrey Barrett, Sonia Goncalves, Ewan Harrison, David K. Jackson, Ian Johnston, Dominic Kwiatkowski, Cordelia Langford, John Sillitoe on behalf of the Wellcome Sanger Institute COVID-19 Surveillance Team |

| | | | |
|---|---|---|---|
| EPI_ISL_3093016, EPI_ISL_3093017 | Lighthouse Lab in Milton Keynes | Wellcome Sanger Institute for the COVID-19 Genomics UK (COG-UK) Consortium | The Lighthouse Lab in Milton Keynes and Alex Alderton, Roberto Amato, Jeffrey Barrett, Sonia Goncalves, Ewan Harrison, David K. Jackson, Ian Johnston, Dominic Kwiatkowski, Cordelia Langford, John Sillitoe on behalf of the Wellcome Sanger Institute COVID-19 Surveillance Team |
| EPI_ISL_3093018 | Lighthouse Lab in Glasgow | Wellcome Sanger Institute for the COVID-19 Genomics UK (COG-UK) Consortium | Harper VanSteenhouse, Yumi Kasai, David Gray, Carol Clugston, Anna Dominiczak and Alex Alderton, Roberto Amato, Jeffrey Barrett, Sonia Goncalves, Ewan Harrison, David K. Jackson, Ian Johnston, Dominic Kwiatkowski, Cordelia Langford, John Sillitoe on behalf of the Wellcome Sanger Institute COVID-19 Surveillance Team |
| EPI_ISL_3093019 | Lighthouse Lab in Alderley Park | Wellcome Sanger Institute for the COVID-19 Genomics UK (COG-UK) Consortium | Jacquelyn Wynn, Mairead Hyland, The Lighthouse Lab in Alderley Park and Alex Alderton, Roberto Amato, Jeffrey Barrett, Sonia Goncalves, Ewan Harrison, David K. Jackson, Ian Johnston, Dominic Kwiatkowski, Cordelia Langford, John Sillitoe on behalf of the Wellcome Sanger Institute COVID-19 Surveillance Team |
| EPI_ISL_3093020 | Lighthouse Lab in Milton Keynes | Wellcome Sanger Institute for the COVID-19 Genomics UK (COG-UK) Consortium | The Lighthouse Lab in Milton Keynes and Alex Alderton, Roberto Amato, Jeffrey Barrett, Sonia Goncalves, Ewan Harrison, David K. Jackson, Ian Johnston, Dominic Kwiatkowski, Cordelia Langford, John Sillitoe on behalf of the Wellcome Sanger Institute COVID-19 Surveillance Team |
| EPI_ISL_3093021, EPI_ISL_3093022, EPI_ISL_3093023 | Lighthouse Lab in Alderley Park | Wellcome Sanger Institute for the COVID-19 Genomics UK (COG-UK) Consortium | Jacquelyn Wynn, Mairead Hyland, The Lighthouse Lab in Alderley Park and Alex Alderton, Roberto Amato, Jeffrey Barrett, Sonia Goncalves, Ewan Harrison, David K. Jackson, Ian Johnston, Dominic Kwiatkowski, Cordelia Langford, John Sillitoe on behalf of the Wellcome Sanger Institute COVID-19 Surveillance Team |
| EPI_ISL_3093024 | Lighthouse Lab in Milton Keynes | Wellcome Sanger Institute for the COVID-19 Genomics UK (COG-UK) Consortium | The Lighthouse Lab in Milton Keynes and Alex Alderton, Roberto Amato, Jeffrey Barrett, Sonia Goncalves, Ewan Harrison, David K. Jackson, Ian Johnston, Dominic Kwiatkowski, Cordelia Langford, John Sillitoe on behalf of the Wellcome Sanger Institute COVID-19 Surveillance Team |
| EPI_ISL_3093025 | Berkshire and Surrey Pathology Services Lighthouse Laboratory | Wellcome Sanger Institute for the COVID-19 Genomics UK (COG-UK) Consortium | Berkshire and Surrey Pathology Services Lighthouse Laboratory and Alex Alderton, Roberto Amato, Jeffrey Barrett, Sonia Goncalves, Ewan Harrison, David K. Jackson, Ian Johnston, Dominic Kwiatkowski, Cordelia Langford, John Sillitoe on behalf of the Wellcome Sanger Institute COVID-19 Surveillance Team |
| EPI_ISL_3093026 | Lighthouse Lab in Alderley Park | Wellcome Sanger Institute for the COVID-19 Genomics UK (COG-UK) Consortium | Jacquelyn Wynn, Mairead Hyland, The Lighthouse Lab in Alderley Park and Alex Alderton, Roberto Amato, Jeffrey Barrett, Sonia Goncalves, Ewan Harrison, David K. Jackson, Ian Johnston, Dominic Kwiatkowski, Cordelia Langford, John Sillitoe on behalf of the Wellcome Sanger Institute COVID-19 Surveillance Team |
| EPI_ISL_3093027 | Lighthouse Lab in Glasgow | Wellcome Sanger Institute for the COVID-19 Genomics UK (COG-UK) Consortium | Harper VanSteenhouse, Yumi Kasai, David Gray, Carol Clugston, Anna Dominiczak and Alex Alderton, Roberto Amato, Jeffrey Barrett, Sonia Goncalves, Ewan Harrison, David K. Jackson, Ian Johnston, Dominic Kwiatkowski, Cordelia Langford, John Sillitoe on behalf of the Wellcome Sanger Institute COVID-19 Surveillance Team |
| EPI_ISL_3093028 | Lighthouse Lab in Milton Keynes | Wellcome Sanger Institute for the COVID-19 Genomics UK (COG-UK) Consortium | The Lighthouse Lab in Milton Keynes and Alex Alderton, Roberto Amato, Jeffrey Barrett, Sonia Goncalves, Ewan Harrison, David K. Jackson, Ian Johnston, Dominic Kwiatkowski, Cordelia Langford, John Sillitoe on behalf of the Wellcome Sanger Institute COVID-19 Surveillance Team |
| EPI_ISL_3093029 | Lighthouse Lab in Glasgow | Wellcome Sanger Institute for the COVID-19 Genomics UK (COG-UK) Consortium | Harper VanSteenhouse, Yumi Kasai, David Gray, Carol Clugston, Anna Dominiczak and Alex Alderton, Roberto Amato, Jeffrey Barrett, Sonia Goncalves, Ewan Harrison, David K. Jackson, Ian Johnston, Dominic Kwiatkowski, Cordelia Langford, John Sillitoe on behalf of the Wellcome Sanger Institute COVID-19 Surveillance Team |
| EPI_ISL_3093030 | Lighthouse Lab in Milton Keynes | Wellcome Sanger Institute for the COVID-19 Genomics UK (COG-UK) Consortium | The Lighthouse Lab in Milton Keynes and Alex Alderton, Roberto Amato, Jeffrey Barrett, Sonia Goncalves, Ewan Harrison, David K. Jackson, Ian Johnston, Dominic Kwiatkowski, Cordelia Langford, John Sillitoe on behalf of the Wellcome Sanger Institute COVID-19 Surveillance Team |
| EPI_ISL_3093031 | Berkshire and Surrey Pathology Services Lighthouse Laboratory | Wellcome Sanger Institute for the COVID-19 Genomics UK (COG-UK) Consortium | Berkshire and Surrey Pathology Services Lighthouse Laboratory and Alex Alderton, Roberto Amato, Jeffrey Barrett, Sonia Goncalves, Ewan Harrison, David K. Jackson, Ian Johnston, Dominic Kwiatkowski, Cordelia Langford, John Sillitoe on behalf of the Wellcome Sanger Institute COVID-19 Surveillance Team |
| EPI_ISL_3093032, EPI_ISL_3093033 | Lighthouse Lab in Alderley Park | Wellcome Sanger Institute for the COVID-19 Genomics UK (COG-UK) Consortium | Jacquelyn Wynn, Mairead Hyland, The Lighthouse Lab in Alderley Park and Alex Alderton, Roberto Amato, Jeffrey Barrett, Sonia Goncalves, Ewan Harrison, David K. Jackson, Ian Johnston, Dominic Kwiatkowski, Cordelia Langford, John Sillitoe on behalf of the Wellcome Sanger Institute COVID-19 Surveillance Team |
| EPI_ISL_3093034 | Lighthouse Lab in Glasgow | Wellcome Sanger Institute for the COVID-19 Genomics UK (COG-UK) Consortium | Harper VanSteenhouse, Yumi Kasai, David Gray, Carol Clugston, Anna Dominiczak and Alex Alderton, Roberto Amato, Jeffrey Barrett, Sonia Goncalves, Ewan Harrison, David K. Jackson, Ian Johnston, Dominic Kwiatkowski, Cordelia Langford, John Sillitoe on behalf of the Wellcome Sanger Institute COVID-19 Surveillance Team |
| EPI_ISL_3093035, EPI_ISL_3093036, EPI_ISL_3093037 | Lighthouse Lab in Alderley Park | Wellcome Sanger Institute for the COVID-19 Genomics UK (COG-UK) Consortium | Jacquelyn Wynn, Mairead Hyland, The Lighthouse Lab in Alderley Park and Alex Alderton, Roberto Amato, Jeffrey Barrett, Sonia Goncalves, Ewan Harrison, David K. Jackson, Ian Johnston, Dominic Kwiatkowski, Cordelia Langford, John Sillitoe on behalf of the Wellcome Sanger Institute COVID-19 Surveillance Team |
| EPI_ISL_3093038 | Lighthouse Lab in Glasgow | Wellcome Sanger Institute for the COVID-19 Genomics UK (COG-UK) Consortium | Harper VanSteenhouse, Yumi Kasai, David Gray, Carol Clugston, Anna Dominiczak and Alex Alderton, Roberto Amato, Jeffrey Barrett, Sonia Goncalves, Ewan Harrison, David K. Jackson, Ian Johnston, Dominic Kwiatkowski, Cordelia Langford, John Sillitoe on behalf of the Wellcome Sanger Institute COVID-19 Surveillance Team |
| EPI_ISL_3093039 | Lighthouse Lab in Alderley Park | Wellcome Sanger Institute for the COVID-19 Genomics UK (COG-UK) Consortium | Jacquelyn Wynn, Mairead Hyland, The Lighthouse Lab in Alderley Park and Alex Alderton, Roberto Amato, Jeffrey Barrett, Sonia Goncalves, Ewan Harrison, David K. Jackson, Ian Johnston, Dominic Kwiatkowski, Cordelia Langford, John Sillitoe on behalf of the Wellcome Sanger Institute COVID-19 Surveillance Team |
| EPI_ISL_3093040, EPI_ISL_3093041 | Lighthouse Lab in Glasgow | Wellcome Sanger Institute for the COVID-19 Genomics UK (COG-UK) Consortium | Harper VanSteenhouse, Yumi Kasai, David Gray, Carol Clugston, Anna Dominiczak and Alex Alderton, Roberto Amato, Jeffrey Barrett, Sonia Goncalves, Ewan Harrison, David K. Jackson, Ian Johnston, Dominic Kwiatkowski, Cordelia Langford, John Sillitoe on behalf of the Wellcome Sanger Institute COVID-19 Surveillance Team |
| EPI_ISL_3093042 | Lighthouse Lab in Alderley Park | Wellcome Sanger Institute for the COVID-19 Genomics UK (COG-UK) Consortium | Jacquelyn Wynn, Mairead Hyland, The Lighthouse Lab in Alderley Park and Alex Alderton, Roberto Amato, Jeffrey Barrett, Sonia Goncalves, Ewan Harrison, David K. Jackson, Ian Johnston, Dominic Kwiatkowski, Cordelia Langford, John Sillitoe on behalf of the Wellcome Sanger Institute COVID-19 Surveillance Team |
| EPI_ISL_3093043 | Berkshire and Surrey Pathology Services Lighthouse Laboratory | Wellcome Sanger Institute for the COVID-19 Genomics UK (COG-UK) Consortium | Berkshire and Surrey Pathology Services Lighthouse Laboratory and Alex Alderton, Roberto Amato, Jeffrey Barrett, Sonia Goncalves, Ewan Harrison, David K. Jackson, Ian Johnston, Dominic Kwiatkowski, Cordelia Langford, John Sillitoe on behalf of the Wellcome Sanger Institute COVID-19 Surveillance Team |
| EPI_ISL_3093044, EPI_ISL_3093045 | Lighthouse Lab in Milton Keynes | Wellcome Sanger Institute for the COVID-19 Genomics UK (COG-UK) Consortium | The Lighthouse Lab in Milton Keynes and Alex Alderton, Roberto Amato, Jeffrey Barrett, Sonia Goncalves, Ewan Harrison, David K. Jackson, Ian Johnston, Dominic Kwiatkowski, Cordelia Langford, John Sillitoe on behalf of the Wellcome Sanger Institute COVID-19 Surveillance Team |
| EPI_ISL_3093046 | Berkshire and Surrey Pathology Services Lighthouse Laboratory | Wellcome Sanger Institute for the COVID-19 Genomics UK (COG-UK) Consortium | Berkshire and Surrey Pathology Services Lighthouse Laboratory and Alex Alderton, Roberto Amato, Jeffrey Barrett, Sonia Goncalves, Ewan Harrison, David K. Jackson, Ian Johnston, Dominic Kwiatkowski, Cordelia Langford, John Sillitoe on behalf of the Wellcome Sanger Institute COVID-19 Surveillance Team |
| EPI_ISL_3093047 | Lighthouse Lab in Milton Keynes | Wellcome Sanger Institute for the COVID-19 Genomics UK (COG-UK) Consortium | The Lighthouse Lab in Milton Keynes and Alex Alderton, Roberto Amato, Jeffrey Barrett, Sonia Goncalves, Ewan Harrison, David K. Jackson, Ian Johnston, Dominic Kwiatkowski, Cordelia Langford, John Sillitoe on behalf of the Wellcome Sanger Institute COVID-19 Surveillance Team |
| EPI_ISL_3093048 | Berkshire and Surrey Pathology Services Lighthouse Laboratory | Wellcome Sanger Institute for the COVID-19 Genomics UK (COG-UK) Consortium | Berkshire and Surrey Pathology Services Lighthouse Laboratory and Alex Alderton, Roberto Amato, Jeffrey Barrett, Sonia Goncalves, Ewan Harrison, David K. Jackson, Ian Johnston, Dominic Kwiatkowski, Cordelia Langford, John Sillitoe on behalf of the Wellcome Sanger Institute COVID-19 Surveillance Team |
| EPI_ISL_3093049 | Lighthouse Lab in Glasgow | Wellcome Sanger Institute for the COVID-19 Genomics UK (COG-UK) Consortium | Harper VanSteenhouse, Yumi Kasai, David Gray, Carol Clugston, Anna Dominiczak and Alex Alderton, Roberto Amato, Jeffrey Barrett, Sonia Goncalves, Ewan Harrison, David K. Jackson, Ian Johnston, Dominic Kwiatkowski, Cordelia Langford, John Sillitoe on behalf of the Wellcome Sanger Institute COVID-19 Surveillance Team |
| EPI_ISL_3093050 | Berkshire and Surrey Pathology Services Lighthouse Laboratory | Wellcome Sanger Institute for the COVID-19 Genomics UK (COG-UK) Consortium | Berkshire and Surrey Pathology Services Lighthouse Laboratory and Alex Alderton, Roberto Amato, Jeffrey Barrett, Sonia Goncalves, Ewan Harrison, David K. Jackson, Ian Johnston, Dominic Kwiatkowski, Cordelia Langford, John Sillitoe on behalf of the Wellcome Sanger Institute COVID-19 Surveillance Team |
| EPI_ISL_3093051, EPI_ISL_3093052, EPI_ISL_3093053 | Lighthouse Lab in Alderley Park | Wellcome Sanger Institute for the COVID-19 Genomics UK (COG-UK) Consortium | Jacquelyn Wynn, Mairead Hyland, The Lighthouse Lab in Alderley Park and Alex Alderton, Roberto Amato, Jeffrey Barrett, Sonia Goncalves, Ewan Harrison, David K. Jackson, Ian Johnston, Dominic Kwiatkowski, Cordelia Langford, John Sillitoe on behalf of the Wellcome Sanger Institute COVID-19 Surveillance Team |
| EPI_ISL_3093054, EPI_ISL_3093055, EPI_ISL_3093056, EPI_ISL_3093057, EPI_ISL_3093058 | Berkshire and Surrey Pathology Services Lighthouse Laboratory | Wellcome Sanger Institute for the COVID-19 Genomics UK (COG-UK) Consortium | Berkshire and Surrey Pathology Services Lighthouse Laboratory and Alex Alderton, Roberto Amato, Jeffrey Barrett, Sonia Goncalves, Ewan Harrison, David K. Jackson, Ian Johnston, Dominic Kwiatkowski, Cordelia Langford, John Sillitoe on behalf of the Wellcome Sanger Institute COVID-19 Surveillance Team |
| EPI_ISL_3093059, EPI_ISL_3093060 | Lighthouse Lab in Alderley Park | Wellcome Sanger Institute for the COVID-19 Genomics UK (COG-UK) Consortium | Jacquelyn Wynn, Mairead Hyland, The Lighthouse Lab in Alderley Park and Alex Alderton, Roberto Amato, Jeffrey Barrett, Sonia Goncalves, Ewan Harrison, David K. Jackson, Ian Johnston, Dominic Kwiatkowski, Cordelia Langford, John Sillitoe on behalf of the Wellcome Sanger Institute COVID-19 Surveillance Team |
| EPI_ISL_3093061 | Lighthouse Lab in Milton Keynes | Wellcome Sanger Institute for the COVID-19 Genomics UK (COG-UK) Consortium | The Lighthouse Lab in Milton Keynes and Alex Alderton, Roberto Amato, Jeffrey Barrett, Sonia Goncalves, Ewan Harrison, David K. Jackson, Ian Johnston, Dominic Kwiatkowski, Cordelia Langford, John Sillitoe on behalf of the Wellcome Sanger Institute COVID-19 Surveillance Team |
| EPI_ISL_3093062 | Lighthouse Lab in Alderley Park | Wellcome Sanger Institute for the COVID-19 Genomics UK (COG-UK) Consortium | Jacquelyn Wynn, Mairead Hyland, The Lighthouse Lab in Alderley Park and Alex Alderton, Roberto Amato, Jeffrey Barrett, Sonia Goncalves, Ewan Harrison, David K. Jackson, Ian Johnston, Dominic Kwiatkowski, Cordelia Langford, John Sillitoe on behalf of the Wellcome Sanger Institute COVID-19 Surveillance Team |
| EPI_ISL_3093063, EPI_ISL_3093064 | Berkshire and Surrey Pathology Services Lighthouse Laboratory | Wellcome Sanger Institute for the COVID-19 Genomics UK (COG-UK) Consortium | Berkshire and Surrey Pathology Services Lighthouse Laboratory and Alex Alderton, Roberto Amato, Jeffrey Barrett, Sonia Goncalves, Ewan Harrison, David K. Jackson, Ian Johnston, Dominic Kwiatkowski, Cordelia Langford, John Sillitoe on behalf of the Wellcome Sanger Institute COVID-19 Surveillance Team |
| EPI_ISL_3093065 | Lighthouse Lab in Glasgow | Wellcome Sanger Institute for the COVID-19 Genomics UK (COG-UK) Consortium | Harper VanSteenhouse, Yumi Kasai, David Gray, Carol Clugston, Anna Dominiczak and Alex Alderton, Roberto Amato, Jeffrey Barrett, Sonia Goncalves, Ewan Harrison, David K. Jackson, Ian Johnston, Dominic Kwiatkowski, Cordelia Langford, John Sillitoe on behalf of the Wellcome Sanger Institute COVID-19 Surveillance Team |
| EPI_ISL_3093066 | Lighthouse Lab in Alderley Park | Wellcome Sanger Institute for the COVID-19 Genomics UK (COG-UK) Consortium | Jacquelyn Wynn, Mairead Hyland, The Lighthouse Lab in Alderley Park and Alex Alderton, Roberto Amato, Jeffrey Barrett, Sonia Goncalves, Ewan Harrison, David K. Jackson, Ian Johnston, Dominic Kwiatkowski, Cordelia Langford, John Sillitoe on behalf of the Wellcome Sanger Institute COVID-19 Surveillance Team |
| EPI_ISL_3093067 | Lighthouse Lab in Milton Keynes | Wellcome Sanger Institute for the COVID-19 Genomics UK (COG-UK) Consortium | The Lighthouse Lab in Milton Keynes and Alex Alderton, Roberto Amato, Jeffrey Barrett, Sonia Goncalves, Ewan Harrison, David K. Jackson, Ian Johnston, Dominic Kwiatkowski, Cordelia Langford, John Sillitoe on behalf of the Wellcome Sanger Institute COVID-19 Surveillance Team |
| EPI_ISL_3093068 | Berkshire and Surrey Pathology Services Lighthouse Laboratory | Wellcome Sanger Institute for the COVID-19 Genomics UK (COG-UK) Consortium | Berkshire and Surrey Pathology Services Lighthouse Laboratory and Alex Alderton, Roberto Amato, Jeffrey Barrett, Sonia Goncalves, Ewan Harrison, David K. Jackson, Ian Johnston, Dominic Kwiatkowski, Cordelia Langford, John Sillitoe on behalf of the Wellcome Sanger Institute COVID-19 Surveillance Team |
| EPI_ISL_3093069 | Lighthouse Lab in Milton Keynes | Wellcome Sanger Institute for the COVID-19 Genomics UK (COG-UK) Consortium | The Lighthouse Lab in Milton Keynes and Alex Alderton, Roberto Amato, Jeffrey Barrett, Sonia Goncalves, Ewan Harrison, David K. Jackson, Ian Johnston, Dominic Kwiatkowski, Cordelia Langford, John Sillitoe on behalf of the Wellcome Sanger Institute COVID-19 Surveillance Team |
| EPI_ISL_3093070, EPI_ISL_3093071 | Berkshire and Surrey Pathology Services Lighthouse Laboratory | Wellcome Sanger Institute for the COVID-19 Genomics UK (COG-UK) Consortium | Berkshire and Surrey Pathology Services Lighthouse Laboratory and Alex Alderton, Roberto Amato, Jeffrey Barrett, Sonia Goncalves, Ewan Harrison, David K. Jackson, Ian Johnston, Dominic Kwiatkowski, Cordelia Langford, John Sillitoe on behalf of the Wellcome Sanger Institute COVID-19 Surveillance Team |
| EPI_ISL_3093072, EPI_ISL_3093073 | Lighthouse Lab in Alderley Park | Wellcome Sanger Institute for the COVID-19 Genomics UK (COG-UK) Consortium | Jacquelyn Wynn, Mairead Hyland, The Lighthouse Lab in Alderley Park and Alex Alderton, Roberto Amato, Jeffrey Barrett, Sonia Goncalves, Ewan Harrison, David K. Jackson, Ian Johnston, Dominic Kwiatkowski, Cordelia Langford, John Sillitoe on behalf of the Wellcome Sanger Institute COVID-19 Surveillance Team |
| EPI_ISL_3093074 | Lighthouse Lab in Glasgow | Wellcome Sanger Institute for the COVID-19 Genomics UK (COG-UK) Consortium | Harper VanSteenhouse, Yumi Kasai, David Gray, Carol Clugston, Anna Dominiczak and Alex Alderton, Roberto Amato, Jeffrey Barrett, Sonia Goncalves, Ewan Harrison, David K. Jackson, Ian Johnston, Dominic Kwiatkowski, Cordelia Langford, John Sillitoe on behalf of the Wellcome Sanger Institute COVID-19 Surveillance Team |
| EPI_ISL_3093075 | Lighthouse Lab in Milton Keynes | Wellcome Sanger Institute for the COVID-19 Genomics UK (COG-UK) Consortium | The Lighthouse Lab in Milton Keynes and Alex Alderton, Roberto Amato, Jeffrey Barrett, Sonia Goncalves, Ewan Harrison, David K. Jackson, Ian Johnston, Dominic Kwiatkowski, Cordelia Langford, John Sillitoe on behalf of the Wellcome Sanger Institute COVID-19 Surveillance Team |
| EPI_ISL_3093076, EPI_ISL_3093077, | Lighthouse Lab in Alderley Park | Wellcome Sanger Institute for the COVID-19 Genomics UK (COG-UK) Consortium | Jacquelyn Wynn, Mairead Hyland, The Lighthouse Lab in Alderley Park and Alex Alderton, Roberto Amato, Jeffrey Barrett, Sonia Goncalves, Ewan Harrison, David K. Jackson, Ian Johnston, Dominic Kwiatkowski, Cordelia Langford, John Sillitoe on behalf of the Wellcome Sanger Institute COVID-19 Surveillance Team |

| | | | |
|---|---|---|---|
| EPI_ISL_3093078, EPI_ISL_3093079, EPI_ISL_3093080 | | | |
| EPI_ISL_3093081 | Berkshire and Surrey Pathology Services Lighthouse Laboratory | Wellcome Sanger Institute for the COVID-19 Genomics UK (COG-UK) Consortium | Berkshire and Surrey Pathology Services Lighthouse Laboratory and Alex Alderton, Roberto Amato, Jeffrey Barrett, Sonia Goncalves, Ewan Harrison, David K. Jackson, Ian Johnston, Dominic Kwiatkowski, Cordelia Langford, John Sillitoe on behalf of the Wellcome Sanger Institute COVID-19 Surveillance Team |
| EPI_ISL_3093082 | Lighthouse Lab in Alderley Park | Wellcome Sanger Institute for the COVID-19 Genomics UK (COG-UK) Consortium | Jacquelyn Wynn, Mairead Hyland, The Lighthouse Lab in Alderley Park and Alex Alderton, Roberto Amato, Jeffrey Barrett, Sonia Goncalves, Ewan Harrison, David K. Jackson, Ian Johnston, Dominic Kwiatkowski, Cordelia Langford, John Sillitoe on behalf of the Wellcome Sanger Institute COVID-19 Surveillance Team |
| EPI_ISL_3093083 | Lighthouse Lab in Milton Keynes | Wellcome Sanger Institute for the COVID-19 Genomics UK (COG-UK) Consortium | The Lighthouse Lab in Milton Keynes and Alex Alderton, Roberto Amato, Jeffrey Barrett, Sonia Goncalves, Ewan Harrison, David K. Jackson, Ian Johnston, Dominic Kwiatkowski, Cordelia Langford, John Sillitoe on behalf of the Wellcome Sanger Institute COVID-19 Surveillance Team |
| EPI_ISL_3093084 | Lighthouse Lab in Glasgow | Wellcome Sanger Institute for the COVID-19 Genomics UK (COG-UK) Consortium | Harper VanSteenhouse, Yumi Kasai, David Gray, Carol Clugston, Anna Dominiczak and Alex Alderton, Roberto Amato, Jeffrey Barrett, Sonia Goncalves, Ewan Harrison, David K. Jackson, Ian Johnston, Dominic Kwiatkowski, Cordelia Langford, John Sillitoe on behalf of the Wellcome Sanger Institute COVID-19 Surveillance Team |
| EPI_ISL_3093085, EPI_ISL_3093086 | Berkshire and Surrey Pathology Services Lighthouse Laboratory | Wellcome Sanger Institute for the COVID-19 Genomics UK (COG-UK) Consortium | Berkshire and Surrey Pathology Services Lighthouse Laboratory and Alex Alderton, Roberto Amato, Jeffrey Barrett, Sonia Goncalves, Ewan Harrison, David K. Jackson, Ian Johnston, Dominic Kwiatkowski, Cordelia Langford, John Sillitoe on behalf of the Wellcome Sanger Institute COVID-19 Surveillance Team |
| EPI_ISL_3093087 | Lighthouse Lab in Alderley Park | Wellcome Sanger Institute for the COVID-19 Genomics UK (COG-UK) Consortium | Jacquelyn Wynn, Mairead Hyland, The Lighthouse Lab in Alderley Park and Alex Alderton, Roberto Amato, Jeffrey Barrett, Sonia Goncalves, Ewan Harrison, David K. Jackson, Ian Johnston, Dominic Kwiatkowski, Cordelia Langford, John Sillitoe on behalf of the Wellcome Sanger Institute COVID-19 Surveillance Team |
| EPI_ISL_3093088 | Lighthouse Lab in Milton Keynes | Wellcome Sanger Institute for the COVID-19 Genomics UK (COG-UK) Consortium | The Lighthouse Lab in Milton Keynes and Alex Alderton, Roberto Amato, Jeffrey Barrett, Sonia Goncalves, Ewan Harrison, David K. Jackson, Ian Johnston, Dominic Kwiatkowski, Cordelia Langford, John Sillitoe on behalf of the Wellcome Sanger Institute COVID-19 Surveillance Team |
| EPI_ISL_3093089, EPI_ISL_3093090, EPI_ISL_3093091 | Lighthouse Lab in Glasgow | Wellcome Sanger Institute for the COVID-19 Genomics UK (COG-UK) Consortium | Harper VanSteenhouse, Yumi Kasai, David Gray, Carol Clugston, Anna Dominiczak and Alex Alderton, Roberto Amato, Jeffrey Barrett, Sonia Goncalves, Ewan Harrison, David K. Jackson, Ian Johnston, Dominic Kwiatkowski, Cordelia Langford, John Sillitoe on behalf of the Wellcome Sanger Institute COVID-19 Surveillance Team |
| EPI_ISL_3093092 | Berkshire and Surrey Pathology Services Lighthouse Laboratory | Wellcome Sanger Institute for the COVID-19 Genomics UK (COG-UK) Consortium | Berkshire and Surrey Pathology Services Lighthouse Laboratory and Alex Alderton, Roberto Amato, Jeffrey Barrett, Sonia Goncalves, Ewan Harrison, David K. Jackson, Ian Johnston, Dominic Kwiatkowski, Cordelia Langford, John Sillitoe on behalf of the Wellcome Sanger Institute COVID-19 Surveillance Team |
| EPI_ISL_3093093 | Lighthouse Lab in Milton Keynes | Wellcome Sanger Institute for the COVID-19 Genomics UK (COG-UK) Consortium | The Lighthouse Lab in Milton Keynes and Alex Alderton, Roberto Amato, Jeffrey Barrett, Sonia Goncalves, Ewan Harrison, David K. Jackson, Ian Johnston, Dominic Kwiatkowski, Cordelia Langford, John Sillitoe on behalf of the Wellcome Sanger Institute COVID-19 Surveillance Team |
| EPI_ISL_3093094 | Lighthouse Lab in Glasgow | Wellcome Sanger Institute for the COVID-19 Genomics UK (COG-UK) Consortium | Harper VanSteenhouse, Yumi Kasai, David Gray, Carol Clugston, Anna Dominiczak and Alex Alderton, Roberto Amato, Jeffrey Barrett, Sonia Goncalves, Ewan Harrison, David K. Jackson, Ian Johnston, Dominic Kwiatkowski, Cordelia Langford, John Sillitoe on behalf of the Wellcome Sanger Institute COVID-19 Surveillance Team |
| EPI_ISL_3093095 | Lighthouse Lab in Alderley Park | Wellcome Sanger Institute for the COVID-19 Genomics UK (COG-UK) Consortium | Jacquelyn Wynn, Mairead Hyland, The Lighthouse Lab in Alderley Park and Alex Alderton, Roberto Amato, Jeffrey Barrett, Sonia Goncalves, Ewan Harrison, David K. Jackson, Ian Johnston, Dominic Kwiatkowski, Cordelia Langford, John Sillitoe on behalf of the Wellcome Sanger Institute COVID-19 Surveillance Team |
| EPI_ISL_3093096 | Berkshire and Surrey Pathology Services Lighthouse Laboratory | Wellcome Sanger Institute for the COVID-19 Genomics UK (COG-UK) Consortium | Berkshire and Surrey Pathology Services Lighthouse Laboratory and Alex Alderton, Roberto Amato, Jeffrey Barrett, Sonia Goncalves, Ewan Harrison, David K. Jackson, Ian Johnston, Dominic Kwiatkowski, Cordelia Langford, John Sillitoe on behalf of the Wellcome Sanger Institute COVID-19 Surveillance Team |
| EPI_ISL_3093097 | Lighthouse Lab in Milton Keynes | Wellcome Sanger Institute for the COVID-19 Genomics UK (COG-UK) Consortium | The Lighthouse Lab in Milton Keynes and Alex Alderton, Roberto Amato, Jeffrey Barrett, Sonia Goncalves, Ewan Harrison, David K. Jackson, Ian Johnston, Dominic Kwiatkowski, Cordelia Langford, John Sillitoe on behalf of the Wellcome Sanger Institute COVID-19 Surveillance Team |
| EPI_ISL_3093098 | Berkshire and Surrey Pathology Services Lighthouse Laboratory | Wellcome Sanger Institute for the COVID-19 Genomics UK (COG-UK) Consortium | Berkshire and Surrey Pathology Services Lighthouse Laboratory and Alex Alderton, Roberto Amato, Jeffrey Barrett, Sonia Goncalves, Ewan Harrison, David K. Jackson, Ian Johnston, Dominic Kwiatkowski, Cordelia Langford, John Sillitoe on behalf of the Wellcome Sanger Institute COVID-19 Surveillance Team |
| EPI_ISL_3093099 | Lighthouse Lab in Glasgow | Wellcome Sanger Institute for the COVID-19 Genomics UK (COG-UK) Consortium | Harper VanSteenhouse, Yumi Kasai, David Gray, Carol Clugston, Anna Dominiczak and Alex Alderton, Roberto Amato, Jeffrey Barrett, Sonia Goncalves, Ewan Harrison, David K. Jackson, Ian Johnston, Dominic Kwiatkowski, Cordelia Langford, John Sillitoe on behalf of the Wellcome Sanger Institute COVID-19 Surveillance Team |
| EPI_ISL_3093100 | Lighthouse Lab in Milton Keynes | Wellcome Sanger Institute for the COVID-19 Genomics UK (COG-UK) Consortium | The Lighthouse Lab in Milton Keynes and Alex Alderton, Roberto Amato, Jeffrey Barrett, Sonia Goncalves, Ewan Harrison, David K. Jackson, Ian Johnston, Dominic Kwiatkowski, Cordelia Langford, John Sillitoe on behalf of the Wellcome Sanger Institute COVID-19 Surveillance Team |
| EPI_ISL_3093101 | Berkshire and Surrey Pathology Services Lighthouse Laboratory | Wellcome Sanger Institute for the COVID-19 Genomics UK (COG-UK) Consortium | Berkshire and Surrey Pathology Services Lighthouse Laboratory and Alex Alderton, Roberto Amato, Jeffrey Barrett, Sonia Goncalves, Ewan Harrison, David K. Jackson, Ian Johnston, Dominic Kwiatkowski, Cordelia Langford, John Sillitoe on behalf of the Wellcome Sanger Institute COVID-19 Surveillance Team |
| EPI_ISL_3093102 | Lighthouse Lab in Milton Keynes | Wellcome Sanger Institute for the COVID-19 Genomics UK (COG-UK) Consortium | The Lighthouse Lab in Milton Keynes and Alex Alderton, Roberto Amato, Jeffrey Barrett, Sonia Goncalves, Ewan Harrison, David K. Jackson, Ian Johnston, Dominic Kwiatkowski, Cordelia Langford, John Sillitoe on behalf of the Wellcome Sanger Institute COVID-19 Surveillance Team |
| EPI_ISL_3093103 | Berkshire and Surrey Pathology Services Lighthouse Laboratory | Wellcome Sanger Institute for the COVID-19 Genomics UK (COG-UK) Consortium | Berkshire and Surrey Pathology Services Lighthouse Laboratory and Alex Alderton, Roberto Amato, Jeffrey Barrett, Sonia Goncalves, Ewan Harrison, David K. Jackson, Ian Johnston, Dominic Kwiatkowski, Cordelia Langford, John Sillitoe on behalf of the Wellcome Sanger Institute COVID-19 Surveillance Team |
| EPI_ISL_3093104 | Lighthouse Lab in Glasgow | Wellcome Sanger Institute for the COVID-19 Genomics UK (COG-UK) Consortium | Harper VanSteenhouse, Yumi Kasai, David Gray, Carol Clugston, Anna Dominiczak and Alex Alderton, Roberto Amato, Jeffrey Barrett, Sonia Goncalves, Ewan Harrison, David K. Jackson, Ian Johnston, Dominic Kwiatkowski, Cordelia Langford, John Sillitoe on behalf of the Wellcome Sanger Institute COVID-19 Surveillance Team |
| EPI_ISL_3093105 | Berkshire and Surrey Pathology Services Lighthouse Laboratory | Wellcome Sanger Institute for the COVID-19 Genomics UK (COG-UK) Consortium | Berkshire and Surrey Pathology Services Lighthouse Laboratory and Alex Alderton, Roberto Amato, Jeffrey Barrett, Sonia Goncalves, Ewan Harrison, David K. Jackson, Ian Johnston, Dominic Kwiatkowski, Cordelia Langford, John Sillitoe on behalf of the Wellcome Sanger Institute COVID-19 Surveillance Team |
| EPI_ISL_3093106 | Lighthouse Lab in Alderley Park | Wellcome Sanger Institute for the COVID-19 Genomics UK (COG-UK) Consortium | Jacquelyn Wynn, Mairead Hyland, The Lighthouse Lab in Alderley Park and Alex Alderton, Roberto Amato, Jeffrey Barrett, Sonia Goncalves, Ewan Harrison, David K. Jackson, Ian Johnston, Dominic Kwiatkowski, Cordelia Langford, John Sillitoe on behalf of the Wellcome Sanger Institute COVID-19 Surveillance Team |
| EPI_ISL_3093107 | Lighthouse Lab in Glasgow | Wellcome Sanger Institute for the COVID-19 Genomics UK (COG-UK) Consortium | Harper VanSteenhouse, Yumi Kasai, David Gray, Carol Clugston, Anna Dominiczak and Alex Alderton, Roberto Amato, Jeffrey Barrett, Sonia Goncalves, Ewan Harrison, David K. Jackson, Ian Johnston, Dominic Kwiatkowski, Cordelia Langford, John Sillitoe on behalf of the Wellcome Sanger Institute COVID-19 Surveillance Team |
| EPI_ISL_3093108 | Lighthouse Lab in Milton Keynes | Wellcome Sanger Institute for the COVID-19 Genomics UK (COG-UK) Consortium | The Lighthouse Lab in Milton Keynes and Alex Alderton, Roberto Amato, Jeffrey Barrett, Sonia Goncalves, Ewan Harrison, David K. Jackson, Ian Johnston, Dominic Kwiatkowski, Cordelia Langford, John Sillitoe on behalf of the Wellcome Sanger Institute COVID-19 Surveillance Team |
| EPI_ISL_3093109, EPI_ISL_3093110 | Lighthouse Lab in Glasgow | Wellcome Sanger Institute for the COVID-19 Genomics UK (COG-UK) Consortium | Harper VanSteenhouse, Yumi Kasai, David Gray, Carol Clugston, Anna Dominiczak and Alex Alderton, Roberto Amato, Jeffrey Barrett, Sonia Goncalves, Ewan Harrison, David K. Jackson, Ian Johnston, Dominic Kwiatkowski, Cordelia Langford, John Sillitoe on behalf of the Wellcome Sanger Institute COVID-19 Surveillance Team |
| EPI_ISL_3093111, EPI_ISL_3093112 | Lighthouse Lab in Milton Keynes | Wellcome Sanger Institute for the COVID-19 Genomics UK (COG-UK) Consortium | The Lighthouse Lab in Milton Keynes and Alex Alderton, Roberto Amato, Jeffrey Barrett, Sonia Goncalves, Ewan Harrison, David K. Jackson, Ian Johnston, Dominic Kwiatkowski, Cordelia Langford, John Sillitoe on behalf of the Wellcome Sanger Institute COVID-19 Surveillance Team |
| EPI_ISL_3093113, EPI_ISL_3093114 | Lighthouse Lab in Alderley Park | Wellcome Sanger Institute for the COVID-19 Genomics UK (COG-UK) Consortium | Jacquelyn Wynn, Mairead Hyland, The Lighthouse Lab in Alderley Park and Alex Alderton, Roberto Amato, Jeffrey Barrett, Sonia Goncalves, Ewan Harrison, David K. Jackson, Ian Johnston, Dominic Kwiatkowski, Cordelia Langford, John Sillitoe on behalf of the Wellcome Sanger Institute COVID-19 Surveillance Team |
| EPI_ISL_3093115, EPI_ISL_3093116 | Lighthouse Lab in Milton Keynes | Wellcome Sanger Institute for the COVID-19 Genomics UK (COG-UK) Consortium | The Lighthouse Lab in Milton Keynes and Alex Alderton, Roberto Amato, Jeffrey Barrett, Sonia Goncalves, Ewan Harrison, David K. Jackson, Ian Johnston, Dominic Kwiatkowski, Cordelia Langford, John Sillitoe on behalf of the Wellcome Sanger Institute COVID-19 Surveillance Team |
| EPI_ISL_3093117 | Lighthouse Lab in Glasgow | Wellcome Sanger Institute for the COVID-19 Genomics UK (COG-UK) Consortium | Harper VanSteenhouse, Yumi Kasai, David Gray, Carol Clugston, Anna Dominiczak and Alex Alderton, Roberto Amato, Jeffrey Barrett, Sonia Goncalves, Ewan Harrison, David K. Jackson, Ian Johnston, Dominic Kwiatkowski, Cordelia Langford, John Sillitoe on behalf of the Wellcome Sanger Institute COVID-19 Surveillance Team |
| EPI_ISL_3093118 | Lighthouse Lab in Milton Keynes | Wellcome Sanger Institute for the COVID-19 Genomics UK (COG-UK) Consortium | The Lighthouse Lab in Milton Keynes and Alex Alderton, Roberto Amato, Jeffrey Barrett, Sonia Goncalves, Ewan Harrison, David K. Jackson, Ian Johnston, Dominic Kwiatkowski, Cordelia Langford, John Sillitoe on behalf of the Wellcome Sanger Institute COVID-19 Surveillance Team |
| EPI_ISL_3093119 | Lighthouse Lab in Alderley Park | Wellcome Sanger Institute for the COVID-19 Genomics UK (COG-UK) Consortium | Jacquelyn Wynn, Mairead Hyland, The Lighthouse Lab in Alderley Park and Alex Alderton, Roberto Amato, Jeffrey Barrett, Sonia Goncalves, Ewan Harrison, David K. Jackson, Ian Johnston, Dominic Kwiatkowski, Cordelia Langford, John Sillitoe on behalf of the Wellcome Sanger Institute COVID-19 Surveillance Team |
| EPI_ISL_3093120 | Berkshire and Surrey Pathology Services Lighthouse Laboratory | Wellcome Sanger Institute for the COVID-19 Genomics UK (COG-UK) Consortium | Berkshire and Surrey Pathology Services Lighthouse Laboratory and Alex Alderton, Roberto Amato, Jeffrey Barrett, Sonia Goncalves, Ewan Harrison, David K. Jackson, Ian Johnston, Dominic Kwiatkowski, Cordelia Langford, John Sillitoe on behalf of the Wellcome Sanger Institute COVID-19 Surveillance Team |
| EPI_ISL_3093121 | Lighthouse Lab in Milton Keynes | Wellcome Sanger Institute for the COVID-19 Genomics UK (COG-UK) Consortium | The Lighthouse Lab in Milton Keynes and Alex Alderton, Roberto Amato, Jeffrey Barrett, Sonia Goncalves, Ewan Harrison, David K. Jackson, Ian Johnston, Dominic Kwiatkowski, Cordelia Langford, John Sillitoe on behalf of the Wellcome Sanger Institute COVID-19 Surveillance Team |
| EPI_ISL_3093122 | Lighthouse Lab in Alderley Park | Wellcome Sanger Institute for the COVID-19 Genomics UK (COG-UK) Consortium | Jacquelyn Wynn, Mairead Hyland, The Lighthouse Lab in Alderley Park and Alex Alderton, Roberto Amato, Jeffrey Barrett, Sonia Goncalves, Ewan Harrison, David K. Jackson, Ian Johnston, Dominic Kwiatkowski, Cordelia Langford, John Sillitoe on behalf of the Wellcome Sanger Institute COVID-19 Surveillance Team |
| EPI_ISL_3093123 | Lighthouse Lab in Glasgow | Wellcome Sanger Institute for the COVID-19 Genomics UK (COG-UK) Consortium | Harper VanSteenhouse, Yumi Kasai, David Gray, Carol Clugston, Anna Dominiczak and Alex Alderton, Roberto Amato, Jeffrey Barrett, Sonia Goncalves, Ewan Harrison, David K. Jackson, Ian Johnston, Dominic Kwiatkowski, Cordelia Langford, John Sillitoe on behalf of the Wellcome Sanger Institute COVID-19 Surveillance Team |
| EPI_ISL_3093124, EPI_ISL_3093125 | Berkshire and Surrey Pathology Services Lighthouse Laboratory | Wellcome Sanger Institute for the COVID-19 Genomics UK (COG-UK) Consortium | Berkshire and Surrey Pathology Services Lighthouse Laboratory and Alex Alderton, Roberto Amato, Jeffrey Barrett, Sonia Goncalves, Ewan Harrison, David K. Jackson, Ian Johnston, Dominic Kwiatkowski, Cordelia Langford, John Sillitoe on behalf of the Wellcome Sanger Institute COVID-19 Surveillance Team |
| EPI_ISL_3093126 | Lighthouse Lab in Alderley Park | Wellcome Sanger Institute for the COVID-19 Genomics UK (COG-UK) Consortium | Jacquelyn Wynn, Mairead Hyland, The Lighthouse Lab in Alderley Park and Alex Alderton, Roberto Amato, Jeffrey Barrett, Sonia Goncalves, Ewan Harrison, David K. Jackson, Ian Johnston, Dominic Kwiatkowski, Cordelia Langford, John Sillitoe on behalf of the Wellcome Sanger Institute COVID-19 Surveillance Team |
| EPI_ISL_3093127, EPI_ISL_3093128, EPI_ISL_3093129 | Berkshire and Surrey Pathology Services Lighthouse Laboratory | Wellcome Sanger Institute for the COVID-19 Genomics UK (COG-UK) Consortium | Berkshire and Surrey Pathology Services Lighthouse Laboratory and Alex Alderton, Roberto Amato, Jeffrey Barrett, Sonia Goncalves, Ewan Harrison, David K. Jackson, Ian Johnston, Dominic Kwiatkowski, Cordelia Langford, John Sillitoe on behalf of the Wellcome Sanger Institute COVID-19 Surveillance Team |
| EPI_ISL_3093130 | Lighthouse Lab in Milton Keynes | Wellcome Sanger Institute for the COVID-19 Genomics UK (COG-UK) Consortium | The Lighthouse Lab in Milton Keynes and Alex Alderton, Roberto Amato, Jeffrey Barrett, Sonia Goncalves, Ewan Harrison, David K. Jackson, Ian Johnston, Dominic Kwiatkowski, Cordelia Langford, John Sillitoe on behalf of the Wellcome Sanger Institute COVID-19 Surveillance Team |
| EPI_ISL_3093131, EPI_ISL_3093132 | Lighthouse Lab in Alderley Park | Wellcome Sanger Institute for the COVID-19 Genomics UK (COG-UK) Consortium | Jacquelyn Wynn, Mairead Hyland, The Lighthouse Lab in Alderley Park and Alex Alderton, Roberto Amato, Jeffrey Barrett, Sonia Goncalves, Ewan Harrison, David K. Jackson, Ian Johnston, Dominic Kwiatkowski, Cordelia Langford, John Sillitoe on behalf of the Wellcome Sanger Institute COVID-19 Surveillance Team |
| EPI_ISL_3093133 | Lighthouse Lab in Milton Keynes | Wellcome Sanger Institute for the COVID-19 Genomics UK (COG-UK) Consortium | The Lighthouse Lab in Milton Keynes and Alex Alderton, Roberto Amato, Jeffrey Barrett, Sonia Goncalves, Ewan Harrison, David K. Jackson, Ian Johnston, Dominic Kwiatkowski, Cordelia Langford, John Sillitoe on behalf of the Wellcome Sanger Institute COVID-19 Surveillance Team |
| EPI_ISL_3093134 | Lighthouse Lab in Glasgow | Wellcome Sanger Institute for the COVID-19 Genomics UK (COG-UK) Consortium | Harper VanSteenhouse, Yumi Kasai, David Gray, Carol Clugston, Anna Dominiczak and Alex Alderton, Roberto Amato, Jeffrey Barrett, Sonia Goncalves, Ewan Harrison, David K. Jackson, Ian Johnston, Dominic Kwiatkowski, Cordelia Langford, John Sillitoe on behalf of the Wellcome Sanger Institute COVID-19 Surveillance Team |

| | | | |
|---|---|---|---|
| EPI_ISL_3093135 | Lighthouse Lab in Milton Keynes | Wellcome Sanger Institute for the COVID-19 Genomics UK (COG-UK) Consortium | The Lighthouse Lab in Milton Keynes and Alex Alderton, Roberto Amato, Jeffrey Barrett, Sonia Goncalves, Ewan Harrison, David K. Jackson, Ian Johnston, Dominic Kwiatkowski, Cordelia Langford, John Sillitoe on behalf of the Wellcome Sanger Institute COVID-19 Surveillance Team |
| EPI_ISL_3093136 | Berkshire and Surrey Pathology Services Lighthouse Laboratory | Wellcome Sanger Institute for the COVID-19 Genomics UK (COG-UK) Consortium | Berkshire and Surrey Pathology Services Lighthouse Laboratory and Alex Alderton, Roberto Amato, Jeffrey Barrett, Sonia Goncalves, Ewan Harrison, David K. Jackson, Ian Johnston, Dominic Kwiatkowski, Cordelia Langford, John Sillitoe on behalf of the Wellcome Sanger Institute COVID-19 Surveillance Team |
| EPI_ISL_3093137 | Lighthouse Lab in Milton Keynes | Wellcome Sanger Institute for the COVID-19 Genomics UK (COG-UK) Consortium | The Lighthouse Lab in Milton Keynes and Alex Alderton, Roberto Amato, Jeffrey Barrett, Sonia Goncalves, Ewan Harrison, David K. Jackson, Ian Johnston, Dominic Kwiatkowski, Cordelia Langford, John Sillitoe on behalf of the Wellcome Sanger Institute COVID-19 Surveillance Team |
| EPI_ISL_3093138 | Lighthouse Lab in Alderley Park | Wellcome Sanger Institute for the COVID-19 Genomics UK (COG-UK) Consortium | Jacquelyn Wynn, Mairead Hyland, The Lighthouse Lab in Alderley Park and Alex Alderton, Roberto Amato, Jeffrey Barrett, Sonia Goncalves, Ewan Harrison, David K. Jackson, Ian Johnston, Dominic Kwiatkowski, Cordelia Langford, John Sillitoe on behalf of the Wellcome Sanger Institute COVID-19 Surveillance Team |
| EPI_ISL_3093139 | Berkshire and Surrey Pathology Services Lighthouse Laboratory | Wellcome Sanger Institute for the COVID-19 Genomics UK (COG-UK) Consortium | Berkshire and Surrey Pathology Services Lighthouse Laboratory and Alex Alderton, Roberto Amato, Jeffrey Barrett, Sonia Goncalves, Ewan Harrison, David K. Jackson, Ian Johnston, Dominic Kwiatkowski, Cordelia Langford, John Sillitoe on behalf of the Wellcome Sanger Institute COVID-19 Surveillance Team |
| EPI_ISL_3093140 | Lighthouse Lab in Glasgow | Wellcome Sanger Institute for the COVID-19 Genomics UK (COG-UK) Consortium | Harper VanSteenhouse, Yumi Kasai, David Gray, Carol Clugston, Anna Dominiczak and Alex Alderton, Roberto Amato, Jeffrey Barrett, Sonia Goncalves, Ewan Harrison, David K. Jackson, Ian Johnston, Dominic Kwiatkowski, Cordelia Langford, John Sillitoe on behalf of the Wellcome Sanger Institute COVID-19 Surveillance Team |
| EPI_ISL_3093141, EPI_ISL_3093142 | Berkshire and Surrey Pathology Services Lighthouse Laboratory | Wellcome Sanger Institute for the COVID-19 Genomics UK (COG-UK) Consortium | Berkshire and Surrey Pathology Services Lighthouse Laboratory and Alex Alderton, Roberto Amato, Jeffrey Barrett, Sonia Goncalves, Ewan Harrison, David K. Jackson, Ian Johnston, Dominic Kwiatkowski, Cordelia Langford, John Sillitoe on behalf of the Wellcome Sanger Institute COVID-19 Surveillance Team |
| EPI_ISL_3093143 | Lighthouse Lab in Glasgow | Wellcome Sanger Institute for the COVID-19 Genomics UK (COG-UK) Consortium | Harper VanSteenhouse, Yumi Kasai, David Gray, Carol Clugston, Anna Dominiczak and Alex Alderton, Roberto Amato, Jeffrey Barrett, Sonia Goncalves, Ewan Harrison, David K. Jackson, Ian Johnston, Dominic Kwiatkowski, Cordelia Langford, John Sillitoe on behalf of the Wellcome Sanger Institute COVID-19 Surveillance Team |
| EPI_ISL_3093144 | Lighthouse Lab in Milton Keynes | Wellcome Sanger Institute for the COVID-19 Genomics UK (COG-UK) Consortium | The Lighthouse Lab in Milton Keynes and Alex Alderton, Roberto Amato, Jeffrey Barrett, Sonia Goncalves, Ewan Harrison, David K. Jackson, Ian Johnston, Dominic Kwiatkowski, Cordelia Langford, John Sillitoe on behalf of the Wellcome Sanger Institute COVID-19 Surveillance Team |
| EPI_ISL_3093145 | Lighthouse Lab in Glasgow | Wellcome Sanger Institute for the COVID-19 Genomics UK (COG-UK) Consortium | Harper VanSteenhouse, Yumi Kasai, David Gray, Carol Clugston, Anna Dominiczak and Alex Alderton, Roberto Amato, Jeffrey Barrett, Sonia Goncalves, Ewan Harrison, David K. Jackson, Ian Johnston, Dominic Kwiatkowski, Cordelia Langford, John Sillitoe on behalf of the Wellcome Sanger Institute COVID-19 Surveillance Team |
| EPI_ISL_3093146 | Berkshire and Surrey Pathology Services Lighthouse Laboratory | Wellcome Sanger Institute for the COVID-19 Genomics UK (COG-UK) Consortium | Berkshire and Surrey Pathology Services Lighthouse Laboratory and Alex Alderton, Roberto Amato, Jeffrey Barrett, Sonia Goncalves, Ewan Harrison, David K. Jackson, Ian Johnston, Dominic Kwiatkowski, Cordelia Langford, John Sillitoe on behalf of the Wellcome Sanger Institute COVID-19 Surveillance Team |
| EPI_ISL_3093147 | Lighthouse Lab in Milton Keynes | Wellcome Sanger Institute for the COVID-19 Genomics UK (COG-UK) Consortium | The Lighthouse Lab in Milton Keynes and Alex Alderton, Roberto Amato, Jeffrey Barrett, Sonia Goncalves, Ewan Harrison, David K. Jackson, Ian Johnston, Dominic Kwiatkowski, Cordelia Langford, John Sillitoe on behalf of the Wellcome Sanger Institute COVID-19 Surveillance Team |
| EPI_ISL_3093148, EPI_ISL_3093149, EPI_ISL_3093150 | Lighthouse Lab in Glasgow | Wellcome Sanger Institute for the COVID-19 Genomics UK (COG-UK) Consortium | Harper VanSteenhouse, Yumi Kasai, David Gray, Carol Clugston, Anna Dominiczak and Alex Alderton, Roberto Amato, Jeffrey Barrett, Sonia Goncalves, Ewan Harrison, David K. Jackson, Ian Johnston, Dominic Kwiatkowski, Cordelia Langford, John Sillitoe on behalf of the Wellcome Sanger Institute COVID-19 Surveillance Team |
| EPI_ISL_3093151, EPI_ISL_3093152 | Berkshire and Surrey Pathology Services Lighthouse Laboratory | Wellcome Sanger Institute for the COVID-19 Genomics UK (COG-UK) Consortium | Berkshire and Surrey Pathology Services Lighthouse Laboratory and Alex Alderton, Roberto Amato, Jeffrey Barrett, Sonia Goncalves, Ewan Harrison, David K. Jackson, Ian Johnston, Dominic Kwiatkowski, Cordelia Langford, John Sillitoe on behalf of the Wellcome Sanger Institute COVID-19 Surveillance Team |
| EPI_ISL_3093153 | Lighthouse Lab in Glasgow | Wellcome Sanger Institute for the COVID-19 Genomics UK (COG-UK) Consortium | Harper VanSteenhouse, Yumi Kasai, David Gray, Carol Clugston, Anna Dominiczak and Alex Alderton, Roberto Amato, Jeffrey Barrett, Sonia Goncalves, Ewan Harrison, David K. Jackson, Ian Johnston, Dominic Kwiatkowski, Cordelia Langford, John Sillitoe on behalf of the Wellcome Sanger Institute COVID-19 Surveillance Team |
| EPI_ISL_3093154, EPI_ISL_3093155 | Lighthouse Lab in Alderley Park | Wellcome Sanger Institute for the COVID-19 Genomics UK (COG-UK) Consortium | Jacquelyn Wynn, Mairead Hyland, The Lighthouse Lab in Alderley Park and Alex Alderton, Roberto Amato, Jeffrey Barrett, Sonia Goncalves, Ewan Harrison, David K. Jackson, Ian Johnston, Dominic Kwiatkowski, Cordelia Langford, John Sillitoe on behalf of the Wellcome Sanger Institute COVID-19 Surveillance Team |
| EPI_ISL_3093156, EPI_ISL_3093157, EPI_ISL_3093158 | Lighthouse Lab in Milton Keynes | Wellcome Sanger Institute for the COVID-19 Genomics UK (COG-UK) Consortium | The Lighthouse Lab in Milton Keynes and Alex Alderton, Roberto Amato, Jeffrey Barrett, Sonia Goncalves, Ewan Harrison, David K. Jackson, Ian Johnston, Dominic Kwiatkowski, Cordelia Langford, John Sillitoe on behalf of the Wellcome Sanger Institute COVID-19 Surveillance Team |
| EPI_ISL_3093159 | Lighthouse Lab in Alderley Park | Wellcome Sanger Institute for the COVID-19 Genomics UK (COG-UK) Consortium | Jacquelyn Wynn, Mairead Hyland, The Lighthouse Lab in Alderley Park and Alex Alderton, Roberto Amato, Jeffrey Barrett, Sonia Goncalves, Ewan Harrison, David K. Jackson, Ian Johnston, Dominic Kwiatkowski, Cordelia Langford, John Sillitoe on behalf of the Wellcome Sanger Institute COVID-19 Surveillance Team |
| EPI_ISL_3093160, EPI_ISL_3093161 | Lighthouse Lab in Milton Keynes | Wellcome Sanger Institute for the COVID-19 Genomics UK (COG-UK) Consortium | The Lighthouse Lab in Milton Keynes and Alex Alderton, Roberto Amato, Jeffrey Barrett, Sonia Goncalves, Ewan Harrison, David K. Jackson, Ian Johnston, Dominic Kwiatkowski, Cordelia Langford, John Sillitoe on behalf of the Wellcome Sanger Institute COVID-19 Surveillance Team |
| EPI_ISL_3093162, EPI_ISL_3093163, EPI_ISL_3093164 | Berkshire and Surrey Pathology Services Lighthouse Laboratory | Wellcome Sanger Institute for the COVID-19 Genomics UK (COG-UK) Consortium | Berkshire and Surrey Pathology Services Lighthouse Laboratory and Alex Alderton, Roberto Amato, Jeffrey Barrett, Sonia Goncalves, Ewan Harrison, David K. Jackson, Ian Johnston, Dominic Kwiatkowski, Cordelia Langford, John Sillitoe on behalf of the Wellcome Sanger Institute COVID-19 Surveillance Team |
| EPI_ISL_3093165, EPI_ISL_3093166 | Lighthouse Lab in Alderley Park | Wellcome Sanger Institute for the COVID-19 Genomics UK (COG-UK) Consortium | Jacquelyn Wynn, Mairead Hyland, The Lighthouse Lab in Alderley Park and Alex Alderton, Roberto Amato, Jeffrey Barrett, Sonia Goncalves, Ewan Harrison, David K. Jackson, Ian Johnston, Dominic Kwiatkowski, Cordelia Langford, John Sillitoe on behalf of the Wellcome Sanger Institute COVID-19 Surveillance Team |
| EPI_ISL_3093167 | Lighthouse Lab in Milton Keynes | Wellcome Sanger Institute for the COVID-19 Genomics UK (COG-UK) Consortium | The Lighthouse Lab in Milton Keynes and Alex Alderton, Roberto Amato, Jeffrey Barrett, Sonia Goncalves, Ewan Harrison, David K. Jackson, Ian Johnston, Dominic Kwiatkowski, Cordelia Langford, John Sillitoe on behalf of the Wellcome Sanger Institute COVID-19 Surveillance Team |
| EPI_ISL_3093168 | Berkshire and Surrey Pathology Services Lighthouse Laboratory | Wellcome Sanger Institute for the COVID-19 Genomics UK (COG-UK) Consortium | Berkshire and Surrey Pathology Services Lighthouse Laboratory and Alex Alderton, Roberto Amato, Jeffrey Barrett, Sonia Goncalves, Ewan Harrison, David K. Jackson, Ian Johnston, Dominic Kwiatkowski, Cordelia Langford, John Sillitoe on behalf of the Wellcome Sanger Institute COVID-19 Surveillance Team |
| EPI_ISL_3093169 | Lighthouse Lab in Glasgow | Wellcome Sanger Institute for the COVID-19 Genomics UK (COG-UK) Consortium | Harper VanSteenhouse, Yumi Kasai, David Gray, Carol Clugston, Anna Dominiczak and Alex Alderton, Roberto Amato, Jeffrey Barrett, Sonia Goncalves, Ewan Harrison, David K. Jackson, Ian Johnston, Dominic Kwiatkowski, Cordelia Langford, John Sillitoe on behalf of the Wellcome Sanger Institute COVID-19 Surveillance Team |
| EPI_ISL_3093170, EPI_ISL_3093171 | Lighthouse Lab in Alderley Park | Wellcome Sanger Institute for the COVID-19 Genomics UK (COG-UK) Consortium | Jacquelyn Wynn, Mairead Hyland, The Lighthouse Lab in Alderley Park and Alex Alderton, Roberto Amato, Jeffrey Barrett, Sonia Goncalves, Ewan Harrison, David K. Jackson, Ian Johnston, Dominic Kwiatkowski, Cordelia Langford, John Sillitoe on behalf of the Wellcome Sanger Institute COVID-19 Surveillance Team |
| EPI_ISL_3093172 | Lighthouse Lab in Glasgow | Wellcome Sanger Institute for the COVID-19 Genomics UK (COG-UK) Consortium | Harper VanSteenhouse, Yumi Kasai, David Gray, Carol Clugston, Anna Dominiczak and Alex Alderton, Roberto Amato, Jeffrey Barrett, Sonia Goncalves, Ewan Harrison, David K. Jackson, Ian Johnston, Dominic Kwiatkowski, Cordelia Langford, John Sillitoe on behalf of the Wellcome Sanger Institute COVID-19 Surveillance Team |
| EPI_ISL_3093173 | Lighthouse Lab in Alderley Park | Wellcome Sanger Institute for the COVID-19 Genomics UK (COG-UK) Consortium | Jacquelyn Wynn, Mairead Hyland, The Lighthouse Lab in Alderley Park and Alex Alderton, Roberto Amato, Jeffrey Barrett, Sonia Goncalves, Ewan Harrison, David K. Jackson, Ian Johnston, Dominic Kwiatkowski, Cordelia Langford, John Sillitoe on behalf of the Wellcome Sanger Institute COVID-19 Surveillance Team |
| EPI_ISL_3093174 | Lighthouse Lab in Milton Keynes | Wellcome Sanger Institute for the COVID-19 Genomics UK (COG-UK) Consortium | The Lighthouse Lab in Milton Keynes and Alex Alderton, Roberto Amato, Jeffrey Barrett, Sonia Goncalves, Ewan Harrison, David K. Jackson, Ian Johnston, Dominic Kwiatkowski, Cordelia Langford, John Sillitoe on behalf of the Wellcome Sanger Institute COVID-19 Surveillance Team |
| EPI_ISL_3093175, EPI_ISL_3093176 | Lighthouse Lab in Glasgow | Wellcome Sanger Institute for the COVID-19 Genomics UK (COG-UK) Consortium | Harper VanSteenhouse, Yumi Kasai, David Gray, Carol Clugston, Anna Dominiczak and Alex Alderton, Roberto Amato, Jeffrey Barrett, Sonia Goncalves, Ewan Harrison, David K. Jackson, Ian Johnston, Dominic Kwiatkowski, Cordelia Langford, John Sillitoe on behalf of the Wellcome Sanger Institute COVID-19 Surveillance Team |
| EPI_ISL_3093177 | Lighthouse Lab in Alderley Park | Wellcome Sanger Institute for the COVID-19 Genomics UK (COG-UK) Consortium | Jacquelyn Wynn, Mairead Hyland, The Lighthouse Lab in Alderley Park and Alex Alderton, Roberto Amato, Jeffrey Barrett, Sonia Goncalves, Ewan Harrison, David K. Jackson, Ian Johnston, Dominic Kwiatkowski, Cordelia Langford, John Sillitoe on behalf of the Wellcome Sanger Institute COVID-19 Surveillance Team |
| EPI_ISL_3093178 | Lighthouse Lab in Glasgow | Wellcome Sanger Institute for the COVID-19 Genomics UK (COG-UK) Consortium | Harper VanSteenhouse, Yumi Kasai, David Gray, Carol Clugston, Anna Dominiczak and Alex Alderton, Roberto Amato, Jeffrey Barrett, Sonia Goncalves, Ewan Harrison, David K. Jackson, Ian Johnston, Dominic Kwiatkowski, Cordelia Langford, John Sillitoe on behalf of the Wellcome Sanger Institute COVID-19 Surveillance Team |
| EPI_ISL_3093179 | Lighthouse Lab in Milton Keynes | Wellcome Sanger Institute for the COVID-19 Genomics UK (COG-UK) Consortium | The Lighthouse Lab in Milton Keynes and Alex Alderton, Roberto Amato, Jeffrey Barrett, Sonia Goncalves, Ewan Harrison, David K. Jackson, Ian Johnston, Dominic Kwiatkowski, Cordelia Langford, John Sillitoe on behalf of the Wellcome Sanger Institute COVID-19 Surveillance Team |
| EPI_ISL_3093180 | Lighthouse Lab in Glasgow | Wellcome Sanger Institute for the COVID-19 Genomics UK (COG-UK) Consortium | Harper VanSteenhouse, Yumi Kasai, David Gray, Carol Clugston, Anna Dominiczak and Alex Alderton, Roberto Amato, Jeffrey Barrett, Sonia Goncalves, Ewan Harrison, David K. Jackson, Ian Johnston, Dominic Kwiatkowski, Cordelia Langford, John Sillitoe on behalf of the Wellcome Sanger Institute COVID-19 Surveillance Team |
| EPI_ISL_3093181 | Berkshire and Surrey Pathology Services Lighthouse Laboratory | Wellcome Sanger Institute for the COVID-19 Genomics UK (COG-UK) Consortium | Berkshire and Surrey Pathology Services Lighthouse Laboratory and Alex Alderton, Roberto Amato, Jeffrey Barrett, Sonia Goncalves, Ewan Harrison, David K. Jackson, Ian Johnston, Dominic Kwiatkowski, Cordelia Langford, John Sillitoe on behalf of the Wellcome Sanger Institute COVID-19 Surveillance Team |
| EPI_ISL_3093182 | Lighthouse Lab in Milton Keynes | Wellcome Sanger Institute for the COVID-19 Genomics UK (COG-UK) Consortium | The Lighthouse Lab in Milton Keynes and Alex Alderton, Roberto Amato, Jeffrey Barrett, Sonia Goncalves, Ewan Harrison, David K. Jackson, Ian Johnston, Dominic Kwiatkowski, Cordelia Langford, John Sillitoe on behalf of the Wellcome Sanger Institute COVID-19 Surveillance Team |
| EPI_ISL_3093183 | Lighthouse Lab in Alderley Park | Wellcome Sanger Institute for the COVID-19 Genomics UK (COG-UK) Consortium | Jacquelyn Wynn, Mairead Hyland, The Lighthouse Lab in Alderley Park and Alex Alderton, Roberto Amato, Jeffrey Barrett, Sonia Goncalves, Ewan Harrison, David K. Jackson, Ian Johnston, Dominic Kwiatkowski, Cordelia Langford, John Sillitoe on behalf of the Wellcome Sanger Institute COVID-19 Surveillance Team |
| EPI_ISL_3093184 | Lighthouse Lab in Glasgow | Wellcome Sanger Institute for the COVID-19 Genomics UK (COG-UK) Consortium | Harper VanSteenhouse, Yumi Kasai, David Gray, Carol Clugston, Anna Dominiczak and Alex Alderton, Roberto Amato, Jeffrey Barrett, Sonia Goncalves, Ewan Harrison, David K. Jackson, Ian Johnston, Dominic Kwiatkowski, Cordelia Langford, John Sillitoe on behalf of the Wellcome Sanger Institute COVID-19 Surveillance Team |
| EPI_ISL_3093185 | Berkshire and Surrey Pathology Services Lighthouse Laboratory | Wellcome Sanger Institute for the COVID-19 Genomics UK (COG-UK) Consortium | Berkshire and Surrey Pathology Services Lighthouse Laboratory and Alex Alderton, Roberto Amato, Jeffrey Barrett, Sonia Goncalves, Ewan Harrison, David K. Jackson, Ian Johnston, Dominic Kwiatkowski, Cordelia Langford, John Sillitoe on behalf of the Wellcome Sanger Institute COVID-19 Surveillance Team |
| EPI_ISL_3093186 | Lighthouse Lab in Glasgow | Wellcome Sanger Institute for the COVID-19 Genomics UK (COG-UK) Consortium | Harper VanSteenhouse, Yumi Kasai, David Gray, Carol Clugston, Anna Dominiczak and Alex Alderton, Roberto Amato, Jeffrey Barrett, Sonia Goncalves, Ewan Harrison, David K. Jackson, Ian Johnston, Dominic Kwiatkowski, Cordelia Langford, John Sillitoe on behalf of the Wellcome Sanger Institute COVID-19 Surveillance Team |
| EPI_ISL_3093187 | Lighthouse Lab in Alderley Park | Wellcome Sanger Institute for the COVID-19 Genomics UK (COG-UK) Consortium | Jacquelyn Wynn, Mairead Hyland, The Lighthouse Lab in Alderley Park and Alex Alderton, Roberto Amato, Jeffrey Barrett, Sonia Goncalves, Ewan Harrison, David K. Jackson, Ian Johnston, Dominic Kwiatkowski, Cordelia Langford, John Sillitoe on behalf of the Wellcome Sanger Institute COVID-19 Surveillance Team |
| EPI_ISL_3093188 | Lighthouse Lab in Glasgow | Wellcome Sanger Institute for the COVID-19 Genomics UK (COG-UK) Consortium | Harper VanSteenhouse, Yumi Kasai, David Gray, Carol Clugston, Anna Dominiczak and Alex Alderton, Roberto Amato, Jeffrey Barrett, Sonia Goncalves, Ewan Harrison, David K. Jackson, Ian Johnston, Dominic Kwiatkowski, Cordelia Langford, John Sillitoe on behalf of the Wellcome Sanger Institute COVID-19 Surveillance Team |
| EPI_ISL_3093189 | Berkshire and Surrey Pathology Services Lighthouse Laboratory | Wellcome Sanger Institute for the COVID-19 Genomics UK (COG-UK) Consortium | Berkshire and Surrey Pathology Services Lighthouse Laboratory and Alex Alderton, Roberto Amato, Jeffrey Barrett, Sonia Goncalves, Ewan Harrison, David K. Jackson, Ian Johnston, Dominic Kwiatkowski, Cordelia Langford, John Sillitoe on behalf of the Wellcome Sanger Institute COVID-19 Surveillance Team |
| EPI_ISL_3093190, EPI_ISL_3093191 | Lighthouse Lab in Alderley Park | Wellcome Sanger Institute for the COVID-19 Genomics UK (COG-UK) Consortium | Jacquelyn Wynn, Mairead Hyland, The Lighthouse Lab in Alderley Park and Alex Alderton, Roberto Amato, Jeffrey Barrett, Sonia Goncalves, Ewan Harrison, David K. Jackson, Ian Johnston, Dominic Kwiatkowski, Cordelia Langford, John Sillitoe on behalf of the Wellcome Sanger Institute COVID-19 Surveillance Team |
| EPI_ISL_3093192 | Lighthouse Lab in Milton Keynes | Wellcome Sanger Institute for the COVID-19 Genomics UK (COG-UK) | The Lighthouse Lab in Milton Keynes and Alex Alderton, Roberto Amato, Jeffrey Barrett, Sonia Goncalves, Ewan Harrison, David K. Jackson, Ian Johnston, Dominic Kwiatkowski, Cordelia Langford, John Sillitoe on behalf of the Wellcome Sanger |

| | | | |
|---|---|---|---|
| | | Consortium | Institute COVID-19 Surveillance Team |
| EPI_ISL_3093193 | Lighthouse Lab in Alderley Park | Wellcome Sanger Institute for the COVID-19 Genomics UK (COG-UK) Consortium | Jacquelyn Wynn, Mairead Hyland, The Lighthouse Lab in Alderley Park and Alex Alderton, Roberto Amato, Jeffrey Barrett, Sonia Goncalves, Ewan Harrison, David K. Jackson, Ian Johnston, Dominic Kwiatkowski, Cordelia Langford, John Sillitoe on behalf of the Wellcome Sanger Institute COVID-19 Surveillance Team |
| EPI_ISL_3093194 | Berkshire and Surrey Pathology Services Lighthouse Laboratory | Wellcome Sanger Institute for the COVID-19 Genomics UK (COG-UK) Consortium | Berkshire and Surrey Pathology Services Lighthouse Laboratory and Alex Alderton, Roberto Amato, Jeffrey Barrett, Sonia Goncalves, Ewan Harrison, David K. Jackson, Ian Johnston, Dominic Kwiatkowski, Cordelia Langford, John Sillitoe on behalf of the Wellcome Sanger Institute COVID-19 Surveillance Team |
| EPI_ISL_3093195 | Lighthouse Lab in Alderley Park | Wellcome Sanger Institute for the COVID-19 Genomics UK (COG-UK) Consortium | Jacquelyn Wynn, Mairead Hyland, The Lighthouse Lab in Alderley Park and Alex Alderton, Roberto Amato, Jeffrey Barrett, Sonia Goncalves, Ewan Harrison, David K. Jackson, Ian Johnston, Dominic Kwiatkowski, Cordelia Langford, John Sillitoe on behalf of the Wellcome Sanger Institute COVID-19 Surveillance Team |
| EPI_ISL_3093196, EPI_ISL_3093197, EPI_ISL_3093198 | Berkshire and Surrey Pathology Services Lighthouse Laboratory | Wellcome Sanger Institute for the COVID-19 Genomics UK (COG-UK) Consortium | Berkshire and Surrey Pathology Services Lighthouse Laboratory and Alex Alderton, Roberto Amato, Jeffrey Barrett, Sonia Goncalves, Ewan Harrison, David K. Jackson, Ian Johnston, Dominic Kwiatkowski, Cordelia Langford, John Sillitoe on behalf of the Wellcome Sanger Institute COVID-19 Surveillance Team |
| EPI_ISL_3093199, EPI_ISL_3093200, EPI_ISL_3093201 | Lighthouse Lab in Alderley Park | Wellcome Sanger Institute for the COVID-19 Genomics UK (COG-UK) Consortium | Jacquelyn Wynn, Mairead Hyland, The Lighthouse Lab in Alderley Park and Alex Alderton, Roberto Amato, Jeffrey Barrett, Sonia Goncalves, Ewan Harrison, David K. Jackson, Ian Johnston, Dominic Kwiatkowski, Cordelia Langford, John Sillitoe on behalf of the Wellcome Sanger Institute COVID-19 Surveillance Team |
| EPI_ISL_3093202 | Lighthouse Laboratory Plymouth | Wellcome Sanger Institute for the COVID-19 Genomics UK (COG-UK) Consortium | Lighthouse Laboratory Plymouth and Alex Alderton, Roberto Amato, Jeffrey Barrett, Sonia Goncalves, Ewan Harrison, David K. Jackson, Ian Johnston, Dominic Kwiatkowski, Cordelia Langford, John Sillitoe on behalf of the Wellcome Sanger Institute COVID-19 Surveillance Team |
| EPI_ISL_3093203, EPI_ISL_3093204, EPI_ISL_3093205, EPI_ISL_3093206, EPI_ISL_3093207, EPI_ISL_3093208 | Lighthouse Lab in Alderley Park | Wellcome Sanger Institute for the COVID-19 Genomics UK (COG-UK) Consortium | Jacquelyn Wynn, Mairead Hyland, The Lighthouse Lab in Alderley Park and Alex Alderton, Roberto Amato, Jeffrey Barrett, Sonia Goncalves, Ewan Harrison, David K. Jackson, Ian Johnston, Dominic Kwiatkowski, Cordelia Langford, John Sillitoe on behalf of the Wellcome Sanger Institute COVID-19 Surveillance Team |
| EPI_ISL_3093209, EPI_ISL_3093210 | Lighthouse Laboratory Plymouth | Wellcome Sanger Institute for the COVID-19 Genomics UK (COG-UK) Consortium | Lighthouse Laboratory Plymouth and Alex Alderton, Roberto Amato, Jeffrey Barrett, Sonia Goncalves, Ewan Harrison, David K. Jackson, Ian Johnston, Dominic Kwiatkowski, Cordelia Langford, John Sillitoe on behalf of the Wellcome Sanger Institute COVID-19 Surveillance Team |
| EPI_ISL_3093211, EPI_ISL_3093212 | Lighthouse Lab in Alderley Park | Wellcome Sanger Institute for the COVID-19 Genomics UK (COG-UK) Consortium | Jacquelyn Wynn, Mairead Hyland, The Lighthouse Lab in Alderley Park and Alex Alderton, Roberto Amato, Jeffrey Barrett, Sonia Goncalves, Ewan Harrison, David K. Jackson, Ian Johnston, Dominic Kwiatkowski, Cordelia Langford, John Sillitoe on behalf of the Wellcome Sanger Institute COVID-19 Surveillance Team |
| EPI_ISL_3093213 | Lighthouse Lab in Glasgow | Wellcome Sanger Institute for the COVID-19 Genomics UK (COG-UK) Consortium | Harper VanSteenhouse, Yumi Kasai, David Gray, Carol Clugston, Anna Dominiczak and Alex Alderton, Roberto Amato, Jeffrey Barrett, Sonia Goncalves, Ewan Harrison, David K. Jackson, Ian Johnston, Dominic Kwiatkowski, Cordelia Langford, John Sillitoe on behalf of the Wellcome Sanger Institute COVID-19 Surveillance Team |
| EPI_ISL_3093214, EPI_ISL_3093215, EPI_ISL_3093216, EPI_ISL_3093217, EPI_ISL_3093218 | Lighthouse Lab in Alderley Park | Wellcome Sanger Institute for the COVID-19 Genomics UK (COG-UK) Consortium | Jacquelyn Wynn, Mairead Hyland, The Lighthouse Lab in Alderley Park and Alex Alderton, Roberto Amato, Jeffrey Barrett, Sonia Goncalves, Ewan Harrison, David K. Jackson, Ian Johnston, Dominic Kwiatkowski, Cordelia Langford, John Sillitoe on behalf of the Wellcome Sanger Institute COVID-19 Surveillance Team |
| EPI_ISL_3093219 | Lighthouse Lab in Glasgow | Wellcome Sanger Institute for the COVID-19 Genomics UK (COG-UK) Consortium | Harper VanSteenhouse, Yumi Kasai, David Gray, Carol Clugston, Anna Dominiczak and Alex Alderton, Roberto Amato, Jeffrey Barrett, Sonia Goncalves, Ewan Harrison, David K. Jackson, Ian Johnston, Dominic Kwiatkowski, Cordelia Langford, John Sillitoe on behalf of the Wellcome Sanger Institute COVID-19 Surveillance Team |
| EPI_ISL_3093220 | Lighthouse Lab in Alderley Park | Wellcome Sanger Institute for the COVID-19 Genomics UK (COG-UK) Consortium | Jacquelyn Wynn, Mairead Hyland, The Lighthouse Lab in Alderley Park and Alex Alderton, Roberto Amato, Jeffrey Barrett, Sonia Goncalves, Ewan Harrison, David K. Jackson, Ian Johnston, Dominic Kwiatkowski, Cordelia Langford, John Sillitoe on behalf of the Wellcome Sanger Institute COVID-19 Surveillance Team |
| EPI_ISL_3093221, EPI_ISL_3093222 | Lighthouse Laboratory Plymouth | Wellcome Sanger Institute for the COVID-19 Genomics UK (COG-UK) Consortium | Lighthouse Laboratory Plymouth and Alex Alderton, Roberto Amato, Jeffrey Barrett, Sonia Goncalves, Ewan Harrison, David K. Jackson, Ian Johnston, Dominic Kwiatkowski, Cordelia Langford, John Sillitoe on behalf of the Wellcome Sanger Institute COVID-19 Surveillance Team |
| EPI_ISL_3093223, EPI_ISL_3093224 | Lighthouse Lab in Alderley Park | Wellcome Sanger Institute for the COVID-19 Genomics UK (COG-UK) Consortium | Jacquelyn Wynn, Mairead Hyland, The Lighthouse Lab in Alderley Park and Alex Alderton, Roberto Amato, Jeffrey Barrett, Sonia Goncalves, Ewan Harrison, David K. Jackson, Ian Johnston, Dominic Kwiatkowski, Cordelia Langford, John Sillitoe on behalf of the Wellcome Sanger Institute COVID-19 Surveillance Team |
| EPI_ISL_3093225 | Lighthouse Laboratory Plymouth | Wellcome Sanger Institute for the COVID-19 Genomics UK (COG-UK) Consortium | Lighthouse Laboratory Plymouth and Alex Alderton, Roberto Amato, Jeffrey Barrett, Sonia Goncalves, Ewan Harrison, David K. Jackson, Ian Johnston, Dominic Kwiatkowski, Cordelia Langford, John Sillitoe on behalf of the Wellcome Sanger Institute COVID-19 Surveillance Team |
| EPI_ISL_3093226 | Lighthouse Lab in Alderley Park | Wellcome Sanger Institute for the COVID-19 Genomics UK (COG-UK) Consortium | Jacquelyn Wynn, Mairead Hyland, The Lighthouse Lab in Alderley Park and Alex Alderton, Roberto Amato, Jeffrey Barrett, Sonia Goncalves, Ewan Harrison, David K. Jackson, Ian Johnston, Dominic Kwiatkowski, Cordelia Langford, John Sillitoe on behalf of the Wellcome Sanger Institute COVID-19 Surveillance Team |
| EPI_ISL_3093227 | Lighthouse Laboratory Plymouth | Wellcome Sanger Institute for the COVID-19 Genomics UK (COG-UK) Consortium | Lighthouse Laboratory Plymouth and Alex Alderton, Roberto Amato, Jeffrey Barrett, Sonia Goncalves, Ewan Harrison, David K. Jackson, Ian Johnston, Dominic Kwiatkowski, Cordelia Langford, John Sillitoe on behalf of the Wellcome Sanger Institute COVID-19 Surveillance Team |
| EPI_ISL_3093228, EPI_ISL_3093229 | Lighthouse Lab in Alderley Park | Wellcome Sanger Institute for the COVID-19 Genomics UK (COG-UK) Consortium | Jacquelyn Wynn, Mairead Hyland, The Lighthouse Lab in Alderley Park and Alex Alderton, Roberto Amato, Jeffrey Barrett, Sonia Goncalves, Ewan Harrison, David K. Jackson, Ian Johnston, Dominic Kwiatkowski, Cordelia Langford, John Sillitoe on behalf of the Wellcome Sanger Institute COVID-19 Surveillance Team |
| EPI_ISL_3093230 | Lighthouse Laboratory Plymouth | Wellcome Sanger Institute for the COVID-19 Genomics UK (COG-UK) Consortium | Lighthouse Laboratory Plymouth and Alex Alderton, Roberto Amato, Jeffrey Barrett, Sonia Goncalves, Ewan Harrison, David K. Jackson, Ian Johnston, Dominic Kwiatkowski, Cordelia Langford, John Sillitoe on behalf of the Wellcome Sanger Institute COVID-19 Surveillance Team |
| EPI_ISL_3093231 | Lighthouse Lab in Glasgow | Wellcome Sanger Institute for the COVID-19 Genomics UK (COG-UK) Consortium | Harper VanSteenhouse, Yumi Kasai, David Gray, Carol Clugston, Anna Dominiczak and Alex Alderton, Roberto Amato, Jeffrey Barrett, Sonia Goncalves, Ewan Harrison, David K. Jackson, Ian Johnston, Dominic Kwiatkowski, Cordelia Langford, John Sillitoe on behalf of the Wellcome Sanger Institute COVID-19 Surveillance Team |
| EPI_ISL_3093232, EPI_ISL_3093233, EPI_ISL_3093234, EPI_ISL_3093235, EPI_ISL_3093236, EPI_ISL_3093237, EPI_ISL_3093238 | Lighthouse Lab in Alderley Park | Wellcome Sanger Institute for the COVID-19 Genomics UK (COG-UK) Consortium | Jacquelyn Wynn, Mairead Hyland, The Lighthouse Lab in Alderley Park and Alex Alderton, Roberto Amato, Jeffrey Barrett, Sonia Goncalves, Ewan Harrison, David K. Jackson, Ian Johnston, Dominic Kwiatkowski, Cordelia Langford, John Sillitoe on behalf of the Wellcome Sanger Institute COVID-19 Surveillance Team |
| EPI_ISL_3093239 | Lighthouse Laboratory Plymouth | Wellcome Sanger Institute for the COVID-19 Genomics UK (COG-UK) Consortium | Lighthouse Laboratory Plymouth and Alex Alderton, Roberto Amato, Jeffrey Barrett, Sonia Goncalves, Ewan Harrison, David K. Jackson, Ian Johnston, Dominic Kwiatkowski, Cordelia Langford, John Sillitoe on behalf of the Wellcome Sanger Institute COVID-19 Surveillance Team |
| EPI_ISL_3093240, EPI_ISL_3093241, EPI_ISL_3093242, EPI_ISL_3093243, EPI_ISL_3093244, EPI_ISL_3093245 | Lighthouse Lab in Alderley Park | Wellcome Sanger Institute for the COVID-19 Genomics UK (COG-UK) Consortium | Jacquelyn Wynn, Mairead Hyland, The Lighthouse Lab in Alderley Park and Alex Alderton, Roberto Amato, Jeffrey Barrett, Sonia Goncalves, Ewan Harrison, David K. Jackson, Ian Johnston, Dominic Kwiatkowski, Cordelia Langford, John Sillitoe on behalf of the Wellcome Sanger Institute COVID-19 Surveillance Team |
| EPI_ISL_3093246 | Lighthouse Lab in Glasgow | Wellcome Sanger Institute for the COVID-19 Genomics UK (COG-UK) Consortium | Harper VanSteenhouse, Yumi Kasai, David Gray, Carol Clugston, Anna Dominiczak and Alex Alderton, Roberto Amato, Jeffrey Barrett, Sonia Goncalves, Ewan Harrison, David K. Jackson, Ian Johnston, Dominic Kwiatkowski, Cordelia Langford, John Sillitoe on behalf of the Wellcome Sanger Institute COVID-19 Surveillance Team |
| EPI_ISL_3093247, EPI_ISL_3093248 | Lighthouse Lab in Alderley Park | Wellcome Sanger Institute for the COVID-19 Genomics UK (COG-UK) Consortium | Jacquelyn Wynn, Mairead Hyland, The Lighthouse Lab in Alderley Park and Alex Alderton, Roberto Amato, Jeffrey Barrett, Sonia Goncalves, Ewan Harrison, David K. Jackson, Ian Johnston, Dominic Kwiatkowski, Cordelia Langford, John Sillitoe on behalf of the Wellcome Sanger Institute COVID-19 Surveillance Team |
| EPI_ISL_3093249 | Lighthouse Lab in Glasgow | Wellcome Sanger Institute for the COVID-19 Genomics UK (COG-UK) Consortium | Harper VanSteenhouse, Yumi Kasai, David Gray, Carol Clugston, Anna Dominiczak and Alex Alderton, Roberto Amato, Jeffrey Barrett, Sonia Goncalves, Ewan Harrison, David K. Jackson, Ian Johnston, Dominic Kwiatkowski, Cordelia Langford, John Sillitoe on behalf of the Wellcome Sanger Institute COVID-19 Surveillance Team |
| EPI_ISL_3093250, EPI_ISL_3093251 | Lighthouse Lab in Alderley Park | Wellcome Sanger Institute for the COVID-19 Genomics UK (COG-UK) Consortium | Jacquelyn Wynn, Mairead Hyland, The Lighthouse Lab in Alderley Park and Alex Alderton, Roberto Amato, Jeffrey Barrett, Sonia Goncalves, Ewan Harrison, David K. Jackson, Ian Johnston, Dominic Kwiatkowski, Cordelia Langford, John Sillitoe on behalf of the Wellcome Sanger Institute COVID-19 Surveillance Team |
| EPI_ISL_3093252, EPI_ISL_3093253 | Lighthouse Laboratory Plymouth | Wellcome Sanger Institute for the COVID-19 Genomics UK (COG-UK) Consortium | Lighthouse Laboratory Plymouth and Alex Alderton, Roberto Amato, Jeffrey Barrett, Sonia Goncalves, Ewan Harrison, David K. Jackson, Ian Johnston, Dominic Kwiatkowski, Cordelia Langford, John Sillitoe on behalf of the Wellcome Sanger Institute COVID-19 Surveillance Team |
| EPI_ISL_3093254, EPI_ISL_3093255 | Lighthouse Lab in Glasgow | Wellcome Sanger Institute for the COVID-19 Genomics UK (COG-UK) Consortium | Harper VanSteenhouse, Yumi Kasai, David Gray, Carol Clugston, Anna Dominiczak and Alex Alderton, Roberto Amato, Jeffrey Barrett, Sonia Goncalves, Ewan Harrison, David K. Jackson, Ian Johnston, Dominic Kwiatkowski, Cordelia Langford, John Sillitoe on behalf of the Wellcome Sanger Institute COVID-19 Surveillance Team |
| EPI_ISL_3093256, EPI_ISL_3093257 | Lighthouse Lab in Alderley Park | Wellcome Sanger Institute for the COVID-19 Genomics UK (COG-UK) Consortium | Jacquelyn Wynn, Mairead Hyland, The Lighthouse Lab in Alderley Park and Alex Alderton, Roberto Amato, Jeffrey Barrett, Sonia Goncalves, Ewan Harrison, David K. Jackson, Ian Johnston, Dominic Kwiatkowski, Cordelia Langford, John Sillitoe on behalf of the Wellcome Sanger Institute COVID-19 Surveillance Team |
| EPI_ISL_3093258 | Lighthouse Lab in Glasgow | Wellcome Sanger Institute for the COVID-19 Genomics UK (COG-UK) Consortium | Harper VanSteenhouse, Yumi Kasai, David Gray, Carol Clugston, Anna Dominiczak and Alex Alderton, Roberto Amato, Jeffrey Barrett, Sonia Goncalves, Ewan Harrison, David K. Jackson, Ian Johnston, Dominic Kwiatkowski, Cordelia Langford, John Sillitoe on behalf of the Wellcome Sanger Institute COVID-19 Surveillance Team |
| EPI_ISL_3093259 | Lighthouse Lab in Alderley Park | Wellcome Sanger Institute for the COVID-19 Genomics UK (COG-UK) Consortium | Jacquelyn Wynn, Mairead Hyland, The Lighthouse Lab in Alderley Park and Alex Alderton, Roberto Amato, Jeffrey Barrett, Sonia Goncalves, Ewan Harrison, David K. Jackson, Ian Johnston, Dominic Kwiatkowski, Cordelia Langford, John Sillitoe on behalf of the Wellcome Sanger Institute COVID-19 Surveillance Team |
| EPI_ISL_3093260 | Lighthouse Lab in Glasgow | Wellcome Sanger Institute for the COVID-19 Genomics UK (COG-UK) Consortium | Harper VanSteenhouse, Yumi Kasai, David Gray, Carol Clugston, Anna Dominiczak and Alex Alderton, Roberto Amato, Jeffrey Barrett, Sonia Goncalves, Ewan Harrison, David K. Jackson, Ian Johnston, Dominic Kwiatkowski, Cordelia Langford, John Sillitoe on behalf of the Wellcome Sanger Institute COVID-19 Surveillance Team |
| EPI_ISL_3093261, EPI_ISL_3093262 | Lighthouse Lab in Alderley Park | Wellcome Sanger Institute for the COVID-19 Genomics UK (COG-UK) Consortium | Jacquelyn Wynn, Mairead Hyland, The Lighthouse Lab in Alderley Park and Alex Alderton, Roberto Amato, Jeffrey Barrett, Sonia Goncalves, Ewan Harrison, David K. Jackson, Ian Johnston, Dominic Kwiatkowski, Cordelia Langford, John Sillitoe on behalf of the Wellcome Sanger Institute COVID-19 Surveillance Team |
| EPI_ISL_3093263 | Lighthouse Lab in Glasgow | Wellcome Sanger Institute for the COVID-19 Genomics UK (COG-UK) Consortium | Harper VanSteenhouse, Yumi Kasai, David Gray, Carol Clugston, Anna Dominiczak and Alex Alderton, Roberto Amato, Jeffrey Barrett, Sonia Goncalves, Ewan Harrison, David K. Jackson, Ian Johnston, Dominic Kwiatkowski, Cordelia Langford, John Sillitoe on behalf of the Wellcome Sanger Institute COVID-19 Surveillance Team |
| EPI_ISL_3093264 | Lighthouse Lab in Alderley Park | Wellcome Sanger Institute for the COVID-19 Genomics UK (COG-UK) Consortium | Jacquelyn Wynn, Mairead Hyland, The Lighthouse Lab in Alderley Park and Alex Alderton, Roberto Amato, Jeffrey Barrett, Sonia Goncalves, Ewan Harrison, David K. Jackson, Ian Johnston, Dominic Kwiatkowski, Cordelia Langford, John Sillitoe on behalf of the Wellcome Sanger Institute COVID-19 Surveillance Team |

| | | | |
|---|---|---|---|
| EPI_ISL_3093265, EPI_ISL_3093266 | Lighthouse Laboratory Plymouth | Wellcome Sanger Institute for the COVID-19 Genomics UK (COG-UK) Consortium | Lighthouse Laboratory Plymouth and Alex Alderton, Roberto Amato, Jeffrey Barrett, Sonia Goncalves, Ewan Harrison, David K. Jackson, Ian Johnston, Dominic Kwiatkowski, Cordelia Langford, John Sillitoe on behalf of the Wellcome Sanger Institute COVID-19 Surveillance Team |
| EPI_ISL_3093267, EPI_ISL_3093268 | Lighthouse Lab in Alderley Park | Wellcome Sanger Institute for the COVID-19 Genomics UK (COG-UK) Consortium | Jacquelyn Wynn, Mairead Hyland, The Lighthouse Lab in Alderley Park and Alex Alderton, Roberto Amato, Jeffrey Barrett, Sonia Goncalves, Ewan Harrison, David K. Jackson, Ian Johnston, Dominic Kwiatkowski, Cordelia Langford, John Sillitoe on behalf of the Wellcome Sanger Institute COVID-19 Surveillance Team |
| EPI_ISL_3093269 | Lighthouse Lab in Glasgow | Wellcome Sanger Institute for the COVID-19 Genomics UK (COG-UK) Consortium | Harper VanSteenhouse, Yumi Kasai, David Gray, Carol Clugston, Anna Dominiczak and Alex Alderton, Roberto Amato, Jeffrey Barrett, Sonia Goncalves, Ewan Harrison, David K. Jackson, Ian Johnston, Dominic Kwiatkowski, Cordelia Langford, John Sillitoe on behalf of the Wellcome Sanger Institute COVID-19 Surveillance Team |
| EPI_ISL_3093270, EPI_ISL_3093271, EPI_ISL_3093272, EPI_ISL_3093273 | Lighthouse Lab in Alderley Park | Wellcome Sanger Institute for the COVID-19 Genomics UK (COG-UK) Consortium | Jacquelyn Wynn, Mairead Hyland, The Lighthouse Lab in Alderley Park and Alex Alderton, Roberto Amato, Jeffrey Barrett, Sonia Goncalves, Ewan Harrison, David K. Jackson, Ian Johnston, Dominic Kwiatkowski, Cordelia Langford, John Sillitoe on behalf of the Wellcome Sanger Institute COVID-19 Surveillance Team |
| EPI_ISL_3093274 | Lighthouse Lab in Glasgow | Wellcome Sanger Institute for the COVID-19 Genomics UK (COG-UK) Consortium | Harper VanSteenhouse, Yumi Kasai, David Gray, Carol Clugston, Anna Dominiczak and Alex Alderton, Roberto Amato, Jeffrey Barrett, Sonia Goncalves, Ewan Harrison, David K. Jackson, Ian Johnston, Dominic Kwiatkowski, Cordelia Langford, John Sillitoe on behalf of the Wellcome Sanger Institute COVID-19 Surveillance Team |
| EPI_ISL_3093275 | Lighthouse Laboratory Plymouth | Wellcome Sanger Institute for the COVID-19 Genomics UK (COG-UK) Consortium | Lighthouse Laboratory Plymouth and Alex Alderton, Roberto Amato, Jeffrey Barrett, Sonia Goncalves, Ewan Harrison, David K. Jackson, Ian Johnston, Dominic Kwiatkowski, Cordelia Langford, John Sillitoe on behalf of the Wellcome Sanger Institute COVID-19 Surveillance Team |
| EPI_ISL_3093276 | Hospital del Oriente de Asturias | Laboratorio de Virología HUCA | Sandoval M, Castelló C, Gómez de Oña J, Boga JA, Rojo S, Alvarez-Arguelles ME, Abreu F, Costales I, Perez-Martínez Z, Martín-Rodríguez G, Coto E, Melón S |
| EPI_ISL_3093277 | Lighthouse Lab in Alderley Park | Wellcome Sanger Institute for the COVID-19 Genomics UK (COG-UK) Consortium | Jacquelyn Wynn, Mairead Hyland, The Lighthouse Lab in Alderley Park and Alex Alderton, Roberto Amato, Jeffrey Barrett, Sonia Goncalves, Ewan Harrison, David K. Jackson, Ian Johnston, Dominic Kwiatkowski, Cordelia Langford, John Sillitoe on behalf of the Wellcome Sanger Institute COVID-19 Surveillance Team |
| EPI_ISL_3093278 | Lighthouse Laboratory Plymouth | Wellcome Sanger Institute for the COVID-19 Genomics UK (COG-UK) Consortium | Lighthouse Laboratory Plymouth and Alex Alderton, Roberto Amato, Jeffrey Barrett, Sonia Goncalves, Ewan Harrison, David K. Jackson, Ian Johnston, Dominic Kwiatkowski, Cordelia Langford, John Sillitoe on behalf of the Wellcome Sanger Institute COVID-19 Surveillance Team |
| EPI_ISL_3093279 | Lighthouse Lab in Alderley Park | Wellcome Sanger Institute for the COVID-19 Genomics UK (COG-UK) Consortium | Jacquelyn Wynn, Mairead Hyland, The Lighthouse Lab in Alderley Park and Alex Alderton, Roberto Amato, Jeffrey Barrett, Sonia Goncalves, Ewan Harrison, David K. Jackson, Ian Johnston, Dominic Kwiatkowski, Cordelia Langford, John Sillitoe on behalf of the Wellcome Sanger Institute COVID-19 Surveillance Team |
| EPI_ISL_3093280 | Lighthouse Laboratory Plymouth | Wellcome Sanger Institute for the COVID-19 Genomics UK (COG-UK) Consortium | Lighthouse Laboratory Plymouth and Alex Alderton, Roberto Amato, Jeffrey Barrett, Sonia Goncalves, Ewan Harrison, David K. Jackson, Ian Johnston, Dominic Kwiatkowski, Cordelia Langford, John Sillitoe on behalf of the Wellcome Sanger Institute COVID-19 Surveillance Team |
| EPI_ISL_3093281, EPI_ISL_3093282 | Lighthouse Lab in Alderley Park | Wellcome Sanger Institute for the COVID-19 Genomics UK (COG-UK) Consortium | Jacquelyn Wynn, Mairead Hyland, The Lighthouse Lab in Alderley Park and Alex Alderton, Roberto Amato, Jeffrey Barrett, Sonia Goncalves, Ewan Harrison, David K. Jackson, Ian Johnston, Dominic Kwiatkowski, Cordelia Langford, John Sillitoe on behalf of the Wellcome Sanger Institute COVID-19 Surveillance Team |
| EPI_ISL_3093283 | Lighthouse Lab in Glasgow | Wellcome Sanger Institute for the COVID-19 Genomics UK (COG-UK) Consortium | Harper VanSteenhouse, Yumi Kasai, David Gray, Carol Clugston, Anna Dominiczak and Alex Alderton, Roberto Amato, Jeffrey Barrett, Sonia Goncalves, Ewan Harrison, David K. Jackson, Ian Johnston, Dominic Kwiatkowski, Cordelia Langford, John Sillitoe on behalf of the Wellcome Sanger Institute COVID-19 Surveillance Team |
| EPI_ISL_3093284 | Lighthouse Laboratory Plymouth | Wellcome Sanger Institute for the COVID-19 Genomics UK (COG-UK) Consortium | Lighthouse Laboratory Plymouth and Alex Alderton, Roberto Amato, Jeffrey Barrett, Sonia Goncalves, Ewan Harrison, David K. Jackson, Ian Johnston, Dominic Kwiatkowski, Cordelia Langford, John Sillitoe on behalf of the Wellcome Sanger Institute COVID-19 Surveillance Team |
| EPI_ISL_3093285 | Lighthouse Lab in Alderley Park | Wellcome Sanger Institute for the COVID-19 Genomics UK (COG-UK) Consortium | Jacquelyn Wynn, Mairead Hyland, The Lighthouse Lab in Alderley Park and Alex Alderton, Roberto Amato, Jeffrey Barrett, Sonia Goncalves, Ewan Harrison, David K. Jackson, Ian Johnston, Dominic Kwiatkowski, Cordelia Langford, John Sillitoe on behalf of the Wellcome Sanger Institute COVID-19 Surveillance Team |
| EPI_ISL_3093286 | Lighthouse Lab in Glasgow | Wellcome Sanger Institute for the COVID-19 Genomics UK (COG-UK) Consortium | Harper VanSteenhouse, Yumi Kasai, David Gray, Carol Clugston, Anna Dominiczak and Alex Alderton, Roberto Amato, Jeffrey Barrett, Sonia Goncalves, Ewan Harrison, David K. Jackson, Ian Johnston, Dominic Kwiatkowski, Cordelia Langford, John Sillitoe on behalf of the Wellcome Sanger Institute COVID-19 Surveillance Team |
| EPI_ISL_3093287, EPI_ISL_3093288, EPI_ISL_3093289, EPI_ISL_3093290, EPI_ISL_3093291, EPI_ISL_3093292, EPI_ISL_3093293, EPI_ISL_3093294 | Lighthouse Lab in Alderley Park | Wellcome Sanger Institute for the COVID-19 Genomics UK (COG-UK) Consortium | Jacquelyn Wynn, Mairead Hyland, The Lighthouse Lab in Alderley Park and Alex Alderton, Roberto Amato, Jeffrey Barrett, Sonia Goncalves, Ewan Harrison, David K. Jackson, Ian Johnston, Dominic Kwiatkowski, Cordelia Langford, John Sillitoe on behalf of the Wellcome Sanger Institute COVID-19 Surveillance Team |
| EPI_ISL_3093295, EPI_ISL_3093296 | Lighthouse Laboratory Plymouth | Wellcome Sanger Institute for the COVID-19 Genomics UK (COG-UK) Consortium | Lighthouse Laboratory Plymouth and Alex Alderton, Roberto Amato, Jeffrey Barrett, Sonia Goncalves, Ewan Harrison, David K. Jackson, Ian Johnston, Dominic Kwiatkowski, Cordelia Langford, John Sillitoe on behalf of the Wellcome Sanger Institute COVID-19 Surveillance Team |
| EPI_ISL_3093297, EPI_ISL_3093298 | Lighthouse Lab in Glasgow | Wellcome Sanger Institute for the COVID-19 Genomics UK (COG-UK) Consortium | Harper VanSteenhouse, Yumi Kasai, David Gray, Carol Clugston, Anna Dominiczak and Alex Alderton, Roberto Amato, Jeffrey Barrett, Sonia Goncalves, Ewan Harrison, David K. Jackson, Ian Johnston, Dominic Kwiatkowski, Cordelia Langford, John Sillitoe on behalf of the Wellcome Sanger Institute COVID-19 Surveillance Team |
| EPI_ISL_3093299 | Lighthouse Lab in Alderley Park | Wellcome Sanger Institute for the COVID-19 Genomics UK (COG-UK) Consortium | Jacquelyn Wynn, Mairead Hyland, The Lighthouse Lab in Alderley Park and Alex Alderton, Roberto Amato, Jeffrey Barrett, Sonia Goncalves, Ewan Harrison, David K. Jackson, Ian Johnston, Dominic Kwiatkowski, Cordelia Langford, John Sillitoe on behalf of the Wellcome Sanger Institute COVID-19 Surveillance Team |
| EPI_ISL_3093300 | Lighthouse Lab in Glasgow | Wellcome Sanger Institute for the COVID-19 Genomics UK (COG-UK) Consortium | Harper VanSteenhouse, Yumi Kasai, David Gray, Carol Clugston, Anna Dominiczak and Alex Alderton, Roberto Amato, Jeffrey Barrett, Sonia Goncalves, Ewan Harrison, David K. Jackson, Ian Johnston, Dominic Kwiatkowski, Cordelia Langford, John Sillitoe on behalf of the Wellcome Sanger Institute COVID-19 Surveillance Team |
| EPI_ISL_3093301 | Lighthouse Lab in Alderley Park | Wellcome Sanger Institute for the COVID-19 Genomics UK (COG-UK) Consortium | Jacquelyn Wynn, Mairead Hyland, The Lighthouse Lab in Alderley Park and Alex Alderton, Roberto Amato, Jeffrey Barrett, Sonia Goncalves, Ewan Harrison, David K. Jackson, Ian Johnston, Dominic Kwiatkowski, Cordelia Langford, John Sillitoe on behalf of the Wellcome Sanger Institute COVID-19 Surveillance Team |
| EPI_ISL_3093302, EPI_ISL_3093303 | Lighthouse Lab in Glasgow | Wellcome Sanger Institute for the COVID-19 Genomics UK (COG-UK) Consortium | Harper VanSteenhouse, Yumi Kasai, David Gray, Carol Clugston, Anna Dominiczak and Alex Alderton, Roberto Amato, Jeffrey Barrett, Sonia Goncalves, Ewan Harrison, David K. Jackson, Ian Johnston, Dominic Kwiatkowski, Cordelia Langford, John Sillitoe on behalf of the Wellcome Sanger Institute COVID-19 Surveillance Team |
| EPI_ISL_3093304, EPI_ISL_3093305, EPI_ISL_3093306 | Lighthouse Lab in Alderley Park | Wellcome Sanger Institute for the COVID-19 Genomics UK (COG-UK) Consortium | Jacquelyn Wynn, Mairead Hyland, The Lighthouse Lab in Alderley Park and Alex Alderton, Roberto Amato, Jeffrey Barrett, Sonia Goncalves, Ewan Harrison, David K. Jackson, Ian Johnston, Dominic Kwiatkowski, Cordelia Langford, John Sillitoe on behalf of the Wellcome Sanger Institute COVID-19 Surveillance Team |
| EPI_ISL_3093307 | Lighthouse Lab in Glasgow | Wellcome Sanger Institute for the COVID-19 Genomics UK (COG-UK) Consortium | Harper VanSteenhouse, Yumi Kasai, David Gray, Carol Clugston, Anna Dominiczak and Alex Alderton, Roberto Amato, Jeffrey Barrett, Sonia Goncalves, Ewan Harrison, David K. Jackson, Ian Johnston, Dominic Kwiatkowski, Cordelia Langford, John Sillitoe on behalf of the Wellcome Sanger Institute COVID-19 Surveillance Team |
| EPI_ISL_3093308 | Lighthouse Laboratory Plymouth | Wellcome Sanger Institute for the COVID-19 Genomics UK (COG-UK) Consortium | Lighthouse Laboratory Plymouth and Alex Alderton, Roberto Amato, Jeffrey Barrett, Sonia Goncalves, Ewan Harrison, David K. Jackson, Ian Johnston, Dominic Kwiatkowski, Cordelia Langford, John Sillitoe on behalf of the Wellcome Sanger Institute COVID-19 Surveillance Team |
| EPI_ISL_3093309, EPI_ISL_3093310 | Lighthouse Lab in Alderley Park | Wellcome Sanger Institute for the COVID-19 Genomics UK (COG-UK) Consortium | Jacquelyn Wynn, Mairead Hyland, The Lighthouse Lab in Alderley Park and Alex Alderton, Roberto Amato, Jeffrey Barrett, Sonia Goncalves, Ewan Harrison, David K. Jackson, Ian Johnston, Dominic Kwiatkowski, Cordelia Langford, John Sillitoe on behalf of the Wellcome Sanger Institute COVID-19 Surveillance Team |
| EPI_ISL_3093311 | Lighthouse Laboratory Plymouth | Wellcome Sanger Institute for the COVID-19 Genomics UK (COG-UK) Consortium | Lighthouse Laboratory Plymouth and Alex Alderton, Roberto Amato, Jeffrey Barrett, Sonia Goncalves, Ewan Harrison, David K. Jackson, Ian Johnston, Dominic Kwiatkowski, Cordelia Langford, John Sillitoe on behalf of the Wellcome Sanger Institute COVID-19 Surveillance Team |
| EPI_ISL_3093312 | Lighthouse Lab in Alderley Park | Wellcome Sanger Institute for the COVID-19 Genomics UK (COG-UK) Consortium | Jacquelyn Wynn, Mairead Hyland, The Lighthouse Lab in Alderley Park and Alex Alderton, Roberto Amato, Jeffrey Barrett, Sonia Goncalves, Ewan Harrison, David K. Jackson, Ian Johnston, Dominic Kwiatkowski, Cordelia Langford, John Sillitoe on behalf of the Wellcome Sanger Institute COVID-19 Surveillance Team |
| EPI_ISL_3093313 | Lighthouse Laboratory Plymouth | Wellcome Sanger Institute for the COVID-19 Genomics UK (COG-UK) Consortium | Lighthouse Laboratory Plymouth and Alex Alderton, Roberto Amato, Jeffrey Barrett, Sonia Goncalves, Ewan Harrison, David K. Jackson, Ian Johnston, Dominic Kwiatkowski, Cordelia Langford, John Sillitoe on behalf of the Wellcome Sanger Institute COVID-19 Surveillance Team |
| EPI_ISL_3093314 | Lighthouse Lab in Alderley Park | Wellcome Sanger Institute for the COVID-19 Genomics UK (COG-UK) Consortium | Jacquelyn Wynn, Mairead Hyland, The Lighthouse Lab in Alderley Park and Alex Alderton, Roberto Amato, Jeffrey Barrett, Sonia Goncalves, Ewan Harrison, David K. Jackson, Ian Johnston, Dominic Kwiatkowski, Cordelia Langford, John Sillitoe on behalf of the Wellcome Sanger Institute COVID-19 Surveillance Team |
| EPI_ISL_3093315 | Lighthouse Laboratory Plymouth | Wellcome Sanger Institute for the COVID-19 Genomics UK (COG-UK) Consortium | Lighthouse Laboratory Plymouth and Alex Alderton, Roberto Amato, Jeffrey Barrett, Sonia Goncalves, Ewan Harrison, David K. Jackson, Ian Johnston, Dominic Kwiatkowski, Cordelia Langford, John Sillitoe on behalf of the Wellcome Sanger Institute COVID-19 Surveillance Team |
| EPI_ISL_3093316, EPI_ISL_3093317 | Lighthouse Lab in Alderley Park | Wellcome Sanger Institute for the COVID-19 Genomics UK (COG-UK) Consortium | Jacquelyn Wynn, Mairead Hyland, The Lighthouse Lab in Alderley Park and Alex Alderton, Roberto Amato, Jeffrey Barrett, Sonia Goncalves, Ewan Harrison, David K. Jackson, Ian Johnston, Dominic Kwiatkowski, Cordelia Langford, John Sillitoe on behalf of the Wellcome Sanger Institute COVID-19 Surveillance Team |
| EPI_ISL_3093318, EPI_ISL_3093319 | Lighthouse Lab in Glasgow | Wellcome Sanger Institute for the COVID-19 Genomics UK (COG-UK) Consortium | Harper VanSteenhouse, Yumi Kasai, David Gray, Carol Clugston, Anna Dominiczak and Alex Alderton, Roberto Amato, Jeffrey Barrett, Sonia Goncalves, Ewan Harrison, David K. Jackson, Ian Johnston, Dominic Kwiatkowski, Cordelia Langford, John Sillitoe on behalf of the Wellcome Sanger Institute COVID-19 Surveillance Team |
| EPI_ISL_3093320 | Lighthouse Lab in Alderley Park | Wellcome Sanger Institute for the COVID-19 Genomics UK (COG-UK) Consortium | Jacquelyn Wynn, Mairead Hyland, The Lighthouse Lab in Alderley Park and Alex Alderton, Roberto Amato, Jeffrey Barrett, Sonia Goncalves, Ewan Harrison, David K. Jackson, Ian Johnston, Dominic Kwiatkowski, Cordelia Langford, John Sillitoe on behalf of the Wellcome Sanger Institute COVID-19 Surveillance Team |
| EPI_ISL_3093321 | Lighthouse Laboratory Plymouth | Wellcome Sanger Institute for the COVID-19 Genomics UK (COG-UK) Consortium | Lighthouse Laboratory Plymouth and Alex Alderton, Roberto Amato, Jeffrey Barrett, Sonia Goncalves, Ewan Harrison, David K. Jackson, Ian Johnston, Dominic Kwiatkowski, Cordelia Langford, John Sillitoe on behalf of the Wellcome Sanger Institute COVID-19 Surveillance Team |
| EPI_ISL_3093322, EPI_ISL_3093323 | Lighthouse Lab in Glasgow | Wellcome Sanger Institute for the COVID-19 Genomics UK (COG-UK) Consortium | Harper VanSteenhouse, Yumi Kasai, David Gray, Carol Clugston, Anna Dominiczak and Alex Alderton, Roberto Amato, Jeffrey Barrett, Sonia Goncalves, Ewan Harrison, David K. Jackson, Ian Johnston, Dominic Kwiatkowski, Cordelia Langford, John Sillitoe on behalf of the Wellcome Sanger Institute COVID-19 Surveillance Team |
| EPI_ISL_3093324 | Lighthouse Lab in Alderley Park | Wellcome Sanger Institute for the COVID-19 Genomics UK (COG-UK) Consortium | Jacquelyn Wynn, Mairead Hyland, The Lighthouse Lab in Alderley Park and Alex Alderton, Roberto Amato, Jeffrey Barrett, Sonia Goncalves, Ewan Harrison, David K. Jackson, Ian Johnston, Dominic Kwiatkowski, Cordelia Langford, John Sillitoe on behalf of the Wellcome Sanger Institute COVID-19 Surveillance Team |
| EPI_ISL_3093325 | Lighthouse Laboratory Plymouth | Wellcome Sanger Institute for the COVID-19 Genomics UK (COG-UK) Consortium | Lighthouse Laboratory Plymouth and Alex Alderton, Roberto Amato, Jeffrey Barrett, Sonia Goncalves, Ewan Harrison, David K. Jackson, Ian Johnston, Dominic Kwiatkowski, Cordelia Langford, John Sillitoe on behalf of the Wellcome Sanger Institute COVID-19 Surveillance Team |
| EPI_ISL_3093326 | Lighthouse Lab in Alderley Park | Wellcome Sanger Institute for the COVID-19 Genomics UK (COG-UK) Consortium | Jacquelyn Wynn, Mairead Hyland, The Lighthouse Lab in Alderley Park and Alex Alderton, Roberto Amato, Jeffrey Barrett, Sonia Goncalves, Ewan Harrison, David K. Jackson, Ian Johnston, Dominic Kwiatkowski, Cordelia Langford, John Sillitoe on behalf of the Wellcome Sanger Institute COVID-19 Surveillance Team |
| EPI_ISL_3093327 | Lighthouse Lab in Glasgow | Wellcome Sanger Institute for the COVID-19 Genomics UK (COG-UK) Consortium | Harper VanSteenhouse, Yumi Kasai, David Gray, Carol Clugston, Anna Dominiczak and Alex Alderton, Roberto Amato, Jeffrey Barrett, Sonia Goncalves, Ewan Harrison, David K. Jackson, Ian Johnston, Dominic Kwiatkowski, Cordelia Langford, John Sillitoe on behalf of the Wellcome Sanger Institute COVID-19 Surveillance Team |
| EPI_ISL_3093328 | Lighthouse Laboratory Plymouth | Wellcome Sanger Institute for the COVID-19 Genomics UK (COG-UK) Consortium | Lighthouse Laboratory Plymouth and Alex Alderton, Roberto Amato, Jeffrey Barrett, Sonia Goncalves, Ewan Harrison, David K. Jackson, Ian Johnston, Dominic Kwiatkowski, Cordelia Langford, John Sillitoe on behalf of the Wellcome Sanger Institute COVID-19 Surveillance Team |

| | | | |
|---|---|---|---|
| EPI_ISL_3093329 | Lighthouse Lab in Glasgow | Wellcome Sanger Institute for the COVID-19 Genomics UK (COG-UK) Consortium | Harper VanSteenhouse, Yumi Kasai, David Gray, Carol Clugston, Anna Dominiczak and Alex Alderton, Roberto Amato, Jeffrey Barrett, Sonia Goncalves, Ewan Harrison, David K. Jackson, Ian Johnston, Dominic Kwiatkowski, Cordelia Langford, John Sillitoe on behalf of the Wellcome Sanger Institute COVID-19 Surveillance Team |
| EPI_ISL_3093330, EPI_ISL_3093331 | Lighthouse Lab in Alderley Park | Wellcome Sanger Institute for the COVID-19 Genomics UK (COG-UK) Consortium | Jacquelyn Wynn, Mairead Hyland, The Lighthouse Lab in Alderley Park and Alex Alderton, Roberto Amato, Jeffrey Barrett, Sonia Goncalves, Ewan Harrison, David K. Jackson, Ian Johnston, Dominic Kwiatkowski, Cordelia Langford, John Sillitoe on behalf of the Wellcome Sanger Institute COVID-19 Surveillance Team |
| EPI_ISL_3093332 | Lighthouse Laboratory Plymouth | Wellcome Sanger Institute for the COVID-19 Genomics UK (COG-UK) Consortium | Lighthouse Laboratory Plymouth and Alex Alderton, Roberto Amato, Jeffrey Barrett, Sonia Goncalves, Ewan Harrison, David K. Jackson, Ian Johnston, Dominic Kwiatkowski, Cordelia Langford, John Sillitoe on behalf of the Wellcome Sanger Institute COVID-19 Surveillance Team |
| EPI_ISL_3093333 | Lighthouse Lab in Alderley Park | Wellcome Sanger Institute for the COVID-19 Genomics UK (COG-UK) Consortium | Jacquelyn Wynn, Mairead Hyland, The Lighthouse Lab in Alderley Park and Alex Alderton, Roberto Amato, Jeffrey Barrett, Sonia Goncalves, Ewan Harrison, David K. Jackson, Ian Johnston, Dominic Kwiatkowski, Cordelia Langford, John Sillitoe on behalf of the Wellcome Sanger Institute COVID-19 Surveillance Team |
| EPI_ISL_3093334 | Lighthouse Lab in Glasgow | Wellcome Sanger Institute for the COVID-19 Genomics UK (COG-UK) Consortium | Harper VanSteenhouse, Yumi Kasai, David Gray, Carol Clugston, Anna Dominiczak and Alex Alderton, Roberto Amato, Jeffrey Barrett, Sonia Goncalves, Ewan Harrison, David K. Jackson, Ian Johnston, Dominic Kwiatkowski, Cordelia Langford, John Sillitoe on behalf of the Wellcome Sanger Institute COVID-19 Surveillance Team |
| EPI_ISL_3093335 | Lighthouse Laboratory Plymouth | Wellcome Sanger Institute for the COVID-19 Genomics UK (COG-UK) Consortium | Lighthouse Laboratory Plymouth and Alex Alderton, Roberto Amato, Jeffrey Barrett, Sonia Goncalves, Ewan Harrison, David K. Jackson, Ian Johnston, Dominic Kwiatkowski, Cordelia Langford, John Sillitoe on behalf of the Wellcome Sanger Institute COVID-19 Surveillance Team |
| EPI_ISL_3093336 | Lighthouse Lab in Glasgow | Wellcome Sanger Institute for the COVID-19 Genomics UK (COG-UK) Consortium | Harper VanSteenhouse, Yumi Kasai, David Gray, Carol Clugston, Anna Dominiczak and Alex Alderton, Roberto Amato, Jeffrey Barrett, Sonia Goncalves, Ewan Harrison, David K. Jackson, Ian Johnston, Dominic Kwiatkowski, Cordelia Langford, John Sillitoe on behalf of the Wellcome Sanger Institute COVID-19 Surveillance Team |
| EPI_ISL_3093337 | Lighthouse Laboratory Plymouth | Wellcome Sanger Institute for the COVID-19 Genomics UK (COG-UK) Consortium | Lighthouse Laboratory Plymouth and Alex Alderton, Roberto Amato, Jeffrey Barrett, Sonia Goncalves, Ewan Harrison, David K. Jackson, Ian Johnston, Dominic Kwiatkowski, Cordelia Langford, John Sillitoe on behalf of the Wellcome Sanger Institute COVID-19 Surveillance Team |
| EPI_ISL_3093338, EPI_ISL_3093339, EPI_ISL_3093340 | Lighthouse Lab in Alderley Park | Wellcome Sanger Institute for the COVID-19 Genomics UK (COG-UK) Consortium | Jacquelyn Wynn, Mairead Hyland, The Lighthouse Lab in Alderley Park and Alex Alderton, Roberto Amato, Jeffrey Barrett, Sonia Goncalves, Ewan Harrison, David K. Jackson, Ian Johnston, Dominic Kwiatkowski, Cordelia Langford, John Sillitoe on behalf of the Wellcome Sanger Institute COVID-19 Surveillance Team |
| EPI_ISL_3093341 | Lighthouse Laboratory Plymouth | Wellcome Sanger Institute for the COVID-19 Genomics UK (COG-UK) Consortium | Lighthouse Laboratory Plymouth and Alex Alderton, Roberto Amato, Jeffrey Barrett, Sonia Goncalves, Ewan Harrison, David K. Jackson, Ian Johnston, Dominic Kwiatkowski, Cordelia Langford, John Sillitoe on behalf of the Wellcome Sanger Institute COVID-19 Surveillance Team |
| EPI_ISL_3093342 | Lighthouse Lab in Alderley Park | Wellcome Sanger Institute for the COVID-19 Genomics UK (COG-UK) Consortium | Jacquelyn Wynn, Mairead Hyland, The Lighthouse Lab in Alderley Park and Alex Alderton, Roberto Amato, Jeffrey Barrett, Sonia Goncalves, Ewan Harrison, David K. Jackson, Ian Johnston, Dominic Kwiatkowski, Cordelia Langford, John Sillitoe on behalf of the Wellcome Sanger Institute COVID-19 Surveillance Team |
| EPI_ISL_3093343 | Lighthouse Lab in Glasgow | Wellcome Sanger Institute for the COVID-19 Genomics UK (COG-UK) Consortium | Harper VanSteenhouse, Yumi Kasai, David Gray, Carol Clugston, Anna Dominiczak and Alex Alderton, Roberto Amato, Jeffrey Barrett, Sonia Goncalves, Ewan Harrison, David K. Jackson, Ian Johnston, Dominic Kwiatkowski, Cordelia Langford, John Sillitoe on behalf of the Wellcome Sanger Institute COVID-19 Surveillance Team |
| EPI_ISL_3093344 | Lighthouse Lab in Alderley Park | Wellcome Sanger Institute for the COVID-19 Genomics UK (COG-UK) Consortium | Jacquelyn Wynn, Mairead Hyland, The Lighthouse Lab in Alderley Park and Alex Alderton, Roberto Amato, Jeffrey Barrett, Sonia Goncalves, Ewan Harrison, David K. Jackson, Ian Johnston, Dominic Kwiatkowski, Cordelia Langford, John Sillitoe on behalf of the Wellcome Sanger Institute COVID-19 Surveillance Team |
| EPI_ISL_3093345 | Lighthouse Laboratory Plymouth | Wellcome Sanger Institute for the COVID-19 Genomics UK (COG-UK) Consortium | Lighthouse Laboratory Plymouth and Alex Alderton, Roberto Amato, Jeffrey Barrett, Sonia Goncalves, Ewan Harrison, David K. Jackson, Ian Johnston, Dominic Kwiatkowski, Cordelia Langford, John Sillitoe on behalf of the Wellcome Sanger Institute COVID-19 Surveillance Team |
| EPI_ISL_3093346, EPI_ISL_3093347, EPI_ISL_3093348 | Lighthouse Lab in Alderley Park | Wellcome Sanger Institute for the COVID-19 Genomics UK (COG-UK) Consortium | Jacquelyn Wynn, Mairead Hyland, The Lighthouse Lab in Alderley Park and Alex Alderton, Roberto Amato, Jeffrey Barrett, Sonia Goncalves, Ewan Harrison, David K. Jackson, Ian Johnston, Dominic Kwiatkowski, Cordelia Langford, John Sillitoe on behalf of the Wellcome Sanger Institute COVID-19 Surveillance Team |
| EPI_ISL_3093349, EPI_ISL_3093350 | Lighthouse Laboratory Plymouth | Wellcome Sanger Institute for the COVID-19 Genomics UK (COG-UK) Consortium | Lighthouse Laboratory Plymouth and Alex Alderton, Roberto Amato, Jeffrey Barrett, Sonia Goncalves, Ewan Harrison, David K. Jackson, Ian Johnston, Dominic Kwiatkowski, Cordelia Langford, John Sillitoe on behalf of the Wellcome Sanger Institute COVID-19 Surveillance Team |
| EPI_ISL_3093351 | Lighthouse Lab in Alderley Park | Wellcome Sanger Institute for the COVID-19 Genomics UK (COG-UK) Consortium | Jacquelyn Wynn, Mairead Hyland, The Lighthouse Lab in Alderley Park and Alex Alderton, Roberto Amato, Jeffrey Barrett, Sonia Goncalves, Ewan Harrison, David K. Jackson, Ian Johnston, Dominic Kwiatkowski, Cordelia Langford, John Sillitoe on behalf of the Wellcome Sanger Institute COVID-19 Surveillance Team |
| EPI_ISL_3093352 | Lighthouse Laboratory Plymouth | Wellcome Sanger Institute for the COVID-19 Genomics UK (COG-UK) Consortium | Lighthouse Laboratory Plymouth and Alex Alderton, Roberto Amato, Jeffrey Barrett, Sonia Goncalves, Ewan Harrison, David K. Jackson, Ian Johnston, Dominic Kwiatkowski, Cordelia Langford, John Sillitoe on behalf of the Wellcome Sanger Institute COVID-19 Surveillance Team |
| EPI_ISL_3093353 | Lighthouse Lab in Alderley Park | Wellcome Sanger Institute for the COVID-19 Genomics UK (COG-UK) Consortium | Jacquelyn Wynn, Mairead Hyland, The Lighthouse Lab in Alderley Park and Alex Alderton, Roberto Amato, Jeffrey Barrett, Sonia Goncalves, Ewan Harrison, David K. Jackson, Ian Johnston, Dominic Kwiatkowski, Cordelia Langford, John Sillitoe on behalf of the Wellcome Sanger Institute COVID-19 Surveillance Team |
| EPI_ISL_3093354 | Lighthouse Lab in Glasgow | Wellcome Sanger Institute for the COVID-19 Genomics UK (COG-UK) Consortium | Harper VanSteenhouse, Yumi Kasai, David Gray, Carol Clugston, Anna Dominiczak and Alex Alderton, Roberto Amato, Jeffrey Barrett, Sonia Goncalves, Ewan Harrison, David K. Jackson, Ian Johnston, Dominic Kwiatkowski, Cordelia Langford, John Sillitoe on behalf of the Wellcome Sanger Institute COVID-19 Surveillance Team |
| EPI_ISL_3093355, EPI_ISL_3093356, EPI_ISL_3093357, EPI_ISL_3093358 | Lighthouse Lab in Alderley Park | Wellcome Sanger Institute for the COVID-19 Genomics UK (COG-UK) Consortium | Jacquelyn Wynn, Mairead Hyland, The Lighthouse Lab in Alderley Park and Alex Alderton, Roberto Amato, Jeffrey Barrett, Sonia Goncalves, Ewan Harrison, David K. Jackson, Ian Johnston, Dominic Kwiatkowski, Cordelia Langford, John Sillitoe on behalf of the Wellcome Sanger Institute COVID-19 Surveillance Team |
| EPI_ISL_3093359 | Lighthouse Laboratory Plymouth | Wellcome Sanger Institute for the COVID-19 Genomics UK (COG-UK) Consortium | Lighthouse Laboratory Plymouth and Alex Alderton, Roberto Amato, Jeffrey Barrett, Sonia Goncalves, Ewan Harrison, David K. Jackson, Ian Johnston, Dominic Kwiatkowski, Cordelia Langford, John Sillitoe on behalf of the Wellcome Sanger Institute COVID-19 Surveillance Team |
| EPI_ISL_3093360, EPI_ISL_3093361, EPI_ISL_3093362, EPI_ISL_3093363, EPI_ISL_3093364 | Lighthouse Lab in Alderley Park | Wellcome Sanger Institute for the COVID-19 Genomics UK (COG-UK) Consortium | Jacquelyn Wynn, Mairead Hyland, The Lighthouse Lab in Alderley Park and Alex Alderton, Roberto Amato, Jeffrey Barrett, Sonia Goncalves, Ewan Harrison, David K. Jackson, Ian Johnston, Dominic Kwiatkowski, Cordelia Langford, John Sillitoe on behalf of the Wellcome Sanger Institute COVID-19 Surveillance Team |
| EPI_ISL_3093365 | Lighthouse Laboratory Plymouth | Wellcome Sanger Institute for the COVID-19 Genomics UK (COG-UK) Consortium | Lighthouse Laboratory Plymouth and Alex Alderton, Roberto Amato, Jeffrey Barrett, Sonia Goncalves, Ewan Harrison, David K. Jackson, Ian Johnston, Dominic Kwiatkowski, Cordelia Langford, John Sillitoe on behalf of the Wellcome Sanger Institute COVID-19 Surveillance Team |
| EPI_ISL_3093366 | Lighthouse Lab in Glasgow | Wellcome Sanger Institute for the COVID-19 Genomics UK (COG-UK) Consortium | Harper VanSteenhouse, Yumi Kasai, David Gray, Carol Clugston, Anna Dominiczak and Alex Alderton, Roberto Amato, Jeffrey Barrett, Sonia Goncalves, Ewan Harrison, David K. Jackson, Ian Johnston, Dominic Kwiatkowski, Cordelia Langford, John Sillitoe on behalf of the Wellcome Sanger Institute COVID-19 Surveillance Team |
| EPI_ISL_3093367 | Lighthouse Lab in Alderley Park | Wellcome Sanger Institute for the COVID-19 Genomics UK (COG-UK) Consortium | Jacquelyn Wynn, Mairead Hyland, The Lighthouse Lab in Alderley Park and Alex Alderton, Roberto Amato, Jeffrey Barrett, Sonia Goncalves, Ewan Harrison, David K. Jackson, Ian Johnston, Dominic Kwiatkowski, Cordelia Langford, John Sillitoe on behalf of the Wellcome Sanger Institute COVID-19 Surveillance Team |
| EPI_ISL_3093368, EPI_ISL_3093369 | Lighthouse Laboratory Plymouth | Wellcome Sanger Institute for the COVID-19 Genomics UK (COG-UK) Consortium | Lighthouse Laboratory Plymouth and Alex Alderton, Roberto Amato, Jeffrey Barrett, Sonia Goncalves, Ewan Harrison, David K. Jackson, Ian Johnston, Dominic Kwiatkowski, Cordelia Langford, John Sillitoe on behalf of the Wellcome Sanger Institute COVID-19 Surveillance Team |
| EPI_ISL_3093370 | Lighthouse Lab in Glasgow | Wellcome Sanger Institute for the COVID-19 Genomics UK (COG-UK) Consortium | Harper VanSteenhouse, Yumi Kasai, David Gray, Carol Clugston, Anna Dominiczak and Alex Alderton, Roberto Amato, Jeffrey Barrett, Sonia Goncalves, Ewan Harrison, David K. Jackson, Ian Johnston, Dominic Kwiatkowski, Cordelia Langford, John Sillitoe on behalf of the Wellcome Sanger Institute COVID-19 Surveillance Team |
| EPI_ISL_3093371, EPI_ISL_3093372 | Lighthouse Lab in Alderley Park | Wellcome Sanger Institute for the COVID-19 Genomics UK (COG-UK) Consortium | Jacquelyn Wynn, Mairead Hyland, The Lighthouse Lab in Alderley Park and Alex Alderton, Roberto Amato, Jeffrey Barrett, Sonia Goncalves, Ewan Harrison, David K. Jackson, Ian Johnston, Dominic Kwiatkowski, Cordelia Langford, John Sillitoe on behalf of the Wellcome Sanger Institute COVID-19 Surveillance Team |
| EPI_ISL_3093373 | Lighthouse Laboratory Plymouth | Wellcome Sanger Institute for the COVID-19 Genomics UK (COG-UK) Consortium | Lighthouse Laboratory Plymouth and Alex Alderton, Roberto Amato, Jeffrey Barrett, Sonia Goncalves, Ewan Harrison, David K. Jackson, Ian Johnston, Dominic Kwiatkowski, Cordelia Langford, John Sillitoe on behalf of the Wellcome Sanger Institute COVID-19 Surveillance Team |
| EPI_ISL_3093374, EPI_ISL_3093375, EPI_ISL_3093376 | Lighthouse Lab in Alderley Park | Wellcome Sanger Institute for the COVID-19 Genomics UK (COG-UK) Consortium | Jacquelyn Wynn, Mairead Hyland, The Lighthouse Lab in Alderley Park and Alex Alderton, Roberto Amato, Jeffrey Barrett, Sonia Goncalves, Ewan Harrison, David K. Jackson, Ian Johnston, Dominic Kwiatkowski, Cordelia Langford, John Sillitoe on behalf of the Wellcome Sanger Institute COVID-19 Surveillance Team |
| EPI_ISL_3093377 | Lighthouse Lab in Glasgow | Wellcome Sanger Institute for the COVID-19 Genomics UK (COG-UK) Consortium | Harper VanSteenhouse, Yumi Kasai, David Gray, Carol Clugston, Anna Dominiczak and Alex Alderton, Roberto Amato, Jeffrey Barrett, Sonia Goncalves, Ewan Harrison, David K. Jackson, Ian Johnston, Dominic Kwiatkowski, Cordelia Langford, John Sillitoe on behalf of the Wellcome Sanger Institute COVID-19 Surveillance Team |
| EPI_ISL_3093378, EPI_ISL_3093379 | Lighthouse Lab in Alderley Park | Wellcome Sanger Institute for the COVID-19 Genomics UK (COG-UK) Consortium | Jacquelyn Wynn, Mairead Hyland, The Lighthouse Lab in Alderley Park and Alex Alderton, Roberto Amato, Jeffrey Barrett, Sonia Goncalves, Ewan Harrison, David K. Jackson, Ian Johnston, Dominic Kwiatkowski, Cordelia Langford, John Sillitoe on behalf of the Wellcome Sanger Institute COVID-19 Surveillance Team |
| EPI_ISL_3093380 | Lighthouse Lab in Glasgow | Wellcome Sanger Institute for the COVID-19 Genomics UK (COG-UK) Consortium | Harper VanSteenhouse, Yumi Kasai, David Gray, Carol Clugston, Anna Dominiczak and Alex Alderton, Roberto Amato, Jeffrey Barrett, Sonia Goncalves, Ewan Harrison, David K. Jackson, Ian Johnston, Dominic Kwiatkowski, Cordelia Langford, John Sillitoe on behalf of the Wellcome Sanger Institute COVID-19 Surveillance Team |
| EPI_ISL_3093381, EPI_ISL_3093382, EPI_ISL_3093383, EPI_ISL_3093384 | Lighthouse Lab in Alderley Park | Wellcome Sanger Institute for the COVID-19 Genomics UK (COG-UK) Consortium | Jacquelyn Wynn, Mairead Hyland, The Lighthouse Lab in Alderley Park and Alex Alderton, Roberto Amato, Jeffrey Barrett, Sonia Goncalves, Ewan Harrison, David K. Jackson, Ian Johnston, Dominic Kwiatkowski, Cordelia Langford, John Sillitoe on behalf of the Wellcome Sanger Institute COVID-19 Surveillance Team |
| EPI_ISL_3093385, EPI_ISL_3093386, EPI_ISL_3093387, EPI_ISL_3093388 | Lighthouse Lab in Glasgow | Wellcome Sanger Institute for the COVID-19 Genomics UK (COG-UK) Consortium | Harper VanSteenhouse, Yumi Kasai, David Gray, Carol Clugston, Anna Dominiczak and Alex Alderton, Roberto Amato, Jeffrey Barrett, Sonia Goncalves, Ewan Harrison, David K. Jackson, Ian Johnston, Dominic Kwiatkowski, Cordelia Langford, John Sillitoe on behalf of the Wellcome Sanger Institute COVID-19 Surveillance Team |
| EPI_ISL_3093389 | Lighthouse Laboratory Plymouth | Wellcome Sanger Institute for the COVID-19 Genomics UK (COG-UK) Consortium | Lighthouse Laboratory Plymouth and Alex Alderton, Roberto Amato, Jeffrey Barrett, Sonia Goncalves, Ewan Harrison, David K. Jackson, Ian Johnston, Dominic Kwiatkowski, Cordelia Langford, John Sillitoe on behalf of the Wellcome Sanger Institute COVID-19 Surveillance Team |
| EPI_ISL_3093390 | Lighthouse Lab in Alderley Park | Wellcome Sanger Institute for the COVID-19 Genomics UK (COG-UK) Consortium | Jacquelyn Wynn, Mairead Hyland, The Lighthouse Lab in Alderley Park and Alex Alderton, Roberto Amato, Jeffrey Barrett, Sonia Goncalves, Ewan Harrison, David K. Jackson, Ian Johnston, Dominic Kwiatkowski, Cordelia Langford, John Sillitoe on behalf of the Wellcome Sanger Institute COVID-19 Surveillance Team |
| EPI_ISL_3093391 | Lighthouse Lab in Glasgow | Wellcome Sanger Institute for the COVID-19 Genomics UK (COG-UK) Consortium | Harper VanSteenhouse, Yumi Kasai, David Gray, Carol Clugston, Anna Dominiczak and Alex Alderton, Roberto Amato, Jeffrey Barrett, Sonia Goncalves, Ewan Harrison, David K. Jackson, Ian Johnston, Dominic Kwiatkowski, Cordelia Langford, John Sillitoe on behalf of the Wellcome Sanger Institute COVID-19 Surveillance Team |
| EPI_ISL_3093392, EPI_ISL_3093393, | Lighthouse Lab in Alderley Park | Wellcome Sanger Institute for the COVID-19 Genomics UK (COG-UK) Consortium | Jacquelyn Wynn, Mairead Hyland, The Lighthouse Lab in Alderley Park and Alex Alderton, Roberto Amato, Jeffrey Barrett, Sonia Goncalves, Ewan Harrison, David K. Jackson, Ian Johnston, Dominic Kwiatkowski, Cordelia Langford, John Sillitoe on behalf of the Wellcome Sanger Institute COVID-19 Surveillance Team |

| | | | |
|---|---|---|---|
| EPI_ISL_3093394 | | | |
| EPI_ISL_3093395 | Lighthouse Laboratory Plymouth | Wellcome Sanger Institute for the COVID-19 Genomics UK (COG-UK) Consortium | Lighthouse Laboratory Plymouth and Alex Alderton, Roberto Amato, Jeffrey Barrett, Sonia Goncalves, Ewan Harrison, David K. Jackson, Ian Johnston, Dominic Kwiatkowski, Cordelia Langford, John Sillitoe on behalf of the Wellcome Sanger Institute COVID-19 Surveillance Team |
| EPI_ISL_3093396, EPI_ISL_3093397, EPI_ISL_3093398 | Lighthouse Lab in Alderley Park | Wellcome Sanger Institute for the COVID-19 Genomics UK (COG-UK) Consortium | Jacquelyn Wynn, Mairead Hyland, The Lighthouse Lab in Alderley Park and Alex Alderton, Roberto Amato, Jeffrey Barrett, Sonia Goncalves, Ewan Harrison, David K. Jackson, Ian Johnston, Dominic Kwiatkowski, Cordelia Langford, John Sillitoe on behalf of the Wellcome Sanger Institute COVID-19 Surveillance Team |
| EPI_ISL_3093399, EPI_ISL_3093400 | Lighthouse Laboratory Plymouth | Wellcome Sanger Institute for the COVID-19 Genomics UK (COG-UK) Consortium | Lighthouse Laboratory Plymouth and Alex Alderton, Roberto Amato, Jeffrey Barrett, Sonia Goncalves, Ewan Harrison, David K. Jackson, Ian Johnston, Dominic Kwiatkowski, Cordelia Langford, John Sillitoe on behalf of the Wellcome Sanger Institute COVID-19 Surveillance Team |
| EPI_ISL_3093401 | Lighthouse Lab in Glasgow | Wellcome Sanger Institute for the COVID-19 Genomics UK (COG-UK) Consortium | Harper VanSteenhouse, Yumi Kasai, David Gray, Carol Clugston, Anna Dominiczak and Alex Alderton, Roberto Amato, Jeffrey Barrett, Sonia Goncalves, Ewan Harrison, David K. Jackson, Ian Johnston, Dominic Kwiatkowski, Cordelia Langford, John Sillitoe on behalf of the Wellcome Sanger Institute COVID-19 Surveillance Team |
| EPI_ISL_3093402, EPI_ISL_3093403 | Lighthouse Laboratory Plymouth | Wellcome Sanger Institute for the COVID-19 Genomics UK (COG-UK) Consortium | Lighthouse Laboratory Plymouth and Alex Alderton, Roberto Amato, Jeffrey Barrett, Sonia Goncalves, Ewan Harrison, David K. Jackson, Ian Johnston, Dominic Kwiatkowski, Cordelia Langford, John Sillitoe on behalf of the Wellcome Sanger Institute COVID-19 Surveillance Team |
| EPI_ISL_3093404, EPI_ISL_3093405 | Lighthouse Lab in Alderley Park | Wellcome Sanger Institute for the COVID-19 Genomics UK (COG-UK) Consortium | Jacquelyn Wynn, Mairead Hyland, The Lighthouse Lab in Alderley Park and Alex Alderton, Roberto Amato, Jeffrey Barrett, Sonia Goncalves, Ewan Harrison, David K. Jackson, Ian Johnston, Dominic Kwiatkowski, Cordelia Langford, John Sillitoe on behalf of the Wellcome Sanger Institute COVID-19 Surveillance Team |
| EPI_ISL_3093406, EPI_ISL_3093407 | Lighthouse Laboratory Plymouth | Wellcome Sanger Institute for the COVID-19 Genomics UK (COG-UK) Consortium | Lighthouse Laboratory Plymouth and Alex Alderton, Roberto Amato, Jeffrey Barrett, Sonia Goncalves, Ewan Harrison, David K. Jackson, Ian Johnston, Dominic Kwiatkowski, Cordelia Langford, John Sillitoe on behalf of the Wellcome Sanger Institute COVID-19 Surveillance Team |
| EPI_ISL_3093408 | Lighthouse Lab in Glasgow | Wellcome Sanger Institute for the COVID-19 Genomics UK (COG-UK) Consortium | Harper VanSteenhouse, Yumi Kasai, David Gray, Carol Clugston, Anna Dominiczak and Alex Alderton, Roberto Amato, Jeffrey Barrett, Sonia Goncalves, Ewan Harrison, David K. Jackson, Ian Johnston, Dominic Kwiatkowski, Cordelia Langford, John Sillitoe on behalf of the Wellcome Sanger Institute COVID-19 Surveillance Team |
| EPI_ISL_3093409 | Lighthouse Laboratory Plymouth | Wellcome Sanger Institute for the COVID-19 Genomics UK (COG-UK) Consortium | Lighthouse Laboratory Plymouth and Alex Alderton, Roberto Amato, Jeffrey Barrett, Sonia Goncalves, Ewan Harrison, David K. Jackson, Ian Johnston, Dominic Kwiatkowski, Cordelia Langford, John Sillitoe on behalf of the Wellcome Sanger Institute COVID-19 Surveillance Team |
| EPI_ISL_3093410 | Lighthouse Lab in Alderley Park | Wellcome Sanger Institute for the COVID-19 Genomics UK (COG-UK) Consortium | Jacquelyn Wynn, Mairead Hyland, The Lighthouse Lab in Alderley Park and Alex Alderton, Roberto Amato, Jeffrey Barrett, Sonia Goncalves, Ewan Harrison, David K. Jackson, Ian Johnston, Dominic Kwiatkowski, Cordelia Langford, John Sillitoe on behalf of the Wellcome Sanger Institute COVID-19 Surveillance Team |
| EPI_ISL_3093411 | Lighthouse Lab in Glasgow | Wellcome Sanger Institute for the COVID-19 Genomics UK (COG-UK) Consortium | Harper VanSteenhouse, Yumi Kasai, David Gray, Carol Clugston, Anna Dominiczak and Alex Alderton, Roberto Amato, Jeffrey Barrett, Sonia Goncalves, Ewan Harrison, David K. Jackson, Ian Johnston, Dominic Kwiatkowski, Cordelia Langford, John Sillitoe on behalf of the Wellcome Sanger Institute COVID-19 Surveillance Team |
| EPI_ISL_3093412, EPI_ISL_3093413 | Lighthouse Lab in Alderley Park | Wellcome Sanger Institute for the COVID-19 Genomics UK (COG-UK) Consortium | Jacquelyn Wynn, Mairead Hyland, The Lighthouse Lab in Alderley Park and Alex Alderton, Roberto Amato, Jeffrey Barrett, Sonia Goncalves, Ewan Harrison, David K. Jackson, Ian Johnston, Dominic Kwiatkowski, Cordelia Langford, John Sillitoe on behalf of the Wellcome Sanger Institute COVID-19 Surveillance Team |
| EPI_ISL_3093414 | Lighthouse Lab in Glasgow | Wellcome Sanger Institute for the COVID-19 Genomics UK (COG-UK) Consortium | Harper VanSteenhouse, Yumi Kasai, David Gray, Carol Clugston, Anna Dominiczak and Alex Alderton, Roberto Amato, Jeffrey Barrett, Sonia Goncalves, Ewan Harrison, David K. Jackson, Ian Johnston, Dominic Kwiatkowski, Cordelia Langford, John Sillitoe on behalf of the Wellcome Sanger Institute COVID-19 Surveillance Team |
| EPI_ISL_3093415 | Lighthouse Lab in Alderley Park | Wellcome Sanger Institute for the COVID-19 Genomics UK (COG-UK) Consortium | Jacquelyn Wynn, Mairead Hyland, The Lighthouse Lab in Alderley Park and Alex Alderton, Roberto Amato, Jeffrey Barrett, Sonia Goncalves, Ewan Harrison, David K. Jackson, Ian Johnston, Dominic Kwiatkowski, Cordelia Langford, John Sillitoe on behalf of the Wellcome Sanger Institute COVID-19 Surveillance Team |
| EPI_ISL_3093416 | Lighthouse Lab in Glasgow | Wellcome Sanger Institute for the COVID-19 Genomics UK (COG-UK) Consortium | Harper VanSteenhouse, Yumi Kasai, David Gray, Carol Clugston, Anna Dominiczak and Alex Alderton, Roberto Amato, Jeffrey Barrett, Sonia Goncalves, Ewan Harrison, David K. Jackson, Ian Johnston, Dominic Kwiatkowski, Cordelia Langford, John Sillitoe on behalf of the Wellcome Sanger Institute COVID-19 Surveillance Team |
| EPI_ISL_3093417 | Lighthouse Lab in Alderley Park | Wellcome Sanger Institute for the COVID-19 Genomics UK (COG-UK) Consortium | Jacquelyn Wynn, Mairead Hyland, The Lighthouse Lab in Alderley Park and Alex Alderton, Roberto Amato, Jeffrey Barrett, Sonia Goncalves, Ewan Harrison, David K. Jackson, Ian Johnston, Dominic Kwiatkowski, Cordelia Langford, John Sillitoe on behalf of the Wellcome Sanger Institute COVID-19 Surveillance Team |
| EPI_ISL_3093418 | Lighthouse Lab in Glasgow | Wellcome Sanger Institute for the COVID-19 Genomics UK (COG-UK) Consortium | Harper VanSteenhouse, Yumi Kasai, David Gray, Carol Clugston, Anna Dominiczak and Alex Alderton, Roberto Amato, Jeffrey Barrett, Sonia Goncalves, Ewan Harrison, David K. Jackson, Ian Johnston, Dominic Kwiatkowski, Cordelia Langford, John Sillitoe on behalf of the Wellcome Sanger Institute COVID-19 Surveillance Team |
| EPI_ISL_3093419, EPI_ISL_3093420, EPI_ISL_3093421, EPI_ISL_3093422 | Lighthouse Lab in Alderley Park | Wellcome Sanger Institute for the COVID-19 Genomics UK (COG-UK) Consortium | Jacquelyn Wynn, Mairead Hyland, The Lighthouse Lab in Alderley Park and Alex Alderton, Roberto Amato, Jeffrey Barrett, Sonia Goncalves, Ewan Harrison, David K. Jackson, Ian Johnston, Dominic Kwiatkowski, Cordelia Langford, John Sillitoe on behalf of the Wellcome Sanger Institute COVID-19 Surveillance Team |
| EPI_ISL_3093423, EPI_ISL_3093424 | Lighthouse Laboratory Plymouth | Wellcome Sanger Institute for the COVID-19 Genomics UK (COG-UK) Consortium | Lighthouse Laboratory Plymouth and Alex Alderton, Roberto Amato, Jeffrey Barrett, Sonia Goncalves, Ewan Harrison, David K. Jackson, Ian Johnston, Dominic Kwiatkowski, Cordelia Langford, John Sillitoe on behalf of the Wellcome Sanger Institute COVID-19 Surveillance Team |
| EPI_ISL_3093425, EPI_ISL_3093426 | Lighthouse Lab in Glasgow | Wellcome Sanger Institute for the COVID-19 Genomics UK (COG-UK) Consortium | Harper VanSteenhouse, Yumi Kasai, David Gray, Carol Clugston, Anna Dominiczak and Alex Alderton, Roberto Amato, Jeffrey Barrett, Sonia Goncalves, Ewan Harrison, David K. Jackson, Ian Johnston, Dominic Kwiatkowski, Cordelia Langford, John Sillitoe on behalf of the Wellcome Sanger Institute COVID-19 Surveillance Team |
| EPI_ISL_3093427 | Lighthouse Lab in Alderley Park | Wellcome Sanger Institute for the COVID-19 Genomics UK (COG-UK) Consortium | Jacquelyn Wynn, Mairead Hyland, The Lighthouse Lab in Alderley Park and Alex Alderton, Roberto Amato, Jeffrey Barrett, Sonia Goncalves, Ewan Harrison, David K. Jackson, Ian Johnston, Dominic Kwiatkowski, Cordelia Langford, John Sillitoe on behalf of the Wellcome Sanger Institute COVID-19 Surveillance Team |
| EPI_ISL_3093428 | Lighthouse Laboratory Plymouth | Wellcome Sanger Institute for the COVID-19 Genomics UK (COG-UK) Consortium | Lighthouse Laboratory Plymouth and Alex Alderton, Roberto Amato, Jeffrey Barrett, Sonia Goncalves, Ewan Harrison, David K. Jackson, Ian Johnston, Dominic Kwiatkowski, Cordelia Langford, John Sillitoe on behalf of the Wellcome Sanger Institute COVID-19 Surveillance Team |
| EPI_ISL_3093429 | Lighthouse Lab in Glasgow | Wellcome Sanger Institute for the COVID-19 Genomics UK (COG-UK) Consortium | Harper VanSteenhouse, Yumi Kasai, David Gray, Carol Clugston, Anna Dominiczak and Alex Alderton, Roberto Amato, Jeffrey Barrett, Sonia Goncalves, Ewan Harrison, David K. Jackson, Ian Johnston, Dominic Kwiatkowski, Cordelia Langford, John Sillitoe on behalf of the Wellcome Sanger Institute COVID-19 Surveillance Team |
| EPI_ISL_3093430 | Lighthouse Lab in Alderley Park | Wellcome Sanger Institute for the COVID-19 Genomics UK (COG-UK) Consortium | Jacquelyn Wynn, Mairead Hyland, The Lighthouse Lab in Alderley Park and Alex Alderton, Roberto Amato, Jeffrey Barrett, Sonia Goncalves, Ewan Harrison, David K. Jackson, Ian Johnston, Dominic Kwiatkowski, Cordelia Langford, John Sillitoe on behalf of the Wellcome Sanger Institute COVID-19 Surveillance Team |
| EPI_ISL_3093431 | Lighthouse Lab in Glasgow | Wellcome Sanger Institute for the COVID-19 Genomics UK (COG-UK) Consortium | Harper VanSteenhouse, Yumi Kasai, David Gray, Carol Clugston, Anna Dominiczak and Alex Alderton, Roberto Amato, Jeffrey Barrett, Sonia Goncalves, Ewan Harrison, David K. Jackson, Ian Johnston, Dominic Kwiatkowski, Cordelia Langford, John Sillitoe on behalf of the Wellcome Sanger Institute COVID-19 Surveillance Team |
| EPI_ISL_3093432 | Lighthouse Laboratory Plymouth | Wellcome Sanger Institute for the COVID-19 Genomics UK (COG-UK) Consortium | Lighthouse Laboratory Plymouth and Alex Alderton, Roberto Amato, Jeffrey Barrett, Sonia Goncalves, Ewan Harrison, David K. Jackson, Ian Johnston, Dominic Kwiatkowski, Cordelia Langford, John Sillitoe on behalf of the Wellcome Sanger Institute COVID-19 Surveillance Team |
| EPI_ISL_3093433 | Lighthouse Lab in Alderley Park | Wellcome Sanger Institute for the COVID-19 Genomics UK (COG-UK) Consortium | Jacquelyn Wynn, Mairead Hyland, The Lighthouse Lab in Alderley Park and Alex Alderton, Roberto Amato, Jeffrey Barrett, Sonia Goncalves, Ewan Harrison, David K. Jackson, Ian Johnston, Dominic Kwiatkowski, Cordelia Langford, John Sillitoe on behalf of the Wellcome Sanger Institute COVID-19 Surveillance Team |
| EPI_ISL_3093434 | Lighthouse Laboratory Plymouth | Wellcome Sanger Institute for the COVID-19 Genomics UK (COG-UK) Consortium | Lighthouse Laboratory Plymouth and Alex Alderton, Roberto Amato, Jeffrey Barrett, Sonia Goncalves, Ewan Harrison, David K. Jackson, Ian Johnston, Dominic Kwiatkowski, Cordelia Langford, John Sillitoe on behalf of the Wellcome Sanger Institute COVID-19 Surveillance Team |
| EPI_ISL_3093435 | Lighthouse Lab in Alderley Park | Wellcome Sanger Institute for the COVID-19 Genomics UK (COG-UK) Consortium | Jacquelyn Wynn, Mairead Hyland, The Lighthouse Lab in Alderley Park and Alex Alderton, Roberto Amato, Jeffrey Barrett, Sonia Goncalves, Ewan Harrison, David K. Jackson, Ian Johnston, Dominic Kwiatkowski, Cordelia Langford, John Sillitoe on behalf of the Wellcome Sanger Institute COVID-19 Surveillance Team |
| EPI_ISL_3093436 | Lighthouse Laboratory Plymouth | Wellcome Sanger Institute for the COVID-19 Genomics UK (COG-UK) Consortium | Lighthouse Laboratory Plymouth and Alex Alderton, Roberto Amato, Jeffrey Barrett, Sonia Goncalves, Ewan Harrison, David K. Jackson, Ian Johnston, Dominic Kwiatkowski, Cordelia Langford, John Sillitoe on behalf of the Wellcome Sanger Institute COVID-19 Surveillance Team |
| EPI_ISL_3093437, EPI_ISL_3093438, EPI_ISL_3093439, EPI_ISL_3093440 | Lighthouse Lab in Alderley Park | Wellcome Sanger Institute for the COVID-19 Genomics UK (COG-UK) Consortium | Jacquelyn Wynn, Mairead Hyland, The Lighthouse Lab in Alderley Park and Alex Alderton, Roberto Amato, Jeffrey Barrett, Sonia Goncalves, Ewan Harrison, David K. Jackson, Ian Johnston, Dominic Kwiatkowski, Cordelia Langford, John Sillitoe on behalf of the Wellcome Sanger Institute COVID-19 Surveillance Team |
| EPI_ISL_3093441 | Lighthouse Laboratory Plymouth | Wellcome Sanger Institute for the COVID-19 Genomics UK (COG-UK) Consortium | Lighthouse Laboratory Plymouth and Alex Alderton, Roberto Amato, Jeffrey Barrett, Sonia Goncalves, Ewan Harrison, David K. Jackson, Ian Johnston, Dominic Kwiatkowski, Cordelia Langford, John Sillitoe on behalf of the Wellcome Sanger Institute COVID-19 Surveillance Team |
| EPI_ISL_3093442 | Lighthouse Lab in Glasgow | Wellcome Sanger Institute for the COVID-19 Genomics UK (COG-UK) Consortium | Harper VanSteenhouse, Yumi Kasai, David Gray, Carol Clugston, Anna Dominiczak and Alex Alderton, Roberto Amato, Jeffrey Barrett, Sonia Goncalves, Ewan Harrison, David K. Jackson, Ian Johnston, Dominic Kwiatkowski, Cordelia Langford, John Sillitoe on behalf of the Wellcome Sanger Institute COVID-19 Surveillance Team |
| EPI_ISL_3093443 | Lighthouse Laboratory Plymouth | Wellcome Sanger Institute for the COVID-19 Genomics UK (COG-UK) Consortium | Lighthouse Laboratory Plymouth and Alex Alderton, Roberto Amato, Jeffrey Barrett, Sonia Goncalves, Ewan Harrison, David K. Jackson, Ian Johnston, Dominic Kwiatkowski, Cordelia Langford, John Sillitoe on behalf of the Wellcome Sanger Institute COVID-19 Surveillance Team |
| EPI_ISL_3093444, EPI_ISL_3093445, EPI_ISL_3093446, EPI_ISL_3093447 | Lighthouse Lab in Alderley Park | Wellcome Sanger Institute for the COVID-19 Genomics UK (COG-UK) Consortium | Jacquelyn Wynn, Mairead Hyland, The Lighthouse Lab in Alderley Park and Alex Alderton, Roberto Amato, Jeffrey Barrett, Sonia Goncalves, Ewan Harrison, David K. Jackson, Ian Johnston, Dominic Kwiatkowski, Cordelia Langford, John Sillitoe on behalf of the Wellcome Sanger Institute COVID-19 Surveillance Team |
| EPI_ISL_3093448 | Lighthouse Lab in Glasgow | Wellcome Sanger Institute for the COVID-19 Genomics UK (COG-UK) Consortium | Harper VanSteenhouse, Yumi Kasai, David Gray, Carol Clugston, Anna Dominiczak and Alex Alderton, Roberto Amato, Jeffrey Barrett, Sonia Goncalves, Ewan Harrison, David K. Jackson, Ian Johnston, Dominic Kwiatkowski, Cordelia Langford, John Sillitoe on behalf of the Wellcome Sanger Institute COVID-19 Surveillance Team |
| EPI_ISL_3093449, EPI_ISL_3093450, EPI_ISL_3093451, EPI_ISL_3093452 | Lighthouse Lab in Alderley Park | Wellcome Sanger Institute for the COVID-19 Genomics UK (COG-UK) Consortium | Jacquelyn Wynn, Mairead Hyland, The Lighthouse Lab in Alderley Park and Alex Alderton, Roberto Amato, Jeffrey Barrett, Sonia Goncalves, Ewan Harrison, David K. Jackson, Ian Johnston, Dominic Kwiatkowski, Cordelia Langford, John Sillitoe on behalf of the Wellcome Sanger Institute COVID-19 Surveillance Team |
| EPI_ISL_3093453 | Lighthouse Laboratory Plymouth | Wellcome Sanger Institute for the COVID-19 Genomics UK (COG-UK) Consortium | Lighthouse Laboratory Plymouth and Alex Alderton, Roberto Amato, Jeffrey Barrett, Sonia Goncalves, Ewan Harrison, David K. Jackson, Ian Johnston, Dominic Kwiatkowski, Cordelia Langford, John Sillitoe on behalf of the Wellcome Sanger Institute COVID-19 Surveillance Team |
| EPI_ISL_3093454 | Lighthouse Lab in Glasgow | Wellcome Sanger Institute for the COVID-19 Genomics UK (COG-UK) Consortium | Harper VanSteenhouse, Yumi Kasai, David Gray, Carol Clugston, Anna Dominiczak and Alex Alderton, Roberto Amato, Jeffrey Barrett, Sonia Goncalves, Ewan Harrison, David K. Jackson, Ian Johnston, Dominic Kwiatkowski, Cordelia Langford, John Sillitoe on behalf of the Wellcome Sanger Institute COVID-19 Surveillance Team |
| EPI_ISL_3093455 | Lighthouse Laboratory Plymouth | Wellcome Sanger Institute for the COVID-19 Genomics UK (COG-UK) Consortium | Lighthouse Laboratory Plymouth and Alex Alderton, Roberto Amato, Jeffrey Barrett, Sonia Goncalves, Ewan Harrison, David K. Jackson, Ian Johnston, Dominic Kwiatkowski, Cordelia Langford, John Sillitoe on behalf of the Wellcome Sanger Institute COVID-19 Surveillance Team |
| EPI_ISL_3093456, | Lighthouse Lab in Alderley Park | Wellcome Sanger Institute for the COVID-19 Genomics UK (COG-UK) | Jacquelyn Wynn, Mairead Hyland, The Lighthouse Lab in Alderley Park and Alex Alderton, Roberto Amato, Jeffrey Barrett, Sonia Goncalves, Ewan Harrison, David K. Jackson, Ian Johnston, Dominic Kwiatkowski, Cordelia Langford, John Sillitoe on |

| | | | |
|---|---|---|---|
| EPI_ISL_3093457 | | Consortium | behalf of the Wellcome Sanger Institute COVID-19 Surveillance Team |
| EPI_ISL_3093458, EPI_ISL_3093459 | Lighthouse Laboratory Plymouth | Wellcome Sanger Institute for the COVID-19 Genomics UK (COG-UK) Consortium | Lighthouse Laboratory Plymouth and Alex Alderton, Roberto Amato, Jeffrey Barrett, Sonia Goncalves, Ewan Harrison, David K. Jackson, Ian Johnston, Dominic Kwiatkowski, Cordelia Langford, John Sillitoe on behalf of the Wellcome Sanger Institute COVID-19 Surveillance Team |
| EPI_ISL_3093460, EPI_ISL_3093461, EPI_ISL_3093462 | Lighthouse Lab in Alderley Park | Wellcome Sanger Institute for the COVID-19 Genomics UK (COG-UK) Consortium | Jacquelyn Wynn, Mairead Hyland, The Lighthouse Lab in Alderley Park and Alex Alderton, Roberto Amato, Jeffrey Barrett, Sonia Goncalves, Ewan Harrison, David K. Jackson, Ian Johnston, Dominic Kwiatkowski, Cordelia Langford, John Sillitoe on behalf of the Wellcome Sanger Institute COVID-19 Surveillance Team |
| EPI_ISL_3093463 | Lighthouse Laboratory Plymouth | Wellcome Sanger Institute for the COVID-19 Genomics UK (COG-UK) Consortium | Lighthouse Laboratory Plymouth and Alex Alderton, Roberto Amato, Jeffrey Barrett, Sonia Goncalves, Ewan Harrison, David K. Jackson, Ian Johnston, Dominic Kwiatkowski, Cordelia Langford, John Sillitoe on behalf of the Wellcome Sanger Institute COVID-19 Surveillance Team |
| EPI_ISL_3093464 | Lighthouse Lab in Alderley Park | Wellcome Sanger Institute for the COVID-19 Genomics UK (COG-UK) Consortium | Jacquelyn Wynn, Mairead Hyland, The Lighthouse Lab in Alderley Park and Alex Alderton, Roberto Amato, Jeffrey Barrett, Sonia Goncalves, Ewan Harrison, David K. Jackson, Ian Johnston, Dominic Kwiatkowski, Cordelia Langford, John Sillitoe on behalf of the Wellcome Sanger Institute COVID-19 Surveillance Team |
| EPI_ISL_3093465 | Lighthouse Lab in Glasgow | Wellcome Sanger Institute for the COVID-19 Genomics UK (COG-UK) Consortium | Harper VanSteenhouse, Yumi Kasai, David Gray, Carol Clugston, Anna Dominiczak and Alex Alderton, Roberto Amato, Jeffrey Barrett, Sonia Goncalves, Ewan Harrison, David K. Jackson, Ian Johnston, Dominic Kwiatkowski, Cordelia Langford, John Sillitoe on behalf of the Wellcome Sanger Institute COVID-19 Surveillance Team |
| EPI_ISL_3093466, EPI_ISL_3093467 | Lighthouse Lab in Alderley Park | Wellcome Sanger Institute for the COVID-19 Genomics UK (COG-UK) Consortium | Jacquelyn Wynn, Mairead Hyland, The Lighthouse Lab in Alderley Park and Alex Alderton, Roberto Amato, Jeffrey Barrett, Sonia Goncalves, Ewan Harrison, David K. Jackson, Ian Johnston, Dominic Kwiatkowski, Cordelia Langford, John Sillitoe on behalf of the Wellcome Sanger Institute COVID-19 Surveillance Team |
| EPI_ISL_3093468 | Lighthouse Lab in Glasgow | Wellcome Sanger Institute for the COVID-19 Genomics UK (COG-UK) Consortium | Harper VanSteenhouse, Yumi Kasai, David Gray, Carol Clugston, Anna Dominiczak and Alex Alderton, Roberto Amato, Jeffrey Barrett, Sonia Goncalves, Ewan Harrison, David K. Jackson, Ian Johnston, Dominic Kwiatkowski, Cordelia Langford, John Sillitoe on behalf of the Wellcome Sanger Institute COVID-19 Surveillance Team |
| EPI_ISL_3093469, EPI_ISL_3093470, EPI_ISL_3093471 | Lighthouse Lab in Alderley Park | Wellcome Sanger Institute for the COVID-19 Genomics UK (COG-UK) Consortium | Jacquelyn Wynn, Mairead Hyland, The Lighthouse Lab in Alderley Park and Alex Alderton, Roberto Amato, Jeffrey Barrett, Sonia Goncalves, Ewan Harrison, David K. Jackson, Ian Johnston, Dominic Kwiatkowski, Cordelia Langford, John Sillitoe on behalf of the Wellcome Sanger Institute COVID-19 Surveillance Team |
| EPI_ISL_3093472 | Lighthouse Laboratory Plymouth | Wellcome Sanger Institute for the COVID-19 Genomics UK (COG-UK) Consortium | Lighthouse Laboratory Plymouth and Alex Alderton, Roberto Amato, Jeffrey Barrett, Sonia Goncalves, Ewan Harrison, David K. Jackson, Ian Johnston, Dominic Kwiatkowski, Cordelia Langford, John Sillitoe on behalf of the Wellcome Sanger Institute COVID-19 Surveillance Team |
| EPI_ISL_3093473, EPI_ISL_3093474 | Lighthouse Lab in Alderley Park | Wellcome Sanger Institute for the COVID-19 Genomics UK (COG-UK) Consortium | Jacquelyn Wynn, Mairead Hyland, The Lighthouse Lab in Alderley Park and Alex Alderton, Roberto Amato, Jeffrey Barrett, Sonia Goncalves, Ewan Harrison, David K. Jackson, Ian Johnston, Dominic Kwiatkowski, Cordelia Langford, John Sillitoe on behalf of the Wellcome Sanger Institute COVID-19 Surveillance Team |
| EPI_ISL_3093475 | Lighthouse Lab in Glasgow | Wellcome Sanger Institute for the COVID-19 Genomics UK (COG-UK) Consortium | Harper VanSteenhouse, Yumi Kasai, David Gray, Carol Clugston, Anna Dominiczak and Alex Alderton, Roberto Amato, Jeffrey Barrett, Sonia Goncalves, Ewan Harrison, David K. Jackson, Ian Johnston, Dominic Kwiatkowski, Cordelia Langford, John Sillitoe on behalf of the Wellcome Sanger Institute COVID-19 Surveillance Team |
| EPI_ISL_3093476 | Lighthouse Laboratory Plymouth | Wellcome Sanger Institute for the COVID-19 Genomics UK (COG-UK) Consortium | Lighthouse Laboratory Plymouth and Alex Alderton, Roberto Amato, Jeffrey Barrett, Sonia Goncalves, Ewan Harrison, David K. Jackson, Ian Johnston, Dominic Kwiatkowski, Cordelia Langford, John Sillitoe on behalf of the Wellcome Sanger Institute COVID-19 Surveillance Team |
| EPI_ISL_3093477 | Lighthouse Lab in Alderley Park | Wellcome Sanger Institute for the COVID-19 Genomics UK (COG-UK) Consortium | Jacquelyn Wynn, Mairead Hyland, The Lighthouse Lab in Alderley Park and Alex Alderton, Roberto Amato, Jeffrey Barrett, Sonia Goncalves, Ewan Harrison, David K. Jackson, Ian Johnston, Dominic Kwiatkowski, Cordelia Langford, John Sillitoe on behalf of the Wellcome Sanger Institute COVID-19 Surveillance Team |
| EPI_ISL_3093478, EPI_ISL_3093479 | Lighthouse Laboratory Plymouth | Wellcome Sanger Institute for the COVID-19 Genomics UK (COG-UK) Consortium | Lighthouse Laboratory Plymouth and Alex Alderton, Roberto Amato, Jeffrey Barrett, Sonia Goncalves, Ewan Harrison, David K. Jackson, Ian Johnston, Dominic Kwiatkowski, Cordelia Langford, John Sillitoe on behalf of the Wellcome Sanger Institute COVID-19 Surveillance Team |
| EPI_ISL_3093480 | Lighthouse Lab in Glasgow | Wellcome Sanger Institute for the COVID-19 Genomics UK (COG-UK) Consortium | Harper VanSteenhouse, Yumi Kasai, David Gray, Carol Clugston, Anna Dominiczak and Alex Alderton, Roberto Amato, Jeffrey Barrett, Sonia Goncalves, Ewan Harrison, David K. Jackson, Ian Johnston, Dominic Kwiatkowski, Cordelia Langford, John Sillitoe on behalf of the Wellcome Sanger Institute COVID-19 Surveillance Team |
| EPI_ISL_3093481, EPI_ISL_3093482, EPI_ISL_3093483 | Lighthouse Lab in Alderley Park | Wellcome Sanger Institute for the COVID-19 Genomics UK (COG-UK) Consortium | Jacquelyn Wynn, Mairead Hyland, The Lighthouse Lab in Alderley Park and Alex Alderton, Roberto Amato, Jeffrey Barrett, Sonia Goncalves, Ewan Harrison, David K. Jackson, Ian Johnston, Dominic Kwiatkowski, Cordelia Langford, John Sillitoe on behalf of the Wellcome Sanger Institute COVID-19 Surveillance Team |
| EPI_ISL_3093484 | Lighthouse Lab in Glasgow | Wellcome Sanger Institute for the COVID-19 Genomics UK (COG-UK) Consortium | Harper VanSteenhouse, Yumi Kasai, David Gray, Carol Clugston, Anna Dominiczak and Alex Alderton, Roberto Amato, Jeffrey Barrett, Sonia Goncalves, Ewan Harrison, David K. Jackson, Ian Johnston, Dominic Kwiatkowski, Cordelia Langford, John Sillitoe on behalf of the Wellcome Sanger Institute COVID-19 Surveillance Team |
| EPI_ISL_3093485, EPI_ISL_3093486 | Lighthouse Laboratory Plymouth | Wellcome Sanger Institute for the COVID-19 Genomics UK (COG-UK) Consortium | Lighthouse Laboratory Plymouth and Alex Alderton, Roberto Amato, Jeffrey Barrett, Sonia Goncalves, Ewan Harrison, David K. Jackson, Ian Johnston, Dominic Kwiatkowski, Cordelia Langford, John Sillitoe on behalf of the Wellcome Sanger Institute COVID-19 Surveillance Team |
| EPI_ISL_3093487, EPI_ISL_3093488 | Lighthouse Lab in Alderley Park | Wellcome Sanger Institute for the COVID-19 Genomics UK (COG-UK) Consortium | Jacquelyn Wynn, Mairead Hyland, The Lighthouse Lab in Alderley Park and Alex Alderton, Roberto Amato, Jeffrey Barrett, Sonia Goncalves, Ewan Harrison, David K. Jackson, Ian Johnston, Dominic Kwiatkowski, Cordelia Langford, John Sillitoe on behalf of the Wellcome Sanger Institute COVID-19 Surveillance Team |
| EPI_ISL_3093489 | Lighthouse Laboratory Plymouth | Wellcome Sanger Institute for the COVID-19 Genomics UK (COG-UK) Consortium | Lighthouse Laboratory Plymouth and Alex Alderton, Roberto Amato, Jeffrey Barrett, Sonia Goncalves, Ewan Harrison, David K. Jackson, Ian Johnston, Dominic Kwiatkowski, Cordelia Langford, John Sillitoe on behalf of the Wellcome Sanger Institute COVID-19 Surveillance Team |
| EPI_ISL_3093490 | Lighthouse Lab in Alderley Park | Wellcome Sanger Institute for the COVID-19 Genomics UK (COG-UK) Consortium | Jacquelyn Wynn, Mairead Hyland, The Lighthouse Lab in Alderley Park and Alex Alderton, Roberto Amato, Jeffrey Barrett, Sonia Goncalves, Ewan Harrison, David K. Jackson, Ian Johnston, Dominic Kwiatkowski, Cordelia Langford, John Sillitoe on behalf of the Wellcome Sanger Institute COVID-19 Surveillance Team |
| EPI_ISL_3093491 | Lighthouse Lab in Glasgow | Wellcome Sanger Institute for the COVID-19 Genomics UK (COG-UK) Consortium | Harper VanSteenhouse, Yumi Kasai, David Gray, Carol Clugston, Anna Dominiczak and Alex Alderton, Roberto Amato, Jeffrey Barrett, Sonia Goncalves, Ewan Harrison, David K. Jackson, Ian Johnston, Dominic Kwiatkowski, Cordelia Langford, John Sillitoe on behalf of the Wellcome Sanger Institute COVID-19 Surveillance Team |
| EPI_ISL_3093492 | Lighthouse Laboratory Plymouth | Wellcome Sanger Institute for the COVID-19 Genomics UK (COG-UK) Consortium | Lighthouse Laboratory Plymouth and Alex Alderton, Roberto Amato, Jeffrey Barrett, Sonia Goncalves, Ewan Harrison, David K. Jackson, Ian Johnston, Dominic Kwiatkowski, Cordelia Langford, John Sillitoe on behalf of the Wellcome Sanger Institute COVID-19 Surveillance Team |
| EPI_ISL_3093493 | Lighthouse Lab in Glasgow | Wellcome Sanger Institute for the COVID-19 Genomics UK (COG-UK) Consortium | Harper VanSteenhouse, Yumi Kasai, David Gray, Carol Clugston, Anna Dominiczak and Alex Alderton, Roberto Amato, Jeffrey Barrett, Sonia Goncalves, Ewan Harrison, David K. Jackson, Ian Johnston, Dominic Kwiatkowski, Cordelia Langford, John Sillitoe on behalf of the Wellcome Sanger Institute COVID-19 Surveillance Team |
| EPI_ISL_3093494 | Lighthouse Laboratory Plymouth | Wellcome Sanger Institute for the COVID-19 Genomics UK (COG-UK) Consortium | Lighthouse Laboratory Plymouth and Alex Alderton, Roberto Amato, Jeffrey Barrett, Sonia Goncalves, Ewan Harrison, David K. Jackson, Ian Johnston, Dominic Kwiatkowski, Cordelia Langford, John Sillitoe on behalf of the Wellcome Sanger Institute COVID-19 Surveillance Team |
| EPI_ISL_3093495 | Lighthouse Lab in Alderley Park | Wellcome Sanger Institute for the COVID-19 Genomics UK (COG-UK) Consortium | Jacquelyn Wynn, Mairead Hyland, The Lighthouse Lab in Alderley Park and Alex Alderton, Roberto Amato, Jeffrey Barrett, Sonia Goncalves, Ewan Harrison, David K. Jackson, Ian Johnston, Dominic Kwiatkowski, Cordelia Langford, John Sillitoe on behalf of the Wellcome Sanger Institute COVID-19 Surveillance Team |
| EPI_ISL_3093496, EPI_ISL_3093497, EPI_ISL_3093498, EPI_ISL_3093499 | Lighthouse Laboratory Plymouth | Wellcome Sanger Institute for the COVID-19 Genomics UK (COG-UK) Consortium | Lighthouse Laboratory Plymouth and Alex Alderton, Roberto Amato, Jeffrey Barrett, Sonia Goncalves, Ewan Harrison, David K. Jackson, Ian Johnston, Dominic Kwiatkowski, Cordelia Langford, John Sillitoe on behalf of the Wellcome Sanger Institute COVID-19 Surveillance Team |
| EPI_ISL_3093500, EPI_ISL_3093501 | Lighthouse Lab in Alderley Park | Wellcome Sanger Institute for the COVID-19 Genomics UK (COG-UK) Consortium | Jacquelyn Wynn, Mairead Hyland, The Lighthouse Lab in Alderley Park and Alex Alderton, Roberto Amato, Jeffrey Barrett, Sonia Goncalves, Ewan Harrison, David K. Jackson, Ian Johnston, Dominic Kwiatkowski, Cordelia Langford, John Sillitoe on behalf of the Wellcome Sanger Institute COVID-19 Surveillance Team |
| EPI_ISL_3093502 | Lighthouse Lab in Glasgow | Wellcome Sanger Institute for the COVID-19 Genomics UK (COG-UK) Consortium | Harper VanSteenhouse, Yumi Kasai, David Gray, Carol Clugston, Anna Dominiczak and Alex Alderton, Roberto Amato, Jeffrey Barrett, Sonia Goncalves, Ewan Harrison, David K. Jackson, Ian Johnston, Dominic Kwiatkowski, Cordelia Langford, John Sillitoe on behalf of the Wellcome Sanger Institute COVID-19 Surveillance Team |
| EPI_ISL_3093503 | Lighthouse Lab in Alderley Park | Wellcome Sanger Institute for the COVID-19 Genomics UK (COG-UK) Consortium | Jacquelyn Wynn, Mairead Hyland, The Lighthouse Lab in Alderley Park and Alex Alderton, Roberto Amato, Jeffrey Barrett, Sonia Goncalves, Ewan Harrison, David K. Jackson, Ian Johnston, Dominic Kwiatkowski, Cordelia Langford, John Sillitoe on behalf of the Wellcome Sanger Institute COVID-19 Surveillance Team |
| EPI_ISL_3093504, EPI_ISL_3093505, EPI_ISL_3093506, EPI_ISL_3093507 | Lighthouse Laboratory Plymouth | Wellcome Sanger Institute for the COVID-19 Genomics UK (COG-UK) Consortium | Lighthouse Laboratory Plymouth and Alex Alderton, Roberto Amato, Jeffrey Barrett, Sonia Goncalves, Ewan Harrison, David K. Jackson, Ian Johnston, Dominic Kwiatkowski, Cordelia Langford, John Sillitoe on behalf of the Wellcome Sanger Institute COVID-19 Surveillance Team |
| EPI_ISL_3093508 | Lighthouse Lab in Alderley Park | Wellcome Sanger Institute for the COVID-19 Genomics UK (COG-UK) Consortium | Jacquelyn Wynn, Mairead Hyland, The Lighthouse Lab in Alderley Park and Alex Alderton, Roberto Amato, Jeffrey Barrett, Sonia Goncalves, Ewan Harrison, David K. Jackson, Ian Johnston, Dominic Kwiatkowski, Cordelia Langford, John Sillitoe on behalf of the Wellcome Sanger Institute COVID-19 Surveillance Team |
| EPI_ISL_3093509, EPI_ISL_3093510 | Lighthouse Laboratory Plymouth | Wellcome Sanger Institute for the COVID-19 Genomics UK (COG-UK) Consortium | Lighthouse Laboratory Plymouth and Alex Alderton, Roberto Amato, Jeffrey Barrett, Sonia Goncalves, Ewan Harrison, David K. Jackson, Ian Johnston, Dominic Kwiatkowski, Cordelia Langford, John Sillitoe on behalf of the Wellcome Sanger Institute COVID-19 Surveillance Team |
| EPI_ISL_3093511 | Lighthouse Lab in Glasgow | Wellcome Sanger Institute for the COVID-19 Genomics UK (COG-UK) Consortium | Harper VanSteenhouse, Yumi Kasai, David Gray, Carol Clugston, Anna Dominiczak and Alex Alderton, Roberto Amato, Jeffrey Barrett, Sonia Goncalves, Ewan Harrison, David K. Jackson, Ian Johnston, Dominic Kwiatkowski, Cordelia Langford, John Sillitoe on behalf of the Wellcome Sanger Institute COVID-19 Surveillance Team |
| EPI_ISL_3093512, EPI_ISL_3093513 | Lighthouse Lab in Alderley Park | Wellcome Sanger Institute for the COVID-19 Genomics UK (COG-UK) Consortium | Jacquelyn Wynn, Mairead Hyland, The Lighthouse Lab in Alderley Park and Alex Alderton, Roberto Amato, Jeffrey Barrett, Sonia Goncalves, Ewan Harrison, David K. Jackson, Ian Johnston, Dominic Kwiatkowski, Cordelia Langford, John Sillitoe on behalf of the Wellcome Sanger Institute COVID-19 Surveillance Team |
| EPI_ISL_3093514 | Lighthouse Laboratory Plymouth | Wellcome Sanger Institute for the COVID-19 Genomics UK (COG-UK) Consortium | Lighthouse Laboratory Plymouth and Alex Alderton, Roberto Amato, Jeffrey Barrett, Sonia Goncalves, Ewan Harrison, David K. Jackson, Ian Johnston, Dominic Kwiatkowski, Cordelia Langford, John Sillitoe on behalf of the Wellcome Sanger Institute COVID-19 Surveillance Team |
| EPI_ISL_3093515 | Lighthouse Lab in Glasgow | Wellcome Sanger Institute for the COVID-19 Genomics UK (COG-UK) Consortium | Harper VanSteenhouse, Yumi Kasai, David Gray, Carol Clugston, Anna Dominiczak and Alex Alderton, Roberto Amato, Jeffrey Barrett, Sonia Goncalves, Ewan Harrison, David K. Jackson, Ian Johnston, Dominic Kwiatkowski, Cordelia Langford, John Sillitoe on behalf of the Wellcome Sanger Institute COVID-19 Surveillance Team |
| EPI_ISL_3093516, EPI_ISL_3093517, EPI_ISL_3093518 | Lighthouse Lab in Alderley Park | Wellcome Sanger Institute for the COVID-19 Genomics UK (COG-UK) Consortium | Jacquelyn Wynn, Mairead Hyland, The Lighthouse Lab in Alderley Park and Alex Alderton, Roberto Amato, Jeffrey Barrett, Sonia Goncalves, Ewan Harrison, David K. Jackson, Ian Johnston, Dominic Kwiatkowski, Cordelia Langford, John Sillitoe on behalf of the Wellcome Sanger Institute COVID-19 Surveillance Team |
| EPI_ISL_3093519, EPI_ISL_3093520, EPI_ISL_3093521, EPI_ISL_3093522 | Berkshire and Surrey Pathology Services Lighthouse Laboratory | Wellcome Sanger Institute for the COVID-19 Genomics UK (COG-UK) Consortium | Berkshire and Surrey Pathology Services Lighthouse Laboratory and Alex Alderton, Roberto Amato, Jeffrey Barrett, Sonia Goncalves, Ewan Harrison, David K. Jackson, Ian Johnston, Dominic Kwiatkowski, Cordelia Langford, John Sillitoe on behalf of the Wellcome Sanger Institute COVID-19 Surveillance Team |
| EPI_ISL_3093523 | Lighthouse Laboratory Plymouth | Wellcome Sanger Institute for the COVID-19 Genomics UK (COG-UK) Consortium | Lighthouse Laboratory Plymouth and Alex Alderton, Roberto Amato, Jeffrey Barrett, Sonia Goncalves, Ewan Harrison, David K. Jackson, Ian Johnston, Dominic Kwiatkowski, Cordelia Langford, John Sillitoe on behalf of the Wellcome Sanger Institute COVID-19 Surveillance Team |
| EPI_ISL_3093524, | Berkshire and Surrey Pathology Services Lighthouse Laboratory | Wellcome Sanger Institute for the COVID-19 Genomics UK (COG-UK) | Berkshire and Surrey Pathology Services Lighthouse Laboratory and Alex Alderton, Roberto Amato, Jeffrey Barrett, Sonia Goncalves, Ewan Harrison, David K. Jackson, Ian Johnston, Dominic Kwiatkowski, Cordelia Langford, John Sillitoe on behalf |

| | | | |
|---|---|---|---|
| EPI_ISL_3093525, EPI_ISL_3093526 | | Consortium | of the Wellcome Sanger Institute COVID-19 Surveillance Team |
| EPI_ISL_3093527 | Lighthouse Laboratory Plymouth | Wellcome Sanger Institute for the COVID-19 Genomics UK (COG-UK) Consortium | Lighthouse Laboratory Plymouth and Alex Alderton, Roberto Amato, Jeffrey Barrett, Sonia Goncalves, Ewan Harrison, David K. Jackson, Ian Johnston, Dominic Kwiatkowski, Cordelia Langford, John Sillitoe on behalf of the Wellcome Sanger Institute COVID-19 Surveillance Team |
| EPI_ISL_3093528 | Berkshire and Surrey Pathology Services Lighthouse Laboratory | Wellcome Sanger Institute for the COVID-19 Genomics UK (COG-UK) Consortium | Berkshire and Surrey Pathology Services Lighthouse Laboratory and Alex Alderton, Roberto Amato, Jeffrey Barrett, Sonia Goncalves, Ewan Harrison, David K. Jackson, Ian Johnston, Dominic Kwiatkowski, Cordelia Langford, John Sillitoe on behalf of the Wellcome Sanger Institute COVID-19 Surveillance Team |
| EPI_ISL_3093529 | Lighthouse Laboratory Plymouth | Wellcome Sanger Institute for the COVID-19 Genomics UK (COG-UK) Consortium | Lighthouse Laboratory Plymouth and Alex Alderton, Roberto Amato, Jeffrey Barrett, Sonia Goncalves, Ewan Harrison, David K. Jackson, Ian Johnston, Dominic Kwiatkowski, Cordelia Langford, John Sillitoe on behalf of the Wellcome Sanger Institute COVID-19 Surveillance Team |
| EPI_ISL_3093530 | Berkshire and Surrey Pathology Services Lighthouse Laboratory | Wellcome Sanger Institute for the COVID-19 Genomics UK (COG-UK) Consortium | Berkshire and Surrey Pathology Services Lighthouse Laboratory and Alex Alderton, Roberto Amato, Jeffrey Barrett, Sonia Goncalves, Ewan Harrison, David K. Jackson, Ian Johnston, Dominic Kwiatkowski, Cordelia Langford, John Sillitoe on behalf of the Wellcome Sanger Institute COVID-19 Surveillance Team |
| EPI_ISL_3093531 | Lighthouse Laboratory Plymouth | Wellcome Sanger Institute for the COVID-19 Genomics UK (COG-UK) Consortium | Lighthouse Laboratory Plymouth and Alex Alderton, Roberto Amato, Jeffrey Barrett, Sonia Goncalves, Ewan Harrison, David K. Jackson, Ian Johnston, Dominic Kwiatkowski, Cordelia Langford, John Sillitoe on behalf of the Wellcome Sanger Institute COVID-19 Surveillance Team |
| EPI_ISL_3093532, EPI_ISL_3093533 | Berkshire and Surrey Pathology Services Lighthouse Laboratory | Wellcome Sanger Institute for the COVID-19 Genomics UK (COG-UK) Consortium | Berkshire and Surrey Pathology Services Lighthouse Laboratory and Alex Alderton, Roberto Amato, Jeffrey Barrett, Sonia Goncalves, Ewan Harrison, David K. Jackson, Ian Johnston, Dominic Kwiatkowski, Cordelia Langford, John Sillitoe on behalf of the Wellcome Sanger Institute COVID-19 Surveillance Team |
| EPI_ISL_3093534, EPI_ISL_3093535 | Lighthouse Laboratory Plymouth | Wellcome Sanger Institute for the COVID-19 Genomics UK (COG-UK) Consortium | Lighthouse Laboratory Plymouth and Alex Alderton, Roberto Amato, Jeffrey Barrett, Sonia Goncalves, Ewan Harrison, David K. Jackson, Ian Johnston, Dominic Kwiatkowski, Cordelia Langford, John Sillitoe on behalf of the Wellcome Sanger Institute COVID-19 Surveillance Team |
| EPI_ISL_3093536 | Berkshire and Surrey Pathology Services Lighthouse Laboratory | Wellcome Sanger Institute for the COVID-19 Genomics UK (COG-UK) Consortium | Berkshire and Surrey Pathology Services Lighthouse Laboratory and Alex Alderton, Roberto Amato, Jeffrey Barrett, Sonia Goncalves, Ewan Harrison, David K. Jackson, Ian Johnston, Dominic Kwiatkowski, Cordelia Langford, John Sillitoe on behalf of the Wellcome Sanger Institute COVID-19 Surveillance Team |
| EPI_ISL_3093537, EPI_ISL_3093538 | Lighthouse Laboratory Plymouth | Wellcome Sanger Institute for the COVID-19 Genomics UK (COG-UK) Consortium | Lighthouse Laboratory Plymouth and Alex Alderton, Roberto Amato, Jeffrey Barrett, Sonia Goncalves, Ewan Harrison, David K. Jackson, Ian Johnston, Dominic Kwiatkowski, Cordelia Langford, John Sillitoe on behalf of the Wellcome Sanger Institute COVID-19 Surveillance Team |
| EPI_ISL_3093539, EPI_ISL_3093540, EPI_ISL_3093541 | Berkshire and Surrey Pathology Services Lighthouse Laboratory | Wellcome Sanger Institute for the COVID-19 Genomics UK (COG-UK) Consortium | Berkshire and Surrey Pathology Services Lighthouse Laboratory and Alex Alderton, Roberto Amato, Jeffrey Barrett, Sonia Goncalves, Ewan Harrison, David K. Jackson, Ian Johnston, Dominic Kwiatkowski, Cordelia Langford, John Sillitoe on behalf of the Wellcome Sanger Institute COVID-19 Surveillance Team |
| EPI_ISL_3093542, EPI_ISL_3093543 | Lighthouse Laboratory Plymouth | Wellcome Sanger Institute for the COVID-19 Genomics UK (COG-UK) Consortium | Lighthouse Laboratory Plymouth and Alex Alderton, Roberto Amato, Jeffrey Barrett, Sonia Goncalves, Ewan Harrison, David K. Jackson, Ian Johnston, Dominic Kwiatkowski, Cordelia Langford, John Sillitoe on behalf of the Wellcome Sanger Institute COVID-19 Surveillance Team |
| EPI_ISL_3093544 | Berkshire and Surrey Pathology Services Lighthouse Laboratory | Wellcome Sanger Institute for the COVID-19 Genomics UK (COG-UK) Consortium | Berkshire and Surrey Pathology Services Lighthouse Laboratory and Alex Alderton, Roberto Amato, Jeffrey Barrett, Sonia Goncalves, Ewan Harrison, David K. Jackson, Ian Johnston, Dominic Kwiatkowski, Cordelia Langford, John Sillitoe on behalf of the Wellcome Sanger Institute COVID-19 Surveillance Team |
| EPI_ISL_3093545, EPI_ISL_3093546 | Lighthouse Laboratory Plymouth | Wellcome Sanger Institute for the COVID-19 Genomics UK (COG-UK) Consortium | Lighthouse Laboratory Plymouth and Alex Alderton, Roberto Amato, Jeffrey Barrett, Sonia Goncalves, Ewan Harrison, David K. Jackson, Ian Johnston, Dominic Kwiatkowski, Cordelia Langford, John Sillitoe on behalf of the Wellcome Sanger Institute COVID-19 Surveillance Team |
| EPI_ISL_3093547 | Berkshire and Surrey Pathology Services Lighthouse Laboratory | Wellcome Sanger Institute for the COVID-19 Genomics UK (COG-UK) Consortium | Berkshire and Surrey Pathology Services Lighthouse Laboratory and Alex Alderton, Roberto Amato, Jeffrey Barrett, Sonia Goncalves, Ewan Harrison, David K. Jackson, Ian Johnston, Dominic Kwiatkowski, Cordelia Langford, John Sillitoe on behalf of the Wellcome Sanger Institute COVID-19 Surveillance Team |
| EPI_ISL_3093548, EPI_ISL_3093549, EPI_ISL_3093550, EPI_ISL_3093551 | Lighthouse Laboratory Plymouth | Wellcome Sanger Institute for the COVID-19 Genomics UK (COG-UK) Consortium | Lighthouse Laboratory Plymouth and Alex Alderton, Roberto Amato, Jeffrey Barrett, Sonia Goncalves, Ewan Harrison, David K. Jackson, Ian Johnston, Dominic Kwiatkowski, Cordelia Langford, John Sillitoe on behalf of the Wellcome Sanger Institute COVID-19 Surveillance Team |
| EPI_ISL_3093552, EPI_ISL_3093553 | Berkshire and Surrey Pathology Services Lighthouse Laboratory | Wellcome Sanger Institute for the COVID-19 Genomics UK (COG-UK) Consortium | Berkshire and Surrey Pathology Services Lighthouse Laboratory and Alex Alderton, Roberto Amato, Jeffrey Barrett, Sonia Goncalves, Ewan Harrison, David K. Jackson, Ian Johnston, Dominic Kwiatkowski, Cordelia Langford, John Sillitoe on behalf of the Wellcome Sanger Institute COVID-19 Surveillance Team |
| EPI_ISL_3093554 | Lighthouse Laboratory Plymouth | Wellcome Sanger Institute for the COVID-19 Genomics UK (COG-UK) Consortium | Lighthouse Laboratory Plymouth and Alex Alderton, Roberto Amato, Jeffrey Barrett, Sonia Goncalves, Ewan Harrison, David K. Jackson, Ian Johnston, Dominic Kwiatkowski, Cordelia Langford, John Sillitoe on behalf of the Wellcome Sanger Institute COVID-19 Surveillance Team |
| EPI_ISL_3093555 | Berkshire and Surrey Pathology Services Lighthouse Laboratory | Wellcome Sanger Institute for the COVID-19 Genomics UK (COG-UK) Consortium | Berkshire and Surrey Pathology Services Lighthouse Laboratory and Alex Alderton, Roberto Amato, Jeffrey Barrett, Sonia Goncalves, Ewan Harrison, David K. Jackson, Ian Johnston, Dominic Kwiatkowski, Cordelia Langford, John Sillitoe on behalf of the Wellcome Sanger Institute COVID-19 Surveillance Team |
| EPI_ISL_3093556, EPI_ISL_3093557 | Lighthouse Laboratory Plymouth | Wellcome Sanger Institute for the COVID-19 Genomics UK (COG-UK) Consortium | Lighthouse Laboratory Plymouth and Alex Alderton, Roberto Amato, Jeffrey Barrett, Sonia Goncalves, Ewan Harrison, David K. Jackson, Ian Johnston, Dominic Kwiatkowski, Cordelia Langford, John Sillitoe on behalf of the Wellcome Sanger Institute COVID-19 Surveillance Team |
| EPI_ISL_3093558 | Berkshire and Surrey Pathology Services Lighthouse Laboratory | Wellcome Sanger Institute for the COVID-19 Genomics UK (COG-UK) Consortium | Berkshire and Surrey Pathology Services Lighthouse Laboratory and Alex Alderton, Roberto Amato, Jeffrey Barrett, Sonia Goncalves, Ewan Harrison, David K. Jackson, Ian Johnston, Dominic Kwiatkowski, Cordelia Langford, John Sillitoe on behalf of the Wellcome Sanger Institute COVID-19 Surveillance Team |
| EPI_ISL_3093559 | Lighthouse Laboratory Plymouth | Wellcome Sanger Institute for the COVID-19 Genomics UK (COG-UK) Consortium | Lighthouse Laboratory Plymouth and Alex Alderton, Roberto Amato, Jeffrey Barrett, Sonia Goncalves, Ewan Harrison, David K. Jackson, Ian Johnston, Dominic Kwiatkowski, Cordelia Langford, John Sillitoe on behalf of the Wellcome Sanger Institute COVID-19 Surveillance Team |
| EPI_ISL_3093560 | Berkshire and Surrey Pathology Services Lighthouse Laboratory | Wellcome Sanger Institute for the COVID-19 Genomics UK (COG-UK) Consortium | Berkshire and Surrey Pathology Services Lighthouse Laboratory and Alex Alderton, Roberto Amato, Jeffrey Barrett, Sonia Goncalves, Ewan Harrison, David K. Jackson, Ian Johnston, Dominic Kwiatkowski, Cordelia Langford, John Sillitoe on behalf of the Wellcome Sanger Institute COVID-19 Surveillance Team |
| EPI_ISL_3093561 | Lighthouse Laboratory Plymouth | Wellcome Sanger Institute for the COVID-19 Genomics UK (COG-UK) Consortium | Lighthouse Laboratory Plymouth and Alex Alderton, Roberto Amato, Jeffrey Barrett, Sonia Goncalves, Ewan Harrison, David K. Jackson, Ian Johnston, Dominic Kwiatkowski, Cordelia Langford, John Sillitoe on behalf of the Wellcome Sanger Institute COVID-19 Surveillance Team |
| EPI_ISL_3093562 | Berkshire and Surrey Pathology Services Lighthouse Laboratory | Wellcome Sanger Institute for the COVID-19 Genomics UK (COG-UK) Consortium | Berkshire and Surrey Pathology Services Lighthouse Laboratory and Alex Alderton, Roberto Amato, Jeffrey Barrett, Sonia Goncalves, Ewan Harrison, David K. Jackson, Ian Johnston, Dominic Kwiatkowski, Cordelia Langford, John Sillitoe on behalf of the Wellcome Sanger Institute COVID-19 Surveillance Team |
| EPI_ISL_3093563, EPI_ISL_3093564, EPI_ISL_3093565 | Lighthouse Laboratory Plymouth | Wellcome Sanger Institute for the COVID-19 Genomics UK (COG-UK) Consortium | Lighthouse Laboratory Plymouth and Alex Alderton, Roberto Amato, Jeffrey Barrett, Sonia Goncalves, Ewan Harrison, David K. Jackson, Ian Johnston, Dominic Kwiatkowski, Cordelia Langford, John Sillitoe on behalf of the Wellcome Sanger Institute COVID-19 Surveillance Team |
| EPI_ISL_3093566 | Berkshire and Surrey Pathology Services Lighthouse Laboratory | Wellcome Sanger Institute for the COVID-19 Genomics UK (COG-UK) Consortium | Berkshire and Surrey Pathology Services Lighthouse Laboratory and Alex Alderton, Roberto Amato, Jeffrey Barrett, Sonia Goncalves, Ewan Harrison, David K. Jackson, Ian Johnston, Dominic Kwiatkowski, Cordelia Langford, John Sillitoe on behalf of the Wellcome Sanger Institute COVID-19 Surveillance Team |
| EPI_ISL_3093567, EPI_ISL_3093568 | Lighthouse Laboratory Plymouth | Wellcome Sanger Institute for the COVID-19 Genomics UK (COG-UK) Consortium | Lighthouse Laboratory Plymouth and Alex Alderton, Roberto Amato, Jeffrey Barrett, Sonia Goncalves, Ewan Harrison, David K. Jackson, Ian Johnston, Dominic Kwiatkowski, Cordelia Langford, John Sillitoe on behalf of the Wellcome Sanger Institute COVID-19 Surveillance Team |
| EPI_ISL_3093569 | Berkshire and Surrey Pathology Services Lighthouse Laboratory | Wellcome Sanger Institute for the COVID-19 Genomics UK (COG-UK) Consortium | Berkshire and Surrey Pathology Services Lighthouse Laboratory and Alex Alderton, Roberto Amato, Jeffrey Barrett, Sonia Goncalves, Ewan Harrison, David K. Jackson, Ian Johnston, Dominic Kwiatkowski, Cordelia Langford, John Sillitoe on behalf of the Wellcome Sanger Institute COVID-19 Surveillance Team |
| EPI_ISL_3093570, EPI_ISL_3093571 | Lighthouse Laboratory Plymouth | Wellcome Sanger Institute for the COVID-19 Genomics UK (COG-UK) Consortium | Lighthouse Laboratory Plymouth and Alex Alderton, Roberto Amato, Jeffrey Barrett, Sonia Goncalves, Ewan Harrison, David K. Jackson, Ian Johnston, Dominic Kwiatkowski, Cordelia Langford, John Sillitoe on behalf of the Wellcome Sanger Institute COVID-19 Surveillance Team |
| EPI_ISL_3093572 | Berkshire and Surrey Pathology Services Lighthouse Laboratory | Wellcome Sanger Institute for the COVID-19 Genomics UK (COG-UK) Consortium | Berkshire and Surrey Pathology Services Lighthouse Laboratory and Alex Alderton, Roberto Amato, Jeffrey Barrett, Sonia Goncalves, Ewan Harrison, David K. Jackson, Ian Johnston, Dominic Kwiatkowski, Cordelia Langford, John Sillitoe on behalf of the Wellcome Sanger Institute COVID-19 Surveillance Team |
| EPI_ISL_3093573 | Lighthouse Laboratory Plymouth | Wellcome Sanger Institute for the COVID-19 Genomics UK (COG-UK) Consortium | Lighthouse Laboratory Plymouth and Alex Alderton, Roberto Amato, Jeffrey Barrett, Sonia Goncalves, Ewan Harrison, David K. Jackson, Ian Johnston, Dominic Kwiatkowski, Cordelia Langford, John Sillitoe on behalf of the Wellcome Sanger Institute COVID-19 Surveillance Team |
| EPI_ISL_3093574 | Berkshire and Surrey Pathology Services Lighthouse Laboratory | Wellcome Sanger Institute for the COVID-19 Genomics UK (COG-UK) Consortium | Berkshire and Surrey Pathology Services Lighthouse Laboratory and Alex Alderton, Roberto Amato, Jeffrey Barrett, Sonia Goncalves, Ewan Harrison, David K. Jackson, Ian Johnston, Dominic Kwiatkowski, Cordelia Langford, John Sillitoe on behalf of the Wellcome Sanger Institute COVID-19 Surveillance Team |
| EPI_ISL_3093575, EPI_ISL_3093576, EPI_ISL_3093577 | Lighthouse Laboratory Plymouth | Wellcome Sanger Institute for the COVID-19 Genomics UK (COG-UK) Consortium | Lighthouse Laboratory Plymouth and Alex Alderton, Roberto Amato, Jeffrey Barrett, Sonia Goncalves, Ewan Harrison, David K. Jackson, Ian Johnston, Dominic Kwiatkowski, Cordelia Langford, John Sillitoe on behalf of the Wellcome Sanger Institute COVID-19 Surveillance Team |
| EPI_ISL_3093578, EPI_ISL_3093579, EPI_ISL_3093580, EPI_ISL_3093581 | Berkshire and Surrey Pathology Services Lighthouse Laboratory | Wellcome Sanger Institute for the COVID-19 Genomics UK (COG-UK) Consortium | Berkshire and Surrey Pathology Services Lighthouse Laboratory and Alex Alderton, Roberto Amato, Jeffrey Barrett, Sonia Goncalves, Ewan Harrison, David K. Jackson, Ian Johnston, Dominic Kwiatkowski, Cordelia Langford, John Sillitoe on behalf of the Wellcome Sanger Institute COVID-19 Surveillance Team |
| EPI_ISL_3093582 | Lighthouse Laboratory Plymouth | Wellcome Sanger Institute for the COVID-19 Genomics UK (COG-UK) Consortium | Lighthouse Laboratory Plymouth and Alex Alderton, Roberto Amato, Jeffrey Barrett, Sonia Goncalves, Ewan Harrison, David K. Jackson, Ian Johnston, Dominic Kwiatkowski, Cordelia Langford, John Sillitoe on behalf of the Wellcome Sanger Institute COVID-19 Surveillance Team |
| EPI_ISL_3093583 | Berkshire and Surrey Pathology Services Lighthouse Laboratory | Wellcome Sanger Institute for the COVID-19 Genomics UK (COG-UK) Consortium | Berkshire and Surrey Pathology Services Lighthouse Laboratory and Alex Alderton, Roberto Amato, Jeffrey Barrett, Sonia Goncalves, Ewan Harrison, David K. Jackson, Ian Johnston, Dominic Kwiatkowski, Cordelia Langford, John Sillitoe on behalf of the Wellcome Sanger Institute COVID-19 Surveillance Team |
| EPI_ISL_3093584 | Lighthouse Laboratory Plymouth | Wellcome Sanger Institute for the COVID-19 Genomics UK (COG-UK) Consortium | Lighthouse Laboratory Plymouth and Alex Alderton, Roberto Amato, Jeffrey Barrett, Sonia Goncalves, Ewan Harrison, David K. Jackson, Ian Johnston, Dominic Kwiatkowski, Cordelia Langford, John Sillitoe on behalf of the Wellcome Sanger Institute COVID-19 Surveillance Team |
| EPI_ISL_3093585 | Berkshire and Surrey Pathology Services Lighthouse Laboratory | Wellcome Sanger Institute for the COVID-19 Genomics UK (COG-UK) Consortium | Berkshire and Surrey Pathology Services Lighthouse Laboratory and Alex Alderton, Roberto Amato, Jeffrey Barrett, Sonia Goncalves, Ewan Harrison, David K. Jackson, Ian Johnston, Dominic Kwiatkowski, Cordelia Langford, John Sillitoe on behalf of the Wellcome Sanger Institute COVID-19 Surveillance Team |
| EPI_ISL_3093586 | Lighthouse Laboratory Plymouth | Wellcome Sanger Institute for the COVID-19 Genomics UK (COG-UK) Consortium | Lighthouse Laboratory Plymouth and Alex Alderton, Roberto Amato, Jeffrey Barrett, Sonia Goncalves, Ewan Harrison, David K. Jackson, Ian Johnston, Dominic Kwiatkowski, Cordelia Langford, John Sillitoe on behalf of the Wellcome Sanger Institute COVID-19 Surveillance Team |
| EPI_ISL_3093587, EPI_ISL_3093588 | Berkshire and Surrey Pathology Services Lighthouse Laboratory | Wellcome Sanger Institute for the COVID-19 Genomics UK (COG-UK) Consortium | Berkshire and Surrey Pathology Services Lighthouse Laboratory and Alex Alderton, Roberto Amato, Jeffrey Barrett, Sonia Goncalves, Ewan Harrison, David K. Jackson, Ian Johnston, Dominic Kwiatkowski, Cordelia Langford, John Sillitoe on behalf of the Wellcome Sanger Institute COVID-19 Surveillance Team |
| EPI_ISL_3093589, EPI_ISL_3093590, EPI_ISL_3093591 | Lighthouse Laboratory Plymouth | Wellcome Sanger Institute for the COVID-19 Genomics UK (COG-UK) Consortium | Lighthouse Laboratory Plymouth and Alex Alderton, Roberto Amato, Jeffrey Barrett, Sonia Goncalves, Ewan Harrison, David K. Jackson, Ian Johnston, Dominic Kwiatkowski, Cordelia Langford, John Sillitoe on behalf of the Wellcome Sanger Institute COVID-19 Surveillance Team |

| | | | |
|---|---|---|---|
| EPI_ISL_3093592, EPI_ISL_3093593, EPI_ISL_3093594, EPI_ISL_3093595 | Berkshire and Surrey Pathology Services Lighthouse Laboratory | Wellcome Sanger Institute for the COVID-19 Genomics UK (COG-UK) Consortium | Berkshire and Surrey Pathology Services Lighthouse Laboratory and Alex Alderton, Roberto Amato, Jeffrey Barrett, Sonia Goncalves, Ewan Harrison, David K. Jackson, Ian Johnston, Dominic Kwiatkowski, Cordelia Langford, John Sillitoe on behalf of the Wellcome Sanger Institute COVID-19 Surveillance Team |
| EPI_ISL_3093596 | Lighthouse Laboratory Plymouth | Wellcome Sanger Institute for the COVID-19 Genomics UK (COG-UK) Consortium | Lighthouse Laboratory Plymouth and Alex Alderton, Roberto Amato, Jeffrey Barrett, Sonia Goncalves, Ewan Harrison, David K. Jackson, Ian Johnston, Dominic Kwiatkowski, Cordelia Langford, John Sillitoe on behalf of the Wellcome Sanger Institute COVID-19 Surveillance Team |
| EPI_ISL_3093597, EPI_ISL_3093598 | Berkshire and Surrey Pathology Services Lighthouse Laboratory | Wellcome Sanger Institute for the COVID-19 Genomics UK (COG-UK) Consortium | Berkshire and Surrey Pathology Services Lighthouse Laboratory and Alex Alderton, Roberto Amato, Jeffrey Barrett, Sonia Goncalves, Ewan Harrison, David K. Jackson, Ian Johnston, Dominic Kwiatkowski, Cordelia Langford, John Sillitoe on behalf of the Wellcome Sanger Institute COVID-19 Surveillance Team |
| EPI_ISL_3093599, EPI_ISL_3093600 | Lighthouse Laboratory Plymouth | Wellcome Sanger Institute for the COVID-19 Genomics UK (COG-UK) Consortium | Lighthouse Laboratory Plymouth and Alex Alderton, Roberto Amato, Jeffrey Barrett, Sonia Goncalves, Ewan Harrison, David K. Jackson, Ian Johnston, Dominic Kwiatkowski, Cordelia Langford, John Sillitoe on behalf of the Wellcome Sanger Institute COVID-19 Surveillance Team |
| EPI_ISL_3093601 | Berkshire and Surrey Pathology Services Lighthouse Laboratory | Wellcome Sanger Institute for the COVID-19 Genomics UK (COG-UK) Consortium | Berkshire and Surrey Pathology Services Lighthouse Laboratory and Alex Alderton, Roberto Amato, Jeffrey Barrett, Sonia Goncalves, Ewan Harrison, David K. Jackson, Ian Johnston, Dominic Kwiatkowski, Cordelia Langford, John Sillitoe on behalf of the Wellcome Sanger Institute COVID-19 Surveillance Team |
| EPI_ISL_3093602, EPI_ISL_3093603, EPI_ISL_3093604, EPI_ISL_3093605 | Lighthouse Laboratory Plymouth | Wellcome Sanger Institute for the COVID-19 Genomics UK (COG-UK) Consortium | Lighthouse Laboratory Plymouth and Alex Alderton, Roberto Amato, Jeffrey Barrett, Sonia Goncalves, Ewan Harrison, David K. Jackson, Ian Johnston, Dominic Kwiatkowski, Cordelia Langford, John Sillitoe on behalf of the Wellcome Sanger Institute COVID-19 Surveillance Team |
| EPI_ISL_3093606 | Berkshire and Surrey Pathology Services Lighthouse Laboratory | Wellcome Sanger Institute for the COVID-19 Genomics UK (COG-UK) Consortium | Berkshire and Surrey Pathology Services Lighthouse Laboratory and Alex Alderton, Roberto Amato, Jeffrey Barrett, Sonia Goncalves, Ewan Harrison, David K. Jackson, Ian Johnston, Dominic Kwiatkowski, Cordelia Langford, John Sillitoe on behalf of the Wellcome Sanger Institute COVID-19 Surveillance Team |
| EPI_ISL_3093607, EPI_ISL_3093608 | Lighthouse Laboratory Plymouth | Wellcome Sanger Institute for the COVID-19 Genomics UK (COG-UK) Consortium | Lighthouse Laboratory Plymouth and Alex Alderton, Roberto Amato, Jeffrey Barrett, Sonia Goncalves, Ewan Harrison, David K. Jackson, Ian Johnston, Dominic Kwiatkowski, Cordelia Langford, John Sillitoe on behalf of the Wellcome Sanger Institute COVID-19 Surveillance Team |
| EPI_ISL_3093609, EPI_ISL_3093610, EPI_ISL_3093611 | Berkshire and Surrey Pathology Services Lighthouse Laboratory | Wellcome Sanger Institute for the COVID-19 Genomics UK (COG-UK) Consortium | Berkshire and Surrey Pathology Services Lighthouse Laboratory and Alex Alderton, Roberto Amato, Jeffrey Barrett, Sonia Goncalves, Ewan Harrison, David K. Jackson, Ian Johnston, Dominic Kwiatkowski, Cordelia Langford, John Sillitoe on behalf of the Wellcome Sanger Institute COVID-19 Surveillance Team |
| EPI_ISL_3093612 | Lighthouse Laboratory Plymouth | Wellcome Sanger Institute for the COVID-19 Genomics UK (COG-UK) Consortium | Lighthouse Laboratory Plymouth and Alex Alderton, Roberto Amato, Jeffrey Barrett, Sonia Goncalves, Ewan Harrison, David K. Jackson, Ian Johnston, Dominic Kwiatkowski, Cordelia Langford, John Sillitoe on behalf of the Wellcome Sanger Institute COVID-19 Surveillance Team |
| EPI_ISL_3093613 | Berkshire and Surrey Pathology Services Lighthouse Laboratory | Wellcome Sanger Institute for the COVID-19 Genomics UK (COG-UK) Consortium | Berkshire and Surrey Pathology Services Lighthouse Laboratory and Alex Alderton, Roberto Amato, Jeffrey Barrett, Sonia Goncalves, Ewan Harrison, David K. Jackson, Ian Johnston, Dominic Kwiatkowski, Cordelia Langford, John Sillitoe on behalf of the Wellcome Sanger Institute COVID-19 Surveillance Team |
| EPI_ISL_3093614, EPI_ISL_3093615, EPI_ISL_3093616 | Lighthouse Laboratory Plymouth | Wellcome Sanger Institute for the COVID-19 Genomics UK (COG-UK) Consortium | Lighthouse Laboratory Plymouth and Alex Alderton, Roberto Amato, Jeffrey Barrett, Sonia Goncalves, Ewan Harrison, David K. Jackson, Ian Johnston, Dominic Kwiatkowski, Cordelia Langford, John Sillitoe on behalf of the Wellcome Sanger Institute COVID-19 Surveillance Team |
| EPI_ISL_3093617, EPI_ISL_3093618, EPI_ISL_3093619 | Berkshire and Surrey Pathology Services Lighthouse Laboratory | Wellcome Sanger Institute for the COVID-19 Genomics UK (COG-UK) Consortium | Berkshire and Surrey Pathology Services Lighthouse Laboratory and Alex Alderton, Roberto Amato, Jeffrey Barrett, Sonia Goncalves, Ewan Harrison, David K. Jackson, Ian Johnston, Dominic Kwiatkowski, Cordelia Langford, John Sillitoe on behalf of the Wellcome Sanger Institute COVID-19 Surveillance Team |
| EPI_ISL_3093620 | Lighthouse Laboratory Plymouth | Wellcome Sanger Institute for the COVID-19 Genomics UK (COG-UK) Consortium | Lighthouse Laboratory Plymouth and Alex Alderton, Roberto Amato, Jeffrey Barrett, Sonia Goncalves, Ewan Harrison, David K. Jackson, Ian Johnston, Dominic Kwiatkowski, Cordelia Langford, John Sillitoe on behalf of the Wellcome Sanger Institute COVID-19 Surveillance Team |
| EPI_ISL_3093621, EPI_ISL_3093622, EPI_ISL_3093623 | Berkshire and Surrey Pathology Services Lighthouse Laboratory | Wellcome Sanger Institute for the COVID-19 Genomics UK (COG-UK) Consortium | Berkshire and Surrey Pathology Services Lighthouse Laboratory and Alex Alderton, Roberto Amato, Jeffrey Barrett, Sonia Goncalves, Ewan Harrison, David K. Jackson, Ian Johnston, Dominic Kwiatkowski, Cordelia Langford, John Sillitoe on behalf of the Wellcome Sanger Institute COVID-19 Surveillance Team |
| EPI_ISL_3093624 | Lighthouse Laboratory Plymouth | Wellcome Sanger Institute for the COVID-19 Genomics UK (COG-UK) Consortium | Lighthouse Laboratory Plymouth and Alex Alderton, Roberto Amato, Jeffrey Barrett, Sonia Goncalves, Ewan Harrison, David K. Jackson, Ian Johnston, Dominic Kwiatkowski, Cordelia Langford, John Sillitoe on behalf of the Wellcome Sanger Institute COVID-19 Surveillance Team |
| EPI_ISL_3093625, EPI_ISL_3093626, EPI_ISL_3093627 | Berkshire and Surrey Pathology Services Lighthouse Laboratory | Wellcome Sanger Institute for the COVID-19 Genomics UK (COG-UK) Consortium | Berkshire and Surrey Pathology Services Lighthouse Laboratory and Alex Alderton, Roberto Amato, Jeffrey Barrett, Sonia Goncalves, Ewan Harrison, David K. Jackson, Ian Johnston, Dominic Kwiatkowski, Cordelia Langford, John Sillitoe on behalf of the Wellcome Sanger Institute COVID-19 Surveillance Team |
| EPI_ISL_3093628, EPI_ISL_3093629 | Lighthouse Laboratory Plymouth | Wellcome Sanger Institute for the COVID-19 Genomics UK (COG-UK) Consortium | Lighthouse Laboratory Plymouth and Alex Alderton, Roberto Amato, Jeffrey Barrett, Sonia Goncalves, Ewan Harrison, David K. Jackson, Ian Johnston, Dominic Kwiatkowski, Cordelia Langford, John Sillitoe on behalf of the Wellcome Sanger Institute COVID-19 Surveillance Team |
| EPI_ISL_3093630 | Berkshire and Surrey Pathology Services Lighthouse Laboratory | Wellcome Sanger Institute for the COVID-19 Genomics UK (COG-UK) Consortium | Berkshire and Surrey Pathology Services Lighthouse Laboratory and Alex Alderton, Roberto Amato, Jeffrey Barrett, Sonia Goncalves, Ewan Harrison, David K. Jackson, Ian Johnston, Dominic Kwiatkowski, Cordelia Langford, John Sillitoe on behalf of the Wellcome Sanger Institute COVID-19 Surveillance Team |
| EPI_ISL_3093631 | Lighthouse Laboratory Plymouth | Wellcome Sanger Institute for the COVID-19 Genomics UK (COG-UK) Consortium | Lighthouse Laboratory Plymouth and Alex Alderton, Roberto Amato, Jeffrey Barrett, Sonia Goncalves, Ewan Harrison, David K. Jackson, Ian Johnston, Dominic Kwiatkowski, Cordelia Langford, John Sillitoe on behalf of the Wellcome Sanger Institute COVID-19 Surveillance Team |
| EPI_ISL_3093632 | Berkshire and Surrey Pathology Services Lighthouse Laboratory | Wellcome Sanger Institute for the COVID-19 Genomics UK (COG-UK) Consortium | Berkshire and Surrey Pathology Services Lighthouse Laboratory and Alex Alderton, Roberto Amato, Jeffrey Barrett, Sonia Goncalves, Ewan Harrison, David K. Jackson, Ian Johnston, Dominic Kwiatkowski, Cordelia Langford, John Sillitoe on behalf of the Wellcome Sanger Institute COVID-19 Surveillance Team |
| EPI_ISL_3093633 | Lighthouse Laboratory Plymouth | Wellcome Sanger Institute for the COVID-19 Genomics UK (COG-UK) Consortium | Lighthouse Laboratory Plymouth and Alex Alderton, Roberto Amato, Jeffrey Barrett, Sonia Goncalves, Ewan Harrison, David K. Jackson, Ian Johnston, Dominic Kwiatkowski, Cordelia Langford, John Sillitoe on behalf of the Wellcome Sanger Institute COVID-19 Surveillance Team |
| EPI_ISL_3093634, EPI_ISL_3093635, EPI_ISL_3093636, EPI_ISL_3093637, EPI_ISL_3093638, EPI_ISL_3093639 | Berkshire and Surrey Pathology Services Lighthouse Laboratory | Wellcome Sanger Institute for the COVID-19 Genomics UK (COG-UK) Consortium | Berkshire and Surrey Pathology Services Lighthouse Laboratory and Alex Alderton, Roberto Amato, Jeffrey Barrett, Sonia Goncalves, Ewan Harrison, David K. Jackson, Ian Johnston, Dominic Kwiatkowski, Cordelia Langford, John Sillitoe on behalf of the Wellcome Sanger Institute COVID-19 Surveillance Team |
| EPI_ISL_3093640, EPI_ISL_3093641, EPI_ISL_3093642, EPI_ISL_3093643 | Lighthouse Laboratory Plymouth | Wellcome Sanger Institute for the COVID-19 Genomics UK (COG-UK) Consortium | Lighthouse Laboratory Plymouth and Alex Alderton, Roberto Amato, Jeffrey Barrett, Sonia Goncalves, Ewan Harrison, David K. Jackson, Ian Johnston, Dominic Kwiatkowski, Cordelia Langford, John Sillitoe on behalf of the Wellcome Sanger Institute COVID-19 Surveillance Team |
| EPI_ISL_3093644 | Berkshire and Surrey Pathology Services Lighthouse Laboratory | Wellcome Sanger Institute for the COVID-19 Genomics UK (COG-UK) Consortium | Berkshire and Surrey Pathology Services Lighthouse Laboratory and Alex Alderton, Roberto Amato, Jeffrey Barrett, Sonia Goncalves, Ewan Harrison, David K. Jackson, Ian Johnston, Dominic Kwiatkowski, Cordelia Langford, John Sillitoe on behalf of the Wellcome Sanger Institute COVID-19 Surveillance Team |
| EPI_ISL_3093645 | Lighthouse Laboratory Plymouth | Wellcome Sanger Institute for the COVID-19 Genomics UK (COG-UK) Consortium | Lighthouse Laboratory Plymouth and Alex Alderton, Roberto Amato, Jeffrey Barrett, Sonia Goncalves, Ewan Harrison, David K. Jackson, Ian Johnston, Dominic Kwiatkowski, Cordelia Langford, John Sillitoe on behalf of the Wellcome Sanger Institute COVID-19 Surveillance Team |
| EPI_ISL_3093646 | Berkshire and Surrey Pathology Services Lighthouse Laboratory | Wellcome Sanger Institute for the COVID-19 Genomics UK (COG-UK) Consortium | Berkshire and Surrey Pathology Services Lighthouse Laboratory and Alex Alderton, Roberto Amato, Jeffrey Barrett, Sonia Goncalves, Ewan Harrison, David K. Jackson, Ian Johnston, Dominic Kwiatkowski, Cordelia Langford, John Sillitoe on behalf of the Wellcome Sanger Institute COVID-19 Surveillance Team |
| EPI_ISL_3093647 | Lighthouse Laboratory Plymouth | Wellcome Sanger Institute for the COVID-19 Genomics UK (COG-UK) Consortium | Lighthouse Laboratory Plymouth and Alex Alderton, Roberto Amato, Jeffrey Barrett, Sonia Goncalves, Ewan Harrison, David K. Jackson, Ian Johnston, Dominic Kwiatkowski, Cordelia Langford, John Sillitoe on behalf of the Wellcome Sanger Institute COVID-19 Surveillance Team |
| EPI_ISL_3093648, EPI_ISL_3093649 | Berkshire and Surrey Pathology Services Lighthouse Laboratory | Wellcome Sanger Institute for the COVID-19 Genomics UK (COG-UK) Consortium | Berkshire and Surrey Pathology Services Lighthouse Laboratory and Alex Alderton, Roberto Amato, Jeffrey Barrett, Sonia Goncalves, Ewan Harrison, David K. Jackson, Ian Johnston, Dominic Kwiatkowski, Cordelia Langford, John Sillitoe on behalf of the Wellcome Sanger Institute COVID-19 Surveillance Team |
| EPI_ISL_3093650 | Lighthouse Laboratory Plymouth | Wellcome Sanger Institute for the COVID-19 Genomics UK (COG-UK) Consortium | Lighthouse Laboratory Plymouth and Alex Alderton, Roberto Amato, Jeffrey Barrett, Sonia Goncalves, Ewan Harrison, David K. Jackson, Ian Johnston, Dominic Kwiatkowski, Cordelia Langford, John Sillitoe on behalf of the Wellcome Sanger Institute COVID-19 Surveillance Team |
| EPI_ISL_3093651 | Berkshire and Surrey Pathology Services Lighthouse Laboratory | Wellcome Sanger Institute for the COVID-19 Genomics UK (COG-UK) Consortium | Berkshire and Surrey Pathology Services Lighthouse Laboratory and Alex Alderton, Roberto Amato, Jeffrey Barrett, Sonia Goncalves, Ewan Harrison, David K. Jackson, Ian Johnston, Dominic Kwiatkowski, Cordelia Langford, John Sillitoe on behalf of the Wellcome Sanger Institute COVID-19 Surveillance Team |
| EPI_ISL_3093652 | Lighthouse Laboratory Plymouth | Wellcome Sanger Institute for the COVID-19 Genomics UK (COG-UK) Consortium | Lighthouse Laboratory Plymouth and Alex Alderton, Roberto Amato, Jeffrey Barrett, Sonia Goncalves, Ewan Harrison, David K. Jackson, Ian Johnston, Dominic Kwiatkowski, Cordelia Langford, John Sillitoe on behalf of the Wellcome Sanger Institute COVID-19 Surveillance Team |
| EPI_ISL_3093653, EPI_ISL_3093654 | Berkshire and Surrey Pathology Services Lighthouse Laboratory | Wellcome Sanger Institute for the COVID-19 Genomics UK (COG-UK) Consortium | Berkshire and Surrey Pathology Services Lighthouse Laboratory and Alex Alderton, Roberto Amato, Jeffrey Barrett, Sonia Goncalves, Ewan Harrison, David K. Jackson, Ian Johnston, Dominic Kwiatkowski, Cordelia Langford, John Sillitoe on behalf of the Wellcome Sanger Institute COVID-19 Surveillance Team |
| EPI_ISL_3093655, EPI_ISL_3093656 | Lighthouse Laboratory Plymouth | Wellcome Sanger Institute for the COVID-19 Genomics UK (COG-UK) Consortium | Lighthouse Laboratory Plymouth and Alex Alderton, Roberto Amato, Jeffrey Barrett, Sonia Goncalves, Ewan Harrison, David K. Jackson, Ian Johnston, Dominic Kwiatkowski, Cordelia Langford, John Sillitoe on behalf of the Wellcome Sanger Institute COVID-19 Surveillance Team |
| EPI_ISL_3093657 | Berkshire and Surrey Pathology Services Lighthouse Laboratory | Wellcome Sanger Institute for the COVID-19 Genomics UK (COG-UK) Consortium | Berkshire and Surrey Pathology Services Lighthouse Laboratory and Alex Alderton, Roberto Amato, Jeffrey Barrett, Sonia Goncalves, Ewan Harrison, David K. Jackson, Ian Johnston, Dominic Kwiatkowski, Cordelia Langford, John Sillitoe on behalf of the Wellcome Sanger Institute COVID-19 Surveillance Team |
| EPI_ISL_3093658, EPI_ISL_3093659 | Lighthouse Laboratory Plymouth | Wellcome Sanger Institute for the COVID-19 Genomics UK (COG-UK) Consortium | Lighthouse Laboratory Plymouth and Alex Alderton, Roberto Amato, Jeffrey Barrett, Sonia Goncalves, Ewan Harrison, David K. Jackson, Ian Johnston, Dominic Kwiatkowski, Cordelia Langford, John Sillitoe on behalf of the Wellcome Sanger Institute COVID-19 Surveillance Team |
| EPI_ISL_3093660 | Berkshire and Surrey Pathology Services Lighthouse Laboratory | Wellcome Sanger Institute for the COVID-19 Genomics UK (COG-UK) Consortium | Berkshire and Surrey Pathology Services Lighthouse Laboratory and Alex Alderton, Roberto Amato, Jeffrey Barrett, Sonia Goncalves, Ewan Harrison, David K. Jackson, Ian Johnston, Dominic Kwiatkowski, Cordelia Langford, John Sillitoe on behalf of the Wellcome Sanger Institute COVID-19 Surveillance Team |
| EPI_ISL_3093661 | Lighthouse Laboratory Plymouth | Wellcome Sanger Institute for the COVID-19 Genomics UK (COG-UK) Consortium | Lighthouse Laboratory Plymouth and Alex Alderton, Roberto Amato, Jeffrey Barrett, Sonia Goncalves, Ewan Harrison, David K. Jackson, Ian Johnston, Dominic Kwiatkowski, Cordelia Langford, John Sillitoe on behalf of the Wellcome Sanger Institute COVID-19 Surveillance Team |
| EPI_ISL_3093662 | Berkshire and Surrey Pathology Services Lighthouse Laboratory | Wellcome Sanger Institute for the COVID-19 Genomics UK (COG-UK) Consortium | Berkshire and Surrey Pathology Services Lighthouse Laboratory and Alex Alderton, Roberto Amato, Jeffrey Barrett, Sonia Goncalves, Ewan Harrison, David K. Jackson, Ian Johnston, Dominic Kwiatkowski, Cordelia Langford, John Sillitoe on behalf of the Wellcome Sanger Institute COVID-19 Surveillance Team |

| | | | |
|---|---|---|---|
| EPI_ISL_3093663 | Lighthouse Laboratory Plymouth | Wellcome Sanger Institute for the COVID-19 Genomics UK (COG-UK) Consortium | Lighthouse Laboratory Plymouth and Alex Alderton, Roberto Amato, Jeffrey Barrett, Sonia Goncalves, Ewan Harrison, David K. Jackson, Ian Johnston, Dominic Kwiatkowski, Cordelia Langford, John Sillitoe on behalf of the Wellcome Sanger Institute COVID-19 Surveillance Team |
| EPI_ISL_3093664, EPI_ISL_3093665, EPI_ISL_3093666, EPI_ISL_3093667 | Berkshire and Surrey Pathology Services Lighthouse Laboratory | Wellcome Sanger Institute for the COVID-19 Genomics UK (COG-UK) Consortium | Berkshire and Surrey Pathology Services Lighthouse Laboratory and Alex Alderton, Roberto Amato, Jeffrey Barrett, Sonia Goncalves, Ewan Harrison, David K. Jackson, Ian Johnston, Dominic Kwiatkowski, Cordelia Langford, John Sillitoe on behalf of the Wellcome Sanger Institute COVID-19 Surveillance Team |
| EPI_ISL_3093668 | Lighthouse Laboratory Plymouth | Wellcome Sanger Institute for the COVID-19 Genomics UK (COG-UK) Consortium | Lighthouse Laboratory Plymouth and Alex Alderton, Roberto Amato, Jeffrey Barrett, Sonia Goncalves, Ewan Harrison, David K. Jackson, Ian Johnston, Dominic Kwiatkowski, Cordelia Langford, John Sillitoe on behalf of the Wellcome Sanger Institute COVID-19 Surveillance Team |
| EPI_ISL_3093669 | Berkshire and Surrey Pathology Services Lighthouse Laboratory | Wellcome Sanger Institute for the COVID-19 Genomics UK (COG-UK) Consortium | Berkshire and Surrey Pathology Services Lighthouse Laboratory and Alex Alderton, Roberto Amato, Jeffrey Barrett, Sonia Goncalves, Ewan Harrison, David K. Jackson, Ian Johnston, Dominic Kwiatkowski, Cordelia Langford, John Sillitoe on behalf of the Wellcome Sanger Institute COVID-19 Surveillance Team |
| EPI_ISL_3093670, EPI_ISL_3093671, EPI_ISL_3093672 | Lighthouse Laboratory Plymouth | Wellcome Sanger Institute for the COVID-19 Genomics UK (COG-UK) Consortium | Lighthouse Laboratory Plymouth and Alex Alderton, Roberto Amato, Jeffrey Barrett, Sonia Goncalves, Ewan Harrison, David K. Jackson, Ian Johnston, Dominic Kwiatkowski, Cordelia Langford, John Sillitoe on behalf of the Wellcome Sanger Institute COVID-19 Surveillance Team |
| EPI_ISL_3093673 | Berkshire and Surrey Pathology Services Lighthouse Laboratory | Wellcome Sanger Institute for the COVID-19 Genomics UK (COG-UK) Consortium | Berkshire and Surrey Pathology Services Lighthouse Laboratory and Alex Alderton, Roberto Amato, Jeffrey Barrett, Sonia Goncalves, Ewan Harrison, David K. Jackson, Ian Johnston, Dominic Kwiatkowski, Cordelia Langford, John Sillitoe on behalf of the Wellcome Sanger Institute COVID-19 Surveillance Team |
| EPI_ISL_3093674, EPI_ISL_3093675, EPI_ISL_3093676 | Lighthouse Laboratory Plymouth | Wellcome Sanger Institute for the COVID-19 Genomics UK (COG-UK) Consortium | Lighthouse Laboratory Plymouth and Alex Alderton, Roberto Amato, Jeffrey Barrett, Sonia Goncalves, Ewan Harrison, David K. Jackson, Ian Johnston, Dominic Kwiatkowski, Cordelia Langford, John Sillitoe on behalf of the Wellcome Sanger Institute COVID-19 Surveillance Team |
| EPI_ISL_3093677 | Berkshire and Surrey Pathology Services Lighthouse Laboratory | Wellcome Sanger Institute for the COVID-19 Genomics UK (COG-UK) Consortium | Berkshire and Surrey Pathology Services Lighthouse Laboratory and Alex Alderton, Roberto Amato, Jeffrey Barrett, Sonia Goncalves, Ewan Harrison, David K. Jackson, Ian Johnston, Dominic Kwiatkowski, Cordelia Langford, John Sillitoe on behalf of the Wellcome Sanger Institute COVID-19 Surveillance Team |
| EPI_ISL_3093678 | Lighthouse Laboratory Plymouth | Wellcome Sanger Institute for the COVID-19 Genomics UK (COG-UK) Consortium | Lighthouse Laboratory Plymouth and Alex Alderton, Roberto Amato, Jeffrey Barrett, Sonia Goncalves, Ewan Harrison, David K. Jackson, Ian Johnston, Dominic Kwiatkowski, Cordelia Langford, John Sillitoe on behalf of the Wellcome Sanger Institute COVID-19 Surveillance Team |
| EPI_ISL_3093679, EPI_ISL_3093680 | Berkshire and Surrey Pathology Services Lighthouse Laboratory | Wellcome Sanger Institute for the COVID-19 Genomics UK (COG-UK) Consortium | Berkshire and Surrey Pathology Services Lighthouse Laboratory and Alex Alderton, Roberto Amato, Jeffrey Barrett, Sonia Goncalves, Ewan Harrison, David K. Jackson, Ian Johnston, Dominic Kwiatkowski, Cordelia Langford, John Sillitoe on behalf of the Wellcome Sanger Institute COVID-19 Surveillance Team |
| EPI_ISL_3093681 | Lighthouse Laboratory Plymouth | Wellcome Sanger Institute for the COVID-19 Genomics UK (COG-UK) Consortium | Lighthouse Laboratory Plymouth and Alex Alderton, Roberto Amato, Jeffrey Barrett, Sonia Goncalves, Ewan Harrison, David K. Jackson, Ian Johnston, Dominic Kwiatkowski, Cordelia Langford, John Sillitoe on behalf of the Wellcome Sanger Institute COVID-19 Surveillance Team |
| EPI_ISL_3093682 | Berkshire and Surrey Pathology Services Lighthouse Laboratory | Wellcome Sanger Institute for the COVID-19 Genomics UK (COG-UK) Consortium | Berkshire and Surrey Pathology Services Lighthouse Laboratory and Alex Alderton, Roberto Amato, Jeffrey Barrett, Sonia Goncalves, Ewan Harrison, David K. Jackson, Ian Johnston, Dominic Kwiatkowski, Cordelia Langford, John Sillitoe on behalf of the Wellcome Sanger Institute COVID-19 Surveillance Team |
| EPI_ISL_3093683, EPI_ISL_3093684, EPI_ISL_3093685 | Lighthouse Laboratory Plymouth | Wellcome Sanger Institute for the COVID-19 Genomics UK (COG-UK) Consortium | Lighthouse Laboratory Plymouth and Alex Alderton, Roberto Amato, Jeffrey Barrett, Sonia Goncalves, Ewan Harrison, David K. Jackson, Ian Johnston, Dominic Kwiatkowski, Cordelia Langford, John Sillitoe on behalf of the Wellcome Sanger Institute COVID-19 Surveillance Team |
| EPI_ISL_3093686, EPI_ISL_3093687, EPI_ISL_3093688, EPI_ISL_3093689 | Berkshire and Surrey Pathology Services Lighthouse Laboratory | Wellcome Sanger Institute for the COVID-19 Genomics UK (COG-UK) Consortium | Berkshire and Surrey Pathology Services Lighthouse Laboratory and Alex Alderton, Roberto Amato, Jeffrey Barrett, Sonia Goncalves, Ewan Harrison, David K. Jackson, Ian Johnston, Dominic Kwiatkowski, Cordelia Langford, John Sillitoe on behalf of the Wellcome Sanger Institute COVID-19 Surveillance Team |
| EPI_ISL_3093690 | Lighthouse Laboratory Plymouth | Wellcome Sanger Institute for the COVID-19 Genomics UK (COG-UK) Consortium | Lighthouse Laboratory Plymouth and Alex Alderton, Roberto Amato, Jeffrey Barrett, Sonia Goncalves, Ewan Harrison, David K. Jackson, Ian Johnston, Dominic Kwiatkowski, Cordelia Langford, John Sillitoe on behalf of the Wellcome Sanger Institute COVID-19 Surveillance Team |
| EPI_ISL_3093691 | Berkshire and Surrey Pathology Services Lighthouse Laboratory | Wellcome Sanger Institute for the COVID-19 Genomics UK (COG-UK) Consortium | Berkshire and Surrey Pathology Services Lighthouse Laboratory and Alex Alderton, Roberto Amato, Jeffrey Barrett, Sonia Goncalves, Ewan Harrison, David K. Jackson, Ian Johnston, Dominic Kwiatkowski, Cordelia Langford, John Sillitoe on behalf of the Wellcome Sanger Institute COVID-19 Surveillance Team |
| EPI_ISL_3093692 | Lighthouse Laboratory Plymouth | Wellcome Sanger Institute for the COVID-19 Genomics UK (COG-UK) Consortium | Lighthouse Laboratory Plymouth and Alex Alderton, Roberto Amato, Jeffrey Barrett, Sonia Goncalves, Ewan Harrison, David K. Jackson, Ian Johnston, Dominic Kwiatkowski, Cordelia Langford, John Sillitoe on behalf of the Wellcome Sanger Institute COVID-19 Surveillance Team |
| EPI_ISL_3093693 | Berkshire and Surrey Pathology Services Lighthouse Laboratory | Wellcome Sanger Institute for the COVID-19 Genomics UK (COG-UK) Consortium | Berkshire and Surrey Pathology Services Lighthouse Laboratory and Alex Alderton, Roberto Amato, Jeffrey Barrett, Sonia Goncalves, Ewan Harrison, David K. Jackson, Ian Johnston, Dominic Kwiatkowski, Cordelia Langford, John Sillitoe on behalf of the Wellcome Sanger Institute COVID-19 Surveillance Team |
| EPI_ISL_3093694 | Lighthouse Laboratory Plymouth | Wellcome Sanger Institute for the COVID-19 Genomics UK (COG-UK) Consortium | Lighthouse Laboratory Plymouth and Alex Alderton, Roberto Amato, Jeffrey Barrett, Sonia Goncalves, Ewan Harrison, David K. Jackson, Ian Johnston, Dominic Kwiatkowski, Cordelia Langford, John Sillitoe on behalf of the Wellcome Sanger Institute COVID-19 Surveillance Team |
| EPI_ISL_3093695, EPI_ISL_3093696 | Berkshire and Surrey Pathology Services Lighthouse Laboratory | Wellcome Sanger Institute for the COVID-19 Genomics UK (COG-UK) Consortium | Berkshire and Surrey Pathology Services Lighthouse Laboratory and Alex Alderton, Roberto Amato, Jeffrey Barrett, Sonia Goncalves, Ewan Harrison, David K. Jackson, Ian Johnston, Dominic Kwiatkowski, Cordelia Langford, John Sillitoe on behalf of the Wellcome Sanger Institute COVID-19 Surveillance Team |
| EPI_ISL_3093697 | Lighthouse Laboratory Plymouth | Wellcome Sanger Institute for the COVID-19 Genomics UK (COG-UK) Consortium | Lighthouse Laboratory Plymouth and Alex Alderton, Roberto Amato, Jeffrey Barrett, Sonia Goncalves, Ewan Harrison, David K. Jackson, Ian Johnston, Dominic Kwiatkowski, Cordelia Langford, John Sillitoe on behalf of the Wellcome Sanger Institute COVID-19 Surveillance Team |
| EPI_ISL_3093698 | Berkshire and Surrey Pathology Services Lighthouse Laboratory | Wellcome Sanger Institute for the COVID-19 Genomics UK (COG-UK) Consortium | Berkshire and Surrey Pathology Services Lighthouse Laboratory and Alex Alderton, Roberto Amato, Jeffrey Barrett, Sonia Goncalves, Ewan Harrison, David K. Jackson, Ian Johnston, Dominic Kwiatkowski, Cordelia Langford, John Sillitoe on behalf of the Wellcome Sanger Institute COVID-19 Surveillance Team |
| EPI_ISL_3093699 | Lighthouse Laboratory Plymouth | Wellcome Sanger Institute for the COVID-19 Genomics UK (COG-UK) Consortium | Lighthouse Laboratory Plymouth and Alex Alderton, Roberto Amato, Jeffrey Barrett, Sonia Goncalves, Ewan Harrison, David K. Jackson, Ian Johnston, Dominic Kwiatkowski, Cordelia Langford, John Sillitoe on behalf of the Wellcome Sanger Institute COVID-19 Surveillance Team |
| EPI_ISL_3093700, EPI_ISL_3093701, EPI_ISL_3093702, EPI_ISL_3093703, EPI_ISL_3093704 | Berkshire and Surrey Pathology Services Lighthouse Laboratory | Wellcome Sanger Institute for the COVID-19 Genomics UK (COG-UK) Consortium | Berkshire and Surrey Pathology Services Lighthouse Laboratory and Alex Alderton, Roberto Amato, Jeffrey Barrett, Sonia Goncalves, Ewan Harrison, David K. Jackson, Ian Johnston, Dominic Kwiatkowski, Cordelia Langford, John Sillitoe on behalf of the Wellcome Sanger Institute COVID-19 Surveillance Team |
| EPI_ISL_3093705 | Lighthouse Laboratory Plymouth | Wellcome Sanger Institute for the COVID-19 Genomics UK (COG-UK) Consortium | Lighthouse Laboratory Plymouth and Alex Alderton, Roberto Amato, Jeffrey Barrett, Sonia Goncalves, Ewan Harrison, David K. Jackson, Ian Johnston, Dominic Kwiatkowski, Cordelia Langford, John Sillitoe on behalf of the Wellcome Sanger Institute COVID-19 Surveillance Team |
| EPI_ISL_3093706 | Berkshire and Surrey Pathology Services Lighthouse Laboratory | Wellcome Sanger Institute for the COVID-19 Genomics UK (COG-UK) Consortium | Berkshire and Surrey Pathology Services Lighthouse Laboratory and Alex Alderton, Roberto Amato, Jeffrey Barrett, Sonia Goncalves, Ewan Harrison, David K. Jackson, Ian Johnston, Dominic Kwiatkowski, Cordelia Langford, John Sillitoe on behalf of the Wellcome Sanger Institute COVID-19 Surveillance Team |
| EPI_ISL_3093707 | Lighthouse Laboratory Plymouth | Wellcome Sanger Institute for the COVID-19 Genomics UK (COG-UK) Consortium | Lighthouse Laboratory Plymouth and Alex Alderton, Roberto Amato, Jeffrey Barrett, Sonia Goncalves, Ewan Harrison, David K. Jackson, Ian Johnston, Dominic Kwiatkowski, Cordelia Langford, John Sillitoe on behalf of the Wellcome Sanger Institute COVID-19 Surveillance Team |
| EPI_ISL_3093708, EPI_ISL_3093709 | Berkshire and Surrey Pathology Services Lighthouse Laboratory | Wellcome Sanger Institute for the COVID-19 Genomics UK (COG-UK) Consortium | Berkshire and Surrey Pathology Services Lighthouse Laboratory and Alex Alderton, Roberto Amato, Jeffrey Barrett, Sonia Goncalves, Ewan Harrison, David K. Jackson, Ian Johnston, Dominic Kwiatkowski, Cordelia Langford, John Sillitoe on behalf of the Wellcome Sanger Institute COVID-19 Surveillance Team |
| EPI_ISL_3093710, EPI_ISL_3093711, EPI_ISL_3093712, EPI_ISL_3093713 | Lighthouse Laboratory Plymouth | Wellcome Sanger Institute for the COVID-19 Genomics UK (COG-UK) Consortium | Lighthouse Laboratory Plymouth and Alex Alderton, Roberto Amato, Jeffrey Barrett, Sonia Goncalves, Ewan Harrison, David K. Jackson, Ian Johnston, Dominic Kwiatkowski, Cordelia Langford, John Sillitoe on behalf of the Wellcome Sanger Institute COVID-19 Surveillance Team |
| EPI_ISL_3093714, EPI_ISL_3093715 | Berkshire and Surrey Pathology Services Lighthouse Laboratory | Wellcome Sanger Institute for the COVID-19 Genomics UK (COG-UK) Consortium | Berkshire and Surrey Pathology Services Lighthouse Laboratory and Alex Alderton, Roberto Amato, Jeffrey Barrett, Sonia Goncalves, Ewan Harrison, David K. Jackson, Ian Johnston, Dominic Kwiatkowski, Cordelia Langford, John Sillitoe on behalf of the Wellcome Sanger Institute COVID-19 Surveillance Team |
| EPI_ISL_3093716, EPI_ISL_3093717 | Lighthouse Laboratory Plymouth | Wellcome Sanger Institute for the COVID-19 Genomics UK (COG-UK) Consortium | Lighthouse Laboratory Plymouth and Alex Alderton, Roberto Amato, Jeffrey Barrett, Sonia Goncalves, Ewan Harrison, David K. Jackson, Ian Johnston, Dominic Kwiatkowski, Cordelia Langford, John Sillitoe on behalf of the Wellcome Sanger Institute COVID-19 Surveillance Team |
| EPI_ISL_3093718 | Berkshire and Surrey Pathology Services Lighthouse Laboratory | Wellcome Sanger Institute for the COVID-19 Genomics UK (COG-UK) Consortium | Berkshire and Surrey Pathology Services Lighthouse Laboratory and Alex Alderton, Roberto Amato, Jeffrey Barrett, Sonia Goncalves, Ewan Harrison, David K. Jackson, Ian Johnston, Dominic Kwiatkowski, Cordelia Langford, John Sillitoe on behalf of the Wellcome Sanger Institute COVID-19 Surveillance Team |
| EPI_ISL_3093719, EPI_ISL_3093720, EPI_ISL_3093721 | Lighthouse Laboratory Plymouth | Wellcome Sanger Institute for the COVID-19 Genomics UK (COG-UK) Consortium | Lighthouse Laboratory Plymouth and Alex Alderton, Roberto Amato, Jeffrey Barrett, Sonia Goncalves, Ewan Harrison, David K. Jackson, Ian Johnston, Dominic Kwiatkowski, Cordelia Langford, John Sillitoe on behalf of the Wellcome Sanger Institute COVID-19 Surveillance Team |
| EPI_ISL_3093722 | Berkshire and Surrey Pathology Services Lighthouse Laboratory | Wellcome Sanger Institute for the COVID-19 Genomics UK (COG-UK) Consortium | Berkshire and Surrey Pathology Services Lighthouse Laboratory and Alex Alderton, Roberto Amato, Jeffrey Barrett, Sonia Goncalves, Ewan Harrison, David K. Jackson, Ian Johnston, Dominic Kwiatkowski, Cordelia Langford, John Sillitoe on behalf of the Wellcome Sanger Institute COVID-19 Surveillance Team |
| EPI_ISL_3093723, EPI_ISL_3093724 | Lighthouse Laboratory Plymouth | Wellcome Sanger Institute for the COVID-19 Genomics UK (COG-UK) Consortium | Lighthouse Laboratory Plymouth and Alex Alderton, Roberto Amato, Jeffrey Barrett, Sonia Goncalves, Ewan Harrison, David K. Jackson, Ian Johnston, Dominic Kwiatkowski, Cordelia Langford, John Sillitoe on behalf of the Wellcome Sanger Institute COVID-19 Surveillance Team |
| EPI_ISL_3093725 | Berkshire and Surrey Pathology Services Lighthouse Laboratory | Wellcome Sanger Institute for the COVID-19 Genomics UK (COG-UK) Consortium | Berkshire and Surrey Pathology Services Lighthouse Laboratory and Alex Alderton, Roberto Amato, Jeffrey Barrett, Sonia Goncalves, Ewan Harrison, David K. Jackson, Ian Johnston, Dominic Kwiatkowski, Cordelia Langford, John Sillitoe on behalf of the Wellcome Sanger Institute COVID-19 Surveillance Team |
| EPI_ISL_3093726, EPI_ISL_3093727 | Lighthouse Laboratory Plymouth | Wellcome Sanger Institute for the COVID-19 Genomics UK (COG-UK) Consortium | Lighthouse Laboratory Plymouth and Alex Alderton, Roberto Amato, Jeffrey Barrett, Sonia Goncalves, Ewan Harrison, David K. Jackson, Ian Johnston, Dominic Kwiatkowski, Cordelia Langford, John Sillitoe on behalf of the Wellcome Sanger Institute COVID-19 Surveillance Team |
| EPI_ISL_3093728 | Berkshire and Surrey Pathology Services Lighthouse Laboratory | Wellcome Sanger Institute for the COVID-19 Genomics UK (COG-UK) Consortium | Berkshire and Surrey Pathology Services Lighthouse Laboratory and Alex Alderton, Roberto Amato, Jeffrey Barrett, Sonia Goncalves, Ewan Harrison, David K. Jackson, Ian Johnston, Dominic Kwiatkowski, Cordelia Langford, John Sillitoe on behalf of the Wellcome Sanger Institute COVID-19 Surveillance Team |
| EPI_ISL_3093729, EPI_ISL_3093730, EPI_ISL_3093731, EPI_ISL_3093732, | Lighthouse Laboratory Plymouth | Wellcome Sanger Institute for the COVID-19 Genomics UK (COG-UK) Consortium | Lighthouse Laboratory Plymouth and Alex Alderton, Roberto Amato, Jeffrey Barrett, Sonia Goncalves, Ewan Harrison, David K. Jackson, Ian Johnston, Dominic Kwiatkowski, Cordelia Langford, John Sillitoe on behalf of the Wellcome Sanger Institute COVID-19 Surveillance Team |

| | | | |
|---|---|---|---|
| EPI_ISL_3093733 | | | |
| EPI_ISL_3093734, EPI_ISL_3093735 | Berkshire and Surrey Pathology Services Lighthouse Laboratory | Wellcome Sanger Institute for the COVID-19 Genomics UK (COG-UK) Consortium | Berkshire and Surrey Pathology Services Lighthouse Laboratory and Alex Alderton, Roberto Amato, Jeffrey Barrett, Sonia Goncalves, Ewan Harrison, David K. Jackson, Ian Johnston, Dominic Kwiatkowski, Cordelia Langford, John Sillitoe on behalf of the Wellcome Sanger Institute COVID-19 Surveillance Team |
| EPI_ISL_3093736 | Lighthouse Laboratory Plymouth | Wellcome Sanger Institute for the COVID-19 Genomics UK (COG-UK) Consortium | Lighthouse Laboratory Plymouth and Alex Alderton, Roberto Amato, Jeffrey Barrett, Sonia Goncalves, Ewan Harrison, David K. Jackson, Ian Johnston, Dominic Kwiatkowski, Cordelia Langford, John Sillitoe on behalf of the Wellcome Sanger Institute COVID-19 Surveillance Team |
| EPI_ISL_3093737, EPI_ISL_3093738, EPI_ISL_3093739 | Berkshire and Surrey Pathology Services Lighthouse Laboratory | Wellcome Sanger Institute for the COVID-19 Genomics UK (COG-UK) Consortium | Berkshire and Surrey Pathology Services Lighthouse Laboratory and Alex Alderton, Roberto Amato, Jeffrey Barrett, Sonia Goncalves, Ewan Harrison, David K. Jackson, Ian Johnston, Dominic Kwiatkowski, Cordelia Langford, John Sillitoe on behalf of the Wellcome Sanger Institute COVID-19 Surveillance Team |
| EPI_ISL_3093740, EPI_ISL_3093741 | Lighthouse Laboratory Plymouth | Wellcome Sanger Institute for the COVID-19 Genomics UK (COG-UK) Consortium | Lighthouse Laboratory Plymouth and Alex Alderton, Roberto Amato, Jeffrey Barrett, Sonia Goncalves, Ewan Harrison, David K. Jackson, Ian Johnston, Dominic Kwiatkowski, Cordelia Langford, John Sillitoe on behalf of the Wellcome Sanger Institute COVID-19 Surveillance Team |
| EPI_ISL_3093742 | Berkshire and Surrey Pathology Services Lighthouse Laboratory | Wellcome Sanger Institute for the COVID-19 Genomics UK (COG-UK) Consortium | Berkshire and Surrey Pathology Services Lighthouse Laboratory and Alex Alderton, Roberto Amato, Jeffrey Barrett, Sonia Goncalves, Ewan Harrison, David K. Jackson, Ian Johnston, Dominic Kwiatkowski, Cordelia Langford, John Sillitoe on behalf of the Wellcome Sanger Institute COVID-19 Surveillance Team |
| EPI_ISL_3093743, EPI_ISL_3093744 | Lighthouse Laboratory Plymouth | Wellcome Sanger Institute for the COVID-19 Genomics UK (COG-UK) Consortium | Lighthouse Laboratory Plymouth and Alex Alderton, Roberto Amato, Jeffrey Barrett, Sonia Goncalves, Ewan Harrison, David K. Jackson, Ian Johnston, Dominic Kwiatkowski, Cordelia Langford, John Sillitoe on behalf of the Wellcome Sanger Institute COVID-19 Surveillance Team |
| EPI_ISL_3093745 | Berkshire and Surrey Pathology Services Lighthouse Laboratory | Wellcome Sanger Institute for the COVID-19 Genomics UK (COG-UK) Consortium | Berkshire and Surrey Pathology Services Lighthouse Laboratory and Alex Alderton, Roberto Amato, Jeffrey Barrett, Sonia Goncalves, Ewan Harrison, David K. Jackson, Ian Johnston, Dominic Kwiatkowski, Cordelia Langford, John Sillitoe on behalf of the Wellcome Sanger Institute COVID-19 Surveillance Team |
| EPI_ISL_3093746 | Lighthouse Laboratory Plymouth | Wellcome Sanger Institute for the COVID-19 Genomics UK (COG-UK) Consortium | Lighthouse Laboratory Plymouth and Alex Alderton, Roberto Amato, Jeffrey Barrett, Sonia Goncalves, Ewan Harrison, David K. Jackson, Ian Johnston, Dominic Kwiatkowski, Cordelia Langford, John Sillitoe on behalf of the Wellcome Sanger Institute COVID-19 Surveillance Team |
| EPI_ISL_3093747, EPI_ISL_3093748, EPI_ISL_3093749, EPI_ISL_3093750, EPI_ISL_3093751 | Berkshire and Surrey Pathology Services Lighthouse Laboratory | Wellcome Sanger Institute for the COVID-19 Genomics UK (COG-UK) Consortium | Berkshire and Surrey Pathology Services Lighthouse Laboratory and Alex Alderton, Roberto Amato, Jeffrey Barrett, Sonia Goncalves, Ewan Harrison, David K. Jackson, Ian Johnston, Dominic Kwiatkowski, Cordelia Langford, John Sillitoe on behalf of the Wellcome Sanger Institute COVID-19 Surveillance Team |
| EPI_ISL_3093752, EPI_ISL_3093753 | Lighthouse Laboratory Plymouth | Wellcome Sanger Institute for the COVID-19 Genomics UK (COG-UK) Consortium | Lighthouse Laboratory Plymouth and Alex Alderton, Roberto Amato, Jeffrey Barrett, Sonia Goncalves, Ewan Harrison, David K. Jackson, Ian Johnston, Dominic Kwiatkowski, Cordelia Langford, John Sillitoe on behalf of the Wellcome Sanger Institute COVID-19 Surveillance Team |
| EPI_ISL_3093754 | Berkshire and Surrey Pathology Services Lighthouse Laboratory | Wellcome Sanger Institute for the COVID-19 Genomics UK (COG-UK) Consortium | Berkshire and Surrey Pathology Services Lighthouse Laboratory and Alex Alderton, Roberto Amato, Jeffrey Barrett, Sonia Goncalves, Ewan Harrison, David K. Jackson, Ian Johnston, Dominic Kwiatkowski, Cordelia Langford, John Sillitoe on behalf of the Wellcome Sanger Institute COVID-19 Surveillance Team |
| EPI_ISL_3093755 | Lighthouse Laboratory Plymouth | Wellcome Sanger Institute for the COVID-19 Genomics UK (COG-UK) Consortium | Lighthouse Laboratory Plymouth and Alex Alderton, Roberto Amato, Jeffrey Barrett, Sonia Goncalves, Ewan Harrison, David K. Jackson, Ian Johnston, Dominic Kwiatkowski, Cordelia Langford, John Sillitoe on behalf of the Wellcome Sanger Institute COVID-19 Surveillance Team |
| EPI_ISL_3093756 | Berkshire and Surrey Pathology Services Lighthouse Laboratory | Wellcome Sanger Institute for the COVID-19 Genomics UK (COG-UK) Consortium | Berkshire and Surrey Pathology Services Lighthouse Laboratory and Alex Alderton, Roberto Amato, Jeffrey Barrett, Sonia Goncalves, Ewan Harrison, David K. Jackson, Ian Johnston, Dominic Kwiatkowski, Cordelia Langford, John Sillitoe on behalf of the Wellcome Sanger Institute COVID-19 Surveillance Team |
| EPI_ISL_3093757, EPI_ISL_3093758 | Lighthouse Laboratory Plymouth | Wellcome Sanger Institute for the COVID-19 Genomics UK (COG-UK) Consortium | Lighthouse Laboratory Plymouth and Alex Alderton, Roberto Amato, Jeffrey Barrett, Sonia Goncalves, Ewan Harrison, David K. Jackson, Ian Johnston, Dominic Kwiatkowski, Cordelia Langford, John Sillitoe on behalf of the Wellcome Sanger Institute COVID-19 Surveillance Team |
| EPI_ISL_3093759 | Berkshire and Surrey Pathology Services Lighthouse Laboratory | Wellcome Sanger Institute for the COVID-19 Genomics UK (COG-UK) Consortium | Berkshire and Surrey Pathology Services Lighthouse Laboratory and Alex Alderton, Roberto Amato, Jeffrey Barrett, Sonia Goncalves, Ewan Harrison, David K. Jackson, Ian Johnston, Dominic Kwiatkowski, Cordelia Langford, John Sillitoe on behalf of the Wellcome Sanger Institute COVID-19 Surveillance Team |
| EPI_ISL_3093760, EPI_ISL_3093761 | Lighthouse Laboratory Plymouth | Wellcome Sanger Institute for the COVID-19 Genomics UK (COG-UK) Consortium | Lighthouse Laboratory Plymouth and Alex Alderton, Roberto Amato, Jeffrey Barrett, Sonia Goncalves, Ewan Harrison, David K. Jackson, Ian Johnston, Dominic Kwiatkowski, Cordelia Langford, John Sillitoe on behalf of the Wellcome Sanger Institute COVID-19 Surveillance Team |
| EPI_ISL_3093762 | Berkshire and Surrey Pathology Services Lighthouse Laboratory | Wellcome Sanger Institute for the COVID-19 Genomics UK (COG-UK) Consortium | Berkshire and Surrey Pathology Services Lighthouse Laboratory and Alex Alderton, Roberto Amato, Jeffrey Barrett, Sonia Goncalves, Ewan Harrison, David K. Jackson, Ian Johnston, Dominic Kwiatkowski, Cordelia Langford, John Sillitoe on behalf of the Wellcome Sanger Institute COVID-19 Surveillance Team |
| EPI_ISL_3093763, EPI_ISL_3093764, EPI_ISL_3093765 | Lighthouse Laboratory Plymouth | Wellcome Sanger Institute for the COVID-19 Genomics UK (COG-UK) Consortium | Lighthouse Laboratory Plymouth and Alex Alderton, Roberto Amato, Jeffrey Barrett, Sonia Goncalves, Ewan Harrison, David K. Jackson, Ian Johnston, Dominic Kwiatkowski, Cordelia Langford, John Sillitoe on behalf of the Wellcome Sanger Institute COVID-19 Surveillance Team |
| EPI_ISL_3093766 | Berkshire and Surrey Pathology Services Lighthouse Laboratory | Wellcome Sanger Institute for the COVID-19 Genomics UK (COG-UK) Consortium | Berkshire and Surrey Pathology Services Lighthouse Laboratory and Alex Alderton, Roberto Amato, Jeffrey Barrett, Sonia Goncalves, Ewan Harrison, David K. Jackson, Ian Johnston, Dominic Kwiatkowski, Cordelia Langford, John Sillitoe on behalf of the Wellcome Sanger Institute COVID-19 Surveillance Team |
| EPI_ISL_3093767, EPI_ISL_3093768 | Lighthouse Laboratory Plymouth | Wellcome Sanger Institute for the COVID-19 Genomics UK (COG-UK) Consortium | Lighthouse Laboratory Plymouth and Alex Alderton, Roberto Amato, Jeffrey Barrett, Sonia Goncalves, Ewan Harrison, David K. Jackson, Ian Johnston, Dominic Kwiatkowski, Cordelia Langford, John Sillitoe on behalf of the Wellcome Sanger Institute COVID-19 Surveillance Team |
| EPI_ISL_3093769, EPI_ISL_3093770 | Berkshire and Surrey Pathology Services Lighthouse Laboratory | Wellcome Sanger Institute for the COVID-19 Genomics UK (COG-UK) Consortium | Berkshire and Surrey Pathology Services Lighthouse Laboratory and Alex Alderton, Roberto Amato, Jeffrey Barrett, Sonia Goncalves, Ewan Harrison, David K. Jackson, Ian Johnston, Dominic Kwiatkowski, Cordelia Langford, John Sillitoe on behalf of the Wellcome Sanger Institute COVID-19 Surveillance Team |
| EPI_ISL_3093771 | Lighthouse Laboratory Plymouth | Wellcome Sanger Institute for the COVID-19 Genomics UK (COG-UK) Consortium | Lighthouse Laboratory Plymouth and Alex Alderton, Roberto Amato, Jeffrey Barrett, Sonia Goncalves, Ewan Harrison, David K. Jackson, Ian Johnston, Dominic Kwiatkowski, Cordelia Langford, John Sillitoe on behalf of the Wellcome Sanger Institute COVID-19 Surveillance Team |
| EPI_ISL_3093772, EPI_ISL_3093773 | Berkshire and Surrey Pathology Services Lighthouse Laboratory | Wellcome Sanger Institute for the COVID-19 Genomics UK (COG-UK) Consortium | Berkshire and Surrey Pathology Services Lighthouse Laboratory and Alex Alderton, Roberto Amato, Jeffrey Barrett, Sonia Goncalves, Ewan Harrison, David K. Jackson, Ian Johnston, Dominic Kwiatkowski, Cordelia Langford, John Sillitoe on behalf of the Wellcome Sanger Institute COVID-19 Surveillance Team |
| EPI_ISL_3093774, EPI_ISL_3093775 | Lighthouse Laboratory Plymouth | Wellcome Sanger Institute for the COVID-19 Genomics UK (COG-UK) Consortium | Lighthouse Laboratory Plymouth and Alex Alderton, Roberto Amato, Jeffrey Barrett, Sonia Goncalves, Ewan Harrison, David K. Jackson, Ian Johnston, Dominic Kwiatkowski, Cordelia Langford, John Sillitoe on behalf of the Wellcome Sanger Institute COVID-19 Surveillance Team |
| EPI_ISL_3093776 | Berkshire and Surrey Pathology Services Lighthouse Laboratory | Wellcome Sanger Institute for the COVID-19 Genomics UK (COG-UK) Consortium | Berkshire and Surrey Pathology Services Lighthouse Laboratory and Alex Alderton, Roberto Amato, Jeffrey Barrett, Sonia Goncalves, Ewan Harrison, David K. Jackson, Ian Johnston, Dominic Kwiatkowski, Cordelia Langford, John Sillitoe on behalf of the Wellcome Sanger Institute COVID-19 Surveillance Team |
| EPI_ISL_3093777, EPI_ISL_3093778 | Lighthouse Laboratory Plymouth | Wellcome Sanger Institute for the COVID-19 Genomics UK (COG-UK) Consortium | Lighthouse Laboratory Plymouth and Alex Alderton, Roberto Amato, Jeffrey Barrett, Sonia Goncalves, Ewan Harrison, David K. Jackson, Ian Johnston, Dominic Kwiatkowski, Cordelia Langford, John Sillitoe on behalf of the Wellcome Sanger Institute COVID-19 Surveillance Team |
| EPI_ISL_3093779 | Berkshire and Surrey Pathology Services Lighthouse Laboratory | Wellcome Sanger Institute for the COVID-19 Genomics UK (COG-UK) Consortium | Berkshire and Surrey Pathology Services Lighthouse Laboratory and Alex Alderton, Roberto Amato, Jeffrey Barrett, Sonia Goncalves, Ewan Harrison, David K. Jackson, Ian Johnston, Dominic Kwiatkowski, Cordelia Langford, John Sillitoe on behalf of the Wellcome Sanger Institute COVID-19 Surveillance Team |
| EPI_ISL_3093780, EPI_ISL_3093781, EPI_ISL_3093782, EPI_ISL_3093783, EPI_ISL_3093784, EPI_ISL_3093785, EPI_ISL_3093786 | Lighthouse Laboratory Plymouth | Wellcome Sanger Institute for the COVID-19 Genomics UK (COG-UK) Consortium | Lighthouse Laboratory Plymouth and Alex Alderton, Roberto Amato, Jeffrey Barrett, Sonia Goncalves, Ewan Harrison, David K. Jackson, Ian Johnston, Dominic Kwiatkowski, Cordelia Langford, John Sillitoe on behalf of the Wellcome Sanger Institute COVID-19 Surveillance Team |
| EPI_ISL_3093787, EPI_ISL_3093788, EPI_ISL_3093789, EPI_ISL_3093790 | Berkshire and Surrey Pathology Services Lighthouse Laboratory | Wellcome Sanger Institute for the COVID-19 Genomics UK (COG-UK) Consortium | Berkshire and Surrey Pathology Services Lighthouse Laboratory and Alex Alderton, Roberto Amato, Jeffrey Barrett, Sonia Goncalves, Ewan Harrison, David K. Jackson, Ian Johnston, Dominic Kwiatkowski, Cordelia Langford, John Sillitoe on behalf of the Wellcome Sanger Institute COVID-19 Surveillance Team |
| EPI_ISL_3093791 | Lighthouse Laboratory Plymouth | Wellcome Sanger Institute for the COVID-19 Genomics UK (COG-UK) Consortium | Lighthouse Laboratory Plymouth and Alex Alderton, Roberto Amato, Jeffrey Barrett, Sonia Goncalves, Ewan Harrison, David K. Jackson, Ian Johnston, Dominic Kwiatkowski, Cordelia Langford, John Sillitoe on behalf of the Wellcome Sanger Institute COVID-19 Surveillance Team |
| EPI_ISL_3093792, EPI_ISL_3093793, EPI_ISL_3093794 | Berkshire and Surrey Pathology Services Lighthouse Laboratory | Wellcome Sanger Institute for the COVID-19 Genomics UK (COG-UK) Consortium | Berkshire and Surrey Pathology Services Lighthouse Laboratory and Alex Alderton, Roberto Amato, Jeffrey Barrett, Sonia Goncalves, Ewan Harrison, David K. Jackson, Ian Johnston, Dominic Kwiatkowski, Cordelia Langford, John Sillitoe on behalf of the Wellcome Sanger Institute COVID-19 Surveillance Team |
| EPI_ISL_3093795 | Lighthouse Laboratory Plymouth | Wellcome Sanger Institute for the COVID-19 Genomics UK (COG-UK) Consortium | Lighthouse Laboratory Plymouth and Alex Alderton, Roberto Amato, Jeffrey Barrett, Sonia Goncalves, Ewan Harrison, David K. Jackson, Ian Johnston, Dominic Kwiatkowski, Cordelia Langford, John Sillitoe on behalf of the Wellcome Sanger Institute COVID-19 Surveillance Team |
| EPI_ISL_3093796 | Berkshire and Surrey Pathology Services Lighthouse Laboratory | Wellcome Sanger Institute for the COVID-19 Genomics UK (COG-UK) Consortium | Berkshire and Surrey Pathology Services Lighthouse Laboratory and Alex Alderton, Roberto Amato, Jeffrey Barrett, Sonia Goncalves, Ewan Harrison, David K. Jackson, Ian Johnston, Dominic Kwiatkowski, Cordelia Langford, John Sillitoe on behalf of the Wellcome Sanger Institute COVID-19 Surveillance Team |
| EPI_ISL_3093797 | Lighthouse Laboratory Plymouth | Wellcome Sanger Institute for the COVID-19 Genomics UK (COG-UK) Consortium | Lighthouse Laboratory Plymouth and Alex Alderton, Roberto Amato, Jeffrey Barrett, Sonia Goncalves, Ewan Harrison, David K. Jackson, Ian Johnston, Dominic Kwiatkowski, Cordelia Langford, John Sillitoe on behalf of the Wellcome Sanger Institute COVID-19 Surveillance Team |
| EPI_ISL_3093798, EPI_ISL_3093799 | Berkshire and Surrey Pathology Services Lighthouse Laboratory | Wellcome Sanger Institute for the COVID-19 Genomics UK (COG-UK) Consortium | Berkshire and Surrey Pathology Services Lighthouse Laboratory and Alex Alderton, Roberto Amato, Jeffrey Barrett, Sonia Goncalves, Ewan Harrison, David K. Jackson, Ian Johnston, Dominic Kwiatkowski, Cordelia Langford, John Sillitoe on behalf of the Wellcome Sanger Institute COVID-19 Surveillance Team |
| EPI_ISL_3093800, EPI_ISL_3093801, EPI_ISL_3093802 | Lighthouse Laboratory Plymouth | Wellcome Sanger Institute for the COVID-19 Genomics UK (COG-UK) Consortium | Lighthouse Laboratory Plymouth and Alex Alderton, Roberto Amato, Jeffrey Barrett, Sonia Goncalves, Ewan Harrison, David K. Jackson, Ian Johnston, Dominic Kwiatkowski, Cordelia Langford, John Sillitoe on behalf of the Wellcome Sanger Institute COVID-19 Surveillance Team |
| EPI_ISL_3093803 | Berkshire and Surrey Pathology Services Lighthouse Laboratory | Wellcome Sanger Institute for the COVID-19 Genomics UK (COG-UK) Consortium | Berkshire and Surrey Pathology Services Lighthouse Laboratory and Alex Alderton, Roberto Amato, Jeffrey Barrett, Sonia Goncalves, Ewan Harrison, David K. Jackson, Ian Johnston, Dominic Kwiatkowski, Cordelia Langford, John Sillitoe on behalf of the Wellcome Sanger Institute COVID-19 Surveillance Team |
| EPI_ISL_3093804, EPI_ISL_3093805, EPI_ISL_3093806, EPI_ISL_3093807 | Lighthouse Laboratory Plymouth | Wellcome Sanger Institute for the COVID-19 Genomics UK (COG-UK) Consortium | Lighthouse Laboratory Plymouth and Alex Alderton, Roberto Amato, Jeffrey Barrett, Sonia Goncalves, Ewan Harrison, David K. Jackson, Ian Johnston, Dominic Kwiatkowski, Cordelia Langford, John Sillitoe on behalf of the Wellcome Sanger Institute COVID-19 Surveillance Team |

| | | | |
|---|---|---|---|
| EPI_ISL_3093808 | Berkshire and Surrey Pathology Services Lighthouse Laboratory | Wellcome Sanger Institute for the COVID-19 Genomics UK (COG-UK) Consortium | Berkshire and Surrey Pathology Services Lighthouse Laboratory and Alex Alderton, Roberto Amato, Jeffrey Barrett, Sonia Goncalves, Ewan Harrison, David K. Jackson, Ian Johnston, Dominic Kwiatkowski, Cordelia Langford, John Sillitoe on behalf of the Wellcome Sanger Institute COVID-19 Surveillance Team |
| EPI_ISL_3093809 | Lighthouse Laboratory Plymouth | Wellcome Sanger Institute for the COVID-19 Genomics UK (COG-UK) Consortium | Lighthouse Laboratory Plymouth and Alex Alderton, Roberto Amato, Jeffrey Barrett, Sonia Goncalves, Ewan Harrison, David K. Jackson, Ian Johnston, Dominic Kwiatkowski, Cordelia Langford, John Sillitoe on behalf of the Wellcome Sanger Institute COVID-19 Surveillance Team |
| EPI_ISL_3093810, EPI_ISL_3093811 | Berkshire and Surrey Pathology Services Lighthouse Laboratory | Wellcome Sanger Institute for the COVID-19 Genomics UK (COG-UK) Consortium | Berkshire and Surrey Pathology Services Lighthouse Laboratory and Alex Alderton, Roberto Amato, Jeffrey Barrett, Sonia Goncalves, Ewan Harrison, David K. Jackson, Ian Johnston, Dominic Kwiatkowski, Cordelia Langford, John Sillitoe on behalf of the Wellcome Sanger Institute COVID-19 Surveillance Team |
| EPI_ISL_3093812, EPI_ISL_3093813 | Lighthouse Laboratory Plymouth | Wellcome Sanger Institute for the COVID-19 Genomics UK (COG-UK) Consortium | Lighthouse Laboratory Plymouth and Alex Alderton, Roberto Amato, Jeffrey Barrett, Sonia Goncalves, Ewan Harrison, David K. Jackson, Ian Johnston, Dominic Kwiatkowski, Cordelia Langford, John Sillitoe on behalf of the Wellcome Sanger Institute COVID-19 Surveillance Team |
| EPI_ISL_3093814 | Berkshire and Surrey Pathology Services Lighthouse Laboratory | Wellcome Sanger Institute for the COVID-19 Genomics UK (COG-UK) Consortium | Berkshire and Surrey Pathology Services Lighthouse Laboratory and Alex Alderton, Roberto Amato, Jeffrey Barrett, Sonia Goncalves, Ewan Harrison, David K. Jackson, Ian Johnston, Dominic Kwiatkowski, Cordelia Langford, John Sillitoe on behalf of the Wellcome Sanger Institute COVID-19 Surveillance Team |
| EPI_ISL_3093815, EPI_ISL_3093816, EPI_ISL_3093817, EPI_ISL_3093818 | Lighthouse Laboratory Plymouth | Wellcome Sanger Institute for the COVID-19 Genomics UK (COG-UK) Consortium | Lighthouse Laboratory Plymouth and Alex Alderton, Roberto Amato, Jeffrey Barrett, Sonia Goncalves, Ewan Harrison, David K. Jackson, Ian Johnston, Dominic Kwiatkowski, Cordelia Langford, John Sillitoe on behalf of the Wellcome Sanger Institute COVID-19 Surveillance Team |
| EPI_ISL_3093819, EPI_ISL_3093820 | Berkshire and Surrey Pathology Services Lighthouse Laboratory | Wellcome Sanger Institute for the COVID-19 Genomics UK (COG-UK) Consortium | Berkshire and Surrey Pathology Services Lighthouse Laboratory and Alex Alderton, Roberto Amato, Jeffrey Barrett, Sonia Goncalves, Ewan Harrison, David K. Jackson, Ian Johnston, Dominic Kwiatkowski, Cordelia Langford, John Sillitoe on behalf of the Wellcome Sanger Institute COVID-19 Surveillance Team |
| EPI_ISL_3093821, EPI_ISL_3093822, EPI_ISL_3093823, EPI_ISL_3093824 | Lighthouse Laboratory Plymouth | Wellcome Sanger Institute for the COVID-19 Genomics UK (COG-UK) Consortium | Lighthouse Laboratory Plymouth and Alex Alderton, Roberto Amato, Jeffrey Barrett, Sonia Goncalves, Ewan Harrison, David K. Jackson, Ian Johnston, Dominic Kwiatkowski, Cordelia Langford, John Sillitoe on behalf of the Wellcome Sanger Institute COVID-19 Surveillance Team |
| EPI_ISL_3093825 | Berkshire and Surrey Pathology Services Lighthouse Laboratory | Wellcome Sanger Institute for the COVID-19 Genomics UK (COG-UK) Consortium | Berkshire and Surrey Pathology Services Lighthouse Laboratory and Alex Alderton, Roberto Amato, Jeffrey Barrett, Sonia Goncalves, Ewan Harrison, David K. Jackson, Ian Johnston, Dominic Kwiatkowski, Cordelia Langford, John Sillitoe on behalf of the Wellcome Sanger Institute COVID-19 Surveillance Team |
| EPI_ISL_3093826 | Lighthouse Laboratory Plymouth | Wellcome Sanger Institute for the COVID-19 Genomics UK (COG-UK) Consortium | Lighthouse Laboratory Plymouth and Alex Alderton, Roberto Amato, Jeffrey Barrett, Sonia Goncalves, Ewan Harrison, David K. Jackson, Ian Johnston, Dominic Kwiatkowski, Cordelia Langford, John Sillitoe on behalf of the Wellcome Sanger Institute COVID-19 Surveillance Team |
| EPI_ISL_3093827, EPI_ISL_3093828 | Berkshire and Surrey Pathology Services Lighthouse Laboratory | Wellcome Sanger Institute for the COVID-19 Genomics UK (COG-UK) Consortium | Berkshire and Surrey Pathology Services Lighthouse Laboratory and Alex Alderton, Roberto Amato, Jeffrey Barrett, Sonia Goncalves, Ewan Harrison, David K. Jackson, Ian Johnston, Dominic Kwiatkowski, Cordelia Langford, John Sillitoe on behalf of the Wellcome Sanger Institute COVID-19 Surveillance Team |
| EPI_ISL_3093829, EPI_ISL_3093830, EPI_ISL_3093831 | Lighthouse Laboratory Plymouth | Wellcome Sanger Institute for the COVID-19 Genomics UK (COG-UK) Consortium | Lighthouse Laboratory Plymouth and Alex Alderton, Roberto Amato, Jeffrey Barrett, Sonia Goncalves, Ewan Harrison, David K. Jackson, Ian Johnston, Dominic Kwiatkowski, Cordelia Langford, John Sillitoe on behalf of the Wellcome Sanger Institute COVID-19 Surveillance Team |
| EPI_ISL_3093832, EPI_ISL_3093833 | Berkshire and Surrey Pathology Services Lighthouse Laboratory | Wellcome Sanger Institute for the COVID-19 Genomics UK (COG-UK) Consortium | Berkshire and Surrey Pathology Services Lighthouse Laboratory and Alex Alderton, Roberto Amato, Jeffrey Barrett, Sonia Goncalves, Ewan Harrison, David K. Jackson, Ian Johnston, Dominic Kwiatkowski, Cordelia Langford, John Sillitoe on behalf of the Wellcome Sanger Institute COVID-19 Surveillance Team |
| EPI_ISL_3093834, EPI_ISL_3093835, EPI_ISL_3093836 | Lighthouse Laboratory Plymouth | Wellcome Sanger Institute for the COVID-19 Genomics UK (COG-UK) Consortium | Lighthouse Laboratory Plymouth and Alex Alderton, Roberto Amato, Jeffrey Barrett, Sonia Goncalves, Ewan Harrison, David K. Jackson, Ian Johnston, Dominic Kwiatkowski, Cordelia Langford, John Sillitoe on behalf of the Wellcome Sanger Institute COVID-19 Surveillance Team |
| EPI_ISL_3093837 | Berkshire and Surrey Pathology Services Lighthouse Laboratory | Wellcome Sanger Institute for the COVID-19 Genomics UK (COG-UK) Consortium | Berkshire and Surrey Pathology Services Lighthouse Laboratory and Alex Alderton, Roberto Amato, Jeffrey Barrett, Sonia Goncalves, Ewan Harrison, David K. Jackson, Ian Johnston, Dominic Kwiatkowski, Cordelia Langford, John Sillitoe on behalf of the Wellcome Sanger Institute COVID-19 Surveillance Team |
| EPI_ISL_3093838, EPI_ISL_3093839, EPI_ISL_3093840, EPI_ISL_3093841 | Lighthouse Laboratory Plymouth | Wellcome Sanger Institute for the COVID-19 Genomics UK (COG-UK) Consortium | Lighthouse Laboratory Plymouth and Alex Alderton, Roberto Amato, Jeffrey Barrett, Sonia Goncalves, Ewan Harrison, David K. Jackson, Ian Johnston, Dominic Kwiatkowski, Cordelia Langford, John Sillitoe on behalf of the Wellcome Sanger Institute COVID-19 Surveillance Team |
| EPI_ISL_3093842, EPI_ISL_3093843 | Lighthouse Lab in Alderley Park | Wellcome Sanger Institute for the COVID-19 Genomics UK (COG-UK) Consortium | Jacquelyn Wynn, Mairead Hyland, The Lighthouse Lab in Alderley Park and Alex Alderton, Roberto Amato, Jeffrey Barrett, Sonia Goncalves, Ewan Harrison, David K. Jackson, Ian Johnston, Dominic Kwiatkowski, Cordelia Langford, John Sillitoe on behalf of the Wellcome Sanger Institute COVID-19 Surveillance Team |
| EPI_ISL_3093844, EPI_ISL_3093845 | Berkshire and Surrey Pathology Services Lighthouse Laboratory | Wellcome Sanger Institute for the COVID-19 Genomics UK (COG-UK) Consortium | Berkshire and Surrey Pathology Services Lighthouse Laboratory and Alex Alderton, Roberto Amato, Jeffrey Barrett, Sonia Goncalves, Ewan Harrison, David K. Jackson, Ian Johnston, Dominic Kwiatkowski, Cordelia Langford, John Sillitoe on behalf of the Wellcome Sanger Institute COVID-19 Surveillance Team |
| EPI_ISL_3093846, EPI_ISL_3093847 | Lighthouse Lab in Alderley Park | Wellcome Sanger Institute for the COVID-19 Genomics UK (COG-UK) Consortium | Jacquelyn Wynn, Mairead Hyland, The Lighthouse Lab in Alderley Park and Alex Alderton, Roberto Amato, Jeffrey Barrett, Sonia Goncalves, Ewan Harrison, David K. Jackson, Ian Johnston, Dominic Kwiatkowski, Cordelia Langford, John Sillitoe on behalf of the Wellcome Sanger Institute COVID-19 Surveillance Team |
| EPI_ISL_3093848 | Berkshire and Surrey Pathology Services Lighthouse Laboratory | Wellcome Sanger Institute for the COVID-19 Genomics UK (COG-UK) Consortium | Berkshire and Surrey Pathology Services Lighthouse Laboratory and Alex Alderton, Roberto Amato, Jeffrey Barrett, Sonia Goncalves, Ewan Harrison, David K. Jackson, Ian Johnston, Dominic Kwiatkowski, Cordelia Langford, John Sillitoe on behalf of the Wellcome Sanger Institute COVID-19 Surveillance Team |
| EPI_ISL_3093849, EPI_ISL_3093850, EPI_ISL_3093851 | Lighthouse Lab in Alderley Park | Wellcome Sanger Institute for the COVID-19 Genomics UK (COG-UK) Consortium | Jacquelyn Wynn, Mairead Hyland, The Lighthouse Lab in Alderley Park and Alex Alderton, Roberto Amato, Jeffrey Barrett, Sonia Goncalves, Ewan Harrison, David K. Jackson, Ian Johnston, Dominic Kwiatkowski, Cordelia Langford, John Sillitoe on behalf of the Wellcome Sanger Institute COVID-19 Surveillance Team |
| EPI_ISL_3093852, EPI_ISL_3093853 | Lighthouse Lab in Glasgow | Wellcome Sanger Institute for the COVID-19 Genomics UK (COG-UK) Consortium | Harper VanSteenhouse, Yumi Kasai, David Gray, Carol Clugston, Anna Dominiczak and Alex Alderton, Roberto Amato, Jeffrey Barrett, Sonia Goncalves, Ewan Harrison, David K. Jackson, Ian Johnston, Dominic Kwiatkowski, Cordelia Langford, John Sillitoe on behalf of the Wellcome Sanger Institute COVID-19 Surveillance Team |
| EPI_ISL_3093854, EPI_ISL_3093855 | Lighthouse Lab in Alderley Park | Wellcome Sanger Institute for the COVID-19 Genomics UK (COG-UK) Consortium | Jacquelyn Wynn, Mairead Hyland, The Lighthouse Lab in Alderley Park and Alex Alderton, Roberto Amato, Jeffrey Barrett, Sonia Goncalves, Ewan Harrison, David K. Jackson, Ian Johnston, Dominic Kwiatkowski, Cordelia Langford, John Sillitoe on behalf of the Wellcome Sanger Institute COVID-19 Surveillance Team |
| EPI_ISL_3093856 | Lighthouse Lab in Glasgow | Wellcome Sanger Institute for the COVID-19 Genomics UK (COG-UK) Consortium | Harper VanSteenhouse, Yumi Kasai, David Gray, Carol Clugston, Anna Dominiczak and Alex Alderton, Roberto Amato, Jeffrey Barrett, Sonia Goncalves, Ewan Harrison, David K. Jackson, Ian Johnston, Dominic Kwiatkowski, Cordelia Langford, John Sillitoe on behalf of the Wellcome Sanger Institute COVID-19 Surveillance Team |
| EPI_ISL_3093857 | Lighthouse Lab in Alderley Park | Wellcome Sanger Institute for the COVID-19 Genomics UK (COG-UK) Consortium | Jacquelyn Wynn, Mairead Hyland, The Lighthouse Lab in Alderley Park and Alex Alderton, Roberto Amato, Jeffrey Barrett, Sonia Goncalves, Ewan Harrison, David K. Jackson, Ian Johnston, Dominic Kwiatkowski, Cordelia Langford, John Sillitoe on behalf of the Wellcome Sanger Institute COVID-19 Surveillance Team |
| EPI_ISL_3093858 | Berkshire and Surrey Pathology Services Lighthouse Laboratory | Wellcome Sanger Institute for the COVID-19 Genomics UK (COG-UK) Consortium | Berkshire and Surrey Pathology Services Lighthouse Laboratory and Alex Alderton, Roberto Amato, Jeffrey Barrett, Sonia Goncalves, Ewan Harrison, David K. Jackson, Ian Johnston, Dominic Kwiatkowski, Cordelia Langford, John Sillitoe on behalf of the Wellcome Sanger Institute COVID-19 Surveillance Team |
| EPI_ISL_3093859 | Lighthouse Lab in Alderley Park | Wellcome Sanger Institute for the COVID-19 Genomics UK (COG-UK) Consortium | Jacquelyn Wynn, Mairead Hyland, The Lighthouse Lab in Alderley Park and Alex Alderton, Roberto Amato, Jeffrey Barrett, Sonia Goncalves, Ewan Harrison, David K. Jackson, Ian Johnston, Dominic Kwiatkowski, Cordelia Langford, John Sillitoe on behalf of the Wellcome Sanger Institute COVID-19 Surveillance Team |
| EPI_ISL_3093860, EPI_ISL_3093861, EPI_ISL_3093862 | Berkshire and Surrey Pathology Services Lighthouse Laboratory | Wellcome Sanger Institute for the COVID-19 Genomics UK (COG-UK) Consortium | Berkshire and Surrey Pathology Services Lighthouse Laboratory and Alex Alderton, Roberto Amato, Jeffrey Barrett, Sonia Goncalves, Ewan Harrison, David K. Jackson, Ian Johnston, Dominic Kwiatkowski, Cordelia Langford, John Sillitoe on behalf of the Wellcome Sanger Institute COVID-19 Surveillance Team |
| EPI_ISL_3093863, EPI_ISL_3093864 | Lighthouse Lab in Glasgow | Wellcome Sanger Institute for the COVID-19 Genomics UK (COG-UK) Consortium | Harper VanSteenhouse, Yumi Kasai, David Gray, Carol Clugston, Anna Dominiczak and Alex Alderton, Roberto Amato, Jeffrey Barrett, Sonia Goncalves, Ewan Harrison, David K. Jackson, Ian Johnston, Dominic Kwiatkowski, Cordelia Langford, John Sillitoe on behalf of the Wellcome Sanger Institute COVID-19 Surveillance Team |
| EPI_ISL_3093865, EPI_ISL_3093866 | Lighthouse Lab in Alderley Park | Wellcome Sanger Institute for the COVID-19 Genomics UK (COG-UK) Consortium | Jacquelyn Wynn, Mairead Hyland, The Lighthouse Lab in Alderley Park and Alex Alderton, Roberto Amato, Jeffrey Barrett, Sonia Goncalves, Ewan Harrison, David K. Jackson, Ian Johnston, Dominic Kwiatkowski, Cordelia Langford, John Sillitoe on behalf of the Wellcome Sanger Institute COVID-19 Surveillance Team |
| EPI_ISL_3093867, EPI_ISL_3093868, EPI_ISL_3093869 | Lighthouse Lab in Glasgow | Wellcome Sanger Institute for the COVID-19 Genomics UK (COG-UK) Consortium | Harper VanSteenhouse, Yumi Kasai, David Gray, Carol Clugston, Anna Dominiczak and Alex Alderton, Roberto Amato, Jeffrey Barrett, Sonia Goncalves, Ewan Harrison, David K. Jackson, Ian Johnston, Dominic Kwiatkowski, Cordelia Langford, John Sillitoe on behalf of the Wellcome Sanger Institute COVID-19 Surveillance Team |
| EPI_ISL_3093870, EPI_ISL_3093871 | Lighthouse Lab in Alderley Park | Wellcome Sanger Institute for the COVID-19 Genomics UK (COG-UK) Consortium | Jacquelyn Wynn, Mairead Hyland, The Lighthouse Lab in Alderley Park and Alex Alderton, Roberto Amato, Jeffrey Barrett, Sonia Goncalves, Ewan Harrison, David K. Jackson, Ian Johnston, Dominic Kwiatkowski, Cordelia Langford, John Sillitoe on behalf of the Wellcome Sanger Institute COVID-19 Surveillance Team |
| EPI_ISL_3093872 | Berkshire and Surrey Pathology Services Lighthouse Laboratory | Wellcome Sanger Institute for the COVID-19 Genomics UK (COG-UK) Consortium | Berkshire and Surrey Pathology Services Lighthouse Laboratory and Alex Alderton, Roberto Amato, Jeffrey Barrett, Sonia Goncalves, Ewan Harrison, David K. Jackson, Ian Johnston, Dominic Kwiatkowski, Cordelia Langford, John Sillitoe on behalf of the Wellcome Sanger Institute COVID-19 Surveillance Team |
| EPI_ISL_3093873 | Lighthouse Lab in Alderley Park | Wellcome Sanger Institute for the COVID-19 Genomics UK (COG-UK) Consortium | Jacquelyn Wynn, Mairead Hyland, The Lighthouse Lab in Alderley Park and Alex Alderton, Roberto Amato, Jeffrey Barrett, Sonia Goncalves, Ewan Harrison, David K. Jackson, Ian Johnston, Dominic Kwiatkowski, Cordelia Langford, John Sillitoe on behalf of the Wellcome Sanger Institute COVID-19 Surveillance Team |
| EPI_ISL_3093874 | Berkshire and Surrey Pathology Services Lighthouse Laboratory | Wellcome Sanger Institute for the COVID-19 Genomics UK (COG-UK) Consortium | Berkshire and Surrey Pathology Services Lighthouse Laboratory and Alex Alderton, Roberto Amato, Jeffrey Barrett, Sonia Goncalves, Ewan Harrison, David K. Jackson, Ian Johnston, Dominic Kwiatkowski, Cordelia Langford, John Sillitoe on behalf of the Wellcome Sanger Institute COVID-19 Surveillance Team |
| EPI_ISL_3093875, EPI_ISL_3093876, EPI_ISL_3093877, EPI_ISL_3093878 | Lighthouse Lab in Alderley Park | Wellcome Sanger Institute for the COVID-19 Genomics UK (COG-UK) Consortium | Jacquelyn Wynn, Mairead Hyland, The Lighthouse Lab in Alderley Park and Alex Alderton, Roberto Amato, Jeffrey Barrett, Sonia Goncalves, Ewan Harrison, David K. Jackson, Ian Johnston, Dominic Kwiatkowski, Cordelia Langford, John Sillitoe on behalf of the Wellcome Sanger Institute COVID-19 Surveillance Team |
| EPI_ISL_3093879 | Lighthouse Lab in Glasgow | Wellcome Sanger Institute for the COVID-19 Genomics UK (COG-UK) Consortium | Harper VanSteenhouse, Yumi Kasai, David Gray, Carol Clugston, Anna Dominiczak and Alex Alderton, Roberto Amato, Jeffrey Barrett, Sonia Goncalves, Ewan Harrison, David K. Jackson, Ian Johnston, Dominic Kwiatkowski, Cordelia Langford, John Sillitoe on behalf of the Wellcome Sanger Institute COVID-19 Surveillance Team |
| EPI_ISL_3093880 | Berkshire and Surrey Pathology Services Lighthouse Laboratory | Wellcome Sanger Institute for the COVID-19 Genomics UK (COG-UK) Consortium | Berkshire and Surrey Pathology Services Lighthouse Laboratory and Alex Alderton, Roberto Amato, Jeffrey Barrett, Sonia Goncalves, Ewan Harrison, David K. Jackson, Ian Johnston, Dominic Kwiatkowski, Cordelia Langford, John Sillitoe on behalf of the Wellcome Sanger Institute COVID-19 Surveillance Team |
| EPI_ISL_3093881 | Lighthouse Lab in Glasgow | Wellcome Sanger Institute for the COVID-19 Genomics UK (COG-UK) Consortium | Harper VanSteenhouse, Yumi Kasai, David Gray, Carol Clugston, Anna Dominiczak and Alex Alderton, Roberto Amato, Jeffrey Barrett, Sonia Goncalves, Ewan Harrison, David K. Jackson, Ian Johnston, Dominic Kwiatkowski, Cordelia Langford, John Sillitoe on behalf of the Wellcome Sanger Institute COVID-19 Surveillance Team |
| EPI_ISL_3093882 | Lighthouse Lab in Alderley Park | Wellcome Sanger Institute for the COVID-19 Genomics UK (COG-UK) Consortium | Jacquelyn Wynn, Mairead Hyland, The Lighthouse Lab in Alderley Park and Alex Alderton, Roberto Amato, Jeffrey Barrett, Sonia Goncalves, Ewan Harrison, David K. Jackson, Ian Johnston, Dominic Kwiatkowski, Cordelia Langford, John Sillitoe on behalf of the Wellcome Sanger Institute COVID-19 Surveillance Team |

| | | | |
|---|---|---|---|
| EPI_ISL_3093883 | Berkshire and Surrey Pathology Services Lighthouse Laboratory | Wellcome Sanger Institute for the COVID-19 Genomics UK (COG-UK) Consortium | Berkshire and Surrey Pathology Services Lighthouse Laboratory and Alex Alderton, Roberto Amato, Jeffrey Barrett, Sonia Goncalves, Ewan Harrison, David K. Jackson, Ian Johnston, Dominic Kwiatkowski, Cordelia Langford, John Sillitoe on behalf of the Wellcome Sanger Institute COVID-19 Surveillance Team |
| EPI_ISL_3093884 | Lighthouse Lab in Alderley Park | Wellcome Sanger Institute for the COVID-19 Genomics UK (COG-UK) Consortium | Jacquelyn Wynn, Mairead Hyland, The Lighthouse Lab in Alderley Park and Alex Alderton, Roberto Amato, Jeffrey Barrett, Sonia Goncalves, Ewan Harrison, David K. Jackson, Ian Johnston, Dominic Kwiatkowski, Cordelia Langford, John Sillitoe on behalf of the Wellcome Sanger Institute COVID-19 Surveillance Team |
| EPI_ISL_3093885 | Lighthouse Lab in Glasgow | Wellcome Sanger Institute for the COVID-19 Genomics UK (COG-UK) Consortium | Harper VanSteenhouse, Yumi Kasai, David Gray, Carol Clugston, Anna Dominiczak and Alex Alderton, Roberto Amato, Jeffrey Barrett, Sonia Goncalves, Ewan Harrison, David K. Jackson, Ian Johnston, Dominic Kwiatkowski, Cordelia Langford, John Sillitoe on behalf of the Wellcome Sanger Institute COVID-19 Surveillance Team |
| EPI_ISL_3093886 | Lighthouse Lab in Alderley Park | Wellcome Sanger Institute for the COVID-19 Genomics UK (COG-UK) Consortium | Jacquelyn Wynn, Mairead Hyland, The Lighthouse Lab in Alderley Park and Alex Alderton, Roberto Amato, Jeffrey Barrett, Sonia Goncalves, Ewan Harrison, David K. Jackson, Ian Johnston, Dominic Kwiatkowski, Cordelia Langford, John Sillitoe on behalf of the Wellcome Sanger Institute COVID-19 Surveillance Team |
| EPI_ISL_3093887, EPI_ISL_3093888 | Lighthouse Lab in Glasgow | Wellcome Sanger Institute for the COVID-19 Genomics UK (COG-UK) Consortium | Harper VanSteenhouse, Yumi Kasai, David Gray, Carol Clugston, Anna Dominiczak and Alex Alderton, Roberto Amato, Jeffrey Barrett, Sonia Goncalves, Ewan Harrison, David K. Jackson, Ian Johnston, Dominic Kwiatkowski, Cordelia Langford, John Sillitoe on behalf of the Wellcome Sanger Institute COVID-19 Surveillance Team |
| EPI_ISL_3093889 | Lighthouse Lab in Alderley Park | Wellcome Sanger Institute for the COVID-19 Genomics UK (COG-UK) Consortium | Jacquelyn Wynn, Mairead Hyland, The Lighthouse Lab in Alderley Park and Alex Alderton, Roberto Amato, Jeffrey Barrett, Sonia Goncalves, Ewan Harrison, David K. Jackson, Ian Johnston, Dominic Kwiatkowski, Cordelia Langford, John Sillitoe on behalf of the Wellcome Sanger Institute COVID-19 Surveillance Team |
| EPI_ISL_3093890 | Lighthouse Lab in Glasgow | Wellcome Sanger Institute for the COVID-19 Genomics UK (COG-UK) Consortium | Harper VanSteenhouse, Yumi Kasai, David Gray, Carol Clugston, Anna Dominiczak and Alex Alderton, Roberto Amato, Jeffrey Barrett, Sonia Goncalves, Ewan Harrison, David K. Jackson, Ian Johnston, Dominic Kwiatkowski, Cordelia Langford, John Sillitoe on behalf of the Wellcome Sanger Institute COVID-19 Surveillance Team |
| EPI_ISL_3093891, EPI_ISL_3093892 | Lighthouse Lab in Alderley Park | Wellcome Sanger Institute for the COVID-19 Genomics UK (COG-UK) Consortium | Jacquelyn Wynn, Mairead Hyland, The Lighthouse Lab in Alderley Park and Alex Alderton, Roberto Amato, Jeffrey Barrett, Sonia Goncalves, Ewan Harrison, David K. Jackson, Ian Johnston, Dominic Kwiatkowski, Cordelia Langford, John Sillitoe on behalf of the Wellcome Sanger Institute COVID-19 Surveillance Team |
| EPI_ISL_3093893 | Berkshire and Surrey Pathology Services Lighthouse Laboratory | Wellcome Sanger Institute for the COVID-19 Genomics UK (COG-UK) Consortium | Berkshire and Surrey Pathology Services Lighthouse Laboratory and Alex Alderton, Roberto Amato, Jeffrey Barrett, Sonia Goncalves, Ewan Harrison, David K. Jackson, Ian Johnston, Dominic Kwiatkowski, Cordelia Langford, John Sillitoe on behalf of the Wellcome Sanger Institute COVID-19 Surveillance Team |
| EPI_ISL_3093894 | Lighthouse Lab in Alderley Park | Wellcome Sanger Institute for the COVID-19 Genomics UK (COG-UK) Consortium | Jacquelyn Wynn, Mairead Hyland, The Lighthouse Lab in Alderley Park and Alex Alderton, Roberto Amato, Jeffrey Barrett, Sonia Goncalves, Ewan Harrison, David K. Jackson, Ian Johnston, Dominic Kwiatkowski, Cordelia Langford, John Sillitoe on behalf of the Wellcome Sanger Institute COVID-19 Surveillance Team |
| EPI_ISL_3093895 | Berkshire and Surrey Pathology Services Lighthouse Laboratory | Wellcome Sanger Institute for the COVID-19 Genomics UK (COG-UK) Consortium | Berkshire and Surrey Pathology Services Lighthouse Laboratory and Alex Alderton, Roberto Amato, Jeffrey Barrett, Sonia Goncalves, Ewan Harrison, David K. Jackson, Ian Johnston, Dominic Kwiatkowski, Cordelia Langford, John Sillitoe on behalf of the Wellcome Sanger Institute COVID-19 Surveillance Team |
| EPI_ISL_3093896, EPI_ISL_3093897 | Lighthouse Lab in Alderley Park | Wellcome Sanger Institute for the COVID-19 Genomics UK (COG-UK) Consortium | Jacquelyn Wynn, Mairead Hyland, The Lighthouse Lab in Alderley Park and Alex Alderton, Roberto Amato, Jeffrey Barrett, Sonia Goncalves, Ewan Harrison, David K. Jackson, Ian Johnston, Dominic Kwiatkowski, Cordelia Langford, John Sillitoe on behalf of the Wellcome Sanger Institute COVID-19 Surveillance Team |
| EPI_ISL_3093898 | Berkshire and Surrey Pathology Services Lighthouse Laboratory | Wellcome Sanger Institute for the COVID-19 Genomics UK (COG-UK) Consortium | Berkshire and Surrey Pathology Services Lighthouse Laboratory and Alex Alderton, Roberto Amato, Jeffrey Barrett, Sonia Goncalves, Ewan Harrison, David K. Jackson, Ian Johnston, Dominic Kwiatkowski, Cordelia Langford, John Sillitoe on behalf of the Wellcome Sanger Institute COVID-19 Surveillance Team |
| EPI_ISL_3093899 | Lighthouse Lab in Alderley Park | Wellcome Sanger Institute for the COVID-19 Genomics UK (COG-UK) Consortium | Jacquelyn Wynn, Mairead Hyland, The Lighthouse Lab in Alderley Park and Alex Alderton, Roberto Amato, Jeffrey Barrett, Sonia Goncalves, Ewan Harrison, David K. Jackson, Ian Johnston, Dominic Kwiatkowski, Cordelia Langford, John Sillitoe on behalf of the Wellcome Sanger Institute COVID-19 Surveillance Team |
| EPI_ISL_3093900 | Lighthouse Lab in Glasgow | Wellcome Sanger Institute for the COVID-19 Genomics UK (COG-UK) Consortium | Harper VanSteenhouse, Yumi Kasai, David Gray, Carol Clugston, Anna Dominiczak and Alex Alderton, Roberto Amato, Jeffrey Barrett, Sonia Goncalves, Ewan Harrison, David K. Jackson, Ian Johnston, Dominic Kwiatkowski, Cordelia Langford, John Sillitoe on behalf of the Wellcome Sanger Institute COVID-19 Surveillance Team |
| EPI_ISL_3093901 | Berkshire and Surrey Pathology Services Lighthouse Laboratory | Wellcome Sanger Institute for the COVID-19 Genomics UK (COG-UK) Consortium | Berkshire and Surrey Pathology Services Lighthouse Laboratory and Alex Alderton, Roberto Amato, Jeffrey Barrett, Sonia Goncalves, Ewan Harrison, David K. Jackson, Ian Johnston, Dominic Kwiatkowski, Cordelia Langford, John Sillitoe on behalf of the Wellcome Sanger Institute COVID-19 Surveillance Team |
| EPI_ISL_3093902 | Lighthouse Lab in Alderley Park | Wellcome Sanger Institute for the COVID-19 Genomics UK (COG-UK) Consortium | Jacquelyn Wynn, Mairead Hyland, The Lighthouse Lab in Alderley Park and Alex Alderton, Roberto Amato, Jeffrey Barrett, Sonia Goncalves, Ewan Harrison, David K. Jackson, Ian Johnston, Dominic Kwiatkowski, Cordelia Langford, John Sillitoe on behalf of the Wellcome Sanger Institute COVID-19 Surveillance Team |
| EPI_ISL_3093903 | Berkshire and Surrey Pathology Services Lighthouse Laboratory | Wellcome Sanger Institute for the COVID-19 Genomics UK (COG-UK) Consortium | Berkshire and Surrey Pathology Services Lighthouse Laboratory and Alex Alderton, Roberto Amato, Jeffrey Barrett, Sonia Goncalves, Ewan Harrison, David K. Jackson, Ian Johnston, Dominic Kwiatkowski, Cordelia Langford, John Sillitoe on behalf of the Wellcome Sanger Institute COVID-19 Surveillance Team |
| EPI_ISL_3093904 | Lighthouse Lab in Glasgow | Wellcome Sanger Institute for the COVID-19 Genomics UK (COG-UK) Consortium | Harper VanSteenhouse, Yumi Kasai, David Gray, Carol Clugston, Anna Dominiczak and Alex Alderton, Roberto Amato, Jeffrey Barrett, Sonia Goncalves, Ewan Harrison, David K. Jackson, Ian Johnston, Dominic Kwiatkowski, Cordelia Langford, John Sillitoe on behalf of the Wellcome Sanger Institute COVID-19 Surveillance Team |
| EPI_ISL_3093905, EPI_ISL_3093906, EPI_ISL_3093907 | Lighthouse Lab in Alderley Park | Wellcome Sanger Institute for the COVID-19 Genomics UK (COG-UK) Consortium | Jacquelyn Wynn, Mairead Hyland, The Lighthouse Lab in Alderley Park and Alex Alderton, Roberto Amato, Jeffrey Barrett, Sonia Goncalves, Ewan Harrison, David K. Jackson, Ian Johnston, Dominic Kwiatkowski, Cordelia Langford, John Sillitoe on behalf of the Wellcome Sanger Institute COVID-19 Surveillance Team |
| EPI_ISL_3093908 | Berkshire and Surrey Pathology Services Lighthouse Laboratory | Wellcome Sanger Institute for the COVID-19 Genomics UK (COG-UK) Consortium | Berkshire and Surrey Pathology Services Lighthouse Laboratory and Alex Alderton, Roberto Amato, Jeffrey Barrett, Sonia Goncalves, Ewan Harrison, David K. Jackson, Ian Johnston, Dominic Kwiatkowski, Cordelia Langford, John Sillitoe on behalf of the Wellcome Sanger Institute COVID-19 Surveillance Team |
| EPI_ISL_3093909 | Lighthouse Lab in Alderley Park | Wellcome Sanger Institute for the COVID-19 Genomics UK (COG-UK) Consortium | Jacquelyn Wynn, Mairead Hyland, The Lighthouse Lab in Alderley Park and Alex Alderton, Roberto Amato, Jeffrey Barrett, Sonia Goncalves, Ewan Harrison, David K. Jackson, Ian Johnston, Dominic Kwiatkowski, Cordelia Langford, John Sillitoe on behalf of the Wellcome Sanger Institute COVID-19 Surveillance Team |
| EPI_ISL_3093910 | Lighthouse Lab in Glasgow | Wellcome Sanger Institute for the COVID-19 Genomics UK (COG-UK) Consortium | Harper VanSteenhouse, Yumi Kasai, David Gray, Carol Clugston, Anna Dominiczak and Alex Alderton, Roberto Amato, Jeffrey Barrett, Sonia Goncalves, Ewan Harrison, David K. Jackson, Ian Johnston, Dominic Kwiatkowski, Cordelia Langford, John Sillitoe on behalf of the Wellcome Sanger Institute COVID-19 Surveillance Team |
| EPI_ISL_3093911 | Berkshire and Surrey Pathology Services Lighthouse Laboratory | Wellcome Sanger Institute for the COVID-19 Genomics UK (COG-UK) Consortium | Berkshire and Surrey Pathology Services Lighthouse Laboratory and Alex Alderton, Roberto Amato, Jeffrey Barrett, Sonia Goncalves, Ewan Harrison, David K. Jackson, Ian Johnston, Dominic Kwiatkowski, Cordelia Langford, John Sillitoe on behalf of the Wellcome Sanger Institute COVID-19 Surveillance Team |
| EPI_ISL_3093912, EPI_ISL_3093913 | Lighthouse Lab in Alderley Park | Wellcome Sanger Institute for the COVID-19 Genomics UK (COG-UK) Consortium | Jacquelyn Wynn, Mairead Hyland, The Lighthouse Lab in Alderley Park and Alex Alderton, Roberto Amato, Jeffrey Barrett, Sonia Goncalves, Ewan Harrison, David K. Jackson, Ian Johnston, Dominic Kwiatkowski, Cordelia Langford, John Sillitoe on behalf of the Wellcome Sanger Institute COVID-19 Surveillance Team |
| EPI_ISL_3093914 | Berkshire and Surrey Pathology Services Lighthouse Laboratory | Wellcome Sanger Institute for the COVID-19 Genomics UK (COG-UK) Consortium | Berkshire and Surrey Pathology Services Lighthouse Laboratory and Alex Alderton, Roberto Amato, Jeffrey Barrett, Sonia Goncalves, Ewan Harrison, David K. Jackson, Ian Johnston, Dominic Kwiatkowski, Cordelia Langford, John Sillitoe on behalf of the Wellcome Sanger Institute COVID-19 Surveillance Team |
| EPI_ISL_3093915, EPI_ISL_3093916, EPI_ISL_3093917 | Lighthouse Lab in Alderley Park | Wellcome Sanger Institute for the COVID-19 Genomics UK (COG-UK) Consortium | Jacquelyn Wynn, Mairead Hyland, The Lighthouse Lab in Alderley Park and Alex Alderton, Roberto Amato, Jeffrey Barrett, Sonia Goncalves, Ewan Harrison, David K. Jackson, Ian Johnston, Dominic Kwiatkowski, Cordelia Langford, John Sillitoe on behalf of the Wellcome Sanger Institute COVID-19 Surveillance Team |
| EPI_ISL_3093918 | Lighthouse Lab in Glasgow | Wellcome Sanger Institute for the COVID-19 Genomics UK (COG-UK) Consortium | Harper VanSteenhouse, Yumi Kasai, David Gray, Carol Clugston, Anna Dominiczak and Alex Alderton, Roberto Amato, Jeffrey Barrett, Sonia Goncalves, Ewan Harrison, David K. Jackson, Ian Johnston, Dominic Kwiatkowski, Cordelia Langford, John Sillitoe on behalf of the Wellcome Sanger Institute COVID-19 Surveillance Team |
| EPI_ISL_3093919, EPI_ISL_3093920, EPI_ISL_3093921 | Lighthouse Lab in Alderley Park | Wellcome Sanger Institute for the COVID-19 Genomics UK (COG-UK) Consortium | Jacquelyn Wynn, Mairead Hyland, The Lighthouse Lab in Alderley Park and Alex Alderton, Roberto Amato, Jeffrey Barrett, Sonia Goncalves, Ewan Harrison, David K. Jackson, Ian Johnston, Dominic Kwiatkowski, Cordelia Langford, John Sillitoe on behalf of the Wellcome Sanger Institute COVID-19 Surveillance Team |
| EPI_ISL_3093922 | Lighthouse Lab in Glasgow | Wellcome Sanger Institute for the COVID-19 Genomics UK (COG-UK) Consortium | Harper VanSteenhouse, Yumi Kasai, David Gray, Carol Clugston, Anna Dominiczak and Alex Alderton, Roberto Amato, Jeffrey Barrett, Sonia Goncalves, Ewan Harrison, David K. Jackson, Ian Johnston, Dominic Kwiatkowski, Cordelia Langford, John Sillitoe on behalf of the Wellcome Sanger Institute COVID-19 Surveillance Team |
| EPI_ISL_3093923 | Lighthouse Lab in Alderley Park | Wellcome Sanger Institute for the COVID-19 Genomics UK (COG-UK) Consortium | Jacquelyn Wynn, Mairead Hyland, The Lighthouse Lab in Alderley Park and Alex Alderton, Roberto Amato, Jeffrey Barrett, Sonia Goncalves, Ewan Harrison, David K. Jackson, Ian Johnston, Dominic Kwiatkowski, Cordelia Langford, John Sillitoe on behalf of the Wellcome Sanger Institute COVID-19 Surveillance Team |
| EPI_ISL_3093924 | Lighthouse Lab in Glasgow | Wellcome Sanger Institute for the COVID-19 Genomics UK (COG-UK) Consortium | Harper VanSteenhouse, Yumi Kasai, David Gray, Carol Clugston, Anna Dominiczak and Alex Alderton, Roberto Amato, Jeffrey Barrett, Sonia Goncalves, Ewan Harrison, David K. Jackson, Ian Johnston, Dominic Kwiatkowski, Cordelia Langford, John Sillitoe on behalf of the Wellcome Sanger Institute COVID-19 Surveillance Team |
| EPI_ISL_3093925, EPI_ISL_3093926 | Berkshire and Surrey Pathology Services Lighthouse Laboratory | Wellcome Sanger Institute for the COVID-19 Genomics UK (COG-UK) Consortium | Berkshire and Surrey Pathology Services Lighthouse Laboratory and Alex Alderton, Roberto Amato, Jeffrey Barrett, Sonia Goncalves, Ewan Harrison, David K. Jackson, Ian Johnston, Dominic Kwiatkowski, Cordelia Langford, John Sillitoe on behalf of the Wellcome Sanger Institute COVID-19 Surveillance Team |
| EPI_ISL_3093927, EPI_ISL_3093928 | Lighthouse Lab in Alderley Park | Wellcome Sanger Institute for the COVID-19 Genomics UK (COG-UK) Consortium | Jacquelyn Wynn, Mairead Hyland, The Lighthouse Lab in Alderley Park and Alex Alderton, Roberto Amato, Jeffrey Barrett, Sonia Goncalves, Ewan Harrison, David K. Jackson, Ian Johnston, Dominic Kwiatkowski, Cordelia Langford, John Sillitoe on behalf of the Wellcome Sanger Institute COVID-19 Surveillance Team |
| EPI_ISL_3093929 | Berkshire and Surrey Pathology Services Lighthouse Laboratory | Wellcome Sanger Institute for the COVID-19 Genomics UK (COG-UK) Consortium | Berkshire and Surrey Pathology Services Lighthouse Laboratory and Alex Alderton, Roberto Amato, Jeffrey Barrett, Sonia Goncalves, Ewan Harrison, David K. Jackson, Ian Johnston, Dominic Kwiatkowski, Cordelia Langford, John Sillitoe on behalf of the Wellcome Sanger Institute COVID-19 Surveillance Team |
| EPI_ISL_3093930, EPI_ISL_3093931 | Lighthouse Lab in Alderley Park | Wellcome Sanger Institute for the COVID-19 Genomics UK (COG-UK) Consortium | Jacquelyn Wynn, Mairead Hyland, The Lighthouse Lab in Alderley Park and Alex Alderton, Roberto Amato, Jeffrey Barrett, Sonia Goncalves, Ewan Harrison, David K. Jackson, Ian Johnston, Dominic Kwiatkowski, Cordelia Langford, John Sillitoe on behalf of the Wellcome Sanger Institute COVID-19 Surveillance Team |
| EPI_ISL_3093932, EPI_ISL_3093933 | Lighthouse Lab in Glasgow | Wellcome Sanger Institute for the COVID-19 Genomics UK (COG-UK) Consortium | Harper VanSteenhouse, Yumi Kasai, David Gray, Carol Clugston, Anna Dominiczak and Alex Alderton, Roberto Amato, Jeffrey Barrett, Sonia Goncalves, Ewan Harrison, David K. Jackson, Ian Johnston, Dominic Kwiatkowski, Cordelia Langford, John Sillitoe on behalf of the Wellcome Sanger Institute COVID-19 Surveillance Team |
| EPI_ISL_3093934, EPI_ISL_3093935 | Lighthouse Lab in Alderley Park | Wellcome Sanger Institute for the COVID-19 Genomics UK (COG-UK) Consortium | Jacquelyn Wynn, Mairead Hyland, The Lighthouse Lab in Alderley Park and Alex Alderton, Roberto Amato, Jeffrey Barrett, Sonia Goncalves, Ewan Harrison, David K. Jackson, Ian Johnston, Dominic Kwiatkowski, Cordelia Langford, John Sillitoe on behalf of the Wellcome Sanger Institute COVID-19 Surveillance Team |
| EPI_ISL_3093936 | Berkshire and Surrey Pathology Services Lighthouse Laboratory | Wellcome Sanger Institute for the COVID-19 Genomics UK (COG-UK) Consortium | Berkshire and Surrey Pathology Services Lighthouse Laboratory and Alex Alderton, Roberto Amato, Jeffrey Barrett, Sonia Goncalves, Ewan Harrison, David K. Jackson, Ian Johnston, Dominic Kwiatkowski, Cordelia Langford, John Sillitoe on behalf of the Wellcome Sanger Institute COVID-19 Surveillance Team |
| EPI_ISL_3093937, EPI_ISL_3093938 | Lighthouse Lab in Alderley Park | Wellcome Sanger Institute for the COVID-19 Genomics UK (COG-UK) Consortium | Jacquelyn Wynn, Mairead Hyland, The Lighthouse Lab in Alderley Park and Alex Alderton, Roberto Amato, Jeffrey Barrett, Sonia Goncalves, Ewan Harrison, David K. Jackson, Ian Johnston, Dominic Kwiatkowski, Cordelia Langford, John Sillitoe on behalf of the Wellcome Sanger Institute COVID-19 Surveillance Team |
| EPI_ISL_3093939 | Berkshire and Surrey Pathology Services Lighthouse Laboratory | Wellcome Sanger Institute for the COVID-19 Genomics UK (COG-UK) Consortium | Berkshire and Surrey Pathology Services Lighthouse Laboratory and Alex Alderton, Roberto Amato, Jeffrey Barrett, Sonia Goncalves, Ewan Harrison, David K. Jackson, Ian Johnston, Dominic Kwiatkowski, Cordelia Langford, John Sillitoe on behalf of the Wellcome Sanger Institute COVID-19 Surveillance Team |
| EPI_ISL_3093940 | Lighthouse Lab in Alderley Park | Wellcome Sanger Institute for the COVID-19 Genomics UK (COG-UK) Consortium | Jacquelyn Wynn, Mairead Hyland, The Lighthouse Lab in Alderley Park and Alex Alderton, Roberto Amato, Jeffrey Barrett, Sonia Goncalves, Ewan Harrison, David K. Jackson, Ian Johnston, Dominic Kwiatkowski, Cordelia Langford, John Sillitoe on behalf of the Wellcome Sanger Institute COVID-19 Surveillance Team |
| EPI_ISL_3093941 | Berkshire and Surrey Pathology Services Lighthouse Laboratory | Wellcome Sanger Institute for the COVID-19 Genomics UK (COG-UK) Consortium | Berkshire and Surrey Pathology Services Lighthouse Laboratory and Alex Alderton, Roberto Amato, Jeffrey Barrett, Sonia Goncalves, Ewan Harrison, David K. Jackson, Ian Johnston, Dominic Kwiatkowski, Cordelia Langford, John Sillitoe on behalf of the Wellcome Sanger Institute COVID-19 Surveillance Team |
| EPI_ISL_3093942 | Lighthouse Lab in Glasgow | Wellcome Sanger Institute for the COVID-19 Genomics UK (COG-UK) Consortium | Harper VanSteenhouse, Yumi Kasai, David Gray, Carol Clugston, Anna Dominiczak and Alex Alderton, Roberto Amato, Jeffrey Barrett, Sonia Goncalves, Ewan Harrison, David K. Jackson, Ian Johnston, Dominic Kwiatkowski, Cordelia Langford, John Sillitoe on behalf of the Wellcome Sanger Institute COVID-19 Surveillance Team |

| | | | |
|---|---|---|---|
| EPI_ISL_3093943 | Lighthouse Lab in Alderley Park | Wellcome Sanger Institute for the COVID-19 Genomics UK (COG-UK) Consortium | Jacquelyn Wynn, Mairead Hyland, The Lighthouse Lab in Alderley Park and Alex Alderton, Roberto Amato, Jeffrey Barrett, Sonia Goncalves, Ewan Harrison, David K. Jackson, Ian Johnston, Dominic Kwiatkowski, Cordelia Langford, John Sillitoe on behalf of the Wellcome Sanger Institute COVID-19 Surveillance Team |
| EPI_ISL_3093944 | Berkshire and Surrey Pathology Services Lighthouse Laboratory | Wellcome Sanger Institute for the COVID-19 Genomics UK (COG-UK) Consortium | Berkshire and Surrey Pathology Services Lighthouse Laboratory and Alex Alderton, Roberto Amato, Jeffrey Barrett, Sonia Goncalves, Ewan Harrison, David K. Jackson, Ian Johnston, Dominic Kwiatkowski, Cordelia Langford, John Sillitoe on behalf of the Wellcome Sanger Institute COVID-19 Surveillance Team |
| EPI_ISL_3093945 | Lighthouse Lab in Glasgow | Wellcome Sanger Institute for the COVID-19 Genomics UK (COG-UK) Consortium | Harper VanSteenhouse, Yumi Kasai, David Gray, Carol Clugston, Anna Dominiczak and Alex Alderton, Roberto Amato, Jeffrey Barrett, Sonia Goncalves, Ewan Harrison, David K. Jackson, Ian Johnston, Dominic Kwiatkowski, Cordelia Langford, John Sillitoe on behalf of the Wellcome Sanger Institute COVID-19 Surveillance Team |
| EPI_ISL_3093946 | Lighthouse Lab in Alderley Park | Wellcome Sanger Institute for the COVID-19 Genomics UK (COG-UK) Consortium | Jacquelyn Wynn, Mairead Hyland, The Lighthouse Lab in Alderley Park and Alex Alderton, Roberto Amato, Jeffrey Barrett, Sonia Goncalves, Ewan Harrison, David K. Jackson, Ian Johnston, Dominic Kwiatkowski, Cordelia Langford, John Sillitoe on behalf of the Wellcome Sanger Institute COVID-19 Surveillance Team |
| EPI_ISL_3093947 | Berkshire and Surrey Pathology Services Lighthouse Laboratory | Wellcome Sanger Institute for the COVID-19 Genomics UK (COG-UK) Consortium | Berkshire and Surrey Pathology Services Lighthouse Laboratory and Alex Alderton, Roberto Amato, Jeffrey Barrett, Sonia Goncalves, Ewan Harrison, David K. Jackson, Ian Johnston, Dominic Kwiatkowski, Cordelia Langford, John Sillitoe on behalf of the Wellcome Sanger Institute COVID-19 Surveillance Team |
| EPI_ISL_3093948, EPI_ISL_3093949 | Lighthouse Lab in Alderley Park | Wellcome Sanger Institute for the COVID-19 Genomics UK (COG-UK) Consortium | Jacquelyn Wynn, Mairead Hyland, The Lighthouse Lab in Alderley Park and Alex Alderton, Roberto Amato, Jeffrey Barrett, Sonia Goncalves, Ewan Harrison, David K. Jackson, Ian Johnston, Dominic Kwiatkowski, Cordelia Langford, John Sillitoe on behalf of the Wellcome Sanger Institute COVID-19 Surveillance Team |
| EPI_ISL_3093950 | Lighthouse Lab in Glasgow | Wellcome Sanger Institute for the COVID-19 Genomics UK (COG-UK) Consortium | Harper VanSteenhouse, Yumi Kasai, David Gray, Carol Clugston, Anna Dominiczak and Alex Alderton, Roberto Amato, Jeffrey Barrett, Sonia Goncalves, Ewan Harrison, David K. Jackson, Ian Johnston, Dominic Kwiatkowski, Cordelia Langford, John Sillitoe on behalf of the Wellcome Sanger Institute COVID-19 Surveillance Team |
| EPI_ISL_3093951, EPI_ISL_3093952 | Berkshire and Surrey Pathology Services Lighthouse Laboratory | Wellcome Sanger Institute for the COVID-19 Genomics UK (COG-UK) Consortium | Berkshire and Surrey Pathology Services Lighthouse Laboratory and Alex Alderton, Roberto Amato, Jeffrey Barrett, Sonia Goncalves, Ewan Harrison, David K. Jackson, Ian Johnston, Dominic Kwiatkowski, Cordelia Langford, John Sillitoe on behalf of the Wellcome Sanger Institute COVID-19 Surveillance Team |
| EPI_ISL_3093953 | Lighthouse Lab in Alderley Park | Wellcome Sanger Institute for the COVID-19 Genomics UK (COG-UK) Consortium | Jacquelyn Wynn, Mairead Hyland, The Lighthouse Lab in Alderley Park and Alex Alderton, Roberto Amato, Jeffrey Barrett, Sonia Goncalves, Ewan Harrison, David K. Jackson, Ian Johnston, Dominic Kwiatkowski, Cordelia Langford, John Sillitoe on behalf of the Wellcome Sanger Institute COVID-19 Surveillance Team |
| EPI_ISL_3093954 | Berkshire and Surrey Pathology Services Lighthouse Laboratory | Wellcome Sanger Institute for the COVID-19 Genomics UK (COG-UK) Consortium | Berkshire and Surrey Pathology Services Lighthouse Laboratory and Alex Alderton, Roberto Amato, Jeffrey Barrett, Sonia Goncalves, Ewan Harrison, David K. Jackson, Ian Johnston, Dominic Kwiatkowski, Cordelia Langford, John Sillitoe on behalf of the Wellcome Sanger Institute COVID-19 Surveillance Team |
| EPI_ISL_3093955, EPI_ISL_3093956, EPI_ISL_3093957, EPI_ISL_3093959 | Lighthouse Lab in Alderley Park | Wellcome Sanger Institute for the COVID-19 Genomics UK (COG-UK) Consortium | Jacquelyn Wynn, Mairead Hyland, The Lighthouse Lab in Alderley Park and Alex Alderton, Roberto Amato, Jeffrey Barrett, Sonia Goncalves, Ewan Harrison, David K. Jackson, Ian Johnston, Dominic Kwiatkowski, Cordelia Langford, John Sillitoe on behalf of the Wellcome Sanger Institute COVID-19 Surveillance Team |
| EPI_ISL_3093960, EPI_ISL_3093962 | Lighthouse Lab in Glasgow | Wellcome Sanger Institute for the COVID-19 Genomics UK (COG-UK) Consortium | Harper VanSteenhouse, Yumi Kasai, David Gray, Carol Clugston, Anna Dominiczak and Alex Alderton, Roberto Amato, Jeffrey Barrett, Sonia Goncalves, Ewan Harrison, David K. Jackson, Ian Johnston, Dominic Kwiatkowski, Cordelia Langford, John Sillitoe on behalf of the Wellcome Sanger Institute COVID-19 Surveillance Team |
| EPI_ISL_3093963 | Lighthouse Lab in Alderley Park | Wellcome Sanger Institute for the COVID-19 Genomics UK (COG-UK) Consortium | Jacquelyn Wynn, Mairead Hyland, The Lighthouse Lab in Alderley Park and Alex Alderton, Roberto Amato, Jeffrey Barrett, Sonia Goncalves, Ewan Harrison, David K. Jackson, Ian Johnston, Dominic Kwiatkowski, Cordelia Langford, John Sillitoe on behalf of the Wellcome Sanger Institute COVID-19 Surveillance Team |
| EPI_ISL_3093964 | Lighthouse Lab in Glasgow | Wellcome Sanger Institute for the COVID-19 Genomics UK (COG-UK) Consortium | Harper VanSteenhouse, Yumi Kasai, David Gray, Carol Clugston, Anna Dominiczak and Alex Alderton, Roberto Amato, Jeffrey Barrett, Sonia Goncalves, Ewan Harrison, David K. Jackson, Ian Johnston, Dominic Kwiatkowski, Cordelia Langford, John Sillitoe on behalf of the Wellcome Sanger Institute COVID-19 Surveillance Team |
| EPI_ISL_3093965 | Lighthouse Lab in Alderley Park | Wellcome Sanger Institute for the COVID-19 Genomics UK (COG-UK) Consortium | Jacquelyn Wynn, Mairead Hyland, The Lighthouse Lab in Alderley Park and Alex Alderton, Roberto Amato, Jeffrey Barrett, Sonia Goncalves, Ewan Harrison, David K. Jackson, Ian Johnston, Dominic Kwiatkowski, Cordelia Langford, John Sillitoe on behalf of the Wellcome Sanger Institute COVID-19 Surveillance Team |
| EPI_ISL_3093966 | Lighthouse Lab in Glasgow | Wellcome Sanger Institute for the COVID-19 Genomics UK (COG-UK) Consortium | Harper VanSteenhouse, Yumi Kasai, David Gray, Carol Clugston, Anna Dominiczak and Alex Alderton, Roberto Amato, Jeffrey Barrett, Sonia Goncalves, Ewan Harrison, David K. Jackson, Ian Johnston, Dominic Kwiatkowski, Cordelia Langford, John Sillitoe on behalf of the Wellcome Sanger Institute COVID-19 Surveillance Team |
| EPI_ISL_3093967 | Lighthouse Lab in Alderley Park | Wellcome Sanger Institute for the COVID-19 Genomics UK (COG-UK) Consortium | Jacquelyn Wynn, Mairead Hyland, The Lighthouse Lab in Alderley Park and Alex Alderton, Roberto Amato, Jeffrey Barrett, Sonia Goncalves, Ewan Harrison, David K. Jackson, Ian Johnston, Dominic Kwiatkowski, Cordelia Langford, John Sillitoe on behalf of the Wellcome Sanger Institute COVID-19 Surveillance Team |
| EPI_ISL_3093968 | Lighthouse Lab in Glasgow | Wellcome Sanger Institute for the COVID-19 Genomics UK (COG-UK) Consortium | Harper VanSteenhouse, Yumi Kasai, David Gray, Carol Clugston, Anna Dominiczak and Alex Alderton, Roberto Amato, Jeffrey Barrett, Sonia Goncalves, Ewan Harrison, David K. Jackson, Ian Johnston, Dominic Kwiatkowski, Cordelia Langford, John Sillitoe on behalf of the Wellcome Sanger Institute COVID-19 Surveillance Team |
| EPI_ISL_3093969 | Berkshire and Surrey Pathology Services Lighthouse Laboratory | Wellcome Sanger Institute for the COVID-19 Genomics UK (COG-UK) Consortium | Berkshire and Surrey Pathology Services Lighthouse Laboratory and Alex Alderton, Roberto Amato, Jeffrey Barrett, Sonia Goncalves, Ewan Harrison, David K. Jackson, Ian Johnston, Dominic Kwiatkowski, Cordelia Langford, John Sillitoe on behalf of the Wellcome Sanger Institute COVID-19 Surveillance Team |
| EPI_ISL_3093970 | Lighthouse Lab in Alderley Park | Wellcome Sanger Institute for the COVID-19 Genomics UK (COG-UK) Consortium | Jacquelyn Wynn, Mairead Hyland, The Lighthouse Lab in Alderley Park and Alex Alderton, Roberto Amato, Jeffrey Barrett, Sonia Goncalves, Ewan Harrison, David K. Jackson, Ian Johnston, Dominic Kwiatkowski, Cordelia Langford, John Sillitoe on behalf of the Wellcome Sanger Institute COVID-19 Surveillance Team |
| EPI_ISL_3093971 | Berkshire and Surrey Pathology Services Lighthouse Laboratory | Wellcome Sanger Institute for the COVID-19 Genomics UK (COG-UK) Consortium | Berkshire and Surrey Pathology Services Lighthouse Laboratory and Alex Alderton, Roberto Amato, Jeffrey Barrett, Sonia Goncalves, Ewan Harrison, David K. Jackson, Ian Johnston, Dominic Kwiatkowski, Cordelia Langford, John Sillitoe on behalf of the Wellcome Sanger Institute COVID-19 Surveillance Team |
| EPI_ISL_3093973 | Lighthouse Lab in Alderley Park | Wellcome Sanger Institute for the COVID-19 Genomics UK (COG-UK) Consortium | Jacquelyn Wynn, Mairead Hyland, The Lighthouse Lab in Alderley Park and Alex Alderton, Roberto Amato, Jeffrey Barrett, Sonia Goncalves, Ewan Harrison, David K. Jackson, Ian Johnston, Dominic Kwiatkowski, Cordelia Langford, John Sillitoe on behalf of the Wellcome Sanger Institute COVID-19 Surveillance Team |
| EPI_ISL_3093974 | Lighthouse Lab in Glasgow | Wellcome Sanger Institute for the COVID-19 Genomics UK (COG-UK) Consortium | Harper VanSteenhouse, Yumi Kasai, David Gray, Carol Clugston, Anna Dominiczak and Alex Alderton, Roberto Amato, Jeffrey Barrett, Sonia Goncalves, Ewan Harrison, David K. Jackson, Ian Johnston, Dominic Kwiatkowski, Cordelia Langford, John Sillitoe on behalf of the Wellcome Sanger Institute COVID-19 Surveillance Team |
| EPI_ISL_3093975, EPI_ISL_3093976 | Lighthouse Lab in Alderley Park | Wellcome Sanger Institute for the COVID-19 Genomics UK (COG-UK) Consortium | Jacquelyn Wynn, Mairead Hyland, The Lighthouse Lab in Alderley Park and Alex Alderton, Roberto Amato, Jeffrey Barrett, Sonia Goncalves, Ewan Harrison, David K. Jackson, Ian Johnston, Dominic Kwiatkowski, Cordelia Langford, John Sillitoe on behalf of the Wellcome Sanger Institute COVID-19 Surveillance Team |
| EPI_ISL_3093978 | Berkshire and Surrey Pathology Services Lighthouse Laboratory | Wellcome Sanger Institute for the COVID-19 Genomics UK (COG-UK) Consortium | Berkshire and Surrey Pathology Services Lighthouse Laboratory and Alex Alderton, Roberto Amato, Jeffrey Barrett, Sonia Goncalves, Ewan Harrison, David K. Jackson, Ian Johnston, Dominic Kwiatkowski, Cordelia Langford, John Sillitoe on behalf of the Wellcome Sanger Institute COVID-19 Surveillance Team |
| EPI_ISL_3093979 | Lighthouse Lab in Glasgow | Wellcome Sanger Institute for the COVID-19 Genomics UK (COG-UK) Consortium | Harper VanSteenhouse, Yumi Kasai, David Gray, Carol Clugston, Anna Dominiczak and Alex Alderton, Roberto Amato, Jeffrey Barrett, Sonia Goncalves, Ewan Harrison, David K. Jackson, Ian Johnston, Dominic Kwiatkowski, Cordelia Langford, John Sillitoe on behalf of the Wellcome Sanger Institute COVID-19 Surveillance Team |
| EPI_ISL_3093981, EPI_ISL_3093982, EPI_ISL_3093983, EPI_ISL_3093985 | Lighthouse Lab in Alderley Park | Wellcome Sanger Institute for the COVID-19 Genomics UK (COG-UK) Consortium | Jacquelyn Wynn, Mairead Hyland, The Lighthouse Lab in Alderley Park and Alex Alderton, Roberto Amato, Jeffrey Barrett, Sonia Goncalves, Ewan Harrison, David K. Jackson, Ian Johnston, Dominic Kwiatkowski, Cordelia Langford, John Sillitoe on behalf of the Wellcome Sanger Institute COVID-19 Surveillance Team |
| EPI_ISL_3093986 | Lighthouse Lab in Glasgow | Wellcome Sanger Institute for the COVID-19 Genomics UK (COG-UK) Consortium | Harper VanSteenhouse, Yumi Kasai, David Gray, Carol Clugston, Anna Dominiczak and Alex Alderton, Roberto Amato, Jeffrey Barrett, Sonia Goncalves, Ewan Harrison, David K. Jackson, Ian Johnston, Dominic Kwiatkowski, Cordelia Langford, John Sillitoe on behalf of the Wellcome Sanger Institute COVID-19 Surveillance Team |
| EPI_ISL_3093987 | Berkshire and Surrey Pathology Services Lighthouse Laboratory | Wellcome Sanger Institute for the COVID-19 Genomics UK (COG-UK) Consortium | Berkshire and Surrey Pathology Services Lighthouse Laboratory and Alex Alderton, Roberto Amato, Jeffrey Barrett, Sonia Goncalves, Ewan Harrison, David K. Jackson, Ian Johnston, Dominic Kwiatkowski, Cordelia Langford, John Sillitoe on behalf of the Wellcome Sanger Institute COVID-19 Surveillance Team |
| EPI_ISL_3093989, EPI_ISL_3093990, EPI_ISL_3093992 | Lighthouse Lab in Alderley Park | Wellcome Sanger Institute for the COVID-19 Genomics UK (COG-UK) Consortium | Jacquelyn Wynn, Mairead Hyland, The Lighthouse Lab in Alderley Park and Alex Alderton, Roberto Amato, Jeffrey Barrett, Sonia Goncalves, Ewan Harrison, David K. Jackson, Ian Johnston, Dominic Kwiatkowski, Cordelia Langford, John Sillitoe on behalf of the Wellcome Sanger Institute COVID-19 Surveillance Team |
| EPI_ISL_3093993, EPI_ISL_3093994 | Berkshire and Surrey Pathology Services Lighthouse Laboratory | Wellcome Sanger Institute for the COVID-19 Genomics UK (COG-UK) Consortium | Berkshire and Surrey Pathology Services Lighthouse Laboratory and Alex Alderton, Roberto Amato, Jeffrey Barrett, Sonia Goncalves, Ewan Harrison, David K. Jackson, Ian Johnston, Dominic Kwiatkowski, Cordelia Langford, John Sillitoe on behalf of the Wellcome Sanger Institute COVID-19 Surveillance Team |
| EPI_ISL_3093995, EPI_ISL_3093997, EPI_ISL_3093998, EPI_ISL_3093999, EPI_ISL_3094001 | Lighthouse Lab in Alderley Park | Wellcome Sanger Institute for the COVID-19 Genomics UK (COG-UK) Consortium | Jacquelyn Wynn, Mairead Hyland, The Lighthouse Lab in Alderley Park and Alex Alderton, Roberto Amato, Jeffrey Barrett, Sonia Goncalves, Ewan Harrison, David K. Jackson, Ian Johnston, Dominic Kwiatkowski, Cordelia Langford, John Sillitoe on behalf of the Wellcome Sanger Institute COVID-19 Surveillance Team |
| EPI_ISL_3094002 | Lighthouse Lab in Glasgow | Wellcome Sanger Institute for the COVID-19 Genomics UK (COG-UK) Consortium | Harper VanSteenhouse, Yumi Kasai, David Gray, Carol Clugston, Anna Dominiczak and Alex Alderton, Roberto Amato, Jeffrey Barrett, Sonia Goncalves, Ewan Harrison, David K. Jackson, Ian Johnston, Dominic Kwiatkowski, Cordelia Langford, John Sillitoe on behalf of the Wellcome Sanger Institute COVID-19 Surveillance Team |
| EPI_ISL_3094003 | Lighthouse Lab in Alderley Park | Wellcome Sanger Institute for the COVID-19 Genomics UK (COG-UK) Consortium | Jacquelyn Wynn, Mairead Hyland, The Lighthouse Lab in Alderley Park and Alex Alderton, Roberto Amato, Jeffrey Barrett, Sonia Goncalves, Ewan Harrison, David K. Jackson, Ian Johnston, Dominic Kwiatkowski, Cordelia Langford, John Sillitoe on behalf of the Wellcome Sanger Institute COVID-19 Surveillance Team |
| EPI_ISL_3094005 | Berkshire and Surrey Pathology Services Lighthouse Laboratory | Wellcome Sanger Institute for the COVID-19 Genomics UK (COG-UK) Consortium | Berkshire and Surrey Pathology Services Lighthouse Laboratory and Alex Alderton, Roberto Amato, Jeffrey Barrett, Sonia Goncalves, Ewan Harrison, David K. Jackson, Ian Johnston, Dominic Kwiatkowski, Cordelia Langford, John Sillitoe on behalf of the Wellcome Sanger Institute COVID-19 Surveillance Team |
| EPI_ISL_3094006, EPI_ISL_3094007 | Lighthouse Lab in Glasgow | Wellcome Sanger Institute for the COVID-19 Genomics UK (COG-UK) Consortium | Harper VanSteenhouse, Yumi Kasai, David Gray, Carol Clugston, Anna Dominiczak and Alex Alderton, Roberto Amato, Jeffrey Barrett, Sonia Goncalves, Ewan Harrison, David K. Jackson, Ian Johnston, Dominic Kwiatkowski, Cordelia Langford, John Sillitoe on behalf of the Wellcome Sanger Institute COVID-19 Surveillance Team |
| EPI_ISL_3094008 | Lighthouse Lab in Alderley Park | Wellcome Sanger Institute for the COVID-19 Genomics UK (COG-UK) Consortium | Jacquelyn Wynn, Mairead Hyland, The Lighthouse Lab in Alderley Park and Alex Alderton, Roberto Amato, Jeffrey Barrett, Sonia Goncalves, Ewan Harrison, David K. Jackson, Ian Johnston, Dominic Kwiatkowski, Cordelia Langford, John Sillitoe on behalf of the Wellcome Sanger Institute COVID-19 Surveillance Team |
| EPI_ISL_3094009, EPI_ISL_3094011 | Berkshire and Surrey Pathology Services Lighthouse Laboratory | Wellcome Sanger Institute for the COVID-19 Genomics UK (COG-UK) Consortium | Berkshire and Surrey Pathology Services Lighthouse Laboratory and Alex Alderton, Roberto Amato, Jeffrey Barrett, Sonia Goncalves, Ewan Harrison, David K. Jackson, Ian Johnston, Dominic Kwiatkowski, Cordelia Langford, John Sillitoe on behalf of the Wellcome Sanger Institute COVID-19 Surveillance Team |
| EPI_ISL_3094012 | Lighthouse Lab in Glasgow | Wellcome Sanger Institute for the COVID-19 Genomics UK (COG-UK) Consortium | Harper VanSteenhouse, Yumi Kasai, David Gray, Carol Clugston, Anna Dominiczak and Alex Alderton, Roberto Amato, Jeffrey Barrett, Sonia Goncalves, Ewan Harrison, David K. Jackson, Ian Johnston, Dominic Kwiatkowski, Cordelia Langford, John Sillitoe on behalf of the Wellcome Sanger Institute COVID-19 Surveillance Team |
| EPI_ISL_3094013, EPI_ISL_3094015, EPI_ISL_3094016 | Lighthouse Lab in Alderley Park | Wellcome Sanger Institute for the COVID-19 Genomics UK (COG-UK) Consortium | Jacquelyn Wynn, Mairead Hyland, The Lighthouse Lab in Alderley Park and Alex Alderton, Roberto Amato, Jeffrey Barrett, Sonia Goncalves, Ewan Harrison, David K. Jackson, Ian Johnston, Dominic Kwiatkowski, Cordelia Langford, John Sillitoe on behalf of the Wellcome Sanger Institute COVID-19 Surveillance Team |
| EPI_ISL_3094017, EPI_ISL_3094019 | Lighthouse Lab in Glasgow | Wellcome Sanger Institute for the COVID-19 Genomics UK (COG-UK) Consortium | Harper VanSteenhouse, Yumi Kasai, David Gray, Carol Clugston, Anna Dominiczak and Alex Alderton, Roberto Amato, Jeffrey Barrett, Sonia Goncalves, Ewan Harrison, David K. Jackson, Ian Johnston, Dominic Kwiatkowski, Cordelia Langford, John Sillitoe on behalf of the Wellcome Sanger Institute COVID-19 Surveillance Team |

| | | | |
|---|---|---|---|
| EPI_ISL_3094020, EPI_ISL_3094021, EPI_ISL_3094023 | Lighthouse Lab in Alderley Park | Wellcome Sanger Institute for the COVID-19 Genomics UK (COG-UK) Consortium | Jacquelyn Wynn, Mairead Hyland, The Lighthouse Lab in Alderley Park and Alex Alderton, Roberto Amato, Jeffrey Barrett, Sonia Goncalves, Ewan Harrison, David K. Jackson, Ian Johnston, Dominic Kwiatkowski, Cordelia Langford, John Sillitoe on behalf of the Wellcome Sanger Institute COVID-19 Surveillance Team |
| EPI_ISL_3094024 | Lighthouse Lab in Glasgow | Wellcome Sanger Institute for the COVID-19 Genomics UK (COG-UK) Consortium | Harper VanSteenhouse, Yumi Kasai, David Gray, Carol Clugston, Anna Dominiczak and Alex Alderton, Roberto Amato, Jeffrey Barrett, Sonia Goncalves, Ewan Harrison, David K. Jackson, Ian Johnston, Dominic Kwiatkowski, Cordelia Langford, John Sillitoe on behalf of the Wellcome Sanger Institute COVID-19 Surveillance Team |
| EPI_ISL_3094025 | Lighthouse Lab in Alderley Park | Wellcome Sanger Institute for the COVID-19 Genomics UK (COG-UK) Consortium | Jacquelyn Wynn, Mairead Hyland, The Lighthouse Lab in Alderley Park and Alex Alderton, Roberto Amato, Jeffrey Barrett, Sonia Goncalves, Ewan Harrison, David K. Jackson, Ian Johnston, Dominic Kwiatkowski, Cordelia Langford, John Sillitoe on behalf of the Wellcome Sanger Institute COVID-19 Surveillance Team |
| EPI_ISL_3094027 | Lighthouse Lab in Glasgow | Wellcome Sanger Institute for the COVID-19 Genomics UK (COG-UK) Consortium | Harper VanSteenhouse, Yumi Kasai, David Gray, Carol Clugston, Anna Dominiczak and Alex Alderton, Roberto Amato, Jeffrey Barrett, Sonia Goncalves, Ewan Harrison, David K. Jackson, Ian Johnston, Dominic Kwiatkowski, Cordelia Langford, John Sillitoe on behalf of the Wellcome Sanger Institute COVID-19 Surveillance Team |
| EPI_ISL_3094028 | Lighthouse Lab in Alderley Park | Wellcome Sanger Institute for the COVID-19 Genomics UK (COG-UK) Consortium | Jacquelyn Wynn, Mairead Hyland, The Lighthouse Lab in Alderley Park and Alex Alderton, Roberto Amato, Jeffrey Barrett, Sonia Goncalves, Ewan Harrison, David K. Jackson, Ian Johnston, Dominic Kwiatkowski, Cordelia Langford, John Sillitoe on behalf of the Wellcome Sanger Institute COVID-19 Surveillance Team |
| EPI_ISL_3094029 | Lighthouse Lab in Glasgow | Wellcome Sanger Institute for the COVID-19 Genomics UK (COG-UK) Consortium | Harper VanSteenhouse, Yumi Kasai, David Gray, Carol Clugston, Anna Dominiczak and Alex Alderton, Roberto Amato, Jeffrey Barrett, Sonia Goncalves, Ewan Harrison, David K. Jackson, Ian Johnston, Dominic Kwiatkowski, Cordelia Langford, John Sillitoe on behalf of the Wellcome Sanger Institute COVID-19 Surveillance Team |
| EPI_ISL_3094030 | Lighthouse Lab in Alderley Park | Wellcome Sanger Institute for the COVID-19 Genomics UK (COG-UK) Consortium | Jacquelyn Wynn, Mairead Hyland, The Lighthouse Lab in Alderley Park and Alex Alderton, Roberto Amato, Jeffrey Barrett, Sonia Goncalves, Ewan Harrison, David K. Jackson, Ian Johnston, Dominic Kwiatkowski, Cordelia Langford, John Sillitoe on behalf of the Wellcome Sanger Institute COVID-19 Surveillance Team |
| EPI_ISL_3094031, EPI_ISL_3094032 | Berkshire and Surrey Pathology Services Lighthouse Laboratory | Wellcome Sanger Institute for the COVID-19 Genomics UK (COG-UK) Consortium | Berkshire and Surrey Pathology Services Lighthouse Laboratory and Alex Alderton, Roberto Amato, Jeffrey Barrett, Sonia Goncalves, Ewan Harrison, David K. Jackson, Ian Johnston, Dominic Kwiatkowski, Cordelia Langford, John Sillitoe on behalf of the Wellcome Sanger Institute COVID-19 Surveillance Team |
| EPI_ISL_3094033 | Lighthouse Lab in Alderley Park | Wellcome Sanger Institute for the COVID-19 Genomics UK (COG-UK) Consortium | Jacquelyn Wynn, Mairead Hyland, The Lighthouse Lab in Alderley Park and Alex Alderton, Roberto Amato, Jeffrey Barrett, Sonia Goncalves, Ewan Harrison, David K. Jackson, Ian Johnston, Dominic Kwiatkowski, Cordelia Langford, John Sillitoe on behalf of the Wellcome Sanger Institute COVID-19 Surveillance Team |
| EPI_ISL_3094034 | Lighthouse Lab in Glasgow | Wellcome Sanger Institute for the COVID-19 Genomics UK (COG-UK) Consortium | Harper VanSteenhouse, Yumi Kasai, David Gray, Carol Clugston, Anna Dominiczak and Alex Alderton, Roberto Amato, Jeffrey Barrett, Sonia Goncalves, Ewan Harrison, David K. Jackson, Ian Johnston, Dominic Kwiatkowski, Cordelia Langford, John Sillitoe on behalf of the Wellcome Sanger Institute COVID-19 Surveillance Team |
| EPI_ISL_3094036 | Lighthouse Lab in Alderley Park | Wellcome Sanger Institute for the COVID-19 Genomics UK (COG-UK) Consortium | Jacquelyn Wynn, Mairead Hyland, The Lighthouse Lab in Alderley Park and Alex Alderton, Roberto Amato, Jeffrey Barrett, Sonia Goncalves, Ewan Harrison, David K. Jackson, Ian Johnston, Dominic Kwiatkowski, Cordelia Langford, John Sillitoe on behalf of the Wellcome Sanger Institute COVID-19 Surveillance Team |
| EPI_ISL_3094037, EPI_ISL_3094039 | Lighthouse Lab in Glasgow | Wellcome Sanger Institute for the COVID-19 Genomics UK (COG-UK) Consortium | Harper VanSteenhouse, Yumi Kasai, David Gray, Carol Clugston, Anna Dominiczak and Alex Alderton, Roberto Amato, Jeffrey Barrett, Sonia Goncalves, Ewan Harrison, David K. Jackson, Ian Johnston, Dominic Kwiatkowski, Cordelia Langford, John Sillitoe on behalf of the Wellcome Sanger Institute COVID-19 Surveillance Team |
| EPI_ISL_3094040 | Lighthouse Lab in Alderley Park | Wellcome Sanger Institute for the COVID-19 Genomics UK (COG-UK) Consortium | Jacquelyn Wynn, Mairead Hyland, The Lighthouse Lab in Alderley Park and Alex Alderton, Roberto Amato, Jeffrey Barrett, Sonia Goncalves, Ewan Harrison, David K. Jackson, Ian Johnston, Dominic Kwiatkowski, Cordelia Langford, John Sillitoe on behalf of the Wellcome Sanger Institute COVID-19 Surveillance Team |
| EPI_ISL_3094041 | Lighthouse Lab in Glasgow | Wellcome Sanger Institute for the COVID-19 Genomics UK (COG-UK) Consortium | Harper VanSteenhouse, Yumi Kasai, David Gray, Carol Clugston, Anna Dominiczak and Alex Alderton, Roberto Amato, Jeffrey Barrett, Sonia Goncalves, Ewan Harrison, David K. Jackson, Ian Johnston, Dominic Kwiatkowski, Cordelia Langford, John Sillitoe on behalf of the Wellcome Sanger Institute COVID-19 Surveillance Team |
| EPI_ISL_3094043, EPI_ISL_3094044, EPI_ISL_3094045 | Lighthouse Lab in Alderley Park | Wellcome Sanger Institute for the COVID-19 Genomics UK (COG-UK) Consortium | Jacquelyn Wynn, Mairead Hyland, The Lighthouse Lab in Alderley Park and Alex Alderton, Roberto Amato, Jeffrey Barrett, Sonia Goncalves, Ewan Harrison, David K. Jackson, Ian Johnston, Dominic Kwiatkowski, Cordelia Langford, John Sillitoe on behalf of the Wellcome Sanger Institute COVID-19 Surveillance Team |
| EPI_ISL_3094046 | Berkshire and Surrey Pathology Services Lighthouse Laboratory | Wellcome Sanger Institute for the COVID-19 Genomics UK (COG-UK) Consortium | Berkshire and Surrey Pathology Services Lighthouse Laboratory and Alex Alderton, Roberto Amato, Jeffrey Barrett, Sonia Goncalves, Ewan Harrison, David K. Jackson, Ian Johnston, Dominic Kwiatkowski, Cordelia Langford, John Sillitoe on behalf of the Wellcome Sanger Institute COVID-19 Surveillance Team |
| EPI_ISL_3094048 | Lighthouse Lab in Alderley Park | Wellcome Sanger Institute for the COVID-19 Genomics UK (COG-UK) Consortium | Jacquelyn Wynn, Mairead Hyland, The Lighthouse Lab in Alderley Park and Alex Alderton, Roberto Amato, Jeffrey Barrett, Sonia Goncalves, Ewan Harrison, David K. Jackson, Ian Johnston, Dominic Kwiatkowski, Cordelia Langford, John Sillitoe on behalf of the Wellcome Sanger Institute COVID-19 Surveillance Team |
| EPI_ISL_3094049 | Lighthouse Lab in Glasgow | Wellcome Sanger Institute for the COVID-19 Genomics UK (COG-UK) Consortium | Harper VanSteenhouse, Yumi Kasai, David Gray, Carol Clugston, Anna Dominiczak and Alex Alderton, Roberto Amato, Jeffrey Barrett, Sonia Goncalves, Ewan Harrison, David K. Jackson, Ian Johnston, Dominic Kwiatkowski, Cordelia Langford, John Sillitoe on behalf of the Wellcome Sanger Institute COVID-19 Surveillance Team |
| EPI_ISL_3094050, EPI_ISL_3094052, EPI_ISL_3094053 | Berkshire and Surrey Pathology Services Lighthouse Laboratory | Wellcome Sanger Institute for the COVID-19 Genomics UK (COG-UK) Consortium | Berkshire and Surrey Pathology Services Lighthouse Laboratory and Alex Alderton, Roberto Amato, Jeffrey Barrett, Sonia Goncalves, Ewan Harrison, David K. Jackson, Ian Johnston, Dominic Kwiatkowski, Cordelia Langford, John Sillitoe on behalf of the Wellcome Sanger Institute COVID-19 Surveillance Team |
| EPI_ISL_3094055 | Lighthouse Lab in Alderley Park | Wellcome Sanger Institute for the COVID-19 Genomics UK (COG-UK) Consortium | Jacquelyn Wynn, Mairead Hyland, The Lighthouse Lab in Alderley Park and Alex Alderton, Roberto Amato, Jeffrey Barrett, Sonia Goncalves, Ewan Harrison, David K. Jackson, Ian Johnston, Dominic Kwiatkowski, Cordelia Langford, John Sillitoe on behalf of the Wellcome Sanger Institute COVID-19 Surveillance Team |
| EPI_ISL_3094056, EPI_ISL_3094058 | Lighthouse Lab in Glasgow | Wellcome Sanger Institute for the COVID-19 Genomics UK (COG-UK) Consortium | Harper VanSteenhouse, Yumi Kasai, David Gray, Carol Clugston, Anna Dominiczak and Alex Alderton, Roberto Amato, Jeffrey Barrett, Sonia Goncalves, Ewan Harrison, David K. Jackson, Ian Johnston, Dominic Kwiatkowski, Cordelia Langford, John Sillitoe on behalf of the Wellcome Sanger Institute COVID-19 Surveillance Team |
| EPI_ISL_3094059 | Lighthouse Lab in Alderley Park | Wellcome Sanger Institute for the COVID-19 Genomics UK (COG-UK) Consortium | Jacquelyn Wynn, Mairead Hyland, The Lighthouse Lab in Alderley Park and Alex Alderton, Roberto Amato, Jeffrey Barrett, Sonia Goncalves, Ewan Harrison, David K. Jackson, Ian Johnston, Dominic Kwiatkowski, Cordelia Langford, John Sillitoe on behalf of the Wellcome Sanger Institute COVID-19 Surveillance Team |
| EPI_ISL_3094060 | Lighthouse Lab in Glasgow | Wellcome Sanger Institute for the COVID-19 Genomics UK (COG-UK) Consortium | Harper VanSteenhouse, Yumi Kasai, David Gray, Carol Clugston, Anna Dominiczak and Alex Alderton, Roberto Amato, Jeffrey Barrett, Sonia Goncalves, Ewan Harrison, David K. Jackson, Ian Johnston, Dominic Kwiatkowski, Cordelia Langford, John Sillitoe on behalf of the Wellcome Sanger Institute COVID-19 Surveillance Team |
| EPI_ISL_3094062, EPI_ISL_3094063, EPI_ISL_3094064 | Lighthouse Lab in Alderley Park | Wellcome Sanger Institute for the COVID-19 Genomics UK (COG-UK) Consortium | Jacquelyn Wynn, Mairead Hyland, The Lighthouse Lab in Alderley Park and Alex Alderton, Roberto Amato, Jeffrey Barrett, Sonia Goncalves, Ewan Harrison, David K. Jackson, Ian Johnston, Dominic Kwiatkowski, Cordelia Langford, John Sillitoe on behalf of the Wellcome Sanger Institute COVID-19 Surveillance Team |
| EPI_ISL_3094065 | Berkshire and Surrey Pathology Services Lighthouse Laboratory | Wellcome Sanger Institute for the COVID-19 Genomics UK (COG-UK) Consortium | Berkshire and Surrey Pathology Services Lighthouse Laboratory and Alex Alderton, Roberto Amato, Jeffrey Barrett, Sonia Goncalves, Ewan Harrison, David K. Jackson, Ian Johnston, Dominic Kwiatkowski, Cordelia Langford, John Sillitoe on behalf of the Wellcome Sanger Institute COVID-19 Surveillance Team |
| EPI_ISL_3094066 | Lighthouse Lab in Alderley Park | Wellcome Sanger Institute for the COVID-19 Genomics UK (COG-UK) Consortium | Jacquelyn Wynn, Mairead Hyland, The Lighthouse Lab in Alderley Park and Alex Alderton, Roberto Amato, Jeffrey Barrett, Sonia Goncalves, Ewan Harrison, David K. Jackson, Ian Johnston, Dominic Kwiatkowski, Cordelia Langford, John Sillitoe on behalf of the Wellcome Sanger Institute COVID-19 Surveillance Team |
| EPI_ISL_3094067 | Berkshire and Surrey Pathology Services Lighthouse Laboratory | Wellcome Sanger Institute for the COVID-19 Genomics UK (COG-UK) Consortium | Berkshire and Surrey Pathology Services Lighthouse Laboratory and Alex Alderton, Roberto Amato, Jeffrey Barrett, Sonia Goncalves, Ewan Harrison, David K. Jackson, Ian Johnston, Dominic Kwiatkowski, Cordelia Langford, John Sillitoe on behalf of the Wellcome Sanger Institute COVID-19 Surveillance Team |
| EPI_ISL_3094069, EPI_ISL_3094070, EPI_ISL_3094071 | Lighthouse Lab in Alderley Park | Wellcome Sanger Institute for the COVID-19 Genomics UK (COG-UK) Consortium | Jacquelyn Wynn, Mairead Hyland, The Lighthouse Lab in Alderley Park and Alex Alderton, Roberto Amato, Jeffrey Barrett, Sonia Goncalves, Ewan Harrison, David K. Jackson, Ian Johnston, Dominic Kwiatkowski, Cordelia Langford, John Sillitoe on behalf of the Wellcome Sanger Institute COVID-19 Surveillance Team |
| EPI_ISL_3094073 | Berkshire and Surrey Pathology Services Lighthouse Laboratory | Wellcome Sanger Institute for the COVID-19 Genomics UK (COG-UK) Consortium | Berkshire and Surrey Pathology Services Lighthouse Laboratory and Alex Alderton, Roberto Amato, Jeffrey Barrett, Sonia Goncalves, Ewan Harrison, David K. Jackson, Ian Johnston, Dominic Kwiatkowski, Cordelia Langford, John Sillitoe on behalf of the Wellcome Sanger Institute COVID-19 Surveillance Team |
| EPI_ISL_3094074 | Lighthouse Lab in Alderley Park | Wellcome Sanger Institute for the COVID-19 Genomics UK (COG-UK) Consortium | Jacquelyn Wynn, Mairead Hyland, The Lighthouse Lab in Alderley Park and Alex Alderton, Roberto Amato, Jeffrey Barrett, Sonia Goncalves, Ewan Harrison, David K. Jackson, Ian Johnston, Dominic Kwiatkowski, Cordelia Langford, John Sillitoe on behalf of the Wellcome Sanger Institute COVID-19 Surveillance Team |
| EPI_ISL_3094075 | Berkshire and Surrey Pathology Services Lighthouse Laboratory | Wellcome Sanger Institute for the COVID-19 Genomics UK (COG-UK) Consortium | Berkshire and Surrey Pathology Services Lighthouse Laboratory and Alex Alderton, Roberto Amato, Jeffrey Barrett, Sonia Goncalves, Ewan Harrison, David K. Jackson, Ian Johnston, Dominic Kwiatkowski, Cordelia Langford, John Sillitoe on behalf of the Wellcome Sanger Institute COVID-19 Surveillance Team |
| EPI_ISL_3094077 | Lighthouse Lab in Alderley Park | Wellcome Sanger Institute for the COVID-19 Genomics UK (COG-UK) Consortium | Jacquelyn Wynn, Mairead Hyland, The Lighthouse Lab in Alderley Park and Alex Alderton, Roberto Amato, Jeffrey Barrett, Sonia Goncalves, Ewan Harrison, David K. Jackson, Ian Johnston, Dominic Kwiatkowski, Cordelia Langford, John Sillitoe on behalf of the Wellcome Sanger Institute COVID-19 Surveillance Team |
| EPI_ISL_3094078 | Berkshire and Surrey Pathology Services Lighthouse Laboratory | Wellcome Sanger Institute for the COVID-19 Genomics UK (COG-UK) Consortium | Berkshire and Surrey Pathology Services Lighthouse Laboratory and Alex Alderton, Roberto Amato, Jeffrey Barrett, Sonia Goncalves, Ewan Harrison, David K. Jackson, Ian Johnston, Dominic Kwiatkowski, Cordelia Langford, John Sillitoe on behalf of the Wellcome Sanger Institute COVID-19 Surveillance Team |
| EPI_ISL_3094080, EPI_ISL_3094081 | Lighthouse Lab in Glasgow | Wellcome Sanger Institute for the COVID-19 Genomics UK (COG-UK) Consortium | Harper VanSteenhouse, Yumi Kasai, David Gray, Carol Clugston, Anna Dominiczak and Alex Alderton, Roberto Amato, Jeffrey Barrett, Sonia Goncalves, Ewan Harrison, David K. Jackson, Ian Johnston, Dominic Kwiatkowski, Cordelia Langford, John Sillitoe on behalf of the Wellcome Sanger Institute COVID-19 Surveillance Team |
| EPI_ISL_3094082 | Lighthouse Lab in Alderley Park | Wellcome Sanger Institute for the COVID-19 Genomics UK (COG-UK) Consortium | Jacquelyn Wynn, Mairead Hyland, The Lighthouse Lab in Alderley Park and Alex Alderton, Roberto Amato, Jeffrey Barrett, Sonia Goncalves, Ewan Harrison, David K. Jackson, Ian Johnston, Dominic Kwiatkowski, Cordelia Langford, John Sillitoe on behalf of the Wellcome Sanger Institute COVID-19 Surveillance Team |
| EPI_ISL_3094084 | Berkshire and Surrey Pathology Services Lighthouse Laboratory | Wellcome Sanger Institute for the COVID-19 Genomics UK (COG-UK) Consortium | Berkshire and Surrey Pathology Services Lighthouse Laboratory and Alex Alderton, Roberto Amato, Jeffrey Barrett, Sonia Goncalves, Ewan Harrison, David K. Jackson, Ian Johnston, Dominic Kwiatkowski, Cordelia Langford, John Sillitoe on behalf of the Wellcome Sanger Institute COVID-19 Surveillance Team |
| EPI_ISL_3094085 | Lighthouse Lab in Glasgow | Wellcome Sanger Institute for the COVID-19 Genomics UK (COG-UK) Consortium | Harper VanSteenhouse, Yumi Kasai, David Gray, Carol Clugston, Anna Dominiczak and Alex Alderton, Roberto Amato, Jeffrey Barrett, Sonia Goncalves, Ewan Harrison, David K. Jackson, Ian Johnston, Dominic Kwiatkowski, Cordelia Langford, John Sillitoe on behalf of the Wellcome Sanger Institute COVID-19 Surveillance Team |
| EPI_ISL_3094086 | Lighthouse Lab in Alderley Park | Wellcome Sanger Institute for the COVID-19 Genomics UK (COG-UK) Consortium | Jacquelyn Wynn, Mairead Hyland, The Lighthouse Lab in Alderley Park and Alex Alderton, Roberto Amato, Jeffrey Barrett, Sonia Goncalves, Ewan Harrison, David K. Jackson, Ian Johnston, Dominic Kwiatkowski, Cordelia Langford, John Sillitoe on behalf of the Wellcome Sanger Institute COVID-19 Surveillance Team |
| EPI_ISL_3094088 | Berkshire and Surrey Pathology Services Lighthouse Laboratory | Wellcome Sanger Institute for the COVID-19 Genomics UK (COG-UK) Consortium | Berkshire and Surrey Pathology Services Lighthouse Laboratory and Alex Alderton, Roberto Amato, Jeffrey Barrett, Sonia Goncalves, Ewan Harrison, David K. Jackson, Ian Johnston, Dominic Kwiatkowski, Cordelia Langford, John Sillitoe on behalf of the Wellcome Sanger Institute COVID-19 Surveillance Team |
| EPI_ISL_3094089 | Lighthouse Lab in Alderley Park | Wellcome Sanger Institute for the COVID-19 Genomics UK (COG-UK) Consortium | Jacquelyn Wynn, Mairead Hyland, The Lighthouse Lab in Alderley Park and Alex Alderton, Roberto Amato, Jeffrey Barrett, Sonia Goncalves, Ewan Harrison, David K. Jackson, Ian Johnston, Dominic Kwiatkowski, Cordelia Langford, John Sillitoe on behalf of the Wellcome Sanger Institute COVID-19 Surveillance Team |
| EPI_ISL_3094091 | Berkshire and Surrey Pathology Services Lighthouse Laboratory | Wellcome Sanger Institute for the COVID-19 Genomics UK (COG-UK) Consortium | Berkshire and Surrey Pathology Services Lighthouse Laboratory and Alex Alderton, Roberto Amato, Jeffrey Barrett, Sonia Goncalves, Ewan Harrison, David K. Jackson, Ian Johnston, Dominic Kwiatkowski, Cordelia Langford, John Sillitoe on behalf of the Wellcome Sanger Institute COVID-19 Surveillance Team |
| EPI_ISL_3094092 | Lighthouse Lab in Alderley Park | Wellcome Sanger Institute for the COVID-19 Genomics UK (COG-UK) Consortium | Jacquelyn Wynn, Mairead Hyland, The Lighthouse Lab in Alderley Park and Alex Alderton, Roberto Amato, Jeffrey Barrett, Sonia Goncalves, Ewan Harrison, David K. Jackson, Ian Johnston, Dominic Kwiatkowski, Cordelia Langford, John Sillitoe on behalf of the Wellcome Sanger Institute COVID-19 Surveillance Team |
| EPI_ISL_3094094 | Lighthouse Lab in Glasgow | Wellcome Sanger Institute for the COVID-19 Genomics UK (COG-UK) Consortium | Harper VanSteenhouse, Yumi Kasai, David Gray, Carol Clugston, Anna Dominiczak and Alex Alderton, Roberto Amato, Jeffrey Barrett, Sonia Goncalves, Ewan Harrison, David K. Jackson, Ian Johnston, Dominic Kwiatkowski, Cordelia Langford, John Sillitoe on behalf of the Wellcome Sanger Institute COVID-19 Surveillance Team |

| | | | |
|---|---|---|---|
| EPI_ISL_3094095 | Lighthouse Lab in Alderley Park | Wellcome Sanger Institute for the COVID-19 Genomics UK (COG-UK) Consortium | Jacquelyn Wynn, Mairead Hyland, The Lighthouse Lab in Alderley Park and Alex Alderton, Roberto Amato, Jeffrey Barrett, Sonia Goncalves, Ewan Harrison, David K. Jackson, Ian Johnston, Dominic Kwiatkowski, Cordelia Langford, John Sillitoe on behalf of the Wellcome Sanger Institute COVID-19 Surveillance Team |
| EPI_ISL_3094097 | Berkshire and Surrey Pathology Services Lighthouse Laboratory | Wellcome Sanger Institute for the COVID-19 Genomics UK (COG-UK) Consortium | Berkshire and Surrey Pathology Services Lighthouse Laboratory and Alex Alderton, Roberto Amato, Jeffrey Barrett, Sonia Goncalves, Ewan Harrison, David K. Jackson, Ian Johnston, Dominic Kwiatkowski, Cordelia Langford, John Sillitoe on behalf of the Wellcome Sanger Institute COVID-19 Surveillance Team |
| EPI_ISL_3094098, EPI_ISL_3094099, EPI_ISL_3094101, EPI_ISL_3094102 | Lighthouse Lab in Alderley Park | Wellcome Sanger Institute for the COVID-19 Genomics UK (COG-UK) Consortium | Jacquelyn Wynn, Mairead Hyland, The Lighthouse Lab in Alderley Park and Alex Alderton, Roberto Amato, Jeffrey Barrett, Sonia Goncalves, Ewan Harrison, David K. Jackson, Ian Johnston, Dominic Kwiatkowski, Cordelia Langford, John Sillitoe on behalf of the Wellcome Sanger Institute COVID-19 Surveillance Team |
| EPI_ISL_3094103, EPI_ISL_3094105 | Lighthouse Lab in Glasgow | Wellcome Sanger Institute for the COVID-19 Genomics UK (COG-UK) Consortium | Harper VanSteenhouse, Yumi Kasai, David Gray, Carol Clugston, Anna Dominiczak and Alex Alderton, Roberto Amato, Jeffrey Barrett, Sonia Goncalves, Ewan Harrison, David K. Jackson, Ian Johnston, Dominic Kwiatkowski, Cordelia Langford, John Sillitoe on behalf of the Wellcome Sanger Institute COVID-19 Surveillance Team |
| EPI_ISL_3094106 | Lighthouse Lab in Alderley Park | Wellcome Sanger Institute for the COVID-19 Genomics UK (COG-UK) Consortium | Jacquelyn Wynn, Mairead Hyland, The Lighthouse Lab in Alderley Park and Alex Alderton, Roberto Amato, Jeffrey Barrett, Sonia Goncalves, Ewan Harrison, David K. Jackson, Ian Johnston, Dominic Kwiatkowski, Cordelia Langford, John Sillitoe on behalf of the Wellcome Sanger Institute COVID-19 Surveillance Team |
| EPI_ISL_3094108 | Lighthouse Lab in Glasgow | Wellcome Sanger Institute for the COVID-19 Genomics UK (COG-UK) Consortium | Harper VanSteenhouse, Yumi Kasai, David Gray, Carol Clugston, Anna Dominiczak and Alex Alderton, Roberto Amato, Jeffrey Barrett, Sonia Goncalves, Ewan Harrison, David K. Jackson, Ian Johnston, Dominic Kwiatkowski, Cordelia Langford, John Sillitoe on behalf of the Wellcome Sanger Institute COVID-19 Surveillance Team |
| EPI_ISL_3094109 | Lighthouse Lab in Alderley Park | Wellcome Sanger Institute for the COVID-19 Genomics UK (COG-UK) Consortium | Jacquelyn Wynn, Mairead Hyland, The Lighthouse Lab in Alderley Park and Alex Alderton, Roberto Amato, Jeffrey Barrett, Sonia Goncalves, Ewan Harrison, David K. Jackson, Ian Johnston, Dominic Kwiatkowski, Cordelia Langford, John Sillitoe on behalf of the Wellcome Sanger Institute COVID-19 Surveillance Team |
| EPI_ISL_3094110 | Lighthouse Lab in Glasgow | Wellcome Sanger Institute for the COVID-19 Genomics UK (COG-UK) Consortium | Harper VanSteenhouse, Yumi Kasai, David Gray, Carol Clugston, Anna Dominiczak and Alex Alderton, Roberto Amato, Jeffrey Barrett, Sonia Goncalves, Ewan Harrison, David K. Jackson, Ian Johnston, Dominic Kwiatkowski, Cordelia Langford, John Sillitoe on behalf of the Wellcome Sanger Institute COVID-19 Surveillance Team |
| EPI_ISL_3094112, EPI_ISL_3094113, EPI_ISL_3094114 | Lighthouse Lab in Alderley Park | Wellcome Sanger Institute for the COVID-19 Genomics UK (COG-UK) Consortium | Jacquelyn Wynn, Mairead Hyland, The Lighthouse Lab in Alderley Park and Alex Alderton, Roberto Amato, Jeffrey Barrett, Sonia Goncalves, Ewan Harrison, David K. Jackson, Ian Johnston, Dominic Kwiatkowski, Cordelia Langford, John Sillitoe on behalf of the Wellcome Sanger Institute COVID-19 Surveillance Team |
| EPI_ISL_3094116 | Lighthouse Lab in Glasgow | Wellcome Sanger Institute for the COVID-19 Genomics UK (COG-UK) Consortium | Harper VanSteenhouse, Yumi Kasai, David Gray, Carol Clugston, Anna Dominiczak and Alex Alderton, Roberto Amato, Jeffrey Barrett, Sonia Goncalves, Ewan Harrison, David K. Jackson, Ian Johnston, Dominic Kwiatkowski, Cordelia Langford, John Sillitoe on behalf of the Wellcome Sanger Institute COVID-19 Surveillance Team |
| EPI_ISL_3094117, EPI_ISL_3094118, EPI_ISL_3094120, EPI_ISL_3094121, EPI_ISL_3094122 | Lighthouse Lab in Alderley Park | Wellcome Sanger Institute for the COVID-19 Genomics UK (COG-UK) Consortium | Jacquelyn Wynn, Mairead Hyland, The Lighthouse Lab in Alderley Park and Alex Alderton, Roberto Amato, Jeffrey Barrett, Sonia Goncalves, Ewan Harrison, David K. Jackson, Ian Johnston, Dominic Kwiatkowski, Cordelia Langford, John Sillitoe on behalf of the Wellcome Sanger Institute COVID-19 Surveillance Team |
| EPI_ISL_3094167 | Lighthouse Lab in Glasgow | Wellcome Sanger Institute for the COVID-19 Genomics UK (COG-UK) Consortium | Harper VanSteenhouse, Yumi Kasai, David Gray, Carol Clugston, Anna Dominiczak and Alex Alderton, Roberto Amato, Jeffrey Barrett, Sonia Goncalves, Ewan Harrison, David K. Jackson, Ian Johnston, Dominic Kwiatkowski, Cordelia Langford, John Sillitoe on behalf of the Wellcome Sanger Institute COVID-19 Surveillance Team |
| EPI_ISL_3094169 | Berkshire and Surrey Pathology Services Lighthouse Laboratory | Wellcome Sanger Institute for the COVID-19 Genomics UK (COG-UK) Consortium | Berkshire and Surrey Pathology Services Lighthouse Laboratory and Alex Alderton, Roberto Amato, Jeffrey Barrett, Sonia Goncalves, Ewan Harrison, David K. Jackson, Ian Johnston, Dominic Kwiatkowski, Cordelia Langford, John Sillitoe on behalf of the Wellcome Sanger Institute COVID-19 Surveillance Team |
| EPI_ISL_3094170 | Lighthouse Lab in Glasgow | Wellcome Sanger Institute for the COVID-19 Genomics UK (COG-UK) Consortium | Harper VanSteenhouse, Yumi Kasai, David Gray, Carol Clugston, Anna Dominiczak and Alex Alderton, Roberto Amato, Jeffrey Barrett, Sonia Goncalves, Ewan Harrison, David K. Jackson, Ian Johnston, Dominic Kwiatkowski, Cordelia Langford, John Sillitoe on behalf of the Wellcome Sanger Institute COVID-19 Surveillance Team |
| EPI_ISL_3094171, EPI_ISL_3094173 | Lighthouse Lab in Alderley Park | Wellcome Sanger Institute for the COVID-19 Genomics UK (COG-UK) Consortium | Jacquelyn Wynn, Mairead Hyland, The Lighthouse Lab in Alderley Park and Alex Alderton, Roberto Amato, Jeffrey Barrett, Sonia Goncalves, Ewan Harrison, David K. Jackson, Ian Johnston, Dominic Kwiatkowski, Cordelia Langford, John Sillitoe on behalf of the Wellcome Sanger Institute COVID-19 Surveillance Team |
| EPI_ISL_3094174 | Berkshire and Surrey Pathology Services Lighthouse Laboratory | Wellcome Sanger Institute for the COVID-19 Genomics UK (COG-UK) Consortium | Berkshire and Surrey Pathology Services Lighthouse Laboratory and Alex Alderton, Roberto Amato, Jeffrey Barrett, Sonia Goncalves, Ewan Harrison, David K. Jackson, Ian Johnston, Dominic Kwiatkowski, Cordelia Langford, John Sillitoe on behalf of the Wellcome Sanger Institute COVID-19 Surveillance Team |
| EPI_ISL_3094176 | Lighthouse Lab in Alderley Park | Wellcome Sanger Institute for the COVID-19 Genomics UK (COG-UK) Consortium | Jacquelyn Wynn, Mairead Hyland, The Lighthouse Lab in Alderley Park and Alex Alderton, Roberto Amato, Jeffrey Barrett, Sonia Goncalves, Ewan Harrison, David K. Jackson, Ian Johnston, Dominic Kwiatkowski, Cordelia Langford, John Sillitoe on behalf of the Wellcome Sanger Institute COVID-19 Surveillance Team |
| EPI_ISL_3094177, EPI_ISL_3094179, EPI_ISL_3094180 | Lighthouse Lab in Glasgow | Wellcome Sanger Institute for the COVID-19 Genomics UK (COG-UK) Consortium | Harper VanSteenhouse, Yumi Kasai, David Gray, Carol Clugston, Anna Dominiczak and Alex Alderton, Roberto Amato, Jeffrey Barrett, Sonia Goncalves, Ewan Harrison, David K. Jackson, Ian Johnston, Dominic Kwiatkowski, Cordelia Langford, John Sillitoe on behalf of the Wellcome Sanger Institute COVID-19 Surveillance Team |
| EPI_ISL_3094181, EPI_ISL_3094183 | Berkshire and Surrey Pathology Services Lighthouse Laboratory | Wellcome Sanger Institute for the COVID-19 Genomics UK (COG-UK) Consortium | Berkshire and Surrey Pathology Services Lighthouse Laboratory and Alex Alderton, Roberto Amato, Jeffrey Barrett, Sonia Goncalves, Ewan Harrison, David K. Jackson, Ian Johnston, Dominic Kwiatkowski, Cordelia Langford, John Sillitoe on behalf of the Wellcome Sanger Institute COVID-19 Surveillance Team |
| EPI_ISL_3094184, EPI_ISL_3094185, EPI_ISL_3094187 | Lighthouse Lab in Alderley Park | Wellcome Sanger Institute for the COVID-19 Genomics UK (COG-UK) Consortium | Jacquelyn Wynn, Mairead Hyland, The Lighthouse Lab in Alderley Park and Alex Alderton, Roberto Amato, Jeffrey Barrett, Sonia Goncalves, Ewan Harrison, David K. Jackson, Ian Johnston, Dominic Kwiatkowski, Cordelia Langford, John Sillitoe on behalf of the Wellcome Sanger Institute COVID-19 Surveillance Team |
| EPI_ISL_3094188, EPI_ISL_3094190, EPI_ISL_3094191 | Lighthouse Lab in Glasgow | Wellcome Sanger Institute for the COVID-19 Genomics UK (COG-UK) Consortium | Harper VanSteenhouse, Yumi Kasai, David Gray, Carol Clugston, Anna Dominiczak and Alex Alderton, Roberto Amato, Jeffrey Barrett, Sonia Goncalves, Ewan Harrison, David K. Jackson, Ian Johnston, Dominic Kwiatkowski, Cordelia Langford, John Sillitoe on behalf of the Wellcome Sanger Institute COVID-19 Surveillance Team |
| EPI_ISL_3094192 | Berkshire and Surrey Pathology Services Lighthouse Laboratory | Wellcome Sanger Institute for the COVID-19 Genomics UK (COG-UK) Consortium | Berkshire and Surrey Pathology Services Lighthouse Laboratory and Alex Alderton, Roberto Amato, Jeffrey Barrett, Sonia Goncalves, Ewan Harrison, David K. Jackson, Ian Johnston, Dominic Kwiatkowski, Cordelia Langford, John Sillitoe on behalf of the Wellcome Sanger Institute COVID-19 Surveillance Team |
| EPI_ISL_3094194 | Lighthouse Lab in Glasgow | Wellcome Sanger Institute for the COVID-19 Genomics UK (COG-UK) Consortium | Harper VanSteenhouse, Yumi Kasai, David Gray, Carol Clugston, Anna Dominiczak and Alex Alderton, Roberto Amato, Jeffrey Barrett, Sonia Goncalves, Ewan Harrison, David K. Jackson, Ian Johnston, Dominic Kwiatkowski, Cordelia Langford, John Sillitoe on behalf of the Wellcome Sanger Institute COVID-19 Surveillance Team |
| EPI_ISL_3094195 | Lighthouse Lab in Alderley Park | Wellcome Sanger Institute for the COVID-19 Genomics UK (COG-UK) Consortium | Jacquelyn Wynn, Mairead Hyland, The Lighthouse Lab in Alderley Park and Alex Alderton, Roberto Amato, Jeffrey Barrett, Sonia Goncalves, Ewan Harrison, David K. Jackson, Ian Johnston, Dominic Kwiatkowski, Cordelia Langford, John Sillitoe on behalf of the Wellcome Sanger Institute COVID-19 Surveillance Team |
| EPI_ISL_3094197 | Berkshire and Surrey Pathology Services Lighthouse Laboratory | Wellcome Sanger Institute for the COVID-19 Genomics UK (COG-UK) Consortium | Berkshire and Surrey Pathology Services Lighthouse Laboratory and Alex Alderton, Roberto Amato, Jeffrey Barrett, Sonia Goncalves, Ewan Harrison, David K. Jackson, Ian Johnston, Dominic Kwiatkowski, Cordelia Langford, John Sillitoe on behalf of the Wellcome Sanger Institute COVID-19 Surveillance Team |
| EPI_ISL_3094198, EPI_ISL_3094199 | Lighthouse Lab in Alderley Park | Wellcome Sanger Institute for the COVID-19 Genomics UK (COG-UK) Consortium | Jacquelyn Wynn, Mairead Hyland, The Lighthouse Lab in Alderley Park and Alex Alderton, Roberto Amato, Jeffrey Barrett, Sonia Goncalves, Ewan Harrison, David K. Jackson, Ian Johnston, Dominic Kwiatkowski, Cordelia Langford, John Sillitoe on behalf of the Wellcome Sanger Institute COVID-19 Surveillance Team |
| EPI_ISL_3094201 | Berkshire and Surrey Pathology Services Lighthouse Laboratory | Wellcome Sanger Institute for the COVID-19 Genomics UK (COG-UK) Consortium | Berkshire and Surrey Pathology Services Lighthouse Laboratory and Alex Alderton, Roberto Amato, Jeffrey Barrett, Sonia Goncalves, Ewan Harrison, David K. Jackson, Ian Johnston, Dominic Kwiatkowski, Cordelia Langford, John Sillitoe on behalf of the Wellcome Sanger Institute COVID-19 Surveillance Team |
| EPI_ISL_3094202 | Lighthouse Lab in Glasgow | Wellcome Sanger Institute for the COVID-19 Genomics UK (COG-UK) Consortium | Harper VanSteenhouse, Yumi Kasai, David Gray, Carol Clugston, Anna Dominiczak and Alex Alderton, Roberto Amato, Jeffrey Barrett, Sonia Goncalves, Ewan Harrison, David K. Jackson, Ian Johnston, Dominic Kwiatkowski, Cordelia Langford, John Sillitoe on behalf of the Wellcome Sanger Institute COVID-19 Surveillance Team |
| EPI_ISL_3094203, EPI_ISL_3094205, EPI_ISL_3094206 | Berkshire and Surrey Pathology Services Lighthouse Laboratory | Wellcome Sanger Institute for the COVID-19 Genomics UK (COG-UK) Consortium | Berkshire and Surrey Pathology Services Lighthouse Laboratory and Alex Alderton, Roberto Amato, Jeffrey Barrett, Sonia Goncalves, Ewan Harrison, David K. Jackson, Ian Johnston, Dominic Kwiatkowski, Cordelia Langford, John Sillitoe on behalf of the Wellcome Sanger Institute COVID-19 Surveillance Team |
| EPI_ISL_3094207 | Lighthouse Lab in Alderley Park | Wellcome Sanger Institute for the COVID-19 Genomics UK (COG-UK) Consortium | Jacquelyn Wynn, Mairead Hyland, The Lighthouse Lab in Alderley Park and Alex Alderton, Roberto Amato, Jeffrey Barrett, Sonia Goncalves, Ewan Harrison, David K. Jackson, Ian Johnston, Dominic Kwiatkowski, Cordelia Langford, John Sillitoe on behalf of the Wellcome Sanger Institute COVID-19 Surveillance Team |
| EPI_ISL_3094209 | Berkshire and Surrey Pathology Services Lighthouse Laboratory | Wellcome Sanger Institute for the COVID-19 Genomics UK (COG-UK) Consortium | Berkshire and Surrey Pathology Services Lighthouse Laboratory and Alex Alderton, Roberto Amato, Jeffrey Barrett, Sonia Goncalves, Ewan Harrison, David K. Jackson, Ian Johnston, Dominic Kwiatkowski, Cordelia Langford, John Sillitoe on behalf of the Wellcome Sanger Institute COVID-19 Surveillance Team |
| EPI_ISL_3094210, EPI_ISL_3094211, EPI_ISL_3094213 | Lighthouse Lab in Alderley Park | Wellcome Sanger Institute for the COVID-19 Genomics UK (COG-UK) Consortium | Jacquelyn Wynn, Mairead Hyland, The Lighthouse Lab in Alderley Park and Alex Alderton, Roberto Amato, Jeffrey Barrett, Sonia Goncalves, Ewan Harrison, David K. Jackson, Ian Johnston, Dominic Kwiatkowski, Cordelia Langford, John Sillitoe on behalf of the Wellcome Sanger Institute COVID-19 Surveillance Team |
| EPI_ISL_3094214, EPI_ISL_3094216 | Lighthouse Lab in Glasgow | Wellcome Sanger Institute for the COVID-19 Genomics UK (COG-UK) Consortium | Harper VanSteenhouse, Yumi Kasai, David Gray, Carol Clugston, Anna Dominiczak and Alex Alderton, Roberto Amato, Jeffrey Barrett, Sonia Goncalves, Ewan Harrison, David K. Jackson, Ian Johnston, Dominic Kwiatkowski, Cordelia Langford, John Sillitoe on behalf of the Wellcome Sanger Institute COVID-19 Surveillance Team |
| EPI_ISL_3094217, EPI_ISL_3094219, EPI_ISL_3094220, EPI_ISL_3094221 | Lighthouse Lab in Alderley Park | Wellcome Sanger Institute for the COVID-19 Genomics UK (COG-UK) Consortium | Jacquelyn Wynn, Mairead Hyland, The Lighthouse Lab in Alderley Park and Alex Alderton, Roberto Amato, Jeffrey Barrett, Sonia Goncalves, Ewan Harrison, David K. Jackson, Ian Johnston, Dominic Kwiatkowski, Cordelia Langford, John Sillitoe on behalf of the Wellcome Sanger Institute COVID-19 Surveillance Team |
| EPI_ISL_3094223 | Lighthouse Lab in Glasgow | Wellcome Sanger Institute for the COVID-19 Genomics UK (COG-UK) Consortium | Harper VanSteenhouse, Yumi Kasai, David Gray, Carol Clugston, Anna Dominiczak and Alex Alderton, Roberto Amato, Jeffrey Barrett, Sonia Goncalves, Ewan Harrison, David K. Jackson, Ian Johnston, Dominic Kwiatkowski, Cordelia Langford, John Sillitoe on behalf of the Wellcome Sanger Institute COVID-19 Surveillance Team |
| EPI_ISL_3094224 | Berkshire and Surrey Pathology Services Lighthouse Laboratory | Wellcome Sanger Institute for the COVID-19 Genomics UK (COG-UK) Consortium | Berkshire and Surrey Pathology Services Lighthouse Laboratory and Alex Alderton, Roberto Amato, Jeffrey Barrett, Sonia Goncalves, Ewan Harrison, David K. Jackson, Ian Johnston, Dominic Kwiatkowski, Cordelia Langford, John Sillitoe on behalf of the Wellcome Sanger Institute COVID-19 Surveillance Team |
| EPI_ISL_3094225 | Lighthouse Lab in Alderley Park | Wellcome Sanger Institute for the COVID-19 Genomics UK (COG-UK) Consortium | Jacquelyn Wynn, Mairead Hyland, The Lighthouse Lab in Alderley Park and Alex Alderton, Roberto Amato, Jeffrey Barrett, Sonia Goncalves, Ewan Harrison, David K. Jackson, Ian Johnston, Dominic Kwiatkowski, Cordelia Langford, John Sillitoe on behalf of the Wellcome Sanger Institute COVID-19 Surveillance Team |
| EPI_ISL_3094227 | Berkshire and Surrey Pathology Services Lighthouse Laboratory | Wellcome Sanger Institute for the COVID-19 Genomics UK (COG-UK) Consortium | Berkshire and Surrey Pathology Services Lighthouse Laboratory and Alex Alderton, Roberto Amato, Jeffrey Barrett, Sonia Goncalves, Ewan Harrison, David K. Jackson, Ian Johnston, Dominic Kwiatkowski, Cordelia Langford, John Sillitoe on behalf of the Wellcome Sanger Institute COVID-19 Surveillance Team |
| EPI_ISL_3094228, EPI_ISL_3094230 | Lighthouse Lab in Alderley Park | Wellcome Sanger Institute for the COVID-19 Genomics UK (COG-UK) Consortium | Jacquelyn Wynn, Mairead Hyland, The Lighthouse Lab in Alderley Park and Alex Alderton, Roberto Amato, Jeffrey Barrett, Sonia Goncalves, Ewan Harrison, David K. Jackson, Ian Johnston, Dominic Kwiatkowski, Cordelia Langford, John Sillitoe on behalf of the Wellcome Sanger Institute COVID-19 Surveillance Team |
| EPI_ISL_3094231 | Lighthouse Lab in Glasgow | Wellcome Sanger Institute for the COVID-19 Genomics UK (COG-UK) Consortium | Harper VanSteenhouse, Yumi Kasai, David Gray, Carol Clugston, Anna Dominiczak and Alex Alderton, Roberto Amato, Jeffrey Barrett, Sonia Goncalves, Ewan Harrison, David K. Jackson, Ian Johnston, Dominic Kwiatkowski, Cordelia Langford, John Sillitoe on behalf of the Wellcome Sanger Institute COVID-19 Surveillance Team |

| | | | |
|---|---|---|---|
| EPI_ISL_3094232 | Berkshire and Surrey Pathology Services Lighthouse Laboratory | Wellcome Sanger Institute for the COVID-19 Genomics UK (COG-UK) Consortium | Berkshire and Surrey Pathology Services Lighthouse Laboratory and Alex Alderton, Roberto Amato, Jeffrey Barrett, Sonia Goncalves, Ewan Harrison, David K. Jackson, Ian Johnston, Dominic Kwiatkowski, Cordelia Langford, John Sillitoe on behalf of the Wellcome Sanger Institute COVID-19 Surveillance Team |
| EPI_ISL_3094234, EPI_ISL_3094235 | Lighthouse Lab in Alderley Park | Wellcome Sanger Institute for the COVID-19 Genomics UK (COG-UK) Consortium | Jacquelyn Wynn, Mairead Hyland, The Lighthouse Lab in Alderley Park and Alex Alderton, Roberto Amato, Jeffrey Barrett, Sonia Goncalves, Ewan Harrison, David K. Jackson, Ian Johnston, Dominic Kwiatkowski, Cordelia Langford, John Sillitoe on behalf of the Wellcome Sanger Institute COVID-19 Surveillance Team |
| EPI_ISL_3094236 | Lighthouse Lab in Glasgow | Wellcome Sanger Institute for the COVID-19 Genomics UK (COG-UK) Consortium | Harper VanSteenhouse, Yumi Kasai, David Gray, Carol Clugston, Anna Dominiczak and Alex Alderton, Roberto Amato, Jeffrey Barrett, Sonia Goncalves, Ewan Harrison, David K. Jackson, Ian Johnston, Dominic Kwiatkowski, Cordelia Langford, John Sillitoe on behalf of the Wellcome Sanger Institute COVID-19 Surveillance Team |
| EPI_ISL_3094238, EPI_ISL_3094239, EPI_ISL_3094241, EPI_ISL_3094242 | Lighthouse Lab in Alderley Park | Wellcome Sanger Institute for the COVID-19 Genomics UK (COG-UK) Consortium | Jacquelyn Wynn, Mairead Hyland, The Lighthouse Lab in Alderley Park and Alex Alderton, Roberto Amato, Jeffrey Barrett, Sonia Goncalves, Ewan Harrison, David K. Jackson, Ian Johnston, Dominic Kwiatkowski, Cordelia Langford, John Sillitoe on behalf of the Wellcome Sanger Institute COVID-19 Surveillance Team |
| EPI_ISL_3094243 | Lighthouse Lab in Glasgow | Wellcome Sanger Institute for the COVID-19 Genomics UK (COG-UK) Consortium | Harper VanSteenhouse, Yumi Kasai, David Gray, Carol Clugston, Anna Dominiczak and Alex Alderton, Roberto Amato, Jeffrey Barrett, Sonia Goncalves, Ewan Harrison, David K. Jackson, Ian Johnston, Dominic Kwiatkowski, Cordelia Langford, John Sillitoe on behalf of the Wellcome Sanger Institute COVID-19 Surveillance Team |
| EPI_ISL_3094245 | Lighthouse Lab in Alderley Park | Wellcome Sanger Institute for the COVID-19 Genomics UK (COG-UK) Consortium | Jacquelyn Wynn, Mairead Hyland, The Lighthouse Lab in Alderley Park and Alex Alderton, Roberto Amato, Jeffrey Barrett, Sonia Goncalves, Ewan Harrison, David K. Jackson, Ian Johnston, Dominic Kwiatkowski, Cordelia Langford, John Sillitoe on behalf of the Wellcome Sanger Institute COVID-19 Surveillance Team |
| EPI_ISL_3094246 | Lighthouse Lab in Glasgow | Wellcome Sanger Institute for the COVID-19 Genomics UK (COG-UK) Consortium | Harper VanSteenhouse, Yumi Kasai, David Gray, Carol Clugston, Anna Dominiczak and Alex Alderton, Roberto Amato, Jeffrey Barrett, Sonia Goncalves, Ewan Harrison, David K. Jackson, Ian Johnston, Dominic Kwiatkowski, Cordelia Langford, John Sillitoe on behalf of the Wellcome Sanger Institute COVID-19 Surveillance Team |
| EPI_ISL_3094247 | Berkshire and Surrey Pathology Services Lighthouse Laboratory | Wellcome Sanger Institute for the COVID-19 Genomics UK (COG-UK) Consortium | Berkshire and Surrey Pathology Services Lighthouse Laboratory and Alex Alderton, Roberto Amato, Jeffrey Barrett, Sonia Goncalves, Ewan Harrison, David K. Jackson, Ian Johnston, Dominic Kwiatkowski, Cordelia Langford, John Sillitoe on behalf of the Wellcome Sanger Institute COVID-19 Surveillance Team |
| EPI_ISL_3094249 | Lighthouse Lab in Alderley Park | Wellcome Sanger Institute for the COVID-19 Genomics UK (COG-UK) Consortium | Jacquelyn Wynn, Mairead Hyland, The Lighthouse Lab in Alderley Park and Alex Alderton, Roberto Amato, Jeffrey Barrett, Sonia Goncalves, Ewan Harrison, David K. Jackson, Ian Johnston, Dominic Kwiatkowski, Cordelia Langford, John Sillitoe on behalf of the Wellcome Sanger Institute COVID-19 Surveillance Team |
| EPI_ISL_3094250 | Berkshire and Surrey Pathology Services Lighthouse Laboratory | Wellcome Sanger Institute for the COVID-19 Genomics UK (COG-UK) Consortium | Berkshire and Surrey Pathology Services Lighthouse Laboratory and Alex Alderton, Roberto Amato, Jeffrey Barrett, Sonia Goncalves, Ewan Harrison, David K. Jackson, Ian Johnston, Dominic Kwiatkowski, Cordelia Langford, John Sillitoe on behalf of the Wellcome Sanger Institute COVID-19 Surveillance Team |
| EPI_ISL_3094252, EPI_ISL_3094253 | Lighthouse Lab in Alderley Park | Wellcome Sanger Institute for the COVID-19 Genomics UK (COG-UK) Consortium | Jacquelyn Wynn, Mairead Hyland, The Lighthouse Lab in Alderley Park and Alex Alderton, Roberto Amato, Jeffrey Barrett, Sonia Goncalves, Ewan Harrison, David K. Jackson, Ian Johnston, Dominic Kwiatkowski, Cordelia Langford, John Sillitoe on behalf of the Wellcome Sanger Institute COVID-19 Surveillance Team |
| EPI_ISL_3094255 | Lighthouse Lab in Glasgow | Wellcome Sanger Institute for the COVID-19 Genomics UK (COG-UK) Consortium | Harper VanSteenhouse, Yumi Kasai, David Gray, Carol Clugston, Anna Dominiczak and Alex Alderton, Roberto Amato, Jeffrey Barrett, Sonia Goncalves, Ewan Harrison, David K. Jackson, Ian Johnston, Dominic Kwiatkowski, Cordelia Langford, John Sillitoe on behalf of the Wellcome Sanger Institute COVID-19 Surveillance Team |
| EPI_ISL_3094256 | Lighthouse Lab in Alderley Park | Wellcome Sanger Institute for the COVID-19 Genomics UK (COG-UK) Consortium | Jacquelyn Wynn, Mairead Hyland, The Lighthouse Lab in Alderley Park and Alex Alderton, Roberto Amato, Jeffrey Barrett, Sonia Goncalves, Ewan Harrison, David K. Jackson, Ian Johnston, Dominic Kwiatkowski, Cordelia Langford, John Sillitoe on behalf of the Wellcome Sanger Institute COVID-19 Surveillance Team |
| EPI_ISL_3094257, EPI_ISL_3094259 | Lighthouse Lab in Glasgow | Wellcome Sanger Institute for the COVID-19 Genomics UK (COG-UK) Consortium | Harper VanSteenhouse, Yumi Kasai, David Gray, Carol Clugston, Anna Dominiczak and Alex Alderton, Roberto Amato, Jeffrey Barrett, Sonia Goncalves, Ewan Harrison, David K. Jackson, Ian Johnston, Dominic Kwiatkowski, Cordelia Langford, John Sillitoe on behalf of the Wellcome Sanger Institute COVID-19 Surveillance Team |
| EPI_ISL_3094260 | Lighthouse Lab in Alderley Park | Wellcome Sanger Institute for the COVID-19 Genomics UK (COG-UK) Consortium | Jacquelyn Wynn, Mairead Hyland, The Lighthouse Lab in Alderley Park and Alex Alderton, Roberto Amato, Jeffrey Barrett, Sonia Goncalves, Ewan Harrison, David K. Jackson, Ian Johnston, Dominic Kwiatkowski, Cordelia Langford, John Sillitoe on behalf of the Wellcome Sanger Institute COVID-19 Surveillance Team |
| EPI_ISL_3094262 | Lighthouse Lab in Glasgow | Wellcome Sanger Institute for the COVID-19 Genomics UK (COG-UK) Consortium | Harper VanSteenhouse, Yumi Kasai, David Gray, Carol Clugston, Anna Dominiczak and Alex Alderton, Roberto Amato, Jeffrey Barrett, Sonia Goncalves, Ewan Harrison, David K. Jackson, Ian Johnston, Dominic Kwiatkowski, Cordelia Langford, John Sillitoe on behalf of the Wellcome Sanger Institute COVID-19 Surveillance Team |
| EPI_ISL_3094263, EPI_ISL_3094265, EPI_ISL_3094266 | Lighthouse Lab in Alderley Park | Wellcome Sanger Institute for the COVID-19 Genomics UK (COG-UK) Consortium | Jacquelyn Wynn, Mairead Hyland, The Lighthouse Lab in Alderley Park and Alex Alderton, Roberto Amato, Jeffrey Barrett, Sonia Goncalves, Ewan Harrison, David K. Jackson, Ian Johnston, Dominic Kwiatkowski, Cordelia Langford, John Sillitoe on behalf of the Wellcome Sanger Institute COVID-19 Surveillance Team |
| EPI_ISL_3094267 | Lighthouse Lab in Glasgow | Wellcome Sanger Institute for the COVID-19 Genomics UK (COG-UK) Consortium | Harper VanSteenhouse, Yumi Kasai, David Gray, Carol Clugston, Anna Dominiczak and Alex Alderton, Roberto Amato, Jeffrey Barrett, Sonia Goncalves, Ewan Harrison, David K. Jackson, Ian Johnston, Dominic Kwiatkowski, Cordelia Langford, John Sillitoe on behalf of the Wellcome Sanger Institute COVID-19 Surveillance Team |
| EPI_ISL_3094269 | Lighthouse Lab in Alderley Park | Wellcome Sanger Institute for the COVID-19 Genomics UK (COG-UK) Consortium | Jacquelyn Wynn, Mairead Hyland, The Lighthouse Lab in Alderley Park and Alex Alderton, Roberto Amato, Jeffrey Barrett, Sonia Goncalves, Ewan Harrison, David K. Jackson, Ian Johnston, Dominic Kwiatkowski, Cordelia Langford, John Sillitoe on behalf of the Wellcome Sanger Institute COVID-19 Surveillance Team |
| EPI_ISL_3094270 | Berkshire and Surrey Pathology Services Lighthouse Laboratory | Wellcome Sanger Institute for the COVID-19 Genomics UK (COG-UK) Consortium | Berkshire and Surrey Pathology Services Lighthouse Laboratory and Alex Alderton, Roberto Amato, Jeffrey Barrett, Sonia Goncalves, Ewan Harrison, David K. Jackson, Ian Johnston, Dominic Kwiatkowski, Cordelia Langford, John Sillitoe on behalf of the Wellcome Sanger Institute COVID-19 Surveillance Team |
| EPI_ISL_3094271 | Lighthouse Lab in Alderley Park | Wellcome Sanger Institute for the COVID-19 Genomics UK (COG-UK) Consortium | Jacquelyn Wynn, Mairead Hyland, The Lighthouse Lab in Alderley Park and Alex Alderton, Roberto Amato, Jeffrey Barrett, Sonia Goncalves, Ewan Harrison, David K. Jackson, Ian Johnston, Dominic Kwiatkowski, Cordelia Langford, John Sillitoe on behalf of the Wellcome Sanger Institute COVID-19 Surveillance Team |
| EPI_ISL_3094273, EPI_ISL_3094274 | Berkshire and Surrey Pathology Services Lighthouse Laboratory | Wellcome Sanger Institute for the COVID-19 Genomics UK (COG-UK) Consortium | Berkshire and Surrey Pathology Services Lighthouse Laboratory and Alex Alderton, Roberto Amato, Jeffrey Barrett, Sonia Goncalves, Ewan Harrison, David K. Jackson, Ian Johnston, Dominic Kwiatkowski, Cordelia Langford, John Sillitoe on behalf of the Wellcome Sanger Institute COVID-19 Surveillance Team |
| EPI_ISL_3094276 | Lighthouse Lab in Glasgow | Wellcome Sanger Institute for the COVID-19 Genomics UK (COG-UK) Consortium | Harper VanSteenhouse, Yumi Kasai, David Gray, Carol Clugston, Anna Dominiczak and Alex Alderton, Roberto Amato, Jeffrey Barrett, Sonia Goncalves, Ewan Harrison, David K. Jackson, Ian Johnston, Dominic Kwiatkowski, Cordelia Langford, John Sillitoe on behalf of the Wellcome Sanger Institute COVID-19 Surveillance Team |
| EPI_ISL_3094277, EPI_ISL_3094279 | Lighthouse Lab in Alderley Park | Wellcome Sanger Institute for the COVID-19 Genomics UK (COG-UK) Consortium | Jacquelyn Wynn, Mairead Hyland, The Lighthouse Lab in Alderley Park and Alex Alderton, Roberto Amato, Jeffrey Barrett, Sonia Goncalves, Ewan Harrison, David K. Jackson, Ian Johnston, Dominic Kwiatkowski, Cordelia Langford, John Sillitoe on behalf of the Wellcome Sanger Institute COVID-19 Surveillance Team |
| EPI_ISL_3094280 | Lighthouse Lab in Glasgow | Wellcome Sanger Institute for the COVID-19 Genomics UK (COG-UK) Consortium | Harper VanSteenhouse, Yumi Kasai, David Gray, Carol Clugston, Anna Dominiczak and Alex Alderton, Roberto Amato, Jeffrey Barrett, Sonia Goncalves, Ewan Harrison, David K. Jackson, Ian Johnston, Dominic Kwiatkowski, Cordelia Langford, John Sillitoe on behalf of the Wellcome Sanger Institute COVID-19 Surveillance Team |
| EPI_ISL_3094281 | Lighthouse Lab in Alderley Park | Wellcome Sanger Institute for the COVID-19 Genomics UK (COG-UK) Consortium | Jacquelyn Wynn, Mairead Hyland, The Lighthouse Lab in Alderley Park and Alex Alderton, Roberto Amato, Jeffrey Barrett, Sonia Goncalves, Ewan Harrison, David K. Jackson, Ian Johnston, Dominic Kwiatkowski, Cordelia Langford, John Sillitoe on behalf of the Wellcome Sanger Institute COVID-19 Surveillance Team |
| EPI_ISL_3094283, EPI_ISL_3094284 | Berkshire and Surrey Pathology Services Lighthouse Laboratory | Wellcome Sanger Institute for the COVID-19 Genomics UK (COG-UK) Consortium | Berkshire and Surrey Pathology Services Lighthouse Laboratory and Alex Alderton, Roberto Amato, Jeffrey Barrett, Sonia Goncalves, Ewan Harrison, David K. Jackson, Ian Johnston, Dominic Kwiatkowski, Cordelia Langford, John Sillitoe on behalf of the Wellcome Sanger Institute COVID-19 Surveillance Team |
| EPI_ISL_3094285, EPI_ISL_3094287 | Lighthouse Lab in Alderley Park | Wellcome Sanger Institute for the COVID-19 Genomics UK (COG-UK) Consortium | Jacquelyn Wynn, Mairead Hyland, The Lighthouse Lab in Alderley Park and Alex Alderton, Roberto Amato, Jeffrey Barrett, Sonia Goncalves, Ewan Harrison, David K. Jackson, Ian Johnston, Dominic Kwiatkowski, Cordelia Langford, John Sillitoe on behalf of the Wellcome Sanger Institute COVID-19 Surveillance Team |
| EPI_ISL_3094288, EPI_ISL_3094290 | Lighthouse Lab in Glasgow | Wellcome Sanger Institute for the COVID-19 Genomics UK (COG-UK) Consortium | Harper VanSteenhouse, Yumi Kasai, David Gray, Carol Clugston, Anna Dominiczak and Alex Alderton, Roberto Amato, Jeffrey Barrett, Sonia Goncalves, Ewan Harrison, David K. Jackson, Ian Johnston, Dominic Kwiatkowski, Cordelia Langford, John Sillitoe on behalf of the Wellcome Sanger Institute COVID-19 Surveillance Team |
| EPI_ISL_3094291 | Berkshire and Surrey Pathology Services Lighthouse Laboratory | Wellcome Sanger Institute for the COVID-19 Genomics UK (COG-UK) Consortium | Berkshire and Surrey Pathology Services Lighthouse Laboratory and Alex Alderton, Roberto Amato, Jeffrey Barrett, Sonia Goncalves, Ewan Harrison, David K. Jackson, Ian Johnston, Dominic Kwiatkowski, Cordelia Langford, John Sillitoe on behalf of the Wellcome Sanger Institute COVID-19 Surveillance Team |
| EPI_ISL_3094292 | Lighthouse Lab in Alderley Park | Wellcome Sanger Institute for the COVID-19 Genomics UK (COG-UK) Consortium | Jacquelyn Wynn, Mairead Hyland, The Lighthouse Lab in Alderley Park and Alex Alderton, Roberto Amato, Jeffrey Barrett, Sonia Goncalves, Ewan Harrison, David K. Jackson, Ian Johnston, Dominic Kwiatkowski, Cordelia Langford, John Sillitoe on behalf of the Wellcome Sanger Institute COVID-19 Surveillance Team |
| EPI_ISL_3094294 | Lighthouse Lab in Glasgow | Wellcome Sanger Institute for the COVID-19 Genomics UK (COG-UK) Consortium | Harper VanSteenhouse, Yumi Kasai, David Gray, Carol Clugston, Anna Dominiczak and Alex Alderton, Roberto Amato, Jeffrey Barrett, Sonia Goncalves, Ewan Harrison, David K. Jackson, Ian Johnston, Dominic Kwiatkowski, Cordelia Langford, John Sillitoe on behalf of the Wellcome Sanger Institute COVID-19 Surveillance Team |
| EPI_ISL_3094295, EPI_ISL_3094296 | Lighthouse Lab in Milton Keynes | Wellcome Sanger Institute for the COVID-19 Genomics UK (COG-UK) Consortium | The Lighthouse Lab in Milton Keynes and Alex Alderton, Roberto Amato, Jeffrey Barrett, Sonia Goncalves, Ewan Harrison, David K. Jackson, Ian Johnston, Dominic Kwiatkowski, Cordelia Langford, John Sillitoe on behalf of the Wellcome Sanger Institute COVID-19 Surveillance Team |
| EPI_ISL_3094298 | Lighthouse Laboratory Plymouth | Wellcome Sanger Institute for the COVID-19 Genomics UK (COG-UK) Consortium | Lighthouse Laboratory Plymouth and Alex Alderton, Roberto Amato, Jeffrey Barrett, Sonia Goncalves, Ewan Harrison, David K. Jackson, Ian Johnston, Dominic Kwiatkowski, Cordelia Langford, John Sillitoe on behalf of the Wellcome Sanger Institute COVID-19 Surveillance Team |
| EPI_ISL_3094299, EPI_ISL_3094300, EPI_ISL_3094302, EPI_ISL_3094303 | Lighthouse Lab in Milton Keynes | Wellcome Sanger Institute for the COVID-19 Genomics UK (COG-UK) Consortium | The Lighthouse Lab in Milton Keynes and Alex Alderton, Roberto Amato, Jeffrey Barrett, Sonia Goncalves, Ewan Harrison, David K. Jackson, Ian Johnston, Dominic Kwiatkowski, Cordelia Langford, John Sillitoe on behalf of the Wellcome Sanger Institute COVID-19 Surveillance Team |
| EPI_ISL_3094304 | Lighthouse Lab in Glasgow | Wellcome Sanger Institute for the COVID-19 Genomics UK (COG-UK) Consortium | Harper VanSteenhouse, Yumi Kasai, David Gray, Carol Clugston, Anna Dominiczak and Alex Alderton, Roberto Amato, Jeffrey Barrett, Sonia Goncalves, Ewan Harrison, David K. Jackson, Ian Johnston, Dominic Kwiatkowski, Cordelia Langford, John Sillitoe on behalf of the Wellcome Sanger Institute COVID-19 Surveillance Team |
| EPI_ISL_3094306, EPI_ISL_3094307 | Lighthouse Lab in Milton Keynes | Wellcome Sanger Institute for the COVID-19 Genomics UK (COG-UK) Consortium | The Lighthouse Lab in Milton Keynes and Alex Alderton, Roberto Amato, Jeffrey Barrett, Sonia Goncalves, Ewan Harrison, David K. Jackson, Ian Johnston, Dominic Kwiatkowski, Cordelia Langford, John Sillitoe on behalf of the Wellcome Sanger Institute COVID-19 Surveillance Team |
| EPI_ISL_3094309 | Lighthouse Lab in Glasgow | Wellcome Sanger Institute for the COVID-19 Genomics UK (COG-UK) Consortium | Harper VanSteenhouse, Yumi Kasai, David Gray, Carol Clugston, Anna Dominiczak and Alex Alderton, Roberto Amato, Jeffrey Barrett, Sonia Goncalves, Ewan Harrison, David K. Jackson, Ian Johnston, Dominic Kwiatkowski, Cordelia Langford, John Sillitoe on behalf of the Wellcome Sanger Institute COVID-19 Surveillance Team |
| EPI_ISL_3094310 | Lighthouse Laboratory Plymouth | Wellcome Sanger Institute for the COVID-19 Genomics UK (COG-UK) Consortium | Lighthouse Laboratory Plymouth and Alex Alderton, Roberto Amato, Jeffrey Barrett, Sonia Goncalves, Ewan Harrison, David K. Jackson, Ian Johnston, Dominic Kwiatkowski, Cordelia Langford, John Sillitoe on behalf of the Wellcome Sanger Institute COVID-19 Surveillance Team |
| EPI_ISL_3094311 | Lighthouse Lab in Glasgow | Wellcome Sanger Institute for the COVID-19 Genomics UK (COG-UK) Consortium | Harper VanSteenhouse, Yumi Kasai, David Gray, Carol Clugston, Anna Dominiczak and Alex Alderton, Roberto Amato, Jeffrey Barrett, Sonia Goncalves, Ewan Harrison, David K. Jackson, Ian Johnston, Dominic Kwiatkowski, Cordelia Langford, John Sillitoe on behalf of the Wellcome Sanger Institute COVID-19 Surveillance Team |
| EPI_ISL_3094313, EPI_ISL_3094314, EPI_ISL_3094315, EPI_ISL_3094317 | Lighthouse Lab in Milton Keynes | Wellcome Sanger Institute for the COVID-19 Genomics UK (COG-UK) Consortium | The Lighthouse Lab in Milton Keynes and Alex Alderton, Roberto Amato, Jeffrey Barrett, Sonia Goncalves, Ewan Harrison, David K. Jackson, Ian Johnston, Dominic Kwiatkowski, Cordelia Langford, John Sillitoe on behalf of the Wellcome Sanger Institute COVID-19 Surveillance Team |
| EPI_ISL_3094318 | Lighthouse Lab in Glasgow | Wellcome Sanger Institute for the COVID-19 Genomics UK (COG-UK) Consortium | Harper VanSteenhouse, Yumi Kasai, David Gray, Carol Clugston, Anna Dominiczak and Alex Alderton, Roberto Amato, Jeffrey Barrett, Sonia Goncalves, Ewan Harrison, David K. Jackson, Ian Johnston, Dominic Kwiatkowski, Cordelia Langford, John Sillitoe on behalf of the Wellcome Sanger Institute COVID-19 Surveillance Team |

| | | | |
|---|---|---|---|
| EPI_ISL_3094319, EPI_ISL_3094321 | Lighthouse Lab in Milton Keynes | Wellcome Sanger Institute for the COVID-19 Genomics UK (COG-UK) Consortium | The Lighthouse Lab in Milton Keynes and Alex Alderton, Roberto Amato, Jeffrey Barrett, Sonia Goncalves, Ewan Harrison, David K. Jackson, Ian Johnston, Dominic Kwiatkowski, Cordelia Langford, John Sillitoe on behalf of the Wellcome Sanger Institute COVID-19 Surveillance Team |
| EPI_ISL_3094322 | Lighthouse Laboratory Plymouth | Wellcome Sanger Institute for the COVID-19 Genomics UK (COG-UK) Consortium | Lighthouse Laboratory Plymouth and Alex Alderton, Roberto Amato, Jeffrey Barrett, Sonia Goncalves, Ewan Harrison, David K. Jackson, Ian Johnston, Dominic Kwiatkowski, Cordelia Langford, John Sillitoe on behalf of the Wellcome Sanger Institute COVID-19 Surveillance Team |
| EPI_ISL_3094324, EPI_ISL_3094325 | Lighthouse Lab in Glasgow | Wellcome Sanger Institute for the COVID-19 Genomics UK (COG-UK) Consortium | Harper VanSteenhouse, Yumi Kasai, David Gray, Carol Clugston, Anna Dominiczak and Alex Alderton, Roberto Amato, Jeffrey Barrett, Sonia Goncalves, Ewan Harrison, David K. Jackson, Ian Johnston, Dominic Kwiatkowski, Cordelia Langford, John Sillitoe on behalf of the Wellcome Sanger Institute COVID-19 Surveillance Team |
| EPI_ISL_3094326, EPI_ISL_3094328, EPI_ISL_3094329, EPI_ISL_3094330, EPI_ISL_3094331 | Lighthouse Lab in Milton Keynes | Wellcome Sanger Institute for the COVID-19 Genomics UK (COG-UK) Consortium | The Lighthouse Lab in Milton Keynes and Alex Alderton, Roberto Amato, Jeffrey Barrett, Sonia Goncalves, Ewan Harrison, David K. Jackson, Ian Johnston, Dominic Kwiatkowski, Cordelia Langford, John Sillitoe on behalf of the Wellcome Sanger Institute COVID-19 Surveillance Team |
| EPI_ISL_3094333 | Lighthouse Lab in Glasgow | Wellcome Sanger Institute for the COVID-19 Genomics UK (COG-UK) Consortium | Harper VanSteenhouse, Yumi Kasai, David Gray, Carol Clugston, Anna Dominiczak and Alex Alderton, Roberto Amato, Jeffrey Barrett, Sonia Goncalves, Ewan Harrison, David K. Jackson, Ian Johnston, Dominic Kwiatkowski, Cordelia Langford, John Sillitoe on behalf of the Wellcome Sanger Institute COVID-19 Surveillance Team |
| EPI_ISL_3094334 | Lighthouse Lab in Milton Keynes | Wellcome Sanger Institute for the COVID-19 Genomics UK (COG-UK) Consortium | The Lighthouse Lab in Milton Keynes and Alex Alderton, Roberto Amato, Jeffrey Barrett, Sonia Goncalves, Ewan Harrison, David K. Jackson, Ian Johnston, Dominic Kwiatkowski, Cordelia Langford, John Sillitoe on behalf of the Wellcome Sanger Institute COVID-19 Surveillance Team |
| EPI_ISL_3094336 | Lighthouse Lab in Glasgow | Wellcome Sanger Institute for the COVID-19 Genomics UK (COG-UK) Consortium | Harper VanSteenhouse, Yumi Kasai, David Gray, Carol Clugston, Anna Dominiczak and Alex Alderton, Roberto Amato, Jeffrey Barrett, Sonia Goncalves, Ewan Harrison, David K. Jackson, Ian Johnston, Dominic Kwiatkowski, Cordelia Langford, John Sillitoe on behalf of the Wellcome Sanger Institute COVID-19 Surveillance Team |
| EPI_ISL_3094337 | Lighthouse Laboratory Plymouth | Wellcome Sanger Institute for the COVID-19 Genomics UK (COG-UK) Consortium | Lighthouse Laboratory Plymouth and Alex Alderton, Roberto Amato, Jeffrey Barrett, Sonia Goncalves, Ewan Harrison, David K. Jackson, Ian Johnston, Dominic Kwiatkowski, Cordelia Langford, John Sillitoe on behalf of the Wellcome Sanger Institute COVID-19 Surveillance Team |
| EPI_ISL_3094338, EPI_ISL_3094340, EPI_ISL_3094341, EPI_ISL_3094343, EPI_ISL_3094344, EPI_ISL_3094345, EPI_ISL_3094347, EPI_ISL_3094348, EPI_ISL_3094350 | Lighthouse Lab in Milton Keynes | Wellcome Sanger Institute for the COVID-19 Genomics UK (COG-UK) Consortium | The Lighthouse Lab in Milton Keynes and Alex Alderton, Roberto Amato, Jeffrey Barrett, Sonia Goncalves, Ewan Harrison, David K. Jackson, Ian Johnston, Dominic Kwiatkowski, Cordelia Langford, John Sillitoe on behalf of the Wellcome Sanger Institute COVID-19 Surveillance Team |
| EPI_ISL_3094351 | Lighthouse Laboratory Plymouth | Wellcome Sanger Institute for the COVID-19 Genomics UK (COG-UK) Consortium | Lighthouse Laboratory Plymouth and Alex Alderton, Roberto Amato, Jeffrey Barrett, Sonia Goncalves, Ewan Harrison, David K. Jackson, Ian Johnston, Dominic Kwiatkowski, Cordelia Langford, John Sillitoe on behalf of the Wellcome Sanger Institute COVID-19 Surveillance Team |
| EPI_ISL_3094352 | Lighthouse Lab in Milton Keynes | Wellcome Sanger Institute for the COVID-19 Genomics UK (COG-UK) Consortium | The Lighthouse Lab in Milton Keynes and Alex Alderton, Roberto Amato, Jeffrey Barrett, Sonia Goncalves, Ewan Harrison, David K. Jackson, Ian Johnston, Dominic Kwiatkowski, Cordelia Langford, John Sillitoe on behalf of the Wellcome Sanger Institute COVID-19 Surveillance Team |
| EPI_ISL_3094354 | Lighthouse Lab in Glasgow | Wellcome Sanger Institute for the COVID-19 Genomics UK (COG-UK) Consortium | Harper VanSteenhouse, Yumi Kasai, David Gray, Carol Clugston, Anna Dominiczak and Alex Alderton, Roberto Amato, Jeffrey Barrett, Sonia Goncalves, Ewan Harrison, David K. Jackson, Ian Johnston, Dominic Kwiatkowski, Cordelia Langford, John Sillitoe on behalf of the Wellcome Sanger Institute COVID-19 Surveillance Team |
| EPI_ISL_3094355, EPI_ISL_3094357, EPI_ISL_3094358, EPI_ISL_3094359, EPI_ISL_3094361, EPI_ISL_3094362, EPI_ISL_3094363, EPI_ISL_3094365 | Lighthouse Lab in Milton Keynes | Wellcome Sanger Institute for the COVID-19 Genomics UK (COG-UK) Consortium | The Lighthouse Lab in Milton Keynes and Alex Alderton, Roberto Amato, Jeffrey Barrett, Sonia Goncalves, Ewan Harrison, David K. Jackson, Ian Johnston, Dominic Kwiatkowski, Cordelia Langford, John Sillitoe on behalf of the Wellcome Sanger Institute COVID-19 Surveillance Team |
| EPI_ISL_3094366 | Lighthouse Lab in Glasgow | Wellcome Sanger Institute for the COVID-19 Genomics UK (COG-UK) Consortium | Harper VanSteenhouse, Yumi Kasai, David Gray, Carol Clugston, Anna Dominiczak and Alex Alderton, Roberto Amato, Jeffrey Barrett, Sonia Goncalves, Ewan Harrison, David K. Jackson, Ian Johnston, Dominic Kwiatkowski, Cordelia Langford, John Sillitoe on behalf of the Wellcome Sanger Institute COVID-19 Surveillance Team |
| EPI_ISL_3094367, EPI_ISL_3094369 | Lighthouse Lab in Milton Keynes | Wellcome Sanger Institute for the COVID-19 Genomics UK (COG-UK) Consortium | The Lighthouse Lab in Milton Keynes and Alex Alderton, Roberto Amato, Jeffrey Barrett, Sonia Goncalves, Ewan Harrison, David K. Jackson, Ian Johnston, Dominic Kwiatkowski, Cordelia Langford, John Sillitoe on behalf of the Wellcome Sanger Institute COVID-19 Surveillance Team |
| EPI_ISL_3094370 | Lighthouse Lab in Glasgow | Wellcome Sanger Institute for the COVID-19 Genomics UK (COG-UK) Consortium | Harper VanSteenhouse, Yumi Kasai, David Gray, Carol Clugston, Anna Dominiczak and Alex Alderton, Roberto Amato, Jeffrey Barrett, Sonia Goncalves, Ewan Harrison, David K. Jackson, Ian Johnston, Dominic Kwiatkowski, Cordelia Langford, John Sillitoe on behalf of the Wellcome Sanger Institute COVID-19 Surveillance Team |
| EPI_ISL_3094372, EPI_ISL_3094373, EPI_ISL_3094374 | Lighthouse Lab in Milton Keynes | Wellcome Sanger Institute for the COVID-19 Genomics UK (COG-UK) Consortium | The Lighthouse Lab in Milton Keynes and Alex Alderton, Roberto Amato, Jeffrey Barrett, Sonia Goncalves, Ewan Harrison, David K. Jackson, Ian Johnston, Dominic Kwiatkowski, Cordelia Langford, John Sillitoe on behalf of the Wellcome Sanger Institute COVID-19 Surveillance Team |
| EPI_ISL_3094376 | Lighthouse Laboratory Plymouth | Wellcome Sanger Institute for the COVID-19 Genomics UK (COG-UK) Consortium | Lighthouse Laboratory Plymouth and Alex Alderton, Roberto Amato, Jeffrey Barrett, Sonia Goncalves, Ewan Harrison, David K. Jackson, Ian Johnston, Dominic Kwiatkowski, Cordelia Langford, John Sillitoe on behalf of the Wellcome Sanger Institute COVID-19 Surveillance Team |
| EPI_ISL_3094377, EPI_ISL_3094379 | Lighthouse Lab in Milton Keynes | Wellcome Sanger Institute for the COVID-19 Genomics UK (COG-UK) Consortium | The Lighthouse Lab in Milton Keynes and Alex Alderton, Roberto Amato, Jeffrey Barrett, Sonia Goncalves, Ewan Harrison, David K. Jackson, Ian Johnston, Dominic Kwiatkowski, Cordelia Langford, John Sillitoe on behalf of the Wellcome Sanger Institute COVID-19 Surveillance Team |
| EPI_ISL_3094380, EPI_ISL_3094381 | Lighthouse Laboratory Plymouth | Wellcome Sanger Institute for the COVID-19 Genomics UK (COG-UK) Consortium | Lighthouse Laboratory Plymouth and Alex Alderton, Roberto Amato, Jeffrey Barrett, Sonia Goncalves, Ewan Harrison, David K. Jackson, Ian Johnston, Dominic Kwiatkowski, Cordelia Langford, John Sillitoe on behalf of the Wellcome Sanger Institute COVID-19 Surveillance Team |
| EPI_ISL_3094383 | Lighthouse Lab in Milton Keynes | Wellcome Sanger Institute for the COVID-19 Genomics UK (COG-UK) Consortium | The Lighthouse Lab in Milton Keynes and Alex Alderton, Roberto Amato, Jeffrey Barrett, Sonia Goncalves, Ewan Harrison, David K. Jackson, Ian Johnston, Dominic Kwiatkowski, Cordelia Langford, John Sillitoe on behalf of the Wellcome Sanger Institute COVID-19 Surveillance Team |
| EPI_ISL_3094384, EPI_ISL_3094386 | Lighthouse Laboratory Plymouth | Wellcome Sanger Institute for the COVID-19 Genomics UK (COG-UK) Consortium | Lighthouse Laboratory Plymouth and Alex Alderton, Roberto Amato, Jeffrey Barrett, Sonia Goncalves, Ewan Harrison, David K. Jackson, Ian Johnston, Dominic Kwiatkowski, Cordelia Langford, John Sillitoe on behalf of the Wellcome Sanger Institute COVID-19 Surveillance Team |
| EPI_ISL_3094387 | Lighthouse Lab in Glasgow | Wellcome Sanger Institute for the COVID-19 Genomics UK (COG-UK) Consortium | Harper VanSteenhouse, Yumi Kasai, David Gray, Carol Clugston, Anna Dominiczak and Alex Alderton, Roberto Amato, Jeffrey Barrett, Sonia Goncalves, Ewan Harrison, David K. Jackson, Ian Johnston, Dominic Kwiatkowski, Cordelia Langford, John Sillitoe on behalf of the Wellcome Sanger Institute COVID-19 Surveillance Team |
| EPI_ISL_3094388 | Lighthouse Laboratory Plymouth | Wellcome Sanger Institute for the COVID-19 Genomics UK (COG-UK) Consortium | Lighthouse Laboratory Plymouth and Alex Alderton, Roberto Amato, Jeffrey Barrett, Sonia Goncalves, Ewan Harrison, David K. Jackson, Ian Johnston, Dominic Kwiatkowski, Cordelia Langford, John Sillitoe on behalf of the Wellcome Sanger Institute COVID-19 Surveillance Team |
| EPI_ISL_3094390 | Lighthouse Lab in Milton Keynes | Wellcome Sanger Institute for the COVID-19 Genomics UK (COG-UK) Consortium | The Lighthouse Lab in Milton Keynes and Alex Alderton, Roberto Amato, Jeffrey Barrett, Sonia Goncalves, Ewan Harrison, David K. Jackson, Ian Johnston, Dominic Kwiatkowski, Cordelia Langford, John Sillitoe on behalf of the Wellcome Sanger Institute COVID-19 Surveillance Team |
| EPI_ISL_3094391 | Lighthouse Laboratory Plymouth | Wellcome Sanger Institute for the COVID-19 Genomics UK (COG-UK) Consortium | Lighthouse Laboratory Plymouth and Alex Alderton, Roberto Amato, Jeffrey Barrett, Sonia Goncalves, Ewan Harrison, David K. Jackson, Ian Johnston, Dominic Kwiatkowski, Cordelia Langford, John Sillitoe on behalf of the Wellcome Sanger Institute COVID-19 Surveillance Team |
| EPI_ISL_3094392 | Lighthouse Lab in Milton Keynes | Wellcome Sanger Institute for the COVID-19 Genomics UK (COG-UK) Consortium | The Lighthouse Lab in Milton Keynes and Alex Alderton, Roberto Amato, Jeffrey Barrett, Sonia Goncalves, Ewan Harrison, David K. Jackson, Ian Johnston, Dominic Kwiatkowski, Cordelia Langford, John Sillitoe on behalf of the Wellcome Sanger Institute COVID-19 Surveillance Team |
| EPI_ISL_3094394 | Lighthouse Lab in Glasgow | Wellcome Sanger Institute for the COVID-19 Genomics UK (COG-UK) Consortium | Harper VanSteenhouse, Yumi Kasai, David Gray, Carol Clugston, Anna Dominiczak and Alex Alderton, Roberto Amato, Jeffrey Barrett, Sonia Goncalves, Ewan Harrison, David K. Jackson, Ian Johnston, Dominic Kwiatkowski, Cordelia Langford, John Sillitoe on behalf of the Wellcome Sanger Institute COVID-19 Surveillance Team |
| EPI_ISL_3094395, EPI_ISL_3094396, EPI_ISL_3094398, EPI_ISL_3094399 | Lighthouse Lab in Milton Keynes | Wellcome Sanger Institute for the COVID-19 Genomics UK (COG-UK) Consortium | The Lighthouse Lab in Milton Keynes and Alex Alderton, Roberto Amato, Jeffrey Barrett, Sonia Goncalves, Ewan Harrison, David K. Jackson, Ian Johnston, Dominic Kwiatkowski, Cordelia Langford, John Sillitoe on behalf of the Wellcome Sanger Institute COVID-19 Surveillance Team |
| EPI_ISL_3094401 | Lighthouse Laboratory Plymouth | Wellcome Sanger Institute for the COVID-19 Genomics UK (COG-UK) Consortium | Lighthouse Laboratory Plymouth and Alex Alderton, Roberto Amato, Jeffrey Barrett, Sonia Goncalves, Ewan Harrison, David K. Jackson, Ian Johnston, Dominic Kwiatkowski, Cordelia Langford, John Sillitoe on behalf of the Wellcome Sanger Institute COVID-19 Surveillance Team |
| EPI_ISL_3094402 | Lighthouse Lab in Glasgow | Wellcome Sanger Institute for the COVID-19 Genomics UK (COG-UK) Consortium | Harper VanSteenhouse, Yumi Kasai, David Gray, Carol Clugston, Anna Dominiczak and Alex Alderton, Roberto Amato, Jeffrey Barrett, Sonia Goncalves, Ewan Harrison, David K. Jackson, Ian Johnston, Dominic Kwiatkowski, Cordelia Langford, John Sillitoe on behalf of the Wellcome Sanger Institute COVID-19 Surveillance Team |
| EPI_ISL_3094403 | Lighthouse Laboratory Plymouth | Wellcome Sanger Institute for the COVID-19 Genomics UK (COG-UK) Consortium | Lighthouse Laboratory Plymouth and Alex Alderton, Roberto Amato, Jeffrey Barrett, Sonia Goncalves, Ewan Harrison, David K. Jackson, Ian Johnston, Dominic Kwiatkowski, Cordelia Langford, John Sillitoe on behalf of the Wellcome Sanger Institute COVID-19 Surveillance Team |
| EPI_ISL_3094405 | Lighthouse Lab in Milton Keynes | Wellcome Sanger Institute for the COVID-19 Genomics UK (COG-UK) Consortium | The Lighthouse Lab in Milton Keynes and Alex Alderton, Roberto Amato, Jeffrey Barrett, Sonia Goncalves, Ewan Harrison, David K. Jackson, Ian Johnston, Dominic Kwiatkowski, Cordelia Langford, John Sillitoe on behalf of the Wellcome Sanger Institute COVID-19 Surveillance Team |
| EPI_ISL_3094406 | Lighthouse Lab in Glasgow | Wellcome Sanger Institute for the COVID-19 Genomics UK (COG-UK) Consortium | Harper VanSteenhouse, Yumi Kasai, David Gray, Carol Clugston, Anna Dominiczak and Alex Alderton, Roberto Amato, Jeffrey Barrett, Sonia Goncalves, Ewan Harrison, David K. Jackson, Ian Johnston, Dominic Kwiatkowski, Cordelia Langford, John Sillitoe on behalf of the Wellcome Sanger Institute COVID-19 Surveillance Team |
| EPI_ISL_3094407, EPI_ISL_3094408, EPI_ISL_3094410 | Lighthouse Lab in Milton Keynes | Wellcome Sanger Institute for the COVID-19 Genomics UK (COG-UK) Consortium | The Lighthouse Lab in Milton Keynes and Alex Alderton, Roberto Amato, Jeffrey Barrett, Sonia Goncalves, Ewan Harrison, David K. Jackson, Ian Johnston, Dominic Kwiatkowski, Cordelia Langford, John Sillitoe on behalf of the Wellcome Sanger Institute COVID-19 Surveillance Team |
| EPI_ISL_3094411, EPI_ISL_3094413 | Lighthouse Laboratory Plymouth | Wellcome Sanger Institute for the COVID-19 Genomics UK (COG-UK) Consortium | Lighthouse Laboratory Plymouth and Alex Alderton, Roberto Amato, Jeffrey Barrett, Sonia Goncalves, Ewan Harrison, David K. Jackson, Ian Johnston, Dominic Kwiatkowski, Cordelia Langford, John Sillitoe on behalf of the Wellcome Sanger Institute COVID-19 Surveillance Team |
| EPI_ISL_3094414 | Lighthouse Lab in Glasgow | Wellcome Sanger Institute for the COVID-19 Genomics UK (COG-UK) | Harper VanSteenhouse, Yumi Kasai, David Gray, Carol Clugston, Anna Dominiczak and Alex Alderton, Roberto Amato, Jeffrey Barrett, Sonia Goncalves, Ewan Harrison, David K. Jackson, Ian Johnston, Dominic Kwiatkowski, Cordelia Langford, |

| | | | |
|---|---|---|---|
| | | Consortium | John Sillitoe on behalf of the Wellcome Sanger Institute COVID-19 Surveillance Team |
| EPI_ISL_3094415 | Lighthouse Lab in Milton Keynes | Wellcome Sanger Institute for the COVID-19 Genomics UK (COG-UK) Consortium | The Lighthouse Lab in Milton Keynes and Alex Alderton, Roberto Amato, Jeffrey Barrett, Sonia Goncalves, Ewan Harrison, David K. Jackson, Ian Johnston, Dominic Kwiatkowski, Cordelia Langford, John Sillitoe on behalf of the Wellcome Sanger Institute COVID-19 Surveillance Team |
| EPI_ISL_3094417 | Lighthouse Laboratory Plymouth | Wellcome Sanger Institute for the COVID-19 Genomics UK (COG-UK) Consortium | Lighthouse Laboratory Plymouth and Alex Alderton, Roberto Amato, Jeffrey Barrett, Sonia Goncalves, Ewan Harrison, David K. Jackson, Ian Johnston, Dominic Kwiatkowski, Cordelia Langford, John Sillitoe on behalf of the Wellcome Sanger Institute COVID-19 Surveillance Team |
| EPI_ISL_3094418 | Lighthouse Lab in Milton Keynes | Wellcome Sanger Institute for the COVID-19 Genomics UK (COG-UK) Consortium | The Lighthouse Lab in Milton Keynes and Alex Alderton, Roberto Amato, Jeffrey Barrett, Sonia Goncalves, Ewan Harrison, David K. Jackson, Ian Johnston, Dominic Kwiatkowski, Cordelia Langford, John Sillitoe on behalf of the Wellcome Sanger Institute COVID-19 Surveillance Team |
| EPI_ISL_3094419, EPI_ISL_3094421 | Lighthouse Laboratory Plymouth | Wellcome Sanger Institute for the COVID-19 Genomics UK (COG-UK) Consortium | Lighthouse Laboratory Plymouth and Alex Alderton, Roberto Amato, Jeffrey Barrett, Sonia Goncalves, Ewan Harrison, David K. Jackson, Ian Johnston, Dominic Kwiatkowski, Cordelia Langford, John Sillitoe on behalf of the Wellcome Sanger Institute COVID-19 Surveillance Team |
| EPI_ISL_3094422, EPI_ISL_3094423, EPI_ISL_3094425 | Lighthouse Lab in Milton Keynes | Wellcome Sanger Institute for the COVID-19 Genomics UK (COG-UK) Consortium | The Lighthouse Lab in Milton Keynes and Alex Alderton, Roberto Amato, Jeffrey Barrett, Sonia Goncalves, Ewan Harrison, David K. Jackson, Ian Johnston, Dominic Kwiatkowski, Cordelia Langford, John Sillitoe on behalf of the Wellcome Sanger Institute COVID-19 Surveillance Team |
| EPI_ISL_3094426 | Lighthouse Laboratory Plymouth | Wellcome Sanger Institute for the COVID-19 Genomics UK (COG-UK) Consortium | Lighthouse Laboratory Plymouth and Alex Alderton, Roberto Amato, Jeffrey Barrett, Sonia Goncalves, Ewan Harrison, David K. Jackson, Ian Johnston, Dominic Kwiatkowski, Cordelia Langford, John Sillitoe on behalf of the Wellcome Sanger Institute COVID-19 Surveillance Team |
| EPI_ISL_3094427 | Lighthouse Lab in Milton Keynes | Wellcome Sanger Institute for the COVID-19 Genomics UK (COG-UK) Consortium | The Lighthouse Lab in Milton Keynes and Alex Alderton, Roberto Amato, Jeffrey Barrett, Sonia Goncalves, Ewan Harrison, David K. Jackson, Ian Johnston, Dominic Kwiatkowski, Cordelia Langford, John Sillitoe on behalf of the Wellcome Sanger Institute COVID-19 Surveillance Team |
| EPI_ISL_3094429, EPI_ISL_3094430 | Lighthouse Laboratory Plymouth | Wellcome Sanger Institute for the COVID-19 Genomics UK (COG-UK) Consortium | Lighthouse Laboratory Plymouth and Alex Alderton, Roberto Amato, Jeffrey Barrett, Sonia Goncalves, Ewan Harrison, David K. Jackson, Ian Johnston, Dominic Kwiatkowski, Cordelia Langford, John Sillitoe on behalf of the Wellcome Sanger Institute COVID-19 Surveillance Team |
| EPI_ISL_3094432 | Lighthouse Lab in Milton Keynes | Wellcome Sanger Institute for the COVID-19 Genomics UK (COG-UK) Consortium | The Lighthouse Lab in Milton Keynes and Alex Alderton, Roberto Amato, Jeffrey Barrett, Sonia Goncalves, Ewan Harrison, David K. Jackson, Ian Johnston, Dominic Kwiatkowski, Cordelia Langford, John Sillitoe on behalf of the Wellcome Sanger Institute COVID-19 Surveillance Team |
| EPI_ISL_3094433, EPI_ISL_3094434, EPI_ISL_3094436 | Lighthouse Laboratory Plymouth | Wellcome Sanger Institute for the COVID-19 Genomics UK (COG-UK) Consortium | Lighthouse Laboratory Plymouth and Alex Alderton, Roberto Amato, Jeffrey Barrett, Sonia Goncalves, Ewan Harrison, David K. Jackson, Ian Johnston, Dominic Kwiatkowski, Cordelia Langford, John Sillitoe on behalf of the Wellcome Sanger Institute COVID-19 Surveillance Team |
| EPI_ISL_3094437 | Lighthouse Lab in Milton Keynes | Wellcome Sanger Institute for the COVID-19 Genomics UK (COG-UK) Consortium | The Lighthouse Lab in Milton Keynes and Alex Alderton, Roberto Amato, Jeffrey Barrett, Sonia Goncalves, Ewan Harrison, David K. Jackson, Ian Johnston, Dominic Kwiatkowski, Cordelia Langford, John Sillitoe on behalf of the Wellcome Sanger Institute COVID-19 Surveillance Team |
| EPI_ISL_3094438, EPI_ISL_3094440 | Lighthouse Lab in Glasgow | Wellcome Sanger Institute for the COVID-19 Genomics UK (COG-UK) Consortium | Harper VanSteenhouse, Yumi Kasai, David Gray, Carol Clugston, Anna Dominiczak and Alex Alderton, Roberto Amato, Jeffrey Barrett, Sonia Goncalves, Ewan Harrison, David K. Jackson, Ian Johnston, Dominic Kwiatkowski, Cordelia Langford, John Sillitoe on behalf of the Wellcome Sanger Institute COVID-19 Surveillance Team |
| EPI_ISL_3094441, EPI_ISL_3094442, EPI_ISL_3094444 | Lighthouse Laboratory Plymouth | Wellcome Sanger Institute for the COVID-19 Genomics UK (COG-UK) Consortium | Lighthouse Laboratory Plymouth and Alex Alderton, Roberto Amato, Jeffrey Barrett, Sonia Goncalves, Ewan Harrison, David K. Jackson, Ian Johnston, Dominic Kwiatkowski, Cordelia Langford, John Sillitoe on behalf of the Wellcome Sanger Institute COVID-19 Surveillance Team |
| EPI_ISL_3094445, EPI_ISL_3094446 | Lighthouse Lab in Milton Keynes | Wellcome Sanger Institute for the COVID-19 Genomics UK (COG-UK) Consortium | The Lighthouse Lab in Milton Keynes and Alex Alderton, Roberto Amato, Jeffrey Barrett, Sonia Goncalves, Ewan Harrison, David K. Jackson, Ian Johnston, Dominic Kwiatkowski, Cordelia Langford, John Sillitoe on behalf of the Wellcome Sanger Institute COVID-19 Surveillance Team |
| EPI_ISL_3094447, EPI_ISL_3094449 | Lighthouse Laboratory Plymouth | Wellcome Sanger Institute for the COVID-19 Genomics UK (COG-UK) Consortium | Lighthouse Laboratory Plymouth and Alex Alderton, Roberto Amato, Jeffrey Barrett, Sonia Goncalves, Ewan Harrison, David K. Jackson, Ian Johnston, Dominic Kwiatkowski, Cordelia Langford, John Sillitoe on behalf of the Wellcome Sanger Institute COVID-19 Surveillance Team |
| EPI_ISL_3094450 | Lighthouse Lab in Glasgow | Wellcome Sanger Institute for the COVID-19 Genomics UK (COG-UK) Consortium | Harper VanSteenhouse, Yumi Kasai, David Gray, Carol Clugston, Anna Dominiczak and Alex Alderton, Roberto Amato, Jeffrey Barrett, Sonia Goncalves, Ewan Harrison, David K. Jackson, Ian Johnston, Dominic Kwiatkowski, Cordelia Langford, John Sillitoe on behalf of the Wellcome Sanger Institute COVID-19 Surveillance Team |
| EPI_ISL_3094451 | Lighthouse Lab in Milton Keynes | Wellcome Sanger Institute for the COVID-19 Genomics UK (COG-UK) Consortium | The Lighthouse Lab in Milton Keynes and Alex Alderton, Roberto Amato, Jeffrey Barrett, Sonia Goncalves, Ewan Harrison, David K. Jackson, Ian Johnston, Dominic Kwiatkowski, Cordelia Langford, John Sillitoe on behalf of the Wellcome Sanger Institute COVID-19 Surveillance Team |
| EPI_ISL_3094453 | Lighthouse Lab in Glasgow | Wellcome Sanger Institute for the COVID-19 Genomics UK (COG-UK) Consortium | Harper VanSteenhouse, Yumi Kasai, David Gray, Carol Clugston, Anna Dominiczak and Alex Alderton, Roberto Amato, Jeffrey Barrett, Sonia Goncalves, Ewan Harrison, David K. Jackson, Ian Johnston, Dominic Kwiatkowski, Cordelia Langford, John Sillitoe on behalf of the Wellcome Sanger Institute COVID-19 Surveillance Team |
| EPI_ISL_3094454 | Lighthouse Lab in Milton Keynes | Wellcome Sanger Institute for the COVID-19 Genomics UK (COG-UK) Consortium | The Lighthouse Lab in Milton Keynes and Alex Alderton, Roberto Amato, Jeffrey Barrett, Sonia Goncalves, Ewan Harrison, David K. Jackson, Ian Johnston, Dominic Kwiatkowski, Cordelia Langford, John Sillitoe on behalf of the Wellcome Sanger Institute COVID-19 Surveillance Team |
| EPI_ISL_3094455 | Lighthouse Laboratory Plymouth | Wellcome Sanger Institute for the COVID-19 Genomics UK (COG-UK) Consortium | Lighthouse Laboratory Plymouth and Alex Alderton, Roberto Amato, Jeffrey Barrett, Sonia Goncalves, Ewan Harrison, David K. Jackson, Ian Johnston, Dominic Kwiatkowski, Cordelia Langford, John Sillitoe on behalf of the Wellcome Sanger Institute COVID-19 Surveillance Team |
| EPI_ISL_3094456, EPI_ISL_3094457 | Lighthouse Lab in Milton Keynes | Wellcome Sanger Institute for the COVID-19 Genomics UK (COG-UK) Consortium | The Lighthouse Lab in Milton Keynes and Alex Alderton, Roberto Amato, Jeffrey Barrett, Sonia Goncalves, Ewan Harrison, David K. Jackson, Ian Johnston, Dominic Kwiatkowski, Cordelia Langford, John Sillitoe on behalf of the Wellcome Sanger Institute COVID-19 Surveillance Team |
| EPI_ISL_3094458 | Lighthouse Lab in Glasgow | Wellcome Sanger Institute for the COVID-19 Genomics UK (COG-UK) Consortium | Harper VanSteenhouse, Yumi Kasai, David Gray, Carol Clugston, Anna Dominiczak and Alex Alderton, Roberto Amato, Jeffrey Barrett, Sonia Goncalves, Ewan Harrison, David K. Jackson, Ian Johnston, Dominic Kwiatkowski, Cordelia Langford, John Sillitoe on behalf of the Wellcome Sanger Institute COVID-19 Surveillance Team |
| EPI_ISL_3094459 | Lighthouse Lab in Milton Keynes | Wellcome Sanger Institute for the COVID-19 Genomics UK (COG-UK) Consortium | The Lighthouse Lab in Milton Keynes and Alex Alderton, Roberto Amato, Jeffrey Barrett, Sonia Goncalves, Ewan Harrison, David K. Jackson, Ian Johnston, Dominic Kwiatkowski, Cordelia Langford, John Sillitoe on behalf of the Wellcome Sanger Institute COVID-19 Surveillance Team |
| EPI_ISL_3094460 | Lighthouse Laboratory Plymouth | Wellcome Sanger Institute for the COVID-19 Genomics UK (COG-UK) Consortium | Lighthouse Laboratory Plymouth and Alex Alderton, Roberto Amato, Jeffrey Barrett, Sonia Goncalves, Ewan Harrison, David K. Jackson, Ian Johnston, Dominic Kwiatkowski, Cordelia Langford, John Sillitoe on behalf of the Wellcome Sanger Institute COVID-19 Surveillance Team |
| EPI_ISL_3094461, EPI_ISL_3094462 | Lighthouse Lab in Milton Keynes | Wellcome Sanger Institute for the COVID-19 Genomics UK (COG-UK) Consortium | The Lighthouse Lab in Milton Keynes and Alex Alderton, Roberto Amato, Jeffrey Barrett, Sonia Goncalves, Ewan Harrison, David K. Jackson, Ian Johnston, Dominic Kwiatkowski, Cordelia Langford, John Sillitoe on behalf of the Wellcome Sanger Institute COVID-19 Surveillance Team |
| EPI_ISL_3094463 | Lighthouse Laboratory Plymouth | Wellcome Sanger Institute for the COVID-19 Genomics UK (COG-UK) Consortium | Lighthouse Laboratory Plymouth and Alex Alderton, Roberto Amato, Jeffrey Barrett, Sonia Goncalves, Ewan Harrison, David K. Jackson, Ian Johnston, Dominic Kwiatkowski, Cordelia Langford, John Sillitoe on behalf of the Wellcome Sanger Institute COVID-19 Surveillance Team |
| EPI_ISL_3094464, EPI_ISL_3094465 | Lighthouse Lab in Milton Keynes | Wellcome Sanger Institute for the COVID-19 Genomics UK (COG-UK) Consortium | The Lighthouse Lab in Milton Keynes and Alex Alderton, Roberto Amato, Jeffrey Barrett, Sonia Goncalves, Ewan Harrison, David K. Jackson, Ian Johnston, Dominic Kwiatkowski, Cordelia Langford, John Sillitoe on behalf of the Wellcome Sanger Institute COVID-19 Surveillance Team |
| EPI_ISL_3094466 | Lighthouse Laboratory Plymouth | Wellcome Sanger Institute for the COVID-19 Genomics UK (COG-UK) Consortium | Lighthouse Laboratory Plymouth and Alex Alderton, Roberto Amato, Jeffrey Barrett, Sonia Goncalves, Ewan Harrison, David K. Jackson, Ian Johnston, Dominic Kwiatkowski, Cordelia Langford, John Sillitoe on behalf of the Wellcome Sanger Institute COVID-19 Surveillance Team |
| EPI_ISL_3094467, EPI_ISL_3094468 | Lighthouse Lab in Milton Keynes | Wellcome Sanger Institute for the COVID-19 Genomics UK (COG-UK) Consortium | The Lighthouse Lab in Milton Keynes and Alex Alderton, Roberto Amato, Jeffrey Barrett, Sonia Goncalves, Ewan Harrison, David K. Jackson, Ian Johnston, Dominic Kwiatkowski, Cordelia Langford, John Sillitoe on behalf of the Wellcome Sanger Institute COVID-19 Surveillance Team |
| EPI_ISL_3094469 | Lighthouse Laboratory Plymouth | Wellcome Sanger Institute for the COVID-19 Genomics UK (COG-UK) Consortium | Lighthouse Laboratory Plymouth and Alex Alderton, Roberto Amato, Jeffrey Barrett, Sonia Goncalves, Ewan Harrison, David K. Jackson, Ian Johnston, Dominic Kwiatkowski, Cordelia Langford, John Sillitoe on behalf of the Wellcome Sanger Institute COVID-19 Surveillance Team |
| EPI_ISL_3094470 | Lighthouse Lab in Milton Keynes | Wellcome Sanger Institute for the COVID-19 Genomics UK (COG-UK) Consortium | The Lighthouse Lab in Milton Keynes and Alex Alderton, Roberto Amato, Jeffrey Barrett, Sonia Goncalves, Ewan Harrison, David K. Jackson, Ian Johnston, Dominic Kwiatkowski, Cordelia Langford, John Sillitoe on behalf of the Wellcome Sanger Institute COVID-19 Surveillance Team |
| EPI_ISL_3094471 | Lighthouse Laboratory Plymouth | Wellcome Sanger Institute for the COVID-19 Genomics UK (COG-UK) Consortium | Lighthouse Laboratory Plymouth and Alex Alderton, Roberto Amato, Jeffrey Barrett, Sonia Goncalves, Ewan Harrison, David K. Jackson, Ian Johnston, Dominic Kwiatkowski, Cordelia Langford, John Sillitoe on behalf of the Wellcome Sanger Institute COVID-19 Surveillance Team |
| EPI_ISL_3094472 | Lighthouse Lab in Glasgow | Wellcome Sanger Institute for the COVID-19 Genomics UK (COG-UK) Consortium | Harper VanSteenhouse, Yumi Kasai, David Gray, Carol Clugston, Anna Dominiczak and Alex Alderton, Roberto Amato, Jeffrey Barrett, Sonia Goncalves, Ewan Harrison, David K. Jackson, Ian Johnston, Dominic Kwiatkowski, Cordelia Langford, John Sillitoe on behalf of the Wellcome Sanger Institute COVID-19 Surveillance Team |
| EPI_ISL_3094473, EPI_ISL_3094474 | Lighthouse Laboratory Plymouth | Wellcome Sanger Institute for the COVID-19 Genomics UK (COG-UK) Consortium | Lighthouse Laboratory Plymouth and Alex Alderton, Roberto Amato, Jeffrey Barrett, Sonia Goncalves, Ewan Harrison, David K. Jackson, Ian Johnston, Dominic Kwiatkowski, Cordelia Langford, John Sillitoe on behalf of the Wellcome Sanger Institute COVID-19 Surveillance Team |
| EPI_ISL_3094475 | Lighthouse Lab in Milton Keynes | Wellcome Sanger Institute for the COVID-19 Genomics UK (COG-UK) Consortium | The Lighthouse Lab in Milton Keynes and Alex Alderton, Roberto Amato, Jeffrey Barrett, Sonia Goncalves, Ewan Harrison, David K. Jackson, Ian Johnston, Dominic Kwiatkowski, Cordelia Langford, John Sillitoe on behalf of the Wellcome Sanger Institute COVID-19 Surveillance Team |
| EPI_ISL_3094476 | Lighthouse Laboratory Plymouth | Wellcome Sanger Institute for the COVID-19 Genomics UK (COG-UK) Consortium | Lighthouse Laboratory Plymouth and Alex Alderton, Roberto Amato, Jeffrey Barrett, Sonia Goncalves, Ewan Harrison, David K. Jackson, Ian Johnston, Dominic Kwiatkowski, Cordelia Langford, John Sillitoe on behalf of the Wellcome Sanger Institute COVID-19 Surveillance Team |
| EPI_ISL_3094477 | Lighthouse Lab in Glasgow | Wellcome Sanger Institute for the COVID-19 Genomics UK (COG-UK) Consortium | Harper VanSteenhouse, Yumi Kasai, David Gray, Carol Clugston, Anna Dominiczak and Alex Alderton, Roberto Amato, Jeffrey Barrett, Sonia Goncalves, Ewan Harrison, David K. Jackson, Ian Johnston, Dominic Kwiatkowski, Cordelia Langford, John Sillitoe on behalf of the Wellcome Sanger Institute COVID-19 Surveillance Team |
| EPI_ISL_3094478, EPI_ISL_3094479, EPI_ISL_3094480, EPI_ISL_3094481, EPI_ISL_3094482, EPI_ISL_3094483, EPI_ISL_3094484 | Lighthouse Lab in Milton Keynes | Wellcome Sanger Institute for the COVID-19 Genomics UK (COG-UK) Consortium | The Lighthouse Lab in Milton Keynes and Alex Alderton, Roberto Amato, Jeffrey Barrett, Sonia Goncalves, Ewan Harrison, David K. Jackson, Ian Johnston, Dominic Kwiatkowski, Cordelia Langford, John Sillitoe on behalf of the Wellcome Sanger Institute COVID-19 Surveillance Team |
| EPI_ISL_3094485, EPI_ISL_3094486 | Lighthouse Lab in Glasgow | Wellcome Sanger Institute for the COVID-19 Genomics UK (COG-UK) Consortium | Harper VanSteenhouse, Yumi Kasai, David Gray, Carol Clugston, Anna Dominiczak and Alex Alderton, Roberto Amato, Jeffrey Barrett, Sonia Goncalves, Ewan Harrison, David K. Jackson, Ian Johnston, Dominic Kwiatkowski, Cordelia Langford, John Sillitoe on behalf of the Wellcome Sanger Institute COVID-19 Surveillance Team |
| EPI_ISL_3094487 | Lighthouse Lab in Milton Keynes | Wellcome Sanger Institute for the COVID-19 Genomics UK (COG-UK) Consortium | The Lighthouse Lab in Milton Keynes and Alex Alderton, Roberto Amato, Jeffrey Barrett, Sonia Goncalves, Ewan Harrison, David K. Jackson, Ian Johnston, Dominic Kwiatkowski, Cordelia Langford, John Sillitoe on behalf of the Wellcome Sanger Institute COVID-19 Surveillance Team |
| EPI_ISL_3094488, EPI_ISL_3094489, EPI_ISL_3094490 | Lighthouse Lab in Glasgow | Wellcome Sanger Institute for the COVID-19 Genomics UK (COG-UK) Consortium | Harper VanSteenhouse, Yumi Kasai, David Gray, Carol Clugston, Anna Dominiczak and Alex Alderton, Roberto Amato, Jeffrey Barrett, Sonia Goncalves, Ewan Harrison, David K. Jackson, Ian Johnston, Dominic Kwiatkowski, Cordelia Langford, John Sillitoe on behalf of the Wellcome Sanger Institute COVID-19 Surveillance Team |

| | | | |
|---|---|---|---|
| EPI_ISL_3094491 | Lighthouse Laboratory Plymouth | Wellcome Sanger Institute for the COVID-19 Genomics UK (COG-UK) Consortium | Lighthouse Laboratory Plymouth and Alex Alderton, Roberto Amato, Jeffrey Barrett, Sonia Goncalves, Ewan Harrison, David K. Jackson, Ian Johnston, Dominic Kwiatkowski, Cordelia Langford, John Sillitoe on behalf of the Wellcome Sanger Institute COVID-19 Surveillance Team |
| EPI_ISL_3094492 | Lighthouse Lab in Glasgow | Wellcome Sanger Institute for the COVID-19 Genomics UK (COG-UK) Consortium | Harper VanSteenhouse, Yumi Kasai, David Gray, Carol Clugston, Anna Dominiczak and Alex Alderton, Roberto Amato, Jeffrey Barrett, Sonia Goncalves, Ewan Harrison, David K. Jackson, Ian Johnston, Dominic Kwiatkowski, Cordelia Langford, John Sillitoe on behalf of the Wellcome Sanger Institute COVID-19 Surveillance Team |
| EPI_ISL_3094493, EPI_ISL_3094494, EPI_ISL_3094495, EPI_ISL_3094496, EPI_ISL_3094497 | Lighthouse Lab in Milton Keynes | Wellcome Sanger Institute for the COVID-19 Genomics UK (COG-UK) Consortium | The Lighthouse Lab in Milton Keynes and Alex Alderton, Roberto Amato, Jeffrey Barrett, Sonia Goncalves, Ewan Harrison, David K. Jackson, Ian Johnston, Dominic Kwiatkowski, Cordelia Langford, John Sillitoe on behalf of the Wellcome Sanger Institute COVID-19 Surveillance Team |
| EPI_ISL_3094498, EPI_ISL_3094499, EPI_ISL_3094500 | Lighthouse Laboratory Plymouth | Wellcome Sanger Institute for the COVID-19 Genomics UK (COG-UK) Consortium | Lighthouse Laboratory Plymouth and Alex Alderton, Roberto Amato, Jeffrey Barrett, Sonia Goncalves, Ewan Harrison, David K. Jackson, Ian Johnston, Dominic Kwiatkowski, Cordelia Langford, John Sillitoe on behalf of the Wellcome Sanger Institute COVID-19 Surveillance Team |
| EPI_ISL_3094501, EPI_ISL_3094502, EPI_ISL_3094503 | Lighthouse Lab in Milton Keynes | Wellcome Sanger Institute for the COVID-19 Genomics UK (COG-UK) Consortium | The Lighthouse Lab in Milton Keynes and Alex Alderton, Roberto Amato, Jeffrey Barrett, Sonia Goncalves, Ewan Harrison, David K. Jackson, Ian Johnston, Dominic Kwiatkowski, Cordelia Langford, John Sillitoe on behalf of the Wellcome Sanger Institute COVID-19 Surveillance Team |
| EPI_ISL_3094504, EPI_ISL_3094505 | Lighthouse Lab in Glasgow | Wellcome Sanger Institute for the COVID-19 Genomics UK (COG-UK) Consortium | Harper VanSteenhouse, Yumi Kasai, David Gray, Carol Clugston, Anna Dominiczak and Alex Alderton, Roberto Amato, Jeffrey Barrett, Sonia Goncalves, Ewan Harrison, David K. Jackson, Ian Johnston, Dominic Kwiatkowski, Cordelia Langford, John Sillitoe on behalf of the Wellcome Sanger Institute COVID-19 Surveillance Team |
| EPI_ISL_3094506 | Lighthouse Lab in Milton Keynes | Wellcome Sanger Institute for the COVID-19 Genomics UK (COG-UK) Consortium | The Lighthouse Lab in Milton Keynes and Alex Alderton, Roberto Amato, Jeffrey Barrett, Sonia Goncalves, Ewan Harrison, David K. Jackson, Ian Johnston, Dominic Kwiatkowski, Cordelia Langford, John Sillitoe on behalf of the Wellcome Sanger Institute COVID-19 Surveillance Team |
| EPI_ISL_3094507 | Lighthouse Lab in Glasgow | Wellcome Sanger Institute for the COVID-19 Genomics UK (COG-UK) Consortium | Harper VanSteenhouse, Yumi Kasai, David Gray, Carol Clugston, Anna Dominiczak and Alex Alderton, Roberto Amato, Jeffrey Barrett, Sonia Goncalves, Ewan Harrison, David K. Jackson, Ian Johnston, Dominic Kwiatkowski, Cordelia Langford, John Sillitoe on behalf of the Wellcome Sanger Institute COVID-19 Surveillance Team |
| EPI_ISL_3094508, EPI_ISL_3094509 | Lighthouse Lab in Milton Keynes | Wellcome Sanger Institute for the COVID-19 Genomics UK (COG-UK) Consortium | The Lighthouse Lab in Milton Keynes and Alex Alderton, Roberto Amato, Jeffrey Barrett, Sonia Goncalves, Ewan Harrison, David K. Jackson, Ian Johnston, Dominic Kwiatkowski, Cordelia Langford, John Sillitoe on behalf of the Wellcome Sanger Institute COVID-19 Surveillance Team |
| EPI_ISL_3094510 | Lighthouse Lab in Glasgow | Wellcome Sanger Institute for the COVID-19 Genomics UK (COG-UK) Consortium | Harper VanSteenhouse, Yumi Kasai, David Gray, Carol Clugston, Anna Dominiczak and Alex Alderton, Roberto Amato, Jeffrey Barrett, Sonia Goncalves, Ewan Harrison, David K. Jackson, Ian Johnston, Dominic Kwiatkowski, Cordelia Langford, John Sillitoe on behalf of the Wellcome Sanger Institute COVID-19 Surveillance Team |
| EPI_ISL_3094511 | Lighthouse Lab in Milton Keynes | Wellcome Sanger Institute for the COVID-19 Genomics UK (COG-UK) Consortium | The Lighthouse Lab in Milton Keynes and Alex Alderton, Roberto Amato, Jeffrey Barrett, Sonia Goncalves, Ewan Harrison, David K. Jackson, Ian Johnston, Dominic Kwiatkowski, Cordelia Langford, John Sillitoe on behalf of the Wellcome Sanger Institute COVID-19 Surveillance Team |
| EPI_ISL_3094512, EPI_ISL_3094513 | Lighthouse Lab in Glasgow | Wellcome Sanger Institute for the COVID-19 Genomics UK (COG-UK) Consortium | Harper VanSteenhouse, Yumi Kasai, David Gray, Carol Clugston, Anna Dominiczak and Alex Alderton, Roberto Amato, Jeffrey Barrett, Sonia Goncalves, Ewan Harrison, David K. Jackson, Ian Johnston, Dominic Kwiatkowski, Cordelia Langford, John Sillitoe on behalf of the Wellcome Sanger Institute COVID-19 Surveillance Team |
| EPI_ISL_3094514, EPI_ISL_3094515 | Lighthouse Lab in Milton Keynes | Wellcome Sanger Institute for the COVID-19 Genomics UK (COG-UK) Consortium | The Lighthouse Lab in Milton Keynes and Alex Alderton, Roberto Amato, Jeffrey Barrett, Sonia Goncalves, Ewan Harrison, David K. Jackson, Ian Johnston, Dominic Kwiatkowski, Cordelia Langford, John Sillitoe on behalf of the Wellcome Sanger Institute COVID-19 Surveillance Team |
| EPI_ISL_3094516 | Lighthouse Laboratory Plymouth | Wellcome Sanger Institute for the COVID-19 Genomics UK (COG-UK) Consortium | Lighthouse Laboratory Plymouth and Alex Alderton, Roberto Amato, Jeffrey Barrett, Sonia Goncalves, Ewan Harrison, David K. Jackson, Ian Johnston, Dominic Kwiatkowski, Cordelia Langford, John Sillitoe on behalf of the Wellcome Sanger Institute COVID-19 Surveillance Team |
| EPI_ISL_3094517, EPI_ISL_3094518 | Lighthouse Lab in Milton Keynes | Wellcome Sanger Institute for the COVID-19 Genomics UK (COG-UK) Consortium | The Lighthouse Lab in Milton Keynes and Alex Alderton, Roberto Amato, Jeffrey Barrett, Sonia Goncalves, Ewan Harrison, David K. Jackson, Ian Johnston, Dominic Kwiatkowski, Cordelia Langford, John Sillitoe on behalf of the Wellcome Sanger Institute COVID-19 Surveillance Team |
| EPI_ISL_3094519, EPI_ISL_3094520, EPI_ISL_3094521 | Lighthouse Lab in Glasgow | Wellcome Sanger Institute for the COVID-19 Genomics UK (COG-UK) Consortium | Harper VanSteenhouse, Yumi Kasai, David Gray, Carol Clugston, Anna Dominiczak and Alex Alderton, Roberto Amato, Jeffrey Barrett, Sonia Goncalves, Ewan Harrison, David K. Jackson, Ian Johnston, Dominic Kwiatkowski, Cordelia Langford, John Sillitoe on behalf of the Wellcome Sanger Institute COVID-19 Surveillance Team |
| EPI_ISL_3094522, EPI_ISL_3094523, EPI_ISL_3094524, EPI_ISL_3094525 | Lighthouse Lab in Milton Keynes | Wellcome Sanger Institute for the COVID-19 Genomics UK (COG-UK) Consortium | The Lighthouse Lab in Milton Keynes and Alex Alderton, Roberto Amato, Jeffrey Barrett, Sonia Goncalves, Ewan Harrison, David K. Jackson, Ian Johnston, Dominic Kwiatkowski, Cordelia Langford, John Sillitoe on behalf of the Wellcome Sanger Institute COVID-19 Surveillance Team |
| EPI_ISL_3094526, EPI_ISL_3094527 | Lighthouse Laboratory Plymouth | Wellcome Sanger Institute for the COVID-19 Genomics UK (COG-UK) Consortium | Lighthouse Laboratory Plymouth and Alex Alderton, Roberto Amato, Jeffrey Barrett, Sonia Goncalves, Ewan Harrison, David K. Jackson, Ian Johnston, Dominic Kwiatkowski, Cordelia Langford, John Sillitoe on behalf of the Wellcome Sanger Institute COVID-19 Surveillance Team |
| EPI_ISL_3094528 | Lighthouse Lab in Glasgow | Wellcome Sanger Institute for the COVID-19 Genomics UK (COG-UK) Consortium | Harper VanSteenhouse, Yumi Kasai, David Gray, Carol Clugston, Anna Dominiczak and Alex Alderton, Roberto Amato, Jeffrey Barrett, Sonia Goncalves, Ewan Harrison, David K. Jackson, Ian Johnston, Dominic Kwiatkowski, Cordelia Langford, John Sillitoe on behalf of the Wellcome Sanger Institute COVID-19 Surveillance Team |
| EPI_ISL_3094529 | Lighthouse Lab in Milton Keynes | Wellcome Sanger Institute for the COVID-19 Genomics UK (COG-UK) Consortium | The Lighthouse Lab in Milton Keynes and Alex Alderton, Roberto Amato, Jeffrey Barrett, Sonia Goncalves, Ewan Harrison, David K. Jackson, Ian Johnston, Dominic Kwiatkowski, Cordelia Langford, John Sillitoe on behalf of the Wellcome Sanger Institute COVID-19 Surveillance Team |
| EPI_ISL_3094530 | Lighthouse Lab in Glasgow | Wellcome Sanger Institute for the COVID-19 Genomics UK (COG-UK) Consortium | Harper VanSteenhouse, Yumi Kasai, David Gray, Carol Clugston, Anna Dominiczak and Alex Alderton, Roberto Amato, Jeffrey Barrett, Sonia Goncalves, Ewan Harrison, David K. Jackson, Ian Johnston, Dominic Kwiatkowski, Cordelia Langford, John Sillitoe on behalf of the Wellcome Sanger Institute COVID-19 Surveillance Team |
| EPI_ISL_3094531 | Lighthouse Lab in Milton Keynes | Wellcome Sanger Institute for the COVID-19 Genomics UK (COG-UK) Consortium | The Lighthouse Lab in Milton Keynes and Alex Alderton, Roberto Amato, Jeffrey Barrett, Sonia Goncalves, Ewan Harrison, David K. Jackson, Ian Johnston, Dominic Kwiatkowski, Cordelia Langford, John Sillitoe on behalf of the Wellcome Sanger Institute COVID-19 Surveillance Team |
| EPI_ISL_3094532 | Lighthouse Lab in Glasgow | Wellcome Sanger Institute for the COVID-19 Genomics UK (COG-UK) Consortium | Harper VanSteenhouse, Yumi Kasai, David Gray, Carol Clugston, Anna Dominiczak and Alex Alderton, Roberto Amato, Jeffrey Barrett, Sonia Goncalves, Ewan Harrison, David K. Jackson, Ian Johnston, Dominic Kwiatkowski, Cordelia Langford, John Sillitoe on behalf of the Wellcome Sanger Institute COVID-19 Surveillance Team |
| EPI_ISL_3094533, EPI_ISL_3094534 | Lighthouse Lab in Milton Keynes | Wellcome Sanger Institute for the COVID-19 Genomics UK (COG-UK) Consortium | The Lighthouse Lab in Milton Keynes and Alex Alderton, Roberto Amato, Jeffrey Barrett, Sonia Goncalves, Ewan Harrison, David K. Jackson, Ian Johnston, Dominic Kwiatkowski, Cordelia Langford, John Sillitoe on behalf of the Wellcome Sanger Institute COVID-19 Surveillance Team |
| EPI_ISL_3094535 | Lighthouse Laboratory Plymouth | Wellcome Sanger Institute for the COVID-19 Genomics UK (COG-UK) Consortium | Lighthouse Laboratory Plymouth and Alex Alderton, Roberto Amato, Jeffrey Barrett, Sonia Goncalves, Ewan Harrison, David K. Jackson, Ian Johnston, Dominic Kwiatkowski, Cordelia Langford, John Sillitoe on behalf of the Wellcome Sanger Institute COVID-19 Surveillance Team |
| EPI_ISL_3094536 | Lighthouse Lab in Milton Keynes | Wellcome Sanger Institute for the COVID-19 Genomics UK (COG-UK) Consortium | The Lighthouse Lab in Milton Keynes and Alex Alderton, Roberto Amato, Jeffrey Barrett, Sonia Goncalves, Ewan Harrison, David K. Jackson, Ian Johnston, Dominic Kwiatkowski, Cordelia Langford, John Sillitoe on behalf of the Wellcome Sanger Institute COVID-19 Surveillance Team |
| EPI_ISL_3094537 | Lighthouse Lab in Glasgow | Wellcome Sanger Institute for the COVID-19 Genomics UK (COG-UK) Consortium | Harper VanSteenhouse, Yumi Kasai, David Gray, Carol Clugston, Anna Dominiczak and Alex Alderton, Roberto Amato, Jeffrey Barrett, Sonia Goncalves, Ewan Harrison, David K. Jackson, Ian Johnston, Dominic Kwiatkowski, Cordelia Langford, John Sillitoe on behalf of the Wellcome Sanger Institute COVID-19 Surveillance Team |
| EPI_ISL_3094538 | Lighthouse Lab in Milton Keynes | Wellcome Sanger Institute for the COVID-19 Genomics UK (COG-UK) Consortium | The Lighthouse Lab in Milton Keynes and Alex Alderton, Roberto Amato, Jeffrey Barrett, Sonia Goncalves, Ewan Harrison, David K. Jackson, Ian Johnston, Dominic Kwiatkowski, Cordelia Langford, John Sillitoe on behalf of the Wellcome Sanger Institute COVID-19 Surveillance Team |
| EPI_ISL_3094539, EPI_ISL_3094540 | Lighthouse Lab in Glasgow | Wellcome Sanger Institute for the COVID-19 Genomics UK (COG-UK) Consortium | Harper VanSteenhouse, Yumi Kasai, David Gray, Carol Clugston, Anna Dominiczak and Alex Alderton, Roberto Amato, Jeffrey Barrett, Sonia Goncalves, Ewan Harrison, David K. Jackson, Ian Johnston, Dominic Kwiatkowski, Cordelia Langford, John Sillitoe on behalf of the Wellcome Sanger Institute COVID-19 Surveillance Team |
| EPI_ISL_3094541 | Lighthouse Laboratory Plymouth | Wellcome Sanger Institute for the COVID-19 Genomics UK (COG-UK) Consortium | Lighthouse Laboratory Plymouth and Alex Alderton, Roberto Amato, Jeffrey Barrett, Sonia Goncalves, Ewan Harrison, David K. Jackson, Ian Johnston, Dominic Kwiatkowski, Cordelia Langford, John Sillitoe on behalf of the Wellcome Sanger Institute COVID-19 Surveillance Team |
| EPI_ISL_3094542, EPI_ISL_3094543, EPI_ISL_3094544 | Lighthouse Lab in Milton Keynes | Wellcome Sanger Institute for the COVID-19 Genomics UK (COG-UK) Consortium | The Lighthouse Lab in Milton Keynes and Alex Alderton, Roberto Amato, Jeffrey Barrett, Sonia Goncalves, Ewan Harrison, David K. Jackson, Ian Johnston, Dominic Kwiatkowski, Cordelia Langford, John Sillitoe on behalf of the Wellcome Sanger Institute COVID-19 Surveillance Team |
| EPI_ISL_3094545 | Lighthouse Lab in Glasgow | Wellcome Sanger Institute for the COVID-19 Genomics UK (COG-UK) Consortium | Harper VanSteenhouse, Yumi Kasai, David Gray, Carol Clugston, Anna Dominiczak and Alex Alderton, Roberto Amato, Jeffrey Barrett, Sonia Goncalves, Ewan Harrison, David K. Jackson, Ian Johnston, Dominic Kwiatkowski, Cordelia Langford, John Sillitoe on behalf of the Wellcome Sanger Institute COVID-19 Surveillance Team |
| EPI_ISL_3094546 | Lighthouse Lab in Milton Keynes | Wellcome Sanger Institute for the COVID-19 Genomics UK (COG-UK) Consortium | The Lighthouse Lab in Milton Keynes and Alex Alderton, Roberto Amato, Jeffrey Barrett, Sonia Goncalves, Ewan Harrison, David K. Jackson, Ian Johnston, Dominic Kwiatkowski, Cordelia Langford, John Sillitoe on behalf of the Wellcome Sanger Institute COVID-19 Surveillance Team |
| EPI_ISL_3094547 | Lighthouse Lab in Glasgow | Wellcome Sanger Institute for the COVID-19 Genomics UK (COG-UK) Consortium | Harper VanSteenhouse, Yumi Kasai, David Gray, Carol Clugston, Anna Dominiczak and Alex Alderton, Roberto Amato, Jeffrey Barrett, Sonia Goncalves, Ewan Harrison, David K. Jackson, Ian Johnston, Dominic Kwiatkowski, Cordelia Langford, John Sillitoe on behalf of the Wellcome Sanger Institute COVID-19 Surveillance Team |
| EPI_ISL_3094548 | Lighthouse Laboratory Plymouth | Wellcome Sanger Institute for the COVID-19 Genomics UK (COG-UK) Consortium | Lighthouse Laboratory Plymouth and Alex Alderton, Roberto Amato, Jeffrey Barrett, Sonia Goncalves, Ewan Harrison, David K. Jackson, Ian Johnston, Dominic Kwiatkowski, Cordelia Langford, John Sillitoe on behalf of the Wellcome Sanger Institute COVID-19 Surveillance Team |
| EPI_ISL_3094549, EPI_ISL_3094550, EPI_ISL_3094551 | Lighthouse Lab in Milton Keynes | Wellcome Sanger Institute for the COVID-19 Genomics UK (COG-UK) Consortium | The Lighthouse Lab in Milton Keynes and Alex Alderton, Roberto Amato, Jeffrey Barrett, Sonia Goncalves, Ewan Harrison, David K. Jackson, Ian Johnston, Dominic Kwiatkowski, Cordelia Langford, John Sillitoe on behalf of the Wellcome Sanger Institute COVID-19 Surveillance Team |
| EPI_ISL_3094552 | Lighthouse Laboratory Plymouth | Wellcome Sanger Institute for the COVID-19 Genomics UK (COG-UK) Consortium | Lighthouse Laboratory Plymouth and Alex Alderton, Roberto Amato, Jeffrey Barrett, Sonia Goncalves, Ewan Harrison, David K. Jackson, Ian Johnston, Dominic Kwiatkowski, Cordelia Langford, John Sillitoe on behalf of the Wellcome Sanger Institute COVID-19 Surveillance Team |
| EPI_ISL_3094553 | Lighthouse Lab in Milton Keynes | Wellcome Sanger Institute for the COVID-19 Genomics UK (COG-UK) Consortium | The Lighthouse Lab in Milton Keynes and Alex Alderton, Roberto Amato, Jeffrey Barrett, Sonia Goncalves, Ewan Harrison, David K. Jackson, Ian Johnston, Dominic Kwiatkowski, Cordelia Langford, John Sillitoe on behalf of the Wellcome Sanger Institute COVID-19 Surveillance Team |
| EPI_ISL_3094554 | Lighthouse Lab in Glasgow | Wellcome Sanger Institute for the COVID-19 Genomics UK (COG-UK) Consortium | Harper VanSteenhouse, Yumi Kasai, David Gray, Carol Clugston, Anna Dominiczak and Alex Alderton, Roberto Amato, Jeffrey Barrett, Sonia Goncalves, Ewan Harrison, David K. Jackson, Ian Johnston, Dominic Kwiatkowski, Cordelia Langford, John Sillitoe on behalf of the Wellcome Sanger Institute COVID-19 Surveillance Team |
| EPI_ISL_3094555 | Lighthouse Lab in Milton Keynes | Wellcome Sanger Institute for the COVID-19 Genomics UK (COG-UK) Consortium | The Lighthouse Lab in Milton Keynes and Alex Alderton, Roberto Amato, Jeffrey Barrett, Sonia Goncalves, Ewan Harrison, David K. Jackson, Ian Johnston, Dominic Kwiatkowski, Cordelia Langford, John Sillitoe on behalf of the Wellcome Sanger Institute COVID-19 Surveillance Team |
| EPI_ISL_3094556 | Lighthouse Laboratory Plymouth | Wellcome Sanger Institute for the COVID-19 Genomics UK (COG-UK) Consortium | Lighthouse Laboratory Plymouth and Alex Alderton, Roberto Amato, Jeffrey Barrett, Sonia Goncalves, Ewan Harrison, David K. Jackson, Ian Johnston, Dominic Kwiatkowski, Cordelia Langford, John Sillitoe on behalf of the Wellcome Sanger Institute COVID-19 Surveillance Team |

| | | | |
|---|---|---|---|
| EPI_ISL_3094557 | Lighthouse Lab in Milton Keynes | Wellcome Sanger Institute for the COVID-19 Genomics UK (COG-UK) Consortium | The Lighthouse Lab in Milton Keynes and Alex Alderton, Roberto Amato, Jeffrey Barrett, Sonia Goncalves, Ewan Harrison, David K. Jackson, Ian Johnston, Dominic Kwiatkowski, Cordelia Langford, John Sillitoe on behalf of the Wellcome Sanger Institute COVID-19 Surveillance Team |
| EPI_ISL_3094558 | Lighthouse Laboratory Plymouth | Wellcome Sanger Institute for the COVID-19 Genomics UK (COG-UK) Consortium | Lighthouse Laboratory Plymouth and Alex Alderton, Roberto Amato, Jeffrey Barrett, Sonia Goncalves, Ewan Harrison, David K. Jackson, Ian Johnston, Dominic Kwiatkowski, Cordelia Langford, John Sillitoe on behalf of the Wellcome Sanger Institute COVID-19 Surveillance Team |
| EPI_ISL_3094559, EPI_ISL_3094560 | Lighthouse Lab in Milton Keynes | Wellcome Sanger Institute for the COVID-19 Genomics UK (COG-UK) Consortium | The Lighthouse Lab in Milton Keynes and Alex Alderton, Roberto Amato, Jeffrey Barrett, Sonia Goncalves, Ewan Harrison, David K. Jackson, Ian Johnston, Dominic Kwiatkowski, Cordelia Langford, John Sillitoe on behalf of the Wellcome Sanger Institute COVID-19 Surveillance Team |
| EPI_ISL_3094561 | Lighthouse Laboratory Plymouth | Wellcome Sanger Institute for the COVID-19 Genomics UK (COG-UK) Consortium | Lighthouse Laboratory Plymouth and Alex Alderton, Roberto Amato, Jeffrey Barrett, Sonia Goncalves, Ewan Harrison, David K. Jackson, Ian Johnston, Dominic Kwiatkowski, Cordelia Langford, John Sillitoe on behalf of the Wellcome Sanger Institute COVID-19 Surveillance Team |
| EPI_ISL_3094562 | Lighthouse Lab in Glasgow | Wellcome Sanger Institute for the COVID-19 Genomics UK (COG-UK) Consortium | Harper VanSteenhouse, Yumi Kasai, David Gray, Carol Clugston, Anna Dominiczak and Alex Alderton, Roberto Amato, Jeffrey Barrett, Sonia Goncalves, Ewan Harrison, David K. Jackson, Ian Johnston, Dominic Kwiatkowski, Cordelia Langford, John Sillitoe on behalf of the Wellcome Sanger Institute COVID-19 Surveillance Team |
| EPI_ISL_3094563, EPI_ISL_3094564 | Lighthouse Lab in Milton Keynes | Wellcome Sanger Institute for the COVID-19 Genomics UK (COG-UK) Consortium | The Lighthouse Lab in Milton Keynes and Alex Alderton, Roberto Amato, Jeffrey Barrett, Sonia Goncalves, Ewan Harrison, David K. Jackson, Ian Johnston, Dominic Kwiatkowski, Cordelia Langford, John Sillitoe on behalf of the Wellcome Sanger Institute COVID-19 Surveillance Team |
| EPI_ISL_3094565, EPI_ISL_3094566, EPI_ISL_3094567 | Lighthouse Laboratory Plymouth | Wellcome Sanger Institute for the COVID-19 Genomics UK (COG-UK) Consortium | Lighthouse Laboratory Plymouth and Alex Alderton, Roberto Amato, Jeffrey Barrett, Sonia Goncalves, Ewan Harrison, David K. Jackson, Ian Johnston, Dominic Kwiatkowski, Cordelia Langford, John Sillitoe on behalf of the Wellcome Sanger Institute COVID-19 Surveillance Team |
| EPI_ISL_3094568 | Lighthouse Lab in Milton Keynes | Wellcome Sanger Institute for the COVID-19 Genomics UK (COG-UK) Consortium | The Lighthouse Lab in Milton Keynes and Alex Alderton, Roberto Amato, Jeffrey Barrett, Sonia Goncalves, Ewan Harrison, David K. Jackson, Ian Johnston, Dominic Kwiatkowski, Cordelia Langford, John Sillitoe on behalf of the Wellcome Sanger Institute COVID-19 Surveillance Team |
| EPI_ISL_3094569, EPI_ISL_3094570 | Lighthouse Lab in Glasgow | Wellcome Sanger Institute for the COVID-19 Genomics UK (COG-UK) Consortium | Harper VanSteenhouse, Yumi Kasai, David Gray, Carol Clugston, Anna Dominiczak and Alex Alderton, Roberto Amato, Jeffrey Barrett, Sonia Goncalves, Ewan Harrison, David K. Jackson, Ian Johnston, Dominic Kwiatkowski, Cordelia Langford, John Sillitoe on behalf of the Wellcome Sanger Institute COVID-19 Surveillance Team |
| EPI_ISL_3094571 | Lighthouse Laboratory Plymouth | Wellcome Sanger Institute for the COVID-19 Genomics UK (COG-UK) Consortium | Lighthouse Laboratory Plymouth and Alex Alderton, Roberto Amato, Jeffrey Barrett, Sonia Goncalves, Ewan Harrison, David K. Jackson, Ian Johnston, Dominic Kwiatkowski, Cordelia Langford, John Sillitoe on behalf of the Wellcome Sanger Institute COVID-19 Surveillance Team |
| EPI_ISL_3094572 | Lighthouse Lab in Milton Keynes | Wellcome Sanger Institute for the COVID-19 Genomics UK (COG-UK) Consortium | The Lighthouse Lab in Milton Keynes and Alex Alderton, Roberto Amato, Jeffrey Barrett, Sonia Goncalves, Ewan Harrison, David K. Jackson, Ian Johnston, Dominic Kwiatkowski, Cordelia Langford, John Sillitoe on behalf of the Wellcome Sanger Institute COVID-19 Surveillance Team |
| EPI_ISL_3094573 | Lighthouse Lab in Glasgow | Wellcome Sanger Institute for the COVID-19 Genomics UK (COG-UK) Consortium | Harper VanSteenhouse, Yumi Kasai, David Gray, Carol Clugston, Anna Dominiczak and Alex Alderton, Roberto Amato, Jeffrey Barrett, Sonia Goncalves, Ewan Harrison, David K. Jackson, Ian Johnston, Dominic Kwiatkowski, Cordelia Langford, John Sillitoe on behalf of the Wellcome Sanger Institute COVID-19 Surveillance Team |
| EPI_ISL_3094574 | Lighthouse Laboratory Plymouth | Wellcome Sanger Institute for the COVID-19 Genomics UK (COG-UK) Consortium | Lighthouse Laboratory Plymouth and Alex Alderton, Roberto Amato, Jeffrey Barrett, Sonia Goncalves, Ewan Harrison, David K. Jackson, Ian Johnston, Dominic Kwiatkowski, Cordelia Langford, John Sillitoe on behalf of the Wellcome Sanger Institute COVID-19 Surveillance Team |
| EPI_ISL_3094575 | Lighthouse Lab in Milton Keynes | Wellcome Sanger Institute for the COVID-19 Genomics UK (COG-UK) Consortium | The Lighthouse Lab in Milton Keynes and Alex Alderton, Roberto Amato, Jeffrey Barrett, Sonia Goncalves, Ewan Harrison, David K. Jackson, Ian Johnston, Dominic Kwiatkowski, Cordelia Langford, John Sillitoe on behalf of the Wellcome Sanger Institute COVID-19 Surveillance Team |
| EPI_ISL_3094576 | Lighthouse Lab in Glasgow | Wellcome Sanger Institute for the COVID-19 Genomics UK (COG-UK) Consortium | Harper VanSteenhouse, Yumi Kasai, David Gray, Carol Clugston, Anna Dominiczak and Alex Alderton, Roberto Amato, Jeffrey Barrett, Sonia Goncalves, Ewan Harrison, David K. Jackson, Ian Johnston, Dominic Kwiatkowski, Cordelia Langford, John Sillitoe on behalf of the Wellcome Sanger Institute COVID-19 Surveillance Team |
| EPI_ISL_3094577 | Lighthouse Laboratory Plymouth | Wellcome Sanger Institute for the COVID-19 Genomics UK (COG-UK) Consortium | Lighthouse Laboratory Plymouth and Alex Alderton, Roberto Amato, Jeffrey Barrett, Sonia Goncalves, Ewan Harrison, David K. Jackson, Ian Johnston, Dominic Kwiatkowski, Cordelia Langford, John Sillitoe on behalf of the Wellcome Sanger Institute COVID-19 Surveillance Team |
| EPI_ISL_3094578 | Lighthouse Lab in Glasgow | Wellcome Sanger Institute for the COVID-19 Genomics UK (COG-UK) Consortium | Harper VanSteenhouse, Yumi Kasai, David Gray, Carol Clugston, Anna Dominiczak and Alex Alderton, Roberto Amato, Jeffrey Barrett, Sonia Goncalves, Ewan Harrison, David K. Jackson, Ian Johnston, Dominic Kwiatkowski, Cordelia Langford, John Sillitoe on behalf of the Wellcome Sanger Institute COVID-19 Surveillance Team |
| EPI_ISL_3094579 | Lighthouse Laboratory Plymouth | Wellcome Sanger Institute for the COVID-19 Genomics UK (COG-UK) Consortium | Lighthouse Laboratory Plymouth and Alex Alderton, Roberto Amato, Jeffrey Barrett, Sonia Goncalves, Ewan Harrison, David K. Jackson, Ian Johnston, Dominic Kwiatkowski, Cordelia Langford, John Sillitoe on behalf of the Wellcome Sanger Institute COVID-19 Surveillance Team |
| EPI_ISL_3094580 | Lighthouse Lab in Milton Keynes | Wellcome Sanger Institute for the COVID-19 Genomics UK (COG-UK) Consortium | The Lighthouse Lab in Milton Keynes and Alex Alderton, Roberto Amato, Jeffrey Barrett, Sonia Goncalves, Ewan Harrison, David K. Jackson, Ian Johnston, Dominic Kwiatkowski, Cordelia Langford, John Sillitoe on behalf of the Wellcome Sanger Institute COVID-19 Surveillance Team |
| EPI_ISL_3094581 | Lighthouse Lab in Glasgow | Wellcome Sanger Institute for the COVID-19 Genomics UK (COG-UK) Consortium | Harper VanSteenhouse, Yumi Kasai, David Gray, Carol Clugston, Anna Dominiczak and Alex Alderton, Roberto Amato, Jeffrey Barrett, Sonia Goncalves, Ewan Harrison, David K. Jackson, Ian Johnston, Dominic Kwiatkowski, Cordelia Langford, John Sillitoe on behalf of the Wellcome Sanger Institute COVID-19 Surveillance Team |
| EPI_ISL_3094582 | Lighthouse Lab in Milton Keynes | Wellcome Sanger Institute for the COVID-19 Genomics UK (COG-UK) Consortium | The Lighthouse Lab in Milton Keynes and Alex Alderton, Roberto Amato, Jeffrey Barrett, Sonia Goncalves, Ewan Harrison, David K. Jackson, Ian Johnston, Dominic Kwiatkowski, Cordelia Langford, John Sillitoe on behalf of the Wellcome Sanger Institute COVID-19 Surveillance Team |
| EPI_ISL_3094583, EPI_ISL_3094584, EPI_ISL_3094585 | Lighthouse Laboratory Plymouth | Wellcome Sanger Institute for the COVID-19 Genomics UK (COG-UK) Consortium | Lighthouse Laboratory Plymouth and Alex Alderton, Roberto Amato, Jeffrey Barrett, Sonia Goncalves, Ewan Harrison, David K. Jackson, Ian Johnston, Dominic Kwiatkowski, Cordelia Langford, John Sillitoe on behalf of the Wellcome Sanger Institute COVID-19 Surveillance Team |
| EPI_ISL_3094586, EPI_ISL_3094587, EPI_ISL_3094588 | Lighthouse Lab in Milton Keynes | Wellcome Sanger Institute for the COVID-19 Genomics UK (COG-UK) Consortium | The Lighthouse Lab in Milton Keynes and Alex Alderton, Roberto Amato, Jeffrey Barrett, Sonia Goncalves, Ewan Harrison, David K. Jackson, Ian Johnston, Dominic Kwiatkowski, Cordelia Langford, John Sillitoe on behalf of the Wellcome Sanger Institute COVID-19 Surveillance Team |
| EPI_ISL_3094589 | Lighthouse Lab in Glasgow | Wellcome Sanger Institute for the COVID-19 Genomics UK (COG-UK) Consortium | Harper VanSteenhouse, Yumi Kasai, David Gray, Carol Clugston, Anna Dominiczak and Alex Alderton, Roberto Amato, Jeffrey Barrett, Sonia Goncalves, Ewan Harrison, David K. Jackson, Ian Johnston, Dominic Kwiatkowski, Cordelia Langford, John Sillitoe on behalf of the Wellcome Sanger Institute COVID-19 Surveillance Team |
| EPI_ISL_3094590, EPI_ISL_3094591 | Lighthouse Laboratory Plymouth | Wellcome Sanger Institute for the COVID-19 Genomics UK (COG-UK) Consortium | Lighthouse Laboratory Plymouth and Alex Alderton, Roberto Amato, Jeffrey Barrett, Sonia Goncalves, Ewan Harrison, David K. Jackson, Ian Johnston, Dominic Kwiatkowski, Cordelia Langford, John Sillitoe on behalf of the Wellcome Sanger Institute COVID-19 Surveillance Team |
| EPI_ISL_3094592, EPI_ISL_3094593 | Lighthouse Lab in Milton Keynes | Wellcome Sanger Institute for the COVID-19 Genomics UK (COG-UK) Consortium | The Lighthouse Lab in Milton Keynes and Alex Alderton, Roberto Amato, Jeffrey Barrett, Sonia Goncalves, Ewan Harrison, David K. Jackson, Ian Johnston, Dominic Kwiatkowski, Cordelia Langford, John Sillitoe on behalf of the Wellcome Sanger Institute COVID-19 Surveillance Team |
| EPI_ISL_3094594, EPI_ISL_3094595 | Lighthouse Lab in Glasgow | Wellcome Sanger Institute for the COVID-19 Genomics UK (COG-UK) Consortium | Harper VanSteenhouse, Yumi Kasai, David Gray, Carol Clugston, Anna Dominiczak and Alex Alderton, Roberto Amato, Jeffrey Barrett, Sonia Goncalves, Ewan Harrison, David K. Jackson, Ian Johnston, Dominic Kwiatkowski, Cordelia Langford, John Sillitoe on behalf of the Wellcome Sanger Institute COVID-19 Surveillance Team |
| EPI_ISL_3094596, EPI_ISL_3094597 | Lighthouse Lab in Milton Keynes | Wellcome Sanger Institute for the COVID-19 Genomics UK (COG-UK) Consortium | The Lighthouse Lab in Milton Keynes and Alex Alderton, Roberto Amato, Jeffrey Barrett, Sonia Goncalves, Ewan Harrison, David K. Jackson, Ian Johnston, Dominic Kwiatkowski, Cordelia Langford, John Sillitoe on behalf of the Wellcome Sanger Institute COVID-19 Surveillance Team |
| EPI_ISL_3094598 | Lighthouse Lab in Glasgow | Wellcome Sanger Institute for the COVID-19 Genomics UK (COG-UK) Consortium | Harper VanSteenhouse, Yumi Kasai, David Gray, Carol Clugston, Anna Dominiczak and Alex Alderton, Roberto Amato, Jeffrey Barrett, Sonia Goncalves, Ewan Harrison, David K. Jackson, Ian Johnston, Dominic Kwiatkowski, Cordelia Langford, John Sillitoe on behalf of the Wellcome Sanger Institute COVID-19 Surveillance Team |
| EPI_ISL_3094599, EPI_ISL_3094600, EPI_ISL_3094601, EPI_ISL_3094602 | Lighthouse Lab in Milton Keynes | Wellcome Sanger Institute for the COVID-19 Genomics UK (COG-UK) Consortium | The Lighthouse Lab in Milton Keynes and Alex Alderton, Roberto Amato, Jeffrey Barrett, Sonia Goncalves, Ewan Harrison, David K. Jackson, Ian Johnston, Dominic Kwiatkowski, Cordelia Langford, John Sillitoe on behalf of the Wellcome Sanger Institute COVID-19 Surveillance Team |
| EPI_ISL_3094603 | Lighthouse Laboratory Plymouth | Wellcome Sanger Institute for the COVID-19 Genomics UK (COG-UK) Consortium | Lighthouse Laboratory Plymouth and Alex Alderton, Roberto Amato, Jeffrey Barrett, Sonia Goncalves, Ewan Harrison, David K. Jackson, Ian Johnston, Dominic Kwiatkowski, Cordelia Langford, John Sillitoe on behalf of the Wellcome Sanger Institute COVID-19 Surveillance Team |
| EPI_ISL_3094604 | Lighthouse Lab in Glasgow | Wellcome Sanger Institute for the COVID-19 Genomics UK (COG-UK) Consortium | Harper VanSteenhouse, Yumi Kasai, David Gray, Carol Clugston, Anna Dominiczak and Alex Alderton, Roberto Amato, Jeffrey Barrett, Sonia Goncalves, Ewan Harrison, David K. Jackson, Ian Johnston, Dominic Kwiatkowski, Cordelia Langford, John Sillitoe on behalf of the Wellcome Sanger Institute COVID-19 Surveillance Team |
| EPI_ISL_3094605, EPI_ISL_3094606, EPI_ISL_3094607 | Lighthouse Lab in Milton Keynes | Wellcome Sanger Institute for the COVID-19 Genomics UK (COG-UK) Consortium | The Lighthouse Lab in Milton Keynes and Alex Alderton, Roberto Amato, Jeffrey Barrett, Sonia Goncalves, Ewan Harrison, David K. Jackson, Ian Johnston, Dominic Kwiatkowski, Cordelia Langford, John Sillitoe on behalf of the Wellcome Sanger Institute COVID-19 Surveillance Team |
| EPI_ISL_3094608 | Lighthouse Lab in Glasgow | Wellcome Sanger Institute for the COVID-19 Genomics UK (COG-UK) Consortium | Harper VanSteenhouse, Yumi Kasai, David Gray, Carol Clugston, Anna Dominiczak and Alex Alderton, Roberto Amato, Jeffrey Barrett, Sonia Goncalves, Ewan Harrison, David K. Jackson, Ian Johnston, Dominic Kwiatkowski, Cordelia Langford, John Sillitoe on behalf of the Wellcome Sanger Institute COVID-19 Surveillance Team |
| EPI_ISL_3094609 | Lighthouse Lab in Milton Keynes | Wellcome Sanger Institute for the COVID-19 Genomics UK (COG-UK) Consortium | The Lighthouse Lab in Milton Keynes and Alex Alderton, Roberto Amato, Jeffrey Barrett, Sonia Goncalves, Ewan Harrison, David K. Jackson, Ian Johnston, Dominic Kwiatkowski, Cordelia Langford, John Sillitoe on behalf of the Wellcome Sanger Institute COVID-19 Surveillance Team |
| EPI_ISL_3094610 | Lighthouse Lab in Glasgow | Wellcome Sanger Institute for the COVID-19 Genomics UK (COG-UK) Consortium | Harper VanSteenhouse, Yumi Kasai, David Gray, Carol Clugston, Anna Dominiczak and Alex Alderton, Roberto Amato, Jeffrey Barrett, Sonia Goncalves, Ewan Harrison, David K. Jackson, Ian Johnston, Dominic Kwiatkowski, Cordelia Langford, John Sillitoe on behalf of the Wellcome Sanger Institute COVID-19 Surveillance Team |
| EPI_ISL_3094611, EPI_ISL_3094612, EPI_ISL_3094613, EPI_ISL_3094614 | Lighthouse Laboratory Plymouth | Wellcome Sanger Institute for the COVID-19 Genomics UK (COG-UK) Consortium | Lighthouse Laboratory Plymouth and Alex Alderton, Roberto Amato, Jeffrey Barrett, Sonia Goncalves, Ewan Harrison, David K. Jackson, Ian Johnston, Dominic Kwiatkowski, Cordelia Langford, John Sillitoe on behalf of the Wellcome Sanger Institute COVID-19 Surveillance Team |
| EPI_ISL_3094615, EPI_ISL_3094616, EPI_ISL_3094617, EPI_ISL_3094618, EPI_ISL_3094619 | Lighthouse Lab in Milton Keynes | Wellcome Sanger Institute for the COVID-19 Genomics UK (COG-UK) Consortium | The Lighthouse Lab in Milton Keynes and Alex Alderton, Roberto Amato, Jeffrey Barrett, Sonia Goncalves, Ewan Harrison, David K. Jackson, Ian Johnston, Dominic Kwiatkowski, Cordelia Langford, John Sillitoe on behalf of the Wellcome Sanger Institute COVID-19 Surveillance Team |
| EPI_ISL_3094620, EPI_ISL_3094621, EPI_ISL_3094622, EPI_ISL_3094623, EPI_ISL_3094624 | Lighthouse Laboratory Plymouth | Wellcome Sanger Institute for the COVID-19 Genomics UK (COG-UK) Consortium | Lighthouse Laboratory Plymouth and Alex Alderton, Roberto Amato, Jeffrey Barrett, Sonia Goncalves, Ewan Harrison, David K. Jackson, Ian Johnston, Dominic Kwiatkowski, Cordelia Langford, John Sillitoe on behalf of the Wellcome Sanger Institute COVID-19 Surveillance Team |
| EPI_ISL_3094625 | Lighthouse Lab in Milton Keynes | Wellcome Sanger Institute for the COVID-19 Genomics UK (COG-UK) | The Lighthouse Lab in Milton Keynes and Alex Alderton, Roberto Amato, Jeffrey Barrett, Sonia Goncalves, Ewan Harrison, David K. Jackson, Ian Johnston, Dominic Kwiatkowski, Cordelia Langford, John Sillitoe on behalf of the Wellcome Sanger |

| | | | |
|---|---|---|---|
| | | Consortium | Institute COVID-19 Surveillance Team |
| EPI_ISL_3094626, EPI_ISL_3094627, EPI_ISL_3094628, EPI_ISL_3094629, EPI_ISL_3094630, EPI_ISL_3094631, EPI_ISL_3094632, EPI_ISL_3094633 | Lighthouse Laboratory Plymouth | Wellcome Sanger Institute for the COVID-19 Genomics UK (COG-UK) Consortium | Lighthouse Laboratory Plymouth and Alex Alderton, Roberto Amato, Jeffrey Barrett, Sonia Goncalves, Ewan Harrison, David K. Jackson, Ian Johnston, Dominic Kwiatkowski, Cordelia Langford, John Sillitoe on behalf of the Wellcome Sanger Institute COVID-19 Surveillance Team |
| EPI_ISL_3094634, EPI_ISL_3094635 | Lighthouse Lab in Milton Keynes | Wellcome Sanger Institute for the COVID-19 Genomics UK (COG-UK) Consortium | The Lighthouse Lab in Milton Keynes and Alex Alderton, Roberto Amato, Jeffrey Barrett, Sonia Goncalves, Ewan Harrison, David K. Jackson, Ian Johnston, Dominic Kwiatkowski, Cordelia Langford, John Sillitoe on behalf of the Wellcome Sanger Institute COVID-19 Surveillance Team |
| EPI_ISL_3094636, EPI_ISL_3094637 | Lighthouse Laboratory Plymouth | Wellcome Sanger Institute for the COVID-19 Genomics UK (COG-UK) Consortium | Lighthouse Laboratory Plymouth and Alex Alderton, Roberto Amato, Jeffrey Barrett, Sonia Goncalves, Ewan Harrison, David K. Jackson, Ian Johnston, Dominic Kwiatkowski, Cordelia Langford, John Sillitoe on behalf of the Wellcome Sanger Institute COVID-19 Surveillance Team |
| EPI_ISL_3094638 | Lighthouse Lab in Milton Keynes | Wellcome Sanger Institute for the COVID-19 Genomics UK (COG-UK) Consortium | The Lighthouse Lab in Milton Keynes and Alex Alderton, Roberto Amato, Jeffrey Barrett, Sonia Goncalves, Ewan Harrison, David K. Jackson, Ian Johnston, Dominic Kwiatkowski, Cordelia Langford, John Sillitoe on behalf of the Wellcome Sanger Institute COVID-19 Surveillance Team |
| EPI_ISL_3094639, EPI_ISL_3094640, EPI_ISL_3094641 | Lighthouse Laboratory Plymouth | Wellcome Sanger Institute for the COVID-19 Genomics UK (COG-UK) Consortium | Lighthouse Laboratory Plymouth and Alex Alderton, Roberto Amato, Jeffrey Barrett, Sonia Goncalves, Ewan Harrison, David K. Jackson, Ian Johnston, Dominic Kwiatkowski, Cordelia Langford, John Sillitoe on behalf of the Wellcome Sanger Institute COVID-19 Surveillance Team |
| EPI_ISL_3094642, EPI_ISL_3094643, EPI_ISL_3094644 | Lighthouse Lab in Milton Keynes | Wellcome Sanger Institute for the COVID-19 Genomics UK (COG-UK) Consortium | The Lighthouse Lab in Milton Keynes and Alex Alderton, Roberto Amato, Jeffrey Barrett, Sonia Goncalves, Ewan Harrison, David K. Jackson, Ian Johnston, Dominic Kwiatkowski, Cordelia Langford, John Sillitoe on behalf of the Wellcome Sanger Institute COVID-19 Surveillance Team |
| EPI_ISL_3094645 | Lighthouse Laboratory Plymouth | Wellcome Sanger Institute for the COVID-19 Genomics UK (COG-UK) Consortium | Lighthouse Laboratory Plymouth and Alex Alderton, Roberto Amato, Jeffrey Barrett, Sonia Goncalves, Ewan Harrison, David K. Jackson, Ian Johnston, Dominic Kwiatkowski, Cordelia Langford, John Sillitoe on behalf of the Wellcome Sanger Institute COVID-19 Surveillance Team |
| EPI_ISL_3094646 | Lighthouse Lab in Milton Keynes | Wellcome Sanger Institute for the COVID-19 Genomics UK (COG-UK) Consortium | The Lighthouse Lab in Milton Keynes and Alex Alderton, Roberto Amato, Jeffrey Barrett, Sonia Goncalves, Ewan Harrison, David K. Jackson, Ian Johnston, Dominic Kwiatkowski, Cordelia Langford, John Sillitoe on behalf of the Wellcome Sanger Institute COVID-19 Surveillance Team |
| EPI_ISL_3094647, EPI_ISL_3094648, EPI_ISL_3094649, EPI_ISL_3094650, EPI_ISL_3094651 | Lighthouse Laboratory Plymouth | Wellcome Sanger Institute for the COVID-19 Genomics UK (COG-UK) Consortium | Lighthouse Laboratory Plymouth and Alex Alderton, Roberto Amato, Jeffrey Barrett, Sonia Goncalves, Ewan Harrison, David K. Jackson, Ian Johnston, Dominic Kwiatkowski, Cordelia Langford, John Sillitoe on behalf of the Wellcome Sanger Institute COVID-19 Surveillance Team |
| EPI_ISL_3094652, EPI_ISL_3094653 | Lighthouse Lab in Milton Keynes | Wellcome Sanger Institute for the COVID-19 Genomics UK (COG-UK) Consortium | The Lighthouse Lab in Milton Keynes and Alex Alderton, Roberto Amato, Jeffrey Barrett, Sonia Goncalves, Ewan Harrison, David K. Jackson, Ian Johnston, Dominic Kwiatkowski, Cordelia Langford, John Sillitoe on behalf of the Wellcome Sanger Institute COVID-19 Surveillance Team |
| EPI_ISL_3094654 | Lighthouse Laboratory Plymouth | Wellcome Sanger Institute for the COVID-19 Genomics UK (COG-UK) Consortium | Lighthouse Laboratory Plymouth and Alex Alderton, Roberto Amato, Jeffrey Barrett, Sonia Goncalves, Ewan Harrison, David K. Jackson, Ian Johnston, Dominic Kwiatkowski, Cordelia Langford, John Sillitoe on behalf of the Wellcome Sanger Institute COVID-19 Surveillance Team |
| EPI_ISL_3094655 | Lighthouse Lab in Milton Keynes | Wellcome Sanger Institute for the COVID-19 Genomics UK (COG-UK) Consortium | The Lighthouse Lab in Milton Keynes and Alex Alderton, Roberto Amato, Jeffrey Barrett, Sonia Goncalves, Ewan Harrison, David K. Jackson, Ian Johnston, Dominic Kwiatkowski, Cordelia Langford, John Sillitoe on behalf of the Wellcome Sanger Institute COVID-19 Surveillance Team |
| EPI_ISL_3094656, EPI_ISL_3094657, EPI_ISL_3094658, EPI_ISL_3094659, EPI_ISL_3094660, EPI_ISL_3094661, EPI_ISL_3094662 | Lighthouse Laboratory Plymouth | Wellcome Sanger Institute for the COVID-19 Genomics UK (COG-UK) Consortium | Lighthouse Laboratory Plymouth and Alex Alderton, Roberto Amato, Jeffrey Barrett, Sonia Goncalves, Ewan Harrison, David K. Jackson, Ian Johnston, Dominic Kwiatkowski, Cordelia Langford, John Sillitoe on behalf of the Wellcome Sanger Institute COVID-19 Surveillance Team |
| EPI_ISL_3094663 | Lighthouse Lab in Milton Keynes | Wellcome Sanger Institute for the COVID-19 Genomics UK (COG-UK) Consortium | The Lighthouse Lab in Milton Keynes and Alex Alderton, Roberto Amato, Jeffrey Barrett, Sonia Goncalves, Ewan Harrison, David K. Jackson, Ian Johnston, Dominic Kwiatkowski, Cordelia Langford, John Sillitoe on behalf of the Wellcome Sanger Institute COVID-19 Surveillance Team |
| EPI_ISL_3094664 | Lighthouse Laboratory Plymouth | Wellcome Sanger Institute for the COVID-19 Genomics UK (COG-UK) Consortium | Lighthouse Laboratory Plymouth and Alex Alderton, Roberto Amato, Jeffrey Barrett, Sonia Goncalves, Ewan Harrison, David K. Jackson, Ian Johnston, Dominic Kwiatkowski, Cordelia Langford, John Sillitoe on behalf of the Wellcome Sanger Institute COVID-19 Surveillance Team |
| EPI_ISL_3094665 | Lighthouse Lab in Milton Keynes | Wellcome Sanger Institute for the COVID-19 Genomics UK (COG-UK) Consortium | The Lighthouse Lab in Milton Keynes and Alex Alderton, Roberto Amato, Jeffrey Barrett, Sonia Goncalves, Ewan Harrison, David K. Jackson, Ian Johnston, Dominic Kwiatkowski, Cordelia Langford, John Sillitoe on behalf of the Wellcome Sanger Institute COVID-19 Surveillance Team |
| EPI_ISL_3094666 | Lighthouse Lab in Glasgow | Wellcome Sanger Institute for the COVID-19 Genomics UK (COG-UK) Consortium | Harper VanSteenhouse, Yumi Kasai, David Gray, Carol Clugston, Anna Dominiczak and Alex Alderton, Roberto Amato, Jeffrey Barrett, Sonia Goncalves, Ewan Harrison, David K. Jackson, Ian Johnston, Dominic Kwiatkowski, Cordelia Langford, John Sillitoe on behalf of the Wellcome Sanger Institute COVID-19 Surveillance Team |
| EPI_ISL_3094667 | Lighthouse Lab in Milton Keynes | Wellcome Sanger Institute for the COVID-19 Genomics UK (COG-UK) Consortium | The Lighthouse Lab in Milton Keynes and Alex Alderton, Roberto Amato, Jeffrey Barrett, Sonia Goncalves, Ewan Harrison, David K. Jackson, Ian Johnston, Dominic Kwiatkowski, Cordelia Langford, John Sillitoe on behalf of the Wellcome Sanger Institute COVID-19 Surveillance Team |
| EPI_ISL_3094668, EPI_ISL_3094669, EPI_ISL_3094670 | Lighthouse Laboratory Plymouth | Wellcome Sanger Institute for the COVID-19 Genomics UK (COG-UK) Consortium | Lighthouse Laboratory Plymouth and Alex Alderton, Roberto Amato, Jeffrey Barrett, Sonia Goncalves, Ewan Harrison, David K. Jackson, Ian Johnston, Dominic Kwiatkowski, Cordelia Langford, John Sillitoe on behalf of the Wellcome Sanger Institute COVID-19 Surveillance Team |
| EPI_ISL_3094671 | Lighthouse Lab in Milton Keynes | Wellcome Sanger Institute for the COVID-19 Genomics UK (COG-UK) Consortium | The Lighthouse Lab in Milton Keynes and Alex Alderton, Roberto Amato, Jeffrey Barrett, Sonia Goncalves, Ewan Harrison, David K. Jackson, Ian Johnston, Dominic Kwiatkowski, Cordelia Langford, John Sillitoe on behalf of the Wellcome Sanger Institute COVID-19 Surveillance Team |
| EPI_ISL_3094672, EPI_ISL_3094673, EPI_ISL_3094674, EPI_ISL_3094675 | Lighthouse Laboratory Plymouth | Wellcome Sanger Institute for the COVID-19 Genomics UK (COG-UK) Consortium | Lighthouse Laboratory Plymouth and Alex Alderton, Roberto Amato, Jeffrey Barrett, Sonia Goncalves, Ewan Harrison, David K. Jackson, Ian Johnston, Dominic Kwiatkowski, Cordelia Langford, John Sillitoe on behalf of the Wellcome Sanger Institute COVID-19 Surveillance Team |
| EPI_ISL_3094676, EPI_ISL_3094677 | Lighthouse Lab in Milton Keynes | Wellcome Sanger Institute for the COVID-19 Genomics UK (COG-UK) Consortium | The Lighthouse Lab in Milton Keynes and Alex Alderton, Roberto Amato, Jeffrey Barrett, Sonia Goncalves, Ewan Harrison, David K. Jackson, Ian Johnston, Dominic Kwiatkowski, Cordelia Langford, John Sillitoe on behalf of the Wellcome Sanger Institute COVID-19 Surveillance Team |
| EPI_ISL_3094678 | Lighthouse Laboratory Plymouth | Wellcome Sanger Institute for the COVID-19 Genomics UK (COG-UK) Consortium | Lighthouse Laboratory Plymouth and Alex Alderton, Roberto Amato, Jeffrey Barrett, Sonia Goncalves, Ewan Harrison, David K. Jackson, Ian Johnston, Dominic Kwiatkowski, Cordelia Langford, John Sillitoe on behalf of the Wellcome Sanger Institute COVID-19 Surveillance Team |
| EPI_ISL_3094679, EPI_ISL_3094680, EPI_ISL_3094681 | Lighthouse Lab in Milton Keynes | Wellcome Sanger Institute for the COVID-19 Genomics UK (COG-UK) Consortium | The Lighthouse Lab in Milton Keynes and Alex Alderton, Roberto Amato, Jeffrey Barrett, Sonia Goncalves, Ewan Harrison, David K. Jackson, Ian Johnston, Dominic Kwiatkowski, Cordelia Langford, John Sillitoe on behalf of the Wellcome Sanger Institute COVID-19 Surveillance Team |
| EPI_ISL_3094682, EPI_ISL_3094683, EPI_ISL_3094684, EPI_ISL_3094685, EPI_ISL_3094686, EPI_ISL_3094687 | Lighthouse Laboratory Plymouth | Wellcome Sanger Institute for the COVID-19 Genomics UK (COG-UK) Consortium | Lighthouse Laboratory Plymouth and Alex Alderton, Roberto Amato, Jeffrey Barrett, Sonia Goncalves, Ewan Harrison, David K. Jackson, Ian Johnston, Dominic Kwiatkowski, Cordelia Langford, John Sillitoe on behalf of the Wellcome Sanger Institute COVID-19 Surveillance Team |
| EPI_ISL_3094688 | Lighthouse Lab in Glasgow | Wellcome Sanger Institute for the COVID-19 Genomics UK (COG-UK) Consortium | Harper VanSteenhouse, Yumi Kasai, David Gray, Carol Clugston, Anna Dominiczak and Alex Alderton, Roberto Amato, Jeffrey Barrett, Sonia Goncalves, Ewan Harrison, David K. Jackson, Ian Johnston, Dominic Kwiatkowski, Cordelia Langford, John Sillitoe on behalf of the Wellcome Sanger Institute COVID-19 Surveillance Team |
| EPI_ISL_3094689, EPI_ISL_3094690, EPI_ISL_3094691 | Lighthouse Laboratory Plymouth | Wellcome Sanger Institute for the COVID-19 Genomics UK (COG-UK) Consortium | Lighthouse Laboratory Plymouth and Alex Alderton, Roberto Amato, Jeffrey Barrett, Sonia Goncalves, Ewan Harrison, David K. Jackson, Ian Johnston, Dominic Kwiatkowski, Cordelia Langford, John Sillitoe on behalf of the Wellcome Sanger Institute COVID-19 Surveillance Team |
| EPI_ISL_3094692 | Lighthouse Lab in Milton Keynes | Wellcome Sanger Institute for the COVID-19 Genomics UK (COG-UK) Consortium | The Lighthouse Lab in Milton Keynes and Alex Alderton, Roberto Amato, Jeffrey Barrett, Sonia Goncalves, Ewan Harrison, David K. Jackson, Ian Johnston, Dominic Kwiatkowski, Cordelia Langford, John Sillitoe on behalf of the Wellcome Sanger Institute COVID-19 Surveillance Team |
| EPI_ISL_3094693, EPI_ISL_3094694, EPI_ISL_3094695 | Lighthouse Laboratory Plymouth | Wellcome Sanger Institute for the COVID-19 Genomics UK (COG-UK) Consortium | Lighthouse Laboratory Plymouth and Alex Alderton, Roberto Amato, Jeffrey Barrett, Sonia Goncalves, Ewan Harrison, David K. Jackson, Ian Johnston, Dominic Kwiatkowski, Cordelia Langford, John Sillitoe on behalf of the Wellcome Sanger Institute COVID-19 Surveillance Team |
| EPI_ISL_3094696 | Lighthouse Lab in Milton Keynes | Wellcome Sanger Institute for the COVID-19 Genomics UK (COG-UK) Consortium | The Lighthouse Lab in Milton Keynes and Alex Alderton, Roberto Amato, Jeffrey Barrett, Sonia Goncalves, Ewan Harrison, David K. Jackson, Ian Johnston, Dominic Kwiatkowski, Cordelia Langford, John Sillitoe on behalf of the Wellcome Sanger Institute COVID-19 Surveillance Team |
| EPI_ISL_3094697, EPI_ISL_3094698, EPI_ISL_3094699, EPI_ISL_3094700, EPI_ISL_3094701 | Lighthouse Laboratory Plymouth | Wellcome Sanger Institute for the COVID-19 Genomics UK (COG-UK) Consortium | Lighthouse Laboratory Plymouth and Alex Alderton, Roberto Amato, Jeffrey Barrett, Sonia Goncalves, Ewan Harrison, David K. Jackson, Ian Johnston, Dominic Kwiatkowski, Cordelia Langford, John Sillitoe on behalf of the Wellcome Sanger Institute COVID-19 Surveillance Team |
| EPI_ISL_3094702, EPI_ISL_3094703 | Lighthouse Lab in Milton Keynes | Wellcome Sanger Institute for the COVID-19 Genomics UK (COG-UK) Consortium | The Lighthouse Lab in Milton Keynes and Alex Alderton, Roberto Amato, Jeffrey Barrett, Sonia Goncalves, Ewan Harrison, David K. Jackson, Ian Johnston, Dominic Kwiatkowski, Cordelia Langford, John Sillitoe on behalf of the Wellcome Sanger Institute COVID-19 Surveillance Team |
| EPI_ISL_3094704, | Lighthouse Laboratory Plymouth | Wellcome Sanger Institute for the COVID-19 Genomics UK (COG-UK) | Lighthouse Laboratory Plymouth and Alex Alderton, Roberto Amato, Jeffrey Barrett, Sonia Goncalves, Ewan Harrison, David K. Jackson, Ian Johnston, Dominic Kwiatkowski, Cordelia Langford, John Sillitoe on behalf of the Wellcome Sanger |

| | | | |
|---|---|---|---|
| EPI_ISL_3094705 | | Consortium | Institute COVID-19 Surveillance Team |
| EPI_ISL_3094706, EPI_ISL_3094707 | Lighthouse Lab in Milton Keynes | Wellcome Sanger Institute for the COVID-19 Genomics UK (COG-UK) Consortium | The Lighthouse Lab in Milton Keynes and Alex Alderton, Roberto Amato, Jeffrey Barrett, Sonia Goncalves, Ewan Harrison, David K. Jackson, Ian Johnston, Dominic Kwiatkowski, Cordelia Langford, John Sillitoe on behalf of the Wellcome Sanger Institute COVID-19 Surveillance Team |
| EPI_ISL_3094708, EPI_ISL_3094709 | Lighthouse Laboratory Plymouth | Wellcome Sanger Institute for the COVID-19 Genomics UK (COG-UK) Consortium | Lighthouse Laboratory Plymouth and Alex Alderton, Roberto Amato, Jeffrey Barrett, Sonia Goncalves, Ewan Harrison, David K. Jackson, Ian Johnston, Dominic Kwiatkowski, Cordelia Langford, John Sillitoe on behalf of the Wellcome Sanger Institute COVID-19 Surveillance Team |
| EPI_ISL_3094710 | Lighthouse Lab in Milton Keynes | Wellcome Sanger Institute for the COVID-19 Genomics UK (COG-UK) Consortium | The Lighthouse Lab in Milton Keynes and Alex Alderton, Roberto Amato, Jeffrey Barrett, Sonia Goncalves, Ewan Harrison, David K. Jackson, Ian Johnston, Dominic Kwiatkowski, Cordelia Langford, John Sillitoe on behalf of the Wellcome Sanger Institute COVID-19 Surveillance Team |
| EPI_ISL_3094711 | Lighthouse Lab in Glasgow | Wellcome Sanger Institute for the COVID-19 Genomics UK (COG-UK) Consortium | Harper VanSteenhouse, Yumi Kasai, David Gray, Carol Clugston, Anna Dominiczak and Alex Alderton, Roberto Amato, Jeffrey Barrett, Sonia Goncalves, Ewan Harrison, David K. Jackson, Ian Johnston, Dominic Kwiatkowski, Cordelia Langford, John Sillitoe on behalf of the Wellcome Sanger Institute COVID-19 Surveillance Team |
| EPI_ISL_3094712, EPI_ISL_3094713 | Lighthouse Laboratory Plymouth | Wellcome Sanger Institute for the COVID-19 Genomics UK (COG-UK) Consortium | Lighthouse Laboratory Plymouth and Alex Alderton, Roberto Amato, Jeffrey Barrett, Sonia Goncalves, Ewan Harrison, David K. Jackson, Ian Johnston, Dominic Kwiatkowski, Cordelia Langford, John Sillitoe on behalf of the Wellcome Sanger Institute COVID-19 Surveillance Team |
| EPI_ISL_3094714 | Lighthouse Lab in Milton Keynes | Wellcome Sanger Institute for the COVID-19 Genomics UK (COG-UK) Consortium | The Lighthouse Lab in Milton Keynes and Alex Alderton, Roberto Amato, Jeffrey Barrett, Sonia Goncalves, Ewan Harrison, David K. Jackson, Ian Johnston, Dominic Kwiatkowski, Cordelia Langford, John Sillitoe on behalf of the Wellcome Sanger Institute COVID-19 Surveillance Team |
| EPI_ISL_3094715, EPI_ISL_3094716, EPI_ISL_3094717, EPI_ISL_3094718, EPI_ISL_3094719, EPI_ISL_3094720, EPI_ISL_3094721, EPI_ISL_3094722 | Lighthouse Laboratory Plymouth | Wellcome Sanger Institute for the COVID-19 Genomics UK (COG-UK) Consortium | Lighthouse Laboratory Plymouth and Alex Alderton, Roberto Amato, Jeffrey Barrett, Sonia Goncalves, Ewan Harrison, David K. Jackson, Ian Johnston, Dominic Kwiatkowski, Cordelia Langford, John Sillitoe on behalf of the Wellcome Sanger Institute COVID-19 Surveillance Team |
| EPI_ISL_3094723, EPI_ISL_3094724 | Lighthouse Lab in Milton Keynes | Wellcome Sanger Institute for the COVID-19 Genomics UK (COG-UK) Consortium | The Lighthouse Lab in Milton Keynes and Alex Alderton, Roberto Amato, Jeffrey Barrett, Sonia Goncalves, Ewan Harrison, David K. Jackson, Ian Johnston, Dominic Kwiatkowski, Cordelia Langford, John Sillitoe on behalf of the Wellcome Sanger Institute COVID-19 Surveillance Team |
| EPI_ISL_3094725 | Lighthouse Laboratory Plymouth | Wellcome Sanger Institute for the COVID-19 Genomics UK (COG-UK) Consortium | Lighthouse Laboratory Plymouth and Alex Alderton, Roberto Amato, Jeffrey Barrett, Sonia Goncalves, Ewan Harrison, David K. Jackson, Ian Johnston, Dominic Kwiatkowski, Cordelia Langford, John Sillitoe on behalf of the Wellcome Sanger Institute COVID-19 Surveillance Team |
| EPI_ISL_3094726 | Lighthouse Lab in Glasgow | Wellcome Sanger Institute for the COVID-19 Genomics UK (COG-UK) Consortium | Harper VanSteenhouse, Yumi Kasai, David Gray, Carol Clugston, Anna Dominiczak and Alex Alderton, Roberto Amato, Jeffrey Barrett, Sonia Goncalves, Ewan Harrison, David K. Jackson, Ian Johnston, Dominic Kwiatkowski, Cordelia Langford, John Sillitoe on behalf of the Wellcome Sanger Institute COVID-19 Surveillance Team |
| EPI_ISL_3094727 | Lighthouse Laboratory Plymouth | Wellcome Sanger Institute for the COVID-19 Genomics UK (COG-UK) Consortium | Lighthouse Laboratory Plymouth and Alex Alderton, Roberto Amato, Jeffrey Barrett, Sonia Goncalves, Ewan Harrison, David K. Jackson, Ian Johnston, Dominic Kwiatkowski, Cordelia Langford, John Sillitoe on behalf of the Wellcome Sanger Institute COVID-19 Surveillance Team |
| EPI_ISL_3094728, EPI_ISL_3094729 | Lighthouse Lab in Milton Keynes | Wellcome Sanger Institute for the COVID-19 Genomics UK (COG-UK) Consortium | The Lighthouse Lab in Milton Keynes and Alex Alderton, Roberto Amato, Jeffrey Barrett, Sonia Goncalves, Ewan Harrison, David K. Jackson, Ian Johnston, Dominic Kwiatkowski, Cordelia Langford, John Sillitoe on behalf of the Wellcome Sanger Institute COVID-19 Surveillance Team |
| EPI_ISL_3094730, EPI_ISL_3094731 | Lighthouse Laboratory Plymouth | Wellcome Sanger Institute for the COVID-19 Genomics UK (COG-UK) Consortium | Lighthouse Laboratory Plymouth and Alex Alderton, Roberto Amato, Jeffrey Barrett, Sonia Goncalves, Ewan Harrison, David K. Jackson, Ian Johnston, Dominic Kwiatkowski, Cordelia Langford, John Sillitoe on behalf of the Wellcome Sanger Institute COVID-19 Surveillance Team |
| EPI_ISL_3094732 | Lighthouse Lab in Milton Keynes | Wellcome Sanger Institute for the COVID-19 Genomics UK (COG-UK) Consortium | The Lighthouse Lab in Milton Keynes and Alex Alderton, Roberto Amato, Jeffrey Barrett, Sonia Goncalves, Ewan Harrison, David K. Jackson, Ian Johnston, Dominic Kwiatkowski, Cordelia Langford, John Sillitoe on behalf of the Wellcome Sanger Institute COVID-19 Surveillance Team |
| EPI_ISL_3094733, EPI_ISL_3094734 | Lighthouse Laboratory Plymouth | Wellcome Sanger Institute for the COVID-19 Genomics UK (COG-UK) Consortium | Lighthouse Laboratory Plymouth and Alex Alderton, Roberto Amato, Jeffrey Barrett, Sonia Goncalves, Ewan Harrison, David K. Jackson, Ian Johnston, Dominic Kwiatkowski, Cordelia Langford, John Sillitoe on behalf of the Wellcome Sanger Institute COVID-19 Surveillance Team |
| EPI_ISL_3094735 | Lighthouse Lab in Milton Keynes | Wellcome Sanger Institute for the COVID-19 Genomics UK (COG-UK) Consortium | The Lighthouse Lab in Milton Keynes and Alex Alderton, Roberto Amato, Jeffrey Barrett, Sonia Goncalves, Ewan Harrison, David K. Jackson, Ian Johnston, Dominic Kwiatkowski, Cordelia Langford, John Sillitoe on behalf of the Wellcome Sanger Institute COVID-19 Surveillance Team |
| EPI_ISL_3094736, EPI_ISL_3094737, EPI_ISL_3094738 | Lighthouse Laboratory Plymouth | Wellcome Sanger Institute for the COVID-19 Genomics UK (COG-UK) Consortium | Lighthouse Laboratory Plymouth and Alex Alderton, Roberto Amato, Jeffrey Barrett, Sonia Goncalves, Ewan Harrison, David K. Jackson, Ian Johnston, Dominic Kwiatkowski, Cordelia Langford, John Sillitoe on behalf of the Wellcome Sanger Institute COVID-19 Surveillance Team |
| EPI_ISL_3094739, EPI_ISL_3094740 | Lighthouse Lab in Glasgow | Wellcome Sanger Institute for the COVID-19 Genomics UK (COG-UK) Consortium | Harper VanSteenhouse, Yumi Kasai, David Gray, Carol Clugston, Anna Dominiczak and Alex Alderton, Roberto Amato, Jeffrey Barrett, Sonia Goncalves, Ewan Harrison, David K. Jackson, Ian Johnston, Dominic Kwiatkowski, Cordelia Langford, John Sillitoe on behalf of the Wellcome Sanger Institute COVID-19 Surveillance Team |
| EPI_ISL_3094741 | Lighthouse Laboratory Plymouth | Wellcome Sanger Institute for the COVID-19 Genomics UK (COG-UK) Consortium | Lighthouse Laboratory Plymouth and Alex Alderton, Roberto Amato, Jeffrey Barrett, Sonia Goncalves, Ewan Harrison, David K. Jackson, Ian Johnston, Dominic Kwiatkowski, Cordelia Langford, John Sillitoe on behalf of the Wellcome Sanger Institute COVID-19 Surveillance Team |
| EPI_ISL_3094742 | Lighthouse Lab in Glasgow | Wellcome Sanger Institute for the COVID-19 Genomics UK (COG-UK) Consortium | Harper VanSteenhouse, Yumi Kasai, David Gray, Carol Clugston, Anna Dominiczak and Alex Alderton, Roberto Amato, Jeffrey Barrett, Sonia Goncalves, Ewan Harrison, David K. Jackson, Ian Johnston, Dominic Kwiatkowski, Cordelia Langford, John Sillitoe on behalf of the Wellcome Sanger Institute COVID-19 Surveillance Team |
| EPI_ISL_3094743, EPI_ISL_3094744, EPI_ISL_3094745, EPI_ISL_3094746 | Lighthouse Laboratory Plymouth | Wellcome Sanger Institute for the COVID-19 Genomics UK (COG-UK) Consortium | Lighthouse Laboratory Plymouth and Alex Alderton, Roberto Amato, Jeffrey Barrett, Sonia Goncalves, Ewan Harrison, David K. Jackson, Ian Johnston, Dominic Kwiatkowski, Cordelia Langford, John Sillitoe on behalf of the Wellcome Sanger Institute COVID-19 Surveillance Team |
| EPI_ISL_3094747 | Lighthouse Lab in Glasgow | Wellcome Sanger Institute for the COVID-19 Genomics UK (COG-UK) Consortium | Harper VanSteenhouse, Yumi Kasai, David Gray, Carol Clugston, Anna Dominiczak and Alex Alderton, Roberto Amato, Jeffrey Barrett, Sonia Goncalves, Ewan Harrison, David K. Jackson, Ian Johnston, Dominic Kwiatkowski, Cordelia Langford, John Sillitoe on behalf of the Wellcome Sanger Institute COVID-19 Surveillance Team |
| EPI_ISL_3094748 | Lighthouse Laboratory Plymouth | Wellcome Sanger Institute for the COVID-19 Genomics UK (COG-UK) Consortium | Lighthouse Laboratory Plymouth and Alex Alderton, Roberto Amato, Jeffrey Barrett, Sonia Goncalves, Ewan Harrison, David K. Jackson, Ian Johnston, Dominic Kwiatkowski, Cordelia Langford, John Sillitoe on behalf of the Wellcome Sanger Institute COVID-19 Surveillance Team |
| EPI_ISL_3094749 | Lighthouse Lab in Milton Keynes | Wellcome Sanger Institute for the COVID-19 Genomics UK (COG-UK) Consortium | The Lighthouse Lab in Milton Keynes and Alex Alderton, Roberto Amato, Jeffrey Barrett, Sonia Goncalves, Ewan Harrison, David K. Jackson, Ian Johnston, Dominic Kwiatkowski, Cordelia Langford, John Sillitoe on behalf of the Wellcome Sanger Institute COVID-19 Surveillance Team |
| EPI_ISL_3094750 | Lighthouse Laboratory Plymouth | Wellcome Sanger Institute for the COVID-19 Genomics UK (COG-UK) Consortium | Lighthouse Laboratory Plymouth and Alex Alderton, Roberto Amato, Jeffrey Barrett, Sonia Goncalves, Ewan Harrison, David K. Jackson, Ian Johnston, Dominic Kwiatkowski, Cordelia Langford, John Sillitoe on behalf of the Wellcome Sanger Institute COVID-19 Surveillance Team |
| EPI_ISL_3094751 | Lighthouse Lab in Milton Keynes | Wellcome Sanger Institute for the COVID-19 Genomics UK (COG-UK) Consortium | The Lighthouse Lab in Milton Keynes and Alex Alderton, Roberto Amato, Jeffrey Barrett, Sonia Goncalves, Ewan Harrison, David K. Jackson, Ian Johnston, Dominic Kwiatkowski, Cordelia Langford, John Sillitoe on behalf of the Wellcome Sanger Institute COVID-19 Surveillance Team |
| EPI_ISL_3094752, EPI_ISL_3094753, EPI_ISL_3094754 | Lighthouse Laboratory Plymouth | Wellcome Sanger Institute for the COVID-19 Genomics UK (COG-UK) Consortium | Lighthouse Laboratory Plymouth and Alex Alderton, Roberto Amato, Jeffrey Barrett, Sonia Goncalves, Ewan Harrison, David K. Jackson, Ian Johnston, Dominic Kwiatkowski, Cordelia Langford, John Sillitoe on behalf of the Wellcome Sanger Institute COVID-19 Surveillance Team |
| EPI_ISL_3094755, EPI_ISL_3094756 | Lighthouse Lab in Milton Keynes | Wellcome Sanger Institute for the COVID-19 Genomics UK (COG-UK) Consortium | The Lighthouse Lab in Milton Keynes and Alex Alderton, Roberto Amato, Jeffrey Barrett, Sonia Goncalves, Ewan Harrison, David K. Jackson, Ian Johnston, Dominic Kwiatkowski, Cordelia Langford, John Sillitoe on behalf of the Wellcome Sanger Institute COVID-19 Surveillance Team |
| EPI_ISL_3094757 | Lighthouse Laboratory Plymouth | Wellcome Sanger Institute for the COVID-19 Genomics UK (COG-UK) Consortium | Lighthouse Laboratory Plymouth and Alex Alderton, Roberto Amato, Jeffrey Barrett, Sonia Goncalves, Ewan Harrison, David K. Jackson, Ian Johnston, Dominic Kwiatkowski, Cordelia Langford, John Sillitoe on behalf of the Wellcome Sanger Institute COVID-19 Surveillance Team |
| EPI_ISL_3094758 | Lighthouse Lab in Glasgow | Wellcome Sanger Institute for the COVID-19 Genomics UK (COG-UK) Consortium | Harper VanSteenhouse, Yumi Kasai, David Gray, Carol Clugston, Anna Dominiczak and Alex Alderton, Roberto Amato, Jeffrey Barrett, Sonia Goncalves, Ewan Harrison, David K. Jackson, Ian Johnston, Dominic Kwiatkowski, Cordelia Langford, John Sillitoe on behalf of the Wellcome Sanger Institute COVID-19 Surveillance Team |
| EPI_ISL_3094759, EPI_ISL_3094760 | Lighthouse Laboratory Plymouth | Wellcome Sanger Institute for the COVID-19 Genomics UK (COG-UK) Consortium | Lighthouse Laboratory Plymouth and Alex Alderton, Roberto Amato, Jeffrey Barrett, Sonia Goncalves, Ewan Harrison, David K. Jackson, Ian Johnston, Dominic Kwiatkowski, Cordelia Langford, John Sillitoe on behalf of the Wellcome Sanger Institute COVID-19 Surveillance Team |
| EPI_ISL_3094761 | Lighthouse Lab in Milton Keynes | Wellcome Sanger Institute for the COVID-19 Genomics UK (COG-UK) Consortium | The Lighthouse Lab in Milton Keynes and Alex Alderton, Roberto Amato, Jeffrey Barrett, Sonia Goncalves, Ewan Harrison, David K. Jackson, Ian Johnston, Dominic Kwiatkowski, Cordelia Langford, John Sillitoe on behalf of the Wellcome Sanger Institute COVID-19 Surveillance Team |
| EPI_ISL_3094762, EPI_ISL_3094763 | Lighthouse Laboratory Plymouth | Wellcome Sanger Institute for the COVID-19 Genomics UK (COG-UK) Consortium | Lighthouse Laboratory Plymouth and Alex Alderton, Roberto Amato, Jeffrey Barrett, Sonia Goncalves, Ewan Harrison, David K. Jackson, Ian Johnston, Dominic Kwiatkowski, Cordelia Langford, John Sillitoe on behalf of the Wellcome Sanger Institute COVID-19 Surveillance Team |
| EPI_ISL_3094764 | Lighthouse Lab in Glasgow | Wellcome Sanger Institute for the COVID-19 Genomics UK (COG-UK) Consortium | Harper VanSteenhouse, Yumi Kasai, David Gray, Carol Clugston, Anna Dominiczak and Alex Alderton, Roberto Amato, Jeffrey Barrett, Sonia Goncalves, Ewan Harrison, David K. Jackson, Ian Johnston, Dominic Kwiatkowski, Cordelia Langford, John Sillitoe on behalf of the Wellcome Sanger Institute COVID-19 Surveillance Team |
| EPI_ISL_3094765 | Lighthouse Lab in Milton Keynes | Wellcome Sanger Institute for the COVID-19 Genomics UK (COG-UK) Consortium | The Lighthouse Lab in Milton Keynes and Alex Alderton, Roberto Amato, Jeffrey Barrett, Sonia Goncalves, Ewan Harrison, David K. Jackson, Ian Johnston, Dominic Kwiatkowski, Cordelia Langford, John Sillitoe on behalf of the Wellcome Sanger Institute COVID-19 Surveillance Team |
| EPI_ISL_3094766, EPI_ISL_3094767 | Lighthouse Laboratory Plymouth | Wellcome Sanger Institute for the COVID-19 Genomics UK (COG-UK) Consortium | Lighthouse Laboratory Plymouth and Alex Alderton, Roberto Amato, Jeffrey Barrett, Sonia Goncalves, Ewan Harrison, David K. Jackson, Ian Johnston, Dominic Kwiatkowski, Cordelia Langford, John Sillitoe on behalf of the Wellcome Sanger Institute COVID-19 Surveillance Team |
| EPI_ISL_3094768, EPI_ISL_3094769 | Lighthouse Lab in Milton Keynes | Wellcome Sanger Institute for the COVID-19 Genomics UK (COG-UK) Consortium | The Lighthouse Lab in Milton Keynes and Alex Alderton, Roberto Amato, Jeffrey Barrett, Sonia Goncalves, Ewan Harrison, David K. Jackson, Ian Johnston, Dominic Kwiatkowski, Cordelia Langford, John Sillitoe on behalf of the Wellcome Sanger Institute COVID-19 Surveillance Team |
| EPI_ISL_3094770, EPI_ISL_3094771, EPI_ISL_3094772, EPI_ISL_3094773, EPI_ISL_3094774, EPI_ISL_3094775, EPI_ISL_3094776, EPI_ISL_3094777 | Lighthouse Laboratory Plymouth | Wellcome Sanger Institute for the COVID-19 Genomics UK (COG-UK) Consortium | Lighthouse Laboratory Plymouth and Alex Alderton, Roberto Amato, Jeffrey Barrett, Sonia Goncalves, Ewan Harrison, David K. Jackson, Ian Johnston, Dominic Kwiatkowski, Cordelia Langford, John Sillitoe on behalf of the Wellcome Sanger Institute COVID-19 Surveillance Team |

| | | | |
|---|---|---|---|
| EPI_ISL_3094778 | Lighthouse Lab in Milton Keynes | Wellcome Sanger Institute for the COVID-19 Genomics UK (COG-UK) Consortium | The Lighthouse Lab in Milton Keynes and Alex Alderton, Roberto Amato, Jeffrey Barrett, Sonia Goncalves, Ewan Harrison, David K. Jackson, Ian Johnston, Dominic Kwiatkowski, Cordelia Langford, John Sillitoe on behalf of the Wellcome Sanger Institute COVID-19 Surveillance Team |
| EPI_ISL_3094779 | Lighthouse Lab in Glasgow | Wellcome Sanger Institute for the COVID-19 Genomics UK (COG-UK) Consortium | Harper VanSteenhouse, Yumi Kasai, David Gray, Carol Clugston, Anna Dominiczak and Alex Alderton, Roberto Amato, Jeffrey Barrett, Sonia Goncalves, Ewan Harrison, David K. Jackson, Ian Johnston, Dominic Kwiatkowski, Cordelia Langford, John Sillitoe on behalf of the Wellcome Sanger Institute COVID-19 Surveillance Team |
| EPI_ISL_3094780, EPI_ISL_3094781 | Lighthouse Lab in Milton Keynes | Wellcome Sanger Institute for the COVID-19 Genomics UK (COG-UK) Consortium | The Lighthouse Lab in Milton Keynes and Alex Alderton, Roberto Amato, Jeffrey Barrett, Sonia Goncalves, Ewan Harrison, David K. Jackson, Ian Johnston, Dominic Kwiatkowski, Cordelia Langford, John Sillitoe on behalf of the Wellcome Sanger Institute COVID-19 Surveillance Team |
| EPI_ISL_3094782 | Lighthouse Laboratory Plymouth | Wellcome Sanger Institute for the COVID-19 Genomics UK (COG-UK) Consortium | Lighthouse Laboratory Plymouth and Alex Alderton, Roberto Amato, Jeffrey Barrett, Sonia Goncalves, Ewan Harrison, David K. Jackson, Ian Johnston, Dominic Kwiatkowski, Cordelia Langford, John Sillitoe on behalf of the Wellcome Sanger Institute COVID-19 Surveillance Team |
| EPI_ISL_3094783 | Lighthouse Lab in Milton Keynes | Wellcome Sanger Institute for the COVID-19 Genomics UK (COG-UK) Consortium | The Lighthouse Lab in Milton Keynes and Alex Alderton, Roberto Amato, Jeffrey Barrett, Sonia Goncalves, Ewan Harrison, David K. Jackson, Ian Johnston, Dominic Kwiatkowski, Cordelia Langford, John Sillitoe on behalf of the Wellcome Sanger Institute COVID-19 Surveillance Team |
| EPI_ISL_3094784 | Lighthouse Laboratory Plymouth | Wellcome Sanger Institute for the COVID-19 Genomics UK (COG-UK) Consortium | Lighthouse Laboratory Plymouth and Alex Alderton, Roberto Amato, Jeffrey Barrett, Sonia Goncalves, Ewan Harrison, David K. Jackson, Ian Johnston, Dominic Kwiatkowski, Cordelia Langford, John Sillitoe on behalf of the Wellcome Sanger Institute COVID-19 Surveillance Team |
| EPI_ISL_3094785, EPI_ISL_3094786, EPI_ISL_3094787 | Lighthouse Lab in Milton Keynes | Wellcome Sanger Institute for the COVID-19 Genomics UK (COG-UK) Consortium | The Lighthouse Lab in Milton Keynes and Alex Alderton, Roberto Amato, Jeffrey Barrett, Sonia Goncalves, Ewan Harrison, David K. Jackson, Ian Johnston, Dominic Kwiatkowski, Cordelia Langford, John Sillitoe on behalf of the Wellcome Sanger Institute COVID-19 Surveillance Team |
| EPI_ISL_3094788, EPI_ISL_3094789 | Lighthouse Laboratory Plymouth | Wellcome Sanger Institute for the COVID-19 Genomics UK (COG-UK) Consortium | Lighthouse Laboratory Plymouth and Alex Alderton, Roberto Amato, Jeffrey Barrett, Sonia Goncalves, Ewan Harrison, David K. Jackson, Ian Johnston, Dominic Kwiatkowski, Cordelia Langford, John Sillitoe on behalf of the Wellcome Sanger Institute COVID-19 Surveillance Team |
| EPI_ISL_3094790 | Lighthouse Lab in Milton Keynes | Wellcome Sanger Institute for the COVID-19 Genomics UK (COG-UK) Consortium | The Lighthouse Lab in Milton Keynes and Alex Alderton, Roberto Amato, Jeffrey Barrett, Sonia Goncalves, Ewan Harrison, David K. Jackson, Ian Johnston, Dominic Kwiatkowski, Cordelia Langford, John Sillitoe on behalf of the Wellcome Sanger Institute COVID-19 Surveillance Team |
| EPI_ISL_3094791, EPI_ISL_3094792 | Lighthouse Laboratory Plymouth | Wellcome Sanger Institute for the COVID-19 Genomics UK (COG-UK) Consortium | Lighthouse Laboratory Plymouth and Alex Alderton, Roberto Amato, Jeffrey Barrett, Sonia Goncalves, Ewan Harrison, David K. Jackson, Ian Johnston, Dominic Kwiatkowski, Cordelia Langford, John Sillitoe on behalf of the Wellcome Sanger Institute COVID-19 Surveillance Team |
| EPI_ISL_3094793 | Lighthouse Lab in Milton Keynes | Wellcome Sanger Institute for the COVID-19 Genomics UK (COG-UK) Consortium | The Lighthouse Lab in Milton Keynes and Alex Alderton, Roberto Amato, Jeffrey Barrett, Sonia Goncalves, Ewan Harrison, David K. Jackson, Ian Johnston, Dominic Kwiatkowski, Cordelia Langford, John Sillitoe on behalf of the Wellcome Sanger Institute COVID-19 Surveillance Team |
| EPI_ISL_3094794 | Lighthouse Lab in Glasgow | Wellcome Sanger Institute for the COVID-19 Genomics UK (COG-UK) Consortium | Harper VanSteenhouse, Yumi Kasai, David Gray, Carol Clugston, Anna Dominiczak and Alex Alderton, Roberto Amato, Jeffrey Barrett, Sonia Goncalves, Ewan Harrison, David K. Jackson, Ian Johnston, Dominic Kwiatkowski, Cordelia Langford, John Sillitoe on behalf of the Wellcome Sanger Institute COVID-19 Surveillance Team |
| EPI_ISL_3094795, EPI_ISL_3094796, EPI_ISL_3094797, EPI_ISL_3094798 | Lighthouse Laboratory Plymouth | Wellcome Sanger Institute for the COVID-19 Genomics UK (COG-UK) Consortium | Lighthouse Laboratory Plymouth and Alex Alderton, Roberto Amato, Jeffrey Barrett, Sonia Goncalves, Ewan Harrison, David K. Jackson, Ian Johnston, Dominic Kwiatkowski, Cordelia Langford, John Sillitoe on behalf of the Wellcome Sanger Institute COVID-19 Surveillance Team |
| EPI_ISL_3094799 | Lighthouse Lab in Milton Keynes | Wellcome Sanger Institute for the COVID-19 Genomics UK (COG-UK) Consortium | The Lighthouse Lab in Milton Keynes and Alex Alderton, Roberto Amato, Jeffrey Barrett, Sonia Goncalves, Ewan Harrison, David K. Jackson, Ian Johnston, Dominic Kwiatkowski, Cordelia Langford, John Sillitoe on behalf of the Wellcome Sanger Institute COVID-19 Surveillance Team |
| EPI_ISL_3094800, EPI_ISL_3094801, EPI_ISL_3094802, EPI_ISL_3094803 | Lighthouse Laboratory Plymouth | Wellcome Sanger Institute for the COVID-19 Genomics UK (COG-UK) Consortium | Lighthouse Laboratory Plymouth and Alex Alderton, Roberto Amato, Jeffrey Barrett, Sonia Goncalves, Ewan Harrison, David K. Jackson, Ian Johnston, Dominic Kwiatkowski, Cordelia Langford, John Sillitoe on behalf of the Wellcome Sanger Institute COVID-19 Surveillance Team |
| EPI_ISL_3094804, EPI_ISL_3094805 | Lighthouse Lab in Milton Keynes | Wellcome Sanger Institute for the COVID-19 Genomics UK (COG-UK) Consortium | The Lighthouse Lab in Milton Keynes and Alex Alderton, Roberto Amato, Jeffrey Barrett, Sonia Goncalves, Ewan Harrison, David K. Jackson, Ian Johnston, Dominic Kwiatkowski, Cordelia Langford, John Sillitoe on behalf of the Wellcome Sanger Institute COVID-19 Surveillance Team |
| EPI_ISL_3094806, EPI_ISL_3094807, EPI_ISL_3094808, EPI_ISL_3094809, EPI_ISL_3094810 | Lighthouse Laboratory Plymouth | Wellcome Sanger Institute for the COVID-19 Genomics UK (COG-UK) Consortium | Lighthouse Laboratory Plymouth and Alex Alderton, Roberto Amato, Jeffrey Barrett, Sonia Goncalves, Ewan Harrison, David K. Jackson, Ian Johnston, Dominic Kwiatkowski, Cordelia Langford, John Sillitoe on behalf of the Wellcome Sanger Institute COVID-19 Surveillance Team |
| EPI_ISL_3094811 | Lighthouse Lab in Milton Keynes | Wellcome Sanger Institute for the COVID-19 Genomics UK (COG-UK) Consortium | The Lighthouse Lab in Milton Keynes and Alex Alderton, Roberto Amato, Jeffrey Barrett, Sonia Goncalves, Ewan Harrison, David K. Jackson, Ian Johnston, Dominic Kwiatkowski, Cordelia Langford, John Sillitoe on behalf of the Wellcome Sanger Institute COVID-19 Surveillance Team |
| EPI_ISL_3094812 | Lighthouse Laboratory Plymouth | Wellcome Sanger Institute for the COVID-19 Genomics UK (COG-UK) Consortium | Lighthouse Laboratory Plymouth and Alex Alderton, Roberto Amato, Jeffrey Barrett, Sonia Goncalves, Ewan Harrison, David K. Jackson, Ian Johnston, Dominic Kwiatkowski, Cordelia Langford, John Sillitoe on behalf of the Wellcome Sanger Institute COVID-19 Surveillance Team |
| EPI_ISL_3094813 | Lighthouse Lab in Milton Keynes | Wellcome Sanger Institute for the COVID-19 Genomics UK (COG-UK) Consortium | The Lighthouse Lab in Milton Keynes and Alex Alderton, Roberto Amato, Jeffrey Barrett, Sonia Goncalves, Ewan Harrison, David K. Jackson, Ian Johnston, Dominic Kwiatkowski, Cordelia Langford, John Sillitoe on behalf of the Wellcome Sanger Institute COVID-19 Surveillance Team |
| EPI_ISL_3094814, EPI_ISL_3094815 | Lighthouse Laboratory Plymouth | Wellcome Sanger Institute for the COVID-19 Genomics UK (COG-UK) Consortium | Lighthouse Laboratory Plymouth and Alex Alderton, Roberto Amato, Jeffrey Barrett, Sonia Goncalves, Ewan Harrison, David K. Jackson, Ian Johnston, Dominic Kwiatkowski, Cordelia Langford, John Sillitoe on behalf of the Wellcome Sanger Institute COVID-19 Surveillance Team |
| EPI_ISL_3094816 | Lighthouse Lab in Milton Keynes | Wellcome Sanger Institute for the COVID-19 Genomics UK (COG-UK) Consortium | The Lighthouse Lab in Milton Keynes and Alex Alderton, Roberto Amato, Jeffrey Barrett, Sonia Goncalves, Ewan Harrison, David K. Jackson, Ian Johnston, Dominic Kwiatkowski, Cordelia Langford, John Sillitoe on behalf of the Wellcome Sanger Institute COVID-19 Surveillance Team |
| EPI_ISL_3094817 | Lighthouse Laboratory Plymouth | Wellcome Sanger Institute for the COVID-19 Genomics UK (COG-UK) Consortium | Lighthouse Laboratory Plymouth and Alex Alderton, Roberto Amato, Jeffrey Barrett, Sonia Goncalves, Ewan Harrison, David K. Jackson, Ian Johnston, Dominic Kwiatkowski, Cordelia Langford, John Sillitoe on behalf of the Wellcome Sanger Institute COVID-19 Surveillance Team |
| EPI_ISL_3094818 | Lighthouse Lab in Glasgow | Wellcome Sanger Institute for the COVID-19 Genomics UK (COG-UK) Consortium | Harper VanSteenhouse, Yumi Kasai, David Gray, Carol Clugston, Anna Dominiczak and Alex Alderton, Roberto Amato, Jeffrey Barrett, Sonia Goncalves, Ewan Harrison, David K. Jackson, Ian Johnston, Dominic Kwiatkowski, Cordelia Langford, John Sillitoe on behalf of the Wellcome Sanger Institute COVID-19 Surveillance Team |
| EPI_ISL_3094819, EPI_ISL_3094820 | Lighthouse Laboratory Plymouth | Wellcome Sanger Institute for the COVID-19 Genomics UK (COG-UK) Consortium | Lighthouse Laboratory Plymouth and Alex Alderton, Roberto Amato, Jeffrey Barrett, Sonia Goncalves, Ewan Harrison, David K. Jackson, Ian Johnston, Dominic Kwiatkowski, Cordelia Langford, John Sillitoe on behalf of the Wellcome Sanger Institute COVID-19 Surveillance Team |
| EPI_ISL_3094821 | Lighthouse Lab in Milton Keynes | Wellcome Sanger Institute for the COVID-19 Genomics UK (COG-UK) Consortium | The Lighthouse Lab in Milton Keynes and Alex Alderton, Roberto Amato, Jeffrey Barrett, Sonia Goncalves, Ewan Harrison, David K. Jackson, Ian Johnston, Dominic Kwiatkowski, Cordelia Langford, John Sillitoe on behalf of the Wellcome Sanger Institute COVID-19 Surveillance Team |
| EPI_ISL_3094822 | Lighthouse Laboratory Plymouth | Wellcome Sanger Institute for the COVID-19 Genomics UK (COG-UK) Consortium | Lighthouse Laboratory Plymouth and Alex Alderton, Roberto Amato, Jeffrey Barrett, Sonia Goncalves, Ewan Harrison, David K. Jackson, Ian Johnston, Dominic Kwiatkowski, Cordelia Langford, John Sillitoe on behalf of the Wellcome Sanger Institute COVID-19 Surveillance Team |
| EPI_ISL_3094823 | Lighthouse Lab in Milton Keynes | Wellcome Sanger Institute for the COVID-19 Genomics UK (COG-UK) Consortium | The Lighthouse Lab in Milton Keynes and Alex Alderton, Roberto Amato, Jeffrey Barrett, Sonia Goncalves, Ewan Harrison, David K. Jackson, Ian Johnston, Dominic Kwiatkowski, Cordelia Langford, John Sillitoe on behalf of the Wellcome Sanger Institute COVID-19 Surveillance Team |
| EPI_ISL_3094824, EPI_ISL_3094825, EPI_ISL_3094826, EPI_ISL_3094827 | Lighthouse Laboratory Plymouth | Wellcome Sanger Institute for the COVID-19 Genomics UK (COG-UK) Consortium | Lighthouse Laboratory Plymouth and Alex Alderton, Roberto Amato, Jeffrey Barrett, Sonia Goncalves, Ewan Harrison, David K. Jackson, Ian Johnston, Dominic Kwiatkowski, Cordelia Langford, John Sillitoe on behalf of the Wellcome Sanger Institute COVID-19 Surveillance Team |
| EPI_ISL_3094828, EPI_ISL_3094829 | Lighthouse Lab in Milton Keynes | Wellcome Sanger Institute for the COVID-19 Genomics UK (COG-UK) Consortium | The Lighthouse Lab in Milton Keynes and Alex Alderton, Roberto Amato, Jeffrey Barrett, Sonia Goncalves, Ewan Harrison, David K. Jackson, Ian Johnston, Dominic Kwiatkowski, Cordelia Langford, John Sillitoe on behalf of the Wellcome Sanger Institute COVID-19 Surveillance Team |
| EPI_ISL_3094830 | Lighthouse Laboratory Plymouth | Wellcome Sanger Institute for the COVID-19 Genomics UK (COG-UK) Consortium | Lighthouse Laboratory Plymouth and Alex Alderton, Roberto Amato, Jeffrey Barrett, Sonia Goncalves, Ewan Harrison, David K. Jackson, Ian Johnston, Dominic Kwiatkowski, Cordelia Langford, John Sillitoe on behalf of the Wellcome Sanger Institute COVID-19 Surveillance Team |
| EPI_ISL_3094831 | Lighthouse Lab in Glasgow | Wellcome Sanger Institute for the COVID-19 Genomics UK (COG-UK) Consortium | Harper VanSteenhouse, Yumi Kasai, David Gray, Carol Clugston, Anna Dominiczak and Alex Alderton, Roberto Amato, Jeffrey Barrett, Sonia Goncalves, Ewan Harrison, David K. Jackson, Ian Johnston, Dominic Kwiatkowski, Cordelia Langford, John Sillitoe on behalf of the Wellcome Sanger Institute COVID-19 Surveillance Team |
| EPI_ISL_3094832 | Lighthouse Lab in Milton Keynes | Wellcome Sanger Institute for the COVID-19 Genomics UK (COG-UK) Consortium | The Lighthouse Lab in Milton Keynes and Alex Alderton, Roberto Amato, Jeffrey Barrett, Sonia Goncalves, Ewan Harrison, David K. Jackson, Ian Johnston, Dominic Kwiatkowski, Cordelia Langford, John Sillitoe on behalf of the Wellcome Sanger Institute COVID-19 Surveillance Team |
| EPI_ISL_3094833 | Lighthouse Laboratory Plymouth | Wellcome Sanger Institute for the COVID-19 Genomics UK (COG-UK) Consortium | Lighthouse Laboratory Plymouth and Alex Alderton, Roberto Amato, Jeffrey Barrett, Sonia Goncalves, Ewan Harrison, David K. Jackson, Ian Johnston, Dominic Kwiatkowski, Cordelia Langford, John Sillitoe on behalf of the Wellcome Sanger Institute COVID-19 Surveillance Team |
| EPI_ISL_3094834 | Lighthouse Lab in Milton Keynes | Wellcome Sanger Institute for the COVID-19 Genomics UK (COG-UK) Consortium | The Lighthouse Lab in Milton Keynes and Alex Alderton, Roberto Amato, Jeffrey Barrett, Sonia Goncalves, Ewan Harrison, David K. Jackson, Ian Johnston, Dominic Kwiatkowski, Cordelia Langford, John Sillitoe on behalf of the Wellcome Sanger Institute COVID-19 Surveillance Team |
| EPI_ISL_3094835, EPI_ISL_3094836, EPI_ISL_3094837, EPI_ISL_3094838 | Lighthouse Laboratory Plymouth | Wellcome Sanger Institute for the COVID-19 Genomics UK (COG-UK) Consortium | Lighthouse Laboratory Plymouth and Alex Alderton, Roberto Amato, Jeffrey Barrett, Sonia Goncalves, Ewan Harrison, David K. Jackson, Ian Johnston, Dominic Kwiatkowski, Cordelia Langford, John Sillitoe on behalf of the Wellcome Sanger Institute COVID-19 Surveillance Team |
| EPI_ISL_3094839 | Lighthouse Lab in Milton Keynes | Wellcome Sanger Institute for the COVID-19 Genomics UK (COG-UK) Consortium | The Lighthouse Lab in Milton Keynes and Alex Alderton, Roberto Amato, Jeffrey Barrett, Sonia Goncalves, Ewan Harrison, David K. Jackson, Ian Johnston, Dominic Kwiatkowski, Cordelia Langford, John Sillitoe on behalf of the Wellcome Sanger Institute COVID-19 Surveillance Team |
| EPI_ISL_3094840, EPI_ISL_3094841, EPI_ISL_3094842 | Lighthouse Laboratory Plymouth | Wellcome Sanger Institute for the COVID-19 Genomics UK (COG-UK) Consortium | Lighthouse Laboratory Plymouth and Alex Alderton, Roberto Amato, Jeffrey Barrett, Sonia Goncalves, Ewan Harrison, David K. Jackson, Ian Johnston, Dominic Kwiatkowski, Cordelia Langford, John Sillitoe on behalf of the Wellcome Sanger Institute COVID-19 Surveillance Team |
| EPI_ISL_3094843, EPI_ISL_3094844 | Lighthouse Lab in Milton Keynes | Wellcome Sanger Institute for the COVID-19 Genomics UK (COG-UK) Consortium | The Lighthouse Lab in Milton Keynes and Alex Alderton, Roberto Amato, Jeffrey Barrett, Sonia Goncalves, Ewan Harrison, David K. Jackson, Ian Johnston, Dominic Kwiatkowski, Cordelia Langford, John Sillitoe on behalf of the Wellcome Sanger Institute COVID-19 Surveillance Team |
| EPI_ISL_3094845 | Lighthouse Laboratory Plymouth | Wellcome Sanger Institute for the COVID-19 Genomics UK (COG-UK) Consortium | Lighthouse Laboratory Plymouth and Alex Alderton, Roberto Amato, Jeffrey Barrett, Sonia Goncalves, Ewan Harrison, David K. Jackson, Ian Johnston, Dominic Kwiatkowski, Cordelia Langford, John Sillitoe on behalf of the Wellcome Sanger Institute COVID-19 Surveillance Team |

| | | | |
|---|---|---|---|
| EPI_ISL_3094846 | Lighthouse Lab in Milton Keynes | Wellcome Sanger Institute for the COVID-19 Genomics UK (COG-UK) Consortium | The Lighthouse Lab in Milton Keynes and Alex Alderton, Roberto Amato, Jeffrey Barrett, Sonia Goncalves, Ewan Harrison, David K. Jackson, Ian Johnston, Dominic Kwiatkowski, Cordelia Langford, John Sillitoe on behalf of the Wellcome Sanger Institute COVID-19 Surveillance Team |
| EPI_ISL_3094847, EPI_ISL_3094848 | Lighthouse Laboratory Plymouth | Wellcome Sanger Institute for the COVID-19 Genomics UK (COG-UK) Consortium | Lighthouse Laboratory Plymouth and Alex Alderton, Roberto Amato, Jeffrey Barrett, Sonia Goncalves, Ewan Harrison, David K. Jackson, Ian Johnston, Dominic Kwiatkowski, Cordelia Langford, John Sillitoe on behalf of the Wellcome Sanger Institute COVID-19 Surveillance Team |
| EPI_ISL_3094849 | Lighthouse Lab in Milton Keynes | Wellcome Sanger Institute for the COVID-19 Genomics UK (COG-UK) Consortium | The Lighthouse Lab in Milton Keynes and Alex Alderton, Roberto Amato, Jeffrey Barrett, Sonia Goncalves, Ewan Harrison, David K. Jackson, Ian Johnston, Dominic Kwiatkowski, Cordelia Langford, John Sillitoe on behalf of the Wellcome Sanger Institute COVID-19 Surveillance Team |
| EPI_ISL_3094850, EPI_ISL_3094851, EPI_ISL_3094852, EPI_ISL_3094853 | Lighthouse Laboratory Plymouth | Wellcome Sanger Institute for the COVID-19 Genomics UK (COG-UK) Consortium | Lighthouse Laboratory Plymouth and Alex Alderton, Roberto Amato, Jeffrey Barrett, Sonia Goncalves, Ewan Harrison, David K. Jackson, Ian Johnston, Dominic Kwiatkowski, Cordelia Langford, John Sillitoe on behalf of the Wellcome Sanger Institute COVID-19 Surveillance Team |
| EPI_ISL_3094854 | Lighthouse Lab in Milton Keynes | Wellcome Sanger Institute for the COVID-19 Genomics UK (COG-UK) Consortium | The Lighthouse Lab in Milton Keynes and Alex Alderton, Roberto Amato, Jeffrey Barrett, Sonia Goncalves, Ewan Harrison, David K. Jackson, Ian Johnston, Dominic Kwiatkowski, Cordelia Langford, John Sillitoe on behalf of the Wellcome Sanger Institute COVID-19 Surveillance Team |
| EPI_ISL_3094855 | Lighthouse Laboratory Plymouth | Wellcome Sanger Institute for the COVID-19 Genomics UK (COG-UK) Consortium | Lighthouse Laboratory Plymouth and Alex Alderton, Roberto Amato, Jeffrey Barrett, Sonia Goncalves, Ewan Harrison, David K. Jackson, Ian Johnston, Dominic Kwiatkowski, Cordelia Langford, John Sillitoe on behalf of the Wellcome Sanger Institute COVID-19 Surveillance Team |
| EPI_ISL_3094856 | Lighthouse Lab in Milton Keynes | Wellcome Sanger Institute for the COVID-19 Genomics UK (COG-UK) Consortium | The Lighthouse Lab in Milton Keynes and Alex Alderton, Roberto Amato, Jeffrey Barrett, Sonia Goncalves, Ewan Harrison, David K. Jackson, Ian Johnston, Dominic Kwiatkowski, Cordelia Langford, John Sillitoe on behalf of the Wellcome Sanger Institute COVID-19 Surveillance Team |
| EPI_ISL_3094857 | Lighthouse Laboratory Plymouth | Wellcome Sanger Institute for the COVID-19 Genomics UK (COG-UK) Consortium | Lighthouse Laboratory Plymouth and Alex Alderton, Roberto Amato, Jeffrey Barrett, Sonia Goncalves, Ewan Harrison, David K. Jackson, Ian Johnston, Dominic Kwiatkowski, Cordelia Langford, John Sillitoe on behalf of the Wellcome Sanger Institute COVID-19 Surveillance Team |
| EPI_ISL_3094858, EPI_ISL_3094859, EPI_ISL_3094860 | Lighthouse Lab in Glasgow | Wellcome Sanger Institute for the COVID-19 Genomics UK (COG-UK) Consortium | Harper VanSteenhouse, Yumi Kasai, David Gray, Carol Clugston, Anna Dominiczak and Alex Alderton, Roberto Amato, Jeffrey Barrett, Sonia Goncalves, Ewan Harrison, David K. Jackson, Ian Johnston, Dominic Kwiatkowski, Cordelia Langford, John Sillitoe on behalf of the Wellcome Sanger Institute COVID-19 Surveillance Team |
| EPI_ISL_3094861 | Lighthouse Lab in Milton Keynes | Wellcome Sanger Institute for the COVID-19 Genomics UK (COG-UK) Consortium | The Lighthouse Lab in Milton Keynes and Alex Alderton, Roberto Amato, Jeffrey Barrett, Sonia Goncalves, Ewan Harrison, David K. Jackson, Ian Johnston, Dominic Kwiatkowski, Cordelia Langford, John Sillitoe on behalf of the Wellcome Sanger Institute COVID-19 Surveillance Team |
| EPI_ISL_3094862 | Lighthouse Lab in Alderley Park | Wellcome Sanger Institute for the COVID-19 Genomics UK (COG-UK) Consortium | Jacquelyn Wynn, Mairead Hyland, The Lighthouse Lab in Alderley Park and Alex Alderton, Roberto Amato, Jeffrey Barrett, Sonia Goncalves, Ewan Harrison, David K. Jackson, Ian Johnston, Dominic Kwiatkowski, Cordelia Langford, John Sillitoe on behalf of the Wellcome Sanger Institute COVID-19 Surveillance Team |
| EPI_ISL_3094863, EPI_ISL_3094864 | Lighthouse Lab in Glasgow | Wellcome Sanger Institute for the COVID-19 Genomics UK (COG-UK) Consortium | Harper VanSteenhouse, Yumi Kasai, David Gray, Carol Clugston, Anna Dominiczak and Alex Alderton, Roberto Amato, Jeffrey Barrett, Sonia Goncalves, Ewan Harrison, David K. Jackson, Ian Johnston, Dominic Kwiatkowski, Cordelia Langford, John Sillitoe on behalf of the Wellcome Sanger Institute COVID-19 Surveillance Team |
| EPI_ISL_3094865, EPI_ISL_3094866 | Lighthouse Lab in Milton Keynes | Wellcome Sanger Institute for the COVID-19 Genomics UK (COG-UK) Consortium | The Lighthouse Lab in Milton Keynes and Alex Alderton, Roberto Amato, Jeffrey Barrett, Sonia Goncalves, Ewan Harrison, David K. Jackson, Ian Johnston, Dominic Kwiatkowski, Cordelia Langford, John Sillitoe on behalf of the Wellcome Sanger Institute COVID-19 Surveillance Team |
| EPI_ISL_3094867 | Lighthouse Lab in Glasgow | Wellcome Sanger Institute for the COVID-19 Genomics UK (COG-UK) Consortium | Harper VanSteenhouse, Yumi Kasai, David Gray, Carol Clugston, Anna Dominiczak and Alex Alderton, Roberto Amato, Jeffrey Barrett, Sonia Goncalves, Ewan Harrison, David K. Jackson, Ian Johnston, Dominic Kwiatkowski, Cordelia Langford, John Sillitoe on behalf of the Wellcome Sanger Institute COVID-19 Surveillance Team |
| EPI_ISL_3094868 | Lighthouse Lab in Alderley Park | Wellcome Sanger Institute for the COVID-19 Genomics UK (COG-UK) Consortium | Jacquelyn Wynn, Mairead Hyland, The Lighthouse Lab in Alderley Park and Alex Alderton, Roberto Amato, Jeffrey Barrett, Sonia Goncalves, Ewan Harrison, David K. Jackson, Ian Johnston, Dominic Kwiatkowski, Cordelia Langford, John Sillitoe on behalf of the Wellcome Sanger Institute COVID-19 Surveillance Team |
| EPI_ISL_3094869 | Lighthouse Lab in Glasgow | Wellcome Sanger Institute for the COVID-19 Genomics UK (COG-UK) Consortium | Harper VanSteenhouse, Yumi Kasai, David Gray, Carol Clugston, Anna Dominiczak and Alex Alderton, Roberto Amato, Jeffrey Barrett, Sonia Goncalves, Ewan Harrison, David K. Jackson, Ian Johnston, Dominic Kwiatkowski, Cordelia Langford, John Sillitoe on behalf of the Wellcome Sanger Institute COVID-19 Surveillance Team |
| EPI_ISL_3094870 | Lighthouse Lab in Alderley Park | Wellcome Sanger Institute for the COVID-19 Genomics UK (COG-UK) Consortium | Jacquelyn Wynn, Mairead Hyland, The Lighthouse Lab in Alderley Park and Alex Alderton, Roberto Amato, Jeffrey Barrett, Sonia Goncalves, Ewan Harrison, David K. Jackson, Ian Johnston, Dominic Kwiatkowski, Cordelia Langford, John Sillitoe on behalf of the Wellcome Sanger Institute COVID-19 Surveillance Team |
| EPI_ISL_3094871 | Lighthouse Lab in Milton Keynes | Wellcome Sanger Institute for the COVID-19 Genomics UK (COG-UK) Consortium | The Lighthouse Lab in Milton Keynes and Alex Alderton, Roberto Amato, Jeffrey Barrett, Sonia Goncalves, Ewan Harrison, David K. Jackson, Ian Johnston, Dominic Kwiatkowski, Cordelia Langford, John Sillitoe on behalf of the Wellcome Sanger Institute COVID-19 Surveillance Team |
| EPI_ISL_3094872, EPI_ISL_3094873 | Lighthouse Lab in Glasgow | Wellcome Sanger Institute for the COVID-19 Genomics UK (COG-UK) Consortium | Harper VanSteenhouse, Yumi Kasai, David Gray, Carol Clugston, Anna Dominiczak and Alex Alderton, Roberto Amato, Jeffrey Barrett, Sonia Goncalves, Ewan Harrison, David K. Jackson, Ian Johnston, Dominic Kwiatkowski, Cordelia Langford, John Sillitoe on behalf of the Wellcome Sanger Institute COVID-19 Surveillance Team |
| EPI_ISL_3094874 | Lighthouse Lab in Milton Keynes | Wellcome Sanger Institute for the COVID-19 Genomics UK (COG-UK) Consortium | The Lighthouse Lab in Milton Keynes and Alex Alderton, Roberto Amato, Jeffrey Barrett, Sonia Goncalves, Ewan Harrison, David K. Jackson, Ian Johnston, Dominic Kwiatkowski, Cordelia Langford, John Sillitoe on behalf of the Wellcome Sanger Institute COVID-19 Surveillance Team |
| EPI_ISL_3094875, EPI_ISL_3094876 | Lighthouse Lab in Glasgow | Wellcome Sanger Institute for the COVID-19 Genomics UK (COG-UK) Consortium | Harper VanSteenhouse, Yumi Kasai, David Gray, Carol Clugston, Anna Dominiczak and Alex Alderton, Roberto Amato, Jeffrey Barrett, Sonia Goncalves, Ewan Harrison, David K. Jackson, Ian Johnston, Dominic Kwiatkowski, Cordelia Langford, John Sillitoe on behalf of the Wellcome Sanger Institute COVID-19 Surveillance Team |
| EPI_ISL_3094877, EPI_ISL_3094878 | Lighthouse Lab in Alderley Park | Wellcome Sanger Institute for the COVID-19 Genomics UK (COG-UK) Consortium | Jacquelyn Wynn, Mairead Hyland, The Lighthouse Lab in Alderley Park and Alex Alderton, Roberto Amato, Jeffrey Barrett, Sonia Goncalves, Ewan Harrison, David K. Jackson, Ian Johnston, Dominic Kwiatkowski, Cordelia Langford, John Sillitoe on behalf of the Wellcome Sanger Institute COVID-19 Surveillance Team |
| EPI_ISL_3094879, EPI_ISL_3094880 | Lighthouse Lab in Milton Keynes | Wellcome Sanger Institute for the COVID-19 Genomics UK (COG-UK) Consortium | The Lighthouse Lab in Milton Keynes and Alex Alderton, Roberto Amato, Jeffrey Barrett, Sonia Goncalves, Ewan Harrison, David K. Jackson, Ian Johnston, Dominic Kwiatkowski, Cordelia Langford, John Sillitoe on behalf of the Wellcome Sanger Institute COVID-19 Surveillance Team |
| EPI_ISL_3094881 | Lighthouse Lab in Glasgow | Wellcome Sanger Institute for the COVID-19 Genomics UK (COG-UK) Consortium | Harper VanSteenhouse, Yumi Kasai, David Gray, Carol Clugston, Anna Dominiczak and Alex Alderton, Roberto Amato, Jeffrey Barrett, Sonia Goncalves, Ewan Harrison, David K. Jackson, Ian Johnston, Dominic Kwiatkowski, Cordelia Langford, John Sillitoe on behalf of the Wellcome Sanger Institute COVID-19 Surveillance Team |
| EPI_ISL_3094882, EPI_ISL_3094883 | Lighthouse Lab in Milton Keynes | Wellcome Sanger Institute for the COVID-19 Genomics UK (COG-UK) Consortium | The Lighthouse Lab in Milton Keynes and Alex Alderton, Roberto Amato, Jeffrey Barrett, Sonia Goncalves, Ewan Harrison, David K. Jackson, Ian Johnston, Dominic Kwiatkowski, Cordelia Langford, John Sillitoe on behalf of the Wellcome Sanger Institute COVID-19 Surveillance Team |
| EPI_ISL_3094884 | Lighthouse Lab in Glasgow | Wellcome Sanger Institute for the COVID-19 Genomics UK (COG-UK) Consortium | Harper VanSteenhouse, Yumi Kasai, David Gray, Carol Clugston, Anna Dominiczak and Alex Alderton, Roberto Amato, Jeffrey Barrett, Sonia Goncalves, Ewan Harrison, David K. Jackson, Ian Johnston, Dominic Kwiatkowski, Cordelia Langford, John Sillitoe on behalf of the Wellcome Sanger Institute COVID-19 Surveillance Team |
| EPI_ISL_3094885 | Lighthouse Lab in Milton Keynes | Wellcome Sanger Institute for the COVID-19 Genomics UK (COG-UK) Consortium | The Lighthouse Lab in Milton Keynes and Alex Alderton, Roberto Amato, Jeffrey Barrett, Sonia Goncalves, Ewan Harrison, David K. Jackson, Ian Johnston, Dominic Kwiatkowski, Cordelia Langford, John Sillitoe on behalf of the Wellcome Sanger Institute COVID-19 Surveillance Team |
| EPI_ISL_3094886, EPI_ISL_3094887 | Lighthouse Lab in Glasgow | Wellcome Sanger Institute for the COVID-19 Genomics UK (COG-UK) Consortium | Harper VanSteenhouse, Yumi Kasai, David Gray, Carol Clugston, Anna Dominiczak and Alex Alderton, Roberto Amato, Jeffrey Barrett, Sonia Goncalves, Ewan Harrison, David K. Jackson, Ian Johnston, Dominic Kwiatkowski, Cordelia Langford, John Sillitoe on behalf of the Wellcome Sanger Institute COVID-19 Surveillance Team |
| EPI_ISL_3094888, EPI_ISL_3094889 | Lighthouse Lab in Alderley Park | Wellcome Sanger Institute for the COVID-19 Genomics UK (COG-UK) Consortium | Jacquelyn Wynn, Mairead Hyland, The Lighthouse Lab in Alderley Park and Alex Alderton, Roberto Amato, Jeffrey Barrett, Sonia Goncalves, Ewan Harrison, David K. Jackson, Ian Johnston, Dominic Kwiatkowski, Cordelia Langford, John Sillitoe on behalf of the Wellcome Sanger Institute COVID-19 Surveillance Team |
| EPI_ISL_3094890, EPI_ISL_3094891 | Lighthouse Lab in Glasgow | Wellcome Sanger Institute for the COVID-19 Genomics UK (COG-UK) Consortium | Harper VanSteenhouse, Yumi Kasai, David Gray, Carol Clugston, Anna Dominiczak and Alex Alderton, Roberto Amato, Jeffrey Barrett, Sonia Goncalves, Ewan Harrison, David K. Jackson, Ian Johnston, Dominic Kwiatkowski, Cordelia Langford, John Sillitoe on behalf of the Wellcome Sanger Institute COVID-19 Surveillance Team |
| EPI_ISL_3094892 | Lighthouse Lab in Alderley Park | Wellcome Sanger Institute for the COVID-19 Genomics UK (COG-UK) Consortium | Jacquelyn Wynn, Mairead Hyland, The Lighthouse Lab in Alderley Park and Alex Alderton, Roberto Amato, Jeffrey Barrett, Sonia Goncalves, Ewan Harrison, David K. Jackson, Ian Johnston, Dominic Kwiatkowski, Cordelia Langford, John Sillitoe on behalf of the Wellcome Sanger Institute COVID-19 Surveillance Team |
| EPI_ISL_3094893 | Lighthouse Lab in Glasgow | Wellcome Sanger Institute for the COVID-19 Genomics UK (COG-UK) Consortium | Harper VanSteenhouse, Yumi Kasai, David Gray, Carol Clugston, Anna Dominiczak and Alex Alderton, Roberto Amato, Jeffrey Barrett, Sonia Goncalves, Ewan Harrison, David K. Jackson, Ian Johnston, Dominic Kwiatkowski, Cordelia Langford, John Sillitoe on behalf of the Wellcome Sanger Institute COVID-19 Surveillance Team |
| EPI_ISL_3094894, EPI_ISL_3094895 | Lighthouse Lab in Alderley Park | Wellcome Sanger Institute for the COVID-19 Genomics UK (COG-UK) Consortium | Jacquelyn Wynn, Mairead Hyland, The Lighthouse Lab in Alderley Park and Alex Alderton, Roberto Amato, Jeffrey Barrett, Sonia Goncalves, Ewan Harrison, David K. Jackson, Ian Johnston, Dominic Kwiatkowski, Cordelia Langford, John Sillitoe on behalf of the Wellcome Sanger Institute COVID-19 Surveillance Team |
| EPI_ISL_3094896 | Lighthouse Lab in Milton Keynes | Wellcome Sanger Institute for the COVID-19 Genomics UK (COG-UK) Consortium | The Lighthouse Lab in Milton Keynes and Alex Alderton, Roberto Amato, Jeffrey Barrett, Sonia Goncalves, Ewan Harrison, David K. Jackson, Ian Johnston, Dominic Kwiatkowski, Cordelia Langford, John Sillitoe on behalf of the Wellcome Sanger Institute COVID-19 Surveillance Team |
| EPI_ISL_3094897 | Lighthouse Lab in Glasgow | Wellcome Sanger Institute for the COVID-19 Genomics UK (COG-UK) Consortium | Harper VanSteenhouse, Yumi Kasai, David Gray, Carol Clugston, Anna Dominiczak and Alex Alderton, Roberto Amato, Jeffrey Barrett, Sonia Goncalves, Ewan Harrison, David K. Jackson, Ian Johnston, Dominic Kwiatkowski, Cordelia Langford, John Sillitoe on behalf of the Wellcome Sanger Institute COVID-19 Surveillance Team |
| EPI_ISL_3094898 | Lighthouse Lab in Milton Keynes | Wellcome Sanger Institute for the COVID-19 Genomics UK (COG-UK) Consortium | The Lighthouse Lab in Milton Keynes and Alex Alderton, Roberto Amato, Jeffrey Barrett, Sonia Goncalves, Ewan Harrison, David K. Jackson, Ian Johnston, Dominic Kwiatkowski, Cordelia Langford, John Sillitoe on behalf of the Wellcome Sanger Institute COVID-19 Surveillance Team |
| EPI_ISL_3094899, EPI_ISL_3094900 | Lighthouse Lab in Alderley Park | Wellcome Sanger Institute for the COVID-19 Genomics UK (COG-UK) Consortium | Jacquelyn Wynn, Mairead Hyland, The Lighthouse Lab in Alderley Park and Alex Alderton, Roberto Amato, Jeffrey Barrett, Sonia Goncalves, Ewan Harrison, David K. Jackson, Ian Johnston, Dominic Kwiatkowski, Cordelia Langford, John Sillitoe on behalf of the Wellcome Sanger Institute COVID-19 Surveillance Team |
| EPI_ISL_3094901, EPI_ISL_3094902 | Lighthouse Lab in Milton Keynes | Wellcome Sanger Institute for the COVID-19 Genomics UK (COG-UK) Consortium | The Lighthouse Lab in Milton Keynes and Alex Alderton, Roberto Amato, Jeffrey Barrett, Sonia Goncalves, Ewan Harrison, David K. Jackson, Ian Johnston, Dominic Kwiatkowski, Cordelia Langford, John Sillitoe on behalf of the Wellcome Sanger Institute COVID-19 Surveillance Team |
| EPI_ISL_3094903 | Lighthouse Lab in Glasgow | Wellcome Sanger Institute for the COVID-19 Genomics UK (COG-UK) Consortium | Harper VanSteenhouse, Yumi Kasai, David Gray, Carol Clugston, Anna Dominiczak and Alex Alderton, Roberto Amato, Jeffrey Barrett, Sonia Goncalves, Ewan Harrison, David K. Jackson, Ian Johnston, Dominic Kwiatkowski, Cordelia Langford, John Sillitoe on behalf of the Wellcome Sanger Institute COVID-19 Surveillance Team |
| EPI_ISL_3094904 | Lighthouse Lab in Milton Keynes | Wellcome Sanger Institute for the COVID-19 Genomics UK (COG-UK) Consortium | The Lighthouse Lab in Milton Keynes and Alex Alderton, Roberto Amato, Jeffrey Barrett, Sonia Goncalves, Ewan Harrison, David K. Jackson, Ian Johnston, Dominic Kwiatkowski, Cordelia Langford, John Sillitoe on behalf of the Wellcome Sanger Institute COVID-19 Surveillance Team |
| EPI_ISL_3094905, EPI_ISL_3094906 | Lighthouse Lab in Glasgow | Wellcome Sanger Institute for the COVID-19 Genomics UK (COG-UK) Consortium | Harper VanSteenhouse, Yumi Kasai, David Gray, Carol Clugston, Anna Dominiczak and Alex Alderton, Roberto Amato, Jeffrey Barrett, Sonia Goncalves, Ewan Harrison, David K. Jackson, Ian Johnston, Dominic Kwiatkowski, Cordelia Langford, John Sillitoe on behalf of the Wellcome Sanger Institute COVID-19 Surveillance Team |
| EPI_ISL_3094907 | Lighthouse Lab in Alderley Park | Wellcome Sanger Institute for the COVID-19 Genomics UK (COG-UK) Consortium | Jacquelyn Wynn, Mairead Hyland, The Lighthouse Lab in Alderley Park and Alex Alderton, Roberto Amato, Jeffrey Barrett, Sonia Goncalves, Ewan Harrison, David K. Jackson, Ian Johnston, Dominic Kwiatkowski, Cordelia Langford, John Sillitoe on behalf of the Wellcome Sanger Institute COVID-19 Surveillance Team |
| EPI_ISL_3094908, EPI_ISL_3094909, EPI_ISL_3094910 | Lighthouse Lab in Glasgow | Wellcome Sanger Institute for the COVID-19 Genomics UK (COG-UK) Consortium | Harper VanSteenhouse, Yumi Kasai, David Gray, Carol Clugston, Anna Dominiczak and Alex Alderton, Roberto Amato, Jeffrey Barrett, Sonia Goncalves, Ewan Harrison, David K. Jackson, Ian Johnston, Dominic Kwiatkowski, Cordelia Langford, John Sillitoe on behalf of the Wellcome Sanger Institute COVID-19 Surveillance Team |
| EPI_ISL_3094911, | Lighthouse Lab in Milton Keynes | Wellcome Sanger Institute for the COVID-19 Genomics UK (COG-UK) | The Lighthouse Lab in Milton Keynes and Alex Alderton, Roberto Amato, Jeffrey Barrett, Sonia Goncalves, Ewan Harrison, David K. Jackson, Ian Johnston, Dominic Kwiatkowski, Cordelia Langford, John Sillitoe on behalf of the Wellcome Sanger |

| | | | |
|---|---|---|---|
| EPI_ISL_3094912 | | Consortium | Institute COVID-19 Surveillance Team |
| EPI_ISL_3094913 | Lighthouse Lab in Alderley Park | Wellcome Sanger Institute for the COVID-19 Genomics UK (COG-UK) Consortium | Jacquelyn Wynn, Mairead Hyland, The Lighthouse Lab in Alderley Park and Alex Alderton, Roberto Amato, Jeffrey Barrett, Sonia Goncalves, Ewan Harrison, David K. Jackson, Ian Johnston, Dominic Kwiatkowski, Cordelia Langford, John Sillitoe on behalf of the Wellcome Sanger Institute COVID-19 Surveillance Team |
| EPI_ISL_3094914, EPI_ISL_3094915 | Lighthouse Lab in Glasgow | Wellcome Sanger Institute for the COVID-19 Genomics UK (COG-UK) Consortium | Harper VanSteenhouse, Yumi Kasai, David Gray, Carol Clugston, Anna Dominiczak and Alex Alderton, Roberto Amato, Jeffrey Barrett, Sonia Goncalves, Ewan Harrison, David K. Jackson, Ian Johnston, Dominic Kwiatkowski, Cordelia Langford, John Sillitoe on behalf of the Wellcome Sanger Institute COVID-19 Surveillance Team |
| EPI_ISL_3094916 | Lighthouse Lab in Milton Keynes | Wellcome Sanger Institute for the COVID-19 Genomics UK (COG-UK) Consortium | The Lighthouse Lab in Milton Keynes and Alex Alderton, Roberto Amato, Jeffrey Barrett, Sonia Goncalves, Ewan Harrison, David K. Jackson, Ian Johnston, Dominic Kwiatkowski, Cordelia Langford, John Sillitoe on behalf of the Wellcome Sanger Institute COVID-19 Surveillance Team |
| EPI_ISL_3094917 | Lighthouse Lab in Alderley Park | Wellcome Sanger Institute for the COVID-19 Genomics UK (COG-UK) Consortium | Jacquelyn Wynn, Mairead Hyland, The Lighthouse Lab in Alderley Park and Alex Alderton, Roberto Amato, Jeffrey Barrett, Sonia Goncalves, Ewan Harrison, David K. Jackson, Ian Johnston, Dominic Kwiatkowski, Cordelia Langford, John Sillitoe on behalf of the Wellcome Sanger Institute COVID-19 Surveillance Team |
| EPI_ISL_3094918 | Lighthouse Lab in Milton Keynes | Wellcome Sanger Institute for the COVID-19 Genomics UK (COG-UK) Consortium | The Lighthouse Lab in Milton Keynes and Alex Alderton, Roberto Amato, Jeffrey Barrett, Sonia Goncalves, Ewan Harrison, David K. Jackson, Ian Johnston, Dominic Kwiatkowski, Cordelia Langford, John Sillitoe on behalf of the Wellcome Sanger Institute COVID-19 Surveillance Team |
| EPI_ISL_3094919, EPI_ISL_3094920 | Lighthouse Lab in Alderley Park | Wellcome Sanger Institute for the COVID-19 Genomics UK (COG-UK) Consortium | Jacquelyn Wynn, Mairead Hyland, The Lighthouse Lab in Alderley Park and Alex Alderton, Roberto Amato, Jeffrey Barrett, Sonia Goncalves, Ewan Harrison, David K. Jackson, Ian Johnston, Dominic Kwiatkowski, Cordelia Langford, John Sillitoe on behalf of the Wellcome Sanger Institute COVID-19 Surveillance Team |
| EPI_ISL_3094921, EPI_ISL_3094922 | Lighthouse Lab in Glasgow | Wellcome Sanger Institute for the COVID-19 Genomics UK (COG-UK) Consortium | Harper VanSteenhouse, Yumi Kasai, David Gray, Carol Clugston, Anna Dominiczak and Alex Alderton, Roberto Amato, Jeffrey Barrett, Sonia Goncalves, Ewan Harrison, David K. Jackson, Ian Johnston, Dominic Kwiatkowski, Cordelia Langford, John Sillitoe on behalf of the Wellcome Sanger Institute COVID-19 Surveillance Team |
| EPI_ISL_3094923 | Lighthouse Lab in Alderley Park | Wellcome Sanger Institute for the COVID-19 Genomics UK (COG-UK) Consortium | Jacquelyn Wynn, Mairead Hyland, The Lighthouse Lab in Alderley Park and Alex Alderton, Roberto Amato, Jeffrey Barrett, Sonia Goncalves, Ewan Harrison, David K. Jackson, Ian Johnston, Dominic Kwiatkowski, Cordelia Langford, John Sillitoe on behalf of the Wellcome Sanger Institute COVID-19 Surveillance Team |
| EPI_ISL_3094924 | Lighthouse Lab in Milton Keynes | Wellcome Sanger Institute for the COVID-19 Genomics UK (COG-UK) Consortium | The Lighthouse Lab in Milton Keynes and Alex Alderton, Roberto Amato, Jeffrey Barrett, Sonia Goncalves, Ewan Harrison, David K. Jackson, Ian Johnston, Dominic Kwiatkowski, Cordelia Langford, John Sillitoe on behalf of the Wellcome Sanger Institute COVID-19 Surveillance Team |
| EPI_ISL_3094925 | Lighthouse Lab in Alderley Park | Wellcome Sanger Institute for the COVID-19 Genomics UK (COG-UK) Consortium | Jacquelyn Wynn, Mairead Hyland, The Lighthouse Lab in Alderley Park and Alex Alderton, Roberto Amato, Jeffrey Barrett, Sonia Goncalves, Ewan Harrison, David K. Jackson, Ian Johnston, Dominic Kwiatkowski, Cordelia Langford, John Sillitoe on behalf of the Wellcome Sanger Institute COVID-19 Surveillance Team |
| EPI_ISL_3094926 | Lighthouse Lab in Glasgow | Wellcome Sanger Institute for the COVID-19 Genomics UK (COG-UK) Consortium | Harper VanSteenhouse, Yumi Kasai, David Gray, Carol Clugston, Anna Dominiczak and Alex Alderton, Roberto Amato, Jeffrey Barrett, Sonia Goncalves, Ewan Harrison, David K. Jackson, Ian Johnston, Dominic Kwiatkowski, Cordelia Langford, John Sillitoe on behalf of the Wellcome Sanger Institute COVID-19 Surveillance Team |
| EPI_ISL_3094927 | Lighthouse Lab in Alderley Park | Wellcome Sanger Institute for the COVID-19 Genomics UK (COG-UK) Consortium | Jacquelyn Wynn, Mairead Hyland, The Lighthouse Lab in Alderley Park and Alex Alderton, Roberto Amato, Jeffrey Barrett, Sonia Goncalves, Ewan Harrison, David K. Jackson, Ian Johnston, Dominic Kwiatkowski, Cordelia Langford, John Sillitoe on behalf of the Wellcome Sanger Institute COVID-19 Surveillance Team |
| EPI_ISL_3094928 | Lighthouse Lab in Glasgow | Wellcome Sanger Institute for the COVID-19 Genomics UK (COG-UK) Consortium | Harper VanSteenhouse, Yumi Kasai, David Gray, Carol Clugston, Anna Dominiczak and Alex Alderton, Roberto Amato, Jeffrey Barrett, Sonia Goncalves, Ewan Harrison, David K. Jackson, Ian Johnston, Dominic Kwiatkowski, Cordelia Langford, John Sillitoe on behalf of the Wellcome Sanger Institute COVID-19 Surveillance Team |
| EPI_ISL_3094929 | Lighthouse Lab in Milton Keynes | Wellcome Sanger Institute for the COVID-19 Genomics UK (COG-UK) Consortium | The Lighthouse Lab in Milton Keynes and Alex Alderton, Roberto Amato, Jeffrey Barrett, Sonia Goncalves, Ewan Harrison, David K. Jackson, Ian Johnston, Dominic Kwiatkowski, Cordelia Langford, John Sillitoe on behalf of the Wellcome Sanger Institute COVID-19 Surveillance Team |
| EPI_ISL_3094930, EPI_ISL_3094931, EPI_ISL_3094932, EPI_ISL_3094933 | Lighthouse Lab in Glasgow | Wellcome Sanger Institute for the COVID-19 Genomics UK (COG-UK) Consortium | Harper VanSteenhouse, Yumi Kasai, David Gray, Carol Clugston, Anna Dominiczak and Alex Alderton, Roberto Amato, Jeffrey Barrett, Sonia Goncalves, Ewan Harrison, David K. Jackson, Ian Johnston, Dominic Kwiatkowski, Cordelia Langford, John Sillitoe on behalf of the Wellcome Sanger Institute COVID-19 Surveillance Team |
| EPI_ISL_3094934, EPI_ISL_3094935 | Lighthouse Lab in Alderley Park | Wellcome Sanger Institute for the COVID-19 Genomics UK (COG-UK) Consortium | Jacquelyn Wynn, Mairead Hyland, The Lighthouse Lab in Alderley Park and Alex Alderton, Roberto Amato, Jeffrey Barrett, Sonia Goncalves, Ewan Harrison, David K. Jackson, Ian Johnston, Dominic Kwiatkowski, Cordelia Langford, John Sillitoe on behalf of the Wellcome Sanger Institute COVID-19 Surveillance Team |
| EPI_ISL_3094936, EPI_ISL_3094937 | Lighthouse Lab in Milton Keynes | Wellcome Sanger Institute for the COVID-19 Genomics UK (COG-UK) Consortium | The Lighthouse Lab in Milton Keynes and Alex Alderton, Roberto Amato, Jeffrey Barrett, Sonia Goncalves, Ewan Harrison, David K. Jackson, Ian Johnston, Dominic Kwiatkowski, Cordelia Langford, John Sillitoe on behalf of the Wellcome Sanger Institute COVID-19 Surveillance Team |
| EPI_ISL_3094938 | Lighthouse Lab in Alderley Park | Wellcome Sanger Institute for the COVID-19 Genomics UK (COG-UK) Consortium | Jacquelyn Wynn, Mairead Hyland, The Lighthouse Lab in Alderley Park and Alex Alderton, Roberto Amato, Jeffrey Barrett, Sonia Goncalves, Ewan Harrison, David K. Jackson, Ian Johnston, Dominic Kwiatkowski, Cordelia Langford, John Sillitoe on behalf of the Wellcome Sanger Institute COVID-19 Surveillance Team |
| EPI_ISL_3094939, EPI_ISL_3094940, EPI_ISL_3094941, EPI_ISL_3094942, EPI_ISL_3094943, EPI_ISL_3094944 | Lighthouse Lab in Glasgow | Wellcome Sanger Institute for the COVID-19 Genomics UK (COG-UK) Consortium | Harper VanSteenhouse, Yumi Kasai, David Gray, Carol Clugston, Anna Dominiczak and Alex Alderton, Roberto Amato, Jeffrey Barrett, Sonia Goncalves, Ewan Harrison, David K. Jackson, Ian Johnston, Dominic Kwiatkowski, Cordelia Langford, John Sillitoe on behalf of the Wellcome Sanger Institute COVID-19 Surveillance Team |
| EPI_ISL_3094945 | Lighthouse Lab in Milton Keynes | Wellcome Sanger Institute for the COVID-19 Genomics UK (COG-UK) Consortium | The Lighthouse Lab in Milton Keynes and Alex Alderton, Roberto Amato, Jeffrey Barrett, Sonia Goncalves, Ewan Harrison, David K. Jackson, Ian Johnston, Dominic Kwiatkowski, Cordelia Langford, John Sillitoe on behalf of the Wellcome Sanger Institute COVID-19 Surveillance Team |
| EPI_ISL_3094946, EPI_ISL_3094947 | Lighthouse Lab in Glasgow | Wellcome Sanger Institute for the COVID-19 Genomics UK (COG-UK) Consortium | Harper VanSteenhouse, Yumi Kasai, David Gray, Carol Clugston, Anna Dominiczak and Alex Alderton, Roberto Amato, Jeffrey Barrett, Sonia Goncalves, Ewan Harrison, David K. Jackson, Ian Johnston, Dominic Kwiatkowski, Cordelia Langford, John Sillitoe on behalf of the Wellcome Sanger Institute COVID-19 Surveillance Team |
| EPI_ISL_3094948 | Lighthouse Lab in Alderley Park | Wellcome Sanger Institute for the COVID-19 Genomics UK (COG-UK) Consortium | Jacquelyn Wynn, Mairead Hyland, The Lighthouse Lab in Alderley Park and Alex Alderton, Roberto Amato, Jeffrey Barrett, Sonia Goncalves, Ewan Harrison, David K. Jackson, Ian Johnston, Dominic Kwiatkowski, Cordelia Langford, John Sillitoe on behalf of the Wellcome Sanger Institute COVID-19 Surveillance Team |
| EPI_ISL_3094949 | Lighthouse Lab in Milton Keynes | Wellcome Sanger Institute for the COVID-19 Genomics UK (COG-UK) Consortium | The Lighthouse Lab in Milton Keynes and Alex Alderton, Roberto Amato, Jeffrey Barrett, Sonia Goncalves, Ewan Harrison, David K. Jackson, Ian Johnston, Dominic Kwiatkowski, Cordelia Langford, John Sillitoe on behalf of the Wellcome Sanger Institute COVID-19 Surveillance Team |
| EPI_ISL_3094950, EPI_ISL_3094951, EPI_ISL_3094952 | Lighthouse Lab in Glasgow | Wellcome Sanger Institute for the COVID-19 Genomics UK (COG-UK) Consortium | Harper VanSteenhouse, Yumi Kasai, David Gray, Carol Clugston, Anna Dominiczak and Alex Alderton, Roberto Amato, Jeffrey Barrett, Sonia Goncalves, Ewan Harrison, David K. Jackson, Ian Johnston, Dominic Kwiatkowski, Cordelia Langford, John Sillitoe on behalf of the Wellcome Sanger Institute COVID-19 Surveillance Team |
| EPI_ISL_3094953 | Lighthouse Lab in Milton Keynes | Wellcome Sanger Institute for the COVID-19 Genomics UK (COG-UK) Consortium | The Lighthouse Lab in Milton Keynes and Alex Alderton, Roberto Amato, Jeffrey Barrett, Sonia Goncalves, Ewan Harrison, David K. Jackson, Ian Johnston, Dominic Kwiatkowski, Cordelia Langford, John Sillitoe on behalf of the Wellcome Sanger Institute COVID-19 Surveillance Team |
| EPI_ISL_3094954, EPI_ISL_3094955 | Lighthouse Lab in Glasgow | Wellcome Sanger Institute for the COVID-19 Genomics UK (COG-UK) Consortium | Harper VanSteenhouse, Yumi Kasai, David Gray, Carol Clugston, Anna Dominiczak and Alex Alderton, Roberto Amato, Jeffrey Barrett, Sonia Goncalves, Ewan Harrison, David K. Jackson, Ian Johnston, Dominic Kwiatkowski, Cordelia Langford, John Sillitoe on behalf of the Wellcome Sanger Institute COVID-19 Surveillance Team |
| EPI_ISL_3094956 | Lighthouse Lab in Milton Keynes | Wellcome Sanger Institute for the COVID-19 Genomics UK (COG-UK) Consortium | The Lighthouse Lab in Milton Keynes and Alex Alderton, Roberto Amato, Jeffrey Barrett, Sonia Goncalves, Ewan Harrison, David K. Jackson, Ian Johnston, Dominic Kwiatkowski, Cordelia Langford, John Sillitoe on behalf of the Wellcome Sanger Institute COVID-19 Surveillance Team |
| EPI_ISL_3094957, EPI_ISL_3094958 | Lighthouse Lab in Glasgow | Wellcome Sanger Institute for the COVID-19 Genomics UK (COG-UK) Consortium | Harper VanSteenhouse, Yumi Kasai, David Gray, Carol Clugston, Anna Dominiczak and Alex Alderton, Roberto Amato, Jeffrey Barrett, Sonia Goncalves, Ewan Harrison, David K. Jackson, Ian Johnston, Dominic Kwiatkowski, Cordelia Langford, John Sillitoe on behalf of the Wellcome Sanger Institute COVID-19 Surveillance Team |
| EPI_ISL_3094959 | Lighthouse Lab in Alderley Park | Wellcome Sanger Institute for the COVID-19 Genomics UK (COG-UK) Consortium | Jacquelyn Wynn, Mairead Hyland, The Lighthouse Lab in Alderley Park and Alex Alderton, Roberto Amato, Jeffrey Barrett, Sonia Goncalves, Ewan Harrison, David K. Jackson, Ian Johnston, Dominic Kwiatkowski, Cordelia Langford, John Sillitoe on behalf of the Wellcome Sanger Institute COVID-19 Surveillance Team |
| EPI_ISL_3094960, EPI_ISL_3094961 | Lighthouse Lab in Milton Keynes | Wellcome Sanger Institute for the COVID-19 Genomics UK (COG-UK) Consortium | The Lighthouse Lab in Milton Keynes and Alex Alderton, Roberto Amato, Jeffrey Barrett, Sonia Goncalves, Ewan Harrison, David K. Jackson, Ian Johnston, Dominic Kwiatkowski, Cordelia Langford, John Sillitoe on behalf of the Wellcome Sanger Institute COVID-19 Surveillance Team |
| EPI_ISL_3094962 | Lighthouse Lab in Alderley Park | Wellcome Sanger Institute for the COVID-19 Genomics UK (COG-UK) Consortium | Jacquelyn Wynn, Mairead Hyland, The Lighthouse Lab in Alderley Park and Alex Alderton, Roberto Amato, Jeffrey Barrett, Sonia Goncalves, Ewan Harrison, David K. Jackson, Ian Johnston, Dominic Kwiatkowski, Cordelia Langford, John Sillitoe on behalf of the Wellcome Sanger Institute COVID-19 Surveillance Team |
| EPI_ISL_3094963 | Lighthouse Lab in Glasgow | Wellcome Sanger Institute for the COVID-19 Genomics UK (COG-UK) Consortium | Harper VanSteenhouse, Yumi Kasai, David Gray, Carol Clugston, Anna Dominiczak and Alex Alderton, Roberto Amato, Jeffrey Barrett, Sonia Goncalves, Ewan Harrison, David K. Jackson, Ian Johnston, Dominic Kwiatkowski, Cordelia Langford, John Sillitoe on behalf of the Wellcome Sanger Institute COVID-19 Surveillance Team |
| EPI_ISL_3094964 | Lighthouse Lab in Alderley Park | Wellcome Sanger Institute for the COVID-19 Genomics UK (COG-UK) Consortium | Jacquelyn Wynn, Mairead Hyland, The Lighthouse Lab in Alderley Park and Alex Alderton, Roberto Amato, Jeffrey Barrett, Sonia Goncalves, Ewan Harrison, David K. Jackson, Ian Johnston, Dominic Kwiatkowski, Cordelia Langford, John Sillitoe on behalf of the Wellcome Sanger Institute COVID-19 Surveillance Team |
| EPI_ISL_3094965 | Lighthouse Lab in Milton Keynes | Wellcome Sanger Institute for the COVID-19 Genomics UK (COG-UK) Consortium | The Lighthouse Lab in Milton Keynes and Alex Alderton, Roberto Amato, Jeffrey Barrett, Sonia Goncalves, Ewan Harrison, David K. Jackson, Ian Johnston, Dominic Kwiatkowski, Cordelia Langford, John Sillitoe on behalf of the Wellcome Sanger Institute COVID-19 Surveillance Team |
| EPI_ISL_3094966, EPI_ISL_3094967 | Lighthouse Lab in Glasgow | Wellcome Sanger Institute for the COVID-19 Genomics UK (COG-UK) Consortium | Harper VanSteenhouse, Yumi Kasai, David Gray, Carol Clugston, Anna Dominiczak and Alex Alderton, Roberto Amato, Jeffrey Barrett, Sonia Goncalves, Ewan Harrison, David K. Jackson, Ian Johnston, Dominic Kwiatkowski, Cordelia Langford, John Sillitoe on behalf of the Wellcome Sanger Institute COVID-19 Surveillance Team |
| EPI_ISL_3094968 | Lighthouse Lab in Alderley Park | Wellcome Sanger Institute for the COVID-19 Genomics UK (COG-UK) Consortium | Jacquelyn Wynn, Mairead Hyland, The Lighthouse Lab in Alderley Park and Alex Alderton, Roberto Amato, Jeffrey Barrett, Sonia Goncalves, Ewan Harrison, David K. Jackson, Ian Johnston, Dominic Kwiatkowski, Cordelia Langford, John Sillitoe on behalf of the Wellcome Sanger Institute COVID-19 Surveillance Team |
| EPI_ISL_3094969, EPI_ISL_3094970 | Lighthouse Lab in Glasgow | Wellcome Sanger Institute for the COVID-19 Genomics UK (COG-UK) Consortium | Harper VanSteenhouse, Yumi Kasai, David Gray, Carol Clugston, Anna Dominiczak and Alex Alderton, Roberto Amato, Jeffrey Barrett, Sonia Goncalves, Ewan Harrison, David K. Jackson, Ian Johnston, Dominic Kwiatkowski, Cordelia Langford, John Sillitoe on behalf of the Wellcome Sanger Institute COVID-19 Surveillance Team |
| EPI_ISL_3094971 | Lighthouse Lab in Alderley Park | Wellcome Sanger Institute for the COVID-19 Genomics UK (COG-UK) Consortium | Jacquelyn Wynn, Mairead Hyland, The Lighthouse Lab in Alderley Park and Alex Alderton, Roberto Amato, Jeffrey Barrett, Sonia Goncalves, Ewan Harrison, David K. Jackson, Ian Johnston, Dominic Kwiatkowski, Cordelia Langford, John Sillitoe on behalf of the Wellcome Sanger Institute COVID-19 Surveillance Team |
| EPI_ISL_3094972, EPI_ISL_3094973, EPI_ISL_3094974 | Lighthouse Lab in Glasgow | Wellcome Sanger Institute for the COVID-19 Genomics UK (COG-UK) Consortium | Harper VanSteenhouse, Yumi Kasai, David Gray, Carol Clugston, Anna Dominiczak and Alex Alderton, Roberto Amato, Jeffrey Barrett, Sonia Goncalves, Ewan Harrison, David K. Jackson, Ian Johnston, Dominic Kwiatkowski, Cordelia Langford, John Sillitoe on behalf of the Wellcome Sanger Institute COVID-19 Surveillance Team |
| EPI_ISL_3094975, EPI_ISL_3094976 | Lighthouse Lab in Milton Keynes | Wellcome Sanger Institute for the COVID-19 Genomics UK (COG-UK) Consortium | The Lighthouse Lab in Milton Keynes and Alex Alderton, Roberto Amato, Jeffrey Barrett, Sonia Goncalves, Ewan Harrison, David K. Jackson, Ian Johnston, Dominic Kwiatkowski, Cordelia Langford, John Sillitoe on behalf of the Wellcome Sanger Institute COVID-19 Surveillance Team |
| EPI_ISL_3094977, EPI_ISL_3094978 | Lighthouse Lab in Glasgow | Wellcome Sanger Institute for the COVID-19 Genomics UK (COG-UK) Consortium | Harper VanSteenhouse, Yumi Kasai, David Gray, Carol Clugston, Anna Dominiczak and Alex Alderton, Roberto Amato, Jeffrey Barrett, Sonia Goncalves, Ewan Harrison, David K. Jackson, Ian Johnston, Dominic Kwiatkowski, Cordelia Langford, John Sillitoe on behalf of the Wellcome Sanger Institute COVID-19 Surveillance Team |
| EPI_ISL_3094979 | Lighthouse Lab in Alderley Park | Wellcome Sanger Institute for the COVID-19 Genomics UK (COG-UK) | Jacquelyn Wynn, Mairead Hyland, The Lighthouse Lab in Alderley Park and Alex Alderton, Roberto Amato, Jeffrey Barrett, Sonia Goncalves, Ewan Harrison, David K. Jackson, Ian Johnston, Dominic Kwiatkowski, Cordelia Langford, John Sillitoe on |

| | | | |
|---|---|---|---|
| | | Consortium | behalf of the Wellcome Sanger Institute COVID-19 Surveillance Team |
| EPI_ISL_3094980, EPI_ISL_3094981, EPI_ISL_3094982 | Lighthouse Lab in Glasgow | Wellcome Sanger Institute for the COVID-19 Genomics UK (COG-UK) Consortium | Harper VanSteenhouse, Yumi Kasai, David Gray, Carol Clugston, Anna Dominiczak and Alex Alderton, Roberto Amato, Jeffrey Barrett, Sonia Goncalves, Ewan Harrison, David K. Jackson, Ian Johnston, Dominic Kwiatkowski, Cordelia Langford, John Sillitoe on behalf of the Wellcome Sanger Institute COVID-19 Surveillance Team |
| EPI_ISL_3094983 | Lighthouse Lab in Alderley Park | Wellcome Sanger Institute for the COVID-19 Genomics UK (COG-UK) Consortium | Jacquelyn Wynn, Mairead Hyland, The Lighthouse Lab in Alderley Park and Alex Alderton, Roberto Amato, Jeffrey Barrett, Sonia Goncalves, Ewan Harrison, David K. Jackson, Ian Johnston, Dominic Kwiatkowski, Cordelia Langford, John Sillitoe on behalf of the Wellcome Sanger Institute COVID-19 Surveillance Team |
| EPI_ISL_3094984 | Lighthouse Lab in Milton Keynes | Wellcome Sanger Institute for the COVID-19 Genomics UK (COG-UK) Consortium | The Lighthouse Lab in Milton Keynes and Alex Alderton, Roberto Amato, Jeffrey Barrett, Sonia Goncalves, Ewan Harrison, David K. Jackson, Ian Johnston, Dominic Kwiatkowski, Cordelia Langford, John Sillitoe on behalf of the Wellcome Sanger Institute COVID-19 Surveillance Team |
| EPI_ISL_3094985 | Lighthouse Lab in Glasgow | Wellcome Sanger Institute for the COVID-19 Genomics UK (COG-UK) Consortium | Harper VanSteenhouse, Yumi Kasai, David Gray, Carol Clugston, Anna Dominiczak and Alex Alderton, Roberto Amato, Jeffrey Barrett, Sonia Goncalves, Ewan Harrison, David K. Jackson, Ian Johnston, Dominic Kwiatkowski, Cordelia Langford, John Sillitoe on behalf of the Wellcome Sanger Institute COVID-19 Surveillance Team |
| EPI_ISL_3094986 | Lighthouse Lab in Alderley Park | Wellcome Sanger Institute for the COVID-19 Genomics UK (COG-UK) Consortium | Jacquelyn Wynn, Mairead Hyland, The Lighthouse Lab in Alderley Park and Alex Alderton, Roberto Amato, Jeffrey Barrett, Sonia Goncalves, Ewan Harrison, David K. Jackson, Ian Johnston, Dominic Kwiatkowski, Cordelia Langford, John Sillitoe on behalf of the Wellcome Sanger Institute COVID-19 Surveillance Team |
| EPI_ISL_3094987 | Lighthouse Lab in Glasgow | Wellcome Sanger Institute for the COVID-19 Genomics UK (COG-UK) Consortium | Harper VanSteenhouse, Yumi Kasai, David Gray, Carol Clugston, Anna Dominiczak and Alex Alderton, Roberto Amato, Jeffrey Barrett, Sonia Goncalves, Ewan Harrison, David K. Jackson, Ian Johnston, Dominic Kwiatkowski, Cordelia Langford, John Sillitoe on behalf of the Wellcome Sanger Institute COVID-19 Surveillance Team |
| EPI_ISL_3094988 | Lighthouse Lab in Alderley Park | Wellcome Sanger Institute for the COVID-19 Genomics UK (COG-UK) Consortium | Jacquelyn Wynn, Mairead Hyland, The Lighthouse Lab in Alderley Park and Alex Alderton, Roberto Amato, Jeffrey Barrett, Sonia Goncalves, Ewan Harrison, David K. Jackson, Ian Johnston, Dominic Kwiatkowski, Cordelia Langford, John Sillitoe on behalf of the Wellcome Sanger Institute COVID-19 Surveillance Team |
| EPI_ISL_3094989 | Lighthouse Lab in Milton Keynes | Wellcome Sanger Institute for the COVID-19 Genomics UK (COG-UK) Consortium | The Lighthouse Lab in Milton Keynes and Alex Alderton, Roberto Amato, Jeffrey Barrett, Sonia Goncalves, Ewan Harrison, David K. Jackson, Ian Johnston, Dominic Kwiatkowski, Cordelia Langford, John Sillitoe on behalf of the Wellcome Sanger Institute COVID-19 Surveillance Team |
| EPI_ISL_3094990 | Lighthouse Lab in Alderley Park | Wellcome Sanger Institute for the COVID-19 Genomics UK (COG-UK) Consortium | Jacquelyn Wynn, Mairead Hyland, The Lighthouse Lab in Alderley Park and Alex Alderton, Roberto Amato, Jeffrey Barrett, Sonia Goncalves, Ewan Harrison, David K. Jackson, Ian Johnston, Dominic Kwiatkowski, Cordelia Langford, John Sillitoe on behalf of the Wellcome Sanger Institute COVID-19 Surveillance Team |
| EPI_ISL_3094991 | Lighthouse Lab in Milton Keynes | Wellcome Sanger Institute for the COVID-19 Genomics UK (COG-UK) Consortium | The Lighthouse Lab in Milton Keynes and Alex Alderton, Roberto Amato, Jeffrey Barrett, Sonia Goncalves, Ewan Harrison, David K. Jackson, Ian Johnston, Dominic Kwiatkowski, Cordelia Langford, John Sillitoe on behalf of the Wellcome Sanger Institute COVID-19 Surveillance Team |
| EPI_ISL_3094992 | Lighthouse Lab in Alderley Park | Wellcome Sanger Institute for the COVID-19 Genomics UK (COG-UK) Consortium | Jacquelyn Wynn, Mairead Hyland, The Lighthouse Lab in Alderley Park and Alex Alderton, Roberto Amato, Jeffrey Barrett, Sonia Goncalves, Ewan Harrison, David K. Jackson, Ian Johnston, Dominic Kwiatkowski, Cordelia Langford, John Sillitoe on behalf of the Wellcome Sanger Institute COVID-19 Surveillance Team |
| EPI_ISL_3094993 | Lighthouse Lab in Milton Keynes | Wellcome Sanger Institute for the COVID-19 Genomics UK (COG-UK) Consortium | The Lighthouse Lab in Milton Keynes and Alex Alderton, Roberto Amato, Jeffrey Barrett, Sonia Goncalves, Ewan Harrison, David K. Jackson, Ian Johnston, Dominic Kwiatkowski, Cordelia Langford, John Sillitoe on behalf of the Wellcome Sanger Institute COVID-19 Surveillance Team |
| EPI_ISL_3094994 | Lighthouse Lab in Alderley Park | Wellcome Sanger Institute for the COVID-19 Genomics UK (COG-UK) Consortium | Jacquelyn Wynn, Mairead Hyland, The Lighthouse Lab in Alderley Park and Alex Alderton, Roberto Amato, Jeffrey Barrett, Sonia Goncalves, Ewan Harrison, David K. Jackson, Ian Johnston, Dominic Kwiatkowski, Cordelia Langford, John Sillitoe on behalf of the Wellcome Sanger Institute COVID-19 Surveillance Team |
| EPI_ISL_3094995, EPI_ISL_3094996 | Lighthouse Lab in Glasgow | Wellcome Sanger Institute for the COVID-19 Genomics UK (COG-UK) Consortium | Harper VanSteenhouse, Yumi Kasai, David Gray, Carol Clugston, Anna Dominiczak and Alex Alderton, Roberto Amato, Jeffrey Barrett, Sonia Goncalves, Ewan Harrison, David K. Jackson, Ian Johnston, Dominic Kwiatkowski, Cordelia Langford, John Sillitoe on behalf of the Wellcome Sanger Institute COVID-19 Surveillance Team |
| EPI_ISL_3094997 | Lighthouse Lab in Milton Keynes | Wellcome Sanger Institute for the COVID-19 Genomics UK (COG-UK) Consortium | The Lighthouse Lab in Milton Keynes and Alex Alderton, Roberto Amato, Jeffrey Barrett, Sonia Goncalves, Ewan Harrison, David K. Jackson, Ian Johnston, Dominic Kwiatkowski, Cordelia Langford, John Sillitoe on behalf of the Wellcome Sanger Institute COVID-19 Surveillance Team |
| EPI_ISL_3094998, EPI_ISL_3094999, EPI_ISL_3095000 | Lighthouse Lab in Glasgow | Wellcome Sanger Institute for the COVID-19 Genomics UK (COG-UK) Consortium | Harper VanSteenhouse, Yumi Kasai, David Gray, Carol Clugston, Anna Dominiczak and Alex Alderton, Roberto Amato, Jeffrey Barrett, Sonia Goncalves, Ewan Harrison, David K. Jackson, Ian Johnston, Dominic Kwiatkowski, Cordelia Langford, John Sillitoe on behalf of the Wellcome Sanger Institute COVID-19 Surveillance Team |
| EPI_ISL_3095001 | Lighthouse Lab in Alderley Park | Wellcome Sanger Institute for the COVID-19 Genomics UK (COG-UK) Consortium | Jacquelyn Wynn, Mairead Hyland, The Lighthouse Lab in Alderley Park and Alex Alderton, Roberto Amato, Jeffrey Barrett, Sonia Goncalves, Ewan Harrison, David K. Jackson, Ian Johnston, Dominic Kwiatkowski, Cordelia Langford, John Sillitoe on behalf of the Wellcome Sanger Institute COVID-19 Surveillance Team |
| EPI_ISL_3095002 | Lighthouse Lab in Milton Keynes | Wellcome Sanger Institute for the COVID-19 Genomics UK (COG-UK) Consortium | The Lighthouse Lab in Milton Keynes and Alex Alderton, Roberto Amato, Jeffrey Barrett, Sonia Goncalves, Ewan Harrison, David K. Jackson, Ian Johnston, Dominic Kwiatkowski, Cordelia Langford, John Sillitoe on behalf of the Wellcome Sanger Institute COVID-19 Surveillance Team |
| EPI_ISL_3095003, EPI_ISL_3095004 | Lighthouse Lab in Alderley Park | Wellcome Sanger Institute for the COVID-19 Genomics UK (COG-UK) Consortium | Jacquelyn Wynn, Mairead Hyland, The Lighthouse Lab in Alderley Park and Alex Alderton, Roberto Amato, Jeffrey Barrett, Sonia Goncalves, Ewan Harrison, David K. Jackson, Ian Johnston, Dominic Kwiatkowski, Cordelia Langford, John Sillitoe on behalf of the Wellcome Sanger Institute COVID-19 Surveillance Team |
| EPI_ISL_3095005 | Lighthouse Lab in Milton Keynes | Wellcome Sanger Institute for the COVID-19 Genomics UK (COG-UK) Consortium | The Lighthouse Lab in Milton Keynes and Alex Alderton, Roberto Amato, Jeffrey Barrett, Sonia Goncalves, Ewan Harrison, David K. Jackson, Ian Johnston, Dominic Kwiatkowski, Cordelia Langford, John Sillitoe on behalf of the Wellcome Sanger Institute COVID-19 Surveillance Team |
| EPI_ISL_3095006 | Lighthouse Lab in Alderley Park | Wellcome Sanger Institute for the COVID-19 Genomics UK (COG-UK) Consortium | Jacquelyn Wynn, Mairead Hyland, The Lighthouse Lab in Alderley Park and Alex Alderton, Roberto Amato, Jeffrey Barrett, Sonia Goncalves, Ewan Harrison, David K. Jackson, Ian Johnston, Dominic Kwiatkowski, Cordelia Langford, John Sillitoe on behalf of the Wellcome Sanger Institute COVID-19 Surveillance Team |
| EPI_ISL_3095007 | Lighthouse Lab in Glasgow | Wellcome Sanger Institute for the COVID-19 Genomics UK (COG-UK) Consortium | Harper VanSteenhouse, Yumi Kasai, David Gray, Carol Clugston, Anna Dominiczak and Alex Alderton, Roberto Amato, Jeffrey Barrett, Sonia Goncalves, Ewan Harrison, David K. Jackson, Ian Johnston, Dominic Kwiatkowski, Cordelia Langford, John Sillitoe on behalf of the Wellcome Sanger Institute COVID-19 Surveillance Team |
| EPI_ISL_3095008 | Lighthouse Lab in Alderley Park | Wellcome Sanger Institute for the COVID-19 Genomics UK (COG-UK) Consortium | Jacquelyn Wynn, Mairead Hyland, The Lighthouse Lab in Alderley Park and Alex Alderton, Roberto Amato, Jeffrey Barrett, Sonia Goncalves, Ewan Harrison, David K. Jackson, Ian Johnston, Dominic Kwiatkowski, Cordelia Langford, John Sillitoe on behalf of the Wellcome Sanger Institute COVID-19 Surveillance Team |
| EPI_ISL_3095009, EPI_ISL_3095010, EPI_ISL_3095011 | Lighthouse Lab in Glasgow | Wellcome Sanger Institute for the COVID-19 Genomics UK (COG-UK) Consortium | Harper VanSteenhouse, Yumi Kasai, David Gray, Carol Clugston, Anna Dominiczak and Alex Alderton, Roberto Amato, Jeffrey Barrett, Sonia Goncalves, Ewan Harrison, David K. Jackson, Ian Johnston, Dominic Kwiatkowski, Cordelia Langford, John Sillitoe on behalf of the Wellcome Sanger Institute COVID-19 Surveillance Team |
| EPI_ISL_3095012 | Lighthouse Lab in Milton Keynes | Wellcome Sanger Institute for the COVID-19 Genomics UK (COG-UK) Consortium | The Lighthouse Lab in Milton Keynes and Alex Alderton, Roberto Amato, Jeffrey Barrett, Sonia Goncalves, Ewan Harrison, David K. Jackson, Ian Johnston, Dominic Kwiatkowski, Cordelia Langford, John Sillitoe on behalf of the Wellcome Sanger Institute COVID-19 Surveillance Team |
| EPI_ISL_3095013 | Lighthouse Lab in Alderley Park | Wellcome Sanger Institute for the COVID-19 Genomics UK (COG-UK) Consortium | Jacquelyn Wynn, Mairead Hyland, The Lighthouse Lab in Alderley Park and Alex Alderton, Roberto Amato, Jeffrey Barrett, Sonia Goncalves, Ewan Harrison, David K. Jackson, Ian Johnston, Dominic Kwiatkowski, Cordelia Langford, John Sillitoe on behalf of the Wellcome Sanger Institute COVID-19 Surveillance Team |
| EPI_ISL_3095014 | Lighthouse Lab in Milton Keynes | Wellcome Sanger Institute for the COVID-19 Genomics UK (COG-UK) Consortium | The Lighthouse Lab in Milton Keynes and Alex Alderton, Roberto Amato, Jeffrey Barrett, Sonia Goncalves, Ewan Harrison, David K. Jackson, Ian Johnston, Dominic Kwiatkowski, Cordelia Langford, John Sillitoe on behalf of the Wellcome Sanger Institute COVID-19 Surveillance Team |
| EPI_ISL_3095015 | Lighthouse Lab in Glasgow | Wellcome Sanger Institute for the COVID-19 Genomics UK (COG-UK) Consortium | Harper VanSteenhouse, Yumi Kasai, David Gray, Carol Clugston, Anna Dominiczak and Alex Alderton, Roberto Amato, Jeffrey Barrett, Sonia Goncalves, Ewan Harrison, David K. Jackson, Ian Johnston, Dominic Kwiatkowski, Cordelia Langford, John Sillitoe on behalf of the Wellcome Sanger Institute COVID-19 Surveillance Team |
| EPI_ISL_3095016 | Lighthouse Lab in Milton Keynes | Wellcome Sanger Institute for the COVID-19 Genomics UK (COG-UK) Consortium | The Lighthouse Lab in Milton Keynes and Alex Alderton, Roberto Amato, Jeffrey Barrett, Sonia Goncalves, Ewan Harrison, David K. Jackson, Ian Johnston, Dominic Kwiatkowski, Cordelia Langford, John Sillitoe on behalf of the Wellcome Sanger Institute COVID-19 Surveillance Team |
| EPI_ISL_3095017 | Lighthouse Lab in Alderley Park | Wellcome Sanger Institute for the COVID-19 Genomics UK (COG-UK) Consortium | Jacquelyn Wynn, Mairead Hyland, The Lighthouse Lab in Alderley Park and Alex Alderton, Roberto Amato, Jeffrey Barrett, Sonia Goncalves, Ewan Harrison, David K. Jackson, Ian Johnston, Dominic Kwiatkowski, Cordelia Langford, John Sillitoe on behalf of the Wellcome Sanger Institute COVID-19 Surveillance Team |
| EPI_ISL_3095018 | Lighthouse Lab in Glasgow | Wellcome Sanger Institute for the COVID-19 Genomics UK (COG-UK) Consortium | Harper VanSteenhouse, Yumi Kasai, David Gray, Carol Clugston, Anna Dominiczak and Alex Alderton, Roberto Amato, Jeffrey Barrett, Sonia Goncalves, Ewan Harrison, David K. Jackson, Ian Johnston, Dominic Kwiatkowski, Cordelia Langford, John Sillitoe on behalf of the Wellcome Sanger Institute COVID-19 Surveillance Team |
| EPI_ISL_3095019, EPI_ISL_3095020 | Lighthouse Lab in Alderley Park | Wellcome Sanger Institute for the COVID-19 Genomics UK (COG-UK) Consortium | Jacquelyn Wynn, Mairead Hyland, The Lighthouse Lab in Alderley Park and Alex Alderton, Roberto Amato, Jeffrey Barrett, Sonia Goncalves, Ewan Harrison, David K. Jackson, Ian Johnston, Dominic Kwiatkowski, Cordelia Langford, John Sillitoe on behalf of the Wellcome Sanger Institute COVID-19 Surveillance Team |
| EPI_ISL_3095021 | Lighthouse Lab in Milton Keynes | Wellcome Sanger Institute for the COVID-19 Genomics UK (COG-UK) Consortium | The Lighthouse Lab in Milton Keynes and Alex Alderton, Roberto Amato, Jeffrey Barrett, Sonia Goncalves, Ewan Harrison, David K. Jackson, Ian Johnston, Dominic Kwiatkowski, Cordelia Langford, John Sillitoe on behalf of the Wellcome Sanger Institute COVID-19 Surveillance Team |
| EPI_ISL_3095022 | Lighthouse Lab in Alderley Park | Wellcome Sanger Institute for the COVID-19 Genomics UK (COG-UK) Consortium | Jacquelyn Wynn, Mairead Hyland, The Lighthouse Lab in Alderley Park and Alex Alderton, Roberto Amato, Jeffrey Barrett, Sonia Goncalves, Ewan Harrison, David K. Jackson, Ian Johnston, Dominic Kwiatkowski, Cordelia Langford, John Sillitoe on behalf of the Wellcome Sanger Institute COVID-19 Surveillance Team |
| EPI_ISL_3095023 | Lighthouse Lab in Glasgow | Wellcome Sanger Institute for the COVID-19 Genomics UK (COG-UK) Consortium | Harper VanSteenhouse, Yumi Kasai, David Gray, Carol Clugston, Anna Dominiczak and Alex Alderton, Roberto Amato, Jeffrey Barrett, Sonia Goncalves, Ewan Harrison, David K. Jackson, Ian Johnston, Dominic Kwiatkowski, Cordelia Langford, John Sillitoe on behalf of the Wellcome Sanger Institute COVID-19 Surveillance Team |
| EPI_ISL_3095024 | Lighthouse Lab in Alderley Park | Wellcome Sanger Institute for the COVID-19 Genomics UK (COG-UK) Consortium | Jacquelyn Wynn, Mairead Hyland, The Lighthouse Lab in Alderley Park and Alex Alderton, Roberto Amato, Jeffrey Barrett, Sonia Goncalves, Ewan Harrison, David K. Jackson, Ian Johnston, Dominic Kwiatkowski, Cordelia Langford, John Sillitoe on behalf of the Wellcome Sanger Institute COVID-19 Surveillance Team |
| EPI_ISL_3095025, EPI_ISL_3095026, EPI_ISL_3095027, EPI_ISL_3095028, EPI_ISL_3095029, EPI_ISL_3095030 | Lighthouse Lab in Glasgow | Wellcome Sanger Institute for the COVID-19 Genomics UK (COG-UK) Consortium | Harper VanSteenhouse, Yumi Kasai, David Gray, Carol Clugston, Anna Dominiczak and Alex Alderton, Roberto Amato, Jeffrey Barrett, Sonia Goncalves, Ewan Harrison, David K. Jackson, Ian Johnston, Dominic Kwiatkowski, Cordelia Langford, John Sillitoe on behalf of the Wellcome Sanger Institute COVID-19 Surveillance Team |
| EPI_ISL_3095031 | Lighthouse Lab in Alderley Park | Wellcome Sanger Institute for the COVID-19 Genomics UK (COG-UK) Consortium | Jacquelyn Wynn, Mairead Hyland, The Lighthouse Lab in Alderley Park and Alex Alderton, Roberto Amato, Jeffrey Barrett, Sonia Goncalves, Ewan Harrison, David K. Jackson, Ian Johnston, Dominic Kwiatkowski, Cordelia Langford, John Sillitoe on behalf of the Wellcome Sanger Institute COVID-19 Surveillance Team |
| EPI_ISL_3095032 | Lighthouse Lab in Milton Keynes | Wellcome Sanger Institute for the COVID-19 Genomics UK (COG-UK) Consortium | The Lighthouse Lab in Milton Keynes and Alex Alderton, Roberto Amato, Jeffrey Barrett, Sonia Goncalves, Ewan Harrison, David K. Jackson, Ian Johnston, Dominic Kwiatkowski, Cordelia Langford, John Sillitoe on behalf of the Wellcome Sanger Institute COVID-19 Surveillance Team |
| EPI_ISL_3095033 | Lighthouse Lab in Alderley Park | Wellcome Sanger Institute for the COVID-19 Genomics UK (COG-UK) Consortium | Jacquelyn Wynn, Mairead Hyland, The Lighthouse Lab in Alderley Park and Alex Alderton, Roberto Amato, Jeffrey Barrett, Sonia Goncalves, Ewan Harrison, David K. Jackson, Ian Johnston, Dominic Kwiatkowski, Cordelia Langford, John Sillitoe on behalf of the Wellcome Sanger Institute COVID-19 Surveillance Team |
| EPI_ISL_3095034 | Lighthouse Lab in Milton Keynes | Wellcome Sanger Institute for the COVID-19 Genomics UK (COG-UK) Consortium | The Lighthouse Lab in Milton Keynes and Alex Alderton, Roberto Amato, Jeffrey Barrett, Sonia Goncalves, Ewan Harrison, David K. Jackson, Ian Johnston, Dominic Kwiatkowski, Cordelia Langford, John Sillitoe on behalf of the Wellcome Sanger Institute COVID-19 Surveillance Team |
| EPI_ISL_3095035 | Lighthouse Lab in Alderley Park | Wellcome Sanger Institute for the COVID-19 Genomics UK (COG-UK) | Jacquelyn Wynn, Mairead Hyland, The Lighthouse Lab in Alderley Park and Alex Alderton, Roberto Amato, Jeffrey Barrett, Sonia Goncalves, Ewan Harrison, David K. Jackson, Ian Johnston, Dominic Kwiatkowski, Cordelia Langford, John Sillitoe on |

| | | | |
|---|---|---|---|
| | | Consortium | behalf of the Wellcome Sanger Institute COVID-19 Surveillance Team |
| EPI_ISL_3095036 | Lighthouse Lab in Milton Keynes | Wellcome Sanger Institute for the COVID-19 Genomics UK (COG-UK) Consortium | The Lighthouse Lab in Milton Keynes and Alex Alderton, Roberto Amato, Jeffrey Barrett, Sonia Goncalves, Ewan Harrison, David K. Jackson, Ian Johnston, Dominic Kwiatkowski, Cordelia Langford, John Sillitoe on behalf of the Wellcome Sanger Institute COVID-19 Surveillance Team |
| EPI_ISL_3095037 | Lighthouse Lab in Alderley Park | Wellcome Sanger Institute for the COVID-19 Genomics UK (COG-UK) Consortium | Jacquelyn Wynn, Mairead Hyland, The Lighthouse Lab in Alderley Park and Alex Alderton, Roberto Amato, Jeffrey Barrett, Sonia Goncalves, Ewan Harrison, David K. Jackson, Ian Johnston, Dominic Kwiatkowski, Cordelia Langford, John Sillitoe on behalf of the Wellcome Sanger Institute COVID-19 Surveillance Team |
| EPI_ISL_3095038 | Lighthouse Lab in Milton Keynes | Wellcome Sanger Institute for the COVID-19 Genomics UK (COG-UK) Consortium | The Lighthouse Lab in Milton Keynes and Alex Alderton, Roberto Amato, Jeffrey Barrett, Sonia Goncalves, Ewan Harrison, David K. Jackson, Ian Johnston, Dominic Kwiatkowski, Cordelia Langford, John Sillitoe on behalf of the Wellcome Sanger Institute COVID-19 Surveillance Team |
| EPI_ISL_3095039 | Lighthouse Lab in Glasgow | Wellcome Sanger Institute for the COVID-19 Genomics UK (COG-UK) Consortium | Harper VanSteenhouse, Yumi Kasai, David Gray, Carol Clugston, Anna Dominiczak and Alex Alderton, Roberto Amato, Jeffrey Barrett, Sonia Goncalves, Ewan Harrison, David K. Jackson, Ian Johnston, Dominic Kwiatkowski, Cordelia Langford, John Sillitoe on behalf of the Wellcome Sanger Institute COVID-19 Surveillance Team |
| EPI_ISL_3095040 | Lighthouse Lab in Milton Keynes | Wellcome Sanger Institute for the COVID-19 Genomics UK (COG-UK) Consortium | The Lighthouse Lab in Milton Keynes and Alex Alderton, Roberto Amato, Jeffrey Barrett, Sonia Goncalves, Ewan Harrison, David K. Jackson, Ian Johnston, Dominic Kwiatkowski, Cordelia Langford, John Sillitoe on behalf of the Wellcome Sanger Institute COVID-19 Surveillance Team |
| EPI_ISL_3095041 | Lighthouse Lab in Glasgow | Wellcome Sanger Institute for the COVID-19 Genomics UK (COG-UK) Consortium | Harper VanSteenhouse, Yumi Kasai, David Gray, Carol Clugston, Anna Dominiczak and Alex Alderton, Roberto Amato, Jeffrey Barrett, Sonia Goncalves, Ewan Harrison, David K. Jackson, Ian Johnston, Dominic Kwiatkowski, Cordelia Langford, John Sillitoe on behalf of the Wellcome Sanger Institute COVID-19 Surveillance Team |
| EPI_ISL_3095042, EPI_ISL_3095043, EPI_ISL_3095044, EPI_ISL_3095045 | Lighthouse Lab in Milton Keynes | Wellcome Sanger Institute for the COVID-19 Genomics UK (COG-UK) Consortium | The Lighthouse Lab in Milton Keynes and Alex Alderton, Roberto Amato, Jeffrey Barrett, Sonia Goncalves, Ewan Harrison, David K. Jackson, Ian Johnston, Dominic Kwiatkowski, Cordelia Langford, John Sillitoe on behalf of the Wellcome Sanger Institute COVID-19 Surveillance Team |
| EPI_ISL_3095046 | Lighthouse Lab in Alderley Park | Wellcome Sanger Institute for the COVID-19 Genomics UK (COG-UK) Consortium | Jacquelyn Wynn, Mairead Hyland, The Lighthouse Lab in Alderley Park and Alex Alderton, Roberto Amato, Jeffrey Barrett, Sonia Goncalves, Ewan Harrison, David K. Jackson, Ian Johnston, Dominic Kwiatkowski, Cordelia Langford, John Sillitoe on behalf of the Wellcome Sanger Institute COVID-19 Surveillance Team |
| EPI_ISL_3095047 | Lighthouse Lab in Glasgow | Wellcome Sanger Institute for the COVID-19 Genomics UK (COG-UK) Consortium | Harper VanSteenhouse, Yumi Kasai, David Gray, Carol Clugston, Anna Dominiczak and Alex Alderton, Roberto Amato, Jeffrey Barrett, Sonia Goncalves, Ewan Harrison, David K. Jackson, Ian Johnston, Dominic Kwiatkowski, Cordelia Langford, John Sillitoe on behalf of the Wellcome Sanger Institute COVID-19 Surveillance Team |
| EPI_ISL_3095048 | Lighthouse Lab in Milton Keynes | Wellcome Sanger Institute for the COVID-19 Genomics UK (COG-UK) Consortium | The Lighthouse Lab in Milton Keynes and Alex Alderton, Roberto Amato, Jeffrey Barrett, Sonia Goncalves, Ewan Harrison, David K. Jackson, Ian Johnston, Dominic Kwiatkowski, Cordelia Langford, John Sillitoe on behalf of the Wellcome Sanger Institute COVID-19 Surveillance Team |
| EPI_ISL_3095049, EPI_ISL_3095050, EPI_ISL_3095051 | Lighthouse Lab in Glasgow | Wellcome Sanger Institute for the COVID-19 Genomics UK (COG-UK) Consortium | Harper VanSteenhouse, Yumi Kasai, David Gray, Carol Clugston, Anna Dominiczak and Alex Alderton, Roberto Amato, Jeffrey Barrett, Sonia Goncalves, Ewan Harrison, David K. Jackson, Ian Johnston, Dominic Kwiatkowski, Cordelia Langford, John Sillitoe on behalf of the Wellcome Sanger Institute COVID-19 Surveillance Team |
| EPI_ISL_3095052, EPI_ISL_3095053 | Lighthouse Lab in Alderley Park | Wellcome Sanger Institute for the COVID-19 Genomics UK (COG-UK) Consortium | Jacquelyn Wynn, Mairead Hyland, The Lighthouse Lab in Alderley Park and Alex Alderton, Roberto Amato, Jeffrey Barrett, Sonia Goncalves, Ewan Harrison, David K. Jackson, Ian Johnston, Dominic Kwiatkowski, Cordelia Langford, John Sillitoe on behalf of the Wellcome Sanger Institute COVID-19 Surveillance Team |
| EPI_ISL_3095054 | Lighthouse Lab in Milton Keynes | Wellcome Sanger Institute for the COVID-19 Genomics UK (COG-UK) Consortium | The Lighthouse Lab in Milton Keynes and Alex Alderton, Roberto Amato, Jeffrey Barrett, Sonia Goncalves, Ewan Harrison, David K. Jackson, Ian Johnston, Dominic Kwiatkowski, Cordelia Langford, John Sillitoe on behalf of the Wellcome Sanger Institute COVID-19 Surveillance Team |
| EPI_ISL_3095055, EPI_ISL_3095056 | Lighthouse Lab in Glasgow | Wellcome Sanger Institute for the COVID-19 Genomics UK (COG-UK) Consortium | Harper VanSteenhouse, Yumi Kasai, David Gray, Carol Clugston, Anna Dominiczak and Alex Alderton, Roberto Amato, Jeffrey Barrett, Sonia Goncalves, Ewan Harrison, David K. Jackson, Ian Johnston, Dominic Kwiatkowski, Cordelia Langford, John Sillitoe on behalf of the Wellcome Sanger Institute COVID-19 Surveillance Team |
| EPI_ISL_3095057 | Lighthouse Lab in Alderley Park | Wellcome Sanger Institute for the COVID-19 Genomics UK (COG-UK) Consortium | Jacquelyn Wynn, Mairead Hyland, The Lighthouse Lab in Alderley Park and Alex Alderton, Roberto Amato, Jeffrey Barrett, Sonia Goncalves, Ewan Harrison, David K. Jackson, Ian Johnston, Dominic Kwiatkowski, Cordelia Langford, John Sillitoe on behalf of the Wellcome Sanger Institute COVID-19 Surveillance Team |
| EPI_ISL_3095058, EPI_ISL_3095059 | Lighthouse Lab in Glasgow | Wellcome Sanger Institute for the COVID-19 Genomics UK (COG-UK) Consortium | Harper VanSteenhouse, Yumi Kasai, David Gray, Carol Clugston, Anna Dominiczak and Alex Alderton, Roberto Amato, Jeffrey Barrett, Sonia Goncalves, Ewan Harrison, David K. Jackson, Ian Johnston, Dominic Kwiatkowski, Cordelia Langford, John Sillitoe on behalf of the Wellcome Sanger Institute COVID-19 Surveillance Team |
| EPI_ISL_3095060 | Lighthouse Lab in Alderley Park | Wellcome Sanger Institute for the COVID-19 Genomics UK (COG-UK) Consortium | Jacquelyn Wynn, Mairead Hyland, The Lighthouse Lab in Alderley Park and Alex Alderton, Roberto Amato, Jeffrey Barrett, Sonia Goncalves, Ewan Harrison, David K. Jackson, Ian Johnston, Dominic Kwiatkowski, Cordelia Langford, John Sillitoe on behalf of the Wellcome Sanger Institute COVID-19 Surveillance Team |
| EPI_ISL_3095061 | Lighthouse Lab in Glasgow | Wellcome Sanger Institute for the COVID-19 Genomics UK (COG-UK) Consortium | Harper VanSteenhouse, Yumi Kasai, David Gray, Carol Clugston, Anna Dominiczak and Alex Alderton, Roberto Amato, Jeffrey Barrett, Sonia Goncalves, Ewan Harrison, David K. Jackson, Ian Johnston, Dominic Kwiatkowski, Cordelia Langford, John Sillitoe on behalf of the Wellcome Sanger Institute COVID-19 Surveillance Team |
| EPI_ISL_3095062 | Lighthouse Lab in Milton Keynes | Wellcome Sanger Institute for the COVID-19 Genomics UK (COG-UK) Consortium | The Lighthouse Lab in Milton Keynes and Alex Alderton, Roberto Amato, Jeffrey Barrett, Sonia Goncalves, Ewan Harrison, David K. Jackson, Ian Johnston, Dominic Kwiatkowski, Cordelia Langford, John Sillitoe on behalf of the Wellcome Sanger Institute COVID-19 Surveillance Team |
| EPI_ISL_3095063 | Lighthouse Lab in Glasgow | Wellcome Sanger Institute for the COVID-19 Genomics UK (COG-UK) Consortium | Harper VanSteenhouse, Yumi Kasai, David Gray, Carol Clugston, Anna Dominiczak and Alex Alderton, Roberto Amato, Jeffrey Barrett, Sonia Goncalves, Ewan Harrison, David K. Jackson, Ian Johnston, Dominic Kwiatkowski, Cordelia Langford, John Sillitoe on behalf of the Wellcome Sanger Institute COVID-19 Surveillance Team |
| EPI_ISL_3095064 | Lighthouse Lab in Milton Keynes | Wellcome Sanger Institute for the COVID-19 Genomics UK (COG-UK) Consortium | The Lighthouse Lab in Milton Keynes and Alex Alderton, Roberto Amato, Jeffrey Barrett, Sonia Goncalves, Ewan Harrison, David K. Jackson, Ian Johnston, Dominic Kwiatkowski, Cordelia Langford, John Sillitoe on behalf of the Wellcome Sanger Institute COVID-19 Surveillance Team |
| EPI_ISL_3095065 | Lighthouse Lab in Glasgow | Wellcome Sanger Institute for the COVID-19 Genomics UK (COG-UK) Consortium | Harper VanSteenhouse, Yumi Kasai, David Gray, Carol Clugston, Anna Dominiczak and Alex Alderton, Roberto Amato, Jeffrey Barrett, Sonia Goncalves, Ewan Harrison, David K. Jackson, Ian Johnston, Dominic Kwiatkowski, Cordelia Langford, John Sillitoe on behalf of the Wellcome Sanger Institute COVID-19 Surveillance Team |
| EPI_ISL_3095066 | Lighthouse Lab in Alderley Park | Wellcome Sanger Institute for the COVID-19 Genomics UK (COG-UK) Consortium | Jacquelyn Wynn, Mairead Hyland, The Lighthouse Lab in Alderley Park and Alex Alderton, Roberto Amato, Jeffrey Barrett, Sonia Goncalves, Ewan Harrison, David K. Jackson, Ian Johnston, Dominic Kwiatkowski, Cordelia Langford, John Sillitoe on behalf of the Wellcome Sanger Institute COVID-19 Surveillance Team |
| EPI_ISL_3095067 | Lighthouse Lab in Glasgow | Wellcome Sanger Institute for the COVID-19 Genomics UK (COG-UK) Consortium | Harper VanSteenhouse, Yumi Kasai, David Gray, Carol Clugston, Anna Dominiczak and Alex Alderton, Roberto Amato, Jeffrey Barrett, Sonia Goncalves, Ewan Harrison, David K. Jackson, Ian Johnston, Dominic Kwiatkowski, Cordelia Langford, John Sillitoe on behalf of the Wellcome Sanger Institute COVID-19 Surveillance Team |
| EPI_ISL_3095068 | Lighthouse Lab in Alderley Park | Wellcome Sanger Institute for the COVID-19 Genomics UK (COG-UK) Consortium | Jacquelyn Wynn, Mairead Hyland, The Lighthouse Lab in Alderley Park and Alex Alderton, Roberto Amato, Jeffrey Barrett, Sonia Goncalves, Ewan Harrison, David K. Jackson, Ian Johnston, Dominic Kwiatkowski, Cordelia Langford, John Sillitoe on behalf of the Wellcome Sanger Institute COVID-19 Surveillance Team |
| EPI_ISL_3095069 | Lighthouse Lab in Milton Keynes | Wellcome Sanger Institute for the COVID-19 Genomics UK (COG-UK) Consortium | The Lighthouse Lab in Milton Keynes and Alex Alderton, Roberto Amato, Jeffrey Barrett, Sonia Goncalves, Ewan Harrison, David K. Jackson, Ian Johnston, Dominic Kwiatkowski, Cordelia Langford, John Sillitoe on behalf of the Wellcome Sanger Institute COVID-19 Surveillance Team |
| EPI_ISL_3095070, EPI_ISL_3095071, EPI_ISL_3095072, EPI_ISL_3095073, EPI_ISL_3095074 | Lighthouse Lab in Glasgow | Wellcome Sanger Institute for the COVID-19 Genomics UK (COG-UK) Consortium | Harper VanSteenhouse, Yumi Kasai, David Gray, Carol Clugston, Anna Dominiczak and Alex Alderton, Roberto Amato, Jeffrey Barrett, Sonia Goncalves, Ewan Harrison, David K. Jackson, Ian Johnston, Dominic Kwiatkowski, Cordelia Langford, John Sillitoe on behalf of the Wellcome Sanger Institute COVID-19 Surveillance Team |
| EPI_ISL_3095075 | Lighthouse Lab in Milton Keynes | Wellcome Sanger Institute for the COVID-19 Genomics UK (COG-UK) Consortium | The Lighthouse Lab in Milton Keynes and Alex Alderton, Roberto Amato, Jeffrey Barrett, Sonia Goncalves, Ewan Harrison, David K. Jackson, Ian Johnston, Dominic Kwiatkowski, Cordelia Langford, John Sillitoe on behalf of the Wellcome Sanger Institute COVID-19 Surveillance Team |
| EPI_ISL_3095076, EPI_ISL_3095077, EPI_ISL_3095078 | Lighthouse Lab in Glasgow | Wellcome Sanger Institute for the COVID-19 Genomics UK (COG-UK) Consortium | Harper VanSteenhouse, Yumi Kasai, David Gray, Carol Clugston, Anna Dominiczak and Alex Alderton, Roberto Amato, Jeffrey Barrett, Sonia Goncalves, Ewan Harrison, David K. Jackson, Ian Johnston, Dominic Kwiatkowski, Cordelia Langford, John Sillitoe on behalf of the Wellcome Sanger Institute COVID-19 Surveillance Team |
| EPI_ISL_3095079 | Lighthouse Lab in Milton Keynes | Wellcome Sanger Institute for the COVID-19 Genomics UK (COG-UK) Consortium | The Lighthouse Lab in Milton Keynes and Alex Alderton, Roberto Amato, Jeffrey Barrett, Sonia Goncalves, Ewan Harrison, David K. Jackson, Ian Johnston, Dominic Kwiatkowski, Cordelia Langford, John Sillitoe on behalf of the Wellcome Sanger Institute COVID-19 Surveillance Team |
| EPI_ISL_3095080 | Lighthouse Lab in Glasgow | Wellcome Sanger Institute for the COVID-19 Genomics UK (COG-UK) Consortium | Harper VanSteenhouse, Yumi Kasai, David Gray, Carol Clugston, Anna Dominiczak and Alex Alderton, Roberto Amato, Jeffrey Barrett, Sonia Goncalves, Ewan Harrison, David K. Jackson, Ian Johnston, Dominic Kwiatkowski, Cordelia Langford, John Sillitoe on behalf of the Wellcome Sanger Institute COVID-19 Surveillance Team |
| EPI_ISL_3095081, EPI_ISL_3095082 | Lighthouse Lab in Alderley Park | Wellcome Sanger Institute for the COVID-19 Genomics UK (COG-UK) Consortium | Jacquelyn Wynn, Mairead Hyland, The Lighthouse Lab in Alderley Park and Alex Alderton, Roberto Amato, Jeffrey Barrett, Sonia Goncalves, Ewan Harrison, David K. Jackson, Ian Johnston, Dominic Kwiatkowski, Cordelia Langford, John Sillitoe on behalf of the Wellcome Sanger Institute COVID-19 Surveillance Team |
| EPI_ISL_3095083 | Lighthouse Lab in Milton Keynes | Wellcome Sanger Institute for the COVID-19 Genomics UK (COG-UK) Consortium | The Lighthouse Lab in Milton Keynes and Alex Alderton, Roberto Amato, Jeffrey Barrett, Sonia Goncalves, Ewan Harrison, David K. Jackson, Ian Johnston, Dominic Kwiatkowski, Cordelia Langford, John Sillitoe on behalf of the Wellcome Sanger Institute COVID-19 Surveillance Team |
| EPI_ISL_3095084 | Lighthouse Lab in Alderley Park | Wellcome Sanger Institute for the COVID-19 Genomics UK (COG-UK) Consortium | Jacquelyn Wynn, Mairead Hyland, The Lighthouse Lab in Alderley Park and Alex Alderton, Roberto Amato, Jeffrey Barrett, Sonia Goncalves, Ewan Harrison, David K. Jackson, Ian Johnston, Dominic Kwiatkowski, Cordelia Langford, John Sillitoe on behalf of the Wellcome Sanger Institute COVID-19 Surveillance Team |
| EPI_ISL_3095085 | Lighthouse Lab in Glasgow | Wellcome Sanger Institute for the COVID-19 Genomics UK (COG-UK) Consortium | Harper VanSteenhouse, Yumi Kasai, David Gray, Carol Clugston, Anna Dominiczak and Alex Alderton, Roberto Amato, Jeffrey Barrett, Sonia Goncalves, Ewan Harrison, David K. Jackson, Ian Johnston, Dominic Kwiatkowski, Cordelia Langford, John Sillitoe on behalf of the Wellcome Sanger Institute COVID-19 Surveillance Team |
| EPI_ISL_3095086, EPI_ISL_3095087 | Lighthouse Lab in Alderley Park | Wellcome Sanger Institute for the COVID-19 Genomics UK (COG-UK) Consortium | Jacquelyn Wynn, Mairead Hyland, The Lighthouse Lab in Alderley Park and Alex Alderton, Roberto Amato, Jeffrey Barrett, Sonia Goncalves, Ewan Harrison, David K. Jackson, Ian Johnston, Dominic Kwiatkowski, Cordelia Langford, John Sillitoe on behalf of the Wellcome Sanger Institute COVID-19 Surveillance Team |
| EPI_ISL_3095088 | Lighthouse Lab in Milton Keynes | Wellcome Sanger Institute for the COVID-19 Genomics UK (COG-UK) Consortium | The Lighthouse Lab in Milton Keynes and Alex Alderton, Roberto Amato, Jeffrey Barrett, Sonia Goncalves, Ewan Harrison, David K. Jackson, Ian Johnston, Dominic Kwiatkowski, Cordelia Langford, John Sillitoe on behalf of the Wellcome Sanger Institute COVID-19 Surveillance Team |
| EPI_ISL_3095089, EPI_ISL_3095090, EPI_ISL_3095091 | Lighthouse Lab in Glasgow | Wellcome Sanger Institute for the COVID-19 Genomics UK (COG-UK) Consortium | Harper VanSteenhouse, Yumi Kasai, David Gray, Carol Clugston, Anna Dominiczak and Alex Alderton, Roberto Amato, Jeffrey Barrett, Sonia Goncalves, Ewan Harrison, David K. Jackson, Ian Johnston, Dominic Kwiatkowski, Cordelia Langford, John Sillitoe on behalf of the Wellcome Sanger Institute COVID-19 Surveillance Team |
| EPI_ISL_3095092 | Lighthouse Lab in Alderley Park | Wellcome Sanger Institute for the COVID-19 Genomics UK (COG-UK) Consortium | Jacquelyn Wynn, Mairead Hyland, The Lighthouse Lab in Alderley Park and Alex Alderton, Roberto Amato, Jeffrey Barrett, Sonia Goncalves, Ewan Harrison, David K. Jackson, Ian Johnston, Dominic Kwiatkowski, Cordelia Langford, John Sillitoe on behalf of the Wellcome Sanger Institute COVID-19 Surveillance Team |
| EPI_ISL_3095093 | Lighthouse Lab in Milton Keynes | Wellcome Sanger Institute for the COVID-19 Genomics UK (COG-UK) Consortium | The Lighthouse Lab in Milton Keynes and Alex Alderton, Roberto Amato, Jeffrey Barrett, Sonia Goncalves, Ewan Harrison, David K. Jackson, Ian Johnston, Dominic Kwiatkowski, Cordelia Langford, John Sillitoe on behalf of the Wellcome Sanger Institute COVID-19 Surveillance Team |
| EPI_ISL_3095094, EPI_ISL_3095095 | Lighthouse Lab in Glasgow | Wellcome Sanger Institute for the COVID-19 Genomics UK (COG-UK) Consortium | Harper VanSteenhouse, Yumi Kasai, David Gray, Carol Clugston, Anna Dominiczak and Alex Alderton, Roberto Amato, Jeffrey Barrett, Sonia Goncalves, Ewan Harrison, David K. Jackson, Ian Johnston, Dominic Kwiatkowski, Cordelia Langford, John Sillitoe on behalf of the Wellcome Sanger Institute COVID-19 Surveillance Team |
| EPI_ISL_3095096 | Lighthouse Lab in Milton Keynes | Wellcome Sanger Institute for the COVID-19 Genomics UK (COG-UK) | The Lighthouse Lab in Milton Keynes and Alex Alderton, Roberto Amato, Jeffrey Barrett, Sonia Goncalves, Ewan Harrison, David K. Jackson, Ian Johnston, Dominic Kwiatkowski, Cordelia Langford, John Sillitoe on behalf of the Wellcome Sanger |

| | | | |
|---|---|---|---|
| | | Consortium | Institute COVID-19 Surveillance Team |
| EPI_ISL_3095097 | Lighthouse Lab in Alderley Park | Wellcome Sanger Institute for the COVID-19 Genomics UK (COG-UK) Consortium | Jacquelyn Wynn, Mairead Hyland, The Lighthouse Lab in Alderley Park and Alex Alderton, Roberto Amato, Jeffrey Barrett, Sonia Goncalves, Ewan Harrison, David K. Jackson, Ian Johnston, Dominic Kwiatkowski, Cordelia Langford, John Sillitoe on behalf of the Wellcome Sanger Institute COVID-19 Surveillance Team |
| EPI_ISL_3095098, EPI_ISL_3095099 | Lighthouse Lab in Glasgow | Wellcome Sanger Institute for the COVID-19 Genomics UK (COG-UK) Consortium | Harper VanSteenhouse, Yumi Kasai, David Gray, Carol Clugston, Anna Dominiczak and Alex Alderton, Roberto Amato, Jeffrey Barrett, Sonia Goncalves, Ewan Harrison, David K. Jackson, Ian Johnston, Dominic Kwiatkowski, Cordelia Langford, John Sillitoe on behalf of the Wellcome Sanger Institute COVID-19 Surveillance Team |
| EPI_ISL_3095100 | Lighthouse Lab in Milton Keynes | Wellcome Sanger Institute for the COVID-19 Genomics UK (COG-UK) Consortium | The Lighthouse Lab in Milton Keynes and Alex Alderton, Roberto Amato, Jeffrey Barrett, Sonia Goncalves, Ewan Harrison, David K. Jackson, Ian Johnston, Dominic Kwiatkowski, Cordelia Langford, John Sillitoe on behalf of the Wellcome Sanger Institute COVID-19 Surveillance Team |
| EPI_ISL_3095101 | Lighthouse Lab in Glasgow | Wellcome Sanger Institute for the COVID-19 Genomics UK (COG-UK) Consortium | Harper VanSteenhouse, Yumi Kasai, David Gray, Carol Clugston, Anna Dominiczak and Alex Alderton, Roberto Amato, Jeffrey Barrett, Sonia Goncalves, Ewan Harrison, David K. Jackson, Ian Johnston, Dominic Kwiatkowski, Cordelia Langford, John Sillitoe on behalf of the Wellcome Sanger Institute COVID-19 Surveillance Team |
| EPI_ISL_3095102 | Lighthouse Lab in Milton Keynes | Wellcome Sanger Institute for the COVID-19 Genomics UK (COG-UK) Consortium | The Lighthouse Lab in Milton Keynes and Alex Alderton, Roberto Amato, Jeffrey Barrett, Sonia Goncalves, Ewan Harrison, David K. Jackson, Ian Johnston, Dominic Kwiatkowski, Cordelia Langford, John Sillitoe on behalf of the Wellcome Sanger Institute COVID-19 Surveillance Team |
| EPI_ISL_3095103 | Lighthouse Lab in Alderley Park | Wellcome Sanger Institute for the COVID-19 Genomics UK (COG-UK) Consortium | Jacquelyn Wynn, Mairead Hyland, The Lighthouse Lab in Alderley Park and Alex Alderton, Roberto Amato, Jeffrey Barrett, Sonia Goncalves, Ewan Harrison, David K. Jackson, Ian Johnston, Dominic Kwiatkowski, Cordelia Langford, John Sillitoe on behalf of the Wellcome Sanger Institute COVID-19 Surveillance Team |
| EPI_ISL_3095104 | Lighthouse Lab in Milton Keynes | Wellcome Sanger Institute for the COVID-19 Genomics UK (COG-UK) Consortium | The Lighthouse Lab in Milton Keynes and Alex Alderton, Roberto Amato, Jeffrey Barrett, Sonia Goncalves, Ewan Harrison, David K. Jackson, Ian Johnston, Dominic Kwiatkowski, Cordelia Langford, John Sillitoe on behalf of the Wellcome Sanger Institute COVID-19 Surveillance Team |
| EPI_ISL_3095105, EPI_ISL_3095106 | Lighthouse Lab in Glasgow | Wellcome Sanger Institute for the COVID-19 Genomics UK (COG-UK) Consortium | Harper VanSteenhouse, Yumi Kasai, David Gray, Carol Clugston, Anna Dominiczak and Alex Alderton, Roberto Amato, Jeffrey Barrett, Sonia Goncalves, Ewan Harrison, David K. Jackson, Ian Johnston, Dominic Kwiatkowski, Cordelia Langford, John Sillitoe on behalf of the Wellcome Sanger Institute COVID-19 Surveillance Team |
| EPI_ISL_3095107 | Lighthouse Lab in Milton Keynes | Wellcome Sanger Institute for the COVID-19 Genomics UK (COG-UK) Consortium | The Lighthouse Lab in Milton Keynes and Alex Alderton, Roberto Amato, Jeffrey Barrett, Sonia Goncalves, Ewan Harrison, David K. Jackson, Ian Johnston, Dominic Kwiatkowski, Cordelia Langford, John Sillitoe on behalf of the Wellcome Sanger Institute COVID-19 Surveillance Team |
| EPI_ISL_3095108 | Lighthouse Lab in Glasgow | Wellcome Sanger Institute for the COVID-19 Genomics UK (COG-UK) Consortium | Harper VanSteenhouse, Yumi Kasai, David Gray, Carol Clugston, Anna Dominiczak and Alex Alderton, Roberto Amato, Jeffrey Barrett, Sonia Goncalves, Ewan Harrison, David K. Jackson, Ian Johnston, Dominic Kwiatkowski, Cordelia Langford, John Sillitoe on behalf of the Wellcome Sanger Institute COVID-19 Surveillance Team |
| EPI_ISL_3095109 | Lighthouse Lab in Alderley Park | Wellcome Sanger Institute for the COVID-19 Genomics UK (COG-UK) Consortium | Jacquelyn Wynn, Mairead Hyland, The Lighthouse Lab in Alderley Park and Alex Alderton, Roberto Amato, Jeffrey Barrett, Sonia Goncalves, Ewan Harrison, David K. Jackson, Ian Johnston, Dominic Kwiatkowski, Cordelia Langford, John Sillitoe on behalf of the Wellcome Sanger Institute COVID-19 Surveillance Team |
| EPI_ISL_3095110 | Lighthouse Lab in Glasgow | Wellcome Sanger Institute for the COVID-19 Genomics UK (COG-UK) Consortium | Harper VanSteenhouse, Yumi Kasai, David Gray, Carol Clugston, Anna Dominiczak and Alex Alderton, Roberto Amato, Jeffrey Barrett, Sonia Goncalves, Ewan Harrison, David K. Jackson, Ian Johnston, Dominic Kwiatkowski, Cordelia Langford, John Sillitoe on behalf of the Wellcome Sanger Institute COVID-19 Surveillance Team |
| EPI_ISL_3095111, EPI_ISL_3095112, EPI_ISL_3095113 | Lighthouse Lab in Alderley Park | Wellcome Sanger Institute for the COVID-19 Genomics UK (COG-UK) Consortium | Jacquelyn Wynn, Mairead Hyland, The Lighthouse Lab in Alderley Park and Alex Alderton, Roberto Amato, Jeffrey Barrett, Sonia Goncalves, Ewan Harrison, David K. Jackson, Ian Johnston, Dominic Kwiatkowski, Cordelia Langford, John Sillitoe on behalf of the Wellcome Sanger Institute COVID-19 Surveillance Team |
| EPI_ISL_3095114 | Lighthouse Lab in Glasgow | Wellcome Sanger Institute for the COVID-19 Genomics UK (COG-UK) Consortium | Harper VanSteenhouse, Yumi Kasai, David Gray, Carol Clugston, Anna Dominiczak and Alex Alderton, Roberto Amato, Jeffrey Barrett, Sonia Goncalves, Ewan Harrison, David K. Jackson, Ian Johnston, Dominic Kwiatkowski, Cordelia Langford, John Sillitoe on behalf of the Wellcome Sanger Institute COVID-19 Surveillance Team |
| EPI_ISL_3095115 | Lighthouse Lab in Alderley Park | Wellcome Sanger Institute for the COVID-19 Genomics UK (COG-UK) Consortium | Jacquelyn Wynn, Mairead Hyland, The Lighthouse Lab in Alderley Park and Alex Alderton, Roberto Amato, Jeffrey Barrett, Sonia Goncalves, Ewan Harrison, David K. Jackson, Ian Johnston, Dominic Kwiatkowski, Cordelia Langford, John Sillitoe on behalf of the Wellcome Sanger Institute COVID-19 Surveillance Team |
| EPI_ISL_3095116, EPI_ISL_3095117 | Lighthouse Lab in Milton Keynes | Wellcome Sanger Institute for the COVID-19 Genomics UK (COG-UK) Consortium | The Lighthouse Lab in Milton Keynes and Alex Alderton, Roberto Amato, Jeffrey Barrett, Sonia Goncalves, Ewan Harrison, David K. Jackson, Ian Johnston, Dominic Kwiatkowski, Cordelia Langford, John Sillitoe on behalf of the Wellcome Sanger Institute COVID-19 Surveillance Team |
| EPI_ISL_3095118 | Lighthouse Lab in Glasgow | Wellcome Sanger Institute for the COVID-19 Genomics UK (COG-UK) Consortium | Harper VanSteenhouse, Yumi Kasai, David Gray, Carol Clugston, Anna Dominiczak and Alex Alderton, Roberto Amato, Jeffrey Barrett, Sonia Goncalves, Ewan Harrison, David K. Jackson, Ian Johnston, Dominic Kwiatkowski, Cordelia Langford, John Sillitoe on behalf of the Wellcome Sanger Institute COVID-19 Surveillance Team |
| EPI_ISL_3095119 | Lighthouse Lab in Milton Keynes | Wellcome Sanger Institute for the COVID-19 Genomics UK (COG-UK) Consortium | The Lighthouse Lab in Milton Keynes and Alex Alderton, Roberto Amato, Jeffrey Barrett, Sonia Goncalves, Ewan Harrison, David K. Jackson, Ian Johnston, Dominic Kwiatkowski, Cordelia Langford, John Sillitoe on behalf of the Wellcome Sanger Institute COVID-19 Surveillance Team |
| EPI_ISL_3095120, EPI_ISL_3095121, EPI_ISL_3095122, EPI_ISL_3095123 | Lighthouse Lab in Glasgow | Wellcome Sanger Institute for the COVID-19 Genomics UK (COG-UK) Consortium | Harper VanSteenhouse, Yumi Kasai, David Gray, Carol Clugston, Anna Dominiczak and Alex Alderton, Roberto Amato, Jeffrey Barrett, Sonia Goncalves, Ewan Harrison, David K. Jackson, Ian Johnston, Dominic Kwiatkowski, Cordelia Langford, John Sillitoe on behalf of the Wellcome Sanger Institute COVID-19 Surveillance Team |
| EPI_ISL_3095124 | Lighthouse Lab in Alderley Park | Wellcome Sanger Institute for the COVID-19 Genomics UK (COG-UK) Consortium | Jacquelyn Wynn, Mairead Hyland, The Lighthouse Lab in Alderley Park and Alex Alderton, Roberto Amato, Jeffrey Barrett, Sonia Goncalves, Ewan Harrison, David K. Jackson, Ian Johnston, Dominic Kwiatkowski, Cordelia Langford, John Sillitoe on behalf of the Wellcome Sanger Institute COVID-19 Surveillance Team |
| EPI_ISL_3095125 | Lighthouse Lab in Glasgow | Wellcome Sanger Institute for the COVID-19 Genomics UK (COG-UK) Consortium | Harper VanSteenhouse, Yumi Kasai, David Gray, Carol Clugston, Anna Dominiczak and Alex Alderton, Roberto Amato, Jeffrey Barrett, Sonia Goncalves, Ewan Harrison, David K. Jackson, Ian Johnston, Dominic Kwiatkowski, Cordelia Langford, John Sillitoe on behalf of the Wellcome Sanger Institute COVID-19 Surveillance Team |
| EPI_ISL_3095126, EPI_ISL_3095127, EPI_ISL_3095128, EPI_ISL_3095129, EPI_ISL_3095130 | Lighthouse Lab in Alderley Park | Wellcome Sanger Institute for the COVID-19 Genomics UK (COG-UK) Consortium | Jacquelyn Wynn, Mairead Hyland, The Lighthouse Lab in Alderley Park and Alex Alderton, Roberto Amato, Jeffrey Barrett, Sonia Goncalves, Ewan Harrison, David K. Jackson, Ian Johnston, Dominic Kwiatkowski, Cordelia Langford, John Sillitoe on behalf of the Wellcome Sanger Institute COVID-19 Surveillance Team |
| EPI_ISL_3095131 | Lighthouse Lab in Glasgow | Wellcome Sanger Institute for the COVID-19 Genomics UK (COG-UK) Consortium | Harper VanSteenhouse, Yumi Kasai, David Gray, Carol Clugston, Anna Dominiczak and Alex Alderton, Roberto Amato, Jeffrey Barrett, Sonia Goncalves, Ewan Harrison, David K. Jackson, Ian Johnston, Dominic Kwiatkowski, Cordelia Langford, John Sillitoe on behalf of the Wellcome Sanger Institute COVID-19 Surveillance Team |
| EPI_ISL_3095132, EPI_ISL_3095133, EPI_ISL_3095134 | Lighthouse Lab in Alderley Park | Wellcome Sanger Institute for the COVID-19 Genomics UK (COG-UK) Consortium | Jacquelyn Wynn, Mairead Hyland, The Lighthouse Lab in Alderley Park and Alex Alderton, Roberto Amato, Jeffrey Barrett, Sonia Goncalves, Ewan Harrison, David K. Jackson, Ian Johnston, Dominic Kwiatkowski, Cordelia Langford, John Sillitoe on behalf of the Wellcome Sanger Institute COVID-19 Surveillance Team |
| EPI_ISL_3095135, EPI_ISL_3095136 | Lighthouse Lab in Glasgow | Wellcome Sanger Institute for the COVID-19 Genomics UK (COG-UK) Consortium | Harper VanSteenhouse, Yumi Kasai, David Gray, Carol Clugston, Anna Dominiczak and Alex Alderton, Roberto Amato, Jeffrey Barrett, Sonia Goncalves, Ewan Harrison, David K. Jackson, Ian Johnston, Dominic Kwiatkowski, Cordelia Langford, John Sillitoe on behalf of the Wellcome Sanger Institute COVID-19 Surveillance Team |
| EPI_ISL_3095137, EPI_ISL_3095138, EPI_ISL_3095139, EPI_ISL_3095140 | Lighthouse Lab in Milton Keynes | Wellcome Sanger Institute for the COVID-19 Genomics UK (COG-UK) Consortium | The Lighthouse Lab in Milton Keynes and Alex Alderton, Roberto Amato, Jeffrey Barrett, Sonia Goncalves, Ewan Harrison, David K. Jackson, Ian Johnston, Dominic Kwiatkowski, Cordelia Langford, John Sillitoe on behalf of the Wellcome Sanger Institute COVID-19 Surveillance Team |
| EPI_ISL_3095141 | Lighthouse Lab in Glasgow | Wellcome Sanger Institute for the COVID-19 Genomics UK (COG-UK) Consortium | Harper VanSteenhouse, Yumi Kasai, David Gray, Carol Clugston, Anna Dominiczak and Alex Alderton, Roberto Amato, Jeffrey Barrett, Sonia Goncalves, Ewan Harrison, David K. Jackson, Ian Johnston, Dominic Kwiatkowski, Cordelia Langford, John Sillitoe on behalf of the Wellcome Sanger Institute COVID-19 Surveillance Team |
| EPI_ISL_3095142, EPI_ISL_3095143, EPI_ISL_3095144, EPI_ISL_3095145, EPI_ISL_3095146, EPI_ISL_3095147, EPI_ISL_3095148, EPI_ISL_3095149, EPI_ISL_3095150 | Lighthouse Lab in Milton Keynes | Wellcome Sanger Institute for the COVID-19 Genomics UK (COG-UK) Consortium | The Lighthouse Lab in Milton Keynes and Alex Alderton, Roberto Amato, Jeffrey Barrett, Sonia Goncalves, Ewan Harrison, David K. Jackson, Ian Johnston, Dominic Kwiatkowski, Cordelia Langford, John Sillitoe on behalf of the Wellcome Sanger Institute COVID-19 Surveillance Team |
| EPI_ISL_3095151 | Lighthouse Lab in Glasgow | Wellcome Sanger Institute for the COVID-19 Genomics UK (COG-UK) Consortium | Harper VanSteenhouse, Yumi Kasai, David Gray, Carol Clugston, Anna Dominiczak and Alex Alderton, Roberto Amato, Jeffrey Barrett, Sonia Goncalves, Ewan Harrison, David K. Jackson, Ian Johnston, Dominic Kwiatkowski, Cordelia Langford, John Sillitoe on behalf of the Wellcome Sanger Institute COVID-19 Surveillance Team |
| EPI_ISL_3095152, EPI_ISL_3095153, EPI_ISL_3095154, EPI_ISL_3095155, EPI_ISL_3095156, EPI_ISL_3095157 | Lighthouse Lab in Milton Keynes | Wellcome Sanger Institute for the COVID-19 Genomics UK (COG-UK) Consortium | The Lighthouse Lab in Milton Keynes and Alex Alderton, Roberto Amato, Jeffrey Barrett, Sonia Goncalves, Ewan Harrison, David K. Jackson, Ian Johnston, Dominic Kwiatkowski, Cordelia Langford, John Sillitoe on behalf of the Wellcome Sanger Institute COVID-19 Surveillance Team |
| EPI_ISL_3095158 | Lighthouse Lab in Glasgow | Wellcome Sanger Institute for the COVID-19 Genomics UK (COG-UK) Consortium | Harper VanSteenhouse, Yumi Kasai, David Gray, Carol Clugston, Anna Dominiczak and Alex Alderton, Roberto Amato, Jeffrey Barrett, Sonia Goncalves, Ewan Harrison, David K. Jackson, Ian Johnston, Dominic Kwiatkowski, Cordelia Langford, John Sillitoe on behalf of the Wellcome Sanger Institute COVID-19 Surveillance Team |
| EPI_ISL_3095159, EPI_ISL_3095160, EPI_ISL_3095161, EPI_ISL_3095162, EPI_ISL_3095163, EPI_ISL_3095164, EPI_ISL_3095165, EPI_ISL_3095166, EPI_ISL_3095167, EPI_ISL_3095168, EPI_ISL_3095169 | | | |
| see above | Lighthouse Lab in Milton Keynes | Wellcome Sanger Institute for the COVID-19 Genomics UK (COG-UK) Consortium | The Lighthouse Lab in Milton Keynes and Alex Alderton, Roberto Amato, Jeffrey Barrett, Sonia Goncalves, Ewan Harrison, David K. Jackson, Ian Johnston, Dominic Kwiatkowski, Cordelia Langford, John Sillitoe on behalf of the Wellcome Sanger Institute COVID-19 Surveillance Team |
| EPI_ISL_3095170, EPI_ISL_3095171 | Lighthouse Lab in Glasgow | Wellcome Sanger Institute for the COVID-19 Genomics UK (COG-UK) Consortium | Harper VanSteenhouse, Yumi Kasai, David Gray, Carol Clugston, Anna Dominiczak and Alex Alderton, Roberto Amato, Jeffrey Barrett, Sonia Goncalves, Ewan Harrison, David K. Jackson, Ian Johnston, Dominic Kwiatkowski, Cordelia Langford, John Sillitoe on behalf of the Wellcome Sanger Institute COVID-19 Surveillance Team |
| EPI_ISL_3095172 | Lighthouse Lab in Milton Keynes | Wellcome Sanger Institute for the COVID-19 Genomics UK (COG-UK) Consortium | The Lighthouse Lab in Milton Keynes and Alex Alderton, Roberto Amato, Jeffrey Barrett, Sonia Goncalves, Ewan Harrison, David K. Jackson, Ian Johnston, Dominic Kwiatkowski, Cordelia Langford, John Sillitoe on behalf of the Wellcome Sanger Institute COVID-19 Surveillance Team |
| EPI_ISL_3095173, EPI_ISL_3095174 | Lighthouse Lab in Glasgow | Wellcome Sanger Institute for the COVID-19 Genomics UK (COG-UK) Consortium | Harper VanSteenhouse, Yumi Kasai, David Gray, Carol Clugston, Anna Dominiczak and Alex Alderton, Roberto Amato, Jeffrey Barrett, Sonia Goncalves, Ewan Harrison, David K. Jackson, Ian Johnston, Dominic Kwiatkowski, Cordelia Langford, John Sillitoe on behalf of the Wellcome Sanger Institute COVID-19 Surveillance Team |
| EPI_ISL_3095175, EPI_ISL_3095176 | Lighthouse Lab in Milton Keynes | Wellcome Sanger Institute for the COVID-19 Genomics UK (COG-UK) Consortium | The Lighthouse Lab in Milton Keynes and Alex Alderton, Roberto Amato, Jeffrey Barrett, Sonia Goncalves, Ewan Harrison, David K. Jackson, Ian Johnston, Dominic Kwiatkowski, Cordelia Langford, John Sillitoe on behalf of the Wellcome Sanger Institute COVID-19 Surveillance Team |

| | | | |
|---|---|---|---|
| EPI_ISL_3095177 | Lighthouse Lab in Glasgow | Wellcome Sanger Institute for the COVID-19 Genomics UK (COG-UK) Consortium | Harper VanSteenhouse, Yumi Kasai, David Gray, Carol Clugston, Anna Dominiczak and Alex Alderton, Roberto Amato, Jeffrey Barrett, Sonia Goncalves, Ewan Harrison, David K. Jackson, Ian Johnston, Dominic Kwiatkowski, Cordelia Langford, John Sillitoe on behalf of the Wellcome Sanger Institute COVID-19 Surveillance Team |
| EPI_ISL_3095178, EPI_ISL_3095179, EPI_ISL_3095180, EPI_ISL_3095181, EPI_ISL_3095182, EPI_ISL_3095183, EPI_ISL_3095184 | Lighthouse Lab in Milton Keynes | Wellcome Sanger Institute for the COVID-19 Genomics UK (COG-UK) Consortium | The Lighthouse Lab in Milton Keynes and Alex Alderton, Roberto Amato, Jeffrey Barrett, Sonia Goncalves, Ewan Harrison, David K. Jackson, Ian Johnston, Dominic Kwiatkowski, Cordelia Langford, John Sillitoe on behalf of the Wellcome Sanger Institute COVID-19 Surveillance Team |
| EPI_ISL_3095185 | Lighthouse Lab in Glasgow | Wellcome Sanger Institute for the COVID-19 Genomics UK (COG-UK) Consortium | Harper VanSteenhouse, Yumi Kasai, David Gray, Carol Clugston, Anna Dominiczak and Alex Alderton, Roberto Amato, Jeffrey Barrett, Sonia Goncalves, Ewan Harrison, David K. Jackson, Ian Johnston, Dominic Kwiatkowski, Cordelia Langford, John Sillitoe on behalf of the Wellcome Sanger Institute COVID-19 Surveillance Team |
| EPI_ISL_3095186, EPI_ISL_3095187, EPI_ISL_3095188 | Lighthouse Lab in Milton Keynes | Wellcome Sanger Institute for the COVID-19 Genomics UK (COG-UK) Consortium | The Lighthouse Lab in Milton Keynes and Alex Alderton, Roberto Amato, Jeffrey Barrett, Sonia Goncalves, Ewan Harrison, David K. Jackson, Ian Johnston, Dominic Kwiatkowski, Cordelia Langford, John Sillitoe on behalf of the Wellcome Sanger Institute COVID-19 Surveillance Team |
| EPI_ISL_3095189 | Lighthouse Lab in Glasgow | Wellcome Sanger Institute for the COVID-19 Genomics UK (COG-UK) Consortium | Harper VanSteenhouse, Yumi Kasai, David Gray, Carol Clugston, Anna Dominiczak and Alex Alderton, Roberto Amato, Jeffrey Barrett, Sonia Goncalves, Ewan Harrison, David K. Jackson, Ian Johnston, Dominic Kwiatkowski, Cordelia Langford, John Sillitoe on behalf of the Wellcome Sanger Institute COVID-19 Surveillance Team |
| EPI_ISL_3095190, EPI_ISL_3095191 | Lighthouse Lab in Milton Keynes | Wellcome Sanger Institute for the COVID-19 Genomics UK (COG-UK) Consortium | The Lighthouse Lab in Milton Keynes and Alex Alderton, Roberto Amato, Jeffrey Barrett, Sonia Goncalves, Ewan Harrison, David K. Jackson, Ian Johnston, Dominic Kwiatkowski, Cordelia Langford, John Sillitoe on behalf of the Wellcome Sanger Institute COVID-19 Surveillance Team |
| EPI_ISL_3095192 | Lighthouse Lab in Glasgow | Wellcome Sanger Institute for the COVID-19 Genomics UK (COG-UK) Consortium | Harper VanSteenhouse, Yumi Kasai, David Gray, Carol Clugston, Anna Dominiczak and Alex Alderton, Roberto Amato, Jeffrey Barrett, Sonia Goncalves, Ewan Harrison, David K. Jackson, Ian Johnston, Dominic Kwiatkowski, Cordelia Langford, John Sillitoe on behalf of the Wellcome Sanger Institute COVID-19 Surveillance Team |
| EPI_ISL_3095193 | Hospital del Oriente de Asturias | Laboratorio de Virología HUCA | Sandoval M, Castelló C, Gómez de Oña J, Boga JA, Rojo S, Alvarez-Arguelles ME, Abreu F, Costales I, Perez-Martínez Z, Martín-Rodríguez G, Coto E, Melón S |
| EPI_ISL_3095194, EPI_ISL_3095195 | Lighthouse Lab in Milton Keynes | Wellcome Sanger Institute for the COVID-19 Genomics UK (COG-UK) Consortium | The Lighthouse Lab in Milton Keynes and Alex Alderton, Roberto Amato, Jeffrey Barrett, Sonia Goncalves, Ewan Harrison, David K. Jackson, Ian Johnston, Dominic Kwiatkowski, Cordelia Langford, John Sillitoe on behalf of the Wellcome Sanger Institute COVID-19 Surveillance Team |
| EPI_ISL_3095196 | Lighthouse Lab in Glasgow | Wellcome Sanger Institute for the COVID-19 Genomics UK (COG-UK) Consortium | Harper VanSteenhouse, Yumi Kasai, David Gray, Carol Clugston, Anna Dominiczak and Alex Alderton, Roberto Amato, Jeffrey Barrett, Sonia Goncalves, Ewan Harrison, David K. Jackson, Ian Johnston, Dominic Kwiatkowski, Cordelia Langford, John Sillitoe on behalf of the Wellcome Sanger Institute COVID-19 Surveillance Team |
| EPI_ISL_3095197, EPI_ISL_3095198, EPI_ISL_3095199, EPI_ISL_3095200, EPI_ISL_3095201 | Lighthouse Lab in Milton Keynes | Wellcome Sanger Institute for the COVID-19 Genomics UK (COG-UK) Consortium | The Lighthouse Lab in Milton Keynes and Alex Alderton, Roberto Amato, Jeffrey Barrett, Sonia Goncalves, Ewan Harrison, David K. Jackson, Ian Johnston, Dominic Kwiatkowski, Cordelia Langford, John Sillitoe on behalf of the Wellcome Sanger Institute COVID-19 Surveillance Team |
| EPI_ISL_3095202 | Lighthouse Lab in Glasgow | Wellcome Sanger Institute for the COVID-19 Genomics UK (COG-UK) Consortium | Harper VanSteenhouse, Yumi Kasai, David Gray, Carol Clugston, Anna Dominiczak and Alex Alderton, Roberto Amato, Jeffrey Barrett, Sonia Goncalves, Ewan Harrison, David K. Jackson, Ian Johnston, Dominic Kwiatkowski, Cordelia Langford, John Sillitoe on behalf of the Wellcome Sanger Institute COVID-19 Surveillance Team |

EPI_ISL_3095203, EPI_ISL_3095204, EPI_ISL_3095205, EPI_ISL_3095206, EPI_ISL_3095207, EPI_ISL_3095208, EPI_ISL_3095209, EPI_ISL_3095210, EPI_ISL_3095211, EPI_ISL_3095212, EPI_ISL_3095214, EPI_ISL_3095215, EPI_ISL_3095216, EPI_ISL_3095217, EPI_ISL_3095218, EPI_ISL_3095219, EPI_ISL_3095220, EPI_ISL_3095221, EPI_ISL_3095222, EPI_ISL_3095223, EPI_ISL_3095224, EPI_ISL_3095225, EPI_ISL_3095226

| | | | |
|---|---|---|---|
| see above | Lighthouse Lab in Milton Keynes | Wellcome Sanger Institute for the COVID-19 Genomics UK (COG-UK) Consortium | The Lighthouse Lab in Milton Keynes and Alex Alderton, Roberto Amato, Jeffrey Barrett, Sonia Goncalves, Ewan Harrison, David K. Jackson, Ian Johnston, Dominic Kwiatkowski, Cordelia Langford, John Sillitoe on behalf of the Wellcome Sanger Institute COVID-19 Surveillance Team |
| EPI_ISL_3095227, EPI_ISL_3095228 | Lighthouse Lab in Glasgow | Wellcome Sanger Institute for the COVID-19 Genomics UK (COG-UK) Consortium | Harper VanSteenhouse, Yumi Kasai, David Gray, Carol Clugston, Anna Dominiczak and Alex Alderton, Roberto Amato, Jeffrey Barrett, Sonia Goncalves, Ewan Harrison, David K. Jackson, Ian Johnston, Dominic Kwiatkowski, Cordelia Langford, John Sillitoe on behalf of the Wellcome Sanger Institute COVID-19 Surveillance Team |
| EPI_ISL_3095229, EPI_ISL_3095230, EPI_ISL_3095231, EPI_ISL_3095232 | Lighthouse Lab in Milton Keynes | Wellcome Sanger Institute for the COVID-19 Genomics UK (COG-UK) Consortium | The Lighthouse Lab in Milton Keynes and Alex Alderton, Roberto Amato, Jeffrey Barrett, Sonia Goncalves, Ewan Harrison, David K. Jackson, Ian Johnston, Dominic Kwiatkowski, Cordelia Langford, John Sillitoe on behalf of the Wellcome Sanger Institute COVID-19 Surveillance Team |
| EPI_ISL_3095233 | Lighthouse Lab in Glasgow | Wellcome Sanger Institute for the COVID-19 Genomics UK (COG-UK) Consortium | Harper VanSteenhouse, Yumi Kasai, David Gray, Carol Clugston, Anna Dominiczak and Alex Alderton, Roberto Amato, Jeffrey Barrett, Sonia Goncalves, Ewan Harrison, David K. Jackson, Ian Johnston, Dominic Kwiatkowski, Cordelia Langford, John Sillitoe on behalf of the Wellcome Sanger Institute COVID-19 Surveillance Team |
| EPI_ISL_3095234 | Lighthouse Lab in Milton Keynes | Wellcome Sanger Institute for the COVID-19 Genomics UK (COG-UK) Consortium | The Lighthouse Lab in Milton Keynes and Alex Alderton, Roberto Amato, Jeffrey Barrett, Sonia Goncalves, Ewan Harrison, David K. Jackson, Ian Johnston, Dominic Kwiatkowski, Cordelia Langford, John Sillitoe on behalf of the Wellcome Sanger Institute COVID-19 Surveillance Team |
| EPI_ISL_3095235 | Lighthouse Lab in Glasgow | Wellcome Sanger Institute for the COVID-19 Genomics UK (COG-UK) Consortium | Harper VanSteenhouse, Yumi Kasai, David Gray, Carol Clugston, Anna Dominiczak and Alex Alderton, Roberto Amato, Jeffrey Barrett, Sonia Goncalves, Ewan Harrison, David K. Jackson, Ian Johnston, Dominic Kwiatkowski, Cordelia Langford, John Sillitoe on behalf of the Wellcome Sanger Institute COVID-19 Surveillance Team |
| EPI_ISL_3095236, EPI_ISL_3095237, EPI_ISL_3095238, EPI_ISL_3095239 | Lighthouse Lab in Milton Keynes | Wellcome Sanger Institute for the COVID-19 Genomics UK (COG-UK) Consortium | The Lighthouse Lab in Milton Keynes and Alex Alderton, Roberto Amato, Jeffrey Barrett, Sonia Goncalves, Ewan Harrison, David K. Jackson, Ian Johnston, Dominic Kwiatkowski, Cordelia Langford, John Sillitoe on behalf of the Wellcome Sanger Institute COVID-19 Surveillance Team |
| EPI_ISL_3095240 | Lighthouse Lab in Glasgow | Wellcome Sanger Institute for the COVID-19 Genomics UK (COG-UK) Consortium | Harper VanSteenhouse, Yumi Kasai, David Gray, Carol Clugston, Anna Dominiczak and Alex Alderton, Roberto Amato, Jeffrey Barrett, Sonia Goncalves, Ewan Harrison, David K. Jackson, Ian Johnston, Dominic Kwiatkowski, Cordelia Langford, John Sillitoe on behalf of the Wellcome Sanger Institute COVID-19 Surveillance Team |
| EPI_ISL_3095241, EPI_ISL_3095242, EPI_ISL_3095243, EPI_ISL_3095244, EPI_ISL_3095245, EPI_ISL_3095246, EPI_ISL_3095247, EPI_ISL_3095248, EPI_ISL_3095249, EPI_ISL_3095250 | Lighthouse Lab in Milton Keynes | Wellcome Sanger Institute for the COVID-19 Genomics UK (COG-UK) Consortium | The Lighthouse Lab in Milton Keynes and Alex Alderton, Roberto Amato, Jeffrey Barrett, Sonia Goncalves, Ewan Harrison, David K. Jackson, Ian Johnston, Dominic Kwiatkowski, Cordelia Langford, John Sillitoe on behalf of the Wellcome Sanger Institute COVID-19 Surveillance Team |
| EPI_ISL_3095251 | Lighthouse Lab in Glasgow | Wellcome Sanger Institute for the COVID-19 Genomics UK (COG-UK) Consortium | Harper VanSteenhouse, Yumi Kasai, David Gray, Carol Clugston, Anna Dominiczak and Alex Alderton, Roberto Amato, Jeffrey Barrett, Sonia Goncalves, Ewan Harrison, David K. Jackson, Ian Johnston, Dominic Kwiatkowski, Cordelia Langford, John Sillitoe on behalf of the Wellcome Sanger Institute COVID-19 Surveillance Team |
| EPI_ISL_3095252, EPI_ISL_3095253 | Lighthouse Lab in Milton Keynes | Wellcome Sanger Institute for the COVID-19 Genomics UK (COG-UK) Consortium | The Lighthouse Lab in Milton Keynes and Alex Alderton, Roberto Amato, Jeffrey Barrett, Sonia Goncalves, Ewan Harrison, David K. Jackson, Ian Johnston, Dominic Kwiatkowski, Cordelia Langford, John Sillitoe on behalf of the Wellcome Sanger Institute COVID-19 Surveillance Team |
| EPI_ISL_3095254 | Lighthouse Lab in Glasgow | Wellcome Sanger Institute for the COVID-19 Genomics UK (COG-UK) Consortium | Harper VanSteenhouse, Yumi Kasai, David Gray, Carol Clugston, Anna Dominiczak and Alex Alderton, Roberto Amato, Jeffrey Barrett, Sonia Goncalves, Ewan Harrison, David K. Jackson, Ian Johnston, Dominic Kwiatkowski, Cordelia Langford, John Sillitoe on behalf of the Wellcome Sanger Institute COVID-19 Surveillance Team |
| EPI_ISL_3095255, EPI_ISL_3095256, EPI_ISL_3095257, EPI_ISL_3095258, EPI_ISL_3095259, EPI_ISL_3095260, EPI_ISL_3095261, EPI_ISL_3095262, EPI_ISL_3095263, EPI_ISL_3095264 | Lighthouse Lab in Milton Keynes | Wellcome Sanger Institute for the COVID-19 Genomics UK (COG-UK) Consortium | The Lighthouse Lab in Milton Keynes and Alex Alderton, Roberto Amato, Jeffrey Barrett, Sonia Goncalves, Ewan Harrison, David K. Jackson, Ian Johnston, Dominic Kwiatkowski, Cordelia Langford, John Sillitoe on behalf of the Wellcome Sanger Institute COVID-19 Surveillance Team |
| EPI_ISL_3095265, EPI_ISL_3095266 | Lighthouse Lab in Glasgow | Wellcome Sanger Institute for the COVID-19 Genomics UK (COG-UK) Consortium | Harper VanSteenhouse, Yumi Kasai, David Gray, Carol Clugston, Anna Dominiczak and Alex Alderton, Roberto Amato, Jeffrey Barrett, Sonia Goncalves, Ewan Harrison, David K. Jackson, Ian Johnston, Dominic Kwiatkowski, Cordelia Langford, John Sillitoe on behalf of the Wellcome Sanger Institute COVID-19 Surveillance Team |
| EPI_ISL_3095267, EPI_ISL_3095268 | Lighthouse Lab in Milton Keynes | Wellcome Sanger Institute for the COVID-19 Genomics UK (COG-UK) Consortium | The Lighthouse Lab in Milton Keynes and Alex Alderton, Roberto Amato, Jeffrey Barrett, Sonia Goncalves, Ewan Harrison, David K. Jackson, Ian Johnston, Dominic Kwiatkowski, Cordelia Langford, John Sillitoe on behalf of the Wellcome Sanger Institute COVID-19 Surveillance Team |
| EPI_ISL_3095269 | Lighthouse Lab in Glasgow | Wellcome Sanger Institute for the COVID-19 Genomics UK (COG-UK) Consortium | Harper VanSteenhouse, Yumi Kasai, David Gray, Carol Clugston, Anna Dominiczak and Alex Alderton, Roberto Amato, Jeffrey Barrett, Sonia Goncalves, Ewan Harrison, David K. Jackson, Ian Johnston, Dominic Kwiatkowski, Cordelia Langford, John Sillitoe on behalf of the Wellcome Sanger Institute COVID-19 Surveillance Team |
| EPI_ISL_3095270 | Lighthouse Lab in Milton Keynes | Wellcome Sanger Institute for the COVID-19 Genomics UK (COG-UK) Consortium | The Lighthouse Lab in Milton Keynes and Alex Alderton, Roberto Amato, Jeffrey Barrett, Sonia Goncalves, Ewan Harrison, David K. Jackson, Ian Johnston, Dominic Kwiatkowski, Cordelia Langford, John Sillitoe on behalf of the Wellcome Sanger Institute COVID-19 Surveillance Team |
| EPI_ISL_3095271 | Lighthouse Lab in Glasgow | Wellcome Sanger Institute for the COVID-19 Genomics UK (COG-UK) Consortium | Harper VanSteenhouse, Yumi Kasai, David Gray, Carol Clugston, Anna Dominiczak and Alex Alderton, Roberto Amato, Jeffrey Barrett, Sonia Goncalves, Ewan Harrison, David K. Jackson, Ian Johnston, Dominic Kwiatkowski, Cordelia Langford, John Sillitoe on behalf of the Wellcome Sanger Institute COVID-19 Surveillance Team |
| EPI_ISL_3095272 | Lighthouse Lab in Milton Keynes | Wellcome Sanger Institute for the COVID-19 Genomics UK (COG-UK) Consortium | The Lighthouse Lab in Milton Keynes and Alex Alderton, Roberto Amato, Jeffrey Barrett, Sonia Goncalves, Ewan Harrison, David K. Jackson, Ian Johnston, Dominic Kwiatkowski, Cordelia Langford, John Sillitoe on behalf of the Wellcome Sanger Institute COVID-19 Surveillance Team |
| EPI_ISL_3095273 | Lighthouse Lab in Glasgow | Wellcome Sanger Institute for the COVID-19 Genomics UK (COG-UK) Consortium | Harper VanSteenhouse, Yumi Kasai, David Gray, Carol Clugston, Anna Dominiczak and Alex Alderton, Roberto Amato, Jeffrey Barrett, Sonia Goncalves, Ewan Harrison, David K. Jackson, Ian Johnston, Dominic Kwiatkowski, Cordelia Langford, John Sillitoe on behalf of the Wellcome Sanger Institute COVID-19 Surveillance Team |
| EPI_ISL_3095274, | Lighthouse Lab in Milton Keynes | Wellcome Sanger Institute for the COVID-19 Genomics UK (COG-UK) | The Lighthouse Lab in Milton Keynes and Alex Alderton, Roberto Amato, Jeffrey Barrett, Sonia Goncalves, Ewan Harrison, David K. Jackson, Ian Johnston, Dominic Kwiatkowski, Cordelia Langford, John Sillitoe on behalf of the Wellcome Sanger |

| | | | |
|---|---|---|---|
| EPI_ISL_3095275, EPI_ISL_3095276, EPI_ISL_3095277 | | Consortium | Institute COVID-19 Surveillance Team |
| EPI_ISL_3095278, EPI_ISL_3095279 | Lighthouse Lab in Glasgow | Wellcome Sanger Institute for the COVID-19 Genomics UK (COG-UK) Consortium | Harper VanSteenhouse, Yumi Kasai, David Gray, Carol Clugston, Anna Dominiczak and Alex Alderton, Roberto Amato, Jeffrey Barrett, Sonia Goncalves, Ewan Harrison, David K. Jackson, Ian Johnston, Dominic Kwiatkowski, Cordelia Langford, John Sillitoe on behalf of the Wellcome Sanger Institute COVID-19 Surveillance Team |
| EPI_ISL_3095280, EPI_ISL_3095281 | Lighthouse Lab in Milton Keynes | Wellcome Sanger Institute for the COVID-19 Genomics UK (COG-UK) Consortium | The Lighthouse Lab in Milton Keynes and Alex Alderton, Roberto Amato, Jeffrey Barrett, Sonia Goncalves, Ewan Harrison, David K. Jackson, Ian Johnston, Dominic Kwiatkowski, Cordelia Langford, John Sillitoe on behalf of the Wellcome Sanger Institute COVID-19 Surveillance Team |
| EPI_ISL_3095282 | Lighthouse Lab in Glasgow | Wellcome Sanger Institute for the COVID-19 Genomics UK (COG-UK) Consortium | Harper VanSteenhouse, Yumi Kasai, David Gray, Carol Clugston, Anna Dominiczak and Alex Alderton, Roberto Amato, Jeffrey Barrett, Sonia Goncalves, Ewan Harrison, David K. Jackson, Ian Johnston, Dominic Kwiatkowski, Cordelia Langford, John Sillitoe on behalf of the Wellcome Sanger Institute COVID-19 Surveillance Team |
| EPI_ISL_3095283 | Lighthouse Lab in Milton Keynes | Wellcome Sanger Institute for the COVID-19 Genomics UK (COG-UK) Consortium | The Lighthouse Lab in Milton Keynes and Alex Alderton, Roberto Amato, Jeffrey Barrett, Sonia Goncalves, Ewan Harrison, David K. Jackson, Ian Johnston, Dominic Kwiatkowski, Cordelia Langford, John Sillitoe on behalf of the Wellcome Sanger Institute COVID-19 Surveillance Team |
| EPI_ISL_3095284 | Lighthouse Lab in Glasgow | Wellcome Sanger Institute for the COVID-19 Genomics UK (COG-UK) Consortium | Harper VanSteenhouse, Yumi Kasai, David Gray, Carol Clugston, Anna Dominiczak and Alex Alderton, Roberto Amato, Jeffrey Barrett, Sonia Goncalves, Ewan Harrison, David K. Jackson, Ian Johnston, Dominic Kwiatkowski, Cordelia Langford, John Sillitoe on behalf of the Wellcome Sanger Institute COVID-19 Surveillance Team |
| EPI_ISL_3095285 | Lighthouse Lab in Milton Keynes | Wellcome Sanger Institute for the COVID-19 Genomics UK (COG-UK) Consortium | The Lighthouse Lab in Milton Keynes and Alex Alderton, Roberto Amato, Jeffrey Barrett, Sonia Goncalves, Ewan Harrison, David K. Jackson, Ian Johnston, Dominic Kwiatkowski, Cordelia Langford, John Sillitoe on behalf of the Wellcome Sanger Institute COVID-19 Surveillance Team |
| EPI_ISL_3095286 | Lighthouse Lab in Glasgow | Wellcome Sanger Institute for the COVID-19 Genomics UK (COG-UK) Consortium | Harper VanSteenhouse, Yumi Kasai, David Gray, Carol Clugston, Anna Dominiczak and Alex Alderton, Roberto Amato, Jeffrey Barrett, Sonia Goncalves, Ewan Harrison, David K. Jackson, Ian Johnston, Dominic Kwiatkowski, Cordelia Langford, John Sillitoe on behalf of the Wellcome Sanger Institute COVID-19 Surveillance Team |
| EPI_ISL_3095287, EPI_ISL_3095288, EPI_ISL_3095289, EPI_ISL_3095290 | Lighthouse Lab in Milton Keynes | Wellcome Sanger Institute for the COVID-19 Genomics UK (COG-UK) Consortium | The Lighthouse Lab in Milton Keynes and Alex Alderton, Roberto Amato, Jeffrey Barrett, Sonia Goncalves, Ewan Harrison, David K. Jackson, Ian Johnston, Dominic Kwiatkowski, Cordelia Langford, John Sillitoe on behalf of the Wellcome Sanger Institute COVID-19 Surveillance Team |
| EPI_ISL_3095291 | Lighthouse Lab in Glasgow | Wellcome Sanger Institute for the COVID-19 Genomics UK (COG-UK) Consortium | Harper VanSteenhouse, Yumi Kasai, David Gray, Carol Clugston, Anna Dominiczak and Alex Alderton, Roberto Amato, Jeffrey Barrett, Sonia Goncalves, Ewan Harrison, David K. Jackson, Ian Johnston, Dominic Kwiatkowski, Cordelia Langford, John Sillitoe on behalf of the Wellcome Sanger Institute COVID-19 Surveillance Team |
| EPI_ISL_3095292, EPI_ISL_3095293, EPI_ISL_3095294 | Lighthouse Lab in Milton Keynes | Wellcome Sanger Institute for the COVID-19 Genomics UK (COG-UK) Consortium | The Lighthouse Lab in Milton Keynes and Alex Alderton, Roberto Amato, Jeffrey Barrett, Sonia Goncalves, Ewan Harrison, David K. Jackson, Ian Johnston, Dominic Kwiatkowski, Cordelia Langford, John Sillitoe on behalf of the Wellcome Sanger Institute COVID-19 Surveillance Team |
| EPI_ISL_3095295 | Lighthouse Lab in Glasgow | Wellcome Sanger Institute for the COVID-19 Genomics UK (COG-UK) Consortium | Harper VanSteenhouse, Yumi Kasai, David Gray, Carol Clugston, Anna Dominiczak and Alex Alderton, Roberto Amato, Jeffrey Barrett, Sonia Goncalves, Ewan Harrison, David K. Jackson, Ian Johnston, Dominic Kwiatkowski, Cordelia Langford, John Sillitoe on behalf of the Wellcome Sanger Institute COVID-19 Surveillance Team |
| EPI_ISL_3095296, EPI_ISL_3095297, EPI_ISL_3095298, EPI_ISL_3095299, EPI_ISL_3095300, EPI_ISL_3095301 | Lighthouse Lab in Milton Keynes | Wellcome Sanger Institute for the COVID-19 Genomics UK (COG-UK) Consortium | The Lighthouse Lab in Milton Keynes and Alex Alderton, Roberto Amato, Jeffrey Barrett, Sonia Goncalves, Ewan Harrison, David K. Jackson, Ian Johnston, Dominic Kwiatkowski, Cordelia Langford, John Sillitoe on behalf of the Wellcome Sanger Institute COVID-19 Surveillance Team |
| EPI_ISL_3095302 | Lighthouse Lab in Glasgow | Wellcome Sanger Institute for the COVID-19 Genomics UK (COG-UK) Consortium | Harper VanSteenhouse, Yumi Kasai, David Gray, Carol Clugston, Anna Dominiczak and Alex Alderton, Roberto Amato, Jeffrey Barrett, Sonia Goncalves, Ewan Harrison, David K. Jackson, Ian Johnston, Dominic Kwiatkowski, Cordelia Langford, John Sillitoe on behalf of the Wellcome Sanger Institute COVID-19 Surveillance Team |
| EPI_ISL_3095303 | Lighthouse Lab in Milton Keynes | Wellcome Sanger Institute for the COVID-19 Genomics UK (COG-UK) Consortium | The Lighthouse Lab in Milton Keynes and Alex Alderton, Roberto Amato, Jeffrey Barrett, Sonia Goncalves, Ewan Harrison, David K. Jackson, Ian Johnston, Dominic Kwiatkowski, Cordelia Langford, John Sillitoe on behalf of the Wellcome Sanger Institute COVID-19 Surveillance Team |
| EPI_ISL_3095304, EPI_ISL_3095305 | Lighthouse Lab in Glasgow | Wellcome Sanger Institute for the COVID-19 Genomics UK (COG-UK) Consortium | Harper VanSteenhouse, Yumi Kasai, David Gray, Carol Clugston, Anna Dominiczak and Alex Alderton, Roberto Amato, Jeffrey Barrett, Sonia Goncalves, Ewan Harrison, David K. Jackson, Ian Johnston, Dominic Kwiatkowski, Cordelia Langford, John Sillitoe on behalf of the Wellcome Sanger Institute COVID-19 Surveillance Team |
| EPI_ISL_3095306 | Lighthouse Lab in Milton Keynes | Wellcome Sanger Institute for the COVID-19 Genomics UK (COG-UK) Consortium | The Lighthouse Lab in Milton Keynes and Alex Alderton, Roberto Amato, Jeffrey Barrett, Sonia Goncalves, Ewan Harrison, David K. Jackson, Ian Johnston, Dominic Kwiatkowski, Cordelia Langford, John Sillitoe on behalf of the Wellcome Sanger Institute COVID-19 Surveillance Team |
| EPI_ISL_3095307 | Lighthouse Lab in Glasgow | Wellcome Sanger Institute for the COVID-19 Genomics UK (COG-UK) Consortium | Harper VanSteenhouse, Yumi Kasai, David Gray, Carol Clugston, Anna Dominiczak and Alex Alderton, Roberto Amato, Jeffrey Barrett, Sonia Goncalves, Ewan Harrison, David K. Jackson, Ian Johnston, Dominic Kwiatkowski, Cordelia Langford, John Sillitoe on behalf of the Wellcome Sanger Institute COVID-19 Surveillance Team |
| EPI_ISL_3095308 | Lighthouse Lab in Milton Keynes | Wellcome Sanger Institute for the COVID-19 Genomics UK (COG-UK) Consortium | The Lighthouse Lab in Milton Keynes and Alex Alderton, Roberto Amato, Jeffrey Barrett, Sonia Goncalves, Ewan Harrison, David K. Jackson, Ian Johnston, Dominic Kwiatkowski, Cordelia Langford, John Sillitoe on behalf of the Wellcome Sanger Institute COVID-19 Surveillance Team |
| EPI_ISL_3095309 | Lighthouse Lab in Glasgow | Wellcome Sanger Institute for the COVID-19 Genomics UK (COG-UK) Consortium | Harper VanSteenhouse, Yumi Kasai, David Gray, Carol Clugston, Anna Dominiczak and Alex Alderton, Roberto Amato, Jeffrey Barrett, Sonia Goncalves, Ewan Harrison, David K. Jackson, Ian Johnston, Dominic Kwiatkowski, Cordelia Langford, John Sillitoe on behalf of the Wellcome Sanger Institute COVID-19 Surveillance Team |
| EPI_ISL_3095310, EPI_ISL_3095311, EPI_ISL_3095312, EPI_ISL_3095313 | Lighthouse Lab in Milton Keynes | Wellcome Sanger Institute for the COVID-19 Genomics UK (COG-UK) Consortium | The Lighthouse Lab in Milton Keynes and Alex Alderton, Roberto Amato, Jeffrey Barrett, Sonia Goncalves, Ewan Harrison, David K. Jackson, Ian Johnston, Dominic Kwiatkowski, Cordelia Langford, John Sillitoe on behalf of the Wellcome Sanger Institute COVID-19 Surveillance Team |
| EPI_ISL_3095314 | Lighthouse Lab in Glasgow | Wellcome Sanger Institute for the COVID-19 Genomics UK (COG-UK) Consortium | Harper VanSteenhouse, Yumi Kasai, David Gray, Carol Clugston, Anna Dominiczak and Alex Alderton, Roberto Amato, Jeffrey Barrett, Sonia Goncalves, Ewan Harrison, David K. Jackson, Ian Johnston, Dominic Kwiatkowski, Cordelia Langford, John Sillitoe on behalf of the Wellcome Sanger Institute COVID-19 Surveillance Team |
| EPI_ISL_3095315, EPI_ISL_3095316, EPI_ISL_3095317, EPI_ISL_3095318, EPI_ISL_3095319, EPI_ISL_3095320, EPI_ISL_3095321, EPI_ISL_3095322, EPI_ISL_3095323 | Lighthouse Lab in Milton Keynes | Wellcome Sanger Institute for the COVID-19 Genomics UK (COG-UK) Consortium | The Lighthouse Lab in Milton Keynes and Alex Alderton, Roberto Amato, Jeffrey Barrett, Sonia Goncalves, Ewan Harrison, David K. Jackson, Ian Johnston, Dominic Kwiatkowski, Cordelia Langford, John Sillitoe on behalf of the Wellcome Sanger Institute COVID-19 Surveillance Team |
| EPI_ISL_3095324 | Lighthouse Lab in Glasgow | Wellcome Sanger Institute for the COVID-19 Genomics UK (COG-UK) Consortium | Harper VanSteenhouse, Yumi Kasai, David Gray, Carol Clugston, Anna Dominiczak and Alex Alderton, Roberto Amato, Jeffrey Barrett, Sonia Goncalves, Ewan Harrison, David K. Jackson, Ian Johnston, Dominic Kwiatkowski, Cordelia Langford, John Sillitoe on behalf of the Wellcome Sanger Institute COVID-19 Surveillance Team |
| EPI_ISL_3095325, EPI_ISL_3095326, EPI_ISL_3095327, EPI_ISL_3095328 | Lighthouse Lab in Milton Keynes | Wellcome Sanger Institute for the COVID-19 Genomics UK (COG-UK) Consortium | The Lighthouse Lab in Milton Keynes and Alex Alderton, Roberto Amato, Jeffrey Barrett, Sonia Goncalves, Ewan Harrison, David K. Jackson, Ian Johnston, Dominic Kwiatkowski, Cordelia Langford, John Sillitoe on behalf of the Wellcome Sanger Institute COVID-19 Surveillance Team |
| EPI_ISL_3095329, EPI_ISL_3095330, EPI_ISL_3095331, EPI_ISL_3095332 | Lighthouse Lab in Glasgow | Wellcome Sanger Institute for the COVID-19 Genomics UK (COG-UK) Consortium | Harper VanSteenhouse, Yumi Kasai, David Gray, Carol Clugston, Anna Dominiczak and Alex Alderton, Roberto Amato, Jeffrey Barrett, Sonia Goncalves, Ewan Harrison, David K. Jackson, Ian Johnston, Dominic Kwiatkowski, Cordelia Langford, John Sillitoe on behalf of the Wellcome Sanger Institute COVID-19 Surveillance Team |
| EPI_ISL_3095333 | Lighthouse Lab in Milton Keynes | Wellcome Sanger Institute for the COVID-19 Genomics UK (COG-UK) Consortium | The Lighthouse Lab in Milton Keynes and Alex Alderton, Roberto Amato, Jeffrey Barrett, Sonia Goncalves, Ewan Harrison, David K. Jackson, Ian Johnston, Dominic Kwiatkowski, Cordelia Langford, John Sillitoe on behalf of the Wellcome Sanger Institute COVID-19 Surveillance Team |
| EPI_ISL_3095334 | Lighthouse Lab in Glasgow | Wellcome Sanger Institute for the COVID-19 Genomics UK (COG-UK) Consortium | Harper VanSteenhouse, Yumi Kasai, David Gray, Carol Clugston, Anna Dominiczak and Alex Alderton, Roberto Amato, Jeffrey Barrett, Sonia Goncalves, Ewan Harrison, David K. Jackson, Ian Johnston, Dominic Kwiatkowski, Cordelia Langford, John Sillitoe on behalf of the Wellcome Sanger Institute COVID-19 Surveillance Team |
| EPI_ISL_3095335, EPI_ISL_3095336, EPI_ISL_3095337, EPI_ISL_3095338, EPI_ISL_3095339, EPI_ISL_3095340, EPI_ISL_3095341, EPI_ISL_3095342, EPI_ISL_3095343, EPI_ISL_3095344 | Lighthouse Lab in Milton Keynes | Wellcome Sanger Institute for the COVID-19 Genomics UK (COG-UK) Consortium | The Lighthouse Lab in Milton Keynes and Alex Alderton, Roberto Amato, Jeffrey Barrett, Sonia Goncalves, Ewan Harrison, David K. Jackson, Ian Johnston, Dominic Kwiatkowski, Cordelia Langford, John Sillitoe on behalf of the Wellcome Sanger Institute COVID-19 Surveillance Team |
| EPI_ISL_3095345 | Lighthouse Lab in Glasgow | Wellcome Sanger Institute for the COVID-19 Genomics UK (COG-UK) Consortium | Harper VanSteenhouse, Yumi Kasai, David Gray, Carol Clugston, Anna Dominiczak and Alex Alderton, Roberto Amato, Jeffrey Barrett, Sonia Goncalves, Ewan Harrison, David K. Jackson, Ian Johnston, Dominic Kwiatkowski, Cordelia Langford, John Sillitoe on behalf of the Wellcome Sanger Institute COVID-19 Surveillance Team |
| EPI_ISL_3095346, EPI_ISL_3095347, EPI_ISL_3095348, EPI_ISL_3095349 | Lighthouse Lab in Milton Keynes | Wellcome Sanger Institute for the COVID-19 Genomics UK (COG-UK) Consortium | The Lighthouse Lab in Milton Keynes and Alex Alderton, Roberto Amato, Jeffrey Barrett, Sonia Goncalves, Ewan Harrison, David K. Jackson, Ian Johnston, Dominic Kwiatkowski, Cordelia Langford, John Sillitoe on behalf of the Wellcome Sanger Institute COVID-19 Surveillance Team |
| EPI_ISL_3095350 | Lighthouse Lab in Glasgow | Wellcome Sanger Institute for the COVID-19 Genomics UK (COG-UK) Consortium | Harper VanSteenhouse, Yumi Kasai, David Gray, Carol Clugston, Anna Dominiczak and Alex Alderton, Roberto Amato, Jeffrey Barrett, Sonia Goncalves, Ewan Harrison, David K. Jackson, Ian Johnston, Dominic Kwiatkowski, Cordelia Langford, John Sillitoe on behalf of the Wellcome Sanger Institute COVID-19 Surveillance Team |

| | | | |
|---|---|---|---|
| EPI_ISL_3095351, EPI_ISL_3095352, EPI_ISL_3095353, EPI_ISL_3095354, EPI_ISL_3095355 | Lighthouse Lab in Milton Keynes | Wellcome Sanger Institute for the COVID-19 Genomics UK (COG-UK) Consortium | The Lighthouse Lab in Milton Keynes and Alex Alderton, Roberto Amato, Jeffrey Barrett, Sonia Goncalves, Ewan Harrison, David K. Jackson, Ian Johnston, Dominic Kwiatkowski, Cordelia Langford, John Sillitoe on behalf of the Wellcome Sanger Institute COVID-19 Surveillance Team |
| EPI_ISL_3095356 | Lighthouse Lab in Glasgow | Wellcome Sanger Institute for the COVID-19 Genomics UK (COG-UK) Consortium | Harper VanSteenhouse, Yumi Kasai, David Gray, Carol Clugston, Anna Dominiczak and Alex Alderton, Roberto Amato, Jeffrey Barrett, Sonia Goncalves, Ewan Harrison, David K. Jackson, Ian Johnston, Dominic Kwiatkowski, Cordelia Langford, John Sillitoe on behalf of the Wellcome Sanger Institute COVID-19 Surveillance Team |
| EPI_ISL_3095357, EPI_ISL_3095358, EPI_ISL_3095359, EPI_ISL_3095360, EPI_ISL_3095361, EPI_ISL_3095362, EPI_ISL_3095363, EPI_ISL_3095364, EPI_ISL_3095365, EPI_ISL_3095366, EPI_ISL_3095367, EPI_ISL_3095368, EPI_ISL_3095369, EPI_ISL_3095370, EPI_ISL_3095371, EPI_ISL_3095372, EPI_ISL_3095373, EPI_ISL_3095374 | | | |
| see above | Lighthouse Lab in Milton Keynes | Wellcome Sanger Institute for the COVID-19 Genomics UK (COG-UK) Consortium | The Lighthouse Lab in Milton Keynes and Alex Alderton, Roberto Amato, Jeffrey Barrett, Sonia Goncalves, Ewan Harrison, David K. Jackson, Ian Johnston, Dominic Kwiatkowski, Cordelia Langford, John Sillitoe on behalf of the Wellcome Sanger Institute COVID-19 Surveillance Team |
| EPI_ISL_3095375, EPI_ISL_3095376 | Lighthouse Lab in Glasgow | Wellcome Sanger Institute for the COVID-19 Genomics UK (COG-UK) Consortium | Harper VanSteenhouse, Yumi Kasai, David Gray, Carol Clugston, Anna Dominiczak and Alex Alderton, Roberto Amato, Jeffrey Barrett, Sonia Goncalves, Ewan Harrison, David K. Jackson, Ian Johnston, Dominic Kwiatkowski, Cordelia Langford, John Sillitoe on behalf of the Wellcome Sanger Institute COVID-19 Surveillance Team |
| EPI_ISL_3095377, EPI_ISL_3095378, EPI_ISL_3095379, EPI_ISL_3095380, EPI_ISL_3095381 | Lighthouse Lab in Milton Keynes | Wellcome Sanger Institute for the COVID-19 Genomics UK (COG-UK) Consortium | The Lighthouse Lab in Milton Keynes and Alex Alderton, Roberto Amato, Jeffrey Barrett, Sonia Goncalves, Ewan Harrison, David K. Jackson, Ian Johnston, Dominic Kwiatkowski, Cordelia Langford, John Sillitoe on behalf of the Wellcome Sanger Institute COVID-19 Surveillance Team |
| EPI_ISL_3095382 | Lighthouse Lab in Glasgow | Wellcome Sanger Institute for the COVID-19 Genomics UK (COG-UK) Consortium | Harper VanSteenhouse, Yumi Kasai, David Gray, Carol Clugston, Anna Dominiczak and Alex Alderton, Roberto Amato, Jeffrey Barrett, Sonia Goncalves, Ewan Harrison, David K. Jackson, Ian Johnston, Dominic Kwiatkowski, Cordelia Langford, John Sillitoe on behalf of the Wellcome Sanger Institute COVID-19 Surveillance Team |
| EPI_ISL_3095383, EPI_ISL_3095384, EPI_ISL_3095385, EPI_ISL_3095386, EPI_ISL_3095387, EPI_ISL_3095388, EPI_ISL_3095389, EPI_ISL_3095390, EPI_ISL_3095391, EPI_ISL_3095392, EPI_ISL_3095393, EPI_ISL_3095394, EPI_ISL_3095395, EPI_ISL_3095396, EPI_ISL_3095397, EPI_ISL_3095398, EPI_ISL_3095399, EPI_ISL_3095400, EPI_ISL_3095401, EPI_ISL_3095402, EPI_ISL_3095403, EPI_ISL_3095404, EPI_ISL_3095405, EPI_ISL_3095406, EPI_ISL_3095407 | | | |
| see above | Lighthouse Lab in Milton Keynes | Wellcome Sanger Institute for the COVID-19 Genomics UK (COG-UK) Consortium | The Lighthouse Lab in Milton Keynes and Alex Alderton, Roberto Amato, Jeffrey Barrett, Sonia Goncalves, Ewan Harrison, David K. Jackson, Ian Johnston, Dominic Kwiatkowski, Cordelia Langford, John Sillitoe on behalf of the Wellcome Sanger Institute COVID-19 Surveillance Team |
| EPI_ISL_3095408 | Lighthouse Lab in Glasgow | Wellcome Sanger Institute for the COVID-19 Genomics UK (COG-UK) Consortium | Harper VanSteenhouse, Yumi Kasai, David Gray, Carol Clugston, Anna Dominiczak and Alex Alderton, Roberto Amato, Jeffrey Barrett, Sonia Goncalves, Ewan Harrison, David K. Jackson, Ian Johnston, Dominic Kwiatkowski, Cordelia Langford, John Sillitoe on behalf of the Wellcome Sanger Institute COVID-19 Surveillance Team |
| EPI_ISL_3095409, EPI_ISL_3095410, EPI_ISL_3095411, EPI_ISL_3095412, EPI_ISL_3095413, EPI_ISL_3095414, EPI_ISL_3095415, EPI_ISL_3095416 | Lighthouse Lab in Milton Keynes | Wellcome Sanger Institute for the COVID-19 Genomics UK (COG-UK) Consortium | The Lighthouse Lab in Milton Keynes and Alex Alderton, Roberto Amato, Jeffrey Barrett, Sonia Goncalves, Ewan Harrison, David K. Jackson, Ian Johnston, Dominic Kwiatkowski, Cordelia Langford, John Sillitoe on behalf of the Wellcome Sanger Institute COVID-19 Surveillance Team |
| EPI_ISL_3095417 | Lighthouse Lab in Glasgow | Wellcome Sanger Institute for the COVID-19 Genomics UK (COG-UK) Consortium | Harper VanSteenhouse, Yumi Kasai, David Gray, Carol Clugston, Anna Dominiczak and Alex Alderton, Roberto Amato, Jeffrey Barrett, Sonia Goncalves, Ewan Harrison, David K. Jackson, Ian Johnston, Dominic Kwiatkowski, Cordelia Langford, John Sillitoe on behalf of the Wellcome Sanger Institute COVID-19 Surveillance Team |
| EPI_ISL_3095418 | Lighthouse Lab in Milton Keynes | Wellcome Sanger Institute for the COVID-19 Genomics UK (COG-UK) Consortium | The Lighthouse Lab in Milton Keynes and Alex Alderton, Roberto Amato, Jeffrey Barrett, Sonia Goncalves, Ewan Harrison, David K. Jackson, Ian Johnston, Dominic Kwiatkowski, Cordelia Langford, John Sillitoe on behalf of the Wellcome Sanger Institute COVID-19 Surveillance Team |
| EPI_ISL_3095419 | Lighthouse Lab in Glasgow | Wellcome Sanger Institute for the COVID-19 Genomics UK (COG-UK) Consortium | Harper VanSteenhouse, Yumi Kasai, David Gray, Carol Clugston, Anna Dominiczak and Alex Alderton, Roberto Amato, Jeffrey Barrett, Sonia Goncalves, Ewan Harrison, David K. Jackson, Ian Johnston, Dominic Kwiatkowski, Cordelia Langford, John Sillitoe on behalf of the Wellcome Sanger Institute COVID-19 Surveillance Team |
| EPI_ISL_3095420 | Lighthouse Lab in Milton Keynes | Wellcome Sanger Institute for the COVID-19 Genomics UK (COG-UK) Consortium | The Lighthouse Lab in Milton Keynes and Alex Alderton, Roberto Amato, Jeffrey Barrett, Sonia Goncalves, Ewan Harrison, David K. Jackson, Ian Johnston, Dominic Kwiatkowski, Cordelia Langford, John Sillitoe on behalf of the Wellcome Sanger Institute COVID-19 Surveillance Team |
| EPI_ISL_3095421 | Lighthouse Lab in Glasgow | Wellcome Sanger Institute for the COVID-19 Genomics UK (COG-UK) Consortium | Harper VanSteenhouse, Yumi Kasai, David Gray, Carol Clugston, Anna Dominiczak and Alex Alderton, Roberto Amato, Jeffrey Barrett, Sonia Goncalves, Ewan Harrison, David K. Jackson, Ian Johnston, Dominic Kwiatkowski, Cordelia Langford, John Sillitoe on behalf of the Wellcome Sanger Institute COVID-19 Surveillance Team |
| EPI_ISL_3095422, EPI_ISL_3095423, EPI_ISL_3095424 | Lighthouse Lab in Milton Keynes | Wellcome Sanger Institute for the COVID-19 Genomics UK (COG-UK) Consortium | The Lighthouse Lab in Milton Keynes and Alex Alderton, Roberto Amato, Jeffrey Barrett, Sonia Goncalves, Ewan Harrison, David K. Jackson, Ian Johnston, Dominic Kwiatkowski, Cordelia Langford, John Sillitoe on behalf of the Wellcome Sanger Institute COVID-19 Surveillance Team |
| EPI_ISL_3095425 | Lighthouse Lab in Glasgow | Wellcome Sanger Institute for the COVID-19 Genomics UK (COG-UK) Consortium | Harper VanSteenhouse, Yumi Kasai, David Gray, Carol Clugston, Anna Dominiczak and Alex Alderton, Roberto Amato, Jeffrey Barrett, Sonia Goncalves, Ewan Harrison, David K. Jackson, Ian Johnston, Dominic Kwiatkowski, Cordelia Langford, John Sillitoe on behalf of the Wellcome Sanger Institute COVID-19 Surveillance Team |
| EPI_ISL_3095426, EPI_ISL_3095427, EPI_ISL_3095428, EPI_ISL_3095429, EPI_ISL_3095430 | Lighthouse Lab in Milton Keynes | Wellcome Sanger Institute for the COVID-19 Genomics UK (COG-UK) Consortium | The Lighthouse Lab in Milton Keynes and Alex Alderton, Roberto Amato, Jeffrey Barrett, Sonia Goncalves, Ewan Harrison, David K. Jackson, Ian Johnston, Dominic Kwiatkowski, Cordelia Langford, John Sillitoe on behalf of the Wellcome Sanger Institute COVID-19 Surveillance Team |
| EPI_ISL_3095431 | Lighthouse Lab in Glasgow | Wellcome Sanger Institute for the COVID-19 Genomics UK (COG-UK) Consortium | Harper VanSteenhouse, Yumi Kasai, David Gray, Carol Clugston, Anna Dominiczak and Alex Alderton, Roberto Amato, Jeffrey Barrett, Sonia Goncalves, Ewan Harrison, David K. Jackson, Ian Johnston, Dominic Kwiatkowski, Cordelia Langford, John Sillitoe on behalf of the Wellcome Sanger Institute COVID-19 Surveillance Team |
| EPI_ISL_3095432 | Lighthouse Lab in Alderley Park | Wellcome Sanger Institute for the COVID-19 Genomics UK (COG-UK) Consortium | Jacquelyn Wynn, Mairead Hyland, The Lighthouse Lab in Alderley Park and Alex Alderton, Roberto Amato, Jeffrey Barrett, Sonia Goncalves, Ewan Harrison, David K. Jackson, Ian Johnston, Dominic Kwiatkowski, Cordelia Langford, John Sillitoe on behalf of the Wellcome Sanger Institute COVID-19 Surveillance Team |
| EPI_ISL_3095433 | Lighthouse Lab in Glasgow | Wellcome Sanger Institute for the COVID-19 Genomics UK (COG-UK) Consortium | Harper VanSteenhouse, Yumi Kasai, David Gray, Carol Clugston, Anna Dominiczak and Alex Alderton, Roberto Amato, Jeffrey Barrett, Sonia Goncalves, Ewan Harrison, David K. Jackson, Ian Johnston, Dominic Kwiatkowski, Cordelia Langford, John Sillitoe on behalf of the Wellcome Sanger Institute COVID-19 Surveillance Team |
| EPI_ISL_3095434, EPI_ISL_3095435, EPI_ISL_3095436 | Lighthouse Lab in Milton Keynes | Wellcome Sanger Institute for the COVID-19 Genomics UK (COG-UK) Consortium | The Lighthouse Lab in Milton Keynes and Alex Alderton, Roberto Amato, Jeffrey Barrett, Sonia Goncalves, Ewan Harrison, David K. Jackson, Ian Johnston, Dominic Kwiatkowski, Cordelia Langford, John Sillitoe on behalf of the Wellcome Sanger Institute COVID-19 Surveillance Team |
| EPI_ISL_3095437, EPI_ISL_3095438, EPI_ISL_3095439 | Lighthouse Lab in Alderley Park | Wellcome Sanger Institute for the COVID-19 Genomics UK (COG-UK) Consortium | Jacquelyn Wynn, Mairead Hyland, The Lighthouse Lab in Alderley Park and Alex Alderton, Roberto Amato, Jeffrey Barrett, Sonia Goncalves, Ewan Harrison, David K. Jackson, Ian Johnston, Dominic Kwiatkowski, Cordelia Langford, John Sillitoe on behalf of the Wellcome Sanger Institute COVID-19 Surveillance Team |
| EPI_ISL_3095440, EPI_ISL_3095441 | Lighthouse Lab in Milton Keynes | Wellcome Sanger Institute for the COVID-19 Genomics UK (COG-UK) Consortium | The Lighthouse Lab in Milton Keynes and Alex Alderton, Roberto Amato, Jeffrey Barrett, Sonia Goncalves, Ewan Harrison, David K. Jackson, Ian Johnston, Dominic Kwiatkowski, Cordelia Langford, John Sillitoe on behalf of the Wellcome Sanger Institute COVID-19 Surveillance Team |
| EPI_ISL_3095442 | Lighthouse Lab in Alderley Park | Wellcome Sanger Institute for the COVID-19 Genomics UK (COG-UK) Consortium | Jacquelyn Wynn, Mairead Hyland, The Lighthouse Lab in Alderley Park and Alex Alderton, Roberto Amato, Jeffrey Barrett, Sonia Goncalves, Ewan Harrison, David K. Jackson, Ian Johnston, Dominic Kwiatkowski, Cordelia Langford, John Sillitoe on behalf of the Wellcome Sanger Institute COVID-19 Surveillance Team |
| EPI_ISL_3095443, EPI_ISL_3095444 | Lighthouse Lab in Milton Keynes | Wellcome Sanger Institute for the COVID-19 Genomics UK (COG-UK) Consortium | The Lighthouse Lab in Milton Keynes and Alex Alderton, Roberto Amato, Jeffrey Barrett, Sonia Goncalves, Ewan Harrison, David K. Jackson, Ian Johnston, Dominic Kwiatkowski, Cordelia Langford, John Sillitoe on behalf of the Wellcome Sanger Institute COVID-19 Surveillance Team |
| EPI_ISL_3095445 | Lighthouse Lab in Glasgow | Wellcome Sanger Institute for the COVID-19 Genomics UK (COG-UK) Consortium | Harper VanSteenhouse, Yumi Kasai, David Gray, Carol Clugston, Anna Dominiczak and Alex Alderton, Roberto Amato, Jeffrey Barrett, Sonia Goncalves, Ewan Harrison, David K. Jackson, Ian Johnston, Dominic Kwiatkowski, Cordelia Langford, John Sillitoe on behalf of the Wellcome Sanger Institute COVID-19 Surveillance Team |
| EPI_ISL_3095446 | Lighthouse Lab in Milton Keynes | Wellcome Sanger Institute for the COVID-19 Genomics UK (COG-UK) Consortium | The Lighthouse Lab in Milton Keynes and Alex Alderton, Roberto Amato, Jeffrey Barrett, Sonia Goncalves, Ewan Harrison, David K. Jackson, Ian Johnston, Dominic Kwiatkowski, Cordelia Langford, John Sillitoe on behalf of the Wellcome Sanger Institute COVID-19 Surveillance Team |
| EPI_ISL_3095447 | Lighthouse Lab in Alderley Park | Wellcome Sanger Institute for the COVID-19 Genomics UK (COG-UK) Consortium | Jacquelyn Wynn, Mairead Hyland, The Lighthouse Lab in Alderley Park and Alex Alderton, Roberto Amato, Jeffrey Barrett, Sonia Goncalves, Ewan Harrison, David K. Jackson, Ian Johnston, Dominic Kwiatkowski, Cordelia Langford, John Sillitoe on behalf of the Wellcome Sanger Institute COVID-19 Surveillance Team |
| EPI_ISL_3095448 | Lighthouse Lab in Milton Keynes | Wellcome Sanger Institute for the COVID-19 Genomics UK (COG-UK) Consortium | The Lighthouse Lab in Milton Keynes and Alex Alderton, Roberto Amato, Jeffrey Barrett, Sonia Goncalves, Ewan Harrison, David K. Jackson, Ian Johnston, Dominic Kwiatkowski, Cordelia Langford, John Sillitoe on behalf of the Wellcome Sanger Institute COVID-19 Surveillance Team |
| EPI_ISL_3095449 | Lighthouse Lab in Glasgow | Wellcome Sanger Institute for the COVID-19 Genomics UK (COG-UK) Consortium | Harper VanSteenhouse, Yumi Kasai, David Gray, Carol Clugston, Anna Dominiczak and Alex Alderton, Roberto Amato, Jeffrey Barrett, Sonia Goncalves, Ewan Harrison, David K. Jackson, Ian Johnston, Dominic Kwiatkowski, Cordelia Langford, John Sillitoe on behalf of the Wellcome Sanger Institute COVID-19 Surveillance Team |
| EPI_ISL_3095450 | Lighthouse Lab in Alderley Park | Wellcome Sanger Institute for the COVID-19 Genomics UK (COG-UK) Consortium | Jacquelyn Wynn, Mairead Hyland, The Lighthouse Lab in Alderley Park and Alex Alderton, Roberto Amato, Jeffrey Barrett, Sonia Goncalves, Ewan Harrison, David K. Jackson, Ian Johnston, Dominic Kwiatkowski, Cordelia Langford, John Sillitoe on behalf of the Wellcome Sanger Institute COVID-19 Surveillance Team |
| EPI_ISL_3095451, EPI_ISL_3095452, EPI_ISL_3095453 | Lighthouse Lab in Milton Keynes | Wellcome Sanger Institute for the COVID-19 Genomics UK (COG-UK) Consortium | The Lighthouse Lab in Milton Keynes and Alex Alderton, Roberto Amato, Jeffrey Barrett, Sonia Goncalves, Ewan Harrison, David K. Jackson, Ian Johnston, Dominic Kwiatkowski, Cordelia Langford, John Sillitoe on behalf of the Wellcome Sanger Institute COVID-19 Surveillance Team |
| EPI_ISL_3095454, EPI_ISL_3095455, EPI_ISL_3095456 | Lighthouse Lab in Alderley Park | Wellcome Sanger Institute for the COVID-19 Genomics UK (COG-UK) Consortium | Jacquelyn Wynn, Mairead Hyland, The Lighthouse Lab in Alderley Park and Alex Alderton, Roberto Amato, Jeffrey Barrett, Sonia Goncalves, Ewan Harrison, David K. Jackson, Ian Johnston, Dominic Kwiatkowski, Cordelia Langford, John Sillitoe on behalf of the Wellcome Sanger Institute COVID-19 Surveillance Team |
| EPI_ISL_3095457 | Lighthouse Lab in Milton Keynes | Wellcome Sanger Institute for the COVID-19 Genomics UK (COG-UK) Consortium | The Lighthouse Lab in Milton Keynes and Alex Alderton, Roberto Amato, Jeffrey Barrett, Sonia Goncalves, Ewan Harrison, David K. Jackson, Ian Johnston, Dominic Kwiatkowski, Cordelia Langford, John Sillitoe on behalf of the Wellcome Sanger Institute COVID-19 Surveillance Team |
| EPI_ISL_3095458 | Lighthouse Lab in Alderley Park | Wellcome Sanger Institute for the COVID-19 Genomics UK (COG-UK) Consortium | Jacquelyn Wynn, Mairead Hyland, The Lighthouse Lab in Alderley Park and Alex Alderton, Roberto Amato, Jeffrey Barrett, Sonia Goncalves, Ewan Harrison, David K. Jackson, Ian Johnston, Dominic Kwiatkowski, Cordelia Langford, John Sillitoe on behalf of the Wellcome Sanger Institute COVID-19 Surveillance Team |

| | | | |
|---|---|---|---|
| EPI_ISL_3095459 | Lighthouse Lab in Milton Keynes | Wellcome Sanger Institute for the COVID-19 Genomics UK (COG-UK) Consortium | The Lighthouse Lab in Milton Keynes and Alex Alderton, Roberto Amato, Jeffrey Barrett, Sonia Goncalves, Ewan Harrison, David K. Jackson, Ian Johnston, Dominic Kwiatkowski, Cordelia Langford, John Sillitoe on behalf of the Wellcome Sanger Institute COVID-19 Surveillance Team |
| EPI_ISL_3095460 | Lighthouse Lab in Alderley Park | Wellcome Sanger Institute for the COVID-19 Genomics UK (COG-UK) Consortium | Jacquelyn Wynn, Mairead Hyland, The Lighthouse Lab in Alderley Park and Alex Alderton, Roberto Amato, Jeffrey Barrett, Sonia Goncalves, Ewan Harrison, David K. Jackson, Ian Johnston, Dominic Kwiatkowski, Cordelia Langford, John Sillitoe on behalf of the Wellcome Sanger Institute COVID-19 Surveillance Team |
| EPI_ISL_3095461, EPI_ISL_3095462 | Lighthouse Lab in Milton Keynes | Wellcome Sanger Institute for the COVID-19 Genomics UK (COG-UK) Consortium | The Lighthouse Lab in Milton Keynes and Alex Alderton, Roberto Amato, Jeffrey Barrett, Sonia Goncalves, Ewan Harrison, David K. Jackson, Ian Johnston, Dominic Kwiatkowski, Cordelia Langford, John Sillitoe on behalf of the Wellcome Sanger Institute COVID-19 Surveillance Team |
| EPI_ISL_3095463 | Lighthouse Lab in Glasgow | Wellcome Sanger Institute for the COVID-19 Genomics UK (COG-UK) Consortium | Harper VanSteenhouse, Yumi Kasai, David Gray, Carol Clugston, Anna Dominiczak and Alex Alderton, Roberto Amato, Jeffrey Barrett, Sonia Goncalves, Ewan Harrison, David K. Jackson, Ian Johnston, Dominic Kwiatkowski, Cordelia Langford, John Sillitoe on behalf of the Wellcome Sanger Institute COVID-19 Surveillance Team |
| EPI_ISL_3095464 | Lighthouse Lab in Alderley Park | Wellcome Sanger Institute for the COVID-19 Genomics UK (COG-UK) Consortium | Jacquelyn Wynn, Mairead Hyland, The Lighthouse Lab in Alderley Park and Alex Alderton, Roberto Amato, Jeffrey Barrett, Sonia Goncalves, Ewan Harrison, David K. Jackson, Ian Johnston, Dominic Kwiatkowski, Cordelia Langford, John Sillitoe on behalf of the Wellcome Sanger Institute COVID-19 Surveillance Team |
| EPI_ISL_3095465 | Lighthouse Lab in Glasgow | Wellcome Sanger Institute for the COVID-19 Genomics UK (COG-UK) Consortium | Harper VanSteenhouse, Yumi Kasai, David Gray, Carol Clugston, Anna Dominiczak and Alex Alderton, Roberto Amato, Jeffrey Barrett, Sonia Goncalves, Ewan Harrison, David K. Jackson, Ian Johnston, Dominic Kwiatkowski, Cordelia Langford, John Sillitoe on behalf of the Wellcome Sanger Institute COVID-19 Surveillance Team |
| EPI_ISL_3095466 | Lighthouse Lab in Alderley Park | Wellcome Sanger Institute for the COVID-19 Genomics UK (COG-UK) Consortium | Jacquelyn Wynn, Mairead Hyland, The Lighthouse Lab in Alderley Park and Alex Alderton, Roberto Amato, Jeffrey Barrett, Sonia Goncalves, Ewan Harrison, David K. Jackson, Ian Johnston, Dominic Kwiatkowski, Cordelia Langford, John Sillitoe on behalf of the Wellcome Sanger Institute COVID-19 Surveillance Team |
| EPI_ISL_3095467 | Lighthouse Lab in Milton Keynes | Wellcome Sanger Institute for the COVID-19 Genomics UK (COG-UK) Consortium | The Lighthouse Lab in Milton Keynes and Alex Alderton, Roberto Amato, Jeffrey Barrett, Sonia Goncalves, Ewan Harrison, David K. Jackson, Ian Johnston, Dominic Kwiatkowski, Cordelia Langford, John Sillitoe on behalf of the Wellcome Sanger Institute COVID-19 Surveillance Team |
| EPI_ISL_3095468 | Lighthouse Lab in Alderley Park | Wellcome Sanger Institute for the COVID-19 Genomics UK (COG-UK) Consortium | Jacquelyn Wynn, Mairead Hyland, The Lighthouse Lab in Alderley Park and Alex Alderton, Roberto Amato, Jeffrey Barrett, Sonia Goncalves, Ewan Harrison, David K. Jackson, Ian Johnston, Dominic Kwiatkowski, Cordelia Langford, John Sillitoe on behalf of the Wellcome Sanger Institute COVID-19 Surveillance Team |
| EPI_ISL_3095469, EPI_ISL_3095470 | Lighthouse Lab in Milton Keynes | Wellcome Sanger Institute for the COVID-19 Genomics UK (COG-UK) Consortium | The Lighthouse Lab in Milton Keynes and Alex Alderton, Roberto Amato, Jeffrey Barrett, Sonia Goncalves, Ewan Harrison, David K. Jackson, Ian Johnston, Dominic Kwiatkowski, Cordelia Langford, John Sillitoe on behalf of the Wellcome Sanger Institute COVID-19 Surveillance Team |
| EPI_ISL_3095471, EPI_ISL_3095472, EPI_ISL_3095473 | Lighthouse Lab in Alderley Park | Wellcome Sanger Institute for the COVID-19 Genomics UK (COG-UK) Consortium | Jacquelyn Wynn, Mairead Hyland, The Lighthouse Lab in Alderley Park and Alex Alderton, Roberto Amato, Jeffrey Barrett, Sonia Goncalves, Ewan Harrison, David K. Jackson, Ian Johnston, Dominic Kwiatkowski, Cordelia Langford, John Sillitoe on behalf of the Wellcome Sanger Institute COVID-19 Surveillance Team |
| EPI_ISL_3095474 | Lighthouse Lab in Milton Keynes | Wellcome Sanger Institute for the COVID-19 Genomics UK (COG-UK) Consortium | The Lighthouse Lab in Milton Keynes and Alex Alderton, Roberto Amato, Jeffrey Barrett, Sonia Goncalves, Ewan Harrison, David K. Jackson, Ian Johnston, Dominic Kwiatkowski, Cordelia Langford, John Sillitoe on behalf of the Wellcome Sanger Institute COVID-19 Surveillance Team |
| EPI_ISL_3095475 | Lighthouse Lab in Glasgow | Wellcome Sanger Institute for the COVID-19 Genomics UK (COG-UK) Consortium | Harper VanSteenhouse, Yumi Kasai, David Gray, Carol Clugston, Anna Dominiczak and Alex Alderton, Roberto Amato, Jeffrey Barrett, Sonia Goncalves, Ewan Harrison, David K. Jackson, Ian Johnston, Dominic Kwiatkowski, Cordelia Langford, John Sillitoe on behalf of the Wellcome Sanger Institute COVID-19 Surveillance Team |
| EPI_ISL_3095476 | Lighthouse Lab in Milton Keynes | Wellcome Sanger Institute for the COVID-19 Genomics UK (COG-UK) Consortium | The Lighthouse Lab in Milton Keynes and Alex Alderton, Roberto Amato, Jeffrey Barrett, Sonia Goncalves, Ewan Harrison, David K. Jackson, Ian Johnston, Dominic Kwiatkowski, Cordelia Langford, John Sillitoe on behalf of the Wellcome Sanger Institute COVID-19 Surveillance Team |
| EPI_ISL_3095477 | Lighthouse Lab in Glasgow | Wellcome Sanger Institute for the COVID-19 Genomics UK (COG-UK) Consortium | Harper VanSteenhouse, Yumi Kasai, David Gray, Carol Clugston, Anna Dominiczak and Alex Alderton, Roberto Amato, Jeffrey Barrett, Sonia Goncalves, Ewan Harrison, David K. Jackson, Ian Johnston, Dominic Kwiatkowski, Cordelia Langford, John Sillitoe on behalf of the Wellcome Sanger Institute COVID-19 Surveillance Team |
| EPI_ISL_3095478, EPI_ISL_3095479, EPI_ISL_3095480 | Lighthouse Lab in Milton Keynes | Wellcome Sanger Institute for the COVID-19 Genomics UK (COG-UK) Consortium | The Lighthouse Lab in Milton Keynes and Alex Alderton, Roberto Amato, Jeffrey Barrett, Sonia Goncalves, Ewan Harrison, David K. Jackson, Ian Johnston, Dominic Kwiatkowski, Cordelia Langford, John Sillitoe on behalf of the Wellcome Sanger Institute COVID-19 Surveillance Team |
| EPI_ISL_3095481 | Lighthouse Lab in Alderley Park | Wellcome Sanger Institute for the COVID-19 Genomics UK (COG-UK) Consortium | Jacquelyn Wynn, Mairead Hyland, The Lighthouse Lab in Alderley Park and Alex Alderton, Roberto Amato, Jeffrey Barrett, Sonia Goncalves, Ewan Harrison, David K. Jackson, Ian Johnston, Dominic Kwiatkowski, Cordelia Langford, John Sillitoe on behalf of the Wellcome Sanger Institute COVID-19 Surveillance Team |
| EPI_ISL_3095482, EPI_ISL_3095483, EPI_ISL_3095484 | Lighthouse Lab in Glasgow | Wellcome Sanger Institute for the COVID-19 Genomics UK (COG-UK) Consortium | Harper VanSteenhouse, Yumi Kasai, David Gray, Carol Clugston, Anna Dominiczak and Alex Alderton, Roberto Amato, Jeffrey Barrett, Sonia Goncalves, Ewan Harrison, David K. Jackson, Ian Johnston, Dominic Kwiatkowski, Cordelia Langford, John Sillitoe on behalf of the Wellcome Sanger Institute COVID-19 Surveillance Team |
| EPI_ISL_3095485, EPI_ISL_3095486 | Lighthouse Lab in Milton Keynes | Wellcome Sanger Institute for the COVID-19 Genomics UK (COG-UK) Consortium | The Lighthouse Lab in Milton Keynes and Alex Alderton, Roberto Amato, Jeffrey Barrett, Sonia Goncalves, Ewan Harrison, David K. Jackson, Ian Johnston, Dominic Kwiatkowski, Cordelia Langford, John Sillitoe on behalf of the Wellcome Sanger Institute COVID-19 Surveillance Team |
| EPI_ISL_3095487, EPI_ISL_3095488 | Lighthouse Lab in Alderley Park | Wellcome Sanger Institute for the COVID-19 Genomics UK (COG-UK) Consortium | Jacquelyn Wynn, Mairead Hyland, The Lighthouse Lab in Alderley Park and Alex Alderton, Roberto Amato, Jeffrey Barrett, Sonia Goncalves, Ewan Harrison, David K. Jackson, Ian Johnston, Dominic Kwiatkowski, Cordelia Langford, John Sillitoe on behalf of the Wellcome Sanger Institute COVID-19 Surveillance Team |
| EPI_ISL_3095489 | Lighthouse Lab in Milton Keynes | Wellcome Sanger Institute for the COVID-19 Genomics UK (COG-UK) Consortium | The Lighthouse Lab in Milton Keynes and Alex Alderton, Roberto Amato, Jeffrey Barrett, Sonia Goncalves, Ewan Harrison, David K. Jackson, Ian Johnston, Dominic Kwiatkowski, Cordelia Langford, John Sillitoe on behalf of the Wellcome Sanger Institute COVID-19 Surveillance Team |
| EPI_ISL_3095490 | Lighthouse Lab in Glasgow | Wellcome Sanger Institute for the COVID-19 Genomics UK (COG-UK) Consortium | Harper VanSteenhouse, Yumi Kasai, David Gray, Carol Clugston, Anna Dominiczak and Alex Alderton, Roberto Amato, Jeffrey Barrett, Sonia Goncalves, Ewan Harrison, David K. Jackson, Ian Johnston, Dominic Kwiatkowski, Cordelia Langford, John Sillitoe on behalf of the Wellcome Sanger Institute COVID-19 Surveillance Team |
| EPI_ISL_3095491 | Lighthouse Lab in Alderley Park | Wellcome Sanger Institute for the COVID-19 Genomics UK (COG-UK) Consortium | Jacquelyn Wynn, Mairead Hyland, The Lighthouse Lab in Alderley Park and Alex Alderton, Roberto Amato, Jeffrey Barrett, Sonia Goncalves, Ewan Harrison, David K. Jackson, Ian Johnston, Dominic Kwiatkowski, Cordelia Langford, John Sillitoe on behalf of the Wellcome Sanger Institute COVID-19 Surveillance Team |
| EPI_ISL_3095492, EPI_ISL_3095493 | Lighthouse Lab in Milton Keynes | Wellcome Sanger Institute for the COVID-19 Genomics UK (COG-UK) Consortium | The Lighthouse Lab in Milton Keynes and Alex Alderton, Roberto Amato, Jeffrey Barrett, Sonia Goncalves, Ewan Harrison, David K. Jackson, Ian Johnston, Dominic Kwiatkowski, Cordelia Langford, John Sillitoe on behalf of the Wellcome Sanger Institute COVID-19 Surveillance Team |
| EPI_ISL_3095494 | Lighthouse Lab in Glasgow | Wellcome Sanger Institute for the COVID-19 Genomics UK (COG-UK) Consortium | Harper VanSteenhouse, Yumi Kasai, David Gray, Carol Clugston, Anna Dominiczak and Alex Alderton, Roberto Amato, Jeffrey Barrett, Sonia Goncalves, Ewan Harrison, David K. Jackson, Ian Johnston, Dominic Kwiatkowski, Cordelia Langford, John Sillitoe on behalf of the Wellcome Sanger Institute COVID-19 Surveillance Team |
| EPI_ISL_3095495, EPI_ISL_3095496 | Lighthouse Lab in Milton Keynes | Wellcome Sanger Institute for the COVID-19 Genomics UK (COG-UK) Consortium | The Lighthouse Lab in Milton Keynes and Alex Alderton, Roberto Amato, Jeffrey Barrett, Sonia Goncalves, Ewan Harrison, David K. Jackson, Ian Johnston, Dominic Kwiatkowski, Cordelia Langford, John Sillitoe on behalf of the Wellcome Sanger Institute COVID-19 Surveillance Team |
| EPI_ISL_3095497 | Lighthouse Lab in Glasgow | Wellcome Sanger Institute for the COVID-19 Genomics UK (COG-UK) Consortium | Harper VanSteenhouse, Yumi Kasai, David Gray, Carol Clugston, Anna Dominiczak and Alex Alderton, Roberto Amato, Jeffrey Barrett, Sonia Goncalves, Ewan Harrison, David K. Jackson, Ian Johnston, Dominic Kwiatkowski, Cordelia Langford, John Sillitoe on behalf of the Wellcome Sanger Institute COVID-19 Surveillance Team |
| EPI_ISL_3095498, EPI_ISL_3095499, EPI_ISL_3095500 | Lighthouse Lab in Milton Keynes | Wellcome Sanger Institute for the COVID-19 Genomics UK (COG-UK) Consortium | The Lighthouse Lab in Milton Keynes and Alex Alderton, Roberto Amato, Jeffrey Barrett, Sonia Goncalves, Ewan Harrison, David K. Jackson, Ian Johnston, Dominic Kwiatkowski, Cordelia Langford, John Sillitoe on behalf of the Wellcome Sanger Institute COVID-19 Surveillance Team |
| EPI_ISL_3095501 | Lighthouse Lab in Alderley Park | Wellcome Sanger Institute for the COVID-19 Genomics UK (COG-UK) Consortium | Jacquelyn Wynn, Mairead Hyland, The Lighthouse Lab in Alderley Park and Alex Alderton, Roberto Amato, Jeffrey Barrett, Sonia Goncalves, Ewan Harrison, David K. Jackson, Ian Johnston, Dominic Kwiatkowski, Cordelia Langford, John Sillitoe on behalf of the Wellcome Sanger Institute COVID-19 Surveillance Team |
| EPI_ISL_3095502 | Lighthouse Lab in Glasgow | Wellcome Sanger Institute for the COVID-19 Genomics UK (COG-UK) Consortium | Harper VanSteenhouse, Yumi Kasai, David Gray, Carol Clugston, Anna Dominiczak and Alex Alderton, Roberto Amato, Jeffrey Barrett, Sonia Goncalves, Ewan Harrison, David K. Jackson, Ian Johnston, Dominic Kwiatkowski, Cordelia Langford, John Sillitoe on behalf of the Wellcome Sanger Institute COVID-19 Surveillance Team |
| EPI_ISL_3095503 | Lighthouse Lab in Alderley Park | Wellcome Sanger Institute for the COVID-19 Genomics UK (COG-UK) Consortium | Jacquelyn Wynn, Mairead Hyland, The Lighthouse Lab in Alderley Park and Alex Alderton, Roberto Amato, Jeffrey Barrett, Sonia Goncalves, Ewan Harrison, David K. Jackson, Ian Johnston, Dominic Kwiatkowski, Cordelia Langford, John Sillitoe on behalf of the Wellcome Sanger Institute COVID-19 Surveillance Team |
| EPI_ISL_3095504 | Lighthouse Lab in Milton Keynes | Wellcome Sanger Institute for the COVID-19 Genomics UK (COG-UK) Consortium | The Lighthouse Lab in Milton Keynes and Alex Alderton, Roberto Amato, Jeffrey Barrett, Sonia Goncalves, Ewan Harrison, David K. Jackson, Ian Johnston, Dominic Kwiatkowski, Cordelia Langford, John Sillitoe on behalf of the Wellcome Sanger Institute COVID-19 Surveillance Team |
| EPI_ISL_3095505 | Lighthouse Lab in Glasgow | Wellcome Sanger Institute for the COVID-19 Genomics UK (COG-UK) Consortium | Harper VanSteenhouse, Yumi Kasai, David Gray, Carol Clugston, Anna Dominiczak and Alex Alderton, Roberto Amato, Jeffrey Barrett, Sonia Goncalves, Ewan Harrison, David K. Jackson, Ian Johnston, Dominic Kwiatkowski, Cordelia Langford, John Sillitoe on behalf of the Wellcome Sanger Institute COVID-19 Surveillance Team |
| EPI_ISL_3095506, EPI_ISL_3095507 | Lighthouse Lab in Alderley Park | Wellcome Sanger Institute for the COVID-19 Genomics UK (COG-UK) Consortium | Jacquelyn Wynn, Mairead Hyland, The Lighthouse Lab in Alderley Park and Alex Alderton, Roberto Amato, Jeffrey Barrett, Sonia Goncalves, Ewan Harrison, David K. Jackson, Ian Johnston, Dominic Kwiatkowski, Cordelia Langford, John Sillitoe on behalf of the Wellcome Sanger Institute COVID-19 Surveillance Team |
| EPI_ISL_3095508 | Lighthouse Lab in Glasgow | Wellcome Sanger Institute for the COVID-19 Genomics UK (COG-UK) Consortium | Harper VanSteenhouse, Yumi Kasai, David Gray, Carol Clugston, Anna Dominiczak and Alex Alderton, Roberto Amato, Jeffrey Barrett, Sonia Goncalves, Ewan Harrison, David K. Jackson, Ian Johnston, Dominic Kwiatkowski, Cordelia Langford, John Sillitoe on behalf of the Wellcome Sanger Institute COVID-19 Surveillance Team |
| EPI_ISL_3095509, EPI_ISL_3095510 | Lighthouse Lab in Milton Keynes | Wellcome Sanger Institute for the COVID-19 Genomics UK (COG-UK) Consortium | The Lighthouse Lab in Milton Keynes and Alex Alderton, Roberto Amato, Jeffrey Barrett, Sonia Goncalves, Ewan Harrison, David K. Jackson, Ian Johnston, Dominic Kwiatkowski, Cordelia Langford, John Sillitoe on behalf of the Wellcome Sanger Institute COVID-19 Surveillance Team |
| EPI_ISL_3095511 | Lighthouse Lab in Alderley Park | Wellcome Sanger Institute for the COVID-19 Genomics UK (COG-UK) Consortium | Jacquelyn Wynn, Mairead Hyland, The Lighthouse Lab in Alderley Park and Alex Alderton, Roberto Amato, Jeffrey Barrett, Sonia Goncalves, Ewan Harrison, David K. Jackson, Ian Johnston, Dominic Kwiatkowski, Cordelia Langford, John Sillitoe on behalf of the Wellcome Sanger Institute COVID-19 Surveillance Team |
| EPI_ISL_3095512 | Lighthouse Lab in Milton Keynes | Wellcome Sanger Institute for the COVID-19 Genomics UK (COG-UK) Consortium | The Lighthouse Lab in Milton Keynes and Alex Alderton, Roberto Amato, Jeffrey Barrett, Sonia Goncalves, Ewan Harrison, David K. Jackson, Ian Johnston, Dominic Kwiatkowski, Cordelia Langford, John Sillitoe on behalf of the Wellcome Sanger Institute COVID-19 Surveillance Team |
| EPI_ISL_3095513, EPI_ISL_3095514 | Lighthouse Lab in Glasgow | Wellcome Sanger Institute for the COVID-19 Genomics UK (COG-UK) Consortium | Harper VanSteenhouse, Yumi Kasai, David Gray, Carol Clugston, Anna Dominiczak and Alex Alderton, Roberto Amato, Jeffrey Barrett, Sonia Goncalves, Ewan Harrison, David K. Jackson, Ian Johnston, Dominic Kwiatkowski, Cordelia Langford, John Sillitoe on behalf of the Wellcome Sanger Institute COVID-19 Surveillance Team |
| EPI_ISL_3095515, EPI_ISL_3095516 | Lighthouse Lab in Milton Keynes | Wellcome Sanger Institute for the COVID-19 Genomics UK (COG-UK) Consortium | The Lighthouse Lab in Milton Keynes and Alex Alderton, Roberto Amato, Jeffrey Barrett, Sonia Goncalves, Ewan Harrison, David K. Jackson, Ian Johnston, Dominic Kwiatkowski, Cordelia Langford, John Sillitoe on behalf of the Wellcome Sanger Institute COVID-19 Surveillance Team |
| EPI_ISL_3095517 | Lighthouse Lab in Alderley Park | Wellcome Sanger Institute for the COVID-19 Genomics UK (COG-UK) Consortium | Jacquelyn Wynn, Mairead Hyland, The Lighthouse Lab in Alderley Park and Alex Alderton, Roberto Amato, Jeffrey Barrett, Sonia Goncalves, Ewan Harrison, David K. Jackson, Ian Johnston, Dominic Kwiatkowski, Cordelia Langford, John Sillitoe on behalf of the Wellcome Sanger Institute COVID-19 Surveillance Team |
| EPI_ISL_3095518 | Lighthouse Lab in Milton Keynes | Wellcome Sanger Institute for the COVID-19 Genomics UK (COG-UK) Consortium | The Lighthouse Lab in Milton Keynes and Alex Alderton, Roberto Amato, Jeffrey Barrett, Sonia Goncalves, Ewan Harrison, David K. Jackson, Ian Johnston, Dominic Kwiatkowski, Cordelia Langford, John Sillitoe on behalf of the Wellcome Sanger Institute COVID-19 Surveillance Team |
| EPI_ISL_3095519, EPI_ISL_3095520 | Lighthouse Lab in Alderley Park | Wellcome Sanger Institute for the COVID-19 Genomics UK (COG-UK) Consortium | Jacquelyn Wynn, Mairead Hyland, The Lighthouse Lab in Alderley Park and Alex Alderton, Roberto Amato, Jeffrey Barrett, Sonia Goncalves, Ewan Harrison, David K. Jackson, Ian Johnston, Dominic Kwiatkowski, Cordelia Langford, John Sillitoe on behalf of the Wellcome Sanger Institute COVID-19 Surveillance Team |

| | | | |
|---|---|---|---|
| EPI_ISL_3095521 | Lighthouse Lab in Milton Keynes | Wellcome Sanger Institute for the COVID-19 Genomics UK (COG-UK) Consortium | The Lighthouse Lab in Milton Keynes and Alex Alderton, Roberto Amato, Jeffrey Barrett, Sonia Goncalves, Ewan Harrison, David K. Jackson, Ian Johnston, Dominic Kwiatkowski, Cordelia Langford, John Sillitoe on behalf of the Wellcome Sanger Institute COVID-19 Surveillance Team |
| EPI_ISL_3095522 | Lighthouse Lab in Alderley Park | Wellcome Sanger Institute for the COVID-19 Genomics UK (COG-UK) Consortium | Jacquelyn Wynn, Mairead Hyland, The Lighthouse Lab in Alderley Park and Alex Alderton, Roberto Amato, Jeffrey Barrett, Sonia Goncalves, Ewan Harrison, David K. Jackson, Ian Johnston, Dominic Kwiatkowski, Cordelia Langford, John Sillitoe on behalf of the Wellcome Sanger Institute COVID-19 Surveillance Team |
| EPI_ISL_3095523, EPI_ISL_3095524 | Lighthouse Lab in Milton Keynes | Wellcome Sanger Institute for the COVID-19 Genomics UK (COG-UK) Consortium | The Lighthouse Lab in Milton Keynes and Alex Alderton, Roberto Amato, Jeffrey Barrett, Sonia Goncalves, Ewan Harrison, David K. Jackson, Ian Johnston, Dominic Kwiatkowski, Cordelia Langford, John Sillitoe on behalf of the Wellcome Sanger Institute COVID-19 Surveillance Team |
| EPI_ISL_3095525 | Lighthouse Lab in Glasgow | Wellcome Sanger Institute for the COVID-19 Genomics UK (COG-UK) Consortium | Harper VanSteenhouse, Yumi Kasai, David Gray, Carol Clugston, Anna Dominiczak and Alex Alderton, Roberto Amato, Jeffrey Barrett, Sonia Goncalves, Ewan Harrison, David K. Jackson, Ian Johnston, Dominic Kwiatkowski, Cordelia Langford, John Sillitoe on behalf of the Wellcome Sanger Institute COVID-19 Surveillance Team |
| EPI_ISL_3095526, EPI_ISL_3095527 | Lighthouse Lab in Milton Keynes | Wellcome Sanger Institute for the COVID-19 Genomics UK (COG-UK) Consortium | The Lighthouse Lab in Milton Keynes and Alex Alderton, Roberto Amato, Jeffrey Barrett, Sonia Goncalves, Ewan Harrison, David K. Jackson, Ian Johnston, Dominic Kwiatkowski, Cordelia Langford, John Sillitoe on behalf of the Wellcome Sanger Institute COVID-19 Surveillance Team |
| EPI_ISL_3095528 | Lighthouse Lab in Glasgow | Wellcome Sanger Institute for the COVID-19 Genomics UK (COG-UK) Consortium | Harper VanSteenhouse, Yumi Kasai, David Gray, Carol Clugston, Anna Dominiczak and Alex Alderton, Roberto Amato, Jeffrey Barrett, Sonia Goncalves, Ewan Harrison, David K. Jackson, Ian Johnston, Dominic Kwiatkowski, Cordelia Langford, John Sillitoe on behalf of the Wellcome Sanger Institute COVID-19 Surveillance Team |
| EPI_ISL_3095529 | Lighthouse Lab in Alderley Park | Wellcome Sanger Institute for the COVID-19 Genomics UK (COG-UK) Consortium | Jacquelyn Wynn, Mairead Hyland, The Lighthouse Lab in Alderley Park and Alex Alderton, Roberto Amato, Jeffrey Barrett, Sonia Goncalves, Ewan Harrison, David K. Jackson, Ian Johnston, Dominic Kwiatkowski, Cordelia Langford, John Sillitoe on behalf of the Wellcome Sanger Institute COVID-19 Surveillance Team |
| EPI_ISL_3095530, EPI_ISL_3095531 | Lighthouse Lab in Milton Keynes | Wellcome Sanger Institute for the COVID-19 Genomics UK (COG-UK) Consortium | The Lighthouse Lab in Milton Keynes and Alex Alderton, Roberto Amato, Jeffrey Barrett, Sonia Goncalves, Ewan Harrison, David K. Jackson, Ian Johnston, Dominic Kwiatkowski, Cordelia Langford, John Sillitoe on behalf of the Wellcome Sanger Institute COVID-19 Surveillance Team |
| EPI_ISL_3095532 | Lighthouse Lab in Alderley Park | Wellcome Sanger Institute for the COVID-19 Genomics UK (COG-UK) Consortium | Jacquelyn Wynn, Mairead Hyland, The Lighthouse Lab in Alderley Park and Alex Alderton, Roberto Amato, Jeffrey Barrett, Sonia Goncalves, Ewan Harrison, David K. Jackson, Ian Johnston, Dominic Kwiatkowski, Cordelia Langford, John Sillitoe on behalf of the Wellcome Sanger Institute COVID-19 Surveillance Team |
| EPI_ISL_3095533 | Lighthouse Lab in Glasgow | Wellcome Sanger Institute for the COVID-19 Genomics UK (COG-UK) Consortium | Harper VanSteenhouse, Yumi Kasai, David Gray, Carol Clugston, Anna Dominiczak and Alex Alderton, Roberto Amato, Jeffrey Barrett, Sonia Goncalves, Ewan Harrison, David K. Jackson, Ian Johnston, Dominic Kwiatkowski, Cordelia Langford, John Sillitoe on behalf of the Wellcome Sanger Institute COVID-19 Surveillance Team |
| EPI_ISL_3095534, EPI_ISL_3095535 | Lighthouse Lab in Milton Keynes | Wellcome Sanger Institute for the COVID-19 Genomics UK (COG-UK) Consortium | The Lighthouse Lab in Milton Keynes and Alex Alderton, Roberto Amato, Jeffrey Barrett, Sonia Goncalves, Ewan Harrison, David K. Jackson, Ian Johnston, Dominic Kwiatkowski, Cordelia Langford, John Sillitoe on behalf of the Wellcome Sanger Institute COVID-19 Surveillance Team |
| EPI_ISL_3095536 | Lighthouse Lab in Alderley Park | Wellcome Sanger Institute for the COVID-19 Genomics UK (COG-UK) Consortium | Jacquelyn Wynn, Mairead Hyland, The Lighthouse Lab in Alderley Park and Alex Alderton, Roberto Amato, Jeffrey Barrett, Sonia Goncalves, Ewan Harrison, David K. Jackson, Ian Johnston, Dominic Kwiatkowski, Cordelia Langford, John Sillitoe on behalf of the Wellcome Sanger Institute COVID-19 Surveillance Team |
| EPI_ISL_3095537, EPI_ISL_3095538 | Lighthouse Lab in Milton Keynes | Wellcome Sanger Institute for the COVID-19 Genomics UK (COG-UK) Consortium | The Lighthouse Lab in Milton Keynes and Alex Alderton, Roberto Amato, Jeffrey Barrett, Sonia Goncalves, Ewan Harrison, David K. Jackson, Ian Johnston, Dominic Kwiatkowski, Cordelia Langford, John Sillitoe on behalf of the Wellcome Sanger Institute COVID-19 Surveillance Team |
| EPI_ISL_3095539, EPI_ISL_3095540 | Lighthouse Lab in Glasgow | Wellcome Sanger Institute for the COVID-19 Genomics UK (COG-UK) Consortium | Harper VanSteenhouse, Yumi Kasai, David Gray, Carol Clugston, Anna Dominiczak and Alex Alderton, Roberto Amato, Jeffrey Barrett, Sonia Goncalves, Ewan Harrison, David K. Jackson, Ian Johnston, Dominic Kwiatkowski, Cordelia Langford, John Sillitoe on behalf of the Wellcome Sanger Institute COVID-19 Surveillance Team |
| EPI_ISL_3095541 | Lighthouse Lab in Alderley Park | Wellcome Sanger Institute for the COVID-19 Genomics UK (COG-UK) Consortium | Jacquelyn Wynn, Mairead Hyland, The Lighthouse Lab in Alderley Park and Alex Alderton, Roberto Amato, Jeffrey Barrett, Sonia Goncalves, Ewan Harrison, David K. Jackson, Ian Johnston, Dominic Kwiatkowski, Cordelia Langford, John Sillitoe on behalf of the Wellcome Sanger Institute COVID-19 Surveillance Team |
| EPI_ISL_3095542, EPI_ISL_3095543 | Lighthouse Lab in Glasgow | Wellcome Sanger Institute for the COVID-19 Genomics UK (COG-UK) Consortium | Harper VanSteenhouse, Yumi Kasai, David Gray, Carol Clugston, Anna Dominiczak and Alex Alderton, Roberto Amato, Jeffrey Barrett, Sonia Goncalves, Ewan Harrison, David K. Jackson, Ian Johnston, Dominic Kwiatkowski, Cordelia Langford, John Sillitoe on behalf of the Wellcome Sanger Institute COVID-19 Surveillance Team |
| EPI_ISL_3095544 | Lighthouse Lab in Alderley Park | Wellcome Sanger Institute for the COVID-19 Genomics UK (COG-UK) Consortium | Jacquelyn Wynn, Mairead Hyland, The Lighthouse Lab in Alderley Park and Alex Alderton, Roberto Amato, Jeffrey Barrett, Sonia Goncalves, Ewan Harrison, David K. Jackson, Ian Johnston, Dominic Kwiatkowski, Cordelia Langford, John Sillitoe on behalf of the Wellcome Sanger Institute COVID-19 Surveillance Team |
| EPI_ISL_3095545, EPI_ISL_3095546, EPI_ISL_3095547 | Lighthouse Lab in Milton Keynes | Wellcome Sanger Institute for the COVID-19 Genomics UK (COG-UK) Consortium | The Lighthouse Lab in Milton Keynes and Alex Alderton, Roberto Amato, Jeffrey Barrett, Sonia Goncalves, Ewan Harrison, David K. Jackson, Ian Johnston, Dominic Kwiatkowski, Cordelia Langford, John Sillitoe on behalf of the Wellcome Sanger Institute COVID-19 Surveillance Team |
| EPI_ISL_3095548 | Lighthouse Lab in Alderley Park | Wellcome Sanger Institute for the COVID-19 Genomics UK (COG-UK) Consortium | Jacquelyn Wynn, Mairead Hyland, The Lighthouse Lab in Alderley Park and Alex Alderton, Roberto Amato, Jeffrey Barrett, Sonia Goncalves, Ewan Harrison, David K. Jackson, Ian Johnston, Dominic Kwiatkowski, Cordelia Langford, John Sillitoe on behalf of the Wellcome Sanger Institute COVID-19 Surveillance Team |
| EPI_ISL_3095549 | Lighthouse Lab in Glasgow | Wellcome Sanger Institute for the COVID-19 Genomics UK (COG-UK) Consortium | Harper VanSteenhouse, Yumi Kasai, David Gray, Carol Clugston, Anna Dominiczak and Alex Alderton, Roberto Amato, Jeffrey Barrett, Sonia Goncalves, Ewan Harrison, David K. Jackson, Ian Johnston, Dominic Kwiatkowski, Cordelia Langford, John Sillitoe on behalf of the Wellcome Sanger Institute COVID-19 Surveillance Team |
| EPI_ISL_3095550, EPI_ISL_3095551, EPI_ISL_3095552, EPI_ISL_3095553, EPI_ISL_3095554, EPI_ISL_3095555 | Lighthouse Lab in Alderley Park | Wellcome Sanger Institute for the COVID-19 Genomics UK (COG-UK) Consortium | Jacquelyn Wynn, Mairead Hyland, The Lighthouse Lab in Alderley Park and Alex Alderton, Roberto Amato, Jeffrey Barrett, Sonia Goncalves, Ewan Harrison, David K. Jackson, Ian Johnston, Dominic Kwiatkowski, Cordelia Langford, John Sillitoe on behalf of the Wellcome Sanger Institute COVID-19 Surveillance Team |
| EPI_ISL_3095556, EPI_ISL_3095557, EPI_ISL_3095558, EPI_ISL_3095559 | Lighthouse Lab in Milton Keynes | Wellcome Sanger Institute for the COVID-19 Genomics UK (COG-UK) Consortium | The Lighthouse Lab in Milton Keynes and Alex Alderton, Roberto Amato, Jeffrey Barrett, Sonia Goncalves, Ewan Harrison, David K. Jackson, Ian Johnston, Dominic Kwiatkowski, Cordelia Langford, John Sillitoe on behalf of the Wellcome Sanger Institute COVID-19 Surveillance Team |
| EPI_ISL_3095560 | Lighthouse Lab in Alderley Park | Wellcome Sanger Institute for the COVID-19 Genomics UK (COG-UK) Consortium | Jacquelyn Wynn, Mairead Hyland, The Lighthouse Lab in Alderley Park and Alex Alderton, Roberto Amato, Jeffrey Barrett, Sonia Goncalves, Ewan Harrison, David K. Jackson, Ian Johnston, Dominic Kwiatkowski, Cordelia Langford, John Sillitoe on behalf of the Wellcome Sanger Institute COVID-19 Surveillance Team |
| EPI_ISL_3095561, EPI_ISL_3095562 | Lighthouse Lab in Glasgow | Wellcome Sanger Institute for the COVID-19 Genomics UK (COG-UK) Consortium | Harper VanSteenhouse, Yumi Kasai, David Gray, Carol Clugston, Anna Dominiczak and Alex Alderton, Roberto Amato, Jeffrey Barrett, Sonia Goncalves, Ewan Harrison, David K. Jackson, Ian Johnston, Dominic Kwiatkowski, Cordelia Langford, John Sillitoe on behalf of the Wellcome Sanger Institute COVID-19 Surveillance Team |
| EPI_ISL_3095563, EPI_ISL_3095564 | Lighthouse Lab in Milton Keynes | Wellcome Sanger Institute for the COVID-19 Genomics UK (COG-UK) Consortium | The Lighthouse Lab in Milton Keynes and Alex Alderton, Roberto Amato, Jeffrey Barrett, Sonia Goncalves, Ewan Harrison, David K. Jackson, Ian Johnston, Dominic Kwiatkowski, Cordelia Langford, John Sillitoe on behalf of the Wellcome Sanger Institute COVID-19 Surveillance Team |
| EPI_ISL_3095565 | Lighthouse Lab in Alderley Park | Wellcome Sanger Institute for the COVID-19 Genomics UK (COG-UK) Consortium | Jacquelyn Wynn, Mairead Hyland, The Lighthouse Lab in Alderley Park and Alex Alderton, Roberto Amato, Jeffrey Barrett, Sonia Goncalves, Ewan Harrison, David K. Jackson, Ian Johnston, Dominic Kwiatkowski, Cordelia Langford, John Sillitoe on behalf of the Wellcome Sanger Institute COVID-19 Surveillance Team |
| EPI_ISL_3095566, EPI_ISL_3095567 | Lighthouse Lab in Milton Keynes | Wellcome Sanger Institute for the COVID-19 Genomics UK (COG-UK) Consortium | The Lighthouse Lab in Milton Keynes and Alex Alderton, Roberto Amato, Jeffrey Barrett, Sonia Goncalves, Ewan Harrison, David K. Jackson, Ian Johnston, Dominic Kwiatkowski, Cordelia Langford, John Sillitoe on behalf of the Wellcome Sanger Institute COVID-19 Surveillance Team |
| EPI_ISL_3095568 | Lighthouse Lab in Glasgow | Wellcome Sanger Institute for the COVID-19 Genomics UK (COG-UK) Consortium | Harper VanSteenhouse, Yumi Kasai, David Gray, Carol Clugston, Anna Dominiczak and Alex Alderton, Roberto Amato, Jeffrey Barrett, Sonia Goncalves, Ewan Harrison, David K. Jackson, Ian Johnston, Dominic Kwiatkowski, Cordelia Langford, John Sillitoe on behalf of the Wellcome Sanger Institute COVID-19 Surveillance Team |
| EPI_ISL_3095569, EPI_ISL_3095570 | Lighthouse Lab in Milton Keynes | Wellcome Sanger Institute for the COVID-19 Genomics UK (COG-UK) Consortium | The Lighthouse Lab in Milton Keynes and Alex Alderton, Roberto Amato, Jeffrey Barrett, Sonia Goncalves, Ewan Harrison, David K. Jackson, Ian Johnston, Dominic Kwiatkowski, Cordelia Langford, John Sillitoe on behalf of the Wellcome Sanger Institute COVID-19 Surveillance Team |
| EPI_ISL_3095571 | Lighthouse Lab in Glasgow | Wellcome Sanger Institute for the COVID-19 Genomics UK (COG-UK) Consortium | Harper VanSteenhouse, Yumi Kasai, David Gray, Carol Clugston, Anna Dominiczak and Alex Alderton, Roberto Amato, Jeffrey Barrett, Sonia Goncalves, Ewan Harrison, David K. Jackson, Ian Johnston, Dominic Kwiatkowski, Cordelia Langford, John Sillitoe on behalf of the Wellcome Sanger Institute COVID-19 Surveillance Team |
| EPI_ISL_3095572 | Lighthouse Lab in Alderley Park | Wellcome Sanger Institute for the COVID-19 Genomics UK (COG-UK) Consortium | Jacquelyn Wynn, Mairead Hyland, The Lighthouse Lab in Alderley Park and Alex Alderton, Roberto Amato, Jeffrey Barrett, Sonia Goncalves, Ewan Harrison, David K. Jackson, Ian Johnston, Dominic Kwiatkowski, Cordelia Langford, John Sillitoe on behalf of the Wellcome Sanger Institute COVID-19 Surveillance Team |
| EPI_ISL_3095573, EPI_ISL_3095574 | Lighthouse Lab in Milton Keynes | Wellcome Sanger Institute for the COVID-19 Genomics UK (COG-UK) Consortium | The Lighthouse Lab in Milton Keynes and Alex Alderton, Roberto Amato, Jeffrey Barrett, Sonia Goncalves, Ewan Harrison, David K. Jackson, Ian Johnston, Dominic Kwiatkowski, Cordelia Langford, John Sillitoe on behalf of the Wellcome Sanger Institute COVID-19 Surveillance Team |
| EPI_ISL_3095575 | Lighthouse Lab in Alderley Park | Wellcome Sanger Institute for the COVID-19 Genomics UK (COG-UK) Consortium | Jacquelyn Wynn, Mairead Hyland, The Lighthouse Lab in Alderley Park and Alex Alderton, Roberto Amato, Jeffrey Barrett, Sonia Goncalves, Ewan Harrison, David K. Jackson, Ian Johnston, Dominic Kwiatkowski, Cordelia Langford, John Sillitoe on behalf of the Wellcome Sanger Institute COVID-19 Surveillance Team |
| EPI_ISL_3095576 | Lighthouse Lab in Milton Keynes | Wellcome Sanger Institute for the COVID-19 Genomics UK (COG-UK) Consortium | The Lighthouse Lab in Milton Keynes and Alex Alderton, Roberto Amato, Jeffrey Barrett, Sonia Goncalves, Ewan Harrison, David K. Jackson, Ian Johnston, Dominic Kwiatkowski, Cordelia Langford, John Sillitoe on behalf of the Wellcome Sanger Institute COVID-19 Surveillance Team |
| EPI_ISL_3095577, EPI_ISL_3095578 | Lighthouse Lab in Glasgow | Wellcome Sanger Institute for the COVID-19 Genomics UK (COG-UK) Consortium | Harper VanSteenhouse, Yumi Kasai, David Gray, Carol Clugston, Anna Dominiczak and Alex Alderton, Roberto Amato, Jeffrey Barrett, Sonia Goncalves, Ewan Harrison, David K. Jackson, Ian Johnston, Dominic Kwiatkowski, Cordelia Langford, John Sillitoe on behalf of the Wellcome Sanger Institute COVID-19 Surveillance Team |
| EPI_ISL_3095579 | Lighthouse Lab in Milton Keynes | Wellcome Sanger Institute for the COVID-19 Genomics UK (COG-UK) Consortium | The Lighthouse Lab in Milton Keynes and Alex Alderton, Roberto Amato, Jeffrey Barrett, Sonia Goncalves, Ewan Harrison, David K. Jackson, Ian Johnston, Dominic Kwiatkowski, Cordelia Langford, John Sillitoe on behalf of the Wellcome Sanger Institute COVID-19 Surveillance Team |
| EPI_ISL_3095580 | Lighthouse Lab in Alderley Park | Wellcome Sanger Institute for the COVID-19 Genomics UK (COG-UK) Consortium | Jacquelyn Wynn, Mairead Hyland, The Lighthouse Lab in Alderley Park and Alex Alderton, Roberto Amato, Jeffrey Barrett, Sonia Goncalves, Ewan Harrison, David K. Jackson, Ian Johnston, Dominic Kwiatkowski, Cordelia Langford, John Sillitoe on behalf of the Wellcome Sanger Institute COVID-19 Surveillance Team |
| EPI_ISL_3095581 | Lighthouse Lab in Milton Keynes | Wellcome Sanger Institute for the COVID-19 Genomics UK (COG-UK) Consortium | The Lighthouse Lab in Milton Keynes and Alex Alderton, Roberto Amato, Jeffrey Barrett, Sonia Goncalves, Ewan Harrison, David K. Jackson, Ian Johnston, Dominic Kwiatkowski, Cordelia Langford, John Sillitoe on behalf of the Wellcome Sanger Institute COVID-19 Surveillance Team |
| EPI_ISL_3095582 | Lighthouse Lab in Alderley Park | Wellcome Sanger Institute for the COVID-19 Genomics UK (COG-UK) Consortium | Jacquelyn Wynn, Mairead Hyland, The Lighthouse Lab in Alderley Park and Alex Alderton, Roberto Amato, Jeffrey Barrett, Sonia Goncalves, Ewan Harrison, David K. Jackson, Ian Johnston, Dominic Kwiatkowski, Cordelia Langford, John Sillitoe on behalf of the Wellcome Sanger Institute COVID-19 Surveillance Team |
| EPI_ISL_3095583 | Lighthouse Lab in Glasgow | Wellcome Sanger Institute for the COVID-19 Genomics UK (COG-UK) Consortium | Harper VanSteenhouse, Yumi Kasai, David Gray, Carol Clugston, Anna Dominiczak and Alex Alderton, Roberto Amato, Jeffrey Barrett, Sonia Goncalves, Ewan Harrison, David K. Jackson, Ian Johnston, Dominic Kwiatkowski, Cordelia Langford, John Sillitoe on behalf of the Wellcome Sanger Institute COVID-19 Surveillance Team |
| EPI_ISL_3095584, EPI_ISL_3095585, EPI_ISL_3095586 | Lighthouse Lab in Alderley Park | Wellcome Sanger Institute for the COVID-19 Genomics UK (COG-UK) Consortium | Jacquelyn Wynn, Mairead Hyland, The Lighthouse Lab in Alderley Park and Alex Alderton, Roberto Amato, Jeffrey Barrett, Sonia Goncalves, Ewan Harrison, David K. Jackson, Ian Johnston, Dominic Kwiatkowski, Cordelia Langford, John Sillitoe on behalf of the Wellcome Sanger Institute COVID-19 Surveillance Team |
| EPI_ISL_3095587, EPI_ISL_3095588 | Lighthouse Lab in Milton Keynes | Wellcome Sanger Institute for the COVID-19 Genomics UK (COG-UK) Consortium | The Lighthouse Lab in Milton Keynes and Alex Alderton, Roberto Amato, Jeffrey Barrett, Sonia Goncalves, Ewan Harrison, David K. Jackson, Ian Johnston, Dominic Kwiatkowski, Cordelia Langford, John Sillitoe on behalf of the Wellcome Sanger Institute COVID-19 Surveillance Team |

| | | | |
|---|---|---|---|
| EPI_ISL_3095589 | Lighthouse Lab in Alderley Park | Wellcome Sanger Institute for the COVID-19 Genomics UK (COG-UK) Consortium | Jacquelyn Wynn, Mairead Hyland, The Lighthouse Lab in Alderley Park and Alex Alderton, Roberto Amato, Jeffrey Barrett, Sonia Goncalves, Ewan Harrison, David K. Jackson, Ian Johnston, Dominic Kwiatkowski, Cordelia Langford, John Sillitoe on behalf of the Wellcome Sanger Institute COVID-19 Surveillance Team |
| EPI_ISL_3095590, EPI_ISL_3095591, EPI_ISL_3095592, EPI_ISL_3095593, EPI_ISL_3095594 | Lighthouse Lab in Milton Keynes | Wellcome Sanger Institute for the COVID-19 Genomics UK (COG-UK) Consortium | The Lighthouse Lab in Milton Keynes and Alex Alderton, Roberto Amato, Jeffrey Barrett, Sonia Goncalves, Ewan Harrison, David K. Jackson, Ian Johnston, Dominic Kwiatkowski, Cordelia Langford, John Sillitoe on behalf of the Wellcome Sanger Institute COVID-19 Surveillance Team |
| EPI_ISL_3095595 | Lighthouse Lab in Alderley Park | Wellcome Sanger Institute for the COVID-19 Genomics UK (COG-UK) Consortium | Jacquelyn Wynn, Mairead Hyland, The Lighthouse Lab in Alderley Park and Alex Alderton, Roberto Amato, Jeffrey Barrett, Sonia Goncalves, Ewan Harrison, David K. Jackson, Ian Johnston, Dominic Kwiatkowski, Cordelia Langford, John Sillitoe on behalf of the Wellcome Sanger Institute COVID-19 Surveillance Team |
| EPI_ISL_3095596, EPI_ISL_3095597, EPI_ISL_3095598 | Lighthouse Lab in Milton Keynes | Wellcome Sanger Institute for the COVID-19 Genomics UK (COG-UK) Consortium | The Lighthouse Lab in Milton Keynes and Alex Alderton, Roberto Amato, Jeffrey Barrett, Sonia Goncalves, Ewan Harrison, David K. Jackson, Ian Johnston, Dominic Kwiatkowski, Cordelia Langford, John Sillitoe on behalf of the Wellcome Sanger Institute COVID-19 Surveillance Team |
| EPI_ISL_3095599 | Lighthouse Lab in Alderley Park | Wellcome Sanger Institute for the COVID-19 Genomics UK (COG-UK) Consortium | Jacquelyn Wynn, Mairead Hyland, The Lighthouse Lab in Alderley Park and Alex Alderton, Roberto Amato, Jeffrey Barrett, Sonia Goncalves, Ewan Harrison, David K. Jackson, Ian Johnston, Dominic Kwiatkowski, Cordelia Langford, John Sillitoe on behalf of the Wellcome Sanger Institute COVID-19 Surveillance Team |
| EPI_ISL_3095600 | Lighthouse Lab in Glasgow | Wellcome Sanger Institute for the COVID-19 Genomics UK (COG-UK) Consortium | Harper VanSteenhouse, Yumi Kasai, David Gray, Carol Clugston, Anna Dominiczak and Alex Alderton, Roberto Amato, Jeffrey Barrett, Sonia Goncalves, Ewan Harrison, David K. Jackson, Ian Johnston, Dominic Kwiatkowski, Cordelia Langford, John Sillitoe on behalf of the Wellcome Sanger Institute COVID-19 Surveillance Team |
| EPI_ISL_3095601 | Lighthouse Lab in Milton Keynes | Wellcome Sanger Institute for the COVID-19 Genomics UK (COG-UK) Consortium | The Lighthouse Lab in Milton Keynes and Alex Alderton, Roberto Amato, Jeffrey Barrett, Sonia Goncalves, Ewan Harrison, David K. Jackson, Ian Johnston, Dominic Kwiatkowski, Cordelia Langford, John Sillitoe on behalf of the Wellcome Sanger Institute COVID-19 Surveillance Team |
| EPI_ISL_3095602 | Lighthouse Lab in Alderley Park | Wellcome Sanger Institute for the COVID-19 Genomics UK (COG-UK) Consortium | Jacquelyn Wynn, Mairead Hyland, The Lighthouse Lab in Alderley Park and Alex Alderton, Roberto Amato, Jeffrey Barrett, Sonia Goncalves, Ewan Harrison, David K. Jackson, Ian Johnston, Dominic Kwiatkowski, Cordelia Langford, John Sillitoe on behalf of the Wellcome Sanger Institute COVID-19 Surveillance Team |
| EPI_ISL_3095603 | Lighthouse Lab in Milton Keynes | Wellcome Sanger Institute for the COVID-19 Genomics UK (COG-UK) Consortium | The Lighthouse Lab in Milton Keynes and Alex Alderton, Roberto Amato, Jeffrey Barrett, Sonia Goncalves, Ewan Harrison, David K. Jackson, Ian Johnston, Dominic Kwiatkowski, Cordelia Langford, John Sillitoe on behalf of the Wellcome Sanger Institute COVID-19 Surveillance Team |
| EPI_ISL_3095604 | Lighthouse Lab in Alderley Park | Wellcome Sanger Institute for the COVID-19 Genomics UK (COG-UK) Consortium | Jacquelyn Wynn, Mairead Hyland, The Lighthouse Lab in Alderley Park and Alex Alderton, Roberto Amato, Jeffrey Barrett, Sonia Goncalves, Ewan Harrison, David K. Jackson, Ian Johnston, Dominic Kwiatkowski, Cordelia Langford, John Sillitoe on behalf of the Wellcome Sanger Institute COVID-19 Surveillance Team |
| EPI_ISL_3095605 | Lighthouse Lab in Milton Keynes | Wellcome Sanger Institute for the COVID-19 Genomics UK (COG-UK) Consortium | The Lighthouse Lab in Milton Keynes and Alex Alderton, Roberto Amato, Jeffrey Barrett, Sonia Goncalves, Ewan Harrison, David K. Jackson, Ian Johnston, Dominic Kwiatkowski, Cordelia Langford, John Sillitoe on behalf of the Wellcome Sanger Institute COVID-19 Surveillance Team |
| EPI_ISL_3095606 | Lighthouse Lab in Glasgow | Wellcome Sanger Institute for the COVID-19 Genomics UK (COG-UK) Consortium | Harper VanSteenhouse, Yumi Kasai, David Gray, Carol Clugston, Anna Dominiczak and Alex Alderton, Roberto Amato, Jeffrey Barrett, Sonia Goncalves, Ewan Harrison, David K. Jackson, Ian Johnston, Dominic Kwiatkowski, Cordelia Langford, John Sillitoe on behalf of the Wellcome Sanger Institute COVID-19 Surveillance Team |
| EPI_ISL_3095607, EPI_ISL_3095608, EPI_ISL_3095609 | Lighthouse Lab in Milton Keynes | Wellcome Sanger Institute for the COVID-19 Genomics UK (COG-UK) Consortium | The Lighthouse Lab in Milton Keynes and Alex Alderton, Roberto Amato, Jeffrey Barrett, Sonia Goncalves, Ewan Harrison, David K. Jackson, Ian Johnston, Dominic Kwiatkowski, Cordelia Langford, John Sillitoe on behalf of the Wellcome Sanger Institute COVID-19 Surveillance Team |
| EPI_ISL_3095610 | Lighthouse Lab in Alderley Park | Wellcome Sanger Institute for the COVID-19 Genomics UK (COG-UK) Consortium | Jacquelyn Wynn, Mairead Hyland, The Lighthouse Lab in Alderley Park and Alex Alderton, Roberto Amato, Jeffrey Barrett, Sonia Goncalves, Ewan Harrison, David K. Jackson, Ian Johnston, Dominic Kwiatkowski, Cordelia Langford, John Sillitoe on behalf of the Wellcome Sanger Institute COVID-19 Surveillance Team |
| EPI_ISL_3095611 | Lighthouse Lab in Milton Keynes | Wellcome Sanger Institute for the COVID-19 Genomics UK (COG-UK) Consortium | The Lighthouse Lab in Milton Keynes and Alex Alderton, Roberto Amato, Jeffrey Barrett, Sonia Goncalves, Ewan Harrison, David K. Jackson, Ian Johnston, Dominic Kwiatkowski, Cordelia Langford, John Sillitoe on behalf of the Wellcome Sanger Institute COVID-19 Surveillance Team |
| EPI_ISL_3095612 | Lighthouse Lab in Alderley Park | Wellcome Sanger Institute for the COVID-19 Genomics UK (COG-UK) Consortium | Jacquelyn Wynn, Mairead Hyland, The Lighthouse Lab in Alderley Park and Alex Alderton, Roberto Amato, Jeffrey Barrett, Sonia Goncalves, Ewan Harrison, David K. Jackson, Ian Johnston, Dominic Kwiatkowski, Cordelia Langford, John Sillitoe on behalf of the Wellcome Sanger Institute COVID-19 Surveillance Team |
| EPI_ISL_3095613, EPI_ISL_3095614, EPI_ISL_3095615, EPI_ISL_3095616, EPI_ISL_3095617 | Lighthouse Lab in Milton Keynes | Wellcome Sanger Institute for the COVID-19 Genomics UK (COG-UK) Consortium | The Lighthouse Lab in Milton Keynes and Alex Alderton, Roberto Amato, Jeffrey Barrett, Sonia Goncalves, Ewan Harrison, David K. Jackson, Ian Johnston, Dominic Kwiatkowski, Cordelia Langford, John Sillitoe on behalf of the Wellcome Sanger Institute COVID-19 Surveillance Team |
| EPI_ISL_3095618 | Lighthouse Lab in Glasgow | Wellcome Sanger Institute for the COVID-19 Genomics UK (COG-UK) Consortium | Harper VanSteenhouse, Yumi Kasai, David Gray, Carol Clugston, Anna Dominiczak and Alex Alderton, Roberto Amato, Jeffrey Barrett, Sonia Goncalves, Ewan Harrison, David K. Jackson, Ian Johnston, Dominic Kwiatkowski, Cordelia Langford, John Sillitoe on behalf of the Wellcome Sanger Institute COVID-19 Surveillance Team |
| EPI_ISL_3095619 | Lighthouse Lab in Milton Keynes | Wellcome Sanger Institute for the COVID-19 Genomics UK (COG-UK) Consortium | The Lighthouse Lab in Milton Keynes and Alex Alderton, Roberto Amato, Jeffrey Barrett, Sonia Goncalves, Ewan Harrison, David K. Jackson, Ian Johnston, Dominic Kwiatkowski, Cordelia Langford, John Sillitoe on behalf of the Wellcome Sanger Institute COVID-19 Surveillance Team |
| EPI_ISL_3095620 | Lighthouse Lab in Glasgow | Wellcome Sanger Institute for the COVID-19 Genomics UK (COG-UK) Consortium | Harper VanSteenhouse, Yumi Kasai, David Gray, Carol Clugston, Anna Dominiczak and Alex Alderton, Roberto Amato, Jeffrey Barrett, Sonia Goncalves, Ewan Harrison, David K. Jackson, Ian Johnston, Dominic Kwiatkowski, Cordelia Langford, John Sillitoe on behalf of the Wellcome Sanger Institute COVID-19 Surveillance Team |
| EPI_ISL_3095621, EPI_ISL_3095622 | Lighthouse Lab in Milton Keynes | Wellcome Sanger Institute for the COVID-19 Genomics UK (COG-UK) Consortium | The Lighthouse Lab in Milton Keynes and Alex Alderton, Roberto Amato, Jeffrey Barrett, Sonia Goncalves, Ewan Harrison, David K. Jackson, Ian Johnston, Dominic Kwiatkowski, Cordelia Langford, John Sillitoe on behalf of the Wellcome Sanger Institute COVID-19 Surveillance Team |
| EPI_ISL_3095623 | Lighthouse Lab in Glasgow | Wellcome Sanger Institute for the COVID-19 Genomics UK (COG-UK) Consortium | Harper VanSteenhouse, Yumi Kasai, David Gray, Carol Clugston, Anna Dominiczak and Alex Alderton, Roberto Amato, Jeffrey Barrett, Sonia Goncalves, Ewan Harrison, David K. Jackson, Ian Johnston, Dominic Kwiatkowski, Cordelia Langford, John Sillitoe on behalf of the Wellcome Sanger Institute COVID-19 Surveillance Team |
| EPI_ISL_3095624, EPI_ISL_3095625 | Lighthouse Lab in Alderley Park | Wellcome Sanger Institute for the COVID-19 Genomics UK (COG-UK) Consortium | Jacquelyn Wynn, Mairead Hyland, The Lighthouse Lab in Alderley Park and Alex Alderton, Roberto Amato, Jeffrey Barrett, Sonia Goncalves, Ewan Harrison, David K. Jackson, Ian Johnston, Dominic Kwiatkowski, Cordelia Langford, John Sillitoe on behalf of the Wellcome Sanger Institute COVID-19 Surveillance Team |
| EPI_ISL_3095626, EPI_ISL_3095627, EPI_ISL_3095628, EPI_ISL_3095629, EPI_ISL_3095630 | Lighthouse Lab in Milton Keynes | Wellcome Sanger Institute for the COVID-19 Genomics UK (COG-UK) Consortium | The Lighthouse Lab in Milton Keynes and Alex Alderton, Roberto Amato, Jeffrey Barrett, Sonia Goncalves, Ewan Harrison, David K. Jackson, Ian Johnston, Dominic Kwiatkowski, Cordelia Langford, John Sillitoe on behalf of the Wellcome Sanger Institute COVID-19 Surveillance Team |
| EPI_ISL_3095631 | Lighthouse Lab in Glasgow | Wellcome Sanger Institute for the COVID-19 Genomics UK (COG-UK) Consortium | Harper VanSteenhouse, Yumi Kasai, David Gray, Carol Clugston, Anna Dominiczak and Alex Alderton, Roberto Amato, Jeffrey Barrett, Sonia Goncalves, Ewan Harrison, David K. Jackson, Ian Johnston, Dominic Kwiatkowski, Cordelia Langford, John Sillitoe on behalf of the Wellcome Sanger Institute COVID-19 Surveillance Team |
| EPI_ISL_3095632, EPI_ISL_3095633, EPI_ISL_3095634 | Lighthouse Lab in Milton Keynes | Wellcome Sanger Institute for the COVID-19 Genomics UK (COG-UK) Consortium | The Lighthouse Lab in Milton Keynes and Alex Alderton, Roberto Amato, Jeffrey Barrett, Sonia Goncalves, Ewan Harrison, David K. Jackson, Ian Johnston, Dominic Kwiatkowski, Cordelia Langford, John Sillitoe on behalf of the Wellcome Sanger Institute COVID-19 Surveillance Team |
| EPI_ISL_3095635 | Lighthouse Lab in Alderley Park | Wellcome Sanger Institute for the COVID-19 Genomics UK (COG-UK) Consortium | Jacquelyn Wynn, Mairead Hyland, The Lighthouse Lab in Alderley Park and Alex Alderton, Roberto Amato, Jeffrey Barrett, Sonia Goncalves, Ewan Harrison, David K. Jackson, Ian Johnston, Dominic Kwiatkowski, Cordelia Langford, John Sillitoe on behalf of the Wellcome Sanger Institute COVID-19 Surveillance Team |
| EPI_ISL_3095636, EPI_ISL_3095637, EPI_ISL_3095638 | Lighthouse Lab in Milton Keynes | Wellcome Sanger Institute for the COVID-19 Genomics UK (COG-UK) Consortium | The Lighthouse Lab in Milton Keynes and Alex Alderton, Roberto Amato, Jeffrey Barrett, Sonia Goncalves, Ewan Harrison, David K. Jackson, Ian Johnston, Dominic Kwiatkowski, Cordelia Langford, John Sillitoe on behalf of the Wellcome Sanger Institute COVID-19 Surveillance Team |
| EPI_ISL_3095639 | Lighthouse Lab in Alderley Park | Wellcome Sanger Institute for the COVID-19 Genomics UK (COG-UK) Consortium | Jacquelyn Wynn, Mairead Hyland, The Lighthouse Lab in Alderley Park and Alex Alderton, Roberto Amato, Jeffrey Barrett, Sonia Goncalves, Ewan Harrison, David K. Jackson, Ian Johnston, Dominic Kwiatkowski, Cordelia Langford, John Sillitoe on behalf of the Wellcome Sanger Institute COVID-19 Surveillance Team |
| EPI_ISL_3095640 | Lighthouse Lab in Milton Keynes | Wellcome Sanger Institute for the COVID-19 Genomics UK (COG-UK) Consortium | The Lighthouse Lab in Milton Keynes and Alex Alderton, Roberto Amato, Jeffrey Barrett, Sonia Goncalves, Ewan Harrison, David K. Jackson, Ian Johnston, Dominic Kwiatkowski, Cordelia Langford, John Sillitoe on behalf of the Wellcome Sanger Institute COVID-19 Surveillance Team |
| EPI_ISL_3095641 | Lighthouse Lab in Glasgow | Wellcome Sanger Institute for the COVID-19 Genomics UK (COG-UK) Consortium | Harper VanSteenhouse, Yumi Kasai, David Gray, Carol Clugston, Anna Dominiczak and Alex Alderton, Roberto Amato, Jeffrey Barrett, Sonia Goncalves, Ewan Harrison, David K. Jackson, Ian Johnston, Dominic Kwiatkowski, Cordelia Langford, John Sillitoe on behalf of the Wellcome Sanger Institute COVID-19 Surveillance Team |
| EPI_ISL_3095642, EPI_ISL_3095643, EPI_ISL_3095644, EPI_ISL_3095645, EPI_ISL_3095646, EPI_ISL_3095647, EPI_ISL_3095648 | Lighthouse Lab in Milton Keynes | Wellcome Sanger Institute for the COVID-19 Genomics UK (COG-UK) Consortium | The Lighthouse Lab in Milton Keynes and Alex Alderton, Roberto Amato, Jeffrey Barrett, Sonia Goncalves, Ewan Harrison, David K. Jackson, Ian Johnston, Dominic Kwiatkowski, Cordelia Langford, John Sillitoe on behalf of the Wellcome Sanger Institute COVID-19 Surveillance Team |
| EPI_ISL_3095649 | Lighthouse Lab in Alderley Park | Wellcome Sanger Institute for the COVID-19 Genomics UK (COG-UK) Consortium | Jacquelyn Wynn, Mairead Hyland, The Lighthouse Lab in Alderley Park and Alex Alderton, Roberto Amato, Jeffrey Barrett, Sonia Goncalves, Ewan Harrison, David K. Jackson, Ian Johnston, Dominic Kwiatkowski, Cordelia Langford, John Sillitoe on behalf of the Wellcome Sanger Institute COVID-19 Surveillance Team |
| EPI_ISL_3095650 | Lighthouse Lab in Milton Keynes | Wellcome Sanger Institute for the COVID-19 Genomics UK (COG-UK) Consortium | The Lighthouse Lab in Milton Keynes and Alex Alderton, Roberto Amato, Jeffrey Barrett, Sonia Goncalves, Ewan Harrison, David K. Jackson, Ian Johnston, Dominic Kwiatkowski, Cordelia Langford, John Sillitoe on behalf of the Wellcome Sanger Institute COVID-19 Surveillance Team |
| EPI_ISL_3095651 | Lighthouse Lab in Alderley Park | Wellcome Sanger Institute for the COVID-19 Genomics UK (COG-UK) Consortium | Jacquelyn Wynn, Mairead Hyland, The Lighthouse Lab in Alderley Park and Alex Alderton, Roberto Amato, Jeffrey Barrett, Sonia Goncalves, Ewan Harrison, David K. Jackson, Ian Johnston, Dominic Kwiatkowski, Cordelia Langford, John Sillitoe on behalf of the Wellcome Sanger Institute COVID-19 Surveillance Team |
| EPI_ISL_3095652 | Lighthouse Lab in Milton Keynes | Wellcome Sanger Institute for the COVID-19 Genomics UK (COG-UK) Consortium | The Lighthouse Lab in Milton Keynes and Alex Alderton, Roberto Amato, Jeffrey Barrett, Sonia Goncalves, Ewan Harrison, David K. Jackson, Ian Johnston, Dominic Kwiatkowski, Cordelia Langford, John Sillitoe on behalf of the Wellcome Sanger Institute COVID-19 Surveillance Team |
| EPI_ISL_3095653 | Lighthouse Lab in Glasgow | Wellcome Sanger Institute for the COVID-19 Genomics UK (COG-UK) Consortium | Harper VanSteenhouse, Yumi Kasai, David Gray, Carol Clugston, Anna Dominiczak and Alex Alderton, Roberto Amato, Jeffrey Barrett, Sonia Goncalves, Ewan Harrison, David K. Jackson, Ian Johnston, Dominic Kwiatkowski, Cordelia Langford, John Sillitoe on behalf of the Wellcome Sanger Institute COVID-19 Surveillance Team |
| EPI_ISL_3095654, | Lighthouse Lab in Alderley Park | Wellcome Sanger Institute for the COVID-19 Genomics UK (COG-UK) | Jacquelyn Wynn, Mairead Hyland, The Lighthouse Lab in Alderley Park and Alex Alderton, Roberto Amato, Jeffrey Barrett, Sonia Goncalves, Ewan Harrison, David K. Jackson, Ian Johnston, Dominic Kwiatkowski, Cordelia Langford, John Sillitoe on |

| | | | |
|---|---|---|---|
| EPI_ISL_3095655 | | Consortium | behalf of the Wellcome Sanger Institute COVID-19 Surveillance Team |
| EPI_ISL_3095656 | Lighthouse Lab in Glasgow | Wellcome Sanger Institute for the COVID-19 Genomics UK (COG-UK) Consortium | Harper VanSteenhouse, Yumi Kasai, David Gray, Carol Clugston, Anna Dominiczak and Alex Alderton, Roberto Amato, Jeffrey Barrett, Sonia Goncalves, Ewan Harrison, David K. Jackson, Ian Johnston, Dominic Kwiatkowski, Cordelia Langford, John Sillitoe on behalf of the Wellcome Sanger Institute COVID-19 Surveillance Team |
| EPI_ISL_3095657, EPI_ISL_3095658 | Lighthouse Lab in Milton Keynes | Wellcome Sanger Institute for the COVID-19 Genomics UK (COG-UK) Consortium | The Lighthouse Lab in Milton Keynes and Alex Alderton, Roberto Amato, Jeffrey Barrett, Sonia Goncalves, Ewan Harrison, David K. Jackson, Ian Johnston, Dominic Kwiatkowski, Cordelia Langford, John Sillitoe on behalf of the Wellcome Sanger Institute COVID-19 Surveillance Team |
| EPI_ISL_3095659 | Lighthouse Lab in Glasgow | Wellcome Sanger Institute for the COVID-19 Genomics UK (COG-UK) Consortium | Harper VanSteenhouse, Yumi Kasai, David Gray, Carol Clugston, Anna Dominiczak and Alex Alderton, Roberto Amato, Jeffrey Barrett, Sonia Goncalves, Ewan Harrison, David K. Jackson, Ian Johnston, Dominic Kwiatkowski, Cordelia Langford, John Sillitoe on behalf of the Wellcome Sanger Institute COVID-19 Surveillance Team |
| EPI_ISL_3095660, EPI_ISL_3095661, EPI_ISL_3095662 | Lighthouse Lab in Milton Keynes | Wellcome Sanger Institute for the COVID-19 Genomics UK (COG-UK) Consortium | The Lighthouse Lab in Milton Keynes and Alex Alderton, Roberto Amato, Jeffrey Barrett, Sonia Goncalves, Ewan Harrison, David K. Jackson, Ian Johnston, Dominic Kwiatkowski, Cordelia Langford, John Sillitoe on behalf of the Wellcome Sanger Institute COVID-19 Surveillance Team |
| EPI_ISL_3095663 | Lighthouse Lab in Alderley Park | Wellcome Sanger Institute for the COVID-19 Genomics UK (COG-UK) Consortium | Jacquelyn Wynn, Mairead Hyland, The Lighthouse Lab in Alderley Park and Alex Alderton, Roberto Amato, Jeffrey Barrett, Sonia Goncalves, Ewan Harrison, David K. Jackson, Ian Johnston, Dominic Kwiatkowski, Cordelia Langford, John Sillitoe on behalf of the Wellcome Sanger Institute COVID-19 Surveillance Team |
| EPI_ISL_3095664, EPI_ISL_3095665, EPI_ISL_3095667 | Lighthouse Lab in Milton Keynes | Wellcome Sanger Institute for the COVID-19 Genomics UK (COG-UK) Consortium | The Lighthouse Lab in Milton Keynes and Alex Alderton, Roberto Amato, Jeffrey Barrett, Sonia Goncalves, Ewan Harrison, David K. Jackson, Ian Johnston, Dominic Kwiatkowski, Cordelia Langford, John Sillitoe on behalf of the Wellcome Sanger Institute COVID-19 Surveillance Team |
| EPI_ISL_3095668 | Lighthouse Lab in Glasgow | Wellcome Sanger Institute for the COVID-19 Genomics UK (COG-UK) Consortium | Harper VanSteenhouse, Yumi Kasai, David Gray, Carol Clugston, Anna Dominiczak and Alex Alderton, Roberto Amato, Jeffrey Barrett, Sonia Goncalves, Ewan Harrison, David K. Jackson, Ian Johnston, Dominic Kwiatkowski, Cordelia Langford, John Sillitoe on behalf of the Wellcome Sanger Institute COVID-19 Surveillance Team |
| EPI_ISL_3095669 | Lighthouse Lab in Milton Keynes | Wellcome Sanger Institute for the COVID-19 Genomics UK (COG-UK) Consortium | The Lighthouse Lab in Milton Keynes and Alex Alderton, Roberto Amato, Jeffrey Barrett, Sonia Goncalves, Ewan Harrison, David K. Jackson, Ian Johnston, Dominic Kwiatkowski, Cordelia Langford, John Sillitoe on behalf of the Wellcome Sanger Institute COVID-19 Surveillance Team |
| EPI_ISL_3095670 | Lighthouse Lab in Alderley Park | Wellcome Sanger Institute for the COVID-19 Genomics UK (COG-UK) Consortium | Jacquelyn Wynn, Mairead Hyland, The Lighthouse Lab in Alderley Park and Alex Alderton, Roberto Amato, Jeffrey Barrett, Sonia Goncalves, Ewan Harrison, David K. Jackson, Ian Johnston, Dominic Kwiatkowski, Cordelia Langford, John Sillitoe on behalf of the Wellcome Sanger Institute COVID-19 Surveillance Team |
| EPI_ISL_3095671 | Lighthouse Lab in Milton Keynes | Wellcome Sanger Institute for the COVID-19 Genomics UK (COG-UK) Consortium | The Lighthouse Lab in Milton Keynes and Alex Alderton, Roberto Amato, Jeffrey Barrett, Sonia Goncalves, Ewan Harrison, David K. Jackson, Ian Johnston, Dominic Kwiatkowski, Cordelia Langford, John Sillitoe on behalf of the Wellcome Sanger Institute COVID-19 Surveillance Team |
| EPI_ISL_3095672 | Lighthouse Lab in Alderley Park | Wellcome Sanger Institute for the COVID-19 Genomics UK (COG-UK) Consortium | Jacquelyn Wynn, Mairead Hyland, The Lighthouse Lab in Alderley Park and Alex Alderton, Roberto Amato, Jeffrey Barrett, Sonia Goncalves, Ewan Harrison, David K. Jackson, Ian Johnston, Dominic Kwiatkowski, Cordelia Langford, John Sillitoe on behalf of the Wellcome Sanger Institute COVID-19 Surveillance Team |
| EPI_ISL_3095673 | Lighthouse Lab in Glasgow | Wellcome Sanger Institute for the COVID-19 Genomics UK (COG-UK) Consortium | Harper VanSteenhouse, Yumi Kasai, David Gray, Carol Clugston, Anna Dominiczak and Alex Alderton, Roberto Amato, Jeffrey Barrett, Sonia Goncalves, Ewan Harrison, David K. Jackson, Ian Johnston, Dominic Kwiatkowski, Cordelia Langford, John Sillitoe on behalf of the Wellcome Sanger Institute COVID-19 Surveillance Team |
| EPI_ISL_3095674 | Lighthouse Lab in Alderley Park | Wellcome Sanger Institute for the COVID-19 Genomics UK (COG-UK) Consortium | Jacquelyn Wynn, Mairead Hyland, The Lighthouse Lab in Alderley Park and Alex Alderton, Roberto Amato, Jeffrey Barrett, Sonia Goncalves, Ewan Harrison, David K. Jackson, Ian Johnston, Dominic Kwiatkowski, Cordelia Langford, John Sillitoe on behalf of the Wellcome Sanger Institute COVID-19 Surveillance Team |
| EPI_ISL_3095675, EPI_ISL_3095676 | Lighthouse Lab in Milton Keynes | Wellcome Sanger Institute for the COVID-19 Genomics UK (COG-UK) Consortium | The Lighthouse Lab in Milton Keynes and Alex Alderton, Roberto Amato, Jeffrey Barrett, Sonia Goncalves, Ewan Harrison, David K. Jackson, Ian Johnston, Dominic Kwiatkowski, Cordelia Langford, John Sillitoe on behalf of the Wellcome Sanger Institute COVID-19 Surveillance Team |
| EPI_ISL_3095677 | Lighthouse Lab in Glasgow | Wellcome Sanger Institute for the COVID-19 Genomics UK (COG-UK) Consortium | Harper VanSteenhouse, Yumi Kasai, David Gray, Carol Clugston, Anna Dominiczak and Alex Alderton, Roberto Amato, Jeffrey Barrett, Sonia Goncalves, Ewan Harrison, David K. Jackson, Ian Johnston, Dominic Kwiatkowski, Cordelia Langford, John Sillitoe on behalf of the Wellcome Sanger Institute COVID-19 Surveillance Team |
| EPI_ISL_3095678, EPI_ISL_3095679, EPI_ISL_3095680, EPI_ISL_3095681 | Lighthouse Lab in Milton Keynes | Wellcome Sanger Institute for the COVID-19 Genomics UK (COG-UK) Consortium | The Lighthouse Lab in Milton Keynes and Alex Alderton, Roberto Amato, Jeffrey Barrett, Sonia Goncalves, Ewan Harrison, David K. Jackson, Ian Johnston, Dominic Kwiatkowski, Cordelia Langford, John Sillitoe on behalf of the Wellcome Sanger Institute COVID-19 Surveillance Team |
| EPI_ISL_3095682, EPI_ISL_3095683 | Lighthouse Lab in Alderley Park | Wellcome Sanger Institute for the COVID-19 Genomics UK (COG-UK) Consortium | Jacquelyn Wynn, Mairead Hyland, The Lighthouse Lab in Alderley Park and Alex Alderton, Roberto Amato, Jeffrey Barrett, Sonia Goncalves, Ewan Harrison, David K. Jackson, Ian Johnston, Dominic Kwiatkowski, Cordelia Langford, John Sillitoe on behalf of the Wellcome Sanger Institute COVID-19 Surveillance Team |
| EPI_ISL_3095684, EPI_ISL_3095685 | Lighthouse Lab in Milton Keynes | Wellcome Sanger Institute for the COVID-19 Genomics UK (COG-UK) Consortium | The Lighthouse Lab in Milton Keynes and Alex Alderton, Roberto Amato, Jeffrey Barrett, Sonia Goncalves, Ewan Harrison, David K. Jackson, Ian Johnston, Dominic Kwiatkowski, Cordelia Langford, John Sillitoe on behalf of the Wellcome Sanger Institute COVID-19 Surveillance Team |
| EPI_ISL_3095686, EPI_ISL_3095687, EPI_ISL_3095688, EPI_ISL_3095689, EPI_ISL_3095690 | Lighthouse Lab in Alderley Park | Wellcome Sanger Institute for the COVID-19 Genomics UK (COG-UK) Consortium | Jacquelyn Wynn, Mairead Hyland, The Lighthouse Lab in Alderley Park and Alex Alderton, Roberto Amato, Jeffrey Barrett, Sonia Goncalves, Ewan Harrison, David K. Jackson, Ian Johnston, Dominic Kwiatkowski, Cordelia Langford, John Sillitoe on behalf of the Wellcome Sanger Institute COVID-19 Surveillance Team |
| EPI_ISL_3095691, EPI_ISL_3095692, EPI_ISL_3095693 | Lighthouse Lab in Milton Keynes | Wellcome Sanger Institute for the COVID-19 Genomics UK (COG-UK) Consortium | The Lighthouse Lab in Milton Keynes and Alex Alderton, Roberto Amato, Jeffrey Barrett, Sonia Goncalves, Ewan Harrison, David K. Jackson, Ian Johnston, Dominic Kwiatkowski, Cordelia Langford, John Sillitoe on behalf of the Wellcome Sanger Institute COVID-19 Surveillance Team |
| EPI_ISL_3095694, EPI_ISL_3095695 | Lighthouse Lab in Glasgow | Wellcome Sanger Institute for the COVID-19 Genomics UK (COG-UK) Consortium | Harper VanSteenhouse, Yumi Kasai, David Gray, Carol Clugston, Anna Dominiczak and Alex Alderton, Roberto Amato, Jeffrey Barrett, Sonia Goncalves, Ewan Harrison, David K. Jackson, Ian Johnston, Dominic Kwiatkowski, Cordelia Langford, John Sillitoe on behalf of the Wellcome Sanger Institute COVID-19 Surveillance Team |
| EPI_ISL_3095696, EPI_ISL_3095697, EPI_ISL_3095698 | Lighthouse Laboratory Plymouth | Wellcome Sanger Institute for the COVID-19 Genomics UK (COG-UK) Consortium | Lighthouse Laboratory Plymouth and Alex Alderton, Roberto Amato, Jeffrey Barrett, Sonia Goncalves, Ewan Harrison, David K. Jackson, Ian Johnston, Dominic Kwiatkowski, Cordelia Langford, John Sillitoe on behalf of the Wellcome Sanger Institute COVID-19 Surveillance Team |
| EPI_ISL_3095699 | Berkshire and Surrey Pathology Services Lighthouse Laboratory | Wellcome Sanger Institute for the COVID-19 Genomics UK (COG-UK) Consortium | Berkshire and Surrey Pathology Services Lighthouse Laboratory and Alex Alderton, Roberto Amato, Jeffrey Barrett, Sonia Goncalves, Ewan Harrison, David K. Jackson, Ian Johnston, Dominic Kwiatkowski, Cordelia Langford, John Sillitoe on behalf of the Wellcome Sanger Institute COVID-19 Surveillance Team |
| EPI_ISL_3095700 | Lighthouse Lab in Glasgow | Wellcome Sanger Institute for the COVID-19 Genomics UK (COG-UK) Consortium | Harper VanSteenhouse, Yumi Kasai, David Gray, Carol Clugston, Anna Dominiczak and Alex Alderton, Roberto Amato, Jeffrey Barrett, Sonia Goncalves, Ewan Harrison, David K. Jackson, Ian Johnston, Dominic Kwiatkowski, Cordelia Langford, John Sillitoe on behalf of the Wellcome Sanger Institute COVID-19 Surveillance Team |
| EPI_ISL_3095701 | Health Services Laboratories | Wellcome Sanger Institute for the COVID-19 Genomics UK (COG-UK) Consortium | Health Services Laboratories and Alex Alderton, Roberto Amato, Jeffrey Barrett, Sonia Goncalves, Ewan Harrison, David K. Jackson, Ian Johnston, Dominic Kwiatkowski, Cordelia Langford, John Sillitoe on behalf of the Wellcome Sanger Institute COVID-19 Surveillance Team |
| EPI_ISL_3095702 | Lighthouse Lab in Glasgow | Wellcome Sanger Institute for the COVID-19 Genomics UK (COG-UK) Consortium | Harper VanSteenhouse, Yumi Kasai, David Gray, Carol Clugston, Anna Dominiczak and Alex Alderton, Roberto Amato, Jeffrey Barrett, Sonia Goncalves, Ewan Harrison, David K. Jackson, Ian Johnston, Dominic Kwiatkowski, Cordelia Langford, John Sillitoe on behalf of the Wellcome Sanger Institute COVID-19 Surveillance Team |
| EPI_ISL_3095703 | Lighthouse Laboratory Plymouth | Wellcome Sanger Institute for the COVID-19 Genomics UK (COG-UK) Consortium | Lighthouse Laboratory Plymouth and Alex Alderton, Roberto Amato, Jeffrey Barrett, Sonia Goncalves, Ewan Harrison, David K. Jackson, Ian Johnston, Dominic Kwiatkowski, Cordelia Langford, John Sillitoe on behalf of the Wellcome Sanger Institute COVID-19 Surveillance Team |
| EPI_ISL_3095704 | Lighthouse Lab in Glasgow | Wellcome Sanger Institute for the COVID-19 Genomics UK (COG-UK) Consortium | Harper VanSteenhouse, Yumi Kasai, David Gray, Carol Clugston, Anna Dominiczak and Alex Alderton, Roberto Amato, Jeffrey Barrett, Sonia Goncalves, Ewan Harrison, David K. Jackson, Ian Johnston, Dominic Kwiatkowski, Cordelia Langford, John Sillitoe on behalf of the Wellcome Sanger Institute COVID-19 Surveillance Team |
| EPI_ISL_3095705, EPI_ISL_3095706 | Lighthouse Laboratory Plymouth | Wellcome Sanger Institute for the COVID-19 Genomics UK (COG-UK) Consortium | Lighthouse Laboratory Plymouth and Alex Alderton, Roberto Amato, Jeffrey Barrett, Sonia Goncalves, Ewan Harrison, David K. Jackson, Ian Johnston, Dominic Kwiatkowski, Cordelia Langford, John Sillitoe on behalf of the Wellcome Sanger Institute COVID-19 Surveillance Team |
| EPI_ISL_3095707, EPI_ISL_3095708, EPI_ISL_3095709, EPI_ISL_3095710, EPI_ISL_3095711, EPI_ISL_3095712 | Lighthouse Lab in Glasgow | Wellcome Sanger Institute for the COVID-19 Genomics UK (COG-UK) Consortium | Harper VanSteenhouse, Yumi Kasai, David Gray, Carol Clugston, Anna Dominiczak and Alex Alderton, Roberto Amato, Jeffrey Barrett, Sonia Goncalves, Ewan Harrison, David K. Jackson, Ian Johnston, Dominic Kwiatkowski, Cordelia Langford, John Sillitoe on behalf of the Wellcome Sanger Institute COVID-19 Surveillance Team |
| EPI_ISL_3095713 | Lighthouse Laboratory Plymouth | Wellcome Sanger Institute for the COVID-19 Genomics UK (COG-UK) Consortium | Lighthouse Laboratory Plymouth and Alex Alderton, Roberto Amato, Jeffrey Barrett, Sonia Goncalves, Ewan Harrison, David K. Jackson, Ian Johnston, Dominic Kwiatkowski, Cordelia Langford, John Sillitoe on behalf of the Wellcome Sanger Institute COVID-19 Surveillance Team |
| EPI_ISL_3095714, EPI_ISL_3095715 | Lighthouse Lab in Glasgow | Wellcome Sanger Institute for the COVID-19 Genomics UK (COG-UK) Consortium | Harper VanSteenhouse, Yumi Kasai, David Gray, Carol Clugston, Anna Dominiczak and Alex Alderton, Roberto Amato, Jeffrey Barrett, Sonia Goncalves, Ewan Harrison, David K. Jackson, Ian Johnston, Dominic Kwiatkowski, Cordelia Langford, John Sillitoe on behalf of the Wellcome Sanger Institute COVID-19 Surveillance Team |
| EPI_ISL_3095716 | Lighthouse Laboratory Plymouth | Wellcome Sanger Institute for the COVID-19 Genomics UK (COG-UK) Consortium | Lighthouse Laboratory Plymouth and Alex Alderton, Roberto Amato, Jeffrey Barrett, Sonia Goncalves, Ewan Harrison, David K. Jackson, Ian Johnston, Dominic Kwiatkowski, Cordelia Langford, John Sillitoe on behalf of the Wellcome Sanger Institute COVID-19 Surveillance Team |
| EPI_ISL_3095717, EPI_ISL_3095718, EPI_ISL_3095719 | Lighthouse Lab in Glasgow | Wellcome Sanger Institute for the COVID-19 Genomics UK (COG-UK) Consortium | Harper VanSteenhouse, Yumi Kasai, David Gray, Carol Clugston, Anna Dominiczak and Alex Alderton, Roberto Amato, Jeffrey Barrett, Sonia Goncalves, Ewan Harrison, David K. Jackson, Ian Johnston, Dominic Kwiatkowski, Cordelia Langford, John Sillitoe on behalf of the Wellcome Sanger Institute COVID-19 Surveillance Team |
| EPI_ISL_3095720 | Berkshire and Surrey Pathology Services Lighthouse Laboratory | Wellcome Sanger Institute for the COVID-19 Genomics UK (COG-UK) Consortium | Berkshire and Surrey Pathology Services Lighthouse Laboratory and Alex Alderton, Roberto Amato, Jeffrey Barrett, Sonia Goncalves, Ewan Harrison, David K. Jackson, Ian Johnston, Dominic Kwiatkowski, Cordelia Langford, John Sillitoe on behalf of the Wellcome Sanger Institute COVID-19 Surveillance Team |
| EPI_ISL_3095721, EPI_ISL_3095722, EPI_ISL_3095723, EPI_ISL_3095724, EPI_ISL_3095725 | Lighthouse Lab in Glasgow | Wellcome Sanger Institute for the COVID-19 Genomics UK (COG-UK) Consortium | Harper VanSteenhouse, Yumi Kasai, David Gray, Carol Clugston, Anna Dominiczak and Alex Alderton, Roberto Amato, Jeffrey Barrett, Sonia Goncalves, Ewan Harrison, David K. Jackson, Ian Johnston, Dominic Kwiatkowski, Cordelia Langford, John Sillitoe on behalf of the Wellcome Sanger Institute COVID-19 Surveillance Team |
| EPI_ISL_3095726 | Lighthouse Laboratory Plymouth | Wellcome Sanger Institute for the COVID-19 Genomics UK (COG-UK) Consortium | Lighthouse Laboratory Plymouth and Alex Alderton, Roberto Amato, Jeffrey Barrett, Sonia Goncalves, Ewan Harrison, David K. Jackson, Ian Johnston, Dominic Kwiatkowski, Cordelia Langford, John Sillitoe on behalf of the Wellcome Sanger Institute COVID-19 Surveillance Team |
| EPI_ISL_3095727 | Lighthouse Lab in Glasgow | Wellcome Sanger Institute for the COVID-19 Genomics UK (COG-UK) | Harper VanSteenhouse, Yumi Kasai, David Gray, Carol Clugston, Anna Dominiczak and Alex Alderton, Roberto Amato, Jeffrey Barrett, Sonia Goncalves, Ewan Harrison, David K. Jackson, Ian Johnston, Dominic Kwiatkowski, Cordelia Langford, |

| | | | |
|---|---|---|---|
| | | Consortium | John Sillitoe on behalf of the Wellcome Sanger Institute COVID-19 Surveillance Team |
| EPI_ISL_3095728 | Berkshire and Surrey Pathology Services Lighthouse Laboratory | Wellcome Sanger Institute for the COVID-19 Genomics UK (COG-UK) Consortium | Berkshire and Surrey Pathology Services Lighthouse Laboratory and Alex Alderton, Roberto Amato, Jeffrey Barrett, Sonia Goncalves, Ewan Harrison, David K. Jackson, Ian Johnston, Dominic Kwiatkowski, Cordelia Langford, John Sillitoe on behalf of the Wellcome Sanger Institute COVID-19 Surveillance Team |
| EPI_ISL_3095729, EPI_ISL_3095730 | Lighthouse Laboratory Plymouth | Wellcome Sanger Institute for the COVID-19 Genomics UK (COG-UK) Consortium | Lighthouse Laboratory Plymouth and Alex Alderton, Roberto Amato, Jeffrey Barrett, Sonia Goncalves, Ewan Harrison, David K. Jackson, Ian Johnston, Dominic Kwiatkowski, Cordelia Langford, John Sillitoe on behalf of the Wellcome Sanger Institute COVID-19 Surveillance Team |
| EPI_ISL_3095731 | Lighthouse Lab in Glasgow | Wellcome Sanger Institute for the COVID-19 Genomics UK (COG-UK) Consortium | Harper VanSteenhouse, Yumi Kasai, David Gray, Carol Clugston, Anna Dominiczak and Alex Alderton, Roberto Amato, Jeffrey Barrett, Sonia Goncalves, Ewan Harrison, David K. Jackson, Ian Johnston, Dominic Kwiatkowski, Cordelia Langford, John Sillitoe on behalf of the Wellcome Sanger Institute COVID-19 Surveillance Team |
| EPI_ISL_3095732 | Berkshire and Surrey Pathology Services Lighthouse Laboratory | Wellcome Sanger Institute for the COVID-19 Genomics UK (COG-UK) Consortium | Berkshire and Surrey Pathology Services Lighthouse Laboratory and Alex Alderton, Roberto Amato, Jeffrey Barrett, Sonia Goncalves, Ewan Harrison, David K. Jackson, Ian Johnston, Dominic Kwiatkowski, Cordelia Langford, John Sillitoe on behalf of the Wellcome Sanger Institute COVID-19 Surveillance Team |
| EPI_ISL_3095733, EPI_ISL_3095734, EPI_ISL_3095735 | Lighthouse Lab in Glasgow | Wellcome Sanger Institute for the COVID-19 Genomics UK (COG-UK) Consortium | Harper VanSteenhouse, Yumi Kasai, David Gray, Carol Clugston, Anna Dominiczak and Alex Alderton, Roberto Amato, Jeffrey Barrett, Sonia Goncalves, Ewan Harrison, David K. Jackson, Ian Johnston, Dominic Kwiatkowski, Cordelia Langford, John Sillitoe on behalf of the Wellcome Sanger Institute COVID-19 Surveillance Team |
| EPI_ISL_3095736 | Berkshire and Surrey Pathology Services Lighthouse Laboratory | Wellcome Sanger Institute for the COVID-19 Genomics UK (COG-UK) Consortium | Berkshire and Surrey Pathology Services Lighthouse Laboratory and Alex Alderton, Roberto Amato, Jeffrey Barrett, Sonia Goncalves, Ewan Harrison, David K. Jackson, Ian Johnston, Dominic Kwiatkowski, Cordelia Langford, John Sillitoe on behalf of the Wellcome Sanger Institute COVID-19 Surveillance Team |
| EPI_ISL_3095737, EPI_ISL_3095738 | Lighthouse Lab in Glasgow | Wellcome Sanger Institute for the COVID-19 Genomics UK (COG-UK) Consortium | Harper VanSteenhouse, Yumi Kasai, David Gray, Carol Clugston, Anna Dominiczak and Alex Alderton, Roberto Amato, Jeffrey Barrett, Sonia Goncalves, Ewan Harrison, David K. Jackson, Ian Johnston, Dominic Kwiatkowski, Cordelia Langford, John Sillitoe on behalf of the Wellcome Sanger Institute COVID-19 Surveillance Team |
| EPI_ISL_3095739 | Lighthouse Laboratory Plymouth | Wellcome Sanger Institute for the COVID-19 Genomics UK (COG-UK) Consortium | Lighthouse Laboratory Plymouth and Alex Alderton, Roberto Amato, Jeffrey Barrett, Sonia Goncalves, Ewan Harrison, David K. Jackson, Ian Johnston, Dominic Kwiatkowski, Cordelia Langford, John Sillitoe on behalf of the Wellcome Sanger Institute COVID-19 Surveillance Team |
| EPI_ISL_3095740 | Berkshire and Surrey Pathology Services Lighthouse Laboratory | Wellcome Sanger Institute for the COVID-19 Genomics UK (COG-UK) Consortium | Berkshire and Surrey Pathology Services Lighthouse Laboratory and Alex Alderton, Roberto Amato, Jeffrey Barrett, Sonia Goncalves, Ewan Harrison, David K. Jackson, Ian Johnston, Dominic Kwiatkowski, Cordelia Langford, John Sillitoe on behalf of the Wellcome Sanger Institute COVID-19 Surveillance Team |
| EPI_ISL_3095741 | Lighthouse Lab in Glasgow | Wellcome Sanger Institute for the COVID-19 Genomics UK (COG-UK) Consortium | Harper VanSteenhouse, Yumi Kasai, David Gray, Carol Clugston, Anna Dominiczak and Alex Alderton, Roberto Amato, Jeffrey Barrett, Sonia Goncalves, Ewan Harrison, David K. Jackson, Ian Johnston, Dominic Kwiatkowski, Cordelia Langford, John Sillitoe on behalf of the Wellcome Sanger Institute COVID-19 Surveillance Team |
| EPI_ISL_3095742 | Berkshire and Surrey Pathology Services Lighthouse Laboratory | Wellcome Sanger Institute for the COVID-19 Genomics UK (COG-UK) Consortium | Berkshire and Surrey Pathology Services Lighthouse Laboratory and Alex Alderton, Roberto Amato, Jeffrey Barrett, Sonia Goncalves, Ewan Harrison, David K. Jackson, Ian Johnston, Dominic Kwiatkowski, Cordelia Langford, John Sillitoe on behalf of the Wellcome Sanger Institute COVID-19 Surveillance Team |
| EPI_ISL_3095743, EPI_ISL_3095744, EPI_ISL_3095745 | Lighthouse Lab in Glasgow | Wellcome Sanger Institute for the COVID-19 Genomics UK (COG-UK) Consortium | Harper VanSteenhouse, Yumi Kasai, David Gray, Carol Clugston, Anna Dominiczak and Alex Alderton, Roberto Amato, Jeffrey Barrett, Sonia Goncalves, Ewan Harrison, David K. Jackson, Ian Johnston, Dominic Kwiatkowski, Cordelia Langford, John Sillitoe on behalf of the Wellcome Sanger Institute COVID-19 Surveillance Team |
| EPI_ISL_3095746 | Lighthouse Laboratory Plymouth | Wellcome Sanger Institute for the COVID-19 Genomics UK (COG-UK) Consortium | Lighthouse Laboratory Plymouth and Alex Alderton, Roberto Amato, Jeffrey Barrett, Sonia Goncalves, Ewan Harrison, David K. Jackson, Ian Johnston, Dominic Kwiatkowski, Cordelia Langford, John Sillitoe on behalf of the Wellcome Sanger Institute COVID-19 Surveillance Team |
| EPI_ISL_3095747 | Lighthouse Lab in Glasgow | Wellcome Sanger Institute for the COVID-19 Genomics UK (COG-UK) Consortium | Harper VanSteenhouse, Yumi Kasai, David Gray, Carol Clugston, Anna Dominiczak and Alex Alderton, Roberto Amato, Jeffrey Barrett, Sonia Goncalves, Ewan Harrison, David K. Jackson, Ian Johnston, Dominic Kwiatkowski, Cordelia Langford, John Sillitoe on behalf of the Wellcome Sanger Institute COVID-19 Surveillance Team |
| EPI_ISL_3095748, EPI_ISL_3095749 | Lighthouse Laboratory Plymouth | Wellcome Sanger Institute for the COVID-19 Genomics UK (COG-UK) Consortium | Lighthouse Laboratory Plymouth and Alex Alderton, Roberto Amato, Jeffrey Barrett, Sonia Goncalves, Ewan Harrison, David K. Jackson, Ian Johnston, Dominic Kwiatkowski, Cordelia Langford, John Sillitoe on behalf of the Wellcome Sanger Institute COVID-19 Surveillance Team |
| EPI_ISL_3095750 | Lighthouse Lab in Glasgow | Wellcome Sanger Institute for the COVID-19 Genomics UK (COG-UK) Consortium | Harper VanSteenhouse, Yumi Kasai, David Gray, Carol Clugston, Anna Dominiczak and Alex Alderton, Roberto Amato, Jeffrey Barrett, Sonia Goncalves, Ewan Harrison, David K. Jackson, Ian Johnston, Dominic Kwiatkowski, Cordelia Langford, John Sillitoe on behalf of the Wellcome Sanger Institute COVID-19 Surveillance Team |
| EPI_ISL_3095751 | Berkshire and Surrey Pathology Services Lighthouse Laboratory | Wellcome Sanger Institute for the COVID-19 Genomics UK (COG-UK) Consortium | Berkshire and Surrey Pathology Services Lighthouse Laboratory and Alex Alderton, Roberto Amato, Jeffrey Barrett, Sonia Goncalves, Ewan Harrison, David K. Jackson, Ian Johnston, Dominic Kwiatkowski, Cordelia Langford, John Sillitoe on behalf of the Wellcome Sanger Institute COVID-19 Surveillance Team |
| EPI_ISL_3095752 | Health Services Laboratories | Wellcome Sanger Institute for the COVID-19 Genomics UK (COG-UK) Consortium | Health Services Laboratories and Alex Alderton, Roberto Amato, Jeffrey Barrett, Sonia Goncalves, Ewan Harrison, David K. Jackson, Ian Johnston, Dominic Kwiatkowski, Cordelia Langford, John Sillitoe on behalf of the Wellcome Sanger Institute COVID-19 Surveillance Team |
| EPI_ISL_3095753, EPI_ISL_3095754 | Lighthouse Laboratory Plymouth | Wellcome Sanger Institute for the COVID-19 Genomics UK (COG-UK) Consortium | Lighthouse Laboratory Plymouth and Alex Alderton, Roberto Amato, Jeffrey Barrett, Sonia Goncalves, Ewan Harrison, David K. Jackson, Ian Johnston, Dominic Kwiatkowski, Cordelia Langford, John Sillitoe on behalf of the Wellcome Sanger Institute COVID-19 Surveillance Team |
| EPI_ISL_3095755 | Berkshire and Surrey Pathology Services Lighthouse Laboratory | Wellcome Sanger Institute for the COVID-19 Genomics UK (COG-UK) Consortium | Berkshire and Surrey Pathology Services Lighthouse Laboratory and Alex Alderton, Roberto Amato, Jeffrey Barrett, Sonia Goncalves, Ewan Harrison, David K. Jackson, Ian Johnston, Dominic Kwiatkowski, Cordelia Langford, John Sillitoe on behalf of the Wellcome Sanger Institute COVID-19 Surveillance Team |
| EPI_ISL_3095756, EPI_ISL_3095757, EPI_ISL_3095758 | Lighthouse Lab in Glasgow | Wellcome Sanger Institute for the COVID-19 Genomics UK (COG-UK) Consortium | Harper VanSteenhouse, Yumi Kasai, David Gray, Carol Clugston, Anna Dominiczak and Alex Alderton, Roberto Amato, Jeffrey Barrett, Sonia Goncalves, Ewan Harrison, David K. Jackson, Ian Johnston, Dominic Kwiatkowski, Cordelia Langford, John Sillitoe on behalf of the Wellcome Sanger Institute COVID-19 Surveillance Team |
| EPI_ISL_3095759 | Lighthouse Laboratory Plymouth | Wellcome Sanger Institute for the COVID-19 Genomics UK (COG-UK) Consortium | Lighthouse Laboratory Plymouth and Alex Alderton, Roberto Amato, Jeffrey Barrett, Sonia Goncalves, Ewan Harrison, David K. Jackson, Ian Johnston, Dominic Kwiatkowski, Cordelia Langford, John Sillitoe on behalf of the Wellcome Sanger Institute COVID-19 Surveillance Team |
| EPI_ISL_3095760, EPI_ISL_3095761 | Lighthouse Lab in Glasgow | Wellcome Sanger Institute for the COVID-19 Genomics UK (COG-UK) Consortium | Harper VanSteenhouse, Yumi Kasai, David Gray, Carol Clugston, Anna Dominiczak and Alex Alderton, Roberto Amato, Jeffrey Barrett, Sonia Goncalves, Ewan Harrison, David K. Jackson, Ian Johnston, Dominic Kwiatkowski, Cordelia Langford, John Sillitoe on behalf of the Wellcome Sanger Institute COVID-19 Surveillance Team |
| EPI_ISL_3095762 | Berkshire and Surrey Pathology Services Lighthouse Laboratory | Wellcome Sanger Institute for the COVID-19 Genomics UK (COG-UK) Consortium | Berkshire and Surrey Pathology Services Lighthouse Laboratory and Alex Alderton, Roberto Amato, Jeffrey Barrett, Sonia Goncalves, Ewan Harrison, David K. Jackson, Ian Johnston, Dominic Kwiatkowski, Cordelia Langford, John Sillitoe on behalf of the Wellcome Sanger Institute COVID-19 Surveillance Team |
| EPI_ISL_3095763, EPI_ISL_3095764 | Lighthouse Lab in Glasgow | Wellcome Sanger Institute for the COVID-19 Genomics UK (COG-UK) Consortium | Harper VanSteenhouse, Yumi Kasai, David Gray, Carol Clugston, Anna Dominiczak and Alex Alderton, Roberto Amato, Jeffrey Barrett, Sonia Goncalves, Ewan Harrison, David K. Jackson, Ian Johnston, Dominic Kwiatkowski, Cordelia Langford, John Sillitoe on behalf of the Wellcome Sanger Institute COVID-19 Surveillance Team |
| EPI_ISL_3095765, EPI_ISL_3095766 | Lighthouse Laboratory Plymouth | Wellcome Sanger Institute for the COVID-19 Genomics UK (COG-UK) Consortium | Lighthouse Laboratory Plymouth and Alex Alderton, Roberto Amato, Jeffrey Barrett, Sonia Goncalves, Ewan Harrison, David K. Jackson, Ian Johnston, Dominic Kwiatkowski, Cordelia Langford, John Sillitoe on behalf of the Wellcome Sanger Institute COVID-19 Surveillance Team |
| EPI_ISL_3095767, EPI_ISL_3095768 | Lighthouse Lab in Glasgow | Wellcome Sanger Institute for the COVID-19 Genomics UK (COG-UK) Consortium | Harper VanSteenhouse, Yumi Kasai, David Gray, Carol Clugston, Anna Dominiczak and Alex Alderton, Roberto Amato, Jeffrey Barrett, Sonia Goncalves, Ewan Harrison, David K. Jackson, Ian Johnston, Dominic Kwiatkowski, Cordelia Langford, John Sillitoe on behalf of the Wellcome Sanger Institute COVID-19 Surveillance Team |
| EPI_ISL_3095769 | Lighthouse Laboratory Plymouth | Wellcome Sanger Institute for the COVID-19 Genomics UK (COG-UK) Consortium | Lighthouse Laboratory Plymouth and Alex Alderton, Roberto Amato, Jeffrey Barrett, Sonia Goncalves, Ewan Harrison, David K. Jackson, Ian Johnston, Dominic Kwiatkowski, Cordelia Langford, John Sillitoe on behalf of the Wellcome Sanger Institute COVID-19 Surveillance Team |
| EPI_ISL_3095770, EPI_ISL_3095771 | Lighthouse Lab in Glasgow | Wellcome Sanger Institute for the COVID-19 Genomics UK (COG-UK) Consortium | Harper VanSteenhouse, Yumi Kasai, David Gray, Carol Clugston, Anna Dominiczak and Alex Alderton, Roberto Amato, Jeffrey Barrett, Sonia Goncalves, Ewan Harrison, David K. Jackson, Ian Johnston, Dominic Kwiatkowski, Cordelia Langford, John Sillitoe on behalf of the Wellcome Sanger Institute COVID-19 Surveillance Team |
| EPI_ISL_3095772 | Lighthouse Laboratory Plymouth | Wellcome Sanger Institute for the COVID-19 Genomics UK (COG-UK) Consortium | Lighthouse Laboratory Plymouth and Alex Alderton, Roberto Amato, Jeffrey Barrett, Sonia Goncalves, Ewan Harrison, David K. Jackson, Ian Johnston, Dominic Kwiatkowski, Cordelia Langford, John Sillitoe on behalf of the Wellcome Sanger Institute COVID-19 Surveillance Team |
| EPI_ISL_3095773 | Lighthouse Lab in Glasgow | Wellcome Sanger Institute for the COVID-19 Genomics UK (COG-UK) Consortium | Harper VanSteenhouse, Yumi Kasai, David Gray, Carol Clugston, Anna Dominiczak and Alex Alderton, Roberto Amato, Jeffrey Barrett, Sonia Goncalves, Ewan Harrison, David K. Jackson, Ian Johnston, Dominic Kwiatkowski, Cordelia Langford, John Sillitoe on behalf of the Wellcome Sanger Institute COVID-19 Surveillance Team |
| EPI_ISL_3095774 | Lighthouse Laboratory Plymouth | Wellcome Sanger Institute for the COVID-19 Genomics UK (COG-UK) Consortium | Lighthouse Laboratory Plymouth and Alex Alderton, Roberto Amato, Jeffrey Barrett, Sonia Goncalves, Ewan Harrison, David K. Jackson, Ian Johnston, Dominic Kwiatkowski, Cordelia Langford, John Sillitoe on behalf of the Wellcome Sanger Institute COVID-19 Surveillance Team |
| EPI_ISL_3095775 | Berkshire and Surrey Pathology Services Lighthouse Laboratory | Wellcome Sanger Institute for the COVID-19 Genomics UK (COG-UK) Consortium | Berkshire and Surrey Pathology Services Lighthouse Laboratory and Alex Alderton, Roberto Amato, Jeffrey Barrett, Sonia Goncalves, Ewan Harrison, David K. Jackson, Ian Johnston, Dominic Kwiatkowski, Cordelia Langford, John Sillitoe on behalf of the Wellcome Sanger Institute COVID-19 Surveillance Team |
| EPI_ISL_3095776, EPI_ISL_3095777, EPI_ISL_3095778, EPI_ISL_3095779 | Lighthouse Lab in Glasgow | Wellcome Sanger Institute for the COVID-19 Genomics UK (COG-UK) Consortium | Harper VanSteenhouse, Yumi Kasai, David Gray, Carol Clugston, Anna Dominiczak and Alex Alderton, Roberto Amato, Jeffrey Barrett, Sonia Goncalves, Ewan Harrison, David K. Jackson, Ian Johnston, Dominic Kwiatkowski, Cordelia Langford, John Sillitoe on behalf of the Wellcome Sanger Institute COVID-19 Surveillance Team |
| EPI_ISL_3095780 | Health Services Laboratories | Wellcome Sanger Institute for the COVID-19 Genomics UK (COG-UK) Consortium | Health Services Laboratories and Alex Alderton, Roberto Amato, Jeffrey Barrett, Sonia Goncalves, Ewan Harrison, David K. Jackson, Ian Johnston, Dominic Kwiatkowski, Cordelia Langford, John Sillitoe on behalf of the Wellcome Sanger Institute COVID-19 Surveillance Team |
| EPI_ISL_3095781, EPI_ISL_3095782 | Lighthouse Laboratory Plymouth | Wellcome Sanger Institute for the COVID-19 Genomics UK (COG-UK) Consortium | Lighthouse Laboratory Plymouth and Alex Alderton, Roberto Amato, Jeffrey Barrett, Sonia Goncalves, Ewan Harrison, David K. Jackson, Ian Johnston, Dominic Kwiatkowski, Cordelia Langford, John Sillitoe on behalf of the Wellcome Sanger Institute COVID-19 Surveillance Team |
| EPI_ISL_3095783, EPI_ISL_3095784 | Berkshire and Surrey Pathology Services Lighthouse Laboratory | Wellcome Sanger Institute for the COVID-19 Genomics UK (COG-UK) Consortium | Berkshire and Surrey Pathology Services Lighthouse Laboratory and Alex Alderton, Roberto Amato, Jeffrey Barrett, Sonia Goncalves, Ewan Harrison, David K. Jackson, Ian Johnston, Dominic Kwiatkowski, Cordelia Langford, John Sillitoe on behalf of the Wellcome Sanger Institute COVID-19 Surveillance Team |
| EPI_ISL_3095785 | Lighthouse Laboratory Plymouth | Wellcome Sanger Institute for the COVID-19 Genomics UK (COG-UK) Consortium | Lighthouse Laboratory Plymouth and Alex Alderton, Roberto Amato, Jeffrey Barrett, Sonia Goncalves, Ewan Harrison, David K. Jackson, Ian Johnston, Dominic Kwiatkowski, Cordelia Langford, John Sillitoe on behalf of the Wellcome Sanger Institute COVID-19 Surveillance Team |
| EPI_ISL_3095786, EPI_ISL_3095787 | Lighthouse Lab in Glasgow | Wellcome Sanger Institute for the COVID-19 Genomics UK (COG-UK) Consortium | Harper VanSteenhouse, Yumi Kasai, David Gray, Carol Clugston, Anna Dominiczak and Alex Alderton, Roberto Amato, Jeffrey Barrett, Sonia Goncalves, Ewan Harrison, David K. Jackson, Ian Johnston, Dominic Kwiatkowski, Cordelia Langford, John Sillitoe on behalf of the Wellcome Sanger Institute COVID-19 Surveillance Team |
| EPI_ISL_3095788 | Berkshire and Surrey Pathology Services Lighthouse Laboratory | Wellcome Sanger Institute for the COVID-19 Genomics UK (COG-UK) Consortium | Berkshire and Surrey Pathology Services Lighthouse Laboratory and Alex Alderton, Roberto Amato, Jeffrey Barrett, Sonia Goncalves, Ewan Harrison, David K. Jackson, Ian Johnston, Dominic Kwiatkowski, Cordelia Langford, John Sillitoe on behalf of the Wellcome Sanger Institute COVID-19 Surveillance Team |
| EPI_ISL_3095789 | Lighthouse Lab in Glasgow | Wellcome Sanger Institute for the COVID-19 Genomics UK (COG-UK) Consortium | Harper VanSteenhouse, Yumi Kasai, David Gray, Carol Clugston, Anna Dominiczak and Alex Alderton, Roberto Amato, Jeffrey Barrett, Sonia Goncalves, Ewan Harrison, David K. Jackson, Ian Johnston, Dominic Kwiatkowski, Cordelia Langford, John Sillitoe on behalf of the Wellcome Sanger Institute COVID-19 Surveillance Team |
| EPI_ISL_3095790 | Berkshire and Surrey Pathology Services Lighthouse Laboratory | Wellcome Sanger Institute for the COVID-19 Genomics UK (COG-UK) Consortium | Berkshire and Surrey Pathology Services Lighthouse Laboratory and Alex Alderton, Roberto Amato, Jeffrey Barrett, Sonia Goncalves, Ewan Harrison, David K. Jackson, Ian Johnston, Dominic Kwiatkowski, Cordelia Langford, John Sillitoe on behalf of the Wellcome Sanger Institute COVID-19 Surveillance Team |
| EPI_ISL_3095791 | Lighthouse Laboratory Plymouth | Wellcome Sanger Institute for the COVID-19 Genomics UK (COG-UK) | Lighthouse Laboratory Plymouth and Alex Alderton, Roberto Amato, Jeffrey Barrett, Sonia Goncalves, Ewan Harrison, David K. Jackson, Ian Johnston, Dominic Kwiatkowski, Cordelia Langford, John Sillitoe on behalf of the Wellcome Sanger |

| | | | |
|---|---|---|---|
| | | Consortium | Institute COVID-19 Surveillance Team |
| EPI_ISL_3095792 | Lighthouse Lab in Glasgow | Wellcome Sanger Institute for the COVID-19 Genomics UK (COG-UK) Consortium | Harper VanSteenhouse, Yumi Kasai, David Gray, Carol Clugston, Anna Dominiczak and Alex Alderton, Roberto Amato, Jeffrey Barrett, Sonia Goncalves, Ewan Harrison, David K. Jackson, Ian Johnston, Dominic Kwiatkowski, Cordelia Langford, John Sillitoe on behalf of the Wellcome Sanger Institute COVID-19 Surveillance Team |
| EPI_ISL_3095793 | Lighthouse Laboratory Plymouth | Wellcome Sanger Institute for the COVID-19 Genomics UK (COG-UK) Consortium | Lighthouse Laboratory Plymouth and Alex Alderton, Roberto Amato, Jeffrey Barrett, Sonia Goncalves, Ewan Harrison, David K. Jackson, Ian Johnston, Dominic Kwiatkowski, Cordelia Langford, John Sillitoe on behalf of the Wellcome Sanger Institute COVID-19 Surveillance Team |
| EPI_ISL_3095794 | Berkshire and Surrey Pathology Services Lighthouse Laboratory | Wellcome Sanger Institute for the COVID-19 Genomics UK (COG-UK) Consortium | Berkshire and Surrey Pathology Services Lighthouse Laboratory and Alex Alderton, Roberto Amato, Jeffrey Barrett, Sonia Goncalves, Ewan Harrison, David K. Jackson, Ian Johnston, Dominic Kwiatkowski, Cordelia Langford, John Sillitoe on behalf of the Wellcome Sanger Institute COVID-19 Surveillance Team |
| EPI_ISL_3095795, EPI_ISL_3095796, EPI_ISL_3095797, EPI_ISL_3095798 | Lighthouse Lab in Glasgow | Wellcome Sanger Institute for the COVID-19 Genomics UK (COG-UK) Consortium | Harper VanSteenhouse, Yumi Kasai, David Gray, Carol Clugston, Anna Dominiczak and Alex Alderton, Roberto Amato, Jeffrey Barrett, Sonia Goncalves, Ewan Harrison, David K. Jackson, Ian Johnston, Dominic Kwiatkowski, Cordelia Langford, John Sillitoe on behalf of the Wellcome Sanger Institute COVID-19 Surveillance Team |
| EPI_ISL_3095799, EPI_ISL_3095800 | Berkshire and Surrey Pathology Services Lighthouse Laboratory | Wellcome Sanger Institute for the COVID-19 Genomics UK (COG-UK) Consortium | Berkshire and Surrey Pathology Services Lighthouse Laboratory and Alex Alderton, Roberto Amato, Jeffrey Barrett, Sonia Goncalves, Ewan Harrison, David K. Jackson, Ian Johnston, Dominic Kwiatkowski, Cordelia Langford, John Sillitoe on behalf of the Wellcome Sanger Institute COVID-19 Surveillance Team |
| EPI_ISL_3095801, EPI_ISL_3095802 | Lighthouse Lab in Glasgow | Wellcome Sanger Institute for the COVID-19 Genomics UK (COG-UK) Consortium | Harper VanSteenhouse, Yumi Kasai, David Gray, Carol Clugston, Anna Dominiczak and Alex Alderton, Roberto Amato, Jeffrey Barrett, Sonia Goncalves, Ewan Harrison, David K. Jackson, Ian Johnston, Dominic Kwiatkowski, Cordelia Langford, John Sillitoe on behalf of the Wellcome Sanger Institute COVID-19 Surveillance Team |
| EPI_ISL_3095803 | Lighthouse Laboratory Plymouth | Wellcome Sanger Institute for the COVID-19 Genomics UK (COG-UK) Consortium | Lighthouse Laboratory Plymouth and Alex Alderton, Roberto Amato, Jeffrey Barrett, Sonia Goncalves, Ewan Harrison, David K. Jackson, Ian Johnston, Dominic Kwiatkowski, Cordelia Langford, John Sillitoe on behalf of the Wellcome Sanger Institute COVID-19 Surveillance Team |
| EPI_ISL_3095804, EPI_ISL_3095805 | Berkshire and Surrey Pathology Services Lighthouse Laboratory | Wellcome Sanger Institute for the COVID-19 Genomics UK (COG-UK) Consortium | Berkshire and Surrey Pathology Services Lighthouse Laboratory and Alex Alderton, Roberto Amato, Jeffrey Barrett, Sonia Goncalves, Ewan Harrison, David K. Jackson, Ian Johnston, Dominic Kwiatkowski, Cordelia Langford, John Sillitoe on behalf of the Wellcome Sanger Institute COVID-19 Surveillance Team |
| EPI_ISL_3095806 | Lighthouse Lab in Glasgow | Wellcome Sanger Institute for the COVID-19 Genomics UK (COG-UK) Consortium | Harper VanSteenhouse, Yumi Kasai, David Gray, Carol Clugston, Anna Dominiczak and Alex Alderton, Roberto Amato, Jeffrey Barrett, Sonia Goncalves, Ewan Harrison, David K. Jackson, Ian Johnston, Dominic Kwiatkowski, Cordelia Langford, John Sillitoe on behalf of the Wellcome Sanger Institute COVID-19 Surveillance Team |
| EPI_ISL_3095807, EPI_ISL_3095808 | Lighthouse Laboratory Plymouth | Wellcome Sanger Institute for the COVID-19 Genomics UK (COG-UK) Consortium | Lighthouse Laboratory Plymouth and Alex Alderton, Roberto Amato, Jeffrey Barrett, Sonia Goncalves, Ewan Harrison, David K. Jackson, Ian Johnston, Dominic Kwiatkowski, Cordelia Langford, John Sillitoe on behalf of the Wellcome Sanger Institute COVID-19 Surveillance Team |
| EPI_ISL_3095809 | Berkshire and Surrey Pathology Services Lighthouse Laboratory | Wellcome Sanger Institute for the COVID-19 Genomics UK (COG-UK) Consortium | Berkshire and Surrey Pathology Services Lighthouse Laboratory and Alex Alderton, Roberto Amato, Jeffrey Barrett, Sonia Goncalves, Ewan Harrison, David K. Jackson, Ian Johnston, Dominic Kwiatkowski, Cordelia Langford, John Sillitoe on behalf of the Wellcome Sanger Institute COVID-19 Surveillance Team |
| EPI_ISL_3095810, EPI_ISL_3095811 | Lighthouse Lab in Glasgow | Wellcome Sanger Institute for the COVID-19 Genomics UK (COG-UK) Consortium | Harper VanSteenhouse, Yumi Kasai, David Gray, Carol Clugston, Anna Dominiczak and Alex Alderton, Roberto Amato, Jeffrey Barrett, Sonia Goncalves, Ewan Harrison, David K. Jackson, Ian Johnston, Dominic Kwiatkowski, Cordelia Langford, John Sillitoe on behalf of the Wellcome Sanger Institute COVID-19 Surveillance Team |
| EPI_ISL_3095812 | Berkshire and Surrey Pathology Services Lighthouse Laboratory | Wellcome Sanger Institute for the COVID-19 Genomics UK (COG-UK) Consortium | Berkshire and Surrey Pathology Services Lighthouse Laboratory and Alex Alderton, Roberto Amato, Jeffrey Barrett, Sonia Goncalves, Ewan Harrison, David K. Jackson, Ian Johnston, Dominic Kwiatkowski, Cordelia Langford, John Sillitoe on behalf of the Wellcome Sanger Institute COVID-19 Surveillance Team |
| EPI_ISL_3095813 | Lighthouse Lab in Glasgow | Wellcome Sanger Institute for the COVID-19 Genomics UK (COG-UK) Consortium | Harper VanSteenhouse, Yumi Kasai, David Gray, Carol Clugston, Anna Dominiczak and Alex Alderton, Roberto Amato, Jeffrey Barrett, Sonia Goncalves, Ewan Harrison, David K. Jackson, Ian Johnston, Dominic Kwiatkowski, Cordelia Langford, John Sillitoe on behalf of the Wellcome Sanger Institute COVID-19 Surveillance Team |
| EPI_ISL_3095814 | Berkshire and Surrey Pathology Services Lighthouse Laboratory | Wellcome Sanger Institute for the COVID-19 Genomics UK (COG-UK) Consortium | Berkshire and Surrey Pathology Services Lighthouse Laboratory and Alex Alderton, Roberto Amato, Jeffrey Barrett, Sonia Goncalves, Ewan Harrison, David K. Jackson, Ian Johnston, Dominic Kwiatkowski, Cordelia Langford, John Sillitoe on behalf of the Wellcome Sanger Institute COVID-19 Surveillance Team |
| EPI_ISL_3095815 | Lighthouse Laboratory Plymouth | Wellcome Sanger Institute for the COVID-19 Genomics UK (COG-UK) Consortium | Lighthouse Laboratory Plymouth and Alex Alderton, Roberto Amato, Jeffrey Barrett, Sonia Goncalves, Ewan Harrison, David K. Jackson, Ian Johnston, Dominic Kwiatkowski, Cordelia Langford, John Sillitoe on behalf of the Wellcome Sanger Institute COVID-19 Surveillance Team |
| EPI_ISL_3095816 | Lighthouse Lab in Glasgow | Wellcome Sanger Institute for the COVID-19 Genomics UK (COG-UK) Consortium | Harper VanSteenhouse, Yumi Kasai, David Gray, Carol Clugston, Anna Dominiczak and Alex Alderton, Roberto Amato, Jeffrey Barrett, Sonia Goncalves, Ewan Harrison, David K. Jackson, Ian Johnston, Dominic Kwiatkowski, Cordelia Langford, John Sillitoe on behalf of the Wellcome Sanger Institute COVID-19 Surveillance Team |
| EPI_ISL_3095817, EPI_ISL_3095818 | Berkshire and Surrey Pathology Services Lighthouse Laboratory | Wellcome Sanger Institute for the COVID-19 Genomics UK (COG-UK) Consortium | Berkshire and Surrey Pathology Services Lighthouse Laboratory and Alex Alderton, Roberto Amato, Jeffrey Barrett, Sonia Goncalves, Ewan Harrison, David K. Jackson, Ian Johnston, Dominic Kwiatkowski, Cordelia Langford, John Sillitoe on behalf of the Wellcome Sanger Institute COVID-19 Surveillance Team |
| EPI_ISL_3095819, EPI_ISL_3095820, EPI_ISL_3095821 | Lighthouse Lab in Glasgow | Wellcome Sanger Institute for the COVID-19 Genomics UK (COG-UK) Consortium | Harper VanSteenhouse, Yumi Kasai, David Gray, Carol Clugston, Anna Dominiczak and Alex Alderton, Roberto Amato, Jeffrey Barrett, Sonia Goncalves, Ewan Harrison, David K. Jackson, Ian Johnston, Dominic Kwiatkowski, Cordelia Langford, John Sillitoe on behalf of the Wellcome Sanger Institute COVID-19 Surveillance Team |
| EPI_ISL_3095822, EPI_ISL_3095823 | Berkshire and Surrey Pathology Services Lighthouse Laboratory | Wellcome Sanger Institute for the COVID-19 Genomics UK (COG-UK) Consortium | Berkshire and Surrey Pathology Services Lighthouse Laboratory and Alex Alderton, Roberto Amato, Jeffrey Barrett, Sonia Goncalves, Ewan Harrison, David K. Jackson, Ian Johnston, Dominic Kwiatkowski, Cordelia Langford, John Sillitoe on behalf of the Wellcome Sanger Institute COVID-19 Surveillance Team |
| EPI_ISL_3095824 | Lighthouse Lab in Glasgow | Wellcome Sanger Institute for the COVID-19 Genomics UK (COG-UK) Consortium | Harper VanSteenhouse, Yumi Kasai, David Gray, Carol Clugston, Anna Dominiczak and Alex Alderton, Roberto Amato, Jeffrey Barrett, Sonia Goncalves, Ewan Harrison, David K. Jackson, Ian Johnston, Dominic Kwiatkowski, Cordelia Langford, John Sillitoe on behalf of the Wellcome Sanger Institute COVID-19 Surveillance Team |
| EPI_ISL_3095825 | Berkshire and Surrey Pathology Services Lighthouse Laboratory | Wellcome Sanger Institute for the COVID-19 Genomics UK (COG-UK) Consortium | Berkshire and Surrey Pathology Services Lighthouse Laboratory and Alex Alderton, Roberto Amato, Jeffrey Barrett, Sonia Goncalves, Ewan Harrison, David K. Jackson, Ian Johnston, Dominic Kwiatkowski, Cordelia Langford, John Sillitoe on behalf of the Wellcome Sanger Institute COVID-19 Surveillance Team |
| EPI_ISL_3095826 | Lighthouse Laboratory Plymouth | Wellcome Sanger Institute for the COVID-19 Genomics UK (COG-UK) Consortium | Lighthouse Laboratory Plymouth and Alex Alderton, Roberto Amato, Jeffrey Barrett, Sonia Goncalves, Ewan Harrison, David K. Jackson, Ian Johnston, Dominic Kwiatkowski, Cordelia Langford, John Sillitoe on behalf of the Wellcome Sanger Institute COVID-19 Surveillance Team |
| EPI_ISL_3095827 | Lighthouse Lab in Glasgow | Wellcome Sanger Institute for the COVID-19 Genomics UK (COG-UK) Consortium | Harper VanSteenhouse, Yumi Kasai, David Gray, Carol Clugston, Anna Dominiczak and Alex Alderton, Roberto Amato, Jeffrey Barrett, Sonia Goncalves, Ewan Harrison, David K. Jackson, Ian Johnston, Dominic Kwiatkowski, Cordelia Langford, John Sillitoe on behalf of the Wellcome Sanger Institute COVID-19 Surveillance Team |
| EPI_ISL_3095828 | Lighthouse Laboratory Plymouth | Wellcome Sanger Institute for the COVID-19 Genomics UK (COG-UK) Consortium | Lighthouse Laboratory Plymouth and Alex Alderton, Roberto Amato, Jeffrey Barrett, Sonia Goncalves, Ewan Harrison, David K. Jackson, Ian Johnston, Dominic Kwiatkowski, Cordelia Langford, John Sillitoe on behalf of the Wellcome Sanger Institute COVID-19 Surveillance Team |
| EPI_ISL_3095829 | Berkshire and Surrey Pathology Services Lighthouse Laboratory | Wellcome Sanger Institute for the COVID-19 Genomics UK (COG-UK) Consortium | Berkshire and Surrey Pathology Services Lighthouse Laboratory and Alex Alderton, Roberto Amato, Jeffrey Barrett, Sonia Goncalves, Ewan Harrison, David K. Jackson, Ian Johnston, Dominic Kwiatkowski, Cordelia Langford, John Sillitoe on behalf of the Wellcome Sanger Institute COVID-19 Surveillance Team |
| EPI_ISL_3095830 | Lighthouse Lab in Glasgow | Wellcome Sanger Institute for the COVID-19 Genomics UK (COG-UK) Consortium | Harper VanSteenhouse, Yumi Kasai, David Gray, Carol Clugston, Anna Dominiczak and Alex Alderton, Roberto Amato, Jeffrey Barrett, Sonia Goncalves, Ewan Harrison, David K. Jackson, Ian Johnston, Dominic Kwiatkowski, Cordelia Langford, John Sillitoe on behalf of the Wellcome Sanger Institute COVID-19 Surveillance Team |
| EPI_ISL_3095831 | Health Services Laboratories | Wellcome Sanger Institute for the COVID-19 Genomics UK (COG-UK) Consortium | Health Services Laboratories and Alex Alderton, Roberto Amato, Jeffrey Barrett, Sonia Goncalves, Ewan Harrison, David K. Jackson, Ian Johnston, Dominic Kwiatkowski, Cordelia Langford, John Sillitoe on behalf of the Wellcome Sanger Institute COVID-19 Surveillance Team |
| EPI_ISL_3095832 | Lighthouse Lab in Glasgow | Wellcome Sanger Institute for the COVID-19 Genomics UK (COG-UK) Consortium | Harper VanSteenhouse, Yumi Kasai, David Gray, Carol Clugston, Anna Dominiczak and Alex Alderton, Roberto Amato, Jeffrey Barrett, Sonia Goncalves, Ewan Harrison, David K. Jackson, Ian Johnston, Dominic Kwiatkowski, Cordelia Langford, John Sillitoe on behalf of the Wellcome Sanger Institute COVID-19 Surveillance Team |
| EPI_ISL_3095833 | Berkshire and Surrey Pathology Services Lighthouse Laboratory | Wellcome Sanger Institute for the COVID-19 Genomics UK (COG-UK) Consortium | Berkshire and Surrey Pathology Services Lighthouse Laboratory and Alex Alderton, Roberto Amato, Jeffrey Barrett, Sonia Goncalves, Ewan Harrison, David K. Jackson, Ian Johnston, Dominic Kwiatkowski, Cordelia Langford, John Sillitoe on behalf of the Wellcome Sanger Institute COVID-19 Surveillance Team |
| EPI_ISL_3095834 | Lighthouse Laboratory Plymouth | Wellcome Sanger Institute for the COVID-19 Genomics UK (COG-UK) Consortium | Lighthouse Laboratory Plymouth and Alex Alderton, Roberto Amato, Jeffrey Barrett, Sonia Goncalves, Ewan Harrison, David K. Jackson, Ian Johnston, Dominic Kwiatkowski, Cordelia Langford, John Sillitoe on behalf of the Wellcome Sanger Institute COVID-19 Surveillance Team |
| EPI_ISL_3095835 | Berkshire and Surrey Pathology Services Lighthouse Laboratory | Wellcome Sanger Institute for the COVID-19 Genomics UK (COG-UK) Consortium | Berkshire and Surrey Pathology Services Lighthouse Laboratory and Alex Alderton, Roberto Amato, Jeffrey Barrett, Sonia Goncalves, Ewan Harrison, David K. Jackson, Ian Johnston, Dominic Kwiatkowski, Cordelia Langford, John Sillitoe on behalf of the Wellcome Sanger Institute COVID-19 Surveillance Team |
| EPI_ISL_3095836, EPI_ISL_3095837 | Lighthouse Lab in Glasgow | Wellcome Sanger Institute for the COVID-19 Genomics UK (COG-UK) Consortium | Harper VanSteenhouse, Yumi Kasai, David Gray, Carol Clugston, Anna Dominiczak and Alex Alderton, Roberto Amato, Jeffrey Barrett, Sonia Goncalves, Ewan Harrison, David K. Jackson, Ian Johnston, Dominic Kwiatkowski, Cordelia Langford, John Sillitoe on behalf of the Wellcome Sanger Institute COVID-19 Surveillance Team |
| EPI_ISL_3095838 | Berkshire and Surrey Pathology Services Lighthouse Laboratory | Wellcome Sanger Institute for the COVID-19 Genomics UK (COG-UK) Consortium | Berkshire and Surrey Pathology Services Lighthouse Laboratory and Alex Alderton, Roberto Amato, Jeffrey Barrett, Sonia Goncalves, Ewan Harrison, David K. Jackson, Ian Johnston, Dominic Kwiatkowski, Cordelia Langford, John Sillitoe on behalf of the Wellcome Sanger Institute COVID-19 Surveillance Team |
| EPI_ISL_3095839 | Lighthouse Laboratory Plymouth | Wellcome Sanger Institute for the COVID-19 Genomics UK (COG-UK) Consortium | Lighthouse Laboratory Plymouth and Alex Alderton, Roberto Amato, Jeffrey Barrett, Sonia Goncalves, Ewan Harrison, David K. Jackson, Ian Johnston, Dominic Kwiatkowski, Cordelia Langford, John Sillitoe on behalf of the Wellcome Sanger Institute COVID-19 Surveillance Team |
| EPI_ISL_3095840 | Berkshire and Surrey Pathology Services Lighthouse Laboratory | Wellcome Sanger Institute for the COVID-19 Genomics UK (COG-UK) Consortium | Berkshire and Surrey Pathology Services Lighthouse Laboratory and Alex Alderton, Roberto Amato, Jeffrey Barrett, Sonia Goncalves, Ewan Harrison, David K. Jackson, Ian Johnston, Dominic Kwiatkowski, Cordelia Langford, John Sillitoe on behalf of the Wellcome Sanger Institute COVID-19 Surveillance Team |
| EPI_ISL_3095841 | Lighthouse Lab in Glasgow | Wellcome Sanger Institute for the COVID-19 Genomics UK (COG-UK) Consortium | Harper VanSteenhouse, Yumi Kasai, David Gray, Carol Clugston, Anna Dominiczak and Alex Alderton, Roberto Amato, Jeffrey Barrett, Sonia Goncalves, Ewan Harrison, David K. Jackson, Ian Johnston, Dominic Kwiatkowski, Cordelia Langford, John Sillitoe on behalf of the Wellcome Sanger Institute COVID-19 Surveillance Team |
| EPI_ISL_3095842 | Berkshire and Surrey Pathology Services Lighthouse Laboratory | Wellcome Sanger Institute for the COVID-19 Genomics UK (COG-UK) Consortium | Berkshire and Surrey Pathology Services Lighthouse Laboratory and Alex Alderton, Roberto Amato, Jeffrey Barrett, Sonia Goncalves, Ewan Harrison, David K. Jackson, Ian Johnston, Dominic Kwiatkowski, Cordelia Langford, John Sillitoe on behalf of the Wellcome Sanger Institute COVID-19 Surveillance Team |
| EPI_ISL_3095843 | Lighthouse Lab in Glasgow | Wellcome Sanger Institute for the COVID-19 Genomics UK (COG-UK) Consortium | Harper VanSteenhouse, Yumi Kasai, David Gray, Carol Clugston, Anna Dominiczak and Alex Alderton, Roberto Amato, Jeffrey Barrett, Sonia Goncalves, Ewan Harrison, David K. Jackson, Ian Johnston, Dominic Kwiatkowski, Cordelia Langford, John Sillitoe on behalf of the Wellcome Sanger Institute COVID-19 Surveillance Team |
| EPI_ISL_3095844 | Berkshire and Surrey Pathology Services Lighthouse Laboratory | Wellcome Sanger Institute for the COVID-19 Genomics UK (COG-UK) Consortium | Berkshire and Surrey Pathology Services Lighthouse Laboratory and Alex Alderton, Roberto Amato, Jeffrey Barrett, Sonia Goncalves, Ewan Harrison, David K. Jackson, Ian Johnston, Dominic Kwiatkowski, Cordelia Langford, John Sillitoe on behalf of the Wellcome Sanger Institute COVID-19 Surveillance Team |
| EPI_ISL_3095845 | Lighthouse Laboratory Plymouth | Wellcome Sanger Institute for the COVID-19 Genomics UK (COG-UK) Consortium | Lighthouse Laboratory Plymouth and Alex Alderton, Roberto Amato, Jeffrey Barrett, Sonia Goncalves, Ewan Harrison, David K. Jackson, Ian Johnston, Dominic Kwiatkowski, Cordelia Langford, John Sillitoe on behalf of the Wellcome Sanger Institute COVID-19 Surveillance Team |
| EPI_ISL_3095846 | Berkshire and Surrey Pathology Services Lighthouse Laboratory | Wellcome Sanger Institute for the COVID-19 Genomics UK (COG-UK) Consortium | Berkshire and Surrey Pathology Services Lighthouse Laboratory and Alex Alderton, Roberto Amato, Jeffrey Barrett, Sonia Goncalves, Ewan Harrison, David K. Jackson, Ian Johnston, Dominic Kwiatkowski, Cordelia Langford, John Sillitoe on behalf of the Wellcome Sanger Institute COVID-19 Surveillance Team |
| EPI_ISL_3095847, EPI_ISL_3095848 | Lighthouse Lab in Glasgow | Wellcome Sanger Institute for the COVID-19 Genomics UK (COG-UK) Consortium | Harper VanSteenhouse, Yumi Kasai, David Gray, Carol Clugston, Anna Dominiczak and Alex Alderton, Roberto Amato, Jeffrey Barrett, Sonia Goncalves, Ewan Harrison, David K. Jackson, Ian Johnston, Dominic Kwiatkowski, Cordelia Langford, John Sillitoe on behalf of the Wellcome Sanger Institute COVID-19 Surveillance Team |

| | | | |
|---|---|---|---|
| EPI_ISL_3095849 | Lighthouse Laboratory Plymouth | Wellcome Sanger Institute for the COVID-19 Genomics UK (COG-UK) Consortium | Lighthouse Laboratory Plymouth and Alex Alderton, Roberto Amato, Jeffrey Barrett, Sonia Goncalves, Ewan Harrison, David K. Jackson, Ian Johnston, Dominic Kwiatkowski, Cordelia Langford, John Sillitoe on behalf of the Wellcome Sanger Institute COVID-19 Surveillance Team |
| EPI_ISL_3095850 | Berkshire and Surrey Pathology Services Lighthouse Laboratory | Wellcome Sanger Institute for the COVID-19 Genomics UK (COG-UK) Consortium | Berkshire and Surrey Pathology Services Lighthouse Laboratory and Alex Alderton, Roberto Amato, Jeffrey Barrett, Sonia Goncalves, Ewan Harrison, David K. Jackson, Ian Johnston, Dominic Kwiatkowski, Cordelia Langford, John Sillitoe on behalf of the Wellcome Sanger Institute COVID-19 Surveillance Team |
| EPI_ISL_3095851, EPI_ISL_3095852 | Lighthouse Lab in Glasgow | Wellcome Sanger Institute for the COVID-19 Genomics UK (COG-UK) Consortium | Harper VanSteenhouse, Yumi Kasai, David Gray, Carol Clugston, Anna Dominiczak and Alex Alderton, Roberto Amato, Jeffrey Barrett, Sonia Goncalves, Ewan Harrison, David K. Jackson, Ian Johnston, Dominic Kwiatkowski, Cordelia Langford, John Sillitoe on behalf of the Wellcome Sanger Institute COVID-19 Surveillance Team |
| EPI_ISL_3095853, EPI_ISL_3095854, EPI_ISL_3095855 | Lighthouse Laboratory Plymouth | Wellcome Sanger Institute for the COVID-19 Genomics UK (COG-UK) Consortium | Lighthouse Laboratory Plymouth and Alex Alderton, Roberto Amato, Jeffrey Barrett, Sonia Goncalves, Ewan Harrison, David K. Jackson, Ian Johnston, Dominic Kwiatkowski, Cordelia Langford, John Sillitoe on behalf of the Wellcome Sanger Institute COVID-19 Surveillance Team |
| EPI_ISL_3095856, EPI_ISL_3095857 | Berkshire and Surrey Pathology Services Lighthouse Laboratory | Wellcome Sanger Institute for the COVID-19 Genomics UK (COG-UK) Consortium | Berkshire and Surrey Pathology Services Lighthouse Laboratory and Alex Alderton, Roberto Amato, Jeffrey Barrett, Sonia Goncalves, Ewan Harrison, David K. Jackson, Ian Johnston, Dominic Kwiatkowski, Cordelia Langford, John Sillitoe on behalf of the Wellcome Sanger Institute COVID-19 Surveillance Team |
| EPI_ISL_3095858, EPI_ISL_3095859, EPI_ISL_3095860, EPI_ISL_3095861, EPI_ISL_3095862, EPI_ISL_3095863 | Lighthouse Lab in Glasgow | Wellcome Sanger Institute for the COVID-19 Genomics UK (COG-UK) Consortium | Harper VanSteenhouse, Yumi Kasai, David Gray, Carol Clugston, Anna Dominiczak and Alex Alderton, Roberto Amato, Jeffrey Barrett, Sonia Goncalves, Ewan Harrison, David K. Jackson, Ian Johnston, Dominic Kwiatkowski, Cordelia Langford, John Sillitoe on behalf of the Wellcome Sanger Institute COVID-19 Surveillance Team |
| EPI_ISL_3095864 | Lighthouse Laboratory Plymouth | Wellcome Sanger Institute for the COVID-19 Genomics UK (COG-UK) Consortium | Lighthouse Laboratory Plymouth and Alex Alderton, Roberto Amato, Jeffrey Barrett, Sonia Goncalves, Ewan Harrison, David K. Jackson, Ian Johnston, Dominic Kwiatkowski, Cordelia Langford, John Sillitoe on behalf of the Wellcome Sanger Institute COVID-19 Surveillance Team |
| EPI_ISL_3095865, EPI_ISL_3095866 | Lighthouse Lab in Glasgow | Wellcome Sanger Institute for the COVID-19 Genomics UK (COG-UK) Consortium | Harper VanSteenhouse, Yumi Kasai, David Gray, Carol Clugston, Anna Dominiczak and Alex Alderton, Roberto Amato, Jeffrey Barrett, Sonia Goncalves, Ewan Harrison, David K. Jackson, Ian Johnston, Dominic Kwiatkowski, Cordelia Langford, John Sillitoe on behalf of the Wellcome Sanger Institute COVID-19 Surveillance Team |
| EPI_ISL_3095867 | Berkshire and Surrey Pathology Services Lighthouse Laboratory | Wellcome Sanger Institute for the COVID-19 Genomics UK (COG-UK) Consortium | Berkshire and Surrey Pathology Services Lighthouse Laboratory and Alex Alderton, Roberto Amato, Jeffrey Barrett, Sonia Goncalves, Ewan Harrison, David K. Jackson, Ian Johnston, Dominic Kwiatkowski, Cordelia Langford, John Sillitoe on behalf of the Wellcome Sanger Institute COVID-19 Surveillance Team |
| EPI_ISL_3095868 | Lighthouse Laboratory Plymouth | Wellcome Sanger Institute for the COVID-19 Genomics UK (COG-UK) Consortium | Lighthouse Laboratory Plymouth and Alex Alderton, Roberto Amato, Jeffrey Barrett, Sonia Goncalves, Ewan Harrison, David K. Jackson, Ian Johnston, Dominic Kwiatkowski, Cordelia Langford, John Sillitoe on behalf of the Wellcome Sanger Institute COVID-19 Surveillance Team |
| EPI_ISL_3095869 | Berkshire and Surrey Pathology Services Lighthouse Laboratory | Wellcome Sanger Institute for the COVID-19 Genomics UK (COG-UK) Consortium | Berkshire and Surrey Pathology Services Lighthouse Laboratory and Alex Alderton, Roberto Amato, Jeffrey Barrett, Sonia Goncalves, Ewan Harrison, David K. Jackson, Ian Johnston, Dominic Kwiatkowski, Cordelia Langford, John Sillitoe on behalf of the Wellcome Sanger Institute COVID-19 Surveillance Team |
| EPI_ISL_3095870, EPI_ISL_3095871, EPI_ISL_3095872, EPI_ISL_3095873, EPI_ISL_3095874, EPI_ISL_3095875 | Lighthouse Lab in Glasgow | Wellcome Sanger Institute for the COVID-19 Genomics UK (COG-UK) Consortium | Harper VanSteenhouse, Yumi Kasai, David Gray, Carol Clugston, Anna Dominiczak and Alex Alderton, Roberto Amato, Jeffrey Barrett, Sonia Goncalves, Ewan Harrison, David K. Jackson, Ian Johnston, Dominic Kwiatkowski, Cordelia Langford, John Sillitoe on behalf of the Wellcome Sanger Institute COVID-19 Surveillance Team |
| EPI_ISL_3095876 | Berkshire and Surrey Pathology Services Lighthouse Laboratory | Wellcome Sanger Institute for the COVID-19 Genomics UK (COG-UK) Consortium | Berkshire and Surrey Pathology Services Lighthouse Laboratory and Alex Alderton, Roberto Amato, Jeffrey Barrett, Sonia Goncalves, Ewan Harrison, David K. Jackson, Ian Johnston, Dominic Kwiatkowski, Cordelia Langford, John Sillitoe on behalf of the Wellcome Sanger Institute COVID-19 Surveillance Team |
| EPI_ISL_3095877 | Lighthouse Lab in Glasgow | Wellcome Sanger Institute for the COVID-19 Genomics UK (COG-UK) Consortium | Harper VanSteenhouse, Yumi Kasai, David Gray, Carol Clugston, Anna Dominiczak and Alex Alderton, Roberto Amato, Jeffrey Barrett, Sonia Goncalves, Ewan Harrison, David K. Jackson, Ian Johnston, Dominic Kwiatkowski, Cordelia Langford, John Sillitoe on behalf of the Wellcome Sanger Institute COVID-19 Surveillance Team |
| EPI_ISL_3095878 | Lighthouse Laboratory Plymouth | Wellcome Sanger Institute for the COVID-19 Genomics UK (COG-UK) Consortium | Lighthouse Laboratory Plymouth and Alex Alderton, Roberto Amato, Jeffrey Barrett, Sonia Goncalves, Ewan Harrison, David K. Jackson, Ian Johnston, Dominic Kwiatkowski, Cordelia Langford, John Sillitoe on behalf of the Wellcome Sanger Institute COVID-19 Surveillance Team |
| EPI_ISL_3095879 | Lighthouse Lab in Glasgow | Wellcome Sanger Institute for the COVID-19 Genomics UK (COG-UK) Consortium | Harper VanSteenhouse, Yumi Kasai, David Gray, Carol Clugston, Anna Dominiczak and Alex Alderton, Roberto Amato, Jeffrey Barrett, Sonia Goncalves, Ewan Harrison, David K. Jackson, Ian Johnston, Dominic Kwiatkowski, Cordelia Langford, John Sillitoe on behalf of the Wellcome Sanger Institute COVID-19 Surveillance Team |
| EPI_ISL_3095880 | Lighthouse Laboratory Plymouth | Wellcome Sanger Institute for the COVID-19 Genomics UK (COG-UK) Consortium | Lighthouse Laboratory Plymouth and Alex Alderton, Roberto Amato, Jeffrey Barrett, Sonia Goncalves, Ewan Harrison, David K. Jackson, Ian Johnston, Dominic Kwiatkowski, Cordelia Langford, John Sillitoe on behalf of the Wellcome Sanger Institute COVID-19 Surveillance Team |
| EPI_ISL_3095881 | Berkshire and Surrey Pathology Services Lighthouse Laboratory | Wellcome Sanger Institute for the COVID-19 Genomics UK (COG-UK) Consortium | Berkshire and Surrey Pathology Services Lighthouse Laboratory and Alex Alderton, Roberto Amato, Jeffrey Barrett, Sonia Goncalves, Ewan Harrison, David K. Jackson, Ian Johnston, Dominic Kwiatkowski, Cordelia Langford, John Sillitoe on behalf of the Wellcome Sanger Institute COVID-19 Surveillance Team |
| EPI_ISL_3095882 | Lighthouse Laboratory Plymouth | Wellcome Sanger Institute for the COVID-19 Genomics UK (COG-UK) Consortium | Lighthouse Laboratory Plymouth and Alex Alderton, Roberto Amato, Jeffrey Barrett, Sonia Goncalves, Ewan Harrison, David K. Jackson, Ian Johnston, Dominic Kwiatkowski, Cordelia Langford, John Sillitoe on behalf of the Wellcome Sanger Institute COVID-19 Surveillance Team |
| EPI_ISL_3095883 | Lighthouse Lab in Glasgow | Wellcome Sanger Institute for the COVID-19 Genomics UK (COG-UK) Consortium | Harper VanSteenhouse, Yumi Kasai, David Gray, Carol Clugston, Anna Dominiczak and Alex Alderton, Roberto Amato, Jeffrey Barrett, Sonia Goncalves, Ewan Harrison, David K. Jackson, Ian Johnston, Dominic Kwiatkowski, Cordelia Langford, John Sillitoe on behalf of the Wellcome Sanger Institute COVID-19 Surveillance Team |
| EPI_ISL_3095884 | Berkshire and Surrey Pathology Services Lighthouse Laboratory | Wellcome Sanger Institute for the COVID-19 Genomics UK (COG-UK) Consortium | Berkshire and Surrey Pathology Services Lighthouse Laboratory and Alex Alderton, Roberto Amato, Jeffrey Barrett, Sonia Goncalves, Ewan Harrison, David K. Jackson, Ian Johnston, Dominic Kwiatkowski, Cordelia Langford, John Sillitoe on behalf of the Wellcome Sanger Institute COVID-19 Surveillance Team |
| EPI_ISL_3095885 | Lighthouse Laboratory Plymouth | Wellcome Sanger Institute for the COVID-19 Genomics UK (COG-UK) Consortium | Lighthouse Laboratory Plymouth and Alex Alderton, Roberto Amato, Jeffrey Barrett, Sonia Goncalves, Ewan Harrison, David K. Jackson, Ian Johnston, Dominic Kwiatkowski, Cordelia Langford, John Sillitoe on behalf of the Wellcome Sanger Institute COVID-19 Surveillance Team |
| EPI_ISL_3095886, EPI_ISL_3095887 | Lighthouse Lab in Glasgow | Wellcome Sanger Institute for the COVID-19 Genomics UK (COG-UK) Consortium | Harper VanSteenhouse, Yumi Kasai, David Gray, Carol Clugston, Anna Dominiczak and Alex Alderton, Roberto Amato, Jeffrey Barrett, Sonia Goncalves, Ewan Harrison, David K. Jackson, Ian Johnston, Dominic Kwiatkowski, Cordelia Langford, John Sillitoe on behalf of the Wellcome Sanger Institute COVID-19 Surveillance Team |
| EPI_ISL_3095888 | Berkshire and Surrey Pathology Services Lighthouse Laboratory | Wellcome Sanger Institute for the COVID-19 Genomics UK (COG-UK) Consortium | Berkshire and Surrey Pathology Services Lighthouse Laboratory and Alex Alderton, Roberto Amato, Jeffrey Barrett, Sonia Goncalves, Ewan Harrison, David K. Jackson, Ian Johnston, Dominic Kwiatkowski, Cordelia Langford, John Sillitoe on behalf of the Wellcome Sanger Institute COVID-19 Surveillance Team |
| EPI_ISL_3095889 | Lighthouse Lab in Glasgow | Wellcome Sanger Institute for the COVID-19 Genomics UK (COG-UK) Consortium | Harper VanSteenhouse, Yumi Kasai, David Gray, Carol Clugston, Anna Dominiczak and Alex Alderton, Roberto Amato, Jeffrey Barrett, Sonia Goncalves, Ewan Harrison, David K. Jackson, Ian Johnston, Dominic Kwiatkowski, Cordelia Langford, John Sillitoe on behalf of the Wellcome Sanger Institute COVID-19 Surveillance Team |
| EPI_ISL_3095890 | Lighthouse Laboratory Plymouth | Wellcome Sanger Institute for the COVID-19 Genomics UK (COG-UK) Consortium | Lighthouse Laboratory Plymouth and Alex Alderton, Roberto Amato, Jeffrey Barrett, Sonia Goncalves, Ewan Harrison, David K. Jackson, Ian Johnston, Dominic Kwiatkowski, Cordelia Langford, John Sillitoe on behalf of the Wellcome Sanger Institute COVID-19 Surveillance Team |
| EPI_ISL_3095891 | Lighthouse Lab in Glasgow | Wellcome Sanger Institute for the COVID-19 Genomics UK (COG-UK) Consortium | Harper VanSteenhouse, Yumi Kasai, David Gray, Carol Clugston, Anna Dominiczak and Alex Alderton, Roberto Amato, Jeffrey Barrett, Sonia Goncalves, Ewan Harrison, David K. Jackson, Ian Johnston, Dominic Kwiatkowski, Cordelia Langford, John Sillitoe on behalf of the Wellcome Sanger Institute COVID-19 Surveillance Team |
| EPI_ISL_3095892 | Lighthouse Laboratory Plymouth | Wellcome Sanger Institute for the COVID-19 Genomics UK (COG-UK) Consortium | Lighthouse Laboratory Plymouth and Alex Alderton, Roberto Amato, Jeffrey Barrett, Sonia Goncalves, Ewan Harrison, David K. Jackson, Ian Johnston, Dominic Kwiatkowski, Cordelia Langford, John Sillitoe on behalf of the Wellcome Sanger Institute COVID-19 Surveillance Team |
| EPI_ISL_3095893, EPI_ISL_3095894, EPI_ISL_3095895 | Lighthouse Lab in Glasgow | Wellcome Sanger Institute for the COVID-19 Genomics UK (COG-UK) Consortium | Harper VanSteenhouse, Yumi Kasai, David Gray, Carol Clugston, Anna Dominiczak and Alex Alderton, Roberto Amato, Jeffrey Barrett, Sonia Goncalves, Ewan Harrison, David K. Jackson, Ian Johnston, Dominic Kwiatkowski, Cordelia Langford, John Sillitoe on behalf of the Wellcome Sanger Institute COVID-19 Surveillance Team |
| EPI_ISL_3095896 | Lighthouse Laboratory Plymouth | Wellcome Sanger Institute for the COVID-19 Genomics UK (COG-UK) Consortium | Lighthouse Laboratory Plymouth and Alex Alderton, Roberto Amato, Jeffrey Barrett, Sonia Goncalves, Ewan Harrison, David K. Jackson, Ian Johnston, Dominic Kwiatkowski, Cordelia Langford, John Sillitoe on behalf of the Wellcome Sanger Institute COVID-19 Surveillance Team |
| EPI_ISL_3095897, EPI_ISL_3095898 | Lighthouse Lab in Glasgow | Wellcome Sanger Institute for the COVID-19 Genomics UK (COG-UK) Consortium | Harper VanSteenhouse, Yumi Kasai, David Gray, Carol Clugston, Anna Dominiczak and Alex Alderton, Roberto Amato, Jeffrey Barrett, Sonia Goncalves, Ewan Harrison, David K. Jackson, Ian Johnston, Dominic Kwiatkowski, Cordelia Langford, John Sillitoe on behalf of the Wellcome Sanger Institute COVID-19 Surveillance Team |
| EPI_ISL_3095899 | Berkshire and Surrey Pathology Services Lighthouse Laboratory | Wellcome Sanger Institute for the COVID-19 Genomics UK (COG-UK) Consortium | Berkshire and Surrey Pathology Services Lighthouse Laboratory and Alex Alderton, Roberto Amato, Jeffrey Barrett, Sonia Goncalves, Ewan Harrison, David K. Jackson, Ian Johnston, Dominic Kwiatkowski, Cordelia Langford, John Sillitoe on behalf of the Wellcome Sanger Institute COVID-19 Surveillance Team |
| EPI_ISL_3095900 | Lighthouse Lab in Glasgow | Wellcome Sanger Institute for the COVID-19 Genomics UK (COG-UK) Consortium | Harper VanSteenhouse, Yumi Kasai, David Gray, Carol Clugston, Anna Dominiczak and Alex Alderton, Roberto Amato, Jeffrey Barrett, Sonia Goncalves, Ewan Harrison, David K. Jackson, Ian Johnston, Dominic Kwiatkowski, Cordelia Langford, John Sillitoe on behalf of the Wellcome Sanger Institute COVID-19 Surveillance Team |
| EPI_ISL_3095901 | Berkshire and Surrey Pathology Services Lighthouse Laboratory | Wellcome Sanger Institute for the COVID-19 Genomics UK (COG-UK) Consortium | Berkshire and Surrey Pathology Services Lighthouse Laboratory and Alex Alderton, Roberto Amato, Jeffrey Barrett, Sonia Goncalves, Ewan Harrison, David K. Jackson, Ian Johnston, Dominic Kwiatkowski, Cordelia Langford, John Sillitoe on behalf of the Wellcome Sanger Institute COVID-19 Surveillance Team |
| EPI_ISL_3095902 | Lighthouse Lab in Glasgow | Wellcome Sanger Institute for the COVID-19 Genomics UK (COG-UK) Consortium | Harper VanSteenhouse, Yumi Kasai, David Gray, Carol Clugston, Anna Dominiczak and Alex Alderton, Roberto Amato, Jeffrey Barrett, Sonia Goncalves, Ewan Harrison, David K. Jackson, Ian Johnston, Dominic Kwiatkowski, Cordelia Langford, John Sillitoe on behalf of the Wellcome Sanger Institute COVID-19 Surveillance Team |
| EPI_ISL_3095903 | Lighthouse Laboratory Plymouth | Wellcome Sanger Institute for the COVID-19 Genomics UK (COG-UK) Consortium | Lighthouse Laboratory Plymouth and Alex Alderton, Roberto Amato, Jeffrey Barrett, Sonia Goncalves, Ewan Harrison, David K. Jackson, Ian Johnston, Dominic Kwiatkowski, Cordelia Langford, John Sillitoe on behalf of the Wellcome Sanger Institute COVID-19 Surveillance Team |
| EPI_ISL_3095904, EPI_ISL_3095905, EPI_ISL_3095906 | Lighthouse Lab in Glasgow | Wellcome Sanger Institute for the COVID-19 Genomics UK (COG-UK) Consortium | Harper VanSteenhouse, Yumi Kasai, David Gray, Carol Clugston, Anna Dominiczak and Alex Alderton, Roberto Amato, Jeffrey Barrett, Sonia Goncalves, Ewan Harrison, David K. Jackson, Ian Johnston, Dominic Kwiatkowski, Cordelia Langford, John Sillitoe on behalf of the Wellcome Sanger Institute COVID-19 Surveillance Team |
| EPI_ISL_3095907, EPI_ISL_3095908, EPI_ISL_3095909, EPI_ISL_3095910 | Lighthouse Laboratory Plymouth | Wellcome Sanger Institute for the COVID-19 Genomics UK (COG-UK) Consortium | Lighthouse Laboratory Plymouth and Alex Alderton, Roberto Amato, Jeffrey Barrett, Sonia Goncalves, Ewan Harrison, David K. Jackson, Ian Johnston, Dominic Kwiatkowski, Cordelia Langford, John Sillitoe on behalf of the Wellcome Sanger Institute COVID-19 Surveillance Team |
| EPI_ISL_3095911 | Lighthouse Lab in Glasgow | Wellcome Sanger Institute for the COVID-19 Genomics UK (COG-UK) Consortium | Harper VanSteenhouse, Yumi Kasai, David Gray, Carol Clugston, Anna Dominiczak and Alex Alderton, Roberto Amato, Jeffrey Barrett, Sonia Goncalves, Ewan Harrison, David K. Jackson, Ian Johnston, Dominic Kwiatkowski, Cordelia Langford, John Sillitoe on behalf of the Wellcome Sanger Institute COVID-19 Surveillance Team |
| EPI_ISL_3095912 | Lighthouse Laboratory Plymouth | Wellcome Sanger Institute for the COVID-19 Genomics UK (COG-UK) | Lighthouse Laboratory Plymouth and Alex Alderton, Roberto Amato, Jeffrey Barrett, Sonia Goncalves, Ewan Harrison, David K. Jackson, Ian Johnston, Dominic Kwiatkowski, Cordelia Langford, John Sillitoe on behalf of the Wellcome Sanger |

| | | | |
|---|---|---|---|
| | | Consortium | Institute COVID-19 Surveillance Team |
| EPI_ISL_3095913 | Berkshire and Surrey Pathology Services Lighthouse Laboratory | Wellcome Sanger Institute for the COVID-19 Genomics UK (COG-UK) Consortium | Berkshire and Surrey Pathology Services Lighthouse Laboratory and Alex Alderton, Roberto Amato, Jeffrey Barrett, Sonia Goncalves, Ewan Harrison, David K. Jackson, Ian Johnston, Dominic Kwiatkowski, Cordelia Langford, John Sillitoe on behalf of the Wellcome Sanger Institute COVID-19 Surveillance Team |
| EPI_ISL_3095914, EPI_ISL_3095915 | Lighthouse Lab in Glasgow | Wellcome Sanger Institute for the COVID-19 Genomics UK (COG-UK) Consortium | Harper VanSteenhouse, Yumi Kasai, David Gray, Carol Clugston, Anna Dominiczak and Alex Alderton, Roberto Amato, Jeffrey Barrett, Sonia Goncalves, Ewan Harrison, David K. Jackson, Ian Johnston, Dominic Kwiatkowski, Cordelia Langford, John Sillitoe on behalf of the Wellcome Sanger Institute COVID-19 Surveillance Team |
| EPI_ISL_3095916 | Berkshire and Surrey Pathology Services Lighthouse Laboratory | Wellcome Sanger Institute for the COVID-19 Genomics UK (COG-UK) Consortium | Berkshire and Surrey Pathology Services Lighthouse Laboratory and Alex Alderton, Roberto Amato, Jeffrey Barrett, Sonia Goncalves, Ewan Harrison, David K. Jackson, Ian Johnston, Dominic Kwiatkowski, Cordelia Langford, John Sillitoe on behalf of the Wellcome Sanger Institute COVID-19 Surveillance Team |
| EPI_ISL_3095917, EPI_ISL_3095918 | Lighthouse Lab in Glasgow | Wellcome Sanger Institute for the COVID-19 Genomics UK (COG-UK) Consortium | Harper VanSteenhouse, Yumi Kasai, David Gray, Carol Clugston, Anna Dominiczak and Alex Alderton, Roberto Amato, Jeffrey Barrett, Sonia Goncalves, Ewan Harrison, David K. Jackson, Ian Johnston, Dominic Kwiatkowski, Cordelia Langford, John Sillitoe on behalf of the Wellcome Sanger Institute COVID-19 Surveillance Team |
| EPI_ISL_3095919, EPI_ISL_3095920 | Lighthouse Laboratory Plymouth | Wellcome Sanger Institute for the COVID-19 Genomics UK (COG-UK) Consortium | Lighthouse Laboratory Plymouth and Alex Alderton, Roberto Amato, Jeffrey Barrett, Sonia Goncalves, Ewan Harrison, David K. Jackson, Ian Johnston, Dominic Kwiatkowski, Cordelia Langford, John Sillitoe on behalf of the Wellcome Sanger Institute COVID-19 Surveillance Team |
| EPI_ISL_3095921 | Berkshire and Surrey Pathology Services Lighthouse Laboratory | Wellcome Sanger Institute for the COVID-19 Genomics UK (COG-UK) Consortium | Berkshire and Surrey Pathology Services Lighthouse Laboratory and Alex Alderton, Roberto Amato, Jeffrey Barrett, Sonia Goncalves, Ewan Harrison, David K. Jackson, Ian Johnston, Dominic Kwiatkowski, Cordelia Langford, John Sillitoe on behalf of the Wellcome Sanger Institute COVID-19 Surveillance Team |
| EPI_ISL_3095922 | Health Services Laboratories | Wellcome Sanger Institute for the COVID-19 Genomics UK (COG-UK) Consortium | Health Services Laboratories and Alex Alderton, Roberto Amato, Jeffrey Barrett, Sonia Goncalves, Ewan Harrison, David K. Jackson, Ian Johnston, Dominic Kwiatkowski, Cordelia Langford, John Sillitoe on behalf of the Wellcome Sanger Institute COVID-19 Surveillance Team |
| EPI_ISL_3095923 | Berkshire and Surrey Pathology Services Lighthouse Laboratory | Wellcome Sanger Institute for the COVID-19 Genomics UK (COG-UK) Consortium | Berkshire and Surrey Pathology Services Lighthouse Laboratory and Alex Alderton, Roberto Amato, Jeffrey Barrett, Sonia Goncalves, Ewan Harrison, David K. Jackson, Ian Johnston, Dominic Kwiatkowski, Cordelia Langford, John Sillitoe on behalf of the Wellcome Sanger Institute COVID-19 Surveillance Team |
| EPI_ISL_3095924 | Lighthouse Lab in Glasgow | Wellcome Sanger Institute for the COVID-19 Genomics UK (COG-UK) Consortium | Harper VanSteenhouse, Yumi Kasai, David Gray, Carol Clugston, Anna Dominiczak and Alex Alderton, Roberto Amato, Jeffrey Barrett, Sonia Goncalves, Ewan Harrison, David K. Jackson, Ian Johnston, Dominic Kwiatkowski, Cordelia Langford, John Sillitoe on behalf of the Wellcome Sanger Institute COVID-19 Surveillance Team |
| EPI_ISL_3095925 | Lighthouse Laboratory Plymouth | Wellcome Sanger Institute for the COVID-19 Genomics UK (COG-UK) Consortium | Lighthouse Laboratory Plymouth and Alex Alderton, Roberto Amato, Jeffrey Barrett, Sonia Goncalves, Ewan Harrison, David K. Jackson, Ian Johnston, Dominic Kwiatkowski, Cordelia Langford, John Sillitoe on behalf of the Wellcome Sanger Institute COVID-19 Surveillance Team |
| EPI_ISL_3095926 | Berkshire and Surrey Pathology Services Lighthouse Laboratory | Wellcome Sanger Institute for the COVID-19 Genomics UK (COG-UK) Consortium | Berkshire and Surrey Pathology Services Lighthouse Laboratory and Alex Alderton, Roberto Amato, Jeffrey Barrett, Sonia Goncalves, Ewan Harrison, David K. Jackson, Ian Johnston, Dominic Kwiatkowski, Cordelia Langford, John Sillitoe on behalf of the Wellcome Sanger Institute COVID-19 Surveillance Team |
| EPI_ISL_3095927 | Lighthouse Laboratory Plymouth | Wellcome Sanger Institute for the COVID-19 Genomics UK (COG-UK) Consortium | Lighthouse Laboratory Plymouth and Alex Alderton, Roberto Amato, Jeffrey Barrett, Sonia Goncalves, Ewan Harrison, David K. Jackson, Ian Johnston, Dominic Kwiatkowski, Cordelia Langford, John Sillitoe on behalf of the Wellcome Sanger Institute COVID-19 Surveillance Team |
| EPI_ISL_3095928, EPI_ISL_3095929, EPI_ISL_3095930 | Lighthouse Lab in Glasgow | Wellcome Sanger Institute for the COVID-19 Genomics UK (COG-UK) Consortium | Harper VanSteenhouse, Yumi Kasai, David Gray, Carol Clugston, Anna Dominiczak and Alex Alderton, Roberto Amato, Jeffrey Barrett, Sonia Goncalves, Ewan Harrison, David K. Jackson, Ian Johnston, Dominic Kwiatkowski, Cordelia Langford, John Sillitoe on behalf of the Wellcome Sanger Institute COVID-19 Surveillance Team |
| EPI_ISL_3095931 | Health Services Laboratories | Wellcome Sanger Institute for the COVID-19 Genomics UK (COG-UK) Consortium | Health Services Laboratories and Alex Alderton, Roberto Amato, Jeffrey Barrett, Sonia Goncalves, Ewan Harrison, David K. Jackson, Ian Johnston, Dominic Kwiatkowski, Cordelia Langford, John Sillitoe on behalf of the Wellcome Sanger Institute COVID-19 Surveillance Team |
| EPI_ISL_3095932, EPI_ISL_3095933, EPI_ISL_3095934 | Lighthouse Lab in Glasgow | Wellcome Sanger Institute for the COVID-19 Genomics UK (COG-UK) Consortium | Harper VanSteenhouse, Yumi Kasai, David Gray, Carol Clugston, Anna Dominiczak and Alex Alderton, Roberto Amato, Jeffrey Barrett, Sonia Goncalves, Ewan Harrison, David K. Jackson, Ian Johnston, Dominic Kwiatkowski, Cordelia Langford, John Sillitoe on behalf of the Wellcome Sanger Institute COVID-19 Surveillance Team |
| EPI_ISL_3095935 | Berkshire and Surrey Pathology Services Lighthouse Laboratory | Wellcome Sanger Institute for the COVID-19 Genomics UK (COG-UK) Consortium | Berkshire and Surrey Pathology Services Lighthouse Laboratory and Alex Alderton, Roberto Amato, Jeffrey Barrett, Sonia Goncalves, Ewan Harrison, David K. Jackson, Ian Johnston, Dominic Kwiatkowski, Cordelia Langford, John Sillitoe on behalf of the Wellcome Sanger Institute COVID-19 Surveillance Team |
| EPI_ISL_3095936, EPI_ISL_3095937, EPI_ISL_3095938, EPI_ISL_3095939 | Lighthouse Lab in Glasgow | Wellcome Sanger Institute for the COVID-19 Genomics UK (COG-UK) Consortium | Harper VanSteenhouse, Yumi Kasai, David Gray, Carol Clugston, Anna Dominiczak and Alex Alderton, Roberto Amato, Jeffrey Barrett, Sonia Goncalves, Ewan Harrison, David K. Jackson, Ian Johnston, Dominic Kwiatkowski, Cordelia Langford, John Sillitoe on behalf of the Wellcome Sanger Institute COVID-19 Surveillance Team |
| EPI_ISL_3095940, EPI_ISL_3095941 | Lighthouse Laboratory Plymouth | Wellcome Sanger Institute for the COVID-19 Genomics UK (COG-UK) Consortium | Lighthouse Laboratory Plymouth and Alex Alderton, Roberto Amato, Jeffrey Barrett, Sonia Goncalves, Ewan Harrison, David K. Jackson, Ian Johnston, Dominic Kwiatkowski, Cordelia Langford, John Sillitoe on behalf of the Wellcome Sanger Institute COVID-19 Surveillance Team |
| EPI_ISL_3095942 | Berkshire and Surrey Pathology Services Lighthouse Laboratory | Wellcome Sanger Institute for the COVID-19 Genomics UK (COG-UK) Consortium | Berkshire and Surrey Pathology Services Lighthouse Laboratory and Alex Alderton, Roberto Amato, Jeffrey Barrett, Sonia Goncalves, Ewan Harrison, David K. Jackson, Ian Johnston, Dominic Kwiatkowski, Cordelia Langford, John Sillitoe on behalf of the Wellcome Sanger Institute COVID-19 Surveillance Team |
| EPI_ISL_3095943, EPI_ISL_3095944, EPI_ISL_3095945, EPI_ISL_3095946 | Lighthouse Lab in Glasgow | Wellcome Sanger Institute for the COVID-19 Genomics UK (COG-UK) Consortium | Harper VanSteenhouse, Yumi Kasai, David Gray, Carol Clugston, Anna Dominiczak and Alex Alderton, Roberto Amato, Jeffrey Barrett, Sonia Goncalves, Ewan Harrison, David K. Jackson, Ian Johnston, Dominic Kwiatkowski, Cordelia Langford, John Sillitoe on behalf of the Wellcome Sanger Institute COVID-19 Surveillance Team |
| EPI_ISL_3095947 | Lighthouse Laboratory Plymouth | Wellcome Sanger Institute for the COVID-19 Genomics UK (COG-UK) Consortium | Lighthouse Laboratory Plymouth and Alex Alderton, Roberto Amato, Jeffrey Barrett, Sonia Goncalves, Ewan Harrison, David K. Jackson, Ian Johnston, Dominic Kwiatkowski, Cordelia Langford, John Sillitoe on behalf of the Wellcome Sanger Institute COVID-19 Surveillance Team |
| EPI_ISL_3095948 | Berkshire and Surrey Pathology Services Lighthouse Laboratory | Wellcome Sanger Institute for the COVID-19 Genomics UK (COG-UK) Consortium | Berkshire and Surrey Pathology Services Lighthouse Laboratory and Alex Alderton, Roberto Amato, Jeffrey Barrett, Sonia Goncalves, Ewan Harrison, David K. Jackson, Ian Johnston, Dominic Kwiatkowski, Cordelia Langford, John Sillitoe on behalf of the Wellcome Sanger Institute COVID-19 Surveillance Team |
| EPI_ISL_3095949, EPI_ISL_3095950, EPI_ISL_3095951, EPI_ISL_3095952, EPI_ISL_3095953 | Lighthouse Lab in Glasgow | Wellcome Sanger Institute for the COVID-19 Genomics UK (COG-UK) Consortium | Harper VanSteenhouse, Yumi Kasai, David Gray, Carol Clugston, Anna Dominiczak and Alex Alderton, Roberto Amato, Jeffrey Barrett, Sonia Goncalves, Ewan Harrison, David K. Jackson, Ian Johnston, Dominic Kwiatkowski, Cordelia Langford, John Sillitoe on behalf of the Wellcome Sanger Institute COVID-19 Surveillance Team |
| EPI_ISL_3095954, EPI_ISL_3095955, EPI_ISL_3095956 | Berkshire and Surrey Pathology Services Lighthouse Laboratory | Wellcome Sanger Institute for the COVID-19 Genomics UK (COG-UK) Consortium | Berkshire and Surrey Pathology Services Lighthouse Laboratory and Alex Alderton, Roberto Amato, Jeffrey Barrett, Sonia Goncalves, Ewan Harrison, David K. Jackson, Ian Johnston, Dominic Kwiatkowski, Cordelia Langford, John Sillitoe on behalf of the Wellcome Sanger Institute COVID-19 Surveillance Team |
| EPI_ISL_3095957 | Lighthouse Laboratory Plymouth | Wellcome Sanger Institute for the COVID-19 Genomics UK (COG-UK) Consortium | Lighthouse Laboratory Plymouth and Alex Alderton, Roberto Amato, Jeffrey Barrett, Sonia Goncalves, Ewan Harrison, David K. Jackson, Ian Johnston, Dominic Kwiatkowski, Cordelia Langford, John Sillitoe on behalf of the Wellcome Sanger Institute COVID-19 Surveillance Team |
| EPI_ISL_3095958, EPI_ISL_3095959 | Lighthouse Lab in Glasgow | Wellcome Sanger Institute for the COVID-19 Genomics UK (COG-UK) Consortium | Harper VanSteenhouse, Yumi Kasai, David Gray, Carol Clugston, Anna Dominiczak and Alex Alderton, Roberto Amato, Jeffrey Barrett, Sonia Goncalves, Ewan Harrison, David K. Jackson, Ian Johnston, Dominic Kwiatkowski, Cordelia Langford, John Sillitoe on behalf of the Wellcome Sanger Institute COVID-19 Surveillance Team |
| EPI_ISL_3095960 | Berkshire and Surrey Pathology Services Lighthouse Laboratory | Wellcome Sanger Institute for the COVID-19 Genomics UK (COG-UK) Consortium | Berkshire and Surrey Pathology Services Lighthouse Laboratory and Alex Alderton, Roberto Amato, Jeffrey Barrett, Sonia Goncalves, Ewan Harrison, David K. Jackson, Ian Johnston, Dominic Kwiatkowski, Cordelia Langford, John Sillitoe on behalf of the Wellcome Sanger Institute COVID-19 Surveillance Team |
| EPI_ISL_3095961, EPI_ISL_3095962 | Lighthouse Lab in Glasgow | Wellcome Sanger Institute for the COVID-19 Genomics UK (COG-UK) Consortium | Harper VanSteenhouse, Yumi Kasai, David Gray, Carol Clugston, Anna Dominiczak and Alex Alderton, Roberto Amato, Jeffrey Barrett, Sonia Goncalves, Ewan Harrison, David K. Jackson, Ian Johnston, Dominic Kwiatkowski, Cordelia Langford, John Sillitoe on behalf of the Wellcome Sanger Institute COVID-19 Surveillance Team |
| EPI_ISL_3095963 | Lighthouse Laboratory Plymouth | Wellcome Sanger Institute for the COVID-19 Genomics UK (COG-UK) Consortium | Lighthouse Laboratory Plymouth and Alex Alderton, Roberto Amato, Jeffrey Barrett, Sonia Goncalves, Ewan Harrison, David K. Jackson, Ian Johnston, Dominic Kwiatkowski, Cordelia Langford, John Sillitoe on behalf of the Wellcome Sanger Institute COVID-19 Surveillance Team |
| EPI_ISL_3095964, EPI_ISL_3095965, EPI_ISL_3095966 | Lighthouse Lab in Glasgow | Wellcome Sanger Institute for the COVID-19 Genomics UK (COG-UK) Consortium | Harper VanSteenhouse, Yumi Kasai, David Gray, Carol Clugston, Anna Dominiczak and Alex Alderton, Roberto Amato, Jeffrey Barrett, Sonia Goncalves, Ewan Harrison, David K. Jackson, Ian Johnston, Dominic Kwiatkowski, Cordelia Langford, John Sillitoe on behalf of the Wellcome Sanger Institute COVID-19 Surveillance Team |
| EPI_ISL_3095967 | Berkshire and Surrey Pathology Services Lighthouse Laboratory | Wellcome Sanger Institute for the COVID-19 Genomics UK (COG-UK) Consortium | Berkshire and Surrey Pathology Services Lighthouse Laboratory and Alex Alderton, Roberto Amato, Jeffrey Barrett, Sonia Goncalves, Ewan Harrison, David K. Jackson, Ian Johnston, Dominic Kwiatkowski, Cordelia Langford, John Sillitoe on behalf of the Wellcome Sanger Institute COVID-19 Surveillance Team |
| EPI_ISL_3095968, EPI_ISL_3095969 | Lighthouse Lab in Glasgow | Wellcome Sanger Institute for the COVID-19 Genomics UK (COG-UK) Consortium | Harper VanSteenhouse, Yumi Kasai, David Gray, Carol Clugston, Anna Dominiczak and Alex Alderton, Roberto Amato, Jeffrey Barrett, Sonia Goncalves, Ewan Harrison, David K. Jackson, Ian Johnston, Dominic Kwiatkowski, Cordelia Langford, John Sillitoe on behalf of the Wellcome Sanger Institute COVID-19 Surveillance Team |
| EPI_ISL_3095970 | Lighthouse Laboratory Plymouth | Wellcome Sanger Institute for the COVID-19 Genomics UK (COG-UK) Consortium | Lighthouse Laboratory Plymouth and Alex Alderton, Roberto Amato, Jeffrey Barrett, Sonia Goncalves, Ewan Harrison, David K. Jackson, Ian Johnston, Dominic Kwiatkowski, Cordelia Langford, John Sillitoe on behalf of the Wellcome Sanger Institute COVID-19 Surveillance Team |
| EPI_ISL_3095971 | Lighthouse Lab in Glasgow | Wellcome Sanger Institute for the COVID-19 Genomics UK (COG-UK) Consortium | Harper VanSteenhouse, Yumi Kasai, David Gray, Carol Clugston, Anna Dominiczak and Alex Alderton, Roberto Amato, Jeffrey Barrett, Sonia Goncalves, Ewan Harrison, David K. Jackson, Ian Johnston, Dominic Kwiatkowski, Cordelia Langford, John Sillitoe on behalf of the Wellcome Sanger Institute COVID-19 Surveillance Team |
| EPI_ISL_3095972 | Berkshire and Surrey Pathology Services Lighthouse Laboratory | Wellcome Sanger Institute for the COVID-19 Genomics UK (COG-UK) Consortium | Berkshire and Surrey Pathology Services Lighthouse Laboratory and Alex Alderton, Roberto Amato, Jeffrey Barrett, Sonia Goncalves, Ewan Harrison, David K. Jackson, Ian Johnston, Dominic Kwiatkowski, Cordelia Langford, John Sillitoe on behalf of the Wellcome Sanger Institute COVID-19 Surveillance Team |
| EPI_ISL_3095973 | Lighthouse Laboratory Plymouth | Wellcome Sanger Institute for the COVID-19 Genomics UK (COG-UK) Consortium | Lighthouse Laboratory Plymouth and Alex Alderton, Roberto Amato, Jeffrey Barrett, Sonia Goncalves, Ewan Harrison, David K. Jackson, Ian Johnston, Dominic Kwiatkowski, Cordelia Langford, John Sillitoe on behalf of the Wellcome Sanger Institute COVID-19 Surveillance Team |
| EPI_ISL_3095974 | Lighthouse Lab in Glasgow | Wellcome Sanger Institute for the COVID-19 Genomics UK (COG-UK) Consortium | Harper VanSteenhouse, Yumi Kasai, David Gray, Carol Clugston, Anna Dominiczak and Alex Alderton, Roberto Amato, Jeffrey Barrett, Sonia Goncalves, Ewan Harrison, David K. Jackson, Ian Johnston, Dominic Kwiatkowski, Cordelia Langford, John Sillitoe on behalf of the Wellcome Sanger Institute COVID-19 Surveillance Team |
| EPI_ISL_3095975 | Berkshire and Surrey Pathology Services Lighthouse Laboratory | Wellcome Sanger Institute for the COVID-19 Genomics UK (COG-UK) Consortium | Berkshire and Surrey Pathology Services Lighthouse Laboratory and Alex Alderton, Roberto Amato, Jeffrey Barrett, Sonia Goncalves, Ewan Harrison, David K. Jackson, Ian Johnston, Dominic Kwiatkowski, Cordelia Langford, John Sillitoe on behalf of the Wellcome Sanger Institute COVID-19 Surveillance Team |
| EPI_ISL_3095976, EPI_ISL_3095977, EPI_ISL_3095978, EPI_ISL_3095979 | Lighthouse Lab in Glasgow | Wellcome Sanger Institute for the COVID-19 Genomics UK (COG-UK) Consortium | Harper VanSteenhouse, Yumi Kasai, David Gray, Carol Clugston, Anna Dominiczak and Alex Alderton, Roberto Amato, Jeffrey Barrett, Sonia Goncalves, Ewan Harrison, David K. Jackson, Ian Johnston, Dominic Kwiatkowski, Cordelia Langford, John Sillitoe on behalf of the Wellcome Sanger Institute COVID-19 Surveillance Team |

| | | | |
|---|---|---|---|
| EPI_ISL_3095980, EPI_ISL_3095981 | Lighthouse Laboratory Plymouth | Wellcome Sanger Institute for the COVID-19 Genomics UK (COG-UK) Consortium | Lighthouse Laboratory Plymouth and Alex Alderton, Roberto Amato, Jeffrey Barrett, Sonia Goncalves, Ewan Harrison, David K. Jackson, Ian Johnston, Dominic Kwiatkowski, Cordelia Langford, John Sillitoe on behalf of the Wellcome Sanger Institute COVID-19 Surveillance Team |
| EPI_ISL_3095982, EPI_ISL_3095983, EPI_ISL_3095984 | Lighthouse Lab in Glasgow | Wellcome Sanger Institute for the COVID-19 Genomics UK (COG-UK) Consortium | Harper VanSteenhouse, Yumi Kasai, David Gray, Carol Clugston, Anna Dominiczak and Alex Alderton, Roberto Amato, Jeffrey Barrett, Sonia Goncalves, Ewan Harrison, David K. Jackson, Ian Johnston, Dominic Kwiatkowski, Cordelia Langford, John Sillitoe on behalf of the Wellcome Sanger Institute COVID-19 Surveillance Team |
| EPI_ISL_3095985 | Berkshire and Surrey Pathology Services Lighthouse Laboratory | Wellcome Sanger Institute for the COVID-19 Genomics UK (COG-UK) Consortium | Berkshire and Surrey Pathology Services Lighthouse Laboratory and Alex Alderton, Roberto Amato, Jeffrey Barrett, Sonia Goncalves, Ewan Harrison, David K. Jackson, Ian Johnston, Dominic Kwiatkowski, Cordelia Langford, John Sillitoe on behalf of the Wellcome Sanger Institute COVID-19 Surveillance Team |
| EPI_ISL_3095986 | Lighthouse Laboratory Plymouth | Wellcome Sanger Institute for the COVID-19 Genomics UK (COG-UK) Consortium | Lighthouse Laboratory Plymouth and Alex Alderton, Roberto Amato, Jeffrey Barrett, Sonia Goncalves, Ewan Harrison, David K. Jackson, Ian Johnston, Dominic Kwiatkowski, Cordelia Langford, John Sillitoe on behalf of the Wellcome Sanger Institute COVID-19 Surveillance Team |
| EPI_ISL_3095987, EPI_ISL_3095988, EPI_ISL_3095989 | Lighthouse Lab in Glasgow | Wellcome Sanger Institute for the COVID-19 Genomics UK (COG-UK) Consortium | Harper VanSteenhouse, Yumi Kasai, David Gray, Carol Clugston, Anna Dominiczak and Alex Alderton, Roberto Amato, Jeffrey Barrett, Sonia Goncalves, Ewan Harrison, David K. Jackson, Ian Johnston, Dominic Kwiatkowski, Cordelia Langford, John Sillitoe on behalf of the Wellcome Sanger Institute COVID-19 Surveillance Team |
| EPI_ISL_3095990 | Berkshire and Surrey Pathology Services Lighthouse Laboratory | Wellcome Sanger Institute for the COVID-19 Genomics UK (COG-UK) Consortium | Berkshire and Surrey Pathology Services Lighthouse Laboratory and Alex Alderton, Roberto Amato, Jeffrey Barrett, Sonia Goncalves, Ewan Harrison, David K. Jackson, Ian Johnston, Dominic Kwiatkowski, Cordelia Langford, John Sillitoe on behalf of the Wellcome Sanger Institute COVID-19 Surveillance Team |
| EPI_ISL_3095991 | Lighthouse Laboratory Plymouth | Wellcome Sanger Institute for the COVID-19 Genomics UK (COG-UK) Consortium | Lighthouse Laboratory Plymouth and Alex Alderton, Roberto Amato, Jeffrey Barrett, Sonia Goncalves, Ewan Harrison, David K. Jackson, Ian Johnston, Dominic Kwiatkowski, Cordelia Langford, John Sillitoe on behalf of the Wellcome Sanger Institute COVID-19 Surveillance Team |
| EPI_ISL_3095992, EPI_ISL_3095993 | Berkshire and Surrey Pathology Services Lighthouse Laboratory | Wellcome Sanger Institute for the COVID-19 Genomics UK (COG-UK) Consortium | Berkshire and Surrey Pathology Services Lighthouse Laboratory and Alex Alderton, Roberto Amato, Jeffrey Barrett, Sonia Goncalves, Ewan Harrison, David K. Jackson, Ian Johnston, Dominic Kwiatkowski, Cordelia Langford, John Sillitoe on behalf of the Wellcome Sanger Institute COVID-19 Surveillance Team |
| EPI_ISL_3095994, EPI_ISL_3095995, EPI_ISL_3095996 | Lighthouse Lab in Glasgow | Wellcome Sanger Institute for the COVID-19 Genomics UK (COG-UK) Consortium | Harper VanSteenhouse, Yumi Kasai, David Gray, Carol Clugston, Anna Dominiczak and Alex Alderton, Roberto Amato, Jeffrey Barrett, Sonia Goncalves, Ewan Harrison, David K. Jackson, Ian Johnston, Dominic Kwiatkowski, Cordelia Langford, John Sillitoe on behalf of the Wellcome Sanger Institute COVID-19 Surveillance Team |
| EPI_ISL_3095997 | Lighthouse Laboratory Plymouth | Wellcome Sanger Institute for the COVID-19 Genomics UK (COG-UK) Consortium | Lighthouse Laboratory Plymouth and Alex Alderton, Roberto Amato, Jeffrey Barrett, Sonia Goncalves, Ewan Harrison, David K. Jackson, Ian Johnston, Dominic Kwiatkowski, Cordelia Langford, John Sillitoe on behalf of the Wellcome Sanger Institute COVID-19 Surveillance Team |
| EPI_ISL_3095998, EPI_ISL_3095999, EPI_ISL_3096000, EPI_ISL_3096001, EPI_ISL_3096002 | Lighthouse Lab in Glasgow | Wellcome Sanger Institute for the COVID-19 Genomics UK (COG-UK) Consortium | Harper VanSteenhouse, Yumi Kasai, David Gray, Carol Clugston, Anna Dominiczak and Alex Alderton, Roberto Amato, Jeffrey Barrett, Sonia Goncalves, Ewan Harrison, David K. Jackson, Ian Johnston, Dominic Kwiatkowski, Cordelia Langford, John Sillitoe on behalf of the Wellcome Sanger Institute COVID-19 Surveillance Team |
| EPI_ISL_3096003, EPI_ISL_3096004, EPI_ISL_3096005, EPI_ISL_3096006, EPI_ISL_3096007 | Lighthouse Laboratory Plymouth | Wellcome Sanger Institute for the COVID-19 Genomics UK (COG-UK) Consortium | Lighthouse Laboratory Plymouth and Alex Alderton, Roberto Amato, Jeffrey Barrett, Sonia Goncalves, Ewan Harrison, David K. Jackson, Ian Johnston, Dominic Kwiatkowski, Cordelia Langford, John Sillitoe on behalf of the Wellcome Sanger Institute COVID-19 Surveillance Team |
| EPI_ISL_3096008, EPI_ISL_3096009 | Lighthouse Lab in Glasgow | Wellcome Sanger Institute for the COVID-19 Genomics UK (COG-UK) Consortium | Harper VanSteenhouse, Yumi Kasai, David Gray, Carol Clugston, Anna Dominiczak and Alex Alderton, Roberto Amato, Jeffrey Barrett, Sonia Goncalves, Ewan Harrison, David K. Jackson, Ian Johnston, Dominic Kwiatkowski, Cordelia Langford, John Sillitoe on behalf of the Wellcome Sanger Institute COVID-19 Surveillance Team |
| EPI_ISL_3096010, EPI_ISL_3096011 | Berkshire and Surrey Pathology Services Lighthouse Laboratory | Wellcome Sanger Institute for the COVID-19 Genomics UK (COG-UK) Consortium | Berkshire and Surrey Pathology Services Lighthouse Laboratory and Alex Alderton, Roberto Amato, Jeffrey Barrett, Sonia Goncalves, Ewan Harrison, David K. Jackson, Ian Johnston, Dominic Kwiatkowski, Cordelia Langford, John Sillitoe on behalf of the Wellcome Sanger Institute COVID-19 Surveillance Team |
| EPI_ISL_3096012, EPI_ISL_3096013 | Lighthouse Lab in Milton Keynes | Wellcome Sanger Institute for the COVID-19 Genomics UK (COG-UK) Consortium | The Lighthouse Lab in Milton Keynes and Alex Alderton, Roberto Amato, Jeffrey Barrett, Sonia Goncalves, Ewan Harrison, David K. Jackson, Ian Johnston, Dominic Kwiatkowski, Cordelia Langford, John Sillitoe on behalf of the Wellcome Sanger Institute COVID-19 Surveillance Team |
| EPI_ISL_3096014 | Berkshire and Surrey Pathology Services Lighthouse Laboratory | Wellcome Sanger Institute for the COVID-19 Genomics UK (COG-UK) Consortium | Berkshire and Surrey Pathology Services Lighthouse Laboratory and Alex Alderton, Roberto Amato, Jeffrey Barrett, Sonia Goncalves, Ewan Harrison, David K. Jackson, Ian Johnston, Dominic Kwiatkowski, Cordelia Langford, John Sillitoe on behalf of the Wellcome Sanger Institute COVID-19 Surveillance Team |
| EPI_ISL_3096015, EPI_ISL_3096016, EPI_ISL_3096017 | Lighthouse Lab in Milton Keynes | Wellcome Sanger Institute for the COVID-19 Genomics UK (COG-UK) Consortium | The Lighthouse Lab in Milton Keynes and Alex Alderton, Roberto Amato, Jeffrey Barrett, Sonia Goncalves, Ewan Harrison, David K. Jackson, Ian Johnston, Dominic Kwiatkowski, Cordelia Langford, John Sillitoe on behalf of the Wellcome Sanger Institute COVID-19 Surveillance Team |
| EPI_ISL_3096018 | Berkshire and Surrey Pathology Services Lighthouse Laboratory | Wellcome Sanger Institute for the COVID-19 Genomics UK (COG-UK) Consortium | Berkshire and Surrey Pathology Services Lighthouse Laboratory and Alex Alderton, Roberto Amato, Jeffrey Barrett, Sonia Goncalves, Ewan Harrison, David K. Jackson, Ian Johnston, Dominic Kwiatkowski, Cordelia Langford, John Sillitoe on behalf of the Wellcome Sanger Institute COVID-19 Surveillance Team |
| EPI_ISL_3096019, EPI_ISL_3096020 | Lighthouse Lab in Milton Keynes | Wellcome Sanger Institute for the COVID-19 Genomics UK (COG-UK) Consortium | The Lighthouse Lab in Milton Keynes and Alex Alderton, Roberto Amato, Jeffrey Barrett, Sonia Goncalves, Ewan Harrison, David K. Jackson, Ian Johnston, Dominic Kwiatkowski, Cordelia Langford, John Sillitoe on behalf of the Wellcome Sanger Institute COVID-19 Surveillance Team |
| EPI_ISL_3096021 | Berkshire and Surrey Pathology Services Lighthouse Laboratory | Wellcome Sanger Institute for the COVID-19 Genomics UK (COG-UK) Consortium | Berkshire and Surrey Pathology Services Lighthouse Laboratory and Alex Alderton, Roberto Amato, Jeffrey Barrett, Sonia Goncalves, Ewan Harrison, David K. Jackson, Ian Johnston, Dominic Kwiatkowski, Cordelia Langford, John Sillitoe on behalf of the Wellcome Sanger Institute COVID-19 Surveillance Team |
| EPI_ISL_3096022 | Lighthouse Lab in Milton Keynes | Wellcome Sanger Institute for the COVID-19 Genomics UK (COG-UK) Consortium | The Lighthouse Lab in Milton Keynes and Alex Alderton, Roberto Amato, Jeffrey Barrett, Sonia Goncalves, Ewan Harrison, David K. Jackson, Ian Johnston, Dominic Kwiatkowski, Cordelia Langford, John Sillitoe on behalf of the Wellcome Sanger Institute COVID-19 Surveillance Team |
| EPI_ISL_3096023 | Berkshire and Surrey Pathology Services Lighthouse Laboratory | Wellcome Sanger Institute for the COVID-19 Genomics UK (COG-UK) Consortium | Berkshire and Surrey Pathology Services Lighthouse Laboratory and Alex Alderton, Roberto Amato, Jeffrey Barrett, Sonia Goncalves, Ewan Harrison, David K. Jackson, Ian Johnston, Dominic Kwiatkowski, Cordelia Langford, John Sillitoe on behalf of the Wellcome Sanger Institute COVID-19 Surveillance Team |
| EPI_ISL_3096024 | Lighthouse Lab in Milton Keynes | Wellcome Sanger Institute for the COVID-19 Genomics UK (COG-UK) Consortium | The Lighthouse Lab in Milton Keynes and Alex Alderton, Roberto Amato, Jeffrey Barrett, Sonia Goncalves, Ewan Harrison, David K. Jackson, Ian Johnston, Dominic Kwiatkowski, Cordelia Langford, John Sillitoe on behalf of the Wellcome Sanger Institute COVID-19 Surveillance Team |
| EPI_ISL_3096025 | Berkshire and Surrey Pathology Services Lighthouse Laboratory | Wellcome Sanger Institute for the COVID-19 Genomics UK (COG-UK) Consortium | Berkshire and Surrey Pathology Services Lighthouse Laboratory and Alex Alderton, Roberto Amato, Jeffrey Barrett, Sonia Goncalves, Ewan Harrison, David K. Jackson, Ian Johnston, Dominic Kwiatkowski, Cordelia Langford, John Sillitoe on behalf of the Wellcome Sanger Institute COVID-19 Surveillance Team |
| EPI_ISL_3096026 | Lighthouse Lab in Milton Keynes | Wellcome Sanger Institute for the COVID-19 Genomics UK (COG-UK) Consortium | The Lighthouse Lab in Milton Keynes and Alex Alderton, Roberto Amato, Jeffrey Barrett, Sonia Goncalves, Ewan Harrison, David K. Jackson, Ian Johnston, Dominic Kwiatkowski, Cordelia Langford, John Sillitoe on behalf of the Wellcome Sanger Institute COVID-19 Surveillance Team |
| EPI_ISL_3096027 | Berkshire and Surrey Pathology Services Lighthouse Laboratory | Wellcome Sanger Institute for the COVID-19 Genomics UK (COG-UK) Consortium | Berkshire and Surrey Pathology Services Lighthouse Laboratory and Alex Alderton, Roberto Amato, Jeffrey Barrett, Sonia Goncalves, Ewan Harrison, David K. Jackson, Ian Johnston, Dominic Kwiatkowski, Cordelia Langford, John Sillitoe on behalf of the Wellcome Sanger Institute COVID-19 Surveillance Team |
| EPI_ISL_3096028 | Lighthouse Lab in Glasgow | Wellcome Sanger Institute for the COVID-19 Genomics UK (COG-UK) Consortium | Harper VanSteenhouse, Yumi Kasai, David Gray, Carol Clugston, Anna Dominiczak and Alex Alderton, Roberto Amato, Jeffrey Barrett, Sonia Goncalves, Ewan Harrison, David K. Jackson, Ian Johnston, Dominic Kwiatkowski, Cordelia Langford, John Sillitoe on behalf of the Wellcome Sanger Institute COVID-19 Surveillance Team |
| EPI_ISL_3096029, EPI_ISL_3096030, EPI_ISL_3096031 | Berkshire and Surrey Pathology Services Lighthouse Laboratory | Wellcome Sanger Institute for the COVID-19 Genomics UK (COG-UK) Consortium | Berkshire and Surrey Pathology Services Lighthouse Laboratory and Alex Alderton, Roberto Amato, Jeffrey Barrett, Sonia Goncalves, Ewan Harrison, David K. Jackson, Ian Johnston, Dominic Kwiatkowski, Cordelia Langford, John Sillitoe on behalf of the Wellcome Sanger Institute COVID-19 Surveillance Team |
| EPI_ISL_3096032 | Lighthouse Lab in Milton Keynes | Wellcome Sanger Institute for the COVID-19 Genomics UK (COG-UK) Consortium | The Lighthouse Lab in Milton Keynes and Alex Alderton, Roberto Amato, Jeffrey Barrett, Sonia Goncalves, Ewan Harrison, David K. Jackson, Ian Johnston, Dominic Kwiatkowski, Cordelia Langford, John Sillitoe on behalf of the Wellcome Sanger Institute COVID-19 Surveillance Team |
| EPI_ISL_3096033 | Berkshire and Surrey Pathology Services Lighthouse Laboratory | Wellcome Sanger Institute for the COVID-19 Genomics UK (COG-UK) Consortium | Berkshire and Surrey Pathology Services Lighthouse Laboratory and Alex Alderton, Roberto Amato, Jeffrey Barrett, Sonia Goncalves, Ewan Harrison, David K. Jackson, Ian Johnston, Dominic Kwiatkowski, Cordelia Langford, John Sillitoe on behalf of the Wellcome Sanger Institute COVID-19 Surveillance Team |
| EPI_ISL_3096034, EPI_ISL_3096035 | Lighthouse Lab in Milton Keynes | Wellcome Sanger Institute for the COVID-19 Genomics UK (COG-UK) Consortium | The Lighthouse Lab in Milton Keynes and Alex Alderton, Roberto Amato, Jeffrey Barrett, Sonia Goncalves, Ewan Harrison, David K. Jackson, Ian Johnston, Dominic Kwiatkowski, Cordelia Langford, John Sillitoe on behalf of the Wellcome Sanger Institute COVID-19 Surveillance Team |
| EPI_ISL_3096036 | Lighthouse Lab in Glasgow | Wellcome Sanger Institute for the COVID-19 Genomics UK (COG-UK) Consortium | Harper VanSteenhouse, Yumi Kasai, David Gray, Carol Clugston, Anna Dominiczak and Alex Alderton, Roberto Amato, Jeffrey Barrett, Sonia Goncalves, Ewan Harrison, David K. Jackson, Ian Johnston, Dominic Kwiatkowski, Cordelia Langford, John Sillitoe on behalf of the Wellcome Sanger Institute COVID-19 Surveillance Team |
| EPI_ISL_3096037 | Lighthouse Lab in Milton Keynes | Wellcome Sanger Institute for the COVID-19 Genomics UK (COG-UK) Consortium | The Lighthouse Lab in Milton Keynes and Alex Alderton, Roberto Amato, Jeffrey Barrett, Sonia Goncalves, Ewan Harrison, David K. Jackson, Ian Johnston, Dominic Kwiatkowski, Cordelia Langford, John Sillitoe on behalf of the Wellcome Sanger Institute COVID-19 Surveillance Team |
| EPI_ISL_3096038 | Berkshire and Surrey Pathology Services Lighthouse Laboratory | Wellcome Sanger Institute for the COVID-19 Genomics UK (COG-UK) Consortium | Berkshire and Surrey Pathology Services Lighthouse Laboratory and Alex Alderton, Roberto Amato, Jeffrey Barrett, Sonia Goncalves, Ewan Harrison, David K. Jackson, Ian Johnston, Dominic Kwiatkowski, Cordelia Langford, John Sillitoe on behalf of the Wellcome Sanger Institute COVID-19 Surveillance Team |
| EPI_ISL_3096039, EPI_ISL_3096040, EPI_ISL_3096041 | Lighthouse Lab in Milton Keynes | Wellcome Sanger Institute for the COVID-19 Genomics UK (COG-UK) Consortium | The Lighthouse Lab in Milton Keynes and Alex Alderton, Roberto Amato, Jeffrey Barrett, Sonia Goncalves, Ewan Harrison, David K. Jackson, Ian Johnston, Dominic Kwiatkowski, Cordelia Langford, John Sillitoe on behalf of the Wellcome Sanger Institute COVID-19 Surveillance Team |
| EPI_ISL_3096042 | Lighthouse Lab in Glasgow | Wellcome Sanger Institute for the COVID-19 Genomics UK (COG-UK) Consortium | Harper VanSteenhouse, Yumi Kasai, David Gray, Carol Clugston, Anna Dominiczak and Alex Alderton, Roberto Amato, Jeffrey Barrett, Sonia Goncalves, Ewan Harrison, David K. Jackson, Ian Johnston, Dominic Kwiatkowski, Cordelia Langford, John Sillitoe on behalf of the Wellcome Sanger Institute COVID-19 Surveillance Team |
| EPI_ISL_3096043 | Lighthouse Lab in Milton Keynes | Wellcome Sanger Institute for the COVID-19 Genomics UK (COG-UK) Consortium | The Lighthouse Lab in Milton Keynes and Alex Alderton, Roberto Amato, Jeffrey Barrett, Sonia Goncalves, Ewan Harrison, David K. Jackson, Ian Johnston, Dominic Kwiatkowski, Cordelia Langford, John Sillitoe on behalf of the Wellcome Sanger Institute COVID-19 Surveillance Team |
| EPI_ISL_3096044 | Lighthouse Lab in Glasgow | Wellcome Sanger Institute for the COVID-19 Genomics UK (COG-UK) Consortium | Harper VanSteenhouse, Yumi Kasai, David Gray, Carol Clugston, Anna Dominiczak and Alex Alderton, Roberto Amato, Jeffrey Barrett, Sonia Goncalves, Ewan Harrison, David K. Jackson, Ian Johnston, Dominic Kwiatkowski, Cordelia Langford, John Sillitoe on behalf of the Wellcome Sanger Institute COVID-19 Surveillance Team |
| EPI_ISL_3096045 | Lighthouse Lab in Milton Keynes | Wellcome Sanger Institute for the COVID-19 Genomics UK (COG-UK) Consortium | The Lighthouse Lab in Milton Keynes and Alex Alderton, Roberto Amato, Jeffrey Barrett, Sonia Goncalves, Ewan Harrison, David K. Jackson, Ian Johnston, Dominic Kwiatkowski, Cordelia Langford, John Sillitoe on behalf of the Wellcome Sanger Institute COVID-19 Surveillance Team |
| EPI_ISL_3096046 | Lighthouse Lab in Glasgow | Wellcome Sanger Institute for the COVID-19 Genomics UK (COG-UK) Consortium | Harper VanSteenhouse, Yumi Kasai, David Gray, Carol Clugston, Anna Dominiczak and Alex Alderton, Roberto Amato, Jeffrey Barrett, Sonia Goncalves, Ewan Harrison, David K. Jackson, Ian Johnston, Dominic Kwiatkowski, Cordelia Langford, John Sillitoe on behalf of the Wellcome Sanger Institute COVID-19 Surveillance Team |

| | | | |
|---|---|---|---|
| EPI_ISL_3096047 | Berkshire and Surrey Pathology Services Lighthouse Laboratory | Wellcome Sanger Institute for the COVID-19 Genomics UK (COG-UK) Consortium | Berkshire and Surrey Pathology Services Lighthouse Laboratory and Alex Alderton, Roberto Amato, Jeffrey Barrett, Sonia Goncalves, Ewan Harrison, David K. Jackson, Ian Johnston, Dominic Kwiatkowski, Cordelia Langford, John Sillitoe on behalf of the Wellcome Sanger Institute COVID-19 Surveillance Team |
| EPI_ISL_3096048 | Lighthouse Lab in Milton Keynes | Wellcome Sanger Institute for the COVID-19 Genomics UK (COG-UK) Consortium | The Lighthouse Lab in Milton Keynes and Alex Alderton, Roberto Amato, Jeffrey Barrett, Sonia Goncalves, Ewan Harrison, David K. Jackson, Ian Johnston, Dominic Kwiatkowski, Cordelia Langford, John Sillitoe on behalf of the Wellcome Sanger Institute COVID-19 Surveillance Team |
| EPI_ISL_3096049 | Berkshire and Surrey Pathology Services Lighthouse Laboratory | Wellcome Sanger Institute for the COVID-19 Genomics UK (COG-UK) Consortium | Berkshire and Surrey Pathology Services Lighthouse Laboratory and Alex Alderton, Roberto Amato, Jeffrey Barrett, Sonia Goncalves, Ewan Harrison, David K. Jackson, Ian Johnston, Dominic Kwiatkowski, Cordelia Langford, John Sillitoe on behalf of the Wellcome Sanger Institute COVID-19 Surveillance Team |
| EPI_ISL_3096050 | Lighthouse Lab in Milton Keynes | Wellcome Sanger Institute for the COVID-19 Genomics UK (COG-UK) Consortium | The Lighthouse Lab in Milton Keynes and Alex Alderton, Roberto Amato, Jeffrey Barrett, Sonia Goncalves, Ewan Harrison, David K. Jackson, Ian Johnston, Dominic Kwiatkowski, Cordelia Langford, John Sillitoe on behalf of the Wellcome Sanger Institute COVID-19 Surveillance Team |
| EPI_ISL_3096051 | Berkshire and Surrey Pathology Services Lighthouse Laboratory | Wellcome Sanger Institute for the COVID-19 Genomics UK (COG-UK) Consortium | Berkshire and Surrey Pathology Services Lighthouse Laboratory and Alex Alderton, Roberto Amato, Jeffrey Barrett, Sonia Goncalves, Ewan Harrison, David K. Jackson, Ian Johnston, Dominic Kwiatkowski, Cordelia Langford, John Sillitoe on behalf of the Wellcome Sanger Institute COVID-19 Surveillance Team |
| EPI_ISL_3096052 | Lighthouse Lab in Milton Keynes | Wellcome Sanger Institute for the COVID-19 Genomics UK (COG-UK) Consortium | The Lighthouse Lab in Milton Keynes and Alex Alderton, Roberto Amato, Jeffrey Barrett, Sonia Goncalves, Ewan Harrison, David K. Jackson, Ian Johnston, Dominic Kwiatkowski, Cordelia Langford, John Sillitoe on behalf of the Wellcome Sanger Institute COVID-19 Surveillance Team |
| EPI_ISL_3096053, EPI_ISL_3096054 | Lighthouse Lab in Glasgow | Wellcome Sanger Institute for the COVID-19 Genomics UK (COG-UK) Consortium | Harper VanSteenhouse, Yumi Kasai, David Gray, Carol Clugston, Anna Dominiczak and Alex Alderton, Roberto Amato, Jeffrey Barrett, Sonia Goncalves, Ewan Harrison, David K. Jackson, Ian Johnston, Dominic Kwiatkowski, Cordelia Langford, John Sillitoe on behalf of the Wellcome Sanger Institute COVID-19 Surveillance Team |
| EPI_ISL_3096055 | Berkshire and Surrey Pathology Services Lighthouse Laboratory | Wellcome Sanger Institute for the COVID-19 Genomics UK (COG-UK) Consortium | Berkshire and Surrey Pathology Services Lighthouse Laboratory and Alex Alderton, Roberto Amato, Jeffrey Barrett, Sonia Goncalves, Ewan Harrison, David K. Jackson, Ian Johnston, Dominic Kwiatkowski, Cordelia Langford, John Sillitoe on behalf of the Wellcome Sanger Institute COVID-19 Surveillance Team |
| EPI_ISL_3096056, EPI_ISL_3096057 | Lighthouse Lab in Milton Keynes | Wellcome Sanger Institute for the COVID-19 Genomics UK (COG-UK) Consortium | The Lighthouse Lab in Milton Keynes and Alex Alderton, Roberto Amato, Jeffrey Barrett, Sonia Goncalves, Ewan Harrison, David K. Jackson, Ian Johnston, Dominic Kwiatkowski, Cordelia Langford, John Sillitoe on behalf of the Wellcome Sanger Institute COVID-19 Surveillance Team |
| EPI_ISL_3096058 | Lighthouse Lab in Glasgow | Wellcome Sanger Institute for the COVID-19 Genomics UK (COG-UK) Consortium | Harper VanSteenhouse, Yumi Kasai, David Gray, Carol Clugston, Anna Dominiczak and Alex Alderton, Roberto Amato, Jeffrey Barrett, Sonia Goncalves, Ewan Harrison, David K. Jackson, Ian Johnston, Dominic Kwiatkowski, Cordelia Langford, John Sillitoe on behalf of the Wellcome Sanger Institute COVID-19 Surveillance Team |
| EPI_ISL_3096059 | Berkshire and Surrey Pathology Services Lighthouse Laboratory | Wellcome Sanger Institute for the COVID-19 Genomics UK (COG-UK) Consortium | Berkshire and Surrey Pathology Services Lighthouse Laboratory and Alex Alderton, Roberto Amato, Jeffrey Barrett, Sonia Goncalves, Ewan Harrison, David K. Jackson, Ian Johnston, Dominic Kwiatkowski, Cordelia Langford, John Sillitoe on behalf of the Wellcome Sanger Institute COVID-19 Surveillance Team |
| EPI_ISL_3096060, EPI_ISL_3096061 | Lighthouse Lab in Milton Keynes | Wellcome Sanger Institute for the COVID-19 Genomics UK (COG-UK) Consortium | The Lighthouse Lab in Milton Keynes and Alex Alderton, Roberto Amato, Jeffrey Barrett, Sonia Goncalves, Ewan Harrison, David K. Jackson, Ian Johnston, Dominic Kwiatkowski, Cordelia Langford, John Sillitoe on behalf of the Wellcome Sanger Institute COVID-19 Surveillance Team |
| EPI_ISL_3096062 | Berkshire and Surrey Pathology Services Lighthouse Laboratory | Wellcome Sanger Institute for the COVID-19 Genomics UK (COG-UK) Consortium | Berkshire and Surrey Pathology Services Lighthouse Laboratory and Alex Alderton, Roberto Amato, Jeffrey Barrett, Sonia Goncalves, Ewan Harrison, David K. Jackson, Ian Johnston, Dominic Kwiatkowski, Cordelia Langford, John Sillitoe on behalf of the Wellcome Sanger Institute COVID-19 Surveillance Team |
| EPI_ISL_3096063, EPI_ISL_3096064 | Lighthouse Lab in Milton Keynes | Wellcome Sanger Institute for the COVID-19 Genomics UK (COG-UK) Consortium | The Lighthouse Lab in Milton Keynes and Alex Alderton, Roberto Amato, Jeffrey Barrett, Sonia Goncalves, Ewan Harrison, David K. Jackson, Ian Johnston, Dominic Kwiatkowski, Cordelia Langford, John Sillitoe on behalf of the Wellcome Sanger Institute COVID-19 Surveillance Team |
| EPI_ISL_3096065 | Lighthouse Lab in Glasgow | Wellcome Sanger Institute for the COVID-19 Genomics UK (COG-UK) Consortium | Harper VanSteenhouse, Yumi Kasai, David Gray, Carol Clugston, Anna Dominiczak and Alex Alderton, Roberto Amato, Jeffrey Barrett, Sonia Goncalves, Ewan Harrison, David K. Jackson, Ian Johnston, Dominic Kwiatkowski, Cordelia Langford, John Sillitoe on behalf of the Wellcome Sanger Institute COVID-19 Surveillance Team |
| EPI_ISL_3096066, EPI_ISL_3096067, EPI_ISL_3096068 | Lighthouse Lab in Milton Keynes | Wellcome Sanger Institute for the COVID-19 Genomics UK (COG-UK) Consortium | The Lighthouse Lab in Milton Keynes and Alex Alderton, Roberto Amato, Jeffrey Barrett, Sonia Goncalves, Ewan Harrison, David K. Jackson, Ian Johnston, Dominic Kwiatkowski, Cordelia Langford, John Sillitoe on behalf of the Wellcome Sanger Institute COVID-19 Surveillance Team |
| EPI_ISL_3096069 | Lighthouse Lab in Glasgow | Wellcome Sanger Institute for the COVID-19 Genomics UK (COG-UK) Consortium | Harper VanSteenhouse, Yumi Kasai, David Gray, Carol Clugston, Anna Dominiczak and Alex Alderton, Roberto Amato, Jeffrey Barrett, Sonia Goncalves, Ewan Harrison, David K. Jackson, Ian Johnston, Dominic Kwiatkowski, Cordelia Langford, John Sillitoe on behalf of the Wellcome Sanger Institute COVID-19 Surveillance Team |
| EPI_ISL_3096070, EPI_ISL_3096071 | Lighthouse Lab in Milton Keynes | Wellcome Sanger Institute for the COVID-19 Genomics UK (COG-UK) Consortium | The Lighthouse Lab in Milton Keynes and Alex Alderton, Roberto Amato, Jeffrey Barrett, Sonia Goncalves, Ewan Harrison, David K. Jackson, Ian Johnston, Dominic Kwiatkowski, Cordelia Langford, John Sillitoe on behalf of the Wellcome Sanger Institute COVID-19 Surveillance Team |
| EPI_ISL_3096072, EPI_ISL_3096073 | Berkshire and Surrey Pathology Services Lighthouse Laboratory | Wellcome Sanger Institute for the COVID-19 Genomics UK (COG-UK) Consortium | Berkshire and Surrey Pathology Services Lighthouse Laboratory and Alex Alderton, Roberto Amato, Jeffrey Barrett, Sonia Goncalves, Ewan Harrison, David K. Jackson, Ian Johnston, Dominic Kwiatkowski, Cordelia Langford, John Sillitoe on behalf of the Wellcome Sanger Institute COVID-19 Surveillance Team |
| EPI_ISL_3096074, EPI_ISL_3096075, EPI_ISL_3096076 | Lighthouse Lab in Milton Keynes | Wellcome Sanger Institute for the COVID-19 Genomics UK (COG-UK) Consortium | The Lighthouse Lab in Milton Keynes and Alex Alderton, Roberto Amato, Jeffrey Barrett, Sonia Goncalves, Ewan Harrison, David K. Jackson, Ian Johnston, Dominic Kwiatkowski, Cordelia Langford, John Sillitoe on behalf of the Wellcome Sanger Institute COVID-19 Surveillance Team |
| EPI_ISL_3096077 | Lighthouse Lab in Glasgow | Wellcome Sanger Institute for the COVID-19 Genomics UK (COG-UK) Consortium | Harper VanSteenhouse, Yumi Kasai, David Gray, Carol Clugston, Anna Dominiczak and Alex Alderton, Roberto Amato, Jeffrey Barrett, Sonia Goncalves, Ewan Harrison, David K. Jackson, Ian Johnston, Dominic Kwiatkowski, Cordelia Langford, John Sillitoe on behalf of the Wellcome Sanger Institute COVID-19 Surveillance Team |
| EPI_ISL_3096078 | Berkshire and Surrey Pathology Services Lighthouse Laboratory | Wellcome Sanger Institute for the COVID-19 Genomics UK (COG-UK) Consortium | Berkshire and Surrey Pathology Services Lighthouse Laboratory and Alex Alderton, Roberto Amato, Jeffrey Barrett, Sonia Goncalves, Ewan Harrison, David K. Jackson, Ian Johnston, Dominic Kwiatkowski, Cordelia Langford, John Sillitoe on behalf of the Wellcome Sanger Institute COVID-19 Surveillance Team |
| EPI_ISL_3096079 | Lighthouse Lab in Glasgow | Wellcome Sanger Institute for the COVID-19 Genomics UK (COG-UK) Consortium | Harper VanSteenhouse, Yumi Kasai, David Gray, Carol Clugston, Anna Dominiczak and Alex Alderton, Roberto Amato, Jeffrey Barrett, Sonia Goncalves, Ewan Harrison, David K. Jackson, Ian Johnston, Dominic Kwiatkowski, Cordelia Langford, John Sillitoe on behalf of the Wellcome Sanger Institute COVID-19 Surveillance Team |
| EPI_ISL_3096080, EPI_ISL_3096081 | Lighthouse Lab in Milton Keynes | Wellcome Sanger Institute for the COVID-19 Genomics UK (COG-UK) Consortium | The Lighthouse Lab in Milton Keynes and Alex Alderton, Roberto Amato, Jeffrey Barrett, Sonia Goncalves, Ewan Harrison, David K. Jackson, Ian Johnston, Dominic Kwiatkowski, Cordelia Langford, John Sillitoe on behalf of the Wellcome Sanger Institute COVID-19 Surveillance Team |
| EPI_ISL_3096082 | Berkshire and Surrey Pathology Services Lighthouse Laboratory | Wellcome Sanger Institute for the COVID-19 Genomics UK (COG-UK) Consortium | Berkshire and Surrey Pathology Services Lighthouse Laboratory and Alex Alderton, Roberto Amato, Jeffrey Barrett, Sonia Goncalves, Ewan Harrison, David K. Jackson, Ian Johnston, Dominic Kwiatkowski, Cordelia Langford, John Sillitoe on behalf of the Wellcome Sanger Institute COVID-19 Surveillance Team |
| EPI_ISL_3096083 | Lighthouse Lab in Glasgow | Wellcome Sanger Institute for the COVID-19 Genomics UK (COG-UK) Consortium | Harper VanSteenhouse, Yumi Kasai, David Gray, Carol Clugston, Anna Dominiczak and Alex Alderton, Roberto Amato, Jeffrey Barrett, Sonia Goncalves, Ewan Harrison, David K. Jackson, Ian Johnston, Dominic Kwiatkowski, Cordelia Langford, John Sillitoe on behalf of the Wellcome Sanger Institute COVID-19 Surveillance Team |
| EPI_ISL_3096084 | Berkshire and Surrey Pathology Services Lighthouse Laboratory | Wellcome Sanger Institute for the COVID-19 Genomics UK (COG-UK) Consortium | Berkshire and Surrey Pathology Services Lighthouse Laboratory and Alex Alderton, Roberto Amato, Jeffrey Barrett, Sonia Goncalves, Ewan Harrison, David K. Jackson, Ian Johnston, Dominic Kwiatkowski, Cordelia Langford, John Sillitoe on behalf of the Wellcome Sanger Institute COVID-19 Surveillance Team |
| EPI_ISL_3096085 | Lighthouse Lab in Milton Keynes | Wellcome Sanger Institute for the COVID-19 Genomics UK (COG-UK) Consortium | The Lighthouse Lab in Milton Keynes and Alex Alderton, Roberto Amato, Jeffrey Barrett, Sonia Goncalves, Ewan Harrison, David K. Jackson, Ian Johnston, Dominic Kwiatkowski, Cordelia Langford, John Sillitoe on behalf of the Wellcome Sanger Institute COVID-19 Surveillance Team |
| EPI_ISL_3096086 | Lighthouse Lab in Glasgow | Wellcome Sanger Institute for the COVID-19 Genomics UK (COG-UK) Consortium | Harper VanSteenhouse, Yumi Kasai, David Gray, Carol Clugston, Anna Dominiczak and Alex Alderton, Roberto Amato, Jeffrey Barrett, Sonia Goncalves, Ewan Harrison, David K. Jackson, Ian Johnston, Dominic Kwiatkowski, Cordelia Langford, John Sillitoe on behalf of the Wellcome Sanger Institute COVID-19 Surveillance Team |
| EPI_ISL_3096087 | Lighthouse Lab in Milton Keynes | Wellcome Sanger Institute for the COVID-19 Genomics UK (COG-UK) Consortium | The Lighthouse Lab in Milton Keynes and Alex Alderton, Roberto Amato, Jeffrey Barrett, Sonia Goncalves, Ewan Harrison, David K. Jackson, Ian Johnston, Dominic Kwiatkowski, Cordelia Langford, John Sillitoe on behalf of the Wellcome Sanger Institute COVID-19 Surveillance Team |
| EPI_ISL_3096088 | Lighthouse Lab in Glasgow | Wellcome Sanger Institute for the COVID-19 Genomics UK (COG-UK) Consortium | Harper VanSteenhouse, Yumi Kasai, David Gray, Carol Clugston, Anna Dominiczak and Alex Alderton, Roberto Amato, Jeffrey Barrett, Sonia Goncalves, Ewan Harrison, David K. Jackson, Ian Johnston, Dominic Kwiatkowski, Cordelia Langford, John Sillitoe on behalf of the Wellcome Sanger Institute COVID-19 Surveillance Team |
| EPI_ISL_3096089 | Lighthouse Lab in Milton Keynes | Wellcome Sanger Institute for the COVID-19 Genomics UK (COG-UK) Consortium | The Lighthouse Lab in Milton Keynes and Alex Alderton, Roberto Amato, Jeffrey Barrett, Sonia Goncalves, Ewan Harrison, David K. Jackson, Ian Johnston, Dominic Kwiatkowski, Cordelia Langford, John Sillitoe on behalf of the Wellcome Sanger Institute COVID-19 Surveillance Team |
| EPI_ISL_3096090 | Lighthouse Lab in Glasgow | Wellcome Sanger Institute for the COVID-19 Genomics UK (COG-UK) Consortium | Harper VanSteenhouse, Yumi Kasai, David Gray, Carol Clugston, Anna Dominiczak and Alex Alderton, Roberto Amato, Jeffrey Barrett, Sonia Goncalves, Ewan Harrison, David K. Jackson, Ian Johnston, Dominic Kwiatkowski, Cordelia Langford, John Sillitoe on behalf of the Wellcome Sanger Institute COVID-19 Surveillance Team |
| EPI_ISL_3096091 | Berkshire and Surrey Pathology Services Lighthouse Laboratory | Wellcome Sanger Institute for the COVID-19 Genomics UK (COG-UK) Consortium | Berkshire and Surrey Pathology Services Lighthouse Laboratory and Alex Alderton, Roberto Amato, Jeffrey Barrett, Sonia Goncalves, Ewan Harrison, David K. Jackson, Ian Johnston, Dominic Kwiatkowski, Cordelia Langford, John Sillitoe on behalf of the Wellcome Sanger Institute COVID-19 Surveillance Team |
| EPI_ISL_3096092, EPI_ISL_3096093 | Lighthouse Lab in Milton Keynes | Wellcome Sanger Institute for the COVID-19 Genomics UK (COG-UK) Consortium | The Lighthouse Lab in Milton Keynes and Alex Alderton, Roberto Amato, Jeffrey Barrett, Sonia Goncalves, Ewan Harrison, David K. Jackson, Ian Johnston, Dominic Kwiatkowski, Cordelia Langford, John Sillitoe on behalf of the Wellcome Sanger Institute COVID-19 Surveillance Team |
| EPI_ISL_3096094 | Lighthouse Lab in Glasgow | Wellcome Sanger Institute for the COVID-19 Genomics UK (COG-UK) Consortium | Harper VanSteenhouse, Yumi Kasai, David Gray, Carol Clugston, Anna Dominiczak and Alex Alderton, Roberto Amato, Jeffrey Barrett, Sonia Goncalves, Ewan Harrison, David K. Jackson, Ian Johnston, Dominic Kwiatkowski, Cordelia Langford, John Sillitoe on behalf of the Wellcome Sanger Institute COVID-19 Surveillance Team |
| EPI_ISL_3096095, EPI_ISL_3096096 | Lighthouse Lab in Milton Keynes | Wellcome Sanger Institute for the COVID-19 Genomics UK (COG-UK) Consortium | The Lighthouse Lab in Milton Keynes and Alex Alderton, Roberto Amato, Jeffrey Barrett, Sonia Goncalves, Ewan Harrison, David K. Jackson, Ian Johnston, Dominic Kwiatkowski, Cordelia Langford, John Sillitoe on behalf of the Wellcome Sanger Institute COVID-19 Surveillance Team |
| EPI_ISL_3096097 | Berkshire and Surrey Pathology Services Lighthouse Laboratory | Wellcome Sanger Institute for the COVID-19 Genomics UK (COG-UK) Consortium | Berkshire and Surrey Pathology Services Lighthouse Laboratory and Alex Alderton, Roberto Amato, Jeffrey Barrett, Sonia Goncalves, Ewan Harrison, David K. Jackson, Ian Johnston, Dominic Kwiatkowski, Cordelia Langford, John Sillitoe on behalf of the Wellcome Sanger Institute COVID-19 Surveillance Team |
| EPI_ISL_3096098 | Lighthouse Lab in Glasgow | Wellcome Sanger Institute for the COVID-19 Genomics UK (COG-UK) Consortium | Harper VanSteenhouse, Yumi Kasai, David Gray, Carol Clugston, Anna Dominiczak and Alex Alderton, Roberto Amato, Jeffrey Barrett, Sonia Goncalves, Ewan Harrison, David K. Jackson, Ian Johnston, Dominic Kwiatkowski, Cordelia Langford, John Sillitoe on behalf of the Wellcome Sanger Institute COVID-19 Surveillance Team |
| EPI_ISL_3096099, EPI_ISL_3096100 | Lighthouse Lab in Milton Keynes | Wellcome Sanger Institute for the COVID-19 Genomics UK (COG-UK) Consortium | The Lighthouse Lab in Milton Keynes and Alex Alderton, Roberto Amato, Jeffrey Barrett, Sonia Goncalves, Ewan Harrison, David K. Jackson, Ian Johnston, Dominic Kwiatkowski, Cordelia Langford, John Sillitoe on behalf of the Wellcome Sanger Institute COVID-19 Surveillance Team |
| EPI_ISL_3096101 | Lighthouse Lab in Glasgow | Wellcome Sanger Institute for the COVID-19 Genomics UK (COG-UK) Consortium | Harper VanSteenhouse, Yumi Kasai, David Gray, Carol Clugston, Anna Dominiczak and Alex Alderton, Roberto Amato, Jeffrey Barrett, Sonia Goncalves, Ewan Harrison, David K. Jackson, Ian Johnston, Dominic Kwiatkowski, Cordelia Langford, John Sillitoe on behalf of the Wellcome Sanger Institute COVID-19 Surveillance Team |
| EPI_ISL_3096102, EPI_ISL_3096104, EPI_ISL_3096105, EPI_ISL_3096106 | Lighthouse Lab in Milton Keynes | Wellcome Sanger Institute for the COVID-19 Genomics UK (COG-UK) Consortium | The Lighthouse Lab in Milton Keynes and Alex Alderton, Roberto Amato, Jeffrey Barrett, Sonia Goncalves, Ewan Harrison, David K. Jackson, Ian Johnston, Dominic Kwiatkowski, Cordelia Langford, John Sillitoe on behalf of the Wellcome Sanger Institute COVID-19 Surveillance Team |
| EPI_ISL_3096107, EPI_ISL_3096108, EPI_ISL_3096109 | Berkshire and Surrey Pathology Services Lighthouse Laboratory | Wellcome Sanger Institute for the COVID-19 Genomics UK (COG-UK) Consortium | Berkshire and Surrey Pathology Services Lighthouse Laboratory and Alex Alderton, Roberto Amato, Jeffrey Barrett, Sonia Goncalves, Ewan Harrison, David K. Jackson, Ian Johnston, Dominic Kwiatkowski, Cordelia Langford, John Sillitoe on behalf of the Wellcome Sanger Institute COVID-19 Surveillance Team |

| | | | |
|---|---|---|---|
| EPI_ISL_3096110 | Lighthouse Lab in Milton Keynes | Wellcome Sanger Institute for the COVID-19 Genomics UK (COG-UK) Consortium | The Lighthouse Lab in Milton Keynes and Alex Alderton, Roberto Amato, Jeffrey Barrett, Sonia Goncalves, Ewan Harrison, David K. Jackson, Ian Johnston, Dominic Kwiatkowski, Cordelia Langford, John Sillitoe on behalf of the Wellcome Sanger Institute COVID-19 Surveillance Team |
| EPI_ISL_3096111 | Lighthouse Lab in Glasgow | Wellcome Sanger Institute for the COVID-19 Genomics UK (COG-UK) Consortium | Harper VanSteenhouse, Yumi Kasai, David Gray, Carol Clugston, Anna Dominiczak and Alex Alderton, Roberto Amato, Jeffrey Barrett, Sonia Goncalves, Ewan Harrison, David K. Jackson, Ian Johnston, Dominic Kwiatkowski, Cordelia Langford, John Sillitoe on behalf of the Wellcome Sanger Institute COVID-19 Surveillance Team |
| EPI_ISL_3096112 | Berkshire and Surrey Pathology Services Lighthouse Laboratory | Wellcome Sanger Institute for the COVID-19 Genomics UK (COG-UK) Consortium | Berkshire and Surrey Pathology Services Lighthouse Laboratory and Alex Alderton, Roberto Amato, Jeffrey Barrett, Sonia Goncalves, Ewan Harrison, David K. Jackson, Ian Johnston, Dominic Kwiatkowski, Cordelia Langford, John Sillitoe on behalf of the Wellcome Sanger Institute COVID-19 Surveillance Team |
| EPI_ISL_3096113 | Lighthouse Lab in Glasgow | Wellcome Sanger Institute for the COVID-19 Genomics UK (COG-UK) Consortium | Harper VanSteenhouse, Yumi Kasai, David Gray, Carol Clugston, Anna Dominiczak and Alex Alderton, Roberto Amato, Jeffrey Barrett, Sonia Goncalves, Ewan Harrison, David K. Jackson, Ian Johnston, Dominic Kwiatkowski, Cordelia Langford, John Sillitoe on behalf of the Wellcome Sanger Institute COVID-19 Surveillance Team |
| EPI_ISL_3096114, EPI_ISL_3096115 | Berkshire and Surrey Pathology Services Lighthouse Laboratory | Wellcome Sanger Institute for the COVID-19 Genomics UK (COG-UK) Consortium | Berkshire and Surrey Pathology Services Lighthouse Laboratory and Alex Alderton, Roberto Amato, Jeffrey Barrett, Sonia Goncalves, Ewan Harrison, David K. Jackson, Ian Johnston, Dominic Kwiatkowski, Cordelia Langford, John Sillitoe on behalf of the Wellcome Sanger Institute COVID-19 Surveillance Team |
| EPI_ISL_3096116 | Lighthouse Lab in Milton Keynes | Wellcome Sanger Institute for the COVID-19 Genomics UK (COG-UK) Consortium | The Lighthouse Lab in Milton Keynes and Alex Alderton, Roberto Amato, Jeffrey Barrett, Sonia Goncalves, Ewan Harrison, David K. Jackson, Ian Johnston, Dominic Kwiatkowski, Cordelia Langford, John Sillitoe on behalf of the Wellcome Sanger Institute COVID-19 Surveillance Team |
| EPI_ISL_3096117 | Berkshire and Surrey Pathology Services Lighthouse Laboratory | Wellcome Sanger Institute for the COVID-19 Genomics UK (COG-UK) Consortium | Berkshire and Surrey Pathology Services Lighthouse Laboratory and Alex Alderton, Roberto Amato, Jeffrey Barrett, Sonia Goncalves, Ewan Harrison, David K. Jackson, Ian Johnston, Dominic Kwiatkowski, Cordelia Langford, John Sillitoe on behalf of the Wellcome Sanger Institute COVID-19 Surveillance Team |
| EPI_ISL_3096118, EPI_ISL_3096119, EPI_ISL_3096120 | Lighthouse Lab in Glasgow | Wellcome Sanger Institute for the COVID-19 Genomics UK (COG-UK) Consortium | Harper VanSteenhouse, Yumi Kasai, David Gray, Carol Clugston, Anna Dominiczak and Alex Alderton, Roberto Amato, Jeffrey Barrett, Sonia Goncalves, Ewan Harrison, David K. Jackson, Ian Johnston, Dominic Kwiatkowski, Cordelia Langford, John Sillitoe on behalf of the Wellcome Sanger Institute COVID-19 Surveillance Team |
| EPI_ISL_3096121, EPI_ISL_3096122 | Lighthouse Lab in Milton Keynes | Wellcome Sanger Institute for the COVID-19 Genomics UK (COG-UK) Consortium | The Lighthouse Lab in Milton Keynes and Alex Alderton, Roberto Amato, Jeffrey Barrett, Sonia Goncalves, Ewan Harrison, David K. Jackson, Ian Johnston, Dominic Kwiatkowski, Cordelia Langford, John Sillitoe on behalf of the Wellcome Sanger Institute COVID-19 Surveillance Team |
| EPI_ISL_3096123 | Lighthouse Lab in Glasgow | Wellcome Sanger Institute for the COVID-19 Genomics UK (COG-UK) Consortium | Harper VanSteenhouse, Yumi Kasai, David Gray, Carol Clugston, Anna Dominiczak and Alex Alderton, Roberto Amato, Jeffrey Barrett, Sonia Goncalves, Ewan Harrison, David K. Jackson, Ian Johnston, Dominic Kwiatkowski, Cordelia Langford, John Sillitoe on behalf of the Wellcome Sanger Institute COVID-19 Surveillance Team |
| EPI_ISL_3096124 | Berkshire and Surrey Pathology Services Lighthouse Laboratory | Wellcome Sanger Institute for the COVID-19 Genomics UK (COG-UK) Consortium | Berkshire and Surrey Pathology Services Lighthouse Laboratory and Alex Alderton, Roberto Amato, Jeffrey Barrett, Sonia Goncalves, Ewan Harrison, David K. Jackson, Ian Johnston, Dominic Kwiatkowski, Cordelia Langford, John Sillitoe on behalf of the Wellcome Sanger Institute COVID-19 Surveillance Team |
| EPI_ISL_3096125 | Lighthouse Lab in Milton Keynes | Wellcome Sanger Institute for the COVID-19 Genomics UK (COG-UK) Consortium | The Lighthouse Lab in Milton Keynes and Alex Alderton, Roberto Amato, Jeffrey Barrett, Sonia Goncalves, Ewan Harrison, David K. Jackson, Ian Johnston, Dominic Kwiatkowski, Cordelia Langford, John Sillitoe on behalf of the Wellcome Sanger Institute COVID-19 Surveillance Team |
| EPI_ISL_3096126, EPI_ISL_3096127 | Lighthouse Lab in Glasgow | Wellcome Sanger Institute for the COVID-19 Genomics UK (COG-UK) Consortium | Harper VanSteenhouse, Yumi Kasai, David Gray, Carol Clugston, Anna Dominiczak and Alex Alderton, Roberto Amato, Jeffrey Barrett, Sonia Goncalves, Ewan Harrison, David K. Jackson, Ian Johnston, Dominic Kwiatkowski, Cordelia Langford, John Sillitoe on behalf of the Wellcome Sanger Institute COVID-19 Surveillance Team |
| EPI_ISL_3096128 | Lighthouse Lab in Milton Keynes | Wellcome Sanger Institute for the COVID-19 Genomics UK (COG-UK) Consortium | The Lighthouse Lab in Milton Keynes and Alex Alderton, Roberto Amato, Jeffrey Barrett, Sonia Goncalves, Ewan Harrison, David K. Jackson, Ian Johnston, Dominic Kwiatkowski, Cordelia Langford, John Sillitoe on behalf of the Wellcome Sanger Institute COVID-19 Surveillance Team |
| EPI_ISL_3096129, EPI_ISL_3096130 | Berkshire and Surrey Pathology Services Lighthouse Laboratory | Wellcome Sanger Institute for the COVID-19 Genomics UK (COG-UK) Consortium | Berkshire and Surrey Pathology Services Lighthouse Laboratory and Alex Alderton, Roberto Amato, Jeffrey Barrett, Sonia Goncalves, Ewan Harrison, David K. Jackson, Ian Johnston, Dominic Kwiatkowski, Cordelia Langford, John Sillitoe on behalf of the Wellcome Sanger Institute COVID-19 Surveillance Team |
| EPI_ISL_3096131 | Lighthouse Lab in Milton Keynes | Wellcome Sanger Institute for the COVID-19 Genomics UK (COG-UK) Consortium | The Lighthouse Lab in Milton Keynes and Alex Alderton, Roberto Amato, Jeffrey Barrett, Sonia Goncalves, Ewan Harrison, David K. Jackson, Ian Johnston, Dominic Kwiatkowski, Cordelia Langford, John Sillitoe on behalf of the Wellcome Sanger Institute COVID-19 Surveillance Team |
| EPI_ISL_3096132 | Berkshire and Surrey Pathology Services Lighthouse Laboratory | Wellcome Sanger Institute for the COVID-19 Genomics UK (COG-UK) Consortium | Berkshire and Surrey Pathology Services Lighthouse Laboratory and Alex Alderton, Roberto Amato, Jeffrey Barrett, Sonia Goncalves, Ewan Harrison, David K. Jackson, Ian Johnston, Dominic Kwiatkowski, Cordelia Langford, John Sillitoe on behalf of the Wellcome Sanger Institute COVID-19 Surveillance Team |
| EPI_ISL_3096133, EPI_ISL_3096134, EPI_ISL_3096135, EPI_ISL_3096136 | Lighthouse Lab in Milton Keynes | Wellcome Sanger Institute for the COVID-19 Genomics UK (COG-UK) Consortium | The Lighthouse Lab in Milton Keynes and Alex Alderton, Roberto Amato, Jeffrey Barrett, Sonia Goncalves, Ewan Harrison, David K. Jackson, Ian Johnston, Dominic Kwiatkowski, Cordelia Langford, John Sillitoe on behalf of the Wellcome Sanger Institute COVID-19 Surveillance Team |
| EPI_ISL_3096137, EPI_ISL_3096138, EPI_ISL_3096139, EPI_ISL_3096140 | Lighthouse Lab in Glasgow | Wellcome Sanger Institute for the COVID-19 Genomics UK (COG-UK) Consortium | Harper VanSteenhouse, Yumi Kasai, David Gray, Carol Clugston, Anna Dominiczak and Alex Alderton, Roberto Amato, Jeffrey Barrett, Sonia Goncalves, Ewan Harrison, David K. Jackson, Ian Johnston, Dominic Kwiatkowski, Cordelia Langford, John Sillitoe on behalf of the Wellcome Sanger Institute COVID-19 Surveillance Team |
| EPI_ISL_3096141 | Lighthouse Lab in Milton Keynes | Wellcome Sanger Institute for the COVID-19 Genomics UK (COG-UK) Consortium | The Lighthouse Lab in Milton Keynes and Alex Alderton, Roberto Amato, Jeffrey Barrett, Sonia Goncalves, Ewan Harrison, David K. Jackson, Ian Johnston, Dominic Kwiatkowski, Cordelia Langford, John Sillitoe on behalf of the Wellcome Sanger Institute COVID-19 Surveillance Team |
| EPI_ISL_3096142, EPI_ISL_3096143 | Lighthouse Lab in Glasgow | Wellcome Sanger Institute for the COVID-19 Genomics UK (COG-UK) Consortium | Harper VanSteenhouse, Yumi Kasai, David Gray, Carol Clugston, Anna Dominiczak and Alex Alderton, Roberto Amato, Jeffrey Barrett, Sonia Goncalves, Ewan Harrison, David K. Jackson, Ian Johnston, Dominic Kwiatkowski, Cordelia Langford, John Sillitoe on behalf of the Wellcome Sanger Institute COVID-19 Surveillance Team |
| EPI_ISL_3096144, EPI_ISL_3096145 | Lighthouse Lab in Milton Keynes | Wellcome Sanger Institute for the COVID-19 Genomics UK (COG-UK) Consortium | The Lighthouse Lab in Milton Keynes and Alex Alderton, Roberto Amato, Jeffrey Barrett, Sonia Goncalves, Ewan Harrison, David K. Jackson, Ian Johnston, Dominic Kwiatkowski, Cordelia Langford, John Sillitoe on behalf of the Wellcome Sanger Institute COVID-19 Surveillance Team |
| EPI_ISL_3096146 | Lighthouse Lab in Glasgow | Wellcome Sanger Institute for the COVID-19 Genomics UK (COG-UK) Consortium | Harper VanSteenhouse, Yumi Kasai, David Gray, Carol Clugston, Anna Dominiczak and Alex Alderton, Roberto Amato, Jeffrey Barrett, Sonia Goncalves, Ewan Harrison, David K. Jackson, Ian Johnston, Dominic Kwiatkowski, Cordelia Langford, John Sillitoe on behalf of the Wellcome Sanger Institute COVID-19 Surveillance Team |
| EPI_ISL_3096147 | Berkshire and Surrey Pathology Services Lighthouse Laboratory | Wellcome Sanger Institute for the COVID-19 Genomics UK (COG-UK) Consortium | Berkshire and Surrey Pathology Services Lighthouse Laboratory and Alex Alderton, Roberto Amato, Jeffrey Barrett, Sonia Goncalves, Ewan Harrison, David K. Jackson, Ian Johnston, Dominic Kwiatkowski, Cordelia Langford, John Sillitoe on behalf of the Wellcome Sanger Institute COVID-19 Surveillance Team |
| EPI_ISL_3096148 | Lighthouse Lab in Milton Keynes | Wellcome Sanger Institute for the COVID-19 Genomics UK (COG-UK) Consortium | The Lighthouse Lab in Milton Keynes and Alex Alderton, Roberto Amato, Jeffrey Barrett, Sonia Goncalves, Ewan Harrison, David K. Jackson, Ian Johnston, Dominic Kwiatkowski, Cordelia Langford, John Sillitoe on behalf of the Wellcome Sanger Institute COVID-19 Surveillance Team |
| EPI_ISL_3096149 | Lighthouse Lab in Glasgow | Wellcome Sanger Institute for the COVID-19 Genomics UK (COG-UK) Consortium | Harper VanSteenhouse, Yumi Kasai, David Gray, Carol Clugston, Anna Dominiczak and Alex Alderton, Roberto Amato, Jeffrey Barrett, Sonia Goncalves, Ewan Harrison, David K. Jackson, Ian Johnston, Dominic Kwiatkowski, Cordelia Langford, John Sillitoe on behalf of the Wellcome Sanger Institute COVID-19 Surveillance Team |
| EPI_ISL_3096150, EPI_ISL_3096151 | Lighthouse Lab in Milton Keynes | Wellcome Sanger Institute for the COVID-19 Genomics UK (COG-UK) Consortium | The Lighthouse Lab in Milton Keynes and Alex Alderton, Roberto Amato, Jeffrey Barrett, Sonia Goncalves, Ewan Harrison, David K. Jackson, Ian Johnston, Dominic Kwiatkowski, Cordelia Langford, John Sillitoe on behalf of the Wellcome Sanger Institute COVID-19 Surveillance Team |
| EPI_ISL_3096152 | Berkshire and Surrey Pathology Services Lighthouse Laboratory | Wellcome Sanger Institute for the COVID-19 Genomics UK (COG-UK) Consortium | Berkshire and Surrey Pathology Services Lighthouse Laboratory and Alex Alderton, Roberto Amato, Jeffrey Barrett, Sonia Goncalves, Ewan Harrison, David K. Jackson, Ian Johnston, Dominic Kwiatkowski, Cordelia Langford, John Sillitoe on behalf of the Wellcome Sanger Institute COVID-19 Surveillance Team |
| EPI_ISL_3096153 | Lighthouse Lab in Glasgow | Wellcome Sanger Institute for the COVID-19 Genomics UK (COG-UK) Consortium | Harper VanSteenhouse, Yumi Kasai, David Gray, Carol Clugston, Anna Dominiczak and Alex Alderton, Roberto Amato, Jeffrey Barrett, Sonia Goncalves, Ewan Harrison, David K. Jackson, Ian Johnston, Dominic Kwiatkowski, Cordelia Langford, John Sillitoe on behalf of the Wellcome Sanger Institute COVID-19 Surveillance Team |
| EPI_ISL_3096154 | Berkshire and Surrey Pathology Services Lighthouse Laboratory | Wellcome Sanger Institute for the COVID-19 Genomics UK (COG-UK) Consortium | Berkshire and Surrey Pathology Services Lighthouse Laboratory and Alex Alderton, Roberto Amato, Jeffrey Barrett, Sonia Goncalves, Ewan Harrison, David K. Jackson, Ian Johnston, Dominic Kwiatkowski, Cordelia Langford, John Sillitoe on behalf of the Wellcome Sanger Institute COVID-19 Surveillance Team |
| EPI_ISL_3096155 | Lighthouse Lab in Milton Keynes | Wellcome Sanger Institute for the COVID-19 Genomics UK (COG-UK) Consortium | The Lighthouse Lab in Milton Keynes and Alex Alderton, Roberto Amato, Jeffrey Barrett, Sonia Goncalves, Ewan Harrison, David K. Jackson, Ian Johnston, Dominic Kwiatkowski, Cordelia Langford, John Sillitoe on behalf of the Wellcome Sanger Institute COVID-19 Surveillance Team |
| EPI_ISL_3096156, EPI_ISL_3096157 | Berkshire and Surrey Pathology Services Lighthouse Laboratory | Wellcome Sanger Institute for the COVID-19 Genomics UK (COG-UK) Consortium | Berkshire and Surrey Pathology Services Lighthouse Laboratory and Alex Alderton, Roberto Amato, Jeffrey Barrett, Sonia Goncalves, Ewan Harrison, David K. Jackson, Ian Johnston, Dominic Kwiatkowski, Cordelia Langford, John Sillitoe on behalf of the Wellcome Sanger Institute COVID-19 Surveillance Team |
| EPI_ISL_3096158, EPI_ISL_3096159, EPI_ISL_3096160, EPI_ISL_3096161, EPI_ISL_3096162, EPI_ISL_3096163, EPI_ISL_3096164, EPI_ISL_3096165 | Lighthouse Lab in Milton Keynes | Wellcome Sanger Institute for the COVID-19 Genomics UK (COG-UK) Consortium | The Lighthouse Lab in Milton Keynes and Alex Alderton, Roberto Amato, Jeffrey Barrett, Sonia Goncalves, Ewan Harrison, David K. Jackson, Ian Johnston, Dominic Kwiatkowski, Cordelia Langford, John Sillitoe on behalf of the Wellcome Sanger Institute COVID-19 Surveillance Team |
| EPI_ISL_3096166 | Lighthouse Lab in Glasgow | Wellcome Sanger Institute for the COVID-19 Genomics UK (COG-UK) Consortium | Harper VanSteenhouse, Yumi Kasai, David Gray, Carol Clugston, Anna Dominiczak and Alex Alderton, Roberto Amato, Jeffrey Barrett, Sonia Goncalves, Ewan Harrison, David K. Jackson, Ian Johnston, Dominic Kwiatkowski, Cordelia Langford, John Sillitoe on behalf of the Wellcome Sanger Institute COVID-19 Surveillance Team |
| EPI_ISL_3096167 | Lighthouse Lab in Milton Keynes | Wellcome Sanger Institute for the COVID-19 Genomics UK (COG-UK) Consortium | The Lighthouse Lab in Milton Keynes and Alex Alderton, Roberto Amato, Jeffrey Barrett, Sonia Goncalves, Ewan Harrison, David K. Jackson, Ian Johnston, Dominic Kwiatkowski, Cordelia Langford, John Sillitoe on behalf of the Wellcome Sanger Institute COVID-19 Surveillance Team |
| EPI_ISL_3096168 | Berkshire and Surrey Pathology Services Lighthouse Laboratory | Wellcome Sanger Institute for the COVID-19 Genomics UK (COG-UK) Consortium | Berkshire and Surrey Pathology Services Lighthouse Laboratory and Alex Alderton, Roberto Amato, Jeffrey Barrett, Sonia Goncalves, Ewan Harrison, David K. Jackson, Ian Johnston, Dominic Kwiatkowski, Cordelia Langford, John Sillitoe on behalf of the Wellcome Sanger Institute COVID-19 Surveillance Team |
| EPI_ISL_3096169, EPI_ISL_3096170 | Lighthouse Lab in Milton Keynes | Wellcome Sanger Institute for the COVID-19 Genomics UK (COG-UK) Consortium | The Lighthouse Lab in Milton Keynes and Alex Alderton, Roberto Amato, Jeffrey Barrett, Sonia Goncalves, Ewan Harrison, David K. Jackson, Ian Johnston, Dominic Kwiatkowski, Cordelia Langford, John Sillitoe on behalf of the Wellcome Sanger Institute COVID-19 Surveillance Team |
| EPI_ISL_3096171 | Berkshire and Surrey Pathology Services Lighthouse Laboratory | Wellcome Sanger Institute for the COVID-19 Genomics UK (COG-UK) Consortium | Berkshire and Surrey Pathology Services Lighthouse Laboratory and Alex Alderton, Roberto Amato, Jeffrey Barrett, Sonia Goncalves, Ewan Harrison, David K. Jackson, Ian Johnston, Dominic Kwiatkowski, Cordelia Langford, John Sillitoe on behalf of the Wellcome Sanger Institute COVID-19 Surveillance Team |
| EPI_ISL_3096172 | Lighthouse Lab in Milton Keynes | Wellcome Sanger Institute for the COVID-19 Genomics UK (COG-UK) Consortium | The Lighthouse Lab in Milton Keynes and Alex Alderton, Roberto Amato, Jeffrey Barrett, Sonia Goncalves, Ewan Harrison, David K. Jackson, Ian Johnston, Dominic Kwiatkowski, Cordelia Langford, John Sillitoe on behalf of the Wellcome Sanger Institute COVID-19 Surveillance Team |
| EPI_ISL_3096173 | Lighthouse Lab in Glasgow | Wellcome Sanger Institute for the COVID-19 Genomics UK (COG-UK) Consortium | Harper VanSteenhouse, Yumi Kasai, David Gray, Carol Clugston, Anna Dominiczak and Alex Alderton, Roberto Amato, Jeffrey Barrett, Sonia Goncalves, Ewan Harrison, David K. Jackson, Ian Johnston, Dominic Kwiatkowski, Cordelia Langford, John Sillitoe on behalf of the Wellcome Sanger Institute COVID-19 Surveillance Team |

| | | | |
|---|---|---|---|
| EPI_ISL_3096174, EPI_ISL_3096175, EPI_ISL_3096176, EPI_ISL_3096177, EPI_ISL_3096178, EPI_ISL_3096179 | Lighthouse Lab in Milton Keynes | Wellcome Sanger Institute for the COVID-19 Genomics UK (COG-UK) Consortium | The Lighthouse Lab in Milton Keynes and Alex Alderton, Roberto Amato, Jeffrey Barrett, Sonia Goncalves, Ewan Harrison, David K. Jackson, Ian Johnston, Dominic Kwiatkowski, Cordelia Langford, John Sillitoe on behalf of the Wellcome Sanger Institute COVID-19 Surveillance Team |
| EPI_ISL_3096180, EPI_ISL_3096181 | Lighthouse Lab in Glasgow | Wellcome Sanger Institute for the COVID-19 Genomics UK (COG-UK) Consortium | Harper VanSteenhouse, Yumi Kasai, David Gray, Carol Clugston, Anna Dominiczak and Alex Alderton, Roberto Amato, Jeffrey Barrett, Sonia Goncalves, Ewan Harrison, David K. Jackson, Ian Johnston, Dominic Kwiatkowski, Cordelia Langford, John Sillitoe on behalf of the Wellcome Sanger Institute COVID-19 Surveillance Team |
| EPI_ISL_3096182 | Berkshire and Surrey Pathology Services Lighthouse Laboratory | Wellcome Sanger Institute for the COVID-19 Genomics UK (COG-UK) Consortium | Berkshire and Surrey Pathology Services Lighthouse Laboratory and Alex Alderton, Roberto Amato, Jeffrey Barrett, Sonia Goncalves, Ewan Harrison, David K. Jackson, Ian Johnston, Dominic Kwiatkowski, Cordelia Langford, John Sillitoe on behalf of the Wellcome Sanger Institute COVID-19 Surveillance Team |
| EPI_ISL_3096183 | Lighthouse Lab in Glasgow | Wellcome Sanger Institute for the COVID-19 Genomics UK (COG-UK) Consortium | Harper VanSteenhouse, Yumi Kasai, David Gray, Carol Clugston, Anna Dominiczak and Alex Alderton, Roberto Amato, Jeffrey Barrett, Sonia Goncalves, Ewan Harrison, David K. Jackson, Ian Johnston, Dominic Kwiatkowski, Cordelia Langford, John Sillitoe on behalf of the Wellcome Sanger Institute COVID-19 Surveillance Team |
| EPI_ISL_3096184, EPI_ISL_3096185 | Lighthouse Lab in Milton Keynes | Wellcome Sanger Institute for the COVID-19 Genomics UK (COG-UK) Consortium | The Lighthouse Lab in Milton Keynes and Alex Alderton, Roberto Amato, Jeffrey Barrett, Sonia Goncalves, Ewan Harrison, David K. Jackson, Ian Johnston, Dominic Kwiatkowski, Cordelia Langford, John Sillitoe on behalf of the Wellcome Sanger Institute COVID-19 Surveillance Team |
| EPI_ISL_3096186 | Lighthouse Lab in Glasgow | Wellcome Sanger Institute for the COVID-19 Genomics UK (COG-UK) Consortium | Harper VanSteenhouse, Yumi Kasai, David Gray, Carol Clugston, Anna Dominiczak and Alex Alderton, Roberto Amato, Jeffrey Barrett, Sonia Goncalves, Ewan Harrison, David K. Jackson, Ian Johnston, Dominic Kwiatkowski, Cordelia Langford, John Sillitoe on behalf of the Wellcome Sanger Institute COVID-19 Surveillance Team |
| EPI_ISL_3096187 | Lighthouse Lab in Milton Keynes | Wellcome Sanger Institute for the COVID-19 Genomics UK (COG-UK) Consortium | The Lighthouse Lab in Milton Keynes and Alex Alderton, Roberto Amato, Jeffrey Barrett, Sonia Goncalves, Ewan Harrison, David K. Jackson, Ian Johnston, Dominic Kwiatkowski, Cordelia Langford, John Sillitoe on behalf of the Wellcome Sanger Institute COVID-19 Surveillance Team |
| EPI_ISL_3096188 | Berkshire and Surrey Pathology Services Lighthouse Laboratory | Wellcome Sanger Institute for the COVID-19 Genomics UK (COG-UK) Consortium | Berkshire and Surrey Pathology Services Lighthouse Laboratory and Alex Alderton, Roberto Amato, Jeffrey Barrett, Sonia Goncalves, Ewan Harrison, David K. Jackson, Ian Johnston, Dominic Kwiatkowski, Cordelia Langford, John Sillitoe on behalf of the Wellcome Sanger Institute COVID-19 Surveillance Team |
| EPI_ISL_3096189 | Lighthouse Lab in Milton Keynes | Wellcome Sanger Institute for the COVID-19 Genomics UK (COG-UK) Consortium | The Lighthouse Lab in Milton Keynes and Alex Alderton, Roberto Amato, Jeffrey Barrett, Sonia Goncalves, Ewan Harrison, David K. Jackson, Ian Johnston, Dominic Kwiatkowski, Cordelia Langford, John Sillitoe on behalf of the Wellcome Sanger Institute COVID-19 Surveillance Team |
| EPI_ISL_3096190 | Berkshire and Surrey Pathology Services Lighthouse Laboratory | Wellcome Sanger Institute for the COVID-19 Genomics UK (COG-UK) Consortium | Berkshire and Surrey Pathology Services Lighthouse Laboratory and Alex Alderton, Roberto Amato, Jeffrey Barrett, Sonia Goncalves, Ewan Harrison, David K. Jackson, Ian Johnston, Dominic Kwiatkowski, Cordelia Langford, John Sillitoe on behalf of the Wellcome Sanger Institute COVID-19 Surveillance Team |
| EPI_ISL_3096191, EPI_ISL_3096192, EPI_ISL_3096193, EPI_ISL_3096194 | Lighthouse Lab in Milton Keynes | Wellcome Sanger Institute for the COVID-19 Genomics UK (COG-UK) Consortium | The Lighthouse Lab in Milton Keynes and Alex Alderton, Roberto Amato, Jeffrey Barrett, Sonia Goncalves, Ewan Harrison, David K. Jackson, Ian Johnston, Dominic Kwiatkowski, Cordelia Langford, John Sillitoe on behalf of the Wellcome Sanger Institute COVID-19 Surveillance Team |
| EPI_ISL_3096195, EPI_ISL_3096196 | Berkshire and Surrey Pathology Services Lighthouse Laboratory | Wellcome Sanger Institute for the COVID-19 Genomics UK (COG-UK) Consortium | Berkshire and Surrey Pathology Services Lighthouse Laboratory and Alex Alderton, Roberto Amato, Jeffrey Barrett, Sonia Goncalves, Ewan Harrison, David K. Jackson, Ian Johnston, Dominic Kwiatkowski, Cordelia Langford, John Sillitoe on behalf of the Wellcome Sanger Institute COVID-19 Surveillance Team |
| EPI_ISL_3096197 | Lighthouse Lab in Milton Keynes | Wellcome Sanger Institute for the COVID-19 Genomics UK (COG-UK) Consortium | The Lighthouse Lab in Milton Keynes and Alex Alderton, Roberto Amato, Jeffrey Barrett, Sonia Goncalves, Ewan Harrison, David K. Jackson, Ian Johnston, Dominic Kwiatkowski, Cordelia Langford, John Sillitoe on behalf of the Wellcome Sanger Institute COVID-19 Surveillance Team |
| EPI_ISL_3096198 | Lighthouse Lab in Glasgow | Wellcome Sanger Institute for the COVID-19 Genomics UK (COG-UK) Consortium | Harper VanSteenhouse, Yumi Kasai, David Gray, Carol Clugston, Anna Dominiczak and Alex Alderton, Roberto Amato, Jeffrey Barrett, Sonia Goncalves, Ewan Harrison, David K. Jackson, Ian Johnston, Dominic Kwiatkowski, Cordelia Langford, John Sillitoe on behalf of the Wellcome Sanger Institute COVID-19 Surveillance Team |
| EPI_ISL_3096199, EPI_ISL_3096200 | Lighthouse Lab in Milton Keynes | Wellcome Sanger Institute for the COVID-19 Genomics UK (COG-UK) Consortium | The Lighthouse Lab in Milton Keynes and Alex Alderton, Roberto Amato, Jeffrey Barrett, Sonia Goncalves, Ewan Harrison, David K. Jackson, Ian Johnston, Dominic Kwiatkowski, Cordelia Langford, John Sillitoe on behalf of the Wellcome Sanger Institute COVID-19 Surveillance Team |
| EPI_ISL_3096201, EPI_ISL_3096202 | Lighthouse Lab in Glasgow | Wellcome Sanger Institute for the COVID-19 Genomics UK (COG-UK) Consortium | Harper VanSteenhouse, Yumi Kasai, David Gray, Carol Clugston, Anna Dominiczak and Alex Alderton, Roberto Amato, Jeffrey Barrett, Sonia Goncalves, Ewan Harrison, David K. Jackson, Ian Johnston, Dominic Kwiatkowski, Cordelia Langford, John Sillitoe on behalf of the Wellcome Sanger Institute COVID-19 Surveillance Team |
| EPI_ISL_3096203 | Lighthouse Lab in Milton Keynes | Wellcome Sanger Institute for the COVID-19 Genomics UK (COG-UK) Consortium | The Lighthouse Lab in Milton Keynes and Alex Alderton, Roberto Amato, Jeffrey Barrett, Sonia Goncalves, Ewan Harrison, David K. Jackson, Ian Johnston, Dominic Kwiatkowski, Cordelia Langford, John Sillitoe on behalf of the Wellcome Sanger Institute COVID-19 Surveillance Team |
| EPI_ISL_3096204 | Berkshire and Surrey Pathology Services Lighthouse Laboratory | Wellcome Sanger Institute for the COVID-19 Genomics UK (COG-UK) Consortium | Berkshire and Surrey Pathology Services Lighthouse Laboratory and Alex Alderton, Roberto Amato, Jeffrey Barrett, Sonia Goncalves, Ewan Harrison, David K. Jackson, Ian Johnston, Dominic Kwiatkowski, Cordelia Langford, John Sillitoe on behalf of the Wellcome Sanger Institute COVID-19 Surveillance Team |
| EPI_ISL_3096205 | Lighthouse Lab in Milton Keynes | Wellcome Sanger Institute for the COVID-19 Genomics UK (COG-UK) Consortium | The Lighthouse Lab in Milton Keynes and Alex Alderton, Roberto Amato, Jeffrey Barrett, Sonia Goncalves, Ewan Harrison, David K. Jackson, Ian Johnston, Dominic Kwiatkowski, Cordelia Langford, John Sillitoe on behalf of the Wellcome Sanger Institute COVID-19 Surveillance Team |
| EPI_ISL_3096206 | Berkshire and Surrey Pathology Services Lighthouse Laboratory | Wellcome Sanger Institute for the COVID-19 Genomics UK (COG-UK) Consortium | Berkshire and Surrey Pathology Services Lighthouse Laboratory and Alex Alderton, Roberto Amato, Jeffrey Barrett, Sonia Goncalves, Ewan Harrison, David K. Jackson, Ian Johnston, Dominic Kwiatkowski, Cordelia Langford, John Sillitoe on behalf of the Wellcome Sanger Institute COVID-19 Surveillance Team |
| EPI_ISL_3096207, EPI_ISL_3096208 | Lighthouse Lab in Milton Keynes | Wellcome Sanger Institute for the COVID-19 Genomics UK (COG-UK) Consortium | The Lighthouse Lab in Milton Keynes and Alex Alderton, Roberto Amato, Jeffrey Barrett, Sonia Goncalves, Ewan Harrison, David K. Jackson, Ian Johnston, Dominic Kwiatkowski, Cordelia Langford, John Sillitoe on behalf of the Wellcome Sanger Institute COVID-19 Surveillance Team |
| EPI_ISL_3096209 | Lighthouse Lab in Glasgow | Wellcome Sanger Institute for the COVID-19 Genomics UK (COG-UK) Consortium | Harper VanSteenhouse, Yumi Kasai, David Gray, Carol Clugston, Anna Dominiczak and Alex Alderton, Roberto Amato, Jeffrey Barrett, Sonia Goncalves, Ewan Harrison, David K. Jackson, Ian Johnston, Dominic Kwiatkowski, Cordelia Langford, John Sillitoe on behalf of the Wellcome Sanger Institute COVID-19 Surveillance Team |
| EPI_ISL_3096210 | Lighthouse Lab in Milton Keynes | Wellcome Sanger Institute for the COVID-19 Genomics UK (COG-UK) Consortium | The Lighthouse Lab in Milton Keynes and Alex Alderton, Roberto Amato, Jeffrey Barrett, Sonia Goncalves, Ewan Harrison, David K. Jackson, Ian Johnston, Dominic Kwiatkowski, Cordelia Langford, John Sillitoe on behalf of the Wellcome Sanger Institute COVID-19 Surveillance Team |
| EPI_ISL_3096211 | Lighthouse Lab in Glasgow | Wellcome Sanger Institute for the COVID-19 Genomics UK (COG-UK) Consortium | Harper VanSteenhouse, Yumi Kasai, David Gray, Carol Clugston, Anna Dominiczak and Alex Alderton, Roberto Amato, Jeffrey Barrett, Sonia Goncalves, Ewan Harrison, David K. Jackson, Ian Johnston, Dominic Kwiatkowski, Cordelia Langford, John Sillitoe on behalf of the Wellcome Sanger Institute COVID-19 Surveillance Team |
| EPI_ISL_3096212, EPI_ISL_3096213 | Berkshire and Surrey Pathology Services Lighthouse Laboratory | Wellcome Sanger Institute for the COVID-19 Genomics UK (COG-UK) Consortium | Berkshire and Surrey Pathology Services Lighthouse Laboratory and Alex Alderton, Roberto Amato, Jeffrey Barrett, Sonia Goncalves, Ewan Harrison, David K. Jackson, Ian Johnston, Dominic Kwiatkowski, Cordelia Langford, John Sillitoe on behalf of the Wellcome Sanger Institute COVID-19 Surveillance Team |
| EPI_ISL_3096214 | Lighthouse Lab in Glasgow | Wellcome Sanger Institute for the COVID-19 Genomics UK (COG-UK) Consortium | Harper VanSteenhouse, Yumi Kasai, David Gray, Carol Clugston, Anna Dominiczak and Alex Alderton, Roberto Amato, Jeffrey Barrett, Sonia Goncalves, Ewan Harrison, David K. Jackson, Ian Johnston, Dominic Kwiatkowski, Cordelia Langford, John Sillitoe on behalf of the Wellcome Sanger Institute COVID-19 Surveillance Team |
| EPI_ISL_3096215, EPI_ISL_3096216 | Lighthouse Lab in Milton Keynes | Wellcome Sanger Institute for the COVID-19 Genomics UK (COG-UK) Consortium | The Lighthouse Lab in Milton Keynes and Alex Alderton, Roberto Amato, Jeffrey Barrett, Sonia Goncalves, Ewan Harrison, David K. Jackson, Ian Johnston, Dominic Kwiatkowski, Cordelia Langford, John Sillitoe on behalf of the Wellcome Sanger Institute COVID-19 Surveillance Team |
| EPI_ISL_3096217 | Berkshire and Surrey Pathology Services Lighthouse Laboratory | Wellcome Sanger Institute for the COVID-19 Genomics UK (COG-UK) Consortium | Berkshire and Surrey Pathology Services Lighthouse Laboratory and Alex Alderton, Roberto Amato, Jeffrey Barrett, Sonia Goncalves, Ewan Harrison, David K. Jackson, Ian Johnston, Dominic Kwiatkowski, Cordelia Langford, John Sillitoe on behalf of the Wellcome Sanger Institute COVID-19 Surveillance Team |
| EPI_ISL_3096218 | Lighthouse Lab in Glasgow | Wellcome Sanger Institute for the COVID-19 Genomics UK (COG-UK) Consortium | Harper VanSteenhouse, Yumi Kasai, David Gray, Carol Clugston, Anna Dominiczak and Alex Alderton, Roberto Amato, Jeffrey Barrett, Sonia Goncalves, Ewan Harrison, David K. Jackson, Ian Johnston, Dominic Kwiatkowski, Cordelia Langford, John Sillitoe on behalf of the Wellcome Sanger Institute COVID-19 Surveillance Team |
| EPI_ISL_3096219, EPI_ISL_3096220, EPI_ISL_3096221, EPI_ISL_3096222 | Lighthouse Lab in Milton Keynes | Wellcome Sanger Institute for the COVID-19 Genomics UK (COG-UK) Consortium | The Lighthouse Lab in Milton Keynes and Alex Alderton, Roberto Amato, Jeffrey Barrett, Sonia Goncalves, Ewan Harrison, David K. Jackson, Ian Johnston, Dominic Kwiatkowski, Cordelia Langford, John Sillitoe on behalf of the Wellcome Sanger Institute COVID-19 Surveillance Team |
| EPI_ISL_3096223 | Lighthouse Lab in Glasgow | Wellcome Sanger Institute for the COVID-19 Genomics UK (COG-UK) Consortium | Harper VanSteenhouse, Yumi Kasai, David Gray, Carol Clugston, Anna Dominiczak and Alex Alderton, Roberto Amato, Jeffrey Barrett, Sonia Goncalves, Ewan Harrison, David K. Jackson, Ian Johnston, Dominic Kwiatkowski, Cordelia Langford, John Sillitoe on behalf of the Wellcome Sanger Institute COVID-19 Surveillance Team |
| EPI_ISL_3096224, EPI_ISL_3096225 | Lighthouse Lab in Milton Keynes | Wellcome Sanger Institute for the COVID-19 Genomics UK (COG-UK) Consortium | The Lighthouse Lab in Milton Keynes and Alex Alderton, Roberto Amato, Jeffrey Barrett, Sonia Goncalves, Ewan Harrison, David K. Jackson, Ian Johnston, Dominic Kwiatkowski, Cordelia Langford, John Sillitoe on behalf of the Wellcome Sanger Institute COVID-19 Surveillance Team |
| EPI_ISL_3096226, EPI_ISL_3096227 | Lighthouse Lab in Glasgow | Wellcome Sanger Institute for the COVID-19 Genomics UK (COG-UK) Consortium | Harper VanSteenhouse, Yumi Kasai, David Gray, Carol Clugston, Anna Dominiczak and Alex Alderton, Roberto Amato, Jeffrey Barrett, Sonia Goncalves, Ewan Harrison, David K. Jackson, Ian Johnston, Dominic Kwiatkowski, Cordelia Langford, John Sillitoe on behalf of the Wellcome Sanger Institute COVID-19 Surveillance Team |
| EPI_ISL_3096228 | Lighthouse Lab in Milton Keynes | Wellcome Sanger Institute for the COVID-19 Genomics UK (COG-UK) Consortium | The Lighthouse Lab in Milton Keynes and Alex Alderton, Roberto Amato, Jeffrey Barrett, Sonia Goncalves, Ewan Harrison, David K. Jackson, Ian Johnston, Dominic Kwiatkowski, Cordelia Langford, John Sillitoe on behalf of the Wellcome Sanger Institute COVID-19 Surveillance Team |
| EPI_ISL_3096229 | Lighthouse Lab in Glasgow | Wellcome Sanger Institute for the COVID-19 Genomics UK (COG-UK) Consortium | Harper VanSteenhouse, Yumi Kasai, David Gray, Carol Clugston, Anna Dominiczak and Alex Alderton, Roberto Amato, Jeffrey Barrett, Sonia Goncalves, Ewan Harrison, David K. Jackson, Ian Johnston, Dominic Kwiatkowski, Cordelia Langford, John Sillitoe on behalf of the Wellcome Sanger Institute COVID-19 Surveillance Team |
| EPI_ISL_3096230 | Berkshire and Surrey Pathology Services Lighthouse Laboratory | Wellcome Sanger Institute for the COVID-19 Genomics UK (COG-UK) Consortium | Berkshire and Surrey Pathology Services Lighthouse Laboratory and Alex Alderton, Roberto Amato, Jeffrey Barrett, Sonia Goncalves, Ewan Harrison, David K. Jackson, Ian Johnston, Dominic Kwiatkowski, Cordelia Langford, John Sillitoe on behalf of the Wellcome Sanger Institute COVID-19 Surveillance Team |
| EPI_ISL_3096231 | Lighthouse Lab in Glasgow | Wellcome Sanger Institute for the COVID-19 Genomics UK (COG-UK) Consortium | Harper VanSteenhouse, Yumi Kasai, David Gray, Carol Clugston, Anna Dominiczak and Alex Alderton, Roberto Amato, Jeffrey Barrett, Sonia Goncalves, Ewan Harrison, David K. Jackson, Ian Johnston, Dominic Kwiatkowski, Cordelia Langford, John Sillitoe on behalf of the Wellcome Sanger Institute COVID-19 Surveillance Team |
| EPI_ISL_3096232 | Lighthouse Lab in Milton Keynes | Wellcome Sanger Institute for the COVID-19 Genomics UK (COG-UK) Consortium | The Lighthouse Lab in Milton Keynes and Alex Alderton, Roberto Amato, Jeffrey Barrett, Sonia Goncalves, Ewan Harrison, David K. Jackson, Ian Johnston, Dominic Kwiatkowski, Cordelia Langford, John Sillitoe on behalf of the Wellcome Sanger Institute COVID-19 Surveillance Team |
| EPI_ISL_3096233 | Lighthouse Lab in Glasgow | Wellcome Sanger Institute for the COVID-19 Genomics UK (COG-UK) Consortium | Harper VanSteenhouse, Yumi Kasai, David Gray, Carol Clugston, Anna Dominiczak and Alex Alderton, Roberto Amato, Jeffrey Barrett, Sonia Goncalves, Ewan Harrison, David K. Jackson, Ian Johnston, Dominic Kwiatkowski, Cordelia Langford, John Sillitoe on behalf of the Wellcome Sanger Institute COVID-19 Surveillance Team |
| EPI_ISL_3096234 | Lighthouse Lab in Milton Keynes | Wellcome Sanger Institute for the COVID-19 Genomics UK (COG-UK) Consortium | The Lighthouse Lab in Milton Keynes and Alex Alderton, Roberto Amato, Jeffrey Barrett, Sonia Goncalves, Ewan Harrison, David K. Jackson, Ian Johnston, Dominic Kwiatkowski, Cordelia Langford, John Sillitoe on behalf of the Wellcome Sanger Institute COVID-19 Surveillance Team |
| EPI_ISL_3096235 | Lighthouse Lab in Glasgow | Wellcome Sanger Institute for the COVID-19 Genomics UK (COG-UK) Consortium | Harper VanSteenhouse, Yumi Kasai, David Gray, Carol Clugston, Anna Dominiczak and Alex Alderton, Roberto Amato, Jeffrey Barrett, Sonia Goncalves, Ewan Harrison, David K. Jackson, Ian Johnston, Dominic Kwiatkowski, Cordelia Langford, John Sillitoe on behalf of the Wellcome Sanger Institute COVID-19 Surveillance Team |
| EPI_ISL_3096236, EPI_ISL_3096237 | Berkshire and Surrey Pathology Services Lighthouse Laboratory | Wellcome Sanger Institute for the COVID-19 Genomics UK (COG-UK) Consortium | Berkshire and Surrey Pathology Services Lighthouse Laboratory and Alex Alderton, Roberto Amato, Jeffrey Barrett, Sonia Goncalves, Ewan Harrison, David K. Jackson, Ian Johnston, Dominic Kwiatkowski, Cordelia Langford, John Sillitoe on behalf of the Wellcome Sanger Institute COVID-19 Surveillance Team |

| | | | |
|---|---|---|---|
| EPI_ISL_3096238 | Lighthouse Lab in Milton Keynes | Wellcome Sanger Institute for the COVID-19 Genomics UK (COG-UK) Consortium | The Lighthouse Lab in Milton Keynes and Alex Alderton, Roberto Amato, Jeffrey Barrett, Sonia Goncalves, Ewan Harrison, David K. Jackson, Ian Johnston, Dominic Kwiatkowski, Cordelia Langford, John Sillitoe on behalf of the Wellcome Sanger Institute COVID-19 Surveillance Team |
| EPI_ISL_3096239 | Lighthouse Lab in Glasgow | Wellcome Sanger Institute for the COVID-19 Genomics UK (COG-UK) Consortium | Harper VanSteenhouse, Yumi Kasai, David Gray, Carol Clugston, Anna Dominiczak and Alex Alderton, Roberto Amato, Jeffrey Barrett, Sonia Goncalves, Ewan Harrison, David K. Jackson, Ian Johnston, Dominic Kwiatkowski, Cordelia Langford, John Sillitoe on behalf of the Wellcome Sanger Institute COVID-19 Surveillance Team |
| EPI_ISL_3096240, EPI_ISL_3096241 | Lighthouse Lab in Milton Keynes | Wellcome Sanger Institute for the COVID-19 Genomics UK (COG-UK) Consortium | The Lighthouse Lab in Milton Keynes and Alex Alderton, Roberto Amato, Jeffrey Barrett, Sonia Goncalves, Ewan Harrison, David K. Jackson, Ian Johnston, Dominic Kwiatkowski, Cordelia Langford, John Sillitoe on behalf of the Wellcome Sanger Institute COVID-19 Surveillance Team |
| EPI_ISL_3096242 | Berkshire and Surrey Pathology Services Lighthouse Laboratory | Wellcome Sanger Institute for the COVID-19 Genomics UK (COG-UK) Consortium | Berkshire and Surrey Pathology Services Lighthouse Laboratory and Alex Alderton, Roberto Amato, Jeffrey Barrett, Sonia Goncalves, Ewan Harrison, David K. Jackson, Ian Johnston, Dominic Kwiatkowski, Cordelia Langford, John Sillitoe on behalf of the Wellcome Sanger Institute COVID-19 Surveillance Team |
| EPI_ISL_3096243 | Lighthouse Lab in Glasgow | Wellcome Sanger Institute for the COVID-19 Genomics UK (COG-UK) Consortium | Harper VanSteenhouse, Yumi Kasai, David Gray, Carol Clugston, Anna Dominiczak and Alex Alderton, Roberto Amato, Jeffrey Barrett, Sonia Goncalves, Ewan Harrison, David K. Jackson, Ian Johnston, Dominic Kwiatkowski, Cordelia Langford, John Sillitoe on behalf of the Wellcome Sanger Institute COVID-19 Surveillance Team |
| EPI_ISL_3096244, EPI_ISL_3096245, EPI_ISL_3096246, EPI_ISL_3096247, EPI_ISL_3096248, EPI_ISL_3096249 | Lighthouse Lab in Milton Keynes | Wellcome Sanger Institute for the COVID-19 Genomics UK (COG-UK) Consortium | The Lighthouse Lab in Milton Keynes and Alex Alderton, Roberto Amato, Jeffrey Barrett, Sonia Goncalves, Ewan Harrison, David K. Jackson, Ian Johnston, Dominic Kwiatkowski, Cordelia Langford, John Sillitoe on behalf of the Wellcome Sanger Institute COVID-19 Surveillance Team |
| EPI_ISL_3096250 | Lighthouse Lab in Glasgow | Wellcome Sanger Institute for the COVID-19 Genomics UK (COG-UK) Consortium | Harper VanSteenhouse, Yumi Kasai, David Gray, Carol Clugston, Anna Dominiczak and Alex Alderton, Roberto Amato, Jeffrey Barrett, Sonia Goncalves, Ewan Harrison, David K. Jackson, Ian Johnston, Dominic Kwiatkowski, Cordelia Langford, John Sillitoe on behalf of the Wellcome Sanger Institute COVID-19 Surveillance Team |
| EPI_ISL_3096251 | Lighthouse Lab in Milton Keynes | Wellcome Sanger Institute for the COVID-19 Genomics UK (COG-UK) Consortium | The Lighthouse Lab in Milton Keynes and Alex Alderton, Roberto Amato, Jeffrey Barrett, Sonia Goncalves, Ewan Harrison, David K. Jackson, Ian Johnston, Dominic Kwiatkowski, Cordelia Langford, John Sillitoe on behalf of the Wellcome Sanger Institute COVID-19 Surveillance Team |
| EPI_ISL_3096252 | Berkshire and Surrey Pathology Services Lighthouse Laboratory | Wellcome Sanger Institute for the COVID-19 Genomics UK (COG-UK) Consortium | Berkshire and Surrey Pathology Services Lighthouse Laboratory and Alex Alderton, Roberto Amato, Jeffrey Barrett, Sonia Goncalves, Ewan Harrison, David K. Jackson, Ian Johnston, Dominic Kwiatkowski, Cordelia Langford, John Sillitoe on behalf of the Wellcome Sanger Institute COVID-19 Surveillance Team |
| EPI_ISL_3096253 | Lighthouse Lab in Glasgow | Wellcome Sanger Institute for the COVID-19 Genomics UK (COG-UK) Consortium | Harper VanSteenhouse, Yumi Kasai, David Gray, Carol Clugston, Anna Dominiczak and Alex Alderton, Roberto Amato, Jeffrey Barrett, Sonia Goncalves, Ewan Harrison, David K. Jackson, Ian Johnston, Dominic Kwiatkowski, Cordelia Langford, John Sillitoe on behalf of the Wellcome Sanger Institute COVID-19 Surveillance Team |
| EPI_ISL_3096254, EPI_ISL_3096255 | Lighthouse Lab in Milton Keynes | Wellcome Sanger Institute for the COVID-19 Genomics UK (COG-UK) Consortium | The Lighthouse Lab in Milton Keynes and Alex Alderton, Roberto Amato, Jeffrey Barrett, Sonia Goncalves, Ewan Harrison, David K. Jackson, Ian Johnston, Dominic Kwiatkowski, Cordelia Langford, John Sillitoe on behalf of the Wellcome Sanger Institute COVID-19 Surveillance Team |
| EPI_ISL_3096256 | Lighthouse Lab in Glasgow | Wellcome Sanger Institute for the COVID-19 Genomics UK (COG-UK) Consortium | Harper VanSteenhouse, Yumi Kasai, David Gray, Carol Clugston, Anna Dominiczak and Alex Alderton, Roberto Amato, Jeffrey Barrett, Sonia Goncalves, Ewan Harrison, David K. Jackson, Ian Johnston, Dominic Kwiatkowski, Cordelia Langford, John Sillitoe on behalf of the Wellcome Sanger Institute COVID-19 Surveillance Team |
| EPI_ISL_3096257, EPI_ISL_3096258, EPI_ISL_3096259, EPI_ISL_3096260, EPI_ISL_3096261 | Lighthouse Lab in Milton Keynes | Wellcome Sanger Institute for the COVID-19 Genomics UK (COG-UK) Consortium | The Lighthouse Lab in Milton Keynes and Alex Alderton, Roberto Amato, Jeffrey Barrett, Sonia Goncalves, Ewan Harrison, David K. Jackson, Ian Johnston, Dominic Kwiatkowski, Cordelia Langford, John Sillitoe on behalf of the Wellcome Sanger Institute COVID-19 Surveillance Team |
| EPI_ISL_3096262 | Lighthouse Lab in Glasgow | Wellcome Sanger Institute for the COVID-19 Genomics UK (COG-UK) Consortium | Harper VanSteenhouse, Yumi Kasai, David Gray, Carol Clugston, Anna Dominiczak and Alex Alderton, Roberto Amato, Jeffrey Barrett, Sonia Goncalves, Ewan Harrison, David K. Jackson, Ian Johnston, Dominic Kwiatkowski, Cordelia Langford, John Sillitoe on behalf of the Wellcome Sanger Institute COVID-19 Surveillance Team |
| EPI_ISL_3096263 | Lighthouse Lab in Milton Keynes | Wellcome Sanger Institute for the COVID-19 Genomics UK (COG-UK) Consortium | The Lighthouse Lab in Milton Keynes and Alex Alderton, Roberto Amato, Jeffrey Barrett, Sonia Goncalves, Ewan Harrison, David K. Jackson, Ian Johnston, Dominic Kwiatkowski, Cordelia Langford, John Sillitoe on behalf of the Wellcome Sanger Institute COVID-19 Surveillance Team |
| EPI_ISL_3096264 | Berkshire and Surrey Pathology Services Lighthouse Laboratory | Wellcome Sanger Institute for the COVID-19 Genomics UK (COG-UK) Consortium | Berkshire and Surrey Pathology Services Lighthouse Laboratory and Alex Alderton, Roberto Amato, Jeffrey Barrett, Sonia Goncalves, Ewan Harrison, David K. Jackson, Ian Johnston, Dominic Kwiatkowski, Cordelia Langford, John Sillitoe on behalf of the Wellcome Sanger Institute COVID-19 Surveillance Team |
| EPI_ISL_3096265 | Lighthouse Lab in Milton Keynes | Wellcome Sanger Institute for the COVID-19 Genomics UK (COG-UK) Consortium | The Lighthouse Lab in Milton Keynes and Alex Alderton, Roberto Amato, Jeffrey Barrett, Sonia Goncalves, Ewan Harrison, David K. Jackson, Ian Johnston, Dominic Kwiatkowski, Cordelia Langford, John Sillitoe on behalf of the Wellcome Sanger Institute COVID-19 Surveillance Team |
| EPI_ISL_3096266 | Lighthouse Lab in Glasgow | Wellcome Sanger Institute for the COVID-19 Genomics UK (COG-UK) Consortium | Harper VanSteenhouse, Yumi Kasai, David Gray, Carol Clugston, Anna Dominiczak and Alex Alderton, Roberto Amato, Jeffrey Barrett, Sonia Goncalves, Ewan Harrison, David K. Jackson, Ian Johnston, Dominic Kwiatkowski, Cordelia Langford, John Sillitoe on behalf of the Wellcome Sanger Institute COVID-19 Surveillance Team |
| EPI_ISL_3096267, EPI_ISL_3096268 | Lighthouse Lab in Milton Keynes | Wellcome Sanger Institute for the COVID-19 Genomics UK (COG-UK) Consortium | The Lighthouse Lab in Milton Keynes and Alex Alderton, Roberto Amato, Jeffrey Barrett, Sonia Goncalves, Ewan Harrison, David K. Jackson, Ian Johnston, Dominic Kwiatkowski, Cordelia Langford, John Sillitoe on behalf of the Wellcome Sanger Institute COVID-19 Surveillance Team |
| EPI_ISL_3096269 | Berkshire and Surrey Pathology Services Lighthouse Laboratory | Wellcome Sanger Institute for the COVID-19 Genomics UK (COG-UK) Consortium | Berkshire and Surrey Pathology Services Lighthouse Laboratory and Alex Alderton, Roberto Amato, Jeffrey Barrett, Sonia Goncalves, Ewan Harrison, David K. Jackson, Ian Johnston, Dominic Kwiatkowski, Cordelia Langford, John Sillitoe on behalf of the Wellcome Sanger Institute COVID-19 Surveillance Team |
| EPI_ISL_3096270 | Lighthouse Lab in Milton Keynes | Wellcome Sanger Institute for the COVID-19 Genomics UK (COG-UK) Consortium | The Lighthouse Lab in Milton Keynes and Alex Alderton, Roberto Amato, Jeffrey Barrett, Sonia Goncalves, Ewan Harrison, David K. Jackson, Ian Johnston, Dominic Kwiatkowski, Cordelia Langford, John Sillitoe on behalf of the Wellcome Sanger Institute COVID-19 Surveillance Team |
| EPI_ISL_3096271, EPI_ISL_3096272 | Lighthouse Lab in Glasgow | Wellcome Sanger Institute for the COVID-19 Genomics UK (COG-UK) Consortium | Harper VanSteenhouse, Yumi Kasai, David Gray, Carol Clugston, Anna Dominiczak and Alex Alderton, Roberto Amato, Jeffrey Barrett, Sonia Goncalves, Ewan Harrison, David K. Jackson, Ian Johnston, Dominic Kwiatkowski, Cordelia Langford, John Sillitoe on behalf of the Wellcome Sanger Institute COVID-19 Surveillance Team |
| EPI_ISL_3096273 | Lighthouse Lab in Milton Keynes | Wellcome Sanger Institute for the COVID-19 Genomics UK (COG-UK) Consortium | The Lighthouse Lab in Milton Keynes and Alex Alderton, Roberto Amato, Jeffrey Barrett, Sonia Goncalves, Ewan Harrison, David K. Jackson, Ian Johnston, Dominic Kwiatkowski, Cordelia Langford, John Sillitoe on behalf of the Wellcome Sanger Institute COVID-19 Surveillance Team |
| EPI_ISL_3096274 | Berkshire and Surrey Pathology Services Lighthouse Laboratory | Wellcome Sanger Institute for the COVID-19 Genomics UK (COG-UK) Consortium | Berkshire and Surrey Pathology Services Lighthouse Laboratory and Alex Alderton, Roberto Amato, Jeffrey Barrett, Sonia Goncalves, Ewan Harrison, David K. Jackson, Ian Johnston, Dominic Kwiatkowski, Cordelia Langford, John Sillitoe on behalf of the Wellcome Sanger Institute COVID-19 Surveillance Team |
| EPI_ISL_3096275, EPI_ISL_3096276 | Lighthouse Lab in Milton Keynes | Wellcome Sanger Institute for the COVID-19 Genomics UK (COG-UK) Consortium | The Lighthouse Lab in Milton Keynes and Alex Alderton, Roberto Amato, Jeffrey Barrett, Sonia Goncalves, Ewan Harrison, David K. Jackson, Ian Johnston, Dominic Kwiatkowski, Cordelia Langford, John Sillitoe on behalf of the Wellcome Sanger Institute COVID-19 Surveillance Team |
| EPI_ISL_3096277 | Berkshire and Surrey Pathology Services Lighthouse Laboratory | Wellcome Sanger Institute for the COVID-19 Genomics UK (COG-UK) Consortium | Berkshire and Surrey Pathology Services Lighthouse Laboratory and Alex Alderton, Roberto Amato, Jeffrey Barrett, Sonia Goncalves, Ewan Harrison, David K. Jackson, Ian Johnston, Dominic Kwiatkowski, Cordelia Langford, John Sillitoe on behalf of the Wellcome Sanger Institute COVID-19 Surveillance Team |
| EPI_ISL_3096278, EPI_ISL_3096279, EPI_ISL_3096280 | Lighthouse Lab in Glasgow | Wellcome Sanger Institute for the COVID-19 Genomics UK (COG-UK) Consortium | Harper VanSteenhouse, Yumi Kasai, David Gray, Carol Clugston, Anna Dominiczak and Alex Alderton, Roberto Amato, Jeffrey Barrett, Sonia Goncalves, Ewan Harrison, David K. Jackson, Ian Johnston, Dominic Kwiatkowski, Cordelia Langford, John Sillitoe on behalf of the Wellcome Sanger Institute COVID-19 Surveillance Team |
| EPI_ISL_3096281 | Lighthouse Lab in Milton Keynes | Wellcome Sanger Institute for the COVID-19 Genomics UK (COG-UK) Consortium | The Lighthouse Lab in Milton Keynes and Alex Alderton, Roberto Amato, Jeffrey Barrett, Sonia Goncalves, Ewan Harrison, David K. Jackson, Ian Johnston, Dominic Kwiatkowski, Cordelia Langford, John Sillitoe on behalf of the Wellcome Sanger Institute COVID-19 Surveillance Team |
| EPI_ISL_3096282 | Berkshire and Surrey Pathology Services Lighthouse Laboratory | Wellcome Sanger Institute for the COVID-19 Genomics UK (COG-UK) Consortium | Berkshire and Surrey Pathology Services Lighthouse Laboratory and Alex Alderton, Roberto Amato, Jeffrey Barrett, Sonia Goncalves, Ewan Harrison, David K. Jackson, Ian Johnston, Dominic Kwiatkowski, Cordelia Langford, John Sillitoe on behalf of the Wellcome Sanger Institute COVID-19 Surveillance Team |
| EPI_ISL_3096283 | Lighthouse Lab in Glasgow | Wellcome Sanger Institute for the COVID-19 Genomics UK (COG-UK) Consortium | Harper VanSteenhouse, Yumi Kasai, David Gray, Carol Clugston, Anna Dominiczak and Alex Alderton, Roberto Amato, Jeffrey Barrett, Sonia Goncalves, Ewan Harrison, David K. Jackson, Ian Johnston, Dominic Kwiatkowski, Cordelia Langford, John Sillitoe on behalf of the Wellcome Sanger Institute COVID-19 Surveillance Team |
| EPI_ISL_3096284, EPI_ISL_3096285 | Berkshire and Surrey Pathology Services Lighthouse Laboratory | Wellcome Sanger Institute for the COVID-19 Genomics UK (COG-UK) Consortium | Berkshire and Surrey Pathology Services Lighthouse Laboratory and Alex Alderton, Roberto Amato, Jeffrey Barrett, Sonia Goncalves, Ewan Harrison, David K. Jackson, Ian Johnston, Dominic Kwiatkowski, Cordelia Langford, John Sillitoe on behalf of the Wellcome Sanger Institute COVID-19 Surveillance Team |
| EPI_ISL_3096286 | Lighthouse Lab in Glasgow | Wellcome Sanger Institute for the COVID-19 Genomics UK (COG-UK) Consortium | Harper VanSteenhouse, Yumi Kasai, David Gray, Carol Clugston, Anna Dominiczak and Alex Alderton, Roberto Amato, Jeffrey Barrett, Sonia Goncalves, Ewan Harrison, David K. Jackson, Ian Johnston, Dominic Kwiatkowski, Cordelia Langford, John Sillitoe on behalf of the Wellcome Sanger Institute COVID-19 Surveillance Team |
| EPI_ISL_3096287, EPI_ISL_3096288 | Lighthouse Lab in Milton Keynes | Wellcome Sanger Institute for the COVID-19 Genomics UK (COG-UK) Consortium | The Lighthouse Lab in Milton Keynes and Alex Alderton, Roberto Amato, Jeffrey Barrett, Sonia Goncalves, Ewan Harrison, David K. Jackson, Ian Johnston, Dominic Kwiatkowski, Cordelia Langford, John Sillitoe on behalf of the Wellcome Sanger Institute COVID-19 Surveillance Team |
| EPI_ISL_3096289, EPI_ISL_3096290 | Berkshire and Surrey Pathology Services Lighthouse Laboratory | Wellcome Sanger Institute for the COVID-19 Genomics UK (COG-UK) Consortium | Berkshire and Surrey Pathology Services Lighthouse Laboratory and Alex Alderton, Roberto Amato, Jeffrey Barrett, Sonia Goncalves, Ewan Harrison, David K. Jackson, Ian Johnston, Dominic Kwiatkowski, Cordelia Langford, John Sillitoe on behalf of the Wellcome Sanger Institute COVID-19 Surveillance Team |
| EPI_ISL_3096291 | Lighthouse Lab in Glasgow | Wellcome Sanger Institute for the COVID-19 Genomics UK (COG-UK) Consortium | Harper VanSteenhouse, Yumi Kasai, David Gray, Carol Clugston, Anna Dominiczak and Alex Alderton, Roberto Amato, Jeffrey Barrett, Sonia Goncalves, Ewan Harrison, David K. Jackson, Ian Johnston, Dominic Kwiatkowski, Cordelia Langford, John Sillitoe on behalf of the Wellcome Sanger Institute COVID-19 Surveillance Team |
| EPI_ISL_3096292 | Lighthouse Lab in Milton Keynes | Wellcome Sanger Institute for the COVID-19 Genomics UK (COG-UK) Consortium | The Lighthouse Lab in Milton Keynes and Alex Alderton, Roberto Amato, Jeffrey Barrett, Sonia Goncalves, Ewan Harrison, David K. Jackson, Ian Johnston, Dominic Kwiatkowski, Cordelia Langford, John Sillitoe on behalf of the Wellcome Sanger Institute COVID-19 Surveillance Team |
| EPI_ISL_3096293 | Berkshire and Surrey Pathology Services Lighthouse Laboratory | Wellcome Sanger Institute for the COVID-19 Genomics UK (COG-UK) Consortium | Berkshire and Surrey Pathology Services Lighthouse Laboratory and Alex Alderton, Roberto Amato, Jeffrey Barrett, Sonia Goncalves, Ewan Harrison, David K. Jackson, Ian Johnston, Dominic Kwiatkowski, Cordelia Langford, John Sillitoe on behalf of the Wellcome Sanger Institute COVID-19 Surveillance Team |
| EPI_ISL_3096294, EPI_ISL_3096295 | Lighthouse Lab in Milton Keynes | Wellcome Sanger Institute for the COVID-19 Genomics UK (COG-UK) Consortium | The Lighthouse Lab in Milton Keynes and Alex Alderton, Roberto Amato, Jeffrey Barrett, Sonia Goncalves, Ewan Harrison, David K. Jackson, Ian Johnston, Dominic Kwiatkowski, Cordelia Langford, John Sillitoe on behalf of the Wellcome Sanger Institute COVID-19 Surveillance Team |
| EPI_ISL_3096296 | Berkshire and Surrey Pathology Services Lighthouse Laboratory | Wellcome Sanger Institute for the COVID-19 Genomics UK (COG-UK) Consortium | Berkshire and Surrey Pathology Services Lighthouse Laboratory and Alex Alderton, Roberto Amato, Jeffrey Barrett, Sonia Goncalves, Ewan Harrison, David K. Jackson, Ian Johnston, Dominic Kwiatkowski, Cordelia Langford, John Sillitoe on behalf of the Wellcome Sanger Institute COVID-19 Surveillance Team |
| EPI_ISL_3096297 | Lighthouse Lab in Glasgow | Wellcome Sanger Institute for the COVID-19 Genomics UK (COG-UK) Consortium | Harper VanSteenhouse, Yumi Kasai, David Gray, Carol Clugston, Anna Dominiczak and Alex Alderton, Roberto Amato, Jeffrey Barrett, Sonia Goncalves, Ewan Harrison, David K. Jackson, Ian Johnston, Dominic Kwiatkowski, Cordelia Langford, John Sillitoe on behalf of the Wellcome Sanger Institute COVID-19 Surveillance Team |
| EPI_ISL_3096298 | Berkshire and Surrey Pathology Services Lighthouse Laboratory | Wellcome Sanger Institute for the COVID-19 Genomics UK (COG-UK) Consortium | Berkshire and Surrey Pathology Services Lighthouse Laboratory and Alex Alderton, Roberto Amato, Jeffrey Barrett, Sonia Goncalves, Ewan Harrison, David K. Jackson, Ian Johnston, Dominic Kwiatkowski, Cordelia Langford, John Sillitoe on behalf of the Wellcome Sanger Institute COVID-19 Surveillance Team |
| EPI_ISL_3096299, | Lighthouse Lab in Glasgow | Wellcome Sanger Institute for the COVID-19 Genomics UK (COG-UK) | Harper VanSteenhouse, Yumi Kasai, David Gray, Carol Clugston, Anna Dominiczak and Alex Alderton, Roberto Amato, Jeffrey Barrett, Sonia Goncalves, Ewan Harrison, David K. Jackson, Ian Johnston, Dominic Kwiatkowski, Cordelia Langford, |

| | | | |
|---|---|---|---|
| EPI_ISL_3096300 | | Consortium | John Sillitoe on behalf of the Wellcome Sanger Institute COVID-19 Surveillance Team |
| EPI_ISL_3096301, EPI_ISL_3096302, EPI_ISL_3096303 | Lighthouse Lab in Milton Keynes | Wellcome Sanger Institute for the COVID-19 Genomics UK (COG-UK) Consortium | The Lighthouse Lab in Milton Keynes and Alex Alderton, Roberto Amato, Jeffrey Barrett, Sonia Goncalves, Ewan Harrison, David K. Jackson, Ian Johnston, Dominic Kwiatkowski, Cordelia Langford, John Sillitoe on behalf of the Wellcome Sanger Institute COVID-19 Surveillance Team |
| EPI_ISL_3096304, EPI_ISL_3096305, EPI_ISL_3096306 | Lighthouse Lab in Glasgow | Wellcome Sanger Institute for the COVID-19 Genomics UK (COG-UK) Consortium | Harper VanSteenhouse, Yumi Kasai, David Gray, Carol Clugston, Anna Dominiczak and Alex Alderton, Roberto Amato, Jeffrey Barrett, Sonia Goncalves, Ewan Harrison, David K. Jackson, Ian Johnston, Dominic Kwiatkowski, Cordelia Langford, John Sillitoe on behalf of the Wellcome Sanger Institute COVID-19 Surveillance Team |
| EPI_ISL_3096307 | Lighthouse Lab in Milton Keynes | Wellcome Sanger Institute for the COVID-19 Genomics UK (COG-UK) Consortium | The Lighthouse Lab in Milton Keynes and Alex Alderton, Roberto Amato, Jeffrey Barrett, Sonia Goncalves, Ewan Harrison, David K. Jackson, Ian Johnston, Dominic Kwiatkowski, Cordelia Langford, John Sillitoe on behalf of the Wellcome Sanger Institute COVID-19 Surveillance Team |
| EPI_ISL_3096308 | Lighthouse Lab in Glasgow | Wellcome Sanger Institute for the COVID-19 Genomics UK (COG-UK) Consortium | Harper VanSteenhouse, Yumi Kasai, David Gray, Carol Clugston, Anna Dominiczak and Alex Alderton, Roberto Amato, Jeffrey Barrett, Sonia Goncalves, Ewan Harrison, David K. Jackson, Ian Johnston, Dominic Kwiatkowski, Cordelia Langford, John Sillitoe on behalf of the Wellcome Sanger Institute COVID-19 Surveillance Team |
| EPI_ISL_3096309, EPI_ISL_3096310 | Lighthouse Lab in Milton Keynes | Wellcome Sanger Institute for the COVID-19 Genomics UK (COG-UK) Consortium | The Lighthouse Lab in Milton Keynes and Alex Alderton, Roberto Amato, Jeffrey Barrett, Sonia Goncalves, Ewan Harrison, David K. Jackson, Ian Johnston, Dominic Kwiatkowski, Cordelia Langford, John Sillitoe on behalf of the Wellcome Sanger Institute COVID-19 Surveillance Team |
| EPI_ISL_3096311 | Lighthouse Lab in Glasgow | Wellcome Sanger Institute for the COVID-19 Genomics UK (COG-UK) Consortium | Harper VanSteenhouse, Yumi Kasai, David Gray, Carol Clugston, Anna Dominiczak and Alex Alderton, Roberto Amato, Jeffrey Barrett, Sonia Goncalves, Ewan Harrison, David K. Jackson, Ian Johnston, Dominic Kwiatkowski, Cordelia Langford, John Sillitoe on behalf of the Wellcome Sanger Institute COVID-19 Surveillance Team |
| EPI_ISL_3096312, EPI_ISL_3096313 | Lighthouse Lab in Milton Keynes | Wellcome Sanger Institute for the COVID-19 Genomics UK (COG-UK) Consortium | The Lighthouse Lab in Milton Keynes and Alex Alderton, Roberto Amato, Jeffrey Barrett, Sonia Goncalves, Ewan Harrison, David K. Jackson, Ian Johnston, Dominic Kwiatkowski, Cordelia Langford, John Sillitoe on behalf of the Wellcome Sanger Institute COVID-19 Surveillance Team |
| EPI_ISL_3096314, EPI_ISL_3096315 | Berkshire and Surrey Pathology Services Lighthouse Laboratory | Wellcome Sanger Institute for the COVID-19 Genomics UK (COG-UK) Consortium | Berkshire and Surrey Pathology Services Lighthouse Laboratory and Alex Alderton, Roberto Amato, Jeffrey Barrett, Sonia Goncalves, Ewan Harrison, David K. Jackson, Ian Johnston, Dominic Kwiatkowski, Cordelia Langford, John Sillitoe on behalf of the Wellcome Sanger Institute COVID-19 Surveillance Team |
| EPI_ISL_3096316, EPI_ISL_3096317 | Lighthouse Lab in Milton Keynes | Wellcome Sanger Institute for the COVID-19 Genomics UK (COG-UK) Consortium | The Lighthouse Lab in Milton Keynes and Alex Alderton, Roberto Amato, Jeffrey Barrett, Sonia Goncalves, Ewan Harrison, David K. Jackson, Ian Johnston, Dominic Kwiatkowski, Cordelia Langford, John Sillitoe on behalf of the Wellcome Sanger Institute COVID-19 Surveillance Team |
| EPI_ISL_3096318 | Lighthouse Lab in Alderley Park | Wellcome Sanger Institute for the COVID-19 Genomics UK (COG-UK) Consortium | Jacquelyn Wynn, Mairead Hyland, The Lighthouse Lab in Alderley Park and Alex Alderton, Roberto Amato, Jeffrey Barrett, Sonia Goncalves, Ewan Harrison, David K. Jackson, Ian Johnston, Dominic Kwiatkowski, Cordelia Langford, John Sillitoe on behalf of the Wellcome Sanger Institute COVID-19 Surveillance Team |
| EPI_ISL_3096319, EPI_ISL_3096320 | Berkshire and Surrey Pathology Services Lighthouse Laboratory | Wellcome Sanger Institute for the COVID-19 Genomics UK (COG-UK) Consortium | Berkshire and Surrey Pathology Services Lighthouse Laboratory and Alex Alderton, Roberto Amato, Jeffrey Barrett, Sonia Goncalves, Ewan Harrison, David K. Jackson, Ian Johnston, Dominic Kwiatkowski, Cordelia Langford, John Sillitoe on behalf of the Wellcome Sanger Institute COVID-19 Surveillance Team |
| EPI_ISL_3096321, EPI_ISL_3096322 | Lighthouse Lab in Alderley Park | Wellcome Sanger Institute for the COVID-19 Genomics UK (COG-UK) Consortium | Jacquelyn Wynn, Mairead Hyland, The Lighthouse Lab in Alderley Park and Alex Alderton, Roberto Amato, Jeffrey Barrett, Sonia Goncalves, Ewan Harrison, David K. Jackson, Ian Johnston, Dominic Kwiatkowski, Cordelia Langford, John Sillitoe on behalf of the Wellcome Sanger Institute COVID-19 Surveillance Team |
| EPI_ISL_3096323, EPI_ISL_3096324, EPI_ISL_3096325, EPI_ISL_3096326 | Berkshire and Surrey Pathology Services Lighthouse Laboratory | Wellcome Sanger Institute for the COVID-19 Genomics UK (COG-UK) Consortium | Berkshire and Surrey Pathology Services Lighthouse Laboratory and Alex Alderton, Roberto Amato, Jeffrey Barrett, Sonia Goncalves, Ewan Harrison, David K. Jackson, Ian Johnston, Dominic Kwiatkowski, Cordelia Langford, John Sillitoe on behalf of the Wellcome Sanger Institute COVID-19 Surveillance Team |
| EPI_ISL_3096327 | Lighthouse Lab in Alderley Park | Wellcome Sanger Institute for the COVID-19 Genomics UK (COG-UK) Consortium | Jacquelyn Wynn, Mairead Hyland, The Lighthouse Lab in Alderley Park and Alex Alderton, Roberto Amato, Jeffrey Barrett, Sonia Goncalves, Ewan Harrison, David K. Jackson, Ian Johnston, Dominic Kwiatkowski, Cordelia Langford, John Sillitoe on behalf of the Wellcome Sanger Institute COVID-19 Surveillance Team |
| EPI_ISL_3096328, EPI_ISL_3096329 | Berkshire and Surrey Pathology Services Lighthouse Laboratory | Wellcome Sanger Institute for the COVID-19 Genomics UK (COG-UK) Consortium | Berkshire and Surrey Pathology Services Lighthouse Laboratory and Alex Alderton, Roberto Amato, Jeffrey Barrett, Sonia Goncalves, Ewan Harrison, David K. Jackson, Ian Johnston, Dominic Kwiatkowski, Cordelia Langford, John Sillitoe on behalf of the Wellcome Sanger Institute COVID-19 Surveillance Team |
| EPI_ISL_3096330, EPI_ISL_3096331, EPI_ISL_3096332 | Lighthouse Lab in Alderley Park | Wellcome Sanger Institute for the COVID-19 Genomics UK (COG-UK) Consortium | Jacquelyn Wynn, Mairead Hyland, The Lighthouse Lab in Alderley Park and Alex Alderton, Roberto Amato, Jeffrey Barrett, Sonia Goncalves, Ewan Harrison, David K. Jackson, Ian Johnston, Dominic Kwiatkowski, Cordelia Langford, John Sillitoe on behalf of the Wellcome Sanger Institute COVID-19 Surveillance Team |
| EPI_ISL_3096333 | Berkshire and Surrey Pathology Services Lighthouse Laboratory | Wellcome Sanger Institute for the COVID-19 Genomics UK (COG-UK) Consortium | Berkshire and Surrey Pathology Services Lighthouse Laboratory and Alex Alderton, Roberto Amato, Jeffrey Barrett, Sonia Goncalves, Ewan Harrison, David K. Jackson, Ian Johnston, Dominic Kwiatkowski, Cordelia Langford, John Sillitoe on behalf of the Wellcome Sanger Institute COVID-19 Surveillance Team |
| EPI_ISL_3096334, EPI_ISL_3096335, EPI_ISL_3096336, EPI_ISL_3096337 | Lighthouse Lab in Alderley Park | Wellcome Sanger Institute for the COVID-19 Genomics UK (COG-UK) Consortium | Jacquelyn Wynn, Mairead Hyland, The Lighthouse Lab in Alderley Park and Alex Alderton, Roberto Amato, Jeffrey Barrett, Sonia Goncalves, Ewan Harrison, David K. Jackson, Ian Johnston, Dominic Kwiatkowski, Cordelia Langford, John Sillitoe on behalf of the Wellcome Sanger Institute COVID-19 Surveillance Team |
| EPI_ISL_3096338 | Berkshire and Surrey Pathology Services Lighthouse Laboratory | Wellcome Sanger Institute for the COVID-19 Genomics UK (COG-UK) Consortium | Berkshire and Surrey Pathology Services Lighthouse Laboratory and Alex Alderton, Roberto Amato, Jeffrey Barrett, Sonia Goncalves, Ewan Harrison, David K. Jackson, Ian Johnston, Dominic Kwiatkowski, Cordelia Langford, John Sillitoe on behalf of the Wellcome Sanger Institute COVID-19 Surveillance Team |
| EPI_ISL_3096339, EPI_ISL_3096340, EPI_ISL_3096341, EPI_ISL_3096342, EPI_ISL_3096343 | Lighthouse Lab in Alderley Park | Wellcome Sanger Institute for the COVID-19 Genomics UK (COG-UK) Consortium | Jacquelyn Wynn, Mairead Hyland, The Lighthouse Lab in Alderley Park and Alex Alderton, Roberto Amato, Jeffrey Barrett, Sonia Goncalves, Ewan Harrison, David K. Jackson, Ian Johnston, Dominic Kwiatkowski, Cordelia Langford, John Sillitoe on behalf of the Wellcome Sanger Institute COVID-19 Surveillance Team |
| EPI_ISL_3096344, EPI_ISL_3096345 | Berkshire and Surrey Pathology Services Lighthouse Laboratory | Wellcome Sanger Institute for the COVID-19 Genomics UK (COG-UK) Consortium | Berkshire and Surrey Pathology Services Lighthouse Laboratory and Alex Alderton, Roberto Amato, Jeffrey Barrett, Sonia Goncalves, Ewan Harrison, David K. Jackson, Ian Johnston, Dominic Kwiatkowski, Cordelia Langford, John Sillitoe on behalf of the Wellcome Sanger Institute COVID-19 Surveillance Team |
| EPI_ISL_3096346, EPI_ISL_3096347, EPI_ISL_3096348, EPI_ISL_3096349, EPI_ISL_3096350 | Lighthouse Lab in Alderley Park | Wellcome Sanger Institute for the COVID-19 Genomics UK (COG-UK) Consortium | Jacquelyn Wynn, Mairead Hyland, The Lighthouse Lab in Alderley Park and Alex Alderton, Roberto Amato, Jeffrey Barrett, Sonia Goncalves, Ewan Harrison, David K. Jackson, Ian Johnston, Dominic Kwiatkowski, Cordelia Langford, John Sillitoe on behalf of the Wellcome Sanger Institute COVID-19 Surveillance Team |
| EPI_ISL_3096351 | Berkshire and Surrey Pathology Services Lighthouse Laboratory | Wellcome Sanger Institute for the COVID-19 Genomics UK (COG-UK) Consortium | Berkshire and Surrey Pathology Services Lighthouse Laboratory and Alex Alderton, Roberto Amato, Jeffrey Barrett, Sonia Goncalves, Ewan Harrison, David K. Jackson, Ian Johnston, Dominic Kwiatkowski, Cordelia Langford, John Sillitoe on behalf of the Wellcome Sanger Institute COVID-19 Surveillance Team |
| EPI_ISL_3096352 | Lighthouse Lab in Alderley Park | Wellcome Sanger Institute for the COVID-19 Genomics UK (COG-UK) Consortium | Jacquelyn Wynn, Mairead Hyland, The Lighthouse Lab in Alderley Park and Alex Alderton, Roberto Amato, Jeffrey Barrett, Sonia Goncalves, Ewan Harrison, David K. Jackson, Ian Johnston, Dominic Kwiatkowski, Cordelia Langford, John Sillitoe on behalf of the Wellcome Sanger Institute COVID-19 Surveillance Team |
| EPI_ISL_3096353 | Berkshire and Surrey Pathology Services Lighthouse Laboratory | Wellcome Sanger Institute for the COVID-19 Genomics UK (COG-UK) Consortium | Berkshire and Surrey Pathology Services Lighthouse Laboratory and Alex Alderton, Roberto Amato, Jeffrey Barrett, Sonia Goncalves, Ewan Harrison, David K. Jackson, Ian Johnston, Dominic Kwiatkowski, Cordelia Langford, John Sillitoe on behalf of the Wellcome Sanger Institute COVID-19 Surveillance Team |
| EPI_ISL_3096354 | Lighthouse Lab in Alderley Park | Wellcome Sanger Institute for the COVID-19 Genomics UK (COG-UK) Consortium | Jacquelyn Wynn, Mairead Hyland, The Lighthouse Lab in Alderley Park and Alex Alderton, Roberto Amato, Jeffrey Barrett, Sonia Goncalves, Ewan Harrison, David K. Jackson, Ian Johnston, Dominic Kwiatkowski, Cordelia Langford, John Sillitoe on behalf of the Wellcome Sanger Institute COVID-19 Surveillance Team |
| EPI_ISL_3096355, EPI_ISL_3096356, EPI_ISL_3096357, EPI_ISL_3096358, EPI_ISL_3096359, EPI_ISL_3096360, EPI_ISL_3096361, EPI_ISL_3096362 | Berkshire and Surrey Pathology Services Lighthouse Laboratory | Wellcome Sanger Institute for the COVID-19 Genomics UK (COG-UK) Consortium | Berkshire and Surrey Pathology Services Lighthouse Laboratory and Alex Alderton, Roberto Amato, Jeffrey Barrett, Sonia Goncalves, Ewan Harrison, David K. Jackson, Ian Johnston, Dominic Kwiatkowski, Cordelia Langford, John Sillitoe on behalf of the Wellcome Sanger Institute COVID-19 Surveillance Team |
| EPI_ISL_3096363 | Lighthouse Lab in Alderley Park | Wellcome Sanger Institute for the COVID-19 Genomics UK (COG-UK) Consortium | Jacquelyn Wynn, Mairead Hyland, The Lighthouse Lab in Alderley Park and Alex Alderton, Roberto Amato, Jeffrey Barrett, Sonia Goncalves, Ewan Harrison, David K. Jackson, Ian Johnston, Dominic Kwiatkowski, Cordelia Langford, John Sillitoe on behalf of the Wellcome Sanger Institute COVID-19 Surveillance Team |
| EPI_ISL_3096364, EPI_ISL_3096365 | Berkshire and Surrey Pathology Services Lighthouse Laboratory | Wellcome Sanger Institute for the COVID-19 Genomics UK (COG-UK) Consortium | Berkshire and Surrey Pathology Services Lighthouse Laboratory and Alex Alderton, Roberto Amato, Jeffrey Barrett, Sonia Goncalves, Ewan Harrison, David K. Jackson, Ian Johnston, Dominic Kwiatkowski, Cordelia Langford, John Sillitoe on behalf of the Wellcome Sanger Institute COVID-19 Surveillance Team |
| EPI_ISL_3096366 | Lighthouse Lab in Alderley Park | Wellcome Sanger Institute for the COVID-19 Genomics UK (COG-UK) Consortium | Jacquelyn Wynn, Mairead Hyland, The Lighthouse Lab in Alderley Park and Alex Alderton, Roberto Amato, Jeffrey Barrett, Sonia Goncalves, Ewan Harrison, David K. Jackson, Ian Johnston, Dominic Kwiatkowski, Cordelia Langford, John Sillitoe on behalf of the Wellcome Sanger Institute COVID-19 Surveillance Team |
| EPI_ISL_3096367 | Berkshire and Surrey Pathology Services Lighthouse Laboratory | Wellcome Sanger Institute for the COVID-19 Genomics UK (COG-UK) Consortium | Berkshire and Surrey Pathology Services Lighthouse Laboratory and Alex Alderton, Roberto Amato, Jeffrey Barrett, Sonia Goncalves, Ewan Harrison, David K. Jackson, Ian Johnston, Dominic Kwiatkowski, Cordelia Langford, John Sillitoe on behalf of the Wellcome Sanger Institute COVID-19 Surveillance Team |
| EPI_ISL_3096368 | Lighthouse Lab in Alderley Park | Wellcome Sanger Institute for the COVID-19 Genomics UK (COG-UK) Consortium | Jacquelyn Wynn, Mairead Hyland, The Lighthouse Lab in Alderley Park and Alex Alderton, Roberto Amato, Jeffrey Barrett, Sonia Goncalves, Ewan Harrison, David K. Jackson, Ian Johnston, Dominic Kwiatkowski, Cordelia Langford, John Sillitoe on behalf of the Wellcome Sanger Institute COVID-19 Surveillance Team |
| EPI_ISL_3096369, EPI_ISL_3096370 | Berkshire and Surrey Pathology Services Lighthouse Laboratory | Wellcome Sanger Institute for the COVID-19 Genomics UK (COG-UK) Consortium | Berkshire and Surrey Pathology Services Lighthouse Laboratory and Alex Alderton, Roberto Amato, Jeffrey Barrett, Sonia Goncalves, Ewan Harrison, David K. Jackson, Ian Johnston, Dominic Kwiatkowski, Cordelia Langford, John Sillitoe on behalf of the Wellcome Sanger Institute COVID-19 Surveillance Team |
| EPI_ISL_3096371, EPI_ISL_3096372 | Lighthouse Lab in Alderley Park | Wellcome Sanger Institute for the COVID-19 Genomics UK (COG-UK) Consortium | Jacquelyn Wynn, Mairead Hyland, The Lighthouse Lab in Alderley Park and Alex Alderton, Roberto Amato, Jeffrey Barrett, Sonia Goncalves, Ewan Harrison, David K. Jackson, Ian Johnston, Dominic Kwiatkowski, Cordelia Langford, John Sillitoe on behalf of the Wellcome Sanger Institute COVID-19 Surveillance Team |
| EPI_ISL_3096373 | Berkshire and Surrey Pathology Services Lighthouse Laboratory | Wellcome Sanger Institute for the COVID-19 Genomics UK (COG-UK) Consortium | Berkshire and Surrey Pathology Services Lighthouse Laboratory and Alex Alderton, Roberto Amato, Jeffrey Barrett, Sonia Goncalves, Ewan Harrison, David K. Jackson, Ian Johnston, Dominic Kwiatkowski, Cordelia Langford, John Sillitoe on behalf of the Wellcome Sanger Institute COVID-19 Surveillance Team |
| EPI_ISL_3096374 | Lighthouse Lab in Alderley Park | Wellcome Sanger Institute for the COVID-19 Genomics UK (COG-UK) Consortium | Jacquelyn Wynn, Mairead Hyland, The Lighthouse Lab in Alderley Park and Alex Alderton, Roberto Amato, Jeffrey Barrett, Sonia Goncalves, Ewan Harrison, David K. Jackson, Ian Johnston, Dominic Kwiatkowski, Cordelia Langford, John Sillitoe on behalf of the Wellcome Sanger Institute COVID-19 Surveillance Team |
| EPI_ISL_3096375, | Berkshire and Surrey Pathology Services Lighthouse Laboratory | Wellcome Sanger Institute for the COVID-19 Genomics UK (COG-UK) | Berkshire and Surrey Pathology Services Lighthouse Laboratory and Alex Alderton, Roberto Amato, Jeffrey Barrett, Sonia Goncalves, Ewan Harrison, David K. Jackson, Ian Johnston, Dominic Kwiatkowski, Cordelia Langford, John Sillitoe on behalf |

| | | | |
|---|---|---|---|
| EPI_ISL_3096376 | | Consortium | of the Wellcome Sanger Institute COVID-19 Surveillance Team |
| EPI_ISL_3096377, EPI_ISL_3096378 | Lighthouse Lab in Alderley Park | Wellcome Sanger Institute for the COVID-19 Genomics UK (COG-UK) Consortium | Jacquelyn Wynn, Mairead Hyland, The Lighthouse Lab in Alderley Park and Alex Alderton, Roberto Amato, Jeffrey Barrett, Sonia Goncalves, Ewan Harrison, David K. Jackson, Ian Johnston, Dominic Kwiatkowski, Cordelia Langford, John Sillitoe on behalf of the Wellcome Sanger Institute COVID-19 Surveillance Team |
| EPI_ISL_3096379 | Berkshire and Surrey Pathology Services Lighthouse Laboratory | Wellcome Sanger Institute for the COVID-19 Genomics UK (COG-UK) Consortium | Berkshire and Surrey Pathology Services Lighthouse Laboratory and Alex Alderton, Roberto Amato, Jeffrey Barrett, Sonia Goncalves, Ewan Harrison, David K. Jackson, Ian Johnston, Dominic Kwiatkowski, Cordelia Langford, John Sillitoe on behalf of the Wellcome Sanger Institute COVID-19 Surveillance Team |
| EPI_ISL_3096380, EPI_ISL_3096381 | Lighthouse Lab in Alderley Park | Wellcome Sanger Institute for the COVID-19 Genomics UK (COG-UK) Consortium | Jacquelyn Wynn, Mairead Hyland, The Lighthouse Lab in Alderley Park and Alex Alderton, Roberto Amato, Jeffrey Barrett, Sonia Goncalves, Ewan Harrison, David K. Jackson, Ian Johnston, Dominic Kwiatkowski, Cordelia Langford, John Sillitoe on behalf of the Wellcome Sanger Institute COVID-19 Surveillance Team |
| EPI_ISL_3096382 | Berkshire and Surrey Pathology Services Lighthouse Laboratory | Wellcome Sanger Institute for the COVID-19 Genomics UK (COG-UK) Consortium | Berkshire and Surrey Pathology Services Lighthouse Laboratory and Alex Alderton, Roberto Amato, Jeffrey Barrett, Sonia Goncalves, Ewan Harrison, David K. Jackson, Ian Johnston, Dominic Kwiatkowski, Cordelia Langford, John Sillitoe on behalf of the Wellcome Sanger Institute COVID-19 Surveillance Team |
| EPI_ISL_3096383 | Lighthouse Lab in Alderley Park | Wellcome Sanger Institute for the COVID-19 Genomics UK (COG-UK) Consortium | Jacquelyn Wynn, Mairead Hyland, The Lighthouse Lab in Alderley Park and Alex Alderton, Roberto Amato, Jeffrey Barrett, Sonia Goncalves, Ewan Harrison, David K. Jackson, Ian Johnston, Dominic Kwiatkowski, Cordelia Langford, John Sillitoe on behalf of the Wellcome Sanger Institute COVID-19 Surveillance Team |
| EPI_ISL_3096384 | Berkshire and Surrey Pathology Services Lighthouse Laboratory | Wellcome Sanger Institute for the COVID-19 Genomics UK (COG-UK) Consortium | Berkshire and Surrey Pathology Services Lighthouse Laboratory and Alex Alderton, Roberto Amato, Jeffrey Barrett, Sonia Goncalves, Ewan Harrison, David K. Jackson, Ian Johnston, Dominic Kwiatkowski, Cordelia Langford, John Sillitoe on behalf of the Wellcome Sanger Institute COVID-19 Surveillance Team |
| EPI_ISL_3096385, EPI_ISL_3096386, EPI_ISL_3096387, EPI_ISL_3096388 | Lighthouse Lab in Alderley Park | Wellcome Sanger Institute for the COVID-19 Genomics UK (COG-UK) Consortium | Jacquelyn Wynn, Mairead Hyland, The Lighthouse Lab in Alderley Park and Alex Alderton, Roberto Amato, Jeffrey Barrett, Sonia Goncalves, Ewan Harrison, David K. Jackson, Ian Johnston, Dominic Kwiatkowski, Cordelia Langford, John Sillitoe on behalf of the Wellcome Sanger Institute COVID-19 Surveillance Team |
| EPI_ISL_3096389, EPI_ISL_3096390 | Berkshire and Surrey Pathology Services Lighthouse Laboratory | Wellcome Sanger Institute for the COVID-19 Genomics UK (COG-UK) Consortium | Berkshire and Surrey Pathology Services Lighthouse Laboratory and Alex Alderton, Roberto Amato, Jeffrey Barrett, Sonia Goncalves, Ewan Harrison, David K. Jackson, Ian Johnston, Dominic Kwiatkowski, Cordelia Langford, John Sillitoe on behalf of the Wellcome Sanger Institute COVID-19 Surveillance Team |
| EPI_ISL_3096391 | Lighthouse Lab in Alderley Park | Wellcome Sanger Institute for the COVID-19 Genomics UK (COG-UK) Consortium | Jacquelyn Wynn, Mairead Hyland, The Lighthouse Lab in Alderley Park and Alex Alderton, Roberto Amato, Jeffrey Barrett, Sonia Goncalves, Ewan Harrison, David K. Jackson, Ian Johnston, Dominic Kwiatkowski, Cordelia Langford, John Sillitoe on behalf of the Wellcome Sanger Institute COVID-19 Surveillance Team |
| EPI_ISL_3096392, EPI_ISL_3096393, EPI_ISL_3096394, EPI_ISL_3096395 | Berkshire and Surrey Pathology Services Lighthouse Laboratory | Wellcome Sanger Institute for the COVID-19 Genomics UK (COG-UK) Consortium | Berkshire and Surrey Pathology Services Lighthouse Laboratory and Alex Alderton, Roberto Amato, Jeffrey Barrett, Sonia Goncalves, Ewan Harrison, David K. Jackson, Ian Johnston, Dominic Kwiatkowski, Cordelia Langford, John Sillitoe on behalf of the Wellcome Sanger Institute COVID-19 Surveillance Team |
| EPI_ISL_3096396 | Lighthouse Lab in Alderley Park | Wellcome Sanger Institute for the COVID-19 Genomics UK (COG-UK) Consortium | Jacquelyn Wynn, Mairead Hyland, The Lighthouse Lab in Alderley Park and Alex Alderton, Roberto Amato, Jeffrey Barrett, Sonia Goncalves, Ewan Harrison, David K. Jackson, Ian Johnston, Dominic Kwiatkowski, Cordelia Langford, John Sillitoe on behalf of the Wellcome Sanger Institute COVID-19 Surveillance Team |
| EPI_ISL_3096397 | Berkshire and Surrey Pathology Services Lighthouse Laboratory | Wellcome Sanger Institute for the COVID-19 Genomics UK (COG-UK) Consortium | Berkshire and Surrey Pathology Services Lighthouse Laboratory and Alex Alderton, Roberto Amato, Jeffrey Barrett, Sonia Goncalves, Ewan Harrison, David K. Jackson, Ian Johnston, Dominic Kwiatkowski, Cordelia Langford, John Sillitoe on behalf of the Wellcome Sanger Institute COVID-19 Surveillance Team |
| EPI_ISL_3096398, EPI_ISL_3096399 | Lighthouse Lab in Alderley Park | Wellcome Sanger Institute for the COVID-19 Genomics UK (COG-UK) Consortium | Jacquelyn Wynn, Mairead Hyland, The Lighthouse Lab in Alderley Park and Alex Alderton, Roberto Amato, Jeffrey Barrett, Sonia Goncalves, Ewan Harrison, David K. Jackson, Ian Johnston, Dominic Kwiatkowski, Cordelia Langford, John Sillitoe on behalf of the Wellcome Sanger Institute COVID-19 Surveillance Team |
| EPI_ISL_3096400, EPI_ISL_3096401, EPI_ISL_3096402 | Berkshire and Surrey Pathology Services Lighthouse Laboratory | Wellcome Sanger Institute for the COVID-19 Genomics UK (COG-UK) Consortium | Berkshire and Surrey Pathology Services Lighthouse Laboratory and Alex Alderton, Roberto Amato, Jeffrey Barrett, Sonia Goncalves, Ewan Harrison, David K. Jackson, Ian Johnston, Dominic Kwiatkowski, Cordelia Langford, John Sillitoe on behalf of the Wellcome Sanger Institute COVID-19 Surveillance Team |
| EPI_ISL_3096403, EPI_ISL_3096404 | Lighthouse Lab in Alderley Park | Wellcome Sanger Institute for the COVID-19 Genomics UK (COG-UK) Consortium | Jacquelyn Wynn, Mairead Hyland, The Lighthouse Lab in Alderley Park and Alex Alderton, Roberto Amato, Jeffrey Barrett, Sonia Goncalves, Ewan Harrison, David K. Jackson, Ian Johnston, Dominic Kwiatkowski, Cordelia Langford, John Sillitoe on behalf of the Wellcome Sanger Institute COVID-19 Surveillance Team |
| EPI_ISL_3096405 | Berkshire and Surrey Pathology Services Lighthouse Laboratory | Wellcome Sanger Institute for the COVID-19 Genomics UK (COG-UK) Consortium | Berkshire and Surrey Pathology Services Lighthouse Laboratory and Alex Alderton, Roberto Amato, Jeffrey Barrett, Sonia Goncalves, Ewan Harrison, David K. Jackson, Ian Johnston, Dominic Kwiatkowski, Cordelia Langford, John Sillitoe on behalf of the Wellcome Sanger Institute COVID-19 Surveillance Team |
| EPI_ISL_3096406 | Lighthouse Lab in Alderley Park | Wellcome Sanger Institute for the COVID-19 Genomics UK (COG-UK) Consortium | Jacquelyn Wynn, Mairead Hyland, The Lighthouse Lab in Alderley Park and Alex Alderton, Roberto Amato, Jeffrey Barrett, Sonia Goncalves, Ewan Harrison, David K. Jackson, Ian Johnston, Dominic Kwiatkowski, Cordelia Langford, John Sillitoe on behalf of the Wellcome Sanger Institute COVID-19 Surveillance Team |
| EPI_ISL_3096407, EPI_ISL_3096408, EPI_ISL_3096409, EPI_ISL_3096410, EPI_ISL_3096411, EPI_ISL_3096412 | Berkshire and Surrey Pathology Services Lighthouse Laboratory | Wellcome Sanger Institute for the COVID-19 Genomics UK (COG-UK) Consortium | Berkshire and Surrey Pathology Services Lighthouse Laboratory and Alex Alderton, Roberto Amato, Jeffrey Barrett, Sonia Goncalves, Ewan Harrison, David K. Jackson, Ian Johnston, Dominic Kwiatkowski, Cordelia Langford, John Sillitoe on behalf of the Wellcome Sanger Institute COVID-19 Surveillance Team |
| EPI_ISL_3096413 | Lighthouse Lab in Alderley Park | Wellcome Sanger Institute for the COVID-19 Genomics UK (COG-UK) Consortium | Jacquelyn Wynn, Mairead Hyland, The Lighthouse Lab in Alderley Park and Alex Alderton, Roberto Amato, Jeffrey Barrett, Sonia Goncalves, Ewan Harrison, David K. Jackson, Ian Johnston, Dominic Kwiatkowski, Cordelia Langford, John Sillitoe on behalf of the Wellcome Sanger Institute COVID-19 Surveillance Team |
| EPI_ISL_3096414, EPI_ISL_3096415, EPI_ISL_3096416 | Berkshire and Surrey Pathology Services Lighthouse Laboratory | Wellcome Sanger Institute for the COVID-19 Genomics UK (COG-UK) Consortium | Berkshire and Surrey Pathology Services Lighthouse Laboratory and Alex Alderton, Roberto Amato, Jeffrey Barrett, Sonia Goncalves, Ewan Harrison, David K. Jackson, Ian Johnston, Dominic Kwiatkowski, Cordelia Langford, John Sillitoe on behalf of the Wellcome Sanger Institute COVID-19 Surveillance Team |
| EPI_ISL_3096417 | Lighthouse Lab in Alderley Park | Wellcome Sanger Institute for the COVID-19 Genomics UK (COG-UK) Consortium | Jacquelyn Wynn, Mairead Hyland, The Lighthouse Lab in Alderley Park and Alex Alderton, Roberto Amato, Jeffrey Barrett, Sonia Goncalves, Ewan Harrison, David K. Jackson, Ian Johnston, Dominic Kwiatkowski, Cordelia Langford, John Sillitoe on behalf of the Wellcome Sanger Institute COVID-19 Surveillance Team |
| EPI_ISL_3096418, EPI_ISL_3096419, EPI_ISL_3096420 | Berkshire and Surrey Pathology Services Lighthouse Laboratory | Wellcome Sanger Institute for the COVID-19 Genomics UK (COG-UK) Consortium | Berkshire and Surrey Pathology Services Lighthouse Laboratory and Alex Alderton, Roberto Amato, Jeffrey Barrett, Sonia Goncalves, Ewan Harrison, David K. Jackson, Ian Johnston, Dominic Kwiatkowski, Cordelia Langford, John Sillitoe on behalf of the Wellcome Sanger Institute COVID-19 Surveillance Team |
| EPI_ISL_3096421, EPI_ISL_3096422, EPI_ISL_3096423 | Lighthouse Lab in Alderley Park | Wellcome Sanger Institute for the COVID-19 Genomics UK (COG-UK) Consortium | Jacquelyn Wynn, Mairead Hyland, The Lighthouse Lab in Alderley Park and Alex Alderton, Roberto Amato, Jeffrey Barrett, Sonia Goncalves, Ewan Harrison, David K. Jackson, Ian Johnston, Dominic Kwiatkowski, Cordelia Langford, John Sillitoe on behalf of the Wellcome Sanger Institute COVID-19 Surveillance Team |
| EPI_ISL_3096424 | Berkshire and Surrey Pathology Services Lighthouse Laboratory | Wellcome Sanger Institute for the COVID-19 Genomics UK (COG-UK) Consortium | Berkshire and Surrey Pathology Services Lighthouse Laboratory and Alex Alderton, Roberto Amato, Jeffrey Barrett, Sonia Goncalves, Ewan Harrison, David K. Jackson, Ian Johnston, Dominic Kwiatkowski, Cordelia Langford, John Sillitoe on behalf of the Wellcome Sanger Institute COVID-19 Surveillance Team |
| EPI_ISL_3096425 | Lighthouse Lab in Alderley Park | Wellcome Sanger Institute for the COVID-19 Genomics UK (COG-UK) Consortium | Jacquelyn Wynn, Mairead Hyland, The Lighthouse Lab in Alderley Park and Alex Alderton, Roberto Amato, Jeffrey Barrett, Sonia Goncalves, Ewan Harrison, David K. Jackson, Ian Johnston, Dominic Kwiatkowski, Cordelia Langford, John Sillitoe on behalf of the Wellcome Sanger Institute COVID-19 Surveillance Team |
| EPI_ISL_3096426 | Berkshire and Surrey Pathology Services Lighthouse Laboratory | Wellcome Sanger Institute for the COVID-19 Genomics UK (COG-UK) Consortium | Berkshire and Surrey Pathology Services Lighthouse Laboratory and Alex Alderton, Roberto Amato, Jeffrey Barrett, Sonia Goncalves, Ewan Harrison, David K. Jackson, Ian Johnston, Dominic Kwiatkowski, Cordelia Langford, John Sillitoe on behalf of the Wellcome Sanger Institute COVID-19 Surveillance Team |
| EPI_ISL_3096427, EPI_ISL_3096428, EPI_ISL_3096429 | Lighthouse Lab in Alderley Park | Wellcome Sanger Institute for the COVID-19 Genomics UK (COG-UK) Consortium | Jacquelyn Wynn, Mairead Hyland, The Lighthouse Lab in Alderley Park and Alex Alderton, Roberto Amato, Jeffrey Barrett, Sonia Goncalves, Ewan Harrison, David K. Jackson, Ian Johnston, Dominic Kwiatkowski, Cordelia Langford, John Sillitoe on behalf of the Wellcome Sanger Institute COVID-19 Surveillance Team |
| EPI_ISL_3096430 | Berkshire and Surrey Pathology Services Lighthouse Laboratory | Wellcome Sanger Institute for the COVID-19 Genomics UK (COG-UK) Consortium | Berkshire and Surrey Pathology Services Lighthouse Laboratory and Alex Alderton, Roberto Amato, Jeffrey Barrett, Sonia Goncalves, Ewan Harrison, David K. Jackson, Ian Johnston, Dominic Kwiatkowski, Cordelia Langford, John Sillitoe on behalf of the Wellcome Sanger Institute COVID-19 Surveillance Team |
| EPI_ISL_3096431 | Lighthouse Lab in Alderley Park | Wellcome Sanger Institute for the COVID-19 Genomics UK (COG-UK) Consortium | Jacquelyn Wynn, Mairead Hyland, The Lighthouse Lab in Alderley Park and Alex Alderton, Roberto Amato, Jeffrey Barrett, Sonia Goncalves, Ewan Harrison, David K. Jackson, Ian Johnston, Dominic Kwiatkowski, Cordelia Langford, John Sillitoe on behalf of the Wellcome Sanger Institute COVID-19 Surveillance Team |
| EPI_ISL_3096432, EPI_ISL_3096433, EPI_ISL_3096434, EPI_ISL_3096435 | Berkshire and Surrey Pathology Services Lighthouse Laboratory | Wellcome Sanger Institute for the COVID-19 Genomics UK (COG-UK) Consortium | Berkshire and Surrey Pathology Services Lighthouse Laboratory and Alex Alderton, Roberto Amato, Jeffrey Barrett, Sonia Goncalves, Ewan Harrison, David K. Jackson, Ian Johnston, Dominic Kwiatkowski, Cordelia Langford, John Sillitoe on behalf of the Wellcome Sanger Institute COVID-19 Surveillance Team |
| EPI_ISL_3096436, EPI_ISL_3096437, EPI_ISL_3096438, EPI_ISL_3096439 | Lighthouse Lab in Alderley Park | Wellcome Sanger Institute for the COVID-19 Genomics UK (COG-UK) Consortium | Jacquelyn Wynn, Mairead Hyland, The Lighthouse Lab in Alderley Park and Alex Alderton, Roberto Amato, Jeffrey Barrett, Sonia Goncalves, Ewan Harrison, David K. Jackson, Ian Johnston, Dominic Kwiatkowski, Cordelia Langford, John Sillitoe on behalf of the Wellcome Sanger Institute COVID-19 Surveillance Team |
| EPI_ISL_3096440, EPI_ISL_3096441 | Berkshire and Surrey Pathology Services Lighthouse Laboratory | Wellcome Sanger Institute for the COVID-19 Genomics UK (COG-UK) Consortium | Berkshire and Surrey Pathology Services Lighthouse Laboratory and Alex Alderton, Roberto Amato, Jeffrey Barrett, Sonia Goncalves, Ewan Harrison, David K. Jackson, Ian Johnston, Dominic Kwiatkowski, Cordelia Langford, John Sillitoe on behalf of the Wellcome Sanger Institute COVID-19 Surveillance Team |
| EPI_ISL_3096442 | Lighthouse Lab in Alderley Park | Wellcome Sanger Institute for the COVID-19 Genomics UK (COG-UK) Consortium | Jacquelyn Wynn, Mairead Hyland, The Lighthouse Lab in Alderley Park and Alex Alderton, Roberto Amato, Jeffrey Barrett, Sonia Goncalves, Ewan Harrison, David K. Jackson, Ian Johnston, Dominic Kwiatkowski, Cordelia Langford, John Sillitoe on behalf of the Wellcome Sanger Institute COVID-19 Surveillance Team |
| EPI_ISL_3096443 | Berkshire and Surrey Pathology Services Lighthouse Laboratory | Wellcome Sanger Institute for the COVID-19 Genomics UK (COG-UK) Consortium | Berkshire and Surrey Pathology Services Lighthouse Laboratory and Alex Alderton, Roberto Amato, Jeffrey Barrett, Sonia Goncalves, Ewan Harrison, David K. Jackson, Ian Johnston, Dominic Kwiatkowski, Cordelia Langford, John Sillitoe on behalf of the Wellcome Sanger Institute COVID-19 Surveillance Team |
| EPI_ISL_3096444, EPI_ISL_3096445, EPI_ISL_3096446 | Lighthouse Lab in Alderley Park | Wellcome Sanger Institute for the COVID-19 Genomics UK (COG-UK) Consortium | Jacquelyn Wynn, Mairead Hyland, The Lighthouse Lab in Alderley Park and Alex Alderton, Roberto Amato, Jeffrey Barrett, Sonia Goncalves, Ewan Harrison, David K. Jackson, Ian Johnston, Dominic Kwiatkowski, Cordelia Langford, John Sillitoe on behalf of the Wellcome Sanger Institute COVID-19 Surveillance Team |
| EPI_ISL_3096447 | Berkshire and Surrey Pathology Services Lighthouse Laboratory | Wellcome Sanger Institute for the COVID-19 Genomics UK (COG-UK) Consortium | Berkshire and Surrey Pathology Services Lighthouse Laboratory and Alex Alderton, Roberto Amato, Jeffrey Barrett, Sonia Goncalves, Ewan Harrison, David K. Jackson, Ian Johnston, Dominic Kwiatkowski, Cordelia Langford, John Sillitoe on behalf of the Wellcome Sanger Institute COVID-19 Surveillance Team |
| EPI_ISL_3096448 | Lighthouse Lab in Alderley Park | Wellcome Sanger Institute for the COVID-19 Genomics UK (COG-UK) Consortium | Jacquelyn Wynn, Mairead Hyland, The Lighthouse Lab in Alderley Park and Alex Alderton, Roberto Amato, Jeffrey Barrett, Sonia Goncalves, Ewan Harrison, David K. Jackson, Ian Johnston, Dominic Kwiatkowski, Cordelia Langford, John Sillitoe on behalf of the Wellcome Sanger Institute COVID-19 Surveillance Team |

| | | | |
|---|---|---|---|
| EPI_ISL_3096449 | Berkshire and Surrey Pathology Services Lighthouse Laboratory | Wellcome Sanger Institute for the COVID-19 Genomics UK (COG-UK) Consortium | Berkshire and Surrey Pathology Services Lighthouse Laboratory and Alex Alderton, Roberto Amato, Jeffrey Barrett, Sonia Goncalves, Ewan Harrison, David K. Jackson, Ian Johnston, Dominic Kwiatkowski, Cordelia Langford, John Sillitoe on behalf of the Wellcome Sanger Institute COVID-19 Surveillance Team |
| EPI_ISL_3096450, EPI_ISL_3096451 | Lighthouse Lab in Alderley Park | Wellcome Sanger Institute for the COVID-19 Genomics UK (COG-UK) Consortium | Jacquelyn Wynn, Mairead Hyland, The Lighthouse Lab in Alderley Park and Alex Alderton, Roberto Amato, Jeffrey Barrett, Sonia Goncalves, Ewan Harrison, David K. Jackson, Ian Johnston, Dominic Kwiatkowski, Cordelia Langford, John Sillitoe on behalf of the Wellcome Sanger Institute COVID-19 Surveillance Team |
| EPI_ISL_3096452, EPI_ISL_3096453 | Berkshire and Surrey Pathology Services Lighthouse Laboratory | Wellcome Sanger Institute for the COVID-19 Genomics UK (COG-UK) Consortium | Berkshire and Surrey Pathology Services Lighthouse Laboratory and Alex Alderton, Roberto Amato, Jeffrey Barrett, Sonia Goncalves, Ewan Harrison, David K. Jackson, Ian Johnston, Dominic Kwiatkowski, Cordelia Langford, John Sillitoe on behalf of the Wellcome Sanger Institute COVID-19 Surveillance Team |
| EPI_ISL_3096454, EPI_ISL_3096455 | Lighthouse Lab in Alderley Park | Wellcome Sanger Institute for the COVID-19 Genomics UK (COG-UK) Consortium | Jacquelyn Wynn, Mairead Hyland, The Lighthouse Lab in Alderley Park and Alex Alderton, Roberto Amato, Jeffrey Barrett, Sonia Goncalves, Ewan Harrison, David K. Jackson, Ian Johnston, Dominic Kwiatkowski, Cordelia Langford, John Sillitoe on behalf of the Wellcome Sanger Institute COVID-19 Surveillance Team |
| EPI_ISL_3096456 | Berkshire and Surrey Pathology Services Lighthouse Laboratory | Wellcome Sanger Institute for the COVID-19 Genomics UK (COG-UK) Consortium | Berkshire and Surrey Pathology Services Lighthouse Laboratory and Alex Alderton, Roberto Amato, Jeffrey Barrett, Sonia Goncalves, Ewan Harrison, David K. Jackson, Ian Johnston, Dominic Kwiatkowski, Cordelia Langford, John Sillitoe on behalf of the Wellcome Sanger Institute COVID-19 Surveillance Team |
| EPI_ISL_3096457, EPI_ISL_3096458 | Lighthouse Lab in Alderley Park | Wellcome Sanger Institute for the COVID-19 Genomics UK (COG-UK) Consortium | Jacquelyn Wynn, Mairead Hyland, The Lighthouse Lab in Alderley Park and Alex Alderton, Roberto Amato, Jeffrey Barrett, Sonia Goncalves, Ewan Harrison, David K. Jackson, Ian Johnston, Dominic Kwiatkowski, Cordelia Langford, John Sillitoe on behalf of the Wellcome Sanger Institute COVID-19 Surveillance Team |
| EPI_ISL_3096459 | Berkshire and Surrey Pathology Services Lighthouse Laboratory | Wellcome Sanger Institute for the COVID-19 Genomics UK (COG-UK) Consortium | Berkshire and Surrey Pathology Services Lighthouse Laboratory and Alex Alderton, Roberto Amato, Jeffrey Barrett, Sonia Goncalves, Ewan Harrison, David K. Jackson, Ian Johnston, Dominic Kwiatkowski, Cordelia Langford, John Sillitoe on behalf of the Wellcome Sanger Institute COVID-19 Surveillance Team |
| EPI_ISL_3096460, EPI_ISL_3096461 | Lighthouse Lab in Alderley Park | Wellcome Sanger Institute for the COVID-19 Genomics UK (COG-UK) Consortium | Jacquelyn Wynn, Mairead Hyland, The Lighthouse Lab in Alderley Park and Alex Alderton, Roberto Amato, Jeffrey Barrett, Sonia Goncalves, Ewan Harrison, David K. Jackson, Ian Johnston, Dominic Kwiatkowski, Cordelia Langford, John Sillitoe on behalf of the Wellcome Sanger Institute COVID-19 Surveillance Team |
| EPI_ISL_3096462, EPI_ISL_3096463 | Berkshire and Surrey Pathology Services Lighthouse Laboratory | Wellcome Sanger Institute for the COVID-19 Genomics UK (COG-UK) Consortium | Berkshire and Surrey Pathology Services Lighthouse Laboratory and Alex Alderton, Roberto Amato, Jeffrey Barrett, Sonia Goncalves, Ewan Harrison, David K. Jackson, Ian Johnston, Dominic Kwiatkowski, Cordelia Langford, John Sillitoe on behalf of the Wellcome Sanger Institute COVID-19 Surveillance Team |
| EPI_ISL_3096464, EPI_ISL_3096465, EPI_ISL_3096466 | Lighthouse Lab in Alderley Park | Wellcome Sanger Institute for the COVID-19 Genomics UK (COG-UK) Consortium | Jacquelyn Wynn, Mairead Hyland, The Lighthouse Lab in Alderley Park and Alex Alderton, Roberto Amato, Jeffrey Barrett, Sonia Goncalves, Ewan Harrison, David K. Jackson, Ian Johnston, Dominic Kwiatkowski, Cordelia Langford, John Sillitoe on behalf of the Wellcome Sanger Institute COVID-19 Surveillance Team |
| EPI_ISL_3096467, EPI_ISL_3096468 | Berkshire and Surrey Pathology Services Lighthouse Laboratory | Wellcome Sanger Institute for the COVID-19 Genomics UK (COG-UK) Consortium | Berkshire and Surrey Pathology Services Lighthouse Laboratory and Alex Alderton, Roberto Amato, Jeffrey Barrett, Sonia Goncalves, Ewan Harrison, David K. Jackson, Ian Johnston, Dominic Kwiatkowski, Cordelia Langford, John Sillitoe on behalf of the Wellcome Sanger Institute COVID-19 Surveillance Team |
| EPI_ISL_3096469, EPI_ISL_3096470 | Lighthouse Lab in Alderley Park | Wellcome Sanger Institute for the COVID-19 Genomics UK (COG-UK) Consortium | Jacquelyn Wynn, Mairead Hyland, The Lighthouse Lab in Alderley Park and Alex Alderton, Roberto Amato, Jeffrey Barrett, Sonia Goncalves, Ewan Harrison, David K. Jackson, Ian Johnston, Dominic Kwiatkowski, Cordelia Langford, John Sillitoe on behalf of the Wellcome Sanger Institute COVID-19 Surveillance Team |
| EPI_ISL_3096471 | Berkshire and Surrey Pathology Services Lighthouse Laboratory | Wellcome Sanger Institute for the COVID-19 Genomics UK (COG-UK) Consortium | Berkshire and Surrey Pathology Services Lighthouse Laboratory and Alex Alderton, Roberto Amato, Jeffrey Barrett, Sonia Goncalves, Ewan Harrison, David K. Jackson, Ian Johnston, Dominic Kwiatkowski, Cordelia Langford, John Sillitoe on behalf of the Wellcome Sanger Institute COVID-19 Surveillance Team |
| EPI_ISL_3096472, EPI_ISL_3096473 | Lighthouse Lab in Alderley Park | Wellcome Sanger Institute for the COVID-19 Genomics UK (COG-UK) Consortium | Jacquelyn Wynn, Mairead Hyland, The Lighthouse Lab in Alderley Park and Alex Alderton, Roberto Amato, Jeffrey Barrett, Sonia Goncalves, Ewan Harrison, David K. Jackson, Ian Johnston, Dominic Kwiatkowski, Cordelia Langford, John Sillitoe on behalf of the Wellcome Sanger Institute COVID-19 Surveillance Team |
| EPI_ISL_3096474, EPI_ISL_3096475 | Berkshire and Surrey Pathology Services Lighthouse Laboratory | Wellcome Sanger Institute for the COVID-19 Genomics UK (COG-UK) Consortium | Berkshire and Surrey Pathology Services Lighthouse Laboratory and Alex Alderton, Roberto Amato, Jeffrey Barrett, Sonia Goncalves, Ewan Harrison, David K. Jackson, Ian Johnston, Dominic Kwiatkowski, Cordelia Langford, John Sillitoe on behalf of the Wellcome Sanger Institute COVID-19 Surveillance Team |
| EPI_ISL_3096476 | Lighthouse Lab in Alderley Park | Wellcome Sanger Institute for the COVID-19 Genomics UK (COG-UK) Consortium | Jacquelyn Wynn, Mairead Hyland, The Lighthouse Lab in Alderley Park and Alex Alderton, Roberto Amato, Jeffrey Barrett, Sonia Goncalves, Ewan Harrison, David K. Jackson, Ian Johnston, Dominic Kwiatkowski, Cordelia Langford, John Sillitoe on behalf of the Wellcome Sanger Institute COVID-19 Surveillance Team |
| EPI_ISL_3096477 | Berkshire and Surrey Pathology Services Lighthouse Laboratory | Wellcome Sanger Institute for the COVID-19 Genomics UK (COG-UK) Consortium | Berkshire and Surrey Pathology Services Lighthouse Laboratory and Alex Alderton, Roberto Amato, Jeffrey Barrett, Sonia Goncalves, Ewan Harrison, David K. Jackson, Ian Johnston, Dominic Kwiatkowski, Cordelia Langford, John Sillitoe on behalf of the Wellcome Sanger Institute COVID-19 Surveillance Team |
| EPI_ISL_3096478 | Lighthouse Lab in Alderley Park | Wellcome Sanger Institute for the COVID-19 Genomics UK (COG-UK) Consortium | Jacquelyn Wynn, Mairead Hyland, The Lighthouse Lab in Alderley Park and Alex Alderton, Roberto Amato, Jeffrey Barrett, Sonia Goncalves, Ewan Harrison, David K. Jackson, Ian Johnston, Dominic Kwiatkowski, Cordelia Langford, John Sillitoe on behalf of the Wellcome Sanger Institute COVID-19 Surveillance Team |
| EPI_ISL_3096479, EPI_ISL_3096480 | Berkshire and Surrey Pathology Services Lighthouse Laboratory | Wellcome Sanger Institute for the COVID-19 Genomics UK (COG-UK) Consortium | Berkshire and Surrey Pathology Services Lighthouse Laboratory and Alex Alderton, Roberto Amato, Jeffrey Barrett, Sonia Goncalves, Ewan Harrison, David K. Jackson, Ian Johnston, Dominic Kwiatkowski, Cordelia Langford, John Sillitoe on behalf of the Wellcome Sanger Institute COVID-19 Surveillance Team |
| EPI_ISL_3096481, EPI_ISL_3096482 | Lighthouse Lab in Alderley Park | Wellcome Sanger Institute for the COVID-19 Genomics UK (COG-UK) Consortium | Jacquelyn Wynn, Mairead Hyland, The Lighthouse Lab in Alderley Park and Alex Alderton, Roberto Amato, Jeffrey Barrett, Sonia Goncalves, Ewan Harrison, David K. Jackson, Ian Johnston, Dominic Kwiatkowski, Cordelia Langford, John Sillitoe on behalf of the Wellcome Sanger Institute COVID-19 Surveillance Team |
| EPI_ISL_3096483, EPI_ISL_3096484, EPI_ISL_3096485 | Berkshire and Surrey Pathology Services Lighthouse Laboratory | Wellcome Sanger Institute for the COVID-19 Genomics UK (COG-UK) Consortium | Berkshire and Surrey Pathology Services Lighthouse Laboratory and Alex Alderton, Roberto Amato, Jeffrey Barrett, Sonia Goncalves, Ewan Harrison, David K. Jackson, Ian Johnston, Dominic Kwiatkowski, Cordelia Langford, John Sillitoe on behalf of the Wellcome Sanger Institute COVID-19 Surveillance Team |
| EPI_ISL_3096486 | Lighthouse Lab in Alderley Park | Wellcome Sanger Institute for the COVID-19 Genomics UK (COG-UK) Consortium | Jacquelyn Wynn, Mairead Hyland, The Lighthouse Lab in Alderley Park and Alex Alderton, Roberto Amato, Jeffrey Barrett, Sonia Goncalves, Ewan Harrison, David K. Jackson, Ian Johnston, Dominic Kwiatkowski, Cordelia Langford, John Sillitoe on behalf of the Wellcome Sanger Institute COVID-19 Surveillance Team |
| EPI_ISL_3096487, EPI_ISL_3096488 | Berkshire and Surrey Pathology Services Lighthouse Laboratory | Wellcome Sanger Institute for the COVID-19 Genomics UK (COG-UK) Consortium | Berkshire and Surrey Pathology Services Lighthouse Laboratory and Alex Alderton, Roberto Amato, Jeffrey Barrett, Sonia Goncalves, Ewan Harrison, David K. Jackson, Ian Johnston, Dominic Kwiatkowski, Cordelia Langford, John Sillitoe on behalf of the Wellcome Sanger Institute COVID-19 Surveillance Team |
| EPI_ISL_3096489 | Lighthouse Lab in Alderley Park | Wellcome Sanger Institute for the COVID-19 Genomics UK (COG-UK) Consortium | Jacquelyn Wynn, Mairead Hyland, The Lighthouse Lab in Alderley Park and Alex Alderton, Roberto Amato, Jeffrey Barrett, Sonia Goncalves, Ewan Harrison, David K. Jackson, Ian Johnston, Dominic Kwiatkowski, Cordelia Langford, John Sillitoe on behalf of the Wellcome Sanger Institute COVID-19 Surveillance Team |
| EPI_ISL_3096490 | Berkshire and Surrey Pathology Services Lighthouse Laboratory | Wellcome Sanger Institute for the COVID-19 Genomics UK (COG-UK) Consortium | Berkshire and Surrey Pathology Services Lighthouse Laboratory and Alex Alderton, Roberto Amato, Jeffrey Barrett, Sonia Goncalves, Ewan Harrison, David K. Jackson, Ian Johnston, Dominic Kwiatkowski, Cordelia Langford, John Sillitoe on behalf of the Wellcome Sanger Institute COVID-19 Surveillance Team |
| EPI_ISL_3096491 | Lighthouse Lab in Alderley Park | Wellcome Sanger Institute for the COVID-19 Genomics UK (COG-UK) Consortium | Jacquelyn Wynn, Mairead Hyland, The Lighthouse Lab in Alderley Park and Alex Alderton, Roberto Amato, Jeffrey Barrett, Sonia Goncalves, Ewan Harrison, David K. Jackson, Ian Johnston, Dominic Kwiatkowski, Cordelia Langford, John Sillitoe on behalf of the Wellcome Sanger Institute COVID-19 Surveillance Team |
| EPI_ISL_3096492, EPI_ISL_3096493, EPI_ISL_3096494 | Berkshire and Surrey Pathology Services Lighthouse Laboratory | Wellcome Sanger Institute for the COVID-19 Genomics UK (COG-UK) Consortium | Berkshire and Surrey Pathology Services Lighthouse Laboratory and Alex Alderton, Roberto Amato, Jeffrey Barrett, Sonia Goncalves, Ewan Harrison, David K. Jackson, Ian Johnston, Dominic Kwiatkowski, Cordelia Langford, John Sillitoe on behalf of the Wellcome Sanger Institute COVID-19 Surveillance Team |
| EPI_ISL_3096495, EPI_ISL_3096496, EPI_ISL_3096497 | Lighthouse Lab in Alderley Park | Wellcome Sanger Institute for the COVID-19 Genomics UK (COG-UK) Consortium | Jacquelyn Wynn, Mairead Hyland, The Lighthouse Lab in Alderley Park and Alex Alderton, Roberto Amato, Jeffrey Barrett, Sonia Goncalves, Ewan Harrison, David K. Jackson, Ian Johnston, Dominic Kwiatkowski, Cordelia Langford, John Sillitoe on behalf of the Wellcome Sanger Institute COVID-19 Surveillance Team |
| EPI_ISL_3096498 | Berkshire and Surrey Pathology Services Lighthouse Laboratory | Wellcome Sanger Institute for the COVID-19 Genomics UK (COG-UK) Consortium | Berkshire and Surrey Pathology Services Lighthouse Laboratory and Alex Alderton, Roberto Amato, Jeffrey Barrett, Sonia Goncalves, Ewan Harrison, David K. Jackson, Ian Johnston, Dominic Kwiatkowski, Cordelia Langford, John Sillitoe on behalf of the Wellcome Sanger Institute COVID-19 Surveillance Team |
| EPI_ISL_3096499 | Lighthouse Lab in Alderley Park | Wellcome Sanger Institute for the COVID-19 Genomics UK (COG-UK) Consortium | Jacquelyn Wynn, Mairead Hyland, The Lighthouse Lab in Alderley Park and Alex Alderton, Roberto Amato, Jeffrey Barrett, Sonia Goncalves, Ewan Harrison, David K. Jackson, Ian Johnston, Dominic Kwiatkowski, Cordelia Langford, John Sillitoe on behalf of the Wellcome Sanger Institute COVID-19 Surveillance Team |
| EPI_ISL_3096500 | Berkshire and Surrey Pathology Services Lighthouse Laboratory | Wellcome Sanger Institute for the COVID-19 Genomics UK (COG-UK) Consortium | Berkshire and Surrey Pathology Services Lighthouse Laboratory and Alex Alderton, Roberto Amato, Jeffrey Barrett, Sonia Goncalves, Ewan Harrison, David K. Jackson, Ian Johnston, Dominic Kwiatkowski, Cordelia Langford, John Sillitoe on behalf of the Wellcome Sanger Institute COVID-19 Surveillance Team |
| EPI_ISL_3096501 | Lighthouse Lab in Alderley Park | Wellcome Sanger Institute for the COVID-19 Genomics UK (COG-UK) Consortium | Jacquelyn Wynn, Mairead Hyland, The Lighthouse Lab in Alderley Park and Alex Alderton, Roberto Amato, Jeffrey Barrett, Sonia Goncalves, Ewan Harrison, David K. Jackson, Ian Johnston, Dominic Kwiatkowski, Cordelia Langford, John Sillitoe on behalf of the Wellcome Sanger Institute COVID-19 Surveillance Team |
| EPI_ISL_3096502, EPI_ISL_3096503, EPI_ISL_3096504, EPI_ISL_3096505, EPI_ISL_3096506 | Berkshire and Surrey Pathology Services Lighthouse Laboratory | Wellcome Sanger Institute for the COVID-19 Genomics UK (COG-UK) Consortium | Berkshire and Surrey Pathology Services Lighthouse Laboratory and Alex Alderton, Roberto Amato, Jeffrey Barrett, Sonia Goncalves, Ewan Harrison, David K. Jackson, Ian Johnston, Dominic Kwiatkowski, Cordelia Langford, John Sillitoe on behalf of the Wellcome Sanger Institute COVID-19 Surveillance Team |
| EPI_ISL_3096507 | Lighthouse Lab in Alderley Park | Wellcome Sanger Institute for the COVID-19 Genomics UK (COG-UK) Consortium | Jacquelyn Wynn, Mairead Hyland, The Lighthouse Lab in Alderley Park and Alex Alderton, Roberto Amato, Jeffrey Barrett, Sonia Goncalves, Ewan Harrison, David K. Jackson, Ian Johnston, Dominic Kwiatkowski, Cordelia Langford, John Sillitoe on behalf of the Wellcome Sanger Institute COVID-19 Surveillance Team |
| EPI_ISL_3096508, EPI_ISL_3096509 | Berkshire and Surrey Pathology Services Lighthouse Laboratory | Wellcome Sanger Institute for the COVID-19 Genomics UK (COG-UK) Consortium | Berkshire and Surrey Pathology Services Lighthouse Laboratory and Alex Alderton, Roberto Amato, Jeffrey Barrett, Sonia Goncalves, Ewan Harrison, David K. Jackson, Ian Johnston, Dominic Kwiatkowski, Cordelia Langford, John Sillitoe on behalf of the Wellcome Sanger Institute COVID-19 Surveillance Team |
| EPI_ISL_3096510, EPI_ISL_3096511, EPI_ISL_3096512, EPI_ISL_3096513, EPI_ISL_3096514, EPI_ISL_3096515, EPI_ISL_3096516, EPI_ISL_3096517, EPI_ISL_3096518, EPI_ISL_3096519, EPI_ISL_3096520, EPI_ISL_3096521, EPI_ISL_3096522, EPI_ISL_3096523, EPI_ISL_3096524 | | | |
| see above | Lighthouse Lab in Alderley Park | Wellcome Sanger Institute for the COVID-19 Genomics UK (COG-UK) Consortium | Jacquelyn Wynn, Mairead Hyland, The Lighthouse Lab in Alderley Park and Alex Alderton, Roberto Amato, Jeffrey Barrett, Sonia Goncalves, Ewan Harrison, David K. Jackson, Ian Johnston, Dominic Kwiatkowski, Cordelia Langford, John Sillitoe on behalf of the Wellcome Sanger Institute COVID-19 Surveillance Team |
| EPI_ISL_3096525 | Berkshire and Surrey Pathology Services Lighthouse Laboratory | Wellcome Sanger Institute for the COVID-19 Genomics UK (COG-UK) Consortium | Berkshire and Surrey Pathology Services Lighthouse Laboratory and Alex Alderton, Roberto Amato, Jeffrey Barrett, Sonia Goncalves, Ewan Harrison, David K. Jackson, Ian Johnston, Dominic Kwiatkowski, Cordelia Langford, John Sillitoe on behalf of the Wellcome Sanger Institute COVID-19 Surveillance Team |
| EPI_ISL_3096526, EPI_ISL_3096527, EPI_ISL_3096528 | Lighthouse Lab in Alderley Park | Wellcome Sanger Institute for the COVID-19 Genomics UK (COG-UK) Consortium | Jacquelyn Wynn, Mairead Hyland, The Lighthouse Lab in Alderley Park and Alex Alderton, Roberto Amato, Jeffrey Barrett, Sonia Goncalves, Ewan Harrison, David K. Jackson, Ian Johnston, Dominic Kwiatkowski, Cordelia Langford, John Sillitoe on behalf of the Wellcome Sanger Institute COVID-19 Surveillance Team |
| EPI_ISL_3096529 | Berkshire and Surrey Pathology Services Lighthouse Laboratory | Wellcome Sanger Institute for the COVID-19 Genomics UK (COG-UK) Consortium | Berkshire and Surrey Pathology Services Lighthouse Laboratory and Alex Alderton, Roberto Amato, Jeffrey Barrett, Sonia Goncalves, Ewan Harrison, David K. Jackson, Ian Johnston, Dominic Kwiatkowski, Cordelia Langford, John Sillitoe on behalf of the Wellcome Sanger Institute COVID-19 Surveillance Team |
| EPI_ISL_3096530 | Lighthouse Lab in Alderley Park | Wellcome Sanger Institute for the COVID-19 Genomics UK (COG-UK) Consortium | Jacquelyn Wynn, Mairead Hyland, The Lighthouse Lab in Alderley Park and Alex Alderton, Roberto Amato, Jeffrey Barrett, Sonia Goncalves, Ewan Harrison, David K. Jackson, Ian Johnston, Dominic Kwiatkowski, Cordelia Langford, John Sillitoe on behalf of the Wellcome Sanger Institute COVID-19 Surveillance Team |
| EPI_ISL_3096531 | Berkshire and Surrey Pathology Services Lighthouse Laboratory | Wellcome Sanger Institute for the COVID-19 Genomics UK (COG-UK) Consortium | Berkshire and Surrey Pathology Services Lighthouse Laboratory and Alex Alderton, Roberto Amato, Jeffrey Barrett, Sonia Goncalves, Ewan Harrison, David K. Jackson, Ian Johnston, Dominic Kwiatkowski, Cordelia Langford, John Sillitoe on behalf of the Wellcome Sanger Institute COVID-19 Surveillance Team |

| | | | |
|---|---|---|---|
| EPI_ISL_3096532 | Lighthouse Lab in Alderley Park | Wellcome Sanger Institute for the COVID-19 Genomics UK (COG-UK) Consortium | Jacquelyn Wynn, Mairead Hyland, The Lighthouse Lab in Alderley Park and Alex Alderton, Roberto Amato, Jeffrey Barrett, Sonia Goncalves, Ewan Harrison, David K. Jackson, Ian Johnston, Dominic Kwiatkowski, Cordelia Langford, John Sillitoe on behalf of the Wellcome Sanger Institute COVID-19 Surveillance Team |
| EPI_ISL_3096533, EPI_ISL_3096534, EPI_ISL_3096535 | Berkshire and Surrey Pathology Services Lighthouse Laboratory | Wellcome Sanger Institute for the COVID-19 Genomics UK (COG-UK) Consortium | Berkshire and Surrey Pathology Services Lighthouse Laboratory and Alex Alderton, Roberto Amato, Jeffrey Barrett, Sonia Goncalves, Ewan Harrison, David K. Jackson, Ian Johnston, Dominic Kwiatkowski, Cordelia Langford, John Sillitoe on behalf of the Wellcome Sanger Institute COVID-19 Surveillance Team |
| EPI_ISL_3096536, EPI_ISL_3096537 | Lighthouse Lab in Alderley Park | Wellcome Sanger Institute for the COVID-19 Genomics UK (COG-UK) Consortium | Jacquelyn Wynn, Mairead Hyland, The Lighthouse Lab in Alderley Park and Alex Alderton, Roberto Amato, Jeffrey Barrett, Sonia Goncalves, Ewan Harrison, David K. Jackson, Ian Johnston, Dominic Kwiatkowski, Cordelia Langford, John Sillitoe on behalf of the Wellcome Sanger Institute COVID-19 Surveillance Team |
| EPI_ISL_3096538 | Berkshire and Surrey Pathology Services Lighthouse Laboratory | Wellcome Sanger Institute for the COVID-19 Genomics UK (COG-UK) Consortium | Berkshire and Surrey Pathology Services Lighthouse Laboratory and Alex Alderton, Roberto Amato, Jeffrey Barrett, Sonia Goncalves, Ewan Harrison, David K. Jackson, Ian Johnston, Dominic Kwiatkowski, Cordelia Langford, John Sillitoe on behalf of the Wellcome Sanger Institute COVID-19 Surveillance Team |
| EPI_ISL_3096539, EPI_ISL_3096540, EPI_ISL_3096541, EPI_ISL_3096542 | Lighthouse Lab in Alderley Park | Wellcome Sanger Institute for the COVID-19 Genomics UK (COG-UK) Consortium | Jacquelyn Wynn, Mairead Hyland, The Lighthouse Lab in Alderley Park and Alex Alderton, Roberto Amato, Jeffrey Barrett, Sonia Goncalves, Ewan Harrison, David K. Jackson, Ian Johnston, Dominic Kwiatkowski, Cordelia Langford, John Sillitoe on behalf of the Wellcome Sanger Institute COVID-19 Surveillance Team |
| EPI_ISL_3096543, EPI_ISL_3096544, EPI_ISL_3096545 | Berkshire and Surrey Pathology Services Lighthouse Laboratory | Wellcome Sanger Institute for the COVID-19 Genomics UK (COG-UK) Consortium | Berkshire and Surrey Pathology Services Lighthouse Laboratory and Alex Alderton, Roberto Amato, Jeffrey Barrett, Sonia Goncalves, Ewan Harrison, David K. Jackson, Ian Johnston, Dominic Kwiatkowski, Cordelia Langford, John Sillitoe on behalf of the Wellcome Sanger Institute COVID-19 Surveillance Team |
| EPI_ISL_3096546 | Lighthouse Lab in Alderley Park | Wellcome Sanger Institute for the COVID-19 Genomics UK (COG-UK) Consortium | Jacquelyn Wynn, Mairead Hyland, The Lighthouse Lab in Alderley Park and Alex Alderton, Roberto Amato, Jeffrey Barrett, Sonia Goncalves, Ewan Harrison, David K. Jackson, Ian Johnston, Dominic Kwiatkowski, Cordelia Langford, John Sillitoe on behalf of the Wellcome Sanger Institute COVID-19 Surveillance Team |
| EPI_ISL_3096547 | Berkshire and Surrey Pathology Services Lighthouse Laboratory | Wellcome Sanger Institute for the COVID-19 Genomics UK (COG-UK) Consortium | Berkshire and Surrey Pathology Services Lighthouse Laboratory and Alex Alderton, Roberto Amato, Jeffrey Barrett, Sonia Goncalves, Ewan Harrison, David K. Jackson, Ian Johnston, Dominic Kwiatkowski, Cordelia Langford, John Sillitoe on behalf of the Wellcome Sanger Institute COVID-19 Surveillance Team |
| EPI_ISL_3096548, EPI_ISL_3096549 | Lighthouse Lab in Alderley Park | Wellcome Sanger Institute for the COVID-19 Genomics UK (COG-UK) Consortium | Jacquelyn Wynn, Mairead Hyland, The Lighthouse Lab in Alderley Park and Alex Alderton, Roberto Amato, Jeffrey Barrett, Sonia Goncalves, Ewan Harrison, David K. Jackson, Ian Johnston, Dominic Kwiatkowski, Cordelia Langford, John Sillitoe on behalf of the Wellcome Sanger Institute COVID-19 Surveillance Team |
| EPI_ISL_3096550, EPI_ISL_3096551 | Berkshire and Surrey Pathology Services Lighthouse Laboratory | Wellcome Sanger Institute for the COVID-19 Genomics UK (COG-UK) Consortium | Berkshire and Surrey Pathology Services Lighthouse Laboratory and Alex Alderton, Roberto Amato, Jeffrey Barrett, Sonia Goncalves, Ewan Harrison, David K. Jackson, Ian Johnston, Dominic Kwiatkowski, Cordelia Langford, John Sillitoe on behalf of the Wellcome Sanger Institute COVID-19 Surveillance Team |
| EPI_ISL_3096552, EPI_ISL_3096553 | Lighthouse Lab in Alderley Park | Wellcome Sanger Institute for the COVID-19 Genomics UK (COG-UK) Consortium | Jacquelyn Wynn, Mairead Hyland, The Lighthouse Lab in Alderley Park and Alex Alderton, Roberto Amato, Jeffrey Barrett, Sonia Goncalves, Ewan Harrison, David K. Jackson, Ian Johnston, Dominic Kwiatkowski, Cordelia Langford, John Sillitoe on behalf of the Wellcome Sanger Institute COVID-19 Surveillance Team |
| EPI_ISL_3096554 | Berkshire and Surrey Pathology Services Lighthouse Laboratory | Wellcome Sanger Institute for the COVID-19 Genomics UK (COG-UK) Consortium | Berkshire and Surrey Pathology Services Lighthouse Laboratory and Alex Alderton, Roberto Amato, Jeffrey Barrett, Sonia Goncalves, Ewan Harrison, David K. Jackson, Ian Johnston, Dominic Kwiatkowski, Cordelia Langford, John Sillitoe on behalf of the Wellcome Sanger Institute COVID-19 Surveillance Team |
| EPI_ISL_3096555, EPI_ISL_3096556 | Lighthouse Lab in Alderley Park | Wellcome Sanger Institute for the COVID-19 Genomics UK (COG-UK) Consortium | Jacquelyn Wynn, Mairead Hyland, The Lighthouse Lab in Alderley Park and Alex Alderton, Roberto Amato, Jeffrey Barrett, Sonia Goncalves, Ewan Harrison, David K. Jackson, Ian Johnston, Dominic Kwiatkowski, Cordelia Langford, John Sillitoe on behalf of the Wellcome Sanger Institute COVID-19 Surveillance Team |
| EPI_ISL_3096557 | Berkshire and Surrey Pathology Services Lighthouse Laboratory | Wellcome Sanger Institute for the COVID-19 Genomics UK (COG-UK) Consortium | Berkshire and Surrey Pathology Services Lighthouse Laboratory and Alex Alderton, Roberto Amato, Jeffrey Barrett, Sonia Goncalves, Ewan Harrison, David K. Jackson, Ian Johnston, Dominic Kwiatkowski, Cordelia Langford, John Sillitoe on behalf of the Wellcome Sanger Institute COVID-19 Surveillance Team |
| EPI_ISL_3096558, EPI_ISL_3096559, EPI_ISL_3096560 | Lighthouse Lab in Alderley Park | Wellcome Sanger Institute for the COVID-19 Genomics UK (COG-UK) Consortium | Jacquelyn Wynn, Mairead Hyland, The Lighthouse Lab in Alderley Park and Alex Alderton, Roberto Amato, Jeffrey Barrett, Sonia Goncalves, Ewan Harrison, David K. Jackson, Ian Johnston, Dominic Kwiatkowski, Cordelia Langford, John Sillitoe on behalf of the Wellcome Sanger Institute COVID-19 Surveillance Team |
| EPI_ISL_3096561 | Berkshire and Surrey Pathology Services Lighthouse Laboratory | Wellcome Sanger Institute for the COVID-19 Genomics UK (COG-UK) Consortium | Berkshire and Surrey Pathology Services Lighthouse Laboratory and Alex Alderton, Roberto Amato, Jeffrey Barrett, Sonia Goncalves, Ewan Harrison, David K. Jackson, Ian Johnston, Dominic Kwiatkowski, Cordelia Langford, John Sillitoe on behalf of the Wellcome Sanger Institute COVID-19 Surveillance Team |
| EPI_ISL_3096562, EPI_ISL_3096563, EPI_ISL_3096564, EPI_ISL_3096565 | Lighthouse Lab in Alderley Park | Wellcome Sanger Institute for the COVID-19 Genomics UK (COG-UK) Consortium | Jacquelyn Wynn, Mairead Hyland, The Lighthouse Lab in Alderley Park and Alex Alderton, Roberto Amato, Jeffrey Barrett, Sonia Goncalves, Ewan Harrison, David K. Jackson, Ian Johnston, Dominic Kwiatkowski, Cordelia Langford, John Sillitoe on behalf of the Wellcome Sanger Institute COVID-19 Surveillance Team |
| EPI_ISL_3096566 | Berkshire and Surrey Pathology Services Lighthouse Laboratory | Wellcome Sanger Institute for the COVID-19 Genomics UK (COG-UK) Consortium | Berkshire and Surrey Pathology Services Lighthouse Laboratory and Alex Alderton, Roberto Amato, Jeffrey Barrett, Sonia Goncalves, Ewan Harrison, David K. Jackson, Ian Johnston, Dominic Kwiatkowski, Cordelia Langford, John Sillitoe on behalf of the Wellcome Sanger Institute COVID-19 Surveillance Team |
| EPI_ISL_3096567 | Lighthouse Lab in Alderley Park | Wellcome Sanger Institute for the COVID-19 Genomics UK (COG-UK) Consortium | Jacquelyn Wynn, Mairead Hyland, The Lighthouse Lab in Alderley Park and Alex Alderton, Roberto Amato, Jeffrey Barrett, Sonia Goncalves, Ewan Harrison, David K. Jackson, Ian Johnston, Dominic Kwiatkowski, Cordelia Langford, John Sillitoe on behalf of the Wellcome Sanger Institute COVID-19 Surveillance Team |
| EPI_ISL_3096568 | Berkshire and Surrey Pathology Services Lighthouse Laboratory | Wellcome Sanger Institute for the COVID-19 Genomics UK (COG-UK) Consortium | Berkshire and Surrey Pathology Services Lighthouse Laboratory and Alex Alderton, Roberto Amato, Jeffrey Barrett, Sonia Goncalves, Ewan Harrison, David K. Jackson, Ian Johnston, Dominic Kwiatkowski, Cordelia Langford, John Sillitoe on behalf of the Wellcome Sanger Institute COVID-19 Surveillance Team |
| EPI_ISL_3096569 | Lighthouse Lab in Alderley Park | Wellcome Sanger Institute for the COVID-19 Genomics UK (COG-UK) Consortium | Jacquelyn Wynn, Mairead Hyland, The Lighthouse Lab in Alderley Park and Alex Alderton, Roberto Amato, Jeffrey Barrett, Sonia Goncalves, Ewan Harrison, David K. Jackson, Ian Johnston, Dominic Kwiatkowski, Cordelia Langford, John Sillitoe on behalf of the Wellcome Sanger Institute COVID-19 Surveillance Team |
| EPI_ISL_3096570 | Berkshire and Surrey Pathology Services Lighthouse Laboratory | Wellcome Sanger Institute for the COVID-19 Genomics UK (COG-UK) Consortium | Berkshire and Surrey Pathology Services Lighthouse Laboratory and Alex Alderton, Roberto Amato, Jeffrey Barrett, Sonia Goncalves, Ewan Harrison, David K. Jackson, Ian Johnston, Dominic Kwiatkowski, Cordelia Langford, John Sillitoe on behalf of the Wellcome Sanger Institute COVID-19 Surveillance Team |
| EPI_ISL_3096571, EPI_ISL_3096572, EPI_ISL_3096573, EPI_ISL_3096574 | Lighthouse Lab in Alderley Park | Wellcome Sanger Institute for the COVID-19 Genomics UK (COG-UK) Consortium | Jacquelyn Wynn, Mairead Hyland, The Lighthouse Lab in Alderley Park and Alex Alderton, Roberto Amato, Jeffrey Barrett, Sonia Goncalves, Ewan Harrison, David K. Jackson, Ian Johnston, Dominic Kwiatkowski, Cordelia Langford, John Sillitoe on behalf of the Wellcome Sanger Institute COVID-19 Surveillance Team |
| EPI_ISL_3096575, EPI_ISL_3096576, EPI_ISL_3096577 | Berkshire and Surrey Pathology Services Lighthouse Laboratory | Wellcome Sanger Institute for the COVID-19 Genomics UK (COG-UK) Consortium | Berkshire and Surrey Pathology Services Lighthouse Laboratory and Alex Alderton, Roberto Amato, Jeffrey Barrett, Sonia Goncalves, Ewan Harrison, David K. Jackson, Ian Johnston, Dominic Kwiatkowski, Cordelia Langford, John Sillitoe on behalf of the Wellcome Sanger Institute COVID-19 Surveillance Team |
| EPI_ISL_3096578 | Lighthouse Lab in Alderley Park | Wellcome Sanger Institute for the COVID-19 Genomics UK (COG-UK) Consortium | Jacquelyn Wynn, Mairead Hyland, The Lighthouse Lab in Alderley Park and Alex Alderton, Roberto Amato, Jeffrey Barrett, Sonia Goncalves, Ewan Harrison, David K. Jackson, Ian Johnston, Dominic Kwiatkowski, Cordelia Langford, John Sillitoe on behalf of the Wellcome Sanger Institute COVID-19 Surveillance Team |
| EPI_ISL_3096579 | Berkshire and Surrey Pathology Services Lighthouse Laboratory | Wellcome Sanger Institute for the COVID-19 Genomics UK (COG-UK) Consortium | Berkshire and Surrey Pathology Services Lighthouse Laboratory and Alex Alderton, Roberto Amato, Jeffrey Barrett, Sonia Goncalves, Ewan Harrison, David K. Jackson, Ian Johnston, Dominic Kwiatkowski, Cordelia Langford, John Sillitoe on behalf of the Wellcome Sanger Institute COVID-19 Surveillance Team |
| EPI_ISL_3096580 | Lighthouse Lab in Alderley Park | Wellcome Sanger Institute for the COVID-19 Genomics UK (COG-UK) Consortium | Jacquelyn Wynn, Mairead Hyland, The Lighthouse Lab in Alderley Park and Alex Alderton, Roberto Amato, Jeffrey Barrett, Sonia Goncalves, Ewan Harrison, David K. Jackson, Ian Johnston, Dominic Kwiatkowski, Cordelia Langford, John Sillitoe on behalf of the Wellcome Sanger Institute COVID-19 Surveillance Team |
| EPI_ISL_3096581, EPI_ISL_3096582, EPI_ISL_3096583 | Berkshire and Surrey Pathology Services Lighthouse Laboratory | Wellcome Sanger Institute for the COVID-19 Genomics UK (COG-UK) Consortium | Berkshire and Surrey Pathology Services Lighthouse Laboratory and Alex Alderton, Roberto Amato, Jeffrey Barrett, Sonia Goncalves, Ewan Harrison, David K. Jackson, Ian Johnston, Dominic Kwiatkowski, Cordelia Langford, John Sillitoe on behalf of the Wellcome Sanger Institute COVID-19 Surveillance Team |
| EPI_ISL_3096584 | Lighthouse Lab in Alderley Park | Wellcome Sanger Institute for the COVID-19 Genomics UK (COG-UK) Consortium | Jacquelyn Wynn, Mairead Hyland, The Lighthouse Lab in Alderley Park and Alex Alderton, Roberto Amato, Jeffrey Barrett, Sonia Goncalves, Ewan Harrison, David K. Jackson, Ian Johnston, Dominic Kwiatkowski, Cordelia Langford, John Sillitoe on behalf of the Wellcome Sanger Institute COVID-19 Surveillance Team |
| EPI_ISL_3096585 | Berkshire and Surrey Pathology Services Lighthouse Laboratory | Wellcome Sanger Institute for the COVID-19 Genomics UK (COG-UK) Consortium | Berkshire and Surrey Pathology Services Lighthouse Laboratory and Alex Alderton, Roberto Amato, Jeffrey Barrett, Sonia Goncalves, Ewan Harrison, David K. Jackson, Ian Johnston, Dominic Kwiatkowski, Cordelia Langford, John Sillitoe on behalf of the Wellcome Sanger Institute COVID-19 Surveillance Team |
| EPI_ISL_3096586 | Lighthouse Lab in Alderley Park | Wellcome Sanger Institute for the COVID-19 Genomics UK (COG-UK) Consortium | Jacquelyn Wynn, Mairead Hyland, The Lighthouse Lab in Alderley Park and Alex Alderton, Roberto Amato, Jeffrey Barrett, Sonia Goncalves, Ewan Harrison, David K. Jackson, Ian Johnston, Dominic Kwiatkowski, Cordelia Langford, John Sillitoe on behalf of the Wellcome Sanger Institute COVID-19 Surveillance Team |
| EPI_ISL_3096587 | Berkshire and Surrey Pathology Services Lighthouse Laboratory | Wellcome Sanger Institute for the COVID-19 Genomics UK (COG-UK) Consortium | Berkshire and Surrey Pathology Services Lighthouse Laboratory and Alex Alderton, Roberto Amato, Jeffrey Barrett, Sonia Goncalves, Ewan Harrison, David K. Jackson, Ian Johnston, Dominic Kwiatkowski, Cordelia Langford, John Sillitoe on behalf of the Wellcome Sanger Institute COVID-19 Surveillance Team |
| EPI_ISL_3096588 | Lighthouse Lab in Alderley Park | Wellcome Sanger Institute for the COVID-19 Genomics UK (COG-UK) Consortium | Jacquelyn Wynn, Mairead Hyland, The Lighthouse Lab in Alderley Park and Alex Alderton, Roberto Amato, Jeffrey Barrett, Sonia Goncalves, Ewan Harrison, David K. Jackson, Ian Johnston, Dominic Kwiatkowski, Cordelia Langford, John Sillitoe on behalf of the Wellcome Sanger Institute COVID-19 Surveillance Team |
| EPI_ISL_3096589 | Berkshire and Surrey Pathology Services Lighthouse Laboratory | Wellcome Sanger Institute for the COVID-19 Genomics UK (COG-UK) Consortium | Berkshire and Surrey Pathology Services Lighthouse Laboratory and Alex Alderton, Roberto Amato, Jeffrey Barrett, Sonia Goncalves, Ewan Harrison, David K. Jackson, Ian Johnston, Dominic Kwiatkowski, Cordelia Langford, John Sillitoe on behalf of the Wellcome Sanger Institute COVID-19 Surveillance Team |
| EPI_ISL_3096590, EPI_ISL_3096591, EPI_ISL_3096592, EPI_ISL_3096593, EPI_ISL_3096594 | Lighthouse Lab in Alderley Park | Wellcome Sanger Institute for the COVID-19 Genomics UK (COG-UK) Consortium | Jacquelyn Wynn, Mairead Hyland, The Lighthouse Lab in Alderley Park and Alex Alderton, Roberto Amato, Jeffrey Barrett, Sonia Goncalves, Ewan Harrison, David K. Jackson, Ian Johnston, Dominic Kwiatkowski, Cordelia Langford, John Sillitoe on behalf of the Wellcome Sanger Institute COVID-19 Surveillance Team |
| EPI_ISL_3096595, EPI_ISL_3096596 | Berkshire and Surrey Pathology Services Lighthouse Laboratory | Wellcome Sanger Institute for the COVID-19 Genomics UK (COG-UK) Consortium | Berkshire and Surrey Pathology Services Lighthouse Laboratory and Alex Alderton, Roberto Amato, Jeffrey Barrett, Sonia Goncalves, Ewan Harrison, David K. Jackson, Ian Johnston, Dominic Kwiatkowski, Cordelia Langford, John Sillitoe on behalf of the Wellcome Sanger Institute COVID-19 Surveillance Team |
| EPI_ISL_3096597, EPI_ISL_3096598, EPI_ISL_3096599, EPI_ISL_3096600 | Lighthouse Lab in Alderley Park | Wellcome Sanger Institute for the COVID-19 Genomics UK (COG-UK) Consortium | Jacquelyn Wynn, Mairead Hyland, The Lighthouse Lab in Alderley Park and Alex Alderton, Roberto Amato, Jeffrey Barrett, Sonia Goncalves, Ewan Harrison, David K. Jackson, Ian Johnston, Dominic Kwiatkowski, Cordelia Langford, John Sillitoe on behalf of the Wellcome Sanger Institute COVID-19 Surveillance Team |
| EPI_ISL_3096601 | Berkshire and Surrey Pathology Services Lighthouse Laboratory | Wellcome Sanger Institute for the COVID-19 Genomics UK (COG-UK) Consortium | Berkshire and Surrey Pathology Services Lighthouse Laboratory and Alex Alderton, Roberto Amato, Jeffrey Barrett, Sonia Goncalves, Ewan Harrison, David K. Jackson, Ian Johnston, Dominic Kwiatkowski, Cordelia Langford, John Sillitoe on behalf of the Wellcome Sanger Institute COVID-19 Surveillance Team |
| EPI_ISL_3096602, | Lighthouse Lab in Alderley Park | Wellcome Sanger Institute for the COVID-19 Genomics UK (COG-UK) | Jacquelyn Wynn, Mairead Hyland, The Lighthouse Lab in Alderley Park and Alex Alderton, Roberto Amato, Jeffrey Barrett, Sonia Goncalves, Ewan Harrison, David K. Jackson, Ian Johnston, Dominic Kwiatkowski, Cordelia Langford, John Sillitoe on |

| | | | |
|---|---|---|---|
| EPI_ISL_3096603, EPI_ISL_3096604 | | Consortium | behalf of the Wellcome Sanger Institute COVID-19 Surveillance Team |
| EPI_ISL_3096605, EPI_ISL_3096606, EPI_ISL_3096607, EPI_ISL_3096608, EPI_ISL_3096609, EPI_ISL_3096610 | Berkshire and Surrey Pathology Services Lighthouse Laboratory | Wellcome Sanger Institute for the COVID-19 Genomics UK (COG-UK) Consortium | Berkshire and Surrey Pathology Services Lighthouse Laboratory and Alex Alderton, Roberto Amato, Jeffrey Barrett, Sonia Goncalves, Ewan Harrison, David K. Jackson, Ian Johnston, Dominic Kwiatkowski, Cordelia Langford, John Sillitoe on behalf of the Wellcome Sanger Institute COVID-19 Surveillance Team |
| EPI_ISL_3096611, EPI_ISL_3096612, EPI_ISL_3096613, EPI_ISL_3096614 | Lighthouse Lab in Alderley Park | Wellcome Sanger Institute for the COVID-19 Genomics UK (COG-UK) Consortium | Jacquelyn Wynn, Mairead Hyland, The Lighthouse Lab in Alderley Park and Alex Alderton, Roberto Amato, Jeffrey Barrett, Sonia Goncalves, Ewan Harrison, David K. Jackson, Ian Johnston, Dominic Kwiatkowski, Cordelia Langford, John Sillitoe on behalf of the Wellcome Sanger Institute COVID-19 Surveillance Team |
| EPI_ISL_3096615, EPI_ISL_3096616, EPI_ISL_3096617 | Berkshire and Surrey Pathology Services Lighthouse Laboratory | Wellcome Sanger Institute for the COVID-19 Genomics UK (COG-UK) Consortium | Berkshire and Surrey Pathology Services Lighthouse Laboratory and Alex Alderton, Roberto Amato, Jeffrey Barrett, Sonia Goncalves, Ewan Harrison, David K. Jackson, Ian Johnston, Dominic Kwiatkowski, Cordelia Langford, John Sillitoe on behalf of the Wellcome Sanger Institute COVID-19 Surveillance Team |
| EPI_ISL_3096618 | Lighthouse Lab in Alderley Park | Wellcome Sanger Institute for the COVID-19 Genomics UK (COG-UK) Consortium | Jacquelyn Wynn, Mairead Hyland, The Lighthouse Lab in Alderley Park and Alex Alderton, Roberto Amato, Jeffrey Barrett, Sonia Goncalves, Ewan Harrison, David K. Jackson, Ian Johnston, Dominic Kwiatkowski, Cordelia Langford, John Sillitoe on behalf of the Wellcome Sanger Institute COVID-19 Surveillance Team |
| EPI_ISL_3096619 | Berkshire and Surrey Pathology Services Lighthouse Laboratory | Wellcome Sanger Institute for the COVID-19 Genomics UK (COG-UK) Consortium | Berkshire and Surrey Pathology Services Lighthouse Laboratory and Alex Alderton, Roberto Amato, Jeffrey Barrett, Sonia Goncalves, Ewan Harrison, David K. Jackson, Ian Johnston, Dominic Kwiatkowski, Cordelia Langford, John Sillitoe on behalf of the Wellcome Sanger Institute COVID-19 Surveillance Team |
| EPI_ISL_3096620 | Lighthouse Lab in Alderley Park | Wellcome Sanger Institute for the COVID-19 Genomics UK (COG-UK) Consortium | Jacquelyn Wynn, Mairead Hyland, The Lighthouse Lab in Alderley Park and Alex Alderton, Roberto Amato, Jeffrey Barrett, Sonia Goncalves, Ewan Harrison, David K. Jackson, Ian Johnston, Dominic Kwiatkowski, Cordelia Langford, John Sillitoe on behalf of the Wellcome Sanger Institute COVID-19 Surveillance Team |
| EPI_ISL_3096621, EPI_ISL_3096622, EPI_ISL_3096623 | Berkshire and Surrey Pathology Services Lighthouse Laboratory | Wellcome Sanger Institute for the COVID-19 Genomics UK (COG-UK) Consortium | Berkshire and Surrey Pathology Services Lighthouse Laboratory and Alex Alderton, Roberto Amato, Jeffrey Barrett, Sonia Goncalves, Ewan Harrison, David K. Jackson, Ian Johnston, Dominic Kwiatkowski, Cordelia Langford, John Sillitoe on behalf of the Wellcome Sanger Institute COVID-19 Surveillance Team |
| EPI_ISL_3096624, EPI_ISL_3096625, EPI_ISL_3096626, EPI_ISL_3096627, EPI_ISL_3096628 | Lighthouse Lab in Alderley Park | Wellcome Sanger Institute for the COVID-19 Genomics UK (COG-UK) Consortium | Jacquelyn Wynn, Mairead Hyland, The Lighthouse Lab in Alderley Park and Alex Alderton, Roberto Amato, Jeffrey Barrett, Sonia Goncalves, Ewan Harrison, David K. Jackson, Ian Johnston, Dominic Kwiatkowski, Cordelia Langford, John Sillitoe on behalf of the Wellcome Sanger Institute COVID-19 Surveillance Team |
| EPI_ISL_3096629, EPI_ISL_3096630, EPI_ISL_3096631 | Berkshire and Surrey Pathology Services Lighthouse Laboratory | Wellcome Sanger Institute for the COVID-19 Genomics UK (COG-UK) Consortium | Berkshire and Surrey Pathology Services Lighthouse Laboratory and Alex Alderton, Roberto Amato, Jeffrey Barrett, Sonia Goncalves, Ewan Harrison, David K. Jackson, Ian Johnston, Dominic Kwiatkowski, Cordelia Langford, John Sillitoe on behalf of the Wellcome Sanger Institute COVID-19 Surveillance Team |
| EPI_ISL_3096632 | Lighthouse Lab in Alderley Park | Wellcome Sanger Institute for the COVID-19 Genomics UK (COG-UK) Consortium | Jacquelyn Wynn, Mairead Hyland, The Lighthouse Lab in Alderley Park and Alex Alderton, Roberto Amato, Jeffrey Barrett, Sonia Goncalves, Ewan Harrison, David K. Jackson, Ian Johnston, Dominic Kwiatkowski, Cordelia Langford, John Sillitoe on behalf of the Wellcome Sanger Institute COVID-19 Surveillance Team |
| EPI_ISL_3096633 | Berkshire and Surrey Pathology Services Lighthouse Laboratory | Wellcome Sanger Institute for the COVID-19 Genomics UK (COG-UK) Consortium | Berkshire and Surrey Pathology Services Lighthouse Laboratory and Alex Alderton, Roberto Amato, Jeffrey Barrett, Sonia Goncalves, Ewan Harrison, David K. Jackson, Ian Johnston, Dominic Kwiatkowski, Cordelia Langford, John Sillitoe on behalf of the Wellcome Sanger Institute COVID-19 Surveillance Team |
| EPI_ISL_3096634 | Lighthouse Lab in Alderley Park | Wellcome Sanger Institute for the COVID-19 Genomics UK (COG-UK) Consortium | Jacquelyn Wynn, Mairead Hyland, The Lighthouse Lab in Alderley Park and Alex Alderton, Roberto Amato, Jeffrey Barrett, Sonia Goncalves, Ewan Harrison, David K. Jackson, Ian Johnston, Dominic Kwiatkowski, Cordelia Langford, John Sillitoe on behalf of the Wellcome Sanger Institute COVID-19 Surveillance Team |
| EPI_ISL_3096635, EPI_ISL_3096636 | Berkshire and Surrey Pathology Services Lighthouse Laboratory | Wellcome Sanger Institute for the COVID-19 Genomics UK (COG-UK) Consortium | Berkshire and Surrey Pathology Services Lighthouse Laboratory and Alex Alderton, Roberto Amato, Jeffrey Barrett, Sonia Goncalves, Ewan Harrison, David K. Jackson, Ian Johnston, Dominic Kwiatkowski, Cordelia Langford, John Sillitoe on behalf of the Wellcome Sanger Institute COVID-19 Surveillance Team |
| EPI_ISL_3096637 | Lighthouse Lab in Milton Keynes | Wellcome Sanger Institute for the COVID-19 Genomics UK (COG-UK) Consortium | The Lighthouse Lab in Milton Keynes and Alex Alderton, Roberto Amato, Jeffrey Barrett, Sonia Goncalves, Ewan Harrison, David K. Jackson, Ian Johnston, Dominic Kwiatkowski, Cordelia Langford, John Sillitoe on behalf of the Wellcome Sanger Institute COVID-19 Surveillance Team |
| EPI_ISL_3096638 | Lighthouse Lab in Alderley Park | Wellcome Sanger Institute for the COVID-19 Genomics UK (COG-UK) Consortium | Jacquelyn Wynn, Mairead Hyland, The Lighthouse Lab in Alderley Park and Alex Alderton, Roberto Amato, Jeffrey Barrett, Sonia Goncalves, Ewan Harrison, David K. Jackson, Ian Johnston, Dominic Kwiatkowski, Cordelia Langford, John Sillitoe on behalf of the Wellcome Sanger Institute COVID-19 Surveillance Team |
| EPI_ISL_3096639 | Berkshire and Surrey Pathology Services Lighthouse Laboratory | Wellcome Sanger Institute for the COVID-19 Genomics UK (COG-UK) Consortium | Berkshire and Surrey Pathology Services Lighthouse Laboratory and Alex Alderton, Roberto Amato, Jeffrey Barrett, Sonia Goncalves, Ewan Harrison, David K. Jackson, Ian Johnston, Dominic Kwiatkowski, Cordelia Langford, John Sillitoe on behalf of the Wellcome Sanger Institute COVID-19 Surveillance Team |
| EPI_ISL_3096640, EPI_ISL_3096641 | Lighthouse Lab in Milton Keynes | Wellcome Sanger Institute for the COVID-19 Genomics UK (COG-UK) Consortium | The Lighthouse Lab in Milton Keynes and Alex Alderton, Roberto Amato, Jeffrey Barrett, Sonia Goncalves, Ewan Harrison, David K. Jackson, Ian Johnston, Dominic Kwiatkowski, Cordelia Langford, John Sillitoe on behalf of the Wellcome Sanger Institute COVID-19 Surveillance Team |
| EPI_ISL_3096642, EPI_ISL_3096643 | Lighthouse Lab in Alderley Park | Wellcome Sanger Institute for the COVID-19 Genomics UK (COG-UK) Consortium | Jacquelyn Wynn, Mairead Hyland, The Lighthouse Lab in Alderley Park and Alex Alderton, Roberto Amato, Jeffrey Barrett, Sonia Goncalves, Ewan Harrison, David K. Jackson, Ian Johnston, Dominic Kwiatkowski, Cordelia Langford, John Sillitoe on behalf of the Wellcome Sanger Institute COVID-19 Surveillance Team |
| EPI_ISL_3096644 | Lighthouse Lab in Milton Keynes | Wellcome Sanger Institute for the COVID-19 Genomics UK (COG-UK) Consortium | The Lighthouse Lab in Milton Keynes and Alex Alderton, Roberto Amato, Jeffrey Barrett, Sonia Goncalves, Ewan Harrison, David K. Jackson, Ian Johnston, Dominic Kwiatkowski, Cordelia Langford, John Sillitoe on behalf of the Wellcome Sanger Institute COVID-19 Surveillance Team |
| EPI_ISL_3096645 | Berkshire and Surrey Pathology Services Lighthouse Laboratory | Wellcome Sanger Institute for the COVID-19 Genomics UK (COG-UK) Consortium | Berkshire and Surrey Pathology Services Lighthouse Laboratory and Alex Alderton, Roberto Amato, Jeffrey Barrett, Sonia Goncalves, Ewan Harrison, David K. Jackson, Ian Johnston, Dominic Kwiatkowski, Cordelia Langford, John Sillitoe on behalf of the Wellcome Sanger Institute COVID-19 Surveillance Team |
| EPI_ISL_3096646 | Lighthouse Lab in Milton Keynes | Wellcome Sanger Institute for the COVID-19 Genomics UK (COG-UK) Consortium | The Lighthouse Lab in Milton Keynes and Alex Alderton, Roberto Amato, Jeffrey Barrett, Sonia Goncalves, Ewan Harrison, David K. Jackson, Ian Johnston, Dominic Kwiatkowski, Cordelia Langford, John Sillitoe on behalf of the Wellcome Sanger Institute COVID-19 Surveillance Team |
| EPI_ISL_3096647 | Berkshire and Surrey Pathology Services Lighthouse Laboratory | Wellcome Sanger Institute for the COVID-19 Genomics UK (COG-UK) Consortium | Berkshire and Surrey Pathology Services Lighthouse Laboratory and Alex Alderton, Roberto Amato, Jeffrey Barrett, Sonia Goncalves, Ewan Harrison, David K. Jackson, Ian Johnston, Dominic Kwiatkowski, Cordelia Langford, John Sillitoe on behalf of the Wellcome Sanger Institute COVID-19 Surveillance Team |
| EPI_ISL_3096648 | Lighthouse Lab in Milton Keynes | Wellcome Sanger Institute for the COVID-19 Genomics UK (COG-UK) Consortium | The Lighthouse Lab in Milton Keynes and Alex Alderton, Roberto Amato, Jeffrey Barrett, Sonia Goncalves, Ewan Harrison, David K. Jackson, Ian Johnston, Dominic Kwiatkowski, Cordelia Langford, John Sillitoe on behalf of the Wellcome Sanger Institute COVID-19 Surveillance Team |
| EPI_ISL_3096649 | Lighthouse Lab in Alderley Park | Wellcome Sanger Institute for the COVID-19 Genomics UK (COG-UK) Consortium | Jacquelyn Wynn, Mairead Hyland, The Lighthouse Lab in Alderley Park and Alex Alderton, Roberto Amato, Jeffrey Barrett, Sonia Goncalves, Ewan Harrison, David K. Jackson, Ian Johnston, Dominic Kwiatkowski, Cordelia Langford, John Sillitoe on behalf of the Wellcome Sanger Institute COVID-19 Surveillance Team |
| EPI_ISL_3096650 | Berkshire and Surrey Pathology Services Lighthouse Laboratory | Wellcome Sanger Institute for the COVID-19 Genomics UK (COG-UK) Consortium | Berkshire and Surrey Pathology Services Lighthouse Laboratory and Alex Alderton, Roberto Amato, Jeffrey Barrett, Sonia Goncalves, Ewan Harrison, David K. Jackson, Ian Johnston, Dominic Kwiatkowski, Cordelia Langford, John Sillitoe on behalf of the Wellcome Sanger Institute COVID-19 Surveillance Team |
| EPI_ISL_3096651 | Lighthouse Lab in Milton Keynes | Wellcome Sanger Institute for the COVID-19 Genomics UK (COG-UK) Consortium | The Lighthouse Lab in Milton Keynes and Alex Alderton, Roberto Amato, Jeffrey Barrett, Sonia Goncalves, Ewan Harrison, David K. Jackson, Ian Johnston, Dominic Kwiatkowski, Cordelia Langford, John Sillitoe on behalf of the Wellcome Sanger Institute COVID-19 Surveillance Team |
| EPI_ISL_3096652, EPI_ISL_3096653 | Berkshire and Surrey Pathology Services Lighthouse Laboratory | Wellcome Sanger Institute for the COVID-19 Genomics UK (COG-UK) Consortium | Berkshire and Surrey Pathology Services Lighthouse Laboratory and Alex Alderton, Roberto Amato, Jeffrey Barrett, Sonia Goncalves, Ewan Harrison, David K. Jackson, Ian Johnston, Dominic Kwiatkowski, Cordelia Langford, John Sillitoe on behalf of the Wellcome Sanger Institute COVID-19 Surveillance Team |
| EPI_ISL_3096654 | Lighthouse Lab in Alderley Park | Wellcome Sanger Institute for the COVID-19 Genomics UK (COG-UK) Consortium | Jacquelyn Wynn, Mairead Hyland, The Lighthouse Lab in Alderley Park and Alex Alderton, Roberto Amato, Jeffrey Barrett, Sonia Goncalves, Ewan Harrison, David K. Jackson, Ian Johnston, Dominic Kwiatkowski, Cordelia Langford, John Sillitoe on behalf of the Wellcome Sanger Institute COVID-19 Surveillance Team |
| EPI_ISL_3096655 | Lighthouse Lab in Milton Keynes | Wellcome Sanger Institute for the COVID-19 Genomics UK (COG-UK) Consortium | The Lighthouse Lab in Milton Keynes and Alex Alderton, Roberto Amato, Jeffrey Barrett, Sonia Goncalves, Ewan Harrison, David K. Jackson, Ian Johnston, Dominic Kwiatkowski, Cordelia Langford, John Sillitoe on behalf of the Wellcome Sanger Institute COVID-19 Surveillance Team |
| EPI_ISL_3096656 | Lighthouse Lab in Alderley Park | Wellcome Sanger Institute for the COVID-19 Genomics UK (COG-UK) Consortium | Jacquelyn Wynn, Mairead Hyland, The Lighthouse Lab in Alderley Park and Alex Alderton, Roberto Amato, Jeffrey Barrett, Sonia Goncalves, Ewan Harrison, David K. Jackson, Ian Johnston, Dominic Kwiatkowski, Cordelia Langford, John Sillitoe on behalf of the Wellcome Sanger Institute COVID-19 Surveillance Team |
| EPI_ISL_3096657, EPI_ISL_3096658, EPI_ISL_3096659 | Lighthouse Lab in Milton Keynes | Wellcome Sanger Institute for the COVID-19 Genomics UK (COG-UK) Consortium | The Lighthouse Lab in Milton Keynes and Alex Alderton, Roberto Amato, Jeffrey Barrett, Sonia Goncalves, Ewan Harrison, David K. Jackson, Ian Johnston, Dominic Kwiatkowski, Cordelia Langford, John Sillitoe on behalf of the Wellcome Sanger Institute COVID-19 Surveillance Team |
| EPI_ISL_3096660 | Berkshire and Surrey Pathology Services Lighthouse Laboratory | Wellcome Sanger Institute for the COVID-19 Genomics UK (COG-UK) Consortium | Berkshire and Surrey Pathology Services Lighthouse Laboratory and Alex Alderton, Roberto Amato, Jeffrey Barrett, Sonia Goncalves, Ewan Harrison, David K. Jackson, Ian Johnston, Dominic Kwiatkowski, Cordelia Langford, John Sillitoe on behalf of the Wellcome Sanger Institute COVID-19 Surveillance Team |
| EPI_ISL_3096661 | Lighthouse Lab in Alderley Park | Wellcome Sanger Institute for the COVID-19 Genomics UK (COG-UK) Consortium | Jacquelyn Wynn, Mairead Hyland, The Lighthouse Lab in Alderley Park and Alex Alderton, Roberto Amato, Jeffrey Barrett, Sonia Goncalves, Ewan Harrison, David K. Jackson, Ian Johnston, Dominic Kwiatkowski, Cordelia Langford, John Sillitoe on behalf of the Wellcome Sanger Institute COVID-19 Surveillance Team |
| EPI_ISL_3096662, EPI_ISL_3096663, EPI_ISL_3096664 | Lighthouse Lab in Milton Keynes | Wellcome Sanger Institute for the COVID-19 Genomics UK (COG-UK) Consortium | The Lighthouse Lab in Milton Keynes and Alex Alderton, Roberto Amato, Jeffrey Barrett, Sonia Goncalves, Ewan Harrison, David K. Jackson, Ian Johnston, Dominic Kwiatkowski, Cordelia Langford, John Sillitoe on behalf of the Wellcome Sanger Institute COVID-19 Surveillance Team |
| EPI_ISL_3096665 | Berkshire and Surrey Pathology Services Lighthouse Laboratory | Wellcome Sanger Institute for the COVID-19 Genomics UK (COG-UK) Consortium | Berkshire and Surrey Pathology Services Lighthouse Laboratory and Alex Alderton, Roberto Amato, Jeffrey Barrett, Sonia Goncalves, Ewan Harrison, David K. Jackson, Ian Johnston, Dominic Kwiatkowski, Cordelia Langford, John Sillitoe on behalf of the Wellcome Sanger Institute COVID-19 Surveillance Team |
| EPI_ISL_3096666, EPI_ISL_3096667, EPI_ISL_3096668 | Lighthouse Lab in Milton Keynes | Wellcome Sanger Institute for the COVID-19 Genomics UK (COG-UK) Consortium | The Lighthouse Lab in Milton Keynes and Alex Alderton, Roberto Amato, Jeffrey Barrett, Sonia Goncalves, Ewan Harrison, David K. Jackson, Ian Johnston, Dominic Kwiatkowski, Cordelia Langford, John Sillitoe on behalf of the Wellcome Sanger Institute COVID-19 Surveillance Team |
| EPI_ISL_3096669 | Berkshire and Surrey Pathology Services Lighthouse Laboratory | Wellcome Sanger Institute for the COVID-19 Genomics UK (COG-UK) Consortium | Berkshire and Surrey Pathology Services Lighthouse Laboratory and Alex Alderton, Roberto Amato, Jeffrey Barrett, Sonia Goncalves, Ewan Harrison, David K. Jackson, Ian Johnston, Dominic Kwiatkowski, Cordelia Langford, John Sillitoe on behalf of the Wellcome Sanger Institute COVID-19 Surveillance Team |
| EPI_ISL_3096670, EPI_ISL_3096671, | Lighthouse Lab in Milton Keynes | Wellcome Sanger Institute for the COVID-19 Genomics UK (COG-UK) Consortium | The Lighthouse Lab in Milton Keynes and Alex Alderton, Roberto Amato, Jeffrey Barrett, Sonia Goncalves, Ewan Harrison, David K. Jackson, Ian Johnston, Dominic Kwiatkowski, Cordelia Langford, John Sillitoe on behalf of the Wellcome Sanger Institute COVID-19 Surveillance Team |

| | | | |
|---|---|---|---|
| EPI_ISL_3096672, EPI_ISL_3096673 | | | |
| EPI_ISL_3096674 | Berkshire and Surrey Pathology Services Lighthouse Laboratory | Wellcome Sanger Institute for the COVID-19 Genomics UK (COG-UK) Consortium | Berkshire and Surrey Pathology Services Lighthouse Laboratory and Alex Alderton, Roberto Amato, Jeffrey Barrett, Sonia Goncalves, Ewan Harrison, David K. Jackson, Ian Johnston, Dominic Kwiatkowski, Cordelia Langford, John Sillitoe on behalf of the Wellcome Sanger Institute COVID-19 Surveillance Team |
| EPI_ISL_3096675, EPI_ISL_3096676 | Lighthouse Lab in Milton Keynes | Wellcome Sanger Institute for the COVID-19 Genomics UK (COG-UK) Consortium | The Lighthouse Lab in Milton Keynes and Alex Alderton, Roberto Amato, Jeffrey Barrett, Sonia Goncalves, Ewan Harrison, David K. Jackson, Ian Johnston, Dominic Kwiatkowski, Cordelia Langford, John Sillitoe on behalf of the Wellcome Sanger Institute COVID-19 Surveillance Team |
| EPI_ISL_3096677 | Berkshire and Surrey Pathology Services Lighthouse Laboratory | Wellcome Sanger Institute for the COVID-19 Genomics UK (COG-UK) Consortium | Berkshire and Surrey Pathology Services Lighthouse Laboratory and Alex Alderton, Roberto Amato, Jeffrey Barrett, Sonia Goncalves, Ewan Harrison, David K. Jackson, Ian Johnston, Dominic Kwiatkowski, Cordelia Langford, John Sillitoe on behalf of the Wellcome Sanger Institute COVID-19 Surveillance Team |
| EPI_ISL_3096678 | Lighthouse Lab in Milton Keynes | Wellcome Sanger Institute for the COVID-19 Genomics UK (COG-UK) Consortium | The Lighthouse Lab in Milton Keynes and Alex Alderton, Roberto Amato, Jeffrey Barrett, Sonia Goncalves, Ewan Harrison, David K. Jackson, Ian Johnston, Dominic Kwiatkowski, Cordelia Langford, John Sillitoe on behalf of the Wellcome Sanger Institute COVID-19 Surveillance Team |
| EPI_ISL_3096679 | Berkshire and Surrey Pathology Services Lighthouse Laboratory | Wellcome Sanger Institute for the COVID-19 Genomics UK (COG-UK) Consortium | Berkshire and Surrey Pathology Services Lighthouse Laboratory and Alex Alderton, Roberto Amato, Jeffrey Barrett, Sonia Goncalves, Ewan Harrison, David K. Jackson, Ian Johnston, Dominic Kwiatkowski, Cordelia Langford, John Sillitoe on behalf of the Wellcome Sanger Institute COVID-19 Surveillance Team |
| EPI_ISL_3096680 | Lighthouse Lab in Milton Keynes | Wellcome Sanger Institute for the COVID-19 Genomics UK (COG-UK) Consortium | The Lighthouse Lab in Milton Keynes and Alex Alderton, Roberto Amato, Jeffrey Barrett, Sonia Goncalves, Ewan Harrison, David K. Jackson, Ian Johnston, Dominic Kwiatkowski, Cordelia Langford, John Sillitoe on behalf of the Wellcome Sanger Institute COVID-19 Surveillance Team |
| EPI_ISL_3096681, EPI_ISL_3096682 | Lighthouse Lab in Alderley Park | Wellcome Sanger Institute for the COVID-19 Genomics UK (COG-UK) Consortium | Jacquelyn Wynn, Mairead Hyland, The Lighthouse Lab in Alderley Park and Alex Alderton, Roberto Amato, Jeffrey Barrett, Sonia Goncalves, Ewan Harrison, David K. Jackson, Ian Johnston, Dominic Kwiatkowski, Cordelia Langford, John Sillitoe on behalf of the Wellcome Sanger Institute COVID-19 Surveillance Team |
| EPI_ISL_3096683 | Lighthouse Lab in Milton Keynes | Wellcome Sanger Institute for the COVID-19 Genomics UK (COG-UK) Consortium | The Lighthouse Lab in Milton Keynes and Alex Alderton, Roberto Amato, Jeffrey Barrett, Sonia Goncalves, Ewan Harrison, David K. Jackson, Ian Johnston, Dominic Kwiatkowski, Cordelia Langford, John Sillitoe on behalf of the Wellcome Sanger Institute COVID-19 Surveillance Team |
| EPI_ISL_3096684, EPI_ISL_3096686, EPI_ISL_3096687 | Berkshire and Surrey Pathology Services Lighthouse Laboratory | Wellcome Sanger Institute for the COVID-19 Genomics UK (COG-UK) Consortium | Berkshire and Surrey Pathology Services Lighthouse Laboratory and Alex Alderton, Roberto Amato, Jeffrey Barrett, Sonia Goncalves, Ewan Harrison, David K. Jackson, Ian Johnston, Dominic Kwiatkowski, Cordelia Langford, John Sillitoe on behalf of the Wellcome Sanger Institute COVID-19 Surveillance Team |
| EPI_ISL_3096688, EPI_ISL_3096689 | Lighthouse Lab in Alderley Park | Wellcome Sanger Institute for the COVID-19 Genomics UK (COG-UK) Consortium | Jacquelyn Wynn, Mairead Hyland, The Lighthouse Lab in Alderley Park and Alex Alderton, Roberto Amato, Jeffrey Barrett, Sonia Goncalves, Ewan Harrison, David K. Jackson, Ian Johnston, Dominic Kwiatkowski, Cordelia Langford, John Sillitoe on behalf of the Wellcome Sanger Institute COVID-19 Surveillance Team |
| EPI_ISL_3096690, EPI_ISL_3096691, EPI_ISL_3096692, EPI_ISL_3096693, EPI_ISL_3096694, EPI_ISL_3096695 | Lighthouse Lab in Milton Keynes | Wellcome Sanger Institute for the COVID-19 Genomics UK (COG-UK) Consortium | The Lighthouse Lab in Milton Keynes and Alex Alderton, Roberto Amato, Jeffrey Barrett, Sonia Goncalves, Ewan Harrison, David K. Jackson, Ian Johnston, Dominic Kwiatkowski, Cordelia Langford, John Sillitoe on behalf of the Wellcome Sanger Institute COVID-19 Surveillance Team |
| EPI_ISL_3096696 | Berkshire and Surrey Pathology Services Lighthouse Laboratory | Wellcome Sanger Institute for the COVID-19 Genomics UK (COG-UK) Consortium | Berkshire and Surrey Pathology Services Lighthouse Laboratory and Alex Alderton, Roberto Amato, Jeffrey Barrett, Sonia Goncalves, Ewan Harrison, David K. Jackson, Ian Johnston, Dominic Kwiatkowski, Cordelia Langford, John Sillitoe on behalf of the Wellcome Sanger Institute COVID-19 Surveillance Team |
| EPI_ISL_3096697, EPI_ISL_3096698 | Lighthouse Lab in Milton Keynes | Wellcome Sanger Institute for the COVID-19 Genomics UK (COG-UK) Consortium | The Lighthouse Lab in Milton Keynes and Alex Alderton, Roberto Amato, Jeffrey Barrett, Sonia Goncalves, Ewan Harrison, David K. Jackson, Ian Johnston, Dominic Kwiatkowski, Cordelia Langford, John Sillitoe on behalf of the Wellcome Sanger Institute COVID-19 Surveillance Team |
| EPI_ISL_3096699 | Berkshire and Surrey Pathology Services Lighthouse Laboratory | Wellcome Sanger Institute for the COVID-19 Genomics UK (COG-UK) Consortium | Berkshire and Surrey Pathology Services Lighthouse Laboratory and Alex Alderton, Roberto Amato, Jeffrey Barrett, Sonia Goncalves, Ewan Harrison, David K. Jackson, Ian Johnston, Dominic Kwiatkowski, Cordelia Langford, John Sillitoe on behalf of the Wellcome Sanger Institute COVID-19 Surveillance Team |
| EPI_ISL_3096700, EPI_ISL_3096701, EPI_ISL_3096702, EPI_ISL_3096703, EPI_ISL_3096704, EPI_ISL_3096705, EPI_ISL_3096706 | Lighthouse Lab in Milton Keynes | Wellcome Sanger Institute for the COVID-19 Genomics UK (COG-UK) Consortium | The Lighthouse Lab in Milton Keynes and Alex Alderton, Roberto Amato, Jeffrey Barrett, Sonia Goncalves, Ewan Harrison, David K. Jackson, Ian Johnston, Dominic Kwiatkowski, Cordelia Langford, John Sillitoe on behalf of the Wellcome Sanger Institute COVID-19 Surveillance Team |
| EPI_ISL_3096707 | Lighthouse Lab in Alderley Park | Wellcome Sanger Institute for the COVID-19 Genomics UK (COG-UK) Consortium | Jacquelyn Wynn, Mairead Hyland, The Lighthouse Lab in Alderley Park and Alex Alderton, Roberto Amato, Jeffrey Barrett, Sonia Goncalves, Ewan Harrison, David K. Jackson, Ian Johnston, Dominic Kwiatkowski, Cordelia Langford, John Sillitoe on behalf of the Wellcome Sanger Institute COVID-19 Surveillance Team |
| EPI_ISL_3096708, EPI_ISL_3096709, EPI_ISL_3096710, EPI_ISL_3096711 | Lighthouse Lab in Milton Keynes | Wellcome Sanger Institute for the COVID-19 Genomics UK (COG-UK) Consortium | The Lighthouse Lab in Milton Keynes and Alex Alderton, Roberto Amato, Jeffrey Barrett, Sonia Goncalves, Ewan Harrison, David K. Jackson, Ian Johnston, Dominic Kwiatkowski, Cordelia Langford, John Sillitoe on behalf of the Wellcome Sanger Institute COVID-19 Surveillance Team |
| EPI_ISL_3096712 | Berkshire and Surrey Pathology Services Lighthouse Laboratory | Wellcome Sanger Institute for the COVID-19 Genomics UK (COG-UK) Consortium | Berkshire and Surrey Pathology Services Lighthouse Laboratory and Alex Alderton, Roberto Amato, Jeffrey Barrett, Sonia Goncalves, Ewan Harrison, David K. Jackson, Ian Johnston, Dominic Kwiatkowski, Cordelia Langford, John Sillitoe on behalf of the Wellcome Sanger Institute COVID-19 Surveillance Team |
| EPI_ISL_3096713 | Lighthouse Lab in Milton Keynes | Wellcome Sanger Institute for the COVID-19 Genomics UK (COG-UK) Consortium | The Lighthouse Lab in Milton Keynes and Alex Alderton, Roberto Amato, Jeffrey Barrett, Sonia Goncalves, Ewan Harrison, David K. Jackson, Ian Johnston, Dominic Kwiatkowski, Cordelia Langford, John Sillitoe on behalf of the Wellcome Sanger Institute COVID-19 Surveillance Team |
| EPI_ISL_3096714 | Lighthouse Lab in Alderley Park | Wellcome Sanger Institute for the COVID-19 Genomics UK (COG-UK) Consortium | Jacquelyn Wynn, Mairead Hyland, The Lighthouse Lab in Alderley Park and Alex Alderton, Roberto Amato, Jeffrey Barrett, Sonia Goncalves, Ewan Harrison, David K. Jackson, Ian Johnston, Dominic Kwiatkowski, Cordelia Langford, John Sillitoe on behalf of the Wellcome Sanger Institute COVID-19 Surveillance Team |
| EPI_ISL_3096715, EPI_ISL_3096716, EPI_ISL_3096717 | Lighthouse Lab in Milton Keynes | Wellcome Sanger Institute for the COVID-19 Genomics UK (COG-UK) Consortium | The Lighthouse Lab in Milton Keynes and Alex Alderton, Roberto Amato, Jeffrey Barrett, Sonia Goncalves, Ewan Harrison, David K. Jackson, Ian Johnston, Dominic Kwiatkowski, Cordelia Langford, John Sillitoe on behalf of the Wellcome Sanger Institute COVID-19 Surveillance Team |
| EPI_ISL_3096718 | Berkshire and Surrey Pathology Services Lighthouse Laboratory | Wellcome Sanger Institute for the COVID-19 Genomics UK (COG-UK) Consortium | Berkshire and Surrey Pathology Services Lighthouse Laboratory and Alex Alderton, Roberto Amato, Jeffrey Barrett, Sonia Goncalves, Ewan Harrison, David K. Jackson, Ian Johnston, Dominic Kwiatkowski, Cordelia Langford, John Sillitoe on behalf of the Wellcome Sanger Institute COVID-19 Surveillance Team |
| EPI_ISL_3096719 | Lighthouse Lab in Milton Keynes | Wellcome Sanger Institute for the COVID-19 Genomics UK (COG-UK) Consortium | The Lighthouse Lab in Milton Keynes and Alex Alderton, Roberto Amato, Jeffrey Barrett, Sonia Goncalves, Ewan Harrison, David K. Jackson, Ian Johnston, Dominic Kwiatkowski, Cordelia Langford, John Sillitoe on behalf of the Wellcome Sanger Institute COVID-19 Surveillance Team |
| EPI_ISL_3096720 | Lighthouse Lab in Alderley Park | Wellcome Sanger Institute for the COVID-19 Genomics UK (COG-UK) Consortium | Jacquelyn Wynn, Mairead Hyland, The Lighthouse Lab in Alderley Park and Alex Alderton, Roberto Amato, Jeffrey Barrett, Sonia Goncalves, Ewan Harrison, David K. Jackson, Ian Johnston, Dominic Kwiatkowski, Cordelia Langford, John Sillitoe on behalf of the Wellcome Sanger Institute COVID-19 Surveillance Team |
| EPI_ISL_3096721, EPI_ISL_3096722, EPI_ISL_3096723 | Lighthouse Lab in Milton Keynes | Wellcome Sanger Institute for the COVID-19 Genomics UK (COG-UK) Consortium | The Lighthouse Lab in Milton Keynes and Alex Alderton, Roberto Amato, Jeffrey Barrett, Sonia Goncalves, Ewan Harrison, David K. Jackson, Ian Johnston, Dominic Kwiatkowski, Cordelia Langford, John Sillitoe on behalf of the Wellcome Sanger Institute COVID-19 Surveillance Team |
| EPI_ISL_3096724 | Lighthouse Lab in Alderley Park | Wellcome Sanger Institute for the COVID-19 Genomics UK (COG-UK) Consortium | Jacquelyn Wynn, Mairead Hyland, The Lighthouse Lab in Alderley Park and Alex Alderton, Roberto Amato, Jeffrey Barrett, Sonia Goncalves, Ewan Harrison, David K. Jackson, Ian Johnston, Dominic Kwiatkowski, Cordelia Langford, John Sillitoe on behalf of the Wellcome Sanger Institute COVID-19 Surveillance Team |
| EPI_ISL_3096725 | Lighthouse Lab in Milton Keynes | Wellcome Sanger Institute for the COVID-19 Genomics UK (COG-UK) Consortium | The Lighthouse Lab in Milton Keynes and Alex Alderton, Roberto Amato, Jeffrey Barrett, Sonia Goncalves, Ewan Harrison, David K. Jackson, Ian Johnston, Dominic Kwiatkowski, Cordelia Langford, John Sillitoe on behalf of the Wellcome Sanger Institute COVID-19 Surveillance Team |
| EPI_ISL_3096726, EPI_ISL_3096727, EPI_ISL_3096728 | Lighthouse Lab in Alderley Park | Wellcome Sanger Institute for the COVID-19 Genomics UK (COG-UK) Consortium | Jacquelyn Wynn, Mairead Hyland, The Lighthouse Lab in Alderley Park and Alex Alderton, Roberto Amato, Jeffrey Barrett, Sonia Goncalves, Ewan Harrison, David K. Jackson, Ian Johnston, Dominic Kwiatkowski, Cordelia Langford, John Sillitoe on behalf of the Wellcome Sanger Institute COVID-19 Surveillance Team |
| EPI_ISL_3096729, EPI_ISL_3096730, EPI_ISL_3096731, EPI_ISL_3096732 | Lighthouse Lab in Milton Keynes | Wellcome Sanger Institute for the COVID-19 Genomics UK (COG-UK) Consortium | The Lighthouse Lab in Milton Keynes and Alex Alderton, Roberto Amato, Jeffrey Barrett, Sonia Goncalves, Ewan Harrison, David K. Jackson, Ian Johnston, Dominic Kwiatkowski, Cordelia Langford, John Sillitoe on behalf of the Wellcome Sanger Institute COVID-19 Surveillance Team |
| EPI_ISL_3096733 | Lighthouse Lab in Alderley Park | Wellcome Sanger Institute for the COVID-19 Genomics UK (COG-UK) Consortium | Jacquelyn Wynn, Mairead Hyland, The Lighthouse Lab in Alderley Park and Alex Alderton, Roberto Amato, Jeffrey Barrett, Sonia Goncalves, Ewan Harrison, David K. Jackson, Ian Johnston, Dominic Kwiatkowski, Cordelia Langford, John Sillitoe on behalf of the Wellcome Sanger Institute COVID-19 Surveillance Team |
| EPI_ISL_3096734, EPI_ISL_3096735 | Lighthouse Lab in Milton Keynes | Wellcome Sanger Institute for the COVID-19 Genomics UK (COG-UK) Consortium | The Lighthouse Lab in Milton Keynes and Alex Alderton, Roberto Amato, Jeffrey Barrett, Sonia Goncalves, Ewan Harrison, David K. Jackson, Ian Johnston, Dominic Kwiatkowski, Cordelia Langford, John Sillitoe on behalf of the Wellcome Sanger Institute COVID-19 Surveillance Team |
| EPI_ISL_3096736 | Berkshire and Surrey Pathology Services Lighthouse Laboratory | Wellcome Sanger Institute for the COVID-19 Genomics UK (COG-UK) Consortium | Berkshire and Surrey Pathology Services Lighthouse Laboratory and Alex Alderton, Roberto Amato, Jeffrey Barrett, Sonia Goncalves, Ewan Harrison, David K. Jackson, Ian Johnston, Dominic Kwiatkowski, Cordelia Langford, John Sillitoe on behalf of the Wellcome Sanger Institute COVID-19 Surveillance Team |
| EPI_ISL_3096737 | Lighthouse Lab in Milton Keynes | Wellcome Sanger Institute for the COVID-19 Genomics UK (COG-UK) Consortium | The Lighthouse Lab in Milton Keynes and Alex Alderton, Roberto Amato, Jeffrey Barrett, Sonia Goncalves, Ewan Harrison, David K. Jackson, Ian Johnston, Dominic Kwiatkowski, Cordelia Langford, John Sillitoe on behalf of the Wellcome Sanger Institute COVID-19 Surveillance Team |
| EPI_ISL_3096738 | Lighthouse Lab in Alderley Park | Wellcome Sanger Institute for the COVID-19 Genomics UK (COG-UK) Consortium | Jacquelyn Wynn, Mairead Hyland, The Lighthouse Lab in Alderley Park and Alex Alderton, Roberto Amato, Jeffrey Barrett, Sonia Goncalves, Ewan Harrison, David K. Jackson, Ian Johnston, Dominic Kwiatkowski, Cordelia Langford, John Sillitoe on behalf of the Wellcome Sanger Institute COVID-19 Surveillance Team |
| EPI_ISL_3096739, EPI_ISL_3096740 | Berkshire and Surrey Pathology Services Lighthouse Laboratory | Wellcome Sanger Institute for the COVID-19 Genomics UK (COG-UK) Consortium | Berkshire and Surrey Pathology Services Lighthouse Laboratory and Alex Alderton, Roberto Amato, Jeffrey Barrett, Sonia Goncalves, Ewan Harrison, David K. Jackson, Ian Johnston, Dominic Kwiatkowski, Cordelia Langford, John Sillitoe on behalf of the Wellcome Sanger Institute COVID-19 Surveillance Team |
| EPI_ISL_3096741 | Lighthouse Lab in Alderley Park | Wellcome Sanger Institute for the COVID-19 Genomics UK (COG-UK) Consortium | Jacquelyn Wynn, Mairead Hyland, The Lighthouse Lab in Alderley Park and Alex Alderton, Roberto Amato, Jeffrey Barrett, Sonia Goncalves, Ewan Harrison, David K. Jackson, Ian Johnston, Dominic Kwiatkowski, Cordelia Langford, John Sillitoe on behalf of the Wellcome Sanger Institute COVID-19 Surveillance Team |
| EPI_ISL_3096742 | Lighthouse Lab in Milton Keynes | Wellcome Sanger Institute for the COVID-19 Genomics UK (COG-UK) Consortium | The Lighthouse Lab in Milton Keynes and Alex Alderton, Roberto Amato, Jeffrey Barrett, Sonia Goncalves, Ewan Harrison, David K. Jackson, Ian Johnston, Dominic Kwiatkowski, Cordelia Langford, John Sillitoe on behalf of the Wellcome Sanger Institute COVID-19 Surveillance Team |

| | | | |
|---|---|---|---|
| EPI_ISL_3096743 | Lighthouse Lab in Alderley Park | Wellcome Sanger Institute for the COVID-19 Genomics UK (COG-UK) Consortium | Jacquelyn Wynn, Mairead Hyland, The Lighthouse Lab in Alderley Park and Alex Alderton, Roberto Amato, Jeffrey Barrett, Sonia Goncalves, Ewan Harrison, David K. Jackson, Ian Johnston, Dominic Kwiatkowski, Cordelia Langford, John Sillitoe on behalf of the Wellcome Sanger Institute COVID-19 Surveillance Team |
| EPI_ISL_3096744, EPI_ISL_3096745, EPI_ISL_3096746 | Lighthouse Lab in Milton Keynes | Wellcome Sanger Institute for the COVID-19 Genomics UK (COG-UK) Consortium | The Lighthouse Lab in Milton Keynes and Alex Alderton, Roberto Amato, Jeffrey Barrett, Sonia Goncalves, Ewan Harrison, David K. Jackson, Ian Johnston, Dominic Kwiatkowski, Cordelia Langford, John Sillitoe on behalf of the Wellcome Sanger Institute COVID-19 Surveillance Team |
| EPI_ISL_3096747 | Berkshire and Surrey Pathology Services Lighthouse Laboratory | Wellcome Sanger Institute for the COVID-19 Genomics UK (COG-UK) Consortium | Berkshire and Surrey Pathology Services Lighthouse Laboratory and Alex Alderton, Roberto Amato, Jeffrey Barrett, Sonia Goncalves, Ewan Harrison, David K. Jackson, Ian Johnston, Dominic Kwiatkowski, Cordelia Langford, John Sillitoe on behalf of the Wellcome Sanger Institute COVID-19 Surveillance Team |
| EPI_ISL_3096748 | Lighthouse Lab in Alderley Park | Wellcome Sanger Institute for the COVID-19 Genomics UK (COG-UK) Consortium | Jacquelyn Wynn, Mairead Hyland, The Lighthouse Lab in Alderley Park and Alex Alderton, Roberto Amato, Jeffrey Barrett, Sonia Goncalves, Ewan Harrison, David K. Jackson, Ian Johnston, Dominic Kwiatkowski, Cordelia Langford, John Sillitoe on behalf of the Wellcome Sanger Institute COVID-19 Surveillance Team |
| EPI_ISL_3096749 | Lighthouse Lab in Milton Keynes | Wellcome Sanger Institute for the COVID-19 Genomics UK (COG-UK) Consortium | The Lighthouse Lab in Milton Keynes and Alex Alderton, Roberto Amato, Jeffrey Barrett, Sonia Goncalves, Ewan Harrison, David K. Jackson, Ian Johnston, Dominic Kwiatkowski, Cordelia Langford, John Sillitoe on behalf of the Wellcome Sanger Institute COVID-19 Surveillance Team |
| EPI_ISL_3096750 | Berkshire and Surrey Pathology Services Lighthouse Laboratory | Wellcome Sanger Institute for the COVID-19 Genomics UK (COG-UK) Consortium | Berkshire and Surrey Pathology Services Lighthouse Laboratory and Alex Alderton, Roberto Amato, Jeffrey Barrett, Sonia Goncalves, Ewan Harrison, David K. Jackson, Ian Johnston, Dominic Kwiatkowski, Cordelia Langford, John Sillitoe on behalf of the Wellcome Sanger Institute COVID-19 Surveillance Team |
| EPI_ISL_3096751 | Lighthouse Lab in Milton Keynes | Wellcome Sanger Institute for the COVID-19 Genomics UK (COG-UK) Consortium | The Lighthouse Lab in Milton Keynes and Alex Alderton, Roberto Amato, Jeffrey Barrett, Sonia Goncalves, Ewan Harrison, David K. Jackson, Ian Johnston, Dominic Kwiatkowski, Cordelia Langford, John Sillitoe on behalf of the Wellcome Sanger Institute COVID-19 Surveillance Team |
| EPI_ISL_3096752 | Berkshire and Surrey Pathology Services Lighthouse Laboratory | Wellcome Sanger Institute for the COVID-19 Genomics UK (COG-UK) Consortium | Berkshire and Surrey Pathology Services Lighthouse Laboratory and Alex Alderton, Roberto Amato, Jeffrey Barrett, Sonia Goncalves, Ewan Harrison, David K. Jackson, Ian Johnston, Dominic Kwiatkowski, Cordelia Langford, John Sillitoe on behalf of the Wellcome Sanger Institute COVID-19 Surveillance Team |
| EPI_ISL_3096753 | Lighthouse Lab in Milton Keynes | Wellcome Sanger Institute for the COVID-19 Genomics UK (COG-UK) Consortium | The Lighthouse Lab in Milton Keynes and Alex Alderton, Roberto Amato, Jeffrey Barrett, Sonia Goncalves, Ewan Harrison, David K. Jackson, Ian Johnston, Dominic Kwiatkowski, Cordelia Langford, John Sillitoe on behalf of the Wellcome Sanger Institute COVID-19 Surveillance Team |
| EPI_ISL_3096754 | Berkshire and Surrey Pathology Services Lighthouse Laboratory | Wellcome Sanger Institute for the COVID-19 Genomics UK (COG-UK) Consortium | Berkshire and Surrey Pathology Services Lighthouse Laboratory and Alex Alderton, Roberto Amato, Jeffrey Barrett, Sonia Goncalves, Ewan Harrison, David K. Jackson, Ian Johnston, Dominic Kwiatkowski, Cordelia Langford, John Sillitoe on behalf of the Wellcome Sanger Institute COVID-19 Surveillance Team |
| EPI_ISL_3096755 | Lighthouse Lab in Milton Keynes | Wellcome Sanger Institute for the COVID-19 Genomics UK (COG-UK) Consortium | The Lighthouse Lab in Milton Keynes and Alex Alderton, Roberto Amato, Jeffrey Barrett, Sonia Goncalves, Ewan Harrison, David K. Jackson, Ian Johnston, Dominic Kwiatkowski, Cordelia Langford, John Sillitoe on behalf of the Wellcome Sanger Institute COVID-19 Surveillance Team |
| EPI_ISL_3096756 | Lighthouse Lab in Alderley Park | Wellcome Sanger Institute for the COVID-19 Genomics UK (COG-UK) Consortium | Jacquelyn Wynn, Mairead Hyland, The Lighthouse Lab in Alderley Park and Alex Alderton, Roberto Amato, Jeffrey Barrett, Sonia Goncalves, Ewan Harrison, David K. Jackson, Ian Johnston, Dominic Kwiatkowski, Cordelia Langford, John Sillitoe on behalf of the Wellcome Sanger Institute COVID-19 Surveillance Team |
| EPI_ISL_3096757 | Lighthouse Lab in Milton Keynes | Wellcome Sanger Institute for the COVID-19 Genomics UK (COG-UK) Consortium | The Lighthouse Lab in Milton Keynes and Alex Alderton, Roberto Amato, Jeffrey Barrett, Sonia Goncalves, Ewan Harrison, David K. Jackson, Ian Johnston, Dominic Kwiatkowski, Cordelia Langford, John Sillitoe on behalf of the Wellcome Sanger Institute COVID-19 Surveillance Team |
| EPI_ISL_3096758 | Berkshire and Surrey Pathology Services Lighthouse Laboratory | Wellcome Sanger Institute for the COVID-19 Genomics UK (COG-UK) Consortium | Berkshire and Surrey Pathology Services Lighthouse Laboratory and Alex Alderton, Roberto Amato, Jeffrey Barrett, Sonia Goncalves, Ewan Harrison, David K. Jackson, Ian Johnston, Dominic Kwiatkowski, Cordelia Langford, John Sillitoe on behalf of the Wellcome Sanger Institute COVID-19 Surveillance Team |
| EPI_ISL_3096759 | Lighthouse Lab in Alderley Park | Wellcome Sanger Institute for the COVID-19 Genomics UK (COG-UK) Consortium | Jacquelyn Wynn, Mairead Hyland, The Lighthouse Lab in Alderley Park and Alex Alderton, Roberto Amato, Jeffrey Barrett, Sonia Goncalves, Ewan Harrison, David K. Jackson, Ian Johnston, Dominic Kwiatkowski, Cordelia Langford, John Sillitoe on behalf of the Wellcome Sanger Institute COVID-19 Surveillance Team |
| EPI_ISL_3096760 | Lighthouse Lab in Milton Keynes | Wellcome Sanger Institute for the COVID-19 Genomics UK (COG-UK) Consortium | The Lighthouse Lab in Milton Keynes and Alex Alderton, Roberto Amato, Jeffrey Barrett, Sonia Goncalves, Ewan Harrison, David K. Jackson, Ian Johnston, Dominic Kwiatkowski, Cordelia Langford, John Sillitoe on behalf of the Wellcome Sanger Institute COVID-19 Surveillance Team |
| EPI_ISL_3096761 | Berkshire and Surrey Pathology Services Lighthouse Laboratory | Wellcome Sanger Institute for the COVID-19 Genomics UK (COG-UK) Consortium | Berkshire and Surrey Pathology Services Lighthouse Laboratory and Alex Alderton, Roberto Amato, Jeffrey Barrett, Sonia Goncalves, Ewan Harrison, David K. Jackson, Ian Johnston, Dominic Kwiatkowski, Cordelia Langford, John Sillitoe on behalf of the Wellcome Sanger Institute COVID-19 Surveillance Team |
| EPI_ISL_3096762, EPI_ISL_3096763 | Lighthouse Lab in Alderley Park | Wellcome Sanger Institute for the COVID-19 Genomics UK (COG-UK) Consortium | Jacquelyn Wynn, Mairead Hyland, The Lighthouse Lab in Alderley Park and Alex Alderton, Roberto Amato, Jeffrey Barrett, Sonia Goncalves, Ewan Harrison, David K. Jackson, Ian Johnston, Dominic Kwiatkowski, Cordelia Langford, John Sillitoe on behalf of the Wellcome Sanger Institute COVID-19 Surveillance Team |
| EPI_ISL_3096764, EPI_ISL_3096765, EPI_ISL_3096766 | Lighthouse Lab in Milton Keynes | Wellcome Sanger Institute for the COVID-19 Genomics UK (COG-UK) Consortium | The Lighthouse Lab in Milton Keynes and Alex Alderton, Roberto Amato, Jeffrey Barrett, Sonia Goncalves, Ewan Harrison, David K. Jackson, Ian Johnston, Dominic Kwiatkowski, Cordelia Langford, John Sillitoe on behalf of the Wellcome Sanger Institute COVID-19 Surveillance Team |
| EPI_ISL_3096767 | Berkshire and Surrey Pathology Services Lighthouse Laboratory | Wellcome Sanger Institute for the COVID-19 Genomics UK (COG-UK) Consortium | Berkshire and Surrey Pathology Services Lighthouse Laboratory and Alex Alderton, Roberto Amato, Jeffrey Barrett, Sonia Goncalves, Ewan Harrison, David K. Jackson, Ian Johnston, Dominic Kwiatkowski, Cordelia Langford, John Sillitoe on behalf of the Wellcome Sanger Institute COVID-19 Surveillance Team |
| EPI_ISL_3096768, EPI_ISL_3096769 | Lighthouse Lab in Milton Keynes | Wellcome Sanger Institute for the COVID-19 Genomics UK (COG-UK) Consortium | The Lighthouse Lab in Milton Keynes and Alex Alderton, Roberto Amato, Jeffrey Barrett, Sonia Goncalves, Ewan Harrison, David K. Jackson, Ian Johnston, Dominic Kwiatkowski, Cordelia Langford, John Sillitoe on behalf of the Wellcome Sanger Institute COVID-19 Surveillance Team |
| EPI_ISL_3096770 | Berkshire and Surrey Pathology Services Lighthouse Laboratory | Wellcome Sanger Institute for the COVID-19 Genomics UK (COG-UK) Consortium | Berkshire and Surrey Pathology Services Lighthouse Laboratory and Alex Alderton, Roberto Amato, Jeffrey Barrett, Sonia Goncalves, Ewan Harrison, David K. Jackson, Ian Johnston, Dominic Kwiatkowski, Cordelia Langford, John Sillitoe on behalf of the Wellcome Sanger Institute COVID-19 Surveillance Team |
| EPI_ISL_3096771 | Lighthouse Lab in Milton Keynes | Wellcome Sanger Institute for the COVID-19 Genomics UK (COG-UK) Consortium | The Lighthouse Lab in Milton Keynes and Alex Alderton, Roberto Amato, Jeffrey Barrett, Sonia Goncalves, Ewan Harrison, David K. Jackson, Ian Johnston, Dominic Kwiatkowski, Cordelia Langford, John Sillitoe on behalf of the Wellcome Sanger Institute COVID-19 Surveillance Team |
| EPI_ISL_3096772 | Berkshire and Surrey Pathology Services Lighthouse Laboratory | Wellcome Sanger Institute for the COVID-19 Genomics UK (COG-UK) Consortium | Berkshire and Surrey Pathology Services Lighthouse Laboratory and Alex Alderton, Roberto Amato, Jeffrey Barrett, Sonia Goncalves, Ewan Harrison, David K. Jackson, Ian Johnston, Dominic Kwiatkowski, Cordelia Langford, John Sillitoe on behalf of the Wellcome Sanger Institute COVID-19 Surveillance Team |
| EPI_ISL_3096773, EPI_ISL_3096774 | Lighthouse Lab in Milton Keynes | Wellcome Sanger Institute for the COVID-19 Genomics UK (COG-UK) Consortium | The Lighthouse Lab in Milton Keynes and Alex Alderton, Roberto Amato, Jeffrey Barrett, Sonia Goncalves, Ewan Harrison, David K. Jackson, Ian Johnston, Dominic Kwiatkowski, Cordelia Langford, John Sillitoe on behalf of the Wellcome Sanger Institute COVID-19 Surveillance Team |
| EPI_ISL_3096775 | Lighthouse Lab in Alderley Park | Wellcome Sanger Institute for the COVID-19 Genomics UK (COG-UK) Consortium | Jacquelyn Wynn, Mairead Hyland, The Lighthouse Lab in Alderley Park and Alex Alderton, Roberto Amato, Jeffrey Barrett, Sonia Goncalves, Ewan Harrison, David K. Jackson, Ian Johnston, Dominic Kwiatkowski, Cordelia Langford, John Sillitoe on behalf of the Wellcome Sanger Institute COVID-19 Surveillance Team |
| EPI_ISL_3096776 | Berkshire and Surrey Pathology Services Lighthouse Laboratory | Wellcome Sanger Institute for the COVID-19 Genomics UK (COG-UK) Consortium | Berkshire and Surrey Pathology Services Lighthouse Laboratory and Alex Alderton, Roberto Amato, Jeffrey Barrett, Sonia Goncalves, Ewan Harrison, David K. Jackson, Ian Johnston, Dominic Kwiatkowski, Cordelia Langford, John Sillitoe on behalf of the Wellcome Sanger Institute COVID-19 Surveillance Team |
| EPI_ISL_3096777, EPI_ISL_3096778 | Lighthouse Lab in Milton Keynes | Wellcome Sanger Institute for the COVID-19 Genomics UK (COG-UK) Consortium | The Lighthouse Lab in Milton Keynes and Alex Alderton, Roberto Amato, Jeffrey Barrett, Sonia Goncalves, Ewan Harrison, David K. Jackson, Ian Johnston, Dominic Kwiatkowski, Cordelia Langford, John Sillitoe on behalf of the Wellcome Sanger Institute COVID-19 Surveillance Team |
| EPI_ISL_3096779 | Lighthouse Lab in Alderley Park | Wellcome Sanger Institute for the COVID-19 Genomics UK (COG-UK) Consortium | Jacquelyn Wynn, Mairead Hyland, The Lighthouse Lab in Alderley Park and Alex Alderton, Roberto Amato, Jeffrey Barrett, Sonia Goncalves, Ewan Harrison, David K. Jackson, Ian Johnston, Dominic Kwiatkowski, Cordelia Langford, John Sillitoe on behalf of the Wellcome Sanger Institute COVID-19 Surveillance Team |
| EPI_ISL_3096780, EPI_ISL_3096781, EPI_ISL_3096782, EPI_ISL_3096783 | Lighthouse Lab in Milton Keynes | Wellcome Sanger Institute for the COVID-19 Genomics UK (COG-UK) Consortium | The Lighthouse Lab in Milton Keynes and Alex Alderton, Roberto Amato, Jeffrey Barrett, Sonia Goncalves, Ewan Harrison, David K. Jackson, Ian Johnston, Dominic Kwiatkowski, Cordelia Langford, John Sillitoe on behalf of the Wellcome Sanger Institute COVID-19 Surveillance Team |
| EPI_ISL_3096784 | Lighthouse Lab in Alderley Park | Wellcome Sanger Institute for the COVID-19 Genomics UK (COG-UK) Consortium | Jacquelyn Wynn, Mairead Hyland, The Lighthouse Lab in Alderley Park and Alex Alderton, Roberto Amato, Jeffrey Barrett, Sonia Goncalves, Ewan Harrison, David K. Jackson, Ian Johnston, Dominic Kwiatkowski, Cordelia Langford, John Sillitoe on behalf of the Wellcome Sanger Institute COVID-19 Surveillance Team |
| EPI_ISL_3096785 | Lighthouse Lab in Milton Keynes | Wellcome Sanger Institute for the COVID-19 Genomics UK (COG-UK) Consortium | The Lighthouse Lab in Milton Keynes and Alex Alderton, Roberto Amato, Jeffrey Barrett, Sonia Goncalves, Ewan Harrison, David K. Jackson, Ian Johnston, Dominic Kwiatkowski, Cordelia Langford, John Sillitoe on behalf of the Wellcome Sanger Institute COVID-19 Surveillance Team |
| EPI_ISL_3096786 | Lighthouse Lab in Alderley Park | Wellcome Sanger Institute for the COVID-19 Genomics UK (COG-UK) Consortium | Jacquelyn Wynn, Mairead Hyland, The Lighthouse Lab in Alderley Park and Alex Alderton, Roberto Amato, Jeffrey Barrett, Sonia Goncalves, Ewan Harrison, David K. Jackson, Ian Johnston, Dominic Kwiatkowski, Cordelia Langford, John Sillitoe on behalf of the Wellcome Sanger Institute COVID-19 Surveillance Team |
| EPI_ISL_3096787, EPI_ISL_3096788 | Lighthouse Lab in Milton Keynes | Wellcome Sanger Institute for the COVID-19 Genomics UK (COG-UK) Consortium | The Lighthouse Lab in Milton Keynes and Alex Alderton, Roberto Amato, Jeffrey Barrett, Sonia Goncalves, Ewan Harrison, David K. Jackson, Ian Johnston, Dominic Kwiatkowski, Cordelia Langford, John Sillitoe on behalf of the Wellcome Sanger Institute COVID-19 Surveillance Team |
| EPI_ISL_3096789, EPI_ISL_3096790, EPI_ISL_3096791 | Lighthouse Lab in Alderley Park | Wellcome Sanger Institute for the COVID-19 Genomics UK (COG-UK) Consortium | Jacquelyn Wynn, Mairead Hyland, The Lighthouse Lab in Alderley Park and Alex Alderton, Roberto Amato, Jeffrey Barrett, Sonia Goncalves, Ewan Harrison, David K. Jackson, Ian Johnston, Dominic Kwiatkowski, Cordelia Langford, John Sillitoe on behalf of the Wellcome Sanger Institute COVID-19 Surveillance Team |
| EPI_ISL_3096792, EPI_ISL_3096793 | Lighthouse Lab in Milton Keynes | Wellcome Sanger Institute for the COVID-19 Genomics UK (COG-UK) Consortium | The Lighthouse Lab in Milton Keynes and Alex Alderton, Roberto Amato, Jeffrey Barrett, Sonia Goncalves, Ewan Harrison, David K. Jackson, Ian Johnston, Dominic Kwiatkowski, Cordelia Langford, John Sillitoe on behalf of the Wellcome Sanger Institute COVID-19 Surveillance Team |
| EPI_ISL_3096794 | Lighthouse Lab in Alderley Park | Wellcome Sanger Institute for the COVID-19 Genomics UK (COG-UK) Consortium | Jacquelyn Wynn, Mairead Hyland, The Lighthouse Lab in Alderley Park and Alex Alderton, Roberto Amato, Jeffrey Barrett, Sonia Goncalves, Ewan Harrison, David K. Jackson, Ian Johnston, Dominic Kwiatkowski, Cordelia Langford, John Sillitoe on behalf of the Wellcome Sanger Institute COVID-19 Surveillance Team |
| EPI_ISL_3096795 | Lighthouse Lab in Milton Keynes | Wellcome Sanger Institute for the COVID-19 Genomics UK (COG-UK) Consortium | The Lighthouse Lab in Milton Keynes and Alex Alderton, Roberto Amato, Jeffrey Barrett, Sonia Goncalves, Ewan Harrison, David K. Jackson, Ian Johnston, Dominic Kwiatkowski, Cordelia Langford, John Sillitoe on behalf of the Wellcome Sanger Institute COVID-19 Surveillance Team |
| EPI_ISL_3096796 | Lighthouse Lab in Alderley Park | Wellcome Sanger Institute for the COVID-19 Genomics UK (COG-UK) Consortium | Jacquelyn Wynn, Mairead Hyland, The Lighthouse Lab in Alderley Park and Alex Alderton, Roberto Amato, Jeffrey Barrett, Sonia Goncalves, Ewan Harrison, David K. Jackson, Ian Johnston, Dominic Kwiatkowski, Cordelia Langford, John Sillitoe on behalf of the Wellcome Sanger Institute COVID-19 Surveillance Team |
| EPI_ISL_3096797, EPI_ISL_3096798, EPI_ISL_3096799 | Lighthouse Lab in Milton Keynes | Wellcome Sanger Institute for the COVID-19 Genomics UK (COG-UK) Consortium | The Lighthouse Lab in Milton Keynes and Alex Alderton, Roberto Amato, Jeffrey Barrett, Sonia Goncalves, Ewan Harrison, David K. Jackson, Ian Johnston, Dominic Kwiatkowski, Cordelia Langford, John Sillitoe on behalf of the Wellcome Sanger Institute COVID-19 Surveillance Team |
| EPI_ISL_3096800, EPI_ISL_3096801, EPI_ISL_3096802 | Lighthouse Lab in Alderley Park | Wellcome Sanger Institute for the COVID-19 Genomics UK (COG-UK) Consortium | Jacquelyn Wynn, Mairead Hyland, The Lighthouse Lab in Alderley Park and Alex Alderton, Roberto Amato, Jeffrey Barrett, Sonia Goncalves, Ewan Harrison, David K. Jackson, Ian Johnston, Dominic Kwiatkowski, Cordelia Langford, John Sillitoe on behalf of the Wellcome Sanger Institute COVID-19 Surveillance Team |
| EPI_ISL_3096803 | Lighthouse Lab in Milton Keynes | Wellcome Sanger Institute for the COVID-19 Genomics UK (COG-UK) Consortium | The Lighthouse Lab in Milton Keynes and Alex Alderton, Roberto Amato, Jeffrey Barrett, Sonia Goncalves, Ewan Harrison, David K. Jackson, Ian Johnston, Dominic Kwiatkowski, Cordelia Langford, John Sillitoe on behalf of the Wellcome Sanger Institute COVID-19 Surveillance Team |

| | | | |
|---|---|---|---|
| EPI_ISL_3096804 | Berkshire and Surrey Pathology Services Lighthouse Laboratory | Wellcome Sanger Institute for the COVID-19 Genomics UK (COG-UK) Consortium | Berkshire and Surrey Pathology Services Lighthouse Laboratory and Alex Alderton, Roberto Amato, Jeffrey Barrett, Sonia Goncalves, Ewan Harrison, David K. Jackson, Ian Johnston, Dominic Kwiatkowski, Cordelia Langford, John Sillitoe on behalf of the Wellcome Sanger Institute COVID-19 Surveillance Team |
| EPI_ISL_3096805, EPI_ISL_3096806 | Lighthouse Lab in Alderley Park | Wellcome Sanger Institute for the COVID-19 Genomics UK (COG-UK) Consortium | Jacquelyn Wynn, Mairead Hyland, The Lighthouse Lab in Alderley Park and Alex Alderton, Roberto Amato, Jeffrey Barrett, Sonia Goncalves, Ewan Harrison, David K. Jackson, Ian Johnston, Dominic Kwiatkowski, Cordelia Langford, John Sillitoe on behalf of the Wellcome Sanger Institute COVID-19 Surveillance Team |
| EPI_ISL_3096807, EPI_ISL_3096808 | Lighthouse Lab in Milton Keynes | Wellcome Sanger Institute for the COVID-19 Genomics UK (COG-UK) Consortium | The Lighthouse Lab in Milton Keynes and Alex Alderton, Roberto Amato, Jeffrey Barrett, Sonia Goncalves, Ewan Harrison, David K. Jackson, Ian Johnston, Dominic Kwiatkowski, Cordelia Langford, John Sillitoe on behalf of the Wellcome Sanger Institute COVID-19 Surveillance Team |
| EPI_ISL_3096809, EPI_ISL_3096810 | Berkshire and Surrey Pathology Services Lighthouse Laboratory | Wellcome Sanger Institute for the COVID-19 Genomics UK (COG-UK) Consortium | Berkshire and Surrey Pathology Services Lighthouse Laboratory and Alex Alderton, Roberto Amato, Jeffrey Barrett, Sonia Goncalves, Ewan Harrison, David K. Jackson, Ian Johnston, Dominic Kwiatkowski, Cordelia Langford, John Sillitoe on behalf of the Wellcome Sanger Institute COVID-19 Surveillance Team |
| EPI_ISL_3096811 | Lighthouse Lab in Alderley Park | Wellcome Sanger Institute for the COVID-19 Genomics UK (COG-UK) Consortium | Jacquelyn Wynn, Mairead Hyland, The Lighthouse Lab in Alderley Park and Alex Alderton, Roberto Amato, Jeffrey Barrett, Sonia Goncalves, Ewan Harrison, David K. Jackson, Ian Johnston, Dominic Kwiatkowski, Cordelia Langford, John Sillitoe on behalf of the Wellcome Sanger Institute COVID-19 Surveillance Team |
| EPI_ISL_3096812 | Berkshire and Surrey Pathology Services Lighthouse Laboratory | Wellcome Sanger Institute for the COVID-19 Genomics UK (COG-UK) Consortium | Berkshire and Surrey Pathology Services Lighthouse Laboratory and Alex Alderton, Roberto Amato, Jeffrey Barrett, Sonia Goncalves, Ewan Harrison, David K. Jackson, Ian Johnston, Dominic Kwiatkowski, Cordelia Langford, John Sillitoe on behalf of the Wellcome Sanger Institute COVID-19 Surveillance Team |
| EPI_ISL_3096813, EPI_ISL_3096814 | Lighthouse Lab in Milton Keynes | Wellcome Sanger Institute for the COVID-19 Genomics UK (COG-UK) Consortium | The Lighthouse Lab in Milton Keynes and Alex Alderton, Roberto Amato, Jeffrey Barrett, Sonia Goncalves, Ewan Harrison, David K. Jackson, Ian Johnston, Dominic Kwiatkowski, Cordelia Langford, John Sillitoe on behalf of the Wellcome Sanger Institute COVID-19 Surveillance Team |
| EPI_ISL_3096815 | Berkshire and Surrey Pathology Services Lighthouse Laboratory | Wellcome Sanger Institute for the COVID-19 Genomics UK (COG-UK) Consortium | Berkshire and Surrey Pathology Services Lighthouse Laboratory and Alex Alderton, Roberto Amato, Jeffrey Barrett, Sonia Goncalves, Ewan Harrison, David K. Jackson, Ian Johnston, Dominic Kwiatkowski, Cordelia Langford, John Sillitoe on behalf of the Wellcome Sanger Institute COVID-19 Surveillance Team |
| EPI_ISL_3096816 | Lighthouse Lab in Milton Keynes | Wellcome Sanger Institute for the COVID-19 Genomics UK (COG-UK) Consortium | The Lighthouse Lab in Milton Keynes and Alex Alderton, Roberto Amato, Jeffrey Barrett, Sonia Goncalves, Ewan Harrison, David K. Jackson, Ian Johnston, Dominic Kwiatkowski, Cordelia Langford, John Sillitoe on behalf of the Wellcome Sanger Institute COVID-19 Surveillance Team |
| EPI_ISL_3096817 | Lighthouse Lab in Alderley Park | Wellcome Sanger Institute for the COVID-19 Genomics UK (COG-UK) Consortium | Jacquelyn Wynn, Mairead Hyland, The Lighthouse Lab in Alderley Park and Alex Alderton, Roberto Amato, Jeffrey Barrett, Sonia Goncalves, Ewan Harrison, David K. Jackson, Ian Johnston, Dominic Kwiatkowski, Cordelia Langford, John Sillitoe on behalf of the Wellcome Sanger Institute COVID-19 Surveillance Team |
| EPI_ISL_3096818 | Lighthouse Lab in Milton Keynes | Wellcome Sanger Institute for the COVID-19 Genomics UK (COG-UK) Consortium | The Lighthouse Lab in Milton Keynes and Alex Alderton, Roberto Amato, Jeffrey Barrett, Sonia Goncalves, Ewan Harrison, David K. Jackson, Ian Johnston, Dominic Kwiatkowski, Cordelia Langford, John Sillitoe on behalf of the Wellcome Sanger Institute COVID-19 Surveillance Team |
| EPI_ISL_3096819 | Lighthouse Lab in Alderley Park | Wellcome Sanger Institute for the COVID-19 Genomics UK (COG-UK) Consortium | Jacquelyn Wynn, Mairead Hyland, The Lighthouse Lab in Alderley Park and Alex Alderton, Roberto Amato, Jeffrey Barrett, Sonia Goncalves, Ewan Harrison, David K. Jackson, Ian Johnston, Dominic Kwiatkowski, Cordelia Langford, John Sillitoe on behalf of the Wellcome Sanger Institute COVID-19 Surveillance Team |
| EPI_ISL_3096820 | Berkshire and Surrey Pathology Services Lighthouse Laboratory | Wellcome Sanger Institute for the COVID-19 Genomics UK (COG-UK) Consortium | Berkshire and Surrey Pathology Services Lighthouse Laboratory and Alex Alderton, Roberto Amato, Jeffrey Barrett, Sonia Goncalves, Ewan Harrison, David K. Jackson, Ian Johnston, Dominic Kwiatkowski, Cordelia Langford, John Sillitoe on behalf of the Wellcome Sanger Institute COVID-19 Surveillance Team |
| EPI_ISL_3096821 | Lighthouse Lab in Milton Keynes | Wellcome Sanger Institute for the COVID-19 Genomics UK (COG-UK) Consortium | The Lighthouse Lab in Milton Keynes and Alex Alderton, Roberto Amato, Jeffrey Barrett, Sonia Goncalves, Ewan Harrison, David K. Jackson, Ian Johnston, Dominic Kwiatkowski, Cordelia Langford, John Sillitoe on behalf of the Wellcome Sanger Institute COVID-19 Surveillance Team |
| EPI_ISL_3096822 | Berkshire and Surrey Pathology Services Lighthouse Laboratory | Wellcome Sanger Institute for the COVID-19 Genomics UK (COG-UK) Consortium | Berkshire and Surrey Pathology Services Lighthouse Laboratory and Alex Alderton, Roberto Amato, Jeffrey Barrett, Sonia Goncalves, Ewan Harrison, David K. Jackson, Ian Johnston, Dominic Kwiatkowski, Cordelia Langford, John Sillitoe on behalf of the Wellcome Sanger Institute COVID-19 Surveillance Team |
| EPI_ISL_3096823, EPI_ISL_3096824 | Lighthouse Lab in Milton Keynes | Wellcome Sanger Institute for the COVID-19 Genomics UK (COG-UK) Consortium | The Lighthouse Lab in Milton Keynes and Alex Alderton, Roberto Amato, Jeffrey Barrett, Sonia Goncalves, Ewan Harrison, David K. Jackson, Ian Johnston, Dominic Kwiatkowski, Cordelia Langford, John Sillitoe on behalf of the Wellcome Sanger Institute COVID-19 Surveillance Team |
| EPI_ISL_3096825 | Berkshire and Surrey Pathology Services Lighthouse Laboratory | Wellcome Sanger Institute for the COVID-19 Genomics UK (COG-UK) Consortium | Berkshire and Surrey Pathology Services Lighthouse Laboratory and Alex Alderton, Roberto Amato, Jeffrey Barrett, Sonia Goncalves, Ewan Harrison, David K. Jackson, Ian Johnston, Dominic Kwiatkowski, Cordelia Langford, John Sillitoe on behalf of the Wellcome Sanger Institute COVID-19 Surveillance Team |
| EPI_ISL_3096826 | Lighthouse Lab in Alderley Park | Wellcome Sanger Institute for the COVID-19 Genomics UK (COG-UK) Consortium | Jacquelyn Wynn, Mairead Hyland, The Lighthouse Lab in Alderley Park and Alex Alderton, Roberto Amato, Jeffrey Barrett, Sonia Goncalves, Ewan Harrison, David K. Jackson, Ian Johnston, Dominic Kwiatkowski, Cordelia Langford, John Sillitoe on behalf of the Wellcome Sanger Institute COVID-19 Surveillance Team |
| EPI_ISL_3096827, EPI_ISL_3096828, EPI_ISL_3096829 | Lighthouse Lab in Milton Keynes | Wellcome Sanger Institute for the COVID-19 Genomics UK (COG-UK) Consortium | The Lighthouse Lab in Milton Keynes and Alex Alderton, Roberto Amato, Jeffrey Barrett, Sonia Goncalves, Ewan Harrison, David K. Jackson, Ian Johnston, Dominic Kwiatkowski, Cordelia Langford, John Sillitoe on behalf of the Wellcome Sanger Institute COVID-19 Surveillance Team |
| EPI_ISL_3096830 | Lighthouse Lab in Alderley Park | Wellcome Sanger Institute for the COVID-19 Genomics UK (COG-UK) Consortium | Jacquelyn Wynn, Mairead Hyland, The Lighthouse Lab in Alderley Park and Alex Alderton, Roberto Amato, Jeffrey Barrett, Sonia Goncalves, Ewan Harrison, David K. Jackson, Ian Johnston, Dominic Kwiatkowski, Cordelia Langford, John Sillitoe on behalf of the Wellcome Sanger Institute COVID-19 Surveillance Team |
| EPI_ISL_3096831 | Lighthouse Lab in Milton Keynes | Wellcome Sanger Institute for the COVID-19 Genomics UK (COG-UK) Consortium | The Lighthouse Lab in Milton Keynes and Alex Alderton, Roberto Amato, Jeffrey Barrett, Sonia Goncalves, Ewan Harrison, David K. Jackson, Ian Johnston, Dominic Kwiatkowski, Cordelia Langford, John Sillitoe on behalf of the Wellcome Sanger Institute COVID-19 Surveillance Team |
| EPI_ISL_3096832 | Lighthouse Lab in Alderley Park | Wellcome Sanger Institute for the COVID-19 Genomics UK (COG-UK) Consortium | Jacquelyn Wynn, Mairead Hyland, The Lighthouse Lab in Alderley Park and Alex Alderton, Roberto Amato, Jeffrey Barrett, Sonia Goncalves, Ewan Harrison, David K. Jackson, Ian Johnston, Dominic Kwiatkowski, Cordelia Langford, John Sillitoe on behalf of the Wellcome Sanger Institute COVID-19 Surveillance Team |
| EPI_ISL_3096833 | Lighthouse Lab in Milton Keynes | Wellcome Sanger Institute for the COVID-19 Genomics UK (COG-UK) Consortium | The Lighthouse Lab in Milton Keynes and Alex Alderton, Roberto Amato, Jeffrey Barrett, Sonia Goncalves, Ewan Harrison, David K. Jackson, Ian Johnston, Dominic Kwiatkowski, Cordelia Langford, John Sillitoe on behalf of the Wellcome Sanger Institute COVID-19 Surveillance Team |
| EPI_ISL_3096834 | Lighthouse Lab in Alderley Park | Wellcome Sanger Institute for the COVID-19 Genomics UK (COG-UK) Consortium | Jacquelyn Wynn, Mairead Hyland, The Lighthouse Lab in Alderley Park and Alex Alderton, Roberto Amato, Jeffrey Barrett, Sonia Goncalves, Ewan Harrison, David K. Jackson, Ian Johnston, Dominic Kwiatkowski, Cordelia Langford, John Sillitoe on behalf of the Wellcome Sanger Institute COVID-19 Surveillance Team |
| EPI_ISL_3096835 | Lighthouse Lab in Milton Keynes | Wellcome Sanger Institute for the COVID-19 Genomics UK (COG-UK) Consortium | The Lighthouse Lab in Milton Keynes and Alex Alderton, Roberto Amato, Jeffrey Barrett, Sonia Goncalves, Ewan Harrison, David K. Jackson, Ian Johnston, Dominic Kwiatkowski, Cordelia Langford, John Sillitoe on behalf of the Wellcome Sanger Institute COVID-19 Surveillance Team |
| EPI_ISL_3096836 | Lighthouse Lab in Alderley Park | Wellcome Sanger Institute for the COVID-19 Genomics UK (COG-UK) Consortium | Jacquelyn Wynn, Mairead Hyland, The Lighthouse Lab in Alderley Park and Alex Alderton, Roberto Amato, Jeffrey Barrett, Sonia Goncalves, Ewan Harrison, David K. Jackson, Ian Johnston, Dominic Kwiatkowski, Cordelia Langford, John Sillitoe on behalf of the Wellcome Sanger Institute COVID-19 Surveillance Team |
| EPI_ISL_3096837 | Berkshire and Surrey Pathology Services Lighthouse Laboratory | Wellcome Sanger Institute for the COVID-19 Genomics UK (COG-UK) Consortium | Berkshire and Surrey Pathology Services Lighthouse Laboratory and Alex Alderton, Roberto Amato, Jeffrey Barrett, Sonia Goncalves, Ewan Harrison, David K. Jackson, Ian Johnston, Dominic Kwiatkowski, Cordelia Langford, John Sillitoe on behalf of the Wellcome Sanger Institute COVID-19 Surveillance Team |
| EPI_ISL_3096838 | Lighthouse Lab in Milton Keynes | Wellcome Sanger Institute for the COVID-19 Genomics UK (COG-UK) Consortium | The Lighthouse Lab in Milton Keynes and Alex Alderton, Roberto Amato, Jeffrey Barrett, Sonia Goncalves, Ewan Harrison, David K. Jackson, Ian Johnston, Dominic Kwiatkowski, Cordelia Langford, John Sillitoe on behalf of the Wellcome Sanger Institute COVID-19 Surveillance Team |
| EPI_ISL_3096839 | Berkshire and Surrey Pathology Services Lighthouse Laboratory | Wellcome Sanger Institute for the COVID-19 Genomics UK (COG-UK) Consortium | Berkshire and Surrey Pathology Services Lighthouse Laboratory and Alex Alderton, Roberto Amato, Jeffrey Barrett, Sonia Goncalves, Ewan Harrison, David K. Jackson, Ian Johnston, Dominic Kwiatkowski, Cordelia Langford, John Sillitoe on behalf of the Wellcome Sanger Institute COVID-19 Surveillance Team |
| EPI_ISL_3096840 | Lighthouse Lab in Milton Keynes | Wellcome Sanger Institute for the COVID-19 Genomics UK (COG-UK) Consortium | The Lighthouse Lab in Milton Keynes and Alex Alderton, Roberto Amato, Jeffrey Barrett, Sonia Goncalves, Ewan Harrison, David K. Jackson, Ian Johnston, Dominic Kwiatkowski, Cordelia Langford, John Sillitoe on behalf of the Wellcome Sanger Institute COVID-19 Surveillance Team |
| EPI_ISL_3096841 | Berkshire and Surrey Pathology Services Lighthouse Laboratory | Wellcome Sanger Institute for the COVID-19 Genomics UK (COG-UK) Consortium | Berkshire and Surrey Pathology Services Lighthouse Laboratory and Alex Alderton, Roberto Amato, Jeffrey Barrett, Sonia Goncalves, Ewan Harrison, David K. Jackson, Ian Johnston, Dominic Kwiatkowski, Cordelia Langford, John Sillitoe on behalf of the Wellcome Sanger Institute COVID-19 Surveillance Team |
| EPI_ISL_3096842, EPI_ISL_3096843 | Lighthouse Lab in Alderley Park | Wellcome Sanger Institute for the COVID-19 Genomics UK (COG-UK) Consortium | Jacquelyn Wynn, Mairead Hyland, The Lighthouse Lab in Alderley Park and Alex Alderton, Roberto Amato, Jeffrey Barrett, Sonia Goncalves, Ewan Harrison, David K. Jackson, Ian Johnston, Dominic Kwiatkowski, Cordelia Langford, John Sillitoe on behalf of the Wellcome Sanger Institute COVID-19 Surveillance Team |
| EPI_ISL_3096844, EPI_ISL_3096845 | Lighthouse Lab in Milton Keynes | Wellcome Sanger Institute for the COVID-19 Genomics UK (COG-UK) Consortium | The Lighthouse Lab in Milton Keynes and Alex Alderton, Roberto Amato, Jeffrey Barrett, Sonia Goncalves, Ewan Harrison, David K. Jackson, Ian Johnston, Dominic Kwiatkowski, Cordelia Langford, John Sillitoe on behalf of the Wellcome Sanger Institute COVID-19 Surveillance Team |
| EPI_ISL_3096846 | Lighthouse Lab in Alderley Park | Wellcome Sanger Institute for the COVID-19 Genomics UK (COG-UK) Consortium | Jacquelyn Wynn, Mairead Hyland, The Lighthouse Lab in Alderley Park and Alex Alderton, Roberto Amato, Jeffrey Barrett, Sonia Goncalves, Ewan Harrison, David K. Jackson, Ian Johnston, Dominic Kwiatkowski, Cordelia Langford, John Sillitoe on behalf of the Wellcome Sanger Institute COVID-19 Surveillance Team |
| EPI_ISL_3096847, EPI_ISL_3096848 | Lighthouse Lab in Milton Keynes | Wellcome Sanger Institute for the COVID-19 Genomics UK (COG-UK) Consortium | The Lighthouse Lab in Milton Keynes and Alex Alderton, Roberto Amato, Jeffrey Barrett, Sonia Goncalves, Ewan Harrison, David K. Jackson, Ian Johnston, Dominic Kwiatkowski, Cordelia Langford, John Sillitoe on behalf of the Wellcome Sanger Institute COVID-19 Surveillance Team |
| EPI_ISL_3096849 | Lighthouse Lab in Alderley Park | Wellcome Sanger Institute for the COVID-19 Genomics UK (COG-UK) Consortium | Jacquelyn Wynn, Mairead Hyland, The Lighthouse Lab in Alderley Park and Alex Alderton, Roberto Amato, Jeffrey Barrett, Sonia Goncalves, Ewan Harrison, David K. Jackson, Ian Johnston, Dominic Kwiatkowski, Cordelia Langford, John Sillitoe on behalf of the Wellcome Sanger Institute COVID-19 Surveillance Team |
| EPI_ISL_3096850, EPI_ISL_3096851 | Lighthouse Lab in Milton Keynes | Wellcome Sanger Institute for the COVID-19 Genomics UK (COG-UK) Consortium | The Lighthouse Lab in Milton Keynes and Alex Alderton, Roberto Amato, Jeffrey Barrett, Sonia Goncalves, Ewan Harrison, David K. Jackson, Ian Johnston, Dominic Kwiatkowski, Cordelia Langford, John Sillitoe on behalf of the Wellcome Sanger Institute COVID-19 Surveillance Team |
| EPI_ISL_3096852, EPI_ISL_3096853 | Berkshire and Surrey Pathology Services Lighthouse Laboratory | Wellcome Sanger Institute for the COVID-19 Genomics UK (COG-UK) Consortium | Berkshire and Surrey Pathology Services Lighthouse Laboratory and Alex Alderton, Roberto Amato, Jeffrey Barrett, Sonia Goncalves, Ewan Harrison, David K. Jackson, Ian Johnston, Dominic Kwiatkowski, Cordelia Langford, John Sillitoe on behalf of the Wellcome Sanger Institute COVID-19 Surveillance Team |
| EPI_ISL_3096854 | Lighthouse Lab in Milton Keynes | Wellcome Sanger Institute for the COVID-19 Genomics UK (COG-UK) Consortium | The Lighthouse Lab in Milton Keynes and Alex Alderton, Roberto Amato, Jeffrey Barrett, Sonia Goncalves, Ewan Harrison, David K. Jackson, Ian Johnston, Dominic Kwiatkowski, Cordelia Langford, John Sillitoe on behalf of the Wellcome Sanger Institute COVID-19 Surveillance Team |
| EPI_ISL_3096855, EPI_ISL_3096856 | Lighthouse Lab in Alderley Park | Wellcome Sanger Institute for the COVID-19 Genomics UK (COG-UK) Consortium | Jacquelyn Wynn, Mairead Hyland, The Lighthouse Lab in Alderley Park and Alex Alderton, Roberto Amato, Jeffrey Barrett, Sonia Goncalves, Ewan Harrison, David K. Jackson, Ian Johnston, Dominic Kwiatkowski, Cordelia Langford, John Sillitoe on behalf of the Wellcome Sanger Institute COVID-19 Surveillance Team |
| EPI_ISL_3096857 | Berkshire and Surrey Pathology Services Lighthouse Laboratory | Wellcome Sanger Institute for the COVID-19 Genomics UK (COG-UK) Consortium | Berkshire and Surrey Pathology Services Lighthouse Laboratory and Alex Alderton, Roberto Amato, Jeffrey Barrett, Sonia Goncalves, Ewan Harrison, David K. Jackson, Ian Johnston, Dominic Kwiatkowski, Cordelia Langford, John Sillitoe on behalf of the Wellcome Sanger Institute COVID-19 Surveillance Team |
| EPI_ISL_3096858 | Lighthouse Lab in Alderley Park | Wellcome Sanger Institute for the COVID-19 Genomics UK (COG-UK) Consortium | Jacquelyn Wynn, Mairead Hyland, The Lighthouse Lab in Alderley Park and Alex Alderton, Roberto Amato, Jeffrey Barrett, Sonia Goncalves, Ewan Harrison, David K. Jackson, Ian Johnston, Dominic Kwiatkowski, Cordelia Langford, John Sillitoe on behalf of the Wellcome Sanger Institute COVID-19 Surveillance Team |
| EPI_ISL_3096859 | Lighthouse Lab in Milton Keynes | Wellcome Sanger Institute for the COVID-19 Genomics UK (COG-UK) Consortium | The Lighthouse Lab in Milton Keynes and Alex Alderton, Roberto Amato, Jeffrey Barrett, Sonia Goncalves, Ewan Harrison, David K. Jackson, Ian Johnston, Dominic Kwiatkowski, Cordelia Langford, John Sillitoe on behalf of the Wellcome Sanger Institute COVID-19 Surveillance Team |
| EPI_ISL_3096860 | Berkshire and Surrey Pathology Services Lighthouse Laboratory | Wellcome Sanger Institute for the COVID-19 Genomics UK (COG-UK) Consortium | Berkshire and Surrey Pathology Services Lighthouse Laboratory and Alex Alderton, Roberto Amato, Jeffrey Barrett, Sonia Goncalves, Ewan Harrison, David K. Jackson, Ian Johnston, Dominic Kwiatkowski, Cordelia Langford, John Sillitoe on behalf of the Wellcome Sanger Institute COVID-19 Surveillance Team |
| EPI_ISL_3096861 | Lighthouse Lab in Milton Keynes | Wellcome Sanger Institute for the COVID-19 Genomics UK (COG-UK) Consortium | The Lighthouse Lab in Milton Keynes and Alex Alderton, Roberto Amato, Jeffrey Barrett, Sonia Goncalves, Ewan Harrison, David K. Jackson, Ian Johnston, Dominic Kwiatkowski, Cordelia Langford, John Sillitoe on behalf of the Wellcome Sanger Institute COVID-19 Surveillance Team |

| | | | |
|---|---|---|---|
| EPI_ISL_3096862 | Lighthouse Lab in Alderley Park | Wellcome Sanger Institute for the COVID-19 Genomics UK (COG-UK) Consortium | Jacquelyn Wynn, Mairead Hyland, The Lighthouse Lab in Alderley Park and Alex Alderton, Roberto Amato, Jeffrey Barrett, Sonia Goncalves, Ewan Harrison, David K. Jackson, Ian Johnston, Dominic Kwiatkowski, Cordelia Langford, John Sillitoe on behalf of the Wellcome Sanger Institute COVID-19 Surveillance Team |
| EPI_ISL_3096863 | Lighthouse Lab in Milton Keynes | Wellcome Sanger Institute for the COVID-19 Genomics UK (COG-UK) Consortium | The Lighthouse Lab in Milton Keynes and Alex Alderton, Roberto Amato, Jeffrey Barrett, Sonia Goncalves, Ewan Harrison, David K. Jackson, Ian Johnston, Dominic Kwiatkowski, Cordelia Langford, John Sillitoe on behalf of the Wellcome Sanger Institute COVID-19 Surveillance Team |
| EPI_ISL_3096864 | Berkshire and Surrey Pathology Services Lighthouse Laboratory | Wellcome Sanger Institute for the COVID-19 Genomics UK (COG-UK) Consortium | Berkshire and Surrey Pathology Services Lighthouse Laboratory and Alex Alderton, Roberto Amato, Jeffrey Barrett, Sonia Goncalves, Ewan Harrison, David K. Jackson, Ian Johnston, Dominic Kwiatkowski, Cordelia Langford, John Sillitoe on behalf of the Wellcome Sanger Institute COVID-19 Surveillance Team |
| EPI_ISL_3096865, EPI_ISL_3096866 | Lighthouse Lab in Alderley Park | Wellcome Sanger Institute for the COVID-19 Genomics UK (COG-UK) Consortium | Jacquelyn Wynn, Mairead Hyland, The Lighthouse Lab in Alderley Park and Alex Alderton, Roberto Amato, Jeffrey Barrett, Sonia Goncalves, Ewan Harrison, David K. Jackson, Ian Johnston, Dominic Kwiatkowski, Cordelia Langford, John Sillitoe on behalf of the Wellcome Sanger Institute COVID-19 Surveillance Team |
| EPI_ISL_3096867 | Lighthouse Lab in Milton Keynes | Wellcome Sanger Institute for the COVID-19 Genomics UK (COG-UK) Consortium | The Lighthouse Lab in Milton Keynes and Alex Alderton, Roberto Amato, Jeffrey Barrett, Sonia Goncalves, Ewan Harrison, David K. Jackson, Ian Johnston, Dominic Kwiatkowski, Cordelia Langford, John Sillitoe on behalf of the Wellcome Sanger Institute COVID-19 Surveillance Team |
| EPI_ISL_3096868, EPI_ISL_3096869, EPI_ISL_3096870, EPI_ISL_3096871 | Berkshire and Surrey Pathology Services Lighthouse Laboratory | Wellcome Sanger Institute for the COVID-19 Genomics UK (COG-UK) Consortium | Berkshire and Surrey Pathology Services Lighthouse Laboratory and Alex Alderton, Roberto Amato, Jeffrey Barrett, Sonia Goncalves, Ewan Harrison, David K. Jackson, Ian Johnston, Dominic Kwiatkowski, Cordelia Langford, John Sillitoe on behalf of the Wellcome Sanger Institute COVID-19 Surveillance Team |
| EPI_ISL_3096872 | Lighthouse Lab in Alderley Park | Wellcome Sanger Institute for the COVID-19 Genomics UK (COG-UK) Consortium | Jacquelyn Wynn, Mairead Hyland, The Lighthouse Lab in Alderley Park and Alex Alderton, Roberto Amato, Jeffrey Barrett, Sonia Goncalves, Ewan Harrison, David K. Jackson, Ian Johnston, Dominic Kwiatkowski, Cordelia Langford, John Sillitoe on behalf of the Wellcome Sanger Institute COVID-19 Surveillance Team |
| EPI_ISL_3096873, EPI_ISL_3096874, EPI_ISL_3096875 | Lighthouse Lab in Milton Keynes | Wellcome Sanger Institute for the COVID-19 Genomics UK (COG-UK) Consortium | The Lighthouse Lab in Milton Keynes and Alex Alderton, Roberto Amato, Jeffrey Barrett, Sonia Goncalves, Ewan Harrison, David K. Jackson, Ian Johnston, Dominic Kwiatkowski, Cordelia Langford, John Sillitoe on behalf of the Wellcome Sanger Institute COVID-19 Surveillance Team |
| EPI_ISL_3096876 | Lighthouse Lab in Alderley Park | Wellcome Sanger Institute for the COVID-19 Genomics UK (COG-UK) Consortium | Jacquelyn Wynn, Mairead Hyland, The Lighthouse Lab in Alderley Park and Alex Alderton, Roberto Amato, Jeffrey Barrett, Sonia Goncalves, Ewan Harrison, David K. Jackson, Ian Johnston, Dominic Kwiatkowski, Cordelia Langford, John Sillitoe on behalf of the Wellcome Sanger Institute COVID-19 Surveillance Team |
| EPI_ISL_3096877 | Lighthouse Lab in Milton Keynes | Wellcome Sanger Institute for the COVID-19 Genomics UK (COG-UK) Consortium | The Lighthouse Lab in Milton Keynes and Alex Alderton, Roberto Amato, Jeffrey Barrett, Sonia Goncalves, Ewan Harrison, David K. Jackson, Ian Johnston, Dominic Kwiatkowski, Cordelia Langford, John Sillitoe on behalf of the Wellcome Sanger Institute COVID-19 Surveillance Team |
| EPI_ISL_3096878, EPI_ISL_3096879, EPI_ISL_3096880 | Lighthouse Lab in Alderley Park | Wellcome Sanger Institute for the COVID-19 Genomics UK (COG-UK) Consortium | Jacquelyn Wynn, Mairead Hyland, The Lighthouse Lab in Alderley Park and Alex Alderton, Roberto Amato, Jeffrey Barrett, Sonia Goncalves, Ewan Harrison, David K. Jackson, Ian Johnston, Dominic Kwiatkowski, Cordelia Langford, John Sillitoe on behalf of the Wellcome Sanger Institute COVID-19 Surveillance Team |
| EPI_ISL_3096881, EPI_ISL_3096882, EPI_ISL_3096883, EPI_ISL_3096884, EPI_ISL_3096885 | Lighthouse Lab in Milton Keynes | Wellcome Sanger Institute for the COVID-19 Genomics UK (COG-UK) Consortium | The Lighthouse Lab in Milton Keynes and Alex Alderton, Roberto Amato, Jeffrey Barrett, Sonia Goncalves, Ewan Harrison, David K. Jackson, Ian Johnston, Dominic Kwiatkowski, Cordelia Langford, John Sillitoe on behalf of the Wellcome Sanger Institute COVID-19 Surveillance Team |
| EPI_ISL_3096886 | Lighthouse Lab in Alderley Park | Wellcome Sanger Institute for the COVID-19 Genomics UK (COG-UK) Consortium | Jacquelyn Wynn, Mairead Hyland, The Lighthouse Lab in Alderley Park and Alex Alderton, Roberto Amato, Jeffrey Barrett, Sonia Goncalves, Ewan Harrison, David K. Jackson, Ian Johnston, Dominic Kwiatkowski, Cordelia Langford, John Sillitoe on behalf of the Wellcome Sanger Institute COVID-19 Surveillance Team |
| EPI_ISL_3096887 | Berkshire and Surrey Pathology Services Lighthouse Laboratory | Wellcome Sanger Institute for the COVID-19 Genomics UK (COG-UK) Consortium | Berkshire and Surrey Pathology Services Lighthouse Laboratory and Alex Alderton, Roberto Amato, Jeffrey Barrett, Sonia Goncalves, Ewan Harrison, David K. Jackson, Ian Johnston, Dominic Kwiatkowski, Cordelia Langford, John Sillitoe on behalf of the Wellcome Sanger Institute COVID-19 Surveillance Team |
| EPI_ISL_3096888, EPI_ISL_3096889 | Lighthouse Lab in Milton Keynes | Wellcome Sanger Institute for the COVID-19 Genomics UK (COG-UK) Consortium | The Lighthouse Lab in Milton Keynes and Alex Alderton, Roberto Amato, Jeffrey Barrett, Sonia Goncalves, Ewan Harrison, David K. Jackson, Ian Johnston, Dominic Kwiatkowski, Cordelia Langford, John Sillitoe on behalf of the Wellcome Sanger Institute COVID-19 Surveillance Team |
| EPI_ISL_3096890, EPI_ISL_3096891 | Berkshire and Surrey Pathology Services Lighthouse Laboratory | Wellcome Sanger Institute for the COVID-19 Genomics UK (COG-UK) Consortium | Berkshire and Surrey Pathology Services Lighthouse Laboratory and Alex Alderton, Roberto Amato, Jeffrey Barrett, Sonia Goncalves, Ewan Harrison, David K. Jackson, Ian Johnston, Dominic Kwiatkowski, Cordelia Langford, John Sillitoe on behalf of the Wellcome Sanger Institute COVID-19 Surveillance Team |
| EPI_ISL_3096892 | Lighthouse Lab in Alderley Park | Wellcome Sanger Institute for the COVID-19 Genomics UK (COG-UK) Consortium | Jacquelyn Wynn, Mairead Hyland, The Lighthouse Lab in Alderley Park and Alex Alderton, Roberto Amato, Jeffrey Barrett, Sonia Goncalves, Ewan Harrison, David K. Jackson, Ian Johnston, Dominic Kwiatkowski, Cordelia Langford, John Sillitoe on behalf of the Wellcome Sanger Institute COVID-19 Surveillance Team |
| EPI_ISL_3096893 | Berkshire and Surrey Pathology Services Lighthouse Laboratory | Wellcome Sanger Institute for the COVID-19 Genomics UK (COG-UK) Consortium | Berkshire and Surrey Pathology Services Lighthouse Laboratory and Alex Alderton, Roberto Amato, Jeffrey Barrett, Sonia Goncalves, Ewan Harrison, David K. Jackson, Ian Johnston, Dominic Kwiatkowski, Cordelia Langford, John Sillitoe on behalf of the Wellcome Sanger Institute COVID-19 Surveillance Team |
| EPI_ISL_3096894, EPI_ISL_3096895 | Lighthouse Lab in Milton Keynes | Wellcome Sanger Institute for the COVID-19 Genomics UK (COG-UK) Consortium | The Lighthouse Lab in Milton Keynes and Alex Alderton, Roberto Amato, Jeffrey Barrett, Sonia Goncalves, Ewan Harrison, David K. Jackson, Ian Johnston, Dominic Kwiatkowski, Cordelia Langford, John Sillitoe on behalf of the Wellcome Sanger Institute COVID-19 Surveillance Team |
| EPI_ISL_3096896 | Lighthouse Lab in Alderley Park | Wellcome Sanger Institute for the COVID-19 Genomics UK (COG-UK) Consortium | Jacquelyn Wynn, Mairead Hyland, The Lighthouse Lab in Alderley Park and Alex Alderton, Roberto Amato, Jeffrey Barrett, Sonia Goncalves, Ewan Harrison, David K. Jackson, Ian Johnston, Dominic Kwiatkowski, Cordelia Langford, John Sillitoe on behalf of the Wellcome Sanger Institute COVID-19 Surveillance Team |
| EPI_ISL_3096897, EPI_ISL_3096898 | Lighthouse Lab in Milton Keynes | Wellcome Sanger Institute for the COVID-19 Genomics UK (COG-UK) Consortium | The Lighthouse Lab in Milton Keynes and Alex Alderton, Roberto Amato, Jeffrey Barrett, Sonia Goncalves, Ewan Harrison, David K. Jackson, Ian Johnston, Dominic Kwiatkowski, Cordelia Langford, John Sillitoe on behalf of the Wellcome Sanger Institute COVID-19 Surveillance Team |
| EPI_ISL_3096899 | Berkshire and Surrey Pathology Services Lighthouse Laboratory | Wellcome Sanger Institute for the COVID-19 Genomics UK (COG-UK) Consortium | Berkshire and Surrey Pathology Services Lighthouse Laboratory and Alex Alderton, Roberto Amato, Jeffrey Barrett, Sonia Goncalves, Ewan Harrison, David K. Jackson, Ian Johnston, Dominic Kwiatkowski, Cordelia Langford, John Sillitoe on behalf of the Wellcome Sanger Institute COVID-19 Surveillance Team |
| EPI_ISL_3096900 | Lighthouse Lab in Alderley Park | Wellcome Sanger Institute for the COVID-19 Genomics UK (COG-UK) Consortium | Jacquelyn Wynn, Mairead Hyland, The Lighthouse Lab in Alderley Park and Alex Alderton, Roberto Amato, Jeffrey Barrett, Sonia Goncalves, Ewan Harrison, David K. Jackson, Ian Johnston, Dominic Kwiatkowski, Cordelia Langford, John Sillitoe on behalf of the Wellcome Sanger Institute COVID-19 Surveillance Team |
| EPI_ISL_3096901 | Berkshire and Surrey Pathology Services Lighthouse Laboratory | Wellcome Sanger Institute for the COVID-19 Genomics UK (COG-UK) Consortium | Berkshire and Surrey Pathology Services Lighthouse Laboratory and Alex Alderton, Roberto Amato, Jeffrey Barrett, Sonia Goncalves, Ewan Harrison, David K. Jackson, Ian Johnston, Dominic Kwiatkowski, Cordelia Langford, John Sillitoe on behalf of the Wellcome Sanger Institute COVID-19 Surveillance Team |
| EPI_ISL_3096902 | Lighthouse Lab in Alderley Park | Wellcome Sanger Institute for the COVID-19 Genomics UK (COG-UK) Consortium | Jacquelyn Wynn, Mairead Hyland, The Lighthouse Lab in Alderley Park and Alex Alderton, Roberto Amato, Jeffrey Barrett, Sonia Goncalves, Ewan Harrison, David K. Jackson, Ian Johnston, Dominic Kwiatkowski, Cordelia Langford, John Sillitoe on behalf of the Wellcome Sanger Institute COVID-19 Surveillance Team |
| EPI_ISL_3096903 | Lighthouse Lab in Milton Keynes | Wellcome Sanger Institute for the COVID-19 Genomics UK (COG-UK) Consortium | The Lighthouse Lab in Milton Keynes and Alex Alderton, Roberto Amato, Jeffrey Barrett, Sonia Goncalves, Ewan Harrison, David K. Jackson, Ian Johnston, Dominic Kwiatkowski, Cordelia Langford, John Sillitoe on behalf of the Wellcome Sanger Institute COVID-19 Surveillance Team |
| EPI_ISL_3096904 | Berkshire and Surrey Pathology Services Lighthouse Laboratory | Wellcome Sanger Institute for the COVID-19 Genomics UK (COG-UK) Consortium | Berkshire and Surrey Pathology Services Lighthouse Laboratory and Alex Alderton, Roberto Amato, Jeffrey Barrett, Sonia Goncalves, Ewan Harrison, David K. Jackson, Ian Johnston, Dominic Kwiatkowski, Cordelia Langford, John Sillitoe on behalf of the Wellcome Sanger Institute COVID-19 Surveillance Team |
| EPI_ISL_3096905, EPI_ISL_3096906 | Lighthouse Lab in Milton Keynes | Wellcome Sanger Institute for the COVID-19 Genomics UK (COG-UK) Consortium | The Lighthouse Lab in Milton Keynes and Alex Alderton, Roberto Amato, Jeffrey Barrett, Sonia Goncalves, Ewan Harrison, David K. Jackson, Ian Johnston, Dominic Kwiatkowski, Cordelia Langford, John Sillitoe on behalf of the Wellcome Sanger Institute COVID-19 Surveillance Team |
| EPI_ISL_3096907, EPI_ISL_3096908 | Lighthouse Lab in Alderley Park | Wellcome Sanger Institute for the COVID-19 Genomics UK (COG-UK) Consortium | Jacquelyn Wynn, Mairead Hyland, The Lighthouse Lab in Alderley Park and Alex Alderton, Roberto Amato, Jeffrey Barrett, Sonia Goncalves, Ewan Harrison, David K. Jackson, Ian Johnston, Dominic Kwiatkowski, Cordelia Langford, John Sillitoe on behalf of the Wellcome Sanger Institute COVID-19 Surveillance Team |
| EPI_ISL_3096909, EPI_ISL_3096910 | Lighthouse Lab in Milton Keynes | Wellcome Sanger Institute for the COVID-19 Genomics UK (COG-UK) Consortium | The Lighthouse Lab in Milton Keynes and Alex Alderton, Roberto Amato, Jeffrey Barrett, Sonia Goncalves, Ewan Harrison, David K. Jackson, Ian Johnston, Dominic Kwiatkowski, Cordelia Langford, John Sillitoe on behalf of the Wellcome Sanger Institute COVID-19 Surveillance Team |
| EPI_ISL_3096911 | Berkshire and Surrey Pathology Services Lighthouse Laboratory | Wellcome Sanger Institute for the COVID-19 Genomics UK (COG-UK) Consortium | Berkshire and Surrey Pathology Services Lighthouse Laboratory and Alex Alderton, Roberto Amato, Jeffrey Barrett, Sonia Goncalves, Ewan Harrison, David K. Jackson, Ian Johnston, Dominic Kwiatkowski, Cordelia Langford, John Sillitoe on behalf of the Wellcome Sanger Institute COVID-19 Surveillance Team |
| EPI_ISL_3096912, EPI_ISL_3096913 | Lighthouse Lab in Milton Keynes | Wellcome Sanger Institute for the COVID-19 Genomics UK (COG-UK) Consortium | The Lighthouse Lab in Milton Keynes and Alex Alderton, Roberto Amato, Jeffrey Barrett, Sonia Goncalves, Ewan Harrison, David K. Jackson, Ian Johnston, Dominic Kwiatkowski, Cordelia Langford, John Sillitoe on behalf of the Wellcome Sanger Institute COVID-19 Surveillance Team |
| EPI_ISL_3096914 | Lighthouse Lab in Alderley Park | Wellcome Sanger Institute for the COVID-19 Genomics UK (COG-UK) Consortium | Jacquelyn Wynn, Mairead Hyland, The Lighthouse Lab in Alderley Park and Alex Alderton, Roberto Amato, Jeffrey Barrett, Sonia Goncalves, Ewan Harrison, David K. Jackson, Ian Johnston, Dominic Kwiatkowski, Cordelia Langford, John Sillitoe on behalf of the Wellcome Sanger Institute COVID-19 Surveillance Team |
| EPI_ISL_3096915 | Berkshire and Surrey Pathology Services Lighthouse Laboratory | Wellcome Sanger Institute for the COVID-19 Genomics UK (COG-UK) Consortium | Berkshire and Surrey Pathology Services Lighthouse Laboratory and Alex Alderton, Roberto Amato, Jeffrey Barrett, Sonia Goncalves, Ewan Harrison, David K. Jackson, Ian Johnston, Dominic Kwiatkowski, Cordelia Langford, John Sillitoe on behalf of the Wellcome Sanger Institute COVID-19 Surveillance Team |
| EPI_ISL_3096916 | Lighthouse Lab in Alderley Park | Wellcome Sanger Institute for the COVID-19 Genomics UK (COG-UK) Consortium | Jacquelyn Wynn, Mairead Hyland, The Lighthouse Lab in Alderley Park and Alex Alderton, Roberto Amato, Jeffrey Barrett, Sonia Goncalves, Ewan Harrison, David K. Jackson, Ian Johnston, Dominic Kwiatkowski, Cordelia Langford, John Sillitoe on behalf of the Wellcome Sanger Institute COVID-19 Surveillance Team |
| EPI_ISL_3096917 | Lighthouse Lab in Milton Keynes | Wellcome Sanger Institute for the COVID-19 Genomics UK (COG-UK) Consortium | The Lighthouse Lab in Milton Keynes and Alex Alderton, Roberto Amato, Jeffrey Barrett, Sonia Goncalves, Ewan Harrison, David K. Jackson, Ian Johnston, Dominic Kwiatkowski, Cordelia Langford, John Sillitoe on behalf of the Wellcome Sanger Institute COVID-19 Surveillance Team |
| EPI_ISL_3096918 | Lighthouse Lab in Alderley Park | Wellcome Sanger Institute for the COVID-19 Genomics UK (COG-UK) Consortium | Jacquelyn Wynn, Mairead Hyland, The Lighthouse Lab in Alderley Park and Alex Alderton, Roberto Amato, Jeffrey Barrett, Sonia Goncalves, Ewan Harrison, David K. Jackson, Ian Johnston, Dominic Kwiatkowski, Cordelia Langford, John Sillitoe on behalf of the Wellcome Sanger Institute COVID-19 Surveillance Team |
| EPI_ISL_3096919, EPI_ISL_3096920 | Lighthouse Lab in Milton Keynes | Wellcome Sanger Institute for the COVID-19 Genomics UK (COG-UK) Consortium | The Lighthouse Lab in Milton Keynes and Alex Alderton, Roberto Amato, Jeffrey Barrett, Sonia Goncalves, Ewan Harrison, David K. Jackson, Ian Johnston, Dominic Kwiatkowski, Cordelia Langford, John Sillitoe on behalf of the Wellcome Sanger Institute COVID-19 Surveillance Team |
| EPI_ISL_3096921 | Lighthouse Lab in Alderley Park | Wellcome Sanger Institute for the COVID-19 Genomics UK (COG-UK) Consortium | Jacquelyn Wynn, Mairead Hyland, The Lighthouse Lab in Alderley Park and Alex Alderton, Roberto Amato, Jeffrey Barrett, Sonia Goncalves, Ewan Harrison, David K. Jackson, Ian Johnston, Dominic Kwiatkowski, Cordelia Langford, John Sillitoe on behalf of the Wellcome Sanger Institute COVID-19 Surveillance Team |
| EPI_ISL_3096922 | Berkshire and Surrey Pathology Services Lighthouse Laboratory | Wellcome Sanger Institute for the COVID-19 Genomics UK (COG-UK) Consortium | Berkshire and Surrey Pathology Services Lighthouse Laboratory and Alex Alderton, Roberto Amato, Jeffrey Barrett, Sonia Goncalves, Ewan Harrison, David K. Jackson, Ian Johnston, Dominic Kwiatkowski, Cordelia Langford, John Sillitoe on behalf of the Wellcome Sanger Institute COVID-19 Surveillance Team |
| EPI_ISL_3096923, EPI_ISL_3096924, EPI_ISL_3096925 | Lighthouse Lab in Milton Keynes | Wellcome Sanger Institute for the COVID-19 Genomics UK (COG-UK) Consortium | The Lighthouse Lab in Milton Keynes and Alex Alderton, Roberto Amato, Jeffrey Barrett, Sonia Goncalves, Ewan Harrison, David K. Jackson, Ian Johnston, Dominic Kwiatkowski, Cordelia Langford, John Sillitoe on behalf of the Wellcome Sanger Institute COVID-19 Surveillance Team |
| EPI_ISL_3096926, | Lighthouse Lab in Alderley Park | Wellcome Sanger Institute for the COVID-19 Genomics UK (COG-UK) | Jacquelyn Wynn, Mairead Hyland, The Lighthouse Lab in Alderley Park and Alex Alderton, Roberto Amato, Jeffrey Barrett, Sonia Goncalves, Ewan Harrison, David K. Jackson, Ian Johnston, Dominic Kwiatkowski, Cordelia Langford, John Sillitoe on |

| | | | |
|---|---|---|---|
| EPI_ISL_3096927 | | Consortium | behalf of the Wellcome Sanger Institute COVID-19 Surveillance Team |
| EPI_ISL_3096928, EPI_ISL_3096929 | Berkshire and Surrey Pathology Services Lighthouse Laboratory | Wellcome Sanger Institute for the COVID-19 Genomics UK (COG-UK) Consortium | Berkshire and Surrey Pathology Services Lighthouse Laboratory and Alex Alderton, Roberto Amato, Jeffrey Barrett, Sonia Goncalves, Ewan Harrison, David K. Jackson, Ian Johnston, Dominic Kwiatkowski, Cordelia Langford, John Sillitoe on behalf of the Wellcome Sanger Institute COVID-19 Surveillance Team |
| EPI_ISL_3096930, EPI_ISL_3096931, EPI_ISL_3096932 | Lighthouse Lab in Milton Keynes | Wellcome Sanger Institute for the COVID-19 Genomics UK (COG-UK) Consortium | The Lighthouse Lab in Milton Keynes and Alex Alderton, Roberto Amato, Jeffrey Barrett, Sonia Goncalves, Ewan Harrison, David K. Jackson, Ian Johnston, Dominic Kwiatkowski, Cordelia Langford, John Sillitoe on behalf of the Wellcome Sanger Institute COVID-19 Surveillance Team |
| EPI_ISL_3096933, EPI_ISL_3096934 | Berkshire and Surrey Pathology Services Lighthouse Laboratory | Wellcome Sanger Institute for the COVID-19 Genomics UK (COG-UK) Consortium | Berkshire and Surrey Pathology Services Lighthouse Laboratory and Alex Alderton, Roberto Amato, Jeffrey Barrett, Sonia Goncalves, Ewan Harrison, David K. Jackson, Ian Johnston, Dominic Kwiatkowski, Cordelia Langford, John Sillitoe on behalf of the Wellcome Sanger Institute COVID-19 Surveillance Team |
| EPI_ISL_3096935, EPI_ISL_3096936 | Lighthouse Lab in Milton Keynes | Wellcome Sanger Institute for the COVID-19 Genomics UK (COG-UK) Consortium | The Lighthouse Lab in Milton Keynes and Alex Alderton, Roberto Amato, Jeffrey Barrett, Sonia Goncalves, Ewan Harrison, David K. Jackson, Ian Johnston, Dominic Kwiatkowski, Cordelia Langford, John Sillitoe on behalf of the Wellcome Sanger Institute COVID-19 Surveillance Team |
| EPI_ISL_3096937, EPI_ISL_3096938 | Lighthouse Lab in Alderley Park | Wellcome Sanger Institute for the COVID-19 Genomics UK (COG-UK) Consortium | Jacquelyn Wynn, Mairead Hyland, The Lighthouse Lab in Alderley Park and Alex Alderton, Roberto Amato, Jeffrey Barrett, Sonia Goncalves, Ewan Harrison, David K. Jackson, Ian Johnston, Dominic Kwiatkowski, Cordelia Langford, John Sillitoe on behalf of the Wellcome Sanger Institute COVID-19 Surveillance Team |
| EPI_ISL_3096939, EPI_ISL_3096940, EPI_ISL_3096941, EPI_ISL_3096942, EPI_ISL_3096943, EPI_ISL_3096944 | Lighthouse Lab in Milton Keynes | Wellcome Sanger Institute for the COVID-19 Genomics UK (COG-UK) Consortium | The Lighthouse Lab in Milton Keynes and Alex Alderton, Roberto Amato, Jeffrey Barrett, Sonia Goncalves, Ewan Harrison, David K. Jackson, Ian Johnston, Dominic Kwiatkowski, Cordelia Langford, John Sillitoe on behalf of the Wellcome Sanger Institute COVID-19 Surveillance Team |
| EPI_ISL_3096945 | Berkshire and Surrey Pathology Services Lighthouse Laboratory | Wellcome Sanger Institute for the COVID-19 Genomics UK (COG-UK) Consortium | Berkshire and Surrey Pathology Services Lighthouse Laboratory and Alex Alderton, Roberto Amato, Jeffrey Barrett, Sonia Goncalves, Ewan Harrison, David K. Jackson, Ian Johnston, Dominic Kwiatkowski, Cordelia Langford, John Sillitoe on behalf of the Wellcome Sanger Institute COVID-19 Surveillance Team |
| EPI_ISL_3096946 | Lighthouse Lab in Milton Keynes | Wellcome Sanger Institute for the COVID-19 Genomics UK (COG-UK) Consortium | The Lighthouse Lab in Milton Keynes and Alex Alderton, Roberto Amato, Jeffrey Barrett, Sonia Goncalves, Ewan Harrison, David K. Jackson, Ian Johnston, Dominic Kwiatkowski, Cordelia Langford, John Sillitoe on behalf of the Wellcome Sanger Institute COVID-19 Surveillance Team |
| EPI_ISL_3096947, EPI_ISL_3096948 | Berkshire and Surrey Pathology Services Lighthouse Laboratory | Wellcome Sanger Institute for the COVID-19 Genomics UK (COG-UK) Consortium | Berkshire and Surrey Pathology Services Lighthouse Laboratory and Alex Alderton, Roberto Amato, Jeffrey Barrett, Sonia Goncalves, Ewan Harrison, David K. Jackson, Ian Johnston, Dominic Kwiatkowski, Cordelia Langford, John Sillitoe on behalf of the Wellcome Sanger Institute COVID-19 Surveillance Team |
| EPI_ISL_3096949 | Lighthouse Lab in Alderley Park | Wellcome Sanger Institute for the COVID-19 Genomics UK (COG-UK) Consortium | Jacquelyn Wynn, Mairead Hyland, The Lighthouse Lab in Alderley Park and Alex Alderton, Roberto Amato, Jeffrey Barrett, Sonia Goncalves, Ewan Harrison, David K. Jackson, Ian Johnston, Dominic Kwiatkowski, Cordelia Langford, John Sillitoe on behalf of the Wellcome Sanger Institute COVID-19 Surveillance Team |
| EPI_ISL_3096950 | Berkshire and Surrey Pathology Services Lighthouse Laboratory | Wellcome Sanger Institute for the COVID-19 Genomics UK (COG-UK) Consortium | Berkshire and Surrey Pathology Services Lighthouse Laboratory and Alex Alderton, Roberto Amato, Jeffrey Barrett, Sonia Goncalves, Ewan Harrison, David K. Jackson, Ian Johnston, Dominic Kwiatkowski, Cordelia Langford, John Sillitoe on behalf of the Wellcome Sanger Institute COVID-19 Surveillance Team |
| EPI_ISL_3096951, EPI_ISL_3096952 | Lighthouse Lab in Alderley Park | Wellcome Sanger Institute for the COVID-19 Genomics UK (COG-UK) Consortium | Jacquelyn Wynn, Mairead Hyland, The Lighthouse Lab in Alderley Park and Alex Alderton, Roberto Amato, Jeffrey Barrett, Sonia Goncalves, Ewan Harrison, David K. Jackson, Ian Johnston, Dominic Kwiatkowski, Cordelia Langford, John Sillitoe on behalf of the Wellcome Sanger Institute COVID-19 Surveillance Team |
| EPI_ISL_3096953 | Lighthouse Lab in Milton Keynes | Wellcome Sanger Institute for the COVID-19 Genomics UK (COG-UK) Consortium | The Lighthouse Lab in Milton Keynes and Alex Alderton, Roberto Amato, Jeffrey Barrett, Sonia Goncalves, Ewan Harrison, David K. Jackson, Ian Johnston, Dominic Kwiatkowski, Cordelia Langford, John Sillitoe on behalf of the Wellcome Sanger Institute COVID-19 Surveillance Team |
| EPI_ISL_3096954, EPI_ISL_3096955 | Health Services Laboratories | Wellcome Sanger Institute for the COVID-19 Genomics UK (COG-UK) Consortium | Health Services Laboratories and Alex Alderton, Roberto Amato, Jeffrey Barrett, Sonia Goncalves, Ewan Harrison, David K. Jackson, Ian Johnston, Dominic Kwiatkowski, Cordelia Langford, John Sillitoe on behalf of the Wellcome Sanger Institute COVID-19 Surveillance Team |
| EPI_ISL_3096956 | Lighthouse Lab in Alderley Park | Wellcome Sanger Institute for the COVID-19 Genomics UK (COG-UK) Consortium | Jacquelyn Wynn, Mairead Hyland, The Lighthouse Lab in Alderley Park and Alex Alderton, Roberto Amato, Jeffrey Barrett, Sonia Goncalves, Ewan Harrison, David K. Jackson, Ian Johnston, Dominic Kwiatkowski, Cordelia Langford, John Sillitoe on behalf of the Wellcome Sanger Institute COVID-19 Surveillance Team |
| EPI_ISL_3096957, EPI_ISL_3096958, EPI_ISL_3096959, EPI_ISL_3096960, EPI_ISL_3096961, EPI_ISL_3096962, EPI_ISL_3096963, EPI_ISL_3096964, EPI_ISL_3096965, EPI_ISL_3096966, EPI_ISL_3096967 | | | |
| see above | Health Services Laboratories | Wellcome Sanger Institute for the COVID-19 Genomics UK (COG-UK) Consortium | Health Services Laboratories and Alex Alderton, Roberto Amato, Jeffrey Barrett, Sonia Goncalves, Ewan Harrison, David K. Jackson, Ian Johnston, Dominic Kwiatkowski, Cordelia Langford, John Sillitoe on behalf of the Wellcome Sanger Institute COVID-19 Surveillance Team |
| EPI_ISL_3096968 | Lighthouse Lab in Alderley Park | Wellcome Sanger Institute for the COVID-19 Genomics UK (COG-UK) Consortium | Jacquelyn Wynn, Mairead Hyland, The Lighthouse Lab in Alderley Park and Alex Alderton, Roberto Amato, Jeffrey Barrett, Sonia Goncalves, Ewan Harrison, David K. Jackson, Ian Johnston, Dominic Kwiatkowski, Cordelia Langford, John Sillitoe on behalf of the Wellcome Sanger Institute COVID-19 Surveillance Team |
| EPI_ISL_3096969, EPI_ISL_3096970, EPI_ISL_3096971, EPI_ISL_3096972, EPI_ISL_3096973, EPI_ISL_3096974 | Health Services Laboratories | Wellcome Sanger Institute for the COVID-19 Genomics UK (COG-UK) Consortium | Health Services Laboratories and Alex Alderton, Roberto Amato, Jeffrey Barrett, Sonia Goncalves, Ewan Harrison, David K. Jackson, Ian Johnston, Dominic Kwiatkowski, Cordelia Langford, John Sillitoe on behalf of the Wellcome Sanger Institute COVID-19 Surveillance Team |
| EPI_ISL_3096975 | Lighthouse Laboratory Plymouth | Wellcome Sanger Institute for the COVID-19 Genomics UK (COG-UK) Consortium | Lighthouse Laboratory Plymouth and Alex Alderton, Roberto Amato, Jeffrey Barrett, Sonia Goncalves, Ewan Harrison, David K. Jackson, Ian Johnston, Dominic Kwiatkowski, Cordelia Langford, John Sillitoe on behalf of the Wellcome Sanger Institute COVID-19 Surveillance Team |
| EPI_ISL_3096976, EPI_ISL_3096977, EPI_ISL_3096978, EPI_ISL_3096979 | Health Services Laboratories | Wellcome Sanger Institute for the COVID-19 Genomics UK (COG-UK) Consortium | Health Services Laboratories and Alex Alderton, Roberto Amato, Jeffrey Barrett, Sonia Goncalves, Ewan Harrison, David K. Jackson, Ian Johnston, Dominic Kwiatkowski, Cordelia Langford, John Sillitoe on behalf of the Wellcome Sanger Institute COVID-19 Surveillance Team |
| EPI_ISL_3096980 | Lighthouse Lab in Alderley Park | Wellcome Sanger Institute for the COVID-19 Genomics UK (COG-UK) Consortium | Jacquelyn Wynn, Mairead Hyland, The Lighthouse Lab in Alderley Park and Alex Alderton, Roberto Amato, Jeffrey Barrett, Sonia Goncalves, Ewan Harrison, David K. Jackson, Ian Johnston, Dominic Kwiatkowski, Cordelia Langford, John Sillitoe on behalf of the Wellcome Sanger Institute COVID-19 Surveillance Team |
| EPI_ISL_3096981, EPI_ISL_3096982 | Health Services Laboratories | Wellcome Sanger Institute for the COVID-19 Genomics UK (COG-UK) Consortium | Health Services Laboratories and Alex Alderton, Roberto Amato, Jeffrey Barrett, Sonia Goncalves, Ewan Harrison, David K. Jackson, Ian Johnston, Dominic Kwiatkowski, Cordelia Langford, John Sillitoe on behalf of the Wellcome Sanger Institute COVID-19 Surveillance Team |
| EPI_ISL_3096983 | Lighthouse Laboratory Plymouth | Wellcome Sanger Institute for the COVID-19 Genomics UK (COG-UK) Consortium | Lighthouse Laboratory Plymouth and Alex Alderton, Roberto Amato, Jeffrey Barrett, Sonia Goncalves, Ewan Harrison, David K. Jackson, Ian Johnston, Dominic Kwiatkowski, Cordelia Langford, John Sillitoe on behalf of the Wellcome Sanger Institute COVID-19 Surveillance Team |
| EPI_ISL_3096984, EPI_ISL_3096985, EPI_ISL_3096986, EPI_ISL_3096987, EPI_ISL_3096988, EPI_ISL_3096989, EPI_ISL_3096990, EPI_ISL_3096991, EPI_ISL_3096992, EPI_ISL_3096993 | Health Services Laboratories | Wellcome Sanger Institute for the COVID-19 Genomics UK (COG-UK) Consortium | Health Services Laboratories and Alex Alderton, Roberto Amato, Jeffrey Barrett, Sonia Goncalves, Ewan Harrison, David K. Jackson, Ian Johnston, Dominic Kwiatkowski, Cordelia Langford, John Sillitoe on behalf of the Wellcome Sanger Institute COVID-19 Surveillance Team |
| EPI_ISL_3096994 | Lighthouse Lab in Milton Keynes | Wellcome Sanger Institute for the COVID-19 Genomics UK (COG-UK) Consortium | The Lighthouse Lab in Milton Keynes and Alex Alderton, Roberto Amato, Jeffrey Barrett, Sonia Goncalves, Ewan Harrison, David K. Jackson, Ian Johnston, Dominic Kwiatkowski, Cordelia Langford, John Sillitoe on behalf of the Wellcome Sanger Institute COVID-19 Surveillance Team |
| EPI_ISL_3096995 | Lighthouse Lab in Glasgow | Wellcome Sanger Institute for the COVID-19 Genomics UK (COG-UK) Consortium | Harper VanSteenhouse, Yumi Kasai, David Gray, Carol Clugston, Anna Dominiczak and Alex Alderton, Roberto Amato, Jeffrey Barrett, Sonia Goncalves, Ewan Harrison, David K. Jackson, Ian Johnston, Dominic Kwiatkowski, Cordelia Langford, John Sillitoe on behalf of the Wellcome Sanger Institute COVID-19 Surveillance Team |
| EPI_ISL_3096996 | Lighthouse Lab in Milton Keynes | Wellcome Sanger Institute for the COVID-19 Genomics UK (COG-UK) Consortium | The Lighthouse Lab in Milton Keynes and Alex Alderton, Roberto Amato, Jeffrey Barrett, Sonia Goncalves, Ewan Harrison, David K. Jackson, Ian Johnston, Dominic Kwiatkowski, Cordelia Langford, John Sillitoe on behalf of the Wellcome Sanger Institute COVID-19 Surveillance Team |
| EPI_ISL_3096997, EPI_ISL_3096998 | Lighthouse Lab in Glasgow | Wellcome Sanger Institute for the COVID-19 Genomics UK (COG-UK) Consortium | Harper VanSteenhouse, Yumi Kasai, David Gray, Carol Clugston, Anna Dominiczak and Alex Alderton, Roberto Amato, Jeffrey Barrett, Sonia Goncalves, Ewan Harrison, David K. Jackson, Ian Johnston, Dominic Kwiatkowski, Cordelia Langford, John Sillitoe on behalf of the Wellcome Sanger Institute COVID-19 Surveillance Team |
| EPI_ISL_3096999, EPI_ISL_3097000 | Lighthouse Lab in Milton Keynes | Wellcome Sanger Institute for the COVID-19 Genomics UK (COG-UK) Consortium | The Lighthouse Lab in Milton Keynes and Alex Alderton, Roberto Amato, Jeffrey Barrett, Sonia Goncalves, Ewan Harrison, David K. Jackson, Ian Johnston, Dominic Kwiatkowski, Cordelia Langford, John Sillitoe on behalf of the Wellcome Sanger Institute COVID-19 Surveillance Team |
| EPI_ISL_3097001 | Lighthouse Lab in Glasgow | Wellcome Sanger Institute for the COVID-19 Genomics UK (COG-UK) Consortium | Harper VanSteenhouse, Yumi Kasai, David Gray, Carol Clugston, Anna Dominiczak and Alex Alderton, Roberto Amato, Jeffrey Barrett, Sonia Goncalves, Ewan Harrison, David K. Jackson, Ian Johnston, Dominic Kwiatkowski, Cordelia Langford, John Sillitoe on behalf of the Wellcome Sanger Institute COVID-19 Surveillance Team |
| EPI_ISL_3097002, EPI_ISL_3097003 | Lighthouse Lab in Milton Keynes | Wellcome Sanger Institute for the COVID-19 Genomics UK (COG-UK) Consortium | The Lighthouse Lab in Milton Keynes and Alex Alderton, Roberto Amato, Jeffrey Barrett, Sonia Goncalves, Ewan Harrison, David K. Jackson, Ian Johnston, Dominic Kwiatkowski, Cordelia Langford, John Sillitoe on behalf of the Wellcome Sanger Institute COVID-19 Surveillance Team |
| EPI_ISL_3097004 | Lighthouse Lab in Glasgow | Wellcome Sanger Institute for the COVID-19 Genomics UK (COG-UK) Consortium | Harper VanSteenhouse, Yumi Kasai, David Gray, Carol Clugston, Anna Dominiczak and Alex Alderton, Roberto Amato, Jeffrey Barrett, Sonia Goncalves, Ewan Harrison, David K. Jackson, Ian Johnston, Dominic Kwiatkowski, Cordelia Langford, John Sillitoe on behalf of the Wellcome Sanger Institute COVID-19 Surveillance Team |
| EPI_ISL_3097005, EPI_ISL_3097006 | Lighthouse Lab in Milton Keynes | Wellcome Sanger Institute for the COVID-19 Genomics UK (COG-UK) Consortium | The Lighthouse Lab in Milton Keynes and Alex Alderton, Roberto Amato, Jeffrey Barrett, Sonia Goncalves, Ewan Harrison, David K. Jackson, Ian Johnston, Dominic Kwiatkowski, Cordelia Langford, John Sillitoe on behalf of the Wellcome Sanger Institute COVID-19 Surveillance Team |
| EPI_ISL_3097007, EPI_ISL_3097008, EPI_ISL_3097009, EPI_ISL_3097010 | Lighthouse Lab in Glasgow | Wellcome Sanger Institute for the COVID-19 Genomics UK (COG-UK) Consortium | Harper VanSteenhouse, Yumi Kasai, David Gray, Carol Clugston, Anna Dominiczak and Alex Alderton, Roberto Amato, Jeffrey Barrett, Sonia Goncalves, Ewan Harrison, David K. Jackson, Ian Johnston, Dominic Kwiatkowski, Cordelia Langford, John Sillitoe on behalf of the Wellcome Sanger Institute COVID-19 Surveillance Team |
| EPI_ISL_3097011, EPI_ISL_3097012 | Lighthouse Lab in Milton Keynes | Wellcome Sanger Institute for the COVID-19 Genomics UK (COG-UK) Consortium | The Lighthouse Lab in Milton Keynes and Alex Alderton, Roberto Amato, Jeffrey Barrett, Sonia Goncalves, Ewan Harrison, David K. Jackson, Ian Johnston, Dominic Kwiatkowski, Cordelia Langford, John Sillitoe on behalf of the Wellcome Sanger Institute COVID-19 Surveillance Team |
| EPI_ISL_3097013, | Lighthouse Lab in Glasgow | Wellcome Sanger Institute for the COVID-19 Genomics UK (COG-UK) | Harper VanSteenhouse, Yumi Kasai, David Gray, Carol Clugston, Anna Dominiczak and Alex Alderton, Roberto Amato, Jeffrey Barrett, Sonia Goncalves, Ewan Harrison, David K. Jackson, Ian Johnston, Dominic Kwiatkowski, Cordelia Langford, |

| | | | |
|---|---|---|---|
| EPI_ISL_3097014 | | Consortium | John Sillitoe on behalf of the Wellcome Sanger Institute COVID-19 Surveillance Team |
| EPI_ISL_3097015, EPI_ISL_3097016 | Lighthouse Lab in Milton Keynes | Wellcome Sanger Institute for the COVID-19 Genomics UK (COG-UK) Consortium | The Lighthouse Lab in Milton Keynes and Alex Alderton, Roberto Amato, Jeffrey Barrett, Sonia Goncalves, Ewan Harrison, David K. Jackson, Ian Johnston, Dominic Kwiatkowski, Cordelia Langford, John Sillitoe on behalf of the Wellcome Sanger Institute COVID-19 Surveillance Team |
| EPI_ISL_3097017, EPI_ISL_3097018 | Lighthouse Lab in Glasgow | Wellcome Sanger Institute for the COVID-19 Genomics UK (COG-UK) Consortium | Harper VanSteenhouse, Yumi Kasai, David Gray, Carol Clugston, Anna Dominiczak and Alex Alderton, Roberto Amato, Jeffrey Barrett, Sonia Goncalves, Ewan Harrison, David K. Jackson, Ian Johnston, Dominic Kwiatkowski, Cordelia Langford, John Sillitoe on behalf of the Wellcome Sanger Institute COVID-19 Surveillance Team |
| EPI_ISL_3097019 | Lighthouse Lab in Milton Keynes | Wellcome Sanger Institute for the COVID-19 Genomics UK (COG-UK) Consortium | The Lighthouse Lab in Milton Keynes and Alex Alderton, Roberto Amato, Jeffrey Barrett, Sonia Goncalves, Ewan Harrison, David K. Jackson, Ian Johnston, Dominic Kwiatkowski, Cordelia Langford, John Sillitoe on behalf of the Wellcome Sanger Institute COVID-19 Surveillance Team |
| EPI_ISL_3097020, EPI_ISL_3097021 | Lighthouse Lab in Glasgow | Wellcome Sanger Institute for the COVID-19 Genomics UK (COG-UK) Consortium | Harper VanSteenhouse, Yumi Kasai, David Gray, Carol Clugston, Anna Dominiczak and Alex Alderton, Roberto Amato, Jeffrey Barrett, Sonia Goncalves, Ewan Harrison, David K. Jackson, Ian Johnston, Dominic Kwiatkowski, Cordelia Langford, John Sillitoe on behalf of the Wellcome Sanger Institute COVID-19 Surveillance Team |
| EPI_ISL_3097022 | Lighthouse Lab in Milton Keynes | Wellcome Sanger Institute for the COVID-19 Genomics UK (COG-UK) Consortium | The Lighthouse Lab in Milton Keynes and Alex Alderton, Roberto Amato, Jeffrey Barrett, Sonia Goncalves, Ewan Harrison, David K. Jackson, Ian Johnston, Dominic Kwiatkowski, Cordelia Langford, John Sillitoe on behalf of the Wellcome Sanger Institute COVID-19 Surveillance Team |
| EPI_ISL_3097023 | Lighthouse Lab in Glasgow | Wellcome Sanger Institute for the COVID-19 Genomics UK (COG-UK) Consortium | Harper VanSteenhouse, Yumi Kasai, David Gray, Carol Clugston, Anna Dominiczak and Alex Alderton, Roberto Amato, Jeffrey Barrett, Sonia Goncalves, Ewan Harrison, David K. Jackson, Ian Johnston, Dominic Kwiatkowski, Cordelia Langford, John Sillitoe on behalf of the Wellcome Sanger Institute COVID-19 Surveillance Team |
| EPI_ISL_3097024 | Lighthouse Lab in Milton Keynes | Wellcome Sanger Institute for the COVID-19 Genomics UK (COG-UK) Consortium | The Lighthouse Lab in Milton Keynes and Alex Alderton, Roberto Amato, Jeffrey Barrett, Sonia Goncalves, Ewan Harrison, David K. Jackson, Ian Johnston, Dominic Kwiatkowski, Cordelia Langford, John Sillitoe on behalf of the Wellcome Sanger Institute COVID-19 Surveillance Team |
| EPI_ISL_3097025, EPI_ISL_3097026 | Lighthouse Lab in Glasgow | Wellcome Sanger Institute for the COVID-19 Genomics UK (COG-UK) Consortium | Harper VanSteenhouse, Yumi Kasai, David Gray, Carol Clugston, Anna Dominiczak and Alex Alderton, Roberto Amato, Jeffrey Barrett, Sonia Goncalves, Ewan Harrison, David K. Jackson, Ian Johnston, Dominic Kwiatkowski, Cordelia Langford, John Sillitoe on behalf of the Wellcome Sanger Institute COVID-19 Surveillance Team |
| EPI_ISL_3097027, EPI_ISL_3097028 | Lighthouse Lab in Milton Keynes | Wellcome Sanger Institute for the COVID-19 Genomics UK (COG-UK) Consortium | The Lighthouse Lab in Milton Keynes and Alex Alderton, Roberto Amato, Jeffrey Barrett, Sonia Goncalves, Ewan Harrison, David K. Jackson, Ian Johnston, Dominic Kwiatkowski, Cordelia Langford, John Sillitoe on behalf of the Wellcome Sanger Institute COVID-19 Surveillance Team |
| EPI_ISL_3097029 | Lighthouse Lab in Glasgow | Wellcome Sanger Institute for the COVID-19 Genomics UK (COG-UK) Consortium | Harper VanSteenhouse, Yumi Kasai, David Gray, Carol Clugston, Anna Dominiczak and Alex Alderton, Roberto Amato, Jeffrey Barrett, Sonia Goncalves, Ewan Harrison, David K. Jackson, Ian Johnston, Dominic Kwiatkowski, Cordelia Langford, John Sillitoe on behalf of the Wellcome Sanger Institute COVID-19 Surveillance Team |
| EPI_ISL_3097030 | Lighthouse Lab in Milton Keynes | Wellcome Sanger Institute for the COVID-19 Genomics UK (COG-UK) Consortium | The Lighthouse Lab in Milton Keynes and Alex Alderton, Roberto Amato, Jeffrey Barrett, Sonia Goncalves, Ewan Harrison, David K. Jackson, Ian Johnston, Dominic Kwiatkowski, Cordelia Langford, John Sillitoe on behalf of the Wellcome Sanger Institute COVID-19 Surveillance Team |
| EPI_ISL_3097031 | Lighthouse Lab in Glasgow | Wellcome Sanger Institute for the COVID-19 Genomics UK (COG-UK) Consortium | Harper VanSteenhouse, Yumi Kasai, David Gray, Carol Clugston, Anna Dominiczak and Alex Alderton, Roberto Amato, Jeffrey Barrett, Sonia Goncalves, Ewan Harrison, David K. Jackson, Ian Johnston, Dominic Kwiatkowski, Cordelia Langford, John Sillitoe on behalf of the Wellcome Sanger Institute COVID-19 Surveillance Team |
| EPI_ISL_3097032, EPI_ISL_3097033 | Lighthouse Lab in Milton Keynes | Wellcome Sanger Institute for the COVID-19 Genomics UK (COG-UK) Consortium | The Lighthouse Lab in Milton Keynes and Alex Alderton, Roberto Amato, Jeffrey Barrett, Sonia Goncalves, Ewan Harrison, David K. Jackson, Ian Johnston, Dominic Kwiatkowski, Cordelia Langford, John Sillitoe on behalf of the Wellcome Sanger Institute COVID-19 Surveillance Team |
| EPI_ISL_3097034, EPI_ISL_3097035 | Lighthouse Lab in Glasgow | Wellcome Sanger Institute for the COVID-19 Genomics UK (COG-UK) Consortium | Harper VanSteenhouse, Yumi Kasai, David Gray, Carol Clugston, Anna Dominiczak and Alex Alderton, Roberto Amato, Jeffrey Barrett, Sonia Goncalves, Ewan Harrison, David K. Jackson, Ian Johnston, Dominic Kwiatkowski, Cordelia Langford, John Sillitoe on behalf of the Wellcome Sanger Institute COVID-19 Surveillance Team |
| EPI_ISL_3097036 | Lighthouse Lab in Milton Keynes | Wellcome Sanger Institute for the COVID-19 Genomics UK (COG-UK) Consortium | The Lighthouse Lab in Milton Keynes and Alex Alderton, Roberto Amato, Jeffrey Barrett, Sonia Goncalves, Ewan Harrison, David K. Jackson, Ian Johnston, Dominic Kwiatkowski, Cordelia Langford, John Sillitoe on behalf of the Wellcome Sanger Institute COVID-19 Surveillance Team |
| EPI_ISL_3097037 | Lighthouse Lab in Glasgow | Wellcome Sanger Institute for the COVID-19 Genomics UK (COG-UK) Consortium | Harper VanSteenhouse, Yumi Kasai, David Gray, Carol Clugston, Anna Dominiczak and Alex Alderton, Roberto Amato, Jeffrey Barrett, Sonia Goncalves, Ewan Harrison, David K. Jackson, Ian Johnston, Dominic Kwiatkowski, Cordelia Langford, John Sillitoe on behalf of the Wellcome Sanger Institute COVID-19 Surveillance Team |
| EPI_ISL_3097038, EPI_ISL_3097039, EPI_ISL_3097040 | Lighthouse Lab in Milton Keynes | Wellcome Sanger Institute for the COVID-19 Genomics UK (COG-UK) Consortium | The Lighthouse Lab in Milton Keynes and Alex Alderton, Roberto Amato, Jeffrey Barrett, Sonia Goncalves, Ewan Harrison, David K. Jackson, Ian Johnston, Dominic Kwiatkowski, Cordelia Langford, John Sillitoe on behalf of the Wellcome Sanger Institute COVID-19 Surveillance Team |
| EPI_ISL_3097041, EPI_ISL_3097042, EPI_ISL_3097043 | Lighthouse Lab in Glasgow | Wellcome Sanger Institute for the COVID-19 Genomics UK (COG-UK) Consortium | Harper VanSteenhouse, Yumi Kasai, David Gray, Carol Clugston, Anna Dominiczak and Alex Alderton, Roberto Amato, Jeffrey Barrett, Sonia Goncalves, Ewan Harrison, David K. Jackson, Ian Johnston, Dominic Kwiatkowski, Cordelia Langford, John Sillitoe on behalf of the Wellcome Sanger Institute COVID-19 Surveillance Team |
| EPI_ISL_3097044, EPI_ISL_3097045 | Lighthouse Lab in Milton Keynes | Wellcome Sanger Institute for the COVID-19 Genomics UK (COG-UK) Consortium | The Lighthouse Lab in Milton Keynes and Alex Alderton, Roberto Amato, Jeffrey Barrett, Sonia Goncalves, Ewan Harrison, David K. Jackson, Ian Johnston, Dominic Kwiatkowski, Cordelia Langford, John Sillitoe on behalf of the Wellcome Sanger Institute COVID-19 Surveillance Team |
| EPI_ISL_3097046, EPI_ISL_3097047 | Lighthouse Lab in Glasgow | Wellcome Sanger Institute for the COVID-19 Genomics UK (COG-UK) Consortium | Harper VanSteenhouse, Yumi Kasai, David Gray, Carol Clugston, Anna Dominiczak and Alex Alderton, Roberto Amato, Jeffrey Barrett, Sonia Goncalves, Ewan Harrison, David K. Jackson, Ian Johnston, Dominic Kwiatkowski, Cordelia Langford, John Sillitoe on behalf of the Wellcome Sanger Institute COVID-19 Surveillance Team |
| EPI_ISL_3097048, EPI_ISL_3097049 | Lighthouse Lab in Milton Keynes | Wellcome Sanger Institute for the COVID-19 Genomics UK (COG-UK) Consortium | The Lighthouse Lab in Milton Keynes and Alex Alderton, Roberto Amato, Jeffrey Barrett, Sonia Goncalves, Ewan Harrison, David K. Jackson, Ian Johnston, Dominic Kwiatkowski, Cordelia Langford, John Sillitoe on behalf of the Wellcome Sanger Institute COVID-19 Surveillance Team |
| EPI_ISL_3097050 | Lighthouse Lab in Glasgow | Wellcome Sanger Institute for the COVID-19 Genomics UK (COG-UK) Consortium | Harper VanSteenhouse, Yumi Kasai, David Gray, Carol Clugston, Anna Dominiczak and Alex Alderton, Roberto Amato, Jeffrey Barrett, Sonia Goncalves, Ewan Harrison, David K. Jackson, Ian Johnston, Dominic Kwiatkowski, Cordelia Langford, John Sillitoe on behalf of the Wellcome Sanger Institute COVID-19 Surveillance Team |
| EPI_ISL_3097051, EPI_ISL_3097052, EPI_ISL_3097053, EPI_ISL_3097054, EPI_ISL_3097055, EPI_ISL_3097056, EPI_ISL_3097057 | Lighthouse Lab in Milton Keynes | Wellcome Sanger Institute for the COVID-19 Genomics UK (COG-UK) Consortium | The Lighthouse Lab in Milton Keynes and Alex Alderton, Roberto Amato, Jeffrey Barrett, Sonia Goncalves, Ewan Harrison, David K. Jackson, Ian Johnston, Dominic Kwiatkowski, Cordelia Langford, John Sillitoe on behalf of the Wellcome Sanger Institute COVID-19 Surveillance Team |
| EPI_ISL_3097058 | Lighthouse Lab in Glasgow | Wellcome Sanger Institute for the COVID-19 Genomics UK (COG-UK) Consortium | Harper VanSteenhouse, Yumi Kasai, David Gray, Carol Clugston, Anna Dominiczak and Alex Alderton, Roberto Amato, Jeffrey Barrett, Sonia Goncalves, Ewan Harrison, David K. Jackson, Ian Johnston, Dominic Kwiatkowski, Cordelia Langford, John Sillitoe on behalf of the Wellcome Sanger Institute COVID-19 Surveillance Team |
| EPI_ISL_3097059 | Lighthouse Lab in Milton Keynes | Wellcome Sanger Institute for the COVID-19 Genomics UK (COG-UK) Consortium | The Lighthouse Lab in Milton Keynes and Alex Alderton, Roberto Amato, Jeffrey Barrett, Sonia Goncalves, Ewan Harrison, David K. Jackson, Ian Johnston, Dominic Kwiatkowski, Cordelia Langford, John Sillitoe on behalf of the Wellcome Sanger Institute COVID-19 Surveillance Team |
| EPI_ISL_3097060 | Lighthouse Lab in Glasgow | Wellcome Sanger Institute for the COVID-19 Genomics UK (COG-UK) Consortium | Harper VanSteenhouse, Yumi Kasai, David Gray, Carol Clugston, Anna Dominiczak and Alex Alderton, Roberto Amato, Jeffrey Barrett, Sonia Goncalves, Ewan Harrison, David K. Jackson, Ian Johnston, Dominic Kwiatkowski, Cordelia Langford, John Sillitoe on behalf of the Wellcome Sanger Institute COVID-19 Surveillance Team |
| EPI_ISL_3097061, EPI_ISL_3097062, EPI_ISL_3097063, EPI_ISL_3097064, EPI_ISL_3097065 | Lighthouse Lab in Milton Keynes | Wellcome Sanger Institute for the COVID-19 Genomics UK (COG-UK) Consortium | The Lighthouse Lab in Milton Keynes and Alex Alderton, Roberto Amato, Jeffrey Barrett, Sonia Goncalves, Ewan Harrison, David K. Jackson, Ian Johnston, Dominic Kwiatkowski, Cordelia Langford, John Sillitoe on behalf of the Wellcome Sanger Institute COVID-19 Surveillance Team |
| EPI_ISL_3097066 | Lighthouse Lab in Glasgow | Wellcome Sanger Institute for the COVID-19 Genomics UK (COG-UK) Consortium | Harper VanSteenhouse, Yumi Kasai, David Gray, Carol Clugston, Anna Dominiczak and Alex Alderton, Roberto Amato, Jeffrey Barrett, Sonia Goncalves, Ewan Harrison, David K. Jackson, Ian Johnston, Dominic Kwiatkowski, Cordelia Langford, John Sillitoe on behalf of the Wellcome Sanger Institute COVID-19 Surveillance Team |
| EPI_ISL_3097067, EPI_ISL_3097068 | Lighthouse Lab in Milton Keynes | Wellcome Sanger Institute for the COVID-19 Genomics UK (COG-UK) Consortium | The Lighthouse Lab in Milton Keynes and Alex Alderton, Roberto Amato, Jeffrey Barrett, Sonia Goncalves, Ewan Harrison, David K. Jackson, Ian Johnston, Dominic Kwiatkowski, Cordelia Langford, John Sillitoe on behalf of the Wellcome Sanger Institute COVID-19 Surveillance Team |
| EPI_ISL_3097069, EPI_ISL_3097070 | Lighthouse Lab in Glasgow | Wellcome Sanger Institute for the COVID-19 Genomics UK (COG-UK) Consortium | Harper VanSteenhouse, Yumi Kasai, David Gray, Carol Clugston, Anna Dominiczak and Alex Alderton, Roberto Amato, Jeffrey Barrett, Sonia Goncalves, Ewan Harrison, David K. Jackson, Ian Johnston, Dominic Kwiatkowski, Cordelia Langford, John Sillitoe on behalf of the Wellcome Sanger Institute COVID-19 Surveillance Team |
| EPI_ISL_3097071, EPI_ISL_3097072 | Lighthouse Lab in Milton Keynes | Wellcome Sanger Institute for the COVID-19 Genomics UK (COG-UK) Consortium | The Lighthouse Lab in Milton Keynes and Alex Alderton, Roberto Amato, Jeffrey Barrett, Sonia Goncalves, Ewan Harrison, David K. Jackson, Ian Johnston, Dominic Kwiatkowski, Cordelia Langford, John Sillitoe on behalf of the Wellcome Sanger Institute COVID-19 Surveillance Team |
| EPI_ISL_3097073 | Lighthouse Lab in Glasgow | Wellcome Sanger Institute for the COVID-19 Genomics UK (COG-UK) Consortium | Harper VanSteenhouse, Yumi Kasai, David Gray, Carol Clugston, Anna Dominiczak and Alex Alderton, Roberto Amato, Jeffrey Barrett, Sonia Goncalves, Ewan Harrison, David K. Jackson, Ian Johnston, Dominic Kwiatkowski, Cordelia Langford, John Sillitoe on behalf of the Wellcome Sanger Institute COVID-19 Surveillance Team |
| EPI_ISL_3097074, EPI_ISL_3097075, EPI_ISL_3097076 | Lighthouse Lab in Milton Keynes | Wellcome Sanger Institute for the COVID-19 Genomics UK (COG-UK) Consortium | The Lighthouse Lab in Milton Keynes and Alex Alderton, Roberto Amato, Jeffrey Barrett, Sonia Goncalves, Ewan Harrison, David K. Jackson, Ian Johnston, Dominic Kwiatkowski, Cordelia Langford, John Sillitoe on behalf of the Wellcome Sanger Institute COVID-19 Surveillance Team |
| EPI_ISL_3097077, EPI_ISL_3097078 | Lighthouse Lab in Glasgow | Wellcome Sanger Institute for the COVID-19 Genomics UK (COG-UK) Consortium | Harper VanSteenhouse, Yumi Kasai, David Gray, Carol Clugston, Anna Dominiczak and Alex Alderton, Roberto Amato, Jeffrey Barrett, Sonia Goncalves, Ewan Harrison, David K. Jackson, Ian Johnston, Dominic Kwiatkowski, Cordelia Langford, John Sillitoe on behalf of the Wellcome Sanger Institute COVID-19 Surveillance Team |
| EPI_ISL_3097079, EPI_ISL_3097080, EPI_ISL_3097081, EPI_ISL_3097082 | Lighthouse Lab in Milton Keynes | Wellcome Sanger Institute for the COVID-19 Genomics UK (COG-UK) Consortium | The Lighthouse Lab in Milton Keynes and Alex Alderton, Roberto Amato, Jeffrey Barrett, Sonia Goncalves, Ewan Harrison, David K. Jackson, Ian Johnston, Dominic Kwiatkowski, Cordelia Langford, John Sillitoe on behalf of the Wellcome Sanger Institute COVID-19 Surveillance Team |
| EPI_ISL_3097083, EPI_ISL_3097084 | Lighthouse Lab in Glasgow | Wellcome Sanger Institute for the COVID-19 Genomics UK (COG-UK) Consortium | Harper VanSteenhouse, Yumi Kasai, David Gray, Carol Clugston, Anna Dominiczak and Alex Alderton, Roberto Amato, Jeffrey Barrett, Sonia Goncalves, Ewan Harrison, David K. Jackson, Ian Johnston, Dominic Kwiatkowski, Cordelia Langford, John Sillitoe on behalf of the Wellcome Sanger Institute COVID-19 Surveillance Team |
| EPI_ISL_3097085 | Lighthouse Lab in Milton Keynes | Wellcome Sanger Institute for the COVID-19 Genomics UK (COG-UK) Consortium | The Lighthouse Lab in Milton Keynes and Alex Alderton, Roberto Amato, Jeffrey Barrett, Sonia Goncalves, Ewan Harrison, David K. Jackson, Ian Johnston, Dominic Kwiatkowski, Cordelia Langford, John Sillitoe on behalf of the Wellcome Sanger Institute COVID-19 Surveillance Team |
| EPI_ISL_3097086, EPI_ISL_3097087 | Lighthouse Lab in Glasgow | Wellcome Sanger Institute for the COVID-19 Genomics UK (COG-UK) Consortium | Harper VanSteenhouse, Yumi Kasai, David Gray, Carol Clugston, Anna Dominiczak and Alex Alderton, Roberto Amato, Jeffrey Barrett, Sonia Goncalves, Ewan Harrison, David K. Jackson, Ian Johnston, Dominic Kwiatkowski, Cordelia Langford, John Sillitoe on behalf of the Wellcome Sanger Institute COVID-19 Surveillance Team |
| EPI_ISL_3097088 | Lighthouse Lab in Milton Keynes | Wellcome Sanger Institute for the COVID-19 Genomics UK (COG-UK) Consortium | The Lighthouse Lab in Milton Keynes and Alex Alderton, Roberto Amato, Jeffrey Barrett, Sonia Goncalves, Ewan Harrison, David K. Jackson, Ian Johnston, Dominic Kwiatkowski, Cordelia Langford, John Sillitoe on behalf of the Wellcome Sanger Institute COVID-19 Surveillance Team |

| | | | |
|---|---|---|---|
| EPI_ISL_3097089 | Lighthouse Lab in Glasgow | Wellcome Sanger Institute for the COVID-19 Genomics UK (COG-UK) Consortium | Harper VanSteenhouse, Yumi Kasai, David Gray, Carol Clugston, Anna Dominiczak and Alex Alderton, Roberto Amato, Jeffrey Barrett, Sonia Goncalves, Ewan Harrison, David K. Jackson, Ian Johnston, Dominic Kwiatkowski, Cordelia Langford, John Sillitoe on behalf of the Wellcome Sanger Institute COVID-19 Surveillance Team |
| EPI_ISL_3097090, EPI_ISL_3097091 | Lighthouse Lab in Milton Keynes | Wellcome Sanger Institute for the COVID-19 Genomics UK (COG-UK) Consortium | The Lighthouse Lab in Milton Keynes and Alex Alderton, Roberto Amato, Jeffrey Barrett, Sonia Goncalves, Ewan Harrison, David K. Jackson, Ian Johnston, Dominic Kwiatkowski, Cordelia Langford, John Sillitoe on behalf of the Wellcome Sanger Institute COVID-19 Surveillance Team |
| EPI_ISL_3097092 | Lighthouse Lab in Glasgow | Wellcome Sanger Institute for the COVID-19 Genomics UK (COG-UK) Consortium | Harper VanSteenhouse, Yumi Kasai, David Gray, Carol Clugston, Anna Dominiczak and Alex Alderton, Roberto Amato, Jeffrey Barrett, Sonia Goncalves, Ewan Harrison, David K. Jackson, Ian Johnston, Dominic Kwiatkowski, Cordelia Langford, John Sillitoe on behalf of the Wellcome Sanger Institute COVID-19 Surveillance Team |
| EPI_ISL_3097093, EPI_ISL_3097094, EPI_ISL_3097095, EPI_ISL_3097096, EPI_ISL_3097097, EPI_ISL_3097098 | Lighthouse Lab in Milton Keynes | Wellcome Sanger Institute for the COVID-19 Genomics UK (COG-UK) Consortium | The Lighthouse Lab in Milton Keynes and Alex Alderton, Roberto Amato, Jeffrey Barrett, Sonia Goncalves, Ewan Harrison, David K. Jackson, Ian Johnston, Dominic Kwiatkowski, Cordelia Langford, John Sillitoe on behalf of the Wellcome Sanger Institute COVID-19 Surveillance Team |
| EPI_ISL_3097099, EPI_ISL_3097100 | Lighthouse Lab in Glasgow | Wellcome Sanger Institute for the COVID-19 Genomics UK (COG-UK) Consortium | Harper VanSteenhouse, Yumi Kasai, David Gray, Carol Clugston, Anna Dominiczak and Alex Alderton, Roberto Amato, Jeffrey Barrett, Sonia Goncalves, Ewan Harrison, David K. Jackson, Ian Johnston, Dominic Kwiatkowski, Cordelia Langford, John Sillitoe on behalf of the Wellcome Sanger Institute COVID-19 Surveillance Team |
| EPI_ISL_3097101, EPI_ISL_3097102 | Lighthouse Lab in Milton Keynes | Wellcome Sanger Institute for the COVID-19 Genomics UK (COG-UK) Consortium | The Lighthouse Lab in Milton Keynes and Alex Alderton, Roberto Amato, Jeffrey Barrett, Sonia Goncalves, Ewan Harrison, David K. Jackson, Ian Johnston, Dominic Kwiatkowski, Cordelia Langford, John Sillitoe on behalf of the Wellcome Sanger Institute COVID-19 Surveillance Team |
| EPI_ISL_3097103, EPI_ISL_3097104, EPI_ISL_3097105 | Lighthouse Lab in Glasgow | Wellcome Sanger Institute for the COVID-19 Genomics UK (COG-UK) Consortium | Harper VanSteenhouse, Yumi Kasai, David Gray, Carol Clugston, Anna Dominiczak and Alex Alderton, Roberto Amato, Jeffrey Barrett, Sonia Goncalves, Ewan Harrison, David K. Jackson, Ian Johnston, Dominic Kwiatkowski, Cordelia Langford, John Sillitoe on behalf of the Wellcome Sanger Institute COVID-19 Surveillance Team |
| EPI_ISL_3097106 | Lighthouse Lab in Milton Keynes | Wellcome Sanger Institute for the COVID-19 Genomics UK (COG-UK) Consortium | The Lighthouse Lab in Milton Keynes and Alex Alderton, Roberto Amato, Jeffrey Barrett, Sonia Goncalves, Ewan Harrison, David K. Jackson, Ian Johnston, Dominic Kwiatkowski, Cordelia Langford, John Sillitoe on behalf of the Wellcome Sanger Institute COVID-19 Surveillance Team |
| EPI_ISL_3097107, EPI_ISL_3097108, EPI_ISL_3097109, EPI_ISL_3097110, EPI_ISL_3097111 | Lighthouse Lab in Glasgow | Wellcome Sanger Institute for the COVID-19 Genomics UK (COG-UK) Consortium | Harper VanSteenhouse, Yumi Kasai, David Gray, Carol Clugston, Anna Dominiczak and Alex Alderton, Roberto Amato, Jeffrey Barrett, Sonia Goncalves, Ewan Harrison, David K. Jackson, Ian Johnston, Dominic Kwiatkowski, Cordelia Langford, John Sillitoe on behalf of the Wellcome Sanger Institute COVID-19 Surveillance Team |
| EPI_ISL_3097112, EPI_ISL_3097113, EPI_ISL_3097114, EPI_ISL_3097115 | Lighthouse Lab in Milton Keynes | Wellcome Sanger Institute for the COVID-19 Genomics UK (COG-UK) Consortium | The Lighthouse Lab in Milton Keynes and Alex Alderton, Roberto Amato, Jeffrey Barrett, Sonia Goncalves, Ewan Harrison, David K. Jackson, Ian Johnston, Dominic Kwiatkowski, Cordelia Langford, John Sillitoe on behalf of the Wellcome Sanger Institute COVID-19 Surveillance Team |
| EPI_ISL_3097116 | Lighthouse Lab in Glasgow | Wellcome Sanger Institute for the COVID-19 Genomics UK (COG-UK) Consortium | Harper VanSteenhouse, Yumi Kasai, David Gray, Carol Clugston, Anna Dominiczak and Alex Alderton, Roberto Amato, Jeffrey Barrett, Sonia Goncalves, Ewan Harrison, David K. Jackson, Ian Johnston, Dominic Kwiatkowski, Cordelia Langford, John Sillitoe on behalf of the Wellcome Sanger Institute COVID-19 Surveillance Team |
| EPI_ISL_3097117 | Lighthouse Lab in Milton Keynes | Wellcome Sanger Institute for the COVID-19 Genomics UK (COG-UK) Consortium | The Lighthouse Lab in Milton Keynes and Alex Alderton, Roberto Amato, Jeffrey Barrett, Sonia Goncalves, Ewan Harrison, David K. Jackson, Ian Johnston, Dominic Kwiatkowski, Cordelia Langford, John Sillitoe on behalf of the Wellcome Sanger Institute COVID-19 Surveillance Team |
| EPI_ISL_3097118, EPI_ISL_3097119 | Lighthouse Lab in Glasgow | Wellcome Sanger Institute for the COVID-19 Genomics UK (COG-UK) Consortium | Harper VanSteenhouse, Yumi Kasai, David Gray, Carol Clugston, Anna Dominiczak and Alex Alderton, Roberto Amato, Jeffrey Barrett, Sonia Goncalves, Ewan Harrison, David K. Jackson, Ian Johnston, Dominic Kwiatkowski, Cordelia Langford, John Sillitoe on behalf of the Wellcome Sanger Institute COVID-19 Surveillance Team |
| EPI_ISL_3097120, EPI_ISL_3097121, EPI_ISL_3097122, EPI_ISL_3097123 | Lighthouse Lab in Milton Keynes | Wellcome Sanger Institute for the COVID-19 Genomics UK (COG-UK) Consortium | The Lighthouse Lab in Milton Keynes and Alex Alderton, Roberto Amato, Jeffrey Barrett, Sonia Goncalves, Ewan Harrison, David K. Jackson, Ian Johnston, Dominic Kwiatkowski, Cordelia Langford, John Sillitoe on behalf of the Wellcome Sanger Institute COVID-19 Surveillance Team |
| EPI_ISL_3097124, EPI_ISL_3097125 | Lighthouse Lab in Glasgow | Wellcome Sanger Institute for the COVID-19 Genomics UK (COG-UK) Consortium | Harper VanSteenhouse, Yumi Kasai, David Gray, Carol Clugston, Anna Dominiczak and Alex Alderton, Roberto Amato, Jeffrey Barrett, Sonia Goncalves, Ewan Harrison, David K. Jackson, Ian Johnston, Dominic Kwiatkowski, Cordelia Langford, John Sillitoe on behalf of the Wellcome Sanger Institute COVID-19 Surveillance Team |
| EPI_ISL_3097126, EPI_ISL_3097127, EPI_ISL_3097128, EPI_ISL_3097129, EPI_ISL_3097130 | Lighthouse Lab in Milton Keynes | Wellcome Sanger Institute for the COVID-19 Genomics UK (COG-UK) Consortium | The Lighthouse Lab in Milton Keynes and Alex Alderton, Roberto Amato, Jeffrey Barrett, Sonia Goncalves, Ewan Harrison, David K. Jackson, Ian Johnston, Dominic Kwiatkowski, Cordelia Langford, John Sillitoe on behalf of the Wellcome Sanger Institute COVID-19 Surveillance Team |
| EPI_ISL_3097131, EPI_ISL_3097132, EPI_ISL_3097133 | Lighthouse Lab in Glasgow | Wellcome Sanger Institute for the COVID-19 Genomics UK (COG-UK) Consortium | Harper VanSteenhouse, Yumi Kasai, David Gray, Carol Clugston, Anna Dominiczak and Alex Alderton, Roberto Amato, Jeffrey Barrett, Sonia Goncalves, Ewan Harrison, David K. Jackson, Ian Johnston, Dominic Kwiatkowski, Cordelia Langford, John Sillitoe on behalf of the Wellcome Sanger Institute COVID-19 Surveillance Team |
| EPI_ISL_3097134 | Lighthouse Lab in Milton Keynes | Wellcome Sanger Institute for the COVID-19 Genomics UK (COG-UK) Consortium | The Lighthouse Lab in Milton Keynes and Alex Alderton, Roberto Amato, Jeffrey Barrett, Sonia Goncalves, Ewan Harrison, David K. Jackson, Ian Johnston, Dominic Kwiatkowski, Cordelia Langford, John Sillitoe on behalf of the Wellcome Sanger Institute COVID-19 Surveillance Team |
| EPI_ISL_3097135, EPI_ISL_3097136 | Lighthouse Lab in Glasgow | Wellcome Sanger Institute for the COVID-19 Genomics UK (COG-UK) Consortium | Harper VanSteenhouse, Yumi Kasai, David Gray, Carol Clugston, Anna Dominiczak and Alex Alderton, Roberto Amato, Jeffrey Barrett, Sonia Goncalves, Ewan Harrison, David K. Jackson, Ian Johnston, Dominic Kwiatkowski, Cordelia Langford, John Sillitoe on behalf of the Wellcome Sanger Institute COVID-19 Surveillance Team |
| EPI_ISL_3097137, EPI_ISL_3097138 | Lighthouse Lab in Milton Keynes | Wellcome Sanger Institute for the COVID-19 Genomics UK (COG-UK) Consortium | The Lighthouse Lab in Milton Keynes and Alex Alderton, Roberto Amato, Jeffrey Barrett, Sonia Goncalves, Ewan Harrison, David K. Jackson, Ian Johnston, Dominic Kwiatkowski, Cordelia Langford, John Sillitoe on behalf of the Wellcome Sanger Institute COVID-19 Surveillance Team |
| EPI_ISL_3097139, EPI_ISL_3097140, EPI_ISL_3097141, EPI_ISL_3097142 | Lighthouse Lab in Glasgow | Wellcome Sanger Institute for the COVID-19 Genomics UK (COG-UK) Consortium | Harper VanSteenhouse, Yumi Kasai, David Gray, Carol Clugston, Anna Dominiczak and Alex Alderton, Roberto Amato, Jeffrey Barrett, Sonia Goncalves, Ewan Harrison, David K. Jackson, Ian Johnston, Dominic Kwiatkowski, Cordelia Langford, John Sillitoe on behalf of the Wellcome Sanger Institute COVID-19 Surveillance Team |
| EPI_ISL_3097143, EPI_ISL_3097144, EPI_ISL_3097145 | Lighthouse Lab in Milton Keynes | Wellcome Sanger Institute for the COVID-19 Genomics UK (COG-UK) Consortium | The Lighthouse Lab in Milton Keynes and Alex Alderton, Roberto Amato, Jeffrey Barrett, Sonia Goncalves, Ewan Harrison, David K. Jackson, Ian Johnston, Dominic Kwiatkowski, Cordelia Langford, John Sillitoe on behalf of the Wellcome Sanger Institute COVID-19 Surveillance Team |
| EPI_ISL_3097146, EPI_ISL_3097147 | Lighthouse Lab in Glasgow | Wellcome Sanger Institute for the COVID-19 Genomics UK (COG-UK) Consortium | Harper VanSteenhouse, Yumi Kasai, David Gray, Carol Clugston, Anna Dominiczak and Alex Alderton, Roberto Amato, Jeffrey Barrett, Sonia Goncalves, Ewan Harrison, David K. Jackson, Ian Johnston, Dominic Kwiatkowski, Cordelia Langford, John Sillitoe on behalf of the Wellcome Sanger Institute COVID-19 Surveillance Team |
| EPI_ISL_3097148, EPI_ISL_3097149, EPI_ISL_3097150, EPI_ISL_3097151 | Lighthouse Lab in Milton Keynes | Wellcome Sanger Institute for the COVID-19 Genomics UK (COG-UK) Consortium | The Lighthouse Lab in Milton Keynes and Alex Alderton, Roberto Amato, Jeffrey Barrett, Sonia Goncalves, Ewan Harrison, David K. Jackson, Ian Johnston, Dominic Kwiatkowski, Cordelia Langford, John Sillitoe on behalf of the Wellcome Sanger Institute COVID-19 Surveillance Team |
| EPI_ISL_3097152 | Lighthouse Lab in Glasgow | Wellcome Sanger Institute for the COVID-19 Genomics UK (COG-UK) Consortium | Harper VanSteenhouse, Yumi Kasai, David Gray, Carol Clugston, Anna Dominiczak and Alex Alderton, Roberto Amato, Jeffrey Barrett, Sonia Goncalves, Ewan Harrison, David K. Jackson, Ian Johnston, Dominic Kwiatkowski, Cordelia Langford, John Sillitoe on behalf of the Wellcome Sanger Institute COVID-19 Surveillance Team |
| EPI_ISL_3097153, EPI_ISL_3097154 | Lighthouse Lab in Milton Keynes | Wellcome Sanger Institute for the COVID-19 Genomics UK (COG-UK) Consortium | The Lighthouse Lab in Milton Keynes and Alex Alderton, Roberto Amato, Jeffrey Barrett, Sonia Goncalves, Ewan Harrison, David K. Jackson, Ian Johnston, Dominic Kwiatkowski, Cordelia Langford, John Sillitoe on behalf of the Wellcome Sanger Institute COVID-19 Surveillance Team |
| EPI_ISL_3097155, EPI_ISL_3097156, EPI_ISL_3097157 | Lighthouse Lab in Glasgow | Wellcome Sanger Institute for the COVID-19 Genomics UK (COG-UK) Consortium | Harper VanSteenhouse, Yumi Kasai, David Gray, Carol Clugston, Anna Dominiczak and Alex Alderton, Roberto Amato, Jeffrey Barrett, Sonia Goncalves, Ewan Harrison, David K. Jackson, Ian Johnston, Dominic Kwiatkowski, Cordelia Langford, John Sillitoe on behalf of the Wellcome Sanger Institute COVID-19 Surveillance Team |
| EPI_ISL_3097158, EPI_ISL_3097159, EPI_ISL_3097160, EPI_ISL_3097161, EPI_ISL_3097162, EPI_ISL_3097163, EPI_ISL_3097164 | Lighthouse Lab in Milton Keynes | Wellcome Sanger Institute for the COVID-19 Genomics UK (COG-UK) Consortium | The Lighthouse Lab in Milton Keynes and Alex Alderton, Roberto Amato, Jeffrey Barrett, Sonia Goncalves, Ewan Harrison, David K. Jackson, Ian Johnston, Dominic Kwiatkowski, Cordelia Langford, John Sillitoe on behalf of the Wellcome Sanger Institute COVID-19 Surveillance Team |
| EPI_ISL_3097165 | Lighthouse Lab in Glasgow | Wellcome Sanger Institute for the COVID-19 Genomics UK (COG-UK) Consortium | Harper VanSteenhouse, Yumi Kasai, David Gray, Carol Clugston, Anna Dominiczak and Alex Alderton, Roberto Amato, Jeffrey Barrett, Sonia Goncalves, Ewan Harrison, David K. Jackson, Ian Johnston, Dominic Kwiatkowski, Cordelia Langford, John Sillitoe on behalf of the Wellcome Sanger Institute COVID-19 Surveillance Team |
| EPI_ISL_3097166 | Lighthouse Lab in Milton Keynes | Wellcome Sanger Institute for the COVID-19 Genomics UK (COG-UK) Consortium | The Lighthouse Lab in Milton Keynes and Alex Alderton, Roberto Amato, Jeffrey Barrett, Sonia Goncalves, Ewan Harrison, David K. Jackson, Ian Johnston, Dominic Kwiatkowski, Cordelia Langford, John Sillitoe on behalf of the Wellcome Sanger Institute COVID-19 Surveillance Team |
| EPI_ISL_3097167 | Lighthouse Lab in Glasgow | Wellcome Sanger Institute for the COVID-19 Genomics UK (COG-UK) Consortium | Harper VanSteenhouse, Yumi Kasai, David Gray, Carol Clugston, Anna Dominiczak and Alex Alderton, Roberto Amato, Jeffrey Barrett, Sonia Goncalves, Ewan Harrison, David K. Jackson, Ian Johnston, Dominic Kwiatkowski, Cordelia Langford, John Sillitoe on behalf of the Wellcome Sanger Institute COVID-19 Surveillance Team |
| EPI_ISL_3097168, EPI_ISL_3097169, EPI_ISL_3097170 | Lighthouse Lab in Milton Keynes | Wellcome Sanger Institute for the COVID-19 Genomics UK (COG-UK) Consortium | The Lighthouse Lab in Milton Keynes and Alex Alderton, Roberto Amato, Jeffrey Barrett, Sonia Goncalves, Ewan Harrison, David K. Jackson, Ian Johnston, Dominic Kwiatkowski, Cordelia Langford, John Sillitoe on behalf of the Wellcome Sanger Institute COVID-19 Surveillance Team |
| EPI_ISL_3097171 | Lighthouse Lab in Glasgow | Wellcome Sanger Institute for the COVID-19 Genomics UK (COG-UK) Consortium | Harper VanSteenhouse, Yumi Kasai, David Gray, Carol Clugston, Anna Dominiczak and Alex Alderton, Roberto Amato, Jeffrey Barrett, Sonia Goncalves, Ewan Harrison, David K. Jackson, Ian Johnston, Dominic Kwiatkowski, Cordelia Langford, John Sillitoe on behalf of the Wellcome Sanger Institute COVID-19 Surveillance Team |
| EPI_ISL_3097172, | Lighthouse Lab in Milton Keynes | Wellcome Sanger Institute for the COVID-19 Genomics UK (COG-UK) | The Lighthouse Lab in Milton Keynes and Alex Alderton, Roberto Amato, Jeffrey Barrett, Sonia Goncalves, Ewan Harrison, David K. Jackson, Ian Johnston, Dominic Kwiatkowski, Cordelia Langford, John Sillitoe on behalf of the Wellcome Sanger |

| | | | |
|---|---|---|---|
| EPI_ISL_3097173, EPI_ISL_3097174, EPI_ISL_3097175, EPI_ISL_3097176, EPI_ISL_3097177, EPI_ISL_3097178, EPI_ISL_3097179 | | Consortium | Institute COVID-19 Surveillance Team |
| EPI_ISL_3097180, EPI_ISL_3097181, EPI_ISL_3097182 | Lighthouse Lab in Glasgow | Wellcome Sanger Institute for the COVID-19 Genomics UK (COG-UK) Consortium | Harper VanSteenhouse, Yumi Kasai, David Gray, Carol Clugston, Anna Dominiczak and Alex Alderton, Roberto Amato, Jeffrey Barrett, Sonia Goncalves, Ewan Harrison, David K. Jackson, Ian Johnston, Dominic Kwiatkowski, Cordelia Langford, John Sillitoe on behalf of the Wellcome Sanger Institute COVID-19 Surveillance Team |
| EPI_ISL_3097183 | Lighthouse Lab in Milton Keynes | Wellcome Sanger Institute for the COVID-19 Genomics UK (COG-UK) Consortium | The Lighthouse Lab in Milton Keynes and Alex Alderton, Roberto Amato, Jeffrey Barrett, Sonia Goncalves, Ewan Harrison, David K. Jackson, Ian Johnston, Dominic Kwiatkowski, Cordelia Langford, John Sillitoe on behalf of the Wellcome Sanger Institute COVID-19 Surveillance Team |
| EPI_ISL_3097184, EPI_ISL_3097185, EPI_ISL_3097186, EPI_ISL_3097187 | Lighthouse Lab in Glasgow | Wellcome Sanger Institute for the COVID-19 Genomics UK (COG-UK) Consortium | Harper VanSteenhouse, Yumi Kasai, David Gray, Carol Clugston, Anna Dominiczak and Alex Alderton, Roberto Amato, Jeffrey Barrett, Sonia Goncalves, Ewan Harrison, David K. Jackson, Ian Johnston, Dominic Kwiatkowski, Cordelia Langford, John Sillitoe on behalf of the Wellcome Sanger Institute COVID-19 Surveillance Team |
| EPI_ISL_3097188 | Lighthouse Lab in Milton Keynes | Wellcome Sanger Institute for the COVID-19 Genomics UK (COG-UK) Consortium | The Lighthouse Lab in Milton Keynes and Alex Alderton, Roberto Amato, Jeffrey Barrett, Sonia Goncalves, Ewan Harrison, David K. Jackson, Ian Johnston, Dominic Kwiatkowski, Cordelia Langford, John Sillitoe on behalf of the Wellcome Sanger Institute COVID-19 Surveillance Team |
| EPI_ISL_3097189, EPI_ISL_3097190 | Lighthouse Lab in Glasgow | Wellcome Sanger Institute for the COVID-19 Genomics UK (COG-UK) Consortium | Harper VanSteenhouse, Yumi Kasai, David Gray, Carol Clugston, Anna Dominiczak and Alex Alderton, Roberto Amato, Jeffrey Barrett, Sonia Goncalves, Ewan Harrison, David K. Jackson, Ian Johnston, Dominic Kwiatkowski, Cordelia Langford, John Sillitoe on behalf of the Wellcome Sanger Institute COVID-19 Surveillance Team |
| EPI_ISL_3097191 | Lighthouse Lab in Milton Keynes | Wellcome Sanger Institute for the COVID-19 Genomics UK (COG-UK) Consortium | The Lighthouse Lab in Milton Keynes and Alex Alderton, Roberto Amato, Jeffrey Barrett, Sonia Goncalves, Ewan Harrison, David K. Jackson, Ian Johnston, Dominic Kwiatkowski, Cordelia Langford, John Sillitoe on behalf of the Wellcome Sanger Institute COVID-19 Surveillance Team |
| EPI_ISL_3097192 | Lighthouse Lab in Glasgow | Wellcome Sanger Institute for the COVID-19 Genomics UK (COG-UK) Consortium | Harper VanSteenhouse, Yumi Kasai, David Gray, Carol Clugston, Anna Dominiczak and Alex Alderton, Roberto Amato, Jeffrey Barrett, Sonia Goncalves, Ewan Harrison, David K. Jackson, Ian Johnston, Dominic Kwiatkowski, Cordelia Langford, John Sillitoe on behalf of the Wellcome Sanger Institute COVID-19 Surveillance Team |
| EPI_ISL_3097193, EPI_ISL_3097194 | Lighthouse Lab in Milton Keynes | Wellcome Sanger Institute for the COVID-19 Genomics UK (COG-UK) Consortium | The Lighthouse Lab in Milton Keynes and Alex Alderton, Roberto Amato, Jeffrey Barrett, Sonia Goncalves, Ewan Harrison, David K. Jackson, Ian Johnston, Dominic Kwiatkowski, Cordelia Langford, John Sillitoe on behalf of the Wellcome Sanger Institute COVID-19 Surveillance Team |
| EPI_ISL_3097195, EPI_ISL_3097196 | Lighthouse Lab in Glasgow | Wellcome Sanger Institute for the COVID-19 Genomics UK (COG-UK) Consortium | Harper VanSteenhouse, Yumi Kasai, David Gray, Carol Clugston, Anna Dominiczak and Alex Alderton, Roberto Amato, Jeffrey Barrett, Sonia Goncalves, Ewan Harrison, David K. Jackson, Ian Johnston, Dominic Kwiatkowski, Cordelia Langford, John Sillitoe on behalf of the Wellcome Sanger Institute COVID-19 Surveillance Team |
| EPI_ISL_3097197, EPI_ISL_3097198 | Lighthouse Lab in Milton Keynes | Wellcome Sanger Institute for the COVID-19 Genomics UK (COG-UK) Consortium | The Lighthouse Lab in Milton Keynes and Alex Alderton, Roberto Amato, Jeffrey Barrett, Sonia Goncalves, Ewan Harrison, David K. Jackson, Ian Johnston, Dominic Kwiatkowski, Cordelia Langford, John Sillitoe on behalf of the Wellcome Sanger Institute COVID-19 Surveillance Team |
| EPI_ISL_3097199, EPI_ISL_3097200, EPI_ISL_3097201 | Lighthouse Lab in Glasgow | Wellcome Sanger Institute for the COVID-19 Genomics UK (COG-UK) Consortium | Harper VanSteenhouse, Yumi Kasai, David Gray, Carol Clugston, Anna Dominiczak and Alex Alderton, Roberto Amato, Jeffrey Barrett, Sonia Goncalves, Ewan Harrison, David K. Jackson, Ian Johnston, Dominic Kwiatkowski, Cordelia Langford, John Sillitoe on behalf of the Wellcome Sanger Institute COVID-19 Surveillance Team |
| EPI_ISL_3097202, EPI_ISL_3097203, EPI_ISL_3097204, EPI_ISL_3097205 | Lighthouse Lab in Milton Keynes | Wellcome Sanger Institute for the COVID-19 Genomics UK (COG-UK) Consortium | The Lighthouse Lab in Milton Keynes and Alex Alderton, Roberto Amato, Jeffrey Barrett, Sonia Goncalves, Ewan Harrison, David K. Jackson, Ian Johnston, Dominic Kwiatkowski, Cordelia Langford, John Sillitoe on behalf of the Wellcome Sanger Institute COVID-19 Surveillance Team |
| EPI_ISL_3097206, EPI_ISL_3097207 | Lighthouse Lab in Glasgow | Wellcome Sanger Institute for the COVID-19 Genomics UK (COG-UK) Consortium | Harper VanSteenhouse, Yumi Kasai, David Gray, Carol Clugston, Anna Dominiczak and Alex Alderton, Roberto Amato, Jeffrey Barrett, Sonia Goncalves, Ewan Harrison, David K. Jackson, Ian Johnston, Dominic Kwiatkowski, Cordelia Langford, John Sillitoe on behalf of the Wellcome Sanger Institute COVID-19 Surveillance Team |
| EPI_ISL_3097208 | Lighthouse Lab in Milton Keynes | Wellcome Sanger Institute for the COVID-19 Genomics UK (COG-UK) Consortium | The Lighthouse Lab in Milton Keynes and Alex Alderton, Roberto Amato, Jeffrey Barrett, Sonia Goncalves, Ewan Harrison, David K. Jackson, Ian Johnston, Dominic Kwiatkowski, Cordelia Langford, John Sillitoe on behalf of the Wellcome Sanger Institute COVID-19 Surveillance Team |
| EPI_ISL_3097209, EPI_ISL_3097210, EPI_ISL_3097211 | Lighthouse Lab in Glasgow | Wellcome Sanger Institute for the COVID-19 Genomics UK (COG-UK) Consortium | Harper VanSteenhouse, Yumi Kasai, David Gray, Carol Clugston, Anna Dominiczak and Alex Alderton, Roberto Amato, Jeffrey Barrett, Sonia Goncalves, Ewan Harrison, David K. Jackson, Ian Johnston, Dominic Kwiatkowski, Cordelia Langford, John Sillitoe on behalf of the Wellcome Sanger Institute COVID-19 Surveillance Team |
| EPI_ISL_3097212, EPI_ISL_3097213, EPI_ISL_3097214, EPI_ISL_3097215 | Lighthouse Lab in Milton Keynes | Wellcome Sanger Institute for the COVID-19 Genomics UK (COG-UK) Consortium | The Lighthouse Lab in Milton Keynes and Alex Alderton, Roberto Amato, Jeffrey Barrett, Sonia Goncalves, Ewan Harrison, David K. Jackson, Ian Johnston, Dominic Kwiatkowski, Cordelia Langford, John Sillitoe on behalf of the Wellcome Sanger Institute COVID-19 Surveillance Team |
| EPI_ISL_3097216, EPI_ISL_3097217 | Lighthouse Lab in Glasgow | Wellcome Sanger Institute for the COVID-19 Genomics UK (COG-UK) Consortium | Harper VanSteenhouse, Yumi Kasai, David Gray, Carol Clugston, Anna Dominiczak and Alex Alderton, Roberto Amato, Jeffrey Barrett, Sonia Goncalves, Ewan Harrison, David K. Jackson, Ian Johnston, Dominic Kwiatkowski, Cordelia Langford, John Sillitoe on behalf of the Wellcome Sanger Institute COVID-19 Surveillance Team |
| EPI_ISL_3097218, EPI_ISL_3097219 | Lighthouse Lab in Milton Keynes | Wellcome Sanger Institute for the COVID-19 Genomics UK (COG-UK) Consortium | The Lighthouse Lab in Milton Keynes and Alex Alderton, Roberto Amato, Jeffrey Barrett, Sonia Goncalves, Ewan Harrison, David K. Jackson, Ian Johnston, Dominic Kwiatkowski, Cordelia Langford, John Sillitoe on behalf of the Wellcome Sanger Institute COVID-19 Surveillance Team |
| EPI_ISL_3097220, EPI_ISL_3097221, EPI_ISL_3097222 | Lighthouse Lab in Glasgow | Wellcome Sanger Institute for the COVID-19 Genomics UK (COG-UK) Consortium | Harper VanSteenhouse, Yumi Kasai, David Gray, Carol Clugston, Anna Dominiczak and Alex Alderton, Roberto Amato, Jeffrey Barrett, Sonia Goncalves, Ewan Harrison, David K. Jackson, Ian Johnston, Dominic Kwiatkowski, Cordelia Langford, John Sillitoe on behalf of the Wellcome Sanger Institute COVID-19 Surveillance Team |
| EPI_ISL_3097223, EPI_ISL_3097224, EPI_ISL_3097225 | Lighthouse Lab in Milton Keynes | Wellcome Sanger Institute for the COVID-19 Genomics UK (COG-UK) Consortium | The Lighthouse Lab in Milton Keynes and Alex Alderton, Roberto Amato, Jeffrey Barrett, Sonia Goncalves, Ewan Harrison, David K. Jackson, Ian Johnston, Dominic Kwiatkowski, Cordelia Langford, John Sillitoe on behalf of the Wellcome Sanger Institute COVID-19 Surveillance Team |
| EPI_ISL_3097226 | Lighthouse Lab in Glasgow | Wellcome Sanger Institute for the COVID-19 Genomics UK (COG-UK) Consortium | Harper VanSteenhouse, Yumi Kasai, David Gray, Carol Clugston, Anna Dominiczak and Alex Alderton, Roberto Amato, Jeffrey Barrett, Sonia Goncalves, Ewan Harrison, David K. Jackson, Ian Johnston, Dominic Kwiatkowski, Cordelia Langford, John Sillitoe on behalf of the Wellcome Sanger Institute COVID-19 Surveillance Team |
| EPI_ISL_3097227, EPI_ISL_3097228, EPI_ISL_3097229 | Lighthouse Lab in Milton Keynes | Wellcome Sanger Institute for the COVID-19 Genomics UK (COG-UK) Consortium | The Lighthouse Lab in Milton Keynes and Alex Alderton, Roberto Amato, Jeffrey Barrett, Sonia Goncalves, Ewan Harrison, David K. Jackson, Ian Johnston, Dominic Kwiatkowski, Cordelia Langford, John Sillitoe on behalf of the Wellcome Sanger Institute COVID-19 Surveillance Team |
| EPI_ISL_3097230 | Lighthouse Lab in Glasgow | Wellcome Sanger Institute for the COVID-19 Genomics UK (COG-UK) Consortium | Harper VanSteenhouse, Yumi Kasai, David Gray, Carol Clugston, Anna Dominiczak and Alex Alderton, Roberto Amato, Jeffrey Barrett, Sonia Goncalves, Ewan Harrison, David K. Jackson, Ian Johnston, Dominic Kwiatkowski, Cordelia Langford, John Sillitoe on behalf of the Wellcome Sanger Institute COVID-19 Surveillance Team |
| EPI_ISL_3097231 | Lighthouse Lab in Milton Keynes | Wellcome Sanger Institute for the COVID-19 Genomics UK (COG-UK) Consortium | The Lighthouse Lab in Milton Keynes and Alex Alderton, Roberto Amato, Jeffrey Barrett, Sonia Goncalves, Ewan Harrison, David K. Jackson, Ian Johnston, Dominic Kwiatkowski, Cordelia Langford, John Sillitoe on behalf of the Wellcome Sanger Institute COVID-19 Surveillance Team |
| EPI_ISL_3097232 | Lighthouse Lab in Glasgow | Wellcome Sanger Institute for the COVID-19 Genomics UK (COG-UK) Consortium | Harper VanSteenhouse, Yumi Kasai, David Gray, Carol Clugston, Anna Dominiczak and Alex Alderton, Roberto Amato, Jeffrey Barrett, Sonia Goncalves, Ewan Harrison, David K. Jackson, Ian Johnston, Dominic Kwiatkowski, Cordelia Langford, John Sillitoe on behalf of the Wellcome Sanger Institute COVID-19 Surveillance Team |
| EPI_ISL_3097233, EPI_ISL_3097234 | Lighthouse Lab in Milton Keynes | Wellcome Sanger Institute for the COVID-19 Genomics UK (COG-UK) Consortium | The Lighthouse Lab in Milton Keynes and Alex Alderton, Roberto Amato, Jeffrey Barrett, Sonia Goncalves, Ewan Harrison, David K. Jackson, Ian Johnston, Dominic Kwiatkowski, Cordelia Langford, John Sillitoe on behalf of the Wellcome Sanger Institute COVID-19 Surveillance Team |
| EPI_ISL_3097235, EPI_ISL_3097236, EPI_ISL_3097237 | Lighthouse Lab in Glasgow | Wellcome Sanger Institute for the COVID-19 Genomics UK (COG-UK) Consortium | Harper VanSteenhouse, Yumi Kasai, David Gray, Carol Clugston, Anna Dominiczak and Alex Alderton, Roberto Amato, Jeffrey Barrett, Sonia Goncalves, Ewan Harrison, David K. Jackson, Ian Johnston, Dominic Kwiatkowski, Cordelia Langford, John Sillitoe on behalf of the Wellcome Sanger Institute COVID-19 Surveillance Team |
| EPI_ISL_3097238 | Lighthouse Lab in Milton Keynes | Wellcome Sanger Institute for the COVID-19 Genomics UK (COG-UK) Consortium | The Lighthouse Lab in Milton Keynes and Alex Alderton, Roberto Amato, Jeffrey Barrett, Sonia Goncalves, Ewan Harrison, David K. Jackson, Ian Johnston, Dominic Kwiatkowski, Cordelia Langford, John Sillitoe on behalf of the Wellcome Sanger Institute COVID-19 Surveillance Team |
| EPI_ISL_3097239, EPI_ISL_3097240 | Lighthouse Lab in Glasgow | Wellcome Sanger Institute for the COVID-19 Genomics UK (COG-UK) Consortium | Harper VanSteenhouse, Yumi Kasai, David Gray, Carol Clugston, Anna Dominiczak and Alex Alderton, Roberto Amato, Jeffrey Barrett, Sonia Goncalves, Ewan Harrison, David K. Jackson, Ian Johnston, Dominic Kwiatkowski, Cordelia Langford, John Sillitoe on behalf of the Wellcome Sanger Institute COVID-19 Surveillance Team |
| EPI_ISL_3097241, EPI_ISL_3097242, EPI_ISL_3097243, EPI_ISL_3097244, EPI_ISL_3097245 | Lighthouse Lab in Milton Keynes | Wellcome Sanger Institute for the COVID-19 Genomics UK (COG-UK) Consortium | The Lighthouse Lab in Milton Keynes and Alex Alderton, Roberto Amato, Jeffrey Barrett, Sonia Goncalves, Ewan Harrison, David K. Jackson, Ian Johnston, Dominic Kwiatkowski, Cordelia Langford, John Sillitoe on behalf of the Wellcome Sanger Institute COVID-19 Surveillance Team |
| EPI_ISL_3097246, EPI_ISL_3097247, EPI_ISL_3097248 | Lighthouse Lab in Glasgow | Wellcome Sanger Institute for the COVID-19 Genomics UK (COG-UK) Consortium | Harper VanSteenhouse, Yumi Kasai, David Gray, Carol Clugston, Anna Dominiczak and Alex Alderton, Roberto Amato, Jeffrey Barrett, Sonia Goncalves, Ewan Harrison, David K. Jackson, Ian Johnston, Dominic Kwiatkowski, Cordelia Langford, John Sillitoe on behalf of the Wellcome Sanger Institute COVID-19 Surveillance Team |
| EPI_ISL_3097249, EPI_ISL_3097250 | Lighthouse Lab in Milton Keynes | Wellcome Sanger Institute for the COVID-19 Genomics UK (COG-UK) Consortium | The Lighthouse Lab in Milton Keynes and Alex Alderton, Roberto Amato, Jeffrey Barrett, Sonia Goncalves, Ewan Harrison, David K. Jackson, Ian Johnston, Dominic Kwiatkowski, Cordelia Langford, John Sillitoe on behalf of the Wellcome Sanger Institute COVID-19 Surveillance Team |
| EPI_ISL_3097251, EPI_ISL_3097252 | Lighthouse Lab in Glasgow | Wellcome Sanger Institute for the COVID-19 Genomics UK (COG-UK) Consortium | Harper VanSteenhouse, Yumi Kasai, David Gray, Carol Clugston, Anna Dominiczak and Alex Alderton, Roberto Amato, Jeffrey Barrett, Sonia Goncalves, Ewan Harrison, David K. Jackson, Ian Johnston, Dominic Kwiatkowski, Cordelia Langford, John Sillitoe on behalf of the Wellcome Sanger Institute COVID-19 Surveillance Team |
| EPI_ISL_3097253, EPI_ISL_3097254, EPI_ISL_3097255 | Lighthouse Lab in Milton Keynes | Wellcome Sanger Institute for the COVID-19 Genomics UK (COG-UK) Consortium | The Lighthouse Lab in Milton Keynes and Alex Alderton, Roberto Amato, Jeffrey Barrett, Sonia Goncalves, Ewan Harrison, David K. Jackson, Ian Johnston, Dominic Kwiatkowski, Cordelia Langford, John Sillitoe on behalf of the Wellcome Sanger Institute COVID-19 Surveillance Team |
| EPI_ISL_3097256 | Lighthouse Lab in Glasgow | Wellcome Sanger Institute for the COVID-19 Genomics UK (COG-UK) Consortium | Harper VanSteenhouse, Yumi Kasai, David Gray, Carol Clugston, Anna Dominiczak and Alex Alderton, Roberto Amato, Jeffrey Barrett, Sonia Goncalves, Ewan Harrison, David K. Jackson, Ian Johnston, Dominic Kwiatkowski, Cordelia Langford, John Sillitoe on behalf of the Wellcome Sanger Institute COVID-19 Surveillance Team |

| | | | |
|---|---|---|---|
| EPI_ISL_3097257 | Lighthouse Lab in Milton Keynes | Wellcome Sanger Institute for the COVID-19 Genomics UK (COG-UK) Consortium | The Lighthouse Lab in Milton Keynes and Alex Alderton, Roberto Amato, Jeffrey Barrett, Sonia Goncalves, Ewan Harrison, David K. Jackson, Ian Johnston, Dominic Kwiatkowski, Cordelia Langford, John Sillitoe on behalf of the Wellcome Sanger Institute COVID-19 Surveillance Team |
| EPI_ISL_3097258 | Lighthouse Lab in Glasgow | Wellcome Sanger Institute for the COVID-19 Genomics UK (COG-UK) Consortium | Harper VanSteenhouse, Yumi Kasai, David Gray, Carol Clugston, Anna Dominiczak and Alex Alderton, Roberto Amato, Jeffrey Barrett, Sonia Goncalves, Ewan Harrison, David K. Jackson, Ian Johnston, Dominic Kwiatkowski, Cordelia Langford, John Sillitoe on behalf of the Wellcome Sanger Institute COVID-19 Surveillance Team |
| EPI_ISL_3097259 | Lighthouse Lab in Milton Keynes | Wellcome Sanger Institute for the COVID-19 Genomics UK (COG-UK) Consortium | The Lighthouse Lab in Milton Keynes and Alex Alderton, Roberto Amato, Jeffrey Barrett, Sonia Goncalves, Ewan Harrison, David K. Jackson, Ian Johnston, Dominic Kwiatkowski, Cordelia Langford, John Sillitoe on behalf of the Wellcome Sanger Institute COVID-19 Surveillance Team |
| EPI_ISL_3097260, EPI_ISL_3097261, EPI_ISL_3097262 | Lighthouse Lab in Glasgow | Wellcome Sanger Institute for the COVID-19 Genomics UK (COG-UK) Consortium | Harper VanSteenhouse, Yumi Kasai, David Gray, Carol Clugston, Anna Dominiczak and Alex Alderton, Roberto Amato, Jeffrey Barrett, Sonia Goncalves, Ewan Harrison, David K. Jackson, Ian Johnston, Dominic Kwiatkowski, Cordelia Langford, John Sillitoe on behalf of the Wellcome Sanger Institute COVID-19 Surveillance Team |
| EPI_ISL_3097263, EPI_ISL_3097264, EPI_ISL_3097265 | Lighthouse Lab in Milton Keynes | Wellcome Sanger Institute for the COVID-19 Genomics UK (COG-UK) Consortium | The Lighthouse Lab in Milton Keynes and Alex Alderton, Roberto Amato, Jeffrey Barrett, Sonia Goncalves, Ewan Harrison, David K. Jackson, Ian Johnston, Dominic Kwiatkowski, Cordelia Langford, John Sillitoe on behalf of the Wellcome Sanger Institute COVID-19 Surveillance Team |
| EPI_ISL_3097266 | Lighthouse Lab in Glasgow | Wellcome Sanger Institute for the COVID-19 Genomics UK (COG-UK) Consortium | Harper VanSteenhouse, Yumi Kasai, David Gray, Carol Clugston, Anna Dominiczak and Alex Alderton, Roberto Amato, Jeffrey Barrett, Sonia Goncalves, Ewan Harrison, David K. Jackson, Ian Johnston, Dominic Kwiatkowski, Cordelia Langford, John Sillitoe on behalf of the Wellcome Sanger Institute COVID-19 Surveillance Team |
| EPI_ISL_3097267 | Lighthouse Lab in Milton Keynes | Wellcome Sanger Institute for the COVID-19 Genomics UK (COG-UK) Consortium | The Lighthouse Lab in Milton Keynes and Alex Alderton, Roberto Amato, Jeffrey Barrett, Sonia Goncalves, Ewan Harrison, David K. Jackson, Ian Johnston, Dominic Kwiatkowski, Cordelia Langford, John Sillitoe on behalf of the Wellcome Sanger Institute COVID-19 Surveillance Team |
| EPI_ISL_3097268 | Lighthouse Lab in Glasgow | Wellcome Sanger Institute for the COVID-19 Genomics UK (COG-UK) Consortium | Harper VanSteenhouse, Yumi Kasai, David Gray, Carol Clugston, Anna Dominiczak and Alex Alderton, Roberto Amato, Jeffrey Barrett, Sonia Goncalves, Ewan Harrison, David K. Jackson, Ian Johnston, Dominic Kwiatkowski, Cordelia Langford, John Sillitoe on behalf of the Wellcome Sanger Institute COVID-19 Surveillance Team |
| EPI_ISL_3097269 | Lighthouse Lab in Milton Keynes | Wellcome Sanger Institute for the COVID-19 Genomics UK (COG-UK) Consortium | The Lighthouse Lab in Milton Keynes and Alex Alderton, Roberto Amato, Jeffrey Barrett, Sonia Goncalves, Ewan Harrison, David K. Jackson, Ian Johnston, Dominic Kwiatkowski, Cordelia Langford, John Sillitoe on behalf of the Wellcome Sanger Institute COVID-19 Surveillance Team |
| EPI_ISL_3097270, EPI_ISL_3097271 | Lighthouse Lab in Glasgow | Wellcome Sanger Institute for the COVID-19 Genomics UK (COG-UK) Consortium | Harper VanSteenhouse, Yumi Kasai, David Gray, Carol Clugston, Anna Dominiczak and Alex Alderton, Roberto Amato, Jeffrey Barrett, Sonia Goncalves, Ewan Harrison, David K. Jackson, Ian Johnston, Dominic Kwiatkowski, Cordelia Langford, John Sillitoe on behalf of the Wellcome Sanger Institute COVID-19 Surveillance Team |
| EPI_ISL_3097272 | Lighthouse Lab in Milton Keynes | Wellcome Sanger Institute for the COVID-19 Genomics UK (COG-UK) Consortium | The Lighthouse Lab in Milton Keynes and Alex Alderton, Roberto Amato, Jeffrey Barrett, Sonia Goncalves, Ewan Harrison, David K. Jackson, Ian Johnston, Dominic Kwiatkowski, Cordelia Langford, John Sillitoe on behalf of the Wellcome Sanger Institute COVID-19 Surveillance Team |
| EPI_ISL_3097273, EPI_ISL_3097274, EPI_ISL_3097275, EPI_ISL_3097276, EPI_ISL_3097277 | Lighthouse Lab in Glasgow | Wellcome Sanger Institute for the COVID-19 Genomics UK (COG-UK) Consortium | Harper VanSteenhouse, Yumi Kasai, David Gray, Carol Clugston, Anna Dominiczak and Alex Alderton, Roberto Amato, Jeffrey Barrett, Sonia Goncalves, Ewan Harrison, David K. Jackson, Ian Johnston, Dominic Kwiatkowski, Cordelia Langford, John Sillitoe on behalf of the Wellcome Sanger Institute COVID-19 Surveillance Team |
| EPI_ISL_3097278, EPI_ISL_3097279 | Lighthouse Lab in Milton Keynes | Wellcome Sanger Institute for the COVID-19 Genomics UK (COG-UK) Consortium | The Lighthouse Lab in Milton Keynes and Alex Alderton, Roberto Amato, Jeffrey Barrett, Sonia Goncalves, Ewan Harrison, David K. Jackson, Ian Johnston, Dominic Kwiatkowski, Cordelia Langford, John Sillitoe on behalf of the Wellcome Sanger Institute COVID-19 Surveillance Team |
| EPI_ISL_3097280 | Lighthouse Lab in Glasgow | Wellcome Sanger Institute for the COVID-19 Genomics UK (COG-UK) Consortium | Harper VanSteenhouse, Yumi Kasai, David Gray, Carol Clugston, Anna Dominiczak and Alex Alderton, Roberto Amato, Jeffrey Barrett, Sonia Goncalves, Ewan Harrison, David K. Jackson, Ian Johnston, Dominic Kwiatkowski, Cordelia Langford, John Sillitoe on behalf of the Wellcome Sanger Institute COVID-19 Surveillance Team |
| EPI_ISL_3097281, EPI_ISL_3097282, EPI_ISL_3097283, EPI_ISL_3097284, EPI_ISL_3097285 | Lighthouse Lab in Milton Keynes | Wellcome Sanger Institute for the COVID-19 Genomics UK (COG-UK) Consortium | The Lighthouse Lab in Milton Keynes and Alex Alderton, Roberto Amato, Jeffrey Barrett, Sonia Goncalves, Ewan Harrison, David K. Jackson, Ian Johnston, Dominic Kwiatkowski, Cordelia Langford, John Sillitoe on behalf of the Wellcome Sanger Institute COVID-19 Surveillance Team |
| EPI_ISL_3097286, EPI_ISL_3097287 | Lighthouse Lab in Glasgow | Wellcome Sanger Institute for the COVID-19 Genomics UK (COG-UK) Consortium | Harper VanSteenhouse, Yumi Kasai, David Gray, Carol Clugston, Anna Dominiczak and Alex Alderton, Roberto Amato, Jeffrey Barrett, Sonia Goncalves, Ewan Harrison, David K. Jackson, Ian Johnston, Dominic Kwiatkowski, Cordelia Langford, John Sillitoe on behalf of the Wellcome Sanger Institute COVID-19 Surveillance Team |
| EPI_ISL_3097288, EPI_ISL_3097289, EPI_ISL_3097290, EPI_ISL_3097291 | Lighthouse Lab in Milton Keynes | Wellcome Sanger Institute for the COVID-19 Genomics UK (COG-UK) Consortium | The Lighthouse Lab in Milton Keynes and Alex Alderton, Roberto Amato, Jeffrey Barrett, Sonia Goncalves, Ewan Harrison, David K. Jackson, Ian Johnston, Dominic Kwiatkowski, Cordelia Langford, John Sillitoe on behalf of the Wellcome Sanger Institute COVID-19 Surveillance Team |
| EPI_ISL_3097292 | Lighthouse Lab in Glasgow | Wellcome Sanger Institute for the COVID-19 Genomics UK (COG-UK) Consortium | Harper VanSteenhouse, Yumi Kasai, David Gray, Carol Clugston, Anna Dominiczak and Alex Alderton, Roberto Amato, Jeffrey Barrett, Sonia Goncalves, Ewan Harrison, David K. Jackson, Ian Johnston, Dominic Kwiatkowski, Cordelia Langford, John Sillitoe on behalf of the Wellcome Sanger Institute COVID-19 Surveillance Team |
| EPI_ISL_3097293 | Berkshire and Surrey Pathology Services Lighthouse Laboratory | Wellcome Sanger Institute for the COVID-19 Genomics UK (COG-UK) Consortium | Berkshire and Surrey Pathology Services Lighthouse Laboratory and Alex Alderton, Roberto Amato, Jeffrey Barrett, Sonia Goncalves, Ewan Harrison, David K. Jackson, Ian Johnston, Dominic Kwiatkowski, Cordelia Langford, John Sillitoe on behalf of the Wellcome Sanger Institute COVID-19 Surveillance Team |
| EPI_ISL_3097294 | Lighthouse Lab in Glasgow | Wellcome Sanger Institute for the COVID-19 Genomics UK (COG-UK) Consortium | Harper VanSteenhouse, Yumi Kasai, David Gray, Carol Clugston, Anna Dominiczak and Alex Alderton, Roberto Amato, Jeffrey Barrett, Sonia Goncalves, Ewan Harrison, David K. Jackson, Ian Johnston, Dominic Kwiatkowski, Cordelia Langford, John Sillitoe on behalf of the Wellcome Sanger Institute COVID-19 Surveillance Team |
| EPI_ISL_3097295, EPI_ISL_3097296 | Berkshire and Surrey Pathology Services Lighthouse Laboratory | Wellcome Sanger Institute for the COVID-19 Genomics UK (COG-UK) Consortium | Berkshire and Surrey Pathology Services Lighthouse Laboratory and Alex Alderton, Roberto Amato, Jeffrey Barrett, Sonia Goncalves, Ewan Harrison, David K. Jackson, Ian Johnston, Dominic Kwiatkowski, Cordelia Langford, John Sillitoe on behalf of the Wellcome Sanger Institute COVID-19 Surveillance Team |
| EPI_ISL_3097297 | Lighthouse Lab in Alderley Park | Wellcome Sanger Institute for the COVID-19 Genomics UK (COG-UK) Consortium | Jacquelyn Wynn, Mairead Hyland, The Lighthouse Lab in Alderley Park and Alex Alderton, Roberto Amato, Jeffrey Barrett, Sonia Goncalves, Ewan Harrison, David K. Jackson, Ian Johnston, Dominic Kwiatkowski, Cordelia Langford, John Sillitoe on behalf of the Wellcome Sanger Institute COVID-19 Surveillance Team |
| EPI_ISL_3097298 | Berkshire and Surrey Pathology Services Lighthouse Laboratory | Wellcome Sanger Institute for the COVID-19 Genomics UK (COG-UK) Consortium | Berkshire and Surrey Pathology Services Lighthouse Laboratory and Alex Alderton, Roberto Amato, Jeffrey Barrett, Sonia Goncalves, Ewan Harrison, David K. Jackson, Ian Johnston, Dominic Kwiatkowski, Cordelia Langford, John Sillitoe on behalf of the Wellcome Sanger Institute COVID-19 Surveillance Team |
| EPI_ISL_3097299 | Lighthouse Lab in Glasgow | Wellcome Sanger Institute for the COVID-19 Genomics UK (COG-UK) Consortium | Harper VanSteenhouse, Yumi Kasai, David Gray, Carol Clugston, Anna Dominiczak and Alex Alderton, Roberto Amato, Jeffrey Barrett, Sonia Goncalves, Ewan Harrison, David K. Jackson, Ian Johnston, Dominic Kwiatkowski, Cordelia Langford, John Sillitoe on behalf of the Wellcome Sanger Institute COVID-19 Surveillance Team |
| EPI_ISL_3097300 | Lighthouse Lab in Alderley Park | Wellcome Sanger Institute for the COVID-19 Genomics UK (COG-UK) Consortium | Jacquelyn Wynn, Mairead Hyland, The Lighthouse Lab in Alderley Park and Alex Alderton, Roberto Amato, Jeffrey Barrett, Sonia Goncalves, Ewan Harrison, David K. Jackson, Ian Johnston, Dominic Kwiatkowski, Cordelia Langford, John Sillitoe on behalf of the Wellcome Sanger Institute COVID-19 Surveillance Team |
| EPI_ISL_3097301 | Berkshire and Surrey Pathology Services Lighthouse Laboratory | Wellcome Sanger Institute for the COVID-19 Genomics UK (COG-UK) Consortium | Berkshire and Surrey Pathology Services Lighthouse Laboratory and Alex Alderton, Roberto Amato, Jeffrey Barrett, Sonia Goncalves, Ewan Harrison, David K. Jackson, Ian Johnston, Dominic Kwiatkowski, Cordelia Langford, John Sillitoe on behalf of the Wellcome Sanger Institute COVID-19 Surveillance Team |
| EPI_ISL_3097302, EPI_ISL_3097303, EPI_ISL_3097304, EPI_ISL_3097305, EPI_ISL_3097306 | Lighthouse Lab in Glasgow | Wellcome Sanger Institute for the COVID-19 Genomics UK (COG-UK) Consortium | Harper VanSteenhouse, Yumi Kasai, David Gray, Carol Clugston, Anna Dominiczak and Alex Alderton, Roberto Amato, Jeffrey Barrett, Sonia Goncalves, Ewan Harrison, David K. Jackson, Ian Johnston, Dominic Kwiatkowski, Cordelia Langford, John Sillitoe on behalf of the Wellcome Sanger Institute COVID-19 Surveillance Team |
| EPI_ISL_3097307 | Berkshire and Surrey Pathology Services Lighthouse Laboratory | Wellcome Sanger Institute for the COVID-19 Genomics UK (COG-UK) Consortium | Berkshire and Surrey Pathology Services Lighthouse Laboratory and Alex Alderton, Roberto Amato, Jeffrey Barrett, Sonia Goncalves, Ewan Harrison, David K. Jackson, Ian Johnston, Dominic Kwiatkowski, Cordelia Langford, John Sillitoe on behalf of the Wellcome Sanger Institute COVID-19 Surveillance Team |
| EPI_ISL_3097308 | Lighthouse Lab in Glasgow | Wellcome Sanger Institute for the COVID-19 Genomics UK (COG-UK) Consortium | Harper VanSteenhouse, Yumi Kasai, David Gray, Carol Clugston, Anna Dominiczak and Alex Alderton, Roberto Amato, Jeffrey Barrett, Sonia Goncalves, Ewan Harrison, David K. Jackson, Ian Johnston, Dominic Kwiatkowski, Cordelia Langford, John Sillitoe on behalf of the Wellcome Sanger Institute COVID-19 Surveillance Team |
| EPI_ISL_3097309, EPI_ISL_3097310 | Berkshire and Surrey Pathology Services Lighthouse Laboratory | Wellcome Sanger Institute for the COVID-19 Genomics UK (COG-UK) Consortium | Berkshire and Surrey Pathology Services Lighthouse Laboratory and Alex Alderton, Roberto Amato, Jeffrey Barrett, Sonia Goncalves, Ewan Harrison, David K. Jackson, Ian Johnston, Dominic Kwiatkowski, Cordelia Langford, John Sillitoe on behalf of the Wellcome Sanger Institute COVID-19 Surveillance Team |
| EPI_ISL_3097311 | Lighthouse Lab in Glasgow | Wellcome Sanger Institute for the COVID-19 Genomics UK (COG-UK) Consortium | Harper VanSteenhouse, Yumi Kasai, David Gray, Carol Clugston, Anna Dominiczak and Alex Alderton, Roberto Amato, Jeffrey Barrett, Sonia Goncalves, Ewan Harrison, David K. Jackson, Ian Johnston, Dominic Kwiatkowski, Cordelia Langford, John Sillitoe on behalf of the Wellcome Sanger Institute COVID-19 Surveillance Team |
| EPI_ISL_3097312 | Berkshire and Surrey Pathology Services Lighthouse Laboratory | Wellcome Sanger Institute for the COVID-19 Genomics UK (COG-UK) Consortium | Berkshire and Surrey Pathology Services Lighthouse Laboratory and Alex Alderton, Roberto Amato, Jeffrey Barrett, Sonia Goncalves, Ewan Harrison, David K. Jackson, Ian Johnston, Dominic Kwiatkowski, Cordelia Langford, John Sillitoe on behalf of the Wellcome Sanger Institute COVID-19 Surveillance Team |
| EPI_ISL_3097313 | Lighthouse Lab in Glasgow | Wellcome Sanger Institute for the COVID-19 Genomics UK (COG-UK) Consortium | Harper VanSteenhouse, Yumi Kasai, David Gray, Carol Clugston, Anna Dominiczak and Alex Alderton, Roberto Amato, Jeffrey Barrett, Sonia Goncalves, Ewan Harrison, David K. Jackson, Ian Johnston, Dominic Kwiatkowski, Cordelia Langford, John Sillitoe on behalf of the Wellcome Sanger Institute COVID-19 Surveillance Team |
| EPI_ISL_3097314 | Berkshire and Surrey Pathology Services Lighthouse Laboratory | Wellcome Sanger Institute for the COVID-19 Genomics UK (COG-UK) Consortium | Berkshire and Surrey Pathology Services Lighthouse Laboratory and Alex Alderton, Roberto Amato, Jeffrey Barrett, Sonia Goncalves, Ewan Harrison, David K. Jackson, Ian Johnston, Dominic Kwiatkowski, Cordelia Langford, John Sillitoe on behalf of the Wellcome Sanger Institute COVID-19 Surveillance Team |
| EPI_ISL_3097315, EPI_ISL_3097316 | Lighthouse Lab in Alderley Park | Wellcome Sanger Institute for the COVID-19 Genomics UK (COG-UK) Consortium | Jacquelyn Wynn, Mairead Hyland, The Lighthouse Lab in Alderley Park and Alex Alderton, Roberto Amato, Jeffrey Barrett, Sonia Goncalves, Ewan Harrison, David K. Jackson, Ian Johnston, Dominic Kwiatkowski, Cordelia Langford, John Sillitoe on behalf of the Wellcome Sanger Institute COVID-19 Surveillance Team |
| EPI_ISL_3097317, EPI_ISL_3097318 | Lighthouse Lab in Glasgow | Wellcome Sanger Institute for the COVID-19 Genomics UK (COG-UK) Consortium | Harper VanSteenhouse, Yumi Kasai, David Gray, Carol Clugston, Anna Dominiczak and Alex Alderton, Roberto Amato, Jeffrey Barrett, Sonia Goncalves, Ewan Harrison, David K. Jackson, Ian Johnston, Dominic Kwiatkowski, Cordelia Langford, John Sillitoe on behalf of the Wellcome Sanger Institute COVID-19 Surveillance Team |
| EPI_ISL_3097319 | Lighthouse Lab in Alderley Park | Wellcome Sanger Institute for the COVID-19 Genomics UK (COG-UK) Consortium | Jacquelyn Wynn, Mairead Hyland, The Lighthouse Lab in Alderley Park and Alex Alderton, Roberto Amato, Jeffrey Barrett, Sonia Goncalves, Ewan Harrison, David K. Jackson, Ian Johnston, Dominic Kwiatkowski, Cordelia Langford, John Sillitoe on behalf of the Wellcome Sanger Institute COVID-19 Surveillance Team |
| EPI_ISL_3097320 | Lighthouse Lab in Glasgow | Wellcome Sanger Institute for the COVID-19 Genomics UK (COG-UK) Consortium | Harper VanSteenhouse, Yumi Kasai, David Gray, Carol Clugston, Anna Dominiczak and Alex Alderton, Roberto Amato, Jeffrey Barrett, Sonia Goncalves, Ewan Harrison, David K. Jackson, Ian Johnston, Dominic Kwiatkowski, Cordelia Langford, John Sillitoe on behalf of the Wellcome Sanger Institute COVID-19 Surveillance Team |
| EPI_ISL_3097321 | Lighthouse Lab in Alderley Park | Wellcome Sanger Institute for the COVID-19 Genomics UK (COG-UK) Consortium | Jacquelyn Wynn, Mairead Hyland, The Lighthouse Lab in Alderley Park and Alex Alderton, Roberto Amato, Jeffrey Barrett, Sonia Goncalves, Ewan Harrison, David K. Jackson, Ian Johnston, Dominic Kwiatkowski, Cordelia Langford, John Sillitoe on behalf of the Wellcome Sanger Institute COVID-19 Surveillance Team |

| | | | |
|---|---|---|---|
| EPI_ISL_3097322 | Lighthouse Lab in Glasgow | Wellcome Sanger Institute for the COVID-19 Genomics UK (COG-UK) Consortium | Harper VanSteenhouse, Yumi Kasai, David Gray, Carol Clugston, Anna Dominiczak and Alex Alderton, Roberto Amato, Jeffrey Barrett, Sonia Goncalves, Ewan Harrison, David K. Jackson, Ian Johnston, Dominic Kwiatkowski, Cordelia Langford, John Sillitoe on behalf of the Wellcome Sanger Institute COVID-19 Surveillance Team |
| EPI_ISL_3097323, EPI_ISL_3097324 | Berkshire and Surrey Pathology Services Lighthouse Laboratory | Wellcome Sanger Institute for the COVID-19 Genomics UK (COG-UK) Consortium | Berkshire and Surrey Pathology Services Lighthouse Laboratory and Alex Alderton, Roberto Amato, Jeffrey Barrett, Sonia Goncalves, Ewan Harrison, David K. Jackson, Ian Johnston, Dominic Kwiatkowski, Cordelia Langford, John Sillitoe on behalf of the Wellcome Sanger Institute COVID-19 Surveillance Team |
| EPI_ISL_3097325 | Lighthouse Lab in Glasgow | Wellcome Sanger Institute for the COVID-19 Genomics UK (COG-UK) Consortium | Harper VanSteenhouse, Yumi Kasai, David Gray, Carol Clugston, Anna Dominiczak and Alex Alderton, Roberto Amato, Jeffrey Barrett, Sonia Goncalves, Ewan Harrison, David K. Jackson, Ian Johnston, Dominic Kwiatkowski, Cordelia Langford, John Sillitoe on behalf of the Wellcome Sanger Institute COVID-19 Surveillance Team |
| EPI_ISL_3097326 | Berkshire and Surrey Pathology Services Lighthouse Laboratory | Wellcome Sanger Institute for the COVID-19 Genomics UK (COG-UK) Consortium | Berkshire and Surrey Pathology Services Lighthouse Laboratory and Alex Alderton, Roberto Amato, Jeffrey Barrett, Sonia Goncalves, Ewan Harrison, David K. Jackson, Ian Johnston, Dominic Kwiatkowski, Cordelia Langford, John Sillitoe on behalf of the Wellcome Sanger Institute COVID-19 Surveillance Team |
| EPI_ISL_3097327 | Lighthouse Lab in Glasgow | Wellcome Sanger Institute for the COVID-19 Genomics UK (COG-UK) Consortium | Harper VanSteenhouse, Yumi Kasai, David Gray, Carol Clugston, Anna Dominiczak and Alex Alderton, Roberto Amato, Jeffrey Barrett, Sonia Goncalves, Ewan Harrison, David K. Jackson, Ian Johnston, Dominic Kwiatkowski, Cordelia Langford, John Sillitoe on behalf of the Wellcome Sanger Institute COVID-19 Surveillance Team |
| EPI_ISL_3097328 | Lighthouse Lab in Alderley Park | Wellcome Sanger Institute for the COVID-19 Genomics UK (COG-UK) Consortium | Jacquelyn Wynn, Mairead Hyland, The Lighthouse Lab in Alderley Park and Alex Alderton, Roberto Amato, Jeffrey Barrett, Sonia Goncalves, Ewan Harrison, David K. Jackson, Ian Johnston, Dominic Kwiatkowski, Cordelia Langford, John Sillitoe on behalf of the Wellcome Sanger Institute COVID-19 Surveillance Team |
| EPI_ISL_3097329, EPI_ISL_3097330, EPI_ISL_3097331, EPI_ISL_3097332, EPI_ISL_3097333 | Lighthouse Lab in Glasgow | Wellcome Sanger Institute for the COVID-19 Genomics UK (COG-UK) Consortium | Harper VanSteenhouse, Yumi Kasai, David Gray, Carol Clugston, Anna Dominiczak and Alex Alderton, Roberto Amato, Jeffrey Barrett, Sonia Goncalves, Ewan Harrison, David K. Jackson, Ian Johnston, Dominic Kwiatkowski, Cordelia Langford, John Sillitoe on behalf of the Wellcome Sanger Institute COVID-19 Surveillance Team |
| EPI_ISL_3097334 | Lighthouse Lab in Alderley Park | Wellcome Sanger Institute for the COVID-19 Genomics UK (COG-UK) Consortium | Jacquelyn Wynn, Mairead Hyland, The Lighthouse Lab in Alderley Park and Alex Alderton, Roberto Amato, Jeffrey Barrett, Sonia Goncalves, Ewan Harrison, David K. Jackson, Ian Johnston, Dominic Kwiatkowski, Cordelia Langford, John Sillitoe on behalf of the Wellcome Sanger Institute COVID-19 Surveillance Team |
| EPI_ISL_3097335 | Lighthouse Lab in Glasgow | Wellcome Sanger Institute for the COVID-19 Genomics UK (COG-UK) Consortium | Harper VanSteenhouse, Yumi Kasai, David Gray, Carol Clugston, Anna Dominiczak and Alex Alderton, Roberto Amato, Jeffrey Barrett, Sonia Goncalves, Ewan Harrison, David K. Jackson, Ian Johnston, Dominic Kwiatkowski, Cordelia Langford, John Sillitoe on behalf of the Wellcome Sanger Institute COVID-19 Surveillance Team |
| EPI_ISL_3097336, EPI_ISL_3097337 | Berkshire and Surrey Pathology Services Lighthouse Laboratory | Wellcome Sanger Institute for the COVID-19 Genomics UK (COG-UK) Consortium | Berkshire and Surrey Pathology Services Lighthouse Laboratory and Alex Alderton, Roberto Amato, Jeffrey Barrett, Sonia Goncalves, Ewan Harrison, David K. Jackson, Ian Johnston, Dominic Kwiatkowski, Cordelia Langford, John Sillitoe on behalf of the Wellcome Sanger Institute COVID-19 Surveillance Team |
| EPI_ISL_3097338 | Lighthouse Lab in Alderley Park | Wellcome Sanger Institute for the COVID-19 Genomics UK (COG-UK) Consortium | Jacquelyn Wynn, Mairead Hyland, The Lighthouse Lab in Alderley Park and Alex Alderton, Roberto Amato, Jeffrey Barrett, Sonia Goncalves, Ewan Harrison, David K. Jackson, Ian Johnston, Dominic Kwiatkowski, Cordelia Langford, John Sillitoe on behalf of the Wellcome Sanger Institute COVID-19 Surveillance Team |
| EPI_ISL_3097339 | Berkshire and Surrey Pathology Services Lighthouse Laboratory | Wellcome Sanger Institute for the COVID-19 Genomics UK (COG-UK) Consortium | Berkshire and Surrey Pathology Services Lighthouse Laboratory and Alex Alderton, Roberto Amato, Jeffrey Barrett, Sonia Goncalves, Ewan Harrison, David K. Jackson, Ian Johnston, Dominic Kwiatkowski, Cordelia Langford, John Sillitoe on behalf of the Wellcome Sanger Institute COVID-19 Surveillance Team |
| EPI_ISL_3097340, EPI_ISL_3097341, EPI_ISL_3097342 | Lighthouse Lab in Glasgow | Wellcome Sanger Institute for the COVID-19 Genomics UK (COG-UK) Consortium | Harper VanSteenhouse, Yumi Kasai, David Gray, Carol Clugston, Anna Dominiczak and Alex Alderton, Roberto Amato, Jeffrey Barrett, Sonia Goncalves, Ewan Harrison, David K. Jackson, Ian Johnston, Dominic Kwiatkowski, Cordelia Langford, John Sillitoe on behalf of the Wellcome Sanger Institute COVID-19 Surveillance Team |
| EPI_ISL_3097343 | Lighthouse Lab in Alderley Park | Wellcome Sanger Institute for the COVID-19 Genomics UK (COG-UK) Consortium | Jacquelyn Wynn, Mairead Hyland, The Lighthouse Lab in Alderley Park and Alex Alderton, Roberto Amato, Jeffrey Barrett, Sonia Goncalves, Ewan Harrison, David K. Jackson, Ian Johnston, Dominic Kwiatkowski, Cordelia Langford, John Sillitoe on behalf of the Wellcome Sanger Institute COVID-19 Surveillance Team |
| EPI_ISL_3097344, EPI_ISL_3097345, EPI_ISL_3097346 | Lighthouse Lab in Glasgow | Wellcome Sanger Institute for the COVID-19 Genomics UK (COG-UK) Consortium | Harper VanSteenhouse, Yumi Kasai, David Gray, Carol Clugston, Anna Dominiczak and Alex Alderton, Roberto Amato, Jeffrey Barrett, Sonia Goncalves, Ewan Harrison, David K. Jackson, Ian Johnston, Dominic Kwiatkowski, Cordelia Langford, John Sillitoe on behalf of the Wellcome Sanger Institute COVID-19 Surveillance Team |
| EPI_ISL_3097347 | Berkshire and Surrey Pathology Services Lighthouse Laboratory | Wellcome Sanger Institute for the COVID-19 Genomics UK (COG-UK) Consortium | Berkshire and Surrey Pathology Services Lighthouse Laboratory and Alex Alderton, Roberto Amato, Jeffrey Barrett, Sonia Goncalves, Ewan Harrison, David K. Jackson, Ian Johnston, Dominic Kwiatkowski, Cordelia Langford, John Sillitoe on behalf of the Wellcome Sanger Institute COVID-19 Surveillance Team |
| EPI_ISL_3097348 | Lighthouse Lab in Alderley Park | Wellcome Sanger Institute for the COVID-19 Genomics UK (COG-UK) Consortium | Jacquelyn Wynn, Mairead Hyland, The Lighthouse Lab in Alderley Park and Alex Alderton, Roberto Amato, Jeffrey Barrett, Sonia Goncalves, Ewan Harrison, David K. Jackson, Ian Johnston, Dominic Kwiatkowski, Cordelia Langford, John Sillitoe on behalf of the Wellcome Sanger Institute COVID-19 Surveillance Team |
| EPI_ISL_3097349 | Berkshire and Surrey Pathology Services Lighthouse Laboratory | Wellcome Sanger Institute for the COVID-19 Genomics UK (COG-UK) Consortium | Berkshire and Surrey Pathology Services Lighthouse Laboratory and Alex Alderton, Roberto Amato, Jeffrey Barrett, Sonia Goncalves, Ewan Harrison, David K. Jackson, Ian Johnston, Dominic Kwiatkowski, Cordelia Langford, John Sillitoe on behalf of the Wellcome Sanger Institute COVID-19 Surveillance Team |
| EPI_ISL_3097350, EPI_ISL_3097351, EPI_ISL_3097352 | Lighthouse Lab in Alderley Park | Wellcome Sanger Institute for the COVID-19 Genomics UK (COG-UK) Consortium | Jacquelyn Wynn, Mairead Hyland, The Lighthouse Lab in Alderley Park and Alex Alderton, Roberto Amato, Jeffrey Barrett, Sonia Goncalves, Ewan Harrison, David K. Jackson, Ian Johnston, Dominic Kwiatkowski, Cordelia Langford, John Sillitoe on behalf of the Wellcome Sanger Institute COVID-19 Surveillance Team |
| EPI_ISL_3097353, EPI_ISL_3097354 | Lighthouse Lab in Glasgow | Wellcome Sanger Institute for the COVID-19 Genomics UK (COG-UK) Consortium | Harper VanSteenhouse, Yumi Kasai, David Gray, Carol Clugston, Anna Dominiczak and Alex Alderton, Roberto Amato, Jeffrey Barrett, Sonia Goncalves, Ewan Harrison, David K. Jackson, Ian Johnston, Dominic Kwiatkowski, Cordelia Langford, John Sillitoe on behalf of the Wellcome Sanger Institute COVID-19 Surveillance Team |
| EPI_ISL_3097355, EPI_ISL_3097356, EPI_ISL_3097357, EPI_ISL_3097358 | Lighthouse Lab in Alderley Park | Wellcome Sanger Institute for the COVID-19 Genomics UK (COG-UK) Consortium | Jacquelyn Wynn, Mairead Hyland, The Lighthouse Lab in Alderley Park and Alex Alderton, Roberto Amato, Jeffrey Barrett, Sonia Goncalves, Ewan Harrison, David K. Jackson, Ian Johnston, Dominic Kwiatkowski, Cordelia Langford, John Sillitoe on behalf of the Wellcome Sanger Institute COVID-19 Surveillance Team |
| EPI_ISL_3097359, EPI_ISL_3097360 | Lighthouse Lab in Glasgow | Wellcome Sanger Institute for the COVID-19 Genomics UK (COG-UK) Consortium | Harper VanSteenhouse, Yumi Kasai, David Gray, Carol Clugston, Anna Dominiczak and Alex Alderton, Roberto Amato, Jeffrey Barrett, Sonia Goncalves, Ewan Harrison, David K. Jackson, Ian Johnston, Dominic Kwiatkowski, Cordelia Langford, John Sillitoe on behalf of the Wellcome Sanger Institute COVID-19 Surveillance Team |
| EPI_ISL_3097361 | Berkshire and Surrey Pathology Services Lighthouse Laboratory | Wellcome Sanger Institute for the COVID-19 Genomics UK (COG-UK) Consortium | Berkshire and Surrey Pathology Services Lighthouse Laboratory and Alex Alderton, Roberto Amato, Jeffrey Barrett, Sonia Goncalves, Ewan Harrison, David K. Jackson, Ian Johnston, Dominic Kwiatkowski, Cordelia Langford, John Sillitoe on behalf of the Wellcome Sanger Institute COVID-19 Surveillance Team |
| EPI_ISL_3097362 | Lighthouse Lab in Alderley Park | Wellcome Sanger Institute for the COVID-19 Genomics UK (COG-UK) Consortium | Jacquelyn Wynn, Mairead Hyland, The Lighthouse Lab in Alderley Park and Alex Alderton, Roberto Amato, Jeffrey Barrett, Sonia Goncalves, Ewan Harrison, David K. Jackson, Ian Johnston, Dominic Kwiatkowski, Cordelia Langford, John Sillitoe on behalf of the Wellcome Sanger Institute COVID-19 Surveillance Team |
| EPI_ISL_3097363, EPI_ISL_3097364 | Lighthouse Lab in Glasgow | Wellcome Sanger Institute for the COVID-19 Genomics UK (COG-UK) Consortium | Harper VanSteenhouse, Yumi Kasai, David Gray, Carol Clugston, Anna Dominiczak and Alex Alderton, Roberto Amato, Jeffrey Barrett, Sonia Goncalves, Ewan Harrison, David K. Jackson, Ian Johnston, Dominic Kwiatkowski, Cordelia Langford, John Sillitoe on behalf of the Wellcome Sanger Institute COVID-19 Surveillance Team |
| EPI_ISL_3097365 | Berkshire and Surrey Pathology Services Lighthouse Laboratory | Wellcome Sanger Institute for the COVID-19 Genomics UK (COG-UK) Consortium | Berkshire and Surrey Pathology Services Lighthouse Laboratory and Alex Alderton, Roberto Amato, Jeffrey Barrett, Sonia Goncalves, Ewan Harrison, David K. Jackson, Ian Johnston, Dominic Kwiatkowski, Cordelia Langford, John Sillitoe on behalf of the Wellcome Sanger Institute COVID-19 Surveillance Team |
| EPI_ISL_3097366 | Lighthouse Lab in Alderley Park | Wellcome Sanger Institute for the COVID-19 Genomics UK (COG-UK) Consortium | Jacquelyn Wynn, Mairead Hyland, The Lighthouse Lab in Alderley Park and Alex Alderton, Roberto Amato, Jeffrey Barrett, Sonia Goncalves, Ewan Harrison, David K. Jackson, Ian Johnston, Dominic Kwiatkowski, Cordelia Langford, John Sillitoe on behalf of the Wellcome Sanger Institute COVID-19 Surveillance Team |
| EPI_ISL_3097367 | Berkshire and Surrey Pathology Services Lighthouse Laboratory | Wellcome Sanger Institute for the COVID-19 Genomics UK (COG-UK) Consortium | Berkshire and Surrey Pathology Services Lighthouse Laboratory and Alex Alderton, Roberto Amato, Jeffrey Barrett, Sonia Goncalves, Ewan Harrison, David K. Jackson, Ian Johnston, Dominic Kwiatkowski, Cordelia Langford, John Sillitoe on behalf of the Wellcome Sanger Institute COVID-19 Surveillance Team |
| EPI_ISL_3097368 | Lighthouse Lab in Glasgow | Wellcome Sanger Institute for the COVID-19 Genomics UK (COG-UK) Consortium | Harper VanSteenhouse, Yumi Kasai, David Gray, Carol Clugston, Anna Dominiczak and Alex Alderton, Roberto Amato, Jeffrey Barrett, Sonia Goncalves, Ewan Harrison, David K. Jackson, Ian Johnston, Dominic Kwiatkowski, Cordelia Langford, John Sillitoe on behalf of the Wellcome Sanger Institute COVID-19 Surveillance Team |
| EPI_ISL_3097369 | Berkshire and Surrey Pathology Services Lighthouse Laboratory | Wellcome Sanger Institute for the COVID-19 Genomics UK (COG-UK) Consortium | Berkshire and Surrey Pathology Services Lighthouse Laboratory and Alex Alderton, Roberto Amato, Jeffrey Barrett, Sonia Goncalves, Ewan Harrison, David K. Jackson, Ian Johnston, Dominic Kwiatkowski, Cordelia Langford, John Sillitoe on behalf of the Wellcome Sanger Institute COVID-19 Surveillance Team |
| EPI_ISL_3097370 | Lighthouse Lab in Glasgow | Wellcome Sanger Institute for the COVID-19 Genomics UK (COG-UK) Consortium | Harper VanSteenhouse, Yumi Kasai, David Gray, Carol Clugston, Anna Dominiczak and Alex Alderton, Roberto Amato, Jeffrey Barrett, Sonia Goncalves, Ewan Harrison, David K. Jackson, Ian Johnston, Dominic Kwiatkowski, Cordelia Langford, John Sillitoe on behalf of the Wellcome Sanger Institute COVID-19 Surveillance Team |
| EPI_ISL_3097371 | Lighthouse Lab in Alderley Park | Wellcome Sanger Institute for the COVID-19 Genomics UK (COG-UK) Consortium | Jacquelyn Wynn, Mairead Hyland, The Lighthouse Lab in Alderley Park and Alex Alderton, Roberto Amato, Jeffrey Barrett, Sonia Goncalves, Ewan Harrison, David K. Jackson, Ian Johnston, Dominic Kwiatkowski, Cordelia Langford, John Sillitoe on behalf of the Wellcome Sanger Institute COVID-19 Surveillance Team |
| EPI_ISL_3097372 | Berkshire and Surrey Pathology Services Lighthouse Laboratory | Wellcome Sanger Institute for the COVID-19 Genomics UK (COG-UK) Consortium | Berkshire and Surrey Pathology Services Lighthouse Laboratory and Alex Alderton, Roberto Amato, Jeffrey Barrett, Sonia Goncalves, Ewan Harrison, David K. Jackson, Ian Johnston, Dominic Kwiatkowski, Cordelia Langford, John Sillitoe on behalf of the Wellcome Sanger Institute COVID-19 Surveillance Team |
| EPI_ISL_3097373 | Lighthouse Lab in Alderley Park | Wellcome Sanger Institute for the COVID-19 Genomics UK (COG-UK) Consortium | Jacquelyn Wynn, Mairead Hyland, The Lighthouse Lab in Alderley Park and Alex Alderton, Roberto Amato, Jeffrey Barrett, Sonia Goncalves, Ewan Harrison, David K. Jackson, Ian Johnston, Dominic Kwiatkowski, Cordelia Langford, John Sillitoe on behalf of the Wellcome Sanger Institute COVID-19 Surveillance Team |
| EPI_ISL_3097374, EPI_ISL_3097375 | Berkshire and Surrey Pathology Services Lighthouse Laboratory | Wellcome Sanger Institute for the COVID-19 Genomics UK (COG-UK) Consortium | Berkshire and Surrey Pathology Services Lighthouse Laboratory and Alex Alderton, Roberto Amato, Jeffrey Barrett, Sonia Goncalves, Ewan Harrison, David K. Jackson, Ian Johnston, Dominic Kwiatkowski, Cordelia Langford, John Sillitoe on behalf of the Wellcome Sanger Institute COVID-19 Surveillance Team |
| EPI_ISL_3097376, EPI_ISL_3097377 | Lighthouse Lab in Alderley Park | Wellcome Sanger Institute for the COVID-19 Genomics UK (COG-UK) Consortium | Jacquelyn Wynn, Mairead Hyland, The Lighthouse Lab in Alderley Park and Alex Alderton, Roberto Amato, Jeffrey Barrett, Sonia Goncalves, Ewan Harrison, David K. Jackson, Ian Johnston, Dominic Kwiatkowski, Cordelia Langford, John Sillitoe on behalf of the Wellcome Sanger Institute COVID-19 Surveillance Team |
| EPI_ISL_3097378 | Lighthouse Lab in Glasgow | Wellcome Sanger Institute for the COVID-19 Genomics UK (COG-UK) Consortium | Harper VanSteenhouse, Yumi Kasai, David Gray, Carol Clugston, Anna Dominiczak and Alex Alderton, Roberto Amato, Jeffrey Barrett, Sonia Goncalves, Ewan Harrison, David K. Jackson, Ian Johnston, Dominic Kwiatkowski, Cordelia Langford, John Sillitoe on behalf of the Wellcome Sanger Institute COVID-19 Surveillance Team |
| EPI_ISL_3097379 | Lighthouse Lab in Alderley Park | Wellcome Sanger Institute for the COVID-19 Genomics UK (COG-UK) Consortium | Jacquelyn Wynn, Mairead Hyland, The Lighthouse Lab in Alderley Park and Alex Alderton, Roberto Amato, Jeffrey Barrett, Sonia Goncalves, Ewan Harrison, David K. Jackson, Ian Johnston, Dominic Kwiatkowski, Cordelia Langford, John Sillitoe on behalf of the Wellcome Sanger Institute COVID-19 Surveillance Team |
| EPI_ISL_3097380, EPI_ISL_3097381 | Lighthouse Lab in Glasgow | Wellcome Sanger Institute for the COVID-19 Genomics UK (COG-UK) Consortium | Harper VanSteenhouse, Yumi Kasai, David Gray, Carol Clugston, Anna Dominiczak and Alex Alderton, Roberto Amato, Jeffrey Barrett, Sonia Goncalves, Ewan Harrison, David K. Jackson, Ian Johnston, Dominic Kwiatkowski, Cordelia Langford, John Sillitoe on behalf of the Wellcome Sanger Institute COVID-19 Surveillance Team |
| EPI_ISL_3097382 | Berkshire and Surrey Pathology Services Lighthouse Laboratory | Wellcome Sanger Institute for the COVID-19 Genomics UK (COG-UK) Consortium | Berkshire and Surrey Pathology Services Lighthouse Laboratory and Alex Alderton, Roberto Amato, Jeffrey Barrett, Sonia Goncalves, Ewan Harrison, David K. Jackson, Ian Johnston, Dominic Kwiatkowski, Cordelia Langford, John Sillitoe on behalf of the Wellcome Sanger Institute COVID-19 Surveillance Team |
| EPI_ISL_3097383, EPI_ISL_3097384 | Lighthouse Lab in Glasgow | Wellcome Sanger Institute for the COVID-19 Genomics UK (COG-UK) Consortium | Harper VanSteenhouse, Yumi Kasai, David Gray, Carol Clugston, Anna Dominiczak and Alex Alderton, Roberto Amato, Jeffrey Barrett, Sonia Goncalves, Ewan Harrison, David K. Jackson, Ian Johnston, Dominic Kwiatkowski, Cordelia Langford, John Sillitoe on behalf of the Wellcome Sanger Institute COVID-19 Surveillance Team |
| EPI_ISL_3097385 | Berkshire and Surrey Pathology Services Lighthouse Laboratory | Wellcome Sanger Institute for the COVID-19 Genomics UK (COG-UK) | Berkshire and Surrey Pathology Services Lighthouse Laboratory and Alex Alderton, Roberto Amato, Jeffrey Barrett, Sonia Goncalves, Ewan Harrison, David K. Jackson, Ian Johnston, Dominic Kwiatkowski, Cordelia Langford, John Sillitoe on behalf |

| | | | |
|---|---|---|---|
| | | Consortium | of the Wellcome Sanger Institute COVID-19 Surveillance Team |
| EPI_ISL_3097386 | Lighthouse Lab in Alderley Park | Wellcome Sanger Institute for the COVID-19 Genomics UK (COG-UK) Consortium | Jacquelyn Wynn, Mairead Hyland, The Lighthouse Lab in Alderley Park and Alex Alderton, Roberto Amato, Jeffrey Barrett, Sonia Goncalves, Ewan Harrison, David K. Jackson, Ian Johnston, Dominic Kwiatkowski, Cordelia Langford, John Sillitoe on behalf of the Wellcome Sanger Institute COVID-19 Surveillance Team |
| EPI_ISL_3097387 | Lighthouse Lab in Glasgow | Wellcome Sanger Institute for the COVID-19 Genomics UK (COG-UK) Consortium | Harper VanSteenhouse, Yumi Kasai, David Gray, Carol Clugston, Anna Dominiczak and Alex Alderton, Roberto Amato, Jeffrey Barrett, Sonia Goncalves, Ewan Harrison, David K. Jackson, Ian Johnston, Dominic Kwiatkowski, Cordelia Langford, John Sillitoe on behalf of the Wellcome Sanger Institute COVID-19 Surveillance Team |
| EPI_ISL_3097388 | Berkshire and Surrey Pathology Services Lighthouse Laboratory | Wellcome Sanger Institute for the COVID-19 Genomics UK (COG-UK) Consortium | Berkshire and Surrey Pathology Services Lighthouse Laboratory and Alex Alderton, Roberto Amato, Jeffrey Barrett, Sonia Goncalves, Ewan Harrison, David K. Jackson, Ian Johnston, Dominic Kwiatkowski, Cordelia Langford, John Sillitoe on behalf of the Wellcome Sanger Institute COVID-19 Surveillance Team |
| EPI_ISL_3097389, EPI_ISL_3097390, EPI_ISL_3097391 | Lighthouse Lab in Glasgow | Wellcome Sanger Institute for the COVID-19 Genomics UK (COG-UK) Consortium | Harper VanSteenhouse, Yumi Kasai, David Gray, Carol Clugston, Anna Dominiczak and Alex Alderton, Roberto Amato, Jeffrey Barrett, Sonia Goncalves, Ewan Harrison, David K. Jackson, Ian Johnston, Dominic Kwiatkowski, Cordelia Langford, John Sillitoe on behalf of the Wellcome Sanger Institute COVID-19 Surveillance Team |
| EPI_ISL_3097392 | Berkshire and Surrey Pathology Services Lighthouse Laboratory | Wellcome Sanger Institute for the COVID-19 Genomics UK (COG-UK) Consortium | Berkshire and Surrey Pathology Services Lighthouse Laboratory and Alex Alderton, Roberto Amato, Jeffrey Barrett, Sonia Goncalves, Ewan Harrison, David K. Jackson, Ian Johnston, Dominic Kwiatkowski, Cordelia Langford, John Sillitoe on behalf of the Wellcome Sanger Institute COVID-19 Surveillance Team |
| EPI_ISL_3097393, EPI_ISL_3097394 | Lighthouse Lab in Glasgow | Wellcome Sanger Institute for the COVID-19 Genomics UK (COG-UK) Consortium | Harper VanSteenhouse, Yumi Kasai, David Gray, Carol Clugston, Anna Dominiczak and Alex Alderton, Roberto Amato, Jeffrey Barrett, Sonia Goncalves, Ewan Harrison, David K. Jackson, Ian Johnston, Dominic Kwiatkowski, Cordelia Langford, John Sillitoe on behalf of the Wellcome Sanger Institute COVID-19 Surveillance Team |
| EPI_ISL_3097395 | Berkshire and Surrey Pathology Services Lighthouse Laboratory | Wellcome Sanger Institute for the COVID-19 Genomics UK (COG-UK) Consortium | Berkshire and Surrey Pathology Services Lighthouse Laboratory and Alex Alderton, Roberto Amato, Jeffrey Barrett, Sonia Goncalves, Ewan Harrison, David K. Jackson, Ian Johnston, Dominic Kwiatkowski, Cordelia Langford, John Sillitoe on behalf of the Wellcome Sanger Institute COVID-19 Surveillance Team |
| EPI_ISL_3097396 | Lighthouse Lab in Alderley Park | Wellcome Sanger Institute for the COVID-19 Genomics UK (COG-UK) Consortium | Jacquelyn Wynn, Mairead Hyland, The Lighthouse Lab in Alderley Park and Alex Alderton, Roberto Amato, Jeffrey Barrett, Sonia Goncalves, Ewan Harrison, David K. Jackson, Ian Johnston, Dominic Kwiatkowski, Cordelia Langford, John Sillitoe on behalf of the Wellcome Sanger Institute COVID-19 Surveillance Team |
| EPI_ISL_3097397 | Berkshire and Surrey Pathology Services Lighthouse Laboratory | Wellcome Sanger Institute for the COVID-19 Genomics UK (COG-UK) Consortium | Berkshire and Surrey Pathology Services Lighthouse Laboratory and Alex Alderton, Roberto Amato, Jeffrey Barrett, Sonia Goncalves, Ewan Harrison, David K. Jackson, Ian Johnston, Dominic Kwiatkowski, Cordelia Langford, John Sillitoe on behalf of the Wellcome Sanger Institute COVID-19 Surveillance Team |
| EPI_ISL_3097398 | Lighthouse Lab in Alderley Park | Wellcome Sanger Institute for the COVID-19 Genomics UK (COG-UK) Consortium | Jacquelyn Wynn, Mairead Hyland, The Lighthouse Lab in Alderley Park and Alex Alderton, Roberto Amato, Jeffrey Barrett, Sonia Goncalves, Ewan Harrison, David K. Jackson, Ian Johnston, Dominic Kwiatkowski, Cordelia Langford, John Sillitoe on behalf of the Wellcome Sanger Institute COVID-19 Surveillance Team |
| EPI_ISL_3097399, EPI_ISL_3097400, EPI_ISL_3097401, EPI_ISL_3097402 | Lighthouse Lab in Glasgow | Wellcome Sanger Institute for the COVID-19 Genomics UK (COG-UK) Consortium | Harper VanSteenhouse, Yumi Kasai, David Gray, Carol Clugston, Anna Dominiczak and Alex Alderton, Roberto Amato, Jeffrey Barrett, Sonia Goncalves, Ewan Harrison, David K. Jackson, Ian Johnston, Dominic Kwiatkowski, Cordelia Langford, John Sillitoe on behalf of the Wellcome Sanger Institute COVID-19 Surveillance Team |
| EPI_ISL_3097403 | Berkshire and Surrey Pathology Services Lighthouse Laboratory | Wellcome Sanger Institute for the COVID-19 Genomics UK (COG-UK) Consortium | Berkshire and Surrey Pathology Services Lighthouse Laboratory and Alex Alderton, Roberto Amato, Jeffrey Barrett, Sonia Goncalves, Ewan Harrison, David K. Jackson, Ian Johnston, Dominic Kwiatkowski, Cordelia Langford, John Sillitoe on behalf of the Wellcome Sanger Institute COVID-19 Surveillance Team |
| EPI_ISL_3097404 | Lighthouse Lab in Alderley Park | Wellcome Sanger Institute for the COVID-19 Genomics UK (COG-UK) Consortium | Jacquelyn Wynn, Mairead Hyland, The Lighthouse Lab in Alderley Park and Alex Alderton, Roberto Amato, Jeffrey Barrett, Sonia Goncalves, Ewan Harrison, David K. Jackson, Ian Johnston, Dominic Kwiatkowski, Cordelia Langford, John Sillitoe on behalf of the Wellcome Sanger Institute COVID-19 Surveillance Team |
| EPI_ISL_3097405 | Berkshire and Surrey Pathology Services Lighthouse Laboratory | Wellcome Sanger Institute for the COVID-19 Genomics UK (COG-UK) Consortium | Berkshire and Surrey Pathology Services Lighthouse Laboratory and Alex Alderton, Roberto Amato, Jeffrey Barrett, Sonia Goncalves, Ewan Harrison, David K. Jackson, Ian Johnston, Dominic Kwiatkowski, Cordelia Langford, John Sillitoe on behalf of the Wellcome Sanger Institute COVID-19 Surveillance Team |
| EPI_ISL_3097406 | Lighthouse Lab in Glasgow | Wellcome Sanger Institute for the COVID-19 Genomics UK (COG-UK) Consortium | Harper VanSteenhouse, Yumi Kasai, David Gray, Carol Clugston, Anna Dominiczak and Alex Alderton, Roberto Amato, Jeffrey Barrett, Sonia Goncalves, Ewan Harrison, David K. Jackson, Ian Johnston, Dominic Kwiatkowski, Cordelia Langford, John Sillitoe on behalf of the Wellcome Sanger Institute COVID-19 Surveillance Team |
| EPI_ISL_3097407, EPI_ISL_3097408 | Berkshire and Surrey Pathology Services Lighthouse Laboratory | Wellcome Sanger Institute for the COVID-19 Genomics UK (COG-UK) Consortium | Berkshire and Surrey Pathology Services Lighthouse Laboratory and Alex Alderton, Roberto Amato, Jeffrey Barrett, Sonia Goncalves, Ewan Harrison, David K. Jackson, Ian Johnston, Dominic Kwiatkowski, Cordelia Langford, John Sillitoe on behalf of the Wellcome Sanger Institute COVID-19 Surveillance Team |
| EPI_ISL_3097409 | Lighthouse Lab in Glasgow | Wellcome Sanger Institute for the COVID-19 Genomics UK (COG-UK) Consortium | Harper VanSteenhouse, Yumi Kasai, David Gray, Carol Clugston, Anna Dominiczak and Alex Alderton, Roberto Amato, Jeffrey Barrett, Sonia Goncalves, Ewan Harrison, David K. Jackson, Ian Johnston, Dominic Kwiatkowski, Cordelia Langford, John Sillitoe on behalf of the Wellcome Sanger Institute COVID-19 Surveillance Team |
| EPI_ISL_3097410 | Berkshire and Surrey Pathology Services Lighthouse Laboratory | Wellcome Sanger Institute for the COVID-19 Genomics UK (COG-UK) Consortium | Berkshire and Surrey Pathology Services Lighthouse Laboratory and Alex Alderton, Roberto Amato, Jeffrey Barrett, Sonia Goncalves, Ewan Harrison, David K. Jackson, Ian Johnston, Dominic Kwiatkowski, Cordelia Langford, John Sillitoe on behalf of the Wellcome Sanger Institute COVID-19 Surveillance Team |
| EPI_ISL_3097411 | Lighthouse Lab in Glasgow | Wellcome Sanger Institute for the COVID-19 Genomics UK (COG-UK) Consortium | Harper VanSteenhouse, Yumi Kasai, David Gray, Carol Clugston, Anna Dominiczak and Alex Alderton, Roberto Amato, Jeffrey Barrett, Sonia Goncalves, Ewan Harrison, David K. Jackson, Ian Johnston, Dominic Kwiatkowski, Cordelia Langford, John Sillitoe on behalf of the Wellcome Sanger Institute COVID-19 Surveillance Team |
| EPI_ISL_3097412, EPI_ISL_3097413 | Lighthouse Lab in Alderley Park | Wellcome Sanger Institute for the COVID-19 Genomics UK (COG-UK) Consortium | Jacquelyn Wynn, Mairead Hyland, The Lighthouse Lab in Alderley Park and Alex Alderton, Roberto Amato, Jeffrey Barrett, Sonia Goncalves, Ewan Harrison, David K. Jackson, Ian Johnston, Dominic Kwiatkowski, Cordelia Langford, John Sillitoe on behalf of the Wellcome Sanger Institute COVID-19 Surveillance Team |
| EPI_ISL_3097414 | Berkshire and Surrey Pathology Services Lighthouse Laboratory | Wellcome Sanger Institute for the COVID-19 Genomics UK (COG-UK) Consortium | Berkshire and Surrey Pathology Services Lighthouse Laboratory and Alex Alderton, Roberto Amato, Jeffrey Barrett, Sonia Goncalves, Ewan Harrison, David K. Jackson, Ian Johnston, Dominic Kwiatkowski, Cordelia Langford, John Sillitoe on behalf of the Wellcome Sanger Institute COVID-19 Surveillance Team |
| EPI_ISL_3097415 | Lighthouse Lab in Glasgow | Wellcome Sanger Institute for the COVID-19 Genomics UK (COG-UK) Consortium | Harper VanSteenhouse, Yumi Kasai, David Gray, Carol Clugston, Anna Dominiczak and Alex Alderton, Roberto Amato, Jeffrey Barrett, Sonia Goncalves, Ewan Harrison, David K. Jackson, Ian Johnston, Dominic Kwiatkowski, Cordelia Langford, John Sillitoe on behalf of the Wellcome Sanger Institute COVID-19 Surveillance Team |
| EPI_ISL_3097416 | Berkshire and Surrey Pathology Services Lighthouse Laboratory | Wellcome Sanger Institute for the COVID-19 Genomics UK (COG-UK) Consortium | Berkshire and Surrey Pathology Services Lighthouse Laboratory and Alex Alderton, Roberto Amato, Jeffrey Barrett, Sonia Goncalves, Ewan Harrison, David K. Jackson, Ian Johnston, Dominic Kwiatkowski, Cordelia Langford, John Sillitoe on behalf of the Wellcome Sanger Institute COVID-19 Surveillance Team |
| EPI_ISL_3097417 | Lighthouse Lab in Alderley Park | Wellcome Sanger Institute for the COVID-19 Genomics UK (COG-UK) Consortium | Jacquelyn Wynn, Mairead Hyland, The Lighthouse Lab in Alderley Park and Alex Alderton, Roberto Amato, Jeffrey Barrett, Sonia Goncalves, Ewan Harrison, David K. Jackson, Ian Johnston, Dominic Kwiatkowski, Cordelia Langford, John Sillitoe on behalf of the Wellcome Sanger Institute COVID-19 Surveillance Team |
| EPI_ISL_3097418 | Lighthouse Lab in Glasgow | Wellcome Sanger Institute for the COVID-19 Genomics UK (COG-UK) Consortium | Harper VanSteenhouse, Yumi Kasai, David Gray, Carol Clugston, Anna Dominiczak and Alex Alderton, Roberto Amato, Jeffrey Barrett, Sonia Goncalves, Ewan Harrison, David K. Jackson, Ian Johnston, Dominic Kwiatkowski, Cordelia Langford, John Sillitoe on behalf of the Wellcome Sanger Institute COVID-19 Surveillance Team |
| EPI_ISL_3097419 | Berkshire and Surrey Pathology Services Lighthouse Laboratory | Wellcome Sanger Institute for the COVID-19 Genomics UK (COG-UK) Consortium | Berkshire and Surrey Pathology Services Lighthouse Laboratory and Alex Alderton, Roberto Amato, Jeffrey Barrett, Sonia Goncalves, Ewan Harrison, David K. Jackson, Ian Johnston, Dominic Kwiatkowski, Cordelia Langford, John Sillitoe on behalf of the Wellcome Sanger Institute COVID-19 Surveillance Team |
| EPI_ISL_3097420 | Lighthouse Lab in Alderley Park | Wellcome Sanger Institute for the COVID-19 Genomics UK (COG-UK) Consortium | Jacquelyn Wynn, Mairead Hyland, The Lighthouse Lab in Alderley Park and Alex Alderton, Roberto Amato, Jeffrey Barrett, Sonia Goncalves, Ewan Harrison, David K. Jackson, Ian Johnston, Dominic Kwiatkowski, Cordelia Langford, John Sillitoe on behalf of the Wellcome Sanger Institute COVID-19 Surveillance Team |
| EPI_ISL_3097421 | Berkshire and Surrey Pathology Services Lighthouse Laboratory | Wellcome Sanger Institute for the COVID-19 Genomics UK (COG-UK) Consortium | Berkshire and Surrey Pathology Services Lighthouse Laboratory and Alex Alderton, Roberto Amato, Jeffrey Barrett, Sonia Goncalves, Ewan Harrison, David K. Jackson, Ian Johnston, Dominic Kwiatkowski, Cordelia Langford, John Sillitoe on behalf of the Wellcome Sanger Institute COVID-19 Surveillance Team |
| EPI_ISL_3097422, EPI_ISL_3097423, EPI_ISL_3097424 | Lighthouse Lab in Glasgow | Wellcome Sanger Institute for the COVID-19 Genomics UK (COG-UK) Consortium | Harper VanSteenhouse, Yumi Kasai, David Gray, Carol Clugston, Anna Dominiczak and Alex Alderton, Roberto Amato, Jeffrey Barrett, Sonia Goncalves, Ewan Harrison, David K. Jackson, Ian Johnston, Dominic Kwiatkowski, Cordelia Langford, John Sillitoe on behalf of the Wellcome Sanger Institute COVID-19 Surveillance Team |
| EPI_ISL_3097425 | Lighthouse Lab in Alderley Park | Wellcome Sanger Institute for the COVID-19 Genomics UK (COG-UK) Consortium | Jacquelyn Wynn, Mairead Hyland, The Lighthouse Lab in Alderley Park and Alex Alderton, Roberto Amato, Jeffrey Barrett, Sonia Goncalves, Ewan Harrison, David K. Jackson, Ian Johnston, Dominic Kwiatkowski, Cordelia Langford, John Sillitoe on behalf of the Wellcome Sanger Institute COVID-19 Surveillance Team |
| EPI_ISL_3097426 | Berkshire and Surrey Pathology Services Lighthouse Laboratory | Wellcome Sanger Institute for the COVID-19 Genomics UK (COG-UK) Consortium | Berkshire and Surrey Pathology Services Lighthouse Laboratory and Alex Alderton, Roberto Amato, Jeffrey Barrett, Sonia Goncalves, Ewan Harrison, David K. Jackson, Ian Johnston, Dominic Kwiatkowski, Cordelia Langford, John Sillitoe on behalf of the Wellcome Sanger Institute COVID-19 Surveillance Team |
| EPI_ISL_3097427, EPI_ISL_3097428 | Lighthouse Lab in Glasgow | Wellcome Sanger Institute for the COVID-19 Genomics UK (COG-UK) Consortium | Harper VanSteenhouse, Yumi Kasai, David Gray, Carol Clugston, Anna Dominiczak and Alex Alderton, Roberto Amato, Jeffrey Barrett, Sonia Goncalves, Ewan Harrison, David K. Jackson, Ian Johnston, Dominic Kwiatkowski, Cordelia Langford, John Sillitoe on behalf of the Wellcome Sanger Institute COVID-19 Surveillance Team |
| EPI_ISL_3097429 | Lighthouse Lab in Alderley Park | Wellcome Sanger Institute for the COVID-19 Genomics UK (COG-UK) Consortium | Jacquelyn Wynn, Mairead Hyland, The Lighthouse Lab in Alderley Park and Alex Alderton, Roberto Amato, Jeffrey Barrett, Sonia Goncalves, Ewan Harrison, David K. Jackson, Ian Johnston, Dominic Kwiatkowski, Cordelia Langford, John Sillitoe on behalf of the Wellcome Sanger Institute COVID-19 Surveillance Team |
| EPI_ISL_3097430, EPI_ISL_3097431 | Lighthouse Lab in Glasgow | Wellcome Sanger Institute for the COVID-19 Genomics UK (COG-UK) Consortium | Harper VanSteenhouse, Yumi Kasai, David Gray, Carol Clugston, Anna Dominiczak and Alex Alderton, Roberto Amato, Jeffrey Barrett, Sonia Goncalves, Ewan Harrison, David K. Jackson, Ian Johnston, Dominic Kwiatkowski, Cordelia Langford, John Sillitoe on behalf of the Wellcome Sanger Institute COVID-19 Surveillance Team |
| EPI_ISL_3097432, EPI_ISL_3097433 | Berkshire and Surrey Pathology Services Lighthouse Laboratory | Wellcome Sanger Institute for the COVID-19 Genomics UK (COG-UK) Consortium | Berkshire and Surrey Pathology Services Lighthouse Laboratory and Alex Alderton, Roberto Amato, Jeffrey Barrett, Sonia Goncalves, Ewan Harrison, David K. Jackson, Ian Johnston, Dominic Kwiatkowski, Cordelia Langford, John Sillitoe on behalf of the Wellcome Sanger Institute COVID-19 Surveillance Team |
| EPI_ISL_3097434 | Lighthouse Lab in Glasgow | Wellcome Sanger Institute for the COVID-19 Genomics UK (COG-UK) Consortium | Harper VanSteenhouse, Yumi Kasai, David Gray, Carol Clugston, Anna Dominiczak and Alex Alderton, Roberto Amato, Jeffrey Barrett, Sonia Goncalves, Ewan Harrison, David K. Jackson, Ian Johnston, Dominic Kwiatkowski, Cordelia Langford, John Sillitoe on behalf of the Wellcome Sanger Institute COVID-19 Surveillance Team |
| EPI_ISL_3097435 | Berkshire and Surrey Pathology Services Lighthouse Laboratory | Wellcome Sanger Institute for the COVID-19 Genomics UK (COG-UK) Consortium | Berkshire and Surrey Pathology Services Lighthouse Laboratory and Alex Alderton, Roberto Amato, Jeffrey Barrett, Sonia Goncalves, Ewan Harrison, David K. Jackson, Ian Johnston, Dominic Kwiatkowski, Cordelia Langford, John Sillitoe on behalf of the Wellcome Sanger Institute COVID-19 Surveillance Team |
| EPI_ISL_3097436, EPI_ISL_3097437, EPI_ISL_3097438 | Lighthouse Lab in Glasgow | Wellcome Sanger Institute for the COVID-19 Genomics UK (COG-UK) Consortium | Harper VanSteenhouse, Yumi Kasai, David Gray, Carol Clugston, Anna Dominiczak and Alex Alderton, Roberto Amato, Jeffrey Barrett, Sonia Goncalves, Ewan Harrison, David K. Jackson, Ian Johnston, Dominic Kwiatkowski, Cordelia Langford, John Sillitoe on behalf of the Wellcome Sanger Institute COVID-19 Surveillance Team |
| EPI_ISL_3097439, EPI_ISL_3097440 | Berkshire and Surrey Pathology Services Lighthouse Laboratory | Wellcome Sanger Institute for the COVID-19 Genomics UK (COG-UK) Consortium | Berkshire and Surrey Pathology Services Lighthouse Laboratory and Alex Alderton, Roberto Amato, Jeffrey Barrett, Sonia Goncalves, Ewan Harrison, David K. Jackson, Ian Johnston, Dominic Kwiatkowski, Cordelia Langford, John Sillitoe on behalf of the Wellcome Sanger Institute COVID-19 Surveillance Team |
| EPI_ISL_3097441 | Lighthouse Lab in Alderley Park | Wellcome Sanger Institute for the COVID-19 Genomics UK (COG-UK) Consortium | Jacquelyn Wynn, Mairead Hyland, The Lighthouse Lab in Alderley Park and Alex Alderton, Roberto Amato, Jeffrey Barrett, Sonia Goncalves, Ewan Harrison, David K. Jackson, Ian Johnston, Dominic Kwiatkowski, Cordelia Langford, John Sillitoe on behalf of the Wellcome Sanger Institute COVID-19 Surveillance Team |
| EPI_ISL_3097442, EPI_ISL_3097443 | Lighthouse Lab in Glasgow | Wellcome Sanger Institute for the COVID-19 Genomics UK (COG-UK) Consortium | Harper VanSteenhouse, Yumi Kasai, David Gray, Carol Clugston, Anna Dominiczak and Alex Alderton, Roberto Amato, Jeffrey Barrett, Sonia Goncalves, Ewan Harrison, David K. Jackson, Ian Johnston, Dominic Kwiatkowski, Cordelia Langford, John Sillitoe on behalf of the Wellcome Sanger Institute COVID-19 Surveillance Team |
| EPI_ISL_3097444 | Lighthouse Lab in Alderley Park | Wellcome Sanger Institute for the COVID-19 Genomics UK (COG-UK) Consortium | Jacquelyn Wynn, Mairead Hyland, The Lighthouse Lab in Alderley Park and Alex Alderton, Roberto Amato, Jeffrey Barrett, Sonia Goncalves, Ewan Harrison, David K. Jackson, Ian Johnston, Dominic Kwiatkowski, Cordelia Langford, John Sillitoe on behalf of the Wellcome Sanger Institute COVID-19 Surveillance Team |
| EPI_ISL_3097445 | Berkshire and Surrey Pathology Services Lighthouse Laboratory | Wellcome Sanger Institute for the COVID-19 Genomics UK (COG-UK) | Berkshire and Surrey Pathology Services Lighthouse Laboratory and Alex Alderton, Roberto Amato, Jeffrey Barrett, Sonia Goncalves, Ewan Harrison, David K. Jackson, Ian Johnston, Dominic Kwiatkowski, Cordelia Langford, John Sillitoe on behalf |

| | | | |
|---|---|---|---|
| | | Consortium | of the Wellcome Sanger Institute COVID-19 Surveillance Team |
| EPI_ISL_3097446 | Lighthouse Lab in Glasgow | Wellcome Sanger Institute for the COVID-19 Genomics UK (COG-UK) Consortium | Harper VanSteenhouse, Yumi Kasai, David Gray, Carol Clugston, Anna Dominiczak and Alex Alderton, Roberto Amato, Jeffrey Barrett, Sonia Goncalves, Ewan Harrison, David K. Jackson, Ian Johnston, Dominic Kwiatkowski, Cordelia Langford, John Sillitoe on behalf of the Wellcome Sanger Institute COVID-19 Surveillance Team |
| EPI_ISL_3097447 | Lighthouse Lab in Alderley Park | Wellcome Sanger Institute for the COVID-19 Genomics UK (COG-UK) Consortium | Jacquelyn Wynn, Mairead Hyland, The Lighthouse Lab in Alderley Park and Alex Alderton, Roberto Amato, Jeffrey Barrett, Sonia Goncalves, Ewan Harrison, David K. Jackson, Ian Johnston, Dominic Kwiatkowski, Cordelia Langford, John Sillitoe on behalf of the Wellcome Sanger Institute COVID-19 Surveillance Team |
| EPI_ISL_3097448 | Lighthouse Lab in Glasgow | Wellcome Sanger Institute for the COVID-19 Genomics UK (COG-UK) Consortium | Harper VanSteenhouse, Yumi Kasai, David Gray, Carol Clugston, Anna Dominiczak and Alex Alderton, Roberto Amato, Jeffrey Barrett, Sonia Goncalves, Ewan Harrison, David K. Jackson, Ian Johnston, Dominic Kwiatkowski, Cordelia Langford, John Sillitoe on behalf of the Wellcome Sanger Institute COVID-19 Surveillance Team |
| EPI_ISL_3097449 | Lighthouse Lab in Alderley Park | Wellcome Sanger Institute for the COVID-19 Genomics UK (COG-UK) Consortium | Jacquelyn Wynn, Mairead Hyland, The Lighthouse Lab in Alderley Park and Alex Alderton, Roberto Amato, Jeffrey Barrett, Sonia Goncalves, Ewan Harrison, David K. Jackson, Ian Johnston, Dominic Kwiatkowski, Cordelia Langford, John Sillitoe on behalf of the Wellcome Sanger Institute COVID-19 Surveillance Team |
| EPI_ISL_3097450 | Lighthouse Lab in Glasgow | Wellcome Sanger Institute for the COVID-19 Genomics UK (COG-UK) Consortium | Harper VanSteenhouse, Yumi Kasai, David Gray, Carol Clugston, Anna Dominiczak and Alex Alderton, Roberto Amato, Jeffrey Barrett, Sonia Goncalves, Ewan Harrison, David K. Jackson, Ian Johnston, Dominic Kwiatkowski, Cordelia Langford, John Sillitoe on behalf of the Wellcome Sanger Institute COVID-19 Surveillance Team |
| EPI_ISL_3097451 | Lighthouse Lab in Alderley Park | Wellcome Sanger Institute for the COVID-19 Genomics UK (COG-UK) Consortium | Jacquelyn Wynn, Mairead Hyland, The Lighthouse Lab in Alderley Park and Alex Alderton, Roberto Amato, Jeffrey Barrett, Sonia Goncalves, Ewan Harrison, David K. Jackson, Ian Johnston, Dominic Kwiatkowski, Cordelia Langford, John Sillitoe on behalf of the Wellcome Sanger Institute COVID-19 Surveillance Team |
| EPI_ISL_3097452 | Berkshire and Surrey Pathology Services Lighthouse Laboratory | Wellcome Sanger Institute for the COVID-19 Genomics UK (COG-UK) Consortium | Berkshire and Surrey Pathology Services Lighthouse Laboratory and Alex Alderton, Roberto Amato, Jeffrey Barrett, Sonia Goncalves, Ewan Harrison, David K. Jackson, Ian Johnston, Dominic Kwiatkowski, Cordelia Langford, John Sillitoe on behalf of the Wellcome Sanger Institute COVID-19 Surveillance Team |
| EPI_ISL_3097453, EPI_ISL_3097454 | Lighthouse Lab in Alderley Park | Wellcome Sanger Institute for the COVID-19 Genomics UK (COG-UK) Consortium | Jacquelyn Wynn, Mairead Hyland, The Lighthouse Lab in Alderley Park and Alex Alderton, Roberto Amato, Jeffrey Barrett, Sonia Goncalves, Ewan Harrison, David K. Jackson, Ian Johnston, Dominic Kwiatkowski, Cordelia Langford, John Sillitoe on behalf of the Wellcome Sanger Institute COVID-19 Surveillance Team |
| EPI_ISL_3097455 | Berkshire and Surrey Pathology Services Lighthouse Laboratory | Wellcome Sanger Institute for the COVID-19 Genomics UK (COG-UK) Consortium | Berkshire and Surrey Pathology Services Lighthouse Laboratory and Alex Alderton, Roberto Amato, Jeffrey Barrett, Sonia Goncalves, Ewan Harrison, David K. Jackson, Ian Johnston, Dominic Kwiatkowski, Cordelia Langford, John Sillitoe on behalf of the Wellcome Sanger Institute COVID-19 Surveillance Team |
| EPI_ISL_3097456 | Lighthouse Lab in Glasgow | Wellcome Sanger Institute for the COVID-19 Genomics UK (COG-UK) Consortium | Harper VanSteenhouse, Yumi Kasai, David Gray, Carol Clugston, Anna Dominiczak and Alex Alderton, Roberto Amato, Jeffrey Barrett, Sonia Goncalves, Ewan Harrison, David K. Jackson, Ian Johnston, Dominic Kwiatkowski, Cordelia Langford, John Sillitoe on behalf of the Wellcome Sanger Institute COVID-19 Surveillance Team |
| EPI_ISL_3097457 | Berkshire and Surrey Pathology Services Lighthouse Laboratory | Wellcome Sanger Institute for the COVID-19 Genomics UK (COG-UK) Consortium | Berkshire and Surrey Pathology Services Lighthouse Laboratory and Alex Alderton, Roberto Amato, Jeffrey Barrett, Sonia Goncalves, Ewan Harrison, David K. Jackson, Ian Johnston, Dominic Kwiatkowski, Cordelia Langford, John Sillitoe on behalf of the Wellcome Sanger Institute COVID-19 Surveillance Team |
| EPI_ISL_3097458 | Lighthouse Lab in Alderley Park | Wellcome Sanger Institute for the COVID-19 Genomics UK (COG-UK) Consortium | Jacquelyn Wynn, Mairead Hyland, The Lighthouse Lab in Alderley Park and Alex Alderton, Roberto Amato, Jeffrey Barrett, Sonia Goncalves, Ewan Harrison, David K. Jackson, Ian Johnston, Dominic Kwiatkowski, Cordelia Langford, John Sillitoe on behalf of the Wellcome Sanger Institute COVID-19 Surveillance Team |
| EPI_ISL_3097459 | Berkshire and Surrey Pathology Services Lighthouse Laboratory | Wellcome Sanger Institute for the COVID-19 Genomics UK (COG-UK) Consortium | Berkshire and Surrey Pathology Services Lighthouse Laboratory and Alex Alderton, Roberto Amato, Jeffrey Barrett, Sonia Goncalves, Ewan Harrison, David K. Jackson, Ian Johnston, Dominic Kwiatkowski, Cordelia Langford, John Sillitoe on behalf of the Wellcome Sanger Institute COVID-19 Surveillance Team |
| EPI_ISL_3097460 | Lighthouse Lab in Glasgow | Wellcome Sanger Institute for the COVID-19 Genomics UK (COG-UK) Consortium | Harper VanSteenhouse, Yumi Kasai, David Gray, Carol Clugston, Anna Dominiczak and Alex Alderton, Roberto Amato, Jeffrey Barrett, Sonia Goncalves, Ewan Harrison, David K. Jackson, Ian Johnston, Dominic Kwiatkowski, Cordelia Langford, John Sillitoe on behalf of the Wellcome Sanger Institute COVID-19 Surveillance Team |
| EPI_ISL_3097461, EPI_ISL_3097462 | Berkshire and Surrey Pathology Services Lighthouse Laboratory | Wellcome Sanger Institute for the COVID-19 Genomics UK (COG-UK) Consortium | Berkshire and Surrey Pathology Services Lighthouse Laboratory and Alex Alderton, Roberto Amato, Jeffrey Barrett, Sonia Goncalves, Ewan Harrison, David K. Jackson, Ian Johnston, Dominic Kwiatkowski, Cordelia Langford, John Sillitoe on behalf of the Wellcome Sanger Institute COVID-19 Surveillance Team |
| EPI_ISL_3097463 | Lighthouse Lab in Glasgow | Wellcome Sanger Institute for the COVID-19 Genomics UK (COG-UK) Consortium | Harper VanSteenhouse, Yumi Kasai, David Gray, Carol Clugston, Anna Dominiczak and Alex Alderton, Roberto Amato, Jeffrey Barrett, Sonia Goncalves, Ewan Harrison, David K. Jackson, Ian Johnston, Dominic Kwiatkowski, Cordelia Langford, John Sillitoe on behalf of the Wellcome Sanger Institute COVID-19 Surveillance Team |
| EPI_ISL_3097464 | Lighthouse Lab in Alderley Park | Wellcome Sanger Institute for the COVID-19 Genomics UK (COG-UK) Consortium | Jacquelyn Wynn, Mairead Hyland, The Lighthouse Lab in Alderley Park and Alex Alderton, Roberto Amato, Jeffrey Barrett, Sonia Goncalves, Ewan Harrison, David K. Jackson, Ian Johnston, Dominic Kwiatkowski, Cordelia Langford, John Sillitoe on behalf of the Wellcome Sanger Institute COVID-19 Surveillance Team |
| EPI_ISL_3097465, EPI_ISL_3097466 | Berkshire and Surrey Pathology Services Lighthouse Laboratory | Wellcome Sanger Institute for the COVID-19 Genomics UK (COG-UK) Consortium | Berkshire and Surrey Pathology Services Lighthouse Laboratory and Alex Alderton, Roberto Amato, Jeffrey Barrett, Sonia Goncalves, Ewan Harrison, David K. Jackson, Ian Johnston, Dominic Kwiatkowski, Cordelia Langford, John Sillitoe on behalf of the Wellcome Sanger Institute COVID-19 Surveillance Team |
| EPI_ISL_3097467 | Lighthouse Lab in Alderley Park | Wellcome Sanger Institute for the COVID-19 Genomics UK (COG-UK) Consortium | Jacquelyn Wynn, Mairead Hyland, The Lighthouse Lab in Alderley Park and Alex Alderton, Roberto Amato, Jeffrey Barrett, Sonia Goncalves, Ewan Harrison, David K. Jackson, Ian Johnston, Dominic Kwiatkowski, Cordelia Langford, John Sillitoe on behalf of the Wellcome Sanger Institute COVID-19 Surveillance Team |
| EPI_ISL_3097468 | Berkshire and Surrey Pathology Services Lighthouse Laboratory | Wellcome Sanger Institute for the COVID-19 Genomics UK (COG-UK) Consortium | Berkshire and Surrey Pathology Services Lighthouse Laboratory and Alex Alderton, Roberto Amato, Jeffrey Barrett, Sonia Goncalves, Ewan Harrison, David K. Jackson, Ian Johnston, Dominic Kwiatkowski, Cordelia Langford, John Sillitoe on behalf of the Wellcome Sanger Institute COVID-19 Surveillance Team |
| EPI_ISL_3097469 | Lighthouse Lab in Glasgow | Wellcome Sanger Institute for the COVID-19 Genomics UK (COG-UK) Consortium | Harper VanSteenhouse, Yumi Kasai, David Gray, Carol Clugston, Anna Dominiczak and Alex Alderton, Roberto Amato, Jeffrey Barrett, Sonia Goncalves, Ewan Harrison, David K. Jackson, Ian Johnston, Dominic Kwiatkowski, Cordelia Langford, John Sillitoe on behalf of the Wellcome Sanger Institute COVID-19 Surveillance Team |
| EPI_ISL_3097470 | Berkshire and Surrey Pathology Services Lighthouse Laboratory | Wellcome Sanger Institute for the COVID-19 Genomics UK (COG-UK) Consortium | Berkshire and Surrey Pathology Services Lighthouse Laboratory and Alex Alderton, Roberto Amato, Jeffrey Barrett, Sonia Goncalves, Ewan Harrison, David K. Jackson, Ian Johnston, Dominic Kwiatkowski, Cordelia Langford, John Sillitoe on behalf of the Wellcome Sanger Institute COVID-19 Surveillance Team |
| EPI_ISL_3097471, EPI_ISL_3097472, EPI_ISL_3097473 | Lighthouse Lab in Alderley Park | Wellcome Sanger Institute for the COVID-19 Genomics UK (COG-UK) Consortium | Jacquelyn Wynn, Mairead Hyland, The Lighthouse Lab in Alderley Park and Alex Alderton, Roberto Amato, Jeffrey Barrett, Sonia Goncalves, Ewan Harrison, David K. Jackson, Ian Johnston, Dominic Kwiatkowski, Cordelia Langford, John Sillitoe on behalf of the Wellcome Sanger Institute COVID-19 Surveillance Team |
| EPI_ISL_3097474 | Lighthouse Lab in Glasgow | Wellcome Sanger Institute for the COVID-19 Genomics UK (COG-UK) Consortium | Harper VanSteenhouse, Yumi Kasai, David Gray, Carol Clugston, Anna Dominiczak and Alex Alderton, Roberto Amato, Jeffrey Barrett, Sonia Goncalves, Ewan Harrison, David K. Jackson, Ian Johnston, Dominic Kwiatkowski, Cordelia Langford, John Sillitoe on behalf of the Wellcome Sanger Institute COVID-19 Surveillance Team |
| EPI_ISL_3097475 | Lighthouse Lab in Alderley Park | Wellcome Sanger Institute for the COVID-19 Genomics UK (COG-UK) Consortium | Jacquelyn Wynn, Mairead Hyland, The Lighthouse Lab in Alderley Park and Alex Alderton, Roberto Amato, Jeffrey Barrett, Sonia Goncalves, Ewan Harrison, David K. Jackson, Ian Johnston, Dominic Kwiatkowski, Cordelia Langford, John Sillitoe on behalf of the Wellcome Sanger Institute COVID-19 Surveillance Team |
| EPI_ISL_3097476 | Lighthouse Lab in Glasgow | Wellcome Sanger Institute for the COVID-19 Genomics UK (COG-UK) Consortium | Harper VanSteenhouse, Yumi Kasai, David Gray, Carol Clugston, Anna Dominiczak and Alex Alderton, Roberto Amato, Jeffrey Barrett, Sonia Goncalves, Ewan Harrison, David K. Jackson, Ian Johnston, Dominic Kwiatkowski, Cordelia Langford, John Sillitoe on behalf of the Wellcome Sanger Institute COVID-19 Surveillance Team |
| EPI_ISL_3097477 | Lighthouse Lab in Alderley Park | Wellcome Sanger Institute for the COVID-19 Genomics UK (COG-UK) Consortium | Jacquelyn Wynn, Mairead Hyland, The Lighthouse Lab in Alderley Park and Alex Alderton, Roberto Amato, Jeffrey Barrett, Sonia Goncalves, Ewan Harrison, David K. Jackson, Ian Johnston, Dominic Kwiatkowski, Cordelia Langford, John Sillitoe on behalf of the Wellcome Sanger Institute COVID-19 Surveillance Team |
| EPI_ISL_3097478, EPI_ISL_3097479, EPI_ISL_3097480, EPI_ISL_3097481 | Lighthouse Lab in Glasgow | Wellcome Sanger Institute for the COVID-19 Genomics UK (COG-UK) Consortium | Harper VanSteenhouse, Yumi Kasai, David Gray, Carol Clugston, Anna Dominiczak and Alex Alderton, Roberto Amato, Jeffrey Barrett, Sonia Goncalves, Ewan Harrison, David K. Jackson, Ian Johnston, Dominic Kwiatkowski, Cordelia Langford, John Sillitoe on behalf of the Wellcome Sanger Institute COVID-19 Surveillance Team |
| EPI_ISL_3097482, EPI_ISL_3097483 | Lighthouse Lab in Alderley Park | Wellcome Sanger Institute for the COVID-19 Genomics UK (COG-UK) Consortium | Jacquelyn Wynn, Mairead Hyland, The Lighthouse Lab in Alderley Park and Alex Alderton, Roberto Amato, Jeffrey Barrett, Sonia Goncalves, Ewan Harrison, David K. Jackson, Ian Johnston, Dominic Kwiatkowski, Cordelia Langford, John Sillitoe on behalf of the Wellcome Sanger Institute COVID-19 Surveillance Team |
| EPI_ISL_3097484, EPI_ISL_3097485, EPI_ISL_3097486 | Lighthouse Lab in Glasgow | Wellcome Sanger Institute for the COVID-19 Genomics UK (COG-UK) Consortium | Harper VanSteenhouse, Yumi Kasai, David Gray, Carol Clugston, Anna Dominiczak and Alex Alderton, Roberto Amato, Jeffrey Barrett, Sonia Goncalves, Ewan Harrison, David K. Jackson, Ian Johnston, Dominic Kwiatkowski, Cordelia Langford, John Sillitoe on behalf of the Wellcome Sanger Institute COVID-19 Surveillance Team |
| EPI_ISL_3097487, EPI_ISL_3097488 | Berkshire and Surrey Pathology Services Lighthouse Laboratory | Wellcome Sanger Institute for the COVID-19 Genomics UK (COG-UK) Consortium | Berkshire and Surrey Pathology Services Lighthouse Laboratory and Alex Alderton, Roberto Amato, Jeffrey Barrett, Sonia Goncalves, Ewan Harrison, David K. Jackson, Ian Johnston, Dominic Kwiatkowski, Cordelia Langford, John Sillitoe on behalf of the Wellcome Sanger Institute COVID-19 Surveillance Team |
| EPI_ISL_3097489, EPI_ISL_3097490 | Lighthouse Lab in Alderley Park | Wellcome Sanger Institute for the COVID-19 Genomics UK (COG-UK) Consortium | Jacquelyn Wynn, Mairead Hyland, The Lighthouse Lab in Alderley Park and Alex Alderton, Roberto Amato, Jeffrey Barrett, Sonia Goncalves, Ewan Harrison, David K. Jackson, Ian Johnston, Dominic Kwiatkowski, Cordelia Langford, John Sillitoe on behalf of the Wellcome Sanger Institute COVID-19 Surveillance Team |
| EPI_ISL_3097491, EPI_ISL_3097492 | Lighthouse Lab in Glasgow | Wellcome Sanger Institute for the COVID-19 Genomics UK (COG-UK) Consortium | Harper VanSteenhouse, Yumi Kasai, David Gray, Carol Clugston, Anna Dominiczak and Alex Alderton, Roberto Amato, Jeffrey Barrett, Sonia Goncalves, Ewan Harrison, David K. Jackson, Ian Johnston, Dominic Kwiatkowski, Cordelia Langford, John Sillitoe on behalf of the Wellcome Sanger Institute COVID-19 Surveillance Team |
| EPI_ISL_3097493 | Lighthouse Lab in Alderley Park | Wellcome Sanger Institute for the COVID-19 Genomics UK (COG-UK) Consortium | Jacquelyn Wynn, Mairead Hyland, The Lighthouse Lab in Alderley Park and Alex Alderton, Roberto Amato, Jeffrey Barrett, Sonia Goncalves, Ewan Harrison, David K. Jackson, Ian Johnston, Dominic Kwiatkowski, Cordelia Langford, John Sillitoe on behalf of the Wellcome Sanger Institute COVID-19 Surveillance Team |
| EPI_ISL_3097494 | Berkshire and Surrey Pathology Services Lighthouse Laboratory | Wellcome Sanger Institute for the COVID-19 Genomics UK (COG-UK) Consortium | Berkshire and Surrey Pathology Services Lighthouse Laboratory and Alex Alderton, Roberto Amato, Jeffrey Barrett, Sonia Goncalves, Ewan Harrison, David K. Jackson, Ian Johnston, Dominic Kwiatkowski, Cordelia Langford, John Sillitoe on behalf of the Wellcome Sanger Institute COVID-19 Surveillance Team |
| EPI_ISL_3097495, EPI_ISL_3097496 | Lighthouse Lab in Glasgow | Wellcome Sanger Institute for the COVID-19 Genomics UK (COG-UK) Consortium | Harper VanSteenhouse, Yumi Kasai, David Gray, Carol Clugston, Anna Dominiczak and Alex Alderton, Roberto Amato, Jeffrey Barrett, Sonia Goncalves, Ewan Harrison, David K. Jackson, Ian Johnston, Dominic Kwiatkowski, Cordelia Langford, John Sillitoe on behalf of the Wellcome Sanger Institute COVID-19 Surveillance Team |
| EPI_ISL_3097497, EPI_ISL_3097498 | Lighthouse Lab in Alderley Park | Wellcome Sanger Institute for the COVID-19 Genomics UK (COG-UK) Consortium | Jacquelyn Wynn, Mairead Hyland, The Lighthouse Lab in Alderley Park and Alex Alderton, Roberto Amato, Jeffrey Barrett, Sonia Goncalves, Ewan Harrison, David K. Jackson, Ian Johnston, Dominic Kwiatkowski, Cordelia Langford, John Sillitoe on behalf of the Wellcome Sanger Institute COVID-19 Surveillance Team |
| EPI_ISL_3097499, EPI_ISL_3097500, EPI_ISL_3097501 | Lighthouse Lab in Glasgow | Wellcome Sanger Institute for the COVID-19 Genomics UK (COG-UK) Consortium | Harper VanSteenhouse, Yumi Kasai, David Gray, Carol Clugston, Anna Dominiczak and Alex Alderton, Roberto Amato, Jeffrey Barrett, Sonia Goncalves, Ewan Harrison, David K. Jackson, Ian Johnston, Dominic Kwiatkowski, Cordelia Langford, John Sillitoe on behalf of the Wellcome Sanger Institute COVID-19 Surveillance Team |
| EPI_ISL_3097502 | Berkshire and Surrey Pathology Services Lighthouse Laboratory | Wellcome Sanger Institute for the COVID-19 Genomics UK (COG-UK) Consortium | Berkshire and Surrey Pathology Services Lighthouse Laboratory and Alex Alderton, Roberto Amato, Jeffrey Barrett, Sonia Goncalves, Ewan Harrison, David K. Jackson, Ian Johnston, Dominic Kwiatkowski, Cordelia Langford, John Sillitoe on behalf of the Wellcome Sanger Institute COVID-19 Surveillance Team |
| EPI_ISL_3097503 | Lighthouse Lab in Glasgow | Wellcome Sanger Institute for the COVID-19 Genomics UK (COG-UK) Consortium | Harper VanSteenhouse, Yumi Kasai, David Gray, Carol Clugston, Anna Dominiczak and Alex Alderton, Roberto Amato, Jeffrey Barrett, Sonia Goncalves, Ewan Harrison, David K. Jackson, Ian Johnston, Dominic Kwiatkowski, Cordelia Langford, John Sillitoe on behalf of the Wellcome Sanger Institute COVID-19 Surveillance Team |
| EPI_ISL_3097504 | Lighthouse Lab in Alderley Park | Wellcome Sanger Institute for the COVID-19 Genomics UK (COG-UK) Consortium | Jacquelyn Wynn, Mairead Hyland, The Lighthouse Lab in Alderley Park and Alex Alderton, Roberto Amato, Jeffrey Barrett, Sonia Goncalves, Ewan Harrison, David K. Jackson, Ian Johnston, Dominic Kwiatkowski, Cordelia Langford, John Sillitoe on behalf of the Wellcome Sanger Institute COVID-19 Surveillance Team |
| EPI_ISL_3097505, EPI_ISL_3097506, EPI_ISL_3097507 | Lighthouse Lab in Glasgow | Wellcome Sanger Institute for the COVID-19 Genomics UK (COG-UK) Consortium | Harper VanSteenhouse, Yumi Kasai, David Gray, Carol Clugston, Anna Dominiczak and Alex Alderton, Roberto Amato, Jeffrey Barrett, Sonia Goncalves, Ewan Harrison, David K. Jackson, Ian Johnston, Dominic Kwiatkowski, Cordelia Langford, John Sillitoe on behalf of the Wellcome Sanger Institute COVID-19 Surveillance Team |

| | | | |
|---|---|---|---|
| EPI_ISL_3097508 | Lighthouse Lab in Alderley Park | Wellcome Sanger Institute for the COVID-19 Genomics UK (COG-UK) Consortium | Jacquelyn Wynn, Mairead Hyland, The Lighthouse Lab in Alderley Park and Alex Alderton, Roberto Amato, Jeffrey Barrett, Sonia Goncalves, Ewan Harrison, David K. Jackson, Ian Johnston, Dominic Kwiatkowski, Cordelia Langford, John Sillitoe on behalf of the Wellcome Sanger Institute COVID-19 Surveillance Team |
| EPI_ISL_3097509 | Lighthouse Lab in Glasgow | Wellcome Sanger Institute for the COVID-19 Genomics UK (COG-UK) Consortium | Harper VanSteenhouse, Yumi Kasai, David Gray, Carol Clugston, Anna Dominiczak and Alex Alderton, Roberto Amato, Jeffrey Barrett, Sonia Goncalves, Ewan Harrison, David K. Jackson, Ian Johnston, Dominic Kwiatkowski, Cordelia Langford, John Sillitoe on behalf of the Wellcome Sanger Institute COVID-19 Surveillance Team |
| EPI_ISL_3097510 | Lighthouse Lab in Alderley Park | Wellcome Sanger Institute for the COVID-19 Genomics UK (COG-UK) Consortium | Jacquelyn Wynn, Mairead Hyland, The Lighthouse Lab in Alderley Park and Alex Alderton, Roberto Amato, Jeffrey Barrett, Sonia Goncalves, Ewan Harrison, David K. Jackson, Ian Johnston, Dominic Kwiatkowski, Cordelia Langford, John Sillitoe on behalf of the Wellcome Sanger Institute COVID-19 Surveillance Team |
| EPI_ISL_3097511 | Lighthouse Lab in Glasgow | Wellcome Sanger Institute for the COVID-19 Genomics UK (COG-UK) Consortium | Harper VanSteenhouse, Yumi Kasai, David Gray, Carol Clugston, Anna Dominiczak and Alex Alderton, Roberto Amato, Jeffrey Barrett, Sonia Goncalves, Ewan Harrison, David K. Jackson, Ian Johnston, Dominic Kwiatkowski, Cordelia Langford, John Sillitoe on behalf of the Wellcome Sanger Institute COVID-19 Surveillance Team |
| EPI_ISL_3097512, EPI_ISL_3097513, EPI_ISL_3097514, EPI_ISL_3097515 | Lighthouse Lab in Alderley Park | Wellcome Sanger Institute for the COVID-19 Genomics UK (COG-UK) Consortium | Jacquelyn Wynn, Mairead Hyland, The Lighthouse Lab in Alderley Park and Alex Alderton, Roberto Amato, Jeffrey Barrett, Sonia Goncalves, Ewan Harrison, David K. Jackson, Ian Johnston, Dominic Kwiatkowski, Cordelia Langford, John Sillitoe on behalf of the Wellcome Sanger Institute COVID-19 Surveillance Team |
| EPI_ISL_3097516 | Lighthouse Lab in Glasgow | Wellcome Sanger Institute for the COVID-19 Genomics UK (COG-UK) Consortium | Harper VanSteenhouse, Yumi Kasai, David Gray, Carol Clugston, Anna Dominiczak and Alex Alderton, Roberto Amato, Jeffrey Barrett, Sonia Goncalves, Ewan Harrison, David K. Jackson, Ian Johnston, Dominic Kwiatkowski, Cordelia Langford, John Sillitoe on behalf of the Wellcome Sanger Institute COVID-19 Surveillance Team |
| EPI_ISL_3097517, EPI_ISL_3097518, EPI_ISL_3097519 | Berkshire and Surrey Pathology Services Lighthouse Laboratory | Wellcome Sanger Institute for the COVID-19 Genomics UK (COG-UK) Consortium | Berkshire and Surrey Pathology Services Lighthouse Laboratory and Alex Alderton, Roberto Amato, Jeffrey Barrett, Sonia Goncalves, Ewan Harrison, David K. Jackson, Ian Johnston, Dominic Kwiatkowski, Cordelia Langford, John Sillitoe on behalf of the Wellcome Sanger Institute COVID-19 Surveillance Team |
| EPI_ISL_3097520, EPI_ISL_3097521 | Lighthouse Lab in Glasgow | Wellcome Sanger Institute for the COVID-19 Genomics UK (COG-UK) Consortium | Harper VanSteenhouse, Yumi Kasai, David Gray, Carol Clugston, Anna Dominiczak and Alex Alderton, Roberto Amato, Jeffrey Barrett, Sonia Goncalves, Ewan Harrison, David K. Jackson, Ian Johnston, Dominic Kwiatkowski, Cordelia Langford, John Sillitoe on behalf of the Wellcome Sanger Institute COVID-19 Surveillance Team |
| EPI_ISL_3097522 | Berkshire and Surrey Pathology Services Lighthouse Laboratory | Wellcome Sanger Institute for the COVID-19 Genomics UK (COG-UK) Consortium | Berkshire and Surrey Pathology Services Lighthouse Laboratory and Alex Alderton, Roberto Amato, Jeffrey Barrett, Sonia Goncalves, Ewan Harrison, David K. Jackson, Ian Johnston, Dominic Kwiatkowski, Cordelia Langford, John Sillitoe on behalf of the Wellcome Sanger Institute COVID-19 Surveillance Team |
| EPI_ISL_3097523 | Lighthouse Lab in Glasgow | Wellcome Sanger Institute for the COVID-19 Genomics UK (COG-UK) Consortium | Harper VanSteenhouse, Yumi Kasai, David Gray, Carol Clugston, Anna Dominiczak and Alex Alderton, Roberto Amato, Jeffrey Barrett, Sonia Goncalves, Ewan Harrison, David K. Jackson, Ian Johnston, Dominic Kwiatkowski, Cordelia Langford, John Sillitoe on behalf of the Wellcome Sanger Institute COVID-19 Surveillance Team |
| EPI_ISL_3097524 | Lighthouse Lab in Alderley Park | Wellcome Sanger Institute for the COVID-19 Genomics UK (COG-UK) Consortium | Jacquelyn Wynn, Mairead Hyland, The Lighthouse Lab in Alderley Park and Alex Alderton, Roberto Amato, Jeffrey Barrett, Sonia Goncalves, Ewan Harrison, David K. Jackson, Ian Johnston, Dominic Kwiatkowski, Cordelia Langford, John Sillitoe on behalf of the Wellcome Sanger Institute COVID-19 Surveillance Team |
| EPI_ISL_3097525 | Berkshire and Surrey Pathology Services Lighthouse Laboratory | Wellcome Sanger Institute for the COVID-19 Genomics UK (COG-UK) Consortium | Berkshire and Surrey Pathology Services Lighthouse Laboratory and Alex Alderton, Roberto Amato, Jeffrey Barrett, Sonia Goncalves, Ewan Harrison, David K. Jackson, Ian Johnston, Dominic Kwiatkowski, Cordelia Langford, John Sillitoe on behalf of the Wellcome Sanger Institute COVID-19 Surveillance Team |
| EPI_ISL_3097526 | Lighthouse Lab in Glasgow | Wellcome Sanger Institute for the COVID-19 Genomics UK (COG-UK) Consortium | Harper VanSteenhouse, Yumi Kasai, David Gray, Carol Clugston, Anna Dominiczak and Alex Alderton, Roberto Amato, Jeffrey Barrett, Sonia Goncalves, Ewan Harrison, David K. Jackson, Ian Johnston, Dominic Kwiatkowski, Cordelia Langford, John Sillitoe on behalf of the Wellcome Sanger Institute COVID-19 Surveillance Team |
| EPI_ISL_3097527 | Lighthouse Lab in Alderley Park | Wellcome Sanger Institute for the COVID-19 Genomics UK (COG-UK) Consortium | Jacquelyn Wynn, Mairead Hyland, The Lighthouse Lab in Alderley Park and Alex Alderton, Roberto Amato, Jeffrey Barrett, Sonia Goncalves, Ewan Harrison, David K. Jackson, Ian Johnston, Dominic Kwiatkowski, Cordelia Langford, John Sillitoe on behalf of the Wellcome Sanger Institute COVID-19 Surveillance Team |
| EPI_ISL_3097528 | Berkshire and Surrey Pathology Services Lighthouse Laboratory | Wellcome Sanger Institute for the COVID-19 Genomics UK (COG-UK) Consortium | Berkshire and Surrey Pathology Services Lighthouse Laboratory and Alex Alderton, Roberto Amato, Jeffrey Barrett, Sonia Goncalves, Ewan Harrison, David K. Jackson, Ian Johnston, Dominic Kwiatkowski, Cordelia Langford, John Sillitoe on behalf of the Wellcome Sanger Institute COVID-19 Surveillance Team |
| EPI_ISL_3097529 | Lighthouse Lab in Glasgow | Wellcome Sanger Institute for the COVID-19 Genomics UK (COG-UK) Consortium | Harper VanSteenhouse, Yumi Kasai, David Gray, Carol Clugston, Anna Dominiczak and Alex Alderton, Roberto Amato, Jeffrey Barrett, Sonia Goncalves, Ewan Harrison, David K. Jackson, Ian Johnston, Dominic Kwiatkowski, Cordelia Langford, John Sillitoe on behalf of the Wellcome Sanger Institute COVID-19 Surveillance Team |
| EPI_ISL_3097530, EPI_ISL_3097531 | Berkshire and Surrey Pathology Services Lighthouse Laboratory | Wellcome Sanger Institute for the COVID-19 Genomics UK (COG-UK) Consortium | Berkshire and Surrey Pathology Services Lighthouse Laboratory and Alex Alderton, Roberto Amato, Jeffrey Barrett, Sonia Goncalves, Ewan Harrison, David K. Jackson, Ian Johnston, Dominic Kwiatkowski, Cordelia Langford, John Sillitoe on behalf of the Wellcome Sanger Institute COVID-19 Surveillance Team |
| EPI_ISL_3097532 | Lighthouse Lab in Alderley Park | Wellcome Sanger Institute for the COVID-19 Genomics UK (COG-UK) Consortium | Jacquelyn Wynn, Mairead Hyland, The Lighthouse Lab in Alderley Park and Alex Alderton, Roberto Amato, Jeffrey Barrett, Sonia Goncalves, Ewan Harrison, David K. Jackson, Ian Johnston, Dominic Kwiatkowski, Cordelia Langford, John Sillitoe on behalf of the Wellcome Sanger Institute COVID-19 Surveillance Team |
| EPI_ISL_3097533 | Lighthouse Lab in Glasgow | Wellcome Sanger Institute for the COVID-19 Genomics UK (COG-UK) Consortium | Harper VanSteenhouse, Yumi Kasai, David Gray, Carol Clugston, Anna Dominiczak and Alex Alderton, Roberto Amato, Jeffrey Barrett, Sonia Goncalves, Ewan Harrison, David K. Jackson, Ian Johnston, Dominic Kwiatkowski, Cordelia Langford, John Sillitoe on behalf of the Wellcome Sanger Institute COVID-19 Surveillance Team |
| EPI_ISL_3097534, EPI_ISL_3097535 | Lighthouse Lab in Alderley Park | Wellcome Sanger Institute for the COVID-19 Genomics UK (COG-UK) Consortium | Jacquelyn Wynn, Mairead Hyland, The Lighthouse Lab in Alderley Park and Alex Alderton, Roberto Amato, Jeffrey Barrett, Sonia Goncalves, Ewan Harrison, David K. Jackson, Ian Johnston, Dominic Kwiatkowski, Cordelia Langford, John Sillitoe on behalf of the Wellcome Sanger Institute COVID-19 Surveillance Team |
| EPI_ISL_3097536 | Berkshire and Surrey Pathology Services Lighthouse Laboratory | Wellcome Sanger Institute for the COVID-19 Genomics UK (COG-UK) Consortium | Berkshire and Surrey Pathology Services Lighthouse Laboratory and Alex Alderton, Roberto Amato, Jeffrey Barrett, Sonia Goncalves, Ewan Harrison, David K. Jackson, Ian Johnston, Dominic Kwiatkowski, Cordelia Langford, John Sillitoe on behalf of the Wellcome Sanger Institute COVID-19 Surveillance Team |
| EPI_ISL_3097537 | Lighthouse Lab in Alderley Park | Wellcome Sanger Institute for the COVID-19 Genomics UK (COG-UK) Consortium | Jacquelyn Wynn, Mairead Hyland, The Lighthouse Lab in Alderley Park and Alex Alderton, Roberto Amato, Jeffrey Barrett, Sonia Goncalves, Ewan Harrison, David K. Jackson, Ian Johnston, Dominic Kwiatkowski, Cordelia Langford, John Sillitoe on behalf of the Wellcome Sanger Institute COVID-19 Surveillance Team |
| EPI_ISL_3097538, EPI_ISL_3097539, EPI_ISL_3097540 | Lighthouse Lab in Glasgow | Wellcome Sanger Institute for the COVID-19 Genomics UK (COG-UK) Consortium | Harper VanSteenhouse, Yumi Kasai, David Gray, Carol Clugston, Anna Dominiczak and Alex Alderton, Roberto Amato, Jeffrey Barrett, Sonia Goncalves, Ewan Harrison, David K. Jackson, Ian Johnston, Dominic Kwiatkowski, Cordelia Langford, John Sillitoe on behalf of the Wellcome Sanger Institute COVID-19 Surveillance Team |
| EPI_ISL_3097541 | Berkshire and Surrey Pathology Services Lighthouse Laboratory | Wellcome Sanger Institute for the COVID-19 Genomics UK (COG-UK) Consortium | Berkshire and Surrey Pathology Services Lighthouse Laboratory and Alex Alderton, Roberto Amato, Jeffrey Barrett, Sonia Goncalves, Ewan Harrison, David K. Jackson, Ian Johnston, Dominic Kwiatkowski, Cordelia Langford, John Sillitoe on behalf of the Wellcome Sanger Institute COVID-19 Surveillance Team |
| EPI_ISL_3097542, EPI_ISL_3097543 | Lighthouse Lab in Alderley Park | Wellcome Sanger Institute for the COVID-19 Genomics UK (COG-UK) Consortium | Jacquelyn Wynn, Mairead Hyland, The Lighthouse Lab in Alderley Park and Alex Alderton, Roberto Amato, Jeffrey Barrett, Sonia Goncalves, Ewan Harrison, David K. Jackson, Ian Johnston, Dominic Kwiatkowski, Cordelia Langford, John Sillitoe on behalf of the Wellcome Sanger Institute COVID-19 Surveillance Team |
| EPI_ISL_3097544 | Lighthouse Lab in Glasgow | Wellcome Sanger Institute for the COVID-19 Genomics UK (COG-UK) Consortium | Harper VanSteenhouse, Yumi Kasai, David Gray, Carol Clugston, Anna Dominiczak and Alex Alderton, Roberto Amato, Jeffrey Barrett, Sonia Goncalves, Ewan Harrison, David K. Jackson, Ian Johnston, Dominic Kwiatkowski, Cordelia Langford, John Sillitoe on behalf of the Wellcome Sanger Institute COVID-19 Surveillance Team |
| EPI_ISL_3097545, EPI_ISL_3097546, EPI_ISL_3097547, EPI_ISL_3097548 | Lighthouse Lab in Alderley Park | Wellcome Sanger Institute for the COVID-19 Genomics UK (COG-UK) Consortium | Jacquelyn Wynn, Mairead Hyland, The Lighthouse Lab in Alderley Park and Alex Alderton, Roberto Amato, Jeffrey Barrett, Sonia Goncalves, Ewan Harrison, David K. Jackson, Ian Johnston, Dominic Kwiatkowski, Cordelia Langford, John Sillitoe on behalf of the Wellcome Sanger Institute COVID-19 Surveillance Team |
| EPI_ISL_3097549 | Berkshire and Surrey Pathology Services Lighthouse Laboratory | Wellcome Sanger Institute for the COVID-19 Genomics UK (COG-UK) Consortium | Berkshire and Surrey Pathology Services Lighthouse Laboratory and Alex Alderton, Roberto Amato, Jeffrey Barrett, Sonia Goncalves, Ewan Harrison, David K. Jackson, Ian Johnston, Dominic Kwiatkowski, Cordelia Langford, John Sillitoe on behalf of the Wellcome Sanger Institute COVID-19 Surveillance Team |
| EPI_ISL_3097550, EPI_ISL_3097551, EPI_ISL_3097552 | Lighthouse Lab in Glasgow | Wellcome Sanger Institute for the COVID-19 Genomics UK (COG-UK) Consortium | Harper VanSteenhouse, Yumi Kasai, David Gray, Carol Clugston, Anna Dominiczak and Alex Alderton, Roberto Amato, Jeffrey Barrett, Sonia Goncalves, Ewan Harrison, David K. Jackson, Ian Johnston, Dominic Kwiatkowski, Cordelia Langford, John Sillitoe on behalf of the Wellcome Sanger Institute COVID-19 Surveillance Team |
| EPI_ISL_3097553 | Berkshire and Surrey Pathology Services Lighthouse Laboratory | Wellcome Sanger Institute for the COVID-19 Genomics UK (COG-UK) Consortium | Berkshire and Surrey Pathology Services Lighthouse Laboratory and Alex Alderton, Roberto Amato, Jeffrey Barrett, Sonia Goncalves, Ewan Harrison, David K. Jackson, Ian Johnston, Dominic Kwiatkowski, Cordelia Langford, John Sillitoe on behalf of the Wellcome Sanger Institute COVID-19 Surveillance Team |
| EPI_ISL_3097554 | Lighthouse Lab in Glasgow | Wellcome Sanger Institute for the COVID-19 Genomics UK (COG-UK) Consortium | Harper VanSteenhouse, Yumi Kasai, David Gray, Carol Clugston, Anna Dominiczak and Alex Alderton, Roberto Amato, Jeffrey Barrett, Sonia Goncalves, Ewan Harrison, David K. Jackson, Ian Johnston, Dominic Kwiatkowski, Cordelia Langford, John Sillitoe on behalf of the Wellcome Sanger Institute COVID-19 Surveillance Team |
| EPI_ISL_3097555 | Berkshire and Surrey Pathology Services Lighthouse Laboratory | Wellcome Sanger Institute for the COVID-19 Genomics UK (COG-UK) Consortium | Berkshire and Surrey Pathology Services Lighthouse Laboratory and Alex Alderton, Roberto Amato, Jeffrey Barrett, Sonia Goncalves, Ewan Harrison, David K. Jackson, Ian Johnston, Dominic Kwiatkowski, Cordelia Langford, John Sillitoe on behalf of the Wellcome Sanger Institute COVID-19 Surveillance Team |
| EPI_ISL_3097556 | Lighthouse Lab in Glasgow | Wellcome Sanger Institute for the COVID-19 Genomics UK (COG-UK) Consortium | Harper VanSteenhouse, Yumi Kasai, David Gray, Carol Clugston, Anna Dominiczak and Alex Alderton, Roberto Amato, Jeffrey Barrett, Sonia Goncalves, Ewan Harrison, David K. Jackson, Ian Johnston, Dominic Kwiatkowski, Cordelia Langford, John Sillitoe on behalf of the Wellcome Sanger Institute COVID-19 Surveillance Team |
| EPI_ISL_3097557, EPI_ISL_3097558, EPI_ISL_3097559, EPI_ISL_3097560, EPI_ISL_3097561, EPI_ISL_3097562 | Berkshire and Surrey Pathology Services Lighthouse Laboratory | Wellcome Sanger Institute for the COVID-19 Genomics UK (COG-UK) Consortium | Berkshire and Surrey Pathology Services Lighthouse Laboratory and Alex Alderton, Roberto Amato, Jeffrey Barrett, Sonia Goncalves, Ewan Harrison, David K. Jackson, Ian Johnston, Dominic Kwiatkowski, Cordelia Langford, John Sillitoe on behalf of the Wellcome Sanger Institute COVID-19 Surveillance Team |
| EPI_ISL_3097563, EPI_ISL_3097564 | Lighthouse Lab in Glasgow | Wellcome Sanger Institute for the COVID-19 Genomics UK (COG-UK) Consortium | Harper VanSteenhouse, Yumi Kasai, David Gray, Carol Clugston, Anna Dominiczak and Alex Alderton, Roberto Amato, Jeffrey Barrett, Sonia Goncalves, Ewan Harrison, David K. Jackson, Ian Johnston, Dominic Kwiatkowski, Cordelia Langford, John Sillitoe on behalf of the Wellcome Sanger Institute COVID-19 Surveillance Team |
| EPI_ISL_3097565 | Lighthouse Lab in Alderley Park | Wellcome Sanger Institute for the COVID-19 Genomics UK (COG-UK) Consortium | Jacquelyn Wynn, Mairead Hyland, The Lighthouse Lab in Alderley Park and Alex Alderton, Roberto Amato, Jeffrey Barrett, Sonia Goncalves, Ewan Harrison, David K. Jackson, Ian Johnston, Dominic Kwiatkowski, Cordelia Langford, John Sillitoe on behalf of the Wellcome Sanger Institute COVID-19 Surveillance Team |
| EPI_ISL_3097566 | Lighthouse Lab in Glasgow | Wellcome Sanger Institute for the COVID-19 Genomics UK (COG-UK) Consortium | Harper VanSteenhouse, Yumi Kasai, David Gray, Carol Clugston, Anna Dominiczak and Alex Alderton, Roberto Amato, Jeffrey Barrett, Sonia Goncalves, Ewan Harrison, David K. Jackson, Ian Johnston, Dominic Kwiatkowski, Cordelia Langford, John Sillitoe on behalf of the Wellcome Sanger Institute COVID-19 Surveillance Team |
| EPI_ISL_3097567 | Berkshire and Surrey Pathology Services Lighthouse Laboratory | Wellcome Sanger Institute for the COVID-19 Genomics UK (COG-UK) Consortium | Berkshire and Surrey Pathology Services Lighthouse Laboratory and Alex Alderton, Roberto Amato, Jeffrey Barrett, Sonia Goncalves, Ewan Harrison, David K. Jackson, Ian Johnston, Dominic Kwiatkowski, Cordelia Langford, John Sillitoe on behalf of the Wellcome Sanger Institute COVID-19 Surveillance Team |
| EPI_ISL_3097568 | Lighthouse Lab in Alderley Park | Wellcome Sanger Institute for the COVID-19 Genomics UK (COG-UK) Consortium | Jacquelyn Wynn, Mairead Hyland, The Lighthouse Lab in Alderley Park and Alex Alderton, Roberto Amato, Jeffrey Barrett, Sonia Goncalves, Ewan Harrison, David K. Jackson, Ian Johnston, Dominic Kwiatkowski, Cordelia Langford, John Sillitoe on behalf of the Wellcome Sanger Institute COVID-19 Surveillance Team |
| EPI_ISL_3097569 | Lighthouse Lab in Glasgow | Wellcome Sanger Institute for the COVID-19 Genomics UK (COG-UK) Consortium | Harper VanSteenhouse, Yumi Kasai, David Gray, Carol Clugston, Anna Dominiczak and Alex Alderton, Roberto Amato, Jeffrey Barrett, Sonia Goncalves, Ewan Harrison, David K. Jackson, Ian Johnston, Dominic Kwiatkowski, Cordelia Langford, John Sillitoe on behalf of the Wellcome Sanger Institute COVID-19 Surveillance Team |
| EPI_ISL_3097570 | Lighthouse Lab in Alderley Park | Wellcome Sanger Institute for the COVID-19 Genomics UK (COG-UK) | Jacquelyn Wynn, Mairead Hyland, The Lighthouse Lab in Alderley Park and Alex Alderton, Roberto Amato, Jeffrey Barrett, Sonia Goncalves, Ewan Harrison, David K. Jackson, Ian Johnston, Dominic Kwiatkowski, Cordelia Langford, John Sillitoe on |

| | | | |
|---|---|---|---|
| | | Consortium | behalf of the Wellcome Sanger Institute COVID-19 Surveillance Team |
| EPI_ISL_3097571 | Lighthouse Lab in Glasgow | Wellcome Sanger Institute for the COVID-19 Genomics UK (COG-UK) Consortium | Harper VanSteenhouse, Yumi Kasai, David Gray, Carol Clugston, Anna Dominiczak and Alex Alderton, Roberto Amato, Jeffrey Barrett, Sonia Goncalves, Ewan Harrison, David K. Jackson, Ian Johnston, Dominic Kwiatkowski, Cordelia Langford, John Sillitoe on behalf of the Wellcome Sanger Institute COVID-19 Surveillance Team |
| EPI_ISL_3097572 | Lighthouse Lab in Alderley Park | Wellcome Sanger Institute for the COVID-19 Genomics UK (COG-UK) Consortium | Jacquelyn Wynn, Mairead Hyland, The Lighthouse Lab in Alderley Park and Alex Alderton, Roberto Amato, Jeffrey Barrett, Sonia Goncalves, Ewan Harrison, David K. Jackson, Ian Johnston, Dominic Kwiatkowski, Cordelia Langford, John Sillitoe on behalf of the Wellcome Sanger Institute COVID-19 Surveillance Team |
| EPI_ISL_3097573 | Lighthouse Lab in Glasgow | Wellcome Sanger Institute for the COVID-19 Genomics UK (COG-UK) Consortium | Harper VanSteenhouse, Yumi Kasai, David Gray, Carol Clugston, Anna Dominiczak and Alex Alderton, Roberto Amato, Jeffrey Barrett, Sonia Goncalves, Ewan Harrison, David K. Jackson, Ian Johnston, Dominic Kwiatkowski, Cordelia Langford, John Sillitoe on behalf of the Wellcome Sanger Institute COVID-19 Surveillance Team |
| EPI_ISL_3097574 | Lighthouse Lab in Alderley Park | Wellcome Sanger Institute for the COVID-19 Genomics UK (COG-UK) Consortium | Jacquelyn Wynn, Mairead Hyland, The Lighthouse Lab in Alderley Park and Alex Alderton, Roberto Amato, Jeffrey Barrett, Sonia Goncalves, Ewan Harrison, David K. Jackson, Ian Johnston, Dominic Kwiatkowski, Cordelia Langford, John Sillitoe on behalf of the Wellcome Sanger Institute COVID-19 Surveillance Team |
| EPI_ISL_3097575, EPI_ISL_3097576 | Lighthouse Lab in Glasgow | Wellcome Sanger Institute for the COVID-19 Genomics UK (COG-UK) Consortium | Harper VanSteenhouse, Yumi Kasai, David Gray, Carol Clugston, Anna Dominiczak and Alex Alderton, Roberto Amato, Jeffrey Barrett, Sonia Goncalves, Ewan Harrison, David K. Jackson, Ian Johnston, Dominic Kwiatkowski, Cordelia Langford, John Sillitoe on behalf of the Wellcome Sanger Institute COVID-19 Surveillance Team |
| EPI_ISL_3097577, EPI_ISL_3097578 | Lighthouse Lab in Alderley Park | Wellcome Sanger Institute for the COVID-19 Genomics UK (COG-UK) Consortium | Jacquelyn Wynn, Mairead Hyland, The Lighthouse Lab in Alderley Park and Alex Alderton, Roberto Amato, Jeffrey Barrett, Sonia Goncalves, Ewan Harrison, David K. Jackson, Ian Johnston, Dominic Kwiatkowski, Cordelia Langford, John Sillitoe on behalf of the Wellcome Sanger Institute COVID-19 Surveillance Team |
| EPI_ISL_3097579 | Lighthouse Lab in Glasgow | Wellcome Sanger Institute for the COVID-19 Genomics UK (COG-UK) Consortium | Harper VanSteenhouse, Yumi Kasai, David Gray, Carol Clugston, Anna Dominiczak and Alex Alderton, Roberto Amato, Jeffrey Barrett, Sonia Goncalves, Ewan Harrison, David K. Jackson, Ian Johnston, Dominic Kwiatkowski, Cordelia Langford, John Sillitoe on behalf of the Wellcome Sanger Institute COVID-19 Surveillance Team |
| EPI_ISL_3097580, EPI_ISL_3097581 | Berkshire and Surrey Pathology Services Lighthouse Laboratory | Wellcome Sanger Institute for the COVID-19 Genomics UK (COG-UK) Consortium | Berkshire and Surrey Pathology Services Lighthouse Laboratory and Alex Alderton, Roberto Amato, Jeffrey Barrett, Sonia Goncalves, Ewan Harrison, David K. Jackson, Ian Johnston, Dominic Kwiatkowski, Cordelia Langford, John Sillitoe on behalf of the Wellcome Sanger Institute COVID-19 Surveillance Team |
| EPI_ISL_3097582, EPI_ISL_3097583 | Lighthouse Lab in Glasgow | Wellcome Sanger Institute for the COVID-19 Genomics UK (COG-UK) Consortium | Harper VanSteenhouse, Yumi Kasai, David Gray, Carol Clugston, Anna Dominiczak and Alex Alderton, Roberto Amato, Jeffrey Barrett, Sonia Goncalves, Ewan Harrison, David K. Jackson, Ian Johnston, Dominic Kwiatkowski, Cordelia Langford, John Sillitoe on behalf of the Wellcome Sanger Institute COVID-19 Surveillance Team |
| EPI_ISL_3097584 | Lighthouse Lab in Alderley Park | Wellcome Sanger Institute for the COVID-19 Genomics UK (COG-UK) Consortium | Jacquelyn Wynn, Mairead Hyland, The Lighthouse Lab in Alderley Park and Alex Alderton, Roberto Amato, Jeffrey Barrett, Sonia Goncalves, Ewan Harrison, David K. Jackson, Ian Johnston, Dominic Kwiatkowski, Cordelia Langford, John Sillitoe on behalf of the Wellcome Sanger Institute COVID-19 Surveillance Team |
| EPI_ISL_3097585 | Lighthouse Lab in Glasgow | Wellcome Sanger Institute for the COVID-19 Genomics UK (COG-UK) Consortium | Harper VanSteenhouse, Yumi Kasai, David Gray, Carol Clugston, Anna Dominiczak and Alex Alderton, Roberto Amato, Jeffrey Barrett, Sonia Goncalves, Ewan Harrison, David K. Jackson, Ian Johnston, Dominic Kwiatkowski, Cordelia Langford, John Sillitoe on behalf of the Wellcome Sanger Institute COVID-19 Surveillance Team |
| EPI_ISL_3097586 | Lighthouse Lab in Alderley Park | Wellcome Sanger Institute for the COVID-19 Genomics UK (COG-UK) Consortium | Jacquelyn Wynn, Mairead Hyland, The Lighthouse Lab in Alderley Park and Alex Alderton, Roberto Amato, Jeffrey Barrett, Sonia Goncalves, Ewan Harrison, David K. Jackson, Ian Johnston, Dominic Kwiatkowski, Cordelia Langford, John Sillitoe on behalf of the Wellcome Sanger Institute COVID-19 Surveillance Team |
| EPI_ISL_3097587, EPI_ISL_3097588, EPI_ISL_3097589 | Lighthouse Lab in Glasgow | Wellcome Sanger Institute for the COVID-19 Genomics UK (COG-UK) Consortium | Harper VanSteenhouse, Yumi Kasai, David Gray, Carol Clugston, Anna Dominiczak and Alex Alderton, Roberto Amato, Jeffrey Barrett, Sonia Goncalves, Ewan Harrison, David K. Jackson, Ian Johnston, Dominic Kwiatkowski, Cordelia Langford, John Sillitoe on behalf of the Wellcome Sanger Institute COVID-19 Surveillance Team |
| EPI_ISL_3097590 | Lighthouse Lab in Alderley Park | Wellcome Sanger Institute for the COVID-19 Genomics UK (COG-UK) Consortium | Jacquelyn Wynn, Mairead Hyland, The Lighthouse Lab in Alderley Park and Alex Alderton, Roberto Amato, Jeffrey Barrett, Sonia Goncalves, Ewan Harrison, David K. Jackson, Ian Johnston, Dominic Kwiatkowski, Cordelia Langford, John Sillitoe on behalf of the Wellcome Sanger Institute COVID-19 Surveillance Team |
| EPI_ISL_3097591 | Lighthouse Lab in Glasgow | Wellcome Sanger Institute for the COVID-19 Genomics UK (COG-UK) Consortium | Harper VanSteenhouse, Yumi Kasai, David Gray, Carol Clugston, Anna Dominiczak and Alex Alderton, Roberto Amato, Jeffrey Barrett, Sonia Goncalves, Ewan Harrison, David K. Jackson, Ian Johnston, Dominic Kwiatkowski, Cordelia Langford, John Sillitoe on behalf of the Wellcome Sanger Institute COVID-19 Surveillance Team |
| EPI_ISL_3097592, EPI_ISL_3097593 | Lighthouse Lab in Alderley Park | Wellcome Sanger Institute for the COVID-19 Genomics UK (COG-UK) Consortium | Jacquelyn Wynn, Mairead Hyland, The Lighthouse Lab in Alderley Park and Alex Alderton, Roberto Amato, Jeffrey Barrett, Sonia Goncalves, Ewan Harrison, David K. Jackson, Ian Johnston, Dominic Kwiatkowski, Cordelia Langford, John Sillitoe on behalf of the Wellcome Sanger Institute COVID-19 Surveillance Team |
| EPI_ISL_3097594 | Lighthouse Lab in Glasgow | Wellcome Sanger Institute for the COVID-19 Genomics UK (COG-UK) Consortium | Harper VanSteenhouse, Yumi Kasai, David Gray, Carol Clugston, Anna Dominiczak and Alex Alderton, Roberto Amato, Jeffrey Barrett, Sonia Goncalves, Ewan Harrison, David K. Jackson, Ian Johnston, Dominic Kwiatkowski, Cordelia Langford, John Sillitoe on behalf of the Wellcome Sanger Institute COVID-19 Surveillance Team |
| EPI_ISL_3097595 | Lighthouse Lab in Alderley Park | Wellcome Sanger Institute for the COVID-19 Genomics UK (COG-UK) Consortium | Jacquelyn Wynn, Mairead Hyland, The Lighthouse Lab in Alderley Park and Alex Alderton, Roberto Amato, Jeffrey Barrett, Sonia Goncalves, Ewan Harrison, David K. Jackson, Ian Johnston, Dominic Kwiatkowski, Cordelia Langford, John Sillitoe on behalf of the Wellcome Sanger Institute COVID-19 Surveillance Team |
| EPI_ISL_3097596, EPI_ISL_3097597, EPI_ISL_3097598 | Lighthouse Lab in Glasgow | Wellcome Sanger Institute for the COVID-19 Genomics UK (COG-UK) Consortium | Harper VanSteenhouse, Yumi Kasai, David Gray, Carol Clugston, Anna Dominiczak and Alex Alderton, Roberto Amato, Jeffrey Barrett, Sonia Goncalves, Ewan Harrison, David K. Jackson, Ian Johnston, Dominic Kwiatkowski, Cordelia Langford, John Sillitoe on behalf of the Wellcome Sanger Institute COVID-19 Surveillance Team |
| EPI_ISL_3097599 | Berkshire and Surrey Pathology Services Lighthouse Laboratory | Wellcome Sanger Institute for the COVID-19 Genomics UK (COG-UK) Consortium | Berkshire and Surrey Pathology Services Lighthouse Laboratory and Alex Alderton, Roberto Amato, Jeffrey Barrett, Sonia Goncalves, Ewan Harrison, David K. Jackson, Ian Johnston, Dominic Kwiatkowski, Cordelia Langford, John Sillitoe on behalf of the Wellcome Sanger Institute COVID-19 Surveillance Team |
| EPI_ISL_3097600 | Lighthouse Lab in Glasgow | Wellcome Sanger Institute for the COVID-19 Genomics UK (COG-UK) Consortium | Harper VanSteenhouse, Yumi Kasai, David Gray, Carol Clugston, Anna Dominiczak and Alex Alderton, Roberto Amato, Jeffrey Barrett, Sonia Goncalves, Ewan Harrison, David K. Jackson, Ian Johnston, Dominic Kwiatkowski, Cordelia Langford, John Sillitoe on behalf of the Wellcome Sanger Institute COVID-19 Surveillance Team |
| EPI_ISL_3097601 | Lighthouse Lab in Alderley Park | Wellcome Sanger Institute for the COVID-19 Genomics UK (COG-UK) Consortium | Jacquelyn Wynn, Mairead Hyland, The Lighthouse Lab in Alderley Park and Alex Alderton, Roberto Amato, Jeffrey Barrett, Sonia Goncalves, Ewan Harrison, David K. Jackson, Ian Johnston, Dominic Kwiatkowski, Cordelia Langford, John Sillitoe on behalf of the Wellcome Sanger Institute COVID-19 Surveillance Team |
| EPI_ISL_3097602 | Berkshire and Surrey Pathology Services Lighthouse Laboratory | Wellcome Sanger Institute for the COVID-19 Genomics UK (COG-UK) Consortium | Berkshire and Surrey Pathology Services Lighthouse Laboratory and Alex Alderton, Roberto Amato, Jeffrey Barrett, Sonia Goncalves, Ewan Harrison, David K. Jackson, Ian Johnston, Dominic Kwiatkowski, Cordelia Langford, John Sillitoe on behalf of the Wellcome Sanger Institute COVID-19 Surveillance Team |
| EPI_ISL_3097603, EPI_ISL_3097604, EPI_ISL_3097605, EPI_ISL_3097606, EPI_ISL_3097607 | Lighthouse Lab in Alderley Park | Wellcome Sanger Institute for the COVID-19 Genomics UK (COG-UK) Consortium | Jacquelyn Wynn, Mairead Hyland, The Lighthouse Lab in Alderley Park and Alex Alderton, Roberto Amato, Jeffrey Barrett, Sonia Goncalves, Ewan Harrison, David K. Jackson, Ian Johnston, Dominic Kwiatkowski, Cordelia Langford, John Sillitoe on behalf of the Wellcome Sanger Institute COVID-19 Surveillance Team |
| EPI_ISL_3097608 | Berkshire and Surrey Pathology Services Lighthouse Laboratory | Wellcome Sanger Institute for the COVID-19 Genomics UK (COG-UK) Consortium | Berkshire and Surrey Pathology Services Lighthouse Laboratory and Alex Alderton, Roberto Amato, Jeffrey Barrett, Sonia Goncalves, Ewan Harrison, David K. Jackson, Ian Johnston, Dominic Kwiatkowski, Cordelia Langford, John Sillitoe on behalf of the Wellcome Sanger Institute COVID-19 Surveillance Team |
| EPI_ISL_3097609 | Lighthouse Lab in Alderley Park | Wellcome Sanger Institute for the COVID-19 Genomics UK (COG-UK) Consortium | Jacquelyn Wynn, Mairead Hyland, The Lighthouse Lab in Alderley Park and Alex Alderton, Roberto Amato, Jeffrey Barrett, Sonia Goncalves, Ewan Harrison, David K. Jackson, Ian Johnston, Dominic Kwiatkowski, Cordelia Langford, John Sillitoe on behalf of the Wellcome Sanger Institute COVID-19 Surveillance Team |
| EPI_ISL_3097610, EPI_ISL_3097611 | Berkshire and Surrey Pathology Services Lighthouse Laboratory | Wellcome Sanger Institute for the COVID-19 Genomics UK (COG-UK) Consortium | Berkshire and Surrey Pathology Services Lighthouse Laboratory and Alex Alderton, Roberto Amato, Jeffrey Barrett, Sonia Goncalves, Ewan Harrison, David K. Jackson, Ian Johnston, Dominic Kwiatkowski, Cordelia Langford, John Sillitoe on behalf of the Wellcome Sanger Institute COVID-19 Surveillance Team |
| EPI_ISL_3097612 | Lighthouse Lab in Alderley Park | Wellcome Sanger Institute for the COVID-19 Genomics UK (COG-UK) Consortium | Jacquelyn Wynn, Mairead Hyland, The Lighthouse Lab in Alderley Park and Alex Alderton, Roberto Amato, Jeffrey Barrett, Sonia Goncalves, Ewan Harrison, David K. Jackson, Ian Johnston, Dominic Kwiatkowski, Cordelia Langford, John Sillitoe on behalf of the Wellcome Sanger Institute COVID-19 Surveillance Team |
| EPI_ISL_3097613 | Lighthouse Lab in Glasgow | Wellcome Sanger Institute for the COVID-19 Genomics UK (COG-UK) Consortium | Harper VanSteenhouse, Yumi Kasai, David Gray, Carol Clugston, Anna Dominiczak and Alex Alderton, Roberto Amato, Jeffrey Barrett, Sonia Goncalves, Ewan Harrison, David K. Jackson, Ian Johnston, Dominic Kwiatkowski, Cordelia Langford, John Sillitoe on behalf of the Wellcome Sanger Institute COVID-19 Surveillance Team |
| EPI_ISL_3097614, EPI_ISL_3097615 | Lighthouse Lab in Alderley Park | Wellcome Sanger Institute for the COVID-19 Genomics UK (COG-UK) Consortium | Jacquelyn Wynn, Mairead Hyland, The Lighthouse Lab in Alderley Park and Alex Alderton, Roberto Amato, Jeffrey Barrett, Sonia Goncalves, Ewan Harrison, David K. Jackson, Ian Johnston, Dominic Kwiatkowski, Cordelia Langford, John Sillitoe on behalf of the Wellcome Sanger Institute COVID-19 Surveillance Team |
| EPI_ISL_3097616 | Berkshire and Surrey Pathology Services Lighthouse Laboratory | Wellcome Sanger Institute for the COVID-19 Genomics UK (COG-UK) Consortium | Berkshire and Surrey Pathology Services Lighthouse Laboratory and Alex Alderton, Roberto Amato, Jeffrey Barrett, Sonia Goncalves, Ewan Harrison, David K. Jackson, Ian Johnston, Dominic Kwiatkowski, Cordelia Langford, John Sillitoe on behalf of the Wellcome Sanger Institute COVID-19 Surveillance Team |
| EPI_ISL_3097617 | Lighthouse Lab in Glasgow | Wellcome Sanger Institute for the COVID-19 Genomics UK (COG-UK) Consortium | Harper VanSteenhouse, Yumi Kasai, David Gray, Carol Clugston, Anna Dominiczak and Alex Alderton, Roberto Amato, Jeffrey Barrett, Sonia Goncalves, Ewan Harrison, David K. Jackson, Ian Johnston, Dominic Kwiatkowski, Cordelia Langford, John Sillitoe on behalf of the Wellcome Sanger Institute COVID-19 Surveillance Team |
| EPI_ISL_3097618, EPI_ISL_3097619 | Lighthouse Lab in Alderley Park | Wellcome Sanger Institute for the COVID-19 Genomics UK (COG-UK) Consortium | Jacquelyn Wynn, Mairead Hyland, The Lighthouse Lab in Alderley Park and Alex Alderton, Roberto Amato, Jeffrey Barrett, Sonia Goncalves, Ewan Harrison, David K. Jackson, Ian Johnston, Dominic Kwiatkowski, Cordelia Langford, John Sillitoe on behalf of the Wellcome Sanger Institute COVID-19 Surveillance Team |
| EPI_ISL_3097620 | Lighthouse Lab in Glasgow | Wellcome Sanger Institute for the COVID-19 Genomics UK (COG-UK) Consortium | Harper VanSteenhouse, Yumi Kasai, David Gray, Carol Clugston, Anna Dominiczak and Alex Alderton, Roberto Amato, Jeffrey Barrett, Sonia Goncalves, Ewan Harrison, David K. Jackson, Ian Johnston, Dominic Kwiatkowski, Cordelia Langford, John Sillitoe on behalf of the Wellcome Sanger Institute COVID-19 Surveillance Team |
| EPI_ISL_3097621 | Lighthouse Lab in Alderley Park | Wellcome Sanger Institute for the COVID-19 Genomics UK (COG-UK) Consortium | Jacquelyn Wynn, Mairead Hyland, The Lighthouse Lab in Alderley Park and Alex Alderton, Roberto Amato, Jeffrey Barrett, Sonia Goncalves, Ewan Harrison, David K. Jackson, Ian Johnston, Dominic Kwiatkowski, Cordelia Langford, John Sillitoe on behalf of the Wellcome Sanger Institute COVID-19 Surveillance Team |
| EPI_ISL_3097622 | Lighthouse Lab in Glasgow | Wellcome Sanger Institute for the COVID-19 Genomics UK (COG-UK) Consortium | Harper VanSteenhouse, Yumi Kasai, David Gray, Carol Clugston, Anna Dominiczak and Alex Alderton, Roberto Amato, Jeffrey Barrett, Sonia Goncalves, Ewan Harrison, David K. Jackson, Ian Johnston, Dominic Kwiatkowski, Cordelia Langford, John Sillitoe on behalf of the Wellcome Sanger Institute COVID-19 Surveillance Team |
| EPI_ISL_3097623 | Berkshire and Surrey Pathology Services Lighthouse Laboratory | Wellcome Sanger Institute for the COVID-19 Genomics UK (COG-UK) Consortium | Berkshire and Surrey Pathology Services Lighthouse Laboratory and Alex Alderton, Roberto Amato, Jeffrey Barrett, Sonia Goncalves, Ewan Harrison, David K. Jackson, Ian Johnston, Dominic Kwiatkowski, Cordelia Langford, John Sillitoe on behalf of the Wellcome Sanger Institute COVID-19 Surveillance Team |
| EPI_ISL_3097624, EPI_ISL_3097625 | Lighthouse Lab in Glasgow | Wellcome Sanger Institute for the COVID-19 Genomics UK (COG-UK) Consortium | Harper VanSteenhouse, Yumi Kasai, David Gray, Carol Clugston, Anna Dominiczak and Alex Alderton, Roberto Amato, Jeffrey Barrett, Sonia Goncalves, Ewan Harrison, David K. Jackson, Ian Johnston, Dominic Kwiatkowski, Cordelia Langford, John Sillitoe on behalf of the Wellcome Sanger Institute COVID-19 Surveillance Team |
| EPI_ISL_3097626 | Lighthouse Lab in Alderley Park | Wellcome Sanger Institute for the COVID-19 Genomics UK (COG-UK) Consortium | Jacquelyn Wynn, Mairead Hyland, The Lighthouse Lab in Alderley Park and Alex Alderton, Roberto Amato, Jeffrey Barrett, Sonia Goncalves, Ewan Harrison, David K. Jackson, Ian Johnston, Dominic Kwiatkowski, Cordelia Langford, John Sillitoe on behalf of the Wellcome Sanger Institute COVID-19 Surveillance Team |
| EPI_ISL_3097627 | Lighthouse Lab in Glasgow | Wellcome Sanger Institute for the COVID-19 Genomics UK (COG-UK) Consortium | Harper VanSteenhouse, Yumi Kasai, David Gray, Carol Clugston, Anna Dominiczak and Alex Alderton, Roberto Amato, Jeffrey Barrett, Sonia Goncalves, Ewan Harrison, David K. Jackson, Ian Johnston, Dominic Kwiatkowski, Cordelia Langford, John Sillitoe on behalf of the Wellcome Sanger Institute COVID-19 Surveillance Team |
| EPI_ISL_3097628 | Lighthouse Lab in Alderley Park | Wellcome Sanger Institute for the COVID-19 Genomics UK (COG-UK) Consortium | Jacquelyn Wynn, Mairead Hyland, The Lighthouse Lab in Alderley Park and Alex Alderton, Roberto Amato, Jeffrey Barrett, Sonia Goncalves, Ewan Harrison, David K. Jackson, Ian Johnston, Dominic Kwiatkowski, Cordelia Langford, John Sillitoe on behalf of the Wellcome Sanger Institute COVID-19 Surveillance Team |
| EPI_ISL_3097629 | Lighthouse Lab in Glasgow | Wellcome Sanger Institute for the COVID-19 Genomics UK (COG-UK) Consortium | Harper VanSteenhouse, Yumi Kasai, David Gray, Carol Clugston, Anna Dominiczak and Alex Alderton, Roberto Amato, Jeffrey Barrett, Sonia Goncalves, Ewan Harrison, David K. Jackson, Ian Johnston, Dominic Kwiatkowski, Cordelia Langford, John Sillitoe on behalf of the Wellcome Sanger Institute COVID-19 Surveillance Team |
| EPI_ISL_3097630 | Berkshire and Surrey Pathology Services Lighthouse Laboratory | Wellcome Sanger Institute for the COVID-19 Genomics UK (COG-UK) | Berkshire and Surrey Pathology Services Lighthouse Laboratory and Alex Alderton, Roberto Amato, Jeffrey Barrett, Sonia Goncalves, Ewan Harrison, David K. Jackson, Ian Johnston, Dominic Kwiatkowski, Cordelia Langford, John Sillitoe on behalf |

| | | | |
|---|---|---|---|
| | | Consortium | of the Wellcome Sanger Institute COVID-19 Surveillance Team |
| EPI_ISL_3097631 | Lighthouse Lab in Alderley Park | Wellcome Sanger Institute for the COVID-19 Genomics UK (COG-UK) Consortium | Jacquelyn Wynn, Mairead Hyland, The Lighthouse Lab in Alderley Park and Alex Alderton, Roberto Amato, Jeffrey Barrett, Sonia Goncalves, Ewan Harrison, David K. Jackson, Ian Johnston, Dominic Kwiatkowski, Cordelia Langford, John Sillitoe on behalf of the Wellcome Sanger Institute COVID-19 Surveillance Team |
| EPI_ISL_3097632 | Berkshire and Surrey Pathology Services Lighthouse Laboratory | Wellcome Sanger Institute for the COVID-19 Genomics UK (COG-UK) Consortium | Berkshire and Surrey Pathology Services Lighthouse Laboratory and Alex Alderton, Roberto Amato, Jeffrey Barrett, Sonia Goncalves, Ewan Harrison, David K. Jackson, Ian Johnston, Dominic Kwiatkowski, Cordelia Langford, John Sillitoe on behalf of the Wellcome Sanger Institute COVID-19 Surveillance Team |
| EPI_ISL_3097633 | Lighthouse Lab in Alderley Park | Wellcome Sanger Institute for the COVID-19 Genomics UK (COG-UK) Consortium | Jacquelyn Wynn, Mairead Hyland, The Lighthouse Lab in Alderley Park and Alex Alderton, Roberto Amato, Jeffrey Barrett, Sonia Goncalves, Ewan Harrison, David K. Jackson, Ian Johnston, Dominic Kwiatkowski, Cordelia Langford, John Sillitoe on behalf of the Wellcome Sanger Institute COVID-19 Surveillance Team |
| EPI_ISL_3097634 | Lighthouse Lab in Glasgow | Wellcome Sanger Institute for the COVID-19 Genomics UK (COG-UK) Consortium | Harper VanSteenhouse, Yumi Kasai, David Gray, Carol Clugston, Anna Dominiczak and Alex Alderton, Roberto Amato, Jeffrey Barrett, Sonia Goncalves, Ewan Harrison, David K. Jackson, Ian Johnston, Dominic Kwiatkowski, Cordelia Langford, John Sillitoe on behalf of the Wellcome Sanger Institute COVID-19 Surveillance Team |
| EPI_ISL_3097635 | Berkshire and Surrey Pathology Services Lighthouse Laboratory | Wellcome Sanger Institute for the COVID-19 Genomics UK (COG-UK) Consortium | Berkshire and Surrey Pathology Services Lighthouse Laboratory and Alex Alderton, Roberto Amato, Jeffrey Barrett, Sonia Goncalves, Ewan Harrison, David K. Jackson, Ian Johnston, Dominic Kwiatkowski, Cordelia Langford, John Sillitoe on behalf of the Wellcome Sanger Institute COVID-19 Surveillance Team |
| EPI_ISL_3097636 | Lighthouse Lab in Glasgow | Wellcome Sanger Institute for the COVID-19 Genomics UK (COG-UK) Consortium | Harper VanSteenhouse, Yumi Kasai, David Gray, Carol Clugston, Anna Dominiczak and Alex Alderton, Roberto Amato, Jeffrey Barrett, Sonia Goncalves, Ewan Harrison, David K. Jackson, Ian Johnston, Dominic Kwiatkowski, Cordelia Langford, John Sillitoe on behalf of the Wellcome Sanger Institute COVID-19 Surveillance Team |
| EPI_ISL_3097637, EPI_ISL_3097638, EPI_ISL_3097639, EPI_ISL_3097640, EPI_ISL_3097641 | Lighthouse Lab in Alderley Park | Wellcome Sanger Institute for the COVID-19 Genomics UK (COG-UK) Consortium | Jacquelyn Wynn, Mairead Hyland, The Lighthouse Lab in Alderley Park and Alex Alderton, Roberto Amato, Jeffrey Barrett, Sonia Goncalves, Ewan Harrison, David K. Jackson, Ian Johnston, Dominic Kwiatkowski, Cordelia Langford, John Sillitoe on behalf of the Wellcome Sanger Institute COVID-19 Surveillance Team |
| EPI_ISL_3097642, EPI_ISL_3097643 | Berkshire and Surrey Pathology Services Lighthouse Laboratory | Wellcome Sanger Institute for the COVID-19 Genomics UK (COG-UK) Consortium | Berkshire and Surrey Pathology Services Lighthouse Laboratory and Alex Alderton, Roberto Amato, Jeffrey Barrett, Sonia Goncalves, Ewan Harrison, David K. Jackson, Ian Johnston, Dominic Kwiatkowski, Cordelia Langford, John Sillitoe on behalf of the Wellcome Sanger Institute COVID-19 Surveillance Team |
| EPI_ISL_3097644, EPI_ISL_3097645 | Lighthouse Lab in Glasgow | Wellcome Sanger Institute for the COVID-19 Genomics UK (COG-UK) Consortium | Harper VanSteenhouse, Yumi Kasai, David Gray, Carol Clugston, Anna Dominiczak and Alex Alderton, Roberto Amato, Jeffrey Barrett, Sonia Goncalves, Ewan Harrison, David K. Jackson, Ian Johnston, Dominic Kwiatkowski, Cordelia Langford, John Sillitoe on behalf of the Wellcome Sanger Institute COVID-19 Surveillance Team |
| EPI_ISL_3097646 | Berkshire and Surrey Pathology Services Lighthouse Laboratory | Wellcome Sanger Institute for the COVID-19 Genomics UK (COG-UK) Consortium | Berkshire and Surrey Pathology Services Lighthouse Laboratory and Alex Alderton, Roberto Amato, Jeffrey Barrett, Sonia Goncalves, Ewan Harrison, David K. Jackson, Ian Johnston, Dominic Kwiatkowski, Cordelia Langford, John Sillitoe on behalf of the Wellcome Sanger Institute COVID-19 Surveillance Team |
| EPI_ISL_3097647 | Lighthouse Lab in Alderley Park | Wellcome Sanger Institute for the COVID-19 Genomics UK (COG-UK) Consortium | Jacquelyn Wynn, Mairead Hyland, The Lighthouse Lab in Alderley Park and Alex Alderton, Roberto Amato, Jeffrey Barrett, Sonia Goncalves, Ewan Harrison, David K. Jackson, Ian Johnston, Dominic Kwiatkowski, Cordelia Langford, John Sillitoe on behalf of the Wellcome Sanger Institute COVID-19 Surveillance Team |
| EPI_ISL_3097648 | Berkshire and Surrey Pathology Services Lighthouse Laboratory | Wellcome Sanger Institute for the COVID-19 Genomics UK (COG-UK) Consortium | Berkshire and Surrey Pathology Services Lighthouse Laboratory and Alex Alderton, Roberto Amato, Jeffrey Barrett, Sonia Goncalves, Ewan Harrison, David K. Jackson, Ian Johnston, Dominic Kwiatkowski, Cordelia Langford, John Sillitoe on behalf of the Wellcome Sanger Institute COVID-19 Surveillance Team |
| EPI_ISL_3097649, EPI_ISL_3097650, EPI_ISL_3097651 | Lighthouse Lab in Alderley Park | Wellcome Sanger Institute for the COVID-19 Genomics UK (COG-UK) Consortium | Jacquelyn Wynn, Mairead Hyland, The Lighthouse Lab in Alderley Park and Alex Alderton, Roberto Amato, Jeffrey Barrett, Sonia Goncalves, Ewan Harrison, David K. Jackson, Ian Johnston, Dominic Kwiatkowski, Cordelia Langford, John Sillitoe on behalf of the Wellcome Sanger Institute COVID-19 Surveillance Team |
| EPI_ISL_3097652 | Berkshire and Surrey Pathology Services Lighthouse Laboratory | Wellcome Sanger Institute for the COVID-19 Genomics UK (COG-UK) Consortium | Berkshire and Surrey Pathology Services Lighthouse Laboratory and Alex Alderton, Roberto Amato, Jeffrey Barrett, Sonia Goncalves, Ewan Harrison, David K. Jackson, Ian Johnston, Dominic Kwiatkowski, Cordelia Langford, John Sillitoe on behalf of the Wellcome Sanger Institute COVID-19 Surveillance Team |
| EPI_ISL_3097653, EPI_ISL_3097654, EPI_ISL_3097655, EPI_ISL_3097656, EPI_ISL_3097657, EPI_ISL_3097658, EPI_ISL_3097659, EPI_ISL_3097660 | Lighthouse Lab in Alderley Park | Wellcome Sanger Institute for the COVID-19 Genomics UK (COG-UK) Consortium | Jacquelyn Wynn, Mairead Hyland, The Lighthouse Lab in Alderley Park and Alex Alderton, Roberto Amato, Jeffrey Barrett, Sonia Goncalves, Ewan Harrison, David K. Jackson, Ian Johnston, Dominic Kwiatkowski, Cordelia Langford, John Sillitoe on behalf of the Wellcome Sanger Institute COVID-19 Surveillance Team |
| EPI_ISL_3097661 | Berkshire and Surrey Pathology Services Lighthouse Laboratory | Wellcome Sanger Institute for the COVID-19 Genomics UK (COG-UK) Consortium | Berkshire and Surrey Pathology Services Lighthouse Laboratory and Alex Alderton, Roberto Amato, Jeffrey Barrett, Sonia Goncalves, Ewan Harrison, David K. Jackson, Ian Johnston, Dominic Kwiatkowski, Cordelia Langford, John Sillitoe on behalf of the Wellcome Sanger Institute COVID-19 Surveillance Team |
| EPI_ISL_3097662, EPI_ISL_3097663, EPI_ISL_3097664 | Lighthouse Lab in Alderley Park | Wellcome Sanger Institute for the COVID-19 Genomics UK (COG-UK) Consortium | Jacquelyn Wynn, Mairead Hyland, The Lighthouse Lab in Alderley Park and Alex Alderton, Roberto Amato, Jeffrey Barrett, Sonia Goncalves, Ewan Harrison, David K. Jackson, Ian Johnston, Dominic Kwiatkowski, Cordelia Langford, John Sillitoe on behalf of the Wellcome Sanger Institute COVID-19 Surveillance Team |
| EPI_ISL_3097665 | Lighthouse Lab in Glasgow | Wellcome Sanger Institute for the COVID-19 Genomics UK (COG-UK) Consortium | Harper VanSteenhouse, Yumi Kasai, David Gray, Carol Clugston, Anna Dominiczak and Alex Alderton, Roberto Amato, Jeffrey Barrett, Sonia Goncalves, Ewan Harrison, David K. Jackson, Ian Johnston, Dominic Kwiatkowski, Cordelia Langford, John Sillitoe on behalf of the Wellcome Sanger Institute COVID-19 Surveillance Team |
| EPI_ISL_3097666, EPI_ISL_3097667, EPI_ISL_3097668, EPI_ISL_3097669, EPI_ISL_3097670 | Lighthouse Lab in Alderley Park | Wellcome Sanger Institute for the COVID-19 Genomics UK (COG-UK) Consortium | Jacquelyn Wynn, Mairead Hyland, The Lighthouse Lab in Alderley Park and Alex Alderton, Roberto Amato, Jeffrey Barrett, Sonia Goncalves, Ewan Harrison, David K. Jackson, Ian Johnston, Dominic Kwiatkowski, Cordelia Langford, John Sillitoe on behalf of the Wellcome Sanger Institute COVID-19 Surveillance Team |
| EPI_ISL_3097671 | Berkshire and Surrey Pathology Services Lighthouse Laboratory | Wellcome Sanger Institute for the COVID-19 Genomics UK (COG-UK) Consortium | Berkshire and Surrey Pathology Services Lighthouse Laboratory and Alex Alderton, Roberto Amato, Jeffrey Barrett, Sonia Goncalves, Ewan Harrison, David K. Jackson, Ian Johnston, Dominic Kwiatkowski, Cordelia Langford, John Sillitoe on behalf of the Wellcome Sanger Institute COVID-19 Surveillance Team |
| EPI_ISL_3097672, EPI_ISL_3097673, EPI_ISL_3097674 | Lighthouse Lab in Glasgow | Wellcome Sanger Institute for the COVID-19 Genomics UK (COG-UK) Consortium | Harper VanSteenhouse, Yumi Kasai, David Gray, Carol Clugston, Anna Dominiczak and Alex Alderton, Roberto Amato, Jeffrey Barrett, Sonia Goncalves, Ewan Harrison, David K. Jackson, Ian Johnston, Dominic Kwiatkowski, Cordelia Langford, John Sillitoe on behalf of the Wellcome Sanger Institute COVID-19 Surveillance Team |
| EPI_ISL_3097675 | Lighthouse Lab in Alderley Park | Wellcome Sanger Institute for the COVID-19 Genomics UK (COG-UK) Consortium | Jacquelyn Wynn, Mairead Hyland, The Lighthouse Lab in Alderley Park and Alex Alderton, Roberto Amato, Jeffrey Barrett, Sonia Goncalves, Ewan Harrison, David K. Jackson, Ian Johnston, Dominic Kwiatkowski, Cordelia Langford, John Sillitoe on behalf of the Wellcome Sanger Institute COVID-19 Surveillance Team |
| EPI_ISL_3097676 | Lighthouse Lab in Glasgow | Wellcome Sanger Institute for the COVID-19 Genomics UK (COG-UK) Consortium | Harper VanSteenhouse, Yumi Kasai, David Gray, Carol Clugston, Anna Dominiczak and Alex Alderton, Roberto Amato, Jeffrey Barrett, Sonia Goncalves, Ewan Harrison, David K. Jackson, Ian Johnston, Dominic Kwiatkowski, Cordelia Langford, John Sillitoe on behalf of the Wellcome Sanger Institute COVID-19 Surveillance Team |
| EPI_ISL_3097677 | Berkshire and Surrey Pathology Services Lighthouse Laboratory | Wellcome Sanger Institute for the COVID-19 Genomics UK (COG-UK) Consortium | Berkshire and Surrey Pathology Services Lighthouse Laboratory and Alex Alderton, Roberto Amato, Jeffrey Barrett, Sonia Goncalves, Ewan Harrison, David K. Jackson, Ian Johnston, Dominic Kwiatkowski, Cordelia Langford, John Sillitoe on behalf of the Wellcome Sanger Institute COVID-19 Surveillance Team |
| EPI_ISL_3097678 | Lighthouse Lab in Glasgow | Wellcome Sanger Institute for the COVID-19 Genomics UK (COG-UK) Consortium | Harper VanSteenhouse, Yumi Kasai, David Gray, Carol Clugston, Anna Dominiczak and Alex Alderton, Roberto Amato, Jeffrey Barrett, Sonia Goncalves, Ewan Harrison, David K. Jackson, Ian Johnston, Dominic Kwiatkowski, Cordelia Langford, John Sillitoe on behalf of the Wellcome Sanger Institute COVID-19 Surveillance Team |
| EPI_ISL_3097679 | Berkshire and Surrey Pathology Services Lighthouse Laboratory | Wellcome Sanger Institute for the COVID-19 Genomics UK (COG-UK) Consortium | Berkshire and Surrey Pathology Services Lighthouse Laboratory and Alex Alderton, Roberto Amato, Jeffrey Barrett, Sonia Goncalves, Ewan Harrison, David K. Jackson, Ian Johnston, Dominic Kwiatkowski, Cordelia Langford, John Sillitoe on behalf of the Wellcome Sanger Institute COVID-19 Surveillance Team |
| EPI_ISL_3097680, EPI_ISL_3097681 | Lighthouse Lab in Alderley Park | Wellcome Sanger Institute for the COVID-19 Genomics UK (COG-UK) Consortium | Jacquelyn Wynn, Mairead Hyland, The Lighthouse Lab in Alderley Park and Alex Alderton, Roberto Amato, Jeffrey Barrett, Sonia Goncalves, Ewan Harrison, David K. Jackson, Ian Johnston, Dominic Kwiatkowski, Cordelia Langford, John Sillitoe on behalf of the Wellcome Sanger Institute COVID-19 Surveillance Team |
| EPI_ISL_3097682 | Lighthouse Lab in Glasgow | Wellcome Sanger Institute for the COVID-19 Genomics UK (COG-UK) Consortium | Harper VanSteenhouse, Yumi Kasai, David Gray, Carol Clugston, Anna Dominiczak and Alex Alderton, Roberto Amato, Jeffrey Barrett, Sonia Goncalves, Ewan Harrison, David K. Jackson, Ian Johnston, Dominic Kwiatkowski, Cordelia Langford, John Sillitoe on behalf of the Wellcome Sanger Institute COVID-19 Surveillance Team |
| EPI_ISL_3097683, EPI_ISL_3097684 | Lighthouse Lab in Alderley Park | Wellcome Sanger Institute for the COVID-19 Genomics UK (COG-UK) Consortium | Jacquelyn Wynn, Mairead Hyland, The Lighthouse Lab in Alderley Park and Alex Alderton, Roberto Amato, Jeffrey Barrett, Sonia Goncalves, Ewan Harrison, David K. Jackson, Ian Johnston, Dominic Kwiatkowski, Cordelia Langford, John Sillitoe on behalf of the Wellcome Sanger Institute COVID-19 Surveillance Team |
| EPI_ISL_3097685 | Berkshire and Surrey Pathology Services Lighthouse Laboratory | Wellcome Sanger Institute for the COVID-19 Genomics UK (COG-UK) Consortium | Berkshire and Surrey Pathology Services Lighthouse Laboratory and Alex Alderton, Roberto Amato, Jeffrey Barrett, Sonia Goncalves, Ewan Harrison, David K. Jackson, Ian Johnston, Dominic Kwiatkowski, Cordelia Langford, John Sillitoe on behalf of the Wellcome Sanger Institute COVID-19 Surveillance Team |
| EPI_ISL_3097686, EPI_ISL_3097687 | Lighthouse Lab in Glasgow | Wellcome Sanger Institute for the COVID-19 Genomics UK (COG-UK) Consortium | Harper VanSteenhouse, Yumi Kasai, David Gray, Carol Clugston, Anna Dominiczak and Alex Alderton, Roberto Amato, Jeffrey Barrett, Sonia Goncalves, Ewan Harrison, David K. Jackson, Ian Johnston, Dominic Kwiatkowski, Cordelia Langford, John Sillitoe on behalf of the Wellcome Sanger Institute COVID-19 Surveillance Team |
| EPI_ISL_3097688 | Berkshire and Surrey Pathology Services Lighthouse Laboratory | Wellcome Sanger Institute for the COVID-19 Genomics UK (COG-UK) Consortium | Berkshire and Surrey Pathology Services Lighthouse Laboratory and Alex Alderton, Roberto Amato, Jeffrey Barrett, Sonia Goncalves, Ewan Harrison, David K. Jackson, Ian Johnston, Dominic Kwiatkowski, Cordelia Langford, John Sillitoe on behalf of the Wellcome Sanger Institute COVID-19 Surveillance Team |
| EPI_ISL_3097689, EPI_ISL_3097690, EPI_ISL_3097691 | Lighthouse Lab in Alderley Park | Wellcome Sanger Institute for the COVID-19 Genomics UK (COG-UK) Consortium | Jacquelyn Wynn, Mairead Hyland, The Lighthouse Lab in Alderley Park and Alex Alderton, Roberto Amato, Jeffrey Barrett, Sonia Goncalves, Ewan Harrison, David K. Jackson, Ian Johnston, Dominic Kwiatkowski, Cordelia Langford, John Sillitoe on behalf of the Wellcome Sanger Institute COVID-19 Surveillance Team |
| EPI_ISL_3097692 | Lighthouse Lab in Glasgow | Wellcome Sanger Institute for the COVID-19 Genomics UK (COG-UK) Consortium | Harper VanSteenhouse, Yumi Kasai, David Gray, Carol Clugston, Anna Dominiczak and Alex Alderton, Roberto Amato, Jeffrey Barrett, Sonia Goncalves, Ewan Harrison, David K. Jackson, Ian Johnston, Dominic Kwiatkowski, Cordelia Langford, John Sillitoe on behalf of the Wellcome Sanger Institute COVID-19 Surveillance Team |
| EPI_ISL_3097693 | Berkshire and Surrey Pathology Services Lighthouse Laboratory | Wellcome Sanger Institute for the COVID-19 Genomics UK (COG-UK) Consortium | Berkshire and Surrey Pathology Services Lighthouse Laboratory and Alex Alderton, Roberto Amato, Jeffrey Barrett, Sonia Goncalves, Ewan Harrison, David K. Jackson, Ian Johnston, Dominic Kwiatkowski, Cordelia Langford, John Sillitoe on behalf of the Wellcome Sanger Institute COVID-19 Surveillance Team |
| EPI_ISL_3097694 | Lighthouse Lab in Glasgow | Wellcome Sanger Institute for the COVID-19 Genomics UK (COG-UK) Consortium | Harper VanSteenhouse, Yumi Kasai, David Gray, Carol Clugston, Anna Dominiczak and Alex Alderton, Roberto Amato, Jeffrey Barrett, Sonia Goncalves, Ewan Harrison, David K. Jackson, Ian Johnston, Dominic Kwiatkowski, Cordelia Langford, John Sillitoe on behalf of the Wellcome Sanger Institute COVID-19 Surveillance Team |
| EPI_ISL_3097695 | Berkshire and Surrey Pathology Services Lighthouse Laboratory | Wellcome Sanger Institute for the COVID-19 Genomics UK (COG-UK) Consortium | Berkshire and Surrey Pathology Services Lighthouse Laboratory and Alex Alderton, Roberto Amato, Jeffrey Barrett, Sonia Goncalves, Ewan Harrison, David K. Jackson, Ian Johnston, Dominic Kwiatkowski, Cordelia Langford, John Sillitoe on behalf of the Wellcome Sanger Institute COVID-19 Surveillance Team |
| EPI_ISL_3097696 | Lighthouse Lab in Alderley Park | Wellcome Sanger Institute for the COVID-19 Genomics UK (COG-UK) Consortium | Jacquelyn Wynn, Mairead Hyland, The Lighthouse Lab in Alderley Park and Alex Alderton, Roberto Amato, Jeffrey Barrett, Sonia Goncalves, Ewan Harrison, David K. Jackson, Ian Johnston, Dominic Kwiatkowski, Cordelia Langford, John Sillitoe on behalf of the Wellcome Sanger Institute COVID-19 Surveillance Team |

| | | | |
|---|---|---|---|
| EPI_ISL_3097697 | Berkshire and Surrey Pathology Services Lighthouse Laboratory | Wellcome Sanger Institute for the COVID-19 Genomics UK (COG-UK) Consortium | Berkshire and Surrey Pathology Services Lighthouse Laboratory and Alex Alderton, Roberto Amato, Jeffrey Barrett, Sonia Goncalves, Ewan Harrison, David K. Jackson, Ian Johnston, Dominic Kwiatkowski, Cordelia Langford, John Sillitoe on behalf of the Wellcome Sanger Institute COVID-19 Surveillance Team |
| EPI_ISL_3097698 | Lighthouse Lab in Glasgow | Wellcome Sanger Institute for the COVID-19 Genomics UK (COG-UK) Consortium | Harper VanSteenhouse, Yumi Kasai, David Gray, Carol Clugston, Anna Dominiczak and Alex Alderton, Roberto Amato, Jeffrey Barrett, Sonia Goncalves, Ewan Harrison, David K. Jackson, Ian Johnston, Dominic Kwiatkowski, Cordelia Langford, John Sillitoe on behalf of the Wellcome Sanger Institute COVID-19 Surveillance Team |
| EPI_ISL_3097699 | Berkshire and Surrey Pathology Services Lighthouse Laboratory | Wellcome Sanger Institute for the COVID-19 Genomics UK (COG-UK) Consortium | Berkshire and Surrey Pathology Services Lighthouse Laboratory and Alex Alderton, Roberto Amato, Jeffrey Barrett, Sonia Goncalves, Ewan Harrison, David K. Jackson, Ian Johnston, Dominic Kwiatkowski, Cordelia Langford, John Sillitoe on behalf of the Wellcome Sanger Institute COVID-19 Surveillance Team |
| EPI_ISL_3097700 | Lighthouse Lab in Alderley Park | Wellcome Sanger Institute for the COVID-19 Genomics UK (COG-UK) Consortium | Jacquelyn Wynn, Mairead Hyland, The Lighthouse Lab in Alderley Park and Alex Alderton, Roberto Amato, Jeffrey Barrett, Sonia Goncalves, Ewan Harrison, David K. Jackson, Ian Johnston, Dominic Kwiatkowski, Cordelia Langford, John Sillitoe on behalf of the Wellcome Sanger Institute COVID-19 Surveillance Team |
| EPI_ISL_3097701 | Berkshire and Surrey Pathology Services Lighthouse Laboratory | Wellcome Sanger Institute for the COVID-19 Genomics UK (COG-UK) Consortium | Berkshire and Surrey Pathology Services Lighthouse Laboratory and Alex Alderton, Roberto Amato, Jeffrey Barrett, Sonia Goncalves, Ewan Harrison, David K. Jackson, Ian Johnston, Dominic Kwiatkowski, Cordelia Langford, John Sillitoe on behalf of the Wellcome Sanger Institute COVID-19 Surveillance Team |
| EPI_ISL_3097702, EPI_ISL_3097703 | Lighthouse Lab in Alderley Park | Wellcome Sanger Institute for the COVID-19 Genomics UK (COG-UK) Consortium | Jacquelyn Wynn, Mairead Hyland, The Lighthouse Lab in Alderley Park and Alex Alderton, Roberto Amato, Jeffrey Barrett, Sonia Goncalves, Ewan Harrison, David K. Jackson, Ian Johnston, Dominic Kwiatkowski, Cordelia Langford, John Sillitoe on behalf of the Wellcome Sanger Institute COVID-19 Surveillance Team |
| EPI_ISL_3097704 | Berkshire and Surrey Pathology Services Lighthouse Laboratory | Wellcome Sanger Institute for the COVID-19 Genomics UK (COG-UK) Consortium | Berkshire and Surrey Pathology Services Lighthouse Laboratory and Alex Alderton, Roberto Amato, Jeffrey Barrett, Sonia Goncalves, Ewan Harrison, David K. Jackson, Ian Johnston, Dominic Kwiatkowski, Cordelia Langford, John Sillitoe on behalf of the Wellcome Sanger Institute COVID-19 Surveillance Team |
| EPI_ISL_3097705, EPI_ISL_3097706 | Lighthouse Lab in Glasgow | Wellcome Sanger Institute for the COVID-19 Genomics UK (COG-UK) Consortium | Harper VanSteenhouse, Yumi Kasai, David Gray, Carol Clugston, Anna Dominiczak and Alex Alderton, Roberto Amato, Jeffrey Barrett, Sonia Goncalves, Ewan Harrison, David K. Jackson, Ian Johnston, Dominic Kwiatkowski, Cordelia Langford, John Sillitoe on behalf of the Wellcome Sanger Institute COVID-19 Surveillance Team |
| EPI_ISL_3097707, EPI_ISL_3097708, EPI_ISL_3097709 | Lighthouse Lab in Alderley Park | Wellcome Sanger Institute for the COVID-19 Genomics UK (COG-UK) Consortium | Jacquelyn Wynn, Mairead Hyland, The Lighthouse Lab in Alderley Park and Alex Alderton, Roberto Amato, Jeffrey Barrett, Sonia Goncalves, Ewan Harrison, David K. Jackson, Ian Johnston, Dominic Kwiatkowski, Cordelia Langford, John Sillitoe on behalf of the Wellcome Sanger Institute COVID-19 Surveillance Team |
| EPI_ISL_3097710 | Lighthouse Lab in Glasgow | Wellcome Sanger Institute for the COVID-19 Genomics UK (COG-UK) Consortium | Harper VanSteenhouse, Yumi Kasai, David Gray, Carol Clugston, Anna Dominiczak and Alex Alderton, Roberto Amato, Jeffrey Barrett, Sonia Goncalves, Ewan Harrison, David K. Jackson, Ian Johnston, Dominic Kwiatkowski, Cordelia Langford, John Sillitoe on behalf of the Wellcome Sanger Institute COVID-19 Surveillance Team |
| EPI_ISL_3097711, EPI_ISL_3097712, EPI_ISL_3097713 | Lighthouse Lab in Alderley Park | Wellcome Sanger Institute for the COVID-19 Genomics UK (COG-UK) Consortium | Jacquelyn Wynn, Mairead Hyland, The Lighthouse Lab in Alderley Park and Alex Alderton, Roberto Amato, Jeffrey Barrett, Sonia Goncalves, Ewan Harrison, David K. Jackson, Ian Johnston, Dominic Kwiatkowski, Cordelia Langford, John Sillitoe on behalf of the Wellcome Sanger Institute COVID-19 Surveillance Team |
| EPI_ISL_3097714 | Berkshire and Surrey Pathology Services Lighthouse Laboratory | Wellcome Sanger Institute for the COVID-19 Genomics UK (COG-UK) Consortium | Berkshire and Surrey Pathology Services Lighthouse Laboratory and Alex Alderton, Roberto Amato, Jeffrey Barrett, Sonia Goncalves, Ewan Harrison, David K. Jackson, Ian Johnston, Dominic Kwiatkowski, Cordelia Langford, John Sillitoe on behalf of the Wellcome Sanger Institute COVID-19 Surveillance Team |
| EPI_ISL_3097715, EPI_ISL_3097716, EPI_ISL_3097717, EPI_ISL_3097718 | Lighthouse Lab in Alderley Park | Wellcome Sanger Institute for the COVID-19 Genomics UK (COG-UK) Consortium | Jacquelyn Wynn, Mairead Hyland, The Lighthouse Lab in Alderley Park and Alex Alderton, Roberto Amato, Jeffrey Barrett, Sonia Goncalves, Ewan Harrison, David K. Jackson, Ian Johnston, Dominic Kwiatkowski, Cordelia Langford, John Sillitoe on behalf of the Wellcome Sanger Institute COVID-19 Surveillance Team |
| EPI_ISL_3097719, EPI_ISL_3097720, EPI_ISL_3097721 | Berkshire and Surrey Pathology Services Lighthouse Laboratory | Wellcome Sanger Institute for the COVID-19 Genomics UK (COG-UK) Consortium | Berkshire and Surrey Pathology Services Lighthouse Laboratory and Alex Alderton, Roberto Amato, Jeffrey Barrett, Sonia Goncalves, Ewan Harrison, David K. Jackson, Ian Johnston, Dominic Kwiatkowski, Cordelia Langford, John Sillitoe on behalf of the Wellcome Sanger Institute COVID-19 Surveillance Team |
| EPI_ISL_3097722, EPI_ISL_3097723 | Lighthouse Lab in Alderley Park | Wellcome Sanger Institute for the COVID-19 Genomics UK (COG-UK) Consortium | Jacquelyn Wynn, Mairead Hyland, The Lighthouse Lab in Alderley Park and Alex Alderton, Roberto Amato, Jeffrey Barrett, Sonia Goncalves, Ewan Harrison, David K. Jackson, Ian Johnston, Dominic Kwiatkowski, Cordelia Langford, John Sillitoe on behalf of the Wellcome Sanger Institute COVID-19 Surveillance Team |
| EPI_ISL_3097724, EPI_ISL_3097725 | Lighthouse Lab in Glasgow | Wellcome Sanger Institute for the COVID-19 Genomics UK (COG-UK) Consortium | Harper VanSteenhouse, Yumi Kasai, David Gray, Carol Clugston, Anna Dominiczak and Alex Alderton, Roberto Amato, Jeffrey Barrett, Sonia Goncalves, Ewan Harrison, David K. Jackson, Ian Johnston, Dominic Kwiatkowski, Cordelia Langford, John Sillitoe on behalf of the Wellcome Sanger Institute COVID-19 Surveillance Team |
| EPI_ISL_3097726 | Berkshire and Surrey Pathology Services Lighthouse Laboratory | Wellcome Sanger Institute for the COVID-19 Genomics UK (COG-UK) Consortium | Berkshire and Surrey Pathology Services Lighthouse Laboratory and Alex Alderton, Roberto Amato, Jeffrey Barrett, Sonia Goncalves, Ewan Harrison, David K. Jackson, Ian Johnston, Dominic Kwiatkowski, Cordelia Langford, John Sillitoe on behalf of the Wellcome Sanger Institute COVID-19 Surveillance Team |
| EPI_ISL_3097727, EPI_ISL_3097728 | Lighthouse Lab in Alderley Park | Wellcome Sanger Institute for the COVID-19 Genomics UK (COG-UK) Consortium | Jacquelyn Wynn, Mairead Hyland, The Lighthouse Lab in Alderley Park and Alex Alderton, Roberto Amato, Jeffrey Barrett, Sonia Goncalves, Ewan Harrison, David K. Jackson, Ian Johnston, Dominic Kwiatkowski, Cordelia Langford, John Sillitoe on behalf of the Wellcome Sanger Institute COVID-19 Surveillance Team |
| EPI_ISL_3097729 | Lighthouse Lab in Glasgow | Wellcome Sanger Institute for the COVID-19 Genomics UK (COG-UK) Consortium | Harper VanSteenhouse, Yumi Kasai, David Gray, Carol Clugston, Anna Dominiczak and Alex Alderton, Roberto Amato, Jeffrey Barrett, Sonia Goncalves, Ewan Harrison, David K. Jackson, Ian Johnston, Dominic Kwiatkowski, Cordelia Langford, John Sillitoe on behalf of the Wellcome Sanger Institute COVID-19 Surveillance Team |
| EPI_ISL_3097730 | Lighthouse Lab in Alderley Park | Wellcome Sanger Institute for the COVID-19 Genomics UK (COG-UK) Consortium | Jacquelyn Wynn, Mairead Hyland, The Lighthouse Lab in Alderley Park and Alex Alderton, Roberto Amato, Jeffrey Barrett, Sonia Goncalves, Ewan Harrison, David K. Jackson, Ian Johnston, Dominic Kwiatkowski, Cordelia Langford, John Sillitoe on behalf of the Wellcome Sanger Institute COVID-19 Surveillance Team |
| EPI_ISL_3097731 | Berkshire and Surrey Pathology Services Lighthouse Laboratory | Wellcome Sanger Institute for the COVID-19 Genomics UK (COG-UK) Consortium | Berkshire and Surrey Pathology Services Lighthouse Laboratory and Alex Alderton, Roberto Amato, Jeffrey Barrett, Sonia Goncalves, Ewan Harrison, David K. Jackson, Ian Johnston, Dominic Kwiatkowski, Cordelia Langford, John Sillitoe on behalf of the Wellcome Sanger Institute COVID-19 Surveillance Team |
| EPI_ISL_3097732 | Lighthouse Lab in Alderley Park | Wellcome Sanger Institute for the COVID-19 Genomics UK (COG-UK) Consortium | Jacquelyn Wynn, Mairead Hyland, The Lighthouse Lab in Alderley Park and Alex Alderton, Roberto Amato, Jeffrey Barrett, Sonia Goncalves, Ewan Harrison, David K. Jackson, Ian Johnston, Dominic Kwiatkowski, Cordelia Langford, John Sillitoe on behalf of the Wellcome Sanger Institute COVID-19 Surveillance Team |
| EPI_ISL_3097733 | Berkshire and Surrey Pathology Services Lighthouse Laboratory | Wellcome Sanger Institute for the COVID-19 Genomics UK (COG-UK) Consortium | Berkshire and Surrey Pathology Services Lighthouse Laboratory and Alex Alderton, Roberto Amato, Jeffrey Barrett, Sonia Goncalves, Ewan Harrison, David K. Jackson, Ian Johnston, Dominic Kwiatkowski, Cordelia Langford, John Sillitoe on behalf of the Wellcome Sanger Institute COVID-19 Surveillance Team |
| EPI_ISL_3097734 | Lighthouse Lab in Glasgow | Wellcome Sanger Institute for the COVID-19 Genomics UK (COG-UK) Consortium | Harper VanSteenhouse, Yumi Kasai, David Gray, Carol Clugston, Anna Dominiczak and Alex Alderton, Roberto Amato, Jeffrey Barrett, Sonia Goncalves, Ewan Harrison, David K. Jackson, Ian Johnston, Dominic Kwiatkowski, Cordelia Langford, John Sillitoe on behalf of the Wellcome Sanger Institute COVID-19 Surveillance Team |
| EPI_ISL_3097735 | Lighthouse Lab in Alderley Park | Wellcome Sanger Institute for the COVID-19 Genomics UK (COG-UK) Consortium | Jacquelyn Wynn, Mairead Hyland, The Lighthouse Lab in Alderley Park and Alex Alderton, Roberto Amato, Jeffrey Barrett, Sonia Goncalves, Ewan Harrison, David K. Jackson, Ian Johnston, Dominic Kwiatkowski, Cordelia Langford, John Sillitoe on behalf of the Wellcome Sanger Institute COVID-19 Surveillance Team |
| EPI_ISL_3097736 | Berkshire and Surrey Pathology Services Lighthouse Laboratory | Wellcome Sanger Institute for the COVID-19 Genomics UK (COG-UK) Consortium | Berkshire and Surrey Pathology Services Lighthouse Laboratory and Alex Alderton, Roberto Amato, Jeffrey Barrett, Sonia Goncalves, Ewan Harrison, David K. Jackson, Ian Johnston, Dominic Kwiatkowski, Cordelia Langford, John Sillitoe on behalf of the Wellcome Sanger Institute COVID-19 Surveillance Team |
| EPI_ISL_3097737, EPI_ISL_3097738 | Lighthouse Lab in Alderley Park | Wellcome Sanger Institute for the COVID-19 Genomics UK (COG-UK) Consortium | Jacquelyn Wynn, Mairead Hyland, The Lighthouse Lab in Alderley Park and Alex Alderton, Roberto Amato, Jeffrey Barrett, Sonia Goncalves, Ewan Harrison, David K. Jackson, Ian Johnston, Dominic Kwiatkowski, Cordelia Langford, John Sillitoe on behalf of the Wellcome Sanger Institute COVID-19 Surveillance Team |
| EPI_ISL_3097739 | Lighthouse Lab in Glasgow | Wellcome Sanger Institute for the COVID-19 Genomics UK (COG-UK) Consortium | Harper VanSteenhouse, Yumi Kasai, David Gray, Carol Clugston, Anna Dominiczak and Alex Alderton, Roberto Amato, Jeffrey Barrett, Sonia Goncalves, Ewan Harrison, David K. Jackson, Ian Johnston, Dominic Kwiatkowski, Cordelia Langford, John Sillitoe on behalf of the Wellcome Sanger Institute COVID-19 Surveillance Team |
| EPI_ISL_3097740, EPI_ISL_3097741 | Lighthouse Lab in Alderley Park | Wellcome Sanger Institute for the COVID-19 Genomics UK (COG-UK) Consortium | Jacquelyn Wynn, Mairead Hyland, The Lighthouse Lab in Alderley Park and Alex Alderton, Roberto Amato, Jeffrey Barrett, Sonia Goncalves, Ewan Harrison, David K. Jackson, Ian Johnston, Dominic Kwiatkowski, Cordelia Langford, John Sillitoe on behalf of the Wellcome Sanger Institute COVID-19 Surveillance Team |
| EPI_ISL_3097742 | Lighthouse Lab in Glasgow | Wellcome Sanger Institute for the COVID-19 Genomics UK (COG-UK) Consortium | Harper VanSteenhouse, Yumi Kasai, David Gray, Carol Clugston, Anna Dominiczak and Alex Alderton, Roberto Amato, Jeffrey Barrett, Sonia Goncalves, Ewan Harrison, David K. Jackson, Ian Johnston, Dominic Kwiatkowski, Cordelia Langford, John Sillitoe on behalf of the Wellcome Sanger Institute COVID-19 Surveillance Team |
| EPI_ISL_3097743 | Berkshire and Surrey Pathology Services Lighthouse Laboratory | Wellcome Sanger Institute for the COVID-19 Genomics UK (COG-UK) Consortium | Berkshire and Surrey Pathology Services Lighthouse Laboratory and Alex Alderton, Roberto Amato, Jeffrey Barrett, Sonia Goncalves, Ewan Harrison, David K. Jackson, Ian Johnston, Dominic Kwiatkowski, Cordelia Langford, John Sillitoe on behalf of the Wellcome Sanger Institute COVID-19 Surveillance Team |
| EPI_ISL_3097744 | Lighthouse Lab in Glasgow | Wellcome Sanger Institute for the COVID-19 Genomics UK (COG-UK) Consortium | Harper VanSteenhouse, Yumi Kasai, David Gray, Carol Clugston, Anna Dominiczak and Alex Alderton, Roberto Amato, Jeffrey Barrett, Sonia Goncalves, Ewan Harrison, David K. Jackson, Ian Johnston, Dominic Kwiatkowski, Cordelia Langford, John Sillitoe on behalf of the Wellcome Sanger Institute COVID-19 Surveillance Team |
| EPI_ISL_3097745 | Lighthouse Lab in Alderley Park | Wellcome Sanger Institute for the COVID-19 Genomics UK (COG-UK) Consortium | Jacquelyn Wynn, Mairead Hyland, The Lighthouse Lab in Alderley Park and Alex Alderton, Roberto Amato, Jeffrey Barrett, Sonia Goncalves, Ewan Harrison, David K. Jackson, Ian Johnston, Dominic Kwiatkowski, Cordelia Langford, John Sillitoe on behalf of the Wellcome Sanger Institute COVID-19 Surveillance Team |
| EPI_ISL_3097746 | Berkshire and Surrey Pathology Services Lighthouse Laboratory | Wellcome Sanger Institute for the COVID-19 Genomics UK (COG-UK) Consortium | Berkshire and Surrey Pathology Services Lighthouse Laboratory and Alex Alderton, Roberto Amato, Jeffrey Barrett, Sonia Goncalves, Ewan Harrison, David K. Jackson, Ian Johnston, Dominic Kwiatkowski, Cordelia Langford, John Sillitoe on behalf of the Wellcome Sanger Institute COVID-19 Surveillance Team |
| EPI_ISL_3097747, EPI_ISL_3097748 | Lighthouse Lab in Glasgow | Wellcome Sanger Institute for the COVID-19 Genomics UK (COG-UK) Consortium | Harper VanSteenhouse, Yumi Kasai, David Gray, Carol Clugston, Anna Dominiczak and Alex Alderton, Roberto Amato, Jeffrey Barrett, Sonia Goncalves, Ewan Harrison, David K. Jackson, Ian Johnston, Dominic Kwiatkowski, Cordelia Langford, John Sillitoe on behalf of the Wellcome Sanger Institute COVID-19 Surveillance Team |
| EPI_ISL_3097749, EPI_ISL_3097750 | Berkshire and Surrey Pathology Services Lighthouse Laboratory | Wellcome Sanger Institute for the COVID-19 Genomics UK (COG-UK) Consortium | Berkshire and Surrey Pathology Services Lighthouse Laboratory and Alex Alderton, Roberto Amato, Jeffrey Barrett, Sonia Goncalves, Ewan Harrison, David K. Jackson, Ian Johnston, Dominic Kwiatkowski, Cordelia Langford, John Sillitoe on behalf of the Wellcome Sanger Institute COVID-19 Surveillance Team |
| EPI_ISL_3097751 | Lighthouse Lab in Alderley Park | Wellcome Sanger Institute for the COVID-19 Genomics UK (COG-UK) Consortium | Jacquelyn Wynn, Mairead Hyland, The Lighthouse Lab in Alderley Park and Alex Alderton, Roberto Amato, Jeffrey Barrett, Sonia Goncalves, Ewan Harrison, David K. Jackson, Ian Johnston, Dominic Kwiatkowski, Cordelia Langford, John Sillitoe on behalf of the Wellcome Sanger Institute COVID-19 Surveillance Team |
| EPI_ISL_3097752 | Lighthouse Lab in Glasgow | Wellcome Sanger Institute for the COVID-19 Genomics UK (COG-UK) Consortium | Harper VanSteenhouse, Yumi Kasai, David Gray, Carol Clugston, Anna Dominiczak and Alex Alderton, Roberto Amato, Jeffrey Barrett, Sonia Goncalves, Ewan Harrison, David K. Jackson, Ian Johnston, Dominic Kwiatkowski, Cordelia Langford, John Sillitoe on behalf of the Wellcome Sanger Institute COVID-19 Surveillance Team |
| EPI_ISL_3097753 | Berkshire and Surrey Pathology Services Lighthouse Laboratory | Wellcome Sanger Institute for the COVID-19 Genomics UK (COG-UK) Consortium | Berkshire and Surrey Pathology Services Lighthouse Laboratory and Alex Alderton, Roberto Amato, Jeffrey Barrett, Sonia Goncalves, Ewan Harrison, David K. Jackson, Ian Johnston, Dominic Kwiatkowski, Cordelia Langford, John Sillitoe on behalf of the Wellcome Sanger Institute COVID-19 Surveillance Team |
| EPI_ISL_3097754, EPI_ISL_3097755 | Lighthouse Lab in Alderley Park | Wellcome Sanger Institute for the COVID-19 Genomics UK (COG-UK) Consortium | Jacquelyn Wynn, Mairead Hyland, The Lighthouse Lab in Alderley Park and Alex Alderton, Roberto Amato, Jeffrey Barrett, Sonia Goncalves, Ewan Harrison, David K. Jackson, Ian Johnston, Dominic Kwiatkowski, Cordelia Langford, John Sillitoe on behalf of the Wellcome Sanger Institute COVID-19 Surveillance Team |
| EPI_ISL_3097756 | Lighthouse Lab in Glasgow | Wellcome Sanger Institute for the COVID-19 Genomics UK (COG-UK) Consortium | Harper VanSteenhouse, Yumi Kasai, David Gray, Carol Clugston, Anna Dominiczak and Alex Alderton, Roberto Amato, Jeffrey Barrett, Sonia Goncalves, Ewan Harrison, David K. Jackson, Ian Johnston, Dominic Kwiatkowski, Cordelia Langford, John Sillitoe on behalf of the Wellcome Sanger Institute COVID-19 Surveillance Team |
| EPI_ISL_3097757 | Lighthouse Lab in Alderley Park | Wellcome Sanger Institute for the COVID-19 Genomics UK (COG-UK) Consortium | Jacquelyn Wynn, Mairead Hyland, The Lighthouse Lab in Alderley Park and Alex Alderton, Roberto Amato, Jeffrey Barrett, Sonia Goncalves, Ewan Harrison, David K. Jackson, Ian Johnston, Dominic Kwiatkowski, Cordelia Langford, John Sillitoe on behalf of the Wellcome Sanger Institute COVID-19 Surveillance Team |
| EPI_ISL_3097758 | Berkshire and Surrey Pathology Services Lighthouse Laboratory | Wellcome Sanger Institute for the COVID-19 Genomics UK (COG-UK) Consortium | Berkshire and Surrey Pathology Services Lighthouse Laboratory and Alex Alderton, Roberto Amato, Jeffrey Barrett, Sonia Goncalves, Ewan Harrison, David K. Jackson, Ian Johnston, Dominic Kwiatkowski, Cordelia Langford, John Sillitoe on behalf of the Wellcome Sanger Institute COVID-19 Surveillance Team |

| | | | |
|---|---|---|---|
| EPI_ISL_3097759, EPI_ISL_3097760 | Lighthouse Lab in Alderley Park | Wellcome Sanger Institute for the COVID-19 Genomics UK (COG-UK) Consortium | Jacquelyn Wynn, Mairead Hyland, The Lighthouse Lab in Alderley Park and Alex Alderton, Roberto Amato, Jeffrey Barrett, Sonia Goncalves, Ewan Harrison, David K. Jackson, Ian Johnston, Dominic Kwiatkowski, Cordelia Langford, John Sillitoe on behalf of the Wellcome Sanger Institute COVID-19 Surveillance Team |
| EPI_ISL_3097761 | Berkshire and Surrey Pathology Services Lighthouse Laboratory | Wellcome Sanger Institute for the COVID-19 Genomics UK (COG-UK) Consortium | Berkshire and Surrey Pathology Services Lighthouse Laboratory and Alex Alderton, Roberto Amato, Jeffrey Barrett, Sonia Goncalves, Ewan Harrison, David K. Jackson, Ian Johnston, Dominic Kwiatkowski, Cordelia Langford, John Sillitoe on behalf of the Wellcome Sanger Institute COVID-19 Surveillance Team |
| EPI_ISL_3097762, EPI_ISL_3097763 | Lighthouse Lab in Alderley Park | Wellcome Sanger Institute for the COVID-19 Genomics UK (COG-UK) Consortium | Jacquelyn Wynn, Mairead Hyland, The Lighthouse Lab in Alderley Park and Alex Alderton, Roberto Amato, Jeffrey Barrett, Sonia Goncalves, Ewan Harrison, David K. Jackson, Ian Johnston, Dominic Kwiatkowski, Cordelia Langford, John Sillitoe on behalf of the Wellcome Sanger Institute COVID-19 Surveillance Team |
| EPI_ISL_3097764 | Berkshire and Surrey Pathology Services Lighthouse Laboratory | Wellcome Sanger Institute for the COVID-19 Genomics UK (COG-UK) Consortium | Berkshire and Surrey Pathology Services Lighthouse Laboratory and Alex Alderton, Roberto Amato, Jeffrey Barrett, Sonia Goncalves, Ewan Harrison, David K. Jackson, Ian Johnston, Dominic Kwiatkowski, Cordelia Langford, John Sillitoe on behalf of the Wellcome Sanger Institute COVID-19 Surveillance Team |
| EPI_ISL_3097765, EPI_ISL_3097766 | Lighthouse Lab in Glasgow | Wellcome Sanger Institute for the COVID-19 Genomics UK (COG-UK) Consortium | Harper VanSteenhouse, Yumi Kasai, David Gray, Carol Clugston, Anna Dominiczak and Alex Alderton, Roberto Amato, Jeffrey Barrett, Sonia Goncalves, Ewan Harrison, David K. Jackson, Ian Johnston, Dominic Kwiatkowski, Cordelia Langford, John Sillitoe on behalf of the Wellcome Sanger Institute COVID-19 Surveillance Team |
| EPI_ISL_3097767 | Lighthouse Lab in Alderley Park | Wellcome Sanger Institute for the COVID-19 Genomics UK (COG-UK) Consortium | Jacquelyn Wynn, Mairead Hyland, The Lighthouse Lab in Alderley Park and Alex Alderton, Roberto Amato, Jeffrey Barrett, Sonia Goncalves, Ewan Harrison, David K. Jackson, Ian Johnston, Dominic Kwiatkowski, Cordelia Langford, John Sillitoe on behalf of the Wellcome Sanger Institute COVID-19 Surveillance Team |
| EPI_ISL_3097768, EPI_ISL_3097769 | Lighthouse Lab in Glasgow | Wellcome Sanger Institute for the COVID-19 Genomics UK (COG-UK) Consortium | Harper VanSteenhouse, Yumi Kasai, David Gray, Carol Clugston, Anna Dominiczak and Alex Alderton, Roberto Amato, Jeffrey Barrett, Sonia Goncalves, Ewan Harrison, David K. Jackson, Ian Johnston, Dominic Kwiatkowski, Cordelia Langford, John Sillitoe on behalf of the Wellcome Sanger Institute COVID-19 Surveillance Team |
| EPI_ISL_3097770 | Lighthouse Lab in Alderley Park | Wellcome Sanger Institute for the COVID-19 Genomics UK (COG-UK) Consortium | Jacquelyn Wynn, Mairead Hyland, The Lighthouse Lab in Alderley Park and Alex Alderton, Roberto Amato, Jeffrey Barrett, Sonia Goncalves, Ewan Harrison, David K. Jackson, Ian Johnston, Dominic Kwiatkowski, Cordelia Langford, John Sillitoe on behalf of the Wellcome Sanger Institute COVID-19 Surveillance Team |
| EPI_ISL_3097771 | Berkshire and Surrey Pathology Services Lighthouse Laboratory | Wellcome Sanger Institute for the COVID-19 Genomics UK (COG-UK) Consortium | Berkshire and Surrey Pathology Services Lighthouse Laboratory and Alex Alderton, Roberto Amato, Jeffrey Barrett, Sonia Goncalves, Ewan Harrison, David K. Jackson, Ian Johnston, Dominic Kwiatkowski, Cordelia Langford, John Sillitoe on behalf of the Wellcome Sanger Institute COVID-19 Surveillance Team |
| EPI_ISL_3097772 | Lighthouse Lab in Alderley Park | Wellcome Sanger Institute for the COVID-19 Genomics UK (COG-UK) Consortium | Jacquelyn Wynn, Mairead Hyland, The Lighthouse Lab in Alderley Park and Alex Alderton, Roberto Amato, Jeffrey Barrett, Sonia Goncalves, Ewan Harrison, David K. Jackson, Ian Johnston, Dominic Kwiatkowski, Cordelia Langford, John Sillitoe on behalf of the Wellcome Sanger Institute COVID-19 Surveillance Team |
| EPI_ISL_3097773, EPI_ISL_3097774 | Lighthouse Lab in Glasgow | Wellcome Sanger Institute for the COVID-19 Genomics UK (COG-UK) Consortium | Harper VanSteenhouse, Yumi Kasai, David Gray, Carol Clugston, Anna Dominiczak and Alex Alderton, Roberto Amato, Jeffrey Barrett, Sonia Goncalves, Ewan Harrison, David K. Jackson, Ian Johnston, Dominic Kwiatkowski, Cordelia Langford, John Sillitoe on behalf of the Wellcome Sanger Institute COVID-19 Surveillance Team |
| EPI_ISL_3097775 | Lighthouse Lab in Alderley Park | Wellcome Sanger Institute for the COVID-19 Genomics UK (COG-UK) Consortium | Jacquelyn Wynn, Mairead Hyland, The Lighthouse Lab in Alderley Park and Alex Alderton, Roberto Amato, Jeffrey Barrett, Sonia Goncalves, Ewan Harrison, David K. Jackson, Ian Johnston, Dominic Kwiatkowski, Cordelia Langford, John Sillitoe on behalf of the Wellcome Sanger Institute COVID-19 Surveillance Team |
| EPI_ISL_3097776, EPI_ISL_3097777 | Berkshire and Surrey Pathology Services Lighthouse Laboratory | Wellcome Sanger Institute for the COVID-19 Genomics UK (COG-UK) Consortium | Berkshire and Surrey Pathology Services Lighthouse Laboratory and Alex Alderton, Roberto Amato, Jeffrey Barrett, Sonia Goncalves, Ewan Harrison, David K. Jackson, Ian Johnston, Dominic Kwiatkowski, Cordelia Langford, John Sillitoe on behalf of the Wellcome Sanger Institute COVID-19 Surveillance Team |
| EPI_ISL_3097778, EPI_ISL_3097779, EPI_ISL_3097780 | Lighthouse Lab in Alderley Park | Wellcome Sanger Institute for the COVID-19 Genomics UK (COG-UK) Consortium | Jacquelyn Wynn, Mairead Hyland, The Lighthouse Lab in Alderley Park and Alex Alderton, Roberto Amato, Jeffrey Barrett, Sonia Goncalves, Ewan Harrison, David K. Jackson, Ian Johnston, Dominic Kwiatkowski, Cordelia Langford, John Sillitoe on behalf of the Wellcome Sanger Institute COVID-19 Surveillance Team |
| EPI_ISL_3097781 | Lighthouse Lab in Glasgow | Wellcome Sanger Institute for the COVID-19 Genomics UK (COG-UK) Consortium | Harper VanSteenhouse, Yumi Kasai, David Gray, Carol Clugston, Anna Dominiczak and Alex Alderton, Roberto Amato, Jeffrey Barrett, Sonia Goncalves, Ewan Harrison, David K. Jackson, Ian Johnston, Dominic Kwiatkowski, Cordelia Langford, John Sillitoe on behalf of the Wellcome Sanger Institute COVID-19 Surveillance Team |
| EPI_ISL_3097782 | Lighthouse Lab in Alderley Park | Wellcome Sanger Institute for the COVID-19 Genomics UK (COG-UK) Consortium | Jacquelyn Wynn, Mairead Hyland, The Lighthouse Lab in Alderley Park and Alex Alderton, Roberto Amato, Jeffrey Barrett, Sonia Goncalves, Ewan Harrison, David K. Jackson, Ian Johnston, Dominic Kwiatkowski, Cordelia Langford, John Sillitoe on behalf of the Wellcome Sanger Institute COVID-19 Surveillance Team |
| EPI_ISL_3097783 | Lighthouse Lab in Glasgow | Wellcome Sanger Institute for the COVID-19 Genomics UK (COG-UK) Consortium | Harper VanSteenhouse, Yumi Kasai, David Gray, Carol Clugston, Anna Dominiczak and Alex Alderton, Roberto Amato, Jeffrey Barrett, Sonia Goncalves, Ewan Harrison, David K. Jackson, Ian Johnston, Dominic Kwiatkowski, Cordelia Langford, John Sillitoe on behalf of the Wellcome Sanger Institute COVID-19 Surveillance Team |
| EPI_ISL_3097784 | Lighthouse Lab in Alderley Park | Wellcome Sanger Institute for the COVID-19 Genomics UK (COG-UK) Consortium | Jacquelyn Wynn, Mairead Hyland, The Lighthouse Lab in Alderley Park and Alex Alderton, Roberto Amato, Jeffrey Barrett, Sonia Goncalves, Ewan Harrison, David K. Jackson, Ian Johnston, Dominic Kwiatkowski, Cordelia Langford, John Sillitoe on behalf of the Wellcome Sanger Institute COVID-19 Surveillance Team |
| EPI_ISL_3097785 | Berkshire and Surrey Pathology Services Lighthouse Laboratory | Wellcome Sanger Institute for the COVID-19 Genomics UK (COG-UK) Consortium | Berkshire and Surrey Pathology Services Lighthouse Laboratory and Alex Alderton, Roberto Amato, Jeffrey Barrett, Sonia Goncalves, Ewan Harrison, David K. Jackson, Ian Johnston, Dominic Kwiatkowski, Cordelia Langford, John Sillitoe on behalf of the Wellcome Sanger Institute COVID-19 Surveillance Team |
| EPI_ISL_3097786, EPI_ISL_3097787, EPI_ISL_3097788 | Lighthouse Lab in Glasgow | Wellcome Sanger Institute for the COVID-19 Genomics UK (COG-UK) Consortium | Harper VanSteenhouse, Yumi Kasai, David Gray, Carol Clugston, Anna Dominiczak and Alex Alderton, Roberto Amato, Jeffrey Barrett, Sonia Goncalves, Ewan Harrison, David K. Jackson, Ian Johnston, Dominic Kwiatkowski, Cordelia Langford, John Sillitoe on behalf of the Wellcome Sanger Institute COVID-19 Surveillance Team |
| EPI_ISL_3097789 | Berkshire and Surrey Pathology Services Lighthouse Laboratory | Wellcome Sanger Institute for the COVID-19 Genomics UK (COG-UK) Consortium | Berkshire and Surrey Pathology Services Lighthouse Laboratory and Alex Alderton, Roberto Amato, Jeffrey Barrett, Sonia Goncalves, Ewan Harrison, David K. Jackson, Ian Johnston, Dominic Kwiatkowski, Cordelia Langford, John Sillitoe on behalf of the Wellcome Sanger Institute COVID-19 Surveillance Team |
| EPI_ISL_3097790 | Lighthouse Lab in Glasgow | Wellcome Sanger Institute for the COVID-19 Genomics UK (COG-UK) Consortium | Harper VanSteenhouse, Yumi Kasai, David Gray, Carol Clugston, Anna Dominiczak and Alex Alderton, Roberto Amato, Jeffrey Barrett, Sonia Goncalves, Ewan Harrison, David K. Jackson, Ian Johnston, Dominic Kwiatkowski, Cordelia Langford, John Sillitoe on behalf of the Wellcome Sanger Institute COVID-19 Surveillance Team |
| EPI_ISL_3097791, EPI_ISL_3097792, EPI_ISL_3097793, EPI_ISL_3097794, EPI_ISL_3097795 | Lighthouse Lab in Alderley Park | Wellcome Sanger Institute for the COVID-19 Genomics UK (COG-UK) Consortium | Jacquelyn Wynn, Mairead Hyland, The Lighthouse Lab in Alderley Park and Alex Alderton, Roberto Amato, Jeffrey Barrett, Sonia Goncalves, Ewan Harrison, David K. Jackson, Ian Johnston, Dominic Kwiatkowski, Cordelia Langford, John Sillitoe on behalf of the Wellcome Sanger Institute COVID-19 Surveillance Team |
| EPI_ISL_3097796 | Berkshire and Surrey Pathology Services Lighthouse Laboratory | Wellcome Sanger Institute for the COVID-19 Genomics UK (COG-UK) Consortium | Berkshire and Surrey Pathology Services Lighthouse Laboratory and Alex Alderton, Roberto Amato, Jeffrey Barrett, Sonia Goncalves, Ewan Harrison, David K. Jackson, Ian Johnston, Dominic Kwiatkowski, Cordelia Langford, John Sillitoe on behalf of the Wellcome Sanger Institute COVID-19 Surveillance Team |
| EPI_ISL_3097797, EPI_ISL_3097798, EPI_ISL_3097799 | Lighthouse Lab in Alderley Park | Wellcome Sanger Institute for the COVID-19 Genomics UK (COG-UK) Consortium | Jacquelyn Wynn, Mairead Hyland, The Lighthouse Lab in Alderley Park and Alex Alderton, Roberto Amato, Jeffrey Barrett, Sonia Goncalves, Ewan Harrison, David K. Jackson, Ian Johnston, Dominic Kwiatkowski, Cordelia Langford, John Sillitoe on behalf of the Wellcome Sanger Institute COVID-19 Surveillance Team |
| EPI_ISL_3097800 | Lighthouse Lab in Glasgow | Wellcome Sanger Institute for the COVID-19 Genomics UK (COG-UK) Consortium | Harper VanSteenhouse, Yumi Kasai, David Gray, Carol Clugston, Anna Dominiczak and Alex Alderton, Roberto Amato, Jeffrey Barrett, Sonia Goncalves, Ewan Harrison, David K. Jackson, Ian Johnston, Dominic Kwiatkowski, Cordelia Langford, John Sillitoe on behalf of the Wellcome Sanger Institute COVID-19 Surveillance Team |
| EPI_ISL_3097801 | Berkshire and Surrey Pathology Services Lighthouse Laboratory | Wellcome Sanger Institute for the COVID-19 Genomics UK (COG-UK) Consortium | Berkshire and Surrey Pathology Services Lighthouse Laboratory and Alex Alderton, Roberto Amato, Jeffrey Barrett, Sonia Goncalves, Ewan Harrison, David K. Jackson, Ian Johnston, Dominic Kwiatkowski, Cordelia Langford, John Sillitoe on behalf of the Wellcome Sanger Institute COVID-19 Surveillance Team |
| EPI_ISL_3097802, EPI_ISL_3097803, EPI_ISL_3097804, EPI_ISL_3097805, EPI_ISL_3097806, EPI_ISL_3097807, EPI_ISL_3097808 | Lighthouse Lab in Alderley Park | Wellcome Sanger Institute for the COVID-19 Genomics UK (COG-UK) Consortium | Jacquelyn Wynn, Mairead Hyland, The Lighthouse Lab in Alderley Park and Alex Alderton, Roberto Amato, Jeffrey Barrett, Sonia Goncalves, Ewan Harrison, David K. Jackson, Ian Johnston, Dominic Kwiatkowski, Cordelia Langford, John Sillitoe on behalf of the Wellcome Sanger Institute COVID-19 Surveillance Team |
| EPI_ISL_3097809, EPI_ISL_3097810 | Berkshire and Surrey Pathology Services Lighthouse Laboratory | Wellcome Sanger Institute for the COVID-19 Genomics UK (COG-UK) Consortium | Berkshire and Surrey Pathology Services Lighthouse Laboratory and Alex Alderton, Roberto Amato, Jeffrey Barrett, Sonia Goncalves, Ewan Harrison, David K. Jackson, Ian Johnston, Dominic Kwiatkowski, Cordelia Langford, John Sillitoe on behalf of the Wellcome Sanger Institute COVID-19 Surveillance Team |
| EPI_ISL_3097811, EPI_ISL_3097812 | Lighthouse Lab in Alderley Park | Wellcome Sanger Institute for the COVID-19 Genomics UK (COG-UK) Consortium | Jacquelyn Wynn, Mairead Hyland, The Lighthouse Lab in Alderley Park and Alex Alderton, Roberto Amato, Jeffrey Barrett, Sonia Goncalves, Ewan Harrison, David K. Jackson, Ian Johnston, Dominic Kwiatkowski, Cordelia Langford, John Sillitoe on behalf of the Wellcome Sanger Institute COVID-19 Surveillance Team |
| EPI_ISL_3097813 | Berkshire and Surrey Pathology Services Lighthouse Laboratory | Wellcome Sanger Institute for the COVID-19 Genomics UK (COG-UK) Consortium | Berkshire and Surrey Pathology Services Lighthouse Laboratory and Alex Alderton, Roberto Amato, Jeffrey Barrett, Sonia Goncalves, Ewan Harrison, David K. Jackson, Ian Johnston, Dominic Kwiatkowski, Cordelia Langford, John Sillitoe on behalf of the Wellcome Sanger Institute COVID-19 Surveillance Team |
| EPI_ISL_3097814 | Lighthouse Lab in Glasgow | Wellcome Sanger Institute for the COVID-19 Genomics UK (COG-UK) Consortium | Harper VanSteenhouse, Yumi Kasai, David Gray, Carol Clugston, Anna Dominiczak and Alex Alderton, Roberto Amato, Jeffrey Barrett, Sonia Goncalves, Ewan Harrison, David K. Jackson, Ian Johnston, Dominic Kwiatkowski, Cordelia Langford, John Sillitoe on behalf of the Wellcome Sanger Institute COVID-19 Surveillance Team |
| EPI_ISL_3097815, EPI_ISL_3097816 | Lighthouse Lab in Alderley Park | Wellcome Sanger Institute for the COVID-19 Genomics UK (COG-UK) Consortium | Jacquelyn Wynn, Mairead Hyland, The Lighthouse Lab in Alderley Park and Alex Alderton, Roberto Amato, Jeffrey Barrett, Sonia Goncalves, Ewan Harrison, David K. Jackson, Ian Johnston, Dominic Kwiatkowski, Cordelia Langford, John Sillitoe on behalf of the Wellcome Sanger Institute COVID-19 Surveillance Team |
| EPI_ISL_3097817 | Berkshire and Surrey Pathology Services Lighthouse Laboratory | Wellcome Sanger Institute for the COVID-19 Genomics UK (COG-UK) Consortium | Berkshire and Surrey Pathology Services Lighthouse Laboratory and Alex Alderton, Roberto Amato, Jeffrey Barrett, Sonia Goncalves, Ewan Harrison, David K. Jackson, Ian Johnston, Dominic Kwiatkowski, Cordelia Langford, John Sillitoe on behalf of the Wellcome Sanger Institute COVID-19 Surveillance Team |
| EPI_ISL_3097818, EPI_ISL_3097819, EPI_ISL_3097820, EPI_ISL_3097821, EPI_ISL_3097822, EPI_ISL_3097823, EPI_ISL_3097824 | Lighthouse Lab in Alderley Park | Wellcome Sanger Institute for the COVID-19 Genomics UK (COG-UK) Consortium | Jacquelyn Wynn, Mairead Hyland, The Lighthouse Lab in Alderley Park and Alex Alderton, Roberto Amato, Jeffrey Barrett, Sonia Goncalves, Ewan Harrison, David K. Jackson, Ian Johnston, Dominic Kwiatkowski, Cordelia Langford, John Sillitoe on behalf of the Wellcome Sanger Institute COVID-19 Surveillance Team |
| EPI_ISL_3097825, EPI_ISL_3097826 | Lighthouse Lab in Glasgow | Wellcome Sanger Institute for the COVID-19 Genomics UK (COG-UK) Consortium | Harper VanSteenhouse, Yumi Kasai, David Gray, Carol Clugston, Anna Dominiczak and Alex Alderton, Roberto Amato, Jeffrey Barrett, Sonia Goncalves, Ewan Harrison, David K. Jackson, Ian Johnston, Dominic Kwiatkowski, Cordelia Langford, John Sillitoe on behalf of the Wellcome Sanger Institute COVID-19 Surveillance Team |
| EPI_ISL_3097827, EPI_ISL_3097828 | Lighthouse Lab in Alderley Park | Wellcome Sanger Institute for the COVID-19 Genomics UK (COG-UK) Consortium | Jacquelyn Wynn, Mairead Hyland, The Lighthouse Lab in Alderley Park and Alex Alderton, Roberto Amato, Jeffrey Barrett, Sonia Goncalves, Ewan Harrison, David K. Jackson, Ian Johnston, Dominic Kwiatkowski, Cordelia Langford, John Sillitoe on behalf of the Wellcome Sanger Institute COVID-19 Surveillance Team |
| EPI_ISL_3097829 | Lighthouse Lab in Glasgow | Wellcome Sanger Institute for the COVID-19 Genomics UK (COG-UK) Consortium | Harper VanSteenhouse, Yumi Kasai, David Gray, Carol Clugston, Anna Dominiczak and Alex Alderton, Roberto Amato, Jeffrey Barrett, Sonia Goncalves, Ewan Harrison, David K. Jackson, Ian Johnston, Dominic Kwiatkowski, Cordelia Langford, John Sillitoe on behalf of the Wellcome Sanger Institute COVID-19 Surveillance Team |
| EPI_ISL_3097830 | Lighthouse Lab in Alderley Park | Wellcome Sanger Institute for the COVID-19 Genomics UK (COG-UK) | Jacquelyn Wynn, Mairead Hyland, The Lighthouse Lab in Alderley Park and Alex Alderton, Roberto Amato, Jeffrey Barrett, Sonia Goncalves, Ewan Harrison, David K. Jackson, Ian Johnston, Dominic Kwiatkowski, Cordelia Langford, John Sillitoe on |

| | | | |
|---|---|---|---|
| | | Consortium | behalf of the Wellcome Sanger Institute COVID-19 Surveillance Team |
| EPI_ISL_3097831 | Berkshire and Surrey Pathology Services Lighthouse Laboratory | Wellcome Sanger Institute for the COVID-19 Genomics UK (COG-UK) Consortium | Berkshire and Surrey Pathology Services Lighthouse Laboratory and Alex Alderton, Roberto Amato, Jeffrey Barrett, Sonia Goncalves, Ewan Harrison, David K. Jackson, Ian Johnston, Dominic Kwiatkowski, Cordelia Langford, John Sillitoe on behalf of the Wellcome Sanger Institute COVID-19 Surveillance Team |
| EPI_ISL_3097832 | Lighthouse Lab in Glasgow | Wellcome Sanger Institute for the COVID-19 Genomics UK (COG-UK) Consortium | Harper VanSteenhouse, Yumi Kasai, David Gray, Carol Clugston, Anna Dominiczak and Alex Alderton, Roberto Amato, Jeffrey Barrett, Sonia Goncalves, Ewan Harrison, David K. Jackson, Ian Johnston, Dominic Kwiatkowski, Cordelia Langford, John Sillitoe on behalf of the Wellcome Sanger Institute COVID-19 Surveillance Team |
| EPI_ISL_3097833, EPI_ISL_3097834 | Lighthouse Lab in Alderley Park | Wellcome Sanger Institute for the COVID-19 Genomics UK (COG-UK) Consortium | Jacquelyn Wynn, Mairead Hyland, The Lighthouse Lab in Alderley Park and Alex Alderton, Roberto Amato, Jeffrey Barrett, Sonia Goncalves, Ewan Harrison, David K. Jackson, Ian Johnston, Dominic Kwiatkowski, Cordelia Langford, John Sillitoe on behalf of the Wellcome Sanger Institute COVID-19 Surveillance Team |
| EPI_ISL_3097835, EPI_ISL_3097836 | Lighthouse Lab in Glasgow | Wellcome Sanger Institute for the COVID-19 Genomics UK (COG-UK) Consortium | Harper VanSteenhouse, Yumi Kasai, David Gray, Carol Clugston, Anna Dominiczak and Alex Alderton, Roberto Amato, Jeffrey Barrett, Sonia Goncalves, Ewan Harrison, David K. Jackson, Ian Johnston, Dominic Kwiatkowski, Cordelia Langford, John Sillitoe on behalf of the Wellcome Sanger Institute COVID-19 Surveillance Team |
| EPI_ISL_3097837, EPI_ISL_3097838 | Lighthouse Lab in Alderley Park | Wellcome Sanger Institute for the COVID-19 Genomics UK (COG-UK) Consortium | Jacquelyn Wynn, Mairead Hyland, The Lighthouse Lab in Alderley Park and Alex Alderton, Roberto Amato, Jeffrey Barrett, Sonia Goncalves, Ewan Harrison, David K. Jackson, Ian Johnston, Dominic Kwiatkowski, Cordelia Langford, John Sillitoe on behalf of the Wellcome Sanger Institute COVID-19 Surveillance Team |
| EPI_ISL_3097839 | Berkshire and Surrey Pathology Services Lighthouse Laboratory | Wellcome Sanger Institute for the COVID-19 Genomics UK (COG-UK) Consortium | Berkshire and Surrey Pathology Services Lighthouse Laboratory and Alex Alderton, Roberto Amato, Jeffrey Barrett, Sonia Goncalves, Ewan Harrison, David K. Jackson, Ian Johnston, Dominic Kwiatkowski, Cordelia Langford, John Sillitoe on behalf of the Wellcome Sanger Institute COVID-19 Surveillance Team |
| EPI_ISL_3097840 | Lighthouse Lab in Glasgow | Wellcome Sanger Institute for the COVID-19 Genomics UK (COG-UK) Consortium | Harper VanSteenhouse, Yumi Kasai, David Gray, Carol Clugston, Anna Dominiczak and Alex Alderton, Roberto Amato, Jeffrey Barrett, Sonia Goncalves, Ewan Harrison, David K. Jackson, Ian Johnston, Dominic Kwiatkowski, Cordelia Langford, John Sillitoe on behalf of the Wellcome Sanger Institute COVID-19 Surveillance Team |
| EPI_ISL_3097841 | Lighthouse Lab in Alderley Park | Wellcome Sanger Institute for the COVID-19 Genomics UK (COG-UK) Consortium | Jacquelyn Wynn, Mairead Hyland, The Lighthouse Lab in Alderley Park and Alex Alderton, Roberto Amato, Jeffrey Barrett, Sonia Goncalves, Ewan Harrison, David K. Jackson, Ian Johnston, Dominic Kwiatkowski, Cordelia Langford, John Sillitoe on behalf of the Wellcome Sanger Institute COVID-19 Surveillance Team |
| EPI_ISL_3097842 | Berkshire and Surrey Pathology Services Lighthouse Laboratory | Wellcome Sanger Institute for the COVID-19 Genomics UK (COG-UK) Consortium | Berkshire and Surrey Pathology Services Lighthouse Laboratory and Alex Alderton, Roberto Amato, Jeffrey Barrett, Sonia Goncalves, Ewan Harrison, David K. Jackson, Ian Johnston, Dominic Kwiatkowski, Cordelia Langford, John Sillitoe on behalf of the Wellcome Sanger Institute COVID-19 Surveillance Team |
| EPI_ISL_3097843, EPI_ISL_3097844 | Lighthouse Lab in Glasgow | Wellcome Sanger Institute for the COVID-19 Genomics UK (COG-UK) Consortium | Harper VanSteenhouse, Yumi Kasai, David Gray, Carol Clugston, Anna Dominiczak and Alex Alderton, Roberto Amato, Jeffrey Barrett, Sonia Goncalves, Ewan Harrison, David K. Jackson, Ian Johnston, Dominic Kwiatkowski, Cordelia Langford, John Sillitoe on behalf of the Wellcome Sanger Institute COVID-19 Surveillance Team |
| EPI_ISL_3097845, EPI_ISL_3097846 | Berkshire and Surrey Pathology Services Lighthouse Laboratory | Wellcome Sanger Institute for the COVID-19 Genomics UK (COG-UK) Consortium | Berkshire and Surrey Pathology Services Lighthouse Laboratory and Alex Alderton, Roberto Amato, Jeffrey Barrett, Sonia Goncalves, Ewan Harrison, David K. Jackson, Ian Johnston, Dominic Kwiatkowski, Cordelia Langford, John Sillitoe on behalf of the Wellcome Sanger Institute COVID-19 Surveillance Team |
| EPI_ISL_3097847, EPI_ISL_3097848, EPI_ISL_3097849 | Lighthouse Lab in Alderley Park | Wellcome Sanger Institute for the COVID-19 Genomics UK (COG-UK) Consortium | Jacquelyn Wynn, Mairead Hyland, The Lighthouse Lab in Alderley Park and Alex Alderton, Roberto Amato, Jeffrey Barrett, Sonia Goncalves, Ewan Harrison, David K. Jackson, Ian Johnston, Dominic Kwiatkowski, Cordelia Langford, John Sillitoe on behalf of the Wellcome Sanger Institute COVID-19 Surveillance Team |
| EPI_ISL_3097850 | Berkshire and Surrey Pathology Services Lighthouse Laboratory | Wellcome Sanger Institute for the COVID-19 Genomics UK (COG-UK) Consortium | Berkshire and Surrey Pathology Services Lighthouse Laboratory and Alex Alderton, Roberto Amato, Jeffrey Barrett, Sonia Goncalves, Ewan Harrison, David K. Jackson, Ian Johnston, Dominic Kwiatkowski, Cordelia Langford, John Sillitoe on behalf of the Wellcome Sanger Institute COVID-19 Surveillance Team |
| EPI_ISL_3097851, EPI_ISL_3097852 | Lighthouse Lab in Alderley Park | Wellcome Sanger Institute for the COVID-19 Genomics UK (COG-UK) Consortium | Jacquelyn Wynn, Mairead Hyland, The Lighthouse Lab in Alderley Park and Alex Alderton, Roberto Amato, Jeffrey Barrett, Sonia Goncalves, Ewan Harrison, David K. Jackson, Ian Johnston, Dominic Kwiatkowski, Cordelia Langford, John Sillitoe on behalf of the Wellcome Sanger Institute COVID-19 Surveillance Team |
| EPI_ISL_3097853 | Lighthouse Lab in Glasgow | Wellcome Sanger Institute for the COVID-19 Genomics UK (COG-UK) Consortium | Harper VanSteenhouse, Yumi Kasai, David Gray, Carol Clugston, Anna Dominiczak and Alex Alderton, Roberto Amato, Jeffrey Barrett, Sonia Goncalves, Ewan Harrison, David K. Jackson, Ian Johnston, Dominic Kwiatkowski, Cordelia Langford, John Sillitoe on behalf of the Wellcome Sanger Institute COVID-19 Surveillance Team |
| EPI_ISL_3097854, EPI_ISL_3097855, EPI_ISL_3097856 | Lighthouse Lab in Alderley Park | Wellcome Sanger Institute for the COVID-19 Genomics UK (COG-UK) Consortium | Jacquelyn Wynn, Mairead Hyland, The Lighthouse Lab in Alderley Park and Alex Alderton, Roberto Amato, Jeffrey Barrett, Sonia Goncalves, Ewan Harrison, David K. Jackson, Ian Johnston, Dominic Kwiatkowski, Cordelia Langford, John Sillitoe on behalf of the Wellcome Sanger Institute COVID-19 Surveillance Team |
| EPI_ISL_3097857 | Lighthouse Lab in Glasgow | Wellcome Sanger Institute for the COVID-19 Genomics UK (COG-UK) Consortium | Harper VanSteenhouse, Yumi Kasai, David Gray, Carol Clugston, Anna Dominiczak and Alex Alderton, Roberto Amato, Jeffrey Barrett, Sonia Goncalves, Ewan Harrison, David K. Jackson, Ian Johnston, Dominic Kwiatkowski, Cordelia Langford, John Sillitoe on behalf of the Wellcome Sanger Institute COVID-19 Surveillance Team |
| EPI_ISL_3097858, EPI_ISL_3097859 | Lighthouse Lab in Alderley Park | Wellcome Sanger Institute for the COVID-19 Genomics UK (COG-UK) Consortium | Jacquelyn Wynn, Mairead Hyland, The Lighthouse Lab in Alderley Park and Alex Alderton, Roberto Amato, Jeffrey Barrett, Sonia Goncalves, Ewan Harrison, David K. Jackson, Ian Johnston, Dominic Kwiatkowski, Cordelia Langford, John Sillitoe on behalf of the Wellcome Sanger Institute COVID-19 Surveillance Team |
| EPI_ISL_3097860 | Berkshire and Surrey Pathology Services Lighthouse Laboratory | Wellcome Sanger Institute for the COVID-19 Genomics UK (COG-UK) Consortium | Berkshire and Surrey Pathology Services Lighthouse Laboratory and Alex Alderton, Roberto Amato, Jeffrey Barrett, Sonia Goncalves, Ewan Harrison, David K. Jackson, Ian Johnston, Dominic Kwiatkowski, Cordelia Langford, John Sillitoe on behalf of the Wellcome Sanger Institute COVID-19 Surveillance Team |
| EPI_ISL_3097861 | Lighthouse Lab in Alderley Park | Wellcome Sanger Institute for the COVID-19 Genomics UK (COG-UK) Consortium | Jacquelyn Wynn, Mairead Hyland, The Lighthouse Lab in Alderley Park and Alex Alderton, Roberto Amato, Jeffrey Barrett, Sonia Goncalves, Ewan Harrison, David K. Jackson, Ian Johnston, Dominic Kwiatkowski, Cordelia Langford, John Sillitoe on behalf of the Wellcome Sanger Institute COVID-19 Surveillance Team |
| EPI_ISL_3097862 | Lighthouse Lab in Glasgow | Wellcome Sanger Institute for the COVID-19 Genomics UK (COG-UK) Consortium | Harper VanSteenhouse, Yumi Kasai, David Gray, Carol Clugston, Anna Dominiczak and Alex Alderton, Roberto Amato, Jeffrey Barrett, Sonia Goncalves, Ewan Harrison, David K. Jackson, Ian Johnston, Dominic Kwiatkowski, Cordelia Langford, John Sillitoe on behalf of the Wellcome Sanger Institute COVID-19 Surveillance Team |
| EPI_ISL_3097863 | Berkshire and Surrey Pathology Services Lighthouse Laboratory | Wellcome Sanger Institute for the COVID-19 Genomics UK (COG-UK) Consortium | Berkshire and Surrey Pathology Services Lighthouse Laboratory and Alex Alderton, Roberto Amato, Jeffrey Barrett, Sonia Goncalves, Ewan Harrison, David K. Jackson, Ian Johnston, Dominic Kwiatkowski, Cordelia Langford, John Sillitoe on behalf of the Wellcome Sanger Institute COVID-19 Surveillance Team |
| EPI_ISL_3097864, EPI_ISL_3097865 | Lighthouse Lab in Alderley Park | Wellcome Sanger Institute for the COVID-19 Genomics UK (COG-UK) Consortium | Jacquelyn Wynn, Mairead Hyland, The Lighthouse Lab in Alderley Park and Alex Alderton, Roberto Amato, Jeffrey Barrett, Sonia Goncalves, Ewan Harrison, David K. Jackson, Ian Johnston, Dominic Kwiatkowski, Cordelia Langford, John Sillitoe on behalf of the Wellcome Sanger Institute COVID-19 Surveillance Team |
| EPI_ISL_3097866 | Berkshire and Surrey Pathology Services Lighthouse Laboratory | Wellcome Sanger Institute for the COVID-19 Genomics UK (COG-UK) Consortium | Berkshire and Surrey Pathology Services Lighthouse Laboratory and Alex Alderton, Roberto Amato, Jeffrey Barrett, Sonia Goncalves, Ewan Harrison, David K. Jackson, Ian Johnston, Dominic Kwiatkowski, Cordelia Langford, John Sillitoe on behalf of the Wellcome Sanger Institute COVID-19 Surveillance Team |
| EPI_ISL_3097867 | Lighthouse Lab in Glasgow | Wellcome Sanger Institute for the COVID-19 Genomics UK (COG-UK) Consortium | Harper VanSteenhouse, Yumi Kasai, David Gray, Carol Clugston, Anna Dominiczak and Alex Alderton, Roberto Amato, Jeffrey Barrett, Sonia Goncalves, Ewan Harrison, David K. Jackson, Ian Johnston, Dominic Kwiatkowski, Cordelia Langford, John Sillitoe on behalf of the Wellcome Sanger Institute COVID-19 Surveillance Team |
| EPI_ISL_3097868 | Berkshire and Surrey Pathology Services Lighthouse Laboratory | Wellcome Sanger Institute for the COVID-19 Genomics UK (COG-UK) Consortium | Berkshire and Surrey Pathology Services Lighthouse Laboratory and Alex Alderton, Roberto Amato, Jeffrey Barrett, Sonia Goncalves, Ewan Harrison, David K. Jackson, Ian Johnston, Dominic Kwiatkowski, Cordelia Langford, John Sillitoe on behalf of the Wellcome Sanger Institute COVID-19 Surveillance Team |
| EPI_ISL_3097869, EPI_ISL_3097870 | Lighthouse Lab in Alderley Park | Wellcome Sanger Institute for the COVID-19 Genomics UK (COG-UK) Consortium | Jacquelyn Wynn, Mairead Hyland, The Lighthouse Lab in Alderley Park and Alex Alderton, Roberto Amato, Jeffrey Barrett, Sonia Goncalves, Ewan Harrison, David K. Jackson, Ian Johnston, Dominic Kwiatkowski, Cordelia Langford, John Sillitoe on behalf of the Wellcome Sanger Institute COVID-19 Surveillance Team |
| EPI_ISL_3097871, EPI_ISL_3097872, EPI_ISL_3097873, EPI_ISL_3097874 | Lighthouse Lab in Glasgow | Wellcome Sanger Institute for the COVID-19 Genomics UK (COG-UK) Consortium | Harper VanSteenhouse, Yumi Kasai, David Gray, Carol Clugston, Anna Dominiczak and Alex Alderton, Roberto Amato, Jeffrey Barrett, Sonia Goncalves, Ewan Harrison, David K. Jackson, Ian Johnston, Dominic Kwiatkowski, Cordelia Langford, John Sillitoe on behalf of the Wellcome Sanger Institute COVID-19 Surveillance Team |
| EPI_ISL_3097875, EPI_ISL_3097876, EPI_ISL_3097877 | Lighthouse Lab in Alderley Park | Wellcome Sanger Institute for the COVID-19 Genomics UK (COG-UK) Consortium | Jacquelyn Wynn, Mairead Hyland, The Lighthouse Lab in Alderley Park and Alex Alderton, Roberto Amato, Jeffrey Barrett, Sonia Goncalves, Ewan Harrison, David K. Jackson, Ian Johnston, Dominic Kwiatkowski, Cordelia Langford, John Sillitoe on behalf of the Wellcome Sanger Institute COVID-19 Surveillance Team |
| EPI_ISL_3097878 | Berkshire and Surrey Pathology Services Lighthouse Laboratory | Wellcome Sanger Institute for the COVID-19 Genomics UK (COG-UK) Consortium | Berkshire and Surrey Pathology Services Lighthouse Laboratory and Alex Alderton, Roberto Amato, Jeffrey Barrett, Sonia Goncalves, Ewan Harrison, David K. Jackson, Ian Johnston, Dominic Kwiatkowski, Cordelia Langford, John Sillitoe on behalf of the Wellcome Sanger Institute COVID-19 Surveillance Team |
| EPI_ISL_3097879 | Lighthouse Lab in Alderley Park | Wellcome Sanger Institute for the COVID-19 Genomics UK (COG-UK) Consortium | Jacquelyn Wynn, Mairead Hyland, The Lighthouse Lab in Alderley Park and Alex Alderton, Roberto Amato, Jeffrey Barrett, Sonia Goncalves, Ewan Harrison, David K. Jackson, Ian Johnston, Dominic Kwiatkowski, Cordelia Langford, John Sillitoe on behalf of the Wellcome Sanger Institute COVID-19 Surveillance Team |
| EPI_ISL_3097880 | Lighthouse Lab in Glasgow | Wellcome Sanger Institute for the COVID-19 Genomics UK (COG-UK) Consortium | Harper VanSteenhouse, Yumi Kasai, David Gray, Carol Clugston, Anna Dominiczak and Alex Alderton, Roberto Amato, Jeffrey Barrett, Sonia Goncalves, Ewan Harrison, David K. Jackson, Ian Johnston, Dominic Kwiatkowski, Cordelia Langford, John Sillitoe on behalf of the Wellcome Sanger Institute COVID-19 Surveillance Team |
| EPI_ISL_3097881, EPI_ISL_3097882 | Berkshire and Surrey Pathology Services Lighthouse Laboratory | Wellcome Sanger Institute for the COVID-19 Genomics UK (COG-UK) Consortium | Berkshire and Surrey Pathology Services Lighthouse Laboratory and Alex Alderton, Roberto Amato, Jeffrey Barrett, Sonia Goncalves, Ewan Harrison, David K. Jackson, Ian Johnston, Dominic Kwiatkowski, Cordelia Langford, John Sillitoe on behalf of the Wellcome Sanger Institute COVID-19 Surveillance Team |
| EPI_ISL_3097883, EPI_ISL_3097884, EPI_ISL_3097885 | Lighthouse Lab in Glasgow | Wellcome Sanger Institute for the COVID-19 Genomics UK (COG-UK) Consortium | Harper VanSteenhouse, Yumi Kasai, David Gray, Carol Clugston, Anna Dominiczak and Alex Alderton, Roberto Amato, Jeffrey Barrett, Sonia Goncalves, Ewan Harrison, David K. Jackson, Ian Johnston, Dominic Kwiatkowski, Cordelia Langford, John Sillitoe on behalf of the Wellcome Sanger Institute COVID-19 Surveillance Team |
| EPI_ISL_3097886 | Lighthouse Lab in Alderley Park | Wellcome Sanger Institute for the COVID-19 Genomics UK (COG-UK) Consortium | Jacquelyn Wynn, Mairead Hyland, The Lighthouse Lab in Alderley Park and Alex Alderton, Roberto Amato, Jeffrey Barrett, Sonia Goncalves, Ewan Harrison, David K. Jackson, Ian Johnston, Dominic Kwiatkowski, Cordelia Langford, John Sillitoe on behalf of the Wellcome Sanger Institute COVID-19 Surveillance Team |
| EPI_ISL_3097887, EPI_ISL_3097888, EPI_ISL_3097889 | Berkshire and Surrey Pathology Services Lighthouse Laboratory | Wellcome Sanger Institute for the COVID-19 Genomics UK (COG-UK) Consortium | Berkshire and Surrey Pathology Services Lighthouse Laboratory and Alex Alderton, Roberto Amato, Jeffrey Barrett, Sonia Goncalves, Ewan Harrison, David K. Jackson, Ian Johnston, Dominic Kwiatkowski, Cordelia Langford, John Sillitoe on behalf of the Wellcome Sanger Institute COVID-19 Surveillance Team |
| EPI_ISL_3097890, EPI_ISL_3097891 | Lighthouse Lab in Alderley Park | Wellcome Sanger Institute for the COVID-19 Genomics UK (COG-UK) Consortium | Jacquelyn Wynn, Mairead Hyland, The Lighthouse Lab in Alderley Park and Alex Alderton, Roberto Amato, Jeffrey Barrett, Sonia Goncalves, Ewan Harrison, David K. Jackson, Ian Johnston, Dominic Kwiatkowski, Cordelia Langford, John Sillitoe on behalf of the Wellcome Sanger Institute COVID-19 Surveillance Team |
| EPI_ISL_3097892 | Lighthouse Lab in Glasgow | Wellcome Sanger Institute for the COVID-19 Genomics UK (COG-UK) Consortium | Harper VanSteenhouse, Yumi Kasai, David Gray, Carol Clugston, Anna Dominiczak and Alex Alderton, Roberto Amato, Jeffrey Barrett, Sonia Goncalves, Ewan Harrison, David K. Jackson, Ian Johnston, Dominic Kwiatkowski, Cordelia Langford, John Sillitoe on behalf of the Wellcome Sanger Institute COVID-19 Surveillance Team |
| EPI_ISL_3097893, EPI_ISL_3097894, EPI_ISL_3097895 | Lighthouse Lab in Alderley Park | Wellcome Sanger Institute for the COVID-19 Genomics UK (COG-UK) Consortium | Jacquelyn Wynn, Mairead Hyland, The Lighthouse Lab in Alderley Park and Alex Alderton, Roberto Amato, Jeffrey Barrett, Sonia Goncalves, Ewan Harrison, David K. Jackson, Ian Johnston, Dominic Kwiatkowski, Cordelia Langford, John Sillitoe on behalf of the Wellcome Sanger Institute COVID-19 Surveillance Team |
| EPI_ISL_3097896 | Lighthouse Lab in Glasgow | Wellcome Sanger Institute for the COVID-19 Genomics UK (COG-UK) Consortium | Harper VanSteenhouse, Yumi Kasai, David Gray, Carol Clugston, Anna Dominiczak and Alex Alderton, Roberto Amato, Jeffrey Barrett, Sonia Goncalves, Ewan Harrison, David K. Jackson, Ian Johnston, Dominic Kwiatkowski, Cordelia Langford, John Sillitoe on behalf of the Wellcome Sanger Institute COVID-19 Surveillance Team |
| EPI_ISL_3097897, EPI_ISL_3097898 | Berkshire and Surrey Pathology Services Lighthouse Laboratory | Wellcome Sanger Institute for the COVID-19 Genomics UK (COG-UK) Consortium | Berkshire and Surrey Pathology Services Lighthouse Laboratory and Alex Alderton, Roberto Amato, Jeffrey Barrett, Sonia Goncalves, Ewan Harrison, David K. Jackson, Ian Johnston, Dominic Kwiatkowski, Cordelia Langford, John Sillitoe on behalf of the Wellcome Sanger Institute COVID-19 Surveillance Team |
| EPI_ISL_3097899 | Lighthouse Lab in Alderley Park | Wellcome Sanger Institute for the COVID-19 Genomics UK (COG-UK) | Jacquelyn Wynn, Mairead Hyland, The Lighthouse Lab in Alderley Park and Alex Alderton, Roberto Amato, Jeffrey Barrett, Sonia Goncalves, Ewan Harrison, David K. Jackson, Ian Johnston, Dominic Kwiatkowski, Cordelia Langford, John Sillitoe on |

| | | | |
|---|---|---|---|
| | | Consortium | behalf of the Wellcome Sanger Institute COVID-19 Surveillance Team |
| EPI_ISL_3097900 | Berkshire and Surrey Pathology Services Lighthouse Laboratory | Wellcome Sanger Institute for the COVID-19 Genomics UK (COG-UK) Consortium | Berkshire and Surrey Pathology Services Lighthouse Laboratory and Alex Alderton, Roberto Amato, Jeffrey Barrett, Sonia Goncalves, Ewan Harrison, David K. Jackson, Ian Johnston, Dominic Kwiatkowski, Cordelia Langford, John Sillitoe on behalf of the Wellcome Sanger Institute COVID-19 Surveillance Team |
| EPI_ISL_3097901 | Lighthouse Lab in Glasgow | Wellcome Sanger Institute for the COVID-19 Genomics UK (COG-UK) Consortium | Harper VanSteenhouse, Yumi Kasai, David Gray, Carol Clugston, Anna Dominiczak and Alex Alderton, Roberto Amato, Jeffrey Barrett, Sonia Goncalves, Ewan Harrison, David K. Jackson, Ian Johnston, Dominic Kwiatkowski, Cordelia Langford, John Sillitoe on behalf of the Wellcome Sanger Institute COVID-19 Surveillance Team |
| EPI_ISL_3097902 | Lighthouse Lab in Alderley Park | Wellcome Sanger Institute for the COVID-19 Genomics UK (COG-UK) Consortium | Jacquelyn Wynn, Mairead Hyland, The Lighthouse Lab in Alderley Park and Alex Alderton, Roberto Amato, Jeffrey Barrett, Sonia Goncalves, Ewan Harrison, David K. Jackson, Ian Johnston, Dominic Kwiatkowski, Cordelia Langford, John Sillitoe on behalf of the Wellcome Sanger Institute COVID-19 Surveillance Team |
| EPI_ISL_3097903 | Lighthouse Lab in Glasgow | Wellcome Sanger Institute for the COVID-19 Genomics UK (COG-UK) Consortium | Harper VanSteenhouse, Yumi Kasai, David Gray, Carol Clugston, Anna Dominiczak and Alex Alderton, Roberto Amato, Jeffrey Barrett, Sonia Goncalves, Ewan Harrison, David K. Jackson, Ian Johnston, Dominic Kwiatkowski, Cordelia Langford, John Sillitoe on behalf of the Wellcome Sanger Institute COVID-19 Surveillance Team |
| EPI_ISL_3097904, EPI_ISL_3097905 | Lighthouse Lab in Alderley Park | Wellcome Sanger Institute for the COVID-19 Genomics UK (COG-UK) Consortium | Jacquelyn Wynn, Mairead Hyland, The Lighthouse Lab in Alderley Park and Alex Alderton, Roberto Amato, Jeffrey Barrett, Sonia Goncalves, Ewan Harrison, David K. Jackson, Ian Johnston, Dominic Kwiatkowski, Cordelia Langford, John Sillitoe on behalf of the Wellcome Sanger Institute COVID-19 Surveillance Team |
| EPI_ISL_3097906, EPI_ISL_3097907, EPI_ISL_3097908 | Lighthouse Lab in Glasgow | Wellcome Sanger Institute for the COVID-19 Genomics UK (COG-UK) Consortium | Harper VanSteenhouse, Yumi Kasai, David Gray, Carol Clugston, Anna Dominiczak and Alex Alderton, Roberto Amato, Jeffrey Barrett, Sonia Goncalves, Ewan Harrison, David K. Jackson, Ian Johnston, Dominic Kwiatkowski, Cordelia Langford, John Sillitoe on behalf of the Wellcome Sanger Institute COVID-19 Surveillance Team |
| EPI_ISL_3097909 | Lighthouse Lab in Alderley Park | Wellcome Sanger Institute for the COVID-19 Genomics UK (COG-UK) Consortium | Jacquelyn Wynn, Mairead Hyland, The Lighthouse Lab in Alderley Park and Alex Alderton, Roberto Amato, Jeffrey Barrett, Sonia Goncalves, Ewan Harrison, David K. Jackson, Ian Johnston, Dominic Kwiatkowski, Cordelia Langford, John Sillitoe on behalf of the Wellcome Sanger Institute COVID-19 Surveillance Team |
| EPI_ISL_3097910 | Lighthouse Lab in Glasgow | Wellcome Sanger Institute for the COVID-19 Genomics UK (COG-UK) Consortium | Harper VanSteenhouse, Yumi Kasai, David Gray, Carol Clugston, Anna Dominiczak and Alex Alderton, Roberto Amato, Jeffrey Barrett, Sonia Goncalves, Ewan Harrison, David K. Jackson, Ian Johnston, Dominic Kwiatkowski, Cordelia Langford, John Sillitoe on behalf of the Wellcome Sanger Institute COVID-19 Surveillance Team |
| EPI_ISL_3097911 | Lighthouse Lab in Alderley Park | Wellcome Sanger Institute for the COVID-19 Genomics UK (COG-UK) Consortium | Jacquelyn Wynn, Mairead Hyland, The Lighthouse Lab in Alderley Park and Alex Alderton, Roberto Amato, Jeffrey Barrett, Sonia Goncalves, Ewan Harrison, David K. Jackson, Ian Johnston, Dominic Kwiatkowski, Cordelia Langford, John Sillitoe on behalf of the Wellcome Sanger Institute COVID-19 Surveillance Team |
| EPI_ISL_3097912, EPI_ISL_3097913 | Lighthouse Lab in Glasgow | Wellcome Sanger Institute for the COVID-19 Genomics UK (COG-UK) Consortium | Harper VanSteenhouse, Yumi Kasai, David Gray, Carol Clugston, Anna Dominiczak and Alex Alderton, Roberto Amato, Jeffrey Barrett, Sonia Goncalves, Ewan Harrison, David K. Jackson, Ian Johnston, Dominic Kwiatkowski, Cordelia Langford, John Sillitoe on behalf of the Wellcome Sanger Institute COVID-19 Surveillance Team |
| EPI_ISL_3097914 | Lighthouse Lab in Alderley Park | Wellcome Sanger Institute for the COVID-19 Genomics UK (COG-UK) Consortium | Jacquelyn Wynn, Mairead Hyland, The Lighthouse Lab in Alderley Park and Alex Alderton, Roberto Amato, Jeffrey Barrett, Sonia Goncalves, Ewan Harrison, David K. Jackson, Ian Johnston, Dominic Kwiatkowski, Cordelia Langford, John Sillitoe on behalf of the Wellcome Sanger Institute COVID-19 Surveillance Team |
| EPI_ISL_3097915 | Lighthouse Lab in Glasgow | Wellcome Sanger Institute for the COVID-19 Genomics UK (COG-UK) Consortium | Harper VanSteenhouse, Yumi Kasai, David Gray, Carol Clugston, Anna Dominiczak and Alex Alderton, Roberto Amato, Jeffrey Barrett, Sonia Goncalves, Ewan Harrison, David K. Jackson, Ian Johnston, Dominic Kwiatkowski, Cordelia Langford, John Sillitoe on behalf of the Wellcome Sanger Institute COVID-19 Surveillance Team |
| EPI_ISL_3097916 | Berkshire and Surrey Pathology Services Lighthouse Laboratory | Wellcome Sanger Institute for the COVID-19 Genomics UK (COG-UK) Consortium | Berkshire and Surrey Pathology Services Lighthouse Laboratory and Alex Alderton, Roberto Amato, Jeffrey Barrett, Sonia Goncalves, Ewan Harrison, David K. Jackson, Ian Johnston, Dominic Kwiatkowski, Cordelia Langford, John Sillitoe on behalf of the Wellcome Sanger Institute COVID-19 Surveillance Team |
| EPI_ISL_3097917, EPI_ISL_3097918 | Lighthouse Lab in Glasgow | Wellcome Sanger Institute for the COVID-19 Genomics UK (COG-UK) Consortium | Harper VanSteenhouse, Yumi Kasai, David Gray, Carol Clugston, Anna Dominiczak and Alex Alderton, Roberto Amato, Jeffrey Barrett, Sonia Goncalves, Ewan Harrison, David K. Jackson, Ian Johnston, Dominic Kwiatkowski, Cordelia Langford, John Sillitoe on behalf of the Wellcome Sanger Institute COVID-19 Surveillance Team |
| EPI_ISL_3097919 | Lighthouse Lab in Alderley Park | Wellcome Sanger Institute for the COVID-19 Genomics UK (COG-UK) Consortium | Jacquelyn Wynn, Mairead Hyland, The Lighthouse Lab in Alderley Park and Alex Alderton, Roberto Amato, Jeffrey Barrett, Sonia Goncalves, Ewan Harrison, David K. Jackson, Ian Johnston, Dominic Kwiatkowski, Cordelia Langford, John Sillitoe on behalf of the Wellcome Sanger Institute COVID-19 Surveillance Team |
| EPI_ISL_3097920 | Berkshire and Surrey Pathology Services Lighthouse Laboratory | Wellcome Sanger Institute for the COVID-19 Genomics UK (COG-UK) Consortium | Berkshire and Surrey Pathology Services Lighthouse Laboratory and Alex Alderton, Roberto Amato, Jeffrey Barrett, Sonia Goncalves, Ewan Harrison, David K. Jackson, Ian Johnston, Dominic Kwiatkowski, Cordelia Langford, John Sillitoe on behalf of the Wellcome Sanger Institute COVID-19 Surveillance Team |
| EPI_ISL_3097921, EPI_ISL_3097922 | Lighthouse Lab in Glasgow | Wellcome Sanger Institute for the COVID-19 Genomics UK (COG-UK) Consortium | Harper VanSteenhouse, Yumi Kasai, David Gray, Carol Clugston, Anna Dominiczak and Alex Alderton, Roberto Amato, Jeffrey Barrett, Sonia Goncalves, Ewan Harrison, David K. Jackson, Ian Johnston, Dominic Kwiatkowski, Cordelia Langford, John Sillitoe on behalf of the Wellcome Sanger Institute COVID-19 Surveillance Team |
| EPI_ISL_3097923 | Berkshire and Surrey Pathology Services Lighthouse Laboratory | Wellcome Sanger Institute for the COVID-19 Genomics UK (COG-UK) Consortium | Berkshire and Surrey Pathology Services Lighthouse Laboratory and Alex Alderton, Roberto Amato, Jeffrey Barrett, Sonia Goncalves, Ewan Harrison, David K. Jackson, Ian Johnston, Dominic Kwiatkowski, Cordelia Langford, John Sillitoe on behalf of the Wellcome Sanger Institute COVID-19 Surveillance Team |
| EPI_ISL_3097924, EPI_ISL_3097925 | Lighthouse Lab in Alderley Park | Wellcome Sanger Institute for the COVID-19 Genomics UK (COG-UK) Consortium | Jacquelyn Wynn, Mairead Hyland, The Lighthouse Lab in Alderley Park and Alex Alderton, Roberto Amato, Jeffrey Barrett, Sonia Goncalves, Ewan Harrison, David K. Jackson, Ian Johnston, Dominic Kwiatkowski, Cordelia Langford, John Sillitoe on behalf of the Wellcome Sanger Institute COVID-19 Surveillance Team |
| EPI_ISL_3097926 | Berkshire and Surrey Pathology Services Lighthouse Laboratory | Wellcome Sanger Institute for the COVID-19 Genomics UK (COG-UK) Consortium | Berkshire and Surrey Pathology Services Lighthouse Laboratory and Alex Alderton, Roberto Amato, Jeffrey Barrett, Sonia Goncalves, Ewan Harrison, David K. Jackson, Ian Johnston, Dominic Kwiatkowski, Cordelia Langford, John Sillitoe on behalf of the Wellcome Sanger Institute COVID-19 Surveillance Team |
| EPI_ISL_3097927, EPI_ISL_3097928 | Lighthouse Lab in Alderley Park | Wellcome Sanger Institute for the COVID-19 Genomics UK (COG-UK) Consortium | Jacquelyn Wynn, Mairead Hyland, The Lighthouse Lab in Alderley Park and Alex Alderton, Roberto Amato, Jeffrey Barrett, Sonia Goncalves, Ewan Harrison, David K. Jackson, Ian Johnston, Dominic Kwiatkowski, Cordelia Langford, John Sillitoe on behalf of the Wellcome Sanger Institute COVID-19 Surveillance Team |
| EPI_ISL_3097929, EPI_ISL_3097930, EPI_ISL_3097931 | Lighthouse Lab in Glasgow | Wellcome Sanger Institute for the COVID-19 Genomics UK (COG-UK) Consortium | Harper VanSteenhouse, Yumi Kasai, David Gray, Carol Clugston, Anna Dominiczak and Alex Alderton, Roberto Amato, Jeffrey Barrett, Sonia Goncalves, Ewan Harrison, David K. Jackson, Ian Johnston, Dominic Kwiatkowski, Cordelia Langford, John Sillitoe on behalf of the Wellcome Sanger Institute COVID-19 Surveillance Team |
| EPI_ISL_3097932 | Lighthouse Lab in Alderley Park | Wellcome Sanger Institute for the COVID-19 Genomics UK (COG-UK) Consortium | Jacquelyn Wynn, Mairead Hyland, The Lighthouse Lab in Alderley Park and Alex Alderton, Roberto Amato, Jeffrey Barrett, Sonia Goncalves, Ewan Harrison, David K. Jackson, Ian Johnston, Dominic Kwiatkowski, Cordelia Langford, John Sillitoe on behalf of the Wellcome Sanger Institute COVID-19 Surveillance Team |
| EPI_ISL_3097933, EPI_ISL_3097934, EPI_ISL_3097935, EPI_ISL_3097936, EPI_ISL_3097937, EPI_ISL_3097938, EPI_ISL_3097939, EPI_ISL_3097940, EPI_ISL_3097941, EPI_ISL_3097942, EPI_ISL_3097943, EPI_ISL_3097944, EPI_ISL_3097945, EPI_ISL_3097946, EPI_ISL_3097947, EPI_ISL_3097948, EPI_ISL_3097949, EPI_ISL_3097950, EPI_ISL_3097951, EPI_ISL_3097952, EPI_ISL_3097953, EPI_ISL_3097954, EPI_ISL_3097955, EPI_ISL_3097956, EPI_ISL_3097957, EPI_ISL_3097958, EPI_ISL_3097959, EPI_ISL_3097960, EPI_ISL_3097961, EPI_ISL_3097962, EPI_ISL_3097963, EPI_ISL_3097964, EPI_ISL_3097965, EPI_ISL_3097966, EPI_ISL_3097967, EPI_ISL_3097968, EPI_ISL_3097969, EPI_ISL_3097970, EPI_ISL_3097971, EPI_ISL_3097972, EPI_ISL_3097973, EPI_ISL_3097974, EPI_ISL_3097975, EPI_ISL_3097976, EPI_ISL_3097977, EPI_ISL_3097978, EPI_ISL_3097979, EPI_ISL_3097980, EPI_ISL_3097981, EPI_ISL_3097982, EPI_ISL_3097983, EPI_ISL_3097984, EPI_ISL_3097985, EPI_ISL_3097986, EPI_ISL_3097987, EPI_ISL_3097988, EPI_ISL_3097989, EPI_ISL_3097990, EPI_ISL_3097991, EPI_ISL_3097992, EPI_ISL_3097993, EPI_ISL_3097994, EPI_ISL_3097995, EPI_ISL_3097996, EPI_ISL_3097997, EPI_ISL_3097998, EPI_ISL_3097999, EPI_ISL_3098000, EPI_ISL_3098001, EPI_ISL_3098002, EPI_ISL_3098003, EPI_ISL_3098004, EPI_ISL_3098005, EPI_ISL_3098006, EPI_ISL_3098007, EPI_ISL_3098008, EPI_ISL_3098009, EPI_ISL_3098010, EPI_ISL_3098011, EPI_ISL_3098012, EPI_ISL_3098013, EPI_ISL_3098014, EPI_ISL_3098015, EPI_ISL_3098016, EPI_ISL_3098017, EPI_ISL_3098018, EPI_ISL_3098019, EPI_ISL_3098020, EPI_ISL_3098022, EPI_ISL_3098023, EPI_ISL_3098024, EPI_ISL_3098025, EPI_ISL_3098026, EPI_ISL_3098027, EPI_ISL_3098028, EPI_ISL_3098030, EPI_ISL_3098031, EPI_ISL_3098032, EPI_ISL_3098033, EPI_ISL_3098035, EPI_ISL_3098036, EPI_ISL_3098037, EPI_ISL_3098039, EPI_ISL_3098040, EPI_ISL_3098041, EPI_ISL_3098043, EPI_ISL_3098044, EPI_ISL_3098045, EPI_ISL_3098047, EPI_ISL_3098048, EPI_ISL_3098049, EPI_ISL_3098050, EPI_ISL_3098052, EPI_ISL_3098053, EPI_ISL_3098055, EPI_ISL_3098056, EPI_ISL_3098057, EPI_ISL_3098058, EPI_ISL_3098060, EPI_ISL_3098061 | | | |
| see above | Lighthouse Lab in Glasgow | Wellcome Sanger Institute for the COVID-19 Genomics UK (COG-UK) Consortium | Harper VanSteenhouse, Yumi Kasai, David Gray, Carol Clugston, Anna Dominiczak and Alex Alderton, Roberto Amato, Jeffrey Barrett, Sonia Goncalves, Ewan Harrison, David K. Jackson, Ian Johnston, Dominic Kwiatkowski, Cordelia Langford, John Sillitoe on behalf of the Wellcome Sanger Institute COVID-19 Surveillance Team |
| EPI_ISL_3098063 | Berkshire and Surrey Pathology Services Lighthouse Laboratory | Wellcome Sanger Institute for the COVID-19 Genomics UK (COG-UK) Consortium | Berkshire and Surrey Pathology Services Lighthouse Laboratory and Alex Alderton, Roberto Amato, Jeffrey Barrett, Sonia Goncalves, Ewan Harrison, David K. Jackson, Ian Johnston, Dominic Kwiatkowski, Cordelia Langford, John Sillitoe on behalf of the Wellcome Sanger Institute COVID-19 Surveillance Team |
| EPI_ISL_3098064 | Lighthouse Lab in Glasgow | Wellcome Sanger Institute for the COVID-19 Genomics UK (COG-UK) Consortium | Harper VanSteenhouse, Yumi Kasai, David Gray, Carol Clugston, Anna Dominiczak and Alex Alderton, Roberto Amato, Jeffrey Barrett, Sonia Goncalves, Ewan Harrison, David K. Jackson, Ian Johnston, Dominic Kwiatkowski, Cordelia Langford, John Sillitoe on behalf of the Wellcome Sanger Institute COVID-19 Surveillance Team |
| EPI_ISL_3098066, EPI_ISL_3098067, EPI_ISL_3098070, EPI_ISL_3098071, EPI_ISL_3098072, EPI_ISL_3098074, EPI_ISL_3098075, EPI_ISL_3098076, EPI_ISL_3098078, EPI_ISL_3098079, EPI_ISL_3098080, EPI_ISL_3098081, EPI_ISL_3098083, EPI_ISL_3098084, EPI_ISL_3098085, EPI_ISL_3098087, EPI_ISL_3098088, EPI_ISL_3098089, EPI_ISL_3098091, EPI_ISL_3098092, EPI_ISL_3098094, EPI_ISL_3098095, EPI_ISL_3098096, EPI_ISL_3098099, EPI_ISL_3098100, EPI_ISL_3098102, EPI_ISL_3098103, EPI_ISL_3098104, EPI_ISL_3098105, EPI_ISL_3098107, EPI_ISL_3098108, EPI_ISL_3098110, EPI_ISL_3098111, EPI_ISL_3098113, EPI_ISL_3098114, EPI_ISL_3098115, EPI_ISL_3098116, EPI_ISL_3098118, EPI_ISL_3098119, EPI_ISL_3098121, EPI_ISL_3098122, EPI_ISL_3098124, EPI_ISL_3098125, EPI_ISL_3098126, EPI_ISL_3098128, EPI_ISL_3098129, EPI_ISL_3098130, EPI_ISL_3098132, EPI_ISL_3098133, EPI_ISL_3098135 | | | |
| see above | Randox Laboratories | Wellcome Sanger Institute for the COVID-19 Genomics UK (COG-UK) Consortium | Randox Laboratories and Alex Alderton, Roberto Amato, Jeffrey Barrett, Sonia Goncalves, Ewan Harrison, David K. Jackson, Ian Johnston, Dominic Kwiatkowski, Cordelia Langford, John Sillitoe on behalf of the Wellcome Sanger Institute COVID-19 Surveillance Team |
| EPI_ISL_3098136, EPI_ISL_3098137, EPI_ISL_3098139, EPI_ISL_3098140, EPI_ISL_3098142, EPI_ISL_3098143, EPI_ISL_3098144, EPI_ISL_3098146, EPI_ISL_3098147, EPI_ISL_3098148, EPI_ISL_3098150, EPI_ISL_3098151, EPI_ISL_3098152, EPI_ISL_3098154, EPI_ISL_3098155, EPI_ISL_3098157, EPI_ISL_3098158 | | | |
| see above | Lighthouse Lab in Glasgow | Wellcome Sanger Institute for the COVID-19 Genomics UK (COG-UK) Consortium | Harper VanSteenhouse, Yumi Kasai, David Gray, Carol Clugston, Anna Dominiczak and Alex Alderton, Roberto Amato, Jeffrey Barrett, Sonia Goncalves, Ewan Harrison, David K. Jackson, Ian Johnston, Dominic Kwiatkowski, Cordelia Langford, John Sillitoe on behalf of the Wellcome Sanger Institute COVID-19 Surveillance Team |
| EPI_ISL_3098159 | Lighthouse Lab in Alderley Park | Wellcome Sanger Institute for the COVID-19 Genomics UK (COG-UK) Consortium | Jacquelyn Wynn, Mairead Hyland, The Lighthouse Lab in Alderley Park and Alex Alderton, Roberto Amato, Jeffrey Barrett, Sonia Goncalves, Ewan Harrison, David K. Jackson, Ian Johnston, Dominic Kwiatkowski, Cordelia Langford, John Sillitoe on behalf of the Wellcome Sanger Institute COVID-19 Surveillance Team |
| EPI_ISL_3098161, EPI_ISL_3098162, EPI_ISL_3098163, EPI_ISL_3098165, EPI_ISL_3098166, EPI_ISL_3098167, EPI_ISL_3098169, EPI_ISL_3098170, EPI_ISL_3098172, EPI_ISL_3098173, EPI_ISL_3098175, EPI_ISL_3098176, EPI_ISL_3098178, EPI_ISL_3098179, EPI_ISL_3098181, EPI_ISL_3098182, EPI_ISL_3098183, EPI_ISL_3098185, EPI_ISL_3098186, EPI_ISL_3098188, EPI_ISL_3098189, EPI_ISL_3098191, EPI_ISL_3098192, EPI_ISL_3098193, EPI_ISL_3098195, EPI_ISL_3098196, EPI_ISL_3098197, EPI_ISL_3098199, EPI_ISL_3098200, EPI_ISL_3098201, EPI_ISL_3098202, EPI_ISL_3098204, EPI_ISL_3098205, EPI_ISL_3098206, EPI_ISL_3098208, EPI_ISL_3098209, EPI_ISL_3098210, EPI_ISL_3098212, EPI_ISL_3098213, EPI_ISL_3098215, EPI_ISL_3098216, EPI_ISL_3098217, EPI_ISL_3098219, EPI_ISL_3098220, EPI_ISL_3098221 | | | |
| see above | Lighthouse Lab in Glasgow | Wellcome Sanger Institute for the COVID-19 Genomics UK (COG-UK) Consortium | Harper VanSteenhouse, Yumi Kasai, David Gray, Carol Clugston, Anna Dominiczak and Alex Alderton, Roberto Amato, Jeffrey Barrett, Sonia Goncalves, Ewan Harrison, David K. Jackson, Ian Johnston, Dominic Kwiatkowski, Cordelia Langford, John Sillitoe on behalf of the Wellcome Sanger Institute COVID-19 Surveillance Team |
| EPI_ISL_3098223 | Randox Laboratories | Wellcome Sanger Institute for the COVID-19 Genomics UK (COG-UK) Consortium | Randox Laboratories and Alex Alderton, Roberto Amato, Jeffrey Barrett, Sonia Goncalves, Ewan Harrison, David K. Jackson, Ian Johnston, Dominic Kwiatkowski, Cordelia Langford, John Sillitoe on behalf of the Wellcome Sanger Institute COVID-19 Surveillance Team |
| EPI_ISL_3098224, EPI_ISL_3098225 | Lighthouse Lab in Glasgow | Wellcome Sanger Institute for the COVID-19 Genomics UK (COG-UK) Consortium | Harper VanSteenhouse, Yumi Kasai, David Gray, Carol Clugston, Anna Dominiczak and Alex Alderton, Roberto Amato, Jeffrey Barrett, Sonia Goncalves, Ewan Harrison, David K. Jackson, Ian Johnston, Dominic Kwiatkowski, Cordelia Langford, John Sillitoe on behalf of the Wellcome Sanger Institute COVID-19 Surveillance Team |
| EPI_ISL_3098226 | Randox Laboratories | Wellcome Sanger Institute for the COVID-19 Genomics UK (COG-UK) Consortium | Randox Laboratories and Alex Alderton, Roberto Amato, Jeffrey Barrett, Sonia Goncalves, Ewan Harrison, David K. Jackson, Ian Johnston, Dominic Kwiatkowski, Cordelia Langford, John Sillitoe on behalf of the Wellcome Sanger Institute COVID-19 Surveillance Team |
| EPI_ISL_3098228 | Lighthouse Lab in Glasgow | Wellcome Sanger Institute for the COVID-19 Genomics UK (COG-UK) Consortium | Harper VanSteenhouse, Yumi Kasai, David Gray, Carol Clugston, Anna Dominiczak and Alex Alderton, Roberto Amato, Jeffrey Barrett, Sonia Goncalves, Ewan Harrison, David K. Jackson, Ian Johnston, Dominic Kwiatkowski, Cordelia Langford, John Sillitoe on behalf of the Wellcome Sanger Institute COVID-19 Surveillance Team |
| EPI_ISL_3098229, EPI_ISL_3098231 | Randox Laboratories | Wellcome Sanger Institute for the COVID-19 Genomics UK (COG-UK) Consortium | Randox Laboratories and Alex Alderton, Roberto Amato, Jeffrey Barrett, Sonia Goncalves, Ewan Harrison, David K. Jackson, Ian Johnston, Dominic Kwiatkowski, Cordelia Langford, John Sillitoe on behalf of the Wellcome Sanger Institute COVID-19 Surveillance Team |
| EPI_ISL_3098232, EPI_ISL_3098234, EPI_ISL_3098235 | Lighthouse Lab in Glasgow | Wellcome Sanger Institute for the COVID-19 Genomics UK (COG-UK) Consortium | Harper VanSteenhouse, Yumi Kasai, David Gray, Carol Clugston, Anna Dominiczak and Alex Alderton, Roberto Amato, Jeffrey Barrett, Sonia Goncalves, Ewan Harrison, David K. Jackson, Ian Johnston, Dominic Kwiatkowski, Cordelia Langford, John Sillitoe on behalf of the Wellcome Sanger Institute COVID-19 Surveillance Team |
| EPI_ISL_3098236, EPI_ISL_3098238, EPI_ISL_3098239 | Randox Laboratories | Wellcome Sanger Institute for the COVID-19 Genomics UK (COG-UK) Consortium | Randox Laboratories and Alex Alderton, Roberto Amato, Jeffrey Barrett, Sonia Goncalves, Ewan Harrison, David K. Jackson, Ian Johnston, Dominic Kwiatkowski, Cordelia Langford, John Sillitoe on behalf of the Wellcome Sanger Institute COVID-19 Surveillance Team |

| | | | |
|---|---|---|---|
| EPI_ISL_3098240 | Lighthouse Lab in Glasgow | Wellcome Sanger Institute for the COVID-19 Genomics UK (COG-UK) Consortium | Harper VanSteenhouse, Yumi Kasai, David Gray, Carol Clugston, Anna Dominiczak and Alex Alderton, Roberto Amato, Jeffrey Barrett, Sonia Goncalves, Ewan Harrison, David K. Jackson, Ian Johnston, Dominic Kwiatkowski, Cordelia Langford, John Sillitoe on behalf of the Wellcome Sanger Institute COVID-19 Surveillance Team |
| EPI_ISL_3098242, EPI_ISL_3098243, EPI_ISL_3098244, EPI_ISL_3098246 | Randox Laboratories | Wellcome Sanger Institute for the COVID-19 Genomics UK (COG-UK) Consortium | Randox Laboratories and Alex Alderton, Roberto Amato, Jeffrey Barrett, Sonia Goncalves, Ewan Harrison, David K. Jackson, Ian Johnston, Dominic Kwiatkowski, Cordelia Langford, John Sillitoe on behalf of the Wellcome Sanger Institute COVID-19 Surveillance Team |
| EPI_ISL_3098247, EPI_ISL_3098248 | Lighthouse Lab in Glasgow | Wellcome Sanger Institute for the COVID-19 Genomics UK (COG-UK) Consortium | Harper VanSteenhouse, Yumi Kasai, David Gray, Carol Clugston, Anna Dominiczak and Alex Alderton, Roberto Amato, Jeffrey Barrett, Sonia Goncalves, Ewan Harrison, David K. Jackson, Ian Johnston, Dominic Kwiatkowski, Cordelia Langford, John Sillitoe on behalf of the Wellcome Sanger Institute COVID-19 Surveillance Team |
| EPI_ISL_3098250, EPI_ISL_3098251, EPI_ISL_3098253 | Randox Laboratories | Wellcome Sanger Institute for the COVID-19 Genomics UK (COG-UK) Consortium | Randox Laboratories and Alex Alderton, Roberto Amato, Jeffrey Barrett, Sonia Goncalves, Ewan Harrison, David K. Jackson, Ian Johnston, Dominic Kwiatkowski, Cordelia Langford, John Sillitoe on behalf of the Wellcome Sanger Institute COVID-19 Surveillance Team |
| EPI_ISL_3098254, EPI_ISL_3098255 | Lighthouse Lab in Glasgow | Wellcome Sanger Institute for the COVID-19 Genomics UK (COG-UK) Consortium | Harper VanSteenhouse, Yumi Kasai, David Gray, Carol Clugston, Anna Dominiczak and Alex Alderton, Roberto Amato, Jeffrey Barrett, Sonia Goncalves, Ewan Harrison, David K. Jackson, Ian Johnston, Dominic Kwiatkowski, Cordelia Langford, John Sillitoe on behalf of the Wellcome Sanger Institute COVID-19 Surveillance Team |
| EPI_ISL_3098257 | Randox Laboratories | Wellcome Sanger Institute for the COVID-19 Genomics UK (COG-UK) Consortium | Randox Laboratories and Alex Alderton, Roberto Amato, Jeffrey Barrett, Sonia Goncalves, Ewan Harrison, David K. Jackson, Ian Johnston, Dominic Kwiatkowski, Cordelia Langford, John Sillitoe on behalf of the Wellcome Sanger Institute COVID-19 Surveillance Team |
| EPI_ISL_3098258, EPI_ISL_3098259, EPI_ISL_3098261, EPI_ISL_3098262 | Lighthouse Lab in Glasgow | Wellcome Sanger Institute for the COVID-19 Genomics UK (COG-UK) Consortium | Harper VanSteenhouse, Yumi Kasai, David Gray, Carol Clugston, Anna Dominiczak and Alex Alderton, Roberto Amato, Jeffrey Barrett, Sonia Goncalves, Ewan Harrison, David K. Jackson, Ian Johnston, Dominic Kwiatkowski, Cordelia Langford, John Sillitoe on behalf of the Wellcome Sanger Institute COVID-19 Surveillance Team |
| EPI_ISL_3098264 | Randox Laboratories | Wellcome Sanger Institute for the COVID-19 Genomics UK (COG-UK) Consortium | Randox Laboratories and Alex Alderton, Roberto Amato, Jeffrey Barrett, Sonia Goncalves, Ewan Harrison, David K. Jackson, Ian Johnston, Dominic Kwiatkowski, Cordelia Langford, John Sillitoe on behalf of the Wellcome Sanger Institute COVID-19 Surveillance Team |
| EPI_ISL_3098265 | Lighthouse Lab in Glasgow | Wellcome Sanger Institute for the COVID-19 Genomics UK (COG-UK) Consortium | Harper VanSteenhouse, Yumi Kasai, David Gray, Carol Clugston, Anna Dominiczak and Alex Alderton, Roberto Amato, Jeffrey Barrett, Sonia Goncalves, Ewan Harrison, David K. Jackson, Ian Johnston, Dominic Kwiatkowski, Cordelia Langford, John Sillitoe on behalf of the Wellcome Sanger Institute COVID-19 Surveillance Team |
| EPI_ISL_3098266 | Randox Laboratories | Wellcome Sanger Institute for the COVID-19 Genomics UK (COG-UK) Consortium | Randox Laboratories and Alex Alderton, Roberto Amato, Jeffrey Barrett, Sonia Goncalves, Ewan Harrison, David K. Jackson, Ian Johnston, Dominic Kwiatkowski, Cordelia Langford, John Sillitoe on behalf of the Wellcome Sanger Institute COVID-19 Surveillance Team |
| EPI_ISL_3098268, EPI_ISL_3098269 | Lighthouse Lab in Glasgow | Wellcome Sanger Institute for the COVID-19 Genomics UK (COG-UK) Consortium | Harper VanSteenhouse, Yumi Kasai, David Gray, Carol Clugston, Anna Dominiczak and Alex Alderton, Roberto Amato, Jeffrey Barrett, Sonia Goncalves, Ewan Harrison, David K. Jackson, Ian Johnston, Dominic Kwiatkowski, Cordelia Langford, John Sillitoe on behalf of the Wellcome Sanger Institute COVID-19 Surveillance Team |
| EPI_ISL_3098270 | Randox Laboratories | Wellcome Sanger Institute for the COVID-19 Genomics UK (COG-UK) Consortium | Randox Laboratories and Alex Alderton, Roberto Amato, Jeffrey Barrett, Sonia Goncalves, Ewan Harrison, David K. Jackson, Ian Johnston, Dominic Kwiatkowski, Cordelia Langford, John Sillitoe on behalf of the Wellcome Sanger Institute COVID-19 Surveillance Team |
| EPI_ISL_3098271, EPI_ISL_3098273, EPI_ISL_3098274, EPI_ISL_3098276, EPI_ISL_3098277, EPI_ISL_3098278, EPI_ISL_3098280, EPI_ISL_3098281, EPI_ISL_3098282, EPI_ISL_3098284, EPI_ISL_3098285 | | | |
| see above | Lighthouse Lab in Glasgow | Wellcome Sanger Institute for the COVID-19 Genomics UK (COG-UK) Consortium | Harper VanSteenhouse, Yumi Kasai, David Gray, Carol Clugston, Anna Dominiczak and Alex Alderton, Roberto Amato, Jeffrey Barrett, Sonia Goncalves, Ewan Harrison, David K. Jackson, Ian Johnston, Dominic Kwiatkowski, Cordelia Langford, John Sillitoe on behalf of the Wellcome Sanger Institute COVID-19 Surveillance Team |
| EPI_ISL_3098286, EPI_ISL_3098288, EPI_ISL_3098289, EPI_ISL_3098290, EPI_ISL_3098292, EPI_ISL_3098293, EPI_ISL_3098294, EPI_ISL_3098296, EPI_ISL_3098297, EPI_ISL_3098298, EPI_ISL_3098300, EPI_ISL_3098301, EPI_ISL_3098302, EPI_ISL_3098303, EPI_ISL_3098305, EPI_ISL_3098306, EPI_ISL_3098307, EPI_ISL_3098309, EPI_ISL_3098310, EPI_ISL_3098313, EPI_ISL_3098314, EPI_ISL_3098316, EPI_ISL_3098317, EPI_ISL_3098318, EPI_ISL_3098320, EPI_ISL_3098368, EPI_ISL_3098370, EPI_ISL_3098371 | | | |
| see above | Randox Laboratories | Wellcome Sanger Institute for the COVID-19 Genomics UK (COG-UK) Consortium | Randox Laboratories and Alex Alderton, Roberto Amato, Jeffrey Barrett, Sonia Goncalves, Ewan Harrison, David K. Jackson, Ian Johnston, Dominic Kwiatkowski, Cordelia Langford, John Sillitoe on behalf of the Wellcome Sanger Institute COVID-19 Surveillance Team |
| EPI_ISL_3098372 | Lighthouse Lab in Milton Keynes | Wellcome Sanger Institute for the COVID-19 Genomics UK (COG-UK) Consortium | The Lighthouse Lab in Milton Keynes and Alex Alderton, Roberto Amato, Jeffrey Barrett, Sonia Goncalves, Ewan Harrison, David K. Jackson, Ian Johnston, Dominic Kwiatkowski, Cordelia Langford, John Sillitoe on behalf of the Wellcome Sanger Institute COVID-19 Surveillance Team |
| EPI_ISL_3098442 | Ochsner Health | Bioinfoexperts, LLC | Amy Feehan, Judy Crabtree, Rebecca Rose, Ben Lain, Chris Huston, Samual Moot, David J. Nolan, Tessa LaFleur, Julia-Garcia-Diaz, Lucio Miele, Susanna L. Lamers |
| EPI_ISL_3098443 | Laboratorio de Virología HUCA | Laboratorio de Virología HUCA | Sandoval M, Castelló C, Gómez de Oña J, Boga JA, Rojo S, Alvarez-Arguelles ME, Abreu F, Costales I, Perez-Martínez Z, Martín-Rodríguez G, Coto E, Melón S |
| EPI_ISL_3098445 | DINAS KESEHATAN DKI JAKARTA | Genomik Solidaritas Indonesia Laboratorium | Normastuti Adhini Tantyo, Dhahlia Agustina Cahyono, Alfin Mohammad Abdillah, Vania Gavrila Wikasa, Carissa Sintca Wijaya, Mohammad Anang Syarifudin, Annisa Muthiah Sukirman, Gracia Felias Enos Korompis, Widyastuti, Dwi Oktavia, Tiranti Vindha, Ngabila Salama, Anuraj Shankar, Meutia Ayuputeri Kumaheri, Ariel Pradipta |
| EPI_ISL_3098447, EPI_ISL_3098455 | Laboratorio de Virología HUCA | Laboratorio de Virología HUCA | Sandoval M, Castelló C, Gómez de Oña J, Boga JA, Rojo S, Alvarez-Arguelles ME, Abreu F, Costales I, Perez-Martínez Z, Martín-Rodríguez G, Coto E, Melón S |
| EPI_ISL_3098458 | Hospital Carmen y Severo Ochoa | Laboratorio de Virología HUCA | Sandoval M, Castelló C, Gómez de Oña J, Boga JA, Rojo S, Alvarez-Arguelles ME, Abreu F, Costales I, Perez-Martínez Z, Martín-Rodríguez G, Coto E, Melón S |
| EPI_ISL_3098460, EPI_ISL_3098463 | Laboratorio de Virología HUCA | Laboratorio de Virología HUCA | Sandoval M, Castelló C, Gómez de Oña J, Boga JA, Rojo S, Alvarez-Arguelles ME, Abreu F, Costales I, Perez-Martínez Z, Martín-Rodríguez G, Coto E, Melón S |
| EPI_ISL_3098470, EPI_ISL_3098471, EPI_ISL_3098472, EPI_ISL_3098473, EPI_ISL_3098581, EPI_ISL_3098582, EPI_ISL_3098583, EPI_ISL_3098586, EPI_ISL_3098636, EPI_ISL_3098637, EPI_ISL_3098638, EPI_ISL_3098676, EPI_ISL_3098677, EPI_ISL_3098678, EPI_ISL_3098679, EPI_ISL_3098680, EPI_ISL_3098681, EPI_ISL_3098682, EPI_ISL_3098684, EPI_ISL_3098685, EPI_ISL_3098686, EPI_ISL_3098687, EPI_ISL_3098688, EPI_ISL_3098689, EPI_ISL_3098690, EPI_ISL_3098691, EPI_ISL_3098692 | | | |
| see above | Johns Hopkins Hospital Department of Pathology | Johns Hopkins Hospital Department of Pathology | C. Paul Morris, Chun Huai Luo, Adannaya Amadi, Matthew Schwartz, Nicholas Gallagher, Heba H. Mostafa |
| EPI_ISL_3098705, EPI_ISL_3098707, EPI_ISL_3098708, EPI_ISL_3098712, EPI_ISL_3098713, EPI_ISL_3098714, EPI_ISL_3098716, EPI_ISL_3098717 | NCSLPH | NCSLPH | Chase K, Miller MC, Greene S, Glover W |
| EPI_ISL_3098728 | TIVOLI Hospital (Rome) | University Campus BioMedico of Rome | Riva E, De Florio L, Fogolari M, Francesconi M, Lintas C, Veralli R, Angeletti S. |
| EPI_ISL_3098729 | ASL Latina | University Campus Bio-Medico of Rome | Riva E., De Florio L., Fogolari M., Francesconi M., Lintas C., Veralli R., Angeletti S. |
| EPI_ISL_3098730 | ASL Latina (LT) | University Campus Bio-Medico of Rome | Riva E., De Florio L., Fogolari M., Francesconi M., Lintas C., Veralli R., Angeletti S. |
| EPI_ISL_3098731, EPI_ISL_3098732, EPI_ISL_3098733, EPI_ISL_3098734 | ASL Latina | University Campus Bio-Medico of Rome | Riva E., De Florio L., Fogolari M., Francesconi M., Lintas C., Veralli R., Angeletti S. |
| EPI_ISL_3098735 | ASL Latina (LT) | University Campus Bio-Medico of Rome | Riva E., De Florio L., Fogolari M., Francesconi M., Lintas C., Veralli R., Angeletti S. |
| EPI_ISL_3098736, EPI_ISL_3098737, EPI_ISL_3098738, EPI_ISL_3098739, EPI_ISL_3098740, EPI_ISL_3098741, EPI_ISL_3098742, EPI_ISL_3098743, EPI_ISL_3098744, EPI_ISL_3098745, EPI_ISL_3098746 | | | |
| see above | Policlinico Tor Vergata Hospital | University Campus Bio-Medico of Rome (UCBM) | Riva E., De Florio L., Fogolari M., Francesconi M., Lintas C., Veralli R., Angeletti S. |
| EPI_ISL_3098747, EPI_ISL_3098748, EPI_ISL_3098751, EPI_ISL_3098752, EPI_ISL_3098753, EPI_ISL_3098754 | Sant'Eugenio Hospital | University Campus Bio-Medico of Rome (UCBM) | Riva E., De Florio L., Fogolari M., Francesconi M., Lintas C., Veralli R., Angeletti S. |
| EPI_ISL_3098759 | DINAS KESEHATAN DKI JAKARTA | Genomik Solidaritas Indonesia Laboratorium | Normastuti Adhini Tantyo, Dhahlia Agustina Cahyono, Alfin Mohammad Abdillah, Vania Gavrila Wikasa, Carissa Sintca Wijaya, Mohammad Anang Syarifudin, Annisa Muthiah Sukirman, Gracia Felias Enos Korompis, Widyastuti, Dwi Oktavia, Tiranti Vindha, Ngabila Salama, Anuraj Shankar, Meutia Ayuputeri Kumaheri, Ariel Pradipta |
| EPI_ISL_3098761 | Ospedale Pediatrico Bambino Gesù (U.O.C. Microbiology and Virology) | University Campus Bio-Medico of Rome (UCBM) | Riva E., De Florio L., Fogolari M., Francesconi M., Lintas C., Veralli R., Angeletti S. |
| EPI_ISL_3098764, EPI_ISL_3098765, EPI_ISL_3098766, EPI_ISL_3098768, EPI_ISL_3098770, EPI_ISL_3098771, EPI_ISL_3098773, EPI_ISL_3098774, EPI_ISL_3098775, EPI_ISL_3098776, EPI_ISL_3098777, EPI_ISL_3098778, EPI_ISL_3098779, EPI_ISL_3098780, EPI_ISL_3098781, EPI_ISL_3098782, EPI_ISL_3098783, EPI_ISL_3098784, EPI_ISL_3098785, EPI_ISL_3098786, EPI_ISL_3098787, EPI_ISL_3098788, EPI_ISL_3098789, EPI_ISL_3098790, EPI_ISL_3098791, EPI_ISL_3098792, EPI_ISL_3098793 | | | |
| see above | IN State Department of Health Laboratory Services | IN State Department of Health Laboratory Services | Cassandra Campion, Jamie Yeadon, Brian Pope, Lixia Liu, Kyle Brownlee, Melissa Hindenlang, Mark Glazier |
| EPI_ISL_3098794 | ASL GENZANO | University Campus Bio-Medico, Rome | Angeletti Silvia, Riva Elisabetta |
| EPI_ISL_3098795, EPI_ISL_3098796, EPI_ISL_3098797, EPI_ISL_3098798, EPI_ISL_3098799, EPI_ISL_3098800 | Diagnostyka. Laboratoria Medyczne. | 1. ViroGenetics - BSL3 Laboratory of Virology, Małopolska Centre of Biotechnology, Jagiellonian University; 2. Diagtron Laboratoria Łukasz Rąbalski | Rabalski L., Gromowski,T., Rabalska-Kupiec M., Swadzba J., Wydmanski,W., Mazur-Panasiuk,N., Kowalski,M., Maciej Kosinski, Natalia Derewonko, Szulc,P., Sylwia Januszczak, Labaj,P.P., Pyrc,K. |
| EPI_ISL_3098802, EPI_ISL_3098808 | ASL GENZANO | University Campus Bio-Medico of Rome | Angeletti Silvia, Riva Elisabetta |
| EPI_ISL_3098809, EPI_ISL_3098810, | Istituto Zooprofilattico Sperimentale della Puglia e della Basilicata | Istituto Zooprofilattico Sperimentale della Puglia e della Basilicata | Parisi A., Bianco A., Capozzi L., Del Sambro L., Simone D., Difato L., Pennuzzi G., Pace L., Galante D. |

| | | | |
|---|---|---|---|
| EPI_ISL_3098811, EPI_ISL_3098812, EPI_ISL_3098813, EPI_ISL_3098814 | | | |
| EPI_ISL_3098815 | Dipartimento di prevenzione Bari/Lagravinese | Istituto Zooprofilattico Sperimentale della Puglia e della Basilicata | Parisi A., Bianco A., Capozzi L., Del Sambro L., Simone D., Difato L., Ridolfi D., Giannico A., Lagravinese D. |
| EPI_ISL_3098816, EPI_ISL_3098817, EPI_ISL_3098818, EPI_ISL_3098819, EPI_ISL_3098820, EPI_ISL_3098821 | Ospedale Vito Fazzi | Istituto Zooprofilattico Sperimentale della Puglia e della Basilicata | Parisi A., Bianco A., Capozzi L., Del Sambro L., Simone D., Difato L., Lobreglio G. |
| EPI_ISL_3098823, EPI_ISL_3098824, EPI_ISL_3098825, EPI_ISL_3098826, EPI_ISL_3098827, EPI_ISL_3098828, EPI_ISL_3098829, EPI_ISL_3098830, EPI_ISL_3098831, EPI_ISL_3098832 | Istituto Zooprofilattico Sperimentale della Puglia e della Basilicata | Istituto Zooprofilattico Sperimentale della Puglia e della Basilicata | Parisi A., Bianco A., Capozzi L., Del Sambro L., Simone D., Difato L., Pennuzzi G., Pace L., Galante D. |
| EPI_ISL_3098833, EPI_ISL_3098834, EPI_ISL_3098835, EPI_ISL_3098836, EPI_ISL_3098837, EPI_ISL_3098838, EPI_ISL_3098839, EPI_ISL_3098840, EPI_ISL_3098841, EPI_ISL_3098842, EPI_ISL_3098844, EPI_ISL_3098845, EPI_ISL_3098846, EPI_ISL_3098847, EPI_ISL_3098850 | | | |
| see above | Presidio di Brindisi Di Summa - Perrino | Istituto Zooprofilattico Sperimentale della Puglia e della Basilicata | Parisi A., Bianco A., Capozzi L., Del Sambro L., Simone D., Difato L., Santoro A., Ridolfi D., Giannico A. |
| EPI_ISL_3098851, EPI_ISL_3098852, EPI_ISL_3098853, EPI_ISL_3098855, EPI_ISL_3098856, EPI_ISL_3098858, EPI_ISL_3098859 | Ospedale Santa Caterina Novella | Istituto Zooprofilattico Sperimentale della Puglia e della Basilicata | Parisi A., Bianco A., Capozzi L., Del Sambro L., Simone D., Difato L., Bruno A. R. |
| EPI_ISL_3098860, EPI_ISL_3098861, EPI_ISL_3098862, EPI_ISL_3098863, EPI_ISL_3098864, EPI_ISL_3098865, EPI_ISL_3098866 | Ospedale Vito Fazzi | Istituto Zooprofilattico Sperimentale della Puglia e della Basilicata | Parisi A., Bianco A., Capozzi L., Del Sambro L., Simone D., Difato L., Lobreglio G. |
| EPI_ISL_3098867, EPI_ISL_3098868, EPI_ISL_3098869, EPI_ISL_3098870, EPI_ISL_3098871, EPI_ISL_3098872 | I.R.C.C.S. "S. De Bellis" - Ente Ospedaliero | Istituto Zooprofilattico Sperimentale della Puglia e della Basilicata | Parisi A., Bianco A., Capozzi L., Del Sambro L., Simone D., Difato L., Notarnicola M., Lippolis A. |
| EPI_ISL_3098873, EPI_ISL_3098875, EPI_ISL_3098876, EPI_ISL_3098877, EPI_ISL_3098878 | Ospedale Santissima Annunziata | Istituto Zooprofilattico Sperimentale della Puglia e della Basilicata | Parisi A., Bianco A., Capozzi L., Del Sambro L., Simone D., Difato L., Ridolfi D., Giannico A, Tampoia M. |
| EPI_ISL_3098879 | ASL GENZANO | Univesrity Campus Bio_Medico of Rome | Angeletti Silvia, Riva Elisabetta |
| EPI_ISL_3098884 | Nehru Zoological Park | CSIR-Centre for Cellular and Molecular Biology | B. Sambasiva Rao, Md Abdul Hakeem, Sofia Banu , Shreekant Verma, Amareshwar Vodapalli ,Ara Sreenivas,B Himasri,Valli Nagalakshmi Undamatla,Payel Mukherjee, Lamuk Zaveri, Onkar Kulkarni,Tulasi Nagabandi, Jandhyala Sai Krishna,Sumedha Avadhanula, Priya Nurkurthy, Vidhyadhari Methuku, Archana Bharadwaj Siva, Karthik Bharadwaj Tallapaka,Rakesh K Mishra, Karthikeyan Vasudevan, Divya Tej Sowpati |
| EPI_ISL_3098886 | ASL GENZANO | University campus Bio-Medico of Rome | Angeletti Silvia, Riva Elisabetta |
| EPI_ISL_3098887, EPI_ISL_3098888, EPI_ISL_3098889, EPI_ISL_3098892, EPI_ISL_3098893, EPI_ISL_3098894, EPI_ISL_3098895, EPI_ISL_3098896, EPI_ISL_3098897, EPI_ISL_3098898, EPI_ISL_3098899, EPI_ISL_3098900, EPI_ISL_3098901, EPI_ISL_3098902, EPI_ISL_3098903, EPI_ISL_3098904, EPI_ISL_3098905, EPI_ISL_3098906, EPI_ISL_3098907, EPI_ISL_3098908, EPI_ISL_3098910, EPI_ISL_3098911, EPI_ISL_3098912, EPI_ISL_3098913, EPI_ISL_3098914, EPI_ISL_3098916, EPI_ISL_3098917, EPI_ISL_3098918, EPI_ISL_3098919, EPI_ISL_3098920, EPI_ISL_3098921, EPI_ISL_3098922, EPI_ISL_3098923, EPI_ISL_3098924, EPI_ISL_3098925, EPI_ISL_3098926, EPI_ISL_3098927, EPI_ISL_3098928, EPI_ISL_3098929, EPI_ISL_3098930, EPI_ISL_3098931, EPI_ISL_3098932, EPI_ISL_3098933, EPI_ISL_3098934, EPI_ISL_3098935, EPI_ISL_3098936, EPI_ISL_3098937, EPI_ISL_3098938 | | | |
| see above | ZOL | Jessa | Berden et al. on behalf of the Jessa_cmdLab |
| EPI_ISL_3098939, EPI_ISL_3098941, EPI_ISL_3098942, EPI_ISL_3098943, EPI_ISL_3098944, EPI_ISL_3098945, EPI_ISL_3098946, EPI_ISL_3098947, EPI_ISL_3098948, EPI_ISL_3098949, EPI_ISL_3098950, EPI_ISL_3098951, EPI_ISL_3098952, EPI_ISL_3098953, EPI_ISL_3098954, EPI_ISL_3098955 | | | |
| see above | Jessa | Jessa | Berden et al. on behalf of the Jessa_cmdLab |
| EPI_ISL_3098956, EPI_ISL_3098957, EPI_ISL_3098982 | ASL GENZANO | University Campus Bio-Medico of Rome | Angeletti Silvia, Riva Elisabetta |
| EPI_ISL_3098985, EPI_ISL_3098986, EPI_ISL_3099021 | OSPEDALE SANDRO PERTINI | University Campus Bio-Medico of Rome | Angeletti Silvia, Riva Elisabetta |
| EPI_ISL_3099196 | Institute of Microbiology, Universidad San Francisco de Quito | Institute of Microbiology, Universidad San Francisco de Quito | Sully Márquez, Juan José Guadalupe, Belén Prado-Vivar, Monica Becerra-Wong, Fernanda Zurita, Erika Muñoz, Bernardo Gutiérrez, Verónica Barragán, Patricio Rojas-Silva, Gabriel Trueba, Michelle Grunauer, Paúl Cárdenas |
| EPI_ISL_3099197, EPI_ISL_3099198, EPI_ISL_3099199, EPI_ISL_3099201, EPI_ISL_3099202, EPI_ISL_3099204, EPI_ISL_3099205, EPI_ISL_3099206, EPI_ISL_3099207, EPI_ISL_3099208, EPI_ISL_3099209, EPI_ISL_3099210, EPI_ISL_3099211, EPI_ISL_3099212, EPI_ISL_3099213, EPI_ISL_3099214, EPI_ISL_3099215, EPI_ISL_3099216, EPI_ISL_3099217, EPI_ISL_3099218, EPI_ISL_3099220, EPI_ISL_3099221, EPI_ISL_3099222, EPI_ISL_3099223, EPI_ISL_3099224, EPI_ISL_3099225, EPI_ISL_3099226, EPI_ISL_3099227, EPI_ISL_3099228, EPI_ISL_3099229, EPI_ISL_3099230, EPI_ISL_3099231, EPI_ISL_3099232, EPI_ISL_3099233, EPI_ISL_3099235, EPI_ISL_3099237, EPI_ISL_3099238, EPI_ISL_3099240, EPI_ISL_3099241, EPI_ISL_3099242, EPI_ISL_3099244, EPI_ISL_3099245, EPI_ISL_3099246, EPI_ISL_3099247, EPI_ISL_3099248, EPI_ISL_3099249, EPI_ISL_3099250, EPI_ISL_3099251, EPI_ISL_3099252, EPI_ISL_3099253, EPI_ISL_3099254, EPI_ISL_3099256, EPI_ISL_3099257, EPI_ISL_3099258, EPI_ISL_3099259, EPI_ISL_3099260, EPI_ISL_3099261, EPI_ISL_3099262, EPI_ISL_3099263, EPI_ISL_3099264, EPI_ISL_3099265, EPI_ISL_3099268, EPI_ISL_3099269, EPI_ISL_3099270, EPI_ISL_3099271, EPI_ISL_3099272, EPI_ISL_3099273, EPI_ISL_3099274, EPI_ISL_3099275, EPI_ISL_3099276, EPI_ISL_3099277, EPI_ISL_3099278, EPI_ISL_3099279, EPI_ISL_3099280, EPI_ISL_3099282, EPI_ISL_3099283, EPI_ISL_3099284, EPI_ISL_3099285, EPI_ISL_3099286, EPI_ISL_3099287, EPI_ISL_3099288, EPI_ISL_3099289, EPI_ISL_3099290, EPI_ISL_3099291, EPI_ISL_3099293, EPI_ISL_3099294, EPI_ISL_3099295, EPI_ISL_3099296, EPI_ISL_3099297, EPI_ISL_3099298, EPI_ISL_3099299, EPI_ISL_3099301, EPI_ISL_3099302, EPI_ISL_3099304, EPI_ISL_3099305, EPI_ISL_3099306, EPI_ISL_3099307, EPI_ISL_3099308, EPI_ISL_3099310, EPI_ISL_3099311, EPI_ISL_3099312, EPI_ISL_3099313, EPI_ISL_3099314, EPI_ISL_3099316, EPI_ISL_3099317, EPI_ISL_3099318, EPI_ISL_3099319, EPI_ISL_3099320, EPI_ISL_3099321, EPI_ISL_3099322, EPI_ISL_3099323, EPI_ISL_3099324, EPI_ISL_3099325, EPI_ISL_3099326, EPI_ISL_3099327, EPI_ISL_3099328, EPI_ISL_3099329, EPI_ISL_3099330, EPI_ISL_3099331, EPI_ISL_3099332, EPI_ISL_3099333, EPI_ISL_3099334, EPI_ISL_3099335, EPI_ISL_3099336, EPI_ISL_3099337, EPI_ISL_3099338, EPI_ISL_3099339, EPI_ISL_3099340, EPI_ISL_3099342, EPI_ISL_3099343, EPI_ISL_3099344, EPI_ISL_3099345, EPI_ISL_3099346, EPI_ISL_3099347, EPI_ISL_3099348, EPI_ISL_3099349, EPI_ISL_3099350, EPI_ISL_3099351, EPI_ISL_3099353, EPI_ISL_3099354, EPI_ISL_3099355, EPI_ISL_3099356, EPI_ISL_3099357, EPI_ISL_3099358, EPI_ISL_3099359, EPI_ISL_3099360, EPI_ISL_3099361, EPI_ISL_3099362, EPI_ISL_3099363, EPI_ISL_3099365, EPI_ISL_3099366, EPI_ISL_3099368, EPI_ISL_3099369, EPI_ISL_3099370, EPI_ISL_3099371, EPI_ISL_3099372, EPI_ISL_3099374, EPI_ISL_3099376, EPI_ISL_3099377, EPI_ISL_3099378, EPI_ISL_3099379, EPI_ISL_3099380, EPI_ISL_3099381, EPI_ISL_3099382, EPI_ISL_3099384 | | | |
| see above | Arkansas Public Health Laboratory, Arkansas Department of Health | University of Minnesota Genomics Center | Daryl M. Gohl, John Garbe, Jaquelyn Kuriger-Laber, Corbin Dirkx |
| EPI_ISL_3099389, EPI_ISL_3099390, EPI_ISL_3099391, EPI_ISL_3099392, EPI_ISL_3099393, EPI_ISL_3099394, EPI_ISL_3099395, EPI_ISL_3099396, EPI_ISL_3099397, EPI_ISL_3099398, EPI_ISL_3099399, EPI_ISL_3099409, EPI_ISL_3099410, EPI_ISL_3099411, EPI_ISL_3099412, EPI_ISL_3099413, EPI_ISL_3099414, EPI_ISL_3099415, EPI_ISL_3099416, EPI_ISL_3099417, EPI_ISL_3099418, EPI_ISL_3099419, EPI_ISL_3099420, EPI_ISL_3099421, EPI_ISL_3099422, EPI_ISL_3099423, EPI_ISL_3099424 | | | |
| see above | Saint Louis County Department of Public Health Environmental Health Laboratory | University of Minnesota Genomics Center | Daryl M. Gohl, John Garbe, Jaquelyn Kuriger-Laber, Corbin Dirkx |
| EPI_ISL_3099437 | Regions Hospital | Minnesota Department of Health, Public Health Laboratory | Alexandra Lorentz, Jacob Garfin, Matt Plumb, and Xiong Wang |
| EPI_ISL_3099439 | North Memorial | Minnesota Department of Health, Public Health Laboratory | Alexandra Lorentz, Jacob Garfin, Matt Plumb, and Xiong Wang |
| EPI_ISL_3099440 | Minnesota Department of Health, Public Health Laboratory | Minnesota Department of Health, Public Health Laboratory | Alexandra Lorentz, Jacob Garfin, Matt Plumb, and Xiong Wang |
| EPI_ISL_3099441, EPI_ISL_3099442 | Regions Hospital | Minnesota Department of Health, Public Health Laboratory | Alexandra Lorentz, Jacob Garfin, Matt Plumb, and Xiong Wang |
| EPI_ISL_3099443, EPI_ISL_3099444, EPI_ISL_3099445 | Minnesota Department of Health, Public Health Laboratory | Minnesota Department of Health, Public Health Laboratory | Alexandra Lorentz, Jacob Garfin, Matt Plumb, and Xiong Wang |
| EPI_ISL_3099447, EPI_ISL_3099449, EPI_ISL_3099450, EPI_ISL_3099451, EPI_ISL_3099452 | Centracare Laboratory Services | Minnesota Department of Health, Public Health Laboratory | Alexandra Lorentz, Jacob Garfin, Matt Plumb, and Xiong Wang |
| EPI_ISL_3099453 | Park Nicollet Methodist Hospital | Minnesota Department of Health, Public Health Laboratory | Alexandra Lorentz, Jacob Garfin, Matt Plumb, and Xiong Wang |
| EPI_ISL_3099455 | Minnesota Department of Health, Public Health Laboratory | Minnesota Department of Health, Public Health Laboratory | Alexandra Lorentz, Jacob Garfin, Matt Plumb, and Xiong Wang |
| EPI_ISL_3099456 | Allina Health Laboratory | Minnesota Department of Health, Public Health Laboratory | Alexandra Lorentz, Jacob Garfin, Matt Plumb, and Xiong Wang |

| | | | |
|---|---|---|---|
| EPI_ISL_3099457, EPI_ISL_3099458 | Minnesota Department of Health, Public Health Laboratory | Minnesota Department of Health, Public Health Laboratory | Alexandra Lorentz, Jacob Garfin, Matt Plumb, and Xiong Wang |
| EPI_ISL_3099459, EPI_ISL_3099460, EPI_ISL_3099461 | Park Nicollet Methodist Hospital | Minnesota Department of Health, Public Health Laboratory | Alexandra Lorentz, Jacob Garfin, Matt Plumb, and Xiong Wang |
| EPI_ISL_3099462, EPI_ISL_3099463, EPI_ISL_3099464, EPI_ISL_3099465, EPI_ISL_3099466, EPI_ISL_3099467 | Minnesota Department of Health, Public Health Laboratory | Minnesota Department of Health, Public Health Laboratory | Alexandra Lorentz, Jacob Garfin, Matt Plumb, and Xiong Wang |
| EPI_ISL_3099468 | Maple Grove Hospital | Minnesota Department of Health, Public Health Laboratory | Alexandra Lorentz, Jacob Garfin, Matt Plumb, and Xiong Wang |
| EPI_ISL_3099469 | M Health Fairview | Minnesota Department of Health, Public Health Laboratory | Alexandra Lorentz, Jacob Garfin, Matt Plumb, and Xiong Wang |
| EPI_ISL_3099470 | Centracare Laboratory Services | Minnesota Department of Health, Public Health Laboratory | Alexandra Lorentz, Jacob Garfin, Matt Plumb, and Xiong Wang |
| EPI_ISL_3099471, EPI_ISL_3099472 | Minnesota Department of Health, Public Health Laboratory | Minnesota Department of Health, Public Health Laboratory | Alexandra Lorentz, Jacob Garfin, Matt Plumb, and Xiong Wang |
| EPI_ISL_3099473, EPI_ISL_3099474, EPI_ISL_3099475, EPI_ISL_3099476, EPI_ISL_3099477 | Park Nicollet Methodist Hospital | Minnesota Department of Health, Public Health Laboratory | Alexandra Lorentz, Jacob Garfin, Matt Plumb, and Xiong Wang |
| EPI_ISL_3099478, EPI_ISL_3099479 | Minnesota Department of Health, Public Health Laboratory | Minnesota Department of Health, Public Health Laboratory | Alexandra Lorentz, Jacob Garfin, Matt Plumb, and Xiong Wang |
| EPI_ISL_3099481, EPI_ISL_3099482, EPI_ISL_3099483 | Regions Hospital | Minnesota Department of Health, Public Health Laboratory | Alexandra Lorentz, Jacob Garfin, Matt Plumb, and Xiong Wang |
| EPI_ISL_3099484, EPI_ISL_3099485, EPI_ISL_3099486, EPI_ISL_3099487, EPI_ISL_3099488, EPI_ISL_3099489, EPI_ISL_3099490, EPI_ISL_3099491, EPI_ISL_3099492, EPI_ISL_3099493, EPI_ISL_3099494, EPI_ISL_3099495, EPI_ISL_3099496, EPI_ISL_3099497, EPI_ISL_3099498, EPI_ISL_3099499, EPI_ISL_3099500, EPI_ISL_3099501, EPI_ISL_3099502, EPI_ISL_3099503, EPI_ISL_3099504, EPI_ISL_3099505, EPI_ISL_3099506, EPI_ISL_3099507, EPI_ISL_3099508, EPI_ISL_3099509, EPI_ISL_3099510, EPI_ISL_3099511, EPI_ISL_3099512, EPI_ISL_3099513, EPI_ISL_3099514, EPI_ISL_3099515, EPI_ISL_3099516, EPI_ISL_3099517, EPI_ISL_3099518, EPI_ISL_3099519, EPI_ISL_3099520, EPI_ISL_3099521, EPI_ISL_3099522, EPI_ISL_3099523, EPI_ISL_3099524 | | | |
| see above | OUCRU | OUCRU | Nguyen Van Vinh Chau, Nghiem My Ngoc, Lam Anh Nguyet, Vo Minh Quang, Dao Bach Khoa, Nguyen Thanh Phong, Nguyen Thi Han Ny, Nguyen Thi Thu Hong, Le Nguyen Truc Nhu, Nguyen Huynh Thanh Truc, Nguyen Thi Ngoc Minh, Mar Choisy, Dinh Nguyen Huy Man, Dinh Thi Bich Ty, Nguyen To Anh, Lam Minh Yen, Le Thi Tam Uyen, Tran Nguyen Hoang Tu, Nguyen Thanh Dung, Le Manh Hung, Nguyen Thanh Truong, Tran Tan Thanh, Guy Thwaites, and Le Van Tan |
| EPI_ISL_3099527 | Allergy, Immunology and Cell Biology Unit (AICBU) | Allergy, Immunology and Cell Biology Unit (AICBU) | Chandima Jeewandara, Deshni Jayathilaka, Dumni Gunasinghe, Tibutius Thanesh, Dinuka Ariyarathne, Ayesha Wijesinghe, Diyanath Ranasinghe, Gathsaurie Neelika Malavige |
| EPI_ISL_3099533, EPI_ISL_3099534, EPI_ISL_3099535, EPI_ISL_3099536, EPI_ISL_3099537, EPI_ISL_3099538, EPI_ISL_3099539, EPI_ISL_3099540 | DeRisi Lab, University of California, San Francisco | DeRisi Lab, University of California, San Francisco | Sabrina Mann, Anthea Mitchell, Matthias Hapte-Selassie, Jamin Liu, Sara Sunshine, Matthew Laurie, Genay Pilarowski, Amy Kistler, Manu Vanaerschot, Patrick Ayscue, Lucy Li, Eric Chow, IDseq Team, James Peng, Carina Marquez, Luis Rubio, Gabriel Chamie, Diane Jones, Jon Jacobo, Susana Rojas, Susy Rojas, Valerie Tulier-Laiwa, Douglas Black, Jackie Martinez, Maya Petersen, Diane Havlir, and Joseph DeRisi |
| EPI_ISL_3099542, EPI_ISL_3099545, EPI_ISL_3099546, EPI_ISL_3099547, EPI_ISL_3099549, EPI_ISL_3099550, EPI_ISL_3099551, EPI_ISL_3099552, EPI_ISL_3099553, EPI_ISL_3099554, EPI_ISL_3099555, EPI_ISL_3099556, EPI_ISL_3099557 | | | |
| see above | Wyoming Public Health Laboratory | Wyoming Public Health Laboratory | Jim Mildenberger, Taylor Fearing, Channing Weber, Ashley Norberg, Chayse Rowley, Marley Goetz, Brian Dominguez, Elliot Thomasson, Sam Britz, Cari Sloma, Robert Petit, and Rob Christensen |
| EPI_ISL_3099577 | CAP Taradell | Banc de Sang i Teixits | Francisco Vidal, Irene Corrales, Nina Borràs, Carlos Hobeich, Natália Comes, Lorena Ramírez, Noemí Gonzalez, Silvia Sauleda, Maria Glòria Soria |
| EPI_ISL_3099578 | CAP Manlleu | Banc de Sang i Teixits | Francisco Vidal, Irene Corrales, Nina Borràs, Carlos Hobeich, Natália Comes, Lorena Ramírez, Noemí Gonzalez, Silvia Sauleda, Maria Glòria Soria |
| EPI_ISL_3099579 | CAP El Remei | Banc de Sang i Teixits | Francisco Vidal, Irene Corrales, Nina Borràs, Carlos Hobeich, Natália Comes, Lorena Ramírez, Noemí Gonzalez, Silvia Sauleda, Maria Glòria Soria |
| EPI_ISL_3099580 | CAP Taradell | Banc de Sang i Teixits | Francisco Vidal, Irene Corrales, Nina Borràs, Carlos Hobeich, Natália Comes, Lorena Ramírez, Noemí Gonzalez, Silvia Sauleda, Maria Glòria Soria |
| EPI_ISL_3099581 | CAP Sant Vicenc de Torello | Banc de Sang i Teixits | Francisco Vidal, Irene Corrales, Nina Borràs, Carlos Hobeich, Natália Comes, Lorena Ramírez, Noemí Gonzalez, Silvia Sauleda, Maria Glòria Soria |
| EPI_ISL_3099582 | CAP Vic Nord | Banc de Sang i Teixits | Francisco Vidal, Irene Corrales, Nina Borràs, Carlos Hobeich, Natália Comes, Lorena Ramírez, Noemí Gonzalez, Silvia Sauleda, Maria Glòria Soria |
| EPI_ISL_3099583, EPI_ISL_3099585 | Fundacio Althaia-Manresa | Banc de Sang i Teixits | Francisco Vidal, Irene Corrales, Nina Borràs, Carlos Hobeich, Natália Comes, Lorena Ramírez, Noemí Gonzalez, Silvia Sauleda, Maria Glòria Soria |
| EPI_ISL_3099586 | CAP Tona | Banc de Sang i Teixits | Francisco Vidal, Irene Corrales, Nina Borràs, Carlos Hobeich, Natália Comes, Lorena Ramírez, Noemí Gonzalez, Silvia Sauleda, Maria Glòria Soria |
| EPI_ISL_3099587 | CAP Torello | Banc de Sang i Teixits | Francisco Vidal, Irene Corrales, Nina Borràs, Carlos Hobeich, Natália Comes, Lorena Ramírez, Noemí Gonzalez, Silvia Sauleda, Maria Glòria Soria |
| EPI_ISL_3099588 | CAP El Remei | Banc de Sang i Teixits | Francisco Vidal, Irene Corrales, Nina Borràs, Carlos Hobeich, Natália Comes, Lorena Ramírez, Noemí Gonzalez, Silvia Sauleda, Maria Glòria Soria |
| EPI_ISL_3099589, EPI_ISL_3099590 | Fundacio Althaia-Manresa | Banc de Sang i Teixits | Francisco Vidal, Irene Corrales, Nina Borràs, Carlos Hobeich, Natália Comes, Lorena Ramírez, Noemí Gonzalez, Silvia Sauleda, Maria Glòria Soria |
| EPI_ISL_3099591 | CAP Torello | Banc de Sang i Teixits | Francisco Vidal, Irene Corrales, Nina Borràs, Carlos Hobeich, Natália Comes, Lorena Ramírez, Noemí Gonzalez, Silvia Sauleda, Maria Glòria Soria |
| EPI_ISL_3099592 | Fundacio Althaia-Manresa | Banc de Sang i Teixits | Francisco Vidal, Irene Corrales, Nina Borràs, Carlos Hobeich, Natália Comes, Lorena Ramírez, Noemí Gonzalez, Silvia Sauleda, Maria Glòria Soria |
| EPI_ISL_3099594 | CAP Tona | Banc de Sang i Teixits | Francisco Vidal, Irene Corrales, Nina Borràs, Carlos Hobeich, Natália Comes, Lorena Ramírez, Noemí Gonzalez, Silvia Sauleda, Maria Glòria Soria |
| EPI_ISL_3099597 | CAP Taradell | Banc de Sang i Teixits | Francisco Vidal, Irene Corrales, Nina Borràs, Carlos Hobeich, Natália Comes, Lorena Ramírez, Noemí Gonzalez, Silvia Sauleda, Maria Glòria Soria |
| EPI_ISL_3099598 | Fundacio Althaia-Manresa | Banc de Sang i Teixits | Francisco Vidal, Irene Corrales, Nina Borràs, Carlos Hobeich, Natália Comes, Lorena Ramírez, Noemí Gonzalez, Silvia Sauleda, Maria Glòria Soria |
| EPI_ISL_3099599 | CAP Tona | Banc de Sang i Teixits | Francisco Vidal, Irene Corrales, Nina Borràs, Carlos Hobeich, Natália Comes, Lorena Ramírez, Noemí Gonzalez, Silvia Sauleda, Maria Glòria Soria |
| EPI_ISL_3099600 | CAP Sant Julia de Vilatorta | Banc de Sang i Teixits | Francisco Vidal, Irene Corrales, Nina Borràs, Carlos Hobeich, Natália Comes, Lorena Ramírez, Noemí Gonzalez, Silvia Sauleda, Maria Glòria Soria |
| EPI_ISL_3099601 | Fundacio Althaia-Manresa | Banc de Sang i Teixits | Francisco Vidal, Irene Corrales, Nina Borràs, Carlos Hobeich, Natália Comes, Lorena Ramírez, Noemí Gonzalez, Silvia Sauleda, Maria Glòria Soria |
| EPI_ISL_3099602 | CAP El Remei | Banc de Sang i Teixits | Francisco Vidal, Irene Corrales, Nina Borràs, Carlos Hobeich, Natália Comes, Lorena Ramírez, Noemí Gonzalez, Silvia Sauleda, Maria Glòria Soria |
| EPI_ISL_3099603 | Fundacio Althaia-Manresa | Banc de Sang i Teixits | Francisco Vidal, Irene Corrales, Nina Borràs, Carlos Hobeich, Natália Comes, Lorena Ramírez, Noemí Gonzalez, Silvia Sauleda, Maria Glòria Soria |
| EPI_ISL_3099605 | CAP Sant Quirze de Besora | Banc de Sang i Teixits | Francisco Vidal, Irene Corrales, Nina Borràs, Carlos Hobeich, Natália Comes, Lorena Ramírez, Noemí Gonzalez, Silvia Sauleda, Maria Glòria Soria |
| EPI_ISL_3099606 | CAP Vic Nord | Banc de Sang i Teixits | Francisco Vidal, Irene Corrales, Nina Borràs, Carlos Hobeich, Natália Comes, Lorena Ramírez, Noemí Gonzalez, Silvia Sauleda, Maria Glòria Soria |
| EPI_ISL_3099607 | NA | Banc de Sang i Teixits | Francisco Vidal, Irene Corrales, Nina Borràs, Carlos Hobeich, Natália Comes, Lorena Ramírez, Noemí Gonzalez, Silvia Sauleda, Maria Glòria Soria |
| EPI_ISL_3099608 | CAP Santa Eugenia de Berga | Banc de Sang i Teixits | Francisco Vidal, Irene Corrales, Nina Borràs, Carlos Hobeich, Natália Comes, Lorena Ramírez, Noemí Gonzalez, Silvia Sauleda, Maria Glòria Soria |
| EPI_ISL_3099609 | CAP Manlleu | Banc de Sang i Teixits | Francisco Vidal, Irene Corrales, Nina Borràs, Carlos Hobeich, Natália Comes, Lorena Ramírez, Noemí Gonzalez, Silvia Sauleda, Maria Glòria Soria |
| EPI_ISL_3099610 | CAP Sant Hipolit de Voltrega | Banc de Sang i Teixits | Francisco Vidal, Irene Corrales, Nina Borràs, Carlos Hobeich, Natália Comes, Lorena Ramírez, Noemí Gonzalez, Silvia Sauleda, Maria Glòria Soria |
| EPI_ISL_3099611, EPI_ISL_3099612 | CAP Tona | Banc de Sang i Teixits | Francisco Vidal, Irene Corrales, Nina Borràs, Carlos Hobeich, Natália Comes, Lorena Ramírez, Noemí Gonzalez, Silvia Sauleda, Maria Glòria Soria |
| EPI_ISL_3099613, EPI_ISL_3099614, EPI_ISL_3099615, EPI_ISL_3099616 | Fundacio Althaia-Manresa | Banc de Sang i Teixits | Francisco Vidal, Irene Corrales, Nina Borràs, Carlos Hobeich, Natália Comes, Lorena Ramírez, Noemí Gonzalez, Silvia Sauleda, Maria Glòria Soria |
| EPI_ISL_3099617 | CAP Montesquiu | Banc de Sang i Teixits | Francisco Vidal, Irene Corrales, Nina Borràs, Carlos Hobeich, Natália Comes, Lorena Ramírez, Noemí Gonzalez, Silvia Sauleda, Maria Glòria Soria |
| EPI_ISL_3099618, EPI_ISL_3099619 | Fundacio Althaia-Manresa | Banc de Sang i Teixits | Francisco Vidal, Irene Corrales, Nina Borràs, Carlos Hobeich, Natália Comes, Lorena Ramírez, Noemí Gonzalez, Silvia Sauleda, Maria Glòria Soria |
| EPI_ISL_3099620 | CAP Tona | Banc de Sang i Teixits | Francisco Vidal, Irene Corrales, Nina Borràs, Carlos Hobeich, Natália Comes, Lorena Ramírez, Noemí Gonzalez, Silvia Sauleda, Maria Glòria Soria |
| EPI_ISL_3099622 | Fundacio Althaia-Manresa | Banc de Sang i Teixits | Francisco Vidal, Irene Corrales, Nina Borràs, Carlos Hobeich, Natália Comes, Lorena Ramírez, Noemí Gonzalez, Silvia Sauleda, Maria Glòria Soria |
| EPI_ISL_3099623 | CAP Manlleu | Banc de Sang i Teixits | Francisco Vidal, Irene Corrales, Nina Borràs, Carlos Hobeich, Natália Comes, Lorena Ramírez, Noemí Gonzalez, Silvia Sauleda, Maria Glòria Soria |
| EPI_ISL_3099624 | Fundacio Althaia-Manresa | Banc de Sang i Teixits | Francisco Vidal, Irene Corrales, Nina Borràs, Carlos Hobeich, Natália Comes, Lorena Ramírez, Noemí Gonzalez, Silvia Sauleda, Maria Glòria Soria |
| EPI_ISL_3099625, EPI_ISL_3099627 | CAP Manlleu | Banc de Sang i Teixits | Francisco Vidal, Irene Corrales, Nina Borràs, Carlos Hobeich, Natália Comes, Lorena Ramírez, Noemí Gonzalez, Silvia Sauleda, Maria Glòria Soria |
| EPI_ISL_3099628 | CAP Sant Hipolit de Voltrega | Banc de Sang i Teixits | Francisco Vidal, Irene Corrales, Nina Borràs, Carlos Hobeich, Natália Comes, Lorena Ramírez, Noemí Gonzalez, Silvia Sauleda, Maria Glòria Soria |
| EPI_ISL_3099629 | CAP Roda de Ter | Banc de Sang i Teixits | Francisco Vidal, Irene Corrales, Nina Borràs, Carlos Hobeich, Natália Comes, Lorena Ramírez, Noemí Gonzalez, Silvia Sauleda, Maria Glòria Soria |

| | | | |
|---|---|---|---|
| EPI_ISL_3099630 | CAP Prats de Llucanes | Banc de Sang i Teixits | Francisco Vidal, Irene Corrales, Nina Borràs, Carlos Hobeich, Natália Comes, Lorena Ramírez, Noemí Gonzalez, Silvia Sauleda, Maria Glòria Soria |
| EPI_ISL_3099631 | CAP Viladrau | Banc de Sang i Teixits | Francisco Vidal, Irene Corrales, Nina Borràs, Carlos Hobeich, Natália Comes, Lorena Ramírez, Noemí Gonzalez, Silvia Sauleda, Maria Glòria Soria |
| EPI_ISL_3099632 | Fundacio Althaia-Manresa | Banc de Sang i Teixits | Francisco Vidal, Irene Corrales, Nina Borràs, Carlos Hobeich, Natália Comes, Lorena Ramírez, Noemí Gonzalez, Silvia Sauleda, Maria Glòria Soria |
| EPI_ISL_3099633 | CAP Viladrau | Banc de Sang i Teixits | Francisco Vidal, Irene Corrales, Nina Borràs, Carlos Hobeich, Natália Comes, Lorena Ramírez, Noemí Gonzalez, Silvia Sauleda, Maria Glòria Soria |
| EPI_ISL_3099634 | CAP Manlleu | Banc de Sang i Teixits | Francisco Vidal, Irene Corrales, Nina Borràs, Carlos Hobeich, Natália Comes, Lorena Ramírez, Noemí Gonzalez, Silvia Sauleda, Maria Glòria Soria |
| EPI_ISL_3099635 | Fundacio Althaia-Manresa | Banc de Sang i Teixits | Francisco Vidal, Irene Corrales, Nina Borràs, Carlos Hobeich, Natália Comes, Lorena Ramírez, Noemí Gonzalez, Silvia Sauleda, Maria Glòria Soria |
| EPI_ISL_3099636 | CAP Vic Nord | Banc de Sang i Teixits | Francisco Vidal, Irene Corrales, Nina Borràs, Carlos Hobeich, Natália Comes, Lorena Ramírez, Noemí Gonzalez, Silvia Sauleda, Maria Glòria Soria |
| EPI_ISL_3099637, EPI_ISL_3099638 | Fundacio Althaia-Manresa | Banc de Sang i Teixits | Francisco Vidal, Irene Corrales, Nina Borràs, Carlos Hobeich, Natália Comes, Lorena Ramírez, Noemí Gonzalez, Silvia Sauleda, Maria Glòria Soria |
| EPI_ISL_3099639 | CAP Sant Quirze de Besora | Banc de Sang i Teixits | Francisco Vidal, Irene Corrales, Nina Borràs, Carlos Hobeich, Natália Comes, Lorena Ramírez, Noemí Gonzalez, Silvia Sauleda, Maria Glòria Soria |
| EPI_ISL_3099640, EPI_ISL_3099641 | CAP Vic Nord | Banc de Sang i Teixits | Francisco Vidal, Irene Corrales, Nina Borràs, Carlos Hobeich, Natália Comes, Lorena Ramírez, Noemí Gonzalez, Silvia Sauleda, Maria Glòria Soria |
| EPI_ISL_3099642, EPI_ISL_3099643 | Fundacio Althaia-Manresa | Banc de Sang i Teixits | Francisco Vidal, Irene Corrales, Nina Borràs, Carlos Hobeich, Natália Comes, Lorena Ramírez, Noemí Gonzalez, Silvia Sauleda, Maria Glòria Soria |
| EPI_ISL_3099644 | CAP Torello | Banc de Sang i Teixits | Francisco Vidal, Irene Corrales, Nina Borràs, Carlos Hobeich, Natália Comes, Lorena Ramírez, Noemí Gonzalez, Silvia Sauleda, Maria Glòria Soria |
| EPI_ISL_3099646 | Fundacio Althaia-Manresa | Banc de Sang i Teixits | Francisco Vidal, Irene Corrales, Nina Borràs, Carlos Hobeich, Natália Comes, Lorena Ramírez, Noemí Gonzalez, Silvia Sauleda, Maria Glòria Soria |
| EPI_ISL_3099647 | CAP Manlleu | Banc de Sang i Teixits | Francisco Vidal, Irene Corrales, Nina Borràs, Carlos Hobeich, Natália Comes, Lorena Ramírez, Noemí Gonzalez, Silvia Sauleda, Maria Glòria Soria |
| EPI_ISL_3099648 | Fundacio Althaia-Manresa | Banc de Sang i Teixits | Francisco Vidal, Irene Corrales, Nina Borràs, Carlos Hobeich, Natália Comes, Lorena Ramírez, Noemí Gonzalez, Silvia Sauleda, Maria Glòria Soria |
| EPI_ISL_3099649 | CAP Manlleu | Banc de Sang i Teixits | Francisco Vidal, Irene Corrales, Nina Borràs, Carlos Hobeich, Natália Comes, Lorena Ramírez, Noemí Gonzalez, Silvia Sauleda, Maria Glòria Soria |
| EPI_ISL_3099650, EPI_ISL_3099651 | Fundacio Althaia-Manresa | Banc de Sang i Teixits | Francisco Vidal, Irene Corrales, Nina Borràs, Carlos Hobeich, Natália Comes, Lorena Ramírez, Noemí Gonzalez, Silvia Sauleda, Maria Glòria Soria |
| EPI_ISL_3099652 | CAP El Remei | Banc de Sang i Teixits | Francisco Vidal, Irene Corrales, Nina Borràs, Carlos Hobeich, Natália Comes, Lorena Ramírez, Noemí Gonzalez, Silvia Sauleda, Maria Glòria Soria |
| EPI_ISL_3099653, EPI_ISL_3099654 | Fundacio Althaia-Manresa | Banc de Sang i Teixits | Francisco Vidal, Irene Corrales, Nina Borràs, Carlos Hobeich, Natália Comes, Lorena Ramírez, Noemí Gonzalez, Silvia Sauleda, Maria Glòria Soria |
| EPI_ISL_3099656 | CAP Torello | Banc de Sang i Teixits | Francisco Vidal, Irene Corrales, Nina Borràs, Carlos Hobeich, Natália Comes, Lorena Ramírez, Noemí Gonzalez, Silvia Sauleda, Maria Glòria Soria |
| EPI_ISL_3099657 | Fundacio Althaia-Manresa | Banc de Sang i Teixits | Francisco Vidal, Irene Corrales, Nina Borràs, Carlos Hobeich, Natália Comes, Lorena Ramírez, Noemí Gonzalez, Silvia Sauleda, Maria Glòria Soria |
| EPI_ISL_3099658 | CAP Olost de Llucanes | Banc de Sang i Teixits | Francisco Vidal, Irene Corrales, Nina Borràs, Carlos Hobeich, Natália Comes, Lorena Ramírez, Noemí Gonzalez, Silvia Sauleda, Maria Glòria Soria |
| EPI_ISL_3099659 | CAP Sant Hipolit de Voltrega | Banc de Sang i Teixits | Francisco Vidal, Irene Corrales, Nina Borràs, Carlos Hobeich, Natália Comes, Lorena Ramírez, Noemí Gonzalez, Silvia Sauleda, Maria Glòria Soria |
| EPI_ISL_3099660 | CAP Tona | Banc de Sang i Teixits | Francisco Vidal, Irene Corrales, Nina Borràs, Carlos Hobeich, Natália Comes, Lorena Ramírez, Noemí Gonzalez, Silvia Sauleda, Maria Glòria Soria |
| EPI_ISL_3099661 | CAP Folgueroles | Banc de Sang i Teixits | Francisco Vidal, Irene Corrales, Nina Borràs, Carlos Hobeich, Natália Comes, Lorena Ramírez, Noemí Gonzalez, Silvia Sauleda, Maria Glòria Soria |
| EPI_ISL_3099662 | Fundacio Althaia-Manresa | Banc de Sang i Teixits | Francisco Vidal, Irene Corrales, Nina Borràs, Carlos Hobeich, Natália Comes, Lorena Ramírez, Noemí Gonzalez, Silvia Sauleda, Maria Glòria Soria |
| EPI_ISL_3099663, EPI_ISL_3099664 | CAP El Remei | Banc de Sang i Teixits | Francisco Vidal, Irene Corrales, Nina Borràs, Carlos Hobeich, Natália Comes, Lorena Ramírez, Noemí Gonzalez, Silvia Sauleda, Maria Glòria Soria |
| EPI_ISL_3099665, EPI_ISL_3099666 | CAP Vic Nord | Banc de Sang i Teixits | Francisco Vidal, Irene Corrales, Nina Borràs, Carlos Hobeich, Natália Comes, Lorena Ramírez, Noemí Gonzalez, Silvia Sauleda, Maria Glòria Soria |
| EPI_ISL_3099667 | Fundacio Althaia-Manresa | Banc de Sang i Teixits | Francisco Vidal, Irene Corrales, Nina Borràs, Carlos Hobeich, Natália Comes, Lorena Ramírez, Noemí Gonzalez, Silvia Sauleda, Maria Glòria Soria |
| EPI_ISL_3099668 | CAP Vic Nord | Banc de Sang i Teixits | Francisco Vidal, Irene Corrales, Nina Borràs, Carlos Hobeich, Natália Comes, Lorena Ramírez, Noemí Gonzalez, Silvia Sauleda, Maria Glòria Soria |
| EPI_ISL_3099669 | CAP Taradell | Banc de Sang i Teixits | Francisco Vidal, Irene Corrales, Nina Borràs, Carlos Hobeich, Natália Comes, Lorena Ramírez, Noemí Gonzalez, Silvia Sauleda, Maria Glòria Soria |
| EPI_ISL_3099670 | CAP Tona | Banc de Sang i Teixits | Francisco Vidal, Irene Corrales, Nina Borràs, Carlos Hobeich, Natália Comes, Lorena Ramírez, Noemí Gonzalez, Silvia Sauleda, Maria Glòria Soria |
| EPI_ISL_3099671, EPI_ISL_3099672 | Fundacio Althaia-Manresa | Banc de Sang i Teixits | Francisco Vidal, Irene Corrales, Nina Borràs, Carlos Hobeich, Natália Comes, Lorena Ramírez, Noemí Gonzalez, Silvia Sauleda, Maria Glòria Soria |
| EPI_ISL_3099674 | Ochsner Health | Bioinfoexperts, LLC | Judy Crabtree, Amy Feehan, Rebecca Rose, David J. Nolan, Tessa LaFleur, Julia-Garcia-Diaz, Lucio Miele, Susanna L. Lamers |
| EPI_ISL_3099675, EPI_ISL_3099676 | Fundacio Althaia-Manresa | Banc de Sang i Teixits | Francisco Vidal, Irene Corrales, Nina Borràs, Carlos Hobeich, Natália Comes, Lorena Ramírez, Noemí Gonzalez, Silvia Sauleda, Maria Glòria Soria |
| EPI_ISL_3099677 | CAP Sant Hipolit de Voltrega | Banc de Sang i Teixits | Francisco Vidal, Irene Corrales, Nina Borràs, Carlos Hobeich, Natália Comes, Lorena Ramírez, Noemí Gonzalez, Silvia Sauleda, Maria Glòria Soria |
| EPI_ISL_3099678 | CAP Les Hortes | Banc de Sang i Teixits | Francisco Vidal, Irene Corrales, Nina Borràs, Carlos Hobeich, Natália Comes, Lorena Ramírez, Noemí Gonzalez, Silvia Sauleda, Maria Glòria Soria |
| EPI_ISL_3099679 | CAP Cotxeres | Banc de Sang i Teixits | Francisco Vidal, Irene Corrales, Nina Borràs, Carlos Hobeich, Natália Comes, Lorena Ramírez, Noemí Gonzalez, Silvia Sauleda, Maria Glòria Soria |
| EPI_ISL_3099680 | CAP Sant Quirze de Besora | Banc de Sang i Teixits | Francisco Vidal, Irene Corrales, Nina Borràs, Carlos Hobeich, Natália Comes, Lorena Ramírez, Noemí Gonzalez, Silvia Sauleda, Maria Glòria Soria |
| EPI_ISL_3099681 | CAP Sagrada Familia | Banc de Sang i Teixits | Francisco Vidal, Irene Corrales, Nina Borràs, Carlos Hobeich, Natália Comes, Lorena Ramírez, Noemí Gonzalez, Silvia Sauleda, Maria Glòria Soria |
| EPI_ISL_3099682 | CAP Sarria | Banc de Sang i Teixits | Francisco Vidal, Irene Corrales, Nina Borràs, Carlos Hobeich, Natália Comes, Lorena Ramírez, Noemí Gonzalez, Silvia Sauleda, Maria Glòria Soria |
| EPI_ISL_3099683 | Fundacio Althaia-Manresa | Banc de Sang i Teixits | Francisco Vidal, Irene Corrales, Nina Borràs, Carlos Hobeich, Natália Comes, Lorena Ramírez, Noemí Gonzalez, Silvia Sauleda, Maria Glòria Soria |
| EPI_ISL_3099684 | CAP Tona | Banc de Sang i Teixits | Francisco Vidal, Irene Corrales, Nina Borràs, Carlos Hobeich, Natália Comes, Lorena Ramírez, Noemí Gonzalez, Silvia Sauleda, Maria Glòria Soria |
| EPI_ISL_3099685 | CAP Sagrada Familia | Banc de Sang i Teixits | Francisco Vidal, Irene Corrales, Nina Borràs, Carlos Hobeich, Natália Comes, Lorena Ramírez, Noemí Gonzalez, Silvia Sauleda, Maria Glòria Soria |
| EPI_ISL_3099686 | Fundacio Althaia-Manresa | Banc de Sang i Teixits | Francisco Vidal, Irene Corrales, Nina Borràs, Carlos Hobeich, Natália Comes, Lorena Ramírez, Noemí Gonzalez, Silvia Sauleda, Maria Glòria Soria |
| EPI_ISL_3099687 | CAP Sant Vicenc de Torello | Banc de Sang i Teixits | Francisco Vidal, Irene Corrales, Nina Borràs, Carlos Hobeich, Natália Comes, Lorena Ramírez, Noemí Gonzalez, Silvia Sauleda, Maria Glòria Soria |
| EPI_ISL_3099688 | Fundacio Althaia-Manresa | Banc de Sang i Teixits | Francisco Vidal, Irene Corrales, Nina Borràs, Carlos Hobeich, Natália Comes, Lorena Ramírez, Noemí Gonzalez, Silvia Sauleda, Maria Glòria Soria |
| EPI_ISL_3099689 | CAP Roger de Flor | Banc de Sang i Teixits | Francisco Vidal, Irene Corrales, Nina Borràs, Carlos Hobeich, Natália Comes, Lorena Ramírez, Noemí Gonzalez, Silvia Sauleda, Maria Glòria Soria |
| EPI_ISL_3099690 | CAP Sagrada Familia | Banc de Sang i Teixits | Francisco Vidal, Irene Corrales, Nina Borràs, Carlos Hobeich, Natália Comes, Lorena Ramírez, Noemí Gonzalez, Silvia Sauleda, Maria Glòria Soria |
| EPI_ISL_3099691 | CAP Les Hortes | Banc de Sang i Teixits | Francisco Vidal, Irene Corrales, Nina Borràs, Carlos Hobeich, Natália Comes, Lorena Ramírez, Noemí Gonzalez, Silvia Sauleda, Maria Glòria Soria |
| EPI_ISL_3099692 | CAP Roger de Flor | Banc de Sang i Teixits | Francisco Vidal, Irene Corrales, Nina Borràs, Carlos Hobeich, Natália Comes, Lorena Ramírez, Noemí Gonzalez, Silvia Sauleda, Maria Glòria Soria |
| EPI_ISL_3099693 | CAP Prats de Llucanes | Banc de Sang i Teixits | Francisco Vidal, Irene Corrales, Nina Borràs, Carlos Hobeich, Natália Comes, Lorena Ramírez, Noemí Gonzalez, Silvia Sauleda, Maria Glòria Soria |
| EPI_ISL_3099694 | CAP Torello | Banc de Sang i Teixits | Francisco Vidal, Irene Corrales, Nina Borràs, Carlos Hobeich, Natália Comes, Lorena Ramírez, Noemí Gonzalez, Silvia Sauleda, Maria Glòria Soria |
| EPI_ISL_3099695 | CAP Roger de Flor | Banc de Sang i Teixits | Francisco Vidal, Irene Corrales, Nina Borràs, Carlos Hobeich, Natália Comes, Lorena Ramírez, Noemí Gonzalez, Silvia Sauleda, Maria Glòria Soria |
| EPI_ISL_3099696 | CAP Sagrada Familia | Banc de Sang i Teixits | Francisco Vidal, Irene Corrales, Nina Borràs, Carlos Hobeich, Natália Comes, Lorena Ramírez, Noemí Gonzalez, Silvia Sauleda, Maria Glòria Soria |
| EPI_ISL_3099697 | CAP Vic Nord | Banc de Sang i Teixits | Francisco Vidal, Irene Corrales, Nina Borràs, Carlos Hobeich, Natália Comes, Lorena Ramírez, Noemí Gonzalez, Silvia Sauleda, Maria Glòria Soria |
| EPI_ISL_3099698 | Fundacio Althaia-Manresa | Banc de Sang i Teixits | Francisco Vidal, Irene Corrales, Nina Borràs, Carlos Hobeich, Natália Comes, Lorena Ramírez, Noemí Gonzalez, Silvia Sauleda, Maria Glòria Soria |
| EPI_ISL_3099699 | CAP El Remei | Banc de Sang i Teixits | Francisco Vidal, Irene Corrales, Nina Borràs, Carlos Hobeich, Natália Comes, Lorena Ramírez, Noemí Gonzalez, Silvia Sauleda, Maria Glòria Soria |
| EPI_ISL_3099700 | Fundacio Althaia-Manresa | Banc de Sang i Teixits | Francisco Vidal, Irene Corrales, Nina Borràs, Carlos Hobeich, Natália Comes, Lorena Ramírez, Noemí Gonzalez, Silvia Sauleda, Maria Glòria Soria |
| EPI_ISL_3099701 | CAP Roda de Ter | Banc de Sang i Teixits | Francisco Vidal, Irene Corrales, Nina Borràs, Carlos Hobeich, Natália Comes, Lorena Ramírez, Noemí Gonzalez, Silvia Sauleda, Maria Glòria Soria |
| EPI_ISL_3099702 | CAP Tona | Banc de Sang i Teixits | Francisco Vidal, Irene Corrales, Nina Borràs, Carlos Hobeich, Natália Comes, Lorena Ramírez, Noemí Gonzalez, Silvia Sauleda, Maria Glòria Soria |
| EPI_ISL_3099703 | CAP Les Hortes | Banc de Sang i Teixits | Francisco Vidal, Irene Corrales, Nina Borràs, Carlos Hobeich, Natália Comes, Lorena Ramírez, Noemí Gonzalez, Silvia Sauleda, Maria Glòria Soria |
| EPI_ISL_3099704 | CAP Cotxeres | Banc de Sang i Teixits | Francisco Vidal, Irene Corrales, Nina Borràs, Carlos Hobeich, Natália Comes, Lorena Ramírez, Noemí Gonzalez, Silvia Sauleda, Maria Glòria Soria |
| EPI_ISL_3099705 | Fundacio Althaia-Manresa | Banc de Sang i Teixits | Francisco Vidal, Irene Corrales, Nina Borràs, Carlos Hobeich, Natália Comes, Lorena Ramírez, Noemí Gonzalez, Silvia Sauleda, Maria Glòria Soria |
| EPI_ISL_3099706 | CAP Les Hortes | Banc de Sang i Teixits | Francisco Vidal, Irene Corrales, Nina Borràs, Carlos Hobeich, Natália Comes, Lorena Ramírez, Noemí Gonzalez, Silvia Sauleda, Maria Glòria Soria |
| EPI_ISL_3099707 | CAP Taradell | Banc de Sang i Teixits | Francisco Vidal, Irene Corrales, Nina Borràs, Carlos Hobeich, Natália Comes, Lorena Ramírez, Noemí Gonzalez, Silvia Sauleda, Maria Glòria Soria |
| EPI_ISL_3099709, EPI_ISL_3099710 | Fundacio Althaia-Manresa | Banc de Sang i Teixits | Francisco Vidal, Irene Corrales, Nina Borràs, Carlos Hobeich, Natália Comes, Lorena Ramírez, Noemí Gonzalez, Silvia Sauleda, Maria Glòria Soria |
| EPI_ISL_3099711 | CAP Roda de Ter | Banc de Sang i Teixits | Francisco Vidal, Irene Corrales, Nina Borràs, Carlos Hobeich, Natália Comes, Lorena Ramírez, Noemí Gonzalez, Silvia Sauleda, Maria Glòria Soria |
| EPI_ISL_3099712 | CAP Les Hortes | Banc de Sang i Teixits | Francisco Vidal, Irene Corrales, Nina Borràs, Carlos Hobeich, Natália Comes, Lorena Ramírez, Noemí Gonzalez, Silvia Sauleda, Maria Glòria Soria |

| | | | |
|---|---|---|---|
| EPI_ISL_3099713, EPI_ISL_3099714 | CAP Roda de Ter | Banc de Sang i Teixits | Francisco Vidal, Irene Corrales, Nina Borràs, Carlos Hobeich, Natália Comes, Lorena Ramírez, Noemí Gonzalez, Silvia Sauleda, Maria Glòria Soria |
| EPI_ISL_3099715 | CAP Les Hortes | Banc de Sang i Teixits | Francisco Vidal, Irene Corrales, Nina Borràs, Carlos Hobeich, Natália Comes, Lorena Ramírez, Noemí Gonzalez, Silvia Sauleda, Maria Glòria Soria |
| EPI_ISL_3099717 | CAP Sarria | Banc de Sang i Teixits | Francisco Vidal, Irene Corrales, Nina Borràs, Carlos Hobeich, Natália Comes, Lorena Ramírez, Noemí Gonzalez, Silvia Sauleda, Maria Glòria Soria |
| EPI_ISL_3099718 | CAP Manlleu | Banc de Sang i Teixits | Francisco Vidal, Irene Corrales, Nina Borràs, Carlos Hobeich, Natália Comes, Lorena Ramírez, Noemí Gonzalez, Silvia Sauleda, Maria Glòria Soria |
| EPI_ISL_3099719 | Fundacio Althaia-Manresa | Banc de Sang i Teixits | Francisco Vidal, Irene Corrales, Nina Borràs, Carlos Hobeich, Natália Comes, Lorena Ramírez, Noemí Gonzalez, Silvia Sauleda, Maria Glòria Soria |
| EPI_ISL_3099720 | CAP Roda de Ter | Banc de Sang i Teixits | Francisco Vidal, Irene Corrales, Nina Borràs, Carlos Hobeich, Natália Comes, Lorena Ramírez, Noemí Gonzalez, Silvia Sauleda, Maria Glòria Soria |
| EPI_ISL_3099721 | CAP Sant Julia de Vilatorta | Banc de Sang i Teixits | Francisco Vidal, Irene Corrales, Nina Borràs, Carlos Hobeich, Natália Comes, Lorena Ramírez, Noemí Gonzalez, Silvia Sauleda, Maria Glòria Soria |
| EPI_ISL_3099722 | Fundacio Althaia-Manresa | Banc de Sang i Teixits | Francisco Vidal, Irene Corrales, Nina Borràs, Carlos Hobeich, Natália Comes, Lorena Ramírez, Noemí Gonzalez, Silvia Sauleda, Maria Glòria Soria |
| EPI_ISL_3099723 | CAP Sant Quirze de Besora | Banc de Sang i Teixits | Francisco Vidal, Irene Corrales, Nina Borràs, Carlos Hobeich, Natália Comes, Lorena Ramírez, Noemí Gonzalez, Silvia Sauleda, Maria Glòria Soria |
| EPI_ISL_3099724, EPI_ISL_3099725, EPI_ISL_3099726 | CAP Sagrada Familia | Banc de Sang i Teixits | Francisco Vidal, Irene Corrales, Nina Borràs, Carlos Hobeich, Natália Comes, Lorena Ramírez, Noemí Gonzalez, Silvia Sauleda, Maria Glòria Soria |
| EPI_ISL_3099727 | Fundacio Althaia-Manresa | Banc de Sang i Teixits | Francisco Vidal, Irene Corrales, Nina Borràs, Carlos Hobeich, Natália Comes, Lorena Ramírez, Noemí Gonzalez, Silvia Sauleda, Maria Glòria Soria |
| EPI_ISL_3099728 | CAP Les Hortes | Banc de Sang i Teixits | Francisco Vidal, Irene Corrales, Nina Borràs, Carlos Hobeich, Natália Comes, Lorena Ramírez, Noemí Gonzalez, Silvia Sauleda, Maria Glòria Soria |
| EPI_ISL_3099729 | CAP Tona | Banc de Sang i Teixits | Francisco Vidal, Irene Corrales, Nina Borràs, Carlos Hobeich, Natália Comes, Lorena Ramírez, Noemí Gonzalez, Silvia Sauleda, Maria Glòria Soria |
| EPI_ISL_3099730 | CAP El Remei | Banc de Sang i Teixits | Francisco Vidal, Irene Corrales, Nina Borràs, Carlos Hobeich, Natália Comes, Lorena Ramírez, Noemí Gonzalez, Silvia Sauleda, Maria Glòria Soria |
| EPI_ISL_3099731 | CAP Roger de Flor | Banc de Sang i Teixits | Francisco Vidal, Irene Corrales, Nina Borràs, Carlos Hobeich, Natália Comes, Lorena Ramírez, Noemí Gonzalez, Silvia Sauleda, Maria Glòria Soria |
| EPI_ISL_3099732 | Fundacio Althaia-Manresa | Banc de Sang i Teixits | Francisco Vidal, Irene Corrales, Nina Borràs, Carlos Hobeich, Natália Comes, Lorena Ramírez, Noemí Gonzalez, Silvia Sauleda, Maria Glòria Soria |
| EPI_ISL_3099733 | CAP Les Hortes | Banc de Sang i Teixits | Francisco Vidal, Irene Corrales, Nina Borràs, Carlos Hobeich, Natália Comes, Lorena Ramírez, Noemí Gonzalez, Silvia Sauleda, Maria Glòria Soria |
| EPI_ISL_3099734 | CAP Vic Nord | Banc de Sang i Teixits | Francisco Vidal, Irene Corrales, Nina Borràs, Carlos Hobeich, Natália Comes, Lorena Ramírez, Noemí Gonzalez, Silvia Sauleda, Maria Glòria Soria |
| EPI_ISL_3099736 | CAP Sant Julia de Vilatorta | Banc de Sang i Teixits | Francisco Vidal, Irene Corrales, Nina Borràs, Carlos Hobeich, Natália Comes, Lorena Ramírez, Noemí Gonzalez, Silvia Sauleda, Maria Glòria Soria |
| EPI_ISL_3099738 | CAP Sarria | Banc de Sang i Teixits | Francisco Vidal, Irene Corrales, Nina Borràs, Carlos Hobeich, Natália Comes, Lorena Ramírez, Noemí Gonzalez, Silvia Sauleda, Maria Glòria Soria |
| EPI_ISL_3099739 | CAP Sant Julia de Vilatorta | Banc de Sang i Teixits | Francisco Vidal, Irene Corrales, Nina Borràs, Carlos Hobeich, Natália Comes, Lorena Ramírez, Noemí Gonzalez, Silvia Sauleda, Maria Glòria Soria |
| EPI_ISL_3099740 | CAP Les Hortes | Banc de Sang i Teixits | Francisco Vidal, Irene Corrales, Nina Borràs, Carlos Hobeich, Natália Comes, Lorena Ramírez, Noemí Gonzalez, Silvia Sauleda, Maria Glòria Soria |
| EPI_ISL_3099741 | CAP Sarria | Banc de Sang i Teixits | Francisco Vidal, Irene Corrales, Nina Borràs, Carlos Hobeich, Natália Comes, Lorena Ramírez, Noemí Gonzalez, Silvia Sauleda, Maria Glòria Soria |
| EPI_ISL_3099742 | CAP Roda de Ter | Banc de Sang i Teixits | Francisco Vidal, Irene Corrales, Nina Borràs, Carlos Hobeich, Natália Comes, Lorena Ramírez, Noemí Gonzalez, Silvia Sauleda, Maria Glòria Soria |
| EPI_ISL_3099743 | CAP Les Hortes | Banc de Sang i Teixits | Francisco Vidal, Irene Corrales, Nina Borràs, Carlos Hobeich, Natália Comes, Lorena Ramírez, Noemí Gonzalez, Silvia Sauleda, Maria Glòria Soria |
| EPI_ISL_3099745 | CAP Manlleu | Banc de Sang i Teixits | Francisco Vidal, Irene Corrales, Nina Borràs, Carlos Hobeich, Natália Comes, Lorena Ramírez, Noemí Gonzalez, Silvia Sauleda, Maria Glòria Soria |
| EPI_ISL_3099746 | CAP Sagrada Familia | Banc de Sang i Teixits | Francisco Vidal, Irene Corrales, Nina Borràs, Carlos Hobeich, Natália Comes, Lorena Ramírez, Noemí Gonzalez, Silvia Sauleda, Maria Glòria Soria |
| EPI_ISL_3099747 | CAP Cotxeres | Banc de Sang i Teixits | Francisco Vidal, Irene Corrales, Nina Borràs, Carlos Hobeich, Natália Comes, Lorena Ramírez, Noemí Gonzalez, Silvia Sauleda, Maria Glòria Soria |
| EPI_ISL_3099748 | CAP Sant Hipolit de Voltrega | Banc de Sang i Teixits | Francisco Vidal, Irene Corrales, Nina Borràs, Carlos Hobeich, Natália Comes, Lorena Ramírez, Noemí Gonzalez, Silvia Sauleda, Maria Glòria Soria |
| EPI_ISL_3099749 | CAP Seva | Banc de Sang i Teixits | Francisco Vidal, Irene Corrales, Nina Borràs, Carlos Hobeich, Natália Comes, Lorena Ramírez, Noemí Gonzalez, Silvia Sauleda, Maria Glòria Soria |
| EPI_ISL_3099750 | CAP Vic Nord | Banc de Sang i Teixits | Francisco Vidal, Irene Corrales, Nina Borràs, Carlos Hobeich, Natália Comes, Lorena Ramírez, Noemí Gonzalez, Silvia Sauleda, Maria Glòria Soria |
| EPI_ISL_3099751 | CAP Sagrada Familia | Banc de Sang i Teixits | Francisco Vidal, Irene Corrales, Nina Borràs, Carlos Hobeich, Natália Comes, Lorena Ramírez, Noemí Gonzalez, Silvia Sauleda, Maria Glòria Soria |
| EPI_ISL_3099753 | Fundacio Althaia-Manresa | Banc de Sang i Teixits | Francisco Vidal, Irene Corrales, Nina Borràs, Carlos Hobeich, Natália Comes, Lorena Ramírez, Noemí Gonzalez, Silvia Sauleda, Maria Glòria Soria |
| EPI_ISL_3099754 | CAP Les Hortes | Banc de Sang i Teixits | Francisco Vidal, Irene Corrales, Nina Borràs, Carlos Hobeich, Natália Comes, Lorena Ramírez, Noemí Gonzalez, Silvia Sauleda, Maria Glòria Soria |
| EPI_ISL_3099755 | CAP Sant Boi de Llucanes | Banc de Sang i Teixits | Francisco Vidal, Irene Corrales, Nina Borràs, Carlos Hobeich, Natália Comes, Lorena Ramírez, Noemí Gonzalez, Silvia Sauleda, Maria Glòria Soria |
| EPI_ISL_3099771, EPI_ISL_3099773, EPI_ISL_3099789 | Wyoming Public Health Laboratory | Wyoming Public Health Laboratory | Jim Mildenberger, Taylor Fearing, Channing Weber, Ashley Norberg, Chayse Rowley, Marley Goetz, Brian Dominguez, Elliot Thomasson, Sam Britz, Cari Sloma, Robert Petit, and Rob Christensen |
| EPI_ISL_3099875, EPI_ISL_3099876, EPI_ISL_3099877, EPI_ISL_3099878, EPI_ISL_3099879, EPI_ISL_3099880, EPI_ISL_3099881, EPI_ISL_3099882, EPI_ISL_3099883, EPI_ISL_3099884, EPI_ISL_3099885, EPI_ISL_3099886, EPI_ISL_3099887, EPI_ISL_3099888, EPI_ISL_3099889 | | | |
| see above | Servicio Microbiología Hospital La Paz | Servicio Microbiología Hospital La Paz | Fernando Lázaro, Rubén Cáceres, Iván Bloise, Jesús Mingorance Cruz, Elie Dahdouh |
| EPI_ISL_3099900, EPI_ISL_3099905, EPI_ISL_3099907, EPI_ISL_3099908, EPI_ISL_3099909, EPI_ISL_3099910, EPI_ISL_3099912, EPI_ISL_3099914, EPI_ISL_3099915, EPI_ISL_3099916, EPI_ISL_3099918, EPI_ISL_3099922, EPI_ISL_3099923, EPI_ISL_3099924, EPI_ISL_3099925, EPI_ISL_3099926, EPI_ISL_3099927, EPI_ISL_3099929, EPI_ISL_3099930 | | | |
| see above | MD PHL | MD PHL | Maryland Department of Health Laboratories Administration |
| EPI_ISL_3099935, EPI_ISL_3099936, EPI_ISL_3099937 | Wichita State University - Molecular Diagnostics Lab | Kansas Health and Environmental Lab | Mike Grose, Katherine Wiggins, Jonathan Barnell, Ben Olsen, and Phil Adam |
| EPI_ISL_3099938, EPI_ISL_3099939, EPI_ISL_3099940, EPI_ISL_3099941, EPI_ISL_3099942, EPI_ISL_3099943, EPI_ISL_3099944, EPI_ISL_3099945, EPI_ISL_3099946, EPI_ISL_3099947, EPI_ISL_3099948, EPI_ISL_3099949, EPI_ISL_3099950, EPI_ISL_3099951, EPI_ISL_3099952, EPI_ISL_3099953, EPI_ISL_3099954, EPI_ISL_3099955, EPI_ISL_3099956, EPI_ISL_3099957, EPI_ISL_3099958, EPI_ISL_3099959, EPI_ISL_3099960, EPI_ISL_3099961, EPI_ISL_3099962, EPI_ISL_3099963, EPI_ISL_3099964, EPI_ISL_3099965, EPI_ISL_3099966 | | | |
| see above | Kansas Health and Environmental Lab | Kansas Health and Environmental Lab | Mike Grose, Katherine Wiggins, Jonathan Barnell, Ben Olsen, and Phil Adam |
| EPI_ISL_3099967, EPI_ISL_3099968, EPI_ISL_3099969, EPI_ISL_3099970, EPI_ISL_3099971, EPI_ISL_3099972, EPI_ISL_3099973, EPI_ISL_3099974, EPI_ISL_3099975, EPI_ISL_3099976, EPI_ISL_3099978, EPI_ISL_3099979, EPI_ISL_3099980, EPI_ISL_3099981, EPI_ISL_3099982, EPI_ISL_3099983, EPI_ISL_3099984, EPI_ISL_3099985, EPI_ISL_3099986, EPI_ISL_3099987, EPI_ISL_3099988, EPI_ISL_3099989, EPI_ISL_3099990, EPI_ISL_3099991, EPI_ISL_3099992, EPI_ISL_3099993, EPI_ISL_3099994, EPI_ISL_3099995, EPI_ISL_3099996, EPI_ISL_3099997, EPI_ISL_3099998, EPI_ISL_3099999, EPI_ISL_3100000, EPI_ISL_3100001, EPI_ISL_3100002, EPI_ISL_3100003, EPI_ISL_3100004, EPI_ISL_3100005, EPI_ISL_3100006, EPI_ISL_3100007 | | | |
| see above | Orange County Public Health Laboratory | Orange County Public Health Laboratory | Angelica Torres, Julia Wolfe |
| EPI_ISL_3100008, EPI_ISL_3100009, EPI_ISL_3100010, EPI_ISL_3100011, EPI_ISL_3100012, EPI_ISL_3100013, EPI_ISL_3100014, EPI_ISL_3100015 | Salzkammergutklinikum Vöcklabruck, Institut für Pathologie | Salzkammergutklinikum Vöcklabruck, Institut für Pathologie | Regina Stitz, Penka Lechner, Senka Rohregger, René Silye |
| EPI_ISL_3100050, EPI_ISL_3100051, EPI_ISL_3100053, EPI_ISL_3100054, EPI_ISL_3100055, EPI_ISL_3100056, EPI_ISL_3100057, EPI_ISL_3100058, EPI_ISL_3100059 | AZ Zeno | AZ Sint Jan | Marijke Reynders, Patrick Descheemaeker, Laurien Hoornaert, Thomas Van Landschoot, Marie Madeleine Chabert-Consen, Jorn Hellemans |
| EPI_ISL_3100060, EPI_ISL_3100061, EPI_ISL_3100062, EPI_ISL_3100063, EPI_ISL_3100064, EPI_ISL_3100065, EPI_ISL_3100066, EPI_ISL_3100067, EPI_ISL_3100068, EPI_ISL_3100069, EPI_ISL_3100070 | | | |
| see above | Center of Advanced Studies and Technology, Molecular Genetics Laboratory | Center of Advanced Studies and Technology, Molecular Genetics Laboratory | Anaclerio Federico, De Fabritiis Simone |
| EPI_ISL_3100071, EPI_ISL_3100072, EPI_ISL_3100074, EPI_ISL_3100075, EPI_ISL_3100078, EPI_ISL_3100079, EPI_ISL_3100081, EPI_ISL_3100082, EPI_ISL_3100083, EPI_ISL_3100084, EPI_ISL_3100086, EPI_ISL_3100087, EPI_ISL_3100088, EPI_ISL_3100089, EPI_ISL_3100090, EPI_ISL_3100091, EPI_ISL_3100092, EPI_ISL_3100093, EPI_ISL_3100094, EPI_ISL_3100095, EPI_ISL_3100096, EPI_ISL_3100097, EPI_ISL_3100098 | | | |
| see above | Klinisch Laboratorium ZNA | Klinisch Laboratorium ZNA | Verstrepen et al. |
| EPI_ISL_3100104, EPI_ISL_3100105, EPI_ISL_3100106, EPI_ISL_3100107, EPI_ISL_3100108, EPI_ISL_3100109, EPI_ISL_3100110, EPI_ISL_3100111, EPI_ISL_3100113, EPI_ISL_3100114, EPI_ISL_3100115, EPI_ISL_3100117, EPI_ISL_3100118, EPI_ISL_3100119, EPI_ISL_3100121, EPI_ISL_3100122 | | | |
| see above | OLVZ Aalst | OLVZ Aalst | Astrid Holderbeke |
| EPI_ISL_3100126 | Tengku Ampuan Afzan Hospital | Institute for Medical Research, Infectious Disease Research Centre, National Institutes of Health, Ministry of Health Malaysia | Suppiah J, Kamel K, Mohd Zawawi Z, Azizan MA, Ramly N, Robert F, Anasir MI, Thayan R |
| EPI_ISL_3100127, EPI_ISL_3100128, EPI_ISL_3100129, EPI_ISL_3100130, EPI_ISL_3100131, EPI_ISL_3100132, EPI_ISL_3100133, EPI_ISL_3100134, EPI_ISL_3100135, EPI_ISL_3100136, EPI_ISL_3100137, EPI_ISL_3100138, EPI_ISL_3100139, EPI_ISL_3100140, EPI_ISL_3100141, EPI_ISL_3100142, EPI_ISL_3100143, EPI_ISL_3100144, EPI_ISL_3100145, EPI_ISL_3100146, EPI_ISL_3100147, EPI_ISL_3100148, EPI_ISL_3100149, EPI_ISL_3100150 | | | |
| see above | Tokyo Metropolitan Institute of Public Health | Tokyo Metropolitan Institute of Public Health | Masaki Hayashi, Takako Yamazaki, Akane Negishi, Hirofumi Miyake, Kenshirou Kuroki, Maya Isogai, Takushi Fujiwara, Mami Nagashima, Jun Suzuki, Kenji Sadamasu |
| EPI_ISL_3100151, | PO AVEZZANO - U.O.C. Laboratorio Analisi | Istituto Zooprofilattico Sperimentale dell'Abruzzo e Molise "G. | Lorusso A, Marcacci M, Di Domenico M, Ancora M, Curini V, Di Lollo Valeria, Mangone I, Rinaldi A, Delli Compagni E, Scialabba S, Caporale M, Di Pasquale A, Cammà C, Puglia I, Calistri P, Savini G |

| | | | |
|---|---|---|---|
| EPI_ISL_3100152 | | Caporale" | |
| EPI_ISL_3100154, EPI_ISL_3100155, EPI_ISL_3100156, EPI_ISL_3100157, EPI_ISL_3100158, EPI_ISL_3100159, EPI_ISL_3100160, EPI_ISL_3100161, EPI_ISL_3100162, EPI_ISL_3100163, EPI_ISL_3100165, EPI_ISL_3100166, EPI_ISL_3100168, EPI_ISL_3100169, EPI_ISL_3100171 | | | |
| see above | OSPEDALE CIVILE TERAMO - CENTRO TRASFUSIONALE | Istituto Zooprofilattico Sperimentale dell'Abruzzo e Molise "G. Caporale" | Lorusso A, Marcacci M, Di Domenico M, Ancora M, Curini V, Di Lollo Valeria, Mangone I, Rinaldi A, Delli Compagni E, Scialabba S, Caporale M, Di Pasquale A, Cammà C, Puglia I, Calistri P, Savini G |
| EPI_ISL_3100172, EPI_ISL_3100173, EPI_ISL_3100174 | Tengku Ampuan Afzan Hospital | Institute for Medical Research, Infectious Disease Research Centre, National Institutes of Health, Ministry of Health Malaysia | Suppiah J, Kamel K, Mohd Zawawi Z, Azizan MA, Ramly N, Robert F, Anasir MI, Thayan R |
| EPI_ISL_3100180 | Sweden/Sodermanland/Unilabs | Unilabs/Eskilstuna/Sweden | Emma Arvidsson |
| EPI_ISL_3100211, EPI_ISL_3100217, EPI_ISL_3100221, EPI_ISL_3100232, EPI_ISL_3100233, EPI_ISL_3100240, EPI_ISL_3100248, EPI_ISL_3100249, EPI_ISL_3100259, EPI_ISL_3100269, EPI_ISL_3100278, EPI_ISL_3100282, EPI_ISL_3100283, EPI_ISL_3100284, EPI_ISL_3100285, EPI_ISL_3100286, EPI_ISL_3100287, EPI_ISL_3100288, EPI_ISL_3100293 | | | |
| see above | Sweden/Vastragotaland/Unilabs | Unilabs/Eskilstuna/Sweden | Emma Arvidsson |
| EPI_ISL_3100298 | Johor Bahru Public Health Laboratory | Institute for Medical Research, Infectious Disease Research Centre, National Institutes of Health, Ministry of Health Malaysia | Suppiah J, Kamel K, Mohd Zawawi Z, Azizan MA, Ramly N, Robert F, Anasir MI, Thayan R |
| EPI_ISL_3100300, EPI_ISL_3100301, EPI_ISL_3100314, EPI_ISL_3100317, EPI_ISL_3100318, EPI_ISL_3100319, EPI_ISL_3100323, EPI_ISL_3100335, EPI_ISL_3100336, EPI_ISL_3100337, EPI_ISL_3100350, EPI_ISL_3100351, EPI_ISL_3100355, EPI_ISL_3100357, EPI_ISL_3100359, EPI_ISL_3100360, EPI_ISL_3100361, EPI_ISL_3100369, EPI_ISL_3100370, EPI_ISL_3100371, EPI_ISL_3100374 | | | |
| see above | Public Health Authority of the Slovak Republic | Laboratory of Genomics and Bioinformatics, Comenius University Science Park | Tatiana Sedláčková, Diana Rusňáková, Miroslav Böhmer, Anna Gičová, Jaroslav Budiš, Tomáš Szemes, Jakub Styk |
| EPI_ISL_3100465 | Johor Bahru Public Health Laboratory | Institute for Medical Research, Infectious Disease Research Centre, National Institutes of Health, Ministry of Health Malaysia | Suppiah J, Kamel K, Mohd Zawawi Z, Azizan MA, Ramly N, Robert F, Anasir MI, Thayan R |
| EPI_ISL_3100511, EPI_ISL_3100512, EPI_ISL_3100513, EPI_ISL_3100514, EPI_ISL_3100515, EPI_ISL_3100516, EPI_ISL_3100517, EPI_ISL_3100518, EPI_ISL_3100519, EPI_ISL_3100520, EPI_ISL_3100521, EPI_ISL_3100522, EPI_ISL_3100524, EPI_ISL_3100525, EPI_ISL_3100526, EPI_ISL_3100527, EPI_ISL_3100528, EPI_ISL_3100529, EPI_ISL_3100530, EPI_ISL_3100531, EPI_ISL_3100532, EPI_ISL_3100533, EPI_ISL_3100534 | | | |
| see above | ASST MONZA | ASST MONZA | Sergio Maria Ivano Malandrin |
| EPI_ISL_3100535 | ASST GOM NIGUARDA | ASST Grande ospedale Metropolitano Niguarda | Alice Nava |
| EPI_ISL_3100536 | ASST MONZA | ASST MONZA | Sergio Maria Ivano Malandrin |
| EPI_ISL_3100537 | ASST GOM NIGUARDA | ASST Grande ospedale Metropolitano Niguarda | Alice Nava |
| EPI_ISL_3100538, EPI_ISL_3100539, EPI_ISL_3100540, EPI_ISL_3100541, EPI_ISL_3100542, EPI_ISL_3100543, EPI_ISL_3100544 | ASST MONZA | ASST MONZA | Sergio Maria Ivano Malandrin |
| EPI_ISL_3100546, EPI_ISL_3100547, EPI_ISL_3100548, EPI_ISL_3100549, EPI_ISL_3100550, EPI_ISL_3100551, EPI_ISL_3100552, EPI_ISL_3100553, EPI_ISL_3100554, EPI_ISL_3100555, EPI_ISL_3100556, EPI_ISL_3100557, EPI_ISL_3100558, EPI_ISL_3100559, EPI_ISL_3100560, EPI_ISL_3100561, EPI_ISL_3100562, EPI_ISL_3100563, EPI_ISL_3100564, EPI_ISL_3100565, EPI_ISL_3100566, EPI_ISL_3100567, EPI_ISL_3100568, EPI_ISL_3100569, EPI_ISL_3100570, EPI_ISL_3100571, EPI_ISL_3100572, EPI_ISL_3100573, EPI_ISL_3100574, EPI_ISL_3100575, EPI_ISL_3100576, EPI_ISL_3100577, EPI_ISL_3100578, EPI_ISL_3100579, EPI_ISL_3100580, EPI_ISL_3100581, EPI_ISL_3100582, EPI_ISL_3100583, EPI_ISL_3100585, EPI_ISL_3100586, EPI_ISL_3100587, EPI_ISL_3100589, EPI_ISL_3100590, EPI_ISL_3100592, EPI_ISL_3100594, EPI_ISL_3100595, EPI_ISL_3100596, EPI_ISL_3100597, EPI_ISL_3100598, EPI_ISL_3100599, EPI_ISL_3100600, EPI_ISL_3100601, EPI_ISL_3100604, EPI_ISL_3100605, EPI_ISL_3100606, EPI_ISL_3100608, EPI_ISL_3100609, EPI_ISL_3100610, EPI_ISL_3100611, EPI_ISL_3100612, EPI_ISL_3100613, EPI_ISL_3100614, EPI_ISL_3100615, EPI_ISL_3100616, EPI_ISL_3100617, EPI_ISL_3100618, EPI_ISL_3100619, EPI_ISL_3100621, EPI_ISL_3100622, EPI_ISL_3100623, EPI_ISL_3100626, EPI_ISL_3100627, EPI_ISL_3100628, EPI_ISL_3100629, EPI_ISL_3100630 | | | |
| see above | Fondazione IRCCS Ca' Granda Ospedale Maggiore Policlinico | Fondazione IRCCS Ca' Granda Ospedale Maggiore Policlinico | Ferruccio Ceriotti, Sara Uceda Renteria |
| EPI_ISL_3100633, EPI_ISL_3100634, EPI_ISL_3100635, EPI_ISL_3100636, EPI_ISL_3100637, EPI_ISL_3100639, EPI_ISL_3100640, EPI_ISL_3100641, EPI_ISL_3100642, EPI_ISL_3100643, EPI_ISL_3100644, EPI_ISL_3100645, EPI_ISL_3100647, EPI_ISL_3100648, EPI_ISL_3100654, EPI_ISL_3100655, EPI_ISL_3100656, EPI_ISL_3100657, EPI_ISL_3100658, EPI_ISL_3100659, EPI_ISL_3100660, EPI_ISL_3100661, EPI_ISL_3100662, EPI_ISL_3100663, EPI_ISL_3100664, EPI_ISL_3100665, EPI_ISL_3100666, EPI_ISL_3100667, EPI_ISL_3100668, EPI_ISL_3100669, EPI_ISL_3100670, EPI_ISL_3100671, EPI_ISL_3100672, EPI_ISL_3100673, EPI_ISL_3100674, EPI_ISL_3100675, EPI_ISL_3100676, EPI_ISL_3100677 | | | |
| see above | Idaho Bureau of Laboratories | Boise VA Medical Center, PALMS | Matthew Burns, Aimee Ceniseros, Christian Loera, Dr. Christopher Ball, Ying Pei, Cheri Lamb McFarlane, James Razor, Emily Bartlette |
| EPI_ISL_3100679, EPI_ISL_3100680 | Johor Bahru Public Health Laboratory | Institute for Medical Research, Infectious Disease Research Centre, National Institutes of Health, Ministry of Health Malaysia | Suppiah J, Kamel K, Mohd Zawawi Z, Azizan MA, Ramly N, Robert F, Anasir MI, Thayan R |
| EPI_ISL_3100681, EPI_ISL_3100682 | Ipoh Public Health Laboratory | Institute for Medical Research, Infectious Disease Research Centre, National Institutes of Health, Ministry of Health Malaysia | Suppiah J, Kamel K, Mohd Zawawi Z, Azizan MA, Ramly N, Robert F, Anasir MI, Thayan R |
| EPI_ISL_3100727, EPI_ISL_3100769, EPI_ISL_3100773, EPI_ISL_3100784, EPI_ISL_3100795, EPI_ISL_3100798, EPI_ISL_3100799, EPI_ISL_3100801, EPI_ISL_3100802, EPI_ISL_3100805, EPI_ISL_3100811, EPI_ISL_3100812, EPI_ISL_3100815, EPI_ISL_3100820, EPI_ISL_3100824, EPI_ISL_3100825, EPI_ISL_3100826, EPI_ISL_3100830, EPI_ISL_3100832, EPI_ISL_3100834, EPI_ISL_3100837, EPI_ISL_3100843, EPI_ISL_3100844, EPI_ISL_3100846, EPI_ISL_3100848, EPI_ISL_3100851, EPI_ISL_3100852, EPI_ISL_3100854, EPI_ISL_3100857, EPI_ISL_3100858, EPI_ISL_3100860, EPI_ISL_3100866, EPI_ISL_3100868, EPI_ISL_3100869, EPI_ISL_3100870, EPI_ISL_3100877, EPI_ISL_3100885, EPI_ISL_3100888, EPI_ISL_3100890, EPI_ISL_3100891, EPI_ISL_3100894, EPI_ISL_3100895, EPI_ISL_3100897, EPI_ISL_3100903, EPI_ISL_3100906, EPI_ISL_3100907, EPI_ISL_3100911, EPI_ISL_3100912, EPI_ISL_3100913, EPI_ISL_3100914, EPI_ISL_3100916, EPI_ISL_3100917, EPI_ISL_3100918, EPI_ISL_3100920, EPI_ISL_3100921, EPI_ISL_3100922, EPI_ISL_3100923, EPI_ISL_3100924, EPI_ISL_3100925, EPI_ISL_3100926, EPI_ISL_3100927, EPI_ISL_3100930, EPI_ISL_3100931, EPI_ISL_3100932, EPI_ISL_3100933, EPI_ISL_3100934, EPI_ISL_3100935, EPI_ISL_3100937, EPI_ISL_3100939, EPI_ISL_3100941, EPI_ISL_3100942, EPI_ISL_3100944, EPI_ISL_3100950, EPI_ISL_3100953, EPI_ISL_3100957, EPI_ISL_3100958, EPI_ISL_3100960, EPI_ISL_3100961, EPI_ISL_3100962, EPI_ISL_3100963, EPI_ISL_3100964, EPI_ISL_3100965, EPI_ISL_3100966, EPI_ISL_3100967, EPI_ISL_3100973, EPI_ISL_3100975, EPI_ISL_3100977, EPI_ISL_3100978, EPI_ISL_3100979, EPI_ISL_3100980 | | | |
| see above | Florida Bureau of Public Health Laboratories | Florida Bureau of Public Health Laboratories | Sarah Schmedes, Jason Blanton |
| EPI_ISL_3100981 | Ipoh Public Health Laboratory | Institute for Medical Research, Infectious Disease Research Centre, National Institutes of Health, Ministry of Health Malaysia | Suppiah J, Kamel K, Mohd Zawawi Z, Azizan MA, Ramly N, Robert F, Anasir MI, Thayan R |
| EPI_ISL_3100982 | Sultanah Bahiyah Hospital, Alor Setar | Institute for Medical Research, Infectious Disease Research Centre, National Institutes of Health, Ministry of Health Malaysia | Suppiah J, Kamel K, Mohd Zawawi Z, Azizan MA, Ramly N, Robert F, Anasir MI, Thayan R |
| EPI_ISL_3100983, EPI_ISL_3100984, EPI_ISL_3100985, EPI_ISL_3100986, EPI_ISL_3100987, EPI_ISL_3100988, EPI_ISL_3100990 | Microbiology and Virology Laboratory, "Policlinico Riuniti, Azienda Ospedaliero Universitaria, Foggia" | Microbiology and Virology Laboratory, "Policlinico Riuniti, Azienda Ospedaliero Universitaria, Foggia" | Fabio Arena, Settimia Altamura, Donatella Cedola, Valeria Delli Carri, Annamaria D'Aprile, Maria Rosaria Lipsi, Loris Micelli, Maria Pia Patrizio, Daniela Pisanelli, Tiziana Rollo, Maurizio Margaglione and Rosella De Nittis. |
| EPI_ISL_3101006 | Kuala Lumpur Hospital | Institute for Medical Research, Infectious Disease Research Centre, National Institutes of Health, Ministry of Health Malaysia | Suppiah J, Kamel K, Mohd Zawawi Z, Azizan MA, Ramly N, Robert F, Anasir MI, Thayan R |
| EPI_ISL_3101007, EPI_ISL_3101008, EPI_ISL_3101009, EPI_ISL_3101010, EPI_ISL_3101011, EPI_ISL_3101012, EPI_ISL_3101013, EPI_ISL_3101014, EPI_ISL_3101015, EPI_ISL_3101016, EPI_ISL_3101017, EPI_ISL_3101018, EPI_ISL_3101019, EPI_ISL_3101020, EPI_ISL_3101022, EPI_ISL_3101023, EPI_ISL_3101024, EPI_ISL_3101025, EPI_ISL_3101026, EPI_ISL_3101027, EPI_ISL_3101028, EPI_ISL_3101029, EPI_ISL_3101030, EPI_ISL_3101031, EPI_ISL_3101032, EPI_ISL_3101033, EPI_ISL_3101034, EPI_ISL_3101035, EPI_ISL_3101036, EPI_ISL_3101037, EPI_ISL_3101038, EPI_ISL_3101039, EPI_ISL_3101040, EPI_ISL_3101041, EPI_ISL_3101042, EPI_ISL_3101043, EPI_ISL_3101044, EPI_ISL_3101045, EPI_ISL_3101046, EPI_ISL_3101047, EPI_ISL_3101051, EPI_ISL_3101053, EPI_ISL_3101054, EPI_ISL_3101055, EPI_ISL_3101057, EPI_ISL_3101058, EPI_ISL_3101059, EPI_ISL_3101060, EPI_ISL_3101061 | | | |
| see above | University of Liège COVID-19 testing center | GIGA Medical Genomics | Keith Durkin, Maria Artesi, Bouchra Boujemla, Nathalie Renotte, Nadine Cambisano, Sébastien Bontems, Cécile Meex, Claire Gourzonès, Olivier Ek, Laurent Gillet, Marie-Pierre Hayette, Vincent Bours |
| EPI_ISL_3101064, EPI_ISL_3101066, EPI_ISL_3101067, EPI_ISL_3101071, EPI_ISL_3101073, EPI_ISL_3101075, EPI_ISL_3101084, EPI_ISL_3101087, EPI_ISL_3101097, EPI_ISL_3101099, EPI_ISL_3101100, EPI_ISL_3101101, EPI_ISL_3101102, EPI_ISL_3101103, EPI_ISL_3101107, EPI_ISL_3101112, EPI_ISL_3101114, EPI_ISL_3101115 | | | |
| see above | Molecular diagnostic laboratory of Federal Budget Institution of Science "Central Research Institute of Epidemiology" of The Federal Service on Customers' Rights Protection and Human Well-being Surveillance | Group of Genomics and Postgenomic Technologies of Central Research Institute of Epidemiology | Samoilov A.E., Kaptelova V.V., Saenko S.S.,Sinitsyn S.O., Korneenko E.V., Smirnova Y.S., Nadtoka M.I., Golubeva A.G., Speranskaya A.S., Tivanova E.V., Shipulina O.Y., Kondrasheva L.Y., Akimkin V.G. |
| EPI_ISL_3101119, EPI_ISL_3101120, EPI_ISL_3101121, EPI_ISL_3101129, EPI_ISL_3101130 | Molecular diagnostic laboratory of Federal Budget Institution of Science "Central Research Institute of Epidemiology" of The Federal Service on Customers' Rights Protection and Human Well-being Surveillance | Group of Genomics and Postgenomic Technologies of Central Research Institute of Epidemiology | Samoilov A.E., Kaptelova V.V., Korneenko E.V., Saenko S.S., Smirnova Y.S., Zuev S.N., Dohoyan A.Y., Nadtoka M.I., Valdohina A.V., Bulanenko V.P., Cherkashina A.S., Golubeva A.G., Zotova M.I., Berlina Y.Y., Solovieva E.D., Shipulina O.Y., Speranskaya A.S., Kondrasheva L.Y., Akimkin V.G. |
| EPI_ISL_3101146, EPI_ISL_3101147, EPI_ISL_3101148, EPI_ISL_3101149, EPI_ISL_3101151, EPI_ISL_3101152, EPI_ISL_3101154 | Molecular diagnostic laboratory of Federal Budget Institution of Science "Central Research Institute of Epidemiology" of The Federal Service on Customers' Rights Protection and Human Well-being Surveillance | Group of Genomics and Postgenomic Technologies of Central Research Institute of Epidemiology | Samoilov A.E., Kaptelova V.V., Korneenko E.V., Sinitsyn S.O., Saenko S.S., Zuev S.N., Nadtoka M.I., Speranskaya A.S., Tivanova E.V., Kondrasheva L.Y.,Shipulina O.Y., Smirnova Y.S., Akimkin V.G. |
| EPI_ISL_3101155 | Molecular diagnostic laboratory of Federal Budget Institution of Science "Central Research Institute of Epidemiology" of The Federal Service on Customers' Rights Protection and Human Well-being Surveillance | Group of Genomics and Postgenomic Technologies of Central Research Institute of Epidemiology | Samoilov A.E., Kaptelova V.V., Korneenko E.V., Saenko S.S., Smirnova Y.S., Zuev S.N., Dohoyan A.Y., Nadtoka M.I., Valdohina A.V., Bulanenko V.P., Cherkashina A.S., Golubeva A.G., Zotova M.I., Berlina Y.Y., Solovieva E.D., Shipulina O.Y., Speranskaya A.S., Kondrasheva L.Y., Akimkin V.G. |
| EPI_ISL_3101156, EPI_ISL_3101157, EPI_ISL_3101158, EPI_ISL_3101159, EPI_ISL_3101160, EPI_ISL_3101161, EPI_ISL_3101162 | Molecular diagnostic laboratory of Federal Budget Institution of Science "Central Research Institute of Epidemiology" of The Federal Service on Customers' Rights Protection and Human Well-being Surveillance | Group of Genomics and Postgenomic Technologies of Central Research Institute of Epidemiology | Samoilov A.E., Kaptelova V.V., Korneenko E.V., Sinitsyn S.O., Saenko S.S., Zuev S.N., Nadtoka M.I., Speranskaya A.S., Tivanova E.V., Kondrasheva L.Y.,Shipulina O.Y., Smirnova Y.S., Akimkin V.G. |

| | | | |
|---|---|---|---|
| EPI_ISL_3101163 | Molecular diagnostic laboratory of Federal Budget Institution of Science "Central Research Institute of Epidemiology" of The Federal Service on Customers' Rights Protection and Human Well-being Surveillance | Group of Genomics and Postgenomic Technologies of Central Research Institute of Epidemiology | Samoilov A.E., Kaptelova V.V., Korneenko E.V., Saenko S.S., Smirnova Y.S., Zuev S.N., Dohoyan A.Y., Nadtoka M.I., Valdohina A.V., Bulanenko V.P., Cherkashina A.S., Golubeva A.G., Zotova M.I., Berlina Y.Y., Solovieva E.D., Shipulina O.Y., Speranskaya A.S., Kondrasheva L.Y., Akimkin V.G. |
| EPI_ISL_3101164, EPI_ISL_3101165, EPI_ISL_3101167, EPI_ISL_3101168, EPI_ISL_3101169, EPI_ISL_3101170, EPI_ISL_3101171, EPI_ISL_3101172, EPI_ISL_3101173, EPI_ISL_3101175, EPI_ISL_3101176, EPI_ISL_3101179 | | | |
| see above | Molecular diagnostic laboratory of Federal Budget Institution of Science "Central Research Institute of Epidemiology" of The Federal Service on Customers' Rights Protection and Human Well-being Surveillance | Group of Genomics and Postgenomic Technologies of Central Research Institute of Epidemiology | Samoilov A.E., Kaptelova V.V., Korneenko E.V., Sinitsyn S.O., Saenko S.S., Zuev S.N., Nadtoka M.I., Speranskaya A.S., Tivanova E.V., Kondrasheva L.Y.,Shipulina O.Y., Smirnova Y.S., Akimkin V.G. |
| EPI_ISL_3101180, EPI_ISL_3101181, EPI_ISL_3101182, EPI_ISL_3101183, EPI_ISL_3101184, EPI_ISL_3101185, EPI_ISL_3101186, EPI_ISL_3101187, EPI_ISL_3101188, EPI_ISL_3101190, EPI_ISL_3101191, EPI_ISL_3101192, EPI_ISL_3101193, EPI_ISL_3101194, EPI_ISL_3101196, EPI_ISL_3101199, EPI_ISL_3101200, EPI_ISL_3101201, EPI_ISL_3101204, EPI_ISL_3101205, EPI_ISL_3101206, EPI_ISL_3101207, EPI_ISL_3101208, EPI_ISL_3101209, EPI_ISL_3101210, EPI_ISL_3101211, EPI_ISL_3101212, EPI_ISL_3101213, EPI_ISL_3101214, EPI_ISL_3101215, EPI_ISL_3101216, EPI_ISL_3101217, EPI_ISL_3101218, EPI_ISL_3101219, EPI_ISL_3101220, EPI_ISL_3101221, EPI_ISL_3101222, EPI_ISL_3101223, EPI_ISL_3101224, EPI_ISL_3101225, EPI_ISL_3101226, EPI_ISL_3101229, EPI_ISL_3101230, EPI_ISL_3101231, EPI_ISL_3101233, EPI_ISL_3101234, EPI_ISL_3101235, EPI_ISL_3101239, EPI_ISL_3101242, EPI_ISL_3101269, EPI_ISL_3101271, EPI_ISL_3101273, EPI_ISL_3101280, EPI_ISL_3101282, EPI_ISL_3101283, EPI_ISL_3101284, EPI_ISL_3101285, EPI_ISL_3101287, EPI_ISL_3101321, EPI_ISL_3101323, EPI_ISL_3101325, EPI_ISL_3101326, EPI_ISL_3101328, EPI_ISL_3101330, EPI_ISL_3101331, EPI_ISL_3101333, EPI_ISL_3101334, EPI_ISL_3101337, EPI_ISL_3101338, EPI_ISL_3101340, EPI_ISL_3101341, EPI_ISL_3101342, EPI_ISL_3101344, EPI_ISL_3101350, EPI_ISL_3101351, EPI_ISL_3101352 | | | |
| see above | Molecular diagnostic laboratory of Federal Budget Institution of Science "Central Research Institute of Epidemiology" of The Federal Service on Customers' Rights Protection and Human Well-being Surveillance | Group of Genomics and Postgenomic Technologies of Central Research Institute of Epidemiology | Samoilov A.E., Kaptelova V.V., Korneenko E.V., Saenko S.S., Smirnova Y.S., Zuev S.N., Dohoyan A.Y., Nadtoka M.I., Valdohina A.V., Bulanenko V.P., Cherkashina A.S., Golubeva A.G., Zotova M.I., Berlina Y.Y., Solovieva E.D., Shipulina O.Y., Speranskaya A.S., Kondrasheva L.Y., Akimkin V.G. |
| EPI_ISL_3101355, EPI_ISL_3101356 | Molecular diagnostic laboratory of Federal Budget Institution of Science "Central Research Institute of Epidemiology" of The Federal Service on Customers' Rights Protection and Human Well-being Surveillance | Group of Genomics and Postgenomic Technologies of Central Research Institute of Epidemiology | Samoilov A.E., Korneenko E.V., Smirnova Y.S., Sinitsyn S.O., Roev G.V., Nadtoka M.I., Buharina A.Y., Shipulina O.Y., Speranskaya A.S., Kondrasheva L.Y., Akimkin V.G. |
| EPI_ISL_3101358, EPI_ISL_3101359 | Molecular diagnostic laboratory of Federal Budget Institution of Science "Central Research Institute of Epidemiology" of The Federal Service on Customers' Rights Protection and Human Well-being Surveillance | Group of Genomics and Postgenomic Technologies of Central Research Institute of Epidemiology | Samoilov A.E., Kaptelova V.V., Korneenko E.V., Saenko S.S., Smirnova Y.S., Zuev S.N., Dohoyan A.Y., Nadtoka M.I., Valdohina A.V., Bulanenko V.P., Cherkashina A.S., Golubeva A.G., Zotova M.I., Berlina Y.Y., Solovieva E.D., Shipulina O.Y., Speranskaya A.S., Kondrasheva L.Y., Akimkin V.G. |
| EPI_ISL_3101361, EPI_ISL_3101362, EPI_ISL_3101363, EPI_ISL_3101364, EPI_ISL_3101365, EPI_ISL_3101366, EPI_ISL_3101367, EPI_ISL_3101368, EPI_ISL_3101369, EPI_ISL_3101370, EPI_ISL_3101371, EPI_ISL_3101372, EPI_ISL_3101373, EPI_ISL_3101374, EPI_ISL_3101375, EPI_ISL_3101376, EPI_ISL_3101377, EPI_ISL_3101378, EPI_ISL_3101379, EPI_ISL_3101380 | | | |
| see above | Molecular diagnostic laboratory of Federal Budget Institution of Science "Central Research Institute of Epidemiology" of The Federal Service on Customers' Rights Protection and Human Well-being Surveillance | Group of Genomics and Postgenomic Technologies of Central Research Institute of Epidemiology | Samoilov A.E., Korneenko E.V., Smirnova Y.S., Sinitsyn S.O., Roev G.V., Nadtoka M.I., Buharina A.Y., Shipulina O.Y., Speranskaya A.S., Kondrasheva L.Y., Akimkin V.G. |
| EPI_ISL_3101382, EPI_ISL_3101383 | Molecular diagnostic laboratory of Federal Budget Institution of Science "Central Research Institute of Epidemiology" of The Federal Service on Customers' Rights Protection and Human Well-being Surveillance | Group of Genomics and Postgenomic Technologies of Central Research Institute of Epidemiology | Samoilov A.E., Korneenko E.V., Smirnova Y.S., Zuev S.N., Dohoyan A.Y., Nadtoka M.I., Valdohina A.V., Bulanenko V.P., Cherkashina A.S., Golubeva A.G., Zotova M.I., Berlina Y.Y., Solovieva E.D., Shipulina O.Y., Speranskaya A.S., Kondrasheva L.Y., Akimkin V.G. |
| EPI_ISL_3101384 | Molecular diagnostic laboratory of Federal Budget Institution of Science "Central Research Institute of Epidemiology" of The Federal Service on Customers' Rights Protection and Human Well-being Surveillance | Group of Genomics and Postgenomic Technologies of Central Research Institute of Epidemiology | Samoilov A.E., Korneenko E.V., Smirnova Y.S., Sinitsyn S.O., Roev G.V., Nadtoka M.I., Buharina A.Y., Shipulina O.Y., Speranskaya A.S., Kondrasheva L.Y., Akimkin V.G. |
| EPI_ISL_3101385 | Molecular diagnostic laboratory of Federal Budget Institution of Science "Central Research Institute of Epidemiology" of The Federal Service on Customers' Rights Protection and Human Well-being Surveillance | Group of Genomics and Postgenomic Technologies of Central Research Institute of Epidemiology | Samoilov A.E., Korneenko E.V., Smirnova Y.S., Zuev S.N., Dohoyan A.Y., Nadtoka M.I., Valdohina A.V., Bulanenko V.P., Cherkashina A.S., Golubeva A.G., Zotova M.I., Berlina Y.Y., Solovieva E.D., Shipulina O.Y., Speranskaya A.S., Kondrasheva L.Y., Akimkin V.G. |
| EPI_ISL_3101387 | SYNLAB | University Hospital Brno, CMBG | Matej Bezdicek, Jan Svaton, Martina Lengerova, Kristyna Dufkova |
| EPI_ISL_3101388, EPI_ISL_3101389 | Cytogeneticka laborator Brno | University Hospital Brno, CMBG | Matej Bezdicek, Jan Svaton, Martina Lengerova, Kristyna Dufkova |
| EPI_ISL_3101390, EPI_ISL_3101391, EPI_ISL_3101392, EPI_ISL_3101393, EPI_ISL_3101394, EPI_ISL_3101395, EPI_ISL_3101396, EPI_ISL_3101397 | University Hospital Brno, OKMI | University Hospital Brno, CMBG | Matej Bezdicek, Jan Svaton, Martina Lengerova, Kristyna Dufkova |
| EPI_ISL_3101398, EPI_ISL_3101399, EPI_ISL_3101400, EPI_ISL_3101401, EPI_ISL_3101402, EPI_ISL_3101403, EPI_ISL_3101404 | Nemocnice Havlickuv Brod | University Hospital Brno, CMBG | Matej Bezdicek, Jan Svaton, Martina Lengerova, Kristyna Dufkova |
| EPI_ISL_3101405, EPI_ISL_3101406, EPI_ISL_3101407, EPI_ISL_3101408, EPI_ISL_3101409, EPI_ISL_3101410, EPI_ISL_3101411, EPI_ISL_3101412, EPI_ISL_3101413, EPI_ISL_3101414, EPI_ISL_3101415, EPI_ISL_3101416, EPI_ISL_3101417 | | | |
| see above | Nemocnice Jihlava | University Hospital Brno, CMBG | Matej Bezdicek, Jan Svaton, Martina Lengerova, Kristyna Dufkova |
| EPI_ISL_3101418, EPI_ISL_3101419, EPI_ISL_3101420, EPI_ISL_3101421, EPI_ISL_3101422, EPI_ISL_3101423, EPI_ISL_3101424 | Cytogeneticka laborator Brno | University Hospital Brno, CMBG | Matej Bezdicek, Jan Svaton, Martina Lengerova, Kristyna Dufkova |
| EPI_ISL_3101425 | Nemocnice Breclav | University Hospital Brno, CMBG | Matej Bezdicek, Jan Svaton, Martina Lengerova, Kristyna Dufkova |
| EPI_ISL_3101426 | IFCOR Brno | University Hospital Brno, CMBG | Matej Bezdicek, Jan Svaton, Martina Lengerova, Kristyna Dufkova |
| EPI_ISL_3101427 | Nemocnice Vyskov | University Hospital Brno, CMBG | Matej Bezdicek, Jan Svaton, Martina Lengerova, Kristyna Dufkova |
| EPI_ISL_3101428 | Nemocnice Boskovice | University Hospital Brno, CMBG | Matej Bezdicek, Jan Svaton, Martina Lengerova, Kristyna Dufkova |
| EPI_ISL_3101429, EPI_ISL_3101430, EPI_ISL_3101431, EPI_ISL_3101432, EPI_ISL_3101435 | St. Anne´s University Hospital Brno | University Hospital Brno, CMBG | Matej Bezdicek, Jan Svaton, Martina Lengerova, Kristyna Dufkova |
| EPI_ISL_3101436, EPI_ISL_3101437, EPI_ISL_3101438, EPI_ISL_3101439, EPI_ISL_3101440, EPI_ISL_3101441, EPI_ISL_3101442 | Nemocnice Trebic | University Hospital Brno, CMBG | Matej Bezdicek, Jan Svaton, Martina Lengerova, Kristyna Dufkova |
| EPI_ISL_3101443 | Cytogeneticka laborator Brno | University Hospital Brno, CMBG | Matej Bezdicek, Jan Svaton, Martina Lengerova, Kristyna Dufkova |
| EPI_ISL_3101445, EPI_ISL_3101446, EPI_ISL_3101447, EPI_ISL_3101448, EPI_ISL_3101449, EPI_ISL_3101450, EPI_ISL_3101451, EPI_ISL_3101452 | St. Anne´s University Hospital Brno | University Hospital Brno, CMBG | Matej Bezdicek, Jan Svaton, Martina Lengerova, Kristyna Dufkova |
| EPI_ISL_3101453, EPI_ISL_3101454 | Cytogeneticka laborator Brno | University Hospital Brno, CMBG | Matej Bezdicek, Jan Svaton, Martina Lengerova, Kristyna Dufkova |
| EPI_ISL_3101455, | Nemocnice Havlickuv Brod | University Hospital Brno, CMBG | Matej Bezdicek, Jan Svaton, Martina Lengerova, Kristyna Dufkova |

| | | | |
|---|---|---|---|
| EPI_ISL_3101456, EPI_ISL_3101458, EPI_ISL_3101459, EPI_ISL_3101460, EPI_ISL_3101461 | | | |
| EPI_ISL_3101462, EPI_ISL_3101463, EPI_ISL_3101464, EPI_ISL_3101465, EPI_ISL_3101466, EPI_ISL_3101467, EPI_ISL_3101469, EPI_ISL_3101470, EPI_ISL_3101471 | Nemocnice Trebic | University Hospital Brno, CMBG | Matej Bezdicek, Jan Svaton, Martina Lengerova, Kristyna Dufkova |
| EPI_ISL_3101476 | TSHEPONG | National Institute for Communicable Diseases of the National Health Laboratory Service | Amoako DG, Everatt J, Scheepers C, Mohale T, Ntuli N, Mahlangu B, Mnguni A, Ismail A, Bhiman JN |
| EPI_ISL_3101477, EPI_ISL_3101478, EPI_ISL_3101479, EPI_ISL_3101480, EPI_ISL_3101481, EPI_ISL_3101482, EPI_ISL_3101483, EPI_ISL_3101484, EPI_ISL_3101485, EPI_ISL_3101486, EPI_ISL_3101487 | | | |
| see above | LEBOWAKGOMO | National Institute for Communicable Diseases of the National Health Laboratory Service | Amoako DG, Everatt J, Scheepers C, Mohale T, Ntuli N, Mahlangu B, Mnguni A, Ismail A, Bhiman JN |
| EPI_ISL_3101489, EPI_ISL_3101490, EPI_ISL_3101491, EPI_ISL_3101492, EPI_ISL_3101493, EPI_ISL_3101494, EPI_ISL_3101496, EPI_ISL_3101497, EPI_ISL_3101498, EPI_ISL_3101499, EPI_ISL_3101500, EPI_ISL_3101501, EPI_ISL_3101502, EPI_ISL_3101503, EPI_ISL_3101504, EPI_ISL_3101506, EPI_ISL_3101507, EPI_ISL_3101509, EPI_ISL_3101510, EPI_ISL_3101511, EPI_ISL_3101512, EPI_ISL_3101513, EPI_ISL_3101514, EPI_ISL_3101515, EPI_ISL_3101516, EPI_ISL_3101517, EPI_ISL_3101518, EPI_ISL_3101519, EPI_ISL_3101520, EPI_ISL_3101521, EPI_ISL_3101523, EPI_ISL_3101524, EPI_ISL_3101525, EPI_ISL_3101526, EPI_ISL_3101527, EPI_ISL_3101528, EPI_ISL_3101529, EPI_ISL_3101530, EPI_ISL_3101531, EPI_ISL_3101532, EPI_ISL_3101533, EPI_ISL_3101534 | | | |
| see above | KOPANONG HOSPITAL | National Institute for Communicable Diseases of the National Health Laboratory Service | Amoako DG, Everatt J, Scheepers C, Mohale T, Ntuli N, Mahlangu B, Mnguni A, Ismail A, Bhiman JN |
| EPI_ISL_3101535, EPI_ISL_3101536, EPI_ISL_3101537, EPI_ISL_3101538, EPI_ISL_3101539, EPI_ISL_3101540, EPI_ISL_3101541, EPI_ISL_3101542, EPI_ISL_3101543, EPI_ISL_3101545, EPI_ISL_3101546, EPI_ISL_3101547, EPI_ISL_3101548, EPI_ISL_3101549, EPI_ISL_3101551, EPI_ISL_3101552, EPI_ISL_3101553, EPI_ISL_3101554, EPI_ISL_3101555, EPI_ISL_3101556, EPI_ISL_3101557 | | | |
| see above | TAMBO MEMORIAL | National Institute for Communicable Diseases of the National Health Laboratory Service | Amoako DG, Everatt J, Scheepers C, Mohale T, Ntuli N, Mahlangu B, Mnguni A, Ismail A, Bhiman JN |
| EPI_ISL_3101558 | TAMBO MEMORIAL | National Institute for Communicable Diseases of the National Health Laboratory Service | Sisonke Team, Amoako DG, Everatt J, Scheepers C, Mohale T, Ntuli N, Mahlangu B, Mnguni A, Ismail A, Bhiman JN |
| EPI_ISL_3101559, EPI_ISL_3101560, EPI_ISL_3101561, EPI_ISL_3101562, EPI_ISL_3101563, EPI_ISL_3101564, EPI_ISL_3101566, EPI_ISL_3101567, EPI_ISL_3101568, EPI_ISL_3101569 | TAMBO MEMORIAL | National Institute for Communicable Diseases of the National Health Laboratory Service | Amoako DG, Everatt J, Scheepers C, Mohale T, Ntuli N, Mahlangu B, Mnguni A, Ismail A, Bhiman JN |
| EPI_ISL_3101571, EPI_ISL_3101572, EPI_ISL_3101573, EPI_ISL_3101574, EPI_ISL_3101575, EPI_ISL_3101577, EPI_ISL_3101580, EPI_ISL_3101582 | POLOKWANE | National Institute for Communicable Diseases of the National Health Laboratory Service | Amoako DG, Everatt J, Scheepers C, Mohale T, Ntuli N, Mahlangu B, Mnguni A, Ismail A, Bhiman JN |
| EPI_ISL_3101583, EPI_ISL_3101584, EPI_ISL_3101585, EPI_ISL_3101586, EPI_ISL_3101588, EPI_ISL_3101589, EPI_ISL_3101590, EPI_ISL_3101591 | Pathcare Vaal | National Institute for Communicable Diseases of the National Health Laboratory Service | Amoako DG, Everatt J, Scheepers C, Mohale T, Ntuli N, Mahlangu B, Mnguni A, Ismail A, Bhiman JN |
| EPI_ISL_3101592, EPI_ISL_3101593, EPI_ISL_3101594, EPI_ISL_3101595, EPI_ISL_3101597, EPI_ISL_3101598, EPI_ISL_3101599, EPI_ISL_3101600, EPI_ISL_3101601, EPI_ISL_3101602, EPI_ISL_3101603, EPI_ISL_3101604, EPI_ISL_3101605, EPI_ISL_3101606, EPI_ISL_3101607, EPI_ISL_3101608, EPI_ISL_3101609, EPI_ISL_3101610, EPI_ISL_3101611, EPI_ISL_3101612, EPI_ISL_3101613, EPI_ISL_3101614, EPI_ISL_3101615, EPI_ISL_3101616, EPI_ISL_3101617 | | | |
| see above | EAST LONDON | National Institute for Communicable Diseases of the National Health Laboratory Service | Amoako DG, Everatt J, Scheepers C, Mohale T, Ntuli N, Mahlangu B, Mnguni A, Ismail A, Bhiman JN |
| EPI_ISL_3101618, EPI_ISL_3101619, EPI_ISL_3101620, EPI_ISL_3101621, EPI_ISL_3101622, EPI_ISL_3101623, EPI_ISL_3101624, EPI_ISL_3101625, EPI_ISL_3101626, EPI_ISL_3101627, EPI_ISL_3101628, EPI_ISL_3101629, EPI_ISL_3101630, EPI_ISL_3101631, EPI_ISL_3101632 | | | |
| see above | MANKWENG | National Institute for Communicable Diseases of the National Health Laboratory Service | Amoako DG, Everatt J, Scheepers C, Mohale T, Ntuli N, Mahlangu B, Mnguni A, Ismail A, Bhiman JN |
| EPI_ISL_3101635, EPI_ISL_3101636, EPI_ISL_3101637, EPI_ISL_3101638, EPI_ISL_3101639, EPI_ISL_3101642, EPI_ISL_3101643, EPI_ISL_3101644, EPI_ISL_3101645, EPI_ISL_3101646, EPI_ISL_3101647, EPI_ISL_3101648, EPI_ISL_3101649, EPI_ISL_3101651, EPI_ISL_3101652, EPI_ISL_3101653, EPI_ISL_3101654, EPI_ISL_3101655 | | | |
| see above | National Institute for Communicable Diseases of the National Health Laboratory Service | National Institute for Communicable Diseases of the National Health Laboratory Service | Amoako DG, Everatt J, Scheepers C, Mohale T, Ntuli N, Mahlangu B, Mnguni A, Ismail A, Bhiman JN |
| EPI_ISL_3101656, EPI_ISL_3101657, EPI_ISL_3101658, EPI_ISL_3101659, EPI_ISL_3101660, EPI_ISL_3101661, EPI_ISL_3101662, EPI_ISL_3101663, EPI_ISL_3101664 | National Institute for Communicable Diseases of the National Health Laboratory Service | National Institute for Communicable Diseases of the National Health Laboratory Service | Sisonke Team, Amoako DG, Everatt J, Scheepers C, Mohale T, Ntuli N, Mahlangu B, Mnguni A, Ismail A, Bhiman JN |
| EPI_ISL_3101984 | ASL GENZANO | University Campus Bio-Medico of Rome (UCBM) | Riva E., De Florio L., Fogolari M., Francesconi M., Lintas C., Veralli R., Angeletti S. |
| EPI_ISL_3101986, EPI_ISL_3101987, EPI_ISL_3101988, EPI_ISL_3101989, EPI_ISL_3101990, EPI_ISL_3101991, EPI_ISL_3101992, EPI_ISL_3101994, EPI_ISL_3101995, EPI_ISL_3101997, EPI_ISL_3101998, EPI_ISL_3101999, EPI_ISL_3102000, EPI_ISL_3102001, EPI_ISL_3102002 | | | |
| see above | Wyoming Public Health Laboratory | Wyoming Public Health Laboratory | Jim Mildenberger, Taylor Fearing, Channing Weber, Ashley Norberg, Chayse Rowley, Marley Goetz, Brian Dominguez, Elliot Thomasson, Sam Britz, Cari Sloma, Robert Petit, and Rob Christensen |
| EPI_ISL_3102003 | Ospedale Sandro Pertini UOC Microbiology and Virology | University Campus Bio-Medico of Rome (UCBM) | Riva E., De Florio L., Fogolari M., Francesconi M., Lintas C., Veralli R., Angeletti S. |
| EPI_ISL_3102004 | OSPEDALE SAN CAMILLO, Roma | University Campus Bio-Medico of Rome | Angeletti Silvia, Riva Elisabetta |
| EPI_ISL_3102005 | Ospedale Sandro Pertini UOC Microbiology and virology | University Campus Bio-Medico of Rome (UCBM) | Riva E., De Florio L., Fogolari M., Francesconi M., Lintas C., Veralli R., Angeletti S |
| EPI_ISL_3102006 | OSPEDALE SAN CAMILLO di Roma | University Campus Bio-Medico di Roma | Angeletti Silvia, Riva Elisabetta |
| EPI_ISL_3102007, EPI_ISL_3102011, EPI_ISL_3102012, EPI_ISL_3102014 | Ospedale Sandro Pertini UOC Microbiology and virology | University Campus Bio-Medico of Rome (UCBM) | Riva E., De Florio L., Fogolari M., Francesconi M., Lintas C., Veralli R., Angeletti S. |
| EPI_ISL_3102015 | IDI-IRCCS di Roma | University Campus Bio-Medico di Roma | Angeletti Silvia, Riva Elisabetta |
| EPI_ISL_3102016 | Azienda Ospedaliera San Camillo Forlanini Rome, Italy | University Campus Bio-Medico of Rome (UCBM | Riva E., De Florio L., Fogolari M., Francesconi M., Lintas C., Veralli R., Angeletti S. |
| EPI_ISL_3102017 | IDI-IRCCS di Roma | University Campus Bio-Medico of Rome | Angeletti Silvia, Riva Elisabetta |
| EPI_ISL_3102018 | Azienda Ospedaliera San Camillo Forlanini Rome Italy | University Campus Bio-Medico of Rome (UCBM) | Riva E., De Florio L., Fogolari M., Francesconi M., Lintas C., Veralli R., Angeletti S |
| EPI_ISL_3102019 | IDI-IRCCS di Roma | University Campus Bio-Medico di Roma | Angeletti Silvia, Riva Elisabetta |
| EPI_ISL_3102020 | Azienda Ospedaliera San Camillo Forlanini Rome Italy | University Campus Bio-Medico of Rome (UCBM) | Riva E., De Florio L., Fogolari M., Francesconi M., Lintas C., Veralli R., Angeletti S. |
| EPI_ISL_3102021 | IDI-IRCCS di Roma | University Campus Bio-Medico di Roma | Angeletti Silvia, riva Elisabetta |
| EPI_ISL_3102022 | OSPEDALE SAN GIOVANNI ADDOLORATA di Roma | University Campus Bio-Medico di Roma | Angeletti Silvia, Riva Elisabetta |
| EPI_ISL_3102023 | Ospedale Sant'Andrea di Roma | University Campus Bio-Medico di Roma | Angeletti Silvia, Riva Elisabetta |
| EPI_ISL_3102025 | Genomik Solidaritas Indonesia Laboratorium | Genomik Solidaritas Indonesia Laboratorium | Himawan Masyhuri, Normastuti Adhini Tantyo, Alfin Muhammad Abdillah, Dhahlia Agustina Cahyono, Anna Christina Brazia, Ahmad Zoebad Foeady, Annisa Muthiah Sukirman, Gracia Felias Enos Korompis, Laely Indrayanti, Louisa Markus, |

| | | | |
|---|---|---|---|
| | | | Anuraj Shankar, Meutia Ayuputeri Kumaheri, Anindya Pradipta Susanto, Ariel Pradipta |
| EPI_ISL_3102026 | UOC Patologia Clinica Spaziani | University Campus Bio-Medico di Roma | Angeletti Silvia, Riva Elisabetta |
| EPI_ISL_3102028 | UOC Patologia Clinica Spaziani | University Campus Bio-Medico di Roma | Angeletti Silvia, riva Elisabetta |
| EPI_ISL_3102030 | Genomik Solidaritas Indonesia Laboratorium | Genomik Solidaritas Indonesia Laboratorium | Himawan Masyhuri, Normastuti Adhini Tantyo, Alfin Muhammad Abdillah, Dhahlia Agustina Cahyono, Anna Christina Brazia, Ahmad Zoebad Foeady, Annisa Muthiah Sukirman, Gracia Felias Enos Korompis, Anuraj Shankar, Meutia Ayuputeri Kumaheri, Ariel Pradipta |
| EPI_ISL_3102031 | IZSLT Roma | University campus Bio-Medico di Roma | Angeletti silvia, Riva Elisabetta |
| EPI_ISL_3102032 | IZSLT Roma | University campus Bio-Medico di Roma | Angeletti Silvia, Riva Elisabetta |
| EPI_ISL_3102033 | IZSLT Roma | University Campus Bio-Medico di Roma | Angeletti Silvia, Riva Elisabetta |
| EPI_ISL_3102034 | Genomik Solidaritas Indonesia Laboratorium | Genomik Solidaritas Indonesia Laboratorium | Neng Hilma Hamidah, Normastuti Adhini Tantyo, Alfin Muhammad Abdillah, Dhahlia Agustina Cahyono, Anna Christina Brazia, Himawan Masyhuri, Ahmad Zoebad Foeady, Annisa Muthiah Sukirman, Gracia Felias Enos Korompis, Anuraj Shankar, Meutia Ayuputeri Kumaheri, Ariel Pradipta |
| EPI_ISL_3102035 | VITERBO ASL Ospedale di Belcolle, 01100 Belcolle VT | University Campus Bio-Medico of Rome (UCBM) | Riva E., De Florio L., Fogolari M., Francesconi M., Lintas C., Veralli R., Angeletti S. |
| EPI_ISL_3102036 | IZSLT Roma | University Campus Bio-Medico di Roma | Angeletti Silvia, Riva Elisabetta |
| EPI_ISL_3102037 | VITERBO ASL Ospedale di Belcolle, 01100 Belcolle VT | University Campus Bio-Medico of Rome (UCBM) | Riva E., De Florio L., Fogolari M., Francesconi M., Lintas C., Veralli R., Angeletti S. |
| EPI_ISL_3102038 | IZSLT Roma | University Campus Bio-Medico di Roma | Angeletti Silvia, riva Elisabetta |
| EPI_ISL_3102040 | VITERBO ASL Ospedale di Belcolle, 01100 Belcolle VT | University Campus Bio-Medico of Rome (UCBM) | Riva E., De Florio L., Fogolari M., Francesconi M., Lintas C., Veralli R., Angeletti S. |
| EPI_ISL_3102041 | Laboratorio di Medicina Genomica Fondazione Santa Lucia | University Campus Bio-Medico di Roma | Angeletti Silvia, Riva Elisabetta |
| EPI_ISL_3102042 | UOC Microbiology and Virology Policlinico Umberto I Rome, Italy | University Campus Bio-Medico of Rome (UCBM) | Riva E., De Florio L., Fogolari M., Francesconi M., Lintas C., Veralli R., Angeletti S. |
| EPI_ISL_3102043 | Laboratorio di Medicina Genomica Fondazione Santa Lucia | University Campus Bio-Medico di Roma | Angeletti Silvia, Riva Elisabetta |
| EPI_ISL_3102044 | Genomik Solidaritas Indonesia Laboratorium | Genomik Solidaritas Indonesia Laboratorium | Neng Hilma Hamidah, Normastuti Adhini Tantyo, Alfin Muhammad Abdillah, Dhahlia Agustina Cahyono, Anna Christina Brazia, Himawan Masyhuri, Ahmad Zoebad Foeady, Annisa Muthiah Sukirman, Gracia Felias Enos Korompis, Anuraj Shankar, Meutia Ayuputeri Kumaheri, Ariel Pradipta |
| EPI_ISL_3102045 | UOC Microbiology and Virology Policlinico Umberto I Rome, Italy | University Campus Bio-Medico of Rome (UCBM) | Riva E., De Florio L., Fogolari M., Francesconi M., Lintas C., Veralli R., Angeletti S. |
| EPI_ISL_3102046 | Laboratorio di Medicina Genomica Fondazione Santa Lucia | University campus Bio-Medico di Roma | Angeletti Silvia, Riva Elisabetta |
| EPI_ISL_3102048 | UOC Microbiology and Virology Policlinico Umberto I Rome, Italy | University Campus Bio-Medico of Rome (UCBM) | Riva E., De Florio L., Fogolari M., Francesconi M., Lintas C., Veralli R., Angeletti S. |
| EPI_ISL_3102049 | ICOT Latina | University Campus Bio-Medico di Roma | Angeletti Silvia, Riva Elisabetta |
| EPI_ISL_3102050, EPI_ISL_3102051 | Laboratory Diagnostic, Veterinary Specialized Institute Kraljevo | Laboratory Diagnostic, Veterinary Specialized Institute Kraljevo | Vidanovic,D., Tesovic,B., Knezevic,A., Jovanovic,T., Jankovic,M., Sekler,M., Banovic Djeri,B., Volkening,J., Afonso,C., Petrovic,T., Djukic,V. |
| EPI_ISL_3102052 | UOC Microbiology and Virology Policlinico Umberto I Rome, Italy | University Campus Bio-Medico of Rome (UCBM) | Riva E., De Florio L., Fogolari M., Francesconi M., Lintas C., Veralli R., Angeletti S. |
| EPI_ISL_3102083, EPI_ISL_3102084, EPI_ISL_3102085, EPI_ISL_3102086, EPI_ISL_3102087, EPI_ISL_3102088, EPI_ISL_3102089, EPI_ISL_3102090, EPI_ISL_3102091, EPI_ISL_3102092, EPI_ISL_3102093, EPI_ISL_3102094, EPI_ISL_3102095, EPI_ISL_3102096, EPI_ISL_3102097, EPI_ISL_3102098, EPI_ISL_3102099 | | | |
| see above | City of Milwaukee Health Department Laboratory | City of Milwaukee Health Department Laboratory | Manjeet Khubbar, Amy Bauer, Jennifer Lentz, Samantha Scott, Sanjib Bhattacharyya |
| EPI_ISL_3102100 | UOC Microbiology and Virology Policlinico Umberto I Rome, Italy | University Campus Bio-Medico of Rome (UCBM) | Riva E., De Florio L., Fogolari M., Francesconi M., Lintas C., Veralli R., Angeletti S. |
| EPI_ISL_3102105 | Translational Health Science and Technology Institute -ESIC medical college and hospital, Faridabad | THSTI Bioassay laboratory | Naseem Ahmed Khan, Heena Shaman, Janmejay Singh, Kamal Pargai, Jigme Wangchuk, Anil Pandey, Akshay Kanakan, Ranjeet Maurya, Janani Srinivasa Vasudevan, Neha Jha, Priyanka Mehta, Rajesh Pandey, Guruprasad R Medigeshi |
| EPI_ISL_3102106, EPI_ISL_3102107, EPI_ISL_3102108, EPI_ISL_3102109, EPI_ISL_3102110, EPI_ISL_3102111 | St James's Hospital, Dublin 8, Ireland. | ELDA biotech | Deirdre M. Broderick, Peter R. Flanagan, Brendan Crowley |
| EPI_ISL_3102136 | Genomik Solidaritas Indonesia Laboratorium | Genomik Solidaritas Indonesia Laboratorium | Himawan Masyhuri, Neng Hilma Hamidah, Normastuti Adhini Tantyo, Alfin Muhammad Abdillah, Dhahlia Agustina Cahyono, Anna Christina Brazia, Ahmad Zoebad Foeady, Annisa Muthiah Sukirman, Gracia Felias Enos Korompis, Anuraj Shankar, Meutia Ayuputeri Kumaheri, Ariel Pradipta |
| EPI_ISL_3102137 | UOC Microbiology and Virology Policlinico Umberto I Rome, Italy | University Campus Bio-Medico of Rome (UCBM) | Riva E., De Florio L., Fogolari M., Francesconi M., Lintas C., Veralli R., Angeletti S. |
| EPI_ISL_3102138 | Ospedale Pediatrico Bambin Gesu' | University Campus Bio-Medico di Roma | Angeletti Silvia, Riva Elisabetta |
| EPI_ISL_3102150 | UOC Microbiology and Virology Policlinico Umberto I Rome, Italy | University Campus Bio-Medico of Rome (UCBM) | Riva E., De Florio L., Fogolari M., Francesconi M., Lintas C., Veralli R., Angeletti S. |
| EPI_ISL_3102151, EPI_ISL_3102152, EPI_ISL_3102153, EPI_ISL_3102155, EPI_ISL_3102157, EPI_ISL_3102158 | TXDSHS | TXDSHS | Rashmi Tuladhar, Bonnie Oh, Jenny Zhang, Maliha Rahman, Mayela Pedrueza, Anita Pokharel, Karen Bobier, Lorraine Rodriguez, Myong Koag, Chun Wang, Rachel Lee, Grace Kubin |
| EPI_ISL_3102159 | Collection of WHO National Influenza Center, N.F. Gamaleya National Research Center for Epidemiology and Microbiology of the Ministry of Health of the Russian Federation | N.F. Gamaleya Research Center for Epidemiology and Microbiology | Olga Voronina, Ekaterina Aksenova, Natalia Ryzhova, Marina Kunda, Kirill Krasnoslobodtsev, Elena Burtseva, Lyudmila Kolobukhina, Svetlana Smetanina, Denis Logunov |
| EPI_ISL_3102546, EPI_ISL_3102547 | Ospedale San Camillo di Roma | University Campus Bio-Medico di Roma | Angeletti Silvia, Riva Elisabetta |
| EPI_ISL_3102548, EPI_ISL_3102549, EPI_ISL_3102550, EPI_ISL_3102551, EPI_ISL_3102552, EPI_ISL_3102553, EPI_ISL_3102554, EPI_ISL_3102555, EPI_ISL_3102556, EPI_ISL_3102557, EPI_ISL_3102558, EPI_ISL_3102559, EPI_ISL_3102560, EPI_ISL_3102561, EPI_ISL_3102562, EPI_ISL_3102563, EPI_ISL_3102564, EPI_ISL_3102565, EPI_ISL_3102566, EPI_ISL_3102567, EPI_ISL_3102568, EPI_ISL_3102569, EPI_ISL_3102570, EPI_ISL_3102571, EPI_ISL_3102572, EPI_ISL_3102573, EPI_ISL_3102574, EPI_ISL_3102575, EPI_ISL_3102576, EPI_ISL_3102577, EPI_ISL_3102578, EPI_ISL_3102579, EPI_ISL_3102580, EPI_ISL_3102581, EPI_ISL_3102582, EPI_ISL_3102584, EPI_ISL_3102585, EPI_ISL_3102586, EPI_ISL_3102587, EPI_ISL_3102588, EPI_ISL_3102589, EPI_ISL_3102590, EPI_ISL_3102591, EPI_ISL_3102592, EPI_ISL_3102593, EPI_ISL_3102594, EPI_ISL_3102595, EPI_ISL_3102596, EPI_ISL_3102597, EPI_ISL_3102598, EPI_ISL_3102599, EPI_ISL_3102600, EPI_ISL_3102601, EPI_ISL_3102603, EPI_ISL_3102604, EPI_ISL_3102605, EPI_ISL_3102606, EPI_ISL_3102607, EPI_ISL_3102608, EPI_ISL_3102609, EPI_ISL_3102610, EPI_ISL_3102611, EPI_ISL_3102612, EPI_ISL_3102613, EPI_ISL_3102614, EPI_ISL_3102615, EPI_ISL_3102616, EPI_ISL_3102617, EPI_ISL_3102618, EPI_ISL_3102619, EPI_ISL_3102620, EPI_ISL_3102621, EPI_ISL_3102622, EPI_ISL_3102623, EPI_ISL_3102624, EPI_ISL_3102625, EPI_ISL_3102626, EPI_ISL_3102627, EPI_ISL_3102628, EPI_ISL_3102629, EPI_ISL_3102630, EPI_ISL_3102631, EPI_ISL_3102632, EPI_ISL_3102633, EPI_ISL_3102634, EPI_ISL_3102635, EPI_ISL_3102636, EPI_ISL_3102637, EPI_ISL_3102638, EPI_ISL_3102639, EPI_ISL_3102640, EPI_ISL_3102641, EPI_ISL_3102642, EPI_ISL_3102643, EPI_ISL_3102644, EPI_ISL_3102645, EPI_ISL_3102646, EPI_ISL_3102647, EPI_ISL_3102648, EPI_ISL_3102649, EPI_ISL_3102650, EPI_ISL_3102651, EPI_ISL_3102653, EPI_ISL_3102654, EPI_ISL_3102655, EPI_ISL_3102656, EPI_ISL_3102657, EPI_ISL_3102658, EPI_ISL_3102659, EPI_ISL_3102660, EPI_ISL_3102661, EPI_ISL_3102662, EPI_ISL_3102663, EPI_ISL_3102664, EPI_ISL_3102665, EPI_ISL_3102666, EPI_ISL_3102667, EPI_ISL_3102668, EPI_ISL_3102669, EPI_ISL_3102670, EPI_ISL_3102671, EPI_ISL_3102672, EPI_ISL_3102673, EPI_ISL_3102674, EPI_ISL_3102675, EPI_ISL_3102676, EPI_ISL_3102677, EPI_ISL_3102678, EPI_ISL_3102679, EPI_ISL_3102680, EPI_ISL_3102681, EPI_ISL_3102682, EPI_ISL_3102683, EPI_ISL_3102684, EPI_ISL_3102685, EPI_ISL_3102686, EPI_ISL_3102687, EPI_ISL_3102688, EPI_ISL_3102689, EPI_ISL_3102691, EPI_ISL_3102692, EPI_ISL_3102693, EPI_ISL_3102694, EPI_ISL_3102696, EPI_ISL_3102697, EPI_ISL_3102698, EPI_ISL_3102699, EPI_ISL_3102700, EPI_ISL_3102701, EPI_ISL_3102702, EPI_ISL_3102703, EPI_ISL_3102704, EPI_ISL_3102705, EPI_ISL_3102706, EPI_ISL_3102707, EPI_ISL_3102708, EPI_ISL_3102709, EPI_ISL_3102710, EPI_ISL_3102711, EPI_ISL_3102712, EPI_ISL_3102713, EPI_ISL_3102714, EPI_ISL_3102717, EPI_ISL_3102718, EPI_ISL_3102719, EPI_ISL_3102720, EPI_ISL_3102721, EPI_ISL_3102722, EPI_ISL_3102723, EPI_ISL_3102724, EPI_ISL_3102725, EPI_ISL_3102726, EPI_ISL_3102727, EPI_ISL_3102728, EPI_ISL_3102729, EPI_ISL_3102730, EPI_ISL_3102731, EPI_ISL_3102732, EPI_ISL_3102733, EPI_ISL_3102734, EPI_ISL_3102735, EPI_ISL_3102736, EPI_ISL_3102737, EPI_ISL_3102738, EPI_ISL_3102739, EPI_ISL_3102740, EPI_ISL_3102741, EPI_ISL_3102742, EPI_ISL_3102743, EPI_ISL_3102744, EPI_ISL_3102745, EPI_ISL_3102746, EPI_ISL_3102747, EPI_ISL_3102748, EPI_ISL_3102749, EPI_ISL_3102750, EPI_ISL_3102751, EPI_ISL_3102752, EPI_ISL_3102753, EPI_ISL_3102755, EPI_ISL_3102756, EPI_ISL_3102757, EPI_ISL_3102758, EPI_ISL_3102759, EPI_ISL_3102760, EPI_ISL_3102761, EPI_ISL_3102762, EPI_ISL_3102763, EPI_ISL_3102764, EPI_ISL_3102765, EPI_ISL_3102766, EPI_ISL_3102767, EPI_ISL_3102768, EPI_ISL_3102769, EPI_ISL_3102770, EPI_ISL_3102771, EPI_ISL_3102772, EPI_ISL_3102773, EPI_ISL_3102774, EPI_ISL_3102775, EPI_ISL_3102776, EPI_ISL_3102777, EPI_ISL_3102778, EPI_ISL_3102779, EPI_ISL_3102780, EPI_ISL_3102784, EPI_ISL_3102785, EPI_ISL_3102786, EPI_ISL_3102787, EPI_ISL_3102788, EPI_ISL_3102789, EPI_ISL_3102790, EPI_ISL_3102791, EPI_ISL_3102792, EPI_ISL_3102793, EPI_ISL_3102794, EPI_ISL_3102795, EPI_ISL_3102796, EPI_ISL_3102797, EPI_ISL_3102798, EPI_ISL_3102799, EPI_ISL_3102800, EPI_ISL_3102801, EPI_ISL_3102802, EPI_ISL_3102803, EPI_ISL_3102804, EPI_ISL_3102805, EPI_ISL_3102806, EPI_ISL_3102807, EPI_ISL_3102808, EPI_ISL_3102809, EPI_ISL_3102810, EPI_ISL_3102811, EPI_ISL_3102812, EPI_ISL_3102814, EPI_ISL_3102815, EPI_ISL_3102816, EPI_ISL_3102817, EPI_ISL_3102818, EPI_ISL_3102819, EPI_ISL_3102820, EPI_ISL_3102821, EPI_ISL_3102822, EPI_ISL_3102823, EPI_ISL_3102824, EPI_ISL_3102825, EPI_ISL_3102826, EPI_ISL_3102827, EPI_ISL_3102829, EPI_ISL_3102830, EPI_ISL_3102831, EPI_ISL_3102832, EPI_ISL_3102833, EPI_ISL_3102834, EPI_ISL_3102835, EPI_ISL_3102836, EPI_ISL_3102837, EPI_ISL_3102838, EPI_ISL_3102839, EPI_ISL_3102840, EPI_ISL_3102841, EPI_ISL_3102842, EPI_ISL_3102843, EPI_ISL_3102844, EPI_ISL_3102845, EPI_ISL_3102846, EPI_ISL_3102847, EPI_ISL_3102848, EPI_ISL_3102849, EPI_ISL_3102850, EPI_ISL_3102851, EPI_ISL_3102853, EPI_ISL_3102854, EPI_ISL_3102855, EPI_ISL_3102856, EPI_ISL_3102857, EPI_ISL_3102858, EPI_ISL_3102859, EPI_ISL_3102860, EPI_ISL_3102861, EPI_ISL_3102862, EPI_ISL_3102863, EPI_ISL_3102864, EPI_ISL_3102865, EPI_ISL_3102866, EPI_ISL_3102867, EPI_ISL_3102868, EPI_ISL_3102869, EPI_ISL_3102870, EPI_ISL_3102871, EPI_ISL_3102872, EPI_ISL_3102873, EPI_ISL_3102874, EPI_ISL_3102875, EPI_ISL_3102876, EPI_ISL_3102877, EPI_ISL_3102878, EPI_ISL_3102879, EPI_ISL_3102880, EPI_ISL_3102881, EPI_ISL_3102882, EPI_ISL_3102883, EPI_ISL_3102884, EPI_ISL_3102885, EPI_ISL_3102886, EPI_ISL_3102887, EPI_ISL_3102890, EPI_ISL_3102891, EPI_ISL_3102892, EPI_ISL_3102893, EPI_ISL_3102894, EPI_ISL_3102896, EPI_ISL_3102897, EPI_ISL_3102898, EPI_ISL_3102899, EPI_ISL_3102900, EPI_ISL_3102901 | | | |
| see above | Nevada State Public Health Laboratory | Nevada State Public Health Laboratory | Andrew Gorzalski, Mark Pandori |
| EPI_ISL_3102902, EPI_ISL_3102903, EPI_ISL_3102904, EPI_ISL_3102905, EPI_ISL_3102906, EPI_ISL_3102907, EPI_ISL_3102908, EPI_ISL_3102909, EPI_ISL_3102910, EPI_ISL_3102911, EPI_ISL_3102912, EPI_ISL_3102913, EPI_ISL_3102914, EPI_ISL_3102915, EPI_ISL_3102916, EPI_ISL_3102917, EPI_ISL_3102918, EPI_ISL_3102919, EPI_ISL_3102920, EPI_ISL_3102921, EPI_ISL_3102922, EPI_ISL_3102923, EPI_ISL_3102924, EPI_ISL_3102925, EPI_ISL_3102926, EPI_ISL_3102927, EPI_ISL_3102928, EPI_ISL_3102929, EPI_ISL_3102930, EPI_ISL_3102931, EPI_ISL_3102932, EPI_ISL_3102933, EPI_ISL_3102934, EPI_ISL_3102935, EPI_ISL_3102936, EPI_ISL_3102937, EPI_ISL_3102938, EPI_ISL_3102939, EPI_ISL_3102940, EPI_ISL_3102941, EPI_ISL_3102942, EPI_ISL_3102943, EPI_ISL_3102944, EPI_ISL_3102945, EPI_ISL_3102946, EPI_ISL_3102947, EPI_ISL_3102948, EPI_ISL_3102949, EPI_ISL_3102950, EPI_ISL_3102951, EPI_ISL_3102952, EPI_ISL_3102953, EPI_ISL_3102954, EPI_ISL_3102955, EPI_ISL_3102956, EPI_ISL_3102957, EPI_ISL_3102958, EPI_ISL_3102959, EPI_ISL_3102960, EPI_ISL_3102961, EPI_ISL_3102962, EPI_ISL_3102963, EPI_ISL_3102964, EPI_ISL_3102965, EPI_ISL_3102966, EPI_ISL_3102967, EPI_ISL_3102968, EPI_ISL_3102969, EPI_ISL_3102970, EPI_ISL_3102971, EPI_ISL_3102972, EPI_ISL_3102973, EPI_ISL_3102974, EPI_ISL_3102975, EPI_ISL_3102976, EPI_ISL_3102977, EPI_ISL_3102978, EPI_ISL_3102979, EPI_ISL_3102980, EPI_ISL_3102981, EPI_ISL_3102982, EPI_ISL_3102983, EPI_ISL_3102984, EPI_ISL_3102985, EPI_ISL_3102986, EPI_ISL_3102987, EPI_ISL_3102988, EPI_ISL_3102989, EPI_ISL_3102990, EPI_ISL_3102991, EPI_ISL_3102992, EPI_ISL_3102993, EPI_ISL_3102994, EPI_ISL_3102995, EPI_ISL_3102996, EPI_ISL_3102997, EPI_ISL_3102998, EPI_ISL_3102999, EPI_ISL_3103000, EPI_ISL_3103001, EPI_ISL_3103002, EPI_ISL_3103003, EPI_ISL_3103004, EPI_ISL_3103005, EPI_ISL_3103006, EPI_ISL_3103007, EPI_ISL_3103008, EPI_ISL_3103009, EPI_ISL_3103010, EPI_ISL_3103011, EPI_ISL_3103012, EPI_ISL_3103013, EPI_ISL_3103014, EPI_ISL_3103015, EPI_ISL_3103016, EPI_ISL_3103017, EPI_ISL_3103018, EPI_ISL_3103019, EPI_ISL_3103020, EPI_ISL_3103021, EPI_ISL_3103022, EPI_ISL_3103023, EPI_ISL_3103024, EPI_ISL_3103025, EPI_ISL_3103026, EPI_ISL_3103027, EPI_ISL_3103028, EPI_ISL_3103029, EPI_ISL_3103030, EPI_ISL_3103031, EPI_ISL_3103032, EPI_ISL_3103033, EPI_ISL_3103034, EPI_ISL_3103035, EPI_ISL_3103036, EPI_ISL_3103037, EPI_ISL_3103038, EPI_ISL_3103039, EPI_ISL_3103040, EPI_ISL_3103041, EPI_ISL_3103042, EPI_ISL_3103043, EPI_ISL_3103044, EPI_ISL_3103046, EPI_ISL_3103047, EPI_ISL_3103048, EPI_ISL_3103049, EPI_ISL_3103050, EPI_ISL_3103051, EPI_ISL_3103052, EPI_ISL_3103053, EPI_ISL_3103054, EPI_ISL_3103055, EPI_ISL_3103056, EPI_ISL_3103057, EPI_ISL_3103058, EPI_ISL_3103059, EPI_ISL_3103060, EPI_ISL_3103061, EPI_ISL_3103062, EPI_ISL_3103063, EPI_ISL_3103064, EPI_ISL_3103065, EPI_ISL_3103066, EPI_ISL_3103067, EPI_ISL_3103068, EPI_ISL_3103069, EPI_ISL_3103070, EPI_ISL_3103071, EPI_ISL_3103072, EPI_ISL_3103073, EPI_ISL_3103074, EPI_ISL_3103075, EPI_ISL_3103076, EPI_ISL_3103077, EPI_ISL_3103078, EPI_ISL_3103079, EPI_ISL_3103080, EPI_ISL_3103081, EPI_ISL_3103082, EPI_ISL_3103083, EPI_ISL_3103084, EPI_ISL_3103085, EPI_ISL_3103086, EPI_ISL_3103087, EPI_ISL_3103088, EPI_ISL_3103089, EPI_ISL_3103090, EPI_ISL_3103091, EPI_ISL_3103092, EPI_ISL_3103093, EPI_ISL_3103094, EPI_ISL_3103095, EPI_ISL_3103096, EPI_ISL_3103097, EPI_ISL_3103098, EPI_ISL_3103099, EPI_ISL_3103100, EPI_ISL_3103101, EPI_ISL_3103102, EPI_ISL_3103103, EPI_ISL_3103104, EPI_ISL_3103105, EPI_ISL_3103106, EPI_ISL_3103107, EPI_ISL_3103108, EPI_ISL_3103109, EPI_ISL_3103110, EPI_ISL_3103111, EPI_ISL_3103112, EPI_ISL_3103113, EPI_ISL_3103114, EPI_ISL_3103115, EPI_ISL_3103116, EPI_ISL_3103117, EPI_ISL_3103118, EPI_ISL_3103119, EPI_ISL_3103120, EPI_ISL_3103121, EPI_ISL_3103122, EPI_ISL_3103123, EPI_ISL_3103124, EPI_ISL_3103125, EPI_ISL_3103126, EPI_ISL_3103127, EPI_ISL_3103128, EPI_ISL_3103129, EPI_ISL_3103130, EPI_ISL_3103131, EPI_ISL_3103132, EPI_ISL_3103133, EPI_ISL_3103134, EPI_ISL_3103135, EPI_ISL_3103136, EPI_ISL_3103137, EPI_ISL_3103138, EPI_ISL_3103139, EPI_ISL_3103140, EPI_ISL_3103141, EPI_ISL_3103142, EPI_ISL_3103143, EPI_ISL_3103144, EPI_ISL_3103145, EPI_ISL_3103146, EPI_ISL_3103147, EPI_ISL_3103148, EPI_ISL_3103149, EPI_ISL_3103150, EPI_ISL_3103151, EPI_ISL_3103152, EPI_ISL_3103153, EPI_ISL_3103154, EPI_ISL_3103155, EPI_ISL_3103156, EPI_ISL_3103157, EPI_ISL_3103158, EPI_ISL_3103159, EPI_ISL_3103160, EPI_ISL_3103161, EPI_ISL_3103162, EPI_ISL_3103163, EPI_ISL_3103164, EPI_ISL_3103165, EPI_ISL_3103166, EPI_ISL_3103168, EPI_ISL_3103169, EPI_ISL_3103170, EPI_ISL_3103171, EPI_ISL_3103172, EPI_ISL_3103173, EPI_ISL_3103174, EPI_ISL_3103175, EPI_ISL_3103176, EPI_ISL_3103177, EPI_ISL_3103178, EPI_ISL_3103179, EPI_ISL_3103180, EPI_ISL_3103181, EPI_ISL_3103182, EPI_ISL_3103183, EPI_ISL_3103184, EPI_ISL_3103185, EPI_ISL_3103186, EPI_ISL_3103187, EPI_ISL_3103188, EPI_ISL_3103189, EPI_ISL_3103190, EPI_ISL_3103191, EPI_ISL_3103192, EPI_ISL_3103193, EPI_ISL_3103194, EPI_ISL_3103195, EPI_ISL_3103196, EPI_ISL_3103197, EPI_ISL_3103198, EPI_ISL_3103199, EPI_ISL_3103200, EPI_ISL_3103201, EPI_ISL_3103202, EPI_ISL_3103203, EPI_ISL_3103204, EPI_ISL_3103205, EPI_ISL_3103206, EPI_ISL_3103207, EPI_ISL_3103209, EPI_ISL_3103210, EPI_ISL_3103211, EPI_ISL_3103212, EPI_ISL_3103213, EPI_ISL_3103214, EPI_ISL_3103215, EPI_ISL_3103216, EPI_ISL_3103217, EPI_ISL_3103218, EPI_ISL_3103219, EPI_ISL_3103220, EPI_ISL_3103221, EPI_ISL_3103222, EPI_ISL_3103223, EPI_ISL_3103224, EPI_ISL_3103225, EPI_ISL_3103226, EPI_ISL_3103227, EPI_ISL_3103228, EPI_ISL_3103229, EPI_ISL_3103230, EPI_ISL_3103231, EPI_ISL_3103232, EPI_ISL_3103233, EPI_ISL_3103234, EPI_ISL_3103235, EPI_ISL_3103236, EPI_ISL_3103237, EPI_ISL_3103238, EPI_ISL_3103239, EPI_ISL_3103240, EPI_ISL_3103241, EPI_ISL_3103242, EPI_ISL_3103243, EPI_ISL_3103244, EPI_ISL_3103245, EPI_ISL_3103246, EPI_ISL_3103247, EPI_ISL_3103248, EPI_ISL_3103249, EPI_ISL_3103250, EPI_ISL_3103251, EPI_ISL_3103252, EPI_ISL_3103253, EPI_ISL_3103254, EPI_ISL_3103255, EPI_ISL_3103256, EPI_ISL_3103257, EPI_ISL_3103258, EPI_ISL_3103259, EPI_ISL_3103260, EPI_ISL_3103261, EPI_ISL_3103262, EPI_ISL_3103263, EPI_ISL_3103264, EPI_ISL_3103265, EPI_ISL_3103266, EPI_ISL_3103267, EPI_ISL_3103268, EPI_ISL_3103269, EPI_ISL_3103270, EPI_ISL_3103271, EPI_ISL_3103272, EPI_ISL_3103273, EPI_ISL_3103274, EPI_ISL_3103275, EPI_ISL_3103276, EPI_ISL_3103277, EPI_ISL_3103278, EPI_ISL_3103279, EPI_ISL_3103280, EPI_ISL_3103281, EPI_ISL_3103282, EPI_ISL_3103283, EPI_ISL_3103284, EPI_ISL_3103285, EPI_ISL_3103286, EPI_ISL_3103287, EPI_ISL_3103288, EPI_ISL_3103289, EPI_ISL_3103290, EPI_ISL_3103291, EPI_ISL_3103292, EPI_ISL_3103293, EPI_ISL_3103294, EPI_ISL_3103295, EPI_ISL_3103296, EPI_ISL_3103297, EPI_ISL_3103298, EPI_ISL_3103299, EPI_ISL_3103300, EPI_ISL_3103301, EPI_ISL_3103302, EPI_ISL_3103303, EPI_ISL_3103304, EPI_ISL_3103305, EPI_ISL_3103306, EPI_ISL_3103307, EPI_ISL_3103308, EPI_ISL_3103309, EPI_ISL_3103310, EPI_ISL_3103311, EPI_ISL_3103312, EPI_ISL_3103313, EPI_ISL_3103314, EPI_ISL_3103315, EPI_ISL_3103316, EPI_ISL_3103317, EPI_ISL_3103318, EPI_ISL_3103319, EPI_ISL_3103320, EPI_ISL_3103321, EPI_ISL_3103322, EPI_ISL_3103323, EPI_ISL_3103324, EPI_ISL_3103325, EPI_ISL_3103326, EPI_ISL_3103327, EPI_ISL_3103328, EPI_ISL_3103329, EPI_ISL_3103331 | | | |
| see above | Department of Bacteria, Parasites and Fungi, Statens Serum Institut, Copenhagen, Denmark | Statens Serum Institut Bioinformatics and Microbial Genomics | Danish Covid-19 Genome Consortium |

| | | | |
|---|---|---|---|
| EPI_ISL_3103333 | Molekylær Medicinsk Afdeling, Aarhus University Hospital, Aarhus, Denmark | Statens Serum Institut Bioinformatics and Microbial Genomics | Danish Covid-19 Genome Consortium |
| EPI_ISL_3103334, EPI_ISL_3103336, EPI_ISL_3103337, EPI_ISL_3103338, EPI_ISL_3103339, EPI_ISL_3103340, EPI_ISL_3103341, EPI_ISL_3103342, EPI_ISL_3103343, EPI_ISL_3103344, EPI_ISL_3103345, EPI_ISL_3103346, EPI_ISL_3103347, EPI_ISL_3103348, EPI_ISL_3103349, EPI_ISL_3103350, EPI_ISL_3103351, EPI_ISL_3103352, EPI_ISL_3103353, EPI_ISL_3103354, EPI_ISL_3103355, EPI_ISL_3103356, EPI_ISL_3103357, EPI_ISL_3103358, EPI_ISL_3103359, EPI_ISL_3103360, EPI_ISL_3103361, EPI_ISL_3103362, EPI_ISL_3103363, EPI_ISL_3103364, EPI_ISL_3103365, EPI_ISL_3103366, EPI_ISL_3103367, EPI_ISL_3103368, EPI_ISL_3103369, EPI_ISL_3103370, EPI_ISL_3103371, EPI_ISL_3103372, EPI_ISL_3103373, EPI_ISL_3103374, EPI_ISL_3103375, EPI_ISL_3103376, EPI_ISL_3103377, EPI_ISL_3103378, EPI_ISL_3103379, EPI_ISL_3103380, EPI_ISL_3103381, EPI_ISL_3103382, EPI_ISL_3103383, EPI_ISL_3103384, EPI_ISL_3103385, EPI_ISL_3103386, EPI_ISL_3103387, EPI_ISL_3103388, EPI_ISL_3103389, EPI_ISL_3103390 | | | |
| see above | Department of Bacteria, Parasites and Fungi, Statens Serum Institut, Copenhagen, Denmark | Statens Serum Institut Bioinformatics and Microbial Genomics | Danish Covid-19 Genome Consortium |
| EPI_ISL_3103391 | Molekylær Medicinsk Afdeling, Aarhus University Hospital, Aarhus, Denmark | Statens Serum Institut Bioinformatics and Microbial Genomics | Danish Covid-19 Genome Consortium |
| EPI_ISL_3103392, EPI_ISL_3103393, EPI_ISL_3103394, EPI_ISL_3103395, EPI_ISL_3103396, EPI_ISL_3103397, EPI_ISL_3103398, EPI_ISL_3103399, EPI_ISL_3103400, EPI_ISL_3103401, EPI_ISL_3103403, EPI_ISL_3103404, EPI_ISL_3103405, EPI_ISL_3103406, EPI_ISL_3103407, EPI_ISL_3103408, EPI_ISL_3103409, EPI_ISL_3103410, EPI_ISL_3103411, EPI_ISL_3103412, EPI_ISL_3103413, EPI_ISL_3103414, EPI_ISL_3103415, EPI_ISL_3103416, EPI_ISL_3103417, EPI_ISL_3103418, EPI_ISL_3103419, EPI_ISL_3103420, EPI_ISL_3103421, EPI_ISL_3103422, EPI_ISL_3103423, EPI_ISL_3103424, EPI_ISL_3103425, EPI_ISL_3103426, EPI_ISL_3103427, EPI_ISL_3103428, EPI_ISL_3103429, EPI_ISL_3103430, EPI_ISL_3103431, EPI_ISL_3103432, EPI_ISL_3103433, EPI_ISL_3103434, EPI_ISL_3103435, EPI_ISL_3103436, EPI_ISL_3103437, EPI_ISL_3103438, EPI_ISL_3103439, EPI_ISL_3103440, EPI_ISL_3103441, EPI_ISL_3103442, EPI_ISL_3103443, EPI_ISL_3103444, EPI_ISL_3103445, EPI_ISL_3103446, EPI_ISL_3103447, EPI_ISL_3103448, EPI_ISL_3103449, EPI_ISL_3103450, EPI_ISL_3103451, EPI_ISL_3103452, EPI_ISL_3103453, EPI_ISL_3103454, EPI_ISL_3103455, EPI_ISL_3103456, EPI_ISL_3103457, EPI_ISL_3103458, EPI_ISL_3103459, EPI_ISL_3103460, EPI_ISL_3103461, EPI_ISL_3103462, EPI_ISL_3103463, EPI_ISL_3103464, EPI_ISL_3103465, EPI_ISL_3103466, EPI_ISL_3103467, EPI_ISL_3103468, EPI_ISL_3103469, EPI_ISL_3103470, EPI_ISL_3103471, EPI_ISL_3103472, EPI_ISL_3103473, EPI_ISL_3103474, EPI_ISL_3103475, EPI_ISL_3103476, EPI_ISL_3103477, EPI_ISL_3103478, EPI_ISL_3103479, EPI_ISL_3103480, EPI_ISL_3103481, EPI_ISL_3103482, EPI_ISL_3103484, EPI_ISL_3103485 | | | |
| see above | Department of Bacteria, Parasites and Fungi, Statens Serum Institut, Copenhagen, Denmark | Statens Serum Institut Bioinformatics and Microbial Genomics | Danish Covid-19 Genome Consortium |
| EPI_ISL_3103486 | Molekylær Medicinsk Afdeling, Aarhus University Hospital, Aarhus, Denmark | Statens Serum Institut Bioinformatics and Microbial Genomics | Danish Covid-19 Genome Consortium |
| EPI_ISL_3103487, EPI_ISL_3103488, EPI_ISL_3103489, EPI_ISL_3103490, EPI_ISL_3103491, EPI_ISL_3103492, EPI_ISL_3103493, EPI_ISL_3103494, EPI_ISL_3103495, EPI_ISL_3103496, EPI_ISL_3103497, EPI_ISL_3103498, EPI_ISL_3103499, EPI_ISL_3103500, EPI_ISL_3103501, EPI_ISL_3103502, EPI_ISL_3103503, EPI_ISL_3103504, EPI_ISL_3103505, EPI_ISL_3103506, EPI_ISL_3103507, EPI_ISL_3103508, EPI_ISL_3103509, EPI_ISL_3103510, EPI_ISL_3103511, EPI_ISL_3103512, EPI_ISL_3103513, EPI_ISL_3103514, EPI_ISL_3103515, EPI_ISL_3103516, EPI_ISL_3103517, EPI_ISL_3103518, EPI_ISL_3103519, EPI_ISL_3103520, EPI_ISL_3103521, EPI_ISL_3103522, EPI_ISL_3103523, EPI_ISL_3103524, EPI_ISL_3103525, EPI_ISL_3103526, EPI_ISL_3103527, EPI_ISL_3103528, EPI_ISL_3103529, EPI_ISL_3103530 | | | |
| see above | Department of Bacteria, Parasites and Fungi, Statens Serum Institut, Copenhagen, Denmark | Statens Serum Institut Bioinformatics and Microbial Genomics | Danish Covid-19 Genome Consortium |
| EPI_ISL_3103531 | Molekylær Medicinsk Afdeling, Aarhus University Hospital, Aarhus, Denmark | Statens Serum Institut Bioinformatics and Microbial Genomics | Danish Covid-19 Genome Consortium |
| EPI_ISL_3103532, EPI_ISL_3103533, EPI_ISL_3103534, EPI_ISL_3103535, EPI_ISL_3103536, EPI_ISL_3103538, EPI_ISL_3103539, EPI_ISL_3103540, EPI_ISL_3103541, EPI_ISL_3103542, EPI_ISL_3103543, EPI_ISL_3103544, EPI_ISL_3103545, EPI_ISL_3103546, EPI_ISL_3103547, EPI_ISL_3103548, EPI_ISL_3103549, EPI_ISL_3103550, EPI_ISL_3103551, EPI_ISL_3103552, EPI_ISL_3103553, EPI_ISL_3103554, EPI_ISL_3103555, EPI_ISL_3103556, EPI_ISL_3103557, EPI_ISL_3103558, EPI_ISL_3103559, EPI_ISL_3103560, EPI_ISL_3103561, EPI_ISL_3103562, EPI_ISL_3103563, EPI_ISL_3103564, EPI_ISL_3103565, EPI_ISL_3103566, EPI_ISL_3103567, EPI_ISL_3103568, EPI_ISL_3103569, EPI_ISL_3103570, EPI_ISL_3103571, EPI_ISL_3103572, EPI_ISL_3103573, EPI_ISL_3103574, EPI_ISL_3103575, EPI_ISL_3103576, EPI_ISL_3103577, EPI_ISL_3103579, EPI_ISL_3103580, EPI_ISL_3103581, EPI_ISL_3103582, EPI_ISL_3103583, EPI_ISL_3103584, EPI_ISL_3103586, EPI_ISL_3103587, EPI_ISL_3103588, EPI_ISL_3103589, EPI_ISL_3103590, EPI_ISL_3103591, EPI_ISL_3103592, EPI_ISL_3103593, EPI_ISL_3103594, EPI_ISL_3103595, EPI_ISL_3103596, EPI_ISL_3103597, EPI_ISL_3103598, EPI_ISL_3103599, EPI_ISL_3103600, EPI_ISL_3103601, EPI_ISL_3103602, EPI_ISL_3103603, EPI_ISL_3103604, EPI_ISL_3103605, EPI_ISL_3103606, EPI_ISL_3103607, EPI_ISL_3103608, EPI_ISL_3103609, EPI_ISL_3103610, EPI_ISL_3103611, EPI_ISL_3103612, EPI_ISL_3103613, EPI_ISL_3103614, EPI_ISL_3103615, EPI_ISL_3103616, EPI_ISL_3103617, EPI_ISL_3103618, EPI_ISL_3103619, EPI_ISL_3103620, EPI_ISL_3103621, EPI_ISL_3103622, EPI_ISL_3103623, EPI_ISL_3103624, EPI_ISL_3103625, EPI_ISL_3103626, EPI_ISL_3103627, EPI_ISL_3103628, EPI_ISL_3103629, EPI_ISL_3103630, EPI_ISL_3103631, EPI_ISL_3103632, EPI_ISL_3103633, EPI_ISL_3103634, EPI_ISL_3103635, EPI_ISL_3103636, EPI_ISL_3103637, EPI_ISL_3103638, EPI_ISL_3103639, EPI_ISL_3103640, EPI_ISL_3103641, EPI_ISL_3103642, EPI_ISL_3103643, EPI_ISL_3103644, EPI_ISL_3103645, EPI_ISL_3103646, EPI_ISL_3103647, EPI_ISL_3103648, EPI_ISL_3103649, EPI_ISL_3103650, EPI_ISL_3103651, EPI_ISL_3103652, EPI_ISL_3103653, EPI_ISL_3103654, EPI_ISL_3103655, EPI_ISL_3103656, EPI_ISL_3103657, EPI_ISL_3103658, EPI_ISL_3103659, EPI_ISL_3103660, EPI_ISL_3103661, EPI_ISL_3103662, EPI_ISL_3103663, EPI_ISL_3103664, EPI_ISL_3103665, EPI_ISL_3103666, EPI_ISL_3103667, EPI_ISL_3103668, EPI_ISL_3103669, EPI_ISL_3103670, EPI_ISL_3103671, EPI_ISL_3103672, EPI_ISL_3103673, EPI_ISL_3103674, EPI_ISL_3103675, EPI_ISL_3103676, EPI_ISL_3103677, EPI_ISL_3103678, EPI_ISL_3103679, EPI_ISL_3103680, EPI_ISL_3103681, EPI_ISL_3103682, EPI_ISL_3103683, EPI_ISL_3103684, EPI_ISL_3103685, EPI_ISL_3103686, EPI_ISL_3103687, EPI_ISL_3103688, EPI_ISL_3103689, EPI_ISL_3103690, EPI_ISL_3103691, EPI_ISL_3103692, EPI_ISL_3103693, EPI_ISL_3103694, EPI_ISL_3103695, EPI_ISL_3103696, EPI_ISL_3103697, EPI_ISL_3103698, EPI_ISL_3103699, EPI_ISL_3103700, EPI_ISL_3103701, EPI_ISL_3103702, EPI_ISL_3103703, EPI_ISL_3103704, EPI_ISL_3103705, EPI_ISL_3103706, EPI_ISL_3103707, EPI_ISL_3103708, EPI_ISL_3103709, EPI_ISL_3103710, EPI_ISL_3103711, EPI_ISL_3103712, EPI_ISL_3103713, EPI_ISL_3103714, EPI_ISL_3103716, EPI_ISL_3103717, EPI_ISL_3103718, EPI_ISL_3103719, EPI_ISL_3103720, EPI_ISL_3103721, EPI_ISL_3103722, EPI_ISL_3103723, EPI_ISL_3103724, EPI_ISL_3103725, EPI_ISL_3103726, EPI_ISL_3103727, EPI_ISL_3103728, EPI_ISL_3103729, EPI_ISL_3103730, EPI_ISL_3103731, EPI_ISL_3103732, EPI_ISL_3103733, EPI_ISL_3103734, EPI_ISL_3103735, EPI_ISL_3103736, EPI_ISL_3103737, EPI_ISL_3103738, EPI_ISL_3103739, EPI_ISL_3103740, EPI_ISL_3103741, EPI_ISL_3103742, EPI_ISL_3103743, EPI_ISL_3103744, EPI_ISL_3103745, EPI_ISL_3103746, EPI_ISL_3103747, EPI_ISL_3103748, EPI_ISL_3103749, EPI_ISL_3103750, EPI_ISL_3103751, EPI_ISL_3103752, EPI_ISL_3103753, EPI_ISL_3103755, EPI_ISL_3103756, EPI_ISL_3103757, EPI_ISL_3103758, EPI_ISL_3103759, EPI_ISL_3103760, EPI_ISL_3103761, EPI_ISL_3103762, EPI_ISL_3103763, EPI_ISL_3103764, EPI_ISL_3103765, EPI_ISL_3103766, EPI_ISL_3103767, EPI_ISL_3103768, EPI_ISL_3103769, EPI_ISL_3103770, EPI_ISL_3103771, EPI_ISL_3103772, EPI_ISL_3103773, EPI_ISL_3103774, EPI_ISL_3103775, EPI_ISL_3103776, EPI_ISL_3103777, EPI_ISL_3103779, EPI_ISL_3103780, EPI_ISL_3103781, EPI_ISL_3103782, EPI_ISL_3103783, EPI_ISL_3103784, EPI_ISL_3103785, EPI_ISL_3103786, EPI_ISL_3103787, EPI_ISL_3103788, EPI_ISL_3103789, EPI_ISL_3103790, EPI_ISL_3103791, EPI_ISL_3103792, EPI_ISL_3103793, EPI_ISL_3103794, EPI_ISL_3103795, EPI_ISL_3103796, EPI_ISL_3103797, EPI_ISL_3103798, EPI_ISL_3103799, EPI_ISL_3103800, EPI_ISL_3103801, EPI_ISL_3103802, EPI_ISL_3103803, EPI_ISL_3103804, EPI_ISL_3103805, EPI_ISL_3103806, EPI_ISL_3103807, EPI_ISL_3103808, EPI_ISL_3103809, EPI_ISL_3103810, EPI_ISL_3103811, EPI_ISL_3103812, EPI_ISL_3103813, EPI_ISL_3103814, EPI_ISL_3103815, EPI_ISL_3103816, EPI_ISL_3103817, EPI_ISL_3103818, EPI_ISL_3103819, EPI_ISL_3103820, EPI_ISL_3103821, EPI_ISL_3103822, EPI_ISL_3103823, EPI_ISL_3103824, EPI_ISL_3103825, EPI_ISL_3103826, EPI_ISL_3103827, EPI_ISL_3103828, EPI_ISL_3103829, EPI_ISL_3103830, EPI_ISL_3103831, EPI_ISL_3103832, EPI_ISL_3103833, EPI_ISL_3103835, EPI_ISL_3103837, EPI_ISL_3103839, EPI_ISL_3103841, EPI_ISL_3103843, EPI_ISL_3103845, EPI_ISL_3103846, EPI_ISL_3103848, EPI_ISL_3103851, EPI_ISL_3103852, EPI_ISL_3103854, EPI_ISL_3103856, EPI_ISL_3103858 | | | |
| see above | Department of Bacteria, Parasites and Fungi, Statens Serum Institut, Copenhagen, Denmark | Statens Serum Institut Bioinformatics and Microbial Genomics | Danish Covid-19 Genome Consortium |
| EPI_ISL_3103859 | Arizona State University | Arizona State University | Ajeet Bains, Peter T. Skidmore, LaRinda A. Holland, Matthew F. Smith, Rabia Maqsood, Nicholas J. Mellor, Joy M. Blain, Valerie Harris, Joshua LaBaer, Vel Murugan, Efrem S. Lim |
| EPI_ISL_3103860, EPI_ISL_3103861, EPI_ISL_3103863, EPI_ISL_3103866, EPI_ISL_3103869, EPI_ISL_3103873, EPI_ISL_3103875, EPI_ISL_3103877, EPI_ISL_3103879, EPI_ISL_3103881, EPI_ISL_3103883, EPI_ISL_3103887, EPI_ISL_3103889, EPI_ISL_3103891, EPI_ISL_3103893, EPI_ISL_3103895, EPI_ISL_3103897, EPI_ISL_3103899, EPI_ISL_3103902, EPI_ISL_3103904, EPI_ISL_3103905, EPI_ISL_3103907 | | | |
| see above | Department of Bacteria, Parasites and Fungi, Statens Serum Institut, Copenhagen, Denmark | Statens Serum Institut Bioinformatics and Microbial Genomics | Danish Covid-19 Genome Consortium |
| EPI_ISL_3103908 | Arizona State University | Arizona State University | Ajeet Bains, Peter T. Skidmore, LaRinda A. Holland, Matthew F. Smith, Rabia Maqsood, Nicholas J. Mellor, Joy M. Blain, Valerie Harris, Joshua LaBaer, Vel Murugan, Efrem S. Lim |
| EPI_ISL_3103909, EPI_ISL_3103911, EPI_ISL_3103913, EPI_ISL_3103915, EPI_ISL_3103917, EPI_ISL_3103919, EPI_ISL_3103921, EPI_ISL_3103923, EPI_ISL_3103924, EPI_ISL_3103926, EPI_ISL_3103928, EPI_ISL_3103930, EPI_ISL_3103932, EPI_ISL_3103934, EPI_ISL_3103935, EPI_ISL_3103937, EPI_ISL_3103939, EPI_ISL_3103941, EPI_ISL_3103943, EPI_ISL_3103945, EPI_ISL_3103947, EPI_ISL_3103949, EPI_ISL_3103951, EPI_ISL_3103953, EPI_ISL_3103955, EPI_ISL_3103956, EPI_ISL_3103958, EPI_ISL_3103960, EPI_ISL_3103962, EPI_ISL_3103964, EPI_ISL_3103969, EPI_ISL_3103971, EPI_ISL_3103972, EPI_ISL_3103974, EPI_ISL_3103976, EPI_ISL_3103978, EPI_ISL_3103979, EPI_ISL_3103981, EPI_ISL_3103983, EPI_ISL_3103985, EPI_ISL_3103987, EPI_ISL_3103989, EPI_ISL_3103991, EPI_ISL_3103993, EPI_ISL_3103995, EPI_ISL_3103996, EPI_ISL_3103998, EPI_ISL_3104000, EPI_ISL_3104002, EPI_ISL_3104004, EPI_ISL_3104006, EPI_ISL_3104007, EPI_ISL_3104009, EPI_ISL_3104011, EPI_ISL_3104013, EPI_ISL_3104015, EPI_ISL_3104017, EPI_ISL_3104019, EPI_ISL_3104021, EPI_ISL_3104022, EPI_ISL_3104024, EPI_ISL_3104026, EPI_ISL_3104028, EPI_ISL_3104030, EPI_ISL_3104032, EPI_ISL_3104033, EPI_ISL_3104035, EPI_ISL_3104037, EPI_ISL_3104039, EPI_ISL_3104041, EPI_ISL_3104043, EPI_ISL_3104045, EPI_ISL_3104046, EPI_ISL_3104048, EPI_ISL_3104050, EPI_ISL_3104052, EPI_ISL_3104054, EPI_ISL_3104055, EPI_ISL_3104057, EPI_ISL_3104059, EPI_ISL_3104061, EPI_ISL_3104062, EPI_ISL_3104063, EPI_ISL_3104064, EPI_ISL_3104065, EPI_ISL_3104066, EPI_ISL_3104067, EPI_ISL_3104068, EPI_ISL_3104069, EPI_ISL_3104070, EPI_ISL_3104071, EPI_ISL_3104072, EPI_ISL_3104073, EPI_ISL_3104074, EPI_ISL_3104075, EPI_ISL_3104076, EPI_ISL_3104077, EPI_ISL_3104078, EPI_ISL_3104079, EPI_ISL_3104080, EPI_ISL_3104081, EPI_ISL_3104082, EPI_ISL_3104083, EPI_ISL_3104084, EPI_ISL_3104085, EPI_ISL_3104086, EPI_ISL_3104087, EPI_ISL_3104088, EPI_ISL_3104089, EPI_ISL_3104090, EPI_ISL_3104091, EPI_ISL_3104092, EPI_ISL_3104093, EPI_ISL_3104094, EPI_ISL_3104095, EPI_ISL_3104096, EPI_ISL_3104097, EPI_ISL_3104098, EPI_ISL_3104099, EPI_ISL_3104100, EPI_ISL_3104101, EPI_ISL_3104102, EPI_ISL_3104103, EPI_ISL_3104104, EPI_ISL_3104105, EPI_ISL_3104106, EPI_ISL_3104107, EPI_ISL_3104108, EPI_ISL_3104109, EPI_ISL_3104110, EPI_ISL_3104111, EPI_ISL_3104112, EPI_ISL_3104113, EPI_ISL_3104114, EPI_ISL_3104115, EPI_ISL_3104116, EPI_ISL_3104117, EPI_ISL_3104118, EPI_ISL_3104119, EPI_ISL_3104120, EPI_ISL_3104121, EPI_ISL_3104122, EPI_ISL_3104123, EPI_ISL_3104124, EPI_ISL_3104125, EPI_ISL_3104126, EPI_ISL_3104127, EPI_ISL_3104128, EPI_ISL_3104129, EPI_ISL_3104130, EPI_ISL_3104131, EPI_ISL_3104132, EPI_ISL_3104133, EPI_ISL_3104134, EPI_ISL_3104135, EPI_ISL_3104136, EPI_ISL_3104137, EPI_ISL_3104138, EPI_ISL_3104139, EPI_ISL_3104141, EPI_ISL_3104142, EPI_ISL_3104143 | | | |
| see above | Department of Bacteria, Parasites and Fungi, Statens Serum Institut, Copenhagen, Denmark | Statens Serum Institut Bioinformatics and Microbial Genomics | Danish Covid-19 Genome Consortium |
| EPI_ISL_3104144 | Molekylær Medicinsk Afdeling, Aarhus University Hospital, Aarhus, Denmark | Statens Serum Institut Bioinformatics and Microbial Genomics | Danish Covid-19 Genome Consortium |
| EPI_ISL_3104145, EPI_ISL_3104146, EPI_ISL_3104147, EPI_ISL_3104148, EPI_ISL_3104149, EPI_ISL_3104150, EPI_ISL_3104151, EPI_ISL_3104152, EPI_ISL_3104153, EPI_ISL_3104154, EPI_ISL_3104155, EPI_ISL_3104156, EPI_ISL_3104157, EPI_ISL_3104158, EPI_ISL_3104159, EPI_ISL_3104160, EPI_ISL_3104161, EPI_ISL_3104162, EPI_ISL_3104163, EPI_ISL_3104164, EPI_ISL_3104165, EPI_ISL_3104166, EPI_ISL_3104167, EPI_ISL_3104168, EPI_ISL_3104169, EPI_ISL_3104170, EPI_ISL_3104171, EPI_ISL_3104172, EPI_ISL_3104173, EPI_ISL_3104174, EPI_ISL_3104175, EPI_ISL_3104176, EPI_ISL_3104177, EPI_ISL_3104178, EPI_ISL_3104179, EPI_ISL_3104180, EPI_ISL_3104181, EPI_ISL_3104182, EPI_ISL_3104183, EPI_ISL_3104184, EPI_ISL_3104185, EPI_ISL_3104186, EPI_ISL_3104187, EPI_ISL_3104188, EPI_ISL_3104189, EPI_ISL_3104190, EPI_ISL_3104191, EPI_ISL_3104192, EPI_ISL_3104193, EPI_ISL_3104194, EPI_ISL_3104195, EPI_ISL_3104196, EPI_ISL_3104197, EPI_ISL_3104199, EPI_ISL_3104200, EPI_ISL_3104201, EPI_ISL_3104202, EPI_ISL_3104203, EPI_ISL_3104204, EPI_ISL_3104205, EPI_ISL_3104206, EPI_ISL_3104207, EPI_ISL_3104208, EPI_ISL_3104209, EPI_ISL_3104210, EPI_ISL_3104211, EPI_ISL_3104212, EPI_ISL_3104213, EPI_ISL_3104214, EPI_ISL_3104215, EPI_ISL_3104216, EPI_ISL_3104217, EPI_ISL_3104218, EPI_ISL_3104219, EPI_ISL_3104220, EPI_ISL_3104221, EPI_ISL_3104222, EPI_ISL_3104223, EPI_ISL_3104224, EPI_ISL_3104225, EPI_ISL_3104226, EPI_ISL_3104227, EPI_ISL_3104228, EPI_ISL_3104229, EPI_ISL_3104230, EPI_ISL_3104231, EPI_ISL_3104232, EPI_ISL_3104233, EPI_ISL_3104234, EPI_ISL_3104235, EPI_ISL_3104236, EPI_ISL_3104237, EPI_ISL_3104238, EPI_ISL_3104239, EPI_ISL_3104240, EPI_ISL_3104241, EPI_ISL_3104242, EPI_ISL_3104243, EPI_ISL_3104244, EPI_ISL_3104245, EPI_ISL_3104246, EPI_ISL_3104247, EPI_ISL_3104248, EPI_ISL_3104249, EPI_ISL_3104250, EPI_ISL_3104251, EPI_ISL_3104252, EPI_ISL_3104253, EPI_ISL_3104254, EPI_ISL_3104255, EPI_ISL_3104256, EPI_ISL_3104257, EPI_ISL_3104258, EPI_ISL_3104259, EPI_ISL_3104260, EPI_ISL_3104261, EPI_ISL_3104262, EPI_ISL_3104263, EPI_ISL_3104264, EPI_ISL_3104265, EPI_ISL_3104266, EPI_ISL_3104267, EPI_ISL_3104268, EPI_ISL_3104269, EPI_ISL_3104270, EPI_ISL_3104271, EPI_ISL_3104272, EPI_ISL_3104273, EPI_ISL_3104274, EPI_ISL_3104275, EPI_ISL_3104276, EPI_ISL_3104277, EPI_ISL_3104278, EPI_ISL_3104279, EPI_ISL_3104280, EPI_ISL_3104281, EPI_ISL_3104282, EPI_ISL_3104283, EPI_ISL_3104284, EPI_ISL_3104285, EPI_ISL_3104286, EPI_ISL_3104287, EPI_ISL_3104288, EPI_ISL_3104289, EPI_ISL_3104290, EPI_ISL_3104291, EPI_ISL_3104292, EPI_ISL_3104293, EPI_ISL_3104294, EPI_ISL_3104295, EPI_ISL_3104296, EPI_ISL_3104297, EPI_ISL_3104298, EPI_ISL_3104299, EPI_ISL_3104300, EPI_ISL_3104301, EPI_ISL_3104302, EPI_ISL_3104303, EPI_ISL_3104304, EPI_ISL_3104305 | | | |
| see above | Department of Bacteria, Parasites and Fungi, Statens Serum Institut, Copenhagen, Denmark | Statens Serum Institut Bioinformatics and Microbial Genomics | Danish Covid-19 Genome Consortium |
| EPI_ISL_3104306 | Department of Clinical Microbiology, Odense University Hospital, Odense, Denmark | Statens Serum Institut Bioinformatics and Microbial Genomics | Danish Covid-19 Genome Consortium |
| EPI_ISL_3104307, EPI_ISL_3104308, EPI_ISL_3104309, EPI_ISL_3104310, EPI_ISL_3104311, EPI_ISL_3104312, EPI_ISL_3104313, EPI_ISL_3104314, EPI_ISL_3104315, EPI_ISL_3104316, EPI_ISL_3104317, EPI_ISL_3104318, EPI_ISL_3104319, EPI_ISL_3104320, EPI_ISL_3104321, EPI_ISL_3104322, EPI_ISL_3104323, EPI_ISL_3104324, EPI_ISL_3104325, EPI_ISL_3104326, EPI_ISL_3104328, EPI_ISL_3104329, EPI_ISL_3104330, EPI_ISL_3104331, EPI_ISL_3104332, EPI_ISL_3104333, EPI_ISL_3104334, EPI_ISL_3104335, EPI_ISL_3104336, EPI_ISL_3104337, EPI_ISL_3104338, EPI_ISL_3104339, EPI_ISL_3104340, EPI_ISL_3104341 | | | |
| see above | Department of Bacteria, Parasites and Fungi, Statens Serum Institut, Copenhagen, Denmark | Statens Serum Institut Bioinformatics and Microbial Genomics | Danish Covid-19 Genome Consortium |
| EPI_ISL_3104342 | Molekylær Medicinsk Afdeling, Aarhus University Hospital, Aarhus, Denmark | Statens Serum Institut Bioinformatics and Microbial Genomics | Danish Covid-19 Genome Consortium |
| EPI_ISL_3104343, EPI_ISL_3104344, EPI_ISL_3104345, EPI_ISL_3104346, EPI_ISL_3104347, EPI_ISL_3104348, EPI_ISL_3104349, EPI_ISL_3104350, EPI_ISL_3104351, EPI_ISL_3104352, EPI_ISL_3104353, EPI_ISL_3104354, EPI_ISL_3104355, EPI_ISL_3104356, EPI_ISL_3104357, EPI_ISL_3104358, EPI_ISL_3104359, EPI_ISL_3104360, EPI_ISL_3104361, EPI_ISL_3104362, EPI_ISL_3104363, EPI_ISL_3104364, EPI_ISL_3104365, EPI_ISL_3104366, EPI_ISL_3104367, EPI_ISL_3104368, EPI_ISL_3104369, EPI_ISL_3104370, EPI_ISL_3104371, EPI_ISL_3104372, EPI_ISL_3104373, EPI_ISL_3104374, EPI_ISL_3104375, EPI_ISL_3104376, EPI_ISL_3104377, EPI_ISL_3104378, EPI_ISL_3104379, EPI_ISL_3104380, EPI_ISL_3104381, EPI_ISL_3104382, EPI_ISL_3104383, EPI_ISL_3104384, EPI_ISL_3104385, EPI_ISL_3104386, EPI_ISL_3104387, EPI_ISL_3104388, EPI_ISL_3104389, EPI_ISL_3104390, EPI_ISL_3104391, EPI_ISL_3104392, EPI_ISL_3104393, EPI_ISL_3104394, EPI_ISL_3104395, EPI_ISL_3104396, EPI_ISL_3104397, EPI_ISL_3104398, EPI_ISL_3104399, EPI_ISL_3104400, EPI_ISL_3104401, EPI_ISL_3104402, EPI_ISL_3104403, EPI_ISL_3104404, EPI_ISL_3104405, EPI_ISL_3104406, EPI_ISL_3104407, EPI_ISL_3104408, EPI_ISL_3104409, EPI_ISL_3104410, EPI_ISL_3104411, EPI_ISL_3104412, EPI_ISL_3104413, EPI_ISL_3104414, EPI_ISL_3104415, EPI_ISL_3104416, EPI_ISL_3104417, EPI_ISL_3104418, EPI_ISL_3104419, EPI_ISL_3104420, EPI_ISL_3104421, EPI_ISL_3104422, EPI_ISL_3104423, EPI_ISL_3104424, EPI_ISL_3104425, EPI_ISL_3104426, EPI_ISL_3104427, EPI_ISL_3104428, EPI_ISL_3104429, EPI_ISL_3104430, EPI_ISL_3104431, EPI_ISL_3104432, EPI_ISL_3104433, EPI_ISL_3104434, EPI_ISL_3104435, EPI_ISL_3104436, EPI_ISL_3104437, EPI_ISL_3104438, EPI_ISL_3104439, EPI_ISL_3104440, EPI_ISL_3104441, EPI_ISL_3104442, EPI_ISL_3104443, EPI_ISL_3104444, EPI_ISL_3104445, EPI_ISL_3104446, EPI_ISL_3104447, EPI_ISL_3104448, EPI_ISL_3104449, EPI_ISL_3104450, EPI_ISL_3104451, EPI_ISL_3104452, EPI_ISL_3104453, EPI_ISL_3104454, EPI_ISL_3104455, EPI_ISL_3104456, EPI_ISL_3104457, EPI_ISL_3104458, EPI_ISL_3104459, EPI_ISL_3104460, EPI_ISL_3104461, EPI_ISL_3104462, EPI_ISL_3104463, EPI_ISL_3104464, EPI_ISL_3104465, EPI_ISL_3104466, EPI_ISL_3104467, EPI_ISL_3104468, EPI_ISL_3104469, EPI_ISL_3104470, EPI_ISL_3104471, EPI_ISL_3104472, EPI_ISL_3104473, EPI_ISL_3104474, EPI_ISL_3104475, EPI_ISL_3104476, EPI_ISL_3104477, EPI_ISL_3104478, EPI_ISL_3104479, EPI_ISL_3104480, EPI_ISL_3104481, EPI_ISL_3104482, EPI_ISL_3104483, EPI_ISL_3104484, EPI_ISL_3104485, EPI_ISL_3104486, EPI_ISL_3104487, EPI_ISL_3104488, EPI_ISL_3104489, EPI_ISL_3104490, EPI_ISL_3104491, EPI_ISL_3104492, EPI_ISL_3104493, EPI_ISL_3104494, EPI_ISL_3104495, EPI_ISL_3104496, EPI_ISL_3104497, EPI_ISL_3104498, EPI_ISL_3104499, EPI_ISL_3104500, EPI_ISL_3104501, EPI_ISL_3104502, EPI_ISL_3104503, EPI_ISL_3104504, EPI_ISL_3104505, EPI_ISL_3104506, EPI_ISL_3104507, EPI_ISL_3104508, EPI_ISL_3104509, EPI_ISL_3104510, EPI_ISL_3104511, EPI_ISL_3104512, EPI_ISL_3104513, EPI_ISL_3104514, EPI_ISL_3104515, EPI_ISL_3104516, EPI_ISL_3104517, EPI_ISL_3104518, EPI_ISL_3104519, EPI_ISL_3104520, EPI_ISL_3104521, EPI_ISL_3104522, EPI_ISL_3104523, EPI_ISL_3104524, EPI_ISL_3104525, EPI_ISL_3104526, EPI_ISL_3104527, EPI_ISL_3104528, EPI_ISL_3104529, EPI_ISL_3104530, EPI_ISL_3104531, EPI_ISL_3104532, EPI_ISL_3104533, EPI_ISL_3104534, EPI_ISL_3104535, EPI_ISL_3104536, EPI_ISL_3104537, EPI_ISL_3104538, EPI_ISL_3104539, EPI_ISL_3104540, EPI_ISL_3104541, EPI_ISL_3104542, EPI_ISL_3104543, EPI_ISL_3104544, EPI_ISL_3104545, EPI_ISL_3104546, EPI_ISL_3104547, EPI_ISL_3104548, EPI_ISL_3104549, EPI_ISL_3104550, EPI_ISL_3104551, EPI_ISL_3104552, EPI_ISL_3104553, EPI_ISL_3104554, EPI_ISL_3104555, EPI_ISL_3104556, EPI_ISL_3104557, EPI_ISL_3104558, EPI_ISL_3104559, EPI_ISL_3104560, EPI_ISL_3104561, EPI_ISL_3104562, EPI_ISL_3104563, EPI_ISL_3104564, EPI_ISL_3104565, EPI_ISL_3104566, EPI_ISL_3104567, EPI_ISL_3104568, EPI_ISL_3104569, EPI_ISL_3104570, EPI_ISL_3104571, EPI_ISL_3104572, EPI_ISL_3104573, EPI_ISL_3104574, EPI_ISL_3104575, EPI_ISL_3104576, EPI_ISL_3104577, EPI_ISL_3104578, EPI_ISL_3104579, EPI_ISL_3104580, EPI_ISL_3104581 | | | |
| see above | Department of Bacteria, Parasites and Fungi, Statens Serum Institut, Copenhagen, Denmark | Statens Serum Institut Bioinformatics and Microbial Genomics | Danish Covid-19 Genome Consortium |
| EPI_ISL_3104582, EPI_ISL_3104583, EPI_ISL_3104585, EPI_ISL_3104586, EPI_ISL_3104587, EPI_ISL_3104588, EPI_ISL_3104589, EPI_ISL_3104590, EPI_ISL_3104591, EPI_ISL_3104592, EPI_ISL_3104593, EPI_ISL_3104594, EPI_ISL_3104595, EPI_ISL_3104596, EPI_ISL_3104597, EPI_ISL_3104598, EPI_ISL_3104599, EPI_ISL_3104600, EPI_ISL_3104601, EPI_ISL_3104602, EPI_ISL_3104603, EPI_ISL_3104604, EPI_ISL_3104607, EPI_ISL_3104608, EPI_ISL_3104609, EPI_ISL_3104610, EPI_ISL_3104611, EPI_ISL_3104612, EPI_ISL_3104613, EPI_ISL_3104614, EPI_ISL_3104615, EPI_ISL_3104616, EPI_ISL_3104617, EPI_ISL_3104618, EPI_ISL_3104619, EPI_ISL_3104620, EPI_ISL_3104621, EPI_ISL_3104622, EPI_ISL_3104623, EPI_ISL_3104624, EPI_ISL_3104625, EPI_ISL_3104626, EPI_ISL_3104627, EPI_ISL_3104628, EPI_ISL_3104629, EPI_ISL_3104630, EPI_ISL_3104631, EPI_ISL_3104633, EPI_ISL_3104634, EPI_ISL_3104635, EPI_ISL_3104636, EPI_ISL_3104637, EPI_ISL_3104638, EPI_ISL_3104639, EPI_ISL_3104640, EPI_ISL_3104641, EPI_ISL_3104642, EPI_ISL_3104643, EPI_ISL_3104644, EPI_ISL_3104645, EPI_ISL_3104646, EPI_ISL_3104647, EPI_ISL_3104648, EPI_ISL_3104649, EPI_ISL_3104651, EPI_ISL_3104652, EPI_ISL_3104653, EPI_ISL_3104656 | | | |
| see above | Alaska State Virology Laboratory | Alaska State Virology Laboratory | Stephanie DeRonde, Elva House, Jacob Zidek, Lisa Smith, Ph.D., Jack Chen, Ph.D. |
| EPI_ISL_3104657 | Instituto de Medicina Tropical & Salud Global Universidad Iberoamericana | Grubaugh Lab - Yale School of Public Health | Joseph Fauver, Mallery Breban, Isabel Ott, Tara Alpert, Mary Petrone, Anderson Brito, Chantal Vogels, Annie Watkins, Chaney Kalinich, Jessica Rothman, Kendall Billig, Tobias Koch, Robert Paulino-Ramirez, Alejandaro Vallejo Degaudenzi, Victor Virgilio Calderon, Elisa Contreras, Esperanza Mendoza, Nathan Grubaugh |
| EPI_ISL_3104658, | Yale Clinical Virology Lab | Grubaugh Lab - Yale School of Public Health | Joseph Fauver, Mallery Breban, Isabel Ott, Tara Alpert, Mary Petrone, Anderson Brito, Chantal Vogels, Annie Watkins, Chaney Kalinich, Jessica Rothman, Kendall Billig, Tobias Koch, Marie L. Landry, Nathan Grubaugh |

| | | | |
|---|---|---|---|
| EPI_ISL_3104660, EPI_ISL_3104661 | | | |
| EPI_ISL_3104662, EPI_ISL_3104665, EPI_ISL_3104667 | Instituto de Medicina Tropical & Salud Global Universidad Iberoamericana | Grubaugh Lab - Yale School of Public Health | Joseph Fauver, Mallery Breban, Isabel Ott, Tara Alpert, Mary Petrone, Anderson Brito, Chantal Vogels, Annie Watkins, Chaney Kalinich, Jessica Rothman, Kendall Billig, Tobias Koch, Robert Paulino-Ramirez, Alejandaro Vallejo Degaudenzi, Victor Virgilio Calderon, Elisa Contreras, Esperanza Mendoza, Nathan Grubaugh |
| EPI_ISL_3104676 | Yale Clinical Virology Lab | Grubaugh Lab - Yale School of Public Health | Joseph Fauver, Mallery Breban, Isabel Ott, Tara Alpert, Mary Petrone, Anderson Brito, Chantal Vogels, Annie Watkins, Chaney Kalinich, Jessica Rothman, Kendall Billig, Tobias Koch, Marie L. Landry, Nathan Grubaugh |
| EPI_ISL_3104677, EPI_ISL_3104679 | Instituto de Medicina Tropical & Salud Global Universidad Iberoamericana | Grubaugh Lab - Yale School of Public Health | Joseph Fauver, Mallery Breban, Isabel Ott, Tara Alpert, Mary Petrone, Anderson Brito, Chantal Vogels, Annie Watkins, Chaney Kalinich, Jessica Rothman, Kendall Billig, Tobias Koch, Robert Paulino-Ramirez, Alejandaro Vallejo Degaudenzi, Victor Virgilio Calderon, Elisa Contreras, Esperanza Mendoza, Nathan Grubaugh |
| EPI_ISL_3104681, EPI_ISL_3104711, EPI_ISL_3104712, EPI_ISL_3104713 | Yale Clinical Virology Lab | Grubaugh Lab - Yale School of Public Health | Joseph Fauver, Mallery Breban, Isabel Ott, Tara Alpert, Mary Petrone, Anderson Brito, Chantal Vogels, Annie Watkins, Chaney Kalinich, Jessica Rothman, Kendall Billig, Tobias Koch, Marie L. Landry, Nathan Grubaugh |
| EPI_ISL_3104714, EPI_ISL_3104715, EPI_ISL_3104716 | UCC (Universidad Central del Caribe) | Grubaugh Lab - Yale School of Public Health | Joseph Fauver, Mallery Breban, Isabel Ott, Tara Alpert, Mary Petrone, Anderson Brito, Chantal Vogels, Annie Watkins, Chaney Kalinich, Jessica Rothman, Kendall Billig, Tobias Koch, Robert Paulino-Ramirez, Alejandaro Vallejo Degaudenzi, Victor Virgilio Calderon, Elisa Contreras, Esperanza Mendoza, Nathan Grubaugh |
| EPI_ISL_3104720, EPI_ISL_3104721 | Yale Clinical Virology Lab | Grubaugh Lab - Yale School of Public Health | Joseph Fauver, Mallery Breban, Isabel Ott, Tara Alpert, Mary Petrone, Anderson Brito, Chantal Vogels, Annie Watkins, Chaney Kalinich, Jessica Rothman, Kendall Billig, Tobias Koch, Marie L. Landry, Nathan Grubaugh |
| EPI_ISL_3104725 | UCC (Universidad Central del Caribe) | Grubaugh Lab - Yale School of Public Health | Joseph Fauver, Mallery Breban, Isabel Ott, Tara Alpert, Mary Petrone, Anderson Brito, Chantal Vogels, Annie Watkins, Chaney Kalinich, Jessica Rothman, Kendall Billig, Tobias Koch, Robert Paulino-Ramirez, Alejandaro Vallejo Degaudenzi, Victor Virgilio Calderon, Elisa Contreras, Esperanza Mendoza, Nathan Grubaugh |
| EPI_ISL_3104726, EPI_ISL_3104727, EPI_ISL_3104728, EPI_ISL_3104729, EPI_ISL_3104730, EPI_ISL_3104731, EPI_ISL_3104732, EPI_ISL_3104733, EPI_ISL_3104734, EPI_ISL_3104735, EPI_ISL_3104736, EPI_ISL_3104737, EPI_ISL_3104738, EPI_ISL_3104739, EPI_ISL_3104740, EPI_ISL_3104741, EPI_ISL_3104742, EPI_ISL_3104743, EPI_ISL_3104744, EPI_ISL_3104745, EPI_ISL_3104746, EPI_ISL_3104747, EPI_ISL_3104748, EPI_ISL_3104749, EPI_ISL_3104750, EPI_ISL_3104751 | | | |
| see above | Yale Clinical Virology Lab | Grubaugh Lab - Yale School of Public Health | Joseph Fauver, Mallery Breban, Isabel Ott, Tara Alpert, Mary Petrone, Anderson Brito, Chantal Vogels, Annie Watkins, Chaney Kalinich, Jessica Rothman, Kendall Billig, Tobias Koch, Marie L. Landry, Nathan Grubaugh |
| EPI_ISL_3104752 | Instituto de Medicina Tropical & Salud Global Universidad Iberoamericana | Grubaugh Lab - Yale School of Public Health | Joseph Fauver, Mallery Breban, Isabel Ott, Tara Alpert, Mary Petrone, Anderson Brito, Chantal Vogels, Annie Watkins, Chaney Kalinich, Jessica Rothman, Kendall Billig, Tobias Koch, Fabiola Cruz-Lopez, David Rivera Aponte, Fabiola Fontanet, Natalie Machargo, Andrea Firpo, Vianca Aponte, Ernesto Vazquez, Patricia Serrano, Tania Mitwalli, Karolane Gonzalez, Nathan Grubaugh |
| EPI_ISL_3104764, EPI_ISL_3104765, EPI_ISL_3104766, EPI_ISL_3104767, EPI_ISL_3104768, EPI_ISL_3104769, EPI_ISL_3104770, EPI_ISL_3104771, EPI_ISL_3104772, EPI_ISL_3104773, EPI_ISL_3104774, EPI_ISL_3104775, EPI_ISL_3104776, EPI_ISL_3104777, EPI_ISL_3104778, EPI_ISL_3104779, EPI_ISL_3104780, EPI_ISL_3104781, EPI_ISL_3104782, EPI_ISL_3104783, EPI_ISL_3104784, EPI_ISL_3104785, EPI_ISL_3104786, EPI_ISL_3104787, EPI_ISL_3104788, EPI_ISL_3104789, EPI_ISL_3104790, EPI_ISL_3104791, EPI_ISL_3104792 | | | |
| see above | UCC (Universidad Central del Caribe) | Grubaugh Lab - Yale School of Public Health | Joseph Fauver, Mallery Breban, Isabel Ott, Tara Alpert, Mary Petrone, Anderson Brito, Chantal Vogels, Annie Watkins, Chaney Kalinich, Jessica Rothman, Kendall Billig, Tobias Koch, Robert Paulino-Ramirez, Alejandaro Vallejo Degaudenzi, Victor Virgilio Calderon, Elisa Contreras, Esperanza Mendoza, Nathan Grubaugh |
| EPI_ISL_3104793 | Yale Pathology Lab | Grubaugh Lab - Yale School of Public Health | Joseph Fauver, Mallery Breban, Isabel Ott, Tara Alpert, Mary Petrone, Anderson Brito, Chantal Vogels, Annie Watkins, Chaney Kalinich, Jessica Rothman, Kendall Billig, Tobias Koch, Jianhui Wang, Chen Liu, Pei Hui, Nathan Grubaugh |
| EPI_ISL_3104800, EPI_ISL_3104801, EPI_ISL_3104802, EPI_ISL_3104803, EPI_ISL_3104804, EPI_ISL_3104805, EPI_ISL_3104806, EPI_ISL_3104808, EPI_ISL_3104809, EPI_ISL_3104810, EPI_ISL_3104811, EPI_ISL_3104812, EPI_ISL_3104813, EPI_ISL_3104814 | | | |
| see above | Department of Microbiology, Yamanashi Institute for Public Health and Environment | Genome Analysis Center, Yamanashi Central Hospital | Yosuke Hirotsu |
| EPI_ISL_3104815, EPI_ISL_3104816, EPI_ISL_3104817, EPI_ISL_3104818 | BIOFAST | Instituto Butantan | Dimas Tadeu Covas, Antonio Jorge Martins, Claudia Renata dos Santos Barros, David Schlesinger, Debora Botequio Moretti, Elaine Cristina Marqueze, Elaine Vieira Santos, Evandra Strazza Rodrigues, Heidge Fukumasu, Jayme Augusto de Souza-Neto, José Salvatore Leister Patané, Luiz Alcantara, Luiz Lehmann Coutinho, Maria Carolina Elias, Maurício Lacerda Nogueira, Rafael dos Santos Bezerra, Raul Machado Neto, Rejane Maria Tommasini Grotto, Ricardo Haddad, Sandra Coccuzzo Sampaio Vessoni, Simone Kashima, Svetoslav Nanev Slavov, Vincent Louis Viala |
| EPI_ISL_3104819 | UBS BELENZINHO / UVIS MOOCA ARICANDUVA | Instituto Butantan | Dimas Tadeu Covas, Antonio Jorge Martins, Claudia Renata dos Santos Barros, David Schlesinger, Debora Botequio Moretti, Elaine Cristina Marqueze, Elaine Vieira Santos, Evandra Strazza Rodrigues, Heidge Fukumasu, Jayme Augusto de Souza-Neto, José Salvatore Leister Patané, Luiz Alcantara, Luiz Lehmann Coutinho, Maria Carolina Elias, Maurício Lacerda Nogueira, Rafael dos Santos Bezerra, Raul Machado Neto, Rejane Maria Tommasini Grotto, Ricardo Haddad, Sandra Coccuzzo Sampaio Vessoni, Simone Kashima, Svetoslav Nanev Slavov, Vincent Louis Viala |
| EPI_ISL_3104820 | INSIDE DIAGNÓSTICOS | Instituto Butantan | Dimas Tadeu Covas, Antonio Jorge Martins, Claudia Renata dos Santos Barros, David Schlesinger, Debora Botequio Moretti, Elaine Cristina Marqueze, Elaine Vieira Santos, Evandra Strazza Rodrigues, Heidge Fukumasu, Jayme Augusto de Souza-Neto, José Salvatore Leister Patané, Luiz Alcantara, Luiz Lehmann Coutinho, Maria Carolina Elias, Maurício Lacerda Nogueira, Rafael dos Santos Bezerra, Raul Machado Neto, Rejane Maria Tommasini Grotto, Ricardo Haddad, Sandra Coccuzzo Sampaio Vessoni, Simone Kashima, Svetoslav Nanev Slavov, Vincent Louis Viala |
| EPI_ISL_3104822 | SECAO CENTRO DE DIAGNOSTICO SECEDI | Instituto Butantan | Dimas Tadeu Covas, Antonio Jorge Martins, Claudia Renata dos Santos Barros, David Schlesinger, Debora Botequio Moretti, Elaine Cristina Marqueze, Elaine Vieira Santos, Evandra Strazza Rodrigues, Heidge Fukumasu, Jayme Augusto de Souza-Neto, José Salvatore Leister Patané, Luiz Alcantara, Luiz Lehmann Coutinho, Maria Carolina Elias, Maurício Lacerda Nogueira, Rafael dos Santos Bezerra, Raul Machado Neto, Rejane Maria Tommasini Grotto, Ricardo Haddad, Sandra Coccuzzo Sampaio Vessoni, Simone Kashima, Svetoslav Nanev Slavov, Vincent Louis Viala |
| EPI_ISL_3104823 | AFIP | Instituto Butantan | Dimas Tadeu Covas, Antonio Jorge Martins, Claudia Renata dos Santos Barros, David Schlesinger, Debora Botequio Moretti, Elaine Cristina Marqueze, Elaine Vieira Santos, Evandra Strazza Rodrigues, Heidge Fukumasu, Jayme Augusto de Souza-Neto, José Salvatore Leister Patané, Luiz Alcantara, Luiz Lehmann Coutinho, Maria Carolina Elias, Maurício Lacerda Nogueira, Rafael dos Santos Bezerra, Raul Machado Neto, Rejane Maria Tommasini Grotto, Ricardo Haddad, Sandra Coccuzzo Sampaio Vessoni, Simone Kashima, Svetoslav Nanev Slavov, Vincent Louis Viala |
| EPI_ISL_3104825, EPI_ISL_3104826 | INSIDE DIAGNÓSTICOS | Instituto Butantan | Dimas Tadeu Covas, Antonio Jorge Martins, Claudia Renata dos Santos Barros, David Schlesinger, Debora Botequio Moretti, Elaine Cristina Marqueze, Elaine Vieira Santos, Evandra Strazza Rodrigues, Heidge Fukumasu, Jayme Augusto de Souza-Neto, José Salvatore Leister Patané, Luiz Alcantara, Luiz Lehmann Coutinho, Maria Carolina Elias, Maurício Lacerda Nogueira, Rafael dos Santos Bezerra, Raul Machado Neto, Rejane Maria Tommasini Grotto, Ricardo Haddad, Sandra Coccuzzo Sampaio Vessoni, Simone Kashima, Svetoslav Nanev Slavov, Vincent Louis Viala |
| EPI_ISL_3104827 | Research Group of Rolf Kaiser, Institute of Virology, Cologne | Center of Medical Microbiology, Virology, and Hospital Hygiene, University of Duesseldorf | Philipp Niklas Ostermann, Elena Timmerberg, Eva Heger, Maximilian Damagnez, Alexander Dilthey, Torsten Houwaart, Lisanna Hülse, Nadine Lübke, Jessica Nicolai, Klaus Pfeffer, Daniel Strelow, Jörg Timm, Andreas Walker, Rolf Kaiser, Heiner Schaal |
| EPI_ISL_3104828, EPI_ISL_3104830, EPI_ISL_3104832, EPI_ISL_3104833 | Northwest Laboratory | Seattle Flu Study | Deborah A. Nickerson, Chris D. Frazar, Jover Lee, Benjamin Pelle, Erica Ryke, Matthew Richardson, Amanda Adler, Elisabeth Brandstetter, Peter D. Han, Kairsten Fay, Misja Ilcisin, Kirsten Lacombe, Thomas R. Sibley, Melissa Truong, Caitlin R. Wolf, Romesh Gautom, Geoff Melly, Brian Hiatt, Philip Dykema, Scott Lindquist, Michael Boeckh, Janet A. Englund, Michael Famulare, Barry R. Lutz, Mark J. Rieder, Lea M. Starita, Matthew Thompson, Helen Y. Chu, Jay Shendure, Trevor Bedford |
| EPI_ISL_3104835 | Atlas Genomics | Seattle Flu Study | Deborah A. Nickerson, Chris D. Frazar, Jover Lee, Benjamin Pelle, Erica Ryke, Matthew Richardson, Amanda Adler, Elisabeth Brandstetter, Peter D. Han, Kairsten Fay, Misja Ilcisin, Kirsten Lacombe, Thomas R. Sibley, Melissa Truong, Caitlin R. Wolf, Romesh Gautom, Geoff Melly, Brian Hiatt, Philip Dykema, Scott Lindquist, Michael Boeckh, Janet A. Englund, Michael Famulare, Barry R. Lutz, Mark J. Rieder, Lea M. Starita, Matthew Thompson, Helen Y. Chu, Jay Shendure, Trevor Bedford |
| EPI_ISL_3104838 | Northwest Laboratory | Seattle Flu Study | Deborah A. Nickerson, Chris D. Frazar, Jover Lee, Benjamin Pelle, Erica Ryke, Matthew Richardson, Amanda Adler, Elisabeth Brandstetter, Peter D. Han, Kairsten Fay, Misja Ilcisin, Kirsten Lacombe, Thomas R. Sibley, Melissa Truong, Caitlin R. Wolf, Romesh Gautom, Geoff Melly, Brian Hiatt, Philip Dykema, Scott Lindquist, Michael Boeckh, Janet A. Englund, Michael Famulare, Barry R. Lutz, Mark J. Rieder, Lea M. Starita, Matthew Thompson, Helen Y. Chu, Jay Shendure, Trevor Bedford |
| EPI_ISL_3104842 | Interpath Laboratory | Seattle Flu Study | Deborah A. Nickerson, Chris D. Frazar, Jover Lee, Benjamin Pelle, Erica Ryke, Matthew Richardson, Amanda Adler, Elisabeth Brandstetter, Peter D. Han, Kairsten Fay, Misja Ilcisin, Kirsten Lacombe, Thomas R. Sibley, Melissa Truong, Caitlin R. Wolf, Romesh Gautom, Geoff Melly, Brian Hiatt, Philip Dykema, Scott Lindquist, Michael Boeckh, Janet A. Englund, Michael Famulare, Barry R. Lutz, Mark J. Rieder, Lea M. Starita, Matthew Thompson, Helen Y. Chu, Jay Shendure, Trevor Bedford |
| EPI_ISL_3104843, EPI_ISL_3104844, EPI_ISL_3104846 | Northwest Laboratory | Seattle Flu Study | Deborah A. Nickerson, Chris D. Frazar, Jover Lee, Benjamin Pelle, Erica Ryke, Matthew Richardson, Amanda Adler, Elisabeth Brandstetter, Peter D. Han, Kairsten Fay, Misja Ilcisin, Kirsten Lacombe, Thomas R. Sibley, Melissa Truong, Caitlin R. Wolf, Romesh Gautom, Geoff Melly, Brian Hiatt, Philip Dykema, Scott Lindquist, Michael Boeckh, Janet A. Englund, Michael Famulare, Barry R. Lutz, Mark J. Rieder, Lea M. Starita, Matthew Thompson, Helen Y. Chu, Jay Shendure, Trevor Bedford |
| EPI_ISL_3104847 | Atlas Genomics | Seattle Flu Study | Deborah A. Nickerson, Chris D. Frazar, Jover Lee, Benjamin Pelle, Erica Ryke, Matthew Richardson, Amanda Adler, Elisabeth Brandstetter, Peter D. Han, Kairsten Fay, Misja Ilcisin, Kirsten Lacombe, Thomas R. Sibley, Melissa Truong, Caitlin R. Wolf, Romesh Gautom, Geoff Melly, Brian Hiatt, Philip Dykema, Scott Lindquist, Michael Boeckh, Janet A. Englund, Michael Famulare, Barry R. Lutz, Mark J. Rieder, Lea M. Starita, Matthew Thompson, Helen Y. Chu, Jay Shendure, Trevor Bedford |
| EPI_ISL_3104848 | Northwest Laboratory | Seattle Flu Study | Deborah A. Nickerson, Chris D. Frazar, Jover Lee, Benjamin Pelle, Erica Ryke, Matthew Richardson, Amanda Adler, Elisabeth Brandstetter, Peter D. Han, Kairsten Fay, Misja Ilcisin, Kirsten Lacombe, Thomas R. Sibley, Melissa Truong, Caitlin R. Wolf, Romesh Gautom, Geoff Melly, Brian Hiatt, Philip Dykema, Scott Lindquist, Michael Boeckh, Janet A. Englund, Michael Famulare, Barry R. Lutz, Mark J. Rieder, Lea M. Starita, Matthew Thompson, Helen Y. Chu, Jay Shendure, Trevor Bedford |
| EPI_ISL_3104855 | Seattle Flu Study | Seattle Flu Study | Deborah A. Nickerson, Chris D. Frazar, Jover Lee, Benjamin Pelle, Erica Ryke, Matthew Richardson, Amanda Adler, Elisabeth Brandstetter, Peter D. Han, Kairsten Fay, Misja Ilcisin, Kirsten Lacombe, Thomas R. Sibley, Melissa Truong, Caitlin R. Wolf, Karen Cowgill, Stephanie Schrag, Jeff Duchin, Michael Boeckh, Janet A. Englund, Michael Famulare, Barry R. Lutz, Mark J. Rieder, Lea M. Starita, Matthew Thompson, Helen Y. Chu, Trevor Bedford, Jay Shendure |
| EPI_ISL_3104857, EPI_ISL_3104858, EPI_ISL_3104862 | Northwest Laboratory | Seattle Flu Study | Deborah A. Nickerson, Chris D. Frazar, Jover Lee, Benjamin Pelle, Erica Ryke, Matthew Richardson, Amanda Adler, Elisabeth Brandstetter, Peter D. Han, Kairsten Fay, Misja Ilcisin, Kirsten Lacombe, Thomas R. Sibley, Melissa Truong, Caitlin R. Wolf, Romesh Gautom, Geoff Melly, Brian Hiatt, Philip Dykema, Scott Lindquist, Michael Boeckh, Janet A. Englund, Michael Famulare, Barry R. Lutz, Mark J. Rieder, Lea M. Starita, Matthew Thompson, Helen Y. Chu, Jay Shendure, Trevor Bedford |
| EPI_ISL_3104864 | Interpath Laboratory | Seattle Flu Study | Deborah A. Nickerson, Chris D. Frazar, Jover Lee, Benjamin Pelle, Erica Ryke, Matthew Richardson, Amanda Adler, Elisabeth Brandstetter, Peter D. Han, Kairsten Fay, Misja Ilcisin, Kirsten Lacombe, Thomas R. Sibley, Melissa Truong, Caitlin R. Wolf, Romesh Gautom, Geoff Melly, Brian Hiatt, Philip Dykema, Scott Lindquist, Michael Boeckh, Janet A. Englund, Michael Famulare, Barry R. Lutz, Mark J. Rieder, Lea M. Starita, Matthew Thompson, Helen Y. Chu, Jay Shendure, Trevor Bedford |
| EPI_ISL_3104866 | Northwest Laboratory | Seattle Flu Study | Deborah A. Nickerson, Chris D. Frazar, Jover Lee, Benjamin Pelle, Erica Ryke, Matthew Richardson, Amanda Adler, Elisabeth Brandstetter, Peter D. Han, Kairsten Fay, Misja Ilcisin, Kirsten Lacombe, Thomas R. Sibley, Melissa Truong, Caitlin R. Wolf, Romesh Gautom, Geoff Melly, Brian Hiatt, Philip Dykema, Scott Lindquist, Michael Boeckh, Janet A. Englund, Michael Famulare, Barry R. Lutz, Mark J. Rieder, Lea M. Starita, Matthew Thompson, Helen Y. Chu, Jay Shendure, Trevor Bedford |
| EPI_ISL_3104868 | Atlas Genomics | Seattle Flu Study | Deborah A. Nickerson, Chris D. Frazar, Jover Lee, Benjamin Pelle, Erica Ryke, Matthew Richardson, Amanda Adler, Elisabeth Brandstetter, Peter D. Han, Kairsten Fay, Misja Ilcisin, Kirsten Lacombe, Thomas R. Sibley, Melissa Truong, Caitlin R. Wolf, Romesh Gautom, Geoff Melly, Brian Hiatt, Philip Dykema, Scott Lindquist, Michael Boeckh, Janet A. Englund, Michael Famulare, Barry R. Lutz, Mark J. Rieder, Lea M. Starita, Matthew Thompson, Helen Y. Chu, Jay Shendure, Trevor Bedford |
| EPI_ISL_3104872 | Northwest Laboratory | Seattle Flu Study | Deborah A. Nickerson, Chris D. Frazar, Jover Lee, Benjamin Pelle, Erica Ryke, Matthew Richardson, Amanda Adler, Elisabeth Brandstetter, Peter D. Han, Kairsten Fay, Misja Ilcisin, Kirsten Lacombe, Thomas R. Sibley, Melissa Truong, Caitlin R. Wolf, Romesh Gautom, Geoff Melly, Brian Hiatt, Philip Dykema, Scott Lindquist, Michael Boeckh, Janet A. Englund, Michael Famulare, Barry R. Lutz, Mark J. Rieder, Lea M. Starita, Matthew Thompson, Helen Y. Chu, Jay Shendure, Trevor Bedford |
| EPI_ISL_3104876, EPI_ISL_3104883 | Seattle Flu Study | Seattle Flu Study | Deborah A. Nickerson, Chris D. Frazar, Jover Lee, Benjamin Pelle, Erica Ryke, Matthew Richardson, Amanda Adler, Elisabeth Brandstetter, Peter D. Han, Kairsten Fay, Misja Ilcisin, Kirsten Lacombe, Thomas R. Sibley, Melissa Truong, Caitlin R. Wolf, Karen Cowgill, Stephanie Schrag, Jeff Duchin, Michael Boeckh, Janet A. Englund, Michael Famulare, Barry R. Lutz, Mark J. Rieder, Lea M. Starita, Matthew Thompson, Helen Y. Chu, Trevor Bedford, Jay Shendure |
| EPI_ISL_3104887 | Northwest Laboratory | Seattle Flu Study | Deborah A. Nickerson, Chris D. Frazar, Jover Lee, Benjamin Pelle, Erica Ryke, Matthew Richardson, Amanda Adler, Elisabeth Brandstetter, Peter D. Han, Kairsten Fay, Misja Ilcisin, Kirsten Lacombe, Thomas R. Sibley, Melissa Truong, Caitlin R. Wolf, Romesh Gautom, Geoff Melly, Brian Hiatt, Philip Dykema, Scott Lindquist, Michael Boeckh, Janet A. Englund, Michael Famulare, Barry R. Lutz, Mark J. Rieder, Lea M. Starita, Matthew Thompson, Helen Y. Chu, Jay Shendure, Trevor Bedford |
| EPI_ISL_3104891 | Incyte Diagnostics Spokane | Seattle Flu Study | Deborah A. Nickerson, Chris D. Frazar, Jover Lee, Benjamin Pelle, Erica Ryke, Matthew Richardson, Amanda Adler, Elisabeth Brandstetter, Peter D. Han, Kairsten Fay, Misja Ilcisin, Kirsten Lacombe, Thomas R. Sibley, Melissa Truong, Caitlin R. Wolf, Romesh Gautom, Geoff Melly, Brian Hiatt, Philip Dykema, Scott Lindquist, Michael Boeckh, Janet A. Englund, Michael Famulare, Barry R. Lutz, Mark J. Rieder, Lea M. Starita, Matthew Thompson, Helen Y. Chu, Jay Shendure, Trevor Bedford |
| EPI_ISL_3104896, EPI_ISL_3104897, EPI_ISL_3104902 | Northwest Laboratory | Seattle Flu Study | Deborah A. Nickerson, Chris D. Frazar, Jover Lee, Benjamin Pelle, Erica Ryke, Matthew Richardson, Amanda Adler, Elisabeth Brandstetter, Peter D. Han, Kairsten Fay, Misja Ilcisin, Kirsten Lacombe, Thomas R. Sibley, Melissa Truong, Caitlin R. Wolf, Romesh Gautom, Geoff Melly, Brian Hiatt, Philip Dykema, Scott Lindquist, Michael Boeckh, Janet A. Englund, Michael Famulare, Barry R. Lutz, Mark J. Rieder, Lea M. Starita, Matthew Thompson, Helen Y. Chu, Jay Shendure, Trevor Bedford |

| | | | |
|---|---|---|---|
| EPI_ISL_3104906, EPI_ISL_3104913 | Atlas Genomics | Seattle Flu Study | Deborah A. Nickerson, Chris D. Frazar, Jover Lee, Benjamin Pelle, Erica Ryke, Matthew Richardson, Amanda Adler, Elisabeth Brandstetter, Peter D. Han, Kairsten Fay, Misja Ilcisin, Kirsten Lacombe, Thomas R. Sibley, Melissa Truong, Caitlin R. Wolf, Romesh Gautom, Geoff Melly, Brian Hiatt, Philip Dykema, Scott Lindquist, Michael Boeckh, Janet A. Englund, Michael Famulare, Barry R. Lutz, Mark J. Rieder, Lea M. Starita, Matthew Thompson, Helen Y. Chu, Jay Shendure, Trevor Bedford |
| EPI_ISL_3104914, EPI_ISL_3104916 | Northwest Laboratory | Seattle Flu Study | Deborah A. Nickerson, Chris D. Frazar, Jover Lee, Benjamin Pelle, Erica Ryke, Matthew Richardson, Amanda Adler, Elisabeth Brandstetter, Peter D. Han, Kairsten Fay, Misja Ilcisin, Kirsten Lacombe, Thomas R. Sibley, Melissa Truong, Caitlin R. Wolf, Romesh Gautom, Geoff Melly, Brian Hiatt, Philip Dykema, Scott Lindquist, Michael Boeckh, Janet A. Englund, Michael Famulare, Barry R. Lutz, Mark J. Rieder, Lea M. Starita, Matthew Thompson, Helen Y. Chu, Jay Shendure, Trevor Bedford |
| EPI_ISL_3104919 | Interpath Laboratory | Seattle Flu Study | Deborah A. Nickerson, Chris D. Frazar, Jover Lee, Benjamin Pelle, Erica Ryke, Matthew Richardson, Amanda Adler, Elisabeth Brandstetter, Peter D. Han, Kairsten Fay, Misja Ilcisin, Kirsten Lacombe, Thomas R. Sibley, Melissa Truong, Caitlin R. Wolf, Romesh Gautom, Geoff Melly, Brian Hiatt, Philip Dykema, Scott Lindquist, Michael Boeckh, Janet A. Englund, Michael Famulare, Barry R. Lutz, Mark J. Rieder, Lea M. Starita, Matthew Thompson, Helen Y. Chu, Jay Shendure, Trevor Bedford |
| EPI_ISL_3104920 | Seattle Flu Study | Seattle Flu Study | Deborah A. Nickerson, Chris D. Frazar, Jover Lee, Benjamin Pelle, Erica Ryke, Matthew Richardson, Amanda Adler, Elisabeth Brandstetter, Peter D. Han, Kairsten Fay, Misja Ilcisin, Kirsten Lacombe, Thomas R. Sibley, Melissa Truong, Caitlin R. Wolf, Michael Boeckh, Janet A. Englund, Michael Famulare, Barry R. Lutz, Mark J. Rieder, Lea M. Starita, Matthew Thompson, Jay Shendure, Trevor Bedford, Helen Y. Chu |
| EPI_ISL_3104923 | Atlas Genomics | Seattle Flu Study | Deborah A. Nickerson, Chris D. Frazar, Jover Lee, Benjamin Pelle, Erica Ryke, Matthew Richardson, Amanda Adler, Elisabeth Brandstetter, Peter D. Han, Kairsten Fay, Misja Ilcisin, Kirsten Lacombe, Thomas R. Sibley, Melissa Truong, Caitlin R. Wolf, Romesh Gautom, Geoff Melly, Brian Hiatt, Philip Dykema, Scott Lindquist, Michael Boeckh, Janet A. Englund, Michael Famulare, Barry R. Lutz, Mark J. Rieder, Lea M. Starita, Matthew Thompson, Helen Y. Chu, Jay Shendure, Trevor Bedford |
| EPI_ISL_3104925 | Northwest Laboratory | Seattle Flu Study | Deborah A. Nickerson, Chris D. Frazar, Jover Lee, Benjamin Pelle, Erica Ryke, Matthew Richardson, Amanda Adler, Elisabeth Brandstetter, Peter D. Han, Kairsten Fay, Misja Ilcisin, Kirsten Lacombe, Thomas R. Sibley, Melissa Truong, Caitlin R. Wolf, Romesh Gautom, Geoff Melly, Brian Hiatt, Philip Dykema, Scott Lindquist, Michael Boeckh, Janet A. Englund, Michael Famulare, Barry R. Lutz, Mark J. Rieder, Lea M. Starita, Matthew Thompson, Helen Y. Chu, Jay Shendure, Trevor Bedford |
| EPI_ISL_3104931 | Overlake Hospital | Seattle Flu Study | Deborah A. Nickerson, Chris D. Frazar, Jover Lee, Benjamin Pelle, Erica Ryke, Matthew Richardson, Amanda Adler, Elisabeth Brandstetter, Peter D. Han, Kairsten Fay, Misja Ilcisin, Kirsten Lacombe, Thomas R. Sibley, Melissa Truong, Caitlin R. Wolf, Romesh Gautom, Geoff Melly, Brian Hiatt, Philip Dykema, Scott Lindquist, Michael Boeckh, Janet A. Englund, Michael Famulare, Barry R. Lutz, Mark J. Rieder, Lea M. Starita, Matthew Thompson, Helen Y. Chu, Jay Shendure, Trevor Bedford |
| EPI_ISL_3104932, EPI_ISL_3104933 | Atlas Genomics | Seattle Flu Study | Deborah A. Nickerson, Chris D. Frazar, Jover Lee, Benjamin Pelle, Erica Ryke, Matthew Richardson, Amanda Adler, Elisabeth Brandstetter, Peter D. Han, Kairsten Fay, Misja Ilcisin, Kirsten Lacombe, Thomas R. Sibley, Melissa Truong, Caitlin R. Wolf, Romesh Gautom, Geoff Melly, Brian Hiatt, Philip Dykema, Scott Lindquist, Michael Boeckh, Janet A. Englund, Michael Famulare, Barry R. Lutz, Mark J. Rieder, Lea M. Starita, Matthew Thompson, Helen Y. Chu, Jay Shendure, Trevor Bedford |
| EPI_ISL_3104936 | Northwest Laboratory | Seattle Flu Study | Deborah A. Nickerson, Chris D. Frazar, Jover Lee, Benjamin Pelle, Erica Ryke, Matthew Richardson, Amanda Adler, Elisabeth Brandstetter, Peter D. Han, Kairsten Fay, Misja Ilcisin, Kirsten Lacombe, Thomas R. Sibley, Melissa Truong, Caitlin R. Wolf, Romesh Gautom, Geoff Melly, Brian Hiatt, Philip Dykema, Scott Lindquist, Michael Boeckh, Janet A. Englund, Michael Famulare, Barry R. Lutz, Mark J. Rieder, Lea M. Starita, Matthew Thompson, Helen Y. Chu, Jay Shendure, Trevor Bedford |
| EPI_ISL_3104937 | Atlas Genomics | Seattle Flu Study | Deborah A. Nickerson, Chris D. Frazar, Jover Lee, Benjamin Pelle, Erica Ryke, Matthew Richardson, Amanda Adler, Elisabeth Brandstetter, Peter D. Han, Kairsten Fay, Misja Ilcisin, Kirsten Lacombe, Thomas R. Sibley, Melissa Truong, Caitlin R. Wolf, Romesh Gautom, Geoff Melly, Brian Hiatt, Philip Dykema, Scott Lindquist, Michael Boeckh, Janet A. Englund, Michael Famulare, Barry R. Lutz, Mark J. Rieder, Lea M. Starita, Matthew Thompson, Helen Y. Chu, Jay Shendure, Trevor Bedford |
| EPI_ISL_3104939, EPI_ISL_3104941 | Northwest Laboratory | Seattle Flu Study | Deborah A. Nickerson, Chris D. Frazar, Jover Lee, Benjamin Pelle, Erica Ryke, Matthew Richardson, Amanda Adler, Elisabeth Brandstetter, Peter D. Han, Kairsten Fay, Misja Ilcisin, Kirsten Lacombe, Thomas R. Sibley, Melissa Truong, Caitlin R. Wolf, Romesh Gautom, Geoff Melly, Brian Hiatt, Philip Dykema, Scott Lindquist, Michael Boeckh, Janet A. Englund, Michael Famulare, Barry R. Lutz, Mark J. Rieder, Lea M. Starita, Matthew Thompson, Helen Y. Chu, Jay Shendure, Trevor Bedford |
| EPI_ISL_3104942 | Seattle Flu Study | Seattle Flu Study | Deborah A. Nickerson, Chris D. Frazar, Jover Lee, Benjamin Pelle, Erica Ryke, Matthew Richardson, Amanda Adler, Elisabeth Brandstetter, Peter D. Han, Kairsten Fay, Misja Ilcisin, Kirsten Lacombe, Thomas R. Sibley, Melissa Truong, Caitlin R. Wolf, Karen Cowgill, Stephanie Schrag, Jeff Duchin, Michael Boeckh, Janet A. Englund, Michael Famulare, Barry R. Lutz, Mark J. Rieder, Lea M. Starita, Matthew Thompson, Helen Y. Chu, Trevor Bedford, Jay Shendure |
| EPI_ISL_3104943, EPI_ISL_3104946 | Incyte Diagnostics Spokane | Seattle Flu Study | Deborah A. Nickerson, Chris D. Frazar, Jover Lee, Benjamin Pelle, Erica Ryke, Matthew Richardson, Amanda Adler, Elisabeth Brandstetter, Peter D. Han, Kairsten Fay, Misja Ilcisin, Kirsten Lacombe, Thomas R. Sibley, Melissa Truong, Caitlin R. Wolf, Romesh Gautom, Geoff Melly, Brian Hiatt, Philip Dykema, Scott Lindquist, Michael Boeckh, Janet A. Englund, Michael Famulare, Barry R. Lutz, Mark J. Rieder, Lea M. Starita, Matthew Thompson, Helen Y. Chu, Jay Shendure, Trevor Bedford |
| EPI_ISL_3104947 | Northwest Laboratory | Seattle Flu Study | Deborah A. Nickerson, Chris D. Frazar, Jover Lee, Benjamin Pelle, Erica Ryke, Matthew Richardson, Amanda Adler, Elisabeth Brandstetter, Peter D. Han, Kairsten Fay, Misja Ilcisin, Kirsten Lacombe, Thomas R. Sibley, Melissa Truong, Caitlin R. Wolf, Romesh Gautom, Geoff Melly, Brian Hiatt, Philip Dykema, Scott Lindquist, Michael Boeckh, Janet A. Englund, Michael Famulare, Barry R. Lutz, Mark J. Rieder, Lea M. Starita, Matthew Thompson, Helen Y. Chu, Jay Shendure, Trevor Bedford |
| EPI_ISL_3104951 | Overlake Hospital | Seattle Flu Study | Deborah A. Nickerson, Chris D. Frazar, Jover Lee, Benjamin Pelle, Erica Ryke, Matthew Richardson, Amanda Adler, Elisabeth Brandstetter, Peter D. Han, Kairsten Fay, Misja Ilcisin, Kirsten Lacombe, Thomas R. Sibley, Melissa Truong, Caitlin R. Wolf, Romesh Gautom, Geoff Melly, Brian Hiatt, Philip Dykema, Scott Lindquist, Michael Boeckh, Janet A. Englund, Michael Famulare, Barry R. Lutz, Mark J. Rieder, Lea M. Starita, Matthew Thompson, Helen Y. Chu, Jay Shendure, Trevor Bedford |
| EPI_ISL_3104953 | Seattle Flu Study | Seattle Flu Study | Deborah A. Nickerson, Chris D. Frazar, Jover Lee, Benjamin Pelle, Erica Ryke, Matthew Richardson, Amanda Adler, Elisabeth Brandstetter, Peter D. Han, Kairsten Fay, Misja Ilcisin, Kirsten Lacombe, Thomas R. Sibley, Melissa Truong, Caitlin R. Wolf, Michael Boeckh, Janet A. Englund, Michael Famulare, Barry R. Lutz, Mark J. Rieder, Lea M. Starita, Matthew Thompson, Jay Shendure, Trevor Bedford, Helen Y. Chu |
| EPI_ISL_3104954 | Atlas Genomics | Seattle Flu Study | Deborah A. Nickerson, Chris D. Frazar, Jover Lee, Benjamin Pelle, Erica Ryke, Matthew Richardson, Amanda Adler, Elisabeth Brandstetter, Peter D. Han, Kairsten Fay, Misja Ilcisin, Kirsten Lacombe, Thomas R. Sibley, Melissa Truong, Caitlin R. Wolf, Romesh Gautom, Geoff Melly, Brian Hiatt, Philip Dykema, Scott Lindquist, Michael Boeckh, Janet A. Englund, Michael Famulare, Barry R. Lutz, Mark J. Rieder, Lea M. Starita, Matthew Thompson, Helen Y. Chu, Jay Shendure, Trevor Bedford |
| EPI_ISL_3104955 | Seattle Flu Study | Seattle Flu Study | Deborah A. Nickerson, Chris D. Frazar, Jover Lee, Benjamin Pelle, Erica Ryke, Matthew Richardson, Amanda Adler, Elisabeth Brandstetter, Peter D. Han, Kairsten Fay, Misja Ilcisin, Kirsten Lacombe, Thomas R. Sibley, Melissa Truong, Caitlin R. Wolf, Karen Cowgill, Stephanie Schrag, Jeff Duchin, Michael Boeckh, Janet A. Englund, Michael Famulare, Barry R. Lutz, Mark J. Rieder, Lea M. Starita, Matthew Thompson, Helen Y. Chu, Trevor Bedford, Jay Shendure |
| EPI_ISL_3104956 | Atlas Genomics | Seattle Flu Study | Deborah A. Nickerson, Chris D. Frazar, Jover Lee, Benjamin Pelle, Erica Ryke, Matthew Richardson, Amanda Adler, Elisabeth Brandstetter, Peter D. Han, Kairsten Fay, Misja Ilcisin, Kirsten Lacombe, Thomas R. Sibley, Melissa Truong, Caitlin R. Wolf, Romesh Gautom, Geoff Melly, Brian Hiatt, Philip Dykema, Scott Lindquist, Michael Boeckh, Janet A. Englund, Michael Famulare, Barry R. Lutz, Mark J. Rieder, Lea M. Starita, Matthew Thompson, Helen Y. Chu, Jay Shendure, Trevor Bedford |
| EPI_ISL_3104958, EPI_ISL_3104960 | Overlake Hospital | Seattle Flu Study | Deborah A. Nickerson, Chris D. Frazar, Jover Lee, Benjamin Pelle, Erica Ryke, Matthew Richardson, Amanda Adler, Elisabeth Brandstetter, Peter D. Han, Kairsten Fay, Misja Ilcisin, Kirsten Lacombe, Thomas R. Sibley, Melissa Truong, Caitlin R. Wolf, Romesh Gautom, Geoff Melly, Brian Hiatt, Philip Dykema, Scott Lindquist, Michael Boeckh, Janet A. Englund, Michael Famulare, Barry R. Lutz, Mark J. Rieder, Lea M. Starita, Matthew Thompson, Helen Y. Chu, Jay Shendure, Trevor Bedford |
| EPI_ISL_3104961 | Interpath Laboratory | Seattle Flu Study | Deborah A. Nickerson, Chris D. Frazar, Jover Lee, Benjamin Pelle, Erica Ryke, Matthew Richardson, Amanda Adler, Elisabeth Brandstetter, Peter D. Han, Kairsten Fay, Misja Ilcisin, Kirsten Lacombe, Thomas R. Sibley, Melissa Truong, Caitlin R. Wolf, Romesh Gautom, Geoff Melly, Brian Hiatt, Philip Dykema, Scott Lindquist, Michael Boeckh, Janet A. Englund, Michael Famulare, Barry R. Lutz, Mark J. Rieder, Lea M. Starita, Matthew Thompson, Helen Y. Chu, Jay Shendure, Trevor Bedford |
| EPI_ISL_3104965 | Atlas Genomics | Seattle Flu Study | Deborah A. Nickerson, Chris D. Frazar, Jover Lee, Benjamin Pelle, Erica Ryke, Matthew Richardson, Amanda Adler, Elisabeth Brandstetter, Peter D. Han, Kairsten Fay, Misja Ilcisin, Kirsten Lacombe, Thomas R. Sibley, Melissa Truong, Caitlin R. Wolf, Romesh Gautom, Geoff Melly, Brian Hiatt, Philip Dykema, Scott Lindquist, Michael Boeckh, Janet A. Englund, Michael Famulare, Barry R. Lutz, Mark J. Rieder, Lea M. Starita, Matthew Thompson, Helen Y. Chu, Jay Shendure, Trevor Bedford |
| EPI_ISL_3104966, EPI_ISL_3104967 | Northwest Laboratory | Seattle Flu Study | Deborah A. Nickerson, Chris D. Frazar, Jover Lee, Benjamin Pelle, Erica Ryke, Matthew Richardson, Amanda Adler, Elisabeth Brandstetter, Peter D. Han, Kairsten Fay, Misja Ilcisin, Kirsten Lacombe, Thomas R. Sibley, Melissa Truong, Caitlin R. Wolf, Romesh Gautom, Geoff Melly, Brian Hiatt, Philip Dykema, Scott Lindquist, Michael Boeckh, Janet A. Englund, Michael Famulare, Barry R. Lutz, Mark J. Rieder, Lea M. Starita, Matthew Thompson, Helen Y. Chu, Jay Shendure, Trevor Bedford |
| EPI_ISL_3104971 | Overlake Hospital | Seattle Flu Study | Deborah A. Nickerson, Chris D. Frazar, Jover Lee, Benjamin Pelle, Erica Ryke, Matthew Richardson, Amanda Adler, Elisabeth Brandstetter, Peter D. Han, Kairsten Fay, Misja Ilcisin, Kirsten Lacombe, Thomas R. Sibley, Melissa Truong, Caitlin R. Wolf, Romesh Gautom, Geoff Melly, Brian Hiatt, Philip Dykema, Scott Lindquist, Michael Boeckh, Janet A. Englund, Michael Famulare, Barry R. Lutz, Mark J. Rieder, Lea M. Starita, Matthew Thompson, Helen Y. Chu, Jay Shendure, Trevor Bedford |
| EPI_ISL_3105022 | Incyte Diagnostics Spokane | Seattle Flu Study | Deborah A. Nickerson, Chris D. Frazar, Jover Lee, Benjamin Pelle, Erica Ryke, Matthew Richardson, Amanda Adler, Elisabeth Brandstetter, Peter D. Han, Kairsten Fay, Misja Ilcisin, Kirsten Lacombe, Thomas R. Sibley, Melissa Truong, Caitlin R. Wolf, Romesh Gautom, Geoff Melly, Brian Hiatt, Philip Dykema, Scott Lindquist, Michael Boeckh, Janet A. Englund, Michael Famulare, Barry R. Lutz, Mark J. Rieder, Lea M. Starita, Matthew Thompson, Helen Y. Chu, Jay Shendure, Trevor Bedford |
| EPI_ISL_3105023 | Atlas Genomics | Seattle Flu Study | Deborah A. Nickerson, Chris D. Frazar, Jover Lee, Benjamin Pelle, Erica Ryke, Matthew Richardson, Amanda Adler, Elisabeth Brandstetter, Peter D. Han, Kairsten Fay, Misja Ilcisin, Kirsten Lacombe, Thomas R. Sibley, Melissa Truong, Caitlin R. Wolf, Romesh Gautom, Geoff Melly, Brian Hiatt, Philip Dykema, Scott Lindquist, Michael Boeckh, Janet A. Englund, Michael Famulare, Barry R. Lutz, Mark J. Rieder, Lea M. Starita, Matthew Thompson, Helen Y. Chu, Jay Shendure, Trevor Bedford |
| EPI_ISL_3105031 | Northwest Laboratory | Seattle Flu Study | Deborah A. Nickerson, Chris D. Frazar, Jover Lee, Benjamin Pelle, Erica Ryke, Matthew Richardson, Amanda Adler, Elisabeth Brandstetter, Peter D. Han, Kairsten Fay, Misja Ilcisin, Kirsten Lacombe, Thomas R. Sibley, Melissa Truong, Caitlin R. Wolf, Romesh Gautom, Geoff Melly, Brian Hiatt, Philip Dykema, Scott Lindquist, Michael Boeckh, Janet A. Englund, Michael Famulare, Barry R. Lutz, Mark J. Rieder, Lea M. Starita, Matthew Thompson, Helen Y. Chu, Jay Shendure, Trevor Bedford |
| EPI_ISL_3105035 | Overlake Hospital | Seattle Flu Study | Deborah A. Nickerson, Chris D. Frazar, Jover Lee, Benjamin Pelle, Erica Ryke, Matthew Richardson, Amanda Adler, Elisabeth Brandstetter, Peter D. Han, Kairsten Fay, Misja Ilcisin, Kirsten Lacombe, Thomas R. Sibley, Melissa Truong, Caitlin R. Wolf, Romesh Gautom, Geoff Melly, Brian Hiatt, Philip Dykema, Scott Lindquist, Michael Boeckh, Janet A. Englund, Michael Famulare, Barry R. Lutz, Mark J. Rieder, Lea M. Starita, Matthew Thompson, Helen Y. Chu, Jay Shendure, Trevor Bedford |
| EPI_ISL_3105041 | Atlas Genomics | Seattle Flu Study | Deborah A. Nickerson, Chris D. Frazar, Jover Lee, Benjamin Pelle, Erica Ryke, Matthew Richardson, Amanda Adler, Elisabeth Brandstetter, Peter D. Han, Kairsten Fay, Misja Ilcisin, Kirsten Lacombe, Thomas R. Sibley, Melissa Truong, Caitlin R. Wolf, Romesh Gautom, Geoff Melly, Brian Hiatt, Philip Dykema, Scott Lindquist, Michael Boeckh, Janet A. Englund, Michael Famulare, Barry R. Lutz, Mark J. Rieder, Lea M. Starita, Matthew Thompson, Helen Y. Chu, Jay Shendure, Trevor Bedford |
| EPI_ISL_3105042, EPI_ISL_3105043 | Northwest Laboratory | Seattle Flu Study | Deborah A. Nickerson, Chris D. Frazar, Jover Lee, Benjamin Pelle, Erica Ryke, Matthew Richardson, Amanda Adler, Elisabeth Brandstetter, Peter D. Han, Kairsten Fay, Misja Ilcisin, Kirsten Lacombe, Thomas R. Sibley, Melissa Truong, Caitlin R. Wolf, Romesh Gautom, Geoff Melly, Brian Hiatt, Philip Dykema, Scott Lindquist, Michael Boeckh, Janet A. Englund, Michael Famulare, Barry R. Lutz, Mark J. Rieder, Lea M. Starita, Matthew Thompson, Helen Y. Chu, Jay Shendure, Trevor Bedford |
| EPI_ISL_3105044 | Atlas Genomics | Seattle Flu Study | Deborah A. Nickerson, Chris D. Frazar, Jover Lee, Benjamin Pelle, Erica Ryke, Matthew Richardson, Amanda Adler, Elisabeth Brandstetter, Peter D. Han, Kairsten Fay, Misja Ilcisin, Kirsten Lacombe, Thomas R. Sibley, Melissa Truong, Caitlin R. Wolf, Romesh Gautom, Geoff Melly, Brian Hiatt, Philip Dykema, Scott Lindquist, Michael Boeckh, Janet A. Englund, Michael Famulare, Barry R. Lutz, Mark J. Rieder, Lea M. Starita, Matthew Thompson, Helen Y. Chu, Jay Shendure, Trevor Bedford |
| EPI_ISL_3105059 | Overlake Hospital | Seattle Flu Study | Deborah A. Nickerson, Chris D. Frazar, Jover Lee, Benjamin Pelle, Erica Ryke, Matthew Richardson, Amanda Adler, Elisabeth Brandstetter, Peter D. Han, Kairsten Fay, Misja Ilcisin, Kirsten Lacombe, Thomas R. Sibley, Melissa Truong, Caitlin R. Wolf, Romesh Gautom, Geoff Melly, Brian Hiatt, Philip Dykema, Scott Lindquist, Michael Boeckh, Janet A. Englund, Michael Famulare, Barry R. Lutz, Mark J. Rieder, Lea M. Starita, Matthew Thompson, Helen Y. Chu, Jay Shendure, Trevor Bedford |
| EPI_ISL_3105060 | Northwest Laboratory | Seattle Flu Study | Deborah A. Nickerson, Chris D. Frazar, Jover Lee, Benjamin Pelle, Erica Ryke, Matthew Richardson, Amanda Adler, Elisabeth Brandstetter, Peter D. Han, Kairsten Fay, Misja Ilcisin, Kirsten Lacombe, Thomas R. Sibley, Melissa Truong, Caitlin R. Wolf, Romesh Gautom, Geoff Melly, Brian Hiatt, Philip Dykema, Scott Lindquist, Michael Boeckh, Janet A. Englund, Michael Famulare, Barry R. Lutz, Mark J. Rieder, Lea M. Starita, Matthew Thompson, Helen Y. Chu, Jay Shendure, Trevor Bedford |
| EPI_ISL_3105061, EPI_ISL_3105062 | Overlake Hospital | Seattle Flu Study | Deborah A. Nickerson, Chris D. Frazar, Jover Lee, Benjamin Pelle, Erica Ryke, Matthew Richardson, Amanda Adler, Elisabeth Brandstetter, Peter D. Han, Kairsten Fay, Misja Ilcisin, Kirsten Lacombe, Thomas R. Sibley, Melissa Truong, Caitlin R. Wolf, Romesh Gautom, Geoff Melly, Brian Hiatt, Philip Dykema, Scott Lindquist, Michael Boeckh, Janet A. Englund, Michael Famulare, Barry R. Lutz, Mark J. Rieder, Lea M. Starita, Matthew Thompson, Helen Y. Chu, Jay Shendure, Trevor Bedford |
| EPI_ISL_3105063 | Incyte Diagnostics Spokane | Seattle Flu Study | Deborah A. Nickerson, Chris D. Frazar, Jover Lee, Benjamin Pelle, Erica Ryke, Matthew Richardson, Amanda Adler, Elisabeth Brandstetter, Peter D. Han, Kairsten Fay, Misja Ilcisin, Kirsten Lacombe, Thomas R. Sibley, Melissa Truong, Caitlin R. Wolf, Romesh Gautom, Geoff Melly, Brian Hiatt, Philip Dykema, Scott Lindquist, Michael Boeckh, Janet A. Englund, Michael Famulare, Barry R. Lutz, Mark J. Rieder, Lea M. Starita, Matthew Thompson, Helen Y. Chu, Jay Shendure, Trevor Bedford |
| EPI_ISL_3105067, EPI_ISL_3105068 | Seattle Flu Study | Seattle Flu Study | Deborah A. Nickerson, Chris D. Frazar, Jover Lee, Benjamin Pelle, Erica Ryke, Matthew Richardson, Amanda Adler, Elisabeth Brandstetter, Peter D. Han, Kairsten Fay, Misja Ilcisin, Kirsten Lacombe, Thomas R. Sibley, Melissa Truong, Caitlin R. Wolf, Michael Boeckh, Janet A. Englund, Michael Famulare, Barry R. Lutz, Mark J. Rieder, Lea M. Starita, Matthew Thompson, Jay Shendure, Trevor Bedford, Helen Y. Chu |
| EPI_ISL_3105069 | Seattle Flu Study | Seattle Flu Study | Deborah A. Nickerson, Chris D. Frazar, Jover Lee, Benjamin Pelle, Erica Ryke, Matthew Richardson, Amanda Adler, Elisabeth Brandstetter, Peter D. Han, Kairsten Fay, Misja Ilcisin, Kirsten Lacombe, Thomas R. Sibley, Melissa Truong, Caitlin R. Wolf, Karen Cowgill, Stephanie Schrag, Jeff Duchin, Michael Boeckh, Janet A. Englund, Michael Famulare, Barry R. Lutz, Mark J. Rieder, Lea M. Starita, Matthew Thompson, Helen Y. Chu, Trevor Bedford, Jay Shendure |
| EPI_ISL_3105070 | Overlake Hospital | Seattle Flu Study | Deborah A. Nickerson, Chris D. Frazar, Jover Lee, Benjamin Pelle, Erica Ryke, Matthew Richardson, Amanda Adler, Elisabeth Brandstetter, Peter D. Han, Kairsten Fay, Misja Ilcisin, Kirsten Lacombe, Thomas R. Sibley, Melissa Truong, Caitlin R. Wolf, Romesh Gautom, Geoff Melly, Brian Hiatt, Philip Dykema, Scott Lindquist, Michael Boeckh, Janet A. Englund, Michael Famulare, Barry R. Lutz, Mark J. Rieder, Lea M. Starita, Matthew Thompson, Helen Y. Chu, Jay Shendure, Trevor Bedford |
| EPI_ISL_3105073, EPI_ISL_3105074, EPI_ISL_3105075, EPI_ISL_3105076, EPI_ISL_3105077 | Incyte Diagnostics Spokane | Seattle Flu Study | Deborah A. Nickerson, Chris D. Frazar, Jover Lee, Benjamin Pelle, Erica Ryke, Matthew Richardson, Amanda Adler, Elisabeth Brandstetter, Peter D. Han, Kairsten Fay, Misja Ilcisin, Kirsten Lacombe, Thomas R. Sibley, Melissa Truong, Caitlin R. Wolf, Romesh Gautom, Geoff Melly, Brian Hiatt, Philip Dykema, Scott Lindquist, Michael Boeckh, Janet A. Englund, Michael Famulare, Barry R. Lutz, Mark J. Rieder, Lea M. Starita, Matthew Thompson, Helen Y. Chu, Jay Shendure, Trevor Bedford |
| EPI_ISL_3105078, EPI_ISL_3105079, EPI_ISL_3105080, EPI_ISL_3105081 | Atlas Genomics | Seattle Flu Study | Deborah A. Nickerson, Chris D. Frazar, Jover Lee, Benjamin Pelle, Erica Ryke, Matthew Richardson, Amanda Adler, Elisabeth Brandstetter, Peter D. Han, Kairsten Fay, Misja Ilcisin, Kirsten Lacombe, Thomas R. Sibley, Melissa Truong, Caitlin R. Wolf, Romesh Gautom, Geoff Melly, Brian Hiatt, Philip Dykema, Scott Lindquist, Michael Boeckh, Janet A. Englund, Michael Famulare, Barry R. Lutz, Mark J. Rieder, Lea M. Starita, Matthew Thompson, Helen Y. Chu, Jay Shendure, Trevor Bedford |
| EPI_ISL_3105082, EPI_ISL_3105083, EPI_ISL_3105084 | Seattle Flu Study | Seattle Flu Study | Deborah A. Nickerson, Chris D. Frazar, Jover Lee, Benjamin Pelle, Erica Ryke, Matthew Richardson, Amanda Adler, Elisabeth Brandstetter, Peter D. Han, Kairsten Fay, Misja Ilcisin, Kirsten Lacombe, Thomas R. Sibley, Melissa Truong, Caitlin R. Wolf, Karen Cowgill, Stephanie Schrag, Jeff Duchin, Michael Boeckh, Janet A. Englund, Michael Famulare, Barry R. Lutz, Mark J. Rieder, Lea M. Starita, Matthew Thompson, Helen Y. Chu, Trevor Bedford, Jay Shendure |
| EPI_ISL_3105085 | Seattle Flu Study | Seattle Flu Study | Deborah A. Nickerson, Chris D. Frazar, Jover Lee, Benjamin Pelle, Erica Ryke, Matthew Richardson, Amanda Adler, Elisabeth Brandstetter, Peter D. Han, Kairsten Fay, Misja Ilcisin, Kirsten Lacombe, Thomas R. Sibley, Melissa Truong, Caitlin R. Wolf, Michael Boeckh, Janet A. Englund, Michael Famulare, Barry R. Lutz, Mark J. Rieder, Lea M. Starita, Matthew Thompson, Jay Shendure, Trevor Bedford, Helen Y. Chu |
| EPI_ISL_3105086, EPI_ISL_3105087, | Seattle Flu Study | Seattle Flu Study | Deborah A. Nickerson, Chris D. Frazar, Jover Lee, Benjamin Pelle, Erica Ryke, Matthew Richardson, Amanda Adler, Elisabeth Brandstetter, Peter D. Han, Kairsten Fay, Misja Ilcisin, Kirsten Lacombe, Thomas R. Sibley, Melissa Truong, Caitlin R. Wolf, Karen Cowgill, Stephanie Schrag, Jeff Duchin, Michael Boeckh, Janet A. Englund, Michael Famulare, Barry R. Lutz, Mark J. Rieder, Lea M. Starita, Matthew Thompson, Helen Y. Chu, Trevor Bedford, Jay Shendure |

| | | | |
|---|---|---|---|
| EPI_ISL_3105088 | | | |
| EPI_ISL_3105089, EPI_ISL_3105090, EPI_ISL_3105091, EPI_ISL_3105092 | Overlake Hospital | Seattle Flu Study | Deborah A. Nickerson, Chris D. Frazar, Jover Lee, Benjamin Pelle, Erica Ryke, Matthew Richardson, Amanda Adler, Elisabeth Brandstetter, Peter D. Han, Kairsten Fay, Misja Ilcisin, Kirsten Lacombe, Thomas R. Sibley, Melissa Truong, Caitlin R. Wolf, Romesh Gautom, Geoff Melly, Brian Hiatt, Philip Dykema, Scott Lindquist, Michael Boeckh, Janet A. Englund, Michael Famulare, Barry R. Lutz, Mark J. Rieder, Lea M. Starita, Matthew Thompson, Helen Y. Chu, Jay Shendure, Trevor Bedford |
| EPI_ISL_3105093, EPI_ISL_3105094, EPI_ISL_3105095, EPI_ISL_3105096, EPI_ISL_3105097, EPI_ISL_3105098, EPI_ISL_3105099, EPI_ISL_3105100, EPI_ISL_3105101, EPI_ISL_3105102, EPI_ISL_3105103, EPI_ISL_3105104, EPI_ISL_3105105, EPI_ISL_3105106, EPI_ISL_3105107, EPI_ISL_3105108 | | | |
| see above | Northwest Laboratory | Seattle Flu Study | Deborah A. Nickerson, Chris D. Frazar, Jover Lee, Benjamin Pelle, Erica Ryke, Matthew Richardson, Amanda Adler, Elisabeth Brandstetter, Peter D. Han, Kairsten Fay, Misja Ilcisin, Kirsten Lacombe, Thomas R. Sibley, Melissa Truong, Caitlin R. Wolf, Romesh Gautom, Geoff Melly, Brian Hiatt, Philip Dykema, Scott Lindquist, Michael Boeckh, Janet A. Englund, Michael Famulare, Barry R. Lutz, Mark J. Rieder, Lea M. Starita, Matthew Thompson, Helen Y. Chu, Jay Shendure, Trevor Bedford |
| EPI_ISL_3105109, EPI_ISL_3105110, EPI_ISL_3105111 | Interpath Laboratory | Seattle Flu Study | Deborah A. Nickerson, Chris D. Frazar, Jover Lee, Benjamin Pelle, Erica Ryke, Matthew Richardson, Amanda Adler, Elisabeth Brandstetter, Peter D. Han, Kairsten Fay, Misja Ilcisin, Kirsten Lacombe, Thomas R. Sibley, Melissa Truong, Caitlin R. Wolf, Romesh Gautom, Geoff Melly, Brian Hiatt, Philip Dykema, Scott Lindquist, Michael Boeckh, Janet A. Englund, Michael Famulare, Barry R. Lutz, Mark J. Rieder, Lea M. Starita, Matthew Thompson, Helen Y. Chu, Jay Shendure, Trevor Bedford |
| EPI_ISL_3105499 | Yale Clinical Virology Lab | Grubaugh Lab - Yale School of Public Health | Joseph Fauver, Mallery Breban, Isabel Ott, Tara Alpert, Mary Petrone, Anderson Brito, Chantal Vogels, Annie Watkins, Chaney Kalinich, Jessica Rothman, Marie L. Landry, Nathan Grubaugh |
| EPI_ISL_3105891 | Fondation Congolaise pour la recherche medicale (FCRM), Francine Ntoumi | Fondation Congolaise pour la Recherche Médicale | Dr Armel Landry Batchi-Bouyou ; Mfoutou Mapanguy Claujens Chastel; Dr. Jean Claude Djontu; Dr. Abel Lissom; Prof. Dr. Thirumalaisamy P. Velavan; Prof. Francine Ntoumi |
| EPI_ISL_3105892 | Lighthouse Lab in Milton Keynes | Wellcome Sanger Institute for the COVID-19 Genomics UK (COG-UK) Consortium | The Lighthouse Lab in Milton Keynes and Alex Alderton, Roberto Amato, Jeffrey Barrett, Sonia Goncalves, Ewan Harrison, David K. Jackson, Ian Johnston, Dominic Kwiatkowski, Cordelia Langford, John Sillitoe on behalf of the Wellcome Sanger Institute COVID-19 Surveillance Team |
| EPI_ISL_3105893 | Lighthouse Lab in Alderley Park | Wellcome Sanger Institute for the COVID-19 Genomics UK (COG-UK) Consortium | Jacquelyn Wynn, Mairead Hyland, The Lighthouse Lab in Alderley Park and Alex Alderton, Roberto Amato, Jeffrey Barrett, Sonia Goncalves, Ewan Harrison, David K. Jackson, Ian Johnston, Dominic Kwiatkowski, Cordelia Langford, John Sillitoe on behalf of the Wellcome Sanger Institute COVID-19 Surveillance Team |
| EPI_ISL_3105894 | Lighthouse Lab in Milton Keynes | Wellcome Sanger Institute for the COVID-19 Genomics UK (COG-UK) Consortium | The Lighthouse Lab in Milton Keynes and Alex Alderton, Roberto Amato, Jeffrey Barrett, Sonia Goncalves, Ewan Harrison, David K. Jackson, Ian Johnston, Dominic Kwiatkowski, Cordelia Langford, John Sillitoe on behalf of the Wellcome Sanger Institute COVID-19 Surveillance Team |
| EPI_ISL_3105895, EPI_ISL_3105896 | Lighthouse Lab in Alderley Park | Wellcome Sanger Institute for the COVID-19 Genomics UK (COG-UK) Consortium | Jacquelyn Wynn, Mairead Hyland, The Lighthouse Lab in Alderley Park and Alex Alderton, Roberto Amato, Jeffrey Barrett, Sonia Goncalves, Ewan Harrison, David K. Jackson, Ian Johnston, Dominic Kwiatkowski, Cordelia Langford, John Sillitoe on behalf of the Wellcome Sanger Institute COVID-19 Surveillance Team |
| EPI_ISL_3105897 | Lighthouse Lab in Glasgow | Wellcome Sanger Institute for the COVID-19 Genomics UK (COG-UK) Consortium | Harper VanSteenhouse, Yumi Kasai, David Gray, Carol Clugston, Anna Dominiczak and Alex Alderton, Roberto Amato, Jeffrey Barrett, Sonia Goncalves, Ewan Harrison, David K. Jackson, Ian Johnston, Dominic Kwiatkowski, Cordelia Langford, John Sillitoe on behalf of the Wellcome Sanger Institute COVID-19 Surveillance Team |
| EPI_ISL_3105898, EPI_ISL_3105899 | Lighthouse Lab in Milton Keynes | Wellcome Sanger Institute for the COVID-19 Genomics UK (COG-UK) Consortium | The Lighthouse Lab in Milton Keynes and Alex Alderton, Roberto Amato, Jeffrey Barrett, Sonia Goncalves, Ewan Harrison, David K. Jackson, Ian Johnston, Dominic Kwiatkowski, Cordelia Langford, John Sillitoe on behalf of the Wellcome Sanger Institute COVID-19 Surveillance Team |
| EPI_ISL_3105900, EPI_ISL_3105901, EPI_ISL_3105902 | Lighthouse Lab in Alderley Park | Wellcome Sanger Institute for the COVID-19 Genomics UK (COG-UK) Consortium | Jacquelyn Wynn, Mairead Hyland, The Lighthouse Lab in Alderley Park and Alex Alderton, Roberto Amato, Jeffrey Barrett, Sonia Goncalves, Ewan Harrison, David K. Jackson, Ian Johnston, Dominic Kwiatkowski, Cordelia Langford, John Sillitoe on behalf of the Wellcome Sanger Institute COVID-19 Surveillance Team |
| EPI_ISL_3105903 | Lighthouse Lab in Milton Keynes | Wellcome Sanger Institute for the COVID-19 Genomics UK (COG-UK) Consortium | The Lighthouse Lab in Milton Keynes and Alex Alderton, Roberto Amato, Jeffrey Barrett, Sonia Goncalves, Ewan Harrison, David K. Jackson, Ian Johnston, Dominic Kwiatkowski, Cordelia Langford, John Sillitoe on behalf of the Wellcome Sanger Institute COVID-19 Surveillance Team |
| EPI_ISL_3105904 | Lighthouse Lab in Glasgow | Wellcome Sanger Institute for the COVID-19 Genomics UK (COG-UK) Consortium | Harper VanSteenhouse, Yumi Kasai, David Gray, Carol Clugston, Anna Dominiczak and Alex Alderton, Roberto Amato, Jeffrey Barrett, Sonia Goncalves, Ewan Harrison, David K. Jackson, Ian Johnston, Dominic Kwiatkowski, Cordelia Langford, John Sillitoe on behalf of the Wellcome Sanger Institute COVID-19 Surveillance Team |
| EPI_ISL_3105905 | Lighthouse Lab in Milton Keynes | Wellcome Sanger Institute for the COVID-19 Genomics UK (COG-UK) Consortium | The Lighthouse Lab in Milton Keynes and Alex Alderton, Roberto Amato, Jeffrey Barrett, Sonia Goncalves, Ewan Harrison, David K. Jackson, Ian Johnston, Dominic Kwiatkowski, Cordelia Langford, John Sillitoe on behalf of the Wellcome Sanger Institute COVID-19 Surveillance Team |
| EPI_ISL_3105906 | Lighthouse Lab in Alderley Park | Wellcome Sanger Institute for the COVID-19 Genomics UK (COG-UK) Consortium | Jacquelyn Wynn, Mairead Hyland, The Lighthouse Lab in Alderley Park and Alex Alderton, Roberto Amato, Jeffrey Barrett, Sonia Goncalves, Ewan Harrison, David K. Jackson, Ian Johnston, Dominic Kwiatkowski, Cordelia Langford, John Sillitoe on behalf of the Wellcome Sanger Institute COVID-19 Surveillance Team |
| EPI_ISL_3105907, EPI_ISL_3105908 | Lighthouse Lab in Milton Keynes | Wellcome Sanger Institute for the COVID-19 Genomics UK (COG-UK) Consortium | The Lighthouse Lab in Milton Keynes and Alex Alderton, Roberto Amato, Jeffrey Barrett, Sonia Goncalves, Ewan Harrison, David K. Jackson, Ian Johnston, Dominic Kwiatkowski, Cordelia Langford, John Sillitoe on behalf of the Wellcome Sanger Institute COVID-19 Surveillance Team |
| EPI_ISL_3105909 | Lighthouse Lab in Alderley Park | Wellcome Sanger Institute for the COVID-19 Genomics UK (COG-UK) Consortium | Jacquelyn Wynn, Mairead Hyland, The Lighthouse Lab in Alderley Park and Alex Alderton, Roberto Amato, Jeffrey Barrett, Sonia Goncalves, Ewan Harrison, David K. Jackson, Ian Johnston, Dominic Kwiatkowski, Cordelia Langford, John Sillitoe on behalf of the Wellcome Sanger Institute COVID-19 Surveillance Team |
| EPI_ISL_3105910 | Lighthouse Lab in Glasgow | Wellcome Sanger Institute for the COVID-19 Genomics UK (COG-UK) Consortium | Harper VanSteenhouse, Yumi Kasai, David Gray, Carol Clugston, Anna Dominiczak and Alex Alderton, Roberto Amato, Jeffrey Barrett, Sonia Goncalves, Ewan Harrison, David K. Jackson, Ian Johnston, Dominic Kwiatkowski, Cordelia Langford, John Sillitoe on behalf of the Wellcome Sanger Institute COVID-19 Surveillance Team |
| EPI_ISL_3105911 | Lighthouse Lab in Alderley Park | Wellcome Sanger Institute for the COVID-19 Genomics UK (COG-UK) Consortium | Jacquelyn Wynn, Mairead Hyland, The Lighthouse Lab in Alderley Park and Alex Alderton, Roberto Amato, Jeffrey Barrett, Sonia Goncalves, Ewan Harrison, David K. Jackson, Ian Johnston, Dominic Kwiatkowski, Cordelia Langford, John Sillitoe on behalf of the Wellcome Sanger Institute COVID-19 Surveillance Team |
| EPI_ISL_3105912, EPI_ISL_3105913 | Lighthouse Lab in Milton Keynes | Wellcome Sanger Institute for the COVID-19 Genomics UK (COG-UK) Consortium | The Lighthouse Lab in Milton Keynes and Alex Alderton, Roberto Amato, Jeffrey Barrett, Sonia Goncalves, Ewan Harrison, David K. Jackson, Ian Johnston, Dominic Kwiatkowski, Cordelia Langford, John Sillitoe on behalf of the Wellcome Sanger Institute COVID-19 Surveillance Team |
| EPI_ISL_3105914 | Lighthouse Lab in Alderley Park | Wellcome Sanger Institute for the COVID-19 Genomics UK (COG-UK) Consortium | Jacquelyn Wynn, Mairead Hyland, The Lighthouse Lab in Alderley Park and Alex Alderton, Roberto Amato, Jeffrey Barrett, Sonia Goncalves, Ewan Harrison, David K. Jackson, Ian Johnston, Dominic Kwiatkowski, Cordelia Langford, John Sillitoe on behalf of the Wellcome Sanger Institute COVID-19 Surveillance Team |
| EPI_ISL_3105915 | Lighthouse Lab in Milton Keynes | Wellcome Sanger Institute for the COVID-19 Genomics UK (COG-UK) Consortium | The Lighthouse Lab in Milton Keynes and Alex Alderton, Roberto Amato, Jeffrey Barrett, Sonia Goncalves, Ewan Harrison, David K. Jackson, Ian Johnston, Dominic Kwiatkowski, Cordelia Langford, John Sillitoe on behalf of the Wellcome Sanger Institute COVID-19 Surveillance Team |
| EPI_ISL_3105916, EPI_ISL_3105917 | Lighthouse Lab in Alderley Park | Wellcome Sanger Institute for the COVID-19 Genomics UK (COG-UK) Consortium | Jacquelyn Wynn, Mairead Hyland, The Lighthouse Lab in Alderley Park and Alex Alderton, Roberto Amato, Jeffrey Barrett, Sonia Goncalves, Ewan Harrison, David K. Jackson, Ian Johnston, Dominic Kwiatkowski, Cordelia Langford, John Sillitoe on behalf of the Wellcome Sanger Institute COVID-19 Surveillance Team |
| EPI_ISL_3105918, EPI_ISL_3105919 | Lighthouse Lab in Milton Keynes | Wellcome Sanger Institute for the COVID-19 Genomics UK (COG-UK) Consortium | The Lighthouse Lab in Milton Keynes and Alex Alderton, Roberto Amato, Jeffrey Barrett, Sonia Goncalves, Ewan Harrison, David K. Jackson, Ian Johnston, Dominic Kwiatkowski, Cordelia Langford, John Sillitoe on behalf of the Wellcome Sanger Institute COVID-19 Surveillance Team |
| EPI_ISL_3105920 | Lighthouse Lab in Alderley Park | Wellcome Sanger Institute for the COVID-19 Genomics UK (COG-UK) Consortium | Jacquelyn Wynn, Mairead Hyland, The Lighthouse Lab in Alderley Park and Alex Alderton, Roberto Amato, Jeffrey Barrett, Sonia Goncalves, Ewan Harrison, David K. Jackson, Ian Johnston, Dominic Kwiatkowski, Cordelia Langford, John Sillitoe on behalf of the Wellcome Sanger Institute COVID-19 Surveillance Team |
| EPI_ISL_3105921 | Lighthouse Lab in Milton Keynes | Wellcome Sanger Institute for the COVID-19 Genomics UK (COG-UK) Consortium | The Lighthouse Lab in Milton Keynes and Alex Alderton, Roberto Amato, Jeffrey Barrett, Sonia Goncalves, Ewan Harrison, David K. Jackson, Ian Johnston, Dominic Kwiatkowski, Cordelia Langford, John Sillitoe on behalf of the Wellcome Sanger Institute COVID-19 Surveillance Team |
| EPI_ISL_3105922 | Lighthouse Lab in Glasgow | Wellcome Sanger Institute for the COVID-19 Genomics UK (COG-UK) Consortium | Harper VanSteenhouse, Yumi Kasai, David Gray, Carol Clugston, Anna Dominiczak and Alex Alderton, Roberto Amato, Jeffrey Barrett, Sonia Goncalves, Ewan Harrison, David K. Jackson, Ian Johnston, Dominic Kwiatkowski, Cordelia Langford, John Sillitoe on behalf of the Wellcome Sanger Institute COVID-19 Surveillance Team |
| EPI_ISL_3105923, EPI_ISL_3105924, EPI_ISL_3105925, EPI_ISL_3105926, EPI_ISL_3105927 | Lighthouse Lab in Milton Keynes | Wellcome Sanger Institute for the COVID-19 Genomics UK (COG-UK) Consortium | The Lighthouse Lab in Milton Keynes and Alex Alderton, Roberto Amato, Jeffrey Barrett, Sonia Goncalves, Ewan Harrison, David K. Jackson, Ian Johnston, Dominic Kwiatkowski, Cordelia Langford, John Sillitoe on behalf of the Wellcome Sanger Institute COVID-19 Surveillance Team |
| EPI_ISL_3105928 | Lighthouse Lab in Alderley Park | Wellcome Sanger Institute for the COVID-19 Genomics UK (COG-UK) Consortium | Jacquelyn Wynn, Mairead Hyland, The Lighthouse Lab in Alderley Park and Alex Alderton, Roberto Amato, Jeffrey Barrett, Sonia Goncalves, Ewan Harrison, David K. Jackson, Ian Johnston, Dominic Kwiatkowski, Cordelia Langford, John Sillitoe on behalf of the Wellcome Sanger Institute COVID-19 Surveillance Team |
| EPI_ISL_3105929, EPI_ISL_3105930 | Lighthouse Lab in Milton Keynes | Wellcome Sanger Institute for the COVID-19 Genomics UK (COG-UK) Consortium | The Lighthouse Lab in Milton Keynes and Alex Alderton, Roberto Amato, Jeffrey Barrett, Sonia Goncalves, Ewan Harrison, David K. Jackson, Ian Johnston, Dominic Kwiatkowski, Cordelia Langford, John Sillitoe on behalf of the Wellcome Sanger Institute COVID-19 Surveillance Team |
| EPI_ISL_3105931 | Lighthouse Lab in Alderley Park | Wellcome Sanger Institute for the COVID-19 Genomics UK (COG-UK) Consortium | Jacquelyn Wynn, Mairead Hyland, The Lighthouse Lab in Alderley Park and Alex Alderton, Roberto Amato, Jeffrey Barrett, Sonia Goncalves, Ewan Harrison, David K. Jackson, Ian Johnston, Dominic Kwiatkowski, Cordelia Langford, John Sillitoe on behalf of the Wellcome Sanger Institute COVID-19 Surveillance Team |
| EPI_ISL_3105932 | Lighthouse Lab in Milton Keynes | Wellcome Sanger Institute for the COVID-19 Genomics UK (COG-UK) Consortium | The Lighthouse Lab in Milton Keynes and Alex Alderton, Roberto Amato, Jeffrey Barrett, Sonia Goncalves, Ewan Harrison, David K. Jackson, Ian Johnston, Dominic Kwiatkowski, Cordelia Langford, John Sillitoe on behalf of the Wellcome Sanger Institute COVID-19 Surveillance Team |
| EPI_ISL_3105933 | Lighthouse Lab in Glasgow | Wellcome Sanger Institute for the COVID-19 Genomics UK (COG-UK) Consortium | Harper VanSteenhouse, Yumi Kasai, David Gray, Carol Clugston, Anna Dominiczak and Alex Alderton, Roberto Amato, Jeffrey Barrett, Sonia Goncalves, Ewan Harrison, David K. Jackson, Ian Johnston, Dominic Kwiatkowski, Cordelia Langford, John Sillitoe on behalf of the Wellcome Sanger Institute COVID-19 Surveillance Team |
| EPI_ISL_3105934, EPI_ISL_3105935 | Lighthouse Lab in Alderley Park | Wellcome Sanger Institute for the COVID-19 Genomics UK (COG-UK) Consortium | Jacquelyn Wynn, Mairead Hyland, The Lighthouse Lab in Alderley Park and Alex Alderton, Roberto Amato, Jeffrey Barrett, Sonia Goncalves, Ewan Harrison, David K. Jackson, Ian Johnston, Dominic Kwiatkowski, Cordelia Langford, John Sillitoe on behalf of the Wellcome Sanger Institute COVID-19 Surveillance Team |
| EPI_ISL_3105936 | Lighthouse Lab in Glasgow | Wellcome Sanger Institute for the COVID-19 Genomics UK (COG-UK) Consortium | Harper VanSteenhouse, Yumi Kasai, David Gray, Carol Clugston, Anna Dominiczak and Alex Alderton, Roberto Amato, Jeffrey Barrett, Sonia Goncalves, Ewan Harrison, David K. Jackson, Ian Johnston, Dominic Kwiatkowski, Cordelia Langford, John Sillitoe on behalf of the Wellcome Sanger Institute COVID-19 Surveillance Team |
| EPI_ISL_3105937 | Lighthouse Lab in Milton Keynes | Wellcome Sanger Institute for the COVID-19 Genomics UK (COG-UK) Consortium | The Lighthouse Lab in Milton Keynes and Alex Alderton, Roberto Amato, Jeffrey Barrett, Sonia Goncalves, Ewan Harrison, David K. Jackson, Ian Johnston, Dominic Kwiatkowski, Cordelia Langford, John Sillitoe on behalf of the Wellcome Sanger Institute COVID-19 Surveillance Team |
| EPI_ISL_3105938 | Lighthouse Lab in Alderley Park | Wellcome Sanger Institute for the COVID-19 Genomics UK (COG-UK) Consortium | Jacquelyn Wynn, Mairead Hyland, The Lighthouse Lab in Alderley Park and Alex Alderton, Roberto Amato, Jeffrey Barrett, Sonia Goncalves, Ewan Harrison, David K. Jackson, Ian Johnston, Dominic Kwiatkowski, Cordelia Langford, John Sillitoe on behalf of the Wellcome Sanger Institute COVID-19 Surveillance Team |
| EPI_ISL_3105939 | Lighthouse Lab in Milton Keynes | Wellcome Sanger Institute for the COVID-19 Genomics UK (COG-UK) Consortium | The Lighthouse Lab in Milton Keynes and Alex Alderton, Roberto Amato, Jeffrey Barrett, Sonia Goncalves, Ewan Harrison, David K. Jackson, Ian Johnston, Dominic Kwiatkowski, Cordelia Langford, John Sillitoe on behalf of the Wellcome Sanger Institute COVID-19 Surveillance Team |
| EPI_ISL_3105940 | Lighthouse Lab in Glasgow | Wellcome Sanger Institute for the COVID-19 Genomics UK (COG-UK) Consortium | Harper VanSteenhouse, Yumi Kasai, David Gray, Carol Clugston, Anna Dominiczak and Alex Alderton, Roberto Amato, Jeffrey Barrett, Sonia Goncalves, Ewan Harrison, David K. Jackson, Ian Johnston, Dominic Kwiatkowski, Cordelia Langford, John Sillitoe on behalf of the Wellcome Sanger Institute COVID-19 Surveillance Team |
| EPI_ISL_3105941 | Lighthouse Lab in Alderley Park | Wellcome Sanger Institute for the COVID-19 Genomics UK (COG-UK) Consortium | Jacquelyn Wynn, Mairead Hyland, The Lighthouse Lab in Alderley Park and Alex Alderton, Roberto Amato, Jeffrey Barrett, Sonia Goncalves, Ewan Harrison, David K. Jackson, Ian Johnston, Dominic Kwiatkowski, Cordelia Langford, John Sillitoe on behalf of the Wellcome Sanger Institute COVID-19 Surveillance Team |
| EPI_ISL_3105942 | Lighthouse Lab in Milton Keynes | Wellcome Sanger Institute for the COVID-19 Genomics UK (COG-UK) | The Lighthouse Lab in Milton Keynes and Alex Alderton, Roberto Amato, Jeffrey Barrett, Sonia Goncalves, Ewan Harrison, David K. Jackson, Ian Johnston, Dominic Kwiatkowski, Cordelia Langford, John Sillitoe on behalf of the Wellcome Sanger |

| | | | |
|---|---|---|---|
| | | Consortium | Institute COVID-19 Surveillance Team |
| EPI_ISL_3105943 | Lighthouse Lab in Alderley Park | Wellcome Sanger Institute for the COVID-19 Genomics UK (COG-UK) Consortium | Jacquelyn Wynn, Mairead Hyland, The Lighthouse Lab in Alderley Park and Alex Alderton, Roberto Amato, Jeffrey Barrett, Sonia Goncalves, Ewan Harrison, David K. Jackson, Ian Johnston, Dominic Kwiatkowski, Cordelia Langford, John Sillitoe on behalf of the Wellcome Sanger Institute COVID-19 Surveillance Team |
| EPI_ISL_3105944, EPI_ISL_3105945, EPI_ISL_3105946, EPI_ISL_3105947 | Lighthouse Lab in Milton Keynes | Wellcome Sanger Institute for the COVID-19 Genomics UK (COG-UK) Consortium | The Lighthouse Lab in Milton Keynes and Alex Alderton, Roberto Amato, Jeffrey Barrett, Sonia Goncalves, Ewan Harrison, David K. Jackson, Ian Johnston, Dominic Kwiatkowski, Cordelia Langford, John Sillitoe on behalf of the Wellcome Sanger Institute COVID-19 Surveillance Team |
| EPI_ISL_3105948 | Lighthouse Lab in Glasgow | Wellcome Sanger Institute for the COVID-19 Genomics UK (COG-UK) Consortium | Harper VanSteenhouse, Yumi Kasai, David Gray, Carol Clugston, Anna Dominiczak and Alex Alderton, Roberto Amato, Jeffrey Barrett, Sonia Goncalves, Ewan Harrison, David K. Jackson, Ian Johnston, Dominic Kwiatkowski, Cordelia Langford, John Sillitoe on behalf of the Wellcome Sanger Institute COVID-19 Surveillance Team |
| EPI_ISL_3105949 | Lighthouse Lab in Alderley Park | Wellcome Sanger Institute for the COVID-19 Genomics UK (COG-UK) Consortium | Jacquelyn Wynn, Mairead Hyland, The Lighthouse Lab in Alderley Park and Alex Alderton, Roberto Amato, Jeffrey Barrett, Sonia Goncalves, Ewan Harrison, David K. Jackson, Ian Johnston, Dominic Kwiatkowski, Cordelia Langford, John Sillitoe on behalf of the Wellcome Sanger Institute COVID-19 Surveillance Team |
| EPI_ISL_3105950, EPI_ISL_3105951 | Lighthouse Lab in Milton Keynes | Wellcome Sanger Institute for the COVID-19 Genomics UK (COG-UK) Consortium | The Lighthouse Lab in Milton Keynes and Alex Alderton, Roberto Amato, Jeffrey Barrett, Sonia Goncalves, Ewan Harrison, David K. Jackson, Ian Johnston, Dominic Kwiatkowski, Cordelia Langford, John Sillitoe on behalf of the Wellcome Sanger Institute COVID-19 Surveillance Team |
| EPI_ISL_3105952 | Lighthouse Lab in Alderley Park | Wellcome Sanger Institute for the COVID-19 Genomics UK (COG-UK) Consortium | Jacquelyn Wynn, Mairead Hyland, The Lighthouse Lab in Alderley Park and Alex Alderton, Roberto Amato, Jeffrey Barrett, Sonia Goncalves, Ewan Harrison, David K. Jackson, Ian Johnston, Dominic Kwiatkowski, Cordelia Langford, John Sillitoe on behalf of the Wellcome Sanger Institute COVID-19 Surveillance Team |
| EPI_ISL_3105953 | Lighthouse Lab in Glasgow | Wellcome Sanger Institute for the COVID-19 Genomics UK (COG-UK) Consortium | Harper VanSteenhouse, Yumi Kasai, David Gray, Carol Clugston, Anna Dominiczak and Alex Alderton, Roberto Amato, Jeffrey Barrett, Sonia Goncalves, Ewan Harrison, David K. Jackson, Ian Johnston, Dominic Kwiatkowski, Cordelia Langford, John Sillitoe on behalf of the Wellcome Sanger Institute COVID-19 Surveillance Team |
| EPI_ISL_3105954, EPI_ISL_3105955 | Lighthouse Lab in Alderley Park | Wellcome Sanger Institute for the COVID-19 Genomics UK (COG-UK) Consortium | Jacquelyn Wynn, Mairead Hyland, The Lighthouse Lab in Alderley Park and Alex Alderton, Roberto Amato, Jeffrey Barrett, Sonia Goncalves, Ewan Harrison, David K. Jackson, Ian Johnston, Dominic Kwiatkowski, Cordelia Langford, John Sillitoe on behalf of the Wellcome Sanger Institute COVID-19 Surveillance Team |
| EPI_ISL_3105956, EPI_ISL_3105957, EPI_ISL_3105958 | Lighthouse Lab in Milton Keynes | Wellcome Sanger Institute for the COVID-19 Genomics UK (COG-UK) Consortium | The Lighthouse Lab in Milton Keynes and Alex Alderton, Roberto Amato, Jeffrey Barrett, Sonia Goncalves, Ewan Harrison, David K. Jackson, Ian Johnston, Dominic Kwiatkowski, Cordelia Langford, John Sillitoe on behalf of the Wellcome Sanger Institute COVID-19 Surveillance Team |
| EPI_ISL_3105959 | Lighthouse Lab in Glasgow | Wellcome Sanger Institute for the COVID-19 Genomics UK (COG-UK) Consortium | Harper VanSteenhouse, Yumi Kasai, David Gray, Carol Clugston, Anna Dominiczak and Alex Alderton, Roberto Amato, Jeffrey Barrett, Sonia Goncalves, Ewan Harrison, David K. Jackson, Ian Johnston, Dominic Kwiatkowski, Cordelia Langford, John Sillitoe on behalf of the Wellcome Sanger Institute COVID-19 Surveillance Team |
| EPI_ISL_3105960 | Lighthouse Lab in Alderley Park | Wellcome Sanger Institute for the COVID-19 Genomics UK (COG-UK) Consortium | Jacquelyn Wynn, Mairead Hyland, The Lighthouse Lab in Alderley Park and Alex Alderton, Roberto Amato, Jeffrey Barrett, Sonia Goncalves, Ewan Harrison, David K. Jackson, Ian Johnston, Dominic Kwiatkowski, Cordelia Langford, John Sillitoe on behalf of the Wellcome Sanger Institute COVID-19 Surveillance Team |
| EPI_ISL_3105961, EPI_ISL_3105962, EPI_ISL_3105963 | Lighthouse Lab in Milton Keynes | Wellcome Sanger Institute for the COVID-19 Genomics UK (COG-UK) Consortium | The Lighthouse Lab in Milton Keynes and Alex Alderton, Roberto Amato, Jeffrey Barrett, Sonia Goncalves, Ewan Harrison, David K. Jackson, Ian Johnston, Dominic Kwiatkowski, Cordelia Langford, John Sillitoe on behalf of the Wellcome Sanger Institute COVID-19 Surveillance Team |
| EPI_ISL_3105964 | Lighthouse Lab in Alderley Park | Wellcome Sanger Institute for the COVID-19 Genomics UK (COG-UK) Consortium | Jacquelyn Wynn, Mairead Hyland, The Lighthouse Lab in Alderley Park and Alex Alderton, Roberto Amato, Jeffrey Barrett, Sonia Goncalves, Ewan Harrison, David K. Jackson, Ian Johnston, Dominic Kwiatkowski, Cordelia Langford, John Sillitoe on behalf of the Wellcome Sanger Institute COVID-19 Surveillance Team |
| EPI_ISL_3105965, EPI_ISL_3105966, EPI_ISL_3105967, EPI_ISL_3105968 | Lighthouse Lab in Milton Keynes | Wellcome Sanger Institute for the COVID-19 Genomics UK (COG-UK) Consortium | The Lighthouse Lab in Milton Keynes and Alex Alderton, Roberto Amato, Jeffrey Barrett, Sonia Goncalves, Ewan Harrison, David K. Jackson, Ian Johnston, Dominic Kwiatkowski, Cordelia Langford, John Sillitoe on behalf of the Wellcome Sanger Institute COVID-19 Surveillance Team |
| EPI_ISL_3105969, EPI_ISL_3105970, EPI_ISL_3105971 | Lighthouse Lab in Glasgow | Wellcome Sanger Institute for the COVID-19 Genomics UK (COG-UK) Consortium | Harper VanSteenhouse, Yumi Kasai, David Gray, Carol Clugston, Anna Dominiczak and Alex Alderton, Roberto Amato, Jeffrey Barrett, Sonia Goncalves, Ewan Harrison, David K. Jackson, Ian Johnston, Dominic Kwiatkowski, Cordelia Langford, John Sillitoe on behalf of the Wellcome Sanger Institute COVID-19 Surveillance Team |
| EPI_ISL_3105972 | Lighthouse Lab in Alderley Park | Wellcome Sanger Institute for the COVID-19 Genomics UK (COG-UK) Consortium | Jacquelyn Wynn, Mairead Hyland, The Lighthouse Lab in Alderley Park and Alex Alderton, Roberto Amato, Jeffrey Barrett, Sonia Goncalves, Ewan Harrison, David K. Jackson, Ian Johnston, Dominic Kwiatkowski, Cordelia Langford, John Sillitoe on behalf of the Wellcome Sanger Institute COVID-19 Surveillance Team |
| EPI_ISL_3105973 | Lighthouse Lab in Glasgow | Wellcome Sanger Institute for the COVID-19 Genomics UK (COG-UK) Consortium | Harper VanSteenhouse, Yumi Kasai, David Gray, Carol Clugston, Anna Dominiczak and Alex Alderton, Roberto Amato, Jeffrey Barrett, Sonia Goncalves, Ewan Harrison, David K. Jackson, Ian Johnston, Dominic Kwiatkowski, Cordelia Langford, John Sillitoe on behalf of the Wellcome Sanger Institute COVID-19 Surveillance Team |
| EPI_ISL_3105974, EPI_ISL_3105975 | Lighthouse Lab in Milton Keynes | Wellcome Sanger Institute for the COVID-19 Genomics UK (COG-UK) Consortium | The Lighthouse Lab in Milton Keynes and Alex Alderton, Roberto Amato, Jeffrey Barrett, Sonia Goncalves, Ewan Harrison, David K. Jackson, Ian Johnston, Dominic Kwiatkowski, Cordelia Langford, John Sillitoe on behalf of the Wellcome Sanger Institute COVID-19 Surveillance Team |
| EPI_ISL_3105976, EPI_ISL_3105977, EPI_ISL_3105978 | Lighthouse Lab in Alderley Park | Wellcome Sanger Institute for the COVID-19 Genomics UK (COG-UK) Consortium | Jacquelyn Wynn, Mairead Hyland, The Lighthouse Lab in Alderley Park and Alex Alderton, Roberto Amato, Jeffrey Barrett, Sonia Goncalves, Ewan Harrison, David K. Jackson, Ian Johnston, Dominic Kwiatkowski, Cordelia Langford, John Sillitoe on behalf of the Wellcome Sanger Institute COVID-19 Surveillance Team |
| EPI_ISL_3105979 | Lighthouse Lab in Glasgow | Wellcome Sanger Institute for the COVID-19 Genomics UK (COG-UK) Consortium | Harper VanSteenhouse, Yumi Kasai, David Gray, Carol Clugston, Anna Dominiczak and Alex Alderton, Roberto Amato, Jeffrey Barrett, Sonia Goncalves, Ewan Harrison, David K. Jackson, Ian Johnston, Dominic Kwiatkowski, Cordelia Langford, John Sillitoe on behalf of the Wellcome Sanger Institute COVID-19 Surveillance Team |
| EPI_ISL_3105980 | Lighthouse Lab in Alderley Park | Wellcome Sanger Institute for the COVID-19 Genomics UK (COG-UK) Consortium | Jacquelyn Wynn, Mairead Hyland, The Lighthouse Lab in Alderley Park and Alex Alderton, Roberto Amato, Jeffrey Barrett, Sonia Goncalves, Ewan Harrison, David K. Jackson, Ian Johnston, Dominic Kwiatkowski, Cordelia Langford, John Sillitoe on behalf of the Wellcome Sanger Institute COVID-19 Surveillance Team |
| EPI_ISL_3105981 | Lighthouse Lab in Glasgow | Wellcome Sanger Institute for the COVID-19 Genomics UK (COG-UK) Consortium | Harper VanSteenhouse, Yumi Kasai, David Gray, Carol Clugston, Anna Dominiczak and Alex Alderton, Roberto Amato, Jeffrey Barrett, Sonia Goncalves, Ewan Harrison, David K. Jackson, Ian Johnston, Dominic Kwiatkowski, Cordelia Langford, John Sillitoe on behalf of the Wellcome Sanger Institute COVID-19 Surveillance Team |
| EPI_ISL_3105982, EPI_ISL_3105983, EPI_ISL_3105984 | Lighthouse Lab in Milton Keynes | Wellcome Sanger Institute for the COVID-19 Genomics UK (COG-UK) Consortium | The Lighthouse Lab in Milton Keynes and Alex Alderton, Roberto Amato, Jeffrey Barrett, Sonia Goncalves, Ewan Harrison, David K. Jackson, Ian Johnston, Dominic Kwiatkowski, Cordelia Langford, John Sillitoe on behalf of the Wellcome Sanger Institute COVID-19 Surveillance Team |
| EPI_ISL_3105985, EPI_ISL_3105986 | Lighthouse Lab in Glasgow | Wellcome Sanger Institute for the COVID-19 Genomics UK (COG-UK) Consortium | Harper VanSteenhouse, Yumi Kasai, David Gray, Carol Clugston, Anna Dominiczak and Alex Alderton, Roberto Amato, Jeffrey Barrett, Sonia Goncalves, Ewan Harrison, David K. Jackson, Ian Johnston, Dominic Kwiatkowski, Cordelia Langford, John Sillitoe on behalf of the Wellcome Sanger Institute COVID-19 Surveillance Team |
| EPI_ISL_3105987 | Lighthouse Lab in Alderley Park | Wellcome Sanger Institute for the COVID-19 Genomics UK (COG-UK) Consortium | Jacquelyn Wynn, Mairead Hyland, The Lighthouse Lab in Alderley Park and Alex Alderton, Roberto Amato, Jeffrey Barrett, Sonia Goncalves, Ewan Harrison, David K. Jackson, Ian Johnston, Dominic Kwiatkowski, Cordelia Langford, John Sillitoe on behalf of the Wellcome Sanger Institute COVID-19 Surveillance Team |
| EPI_ISL_3105988 | Lighthouse Lab in Glasgow | Wellcome Sanger Institute for the COVID-19 Genomics UK (COG-UK) Consortium | Harper VanSteenhouse, Yumi Kasai, David Gray, Carol Clugston, Anna Dominiczak and Alex Alderton, Roberto Amato, Jeffrey Barrett, Sonia Goncalves, Ewan Harrison, David K. Jackson, Ian Johnston, Dominic Kwiatkowski, Cordelia Langford, John Sillitoe on behalf of the Wellcome Sanger Institute COVID-19 Surveillance Team |
| EPI_ISL_3105989 | Lighthouse Lab in Alderley Park | Wellcome Sanger Institute for the COVID-19 Genomics UK (COG-UK) Consortium | Jacquelyn Wynn, Mairead Hyland, The Lighthouse Lab in Alderley Park and Alex Alderton, Roberto Amato, Jeffrey Barrett, Sonia Goncalves, Ewan Harrison, David K. Jackson, Ian Johnston, Dominic Kwiatkowski, Cordelia Langford, John Sillitoe on behalf of the Wellcome Sanger Institute COVID-19 Surveillance Team |
| EPI_ISL_3105990, EPI_ISL_3105991, EPI_ISL_3105992, EPI_ISL_3105993 | Lighthouse Lab in Milton Keynes | Wellcome Sanger Institute for the COVID-19 Genomics UK (COG-UK) Consortium | The Lighthouse Lab in Milton Keynes and Alex Alderton, Roberto Amato, Jeffrey Barrett, Sonia Goncalves, Ewan Harrison, David K. Jackson, Ian Johnston, Dominic Kwiatkowski, Cordelia Langford, John Sillitoe on behalf of the Wellcome Sanger Institute COVID-19 Surveillance Team |
| EPI_ISL_3105994 | Lighthouse Lab in Glasgow | Wellcome Sanger Institute for the COVID-19 Genomics UK (COG-UK) Consortium | Harper VanSteenhouse, Yumi Kasai, David Gray, Carol Clugston, Anna Dominiczak and Alex Alderton, Roberto Amato, Jeffrey Barrett, Sonia Goncalves, Ewan Harrison, David K. Jackson, Ian Johnston, Dominic Kwiatkowski, Cordelia Langford, John Sillitoe on behalf of the Wellcome Sanger Institute COVID-19 Surveillance Team |
| EPI_ISL_3105995 | Lighthouse Lab in Alderley Park | Wellcome Sanger Institute for the COVID-19 Genomics UK (COG-UK) Consortium | Jacquelyn Wynn, Mairead Hyland, The Lighthouse Lab in Alderley Park and Alex Alderton, Roberto Amato, Jeffrey Barrett, Sonia Goncalves, Ewan Harrison, David K. Jackson, Ian Johnston, Dominic Kwiatkowski, Cordelia Langford, John Sillitoe on behalf of the Wellcome Sanger Institute COVID-19 Surveillance Team |
| EPI_ISL_3105996 | Lighthouse Lab in Glasgow | Wellcome Sanger Institute for the COVID-19 Genomics UK (COG-UK) Consortium | Harper VanSteenhouse, Yumi Kasai, David Gray, Carol Clugston, Anna Dominiczak and Alex Alderton, Roberto Amato, Jeffrey Barrett, Sonia Goncalves, Ewan Harrison, David K. Jackson, Ian Johnston, Dominic Kwiatkowski, Cordelia Langford, John Sillitoe on behalf of the Wellcome Sanger Institute COVID-19 Surveillance Team |
| EPI_ISL_3105997 | Lighthouse Lab in Alderley Park | Wellcome Sanger Institute for the COVID-19 Genomics UK (COG-UK) Consortium | Jacquelyn Wynn, Mairead Hyland, The Lighthouse Lab in Alderley Park and Alex Alderton, Roberto Amato, Jeffrey Barrett, Sonia Goncalves, Ewan Harrison, David K. Jackson, Ian Johnston, Dominic Kwiatkowski, Cordelia Langford, John Sillitoe on behalf of the Wellcome Sanger Institute COVID-19 Surveillance Team |
| EPI_ISL_3105998, EPI_ISL_3105999 | Lighthouse Lab in Glasgow | Wellcome Sanger Institute for the COVID-19 Genomics UK (COG-UK) Consortium | Harper VanSteenhouse, Yumi Kasai, David Gray, Carol Clugston, Anna Dominiczak and Alex Alderton, Roberto Amato, Jeffrey Barrett, Sonia Goncalves, Ewan Harrison, David K. Jackson, Ian Johnston, Dominic Kwiatkowski, Cordelia Langford, John Sillitoe on behalf of the Wellcome Sanger Institute COVID-19 Surveillance Team |
| EPI_ISL_3106000, EPI_ISL_3106001 | Lighthouse Lab in Alderley Park | Wellcome Sanger Institute for the COVID-19 Genomics UK (COG-UK) Consortium | Jacquelyn Wynn, Mairead Hyland, The Lighthouse Lab in Alderley Park and Alex Alderton, Roberto Amato, Jeffrey Barrett, Sonia Goncalves, Ewan Harrison, David K. Jackson, Ian Johnston, Dominic Kwiatkowski, Cordelia Langford, John Sillitoe on behalf of the Wellcome Sanger Institute COVID-19 Surveillance Team |
| EPI_ISL_3106002 | Lighthouse Lab in Milton Keynes | Wellcome Sanger Institute for the COVID-19 Genomics UK (COG-UK) Consortium | The Lighthouse Lab in Milton Keynes and Alex Alderton, Roberto Amato, Jeffrey Barrett, Sonia Goncalves, Ewan Harrison, David K. Jackson, Ian Johnston, Dominic Kwiatkowski, Cordelia Langford, John Sillitoe on behalf of the Wellcome Sanger Institute COVID-19 Surveillance Team |
| EPI_ISL_3106003 | Lighthouse Lab in Alderley Park | Wellcome Sanger Institute for the COVID-19 Genomics UK (COG-UK) Consortium | Jacquelyn Wynn, Mairead Hyland, The Lighthouse Lab in Alderley Park and Alex Alderton, Roberto Amato, Jeffrey Barrett, Sonia Goncalves, Ewan Harrison, David K. Jackson, Ian Johnston, Dominic Kwiatkowski, Cordelia Langford, John Sillitoe on behalf of the Wellcome Sanger Institute COVID-19 Surveillance Team |
| EPI_ISL_3106004 | Lighthouse Lab in Milton Keynes | Wellcome Sanger Institute for the COVID-19 Genomics UK (COG-UK) Consortium | The Lighthouse Lab in Milton Keynes and Alex Alderton, Roberto Amato, Jeffrey Barrett, Sonia Goncalves, Ewan Harrison, David K. Jackson, Ian Johnston, Dominic Kwiatkowski, Cordelia Langford, John Sillitoe on behalf of the Wellcome Sanger Institute COVID-19 Surveillance Team |
| EPI_ISL_3106005 | Lighthouse Lab in Glasgow | Wellcome Sanger Institute for the COVID-19 Genomics UK (COG-UK) Consortium | Harper VanSteenhouse, Yumi Kasai, David Gray, Carol Clugston, Anna Dominiczak and Alex Alderton, Roberto Amato, Jeffrey Barrett, Sonia Goncalves, Ewan Harrison, David K. Jackson, Ian Johnston, Dominic Kwiatkowski, Cordelia Langford, John Sillitoe on behalf of the Wellcome Sanger Institute COVID-19 Surveillance Team |
| EPI_ISL_3106006 | Lighthouse Lab in Milton Keynes | Wellcome Sanger Institute for the COVID-19 Genomics UK (COG-UK) Consortium | The Lighthouse Lab in Milton Keynes and Alex Alderton, Roberto Amato, Jeffrey Barrett, Sonia Goncalves, Ewan Harrison, David K. Jackson, Ian Johnston, Dominic Kwiatkowski, Cordelia Langford, John Sillitoe on behalf of the Wellcome Sanger Institute COVID-19 Surveillance Team |
| EPI_ISL_3106007, EPI_ISL_3106008, | Lighthouse Lab in Glasgow | Wellcome Sanger Institute for the COVID-19 Genomics UK (COG-UK) Consortium | Harper VanSteenhouse, Yumi Kasai, David Gray, Carol Clugston, Anna Dominiczak and Alex Alderton, Roberto Amato, Jeffrey Barrett, Sonia Goncalves, Ewan Harrison, David K. Jackson, Ian Johnston, Dominic Kwiatkowski, Cordelia Langford, John Sillitoe on behalf of the Wellcome Sanger Institute COVID-19 Surveillance Team |

| | | | |
|---|---|---|---|
| EPI_ISL_3106009 | | | |
| EPI_ISL_3106010, EPI_ISL_3106011 | Lighthouse Lab in Milton Keynes | Wellcome Sanger Institute for the COVID-19 Genomics UK (COG-UK) Consortium | The Lighthouse Lab in Milton Keynes and Alex Alderton, Roberto Amato, Jeffrey Barrett, Sonia Goncalves, Ewan Harrison, David K. Jackson, Ian Johnston, Dominic Kwiatkowski, Cordelia Langford, John Sillitoe on behalf of the Wellcome Sanger Institute COVID-19 Surveillance Team |
| EPI_ISL_3106012 | Lighthouse Lab in Glasgow | Wellcome Sanger Institute for the COVID-19 Genomics UK (COG-UK) Consortium | Harper VanSteenhouse, Yumi Kasai, David Gray, Carol Clugston, Anna Dominiczak and Alex Alderton, Roberto Amato, Jeffrey Barrett, Sonia Goncalves, Ewan Harrison, David K. Jackson, Ian Johnston, Dominic Kwiatkowski, Cordelia Langford, John Sillitoe on behalf of the Wellcome Sanger Institute COVID-19 Surveillance Team |
| EPI_ISL_3106013 | Lighthouse Lab in Alderley Park | Wellcome Sanger Institute for the COVID-19 Genomics UK (COG-UK) Consortium | Jacquelyn Wynn, Mairead Hyland, The Lighthouse Lab in Alderley Park and Alex Alderton, Roberto Amato, Jeffrey Barrett, Sonia Goncalves, Ewan Harrison, David K. Jackson, Ian Johnston, Dominic Kwiatkowski, Cordelia Langford, John Sillitoe on behalf of the Wellcome Sanger Institute COVID-19 Surveillance Team |
| EPI_ISL_3106014 | Lighthouse Lab in Milton Keynes | Wellcome Sanger Institute for the COVID-19 Genomics UK (COG-UK) Consortium | The Lighthouse Lab in Milton Keynes and Alex Alderton, Roberto Amato, Jeffrey Barrett, Sonia Goncalves, Ewan Harrison, David K. Jackson, Ian Johnston, Dominic Kwiatkowski, Cordelia Langford, John Sillitoe on behalf of the Wellcome Sanger Institute COVID-19 Surveillance Team |
| EPI_ISL_3106015 | Lighthouse Lab in Alderley Park | Wellcome Sanger Institute for the COVID-19 Genomics UK (COG-UK) Consortium | Jacquelyn Wynn, Mairead Hyland, The Lighthouse Lab in Alderley Park and Alex Alderton, Roberto Amato, Jeffrey Barrett, Sonia Goncalves, Ewan Harrison, David K. Jackson, Ian Johnston, Dominic Kwiatkowski, Cordelia Langford, John Sillitoe on behalf of the Wellcome Sanger Institute COVID-19 Surveillance Team |
| EPI_ISL_3106016, EPI_ISL_3106017, EPI_ISL_3106018 | Lighthouse Lab in Milton Keynes | Wellcome Sanger Institute for the COVID-19 Genomics UK (COG-UK) Consortium | The Lighthouse Lab in Milton Keynes and Alex Alderton, Roberto Amato, Jeffrey Barrett, Sonia Goncalves, Ewan Harrison, David K. Jackson, Ian Johnston, Dominic Kwiatkowski, Cordelia Langford, John Sillitoe on behalf of the Wellcome Sanger Institute COVID-19 Surveillance Team |
| EPI_ISL_3106019 | Lighthouse Lab in Glasgow | Wellcome Sanger Institute for the COVID-19 Genomics UK (COG-UK) Consortium | Harper VanSteenhouse, Yumi Kasai, David Gray, Carol Clugston, Anna Dominiczak and Alex Alderton, Roberto Amato, Jeffrey Barrett, Sonia Goncalves, Ewan Harrison, David K. Jackson, Ian Johnston, Dominic Kwiatkowski, Cordelia Langford, John Sillitoe on behalf of the Wellcome Sanger Institute COVID-19 Surveillance Team |
| EPI_ISL_3106020 | Lighthouse Lab in Milton Keynes | Wellcome Sanger Institute for the COVID-19 Genomics UK (COG-UK) Consortium | The Lighthouse Lab in Milton Keynes and Alex Alderton, Roberto Amato, Jeffrey Barrett, Sonia Goncalves, Ewan Harrison, David K. Jackson, Ian Johnston, Dominic Kwiatkowski, Cordelia Langford, John Sillitoe on behalf of the Wellcome Sanger Institute COVID-19 Surveillance Team |
| EPI_ISL_3106021, EPI_ISL_3106022 | Lighthouse Lab in Alderley Park | Wellcome Sanger Institute for the COVID-19 Genomics UK (COG-UK) Consortium | Jacquelyn Wynn, Mairead Hyland, The Lighthouse Lab in Alderley Park and Alex Alderton, Roberto Amato, Jeffrey Barrett, Sonia Goncalves, Ewan Harrison, David K. Jackson, Ian Johnston, Dominic Kwiatkowski, Cordelia Langford, John Sillitoe on behalf of the Wellcome Sanger Institute COVID-19 Surveillance Team |
| EPI_ISL_3106023, EPI_ISL_3106024 | Lighthouse Lab in Milton Keynes | Wellcome Sanger Institute for the COVID-19 Genomics UK (COG-UK) Consortium | The Lighthouse Lab in Milton Keynes and Alex Alderton, Roberto Amato, Jeffrey Barrett, Sonia Goncalves, Ewan Harrison, David K. Jackson, Ian Johnston, Dominic Kwiatkowski, Cordelia Langford, John Sillitoe on behalf of the Wellcome Sanger Institute COVID-19 Surveillance Team |
| EPI_ISL_3106025 | Lighthouse Lab in Glasgow | Wellcome Sanger Institute for the COVID-19 Genomics UK (COG-UK) Consortium | Harper VanSteenhouse, Yumi Kasai, David Gray, Carol Clugston, Anna Dominiczak and Alex Alderton, Roberto Amato, Jeffrey Barrett, Sonia Goncalves, Ewan Harrison, David K. Jackson, Ian Johnston, Dominic Kwiatkowski, Cordelia Langford, John Sillitoe on behalf of the Wellcome Sanger Institute COVID-19 Surveillance Team |
| EPI_ISL_3106026 | Lighthouse Lab in Alderley Park | Wellcome Sanger Institute for the COVID-19 Genomics UK (COG-UK) Consortium | Jacquelyn Wynn, Mairead Hyland, The Lighthouse Lab in Alderley Park and Alex Alderton, Roberto Amato, Jeffrey Barrett, Sonia Goncalves, Ewan Harrison, David K. Jackson, Ian Johnston, Dominic Kwiatkowski, Cordelia Langford, John Sillitoe on behalf of the Wellcome Sanger Institute COVID-19 Surveillance Team |
| EPI_ISL_3106027 | Lighthouse Lab in Milton Keynes | Wellcome Sanger Institute for the COVID-19 Genomics UK (COG-UK) Consortium | The Lighthouse Lab in Milton Keynes and Alex Alderton, Roberto Amato, Jeffrey Barrett, Sonia Goncalves, Ewan Harrison, David K. Jackson, Ian Johnston, Dominic Kwiatkowski, Cordelia Langford, John Sillitoe on behalf of the Wellcome Sanger Institute COVID-19 Surveillance Team |
| EPI_ISL_3106028 | Lighthouse Lab in Alderley Park | Wellcome Sanger Institute for the COVID-19 Genomics UK (COG-UK) Consortium | Jacquelyn Wynn, Mairead Hyland, The Lighthouse Lab in Alderley Park and Alex Alderton, Roberto Amato, Jeffrey Barrett, Sonia Goncalves, Ewan Harrison, David K. Jackson, Ian Johnston, Dominic Kwiatkowski, Cordelia Langford, John Sillitoe on behalf of the Wellcome Sanger Institute COVID-19 Surveillance Team |
| EPI_ISL_3106029, EPI_ISL_3106030, EPI_ISL_3106031, EPI_ISL_3106032, EPI_ISL_3106033 | Lighthouse Lab in Milton Keynes | Wellcome Sanger Institute for the COVID-19 Genomics UK (COG-UK) Consortium | The Lighthouse Lab in Milton Keynes and Alex Alderton, Roberto Amato, Jeffrey Barrett, Sonia Goncalves, Ewan Harrison, David K. Jackson, Ian Johnston, Dominic Kwiatkowski, Cordelia Langford, John Sillitoe on behalf of the Wellcome Sanger Institute COVID-19 Surveillance Team |
| EPI_ISL_3106034 | Lighthouse Lab in Alderley Park | Wellcome Sanger Institute for the COVID-19 Genomics UK (COG-UK) Consortium | Jacquelyn Wynn, Mairead Hyland, The Lighthouse Lab in Alderley Park and Alex Alderton, Roberto Amato, Jeffrey Barrett, Sonia Goncalves, Ewan Harrison, David K. Jackson, Ian Johnston, Dominic Kwiatkowski, Cordelia Langford, John Sillitoe on behalf of the Wellcome Sanger Institute COVID-19 Surveillance Team |
| EPI_ISL_3106035 | Lighthouse Lab in Glasgow | Wellcome Sanger Institute for the COVID-19 Genomics UK (COG-UK) Consortium | Harper VanSteenhouse, Yumi Kasai, David Gray, Carol Clugston, Anna Dominiczak and Alex Alderton, Roberto Amato, Jeffrey Barrett, Sonia Goncalves, Ewan Harrison, David K. Jackson, Ian Johnston, Dominic Kwiatkowski, Cordelia Langford, John Sillitoe on behalf of the Wellcome Sanger Institute COVID-19 Surveillance Team |
| EPI_ISL_3106036 | Lighthouse Lab in Milton Keynes | Wellcome Sanger Institute for the COVID-19 Genomics UK (COG-UK) Consortium | The Lighthouse Lab in Milton Keynes and Alex Alderton, Roberto Amato, Jeffrey Barrett, Sonia Goncalves, Ewan Harrison, David K. Jackson, Ian Johnston, Dominic Kwiatkowski, Cordelia Langford, John Sillitoe on behalf of the Wellcome Sanger Institute COVID-19 Surveillance Team |
| EPI_ISL_3106037 | Lighthouse Lab in Glasgow | Wellcome Sanger Institute for the COVID-19 Genomics UK (COG-UK) Consortium | Harper VanSteenhouse, Yumi Kasai, David Gray, Carol Clugston, Anna Dominiczak and Alex Alderton, Roberto Amato, Jeffrey Barrett, Sonia Goncalves, Ewan Harrison, David K. Jackson, Ian Johnston, Dominic Kwiatkowski, Cordelia Langford, John Sillitoe on behalf of the Wellcome Sanger Institute COVID-19 Surveillance Team |
| EPI_ISL_3106038 | Lighthouse Lab in Milton Keynes | Wellcome Sanger Institute for the COVID-19 Genomics UK (COG-UK) Consortium | The Lighthouse Lab in Milton Keynes and Alex Alderton, Roberto Amato, Jeffrey Barrett, Sonia Goncalves, Ewan Harrison, David K. Jackson, Ian Johnston, Dominic Kwiatkowski, Cordelia Langford, John Sillitoe on behalf of the Wellcome Sanger Institute COVID-19 Surveillance Team |
| EPI_ISL_3106039 | Lighthouse Lab in Glasgow | Wellcome Sanger Institute for the COVID-19 Genomics UK (COG-UK) Consortium | Harper VanSteenhouse, Yumi Kasai, David Gray, Carol Clugston, Anna Dominiczak and Alex Alderton, Roberto Amato, Jeffrey Barrett, Sonia Goncalves, Ewan Harrison, David K. Jackson, Ian Johnston, Dominic Kwiatkowski, Cordelia Langford, John Sillitoe on behalf of the Wellcome Sanger Institute COVID-19 Surveillance Team |
| EPI_ISL_3106040 | Lighthouse Lab in Alderley Park | Wellcome Sanger Institute for the COVID-19 Genomics UK (COG-UK) Consortium | Jacquelyn Wynn, Mairead Hyland, The Lighthouse Lab in Alderley Park and Alex Alderton, Roberto Amato, Jeffrey Barrett, Sonia Goncalves, Ewan Harrison, David K. Jackson, Ian Johnston, Dominic Kwiatkowski, Cordelia Langford, John Sillitoe on behalf of the Wellcome Sanger Institute COVID-19 Surveillance Team |
| EPI_ISL_3106041, EPI_ISL_3106042, EPI_ISL_3106043 | Lighthouse Lab in Milton Keynes | Wellcome Sanger Institute for the COVID-19 Genomics UK (COG-UK) Consortium | The Lighthouse Lab in Milton Keynes and Alex Alderton, Roberto Amato, Jeffrey Barrett, Sonia Goncalves, Ewan Harrison, David K. Jackson, Ian Johnston, Dominic Kwiatkowski, Cordelia Langford, John Sillitoe on behalf of the Wellcome Sanger Institute COVID-19 Surveillance Team |
| EPI_ISL_3106044 | Lighthouse Lab in Glasgow | Wellcome Sanger Institute for the COVID-19 Genomics UK (COG-UK) Consortium | Harper VanSteenhouse, Yumi Kasai, David Gray, Carol Clugston, Anna Dominiczak and Alex Alderton, Roberto Amato, Jeffrey Barrett, Sonia Goncalves, Ewan Harrison, David K. Jackson, Ian Johnston, Dominic Kwiatkowski, Cordelia Langford, John Sillitoe on behalf of the Wellcome Sanger Institute COVID-19 Surveillance Team |
| EPI_ISL_3106045 | Lighthouse Lab in Milton Keynes | Wellcome Sanger Institute for the COVID-19 Genomics UK (COG-UK) Consortium | The Lighthouse Lab in Milton Keynes and Alex Alderton, Roberto Amato, Jeffrey Barrett, Sonia Goncalves, Ewan Harrison, David K. Jackson, Ian Johnston, Dominic Kwiatkowski, Cordelia Langford, John Sillitoe on behalf of the Wellcome Sanger Institute COVID-19 Surveillance Team |
| EPI_ISL_3106046 | Lighthouse Lab in Alderley Park | Wellcome Sanger Institute for the COVID-19 Genomics UK (COG-UK) Consortium | Jacquelyn Wynn, Mairead Hyland, The Lighthouse Lab in Alderley Park and Alex Alderton, Roberto Amato, Jeffrey Barrett, Sonia Goncalves, Ewan Harrison, David K. Jackson, Ian Johnston, Dominic Kwiatkowski, Cordelia Langford, John Sillitoe on behalf of the Wellcome Sanger Institute COVID-19 Surveillance Team |
| EPI_ISL_3106047 | Lighthouse Lab in Milton Keynes | Wellcome Sanger Institute for the COVID-19 Genomics UK (COG-UK) Consortium | The Lighthouse Lab in Milton Keynes and Alex Alderton, Roberto Amato, Jeffrey Barrett, Sonia Goncalves, Ewan Harrison, David K. Jackson, Ian Johnston, Dominic Kwiatkowski, Cordelia Langford, John Sillitoe on behalf of the Wellcome Sanger Institute COVID-19 Surveillance Team |
| EPI_ISL_3106048 | Lighthouse Lab in Glasgow | Wellcome Sanger Institute for the COVID-19 Genomics UK (COG-UK) Consortium | Harper VanSteenhouse, Yumi Kasai, David Gray, Carol Clugston, Anna Dominiczak and Alex Alderton, Roberto Amato, Jeffrey Barrett, Sonia Goncalves, Ewan Harrison, David K. Jackson, Ian Johnston, Dominic Kwiatkowski, Cordelia Langford, John Sillitoe on behalf of the Wellcome Sanger Institute COVID-19 Surveillance Team |
| EPI_ISL_3106049, EPI_ISL_3106050 | Lighthouse Lab in Alderley Park | Wellcome Sanger Institute for the COVID-19 Genomics UK (COG-UK) Consortium | Jacquelyn Wynn, Mairead Hyland, The Lighthouse Lab in Alderley Park and Alex Alderton, Roberto Amato, Jeffrey Barrett, Sonia Goncalves, Ewan Harrison, David K. Jackson, Ian Johnston, Dominic Kwiatkowski, Cordelia Langford, John Sillitoe on behalf of the Wellcome Sanger Institute COVID-19 Surveillance Team |
| EPI_ISL_3106051 | Lighthouse Lab in Glasgow | Wellcome Sanger Institute for the COVID-19 Genomics UK (COG-UK) Consortium | Harper VanSteenhouse, Yumi Kasai, David Gray, Carol Clugston, Anna Dominiczak and Alex Alderton, Roberto Amato, Jeffrey Barrett, Sonia Goncalves, Ewan Harrison, David K. Jackson, Ian Johnston, Dominic Kwiatkowski, Cordelia Langford, John Sillitoe on behalf of the Wellcome Sanger Institute COVID-19 Surveillance Team |
| EPI_ISL_3106052, EPI_ISL_3106053 | Lighthouse Lab in Milton Keynes | Wellcome Sanger Institute for the COVID-19 Genomics UK (COG-UK) Consortium | The Lighthouse Lab in Milton Keynes and Alex Alderton, Roberto Amato, Jeffrey Barrett, Sonia Goncalves, Ewan Harrison, David K. Jackson, Ian Johnston, Dominic Kwiatkowski, Cordelia Langford, John Sillitoe on behalf of the Wellcome Sanger Institute COVID-19 Surveillance Team |
| EPI_ISL_3106054 | Lighthouse Lab in Alderley Park | Wellcome Sanger Institute for the COVID-19 Genomics UK (COG-UK) Consortium | Jacquelyn Wynn, Mairead Hyland, The Lighthouse Lab in Alderley Park and Alex Alderton, Roberto Amato, Jeffrey Barrett, Sonia Goncalves, Ewan Harrison, David K. Jackson, Ian Johnston, Dominic Kwiatkowski, Cordelia Langford, John Sillitoe on behalf of the Wellcome Sanger Institute COVID-19 Surveillance Team |
| EPI_ISL_3106055 | Lighthouse Lab in Milton Keynes | Wellcome Sanger Institute for the COVID-19 Genomics UK (COG-UK) Consortium | The Lighthouse Lab in Milton Keynes and Alex Alderton, Roberto Amato, Jeffrey Barrett, Sonia Goncalves, Ewan Harrison, David K. Jackson, Ian Johnston, Dominic Kwiatkowski, Cordelia Langford, John Sillitoe on behalf of the Wellcome Sanger Institute COVID-19 Surveillance Team |
| EPI_ISL_3106056 | Lighthouse Lab in Alderley Park | Wellcome Sanger Institute for the COVID-19 Genomics UK (COG-UK) Consortium | Jacquelyn Wynn, Mairead Hyland, The Lighthouse Lab in Alderley Park and Alex Alderton, Roberto Amato, Jeffrey Barrett, Sonia Goncalves, Ewan Harrison, David K. Jackson, Ian Johnston, Dominic Kwiatkowski, Cordelia Langford, John Sillitoe on behalf of the Wellcome Sanger Institute COVID-19 Surveillance Team |
| EPI_ISL_3106057 | Lighthouse Lab in Glasgow | Wellcome Sanger Institute for the COVID-19 Genomics UK (COG-UK) Consortium | Harper VanSteenhouse, Yumi Kasai, David Gray, Carol Clugston, Anna Dominiczak and Alex Alderton, Roberto Amato, Jeffrey Barrett, Sonia Goncalves, Ewan Harrison, David K. Jackson, Ian Johnston, Dominic Kwiatkowski, Cordelia Langford, John Sillitoe on behalf of the Wellcome Sanger Institute COVID-19 Surveillance Team |
| EPI_ISL_3106058, EPI_ISL_3106059, EPI_ISL_3106060, EPI_ISL_3106061, EPI_ISL_3106062, EPI_ISL_3106063, EPI_ISL_3106064, EPI_ISL_3106065 | Lighthouse Lab in Milton Keynes | Wellcome Sanger Institute for the COVID-19 Genomics UK (COG-UK) Consortium | The Lighthouse Lab in Milton Keynes and Alex Alderton, Roberto Amato, Jeffrey Barrett, Sonia Goncalves, Ewan Harrison, David K. Jackson, Ian Johnston, Dominic Kwiatkowski, Cordelia Langford, John Sillitoe on behalf of the Wellcome Sanger Institute COVID-19 Surveillance Team |
| EPI_ISL_3106066 | Lighthouse Lab in Glasgow | Wellcome Sanger Institute for the COVID-19 Genomics UK (COG-UK) Consortium | Harper VanSteenhouse, Yumi Kasai, David Gray, Carol Clugston, Anna Dominiczak and Alex Alderton, Roberto Amato, Jeffrey Barrett, Sonia Goncalves, Ewan Harrison, David K. Jackson, Ian Johnston, Dominic Kwiatkowski, Cordelia Langford, John Sillitoe on behalf of the Wellcome Sanger Institute COVID-19 Surveillance Team |
| EPI_ISL_3106067, EPI_ISL_3106068 | Lighthouse Lab in Milton Keynes | Wellcome Sanger Institute for the COVID-19 Genomics UK (COG-UK) Consortium | The Lighthouse Lab in Milton Keynes and Alex Alderton, Roberto Amato, Jeffrey Barrett, Sonia Goncalves, Ewan Harrison, David K. Jackson, Ian Johnston, Dominic Kwiatkowski, Cordelia Langford, John Sillitoe on behalf of the Wellcome Sanger Institute COVID-19 Surveillance Team |
| EPI_ISL_3106069 | Lighthouse Lab in Alderley Park | Wellcome Sanger Institute for the COVID-19 Genomics UK (COG-UK) Consortium | Jacquelyn Wynn, Mairead Hyland, The Lighthouse Lab in Alderley Park and Alex Alderton, Roberto Amato, Jeffrey Barrett, Sonia Goncalves, Ewan Harrison, David K. Jackson, Ian Johnston, Dominic Kwiatkowski, Cordelia Langford, John Sillitoe on behalf of the Wellcome Sanger Institute COVID-19 Surveillance Team |
| EPI_ISL_3106070 | Lighthouse Lab in Milton Keynes | Wellcome Sanger Institute for the COVID-19 Genomics UK (COG-UK) Consortium | The Lighthouse Lab in Milton Keynes and Alex Alderton, Roberto Amato, Jeffrey Barrett, Sonia Goncalves, Ewan Harrison, David K. Jackson, Ian Johnston, Dominic Kwiatkowski, Cordelia Langford, John Sillitoe on behalf of the Wellcome Sanger Institute COVID-19 Surveillance Team |

| | | | |
|---|---|---|---|
| EPI_ISL_3106071 | Lighthouse Lab in Glasgow | Wellcome Sanger Institute for the COVID-19 Genomics UK (COG-UK) Consortium | Harper VanSteenhouse, Yumi Kasai, David Gray, Carol Clugston, Anna Dominiczak and Alex Alderton, Roberto Amato, Jeffrey Barrett, Sonia Goncalves, Ewan Harrison, David K. Jackson, Ian Johnston, Dominic Kwiatkowski, Cordelia Langford, John Sillitoe on behalf of the Wellcome Sanger Institute COVID-19 Surveillance Team |
| EPI_ISL_3106072, EPI_ISL_3106073 | Lighthouse Lab in Milton Keynes | Wellcome Sanger Institute for the COVID-19 Genomics UK (COG-UK) Consortium | The Lighthouse Lab in Milton Keynes and Alex Alderton, Roberto Amato, Jeffrey Barrett, Sonia Goncalves, Ewan Harrison, David K. Jackson, Ian Johnston, Dominic Kwiatkowski, Cordelia Langford, John Sillitoe on behalf of the Wellcome Sanger Institute COVID-19 Surveillance Team |
| EPI_ISL_3106074 | Lighthouse Lab in Glasgow | Wellcome Sanger Institute for the COVID-19 Genomics UK (COG-UK) Consortium | Harper VanSteenhouse, Yumi Kasai, David Gray, Carol Clugston, Anna Dominiczak and Alex Alderton, Roberto Amato, Jeffrey Barrett, Sonia Goncalves, Ewan Harrison, David K. Jackson, Ian Johnston, Dominic Kwiatkowski, Cordelia Langford, John Sillitoe on behalf of the Wellcome Sanger Institute COVID-19 Surveillance Team |
| EPI_ISL_3106075 | Lighthouse Lab in Milton Keynes | Wellcome Sanger Institute for the COVID-19 Genomics UK (COG-UK) Consortium | The Lighthouse Lab in Milton Keynes and Alex Alderton, Roberto Amato, Jeffrey Barrett, Sonia Goncalves, Ewan Harrison, David K. Jackson, Ian Johnston, Dominic Kwiatkowski, Cordelia Langford, John Sillitoe on behalf of the Wellcome Sanger Institute COVID-19 Surveillance Team |
| EPI_ISL_3106076, EPI_ISL_3106077 | Lighthouse Lab in Alderley Park | Wellcome Sanger Institute for the COVID-19 Genomics UK (COG-UK) Consortium | Jacquelyn Wynn, Mairead Hyland, The Lighthouse Lab in Alderley Park and Alex Alderton, Roberto Amato, Jeffrey Barrett, Sonia Goncalves, Ewan Harrison, David K. Jackson, Ian Johnston, Dominic Kwiatkowski, Cordelia Langford, John Sillitoe on behalf of the Wellcome Sanger Institute COVID-19 Surveillance Team |
| EPI_ISL_3106078 | Lighthouse Lab in Milton Keynes | Wellcome Sanger Institute for the COVID-19 Genomics UK (COG-UK) Consortium | The Lighthouse Lab in Milton Keynes and Alex Alderton, Roberto Amato, Jeffrey Barrett, Sonia Goncalves, Ewan Harrison, David K. Jackson, Ian Johnston, Dominic Kwiatkowski, Cordelia Langford, John Sillitoe on behalf of the Wellcome Sanger Institute COVID-19 Surveillance Team |
| EPI_ISL_3106079 | Lighthouse Lab in Alderley Park | Wellcome Sanger Institute for the COVID-19 Genomics UK (COG-UK) Consortium | Jacquelyn Wynn, Mairead Hyland, The Lighthouse Lab in Alderley Park and Alex Alderton, Roberto Amato, Jeffrey Barrett, Sonia Goncalves, Ewan Harrison, David K. Jackson, Ian Johnston, Dominic Kwiatkowski, Cordelia Langford, John Sillitoe on behalf of the Wellcome Sanger Institute COVID-19 Surveillance Team |
| EPI_ISL_3106080, EPI_ISL_3106081, EPI_ISL_3106082 | Lighthouse Lab in Milton Keynes | Wellcome Sanger Institute for the COVID-19 Genomics UK (COG-UK) Consortium | The Lighthouse Lab in Milton Keynes and Alex Alderton, Roberto Amato, Jeffrey Barrett, Sonia Goncalves, Ewan Harrison, David K. Jackson, Ian Johnston, Dominic Kwiatkowski, Cordelia Langford, John Sillitoe on behalf of the Wellcome Sanger Institute COVID-19 Surveillance Team |
| EPI_ISL_3106083 | Lighthouse Lab in Alderley Park | Wellcome Sanger Institute for the COVID-19 Genomics UK (COG-UK) Consortium | Jacquelyn Wynn, Mairead Hyland, The Lighthouse Lab in Alderley Park and Alex Alderton, Roberto Amato, Jeffrey Barrett, Sonia Goncalves, Ewan Harrison, David K. Jackson, Ian Johnston, Dominic Kwiatkowski, Cordelia Langford, John Sillitoe on behalf of the Wellcome Sanger Institute COVID-19 Surveillance Team |
| EPI_ISL_3106084 | Lighthouse Lab in Glasgow | Wellcome Sanger Institute for the COVID-19 Genomics UK (COG-UK) Consortium | Harper VanSteenhouse, Yumi Kasai, David Gray, Carol Clugston, Anna Dominiczak and Alex Alderton, Roberto Amato, Jeffrey Barrett, Sonia Goncalves, Ewan Harrison, David K. Jackson, Ian Johnston, Dominic Kwiatkowski, Cordelia Langford, John Sillitoe on behalf of the Wellcome Sanger Institute COVID-19 Surveillance Team |
| EPI_ISL_3106085, EPI_ISL_3106086, EPI_ISL_3106087, EPI_ISL_3106088, EPI_ISL_3106089 | Lighthouse Lab in Milton Keynes | Wellcome Sanger Institute for the COVID-19 Genomics UK (COG-UK) Consortium | The Lighthouse Lab in Milton Keynes and Alex Alderton, Roberto Amato, Jeffrey Barrett, Sonia Goncalves, Ewan Harrison, David K. Jackson, Ian Johnston, Dominic Kwiatkowski, Cordelia Langford, John Sillitoe on behalf of the Wellcome Sanger Institute COVID-19 Surveillance Team |
| EPI_ISL_3106090 | Lighthouse Lab in Alderley Park | Wellcome Sanger Institute for the COVID-19 Genomics UK (COG-UK) Consortium | Jacquelyn Wynn, Mairead Hyland, The Lighthouse Lab in Alderley Park and Alex Alderton, Roberto Amato, Jeffrey Barrett, Sonia Goncalves, Ewan Harrison, David K. Jackson, Ian Johnston, Dominic Kwiatkowski, Cordelia Langford, John Sillitoe on behalf of the Wellcome Sanger Institute COVID-19 Surveillance Team |
| EPI_ISL_3106091, EPI_ISL_3106092, EPI_ISL_3106093, EPI_ISL_3106094 | Lighthouse Lab in Milton Keynes | Wellcome Sanger Institute for the COVID-19 Genomics UK (COG-UK) Consortium | The Lighthouse Lab in Milton Keynes and Alex Alderton, Roberto Amato, Jeffrey Barrett, Sonia Goncalves, Ewan Harrison, David K. Jackson, Ian Johnston, Dominic Kwiatkowski, Cordelia Langford, John Sillitoe on behalf of the Wellcome Sanger Institute COVID-19 Surveillance Team |
| EPI_ISL_3106095, EPI_ISL_3106096 | Lighthouse Lab in Alderley Park | Wellcome Sanger Institute for the COVID-19 Genomics UK (COG-UK) Consortium | Jacquelyn Wynn, Mairead Hyland, The Lighthouse Lab in Alderley Park and Alex Alderton, Roberto Amato, Jeffrey Barrett, Sonia Goncalves, Ewan Harrison, David K. Jackson, Ian Johnston, Dominic Kwiatkowski, Cordelia Langford, John Sillitoe on behalf of the Wellcome Sanger Institute COVID-19 Surveillance Team |
| EPI_ISL_3106097 | Lighthouse Lab in Milton Keynes | Wellcome Sanger Institute for the COVID-19 Genomics UK (COG-UK) Consortium | The Lighthouse Lab in Milton Keynes and Alex Alderton, Roberto Amato, Jeffrey Barrett, Sonia Goncalves, Ewan Harrison, David K. Jackson, Ian Johnston, Dominic Kwiatkowski, Cordelia Langford, John Sillitoe on behalf of the Wellcome Sanger Institute COVID-19 Surveillance Team |
| EPI_ISL_3106098 | Lighthouse Lab in Alderley Park | Wellcome Sanger Institute for the COVID-19 Genomics UK (COG-UK) Consortium | Jacquelyn Wynn, Mairead Hyland, The Lighthouse Lab in Alderley Park and Alex Alderton, Roberto Amato, Jeffrey Barrett, Sonia Goncalves, Ewan Harrison, David K. Jackson, Ian Johnston, Dominic Kwiatkowski, Cordelia Langford, John Sillitoe on behalf of the Wellcome Sanger Institute COVID-19 Surveillance Team |
| EPI_ISL_3106099 | Lighthouse Lab in Glasgow | Wellcome Sanger Institute for the COVID-19 Genomics UK (COG-UK) Consortium | Harper VanSteenhouse, Yumi Kasai, David Gray, Carol Clugston, Anna Dominiczak and Alex Alderton, Roberto Amato, Jeffrey Barrett, Sonia Goncalves, Ewan Harrison, David K. Jackson, Ian Johnston, Dominic Kwiatkowski, Cordelia Langford, John Sillitoe on behalf of the Wellcome Sanger Institute COVID-19 Surveillance Team |
| EPI_ISL_3106100 | Lighthouse Lab in Milton Keynes | Wellcome Sanger Institute for the COVID-19 Genomics UK (COG-UK) Consortium | The Lighthouse Lab in Milton Keynes and Alex Alderton, Roberto Amato, Jeffrey Barrett, Sonia Goncalves, Ewan Harrison, David K. Jackson, Ian Johnston, Dominic Kwiatkowski, Cordelia Langford, John Sillitoe on behalf of the Wellcome Sanger Institute COVID-19 Surveillance Team |
| EPI_ISL_3106101 | Lighthouse Lab in Glasgow | Wellcome Sanger Institute for the COVID-19 Genomics UK (COG-UK) Consortium | Harper VanSteenhouse, Yumi Kasai, David Gray, Carol Clugston, Anna Dominiczak and Alex Alderton, Roberto Amato, Jeffrey Barrett, Sonia Goncalves, Ewan Harrison, David K. Jackson, Ian Johnston, Dominic Kwiatkowski, Cordelia Langford, John Sillitoe on behalf of the Wellcome Sanger Institute COVID-19 Surveillance Team |
| EPI_ISL_3106102 | Lighthouse Lab in Milton Keynes | Wellcome Sanger Institute for the COVID-19 Genomics UK (COG-UK) Consortium | The Lighthouse Lab in Milton Keynes and Alex Alderton, Roberto Amato, Jeffrey Barrett, Sonia Goncalves, Ewan Harrison, David K. Jackson, Ian Johnston, Dominic Kwiatkowski, Cordelia Langford, John Sillitoe on behalf of the Wellcome Sanger Institute COVID-19 Surveillance Team |
| EPI_ISL_3106103 | Lighthouse Lab in Alderley Park | Wellcome Sanger Institute for the COVID-19 Genomics UK (COG-UK) Consortium | Jacquelyn Wynn, Mairead Hyland, The Lighthouse Lab in Alderley Park and Alex Alderton, Roberto Amato, Jeffrey Barrett, Sonia Goncalves, Ewan Harrison, David K. Jackson, Ian Johnston, Dominic Kwiatkowski, Cordelia Langford, John Sillitoe on behalf of the Wellcome Sanger Institute COVID-19 Surveillance Team |
| EPI_ISL_3106104 | Lighthouse Lab in Milton Keynes | Wellcome Sanger Institute for the COVID-19 Genomics UK (COG-UK) Consortium | The Lighthouse Lab in Milton Keynes and Alex Alderton, Roberto Amato, Jeffrey Barrett, Sonia Goncalves, Ewan Harrison, David K. Jackson, Ian Johnston, Dominic Kwiatkowski, Cordelia Langford, John Sillitoe on behalf of the Wellcome Sanger Institute COVID-19 Surveillance Team |
| EPI_ISL_3106105, EPI_ISL_3106106, EPI_ISL_3106107, EPI_ISL_3106108 | Lighthouse Lab in Alderley Park | Wellcome Sanger Institute for the COVID-19 Genomics UK (COG-UK) Consortium | Jacquelyn Wynn, Mairead Hyland, The Lighthouse Lab in Alderley Park and Alex Alderton, Roberto Amato, Jeffrey Barrett, Sonia Goncalves, Ewan Harrison, David K. Jackson, Ian Johnston, Dominic Kwiatkowski, Cordelia Langford, John Sillitoe on behalf of the Wellcome Sanger Institute COVID-19 Surveillance Team |
| EPI_ISL_3106109 | Lighthouse Lab in Milton Keynes | Wellcome Sanger Institute for the COVID-19 Genomics UK (COG-UK) Consortium | The Lighthouse Lab in Milton Keynes and Alex Alderton, Roberto Amato, Jeffrey Barrett, Sonia Goncalves, Ewan Harrison, David K. Jackson, Ian Johnston, Dominic Kwiatkowski, Cordelia Langford, John Sillitoe on behalf of the Wellcome Sanger Institute COVID-19 Surveillance Team |
| EPI_ISL_3106110 | Lighthouse Lab in Alderley Park | Wellcome Sanger Institute for the COVID-19 Genomics UK (COG-UK) Consortium | Jacquelyn Wynn, Mairead Hyland, The Lighthouse Lab in Alderley Park and Alex Alderton, Roberto Amato, Jeffrey Barrett, Sonia Goncalves, Ewan Harrison, David K. Jackson, Ian Johnston, Dominic Kwiatkowski, Cordelia Langford, John Sillitoe on behalf of the Wellcome Sanger Institute COVID-19 Surveillance Team |
| EPI_ISL_3106111 | Lighthouse Lab in Glasgow | Wellcome Sanger Institute for the COVID-19 Genomics UK (COG-UK) Consortium | Harper VanSteenhouse, Yumi Kasai, David Gray, Carol Clugston, Anna Dominiczak and Alex Alderton, Roberto Amato, Jeffrey Barrett, Sonia Goncalves, Ewan Harrison, David K. Jackson, Ian Johnston, Dominic Kwiatkowski, Cordelia Langford, John Sillitoe on behalf of the Wellcome Sanger Institute COVID-19 Surveillance Team |
| EPI_ISL_3106112, EPI_ISL_3106113, EPI_ISL_3106114, EPI_ISL_3106115 | Lighthouse Lab in Milton Keynes | Wellcome Sanger Institute for the COVID-19 Genomics UK (COG-UK) Consortium | The Lighthouse Lab in Milton Keynes and Alex Alderton, Roberto Amato, Jeffrey Barrett, Sonia Goncalves, Ewan Harrison, David K. Jackson, Ian Johnston, Dominic Kwiatkowski, Cordelia Langford, John Sillitoe on behalf of the Wellcome Sanger Institute COVID-19 Surveillance Team |
| EPI_ISL_3106116 | Lighthouse Lab in Glasgow | Wellcome Sanger Institute for the COVID-19 Genomics UK (COG-UK) Consortium | Harper VanSteenhouse, Yumi Kasai, David Gray, Carol Clugston, Anna Dominiczak and Alex Alderton, Roberto Amato, Jeffrey Barrett, Sonia Goncalves, Ewan Harrison, David K. Jackson, Ian Johnston, Dominic Kwiatkowski, Cordelia Langford, John Sillitoe on behalf of the Wellcome Sanger Institute COVID-19 Surveillance Team |
| EPI_ISL_3106117, EPI_ISL_3106118, EPI_ISL_3106119, EPI_ISL_3106120, EPI_ISL_3106121, EPI_ISL_3106122, EPI_ISL_3106123 | Lighthouse Lab in Milton Keynes | Wellcome Sanger Institute for the COVID-19 Genomics UK (COG-UK) Consortium | The Lighthouse Lab in Milton Keynes and Alex Alderton, Roberto Amato, Jeffrey Barrett, Sonia Goncalves, Ewan Harrison, David K. Jackson, Ian Johnston, Dominic Kwiatkowski, Cordelia Langford, John Sillitoe on behalf of the Wellcome Sanger Institute COVID-19 Surveillance Team |
| EPI_ISL_3106124 | Lighthouse Lab in Alderley Park | Wellcome Sanger Institute for the COVID-19 Genomics UK (COG-UK) Consortium | Jacquelyn Wynn, Mairead Hyland, The Lighthouse Lab in Alderley Park and Alex Alderton, Roberto Amato, Jeffrey Barrett, Sonia Goncalves, Ewan Harrison, David K. Jackson, Ian Johnston, Dominic Kwiatkowski, Cordelia Langford, John Sillitoe on behalf of the Wellcome Sanger Institute COVID-19 Surveillance Team |
| EPI_ISL_3106125 | Lighthouse Lab in Milton Keynes | Wellcome Sanger Institute for the COVID-19 Genomics UK (COG-UK) Consortium | The Lighthouse Lab in Milton Keynes and Alex Alderton, Roberto Amato, Jeffrey Barrett, Sonia Goncalves, Ewan Harrison, David K. Jackson, Ian Johnston, Dominic Kwiatkowski, Cordelia Langford, John Sillitoe on behalf of the Wellcome Sanger Institute COVID-19 Surveillance Team |
| EPI_ISL_3106126 | Lighthouse Lab in Glasgow | Wellcome Sanger Institute for the COVID-19 Genomics UK (COG-UK) Consortium | Harper VanSteenhouse, Yumi Kasai, David Gray, Carol Clugston, Anna Dominiczak and Alex Alderton, Roberto Amato, Jeffrey Barrett, Sonia Goncalves, Ewan Harrison, David K. Jackson, Ian Johnston, Dominic Kwiatkowski, Cordelia Langford, John Sillitoe on behalf of the Wellcome Sanger Institute COVID-19 Surveillance Team |
| EPI_ISL_3106127, EPI_ISL_3106128, EPI_ISL_3106129 | Lighthouse Lab in Milton Keynes | Wellcome Sanger Institute for the COVID-19 Genomics UK (COG-UK) Consortium | The Lighthouse Lab in Milton Keynes and Alex Alderton, Roberto Amato, Jeffrey Barrett, Sonia Goncalves, Ewan Harrison, David K. Jackson, Ian Johnston, Dominic Kwiatkowski, Cordelia Langford, John Sillitoe on behalf of the Wellcome Sanger Institute COVID-19 Surveillance Team |
| EPI_ISL_3106130, EPI_ISL_3106131, EPI_ISL_3106132 | Lighthouse Lab in Alderley Park | Wellcome Sanger Institute for the COVID-19 Genomics UK (COG-UK) Consortium | Jacquelyn Wynn, Mairead Hyland, The Lighthouse Lab in Alderley Park and Alex Alderton, Roberto Amato, Jeffrey Barrett, Sonia Goncalves, Ewan Harrison, David K. Jackson, Ian Johnston, Dominic Kwiatkowski, Cordelia Langford, John Sillitoe on behalf of the Wellcome Sanger Institute COVID-19 Surveillance Team |
| EPI_ISL_3106133 | Lighthouse Lab in Milton Keynes | Wellcome Sanger Institute for the COVID-19 Genomics UK (COG-UK) Consortium | The Lighthouse Lab in Milton Keynes and Alex Alderton, Roberto Amato, Jeffrey Barrett, Sonia Goncalves, Ewan Harrison, David K. Jackson, Ian Johnston, Dominic Kwiatkowski, Cordelia Langford, John Sillitoe on behalf of the Wellcome Sanger Institute COVID-19 Surveillance Team |
| EPI_ISL_3106134, EPI_ISL_3106135 | Lighthouse Lab in Alderley Park | Wellcome Sanger Institute for the COVID-19 Genomics UK (COG-UK) Consortium | Jacquelyn Wynn, Mairead Hyland, The Lighthouse Lab in Alderley Park and Alex Alderton, Roberto Amato, Jeffrey Barrett, Sonia Goncalves, Ewan Harrison, David K. Jackson, Ian Johnston, Dominic Kwiatkowski, Cordelia Langford, John Sillitoe on behalf of the Wellcome Sanger Institute COVID-19 Surveillance Team |
| EPI_ISL_3106136, EPI_ISL_3106137 | Lighthouse Lab in Milton Keynes | Wellcome Sanger Institute for the COVID-19 Genomics UK (COG-UK) Consortium | The Lighthouse Lab in Milton Keynes and Alex Alderton, Roberto Amato, Jeffrey Barrett, Sonia Goncalves, Ewan Harrison, David K. Jackson, Ian Johnston, Dominic Kwiatkowski, Cordelia Langford, John Sillitoe on behalf of the Wellcome Sanger Institute COVID-19 Surveillance Team |
| EPI_ISL_3106138 | Lighthouse Lab in Glasgow | Wellcome Sanger Institute for the COVID-19 Genomics UK (COG-UK) Consortium | Harper VanSteenhouse, Yumi Kasai, David Gray, Carol Clugston, Anna Dominiczak and Alex Alderton, Roberto Amato, Jeffrey Barrett, Sonia Goncalves, Ewan Harrison, David K. Jackson, Ian Johnston, Dominic Kwiatkowski, Cordelia Langford, John Sillitoe on behalf of the Wellcome Sanger Institute COVID-19 Surveillance Team |

| | | | |
|---|---|---|---|
| EPI_ISL_3106139, EPI_ISL_3106140 | Lighthouse Lab in Milton Keynes | Wellcome Sanger Institute for the COVID-19 Genomics UK (COG-UK) Consortium | The Lighthouse Lab in Milton Keynes and Alex Alderton, Roberto Amato, Jeffrey Barrett, Sonia Goncalves, Ewan Harrison, David K. Jackson, Ian Johnston, Dominic Kwiatkowski, Cordelia Langford, John Sillitoe on behalf of the Wellcome Sanger Institute COVID-19 Surveillance Team |
| EPI_ISL_3106141 | Lighthouse Lab in Alderley Park | Wellcome Sanger Institute for the COVID-19 Genomics UK (COG-UK) Consortium | Jacquelyn Wynn, Mairead Hyland, The Lighthouse Lab in Alderley Park and Alex Alderton, Roberto Amato, Jeffrey Barrett, Sonia Goncalves, Ewan Harrison, David K. Jackson, Ian Johnston, Dominic Kwiatkowski, Cordelia Langford, John Sillitoe on behalf of the Wellcome Sanger Institute COVID-19 Surveillance Team |
| EPI_ISL_3106142, EPI_ISL_3106143 | Lighthouse Lab in Milton Keynes | Wellcome Sanger Institute for the COVID-19 Genomics UK (COG-UK) Consortium | The Lighthouse Lab in Milton Keynes and Alex Alderton, Roberto Amato, Jeffrey Barrett, Sonia Goncalves, Ewan Harrison, David K. Jackson, Ian Johnston, Dominic Kwiatkowski, Cordelia Langford, John Sillitoe on behalf of the Wellcome Sanger Institute COVID-19 Surveillance Team |
| EPI_ISL_3106144 | Lighthouse Lab in Alderley Park | Wellcome Sanger Institute for the COVID-19 Genomics UK (COG-UK) Consortium | Jacquelyn Wynn, Mairead Hyland, The Lighthouse Lab in Alderley Park and Alex Alderton, Roberto Amato, Jeffrey Barrett, Sonia Goncalves, Ewan Harrison, David K. Jackson, Ian Johnston, Dominic Kwiatkowski, Cordelia Langford, John Sillitoe on behalf of the Wellcome Sanger Institute COVID-19 Surveillance Team |
| EPI_ISL_3106145, EPI_ISL_3106146, EPI_ISL_3106147, EPI_ISL_3106148 | Lighthouse Lab in Milton Keynes | Wellcome Sanger Institute for the COVID-19 Genomics UK (COG-UK) Consortium | The Lighthouse Lab in Milton Keynes and Alex Alderton, Roberto Amato, Jeffrey Barrett, Sonia Goncalves, Ewan Harrison, David K. Jackson, Ian Johnston, Dominic Kwiatkowski, Cordelia Langford, John Sillitoe on behalf of the Wellcome Sanger Institute COVID-19 Surveillance Team |
| EPI_ISL_3106149 | Lighthouse Lab in Glasgow | Wellcome Sanger Institute for the COVID-19 Genomics UK (COG-UK) Consortium | Harper VanSteenhouse, Yumi Kasai, David Gray, Carol Clugston, Anna Dominiczak and Alex Alderton, Roberto Amato, Jeffrey Barrett, Sonia Goncalves, Ewan Harrison, David K. Jackson, Ian Johnston, Dominic Kwiatkowski, Cordelia Langford, John Sillitoe on behalf of the Wellcome Sanger Institute COVID-19 Surveillance Team |
| EPI_ISL_3106150 | Lighthouse Lab in Alderley Park | Wellcome Sanger Institute for the COVID-19 Genomics UK (COG-UK) Consortium | Jacquelyn Wynn, Mairead Hyland, The Lighthouse Lab in Alderley Park and Alex Alderton, Roberto Amato, Jeffrey Barrett, Sonia Goncalves, Ewan Harrison, David K. Jackson, Ian Johnston, Dominic Kwiatkowski, Cordelia Langford, John Sillitoe on behalf of the Wellcome Sanger Institute COVID-19 Surveillance Team |
| EPI_ISL_3106151 | Lighthouse Lab in Glasgow | Wellcome Sanger Institute for the COVID-19 Genomics UK (COG-UK) Consortium | Harper VanSteenhouse, Yumi Kasai, David Gray, Carol Clugston, Anna Dominiczak and Alex Alderton, Roberto Amato, Jeffrey Barrett, Sonia Goncalves, Ewan Harrison, David K. Jackson, Ian Johnston, Dominic Kwiatkowski, Cordelia Langford, John Sillitoe on behalf of the Wellcome Sanger Institute COVID-19 Surveillance Team |
| EPI_ISL_3106152 | Lighthouse Lab in Alderley Park | Wellcome Sanger Institute for the COVID-19 Genomics UK (COG-UK) Consortium | Jacquelyn Wynn, Mairead Hyland, The Lighthouse Lab in Alderley Park and Alex Alderton, Roberto Amato, Jeffrey Barrett, Sonia Goncalves, Ewan Harrison, David K. Jackson, Ian Johnston, Dominic Kwiatkowski, Cordelia Langford, John Sillitoe on behalf of the Wellcome Sanger Institute COVID-19 Surveillance Team |
| EPI_ISL_3106153 | Lighthouse Lab in Milton Keynes | Wellcome Sanger Institute for the COVID-19 Genomics UK (COG-UK) Consortium | The Lighthouse Lab in Milton Keynes and Alex Alderton, Roberto Amato, Jeffrey Barrett, Sonia Goncalves, Ewan Harrison, David K. Jackson, Ian Johnston, Dominic Kwiatkowski, Cordelia Langford, John Sillitoe on behalf of the Wellcome Sanger Institute COVID-19 Surveillance Team |
| EPI_ISL_3106154, EPI_ISL_3106155, EPI_ISL_3106156 | Lighthouse Lab in Glasgow | Wellcome Sanger Institute for the COVID-19 Genomics UK (COG-UK) Consortium | Harper VanSteenhouse, Yumi Kasai, David Gray, Carol Clugston, Anna Dominiczak and Alex Alderton, Roberto Amato, Jeffrey Barrett, Sonia Goncalves, Ewan Harrison, David K. Jackson, Ian Johnston, Dominic Kwiatkowski, Cordelia Langford, John Sillitoe on behalf of the Wellcome Sanger Institute COVID-19 Surveillance Team |
| EPI_ISL_3106157 | Lighthouse Lab in Alderley Park | Wellcome Sanger Institute for the COVID-19 Genomics UK (COG-UK) Consortium | Jacquelyn Wynn, Mairead Hyland, The Lighthouse Lab in Alderley Park and Alex Alderton, Roberto Amato, Jeffrey Barrett, Sonia Goncalves, Ewan Harrison, David K. Jackson, Ian Johnston, Dominic Kwiatkowski, Cordelia Langford, John Sillitoe on behalf of the Wellcome Sanger Institute COVID-19 Surveillance Team |
| EPI_ISL_3106158, EPI_ISL_3106159 | Lighthouse Lab in Milton Keynes | Wellcome Sanger Institute for the COVID-19 Genomics UK (COG-UK) Consortium | The Lighthouse Lab in Milton Keynes and Alex Alderton, Roberto Amato, Jeffrey Barrett, Sonia Goncalves, Ewan Harrison, David K. Jackson, Ian Johnston, Dominic Kwiatkowski, Cordelia Langford, John Sillitoe on behalf of the Wellcome Sanger Institute COVID-19 Surveillance Team |
| EPI_ISL_3106160 | Lighthouse Lab in Alderley Park | Wellcome Sanger Institute for the COVID-19 Genomics UK (COG-UK) Consortium | Jacquelyn Wynn, Mairead Hyland, The Lighthouse Lab in Alderley Park and Alex Alderton, Roberto Amato, Jeffrey Barrett, Sonia Goncalves, Ewan Harrison, David K. Jackson, Ian Johnston, Dominic Kwiatkowski, Cordelia Langford, John Sillitoe on behalf of the Wellcome Sanger Institute COVID-19 Surveillance Team |
| EPI_ISL_3106161 | Lighthouse Lab in Milton Keynes | Wellcome Sanger Institute for the COVID-19 Genomics UK (COG-UK) Consortium | The Lighthouse Lab in Milton Keynes and Alex Alderton, Roberto Amato, Jeffrey Barrett, Sonia Goncalves, Ewan Harrison, David K. Jackson, Ian Johnston, Dominic Kwiatkowski, Cordelia Langford, John Sillitoe on behalf of the Wellcome Sanger Institute COVID-19 Surveillance Team |
| EPI_ISL_3106162 | Lighthouse Lab in Alderley Park | Wellcome Sanger Institute for the COVID-19 Genomics UK (COG-UK) Consortium | Jacquelyn Wynn, Mairead Hyland, The Lighthouse Lab in Alderley Park and Alex Alderton, Roberto Amato, Jeffrey Barrett, Sonia Goncalves, Ewan Harrison, David K. Jackson, Ian Johnston, Dominic Kwiatkowski, Cordelia Langford, John Sillitoe on behalf of the Wellcome Sanger Institute COVID-19 Surveillance Team |
| EPI_ISL_3106163, EPI_ISL_3106164, EPI_ISL_3106165, EPI_ISL_3106166, EPI_ISL_3106167 | Lighthouse Lab in Milton Keynes | Wellcome Sanger Institute for the COVID-19 Genomics UK (COG-UK) Consortium | The Lighthouse Lab in Milton Keynes and Alex Alderton, Roberto Amato, Jeffrey Barrett, Sonia Goncalves, Ewan Harrison, David K. Jackson, Ian Johnston, Dominic Kwiatkowski, Cordelia Langford, John Sillitoe on behalf of the Wellcome Sanger Institute COVID-19 Surveillance Team |
| EPI_ISL_3106168 | Lighthouse Lab in Glasgow | Wellcome Sanger Institute for the COVID-19 Genomics UK (COG-UK) Consortium | Harper VanSteenhouse, Yumi Kasai, David Gray, Carol Clugston, Anna Dominiczak and Alex Alderton, Roberto Amato, Jeffrey Barrett, Sonia Goncalves, Ewan Harrison, David K. Jackson, Ian Johnston, Dominic Kwiatkowski, Cordelia Langford, John Sillitoe on behalf of the Wellcome Sanger Institute COVID-19 Surveillance Team |
| EPI_ISL_3106169 | Lighthouse Lab in Alderley Park | Wellcome Sanger Institute for the COVID-19 Genomics UK (COG-UK) Consortium | Jacquelyn Wynn, Mairead Hyland, The Lighthouse Lab in Alderley Park and Alex Alderton, Roberto Amato, Jeffrey Barrett, Sonia Goncalves, Ewan Harrison, David K. Jackson, Ian Johnston, Dominic Kwiatkowski, Cordelia Langford, John Sillitoe on behalf of the Wellcome Sanger Institute COVID-19 Surveillance Team |
| EPI_ISL_3106170, EPI_ISL_3106171, EPI_ISL_3106172, EPI_ISL_3106173, EPI_ISL_3106174 | Lighthouse Lab in Milton Keynes | Wellcome Sanger Institute for the COVID-19 Genomics UK (COG-UK) Consortium | The Lighthouse Lab in Milton Keynes and Alex Alderton, Roberto Amato, Jeffrey Barrett, Sonia Goncalves, Ewan Harrison, David K. Jackson, Ian Johnston, Dominic Kwiatkowski, Cordelia Langford, John Sillitoe on behalf of the Wellcome Sanger Institute COVID-19 Surveillance Team |
| EPI_ISL_3106175 | Lighthouse Lab in Glasgow | Wellcome Sanger Institute for the COVID-19 Genomics UK (COG-UK) Consortium | Harper VanSteenhouse, Yumi Kasai, David Gray, Carol Clugston, Anna Dominiczak and Alex Alderton, Roberto Amato, Jeffrey Barrett, Sonia Goncalves, Ewan Harrison, David K. Jackson, Ian Johnston, Dominic Kwiatkowski, Cordelia Langford, John Sillitoe on behalf of the Wellcome Sanger Institute COVID-19 Surveillance Team |
| EPI_ISL_3106176, EPI_ISL_3106177 | Lighthouse Lab in Milton Keynes | Wellcome Sanger Institute for the COVID-19 Genomics UK (COG-UK) Consortium | The Lighthouse Lab in Milton Keynes and Alex Alderton, Roberto Amato, Jeffrey Barrett, Sonia Goncalves, Ewan Harrison, David K. Jackson, Ian Johnston, Dominic Kwiatkowski, Cordelia Langford, John Sillitoe on behalf of the Wellcome Sanger Institute COVID-19 Surveillance Team |
| EPI_ISL_3106178 | Lighthouse Lab in Alderley Park | Wellcome Sanger Institute for the COVID-19 Genomics UK (COG-UK) Consortium | Jacquelyn Wynn, Mairead Hyland, The Lighthouse Lab in Alderley Park and Alex Alderton, Roberto Amato, Jeffrey Barrett, Sonia Goncalves, Ewan Harrison, David K. Jackson, Ian Johnston, Dominic Kwiatkowski, Cordelia Langford, John Sillitoe on behalf of the Wellcome Sanger Institute COVID-19 Surveillance Team |
| EPI_ISL_3106179, EPI_ISL_3106180 | Lighthouse Lab in Milton Keynes | Wellcome Sanger Institute for the COVID-19 Genomics UK (COG-UK) Consortium | The Lighthouse Lab in Milton Keynes and Alex Alderton, Roberto Amato, Jeffrey Barrett, Sonia Goncalves, Ewan Harrison, David K. Jackson, Ian Johnston, Dominic Kwiatkowski, Cordelia Langford, John Sillitoe on behalf of the Wellcome Sanger Institute COVID-19 Surveillance Team |
| EPI_ISL_3106181 | Lighthouse Lab in Glasgow | Wellcome Sanger Institute for the COVID-19 Genomics UK (COG-UK) Consortium | Harper VanSteenhouse, Yumi Kasai, David Gray, Carol Clugston, Anna Dominiczak and Alex Alderton, Roberto Amato, Jeffrey Barrett, Sonia Goncalves, Ewan Harrison, David K. Jackson, Ian Johnston, Dominic Kwiatkowski, Cordelia Langford, John Sillitoe on behalf of the Wellcome Sanger Institute COVID-19 Surveillance Team |
| EPI_ISL_3106182 | Lighthouse Lab in Alderley Park | Wellcome Sanger Institute for the COVID-19 Genomics UK (COG-UK) Consortium | Jacquelyn Wynn, Mairead Hyland, The Lighthouse Lab in Alderley Park and Alex Alderton, Roberto Amato, Jeffrey Barrett, Sonia Goncalves, Ewan Harrison, David K. Jackson, Ian Johnston, Dominic Kwiatkowski, Cordelia Langford, John Sillitoe on behalf of the Wellcome Sanger Institute COVID-19 Surveillance Team |
| EPI_ISL_3106183, EPI_ISL_3106184, EPI_ISL_3106185, EPI_ISL_3106186, EPI_ISL_3106187 | Lighthouse Lab in Glasgow | Wellcome Sanger Institute for the COVID-19 Genomics UK (COG-UK) Consortium | Harper VanSteenhouse, Yumi Kasai, David Gray, Carol Clugston, Anna Dominiczak and Alex Alderton, Roberto Amato, Jeffrey Barrett, Sonia Goncalves, Ewan Harrison, David K. Jackson, Ian Johnston, Dominic Kwiatkowski, Cordelia Langford, John Sillitoe on behalf of the Wellcome Sanger Institute COVID-19 Surveillance Team |
| EPI_ISL_3106188 | Lighthouse Lab in Alderley Park | Wellcome Sanger Institute for the COVID-19 Genomics UK (COG-UK) Consortium | Jacquelyn Wynn, Mairead Hyland, The Lighthouse Lab in Alderley Park and Alex Alderton, Roberto Amato, Jeffrey Barrett, Sonia Goncalves, Ewan Harrison, David K. Jackson, Ian Johnston, Dominic Kwiatkowski, Cordelia Langford, John Sillitoe on behalf of the Wellcome Sanger Institute COVID-19 Surveillance Team |
| EPI_ISL_3106189 | Lighthouse Lab in Glasgow | Wellcome Sanger Institute for the COVID-19 Genomics UK (COG-UK) Consortium | Harper VanSteenhouse, Yumi Kasai, David Gray, Carol Clugston, Anna Dominiczak and Alex Alderton, Roberto Amato, Jeffrey Barrett, Sonia Goncalves, Ewan Harrison, David K. Jackson, Ian Johnston, Dominic Kwiatkowski, Cordelia Langford, John Sillitoe on behalf of the Wellcome Sanger Institute COVID-19 Surveillance Team |
| EPI_ISL_3106190 | Lighthouse Lab in Alderley Park | Wellcome Sanger Institute for the COVID-19 Genomics UK (COG-UK) Consortium | Jacquelyn Wynn, Mairead Hyland, The Lighthouse Lab in Alderley Park and Alex Alderton, Roberto Amato, Jeffrey Barrett, Sonia Goncalves, Ewan Harrison, David K. Jackson, Ian Johnston, Dominic Kwiatkowski, Cordelia Langford, John Sillitoe on behalf of the Wellcome Sanger Institute COVID-19 Surveillance Team |
| EPI_ISL_3106191, EPI_ISL_3106192 | Lighthouse Lab in Glasgow | Wellcome Sanger Institute for the COVID-19 Genomics UK (COG-UK) Consortium | Harper VanSteenhouse, Yumi Kasai, David Gray, Carol Clugston, Anna Dominiczak and Alex Alderton, Roberto Amato, Jeffrey Barrett, Sonia Goncalves, Ewan Harrison, David K. Jackson, Ian Johnston, Dominic Kwiatkowski, Cordelia Langford, John Sillitoe on behalf of the Wellcome Sanger Institute COVID-19 Surveillance Team |
| EPI_ISL_3106193, EPI_ISL_3106194 | Lighthouse Lab in Alderley Park | Wellcome Sanger Institute for the COVID-19 Genomics UK (COG-UK) Consortium | Jacquelyn Wynn, Mairead Hyland, The Lighthouse Lab in Alderley Park and Alex Alderton, Roberto Amato, Jeffrey Barrett, Sonia Goncalves, Ewan Harrison, David K. Jackson, Ian Johnston, Dominic Kwiatkowski, Cordelia Langford, John Sillitoe on behalf of the Wellcome Sanger Institute COVID-19 Surveillance Team |
| EPI_ISL_3106195, EPI_ISL_3106196, EPI_ISL_3106197, EPI_ISL_3106198, EPI_ISL_3106199, EPI_ISL_3106200 | Lighthouse Lab in Glasgow | Wellcome Sanger Institute for the COVID-19 Genomics UK (COG-UK) Consortium | Harper VanSteenhouse, Yumi Kasai, David Gray, Carol Clugston, Anna Dominiczak and Alex Alderton, Roberto Amato, Jeffrey Barrett, Sonia Goncalves, Ewan Harrison, David K. Jackson, Ian Johnston, Dominic Kwiatkowski, Cordelia Langford, John Sillitoe on behalf of the Wellcome Sanger Institute COVID-19 Surveillance Team |
| EPI_ISL_3106201 | Lighthouse Lab in Alderley Park | Wellcome Sanger Institute for the COVID-19 Genomics UK (COG-UK) Consortium | Jacquelyn Wynn, Mairead Hyland, The Lighthouse Lab in Alderley Park and Alex Alderton, Roberto Amato, Jeffrey Barrett, Sonia Goncalves, Ewan Harrison, David K. Jackson, Ian Johnston, Dominic Kwiatkowski, Cordelia Langford, John Sillitoe on behalf of the Wellcome Sanger Institute COVID-19 Surveillance Team |
| EPI_ISL_3106202, EPI_ISL_3106203 | Lighthouse Lab in Glasgow | Wellcome Sanger Institute for the COVID-19 Genomics UK (COG-UK) Consortium | Harper VanSteenhouse, Yumi Kasai, David Gray, Carol Clugston, Anna Dominiczak and Alex Alderton, Roberto Amato, Jeffrey Barrett, Sonia Goncalves, Ewan Harrison, David K. Jackson, Ian Johnston, Dominic Kwiatkowski, Cordelia Langford, John Sillitoe on behalf of the Wellcome Sanger Institute COVID-19 Surveillance Team |
| EPI_ISL_3106204 | Lighthouse Lab in Alderley Park | Wellcome Sanger Institute for the COVID-19 Genomics UK (COG-UK) Consortium | Jacquelyn Wynn, Mairead Hyland, The Lighthouse Lab in Alderley Park and Alex Alderton, Roberto Amato, Jeffrey Barrett, Sonia Goncalves, Ewan Harrison, David K. Jackson, Ian Johnston, Dominic Kwiatkowski, Cordelia Langford, John Sillitoe on behalf of the Wellcome Sanger Institute COVID-19 Surveillance Team |
| EPI_ISL_3106205, EPI_ISL_3106206 | Lighthouse Lab in Glasgow | Wellcome Sanger Institute for the COVID-19 Genomics UK (COG-UK) Consortium | Harper VanSteenhouse, Yumi Kasai, David Gray, Carol Clugston, Anna Dominiczak and Alex Alderton, Roberto Amato, Jeffrey Barrett, Sonia Goncalves, Ewan Harrison, David K. Jackson, Ian Johnston, Dominic Kwiatkowski, Cordelia Langford, John Sillitoe on behalf of the Wellcome Sanger Institute COVID-19 Surveillance Team |
| EPI_ISL_3106207 | Lighthouse Lab in Alderley Park | Wellcome Sanger Institute for the COVID-19 Genomics UK (COG-UK) Consortium | Jacquelyn Wynn, Mairead Hyland, The Lighthouse Lab in Alderley Park and Alex Alderton, Roberto Amato, Jeffrey Barrett, Sonia Goncalves, Ewan Harrison, David K. Jackson, Ian Johnston, Dominic Kwiatkowski, Cordelia Langford, John Sillitoe on behalf of the Wellcome Sanger Institute COVID-19 Surveillance Team |
| EPI_ISL_3106208, | Lighthouse Lab in Glasgow | Wellcome Sanger Institute for the COVID-19 Genomics UK (COG-UK) | Harper VanSteenhouse, Yumi Kasai, David Gray, Carol Clugston, Anna Dominiczak and Alex Alderton, Roberto Amato, Jeffrey Barrett, Sonia Goncalves, Ewan Harrison, David K. Jackson, Ian Johnston, Dominic Kwiatkowski, Cordelia Langford, |

| | | | |
|---|---|---|---|
| EPI_ISL_3106209, EPI_ISL_3106210, EPI_ISL_3106211, EPI_ISL_3106212, EPI_ISL_3106213, EPI_ISL_3106214, EPI_ISL_3106215 | | Consortium | John Sillitoe on behalf of the Wellcome Sanger Institute COVID-19 Surveillance Team |
| EPI_ISL_3106216 | Lighthouse Lab in Alderley Park | Wellcome Sanger Institute for the COVID-19 Genomics UK (COG-UK) Consortium | Jacquelyn Wynn, Mairead Hyland, The Lighthouse Lab in Alderley Park and Alex Alderton, Roberto Amato, Jeffrey Barrett, Sonia Goncalves, Ewan Harrison, David K. Jackson, Ian Johnston, Dominic Kwiatkowski, Cordelia Langford, John Sillitoe on behalf of the Wellcome Sanger Institute COVID-19 Surveillance Team |
| EPI_ISL_3106217 | Lighthouse Lab in Glasgow | Wellcome Sanger Institute for the COVID-19 Genomics UK (COG-UK) Consortium | Harper VanSteenhouse, Yumi Kasai, David Gray, Carol Clugston, Anna Dominiczak and Alex Alderton, Roberto Amato, Jeffrey Barrett, Sonia Goncalves, Ewan Harrison, David K. Jackson, Ian Johnston, Dominic Kwiatkowski, Cordelia Langford, John Sillitoe on behalf of the Wellcome Sanger Institute COVID-19 Surveillance Team |
| EPI_ISL_3106218 | Lighthouse Lab in Alderley Park | Wellcome Sanger Institute for the COVID-19 Genomics UK (COG-UK) Consortium | Jacquelyn Wynn, Mairead Hyland, The Lighthouse Lab in Alderley Park and Alex Alderton, Roberto Amato, Jeffrey Barrett, Sonia Goncalves, Ewan Harrison, David K. Jackson, Ian Johnston, Dominic Kwiatkowski, Cordelia Langford, John Sillitoe on behalf of the Wellcome Sanger Institute COVID-19 Surveillance Team |
| EPI_ISL_3106219, EPI_ISL_3106220, EPI_ISL_3106221, EPI_ISL_3106222, EPI_ISL_3106223, EPI_ISL_3106224, EPI_ISL_3106225, EPI_ISL_3106226, EPI_ISL_3106227, EPI_ISL_3106228, EPI_ISL_3106229, EPI_ISL_3106230, EPI_ISL_3106231 | | | |
| see above | Lighthouse Lab in Glasgow | Wellcome Sanger Institute for the COVID-19 Genomics UK (COG-UK) Consortium | Harper VanSteenhouse, Yumi Kasai, David Gray, Carol Clugston, Anna Dominiczak and Alex Alderton, Roberto Amato, Jeffrey Barrett, Sonia Goncalves, Ewan Harrison, David K. Jackson, Ian Johnston, Dominic Kwiatkowski, Cordelia Langford, John Sillitoe on behalf of the Wellcome Sanger Institute COVID-19 Surveillance Team |
| EPI_ISL_3106232 | Lighthouse Lab in Alderley Park | Wellcome Sanger Institute for the COVID-19 Genomics UK (COG-UK) Consortium | Jacquelyn Wynn, Mairead Hyland, The Lighthouse Lab in Alderley Park and Alex Alderton, Roberto Amato, Jeffrey Barrett, Sonia Goncalves, Ewan Harrison, David K. Jackson, Ian Johnston, Dominic Kwiatkowski, Cordelia Langford, John Sillitoe on behalf of the Wellcome Sanger Institute COVID-19 Surveillance Team |
| EPI_ISL_3106233, EPI_ISL_3106234 | Lighthouse Lab in Glasgow | Wellcome Sanger Institute for the COVID-19 Genomics UK (COG-UK) Consortium | Harper VanSteenhouse, Yumi Kasai, David Gray, Carol Clugston, Anna Dominiczak and Alex Alderton, Roberto Amato, Jeffrey Barrett, Sonia Goncalves, Ewan Harrison, David K. Jackson, Ian Johnston, Dominic Kwiatkowski, Cordelia Langford, John Sillitoe on behalf of the Wellcome Sanger Institute COVID-19 Surveillance Team |
| EPI_ISL_3106235, EPI_ISL_3106236 | Lighthouse Lab in Alderley Park | Wellcome Sanger Institute for the COVID-19 Genomics UK (COG-UK) Consortium | Jacquelyn Wynn, Mairead Hyland, The Lighthouse Lab in Alderley Park and Alex Alderton, Roberto Amato, Jeffrey Barrett, Sonia Goncalves, Ewan Harrison, David K. Jackson, Ian Johnston, Dominic Kwiatkowski, Cordelia Langford, John Sillitoe on behalf of the Wellcome Sanger Institute COVID-19 Surveillance Team |
| EPI_ISL_3106237 | Lighthouse Lab in Glasgow | Wellcome Sanger Institute for the COVID-19 Genomics UK (COG-UK) Consortium | Harper VanSteenhouse, Yumi Kasai, David Gray, Carol Clugston, Anna Dominiczak and Alex Alderton, Roberto Amato, Jeffrey Barrett, Sonia Goncalves, Ewan Harrison, David K. Jackson, Ian Johnston, Dominic Kwiatkowski, Cordelia Langford, John Sillitoe on behalf of the Wellcome Sanger Institute COVID-19 Surveillance Team |
| EPI_ISL_3106238, EPI_ISL_3106239 | Lighthouse Lab in Alderley Park | Wellcome Sanger Institute for the COVID-19 Genomics UK (COG-UK) Consortium | Jacquelyn Wynn, Mairead Hyland, The Lighthouse Lab in Alderley Park and Alex Alderton, Roberto Amato, Jeffrey Barrett, Sonia Goncalves, Ewan Harrison, David K. Jackson, Ian Johnston, Dominic Kwiatkowski, Cordelia Langford, John Sillitoe on behalf of the Wellcome Sanger Institute COVID-19 Surveillance Team |
| EPI_ISL_3106240, EPI_ISL_3106241 | Lighthouse Lab in Glasgow | Wellcome Sanger Institute for the COVID-19 Genomics UK (COG-UK) Consortium | Harper VanSteenhouse, Yumi Kasai, David Gray, Carol Clugston, Anna Dominiczak and Alex Alderton, Roberto Amato, Jeffrey Barrett, Sonia Goncalves, Ewan Harrison, David K. Jackson, Ian Johnston, Dominic Kwiatkowski, Cordelia Langford, John Sillitoe on behalf of the Wellcome Sanger Institute COVID-19 Surveillance Team |
| EPI_ISL_3106242 | Lighthouse Lab in Alderley Park | Wellcome Sanger Institute for the COVID-19 Genomics UK (COG-UK) Consortium | Jacquelyn Wynn, Mairead Hyland, The Lighthouse Lab in Alderley Park and Alex Alderton, Roberto Amato, Jeffrey Barrett, Sonia Goncalves, Ewan Harrison, David K. Jackson, Ian Johnston, Dominic Kwiatkowski, Cordelia Langford, John Sillitoe on behalf of the Wellcome Sanger Institute COVID-19 Surveillance Team |
| EPI_ISL_3106243 | Lighthouse Lab in Glasgow | Wellcome Sanger Institute for the COVID-19 Genomics UK (COG-UK) Consortium | Harper VanSteenhouse, Yumi Kasai, David Gray, Carol Clugston, Anna Dominiczak and Alex Alderton, Roberto Amato, Jeffrey Barrett, Sonia Goncalves, Ewan Harrison, David K. Jackson, Ian Johnston, Dominic Kwiatkowski, Cordelia Langford, John Sillitoe on behalf of the Wellcome Sanger Institute COVID-19 Surveillance Team |
| EPI_ISL_3106244 | Lighthouse Lab in Alderley Park | Wellcome Sanger Institute for the COVID-19 Genomics UK (COG-UK) Consortium | Jacquelyn Wynn, Mairead Hyland, The Lighthouse Lab in Alderley Park and Alex Alderton, Roberto Amato, Jeffrey Barrett, Sonia Goncalves, Ewan Harrison, David K. Jackson, Ian Johnston, Dominic Kwiatkowski, Cordelia Langford, John Sillitoe on behalf of the Wellcome Sanger Institute COVID-19 Surveillance Team |
| EPI_ISL_3106245 | Lighthouse Lab in Glasgow | Wellcome Sanger Institute for the COVID-19 Genomics UK (COG-UK) Consortium | Harper VanSteenhouse, Yumi Kasai, David Gray, Carol Clugston, Anna Dominiczak and Alex Alderton, Roberto Amato, Jeffrey Barrett, Sonia Goncalves, Ewan Harrison, David K. Jackson, Ian Johnston, Dominic Kwiatkowski, Cordelia Langford, John Sillitoe on behalf of the Wellcome Sanger Institute COVID-19 Surveillance Team |
| EPI_ISL_3106246 | Lighthouse Lab in Alderley Park | Wellcome Sanger Institute for the COVID-19 Genomics UK (COG-UK) Consortium | Jacquelyn Wynn, Mairead Hyland, The Lighthouse Lab in Alderley Park and Alex Alderton, Roberto Amato, Jeffrey Barrett, Sonia Goncalves, Ewan Harrison, David K. Jackson, Ian Johnston, Dominic Kwiatkowski, Cordelia Langford, John Sillitoe on behalf of the Wellcome Sanger Institute COVID-19 Surveillance Team |
| EPI_ISL_3106247 | Lighthouse Lab in Glasgow | Wellcome Sanger Institute for the COVID-19 Genomics UK (COG-UK) Consortium | Harper VanSteenhouse, Yumi Kasai, David Gray, Carol Clugston, Anna Dominiczak and Alex Alderton, Roberto Amato, Jeffrey Barrett, Sonia Goncalves, Ewan Harrison, David K. Jackson, Ian Johnston, Dominic Kwiatkowski, Cordelia Langford, John Sillitoe on behalf of the Wellcome Sanger Institute COVID-19 Surveillance Team |
| EPI_ISL_3106248 | Lighthouse Lab in Alderley Park | Wellcome Sanger Institute for the COVID-19 Genomics UK (COG-UK) Consortium | Jacquelyn Wynn, Mairead Hyland, The Lighthouse Lab in Alderley Park and Alex Alderton, Roberto Amato, Jeffrey Barrett, Sonia Goncalves, Ewan Harrison, David K. Jackson, Ian Johnston, Dominic Kwiatkowski, Cordelia Langford, John Sillitoe on behalf of the Wellcome Sanger Institute COVID-19 Surveillance Team |
| EPI_ISL_3106249, EPI_ISL_3106250, EPI_ISL_3106251, EPI_ISL_3106252, EPI_ISL_3106253, EPI_ISL_3106254, EPI_ISL_3106255, EPI_ISL_3106256, EPI_ISL_3106257, EPI_ISL_3106258, EPI_ISL_3106259, EPI_ISL_3106260 | | | |
| see above | Lighthouse Lab in Glasgow | Wellcome Sanger Institute for the COVID-19 Genomics UK (COG-UK) Consortium | Harper VanSteenhouse, Yumi Kasai, David Gray, Carol Clugston, Anna Dominiczak and Alex Alderton, Roberto Amato, Jeffrey Barrett, Sonia Goncalves, Ewan Harrison, David K. Jackson, Ian Johnston, Dominic Kwiatkowski, Cordelia Langford, John Sillitoe on behalf of the Wellcome Sanger Institute COVID-19 Surveillance Team |
| EPI_ISL_3106261 | Lighthouse Lab in Alderley Park | Wellcome Sanger Institute for the COVID-19 Genomics UK (COG-UK) Consortium | Jacquelyn Wynn, Mairead Hyland, The Lighthouse Lab in Alderley Park and Alex Alderton, Roberto Amato, Jeffrey Barrett, Sonia Goncalves, Ewan Harrison, David K. Jackson, Ian Johnston, Dominic Kwiatkowski, Cordelia Langford, John Sillitoe on behalf of the Wellcome Sanger Institute COVID-19 Surveillance Team |
| EPI_ISL_3106262, EPI_ISL_3106263 | Lighthouse Lab in Glasgow | Wellcome Sanger Institute for the COVID-19 Genomics UK (COG-UK) Consortium | Harper VanSteenhouse, Yumi Kasai, David Gray, Carol Clugston, Anna Dominiczak and Alex Alderton, Roberto Amato, Jeffrey Barrett, Sonia Goncalves, Ewan Harrison, David K. Jackson, Ian Johnston, Dominic Kwiatkowski, Cordelia Langford, John Sillitoe on behalf of the Wellcome Sanger Institute COVID-19 Surveillance Team |
| EPI_ISL_3106264 | Lighthouse Lab in Alderley Park | Wellcome Sanger Institute for the COVID-19 Genomics UK (COG-UK) Consortium | Jacquelyn Wynn, Mairead Hyland, The Lighthouse Lab in Alderley Park and Alex Alderton, Roberto Amato, Jeffrey Barrett, Sonia Goncalves, Ewan Harrison, David K. Jackson, Ian Johnston, Dominic Kwiatkowski, Cordelia Langford, John Sillitoe on behalf of the Wellcome Sanger Institute COVID-19 Surveillance Team |
| EPI_ISL_3106265, EPI_ISL_3106266, EPI_ISL_3106267 | Lighthouse Lab in Glasgow | Wellcome Sanger Institute for the COVID-19 Genomics UK (COG-UK) Consortium | Harper VanSteenhouse, Yumi Kasai, David Gray, Carol Clugston, Anna Dominiczak and Alex Alderton, Roberto Amato, Jeffrey Barrett, Sonia Goncalves, Ewan Harrison, David K. Jackson, Ian Johnston, Dominic Kwiatkowski, Cordelia Langford, John Sillitoe on behalf of the Wellcome Sanger Institute COVID-19 Surveillance Team |
| EPI_ISL_3106268 | Lighthouse Lab in Alderley Park | Wellcome Sanger Institute for the COVID-19 Genomics UK (COG-UK) Consortium | Jacquelyn Wynn, Mairead Hyland, The Lighthouse Lab in Alderley Park and Alex Alderton, Roberto Amato, Jeffrey Barrett, Sonia Goncalves, Ewan Harrison, David K. Jackson, Ian Johnston, Dominic Kwiatkowski, Cordelia Langford, John Sillitoe on behalf of the Wellcome Sanger Institute COVID-19 Surveillance Team |
| EPI_ISL_3106269, EPI_ISL_3106270, EPI_ISL_3106271 | Lighthouse Lab in Glasgow | Wellcome Sanger Institute for the COVID-19 Genomics UK (COG-UK) Consortium | Harper VanSteenhouse, Yumi Kasai, David Gray, Carol Clugston, Anna Dominiczak and Alex Alderton, Roberto Amato, Jeffrey Barrett, Sonia Goncalves, Ewan Harrison, David K. Jackson, Ian Johnston, Dominic Kwiatkowski, Cordelia Langford, John Sillitoe on behalf of the Wellcome Sanger Institute COVID-19 Surveillance Team |
| EPI_ISL_3106272 | Lighthouse Lab in Alderley Park | Wellcome Sanger Institute for the COVID-19 Genomics UK (COG-UK) Consortium | Jacquelyn Wynn, Mairead Hyland, The Lighthouse Lab in Alderley Park and Alex Alderton, Roberto Amato, Jeffrey Barrett, Sonia Goncalves, Ewan Harrison, David K. Jackson, Ian Johnston, Dominic Kwiatkowski, Cordelia Langford, John Sillitoe on behalf of the Wellcome Sanger Institute COVID-19 Surveillance Team |
| EPI_ISL_3106273, EPI_ISL_3106274 | Lighthouse Lab in Glasgow | Wellcome Sanger Institute for the COVID-19 Genomics UK (COG-UK) Consortium | Harper VanSteenhouse, Yumi Kasai, David Gray, Carol Clugston, Anna Dominiczak and Alex Alderton, Roberto Amato, Jeffrey Barrett, Sonia Goncalves, Ewan Harrison, David K. Jackson, Ian Johnston, Dominic Kwiatkowski, Cordelia Langford, John Sillitoe on behalf of the Wellcome Sanger Institute COVID-19 Surveillance Team |
| EPI_ISL_3106275 | Lighthouse Lab in Alderley Park | Wellcome Sanger Institute for the COVID-19 Genomics UK (COG-UK) Consortium | Jacquelyn Wynn, Mairead Hyland, The Lighthouse Lab in Alderley Park and Alex Alderton, Roberto Amato, Jeffrey Barrett, Sonia Goncalves, Ewan Harrison, David K. Jackson, Ian Johnston, Dominic Kwiatkowski, Cordelia Langford, John Sillitoe on behalf of the Wellcome Sanger Institute COVID-19 Surveillance Team |
| EPI_ISL_3106276, EPI_ISL_3106277, EPI_ISL_3106278 | Lighthouse Lab in Glasgow | Wellcome Sanger Institute for the COVID-19 Genomics UK (COG-UK) Consortium | Harper VanSteenhouse, Yumi Kasai, David Gray, Carol Clugston, Anna Dominiczak and Alex Alderton, Roberto Amato, Jeffrey Barrett, Sonia Goncalves, Ewan Harrison, David K. Jackson, Ian Johnston, Dominic Kwiatkowski, Cordelia Langford, John Sillitoe on behalf of the Wellcome Sanger Institute COVID-19 Surveillance Team |
| EPI_ISL_3106279, EPI_ISL_3106280, EPI_ISL_3106281 | Lighthouse Lab in Alderley Park | Wellcome Sanger Institute for the COVID-19 Genomics UK (COG-UK) Consortium | Jacquelyn Wynn, Mairead Hyland, The Lighthouse Lab in Alderley Park and Alex Alderton, Roberto Amato, Jeffrey Barrett, Sonia Goncalves, Ewan Harrison, David K. Jackson, Ian Johnston, Dominic Kwiatkowski, Cordelia Langford, John Sillitoe on behalf of the Wellcome Sanger Institute COVID-19 Surveillance Team |
| EPI_ISL_3106282, EPI_ISL_3106283, EPI_ISL_3106284, EPI_ISL_3106285, EPI_ISL_3106286, EPI_ISL_3106287, EPI_ISL_3106288, EPI_ISL_3106289, EPI_ISL_3106290, EPI_ISL_3106291 | Lighthouse Lab in Glasgow | Wellcome Sanger Institute for the COVID-19 Genomics UK (COG-UK) Consortium | Harper VanSteenhouse, Yumi Kasai, David Gray, Carol Clugston, Anna Dominiczak and Alex Alderton, Roberto Amato, Jeffrey Barrett, Sonia Goncalves, Ewan Harrison, David K. Jackson, Ian Johnston, Dominic Kwiatkowski, Cordelia Langford, John Sillitoe on behalf of the Wellcome Sanger Institute COVID-19 Surveillance Team |
| EPI_ISL_3106292, EPI_ISL_3106293 | Lighthouse Lab in Alderley Park | Wellcome Sanger Institute for the COVID-19 Genomics UK (COG-UK) Consortium | Jacquelyn Wynn, Mairead Hyland, The Lighthouse Lab in Alderley Park and Alex Alderton, Roberto Amato, Jeffrey Barrett, Sonia Goncalves, Ewan Harrison, David K. Jackson, Ian Johnston, Dominic Kwiatkowski, Cordelia Langford, John Sillitoe on behalf of the Wellcome Sanger Institute COVID-19 Surveillance Team |
| EPI_ISL_3106294 | Lighthouse Lab in Glasgow | Wellcome Sanger Institute for the COVID-19 Genomics UK (COG-UK) Consortium | Harper VanSteenhouse, Yumi Kasai, David Gray, Carol Clugston, Anna Dominiczak and Alex Alderton, Roberto Amato, Jeffrey Barrett, Sonia Goncalves, Ewan Harrison, David K. Jackson, Ian Johnston, Dominic Kwiatkowski, Cordelia Langford, John Sillitoe on behalf of the Wellcome Sanger Institute COVID-19 Surveillance Team |
| EPI_ISL_3106295 | Lighthouse Lab in Alderley Park | Wellcome Sanger Institute for the COVID-19 Genomics UK (COG-UK) Consortium | Jacquelyn Wynn, Mairead Hyland, The Lighthouse Lab in Alderley Park and Alex Alderton, Roberto Amato, Jeffrey Barrett, Sonia Goncalves, Ewan Harrison, David K. Jackson, Ian Johnston, Dominic Kwiatkowski, Cordelia Langford, John Sillitoe on behalf of the Wellcome Sanger Institute COVID-19 Surveillance Team |
| EPI_ISL_3106296, EPI_ISL_3106297, EPI_ISL_3106298, EPI_ISL_3106299, EPI_ISL_3106300, EPI_ISL_3106301, EPI_ISL_3106302 | Lighthouse Lab in Glasgow | Wellcome Sanger Institute for the COVID-19 Genomics UK (COG-UK) Consortium | Harper VanSteenhouse, Yumi Kasai, David Gray, Carol Clugston, Anna Dominiczak and Alex Alderton, Roberto Amato, Jeffrey Barrett, Sonia Goncalves, Ewan Harrison, David K. Jackson, Ian Johnston, Dominic Kwiatkowski, Cordelia Langford, John Sillitoe on behalf of the Wellcome Sanger Institute COVID-19 Surveillance Team |

| | | | |
|---|---|---|---|
| EPI_ISL_3106303, EPI_ISL_3106304 | Lighthouse Lab in Alderley Park | Wellcome Sanger Institute for the COVID-19 Genomics UK (COG-UK) Consortium | Jacquelyn Wynn, Mairead Hyland, The Lighthouse Lab in Alderley Park and Alex Alderton, Roberto Amato, Jeffrey Barrett, Sonia Goncalves, Ewan Harrison, David K. Jackson, Ian Johnston, Dominic Kwiatkowski, Cordelia Langford, John Sillitoe on behalf of the Wellcome Sanger Institute COVID-19 Surveillance Team |
| EPI_ISL_3106305, EPI_ISL_3106306, EPI_ISL_3106307, EPI_ISL_3106308, EPI_ISL_3106309, EPI_ISL_3106310, EPI_ISL_3106311, EPI_ISL_3106312, EPI_ISL_3106313 | Lighthouse Lab in Glasgow | Wellcome Sanger Institute for the COVID-19 Genomics UK (COG-UK) Consortium | Harper VanSteenhouse, Yumi Kasai, David Gray, Carol Clugston, Anna Dominiczak and Alex Alderton, Roberto Amato, Jeffrey Barrett, Sonia Goncalves, Ewan Harrison, David K. Jackson, Ian Johnston, Dominic Kwiatkowski, Cordelia Langford, John Sillitoe on behalf of the Wellcome Sanger Institute COVID-19 Surveillance Team |
| EPI_ISL_3106314, EPI_ISL_3106315 | Lighthouse Lab in Alderley Park | Wellcome Sanger Institute for the COVID-19 Genomics UK (COG-UK) Consortium | Jacquelyn Wynn, Mairead Hyland, The Lighthouse Lab in Alderley Park and Alex Alderton, Roberto Amato, Jeffrey Barrett, Sonia Goncalves, Ewan Harrison, David K. Jackson, Ian Johnston, Dominic Kwiatkowski, Cordelia Langford, John Sillitoe on behalf of the Wellcome Sanger Institute COVID-19 Surveillance Team |
| EPI_ISL_3106316, EPI_ISL_3106317, EPI_ISL_3106318, EPI_ISL_3106319, EPI_ISL_3106320, EPI_ISL_3106321, EPI_ISL_3106322, EPI_ISL_3106323 | Lighthouse Lab in Glasgow | Wellcome Sanger Institute for the COVID-19 Genomics UK (COG-UK) Consortium | Harper VanSteenhouse, Yumi Kasai, David Gray, Carol Clugston, Anna Dominiczak and Alex Alderton, Roberto Amato, Jeffrey Barrett, Sonia Goncalves, Ewan Harrison, David K. Jackson, Ian Johnston, Dominic Kwiatkowski, Cordelia Langford, John Sillitoe on behalf of the Wellcome Sanger Institute COVID-19 Surveillance Team |
| EPI_ISL_3106324, EPI_ISL_3106325 | Lighthouse Lab in Alderley Park | Wellcome Sanger Institute for the COVID-19 Genomics UK (COG-UK) Consortium | Jacquelyn Wynn, Mairead Hyland, The Lighthouse Lab in Alderley Park and Alex Alderton, Roberto Amato, Jeffrey Barrett, Sonia Goncalves, Ewan Harrison, David K. Jackson, Ian Johnston, Dominic Kwiatkowski, Cordelia Langford, John Sillitoe on behalf of the Wellcome Sanger Institute COVID-19 Surveillance Team |
| EPI_ISL_3106326, EPI_ISL_3106327 | Lighthouse Lab in Glasgow | Wellcome Sanger Institute for the COVID-19 Genomics UK (COG-UK) Consortium | Harper VanSteenhouse, Yumi Kasai, David Gray, Carol Clugston, Anna Dominiczak and Alex Alderton, Roberto Amato, Jeffrey Barrett, Sonia Goncalves, Ewan Harrison, David K. Jackson, Ian Johnston, Dominic Kwiatkowski, Cordelia Langford, John Sillitoe on behalf of the Wellcome Sanger Institute COVID-19 Surveillance Team |
| EPI_ISL_3106328, EPI_ISL_3106329, EPI_ISL_3106330 | Lighthouse Lab in Alderley Park | Wellcome Sanger Institute for the COVID-19 Genomics UK (COG-UK) Consortium | Jacquelyn Wynn, Mairead Hyland, The Lighthouse Lab in Alderley Park and Alex Alderton, Roberto Amato, Jeffrey Barrett, Sonia Goncalves, Ewan Harrison, David K. Jackson, Ian Johnston, Dominic Kwiatkowski, Cordelia Langford, John Sillitoe on behalf of the Wellcome Sanger Institute COVID-19 Surveillance Team |
| EPI_ISL_3106331, EPI_ISL_3106332, EPI_ISL_3106333, EPI_ISL_3106334, EPI_ISL_3106335, EPI_ISL_3106336, EPI_ISL_3106337, EPI_ISL_3106338, EPI_ISL_3106339, EPI_ISL_3106340, EPI_ISL_3106341, EPI_ISL_3106342, EPI_ISL_3106343, EPI_ISL_3106344 | | | |
| see above | Lighthouse Lab in Glasgow | Wellcome Sanger Institute for the COVID-19 Genomics UK (COG-UK) Consortium | Harper VanSteenhouse, Yumi Kasai, David Gray, Carol Clugston, Anna Dominiczak and Alex Alderton, Roberto Amato, Jeffrey Barrett, Sonia Goncalves, Ewan Harrison, David K. Jackson, Ian Johnston, Dominic Kwiatkowski, Cordelia Langford, John Sillitoe on behalf of the Wellcome Sanger Institute COVID-19 Surveillance Team |
| EPI_ISL_3106345, EPI_ISL_3106346 | Lighthouse Lab in Alderley Park | Wellcome Sanger Institute for the COVID-19 Genomics UK (COG-UK) Consortium | Jacquelyn Wynn, Mairead Hyland, The Lighthouse Lab in Alderley Park and Alex Alderton, Roberto Amato, Jeffrey Barrett, Sonia Goncalves, Ewan Harrison, David K. Jackson, Ian Johnston, Dominic Kwiatkowski, Cordelia Langford, John Sillitoe on behalf of the Wellcome Sanger Institute COVID-19 Surveillance Team |
| EPI_ISL_3106347, EPI_ISL_3106348 | Lighthouse Lab in Glasgow | Wellcome Sanger Institute for the COVID-19 Genomics UK (COG-UK) Consortium | Harper VanSteenhouse, Yumi Kasai, David Gray, Carol Clugston, Anna Dominiczak and Alex Alderton, Roberto Amato, Jeffrey Barrett, Sonia Goncalves, Ewan Harrison, David K. Jackson, Ian Johnston, Dominic Kwiatkowski, Cordelia Langford, John Sillitoe on behalf of the Wellcome Sanger Institute COVID-19 Surveillance Team |
| EPI_ISL_3106349 | Lighthouse Lab in Alderley Park | Wellcome Sanger Institute for the COVID-19 Genomics UK (COG-UK) Consortium | Jacquelyn Wynn, Mairead Hyland, The Lighthouse Lab in Alderley Park and Alex Alderton, Roberto Amato, Jeffrey Barrett, Sonia Goncalves, Ewan Harrison, David K. Jackson, Ian Johnston, Dominic Kwiatkowski, Cordelia Langford, John Sillitoe on behalf of the Wellcome Sanger Institute COVID-19 Surveillance Team |
| EPI_ISL_3106350, EPI_ISL_3106351 | Lighthouse Lab in Glasgow | Wellcome Sanger Institute for the COVID-19 Genomics UK (COG-UK) Consortium | Harper VanSteenhouse, Yumi Kasai, David Gray, Carol Clugston, Anna Dominiczak and Alex Alderton, Roberto Amato, Jeffrey Barrett, Sonia Goncalves, Ewan Harrison, David K. Jackson, Ian Johnston, Dominic Kwiatkowski, Cordelia Langford, John Sillitoe on behalf of the Wellcome Sanger Institute COVID-19 Surveillance Team |
| EPI_ISL_3106352 | Lighthouse Lab in Alderley Park | Wellcome Sanger Institute for the COVID-19 Genomics UK (COG-UK) Consortium | Jacquelyn Wynn, Mairead Hyland, The Lighthouse Lab in Alderley Park and Alex Alderton, Roberto Amato, Jeffrey Barrett, Sonia Goncalves, Ewan Harrison, David K. Jackson, Ian Johnston, Dominic Kwiatkowski, Cordelia Langford, John Sillitoe on behalf of the Wellcome Sanger Institute COVID-19 Surveillance Team |
| EPI_ISL_3106353, EPI_ISL_3106354, EPI_ISL_3106355 | Lighthouse Lab in Glasgow | Wellcome Sanger Institute for the COVID-19 Genomics UK (COG-UK) Consortium | Harper VanSteenhouse, Yumi Kasai, David Gray, Carol Clugston, Anna Dominiczak and Alex Alderton, Roberto Amato, Jeffrey Barrett, Sonia Goncalves, Ewan Harrison, David K. Jackson, Ian Johnston, Dominic Kwiatkowski, Cordelia Langford, John Sillitoe on behalf of the Wellcome Sanger Institute COVID-19 Surveillance Team |
| EPI_ISL_3106356, EPI_ISL_3106357 | Lighthouse Lab in Alderley Park | Wellcome Sanger Institute for the COVID-19 Genomics UK (COG-UK) Consortium | Jacquelyn Wynn, Mairead Hyland, The Lighthouse Lab in Alderley Park and Alex Alderton, Roberto Amato, Jeffrey Barrett, Sonia Goncalves, Ewan Harrison, David K. Jackson, Ian Johnston, Dominic Kwiatkowski, Cordelia Langford, John Sillitoe on behalf of the Wellcome Sanger Institute COVID-19 Surveillance Team |
| EPI_ISL_3106358 | Lighthouse Lab in Glasgow | Wellcome Sanger Institute for the COVID-19 Genomics UK (COG-UK) Consortium | Harper VanSteenhouse, Yumi Kasai, David Gray, Carol Clugston, Anna Dominiczak and Alex Alderton, Roberto Amato, Jeffrey Barrett, Sonia Goncalves, Ewan Harrison, David K. Jackson, Ian Johnston, Dominic Kwiatkowski, Cordelia Langford, John Sillitoe on behalf of the Wellcome Sanger Institute COVID-19 Surveillance Team |
| EPI_ISL_3106359, EPI_ISL_3106360, EPI_ISL_3106361 | Lighthouse Lab in Alderley Park | Wellcome Sanger Institute for the COVID-19 Genomics UK (COG-UK) Consortium | Jacquelyn Wynn, Mairead Hyland, The Lighthouse Lab in Alderley Park and Alex Alderton, Roberto Amato, Jeffrey Barrett, Sonia Goncalves, Ewan Harrison, David K. Jackson, Ian Johnston, Dominic Kwiatkowski, Cordelia Langford, John Sillitoe on behalf of the Wellcome Sanger Institute COVID-19 Surveillance Team |
| EPI_ISL_3106362, EPI_ISL_3106363, EPI_ISL_3106364, EPI_ISL_3106365, EPI_ISL_3106366 | Lighthouse Lab in Glasgow | Wellcome Sanger Institute for the COVID-19 Genomics UK (COG-UK) Consortium | Harper VanSteenhouse, Yumi Kasai, David Gray, Carol Clugston, Anna Dominiczak and Alex Alderton, Roberto Amato, Jeffrey Barrett, Sonia Goncalves, Ewan Harrison, David K. Jackson, Ian Johnston, Dominic Kwiatkowski, Cordelia Langford, John Sillitoe on behalf of the Wellcome Sanger Institute COVID-19 Surveillance Team |
| EPI_ISL_3106367, EPI_ISL_3106368, EPI_ISL_3106369 | Lighthouse Lab in Alderley Park | Wellcome Sanger Institute for the COVID-19 Genomics UK (COG-UK) Consortium | Jacquelyn Wynn, Mairead Hyland, The Lighthouse Lab in Alderley Park and Alex Alderton, Roberto Amato, Jeffrey Barrett, Sonia Goncalves, Ewan Harrison, David K. Jackson, Ian Johnston, Dominic Kwiatkowski, Cordelia Langford, John Sillitoe on behalf of the Wellcome Sanger Institute COVID-19 Surveillance Team |
| EPI_ISL_3106370 | Lighthouse Lab in Glasgow | Wellcome Sanger Institute for the COVID-19 Genomics UK (COG-UK) Consortium | Harper VanSteenhouse, Yumi Kasai, David Gray, Carol Clugston, Anna Dominiczak and Alex Alderton, Roberto Amato, Jeffrey Barrett, Sonia Goncalves, Ewan Harrison, David K. Jackson, Ian Johnston, Dominic Kwiatkowski, Cordelia Langford, John Sillitoe on behalf of the Wellcome Sanger Institute COVID-19 Surveillance Team |
| EPI_ISL_3106371 | Lighthouse Lab in Alderley Park | Wellcome Sanger Institute for the COVID-19 Genomics UK (COG-UK) Consortium | Jacquelyn Wynn, Mairead Hyland, The Lighthouse Lab in Alderley Park and Alex Alderton, Roberto Amato, Jeffrey Barrett, Sonia Goncalves, Ewan Harrison, David K. Jackson, Ian Johnston, Dominic Kwiatkowski, Cordelia Langford, John Sillitoe on behalf of the Wellcome Sanger Institute COVID-19 Surveillance Team |
| EPI_ISL_3106372, EPI_ISL_3106373, EPI_ISL_3106374, EPI_ISL_3106375, EPI_ISL_3106376, EPI_ISL_3106377, EPI_ISL_3106378 | Lighthouse Lab in Glasgow | Wellcome Sanger Institute for the COVID-19 Genomics UK (COG-UK) Consortium | Harper VanSteenhouse, Yumi Kasai, David Gray, Carol Clugston, Anna Dominiczak and Alex Alderton, Roberto Amato, Jeffrey Barrett, Sonia Goncalves, Ewan Harrison, David K. Jackson, Ian Johnston, Dominic Kwiatkowski, Cordelia Langford, John Sillitoe on behalf of the Wellcome Sanger Institute COVID-19 Surveillance Team |
| EPI_ISL_3106379, EPI_ISL_3106380, EPI_ISL_3106381 | Lighthouse Lab in Alderley Park | Wellcome Sanger Institute for the COVID-19 Genomics UK (COG-UK) Consortium | Jacquelyn Wynn, Mairead Hyland, The Lighthouse Lab in Alderley Park and Alex Alderton, Roberto Amato, Jeffrey Barrett, Sonia Goncalves, Ewan Harrison, David K. Jackson, Ian Johnston, Dominic Kwiatkowski, Cordelia Langford, John Sillitoe on behalf of the Wellcome Sanger Institute COVID-19 Surveillance Team |
| EPI_ISL_3106382 | Lighthouse Lab in Glasgow | Wellcome Sanger Institute for the COVID-19 Genomics UK (COG-UK) Consortium | Harper VanSteenhouse, Yumi Kasai, David Gray, Carol Clugston, Anna Dominiczak and Alex Alderton, Roberto Amato, Jeffrey Barrett, Sonia Goncalves, Ewan Harrison, David K. Jackson, Ian Johnston, Dominic Kwiatkowski, Cordelia Langford, John Sillitoe on behalf of the Wellcome Sanger Institute COVID-19 Surveillance Team |
| EPI_ISL_3106383 | Lighthouse Lab in Alderley Park | Wellcome Sanger Institute for the COVID-19 Genomics UK (COG-UK) Consortium | Jacquelyn Wynn, Mairead Hyland, The Lighthouse Lab in Alderley Park and Alex Alderton, Roberto Amato, Jeffrey Barrett, Sonia Goncalves, Ewan Harrison, David K. Jackson, Ian Johnston, Dominic Kwiatkowski, Cordelia Langford, John Sillitoe on behalf of the Wellcome Sanger Institute COVID-19 Surveillance Team |
| EPI_ISL_3106384, EPI_ISL_3106385, EPI_ISL_3106386 | Lighthouse Lab in Glasgow | Wellcome Sanger Institute for the COVID-19 Genomics UK (COG-UK) Consortium | Harper VanSteenhouse, Yumi Kasai, David Gray, Carol Clugston, Anna Dominiczak and Alex Alderton, Roberto Amato, Jeffrey Barrett, Sonia Goncalves, Ewan Harrison, David K. Jackson, Ian Johnston, Dominic Kwiatkowski, Cordelia Langford, John Sillitoe on behalf of the Wellcome Sanger Institute COVID-19 Surveillance Team |
| EPI_ISL_3106387, EPI_ISL_3106388, EPI_ISL_3106389 | Lighthouse Lab in Alderley Park | Wellcome Sanger Institute for the COVID-19 Genomics UK (COG-UK) Consortium | Jacquelyn Wynn, Mairead Hyland, The Lighthouse Lab in Alderley Park and Alex Alderton, Roberto Amato, Jeffrey Barrett, Sonia Goncalves, Ewan Harrison, David K. Jackson, Ian Johnston, Dominic Kwiatkowski, Cordelia Langford, John Sillitoe on behalf of the Wellcome Sanger Institute COVID-19 Surveillance Team |
| EPI_ISL_3106390 | Lighthouse Lab in Glasgow | Wellcome Sanger Institute for the COVID-19 Genomics UK (COG-UK) Consortium | Harper VanSteenhouse, Yumi Kasai, David Gray, Carol Clugston, Anna Dominiczak and Alex Alderton, Roberto Amato, Jeffrey Barrett, Sonia Goncalves, Ewan Harrison, David K. Jackson, Ian Johnston, Dominic Kwiatkowski, Cordelia Langford, John Sillitoe on behalf of the Wellcome Sanger Institute COVID-19 Surveillance Team |
| EPI_ISL_3106391 | Lighthouse Lab in Alderley Park | Wellcome Sanger Institute for the COVID-19 Genomics UK (COG-UK) Consortium | Jacquelyn Wynn, Mairead Hyland, The Lighthouse Lab in Alderley Park and Alex Alderton, Roberto Amato, Jeffrey Barrett, Sonia Goncalves, Ewan Harrison, David K. Jackson, Ian Johnston, Dominic Kwiatkowski, Cordelia Langford, John Sillitoe on behalf of the Wellcome Sanger Institute COVID-19 Surveillance Team |
| EPI_ISL_3106392, EPI_ISL_3106393 | Lighthouse Lab in Glasgow | Wellcome Sanger Institute for the COVID-19 Genomics UK (COG-UK) Consortium | Harper VanSteenhouse, Yumi Kasai, David Gray, Carol Clugston, Anna Dominiczak and Alex Alderton, Roberto Amato, Jeffrey Barrett, Sonia Goncalves, Ewan Harrison, David K. Jackson, Ian Johnston, Dominic Kwiatkowski, Cordelia Langford, John Sillitoe on behalf of the Wellcome Sanger Institute COVID-19 Surveillance Team |
| EPI_ISL_3106394 | Lighthouse Lab in Alderley Park | Wellcome Sanger Institute for the COVID-19 Genomics UK (COG-UK) Consortium | Jacquelyn Wynn, Mairead Hyland, The Lighthouse Lab in Alderley Park and Alex Alderton, Roberto Amato, Jeffrey Barrett, Sonia Goncalves, Ewan Harrison, David K. Jackson, Ian Johnston, Dominic Kwiatkowski, Cordelia Langford, John Sillitoe on behalf of the Wellcome Sanger Institute COVID-19 Surveillance Team |
| EPI_ISL_3106395, EPI_ISL_3106396 | Lighthouse Lab in Glasgow | Wellcome Sanger Institute for the COVID-19 Genomics UK (COG-UK) Consortium | Harper VanSteenhouse, Yumi Kasai, David Gray, Carol Clugston, Anna Dominiczak and Alex Alderton, Roberto Amato, Jeffrey Barrett, Sonia Goncalves, Ewan Harrison, David K. Jackson, Ian Johnston, Dominic Kwiatkowski, Cordelia Langford, John Sillitoe on behalf of the Wellcome Sanger Institute COVID-19 Surveillance Team |
| EPI_ISL_3106397 | Lighthouse Lab in Alderley Park | Wellcome Sanger Institute for the COVID-19 Genomics UK (COG-UK) | Jacquelyn Wynn, Mairead Hyland, The Lighthouse Lab in Alderley Park and Alex Alderton, Roberto Amato, Jeffrey Barrett, Sonia Goncalves, Ewan Harrison, David K. Jackson, Ian Johnston, Dominic Kwiatkowski, Cordelia Langford, John Sillitoe on |

| | | | |
|---|---|---|---|
| | | Consortium | behalf of the Wellcome Sanger Institute COVID-19 Surveillance Team |
| EPI_ISL_3106398 | Lighthouse Lab in Glasgow | Wellcome Sanger Institute for the COVID-19 Genomics UK (COG-UK) Consortium | Harper VanSteenhouse, Yumi Kasai, David Gray, Carol Clugston, Anna Dominiczak and Alex Alderton, Roberto Amato, Jeffrey Barrett, Sonia Goncalves, Ewan Harrison, David K. Jackson, Ian Johnston, Dominic Kwiatkowski, Cordelia Langford, John Sillitoe on behalf of the Wellcome Sanger Institute COVID-19 Surveillance Team |
| EPI_ISL_3106399 | Lighthouse Lab in Alderley Park | Wellcome Sanger Institute for the COVID-19 Genomics UK (COG-UK) Consortium | Jacquelyn Wynn, Mairead Hyland, The Lighthouse Lab in Alderley Park and Alex Alderton, Roberto Amato, Jeffrey Barrett, Sonia Goncalves, Ewan Harrison, David K. Jackson, Ian Johnston, Dominic Kwiatkowski, Cordelia Langford, John Sillitoe on behalf of the Wellcome Sanger Institute COVID-19 Surveillance Team |
| EPI_ISL_3106400, EPI_ISL_3106401 | Lighthouse Lab in Glasgow | Wellcome Sanger Institute for the COVID-19 Genomics UK (COG-UK) Consortium | Harper VanSteenhouse, Yumi Kasai, David Gray, Carol Clugston, Anna Dominiczak and Alex Alderton, Roberto Amato, Jeffrey Barrett, Sonia Goncalves, Ewan Harrison, David K. Jackson, Ian Johnston, Dominic Kwiatkowski, Cordelia Langford, John Sillitoe on behalf of the Wellcome Sanger Institute COVID-19 Surveillance Team |
| EPI_ISL_3106402 | Lighthouse Lab in Alderley Park | Wellcome Sanger Institute for the COVID-19 Genomics UK (COG-UK) Consortium | Jacquelyn Wynn, Mairead Hyland, The Lighthouse Lab in Alderley Park and Alex Alderton, Roberto Amato, Jeffrey Barrett, Sonia Goncalves, Ewan Harrison, David K. Jackson, Ian Johnston, Dominic Kwiatkowski, Cordelia Langford, John Sillitoe on behalf of the Wellcome Sanger Institute COVID-19 Surveillance Team |
| EPI_ISL_3106403, EPI_ISL_3106404, EPI_ISL_3106405, EPI_ISL_3106406, EPI_ISL_3106407, EPI_ISL_3106408 | Lighthouse Lab in Glasgow | Wellcome Sanger Institute for the COVID-19 Genomics UK (COG-UK) Consortium | Harper VanSteenhouse, Yumi Kasai, David Gray, Carol Clugston, Anna Dominiczak and Alex Alderton, Roberto Amato, Jeffrey Barrett, Sonia Goncalves, Ewan Harrison, David K. Jackson, Ian Johnston, Dominic Kwiatkowski, Cordelia Langford, John Sillitoe on behalf of the Wellcome Sanger Institute COVID-19 Surveillance Team |
| EPI_ISL_3106409 | Lighthouse Lab in Alderley Park | Wellcome Sanger Institute for the COVID-19 Genomics UK (COG-UK) Consortium | Jacquelyn Wynn, Mairead Hyland, The Lighthouse Lab in Alderley Park and Alex Alderton, Roberto Amato, Jeffrey Barrett, Sonia Goncalves, Ewan Harrison, David K. Jackson, Ian Johnston, Dominic Kwiatkowski, Cordelia Langford, John Sillitoe on behalf of the Wellcome Sanger Institute COVID-19 Surveillance Team |
| EPI_ISL_3106410, EPI_ISL_3106411 | Lighthouse Lab in Glasgow | Wellcome Sanger Institute for the COVID-19 Genomics UK (COG-UK) Consortium | Harper VanSteenhouse, Yumi Kasai, David Gray, Carol Clugston, Anna Dominiczak and Alex Alderton, Roberto Amato, Jeffrey Barrett, Sonia Goncalves, Ewan Harrison, David K. Jackson, Ian Johnston, Dominic Kwiatkowski, Cordelia Langford, John Sillitoe on behalf of the Wellcome Sanger Institute COVID-19 Surveillance Team |
| EPI_ISL_3106412 | Lighthouse Lab in Alderley Park | Wellcome Sanger Institute for the COVID-19 Genomics UK (COG-UK) Consortium | Jacquelyn Wynn, Mairead Hyland, The Lighthouse Lab in Alderley Park and Alex Alderton, Roberto Amato, Jeffrey Barrett, Sonia Goncalves, Ewan Harrison, David K. Jackson, Ian Johnston, Dominic Kwiatkowski, Cordelia Langford, John Sillitoe on behalf of the Wellcome Sanger Institute COVID-19 Surveillance Team |
| EPI_ISL_3106413 | Lighthouse Lab in Glasgow | Wellcome Sanger Institute for the COVID-19 Genomics UK (COG-UK) Consortium | Harper VanSteenhouse, Yumi Kasai, David Gray, Carol Clugston, Anna Dominiczak and Alex Alderton, Roberto Amato, Jeffrey Barrett, Sonia Goncalves, Ewan Harrison, David K. Jackson, Ian Johnston, Dominic Kwiatkowski, Cordelia Langford, John Sillitoe on behalf of the Wellcome Sanger Institute COVID-19 Surveillance Team |
| EPI_ISL_3106414 | Lighthouse Lab in Alderley Park | Wellcome Sanger Institute for the COVID-19 Genomics UK (COG-UK) Consortium | Jacquelyn Wynn, Mairead Hyland, The Lighthouse Lab in Alderley Park and Alex Alderton, Roberto Amato, Jeffrey Barrett, Sonia Goncalves, Ewan Harrison, David K. Jackson, Ian Johnston, Dominic Kwiatkowski, Cordelia Langford, John Sillitoe on behalf of the Wellcome Sanger Institute COVID-19 Surveillance Team |
| EPI_ISL_3106415, EPI_ISL_3106416, EPI_ISL_3106417, EPI_ISL_3106418 | Lighthouse Lab in Glasgow | Wellcome Sanger Institute for the COVID-19 Genomics UK (COG-UK) Consortium | Harper VanSteenhouse, Yumi Kasai, David Gray, Carol Clugston, Anna Dominiczak and Alex Alderton, Roberto Amato, Jeffrey Barrett, Sonia Goncalves, Ewan Harrison, David K. Jackson, Ian Johnston, Dominic Kwiatkowski, Cordelia Langford, John Sillitoe on behalf of the Wellcome Sanger Institute COVID-19 Surveillance Team |
| EPI_ISL_3106419 | Lighthouse Lab in Alderley Park | Wellcome Sanger Institute for the COVID-19 Genomics UK (COG-UK) Consortium | Jacquelyn Wynn, Mairead Hyland, The Lighthouse Lab in Alderley Park and Alex Alderton, Roberto Amato, Jeffrey Barrett, Sonia Goncalves, Ewan Harrison, David K. Jackson, Ian Johnston, Dominic Kwiatkowski, Cordelia Langford, John Sillitoe on behalf of the Wellcome Sanger Institute COVID-19 Surveillance Team |
| EPI_ISL_3106420, EPI_ISL_3106421 | Lighthouse Lab in Glasgow | Wellcome Sanger Institute for the COVID-19 Genomics UK (COG-UK) Consortium | Harper VanSteenhouse, Yumi Kasai, David Gray, Carol Clugston, Anna Dominiczak and Alex Alderton, Roberto Amato, Jeffrey Barrett, Sonia Goncalves, Ewan Harrison, David K. Jackson, Ian Johnston, Dominic Kwiatkowski, Cordelia Langford, John Sillitoe on behalf of the Wellcome Sanger Institute COVID-19 Surveillance Team |
| EPI_ISL_3106422 | Lighthouse Lab in Alderley Park | Wellcome Sanger Institute for the COVID-19 Genomics UK (COG-UK) Consortium | Jacquelyn Wynn, Mairead Hyland, The Lighthouse Lab in Alderley Park and Alex Alderton, Roberto Amato, Jeffrey Barrett, Sonia Goncalves, Ewan Harrison, David K. Jackson, Ian Johnston, Dominic Kwiatkowski, Cordelia Langford, John Sillitoe on behalf of the Wellcome Sanger Institute COVID-19 Surveillance Team |
| EPI_ISL_3106423, EPI_ISL_3106424 | Lighthouse Lab in Glasgow | Wellcome Sanger Institute for the COVID-19 Genomics UK (COG-UK) Consortium | Harper VanSteenhouse, Yumi Kasai, David Gray, Carol Clugston, Anna Dominiczak and Alex Alderton, Roberto Amato, Jeffrey Barrett, Sonia Goncalves, Ewan Harrison, David K. Jackson, Ian Johnston, Dominic Kwiatkowski, Cordelia Langford, John Sillitoe on behalf of the Wellcome Sanger Institute COVID-19 Surveillance Team |
| EPI_ISL_3106425 | Lighthouse Lab in Alderley Park | Wellcome Sanger Institute for the COVID-19 Genomics UK (COG-UK) Consortium | Jacquelyn Wynn, Mairead Hyland, The Lighthouse Lab in Alderley Park and Alex Alderton, Roberto Amato, Jeffrey Barrett, Sonia Goncalves, Ewan Harrison, David K. Jackson, Ian Johnston, Dominic Kwiatkowski, Cordelia Langford, John Sillitoe on behalf of the Wellcome Sanger Institute COVID-19 Surveillance Team |
| EPI_ISL_3106426 | Lighthouse Lab in Glasgow | Wellcome Sanger Institute for the COVID-19 Genomics UK (COG-UK) Consortium | Harper VanSteenhouse, Yumi Kasai, David Gray, Carol Clugston, Anna Dominiczak and Alex Alderton, Roberto Amato, Jeffrey Barrett, Sonia Goncalves, Ewan Harrison, David K. Jackson, Ian Johnston, Dominic Kwiatkowski, Cordelia Langford, John Sillitoe on behalf of the Wellcome Sanger Institute COVID-19 Surveillance Team |
| EPI_ISL_3106427 | Lighthouse Lab in Alderley Park | Wellcome Sanger Institute for the COVID-19 Genomics UK (COG-UK) Consortium | Jacquelyn Wynn, Mairead Hyland, The Lighthouse Lab in Alderley Park and Alex Alderton, Roberto Amato, Jeffrey Barrett, Sonia Goncalves, Ewan Harrison, David K. Jackson, Ian Johnston, Dominic Kwiatkowski, Cordelia Langford, John Sillitoe on behalf of the Wellcome Sanger Institute COVID-19 Surveillance Team |
| EPI_ISL_3106428, EPI_ISL_3106429, EPI_ISL_3106430, EPI_ISL_3106431 | Lighthouse Lab in Glasgow | Wellcome Sanger Institute for the COVID-19 Genomics UK (COG-UK) Consortium | Harper VanSteenhouse, Yumi Kasai, David Gray, Carol Clugston, Anna Dominiczak and Alex Alderton, Roberto Amato, Jeffrey Barrett, Sonia Goncalves, Ewan Harrison, David K. Jackson, Ian Johnston, Dominic Kwiatkowski, Cordelia Langford, John Sillitoe on behalf of the Wellcome Sanger Institute COVID-19 Surveillance Team |
| EPI_ISL_3106432, EPI_ISL_3106433, EPI_ISL_3106434 | Lighthouse Lab in Alderley Park | Wellcome Sanger Institute for the COVID-19 Genomics UK (COG-UK) Consortium | Jacquelyn Wynn, Mairead Hyland, The Lighthouse Lab in Alderley Park and Alex Alderton, Roberto Amato, Jeffrey Barrett, Sonia Goncalves, Ewan Harrison, David K. Jackson, Ian Johnston, Dominic Kwiatkowski, Cordelia Langford, John Sillitoe on behalf of the Wellcome Sanger Institute COVID-19 Surveillance Team |
| EPI_ISL_3106435 | Lighthouse Lab in Glasgow | Wellcome Sanger Institute for the COVID-19 Genomics UK (COG-UK) Consortium | Harper VanSteenhouse, Yumi Kasai, David Gray, Carol Clugston, Anna Dominiczak and Alex Alderton, Roberto Amato, Jeffrey Barrett, Sonia Goncalves, Ewan Harrison, David K. Jackson, Ian Johnston, Dominic Kwiatkowski, Cordelia Langford, John Sillitoe on behalf of the Wellcome Sanger Institute COVID-19 Surveillance Team |
| EPI_ISL_3106436, EPI_ISL_3106437, EPI_ISL_3106438 | Lighthouse Lab in Alderley Park | Wellcome Sanger Institute for the COVID-19 Genomics UK (COG-UK) Consortium | Jacquelyn Wynn, Mairead Hyland, The Lighthouse Lab in Alderley Park and Alex Alderton, Roberto Amato, Jeffrey Barrett, Sonia Goncalves, Ewan Harrison, David K. Jackson, Ian Johnston, Dominic Kwiatkowski, Cordelia Langford, John Sillitoe on behalf of the Wellcome Sanger Institute COVID-19 Surveillance Team |
| EPI_ISL_3106439, EPI_ISL_3106440, EPI_ISL_3106441 | Lighthouse Lab in Glasgow | Wellcome Sanger Institute for the COVID-19 Genomics UK (COG-UK) Consortium | Harper VanSteenhouse, Yumi Kasai, David Gray, Carol Clugston, Anna Dominiczak and Alex Alderton, Roberto Amato, Jeffrey Barrett, Sonia Goncalves, Ewan Harrison, David K. Jackson, Ian Johnston, Dominic Kwiatkowski, Cordelia Langford, John Sillitoe on behalf of the Wellcome Sanger Institute COVID-19 Surveillance Team |
| EPI_ISL_3106442, EPI_ISL_3106443, EPI_ISL_3106444, EPI_ISL_3106445 | Lighthouse Lab in Alderley Park | Wellcome Sanger Institute for the COVID-19 Genomics UK (COG-UK) Consortium | Jacquelyn Wynn, Mairead Hyland, The Lighthouse Lab in Alderley Park and Alex Alderton, Roberto Amato, Jeffrey Barrett, Sonia Goncalves, Ewan Harrison, David K. Jackson, Ian Johnston, Dominic Kwiatkowski, Cordelia Langford, John Sillitoe on behalf of the Wellcome Sanger Institute COVID-19 Surveillance Team |
| EPI_ISL_3106446 | Lighthouse Lab in Glasgow | Wellcome Sanger Institute for the COVID-19 Genomics UK (COG-UK) Consortium | Harper VanSteenhouse, Yumi Kasai, David Gray, Carol Clugston, Anna Dominiczak and Alex Alderton, Roberto Amato, Jeffrey Barrett, Sonia Goncalves, Ewan Harrison, David K. Jackson, Ian Johnston, Dominic Kwiatkowski, Cordelia Langford, John Sillitoe on behalf of the Wellcome Sanger Institute COVID-19 Surveillance Team |
| EPI_ISL_3106447 | Lighthouse Lab in Alderley Park | Wellcome Sanger Institute for the COVID-19 Genomics UK (COG-UK) Consortium | Jacquelyn Wynn, Mairead Hyland, The Lighthouse Lab in Alderley Park and Alex Alderton, Roberto Amato, Jeffrey Barrett, Sonia Goncalves, Ewan Harrison, David K. Jackson, Ian Johnston, Dominic Kwiatkowski, Cordelia Langford, John Sillitoe on behalf of the Wellcome Sanger Institute COVID-19 Surveillance Team |
| EPI_ISL_3106448 | Lighthouse Lab in Glasgow | Wellcome Sanger Institute for the COVID-19 Genomics UK (COG-UK) Consortium | Harper VanSteenhouse, Yumi Kasai, David Gray, Carol Clugston, Anna Dominiczak and Alex Alderton, Roberto Amato, Jeffrey Barrett, Sonia Goncalves, Ewan Harrison, David K. Jackson, Ian Johnston, Dominic Kwiatkowski, Cordelia Langford, John Sillitoe on behalf of the Wellcome Sanger Institute COVID-19 Surveillance Team |
| EPI_ISL_3106449, EPI_ISL_3106450 | Lighthouse Lab in Alderley Park | Wellcome Sanger Institute for the COVID-19 Genomics UK (COG-UK) Consortium | Jacquelyn Wynn, Mairead Hyland, The Lighthouse Lab in Alderley Park and Alex Alderton, Roberto Amato, Jeffrey Barrett, Sonia Goncalves, Ewan Harrison, David K. Jackson, Ian Johnston, Dominic Kwiatkowski, Cordelia Langford, John Sillitoe on behalf of the Wellcome Sanger Institute COVID-19 Surveillance Team |
| EPI_ISL_3106451 | Lighthouse Lab in Glasgow | Wellcome Sanger Institute for the COVID-19 Genomics UK (COG-UK) Consortium | Harper VanSteenhouse, Yumi Kasai, David Gray, Carol Clugston, Anna Dominiczak and Alex Alderton, Roberto Amato, Jeffrey Barrett, Sonia Goncalves, Ewan Harrison, David K. Jackson, Ian Johnston, Dominic Kwiatkowski, Cordelia Langford, John Sillitoe on behalf of the Wellcome Sanger Institute COVID-19 Surveillance Team |
| EPI_ISL_3106452 | Lighthouse Lab in Alderley Park | Wellcome Sanger Institute for the COVID-19 Genomics UK (COG-UK) Consortium | Jacquelyn Wynn, Mairead Hyland, The Lighthouse Lab in Alderley Park and Alex Alderton, Roberto Amato, Jeffrey Barrett, Sonia Goncalves, Ewan Harrison, David K. Jackson, Ian Johnston, Dominic Kwiatkowski, Cordelia Langford, John Sillitoe on behalf of the Wellcome Sanger Institute COVID-19 Surveillance Team |
| EPI_ISL_3106453 | Lighthouse Lab in Glasgow | Wellcome Sanger Institute for the COVID-19 Genomics UK (COG-UK) Consortium | Harper VanSteenhouse, Yumi Kasai, David Gray, Carol Clugston, Anna Dominiczak and Alex Alderton, Roberto Amato, Jeffrey Barrett, Sonia Goncalves, Ewan Harrison, David K. Jackson, Ian Johnston, Dominic Kwiatkowski, Cordelia Langford, John Sillitoe on behalf of the Wellcome Sanger Institute COVID-19 Surveillance Team |
| EPI_ISL_3106454, EPI_ISL_3106455, EPI_ISL_3106456 | Lighthouse Lab in Alderley Park | Wellcome Sanger Institute for the COVID-19 Genomics UK (COG-UK) Consortium | Jacquelyn Wynn, Mairead Hyland, The Lighthouse Lab in Alderley Park and Alex Alderton, Roberto Amato, Jeffrey Barrett, Sonia Goncalves, Ewan Harrison, David K. Jackson, Ian Johnston, Dominic Kwiatkowski, Cordelia Langford, John Sillitoe on behalf of the Wellcome Sanger Institute COVID-19 Surveillance Team |
| EPI_ISL_3106457 | Lighthouse Lab in Glasgow | Wellcome Sanger Institute for the COVID-19 Genomics UK (COG-UK) Consortium | Harper VanSteenhouse, Yumi Kasai, David Gray, Carol Clugston, Anna Dominiczak and Alex Alderton, Roberto Amato, Jeffrey Barrett, Sonia Goncalves, Ewan Harrison, David K. Jackson, Ian Johnston, Dominic Kwiatkowski, Cordelia Langford, John Sillitoe on behalf of the Wellcome Sanger Institute COVID-19 Surveillance Team |
| EPI_ISL_3106458 | Lighthouse Lab in Alderley Park | Wellcome Sanger Institute for the COVID-19 Genomics UK (COG-UK) Consortium | Jacquelyn Wynn, Mairead Hyland, The Lighthouse Lab in Alderley Park and Alex Alderton, Roberto Amato, Jeffrey Barrett, Sonia Goncalves, Ewan Harrison, David K. Jackson, Ian Johnston, Dominic Kwiatkowski, Cordelia Langford, John Sillitoe on behalf of the Wellcome Sanger Institute COVID-19 Surveillance Team |
| EPI_ISL_3106459, EPI_ISL_3106460 | Lighthouse Lab in Glasgow | Wellcome Sanger Institute for the COVID-19 Genomics UK (COG-UK) Consortium | Harper VanSteenhouse, Yumi Kasai, David Gray, Carol Clugston, Anna Dominiczak and Alex Alderton, Roberto Amato, Jeffrey Barrett, Sonia Goncalves, Ewan Harrison, David K. Jackson, Ian Johnston, Dominic Kwiatkowski, Cordelia Langford, John Sillitoe on behalf of the Wellcome Sanger Institute COVID-19 Surveillance Team |
| EPI_ISL_3106461 | Lighthouse Lab in Milton Keynes | Wellcome Sanger Institute for the COVID-19 Genomics UK (COG-UK) Consortium | The Lighthouse Lab in Milton Keynes and Alex Alderton, Roberto Amato, Jeffrey Barrett, Sonia Goncalves, Ewan Harrison, David K. Jackson, Ian Johnston, Dominic Kwiatkowski, Cordelia Langford, John Sillitoe on behalf of the Wellcome Sanger Institute COVID-19 Surveillance Team |
| EPI_ISL_3106462, EPI_ISL_3106463 | Lighthouse Lab in Glasgow | Wellcome Sanger Institute for the COVID-19 Genomics UK (COG-UK) Consortium | Harper VanSteenhouse, Yumi Kasai, David Gray, Carol Clugston, Anna Dominiczak and Alex Alderton, Roberto Amato, Jeffrey Barrett, Sonia Goncalves, Ewan Harrison, David K. Jackson, Ian Johnston, Dominic Kwiatkowski, Cordelia Langford, John Sillitoe on behalf of the Wellcome Sanger Institute COVID-19 Surveillance Team |
| EPI_ISL_3106464, EPI_ISL_3106465 | Lighthouse Lab in Milton Keynes | Wellcome Sanger Institute for the COVID-19 Genomics UK (COG-UK) Consortium | The Lighthouse Lab in Milton Keynes and Alex Alderton, Roberto Amato, Jeffrey Barrett, Sonia Goncalves, Ewan Harrison, David K. Jackson, Ian Johnston, Dominic Kwiatkowski, Cordelia Langford, John Sillitoe on behalf of the Wellcome Sanger Institute COVID-19 Surveillance Team |
| EPI_ISL_3106466 | Lighthouse Lab in Glasgow | Wellcome Sanger Institute for the COVID-19 Genomics UK (COG-UK) Consortium | Harper VanSteenhouse, Yumi Kasai, David Gray, Carol Clugston, Anna Dominiczak and Alex Alderton, Roberto Amato, Jeffrey Barrett, Sonia Goncalves, Ewan Harrison, David K. Jackson, Ian Johnston, Dominic Kwiatkowski, Cordelia Langford, John Sillitoe on behalf of the Wellcome Sanger Institute COVID-19 Surveillance Team |

| | | | |
|---|---|---|---|
| EPI_ISL_3106467, EPI_ISL_3106468, EPI_ISL_3106469, EPI_ISL_3106470 | Lighthouse Lab in Alderley Park | Wellcome Sanger Institute for the COVID-19 Genomics UK (COG-UK) Consortium | Jacquelyn Wynn, Mairead Hyland, The Lighthouse Lab in Alderley Park and Alex Alderton, Roberto Amato, Jeffrey Barrett, Sonia Goncalves, Ewan Harrison, David K. Jackson, Ian Johnston, Dominic Kwiatkowski, Cordelia Langford, John Sillitoe on behalf of the Wellcome Sanger Institute COVID-19 Surveillance Team |
| EPI_ISL_3106471, EPI_ISL_3106472, EPI_ISL_3106473 | Lighthouse Lab in Milton Keynes | Wellcome Sanger Institute for the COVID-19 Genomics UK (COG-UK) Consortium | The Lighthouse Lab in Milton Keynes and Alex Alderton, Roberto Amato, Jeffrey Barrett, Sonia Goncalves, Ewan Harrison, David K. Jackson, Ian Johnston, Dominic Kwiatkowski, Cordelia Langford, John Sillitoe on behalf of the Wellcome Sanger Institute COVID-19 Surveillance Team |
| EPI_ISL_3106474, EPI_ISL_3106475 | Lighthouse Lab in Alderley Park | Wellcome Sanger Institute for the COVID-19 Genomics UK (COG-UK) Consortium | Jacquelyn Wynn, Mairead Hyland, The Lighthouse Lab in Alderley Park and Alex Alderton, Roberto Amato, Jeffrey Barrett, Sonia Goncalves, Ewan Harrison, David K. Jackson, Ian Johnston, Dominic Kwiatkowski, Cordelia Langford, John Sillitoe on behalf of the Wellcome Sanger Institute COVID-19 Surveillance Team |
| EPI_ISL_3106476, EPI_ISL_3106477 | Lighthouse Lab in Milton Keynes | Wellcome Sanger Institute for the COVID-19 Genomics UK (COG-UK) Consortium | The Lighthouse Lab in Milton Keynes and Alex Alderton, Roberto Amato, Jeffrey Barrett, Sonia Goncalves, Ewan Harrison, David K. Jackson, Ian Johnston, Dominic Kwiatkowski, Cordelia Langford, John Sillitoe on behalf of the Wellcome Sanger Institute COVID-19 Surveillance Team |
| EPI_ISL_3106478, EPI_ISL_3106479, EPI_ISL_3106480, EPI_ISL_3106481, EPI_ISL_3106482 | Lighthouse Lab in Alderley Park | Wellcome Sanger Institute for the COVID-19 Genomics UK (COG-UK) Consortium | Jacquelyn Wynn, Mairead Hyland, The Lighthouse Lab in Alderley Park and Alex Alderton, Roberto Amato, Jeffrey Barrett, Sonia Goncalves, Ewan Harrison, David K. Jackson, Ian Johnston, Dominic Kwiatkowski, Cordelia Langford, John Sillitoe on behalf of the Wellcome Sanger Institute COVID-19 Surveillance Team |
| EPI_ISL_3106483 | Lighthouse Lab in Milton Keynes | Wellcome Sanger Institute for the COVID-19 Genomics UK (COG-UK) Consortium | The Lighthouse Lab in Milton Keynes and Alex Alderton, Roberto Amato, Jeffrey Barrett, Sonia Goncalves, Ewan Harrison, David K. Jackson, Ian Johnston, Dominic Kwiatkowski, Cordelia Langford, John Sillitoe on behalf of the Wellcome Sanger Institute COVID-19 Surveillance Team |
| EPI_ISL_3106484, EPI_ISL_3106485 | Lighthouse Lab in Alderley Park | Wellcome Sanger Institute for the COVID-19 Genomics UK (COG-UK) Consortium | Jacquelyn Wynn, Mairead Hyland, The Lighthouse Lab in Alderley Park and Alex Alderton, Roberto Amato, Jeffrey Barrett, Sonia Goncalves, Ewan Harrison, David K. Jackson, Ian Johnston, Dominic Kwiatkowski, Cordelia Langford, John Sillitoe on behalf of the Wellcome Sanger Institute COVID-19 Surveillance Team |
| EPI_ISL_3106486, EPI_ISL_3106487, EPI_ISL_3106488, EPI_ISL_3106489 | Lighthouse Lab in Milton Keynes | Wellcome Sanger Institute for the COVID-19 Genomics UK (COG-UK) Consortium | The Lighthouse Lab in Milton Keynes and Alex Alderton, Roberto Amato, Jeffrey Barrett, Sonia Goncalves, Ewan Harrison, David K. Jackson, Ian Johnston, Dominic Kwiatkowski, Cordelia Langford, John Sillitoe on behalf of the Wellcome Sanger Institute COVID-19 Surveillance Team |
| EPI_ISL_3106490, EPI_ISL_3106491 | Lighthouse Lab in Glasgow | Wellcome Sanger Institute for the COVID-19 Genomics UK (COG-UK) Consortium | Harper VanSteenhouse, Yumi Kasai, David Gray, Carol Clugston, Anna Dominiczak and Alex Alderton, Roberto Amato, Jeffrey Barrett, Sonia Goncalves, Ewan Harrison, David K. Jackson, Ian Johnston, Dominic Kwiatkowski, Cordelia Langford, John Sillitoe on behalf of the Wellcome Sanger Institute COVID-19 Surveillance Team |
| EPI_ISL_3106492 | Lighthouse Lab in Milton Keynes | Wellcome Sanger Institute for the COVID-19 Genomics UK (COG-UK) Consortium | The Lighthouse Lab in Milton Keynes and Alex Alderton, Roberto Amato, Jeffrey Barrett, Sonia Goncalves, Ewan Harrison, David K. Jackson, Ian Johnston, Dominic Kwiatkowski, Cordelia Langford, John Sillitoe on behalf of the Wellcome Sanger Institute COVID-19 Surveillance Team |
| EPI_ISL_3106493 | Lighthouse Lab in Alderley Park | Wellcome Sanger Institute for the COVID-19 Genomics UK (COG-UK) Consortium | Jacquelyn Wynn, Mairead Hyland, The Lighthouse Lab in Alderley Park and Alex Alderton, Roberto Amato, Jeffrey Barrett, Sonia Goncalves, Ewan Harrison, David K. Jackson, Ian Johnston, Dominic Kwiatkowski, Cordelia Langford, John Sillitoe on behalf of the Wellcome Sanger Institute COVID-19 Surveillance Team |
| EPI_ISL_3106494 | Lighthouse Lab in Milton Keynes | Wellcome Sanger Institute for the COVID-19 Genomics UK (COG-UK) Consortium | The Lighthouse Lab in Milton Keynes and Alex Alderton, Roberto Amato, Jeffrey Barrett, Sonia Goncalves, Ewan Harrison, David K. Jackson, Ian Johnston, Dominic Kwiatkowski, Cordelia Langford, John Sillitoe on behalf of the Wellcome Sanger Institute COVID-19 Surveillance Team |
| EPI_ISL_3106495, EPI_ISL_3106496 | Lighthouse Lab in Alderley Park | Wellcome Sanger Institute for the COVID-19 Genomics UK (COG-UK) Consortium | Jacquelyn Wynn, Mairead Hyland, The Lighthouse Lab in Alderley Park and Alex Alderton, Roberto Amato, Jeffrey Barrett, Sonia Goncalves, Ewan Harrison, David K. Jackson, Ian Johnston, Dominic Kwiatkowski, Cordelia Langford, John Sillitoe on behalf of the Wellcome Sanger Institute COVID-19 Surveillance Team |
| EPI_ISL_3106497, EPI_ISL_3106498 | Lighthouse Lab in Milton Keynes | Wellcome Sanger Institute for the COVID-19 Genomics UK (COG-UK) Consortium | The Lighthouse Lab in Milton Keynes and Alex Alderton, Roberto Amato, Jeffrey Barrett, Sonia Goncalves, Ewan Harrison, David K. Jackson, Ian Johnston, Dominic Kwiatkowski, Cordelia Langford, John Sillitoe on behalf of the Wellcome Sanger Institute COVID-19 Surveillance Team |
| EPI_ISL_3106499 | Lighthouse Lab in Alderley Park | Wellcome Sanger Institute for the COVID-19 Genomics UK (COG-UK) Consortium | Jacquelyn Wynn, Mairead Hyland, The Lighthouse Lab in Alderley Park and Alex Alderton, Roberto Amato, Jeffrey Barrett, Sonia Goncalves, Ewan Harrison, David K. Jackson, Ian Johnston, Dominic Kwiatkowski, Cordelia Langford, John Sillitoe on behalf of the Wellcome Sanger Institute COVID-19 Surveillance Team |
| EPI_ISL_3106500, EPI_ISL_3106501 | Lighthouse Lab in Milton Keynes | Wellcome Sanger Institute for the COVID-19 Genomics UK (COG-UK) Consortium | The Lighthouse Lab in Milton Keynes and Alex Alderton, Roberto Amato, Jeffrey Barrett, Sonia Goncalves, Ewan Harrison, David K. Jackson, Ian Johnston, Dominic Kwiatkowski, Cordelia Langford, John Sillitoe on behalf of the Wellcome Sanger Institute COVID-19 Surveillance Team |
| EPI_ISL_3106502 | Lighthouse Lab in Alderley Park | Wellcome Sanger Institute for the COVID-19 Genomics UK (COG-UK) Consortium | Jacquelyn Wynn, Mairead Hyland, The Lighthouse Lab in Alderley Park and Alex Alderton, Roberto Amato, Jeffrey Barrett, Sonia Goncalves, Ewan Harrison, David K. Jackson, Ian Johnston, Dominic Kwiatkowski, Cordelia Langford, John Sillitoe on behalf of the Wellcome Sanger Institute COVID-19 Surveillance Team |
| EPI_ISL_3106503 | Lighthouse Lab in Milton Keynes | Wellcome Sanger Institute for the COVID-19 Genomics UK (COG-UK) Consortium | The Lighthouse Lab in Milton Keynes and Alex Alderton, Roberto Amato, Jeffrey Barrett, Sonia Goncalves, Ewan Harrison, David K. Jackson, Ian Johnston, Dominic Kwiatkowski, Cordelia Langford, John Sillitoe on behalf of the Wellcome Sanger Institute COVID-19 Surveillance Team |
| EPI_ISL_3106504, EPI_ISL_3106505, EPI_ISL_3106506 | Lighthouse Lab in Alderley Park | Wellcome Sanger Institute for the COVID-19 Genomics UK (COG-UK) Consortium | Jacquelyn Wynn, Mairead Hyland, The Lighthouse Lab in Alderley Park and Alex Alderton, Roberto Amato, Jeffrey Barrett, Sonia Goncalves, Ewan Harrison, David K. Jackson, Ian Johnston, Dominic Kwiatkowski, Cordelia Langford, John Sillitoe on behalf of the Wellcome Sanger Institute COVID-19 Surveillance Team |
| EPI_ISL_3106507 | Lighthouse Lab in Glasgow | Wellcome Sanger Institute for the COVID-19 Genomics UK (COG-UK) Consortium | Harper VanSteenhouse, Yumi Kasai, David Gray, Carol Clugston, Anna Dominiczak and Alex Alderton, Roberto Amato, Jeffrey Barrett, Sonia Goncalves, Ewan Harrison, David K. Jackson, Ian Johnston, Dominic Kwiatkowski, Cordelia Langford, John Sillitoe on behalf of the Wellcome Sanger Institute COVID-19 Surveillance Team |
| EPI_ISL_3106508, EPI_ISL_3106509 | Lighthouse Lab in Milton Keynes | Wellcome Sanger Institute for the COVID-19 Genomics UK (COG-UK) Consortium | The Lighthouse Lab in Milton Keynes and Alex Alderton, Roberto Amato, Jeffrey Barrett, Sonia Goncalves, Ewan Harrison, David K. Jackson, Ian Johnston, Dominic Kwiatkowski, Cordelia Langford, John Sillitoe on behalf of the Wellcome Sanger Institute COVID-19 Surveillance Team |
| EPI_ISL_3106510 | Lighthouse Lab in Alderley Park | Wellcome Sanger Institute for the COVID-19 Genomics UK (COG-UK) Consortium | Jacquelyn Wynn, Mairead Hyland, The Lighthouse Lab in Alderley Park and Alex Alderton, Roberto Amato, Jeffrey Barrett, Sonia Goncalves, Ewan Harrison, David K. Jackson, Ian Johnston, Dominic Kwiatkowski, Cordelia Langford, John Sillitoe on behalf of the Wellcome Sanger Institute COVID-19 Surveillance Team |
| EPI_ISL_3106511, EPI_ISL_3106512 | Lighthouse Lab in Milton Keynes | Wellcome Sanger Institute for the COVID-19 Genomics UK (COG-UK) Consortium | The Lighthouse Lab in Milton Keynes and Alex Alderton, Roberto Amato, Jeffrey Barrett, Sonia Goncalves, Ewan Harrison, David K. Jackson, Ian Johnston, Dominic Kwiatkowski, Cordelia Langford, John Sillitoe on behalf of the Wellcome Sanger Institute COVID-19 Surveillance Team |
| EPI_ISL_3106513, EPI_ISL_3106514 | Lighthouse Lab in Glasgow | Wellcome Sanger Institute for the COVID-19 Genomics UK (COG-UK) Consortium | Harper VanSteenhouse, Yumi Kasai, David Gray, Carol Clugston, Anna Dominiczak and Alex Alderton, Roberto Amato, Jeffrey Barrett, Sonia Goncalves, Ewan Harrison, David K. Jackson, Ian Johnston, Dominic Kwiatkowski, Cordelia Langford, John Sillitoe on behalf of the Wellcome Sanger Institute COVID-19 Surveillance Team |
| EPI_ISL_3106515 | Lighthouse Lab in Milton Keynes | Wellcome Sanger Institute for the COVID-19 Genomics UK (COG-UK) Consortium | The Lighthouse Lab in Milton Keynes and Alex Alderton, Roberto Amato, Jeffrey Barrett, Sonia Goncalves, Ewan Harrison, David K. Jackson, Ian Johnston, Dominic Kwiatkowski, Cordelia Langford, John Sillitoe on behalf of the Wellcome Sanger Institute COVID-19 Surveillance Team |
| EPI_ISL_3106516 | Lighthouse Lab in Glasgow | Wellcome Sanger Institute for the COVID-19 Genomics UK (COG-UK) Consortium | Harper VanSteenhouse, Yumi Kasai, David Gray, Carol Clugston, Anna Dominiczak and Alex Alderton, Roberto Amato, Jeffrey Barrett, Sonia Goncalves, Ewan Harrison, David K. Jackson, Ian Johnston, Dominic Kwiatkowski, Cordelia Langford, John Sillitoe on behalf of the Wellcome Sanger Institute COVID-19 Surveillance Team |
| EPI_ISL_3106517, EPI_ISL_3106518, EPI_ISL_3106519 | Lighthouse Lab in Alderley Park | Wellcome Sanger Institute for the COVID-19 Genomics UK (COG-UK) Consortium | Jacquelyn Wynn, Mairead Hyland, The Lighthouse Lab in Alderley Park and Alex Alderton, Roberto Amato, Jeffrey Barrett, Sonia Goncalves, Ewan Harrison, David K. Jackson, Ian Johnston, Dominic Kwiatkowski, Cordelia Langford, John Sillitoe on behalf of the Wellcome Sanger Institute COVID-19 Surveillance Team |
| EPI_ISL_3106520, EPI_ISL_3106521, EPI_ISL_3106522, EPI_ISL_3106523 | Lighthouse Lab in Glasgow | Wellcome Sanger Institute for the COVID-19 Genomics UK (COG-UK) Consortium | Harper VanSteenhouse, Yumi Kasai, David Gray, Carol Clugston, Anna Dominiczak and Alex Alderton, Roberto Amato, Jeffrey Barrett, Sonia Goncalves, Ewan Harrison, David K. Jackson, Ian Johnston, Dominic Kwiatkowski, Cordelia Langford, John Sillitoe on behalf of the Wellcome Sanger Institute COVID-19 Surveillance Team |
| EPI_ISL_3106524 | Lighthouse Lab in Alderley Park | Wellcome Sanger Institute for the COVID-19 Genomics UK (COG-UK) Consortium | Jacquelyn Wynn, Mairead Hyland, The Lighthouse Lab in Alderley Park and Alex Alderton, Roberto Amato, Jeffrey Barrett, Sonia Goncalves, Ewan Harrison, David K. Jackson, Ian Johnston, Dominic Kwiatkowski, Cordelia Langford, John Sillitoe on behalf of the Wellcome Sanger Institute COVID-19 Surveillance Team |
| EPI_ISL_3106525 | Lighthouse Lab in Milton Keynes | Wellcome Sanger Institute for the COVID-19 Genomics UK (COG-UK) Consortium | The Lighthouse Lab in Milton Keynes and Alex Alderton, Roberto Amato, Jeffrey Barrett, Sonia Goncalves, Ewan Harrison, David K. Jackson, Ian Johnston, Dominic Kwiatkowski, Cordelia Langford, John Sillitoe on behalf of the Wellcome Sanger Institute COVID-19 Surveillance Team |
| EPI_ISL_3106526 | Lighthouse Lab in Alderley Park | Wellcome Sanger Institute for the COVID-19 Genomics UK (COG-UK) Consortium | Jacquelyn Wynn, Mairead Hyland, The Lighthouse Lab in Alderley Park and Alex Alderton, Roberto Amato, Jeffrey Barrett, Sonia Goncalves, Ewan Harrison, David K. Jackson, Ian Johnston, Dominic Kwiatkowski, Cordelia Langford, John Sillitoe on behalf of the Wellcome Sanger Institute COVID-19 Surveillance Team |
| EPI_ISL_3106527 | Lighthouse Lab in Glasgow | Wellcome Sanger Institute for the COVID-19 Genomics UK (COG-UK) Consortium | Harper VanSteenhouse, Yumi Kasai, David Gray, Carol Clugston, Anna Dominiczak and Alex Alderton, Roberto Amato, Jeffrey Barrett, Sonia Goncalves, Ewan Harrison, David K. Jackson, Ian Johnston, Dominic Kwiatkowski, Cordelia Langford, John Sillitoe on behalf of the Wellcome Sanger Institute COVID-19 Surveillance Team |
| EPI_ISL_3106528, EPI_ISL_3106529 | Lighthouse Lab in Milton Keynes | Wellcome Sanger Institute for the COVID-19 Genomics UK (COG-UK) Consortium | The Lighthouse Lab in Milton Keynes and Alex Alderton, Roberto Amato, Jeffrey Barrett, Sonia Goncalves, Ewan Harrison, David K. Jackson, Ian Johnston, Dominic Kwiatkowski, Cordelia Langford, John Sillitoe on behalf of the Wellcome Sanger Institute COVID-19 Surveillance Team |
| EPI_ISL_3106530 | Lighthouse Lab in Alderley Park | Wellcome Sanger Institute for the COVID-19 Genomics UK (COG-UK) Consortium | Jacquelyn Wynn, Mairead Hyland, The Lighthouse Lab in Alderley Park and Alex Alderton, Roberto Amato, Jeffrey Barrett, Sonia Goncalves, Ewan Harrison, David K. Jackson, Ian Johnston, Dominic Kwiatkowski, Cordelia Langford, John Sillitoe on behalf of the Wellcome Sanger Institute COVID-19 Surveillance Team |
| EPI_ISL_3106531, EPI_ISL_3106532 | Lighthouse Lab in Glasgow | Wellcome Sanger Institute for the COVID-19 Genomics UK (COG-UK) Consortium | Harper VanSteenhouse, Yumi Kasai, David Gray, Carol Clugston, Anna Dominiczak and Alex Alderton, Roberto Amato, Jeffrey Barrett, Sonia Goncalves, Ewan Harrison, David K. Jackson, Ian Johnston, Dominic Kwiatkowski, Cordelia Langford, John Sillitoe on behalf of the Wellcome Sanger Institute COVID-19 Surveillance Team |
| EPI_ISL_3106533, EPI_ISL_3106534, EPI_ISL_3106535 | Lighthouse Lab in Milton Keynes | Wellcome Sanger Institute for the COVID-19 Genomics UK (COG-UK) Consortium | The Lighthouse Lab in Milton Keynes and Alex Alderton, Roberto Amato, Jeffrey Barrett, Sonia Goncalves, Ewan Harrison, David K. Jackson, Ian Johnston, Dominic Kwiatkowski, Cordelia Langford, John Sillitoe on behalf of the Wellcome Sanger Institute COVID-19 Surveillance Team |
| EPI_ISL_3106536 | Lighthouse Lab in Alderley Park | Wellcome Sanger Institute for the COVID-19 Genomics UK (COG-UK) Consortium | Jacquelyn Wynn, Mairead Hyland, The Lighthouse Lab in Alderley Park and Alex Alderton, Roberto Amato, Jeffrey Barrett, Sonia Goncalves, Ewan Harrison, David K. Jackson, Ian Johnston, Dominic Kwiatkowski, Cordelia Langford, John Sillitoe on behalf of the Wellcome Sanger Institute COVID-19 Surveillance Team |
| EPI_ISL_3106537 | Lighthouse Lab in Milton Keynes | Wellcome Sanger Institute for the COVID-19 Genomics UK (COG-UK) Consortium | The Lighthouse Lab in Milton Keynes and Alex Alderton, Roberto Amato, Jeffrey Barrett, Sonia Goncalves, Ewan Harrison, David K. Jackson, Ian Johnston, Dominic Kwiatkowski, Cordelia Langford, John Sillitoe on behalf of the Wellcome Sanger Institute COVID-19 Surveillance Team |
| EPI_ISL_3106538, EPI_ISL_3106539 | Lighthouse Lab in Alderley Park | Wellcome Sanger Institute for the COVID-19 Genomics UK (COG-UK) Consortium | Jacquelyn Wynn, Mairead Hyland, The Lighthouse Lab in Alderley Park and Alex Alderton, Roberto Amato, Jeffrey Barrett, Sonia Goncalves, Ewan Harrison, David K. Jackson, Ian Johnston, Dominic Kwiatkowski, Cordelia Langford, John Sillitoe on behalf of the Wellcome Sanger Institute COVID-19 Surveillance Team |
| EPI_ISL_3106540, | Lighthouse Lab in Milton Keynes | Wellcome Sanger Institute for the COVID-19 Genomics UK (COG-UK) | The Lighthouse Lab in Milton Keynes and Alex Alderton, Roberto Amato, Jeffrey Barrett, Sonia Goncalves, Ewan Harrison, David K. Jackson, Ian Johnston, Dominic Kwiatkowski, Cordelia Langford, John Sillitoe on behalf of the Wellcome Sanger |

| | | | |
|---|---|---|---|
| EPI_ISL_3106541 | | Consortium | Institute COVID-19 Surveillance Team |
| EPI_ISL_3106542 | Lighthouse Lab in Alderley Park | Wellcome Sanger Institute for the COVID-19 Genomics UK (COG-UK) Consortium | Jacquelyn Wynn, Mairead Hyland, The Lighthouse Lab in Alderley Park and Alex Alderton, Roberto Amato, Jeffrey Barrett, Sonia Goncalves, Ewan Harrison, David K. Jackson, Ian Johnston, Dominic Kwiatkowski, Cordelia Langford, John Sillitoe on behalf of the Wellcome Sanger Institute COVID-19 Surveillance Team |
| EPI_ISL_3106543, EPI_ISL_3106544, EPI_ISL_3106545 | Lighthouse Lab in Milton Keynes | Wellcome Sanger Institute for the COVID-19 Genomics UK (COG-UK) Consortium | The Lighthouse Lab in Milton Keynes and Alex Alderton, Roberto Amato, Jeffrey Barrett, Sonia Goncalves, Ewan Harrison, David K. Jackson, Ian Johnston, Dominic Kwiatkowski, Cordelia Langford, John Sillitoe on behalf of the Wellcome Sanger Institute COVID-19 Surveillance Team |
| EPI_ISL_3106546 | Lighthouse Lab in Glasgow | Wellcome Sanger Institute for the COVID-19 Genomics UK (COG-UK) Consortium | Harper VanSteenhouse, Yumi Kasai, David Gray, Carol Clugston, Anna Dominiczak and Alex Alderton, Roberto Amato, Jeffrey Barrett, Sonia Goncalves, Ewan Harrison, David K. Jackson, Ian Johnston, Dominic Kwiatkowski, Cordelia Langford, John Sillitoe on behalf of the Wellcome Sanger Institute COVID-19 Surveillance Team |
| EPI_ISL_3106547 | Lighthouse Lab in Alderley Park | Wellcome Sanger Institute for the COVID-19 Genomics UK (COG-UK) Consortium | Jacquelyn Wynn, Mairead Hyland, The Lighthouse Lab in Alderley Park and Alex Alderton, Roberto Amato, Jeffrey Barrett, Sonia Goncalves, Ewan Harrison, David K. Jackson, Ian Johnston, Dominic Kwiatkowski, Cordelia Langford, John Sillitoe on behalf of the Wellcome Sanger Institute COVID-19 Surveillance Team |
| EPI_ISL_3106548, EPI_ISL_3106549, EPI_ISL_3106550 | Lighthouse Lab in Milton Keynes | Wellcome Sanger Institute for the COVID-19 Genomics UK (COG-UK) Consortium | The Lighthouse Lab in Milton Keynes and Alex Alderton, Roberto Amato, Jeffrey Barrett, Sonia Goncalves, Ewan Harrison, David K. Jackson, Ian Johnston, Dominic Kwiatkowski, Cordelia Langford, John Sillitoe on behalf of the Wellcome Sanger Institute COVID-19 Surveillance Team |
| EPI_ISL_3106551 | Lighthouse Lab in Alderley Park | Wellcome Sanger Institute for the COVID-19 Genomics UK (COG-UK) Consortium | Jacquelyn Wynn, Mairead Hyland, The Lighthouse Lab in Alderley Park and Alex Alderton, Roberto Amato, Jeffrey Barrett, Sonia Goncalves, Ewan Harrison, David K. Jackson, Ian Johnston, Dominic Kwiatkowski, Cordelia Langford, John Sillitoe on behalf of the Wellcome Sanger Institute COVID-19 Surveillance Team |
| EPI_ISL_3106552, EPI_ISL_3106553 | Lighthouse Lab in Milton Keynes | Wellcome Sanger Institute for the COVID-19 Genomics UK (COG-UK) Consortium | The Lighthouse Lab in Milton Keynes and Alex Alderton, Roberto Amato, Jeffrey Barrett, Sonia Goncalves, Ewan Harrison, David K. Jackson, Ian Johnston, Dominic Kwiatkowski, Cordelia Langford, John Sillitoe on behalf of the Wellcome Sanger Institute COVID-19 Surveillance Team |
| EPI_ISL_3106554 | Lighthouse Lab in Glasgow | Wellcome Sanger Institute for the COVID-19 Genomics UK (COG-UK) Consortium | Harper VanSteenhouse, Yumi Kasai, David Gray, Carol Clugston, Anna Dominiczak and Alex Alderton, Roberto Amato, Jeffrey Barrett, Sonia Goncalves, Ewan Harrison, David K. Jackson, Ian Johnston, Dominic Kwiatkowski, Cordelia Langford, John Sillitoe on behalf of the Wellcome Sanger Institute COVID-19 Surveillance Team |
| EPI_ISL_3106555 | Lighthouse Lab in Milton Keynes | Wellcome Sanger Institute for the COVID-19 Genomics UK (COG-UK) Consortium | The Lighthouse Lab in Milton Keynes and Alex Alderton, Roberto Amato, Jeffrey Barrett, Sonia Goncalves, Ewan Harrison, David K. Jackson, Ian Johnston, Dominic Kwiatkowski, Cordelia Langford, John Sillitoe on behalf of the Wellcome Sanger Institute COVID-19 Surveillance Team |
| EPI_ISL_3106556 | Lighthouse Lab in Alderley Park | Wellcome Sanger Institute for the COVID-19 Genomics UK (COG-UK) Consortium | Jacquelyn Wynn, Mairead Hyland, The Lighthouse Lab in Alderley Park and Alex Alderton, Roberto Amato, Jeffrey Barrett, Sonia Goncalves, Ewan Harrison, David K. Jackson, Ian Johnston, Dominic Kwiatkowski, Cordelia Langford, John Sillitoe on behalf of the Wellcome Sanger Institute COVID-19 Surveillance Team |
| EPI_ISL_3106557, EPI_ISL_3106558, EPI_ISL_3106559, EPI_ISL_3106560, EPI_ISL_3106561 | Lighthouse Lab in Milton Keynes | Wellcome Sanger Institute for the COVID-19 Genomics UK (COG-UK) Consortium | The Lighthouse Lab in Milton Keynes and Alex Alderton, Roberto Amato, Jeffrey Barrett, Sonia Goncalves, Ewan Harrison, David K. Jackson, Ian Johnston, Dominic Kwiatkowski, Cordelia Langford, John Sillitoe on behalf of the Wellcome Sanger Institute COVID-19 Surveillance Team |
| EPI_ISL_3106562, EPI_ISL_3106563, EPI_ISL_3106564 | Lighthouse Lab in Glasgow | Wellcome Sanger Institute for the COVID-19 Genomics UK (COG-UK) Consortium | Harper VanSteenhouse, Yumi Kasai, David Gray, Carol Clugston, Anna Dominiczak and Alex Alderton, Roberto Amato, Jeffrey Barrett, Sonia Goncalves, Ewan Harrison, David K. Jackson, Ian Johnston, Dominic Kwiatkowski, Cordelia Langford, John Sillitoe on behalf of the Wellcome Sanger Institute COVID-19 Surveillance Team |
| EPI_ISL_3106565, EPI_ISL_3106566, EPI_ISL_3106567 | Lighthouse Lab in Milton Keynes | Wellcome Sanger Institute for the COVID-19 Genomics UK (COG-UK) Consortium | The Lighthouse Lab in Milton Keynes and Alex Alderton, Roberto Amato, Jeffrey Barrett, Sonia Goncalves, Ewan Harrison, David K. Jackson, Ian Johnston, Dominic Kwiatkowski, Cordelia Langford, John Sillitoe on behalf of the Wellcome Sanger Institute COVID-19 Surveillance Team |
| EPI_ISL_3106568 | Lighthouse Lab in Alderley Park | Wellcome Sanger Institute for the COVID-19 Genomics UK (COG-UK) Consortium | Jacquelyn Wynn, Mairead Hyland, The Lighthouse Lab in Alderley Park and Alex Alderton, Roberto Amato, Jeffrey Barrett, Sonia Goncalves, Ewan Harrison, David K. Jackson, Ian Johnston, Dominic Kwiatkowski, Cordelia Langford, John Sillitoe on behalf of the Wellcome Sanger Institute COVID-19 Surveillance Team |
| EPI_ISL_3106569 | Lighthouse Lab in Glasgow | Wellcome Sanger Institute for the COVID-19 Genomics UK (COG-UK) Consortium | Harper VanSteenhouse, Yumi Kasai, David Gray, Carol Clugston, Anna Dominiczak and Alex Alderton, Roberto Amato, Jeffrey Barrett, Sonia Goncalves, Ewan Harrison, David K. Jackson, Ian Johnston, Dominic Kwiatkowski, Cordelia Langford, John Sillitoe on behalf of the Wellcome Sanger Institute COVID-19 Surveillance Team |
| EPI_ISL_3106570 | Lighthouse Lab in Milton Keynes | Wellcome Sanger Institute for the COVID-19 Genomics UK (COG-UK) Consortium | The Lighthouse Lab in Milton Keynes and Alex Alderton, Roberto Amato, Jeffrey Barrett, Sonia Goncalves, Ewan Harrison, David K. Jackson, Ian Johnston, Dominic Kwiatkowski, Cordelia Langford, John Sillitoe on behalf of the Wellcome Sanger Institute COVID-19 Surveillance Team |
| EPI_ISL_3106571 | Lighthouse Lab in Glasgow | Wellcome Sanger Institute for the COVID-19 Genomics UK (COG-UK) Consortium | Harper VanSteenhouse, Yumi Kasai, David Gray, Carol Clugston, Anna Dominiczak and Alex Alderton, Roberto Amato, Jeffrey Barrett, Sonia Goncalves, Ewan Harrison, David K. Jackson, Ian Johnston, Dominic Kwiatkowski, Cordelia Langford, John Sillitoe on behalf of the Wellcome Sanger Institute COVID-19 Surveillance Team |
| EPI_ISL_3106572 | Lighthouse Lab in Milton Keynes | Wellcome Sanger Institute for the COVID-19 Genomics UK (COG-UK) Consortium | The Lighthouse Lab in Milton Keynes and Alex Alderton, Roberto Amato, Jeffrey Barrett, Sonia Goncalves, Ewan Harrison, David K. Jackson, Ian Johnston, Dominic Kwiatkowski, Cordelia Langford, John Sillitoe on behalf of the Wellcome Sanger Institute COVID-19 Surveillance Team |
| EPI_ISL_3106573 | Lighthouse Lab in Alderley Park | Wellcome Sanger Institute for the COVID-19 Genomics UK (COG-UK) Consortium | Jacquelyn Wynn, Mairead Hyland, The Lighthouse Lab in Alderley Park and Alex Alderton, Roberto Amato, Jeffrey Barrett, Sonia Goncalves, Ewan Harrison, David K. Jackson, Ian Johnston, Dominic Kwiatkowski, Cordelia Langford, John Sillitoe on behalf of the Wellcome Sanger Institute COVID-19 Surveillance Team |
| EPI_ISL_3106574 | Lighthouse Lab in Milton Keynes | Wellcome Sanger Institute for the COVID-19 Genomics UK (COG-UK) Consortium | The Lighthouse Lab in Milton Keynes and Alex Alderton, Roberto Amato, Jeffrey Barrett, Sonia Goncalves, Ewan Harrison, David K. Jackson, Ian Johnston, Dominic Kwiatkowski, Cordelia Langford, John Sillitoe on behalf of the Wellcome Sanger Institute COVID-19 Surveillance Team |
| EPI_ISL_3106575, EPI_ISL_3106576 | Lighthouse Lab in Glasgow | Wellcome Sanger Institute for the COVID-19 Genomics UK (COG-UK) Consortium | Harper VanSteenhouse, Yumi Kasai, David Gray, Carol Clugston, Anna Dominiczak and Alex Alderton, Roberto Amato, Jeffrey Barrett, Sonia Goncalves, Ewan Harrison, David K. Jackson, Ian Johnston, Dominic Kwiatkowski, Cordelia Langford, John Sillitoe on behalf of the Wellcome Sanger Institute COVID-19 Surveillance Team |
| EPI_ISL_3106577, EPI_ISL_3106578 | Lighthouse Lab in Milton Keynes | Wellcome Sanger Institute for the COVID-19 Genomics UK (COG-UK) Consortium | The Lighthouse Lab in Milton Keynes and Alex Alderton, Roberto Amato, Jeffrey Barrett, Sonia Goncalves, Ewan Harrison, David K. Jackson, Ian Johnston, Dominic Kwiatkowski, Cordelia Langford, John Sillitoe on behalf of the Wellcome Sanger Institute COVID-19 Surveillance Team |
| EPI_ISL_3106579 | Lighthouse Lab in Alderley Park | Wellcome Sanger Institute for the COVID-19 Genomics UK (COG-UK) Consortium | Jacquelyn Wynn, Mairead Hyland, The Lighthouse Lab in Alderley Park and Alex Alderton, Roberto Amato, Jeffrey Barrett, Sonia Goncalves, Ewan Harrison, David K. Jackson, Ian Johnston, Dominic Kwiatkowski, Cordelia Langford, John Sillitoe on behalf of the Wellcome Sanger Institute COVID-19 Surveillance Team |
| EPI_ISL_3106580 | Lighthouse Lab in Milton Keynes | Wellcome Sanger Institute for the COVID-19 Genomics UK (COG-UK) Consortium | The Lighthouse Lab in Milton Keynes and Alex Alderton, Roberto Amato, Jeffrey Barrett, Sonia Goncalves, Ewan Harrison, David K. Jackson, Ian Johnston, Dominic Kwiatkowski, Cordelia Langford, John Sillitoe on behalf of the Wellcome Sanger Institute COVID-19 Surveillance Team |
| EPI_ISL_3106581 | Lighthouse Lab in Glasgow | Wellcome Sanger Institute for the COVID-19 Genomics UK (COG-UK) Consortium | Harper VanSteenhouse, Yumi Kasai, David Gray, Carol Clugston, Anna Dominiczak and Alex Alderton, Roberto Amato, Jeffrey Barrett, Sonia Goncalves, Ewan Harrison, David K. Jackson, Ian Johnston, Dominic Kwiatkowski, Cordelia Langford, John Sillitoe on behalf of the Wellcome Sanger Institute COVID-19 Surveillance Team |
| EPI_ISL_3106582 | Lighthouse Lab in Alderley Park | Wellcome Sanger Institute for the COVID-19 Genomics UK (COG-UK) Consortium | Jacquelyn Wynn, Mairead Hyland, The Lighthouse Lab in Alderley Park and Alex Alderton, Roberto Amato, Jeffrey Barrett, Sonia Goncalves, Ewan Harrison, David K. Jackson, Ian Johnston, Dominic Kwiatkowski, Cordelia Langford, John Sillitoe on behalf of the Wellcome Sanger Institute COVID-19 Surveillance Team |
| EPI_ISL_3106583, EPI_ISL_3106584 | Lighthouse Lab in Milton Keynes | Wellcome Sanger Institute for the COVID-19 Genomics UK (COG-UK) Consortium | The Lighthouse Lab in Milton Keynes and Alex Alderton, Roberto Amato, Jeffrey Barrett, Sonia Goncalves, Ewan Harrison, David K. Jackson, Ian Johnston, Dominic Kwiatkowski, Cordelia Langford, John Sillitoe on behalf of the Wellcome Sanger Institute COVID-19 Surveillance Team |
| EPI_ISL_3106585 | Lighthouse Lab in Glasgow | Wellcome Sanger Institute for the COVID-19 Genomics UK (COG-UK) Consortium | Harper VanSteenhouse, Yumi Kasai, David Gray, Carol Clugston, Anna Dominiczak and Alex Alderton, Roberto Amato, Jeffrey Barrett, Sonia Goncalves, Ewan Harrison, David K. Jackson, Ian Johnston, Dominic Kwiatkowski, Cordelia Langford, John Sillitoe on behalf of the Wellcome Sanger Institute COVID-19 Surveillance Team |
| EPI_ISL_3106586, EPI_ISL_3106587, EPI_ISL_3106588 | Lighthouse Lab in Milton Keynes | Wellcome Sanger Institute for the COVID-19 Genomics UK (COG-UK) Consortium | The Lighthouse Lab in Milton Keynes and Alex Alderton, Roberto Amato, Jeffrey Barrett, Sonia Goncalves, Ewan Harrison, David K. Jackson, Ian Johnston, Dominic Kwiatkowski, Cordelia Langford, John Sillitoe on behalf of the Wellcome Sanger Institute COVID-19 Surveillance Team |
| EPI_ISL_3106589 | Lighthouse Lab in Glasgow | Wellcome Sanger Institute for the COVID-19 Genomics UK (COG-UK) Consortium | Harper VanSteenhouse, Yumi Kasai, David Gray, Carol Clugston, Anna Dominiczak and Alex Alderton, Roberto Amato, Jeffrey Barrett, Sonia Goncalves, Ewan Harrison, David K. Jackson, Ian Johnston, Dominic Kwiatkowski, Cordelia Langford, John Sillitoe on behalf of the Wellcome Sanger Institute COVID-19 Surveillance Team |
| EPI_ISL_3106590 | Lighthouse Lab in Milton Keynes | Wellcome Sanger Institute for the COVID-19 Genomics UK (COG-UK) Consortium | The Lighthouse Lab in Milton Keynes and Alex Alderton, Roberto Amato, Jeffrey Barrett, Sonia Goncalves, Ewan Harrison, David K. Jackson, Ian Johnston, Dominic Kwiatkowski, Cordelia Langford, John Sillitoe on behalf of the Wellcome Sanger Institute COVID-19 Surveillance Team |
| EPI_ISL_3106591 | Lighthouse Lab in Alderley Park | Wellcome Sanger Institute for the COVID-19 Genomics UK (COG-UK) Consortium | Jacquelyn Wynn, Mairead Hyland, The Lighthouse Lab in Alderley Park and Alex Alderton, Roberto Amato, Jeffrey Barrett, Sonia Goncalves, Ewan Harrison, David K. Jackson, Ian Johnston, Dominic Kwiatkowski, Cordelia Langford, John Sillitoe on behalf of the Wellcome Sanger Institute COVID-19 Surveillance Team |
| EPI_ISL_3106592 | Lighthouse Lab in Milton Keynes | Wellcome Sanger Institute for the COVID-19 Genomics UK (COG-UK) Consortium | The Lighthouse Lab in Milton Keynes and Alex Alderton, Roberto Amato, Jeffrey Barrett, Sonia Goncalves, Ewan Harrison, David K. Jackson, Ian Johnston, Dominic Kwiatkowski, Cordelia Langford, John Sillitoe on behalf of the Wellcome Sanger Institute COVID-19 Surveillance Team |
| EPI_ISL_3106593 | Lighthouse Lab in Glasgow | Wellcome Sanger Institute for the COVID-19 Genomics UK (COG-UK) Consortium | Harper VanSteenhouse, Yumi Kasai, David Gray, Carol Clugston, Anna Dominiczak and Alex Alderton, Roberto Amato, Jeffrey Barrett, Sonia Goncalves, Ewan Harrison, David K. Jackson, Ian Johnston, Dominic Kwiatkowski, Cordelia Langford, John Sillitoe on behalf of the Wellcome Sanger Institute COVID-19 Surveillance Team |
| EPI_ISL_3106594 | Lighthouse Lab in Milton Keynes | Wellcome Sanger Institute for the COVID-19 Genomics UK (COG-UK) Consortium | The Lighthouse Lab in Milton Keynes and Alex Alderton, Roberto Amato, Jeffrey Barrett, Sonia Goncalves, Ewan Harrison, David K. Jackson, Ian Johnston, Dominic Kwiatkowski, Cordelia Langford, John Sillitoe on behalf of the Wellcome Sanger Institute COVID-19 Surveillance Team |
| EPI_ISL_3106595, EPI_ISL_3106596 | Lighthouse Lab in Glasgow | Wellcome Sanger Institute for the COVID-19 Genomics UK (COG-UK) Consortium | Harper VanSteenhouse, Yumi Kasai, David Gray, Carol Clugston, Anna Dominiczak and Alex Alderton, Roberto Amato, Jeffrey Barrett, Sonia Goncalves, Ewan Harrison, David K. Jackson, Ian Johnston, Dominic Kwiatkowski, Cordelia Langford, John Sillitoe on behalf of the Wellcome Sanger Institute COVID-19 Surveillance Team |
| EPI_ISL_3106597, EPI_ISL_3106598, EPI_ISL_3106599, EPI_ISL_3106600, EPI_ISL_3106601 | Lighthouse Lab in Milton Keynes | Wellcome Sanger Institute for the COVID-19 Genomics UK (COG-UK) Consortium | The Lighthouse Lab in Milton Keynes and Alex Alderton, Roberto Amato, Jeffrey Barrett, Sonia Goncalves, Ewan Harrison, David K. Jackson, Ian Johnston, Dominic Kwiatkowski, Cordelia Langford, John Sillitoe on behalf of the Wellcome Sanger Institute COVID-19 Surveillance Team |
| EPI_ISL_3106602, EPI_ISL_3106603 | Lighthouse Lab in Alderley Park | Wellcome Sanger Institute for the COVID-19 Genomics UK (COG-UK) Consortium | Jacquelyn Wynn, Mairead Hyland, The Lighthouse Lab in Alderley Park and Alex Alderton, Roberto Amato, Jeffrey Barrett, Sonia Goncalves, Ewan Harrison, David K. Jackson, Ian Johnston, Dominic Kwiatkowski, Cordelia Langford, John Sillitoe on behalf of the Wellcome Sanger Institute COVID-19 Surveillance Team |
| EPI_ISL_3106604 | Lighthouse Lab in Glasgow | Wellcome Sanger Institute for the COVID-19 Genomics UK (COG-UK) Consortium | Harper VanSteenhouse, Yumi Kasai, David Gray, Carol Clugston, Anna Dominiczak and Alex Alderton, Roberto Amato, Jeffrey Barrett, Sonia Goncalves, Ewan Harrison, David K. Jackson, Ian Johnston, Dominic Kwiatkowski, Cordelia Langford, John Sillitoe on behalf of the Wellcome Sanger Institute COVID-19 Surveillance Team |
| EPI_ISL_3106605 | Lighthouse Lab in Milton Keynes | Wellcome Sanger Institute for the COVID-19 Genomics UK (COG-UK) Consortium | The Lighthouse Lab in Milton Keynes and Alex Alderton, Roberto Amato, Jeffrey Barrett, Sonia Goncalves, Ewan Harrison, David K. Jackson, Ian Johnston, Dominic Kwiatkowski, Cordelia Langford, John Sillitoe on behalf of the Wellcome Sanger Institute COVID-19 Surveillance Team |

| | | | |
|---|---|---|---|
| EPI_ISL_3106606 | Lighthouse Lab in Glasgow | Wellcome Sanger Institute for the COVID-19 Genomics UK (COG-UK) Consortium | Harper VanSteenhouse, Yumi Kasai, David Gray, Carol Clugston, Anna Dominiczak and Alex Alderton, Roberto Amato, Jeffrey Barrett, Sonia Goncalves, Ewan Harrison, David K. Jackson, Ian Johnston, Dominic Kwiatkowski, Cordelia Langford, John Sillitoe on behalf of the Wellcome Sanger Institute COVID-19 Surveillance Team |
| EPI_ISL_3106607, EPI_ISL_3106608, EPI_ISL_3106609 | Lighthouse Lab in Milton Keynes | Wellcome Sanger Institute for the COVID-19 Genomics UK (COG-UK) Consortium | The Lighthouse Lab in Milton Keynes and Alex Alderton, Roberto Amato, Jeffrey Barrett, Sonia Goncalves, Ewan Harrison, David K. Jackson, Ian Johnston, Dominic Kwiatkowski, Cordelia Langford, John Sillitoe on behalf of the Wellcome Sanger Institute COVID-19 Surveillance Team |
| EPI_ISL_3106610 | Lighthouse Lab in Glasgow | Wellcome Sanger Institute for the COVID-19 Genomics UK (COG-UK) Consortium | Harper VanSteenhouse, Yumi Kasai, David Gray, Carol Clugston, Anna Dominiczak and Alex Alderton, Roberto Amato, Jeffrey Barrett, Sonia Goncalves, Ewan Harrison, David K. Jackson, Ian Johnston, Dominic Kwiatkowski, Cordelia Langford, John Sillitoe on behalf of the Wellcome Sanger Institute COVID-19 Surveillance Team |
| EPI_ISL_3106611, EPI_ISL_3106612, EPI_ISL_3106613, EPI_ISL_3106614 | Lighthouse Lab in Milton Keynes | Wellcome Sanger Institute for the COVID-19 Genomics UK (COG-UK) Consortium | The Lighthouse Lab in Milton Keynes and Alex Alderton, Roberto Amato, Jeffrey Barrett, Sonia Goncalves, Ewan Harrison, David K. Jackson, Ian Johnston, Dominic Kwiatkowski, Cordelia Langford, John Sillitoe on behalf of the Wellcome Sanger Institute COVID-19 Surveillance Team |
| EPI_ISL_3106615, EPI_ISL_3106616 | Lighthouse Lab in Glasgow | Wellcome Sanger Institute for the COVID-19 Genomics UK (COG-UK) Consortium | Harper VanSteenhouse, Yumi Kasai, David Gray, Carol Clugston, Anna Dominiczak and Alex Alderton, Roberto Amato, Jeffrey Barrett, Sonia Goncalves, Ewan Harrison, David K. Jackson, Ian Johnston, Dominic Kwiatkowski, Cordelia Langford, John Sillitoe on behalf of the Wellcome Sanger Institute COVID-19 Surveillance Team |
| EPI_ISL_3106617, EPI_ISL_3106618 | Lighthouse Lab in Milton Keynes | Wellcome Sanger Institute for the COVID-19 Genomics UK (COG-UK) Consortium | The Lighthouse Lab in Milton Keynes and Alex Alderton, Roberto Amato, Jeffrey Barrett, Sonia Goncalves, Ewan Harrison, David K. Jackson, Ian Johnston, Dominic Kwiatkowski, Cordelia Langford, John Sillitoe on behalf of the Wellcome Sanger Institute COVID-19 Surveillance Team |
| EPI_ISL_3106619 | Lighthouse Lab in Alderley Park | Wellcome Sanger Institute for the COVID-19 Genomics UK (COG-UK) Consortium | Jacquelyn Wynn, Mairead Hyland, The Lighthouse Lab in Alderley Park and Alex Alderton, Roberto Amato, Jeffrey Barrett, Sonia Goncalves, Ewan Harrison, David K. Jackson, Ian Johnston, Dominic Kwiatkowski, Cordelia Langford, John Sillitoe on behalf of the Wellcome Sanger Institute COVID-19 Surveillance Team |
| EPI_ISL_3106620 | Lighthouse Lab in Glasgow | Wellcome Sanger Institute for the COVID-19 Genomics UK (COG-UK) Consortium | Harper VanSteenhouse, Yumi Kasai, David Gray, Carol Clugston, Anna Dominiczak and Alex Alderton, Roberto Amato, Jeffrey Barrett, Sonia Goncalves, Ewan Harrison, David K. Jackson, Ian Johnston, Dominic Kwiatkowski, Cordelia Langford, John Sillitoe on behalf of the Wellcome Sanger Institute COVID-19 Surveillance Team |
| EPI_ISL_3106621, EPI_ISL_3106622 | Lighthouse Lab in Milton Keynes | Wellcome Sanger Institute for the COVID-19 Genomics UK (COG-UK) Consortium | The Lighthouse Lab in Milton Keynes and Alex Alderton, Roberto Amato, Jeffrey Barrett, Sonia Goncalves, Ewan Harrison, David K. Jackson, Ian Johnston, Dominic Kwiatkowski, Cordelia Langford, John Sillitoe on behalf of the Wellcome Sanger Institute COVID-19 Surveillance Team |
| EPI_ISL_3106623 | Lighthouse Lab in Alderley Park | Wellcome Sanger Institute for the COVID-19 Genomics UK (COG-UK) Consortium | Jacquelyn Wynn, Mairead Hyland, The Lighthouse Lab in Alderley Park and Alex Alderton, Roberto Amato, Jeffrey Barrett, Sonia Goncalves, Ewan Harrison, David K. Jackson, Ian Johnston, Dominic Kwiatkowski, Cordelia Langford, John Sillitoe on behalf of the Wellcome Sanger Institute COVID-19 Surveillance Team |
| EPI_ISL_3106624, EPI_ISL_3106625 | Lighthouse Lab in Glasgow | Wellcome Sanger Institute for the COVID-19 Genomics UK (COG-UK) Consortium | Harper VanSteenhouse, Yumi Kasai, David Gray, Carol Clugston, Anna Dominiczak and Alex Alderton, Roberto Amato, Jeffrey Barrett, Sonia Goncalves, Ewan Harrison, David K. Jackson, Ian Johnston, Dominic Kwiatkowski, Cordelia Langford, John Sillitoe on behalf of the Wellcome Sanger Institute COVID-19 Surveillance Team |
| EPI_ISL_3106626 | Lighthouse Lab in Alderley Park | Wellcome Sanger Institute for the COVID-19 Genomics UK (COG-UK) Consortium | Jacquelyn Wynn, Mairead Hyland, The Lighthouse Lab in Alderley Park and Alex Alderton, Roberto Amato, Jeffrey Barrett, Sonia Goncalves, Ewan Harrison, David K. Jackson, Ian Johnston, Dominic Kwiatkowski, Cordelia Langford, John Sillitoe on behalf of the Wellcome Sanger Institute COVID-19 Surveillance Team |
| EPI_ISL_3106627 | Lighthouse Lab in Milton Keynes | Wellcome Sanger Institute for the COVID-19 Genomics UK (COG-UK) Consortium | The Lighthouse Lab in Milton Keynes and Alex Alderton, Roberto Amato, Jeffrey Barrett, Sonia Goncalves, Ewan Harrison, David K. Jackson, Ian Johnston, Dominic Kwiatkowski, Cordelia Langford, John Sillitoe on behalf of the Wellcome Sanger Institute COVID-19 Surveillance Team |
| EPI_ISL_3106628 | Lighthouse Lab in Alderley Park | Wellcome Sanger Institute for the COVID-19 Genomics UK (COG-UK) Consortium | Jacquelyn Wynn, Mairead Hyland, The Lighthouse Lab in Alderley Park and Alex Alderton, Roberto Amato, Jeffrey Barrett, Sonia Goncalves, Ewan Harrison, David K. Jackson, Ian Johnston, Dominic Kwiatkowski, Cordelia Langford, John Sillitoe on behalf of the Wellcome Sanger Institute COVID-19 Surveillance Team |
| EPI_ISL_3106629 | Lighthouse Lab in Milton Keynes | Wellcome Sanger Institute for the COVID-19 Genomics UK (COG-UK) Consortium | The Lighthouse Lab in Milton Keynes and Alex Alderton, Roberto Amato, Jeffrey Barrett, Sonia Goncalves, Ewan Harrison, David K. Jackson, Ian Johnston, Dominic Kwiatkowski, Cordelia Langford, John Sillitoe on behalf of the Wellcome Sanger Institute COVID-19 Surveillance Team |
| EPI_ISL_3106630, EPI_ISL_3106631 | Lighthouse Lab in Alderley Park | Wellcome Sanger Institute for the COVID-19 Genomics UK (COG-UK) Consortium | Jacquelyn Wynn, Mairead Hyland, The Lighthouse Lab in Alderley Park and Alex Alderton, Roberto Amato, Jeffrey Barrett, Sonia Goncalves, Ewan Harrison, David K. Jackson, Ian Johnston, Dominic Kwiatkowski, Cordelia Langford, John Sillitoe on behalf of the Wellcome Sanger Institute COVID-19 Surveillance Team |
| EPI_ISL_3106632 | Lighthouse Lab in Milton Keynes | Wellcome Sanger Institute for the COVID-19 Genomics UK (COG-UK) Consortium | The Lighthouse Lab in Milton Keynes and Alex Alderton, Roberto Amato, Jeffrey Barrett, Sonia Goncalves, Ewan Harrison, David K. Jackson, Ian Johnston, Dominic Kwiatkowski, Cordelia Langford, John Sillitoe on behalf of the Wellcome Sanger Institute COVID-19 Surveillance Team |
| EPI_ISL_3106633 | Lighthouse Lab in Glasgow | Wellcome Sanger Institute for the COVID-19 Genomics UK (COG-UK) Consortium | Harper VanSteenhouse, Yumi Kasai, David Gray, Carol Clugston, Anna Dominiczak and Alex Alderton, Roberto Amato, Jeffrey Barrett, Sonia Goncalves, Ewan Harrison, David K. Jackson, Ian Johnston, Dominic Kwiatkowski, Cordelia Langford, John Sillitoe on behalf of the Wellcome Sanger Institute COVID-19 Surveillance Team |
| EPI_ISL_3106634, EPI_ISL_3106635 | Lighthouse Lab in Milton Keynes | Wellcome Sanger Institute for the COVID-19 Genomics UK (COG-UK) Consortium | The Lighthouse Lab in Milton Keynes and Alex Alderton, Roberto Amato, Jeffrey Barrett, Sonia Goncalves, Ewan Harrison, David K. Jackson, Ian Johnston, Dominic Kwiatkowski, Cordelia Langford, John Sillitoe on behalf of the Wellcome Sanger Institute COVID-19 Surveillance Team |
| EPI_ISL_3106636 | Lighthouse Lab in Glasgow | Wellcome Sanger Institute for the COVID-19 Genomics UK (COG-UK) Consortium | Harper VanSteenhouse, Yumi Kasai, David Gray, Carol Clugston, Anna Dominiczak and Alex Alderton, Roberto Amato, Jeffrey Barrett, Sonia Goncalves, Ewan Harrison, David K. Jackson, Ian Johnston, Dominic Kwiatkowski, Cordelia Langford, John Sillitoe on behalf of the Wellcome Sanger Institute COVID-19 Surveillance Team |
| EPI_ISL_3106637 | Lighthouse Lab in Milton Keynes | Wellcome Sanger Institute for the COVID-19 Genomics UK (COG-UK) Consortium | The Lighthouse Lab in Milton Keynes and Alex Alderton, Roberto Amato, Jeffrey Barrett, Sonia Goncalves, Ewan Harrison, David K. Jackson, Ian Johnston, Dominic Kwiatkowski, Cordelia Langford, John Sillitoe on behalf of the Wellcome Sanger Institute COVID-19 Surveillance Team |
| EPI_ISL_3106638 | Lighthouse Lab in Glasgow | Wellcome Sanger Institute for the COVID-19 Genomics UK (COG-UK) Consortium | Harper VanSteenhouse, Yumi Kasai, David Gray, Carol Clugston, Anna Dominiczak and Alex Alderton, Roberto Amato, Jeffrey Barrett, Sonia Goncalves, Ewan Harrison, David K. Jackson, Ian Johnston, Dominic Kwiatkowski, Cordelia Langford, John Sillitoe on behalf of the Wellcome Sanger Institute COVID-19 Surveillance Team |
| EPI_ISL_3106639, EPI_ISL_3106640 | Lighthouse Lab in Milton Keynes | Wellcome Sanger Institute for the COVID-19 Genomics UK (COG-UK) Consortium | The Lighthouse Lab in Milton Keynes and Alex Alderton, Roberto Amato, Jeffrey Barrett, Sonia Goncalves, Ewan Harrison, David K. Jackson, Ian Johnston, Dominic Kwiatkowski, Cordelia Langford, John Sillitoe on behalf of the Wellcome Sanger Institute COVID-19 Surveillance Team |
| EPI_ISL_3106641 | Lighthouse Lab in Alderley Park | Wellcome Sanger Institute for the COVID-19 Genomics UK (COG-UK) Consortium | Jacquelyn Wynn, Mairead Hyland, The Lighthouse Lab in Alderley Park and Alex Alderton, Roberto Amato, Jeffrey Barrett, Sonia Goncalves, Ewan Harrison, David K. Jackson, Ian Johnston, Dominic Kwiatkowski, Cordelia Langford, John Sillitoe on behalf of the Wellcome Sanger Institute COVID-19 Surveillance Team |
| EPI_ISL_3106642, EPI_ISL_3106643 | Lighthouse Lab in Milton Keynes | Wellcome Sanger Institute for the COVID-19 Genomics UK (COG-UK) Consortium | The Lighthouse Lab in Milton Keynes and Alex Alderton, Roberto Amato, Jeffrey Barrett, Sonia Goncalves, Ewan Harrison, David K. Jackson, Ian Johnston, Dominic Kwiatkowski, Cordelia Langford, John Sillitoe on behalf of the Wellcome Sanger Institute COVID-19 Surveillance Team |
| EPI_ISL_3106644 | Lighthouse Lab in Alderley Park | Wellcome Sanger Institute for the COVID-19 Genomics UK (COG-UK) Consortium | Jacquelyn Wynn, Mairead Hyland, The Lighthouse Lab in Alderley Park and Alex Alderton, Roberto Amato, Jeffrey Barrett, Sonia Goncalves, Ewan Harrison, David K. Jackson, Ian Johnston, Dominic Kwiatkowski, Cordelia Langford, John Sillitoe on behalf of the Wellcome Sanger Institute COVID-19 Surveillance Team |
| EPI_ISL_3106645, EPI_ISL_3106646, EPI_ISL_3106647, EPI_ISL_3106648, EPI_ISL_3106649, EPI_ISL_3106650, EPI_ISL_3106651 | Lighthouse Lab in Milton Keynes | Wellcome Sanger Institute for the COVID-19 Genomics UK (COG-UK) Consortium | The Lighthouse Lab in Milton Keynes and Alex Alderton, Roberto Amato, Jeffrey Barrett, Sonia Goncalves, Ewan Harrison, David K. Jackson, Ian Johnston, Dominic Kwiatkowski, Cordelia Langford, John Sillitoe on behalf of the Wellcome Sanger Institute COVID-19 Surveillance Team |
| EPI_ISL_3106652 | Lighthouse Lab in Alderley Park | Wellcome Sanger Institute for the COVID-19 Genomics UK (COG-UK) Consortium | Jacquelyn Wynn, Mairead Hyland, The Lighthouse Lab in Alderley Park and Alex Alderton, Roberto Amato, Jeffrey Barrett, Sonia Goncalves, Ewan Harrison, David K. Jackson, Ian Johnston, Dominic Kwiatkowski, Cordelia Langford, John Sillitoe on behalf of the Wellcome Sanger Institute COVID-19 Surveillance Team |
| EPI_ISL_3106653 | Lighthouse Lab in Milton Keynes | Wellcome Sanger Institute for the COVID-19 Genomics UK (COG-UK) Consortium | The Lighthouse Lab in Milton Keynes and Alex Alderton, Roberto Amato, Jeffrey Barrett, Sonia Goncalves, Ewan Harrison, David K. Jackson, Ian Johnston, Dominic Kwiatkowski, Cordelia Langford, John Sillitoe on behalf of the Wellcome Sanger Institute COVID-19 Surveillance Team |
| EPI_ISL_3106654, EPI_ISL_3106655, EPI_ISL_3106656 | Lighthouse Lab in Alderley Park | Wellcome Sanger Institute for the COVID-19 Genomics UK (COG-UK) Consortium | Jacquelyn Wynn, Mairead Hyland, The Lighthouse Lab in Alderley Park and Alex Alderton, Roberto Amato, Jeffrey Barrett, Sonia Goncalves, Ewan Harrison, David K. Jackson, Ian Johnston, Dominic Kwiatkowski, Cordelia Langford, John Sillitoe on behalf of the Wellcome Sanger Institute COVID-19 Surveillance Team |
| EPI_ISL_3106657, EPI_ISL_3106658, EPI_ISL_3106659 | Lighthouse Lab in Milton Keynes | Wellcome Sanger Institute for the COVID-19 Genomics UK (COG-UK) Consortium | The Lighthouse Lab in Milton Keynes and Alex Alderton, Roberto Amato, Jeffrey Barrett, Sonia Goncalves, Ewan Harrison, David K. Jackson, Ian Johnston, Dominic Kwiatkowski, Cordelia Langford, John Sillitoe on behalf of the Wellcome Sanger Institute COVID-19 Surveillance Team |
| EPI_ISL_3106660, EPI_ISL_3106661 | Lighthouse Lab in Alderley Park | Wellcome Sanger Institute for the COVID-19 Genomics UK (COG-UK) Consortium | Jacquelyn Wynn, Mairead Hyland, The Lighthouse Lab in Alderley Park and Alex Alderton, Roberto Amato, Jeffrey Barrett, Sonia Goncalves, Ewan Harrison, David K. Jackson, Ian Johnston, Dominic Kwiatkowski, Cordelia Langford, John Sillitoe on behalf of the Wellcome Sanger Institute COVID-19 Surveillance Team |
| EPI_ISL_3106662, EPI_ISL_3106663, EPI_ISL_3106664, EPI_ISL_3106665 | Lighthouse Lab in Milton Keynes | Wellcome Sanger Institute for the COVID-19 Genomics UK (COG-UK) Consortium | The Lighthouse Lab in Milton Keynes and Alex Alderton, Roberto Amato, Jeffrey Barrett, Sonia Goncalves, Ewan Harrison, David K. Jackson, Ian Johnston, Dominic Kwiatkowski, Cordelia Langford, John Sillitoe on behalf of the Wellcome Sanger Institute COVID-19 Surveillance Team |
| EPI_ISL_3106666, EPI_ISL_3106667 | Lighthouse Lab in Alderley Park | Wellcome Sanger Institute for the COVID-19 Genomics UK (COG-UK) Consortium | Jacquelyn Wynn, Mairead Hyland, The Lighthouse Lab in Alderley Park and Alex Alderton, Roberto Amato, Jeffrey Barrett, Sonia Goncalves, Ewan Harrison, David K. Jackson, Ian Johnston, Dominic Kwiatkowski, Cordelia Langford, John Sillitoe on behalf of the Wellcome Sanger Institute COVID-19 Surveillance Team |
| EPI_ISL_3106668 | Lighthouse Lab in Glasgow | Wellcome Sanger Institute for the COVID-19 Genomics UK (COG-UK) Consortium | Harper VanSteenhouse, Yumi Kasai, David Gray, Carol Clugston, Anna Dominiczak and Alex Alderton, Roberto Amato, Jeffrey Barrett, Sonia Goncalves, Ewan Harrison, David K. Jackson, Ian Johnston, Dominic Kwiatkowski, Cordelia Langford, John Sillitoe on behalf of the Wellcome Sanger Institute COVID-19 Surveillance Team |
| EPI_ISL_3106669, EPI_ISL_3106670, EPI_ISL_3106671, EPI_ISL_3106672 | Lighthouse Lab in Milton Keynes | Wellcome Sanger Institute for the COVID-19 Genomics UK (COG-UK) Consortium | The Lighthouse Lab in Milton Keynes and Alex Alderton, Roberto Amato, Jeffrey Barrett, Sonia Goncalves, Ewan Harrison, David K. Jackson, Ian Johnston, Dominic Kwiatkowski, Cordelia Langford, John Sillitoe on behalf of the Wellcome Sanger Institute COVID-19 Surveillance Team |
| EPI_ISL_3106673, EPI_ISL_3106674 | Lighthouse Lab in Alderley Park | Wellcome Sanger Institute for the COVID-19 Genomics UK (COG-UK) Consortium | Jacquelyn Wynn, Mairead Hyland, The Lighthouse Lab in Alderley Park and Alex Alderton, Roberto Amato, Jeffrey Barrett, Sonia Goncalves, Ewan Harrison, David K. Jackson, Ian Johnston, Dominic Kwiatkowski, Cordelia Langford, John Sillitoe on behalf of the Wellcome Sanger Institute COVID-19 Surveillance Team |
| EPI_ISL_3106675 | Lighthouse Lab in Milton Keynes | Wellcome Sanger Institute for the COVID-19 Genomics UK (COG-UK) Consortium | The Lighthouse Lab in Milton Keynes and Alex Alderton, Roberto Amato, Jeffrey Barrett, Sonia Goncalves, Ewan Harrison, David K. Jackson, Ian Johnston, Dominic Kwiatkowski, Cordelia Langford, John Sillitoe on behalf of the Wellcome Sanger Institute COVID-19 Surveillance Team |
| EPI_ISL_3106676, EPI_ISL_3106677, EPI_ISL_3106678, | Lighthouse Lab in Alderley Park | Wellcome Sanger Institute for the COVID-19 Genomics UK (COG-UK) Consortium | Jacquelyn Wynn, Mairead Hyland, The Lighthouse Lab in Alderley Park and Alex Alderton, Roberto Amato, Jeffrey Barrett, Sonia Goncalves, Ewan Harrison, David K. Jackson, Ian Johnston, Dominic Kwiatkowski, Cordelia Langford, John Sillitoe on behalf of the Wellcome Sanger Institute COVID-19 Surveillance Team |

| | | | |
|---|---|---|---|
| EPI_ISL_3106679, EPI_ISL_3106680 | | | |
| EPI_ISL_3106681 | Lighthouse Lab in Milton Keynes | Wellcome Sanger Institute for the COVID-19 Genomics UK (COG-UK) Consortium | The Lighthouse Lab in Milton Keynes and Alex Alderton, Roberto Amato, Jeffrey Barrett, Sonia Goncalves, Ewan Harrison, David K. Jackson, Ian Johnston, Dominic Kwiatkowski, Cordelia Langford, John Sillitoe on behalf of the Wellcome Sanger Institute COVID-19 Surveillance Team |
| EPI_ISL_3106682 | Lighthouse Lab in Glasgow | Wellcome Sanger Institute for the COVID-19 Genomics UK (COG-UK) Consortium | Harper VanSteenhouse, Yumi Kasai, David Gray, Carol Clugston, Anna Dominiczak and Alex Alderton, Roberto Amato, Jeffrey Barrett, Sonia Goncalves, Ewan Harrison, David K. Jackson, Ian Johnston, Dominic Kwiatkowski, Cordelia Langford, John Sillitoe on behalf of the Wellcome Sanger Institute COVID-19 Surveillance Team |
| EPI_ISL_3106683, EPI_ISL_3106684 | Lighthouse Lab in Milton Keynes | Wellcome Sanger Institute for the COVID-19 Genomics UK (COG-UK) Consortium | The Lighthouse Lab in Milton Keynes and Alex Alderton, Roberto Amato, Jeffrey Barrett, Sonia Goncalves, Ewan Harrison, David K. Jackson, Ian Johnston, Dominic Kwiatkowski, Cordelia Langford, John Sillitoe on behalf of the Wellcome Sanger Institute COVID-19 Surveillance Team |
| EPI_ISL_3106685 | Lighthouse Lab in Alderley Park | Wellcome Sanger Institute for the COVID-19 Genomics UK (COG-UK) Consortium | Jacquelyn Wynn, Mairead Hyland, The Lighthouse Lab in Alderley Park and Alex Alderton, Roberto Amato, Jeffrey Barrett, Sonia Goncalves, Ewan Harrison, David K. Jackson, Ian Johnston, Dominic Kwiatkowski, Cordelia Langford, John Sillitoe on behalf of the Wellcome Sanger Institute COVID-19 Surveillance Team |
| EPI_ISL_3106686 | Lighthouse Lab in Milton Keynes | Wellcome Sanger Institute for the COVID-19 Genomics UK (COG-UK) Consortium | The Lighthouse Lab in Milton Keynes and Alex Alderton, Roberto Amato, Jeffrey Barrett, Sonia Goncalves, Ewan Harrison, David K. Jackson, Ian Johnston, Dominic Kwiatkowski, Cordelia Langford, John Sillitoe on behalf of the Wellcome Sanger Institute COVID-19 Surveillance Team |
| EPI_ISL_3106687 | Lighthouse Lab in Glasgow | Wellcome Sanger Institute for the COVID-19 Genomics UK (COG-UK) Consortium | Harper VanSteenhouse, Yumi Kasai, David Gray, Carol Clugston, Anna Dominiczak and Alex Alderton, Roberto Amato, Jeffrey Barrett, Sonia Goncalves, Ewan Harrison, David K. Jackson, Ian Johnston, Dominic Kwiatkowski, Cordelia Langford, John Sillitoe on behalf of the Wellcome Sanger Institute COVID-19 Surveillance Team |
| EPI_ISL_3106688, EPI_ISL_3106689 | Lighthouse Lab in Milton Keynes | Wellcome Sanger Institute for the COVID-19 Genomics UK (COG-UK) Consortium | The Lighthouse Lab in Milton Keynes and Alex Alderton, Roberto Amato, Jeffrey Barrett, Sonia Goncalves, Ewan Harrison, David K. Jackson, Ian Johnston, Dominic Kwiatkowski, Cordelia Langford, John Sillitoe on behalf of the Wellcome Sanger Institute COVID-19 Surveillance Team |
| EPI_ISL_3106690 | Lighthouse Lab in Alderley Park | Wellcome Sanger Institute for the COVID-19 Genomics UK (COG-UK) Consortium | Jacquelyn Wynn, Mairead Hyland, The Lighthouse Lab in Alderley Park and Alex Alderton, Roberto Amato, Jeffrey Barrett, Sonia Goncalves, Ewan Harrison, David K. Jackson, Ian Johnston, Dominic Kwiatkowski, Cordelia Langford, John Sillitoe on behalf of the Wellcome Sanger Institute COVID-19 Surveillance Team |
| EPI_ISL_3106691 | Lighthouse Lab in Milton Keynes | Wellcome Sanger Institute for the COVID-19 Genomics UK (COG-UK) Consortium | The Lighthouse Lab in Milton Keynes and Alex Alderton, Roberto Amato, Jeffrey Barrett, Sonia Goncalves, Ewan Harrison, David K. Jackson, Ian Johnston, Dominic Kwiatkowski, Cordelia Langford, John Sillitoe on behalf of the Wellcome Sanger Institute COVID-19 Surveillance Team |
| EPI_ISL_3106692 | Lighthouse Lab in Alderley Park | Wellcome Sanger Institute for the COVID-19 Genomics UK (COG-UK) Consortium | Jacquelyn Wynn, Mairead Hyland, The Lighthouse Lab in Alderley Park and Alex Alderton, Roberto Amato, Jeffrey Barrett, Sonia Goncalves, Ewan Harrison, David K. Jackson, Ian Johnston, Dominic Kwiatkowski, Cordelia Langford, John Sillitoe on behalf of the Wellcome Sanger Institute COVID-19 Surveillance Team |
| EPI_ISL_3106693, EPI_ISL_3106694 | Lighthouse Lab in Milton Keynes | Wellcome Sanger Institute for the COVID-19 Genomics UK (COG-UK) Consortium | The Lighthouse Lab in Milton Keynes and Alex Alderton, Roberto Amato, Jeffrey Barrett, Sonia Goncalves, Ewan Harrison, David K. Jackson, Ian Johnston, Dominic Kwiatkowski, Cordelia Langford, John Sillitoe on behalf of the Wellcome Sanger Institute COVID-19 Surveillance Team |
| EPI_ISL_3106695, EPI_ISL_3106696, EPI_ISL_3106697, EPI_ISL_3106698 | Lighthouse Lab in Glasgow | Wellcome Sanger Institute for the COVID-19 Genomics UK (COG-UK) Consortium | Harper VanSteenhouse, Yumi Kasai, David Gray, Carol Clugston, Anna Dominiczak and Alex Alderton, Roberto Amato, Jeffrey Barrett, Sonia Goncalves, Ewan Harrison, David K. Jackson, Ian Johnston, Dominic Kwiatkowski, Cordelia Langford, John Sillitoe on behalf of the Wellcome Sanger Institute COVID-19 Surveillance Team |
| EPI_ISL_3106699 | Lighthouse Lab in Milton Keynes | Wellcome Sanger Institute for the COVID-19 Genomics UK (COG-UK) Consortium | The Lighthouse Lab in Milton Keynes and Alex Alderton, Roberto Amato, Jeffrey Barrett, Sonia Goncalves, Ewan Harrison, David K. Jackson, Ian Johnston, Dominic Kwiatkowski, Cordelia Langford, John Sillitoe on behalf of the Wellcome Sanger Institute COVID-19 Surveillance Team |
| EPI_ISL_3106700 | Lighthouse Lab in Alderley Park | Wellcome Sanger Institute for the COVID-19 Genomics UK (COG-UK) Consortium | Jacquelyn Wynn, Mairead Hyland, The Lighthouse Lab in Alderley Park and Alex Alderton, Roberto Amato, Jeffrey Barrett, Sonia Goncalves, Ewan Harrison, David K. Jackson, Ian Johnston, Dominic Kwiatkowski, Cordelia Langford, John Sillitoe on behalf of the Wellcome Sanger Institute COVID-19 Surveillance Team |
| EPI_ISL_3106701, EPI_ISL_3106702, EPI_ISL_3106703, EPI_ISL_3106704 | Lighthouse Lab in Milton Keynes | Wellcome Sanger Institute for the COVID-19 Genomics UK (COG-UK) Consortium | The Lighthouse Lab in Milton Keynes and Alex Alderton, Roberto Amato, Jeffrey Barrett, Sonia Goncalves, Ewan Harrison, David K. Jackson, Ian Johnston, Dominic Kwiatkowski, Cordelia Langford, John Sillitoe on behalf of the Wellcome Sanger Institute COVID-19 Surveillance Team |
| EPI_ISL_3106705 | Lighthouse Lab in Alderley Park | Wellcome Sanger Institute for the COVID-19 Genomics UK (COG-UK) Consortium | Jacquelyn Wynn, Mairead Hyland, The Lighthouse Lab in Alderley Park and Alex Alderton, Roberto Amato, Jeffrey Barrett, Sonia Goncalves, Ewan Harrison, David K. Jackson, Ian Johnston, Dominic Kwiatkowski, Cordelia Langford, John Sillitoe on behalf of the Wellcome Sanger Institute COVID-19 Surveillance Team |
| EPI_ISL_3106706, EPI_ISL_3106707 | Lighthouse Lab in Milton Keynes | Wellcome Sanger Institute for the COVID-19 Genomics UK (COG-UK) Consortium | The Lighthouse Lab in Milton Keynes and Alex Alderton, Roberto Amato, Jeffrey Barrett, Sonia Goncalves, Ewan Harrison, David K. Jackson, Ian Johnston, Dominic Kwiatkowski, Cordelia Langford, John Sillitoe on behalf of the Wellcome Sanger Institute COVID-19 Surveillance Team |
| EPI_ISL_3106708 | Lighthouse Lab in Alderley Park | Wellcome Sanger Institute for the COVID-19 Genomics UK (COG-UK) Consortium | Jacquelyn Wynn, Mairead Hyland, The Lighthouse Lab in Alderley Park and Alex Alderton, Roberto Amato, Jeffrey Barrett, Sonia Goncalves, Ewan Harrison, David K. Jackson, Ian Johnston, Dominic Kwiatkowski, Cordelia Langford, John Sillitoe on behalf of the Wellcome Sanger Institute COVID-19 Surveillance Team |
| EPI_ISL_3106709, EPI_ISL_3106710, EPI_ISL_3106711, EPI_ISL_3106712, EPI_ISL_3106713, EPI_ISL_3106714 | Lighthouse Lab in Milton Keynes | Wellcome Sanger Institute for the COVID-19 Genomics UK (COG-UK) Consortium | The Lighthouse Lab in Milton Keynes and Alex Alderton, Roberto Amato, Jeffrey Barrett, Sonia Goncalves, Ewan Harrison, David K. Jackson, Ian Johnston, Dominic Kwiatkowski, Cordelia Langford, John Sillitoe on behalf of the Wellcome Sanger Institute COVID-19 Surveillance Team |
| EPI_ISL_3106715 | Lighthouse Lab in Glasgow | Wellcome Sanger Institute for the COVID-19 Genomics UK (COG-UK) Consortium | Harper VanSteenhouse, Yumi Kasai, David Gray, Carol Clugston, Anna Dominiczak and Alex Alderton, Roberto Amato, Jeffrey Barrett, Sonia Goncalves, Ewan Harrison, David K. Jackson, Ian Johnston, Dominic Kwiatkowski, Cordelia Langford, John Sillitoe on behalf of the Wellcome Sanger Institute COVID-19 Surveillance Team |
| EPI_ISL_3106716 | Lighthouse Lab in Milton Keynes | Wellcome Sanger Institute for the COVID-19 Genomics UK (COG-UK) Consortium | The Lighthouse Lab in Milton Keynes and Alex Alderton, Roberto Amato, Jeffrey Barrett, Sonia Goncalves, Ewan Harrison, David K. Jackson, Ian Johnston, Dominic Kwiatkowski, Cordelia Langford, John Sillitoe on behalf of the Wellcome Sanger Institute COVID-19 Surveillance Team |
| EPI_ISL_3106717 | Lighthouse Lab in Alderley Park | Wellcome Sanger Institute for the COVID-19 Genomics UK (COG-UK) Consortium | Jacquelyn Wynn, Mairead Hyland, The Lighthouse Lab in Alderley Park and Alex Alderton, Roberto Amato, Jeffrey Barrett, Sonia Goncalves, Ewan Harrison, David K. Jackson, Ian Johnston, Dominic Kwiatkowski, Cordelia Langford, John Sillitoe on behalf of the Wellcome Sanger Institute COVID-19 Surveillance Team |
| EPI_ISL_3106718 | Lighthouse Lab in Glasgow | Wellcome Sanger Institute for the COVID-19 Genomics UK (COG-UK) Consortium | Harper VanSteenhouse, Yumi Kasai, David Gray, Carol Clugston, Anna Dominiczak and Alex Alderton, Roberto Amato, Jeffrey Barrett, Sonia Goncalves, Ewan Harrison, David K. Jackson, Ian Johnston, Dominic Kwiatkowski, Cordelia Langford, John Sillitoe on behalf of the Wellcome Sanger Institute COVID-19 Surveillance Team |
| EPI_ISL_3106719 | Lighthouse Lab in Milton Keynes | Wellcome Sanger Institute for the COVID-19 Genomics UK (COG-UK) Consortium | The Lighthouse Lab in Milton Keynes and Alex Alderton, Roberto Amato, Jeffrey Barrett, Sonia Goncalves, Ewan Harrison, David K. Jackson, Ian Johnston, Dominic Kwiatkowski, Cordelia Langford, John Sillitoe on behalf of the Wellcome Sanger Institute COVID-19 Surveillance Team |
| EPI_ISL_3106720 | Lighthouse Lab in Alderley Park | Wellcome Sanger Institute for the COVID-19 Genomics UK (COG-UK) Consortium | Jacquelyn Wynn, Mairead Hyland, The Lighthouse Lab in Alderley Park and Alex Alderton, Roberto Amato, Jeffrey Barrett, Sonia Goncalves, Ewan Harrison, David K. Jackson, Ian Johnston, Dominic Kwiatkowski, Cordelia Langford, John Sillitoe on behalf of the Wellcome Sanger Institute COVID-19 Surveillance Team |
| EPI_ISL_3106721 | Lighthouse Lab in Glasgow | Wellcome Sanger Institute for the COVID-19 Genomics UK (COG-UK) Consortium | Harper VanSteenhouse, Yumi Kasai, David Gray, Carol Clugston, Anna Dominiczak and Alex Alderton, Roberto Amato, Jeffrey Barrett, Sonia Goncalves, Ewan Harrison, David K. Jackson, Ian Johnston, Dominic Kwiatkowski, Cordelia Langford, John Sillitoe on behalf of the Wellcome Sanger Institute COVID-19 Surveillance Team |
| EPI_ISL_3106722 | Lighthouse Lab in Milton Keynes | Wellcome Sanger Institute for the COVID-19 Genomics UK (COG-UK) Consortium | The Lighthouse Lab in Milton Keynes and Alex Alderton, Roberto Amato, Jeffrey Barrett, Sonia Goncalves, Ewan Harrison, David K. Jackson, Ian Johnston, Dominic Kwiatkowski, Cordelia Langford, John Sillitoe on behalf of the Wellcome Sanger Institute COVID-19 Surveillance Team |
| EPI_ISL_3106723, EPI_ISL_3106724 | Lighthouse Lab in Glasgow | Wellcome Sanger Institute for the COVID-19 Genomics UK (COG-UK) Consortium | Harper VanSteenhouse, Yumi Kasai, David Gray, Carol Clugston, Anna Dominiczak and Alex Alderton, Roberto Amato, Jeffrey Barrett, Sonia Goncalves, Ewan Harrison, David K. Jackson, Ian Johnston, Dominic Kwiatkowski, Cordelia Langford, John Sillitoe on behalf of the Wellcome Sanger Institute COVID-19 Surveillance Team |
| EPI_ISL_3106725, EPI_ISL_3106726 | Lighthouse Lab in Milton Keynes | Wellcome Sanger Institute for the COVID-19 Genomics UK (COG-UK) Consortium | The Lighthouse Lab in Milton Keynes and Alex Alderton, Roberto Amato, Jeffrey Barrett, Sonia Goncalves, Ewan Harrison, David K. Jackson, Ian Johnston, Dominic Kwiatkowski, Cordelia Langford, John Sillitoe on behalf of the Wellcome Sanger Institute COVID-19 Surveillance Team |
| EPI_ISL_3106727 | Lighthouse Lab in Glasgow | Wellcome Sanger Institute for the COVID-19 Genomics UK (COG-UK) Consortium | Harper VanSteenhouse, Yumi Kasai, David Gray, Carol Clugston, Anna Dominiczak and Alex Alderton, Roberto Amato, Jeffrey Barrett, Sonia Goncalves, Ewan Harrison, David K. Jackson, Ian Johnston, Dominic Kwiatkowski, Cordelia Langford, John Sillitoe on behalf of the Wellcome Sanger Institute COVID-19 Surveillance Team |
| EPI_ISL_3106728 | Lighthouse Lab in Alderley Park | Wellcome Sanger Institute for the COVID-19 Genomics UK (COG-UK) Consortium | Jacquelyn Wynn, Mairead Hyland, The Lighthouse Lab in Alderley Park and Alex Alderton, Roberto Amato, Jeffrey Barrett, Sonia Goncalves, Ewan Harrison, David K. Jackson, Ian Johnston, Dominic Kwiatkowski, Cordelia Langford, John Sillitoe on behalf of the Wellcome Sanger Institute COVID-19 Surveillance Team |
| EPI_ISL_3106729, EPI_ISL_3106730, EPI_ISL_3106731 | Lighthouse Lab in Milton Keynes | Wellcome Sanger Institute for the COVID-19 Genomics UK (COG-UK) Consortium | The Lighthouse Lab in Milton Keynes and Alex Alderton, Roberto Amato, Jeffrey Barrett, Sonia Goncalves, Ewan Harrison, David K. Jackson, Ian Johnston, Dominic Kwiatkowski, Cordelia Langford, John Sillitoe on behalf of the Wellcome Sanger Institute COVID-19 Surveillance Team |
| EPI_ISL_3106732, EPI_ISL_3106733, EPI_ISL_3106734 | Lighthouse Lab in Alderley Park | Wellcome Sanger Institute for the COVID-19 Genomics UK (COG-UK) Consortium | Jacquelyn Wynn, Mairead Hyland, The Lighthouse Lab in Alderley Park and Alex Alderton, Roberto Amato, Jeffrey Barrett, Sonia Goncalves, Ewan Harrison, David K. Jackson, Ian Johnston, Dominic Kwiatkowski, Cordelia Langford, John Sillitoe on behalf of the Wellcome Sanger Institute COVID-19 Surveillance Team |
| EPI_ISL_3106735 | Lighthouse Lab in Milton Keynes | Wellcome Sanger Institute for the COVID-19 Genomics UK (COG-UK) Consortium | The Lighthouse Lab in Milton Keynes and Alex Alderton, Roberto Amato, Jeffrey Barrett, Sonia Goncalves, Ewan Harrison, David K. Jackson, Ian Johnston, Dominic Kwiatkowski, Cordelia Langford, John Sillitoe on behalf of the Wellcome Sanger Institute COVID-19 Surveillance Team |
| EPI_ISL_3106736 | Lighthouse Lab in Alderley Park | Wellcome Sanger Institute for the COVID-19 Genomics UK (COG-UK) Consortium | Jacquelyn Wynn, Mairead Hyland, The Lighthouse Lab in Alderley Park and Alex Alderton, Roberto Amato, Jeffrey Barrett, Sonia Goncalves, Ewan Harrison, David K. Jackson, Ian Johnston, Dominic Kwiatkowski, Cordelia Langford, John Sillitoe on behalf of the Wellcome Sanger Institute COVID-19 Surveillance Team |
| EPI_ISL_3106737, EPI_ISL_3106738, EPI_ISL_3106739 | Lighthouse Lab in Milton Keynes | Wellcome Sanger Institute for the COVID-19 Genomics UK (COG-UK) Consortium | The Lighthouse Lab in Milton Keynes and Alex Alderton, Roberto Amato, Jeffrey Barrett, Sonia Goncalves, Ewan Harrison, David K. Jackson, Ian Johnston, Dominic Kwiatkowski, Cordelia Langford, John Sillitoe on behalf of the Wellcome Sanger Institute COVID-19 Surveillance Team |
| EPI_ISL_3106740 | Lighthouse Lab in Glasgow | Wellcome Sanger Institute for the COVID-19 Genomics UK (COG-UK) Consortium | Harper VanSteenhouse, Yumi Kasai, David Gray, Carol Clugston, Anna Dominiczak and Alex Alderton, Roberto Amato, Jeffrey Barrett, Sonia Goncalves, Ewan Harrison, David K. Jackson, Ian Johnston, Dominic Kwiatkowski, Cordelia Langford, John Sillitoe on behalf of the Wellcome Sanger Institute COVID-19 Surveillance Team |
| EPI_ISL_3106741, EPI_ISL_3106742, EPI_ISL_3106743, EPI_ISL_3106744 | Lighthouse Lab in Milton Keynes | Wellcome Sanger Institute for the COVID-19 Genomics UK (COG-UK) Consortium | The Lighthouse Lab in Milton Keynes and Alex Alderton, Roberto Amato, Jeffrey Barrett, Sonia Goncalves, Ewan Harrison, David K. Jackson, Ian Johnston, Dominic Kwiatkowski, Cordelia Langford, John Sillitoe on behalf of the Wellcome Sanger Institute COVID-19 Surveillance Team |
| EPI_ISL_3106745 | Lighthouse Lab in Alderley Park | Wellcome Sanger Institute for the COVID-19 Genomics UK (COG-UK) Consortium | Jacquelyn Wynn, Mairead Hyland, The Lighthouse Lab in Alderley Park and Alex Alderton, Roberto Amato, Jeffrey Barrett, Sonia Goncalves, Ewan Harrison, David K. Jackson, Ian Johnston, Dominic Kwiatkowski, Cordelia Langford, John Sillitoe on behalf of the Wellcome Sanger Institute COVID-19 Surveillance Team |

| | | | |
|---|---|---|---|
| EPI_ISL_3106746 | Lighthouse Lab in Milton Keynes | Wellcome Sanger Institute for the COVID-19 Genomics UK (COG-UK) Consortium | The Lighthouse Lab in Milton Keynes and Alex Alderton, Roberto Amato, Jeffrey Barrett, Sonia Goncalves, Ewan Harrison, David K. Jackson, Ian Johnston, Dominic Kwiatkowski, Cordelia Langford, John Sillitoe on behalf of the Wellcome Sanger Institute COVID-19 Surveillance Team |
| EPI_ISL_3106747 | Lighthouse Lab in Glasgow | Wellcome Sanger Institute for the COVID-19 Genomics UK (COG-UK) Consortium | Harper VanSteenhouse, Yumi Kasai, David Gray, Carol Clugston, Anna Dominiczak and Alex Alderton, Roberto Amato, Jeffrey Barrett, Sonia Goncalves, Ewan Harrison, David K. Jackson, Ian Johnston, Dominic Kwiatkowski, Cordelia Langford, John Sillitoe on behalf of the Wellcome Sanger Institute COVID-19 Surveillance Team |
| EPI_ISL_3106748 | Lighthouse Lab in Alderley Park | Wellcome Sanger Institute for the COVID-19 Genomics UK (COG-UK) Consortium | Jacquelyn Wynn, Mairead Hyland, The Lighthouse Lab in Alderley Park and Alex Alderton, Roberto Amato, Jeffrey Barrett, Sonia Goncalves, Ewan Harrison, David K. Jackson, Ian Johnston, Dominic Kwiatkowski, Cordelia Langford, John Sillitoe on behalf of the Wellcome Sanger Institute COVID-19 Surveillance Team |
| EPI_ISL_3106749 | Lighthouse Lab in Glasgow | Wellcome Sanger Institute for the COVID-19 Genomics UK (COG-UK) Consortium | Harper VanSteenhouse, Yumi Kasai, David Gray, Carol Clugston, Anna Dominiczak and Alex Alderton, Roberto Amato, Jeffrey Barrett, Sonia Goncalves, Ewan Harrison, David K. Jackson, Ian Johnston, Dominic Kwiatkowski, Cordelia Langford, John Sillitoe on behalf of the Wellcome Sanger Institute COVID-19 Surveillance Team |
| EPI_ISL_3106750 | Lighthouse Lab in Milton Keynes | Wellcome Sanger Institute for the COVID-19 Genomics UK (COG-UK) Consortium | The Lighthouse Lab in Milton Keynes and Alex Alderton, Roberto Amato, Jeffrey Barrett, Sonia Goncalves, Ewan Harrison, David K. Jackson, Ian Johnston, Dominic Kwiatkowski, Cordelia Langford, John Sillitoe on behalf of the Wellcome Sanger Institute COVID-19 Surveillance Team |
| EPI_ISL_3106751 | Lighthouse Lab in Glasgow | Wellcome Sanger Institute for the COVID-19 Genomics UK (COG-UK) Consortium | Harper VanSteenhouse, Yumi Kasai, David Gray, Carol Clugston, Anna Dominiczak and Alex Alderton, Roberto Amato, Jeffrey Barrett, Sonia Goncalves, Ewan Harrison, David K. Jackson, Ian Johnston, Dominic Kwiatkowski, Cordelia Langford, John Sillitoe on behalf of the Wellcome Sanger Institute COVID-19 Surveillance Team |
| EPI_ISL_3106752 | Lighthouse Lab in Alderley Park | Wellcome Sanger Institute for the COVID-19 Genomics UK (COG-UK) Consortium | Jacquelyn Wynn, Mairead Hyland, The Lighthouse Lab in Alderley Park and Alex Alderton, Roberto Amato, Jeffrey Barrett, Sonia Goncalves, Ewan Harrison, David K. Jackson, Ian Johnston, Dominic Kwiatkowski, Cordelia Langford, John Sillitoe on behalf of the Wellcome Sanger Institute COVID-19 Surveillance Team |
| EPI_ISL_3106753 | Lighthouse Lab in Glasgow | Wellcome Sanger Institute for the COVID-19 Genomics UK (COG-UK) Consortium | Harper VanSteenhouse, Yumi Kasai, David Gray, Carol Clugston, Anna Dominiczak and Alex Alderton, Roberto Amato, Jeffrey Barrett, Sonia Goncalves, Ewan Harrison, David K. Jackson, Ian Johnston, Dominic Kwiatkowski, Cordelia Langford, John Sillitoe on behalf of the Wellcome Sanger Institute COVID-19 Surveillance Team |
| EPI_ISL_3106754 | Lighthouse Lab in Milton Keynes | Wellcome Sanger Institute for the COVID-19 Genomics UK (COG-UK) Consortium | The Lighthouse Lab in Milton Keynes and Alex Alderton, Roberto Amato, Jeffrey Barrett, Sonia Goncalves, Ewan Harrison, David K. Jackson, Ian Johnston, Dominic Kwiatkowski, Cordelia Langford, John Sillitoe on behalf of the Wellcome Sanger Institute COVID-19 Surveillance Team |
| EPI_ISL_3106755 | Lighthouse Lab in Glasgow | Wellcome Sanger Institute for the COVID-19 Genomics UK (COG-UK) Consortium | Harper VanSteenhouse, Yumi Kasai, David Gray, Carol Clugston, Anna Dominiczak and Alex Alderton, Roberto Amato, Jeffrey Barrett, Sonia Goncalves, Ewan Harrison, David K. Jackson, Ian Johnston, Dominic Kwiatkowski, Cordelia Langford, John Sillitoe on behalf of the Wellcome Sanger Institute COVID-19 Surveillance Team |
| EPI_ISL_3106756 | Lighthouse Lab in Milton Keynes | Wellcome Sanger Institute for the COVID-19 Genomics UK (COG-UK) Consortium | The Lighthouse Lab in Milton Keynes and Alex Alderton, Roberto Amato, Jeffrey Barrett, Sonia Goncalves, Ewan Harrison, David K. Jackson, Ian Johnston, Dominic Kwiatkowski, Cordelia Langford, John Sillitoe on behalf of the Wellcome Sanger Institute COVID-19 Surveillance Team |
| EPI_ISL_3106757, EPI_ISL_3106758, EPI_ISL_3106759 | Lighthouse Lab in Alderley Park | Wellcome Sanger Institute for the COVID-19 Genomics UK (COG-UK) Consortium | Jacquelyn Wynn, Mairead Hyland, The Lighthouse Lab in Alderley Park and Alex Alderton, Roberto Amato, Jeffrey Barrett, Sonia Goncalves, Ewan Harrison, David K. Jackson, Ian Johnston, Dominic Kwiatkowski, Cordelia Langford, John Sillitoe on behalf of the Wellcome Sanger Institute COVID-19 Surveillance Team |
| EPI_ISL_3106760 | Lighthouse Lab in Glasgow | Wellcome Sanger Institute for the COVID-19 Genomics UK (COG-UK) Consortium | Harper VanSteenhouse, Yumi Kasai, David Gray, Carol Clugston, Anna Dominiczak and Alex Alderton, Roberto Amato, Jeffrey Barrett, Sonia Goncalves, Ewan Harrison, David K. Jackson, Ian Johnston, Dominic Kwiatkowski, Cordelia Langford, John Sillitoe on behalf of the Wellcome Sanger Institute COVID-19 Surveillance Team |
| EPI_ISL_3106761 | Lighthouse Lab in Milton Keynes | Wellcome Sanger Institute for the COVID-19 Genomics UK (COG-UK) Consortium | The Lighthouse Lab in Milton Keynes and Alex Alderton, Roberto Amato, Jeffrey Barrett, Sonia Goncalves, Ewan Harrison, David K. Jackson, Ian Johnston, Dominic Kwiatkowski, Cordelia Langford, John Sillitoe on behalf of the Wellcome Sanger Institute COVID-19 Surveillance Team |
| EPI_ISL_3106762 | Lighthouse Lab in Glasgow | Wellcome Sanger Institute for the COVID-19 Genomics UK (COG-UK) Consortium | Harper VanSteenhouse, Yumi Kasai, David Gray, Carol Clugston, Anna Dominiczak and Alex Alderton, Roberto Amato, Jeffrey Barrett, Sonia Goncalves, Ewan Harrison, David K. Jackson, Ian Johnston, Dominic Kwiatkowski, Cordelia Langford, John Sillitoe on behalf of the Wellcome Sanger Institute COVID-19 Surveillance Team |
| EPI_ISL_3106763, EPI_ISL_3106764 | Lighthouse Lab in Milton Keynes | Wellcome Sanger Institute for the COVID-19 Genomics UK (COG-UK) Consortium | The Lighthouse Lab in Milton Keynes and Alex Alderton, Roberto Amato, Jeffrey Barrett, Sonia Goncalves, Ewan Harrison, David K. Jackson, Ian Johnston, Dominic Kwiatkowski, Cordelia Langford, John Sillitoe on behalf of the Wellcome Sanger Institute COVID-19 Surveillance Team |
| EPI_ISL_3106765, EPI_ISL_3106766, EPI_ISL_3106767 | Lighthouse Lab in Glasgow | Wellcome Sanger Institute for the COVID-19 Genomics UK (COG-UK) Consortium | Harper VanSteenhouse, Yumi Kasai, David Gray, Carol Clugston, Anna Dominiczak and Alex Alderton, Roberto Amato, Jeffrey Barrett, Sonia Goncalves, Ewan Harrison, David K. Jackson, Ian Johnston, Dominic Kwiatkowski, Cordelia Langford, John Sillitoe on behalf of the Wellcome Sanger Institute COVID-19 Surveillance Team |
| EPI_ISL_3106768, EPI_ISL_3106769, EPI_ISL_3106770, EPI_ISL_3106771, EPI_ISL_3106772, EPI_ISL_3106773 | Lighthouse Lab in Alderley Park | Wellcome Sanger Institute for the COVID-19 Genomics UK (COG-UK) Consortium | Jacquelyn Wynn, Mairead Hyland, The Lighthouse Lab in Alderley Park and Alex Alderton, Roberto Amato, Jeffrey Barrett, Sonia Goncalves, Ewan Harrison, David K. Jackson, Ian Johnston, Dominic Kwiatkowski, Cordelia Langford, John Sillitoe on behalf of the Wellcome Sanger Institute COVID-19 Surveillance Team |
| EPI_ISL_3106774 | Lighthouse Lab in Glasgow | Wellcome Sanger Institute for the COVID-19 Genomics UK (COG-UK) Consortium | Harper VanSteenhouse, Yumi Kasai, David Gray, Carol Clugston, Anna Dominiczak and Alex Alderton, Roberto Amato, Jeffrey Barrett, Sonia Goncalves, Ewan Harrison, David K. Jackson, Ian Johnston, Dominic Kwiatkowski, Cordelia Langford, John Sillitoe on behalf of the Wellcome Sanger Institute COVID-19 Surveillance Team |
| EPI_ISL_3106775, EPI_ISL_3106776 | Lighthouse Lab in Alderley Park | Wellcome Sanger Institute for the COVID-19 Genomics UK (COG-UK) Consortium | Jacquelyn Wynn, Mairead Hyland, The Lighthouse Lab in Alderley Park and Alex Alderton, Roberto Amato, Jeffrey Barrett, Sonia Goncalves, Ewan Harrison, David K. Jackson, Ian Johnston, Dominic Kwiatkowski, Cordelia Langford, John Sillitoe on behalf of the Wellcome Sanger Institute COVID-19 Surveillance Team |
| EPI_ISL_3106777, EPI_ISL_3106778, EPI_ISL_3106779 | Lighthouse Lab in Glasgow | Wellcome Sanger Institute for the COVID-19 Genomics UK (COG-UK) Consortium | Harper VanSteenhouse, Yumi Kasai, David Gray, Carol Clugston, Anna Dominiczak and Alex Alderton, Roberto Amato, Jeffrey Barrett, Sonia Goncalves, Ewan Harrison, David K. Jackson, Ian Johnston, Dominic Kwiatkowski, Cordelia Langford, John Sillitoe on behalf of the Wellcome Sanger Institute COVID-19 Surveillance Team |
| EPI_ISL_3106780 | Lighthouse Lab in Alderley Park | Wellcome Sanger Institute for the COVID-19 Genomics UK (COG-UK) Consortium | Jacquelyn Wynn, Mairead Hyland, The Lighthouse Lab in Alderley Park and Alex Alderton, Roberto Amato, Jeffrey Barrett, Sonia Goncalves, Ewan Harrison, David K. Jackson, Ian Johnston, Dominic Kwiatkowski, Cordelia Langford, John Sillitoe on behalf of the Wellcome Sanger Institute COVID-19 Surveillance Team |
| EPI_ISL_3106781, EPI_ISL_3106782 | Lighthouse Lab in Glasgow | Wellcome Sanger Institute for the COVID-19 Genomics UK (COG-UK) Consortium | Harper VanSteenhouse, Yumi Kasai, David Gray, Carol Clugston, Anna Dominiczak and Alex Alderton, Roberto Amato, Jeffrey Barrett, Sonia Goncalves, Ewan Harrison, David K. Jackson, Ian Johnston, Dominic Kwiatkowski, Cordelia Langford, John Sillitoe on behalf of the Wellcome Sanger Institute COVID-19 Surveillance Team |
| EPI_ISL_3106783, EPI_ISL_3106784, EPI_ISL_3106785 | Lighthouse Lab in Alderley Park | Wellcome Sanger Institute for the COVID-19 Genomics UK (COG-UK) Consortium | Jacquelyn Wynn, Mairead Hyland, The Lighthouse Lab in Alderley Park and Alex Alderton, Roberto Amato, Jeffrey Barrett, Sonia Goncalves, Ewan Harrison, David K. Jackson, Ian Johnston, Dominic Kwiatkowski, Cordelia Langford, John Sillitoe on behalf of the Wellcome Sanger Institute COVID-19 Surveillance Team |
| EPI_ISL_3106786 | Lighthouse Lab in Glasgow | Wellcome Sanger Institute for the COVID-19 Genomics UK (COG-UK) Consortium | Harper VanSteenhouse, Yumi Kasai, David Gray, Carol Clugston, Anna Dominiczak and Alex Alderton, Roberto Amato, Jeffrey Barrett, Sonia Goncalves, Ewan Harrison, David K. Jackson, Ian Johnston, Dominic Kwiatkowski, Cordelia Langford, John Sillitoe on behalf of the Wellcome Sanger Institute COVID-19 Surveillance Team |
| EPI_ISL_3106787, EPI_ISL_3106788, EPI_ISL_3106789 | Lighthouse Lab in Alderley Park | Wellcome Sanger Institute for the COVID-19 Genomics UK (COG-UK) Consortium | Jacquelyn Wynn, Mairead Hyland, The Lighthouse Lab in Alderley Park and Alex Alderton, Roberto Amato, Jeffrey Barrett, Sonia Goncalves, Ewan Harrison, David K. Jackson, Ian Johnston, Dominic Kwiatkowski, Cordelia Langford, John Sillitoe on behalf of the Wellcome Sanger Institute COVID-19 Surveillance Team |
| EPI_ISL_3106790 | Lighthouse Lab in Glasgow | Wellcome Sanger Institute for the COVID-19 Genomics UK (COG-UK) Consortium | Harper VanSteenhouse, Yumi Kasai, David Gray, Carol Clugston, Anna Dominiczak and Alex Alderton, Roberto Amato, Jeffrey Barrett, Sonia Goncalves, Ewan Harrison, David K. Jackson, Ian Johnston, Dominic Kwiatkowski, Cordelia Langford, John Sillitoe on behalf of the Wellcome Sanger Institute COVID-19 Surveillance Team |
| EPI_ISL_3106791, EPI_ISL_3106792 | Lighthouse Lab in Alderley Park | Wellcome Sanger Institute for the COVID-19 Genomics UK (COG-UK) Consortium | Jacquelyn Wynn, Mairead Hyland, The Lighthouse Lab in Alderley Park and Alex Alderton, Roberto Amato, Jeffrey Barrett, Sonia Goncalves, Ewan Harrison, David K. Jackson, Ian Johnston, Dominic Kwiatkowski, Cordelia Langford, John Sillitoe on behalf of the Wellcome Sanger Institute COVID-19 Surveillance Team |
| EPI_ISL_3106793 | Lighthouse Lab in Glasgow | Wellcome Sanger Institute for the COVID-19 Genomics UK (COG-UK) Consortium | Harper VanSteenhouse, Yumi Kasai, David Gray, Carol Clugston, Anna Dominiczak and Alex Alderton, Roberto Amato, Jeffrey Barrett, Sonia Goncalves, Ewan Harrison, David K. Jackson, Ian Johnston, Dominic Kwiatkowski, Cordelia Langford, John Sillitoe on behalf of the Wellcome Sanger Institute COVID-19 Surveillance Team |
| EPI_ISL_3106794, EPI_ISL_3106795, EPI_ISL_3106796 | Lighthouse Lab in Alderley Park | Wellcome Sanger Institute for the COVID-19 Genomics UK (COG-UK) Consortium | Jacquelyn Wynn, Mairead Hyland, The Lighthouse Lab in Alderley Park and Alex Alderton, Roberto Amato, Jeffrey Barrett, Sonia Goncalves, Ewan Harrison, David K. Jackson, Ian Johnston, Dominic Kwiatkowski, Cordelia Langford, John Sillitoe on behalf of the Wellcome Sanger Institute COVID-19 Surveillance Team |
| EPI_ISL_3106797 | Lighthouse Lab in Glasgow | Wellcome Sanger Institute for the COVID-19 Genomics UK (COG-UK) Consortium | Harper VanSteenhouse, Yumi Kasai, David Gray, Carol Clugston, Anna Dominiczak and Alex Alderton, Roberto Amato, Jeffrey Barrett, Sonia Goncalves, Ewan Harrison, David K. Jackson, Ian Johnston, Dominic Kwiatkowski, Cordelia Langford, John Sillitoe on behalf of the Wellcome Sanger Institute COVID-19 Surveillance Team |
| EPI_ISL_3106798 | Lighthouse Lab in Alderley Park | Wellcome Sanger Institute for the COVID-19 Genomics UK (COG-UK) Consortium | Jacquelyn Wynn, Mairead Hyland, The Lighthouse Lab in Alderley Park and Alex Alderton, Roberto Amato, Jeffrey Barrett, Sonia Goncalves, Ewan Harrison, David K. Jackson, Ian Johnston, Dominic Kwiatkowski, Cordelia Langford, John Sillitoe on behalf of the Wellcome Sanger Institute COVID-19 Surveillance Team |
| EPI_ISL_3106799, EPI_ISL_3106800, EPI_ISL_3106801 | Lighthouse Lab in Glasgow | Wellcome Sanger Institute for the COVID-19 Genomics UK (COG-UK) Consortium | Harper VanSteenhouse, Yumi Kasai, David Gray, Carol Clugston, Anna Dominiczak and Alex Alderton, Roberto Amato, Jeffrey Barrett, Sonia Goncalves, Ewan Harrison, David K. Jackson, Ian Johnston, Dominic Kwiatkowski, Cordelia Langford, John Sillitoe on behalf of the Wellcome Sanger Institute COVID-19 Surveillance Team |
| EPI_ISL_3106802 | Lighthouse Lab in Alderley Park | Wellcome Sanger Institute for the COVID-19 Genomics UK (COG-UK) Consortium | Jacquelyn Wynn, Mairead Hyland, The Lighthouse Lab in Alderley Park and Alex Alderton, Roberto Amato, Jeffrey Barrett, Sonia Goncalves, Ewan Harrison, David K. Jackson, Ian Johnston, Dominic Kwiatkowski, Cordelia Langford, John Sillitoe on behalf of the Wellcome Sanger Institute COVID-19 Surveillance Team |
| EPI_ISL_3106803 | Lighthouse Lab in Glasgow | Wellcome Sanger Institute for the COVID-19 Genomics UK (COG-UK) Consortium | Harper VanSteenhouse, Yumi Kasai, David Gray, Carol Clugston, Anna Dominiczak and Alex Alderton, Roberto Amato, Jeffrey Barrett, Sonia Goncalves, Ewan Harrison, David K. Jackson, Ian Johnston, Dominic Kwiatkowski, Cordelia Langford, John Sillitoe on behalf of the Wellcome Sanger Institute COVID-19 Surveillance Team |
| EPI_ISL_3106804 | Lighthouse Lab in Alderley Park | Wellcome Sanger Institute for the COVID-19 Genomics UK (COG-UK) Consortium | Jacquelyn Wynn, Mairead Hyland, The Lighthouse Lab in Alderley Park and Alex Alderton, Roberto Amato, Jeffrey Barrett, Sonia Goncalves, Ewan Harrison, David K. Jackson, Ian Johnston, Dominic Kwiatkowski, Cordelia Langford, John Sillitoe on behalf of the Wellcome Sanger Institute COVID-19 Surveillance Team |
| EPI_ISL_3106805, EPI_ISL_3106806, EPI_ISL_3106807 | Lighthouse Lab in Glasgow | Wellcome Sanger Institute for the COVID-19 Genomics UK (COG-UK) Consortium | Harper VanSteenhouse, Yumi Kasai, David Gray, Carol Clugston, Anna Dominiczak and Alex Alderton, Roberto Amato, Jeffrey Barrett, Sonia Goncalves, Ewan Harrison, David K. Jackson, Ian Johnston, Dominic Kwiatkowski, Cordelia Langford, John Sillitoe on behalf of the Wellcome Sanger Institute COVID-19 Surveillance Team |
| EPI_ISL_3106808 | Lighthouse Lab in Alderley Park | Wellcome Sanger Institute for the COVID-19 Genomics UK (COG-UK) Consortium | Jacquelyn Wynn, Mairead Hyland, The Lighthouse Lab in Alderley Park and Alex Alderton, Roberto Amato, Jeffrey Barrett, Sonia Goncalves, Ewan Harrison, David K. Jackson, Ian Johnston, Dominic Kwiatkowski, Cordelia Langford, John Sillitoe on behalf of the Wellcome Sanger Institute COVID-19 Surveillance Team |
| EPI_ISL_3106809 | Lighthouse Lab in Glasgow | Wellcome Sanger Institute for the COVID-19 Genomics UK (COG-UK) Consortium | Harper VanSteenhouse, Yumi Kasai, David Gray, Carol Clugston, Anna Dominiczak and Alex Alderton, Roberto Amato, Jeffrey Barrett, Sonia Goncalves, Ewan Harrison, David K. Jackson, Ian Johnston, Dominic Kwiatkowski, Cordelia Langford, John Sillitoe on behalf of the Wellcome Sanger Institute COVID-19 Surveillance Team |
| EPI_ISL_3106810 | Lighthouse Lab in Alderley Park | Wellcome Sanger Institute for the COVID-19 Genomics UK (COG-UK) Consortium | Jacquelyn Wynn, Mairead Hyland, The Lighthouse Lab in Alderley Park and Alex Alderton, Roberto Amato, Jeffrey Barrett, Sonia Goncalves, Ewan Harrison, David K. Jackson, Ian Johnston, Dominic Kwiatkowski, Cordelia Langford, John Sillitoe on behalf of the Wellcome Sanger Institute COVID-19 Surveillance Team |
| EPI_ISL_3106811 | Lighthouse Lab in Glasgow | Wellcome Sanger Institute for the COVID-19 Genomics UK (COG-UK) | Harper VanSteenhouse, Yumi Kasai, David Gray, Carol Clugston, Anna Dominiczak and Alex Alderton, Roberto Amato, Jeffrey Barrett, Sonia Goncalves, Ewan Harrison, David K. Jackson, Ian Johnston, Dominic Kwiatkowski, Cordelia Langford, |

| | | | |
|---|---|---|---|
| | | Consortium | John Sillitoe on behalf of the Wellcome Sanger Institute COVID-19 Surveillance Team |
| EPI_ISL_3106812 | Lighthouse Lab in Alderley Park | Wellcome Sanger Institute for the COVID-19 Genomics UK (COG-UK) Consortium | Jacquelyn Wynn, Mairead Hyland, The Lighthouse Lab in Alderley Park and Alex Alderton, Roberto Amato, Jeffrey Barrett, Sonia Goncalves, Ewan Harrison, David K. Jackson, Ian Johnston, Dominic Kwiatkowski, Cordelia Langford, John Sillitoe on behalf of the Wellcome Sanger Institute COVID-19 Surveillance Team |
| EPI_ISL_3106813 | Lighthouse Lab in Glasgow | Wellcome Sanger Institute for the COVID-19 Genomics UK (COG-UK) Consortium | Harper VanSteenhouse, Yumi Kasai, David Gray, Carol Clugston, Anna Dominiczak and Alex Alderton, Roberto Amato, Jeffrey Barrett, Sonia Goncalves, Ewan Harrison, David K. Jackson, Ian Johnston, Dominic Kwiatkowski, Cordelia Langford, John Sillitoe on behalf of the Wellcome Sanger Institute COVID-19 Surveillance Team |
| EPI_ISL_3106814 | Lighthouse Lab in Alderley Park | Wellcome Sanger Institute for the COVID-19 Genomics UK (COG-UK) Consortium | Jacquelyn Wynn, Mairead Hyland, The Lighthouse Lab in Alderley Park and Alex Alderton, Roberto Amato, Jeffrey Barrett, Sonia Goncalves, Ewan Harrison, David K. Jackson, Ian Johnston, Dominic Kwiatkowski, Cordelia Langford, John Sillitoe on behalf of the Wellcome Sanger Institute COVID-19 Surveillance Team |
| EPI_ISL_3106815 | Lighthouse Lab in Glasgow | Wellcome Sanger Institute for the COVID-19 Genomics UK (COG-UK) Consortium | Harper VanSteenhouse, Yumi Kasai, David Gray, Carol Clugston, Anna Dominiczak and Alex Alderton, Roberto Amato, Jeffrey Barrett, Sonia Goncalves, Ewan Harrison, David K. Jackson, Ian Johnston, Dominic Kwiatkowski, Cordelia Langford, John Sillitoe on behalf of the Wellcome Sanger Institute COVID-19 Surveillance Team |
| EPI_ISL_3106816, EPI_ISL_3106817, EPI_ISL_3106818 | Lighthouse Lab in Alderley Park | Wellcome Sanger Institute for the COVID-19 Genomics UK (COG-UK) Consortium | Jacquelyn Wynn, Mairead Hyland, The Lighthouse Lab in Alderley Park and Alex Alderton, Roberto Amato, Jeffrey Barrett, Sonia Goncalves, Ewan Harrison, David K. Jackson, Ian Johnston, Dominic Kwiatkowski, Cordelia Langford, John Sillitoe on behalf of the Wellcome Sanger Institute COVID-19 Surveillance Team |
| EPI_ISL_3106819, EPI_ISL_3106820 | Lighthouse Lab in Glasgow | Wellcome Sanger Institute for the COVID-19 Genomics UK (COG-UK) Consortium | Harper VanSteenhouse, Yumi Kasai, David Gray, Carol Clugston, Anna Dominiczak and Alex Alderton, Roberto Amato, Jeffrey Barrett, Sonia Goncalves, Ewan Harrison, David K. Jackson, Ian Johnston, Dominic Kwiatkowski, Cordelia Langford, John Sillitoe on behalf of the Wellcome Sanger Institute COVID-19 Surveillance Team |
| EPI_ISL_3106821, EPI_ISL_3106822, EPI_ISL_3106823 | Lighthouse Lab in Alderley Park | Wellcome Sanger Institute for the COVID-19 Genomics UK (COG-UK) Consortium | Jacquelyn Wynn, Mairead Hyland, The Lighthouse Lab in Alderley Park and Alex Alderton, Roberto Amato, Jeffrey Barrett, Sonia Goncalves, Ewan Harrison, David K. Jackson, Ian Johnston, Dominic Kwiatkowski, Cordelia Langford, John Sillitoe on behalf of the Wellcome Sanger Institute COVID-19 Surveillance Team |
| EPI_ISL_3106824, EPI_ISL_3106825 | Lighthouse Lab in Glasgow | Wellcome Sanger Institute for the COVID-19 Genomics UK (COG-UK) Consortium | Harper VanSteenhouse, Yumi Kasai, David Gray, Carol Clugston, Anna Dominiczak and Alex Alderton, Roberto Amato, Jeffrey Barrett, Sonia Goncalves, Ewan Harrison, David K. Jackson, Ian Johnston, Dominic Kwiatkowski, Cordelia Langford, John Sillitoe on behalf of the Wellcome Sanger Institute COVID-19 Surveillance Team |
| EPI_ISL_3106826 | Lighthouse Lab in Alderley Park | Wellcome Sanger Institute for the COVID-19 Genomics UK (COG-UK) Consortium | Jacquelyn Wynn, Mairead Hyland, The Lighthouse Lab in Alderley Park and Alex Alderton, Roberto Amato, Jeffrey Barrett, Sonia Goncalves, Ewan Harrison, David K. Jackson, Ian Johnston, Dominic Kwiatkowski, Cordelia Langford, John Sillitoe on behalf of the Wellcome Sanger Institute COVID-19 Surveillance Team |
| EPI_ISL_3106827, EPI_ISL_3106828, EPI_ISL_3106829, EPI_ISL_3106830, EPI_ISL_3106831, EPI_ISL_3106832 | Lighthouse Lab in Glasgow | Wellcome Sanger Institute for the COVID-19 Genomics UK (COG-UK) Consortium | Harper VanSteenhouse, Yumi Kasai, David Gray, Carol Clugston, Anna Dominiczak and Alex Alderton, Roberto Amato, Jeffrey Barrett, Sonia Goncalves, Ewan Harrison, David K. Jackson, Ian Johnston, Dominic Kwiatkowski, Cordelia Langford, John Sillitoe on behalf of the Wellcome Sanger Institute COVID-19 Surveillance Team |
| EPI_ISL_3106833, EPI_ISL_3106834, EPI_ISL_3106835 | Lighthouse Lab in Alderley Park | Wellcome Sanger Institute for the COVID-19 Genomics UK (COG-UK) Consortium | Jacquelyn Wynn, Mairead Hyland, The Lighthouse Lab in Alderley Park and Alex Alderton, Roberto Amato, Jeffrey Barrett, Sonia Goncalves, Ewan Harrison, David K. Jackson, Ian Johnston, Dominic Kwiatkowski, Cordelia Langford, John Sillitoe on behalf of the Wellcome Sanger Institute COVID-19 Surveillance Team |
| EPI_ISL_3106836 | Lighthouse Lab in Glasgow | Wellcome Sanger Institute for the COVID-19 Genomics UK (COG-UK) Consortium | Harper VanSteenhouse, Yumi Kasai, David Gray, Carol Clugston, Anna Dominiczak and Alex Alderton, Roberto Amato, Jeffrey Barrett, Sonia Goncalves, Ewan Harrison, David K. Jackson, Ian Johnston, Dominic Kwiatkowski, Cordelia Langford, John Sillitoe on behalf of the Wellcome Sanger Institute COVID-19 Surveillance Team |
| EPI_ISL_3106837, EPI_ISL_3106838 | Lighthouse Lab in Alderley Park | Wellcome Sanger Institute for the COVID-19 Genomics UK (COG-UK) Consortium | Jacquelyn Wynn, Mairead Hyland, The Lighthouse Lab in Alderley Park and Alex Alderton, Roberto Amato, Jeffrey Barrett, Sonia Goncalves, Ewan Harrison, David K. Jackson, Ian Johnston, Dominic Kwiatkowski, Cordelia Langford, John Sillitoe on behalf of the Wellcome Sanger Institute COVID-19 Surveillance Team |
| EPI_ISL_3106839 | Lighthouse Lab in Glasgow | Wellcome Sanger Institute for the COVID-19 Genomics UK (COG-UK) Consortium | Harper VanSteenhouse, Yumi Kasai, David Gray, Carol Clugston, Anna Dominiczak and Alex Alderton, Roberto Amato, Jeffrey Barrett, Sonia Goncalves, Ewan Harrison, David K. Jackson, Ian Johnston, Dominic Kwiatkowski, Cordelia Langford, John Sillitoe on behalf of the Wellcome Sanger Institute COVID-19 Surveillance Team |
| EPI_ISL_3106840 | Lighthouse Lab in Alderley Park | Wellcome Sanger Institute for the COVID-19 Genomics UK (COG-UK) Consortium | Jacquelyn Wynn, Mairead Hyland, The Lighthouse Lab in Alderley Park and Alex Alderton, Roberto Amato, Jeffrey Barrett, Sonia Goncalves, Ewan Harrison, David K. Jackson, Ian Johnston, Dominic Kwiatkowski, Cordelia Langford, John Sillitoe on behalf of the Wellcome Sanger Institute COVID-19 Surveillance Team |
| EPI_ISL_3106841 | Lighthouse Lab in Glasgow | Wellcome Sanger Institute for the COVID-19 Genomics UK (COG-UK) Consortium | Harper VanSteenhouse, Yumi Kasai, David Gray, Carol Clugston, Anna Dominiczak and Alex Alderton, Roberto Amato, Jeffrey Barrett, Sonia Goncalves, Ewan Harrison, David K. Jackson, Ian Johnston, Dominic Kwiatkowski, Cordelia Langford, John Sillitoe on behalf of the Wellcome Sanger Institute COVID-19 Surveillance Team |
| EPI_ISL_3106842, EPI_ISL_3106843 | Lighthouse Lab in Alderley Park | Wellcome Sanger Institute for the COVID-19 Genomics UK (COG-UK) Consortium | Jacquelyn Wynn, Mairead Hyland, The Lighthouse Lab in Alderley Park and Alex Alderton, Roberto Amato, Jeffrey Barrett, Sonia Goncalves, Ewan Harrison, David K. Jackson, Ian Johnston, Dominic Kwiatkowski, Cordelia Langford, John Sillitoe on behalf of the Wellcome Sanger Institute COVID-19 Surveillance Team |
| EPI_ISL_3106844 | Lighthouse Lab in Glasgow | Wellcome Sanger Institute for the COVID-19 Genomics UK (COG-UK) Consortium | Harper VanSteenhouse, Yumi Kasai, David Gray, Carol Clugston, Anna Dominiczak and Alex Alderton, Roberto Amato, Jeffrey Barrett, Sonia Goncalves, Ewan Harrison, David K. Jackson, Ian Johnston, Dominic Kwiatkowski, Cordelia Langford, John Sillitoe on behalf of the Wellcome Sanger Institute COVID-19 Surveillance Team |
| EPI_ISL_3106845 | Lighthouse Lab in Alderley Park | Wellcome Sanger Institute for the COVID-19 Genomics UK (COG-UK) Consortium | Jacquelyn Wynn, Mairead Hyland, The Lighthouse Lab in Alderley Park and Alex Alderton, Roberto Amato, Jeffrey Barrett, Sonia Goncalves, Ewan Harrison, David K. Jackson, Ian Johnston, Dominic Kwiatkowski, Cordelia Langford, John Sillitoe on behalf of the Wellcome Sanger Institute COVID-19 Surveillance Team |
| EPI_ISL_3106846, EPI_ISL_3106847, EPI_ISL_3106848 | Lighthouse Lab in Glasgow | Wellcome Sanger Institute for the COVID-19 Genomics UK (COG-UK) Consortium | Harper VanSteenhouse, Yumi Kasai, David Gray, Carol Clugston, Anna Dominiczak and Alex Alderton, Roberto Amato, Jeffrey Barrett, Sonia Goncalves, Ewan Harrison, David K. Jackson, Ian Johnston, Dominic Kwiatkowski, Cordelia Langford, John Sillitoe on behalf of the Wellcome Sanger Institute COVID-19 Surveillance Team |
| EPI_ISL_3106849 | Lighthouse Lab in Alderley Park | Wellcome Sanger Institute for the COVID-19 Genomics UK (COG-UK) Consortium | Jacquelyn Wynn, Mairead Hyland, The Lighthouse Lab in Alderley Park and Alex Alderton, Roberto Amato, Jeffrey Barrett, Sonia Goncalves, Ewan Harrison, David K. Jackson, Ian Johnston, Dominic Kwiatkowski, Cordelia Langford, John Sillitoe on behalf of the Wellcome Sanger Institute COVID-19 Surveillance Team |
| EPI_ISL_3106850 | Lighthouse Lab in Glasgow | Wellcome Sanger Institute for the COVID-19 Genomics UK (COG-UK) Consortium | Harper VanSteenhouse, Yumi Kasai, David Gray, Carol Clugston, Anna Dominiczak and Alex Alderton, Roberto Amato, Jeffrey Barrett, Sonia Goncalves, Ewan Harrison, David K. Jackson, Ian Johnston, Dominic Kwiatkowski, Cordelia Langford, John Sillitoe on behalf of the Wellcome Sanger Institute COVID-19 Surveillance Team |
| EPI_ISL_3106851, EPI_ISL_3106852, EPI_ISL_3106853, EPI_ISL_3106854 | Lighthouse Lab in Alderley Park | Wellcome Sanger Institute for the COVID-19 Genomics UK (COG-UK) Consortium | Jacquelyn Wynn, Mairead Hyland, The Lighthouse Lab in Alderley Park and Alex Alderton, Roberto Amato, Jeffrey Barrett, Sonia Goncalves, Ewan Harrison, David K. Jackson, Ian Johnston, Dominic Kwiatkowski, Cordelia Langford, John Sillitoe on behalf of the Wellcome Sanger Institute COVID-19 Surveillance Team |
| EPI_ISL_3106855, EPI_ISL_3106856 | Lighthouse Lab in Glasgow | Wellcome Sanger Institute for the COVID-19 Genomics UK (COG-UK) Consortium | Harper VanSteenhouse, Yumi Kasai, David Gray, Carol Clugston, Anna Dominiczak and Alex Alderton, Roberto Amato, Jeffrey Barrett, Sonia Goncalves, Ewan Harrison, David K. Jackson, Ian Johnston, Dominic Kwiatkowski, Cordelia Langford, John Sillitoe on behalf of the Wellcome Sanger Institute COVID-19 Surveillance Team |
| EPI_ISL_3106857, EPI_ISL_3106858, EPI_ISL_3106859 | Lighthouse Lab in Alderley Park | Wellcome Sanger Institute for the COVID-19 Genomics UK (COG-UK) Consortium | Jacquelyn Wynn, Mairead Hyland, The Lighthouse Lab in Alderley Park and Alex Alderton, Roberto Amato, Jeffrey Barrett, Sonia Goncalves, Ewan Harrison, David K. Jackson, Ian Johnston, Dominic Kwiatkowski, Cordelia Langford, John Sillitoe on behalf of the Wellcome Sanger Institute COVID-19 Surveillance Team |
| EPI_ISL_3106860, EPI_ISL_3106861, EPI_ISL_3106862, EPI_ISL_3106863 | Lighthouse Lab in Glasgow | Wellcome Sanger Institute for the COVID-19 Genomics UK (COG-UK) Consortium | Harper VanSteenhouse, Yumi Kasai, David Gray, Carol Clugston, Anna Dominiczak and Alex Alderton, Roberto Amato, Jeffrey Barrett, Sonia Goncalves, Ewan Harrison, David K. Jackson, Ian Johnston, Dominic Kwiatkowski, Cordelia Langford, John Sillitoe on behalf of the Wellcome Sanger Institute COVID-19 Surveillance Team |
| EPI_ISL_3106864 | Lighthouse Lab in Alderley Park | Wellcome Sanger Institute for the COVID-19 Genomics UK (COG-UK) Consortium | Jacquelyn Wynn, Mairead Hyland, The Lighthouse Lab in Alderley Park and Alex Alderton, Roberto Amato, Jeffrey Barrett, Sonia Goncalves, Ewan Harrison, David K. Jackson, Ian Johnston, Dominic Kwiatkowski, Cordelia Langford, John Sillitoe on behalf of the Wellcome Sanger Institute COVID-19 Surveillance Team |
| EPI_ISL_3106865 | Lighthouse Lab in Glasgow | Wellcome Sanger Institute for the COVID-19 Genomics UK (COG-UK) Consortium | Harper VanSteenhouse, Yumi Kasai, David Gray, Carol Clugston, Anna Dominiczak and Alex Alderton, Roberto Amato, Jeffrey Barrett, Sonia Goncalves, Ewan Harrison, David K. Jackson, Ian Johnston, Dominic Kwiatkowski, Cordelia Langford, John Sillitoe on behalf of the Wellcome Sanger Institute COVID-19 Surveillance Team |
| EPI_ISL_3106866, EPI_ISL_3106867 | Lighthouse Lab in Alderley Park | Wellcome Sanger Institute for the COVID-19 Genomics UK (COG-UK) Consortium | Jacquelyn Wynn, Mairead Hyland, The Lighthouse Lab in Alderley Park and Alex Alderton, Roberto Amato, Jeffrey Barrett, Sonia Goncalves, Ewan Harrison, David K. Jackson, Ian Johnston, Dominic Kwiatkowski, Cordelia Langford, John Sillitoe on behalf of the Wellcome Sanger Institute COVID-19 Surveillance Team |
| EPI_ISL_3106868 | Lighthouse Lab in Glasgow | Wellcome Sanger Institute for the COVID-19 Genomics UK (COG-UK) Consortium | Harper VanSteenhouse, Yumi Kasai, David Gray, Carol Clugston, Anna Dominiczak and Alex Alderton, Roberto Amato, Jeffrey Barrett, Sonia Goncalves, Ewan Harrison, David K. Jackson, Ian Johnston, Dominic Kwiatkowski, Cordelia Langford, John Sillitoe on behalf of the Wellcome Sanger Institute COVID-19 Surveillance Team |
| EPI_ISL_3106869 | Lighthouse Lab in Alderley Park | Wellcome Sanger Institute for the COVID-19 Genomics UK (COG-UK) Consortium | Jacquelyn Wynn, Mairead Hyland, The Lighthouse Lab in Alderley Park and Alex Alderton, Roberto Amato, Jeffrey Barrett, Sonia Goncalves, Ewan Harrison, David K. Jackson, Ian Johnston, Dominic Kwiatkowski, Cordelia Langford, John Sillitoe on behalf of the Wellcome Sanger Institute COVID-19 Surveillance Team |
| EPI_ISL_3106870 | Lighthouse Lab in Glasgow | Wellcome Sanger Institute for the COVID-19 Genomics UK (COG-UK) Consortium | Harper VanSteenhouse, Yumi Kasai, David Gray, Carol Clugston, Anna Dominiczak and Alex Alderton, Roberto Amato, Jeffrey Barrett, Sonia Goncalves, Ewan Harrison, David K. Jackson, Ian Johnston, Dominic Kwiatkowski, Cordelia Langford, John Sillitoe on behalf of the Wellcome Sanger Institute COVID-19 Surveillance Team |
| EPI_ISL_3106871, EPI_ISL_3106872, EPI_ISL_3106873, EPI_ISL_3106874, EPI_ISL_3106875, EPI_ISL_3106876 | Lighthouse Lab in Alderley Park | Wellcome Sanger Institute for the COVID-19 Genomics UK (COG-UK) Consortium | Jacquelyn Wynn, Mairead Hyland, The Lighthouse Lab in Alderley Park and Alex Alderton, Roberto Amato, Jeffrey Barrett, Sonia Goncalves, Ewan Harrison, David K. Jackson, Ian Johnston, Dominic Kwiatkowski, Cordelia Langford, John Sillitoe on behalf of the Wellcome Sanger Institute COVID-19 Surveillance Team |
| EPI_ISL_3106877 | Lighthouse Lab in Glasgow | Wellcome Sanger Institute for the COVID-19 Genomics UK (COG-UK) Consortium | Harper VanSteenhouse, Yumi Kasai, David Gray, Carol Clugston, Anna Dominiczak and Alex Alderton, Roberto Amato, Jeffrey Barrett, Sonia Goncalves, Ewan Harrison, David K. Jackson, Ian Johnston, Dominic Kwiatkowski, Cordelia Langford, John Sillitoe on behalf of the Wellcome Sanger Institute COVID-19 Surveillance Team |
| EPI_ISL_3106878, EPI_ISL_3106879, EPI_ISL_3106880, EPI_ISL_3106881 | Lighthouse Lab in Alderley Park | Wellcome Sanger Institute for the COVID-19 Genomics UK (COG-UK) Consortium | Jacquelyn Wynn, Mairead Hyland, The Lighthouse Lab in Alderley Park and Alex Alderton, Roberto Amato, Jeffrey Barrett, Sonia Goncalves, Ewan Harrison, David K. Jackson, Ian Johnston, Dominic Kwiatkowski, Cordelia Langford, John Sillitoe on behalf of the Wellcome Sanger Institute COVID-19 Surveillance Team |
| EPI_ISL_3106882, EPI_ISL_3106883 | Lighthouse Lab in Glasgow | Wellcome Sanger Institute for the COVID-19 Genomics UK (COG-UK) Consortium | Harper VanSteenhouse, Yumi Kasai, David Gray, Carol Clugston, Anna Dominiczak and Alex Alderton, Roberto Amato, Jeffrey Barrett, Sonia Goncalves, Ewan Harrison, David K. Jackson, Ian Johnston, Dominic Kwiatkowski, Cordelia Langford, John Sillitoe on behalf of the Wellcome Sanger Institute COVID-19 Surveillance Team |
| EPI_ISL_3106884, | Lighthouse Lab in Alderley Park | Wellcome Sanger Institute for the COVID-19 Genomics UK (COG-UK) | Jacquelyn Wynn, Mairead Hyland, The Lighthouse Lab in Alderley Park and Alex Alderton, Roberto Amato, Jeffrey Barrett, Sonia Goncalves, Ewan Harrison, David K. Jackson, Ian Johnston, Dominic Kwiatkowski, Cordelia Langford, John Sillitoe on |

| | | | |
|---|---|---|---|
| EPI_ISL_3106885, EPI_ISL_3106886, EPI_ISL_3106887, EPI_ISL_3106888 | | Consortium | behalf of the Wellcome Sanger Institute COVID-19 Surveillance Team |
| EPI_ISL_3106889 | Lighthouse Lab in Glasgow | Wellcome Sanger Institute for the COVID-19 Genomics UK (COG-UK) Consortium | Harper VanSteenhouse, Yumi Kasai, David Gray, Carol Clugston, Anna Dominiczak and Alex Alderton, Roberto Amato, Jeffrey Barrett, Sonia Goncalves, Ewan Harrison, David K. Jackson, Ian Johnston, Dominic Kwiatkowski, Cordelia Langford, John Sillitoe on behalf of the Wellcome Sanger Institute COVID-19 Surveillance Team |
| EPI_ISL_3106890 | Lighthouse Lab in Alderley Park | Wellcome Sanger Institute for the COVID-19 Genomics UK (COG-UK) Consortium | Jacquelyn Wynn, Mairead Hyland, The Lighthouse Lab in Alderley Park and Alex Alderton, Roberto Amato, Jeffrey Barrett, Sonia Goncalves, Ewan Harrison, David K. Jackson, Ian Johnston, Dominic Kwiatkowski, Cordelia Langford, John Sillitoe on behalf of the Wellcome Sanger Institute COVID-19 Surveillance Team |
| EPI_ISL_3106891 | Lighthouse Lab in Glasgow | Wellcome Sanger Institute for the COVID-19 Genomics UK (COG-UK) Consortium | Harper VanSteenhouse, Yumi Kasai, David Gray, Carol Clugston, Anna Dominiczak and Alex Alderton, Roberto Amato, Jeffrey Barrett, Sonia Goncalves, Ewan Harrison, David K. Jackson, Ian Johnston, Dominic Kwiatkowski, Cordelia Langford, John Sillitoe on behalf of the Wellcome Sanger Institute COVID-19 Surveillance Team |
| EPI_ISL_3106892 | Lighthouse Lab in Alderley Park | Wellcome Sanger Institute for the COVID-19 Genomics UK (COG-UK) Consortium | Jacquelyn Wynn, Mairead Hyland, The Lighthouse Lab in Alderley Park and Alex Alderton, Roberto Amato, Jeffrey Barrett, Sonia Goncalves, Ewan Harrison, David K. Jackson, Ian Johnston, Dominic Kwiatkowski, Cordelia Langford, John Sillitoe on behalf of the Wellcome Sanger Institute COVID-19 Surveillance Team |
| EPI_ISL_3106893 | Lighthouse Lab in Glasgow | Wellcome Sanger Institute for the COVID-19 Genomics UK (COG-UK) Consortium | Harper VanSteenhouse, Yumi Kasai, David Gray, Carol Clugston, Anna Dominiczak and Alex Alderton, Roberto Amato, Jeffrey Barrett, Sonia Goncalves, Ewan Harrison, David K. Jackson, Ian Johnston, Dominic Kwiatkowski, Cordelia Langford, John Sillitoe on behalf of the Wellcome Sanger Institute COVID-19 Surveillance Team |
| EPI_ISL_3106894 | Lighthouse Lab in Alderley Park | Wellcome Sanger Institute for the COVID-19 Genomics UK (COG-UK) Consortium | Jacquelyn Wynn, Mairead Hyland, The Lighthouse Lab in Alderley Park and Alex Alderton, Roberto Amato, Jeffrey Barrett, Sonia Goncalves, Ewan Harrison, David K. Jackson, Ian Johnston, Dominic Kwiatkowski, Cordelia Langford, John Sillitoe on behalf of the Wellcome Sanger Institute COVID-19 Surveillance Team |
| EPI_ISL_3106895, EPI_ISL_3106896 | Lighthouse Lab in Glasgow | Wellcome Sanger Institute for the COVID-19 Genomics UK (COG-UK) Consortium | Harper VanSteenhouse, Yumi Kasai, David Gray, Carol Clugston, Anna Dominiczak and Alex Alderton, Roberto Amato, Jeffrey Barrett, Sonia Goncalves, Ewan Harrison, David K. Jackson, Ian Johnston, Dominic Kwiatkowski, Cordelia Langford, John Sillitoe on behalf of the Wellcome Sanger Institute COVID-19 Surveillance Team |
| EPI_ISL_3106897, EPI_ISL_3106898, EPI_ISL_3106899, EPI_ISL_3106900, EPI_ISL_3106901, EPI_ISL_3106902 | Lighthouse Lab in Alderley Park | Wellcome Sanger Institute for the COVID-19 Genomics UK (COG-UK) Consortium | Jacquelyn Wynn, Mairead Hyland, The Lighthouse Lab in Alderley Park and Alex Alderton, Roberto Amato, Jeffrey Barrett, Sonia Goncalves, Ewan Harrison, David K. Jackson, Ian Johnston, Dominic Kwiatkowski, Cordelia Langford, John Sillitoe on behalf of the Wellcome Sanger Institute COVID-19 Surveillance Team |
| EPI_ISL_3106903 | Lighthouse Lab in Glasgow | Wellcome Sanger Institute for the COVID-19 Genomics UK (COG-UK) Consortium | Harper VanSteenhouse, Yumi Kasai, David Gray, Carol Clugston, Anna Dominiczak and Alex Alderton, Roberto Amato, Jeffrey Barrett, Sonia Goncalves, Ewan Harrison, David K. Jackson, Ian Johnston, Dominic Kwiatkowski, Cordelia Langford, John Sillitoe on behalf of the Wellcome Sanger Institute COVID-19 Surveillance Team |
| EPI_ISL_3106904 | Lighthouse Lab in Alderley Park | Wellcome Sanger Institute for the COVID-19 Genomics UK (COG-UK) Consortium | Jacquelyn Wynn, Mairead Hyland, The Lighthouse Lab in Alderley Park and Alex Alderton, Roberto Amato, Jeffrey Barrett, Sonia Goncalves, Ewan Harrison, David K. Jackson, Ian Johnston, Dominic Kwiatkowski, Cordelia Langford, John Sillitoe on behalf of the Wellcome Sanger Institute COVID-19 Surveillance Team |
| EPI_ISL_3106905 | Lighthouse Lab in Glasgow | Wellcome Sanger Institute for the COVID-19 Genomics UK (COG-UK) Consortium | Harper VanSteenhouse, Yumi Kasai, David Gray, Carol Clugston, Anna Dominiczak and Alex Alderton, Roberto Amato, Jeffrey Barrett, Sonia Goncalves, Ewan Harrison, David K. Jackson, Ian Johnston, Dominic Kwiatkowski, Cordelia Langford, John Sillitoe on behalf of the Wellcome Sanger Institute COVID-19 Surveillance Team |
| EPI_ISL_3106906 | Lighthouse Lab in Alderley Park | Wellcome Sanger Institute for the COVID-19 Genomics UK (COG-UK) Consortium | Jacquelyn Wynn, Mairead Hyland, The Lighthouse Lab in Alderley Park and Alex Alderton, Roberto Amato, Jeffrey Barrett, Sonia Goncalves, Ewan Harrison, David K. Jackson, Ian Johnston, Dominic Kwiatkowski, Cordelia Langford, John Sillitoe on behalf of the Wellcome Sanger Institute COVID-19 Surveillance Team |
| EPI_ISL_3106907, EPI_ISL_3106908 | Lighthouse Lab in Glasgow | Wellcome Sanger Institute for the COVID-19 Genomics UK (COG-UK) Consortium | Harper VanSteenhouse, Yumi Kasai, David Gray, Carol Clugston, Anna Dominiczak and Alex Alderton, Roberto Amato, Jeffrey Barrett, Sonia Goncalves, Ewan Harrison, David K. Jackson, Ian Johnston, Dominic Kwiatkowski, Cordelia Langford, John Sillitoe on behalf of the Wellcome Sanger Institute COVID-19 Surveillance Team |
| EPI_ISL_3106909, EPI_ISL_3106910 | Lighthouse Lab in Alderley Park | Wellcome Sanger Institute for the COVID-19 Genomics UK (COG-UK) Consortium | Jacquelyn Wynn, Mairead Hyland, The Lighthouse Lab in Alderley Park and Alex Alderton, Roberto Amato, Jeffrey Barrett, Sonia Goncalves, Ewan Harrison, David K. Jackson, Ian Johnston, Dominic Kwiatkowski, Cordelia Langford, John Sillitoe on behalf of the Wellcome Sanger Institute COVID-19 Surveillance Team |
| EPI_ISL_3106911, EPI_ISL_3106912 | Lighthouse Lab in Glasgow | Wellcome Sanger Institute for the COVID-19 Genomics UK (COG-UK) Consortium | Harper VanSteenhouse, Yumi Kasai, David Gray, Carol Clugston, Anna Dominiczak and Alex Alderton, Roberto Amato, Jeffrey Barrett, Sonia Goncalves, Ewan Harrison, David K. Jackson, Ian Johnston, Dominic Kwiatkowski, Cordelia Langford, John Sillitoe on behalf of the Wellcome Sanger Institute COVID-19 Surveillance Team |
| EPI_ISL_3106913 | Lighthouse Lab in Alderley Park | Wellcome Sanger Institute for the COVID-19 Genomics UK (COG-UK) Consortium | Jacquelyn Wynn, Mairead Hyland, The Lighthouse Lab in Alderley Park and Alex Alderton, Roberto Amato, Jeffrey Barrett, Sonia Goncalves, Ewan Harrison, David K. Jackson, Ian Johnston, Dominic Kwiatkowski, Cordelia Langford, John Sillitoe on behalf of the Wellcome Sanger Institute COVID-19 Surveillance Team |
| EPI_ISL_3106914, EPI_ISL_3106915, EPI_ISL_3106916 | Lighthouse Lab in Glasgow | Wellcome Sanger Institute for the COVID-19 Genomics UK (COG-UK) Consortium | Harper VanSteenhouse, Yumi Kasai, David Gray, Carol Clugston, Anna Dominiczak and Alex Alderton, Roberto Amato, Jeffrey Barrett, Sonia Goncalves, Ewan Harrison, David K. Jackson, Ian Johnston, Dominic Kwiatkowski, Cordelia Langford, John Sillitoe on behalf of the Wellcome Sanger Institute COVID-19 Surveillance Team |
| EPI_ISL_3106917 | Lighthouse Lab in Alderley Park | Wellcome Sanger Institute for the COVID-19 Genomics UK (COG-UK) Consortium | Jacquelyn Wynn, Mairead Hyland, The Lighthouse Lab in Alderley Park and Alex Alderton, Roberto Amato, Jeffrey Barrett, Sonia Goncalves, Ewan Harrison, David K. Jackson, Ian Johnston, Dominic Kwiatkowski, Cordelia Langford, John Sillitoe on behalf of the Wellcome Sanger Institute COVID-19 Surveillance Team |
| EPI_ISL_3106918 | Lighthouse Lab in Glasgow | Wellcome Sanger Institute for the COVID-19 Genomics UK (COG-UK) Consortium | Harper VanSteenhouse, Yumi Kasai, David Gray, Carol Clugston, Anna Dominiczak and Alex Alderton, Roberto Amato, Jeffrey Barrett, Sonia Goncalves, Ewan Harrison, David K. Jackson, Ian Johnston, Dominic Kwiatkowski, Cordelia Langford, John Sillitoe on behalf of the Wellcome Sanger Institute COVID-19 Surveillance Team |
| EPI_ISL_3106919 | Lighthouse Lab in Alderley Park | Wellcome Sanger Institute for the COVID-19 Genomics UK (COG-UK) Consortium | Jacquelyn Wynn, Mairead Hyland, The Lighthouse Lab in Alderley Park and Alex Alderton, Roberto Amato, Jeffrey Barrett, Sonia Goncalves, Ewan Harrison, David K. Jackson, Ian Johnston, Dominic Kwiatkowski, Cordelia Langford, John Sillitoe on behalf of the Wellcome Sanger Institute COVID-19 Surveillance Team |
| EPI_ISL_3106920 | Lighthouse Lab in Glasgow | Wellcome Sanger Institute for the COVID-19 Genomics UK (COG-UK) Consortium | Harper VanSteenhouse, Yumi Kasai, David Gray, Carol Clugston, Anna Dominiczak and Alex Alderton, Roberto Amato, Jeffrey Barrett, Sonia Goncalves, Ewan Harrison, David K. Jackson, Ian Johnston, Dominic Kwiatkowski, Cordelia Langford, John Sillitoe on behalf of the Wellcome Sanger Institute COVID-19 Surveillance Team |
| EPI_ISL_3106921, EPI_ISL_3106922 | Lighthouse Lab in Alderley Park | Wellcome Sanger Institute for the COVID-19 Genomics UK (COG-UK) Consortium | Jacquelyn Wynn, Mairead Hyland, The Lighthouse Lab in Alderley Park and Alex Alderton, Roberto Amato, Jeffrey Barrett, Sonia Goncalves, Ewan Harrison, David K. Jackson, Ian Johnston, Dominic Kwiatkowski, Cordelia Langford, John Sillitoe on behalf of the Wellcome Sanger Institute COVID-19 Surveillance Team |
| EPI_ISL_3106923 | Lighthouse Lab in Glasgow | Wellcome Sanger Institute for the COVID-19 Genomics UK (COG-UK) Consortium | Harper VanSteenhouse, Yumi Kasai, David Gray, Carol Clugston, Anna Dominiczak and Alex Alderton, Roberto Amato, Jeffrey Barrett, Sonia Goncalves, Ewan Harrison, David K. Jackson, Ian Johnston, Dominic Kwiatkowski, Cordelia Langford, John Sillitoe on behalf of the Wellcome Sanger Institute COVID-19 Surveillance Team |
| EPI_ISL_3106924 | Lighthouse Lab in Alderley Park | Wellcome Sanger Institute for the COVID-19 Genomics UK (COG-UK) Consortium | Jacquelyn Wynn, Mairead Hyland, The Lighthouse Lab in Alderley Park and Alex Alderton, Roberto Amato, Jeffrey Barrett, Sonia Goncalves, Ewan Harrison, David K. Jackson, Ian Johnston, Dominic Kwiatkowski, Cordelia Langford, John Sillitoe on behalf of the Wellcome Sanger Institute COVID-19 Surveillance Team |
| EPI_ISL_3106925, EPI_ISL_3106926 | Lighthouse Lab in Glasgow | Wellcome Sanger Institute for the COVID-19 Genomics UK (COG-UK) Consortium | Harper VanSteenhouse, Yumi Kasai, David Gray, Carol Clugston, Anna Dominiczak and Alex Alderton, Roberto Amato, Jeffrey Barrett, Sonia Goncalves, Ewan Harrison, David K. Jackson, Ian Johnston, Dominic Kwiatkowski, Cordelia Langford, John Sillitoe on behalf of the Wellcome Sanger Institute COVID-19 Surveillance Team |
| EPI_ISL_3106927, EPI_ISL_3106928 | Lighthouse Lab in Alderley Park | Wellcome Sanger Institute for the COVID-19 Genomics UK (COG-UK) Consortium | Jacquelyn Wynn, Mairead Hyland, The Lighthouse Lab in Alderley Park and Alex Alderton, Roberto Amato, Jeffrey Barrett, Sonia Goncalves, Ewan Harrison, David K. Jackson, Ian Johnston, Dominic Kwiatkowski, Cordelia Langford, John Sillitoe on behalf of the Wellcome Sanger Institute COVID-19 Surveillance Team |
| EPI_ISL_3106929 | Lighthouse Lab in Glasgow | Wellcome Sanger Institute for the COVID-19 Genomics UK (COG-UK) Consortium | Harper VanSteenhouse, Yumi Kasai, David Gray, Carol Clugston, Anna Dominiczak and Alex Alderton, Roberto Amato, Jeffrey Barrett, Sonia Goncalves, Ewan Harrison, David K. Jackson, Ian Johnston, Dominic Kwiatkowski, Cordelia Langford, John Sillitoe on behalf of the Wellcome Sanger Institute COVID-19 Surveillance Team |
| EPI_ISL_3106930 | Lighthouse Lab in Alderley Park | Wellcome Sanger Institute for the COVID-19 Genomics UK (COG-UK) Consortium | Jacquelyn Wynn, Mairead Hyland, The Lighthouse Lab in Alderley Park and Alex Alderton, Roberto Amato, Jeffrey Barrett, Sonia Goncalves, Ewan Harrison, David K. Jackson, Ian Johnston, Dominic Kwiatkowski, Cordelia Langford, John Sillitoe on behalf of the Wellcome Sanger Institute COVID-19 Surveillance Team |
| EPI_ISL_3106931, EPI_ISL_3106932, EPI_ISL_3106933 | Lighthouse Lab in Glasgow | Wellcome Sanger Institute for the COVID-19 Genomics UK (COG-UK) Consortium | Harper VanSteenhouse, Yumi Kasai, David Gray, Carol Clugston, Anna Dominiczak and Alex Alderton, Roberto Amato, Jeffrey Barrett, Sonia Goncalves, Ewan Harrison, David K. Jackson, Ian Johnston, Dominic Kwiatkowski, Cordelia Langford, John Sillitoe on behalf of the Wellcome Sanger Institute COVID-19 Surveillance Team |
| EPI_ISL_3106934 | Lighthouse Lab in Alderley Park | Wellcome Sanger Institute for the COVID-19 Genomics UK (COG-UK) Consortium | Jacquelyn Wynn, Mairead Hyland, The Lighthouse Lab in Alderley Park and Alex Alderton, Roberto Amato, Jeffrey Barrett, Sonia Goncalves, Ewan Harrison, David K. Jackson, Ian Johnston, Dominic Kwiatkowski, Cordelia Langford, John Sillitoe on behalf of the Wellcome Sanger Institute COVID-19 Surveillance Team |
| EPI_ISL_3106935 | Lighthouse Lab in Glasgow | Wellcome Sanger Institute for the COVID-19 Genomics UK (COG-UK) Consortium | Harper VanSteenhouse, Yumi Kasai, David Gray, Carol Clugston, Anna Dominiczak and Alex Alderton, Roberto Amato, Jeffrey Barrett, Sonia Goncalves, Ewan Harrison, David K. Jackson, Ian Johnston, Dominic Kwiatkowski, Cordelia Langford, John Sillitoe on behalf of the Wellcome Sanger Institute COVID-19 Surveillance Team |
| EPI_ISL_3106936, EPI_ISL_3106937 | Lighthouse Lab in Alderley Park | Wellcome Sanger Institute for the COVID-19 Genomics UK (COG-UK) Consortium | Jacquelyn Wynn, Mairead Hyland, The Lighthouse Lab in Alderley Park and Alex Alderton, Roberto Amato, Jeffrey Barrett, Sonia Goncalves, Ewan Harrison, David K. Jackson, Ian Johnston, Dominic Kwiatkowski, Cordelia Langford, John Sillitoe on behalf of the Wellcome Sanger Institute COVID-19 Surveillance Team |
| EPI_ISL_3106938, EPI_ISL_3106939 | Lighthouse Lab in Glasgow | Wellcome Sanger Institute for the COVID-19 Genomics UK (COG-UK) Consortium | Harper VanSteenhouse, Yumi Kasai, David Gray, Carol Clugston, Anna Dominiczak and Alex Alderton, Roberto Amato, Jeffrey Barrett, Sonia Goncalves, Ewan Harrison, David K. Jackson, Ian Johnston, Dominic Kwiatkowski, Cordelia Langford, John Sillitoe on behalf of the Wellcome Sanger Institute COVID-19 Surveillance Team |
| EPI_ISL_3106940 | Lighthouse Lab in Alderley Park | Wellcome Sanger Institute for the COVID-19 Genomics UK (COG-UK) Consortium | Jacquelyn Wynn, Mairead Hyland, The Lighthouse Lab in Alderley Park and Alex Alderton, Roberto Amato, Jeffrey Barrett, Sonia Goncalves, Ewan Harrison, David K. Jackson, Ian Johnston, Dominic Kwiatkowski, Cordelia Langford, John Sillitoe on behalf of the Wellcome Sanger Institute COVID-19 Surveillance Team |
| EPI_ISL_3106941, EPI_ISL_3106942, EPI_ISL_3106943 | Lighthouse Lab in Glasgow | Wellcome Sanger Institute for the COVID-19 Genomics UK (COG-UK) Consortium | Harper VanSteenhouse, Yumi Kasai, David Gray, Carol Clugston, Anna Dominiczak and Alex Alderton, Roberto Amato, Jeffrey Barrett, Sonia Goncalves, Ewan Harrison, David K. Jackson, Ian Johnston, Dominic Kwiatkowski, Cordelia Langford, John Sillitoe on behalf of the Wellcome Sanger Institute COVID-19 Surveillance Team |
| EPI_ISL_3106944, EPI_ISL_3106945, EPI_ISL_3106946, EPI_ISL_3106947, EPI_ISL_3106948 | Lighthouse Lab in Alderley Park | Wellcome Sanger Institute for the COVID-19 Genomics UK (COG-UK) Consortium | Jacquelyn Wynn, Mairead Hyland, The Lighthouse Lab in Alderley Park and Alex Alderton, Roberto Amato, Jeffrey Barrett, Sonia Goncalves, Ewan Harrison, David K. Jackson, Ian Johnston, Dominic Kwiatkowski, Cordelia Langford, John Sillitoe on behalf of the Wellcome Sanger Institute COVID-19 Surveillance Team |
| EPI_ISL_3106949, EPI_ISL_3106950 | Lighthouse Lab in Glasgow | Wellcome Sanger Institute for the COVID-19 Genomics UK (COG-UK) Consortium | Harper VanSteenhouse, Yumi Kasai, David Gray, Carol Clugston, Anna Dominiczak and Alex Alderton, Roberto Amato, Jeffrey Barrett, Sonia Goncalves, Ewan Harrison, David K. Jackson, Ian Johnston, Dominic Kwiatkowski, Cordelia Langford, John Sillitoe on behalf of the Wellcome Sanger Institute COVID-19 Surveillance Team |
| EPI_ISL_3106951, EPI_ISL_3106952, EPI_ISL_3106953, EPI_ISL_3106954, EPI_ISL_3106955 | Lighthouse Lab in Alderley Park | Wellcome Sanger Institute for the COVID-19 Genomics UK (COG-UK) Consortium | Jacquelyn Wynn, Mairead Hyland, The Lighthouse Lab in Alderley Park and Alex Alderton, Roberto Amato, Jeffrey Barrett, Sonia Goncalves, Ewan Harrison, David K. Jackson, Ian Johnston, Dominic Kwiatkowski, Cordelia Langford, John Sillitoe on behalf of the Wellcome Sanger Institute COVID-19 Surveillance Team |

| | | | |
|---|---|---|---|
| EPI_ISL_3106956 | Lighthouse Lab in Glasgow | Wellcome Sanger Institute for the COVID-19 Genomics UK (COG-UK) Consortium | Harper VanSteenhouse, Yumi Kasai, David Gray, Carol Clugston, Anna Dominiczak and Alex Alderton, Roberto Amato, Jeffrey Barrett, Sonia Goncalves, Ewan Harrison, David K. Jackson, Ian Johnston, Dominic Kwiatkowski, Cordelia Langford, John Sillitoe on behalf of the Wellcome Sanger Institute COVID-19 Surveillance Team |
| EPI_ISL_3106957, EPI_ISL_3106958 | Lighthouse Lab in Alderley Park | Wellcome Sanger Institute for the COVID-19 Genomics UK (COG-UK) Consortium | Jacquelyn Wynn, Mairead Hyland, The Lighthouse Lab in Alderley Park and Alex Alderton, Roberto Amato, Jeffrey Barrett, Sonia Goncalves, Ewan Harrison, David K. Jackson, Ian Johnston, Dominic Kwiatkowski, Cordelia Langford, John Sillitoe on behalf of the Wellcome Sanger Institute COVID-19 Surveillance Team |
| EPI_ISL_3106959, EPI_ISL_3106960 | Lighthouse Lab in Glasgow | Wellcome Sanger Institute for the COVID-19 Genomics UK (COG-UK) Consortium | Harper VanSteenhouse, Yumi Kasai, David Gray, Carol Clugston, Anna Dominiczak and Alex Alderton, Roberto Amato, Jeffrey Barrett, Sonia Goncalves, Ewan Harrison, David K. Jackson, Ian Johnston, Dominic Kwiatkowski, Cordelia Langford, John Sillitoe on behalf of the Wellcome Sanger Institute COVID-19 Surveillance Team |
| EPI_ISL_3106961, EPI_ISL_3106962, EPI_ISL_3106963, EPI_ISL_3106964, EPI_ISL_3106965, EPI_ISL_3106966, EPI_ISL_3106967 | Lighthouse Lab in Alderley Park | Wellcome Sanger Institute for the COVID-19 Genomics UK (COG-UK) Consortium | Jacquelyn Wynn, Mairead Hyland, The Lighthouse Lab in Alderley Park and Alex Alderton, Roberto Amato, Jeffrey Barrett, Sonia Goncalves, Ewan Harrison, David K. Jackson, Ian Johnston, Dominic Kwiatkowski, Cordelia Langford, John Sillitoe on behalf of the Wellcome Sanger Institute COVID-19 Surveillance Team |
| EPI_ISL_3106968, EPI_ISL_3106969, EPI_ISL_3106970 | Lighthouse Lab in Glasgow | Wellcome Sanger Institute for the COVID-19 Genomics UK (COG-UK) Consortium | Harper VanSteenhouse, Yumi Kasai, David Gray, Carol Clugston, Anna Dominiczak and Alex Alderton, Roberto Amato, Jeffrey Barrett, Sonia Goncalves, Ewan Harrison, David K. Jackson, Ian Johnston, Dominic Kwiatkowski, Cordelia Langford, John Sillitoe on behalf of the Wellcome Sanger Institute COVID-19 Surveillance Team |
| EPI_ISL_3106971, EPI_ISL_3106972 | Lighthouse Lab in Alderley Park | Wellcome Sanger Institute for the COVID-19 Genomics UK (COG-UK) Consortium | Jacquelyn Wynn, Mairead Hyland, The Lighthouse Lab in Alderley Park and Alex Alderton, Roberto Amato, Jeffrey Barrett, Sonia Goncalves, Ewan Harrison, David K. Jackson, Ian Johnston, Dominic Kwiatkowski, Cordelia Langford, John Sillitoe on behalf of the Wellcome Sanger Institute COVID-19 Surveillance Team |
| EPI_ISL_3106973 | Lighthouse Lab in Glasgow | Wellcome Sanger Institute for the COVID-19 Genomics UK (COG-UK) Consortium | Harper VanSteenhouse, Yumi Kasai, David Gray, Carol Clugston, Anna Dominiczak and Alex Alderton, Roberto Amato, Jeffrey Barrett, Sonia Goncalves, Ewan Harrison, David K. Jackson, Ian Johnston, Dominic Kwiatkowski, Cordelia Langford, John Sillitoe on behalf of the Wellcome Sanger Institute COVID-19 Surveillance Team |
| EPI_ISL_3106974 | Lighthouse Lab in Alderley Park | Wellcome Sanger Institute for the COVID-19 Genomics UK (COG-UK) Consortium | Jacquelyn Wynn, Mairead Hyland, The Lighthouse Lab in Alderley Park and Alex Alderton, Roberto Amato, Jeffrey Barrett, Sonia Goncalves, Ewan Harrison, David K. Jackson, Ian Johnston, Dominic Kwiatkowski, Cordelia Langford, John Sillitoe on behalf of the Wellcome Sanger Institute COVID-19 Surveillance Team |
| EPI_ISL_3106975 | Lighthouse Lab in Glasgow | Wellcome Sanger Institute for the COVID-19 Genomics UK (COG-UK) Consortium | Harper VanSteenhouse, Yumi Kasai, David Gray, Carol Clugston, Anna Dominiczak and Alex Alderton, Roberto Amato, Jeffrey Barrett, Sonia Goncalves, Ewan Harrison, David K. Jackson, Ian Johnston, Dominic Kwiatkowski, Cordelia Langford, John Sillitoe on behalf of the Wellcome Sanger Institute COVID-19 Surveillance Team |
| EPI_ISL_3106976 | Lighthouse Lab in Alderley Park | Wellcome Sanger Institute for the COVID-19 Genomics UK (COG-UK) Consortium | Jacquelyn Wynn, Mairead Hyland, The Lighthouse Lab in Alderley Park and Alex Alderton, Roberto Amato, Jeffrey Barrett, Sonia Goncalves, Ewan Harrison, David K. Jackson, Ian Johnston, Dominic Kwiatkowski, Cordelia Langford, John Sillitoe on behalf of the Wellcome Sanger Institute COVID-19 Surveillance Team |
| EPI_ISL_3106977 | Lighthouse Lab in Glasgow | Wellcome Sanger Institute for the COVID-19 Genomics UK (COG-UK) Consortium | Harper VanSteenhouse, Yumi Kasai, David Gray, Carol Clugston, Anna Dominiczak and Alex Alderton, Roberto Amato, Jeffrey Barrett, Sonia Goncalves, Ewan Harrison, David K. Jackson, Ian Johnston, Dominic Kwiatkowski, Cordelia Langford, John Sillitoe on behalf of the Wellcome Sanger Institute COVID-19 Surveillance Team |
| EPI_ISL_3106978 | Lighthouse Lab in Alderley Park | Wellcome Sanger Institute for the COVID-19 Genomics UK (COG-UK) Consortium | Jacquelyn Wynn, Mairead Hyland, The Lighthouse Lab in Alderley Park and Alex Alderton, Roberto Amato, Jeffrey Barrett, Sonia Goncalves, Ewan Harrison, David K. Jackson, Ian Johnston, Dominic Kwiatkowski, Cordelia Langford, John Sillitoe on behalf of the Wellcome Sanger Institute COVID-19 Surveillance Team |
| EPI_ISL_3106979, EPI_ISL_3106980 | Lighthouse Lab in Glasgow | Wellcome Sanger Institute for the COVID-19 Genomics UK (COG-UK) Consortium | Harper VanSteenhouse, Yumi Kasai, David Gray, Carol Clugston, Anna Dominiczak and Alex Alderton, Roberto Amato, Jeffrey Barrett, Sonia Goncalves, Ewan Harrison, David K. Jackson, Ian Johnston, Dominic Kwiatkowski, Cordelia Langford, John Sillitoe on behalf of the Wellcome Sanger Institute COVID-19 Surveillance Team |
| EPI_ISL_3106981, EPI_ISL_3106982, EPI_ISL_3106983, EPI_ISL_3106984, EPI_ISL_3106985, EPI_ISL_3106986, EPI_ISL_3106987 | Lighthouse Lab in Alderley Park | Wellcome Sanger Institute for the COVID-19 Genomics UK (COG-UK) Consortium | Jacquelyn Wynn, Mairead Hyland, The Lighthouse Lab in Alderley Park and Alex Alderton, Roberto Amato, Jeffrey Barrett, Sonia Goncalves, Ewan Harrison, David K. Jackson, Ian Johnston, Dominic Kwiatkowski, Cordelia Langford, John Sillitoe on behalf of the Wellcome Sanger Institute COVID-19 Surveillance Team |
| EPI_ISL_3106988 | Lighthouse Lab in Glasgow | Wellcome Sanger Institute for the COVID-19 Genomics UK (COG-UK) Consortium | Harper VanSteenhouse, Yumi Kasai, David Gray, Carol Clugston, Anna Dominiczak and Alex Alderton, Roberto Amato, Jeffrey Barrett, Sonia Goncalves, Ewan Harrison, David K. Jackson, Ian Johnston, Dominic Kwiatkowski, Cordelia Langford, John Sillitoe on behalf of the Wellcome Sanger Institute COVID-19 Surveillance Team |
| EPI_ISL_3106989 | Lighthouse Lab in Alderley Park | Wellcome Sanger Institute for the COVID-19 Genomics UK (COG-UK) Consortium | Jacquelyn Wynn, Mairead Hyland, The Lighthouse Lab in Alderley Park and Alex Alderton, Roberto Amato, Jeffrey Barrett, Sonia Goncalves, Ewan Harrison, David K. Jackson, Ian Johnston, Dominic Kwiatkowski, Cordelia Langford, John Sillitoe on behalf of the Wellcome Sanger Institute COVID-19 Surveillance Team |
| EPI_ISL_3106990 | Lighthouse Lab in Glasgow | Wellcome Sanger Institute for the COVID-19 Genomics UK (COG-UK) Consortium | Harper VanSteenhouse, Yumi Kasai, David Gray, Carol Clugston, Anna Dominiczak and Alex Alderton, Roberto Amato, Jeffrey Barrett, Sonia Goncalves, Ewan Harrison, David K. Jackson, Ian Johnston, Dominic Kwiatkowski, Cordelia Langford, John Sillitoe on behalf of the Wellcome Sanger Institute COVID-19 Surveillance Team |
| EPI_ISL_3106991 | Lighthouse Lab in Alderley Park | Wellcome Sanger Institute for the COVID-19 Genomics UK (COG-UK) Consortium | Jacquelyn Wynn, Mairead Hyland, The Lighthouse Lab in Alderley Park and Alex Alderton, Roberto Amato, Jeffrey Barrett, Sonia Goncalves, Ewan Harrison, David K. Jackson, Ian Johnston, Dominic Kwiatkowski, Cordelia Langford, John Sillitoe on behalf of the Wellcome Sanger Institute COVID-19 Surveillance Team |
| EPI_ISL_3106992 | Lighthouse Lab in Glasgow | Wellcome Sanger Institute for the COVID-19 Genomics UK (COG-UK) Consortium | Harper VanSteenhouse, Yumi Kasai, David Gray, Carol Clugston, Anna Dominiczak and Alex Alderton, Roberto Amato, Jeffrey Barrett, Sonia Goncalves, Ewan Harrison, David K. Jackson, Ian Johnston, Dominic Kwiatkowski, Cordelia Langford, John Sillitoe on behalf of the Wellcome Sanger Institute COVID-19 Surveillance Team |
| EPI_ISL_3106993 | Lighthouse Lab in Alderley Park | Wellcome Sanger Institute for the COVID-19 Genomics UK (COG-UK) Consortium | Jacquelyn Wynn, Mairead Hyland, The Lighthouse Lab in Alderley Park and Alex Alderton, Roberto Amato, Jeffrey Barrett, Sonia Goncalves, Ewan Harrison, David K. Jackson, Ian Johnston, Dominic Kwiatkowski, Cordelia Langford, John Sillitoe on behalf of the Wellcome Sanger Institute COVID-19 Surveillance Team |
| EPI_ISL_3106994 | Lighthouse Lab in Glasgow | Wellcome Sanger Institute for the COVID-19 Genomics UK (COG-UK) Consortium | Harper VanSteenhouse, Yumi Kasai, David Gray, Carol Clugston, Anna Dominiczak and Alex Alderton, Roberto Amato, Jeffrey Barrett, Sonia Goncalves, Ewan Harrison, David K. Jackson, Ian Johnston, Dominic Kwiatkowski, Cordelia Langford, John Sillitoe on behalf of the Wellcome Sanger Institute COVID-19 Surveillance Team |
| EPI_ISL_3106995, EPI_ISL_3106996, EPI_ISL_3106997 | Lighthouse Lab in Alderley Park | Wellcome Sanger Institute for the COVID-19 Genomics UK (COG-UK) Consortium | Jacquelyn Wynn, Mairead Hyland, The Lighthouse Lab in Alderley Park and Alex Alderton, Roberto Amato, Jeffrey Barrett, Sonia Goncalves, Ewan Harrison, David K. Jackson, Ian Johnston, Dominic Kwiatkowski, Cordelia Langford, John Sillitoe on behalf of the Wellcome Sanger Institute COVID-19 Surveillance Team |
| EPI_ISL_3106998 | Lighthouse Lab in Glasgow | Wellcome Sanger Institute for the COVID-19 Genomics UK (COG-UK) Consortium | Harper VanSteenhouse, Yumi Kasai, David Gray, Carol Clugston, Anna Dominiczak and Alex Alderton, Roberto Amato, Jeffrey Barrett, Sonia Goncalves, Ewan Harrison, David K. Jackson, Ian Johnston, Dominic Kwiatkowski, Cordelia Langford, John Sillitoe on behalf of the Wellcome Sanger Institute COVID-19 Surveillance Team |
| EPI_ISL_3106999 | Lighthouse Lab in Alderley Park | Wellcome Sanger Institute for the COVID-19 Genomics UK (COG-UK) Consortium | Jacquelyn Wynn, Mairead Hyland, The Lighthouse Lab in Alderley Park and Alex Alderton, Roberto Amato, Jeffrey Barrett, Sonia Goncalves, Ewan Harrison, David K. Jackson, Ian Johnston, Dominic Kwiatkowski, Cordelia Langford, John Sillitoe on behalf of the Wellcome Sanger Institute COVID-19 Surveillance Team |
| EPI_ISL_3107000, EPI_ISL_3107001 | Lighthouse Lab in Glasgow | Wellcome Sanger Institute for the COVID-19 Genomics UK (COG-UK) Consortium | Harper VanSteenhouse, Yumi Kasai, David Gray, Carol Clugston, Anna Dominiczak and Alex Alderton, Roberto Amato, Jeffrey Barrett, Sonia Goncalves, Ewan Harrison, David K. Jackson, Ian Johnston, Dominic Kwiatkowski, Cordelia Langford, John Sillitoe on behalf of the Wellcome Sanger Institute COVID-19 Surveillance Team |
| EPI_ISL_3107002 | Lighthouse Lab in Alderley Park | Wellcome Sanger Institute for the COVID-19 Genomics UK (COG-UK) Consortium | Jacquelyn Wynn, Mairead Hyland, The Lighthouse Lab in Alderley Park and Alex Alderton, Roberto Amato, Jeffrey Barrett, Sonia Goncalves, Ewan Harrison, David K. Jackson, Ian Johnston, Dominic Kwiatkowski, Cordelia Langford, John Sillitoe on behalf of the Wellcome Sanger Institute COVID-19 Surveillance Team |
| EPI_ISL_3107003, EPI_ISL_3107004, EPI_ISL_3107005 | Lighthouse Lab in Glasgow | Wellcome Sanger Institute for the COVID-19 Genomics UK (COG-UK) Consortium | Harper VanSteenhouse, Yumi Kasai, David Gray, Carol Clugston, Anna Dominiczak and Alex Alderton, Roberto Amato, Jeffrey Barrett, Sonia Goncalves, Ewan Harrison, David K. Jackson, Ian Johnston, Dominic Kwiatkowski, Cordelia Langford, John Sillitoe on behalf of the Wellcome Sanger Institute COVID-19 Surveillance Team |
| EPI_ISL_3107006 | Lighthouse Lab in Alderley Park | Wellcome Sanger Institute for the COVID-19 Genomics UK (COG-UK) Consortium | Jacquelyn Wynn, Mairead Hyland, The Lighthouse Lab in Alderley Park and Alex Alderton, Roberto Amato, Jeffrey Barrett, Sonia Goncalves, Ewan Harrison, David K. Jackson, Ian Johnston, Dominic Kwiatkowski, Cordelia Langford, John Sillitoe on behalf of the Wellcome Sanger Institute COVID-19 Surveillance Team |
| EPI_ISL_3107007, EPI_ISL_3107008 | Lighthouse Lab in Glasgow | Wellcome Sanger Institute for the COVID-19 Genomics UK (COG-UK) Consortium | Harper VanSteenhouse, Yumi Kasai, David Gray, Carol Clugston, Anna Dominiczak and Alex Alderton, Roberto Amato, Jeffrey Barrett, Sonia Goncalves, Ewan Harrison, David K. Jackson, Ian Johnston, Dominic Kwiatkowski, Cordelia Langford, John Sillitoe on behalf of the Wellcome Sanger Institute COVID-19 Surveillance Team |
| EPI_ISL_3107009, EPI_ISL_3107010 | Lighthouse Lab in Alderley Park | Wellcome Sanger Institute for the COVID-19 Genomics UK (COG-UK) Consortium | Jacquelyn Wynn, Mairead Hyland, The Lighthouse Lab in Alderley Park and Alex Alderton, Roberto Amato, Jeffrey Barrett, Sonia Goncalves, Ewan Harrison, David K. Jackson, Ian Johnston, Dominic Kwiatkowski, Cordelia Langford, John Sillitoe on behalf of the Wellcome Sanger Institute COVID-19 Surveillance Team |
| EPI_ISL_3107011 | Lighthouse Lab in Glasgow | Wellcome Sanger Institute for the COVID-19 Genomics UK (COG-UK) Consortium | Harper VanSteenhouse, Yumi Kasai, David Gray, Carol Clugston, Anna Dominiczak and Alex Alderton, Roberto Amato, Jeffrey Barrett, Sonia Goncalves, Ewan Harrison, David K. Jackson, Ian Johnston, Dominic Kwiatkowski, Cordelia Langford, John Sillitoe on behalf of the Wellcome Sanger Institute COVID-19 Surveillance Team |
| EPI_ISL_3107012, EPI_ISL_3107013 | Lighthouse Lab in Alderley Park | Wellcome Sanger Institute for the COVID-19 Genomics UK (COG-UK) Consortium | Jacquelyn Wynn, Mairead Hyland, The Lighthouse Lab in Alderley Park and Alex Alderton, Roberto Amato, Jeffrey Barrett, Sonia Goncalves, Ewan Harrison, David K. Jackson, Ian Johnston, Dominic Kwiatkowski, Cordelia Langford, John Sillitoe on behalf of the Wellcome Sanger Institute COVID-19 Surveillance Team |
| EPI_ISL_3107014 | Lighthouse Lab in Glasgow | Wellcome Sanger Institute for the COVID-19 Genomics UK (COG-UK) Consortium | Harper VanSteenhouse, Yumi Kasai, David Gray, Carol Clugston, Anna Dominiczak and Alex Alderton, Roberto Amato, Jeffrey Barrett, Sonia Goncalves, Ewan Harrison, David K. Jackson, Ian Johnston, Dominic Kwiatkowski, Cordelia Langford, John Sillitoe on behalf of the Wellcome Sanger Institute COVID-19 Surveillance Team |
| EPI_ISL_3107015, EPI_ISL_3107016, EPI_ISL_3107017, EPI_ISL_3107018, EPI_ISL_3107019 | Lighthouse Lab in Alderley Park | Wellcome Sanger Institute for the COVID-19 Genomics UK (COG-UK) Consortium | Jacquelyn Wynn, Mairead Hyland, The Lighthouse Lab in Alderley Park and Alex Alderton, Roberto Amato, Jeffrey Barrett, Sonia Goncalves, Ewan Harrison, David K. Jackson, Ian Johnston, Dominic Kwiatkowski, Cordelia Langford, John Sillitoe on behalf of the Wellcome Sanger Institute COVID-19 Surveillance Team |
| EPI_ISL_3107020 | Lighthouse Lab in Glasgow | Wellcome Sanger Institute for the COVID-19 Genomics UK (COG-UK) Consortium | Harper VanSteenhouse, Yumi Kasai, David Gray, Carol Clugston, Anna Dominiczak and Alex Alderton, Roberto Amato, Jeffrey Barrett, Sonia Goncalves, Ewan Harrison, David K. Jackson, Ian Johnston, Dominic Kwiatkowski, Cordelia Langford, John Sillitoe on behalf of the Wellcome Sanger Institute COVID-19 Surveillance Team |
| EPI_ISL_3107021, EPI_ISL_3107022 | Lighthouse Lab in Alderley Park | Wellcome Sanger Institute for the COVID-19 Genomics UK (COG-UK) Consortium | Jacquelyn Wynn, Mairead Hyland, The Lighthouse Lab in Alderley Park and Alex Alderton, Roberto Amato, Jeffrey Barrett, Sonia Goncalves, Ewan Harrison, David K. Jackson, Ian Johnston, Dominic Kwiatkowski, Cordelia Langford, John Sillitoe on behalf of the Wellcome Sanger Institute COVID-19 Surveillance Team |
| EPI_ISL_3107023, EPI_ISL_3107024, EPI_ISL_3107025 | Lighthouse Lab in Glasgow | Wellcome Sanger Institute for the COVID-19 Genomics UK (COG-UK) Consortium | Harper VanSteenhouse, Yumi Kasai, David Gray, Carol Clugston, Anna Dominiczak and Alex Alderton, Roberto Amato, Jeffrey Barrett, Sonia Goncalves, Ewan Harrison, David K. Jackson, Ian Johnston, Dominic Kwiatkowski, Cordelia Langford, John Sillitoe on behalf of the Wellcome Sanger Institute COVID-19 Surveillance Team |
| EPI_ISL_3107026 | Lighthouse Lab in Alderley Park | Wellcome Sanger Institute for the COVID-19 Genomics UK (COG-UK) Consortium | Jacquelyn Wynn, Mairead Hyland, The Lighthouse Lab in Alderley Park and Alex Alderton, Roberto Amato, Jeffrey Barrett, Sonia Goncalves, Ewan Harrison, David K. Jackson, Ian Johnston, Dominic Kwiatkowski, Cordelia Langford, John Sillitoe on behalf of the Wellcome Sanger Institute COVID-19 Surveillance Team |

| | | | |
|---|---|---|---|
| EPI_ISL_3107027, EPI_ISL_3107028 | Lighthouse Lab in Glasgow | Wellcome Sanger Institute for the COVID-19 Genomics UK (COG-UK) Consortium | Harper VanSteenhouse, Yumi Kasai, David Gray, Carol Clugston, Anna Dominiczak and Alex Alderton, Roberto Amato, Jeffrey Barrett, Sonia Goncalves, Ewan Harrison, David K. Jackson, Ian Johnston, Dominic Kwiatkowski, Cordelia Langford, John Sillitoe on behalf of the Wellcome Sanger Institute COVID-19 Surveillance Team |
| EPI_ISL_3107029 | Lighthouse Lab in Alderley Park | Wellcome Sanger Institute for the COVID-19 Genomics UK (COG-UK) Consortium | Jacquelyn Wynn, Mairead Hyland, The Lighthouse Lab in Alderley Park and Alex Alderton, Roberto Amato, Jeffrey Barrett, Sonia Goncalves, Ewan Harrison, David K. Jackson, Ian Johnston, Dominic Kwiatkowski, Cordelia Langford, John Sillitoe on behalf of the Wellcome Sanger Institute COVID-19 Surveillance Team |
| EPI_ISL_3107030, EPI_ISL_3107031 | Lighthouse Lab in Glasgow | Wellcome Sanger Institute for the COVID-19 Genomics UK (COG-UK) Consortium | Harper VanSteenhouse, Yumi Kasai, David Gray, Carol Clugston, Anna Dominiczak and Alex Alderton, Roberto Amato, Jeffrey Barrett, Sonia Goncalves, Ewan Harrison, David K. Jackson, Ian Johnston, Dominic Kwiatkowski, Cordelia Langford, John Sillitoe on behalf of the Wellcome Sanger Institute COVID-19 Surveillance Team |
| EPI_ISL_3107032, EPI_ISL_3107033, EPI_ISL_3107034 | Lighthouse Lab in Alderley Park | Wellcome Sanger Institute for the COVID-19 Genomics UK (COG-UK) Consortium | Jacquelyn Wynn, Mairead Hyland, The Lighthouse Lab in Alderley Park and Alex Alderton, Roberto Amato, Jeffrey Barrett, Sonia Goncalves, Ewan Harrison, David K. Jackson, Ian Johnston, Dominic Kwiatkowski, Cordelia Langford, John Sillitoe on behalf of the Wellcome Sanger Institute COVID-19 Surveillance Team |
| EPI_ISL_3107035 | Lighthouse Lab in Glasgow | Wellcome Sanger Institute for the COVID-19 Genomics UK (COG-UK) Consortium | Harper VanSteenhouse, Yumi Kasai, David Gray, Carol Clugston, Anna Dominiczak and Alex Alderton, Roberto Amato, Jeffrey Barrett, Sonia Goncalves, Ewan Harrison, David K. Jackson, Ian Johnston, Dominic Kwiatkowski, Cordelia Langford, John Sillitoe on behalf of the Wellcome Sanger Institute COVID-19 Surveillance Team |
| EPI_ISL_3107036, EPI_ISL_3107037, EPI_ISL_3107038 | Lighthouse Lab in Alderley Park | Wellcome Sanger Institute for the COVID-19 Genomics UK (COG-UK) Consortium | Jacquelyn Wynn, Mairead Hyland, The Lighthouse Lab in Alderley Park and Alex Alderton, Roberto Amato, Jeffrey Barrett, Sonia Goncalves, Ewan Harrison, David K. Jackson, Ian Johnston, Dominic Kwiatkowski, Cordelia Langford, John Sillitoe on behalf of the Wellcome Sanger Institute COVID-19 Surveillance Team |
| EPI_ISL_3107039 | Lighthouse Lab in Glasgow | Wellcome Sanger Institute for the COVID-19 Genomics UK (COG-UK) Consortium | Harper VanSteenhouse, Yumi Kasai, David Gray, Carol Clugston, Anna Dominiczak and Alex Alderton, Roberto Amato, Jeffrey Barrett, Sonia Goncalves, Ewan Harrison, David K. Jackson, Ian Johnston, Dominic Kwiatkowski, Cordelia Langford, John Sillitoe on behalf of the Wellcome Sanger Institute COVID-19 Surveillance Team |
| EPI_ISL_3107040, EPI_ISL_3107041, EPI_ISL_3107042 | Lighthouse Lab in Alderley Park | Wellcome Sanger Institute for the COVID-19 Genomics UK (COG-UK) Consortium | Jacquelyn Wynn, Mairead Hyland, The Lighthouse Lab in Alderley Park and Alex Alderton, Roberto Amato, Jeffrey Barrett, Sonia Goncalves, Ewan Harrison, David K. Jackson, Ian Johnston, Dominic Kwiatkowski, Cordelia Langford, John Sillitoe on behalf of the Wellcome Sanger Institute COVID-19 Surveillance Team |
| EPI_ISL_3107043 | Lighthouse Lab in Glasgow | Wellcome Sanger Institute for the COVID-19 Genomics UK (COG-UK) Consortium | Harper VanSteenhouse, Yumi Kasai, David Gray, Carol Clugston, Anna Dominiczak and Alex Alderton, Roberto Amato, Jeffrey Barrett, Sonia Goncalves, Ewan Harrison, David K. Jackson, Ian Johnston, Dominic Kwiatkowski, Cordelia Langford, John Sillitoe on behalf of the Wellcome Sanger Institute COVID-19 Surveillance Team |
| EPI_ISL_3107044 | Lighthouse Lab in Alderley Park | Wellcome Sanger Institute for the COVID-19 Genomics UK (COG-UK) Consortium | Jacquelyn Wynn, Mairead Hyland, The Lighthouse Lab in Alderley Park and Alex Alderton, Roberto Amato, Jeffrey Barrett, Sonia Goncalves, Ewan Harrison, David K. Jackson, Ian Johnston, Dominic Kwiatkowski, Cordelia Langford, John Sillitoe on behalf of the Wellcome Sanger Institute COVID-19 Surveillance Team |
| EPI_ISL_3107045, EPI_ISL_3107046 | Lighthouse Lab in Glasgow | Wellcome Sanger Institute for the COVID-19 Genomics UK (COG-UK) Consortium | Harper VanSteenhouse, Yumi Kasai, David Gray, Carol Clugston, Anna Dominiczak and Alex Alderton, Roberto Amato, Jeffrey Barrett, Sonia Goncalves, Ewan Harrison, David K. Jackson, Ian Johnston, Dominic Kwiatkowski, Cordelia Langford, John Sillitoe on behalf of the Wellcome Sanger Institute COVID-19 Surveillance Team |
| EPI_ISL_3107047, EPI_ISL_3107048 | Lighthouse Lab in Alderley Park | Wellcome Sanger Institute for the COVID-19 Genomics UK (COG-UK) Consortium | Jacquelyn Wynn, Mairead Hyland, The Lighthouse Lab in Alderley Park and Alex Alderton, Roberto Amato, Jeffrey Barrett, Sonia Goncalves, Ewan Harrison, David K. Jackson, Ian Johnston, Dominic Kwiatkowski, Cordelia Langford, John Sillitoe on behalf of the Wellcome Sanger Institute COVID-19 Surveillance Team |
| EPI_ISL_3107049, EPI_ISL_3107050 | Berkshire and Surrey Pathology Services Lighthouse Laboratory | Wellcome Sanger Institute for the COVID-19 Genomics UK (COG-UK) Consortium | Berkshire and Surrey Pathology Services Lighthouse Laboratory and Alex Alderton, Roberto Amato, Jeffrey Barrett, Sonia Goncalves, Ewan Harrison, David K. Jackson, Ian Johnston, Dominic Kwiatkowski, Cordelia Langford, John Sillitoe on behalf of the Wellcome Sanger Institute COVID-19 Surveillance Team |
| EPI_ISL_3107051 | Lighthouse Lab in Alderley Park | Wellcome Sanger Institute for the COVID-19 Genomics UK (COG-UK) Consortium | Jacquelyn Wynn, Mairead Hyland, The Lighthouse Lab in Alderley Park and Alex Alderton, Roberto Amato, Jeffrey Barrett, Sonia Goncalves, Ewan Harrison, David K. Jackson, Ian Johnston, Dominic Kwiatkowski, Cordelia Langford, John Sillitoe on behalf of the Wellcome Sanger Institute COVID-19 Surveillance Team |
| EPI_ISL_3107052, EPI_ISL_3107053 | Berkshire and Surrey Pathology Services Lighthouse Laboratory | Wellcome Sanger Institute for the COVID-19 Genomics UK (COG-UK) Consortium | Berkshire and Surrey Pathology Services Lighthouse Laboratory and Alex Alderton, Roberto Amato, Jeffrey Barrett, Sonia Goncalves, Ewan Harrison, David K. Jackson, Ian Johnston, Dominic Kwiatkowski, Cordelia Langford, John Sillitoe on behalf of the Wellcome Sanger Institute COVID-19 Surveillance Team |
| EPI_ISL_3107054 | Lighthouse Lab in Alderley Park | Wellcome Sanger Institute for the COVID-19 Genomics UK (COG-UK) Consortium | Jacquelyn Wynn, Mairead Hyland, The Lighthouse Lab in Alderley Park and Alex Alderton, Roberto Amato, Jeffrey Barrett, Sonia Goncalves, Ewan Harrison, David K. Jackson, Ian Johnston, Dominic Kwiatkowski, Cordelia Langford, John Sillitoe on behalf of the Wellcome Sanger Institute COVID-19 Surveillance Team |
| EPI_ISL_3107055, EPI_ISL_3107056, EPI_ISL_3107057, EPI_ISL_3107058, EPI_ISL_3107059, EPI_ISL_3107060, EPI_ISL_3107061 | Berkshire and Surrey Pathology Services Lighthouse Laboratory | Wellcome Sanger Institute for the COVID-19 Genomics UK (COG-UK) Consortium | Berkshire and Surrey Pathology Services Lighthouse Laboratory and Alex Alderton, Roberto Amato, Jeffrey Barrett, Sonia Goncalves, Ewan Harrison, David K. Jackson, Ian Johnston, Dominic Kwiatkowski, Cordelia Langford, John Sillitoe on behalf of the Wellcome Sanger Institute COVID-19 Surveillance Team |
| EPI_ISL_3107062 | Lighthouse Lab in Alderley Park | Wellcome Sanger Institute for the COVID-19 Genomics UK (COG-UK) Consortium | Jacquelyn Wynn, Mairead Hyland, The Lighthouse Lab in Alderley Park and Alex Alderton, Roberto Amato, Jeffrey Barrett, Sonia Goncalves, Ewan Harrison, David K. Jackson, Ian Johnston, Dominic Kwiatkowski, Cordelia Langford, John Sillitoe on behalf of the Wellcome Sanger Institute COVID-19 Surveillance Team |
| EPI_ISL_3107063 | Berkshire and Surrey Pathology Services Lighthouse Laboratory | Wellcome Sanger Institute for the COVID-19 Genomics UK (COG-UK) Consortium | Berkshire and Surrey Pathology Services Lighthouse Laboratory and Alex Alderton, Roberto Amato, Jeffrey Barrett, Sonia Goncalves, Ewan Harrison, David K. Jackson, Ian Johnston, Dominic Kwiatkowski, Cordelia Langford, John Sillitoe on behalf of the Wellcome Sanger Institute COVID-19 Surveillance Team |
| EPI_ISL_3107064, EPI_ISL_3107065 | Lighthouse Lab in Alderley Park | Wellcome Sanger Institute for the COVID-19 Genomics UK (COG-UK) Consortium | Jacquelyn Wynn, Mairead Hyland, The Lighthouse Lab in Alderley Park and Alex Alderton, Roberto Amato, Jeffrey Barrett, Sonia Goncalves, Ewan Harrison, David K. Jackson, Ian Johnston, Dominic Kwiatkowski, Cordelia Langford, John Sillitoe on behalf of the Wellcome Sanger Institute COVID-19 Surveillance Team |
| EPI_ISL_3107066 | Berkshire and Surrey Pathology Services Lighthouse Laboratory | Wellcome Sanger Institute for the COVID-19 Genomics UK (COG-UK) Consortium | Berkshire and Surrey Pathology Services Lighthouse Laboratory and Alex Alderton, Roberto Amato, Jeffrey Barrett, Sonia Goncalves, Ewan Harrison, David K. Jackson, Ian Johnston, Dominic Kwiatkowski, Cordelia Langford, John Sillitoe on behalf of the Wellcome Sanger Institute COVID-19 Surveillance Team |
| EPI_ISL_3107067, EPI_ISL_3107068 | Lighthouse Lab in Alderley Park | Wellcome Sanger Institute for the COVID-19 Genomics UK (COG-UK) Consortium | Jacquelyn Wynn, Mairead Hyland, The Lighthouse Lab in Alderley Park and Alex Alderton, Roberto Amato, Jeffrey Barrett, Sonia Goncalves, Ewan Harrison, David K. Jackson, Ian Johnston, Dominic Kwiatkowski, Cordelia Langford, John Sillitoe on behalf of the Wellcome Sanger Institute COVID-19 Surveillance Team |
| EPI_ISL_3107069 | Berkshire and Surrey Pathology Services Lighthouse Laboratory | Wellcome Sanger Institute for the COVID-19 Genomics UK (COG-UK) Consortium | Berkshire and Surrey Pathology Services Lighthouse Laboratory and Alex Alderton, Roberto Amato, Jeffrey Barrett, Sonia Goncalves, Ewan Harrison, David K. Jackson, Ian Johnston, Dominic Kwiatkowski, Cordelia Langford, John Sillitoe on behalf of the Wellcome Sanger Institute COVID-19 Surveillance Team |
| EPI_ISL_3107070, EPI_ISL_3107071, EPI_ISL_3107072 | Lighthouse Lab in Milton Keynes | Wellcome Sanger Institute for the COVID-19 Genomics UK (COG-UK) Consortium | The Lighthouse Lab in Milton Keynes and Alex Alderton, Roberto Amato, Jeffrey Barrett, Sonia Goncalves, Ewan Harrison, David K. Jackson, Ian Johnston, Dominic Kwiatkowski, Cordelia Langford, John Sillitoe on behalf of the Wellcome Sanger Institute COVID-19 Surveillance Team |
| EPI_ISL_3107073 | Lighthouse Lab in Alderley Park | Wellcome Sanger Institute for the COVID-19 Genomics UK (COG-UK) Consortium | Jacquelyn Wynn, Mairead Hyland, The Lighthouse Lab in Alderley Park and Alex Alderton, Roberto Amato, Jeffrey Barrett, Sonia Goncalves, Ewan Harrison, David K. Jackson, Ian Johnston, Dominic Kwiatkowski, Cordelia Langford, John Sillitoe on behalf of the Wellcome Sanger Institute COVID-19 Surveillance Team |
| EPI_ISL_3107074 | Berkshire and Surrey Pathology Services Lighthouse Laboratory | Wellcome Sanger Institute for the COVID-19 Genomics UK (COG-UK) Consortium | Berkshire and Surrey Pathology Services Lighthouse Laboratory and Alex Alderton, Roberto Amato, Jeffrey Barrett, Sonia Goncalves, Ewan Harrison, David K. Jackson, Ian Johnston, Dominic Kwiatkowski, Cordelia Langford, John Sillitoe on behalf of the Wellcome Sanger Institute COVID-19 Surveillance Team |
| EPI_ISL_3107075, EPI_ISL_3107076 | Lighthouse Lab in Alderley Park | Wellcome Sanger Institute for the COVID-19 Genomics UK (COG-UK) Consortium | Jacquelyn Wynn, Mairead Hyland, The Lighthouse Lab in Alderley Park and Alex Alderton, Roberto Amato, Jeffrey Barrett, Sonia Goncalves, Ewan Harrison, David K. Jackson, Ian Johnston, Dominic Kwiatkowski, Cordelia Langford, John Sillitoe on behalf of the Wellcome Sanger Institute COVID-19 Surveillance Team |
| EPI_ISL_3107077, EPI_ISL_3107078 | Berkshire and Surrey Pathology Services Lighthouse Laboratory | Wellcome Sanger Institute for the COVID-19 Genomics UK (COG-UK) Consortium | Berkshire and Surrey Pathology Services Lighthouse Laboratory and Alex Alderton, Roberto Amato, Jeffrey Barrett, Sonia Goncalves, Ewan Harrison, David K. Jackson, Ian Johnston, Dominic Kwiatkowski, Cordelia Langford, John Sillitoe on behalf of the Wellcome Sanger Institute COVID-19 Surveillance Team |
| EPI_ISL_3107079 | Lighthouse Lab in Alderley Park | Wellcome Sanger Institute for the COVID-19 Genomics UK (COG-UK) Consortium | Jacquelyn Wynn, Mairead Hyland, The Lighthouse Lab in Alderley Park and Alex Alderton, Roberto Amato, Jeffrey Barrett, Sonia Goncalves, Ewan Harrison, David K. Jackson, Ian Johnston, Dominic Kwiatkowski, Cordelia Langford, John Sillitoe on behalf of the Wellcome Sanger Institute COVID-19 Surveillance Team |
| EPI_ISL_3107080 | Berkshire and Surrey Pathology Services Lighthouse Laboratory | Wellcome Sanger Institute for the COVID-19 Genomics UK (COG-UK) Consortium | Berkshire and Surrey Pathology Services Lighthouse Laboratory and Alex Alderton, Roberto Amato, Jeffrey Barrett, Sonia Goncalves, Ewan Harrison, David K. Jackson, Ian Johnston, Dominic Kwiatkowski, Cordelia Langford, John Sillitoe on behalf of the Wellcome Sanger Institute COVID-19 Surveillance Team |
| EPI_ISL_3107081 | Lighthouse Lab in Alderley Park | Wellcome Sanger Institute for the COVID-19 Genomics UK (COG-UK) Consortium | Jacquelyn Wynn, Mairead Hyland, The Lighthouse Lab in Alderley Park and Alex Alderton, Roberto Amato, Jeffrey Barrett, Sonia Goncalves, Ewan Harrison, David K. Jackson, Ian Johnston, Dominic Kwiatkowski, Cordelia Langford, John Sillitoe on behalf of the Wellcome Sanger Institute COVID-19 Surveillance Team |
| EPI_ISL_3107082, EPI_ISL_3107083 | Berkshire and Surrey Pathology Services Lighthouse Laboratory | Wellcome Sanger Institute for the COVID-19 Genomics UK (COG-UK) Consortium | Berkshire and Surrey Pathology Services Lighthouse Laboratory and Alex Alderton, Roberto Amato, Jeffrey Barrett, Sonia Goncalves, Ewan Harrison, David K. Jackson, Ian Johnston, Dominic Kwiatkowski, Cordelia Langford, John Sillitoe on behalf of the Wellcome Sanger Institute COVID-19 Surveillance Team |
| EPI_ISL_3107084 | Lighthouse Lab in Milton Keynes | Wellcome Sanger Institute for the COVID-19 Genomics UK (COG-UK) Consortium | The Lighthouse Lab in Milton Keynes and Alex Alderton, Roberto Amato, Jeffrey Barrett, Sonia Goncalves, Ewan Harrison, David K. Jackson, Ian Johnston, Dominic Kwiatkowski, Cordelia Langford, John Sillitoe on behalf of the Wellcome Sanger Institute COVID-19 Surveillance Team |
| EPI_ISL_3107085 | Berkshire and Surrey Pathology Services Lighthouse Laboratory | Wellcome Sanger Institute for the COVID-19 Genomics UK (COG-UK) Consortium | Berkshire and Surrey Pathology Services Lighthouse Laboratory and Alex Alderton, Roberto Amato, Jeffrey Barrett, Sonia Goncalves, Ewan Harrison, David K. Jackson, Ian Johnston, Dominic Kwiatkowski, Cordelia Langford, John Sillitoe on behalf of the Wellcome Sanger Institute COVID-19 Surveillance Team |
| EPI_ISL_3107086 | Lighthouse Lab in Alderley Park | Wellcome Sanger Institute for the COVID-19 Genomics UK (COG-UK) Consortium | Jacquelyn Wynn, Mairead Hyland, The Lighthouse Lab in Alderley Park and Alex Alderton, Roberto Amato, Jeffrey Barrett, Sonia Goncalves, Ewan Harrison, David K. Jackson, Ian Johnston, Dominic Kwiatkowski, Cordelia Langford, John Sillitoe on behalf of the Wellcome Sanger Institute COVID-19 Surveillance Team |
| EPI_ISL_3107087 | Berkshire and Surrey Pathology Services Lighthouse Laboratory | Wellcome Sanger Institute for the COVID-19 Genomics UK (COG-UK) Consortium | Berkshire and Surrey Pathology Services Lighthouse Laboratory and Alex Alderton, Roberto Amato, Jeffrey Barrett, Sonia Goncalves, Ewan Harrison, David K. Jackson, Ian Johnston, Dominic Kwiatkowski, Cordelia Langford, John Sillitoe on behalf of the Wellcome Sanger Institute COVID-19 Surveillance Team |
| EPI_ISL_3107088, EPI_ISL_3107089, EPI_ISL_3107090 | Lighthouse Lab in Milton Keynes | Wellcome Sanger Institute for the COVID-19 Genomics UK (COG-UK) Consortium | The Lighthouse Lab in Milton Keynes and Alex Alderton, Roberto Amato, Jeffrey Barrett, Sonia Goncalves, Ewan Harrison, David K. Jackson, Ian Johnston, Dominic Kwiatkowski, Cordelia Langford, John Sillitoe on behalf of the Wellcome Sanger Institute COVID-19 Surveillance Team |
| EPI_ISL_3107091 | Lighthouse Lab in Alderley Park | Wellcome Sanger Institute for the COVID-19 Genomics UK (COG-UK) Consortium | Jacquelyn Wynn, Mairead Hyland, The Lighthouse Lab in Alderley Park and Alex Alderton, Roberto Amato, Jeffrey Barrett, Sonia Goncalves, Ewan Harrison, David K. Jackson, Ian Johnston, Dominic Kwiatkowski, Cordelia Langford, John Sillitoe on behalf of the Wellcome Sanger Institute COVID-19 Surveillance Team |
| EPI_ISL_3107092 | Lighthouse Lab in Milton Keynes | Wellcome Sanger Institute for the COVID-19 Genomics UK (COG-UK) Consortium | The Lighthouse Lab in Milton Keynes and Alex Alderton, Roberto Amato, Jeffrey Barrett, Sonia Goncalves, Ewan Harrison, David K. Jackson, Ian Johnston, Dominic Kwiatkowski, Cordelia Langford, John Sillitoe on behalf of the Wellcome Sanger Institute COVID-19 Surveillance Team |
| EPI_ISL_3107093, EPI_ISL_3107094 | Berkshire and Surrey Pathology Services Lighthouse Laboratory | Wellcome Sanger Institute for the COVID-19 Genomics UK (COG-UK) Consortium | Berkshire and Surrey Pathology Services Lighthouse Laboratory and Alex Alderton, Roberto Amato, Jeffrey Barrett, Sonia Goncalves, Ewan Harrison, David K. Jackson, Ian Johnston, Dominic Kwiatkowski, Cordelia Langford, John Sillitoe on behalf of the Wellcome Sanger Institute COVID-19 Surveillance Team |
| EPI_ISL_3107095 | Lighthouse Lab in Alderley Park | Wellcome Sanger Institute for the COVID-19 Genomics UK (COG-UK) Consortium | Jacquelyn Wynn, Mairead Hyland, The Lighthouse Lab in Alderley Park and Alex Alderton, Roberto Amato, Jeffrey Barrett, Sonia Goncalves, Ewan Harrison, David K. Jackson, Ian Johnston, Dominic Kwiatkowski, Cordelia Langford, John Sillitoe on behalf of the Wellcome Sanger Institute COVID-19 Surveillance Team |

| | | | |
|---|---|---|---|
| EPI_ISL_3107096 | Berkshire and Surrey Pathology Services Lighthouse Laboratory | Wellcome Sanger Institute for the COVID-19 Genomics UK (COG-UK) Consortium | Berkshire and Surrey Pathology Services Lighthouse Laboratory and Alex Alderton, Roberto Amato, Jeffrey Barrett, Sonia Goncalves, Ewan Harrison, David K. Jackson, Ian Johnston, Dominic Kwiatkowski, Cordelia Langford, John Sillitoe on behalf of the Wellcome Sanger Institute COVID-19 Surveillance Team |
| EPI_ISL_3107097 | Lighthouse Lab in Alderley Park | Wellcome Sanger Institute for the COVID-19 Genomics UK (COG-UK) Consortium | Jacquelyn Wynn, Mairead Hyland, The Lighthouse Lab in Alderley Park and Alex Alderton, Roberto Amato, Jeffrey Barrett, Sonia Goncalves, Ewan Harrison, David K. Jackson, Ian Johnston, Dominic Kwiatkowski, Cordelia Langford, John Sillitoe on behalf of the Wellcome Sanger Institute COVID-19 Surveillance Team |
| EPI_ISL_3107098 | Berkshire and Surrey Pathology Services Lighthouse Laboratory | Wellcome Sanger Institute for the COVID-19 Genomics UK (COG-UK) Consortium | Berkshire and Surrey Pathology Services Lighthouse Laboratory and Alex Alderton, Roberto Amato, Jeffrey Barrett, Sonia Goncalves, Ewan Harrison, David K. Jackson, Ian Johnston, Dominic Kwiatkowski, Cordelia Langford, John Sillitoe on behalf of the Wellcome Sanger Institute COVID-19 Surveillance Team |
| EPI_ISL_3107099 | Lighthouse Lab in Alderley Park | Wellcome Sanger Institute for the COVID-19 Genomics UK (COG-UK) Consortium | Jacquelyn Wynn, Mairead Hyland, The Lighthouse Lab in Alderley Park and Alex Alderton, Roberto Amato, Jeffrey Barrett, Sonia Goncalves, Ewan Harrison, David K. Jackson, Ian Johnston, Dominic Kwiatkowski, Cordelia Langford, John Sillitoe on behalf of the Wellcome Sanger Institute COVID-19 Surveillance Team |
| EPI_ISL_3107100 | Berkshire and Surrey Pathology Services Lighthouse Laboratory | Wellcome Sanger Institute for the COVID-19 Genomics UK (COG-UK) Consortium | Berkshire and Surrey Pathology Services Lighthouse Laboratory and Alex Alderton, Roberto Amato, Jeffrey Barrett, Sonia Goncalves, Ewan Harrison, David K. Jackson, Ian Johnston, Dominic Kwiatkowski, Cordelia Langford, John Sillitoe on behalf of the Wellcome Sanger Institute COVID-19 Surveillance Team |
| EPI_ISL_3107101 | Lighthouse Lab in Alderley Park | Wellcome Sanger Institute for the COVID-19 Genomics UK (COG-UK) Consortium | Jacquelyn Wynn, Mairead Hyland, The Lighthouse Lab in Alderley Park and Alex Alderton, Roberto Amato, Jeffrey Barrett, Sonia Goncalves, Ewan Harrison, David K. Jackson, Ian Johnston, Dominic Kwiatkowski, Cordelia Langford, John Sillitoe on behalf of the Wellcome Sanger Institute COVID-19 Surveillance Team |
| EPI_ISL_3107102 | Berkshire and Surrey Pathology Services Lighthouse Laboratory | Wellcome Sanger Institute for the COVID-19 Genomics UK (COG-UK) Consortium | Berkshire and Surrey Pathology Services Lighthouse Laboratory and Alex Alderton, Roberto Amato, Jeffrey Barrett, Sonia Goncalves, Ewan Harrison, David K. Jackson, Ian Johnston, Dominic Kwiatkowski, Cordelia Langford, John Sillitoe on behalf of the Wellcome Sanger Institute COVID-19 Surveillance Team |
| EPI_ISL_3107103, EPI_ISL_3107104 | Lighthouse Lab in Milton Keynes | Wellcome Sanger Institute for the COVID-19 Genomics UK (COG-UK) Consortium | The Lighthouse Lab in Milton Keynes and Alex Alderton, Roberto Amato, Jeffrey Barrett, Sonia Goncalves, Ewan Harrison, David K. Jackson, Ian Johnston, Dominic Kwiatkowski, Cordelia Langford, John Sillitoe on behalf of the Wellcome Sanger Institute COVID-19 Surveillance Team |
| EPI_ISL_3107105 | Berkshire and Surrey Pathology Services Lighthouse Laboratory | Wellcome Sanger Institute for the COVID-19 Genomics UK (COG-UK) Consortium | Berkshire and Surrey Pathology Services Lighthouse Laboratory and Alex Alderton, Roberto Amato, Jeffrey Barrett, Sonia Goncalves, Ewan Harrison, David K. Jackson, Ian Johnston, Dominic Kwiatkowski, Cordelia Langford, John Sillitoe on behalf of the Wellcome Sanger Institute COVID-19 Surveillance Team |
| EPI_ISL_3107106, EPI_ISL_3107107, EPI_ISL_3107108 | Lighthouse Lab in Alderley Park | Wellcome Sanger Institute for the COVID-19 Genomics UK (COG-UK) Consortium | Jacquelyn Wynn, Mairead Hyland, The Lighthouse Lab in Alderley Park and Alex Alderton, Roberto Amato, Jeffrey Barrett, Sonia Goncalves, Ewan Harrison, David K. Jackson, Ian Johnston, Dominic Kwiatkowski, Cordelia Langford, John Sillitoe on behalf of the Wellcome Sanger Institute COVID-19 Surveillance Team |
| EPI_ISL_3107109 | Berkshire and Surrey Pathology Services Lighthouse Laboratory | Wellcome Sanger Institute for the COVID-19 Genomics UK (COG-UK) Consortium | Berkshire and Surrey Pathology Services Lighthouse Laboratory and Alex Alderton, Roberto Amato, Jeffrey Barrett, Sonia Goncalves, Ewan Harrison, David K. Jackson, Ian Johnston, Dominic Kwiatkowski, Cordelia Langford, John Sillitoe on behalf of the Wellcome Sanger Institute COVID-19 Surveillance Team |
| EPI_ISL_3107110 | Lighthouse Lab in Alderley Park | Wellcome Sanger Institute for the COVID-19 Genomics UK (COG-UK) Consortium | Jacquelyn Wynn, Mairead Hyland, The Lighthouse Lab in Alderley Park and Alex Alderton, Roberto Amato, Jeffrey Barrett, Sonia Goncalves, Ewan Harrison, David K. Jackson, Ian Johnston, Dominic Kwiatkowski, Cordelia Langford, John Sillitoe on behalf of the Wellcome Sanger Institute COVID-19 Surveillance Team |
| EPI_ISL_3107111 | Lighthouse Lab in Milton Keynes | Wellcome Sanger Institute for the COVID-19 Genomics UK (COG-UK) Consortium | The Lighthouse Lab in Milton Keynes and Alex Alderton, Roberto Amato, Jeffrey Barrett, Sonia Goncalves, Ewan Harrison, David K. Jackson, Ian Johnston, Dominic Kwiatkowski, Cordelia Langford, John Sillitoe on behalf of the Wellcome Sanger Institute COVID-19 Surveillance Team |
| EPI_ISL_3107112 | Lighthouse Lab in Alderley Park | Wellcome Sanger Institute for the COVID-19 Genomics UK (COG-UK) Consortium | Jacquelyn Wynn, Mairead Hyland, The Lighthouse Lab in Alderley Park and Alex Alderton, Roberto Amato, Jeffrey Barrett, Sonia Goncalves, Ewan Harrison, David K. Jackson, Ian Johnston, Dominic Kwiatkowski, Cordelia Langford, John Sillitoe on behalf of the Wellcome Sanger Institute COVID-19 Surveillance Team |
| EPI_ISL_3107113, EPI_ISL_3107114 | Berkshire and Surrey Pathology Services Lighthouse Laboratory | Wellcome Sanger Institute for the COVID-19 Genomics UK (COG-UK) Consortium | Berkshire and Surrey Pathology Services Lighthouse Laboratory and Alex Alderton, Roberto Amato, Jeffrey Barrett, Sonia Goncalves, Ewan Harrison, David K. Jackson, Ian Johnston, Dominic Kwiatkowski, Cordelia Langford, John Sillitoe on behalf of the Wellcome Sanger Institute COVID-19 Surveillance Team |
| EPI_ISL_3107115 | Lighthouse Lab in Milton Keynes | Wellcome Sanger Institute for the COVID-19 Genomics UK (COG-UK) Consortium | The Lighthouse Lab in Milton Keynes and Alex Alderton, Roberto Amato, Jeffrey Barrett, Sonia Goncalves, Ewan Harrison, David K. Jackson, Ian Johnston, Dominic Kwiatkowski, Cordelia Langford, John Sillitoe on behalf of the Wellcome Sanger Institute COVID-19 Surveillance Team |
| EPI_ISL_3107116, EPI_ISL_3107117 | Berkshire and Surrey Pathology Services Lighthouse Laboratory | Wellcome Sanger Institute for the COVID-19 Genomics UK (COG-UK) Consortium | Berkshire and Surrey Pathology Services Lighthouse Laboratory and Alex Alderton, Roberto Amato, Jeffrey Barrett, Sonia Goncalves, Ewan Harrison, David K. Jackson, Ian Johnston, Dominic Kwiatkowski, Cordelia Langford, John Sillitoe on behalf of the Wellcome Sanger Institute COVID-19 Surveillance Team |
| EPI_ISL_3107118 | Lighthouse Lab in Milton Keynes | Wellcome Sanger Institute for the COVID-19 Genomics UK (COG-UK) Consortium | The Lighthouse Lab in Milton Keynes and Alex Alderton, Roberto Amato, Jeffrey Barrett, Sonia Goncalves, Ewan Harrison, David K. Jackson, Ian Johnston, Dominic Kwiatkowski, Cordelia Langford, John Sillitoe on behalf of the Wellcome Sanger Institute COVID-19 Surveillance Team |
| EPI_ISL_3107119, EPI_ISL_3107120 | Lighthouse Lab in Alderley Park | Wellcome Sanger Institute for the COVID-19 Genomics UK (COG-UK) Consortium | Jacquelyn Wynn, Mairead Hyland, The Lighthouse Lab in Alderley Park and Alex Alderton, Roberto Amato, Jeffrey Barrett, Sonia Goncalves, Ewan Harrison, David K. Jackson, Ian Johnston, Dominic Kwiatkowski, Cordelia Langford, John Sillitoe on behalf of the Wellcome Sanger Institute COVID-19 Surveillance Team |
| EPI_ISL_3107121, EPI_ISL_3107122, EPI_ISL_3107123 | Berkshire and Surrey Pathology Services Lighthouse Laboratory | Wellcome Sanger Institute for the COVID-19 Genomics UK (COG-UK) Consortium | Berkshire and Surrey Pathology Services Lighthouse Laboratory and Alex Alderton, Roberto Amato, Jeffrey Barrett, Sonia Goncalves, Ewan Harrison, David K. Jackson, Ian Johnston, Dominic Kwiatkowski, Cordelia Langford, John Sillitoe on behalf of the Wellcome Sanger Institute COVID-19 Surveillance Team |
| EPI_ISL_3107124 | Lighthouse Lab in Milton Keynes | Wellcome Sanger Institute for the COVID-19 Genomics UK (COG-UK) Consortium | The Lighthouse Lab in Milton Keynes and Alex Alderton, Roberto Amato, Jeffrey Barrett, Sonia Goncalves, Ewan Harrison, David K. Jackson, Ian Johnston, Dominic Kwiatkowski, Cordelia Langford, John Sillitoe on behalf of the Wellcome Sanger Institute COVID-19 Surveillance Team |
| EPI_ISL_3107125 | Berkshire and Surrey Pathology Services Lighthouse Laboratory | Wellcome Sanger Institute for the COVID-19 Genomics UK (COG-UK) Consortium | Berkshire and Surrey Pathology Services Lighthouse Laboratory and Alex Alderton, Roberto Amato, Jeffrey Barrett, Sonia Goncalves, Ewan Harrison, David K. Jackson, Ian Johnston, Dominic Kwiatkowski, Cordelia Langford, John Sillitoe on behalf of the Wellcome Sanger Institute COVID-19 Surveillance Team |
| EPI_ISL_3107126 | Lighthouse Lab in Milton Keynes | Wellcome Sanger Institute for the COVID-19 Genomics UK (COG-UK) Consortium | The Lighthouse Lab in Milton Keynes and Alex Alderton, Roberto Amato, Jeffrey Barrett, Sonia Goncalves, Ewan Harrison, David K. Jackson, Ian Johnston, Dominic Kwiatkowski, Cordelia Langford, John Sillitoe on behalf of the Wellcome Sanger Institute COVID-19 Surveillance Team |
| EPI_ISL_3107127, EPI_ISL_3107128 | Berkshire and Surrey Pathology Services Lighthouse Laboratory | Wellcome Sanger Institute for the COVID-19 Genomics UK (COG-UK) Consortium | Berkshire and Surrey Pathology Services Lighthouse Laboratory and Alex Alderton, Roberto Amato, Jeffrey Barrett, Sonia Goncalves, Ewan Harrison, David K. Jackson, Ian Johnston, Dominic Kwiatkowski, Cordelia Langford, John Sillitoe on behalf of the Wellcome Sanger Institute COVID-19 Surveillance Team |
| EPI_ISL_3107129 | Lighthouse Lab in Milton Keynes | Wellcome Sanger Institute for the COVID-19 Genomics UK (COG-UK) Consortium | The Lighthouse Lab in Milton Keynes and Alex Alderton, Roberto Amato, Jeffrey Barrett, Sonia Goncalves, Ewan Harrison, David K. Jackson, Ian Johnston, Dominic Kwiatkowski, Cordelia Langford, John Sillitoe on behalf of the Wellcome Sanger Institute COVID-19 Surveillance Team |
| EPI_ISL_3107130 | Berkshire and Surrey Pathology Services Lighthouse Laboratory | Wellcome Sanger Institute for the COVID-19 Genomics UK (COG-UK) Consortium | Berkshire and Surrey Pathology Services Lighthouse Laboratory and Alex Alderton, Roberto Amato, Jeffrey Barrett, Sonia Goncalves, Ewan Harrison, David K. Jackson, Ian Johnston, Dominic Kwiatkowski, Cordelia Langford, John Sillitoe on behalf of the Wellcome Sanger Institute COVID-19 Surveillance Team |
| EPI_ISL_3107131 | Lighthouse Lab in Milton Keynes | Wellcome Sanger Institute for the COVID-19 Genomics UK (COG-UK) Consortium | The Lighthouse Lab in Milton Keynes and Alex Alderton, Roberto Amato, Jeffrey Barrett, Sonia Goncalves, Ewan Harrison, David K. Jackson, Ian Johnston, Dominic Kwiatkowski, Cordelia Langford, John Sillitoe on behalf of the Wellcome Sanger Institute COVID-19 Surveillance Team |
| EPI_ISL_3107132 | Berkshire and Surrey Pathology Services Lighthouse Laboratory | Wellcome Sanger Institute for the COVID-19 Genomics UK (COG-UK) Consortium | Berkshire and Surrey Pathology Services Lighthouse Laboratory and Alex Alderton, Roberto Amato, Jeffrey Barrett, Sonia Goncalves, Ewan Harrison, David K. Jackson, Ian Johnston, Dominic Kwiatkowski, Cordelia Langford, John Sillitoe on behalf of the Wellcome Sanger Institute COVID-19 Surveillance Team |
| EPI_ISL_3107133 | Lighthouse Lab in Milton Keynes | Wellcome Sanger Institute for the COVID-19 Genomics UK (COG-UK) Consortium | The Lighthouse Lab in Milton Keynes and Alex Alderton, Roberto Amato, Jeffrey Barrett, Sonia Goncalves, Ewan Harrison, David K. Jackson, Ian Johnston, Dominic Kwiatkowski, Cordelia Langford, John Sillitoe on behalf of the Wellcome Sanger Institute COVID-19 Surveillance Team |
| EPI_ISL_3107134 | Lighthouse Lab in Alderley Park | Wellcome Sanger Institute for the COVID-19 Genomics UK (COG-UK) Consortium | Jacquelyn Wynn, Mairead Hyland, The Lighthouse Lab in Alderley Park and Alex Alderton, Roberto Amato, Jeffrey Barrett, Sonia Goncalves, Ewan Harrison, David K. Jackson, Ian Johnston, Dominic Kwiatkowski, Cordelia Langford, John Sillitoe on behalf of the Wellcome Sanger Institute COVID-19 Surveillance Team |
| EPI_ISL_3107135 | Berkshire and Surrey Pathology Services Lighthouse Laboratory | Wellcome Sanger Institute for the COVID-19 Genomics UK (COG-UK) Consortium | Berkshire and Surrey Pathology Services Lighthouse Laboratory and Alex Alderton, Roberto Amato, Jeffrey Barrett, Sonia Goncalves, Ewan Harrison, David K. Jackson, Ian Johnston, Dominic Kwiatkowski, Cordelia Langford, John Sillitoe on behalf of the Wellcome Sanger Institute COVID-19 Surveillance Team |
| EPI_ISL_3107136, EPI_ISL_3107137 | Lighthouse Lab in Alderley Park | Wellcome Sanger Institute for the COVID-19 Genomics UK (COG-UK) Consortium | Jacquelyn Wynn, Mairead Hyland, The Lighthouse Lab in Alderley Park and Alex Alderton, Roberto Amato, Jeffrey Barrett, Sonia Goncalves, Ewan Harrison, David K. Jackson, Ian Johnston, Dominic Kwiatkowski, Cordelia Langford, John Sillitoe on behalf of the Wellcome Sanger Institute COVID-19 Surveillance Team |
| EPI_ISL_3107138, EPI_ISL_3107139, EPI_ISL_3107140 | Berkshire and Surrey Pathology Services Lighthouse Laboratory | Wellcome Sanger Institute for the COVID-19 Genomics UK (COG-UK) Consortium | Berkshire and Surrey Pathology Services Lighthouse Laboratory and Alex Alderton, Roberto Amato, Jeffrey Barrett, Sonia Goncalves, Ewan Harrison, David K. Jackson, Ian Johnston, Dominic Kwiatkowski, Cordelia Langford, John Sillitoe on behalf of the Wellcome Sanger Institute COVID-19 Surveillance Team |
| EPI_ISL_3107141 | Lighthouse Lab in Milton Keynes | Wellcome Sanger Institute for the COVID-19 Genomics UK (COG-UK) Consortium | The Lighthouse Lab in Milton Keynes and Alex Alderton, Roberto Amato, Jeffrey Barrett, Sonia Goncalves, Ewan Harrison, David K. Jackson, Ian Johnston, Dominic Kwiatkowski, Cordelia Langford, John Sillitoe on behalf of the Wellcome Sanger Institute COVID-19 Surveillance Team |
| EPI_ISL_3107142, EPI_ISL_3107143 | Berkshire and Surrey Pathology Services Lighthouse Laboratory | Wellcome Sanger Institute for the COVID-19 Genomics UK (COG-UK) Consortium | Berkshire and Surrey Pathology Services Lighthouse Laboratory and Alex Alderton, Roberto Amato, Jeffrey Barrett, Sonia Goncalves, Ewan Harrison, David K. Jackson, Ian Johnston, Dominic Kwiatkowski, Cordelia Langford, John Sillitoe on behalf of the Wellcome Sanger Institute COVID-19 Surveillance Team |
| EPI_ISL_3107144, EPI_ISL_3107145 | Lighthouse Lab in Milton Keynes | Wellcome Sanger Institute for the COVID-19 Genomics UK (COG-UK) Consortium | The Lighthouse Lab in Milton Keynes and Alex Alderton, Roberto Amato, Jeffrey Barrett, Sonia Goncalves, Ewan Harrison, David K. Jackson, Ian Johnston, Dominic Kwiatkowski, Cordelia Langford, John Sillitoe on behalf of the Wellcome Sanger Institute COVID-19 Surveillance Team |
| EPI_ISL_3107146 | Berkshire and Surrey Pathology Services Lighthouse Laboratory | Wellcome Sanger Institute for the COVID-19 Genomics UK (COG-UK) Consortium | Berkshire and Surrey Pathology Services Lighthouse Laboratory and Alex Alderton, Roberto Amato, Jeffrey Barrett, Sonia Goncalves, Ewan Harrison, David K. Jackson, Ian Johnston, Dominic Kwiatkowski, Cordelia Langford, John Sillitoe on behalf of the Wellcome Sanger Institute COVID-19 Surveillance Team |
| EPI_ISL_3107147 | Lighthouse Lab in Alderley Park | Wellcome Sanger Institute for the COVID-19 Genomics UK (COG-UK) Consortium | Jacquelyn Wynn, Mairead Hyland, The Lighthouse Lab in Alderley Park and Alex Alderton, Roberto Amato, Jeffrey Barrett, Sonia Goncalves, Ewan Harrison, David K. Jackson, Ian Johnston, Dominic Kwiatkowski, Cordelia Langford, John Sillitoe on behalf of the Wellcome Sanger Institute COVID-19 Surveillance Team |
| EPI_ISL_3107148 | Berkshire and Surrey Pathology Services Lighthouse Laboratory | Wellcome Sanger Institute for the COVID-19 Genomics UK (COG-UK) Consortium | Berkshire and Surrey Pathology Services Lighthouse Laboratory and Alex Alderton, Roberto Amato, Jeffrey Barrett, Sonia Goncalves, Ewan Harrison, David K. Jackson, Ian Johnston, Dominic Kwiatkowski, Cordelia Langford, John Sillitoe on behalf of the Wellcome Sanger Institute COVID-19 Surveillance Team |
| EPI_ISL_3107149, EPI_ISL_3107150, EPI_ISL_3107151 | Lighthouse Lab in Alderley Park | Wellcome Sanger Institute for the COVID-19 Genomics UK (COG-UK) Consortium | Jacquelyn Wynn, Mairead Hyland, The Lighthouse Lab in Alderley Park and Alex Alderton, Roberto Amato, Jeffrey Barrett, Sonia Goncalves, Ewan Harrison, David K. Jackson, Ian Johnston, Dominic Kwiatkowski, Cordelia Langford, John Sillitoe on behalf of the Wellcome Sanger Institute COVID-19 Surveillance Team |
| EPI_ISL_3107152 | Berkshire and Surrey Pathology Services Lighthouse Laboratory | Wellcome Sanger Institute for the COVID-19 Genomics UK (COG-UK) Consortium | Berkshire and Surrey Pathology Services Lighthouse Laboratory and Alex Alderton, Roberto Amato, Jeffrey Barrett, Sonia Goncalves, Ewan Harrison, David K. Jackson, Ian Johnston, Dominic Kwiatkowski, Cordelia Langford, John Sillitoe on behalf of the Wellcome Sanger Institute COVID-19 Surveillance Team |
| EPI_ISL_3107153 | Lighthouse Lab in Alderley Park | Wellcome Sanger Institute for the COVID-19 Genomics UK (COG-UK) Consortium | Jacquelyn Wynn, Mairead Hyland, The Lighthouse Lab in Alderley Park and Alex Alderton, Roberto Amato, Jeffrey Barrett, Sonia Goncalves, Ewan Harrison, David K. Jackson, Ian Johnston, Dominic Kwiatkowski, Cordelia Langford, John Sillitoe on behalf of the Wellcome Sanger Institute COVID-19 Surveillance Team |
| EPI_ISL_3107154 | Berkshire and Surrey Pathology Services Lighthouse Laboratory | Wellcome Sanger Institute for the COVID-19 Genomics UK (COG-UK) Consortium | Berkshire and Surrey Pathology Services Lighthouse Laboratory and Alex Alderton, Roberto Amato, Jeffrey Barrett, Sonia Goncalves, Ewan Harrison, David K. Jackson, Ian Johnston, Dominic Kwiatkowski, Cordelia Langford, John Sillitoe on behalf of the Wellcome Sanger Institute COVID-19 Surveillance Team |

| | | | |
|---|---|---|---|
| EPI_ISL_3107155 | Lighthouse Lab in Milton Keynes | Wellcome Sanger Institute for the COVID-19 Genomics UK (COG-UK) Consortium | The Lighthouse Lab in Milton Keynes and Alex Alderton, Roberto Amato, Jeffrey Barrett, Sonia Goncalves, Ewan Harrison, David K. Jackson, Ian Johnston, Dominic Kwiatkowski, Cordelia Langford, John Sillitoe on behalf of the Wellcome Sanger Institute COVID-19 Surveillance Team |
| EPI_ISL_3107156, EPI_ISL_3107157 | Berkshire and Surrey Pathology Services Lighthouse Laboratory | Wellcome Sanger Institute for the COVID-19 Genomics UK (COG-UK) Consortium | Berkshire and Surrey Pathology Services Lighthouse Laboratory and Alex Alderton, Roberto Amato, Jeffrey Barrett, Sonia Goncalves, Ewan Harrison, David K. Jackson, Ian Johnston, Dominic Kwiatkowski, Cordelia Langford, John Sillitoe on behalf of the Wellcome Sanger Institute COVID-19 Surveillance Team |
| EPI_ISL_3107158 | Lighthouse Lab in Alderley Park | Wellcome Sanger Institute for the COVID-19 Genomics UK (COG-UK) Consortium | Jacquelyn Wynn, Mairead Hyland, The Lighthouse Lab in Alderley Park and Alex Alderton, Roberto Amato, Jeffrey Barrett, Sonia Goncalves, Ewan Harrison, David K. Jackson, Ian Johnston, Dominic Kwiatkowski, Cordelia Langford, John Sillitoe on behalf of the Wellcome Sanger Institute COVID-19 Surveillance Team |
| EPI_ISL_3107159 | Lighthouse Lab in Milton Keynes | Wellcome Sanger Institute for the COVID-19 Genomics UK (COG-UK) Consortium | The Lighthouse Lab in Milton Keynes and Alex Alderton, Roberto Amato, Jeffrey Barrett, Sonia Goncalves, Ewan Harrison, David K. Jackson, Ian Johnston, Dominic Kwiatkowski, Cordelia Langford, John Sillitoe on behalf of the Wellcome Sanger Institute COVID-19 Surveillance Team |
| EPI_ISL_3107160, EPI_ISL_3107161 | Berkshire and Surrey Pathology Services Lighthouse Laboratory | Wellcome Sanger Institute for the COVID-19 Genomics UK (COG-UK) Consortium | Berkshire and Surrey Pathology Services Lighthouse Laboratory and Alex Alderton, Roberto Amato, Jeffrey Barrett, Sonia Goncalves, Ewan Harrison, David K. Jackson, Ian Johnston, Dominic Kwiatkowski, Cordelia Langford, John Sillitoe on behalf of the Wellcome Sanger Institute COVID-19 Surveillance Team |
| EPI_ISL_3107162, EPI_ISL_3107163 | Lighthouse Lab in Milton Keynes | Wellcome Sanger Institute for the COVID-19 Genomics UK (COG-UK) Consortium | The Lighthouse Lab in Milton Keynes and Alex Alderton, Roberto Amato, Jeffrey Barrett, Sonia Goncalves, Ewan Harrison, David K. Jackson, Ian Johnston, Dominic Kwiatkowski, Cordelia Langford, John Sillitoe on behalf of the Wellcome Sanger Institute COVID-19 Surveillance Team |
| EPI_ISL_3107164 | Berkshire and Surrey Pathology Services Lighthouse Laboratory | Wellcome Sanger Institute for the COVID-19 Genomics UK (COG-UK) Consortium | Berkshire and Surrey Pathology Services Lighthouse Laboratory and Alex Alderton, Roberto Amato, Jeffrey Barrett, Sonia Goncalves, Ewan Harrison, David K. Jackson, Ian Johnston, Dominic Kwiatkowski, Cordelia Langford, John Sillitoe on behalf of the Wellcome Sanger Institute COVID-19 Surveillance Team |
| EPI_ISL_3107165 | Lighthouse Lab in Alderley Park | Wellcome Sanger Institute for the COVID-19 Genomics UK (COG-UK) Consortium | Jacquelyn Wynn, Mairead Hyland, The Lighthouse Lab in Alderley Park and Alex Alderton, Roberto Amato, Jeffrey Barrett, Sonia Goncalves, Ewan Harrison, David K. Jackson, Ian Johnston, Dominic Kwiatkowski, Cordelia Langford, John Sillitoe on behalf of the Wellcome Sanger Institute COVID-19 Surveillance Team |
| EPI_ISL_3107166, EPI_ISL_3107167 | Berkshire and Surrey Pathology Services Lighthouse Laboratory | Wellcome Sanger Institute for the COVID-19 Genomics UK (COG-UK) Consortium | Berkshire and Surrey Pathology Services Lighthouse Laboratory and Alex Alderton, Roberto Amato, Jeffrey Barrett, Sonia Goncalves, Ewan Harrison, David K. Jackson, Ian Johnston, Dominic Kwiatkowski, Cordelia Langford, John Sillitoe on behalf of the Wellcome Sanger Institute COVID-19 Surveillance Team |
| EPI_ISL_3107168, EPI_ISL_3107169 | Lighthouse Lab in Alderley Park | Wellcome Sanger Institute for the COVID-19 Genomics UK (COG-UK) Consortium | Jacquelyn Wynn, Mairead Hyland, The Lighthouse Lab in Alderley Park and Alex Alderton, Roberto Amato, Jeffrey Barrett, Sonia Goncalves, Ewan Harrison, David K. Jackson, Ian Johnston, Dominic Kwiatkowski, Cordelia Langford, John Sillitoe on behalf of the Wellcome Sanger Institute COVID-19 Surveillance Team |
| EPI_ISL_3107170, EPI_ISL_3107171 | Berkshire and Surrey Pathology Services Lighthouse Laboratory | Wellcome Sanger Institute for the COVID-19 Genomics UK (COG-UK) Consortium | Berkshire and Surrey Pathology Services Lighthouse Laboratory and Alex Alderton, Roberto Amato, Jeffrey Barrett, Sonia Goncalves, Ewan Harrison, David K. Jackson, Ian Johnston, Dominic Kwiatkowski, Cordelia Langford, John Sillitoe on behalf of the Wellcome Sanger Institute COVID-19 Surveillance Team |
| EPI_ISL_3107172 | Lighthouse Lab in Alderley Park | Wellcome Sanger Institute for the COVID-19 Genomics UK (COG-UK) Consortium | Jacquelyn Wynn, Mairead Hyland, The Lighthouse Lab in Alderley Park and Alex Alderton, Roberto Amato, Jeffrey Barrett, Sonia Goncalves, Ewan Harrison, David K. Jackson, Ian Johnston, Dominic Kwiatkowski, Cordelia Langford, John Sillitoe on behalf of the Wellcome Sanger Institute COVID-19 Surveillance Team |
| EPI_ISL_3107173 | Berkshire and Surrey Pathology Services Lighthouse Laboratory | Wellcome Sanger Institute for the COVID-19 Genomics UK (COG-UK) Consortium | Berkshire and Surrey Pathology Services Lighthouse Laboratory and Alex Alderton, Roberto Amato, Jeffrey Barrett, Sonia Goncalves, Ewan Harrison, David K. Jackson, Ian Johnston, Dominic Kwiatkowski, Cordelia Langford, John Sillitoe on behalf of the Wellcome Sanger Institute COVID-19 Surveillance Team |
| EPI_ISL_3107174 | Lighthouse Lab in Alderley Park | Wellcome Sanger Institute for the COVID-19 Genomics UK (COG-UK) Consortium | Jacquelyn Wynn, Mairead Hyland, The Lighthouse Lab in Alderley Park and Alex Alderton, Roberto Amato, Jeffrey Barrett, Sonia Goncalves, Ewan Harrison, David K. Jackson, Ian Johnston, Dominic Kwiatkowski, Cordelia Langford, John Sillitoe on behalf of the Wellcome Sanger Institute COVID-19 Surveillance Team |
| EPI_ISL_3107175, EPI_ISL_3107176, EPI_ISL_3107177 | Lighthouse Lab in Milton Keynes | Wellcome Sanger Institute for the COVID-19 Genomics UK (COG-UK) Consortium | The Lighthouse Lab in Milton Keynes and Alex Alderton, Roberto Amato, Jeffrey Barrett, Sonia Goncalves, Ewan Harrison, David K. Jackson, Ian Johnston, Dominic Kwiatkowski, Cordelia Langford, John Sillitoe on behalf of the Wellcome Sanger Institute COVID-19 Surveillance Team |
| EPI_ISL_3107178, EPI_ISL_3107179 | Berkshire and Surrey Pathology Services Lighthouse Laboratory | Wellcome Sanger Institute for the COVID-19 Genomics UK (COG-UK) Consortium | Berkshire and Surrey Pathology Services Lighthouse Laboratory and Alex Alderton, Roberto Amato, Jeffrey Barrett, Sonia Goncalves, Ewan Harrison, David K. Jackson, Ian Johnston, Dominic Kwiatkowski, Cordelia Langford, John Sillitoe on behalf of the Wellcome Sanger Institute COVID-19 Surveillance Team |
| EPI_ISL_3107180 | Lighthouse Lab in Milton Keynes | Wellcome Sanger Institute for the COVID-19 Genomics UK (COG-UK) Consortium | The Lighthouse Lab in Milton Keynes and Alex Alderton, Roberto Amato, Jeffrey Barrett, Sonia Goncalves, Ewan Harrison, David K. Jackson, Ian Johnston, Dominic Kwiatkowski, Cordelia Langford, John Sillitoe on behalf of the Wellcome Sanger Institute COVID-19 Surveillance Team |
| EPI_ISL_3107181, EPI_ISL_3107182, EPI_ISL_3107183 | Berkshire and Surrey Pathology Services Lighthouse Laboratory | Wellcome Sanger Institute for the COVID-19 Genomics UK (COG-UK) Consortium | Berkshire and Surrey Pathology Services Lighthouse Laboratory and Alex Alderton, Roberto Amato, Jeffrey Barrett, Sonia Goncalves, Ewan Harrison, David K. Jackson, Ian Johnston, Dominic Kwiatkowski, Cordelia Langford, John Sillitoe on behalf of the Wellcome Sanger Institute COVID-19 Surveillance Team |
| EPI_ISL_3107184 | Lighthouse Lab in Milton Keynes | Wellcome Sanger Institute for the COVID-19 Genomics UK (COG-UK) Consortium | The Lighthouse Lab in Milton Keynes and Alex Alderton, Roberto Amato, Jeffrey Barrett, Sonia Goncalves, Ewan Harrison, David K. Jackson, Ian Johnston, Dominic Kwiatkowski, Cordelia Langford, John Sillitoe on behalf of the Wellcome Sanger Institute COVID-19 Surveillance Team |
| EPI_ISL_3107185 | Berkshire and Surrey Pathology Services Lighthouse Laboratory | Wellcome Sanger Institute for the COVID-19 Genomics UK (COG-UK) Consortium | Berkshire and Surrey Pathology Services Lighthouse Laboratory and Alex Alderton, Roberto Amato, Jeffrey Barrett, Sonia Goncalves, Ewan Harrison, David K. Jackson, Ian Johnston, Dominic Kwiatkowski, Cordelia Langford, John Sillitoe on behalf of the Wellcome Sanger Institute COVID-19 Surveillance Team |
| EPI_ISL_3107186 | Lighthouse Lab in Milton Keynes | Wellcome Sanger Institute for the COVID-19 Genomics UK (COG-UK) Consortium | The Lighthouse Lab in Milton Keynes and Alex Alderton, Roberto Amato, Jeffrey Barrett, Sonia Goncalves, Ewan Harrison, David K. Jackson, Ian Johnston, Dominic Kwiatkowski, Cordelia Langford, John Sillitoe on behalf of the Wellcome Sanger Institute COVID-19 Surveillance Team |
| EPI_ISL_3107187 | Berkshire and Surrey Pathology Services Lighthouse Laboratory | Wellcome Sanger Institute for the COVID-19 Genomics UK (COG-UK) Consortium | Berkshire and Surrey Pathology Services Lighthouse Laboratory and Alex Alderton, Roberto Amato, Jeffrey Barrett, Sonia Goncalves, Ewan Harrison, David K. Jackson, Ian Johnston, Dominic Kwiatkowski, Cordelia Langford, John Sillitoe on behalf of the Wellcome Sanger Institute COVID-19 Surveillance Team |
| EPI_ISL_3107188, EPI_ISL_3107189 | Lighthouse Lab in Milton Keynes | Wellcome Sanger Institute for the COVID-19 Genomics UK (COG-UK) Consortium | The Lighthouse Lab in Milton Keynes and Alex Alderton, Roberto Amato, Jeffrey Barrett, Sonia Goncalves, Ewan Harrison, David K. Jackson, Ian Johnston, Dominic Kwiatkowski, Cordelia Langford, John Sillitoe on behalf of the Wellcome Sanger Institute COVID-19 Surveillance Team |
| EPI_ISL_3107190, EPI_ISL_3107191 | Berkshire and Surrey Pathology Services Lighthouse Laboratory | Wellcome Sanger Institute for the COVID-19 Genomics UK (COG-UK) Consortium | Berkshire and Surrey Pathology Services Lighthouse Laboratory and Alex Alderton, Roberto Amato, Jeffrey Barrett, Sonia Goncalves, Ewan Harrison, David K. Jackson, Ian Johnston, Dominic Kwiatkowski, Cordelia Langford, John Sillitoe on behalf of the Wellcome Sanger Institute COVID-19 Surveillance Team |
| EPI_ISL_3107192 | Lighthouse Lab in Milton Keynes | Wellcome Sanger Institute for the COVID-19 Genomics UK (COG-UK) Consortium | The Lighthouse Lab in Milton Keynes and Alex Alderton, Roberto Amato, Jeffrey Barrett, Sonia Goncalves, Ewan Harrison, David K. Jackson, Ian Johnston, Dominic Kwiatkowski, Cordelia Langford, John Sillitoe on behalf of the Wellcome Sanger Institute COVID-19 Surveillance Team |
| EPI_ISL_3107193 | Berkshire and Surrey Pathology Services Lighthouse Laboratory | Wellcome Sanger Institute for the COVID-19 Genomics UK (COG-UK) Consortium | Berkshire and Surrey Pathology Services Lighthouse Laboratory and Alex Alderton, Roberto Amato, Jeffrey Barrett, Sonia Goncalves, Ewan Harrison, David K. Jackson, Ian Johnston, Dominic Kwiatkowski, Cordelia Langford, John Sillitoe on behalf of the Wellcome Sanger Institute COVID-19 Surveillance Team |
| EPI_ISL_3107194 | Lighthouse Lab in Alderley Park | Wellcome Sanger Institute for the COVID-19 Genomics UK (COG-UK) Consortium | Jacquelyn Wynn, Mairead Hyland, The Lighthouse Lab in Alderley Park and Alex Alderton, Roberto Amato, Jeffrey Barrett, Sonia Goncalves, Ewan Harrison, David K. Jackson, Ian Johnston, Dominic Kwiatkowski, Cordelia Langford, John Sillitoe on behalf of the Wellcome Sanger Institute COVID-19 Surveillance Team |
| EPI_ISL_3107195, EPI_ISL_3107196, EPI_ISL_3107197, EPI_ISL_3107198 | Berkshire and Surrey Pathology Services Lighthouse Laboratory | Wellcome Sanger Institute for the COVID-19 Genomics UK (COG-UK) Consortium | Berkshire and Surrey Pathology Services Lighthouse Laboratory and Alex Alderton, Roberto Amato, Jeffrey Barrett, Sonia Goncalves, Ewan Harrison, David K. Jackson, Ian Johnston, Dominic Kwiatkowski, Cordelia Langford, John Sillitoe on behalf of the Wellcome Sanger Institute COVID-19 Surveillance Team |
| EPI_ISL_3107199 | Lighthouse Lab in Milton Keynes | Wellcome Sanger Institute for the COVID-19 Genomics UK (COG-UK) Consortium | The Lighthouse Lab in Milton Keynes and Alex Alderton, Roberto Amato, Jeffrey Barrett, Sonia Goncalves, Ewan Harrison, David K. Jackson, Ian Johnston, Dominic Kwiatkowski, Cordelia Langford, John Sillitoe on behalf of the Wellcome Sanger Institute COVID-19 Surveillance Team |
| EPI_ISL_3107200 | Berkshire and Surrey Pathology Services Lighthouse Laboratory | Wellcome Sanger Institute for the COVID-19 Genomics UK (COG-UK) Consortium | Berkshire and Surrey Pathology Services Lighthouse Laboratory and Alex Alderton, Roberto Amato, Jeffrey Barrett, Sonia Goncalves, Ewan Harrison, David K. Jackson, Ian Johnston, Dominic Kwiatkowski, Cordelia Langford, John Sillitoe on behalf of the Wellcome Sanger Institute COVID-19 Surveillance Team |
| EPI_ISL_3107201 | Lighthouse Lab in Alderley Park | Wellcome Sanger Institute for the COVID-19 Genomics UK (COG-UK) Consortium | Jacquelyn Wynn, Mairead Hyland, The Lighthouse Lab in Alderley Park and Alex Alderton, Roberto Amato, Jeffrey Barrett, Sonia Goncalves, Ewan Harrison, David K. Jackson, Ian Johnston, Dominic Kwiatkowski, Cordelia Langford, John Sillitoe on behalf of the Wellcome Sanger Institute COVID-19 Surveillance Team |
| EPI_ISL_3107202 | Lighthouse Lab in Milton Keynes | Wellcome Sanger Institute for the COVID-19 Genomics UK (COG-UK) Consortium | The Lighthouse Lab in Milton Keynes and Alex Alderton, Roberto Amato, Jeffrey Barrett, Sonia Goncalves, Ewan Harrison, David K. Jackson, Ian Johnston, Dominic Kwiatkowski, Cordelia Langford, John Sillitoe on behalf of the Wellcome Sanger Institute COVID-19 Surveillance Team |
| EPI_ISL_3107203 | Berkshire and Surrey Pathology Services Lighthouse Laboratory | Wellcome Sanger Institute for the COVID-19 Genomics UK (COG-UK) Consortium | Berkshire and Surrey Pathology Services Lighthouse Laboratory and Alex Alderton, Roberto Amato, Jeffrey Barrett, Sonia Goncalves, Ewan Harrison, David K. Jackson, Ian Johnston, Dominic Kwiatkowski, Cordelia Langford, John Sillitoe on behalf of the Wellcome Sanger Institute COVID-19 Surveillance Team |
| EPI_ISL_3107204 | Lighthouse Lab in Milton Keynes | Wellcome Sanger Institute for the COVID-19 Genomics UK (COG-UK) Consortium | The Lighthouse Lab in Milton Keynes and Alex Alderton, Roberto Amato, Jeffrey Barrett, Sonia Goncalves, Ewan Harrison, David K. Jackson, Ian Johnston, Dominic Kwiatkowski, Cordelia Langford, John Sillitoe on behalf of the Wellcome Sanger Institute COVID-19 Surveillance Team |
| EPI_ISL_3107205, EPI_ISL_3107206 | Lighthouse Lab in Alderley Park | Wellcome Sanger Institute for the COVID-19 Genomics UK (COG-UK) Consortium | Jacquelyn Wynn, Mairead Hyland, The Lighthouse Lab in Alderley Park and Alex Alderton, Roberto Amato, Jeffrey Barrett, Sonia Goncalves, Ewan Harrison, David K. Jackson, Ian Johnston, Dominic Kwiatkowski, Cordelia Langford, John Sillitoe on behalf of the Wellcome Sanger Institute COVID-19 Surveillance Team |
| EPI_ISL_3107207, EPI_ISL_3107208 | Berkshire and Surrey Pathology Services Lighthouse Laboratory | Wellcome Sanger Institute for the COVID-19 Genomics UK (COG-UK) Consortium | Berkshire and Surrey Pathology Services Lighthouse Laboratory and Alex Alderton, Roberto Amato, Jeffrey Barrett, Sonia Goncalves, Ewan Harrison, David K. Jackson, Ian Johnston, Dominic Kwiatkowski, Cordelia Langford, John Sillitoe on behalf of the Wellcome Sanger Institute COVID-19 Surveillance Team |
| EPI_ISL_3107209, EPI_ISL_3107210 | Lighthouse Lab in Alderley Park | Wellcome Sanger Institute for the COVID-19 Genomics UK (COG-UK) Consortium | Jacquelyn Wynn, Mairead Hyland, The Lighthouse Lab in Alderley Park and Alex Alderton, Roberto Amato, Jeffrey Barrett, Sonia Goncalves, Ewan Harrison, David K. Jackson, Ian Johnston, Dominic Kwiatkowski, Cordelia Langford, John Sillitoe on behalf of the Wellcome Sanger Institute COVID-19 Surveillance Team |
| EPI_ISL_3107211 | Lighthouse Lab in Milton Keynes | Wellcome Sanger Institute for the COVID-19 Genomics UK (COG-UK) Consortium | The Lighthouse Lab in Milton Keynes and Alex Alderton, Roberto Amato, Jeffrey Barrett, Sonia Goncalves, Ewan Harrison, David K. Jackson, Ian Johnston, Dominic Kwiatkowski, Cordelia Langford, John Sillitoe on behalf of the Wellcome Sanger Institute COVID-19 Surveillance Team |
| EPI_ISL_3107212, EPI_ISL_3107213 | Berkshire and Surrey Pathology Services Lighthouse Laboratory | Wellcome Sanger Institute for the COVID-19 Genomics UK (COG-UK) Consortium | Berkshire and Surrey Pathology Services Lighthouse Laboratory and Alex Alderton, Roberto Amato, Jeffrey Barrett, Sonia Goncalves, Ewan Harrison, David K. Jackson, Ian Johnston, Dominic Kwiatkowski, Cordelia Langford, John Sillitoe on behalf of the Wellcome Sanger Institute COVID-19 Surveillance Team |
| EPI_ISL_3107214, EPI_ISL_3107215 | Lighthouse Lab in Milton Keynes | Wellcome Sanger Institute for the COVID-19 Genomics UK (COG-UK) Consortium | The Lighthouse Lab in Milton Keynes and Alex Alderton, Roberto Amato, Jeffrey Barrett, Sonia Goncalves, Ewan Harrison, David K. Jackson, Ian Johnston, Dominic Kwiatkowski, Cordelia Langford, John Sillitoe on behalf of the Wellcome Sanger Institute COVID-19 Surveillance Team |
| EPI_ISL_3107216 | Berkshire and Surrey Pathology Services Lighthouse Laboratory | Wellcome Sanger Institute for the COVID-19 Genomics UK (COG-UK) Consortium | Berkshire and Surrey Pathology Services Lighthouse Laboratory and Alex Alderton, Roberto Amato, Jeffrey Barrett, Sonia Goncalves, Ewan Harrison, David K. Jackson, Ian Johnston, Dominic Kwiatkowski, Cordelia Langford, John Sillitoe on behalf of the Wellcome Sanger Institute COVID-19 Surveillance Team |
| EPI_ISL_3107217, EPI_ISL_3107218, EPI_ISL_3107219 | Lighthouse Lab in Milton Keynes | Wellcome Sanger Institute for the COVID-19 Genomics UK (COG-UK) Consortium | The Lighthouse Lab in Milton Keynes and Alex Alderton, Roberto Amato, Jeffrey Barrett, Sonia Goncalves, Ewan Harrison, David K. Jackson, Ian Johnston, Dominic Kwiatkowski, Cordelia Langford, John Sillitoe on behalf of the Wellcome Sanger Institute COVID-19 Surveillance Team |
| EPI_ISL_3107220 | Lighthouse Lab in Alderley Park | Wellcome Sanger Institute for the COVID-19 Genomics UK (COG-UK) Consortium | Jacquelyn Wynn, Mairead Hyland, The Lighthouse Lab in Alderley Park and Alex Alderton, Roberto Amato, Jeffrey Barrett, Sonia Goncalves, Ewan Harrison, David K. Jackson, Ian Johnston, Dominic Kwiatkowski, Cordelia Langford, John Sillitoe on behalf of the Wellcome Sanger Institute COVID-19 Surveillance Team |

| | | | |
|---|---|---|---|
| EPI_ISL_3107221 | Lighthouse Lab in Milton Keynes | Wellcome Sanger Institute for the COVID-19 Genomics UK (COG-UK) Consortium | The Lighthouse Lab in Milton Keynes and Alex Alderton, Roberto Amato, Jeffrey Barrett, Sonia Goncalves, Ewan Harrison, David K. Jackson, Ian Johnston, Dominic Kwiatkowski, Cordelia Langford, John Sillitoe on behalf of the Wellcome Sanger Institute COVID-19 Surveillance Team |
| EPI_ISL_3107222 | Lighthouse Lab in Alderley Park | Wellcome Sanger Institute for the COVID-19 Genomics UK (COG-UK) Consortium | Jacquelyn Wynn, Mairead Hyland, The Lighthouse Lab in Alderley Park and Alex Alderton, Roberto Amato, Jeffrey Barrett, Sonia Goncalves, Ewan Harrison, David K. Jackson, Ian Johnston, Dominic Kwiatkowski, Cordelia Langford, John Sillitoe on behalf of the Wellcome Sanger Institute COVID-19 Surveillance Team |
| EPI_ISL_3107223 | Lighthouse Lab in Milton Keynes | Wellcome Sanger Institute for the COVID-19 Genomics UK (COG-UK) Consortium | The Lighthouse Lab in Milton Keynes and Alex Alderton, Roberto Amato, Jeffrey Barrett, Sonia Goncalves, Ewan Harrison, David K. Jackson, Ian Johnston, Dominic Kwiatkowski, Cordelia Langford, John Sillitoe on behalf of the Wellcome Sanger Institute COVID-19 Surveillance Team |
| EPI_ISL_3107224 | Lighthouse Lab in Alderley Park | Wellcome Sanger Institute for the COVID-19 Genomics UK (COG-UK) Consortium | Jacquelyn Wynn, Mairead Hyland, The Lighthouse Lab in Alderley Park and Alex Alderton, Roberto Amato, Jeffrey Barrett, Sonia Goncalves, Ewan Harrison, David K. Jackson, Ian Johnston, Dominic Kwiatkowski, Cordelia Langford, John Sillitoe on behalf of the Wellcome Sanger Institute COVID-19 Surveillance Team |
| EPI_ISL_3107225, EPI_ISL_3107226 | Berkshire and Surrey Pathology Services Lighthouse Laboratory | Wellcome Sanger Institute for the COVID-19 Genomics UK (COG-UK) Consortium | Berkshire and Surrey Pathology Services Lighthouse Laboratory and Alex Alderton, Roberto Amato, Jeffrey Barrett, Sonia Goncalves, Ewan Harrison, David K. Jackson, Ian Johnston, Dominic Kwiatkowski, Cordelia Langford, John Sillitoe on behalf of the Wellcome Sanger Institute COVID-19 Surveillance Team |
| EPI_ISL_3107227, EPI_ISL_3107228 | Lighthouse Lab in Alderley Park | Wellcome Sanger Institute for the COVID-19 Genomics UK (COG-UK) Consortium | Jacquelyn Wynn, Mairead Hyland, The Lighthouse Lab in Alderley Park and Alex Alderton, Roberto Amato, Jeffrey Barrett, Sonia Goncalves, Ewan Harrison, David K. Jackson, Ian Johnston, Dominic Kwiatkowski, Cordelia Langford, John Sillitoe on behalf of the Wellcome Sanger Institute COVID-19 Surveillance Team |
| EPI_ISL_3107229 | Berkshire and Surrey Pathology Services Lighthouse Laboratory | Wellcome Sanger Institute for the COVID-19 Genomics UK (COG-UK) Consortium | Berkshire and Surrey Pathology Services Lighthouse Laboratory and Alex Alderton, Roberto Amato, Jeffrey Barrett, Sonia Goncalves, Ewan Harrison, David K. Jackson, Ian Johnston, Dominic Kwiatkowski, Cordelia Langford, John Sillitoe on behalf of the Wellcome Sanger Institute COVID-19 Surveillance Team |
| EPI_ISL_3107230 | Lighthouse Lab in Milton Keynes | Wellcome Sanger Institute for the COVID-19 Genomics UK (COG-UK) Consortium | The Lighthouse Lab in Milton Keynes and Alex Alderton, Roberto Amato, Jeffrey Barrett, Sonia Goncalves, Ewan Harrison, David K. Jackson, Ian Johnston, Dominic Kwiatkowski, Cordelia Langford, John Sillitoe on behalf of the Wellcome Sanger Institute COVID-19 Surveillance Team |
| EPI_ISL_3107231 | Lighthouse Lab in Alderley Park | Wellcome Sanger Institute for the COVID-19 Genomics UK (COG-UK) Consortium | Jacquelyn Wynn, Mairead Hyland, The Lighthouse Lab in Alderley Park and Alex Alderton, Roberto Amato, Jeffrey Barrett, Sonia Goncalves, Ewan Harrison, David K. Jackson, Ian Johnston, Dominic Kwiatkowski, Cordelia Langford, John Sillitoe on behalf of the Wellcome Sanger Institute COVID-19 Surveillance Team |
| EPI_ISL_3107232, EPI_ISL_3107233 | Lighthouse Lab in Milton Keynes | Wellcome Sanger Institute for the COVID-19 Genomics UK (COG-UK) Consortium | The Lighthouse Lab in Milton Keynes and Alex Alderton, Roberto Amato, Jeffrey Barrett, Sonia Goncalves, Ewan Harrison, David K. Jackson, Ian Johnston, Dominic Kwiatkowski, Cordelia Langford, John Sillitoe on behalf of the Wellcome Sanger Institute COVID-19 Surveillance Team |
| EPI_ISL_3107234, EPI_ISL_3107235 | Lighthouse Lab in Alderley Park | Wellcome Sanger Institute for the COVID-19 Genomics UK (COG-UK) Consortium | Jacquelyn Wynn, Mairead Hyland, The Lighthouse Lab in Alderley Park and Alex Alderton, Roberto Amato, Jeffrey Barrett, Sonia Goncalves, Ewan Harrison, David K. Jackson, Ian Johnston, Dominic Kwiatkowski, Cordelia Langford, John Sillitoe on behalf of the Wellcome Sanger Institute COVID-19 Surveillance Team |
| EPI_ISL_3107236 | Berkshire and Surrey Pathology Services Lighthouse Laboratory | Wellcome Sanger Institute for the COVID-19 Genomics UK (COG-UK) Consortium | Berkshire and Surrey Pathology Services Lighthouse Laboratory and Alex Alderton, Roberto Amato, Jeffrey Barrett, Sonia Goncalves, Ewan Harrison, David K. Jackson, Ian Johnston, Dominic Kwiatkowski, Cordelia Langford, John Sillitoe on behalf of the Wellcome Sanger Institute COVID-19 Surveillance Team |
| EPI_ISL_3107237 | Lighthouse Lab in Milton Keynes | Wellcome Sanger Institute for the COVID-19 Genomics UK (COG-UK) Consortium | The Lighthouse Lab in Milton Keynes and Alex Alderton, Roberto Amato, Jeffrey Barrett, Sonia Goncalves, Ewan Harrison, David K. Jackson, Ian Johnston, Dominic Kwiatkowski, Cordelia Langford, John Sillitoe on behalf of the Wellcome Sanger Institute COVID-19 Surveillance Team |
| EPI_ISL_3107238 | Lighthouse Lab in Alderley Park | Wellcome Sanger Institute for the COVID-19 Genomics UK (COG-UK) Consortium | Jacquelyn Wynn, Mairead Hyland, The Lighthouse Lab in Alderley Park and Alex Alderton, Roberto Amato, Jeffrey Barrett, Sonia Goncalves, Ewan Harrison, David K. Jackson, Ian Johnston, Dominic Kwiatkowski, Cordelia Langford, John Sillitoe on behalf of the Wellcome Sanger Institute COVID-19 Surveillance Team |
| EPI_ISL_3107239, EPI_ISL_3107240, EPI_ISL_3107241 | Berkshire and Surrey Pathology Services Lighthouse Laboratory | Wellcome Sanger Institute for the COVID-19 Genomics UK (COG-UK) Consortium | Berkshire and Surrey Pathology Services Lighthouse Laboratory and Alex Alderton, Roberto Amato, Jeffrey Barrett, Sonia Goncalves, Ewan Harrison, David K. Jackson, Ian Johnston, Dominic Kwiatkowski, Cordelia Langford, John Sillitoe on behalf of the Wellcome Sanger Institute COVID-19 Surveillance Team |
| EPI_ISL_3107242 | Lighthouse Lab in Alderley Park | Wellcome Sanger Institute for the COVID-19 Genomics UK (COG-UK) Consortium | Jacquelyn Wynn, Mairead Hyland, The Lighthouse Lab in Alderley Park and Alex Alderton, Roberto Amato, Jeffrey Barrett, Sonia Goncalves, Ewan Harrison, David K. Jackson, Ian Johnston, Dominic Kwiatkowski, Cordelia Langford, John Sillitoe on behalf of the Wellcome Sanger Institute COVID-19 Surveillance Team |
| EPI_ISL_3107243 | Berkshire and Surrey Pathology Services Lighthouse Laboratory | Wellcome Sanger Institute for the COVID-19 Genomics UK (COG-UK) Consortium | Berkshire and Surrey Pathology Services Lighthouse Laboratory and Alex Alderton, Roberto Amato, Jeffrey Barrett, Sonia Goncalves, Ewan Harrison, David K. Jackson, Ian Johnston, Dominic Kwiatkowski, Cordelia Langford, John Sillitoe on behalf of the Wellcome Sanger Institute COVID-19 Surveillance Team |
| EPI_ISL_3107244 | Lighthouse Lab in Alderley Park | Wellcome Sanger Institute for the COVID-19 Genomics UK (COG-UK) Consortium | Jacquelyn Wynn, Mairead Hyland, The Lighthouse Lab in Alderley Park and Alex Alderton, Roberto Amato, Jeffrey Barrett, Sonia Goncalves, Ewan Harrison, David K. Jackson, Ian Johnston, Dominic Kwiatkowski, Cordelia Langford, John Sillitoe on behalf of the Wellcome Sanger Institute COVID-19 Surveillance Team |
| EPI_ISL_3107245 | Berkshire and Surrey Pathology Services Lighthouse Laboratory | Wellcome Sanger Institute for the COVID-19 Genomics UK (COG-UK) Consortium | Berkshire and Surrey Pathology Services Lighthouse Laboratory and Alex Alderton, Roberto Amato, Jeffrey Barrett, Sonia Goncalves, Ewan Harrison, David K. Jackson, Ian Johnston, Dominic Kwiatkowski, Cordelia Langford, John Sillitoe on behalf of the Wellcome Sanger Institute COVID-19 Surveillance Team |
| EPI_ISL_3107246 | Lighthouse Lab in Milton Keynes | Wellcome Sanger Institute for the COVID-19 Genomics UK (COG-UK) Consortium | The Lighthouse Lab in Milton Keynes and Alex Alderton, Roberto Amato, Jeffrey Barrett, Sonia Goncalves, Ewan Harrison, David K. Jackson, Ian Johnston, Dominic Kwiatkowski, Cordelia Langford, John Sillitoe on behalf of the Wellcome Sanger Institute COVID-19 Surveillance Team |
| EPI_ISL_3107247, EPI_ISL_3107248 | Berkshire and Surrey Pathology Services Lighthouse Laboratory | Wellcome Sanger Institute for the COVID-19 Genomics UK (COG-UK) Consortium | Berkshire and Surrey Pathology Services Lighthouse Laboratory and Alex Alderton, Roberto Amato, Jeffrey Barrett, Sonia Goncalves, Ewan Harrison, David K. Jackson, Ian Johnston, Dominic Kwiatkowski, Cordelia Langford, John Sillitoe on behalf of the Wellcome Sanger Institute COVID-19 Surveillance Team |
| EPI_ISL_3107249 | Lighthouse Lab in Milton Keynes | Wellcome Sanger Institute for the COVID-19 Genomics UK (COG-UK) Consortium | The Lighthouse Lab in Milton Keynes and Alex Alderton, Roberto Amato, Jeffrey Barrett, Sonia Goncalves, Ewan Harrison, David K. Jackson, Ian Johnston, Dominic Kwiatkowski, Cordelia Langford, John Sillitoe on behalf of the Wellcome Sanger Institute COVID-19 Surveillance Team |
| EPI_ISL_3107250 | Lighthouse Lab in Alderley Park | Wellcome Sanger Institute for the COVID-19 Genomics UK (COG-UK) Consortium | Jacquelyn Wynn, Mairead Hyland, The Lighthouse Lab in Alderley Park and Alex Alderton, Roberto Amato, Jeffrey Barrett, Sonia Goncalves, Ewan Harrison, David K. Jackson, Ian Johnston, Dominic Kwiatkowski, Cordelia Langford, John Sillitoe on behalf of the Wellcome Sanger Institute COVID-19 Surveillance Team |
| EPI_ISL_3107251 | Berkshire and Surrey Pathology Services Lighthouse Laboratory | Wellcome Sanger Institute for the COVID-19 Genomics UK (COG-UK) Consortium | Berkshire and Surrey Pathology Services Lighthouse Laboratory and Alex Alderton, Roberto Amato, Jeffrey Barrett, Sonia Goncalves, Ewan Harrison, David K. Jackson, Ian Johnston, Dominic Kwiatkowski, Cordelia Langford, John Sillitoe on behalf of the Wellcome Sanger Institute COVID-19 Surveillance Team |
| EPI_ISL_3107252 | Lighthouse Lab in Alderley Park | Wellcome Sanger Institute for the COVID-19 Genomics UK (COG-UK) Consortium | Jacquelyn Wynn, Mairead Hyland, The Lighthouse Lab in Alderley Park and Alex Alderton, Roberto Amato, Jeffrey Barrett, Sonia Goncalves, Ewan Harrison, David K. Jackson, Ian Johnston, Dominic Kwiatkowski, Cordelia Langford, John Sillitoe on behalf of the Wellcome Sanger Institute COVID-19 Surveillance Team |
| EPI_ISL_3107253 | Lighthouse Lab in Milton Keynes | Wellcome Sanger Institute for the COVID-19 Genomics UK (COG-UK) Consortium | The Lighthouse Lab in Milton Keynes and Alex Alderton, Roberto Amato, Jeffrey Barrett, Sonia Goncalves, Ewan Harrison, David K. Jackson, Ian Johnston, Dominic Kwiatkowski, Cordelia Langford, John Sillitoe on behalf of the Wellcome Sanger Institute COVID-19 Surveillance Team |
| EPI_ISL_3107254 | Lighthouse Lab in Alderley Park | Wellcome Sanger Institute for the COVID-19 Genomics UK (COG-UK) Consortium | Jacquelyn Wynn, Mairead Hyland, The Lighthouse Lab in Alderley Park and Alex Alderton, Roberto Amato, Jeffrey Barrett, Sonia Goncalves, Ewan Harrison, David K. Jackson, Ian Johnston, Dominic Kwiatkowski, Cordelia Langford, John Sillitoe on behalf of the Wellcome Sanger Institute COVID-19 Surveillance Team |
| EPI_ISL_3107255 | Berkshire and Surrey Pathology Services Lighthouse Laboratory | Wellcome Sanger Institute for the COVID-19 Genomics UK (COG-UK) Consortium | Berkshire and Surrey Pathology Services Lighthouse Laboratory and Alex Alderton, Roberto Amato, Jeffrey Barrett, Sonia Goncalves, Ewan Harrison, David K. Jackson, Ian Johnston, Dominic Kwiatkowski, Cordelia Langford, John Sillitoe on behalf of the Wellcome Sanger Institute COVID-19 Surveillance Team |
| EPI_ISL_3107256, EPI_ISL_3107257 | Lighthouse Lab in Alderley Park | Wellcome Sanger Institute for the COVID-19 Genomics UK (COG-UK) Consortium | Jacquelyn Wynn, Mairead Hyland, The Lighthouse Lab in Alderley Park and Alex Alderton, Roberto Amato, Jeffrey Barrett, Sonia Goncalves, Ewan Harrison, David K. Jackson, Ian Johnston, Dominic Kwiatkowski, Cordelia Langford, John Sillitoe on behalf of the Wellcome Sanger Institute COVID-19 Surveillance Team |
| EPI_ISL_3107258, EPI_ISL_3107259, EPI_ISL_3107260 | Berkshire and Surrey Pathology Services Lighthouse Laboratory | Wellcome Sanger Institute for the COVID-19 Genomics UK (COG-UK) Consortium | Berkshire and Surrey Pathology Services Lighthouse Laboratory and Alex Alderton, Roberto Amato, Jeffrey Barrett, Sonia Goncalves, Ewan Harrison, David K. Jackson, Ian Johnston, Dominic Kwiatkowski, Cordelia Langford, John Sillitoe on behalf of the Wellcome Sanger Institute COVID-19 Surveillance Team |
| EPI_ISL_3107261 | Lighthouse Lab in Milton Keynes | Wellcome Sanger Institute for the COVID-19 Genomics UK (COG-UK) Consortium | The Lighthouse Lab in Milton Keynes and Alex Alderton, Roberto Amato, Jeffrey Barrett, Sonia Goncalves, Ewan Harrison, David K. Jackson, Ian Johnston, Dominic Kwiatkowski, Cordelia Langford, John Sillitoe on behalf of the Wellcome Sanger Institute COVID-19 Surveillance Team |
| EPI_ISL_3107262 | Berkshire and Surrey Pathology Services Lighthouse Laboratory | Wellcome Sanger Institute for the COVID-19 Genomics UK (COG-UK) Consortium | Berkshire and Surrey Pathology Services Lighthouse Laboratory and Alex Alderton, Roberto Amato, Jeffrey Barrett, Sonia Goncalves, Ewan Harrison, David K. Jackson, Ian Johnston, Dominic Kwiatkowski, Cordelia Langford, John Sillitoe on behalf of the Wellcome Sanger Institute COVID-19 Surveillance Team |
| EPI_ISL_3107263, EPI_ISL_3107264 | Lighthouse Lab in Alderley Park | Wellcome Sanger Institute for the COVID-19 Genomics UK (COG-UK) Consortium | Jacquelyn Wynn, Mairead Hyland, The Lighthouse Lab in Alderley Park and Alex Alderton, Roberto Amato, Jeffrey Barrett, Sonia Goncalves, Ewan Harrison, David K. Jackson, Ian Johnston, Dominic Kwiatkowski, Cordelia Langford, John Sillitoe on behalf of the Wellcome Sanger Institute COVID-19 Surveillance Team |
| EPI_ISL_3107265 | Lighthouse Lab in Milton Keynes | Wellcome Sanger Institute for the COVID-19 Genomics UK (COG-UK) Consortium | The Lighthouse Lab in Milton Keynes and Alex Alderton, Roberto Amato, Jeffrey Barrett, Sonia Goncalves, Ewan Harrison, David K. Jackson, Ian Johnston, Dominic Kwiatkowski, Cordelia Langford, John Sillitoe on behalf of the Wellcome Sanger Institute COVID-19 Surveillance Team |
| EPI_ISL_3107266 | Berkshire and Surrey Pathology Services Lighthouse Laboratory | Wellcome Sanger Institute for the COVID-19 Genomics UK (COG-UK) Consortium | Berkshire and Surrey Pathology Services Lighthouse Laboratory and Alex Alderton, Roberto Amato, Jeffrey Barrett, Sonia Goncalves, Ewan Harrison, David K. Jackson, Ian Johnston, Dominic Kwiatkowski, Cordelia Langford, John Sillitoe on behalf of the Wellcome Sanger Institute COVID-19 Surveillance Team |
| EPI_ISL_3107267 | Lighthouse Lab in Alderley Park | Wellcome Sanger Institute for the COVID-19 Genomics UK (COG-UK) Consortium | Jacquelyn Wynn, Mairead Hyland, The Lighthouse Lab in Alderley Park and Alex Alderton, Roberto Amato, Jeffrey Barrett, Sonia Goncalves, Ewan Harrison, David K. Jackson, Ian Johnston, Dominic Kwiatkowski, Cordelia Langford, John Sillitoe on behalf of the Wellcome Sanger Institute COVID-19 Surveillance Team |
| EPI_ISL_3107268, EPI_ISL_3107269 | Berkshire and Surrey Pathology Services Lighthouse Laboratory | Wellcome Sanger Institute for the COVID-19 Genomics UK (COG-UK) Consortium | Berkshire and Surrey Pathology Services Lighthouse Laboratory and Alex Alderton, Roberto Amato, Jeffrey Barrett, Sonia Goncalves, Ewan Harrison, David K. Jackson, Ian Johnston, Dominic Kwiatkowski, Cordelia Langford, John Sillitoe on behalf of the Wellcome Sanger Institute COVID-19 Surveillance Team |
| EPI_ISL_3107270 | Lighthouse Lab in Alderley Park | Wellcome Sanger Institute for the COVID-19 Genomics UK (COG-UK) Consortium | Jacquelyn Wynn, Mairead Hyland, The Lighthouse Lab in Alderley Park and Alex Alderton, Roberto Amato, Jeffrey Barrett, Sonia Goncalves, Ewan Harrison, David K. Jackson, Ian Johnston, Dominic Kwiatkowski, Cordelia Langford, John Sillitoe on behalf of the Wellcome Sanger Institute COVID-19 Surveillance Team |
| EPI_ISL_3107271, EPI_ISL_3107272 | Berkshire and Surrey Pathology Services Lighthouse Laboratory | Wellcome Sanger Institute for the COVID-19 Genomics UK (COG-UK) Consortium | Berkshire and Surrey Pathology Services Lighthouse Laboratory and Alex Alderton, Roberto Amato, Jeffrey Barrett, Sonia Goncalves, Ewan Harrison, David K. Jackson, Ian Johnston, Dominic Kwiatkowski, Cordelia Langford, John Sillitoe on behalf of the Wellcome Sanger Institute COVID-19 Surveillance Team |
| EPI_ISL_3107273 | Lighthouse Lab in Milton Keynes | Wellcome Sanger Institute for the COVID-19 Genomics UK (COG-UK) Consortium | The Lighthouse Lab in Milton Keynes and Alex Alderton, Roberto Amato, Jeffrey Barrett, Sonia Goncalves, Ewan Harrison, David K. Jackson, Ian Johnston, Dominic Kwiatkowski, Cordelia Langford, John Sillitoe on behalf of the Wellcome Sanger Institute COVID-19 Surveillance Team |
| EPI_ISL_3107274 | Lighthouse Lab in Alderley Park | Wellcome Sanger Institute for the COVID-19 Genomics UK (COG-UK) Consortium | Jacquelyn Wynn, Mairead Hyland, The Lighthouse Lab in Alderley Park and Alex Alderton, Roberto Amato, Jeffrey Barrett, Sonia Goncalves, Ewan Harrison, David K. Jackson, Ian Johnston, Dominic Kwiatkowski, Cordelia Langford, John Sillitoe on behalf of the Wellcome Sanger Institute COVID-19 Surveillance Team |
| EPI_ISL_3107275 | Berkshire and Surrey Pathology Services Lighthouse Laboratory | Wellcome Sanger Institute for the COVID-19 Genomics UK (COG-UK) Consortium | Berkshire and Surrey Pathology Services Lighthouse Laboratory and Alex Alderton, Roberto Amato, Jeffrey Barrett, Sonia Goncalves, Ewan Harrison, David K. Jackson, Ian Johnston, Dominic Kwiatkowski, Cordelia Langford, John Sillitoe on behalf of the Wellcome Sanger Institute COVID-19 Surveillance Team |
| EPI_ISL_3107276 | Lighthouse Lab in Alderley Park | Wellcome Sanger Institute for the COVID-19 Genomics UK (COG-UK) Consortium | Jacquelyn Wynn, Mairead Hyland, The Lighthouse Lab in Alderley Park and Alex Alderton, Roberto Amato, Jeffrey Barrett, Sonia Goncalves, Ewan Harrison, David K. Jackson, Ian Johnston, Dominic Kwiatkowski, Cordelia Langford, John Sillitoe on behalf of the Wellcome Sanger Institute COVID-19 Surveillance Team |
| EPI_ISL_3107277 | Lighthouse Lab in Milton Keynes | Wellcome Sanger Institute for the COVID-19 Genomics UK (COG-UK) Consortium | The Lighthouse Lab in Milton Keynes and Alex Alderton, Roberto Amato, Jeffrey Barrett, Sonia Goncalves, Ewan Harrison, David K. Jackson, Ian Johnston, Dominic Kwiatkowski, Cordelia Langford, John Sillitoe on behalf of the Wellcome Sanger Institute COVID-19 Surveillance Team |

| | | | |
|---|---|---|---|
| EPI_ISL_3107278, EPI_ISL_3107279 | Berkshire and Surrey Pathology Services Lighthouse Laboratory | Wellcome Sanger Institute for the COVID-19 Genomics UK (COG-UK) Consortium | Berkshire and Surrey Pathology Services Lighthouse Laboratory and Alex Alderton, Roberto Amato, Jeffrey Barrett, Sonia Goncalves, Ewan Harrison, David K. Jackson, Ian Johnston, Dominic Kwiatkowski, Cordelia Langford, John Sillitoe on behalf of the Wellcome Sanger Institute COVID-19 Surveillance Team |
| EPI_ISL_3107280 | Lighthouse Lab in Milton Keynes | Wellcome Sanger Institute for the COVID-19 Genomics UK (COG-UK) Consortium | The Lighthouse Lab in Milton Keynes and Alex Alderton, Roberto Amato, Jeffrey Barrett, Sonia Goncalves, Ewan Harrison, David K. Jackson, Ian Johnston, Dominic Kwiatkowski, Cordelia Langford, John Sillitoe on behalf of the Wellcome Sanger Institute COVID-19 Surveillance Team |
| EPI_ISL_3107281, EPI_ISL_3107282 | Berkshire and Surrey Pathology Services Lighthouse Laboratory | Wellcome Sanger Institute for the COVID-19 Genomics UK (COG-UK) Consortium | Berkshire and Surrey Pathology Services Lighthouse Laboratory and Alex Alderton, Roberto Amato, Jeffrey Barrett, Sonia Goncalves, Ewan Harrison, David K. Jackson, Ian Johnston, Dominic Kwiatkowski, Cordelia Langford, John Sillitoe on behalf of the Wellcome Sanger Institute COVID-19 Surveillance Team |
| EPI_ISL_3107283 | Lighthouse Lab in Milton Keynes | Wellcome Sanger Institute for the COVID-19 Genomics UK (COG-UK) Consortium | The Lighthouse Lab in Milton Keynes and Alex Alderton, Roberto Amato, Jeffrey Barrett, Sonia Goncalves, Ewan Harrison, David K. Jackson, Ian Johnston, Dominic Kwiatkowski, Cordelia Langford, John Sillitoe on behalf of the Wellcome Sanger Institute COVID-19 Surveillance Team |
| EPI_ISL_3107284, EPI_ISL_3107285, EPI_ISL_3107286, EPI_ISL_3107287 | Berkshire and Surrey Pathology Services Lighthouse Laboratory | Wellcome Sanger Institute for the COVID-19 Genomics UK (COG-UK) Consortium | Berkshire and Surrey Pathology Services Lighthouse Laboratory and Alex Alderton, Roberto Amato, Jeffrey Barrett, Sonia Goncalves, Ewan Harrison, David K. Jackson, Ian Johnston, Dominic Kwiatkowski, Cordelia Langford, John Sillitoe on behalf of the Wellcome Sanger Institute COVID-19 Surveillance Team |
| EPI_ISL_3107288 | Lighthouse Lab in Alderley Park | Wellcome Sanger Institute for the COVID-19 Genomics UK (COG-UK) Consortium | Jacquelyn Wynn, Mairead Hyland, The Lighthouse Lab in Alderley Park and Alex Alderton, Roberto Amato, Jeffrey Barrett, Sonia Goncalves, Ewan Harrison, David K. Jackson, Ian Johnston, Dominic Kwiatkowski, Cordelia Langford, John Sillitoe on behalf of the Wellcome Sanger Institute COVID-19 Surveillance Team |
| EPI_ISL_3107289 | Lighthouse Lab in Milton Keynes | Wellcome Sanger Institute for the COVID-19 Genomics UK (COG-UK) Consortium | The Lighthouse Lab in Milton Keynes and Alex Alderton, Roberto Amato, Jeffrey Barrett, Sonia Goncalves, Ewan Harrison, David K. Jackson, Ian Johnston, Dominic Kwiatkowski, Cordelia Langford, John Sillitoe on behalf of the Wellcome Sanger Institute COVID-19 Surveillance Team |
| EPI_ISL_3107290 | Berkshire and Surrey Pathology Services Lighthouse Laboratory | Wellcome Sanger Institute for the COVID-19 Genomics UK (COG-UK) Consortium | Berkshire and Surrey Pathology Services Lighthouse Laboratory and Alex Alderton, Roberto Amato, Jeffrey Barrett, Sonia Goncalves, Ewan Harrison, David K. Jackson, Ian Johnston, Dominic Kwiatkowski, Cordelia Langford, John Sillitoe on behalf of the Wellcome Sanger Institute COVID-19 Surveillance Team |
| EPI_ISL_3107291 | Lighthouse Lab in Milton Keynes | Wellcome Sanger Institute for the COVID-19 Genomics UK (COG-UK) Consortium | The Lighthouse Lab in Milton Keynes and Alex Alderton, Roberto Amato, Jeffrey Barrett, Sonia Goncalves, Ewan Harrison, David K. Jackson, Ian Johnston, Dominic Kwiatkowski, Cordelia Langford, John Sillitoe on behalf of the Wellcome Sanger Institute COVID-19 Surveillance Team |
| EPI_ISL_3107292 | Berkshire and Surrey Pathology Services Lighthouse Laboratory | Wellcome Sanger Institute for the COVID-19 Genomics UK (COG-UK) Consortium | Berkshire and Surrey Pathology Services Lighthouse Laboratory and Alex Alderton, Roberto Amato, Jeffrey Barrett, Sonia Goncalves, Ewan Harrison, David K. Jackson, Ian Johnston, Dominic Kwiatkowski, Cordelia Langford, John Sillitoe on behalf of the Wellcome Sanger Institute COVID-19 Surveillance Team |
| EPI_ISL_3107293 | Lighthouse Lab in Milton Keynes | Wellcome Sanger Institute for the COVID-19 Genomics UK (COG-UK) Consortium | The Lighthouse Lab in Milton Keynes and Alex Alderton, Roberto Amato, Jeffrey Barrett, Sonia Goncalves, Ewan Harrison, David K. Jackson, Ian Johnston, Dominic Kwiatkowski, Cordelia Langford, John Sillitoe on behalf of the Wellcome Sanger Institute COVID-19 Surveillance Team |
| EPI_ISL_3107294, EPI_ISL_3107295, EPI_ISL_3107296, EPI_ISL_3107297, EPI_ISL_3107298 | Berkshire and Surrey Pathology Services Lighthouse Laboratory | Wellcome Sanger Institute for the COVID-19 Genomics UK (COG-UK) Consortium | Berkshire and Surrey Pathology Services Lighthouse Laboratory and Alex Alderton, Roberto Amato, Jeffrey Barrett, Sonia Goncalves, Ewan Harrison, David K. Jackson, Ian Johnston, Dominic Kwiatkowski, Cordelia Langford, John Sillitoe on behalf of the Wellcome Sanger Institute COVID-19 Surveillance Team |
| EPI_ISL_3107299, EPI_ISL_3107300 | Lighthouse Lab in Alderley Park | Wellcome Sanger Institute for the COVID-19 Genomics UK (COG-UK) Consortium | Jacquelyn Wynn, Mairead Hyland, The Lighthouse Lab in Alderley Park and Alex Alderton, Roberto Amato, Jeffrey Barrett, Sonia Goncalves, Ewan Harrison, David K. Jackson, Ian Johnston, Dominic Kwiatkowski, Cordelia Langford, John Sillitoe on behalf of the Wellcome Sanger Institute COVID-19 Surveillance Team |
| EPI_ISL_3107301, EPI_ISL_3107302 | Berkshire and Surrey Pathology Services Lighthouse Laboratory | Wellcome Sanger Institute for the COVID-19 Genomics UK (COG-UK) Consortium | Berkshire and Surrey Pathology Services Lighthouse Laboratory and Alex Alderton, Roberto Amato, Jeffrey Barrett, Sonia Goncalves, Ewan Harrison, David K. Jackson, Ian Johnston, Dominic Kwiatkowski, Cordelia Langford, John Sillitoe on behalf of the Wellcome Sanger Institute COVID-19 Surveillance Team |
| EPI_ISL_3107303 | Lighthouse Lab in Milton Keynes | Wellcome Sanger Institute for the COVID-19 Genomics UK (COG-UK) Consortium | The Lighthouse Lab in Milton Keynes and Alex Alderton, Roberto Amato, Jeffrey Barrett, Sonia Goncalves, Ewan Harrison, David K. Jackson, Ian Johnston, Dominic Kwiatkowski, Cordelia Langford, John Sillitoe on behalf of the Wellcome Sanger Institute COVID-19 Surveillance Team |
| EPI_ISL_3107304 | Berkshire and Surrey Pathology Services Lighthouse Laboratory | Wellcome Sanger Institute for the COVID-19 Genomics UK (COG-UK) Consortium | Berkshire and Surrey Pathology Services Lighthouse Laboratory and Alex Alderton, Roberto Amato, Jeffrey Barrett, Sonia Goncalves, Ewan Harrison, David K. Jackson, Ian Johnston, Dominic Kwiatkowski, Cordelia Langford, John Sillitoe on behalf of the Wellcome Sanger Institute COVID-19 Surveillance Team |
| EPI_ISL_3107305 | Lighthouse Lab in Alderley Park | Wellcome Sanger Institute for the COVID-19 Genomics UK (COG-UK) Consortium | Jacquelyn Wynn, Mairead Hyland, The Lighthouse Lab in Alderley Park and Alex Alderton, Roberto Amato, Jeffrey Barrett, Sonia Goncalves, Ewan Harrison, David K. Jackson, Ian Johnston, Dominic Kwiatkowski, Cordelia Langford, John Sillitoe on behalf of the Wellcome Sanger Institute COVID-19 Surveillance Team |
| EPI_ISL_3107306 | Berkshire and Surrey Pathology Services Lighthouse Laboratory | Wellcome Sanger Institute for the COVID-19 Genomics UK (COG-UK) Consortium | Berkshire and Surrey Pathology Services Lighthouse Laboratory and Alex Alderton, Roberto Amato, Jeffrey Barrett, Sonia Goncalves, Ewan Harrison, David K. Jackson, Ian Johnston, Dominic Kwiatkowski, Cordelia Langford, John Sillitoe on behalf of the Wellcome Sanger Institute COVID-19 Surveillance Team |
| EPI_ISL_3107307, EPI_ISL_3107308 | Lighthouse Lab in Milton Keynes | Wellcome Sanger Institute for the COVID-19 Genomics UK (COG-UK) Consortium | The Lighthouse Lab in Milton Keynes and Alex Alderton, Roberto Amato, Jeffrey Barrett, Sonia Goncalves, Ewan Harrison, David K. Jackson, Ian Johnston, Dominic Kwiatkowski, Cordelia Langford, John Sillitoe on behalf of the Wellcome Sanger Institute COVID-19 Surveillance Team |
| EPI_ISL_3107309 | Lighthouse Lab in Alderley Park | Wellcome Sanger Institute for the COVID-19 Genomics UK (COG-UK) Consortium | Jacquelyn Wynn, Mairead Hyland, The Lighthouse Lab in Alderley Park and Alex Alderton, Roberto Amato, Jeffrey Barrett, Sonia Goncalves, Ewan Harrison, David K. Jackson, Ian Johnston, Dominic Kwiatkowski, Cordelia Langford, John Sillitoe on behalf of the Wellcome Sanger Institute COVID-19 Surveillance Team |
| EPI_ISL_3107310 | Lighthouse Lab in Milton Keynes | Wellcome Sanger Institute for the COVID-19 Genomics UK (COG-UK) Consortium | The Lighthouse Lab in Milton Keynes and Alex Alderton, Roberto Amato, Jeffrey Barrett, Sonia Goncalves, Ewan Harrison, David K. Jackson, Ian Johnston, Dominic Kwiatkowski, Cordelia Langford, John Sillitoe on behalf of the Wellcome Sanger Institute COVID-19 Surveillance Team |
| EPI_ISL_3107311 | Berkshire and Surrey Pathology Services Lighthouse Laboratory | Wellcome Sanger Institute for the COVID-19 Genomics UK (COG-UK) Consortium | Berkshire and Surrey Pathology Services Lighthouse Laboratory and Alex Alderton, Roberto Amato, Jeffrey Barrett, Sonia Goncalves, Ewan Harrison, David K. Jackson, Ian Johnston, Dominic Kwiatkowski, Cordelia Langford, John Sillitoe on behalf of the Wellcome Sanger Institute COVID-19 Surveillance Team |
| EPI_ISL_3107312 | Lighthouse Lab in Alderley Park | Wellcome Sanger Institute for the COVID-19 Genomics UK (COG-UK) Consortium | Jacquelyn Wynn, Mairead Hyland, The Lighthouse Lab in Alderley Park and Alex Alderton, Roberto Amato, Jeffrey Barrett, Sonia Goncalves, Ewan Harrison, David K. Jackson, Ian Johnston, Dominic Kwiatkowski, Cordelia Langford, John Sillitoe on behalf of the Wellcome Sanger Institute COVID-19 Surveillance Team |
| EPI_ISL_3107313, EPI_ISL_3107314 | Berkshire and Surrey Pathology Services Lighthouse Laboratory | Wellcome Sanger Institute for the COVID-19 Genomics UK (COG-UK) Consortium | Berkshire and Surrey Pathology Services Lighthouse Laboratory and Alex Alderton, Roberto Amato, Jeffrey Barrett, Sonia Goncalves, Ewan Harrison, David K. Jackson, Ian Johnston, Dominic Kwiatkowski, Cordelia Langford, John Sillitoe on behalf of the Wellcome Sanger Institute COVID-19 Surveillance Team |
| EPI_ISL_3107315, EPI_ISL_3107316, EPI_ISL_3107317, EPI_ISL_3107318 | Lighthouse Lab in Alderley Park | Wellcome Sanger Institute for the COVID-19 Genomics UK (COG-UK) Consortium | Jacquelyn Wynn, Mairead Hyland, The Lighthouse Lab in Alderley Park and Alex Alderton, Roberto Amato, Jeffrey Barrett, Sonia Goncalves, Ewan Harrison, David K. Jackson, Ian Johnston, Dominic Kwiatkowski, Cordelia Langford, John Sillitoe on behalf of the Wellcome Sanger Institute COVID-19 Surveillance Team |
| EPI_ISL_3107319 | Lighthouse Lab in Milton Keynes | Wellcome Sanger Institute for the COVID-19 Genomics UK (COG-UK) Consortium | The Lighthouse Lab in Milton Keynes and Alex Alderton, Roberto Amato, Jeffrey Barrett, Sonia Goncalves, Ewan Harrison, David K. Jackson, Ian Johnston, Dominic Kwiatkowski, Cordelia Langford, John Sillitoe on behalf of the Wellcome Sanger Institute COVID-19 Surveillance Team |
| EPI_ISL_3107320 | Berkshire and Surrey Pathology Services Lighthouse Laboratory | Wellcome Sanger Institute for the COVID-19 Genomics UK (COG-UK) Consortium | Berkshire and Surrey Pathology Services Lighthouse Laboratory and Alex Alderton, Roberto Amato, Jeffrey Barrett, Sonia Goncalves, Ewan Harrison, David K. Jackson, Ian Johnston, Dominic Kwiatkowski, Cordelia Langford, John Sillitoe on behalf of the Wellcome Sanger Institute COVID-19 Surveillance Team |
| EPI_ISL_3107321 | Lighthouse Lab in Alderley Park | Wellcome Sanger Institute for the COVID-19 Genomics UK (COG-UK) Consortium | Jacquelyn Wynn, Mairead Hyland, The Lighthouse Lab in Alderley Park and Alex Alderton, Roberto Amato, Jeffrey Barrett, Sonia Goncalves, Ewan Harrison, David K. Jackson, Ian Johnston, Dominic Kwiatkowski, Cordelia Langford, John Sillitoe on behalf of the Wellcome Sanger Institute COVID-19 Surveillance Team |
| EPI_ISL_3107322 | Lighthouse Lab in Milton Keynes | Wellcome Sanger Institute for the COVID-19 Genomics UK (COG-UK) Consortium | The Lighthouse Lab in Milton Keynes and Alex Alderton, Roberto Amato, Jeffrey Barrett, Sonia Goncalves, Ewan Harrison, David K. Jackson, Ian Johnston, Dominic Kwiatkowski, Cordelia Langford, John Sillitoe on behalf of the Wellcome Sanger Institute COVID-19 Surveillance Team |
| EPI_ISL_3107323 | Berkshire and Surrey Pathology Services Lighthouse Laboratory | Wellcome Sanger Institute for the COVID-19 Genomics UK (COG-UK) Consortium | Berkshire and Surrey Pathology Services Lighthouse Laboratory and Alex Alderton, Roberto Amato, Jeffrey Barrett, Sonia Goncalves, Ewan Harrison, David K. Jackson, Ian Johnston, Dominic Kwiatkowski, Cordelia Langford, John Sillitoe on behalf of the Wellcome Sanger Institute COVID-19 Surveillance Team |
| EPI_ISL_3107324, EPI_ISL_3107325, EPI_ISL_3107326, EPI_ISL_3107327, EPI_ISL_3107328, EPI_ISL_3107329, EPI_ISL_3107330 | Lighthouse Lab in Alderley Park | Wellcome Sanger Institute for the COVID-19 Genomics UK (COG-UK) Consortium | Jacquelyn Wynn, Mairead Hyland, The Lighthouse Lab in Alderley Park and Alex Alderton, Roberto Amato, Jeffrey Barrett, Sonia Goncalves, Ewan Harrison, David K. Jackson, Ian Johnston, Dominic Kwiatkowski, Cordelia Langford, John Sillitoe on behalf of the Wellcome Sanger Institute COVID-19 Surveillance Team |
| EPI_ISL_3107331 | Lighthouse Lab in Milton Keynes | Wellcome Sanger Institute for the COVID-19 Genomics UK (COG-UK) Consortium | The Lighthouse Lab in Milton Keynes and Alex Alderton, Roberto Amato, Jeffrey Barrett, Sonia Goncalves, Ewan Harrison, David K. Jackson, Ian Johnston, Dominic Kwiatkowski, Cordelia Langford, John Sillitoe on behalf of the Wellcome Sanger Institute COVID-19 Surveillance Team |
| EPI_ISL_3107332 | Lighthouse Lab in Alderley Park | Wellcome Sanger Institute for the COVID-19 Genomics UK (COG-UK) Consortium | Jacquelyn Wynn, Mairead Hyland, The Lighthouse Lab in Alderley Park and Alex Alderton, Roberto Amato, Jeffrey Barrett, Sonia Goncalves, Ewan Harrison, David K. Jackson, Ian Johnston, Dominic Kwiatkowski, Cordelia Langford, John Sillitoe on behalf of the Wellcome Sanger Institute COVID-19 Surveillance Team |
| EPI_ISL_3107333 | Lighthouse Lab in Milton Keynes | Wellcome Sanger Institute for the COVID-19 Genomics UK (COG-UK) Consortium | The Lighthouse Lab in Milton Keynes and Alex Alderton, Roberto Amato, Jeffrey Barrett, Sonia Goncalves, Ewan Harrison, David K. Jackson, Ian Johnston, Dominic Kwiatkowski, Cordelia Langford, John Sillitoe on behalf of the Wellcome Sanger Institute COVID-19 Surveillance Team |
| EPI_ISL_3107334, EPI_ISL_3107335 | Lighthouse Lab in Alderley Park | Wellcome Sanger Institute for the COVID-19 Genomics UK (COG-UK) Consortium | Jacquelyn Wynn, Mairead Hyland, The Lighthouse Lab in Alderley Park and Alex Alderton, Roberto Amato, Jeffrey Barrett, Sonia Goncalves, Ewan Harrison, David K. Jackson, Ian Johnston, Dominic Kwiatkowski, Cordelia Langford, John Sillitoe on behalf of the Wellcome Sanger Institute COVID-19 Surveillance Team |
| EPI_ISL_3107336 | Lighthouse Lab in Milton Keynes | Wellcome Sanger Institute for the COVID-19 Genomics UK (COG-UK) Consortium | The Lighthouse Lab in Milton Keynes and Alex Alderton, Roberto Amato, Jeffrey Barrett, Sonia Goncalves, Ewan Harrison, David K. Jackson, Ian Johnston, Dominic Kwiatkowski, Cordelia Langford, John Sillitoe on behalf of the Wellcome Sanger Institute COVID-19 Surveillance Team |
| EPI_ISL_3107337, EPI_ISL_3107338 | Lighthouse Lab in Alderley Park | Wellcome Sanger Institute for the COVID-19 Genomics UK (COG-UK) Consortium | Jacquelyn Wynn, Mairead Hyland, The Lighthouse Lab in Alderley Park and Alex Alderton, Roberto Amato, Jeffrey Barrett, Sonia Goncalves, Ewan Harrison, David K. Jackson, Ian Johnston, Dominic Kwiatkowski, Cordelia Langford, John Sillitoe on behalf of the Wellcome Sanger Institute COVID-19 Surveillance Team |
| EPI_ISL_3107339 | Lighthouse Lab in Milton Keynes | Wellcome Sanger Institute for the COVID-19 Genomics UK (COG-UK) Consortium | The Lighthouse Lab in Milton Keynes and Alex Alderton, Roberto Amato, Jeffrey Barrett, Sonia Goncalves, Ewan Harrison, David K. Jackson, Ian Johnston, Dominic Kwiatkowski, Cordelia Langford, John Sillitoe on behalf of the Wellcome Sanger Institute COVID-19 Surveillance Team |
| EPI_ISL_3107340 | Lighthouse Lab in Alderley Park | Wellcome Sanger Institute for the COVID-19 Genomics UK (COG-UK) Consortium | Jacquelyn Wynn, Mairead Hyland, The Lighthouse Lab in Alderley Park and Alex Alderton, Roberto Amato, Jeffrey Barrett, Sonia Goncalves, Ewan Harrison, David K. Jackson, Ian Johnston, Dominic Kwiatkowski, Cordelia Langford, John Sillitoe on behalf of the Wellcome Sanger Institute COVID-19 Surveillance Team |
| EPI_ISL_3107341 | Lighthouse Lab in Milton Keynes | Wellcome Sanger Institute for the COVID-19 Genomics UK (COG-UK) Consortium | The Lighthouse Lab in Milton Keynes and Alex Alderton, Roberto Amato, Jeffrey Barrett, Sonia Goncalves, Ewan Harrison, David K. Jackson, Ian Johnston, Dominic Kwiatkowski, Cordelia Langford, John Sillitoe on behalf of the Wellcome Sanger Institute COVID-19 Surveillance Team |

| | | | |
|---|---|---|---|
| EPI_ISL_3107342, EPI_ISL_3107343 | Lighthouse Lab in Alderley Park | Wellcome Sanger Institute for the COVID-19 Genomics UK (COG-UK) Consortium | Jacquelyn Wynn, Mairead Hyland, The Lighthouse Lab in Alderley Park and Alex Alderton, Roberto Amato, Jeffrey Barrett, Sonia Goncalves, Ewan Harrison, David K. Jackson, Ian Johnston, Dominic Kwiatkowski, Cordelia Langford, John Sillitoe on behalf of the Wellcome Sanger Institute COVID-19 Surveillance Team |
| EPI_ISL_3107344 | Lighthouse Lab in Milton Keynes | Wellcome Sanger Institute for the COVID-19 Genomics UK (COG-UK) Consortium | The Lighthouse Lab in Milton Keynes and Alex Alderton, Roberto Amato, Jeffrey Barrett, Sonia Goncalves, Ewan Harrison, David K. Jackson, Ian Johnston, Dominic Kwiatkowski, Cordelia Langford, John Sillitoe on behalf of the Wellcome Sanger Institute COVID-19 Surveillance Team |
| EPI_ISL_3107345, EPI_ISL_3107346, EPI_ISL_3107347, EPI_ISL_3107348, EPI_ISL_3107349, EPI_ISL_3107350 | Lighthouse Lab in Alderley Park | Wellcome Sanger Institute for the COVID-19 Genomics UK (COG-UK) Consortium | Jacquelyn Wynn, Mairead Hyland, The Lighthouse Lab in Alderley Park and Alex Alderton, Roberto Amato, Jeffrey Barrett, Sonia Goncalves, Ewan Harrison, David K. Jackson, Ian Johnston, Dominic Kwiatkowski, Cordelia Langford, John Sillitoe on behalf of the Wellcome Sanger Institute COVID-19 Surveillance Team |
| EPI_ISL_3107351 | Lighthouse Lab in Milton Keynes | Wellcome Sanger Institute for the COVID-19 Genomics UK (COG-UK) Consortium | The Lighthouse Lab in Milton Keynes and Alex Alderton, Roberto Amato, Jeffrey Barrett, Sonia Goncalves, Ewan Harrison, David K. Jackson, Ian Johnston, Dominic Kwiatkowski, Cordelia Langford, John Sillitoe on behalf of the Wellcome Sanger Institute COVID-19 Surveillance Team |
| EPI_ISL_3107352, EPI_ISL_3107353 | Lighthouse Lab in Alderley Park | Wellcome Sanger Institute for the COVID-19 Genomics UK (COG-UK) Consortium | Jacquelyn Wynn, Mairead Hyland, The Lighthouse Lab in Alderley Park and Alex Alderton, Roberto Amato, Jeffrey Barrett, Sonia Goncalves, Ewan Harrison, David K. Jackson, Ian Johnston, Dominic Kwiatkowski, Cordelia Langford, John Sillitoe on behalf of the Wellcome Sanger Institute COVID-19 Surveillance Team |
| EPI_ISL_3107354, EPI_ISL_3107355 | Lighthouse Lab in Milton Keynes | Wellcome Sanger Institute for the COVID-19 Genomics UK (COG-UK) Consortium | The Lighthouse Lab in Milton Keynes and Alex Alderton, Roberto Amato, Jeffrey Barrett, Sonia Goncalves, Ewan Harrison, David K. Jackson, Ian Johnston, Dominic Kwiatkowski, Cordelia Langford, John Sillitoe on behalf of the Wellcome Sanger Institute COVID-19 Surveillance Team |
| EPI_ISL_3107356, EPI_ISL_3107357, EPI_ISL_3107358 | Lighthouse Lab in Alderley Park | Wellcome Sanger Institute for the COVID-19 Genomics UK (COG-UK) Consortium | Jacquelyn Wynn, Mairead Hyland, The Lighthouse Lab in Alderley Park and Alex Alderton, Roberto Amato, Jeffrey Barrett, Sonia Goncalves, Ewan Harrison, David K. Jackson, Ian Johnston, Dominic Kwiatkowski, Cordelia Langford, John Sillitoe on behalf of the Wellcome Sanger Institute COVID-19 Surveillance Team |
| EPI_ISL_3107359 | Lighthouse Lab in Milton Keynes | Wellcome Sanger Institute for the COVID-19 Genomics UK (COG-UK) Consortium | The Lighthouse Lab in Milton Keynes and Alex Alderton, Roberto Amato, Jeffrey Barrett, Sonia Goncalves, Ewan Harrison, David K. Jackson, Ian Johnston, Dominic Kwiatkowski, Cordelia Langford, John Sillitoe on behalf of the Wellcome Sanger Institute COVID-19 Surveillance Team |
| EPI_ISL_3107360, EPI_ISL_3107361, EPI_ISL_3107362 | Lighthouse Lab in Alderley Park | Wellcome Sanger Institute for the COVID-19 Genomics UK (COG-UK) Consortium | Jacquelyn Wynn, Mairead Hyland, The Lighthouse Lab in Alderley Park and Alex Alderton, Roberto Amato, Jeffrey Barrett, Sonia Goncalves, Ewan Harrison, David K. Jackson, Ian Johnston, Dominic Kwiatkowski, Cordelia Langford, John Sillitoe on behalf of the Wellcome Sanger Institute COVID-19 Surveillance Team |
| EPI_ISL_3107363, EPI_ISL_3107364, EPI_ISL_3107365, EPI_ISL_3107366 | Lighthouse Lab in Milton Keynes | Wellcome Sanger Institute for the COVID-19 Genomics UK (COG-UK) Consortium | The Lighthouse Lab in Milton Keynes and Alex Alderton, Roberto Amato, Jeffrey Barrett, Sonia Goncalves, Ewan Harrison, David K. Jackson, Ian Johnston, Dominic Kwiatkowski, Cordelia Langford, John Sillitoe on behalf of the Wellcome Sanger Institute COVID-19 Surveillance Team |
| EPI_ISL_3107367, EPI_ISL_3107368, EPI_ISL_3107369, EPI_ISL_3107370, EPI_ISL_3107371, EPI_ISL_3107372, EPI_ISL_3107373, EPI_ISL_3107374 | Lighthouse Lab in Alderley Park | Wellcome Sanger Institute for the COVID-19 Genomics UK (COG-UK) Consortium | Jacquelyn Wynn, Mairead Hyland, The Lighthouse Lab in Alderley Park and Alex Alderton, Roberto Amato, Jeffrey Barrett, Sonia Goncalves, Ewan Harrison, David K. Jackson, Ian Johnston, Dominic Kwiatkowski, Cordelia Langford, John Sillitoe on behalf of the Wellcome Sanger Institute COVID-19 Surveillance Team |
| EPI_ISL_3107375 | Lighthouse Lab in Milton Keynes | Wellcome Sanger Institute for the COVID-19 Genomics UK (COG-UK) Consortium | The Lighthouse Lab in Milton Keynes and Alex Alderton, Roberto Amato, Jeffrey Barrett, Sonia Goncalves, Ewan Harrison, David K. Jackson, Ian Johnston, Dominic Kwiatkowski, Cordelia Langford, John Sillitoe on behalf of the Wellcome Sanger Institute COVID-19 Surveillance Team |
| EPI_ISL_3107376, EPI_ISL_3107377, EPI_ISL_3107378 | Lighthouse Lab in Alderley Park | Wellcome Sanger Institute for the COVID-19 Genomics UK (COG-UK) Consortium | Jacquelyn Wynn, Mairead Hyland, The Lighthouse Lab in Alderley Park and Alex Alderton, Roberto Amato, Jeffrey Barrett, Sonia Goncalves, Ewan Harrison, David K. Jackson, Ian Johnston, Dominic Kwiatkowski, Cordelia Langford, John Sillitoe on behalf of the Wellcome Sanger Institute COVID-19 Surveillance Team |
| EPI_ISL_3107379 | Lighthouse Lab in Milton Keynes | Wellcome Sanger Institute for the COVID-19 Genomics UK (COG-UK) Consortium | The Lighthouse Lab in Milton Keynes and Alex Alderton, Roberto Amato, Jeffrey Barrett, Sonia Goncalves, Ewan Harrison, David K. Jackson, Ian Johnston, Dominic Kwiatkowski, Cordelia Langford, John Sillitoe on behalf of the Wellcome Sanger Institute COVID-19 Surveillance Team |
| EPI_ISL_3107380, EPI_ISL_3107381 | Lighthouse Lab in Alderley Park | Wellcome Sanger Institute for the COVID-19 Genomics UK (COG-UK) Consortium | Jacquelyn Wynn, Mairead Hyland, The Lighthouse Lab in Alderley Park and Alex Alderton, Roberto Amato, Jeffrey Barrett, Sonia Goncalves, Ewan Harrison, David K. Jackson, Ian Johnston, Dominic Kwiatkowski, Cordelia Langford, John Sillitoe on behalf of the Wellcome Sanger Institute COVID-19 Surveillance Team |
| EPI_ISL_3107382 | Lighthouse Lab in Milton Keynes | Wellcome Sanger Institute for the COVID-19 Genomics UK (COG-UK) Consortium | The Lighthouse Lab in Milton Keynes and Alex Alderton, Roberto Amato, Jeffrey Barrett, Sonia Goncalves, Ewan Harrison, David K. Jackson, Ian Johnston, Dominic Kwiatkowski, Cordelia Langford, John Sillitoe on behalf of the Wellcome Sanger Institute COVID-19 Surveillance Team |
| EPI_ISL_3107383, EPI_ISL_3107384, EPI_ISL_3107385, EPI_ISL_3107386, EPI_ISL_3107387, EPI_ISL_3107388, EPI_ISL_3107389, EPI_ISL_3107390, EPI_ISL_3107391, EPI_ISL_3107392, EPI_ISL_3107393, EPI_ISL_3107394, EPI_ISL_3107395 | | | |
| see above | Lighthouse Lab in Alderley Park | Wellcome Sanger Institute for the COVID-19 Genomics UK (COG-UK) Consortium | Jacquelyn Wynn, Mairead Hyland, The Lighthouse Lab in Alderley Park and Alex Alderton, Roberto Amato, Jeffrey Barrett, Sonia Goncalves, Ewan Harrison, David K. Jackson, Ian Johnston, Dominic Kwiatkowski, Cordelia Langford, John Sillitoe on behalf of the Wellcome Sanger Institute COVID-19 Surveillance Team |
| EPI_ISL_3107396 | Lighthouse Lab in Milton Keynes | Wellcome Sanger Institute for the COVID-19 Genomics UK (COG-UK) Consortium | The Lighthouse Lab in Milton Keynes and Alex Alderton, Roberto Amato, Jeffrey Barrett, Sonia Goncalves, Ewan Harrison, David K. Jackson, Ian Johnston, Dominic Kwiatkowski, Cordelia Langford, John Sillitoe on behalf of the Wellcome Sanger Institute COVID-19 Surveillance Team |
| EPI_ISL_3107397 | Lighthouse Lab in Alderley Park | Wellcome Sanger Institute for the COVID-19 Genomics UK (COG-UK) Consortium | Jacquelyn Wynn, Mairead Hyland, The Lighthouse Lab in Alderley Park and Alex Alderton, Roberto Amato, Jeffrey Barrett, Sonia Goncalves, Ewan Harrison, David K. Jackson, Ian Johnston, Dominic Kwiatkowski, Cordelia Langford, John Sillitoe on behalf of the Wellcome Sanger Institute COVID-19 Surveillance Team |
| EPI_ISL_3107398 | Lighthouse Lab in Milton Keynes | Wellcome Sanger Institute for the COVID-19 Genomics UK (COG-UK) Consortium | The Lighthouse Lab in Milton Keynes and Alex Alderton, Roberto Amato, Jeffrey Barrett, Sonia Goncalves, Ewan Harrison, David K. Jackson, Ian Johnston, Dominic Kwiatkowski, Cordelia Langford, John Sillitoe on behalf of the Wellcome Sanger Institute COVID-19 Surveillance Team |
| EPI_ISL_3107399, EPI_ISL_3107400, EPI_ISL_3107401 | Lighthouse Lab in Alderley Park | Wellcome Sanger Institute for the COVID-19 Genomics UK (COG-UK) Consortium | Jacquelyn Wynn, Mairead Hyland, The Lighthouse Lab in Alderley Park and Alex Alderton, Roberto Amato, Jeffrey Barrett, Sonia Goncalves, Ewan Harrison, David K. Jackson, Ian Johnston, Dominic Kwiatkowski, Cordelia Langford, John Sillitoe on behalf of the Wellcome Sanger Institute COVID-19 Surveillance Team |
| EPI_ISL_3107402 | Lighthouse Lab in Milton Keynes | Wellcome Sanger Institute for the COVID-19 Genomics UK (COG-UK) Consortium | The Lighthouse Lab in Milton Keynes and Alex Alderton, Roberto Amato, Jeffrey Barrett, Sonia Goncalves, Ewan Harrison, David K. Jackson, Ian Johnston, Dominic Kwiatkowski, Cordelia Langford, John Sillitoe on behalf of the Wellcome Sanger Institute COVID-19 Surveillance Team |
| EPI_ISL_3107403, EPI_ISL_3107404, EPI_ISL_3107405, EPI_ISL_3107406, EPI_ISL_3107407, EPI_ISL_3107408, EPI_ISL_3107409, EPI_ISL_3107410, EPI_ISL_3107411, EPI_ISL_3107412, EPI_ISL_3107413, EPI_ISL_3107414, EPI_ISL_3107415, EPI_ISL_3107416 | | | |
| see above | Lighthouse Lab in Alderley Park | Wellcome Sanger Institute for the COVID-19 Genomics UK (COG-UK) Consortium | Jacquelyn Wynn, Mairead Hyland, The Lighthouse Lab in Alderley Park and Alex Alderton, Roberto Amato, Jeffrey Barrett, Sonia Goncalves, Ewan Harrison, David K. Jackson, Ian Johnston, Dominic Kwiatkowski, Cordelia Langford, John Sillitoe on behalf of the Wellcome Sanger Institute COVID-19 Surveillance Team |
| EPI_ISL_3107417 | Lighthouse Lab in Milton Keynes | Wellcome Sanger Institute for the COVID-19 Genomics UK (COG-UK) Consortium | The Lighthouse Lab in Milton Keynes and Alex Alderton, Roberto Amato, Jeffrey Barrett, Sonia Goncalves, Ewan Harrison, David K. Jackson, Ian Johnston, Dominic Kwiatkowski, Cordelia Langford, John Sillitoe on behalf of the Wellcome Sanger Institute COVID-19 Surveillance Team |
| EPI_ISL_3107418 | Lighthouse Lab in Alderley Park | Wellcome Sanger Institute for the COVID-19 Genomics UK (COG-UK) Consortium | Jacquelyn Wynn, Mairead Hyland, The Lighthouse Lab in Alderley Park and Alex Alderton, Roberto Amato, Jeffrey Barrett, Sonia Goncalves, Ewan Harrison, David K. Jackson, Ian Johnston, Dominic Kwiatkowski, Cordelia Langford, John Sillitoe on behalf of the Wellcome Sanger Institute COVID-19 Surveillance Team |
| EPI_ISL_3107419 | Lighthouse Lab in Milton Keynes | Wellcome Sanger Institute for the COVID-19 Genomics UK (COG-UK) Consortium | The Lighthouse Lab in Milton Keynes and Alex Alderton, Roberto Amato, Jeffrey Barrett, Sonia Goncalves, Ewan Harrison, David K. Jackson, Ian Johnston, Dominic Kwiatkowski, Cordelia Langford, John Sillitoe on behalf of the Wellcome Sanger Institute COVID-19 Surveillance Team |
| EPI_ISL_3107420, EPI_ISL_3107421, EPI_ISL_3107422 | Lighthouse Lab in Alderley Park | Wellcome Sanger Institute for the COVID-19 Genomics UK (COG-UK) Consortium | Jacquelyn Wynn, Mairead Hyland, The Lighthouse Lab in Alderley Park and Alex Alderton, Roberto Amato, Jeffrey Barrett, Sonia Goncalves, Ewan Harrison, David K. Jackson, Ian Johnston, Dominic Kwiatkowski, Cordelia Langford, John Sillitoe on behalf of the Wellcome Sanger Institute COVID-19 Surveillance Team |
| EPI_ISL_3107423 | Lighthouse Lab in Milton Keynes | Wellcome Sanger Institute for the COVID-19 Genomics UK (COG-UK) Consortium | The Lighthouse Lab in Milton Keynes and Alex Alderton, Roberto Amato, Jeffrey Barrett, Sonia Goncalves, Ewan Harrison, David K. Jackson, Ian Johnston, Dominic Kwiatkowski, Cordelia Langford, John Sillitoe on behalf of the Wellcome Sanger Institute COVID-19 Surveillance Team |
| EPI_ISL_3107424, EPI_ISL_3107425, EPI_ISL_3107426 | Lighthouse Lab in Alderley Park | Wellcome Sanger Institute for the COVID-19 Genomics UK (COG-UK) Consortium | Jacquelyn Wynn, Mairead Hyland, The Lighthouse Lab in Alderley Park and Alex Alderton, Roberto Amato, Jeffrey Barrett, Sonia Goncalves, Ewan Harrison, David K. Jackson, Ian Johnston, Dominic Kwiatkowski, Cordelia Langford, John Sillitoe on behalf of the Wellcome Sanger Institute COVID-19 Surveillance Team |
| EPI_ISL_3107427 | Lighthouse Lab in Milton Keynes | Wellcome Sanger Institute for the COVID-19 Genomics UK (COG-UK) Consortium | The Lighthouse Lab in Milton Keynes and Alex Alderton, Roberto Amato, Jeffrey Barrett, Sonia Goncalves, Ewan Harrison, David K. Jackson, Ian Johnston, Dominic Kwiatkowski, Cordelia Langford, John Sillitoe on behalf of the Wellcome Sanger Institute COVID-19 Surveillance Team |
| EPI_ISL_3107428, EPI_ISL_3107429, EPI_ISL_3107430, EPI_ISL_3107431, EPI_ISL_3107432 | Lighthouse Lab in Alderley Park | Wellcome Sanger Institute for the COVID-19 Genomics UK (COG-UK) Consortium | Jacquelyn Wynn, Mairead Hyland, The Lighthouse Lab in Alderley Park and Alex Alderton, Roberto Amato, Jeffrey Barrett, Sonia Goncalves, Ewan Harrison, David K. Jackson, Ian Johnston, Dominic Kwiatkowski, Cordelia Langford, John Sillitoe on behalf of the Wellcome Sanger Institute COVID-19 Surveillance Team |
| EPI_ISL_3107433 | Lighthouse Lab in Milton Keynes | Wellcome Sanger Institute for the COVID-19 Genomics UK (COG-UK) Consortium | The Lighthouse Lab in Milton Keynes and Alex Alderton, Roberto Amato, Jeffrey Barrett, Sonia Goncalves, Ewan Harrison, David K. Jackson, Ian Johnston, Dominic Kwiatkowski, Cordelia Langford, John Sillitoe on behalf of the Wellcome Sanger Institute COVID-19 Surveillance Team |
| EPI_ISL_3107434, EPI_ISL_3107435, EPI_ISL_3107436 | Lighthouse Lab in Alderley Park | Wellcome Sanger Institute for the COVID-19 Genomics UK (COG-UK) Consortium | Jacquelyn Wynn, Mairead Hyland, The Lighthouse Lab in Alderley Park and Alex Alderton, Roberto Amato, Jeffrey Barrett, Sonia Goncalves, Ewan Harrison, David K. Jackson, Ian Johnston, Dominic Kwiatkowski, Cordelia Langford, John Sillitoe on behalf of the Wellcome Sanger Institute COVID-19 Surveillance Team |
| EPI_ISL_3107437 | Lighthouse Lab in Milton Keynes | Wellcome Sanger Institute for the COVID-19 Genomics UK (COG-UK) Consortium | The Lighthouse Lab in Milton Keynes and Alex Alderton, Roberto Amato, Jeffrey Barrett, Sonia Goncalves, Ewan Harrison, David K. Jackson, Ian Johnston, Dominic Kwiatkowski, Cordelia Langford, John Sillitoe on behalf of the Wellcome Sanger Institute COVID-19 Surveillance Team |
| EPI_ISL_3107438, EPI_ISL_3107439 | Lighthouse Lab in Alderley Park | Wellcome Sanger Institute for the COVID-19 Genomics UK (COG-UK) Consortium | Jacquelyn Wynn, Mairead Hyland, The Lighthouse Lab in Alderley Park and Alex Alderton, Roberto Amato, Jeffrey Barrett, Sonia Goncalves, Ewan Harrison, David K. Jackson, Ian Johnston, Dominic Kwiatkowski, Cordelia Langford, John Sillitoe on behalf of the Wellcome Sanger Institute COVID-19 Surveillance Team |

| | | | |
|---|---|---|---|
| EPI_ISL_3107440, EPI_ISL_3107441 | Lighthouse Lab in Milton Keynes | Wellcome Sanger Institute for the COVID-19 Genomics UK (COG-UK) Consortium | The Lighthouse Lab in Milton Keynes and Alex Alderton, Roberto Amato, Jeffrey Barrett, Sonia Goncalves, Ewan Harrison, David K. Jackson, Ian Johnston, Dominic Kwiatkowski, Cordelia Langford, John Sillitoe on behalf of the Wellcome Sanger Institute COVID-19 Surveillance Team |
| EPI_ISL_3107442, EPI_ISL_3107443, EPI_ISL_3107444, EPI_ISL_3107445, EPI_ISL_3107446 | Lighthouse Lab in Alderley Park | Wellcome Sanger Institute for the COVID-19 Genomics UK (COG-UK) Consortium | Jacquelyn Wynn, Mairead Hyland, The Lighthouse Lab in Alderley Park and Alex Alderton, Roberto Amato, Jeffrey Barrett, Sonia Goncalves, Ewan Harrison, David K. Jackson, Ian Johnston, Dominic Kwiatkowski, Cordelia Langford, John Sillitoe on behalf of the Wellcome Sanger Institute COVID-19 Surveillance Team |
| EPI_ISL_3107447 | Lighthouse Lab in Milton Keynes | Wellcome Sanger Institute for the COVID-19 Genomics UK (COG-UK) Consortium | The Lighthouse Lab in Milton Keynes and Alex Alderton, Roberto Amato, Jeffrey Barrett, Sonia Goncalves, Ewan Harrison, David K. Jackson, Ian Johnston, Dominic Kwiatkowski, Cordelia Langford, John Sillitoe on behalf of the Wellcome Sanger Institute COVID-19 Surveillance Team |
| EPI_ISL_3107448 | Lighthouse Lab in Alderley Park | Wellcome Sanger Institute for the COVID-19 Genomics UK (COG-UK) Consortium | Jacquelyn Wynn, Mairead Hyland, The Lighthouse Lab in Alderley Park and Alex Alderton, Roberto Amato, Jeffrey Barrett, Sonia Goncalves, Ewan Harrison, David K. Jackson, Ian Johnston, Dominic Kwiatkowski, Cordelia Langford, John Sillitoe on behalf of the Wellcome Sanger Institute COVID-19 Surveillance Team |
| EPI_ISL_3107449, EPI_ISL_3107450 | Lighthouse Lab in Milton Keynes | Wellcome Sanger Institute for the COVID-19 Genomics UK (COG-UK) Consortium | The Lighthouse Lab in Milton Keynes and Alex Alderton, Roberto Amato, Jeffrey Barrett, Sonia Goncalves, Ewan Harrison, David K. Jackson, Ian Johnston, Dominic Kwiatkowski, Cordelia Langford, John Sillitoe on behalf of the Wellcome Sanger Institute COVID-19 Surveillance Team |
| EPI_ISL_3107451, EPI_ISL_3107452, EPI_ISL_3107453, EPI_ISL_3107454 | Lighthouse Lab in Alderley Park | Wellcome Sanger Institute for the COVID-19 Genomics UK (COG-UK) Consortium | Jacquelyn Wynn, Mairead Hyland, The Lighthouse Lab in Alderley Park and Alex Alderton, Roberto Amato, Jeffrey Barrett, Sonia Goncalves, Ewan Harrison, David K. Jackson, Ian Johnston, Dominic Kwiatkowski, Cordelia Langford, John Sillitoe on behalf of the Wellcome Sanger Institute COVID-19 Surveillance Team |
| EPI_ISL_3107455 | Lighthouse Lab in Milton Keynes | Wellcome Sanger Institute for the COVID-19 Genomics UK (COG-UK) Consortium | The Lighthouse Lab in Milton Keynes and Alex Alderton, Roberto Amato, Jeffrey Barrett, Sonia Goncalves, Ewan Harrison, David K. Jackson, Ian Johnston, Dominic Kwiatkowski, Cordelia Langford, John Sillitoe on behalf of the Wellcome Sanger Institute COVID-19 Surveillance Team |
| EPI_ISL_3107456, EPI_ISL_3107457, EPI_ISL_3107458, EPI_ISL_3107459, EPI_ISL_3107460 | Lighthouse Lab in Alderley Park | Wellcome Sanger Institute for the COVID-19 Genomics UK (COG-UK) Consortium | Jacquelyn Wynn, Mairead Hyland, The Lighthouse Lab in Alderley Park and Alex Alderton, Roberto Amato, Jeffrey Barrett, Sonia Goncalves, Ewan Harrison, David K. Jackson, Ian Johnston, Dominic Kwiatkowski, Cordelia Langford, John Sillitoe on behalf of the Wellcome Sanger Institute COVID-19 Surveillance Team |
| EPI_ISL_3107461 | Lighthouse Lab in Milton Keynes | Wellcome Sanger Institute for the COVID-19 Genomics UK (COG-UK) Consortium | The Lighthouse Lab in Milton Keynes and Alex Alderton, Roberto Amato, Jeffrey Barrett, Sonia Goncalves, Ewan Harrison, David K. Jackson, Ian Johnston, Dominic Kwiatkowski, Cordelia Langford, John Sillitoe on behalf of the Wellcome Sanger Institute COVID-19 Surveillance Team |
| EPI_ISL_3107462, EPI_ISL_3107463, EPI_ISL_3107464 | Lighthouse Lab in Alderley Park | Wellcome Sanger Institute for the COVID-19 Genomics UK (COG-UK) Consortium | Jacquelyn Wynn, Mairead Hyland, The Lighthouse Lab in Alderley Park and Alex Alderton, Roberto Amato, Jeffrey Barrett, Sonia Goncalves, Ewan Harrison, David K. Jackson, Ian Johnston, Dominic Kwiatkowski, Cordelia Langford, John Sillitoe on behalf of the Wellcome Sanger Institute COVID-19 Surveillance Team |
| EPI_ISL_3107465, EPI_ISL_3107466 | Lighthouse Lab in Milton Keynes | Wellcome Sanger Institute for the COVID-19 Genomics UK (COG-UK) Consortium | The Lighthouse Lab in Milton Keynes and Alex Alderton, Roberto Amato, Jeffrey Barrett, Sonia Goncalves, Ewan Harrison, David K. Jackson, Ian Johnston, Dominic Kwiatkowski, Cordelia Langford, John Sillitoe on behalf of the Wellcome Sanger Institute COVID-19 Surveillance Team |
| EPI_ISL_3107467, EPI_ISL_3107468, EPI_ISL_3107469 | Lighthouse Lab in Alderley Park | Wellcome Sanger Institute for the COVID-19 Genomics UK (COG-UK) Consortium | Jacquelyn Wynn, Mairead Hyland, The Lighthouse Lab in Alderley Park and Alex Alderton, Roberto Amato, Jeffrey Barrett, Sonia Goncalves, Ewan Harrison, David K. Jackson, Ian Johnston, Dominic Kwiatkowski, Cordelia Langford, John Sillitoe on behalf of the Wellcome Sanger Institute COVID-19 Surveillance Team |
| EPI_ISL_3107470, EPI_ISL_3107471 | Lighthouse Lab in Milton Keynes | Wellcome Sanger Institute for the COVID-19 Genomics UK (COG-UK) Consortium | The Lighthouse Lab in Milton Keynes and Alex Alderton, Roberto Amato, Jeffrey Barrett, Sonia Goncalves, Ewan Harrison, David K. Jackson, Ian Johnston, Dominic Kwiatkowski, Cordelia Langford, John Sillitoe on behalf of the Wellcome Sanger Institute COVID-19 Surveillance Team |
| EPI_ISL_3107472 | Lighthouse Lab in Alderley Park | Wellcome Sanger Institute for the COVID-19 Genomics UK (COG-UK) Consortium | Jacquelyn Wynn, Mairead Hyland, The Lighthouse Lab in Alderley Park and Alex Alderton, Roberto Amato, Jeffrey Barrett, Sonia Goncalves, Ewan Harrison, David K. Jackson, Ian Johnston, Dominic Kwiatkowski, Cordelia Langford, John Sillitoe on behalf of the Wellcome Sanger Institute COVID-19 Surveillance Team |
| EPI_ISL_3107473, EPI_ISL_3107474, EPI_ISL_3107475 | Lighthouse Lab in Milton Keynes | Wellcome Sanger Institute for the COVID-19 Genomics UK (COG-UK) Consortium | The Lighthouse Lab in Milton Keynes and Alex Alderton, Roberto Amato, Jeffrey Barrett, Sonia Goncalves, Ewan Harrison, David K. Jackson, Ian Johnston, Dominic Kwiatkowski, Cordelia Langford, John Sillitoe on behalf of the Wellcome Sanger Institute COVID-19 Surveillance Team |
| EPI_ISL_3107476, EPI_ISL_3107477, EPI_ISL_3107478, EPI_ISL_3107479, EPI_ISL_3107480, EPI_ISL_3107481, EPI_ISL_3107482, EPI_ISL_3107483, EPI_ISL_3107484, EPI_ISL_3107485 | Lighthouse Lab in Alderley Park | Wellcome Sanger Institute for the COVID-19 Genomics UK (COG-UK) Consortium | Jacquelyn Wynn, Mairead Hyland, The Lighthouse Lab in Alderley Park and Alex Alderton, Roberto Amato, Jeffrey Barrett, Sonia Goncalves, Ewan Harrison, David K. Jackson, Ian Johnston, Dominic Kwiatkowski, Cordelia Langford, John Sillitoe on behalf of the Wellcome Sanger Institute COVID-19 Surveillance Team |
| EPI_ISL_3107486 | Lighthouse Lab in Milton Keynes | Wellcome Sanger Institute for the COVID-19 Genomics UK (COG-UK) Consortium | The Lighthouse Lab in Milton Keynes and Alex Alderton, Roberto Amato, Jeffrey Barrett, Sonia Goncalves, Ewan Harrison, David K. Jackson, Ian Johnston, Dominic Kwiatkowski, Cordelia Langford, John Sillitoe on behalf of the Wellcome Sanger Institute COVID-19 Surveillance Team |
| EPI_ISL_3107487, EPI_ISL_3107488, EPI_ISL_3107489, EPI_ISL_3107490, EPI_ISL_3107491, EPI_ISL_3107492, EPI_ISL_3107493, EPI_ISL_3107494 | Lighthouse Lab in Alderley Park | Wellcome Sanger Institute for the COVID-19 Genomics UK (COG-UK) Consortium | Jacquelyn Wynn, Mairead Hyland, The Lighthouse Lab in Alderley Park and Alex Alderton, Roberto Amato, Jeffrey Barrett, Sonia Goncalves, Ewan Harrison, David K. Jackson, Ian Johnston, Dominic Kwiatkowski, Cordelia Langford, John Sillitoe on behalf of the Wellcome Sanger Institute COVID-19 Surveillance Team |
| EPI_ISL_3107495 | Lighthouse Lab in Milton Keynes | Wellcome Sanger Institute for the COVID-19 Genomics UK (COG-UK) Consortium | The Lighthouse Lab in Milton Keynes and Alex Alderton, Roberto Amato, Jeffrey Barrett, Sonia Goncalves, Ewan Harrison, David K. Jackson, Ian Johnston, Dominic Kwiatkowski, Cordelia Langford, John Sillitoe on behalf of the Wellcome Sanger Institute COVID-19 Surveillance Team |
| EPI_ISL_3107496, EPI_ISL_3107497, EPI_ISL_3107498 | Lighthouse Lab in Alderley Park | Wellcome Sanger Institute for the COVID-19 Genomics UK (COG-UK) Consortium | Jacquelyn Wynn, Mairead Hyland, The Lighthouse Lab in Alderley Park and Alex Alderton, Roberto Amato, Jeffrey Barrett, Sonia Goncalves, Ewan Harrison, David K. Jackson, Ian Johnston, Dominic Kwiatkowski, Cordelia Langford, John Sillitoe on behalf of the Wellcome Sanger Institute COVID-19 Surveillance Team |
| EPI_ISL_3107499 | Lighthouse Lab in Milton Keynes | Wellcome Sanger Institute for the COVID-19 Genomics UK (COG-UK) Consortium | The Lighthouse Lab in Milton Keynes and Alex Alderton, Roberto Amato, Jeffrey Barrett, Sonia Goncalves, Ewan Harrison, David K. Jackson, Ian Johnston, Dominic Kwiatkowski, Cordelia Langford, John Sillitoe on behalf of the Wellcome Sanger Institute COVID-19 Surveillance Team |
| EPI_ISL_3107500, EPI_ISL_3107501 | Lighthouse Lab in Alderley Park | Wellcome Sanger Institute for the COVID-19 Genomics UK (COG-UK) Consortium | Jacquelyn Wynn, Mairead Hyland, The Lighthouse Lab in Alderley Park and Alex Alderton, Roberto Amato, Jeffrey Barrett, Sonia Goncalves, Ewan Harrison, David K. Jackson, Ian Johnston, Dominic Kwiatkowski, Cordelia Langford, John Sillitoe on behalf of the Wellcome Sanger Institute COVID-19 Surveillance Team |
| EPI_ISL_3107502 | Lighthouse Lab in Milton Keynes | Wellcome Sanger Institute for the COVID-19 Genomics UK (COG-UK) Consortium | The Lighthouse Lab in Milton Keynes and Alex Alderton, Roberto Amato, Jeffrey Barrett, Sonia Goncalves, Ewan Harrison, David K. Jackson, Ian Johnston, Dominic Kwiatkowski, Cordelia Langford, John Sillitoe on behalf of the Wellcome Sanger Institute COVID-19 Surveillance Team |
| EPI_ISL_3107503 | Lighthouse Lab in Alderley Park | Wellcome Sanger Institute for the COVID-19 Genomics UK (COG-UK) Consortium | Jacquelyn Wynn, Mairead Hyland, The Lighthouse Lab in Alderley Park and Alex Alderton, Roberto Amato, Jeffrey Barrett, Sonia Goncalves, Ewan Harrison, David K. Jackson, Ian Johnston, Dominic Kwiatkowski, Cordelia Langford, John Sillitoe on behalf of the Wellcome Sanger Institute COVID-19 Surveillance Team |
| EPI_ISL_3107504 | Lighthouse Lab in Milton Keynes | Wellcome Sanger Institute for the COVID-19 Genomics UK (COG-UK) Consortium | The Lighthouse Lab in Milton Keynes and Alex Alderton, Roberto Amato, Jeffrey Barrett, Sonia Goncalves, Ewan Harrison, David K. Jackson, Ian Johnston, Dominic Kwiatkowski, Cordelia Langford, John Sillitoe on behalf of the Wellcome Sanger Institute COVID-19 Surveillance Team |
| EPI_ISL_3107505, EPI_ISL_3107506, EPI_ISL_3107507 | Lighthouse Lab in Alderley Park | Wellcome Sanger Institute for the COVID-19 Genomics UK (COG-UK) Consortium | Jacquelyn Wynn, Mairead Hyland, The Lighthouse Lab in Alderley Park and Alex Alderton, Roberto Amato, Jeffrey Barrett, Sonia Goncalves, Ewan Harrison, David K. Jackson, Ian Johnston, Dominic Kwiatkowski, Cordelia Langford, John Sillitoe on behalf of the Wellcome Sanger Institute COVID-19 Surveillance Team |
| EPI_ISL_3107508, EPI_ISL_3107509 | Lighthouse Lab in Milton Keynes | Wellcome Sanger Institute for the COVID-19 Genomics UK (COG-UK) Consortium | The Lighthouse Lab in Milton Keynes and Alex Alderton, Roberto Amato, Jeffrey Barrett, Sonia Goncalves, Ewan Harrison, David K. Jackson, Ian Johnston, Dominic Kwiatkowski, Cordelia Langford, John Sillitoe on behalf of the Wellcome Sanger Institute COVID-19 Surveillance Team |
| EPI_ISL_3107510, EPI_ISL_3107511 | Lighthouse Lab in Alderley Park | Wellcome Sanger Institute for the COVID-19 Genomics UK (COG-UK) Consortium | Jacquelyn Wynn, Mairead Hyland, The Lighthouse Lab in Alderley Park and Alex Alderton, Roberto Amato, Jeffrey Barrett, Sonia Goncalves, Ewan Harrison, David K. Jackson, Ian Johnston, Dominic Kwiatkowski, Cordelia Langford, John Sillitoe on behalf of the Wellcome Sanger Institute COVID-19 Surveillance Team |
| EPI_ISL_3107512, EPI_ISL_3107513 | Lighthouse Lab in Milton Keynes | Wellcome Sanger Institute for the COVID-19 Genomics UK (COG-UK) Consortium | The Lighthouse Lab in Milton Keynes and Alex Alderton, Roberto Amato, Jeffrey Barrett, Sonia Goncalves, Ewan Harrison, David K. Jackson, Ian Johnston, Dominic Kwiatkowski, Cordelia Langford, John Sillitoe on behalf of the Wellcome Sanger Institute COVID-19 Surveillance Team |
| EPI_ISL_3107514, EPI_ISL_3107515, EPI_ISL_3107516, EPI_ISL_3107517, EPI_ISL_3107518, EPI_ISL_3107519 | Lighthouse Lab in Alderley Park | Wellcome Sanger Institute for the COVID-19 Genomics UK (COG-UK) Consortium | Jacquelyn Wynn, Mairead Hyland, The Lighthouse Lab in Alderley Park and Alex Alderton, Roberto Amato, Jeffrey Barrett, Sonia Goncalves, Ewan Harrison, David K. Jackson, Ian Johnston, Dominic Kwiatkowski, Cordelia Langford, John Sillitoe on behalf of the Wellcome Sanger Institute COVID-19 Surveillance Team |
| EPI_ISL_3107520, EPI_ISL_3107521 | Lighthouse Lab in Milton Keynes | Wellcome Sanger Institute for the COVID-19 Genomics UK (COG-UK) Consortium | The Lighthouse Lab in Milton Keynes and Alex Alderton, Roberto Amato, Jeffrey Barrett, Sonia Goncalves, Ewan Harrison, David K. Jackson, Ian Johnston, Dominic Kwiatkowski, Cordelia Langford, John Sillitoe on behalf of the Wellcome Sanger Institute COVID-19 Surveillance Team |
| EPI_ISL_3107522, EPI_ISL_3107523 | Lighthouse Lab in Alderley Park | Wellcome Sanger Institute for the COVID-19 Genomics UK (COG-UK) Consortium | Jacquelyn Wynn, Mairead Hyland, The Lighthouse Lab in Alderley Park and Alex Alderton, Roberto Amato, Jeffrey Barrett, Sonia Goncalves, Ewan Harrison, David K. Jackson, Ian Johnston, Dominic Kwiatkowski, Cordelia Langford, John Sillitoe on behalf of the Wellcome Sanger Institute COVID-19 Surveillance Team |
| EPI_ISL_3107524 | Lighthouse Lab in Milton Keynes | Wellcome Sanger Institute for the COVID-19 Genomics UK (COG-UK) | The Lighthouse Lab in Milton Keynes and Alex Alderton, Roberto Amato, Jeffrey Barrett, Sonia Goncalves, Ewan Harrison, David K. Jackson, Ian Johnston, Dominic Kwiatkowski, Cordelia Langford, John Sillitoe on behalf of the Wellcome Sanger |

| | | | |
|---|---|---|---|
| | | Consortium | Institute COVID-19 Surveillance Team |
| EPI_ISL_3107525, EPI_ISL_3107526, EPI_ISL_3107527, EPI_ISL_3107528 | Lighthouse Lab in Alderley Park | Wellcome Sanger Institute for the COVID-19 Genomics UK (COG-UK) Consortium | Jacquelyn Wynn, Mairead Hyland, The Lighthouse Lab in Alderley Park and Alex Alderton, Roberto Amato, Jeffrey Barrett, Sonia Goncalves, Ewan Harrison, David K. Jackson, Ian Johnston, Dominic Kwiatkowski, Cordelia Langford, John Sillitoe on behalf of the Wellcome Sanger Institute COVID-19 Surveillance Team |
| EPI_ISL_3107529, EPI_ISL_3107530 | Lighthouse Lab in Milton Keynes | Wellcome Sanger Institute for the COVID-19 Genomics UK (COG-UK) Consortium | The Lighthouse Lab in Milton Keynes and Alex Alderton, Roberto Amato, Jeffrey Barrett, Sonia Goncalves, Ewan Harrison, David K. Jackson, Ian Johnston, Dominic Kwiatkowski, Cordelia Langford, John Sillitoe on behalf of the Wellcome Sanger Institute COVID-19 Surveillance Team |
| EPI_ISL_3107531, EPI_ISL_3107532, EPI_ISL_3107533 | Lighthouse Lab in Alderley Park | Wellcome Sanger Institute for the COVID-19 Genomics UK (COG-UK) Consortium | Jacquelyn Wynn, Mairead Hyland, The Lighthouse Lab in Alderley Park and Alex Alderton, Roberto Amato, Jeffrey Barrett, Sonia Goncalves, Ewan Harrison, David K. Jackson, Ian Johnston, Dominic Kwiatkowski, Cordelia Langford, John Sillitoe on behalf of the Wellcome Sanger Institute COVID-19 Surveillance Team |
| EPI_ISL_3107534 | Lighthouse Lab in Milton Keynes | Wellcome Sanger Institute for the COVID-19 Genomics UK (COG-UK) Consortium | The Lighthouse Lab in Milton Keynes and Alex Alderton, Roberto Amato, Jeffrey Barrett, Sonia Goncalves, Ewan Harrison, David K. Jackson, Ian Johnston, Dominic Kwiatkowski, Cordelia Langford, John Sillitoe on behalf of the Wellcome Sanger Institute COVID-19 Surveillance Team |
| EPI_ISL_3107535, EPI_ISL_3107536, EPI_ISL_3107537 | Lighthouse Lab in Alderley Park | Wellcome Sanger Institute for the COVID-19 Genomics UK (COG-UK) Consortium | Jacquelyn Wynn, Mairead Hyland, The Lighthouse Lab in Alderley Park and Alex Alderton, Roberto Amato, Jeffrey Barrett, Sonia Goncalves, Ewan Harrison, David K. Jackson, Ian Johnston, Dominic Kwiatkowski, Cordelia Langford, John Sillitoe on behalf of the Wellcome Sanger Institute COVID-19 Surveillance Team |
| EPI_ISL_3107538 | Lighthouse Lab in Milton Keynes | Wellcome Sanger Institute for the COVID-19 Genomics UK (COG-UK) Consortium | The Lighthouse Lab in Milton Keynes and Alex Alderton, Roberto Amato, Jeffrey Barrett, Sonia Goncalves, Ewan Harrison, David K. Jackson, Ian Johnston, Dominic Kwiatkowski, Cordelia Langford, John Sillitoe on behalf of the Wellcome Sanger Institute COVID-19 Surveillance Team |
| EPI_ISL_3107539, EPI_ISL_3107540, EPI_ISL_3107541, EPI_ISL_3107542, EPI_ISL_3107543, EPI_ISL_3107544 | Lighthouse Lab in Alderley Park | Wellcome Sanger Institute for the COVID-19 Genomics UK (COG-UK) Consortium | Jacquelyn Wynn, Mairead Hyland, The Lighthouse Lab in Alderley Park and Alex Alderton, Roberto Amato, Jeffrey Barrett, Sonia Goncalves, Ewan Harrison, David K. Jackson, Ian Johnston, Dominic Kwiatkowski, Cordelia Langford, John Sillitoe on behalf of the Wellcome Sanger Institute COVID-19 Surveillance Team |
| EPI_ISL_3107545 | Lighthouse Lab in Milton Keynes | Wellcome Sanger Institute for the COVID-19 Genomics UK (COG-UK) Consortium | The Lighthouse Lab in Milton Keynes and Alex Alderton, Roberto Amato, Jeffrey Barrett, Sonia Goncalves, Ewan Harrison, David K. Jackson, Ian Johnston, Dominic Kwiatkowski, Cordelia Langford, John Sillitoe on behalf of the Wellcome Sanger Institute COVID-19 Surveillance Team |
| EPI_ISL_3107546, EPI_ISL_3107547, EPI_ISL_3107548, EPI_ISL_3107549, EPI_ISL_3107550, EPI_ISL_3107551, EPI_ISL_3107552, EPI_ISL_3107553 | Lighthouse Lab in Alderley Park | Wellcome Sanger Institute for the COVID-19 Genomics UK (COG-UK) Consortium | Jacquelyn Wynn, Mairead Hyland, The Lighthouse Lab in Alderley Park and Alex Alderton, Roberto Amato, Jeffrey Barrett, Sonia Goncalves, Ewan Harrison, David K. Jackson, Ian Johnston, Dominic Kwiatkowski, Cordelia Langford, John Sillitoe on behalf of the Wellcome Sanger Institute COVID-19 Surveillance Team |
| EPI_ISL_3107554 | Lighthouse Lab in Milton Keynes | Wellcome Sanger Institute for the COVID-19 Genomics UK (COG-UK) Consortium | The Lighthouse Lab in Milton Keynes and Alex Alderton, Roberto Amato, Jeffrey Barrett, Sonia Goncalves, Ewan Harrison, David K. Jackson, Ian Johnston, Dominic Kwiatkowski, Cordelia Langford, John Sillitoe on behalf of the Wellcome Sanger Institute COVID-19 Surveillance Team |
| EPI_ISL_3107555, EPI_ISL_3107556, EPI_ISL_3107557, EPI_ISL_3107558, EPI_ISL_3107559 | Lighthouse Lab in Alderley Park | Wellcome Sanger Institute for the COVID-19 Genomics UK (COG-UK) Consortium | Jacquelyn Wynn, Mairead Hyland, The Lighthouse Lab in Alderley Park and Alex Alderton, Roberto Amato, Jeffrey Barrett, Sonia Goncalves, Ewan Harrison, David K. Jackson, Ian Johnston, Dominic Kwiatkowski, Cordelia Langford, John Sillitoe on behalf of the Wellcome Sanger Institute COVID-19 Surveillance Team |
| EPI_ISL_3107560 | Lighthouse Lab in Milton Keynes | Wellcome Sanger Institute for the COVID-19 Genomics UK (COG-UK) Consortium | The Lighthouse Lab in Milton Keynes and Alex Alderton, Roberto Amato, Jeffrey Barrett, Sonia Goncalves, Ewan Harrison, David K. Jackson, Ian Johnston, Dominic Kwiatkowski, Cordelia Langford, John Sillitoe on behalf of the Wellcome Sanger Institute COVID-19 Surveillance Team |
| EPI_ISL_3107561 | Lighthouse Lab in Alderley Park | Wellcome Sanger Institute for the COVID-19 Genomics UK (COG-UK) Consortium | Jacquelyn Wynn, Mairead Hyland, The Lighthouse Lab in Alderley Park and Alex Alderton, Roberto Amato, Jeffrey Barrett, Sonia Goncalves, Ewan Harrison, David K. Jackson, Ian Johnston, Dominic Kwiatkowski, Cordelia Langford, John Sillitoe on behalf of the Wellcome Sanger Institute COVID-19 Surveillance Team |
| EPI_ISL_3107562 | Lighthouse Lab in Milton Keynes | Wellcome Sanger Institute for the COVID-19 Genomics UK (COG-UK) Consortium | The Lighthouse Lab in Milton Keynes and Alex Alderton, Roberto Amato, Jeffrey Barrett, Sonia Goncalves, Ewan Harrison, David K. Jackson, Ian Johnston, Dominic Kwiatkowski, Cordelia Langford, John Sillitoe on behalf of the Wellcome Sanger Institute COVID-19 Surveillance Team |
| EPI_ISL_3107563 | Lighthouse Lab in Alderley Park | Wellcome Sanger Institute for the COVID-19 Genomics UK (COG-UK) Consortium | Jacquelyn Wynn, Mairead Hyland, The Lighthouse Lab in Alderley Park and Alex Alderton, Roberto Amato, Jeffrey Barrett, Sonia Goncalves, Ewan Harrison, David K. Jackson, Ian Johnston, Dominic Kwiatkowski, Cordelia Langford, John Sillitoe on behalf of the Wellcome Sanger Institute COVID-19 Surveillance Team |
| EPI_ISL_3107564, EPI_ISL_3107565 | Lighthouse Lab in Milton Keynes | Wellcome Sanger Institute for the COVID-19 Genomics UK (COG-UK) Consortium | The Lighthouse Lab in Milton Keynes and Alex Alderton, Roberto Amato, Jeffrey Barrett, Sonia Goncalves, Ewan Harrison, David K. Jackson, Ian Johnston, Dominic Kwiatkowski, Cordelia Langford, John Sillitoe on behalf of the Wellcome Sanger Institute COVID-19 Surveillance Team |
| EPI_ISL_3107566, EPI_ISL_3107567, EPI_ISL_3107568, EPI_ISL_3107569 | Lighthouse Lab in Alderley Park | Wellcome Sanger Institute for the COVID-19 Genomics UK (COG-UK) Consortium | Jacquelyn Wynn, Mairead Hyland, The Lighthouse Lab in Alderley Park and Alex Alderton, Roberto Amato, Jeffrey Barrett, Sonia Goncalves, Ewan Harrison, David K. Jackson, Ian Johnston, Dominic Kwiatkowski, Cordelia Langford, John Sillitoe on behalf of the Wellcome Sanger Institute COVID-19 Surveillance Team |
| EPI_ISL_3107570 | Lighthouse Lab in Milton Keynes | Wellcome Sanger Institute for the COVID-19 Genomics UK (COG-UK) Consortium | The Lighthouse Lab in Milton Keynes and Alex Alderton, Roberto Amato, Jeffrey Barrett, Sonia Goncalves, Ewan Harrison, David K. Jackson, Ian Johnston, Dominic Kwiatkowski, Cordelia Langford, John Sillitoe on behalf of the Wellcome Sanger Institute COVID-19 Surveillance Team |
| EPI_ISL_3107571 | Lighthouse Lab in Alderley Park | Wellcome Sanger Institute for the COVID-19 Genomics UK (COG-UK) Consortium | Jacquelyn Wynn, Mairead Hyland, The Lighthouse Lab in Alderley Park and Alex Alderton, Roberto Amato, Jeffrey Barrett, Sonia Goncalves, Ewan Harrison, David K. Jackson, Ian Johnston, Dominic Kwiatkowski, Cordelia Langford, John Sillitoe on behalf of the Wellcome Sanger Institute COVID-19 Surveillance Team |
| EPI_ISL_3107572 | Lighthouse Lab in Milton Keynes | Wellcome Sanger Institute for the COVID-19 Genomics UK (COG-UK) Consortium | The Lighthouse Lab in Milton Keynes and Alex Alderton, Roberto Amato, Jeffrey Barrett, Sonia Goncalves, Ewan Harrison, David K. Jackson, Ian Johnston, Dominic Kwiatkowski, Cordelia Langford, John Sillitoe on behalf of the Wellcome Sanger Institute COVID-19 Surveillance Team |
| EPI_ISL_3107573, EPI_ISL_3107574 | Lighthouse Lab in Alderley Park | Wellcome Sanger Institute for the COVID-19 Genomics UK (COG-UK) Consortium | Jacquelyn Wynn, Mairead Hyland, The Lighthouse Lab in Alderley Park and Alex Alderton, Roberto Amato, Jeffrey Barrett, Sonia Goncalves, Ewan Harrison, David K. Jackson, Ian Johnston, Dominic Kwiatkowski, Cordelia Langford, John Sillitoe on behalf of the Wellcome Sanger Institute COVID-19 Surveillance Team |
| EPI_ISL_3107575, EPI_ISL_3107576 | Lighthouse Lab in Milton Keynes | Wellcome Sanger Institute for the COVID-19 Genomics UK (COG-UK) Consortium | The Lighthouse Lab in Milton Keynes and Alex Alderton, Roberto Amato, Jeffrey Barrett, Sonia Goncalves, Ewan Harrison, David K. Jackson, Ian Johnston, Dominic Kwiatkowski, Cordelia Langford, John Sillitoe on behalf of the Wellcome Sanger Institute COVID-19 Surveillance Team |
| EPI_ISL_3107577, EPI_ISL_3107578 | Lighthouse Lab in Alderley Park | Wellcome Sanger Institute for the COVID-19 Genomics UK (COG-UK) Consortium | Jacquelyn Wynn, Mairead Hyland, The Lighthouse Lab in Alderley Park and Alex Alderton, Roberto Amato, Jeffrey Barrett, Sonia Goncalves, Ewan Harrison, David K. Jackson, Ian Johnston, Dominic Kwiatkowski, Cordelia Langford, John Sillitoe on behalf of the Wellcome Sanger Institute COVID-19 Surveillance Team |
| EPI_ISL_3107579 | Lighthouse Lab in Milton Keynes | Wellcome Sanger Institute for the COVID-19 Genomics UK (COG-UK) Consortium | The Lighthouse Lab in Milton Keynes and Alex Alderton, Roberto Amato, Jeffrey Barrett, Sonia Goncalves, Ewan Harrison, David K. Jackson, Ian Johnston, Dominic Kwiatkowski, Cordelia Langford, John Sillitoe on behalf of the Wellcome Sanger Institute COVID-19 Surveillance Team |
| EPI_ISL_3107580 | Lighthouse Lab in Alderley Park | Wellcome Sanger Institute for the COVID-19 Genomics UK (COG-UK) Consortium | Jacquelyn Wynn, Mairead Hyland, The Lighthouse Lab in Alderley Park and Alex Alderton, Roberto Amato, Jeffrey Barrett, Sonia Goncalves, Ewan Harrison, David K. Jackson, Ian Johnston, Dominic Kwiatkowski, Cordelia Langford, John Sillitoe on behalf of the Wellcome Sanger Institute COVID-19 Surveillance Team |
| EPI_ISL_3107581, EPI_ISL_3107582 | Lighthouse Lab in Milton Keynes | Wellcome Sanger Institute for the COVID-19 Genomics UK (COG-UK) Consortium | The Lighthouse Lab in Milton Keynes and Alex Alderton, Roberto Amato, Jeffrey Barrett, Sonia Goncalves, Ewan Harrison, David K. Jackson, Ian Johnston, Dominic Kwiatkowski, Cordelia Langford, John Sillitoe on behalf of the Wellcome Sanger Institute COVID-19 Surveillance Team |
| EPI_ISL_3107583, EPI_ISL_3107584 | Lighthouse Lab in Alderley Park | Wellcome Sanger Institute for the COVID-19 Genomics UK (COG-UK) Consortium | Jacquelyn Wynn, Mairead Hyland, The Lighthouse Lab in Alderley Park and Alex Alderton, Roberto Amato, Jeffrey Barrett, Sonia Goncalves, Ewan Harrison, David K. Jackson, Ian Johnston, Dominic Kwiatkowski, Cordelia Langford, John Sillitoe on behalf of the Wellcome Sanger Institute COVID-19 Surveillance Team |
| EPI_ISL_3107585 | Lighthouse Lab in Milton Keynes | Wellcome Sanger Institute for the COVID-19 Genomics UK (COG-UK) Consortium | The Lighthouse Lab in Milton Keynes and Alex Alderton, Roberto Amato, Jeffrey Barrett, Sonia Goncalves, Ewan Harrison, David K. Jackson, Ian Johnston, Dominic Kwiatkowski, Cordelia Langford, John Sillitoe on behalf of the Wellcome Sanger Institute COVID-19 Surveillance Team |
| EPI_ISL_3107586, EPI_ISL_3107587 | Lighthouse Lab in Alderley Park | Wellcome Sanger Institute for the COVID-19 Genomics UK (COG-UK) Consortium | Jacquelyn Wynn, Mairead Hyland, The Lighthouse Lab in Alderley Park and Alex Alderton, Roberto Amato, Jeffrey Barrett, Sonia Goncalves, Ewan Harrison, David K. Jackson, Ian Johnston, Dominic Kwiatkowski, Cordelia Langford, John Sillitoe on behalf of the Wellcome Sanger Institute COVID-19 Surveillance Team |
| EPI_ISL_3107588 | Lighthouse Lab in Milton Keynes | Wellcome Sanger Institute for the COVID-19 Genomics UK (COG-UK) Consortium | The Lighthouse Lab in Milton Keynes and Alex Alderton, Roberto Amato, Jeffrey Barrett, Sonia Goncalves, Ewan Harrison, David K. Jackson, Ian Johnston, Dominic Kwiatkowski, Cordelia Langford, John Sillitoe on behalf of the Wellcome Sanger Institute COVID-19 Surveillance Team |
| EPI_ISL_3107589, EPI_ISL_3107590, EPI_ISL_3107591, EPI_ISL_3107592, EPI_ISL_3107593, EPI_ISL_3107594, EPI_ISL_3107595, EPI_ISL_3107596, EPI_ISL_3107597 | Lighthouse Lab in Alderley Park | Wellcome Sanger Institute for the COVID-19 Genomics UK (COG-UK) Consortium | Jacquelyn Wynn, Mairead Hyland, The Lighthouse Lab in Alderley Park and Alex Alderton, Roberto Amato, Jeffrey Barrett, Sonia Goncalves, Ewan Harrison, David K. Jackson, Ian Johnston, Dominic Kwiatkowski, Cordelia Langford, John Sillitoe on behalf of the Wellcome Sanger Institute COVID-19 Surveillance Team |
| EPI_ISL_3107598 | Lighthouse Lab in Milton Keynes | Wellcome Sanger Institute for the COVID-19 Genomics UK (COG-UK) Consortium | The Lighthouse Lab in Milton Keynes and Alex Alderton, Roberto Amato, Jeffrey Barrett, Sonia Goncalves, Ewan Harrison, David K. Jackson, Ian Johnston, Dominic Kwiatkowski, Cordelia Langford, John Sillitoe on behalf of the Wellcome Sanger Institute COVID-19 Surveillance Team |
| EPI_ISL_3107599, EPI_ISL_3107600, EPI_ISL_3107601 | Lighthouse Lab in Alderley Park | Wellcome Sanger Institute for the COVID-19 Genomics UK (COG-UK) Consortium | Jacquelyn Wynn, Mairead Hyland, The Lighthouse Lab in Alderley Park and Alex Alderton, Roberto Amato, Jeffrey Barrett, Sonia Goncalves, Ewan Harrison, David K. Jackson, Ian Johnston, Dominic Kwiatkowski, Cordelia Langford, John Sillitoe on behalf of the Wellcome Sanger Institute COVID-19 Surveillance Team |
| EPI_ISL_3107602 | Lighthouse Lab in Milton Keynes | Wellcome Sanger Institute for the COVID-19 Genomics UK (COG-UK) | The Lighthouse Lab in Milton Keynes and Alex Alderton, Roberto Amato, Jeffrey Barrett, Sonia Goncalves, Ewan Harrison, David K. Jackson, Ian Johnston, Dominic Kwiatkowski, Cordelia Langford, John Sillitoe on behalf of the Wellcome Sanger |

| | | | |
|---|---|---|---|
| | | Consortium | Institute COVID-19 Surveillance Team |
| EPI_ISL_3107603, EPI_ISL_3107604, EPI_ISL_3107605, EPI_ISL_3107606, EPI_ISL_3107607, EPI_ISL_3107608, EPI_ISL_3107609, EPI_ISL_3107610 | Lighthouse Lab in Alderley Park | Wellcome Sanger Institute for the COVID-19 Genomics UK (COG-UK) Consortium | Jacquelyn Wynn, Mairead Hyland, The Lighthouse Lab in Alderley Park and Alex Alderton, Roberto Amato, Jeffrey Barrett, Sonia Goncalves, Ewan Harrison, David K. Jackson, Ian Johnston, Dominic Kwiatkowski, Cordelia Langford, John Sillitoe on behalf of the Wellcome Sanger Institute COVID-19 Surveillance Team |
| EPI_ISL_3107611, EPI_ISL_3107612 | Lighthouse Lab in Milton Keynes | Wellcome Sanger Institute for the COVID-19 Genomics UK (COG-UK) Consortium | The Lighthouse Lab in Milton Keynes and Alex Alderton, Roberto Amato, Jeffrey Barrett, Sonia Goncalves, Ewan Harrison, David K. Jackson, Ian Johnston, Dominic Kwiatkowski, Cordelia Langford, John Sillitoe on behalf of the Wellcome Sanger Institute COVID-19 Surveillance Team |
| EPI_ISL_3107613, EPI_ISL_3107614, EPI_ISL_3107615, EPI_ISL_3107616, EPI_ISL_3107617, EPI_ISL_3107618, EPI_ISL_3107619, EPI_ISL_3107620, EPI_ISL_3107621, EPI_ISL_3107622, EPI_ISL_3107623, EPI_ISL_3107624, EPI_ISL_3107625, EPI_ISL_3107626, EPI_ISL_3107627, EPI_ISL_3107628, EPI_ISL_3107629 | | | |
| see above | Lighthouse Lab in Alderley Park | Wellcome Sanger Institute for the COVID-19 Genomics UK (COG-UK) Consortium | Jacquelyn Wynn, Mairead Hyland, The Lighthouse Lab in Alderley Park and Alex Alderton, Roberto Amato, Jeffrey Barrett, Sonia Goncalves, Ewan Harrison, David K. Jackson, Ian Johnston, Dominic Kwiatkowski, Cordelia Langford, John Sillitoe on behalf of the Wellcome Sanger Institute COVID-19 Surveillance Team |
| EPI_ISL_3107630, EPI_ISL_3107631, EPI_ISL_3107632 | Lighthouse Lab in Milton Keynes | Wellcome Sanger Institute for the COVID-19 Genomics UK (COG-UK) Consortium | The Lighthouse Lab in Milton Keynes and Alex Alderton, Roberto Amato, Jeffrey Barrett, Sonia Goncalves, Ewan Harrison, David K. Jackson, Ian Johnston, Dominic Kwiatkowski, Cordelia Langford, John Sillitoe on behalf of the Wellcome Sanger Institute COVID-19 Surveillance Team |
| EPI_ISL_3107633, EPI_ISL_3107634, EPI_ISL_3107635, EPI_ISL_3107636, EPI_ISL_3107637, EPI_ISL_3107638, EPI_ISL_3107639, EPI_ISL_3107640, EPI_ISL_3107641, EPI_ISL_3107642, EPI_ISL_3107643 | | | |
| see above | Lighthouse Lab in Alderley Park | Wellcome Sanger Institute for the COVID-19 Genomics UK (COG-UK) Consortium | Jacquelyn Wynn, Mairead Hyland, The Lighthouse Lab in Alderley Park and Alex Alderton, Roberto Amato, Jeffrey Barrett, Sonia Goncalves, Ewan Harrison, David K. Jackson, Ian Johnston, Dominic Kwiatkowski, Cordelia Langford, John Sillitoe on behalf of the Wellcome Sanger Institute COVID-19 Surveillance Team |
| EPI_ISL_3107644 | Lighthouse Lab in Milton Keynes | Wellcome Sanger Institute for the COVID-19 Genomics UK (COG-UK) Consortium | The Lighthouse Lab in Milton Keynes and Alex Alderton, Roberto Amato, Jeffrey Barrett, Sonia Goncalves, Ewan Harrison, David K. Jackson, Ian Johnston, Dominic Kwiatkowski, Cordelia Langford, John Sillitoe on behalf of the Wellcome Sanger Institute COVID-19 Surveillance Team |
| EPI_ISL_3107645 | Lighthouse Lab in Alderley Park | Wellcome Sanger Institute for the COVID-19 Genomics UK (COG-UK) Consortium | Jacquelyn Wynn, Mairead Hyland, The Lighthouse Lab in Alderley Park and Alex Alderton, Roberto Amato, Jeffrey Barrett, Sonia Goncalves, Ewan Harrison, David K. Jackson, Ian Johnston, Dominic Kwiatkowski, Cordelia Langford, John Sillitoe on behalf of the Wellcome Sanger Institute COVID-19 Surveillance Team |
| EPI_ISL_3107646, EPI_ISL_3107647 | Lighthouse Lab in Milton Keynes | Wellcome Sanger Institute for the COVID-19 Genomics UK (COG-UK) Consortium | The Lighthouse Lab in Milton Keynes and Alex Alderton, Roberto Amato, Jeffrey Barrett, Sonia Goncalves, Ewan Harrison, David K. Jackson, Ian Johnston, Dominic Kwiatkowski, Cordelia Langford, John Sillitoe on behalf of the Wellcome Sanger Institute COVID-19 Surveillance Team |
| EPI_ISL_3107648, EPI_ISL_3107649, EPI_ISL_3107650 | Lighthouse Lab in Alderley Park | Wellcome Sanger Institute for the COVID-19 Genomics UK (COG-UK) Consortium | Jacquelyn Wynn, Mairead Hyland, The Lighthouse Lab in Alderley Park and Alex Alderton, Roberto Amato, Jeffrey Barrett, Sonia Goncalves, Ewan Harrison, David K. Jackson, Ian Johnston, Dominic Kwiatkowski, Cordelia Langford, John Sillitoe on behalf of the Wellcome Sanger Institute COVID-19 Surveillance Team |
| EPI_ISL_3107651 | Lighthouse Lab in Milton Keynes | Wellcome Sanger Institute for the COVID-19 Genomics UK (COG-UK) Consortium | The Lighthouse Lab in Milton Keynes and Alex Alderton, Roberto Amato, Jeffrey Barrett, Sonia Goncalves, Ewan Harrison, David K. Jackson, Ian Johnston, Dominic Kwiatkowski, Cordelia Langford, John Sillitoe on behalf of the Wellcome Sanger Institute COVID-19 Surveillance Team |
| EPI_ISL_3107652, EPI_ISL_3107653, EPI_ISL_3107654 | Lighthouse Lab in Alderley Park | Wellcome Sanger Institute for the COVID-19 Genomics UK (COG-UK) Consortium | Jacquelyn Wynn, Mairead Hyland, The Lighthouse Lab in Alderley Park and Alex Alderton, Roberto Amato, Jeffrey Barrett, Sonia Goncalves, Ewan Harrison, David K. Jackson, Ian Johnston, Dominic Kwiatkowski, Cordelia Langford, John Sillitoe on behalf of the Wellcome Sanger Institute COVID-19 Surveillance Team |
| EPI_ISL_3107655 | Lighthouse Lab in Milton Keynes | Wellcome Sanger Institute for the COVID-19 Genomics UK (COG-UK) Consortium | The Lighthouse Lab in Milton Keynes and Alex Alderton, Roberto Amato, Jeffrey Barrett, Sonia Goncalves, Ewan Harrison, David K. Jackson, Ian Johnston, Dominic Kwiatkowski, Cordelia Langford, John Sillitoe on behalf of the Wellcome Sanger Institute COVID-19 Surveillance Team |
| EPI_ISL_3107656 | Lighthouse Lab in Alderley Park | Wellcome Sanger Institute for the COVID-19 Genomics UK (COG-UK) Consortium | Jacquelyn Wynn, Mairead Hyland, The Lighthouse Lab in Alderley Park and Alex Alderton, Roberto Amato, Jeffrey Barrett, Sonia Goncalves, Ewan Harrison, David K. Jackson, Ian Johnston, Dominic Kwiatkowski, Cordelia Langford, John Sillitoe on behalf of the Wellcome Sanger Institute COVID-19 Surveillance Team |
| EPI_ISL_3107657 | Lighthouse Lab in Milton Keynes | Wellcome Sanger Institute for the COVID-19 Genomics UK (COG-UK) Consortium | The Lighthouse Lab in Milton Keynes and Alex Alderton, Roberto Amato, Jeffrey Barrett, Sonia Goncalves, Ewan Harrison, David K. Jackson, Ian Johnston, Dominic Kwiatkowski, Cordelia Langford, John Sillitoe on behalf of the Wellcome Sanger Institute COVID-19 Surveillance Team |
| EPI_ISL_3107658, EPI_ISL_3107659 | Lighthouse Lab in Alderley Park | Wellcome Sanger Institute for the COVID-19 Genomics UK (COG-UK) Consortium | Jacquelyn Wynn, Mairead Hyland, The Lighthouse Lab in Alderley Park and Alex Alderton, Roberto Amato, Jeffrey Barrett, Sonia Goncalves, Ewan Harrison, David K. Jackson, Ian Johnston, Dominic Kwiatkowski, Cordelia Langford, John Sillitoe on behalf of the Wellcome Sanger Institute COVID-19 Surveillance Team |
| EPI_ISL_3107660 | Lighthouse Lab in Glasgow | Wellcome Sanger Institute for the COVID-19 Genomics UK (COG-UK) Consortium | Harper VanSteenhouse, Yumi Kasai, David Gray, Carol Clugston, Anna Dominiczak and Alex Alderton, Roberto Amato, Jeffrey Barrett, Sonia Goncalves, Ewan Harrison, David K. Jackson, Ian Johnston, Dominic Kwiatkowski, Cordelia Langford, John Sillitoe on behalf of the Wellcome Sanger Institute COVID-19 Surveillance Team |
| EPI_ISL_3107661 | Lighthouse Lab in Milton Keynes | Wellcome Sanger Institute for the COVID-19 Genomics UK (COG-UK) Consortium | The Lighthouse Lab in Milton Keynes and Alex Alderton, Roberto Amato, Jeffrey Barrett, Sonia Goncalves, Ewan Harrison, David K. Jackson, Ian Johnston, Dominic Kwiatkowski, Cordelia Langford, John Sillitoe on behalf of the Wellcome Sanger Institute COVID-19 Surveillance Team |
| EPI_ISL_3107662 | Lighthouse Lab in Alderley Park | Wellcome Sanger Institute for the COVID-19 Genomics UK (COG-UK) Consortium | Jacquelyn Wynn, Mairead Hyland, The Lighthouse Lab in Alderley Park and Alex Alderton, Roberto Amato, Jeffrey Barrett, Sonia Goncalves, Ewan Harrison, David K. Jackson, Ian Johnston, Dominic Kwiatkowski, Cordelia Langford, John Sillitoe on behalf of the Wellcome Sanger Institute COVID-19 Surveillance Team |
| EPI_ISL_3107663 | Lighthouse Laboratory Plymouth | Wellcome Sanger Institute for the COVID-19 Genomics UK (COG-UK) Consortium | Lighthouse Laboratory Plymouth and Alex Alderton, Roberto Amato, Jeffrey Barrett, Sonia Goncalves, Ewan Harrison, David K. Jackson, Ian Johnston, Dominic Kwiatkowski, Cordelia Langford, John Sillitoe on behalf of the Wellcome Sanger Institute COVID-19 Surveillance Team |
| EPI_ISL_3107664 | Lighthouse Lab in Alderley Park | Wellcome Sanger Institute for the COVID-19 Genomics UK (COG-UK) Consortium | Jacquelyn Wynn, Mairead Hyland, The Lighthouse Lab in Alderley Park and Alex Alderton, Roberto Amato, Jeffrey Barrett, Sonia Goncalves, Ewan Harrison, David K. Jackson, Ian Johnston, Dominic Kwiatkowski, Cordelia Langford, John Sillitoe on behalf of the Wellcome Sanger Institute COVID-19 Surveillance Team |
| EPI_ISL_3107665 | Lighthouse Lab in Milton Keynes | Wellcome Sanger Institute for the COVID-19 Genomics UK (COG-UK) Consortium | The Lighthouse Lab in Milton Keynes and Alex Alderton, Roberto Amato, Jeffrey Barrett, Sonia Goncalves, Ewan Harrison, David K. Jackson, Ian Johnston, Dominic Kwiatkowski, Cordelia Langford, John Sillitoe on behalf of the Wellcome Sanger Institute COVID-19 Surveillance Team |
| EPI_ISL_3107666 | Lighthouse Lab in Alderley Park | Wellcome Sanger Institute for the COVID-19 Genomics UK (COG-UK) Consortium | Jacquelyn Wynn, Mairead Hyland, The Lighthouse Lab in Alderley Park and Alex Alderton, Roberto Amato, Jeffrey Barrett, Sonia Goncalves, Ewan Harrison, David K. Jackson, Ian Johnston, Dominic Kwiatkowski, Cordelia Langford, John Sillitoe on behalf of the Wellcome Sanger Institute COVID-19 Surveillance Team |
| EPI_ISL_3107667 | Lighthouse Lab in Glasgow | Wellcome Sanger Institute for the COVID-19 Genomics UK (COG-UK) Consortium | Harper VanSteenhouse, Yumi Kasai, David Gray, Carol Clugston, Anna Dominiczak and Alex Alderton, Roberto Amato, Jeffrey Barrett, Sonia Goncalves, Ewan Harrison, David K. Jackson, Ian Johnston, Dominic Kwiatkowski, Cordelia Langford, John Sillitoe on behalf of the Wellcome Sanger Institute COVID-19 Surveillance Team |
| EPI_ISL_3107668 | Lighthouse Lab in Milton Keynes | Wellcome Sanger Institute for the COVID-19 Genomics UK (COG-UK) Consortium | The Lighthouse Lab in Milton Keynes and Alex Alderton, Roberto Amato, Jeffrey Barrett, Sonia Goncalves, Ewan Harrison, David K. Jackson, Ian Johnston, Dominic Kwiatkowski, Cordelia Langford, John Sillitoe on behalf of the Wellcome Sanger Institute COVID-19 Surveillance Team |
| EPI_ISL_3107669 | Lighthouse Laboratory Plymouth | Wellcome Sanger Institute for the COVID-19 Genomics UK (COG-UK) Consortium | Lighthouse Laboratory Plymouth and Alex Alderton, Roberto Amato, Jeffrey Barrett, Sonia Goncalves, Ewan Harrison, David K. Jackson, Ian Johnston, Dominic Kwiatkowski, Cordelia Langford, John Sillitoe on behalf of the Wellcome Sanger Institute COVID-19 Surveillance Team |
| EPI_ISL_3107670 | Lighthouse Lab in Glasgow | Wellcome Sanger Institute for the COVID-19 Genomics UK (COG-UK) Consortium | Harper VanSteenhouse, Yumi Kasai, David Gray, Carol Clugston, Anna Dominiczak and Alex Alderton, Roberto Amato, Jeffrey Barrett, Sonia Goncalves, Ewan Harrison, David K. Jackson, Ian Johnston, Dominic Kwiatkowski, Cordelia Langford, John Sillitoe on behalf of the Wellcome Sanger Institute COVID-19 Surveillance Team |
| EPI_ISL_3107671, EPI_ISL_3107672 | Lighthouse Lab in Milton Keynes | Wellcome Sanger Institute for the COVID-19 Genomics UK (COG-UK) Consortium | The Lighthouse Lab in Milton Keynes and Alex Alderton, Roberto Amato, Jeffrey Barrett, Sonia Goncalves, Ewan Harrison, David K. Jackson, Ian Johnston, Dominic Kwiatkowski, Cordelia Langford, John Sillitoe on behalf of the Wellcome Sanger Institute COVID-19 Surveillance Team |
| EPI_ISL_3107673 | Lighthouse Lab in Alderley Park | Wellcome Sanger Institute for the COVID-19 Genomics UK (COG-UK) Consortium | Jacquelyn Wynn, Mairead Hyland, The Lighthouse Lab in Alderley Park and Alex Alderton, Roberto Amato, Jeffrey Barrett, Sonia Goncalves, Ewan Harrison, David K. Jackson, Ian Johnston, Dominic Kwiatkowski, Cordelia Langford, John Sillitoe on behalf of the Wellcome Sanger Institute COVID-19 Surveillance Team |
| EPI_ISL_3107674 | Lighthouse Lab in Glasgow | Wellcome Sanger Institute for the COVID-19 Genomics UK (COG-UK) Consortium | Harper VanSteenhouse, Yumi Kasai, David Gray, Carol Clugston, Anna Dominiczak and Alex Alderton, Roberto Amato, Jeffrey Barrett, Sonia Goncalves, Ewan Harrison, David K. Jackson, Ian Johnston, Dominic Kwiatkowski, Cordelia Langford, John Sillitoe on behalf of the Wellcome Sanger Institute COVID-19 Surveillance Team |
| EPI_ISL_3107675 | Lighthouse Laboratory Plymouth | Wellcome Sanger Institute for the COVID-19 Genomics UK (COG-UK) Consortium | Lighthouse Laboratory Plymouth and Alex Alderton, Roberto Amato, Jeffrey Barrett, Sonia Goncalves, Ewan Harrison, David K. Jackson, Ian Johnston, Dominic Kwiatkowski, Cordelia Langford, John Sillitoe on behalf of the Wellcome Sanger Institute COVID-19 Surveillance Team |
| EPI_ISL_3107676 | Lighthouse Lab in Alderley Park | Wellcome Sanger Institute for the COVID-19 Genomics UK (COG-UK) Consortium | Jacquelyn Wynn, Mairead Hyland, The Lighthouse Lab in Alderley Park and Alex Alderton, Roberto Amato, Jeffrey Barrett, Sonia Goncalves, Ewan Harrison, David K. Jackson, Ian Johnston, Dominic Kwiatkowski, Cordelia Langford, John Sillitoe on behalf of the Wellcome Sanger Institute COVID-19 Surveillance Team |
| EPI_ISL_3107677 | Lighthouse Lab in Milton Keynes | Wellcome Sanger Institute for the COVID-19 Genomics UK (COG-UK) Consortium | The Lighthouse Lab in Milton Keynes and Alex Alderton, Roberto Amato, Jeffrey Barrett, Sonia Goncalves, Ewan Harrison, David K. Jackson, Ian Johnston, Dominic Kwiatkowski, Cordelia Langford, John Sillitoe on behalf of the Wellcome Sanger Institute COVID-19 Surveillance Team |
| EPI_ISL_3107678 | Lighthouse Laboratory Plymouth | Wellcome Sanger Institute for the COVID-19 Genomics UK (COG-UK) Consortium | Lighthouse Laboratory Plymouth and Alex Alderton, Roberto Amato, Jeffrey Barrett, Sonia Goncalves, Ewan Harrison, David K. Jackson, Ian Johnston, Dominic Kwiatkowski, Cordelia Langford, John Sillitoe on behalf of the Wellcome Sanger Institute COVID-19 Surveillance Team |
| EPI_ISL_3107679 | Lighthouse Lab in Glasgow | Wellcome Sanger Institute for the COVID-19 Genomics UK (COG-UK) Consortium | Harper VanSteenhouse, Yumi Kasai, David Gray, Carol Clugston, Anna Dominiczak and Alex Alderton, Roberto Amato, Jeffrey Barrett, Sonia Goncalves, Ewan Harrison, David K. Jackson, Ian Johnston, Dominic Kwiatkowski, Cordelia Langford, John Sillitoe on behalf of the Wellcome Sanger Institute COVID-19 Surveillance Team |
| EPI_ISL_3107680, EPI_ISL_3107681, EPI_ISL_3107682, EPI_ISL_3107683 | Lighthouse Lab in Alderley Park | Wellcome Sanger Institute for the COVID-19 Genomics UK (COG-UK) Consortium | Jacquelyn Wynn, Mairead Hyland, The Lighthouse Lab in Alderley Park and Alex Alderton, Roberto Amato, Jeffrey Barrett, Sonia Goncalves, Ewan Harrison, David K. Jackson, Ian Johnston, Dominic Kwiatkowski, Cordelia Langford, John Sillitoe on behalf of the Wellcome Sanger Institute COVID-19 Surveillance Team |
| EPI_ISL_3107684, EPI_ISL_3107685 | Lighthouse Laboratory Plymouth | Wellcome Sanger Institute for the COVID-19 Genomics UK (COG-UK) Consortium | Lighthouse Laboratory Plymouth and Alex Alderton, Roberto Amato, Jeffrey Barrett, Sonia Goncalves, Ewan Harrison, David K. Jackson, Ian Johnston, Dominic Kwiatkowski, Cordelia Langford, John Sillitoe on behalf of the Wellcome Sanger Institute COVID-19 Surveillance Team |
| EPI_ISL_3107686 | Lighthouse Lab in Alderley Park | Wellcome Sanger Institute for the COVID-19 Genomics UK (COG-UK) Consortium | Jacquelyn Wynn, Mairead Hyland, The Lighthouse Lab in Alderley Park and Alex Alderton, Roberto Amato, Jeffrey Barrett, Sonia Goncalves, Ewan Harrison, David K. Jackson, Ian Johnston, Dominic Kwiatkowski, Cordelia Langford, John Sillitoe on behalf of the Wellcome Sanger Institute COVID-19 Surveillance Team |
| EPI_ISL_3107687 | Lighthouse Laboratory Plymouth | Wellcome Sanger Institute for the COVID-19 Genomics UK (COG-UK) Consortium | Lighthouse Laboratory Plymouth and Alex Alderton, Roberto Amato, Jeffrey Barrett, Sonia Goncalves, Ewan Harrison, David K. Jackson, Ian Johnston, Dominic Kwiatkowski, Cordelia Langford, John Sillitoe on behalf of the Wellcome Sanger Institute COVID-19 Surveillance Team |
| EPI_ISL_3107688 | National Center of Infectious and Parasitic Diseases | National Center of Infectious and Parasitic Diseases | Alexiev et al |

| | | | |
|---|---|---|---|
| EPI_ISL_3107689, EPI_ISL_3107691 | Lighthouse Lab in Alderley Park | Wellcome Sanger Institute for the COVID-19 Genomics UK (COG-UK) Consortium | Jacquelyn Wynn, Mairead Hyland, The Lighthouse Lab in Alderley Park and Alex Alderton, Roberto Amato, Jeffrey Barrett, Sonia Goncalves, Ewan Harrison, David K. Jackson, Ian Johnston, Dominic Kwiatkowski, Cordelia Langford, John Sillitoe on behalf of the Wellcome Sanger Institute COVID-19 Surveillance Team |
| EPI_ISL_3107692 | Lighthouse Laboratory Plymouth | Wellcome Sanger Institute for the COVID-19 Genomics UK (COG-UK) Consortium | Lighthouse Laboratory Plymouth and Alex Alderton, Roberto Amato, Jeffrey Barrett, Sonia Goncalves, Ewan Harrison, David K. Jackson, Ian Johnston, Dominic Kwiatkowski, Cordelia Langford, John Sillitoe on behalf of the Wellcome Sanger Institute COVID-19 Surveillance Team |
| EPI_ISL_3107693 | Lighthouse Lab in Milton Keynes | Wellcome Sanger Institute for the COVID-19 Genomics UK (COG-UK) Consortium | The Lighthouse Lab in Milton Keynes and Alex Alderton, Roberto Amato, Jeffrey Barrett, Sonia Goncalves, Ewan Harrison, David K. Jackson, Ian Johnston, Dominic Kwiatkowski, Cordelia Langford, John Sillitoe on behalf of the Wellcome Sanger Institute COVID-19 Surveillance Team |
| EPI_ISL_3107694 | Lighthouse Lab in Glasgow | Wellcome Sanger Institute for the COVID-19 Genomics UK (COG-UK) Consortium | Harper VanSteenhouse, Yumi Kasai, David Gray, Carol Clugston, Anna Dominiczak and Alex Alderton, Roberto Amato, Jeffrey Barrett, Sonia Goncalves, Ewan Harrison, David K. Jackson, Ian Johnston, Dominic Kwiatkowski, Cordelia Langford, John Sillitoe on behalf of the Wellcome Sanger Institute COVID-19 Surveillance Team |
| EPI_ISL_3107695, EPI_ISL_3107696, EPI_ISL_3107697, EPI_ISL_3107698 | Lighthouse Lab in Milton Keynes | Wellcome Sanger Institute for the COVID-19 Genomics UK (COG-UK) Consortium | The Lighthouse Lab in Milton Keynes and Alex Alderton, Roberto Amato, Jeffrey Barrett, Sonia Goncalves, Ewan Harrison, David K. Jackson, Ian Johnston, Dominic Kwiatkowski, Cordelia Langford, John Sillitoe on behalf of the Wellcome Sanger Institute COVID-19 Surveillance Team |
| EPI_ISL_3107699 | Lighthouse Lab in Glasgow | Wellcome Sanger Institute for the COVID-19 Genomics UK (COG-UK) Consortium | Harper VanSteenhouse, Yumi Kasai, David Gray, Carol Clugston, Anna Dominiczak and Alex Alderton, Roberto Amato, Jeffrey Barrett, Sonia Goncalves, Ewan Harrison, David K. Jackson, Ian Johnston, Dominic Kwiatkowski, Cordelia Langford, John Sillitoe on behalf of the Wellcome Sanger Institute COVID-19 Surveillance Team |
| EPI_ISL_3107700, EPI_ISL_3107701 | Lighthouse Lab in Alderley Park | Wellcome Sanger Institute for the COVID-19 Genomics UK (COG-UK) Consortium | Jacquelyn Wynn, Mairead Hyland, The Lighthouse Lab in Alderley Park and Alex Alderton, Roberto Amato, Jeffrey Barrett, Sonia Goncalves, Ewan Harrison, David K. Jackson, Ian Johnston, Dominic Kwiatkowski, Cordelia Langford, John Sillitoe on behalf of the Wellcome Sanger Institute COVID-19 Surveillance Team |
| EPI_ISL_3107702, EPI_ISL_3107703, EPI_ISL_3107704, EPI_ISL_3107705, EPI_ISL_3107706 | Lighthouse Laboratory Plymouth | Wellcome Sanger Institute for the COVID-19 Genomics UK (COG-UK) Consortium | Lighthouse Laboratory Plymouth and Alex Alderton, Roberto Amato, Jeffrey Barrett, Sonia Goncalves, Ewan Harrison, David K. Jackson, Ian Johnston, Dominic Kwiatkowski, Cordelia Langford, John Sillitoe on behalf of the Wellcome Sanger Institute COVID-19 Surveillance Team |
| EPI_ISL_3107707 | Lighthouse Lab in Alderley Park | Wellcome Sanger Institute for the COVID-19 Genomics UK (COG-UK) Consortium | Jacquelyn Wynn, Mairead Hyland, The Lighthouse Lab in Alderley Park and Alex Alderton, Roberto Amato, Jeffrey Barrett, Sonia Goncalves, Ewan Harrison, David K. Jackson, Ian Johnston, Dominic Kwiatkowski, Cordelia Langford, John Sillitoe on behalf of the Wellcome Sanger Institute COVID-19 Surveillance Team |
| EPI_ISL_3107708 | Lighthouse Lab in Milton Keynes | Wellcome Sanger Institute for the COVID-19 Genomics UK (COG-UK) Consortium | The Lighthouse Lab in Milton Keynes and Alex Alderton, Roberto Amato, Jeffrey Barrett, Sonia Goncalves, Ewan Harrison, David K. Jackson, Ian Johnston, Dominic Kwiatkowski, Cordelia Langford, John Sillitoe on behalf of the Wellcome Sanger Institute COVID-19 Surveillance Team |
| EPI_ISL_3107709 | Lighthouse Lab in Glasgow | Wellcome Sanger Institute for the COVID-19 Genomics UK (COG-UK) Consortium | Harper VanSteenhouse, Yumi Kasai, David Gray, Carol Clugston, Anna Dominiczak and Alex Alderton, Roberto Amato, Jeffrey Barrett, Sonia Goncalves, Ewan Harrison, David K. Jackson, Ian Johnston, Dominic Kwiatkowski, Cordelia Langford, John Sillitoe on behalf of the Wellcome Sanger Institute COVID-19 Surveillance Team |
| EPI_ISL_3107710 | Lighthouse Laboratory Plymouth | Wellcome Sanger Institute for the COVID-19 Genomics UK (COG-UK) Consortium | Lighthouse Laboratory Plymouth and Alex Alderton, Roberto Amato, Jeffrey Barrett, Sonia Goncalves, Ewan Harrison, David K. Jackson, Ian Johnston, Dominic Kwiatkowski, Cordelia Langford, John Sillitoe on behalf of the Wellcome Sanger Institute COVID-19 Surveillance Team |
| EPI_ISL_3107711 | Lighthouse Lab in Glasgow | Wellcome Sanger Institute for the COVID-19 Genomics UK (COG-UK) Consortium | Harper VanSteenhouse, Yumi Kasai, David Gray, Carol Clugston, Anna Dominiczak and Alex Alderton, Roberto Amato, Jeffrey Barrett, Sonia Goncalves, Ewan Harrison, David K. Jackson, Ian Johnston, Dominic Kwiatkowski, Cordelia Langford, John Sillitoe on behalf of the Wellcome Sanger Institute COVID-19 Surveillance Team |
| EPI_ISL_3107712, EPI_ISL_3107713 | Lighthouse Lab in Milton Keynes | Wellcome Sanger Institute for the COVID-19 Genomics UK (COG-UK) Consortium | The Lighthouse Lab in Milton Keynes and Alex Alderton, Roberto Amato, Jeffrey Barrett, Sonia Goncalves, Ewan Harrison, David K. Jackson, Ian Johnston, Dominic Kwiatkowski, Cordelia Langford, John Sillitoe on behalf of the Wellcome Sanger Institute COVID-19 Surveillance Team |
| EPI_ISL_3107714 | Lighthouse Lab in Alderley Park | Wellcome Sanger Institute for the COVID-19 Genomics UK (COG-UK) Consortium | Jacquelyn Wynn, Mairead Hyland, The Lighthouse Lab in Alderley Park and Alex Alderton, Roberto Amato, Jeffrey Barrett, Sonia Goncalves, Ewan Harrison, David K. Jackson, Ian Johnston, Dominic Kwiatkowski, Cordelia Langford, John Sillitoe on behalf of the Wellcome Sanger Institute COVID-19 Surveillance Team |
| EPI_ISL_3107715 | Lighthouse Laboratory Plymouth | Wellcome Sanger Institute for the COVID-19 Genomics UK (COG-UK) Consortium | Lighthouse Laboratory Plymouth and Alex Alderton, Roberto Amato, Jeffrey Barrett, Sonia Goncalves, Ewan Harrison, David K. Jackson, Ian Johnston, Dominic Kwiatkowski, Cordelia Langford, John Sillitoe on behalf of the Wellcome Sanger Institute COVID-19 Surveillance Team |
| EPI_ISL_3107716 | Lighthouse Lab in Alderley Park | Wellcome Sanger Institute for the COVID-19 Genomics UK (COG-UK) Consortium | Jacquelyn Wynn, Mairead Hyland, The Lighthouse Lab in Alderley Park and Alex Alderton, Roberto Amato, Jeffrey Barrett, Sonia Goncalves, Ewan Harrison, David K. Jackson, Ian Johnston, Dominic Kwiatkowski, Cordelia Langford, John Sillitoe on behalf of the Wellcome Sanger Institute COVID-19 Surveillance Team |
| EPI_ISL_3107717 | Lighthouse Lab in Glasgow | Wellcome Sanger Institute for the COVID-19 Genomics UK (COG-UK) Consortium | Harper VanSteenhouse, Yumi Kasai, David Gray, Carol Clugston, Anna Dominiczak and Alex Alderton, Roberto Amato, Jeffrey Barrett, Sonia Goncalves, Ewan Harrison, David K. Jackson, Ian Johnston, Dominic Kwiatkowski, Cordelia Langford, John Sillitoe on behalf of the Wellcome Sanger Institute COVID-19 Surveillance Team |
| EPI_ISL_3107718 | Lighthouse Laboratory Plymouth | Wellcome Sanger Institute for the COVID-19 Genomics UK (COG-UK) Consortium | Lighthouse Laboratory Plymouth and Alex Alderton, Roberto Amato, Jeffrey Barrett, Sonia Goncalves, Ewan Harrison, David K. Jackson, Ian Johnston, Dominic Kwiatkowski, Cordelia Langford, John Sillitoe on behalf of the Wellcome Sanger Institute COVID-19 Surveillance Team |
| EPI_ISL_3107719 | Lighthouse Lab in Milton Keynes | Wellcome Sanger Institute for the COVID-19 Genomics UK (COG-UK) Consortium | The Lighthouse Lab in Milton Keynes and Alex Alderton, Roberto Amato, Jeffrey Barrett, Sonia Goncalves, Ewan Harrison, David K. Jackson, Ian Johnston, Dominic Kwiatkowski, Cordelia Langford, John Sillitoe on behalf of the Wellcome Sanger Institute COVID-19 Surveillance Team |
| EPI_ISL_3107720 | Lighthouse Laboratory Plymouth | Wellcome Sanger Institute for the COVID-19 Genomics UK (COG-UK) Consortium | Lighthouse Laboratory Plymouth and Alex Alderton, Roberto Amato, Jeffrey Barrett, Sonia Goncalves, Ewan Harrison, David K. Jackson, Ian Johnston, Dominic Kwiatkowski, Cordelia Langford, John Sillitoe on behalf of the Wellcome Sanger Institute COVID-19 Surveillance Team |
| EPI_ISL_3107721 | Lighthouse Lab in Glasgow | Wellcome Sanger Institute for the COVID-19 Genomics UK (COG-UK) Consortium | Harper VanSteenhouse, Yumi Kasai, David Gray, Carol Clugston, Anna Dominiczak and Alex Alderton, Roberto Amato, Jeffrey Barrett, Sonia Goncalves, Ewan Harrison, David K. Jackson, Ian Johnston, Dominic Kwiatkowski, Cordelia Langford, John Sillitoe on behalf of the Wellcome Sanger Institute COVID-19 Surveillance Team |
| EPI_ISL_3107722, EPI_ISL_3107723 | Lighthouse Lab in Alderley Park | Wellcome Sanger Institute for the COVID-19 Genomics UK (COG-UK) Consortium | Jacquelyn Wynn, Mairead Hyland, The Lighthouse Lab in Alderley Park and Alex Alderton, Roberto Amato, Jeffrey Barrett, Sonia Goncalves, Ewan Harrison, David K. Jackson, Ian Johnston, Dominic Kwiatkowski, Cordelia Langford, John Sillitoe on behalf of the Wellcome Sanger Institute COVID-19 Surveillance Team |
| EPI_ISL_3107724, EPI_ISL_3107725, EPI_ISL_3107726, EPI_ISL_3107727 | Lighthouse Laboratory Plymouth | Wellcome Sanger Institute for the COVID-19 Genomics UK (COG-UK) Consortium | Lighthouse Laboratory Plymouth and Alex Alderton, Roberto Amato, Jeffrey Barrett, Sonia Goncalves, Ewan Harrison, David K. Jackson, Ian Johnston, Dominic Kwiatkowski, Cordelia Langford, John Sillitoe on behalf of the Wellcome Sanger Institute COVID-19 Surveillance Team |
| EPI_ISL_3107728, EPI_ISL_3107729, EPI_ISL_3107730, EPI_ISL_3107731 | Lighthouse Lab in Alderley Park | Wellcome Sanger Institute for the COVID-19 Genomics UK (COG-UK) Consortium | Jacquelyn Wynn, Mairead Hyland, The Lighthouse Lab in Alderley Park and Alex Alderton, Roberto Amato, Jeffrey Barrett, Sonia Goncalves, Ewan Harrison, David K. Jackson, Ian Johnston, Dominic Kwiatkowski, Cordelia Langford, John Sillitoe on behalf of the Wellcome Sanger Institute COVID-19 Surveillance Team |
| EPI_ISL_3107732 | Lighthouse Laboratory Plymouth | Wellcome Sanger Institute for the COVID-19 Genomics UK (COG-UK) Consortium | Lighthouse Laboratory Plymouth and Alex Alderton, Roberto Amato, Jeffrey Barrett, Sonia Goncalves, Ewan Harrison, David K. Jackson, Ian Johnston, Dominic Kwiatkowski, Cordelia Langford, John Sillitoe on behalf of the Wellcome Sanger Institute COVID-19 Surveillance Team |
| EPI_ISL_3107733, EPI_ISL_3107734, EPI_ISL_3107735 | Lighthouse Lab in Alderley Park | Wellcome Sanger Institute for the COVID-19 Genomics UK (COG-UK) Consortium | Jacquelyn Wynn, Mairead Hyland, The Lighthouse Lab in Alderley Park and Alex Alderton, Roberto Amato, Jeffrey Barrett, Sonia Goncalves, Ewan Harrison, David K. Jackson, Ian Johnston, Dominic Kwiatkowski, Cordelia Langford, John Sillitoe on behalf of the Wellcome Sanger Institute COVID-19 Surveillance Team |
| EPI_ISL_3107736 | Lighthouse Laboratory Plymouth | Wellcome Sanger Institute for the COVID-19 Genomics UK (COG-UK) Consortium | Lighthouse Laboratory Plymouth and Alex Alderton, Roberto Amato, Jeffrey Barrett, Sonia Goncalves, Ewan Harrison, David K. Jackson, Ian Johnston, Dominic Kwiatkowski, Cordelia Langford, John Sillitoe on behalf of the Wellcome Sanger Institute COVID-19 Surveillance Team |
| EPI_ISL_3107737 | Lighthouse Lab in Glasgow | Wellcome Sanger Institute for the COVID-19 Genomics UK (COG-UK) Consortium | Harper VanSteenhouse, Yumi Kasai, David Gray, Carol Clugston, Anna Dominiczak and Alex Alderton, Roberto Amato, Jeffrey Barrett, Sonia Goncalves, Ewan Harrison, David K. Jackson, Ian Johnston, Dominic Kwiatkowski, Cordelia Langford, John Sillitoe on behalf of the Wellcome Sanger Institute COVID-19 Surveillance Team |
| EPI_ISL_3107738 | Lighthouse Lab in Alderley Park | Wellcome Sanger Institute for the COVID-19 Genomics UK (COG-UK) Consortium | Jacquelyn Wynn, Mairead Hyland, The Lighthouse Lab in Alderley Park and Alex Alderton, Roberto Amato, Jeffrey Barrett, Sonia Goncalves, Ewan Harrison, David K. Jackson, Ian Johnston, Dominic Kwiatkowski, Cordelia Langford, John Sillitoe on behalf of the Wellcome Sanger Institute COVID-19 Surveillance Team |
| EPI_ISL_3107739 | Lighthouse Lab in Milton Keynes | Wellcome Sanger Institute for the COVID-19 Genomics UK (COG-UK) Consortium | The Lighthouse Lab in Milton Keynes and Alex Alderton, Roberto Amato, Jeffrey Barrett, Sonia Goncalves, Ewan Harrison, David K. Jackson, Ian Johnston, Dominic Kwiatkowski, Cordelia Langford, John Sillitoe on behalf of the Wellcome Sanger Institute COVID-19 Surveillance Team |
| EPI_ISL_3107740, EPI_ISL_3107741 | Lighthouse Laboratory Plymouth | Wellcome Sanger Institute for the COVID-19 Genomics UK (COG-UK) Consortium | Lighthouse Laboratory Plymouth and Alex Alderton, Roberto Amato, Jeffrey Barrett, Sonia Goncalves, Ewan Harrison, David K. Jackson, Ian Johnston, Dominic Kwiatkowski, Cordelia Langford, John Sillitoe on behalf of the Wellcome Sanger Institute COVID-19 Surveillance Team |
| EPI_ISL_3107742, EPI_ISL_3107743, EPI_ISL_3107744, EPI_ISL_3107745 | Lighthouse Lab in Alderley Park | Wellcome Sanger Institute for the COVID-19 Genomics UK (COG-UK) Consortium | Jacquelyn Wynn, Mairead Hyland, The Lighthouse Lab in Alderley Park and Alex Alderton, Roberto Amato, Jeffrey Barrett, Sonia Goncalves, Ewan Harrison, David K. Jackson, Ian Johnston, Dominic Kwiatkowski, Cordelia Langford, John Sillitoe on behalf of the Wellcome Sanger Institute COVID-19 Surveillance Team |
| EPI_ISL_3107746 | Lighthouse Lab in Milton Keynes | Wellcome Sanger Institute for the COVID-19 Genomics UK (COG-UK) Consortium | The Lighthouse Lab in Milton Keynes and Alex Alderton, Roberto Amato, Jeffrey Barrett, Sonia Goncalves, Ewan Harrison, David K. Jackson, Ian Johnston, Dominic Kwiatkowski, Cordelia Langford, John Sillitoe on behalf of the Wellcome Sanger Institute COVID-19 Surveillance Team |
| EPI_ISL_3107747, EPI_ISL_3107748 | Lighthouse Lab in Alderley Park | Wellcome Sanger Institute for the COVID-19 Genomics UK (COG-UK) Consortium | Jacquelyn Wynn, Mairead Hyland, The Lighthouse Lab in Alderley Park and Alex Alderton, Roberto Amato, Jeffrey Barrett, Sonia Goncalves, Ewan Harrison, David K. Jackson, Ian Johnston, Dominic Kwiatkowski, Cordelia Langford, John Sillitoe on behalf of the Wellcome Sanger Institute COVID-19 Surveillance Team |
| EPI_ISL_3107749, EPI_ISL_3107750 | Lighthouse Laboratory Plymouth | Wellcome Sanger Institute for the COVID-19 Genomics UK (COG-UK) Consortium | Lighthouse Laboratory Plymouth and Alex Alderton, Roberto Amato, Jeffrey Barrett, Sonia Goncalves, Ewan Harrison, David K. Jackson, Ian Johnston, Dominic Kwiatkowski, Cordelia Langford, John Sillitoe on behalf of the Wellcome Sanger Institute COVID-19 Surveillance Team |
| EPI_ISL_3107751, EPI_ISL_3107752 | Lighthouse Lab in Alderley Park | Wellcome Sanger Institute for the COVID-19 Genomics UK (COG-UK) Consortium | Jacquelyn Wynn, Mairead Hyland, The Lighthouse Lab in Alderley Park and Alex Alderton, Roberto Amato, Jeffrey Barrett, Sonia Goncalves, Ewan Harrison, David K. Jackson, Ian Johnston, Dominic Kwiatkowski, Cordelia Langford, John Sillitoe on behalf of the Wellcome Sanger Institute COVID-19 Surveillance Team |
| EPI_ISL_3107753 | Lighthouse Lab in Milton Keynes | Wellcome Sanger Institute for the COVID-19 Genomics UK (COG-UK) Consortium | The Lighthouse Lab in Milton Keynes and Alex Alderton, Roberto Amato, Jeffrey Barrett, Sonia Goncalves, Ewan Harrison, David K. Jackson, Ian Johnston, Dominic Kwiatkowski, Cordelia Langford, John Sillitoe on behalf of the Wellcome Sanger Institute COVID-19 Surveillance Team |
| EPI_ISL_3107754 | Lighthouse Laboratory Plymouth | Wellcome Sanger Institute for the COVID-19 Genomics UK (COG-UK) Consortium | Lighthouse Laboratory Plymouth and Alex Alderton, Roberto Amato, Jeffrey Barrett, Sonia Goncalves, Ewan Harrison, David K. Jackson, Ian Johnston, Dominic Kwiatkowski, Cordelia Langford, John Sillitoe on behalf of the Wellcome Sanger Institute COVID-19 Surveillance Team |
| EPI_ISL_3107755 | Lighthouse Lab in Glasgow | Wellcome Sanger Institute for the COVID-19 Genomics UK (COG-UK) Consortium | Harper VanSteenhouse, Yumi Kasai, David Gray, Carol Clugston, Anna Dominiczak and Alex Alderton, Roberto Amato, Jeffrey Barrett, Sonia Goncalves, Ewan Harrison, David K. Jackson, Ian Johnston, Dominic Kwiatkowski, Cordelia Langford, John Sillitoe on behalf of the Wellcome Sanger Institute COVID-19 Surveillance Team |
| EPI_ISL_3107756 | Lighthouse Lab in Alderley Park | Wellcome Sanger Institute for the COVID-19 Genomics UK (COG-UK) | Jacquelyn Wynn, Mairead Hyland, The Lighthouse Lab in Alderley Park and Alex Alderton, Roberto Amato, Jeffrey Barrett, Sonia Goncalves, Ewan Harrison, David K. Jackson, Ian Johnston, Dominic Kwiatkowski, Cordelia Langford, John Sillitoe on |

| | | | |
|---|---|---|---|
| | | Consortium | behalf of the Wellcome Sanger Institute COVID-19 Surveillance Team |
| EPI_ISL_3107757 | Lighthouse Laboratory Plymouth | Wellcome Sanger Institute for the COVID-19 Genomics UK (COG-UK) Consortium | Lighthouse Laboratory Plymouth and Alex Alderton, Roberto Amato, Jeffrey Barrett, Sonia Goncalves, Ewan Harrison, David K. Jackson, Ian Johnston, Dominic Kwiatkowski, Cordelia Langford, John Sillitoe on behalf of the Wellcome Sanger Institute COVID-19 Surveillance Team |
| EPI_ISL_3107758 | Lighthouse Lab in Alderley Park | Wellcome Sanger Institute for the COVID-19 Genomics UK (COG-UK) Consortium | Jacquelyn Wynn, Mairead Hyland, The Lighthouse Lab in Alderley Park and Alex Alderton, Roberto Amato, Jeffrey Barrett, Sonia Goncalves, Ewan Harrison, David K. Jackson, Ian Johnston, Dominic Kwiatkowski, Cordelia Langford, John Sillitoe on behalf of the Wellcome Sanger Institute COVID-19 Surveillance Team |
| EPI_ISL_3107759, EPI_ISL_3107760, EPI_ISL_3107761, EPI_ISL_3107762, EPI_ISL_3107763 | Lighthouse Laboratory Plymouth | Wellcome Sanger Institute for the COVID-19 Genomics UK (COG-UK) Consortium | Lighthouse Laboratory Plymouth and Alex Alderton, Roberto Amato, Jeffrey Barrett, Sonia Goncalves, Ewan Harrison, David K. Jackson, Ian Johnston, Dominic Kwiatkowski, Cordelia Langford, John Sillitoe on behalf of the Wellcome Sanger Institute COVID-19 Surveillance Team |
| EPI_ISL_3107764, EPI_ISL_3107765 | Lighthouse Lab in Milton Keynes | Wellcome Sanger Institute for the COVID-19 Genomics UK (COG-UK) Consortium | The Lighthouse Lab in Milton Keynes and Alex Alderton, Roberto Amato, Jeffrey Barrett, Sonia Goncalves, Ewan Harrison, David K. Jackson, Ian Johnston, Dominic Kwiatkowski, Cordelia Langford, John Sillitoe on behalf of the Wellcome Sanger Institute COVID-19 Surveillance Team |
| EPI_ISL_3107766, EPI_ISL_3107767 | Lighthouse Lab in Alderley Park | Wellcome Sanger Institute for the COVID-19 Genomics UK (COG-UK) Consortium | Jacquelyn Wynn, Mairead Hyland, The Lighthouse Lab in Alderley Park and Alex Alderton, Roberto Amato, Jeffrey Barrett, Sonia Goncalves, Ewan Harrison, David K. Jackson, Ian Johnston, Dominic Kwiatkowski, Cordelia Langford, John Sillitoe on behalf of the Wellcome Sanger Institute COVID-19 Surveillance Team |
| EPI_ISL_3107768, EPI_ISL_3107769, EPI_ISL_3107770 | Lighthouse Laboratory Plymouth | Wellcome Sanger Institute for the COVID-19 Genomics UK (COG-UK) Consortium | Lighthouse Laboratory Plymouth and Alex Alderton, Roberto Amato, Jeffrey Barrett, Sonia Goncalves, Ewan Harrison, David K. Jackson, Ian Johnston, Dominic Kwiatkowski, Cordelia Langford, John Sillitoe on behalf of the Wellcome Sanger Institute COVID-19 Surveillance Team |
| EPI_ISL_3107771, EPI_ISL_3107772 | Lighthouse Lab in Alderley Park | Wellcome Sanger Institute for the COVID-19 Genomics UK (COG-UK) Consortium | Jacquelyn Wynn, Mairead Hyland, The Lighthouse Lab in Alderley Park and Alex Alderton, Roberto Amato, Jeffrey Barrett, Sonia Goncalves, Ewan Harrison, David K. Jackson, Ian Johnston, Dominic Kwiatkowski, Cordelia Langford, John Sillitoe on behalf of the Wellcome Sanger Institute COVID-19 Surveillance Team |
| EPI_ISL_3107773 | Lighthouse Lab in Milton Keynes | Wellcome Sanger Institute for the COVID-19 Genomics UK (COG-UK) Consortium | The Lighthouse Lab in Milton Keynes and Alex Alderton, Roberto Amato, Jeffrey Barrett, Sonia Goncalves, Ewan Harrison, David K. Jackson, Ian Johnston, Dominic Kwiatkowski, Cordelia Langford, John Sillitoe on behalf of the Wellcome Sanger Institute COVID-19 Surveillance Team |
| EPI_ISL_3107774 | Lighthouse Lab in Alderley Park | Wellcome Sanger Institute for the COVID-19 Genomics UK (COG-UK) Consortium | Jacquelyn Wynn, Mairead Hyland, The Lighthouse Lab in Alderley Park and Alex Alderton, Roberto Amato, Jeffrey Barrett, Sonia Goncalves, Ewan Harrison, David K. Jackson, Ian Johnston, Dominic Kwiatkowski, Cordelia Langford, John Sillitoe on behalf of the Wellcome Sanger Institute COVID-19 Surveillance Team |
| EPI_ISL_3107775 | Lighthouse Laboratory Plymouth | Wellcome Sanger Institute for the COVID-19 Genomics UK (COG-UK) Consortium | Lighthouse Laboratory Plymouth and Alex Alderton, Roberto Amato, Jeffrey Barrett, Sonia Goncalves, Ewan Harrison, David K. Jackson, Ian Johnston, Dominic Kwiatkowski, Cordelia Langford, John Sillitoe on behalf of the Wellcome Sanger Institute COVID-19 Surveillance Team |
| EPI_ISL_3107776 | Lighthouse Lab in Glasgow | Wellcome Sanger Institute for the COVID-19 Genomics UK (COG-UK) Consortium | Harper VanSteenhouse, Yumi Kasai, David Gray, Carol Clugston, Anna Dominiczak and Alex Alderton, Roberto Amato, Jeffrey Barrett, Sonia Goncalves, Ewan Harrison, David K. Jackson, Ian Johnston, Dominic Kwiatkowski, Cordelia Langford, John Sillitoe on behalf of the Wellcome Sanger Institute COVID-19 Surveillance Team |
| EPI_ISL_3107777 | Lighthouse Lab in Milton Keynes | Wellcome Sanger Institute for the COVID-19 Genomics UK (COG-UK) Consortium | The Lighthouse Lab in Milton Keynes and Alex Alderton, Roberto Amato, Jeffrey Barrett, Sonia Goncalves, Ewan Harrison, David K. Jackson, Ian Johnston, Dominic Kwiatkowski, Cordelia Langford, John Sillitoe on behalf of the Wellcome Sanger Institute COVID-19 Surveillance Team |
| EPI_ISL_3107778 | Lighthouse Laboratory Plymouth | Wellcome Sanger Institute for the COVID-19 Genomics UK (COG-UK) Consortium | Lighthouse Laboratory Plymouth and Alex Alderton, Roberto Amato, Jeffrey Barrett, Sonia Goncalves, Ewan Harrison, David K. Jackson, Ian Johnston, Dominic Kwiatkowski, Cordelia Langford, John Sillitoe on behalf of the Wellcome Sanger Institute COVID-19 Surveillance Team |
| EPI_ISL_3107779 | Lighthouse Lab in Alderley Park | Wellcome Sanger Institute for the COVID-19 Genomics UK (COG-UK) Consortium | Jacquelyn Wynn, Mairead Hyland, The Lighthouse Lab in Alderley Park and Alex Alderton, Roberto Amato, Jeffrey Barrett, Sonia Goncalves, Ewan Harrison, David K. Jackson, Ian Johnston, Dominic Kwiatkowski, Cordelia Langford, John Sillitoe on behalf of the Wellcome Sanger Institute COVID-19 Surveillance Team |
| EPI_ISL_3107780 | Lighthouse Lab in Glasgow | Wellcome Sanger Institute for the COVID-19 Genomics UK (COG-UK) Consortium | Harper VanSteenhouse, Yumi Kasai, David Gray, Carol Clugston, Anna Dominiczak and Alex Alderton, Roberto Amato, Jeffrey Barrett, Sonia Goncalves, Ewan Harrison, David K. Jackson, Ian Johnston, Dominic Kwiatkowski, Cordelia Langford, John Sillitoe on behalf of the Wellcome Sanger Institute COVID-19 Surveillance Team |
| EPI_ISL_3107781 | Lighthouse Lab in Milton Keynes | Wellcome Sanger Institute for the COVID-19 Genomics UK (COG-UK) Consortium | The Lighthouse Lab in Milton Keynes and Alex Alderton, Roberto Amato, Jeffrey Barrett, Sonia Goncalves, Ewan Harrison, David K. Jackson, Ian Johnston, Dominic Kwiatkowski, Cordelia Langford, John Sillitoe on behalf of the Wellcome Sanger Institute COVID-19 Surveillance Team |
| EPI_ISL_3107782 | Lighthouse Lab in Alderley Park | Wellcome Sanger Institute for the COVID-19 Genomics UK (COG-UK) Consortium | Jacquelyn Wynn, Mairead Hyland, The Lighthouse Lab in Alderley Park and Alex Alderton, Roberto Amato, Jeffrey Barrett, Sonia Goncalves, Ewan Harrison, David K. Jackson, Ian Johnston, Dominic Kwiatkowski, Cordelia Langford, John Sillitoe on behalf of the Wellcome Sanger Institute COVID-19 Surveillance Team |
| EPI_ISL_3107783 | Lighthouse Laboratory Plymouth | Wellcome Sanger Institute for the COVID-19 Genomics UK (COG-UK) Consortium | Lighthouse Laboratory Plymouth and Alex Alderton, Roberto Amato, Jeffrey Barrett, Sonia Goncalves, Ewan Harrison, David K. Jackson, Ian Johnston, Dominic Kwiatkowski, Cordelia Langford, John Sillitoe on behalf of the Wellcome Sanger Institute COVID-19 Surveillance Team |
| EPI_ISL_3107784 | Lighthouse Lab in Glasgow | Wellcome Sanger Institute for the COVID-19 Genomics UK (COG-UK) Consortium | Harper VanSteenhouse, Yumi Kasai, David Gray, Carol Clugston, Anna Dominiczak and Alex Alderton, Roberto Amato, Jeffrey Barrett, Sonia Goncalves, Ewan Harrison, David K. Jackson, Ian Johnston, Dominic Kwiatkowski, Cordelia Langford, John Sillitoe on behalf of the Wellcome Sanger Institute COVID-19 Surveillance Team |
| EPI_ISL_3107785 | Lighthouse Lab in Alderley Park | Wellcome Sanger Institute for the COVID-19 Genomics UK (COG-UK) Consortium | Jacquelyn Wynn, Mairead Hyland, The Lighthouse Lab in Alderley Park and Alex Alderton, Roberto Amato, Jeffrey Barrett, Sonia Goncalves, Ewan Harrison, David K. Jackson, Ian Johnston, Dominic Kwiatkowski, Cordelia Langford, John Sillitoe on behalf of the Wellcome Sanger Institute COVID-19 Surveillance Team |
| EPI_ISL_3107786 | Lighthouse Laboratory Plymouth | Wellcome Sanger Institute for the COVID-19 Genomics UK (COG-UK) Consortium | Lighthouse Laboratory Plymouth and Alex Alderton, Roberto Amato, Jeffrey Barrett, Sonia Goncalves, Ewan Harrison, David K. Jackson, Ian Johnston, Dominic Kwiatkowski, Cordelia Langford, John Sillitoe on behalf of the Wellcome Sanger Institute COVID-19 Surveillance Team |
| EPI_ISL_3107787 | Lighthouse Lab in Milton Keynes | Wellcome Sanger Institute for the COVID-19 Genomics UK (COG-UK) Consortium | The Lighthouse Lab in Milton Keynes and Alex Alderton, Roberto Amato, Jeffrey Barrett, Sonia Goncalves, Ewan Harrison, David K. Jackson, Ian Johnston, Dominic Kwiatkowski, Cordelia Langford, John Sillitoe on behalf of the Wellcome Sanger Institute COVID-19 Surveillance Team |
| EPI_ISL_3107788, EPI_ISL_3107789, EPI_ISL_3107790 | Lighthouse Laboratory Plymouth | Wellcome Sanger Institute for the COVID-19 Genomics UK (COG-UK) Consortium | Lighthouse Laboratory Plymouth and Alex Alderton, Roberto Amato, Jeffrey Barrett, Sonia Goncalves, Ewan Harrison, David K. Jackson, Ian Johnston, Dominic Kwiatkowski, Cordelia Langford, John Sillitoe on behalf of the Wellcome Sanger Institute COVID-19 Surveillance Team |
| EPI_ISL_3107791 | Lighthouse Lab in Glasgow | Wellcome Sanger Institute for the COVID-19 Genomics UK (COG-UK) Consortium | Harper VanSteenhouse, Yumi Kasai, David Gray, Carol Clugston, Anna Dominiczak and Alex Alderton, Roberto Amato, Jeffrey Barrett, Sonia Goncalves, Ewan Harrison, David K. Jackson, Ian Johnston, Dominic Kwiatkowski, Cordelia Langford, John Sillitoe on behalf of the Wellcome Sanger Institute COVID-19 Surveillance Team |
| EPI_ISL_3107792 | Lighthouse Lab in Alderley Park | Wellcome Sanger Institute for the COVID-19 Genomics UK (COG-UK) Consortium | Jacquelyn Wynn, Mairead Hyland, The Lighthouse Lab in Alderley Park and Alex Alderton, Roberto Amato, Jeffrey Barrett, Sonia Goncalves, Ewan Harrison, David K. Jackson, Ian Johnston, Dominic Kwiatkowski, Cordelia Langford, John Sillitoe on behalf of the Wellcome Sanger Institute COVID-19 Surveillance Team |
| EPI_ISL_3107793, EPI_ISL_3107794, EPI_ISL_3107795, EPI_ISL_3107796 | Lighthouse Laboratory Plymouth | Wellcome Sanger Institute for the COVID-19 Genomics UK (COG-UK) Consortium | Lighthouse Laboratory Plymouth and Alex Alderton, Roberto Amato, Jeffrey Barrett, Sonia Goncalves, Ewan Harrison, David K. Jackson, Ian Johnston, Dominic Kwiatkowski, Cordelia Langford, John Sillitoe on behalf of the Wellcome Sanger Institute COVID-19 Surveillance Team |
| EPI_ISL_3107797, EPI_ISL_3107798 | Lighthouse Lab in Alderley Park | Wellcome Sanger Institute for the COVID-19 Genomics UK (COG-UK) Consortium | Jacquelyn Wynn, Mairead Hyland, The Lighthouse Lab in Alderley Park and Alex Alderton, Roberto Amato, Jeffrey Barrett, Sonia Goncalves, Ewan Harrison, David K. Jackson, Ian Johnston, Dominic Kwiatkowski, Cordelia Langford, John Sillitoe on behalf of the Wellcome Sanger Institute COVID-19 Surveillance Team |
| EPI_ISL_3107799, EPI_ISL_3107800 | Lighthouse Lab in Milton Keynes | Wellcome Sanger Institute for the COVID-19 Genomics UK (COG-UK) Consortium | The Lighthouse Lab in Milton Keynes and Alex Alderton, Roberto Amato, Jeffrey Barrett, Sonia Goncalves, Ewan Harrison, David K. Jackson, Ian Johnston, Dominic Kwiatkowski, Cordelia Langford, John Sillitoe on behalf of the Wellcome Sanger Institute COVID-19 Surveillance Team |
| EPI_ISL_3107801 | Lighthouse Laboratory Plymouth | Wellcome Sanger Institute for the COVID-19 Genomics UK (COG-UK) Consortium | Lighthouse Laboratory Plymouth and Alex Alderton, Roberto Amato, Jeffrey Barrett, Sonia Goncalves, Ewan Harrison, David K. Jackson, Ian Johnston, Dominic Kwiatkowski, Cordelia Langford, John Sillitoe on behalf of the Wellcome Sanger Institute COVID-19 Surveillance Team |
| EPI_ISL_3107802, EPI_ISL_3107803 | Lighthouse Lab in Milton Keynes | Wellcome Sanger Institute for the COVID-19 Genomics UK (COG-UK) Consortium | The Lighthouse Lab in Milton Keynes and Alex Alderton, Roberto Amato, Jeffrey Barrett, Sonia Goncalves, Ewan Harrison, David K. Jackson, Ian Johnston, Dominic Kwiatkowski, Cordelia Langford, John Sillitoe on behalf of the Wellcome Sanger Institute COVID-19 Surveillance Team |
| EPI_ISL_3107804 | Lighthouse Lab in Alderley Park | Wellcome Sanger Institute for the COVID-19 Genomics UK (COG-UK) Consortium | Jacquelyn Wynn, Mairead Hyland, The Lighthouse Lab in Alderley Park and Alex Alderton, Roberto Amato, Jeffrey Barrett, Sonia Goncalves, Ewan Harrison, David K. Jackson, Ian Johnston, Dominic Kwiatkowski, Cordelia Langford, John Sillitoe on behalf of the Wellcome Sanger Institute COVID-19 Surveillance Team |
| EPI_ISL_3107805 | Lighthouse Laboratory Plymouth | Wellcome Sanger Institute for the COVID-19 Genomics UK (COG-UK) Consortium | Lighthouse Laboratory Plymouth and Alex Alderton, Roberto Amato, Jeffrey Barrett, Sonia Goncalves, Ewan Harrison, David K. Jackson, Ian Johnston, Dominic Kwiatkowski, Cordelia Langford, John Sillitoe on behalf of the Wellcome Sanger Institute COVID-19 Surveillance Team |
| EPI_ISL_3107806 | Lighthouse Lab in Glasgow | Wellcome Sanger Institute for the COVID-19 Genomics UK (COG-UK) Consortium | Harper VanSteenhouse, Yumi Kasai, David Gray, Carol Clugston, Anna Dominiczak and Alex Alderton, Roberto Amato, Jeffrey Barrett, Sonia Goncalves, Ewan Harrison, David K. Jackson, Ian Johnston, Dominic Kwiatkowski, Cordelia Langford, John Sillitoe on behalf of the Wellcome Sanger Institute COVID-19 Surveillance Team |
| EPI_ISL_3107807 | Lighthouse Laboratory Plymouth | Wellcome Sanger Institute for the COVID-19 Genomics UK (COG-UK) Consortium | Lighthouse Laboratory Plymouth and Alex Alderton, Roberto Amato, Jeffrey Barrett, Sonia Goncalves, Ewan Harrison, David K. Jackson, Ian Johnston, Dominic Kwiatkowski, Cordelia Langford, John Sillitoe on behalf of the Wellcome Sanger Institute COVID-19 Surveillance Team |
| EPI_ISL_3107808 | Lighthouse Lab in Milton Keynes | Wellcome Sanger Institute for the COVID-19 Genomics UK (COG-UK) Consortium | The Lighthouse Lab in Milton Keynes and Alex Alderton, Roberto Amato, Jeffrey Barrett, Sonia Goncalves, Ewan Harrison, David K. Jackson, Ian Johnston, Dominic Kwiatkowski, Cordelia Langford, John Sillitoe on behalf of the Wellcome Sanger Institute COVID-19 Surveillance Team |
| EPI_ISL_3107809 | Lighthouse Lab in Alderley Park | Wellcome Sanger Institute for the COVID-19 Genomics UK (COG-UK) Consortium | Jacquelyn Wynn, Mairead Hyland, The Lighthouse Lab in Alderley Park and Alex Alderton, Roberto Amato, Jeffrey Barrett, Sonia Goncalves, Ewan Harrison, David K. Jackson, Ian Johnston, Dominic Kwiatkowski, Cordelia Langford, John Sillitoe on behalf of the Wellcome Sanger Institute COVID-19 Surveillance Team |
| EPI_ISL_3107810, EPI_ISL_3107811 | Lighthouse Laboratory Plymouth | Wellcome Sanger Institute for the COVID-19 Genomics UK (COG-UK) Consortium | Lighthouse Laboratory Plymouth and Alex Alderton, Roberto Amato, Jeffrey Barrett, Sonia Goncalves, Ewan Harrison, David K. Jackson, Ian Johnston, Dominic Kwiatkowski, Cordelia Langford, John Sillitoe on behalf of the Wellcome Sanger Institute COVID-19 Surveillance Team |
| EPI_ISL_3107812 | Lighthouse Lab in Alderley Park | Wellcome Sanger Institute for the COVID-19 Genomics UK (COG-UK) Consortium | Jacquelyn Wynn, Mairead Hyland, The Lighthouse Lab in Alderley Park and Alex Alderton, Roberto Amato, Jeffrey Barrett, Sonia Goncalves, Ewan Harrison, David K. Jackson, Ian Johnston, Dominic Kwiatkowski, Cordelia Langford, John Sillitoe on behalf of the Wellcome Sanger Institute COVID-19 Surveillance Team |
| EPI_ISL_3107813 | Lighthouse Lab in Milton Keynes | Wellcome Sanger Institute for the COVID-19 Genomics UK (COG-UK) Consortium | The Lighthouse Lab in Milton Keynes and Alex Alderton, Roberto Amato, Jeffrey Barrett, Sonia Goncalves, Ewan Harrison, David K. Jackson, Ian Johnston, Dominic Kwiatkowski, Cordelia Langford, John Sillitoe on behalf of the Wellcome Sanger Institute COVID-19 Surveillance Team |
| EPI_ISL_3107814 | Lighthouse Laboratory Plymouth | Wellcome Sanger Institute for the COVID-19 Genomics UK (COG-UK) Consortium | Lighthouse Laboratory Plymouth and Alex Alderton, Roberto Amato, Jeffrey Barrett, Sonia Goncalves, Ewan Harrison, David K. Jackson, Ian Johnston, Dominic Kwiatkowski, Cordelia Langford, John Sillitoe on behalf of the Wellcome Sanger Institute COVID-19 Surveillance Team |
| EPI_ISL_3107815 | Lighthouse Lab in Milton Keynes | Wellcome Sanger Institute for the COVID-19 Genomics UK (COG-UK) Consortium | The Lighthouse Lab in Milton Keynes and Alex Alderton, Roberto Amato, Jeffrey Barrett, Sonia Goncalves, Ewan Harrison, David K. Jackson, Ian Johnston, Dominic Kwiatkowski, Cordelia Langford, John Sillitoe on behalf of the Wellcome Sanger Institute COVID-19 Surveillance Team |
| EPI_ISL_3107816, | Lighthouse Lab in Alderley Park | Wellcome Sanger Institute for the COVID-19 Genomics UK (COG-UK) | Jacquelyn Wynn, Mairead Hyland, The Lighthouse Lab in Alderley Park and Alex Alderton, Roberto Amato, Jeffrey Barrett, Sonia Goncalves, Ewan Harrison, David K. Jackson, Ian Johnston, Dominic Kwiatkowski, Cordelia Langford, John Sillitoe on |

| | | | |
|---|---|---|---|
| EPI_ISL_3107817 | | Consortium | behalf of the Wellcome Sanger Institute COVID-19 Surveillance Team |
| EPI_ISL_3107818, EPI_ISL_3107819 | Lighthouse Lab in Milton Keynes | Wellcome Sanger Institute for the COVID-19 Genomics UK (COG-UK) Consortium | The Lighthouse Lab in Milton Keynes and Alex Alderton, Roberto Amato, Jeffrey Barrett, Sonia Goncalves, Ewan Harrison, David K. Jackson, Ian Johnston, Dominic Kwiatkowski, Cordelia Langford, John Sillitoe on behalf of the Wellcome Sanger Institute COVID-19 Surveillance Team |
| EPI_ISL_3107820 | Lighthouse Lab in Alderley Park | Wellcome Sanger Institute for the COVID-19 Genomics UK (COG-UK) Consortium | Jacquelyn Wynn, Mairead Hyland, The Lighthouse Lab in Alderley Park and Alex Alderton, Roberto Amato, Jeffrey Barrett, Sonia Goncalves, Ewan Harrison, David K. Jackson, Ian Johnston, Dominic Kwiatkowski, Cordelia Langford, John Sillitoe on behalf of the Wellcome Sanger Institute COVID-19 Surveillance Team |
| EPI_ISL_3107821 | Lighthouse Laboratory Plymouth | Wellcome Sanger Institute for the COVID-19 Genomics UK (COG-UK) Consortium | Lighthouse Laboratory Plymouth and Alex Alderton, Roberto Amato, Jeffrey Barrett, Sonia Goncalves, Ewan Harrison, David K. Jackson, Ian Johnston, Dominic Kwiatkowski, Cordelia Langford, John Sillitoe on behalf of the Wellcome Sanger Institute COVID-19 Surveillance Team |
| EPI_ISL_3107822 | Lighthouse Lab in Milton Keynes | Wellcome Sanger Institute for the COVID-19 Genomics UK (COG-UK) Consortium | The Lighthouse Lab in Milton Keynes and Alex Alderton, Roberto Amato, Jeffrey Barrett, Sonia Goncalves, Ewan Harrison, David K. Jackson, Ian Johnston, Dominic Kwiatkowski, Cordelia Langford, John Sillitoe on behalf of the Wellcome Sanger Institute COVID-19 Surveillance Team |
| EPI_ISL_3107823 | Lighthouse Lab in Alderley Park | Wellcome Sanger Institute for the COVID-19 Genomics UK (COG-UK) Consortium | Jacquelyn Wynn, Mairead Hyland, The Lighthouse Lab in Alderley Park and Alex Alderton, Roberto Amato, Jeffrey Barrett, Sonia Goncalves, Ewan Harrison, David K. Jackson, Ian Johnston, Dominic Kwiatkowski, Cordelia Langford, John Sillitoe on behalf of the Wellcome Sanger Institute COVID-19 Surveillance Team |
| EPI_ISL_3107824 | Lighthouse Lab in Milton Keynes | Wellcome Sanger Institute for the COVID-19 Genomics UK (COG-UK) Consortium | The Lighthouse Lab in Milton Keynes and Alex Alderton, Roberto Amato, Jeffrey Barrett, Sonia Goncalves, Ewan Harrison, David K. Jackson, Ian Johnston, Dominic Kwiatkowski, Cordelia Langford, John Sillitoe on behalf of the Wellcome Sanger Institute COVID-19 Surveillance Team |
| EPI_ISL_3107825, EPI_ISL_3107826, EPI_ISL_3107827, EPI_ISL_3107828 | Lighthouse Laboratory Plymouth | Wellcome Sanger Institute for the COVID-19 Genomics UK (COG-UK) Consortium | Lighthouse Laboratory Plymouth and Alex Alderton, Roberto Amato, Jeffrey Barrett, Sonia Goncalves, Ewan Harrison, David K. Jackson, Ian Johnston, Dominic Kwiatkowski, Cordelia Langford, John Sillitoe on behalf of the Wellcome Sanger Institute COVID-19 Surveillance Team |
| EPI_ISL_3107829 | Lighthouse Lab in Alderley Park | Wellcome Sanger Institute for the COVID-19 Genomics UK (COG-UK) Consortium | Jacquelyn Wynn, Mairead Hyland, The Lighthouse Lab in Alderley Park and Alex Alderton, Roberto Amato, Jeffrey Barrett, Sonia Goncalves, Ewan Harrison, David K. Jackson, Ian Johnston, Dominic Kwiatkowski, Cordelia Langford, John Sillitoe on behalf of the Wellcome Sanger Institute COVID-19 Surveillance Team |
| EPI_ISL_3107830, EPI_ISL_3107831 | Lighthouse Laboratory Plymouth | Wellcome Sanger Institute for the COVID-19 Genomics UK (COG-UK) Consortium | Lighthouse Laboratory Plymouth and Alex Alderton, Roberto Amato, Jeffrey Barrett, Sonia Goncalves, Ewan Harrison, David K. Jackson, Ian Johnston, Dominic Kwiatkowski, Cordelia Langford, John Sillitoe on behalf of the Wellcome Sanger Institute COVID-19 Surveillance Team |
| EPI_ISL_3107832, EPI_ISL_3107833 | Lighthouse Lab in Alderley Park | Wellcome Sanger Institute for the COVID-19 Genomics UK (COG-UK) Consortium | Jacquelyn Wynn, Mairead Hyland, The Lighthouse Lab in Alderley Park and Alex Alderton, Roberto Amato, Jeffrey Barrett, Sonia Goncalves, Ewan Harrison, David K. Jackson, Ian Johnston, Dominic Kwiatkowski, Cordelia Langford, John Sillitoe on behalf of the Wellcome Sanger Institute COVID-19 Surveillance Team |
| EPI_ISL_3107834 | Lighthouse Laboratory Plymouth | Wellcome Sanger Institute for the COVID-19 Genomics UK (COG-UK) Consortium | Lighthouse Laboratory Plymouth and Alex Alderton, Roberto Amato, Jeffrey Barrett, Sonia Goncalves, Ewan Harrison, David K. Jackson, Ian Johnston, Dominic Kwiatkowski, Cordelia Langford, John Sillitoe on behalf of the Wellcome Sanger Institute COVID-19 Surveillance Team |
| EPI_ISL_3107835, EPI_ISL_3107836 | Lighthouse Lab in Milton Keynes | Wellcome Sanger Institute for the COVID-19 Genomics UK (COG-UK) Consortium | The Lighthouse Lab in Milton Keynes and Alex Alderton, Roberto Amato, Jeffrey Barrett, Sonia Goncalves, Ewan Harrison, David K. Jackson, Ian Johnston, Dominic Kwiatkowski, Cordelia Langford, John Sillitoe on behalf of the Wellcome Sanger Institute COVID-19 Surveillance Team |
| EPI_ISL_3107837, EPI_ISL_3107838 | Lighthouse Lab in Alderley Park | Wellcome Sanger Institute for the COVID-19 Genomics UK (COG-UK) Consortium | Jacquelyn Wynn, Mairead Hyland, The Lighthouse Lab in Alderley Park and Alex Alderton, Roberto Amato, Jeffrey Barrett, Sonia Goncalves, Ewan Harrison, David K. Jackson, Ian Johnston, Dominic Kwiatkowski, Cordelia Langford, John Sillitoe on behalf of the Wellcome Sanger Institute COVID-19 Surveillance Team |
| EPI_ISL_3107839 | Lighthouse Lab in Milton Keynes | Wellcome Sanger Institute for the COVID-19 Genomics UK (COG-UK) Consortium | The Lighthouse Lab in Milton Keynes and Alex Alderton, Roberto Amato, Jeffrey Barrett, Sonia Goncalves, Ewan Harrison, David K. Jackson, Ian Johnston, Dominic Kwiatkowski, Cordelia Langford, John Sillitoe on behalf of the Wellcome Sanger Institute COVID-19 Surveillance Team |
| EPI_ISL_3107840 | Lighthouse Lab in Glasgow | Wellcome Sanger Institute for the COVID-19 Genomics UK (COG-UK) Consortium | Harper VanSteenhouse, Yumi Kasai, David Gray, Carol Clugston, Anna Dominiczak and Alex Alderton, Roberto Amato, Jeffrey Barrett, Sonia Goncalves, Ewan Harrison, David K. Jackson, Ian Johnston, Dominic Kwiatkowski, Cordelia Langford, John Sillitoe on behalf of the Wellcome Sanger Institute COVID-19 Surveillance Team |
| EPI_ISL_3107841 | Lighthouse Lab in Milton Keynes | Wellcome Sanger Institute for the COVID-19 Genomics UK (COG-UK) Consortium | The Lighthouse Lab in Milton Keynes and Alex Alderton, Roberto Amato, Jeffrey Barrett, Sonia Goncalves, Ewan Harrison, David K. Jackson, Ian Johnston, Dominic Kwiatkowski, Cordelia Langford, John Sillitoe on behalf of the Wellcome Sanger Institute COVID-19 Surveillance Team |
| EPI_ISL_3107842 | Lighthouse Lab in Alderley Park | Wellcome Sanger Institute for the COVID-19 Genomics UK (COG-UK) Consortium | Jacquelyn Wynn, Mairead Hyland, The Lighthouse Lab in Alderley Park and Alex Alderton, Roberto Amato, Jeffrey Barrett, Sonia Goncalves, Ewan Harrison, David K. Jackson, Ian Johnston, Dominic Kwiatkowski, Cordelia Langford, John Sillitoe on behalf of the Wellcome Sanger Institute COVID-19 Surveillance Team |
| EPI_ISL_3107843, EPI_ISL_3107844 | Lighthouse Lab in Glasgow | Wellcome Sanger Institute for the COVID-19 Genomics UK (COG-UK) Consortium | Harper VanSteenhouse, Yumi Kasai, David Gray, Carol Clugston, Anna Dominiczak and Alex Alderton, Roberto Amato, Jeffrey Barrett, Sonia Goncalves, Ewan Harrison, David K. Jackson, Ian Johnston, Dominic Kwiatkowski, Cordelia Langford, John Sillitoe on behalf of the Wellcome Sanger Institute COVID-19 Surveillance Team |
| EPI_ISL_3107845 | Lighthouse Laboratory Plymouth | Wellcome Sanger Institute for the COVID-19 Genomics UK (COG-UK) Consortium | Lighthouse Laboratory Plymouth and Alex Alderton, Roberto Amato, Jeffrey Barrett, Sonia Goncalves, Ewan Harrison, David K. Jackson, Ian Johnston, Dominic Kwiatkowski, Cordelia Langford, John Sillitoe on behalf of the Wellcome Sanger Institute COVID-19 Surveillance Team |
| EPI_ISL_3107846 | Lighthouse Lab in Alderley Park | Wellcome Sanger Institute for the COVID-19 Genomics UK (COG-UK) Consortium | Jacquelyn Wynn, Mairead Hyland, The Lighthouse Lab in Alderley Park and Alex Alderton, Roberto Amato, Jeffrey Barrett, Sonia Goncalves, Ewan Harrison, David K. Jackson, Ian Johnston, Dominic Kwiatkowski, Cordelia Langford, John Sillitoe on behalf of the Wellcome Sanger Institute COVID-19 Surveillance Team |
| EPI_ISL_3107847 | Lighthouse Lab in Milton Keynes | Wellcome Sanger Institute for the COVID-19 Genomics UK (COG-UK) Consortium | The Lighthouse Lab in Milton Keynes and Alex Alderton, Roberto Amato, Jeffrey Barrett, Sonia Goncalves, Ewan Harrison, David K. Jackson, Ian Johnston, Dominic Kwiatkowski, Cordelia Langford, John Sillitoe on behalf of the Wellcome Sanger Institute COVID-19 Surveillance Team |
| EPI_ISL_3107848 | Lighthouse Lab in Glasgow | Wellcome Sanger Institute for the COVID-19 Genomics UK (COG-UK) Consortium | Harper VanSteenhouse, Yumi Kasai, David Gray, Carol Clugston, Anna Dominiczak and Alex Alderton, Roberto Amato, Jeffrey Barrett, Sonia Goncalves, Ewan Harrison, David K. Jackson, Ian Johnston, Dominic Kwiatkowski, Cordelia Langford, John Sillitoe on behalf of the Wellcome Sanger Institute COVID-19 Surveillance Team |
| EPI_ISL_3107849 | Lighthouse Lab in Alderley Park | Wellcome Sanger Institute for the COVID-19 Genomics UK (COG-UK) Consortium | Jacquelyn Wynn, Mairead Hyland, The Lighthouse Lab in Alderley Park and Alex Alderton, Roberto Amato, Jeffrey Barrett, Sonia Goncalves, Ewan Harrison, David K. Jackson, Ian Johnston, Dominic Kwiatkowski, Cordelia Langford, John Sillitoe on behalf of the Wellcome Sanger Institute COVID-19 Surveillance Team |
| EPI_ISL_3107850 | Lighthouse Laboratory Plymouth | Wellcome Sanger Institute for the COVID-19 Genomics UK (COG-UK) Consortium | Lighthouse Laboratory Plymouth and Alex Alderton, Roberto Amato, Jeffrey Barrett, Sonia Goncalves, Ewan Harrison, David K. Jackson, Ian Johnston, Dominic Kwiatkowski, Cordelia Langford, John Sillitoe on behalf of the Wellcome Sanger Institute COVID-19 Surveillance Team |
| EPI_ISL_3107851 | Lighthouse Lab in Alderley Park | Wellcome Sanger Institute for the COVID-19 Genomics UK (COG-UK) Consortium | Jacquelyn Wynn, Mairead Hyland, The Lighthouse Lab in Alderley Park and Alex Alderton, Roberto Amato, Jeffrey Barrett, Sonia Goncalves, Ewan Harrison, David K. Jackson, Ian Johnston, Dominic Kwiatkowski, Cordelia Langford, John Sillitoe on behalf of the Wellcome Sanger Institute COVID-19 Surveillance Team |
| EPI_ISL_3107852 | Lighthouse Lab in Glasgow | Wellcome Sanger Institute for the COVID-19 Genomics UK (COG-UK) Consortium | Harper VanSteenhouse, Yumi Kasai, David Gray, Carol Clugston, Anna Dominiczak and Alex Alderton, Roberto Amato, Jeffrey Barrett, Sonia Goncalves, Ewan Harrison, David K. Jackson, Ian Johnston, Dominic Kwiatkowski, Cordelia Langford, John Sillitoe on behalf of the Wellcome Sanger Institute COVID-19 Surveillance Team |
| EPI_ISL_3107853, EPI_ISL_3107854 | Lighthouse Lab in Alderley Park | Wellcome Sanger Institute for the COVID-19 Genomics UK (COG-UK) Consortium | Jacquelyn Wynn, Mairead Hyland, The Lighthouse Lab in Alderley Park and Alex Alderton, Roberto Amato, Jeffrey Barrett, Sonia Goncalves, Ewan Harrison, David K. Jackson, Ian Johnston, Dominic Kwiatkowski, Cordelia Langford, John Sillitoe on behalf of the Wellcome Sanger Institute COVID-19 Surveillance Team |
| EPI_ISL_3107855 | Lighthouse Lab in Milton Keynes | Wellcome Sanger Institute for the COVID-19 Genomics UK (COG-UK) Consortium | The Lighthouse Lab in Milton Keynes and Alex Alderton, Roberto Amato, Jeffrey Barrett, Sonia Goncalves, Ewan Harrison, David K. Jackson, Ian Johnston, Dominic Kwiatkowski, Cordelia Langford, John Sillitoe on behalf of the Wellcome Sanger Institute COVID-19 Surveillance Team |
| EPI_ISL_3107856, EPI_ISL_3107857 | Lighthouse Lab in Alderley Park | Wellcome Sanger Institute for the COVID-19 Genomics UK (COG-UK) Consortium | Jacquelyn Wynn, Mairead Hyland, The Lighthouse Lab in Alderley Park and Alex Alderton, Roberto Amato, Jeffrey Barrett, Sonia Goncalves, Ewan Harrison, David K. Jackson, Ian Johnston, Dominic Kwiatkowski, Cordelia Langford, John Sillitoe on behalf of the Wellcome Sanger Institute COVID-19 Surveillance Team |
| EPI_ISL_3107858 | Lighthouse Laboratory Plymouth | Wellcome Sanger Institute for the COVID-19 Genomics UK (COG-UK) Consortium | Lighthouse Laboratory Plymouth and Alex Alderton, Roberto Amato, Jeffrey Barrett, Sonia Goncalves, Ewan Harrison, David K. Jackson, Ian Johnston, Dominic Kwiatkowski, Cordelia Langford, John Sillitoe on behalf of the Wellcome Sanger Institute COVID-19 Surveillance Team |
| EPI_ISL_3107859, EPI_ISL_3107860 | Lighthouse Lab in Alderley Park | Wellcome Sanger Institute for the COVID-19 Genomics UK (COG-UK) Consortium | Jacquelyn Wynn, Mairead Hyland, The Lighthouse Lab in Alderley Park and Alex Alderton, Roberto Amato, Jeffrey Barrett, Sonia Goncalves, Ewan Harrison, David K. Jackson, Ian Johnston, Dominic Kwiatkowski, Cordelia Langford, John Sillitoe on behalf of the Wellcome Sanger Institute COVID-19 Surveillance Team |
| EPI_ISL_3107861 | Lighthouse Lab in Glasgow | Wellcome Sanger Institute for the COVID-19 Genomics UK (COG-UK) Consortium | Harper VanSteenhouse, Yumi Kasai, David Gray, Carol Clugston, Anna Dominiczak and Alex Alderton, Roberto Amato, Jeffrey Barrett, Sonia Goncalves, Ewan Harrison, David K. Jackson, Ian Johnston, Dominic Kwiatkowski, Cordelia Langford, John Sillitoe on behalf of the Wellcome Sanger Institute COVID-19 Surveillance Team |
| EPI_ISL_3107862 | Lighthouse Lab in Alderley Park | Wellcome Sanger Institute for the COVID-19 Genomics UK (COG-UK) Consortium | Jacquelyn Wynn, Mairead Hyland, The Lighthouse Lab in Alderley Park and Alex Alderton, Roberto Amato, Jeffrey Barrett, Sonia Goncalves, Ewan Harrison, David K. Jackson, Ian Johnston, Dominic Kwiatkowski, Cordelia Langford, John Sillitoe on behalf of the Wellcome Sanger Institute COVID-19 Surveillance Team |
| EPI_ISL_3107863 | Lighthouse Lab in Glasgow | Wellcome Sanger Institute for the COVID-19 Genomics UK (COG-UK) Consortium | Harper VanSteenhouse, Yumi Kasai, David Gray, Carol Clugston, Anna Dominiczak and Alex Alderton, Roberto Amato, Jeffrey Barrett, Sonia Goncalves, Ewan Harrison, David K. Jackson, Ian Johnston, Dominic Kwiatkowski, Cordelia Langford, John Sillitoe on behalf of the Wellcome Sanger Institute COVID-19 Surveillance Team |
| EPI_ISL_3107864 | Lighthouse Lab in Alderley Park | Wellcome Sanger Institute for the COVID-19 Genomics UK (COG-UK) Consortium | Jacquelyn Wynn, Mairead Hyland, The Lighthouse Lab in Alderley Park and Alex Alderton, Roberto Amato, Jeffrey Barrett, Sonia Goncalves, Ewan Harrison, David K. Jackson, Ian Johnston, Dominic Kwiatkowski, Cordelia Langford, John Sillitoe on behalf of the Wellcome Sanger Institute COVID-19 Surveillance Team |
| EPI_ISL_3107865 | Lighthouse Lab in Milton Keynes | Wellcome Sanger Institute for the COVID-19 Genomics UK (COG-UK) Consortium | The Lighthouse Lab in Milton Keynes and Alex Alderton, Roberto Amato, Jeffrey Barrett, Sonia Goncalves, Ewan Harrison, David K. Jackson, Ian Johnston, Dominic Kwiatkowski, Cordelia Langford, John Sillitoe on behalf of the Wellcome Sanger Institute COVID-19 Surveillance Team |
| EPI_ISL_3107866 | Lighthouse Laboratory Plymouth | Wellcome Sanger Institute for the COVID-19 Genomics UK (COG-UK) Consortium | Lighthouse Laboratory Plymouth and Alex Alderton, Roberto Amato, Jeffrey Barrett, Sonia Goncalves, Ewan Harrison, David K. Jackson, Ian Johnston, Dominic Kwiatkowski, Cordelia Langford, John Sillitoe on behalf of the Wellcome Sanger Institute COVID-19 Surveillance Team |
| EPI_ISL_3107867 | Lighthouse Lab in Milton Keynes | Wellcome Sanger Institute for the COVID-19 Genomics UK (COG-UK) Consortium | The Lighthouse Lab in Milton Keynes and Alex Alderton, Roberto Amato, Jeffrey Barrett, Sonia Goncalves, Ewan Harrison, David K. Jackson, Ian Johnston, Dominic Kwiatkowski, Cordelia Langford, John Sillitoe on behalf of the Wellcome Sanger Institute COVID-19 Surveillance Team |
| EPI_ISL_3107868, EPI_ISL_3107869 | Lighthouse Laboratory Plymouth | Wellcome Sanger Institute for the COVID-19 Genomics UK (COG-UK) Consortium | Lighthouse Laboratory Plymouth and Alex Alderton, Roberto Amato, Jeffrey Barrett, Sonia Goncalves, Ewan Harrison, David K. Jackson, Ian Johnston, Dominic Kwiatkowski, Cordelia Langford, John Sillitoe on behalf of the Wellcome Sanger Institute COVID-19 Surveillance Team |
| EPI_ISL_3107870 | Lighthouse Lab in Glasgow | Wellcome Sanger Institute for the COVID-19 Genomics UK (COG-UK) Consortium | Harper VanSteenhouse, Yumi Kasai, David Gray, Carol Clugston, Anna Dominiczak and Alex Alderton, Roberto Amato, Jeffrey Barrett, Sonia Goncalves, Ewan Harrison, David K. Jackson, Ian Johnston, Dominic Kwiatkowski, Cordelia Langford, John Sillitoe on behalf of the Wellcome Sanger Institute COVID-19 Surveillance Team |
| EPI_ISL_3107871, EPI_ISL_3107872 | Lighthouse Laboratory Plymouth | Wellcome Sanger Institute for the COVID-19 Genomics UK (COG-UK) Consortium | Lighthouse Laboratory Plymouth and Alex Alderton, Roberto Amato, Jeffrey Barrett, Sonia Goncalves, Ewan Harrison, David K. Jackson, Ian Johnston, Dominic Kwiatkowski, Cordelia Langford, John Sillitoe on behalf of the Wellcome Sanger Institute COVID-19 Surveillance Team |
| EPI_ISL_3107873 | Lighthouse Lab in Milton Keynes | Wellcome Sanger Institute for the COVID-19 Genomics UK (COG-UK) Consortium | The Lighthouse Lab in Milton Keynes and Alex Alderton, Roberto Amato, Jeffrey Barrett, Sonia Goncalves, Ewan Harrison, David K. Jackson, Ian Johnston, Dominic Kwiatkowski, Cordelia Langford, John Sillitoe on behalf of the Wellcome Sanger Institute COVID-19 Surveillance Team |
| EPI_ISL_3107874 | Lighthouse Laboratory Plymouth | Wellcome Sanger Institute for the COVID-19 Genomics UK (COG-UK) Consortium | Lighthouse Laboratory Plymouth and Alex Alderton, Roberto Amato, Jeffrey Barrett, Sonia Goncalves, Ewan Harrison, David K. Jackson, Ian Johnston, Dominic Kwiatkowski, Cordelia Langford, John Sillitoe on behalf of the Wellcome Sanger Institute COVID-19 Surveillance Team |
| EPI_ISL_3107875 | Lighthouse Lab in Alderley Park | Wellcome Sanger Institute for the COVID-19 Genomics UK (COG-UK) | Jacquelyn Wynn, Mairead Hyland, The Lighthouse Lab in Alderley Park and Alex Alderton, Roberto Amato, Jeffrey Barrett, Sonia Goncalves, Ewan Harrison, David K. Jackson, Ian Johnston, Dominic Kwiatkowski, Cordelia Langford, John Sillitoe on |

| | | | |
|---|---|---|---|
| | | Consortium | behalf of the Wellcome Sanger Institute COVID-19 Surveillance Team |
| EPI_ISL_3107876 | Lighthouse Laboratory Plymouth | Wellcome Sanger Institute for the COVID-19 Genomics UK (COG-UK) Consortium | Lighthouse Laboratory Plymouth and Alex Alderton, Roberto Amato, Jeffrey Barrett, Sonia Goncalves, Ewan Harrison, David K. Jackson, Ian Johnston, Dominic Kwiatkowski, Cordelia Langford, John Sillitoe on behalf of the Wellcome Sanger Institute COVID-19 Surveillance Team |
| EPI_ISL_3107877, EPI_ISL_3107878 | Lighthouse Lab in Alderley Park | Wellcome Sanger Institute for the COVID-19 Genomics UK (COG-UK) Consortium | Jacquelyn Wynn, Mairead Hyland, The Lighthouse Lab in Alderley Park and Alex Alderton, Roberto Amato, Jeffrey Barrett, Sonia Goncalves, Ewan Harrison, David K. Jackson, Ian Johnston, Dominic Kwiatkowski, Cordelia Langford, John Sillitoe on behalf of the Wellcome Sanger Institute COVID-19 Surveillance Team |
| EPI_ISL_3107879 | Lighthouse Lab in Milton Keynes | Wellcome Sanger Institute for the COVID-19 Genomics UK (COG-UK) Consortium | The Lighthouse Lab in Milton Keynes and Alex Alderton, Roberto Amato, Jeffrey Barrett, Sonia Goncalves, Ewan Harrison, David K. Jackson, Ian Johnston, Dominic Kwiatkowski, Cordelia Langford, John Sillitoe on behalf of the Wellcome Sanger Institute COVID-19 Surveillance Team |
| EPI_ISL_3107880 | Lighthouse Lab in Glasgow | Wellcome Sanger Institute for the COVID-19 Genomics UK (COG-UK) Consortium | Harper VanSteenhouse, Yumi Kasai, David Gray, Carol Clugston, Anna Dominiczak and Alex Alderton, Roberto Amato, Jeffrey Barrett, Sonia Goncalves, Ewan Harrison, David K. Jackson, Ian Johnston, Dominic Kwiatkowski, Cordelia Langford, John Sillitoe on behalf of the Wellcome Sanger Institute COVID-19 Surveillance Team |
| EPI_ISL_3107881 | Lighthouse Lab in Milton Keynes | Wellcome Sanger Institute for the COVID-19 Genomics UK (COG-UK) Consortium | The Lighthouse Lab in Milton Keynes and Alex Alderton, Roberto Amato, Jeffrey Barrett, Sonia Goncalves, Ewan Harrison, David K. Jackson, Ian Johnston, Dominic Kwiatkowski, Cordelia Langford, John Sillitoe on behalf of the Wellcome Sanger Institute COVID-19 Surveillance Team |
| EPI_ISL_3107882 | Lighthouse Laboratory Plymouth | Wellcome Sanger Institute for the COVID-19 Genomics UK (COG-UK) Consortium | Lighthouse Laboratory Plymouth and Alex Alderton, Roberto Amato, Jeffrey Barrett, Sonia Goncalves, Ewan Harrison, David K. Jackson, Ian Johnston, Dominic Kwiatkowski, Cordelia Langford, John Sillitoe on behalf of the Wellcome Sanger Institute COVID-19 Surveillance Team |
| EPI_ISL_3107883 | Lighthouse Lab in Glasgow | Wellcome Sanger Institute for the COVID-19 Genomics UK (COG-UK) Consortium | Harper VanSteenhouse, Yumi Kasai, David Gray, Carol Clugston, Anna Dominiczak and Alex Alderton, Roberto Amato, Jeffrey Barrett, Sonia Goncalves, Ewan Harrison, David K. Jackson, Ian Johnston, Dominic Kwiatkowski, Cordelia Langford, John Sillitoe on behalf of the Wellcome Sanger Institute COVID-19 Surveillance Team |
| EPI_ISL_3107884 | Lighthouse Lab in Milton Keynes | Wellcome Sanger Institute for the COVID-19 Genomics UK (COG-UK) Consortium | The Lighthouse Lab in Milton Keynes and Alex Alderton, Roberto Amato, Jeffrey Barrett, Sonia Goncalves, Ewan Harrison, David K. Jackson, Ian Johnston, Dominic Kwiatkowski, Cordelia Langford, John Sillitoe on behalf of the Wellcome Sanger Institute COVID-19 Surveillance Team |
| EPI_ISL_3107885 | Lighthouse Laboratory Plymouth | Wellcome Sanger Institute for the COVID-19 Genomics UK (COG-UK) Consortium | Lighthouse Laboratory Plymouth and Alex Alderton, Roberto Amato, Jeffrey Barrett, Sonia Goncalves, Ewan Harrison, David K. Jackson, Ian Johnston, Dominic Kwiatkowski, Cordelia Langford, John Sillitoe on behalf of the Wellcome Sanger Institute COVID-19 Surveillance Team |
| EPI_ISL_3107886, EPI_ISL_3107887 | Lighthouse Lab in Alderley Park | Wellcome Sanger Institute for the COVID-19 Genomics UK (COG-UK) Consortium | Jacquelyn Wynn, Mairead Hyland, The Lighthouse Lab in Alderley Park and Alex Alderton, Roberto Amato, Jeffrey Barrett, Sonia Goncalves, Ewan Harrison, David K. Jackson, Ian Johnston, Dominic Kwiatkowski, Cordelia Langford, John Sillitoe on behalf of the Wellcome Sanger Institute COVID-19 Surveillance Team |
| EPI_ISL_3107888 | Lighthouse Lab in Glasgow | Wellcome Sanger Institute for the COVID-19 Genomics UK (COG-UK) Consortium | Harper VanSteenhouse, Yumi Kasai, David Gray, Carol Clugston, Anna Dominiczak and Alex Alderton, Roberto Amato, Jeffrey Barrett, Sonia Goncalves, Ewan Harrison, David K. Jackson, Ian Johnston, Dominic Kwiatkowski, Cordelia Langford, John Sillitoe on behalf of the Wellcome Sanger Institute COVID-19 Surveillance Team |
| EPI_ISL_3107889 | Lighthouse Lab in Milton Keynes | Wellcome Sanger Institute for the COVID-19 Genomics UK (COG-UK) Consortium | The Lighthouse Lab in Milton Keynes and Alex Alderton, Roberto Amato, Jeffrey Barrett, Sonia Goncalves, Ewan Harrison, David K. Jackson, Ian Johnston, Dominic Kwiatkowski, Cordelia Langford, John Sillitoe on behalf of the Wellcome Sanger Institute COVID-19 Surveillance Team |
| EPI_ISL_3107890 | Lighthouse Laboratory Plymouth | Wellcome Sanger Institute for the COVID-19 Genomics UK (COG-UK) Consortium | Lighthouse Laboratory Plymouth and Alex Alderton, Roberto Amato, Jeffrey Barrett, Sonia Goncalves, Ewan Harrison, David K. Jackson, Ian Johnston, Dominic Kwiatkowski, Cordelia Langford, John Sillitoe on behalf of the Wellcome Sanger Institute COVID-19 Surveillance Team |
| EPI_ISL_3107891 | Lighthouse Lab in Alderley Park | Wellcome Sanger Institute for the COVID-19 Genomics UK (COG-UK) Consortium | Jacquelyn Wynn, Mairead Hyland, The Lighthouse Lab in Alderley Park and Alex Alderton, Roberto Amato, Jeffrey Barrett, Sonia Goncalves, Ewan Harrison, David K. Jackson, Ian Johnston, Dominic Kwiatkowski, Cordelia Langford, John Sillitoe on behalf of the Wellcome Sanger Institute COVID-19 Surveillance Team |
| EPI_ISL_3107892 | Lighthouse Laboratory Plymouth | Wellcome Sanger Institute for the COVID-19 Genomics UK (COG-UK) Consortium | Lighthouse Laboratory Plymouth and Alex Alderton, Roberto Amato, Jeffrey Barrett, Sonia Goncalves, Ewan Harrison, David K. Jackson, Ian Johnston, Dominic Kwiatkowski, Cordelia Langford, John Sillitoe on behalf of the Wellcome Sanger Institute COVID-19 Surveillance Team |
| EPI_ISL_3107893, EPI_ISL_3107894 | Lighthouse Lab in Glasgow | Wellcome Sanger Institute for the COVID-19 Genomics UK (COG-UK) Consortium | Harper VanSteenhouse, Yumi Kasai, David Gray, Carol Clugston, Anna Dominiczak and Alex Alderton, Roberto Amato, Jeffrey Barrett, Sonia Goncalves, Ewan Harrison, David K. Jackson, Ian Johnston, Dominic Kwiatkowski, Cordelia Langford, John Sillitoe on behalf of the Wellcome Sanger Institute COVID-19 Surveillance Team |
| EPI_ISL_3107895, EPI_ISL_3107896 | Lighthouse Lab in Milton Keynes | Wellcome Sanger Institute for the COVID-19 Genomics UK (COG-UK) Consortium | The Lighthouse Lab in Milton Keynes and Alex Alderton, Roberto Amato, Jeffrey Barrett, Sonia Goncalves, Ewan Harrison, David K. Jackson, Ian Johnston, Dominic Kwiatkowski, Cordelia Langford, John Sillitoe on behalf of the Wellcome Sanger Institute COVID-19 Surveillance Team |
| EPI_ISL_3107897 | Lighthouse Lab in Alderley Park | Wellcome Sanger Institute for the COVID-19 Genomics UK (COG-UK) Consortium | Jacquelyn Wynn, Mairead Hyland, The Lighthouse Lab in Alderley Park and Alex Alderton, Roberto Amato, Jeffrey Barrett, Sonia Goncalves, Ewan Harrison, David K. Jackson, Ian Johnston, Dominic Kwiatkowski, Cordelia Langford, John Sillitoe on behalf of the Wellcome Sanger Institute COVID-19 Surveillance Team |
| EPI_ISL_3107898, EPI_ISL_3107899 | Lighthouse Laboratory Plymouth | Wellcome Sanger Institute for the COVID-19 Genomics UK (COG-UK) Consortium | Lighthouse Laboratory Plymouth and Alex Alderton, Roberto Amato, Jeffrey Barrett, Sonia Goncalves, Ewan Harrison, David K. Jackson, Ian Johnston, Dominic Kwiatkowski, Cordelia Langford, John Sillitoe on behalf of the Wellcome Sanger Institute COVID-19 Surveillance Team |
| EPI_ISL_3107900, EPI_ISL_3107901 | Lighthouse Lab in Milton Keynes | Wellcome Sanger Institute for the COVID-19 Genomics UK (COG-UK) Consortium | The Lighthouse Lab in Milton Keynes and Alex Alderton, Roberto Amato, Jeffrey Barrett, Sonia Goncalves, Ewan Harrison, David K. Jackson, Ian Johnston, Dominic Kwiatkowski, Cordelia Langford, John Sillitoe on behalf of the Wellcome Sanger Institute COVID-19 Surveillance Team |
| EPI_ISL_3107902 | Lighthouse Lab in Alderley Park | Wellcome Sanger Institute for the COVID-19 Genomics UK (COG-UK) Consortium | Jacquelyn Wynn, Mairead Hyland, The Lighthouse Lab in Alderley Park and Alex Alderton, Roberto Amato, Jeffrey Barrett, Sonia Goncalves, Ewan Harrison, David K. Jackson, Ian Johnston, Dominic Kwiatkowski, Cordelia Langford, John Sillitoe on behalf of the Wellcome Sanger Institute COVID-19 Surveillance Team |
| EPI_ISL_3107903, EPI_ISL_3107904 | Lighthouse Lab in Milton Keynes | Wellcome Sanger Institute for the COVID-19 Genomics UK (COG-UK) Consortium | The Lighthouse Lab in Milton Keynes and Alex Alderton, Roberto Amato, Jeffrey Barrett, Sonia Goncalves, Ewan Harrison, David K. Jackson, Ian Johnston, Dominic Kwiatkowski, Cordelia Langford, John Sillitoe on behalf of the Wellcome Sanger Institute COVID-19 Surveillance Team |
| EPI_ISL_3107905, EPI_ISL_3107906 | Lighthouse Lab in Alderley Park | Wellcome Sanger Institute for the COVID-19 Genomics UK (COG-UK) Consortium | Jacquelyn Wynn, Mairead Hyland, The Lighthouse Lab in Alderley Park and Alex Alderton, Roberto Amato, Jeffrey Barrett, Sonia Goncalves, Ewan Harrison, David K. Jackson, Ian Johnston, Dominic Kwiatkowski, Cordelia Langford, John Sillitoe on behalf of the Wellcome Sanger Institute COVID-19 Surveillance Team |
| EPI_ISL_3107907 | Lighthouse Laboratory Plymouth | Wellcome Sanger Institute for the COVID-19 Genomics UK (COG-UK) Consortium | Lighthouse Laboratory Plymouth and Alex Alderton, Roberto Amato, Jeffrey Barrett, Sonia Goncalves, Ewan Harrison, David K. Jackson, Ian Johnston, Dominic Kwiatkowski, Cordelia Langford, John Sillitoe on behalf of the Wellcome Sanger Institute COVID-19 Surveillance Team |
| EPI_ISL_3107908 | Lighthouse Lab in Glasgow | Wellcome Sanger Institute for the COVID-19 Genomics UK (COG-UK) Consortium | Harper VanSteenhouse, Yumi Kasai, David Gray, Carol Clugston, Anna Dominiczak and Alex Alderton, Roberto Amato, Jeffrey Barrett, Sonia Goncalves, Ewan Harrison, David K. Jackson, Ian Johnston, Dominic Kwiatkowski, Cordelia Langford, John Sillitoe on behalf of the Wellcome Sanger Institute COVID-19 Surveillance Team |
| EPI_ISL_3107909 | Lighthouse Lab in Milton Keynes | Wellcome Sanger Institute for the COVID-19 Genomics UK (COG-UK) Consortium | The Lighthouse Lab in Milton Keynes and Alex Alderton, Roberto Amato, Jeffrey Barrett, Sonia Goncalves, Ewan Harrison, David K. Jackson, Ian Johnston, Dominic Kwiatkowski, Cordelia Langford, John Sillitoe on behalf of the Wellcome Sanger Institute COVID-19 Surveillance Team |
| EPI_ISL_3107910 | Lighthouse Laboratory Plymouth | Wellcome Sanger Institute for the COVID-19 Genomics UK (COG-UK) Consortium | Lighthouse Laboratory Plymouth and Alex Alderton, Roberto Amato, Jeffrey Barrett, Sonia Goncalves, Ewan Harrison, David K. Jackson, Ian Johnston, Dominic Kwiatkowski, Cordelia Langford, John Sillitoe on behalf of the Wellcome Sanger Institute COVID-19 Surveillance Team |
| EPI_ISL_3107911 | Lighthouse Lab in Glasgow | Wellcome Sanger Institute for the COVID-19 Genomics UK (COG-UK) Consortium | Harper VanSteenhouse, Yumi Kasai, David Gray, Carol Clugston, Anna Dominiczak and Alex Alderton, Roberto Amato, Jeffrey Barrett, Sonia Goncalves, Ewan Harrison, David K. Jackson, Ian Johnston, Dominic Kwiatkowski, Cordelia Langford, John Sillitoe on behalf of the Wellcome Sanger Institute COVID-19 Surveillance Team |
| EPI_ISL_3107912 | Lighthouse Lab in Alderley Park | Wellcome Sanger Institute for the COVID-19 Genomics UK (COG-UK) Consortium | Jacquelyn Wynn, Mairead Hyland, The Lighthouse Lab in Alderley Park and Alex Alderton, Roberto Amato, Jeffrey Barrett, Sonia Goncalves, Ewan Harrison, David K. Jackson, Ian Johnston, Dominic Kwiatkowski, Cordelia Langford, John Sillitoe on behalf of the Wellcome Sanger Institute COVID-19 Surveillance Team |
| EPI_ISL_3107913, EPI_ISL_3107914, EPI_ISL_3107915 | Lighthouse Lab in Milton Keynes | Wellcome Sanger Institute for the COVID-19 Genomics UK (COG-UK) Consortium | The Lighthouse Lab in Milton Keynes and Alex Alderton, Roberto Amato, Jeffrey Barrett, Sonia Goncalves, Ewan Harrison, David K. Jackson, Ian Johnston, Dominic Kwiatkowski, Cordelia Langford, John Sillitoe on behalf of the Wellcome Sanger Institute COVID-19 Surveillance Team |
| EPI_ISL_3107916 | Lighthouse Laboratory Plymouth | Wellcome Sanger Institute for the COVID-19 Genomics UK (COG-UK) Consortium | Lighthouse Laboratory Plymouth and Alex Alderton, Roberto Amato, Jeffrey Barrett, Sonia Goncalves, Ewan Harrison, David K. Jackson, Ian Johnston, Dominic Kwiatkowski, Cordelia Langford, John Sillitoe on behalf of the Wellcome Sanger Institute COVID-19 Surveillance Team |
| EPI_ISL_3107917 | Lighthouse Lab in Glasgow | Wellcome Sanger Institute for the COVID-19 Genomics UK (COG-UK) Consortium | Harper VanSteenhouse, Yumi Kasai, David Gray, Carol Clugston, Anna Dominiczak and Alex Alderton, Roberto Amato, Jeffrey Barrett, Sonia Goncalves, Ewan Harrison, David K. Jackson, Ian Johnston, Dominic Kwiatkowski, Cordelia Langford, John Sillitoe on behalf of the Wellcome Sanger Institute COVID-19 Surveillance Team |
| EPI_ISL_3107918, EPI_ISL_3107919 | Lighthouse Lab in Milton Keynes | Wellcome Sanger Institute for the COVID-19 Genomics UK (COG-UK) Consortium | The Lighthouse Lab in Milton Keynes and Alex Alderton, Roberto Amato, Jeffrey Barrett, Sonia Goncalves, Ewan Harrison, David K. Jackson, Ian Johnston, Dominic Kwiatkowski, Cordelia Langford, John Sillitoe on behalf of the Wellcome Sanger Institute COVID-19 Surveillance Team |
| EPI_ISL_3107920 | Lighthouse Lab in Alderley Park | Wellcome Sanger Institute for the COVID-19 Genomics UK (COG-UK) Consortium | Jacquelyn Wynn, Mairead Hyland, The Lighthouse Lab in Alderley Park and Alex Alderton, Roberto Amato, Jeffrey Barrett, Sonia Goncalves, Ewan Harrison, David K. Jackson, Ian Johnston, Dominic Kwiatkowski, Cordelia Langford, John Sillitoe on behalf of the Wellcome Sanger Institute COVID-19 Surveillance Team |
| EPI_ISL_3107921 | Lighthouse Lab in Glasgow | Wellcome Sanger Institute for the COVID-19 Genomics UK (COG-UK) Consortium | Harper VanSteenhouse, Yumi Kasai, David Gray, Carol Clugston, Anna Dominiczak and Alex Alderton, Roberto Amato, Jeffrey Barrett, Sonia Goncalves, Ewan Harrison, David K. Jackson, Ian Johnston, Dominic Kwiatkowski, Cordelia Langford, John Sillitoe on behalf of the Wellcome Sanger Institute COVID-19 Surveillance Team |
| EPI_ISL_3107922, EPI_ISL_3107923 | Lighthouse Lab in Milton Keynes | Wellcome Sanger Institute for the COVID-19 Genomics UK (COG-UK) Consortium | The Lighthouse Lab in Milton Keynes and Alex Alderton, Roberto Amato, Jeffrey Barrett, Sonia Goncalves, Ewan Harrison, David K. Jackson, Ian Johnston, Dominic Kwiatkowski, Cordelia Langford, John Sillitoe on behalf of the Wellcome Sanger Institute COVID-19 Surveillance Team |
| EPI_ISL_3107924, EPI_ISL_3107925, EPI_ISL_3107926 | Lighthouse Lab in Alderley Park | Wellcome Sanger Institute for the COVID-19 Genomics UK (COG-UK) Consortium | Jacquelyn Wynn, Mairead Hyland, The Lighthouse Lab in Alderley Park and Alex Alderton, Roberto Amato, Jeffrey Barrett, Sonia Goncalves, Ewan Harrison, David K. Jackson, Ian Johnston, Dominic Kwiatkowski, Cordelia Langford, John Sillitoe on behalf of the Wellcome Sanger Institute COVID-19 Surveillance Team |
| EPI_ISL_3107927, EPI_ISL_3107928, EPI_ISL_3107929 | Lighthouse Laboratory Plymouth | Wellcome Sanger Institute for the COVID-19 Genomics UK (COG-UK) Consortium | Lighthouse Laboratory Plymouth and Alex Alderton, Roberto Amato, Jeffrey Barrett, Sonia Goncalves, Ewan Harrison, David K. Jackson, Ian Johnston, Dominic Kwiatkowski, Cordelia Langford, John Sillitoe on behalf of the Wellcome Sanger Institute COVID-19 Surveillance Team |
| EPI_ISL_3107930 | Lighthouse Lab in Alderley Park | Wellcome Sanger Institute for the COVID-19 Genomics UK (COG-UK) Consortium | Jacquelyn Wynn, Mairead Hyland, The Lighthouse Lab in Alderley Park and Alex Alderton, Roberto Amato, Jeffrey Barrett, Sonia Goncalves, Ewan Harrison, David K. Jackson, Ian Johnston, Dominic Kwiatkowski, Cordelia Langford, John Sillitoe on behalf of the Wellcome Sanger Institute COVID-19 Surveillance Team |
| EPI_ISL_3107931, EPI_ISL_3107932, EPI_ISL_3107933 | Lighthouse Lab in Milton Keynes | Wellcome Sanger Institute for the COVID-19 Genomics UK (COG-UK) Consortium | The Lighthouse Lab in Milton Keynes and Alex Alderton, Roberto Amato, Jeffrey Barrett, Sonia Goncalves, Ewan Harrison, David K. Jackson, Ian Johnston, Dominic Kwiatkowski, Cordelia Langford, John Sillitoe on behalf of the Wellcome Sanger Institute COVID-19 Surveillance Team |
| EPI_ISL_3107934 | Lighthouse Laboratory Plymouth | Wellcome Sanger Institute for the COVID-19 Genomics UK (COG-UK) Consortium | Lighthouse Laboratory Plymouth and Alex Alderton, Roberto Amato, Jeffrey Barrett, Sonia Goncalves, Ewan Harrison, David K. Jackson, Ian Johnston, Dominic Kwiatkowski, Cordelia Langford, John Sillitoe on behalf of the Wellcome Sanger Institute COVID-19 Surveillance Team |
| EPI_ISL_3107935 | Lighthouse Lab in Alderley Park | Wellcome Sanger Institute for the COVID-19 Genomics UK (COG-UK) Consortium | Jacquelyn Wynn, Mairead Hyland, The Lighthouse Lab in Alderley Park and Alex Alderton, Roberto Amato, Jeffrey Barrett, Sonia Goncalves, Ewan Harrison, David K. Jackson, Ian Johnston, Dominic Kwiatkowski, Cordelia Langford, John Sillitoe on behalf of the Wellcome Sanger Institute COVID-19 Surveillance Team |
| EPI_ISL_3107936 | Lighthouse Laboratory Plymouth | Wellcome Sanger Institute for the COVID-19 Genomics UK (COG-UK) Consortium | Lighthouse Laboratory Plymouth and Alex Alderton, Roberto Amato, Jeffrey Barrett, Sonia Goncalves, Ewan Harrison, David K. Jackson, Ian Johnston, Dominic Kwiatkowski, Cordelia Langford, John Sillitoe on behalf of the Wellcome Sanger Institute COVID-19 Surveillance Team |

| | | | |
|---|---|---|---|
| EPI_ISL_3107937, EPI_ISL_3107938, EPI_ISL_3107939, EPI_ISL_3107940 | Lighthouse Lab in Alderley Park | Wellcome Sanger Institute for the COVID-19 Genomics UK (COG-UK) Consortium | Jacquelyn Wynn, Mairead Hyland, The Lighthouse Lab in Alderley Park and Alex Alderton, Roberto Amato, Jeffrey Barrett, Sonia Goncalves, Ewan Harrison, David K. Jackson, Ian Johnston, Dominic Kwiatkowski, Cordelia Langford, John Sillitoe on behalf of the Wellcome Sanger Institute COVID-19 Surveillance Team |
| EPI_ISL_3107941, EPI_ISL_3107942 | Lighthouse Lab in Glasgow | Wellcome Sanger Institute for the COVID-19 Genomics UK (COG-UK) Consortium | Harper VanSteenhouse, Yumi Kasai, David Gray, Carol Clugston, Anna Dominiczak and Alex Alderton, Roberto Amato, Jeffrey Barrett, Sonia Goncalves, Ewan Harrison, David K. Jackson, Ian Johnston, Dominic Kwiatkowski, Cordelia Langford, John Sillitoe on behalf of the Wellcome Sanger Institute COVID-19 Surveillance Team |
| EPI_ISL_3107943 | Lighthouse Lab in Alderley Park | Wellcome Sanger Institute for the COVID-19 Genomics UK (COG-UK) Consortium | Jacquelyn Wynn, Mairead Hyland, The Lighthouse Lab in Alderley Park and Alex Alderton, Roberto Amato, Jeffrey Barrett, Sonia Goncalves, Ewan Harrison, David K. Jackson, Ian Johnston, Dominic Kwiatkowski, Cordelia Langford, John Sillitoe on behalf of the Wellcome Sanger Institute COVID-19 Surveillance Team |
| EPI_ISL_3107944 | Lighthouse Lab in Glasgow | Wellcome Sanger Institute for the COVID-19 Genomics UK (COG-UK) Consortium | Harper VanSteenhouse, Yumi Kasai, David Gray, Carol Clugston, Anna Dominiczak and Alex Alderton, Roberto Amato, Jeffrey Barrett, Sonia Goncalves, Ewan Harrison, David K. Jackson, Ian Johnston, Dominic Kwiatkowski, Cordelia Langford, John Sillitoe on behalf of the Wellcome Sanger Institute COVID-19 Surveillance Team |
| EPI_ISL_3107945 | Lighthouse Lab in Alderley Park | Wellcome Sanger Institute for the COVID-19 Genomics UK (COG-UK) Consortium | Jacquelyn Wynn, Mairead Hyland, The Lighthouse Lab in Alderley Park and Alex Alderton, Roberto Amato, Jeffrey Barrett, Sonia Goncalves, Ewan Harrison, David K. Jackson, Ian Johnston, Dominic Kwiatkowski, Cordelia Langford, John Sillitoe on behalf of the Wellcome Sanger Institute COVID-19 Surveillance Team |
| EPI_ISL_3107946, EPI_ISL_3107947 | Lighthouse Lab in Glasgow | Wellcome Sanger Institute for the COVID-19 Genomics UK (COG-UK) Consortium | Harper VanSteenhouse, Yumi Kasai, David Gray, Carol Clugston, Anna Dominiczak and Alex Alderton, Roberto Amato, Jeffrey Barrett, Sonia Goncalves, Ewan Harrison, David K. Jackson, Ian Johnston, Dominic Kwiatkowski, Cordelia Langford, John Sillitoe on behalf of the Wellcome Sanger Institute COVID-19 Surveillance Team |
| EPI_ISL_3107948, EPI_ISL_3107949, EPI_ISL_3107950, EPI_ISL_3107951, EPI_ISL_3107952, EPI_ISL_3107953, EPI_ISL_3107954 | Lighthouse Lab in Alderley Park | Wellcome Sanger Institute for the COVID-19 Genomics UK (COG-UK) Consortium | Jacquelyn Wynn, Mairead Hyland, The Lighthouse Lab in Alderley Park and Alex Alderton, Roberto Amato, Jeffrey Barrett, Sonia Goncalves, Ewan Harrison, David K. Jackson, Ian Johnston, Dominic Kwiatkowski, Cordelia Langford, John Sillitoe on behalf of the Wellcome Sanger Institute COVID-19 Surveillance Team |
| EPI_ISL_3107955 | Lighthouse Lab in Glasgow | Wellcome Sanger Institute for the COVID-19 Genomics UK (COG-UK) Consortium | Harper VanSteenhouse, Yumi Kasai, David Gray, Carol Clugston, Anna Dominiczak and Alex Alderton, Roberto Amato, Jeffrey Barrett, Sonia Goncalves, Ewan Harrison, David K. Jackson, Ian Johnston, Dominic Kwiatkowski, Cordelia Langford, John Sillitoe on behalf of the Wellcome Sanger Institute COVID-19 Surveillance Team |
| EPI_ISL_3107956, EPI_ISL_3107957, EPI_ISL_3107958, EPI_ISL_3107959, EPI_ISL_3107960, EPI_ISL_3107961, EPI_ISL_3107962, EPI_ISL_3107963, EPI_ISL_3107964, EPI_ISL_3107965 | Lighthouse Lab in Alderley Park | Wellcome Sanger Institute for the COVID-19 Genomics UK (COG-UK) Consortium | Jacquelyn Wynn, Mairead Hyland, The Lighthouse Lab in Alderley Park and Alex Alderton, Roberto Amato, Jeffrey Barrett, Sonia Goncalves, Ewan Harrison, David K. Jackson, Ian Johnston, Dominic Kwiatkowski, Cordelia Langford, John Sillitoe on behalf of the Wellcome Sanger Institute COVID-19 Surveillance Team |
| EPI_ISL_3107966 | Lighthouse Lab in Glasgow | Wellcome Sanger Institute for the COVID-19 Genomics UK (COG-UK) Consortium | Harper VanSteenhouse, Yumi Kasai, David Gray, Carol Clugston, Anna Dominiczak and Alex Alderton, Roberto Amato, Jeffrey Barrett, Sonia Goncalves, Ewan Harrison, David K. Jackson, Ian Johnston, Dominic Kwiatkowski, Cordelia Langford, John Sillitoe on behalf of the Wellcome Sanger Institute COVID-19 Surveillance Team |
| EPI_ISL_3107967, EPI_ISL_3107968, EPI_ISL_3107969 | Lighthouse Lab in Alderley Park | Wellcome Sanger Institute for the COVID-19 Genomics UK (COG-UK) Consortium | Jacquelyn Wynn, Mairead Hyland, The Lighthouse Lab in Alderley Park and Alex Alderton, Roberto Amato, Jeffrey Barrett, Sonia Goncalves, Ewan Harrison, David K. Jackson, Ian Johnston, Dominic Kwiatkowski, Cordelia Langford, John Sillitoe on behalf of the Wellcome Sanger Institute COVID-19 Surveillance Team |
| EPI_ISL_3107970, EPI_ISL_3107971 | Lighthouse Lab in Glasgow | Wellcome Sanger Institute for the COVID-19 Genomics UK (COG-UK) Consortium | Harper VanSteenhouse, Yumi Kasai, David Gray, Carol Clugston, Anna Dominiczak and Alex Alderton, Roberto Amato, Jeffrey Barrett, Sonia Goncalves, Ewan Harrison, David K. Jackson, Ian Johnston, Dominic Kwiatkowski, Cordelia Langford, John Sillitoe on behalf of the Wellcome Sanger Institute COVID-19 Surveillance Team |
| EPI_ISL_3107972 | Lighthouse Lab in Alderley Park | Wellcome Sanger Institute for the COVID-19 Genomics UK (COG-UK) Consortium | Jacquelyn Wynn, Mairead Hyland, The Lighthouse Lab in Alderley Park and Alex Alderton, Roberto Amato, Jeffrey Barrett, Sonia Goncalves, Ewan Harrison, David K. Jackson, Ian Johnston, Dominic Kwiatkowski, Cordelia Langford, John Sillitoe on behalf of the Wellcome Sanger Institute COVID-19 Surveillance Team |
| EPI_ISL_3107973, EPI_ISL_3107974, EPI_ISL_3107975, EPI_ISL_3107976 | Lighthouse Lab in Glasgow | Wellcome Sanger Institute for the COVID-19 Genomics UK (COG-UK) Consortium | Harper VanSteenhouse, Yumi Kasai, David Gray, Carol Clugston, Anna Dominiczak and Alex Alderton, Roberto Amato, Jeffrey Barrett, Sonia Goncalves, Ewan Harrison, David K. Jackson, Ian Johnston, Dominic Kwiatkowski, Cordelia Langford, John Sillitoe on behalf of the Wellcome Sanger Institute COVID-19 Surveillance Team |
| EPI_ISL_3107977, EPI_ISL_3107978, EPI_ISL_3107979, EPI_ISL_3107980, EPI_ISL_3107981, EPI_ISL_3107982, EPI_ISL_3107983 | Lighthouse Lab in Alderley Park | Wellcome Sanger Institute for the COVID-19 Genomics UK (COG-UK) Consortium | Jacquelyn Wynn, Mairead Hyland, The Lighthouse Lab in Alderley Park and Alex Alderton, Roberto Amato, Jeffrey Barrett, Sonia Goncalves, Ewan Harrison, David K. Jackson, Ian Johnston, Dominic Kwiatkowski, Cordelia Langford, John Sillitoe on behalf of the Wellcome Sanger Institute COVID-19 Surveillance Team |
| EPI_ISL_3107984, EPI_ISL_3107985, EPI_ISL_3107986 | Lighthouse Lab in Glasgow | Wellcome Sanger Institute for the COVID-19 Genomics UK (COG-UK) Consortium | Harper VanSteenhouse, Yumi Kasai, David Gray, Carol Clugston, Anna Dominiczak and Alex Alderton, Roberto Amato, Jeffrey Barrett, Sonia Goncalves, Ewan Harrison, David K. Jackson, Ian Johnston, Dominic Kwiatkowski, Cordelia Langford, John Sillitoe on behalf of the Wellcome Sanger Institute COVID-19 Surveillance Team |
| EPI_ISL_3107987, EPI_ISL_3107988, EPI_ISL_3107989 | Lighthouse Lab in Alderley Park | Wellcome Sanger Institute for the COVID-19 Genomics UK (COG-UK) Consortium | Jacquelyn Wynn, Mairead Hyland, The Lighthouse Lab in Alderley Park and Alex Alderton, Roberto Amato, Jeffrey Barrett, Sonia Goncalves, Ewan Harrison, David K. Jackson, Ian Johnston, Dominic Kwiatkowski, Cordelia Langford, John Sillitoe on behalf of the Wellcome Sanger Institute COVID-19 Surveillance Team |
| EPI_ISL_3107990 | Lighthouse Lab in Glasgow | Wellcome Sanger Institute for the COVID-19 Genomics UK (COG-UK) Consortium | Harper VanSteenhouse, Yumi Kasai, David Gray, Carol Clugston, Anna Dominiczak and Alex Alderton, Roberto Amato, Jeffrey Barrett, Sonia Goncalves, Ewan Harrison, David K. Jackson, Ian Johnston, Dominic Kwiatkowski, Cordelia Langford, John Sillitoe on behalf of the Wellcome Sanger Institute COVID-19 Surveillance Team |
| EPI_ISL_3107991 | Lighthouse Lab in Alderley Park | Wellcome Sanger Institute for the COVID-19 Genomics UK (COG-UK) Consortium | Jacquelyn Wynn, Mairead Hyland, The Lighthouse Lab in Alderley Park and Alex Alderton, Roberto Amato, Jeffrey Barrett, Sonia Goncalves, Ewan Harrison, David K. Jackson, Ian Johnston, Dominic Kwiatkowski, Cordelia Langford, John Sillitoe on behalf of the Wellcome Sanger Institute COVID-19 Surveillance Team |
| EPI_ISL_3107992 | Lighthouse Lab in Glasgow | Wellcome Sanger Institute for the COVID-19 Genomics UK (COG-UK) Consortium | Harper VanSteenhouse, Yumi Kasai, David Gray, Carol Clugston, Anna Dominiczak and Alex Alderton, Roberto Amato, Jeffrey Barrett, Sonia Goncalves, Ewan Harrison, David K. Jackson, Ian Johnston, Dominic Kwiatkowski, Cordelia Langford, John Sillitoe on behalf of the Wellcome Sanger Institute COVID-19 Surveillance Team |
| EPI_ISL_3107993, EPI_ISL_3107994, EPI_ISL_3107995 | Lighthouse Lab in Alderley Park | Wellcome Sanger Institute for the COVID-19 Genomics UK (COG-UK) Consortium | Jacquelyn Wynn, Mairead Hyland, The Lighthouse Lab in Alderley Park and Alex Alderton, Roberto Amato, Jeffrey Barrett, Sonia Goncalves, Ewan Harrison, David K. Jackson, Ian Johnston, Dominic Kwiatkowski, Cordelia Langford, John Sillitoe on behalf of the Wellcome Sanger Institute COVID-19 Surveillance Team |
| EPI_ISL_3107996 | Lighthouse Lab in Glasgow | Wellcome Sanger Institute for the COVID-19 Genomics UK (COG-UK) Consortium | Harper VanSteenhouse, Yumi Kasai, David Gray, Carol Clugston, Anna Dominiczak and Alex Alderton, Roberto Amato, Jeffrey Barrett, Sonia Goncalves, Ewan Harrison, David K. Jackson, Ian Johnston, Dominic Kwiatkowski, Cordelia Langford, John Sillitoe on behalf of the Wellcome Sanger Institute COVID-19 Surveillance Team |
| EPI_ISL_3107997, EPI_ISL_3107998, EPI_ISL_3107999, EPI_ISL_3108000, EPI_ISL_3108001, EPI_ISL_3108002, EPI_ISL_3108003, EPI_ISL_3108004, EPI_ISL_3108005, EPI_ISL_3108006, EPI_ISL_3108007, EPI_ISL_3108008, EPI_ISL_3108009 | | | |
| see above | Lighthouse Lab in Alderley Park | Wellcome Sanger Institute for the COVID-19 Genomics UK (COG-UK) Consortium | Jacquelyn Wynn, Mairead Hyland, The Lighthouse Lab in Alderley Park and Alex Alderton, Roberto Amato, Jeffrey Barrett, Sonia Goncalves, Ewan Harrison, David K. Jackson, Ian Johnston, Dominic Kwiatkowski, Cordelia Langford, John Sillitoe on behalf of the Wellcome Sanger Institute COVID-19 Surveillance Team |
| EPI_ISL_3108010 | Lighthouse Lab in Glasgow | Wellcome Sanger Institute for the COVID-19 Genomics UK (COG-UK) Consortium | Harper VanSteenhouse, Yumi Kasai, David Gray, Carol Clugston, Anna Dominiczak and Alex Alderton, Roberto Amato, Jeffrey Barrett, Sonia Goncalves, Ewan Harrison, David K. Jackson, Ian Johnston, Dominic Kwiatkowski, Cordelia Langford, John Sillitoe on behalf of the Wellcome Sanger Institute COVID-19 Surveillance Team |
| EPI_ISL_3108011, EPI_ISL_3108012, EPI_ISL_3108013, EPI_ISL_3108014, EPI_ISL_3108015, EPI_ISL_3108016, EPI_ISL_3108017, EPI_ISL_3108018 | Lighthouse Lab in Alderley Park | Wellcome Sanger Institute for the COVID-19 Genomics UK (COG-UK) Consortium | Jacquelyn Wynn, Mairead Hyland, The Lighthouse Lab in Alderley Park and Alex Alderton, Roberto Amato, Jeffrey Barrett, Sonia Goncalves, Ewan Harrison, David K. Jackson, Ian Johnston, Dominic Kwiatkowski, Cordelia Langford, John Sillitoe on behalf of the Wellcome Sanger Institute COVID-19 Surveillance Team |
| EPI_ISL_3108019 | Lighthouse Lab in Glasgow | Wellcome Sanger Institute for the COVID-19 Genomics UK (COG-UK) Consortium | Harper VanSteenhouse, Yumi Kasai, David Gray, Carol Clugston, Anna Dominiczak and Alex Alderton, Roberto Amato, Jeffrey Barrett, Sonia Goncalves, Ewan Harrison, David K. Jackson, Ian Johnston, Dominic Kwiatkowski, Cordelia Langford, John Sillitoe on behalf of the Wellcome Sanger Institute COVID-19 Surveillance Team |
| EPI_ISL_3108020, EPI_ISL_3108021, EPI_ISL_3108022, EPI_ISL_3108023, EPI_ISL_3108024, EPI_ISL_3108025, EPI_ISL_3108026, EPI_ISL_3108027, EPI_ISL_3108028 | Lighthouse Lab in Alderley Park | Wellcome Sanger Institute for the COVID-19 Genomics UK (COG-UK) Consortium | Jacquelyn Wynn, Mairead Hyland, The Lighthouse Lab in Alderley Park and Alex Alderton, Roberto Amato, Jeffrey Barrett, Sonia Goncalves, Ewan Harrison, David K. Jackson, Ian Johnston, Dominic Kwiatkowski, Cordelia Langford, John Sillitoe on behalf of the Wellcome Sanger Institute COVID-19 Surveillance Team |
| EPI_ISL_3108029 | Lighthouse Lab in Glasgow | Wellcome Sanger Institute for the COVID-19 Genomics UK (COG-UK) Consortium | Harper VanSteenhouse, Yumi Kasai, David Gray, Carol Clugston, Anna Dominiczak and Alex Alderton, Roberto Amato, Jeffrey Barrett, Sonia Goncalves, Ewan Harrison, David K. Jackson, Ian Johnston, Dominic Kwiatkowski, Cordelia Langford, John Sillitoe on behalf of the Wellcome Sanger Institute COVID-19 Surveillance Team |
| EPI_ISL_3108030, | Lighthouse Lab in Alderley Park | Wellcome Sanger Institute for the COVID-19 Genomics UK (COG-UK) | Jacquelyn Wynn, Mairead Hyland, The Lighthouse Lab in Alderley Park and Alex Alderton, Roberto Amato, Jeffrey Barrett, Sonia Goncalves, Ewan Harrison, David K. Jackson, Ian Johnston, Dominic Kwiatkowski, Cordelia Langford, John Sillitoe on |

| | | | |
|---|---|---|---|
| EPI_ISL_3108031, EPI_ISL_3108032 | | Consortium | behalf of the Wellcome Sanger Institute COVID-19 Surveillance Team |
| EPI_ISL_3108033 | Lighthouse Lab in Glasgow | Wellcome Sanger Institute for the COVID-19 Genomics UK (COG-UK) Consortium | Harper VanSteenhouse, Yumi Kasai, David Gray, Carol Clugston, Anna Dominiczak and Alex Alderton, Roberto Amato, Jeffrey Barrett, Sonia Goncalves, Ewan Harrison, David K. Jackson, Ian Johnston, Dominic Kwiatkowski, Cordelia Langford, John Sillitoe on behalf of the Wellcome Sanger Institute COVID-19 Surveillance Team |
| EPI_ISL_3108034 | Lighthouse Lab in Alderley Park | Wellcome Sanger Institute for the COVID-19 Genomics UK (COG-UK) Consortium | Jacquelyn Wynn, Mairead Hyland, The Lighthouse Lab in Alderley Park and Alex Alderton, Roberto Amato, Jeffrey Barrett, Sonia Goncalves, Ewan Harrison, David K. Jackson, Ian Johnston, Dominic Kwiatkowski, Cordelia Langford, John Sillitoe on behalf of the Wellcome Sanger Institute COVID-19 Surveillance Team |
| EPI_ISL_3108035 | Lighthouse Lab in Glasgow | Wellcome Sanger Institute for the COVID-19 Genomics UK (COG-UK) Consortium | Harper VanSteenhouse, Yumi Kasai, David Gray, Carol Clugston, Anna Dominiczak and Alex Alderton, Roberto Amato, Jeffrey Barrett, Sonia Goncalves, Ewan Harrison, David K. Jackson, Ian Johnston, Dominic Kwiatkowski, Cordelia Langford, John Sillitoe on behalf of the Wellcome Sanger Institute COVID-19 Surveillance Team |
| EPI_ISL_3108036, EPI_ISL_3108037, EPI_ISL_3108038, EPI_ISL_3108039, EPI_ISL_3108040, EPI_ISL_3108041, EPI_ISL_3108042, EPI_ISL_3108043, EPI_ISL_3108044, EPI_ISL_3108045, EPI_ISL_3108046 | | | |
| see above | Lighthouse Lab in Alderley Park | Wellcome Sanger Institute for the COVID-19 Genomics UK (COG-UK) Consortium | Jacquelyn Wynn, Mairead Hyland, The Lighthouse Lab in Alderley Park and Alex Alderton, Roberto Amato, Jeffrey Barrett, Sonia Goncalves, Ewan Harrison, David K. Jackson, Ian Johnston, Dominic Kwiatkowski, Cordelia Langford, John Sillitoe on behalf of the Wellcome Sanger Institute COVID-19 Surveillance Team |
| EPI_ISL_3108047, EPI_ISL_3108048 | Lighthouse Lab in Glasgow | Wellcome Sanger Institute for the COVID-19 Genomics UK (COG-UK) Consortium | Harper VanSteenhouse, Yumi Kasai, David Gray, Carol Clugston, Anna Dominiczak and Alex Alderton, Roberto Amato, Jeffrey Barrett, Sonia Goncalves, Ewan Harrison, David K. Jackson, Ian Johnston, Dominic Kwiatkowski, Cordelia Langford, John Sillitoe on behalf of the Wellcome Sanger Institute COVID-19 Surveillance Team |
| EPI_ISL_3108049, EPI_ISL_3108050, EPI_ISL_3108051, EPI_ISL_3108052, EPI_ISL_3108053, EPI_ISL_3108054, EPI_ISL_3108055, EPI_ISL_3108056 | Lighthouse Lab in Alderley Park | Wellcome Sanger Institute for the COVID-19 Genomics UK (COG-UK) Consortium | Jacquelyn Wynn, Mairead Hyland, The Lighthouse Lab in Alderley Park and Alex Alderton, Roberto Amato, Jeffrey Barrett, Sonia Goncalves, Ewan Harrison, David K. Jackson, Ian Johnston, Dominic Kwiatkowski, Cordelia Langford, John Sillitoe on behalf of the Wellcome Sanger Institute COVID-19 Surveillance Team |
| EPI_ISL_3108057, EPI_ISL_3108058 | Lighthouse Lab in Glasgow | Wellcome Sanger Institute for the COVID-19 Genomics UK (COG-UK) Consortium | Harper VanSteenhouse, Yumi Kasai, David Gray, Carol Clugston, Anna Dominiczak and Alex Alderton, Roberto Amato, Jeffrey Barrett, Sonia Goncalves, Ewan Harrison, David K. Jackson, Ian Johnston, Dominic Kwiatkowski, Cordelia Langford, John Sillitoe on behalf of the Wellcome Sanger Institute COVID-19 Surveillance Team |
| EPI_ISL_3108059 | Lighthouse Lab in Alderley Park | Wellcome Sanger Institute for the COVID-19 Genomics UK (COG-UK) Consortium | Jacquelyn Wynn, Mairead Hyland, The Lighthouse Lab in Alderley Park and Alex Alderton, Roberto Amato, Jeffrey Barrett, Sonia Goncalves, Ewan Harrison, David K. Jackson, Ian Johnston, Dominic Kwiatkowski, Cordelia Langford, John Sillitoe on behalf of the Wellcome Sanger Institute COVID-19 Surveillance Team |
| EPI_ISL_3108060 | Lighthouse Lab in Glasgow | Wellcome Sanger Institute for the COVID-19 Genomics UK (COG-UK) Consortium | Harper VanSteenhouse, Yumi Kasai, David Gray, Carol Clugston, Anna Dominiczak and Alex Alderton, Roberto Amato, Jeffrey Barrett, Sonia Goncalves, Ewan Harrison, David K. Jackson, Ian Johnston, Dominic Kwiatkowski, Cordelia Langford, John Sillitoe on behalf of the Wellcome Sanger Institute COVID-19 Surveillance Team |
| EPI_ISL_3108061, EPI_ISL_3108062 | Lighthouse Lab in Alderley Park | Wellcome Sanger Institute for the COVID-19 Genomics UK (COG-UK) Consortium | Jacquelyn Wynn, Mairead Hyland, The Lighthouse Lab in Alderley Park and Alex Alderton, Roberto Amato, Jeffrey Barrett, Sonia Goncalves, Ewan Harrison, David K. Jackson, Ian Johnston, Dominic Kwiatkowski, Cordelia Langford, John Sillitoe on behalf of the Wellcome Sanger Institute COVID-19 Surveillance Team |
| EPI_ISL_3108063 | Lighthouse Lab in Glasgow | Wellcome Sanger Institute for the COVID-19 Genomics UK (COG-UK) Consortium | Harper VanSteenhouse, Yumi Kasai, David Gray, Carol Clugston, Anna Dominiczak and Alex Alderton, Roberto Amato, Jeffrey Barrett, Sonia Goncalves, Ewan Harrison, David K. Jackson, Ian Johnston, Dominic Kwiatkowski, Cordelia Langford, John Sillitoe on behalf of the Wellcome Sanger Institute COVID-19 Surveillance Team |
| EPI_ISL_3108064, EPI_ISL_3108065 | Lighthouse Lab in Alderley Park | Wellcome Sanger Institute for the COVID-19 Genomics UK (COG-UK) Consortium | Jacquelyn Wynn, Mairead Hyland, The Lighthouse Lab in Alderley Park and Alex Alderton, Roberto Amato, Jeffrey Barrett, Sonia Goncalves, Ewan Harrison, David K. Jackson, Ian Johnston, Dominic Kwiatkowski, Cordelia Langford, John Sillitoe on behalf of the Wellcome Sanger Institute COVID-19 Surveillance Team |
| EPI_ISL_3108066 | Lighthouse Lab in Glasgow | Wellcome Sanger Institute for the COVID-19 Genomics UK (COG-UK) Consortium | Harper VanSteenhouse, Yumi Kasai, David Gray, Carol Clugston, Anna Dominiczak and Alex Alderton, Roberto Amato, Jeffrey Barrett, Sonia Goncalves, Ewan Harrison, David K. Jackson, Ian Johnston, Dominic Kwiatkowski, Cordelia Langford, John Sillitoe on behalf of the Wellcome Sanger Institute COVID-19 Surveillance Team |
| EPI_ISL_3108067, EPI_ISL_3108068, EPI_ISL_3108069, EPI_ISL_3108070, EPI_ISL_3108071 | Lighthouse Lab in Alderley Park | Wellcome Sanger Institute for the COVID-19 Genomics UK (COG-UK) Consortium | Jacquelyn Wynn, Mairead Hyland, The Lighthouse Lab in Alderley Park and Alex Alderton, Roberto Amato, Jeffrey Barrett, Sonia Goncalves, Ewan Harrison, David K. Jackson, Ian Johnston, Dominic Kwiatkowski, Cordelia Langford, John Sillitoe on behalf of the Wellcome Sanger Institute COVID-19 Surveillance Team |
| EPI_ISL_3108072, EPI_ISL_3108073 | Lighthouse Lab in Glasgow | Wellcome Sanger Institute for the COVID-19 Genomics UK (COG-UK) Consortium | Harper VanSteenhouse, Yumi Kasai, David Gray, Carol Clugston, Anna Dominiczak and Alex Alderton, Roberto Amato, Jeffrey Barrett, Sonia Goncalves, Ewan Harrison, David K. Jackson, Ian Johnston, Dominic Kwiatkowski, Cordelia Langford, John Sillitoe on behalf of the Wellcome Sanger Institute COVID-19 Surveillance Team |
| EPI_ISL_3108074, EPI_ISL_3108075, EPI_ISL_3108076, EPI_ISL_3108077 | Lighthouse Lab in Alderley Park | Wellcome Sanger Institute for the COVID-19 Genomics UK (COG-UK) Consortium | Jacquelyn Wynn, Mairead Hyland, The Lighthouse Lab in Alderley Park and Alex Alderton, Roberto Amato, Jeffrey Barrett, Sonia Goncalves, Ewan Harrison, David K. Jackson, Ian Johnston, Dominic Kwiatkowski, Cordelia Langford, John Sillitoe on behalf of the Wellcome Sanger Institute COVID-19 Surveillance Team |
| EPI_ISL_3108078 | Lighthouse Lab in Glasgow | Wellcome Sanger Institute for the COVID-19 Genomics UK (COG-UK) Consortium | Harper VanSteenhouse, Yumi Kasai, David Gray, Carol Clugston, Anna Dominiczak and Alex Alderton, Roberto Amato, Jeffrey Barrett, Sonia Goncalves, Ewan Harrison, David K. Jackson, Ian Johnston, Dominic Kwiatkowski, Cordelia Langford, John Sillitoe on behalf of the Wellcome Sanger Institute COVID-19 Surveillance Team |
| EPI_ISL_3108079, EPI_ISL_3108080, EPI_ISL_3108081, EPI_ISL_3108082 | Lighthouse Lab in Alderley Park | Wellcome Sanger Institute for the COVID-19 Genomics UK (COG-UK) Consortium | Jacquelyn Wynn, Mairead Hyland, The Lighthouse Lab in Alderley Park and Alex Alderton, Roberto Amato, Jeffrey Barrett, Sonia Goncalves, Ewan Harrison, David K. Jackson, Ian Johnston, Dominic Kwiatkowski, Cordelia Langford, John Sillitoe on behalf of the Wellcome Sanger Institute COVID-19 Surveillance Team |
| EPI_ISL_3108083 | Lighthouse Lab in Glasgow | Wellcome Sanger Institute for the COVID-19 Genomics UK (COG-UK) Consortium | Harper VanSteenhouse, Yumi Kasai, David Gray, Carol Clugston, Anna Dominiczak and Alex Alderton, Roberto Amato, Jeffrey Barrett, Sonia Goncalves, Ewan Harrison, David K. Jackson, Ian Johnston, Dominic Kwiatkowski, Cordelia Langford, John Sillitoe on behalf of the Wellcome Sanger Institute COVID-19 Surveillance Team |
| EPI_ISL_3108084, EPI_ISL_3108085 | Lighthouse Lab in Alderley Park | Wellcome Sanger Institute for the COVID-19 Genomics UK (COG-UK) Consortium | Jacquelyn Wynn, Mairead Hyland, The Lighthouse Lab in Alderley Park and Alex Alderton, Roberto Amato, Jeffrey Barrett, Sonia Goncalves, Ewan Harrison, David K. Jackson, Ian Johnston, Dominic Kwiatkowski, Cordelia Langford, John Sillitoe on behalf of the Wellcome Sanger Institute COVID-19 Surveillance Team |
| EPI_ISL_3108086 | Lighthouse Lab in Glasgow | Wellcome Sanger Institute for the COVID-19 Genomics UK (COG-UK) Consortium | Harper VanSteenhouse, Yumi Kasai, David Gray, Carol Clugston, Anna Dominiczak and Alex Alderton, Roberto Amato, Jeffrey Barrett, Sonia Goncalves, Ewan Harrison, David K. Jackson, Ian Johnston, Dominic Kwiatkowski, Cordelia Langford, John Sillitoe on behalf of the Wellcome Sanger Institute COVID-19 Surveillance Team |
| EPI_ISL_3108087 | Lighthouse Lab in Alderley Park | Wellcome Sanger Institute for the COVID-19 Genomics UK (COG-UK) Consortium | Jacquelyn Wynn, Mairead Hyland, The Lighthouse Lab in Alderley Park and Alex Alderton, Roberto Amato, Jeffrey Barrett, Sonia Goncalves, Ewan Harrison, David K. Jackson, Ian Johnston, Dominic Kwiatkowski, Cordelia Langford, John Sillitoe on behalf of the Wellcome Sanger Institute COVID-19 Surveillance Team |
| EPI_ISL_3108088 | Lighthouse Lab in Glasgow | Wellcome Sanger Institute for the COVID-19 Genomics UK (COG-UK) Consortium | Harper VanSteenhouse, Yumi Kasai, David Gray, Carol Clugston, Anna Dominiczak and Alex Alderton, Roberto Amato, Jeffrey Barrett, Sonia Goncalves, Ewan Harrison, David K. Jackson, Ian Johnston, Dominic Kwiatkowski, Cordelia Langford, John Sillitoe on behalf of the Wellcome Sanger Institute COVID-19 Surveillance Team |
| EPI_ISL_3108089, EPI_ISL_3108090, EPI_ISL_3108091 | Lighthouse Lab in Alderley Park | Wellcome Sanger Institute for the COVID-19 Genomics UK (COG-UK) Consortium | Jacquelyn Wynn, Mairead Hyland, The Lighthouse Lab in Alderley Park and Alex Alderton, Roberto Amato, Jeffrey Barrett, Sonia Goncalves, Ewan Harrison, David K. Jackson, Ian Johnston, Dominic Kwiatkowski, Cordelia Langford, John Sillitoe on behalf of the Wellcome Sanger Institute COVID-19 Surveillance Team |
| EPI_ISL_3108092 | Lighthouse Lab in Glasgow | Wellcome Sanger Institute for the COVID-19 Genomics UK (COG-UK) Consortium | Harper VanSteenhouse, Yumi Kasai, David Gray, Carol Clugston, Anna Dominiczak and Alex Alderton, Roberto Amato, Jeffrey Barrett, Sonia Goncalves, Ewan Harrison, David K. Jackson, Ian Johnston, Dominic Kwiatkowski, Cordelia Langford, John Sillitoe on behalf of the Wellcome Sanger Institute COVID-19 Surveillance Team |
| EPI_ISL_3108093 | Lighthouse Lab in Alderley Park | Wellcome Sanger Institute for the COVID-19 Genomics UK (COG-UK) Consortium | Jacquelyn Wynn, Mairead Hyland, The Lighthouse Lab in Alderley Park and Alex Alderton, Roberto Amato, Jeffrey Barrett, Sonia Goncalves, Ewan Harrison, David K. Jackson, Ian Johnston, Dominic Kwiatkowski, Cordelia Langford, John Sillitoe on behalf of the Wellcome Sanger Institute COVID-19 Surveillance Team |
| EPI_ISL_3108094 | Lighthouse Lab in Glasgow | Wellcome Sanger Institute for the COVID-19 Genomics UK (COG-UK) Consortium | Harper VanSteenhouse, Yumi Kasai, David Gray, Carol Clugston, Anna Dominiczak and Alex Alderton, Roberto Amato, Jeffrey Barrett, Sonia Goncalves, Ewan Harrison, David K. Jackson, Ian Johnston, Dominic Kwiatkowski, Cordelia Langford, John Sillitoe on behalf of the Wellcome Sanger Institute COVID-19 Surveillance Team |
| EPI_ISL_3108095, EPI_ISL_3108096, EPI_ISL_3108097, EPI_ISL_3108098, EPI_ISL_3108099, EPI_ISL_3108100, EPI_ISL_3108101, EPI_ISL_3108102, EPI_ISL_3108103, EPI_ISL_3108104 | Lighthouse Lab in Alderley Park | Wellcome Sanger Institute for the COVID-19 Genomics UK (COG-UK) Consortium | Jacquelyn Wynn, Mairead Hyland, The Lighthouse Lab in Alderley Park and Alex Alderton, Roberto Amato, Jeffrey Barrett, Sonia Goncalves, Ewan Harrison, David K. Jackson, Ian Johnston, Dominic Kwiatkowski, Cordelia Langford, John Sillitoe on behalf of the Wellcome Sanger Institute COVID-19 Surveillance Team |
| EPI_ISL_3108105 | Lighthouse Lab in Glasgow | Wellcome Sanger Institute for the COVID-19 Genomics UK (COG-UK) Consortium | Harper VanSteenhouse, Yumi Kasai, David Gray, Carol Clugston, Anna Dominiczak and Alex Alderton, Roberto Amato, Jeffrey Barrett, Sonia Goncalves, Ewan Harrison, David K. Jackson, Ian Johnston, Dominic Kwiatkowski, Cordelia Langford, John Sillitoe on behalf of the Wellcome Sanger Institute COVID-19 Surveillance Team |
| EPI_ISL_3108106 | Lighthouse Lab in Alderley Park | Wellcome Sanger Institute for the COVID-19 Genomics UK (COG-UK) Consortium | Jacquelyn Wynn, Mairead Hyland, The Lighthouse Lab in Alderley Park and Alex Alderton, Roberto Amato, Jeffrey Barrett, Sonia Goncalves, Ewan Harrison, David K. Jackson, Ian Johnston, Dominic Kwiatkowski, Cordelia Langford, John Sillitoe on behalf of the Wellcome Sanger Institute COVID-19 Surveillance Team |
| EPI_ISL_3108107 | Lighthouse Lab in Glasgow | Wellcome Sanger Institute for the COVID-19 Genomics UK (COG-UK) Consortium | Harper VanSteenhouse, Yumi Kasai, David Gray, Carol Clugston, Anna Dominiczak and Alex Alderton, Roberto Amato, Jeffrey Barrett, Sonia Goncalves, Ewan Harrison, David K. Jackson, Ian Johnston, Dominic Kwiatkowski, Cordelia Langford, John Sillitoe on behalf of the Wellcome Sanger Institute COVID-19 Surveillance Team |
| EPI_ISL_3108108, EPI_ISL_3108109, EPI_ISL_3108110, EPI_ISL_3108111, EPI_ISL_3108112, EPI_ISL_3108113 | Lighthouse Lab in Alderley Park | Wellcome Sanger Institute for the COVID-19 Genomics UK (COG-UK) Consortium | Jacquelyn Wynn, Mairead Hyland, The Lighthouse Lab in Alderley Park and Alex Alderton, Roberto Amato, Jeffrey Barrett, Sonia Goncalves, Ewan Harrison, David K. Jackson, Ian Johnston, Dominic Kwiatkowski, Cordelia Langford, John Sillitoe on behalf of the Wellcome Sanger Institute COVID-19 Surveillance Team |
| EPI_ISL_3108114 | Lighthouse Lab in Glasgow | Wellcome Sanger Institute for the COVID-19 Genomics UK (COG-UK) Consortium | Harper VanSteenhouse, Yumi Kasai, David Gray, Carol Clugston, Anna Dominiczak and Alex Alderton, Roberto Amato, Jeffrey Barrett, Sonia Goncalves, Ewan Harrison, David K. Jackson, Ian Johnston, Dominic Kwiatkowski, Cordelia Langford, John Sillitoe on behalf of the Wellcome Sanger Institute COVID-19 Surveillance Team |

| | | | |
|---|---|---|---|
| EPI_ISL_3108115 | Lighthouse Lab in Alderley Park | Wellcome Sanger Institute for the COVID-19 Genomics UK (COG-UK) Consortium | Jacquelyn Wynn, Mairead Hyland, The Lighthouse Lab in Alderley Park and Alex Alderton, Roberto Amato, Jeffrey Barrett, Sonia Goncalves, Ewan Harrison, David K. Jackson, Ian Johnston, Dominic Kwiatkowski, Cordelia Langford, John Sillitoe on behalf of the Wellcome Sanger Institute COVID-19 Surveillance Team |
| EPI_ISL_3108116 | Lighthouse Lab in Glasgow | Wellcome Sanger Institute for the COVID-19 Genomics UK (COG-UK) Consortium | Harper VanSteenhouse, Yumi Kasai, David Gray, Carol Clugston, Anna Dominiczak and Alex Alderton, Roberto Amato, Jeffrey Barrett, Sonia Goncalves, Ewan Harrison, David K. Jackson, Ian Johnston, Dominic Kwiatkowski, Cordelia Langford, John Sillitoe on behalf of the Wellcome Sanger Institute COVID-19 Surveillance Team |
| EPI_ISL_3108117 | Lighthouse Lab in Alderley Park | Wellcome Sanger Institute for the COVID-19 Genomics UK (COG-UK) Consortium | Jacquelyn Wynn, Mairead Hyland, The Lighthouse Lab in Alderley Park and Alex Alderton, Roberto Amato, Jeffrey Barrett, Sonia Goncalves, Ewan Harrison, David K. Jackson, Ian Johnston, Dominic Kwiatkowski, Cordelia Langford, John Sillitoe on behalf of the Wellcome Sanger Institute COVID-19 Surveillance Team |
| EPI_ISL_3108118, EPI_ISL_3108119 | Lighthouse Lab in Glasgow | Wellcome Sanger Institute for the COVID-19 Genomics UK (COG-UK) Consortium | Harper VanSteenhouse, Yumi Kasai, David Gray, Carol Clugston, Anna Dominiczak and Alex Alderton, Roberto Amato, Jeffrey Barrett, Sonia Goncalves, Ewan Harrison, David K. Jackson, Ian Johnston, Dominic Kwiatkowski, Cordelia Langford, John Sillitoe on behalf of the Wellcome Sanger Institute COVID-19 Surveillance Team |
| EPI_ISL_3108120, EPI_ISL_3108121, EPI_ISL_3108122, EPI_ISL_3108123, EPI_ISL_3108124, EPI_ISL_3108125 | Lighthouse Lab in Alderley Park | Wellcome Sanger Institute for the COVID-19 Genomics UK (COG-UK) Consortium | Jacquelyn Wynn, Mairead Hyland, The Lighthouse Lab in Alderley Park and Alex Alderton, Roberto Amato, Jeffrey Barrett, Sonia Goncalves, Ewan Harrison, David K. Jackson, Ian Johnston, Dominic Kwiatkowski, Cordelia Langford, John Sillitoe on behalf of the Wellcome Sanger Institute COVID-19 Surveillance Team |
| EPI_ISL_3108126 | Lighthouse Lab in Glasgow | Wellcome Sanger Institute for the COVID-19 Genomics UK (COG-UK) Consortium | Harper VanSteenhouse, Yumi Kasai, David Gray, Carol Clugston, Anna Dominiczak and Alex Alderton, Roberto Amato, Jeffrey Barrett, Sonia Goncalves, Ewan Harrison, David K. Jackson, Ian Johnston, Dominic Kwiatkowski, Cordelia Langford, John Sillitoe on behalf of the Wellcome Sanger Institute COVID-19 Surveillance Team |
| EPI_ISL_3108127, EPI_ISL_3108128, EPI_ISL_3108129, EPI_ISL_3108130, EPI_ISL_3108131, EPI_ISL_3108132 | Lighthouse Lab in Alderley Park | Wellcome Sanger Institute for the COVID-19 Genomics UK (COG-UK) Consortium | Jacquelyn Wynn, Mairead Hyland, The Lighthouse Lab in Alderley Park and Alex Alderton, Roberto Amato, Jeffrey Barrett, Sonia Goncalves, Ewan Harrison, David K. Jackson, Ian Johnston, Dominic Kwiatkowski, Cordelia Langford, John Sillitoe on behalf of the Wellcome Sanger Institute COVID-19 Surveillance Team |
| EPI_ISL_3108133, EPI_ISL_3108134 | Lighthouse Lab in Glasgow | Wellcome Sanger Institute for the COVID-19 Genomics UK (COG-UK) Consortium | Harper VanSteenhouse, Yumi Kasai, David Gray, Carol Clugston, Anna Dominiczak and Alex Alderton, Roberto Amato, Jeffrey Barrett, Sonia Goncalves, Ewan Harrison, David K. Jackson, Ian Johnston, Dominic Kwiatkowski, Cordelia Langford, John Sillitoe on behalf of the Wellcome Sanger Institute COVID-19 Surveillance Team |
| EPI_ISL_3108135, EPI_ISL_3108136 | Lighthouse Lab in Alderley Park | Wellcome Sanger Institute for the COVID-19 Genomics UK (COG-UK) Consortium | Jacquelyn Wynn, Mairead Hyland, The Lighthouse Lab in Alderley Park and Alex Alderton, Roberto Amato, Jeffrey Barrett, Sonia Goncalves, Ewan Harrison, David K. Jackson, Ian Johnston, Dominic Kwiatkowski, Cordelia Langford, John Sillitoe on behalf of the Wellcome Sanger Institute COVID-19 Surveillance Team |
| EPI_ISL_3108137 | Lighthouse Lab in Glasgow | Wellcome Sanger Institute for the COVID-19 Genomics UK (COG-UK) Consortium | Harper VanSteenhouse, Yumi Kasai, David Gray, Carol Clugston, Anna Dominiczak and Alex Alderton, Roberto Amato, Jeffrey Barrett, Sonia Goncalves, Ewan Harrison, David K. Jackson, Ian Johnston, Dominic Kwiatkowski, Cordelia Langford, John Sillitoe on behalf of the Wellcome Sanger Institute COVID-19 Surveillance Team |
| EPI_ISL_3108138 | Lighthouse Lab in Alderley Park | Wellcome Sanger Institute for the COVID-19 Genomics UK (COG-UK) Consortium | Jacquelyn Wynn, Mairead Hyland, The Lighthouse Lab in Alderley Park and Alex Alderton, Roberto Amato, Jeffrey Barrett, Sonia Goncalves, Ewan Harrison, David K. Jackson, Ian Johnston, Dominic Kwiatkowski, Cordelia Langford, John Sillitoe on behalf of the Wellcome Sanger Institute COVID-19 Surveillance Team |
| EPI_ISL_3108139 | Lighthouse Lab in Glasgow | Wellcome Sanger Institute for the COVID-19 Genomics UK (COG-UK) Consortium | Harper VanSteenhouse, Yumi Kasai, David Gray, Carol Clugston, Anna Dominiczak and Alex Alderton, Roberto Amato, Jeffrey Barrett, Sonia Goncalves, Ewan Harrison, David K. Jackson, Ian Johnston, Dominic Kwiatkowski, Cordelia Langford, John Sillitoe on behalf of the Wellcome Sanger Institute COVID-19 Surveillance Team |
| EPI_ISL_3108140, EPI_ISL_3108141, EPI_ISL_3108142, EPI_ISL_3108143, EPI_ISL_3108144, EPI_ISL_3108145, EPI_ISL_3108146, EPI_ISL_3108147, EPI_ISL_3108148, EPI_ISL_3108149, EPI_ISL_3108150, EPI_ISL_3108151, EPI_ISL_3108152, EPI_ISL_3108153, EPI_ISL_3108154, EPI_ISL_3108155, EPI_ISL_3108156 | | | |
| see above | Lighthouse Lab in Alderley Park | Wellcome Sanger Institute for the COVID-19 Genomics UK (COG-UK) Consortium | Jacquelyn Wynn, Mairead Hyland, The Lighthouse Lab in Alderley Park and Alex Alderton, Roberto Amato, Jeffrey Barrett, Sonia Goncalves, Ewan Harrison, David K. Jackson, Ian Johnston, Dominic Kwiatkowski, Cordelia Langford, John Sillitoe on behalf of the Wellcome Sanger Institute COVID-19 Surveillance Team |
| EPI_ISL_3108157 | Lighthouse Lab in Glasgow | Wellcome Sanger Institute for the COVID-19 Genomics UK (COG-UK) Consortium | Harper VanSteenhouse, Yumi Kasai, David Gray, Carol Clugston, Anna Dominiczak and Alex Alderton, Roberto Amato, Jeffrey Barrett, Sonia Goncalves, Ewan Harrison, David K. Jackson, Ian Johnston, Dominic Kwiatkowski, Cordelia Langford, John Sillitoe on behalf of the Wellcome Sanger Institute COVID-19 Surveillance Team |
| EPI_ISL_3108158, EPI_ISL_3108159, EPI_ISL_3108160, EPI_ISL_3108161, EPI_ISL_3108162, EPI_ISL_3108163 | Lighthouse Lab in Alderley Park | Wellcome Sanger Institute for the COVID-19 Genomics UK (COG-UK) Consortium | Jacquelyn Wynn, Mairead Hyland, The Lighthouse Lab in Alderley Park and Alex Alderton, Roberto Amato, Jeffrey Barrett, Sonia Goncalves, Ewan Harrison, David K. Jackson, Ian Johnston, Dominic Kwiatkowski, Cordelia Langford, John Sillitoe on behalf of the Wellcome Sanger Institute COVID-19 Surveillance Team |
| EPI_ISL_3108164, EPI_ISL_3108165 | Lighthouse Lab in Glasgow | Wellcome Sanger Institute for the COVID-19 Genomics UK (COG-UK) Consortium | Harper VanSteenhouse, Yumi Kasai, David Gray, Carol Clugston, Anna Dominiczak and Alex Alderton, Roberto Amato, Jeffrey Barrett, Sonia Goncalves, Ewan Harrison, David K. Jackson, Ian Johnston, Dominic Kwiatkowski, Cordelia Langford, John Sillitoe on behalf of the Wellcome Sanger Institute COVID-19 Surveillance Team |
| EPI_ISL_3108166, EPI_ISL_3108167 | Lighthouse Lab in Alderley Park | Wellcome Sanger Institute for the COVID-19 Genomics UK (COG-UK) Consortium | Jacquelyn Wynn, Mairead Hyland, The Lighthouse Lab in Alderley Park and Alex Alderton, Roberto Amato, Jeffrey Barrett, Sonia Goncalves, Ewan Harrison, David K. Jackson, Ian Johnston, Dominic Kwiatkowski, Cordelia Langford, John Sillitoe on behalf of the Wellcome Sanger Institute COVID-19 Surveillance Team |
| EPI_ISL_3108168, EPI_ISL_3108169 | Lighthouse Lab in Glasgow | Wellcome Sanger Institute for the COVID-19 Genomics UK (COG-UK) Consortium | Harper VanSteenhouse, Yumi Kasai, David Gray, Carol Clugston, Anna Dominiczak and Alex Alderton, Roberto Amato, Jeffrey Barrett, Sonia Goncalves, Ewan Harrison, David K. Jackson, Ian Johnston, Dominic Kwiatkowski, Cordelia Langford, John Sillitoe on behalf of the Wellcome Sanger Institute COVID-19 Surveillance Team |
| EPI_ISL_3108170 | Lighthouse Lab in Alderley Park | Wellcome Sanger Institute for the COVID-19 Genomics UK (COG-UK) Consortium | Jacquelyn Wynn, Mairead Hyland, The Lighthouse Lab in Alderley Park and Alex Alderton, Roberto Amato, Jeffrey Barrett, Sonia Goncalves, Ewan Harrison, David K. Jackson, Ian Johnston, Dominic Kwiatkowski, Cordelia Langford, John Sillitoe on behalf of the Wellcome Sanger Institute COVID-19 Surveillance Team |
| EPI_ISL_3108171 | Lighthouse Lab in Glasgow | Wellcome Sanger Institute for the COVID-19 Genomics UK (COG-UK) Consortium | Harper VanSteenhouse, Yumi Kasai, David Gray, Carol Clugston, Anna Dominiczak and Alex Alderton, Roberto Amato, Jeffrey Barrett, Sonia Goncalves, Ewan Harrison, David K. Jackson, Ian Johnston, Dominic Kwiatkowski, Cordelia Langford, John Sillitoe on behalf of the Wellcome Sanger Institute COVID-19 Surveillance Team |
| EPI_ISL_3108172, EPI_ISL_3108173, EPI_ISL_3108174 | Lighthouse Lab in Alderley Park | Wellcome Sanger Institute for the COVID-19 Genomics UK (COG-UK) Consortium | Jacquelyn Wynn, Mairead Hyland, The Lighthouse Lab in Alderley Park and Alex Alderton, Roberto Amato, Jeffrey Barrett, Sonia Goncalves, Ewan Harrison, David K. Jackson, Ian Johnston, Dominic Kwiatkowski, Cordelia Langford, John Sillitoe on behalf of the Wellcome Sanger Institute COVID-19 Surveillance Team |
| EPI_ISL_3108175, EPI_ISL_3108176 | Lighthouse Lab in Glasgow | Wellcome Sanger Institute for the COVID-19 Genomics UK (COG-UK) Consortium | Harper VanSteenhouse, Yumi Kasai, David Gray, Carol Clugston, Anna Dominiczak and Alex Alderton, Roberto Amato, Jeffrey Barrett, Sonia Goncalves, Ewan Harrison, David K. Jackson, Ian Johnston, Dominic Kwiatkowski, Cordelia Langford, John Sillitoe on behalf of the Wellcome Sanger Institute COVID-19 Surveillance Team |
| EPI_ISL_3108177 | Lighthouse Lab in Alderley Park | Wellcome Sanger Institute for the COVID-19 Genomics UK (COG-UK) Consortium | Jacquelyn Wynn, Mairead Hyland, The Lighthouse Lab in Alderley Park and Alex Alderton, Roberto Amato, Jeffrey Barrett, Sonia Goncalves, Ewan Harrison, David K. Jackson, Ian Johnston, Dominic Kwiatkowski, Cordelia Langford, John Sillitoe on behalf of the Wellcome Sanger Institute COVID-19 Surveillance Team |
| EPI_ISL_3108178 | Lighthouse Lab in Glasgow | Wellcome Sanger Institute for the COVID-19 Genomics UK (COG-UK) Consortium | Harper VanSteenhouse, Yumi Kasai, David Gray, Carol Clugston, Anna Dominiczak and Alex Alderton, Roberto Amato, Jeffrey Barrett, Sonia Goncalves, Ewan Harrison, David K. Jackson, Ian Johnston, Dominic Kwiatkowski, Cordelia Langford, John Sillitoe on behalf of the Wellcome Sanger Institute COVID-19 Surveillance Team |
| EPI_ISL_3108179, EPI_ISL_3108180 | Lighthouse Lab in Alderley Park | Wellcome Sanger Institute for the COVID-19 Genomics UK (COG-UK) Consortium | Jacquelyn Wynn, Mairead Hyland, The Lighthouse Lab in Alderley Park and Alex Alderton, Roberto Amato, Jeffrey Barrett, Sonia Goncalves, Ewan Harrison, David K. Jackson, Ian Johnston, Dominic Kwiatkowski, Cordelia Langford, John Sillitoe on behalf of the Wellcome Sanger Institute COVID-19 Surveillance Team |
| EPI_ISL_3108181 | Lighthouse Lab in Glasgow | Wellcome Sanger Institute for the COVID-19 Genomics UK (COG-UK) Consortium | Harper VanSteenhouse, Yumi Kasai, David Gray, Carol Clugston, Anna Dominiczak and Alex Alderton, Roberto Amato, Jeffrey Barrett, Sonia Goncalves, Ewan Harrison, David K. Jackson, Ian Johnston, Dominic Kwiatkowski, Cordelia Langford, John Sillitoe on behalf of the Wellcome Sanger Institute COVID-19 Surveillance Team |
| EPI_ISL_3108182 | Lighthouse Lab in Alderley Park | Wellcome Sanger Institute for the COVID-19 Genomics UK (COG-UK) Consortium | Jacquelyn Wynn, Mairead Hyland, The Lighthouse Lab in Alderley Park and Alex Alderton, Roberto Amato, Jeffrey Barrett, Sonia Goncalves, Ewan Harrison, David K. Jackson, Ian Johnston, Dominic Kwiatkowski, Cordelia Langford, John Sillitoe on behalf of the Wellcome Sanger Institute COVID-19 Surveillance Team |
| EPI_ISL_3108183 | Lighthouse Lab in Glasgow | Wellcome Sanger Institute for the COVID-19 Genomics UK (COG-UK) Consortium | Harper VanSteenhouse, Yumi Kasai, David Gray, Carol Clugston, Anna Dominiczak and Alex Alderton, Roberto Amato, Jeffrey Barrett, Sonia Goncalves, Ewan Harrison, David K. Jackson, Ian Johnston, Dominic Kwiatkowski, Cordelia Langford, John Sillitoe on behalf of the Wellcome Sanger Institute COVID-19 Surveillance Team |
| EPI_ISL_3108184, EPI_ISL_3108185, EPI_ISL_3108186, EPI_ISL_3108187, EPI_ISL_3108188, EPI_ISL_3108189, EPI_ISL_3108190 | Lighthouse Lab in Alderley Park | Wellcome Sanger Institute for the COVID-19 Genomics UK (COG-UK) Consortium | Jacquelyn Wynn, Mairead Hyland, The Lighthouse Lab in Alderley Park and Alex Alderton, Roberto Amato, Jeffrey Barrett, Sonia Goncalves, Ewan Harrison, David K. Jackson, Ian Johnston, Dominic Kwiatkowski, Cordelia Langford, John Sillitoe on behalf of the Wellcome Sanger Institute COVID-19 Surveillance Team |
| EPI_ISL_3108191, EPI_ISL_3108192 | Lighthouse Lab in Glasgow | Wellcome Sanger Institute for the COVID-19 Genomics UK (COG-UK) Consortium | Harper VanSteenhouse, Yumi Kasai, David Gray, Carol Clugston, Anna Dominiczak and Alex Alderton, Roberto Amato, Jeffrey Barrett, Sonia Goncalves, Ewan Harrison, David K. Jackson, Ian Johnston, Dominic Kwiatkowski, Cordelia Langford, John Sillitoe on behalf of the Wellcome Sanger Institute COVID-19 Surveillance Team |
| EPI_ISL_3108193, EPI_ISL_3108194, EPI_ISL_3108195 | Lighthouse Lab in Alderley Park | Wellcome Sanger Institute for the COVID-19 Genomics UK (COG-UK) Consortium | Jacquelyn Wynn, Mairead Hyland, The Lighthouse Lab in Alderley Park and Alex Alderton, Roberto Amato, Jeffrey Barrett, Sonia Goncalves, Ewan Harrison, David K. Jackson, Ian Johnston, Dominic Kwiatkowski, Cordelia Langford, John Sillitoe on behalf of the Wellcome Sanger Institute COVID-19 Surveillance Team |
| EPI_ISL_3108196 | Lighthouse Lab in Glasgow | Wellcome Sanger Institute for the COVID-19 Genomics UK (COG-UK) Consortium | Harper VanSteenhouse, Yumi Kasai, David Gray, Carol Clugston, Anna Dominiczak and Alex Alderton, Roberto Amato, Jeffrey Barrett, Sonia Goncalves, Ewan Harrison, David K. Jackson, Ian Johnston, Dominic Kwiatkowski, Cordelia Langford, John Sillitoe on behalf of the Wellcome Sanger Institute COVID-19 Surveillance Team |
| EPI_ISL_3108197, EPI_ISL_3108198, EPI_ISL_3108199, EPI_ISL_3108200, EPI_ISL_3108201, EPI_ISL_3108202, EPI_ISL_3108203, EPI_ISL_3108204, EPI_ISL_3108205, EPI_ISL_3108206, EPI_ISL_3108207, EPI_ISL_3108208, EPI_ISL_3108209, EPI_ISL_3108210, EPI_ISL_3108211 | | | |
| see above | Lighthouse Lab in Alderley Park | Wellcome Sanger Institute for the COVID-19 Genomics UK (COG-UK) Consortium | Jacquelyn Wynn, Mairead Hyland, The Lighthouse Lab in Alderley Park and Alex Alderton, Roberto Amato, Jeffrey Barrett, Sonia Goncalves, Ewan Harrison, David K. Jackson, Ian Johnston, Dominic Kwiatkowski, Cordelia Langford, John Sillitoe on behalf of the Wellcome Sanger Institute COVID-19 Surveillance Team |
| EPI_ISL_3108212 | Lighthouse Lab in Glasgow | Wellcome Sanger Institute for the COVID-19 Genomics UK (COG-UK) Consortium | Harper VanSteenhouse, Yumi Kasai, David Gray, Carol Clugston, Anna Dominiczak and Alex Alderton, Roberto Amato, Jeffrey Barrett, Sonia Goncalves, Ewan Harrison, David K. Jackson, Ian Johnston, Dominic Kwiatkowski, Cordelia Langford, John Sillitoe on behalf of the Wellcome Sanger Institute COVID-19 Surveillance Team |
| EPI_ISL_3108213, EPI_ISL_3108214 | Lighthouse Lab in Alderley Park | Wellcome Sanger Institute for the COVID-19 Genomics UK (COG-UK) Consortium | Jacquelyn Wynn, Mairead Hyland, The Lighthouse Lab in Alderley Park and Alex Alderton, Roberto Amato, Jeffrey Barrett, Sonia Goncalves, Ewan Harrison, David K. Jackson, Ian Johnston, Dominic Kwiatkowski, Cordelia Langford, John Sillitoe on behalf of the Wellcome Sanger Institute COVID-19 Surveillance Team |
| EPI_ISL_3108215 | Lighthouse Lab in Glasgow | Wellcome Sanger Institute for the COVID-19 Genomics UK (COG-UK) Consortium | Harper VanSteenhouse, Yumi Kasai, David Gray, Carol Clugston, Anna Dominiczak and Alex Alderton, Roberto Amato, Jeffrey Barrett, Sonia Goncalves, Ewan Harrison, David K. Jackson, Ian Johnston, Dominic Kwiatkowski, Cordelia Langford, John Sillitoe on behalf of the Wellcome Sanger Institute COVID-19 Surveillance Team |

| | | | |
|---|---|---|---|
| EPI_ISL_3108216, EPI_ISL_3108217 | Lighthouse Lab in Alderley Park | Wellcome Sanger Institute for the COVID-19 Genomics UK (COG-UK) Consortium | Jacquelyn Wynn, Mairead Hyland, The Lighthouse Lab in Alderley Park and Alex Alderton, Roberto Amato, Jeffrey Barrett, Sonia Goncalves, Ewan Harrison, David K. Jackson, Ian Johnston, Dominic Kwiatkowski, Cordelia Langford, John Sillitoe on behalf of the Wellcome Sanger Institute COVID-19 Surveillance Team |
| EPI_ISL_3108218 | Lighthouse Lab in Glasgow | Wellcome Sanger Institute for the COVID-19 Genomics UK (COG-UK) Consortium | Harper VanSteenhouse, Yumi Kasai, David Gray, Carol Clugston, Anna Dominiczak and Alex Alderton, Roberto Amato, Jeffrey Barrett, Sonia Goncalves, Ewan Harrison, David K. Jackson, Ian Johnston, Dominic Kwiatkowski, Cordelia Langford, John Sillitoe on behalf of the Wellcome Sanger Institute COVID-19 Surveillance Team |
| EPI_ISL_3108219 | Lighthouse Lab in Alderley Park | Wellcome Sanger Institute for the COVID-19 Genomics UK (COG-UK) Consortium | Jacquelyn Wynn, Mairead Hyland, The Lighthouse Lab in Alderley Park and Alex Alderton, Roberto Amato, Jeffrey Barrett, Sonia Goncalves, Ewan Harrison, David K. Jackson, Ian Johnston, Dominic Kwiatkowski, Cordelia Langford, John Sillitoe on behalf of the Wellcome Sanger Institute COVID-19 Surveillance Team |
| EPI_ISL_3108220 | Lighthouse Lab in Glasgow | Wellcome Sanger Institute for the COVID-19 Genomics UK (COG-UK) Consortium | Harper VanSteenhouse, Yumi Kasai, David Gray, Carol Clugston, Anna Dominiczak and Alex Alderton, Roberto Amato, Jeffrey Barrett, Sonia Goncalves, Ewan Harrison, David K. Jackson, Ian Johnston, Dominic Kwiatkowski, Cordelia Langford, John Sillitoe on behalf of the Wellcome Sanger Institute COVID-19 Surveillance Team |
| EPI_ISL_3108221 | Lighthouse Lab in Alderley Park | Wellcome Sanger Institute for the COVID-19 Genomics UK (COG-UK) Consortium | Jacquelyn Wynn, Mairead Hyland, The Lighthouse Lab in Alderley Park and Alex Alderton, Roberto Amato, Jeffrey Barrett, Sonia Goncalves, Ewan Harrison, David K. Jackson, Ian Johnston, Dominic Kwiatkowski, Cordelia Langford, John Sillitoe on behalf of the Wellcome Sanger Institute COVID-19 Surveillance Team |
| EPI_ISL_3108222 | Lighthouse Lab in Glasgow | Wellcome Sanger Institute for the COVID-19 Genomics UK (COG-UK) Consortium | Harper VanSteenhouse, Yumi Kasai, David Gray, Carol Clugston, Anna Dominiczak and Alex Alderton, Roberto Amato, Jeffrey Barrett, Sonia Goncalves, Ewan Harrison, David K. Jackson, Ian Johnston, Dominic Kwiatkowski, Cordelia Langford, John Sillitoe on behalf of the Wellcome Sanger Institute COVID-19 Surveillance Team |
| EPI_ISL_3108223, EPI_ISL_3108224, EPI_ISL_3108225, EPI_ISL_3108226, EPI_ISL_3108227, EPI_ISL_3108228, EPI_ISL_3108229, EPI_ISL_3108230, EPI_ISL_3108231, EPI_ISL_3108232 | Lighthouse Lab in Alderley Park | Wellcome Sanger Institute for the COVID-19 Genomics UK (COG-UK) Consortium | Jacquelyn Wynn, Mairead Hyland, The Lighthouse Lab in Alderley Park and Alex Alderton, Roberto Amato, Jeffrey Barrett, Sonia Goncalves, Ewan Harrison, David K. Jackson, Ian Johnston, Dominic Kwiatkowski, Cordelia Langford, John Sillitoe on behalf of the Wellcome Sanger Institute COVID-19 Surveillance Team |
| EPI_ISL_3108233 | Lighthouse Lab in Glasgow | Wellcome Sanger Institute for the COVID-19 Genomics UK (COG-UK) Consortium | Harper VanSteenhouse, Yumi Kasai, David Gray, Carol Clugston, Anna Dominiczak and Alex Alderton, Roberto Amato, Jeffrey Barrett, Sonia Goncalves, Ewan Harrison, David K. Jackson, Ian Johnston, Dominic Kwiatkowski, Cordelia Langford, John Sillitoe on behalf of the Wellcome Sanger Institute COVID-19 Surveillance Team |
| EPI_ISL_3108234 | Lighthouse Lab in Alderley Park | Wellcome Sanger Institute for the COVID-19 Genomics UK (COG-UK) Consortium | Jacquelyn Wynn, Mairead Hyland, The Lighthouse Lab in Alderley Park and Alex Alderton, Roberto Amato, Jeffrey Barrett, Sonia Goncalves, Ewan Harrison, David K. Jackson, Ian Johnston, Dominic Kwiatkowski, Cordelia Langford, John Sillitoe on behalf of the Wellcome Sanger Institute COVID-19 Surveillance Team |
| EPI_ISL_3108235 | Lighthouse Lab in Glasgow | Wellcome Sanger Institute for the COVID-19 Genomics UK (COG-UK) Consortium | Harper VanSteenhouse, Yumi Kasai, David Gray, Carol Clugston, Anna Dominiczak and Alex Alderton, Roberto Amato, Jeffrey Barrett, Sonia Goncalves, Ewan Harrison, David K. Jackson, Ian Johnston, Dominic Kwiatkowski, Cordelia Langford, John Sillitoe on behalf of the Wellcome Sanger Institute COVID-19 Surveillance Team |
| EPI_ISL_3108236 | Lighthouse Lab in Alderley Park | Wellcome Sanger Institute for the COVID-19 Genomics UK (COG-UK) Consortium | Jacquelyn Wynn, Mairead Hyland, The Lighthouse Lab in Alderley Park and Alex Alderton, Roberto Amato, Jeffrey Barrett, Sonia Goncalves, Ewan Harrison, David K. Jackson, Ian Johnston, Dominic Kwiatkowski, Cordelia Langford, John Sillitoe on behalf of the Wellcome Sanger Institute COVID-19 Surveillance Team |
| EPI_ISL_3108237 | Lighthouse Lab in Glasgow | Wellcome Sanger Institute for the COVID-19 Genomics UK (COG-UK) Consortium | Harper VanSteenhouse, Yumi Kasai, David Gray, Carol Clugston, Anna Dominiczak and Alex Alderton, Roberto Amato, Jeffrey Barrett, Sonia Goncalves, Ewan Harrison, David K. Jackson, Ian Johnston, Dominic Kwiatkowski, Cordelia Langford, John Sillitoe on behalf of the Wellcome Sanger Institute COVID-19 Surveillance Team |
| EPI_ISL_3108238 | Lighthouse Lab in Alderley Park | Wellcome Sanger Institute for the COVID-19 Genomics UK (COG-UK) Consortium | Jacquelyn Wynn, Mairead Hyland, The Lighthouse Lab in Alderley Park and Alex Alderton, Roberto Amato, Jeffrey Barrett, Sonia Goncalves, Ewan Harrison, David K. Jackson, Ian Johnston, Dominic Kwiatkowski, Cordelia Langford, John Sillitoe on behalf of the Wellcome Sanger Institute COVID-19 Surveillance Team |
| EPI_ISL_3108239 | Lighthouse Lab in Glasgow | Wellcome Sanger Institute for the COVID-19 Genomics UK (COG-UK) Consortium | Harper VanSteenhouse, Yumi Kasai, David Gray, Carol Clugston, Anna Dominiczak and Alex Alderton, Roberto Amato, Jeffrey Barrett, Sonia Goncalves, Ewan Harrison, David K. Jackson, Ian Johnston, Dominic Kwiatkowski, Cordelia Langford, John Sillitoe on behalf of the Wellcome Sanger Institute COVID-19 Surveillance Team |
| EPI_ISL_3108240 | Lighthouse Lab in Alderley Park | Wellcome Sanger Institute for the COVID-19 Genomics UK (COG-UK) Consortium | Jacquelyn Wynn, Mairead Hyland, The Lighthouse Lab in Alderley Park and Alex Alderton, Roberto Amato, Jeffrey Barrett, Sonia Goncalves, Ewan Harrison, David K. Jackson, Ian Johnston, Dominic Kwiatkowski, Cordelia Langford, John Sillitoe on behalf of the Wellcome Sanger Institute COVID-19 Surveillance Team |
| EPI_ISL_3108241 | Lighthouse Lab in Glasgow | Wellcome Sanger Institute for the COVID-19 Genomics UK (COG-UK) Consortium | Harper VanSteenhouse, Yumi Kasai, David Gray, Carol Clugston, Anna Dominiczak and Alex Alderton, Roberto Amato, Jeffrey Barrett, Sonia Goncalves, Ewan Harrison, David K. Jackson, Ian Johnston, Dominic Kwiatkowski, Cordelia Langford, John Sillitoe on behalf of the Wellcome Sanger Institute COVID-19 Surveillance Team |
| EPI_ISL_3108242 | Lighthouse Lab in Alderley Park | Wellcome Sanger Institute for the COVID-19 Genomics UK (COG-UK) Consortium | Jacquelyn Wynn, Mairead Hyland, The Lighthouse Lab in Alderley Park and Alex Alderton, Roberto Amato, Jeffrey Barrett, Sonia Goncalves, Ewan Harrison, David K. Jackson, Ian Johnston, Dominic Kwiatkowski, Cordelia Langford, John Sillitoe on behalf of the Wellcome Sanger Institute COVID-19 Surveillance Team |
| EPI_ISL_3108243, EPI_ISL_3108244, EPI_ISL_3108245 | Lighthouse Lab in Glasgow | Wellcome Sanger Institute for the COVID-19 Genomics UK (COG-UK) Consortium | Harper VanSteenhouse, Yumi Kasai, David Gray, Carol Clugston, Anna Dominiczak and Alex Alderton, Roberto Amato, Jeffrey Barrett, Sonia Goncalves, Ewan Harrison, David K. Jackson, Ian Johnston, Dominic Kwiatkowski, Cordelia Langford, John Sillitoe on behalf of the Wellcome Sanger Institute COVID-19 Surveillance Team |
| EPI_ISL_3108246, EPI_ISL_3108247 | Lighthouse Lab in Alderley Park | Wellcome Sanger Institute for the COVID-19 Genomics UK (COG-UK) Consortium | Jacquelyn Wynn, Mairead Hyland, The Lighthouse Lab in Alderley Park and Alex Alderton, Roberto Amato, Jeffrey Barrett, Sonia Goncalves, Ewan Harrison, David K. Jackson, Ian Johnston, Dominic Kwiatkowski, Cordelia Langford, John Sillitoe on behalf of the Wellcome Sanger Institute COVID-19 Surveillance Team |
| EPI_ISL_3108248, EPI_ISL_3108249 | Lighthouse Lab in Glasgow | Wellcome Sanger Institute for the COVID-19 Genomics UK (COG-UK) Consortium | Harper VanSteenhouse, Yumi Kasai, David Gray, Carol Clugston, Anna Dominiczak and Alex Alderton, Roberto Amato, Jeffrey Barrett, Sonia Goncalves, Ewan Harrison, David K. Jackson, Ian Johnston, Dominic Kwiatkowski, Cordelia Langford, John Sillitoe on behalf of the Wellcome Sanger Institute COVID-19 Surveillance Team |
| EPI_ISL_3108250 | Lighthouse Lab in Milton Keynes | Wellcome Sanger Institute for the COVID-19 Genomics UK (COG-UK) Consortium | The Lighthouse Lab in Milton Keynes and Alex Alderton, Roberto Amato, Jeffrey Barrett, Sonia Goncalves, Ewan Harrison, David K. Jackson, Ian Johnston, Dominic Kwiatkowski, Cordelia Langford, John Sillitoe on behalf of the Wellcome Sanger Institute COVID-19 Surveillance Team |
| EPI_ISL_3108251 | Lighthouse Lab in Alderley Park | Wellcome Sanger Institute for the COVID-19 Genomics UK (COG-UK) Consortium | Jacquelyn Wynn, Mairead Hyland, The Lighthouse Lab in Alderley Park and Alex Alderton, Roberto Amato, Jeffrey Barrett, Sonia Goncalves, Ewan Harrison, David K. Jackson, Ian Johnston, Dominic Kwiatkowski, Cordelia Langford, John Sillitoe on behalf of the Wellcome Sanger Institute COVID-19 Surveillance Team |
| EPI_ISL_3108252, EPI_ISL_3108253, EPI_ISL_3108254 | Lighthouse Lab in Milton Keynes | Wellcome Sanger Institute for the COVID-19 Genomics UK (COG-UK) Consortium | The Lighthouse Lab in Milton Keynes and Alex Alderton, Roberto Amato, Jeffrey Barrett, Sonia Goncalves, Ewan Harrison, David K. Jackson, Ian Johnston, Dominic Kwiatkowski, Cordelia Langford, John Sillitoe on behalf of the Wellcome Sanger Institute COVID-19 Surveillance Team |
| EPI_ISL_3108255, EPI_ISL_3108256, EPI_ISL_3108257 | Lighthouse Lab in Glasgow | Wellcome Sanger Institute for the COVID-19 Genomics UK (COG-UK) Consortium | Harper VanSteenhouse, Yumi Kasai, David Gray, Carol Clugston, Anna Dominiczak and Alex Alderton, Roberto Amato, Jeffrey Barrett, Sonia Goncalves, Ewan Harrison, David K. Jackson, Ian Johnston, Dominic Kwiatkowski, Cordelia Langford, John Sillitoe on behalf of the Wellcome Sanger Institute COVID-19 Surveillance Team |
| EPI_ISL_3108258, EPI_ISL_3108259 | Lighthouse Lab in Milton Keynes | Wellcome Sanger Institute for the COVID-19 Genomics UK (COG-UK) Consortium | The Lighthouse Lab in Milton Keynes and Alex Alderton, Roberto Amato, Jeffrey Barrett, Sonia Goncalves, Ewan Harrison, David K. Jackson, Ian Johnston, Dominic Kwiatkowski, Cordelia Langford, John Sillitoe on behalf of the Wellcome Sanger Institute COVID-19 Surveillance Team |
| EPI_ISL_3108260 | Lighthouse Lab in Glasgow | Wellcome Sanger Institute for the COVID-19 Genomics UK (COG-UK) Consortium | Harper VanSteenhouse, Yumi Kasai, David Gray, Carol Clugston, Anna Dominiczak and Alex Alderton, Roberto Amato, Jeffrey Barrett, Sonia Goncalves, Ewan Harrison, David K. Jackson, Ian Johnston, Dominic Kwiatkowski, Cordelia Langford, John Sillitoe on behalf of the Wellcome Sanger Institute COVID-19 Surveillance Team |
| EPI_ISL_3108261 | Lighthouse Lab in Milton Keynes | Wellcome Sanger Institute for the COVID-19 Genomics UK (COG-UK) Consortium | The Lighthouse Lab in Milton Keynes and Alex Alderton, Roberto Amato, Jeffrey Barrett, Sonia Goncalves, Ewan Harrison, David K. Jackson, Ian Johnston, Dominic Kwiatkowski, Cordelia Langford, John Sillitoe on behalf of the Wellcome Sanger Institute COVID-19 Surveillance Team |
| EPI_ISL_3108262, EPI_ISL_3108263 | Lighthouse Lab in Alderley Park | Wellcome Sanger Institute for the COVID-19 Genomics UK (COG-UK) Consortium | Jacquelyn Wynn, Mairead Hyland, The Lighthouse Lab in Alderley Park and Alex Alderton, Roberto Amato, Jeffrey Barrett, Sonia Goncalves, Ewan Harrison, David K. Jackson, Ian Johnston, Dominic Kwiatkowski, Cordelia Langford, John Sillitoe on behalf of the Wellcome Sanger Institute COVID-19 Surveillance Team |
| EPI_ISL_3108264 | Lighthouse Lab in Glasgow | Wellcome Sanger Institute for the COVID-19 Genomics UK (COG-UK) Consortium | Harper VanSteenhouse, Yumi Kasai, David Gray, Carol Clugston, Anna Dominiczak and Alex Alderton, Roberto Amato, Jeffrey Barrett, Sonia Goncalves, Ewan Harrison, David K. Jackson, Ian Johnston, Dominic Kwiatkowski, Cordelia Langford, John Sillitoe on behalf of the Wellcome Sanger Institute COVID-19 Surveillance Team |
| EPI_ISL_3108265, EPI_ISL_3108266, EPI_ISL_3108267 | Lighthouse Lab in Milton Keynes | Wellcome Sanger Institute for the COVID-19 Genomics UK (COG-UK) Consortium | The Lighthouse Lab in Milton Keynes and Alex Alderton, Roberto Amato, Jeffrey Barrett, Sonia Goncalves, Ewan Harrison, David K. Jackson, Ian Johnston, Dominic Kwiatkowski, Cordelia Langford, John Sillitoe on behalf of the Wellcome Sanger Institute COVID-19 Surveillance Team |
| EPI_ISL_3108268 | Lighthouse Lab in Glasgow | Wellcome Sanger Institute for the COVID-19 Genomics UK (COG-UK) Consortium | Harper VanSteenhouse, Yumi Kasai, David Gray, Carol Clugston, Anna Dominiczak and Alex Alderton, Roberto Amato, Jeffrey Barrett, Sonia Goncalves, Ewan Harrison, David K. Jackson, Ian Johnston, Dominic Kwiatkowski, Cordelia Langford, John Sillitoe on behalf of the Wellcome Sanger Institute COVID-19 Surveillance Team |
| EPI_ISL_3108269, EPI_ISL_3108270 | Lighthouse Lab in Alderley Park | Wellcome Sanger Institute for the COVID-19 Genomics UK (COG-UK) Consortium | Jacquelyn Wynn, Mairead Hyland, The Lighthouse Lab in Alderley Park and Alex Alderton, Roberto Amato, Jeffrey Barrett, Sonia Goncalves, Ewan Harrison, David K. Jackson, Ian Johnston, Dominic Kwiatkowski, Cordelia Langford, John Sillitoe on behalf of the Wellcome Sanger Institute COVID-19 Surveillance Team |
| EPI_ISL_3108271, EPI_ISL_3108272 | Lighthouse Lab in Glasgow | Wellcome Sanger Institute for the COVID-19 Genomics UK (COG-UK) Consortium | Harper VanSteenhouse, Yumi Kasai, David Gray, Carol Clugston, Anna Dominiczak and Alex Alderton, Roberto Amato, Jeffrey Barrett, Sonia Goncalves, Ewan Harrison, David K. Jackson, Ian Johnston, Dominic Kwiatkowski, Cordelia Langford, John Sillitoe on behalf of the Wellcome Sanger Institute COVID-19 Surveillance Team |
| EPI_ISL_3108273 | Lighthouse Lab in Milton Keynes | Wellcome Sanger Institute for the COVID-19 Genomics UK (COG-UK) Consortium | The Lighthouse Lab in Milton Keynes and Alex Alderton, Roberto Amato, Jeffrey Barrett, Sonia Goncalves, Ewan Harrison, David K. Jackson, Ian Johnston, Dominic Kwiatkowski, Cordelia Langford, John Sillitoe on behalf of the Wellcome Sanger Institute COVID-19 Surveillance Team |
| EPI_ISL_3108274, EPI_ISL_3108275 | Lighthouse Lab in Alderley Park | Wellcome Sanger Institute for the COVID-19 Genomics UK (COG-UK) Consortium | Jacquelyn Wynn, Mairead Hyland, The Lighthouse Lab in Alderley Park and Alex Alderton, Roberto Amato, Jeffrey Barrett, Sonia Goncalves, Ewan Harrison, David K. Jackson, Ian Johnston, Dominic Kwiatkowski, Cordelia Langford, John Sillitoe on behalf of the Wellcome Sanger Institute COVID-19 Surveillance Team |
| EPI_ISL_3108276, EPI_ISL_3108277, EPI_ISL_3108278, EPI_ISL_3108279 | Lighthouse Lab in Milton Keynes | Wellcome Sanger Institute for the COVID-19 Genomics UK (COG-UK) Consortium | The Lighthouse Lab in Milton Keynes and Alex Alderton, Roberto Amato, Jeffrey Barrett, Sonia Goncalves, Ewan Harrison, David K. Jackson, Ian Johnston, Dominic Kwiatkowski, Cordelia Langford, John Sillitoe on behalf of the Wellcome Sanger Institute COVID-19 Surveillance Team |
| EPI_ISL_3108280, EPI_ISL_3108281 | Lighthouse Lab in Alderley Park | Wellcome Sanger Institute for the COVID-19 Genomics UK (COG-UK) Consortium | Jacquelyn Wynn, Mairead Hyland, The Lighthouse Lab in Alderley Park and Alex Alderton, Roberto Amato, Jeffrey Barrett, Sonia Goncalves, Ewan Harrison, David K. Jackson, Ian Johnston, Dominic Kwiatkowski, Cordelia Langford, John Sillitoe on behalf of the Wellcome Sanger Institute COVID-19 Surveillance Team |
| EPI_ISL_3108282 | Lighthouse Lab in Milton Keynes | Wellcome Sanger Institute for the COVID-19 Genomics UK (COG-UK) Consortium | The Lighthouse Lab in Milton Keynes and Alex Alderton, Roberto Amato, Jeffrey Barrett, Sonia Goncalves, Ewan Harrison, David K. Jackson, Ian Johnston, Dominic Kwiatkowski, Cordelia Langford, John Sillitoe on behalf of the Wellcome Sanger Institute COVID-19 Surveillance Team |
| EPI_ISL_3108283, EPI_ISL_3108284 | Lighthouse Lab in Glasgow | Wellcome Sanger Institute for the COVID-19 Genomics UK (COG-UK) Consortium | Harper VanSteenhouse, Yumi Kasai, David Gray, Carol Clugston, Anna Dominiczak and Alex Alderton, Roberto Amato, Jeffrey Barrett, Sonia Goncalves, Ewan Harrison, David K. Jackson, Ian Johnston, Dominic Kwiatkowski, Cordelia Langford, John Sillitoe on behalf of the Wellcome Sanger Institute COVID-19 Surveillance Team |
| EPI_ISL_3108285, | Lighthouse Lab in Alderley Park | Wellcome Sanger Institute for the COVID-19 Genomics UK (COG-UK) | Jacquelyn Wynn, Mairead Hyland, The Lighthouse Lab in Alderley Park and Alex Alderton, Roberto Amato, Jeffrey Barrett, Sonia Goncalves, Ewan Harrison, David K. Jackson, Ian Johnston, Dominic Kwiatkowski, Cordelia Langford, John Sillitoe on |

| | | | |
|---|---|---|---|
| EPI_ISL_3108286, EPI_ISL_3108287, EPI_ISL_3108288, EPI_ISL_3108289, EPI_ISL_3108290 | | Consortium | behalf of the Wellcome Sanger Institute COVID-19 Surveillance Team |
| EPI_ISL_3108291 | Lighthouse Lab in Milton Keynes | Wellcome Sanger Institute for the COVID-19 Genomics UK (COG-UK) Consortium | The Lighthouse Lab in Milton Keynes and Alex Alderton, Roberto Amato, Jeffrey Barrett, Sonia Goncalves, Ewan Harrison, David K. Jackson, Ian Johnston, Dominic Kwiatkowski, Cordelia Langford, John Sillitoe on behalf of the Wellcome Sanger Institute COVID-19 Surveillance Team |
| EPI_ISL_3108292, EPI_ISL_3108293, EPI_ISL_3108294, EPI_ISL_3108295 | Lighthouse Lab in Alderley Park | Wellcome Sanger Institute for the COVID-19 Genomics UK (COG-UK) Consortium | Jacquelyn Wynn, Mairead Hyland, The Lighthouse Lab in Alderley Park and Alex Alderton, Roberto Amato, Jeffrey Barrett, Sonia Goncalves, Ewan Harrison, David K. Jackson, Ian Johnston, Dominic Kwiatkowski, Cordelia Langford, John Sillitoe on behalf of the Wellcome Sanger Institute COVID-19 Surveillance Team |
| EPI_ISL_3108296 | Lighthouse Lab in Milton Keynes | Wellcome Sanger Institute for the COVID-19 Genomics UK (COG-UK) Consortium | The Lighthouse Lab in Milton Keynes and Alex Alderton, Roberto Amato, Jeffrey Barrett, Sonia Goncalves, Ewan Harrison, David K. Jackson, Ian Johnston, Dominic Kwiatkowski, Cordelia Langford, John Sillitoe on behalf of the Wellcome Sanger Institute COVID-19 Surveillance Team |
| EPI_ISL_3108297, EPI_ISL_3108298 | Lighthouse Lab in Alderley Park | Wellcome Sanger Institute for the COVID-19 Genomics UK (COG-UK) Consortium | Jacquelyn Wynn, Mairead Hyland, The Lighthouse Lab in Alderley Park and Alex Alderton, Roberto Amato, Jeffrey Barrett, Sonia Goncalves, Ewan Harrison, David K. Jackson, Ian Johnston, Dominic Kwiatkowski, Cordelia Langford, John Sillitoe on behalf of the Wellcome Sanger Institute COVID-19 Surveillance Team |
| EPI_ISL_3108299 | Lighthouse Lab in Milton Keynes | Wellcome Sanger Institute for the COVID-19 Genomics UK (COG-UK) Consortium | The Lighthouse Lab in Milton Keynes and Alex Alderton, Roberto Amato, Jeffrey Barrett, Sonia Goncalves, Ewan Harrison, David K. Jackson, Ian Johnston, Dominic Kwiatkowski, Cordelia Langford, John Sillitoe on behalf of the Wellcome Sanger Institute COVID-19 Surveillance Team |
| EPI_ISL_3108300, EPI_ISL_3108301 | Lighthouse Lab in Alderley Park | Wellcome Sanger Institute for the COVID-19 Genomics UK (COG-UK) Consortium | Jacquelyn Wynn, Mairead Hyland, The Lighthouse Lab in Alderley Park and Alex Alderton, Roberto Amato, Jeffrey Barrett, Sonia Goncalves, Ewan Harrison, David K. Jackson, Ian Johnston, Dominic Kwiatkowski, Cordelia Langford, John Sillitoe on behalf of the Wellcome Sanger Institute COVID-19 Surveillance Team |
| EPI_ISL_3108302 | Lighthouse Lab in Glasgow | Wellcome Sanger Institute for the COVID-19 Genomics UK (COG-UK) Consortium | Harper VanSteenhouse, Yumi Kasai, David Gray, Carol Clugston, Anna Dominiczak and Alex Alderton, Roberto Amato, Jeffrey Barrett, Sonia Goncalves, Ewan Harrison, David K. Jackson, Ian Johnston, Dominic Kwiatkowski, Cordelia Langford, John Sillitoe on behalf of the Wellcome Sanger Institute COVID-19 Surveillance Team |
| EPI_ISL_3108303 | Lighthouse Lab in Milton Keynes | Wellcome Sanger Institute for the COVID-19 Genomics UK (COG-UK) Consortium | The Lighthouse Lab in Milton Keynes and Alex Alderton, Roberto Amato, Jeffrey Barrett, Sonia Goncalves, Ewan Harrison, David K. Jackson, Ian Johnston, Dominic Kwiatkowski, Cordelia Langford, John Sillitoe on behalf of the Wellcome Sanger Institute COVID-19 Surveillance Team |
| EPI_ISL_3108304, EPI_ISL_3108305 | Lighthouse Lab in Alderley Park | Wellcome Sanger Institute for the COVID-19 Genomics UK (COG-UK) Consortium | Jacquelyn Wynn, Mairead Hyland, The Lighthouse Lab in Alderley Park and Alex Alderton, Roberto Amato, Jeffrey Barrett, Sonia Goncalves, Ewan Harrison, David K. Jackson, Ian Johnston, Dominic Kwiatkowski, Cordelia Langford, John Sillitoe on behalf of the Wellcome Sanger Institute COVID-19 Surveillance Team |
| EPI_ISL_3108306 | Lighthouse Lab in Glasgow | Wellcome Sanger Institute for the COVID-19 Genomics UK (COG-UK) Consortium | Harper VanSteenhouse, Yumi Kasai, David Gray, Carol Clugston, Anna Dominiczak and Alex Alderton, Roberto Amato, Jeffrey Barrett, Sonia Goncalves, Ewan Harrison, David K. Jackson, Ian Johnston, Dominic Kwiatkowski, Cordelia Langford, John Sillitoe on behalf of the Wellcome Sanger Institute COVID-19 Surveillance Team |
| EPI_ISL_3108307, EPI_ISL_3108308 | Lighthouse Lab in Alderley Park | Wellcome Sanger Institute for the COVID-19 Genomics UK (COG-UK) Consortium | Jacquelyn Wynn, Mairead Hyland, The Lighthouse Lab in Alderley Park and Alex Alderton, Roberto Amato, Jeffrey Barrett, Sonia Goncalves, Ewan Harrison, David K. Jackson, Ian Johnston, Dominic Kwiatkowski, Cordelia Langford, John Sillitoe on behalf of the Wellcome Sanger Institute COVID-19 Surveillance Team |
| EPI_ISL_3108309, EPI_ISL_3108310, EPI_ISL_3108311 | Lighthouse Lab in Glasgow | Wellcome Sanger Institute for the COVID-19 Genomics UK (COG-UK) Consortium | Harper VanSteenhouse, Yumi Kasai, David Gray, Carol Clugston, Anna Dominiczak and Alex Alderton, Roberto Amato, Jeffrey Barrett, Sonia Goncalves, Ewan Harrison, David K. Jackson, Ian Johnston, Dominic Kwiatkowski, Cordelia Langford, John Sillitoe on behalf of the Wellcome Sanger Institute COVID-19 Surveillance Team |
| EPI_ISL_3108312 | Lighthouse Lab in Milton Keynes | Wellcome Sanger Institute for the COVID-19 Genomics UK (COG-UK) Consortium | The Lighthouse Lab in Milton Keynes and Alex Alderton, Roberto Amato, Jeffrey Barrett, Sonia Goncalves, Ewan Harrison, David K. Jackson, Ian Johnston, Dominic Kwiatkowski, Cordelia Langford, John Sillitoe on behalf of the Wellcome Sanger Institute COVID-19 Surveillance Team |
| EPI_ISL_3108313 | Lighthouse Lab in Alderley Park | Wellcome Sanger Institute for the COVID-19 Genomics UK (COG-UK) Consortium | Jacquelyn Wynn, Mairead Hyland, The Lighthouse Lab in Alderley Park and Alex Alderton, Roberto Amato, Jeffrey Barrett, Sonia Goncalves, Ewan Harrison, David K. Jackson, Ian Johnston, Dominic Kwiatkowski, Cordelia Langford, John Sillitoe on behalf of the Wellcome Sanger Institute COVID-19 Surveillance Team |
| EPI_ISL_3108314, EPI_ISL_3108315 | Lighthouse Lab in Glasgow | Wellcome Sanger Institute for the COVID-19 Genomics UK (COG-UK) Consortium | Harper VanSteenhouse, Yumi Kasai, David Gray, Carol Clugston, Anna Dominiczak and Alex Alderton, Roberto Amato, Jeffrey Barrett, Sonia Goncalves, Ewan Harrison, David K. Jackson, Ian Johnston, Dominic Kwiatkowski, Cordelia Langford, John Sillitoe on behalf of the Wellcome Sanger Institute COVID-19 Surveillance Team |
| EPI_ISL_3108316 | Lighthouse Lab in Alderley Park | Wellcome Sanger Institute for the COVID-19 Genomics UK (COG-UK) Consortium | Jacquelyn Wynn, Mairead Hyland, The Lighthouse Lab in Alderley Park and Alex Alderton, Roberto Amato, Jeffrey Barrett, Sonia Goncalves, Ewan Harrison, David K. Jackson, Ian Johnston, Dominic Kwiatkowski, Cordelia Langford, John Sillitoe on behalf of the Wellcome Sanger Institute COVID-19 Surveillance Team |
| EPI_ISL_3108317, EPI_ISL_3108318, EPI_ISL_3108319 | Lighthouse Lab in Milton Keynes | Wellcome Sanger Institute for the COVID-19 Genomics UK (COG-UK) Consortium | The Lighthouse Lab in Milton Keynes and Alex Alderton, Roberto Amato, Jeffrey Barrett, Sonia Goncalves, Ewan Harrison, David K. Jackson, Ian Johnston, Dominic Kwiatkowski, Cordelia Langford, John Sillitoe on behalf of the Wellcome Sanger Institute COVID-19 Surveillance Team |
| EPI_ISL_3108320 | Lighthouse Lab in Alderley Park | Wellcome Sanger Institute for the COVID-19 Genomics UK (COG-UK) Consortium | Jacquelyn Wynn, Mairead Hyland, The Lighthouse Lab in Alderley Park and Alex Alderton, Roberto Amato, Jeffrey Barrett, Sonia Goncalves, Ewan Harrison, David K. Jackson, Ian Johnston, Dominic Kwiatkowski, Cordelia Langford, John Sillitoe on behalf of the Wellcome Sanger Institute COVID-19 Surveillance Team |
| EPI_ISL_3108321 | Lighthouse Lab in Glasgow | Wellcome Sanger Institute for the COVID-19 Genomics UK (COG-UK) Consortium | Harper VanSteenhouse, Yumi Kasai, David Gray, Carol Clugston, Anna Dominiczak and Alex Alderton, Roberto Amato, Jeffrey Barrett, Sonia Goncalves, Ewan Harrison, David K. Jackson, Ian Johnston, Dominic Kwiatkowski, Cordelia Langford, John Sillitoe on behalf of the Wellcome Sanger Institute COVID-19 Surveillance Team |
| EPI_ISL_3108322, EPI_ISL_3108323, EPI_ISL_3108324, EPI_ISL_3108325 | Lighthouse Lab in Alderley Park | Wellcome Sanger Institute for the COVID-19 Genomics UK (COG-UK) Consortium | Jacquelyn Wynn, Mairead Hyland, The Lighthouse Lab in Alderley Park and Alex Alderton, Roberto Amato, Jeffrey Barrett, Sonia Goncalves, Ewan Harrison, David K. Jackson, Ian Johnston, Dominic Kwiatkowski, Cordelia Langford, John Sillitoe on behalf of the Wellcome Sanger Institute COVID-19 Surveillance Team |
| EPI_ISL_3108326 | Lighthouse Lab in Glasgow | Wellcome Sanger Institute for the COVID-19 Genomics UK (COG-UK) Consortium | Harper VanSteenhouse, Yumi Kasai, David Gray, Carol Clugston, Anna Dominiczak and Alex Alderton, Roberto Amato, Jeffrey Barrett, Sonia Goncalves, Ewan Harrison, David K. Jackson, Ian Johnston, Dominic Kwiatkowski, Cordelia Langford, John Sillitoe on behalf of the Wellcome Sanger Institute COVID-19 Surveillance Team |
| EPI_ISL_3108327 | Lighthouse Lab in Milton Keynes | Wellcome Sanger Institute for the COVID-19 Genomics UK (COG-UK) Consortium | The Lighthouse Lab in Milton Keynes and Alex Alderton, Roberto Amato, Jeffrey Barrett, Sonia Goncalves, Ewan Harrison, David K. Jackson, Ian Johnston, Dominic Kwiatkowski, Cordelia Langford, John Sillitoe on behalf of the Wellcome Sanger Institute COVID-19 Surveillance Team |
| EPI_ISL_3108328 | Lighthouse Lab in Alderley Park | Wellcome Sanger Institute for the COVID-19 Genomics UK (COG-UK) Consortium | Jacquelyn Wynn, Mairead Hyland, The Lighthouse Lab in Alderley Park and Alex Alderton, Roberto Amato, Jeffrey Barrett, Sonia Goncalves, Ewan Harrison, David K. Jackson, Ian Johnston, Dominic Kwiatkowski, Cordelia Langford, John Sillitoe on behalf of the Wellcome Sanger Institute COVID-19 Surveillance Team |
| EPI_ISL_3108329 | Lighthouse Lab in Milton Keynes | Wellcome Sanger Institute for the COVID-19 Genomics UK (COG-UK) Consortium | The Lighthouse Lab in Milton Keynes and Alex Alderton, Roberto Amato, Jeffrey Barrett, Sonia Goncalves, Ewan Harrison, David K. Jackson, Ian Johnston, Dominic Kwiatkowski, Cordelia Langford, John Sillitoe on behalf of the Wellcome Sanger Institute COVID-19 Surveillance Team |
| EPI_ISL_3108330, EPI_ISL_3108331 | Lighthouse Lab in Glasgow | Wellcome Sanger Institute for the COVID-19 Genomics UK (COG-UK) Consortium | Harper VanSteenhouse, Yumi Kasai, David Gray, Carol Clugston, Anna Dominiczak and Alex Alderton, Roberto Amato, Jeffrey Barrett, Sonia Goncalves, Ewan Harrison, David K. Jackson, Ian Johnston, Dominic Kwiatkowski, Cordelia Langford, John Sillitoe on behalf of the Wellcome Sanger Institute COVID-19 Surveillance Team |
| EPI_ISL_3108332 | Lighthouse Lab in Milton Keynes | Wellcome Sanger Institute for the COVID-19 Genomics UK (COG-UK) Consortium | The Lighthouse Lab in Milton Keynes and Alex Alderton, Roberto Amato, Jeffrey Barrett, Sonia Goncalves, Ewan Harrison, David K. Jackson, Ian Johnston, Dominic Kwiatkowski, Cordelia Langford, John Sillitoe on behalf of the Wellcome Sanger Institute COVID-19 Surveillance Team |
| EPI_ISL_3108333, EPI_ISL_3108334, EPI_ISL_3108335, EPI_ISL_3108336 | Lighthouse Lab in Alderley Park | Wellcome Sanger Institute for the COVID-19 Genomics UK (COG-UK) Consortium | Jacquelyn Wynn, Mairead Hyland, The Lighthouse Lab in Alderley Park and Alex Alderton, Roberto Amato, Jeffrey Barrett, Sonia Goncalves, Ewan Harrison, David K. Jackson, Ian Johnston, Dominic Kwiatkowski, Cordelia Langford, John Sillitoe on behalf of the Wellcome Sanger Institute COVID-19 Surveillance Team |
| EPI_ISL_3108337 | Lighthouse Lab in Milton Keynes | Wellcome Sanger Institute for the COVID-19 Genomics UK (COG-UK) Consortium | The Lighthouse Lab in Milton Keynes and Alex Alderton, Roberto Amato, Jeffrey Barrett, Sonia Goncalves, Ewan Harrison, David K. Jackson, Ian Johnston, Dominic Kwiatkowski, Cordelia Langford, John Sillitoe on behalf of the Wellcome Sanger Institute COVID-19 Surveillance Team |
| EPI_ISL_3108338, EPI_ISL_3108339 | Lighthouse Lab in Alderley Park | Wellcome Sanger Institute for the COVID-19 Genomics UK (COG-UK) Consortium | Jacquelyn Wynn, Mairead Hyland, The Lighthouse Lab in Alderley Park and Alex Alderton, Roberto Amato, Jeffrey Barrett, Sonia Goncalves, Ewan Harrison, David K. Jackson, Ian Johnston, Dominic Kwiatkowski, Cordelia Langford, John Sillitoe on behalf of the Wellcome Sanger Institute COVID-19 Surveillance Team |
| EPI_ISL_3108340, EPI_ISL_3108341 | Lighthouse Lab in Milton Keynes | Wellcome Sanger Institute for the COVID-19 Genomics UK (COG-UK) Consortium | The Lighthouse Lab in Milton Keynes and Alex Alderton, Roberto Amato, Jeffrey Barrett, Sonia Goncalves, Ewan Harrison, David K. Jackson, Ian Johnston, Dominic Kwiatkowski, Cordelia Langford, John Sillitoe on behalf of the Wellcome Sanger Institute COVID-19 Surveillance Team |
| EPI_ISL_3108342 | Lighthouse Lab in Glasgow | Wellcome Sanger Institute for the COVID-19 Genomics UK (COG-UK) Consortium | Harper VanSteenhouse, Yumi Kasai, David Gray, Carol Clugston, Anna Dominiczak and Alex Alderton, Roberto Amato, Jeffrey Barrett, Sonia Goncalves, Ewan Harrison, David K. Jackson, Ian Johnston, Dominic Kwiatkowski, Cordelia Langford, John Sillitoe on behalf of the Wellcome Sanger Institute COVID-19 Surveillance Team |
| EPI_ISL_3108343 | Lighthouse Lab in Milton Keynes | Wellcome Sanger Institute for the COVID-19 Genomics UK (COG-UK) Consortium | The Lighthouse Lab in Milton Keynes and Alex Alderton, Roberto Amato, Jeffrey Barrett, Sonia Goncalves, Ewan Harrison, David K. Jackson, Ian Johnston, Dominic Kwiatkowski, Cordelia Langford, John Sillitoe on behalf of the Wellcome Sanger Institute COVID-19 Surveillance Team |
| EPI_ISL_3108344, EPI_ISL_3108345 | Lighthouse Lab in Glasgow | Wellcome Sanger Institute for the COVID-19 Genomics UK (COG-UK) Consortium | Harper VanSteenhouse, Yumi Kasai, David Gray, Carol Clugston, Anna Dominiczak and Alex Alderton, Roberto Amato, Jeffrey Barrett, Sonia Goncalves, Ewan Harrison, David K. Jackson, Ian Johnston, Dominic Kwiatkowski, Cordelia Langford, John Sillitoe on behalf of the Wellcome Sanger Institute COVID-19 Surveillance Team |
| EPI_ISL_3108346 | Lighthouse Lab in Milton Keynes | Wellcome Sanger Institute for the COVID-19 Genomics UK (COG-UK) Consortium | The Lighthouse Lab in Milton Keynes and Alex Alderton, Roberto Amato, Jeffrey Barrett, Sonia Goncalves, Ewan Harrison, David K. Jackson, Ian Johnston, Dominic Kwiatkowski, Cordelia Langford, John Sillitoe on behalf of the Wellcome Sanger Institute COVID-19 Surveillance Team |
| EPI_ISL_3108347, EPI_ISL_3108348, EPI_ISL_3108349 | Lighthouse Lab in Alderley Park | Wellcome Sanger Institute for the COVID-19 Genomics UK (COG-UK) Consortium | Jacquelyn Wynn, Mairead Hyland, The Lighthouse Lab in Alderley Park and Alex Alderton, Roberto Amato, Jeffrey Barrett, Sonia Goncalves, Ewan Harrison, David K. Jackson, Ian Johnston, Dominic Kwiatkowski, Cordelia Langford, John Sillitoe on behalf of the Wellcome Sanger Institute COVID-19 Surveillance Team |
| EPI_ISL_3108350 | Lighthouse Lab in Milton Keynes | Wellcome Sanger Institute for the COVID-19 Genomics UK (COG-UK) Consortium | The Lighthouse Lab in Milton Keynes and Alex Alderton, Roberto Amato, Jeffrey Barrett, Sonia Goncalves, Ewan Harrison, David K. Jackson, Ian Johnston, Dominic Kwiatkowski, Cordelia Langford, John Sillitoe on behalf of the Wellcome Sanger Institute COVID-19 Surveillance Team |
| EPI_ISL_3108351, EPI_ISL_3108352 | Lighthouse Lab in Alderley Park | Wellcome Sanger Institute for the COVID-19 Genomics UK (COG-UK) Consortium | Jacquelyn Wynn, Mairead Hyland, The Lighthouse Lab in Alderley Park and Alex Alderton, Roberto Amato, Jeffrey Barrett, Sonia Goncalves, Ewan Harrison, David K. Jackson, Ian Johnston, Dominic Kwiatkowski, Cordelia Langford, John Sillitoe on behalf of the Wellcome Sanger Institute COVID-19 Surveillance Team |
| EPI_ISL_3108353 | Lighthouse Lab in Glasgow | Wellcome Sanger Institute for the COVID-19 Genomics UK (COG-UK) Consortium | Harper VanSteenhouse, Yumi Kasai, David Gray, Carol Clugston, Anna Dominiczak and Alex Alderton, Roberto Amato, Jeffrey Barrett, Sonia Goncalves, Ewan Harrison, David K. Jackson, Ian Johnston, Dominic Kwiatkowski, Cordelia Langford, John Sillitoe on behalf of the Wellcome Sanger Institute COVID-19 Surveillance Team |
| EPI_ISL_3108354 | Lighthouse Lab in Alderley Park | Wellcome Sanger Institute for the COVID-19 Genomics UK (COG-UK) Consortium | Jacquelyn Wynn, Mairead Hyland, The Lighthouse Lab in Alderley Park and Alex Alderton, Roberto Amato, Jeffrey Barrett, Sonia Goncalves, Ewan Harrison, David K. Jackson, Ian Johnston, Dominic Kwiatkowski, Cordelia Langford, John Sillitoe on behalf of the Wellcome Sanger Institute COVID-19 Surveillance Team |

| | | | |
|---|---|---|---|
| EPI_ISL_3108355 | Lighthouse Lab in Milton Keynes | Wellcome Sanger Institute for the COVID-19 Genomics UK (COG-UK) Consortium | The Lighthouse Lab in Milton Keynes and Alex Alderton, Roberto Amato, Jeffrey Barrett, Sonia Goncalves, Ewan Harrison, David K. Jackson, Ian Johnston, Dominic Kwiatkowski, Cordelia Langford, John Sillitoe on behalf of the Wellcome Sanger Institute COVID-19 Surveillance Team |
| EPI_ISL_3108356, EPI_ISL_3108357 | Lighthouse Lab in Alderley Park | Wellcome Sanger Institute for the COVID-19 Genomics UK (COG-UK) Consortium | Jacquelyn Wynn, Mairead Hyland, The Lighthouse Lab in Alderley Park and Alex Alderton, Roberto Amato, Jeffrey Barrett, Sonia Goncalves, Ewan Harrison, David K. Jackson, Ian Johnston, Dominic Kwiatkowski, Cordelia Langford, John Sillitoe on behalf of the Wellcome Sanger Institute COVID-19 Surveillance Team |
| EPI_ISL_3108358 | Lighthouse Lab in Milton Keynes | Wellcome Sanger Institute for the COVID-19 Genomics UK (COG-UK) Consortium | The Lighthouse Lab in Milton Keynes and Alex Alderton, Roberto Amato, Jeffrey Barrett, Sonia Goncalves, Ewan Harrison, David K. Jackson, Ian Johnston, Dominic Kwiatkowski, Cordelia Langford, John Sillitoe on behalf of the Wellcome Sanger Institute COVID-19 Surveillance Team |
| EPI_ISL_3108359, EPI_ISL_3108360 | Lighthouse Lab in Alderley Park | Wellcome Sanger Institute for the COVID-19 Genomics UK (COG-UK) Consortium | Jacquelyn Wynn, Mairead Hyland, The Lighthouse Lab in Alderley Park and Alex Alderton, Roberto Amato, Jeffrey Barrett, Sonia Goncalves, Ewan Harrison, David K. Jackson, Ian Johnston, Dominic Kwiatkowski, Cordelia Langford, John Sillitoe on behalf of the Wellcome Sanger Institute COVID-19 Surveillance Team |
| EPI_ISL_3108361 | Lighthouse Lab in Milton Keynes | Wellcome Sanger Institute for the COVID-19 Genomics UK (COG-UK) Consortium | The Lighthouse Lab in Milton Keynes and Alex Alderton, Roberto Amato, Jeffrey Barrett, Sonia Goncalves, Ewan Harrison, David K. Jackson, Ian Johnston, Dominic Kwiatkowski, Cordelia Langford, John Sillitoe on behalf of the Wellcome Sanger Institute COVID-19 Surveillance Team |
| EPI_ISL_3108362 | Lighthouse Lab in Alderley Park | Wellcome Sanger Institute for the COVID-19 Genomics UK (COG-UK) Consortium | Jacquelyn Wynn, Mairead Hyland, The Lighthouse Lab in Alderley Park and Alex Alderton, Roberto Amato, Jeffrey Barrett, Sonia Goncalves, Ewan Harrison, David K. Jackson, Ian Johnston, Dominic Kwiatkowski, Cordelia Langford, John Sillitoe on behalf of the Wellcome Sanger Institute COVID-19 Surveillance Team |
| EPI_ISL_3108363, EPI_ISL_3108364 | Lighthouse Lab in Glasgow | Wellcome Sanger Institute for the COVID-19 Genomics UK (COG-UK) Consortium | Harper VanSteenhouse, Yumi Kasai, David Gray, Carol Clugston, Anna Dominiczak and Alex Alderton, Roberto Amato, Jeffrey Barrett, Sonia Goncalves, Ewan Harrison, David K. Jackson, Ian Johnston, Dominic Kwiatkowski, Cordelia Langford, John Sillitoe on behalf of the Wellcome Sanger Institute COVID-19 Surveillance Team |
| EPI_ISL_3108365, EPI_ISL_3108366 | Lighthouse Lab in Alderley Park | Wellcome Sanger Institute for the COVID-19 Genomics UK (COG-UK) Consortium | Jacquelyn Wynn, Mairead Hyland, The Lighthouse Lab in Alderley Park and Alex Alderton, Roberto Amato, Jeffrey Barrett, Sonia Goncalves, Ewan Harrison, David K. Jackson, Ian Johnston, Dominic Kwiatkowski, Cordelia Langford, John Sillitoe on behalf of the Wellcome Sanger Institute COVID-19 Surveillance Team |
| EPI_ISL_3108367 | Lighthouse Lab in Milton Keynes | Wellcome Sanger Institute for the COVID-19 Genomics UK (COG-UK) Consortium | The Lighthouse Lab in Milton Keynes and Alex Alderton, Roberto Amato, Jeffrey Barrett, Sonia Goncalves, Ewan Harrison, David K. Jackson, Ian Johnston, Dominic Kwiatkowski, Cordelia Langford, John Sillitoe on behalf of the Wellcome Sanger Institute COVID-19 Surveillance Team |
| EPI_ISL_3108368, EPI_ISL_3108369, EPI_ISL_3108370, EPI_ISL_3108371, EPI_ISL_3108372, EPI_ISL_3108373, EPI_ISL_3108374, EPI_ISL_3108375, EPI_ISL_3108376, EPI_ISL_3108377, EPI_ISL_3108378 | | | |
| see above | Lighthouse Lab in Alderley Park | Wellcome Sanger Institute for the COVID-19 Genomics UK (COG-UK) Consortium | Jacquelyn Wynn, Mairead Hyland, The Lighthouse Lab in Alderley Park and Alex Alderton, Roberto Amato, Jeffrey Barrett, Sonia Goncalves, Ewan Harrison, David K. Jackson, Ian Johnston, Dominic Kwiatkowski, Cordelia Langford, John Sillitoe on behalf of the Wellcome Sanger Institute COVID-19 Surveillance Team |
| EPI_ISL_3108379 | Lighthouse Lab in Glasgow | Wellcome Sanger Institute for the COVID-19 Genomics UK (COG-UK) Consortium | Harper VanSteenhouse, Yumi Kasai, David Gray, Carol Clugston, Anna Dominiczak and Alex Alderton, Roberto Amato, Jeffrey Barrett, Sonia Goncalves, Ewan Harrison, David K. Jackson, Ian Johnston, Dominic Kwiatkowski, Cordelia Langford, John Sillitoe on behalf of the Wellcome Sanger Institute COVID-19 Surveillance Team |
| EPI_ISL_3108380, EPI_ISL_3108381, EPI_ISL_3108382 | Lighthouse Lab in Alderley Park | Wellcome Sanger Institute for the COVID-19 Genomics UK (COG-UK) Consortium | Jacquelyn Wynn, Mairead Hyland, The Lighthouse Lab in Alderley Park and Alex Alderton, Roberto Amato, Jeffrey Barrett, Sonia Goncalves, Ewan Harrison, David K. Jackson, Ian Johnston, Dominic Kwiatkowski, Cordelia Langford, John Sillitoe on behalf of the Wellcome Sanger Institute COVID-19 Surveillance Team |
| EPI_ISL_3108383 | Lighthouse Lab in Milton Keynes | Wellcome Sanger Institute for the COVID-19 Genomics UK (COG-UK) Consortium | The Lighthouse Lab in Milton Keynes and Alex Alderton, Roberto Amato, Jeffrey Barrett, Sonia Goncalves, Ewan Harrison, David K. Jackson, Ian Johnston, Dominic Kwiatkowski, Cordelia Langford, John Sillitoe on behalf of the Wellcome Sanger Institute COVID-19 Surveillance Team |
| EPI_ISL_3108384, EPI_ISL_3108385 | Lighthouse Lab in Alderley Park | Wellcome Sanger Institute for the COVID-19 Genomics UK (COG-UK) Consortium | Jacquelyn Wynn, Mairead Hyland, The Lighthouse Lab in Alderley Park and Alex Alderton, Roberto Amato, Jeffrey Barrett, Sonia Goncalves, Ewan Harrison, David K. Jackson, Ian Johnston, Dominic Kwiatkowski, Cordelia Langford, John Sillitoe on behalf of the Wellcome Sanger Institute COVID-19 Surveillance Team |
| EPI_ISL_3108386, EPI_ISL_3108387, EPI_ISL_3108388 | Lighthouse Lab in Milton Keynes | Wellcome Sanger Institute for the COVID-19 Genomics UK (COG-UK) Consortium | The Lighthouse Lab in Milton Keynes and Alex Alderton, Roberto Amato, Jeffrey Barrett, Sonia Goncalves, Ewan Harrison, David K. Jackson, Ian Johnston, Dominic Kwiatkowski, Cordelia Langford, John Sillitoe on behalf of the Wellcome Sanger Institute COVID-19 Surveillance Team |
| EPI_ISL_3108389 | Lighthouse Lab in Glasgow | Wellcome Sanger Institute for the COVID-19 Genomics UK (COG-UK) Consortium | Harper VanSteenhouse, Yumi Kasai, David Gray, Carol Clugston, Anna Dominiczak and Alex Alderton, Roberto Amato, Jeffrey Barrett, Sonia Goncalves, Ewan Harrison, David K. Jackson, Ian Johnston, Dominic Kwiatkowski, Cordelia Langford, John Sillitoe on behalf of the Wellcome Sanger Institute COVID-19 Surveillance Team |
| EPI_ISL_3108390 | Lighthouse Lab in Milton Keynes | Wellcome Sanger Institute for the COVID-19 Genomics UK (COG-UK) Consortium | The Lighthouse Lab in Milton Keynes and Alex Alderton, Roberto Amato, Jeffrey Barrett, Sonia Goncalves, Ewan Harrison, David K. Jackson, Ian Johnston, Dominic Kwiatkowski, Cordelia Langford, John Sillitoe on behalf of the Wellcome Sanger Institute COVID-19 Surveillance Team |
| EPI_ISL_3108391, EPI_ISL_3108392, EPI_ISL_3108393, EPI_ISL_3108394, EPI_ISL_3108395, EPI_ISL_3108396, EPI_ISL_3108397 | Lighthouse Lab in Alderley Park | Wellcome Sanger Institute for the COVID-19 Genomics UK (COG-UK) Consortium | Jacquelyn Wynn, Mairead Hyland, The Lighthouse Lab in Alderley Park and Alex Alderton, Roberto Amato, Jeffrey Barrett, Sonia Goncalves, Ewan Harrison, David K. Jackson, Ian Johnston, Dominic Kwiatkowski, Cordelia Langford, John Sillitoe on behalf of the Wellcome Sanger Institute COVID-19 Surveillance Team |
| EPI_ISL_3108398 | Lighthouse Lab in Glasgow | Wellcome Sanger Institute for the COVID-19 Genomics UK (COG-UK) Consortium | Harper VanSteenhouse, Yumi Kasai, David Gray, Carol Clugston, Anna Dominiczak and Alex Alderton, Roberto Amato, Jeffrey Barrett, Sonia Goncalves, Ewan Harrison, David K. Jackson, Ian Johnston, Dominic Kwiatkowski, Cordelia Langford, John Sillitoe on behalf of the Wellcome Sanger Institute COVID-19 Surveillance Team |
| EPI_ISL_3108399 | Lighthouse Lab in Milton Keynes | Wellcome Sanger Institute for the COVID-19 Genomics UK (COG-UK) Consortium | The Lighthouse Lab in Milton Keynes and Alex Alderton, Roberto Amato, Jeffrey Barrett, Sonia Goncalves, Ewan Harrison, David K. Jackson, Ian Johnston, Dominic Kwiatkowski, Cordelia Langford, John Sillitoe on behalf of the Wellcome Sanger Institute COVID-19 Surveillance Team |
| EPI_ISL_3108400, EPI_ISL_3108401, EPI_ISL_3108402, EPI_ISL_3108403, EPI_ISL_3108404, EPI_ISL_3108405 | Lighthouse Lab in Alderley Park | Wellcome Sanger Institute for the COVID-19 Genomics UK (COG-UK) Consortium | Jacquelyn Wynn, Mairead Hyland, The Lighthouse Lab in Alderley Park and Alex Alderton, Roberto Amato, Jeffrey Barrett, Sonia Goncalves, Ewan Harrison, David K. Jackson, Ian Johnston, Dominic Kwiatkowski, Cordelia Langford, John Sillitoe on behalf of the Wellcome Sanger Institute COVID-19 Surveillance Team |
| EPI_ISL_3108406, EPI_ISL_3108407, EPI_ISL_3108408 | Lighthouse Lab in Milton Keynes | Wellcome Sanger Institute for the COVID-19 Genomics UK (COG-UK) Consortium | The Lighthouse Lab in Milton Keynes and Alex Alderton, Roberto Amato, Jeffrey Barrett, Sonia Goncalves, Ewan Harrison, David K. Jackson, Ian Johnston, Dominic Kwiatkowski, Cordelia Langford, John Sillitoe on behalf of the Wellcome Sanger Institute COVID-19 Surveillance Team |
| EPI_ISL_3108409, EPI_ISL_3108410, EPI_ISL_3108411 | Lighthouse Lab in Alderley Park | Wellcome Sanger Institute for the COVID-19 Genomics UK (COG-UK) Consortium | Jacquelyn Wynn, Mairead Hyland, The Lighthouse Lab in Alderley Park and Alex Alderton, Roberto Amato, Jeffrey Barrett, Sonia Goncalves, Ewan Harrison, David K. Jackson, Ian Johnston, Dominic Kwiatkowski, Cordelia Langford, John Sillitoe on behalf of the Wellcome Sanger Institute COVID-19 Surveillance Team |
| EPI_ISL_3108412 | Lighthouse Lab in Glasgow | Wellcome Sanger Institute for the COVID-19 Genomics UK (COG-UK) Consortium | Harper VanSteenhouse, Yumi Kasai, David Gray, Carol Clugston, Anna Dominiczak and Alex Alderton, Roberto Amato, Jeffrey Barrett, Sonia Goncalves, Ewan Harrison, David K. Jackson, Ian Johnston, Dominic Kwiatkowski, Cordelia Langford, John Sillitoe on behalf of the Wellcome Sanger Institute COVID-19 Surveillance Team |
| EPI_ISL_3108413 | Lighthouse Lab in Alderley Park | Wellcome Sanger Institute for the COVID-19 Genomics UK (COG-UK) Consortium | Jacquelyn Wynn, Mairead Hyland, The Lighthouse Lab in Alderley Park and Alex Alderton, Roberto Amato, Jeffrey Barrett, Sonia Goncalves, Ewan Harrison, David K. Jackson, Ian Johnston, Dominic Kwiatkowski, Cordelia Langford, John Sillitoe on behalf of the Wellcome Sanger Institute COVID-19 Surveillance Team |
| EPI_ISL_3108414 | Lighthouse Lab in Milton Keynes | Wellcome Sanger Institute for the COVID-19 Genomics UK (COG-UK) Consortium | The Lighthouse Lab in Milton Keynes and Alex Alderton, Roberto Amato, Jeffrey Barrett, Sonia Goncalves, Ewan Harrison, David K. Jackson, Ian Johnston, Dominic Kwiatkowski, Cordelia Langford, John Sillitoe on behalf of the Wellcome Sanger Institute COVID-19 Surveillance Team |
| EPI_ISL_3108415, EPI_ISL_3108416, EPI_ISL_3108417, EPI_ISL_3108418 | Lighthouse Lab in Alderley Park | Wellcome Sanger Institute for the COVID-19 Genomics UK (COG-UK) Consortium | Jacquelyn Wynn, Mairead Hyland, The Lighthouse Lab in Alderley Park and Alex Alderton, Roberto Amato, Jeffrey Barrett, Sonia Goncalves, Ewan Harrison, David K. Jackson, Ian Johnston, Dominic Kwiatkowski, Cordelia Langford, John Sillitoe on behalf of the Wellcome Sanger Institute COVID-19 Surveillance Team |
| EPI_ISL_3108419 | Lighthouse Lab in Milton Keynes | Wellcome Sanger Institute for the COVID-19 Genomics UK (COG-UK) Consortium | The Lighthouse Lab in Milton Keynes and Alex Alderton, Roberto Amato, Jeffrey Barrett, Sonia Goncalves, Ewan Harrison, David K. Jackson, Ian Johnston, Dominic Kwiatkowski, Cordelia Langford, John Sillitoe on behalf of the Wellcome Sanger Institute COVID-19 Surveillance Team |
| EPI_ISL_3108420 | Lighthouse Lab in Alderley Park | Wellcome Sanger Institute for the COVID-19 Genomics UK (COG-UK) Consortium | Jacquelyn Wynn, Mairead Hyland, The Lighthouse Lab in Alderley Park and Alex Alderton, Roberto Amato, Jeffrey Barrett, Sonia Goncalves, Ewan Harrison, David K. Jackson, Ian Johnston, Dominic Kwiatkowski, Cordelia Langford, John Sillitoe on behalf of the Wellcome Sanger Institute COVID-19 Surveillance Team |
| EPI_ISL_3108421 | Lighthouse Lab in Glasgow | Wellcome Sanger Institute for the COVID-19 Genomics UK (COG-UK) Consortium | Harper VanSteenhouse, Yumi Kasai, David Gray, Carol Clugston, Anna Dominiczak and Alex Alderton, Roberto Amato, Jeffrey Barrett, Sonia Goncalves, Ewan Harrison, David K. Jackson, Ian Johnston, Dominic Kwiatkowski, Cordelia Langford, John Sillitoe on behalf of the Wellcome Sanger Institute COVID-19 Surveillance Team |
| EPI_ISL_3108422 | Lighthouse Lab in Alderley Park | Wellcome Sanger Institute for the COVID-19 Genomics UK (COG-UK) Consortium | Jacquelyn Wynn, Mairead Hyland, The Lighthouse Lab in Alderley Park and Alex Alderton, Roberto Amato, Jeffrey Barrett, Sonia Goncalves, Ewan Harrison, David K. Jackson, Ian Johnston, Dominic Kwiatkowski, Cordelia Langford, John Sillitoe on behalf of the Wellcome Sanger Institute COVID-19 Surveillance Team |
| EPI_ISL_3108423 | Lighthouse Lab in Milton Keynes | Wellcome Sanger Institute for the COVID-19 Genomics UK (COG-UK) Consortium | The Lighthouse Lab in Milton Keynes and Alex Alderton, Roberto Amato, Jeffrey Barrett, Sonia Goncalves, Ewan Harrison, David K. Jackson, Ian Johnston, Dominic Kwiatkowski, Cordelia Langford, John Sillitoe on behalf of the Wellcome Sanger Institute COVID-19 Surveillance Team |
| EPI_ISL_3108424 | Lighthouse Lab in Glasgow | Wellcome Sanger Institute for the COVID-19 Genomics UK (COG-UK) Consortium | Harper VanSteenhouse, Yumi Kasai, David Gray, Carol Clugston, Anna Dominiczak and Alex Alderton, Roberto Amato, Jeffrey Barrett, Sonia Goncalves, Ewan Harrison, David K. Jackson, Ian Johnston, Dominic Kwiatkowski, Cordelia Langford, John Sillitoe on behalf of the Wellcome Sanger Institute COVID-19 Surveillance Team |
| EPI_ISL_3108425 | Lighthouse Lab in Milton Keynes | Wellcome Sanger Institute for the COVID-19 Genomics UK (COG-UK) Consortium | The Lighthouse Lab in Milton Keynes and Alex Alderton, Roberto Amato, Jeffrey Barrett, Sonia Goncalves, Ewan Harrison, David K. Jackson, Ian Johnston, Dominic Kwiatkowski, Cordelia Langford, John Sillitoe on behalf of the Wellcome Sanger Institute COVID-19 Surveillance Team |
| EPI_ISL_3108426 | Lighthouse Lab in Alderley Park | Wellcome Sanger Institute for the COVID-19 Genomics UK (COG-UK) Consortium | Jacquelyn Wynn, Mairead Hyland, The Lighthouse Lab in Alderley Park and Alex Alderton, Roberto Amato, Jeffrey Barrett, Sonia Goncalves, Ewan Harrison, David K. Jackson, Ian Johnston, Dominic Kwiatkowski, Cordelia Langford, John Sillitoe on behalf of the Wellcome Sanger Institute COVID-19 Surveillance Team |
| EPI_ISL_3108427 | Lighthouse Lab in Milton Keynes | Wellcome Sanger Institute for the COVID-19 Genomics UK (COG-UK) Consortium | The Lighthouse Lab in Milton Keynes and Alex Alderton, Roberto Amato, Jeffrey Barrett, Sonia Goncalves, Ewan Harrison, David K. Jackson, Ian Johnston, Dominic Kwiatkowski, Cordelia Langford, John Sillitoe on behalf of the Wellcome Sanger Institute COVID-19 Surveillance Team |
| EPI_ISL_3108428 | Lighthouse Lab in Alderley Park | Wellcome Sanger Institute for the COVID-19 Genomics UK (COG-UK) Consortium | Jacquelyn Wynn, Mairead Hyland, The Lighthouse Lab in Alderley Park and Alex Alderton, Roberto Amato, Jeffrey Barrett, Sonia Goncalves, Ewan Harrison, David K. Jackson, Ian Johnston, Dominic Kwiatkowski, Cordelia Langford, John Sillitoe on behalf of the Wellcome Sanger Institute COVID-19 Surveillance Team |
| EPI_ISL_3108429 | Lighthouse Lab in Glasgow | Wellcome Sanger Institute for the COVID-19 Genomics UK (COG-UK) Consortium | Harper VanSteenhouse, Yumi Kasai, David Gray, Carol Clugston, Anna Dominiczak and Alex Alderton, Roberto Amato, Jeffrey Barrett, Sonia Goncalves, Ewan Harrison, David K. Jackson, Ian Johnston, Dominic Kwiatkowski, Cordelia Langford, John Sillitoe on behalf of the Wellcome Sanger Institute COVID-19 Surveillance Team |

| | | | |
|---|---|---|---|
| EPI_ISL_3108430 | Lighthouse Lab in Alderley Park | Wellcome Sanger Institute for the COVID-19 Genomics UK (COG-UK) Consortium | Jacquelyn Wynn, Mairead Hyland, The Lighthouse Lab in Alderley Park and Alex Alderton, Roberto Amato, Jeffrey Barrett, Sonia Goncalves, Ewan Harrison, David K. Jackson, Ian Johnston, Dominic Kwiatkowski, Cordelia Langford, John Sillitoe on behalf of the Wellcome Sanger Institute COVID-19 Surveillance Team |
| EPI_ISL_3108431 | Lighthouse Lab in Milton Keynes | Wellcome Sanger Institute for the COVID-19 Genomics UK (COG-UK) Consortium | The Lighthouse Lab in Milton Keynes and Alex Alderton, Roberto Amato, Jeffrey Barrett, Sonia Goncalves, Ewan Harrison, David K. Jackson, Ian Johnston, Dominic Kwiatkowski, Cordelia Langford, John Sillitoe on behalf of the Wellcome Sanger Institute COVID-19 Surveillance Team |
| EPI_ISL_3108432 | Lighthouse Lab in Alderley Park | Wellcome Sanger Institute for the COVID-19 Genomics UK (COG-UK) Consortium | Jacquelyn Wynn, Mairead Hyland, The Lighthouse Lab in Alderley Park and Alex Alderton, Roberto Amato, Jeffrey Barrett, Sonia Goncalves, Ewan Harrison, David K. Jackson, Ian Johnston, Dominic Kwiatkowski, Cordelia Langford, John Sillitoe on behalf of the Wellcome Sanger Institute COVID-19 Surveillance Team |
| EPI_ISL_3108433 | Lighthouse Lab in Glasgow | Wellcome Sanger Institute for the COVID-19 Genomics UK (COG-UK) Consortium | Harper VanSteenhouse, Yumi Kasai, David Gray, Carol Clugston, Anna Dominiczak and Alex Alderton, Roberto Amato, Jeffrey Barrett, Sonia Goncalves, Ewan Harrison, David K. Jackson, Ian Johnston, Dominic Kwiatkowski, Cordelia Langford, John Sillitoe on behalf of the Wellcome Sanger Institute COVID-19 Surveillance Team |
| EPI_ISL_3108434, EPI_ISL_3108435 | Lighthouse Lab in Milton Keynes | Wellcome Sanger Institute for the COVID-19 Genomics UK (COG-UK) Consortium | The Lighthouse Lab in Milton Keynes and Alex Alderton, Roberto Amato, Jeffrey Barrett, Sonia Goncalves, Ewan Harrison, David K. Jackson, Ian Johnston, Dominic Kwiatkowski, Cordelia Langford, John Sillitoe on behalf of the Wellcome Sanger Institute COVID-19 Surveillance Team |
| EPI_ISL_3108436, EPI_ISL_3108437 | Lighthouse Lab in Alderley Park | Wellcome Sanger Institute for the COVID-19 Genomics UK (COG-UK) Consortium | Jacquelyn Wynn, Mairead Hyland, The Lighthouse Lab in Alderley Park and Alex Alderton, Roberto Amato, Jeffrey Barrett, Sonia Goncalves, Ewan Harrison, David K. Jackson, Ian Johnston, Dominic Kwiatkowski, Cordelia Langford, John Sillitoe on behalf of the Wellcome Sanger Institute COVID-19 Surveillance Team |
| EPI_ISL_3108438, EPI_ISL_3108439 | Lighthouse Lab in Milton Keynes | Wellcome Sanger Institute for the COVID-19 Genomics UK (COG-UK) Consortium | The Lighthouse Lab in Milton Keynes and Alex Alderton, Roberto Amato, Jeffrey Barrett, Sonia Goncalves, Ewan Harrison, David K. Jackson, Ian Johnston, Dominic Kwiatkowski, Cordelia Langford, John Sillitoe on behalf of the Wellcome Sanger Institute COVID-19 Surveillance Team |
| EPI_ISL_3108440 | Lighthouse Lab in Alderley Park | Wellcome Sanger Institute for the COVID-19 Genomics UK (COG-UK) Consortium | Jacquelyn Wynn, Mairead Hyland, The Lighthouse Lab in Alderley Park and Alex Alderton, Roberto Amato, Jeffrey Barrett, Sonia Goncalves, Ewan Harrison, David K. Jackson, Ian Johnston, Dominic Kwiatkowski, Cordelia Langford, John Sillitoe on behalf of the Wellcome Sanger Institute COVID-19 Surveillance Team |
| EPI_ISL_3108441 | Lighthouse Lab in Glasgow | Wellcome Sanger Institute for the COVID-19 Genomics UK (COG-UK) Consortium | Harper VanSteenhouse, Yumi Kasai, David Gray, Carol Clugston, Anna Dominiczak and Alex Alderton, Roberto Amato, Jeffrey Barrett, Sonia Goncalves, Ewan Harrison, David K. Jackson, Ian Johnston, Dominic Kwiatkowski, Cordelia Langford, John Sillitoe on behalf of the Wellcome Sanger Institute COVID-19 Surveillance Team |
| EPI_ISL_3108442, EPI_ISL_3108443 | Lighthouse Lab in Alderley Park | Wellcome Sanger Institute for the COVID-19 Genomics UK (COG-UK) Consortium | Jacquelyn Wynn, Mairead Hyland, The Lighthouse Lab in Alderley Park and Alex Alderton, Roberto Amato, Jeffrey Barrett, Sonia Goncalves, Ewan Harrison, David K. Jackson, Ian Johnston, Dominic Kwiatkowski, Cordelia Langford, John Sillitoe on behalf of the Wellcome Sanger Institute COVID-19 Surveillance Team |
| EPI_ISL_3108444 | Lighthouse Lab in Milton Keynes | Wellcome Sanger Institute for the COVID-19 Genomics UK (COG-UK) Consortium | The Lighthouse Lab in Milton Keynes and Alex Alderton, Roberto Amato, Jeffrey Barrett, Sonia Goncalves, Ewan Harrison, David K. Jackson, Ian Johnston, Dominic Kwiatkowski, Cordelia Langford, John Sillitoe on behalf of the Wellcome Sanger Institute COVID-19 Surveillance Team |
| EPI_ISL_3108445, EPI_ISL_3108446 | Lighthouse Lab in Alderley Park | Wellcome Sanger Institute for the COVID-19 Genomics UK (COG-UK) Consortium | Jacquelyn Wynn, Mairead Hyland, The Lighthouse Lab in Alderley Park and Alex Alderton, Roberto Amato, Jeffrey Barrett, Sonia Goncalves, Ewan Harrison, David K. Jackson, Ian Johnston, Dominic Kwiatkowski, Cordelia Langford, John Sillitoe on behalf of the Wellcome Sanger Institute COVID-19 Surveillance Team |
| EPI_ISL_3108447 | Lighthouse Lab in Milton Keynes | Wellcome Sanger Institute for the COVID-19 Genomics UK (COG-UK) Consortium | The Lighthouse Lab in Milton Keynes and Alex Alderton, Roberto Amato, Jeffrey Barrett, Sonia Goncalves, Ewan Harrison, David K. Jackson, Ian Johnston, Dominic Kwiatkowski, Cordelia Langford, John Sillitoe on behalf of the Wellcome Sanger Institute COVID-19 Surveillance Team |
| EPI_ISL_3108448 | Lighthouse Lab in Glasgow | Wellcome Sanger Institute for the COVID-19 Genomics UK (COG-UK) Consortium | Harper VanSteenhouse, Yumi Kasai, David Gray, Carol Clugston, Anna Dominiczak and Alex Alderton, Roberto Amato, Jeffrey Barrett, Sonia Goncalves, Ewan Harrison, David K. Jackson, Ian Johnston, Dominic Kwiatkowski, Cordelia Langford, John Sillitoe on behalf of the Wellcome Sanger Institute COVID-19 Surveillance Team |
| EPI_ISL_3108449 | Lighthouse Lab in Milton Keynes | Wellcome Sanger Institute for the COVID-19 Genomics UK (COG-UK) Consortium | The Lighthouse Lab in Milton Keynes and Alex Alderton, Roberto Amato, Jeffrey Barrett, Sonia Goncalves, Ewan Harrison, David K. Jackson, Ian Johnston, Dominic Kwiatkowski, Cordelia Langford, John Sillitoe on behalf of the Wellcome Sanger Institute COVID-19 Surveillance Team |
| EPI_ISL_3108450, EPI_ISL_3108451, EPI_ISL_3108452, EPI_ISL_3108453, EPI_ISL_3108454, EPI_ISL_3108455, EPI_ISL_3108456, EPI_ISL_3108457 | Lighthouse Lab in Alderley Park | Wellcome Sanger Institute for the COVID-19 Genomics UK (COG-UK) Consortium | Jacquelyn Wynn, Mairead Hyland, The Lighthouse Lab in Alderley Park and Alex Alderton, Roberto Amato, Jeffrey Barrett, Sonia Goncalves, Ewan Harrison, David K. Jackson, Ian Johnston, Dominic Kwiatkowski, Cordelia Langford, John Sillitoe on behalf of the Wellcome Sanger Institute COVID-19 Surveillance Team |
| EPI_ISL_3108458 | Lighthouse Lab in Milton Keynes | Wellcome Sanger Institute for the COVID-19 Genomics UK (COG-UK) Consortium | The Lighthouse Lab in Milton Keynes and Alex Alderton, Roberto Amato, Jeffrey Barrett, Sonia Goncalves, Ewan Harrison, David K. Jackson, Ian Johnston, Dominic Kwiatkowski, Cordelia Langford, John Sillitoe on behalf of the Wellcome Sanger Institute COVID-19 Surveillance Team |
| EPI_ISL_3108459 | Lighthouse Lab in Glasgow | Wellcome Sanger Institute for the COVID-19 Genomics UK (COG-UK) Consortium | Harper VanSteenhouse, Yumi Kasai, David Gray, Carol Clugston, Anna Dominiczak and Alex Alderton, Roberto Amato, Jeffrey Barrett, Sonia Goncalves, Ewan Harrison, David K. Jackson, Ian Johnston, Dominic Kwiatkowski, Cordelia Langford, John Sillitoe on behalf of the Wellcome Sanger Institute COVID-19 Surveillance Team |
| EPI_ISL_3108460 | Lighthouse Lab in Milton Keynes | Wellcome Sanger Institute for the COVID-19 Genomics UK (COG-UK) Consortium | The Lighthouse Lab in Milton Keynes and Alex Alderton, Roberto Amato, Jeffrey Barrett, Sonia Goncalves, Ewan Harrison, David K. Jackson, Ian Johnston, Dominic Kwiatkowski, Cordelia Langford, John Sillitoe on behalf of the Wellcome Sanger Institute COVID-19 Surveillance Team |
| EPI_ISL_3108461, EPI_ISL_3108462, EPI_ISL_3108463 | Lighthouse Lab in Alderley Park | Wellcome Sanger Institute for the COVID-19 Genomics UK (COG-UK) Consortium | Jacquelyn Wynn, Mairead Hyland, The Lighthouse Lab in Alderley Park and Alex Alderton, Roberto Amato, Jeffrey Barrett, Sonia Goncalves, Ewan Harrison, David K. Jackson, Ian Johnston, Dominic Kwiatkowski, Cordelia Langford, John Sillitoe on behalf of the Wellcome Sanger Institute COVID-19 Surveillance Team |
| EPI_ISL_3108464 | Lighthouse Lab in Milton Keynes | Wellcome Sanger Institute for the COVID-19 Genomics UK (COG-UK) Consortium | The Lighthouse Lab in Milton Keynes and Alex Alderton, Roberto Amato, Jeffrey Barrett, Sonia Goncalves, Ewan Harrison, David K. Jackson, Ian Johnston, Dominic Kwiatkowski, Cordelia Langford, John Sillitoe on behalf of the Wellcome Sanger Institute COVID-19 Surveillance Team |
| EPI_ISL_3108465 | Lighthouse Lab in Alderley Park | Wellcome Sanger Institute for the COVID-19 Genomics UK (COG-UK) Consortium | Jacquelyn Wynn, Mairead Hyland, The Lighthouse Lab in Alderley Park and Alex Alderton, Roberto Amato, Jeffrey Barrett, Sonia Goncalves, Ewan Harrison, David K. Jackson, Ian Johnston, Dominic Kwiatkowski, Cordelia Langford, John Sillitoe on behalf of the Wellcome Sanger Institute COVID-19 Surveillance Team |
| EPI_ISL_3108466 | Lighthouse Lab in Glasgow | Wellcome Sanger Institute for the COVID-19 Genomics UK (COG-UK) Consortium | Harper VanSteenhouse, Yumi Kasai, David Gray, Carol Clugston, Anna Dominiczak and Alex Alderton, Roberto Amato, Jeffrey Barrett, Sonia Goncalves, Ewan Harrison, David K. Jackson, Ian Johnston, Dominic Kwiatkowski, Cordelia Langford, John Sillitoe on behalf of the Wellcome Sanger Institute COVID-19 Surveillance Team |
| EPI_ISL_3108467 | Lighthouse Lab in Alderley Park | Wellcome Sanger Institute for the COVID-19 Genomics UK (COG-UK) Consortium | Jacquelyn Wynn, Mairead Hyland, The Lighthouse Lab in Alderley Park and Alex Alderton, Roberto Amato, Jeffrey Barrett, Sonia Goncalves, Ewan Harrison, David K. Jackson, Ian Johnston, Dominic Kwiatkowski, Cordelia Langford, John Sillitoe on behalf of the Wellcome Sanger Institute COVID-19 Surveillance Team |
| EPI_ISL_3108468 | Lighthouse Lab in Milton Keynes | Wellcome Sanger Institute for the COVID-19 Genomics UK (COG-UK) Consortium | The Lighthouse Lab in Milton Keynes and Alex Alderton, Roberto Amato, Jeffrey Barrett, Sonia Goncalves, Ewan Harrison, David K. Jackson, Ian Johnston, Dominic Kwiatkowski, Cordelia Langford, John Sillitoe on behalf of the Wellcome Sanger Institute COVID-19 Surveillance Team |
| EPI_ISL_3108469, EPI_ISL_3108470, EPI_ISL_3108471, EPI_ISL_3108472, EPI_ISL_3108473, EPI_ISL_3108474, EPI_ISL_3108475 | Lighthouse Lab in Alderley Park | Wellcome Sanger Institute for the COVID-19 Genomics UK (COG-UK) Consortium | Jacquelyn Wynn, Mairead Hyland, The Lighthouse Lab in Alderley Park and Alex Alderton, Roberto Amato, Jeffrey Barrett, Sonia Goncalves, Ewan Harrison, David K. Jackson, Ian Johnston, Dominic Kwiatkowski, Cordelia Langford, John Sillitoe on behalf of the Wellcome Sanger Institute COVID-19 Surveillance Team |
| EPI_ISL_3108476, EPI_ISL_3108477 | Lighthouse Lab in Milton Keynes | Wellcome Sanger Institute for the COVID-19 Genomics UK (COG-UK) Consortium | The Lighthouse Lab in Milton Keynes and Alex Alderton, Roberto Amato, Jeffrey Barrett, Sonia Goncalves, Ewan Harrison, David K. Jackson, Ian Johnston, Dominic Kwiatkowski, Cordelia Langford, John Sillitoe on behalf of the Wellcome Sanger Institute COVID-19 Surveillance Team |
| EPI_ISL_3108478, EPI_ISL_3108479, EPI_ISL_3108480, EPI_ISL_3108481, EPI_ISL_3108482 | Lighthouse Lab in Alderley Park | Wellcome Sanger Institute for the COVID-19 Genomics UK (COG-UK) Consortium | Jacquelyn Wynn, Mairead Hyland, The Lighthouse Lab in Alderley Park and Alex Alderton, Roberto Amato, Jeffrey Barrett, Sonia Goncalves, Ewan Harrison, David K. Jackson, Ian Johnston, Dominic Kwiatkowski, Cordelia Langford, John Sillitoe on behalf of the Wellcome Sanger Institute COVID-19 Surveillance Team |
| EPI_ISL_3108483 | Lighthouse Lab in Milton Keynes | Wellcome Sanger Institute for the COVID-19 Genomics UK (COG-UK) Consortium | The Lighthouse Lab in Milton Keynes and Alex Alderton, Roberto Amato, Jeffrey Barrett, Sonia Goncalves, Ewan Harrison, David K. Jackson, Ian Johnston, Dominic Kwiatkowski, Cordelia Langford, John Sillitoe on behalf of the Wellcome Sanger Institute COVID-19 Surveillance Team |
| EPI_ISL_3108484, EPI_ISL_3108485 | Lighthouse Lab in Glasgow | Wellcome Sanger Institute for the COVID-19 Genomics UK (COG-UK) Consortium | Harper VanSteenhouse, Yumi Kasai, David Gray, Carol Clugston, Anna Dominiczak and Alex Alderton, Roberto Amato, Jeffrey Barrett, Sonia Goncalves, Ewan Harrison, David K. Jackson, Ian Johnston, Dominic Kwiatkowski, Cordelia Langford, John Sillitoe on behalf of the Wellcome Sanger Institute COVID-19 Surveillance Team |
| EPI_ISL_3108486 | Lighthouse Lab in Milton Keynes | Wellcome Sanger Institute for the COVID-19 Genomics UK (COG-UK) Consortium | The Lighthouse Lab in Milton Keynes and Alex Alderton, Roberto Amato, Jeffrey Barrett, Sonia Goncalves, Ewan Harrison, David K. Jackson, Ian Johnston, Dominic Kwiatkowski, Cordelia Langford, John Sillitoe on behalf of the Wellcome Sanger Institute COVID-19 Surveillance Team |
| EPI_ISL_3108487 | Lighthouse Lab in Glasgow | Wellcome Sanger Institute for the COVID-19 Genomics UK (COG-UK) Consortium | Harper VanSteenhouse, Yumi Kasai, David Gray, Carol Clugston, Anna Dominiczak and Alex Alderton, Roberto Amato, Jeffrey Barrett, Sonia Goncalves, Ewan Harrison, David K. Jackson, Ian Johnston, Dominic Kwiatkowski, Cordelia Langford, John Sillitoe on behalf of the Wellcome Sanger Institute COVID-19 Surveillance Team |
| EPI_ISL_3108488 | Lighthouse Lab in Alderley Park | Wellcome Sanger Institute for the COVID-19 Genomics UK (COG-UK) Consortium | Jacquelyn Wynn, Mairead Hyland, The Lighthouse Lab in Alderley Park and Alex Alderton, Roberto Amato, Jeffrey Barrett, Sonia Goncalves, Ewan Harrison, David K. Jackson, Ian Johnston, Dominic Kwiatkowski, Cordelia Langford, John Sillitoe on behalf of the Wellcome Sanger Institute COVID-19 Surveillance Team |
| EPI_ISL_3108489, EPI_ISL_3108490 | Lighthouse Lab in Glasgow | Wellcome Sanger Institute for the COVID-19 Genomics UK (COG-UK) Consortium | Harper VanSteenhouse, Yumi Kasai, David Gray, Carol Clugston, Anna Dominiczak and Alex Alderton, Roberto Amato, Jeffrey Barrett, Sonia Goncalves, Ewan Harrison, David K. Jackson, Ian Johnston, Dominic Kwiatkowski, Cordelia Langford, John Sillitoe on behalf of the Wellcome Sanger Institute COVID-19 Surveillance Team |
| EPI_ISL_3108491, EPI_ISL_3108492, EPI_ISL_3108493 | Lighthouse Lab in Alderley Park | Wellcome Sanger Institute for the COVID-19 Genomics UK (COG-UK) Consortium | Jacquelyn Wynn, Mairead Hyland, The Lighthouse Lab in Alderley Park and Alex Alderton, Roberto Amato, Jeffrey Barrett, Sonia Goncalves, Ewan Harrison, David K. Jackson, Ian Johnston, Dominic Kwiatkowski, Cordelia Langford, John Sillitoe on behalf of the Wellcome Sanger Institute COVID-19 Surveillance Team |
| EPI_ISL_3108494 | Lighthouse Lab in Milton Keynes | Wellcome Sanger Institute for the COVID-19 Genomics UK (COG-UK) Consortium | The Lighthouse Lab in Milton Keynes and Alex Alderton, Roberto Amato, Jeffrey Barrett, Sonia Goncalves, Ewan Harrison, David K. Jackson, Ian Johnston, Dominic Kwiatkowski, Cordelia Langford, John Sillitoe on behalf of the Wellcome Sanger Institute COVID-19 Surveillance Team |
| EPI_ISL_3108495 | Lighthouse Lab in Alderley Park | Wellcome Sanger Institute for the COVID-19 Genomics UK (COG-UK) Consortium | Jacquelyn Wynn, Mairead Hyland, The Lighthouse Lab in Alderley Park and Alex Alderton, Roberto Amato, Jeffrey Barrett, Sonia Goncalves, Ewan Harrison, David K. Jackson, Ian Johnston, Dominic Kwiatkowski, Cordelia Langford, John Sillitoe on behalf of the Wellcome Sanger Institute COVID-19 Surveillance Team |
| EPI_ISL_3108496 | Lighthouse Lab in Milton Keynes | Wellcome Sanger Institute for the COVID-19 Genomics UK (COG-UK) Consortium | The Lighthouse Lab in Milton Keynes and Alex Alderton, Roberto Amato, Jeffrey Barrett, Sonia Goncalves, Ewan Harrison, David K. Jackson, Ian Johnston, Dominic Kwiatkowski, Cordelia Langford, John Sillitoe on behalf of the Wellcome Sanger Institute COVID-19 Surveillance Team |
| EPI_ISL_3108497, | Lighthouse Lab in Alderley Park | Wellcome Sanger Institute for the COVID-19 Genomics UK (COG-UK) | Jacquelyn Wynn, Mairead Hyland, The Lighthouse Lab in Alderley Park and Alex Alderton, Roberto Amato, Jeffrey Barrett, Sonia Goncalves, Ewan Harrison, David K. Jackson, Ian Johnston, Dominic Kwiatkowski, Cordelia Langford, John Sillitoe on |

| | | | |
|---|---|---|---|
| EPI_ISL_3108498 | | Consortium | behalf of the Wellcome Sanger Institute COVID-19 Surveillance Team |
| EPI_ISL_3108499 | Lighthouse Lab in Glasgow | Wellcome Sanger Institute for the COVID-19 Genomics UK (COG-UK) Consortium | Harper VanSteenhouse, Yumi Kasai, David Gray, Carol Clugston, Anna Dominiczak and Alex Alderton, Roberto Amato, Jeffrey Barrett, Sonia Goncalves, Ewan Harrison, David K. Jackson, Ian Johnston, Dominic Kwiatkowski, Cordelia Langford, John Sillitoe on behalf of the Wellcome Sanger Institute COVID-19 Surveillance Team |
| EPI_ISL_3108500, EPI_ISL_3108501, EPI_ISL_3108502, EPI_ISL_3108503, EPI_ISL_3108504 | Lighthouse Lab in Alderley Park | Wellcome Sanger Institute for the COVID-19 Genomics UK (COG-UK) Consortium | Jacquelyn Wynn, Mairead Hyland, The Lighthouse Lab in Alderley Park and Alex Alderton, Roberto Amato, Jeffrey Barrett, Sonia Goncalves, Ewan Harrison, David K. Jackson, Ian Johnston, Dominic Kwiatkowski, Cordelia Langford, John Sillitoe on behalf of the Wellcome Sanger Institute COVID-19 Surveillance Team |
| EPI_ISL_3108505 | Lighthouse Lab in Milton Keynes | Wellcome Sanger Institute for the COVID-19 Genomics UK (COG-UK) Consortium | The Lighthouse Lab in Milton Keynes and Alex Alderton, Roberto Amato, Jeffrey Barrett, Sonia Goncalves, Ewan Harrison, David K. Jackson, Ian Johnston, Dominic Kwiatkowski, Cordelia Langford, John Sillitoe on behalf of the Wellcome Sanger Institute COVID-19 Surveillance Team |
| EPI_ISL_3108506 | Lighthouse Lab in Glasgow | Wellcome Sanger Institute for the COVID-19 Genomics UK (COG-UK) Consortium | Harper VanSteenhouse, Yumi Kasai, David Gray, Carol Clugston, Anna Dominiczak and Alex Alderton, Roberto Amato, Jeffrey Barrett, Sonia Goncalves, Ewan Harrison, David K. Jackson, Ian Johnston, Dominic Kwiatkowski, Cordelia Langford, John Sillitoe on behalf of the Wellcome Sanger Institute COVID-19 Surveillance Team |
| EPI_ISL_3108507, EPI_ISL_3108508, EPI_ISL_3108509 | Lighthouse Lab in Alderley Park | Wellcome Sanger Institute for the COVID-19 Genomics UK (COG-UK) Consortium | Jacquelyn Wynn, Mairead Hyland, The Lighthouse Lab in Alderley Park and Alex Alderton, Roberto Amato, Jeffrey Barrett, Sonia Goncalves, Ewan Harrison, David K. Jackson, Ian Johnston, Dominic Kwiatkowski, Cordelia Langford, John Sillitoe on behalf of the Wellcome Sanger Institute COVID-19 Surveillance Team |
| EPI_ISL_3108510, EPI_ISL_3108511 | Lighthouse Lab in Milton Keynes | Wellcome Sanger Institute for the COVID-19 Genomics UK (COG-UK) Consortium | The Lighthouse Lab in Milton Keynes and Alex Alderton, Roberto Amato, Jeffrey Barrett, Sonia Goncalves, Ewan Harrison, David K. Jackson, Ian Johnston, Dominic Kwiatkowski, Cordelia Langford, John Sillitoe on behalf of the Wellcome Sanger Institute COVID-19 Surveillance Team |
| EPI_ISL_3108512, EPI_ISL_3108513 | Lighthouse Lab in Alderley Park | Wellcome Sanger Institute for the COVID-19 Genomics UK (COG-UK) Consortium | Jacquelyn Wynn, Mairead Hyland, The Lighthouse Lab in Alderley Park and Alex Alderton, Roberto Amato, Jeffrey Barrett, Sonia Goncalves, Ewan Harrison, David K. Jackson, Ian Johnston, Dominic Kwiatkowski, Cordelia Langford, John Sillitoe on behalf of the Wellcome Sanger Institute COVID-19 Surveillance Team |
| EPI_ISL_3108514 | Lighthouse Lab in Glasgow | Wellcome Sanger Institute for the COVID-19 Genomics UK (COG-UK) Consortium | Harper VanSteenhouse, Yumi Kasai, David Gray, Carol Clugston, Anna Dominiczak and Alex Alderton, Roberto Amato, Jeffrey Barrett, Sonia Goncalves, Ewan Harrison, David K. Jackson, Ian Johnston, Dominic Kwiatkowski, Cordelia Langford, John Sillitoe on behalf of the Wellcome Sanger Institute COVID-19 Surveillance Team |
| EPI_ISL_3108515 | Lighthouse Lab in Milton Keynes | Wellcome Sanger Institute for the COVID-19 Genomics UK (COG-UK) Consortium | The Lighthouse Lab in Milton Keynes and Alex Alderton, Roberto Amato, Jeffrey Barrett, Sonia Goncalves, Ewan Harrison, David K. Jackson, Ian Johnston, Dominic Kwiatkowski, Cordelia Langford, John Sillitoe on behalf of the Wellcome Sanger Institute COVID-19 Surveillance Team |
| EPI_ISL_3108516 | Lighthouse Lab in Alderley Park | Wellcome Sanger Institute for the COVID-19 Genomics UK (COG-UK) Consortium | Jacquelyn Wynn, Mairead Hyland, The Lighthouse Lab in Alderley Park and Alex Alderton, Roberto Amato, Jeffrey Barrett, Sonia Goncalves, Ewan Harrison, David K. Jackson, Ian Johnston, Dominic Kwiatkowski, Cordelia Langford, John Sillitoe on behalf of the Wellcome Sanger Institute COVID-19 Surveillance Team |
| EPI_ISL_3108517 | Lighthouse Lab in Milton Keynes | Wellcome Sanger Institute for the COVID-19 Genomics UK (COG-UK) Consortium | The Lighthouse Lab in Milton Keynes and Alex Alderton, Roberto Amato, Jeffrey Barrett, Sonia Goncalves, Ewan Harrison, David K. Jackson, Ian Johnston, Dominic Kwiatkowski, Cordelia Langford, John Sillitoe on behalf of the Wellcome Sanger Institute COVID-19 Surveillance Team |
| EPI_ISL_3108518 | Lighthouse Lab in Glasgow | Wellcome Sanger Institute for the COVID-19 Genomics UK (COG-UK) Consortium | Harper VanSteenhouse, Yumi Kasai, David Gray, Carol Clugston, Anna Dominiczak and Alex Alderton, Roberto Amato, Jeffrey Barrett, Sonia Goncalves, Ewan Harrison, David K. Jackson, Ian Johnston, Dominic Kwiatkowski, Cordelia Langford, John Sillitoe on behalf of the Wellcome Sanger Institute COVID-19 Surveillance Team |
| EPI_ISL_3108519, EPI_ISL_3108520, EPI_ISL_3108521, EPI_ISL_3108522, EPI_ISL_3108523 | Lighthouse Lab in Alderley Park | Wellcome Sanger Institute for the COVID-19 Genomics UK (COG-UK) Consortium | Jacquelyn Wynn, Mairead Hyland, The Lighthouse Lab in Alderley Park and Alex Alderton, Roberto Amato, Jeffrey Barrett, Sonia Goncalves, Ewan Harrison, David K. Jackson, Ian Johnston, Dominic Kwiatkowski, Cordelia Langford, John Sillitoe on behalf of the Wellcome Sanger Institute COVID-19 Surveillance Team |
| EPI_ISL_3108524 | Lighthouse Lab in Milton Keynes | Wellcome Sanger Institute for the COVID-19 Genomics UK (COG-UK) Consortium | The Lighthouse Lab in Milton Keynes and Alex Alderton, Roberto Amato, Jeffrey Barrett, Sonia Goncalves, Ewan Harrison, David K. Jackson, Ian Johnston, Dominic Kwiatkowski, Cordelia Langford, John Sillitoe on behalf of the Wellcome Sanger Institute COVID-19 Surveillance Team |
| EPI_ISL_3108525 | Lighthouse Lab in Glasgow | Wellcome Sanger Institute for the COVID-19 Genomics UK (COG-UK) Consortium | Harper VanSteenhouse, Yumi Kasai, David Gray, Carol Clugston, Anna Dominiczak and Alex Alderton, Roberto Amato, Jeffrey Barrett, Sonia Goncalves, Ewan Harrison, David K. Jackson, Ian Johnston, Dominic Kwiatkowski, Cordelia Langford, John Sillitoe on behalf of the Wellcome Sanger Institute COVID-19 Surveillance Team |
| EPI_ISL_3108526, EPI_ISL_3108527, EPI_ISL_3108528 | Lighthouse Lab in Alderley Park | Wellcome Sanger Institute for the COVID-19 Genomics UK (COG-UK) Consortium | Jacquelyn Wynn, Mairead Hyland, The Lighthouse Lab in Alderley Park and Alex Alderton, Roberto Amato, Jeffrey Barrett, Sonia Goncalves, Ewan Harrison, David K. Jackson, Ian Johnston, Dominic Kwiatkowski, Cordelia Langford, John Sillitoe on behalf of the Wellcome Sanger Institute COVID-19 Surveillance Team |
| EPI_ISL_3108529 | Lighthouse Lab in Glasgow | Wellcome Sanger Institute for the COVID-19 Genomics UK (COG-UK) Consortium | Harper VanSteenhouse, Yumi Kasai, David Gray, Carol Clugston, Anna Dominiczak and Alex Alderton, Roberto Amato, Jeffrey Barrett, Sonia Goncalves, Ewan Harrison, David K. Jackson, Ian Johnston, Dominic Kwiatkowski, Cordelia Langford, John Sillitoe on behalf of the Wellcome Sanger Institute COVID-19 Surveillance Team |
| EPI_ISL_3108530 | Lighthouse Lab in Alderley Park | Wellcome Sanger Institute for the COVID-19 Genomics UK (COG-UK) Consortium | Jacquelyn Wynn, Mairead Hyland, The Lighthouse Lab in Alderley Park and Alex Alderton, Roberto Amato, Jeffrey Barrett, Sonia Goncalves, Ewan Harrison, David K. Jackson, Ian Johnston, Dominic Kwiatkowski, Cordelia Langford, John Sillitoe on behalf of the Wellcome Sanger Institute COVID-19 Surveillance Team |
| EPI_ISL_3108531, EPI_ISL_3108532 | Lighthouse Lab in Milton Keynes | Wellcome Sanger Institute for the COVID-19 Genomics UK (COG-UK) Consortium | The Lighthouse Lab in Milton Keynes and Alex Alderton, Roberto Amato, Jeffrey Barrett, Sonia Goncalves, Ewan Harrison, David K. Jackson, Ian Johnston, Dominic Kwiatkowski, Cordelia Langford, John Sillitoe on behalf of the Wellcome Sanger Institute COVID-19 Surveillance Team |
| EPI_ISL_3108533 | Lighthouse Lab in Glasgow | Wellcome Sanger Institute for the COVID-19 Genomics UK (COG-UK) Consortium | Harper VanSteenhouse, Yumi Kasai, David Gray, Carol Clugston, Anna Dominiczak and Alex Alderton, Roberto Amato, Jeffrey Barrett, Sonia Goncalves, Ewan Harrison, David K. Jackson, Ian Johnston, Dominic Kwiatkowski, Cordelia Langford, John Sillitoe on behalf of the Wellcome Sanger Institute COVID-19 Surveillance Team |
| EPI_ISL_3108534, EPI_ISL_3108535 | Lighthouse Lab in Alderley Park | Wellcome Sanger Institute for the COVID-19 Genomics UK (COG-UK) Consortium | Jacquelyn Wynn, Mairead Hyland, The Lighthouse Lab in Alderley Park and Alex Alderton, Roberto Amato, Jeffrey Barrett, Sonia Goncalves, Ewan Harrison, David K. Jackson, Ian Johnston, Dominic Kwiatkowski, Cordelia Langford, John Sillitoe on behalf of the Wellcome Sanger Institute COVID-19 Surveillance Team |
| EPI_ISL_3108536 | Lighthouse Lab in Glasgow | Wellcome Sanger Institute for the COVID-19 Genomics UK (COG-UK) Consortium | Harper VanSteenhouse, Yumi Kasai, David Gray, Carol Clugston, Anna Dominiczak and Alex Alderton, Roberto Amato, Jeffrey Barrett, Sonia Goncalves, Ewan Harrison, David K. Jackson, Ian Johnston, Dominic Kwiatkowski, Cordelia Langford, John Sillitoe on behalf of the Wellcome Sanger Institute COVID-19 Surveillance Team |
| EPI_ISL_3108537, EPI_ISL_3108538, EPI_ISL_3108539 | Lighthouse Lab in Alderley Park | Wellcome Sanger Institute for the COVID-19 Genomics UK (COG-UK) Consortium | Jacquelyn Wynn, Mairead Hyland, The Lighthouse Lab in Alderley Park and Alex Alderton, Roberto Amato, Jeffrey Barrett, Sonia Goncalves, Ewan Harrison, David K. Jackson, Ian Johnston, Dominic Kwiatkowski, Cordelia Langford, John Sillitoe on behalf of the Wellcome Sanger Institute COVID-19 Surveillance Team |
| EPI_ISL_3108540 | Lighthouse Lab in Milton Keynes | Wellcome Sanger Institute for the COVID-19 Genomics UK (COG-UK) Consortium | The Lighthouse Lab in Milton Keynes and Alex Alderton, Roberto Amato, Jeffrey Barrett, Sonia Goncalves, Ewan Harrison, David K. Jackson, Ian Johnston, Dominic Kwiatkowski, Cordelia Langford, John Sillitoe on behalf of the Wellcome Sanger Institute COVID-19 Surveillance Team |
| EPI_ISL_3108541, EPI_ISL_3108542 | Lighthouse Lab in Alderley Park | Wellcome Sanger Institute for the COVID-19 Genomics UK (COG-UK) Consortium | Jacquelyn Wynn, Mairead Hyland, The Lighthouse Lab in Alderley Park and Alex Alderton, Roberto Amato, Jeffrey Barrett, Sonia Goncalves, Ewan Harrison, David K. Jackson, Ian Johnston, Dominic Kwiatkowski, Cordelia Langford, John Sillitoe on behalf of the Wellcome Sanger Institute COVID-19 Surveillance Team |
| EPI_ISL_3108543 | Lighthouse Lab in Glasgow | Wellcome Sanger Institute for the COVID-19 Genomics UK (COG-UK) Consortium | Harper VanSteenhouse, Yumi Kasai, David Gray, Carol Clugston, Anna Dominiczak and Alex Alderton, Roberto Amato, Jeffrey Barrett, Sonia Goncalves, Ewan Harrison, David K. Jackson, Ian Johnston, Dominic Kwiatkowski, Cordelia Langford, John Sillitoe on behalf of the Wellcome Sanger Institute COVID-19 Surveillance Team |
| EPI_ISL_3108544 | Lighthouse Lab in Milton Keynes | Wellcome Sanger Institute for the COVID-19 Genomics UK (COG-UK) Consortium | The Lighthouse Lab in Milton Keynes and Alex Alderton, Roberto Amato, Jeffrey Barrett, Sonia Goncalves, Ewan Harrison, David K. Jackson, Ian Johnston, Dominic Kwiatkowski, Cordelia Langford, John Sillitoe on behalf of the Wellcome Sanger Institute COVID-19 Surveillance Team |
| EPI_ISL_3108545, EPI_ISL_3108546, EPI_ISL_3108547 | Lighthouse Lab in Alderley Park | Wellcome Sanger Institute for the COVID-19 Genomics UK (COG-UK) Consortium | Jacquelyn Wynn, Mairead Hyland, The Lighthouse Lab in Alderley Park and Alex Alderton, Roberto Amato, Jeffrey Barrett, Sonia Goncalves, Ewan Harrison, David K. Jackson, Ian Johnston, Dominic Kwiatkowski, Cordelia Langford, John Sillitoe on behalf of the Wellcome Sanger Institute COVID-19 Surveillance Team |
| EPI_ISL_3108548, EPI_ISL_3108549 | Berkshire and Surrey Pathology Services Lighthouse Laboratory | Wellcome Sanger Institute for the COVID-19 Genomics UK (COG-UK) Consortium | Berkshire and Surrey Pathology Services Lighthouse Laboratory and Alex Alderton, Roberto Amato, Jeffrey Barrett, Sonia Goncalves, Ewan Harrison, David K. Jackson, Ian Johnston, Dominic Kwiatkowski, Cordelia Langford, John Sillitoe on behalf of the Wellcome Sanger Institute COVID-19 Surveillance Team |
| EPI_ISL_3108550 | Lighthouse Lab in Milton Keynes | Wellcome Sanger Institute for the COVID-19 Genomics UK (COG-UK) Consortium | The Lighthouse Lab in Milton Keynes and Alex Alderton, Roberto Amato, Jeffrey Barrett, Sonia Goncalves, Ewan Harrison, David K. Jackson, Ian Johnston, Dominic Kwiatkowski, Cordelia Langford, John Sillitoe on behalf of the Wellcome Sanger Institute COVID-19 Surveillance Team |
| EPI_ISL_3108551 | Lighthouse Lab in Alderley Park | Wellcome Sanger Institute for the COVID-19 Genomics UK (COG-UK) Consortium | Jacquelyn Wynn, Mairead Hyland, The Lighthouse Lab in Alderley Park and Alex Alderton, Roberto Amato, Jeffrey Barrett, Sonia Goncalves, Ewan Harrison, David K. Jackson, Ian Johnston, Dominic Kwiatkowski, Cordelia Langford, John Sillitoe on behalf of the Wellcome Sanger Institute COVID-19 Surveillance Team |
| EPI_ISL_3108552, EPI_ISL_3108553, EPI_ISL_3108554, EPI_ISL_3108555 | Lighthouse Lab in Milton Keynes | Wellcome Sanger Institute for the COVID-19 Genomics UK (COG-UK) Consortium | The Lighthouse Lab in Milton Keynes and Alex Alderton, Roberto Amato, Jeffrey Barrett, Sonia Goncalves, Ewan Harrison, David K. Jackson, Ian Johnston, Dominic Kwiatkowski, Cordelia Langford, John Sillitoe on behalf of the Wellcome Sanger Institute COVID-19 Surveillance Team |
| EPI_ISL_3108556 | Berkshire and Surrey Pathology Services Lighthouse Laboratory | Wellcome Sanger Institute for the COVID-19 Genomics UK (COG-UK) Consortium | Berkshire and Surrey Pathology Services Lighthouse Laboratory and Alex Alderton, Roberto Amato, Jeffrey Barrett, Sonia Goncalves, Ewan Harrison, David K. Jackson, Ian Johnston, Dominic Kwiatkowski, Cordelia Langford, John Sillitoe on behalf of the Wellcome Sanger Institute COVID-19 Surveillance Team |
| EPI_ISL_3108557 | Lighthouse Lab in Milton Keynes | Wellcome Sanger Institute for the COVID-19 Genomics UK (COG-UK) Consortium | The Lighthouse Lab in Milton Keynes and Alex Alderton, Roberto Amato, Jeffrey Barrett, Sonia Goncalves, Ewan Harrison, David K. Jackson, Ian Johnston, Dominic Kwiatkowski, Cordelia Langford, John Sillitoe on behalf of the Wellcome Sanger Institute COVID-19 Surveillance Team |
| EPI_ISL_3108558 | Lighthouse Lab in Alderley Park | Wellcome Sanger Institute for the COVID-19 Genomics UK (COG-UK) Consortium | Jacquelyn Wynn, Mairead Hyland, The Lighthouse Lab in Alderley Park and Alex Alderton, Roberto Amato, Jeffrey Barrett, Sonia Goncalves, Ewan Harrison, David K. Jackson, Ian Johnston, Dominic Kwiatkowski, Cordelia Langford, John Sillitoe on behalf of the Wellcome Sanger Institute COVID-19 Surveillance Team |
| EPI_ISL_3108559 | Berkshire and Surrey Pathology Services Lighthouse Laboratory | Wellcome Sanger Institute for the COVID-19 Genomics UK (COG-UK) Consortium | Berkshire and Surrey Pathology Services Lighthouse Laboratory and Alex Alderton, Roberto Amato, Jeffrey Barrett, Sonia Goncalves, Ewan Harrison, David K. Jackson, Ian Johnston, Dominic Kwiatkowski, Cordelia Langford, John Sillitoe on behalf of the Wellcome Sanger Institute COVID-19 Surveillance Team |
| EPI_ISL_3108560, EPI_ISL_3108561, EPI_ISL_3108562, EPI_ISL_3108563 | Lighthouse Lab in Milton Keynes | Wellcome Sanger Institute for the COVID-19 Genomics UK (COG-UK) Consortium | The Lighthouse Lab in Milton Keynes and Alex Alderton, Roberto Amato, Jeffrey Barrett, Sonia Goncalves, Ewan Harrison, David K. Jackson, Ian Johnston, Dominic Kwiatkowski, Cordelia Langford, John Sillitoe on behalf of the Wellcome Sanger Institute COVID-19 Surveillance Team |
| EPI_ISL_3108564 | Berkshire and Surrey Pathology Services Lighthouse Laboratory | Wellcome Sanger Institute for the COVID-19 Genomics UK (COG-UK) Consortium | Berkshire and Surrey Pathology Services Lighthouse Laboratory and Alex Alderton, Roberto Amato, Jeffrey Barrett, Sonia Goncalves, Ewan Harrison, David K. Jackson, Ian Johnston, Dominic Kwiatkowski, Cordelia Langford, John Sillitoe on behalf of the Wellcome Sanger Institute COVID-19 Surveillance Team |
| EPI_ISL_3108565, | Lighthouse Lab in Milton Keynes | Wellcome Sanger Institute for the COVID-19 Genomics UK (COG-UK) | The Lighthouse Lab in Milton Keynes and Alex Alderton, Roberto Amato, Jeffrey Barrett, Sonia Goncalves, Ewan Harrison, David K. Jackson, Ian Johnston, Dominic Kwiatkowski, Cordelia Langford, John Sillitoe on behalf of the Wellcome Sanger |

| | | | |
|---|---|---|---|
| EPI_ISL_3108566, EPI_ISL_3108567, EPI_ISL_3108568 | | Consortium | Institute COVID-19 Surveillance Team |
| EPI_ISL_3108569 | Lighthouse Lab in Alderley Park | Wellcome Sanger Institute for the COVID-19 Genomics UK (COG-UK) Consortium | Jacquelyn Wynn, Mairead Hyland, The Lighthouse Lab in Alderley Park and Alex Alderton, Roberto Amato, Jeffrey Barrett, Sonia Goncalves, Ewan Harrison, David K. Jackson, Ian Johnston, Dominic Kwiatkowski, Cordelia Langford, John Sillitoe on behalf of the Wellcome Sanger Institute COVID-19 Surveillance Team |
| EPI_ISL_3108570 | Lighthouse Lab in Milton Keynes | Wellcome Sanger Institute for the COVID-19 Genomics UK (COG-UK) Consortium | The Lighthouse Lab in Milton Keynes and Alex Alderton, Roberto Amato, Jeffrey Barrett, Sonia Goncalves, Ewan Harrison, David K. Jackson, Ian Johnston, Dominic Kwiatkowski, Cordelia Langford, John Sillitoe on behalf of the Wellcome Sanger Institute COVID-19 Surveillance Team |
| EPI_ISL_3108571 | Berkshire and Surrey Pathology Services Lighthouse Laboratory | Wellcome Sanger Institute for the COVID-19 Genomics UK (COG-UK) Consortium | Berkshire and Surrey Pathology Services Lighthouse Laboratory and Alex Alderton, Roberto Amato, Jeffrey Barrett, Sonia Goncalves, Ewan Harrison, David K. Jackson, Ian Johnston, Dominic Kwiatkowski, Cordelia Langford, John Sillitoe on behalf of the Wellcome Sanger Institute COVID-19 Surveillance Team |
| EPI_ISL_3108572, EPI_ISL_3108573, EPI_ISL_3108574 | Lighthouse Lab in Milton Keynes | Wellcome Sanger Institute for the COVID-19 Genomics UK (COG-UK) Consortium | The Lighthouse Lab in Milton Keynes and Alex Alderton, Roberto Amato, Jeffrey Barrett, Sonia Goncalves, Ewan Harrison, David K. Jackson, Ian Johnston, Dominic Kwiatkowski, Cordelia Langford, John Sillitoe on behalf of the Wellcome Sanger Institute COVID-19 Surveillance Team |
| EPI_ISL_3108575 | Berkshire and Surrey Pathology Services Lighthouse Laboratory | Wellcome Sanger Institute for the COVID-19 Genomics UK (COG-UK) Consortium | Berkshire and Surrey Pathology Services Lighthouse Laboratory and Alex Alderton, Roberto Amato, Jeffrey Barrett, Sonia Goncalves, Ewan Harrison, David K. Jackson, Ian Johnston, Dominic Kwiatkowski, Cordelia Langford, John Sillitoe on behalf of the Wellcome Sanger Institute COVID-19 Surveillance Team |
| EPI_ISL_3108576 | Lighthouse Lab in Milton Keynes | Wellcome Sanger Institute for the COVID-19 Genomics UK (COG-UK) Consortium | The Lighthouse Lab in Milton Keynes and Alex Alderton, Roberto Amato, Jeffrey Barrett, Sonia Goncalves, Ewan Harrison, David K. Jackson, Ian Johnston, Dominic Kwiatkowski, Cordelia Langford, John Sillitoe on behalf of the Wellcome Sanger Institute COVID-19 Surveillance Team |
| EPI_ISL_3108577 | Lighthouse Lab in Alderley Park | Wellcome Sanger Institute for the COVID-19 Genomics UK (COG-UK) Consortium | Jacquelyn Wynn, Mairead Hyland, The Lighthouse Lab in Alderley Park and Alex Alderton, Roberto Amato, Jeffrey Barrett, Sonia Goncalves, Ewan Harrison, David K. Jackson, Ian Johnston, Dominic Kwiatkowski, Cordelia Langford, John Sillitoe on behalf of the Wellcome Sanger Institute COVID-19 Surveillance Team |
| EPI_ISL_3108578 | Berkshire and Surrey Pathology Services Lighthouse Laboratory | Wellcome Sanger Institute for the COVID-19 Genomics UK (COG-UK) Consortium | Berkshire and Surrey Pathology Services Lighthouse Laboratory and Alex Alderton, Roberto Amato, Jeffrey Barrett, Sonia Goncalves, Ewan Harrison, David K. Jackson, Ian Johnston, Dominic Kwiatkowski, Cordelia Langford, John Sillitoe on behalf of the Wellcome Sanger Institute COVID-19 Surveillance Team |
| EPI_ISL_3108579 | Lighthouse Lab in Alderley Park | Wellcome Sanger Institute for the COVID-19 Genomics UK (COG-UK) Consortium | Jacquelyn Wynn, Mairead Hyland, The Lighthouse Lab in Alderley Park and Alex Alderton, Roberto Amato, Jeffrey Barrett, Sonia Goncalves, Ewan Harrison, David K. Jackson, Ian Johnston, Dominic Kwiatkowski, Cordelia Langford, John Sillitoe on behalf of the Wellcome Sanger Institute COVID-19 Surveillance Team |
| EPI_ISL_3108580, EPI_ISL_3108581, EPI_ISL_3108582, EPI_ISL_3108583, EPI_ISL_3108584, EPI_ISL_3108585 | Lighthouse Lab in Milton Keynes | Wellcome Sanger Institute for the COVID-19 Genomics UK (COG-UK) Consortium | The Lighthouse Lab in Milton Keynes and Alex Alderton, Roberto Amato, Jeffrey Barrett, Sonia Goncalves, Ewan Harrison, David K. Jackson, Ian Johnston, Dominic Kwiatkowski, Cordelia Langford, John Sillitoe on behalf of the Wellcome Sanger Institute COVID-19 Surveillance Team |
| EPI_ISL_3108586 | Lighthouse Lab in Alderley Park | Wellcome Sanger Institute for the COVID-19 Genomics UK (COG-UK) Consortium | Jacquelyn Wynn, Mairead Hyland, The Lighthouse Lab in Alderley Park and Alex Alderton, Roberto Amato, Jeffrey Barrett, Sonia Goncalves, Ewan Harrison, David K. Jackson, Ian Johnston, Dominic Kwiatkowski, Cordelia Langford, John Sillitoe on behalf of the Wellcome Sanger Institute COVID-19 Surveillance Team |
| EPI_ISL_3108587 | Lighthouse Lab in Milton Keynes | Wellcome Sanger Institute for the COVID-19 Genomics UK (COG-UK) Consortium | The Lighthouse Lab in Milton Keynes and Alex Alderton, Roberto Amato, Jeffrey Barrett, Sonia Goncalves, Ewan Harrison, David K. Jackson, Ian Johnston, Dominic Kwiatkowski, Cordelia Langford, John Sillitoe on behalf of the Wellcome Sanger Institute COVID-19 Surveillance Team |
| EPI_ISL_3108588 | Lighthouse Lab in Alderley Park | Wellcome Sanger Institute for the COVID-19 Genomics UK (COG-UK) Consortium | Jacquelyn Wynn, Mairead Hyland, The Lighthouse Lab in Alderley Park and Alex Alderton, Roberto Amato, Jeffrey Barrett, Sonia Goncalves, Ewan Harrison, David K. Jackson, Ian Johnston, Dominic Kwiatkowski, Cordelia Langford, John Sillitoe on behalf of the Wellcome Sanger Institute COVID-19 Surveillance Team |
| EPI_ISL_3108589, EPI_ISL_3108590 | Berkshire and Surrey Pathology Services Lighthouse Laboratory | Wellcome Sanger Institute for the COVID-19 Genomics UK (COG-UK) Consortium | Berkshire and Surrey Pathology Services Lighthouse Laboratory and Alex Alderton, Roberto Amato, Jeffrey Barrett, Sonia Goncalves, Ewan Harrison, David K. Jackson, Ian Johnston, Dominic Kwiatkowski, Cordelia Langford, John Sillitoe on behalf of the Wellcome Sanger Institute COVID-19 Surveillance Team |
| EPI_ISL_3108591, EPI_ISL_3108592, EPI_ISL_3108593, EPI_ISL_3108594, EPI_ISL_3108595 | Lighthouse Lab in Milton Keynes | Wellcome Sanger Institute for the COVID-19 Genomics UK (COG-UK) Consortium | The Lighthouse Lab in Milton Keynes and Alex Alderton, Roberto Amato, Jeffrey Barrett, Sonia Goncalves, Ewan Harrison, David K. Jackson, Ian Johnston, Dominic Kwiatkowski, Cordelia Langford, John Sillitoe on behalf of the Wellcome Sanger Institute COVID-19 Surveillance Team |
| EPI_ISL_3108596, EPI_ISL_3108597 | Berkshire and Surrey Pathology Services Lighthouse Laboratory | Wellcome Sanger Institute for the COVID-19 Genomics UK (COG-UK) Consortium | Berkshire and Surrey Pathology Services Lighthouse Laboratory and Alex Alderton, Roberto Amato, Jeffrey Barrett, Sonia Goncalves, Ewan Harrison, David K. Jackson, Ian Johnston, Dominic Kwiatkowski, Cordelia Langford, John Sillitoe on behalf of the Wellcome Sanger Institute COVID-19 Surveillance Team |
| EPI_ISL_3108598 | Lighthouse Lab in Alderley Park | Wellcome Sanger Institute for the COVID-19 Genomics UK (COG-UK) Consortium | Jacquelyn Wynn, Mairead Hyland, The Lighthouse Lab in Alderley Park and Alex Alderton, Roberto Amato, Jeffrey Barrett, Sonia Goncalves, Ewan Harrison, David K. Jackson, Ian Johnston, Dominic Kwiatkowski, Cordelia Langford, John Sillitoe on behalf of the Wellcome Sanger Institute COVID-19 Surveillance Team |
| EPI_ISL_3108599, EPI_ISL_3108600 | Lighthouse Lab in Milton Keynes | Wellcome Sanger Institute for the COVID-19 Genomics UK (COG-UK) Consortium | The Lighthouse Lab in Milton Keynes and Alex Alderton, Roberto Amato, Jeffrey Barrett, Sonia Goncalves, Ewan Harrison, David K. Jackson, Ian Johnston, Dominic Kwiatkowski, Cordelia Langford, John Sillitoe on behalf of the Wellcome Sanger Institute COVID-19 Surveillance Team |
| EPI_ISL_3108601 | Berkshire and Surrey Pathology Services Lighthouse Laboratory | Wellcome Sanger Institute for the COVID-19 Genomics UK (COG-UK) Consortium | Berkshire and Surrey Pathology Services Lighthouse Laboratory and Alex Alderton, Roberto Amato, Jeffrey Barrett, Sonia Goncalves, Ewan Harrison, David K. Jackson, Ian Johnston, Dominic Kwiatkowski, Cordelia Langford, John Sillitoe on behalf of the Wellcome Sanger Institute COVID-19 Surveillance Team |
| EPI_ISL_3108602, EPI_ISL_3108603, EPI_ISL_3108604 | Lighthouse Lab in Alderley Park | Wellcome Sanger Institute for the COVID-19 Genomics UK (COG-UK) Consortium | Jacquelyn Wynn, Mairead Hyland, The Lighthouse Lab in Alderley Park and Alex Alderton, Roberto Amato, Jeffrey Barrett, Sonia Goncalves, Ewan Harrison, David K. Jackson, Ian Johnston, Dominic Kwiatkowski, Cordelia Langford, John Sillitoe on behalf of the Wellcome Sanger Institute COVID-19 Surveillance Team |
| EPI_ISL_3108605 | Lighthouse Lab in Milton Keynes | Wellcome Sanger Institute for the COVID-19 Genomics UK (COG-UK) Consortium | The Lighthouse Lab in Milton Keynes and Alex Alderton, Roberto Amato, Jeffrey Barrett, Sonia Goncalves, Ewan Harrison, David K. Jackson, Ian Johnston, Dominic Kwiatkowski, Cordelia Langford, John Sillitoe on behalf of the Wellcome Sanger Institute COVID-19 Surveillance Team |
| EPI_ISL_3108606 | Lighthouse Lab in Alderley Park | Wellcome Sanger Institute for the COVID-19 Genomics UK (COG-UK) Consortium | Jacquelyn Wynn, Mairead Hyland, The Lighthouse Lab in Alderley Park and Alex Alderton, Roberto Amato, Jeffrey Barrett, Sonia Goncalves, Ewan Harrison, David K. Jackson, Ian Johnston, Dominic Kwiatkowski, Cordelia Langford, John Sillitoe on behalf of the Wellcome Sanger Institute COVID-19 Surveillance Team |
| EPI_ISL_3108607 | Lighthouse Lab in Milton Keynes | Wellcome Sanger Institute for the COVID-19 Genomics UK (COG-UK) Consortium | The Lighthouse Lab in Milton Keynes and Alex Alderton, Roberto Amato, Jeffrey Barrett, Sonia Goncalves, Ewan Harrison, David K. Jackson, Ian Johnston, Dominic Kwiatkowski, Cordelia Langford, John Sillitoe on behalf of the Wellcome Sanger Institute COVID-19 Surveillance Team |
| EPI_ISL_3108608, EPI_ISL_3108609 | Berkshire and Surrey Pathology Services Lighthouse Laboratory | Wellcome Sanger Institute for the COVID-19 Genomics UK (COG-UK) Consortium | Berkshire and Surrey Pathology Services Lighthouse Laboratory and Alex Alderton, Roberto Amato, Jeffrey Barrett, Sonia Goncalves, Ewan Harrison, David K. Jackson, Ian Johnston, Dominic Kwiatkowski, Cordelia Langford, John Sillitoe on behalf of the Wellcome Sanger Institute COVID-19 Surveillance Team |
| EPI_ISL_3108610, EPI_ISL_3108611, EPI_ISL_3108612, EPI_ISL_3108613 | Lighthouse Lab in Milton Keynes | Wellcome Sanger Institute for the COVID-19 Genomics UK (COG-UK) Consortium | The Lighthouse Lab in Milton Keynes and Alex Alderton, Roberto Amato, Jeffrey Barrett, Sonia Goncalves, Ewan Harrison, David K. Jackson, Ian Johnston, Dominic Kwiatkowski, Cordelia Langford, John Sillitoe on behalf of the Wellcome Sanger Institute COVID-19 Surveillance Team |
| EPI_ISL_3108614 | Lighthouse Lab in Alderley Park | Wellcome Sanger Institute for the COVID-19 Genomics UK (COG-UK) Consortium | Jacquelyn Wynn, Mairead Hyland, The Lighthouse Lab in Alderley Park and Alex Alderton, Roberto Amato, Jeffrey Barrett, Sonia Goncalves, Ewan Harrison, David K. Jackson, Ian Johnston, Dominic Kwiatkowski, Cordelia Langford, John Sillitoe on behalf of the Wellcome Sanger Institute COVID-19 Surveillance Team |
| EPI_ISL_3108615 | Lighthouse Lab in Milton Keynes | Wellcome Sanger Institute for the COVID-19 Genomics UK (COG-UK) Consortium | The Lighthouse Lab in Milton Keynes and Alex Alderton, Roberto Amato, Jeffrey Barrett, Sonia Goncalves, Ewan Harrison, David K. Jackson, Ian Johnston, Dominic Kwiatkowski, Cordelia Langford, John Sillitoe on behalf of the Wellcome Sanger Institute COVID-19 Surveillance Team |
| EPI_ISL_3108616 | Lighthouse Lab in Alderley Park | Wellcome Sanger Institute for the COVID-19 Genomics UK (COG-UK) Consortium | Jacquelyn Wynn, Mairead Hyland, The Lighthouse Lab in Alderley Park and Alex Alderton, Roberto Amato, Jeffrey Barrett, Sonia Goncalves, Ewan Harrison, David K. Jackson, Ian Johnston, Dominic Kwiatkowski, Cordelia Langford, John Sillitoe on behalf of the Wellcome Sanger Institute COVID-19 Surveillance Team |
| EPI_ISL_3108617, EPI_ISL_3108618 | Lighthouse Lab in Milton Keynes | Wellcome Sanger Institute for the COVID-19 Genomics UK (COG-UK) Consortium | The Lighthouse Lab in Milton Keynes and Alex Alderton, Roberto Amato, Jeffrey Barrett, Sonia Goncalves, Ewan Harrison, David K. Jackson, Ian Johnston, Dominic Kwiatkowski, Cordelia Langford, John Sillitoe on behalf of the Wellcome Sanger Institute COVID-19 Surveillance Team |
| EPI_ISL_3108619, EPI_ISL_3108620, EPI_ISL_3108621 | Lighthouse Lab in Alderley Park | Wellcome Sanger Institute for the COVID-19 Genomics UK (COG-UK) Consortium | Jacquelyn Wynn, Mairead Hyland, The Lighthouse Lab in Alderley Park and Alex Alderton, Roberto Amato, Jeffrey Barrett, Sonia Goncalves, Ewan Harrison, David K. Jackson, Ian Johnston, Dominic Kwiatkowski, Cordelia Langford, John Sillitoe on behalf of the Wellcome Sanger Institute COVID-19 Surveillance Team |
| EPI_ISL_3108622 | Lighthouse Lab in Milton Keynes | Wellcome Sanger Institute for the COVID-19 Genomics UK (COG-UK) Consortium | The Lighthouse Lab in Milton Keynes and Alex Alderton, Roberto Amato, Jeffrey Barrett, Sonia Goncalves, Ewan Harrison, David K. Jackson, Ian Johnston, Dominic Kwiatkowski, Cordelia Langford, John Sillitoe on behalf of the Wellcome Sanger Institute COVID-19 Surveillance Team |
| EPI_ISL_3108623 | Berkshire and Surrey Pathology Services Lighthouse Laboratory | Wellcome Sanger Institute for the COVID-19 Genomics UK (COG-UK) Consortium | Berkshire and Surrey Pathology Services Lighthouse Laboratory and Alex Alderton, Roberto Amato, Jeffrey Barrett, Sonia Goncalves, Ewan Harrison, David K. Jackson, Ian Johnston, Dominic Kwiatkowski, Cordelia Langford, John Sillitoe on behalf of the Wellcome Sanger Institute COVID-19 Surveillance Team |
| EPI_ISL_3108624 | Lighthouse Lab in Milton Keynes | Wellcome Sanger Institute for the COVID-19 Genomics UK (COG-UK) Consortium | The Lighthouse Lab in Milton Keynes and Alex Alderton, Roberto Amato, Jeffrey Barrett, Sonia Goncalves, Ewan Harrison, David K. Jackson, Ian Johnston, Dominic Kwiatkowski, Cordelia Langford, John Sillitoe on behalf of the Wellcome Sanger Institute COVID-19 Surveillance Team |
| EPI_ISL_3108625, EPI_ISL_3108626, EPI_ISL_3108627 | Lighthouse Lab in Alderley Park | Wellcome Sanger Institute for the COVID-19 Genomics UK (COG-UK) Consortium | Jacquelyn Wynn, Mairead Hyland, The Lighthouse Lab in Alderley Park and Alex Alderton, Roberto Amato, Jeffrey Barrett, Sonia Goncalves, Ewan Harrison, David K. Jackson, Ian Johnston, Dominic Kwiatkowski, Cordelia Langford, John Sillitoe on behalf of the Wellcome Sanger Institute COVID-19 Surveillance Team |
| EPI_ISL_3108628 | Lighthouse Lab in Milton Keynes | Wellcome Sanger Institute for the COVID-19 Genomics UK (COG-UK) Consortium | The Lighthouse Lab in Milton Keynes and Alex Alderton, Roberto Amato, Jeffrey Barrett, Sonia Goncalves, Ewan Harrison, David K. Jackson, Ian Johnston, Dominic Kwiatkowski, Cordelia Langford, John Sillitoe on behalf of the Wellcome Sanger Institute COVID-19 Surveillance Team |
| EPI_ISL_3108629 | Lighthouse Lab in Alderley Park | Wellcome Sanger Institute for the COVID-19 Genomics UK (COG-UK) Consortium | Jacquelyn Wynn, Mairead Hyland, The Lighthouse Lab in Alderley Park and Alex Alderton, Roberto Amato, Jeffrey Barrett, Sonia Goncalves, Ewan Harrison, David K. Jackson, Ian Johnston, Dominic Kwiatkowski, Cordelia Langford, John Sillitoe on behalf of the Wellcome Sanger Institute COVID-19 Surveillance Team |
| EPI_ISL_3108630 | Lighthouse Lab in Milton Keynes | Wellcome Sanger Institute for the COVID-19 Genomics UK (COG-UK) Consortium | The Lighthouse Lab in Milton Keynes and Alex Alderton, Roberto Amato, Jeffrey Barrett, Sonia Goncalves, Ewan Harrison, David K. Jackson, Ian Johnston, Dominic Kwiatkowski, Cordelia Langford, John Sillitoe on behalf of the Wellcome Sanger Institute COVID-19 Surveillance Team |
| EPI_ISL_3108631 | Berkshire and Surrey Pathology Services Lighthouse Laboratory | Wellcome Sanger Institute for the COVID-19 Genomics UK (COG-UK) Consortium | Berkshire and Surrey Pathology Services Lighthouse Laboratory and Alex Alderton, Roberto Amato, Jeffrey Barrett, Sonia Goncalves, Ewan Harrison, David K. Jackson, Ian Johnston, Dominic Kwiatkowski, Cordelia Langford, John Sillitoe on behalf of the Wellcome Sanger Institute COVID-19 Surveillance Team |
| EPI_ISL_3108632 | Lighthouse Lab in Milton Keynes | Wellcome Sanger Institute for the COVID-19 Genomics UK (COG-UK) | The Lighthouse Lab in Milton Keynes and Alex Alderton, Roberto Amato, Jeffrey Barrett, Sonia Goncalves, Ewan Harrison, David K. Jackson, Ian Johnston, Dominic Kwiatkowski, Cordelia Langford, John Sillitoe on behalf of the Wellcome Sanger |

| | | | |
|---|---|---|---|
| | | Consortium | Institute COVID-19 Surveillance Team |
| EPI_ISL_3108633 | Berkshire and Surrey Pathology Services Lighthouse Laboratory | Wellcome Sanger Institute for the COVID-19 Genomics UK (COG-UK) Consortium | Berkshire and Surrey Pathology Services Lighthouse Laboratory and Alex Alderton, Roberto Amato, Jeffrey Barrett, Sonia Goncalves, Ewan Harrison, David K. Jackson, Ian Johnston, Dominic Kwiatkowski, Cordelia Langford, John Sillitoe on behalf of the Wellcome Sanger Institute COVID-19 Surveillance Team |
| EPI_ISL_3108634 | Lighthouse Lab in Alderley Park | Wellcome Sanger Institute for the COVID-19 Genomics UK (COG-UK) Consortium | Jacquelyn Wynn, Mairead Hyland, The Lighthouse Lab in Alderley Park and Alex Alderton, Roberto Amato, Jeffrey Barrett, Sonia Goncalves, Ewan Harrison, David K. Jackson, Ian Johnston, Dominic Kwiatkowski, Cordelia Langford, John Sillitoe on behalf of the Wellcome Sanger Institute COVID-19 Surveillance Team |
| EPI_ISL_3108635, EPI_ISL_3108636, EPI_ISL_3108637 | Lighthouse Lab in Milton Keynes | Wellcome Sanger Institute for the COVID-19 Genomics UK (COG-UK) Consortium | The Lighthouse Lab in Milton Keynes and Alex Alderton, Roberto Amato, Jeffrey Barrett, Sonia Goncalves, Ewan Harrison, David K. Jackson, Ian Johnston, Dominic Kwiatkowski, Cordelia Langford, John Sillitoe on behalf of the Wellcome Sanger Institute COVID-19 Surveillance Team |
| EPI_ISL_3108638, EPI_ISL_3108639 | Lighthouse Lab in Alderley Park | Wellcome Sanger Institute for the COVID-19 Genomics UK (COG-UK) Consortium | Jacquelyn Wynn, Mairead Hyland, The Lighthouse Lab in Alderley Park and Alex Alderton, Roberto Amato, Jeffrey Barrett, Sonia Goncalves, Ewan Harrison, David K. Jackson, Ian Johnston, Dominic Kwiatkowski, Cordelia Langford, John Sillitoe on behalf of the Wellcome Sanger Institute COVID-19 Surveillance Team |
| EPI_ISL_3108640, EPI_ISL_3108641, EPI_ISL_3108642, EPI_ISL_3108643, EPI_ISL_3108644 | Lighthouse Lab in Milton Keynes | Wellcome Sanger Institute for the COVID-19 Genomics UK (COG-UK) Consortium | The Lighthouse Lab in Milton Keynes and Alex Alderton, Roberto Amato, Jeffrey Barrett, Sonia Goncalves, Ewan Harrison, David K. Jackson, Ian Johnston, Dominic Kwiatkowski, Cordelia Langford, John Sillitoe on behalf of the Wellcome Sanger Institute COVID-19 Surveillance Team |
| EPI_ISL_3108645, EPI_ISL_3108646 | Lighthouse Lab in Alderley Park | Wellcome Sanger Institute for the COVID-19 Genomics UK (COG-UK) Consortium | Jacquelyn Wynn, Mairead Hyland, The Lighthouse Lab in Alderley Park and Alex Alderton, Roberto Amato, Jeffrey Barrett, Sonia Goncalves, Ewan Harrison, David K. Jackson, Ian Johnston, Dominic Kwiatkowski, Cordelia Langford, John Sillitoe on behalf of the Wellcome Sanger Institute COVID-19 Surveillance Team |
| EPI_ISL_3108647 | Lighthouse Lab in Milton Keynes | Wellcome Sanger Institute for the COVID-19 Genomics UK (COG-UK) Consortium | The Lighthouse Lab in Milton Keynes and Alex Alderton, Roberto Amato, Jeffrey Barrett, Sonia Goncalves, Ewan Harrison, David K. Jackson, Ian Johnston, Dominic Kwiatkowski, Cordelia Langford, John Sillitoe on behalf of the Wellcome Sanger Institute COVID-19 Surveillance Team |
| EPI_ISL_3108648 | Lighthouse Lab in Alderley Park | Wellcome Sanger Institute for the COVID-19 Genomics UK (COG-UK) Consortium | Jacquelyn Wynn, Mairead Hyland, The Lighthouse Lab in Alderley Park and Alex Alderton, Roberto Amato, Jeffrey Barrett, Sonia Goncalves, Ewan Harrison, David K. Jackson, Ian Johnston, Dominic Kwiatkowski, Cordelia Langford, John Sillitoe on behalf of the Wellcome Sanger Institute COVID-19 Surveillance Team |
| EPI_ISL_3108649 | Lighthouse Lab in Milton Keynes | Wellcome Sanger Institute for the COVID-19 Genomics UK (COG-UK) Consortium | The Lighthouse Lab in Milton Keynes and Alex Alderton, Roberto Amato, Jeffrey Barrett, Sonia Goncalves, Ewan Harrison, David K. Jackson, Ian Johnston, Dominic Kwiatkowski, Cordelia Langford, John Sillitoe on behalf of the Wellcome Sanger Institute COVID-19 Surveillance Team |
| EPI_ISL_3108650 | Berkshire and Surrey Pathology Services Lighthouse Laboratory | Wellcome Sanger Institute for the COVID-19 Genomics UK (COG-UK) Consortium | Berkshire and Surrey Pathology Services Lighthouse Laboratory and Alex Alderton, Roberto Amato, Jeffrey Barrett, Sonia Goncalves, Ewan Harrison, David K. Jackson, Ian Johnston, Dominic Kwiatkowski, Cordelia Langford, John Sillitoe on behalf of the Wellcome Sanger Institute COVID-19 Surveillance Team |
| EPI_ISL_3108651 | Lighthouse Lab in Alderley Park | Wellcome Sanger Institute for the COVID-19 Genomics UK (COG-UK) Consortium | Jacquelyn Wynn, Mairead Hyland, The Lighthouse Lab in Alderley Park and Alex Alderton, Roberto Amato, Jeffrey Barrett, Sonia Goncalves, Ewan Harrison, David K. Jackson, Ian Johnston, Dominic Kwiatkowski, Cordelia Langford, John Sillitoe on behalf of the Wellcome Sanger Institute COVID-19 Surveillance Team |
| EPI_ISL_3108652 | Lighthouse Lab in Milton Keynes | Wellcome Sanger Institute for the COVID-19 Genomics UK (COG-UK) Consortium | The Lighthouse Lab in Milton Keynes and Alex Alderton, Roberto Amato, Jeffrey Barrett, Sonia Goncalves, Ewan Harrison, David K. Jackson, Ian Johnston, Dominic Kwiatkowski, Cordelia Langford, John Sillitoe on behalf of the Wellcome Sanger Institute COVID-19 Surveillance Team |
| EPI_ISL_3108653, EPI_ISL_3108654, EPI_ISL_3108655 | Berkshire and Surrey Pathology Services Lighthouse Laboratory | Wellcome Sanger Institute for the COVID-19 Genomics UK (COG-UK) Consortium | Berkshire and Surrey Pathology Services Lighthouse Laboratory and Alex Alderton, Roberto Amato, Jeffrey Barrett, Sonia Goncalves, Ewan Harrison, David K. Jackson, Ian Johnston, Dominic Kwiatkowski, Cordelia Langford, John Sillitoe on behalf of the Wellcome Sanger Institute COVID-19 Surveillance Team |
| EPI_ISL_3108656 | Lighthouse Lab in Milton Keynes | Wellcome Sanger Institute for the COVID-19 Genomics UK (COG-UK) Consortium | The Lighthouse Lab in Milton Keynes and Alex Alderton, Roberto Amato, Jeffrey Barrett, Sonia Goncalves, Ewan Harrison, David K. Jackson, Ian Johnston, Dominic Kwiatkowski, Cordelia Langford, John Sillitoe on behalf of the Wellcome Sanger Institute COVID-19 Surveillance Team |
| EPI_ISL_3108657 | Lighthouse Lab in Alderley Park | Wellcome Sanger Institute for the COVID-19 Genomics UK (COG-UK) Consortium | Jacquelyn Wynn, Mairead Hyland, The Lighthouse Lab in Alderley Park and Alex Alderton, Roberto Amato, Jeffrey Barrett, Sonia Goncalves, Ewan Harrison, David K. Jackson, Ian Johnston, Dominic Kwiatkowski, Cordelia Langford, John Sillitoe on behalf of the Wellcome Sanger Institute COVID-19 Surveillance Team |
| EPI_ISL_3108658, EPI_ISL_3108659 | Lighthouse Lab in Milton Keynes | Wellcome Sanger Institute for the COVID-19 Genomics UK (COG-UK) Consortium | The Lighthouse Lab in Milton Keynes and Alex Alderton, Roberto Amato, Jeffrey Barrett, Sonia Goncalves, Ewan Harrison, David K. Jackson, Ian Johnston, Dominic Kwiatkowski, Cordelia Langford, John Sillitoe on behalf of the Wellcome Sanger Institute COVID-19 Surveillance Team |
| EPI_ISL_3108660, EPI_ISL_3108661 | Berkshire and Surrey Pathology Services Lighthouse Laboratory | Wellcome Sanger Institute for the COVID-19 Genomics UK (COG-UK) Consortium | Berkshire and Surrey Pathology Services Lighthouse Laboratory and Alex Alderton, Roberto Amato, Jeffrey Barrett, Sonia Goncalves, Ewan Harrison, David K. Jackson, Ian Johnston, Dominic Kwiatkowski, Cordelia Langford, John Sillitoe on behalf of the Wellcome Sanger Institute COVID-19 Surveillance Team |
| EPI_ISL_3108662, EPI_ISL_3108663, EPI_ISL_3108664, EPI_ISL_3108665, EPI_ISL_3108666, EPI_ISL_3108667, EPI_ISL_3108668, EPI_ISL_3108669, EPI_ISL_3108670 | Lighthouse Lab in Milton Keynes | Wellcome Sanger Institute for the COVID-19 Genomics UK (COG-UK) Consortium | The Lighthouse Lab in Milton Keynes and Alex Alderton, Roberto Amato, Jeffrey Barrett, Sonia Goncalves, Ewan Harrison, David K. Jackson, Ian Johnston, Dominic Kwiatkowski, Cordelia Langford, John Sillitoe on behalf of the Wellcome Sanger Institute COVID-19 Surveillance Team |
| EPI_ISL_3108671, EPI_ISL_3108672 | Berkshire and Surrey Pathology Services Lighthouse Laboratory | Wellcome Sanger Institute for the COVID-19 Genomics UK (COG-UK) Consortium | Berkshire and Surrey Pathology Services Lighthouse Laboratory and Alex Alderton, Roberto Amato, Jeffrey Barrett, Sonia Goncalves, Ewan Harrison, David K. Jackson, Ian Johnston, Dominic Kwiatkowski, Cordelia Langford, John Sillitoe on behalf of the Wellcome Sanger Institute COVID-19 Surveillance Team |
| EPI_ISL_3108673, EPI_ISL_3108674 | Lighthouse Lab in Alderley Park | Wellcome Sanger Institute for the COVID-19 Genomics UK (COG-UK) Consortium | Jacquelyn Wynn, Mairead Hyland, The Lighthouse Lab in Alderley Park and Alex Alderton, Roberto Amato, Jeffrey Barrett, Sonia Goncalves, Ewan Harrison, David K. Jackson, Ian Johnston, Dominic Kwiatkowski, Cordelia Langford, John Sillitoe on behalf of the Wellcome Sanger Institute COVID-19 Surveillance Team |
| EPI_ISL_3108675 | Berkshire and Surrey Pathology Services Lighthouse Laboratory | Wellcome Sanger Institute for the COVID-19 Genomics UK (COG-UK) Consortium | Berkshire and Surrey Pathology Services Lighthouse Laboratory and Alex Alderton, Roberto Amato, Jeffrey Barrett, Sonia Goncalves, Ewan Harrison, David K. Jackson, Ian Johnston, Dominic Kwiatkowski, Cordelia Langford, John Sillitoe on behalf of the Wellcome Sanger Institute COVID-19 Surveillance Team |
| EPI_ISL_3108677 | Lighthouse Lab in Milton Keynes | Wellcome Sanger Institute for the COVID-19 Genomics UK (COG-UK) Consortium | The Lighthouse Lab in Milton Keynes and Alex Alderton, Roberto Amato, Jeffrey Barrett, Sonia Goncalves, Ewan Harrison, David K. Jackson, Ian Johnston, Dominic Kwiatkowski, Cordelia Langford, John Sillitoe on behalf of the Wellcome Sanger Institute COVID-19 Surveillance Team |
| EPI_ISL_3108678 | Lighthouse Lab in Alderley Park | Wellcome Sanger Institute for the COVID-19 Genomics UK (COG-UK) Consortium | Jacquelyn Wynn, Mairead Hyland, The Lighthouse Lab in Alderley Park and Alex Alderton, Roberto Amato, Jeffrey Barrett, Sonia Goncalves, Ewan Harrison, David K. Jackson, Ian Johnston, Dominic Kwiatkowski, Cordelia Langford, John Sillitoe on behalf of the Wellcome Sanger Institute COVID-19 Surveillance Team |
| EPI_ISL_3108679, EPI_ISL_3108680, EPI_ISL_3108681, EPI_ISL_3108682, EPI_ISL_3108683, EPI_ISL_3108684 | Lighthouse Lab in Milton Keynes | Wellcome Sanger Institute for the COVID-19 Genomics UK (COG-UK) Consortium | The Lighthouse Lab in Milton Keynes and Alex Alderton, Roberto Amato, Jeffrey Barrett, Sonia Goncalves, Ewan Harrison, David K. Jackson, Ian Johnston, Dominic Kwiatkowski, Cordelia Langford, John Sillitoe on behalf of the Wellcome Sanger Institute COVID-19 Surveillance Team |
| EPI_ISL_3108685, EPI_ISL_3108686 | Lighthouse Lab in Alderley Park | Wellcome Sanger Institute for the COVID-19 Genomics UK (COG-UK) Consortium | Jacquelyn Wynn, Mairead Hyland, The Lighthouse Lab in Alderley Park and Alex Alderton, Roberto Amato, Jeffrey Barrett, Sonia Goncalves, Ewan Harrison, David K. Jackson, Ian Johnston, Dominic Kwiatkowski, Cordelia Langford, John Sillitoe on behalf of the Wellcome Sanger Institute COVID-19 Surveillance Team |
| EPI_ISL_3108687 | Berkshire and Surrey Pathology Services Lighthouse Laboratory | Wellcome Sanger Institute for the COVID-19 Genomics UK (COG-UK) Consortium | Berkshire and Surrey Pathology Services Lighthouse Laboratory and Alex Alderton, Roberto Amato, Jeffrey Barrett, Sonia Goncalves, Ewan Harrison, David K. Jackson, Ian Johnston, Dominic Kwiatkowski, Cordelia Langford, John Sillitoe on behalf of the Wellcome Sanger Institute COVID-19 Surveillance Team |
| EPI_ISL_3108688, EPI_ISL_3108689 | Lighthouse Lab in Milton Keynes | Wellcome Sanger Institute for the COVID-19 Genomics UK (COG-UK) Consortium | The Lighthouse Lab in Milton Keynes and Alex Alderton, Roberto Amato, Jeffrey Barrett, Sonia Goncalves, Ewan Harrison, David K. Jackson, Ian Johnston, Dominic Kwiatkowski, Cordelia Langford, John Sillitoe on behalf of the Wellcome Sanger Institute COVID-19 Surveillance Team |
| EPI_ISL_3108690 | Berkshire and Surrey Pathology Services Lighthouse Laboratory | Wellcome Sanger Institute for the COVID-19 Genomics UK (COG-UK) Consortium | Berkshire and Surrey Pathology Services Lighthouse Laboratory and Alex Alderton, Roberto Amato, Jeffrey Barrett, Sonia Goncalves, Ewan Harrison, David K. Jackson, Ian Johnston, Dominic Kwiatkowski, Cordelia Langford, John Sillitoe on behalf of the Wellcome Sanger Institute COVID-19 Surveillance Team |
| EPI_ISL_3108691 | Lighthouse Lab in Milton Keynes | Wellcome Sanger Institute for the COVID-19 Genomics UK (COG-UK) Consortium | The Lighthouse Lab in Milton Keynes and Alex Alderton, Roberto Amato, Jeffrey Barrett, Sonia Goncalves, Ewan Harrison, David K. Jackson, Ian Johnston, Dominic Kwiatkowski, Cordelia Langford, John Sillitoe on behalf of the Wellcome Sanger Institute COVID-19 Surveillance Team |
| EPI_ISL_3108692 | Lighthouse Lab in Alderley Park | Wellcome Sanger Institute for the COVID-19 Genomics UK (COG-UK) Consortium | Jacquelyn Wynn, Mairead Hyland, The Lighthouse Lab in Alderley Park and Alex Alderton, Roberto Amato, Jeffrey Barrett, Sonia Goncalves, Ewan Harrison, David K. Jackson, Ian Johnston, Dominic Kwiatkowski, Cordelia Langford, John Sillitoe on behalf of the Wellcome Sanger Institute COVID-19 Surveillance Team |
| EPI_ISL_3108693 | Lighthouse Lab in Milton Keynes | Wellcome Sanger Institute for the COVID-19 Genomics UK (COG-UK) Consortium | The Lighthouse Lab in Milton Keynes and Alex Alderton, Roberto Amato, Jeffrey Barrett, Sonia Goncalves, Ewan Harrison, David K. Jackson, Ian Johnston, Dominic Kwiatkowski, Cordelia Langford, John Sillitoe on behalf of the Wellcome Sanger Institute COVID-19 Surveillance Team |
| EPI_ISL_3108694 | Lighthouse Lab in Alderley Park | Wellcome Sanger Institute for the COVID-19 Genomics UK (COG-UK) Consortium | Jacquelyn Wynn, Mairead Hyland, The Lighthouse Lab in Alderley Park and Alex Alderton, Roberto Amato, Jeffrey Barrett, Sonia Goncalves, Ewan Harrison, David K. Jackson, Ian Johnston, Dominic Kwiatkowski, Cordelia Langford, John Sillitoe on behalf of the Wellcome Sanger Institute COVID-19 Surveillance Team |
| EPI_ISL_3108695, EPI_ISL_3108696, EPI_ISL_3108697 | Lighthouse Lab in Milton Keynes | Wellcome Sanger Institute for the COVID-19 Genomics UK (COG-UK) Consortium | The Lighthouse Lab in Milton Keynes and Alex Alderton, Roberto Amato, Jeffrey Barrett, Sonia Goncalves, Ewan Harrison, David K. Jackson, Ian Johnston, Dominic Kwiatkowski, Cordelia Langford, John Sillitoe on behalf of the Wellcome Sanger Institute COVID-19 Surveillance Team |
| EPI_ISL_3108698 | Berkshire and Surrey Pathology Services Lighthouse Laboratory | Wellcome Sanger Institute for the COVID-19 Genomics UK (COG-UK) Consortium | Berkshire and Surrey Pathology Services Lighthouse Laboratory and Alex Alderton, Roberto Amato, Jeffrey Barrett, Sonia Goncalves, Ewan Harrison, David K. Jackson, Ian Johnston, Dominic Kwiatkowski, Cordelia Langford, John Sillitoe on behalf of the Wellcome Sanger Institute COVID-19 Surveillance Team |
| EPI_ISL_3108699 | Lighthouse Lab in Alderley Park | Wellcome Sanger Institute for the COVID-19 Genomics UK (COG-UK) Consortium | Jacquelyn Wynn, Mairead Hyland, The Lighthouse Lab in Alderley Park and Alex Alderton, Roberto Amato, Jeffrey Barrett, Sonia Goncalves, Ewan Harrison, David K. Jackson, Ian Johnston, Dominic Kwiatkowski, Cordelia Langford, John Sillitoe on behalf of the Wellcome Sanger Institute COVID-19 Surveillance Team |
| EPI_ISL_3108700, EPI_ISL_3108701, EPI_ISL_3108702, EPI_ISL_3108703 | Lighthouse Lab in Milton Keynes | Wellcome Sanger Institute for the COVID-19 Genomics UK (COG-UK) Consortium | The Lighthouse Lab in Milton Keynes and Alex Alderton, Roberto Amato, Jeffrey Barrett, Sonia Goncalves, Ewan Harrison, David K. Jackson, Ian Johnston, Dominic Kwiatkowski, Cordelia Langford, John Sillitoe on behalf of the Wellcome Sanger Institute COVID-19 Surveillance Team |
| EPI_ISL_3108704 | Berkshire and Surrey Pathology Services Lighthouse Laboratory | Wellcome Sanger Institute for the COVID-19 Genomics UK (COG-UK) Consortium | Berkshire and Surrey Pathology Services Lighthouse Laboratory and Alex Alderton, Roberto Amato, Jeffrey Barrett, Sonia Goncalves, Ewan Harrison, David K. Jackson, Ian Johnston, Dominic Kwiatkowski, Cordelia Langford, John Sillitoe on behalf of the Wellcome Sanger Institute COVID-19 Surveillance Team |

| | | | |
|---|---|---|---|
| EPI_ISL_3108705, EPI_ISL_3108706 | Lighthouse Lab in Milton Keynes | Wellcome Sanger Institute for the COVID-19 Genomics UK (COG-UK) Consortium | The Lighthouse Lab in Milton Keynes and Alex Alderton, Roberto Amato, Jeffrey Barrett, Sonia Goncalves, Ewan Harrison, David K. Jackson, Ian Johnston, Dominic Kwiatkowski, Cordelia Langford, John Sillitoe on behalf of the Wellcome Sanger Institute COVID-19 Surveillance Team |
| EPI_ISL_3108707 | Berkshire and Surrey Pathology Services Lighthouse Laboratory | Wellcome Sanger Institute for the COVID-19 Genomics UK (COG-UK) Consortium | Berkshire and Surrey Pathology Services Lighthouse Laboratory and Alex Alderton, Roberto Amato, Jeffrey Barrett, Sonia Goncalves, Ewan Harrison, David K. Jackson, Ian Johnston, Dominic Kwiatkowski, Cordelia Langford, John Sillitoe on behalf of the Wellcome Sanger Institute COVID-19 Surveillance Team |
| EPI_ISL_3108708 | Lighthouse Lab in Milton Keynes | Wellcome Sanger Institute for the COVID-19 Genomics UK (COG-UK) Consortium | The Lighthouse Lab in Milton Keynes and Alex Alderton, Roberto Amato, Jeffrey Barrett, Sonia Goncalves, Ewan Harrison, David K. Jackson, Ian Johnston, Dominic Kwiatkowski, Cordelia Langford, John Sillitoe on behalf of the Wellcome Sanger Institute COVID-19 Surveillance Team |
| EPI_ISL_3108709 | Berkshire and Surrey Pathology Services Lighthouse Laboratory | Wellcome Sanger Institute for the COVID-19 Genomics UK (COG-UK) Consortium | Berkshire and Surrey Pathology Services Lighthouse Laboratory and Alex Alderton, Roberto Amato, Jeffrey Barrett, Sonia Goncalves, Ewan Harrison, David K. Jackson, Ian Johnston, Dominic Kwiatkowski, Cordelia Langford, John Sillitoe on behalf of the Wellcome Sanger Institute COVID-19 Surveillance Team |
| EPI_ISL_3108710, EPI_ISL_3108711, EPI_ISL_3108712, EPI_ISL_3108713, EPI_ISL_3108714, EPI_ISL_3108715, EPI_ISL_3108716, EPI_ISL_3108717 | Lighthouse Lab in Milton Keynes | Wellcome Sanger Institute for the COVID-19 Genomics UK (COG-UK) Consortium | The Lighthouse Lab in Milton Keynes and Alex Alderton, Roberto Amato, Jeffrey Barrett, Sonia Goncalves, Ewan Harrison, David K. Jackson, Ian Johnston, Dominic Kwiatkowski, Cordelia Langford, John Sillitoe on behalf of the Wellcome Sanger Institute COVID-19 Surveillance Team |
| EPI_ISL_3108718 | Lighthouse Lab in Alderley Park | Wellcome Sanger Institute for the COVID-19 Genomics UK (COG-UK) Consortium | Jacquelyn Wynn, Mairead Hyland, The Lighthouse Lab in Alderley Park and Alex Alderton, Roberto Amato, Jeffrey Barrett, Sonia Goncalves, Ewan Harrison, David K. Jackson, Ian Johnston, Dominic Kwiatkowski, Cordelia Langford, John Sillitoe on behalf of the Wellcome Sanger Institute COVID-19 Surveillance Team |
| EPI_ISL_3108719 | Berkshire and Surrey Pathology Services Lighthouse Laboratory | Wellcome Sanger Institute for the COVID-19 Genomics UK (COG-UK) Consortium | Berkshire and Surrey Pathology Services Lighthouse Laboratory and Alex Alderton, Roberto Amato, Jeffrey Barrett, Sonia Goncalves, Ewan Harrison, David K. Jackson, Ian Johnston, Dominic Kwiatkowski, Cordelia Langford, John Sillitoe on behalf of the Wellcome Sanger Institute COVID-19 Surveillance Team |
| EPI_ISL_3108720 | Lighthouse Lab in Alderley Park | Wellcome Sanger Institute for the COVID-19 Genomics UK (COG-UK) Consortium | Jacquelyn Wynn, Mairead Hyland, The Lighthouse Lab in Alderley Park and Alex Alderton, Roberto Amato, Jeffrey Barrett, Sonia Goncalves, Ewan Harrison, David K. Jackson, Ian Johnston, Dominic Kwiatkowski, Cordelia Langford, John Sillitoe on behalf of the Wellcome Sanger Institute COVID-19 Surveillance Team |
| EPI_ISL_3108721, EPI_ISL_3108722, EPI_ISL_3108723 | Lighthouse Lab in Milton Keynes | Wellcome Sanger Institute for the COVID-19 Genomics UK (COG-UK) Consortium | The Lighthouse Lab in Milton Keynes and Alex Alderton, Roberto Amato, Jeffrey Barrett, Sonia Goncalves, Ewan Harrison, David K. Jackson, Ian Johnston, Dominic Kwiatkowski, Cordelia Langford, John Sillitoe on behalf of the Wellcome Sanger Institute COVID-19 Surveillance Team |
| EPI_ISL_3108724 | Lighthouse Lab in Alderley Park | Wellcome Sanger Institute for the COVID-19 Genomics UK (COG-UK) Consortium | Jacquelyn Wynn, Mairead Hyland, The Lighthouse Lab in Alderley Park and Alex Alderton, Roberto Amato, Jeffrey Barrett, Sonia Goncalves, Ewan Harrison, David K. Jackson, Ian Johnston, Dominic Kwiatkowski, Cordelia Langford, John Sillitoe on behalf of the Wellcome Sanger Institute COVID-19 Surveillance Team |
| EPI_ISL_3108725, EPI_ISL_3108726, EPI_ISL_3108727 | Lighthouse Lab in Milton Keynes | Wellcome Sanger Institute for the COVID-19 Genomics UK (COG-UK) Consortium | The Lighthouse Lab in Milton Keynes and Alex Alderton, Roberto Amato, Jeffrey Barrett, Sonia Goncalves, Ewan Harrison, David K. Jackson, Ian Johnston, Dominic Kwiatkowski, Cordelia Langford, John Sillitoe on behalf of the Wellcome Sanger Institute COVID-19 Surveillance Team |
| EPI_ISL_3108728 | Berkshire and Surrey Pathology Services Lighthouse Laboratory | Wellcome Sanger Institute for the COVID-19 Genomics UK (COG-UK) Consortium | Berkshire and Surrey Pathology Services Lighthouse Laboratory and Alex Alderton, Roberto Amato, Jeffrey Barrett, Sonia Goncalves, Ewan Harrison, David K. Jackson, Ian Johnston, Dominic Kwiatkowski, Cordelia Langford, John Sillitoe on behalf of the Wellcome Sanger Institute COVID-19 Surveillance Team |
| EPI_ISL_3108729 | Lighthouse Lab in Milton Keynes | Wellcome Sanger Institute for the COVID-19 Genomics UK (COG-UK) Consortium | The Lighthouse Lab in Milton Keynes and Alex Alderton, Roberto Amato, Jeffrey Barrett, Sonia Goncalves, Ewan Harrison, David K. Jackson, Ian Johnston, Dominic Kwiatkowski, Cordelia Langford, John Sillitoe on behalf of the Wellcome Sanger Institute COVID-19 Surveillance Team |
| EPI_ISL_3108730, EPI_ISL_3108731 | Lighthouse Lab in Alderley Park | Wellcome Sanger Institute for the COVID-19 Genomics UK (COG-UK) Consortium | Jacquelyn Wynn, Mairead Hyland, The Lighthouse Lab in Alderley Park and Alex Alderton, Roberto Amato, Jeffrey Barrett, Sonia Goncalves, Ewan Harrison, David K. Jackson, Ian Johnston, Dominic Kwiatkowski, Cordelia Langford, John Sillitoe on behalf of the Wellcome Sanger Institute COVID-19 Surveillance Team |
| EPI_ISL_3108732 | Lighthouse Lab in Milton Keynes | Wellcome Sanger Institute for the COVID-19 Genomics UK (COG-UK) Consortium | The Lighthouse Lab in Milton Keynes and Alex Alderton, Roberto Amato, Jeffrey Barrett, Sonia Goncalves, Ewan Harrison, David K. Jackson, Ian Johnston, Dominic Kwiatkowski, Cordelia Langford, John Sillitoe on behalf of the Wellcome Sanger Institute COVID-19 Surveillance Team |
| EPI_ISL_3108733 | Lighthouse Lab in Alderley Park | Wellcome Sanger Institute for the COVID-19 Genomics UK (COG-UK) Consortium | Jacquelyn Wynn, Mairead Hyland, The Lighthouse Lab in Alderley Park and Alex Alderton, Roberto Amato, Jeffrey Barrett, Sonia Goncalves, Ewan Harrison, David K. Jackson, Ian Johnston, Dominic Kwiatkowski, Cordelia Langford, John Sillitoe on behalf of the Wellcome Sanger Institute COVID-19 Surveillance Team |
| EPI_ISL_3108734, EPI_ISL_3108735 | Berkshire and Surrey Pathology Services Lighthouse Laboratory | Wellcome Sanger Institute for the COVID-19 Genomics UK (COG-UK) Consortium | Berkshire and Surrey Pathology Services Lighthouse Laboratory and Alex Alderton, Roberto Amato, Jeffrey Barrett, Sonia Goncalves, Ewan Harrison, David K. Jackson, Ian Johnston, Dominic Kwiatkowski, Cordelia Langford, John Sillitoe on behalf of the Wellcome Sanger Institute COVID-19 Surveillance Team |
| EPI_ISL_3108736 | Lighthouse Lab in Alderley Park | Wellcome Sanger Institute for the COVID-19 Genomics UK (COG-UK) Consortium | Jacquelyn Wynn, Mairead Hyland, The Lighthouse Lab in Alderley Park and Alex Alderton, Roberto Amato, Jeffrey Barrett, Sonia Goncalves, Ewan Harrison, David K. Jackson, Ian Johnston, Dominic Kwiatkowski, Cordelia Langford, John Sillitoe on behalf of the Wellcome Sanger Institute COVID-19 Surveillance Team |
| EPI_ISL_3108737 | Lighthouse Lab in Milton Keynes | Wellcome Sanger Institute for the COVID-19 Genomics UK (COG-UK) Consortium | The Lighthouse Lab in Milton Keynes and Alex Alderton, Roberto Amato, Jeffrey Barrett, Sonia Goncalves, Ewan Harrison, David K. Jackson, Ian Johnston, Dominic Kwiatkowski, Cordelia Langford, John Sillitoe on behalf of the Wellcome Sanger Institute COVID-19 Surveillance Team |
| EPI_ISL_3108738 | Berkshire and Surrey Pathology Services Lighthouse Laboratory | Wellcome Sanger Institute for the COVID-19 Genomics UK (COG-UK) Consortium | Berkshire and Surrey Pathology Services Lighthouse Laboratory and Alex Alderton, Roberto Amato, Jeffrey Barrett, Sonia Goncalves, Ewan Harrison, David K. Jackson, Ian Johnston, Dominic Kwiatkowski, Cordelia Langford, John Sillitoe on behalf of the Wellcome Sanger Institute COVID-19 Surveillance Team |
| EPI_ISL_3108739, EPI_ISL_3108740 | Lighthouse Lab in Milton Keynes | Wellcome Sanger Institute for the COVID-19 Genomics UK (COG-UK) Consortium | The Lighthouse Lab in Milton Keynes and Alex Alderton, Roberto Amato, Jeffrey Barrett, Sonia Goncalves, Ewan Harrison, David K. Jackson, Ian Johnston, Dominic Kwiatkowski, Cordelia Langford, John Sillitoe on behalf of the Wellcome Sanger Institute COVID-19 Surveillance Team |
| EPI_ISL_3108741 | Lighthouse Lab in Alderley Park | Wellcome Sanger Institute for the COVID-19 Genomics UK (COG-UK) Consortium | Jacquelyn Wynn, Mairead Hyland, The Lighthouse Lab in Alderley Park and Alex Alderton, Roberto Amato, Jeffrey Barrett, Sonia Goncalves, Ewan Harrison, David K. Jackson, Ian Johnston, Dominic Kwiatkowski, Cordelia Langford, John Sillitoe on behalf of the Wellcome Sanger Institute COVID-19 Surveillance Team |
| EPI_ISL_3108742 | Berkshire and Surrey Pathology Services Lighthouse Laboratory | Wellcome Sanger Institute for the COVID-19 Genomics UK (COG-UK) Consortium | Berkshire and Surrey Pathology Services Lighthouse Laboratory and Alex Alderton, Roberto Amato, Jeffrey Barrett, Sonia Goncalves, Ewan Harrison, David K. Jackson, Ian Johnston, Dominic Kwiatkowski, Cordelia Langford, John Sillitoe on behalf of the Wellcome Sanger Institute COVID-19 Surveillance Team |
| EPI_ISL_3108743 | Lighthouse Lab in Milton Keynes | Wellcome Sanger Institute for the COVID-19 Genomics UK (COG-UK) Consortium | The Lighthouse Lab in Milton Keynes and Alex Alderton, Roberto Amato, Jeffrey Barrett, Sonia Goncalves, Ewan Harrison, David K. Jackson, Ian Johnston, Dominic Kwiatkowski, Cordelia Langford, John Sillitoe on behalf of the Wellcome Sanger Institute COVID-19 Surveillance Team |
| EPI_ISL_3108744 | Berkshire and Surrey Pathology Services Lighthouse Laboratory | Wellcome Sanger Institute for the COVID-19 Genomics UK (COG-UK) Consortium | Berkshire and Surrey Pathology Services Lighthouse Laboratory and Alex Alderton, Roberto Amato, Jeffrey Barrett, Sonia Goncalves, Ewan Harrison, David K. Jackson, Ian Johnston, Dominic Kwiatkowski, Cordelia Langford, John Sillitoe on behalf of the Wellcome Sanger Institute COVID-19 Surveillance Team |
| EPI_ISL_3108745 | Lighthouse Lab in Milton Keynes | Wellcome Sanger Institute for the COVID-19 Genomics UK (COG-UK) Consortium | The Lighthouse Lab in Milton Keynes and Alex Alderton, Roberto Amato, Jeffrey Barrett, Sonia Goncalves, Ewan Harrison, David K. Jackson, Ian Johnston, Dominic Kwiatkowski, Cordelia Langford, John Sillitoe on behalf of the Wellcome Sanger Institute COVID-19 Surveillance Team |
| EPI_ISL_3108746, EPI_ISL_3108747 | Berkshire and Surrey Pathology Services Lighthouse Laboratory | Wellcome Sanger Institute for the COVID-19 Genomics UK (COG-UK) Consortium | Berkshire and Surrey Pathology Services Lighthouse Laboratory and Alex Alderton, Roberto Amato, Jeffrey Barrett, Sonia Goncalves, Ewan Harrison, David K. Jackson, Ian Johnston, Dominic Kwiatkowski, Cordelia Langford, John Sillitoe on behalf of the Wellcome Sanger Institute COVID-19 Surveillance Team |
| EPI_ISL_3108748 | Lighthouse Lab in Alderley Park | Wellcome Sanger Institute for the COVID-19 Genomics UK (COG-UK) Consortium | Jacquelyn Wynn, Mairead Hyland, The Lighthouse Lab in Alderley Park and Alex Alderton, Roberto Amato, Jeffrey Barrett, Sonia Goncalves, Ewan Harrison, David K. Jackson, Ian Johnston, Dominic Kwiatkowski, Cordelia Langford, John Sillitoe on behalf of the Wellcome Sanger Institute COVID-19 Surveillance Team |
| EPI_ISL_3108749, EPI_ISL_3108750 | Berkshire and Surrey Pathology Services Lighthouse Laboratory | Wellcome Sanger Institute for the COVID-19 Genomics UK (COG-UK) Consortium | Berkshire and Surrey Pathology Services Lighthouse Laboratory and Alex Alderton, Roberto Amato, Jeffrey Barrett, Sonia Goncalves, Ewan Harrison, David K. Jackson, Ian Johnston, Dominic Kwiatkowski, Cordelia Langford, John Sillitoe on behalf of the Wellcome Sanger Institute COVID-19 Surveillance Team |
| EPI_ISL_3108751 | Lighthouse Lab in Alderley Park | Wellcome Sanger Institute for the COVID-19 Genomics UK (COG-UK) Consortium | Jacquelyn Wynn, Mairead Hyland, The Lighthouse Lab in Alderley Park and Alex Alderton, Roberto Amato, Jeffrey Barrett, Sonia Goncalves, Ewan Harrison, David K. Jackson, Ian Johnston, Dominic Kwiatkowski, Cordelia Langford, John Sillitoe on behalf of the Wellcome Sanger Institute COVID-19 Surveillance Team |
| EPI_ISL_3108752, EPI_ISL_3108753 | Lighthouse Lab in Milton Keynes | Wellcome Sanger Institute for the COVID-19 Genomics UK (COG-UK) Consortium | The Lighthouse Lab in Milton Keynes and Alex Alderton, Roberto Amato, Jeffrey Barrett, Sonia Goncalves, Ewan Harrison, David K. Jackson, Ian Johnston, Dominic Kwiatkowski, Cordelia Langford, John Sillitoe on behalf of the Wellcome Sanger Institute COVID-19 Surveillance Team |
| EPI_ISL_3108754 | Lighthouse Lab in Alderley Park | Wellcome Sanger Institute for the COVID-19 Genomics UK (COG-UK) Consortium | Jacquelyn Wynn, Mairead Hyland, The Lighthouse Lab in Alderley Park and Alex Alderton, Roberto Amato, Jeffrey Barrett, Sonia Goncalves, Ewan Harrison, David K. Jackson, Ian Johnston, Dominic Kwiatkowski, Cordelia Langford, John Sillitoe on behalf of the Wellcome Sanger Institute COVID-19 Surveillance Team |
| EPI_ISL_3108755, EPI_ISL_3108756, EPI_ISL_3108757, EPI_ISL_3108758 | Lighthouse Lab in Milton Keynes | Wellcome Sanger Institute for the COVID-19 Genomics UK (COG-UK) Consortium | The Lighthouse Lab in Milton Keynes and Alex Alderton, Roberto Amato, Jeffrey Barrett, Sonia Goncalves, Ewan Harrison, David K. Jackson, Ian Johnston, Dominic Kwiatkowski, Cordelia Langford, John Sillitoe on behalf of the Wellcome Sanger Institute COVID-19 Surveillance Team |
| EPI_ISL_3108759 | Lighthouse Lab in Alderley Park | Wellcome Sanger Institute for the COVID-19 Genomics UK (COG-UK) Consortium | Jacquelyn Wynn, Mairead Hyland, The Lighthouse Lab in Alderley Park and Alex Alderton, Roberto Amato, Jeffrey Barrett, Sonia Goncalves, Ewan Harrison, David K. Jackson, Ian Johnston, Dominic Kwiatkowski, Cordelia Langford, John Sillitoe on behalf of the Wellcome Sanger Institute COVID-19 Surveillance Team |
| EPI_ISL_3108760, EPI_ISL_3108761, EPI_ISL_3108762 | Lighthouse Lab in Milton Keynes | Wellcome Sanger Institute for the COVID-19 Genomics UK (COG-UK) Consortium | The Lighthouse Lab in Milton Keynes and Alex Alderton, Roberto Amato, Jeffrey Barrett, Sonia Goncalves, Ewan Harrison, David K. Jackson, Ian Johnston, Dominic Kwiatkowski, Cordelia Langford, John Sillitoe on behalf of the Wellcome Sanger Institute COVID-19 Surveillance Team |
| EPI_ISL_3108763 | Berkshire and Surrey Pathology Services Lighthouse Laboratory | Wellcome Sanger Institute for the COVID-19 Genomics UK (COG-UK) Consortium | Berkshire and Surrey Pathology Services Lighthouse Laboratory and Alex Alderton, Roberto Amato, Jeffrey Barrett, Sonia Goncalves, Ewan Harrison, David K. Jackson, Ian Johnston, Dominic Kwiatkowski, Cordelia Langford, John Sillitoe on behalf of the Wellcome Sanger Institute COVID-19 Surveillance Team |
| EPI_ISL_3108764, EPI_ISL_3108765, EPI_ISL_3108766 | Lighthouse Lab in Milton Keynes | Wellcome Sanger Institute for the COVID-19 Genomics UK (COG-UK) Consortium | The Lighthouse Lab in Milton Keynes and Alex Alderton, Roberto Amato, Jeffrey Barrett, Sonia Goncalves, Ewan Harrison, David K. Jackson, Ian Johnston, Dominic Kwiatkowski, Cordelia Langford, John Sillitoe on behalf of the Wellcome Sanger Institute COVID-19 Surveillance Team |
| EPI_ISL_3108767 | Berkshire and Surrey Pathology Services Lighthouse Laboratory | Wellcome Sanger Institute for the COVID-19 Genomics UK (COG-UK) Consortium | Berkshire and Surrey Pathology Services Lighthouse Laboratory and Alex Alderton, Roberto Amato, Jeffrey Barrett, Sonia Goncalves, Ewan Harrison, David K. Jackson, Ian Johnston, Dominic Kwiatkowski, Cordelia Langford, John Sillitoe on behalf of the Wellcome Sanger Institute COVID-19 Surveillance Team |
| EPI_ISL_3108768 | Lighthouse Lab in Milton Keynes | Wellcome Sanger Institute for the COVID-19 Genomics UK (COG-UK) Consortium | The Lighthouse Lab in Milton Keynes and Alex Alderton, Roberto Amato, Jeffrey Barrett, Sonia Goncalves, Ewan Harrison, David K. Jackson, Ian Johnston, Dominic Kwiatkowski, Cordelia Langford, John Sillitoe on behalf of the Wellcome Sanger Institute COVID-19 Surveillance Team |
| EPI_ISL_3108769 | Berkshire and Surrey Pathology Services Lighthouse Laboratory | Wellcome Sanger Institute for the COVID-19 Genomics UK (COG-UK) Consortium | Berkshire and Surrey Pathology Services Lighthouse Laboratory and Alex Alderton, Roberto Amato, Jeffrey Barrett, Sonia Goncalves, Ewan Harrison, David K. Jackson, Ian Johnston, Dominic Kwiatkowski, Cordelia Langford, John Sillitoe on behalf of the Wellcome Sanger Institute COVID-19 Surveillance Team |

| | | | |
|---|---|---|---|
| EPI_ISL_3108770 | Lighthouse Lab in Milton Keynes | Wellcome Sanger Institute for the COVID-19 Genomics UK (COG-UK) Consortium | The Lighthouse Lab in Milton Keynes and Alex Alderton, Roberto Amato, Jeffrey Barrett, Sonia Goncalves, Ewan Harrison, David K. Jackson, Ian Johnston, Dominic Kwiatkowski, Cordelia Langford, John Sillitoe on behalf of the Wellcome Sanger Institute COVID-19 Surveillance Team |
| EPI_ISL_3108771 | Berkshire and Surrey Pathology Services Lighthouse Laboratory | Wellcome Sanger Institute for the COVID-19 Genomics UK (COG-UK) Consortium | Berkshire and Surrey Pathology Services Lighthouse Laboratory and Alex Alderton, Roberto Amato, Jeffrey Barrett, Sonia Goncalves, Ewan Harrison, David K. Jackson, Ian Johnston, Dominic Kwiatkowski, Cordelia Langford, John Sillitoe on behalf of the Wellcome Sanger Institute COVID-19 Surveillance Team |
| EPI_ISL_3108772 | Lighthouse Lab in Milton Keynes | Wellcome Sanger Institute for the COVID-19 Genomics UK (COG-UK) Consortium | The Lighthouse Lab in Milton Keynes and Alex Alderton, Roberto Amato, Jeffrey Barrett, Sonia Goncalves, Ewan Harrison, David K. Jackson, Ian Johnston, Dominic Kwiatkowski, Cordelia Langford, John Sillitoe on behalf of the Wellcome Sanger Institute COVID-19 Surveillance Team |
| EPI_ISL_3108773 | Lighthouse Lab in Alderley Park | Wellcome Sanger Institute for the COVID-19 Genomics UK (COG-UK) Consortium | Jacquelyn Wynn, Mairead Hyland, The Lighthouse Lab in Alderley Park and Alex Alderton, Roberto Amato, Jeffrey Barrett, Sonia Goncalves, Ewan Harrison, David K. Jackson, Ian Johnston, Dominic Kwiatkowski, Cordelia Langford, John Sillitoe on behalf of the Wellcome Sanger Institute COVID-19 Surveillance Team |
| EPI_ISL_3108774, EPI_ISL_3108775 | Berkshire and Surrey Pathology Services Lighthouse Laboratory | Wellcome Sanger Institute for the COVID-19 Genomics UK (COG-UK) Consortium | Berkshire and Surrey Pathology Services Lighthouse Laboratory and Alex Alderton, Roberto Amato, Jeffrey Barrett, Sonia Goncalves, Ewan Harrison, David K. Jackson, Ian Johnston, Dominic Kwiatkowski, Cordelia Langford, John Sillitoe on behalf of the Wellcome Sanger Institute COVID-19 Surveillance Team |
| EPI_ISL_3108776 | Lighthouse Lab in Milton Keynes | Wellcome Sanger Institute for the COVID-19 Genomics UK (COG-UK) Consortium | The Lighthouse Lab in Milton Keynes and Alex Alderton, Roberto Amato, Jeffrey Barrett, Sonia Goncalves, Ewan Harrison, David K. Jackson, Ian Johnston, Dominic Kwiatkowski, Cordelia Langford, John Sillitoe on behalf of the Wellcome Sanger Institute COVID-19 Surveillance Team |
| EPI_ISL_3108777 | Berkshire and Surrey Pathology Services Lighthouse Laboratory | Wellcome Sanger Institute for the COVID-19 Genomics UK (COG-UK) Consortium | Berkshire and Surrey Pathology Services Lighthouse Laboratory and Alex Alderton, Roberto Amato, Jeffrey Barrett, Sonia Goncalves, Ewan Harrison, David K. Jackson, Ian Johnston, Dominic Kwiatkowski, Cordelia Langford, John Sillitoe on behalf of the Wellcome Sanger Institute COVID-19 Surveillance Team |
| EPI_ISL_3108778, EPI_ISL_3108779 | Lighthouse Lab in Milton Keynes | Wellcome Sanger Institute for the COVID-19 Genomics UK (COG-UK) Consortium | The Lighthouse Lab in Milton Keynes and Alex Alderton, Roberto Amato, Jeffrey Barrett, Sonia Goncalves, Ewan Harrison, David K. Jackson, Ian Johnston, Dominic Kwiatkowski, Cordelia Langford, John Sillitoe on behalf of the Wellcome Sanger Institute COVID-19 Surveillance Team |
| EPI_ISL_3108780, EPI_ISL_3108781, EPI_ISL_3108782 | Lighthouse Lab in Alderley Park | Wellcome Sanger Institute for the COVID-19 Genomics UK (COG-UK) Consortium | Jacquelyn Wynn, Mairead Hyland, The Lighthouse Lab in Alderley Park and Alex Alderton, Roberto Amato, Jeffrey Barrett, Sonia Goncalves, Ewan Harrison, David K. Jackson, Ian Johnston, Dominic Kwiatkowski, Cordelia Langford, John Sillitoe on behalf of the Wellcome Sanger Institute COVID-19 Surveillance Team |
| EPI_ISL_3108783 | Berkshire and Surrey Pathology Services Lighthouse Laboratory | Wellcome Sanger Institute for the COVID-19 Genomics UK (COG-UK) Consortium | Berkshire and Surrey Pathology Services Lighthouse Laboratory and Alex Alderton, Roberto Amato, Jeffrey Barrett, Sonia Goncalves, Ewan Harrison, David K. Jackson, Ian Johnston, Dominic Kwiatkowski, Cordelia Langford, John Sillitoe on behalf of the Wellcome Sanger Institute COVID-19 Surveillance Team |
| EPI_ISL_3108784, EPI_ISL_3108785, EPI_ISL_3108786, EPI_ISL_3108787, EPI_ISL_3108788 | Lighthouse Lab in Milton Keynes | Wellcome Sanger Institute for the COVID-19 Genomics UK (COG-UK) Consortium | The Lighthouse Lab in Milton Keynes and Alex Alderton, Roberto Amato, Jeffrey Barrett, Sonia Goncalves, Ewan Harrison, David K. Jackson, Ian Johnston, Dominic Kwiatkowski, Cordelia Langford, John Sillitoe on behalf of the Wellcome Sanger Institute COVID-19 Surveillance Team |
| EPI_ISL_3108789 | Berkshire and Surrey Pathology Services Lighthouse Laboratory | Wellcome Sanger Institute for the COVID-19 Genomics UK (COG-UK) Consortium | Berkshire and Surrey Pathology Services Lighthouse Laboratory and Alex Alderton, Roberto Amato, Jeffrey Barrett, Sonia Goncalves, Ewan Harrison, David K. Jackson, Ian Johnston, Dominic Kwiatkowski, Cordelia Langford, John Sillitoe on behalf of the Wellcome Sanger Institute COVID-19 Surveillance Team |
| EPI_ISL_3108790 | Lighthouse Lab in Milton Keynes | Wellcome Sanger Institute for the COVID-19 Genomics UK (COG-UK) Consortium | The Lighthouse Lab in Milton Keynes and Alex Alderton, Roberto Amato, Jeffrey Barrett, Sonia Goncalves, Ewan Harrison, David K. Jackson, Ian Johnston, Dominic Kwiatkowski, Cordelia Langford, John Sillitoe on behalf of the Wellcome Sanger Institute COVID-19 Surveillance Team |
| EPI_ISL_3108791 | Berkshire and Surrey Pathology Services Lighthouse Laboratory | Wellcome Sanger Institute for the COVID-19 Genomics UK (COG-UK) Consortium | Berkshire and Surrey Pathology Services Lighthouse Laboratory and Alex Alderton, Roberto Amato, Jeffrey Barrett, Sonia Goncalves, Ewan Harrison, David K. Jackson, Ian Johnston, Dominic Kwiatkowski, Cordelia Langford, John Sillitoe on behalf of the Wellcome Sanger Institute COVID-19 Surveillance Team |
| EPI_ISL_3108792 | Lighthouse Lab in Alderley Park | Wellcome Sanger Institute for the COVID-19 Genomics UK (COG-UK) Consortium | Jacquelyn Wynn, Mairead Hyland, The Lighthouse Lab in Alderley Park and Alex Alderton, Roberto Amato, Jeffrey Barrett, Sonia Goncalves, Ewan Harrison, David K. Jackson, Ian Johnston, Dominic Kwiatkowski, Cordelia Langford, John Sillitoe on behalf of the Wellcome Sanger Institute COVID-19 Surveillance Team |
| EPI_ISL_3108793 | Berkshire and Surrey Pathology Services Lighthouse Laboratory | Wellcome Sanger Institute for the COVID-19 Genomics UK (COG-UK) Consortium | Berkshire and Surrey Pathology Services Lighthouse Laboratory and Alex Alderton, Roberto Amato, Jeffrey Barrett, Sonia Goncalves, Ewan Harrison, David K. Jackson, Ian Johnston, Dominic Kwiatkowski, Cordelia Langford, John Sillitoe on behalf of the Wellcome Sanger Institute COVID-19 Surveillance Team |
| EPI_ISL_3108794 | Lighthouse Lab in Alderley Park | Wellcome Sanger Institute for the COVID-19 Genomics UK (COG-UK) Consortium | Jacquelyn Wynn, Mairead Hyland, The Lighthouse Lab in Alderley Park and Alex Alderton, Roberto Amato, Jeffrey Barrett, Sonia Goncalves, Ewan Harrison, David K. Jackson, Ian Johnston, Dominic Kwiatkowski, Cordelia Langford, John Sillitoe on behalf of the Wellcome Sanger Institute COVID-19 Surveillance Team |
| EPI_ISL_3108795 | Berkshire and Surrey Pathology Services Lighthouse Laboratory | Wellcome Sanger Institute for the COVID-19 Genomics UK (COG-UK) Consortium | Berkshire and Surrey Pathology Services Lighthouse Laboratory and Alex Alderton, Roberto Amato, Jeffrey Barrett, Sonia Goncalves, Ewan Harrison, David K. Jackson, Ian Johnston, Dominic Kwiatkowski, Cordelia Langford, John Sillitoe on behalf of the Wellcome Sanger Institute COVID-19 Surveillance Team |
| EPI_ISL_3108796, EPI_ISL_3108797, EPI_ISL_3108798 | Lighthouse Lab in Milton Keynes | Wellcome Sanger Institute for the COVID-19 Genomics UK (COG-UK) Consortium | The Lighthouse Lab in Milton Keynes and Alex Alderton, Roberto Amato, Jeffrey Barrett, Sonia Goncalves, Ewan Harrison, David K. Jackson, Ian Johnston, Dominic Kwiatkowski, Cordelia Langford, John Sillitoe on behalf of the Wellcome Sanger Institute COVID-19 Surveillance Team |
| EPI_ISL_3108799, EPI_ISL_3108800 | Berkshire and Surrey Pathology Services Lighthouse Laboratory | Wellcome Sanger Institute for the COVID-19 Genomics UK (COG-UK) Consortium | Berkshire and Surrey Pathology Services Lighthouse Laboratory and Alex Alderton, Roberto Amato, Jeffrey Barrett, Sonia Goncalves, Ewan Harrison, David K. Jackson, Ian Johnston, Dominic Kwiatkowski, Cordelia Langford, John Sillitoe on behalf of the Wellcome Sanger Institute COVID-19 Surveillance Team |
| EPI_ISL_3108801 | Lighthouse Lab in Milton Keynes | Wellcome Sanger Institute for the COVID-19 Genomics UK (COG-UK) Consortium | The Lighthouse Lab in Milton Keynes and Alex Alderton, Roberto Amato, Jeffrey Barrett, Sonia Goncalves, Ewan Harrison, David K. Jackson, Ian Johnston, Dominic Kwiatkowski, Cordelia Langford, John Sillitoe on behalf of the Wellcome Sanger Institute COVID-19 Surveillance Team |
| EPI_ISL_3108802 | Berkshire and Surrey Pathology Services Lighthouse Laboratory | Wellcome Sanger Institute for the COVID-19 Genomics UK (COG-UK) Consortium | Berkshire and Surrey Pathology Services Lighthouse Laboratory and Alex Alderton, Roberto Amato, Jeffrey Barrett, Sonia Goncalves, Ewan Harrison, David K. Jackson, Ian Johnston, Dominic Kwiatkowski, Cordelia Langford, John Sillitoe on behalf of the Wellcome Sanger Institute COVID-19 Surveillance Team |
| EPI_ISL_3108803 | Lighthouse Lab in Alderley Park | Wellcome Sanger Institute for the COVID-19 Genomics UK (COG-UK) Consortium | Jacquelyn Wynn, Mairead Hyland, The Lighthouse Lab in Alderley Park and Alex Alderton, Roberto Amato, Jeffrey Barrett, Sonia Goncalves, Ewan Harrison, David K. Jackson, Ian Johnston, Dominic Kwiatkowski, Cordelia Langford, John Sillitoe on behalf of the Wellcome Sanger Institute COVID-19 Surveillance Team |
| EPI_ISL_3108804 | Berkshire and Surrey Pathology Services Lighthouse Laboratory | Wellcome Sanger Institute for the COVID-19 Genomics UK (COG-UK) Consortium | Berkshire and Surrey Pathology Services Lighthouse Laboratory and Alex Alderton, Roberto Amato, Jeffrey Barrett, Sonia Goncalves, Ewan Harrison, David K. Jackson, Ian Johnston, Dominic Kwiatkowski, Cordelia Langford, John Sillitoe on behalf of the Wellcome Sanger Institute COVID-19 Surveillance Team |
| EPI_ISL_3108805 | Lighthouse Lab in Alderley Park | Wellcome Sanger Institute for the COVID-19 Genomics UK (COG-UK) Consortium | Jacquelyn Wynn, Mairead Hyland, The Lighthouse Lab in Alderley Park and Alex Alderton, Roberto Amato, Jeffrey Barrett, Sonia Goncalves, Ewan Harrison, David K. Jackson, Ian Johnston, Dominic Kwiatkowski, Cordelia Langford, John Sillitoe on behalf of the Wellcome Sanger Institute COVID-19 Surveillance Team |
| EPI_ISL_3108806 | Lighthouse Lab in Milton Keynes | Wellcome Sanger Institute for the COVID-19 Genomics UK (COG-UK) Consortium | The Lighthouse Lab in Milton Keynes and Alex Alderton, Roberto Amato, Jeffrey Barrett, Sonia Goncalves, Ewan Harrison, David K. Jackson, Ian Johnston, Dominic Kwiatkowski, Cordelia Langford, John Sillitoe on behalf of the Wellcome Sanger Institute COVID-19 Surveillance Team |
| EPI_ISL_3108807, EPI_ISL_3108808 | Lighthouse Lab in Alderley Park | Wellcome Sanger Institute for the COVID-19 Genomics UK (COG-UK) Consortium | Jacquelyn Wynn, Mairead Hyland, The Lighthouse Lab in Alderley Park and Alex Alderton, Roberto Amato, Jeffrey Barrett, Sonia Goncalves, Ewan Harrison, David K. Jackson, Ian Johnston, Dominic Kwiatkowski, Cordelia Langford, John Sillitoe on behalf of the Wellcome Sanger Institute COVID-19 Surveillance Team |
| EPI_ISL_3108809 | Berkshire and Surrey Pathology Services Lighthouse Laboratory | Wellcome Sanger Institute for the COVID-19 Genomics UK (COG-UK) Consortium | Berkshire and Surrey Pathology Services Lighthouse Laboratory and Alex Alderton, Roberto Amato, Jeffrey Barrett, Sonia Goncalves, Ewan Harrison, David K. Jackson, Ian Johnston, Dominic Kwiatkowski, Cordelia Langford, John Sillitoe on behalf of the Wellcome Sanger Institute COVID-19 Surveillance Team |
| EPI_ISL_3108810 | Lighthouse Lab in Alderley Park | Wellcome Sanger Institute for the COVID-19 Genomics UK (COG-UK) Consortium | Jacquelyn Wynn, Mairead Hyland, The Lighthouse Lab in Alderley Park and Alex Alderton, Roberto Amato, Jeffrey Barrett, Sonia Goncalves, Ewan Harrison, David K. Jackson, Ian Johnston, Dominic Kwiatkowski, Cordelia Langford, John Sillitoe on behalf of the Wellcome Sanger Institute COVID-19 Surveillance Team |
| EPI_ISL_3108811, EPI_ISL_3108812 | Lighthouse Lab in Milton Keynes | Wellcome Sanger Institute for the COVID-19 Genomics UK (COG-UK) Consortium | The Lighthouse Lab in Milton Keynes and Alex Alderton, Roberto Amato, Jeffrey Barrett, Sonia Goncalves, Ewan Harrison, David K. Jackson, Ian Johnston, Dominic Kwiatkowski, Cordelia Langford, John Sillitoe on behalf of the Wellcome Sanger Institute COVID-19 Surveillance Team |
| EPI_ISL_3108813 | Lighthouse Lab in Alderley Park | Wellcome Sanger Institute for the COVID-19 Genomics UK (COG-UK) Consortium | Jacquelyn Wynn, Mairead Hyland, The Lighthouse Lab in Alderley Park and Alex Alderton, Roberto Amato, Jeffrey Barrett, Sonia Goncalves, Ewan Harrison, David K. Jackson, Ian Johnston, Dominic Kwiatkowski, Cordelia Langford, John Sillitoe on behalf of the Wellcome Sanger Institute COVID-19 Surveillance Team |
| EPI_ISL_3108814 | Berkshire and Surrey Pathology Services Lighthouse Laboratory | Wellcome Sanger Institute for the COVID-19 Genomics UK (COG-UK) Consortium | Berkshire and Surrey Pathology Services Lighthouse Laboratory and Alex Alderton, Roberto Amato, Jeffrey Barrett, Sonia Goncalves, Ewan Harrison, David K. Jackson, Ian Johnston, Dominic Kwiatkowski, Cordelia Langford, John Sillitoe on behalf of the Wellcome Sanger Institute COVID-19 Surveillance Team |
| EPI_ISL_3108815 | Lighthouse Lab in Milton Keynes | Wellcome Sanger Institute for the COVID-19 Genomics UK (COG-UK) Consortium | The Lighthouse Lab in Milton Keynes and Alex Alderton, Roberto Amato, Jeffrey Barrett, Sonia Goncalves, Ewan Harrison, David K. Jackson, Ian Johnston, Dominic Kwiatkowski, Cordelia Langford, John Sillitoe on behalf of the Wellcome Sanger Institute COVID-19 Surveillance Team |
| EPI_ISL_3108816 | Lighthouse Lab in Alderley Park | Wellcome Sanger Institute for the COVID-19 Genomics UK (COG-UK) Consortium | Jacquelyn Wynn, Mairead Hyland, The Lighthouse Lab in Alderley Park and Alex Alderton, Roberto Amato, Jeffrey Barrett, Sonia Goncalves, Ewan Harrison, David K. Jackson, Ian Johnston, Dominic Kwiatkowski, Cordelia Langford, John Sillitoe on behalf of the Wellcome Sanger Institute COVID-19 Surveillance Team |
| EPI_ISL_3108817 | Lighthouse Lab in Milton Keynes | Wellcome Sanger Institute for the COVID-19 Genomics UK (COG-UK) Consortium | The Lighthouse Lab in Milton Keynes and Alex Alderton, Roberto Amato, Jeffrey Barrett, Sonia Goncalves, Ewan Harrison, David K. Jackson, Ian Johnston, Dominic Kwiatkowski, Cordelia Langford, John Sillitoe on behalf of the Wellcome Sanger Institute COVID-19 Surveillance Team |
| EPI_ISL_3108818 | Lighthouse Lab in Alderley Park | Wellcome Sanger Institute for the COVID-19 Genomics UK (COG-UK) Consortium | Jacquelyn Wynn, Mairead Hyland, The Lighthouse Lab in Alderley Park and Alex Alderton, Roberto Amato, Jeffrey Barrett, Sonia Goncalves, Ewan Harrison, David K. Jackson, Ian Johnston, Dominic Kwiatkowski, Cordelia Langford, John Sillitoe on behalf of the Wellcome Sanger Institute COVID-19 Surveillance Team |
| EPI_ISL_3108819, EPI_ISL_3108820 | Lighthouse Lab in Milton Keynes | Wellcome Sanger Institute for the COVID-19 Genomics UK (COG-UK) Consortium | The Lighthouse Lab in Milton Keynes and Alex Alderton, Roberto Amato, Jeffrey Barrett, Sonia Goncalves, Ewan Harrison, David K. Jackson, Ian Johnston, Dominic Kwiatkowski, Cordelia Langford, John Sillitoe on behalf of the Wellcome Sanger Institute COVID-19 Surveillance Team |
| EPI_ISL_3108821 | Lighthouse Lab in Alderley Park | Wellcome Sanger Institute for the COVID-19 Genomics UK (COG-UK) Consortium | Jacquelyn Wynn, Mairead Hyland, The Lighthouse Lab in Alderley Park and Alex Alderton, Roberto Amato, Jeffrey Barrett, Sonia Goncalves, Ewan Harrison, David K. Jackson, Ian Johnston, Dominic Kwiatkowski, Cordelia Langford, John Sillitoe on behalf of the Wellcome Sanger Institute COVID-19 Surveillance Team |
| EPI_ISL_3108822 | Berkshire and Surrey Pathology Services Lighthouse Laboratory | Wellcome Sanger Institute for the COVID-19 Genomics UK (COG-UK) Consortium | Berkshire and Surrey Pathology Services Lighthouse Laboratory and Alex Alderton, Roberto Amato, Jeffrey Barrett, Sonia Goncalves, Ewan Harrison, David K. Jackson, Ian Johnston, Dominic Kwiatkowski, Cordelia Langford, John Sillitoe on behalf of the Wellcome Sanger Institute COVID-19 Surveillance Team |
| EPI_ISL_3108823 | Lighthouse Lab in Milton Keynes | Wellcome Sanger Institute for the COVID-19 Genomics UK (COG-UK) Consortium | The Lighthouse Lab in Milton Keynes and Alex Alderton, Roberto Amato, Jeffrey Barrett, Sonia Goncalves, Ewan Harrison, David K. Jackson, Ian Johnston, Dominic Kwiatkowski, Cordelia Langford, John Sillitoe on behalf of the Wellcome Sanger Institute COVID-19 Surveillance Team |
| EPI_ISL_3108824, EPI_ISL_3108825, EPI_ISL_3108826 | Lighthouse Lab in Alderley Park | Wellcome Sanger Institute for the COVID-19 Genomics UK (COG-UK) Consortium | Jacquelyn Wynn, Mairead Hyland, The Lighthouse Lab in Alderley Park and Alex Alderton, Roberto Amato, Jeffrey Barrett, Sonia Goncalves, Ewan Harrison, David K. Jackson, Ian Johnston, Dominic Kwiatkowski, Cordelia Langford, John Sillitoe on behalf of the Wellcome Sanger Institute COVID-19 Surveillance Team |
| EPI_ISL_3108827, EPI_ISL_3108828, EPI_ISL_3108829, EPI_ISL_3108830 | Lighthouse Lab in Milton Keynes | Wellcome Sanger Institute for the COVID-19 Genomics UK (COG-UK) Consortium | The Lighthouse Lab in Milton Keynes and Alex Alderton, Roberto Amato, Jeffrey Barrett, Sonia Goncalves, Ewan Harrison, David K. Jackson, Ian Johnston, Dominic Kwiatkowski, Cordelia Langford, John Sillitoe on behalf of the Wellcome Sanger Institute COVID-19 Surveillance Team |

| | | | |
|---|---|---|---|
| EPI_ISL_3108831 | Lighthouse Lab in Alderley Park | Wellcome Sanger Institute for the COVID-19 Genomics UK (COG-UK) Consortium | Jacquelyn Wynn, Mairead Hyland, The Lighthouse Lab in Alderley Park and Alex Alderton, Roberto Amato, Jeffrey Barrett, Sonia Goncalves, Ewan Harrison, David K. Jackson, Ian Johnston, Dominic Kwiatkowski, Cordelia Langford, John Sillitoe on behalf of the Wellcome Sanger Institute COVID-19 Surveillance Team |
| EPI_ISL_3108832, EPI_ISL_3108833, EPI_ISL_3108834, EPI_ISL_3108835, EPI_ISL_3108836, EPI_ISL_3108837 | Lighthouse Lab in Milton Keynes | Wellcome Sanger Institute for the COVID-19 Genomics UK (COG-UK) Consortium | The Lighthouse Lab in Milton Keynes and Alex Alderton, Roberto Amato, Jeffrey Barrett, Sonia Goncalves, Ewan Harrison, David K. Jackson, Ian Johnston, Dominic Kwiatkowski, Cordelia Langford, John Sillitoe on behalf of the Wellcome Sanger Institute COVID-19 Surveillance Team |
| EPI_ISL_3108838, EPI_ISL_3108839 | Berkshire and Surrey Pathology Services Lighthouse Laboratory | Wellcome Sanger Institute for the COVID-19 Genomics UK (COG-UK) Consortium | Berkshire and Surrey Pathology Services Lighthouse Laboratory and Alex Alderton, Roberto Amato, Jeffrey Barrett, Sonia Goncalves, Ewan Harrison, David K. Jackson, Ian Johnston, Dominic Kwiatkowski, Cordelia Langford, John Sillitoe on behalf of the Wellcome Sanger Institute COVID-19 Surveillance Team |
| EPI_ISL_3108840 | Lighthouse Lab in Milton Keynes | Wellcome Sanger Institute for the COVID-19 Genomics UK (COG-UK) Consortium | The Lighthouse Lab in Milton Keynes and Alex Alderton, Roberto Amato, Jeffrey Barrett, Sonia Goncalves, Ewan Harrison, David K. Jackson, Ian Johnston, Dominic Kwiatkowski, Cordelia Langford, John Sillitoe on behalf of the Wellcome Sanger Institute COVID-19 Surveillance Team |
| EPI_ISL_3108841 | Lighthouse Lab in Alderley Park | Wellcome Sanger Institute for the COVID-19 Genomics UK (COG-UK) Consortium | Jacquelyn Wynn, Mairead Hyland, The Lighthouse Lab in Alderley Park and Alex Alderton, Roberto Amato, Jeffrey Barrett, Sonia Goncalves, Ewan Harrison, David K. Jackson, Ian Johnston, Dominic Kwiatkowski, Cordelia Langford, John Sillitoe on behalf of the Wellcome Sanger Institute COVID-19 Surveillance Team |
| EPI_ISL_3108842 | Berkshire and Surrey Pathology Services Lighthouse Laboratory | Wellcome Sanger Institute for the COVID-19 Genomics UK (COG-UK) Consortium | Berkshire and Surrey Pathology Services Lighthouse Laboratory and Alex Alderton, Roberto Amato, Jeffrey Barrett, Sonia Goncalves, Ewan Harrison, David K. Jackson, Ian Johnston, Dominic Kwiatkowski, Cordelia Langford, John Sillitoe on behalf of the Wellcome Sanger Institute COVID-19 Surveillance Team |
| EPI_ISL_3108843 | Lighthouse Lab in Alderley Park | Wellcome Sanger Institute for the COVID-19 Genomics UK (COG-UK) Consortium | Jacquelyn Wynn, Mairead Hyland, The Lighthouse Lab in Alderley Park and Alex Alderton, Roberto Amato, Jeffrey Barrett, Sonia Goncalves, Ewan Harrison, David K. Jackson, Ian Johnston, Dominic Kwiatkowski, Cordelia Langford, John Sillitoe on behalf of the Wellcome Sanger Institute COVID-19 Surveillance Team |
| EPI_ISL_3108844 | Berkshire and Surrey Pathology Services Lighthouse Laboratory | Wellcome Sanger Institute for the COVID-19 Genomics UK (COG-UK) Consortium | Berkshire and Surrey Pathology Services Lighthouse Laboratory and Alex Alderton, Roberto Amato, Jeffrey Barrett, Sonia Goncalves, Ewan Harrison, David K. Jackson, Ian Johnston, Dominic Kwiatkowski, Cordelia Langford, John Sillitoe on behalf of the Wellcome Sanger Institute COVID-19 Surveillance Team |
| EPI_ISL_3108845, EPI_ISL_3108846, EPI_ISL_3108847, EPI_ISL_3108848, EPI_ISL_3108849, EPI_ISL_3108850, EPI_ISL_3108851, EPI_ISL_3108852, EPI_ISL_3108853, EPI_ISL_3108854, EPI_ISL_3108855 | | | |
| see above | Lighthouse Lab in Milton Keynes | Wellcome Sanger Institute for the COVID-19 Genomics UK (COG-UK) Consortium | The Lighthouse Lab in Milton Keynes and Alex Alderton, Roberto Amato, Jeffrey Barrett, Sonia Goncalves, Ewan Harrison, David K. Jackson, Ian Johnston, Dominic Kwiatkowski, Cordelia Langford, John Sillitoe on behalf of the Wellcome Sanger Institute COVID-19 Surveillance Team |
| EPI_ISL_3108856 | Lighthouse Lab in Alderley Park | Wellcome Sanger Institute for the COVID-19 Genomics UK (COG-UK) Consortium | Jacquelyn Wynn, Mairead Hyland, The Lighthouse Lab in Alderley Park and Alex Alderton, Roberto Amato, Jeffrey Barrett, Sonia Goncalves, Ewan Harrison, David K. Jackson, Ian Johnston, Dominic Kwiatkowski, Cordelia Langford, John Sillitoe on behalf of the Wellcome Sanger Institute COVID-19 Surveillance Team |
| EPI_ISL_3108857 | Lighthouse Lab in Milton Keynes | Wellcome Sanger Institute for the COVID-19 Genomics UK (COG-UK) Consortium | The Lighthouse Lab in Milton Keynes and Alex Alderton, Roberto Amato, Jeffrey Barrett, Sonia Goncalves, Ewan Harrison, David K. Jackson, Ian Johnston, Dominic Kwiatkowski, Cordelia Langford, John Sillitoe on behalf of the Wellcome Sanger Institute COVID-19 Surveillance Team |
| EPI_ISL_3108858, EPI_ISL_3108859 | Lighthouse Lab in Alderley Park | Wellcome Sanger Institute for the COVID-19 Genomics UK (COG-UK) Consortium | Jacquelyn Wynn, Mairead Hyland, The Lighthouse Lab in Alderley Park and Alex Alderton, Roberto Amato, Jeffrey Barrett, Sonia Goncalves, Ewan Harrison, David K. Jackson, Ian Johnston, Dominic Kwiatkowski, Cordelia Langford, John Sillitoe on behalf of the Wellcome Sanger Institute COVID-19 Surveillance Team |
| EPI_ISL_3108860 | Lighthouse Lab in Milton Keynes | Wellcome Sanger Institute for the COVID-19 Genomics UK (COG-UK) Consortium | The Lighthouse Lab in Milton Keynes and Alex Alderton, Roberto Amato, Jeffrey Barrett, Sonia Goncalves, Ewan Harrison, David K. Jackson, Ian Johnston, Dominic Kwiatkowski, Cordelia Langford, John Sillitoe on behalf of the Wellcome Sanger Institute COVID-19 Surveillance Team |
| EPI_ISL_3108861 | Lighthouse Lab in Glasgow | Wellcome Sanger Institute for the COVID-19 Genomics UK (COG-UK) Consortium | Harper VanSteenhouse, Yumi Kasai, David Gray, Carol Clugston, Anna Dominiczak and Alex Alderton, Roberto Amato, Jeffrey Barrett, Sonia Goncalves, Ewan Harrison, David K. Jackson, Ian Johnston, Dominic Kwiatkowski, Cordelia Langford, John Sillitoe on behalf of the Wellcome Sanger Institute COVID-19 Surveillance Team |
| EPI_ISL_3108862, EPI_ISL_3108863, EPI_ISL_3108864 | Lighthouse Lab in Alderley Park | Wellcome Sanger Institute for the COVID-19 Genomics UK (COG-UK) Consortium | Jacquelyn Wynn, Mairead Hyland, The Lighthouse Lab in Alderley Park and Alex Alderton, Roberto Amato, Jeffrey Barrett, Sonia Goncalves, Ewan Harrison, David K. Jackson, Ian Johnston, Dominic Kwiatkowski, Cordelia Langford, John Sillitoe on behalf of the Wellcome Sanger Institute COVID-19 Surveillance Team |
| EPI_ISL_3108865 | Lighthouse Lab in Milton Keynes | Wellcome Sanger Institute for the COVID-19 Genomics UK (COG-UK) Consortium | The Lighthouse Lab in Milton Keynes and Alex Alderton, Roberto Amato, Jeffrey Barrett, Sonia Goncalves, Ewan Harrison, David K. Jackson, Ian Johnston, Dominic Kwiatkowski, Cordelia Langford, John Sillitoe on behalf of the Wellcome Sanger Institute COVID-19 Surveillance Team |
| EPI_ISL_3108866 | Lighthouse Lab in Glasgow | Wellcome Sanger Institute for the COVID-19 Genomics UK (COG-UK) Consortium | Harper VanSteenhouse, Yumi Kasai, David Gray, Carol Clugston, Anna Dominiczak and Alex Alderton, Roberto Amato, Jeffrey Barrett, Sonia Goncalves, Ewan Harrison, David K. Jackson, Ian Johnston, Dominic Kwiatkowski, Cordelia Langford, John Sillitoe on behalf of the Wellcome Sanger Institute COVID-19 Surveillance Team |
| EPI_ISL_3108867, EPI_ISL_3108868 | Lighthouse Lab in Milton Keynes | Wellcome Sanger Institute for the COVID-19 Genomics UK (COG-UK) Consortium | The Lighthouse Lab in Milton Keynes and Alex Alderton, Roberto Amato, Jeffrey Barrett, Sonia Goncalves, Ewan Harrison, David K. Jackson, Ian Johnston, Dominic Kwiatkowski, Cordelia Langford, John Sillitoe on behalf of the Wellcome Sanger Institute COVID-19 Surveillance Team |
| EPI_ISL_3108869 | Lighthouse Lab in Glasgow | Wellcome Sanger Institute for the COVID-19 Genomics UK (COG-UK) Consortium | Harper VanSteenhouse, Yumi Kasai, David Gray, Carol Clugston, Anna Dominiczak and Alex Alderton, Roberto Amato, Jeffrey Barrett, Sonia Goncalves, Ewan Harrison, David K. Jackson, Ian Johnston, Dominic Kwiatkowski, Cordelia Langford, John Sillitoe on behalf of the Wellcome Sanger Institute COVID-19 Surveillance Team |
| EPI_ISL_3108870 | Lighthouse Lab in Alderley Park | Wellcome Sanger Institute for the COVID-19 Genomics UK (COG-UK) Consortium | Jacquelyn Wynn, Mairead Hyland, The Lighthouse Lab in Alderley Park and Alex Alderton, Roberto Amato, Jeffrey Barrett, Sonia Goncalves, Ewan Harrison, David K. Jackson, Ian Johnston, Dominic Kwiatkowski, Cordelia Langford, John Sillitoe on behalf of the Wellcome Sanger Institute COVID-19 Surveillance Team |
| EPI_ISL_3108871 | Lighthouse Lab in Glasgow | Wellcome Sanger Institute for the COVID-19 Genomics UK (COG-UK) Consortium | Harper VanSteenhouse, Yumi Kasai, David Gray, Carol Clugston, Anna Dominiczak and Alex Alderton, Roberto Amato, Jeffrey Barrett, Sonia Goncalves, Ewan Harrison, David K. Jackson, Ian Johnston, Dominic Kwiatkowski, Cordelia Langford, John Sillitoe on behalf of the Wellcome Sanger Institute COVID-19 Surveillance Team |
| EPI_ISL_3108872, EPI_ISL_3108873, EPI_ISL_3108874, EPI_ISL_3108875 | Lighthouse Lab in Milton Keynes | Wellcome Sanger Institute for the COVID-19 Genomics UK (COG-UK) Consortium | The Lighthouse Lab in Milton Keynes and Alex Alderton, Roberto Amato, Jeffrey Barrett, Sonia Goncalves, Ewan Harrison, David K. Jackson, Ian Johnston, Dominic Kwiatkowski, Cordelia Langford, John Sillitoe on behalf of the Wellcome Sanger Institute COVID-19 Surveillance Team |
| EPI_ISL_3108876 | Lighthouse Lab in Alderley Park | Wellcome Sanger Institute for the COVID-19 Genomics UK (COG-UK) Consortium | Jacquelyn Wynn, Mairead Hyland, The Lighthouse Lab in Alderley Park and Alex Alderton, Roberto Amato, Jeffrey Barrett, Sonia Goncalves, Ewan Harrison, David K. Jackson, Ian Johnston, Dominic Kwiatkowski, Cordelia Langford, John Sillitoe on behalf of the Wellcome Sanger Institute COVID-19 Surveillance Team |
| EPI_ISL_3108877, EPI_ISL_3108878, EPI_ISL_3108879 | Lighthouse Lab in Milton Keynes | Wellcome Sanger Institute for the COVID-19 Genomics UK (COG-UK) Consortium | The Lighthouse Lab in Milton Keynes and Alex Alderton, Roberto Amato, Jeffrey Barrett, Sonia Goncalves, Ewan Harrison, David K. Jackson, Ian Johnston, Dominic Kwiatkowski, Cordelia Langford, John Sillitoe on behalf of the Wellcome Sanger Institute COVID-19 Surveillance Team |
| EPI_ISL_3108880 | Lighthouse Lab in Glasgow | Wellcome Sanger Institute for the COVID-19 Genomics UK (COG-UK) Consortium | Harper VanSteenhouse, Yumi Kasai, David Gray, Carol Clugston, Anna Dominiczak and Alex Alderton, Roberto Amato, Jeffrey Barrett, Sonia Goncalves, Ewan Harrison, David K. Jackson, Ian Johnston, Dominic Kwiatkowski, Cordelia Langford, John Sillitoe on behalf of the Wellcome Sanger Institute COVID-19 Surveillance Team |
| EPI_ISL_3108881, EPI_ISL_3108882, EPI_ISL_3108883 | Lighthouse Lab in Alderley Park | Wellcome Sanger Institute for the COVID-19 Genomics UK (COG-UK) Consortium | Jacquelyn Wynn, Mairead Hyland, The Lighthouse Lab in Alderley Park and Alex Alderton, Roberto Amato, Jeffrey Barrett, Sonia Goncalves, Ewan Harrison, David K. Jackson, Ian Johnston, Dominic Kwiatkowski, Cordelia Langford, John Sillitoe on behalf of the Wellcome Sanger Institute COVID-19 Surveillance Team |
| EPI_ISL_3108884 | Lighthouse Lab in Milton Keynes | Wellcome Sanger Institute for the COVID-19 Genomics UK (COG-UK) Consortium | The Lighthouse Lab in Milton Keynes and Alex Alderton, Roberto Amato, Jeffrey Barrett, Sonia Goncalves, Ewan Harrison, David K. Jackson, Ian Johnston, Dominic Kwiatkowski, Cordelia Langford, John Sillitoe on behalf of the Wellcome Sanger Institute COVID-19 Surveillance Team |
| EPI_ISL_3108885 | Lighthouse Lab in Glasgow | Wellcome Sanger Institute for the COVID-19 Genomics UK (COG-UK) Consortium | Harper VanSteenhouse, Yumi Kasai, David Gray, Carol Clugston, Anna Dominiczak and Alex Alderton, Roberto Amato, Jeffrey Barrett, Sonia Goncalves, Ewan Harrison, David K. Jackson, Ian Johnston, Dominic Kwiatkowski, Cordelia Langford, John Sillitoe on behalf of the Wellcome Sanger Institute COVID-19 Surveillance Team |
| EPI_ISL_3108886 | Lighthouse Lab in Milton Keynes | Wellcome Sanger Institute for the COVID-19 Genomics UK (COG-UK) Consortium | The Lighthouse Lab in Milton Keynes and Alex Alderton, Roberto Amato, Jeffrey Barrett, Sonia Goncalves, Ewan Harrison, David K. Jackson, Ian Johnston, Dominic Kwiatkowski, Cordelia Langford, John Sillitoe on behalf of the Wellcome Sanger Institute COVID-19 Surveillance Team |
| EPI_ISL_3108887 | Lighthouse Lab in Glasgow | Wellcome Sanger Institute for the COVID-19 Genomics UK (COG-UK) Consortium | Harper VanSteenhouse, Yumi Kasai, David Gray, Carol Clugston, Anna Dominiczak and Alex Alderton, Roberto Amato, Jeffrey Barrett, Sonia Goncalves, Ewan Harrison, David K. Jackson, Ian Johnston, Dominic Kwiatkowski, Cordelia Langford, John Sillitoe on behalf of the Wellcome Sanger Institute COVID-19 Surveillance Team |
| EPI_ISL_3108888, EPI_ISL_3108889, EPI_ISL_3108890 | Lighthouse Lab in Milton Keynes | Wellcome Sanger Institute for the COVID-19 Genomics UK (COG-UK) Consortium | The Lighthouse Lab in Milton Keynes and Alex Alderton, Roberto Amato, Jeffrey Barrett, Sonia Goncalves, Ewan Harrison, David K. Jackson, Ian Johnston, Dominic Kwiatkowski, Cordelia Langford, John Sillitoe on behalf of the Wellcome Sanger Institute COVID-19 Surveillance Team |
| EPI_ISL_3108891, EPI_ISL_3108892 | Lighthouse Lab in Glasgow | Wellcome Sanger Institute for the COVID-19 Genomics UK (COG-UK) Consortium | Harper VanSteenhouse, Yumi Kasai, David Gray, Carol Clugston, Anna Dominiczak and Alex Alderton, Roberto Amato, Jeffrey Barrett, Sonia Goncalves, Ewan Harrison, David K. Jackson, Ian Johnston, Dominic Kwiatkowski, Cordelia Langford, John Sillitoe on behalf of the Wellcome Sanger Institute COVID-19 Surveillance Team |
| EPI_ISL_3108893 | Lighthouse Lab in Milton Keynes | Wellcome Sanger Institute for the COVID-19 Genomics UK (COG-UK) Consortium | The Lighthouse Lab in Milton Keynes and Alex Alderton, Roberto Amato, Jeffrey Barrett, Sonia Goncalves, Ewan Harrison, David K. Jackson, Ian Johnston, Dominic Kwiatkowski, Cordelia Langford, John Sillitoe on behalf of the Wellcome Sanger Institute COVID-19 Surveillance Team |
| EPI_ISL_3108894 | Lighthouse Lab in Glasgow | Wellcome Sanger Institute for the COVID-19 Genomics UK (COG-UK) Consortium | Harper VanSteenhouse, Yumi Kasai, David Gray, Carol Clugston, Anna Dominiczak and Alex Alderton, Roberto Amato, Jeffrey Barrett, Sonia Goncalves, Ewan Harrison, David K. Jackson, Ian Johnston, Dominic Kwiatkowski, Cordelia Langford, John Sillitoe on behalf of the Wellcome Sanger Institute COVID-19 Surveillance Team |
| EPI_ISL_3108895 | Lighthouse Lab in Alderley Park | Wellcome Sanger Institute for the COVID-19 Genomics UK (COG-UK) Consortium | Jacquelyn Wynn, Mairead Hyland, The Lighthouse Lab in Alderley Park and Alex Alderton, Roberto Amato, Jeffrey Barrett, Sonia Goncalves, Ewan Harrison, David K. Jackson, Ian Johnston, Dominic Kwiatkowski, Cordelia Langford, John Sillitoe on behalf of the Wellcome Sanger Institute COVID-19 Surveillance Team |
| EPI_ISL_3108896 | Lighthouse Lab in Milton Keynes | Wellcome Sanger Institute for the COVID-19 Genomics UK (COG-UK) Consortium | The Lighthouse Lab in Milton Keynes and Alex Alderton, Roberto Amato, Jeffrey Barrett, Sonia Goncalves, Ewan Harrison, David K. Jackson, Ian Johnston, Dominic Kwiatkowski, Cordelia Langford, John Sillitoe on behalf of the Wellcome Sanger Institute COVID-19 Surveillance Team |
| EPI_ISL_3108897 | Lighthouse Lab in Alderley Park | Wellcome Sanger Institute for the COVID-19 Genomics UK (COG-UK) Consortium | Jacquelyn Wynn, Mairead Hyland, The Lighthouse Lab in Alderley Park and Alex Alderton, Roberto Amato, Jeffrey Barrett, Sonia Goncalves, Ewan Harrison, David K. Jackson, Ian Johnston, Dominic Kwiatkowski, Cordelia Langford, John Sillitoe on behalf of the Wellcome Sanger Institute COVID-19 Surveillance Team |
| EPI_ISL_3108898, EPI_ISL_3108899, EPI_ISL_3108900, EPI_ISL_3108901, EPI_ISL_3108902, EPI_ISL_3108903, EPI_ISL_3108904 | Lighthouse Lab in Milton Keynes | Wellcome Sanger Institute for the COVID-19 Genomics UK (COG-UK) Consortium | The Lighthouse Lab in Milton Keynes and Alex Alderton, Roberto Amato, Jeffrey Barrett, Sonia Goncalves, Ewan Harrison, David K. Jackson, Ian Johnston, Dominic Kwiatkowski, Cordelia Langford, John Sillitoe on behalf of the Wellcome Sanger Institute COVID-19 Surveillance Team |
| EPI_ISL_3108905, | Lighthouse Lab in Alderley Park | Wellcome Sanger Institute for the COVID-19 Genomics UK (COG-UK) | Jacquelyn Wynn, Mairead Hyland, The Lighthouse Lab in Alderley Park and Alex Alderton, Roberto Amato, Jeffrey Barrett, Sonia Goncalves, Ewan Harrison, David K. Jackson, Ian Johnston, Dominic Kwiatkowski, Cordelia Langford, John Sillitoe on |

| | | | |
|---|---|---|---|
| EPI_ISL_3108906, EPI_ISL_3108907 | | Consortium | behalf of the Wellcome Sanger Institute COVID-19 Surveillance Team |
| EPI_ISL_3108908 | Lighthouse Lab in Glasgow | Wellcome Sanger Institute for the COVID-19 Genomics UK (COG-UK) Consortium | Harper VanSteenhouse, Yumi Kasai, David Gray, Carol Clugston, Anna Dominiczak and Alex Alderton, Roberto Amato, Jeffrey Barrett, Sonia Goncalves, Ewan Harrison, David K. Jackson, Ian Johnston, Dominic Kwiatkowski, Cordelia Langford, John Sillitoe on behalf of the Wellcome Sanger Institute COVID-19 Surveillance Team |
| EPI_ISL_3108909, EPI_ISL_3108910, EPI_ISL_3108911, EPI_ISL_3108912 | Lighthouse Lab in Milton Keynes | Wellcome Sanger Institute for the COVID-19 Genomics UK (COG-UK) Consortium | The Lighthouse Lab in Milton Keynes and Alex Alderton, Roberto Amato, Jeffrey Barrett, Sonia Goncalves, Ewan Harrison, David K. Jackson, Ian Johnston, Dominic Kwiatkowski, Cordelia Langford, John Sillitoe on behalf of the Wellcome Sanger Institute COVID-19 Surveillance Team |
| EPI_ISL_3108913 | Lighthouse Lab in Alderley Park | Wellcome Sanger Institute for the COVID-19 Genomics UK (COG-UK) Consortium | Jacquelyn Wynn, Mairead Hyland, The Lighthouse Lab in Alderley Park and Alex Alderton, Roberto Amato, Jeffrey Barrett, Sonia Goncalves, Ewan Harrison, David K. Jackson, Ian Johnston, Dominic Kwiatkowski, Cordelia Langford, John Sillitoe on behalf of the Wellcome Sanger Institute COVID-19 Surveillance Team |
| EPI_ISL_3108914, EPI_ISL_3108915, EPI_ISL_3108916 | Lighthouse Lab in Milton Keynes | Wellcome Sanger Institute for the COVID-19 Genomics UK (COG-UK) Consortium | The Lighthouse Lab in Milton Keynes and Alex Alderton, Roberto Amato, Jeffrey Barrett, Sonia Goncalves, Ewan Harrison, David K. Jackson, Ian Johnston, Dominic Kwiatkowski, Cordelia Langford, John Sillitoe on behalf of the Wellcome Sanger Institute COVID-19 Surveillance Team |
| EPI_ISL_3108917 | Lighthouse Lab in Alderley Park | Wellcome Sanger Institute for the COVID-19 Genomics UK (COG-UK) Consortium | Jacquelyn Wynn, Mairead Hyland, The Lighthouse Lab in Alderley Park and Alex Alderton, Roberto Amato, Jeffrey Barrett, Sonia Goncalves, Ewan Harrison, David K. Jackson, Ian Johnston, Dominic Kwiatkowski, Cordelia Langford, John Sillitoe on behalf of the Wellcome Sanger Institute COVID-19 Surveillance Team |
| EPI_ISL_3108918 | Lighthouse Lab in Milton Keynes | Wellcome Sanger Institute for the COVID-19 Genomics UK (COG-UK) Consortium | The Lighthouse Lab in Milton Keynes and Alex Alderton, Roberto Amato, Jeffrey Barrett, Sonia Goncalves, Ewan Harrison, David K. Jackson, Ian Johnston, Dominic Kwiatkowski, Cordelia Langford, John Sillitoe on behalf of the Wellcome Sanger Institute COVID-19 Surveillance Team |
| EPI_ISL_3108919 | Lighthouse Lab in Alderley Park | Wellcome Sanger Institute for the COVID-19 Genomics UK (COG-UK) Consortium | Jacquelyn Wynn, Mairead Hyland, The Lighthouse Lab in Alderley Park and Alex Alderton, Roberto Amato, Jeffrey Barrett, Sonia Goncalves, Ewan Harrison, David K. Jackson, Ian Johnston, Dominic Kwiatkowski, Cordelia Langford, John Sillitoe on behalf of the Wellcome Sanger Institute COVID-19 Surveillance Team |
| EPI_ISL_3108920 | Lighthouse Lab in Milton Keynes | Wellcome Sanger Institute for the COVID-19 Genomics UK (COG-UK) Consortium | The Lighthouse Lab in Milton Keynes and Alex Alderton, Roberto Amato, Jeffrey Barrett, Sonia Goncalves, Ewan Harrison, David K. Jackson, Ian Johnston, Dominic Kwiatkowski, Cordelia Langford, John Sillitoe on behalf of the Wellcome Sanger Institute COVID-19 Surveillance Team |
| EPI_ISL_3108921 | Lighthouse Lab in Alderley Park | Wellcome Sanger Institute for the COVID-19 Genomics UK (COG-UK) Consortium | Jacquelyn Wynn, Mairead Hyland, The Lighthouse Lab in Alderley Park and Alex Alderton, Roberto Amato, Jeffrey Barrett, Sonia Goncalves, Ewan Harrison, David K. Jackson, Ian Johnston, Dominic Kwiatkowski, Cordelia Langford, John Sillitoe on behalf of the Wellcome Sanger Institute COVID-19 Surveillance Team |
| EPI_ISL_3108922 | Lighthouse Lab in Milton Keynes | Wellcome Sanger Institute for the COVID-19 Genomics UK (COG-UK) Consortium | The Lighthouse Lab in Milton Keynes and Alex Alderton, Roberto Amato, Jeffrey Barrett, Sonia Goncalves, Ewan Harrison, David K. Jackson, Ian Johnston, Dominic Kwiatkowski, Cordelia Langford, John Sillitoe on behalf of the Wellcome Sanger Institute COVID-19 Surveillance Team |
| EPI_ISL_3108923, EPI_ISL_3108924 | Lighthouse Lab in Alderley Park | Wellcome Sanger Institute for the COVID-19 Genomics UK (COG-UK) Consortium | Jacquelyn Wynn, Mairead Hyland, The Lighthouse Lab in Alderley Park and Alex Alderton, Roberto Amato, Jeffrey Barrett, Sonia Goncalves, Ewan Harrison, David K. Jackson, Ian Johnston, Dominic Kwiatkowski, Cordelia Langford, John Sillitoe on behalf of the Wellcome Sanger Institute COVID-19 Surveillance Team |
| EPI_ISL_3108925, EPI_ISL_3108926, EPI_ISL_3108927 | Lighthouse Lab in Milton Keynes | Wellcome Sanger Institute for the COVID-19 Genomics UK (COG-UK) Consortium | The Lighthouse Lab in Milton Keynes and Alex Alderton, Roberto Amato, Jeffrey Barrett, Sonia Goncalves, Ewan Harrison, David K. Jackson, Ian Johnston, Dominic Kwiatkowski, Cordelia Langford, John Sillitoe on behalf of the Wellcome Sanger Institute COVID-19 Surveillance Team |
| EPI_ISL_3108928, EPI_ISL_3108929 | Lighthouse Lab in Alderley Park | Wellcome Sanger Institute for the COVID-19 Genomics UK (COG-UK) Consortium | Jacquelyn Wynn, Mairead Hyland, The Lighthouse Lab in Alderley Park and Alex Alderton, Roberto Amato, Jeffrey Barrett, Sonia Goncalves, Ewan Harrison, David K. Jackson, Ian Johnston, Dominic Kwiatkowski, Cordelia Langford, John Sillitoe on behalf of the Wellcome Sanger Institute COVID-19 Surveillance Team |
| EPI_ISL_3108930, EPI_ISL_3108931, EPI_ISL_3108932, EPI_ISL_3108933 | Lighthouse Lab in Milton Keynes | Wellcome Sanger Institute for the COVID-19 Genomics UK (COG-UK) Consortium | The Lighthouse Lab in Milton Keynes and Alex Alderton, Roberto Amato, Jeffrey Barrett, Sonia Goncalves, Ewan Harrison, David K. Jackson, Ian Johnston, Dominic Kwiatkowski, Cordelia Langford, John Sillitoe on behalf of the Wellcome Sanger Institute COVID-19 Surveillance Team |
| EPI_ISL_3108934 | Lighthouse Lab in Alderley Park | Wellcome Sanger Institute for the COVID-19 Genomics UK (COG-UK) Consortium | Jacquelyn Wynn, Mairead Hyland, The Lighthouse Lab in Alderley Park and Alex Alderton, Roberto Amato, Jeffrey Barrett, Sonia Goncalves, Ewan Harrison, David K. Jackson, Ian Johnston, Dominic Kwiatkowski, Cordelia Langford, John Sillitoe on behalf of the Wellcome Sanger Institute COVID-19 Surveillance Team |
| EPI_ISL_3108935 | Lighthouse Lab in Glasgow | Wellcome Sanger Institute for the COVID-19 Genomics UK (COG-UK) Consortium | Harper VanSteenhouse, Yumi Kasai, David Gray, Carol Clugston, Anna Dominiczak and Alex Alderton, Roberto Amato, Jeffrey Barrett, Sonia Goncalves, Ewan Harrison, David K. Jackson, Ian Johnston, Dominic Kwiatkowski, Cordelia Langford, John Sillitoe on behalf of the Wellcome Sanger Institute COVID-19 Surveillance Team |
| EPI_ISL_3108936 | Lighthouse Lab in Alderley Park | Wellcome Sanger Institute for the COVID-19 Genomics UK (COG-UK) Consortium | Jacquelyn Wynn, Mairead Hyland, The Lighthouse Lab in Alderley Park and Alex Alderton, Roberto Amato, Jeffrey Barrett, Sonia Goncalves, Ewan Harrison, David K. Jackson, Ian Johnston, Dominic Kwiatkowski, Cordelia Langford, John Sillitoe on behalf of the Wellcome Sanger Institute COVID-19 Surveillance Team |
| EPI_ISL_3108937, EPI_ISL_3108938 | Lighthouse Lab in Milton Keynes | Wellcome Sanger Institute for the COVID-19 Genomics UK (COG-UK) Consortium | The Lighthouse Lab in Milton Keynes and Alex Alderton, Roberto Amato, Jeffrey Barrett, Sonia Goncalves, Ewan Harrison, David K. Jackson, Ian Johnston, Dominic Kwiatkowski, Cordelia Langford, John Sillitoe on behalf of the Wellcome Sanger Institute COVID-19 Surveillance Team |
| EPI_ISL_3108939 | Lighthouse Lab in Glasgow | Wellcome Sanger Institute for the COVID-19 Genomics UK (COG-UK) Consortium | Harper VanSteenhouse, Yumi Kasai, David Gray, Carol Clugston, Anna Dominiczak and Alex Alderton, Roberto Amato, Jeffrey Barrett, Sonia Goncalves, Ewan Harrison, David K. Jackson, Ian Johnston, Dominic Kwiatkowski, Cordelia Langford, John Sillitoe on behalf of the Wellcome Sanger Institute COVID-19 Surveillance Team |
| EPI_ISL_3108940 | Lighthouse Lab in Alderley Park | Wellcome Sanger Institute for the COVID-19 Genomics UK (COG-UK) Consortium | Jacquelyn Wynn, Mairead Hyland, The Lighthouse Lab in Alderley Park and Alex Alderton, Roberto Amato, Jeffrey Barrett, Sonia Goncalves, Ewan Harrison, David K. Jackson, Ian Johnston, Dominic Kwiatkowski, Cordelia Langford, John Sillitoe on behalf of the Wellcome Sanger Institute COVID-19 Surveillance Team |
| EPI_ISL_3108941 | Lighthouse Lab in Milton Keynes | Wellcome Sanger Institute for the COVID-19 Genomics UK (COG-UK) Consortium | The Lighthouse Lab in Milton Keynes and Alex Alderton, Roberto Amato, Jeffrey Barrett, Sonia Goncalves, Ewan Harrison, David K. Jackson, Ian Johnston, Dominic Kwiatkowski, Cordelia Langford, John Sillitoe on behalf of the Wellcome Sanger Institute COVID-19 Surveillance Team |
| EPI_ISL_3108942 | Lighthouse Lab in Alderley Park | Wellcome Sanger Institute for the COVID-19 Genomics UK (COG-UK) Consortium | Jacquelyn Wynn, Mairead Hyland, The Lighthouse Lab in Alderley Park and Alex Alderton, Roberto Amato, Jeffrey Barrett, Sonia Goncalves, Ewan Harrison, David K. Jackson, Ian Johnston, Dominic Kwiatkowski, Cordelia Langford, John Sillitoe on behalf of the Wellcome Sanger Institute COVID-19 Surveillance Team |
| EPI_ISL_3108943 | Lighthouse Lab in Milton Keynes | Wellcome Sanger Institute for the COVID-19 Genomics UK (COG-UK) Consortium | The Lighthouse Lab in Milton Keynes and Alex Alderton, Roberto Amato, Jeffrey Barrett, Sonia Goncalves, Ewan Harrison, David K. Jackson, Ian Johnston, Dominic Kwiatkowski, Cordelia Langford, John Sillitoe on behalf of the Wellcome Sanger Institute COVID-19 Surveillance Team |
| EPI_ISL_3108944 | Lighthouse Lab in Alderley Park | Wellcome Sanger Institute for the COVID-19 Genomics UK (COG-UK) Consortium | Jacquelyn Wynn, Mairead Hyland, The Lighthouse Lab in Alderley Park and Alex Alderton, Roberto Amato, Jeffrey Barrett, Sonia Goncalves, Ewan Harrison, David K. Jackson, Ian Johnston, Dominic Kwiatkowski, Cordelia Langford, John Sillitoe on behalf of the Wellcome Sanger Institute COVID-19 Surveillance Team |
| EPI_ISL_3108945 | Lighthouse Lab in Glasgow | Wellcome Sanger Institute for the COVID-19 Genomics UK (COG-UK) Consortium | Harper VanSteenhouse, Yumi Kasai, David Gray, Carol Clugston, Anna Dominiczak and Alex Alderton, Roberto Amato, Jeffrey Barrett, Sonia Goncalves, Ewan Harrison, David K. Jackson, Ian Johnston, Dominic Kwiatkowski, Cordelia Langford, John Sillitoe on behalf of the Wellcome Sanger Institute COVID-19 Surveillance Team |
| EPI_ISL_3108946, EPI_ISL_3108947 | Lighthouse Lab in Milton Keynes | Wellcome Sanger Institute for the COVID-19 Genomics UK (COG-UK) Consortium | The Lighthouse Lab in Milton Keynes and Alex Alderton, Roberto Amato, Jeffrey Barrett, Sonia Goncalves, Ewan Harrison, David K. Jackson, Ian Johnston, Dominic Kwiatkowski, Cordelia Langford, John Sillitoe on behalf of the Wellcome Sanger Institute COVID-19 Surveillance Team |
| EPI_ISL_3108948 | Lighthouse Lab in Alderley Park | Wellcome Sanger Institute for the COVID-19 Genomics UK (COG-UK) Consortium | Jacquelyn Wynn, Mairead Hyland, The Lighthouse Lab in Alderley Park and Alex Alderton, Roberto Amato, Jeffrey Barrett, Sonia Goncalves, Ewan Harrison, David K. Jackson, Ian Johnston, Dominic Kwiatkowski, Cordelia Langford, John Sillitoe on behalf of the Wellcome Sanger Institute COVID-19 Surveillance Team |
| EPI_ISL_3108949 | Lighthouse Lab in Milton Keynes | Wellcome Sanger Institute for the COVID-19 Genomics UK (COG-UK) Consortium | The Lighthouse Lab in Milton Keynes and Alex Alderton, Roberto Amato, Jeffrey Barrett, Sonia Goncalves, Ewan Harrison, David K. Jackson, Ian Johnston, Dominic Kwiatkowski, Cordelia Langford, John Sillitoe on behalf of the Wellcome Sanger Institute COVID-19 Surveillance Team |
| EPI_ISL_3108950 | Lighthouse Lab in Alderley Park | Wellcome Sanger Institute for the COVID-19 Genomics UK (COG-UK) Consortium | Jacquelyn Wynn, Mairead Hyland, The Lighthouse Lab in Alderley Park and Alex Alderton, Roberto Amato, Jeffrey Barrett, Sonia Goncalves, Ewan Harrison, David K. Jackson, Ian Johnston, Dominic Kwiatkowski, Cordelia Langford, John Sillitoe on behalf of the Wellcome Sanger Institute COVID-19 Surveillance Team |
| EPI_ISL_3108951 | Lighthouse Lab in Milton Keynes | Wellcome Sanger Institute for the COVID-19 Genomics UK (COG-UK) Consortium | The Lighthouse Lab in Milton Keynes and Alex Alderton, Roberto Amato, Jeffrey Barrett, Sonia Goncalves, Ewan Harrison, David K. Jackson, Ian Johnston, Dominic Kwiatkowski, Cordelia Langford, John Sillitoe on behalf of the Wellcome Sanger Institute COVID-19 Surveillance Team |
| EPI_ISL_3108952 | Lighthouse Lab in Alderley Park | Wellcome Sanger Institute for the COVID-19 Genomics UK (COG-UK) Consortium | Jacquelyn Wynn, Mairead Hyland, The Lighthouse Lab in Alderley Park and Alex Alderton, Roberto Amato, Jeffrey Barrett, Sonia Goncalves, Ewan Harrison, David K. Jackson, Ian Johnston, Dominic Kwiatkowski, Cordelia Langford, John Sillitoe on behalf of the Wellcome Sanger Institute COVID-19 Surveillance Team |
| EPI_ISL_3108953 | Lighthouse Lab in Milton Keynes | Wellcome Sanger Institute for the COVID-19 Genomics UK (COG-UK) Consortium | The Lighthouse Lab in Milton Keynes and Alex Alderton, Roberto Amato, Jeffrey Barrett, Sonia Goncalves, Ewan Harrison, David K. Jackson, Ian Johnston, Dominic Kwiatkowski, Cordelia Langford, John Sillitoe on behalf of the Wellcome Sanger Institute COVID-19 Surveillance Team |
| EPI_ISL_3108954 | Lighthouse Lab in Alderley Park | Wellcome Sanger Institute for the COVID-19 Genomics UK (COG-UK) Consortium | Jacquelyn Wynn, Mairead Hyland, The Lighthouse Lab in Alderley Park and Alex Alderton, Roberto Amato, Jeffrey Barrett, Sonia Goncalves, Ewan Harrison, David K. Jackson, Ian Johnston, Dominic Kwiatkowski, Cordelia Langford, John Sillitoe on behalf of the Wellcome Sanger Institute COVID-19 Surveillance Team |
| EPI_ISL_3108955, EPI_ISL_3108956, EPI_ISL_3108957, EPI_ISL_3108958, EPI_ISL_3108959, EPI_ISL_3108960, EPI_ISL_3108961, EPI_ISL_3108962, EPI_ISL_3108963, EPI_ISL_3108964 | Lighthouse Lab in Milton Keynes | Wellcome Sanger Institute for the COVID-19 Genomics UK (COG-UK) Consortium | The Lighthouse Lab in Milton Keynes and Alex Alderton, Roberto Amato, Jeffrey Barrett, Sonia Goncalves, Ewan Harrison, David K. Jackson, Ian Johnston, Dominic Kwiatkowski, Cordelia Langford, John Sillitoe on behalf of the Wellcome Sanger Institute COVID-19 Surveillance Team |
| EPI_ISL_3108965 | Lighthouse Lab in Alderley Park | Wellcome Sanger Institute for the COVID-19 Genomics UK (COG-UK) Consortium | Jacquelyn Wynn, Mairead Hyland, The Lighthouse Lab in Alderley Park and Alex Alderton, Roberto Amato, Jeffrey Barrett, Sonia Goncalves, Ewan Harrison, David K. Jackson, Ian Johnston, Dominic Kwiatkowski, Cordelia Langford, John Sillitoe on behalf of the Wellcome Sanger Institute COVID-19 Surveillance Team |
| EPI_ISL_3108966 | Lighthouse Lab in Milton Keynes | Wellcome Sanger Institute for the COVID-19 Genomics UK (COG-UK) Consortium | The Lighthouse Lab in Milton Keynes and Alex Alderton, Roberto Amato, Jeffrey Barrett, Sonia Goncalves, Ewan Harrison, David K. Jackson, Ian Johnston, Dominic Kwiatkowski, Cordelia Langford, John Sillitoe on behalf of the Wellcome Sanger Institute COVID-19 Surveillance Team |
| EPI_ISL_3108967 | Lighthouse Lab in Alderley Park | Wellcome Sanger Institute for the COVID-19 Genomics UK (COG-UK) Consortium | Jacquelyn Wynn, Mairead Hyland, The Lighthouse Lab in Alderley Park and Alex Alderton, Roberto Amato, Jeffrey Barrett, Sonia Goncalves, Ewan Harrison, David K. Jackson, Ian Johnston, Dominic Kwiatkowski, Cordelia Langford, John Sillitoe on behalf of the Wellcome Sanger Institute COVID-19 Surveillance Team |
| EPI_ISL_3108968, EPI_ISL_3108969, EPI_ISL_3108970 | Lighthouse Lab in Milton Keynes | Wellcome Sanger Institute for the COVID-19 Genomics UK (COG-UK) Consortium | The Lighthouse Lab in Milton Keynes and Alex Alderton, Roberto Amato, Jeffrey Barrett, Sonia Goncalves, Ewan Harrison, David K. Jackson, Ian Johnston, Dominic Kwiatkowski, Cordelia Langford, John Sillitoe on behalf of the Wellcome Sanger Institute COVID-19 Surveillance Team |

| | | | |
|---|---|---|---|
| EPI_ISL_3108971, EPI_ISL_3108972 | Lighthouse Lab in Alderley Park | Wellcome Sanger Institute for the COVID-19 Genomics UK (COG-UK) Consortium | Jacquelyn Wynn, Mairead Hyland, The Lighthouse Lab in Alderley Park and Alex Alderton, Roberto Amato, Jeffrey Barrett, Sonia Goncalves, Ewan Harrison, David K. Jackson, Ian Johnston, Dominic Kwiatkowski, Cordelia Langford, John Sillitoe on behalf of the Wellcome Sanger Institute COVID-19 Surveillance Team |
| EPI_ISL_3108973, EPI_ISL_3108974 | Lighthouse Lab in Milton Keynes | Wellcome Sanger Institute for the COVID-19 Genomics UK (COG-UK) Consortium | The Lighthouse Lab in Milton Keynes and Alex Alderton, Roberto Amato, Jeffrey Barrett, Sonia Goncalves, Ewan Harrison, David K. Jackson, Ian Johnston, Dominic Kwiatkowski, Cordelia Langford, John Sillitoe on behalf of the Wellcome Sanger Institute COVID-19 Surveillance Team |
| EPI_ISL_3108975, EPI_ISL_3108976, EPI_ISL_3108977 | Lighthouse Lab in Alderley Park | Wellcome Sanger Institute for the COVID-19 Genomics UK (COG-UK) Consortium | Jacquelyn Wynn, Mairead Hyland, The Lighthouse Lab in Alderley Park and Alex Alderton, Roberto Amato, Jeffrey Barrett, Sonia Goncalves, Ewan Harrison, David K. Jackson, Ian Johnston, Dominic Kwiatkowski, Cordelia Langford, John Sillitoe on behalf of the Wellcome Sanger Institute COVID-19 Surveillance Team |
| EPI_ISL_3108978 | Lighthouse Lab in Glasgow | Wellcome Sanger Institute for the COVID-19 Genomics UK (COG-UK) Consortium | Harper VanSteenhouse, Yumi Kasai, David Gray, Carol Clugston, Anna Dominiczak and Alex Alderton, Roberto Amato, Jeffrey Barrett, Sonia Goncalves, Ewan Harrison, David K. Jackson, Ian Johnston, Dominic Kwiatkowski, Cordelia Langford, John Sillitoe on behalf of the Wellcome Sanger Institute COVID-19 Surveillance Team |
| EPI_ISL_3108979, EPI_ISL_3108980 | Lighthouse Lab in Milton Keynes | Wellcome Sanger Institute for the COVID-19 Genomics UK (COG-UK) Consortium | The Lighthouse Lab in Milton Keynes and Alex Alderton, Roberto Amato, Jeffrey Barrett, Sonia Goncalves, Ewan Harrison, David K. Jackson, Ian Johnston, Dominic Kwiatkowski, Cordelia Langford, John Sillitoe on behalf of the Wellcome Sanger Institute COVID-19 Surveillance Team |
| EPI_ISL_3108981, EPI_ISL_3108982 | Lighthouse Lab in Alderley Park | Wellcome Sanger Institute for the COVID-19 Genomics UK (COG-UK) Consortium | Jacquelyn Wynn, Mairead Hyland, The Lighthouse Lab in Alderley Park and Alex Alderton, Roberto Amato, Jeffrey Barrett, Sonia Goncalves, Ewan Harrison, David K. Jackson, Ian Johnston, Dominic Kwiatkowski, Cordelia Langford, John Sillitoe on behalf of the Wellcome Sanger Institute COVID-19 Surveillance Team |
| EPI_ISL_3108983 | Lighthouse Lab in Milton Keynes | Wellcome Sanger Institute for the COVID-19 Genomics UK (COG-UK) Consortium | The Lighthouse Lab in Milton Keynes and Alex Alderton, Roberto Amato, Jeffrey Barrett, Sonia Goncalves, Ewan Harrison, David K. Jackson, Ian Johnston, Dominic Kwiatkowski, Cordelia Langford, John Sillitoe on behalf of the Wellcome Sanger Institute COVID-19 Surveillance Team |
| EPI_ISL_3108984, EPI_ISL_3108985 | Lighthouse Lab in Alderley Park | Wellcome Sanger Institute for the COVID-19 Genomics UK (COG-UK) Consortium | Jacquelyn Wynn, Mairead Hyland, The Lighthouse Lab in Alderley Park and Alex Alderton, Roberto Amato, Jeffrey Barrett, Sonia Goncalves, Ewan Harrison, David K. Jackson, Ian Johnston, Dominic Kwiatkowski, Cordelia Langford, John Sillitoe on behalf of the Wellcome Sanger Institute COVID-19 Surveillance Team |
| EPI_ISL_3108986 | Lighthouse Lab in Milton Keynes | Wellcome Sanger Institute for the COVID-19 Genomics UK (COG-UK) Consortium | The Lighthouse Lab in Milton Keynes and Alex Alderton, Roberto Amato, Jeffrey Barrett, Sonia Goncalves, Ewan Harrison, David K. Jackson, Ian Johnston, Dominic Kwiatkowski, Cordelia Langford, John Sillitoe on behalf of the Wellcome Sanger Institute COVID-19 Surveillance Team |
| EPI_ISL_3108987, EPI_ISL_3108988 | Lighthouse Lab in Glasgow | Wellcome Sanger Institute for the COVID-19 Genomics UK (COG-UK) Consortium | Harper VanSteenhouse, Yumi Kasai, David Gray, Carol Clugston, Anna Dominiczak and Alex Alderton, Roberto Amato, Jeffrey Barrett, Sonia Goncalves, Ewan Harrison, David K. Jackson, Ian Johnston, Dominic Kwiatkowski, Cordelia Langford, John Sillitoe on behalf of the Wellcome Sanger Institute COVID-19 Surveillance Team |
| EPI_ISL_3108989 | Lighthouse Lab in Milton Keynes | Wellcome Sanger Institute for the COVID-19 Genomics UK (COG-UK) Consortium | The Lighthouse Lab in Milton Keynes and Alex Alderton, Roberto Amato, Jeffrey Barrett, Sonia Goncalves, Ewan Harrison, David K. Jackson, Ian Johnston, Dominic Kwiatkowski, Cordelia Langford, John Sillitoe on behalf of the Wellcome Sanger Institute COVID-19 Surveillance Team |
| EPI_ISL_3108990 | Lighthouse Lab in Alderley Park | Wellcome Sanger Institute for the COVID-19 Genomics UK (COG-UK) Consortium | Jacquelyn Wynn, Mairead Hyland, The Lighthouse Lab in Alderley Park and Alex Alderton, Roberto Amato, Jeffrey Barrett, Sonia Goncalves, Ewan Harrison, David K. Jackson, Ian Johnston, Dominic Kwiatkowski, Cordelia Langford, John Sillitoe on behalf of the Wellcome Sanger Institute COVID-19 Surveillance Team |
| EPI_ISL_3108991, EPI_ISL_3108992, EPI_ISL_3108993, EPI_ISL_3108994, EPI_ISL_3108995, EPI_ISL_3108996 | Lighthouse Lab in Milton Keynes | Wellcome Sanger Institute for the COVID-19 Genomics UK (COG-UK) Consortium | The Lighthouse Lab in Milton Keynes and Alex Alderton, Roberto Amato, Jeffrey Barrett, Sonia Goncalves, Ewan Harrison, David K. Jackson, Ian Johnston, Dominic Kwiatkowski, Cordelia Langford, John Sillitoe on behalf of the Wellcome Sanger Institute COVID-19 Surveillance Team |
| EPI_ISL_3108997 | Lighthouse Lab in Glasgow | Wellcome Sanger Institute for the COVID-19 Genomics UK (COG-UK) Consortium | Harper VanSteenhouse, Yumi Kasai, David Gray, Carol Clugston, Anna Dominiczak and Alex Alderton, Roberto Amato, Jeffrey Barrett, Sonia Goncalves, Ewan Harrison, David K. Jackson, Ian Johnston, Dominic Kwiatkowski, Cordelia Langford, John Sillitoe on behalf of the Wellcome Sanger Institute COVID-19 Surveillance Team |
| EPI_ISL_3108998 | Lighthouse Lab in Alderley Park | Wellcome Sanger Institute for the COVID-19 Genomics UK (COG-UK) Consortium | Jacquelyn Wynn, Mairead Hyland, The Lighthouse Lab in Alderley Park and Alex Alderton, Roberto Amato, Jeffrey Barrett, Sonia Goncalves, Ewan Harrison, David K. Jackson, Ian Johnston, Dominic Kwiatkowski, Cordelia Langford, John Sillitoe on behalf of the Wellcome Sanger Institute COVID-19 Surveillance Team |
| EPI_ISL_3108999, EPI_ISL_3109000 | Lighthouse Lab in Milton Keynes | Wellcome Sanger Institute for the COVID-19 Genomics UK (COG-UK) Consortium | The Lighthouse Lab in Milton Keynes and Alex Alderton, Roberto Amato, Jeffrey Barrett, Sonia Goncalves, Ewan Harrison, David K. Jackson, Ian Johnston, Dominic Kwiatkowski, Cordelia Langford, John Sillitoe on behalf of the Wellcome Sanger Institute COVID-19 Surveillance Team |
| EPI_ISL_3109001 | Lighthouse Lab in Alderley Park | Wellcome Sanger Institute for the COVID-19 Genomics UK (COG-UK) Consortium | Jacquelyn Wynn, Mairead Hyland, The Lighthouse Lab in Alderley Park and Alex Alderton, Roberto Amato, Jeffrey Barrett, Sonia Goncalves, Ewan Harrison, David K. Jackson, Ian Johnston, Dominic Kwiatkowski, Cordelia Langford, John Sillitoe on behalf of the Wellcome Sanger Institute COVID-19 Surveillance Team |
| EPI_ISL_3109002 | Lighthouse Lab in Milton Keynes | Wellcome Sanger Institute for the COVID-19 Genomics UK (COG-UK) Consortium | The Lighthouse Lab in Milton Keynes and Alex Alderton, Roberto Amato, Jeffrey Barrett, Sonia Goncalves, Ewan Harrison, David K. Jackson, Ian Johnston, Dominic Kwiatkowski, Cordelia Langford, John Sillitoe on behalf of the Wellcome Sanger Institute COVID-19 Surveillance Team |
| EPI_ISL_3109003 | Lighthouse Lab in Glasgow | Wellcome Sanger Institute for the COVID-19 Genomics UK (COG-UK) Consortium | Harper VanSteenhouse, Yumi Kasai, David Gray, Carol Clugston, Anna Dominiczak and Alex Alderton, Roberto Amato, Jeffrey Barrett, Sonia Goncalves, Ewan Harrison, David K. Jackson, Ian Johnston, Dominic Kwiatkowski, Cordelia Langford, John Sillitoe on behalf of the Wellcome Sanger Institute COVID-19 Surveillance Team |
| EPI_ISL_3109004 | Lighthouse Lab in Alderley Park | Wellcome Sanger Institute for the COVID-19 Genomics UK (COG-UK) Consortium | Jacquelyn Wynn, Mairead Hyland, The Lighthouse Lab in Alderley Park and Alex Alderton, Roberto Amato, Jeffrey Barrett, Sonia Goncalves, Ewan Harrison, David K. Jackson, Ian Johnston, Dominic Kwiatkowski, Cordelia Langford, John Sillitoe on behalf of the Wellcome Sanger Institute COVID-19 Surveillance Team |
| EPI_ISL_3109005 | Lighthouse Lab in Milton Keynes | Wellcome Sanger Institute for the COVID-19 Genomics UK (COG-UK) Consortium | The Lighthouse Lab in Milton Keynes and Alex Alderton, Roberto Amato, Jeffrey Barrett, Sonia Goncalves, Ewan Harrison, David K. Jackson, Ian Johnston, Dominic Kwiatkowski, Cordelia Langford, John Sillitoe on behalf of the Wellcome Sanger Institute COVID-19 Surveillance Team |
| EPI_ISL_3109006 | Lighthouse Lab in Alderley Park | Wellcome Sanger Institute for the COVID-19 Genomics UK (COG-UK) Consortium | Jacquelyn Wynn, Mairead Hyland, The Lighthouse Lab in Alderley Park and Alex Alderton, Roberto Amato, Jeffrey Barrett, Sonia Goncalves, Ewan Harrison, David K. Jackson, Ian Johnston, Dominic Kwiatkowski, Cordelia Langford, John Sillitoe on behalf of the Wellcome Sanger Institute COVID-19 Surveillance Team |
| EPI_ISL_3109007, EPI_ISL_3109008, EPI_ISL_3109009, EPI_ISL_3109010 | Lighthouse Lab in Milton Keynes | Wellcome Sanger Institute for the COVID-19 Genomics UK (COG-UK) Consortium | The Lighthouse Lab in Milton Keynes and Alex Alderton, Roberto Amato, Jeffrey Barrett, Sonia Goncalves, Ewan Harrison, David K. Jackson, Ian Johnston, Dominic Kwiatkowski, Cordelia Langford, John Sillitoe on behalf of the Wellcome Sanger Institute COVID-19 Surveillance Team |
| EPI_ISL_3109011 | Lighthouse Lab in Alderley Park | Wellcome Sanger Institute for the COVID-19 Genomics UK (COG-UK) Consortium | Jacquelyn Wynn, Mairead Hyland, The Lighthouse Lab in Alderley Park and Alex Alderton, Roberto Amato, Jeffrey Barrett, Sonia Goncalves, Ewan Harrison, David K. Jackson, Ian Johnston, Dominic Kwiatkowski, Cordelia Langford, John Sillitoe on behalf of the Wellcome Sanger Institute COVID-19 Surveillance Team |
| EPI_ISL_3109012 | Lighthouse Lab in Glasgow | Wellcome Sanger Institute for the COVID-19 Genomics UK (COG-UK) Consortium | Harper VanSteenhouse, Yumi Kasai, David Gray, Carol Clugston, Anna Dominiczak and Alex Alderton, Roberto Amato, Jeffrey Barrett, Sonia Goncalves, Ewan Harrison, David K. Jackson, Ian Johnston, Dominic Kwiatkowski, Cordelia Langford, John Sillitoe on behalf of the Wellcome Sanger Institute COVID-19 Surveillance Team |
| EPI_ISL_3109013, EPI_ISL_3109014, EPI_ISL_3109015 | Lighthouse Lab in Milton Keynes | Wellcome Sanger Institute for the COVID-19 Genomics UK (COG-UK) Consortium | The Lighthouse Lab in Milton Keynes and Alex Alderton, Roberto Amato, Jeffrey Barrett, Sonia Goncalves, Ewan Harrison, David K. Jackson, Ian Johnston, Dominic Kwiatkowski, Cordelia Langford, John Sillitoe on behalf of the Wellcome Sanger Institute COVID-19 Surveillance Team |
| EPI_ISL_3109016 | Lighthouse Lab in Glasgow | Wellcome Sanger Institute for the COVID-19 Genomics UK (COG-UK) Consortium | Harper VanSteenhouse, Yumi Kasai, David Gray, Carol Clugston, Anna Dominiczak and Alex Alderton, Roberto Amato, Jeffrey Barrett, Sonia Goncalves, Ewan Harrison, David K. Jackson, Ian Johnston, Dominic Kwiatkowski, Cordelia Langford, John Sillitoe on behalf of the Wellcome Sanger Institute COVID-19 Surveillance Team |
| EPI_ISL_3109017, EPI_ISL_3109018, EPI_ISL_3109019, EPI_ISL_3109020 | Lighthouse Lab in Milton Keynes | Wellcome Sanger Institute for the COVID-19 Genomics UK (COG-UK) Consortium | The Lighthouse Lab in Milton Keynes and Alex Alderton, Roberto Amato, Jeffrey Barrett, Sonia Goncalves, Ewan Harrison, David K. Jackson, Ian Johnston, Dominic Kwiatkowski, Cordelia Langford, John Sillitoe on behalf of the Wellcome Sanger Institute COVID-19 Surveillance Team |
| EPI_ISL_3109021 | Lighthouse Lab in Glasgow | Wellcome Sanger Institute for the COVID-19 Genomics UK (COG-UK) Consortium | Harper VanSteenhouse, Yumi Kasai, David Gray, Carol Clugston, Anna Dominiczak and Alex Alderton, Roberto Amato, Jeffrey Barrett, Sonia Goncalves, Ewan Harrison, David K. Jackson, Ian Johnston, Dominic Kwiatkowski, Cordelia Langford, John Sillitoe on behalf of the Wellcome Sanger Institute COVID-19 Surveillance Team |
| EPI_ISL_3109022, EPI_ISL_3109023 | Lighthouse Lab in Milton Keynes | Wellcome Sanger Institute for the COVID-19 Genomics UK (COG-UK) Consortium | The Lighthouse Lab in Milton Keynes and Alex Alderton, Roberto Amato, Jeffrey Barrett, Sonia Goncalves, Ewan Harrison, David K. Jackson, Ian Johnston, Dominic Kwiatkowski, Cordelia Langford, John Sillitoe on behalf of the Wellcome Sanger Institute COVID-19 Surveillance Team |
| EPI_ISL_3109024 | Lighthouse Lab in Alderley Park | Wellcome Sanger Institute for the COVID-19 Genomics UK (COG-UK) Consortium | Jacquelyn Wynn, Mairead Hyland, The Lighthouse Lab in Alderley Park and Alex Alderton, Roberto Amato, Jeffrey Barrett, Sonia Goncalves, Ewan Harrison, David K. Jackson, Ian Johnston, Dominic Kwiatkowski, Cordelia Langford, John Sillitoe on behalf of the Wellcome Sanger Institute COVID-19 Surveillance Team |
| EPI_ISL_3109025 | Lighthouse Lab in Glasgow | Wellcome Sanger Institute for the COVID-19 Genomics UK (COG-UK) Consortium | Harper VanSteenhouse, Yumi Kasai, David Gray, Carol Clugston, Anna Dominiczak and Alex Alderton, Roberto Amato, Jeffrey Barrett, Sonia Goncalves, Ewan Harrison, David K. Jackson, Ian Johnston, Dominic Kwiatkowski, Cordelia Langford, John Sillitoe on behalf of the Wellcome Sanger Institute COVID-19 Surveillance Team |
| EPI_ISL_3109026, EPI_ISL_3109027, EPI_ISL_3109028 | Lighthouse Lab in Milton Keynes | Wellcome Sanger Institute for the COVID-19 Genomics UK (COG-UK) Consortium | The Lighthouse Lab in Milton Keynes and Alex Alderton, Roberto Amato, Jeffrey Barrett, Sonia Goncalves, Ewan Harrison, David K. Jackson, Ian Johnston, Dominic Kwiatkowski, Cordelia Langford, John Sillitoe on behalf of the Wellcome Sanger Institute COVID-19 Surveillance Team |
| EPI_ISL_3109029 | Lighthouse Lab in Alderley Park | Wellcome Sanger Institute for the COVID-19 Genomics UK (COG-UK) Consortium | Jacquelyn Wynn, Mairead Hyland, The Lighthouse Lab in Alderley Park and Alex Alderton, Roberto Amato, Jeffrey Barrett, Sonia Goncalves, Ewan Harrison, David K. Jackson, Ian Johnston, Dominic Kwiatkowski, Cordelia Langford, John Sillitoe on behalf of the Wellcome Sanger Institute COVID-19 Surveillance Team |
| EPI_ISL_3109030 | Lighthouse Lab in Milton Keynes | Wellcome Sanger Institute for the COVID-19 Genomics UK (COG-UK) Consortium | The Lighthouse Lab in Milton Keynes and Alex Alderton, Roberto Amato, Jeffrey Barrett, Sonia Goncalves, Ewan Harrison, David K. Jackson, Ian Johnston, Dominic Kwiatkowski, Cordelia Langford, John Sillitoe on behalf of the Wellcome Sanger Institute COVID-19 Surveillance Team |
| EPI_ISL_3109031 | Lighthouse Lab in Alderley Park | Wellcome Sanger Institute for the COVID-19 Genomics UK (COG-UK) Consortium | Jacquelyn Wynn, Mairead Hyland, The Lighthouse Lab in Alderley Park and Alex Alderton, Roberto Amato, Jeffrey Barrett, Sonia Goncalves, Ewan Harrison, David K. Jackson, Ian Johnston, Dominic Kwiatkowski, Cordelia Langford, John Sillitoe on behalf of the Wellcome Sanger Institute COVID-19 Surveillance Team |
| EPI_ISL_3109032, EPI_ISL_3109033, EPI_ISL_3109034, EPI_ISL_3109035 | Lighthouse Lab in Milton Keynes | Wellcome Sanger Institute for the COVID-19 Genomics UK (COG-UK) Consortium | The Lighthouse Lab in Milton Keynes and Alex Alderton, Roberto Amato, Jeffrey Barrett, Sonia Goncalves, Ewan Harrison, David K. Jackson, Ian Johnston, Dominic Kwiatkowski, Cordelia Langford, John Sillitoe on behalf of the Wellcome Sanger Institute COVID-19 Surveillance Team |
| EPI_ISL_3109036 | Lighthouse Lab in Glasgow | Wellcome Sanger Institute for the COVID-19 Genomics UK (COG-UK) Consortium | Harper VanSteenhouse, Yumi Kasai, David Gray, Carol Clugston, Anna Dominiczak and Alex Alderton, Roberto Amato, Jeffrey Barrett, Sonia Goncalves, Ewan Harrison, David K. Jackson, Ian Johnston, Dominic Kwiatkowski, Cordelia Langford, John Sillitoe on behalf of the Wellcome Sanger Institute COVID-19 Surveillance Team |
| EPI_ISL_3109037 | Lighthouse Lab in Alderley Park | Wellcome Sanger Institute for the COVID-19 Genomics UK (COG-UK) Consortium | Jacquelyn Wynn, Mairead Hyland, The Lighthouse Lab in Alderley Park and Alex Alderton, Roberto Amato, Jeffrey Barrett, Sonia Goncalves, Ewan Harrison, David K. Jackson, Ian Johnston, Dominic Kwiatkowski, Cordelia Langford, John Sillitoe on behalf of the Wellcome Sanger Institute COVID-19 Surveillance Team |
| EPI_ISL_3109038, EPI_ISL_3109039 | Lighthouse Lab in Glasgow | Wellcome Sanger Institute for the COVID-19 Genomics UK (COG-UK) Consortium | Harper VanSteenhouse, Yumi Kasai, David Gray, Carol Clugston, Anna Dominiczak and Alex Alderton, Roberto Amato, Jeffrey Barrett, Sonia Goncalves, Ewan Harrison, David K. Jackson, Ian Johnston, Dominic Kwiatkowski, Cordelia Langford, John Sillitoe on behalf of the Wellcome Sanger Institute COVID-19 Surveillance Team |

| | | | |
|---|---|---|---|
| EPI_ISL_3109040, EPI_ISL_3109041, EPI_ISL_3109042, EPI_ISL_3109043 | Lighthouse Lab in Alderley Park | Wellcome Sanger Institute for the COVID-19 Genomics UK (COG-UK) Consortium | Jacquelyn Wynn, Mairead Hyland, The Lighthouse Lab in Alderley Park and Alex Alderton, Roberto Amato, Jeffrey Barrett, Sonia Goncalves, Ewan Harrison, David K. Jackson, Ian Johnston, Dominic Kwiatkowski, Cordelia Langford, John Sillitoe on behalf of the Wellcome Sanger Institute COVID-19 Surveillance Team |
| EPI_ISL_3109044, EPI_ISL_3109045 | Lighthouse Lab in Milton Keynes | Wellcome Sanger Institute for the COVID-19 Genomics UK (COG-UK) Consortium | The Lighthouse Lab in Milton Keynes and Alex Alderton, Roberto Amato, Jeffrey Barrett, Sonia Goncalves, Ewan Harrison, David K. Jackson, Ian Johnston, Dominic Kwiatkowski, Cordelia Langford, John Sillitoe on behalf of the Wellcome Sanger Institute COVID-19 Surveillance Team |
| EPI_ISL_3109046 | Lighthouse Lab in Alderley Park | Wellcome Sanger Institute for the COVID-19 Genomics UK (COG-UK) Consortium | Jacquelyn Wynn, Mairead Hyland, The Lighthouse Lab in Alderley Park and Alex Alderton, Roberto Amato, Jeffrey Barrett, Sonia Goncalves, Ewan Harrison, David K. Jackson, Ian Johnston, Dominic Kwiatkowski, Cordelia Langford, John Sillitoe on behalf of the Wellcome Sanger Institute COVID-19 Surveillance Team |
| EPI_ISL_3109047 | Lighthouse Lab in Milton Keynes | Wellcome Sanger Institute for the COVID-19 Genomics UK (COG-UK) Consortium | The Lighthouse Lab in Milton Keynes and Alex Alderton, Roberto Amato, Jeffrey Barrett, Sonia Goncalves, Ewan Harrison, David K. Jackson, Ian Johnston, Dominic Kwiatkowski, Cordelia Langford, John Sillitoe on behalf of the Wellcome Sanger Institute COVID-19 Surveillance Team |
| EPI_ISL_3109048, EPI_ISL_3109049, EPI_ISL_3109050, EPI_ISL_3109051 | Lighthouse Lab in Alderley Park | Wellcome Sanger Institute for the COVID-19 Genomics UK (COG-UK) Consortium | Jacquelyn Wynn, Mairead Hyland, The Lighthouse Lab in Alderley Park and Alex Alderton, Roberto Amato, Jeffrey Barrett, Sonia Goncalves, Ewan Harrison, David K. Jackson, Ian Johnston, Dominic Kwiatkowski, Cordelia Langford, John Sillitoe on behalf of the Wellcome Sanger Institute COVID-19 Surveillance Team |
| EPI_ISL_3109052, EPI_ISL_3109053 | Lighthouse Lab in Milton Keynes | Wellcome Sanger Institute for the COVID-19 Genomics UK (COG-UK) Consortium | The Lighthouse Lab in Milton Keynes and Alex Alderton, Roberto Amato, Jeffrey Barrett, Sonia Goncalves, Ewan Harrison, David K. Jackson, Ian Johnston, Dominic Kwiatkowski, Cordelia Langford, John Sillitoe on behalf of the Wellcome Sanger Institute COVID-19 Surveillance Team |
| EPI_ISL_3109054 | Lighthouse Lab in Alderley Park | Wellcome Sanger Institute for the COVID-19 Genomics UK (COG-UK) Consortium | Jacquelyn Wynn, Mairead Hyland, The Lighthouse Lab in Alderley Park and Alex Alderton, Roberto Amato, Jeffrey Barrett, Sonia Goncalves, Ewan Harrison, David K. Jackson, Ian Johnston, Dominic Kwiatkowski, Cordelia Langford, John Sillitoe on behalf of the Wellcome Sanger Institute COVID-19 Surveillance Team |
| EPI_ISL_3109055 | Lighthouse Lab in Milton Keynes | Wellcome Sanger Institute for the COVID-19 Genomics UK (COG-UK) Consortium | The Lighthouse Lab in Milton Keynes and Alex Alderton, Roberto Amato, Jeffrey Barrett, Sonia Goncalves, Ewan Harrison, David K. Jackson, Ian Johnston, Dominic Kwiatkowski, Cordelia Langford, John Sillitoe on behalf of the Wellcome Sanger Institute COVID-19 Surveillance Team |
| EPI_ISL_3109056 | Lighthouse Lab in Glasgow | Wellcome Sanger Institute for the COVID-19 Genomics UK (COG-UK) Consortium | Harper VanSteenhouse, Yumi Kasai, David Gray, Carol Clugston, Anna Dominiczak and Alex Alderton, Roberto Amato, Jeffrey Barrett, Sonia Goncalves, Ewan Harrison, David K. Jackson, Ian Johnston, Dominic Kwiatkowski, Cordelia Langford, John Sillitoe on behalf of the Wellcome Sanger Institute COVID-19 Surveillance Team |
| EPI_ISL_3109057, EPI_ISL_3109058, EPI_ISL_3109059, EPI_ISL_3109060, EPI_ISL_3109061, EPI_ISL_3109062 | Lighthouse Lab in Milton Keynes | Wellcome Sanger Institute for the COVID-19 Genomics UK (COG-UK) Consortium | The Lighthouse Lab in Milton Keynes and Alex Alderton, Roberto Amato, Jeffrey Barrett, Sonia Goncalves, Ewan Harrison, David K. Jackson, Ian Johnston, Dominic Kwiatkowski, Cordelia Langford, John Sillitoe on behalf of the Wellcome Sanger Institute COVID-19 Surveillance Team |
| EPI_ISL_3109063 | Lighthouse Lab in Glasgow | Wellcome Sanger Institute for the COVID-19 Genomics UK (COG-UK) Consortium | Harper VanSteenhouse, Yumi Kasai, David Gray, Carol Clugston, Anna Dominiczak and Alex Alderton, Roberto Amato, Jeffrey Barrett, Sonia Goncalves, Ewan Harrison, David K. Jackson, Ian Johnston, Dominic Kwiatkowski, Cordelia Langford, John Sillitoe on behalf of the Wellcome Sanger Institute COVID-19 Surveillance Team |
| EPI_ISL_3109064, EPI_ISL_3109065 | Lighthouse Lab in Milton Keynes | Wellcome Sanger Institute for the COVID-19 Genomics UK (COG-UK) Consortium | The Lighthouse Lab in Milton Keynes and Alex Alderton, Roberto Amato, Jeffrey Barrett, Sonia Goncalves, Ewan Harrison, David K. Jackson, Ian Johnston, Dominic Kwiatkowski, Cordelia Langford, John Sillitoe on behalf of the Wellcome Sanger Institute COVID-19 Surveillance Team |
| EPI_ISL_3109066 | Lighthouse Lab in Glasgow | Wellcome Sanger Institute for the COVID-19 Genomics UK (COG-UK) Consortium | Harper VanSteenhouse, Yumi Kasai, David Gray, Carol Clugston, Anna Dominiczak and Alex Alderton, Roberto Amato, Jeffrey Barrett, Sonia Goncalves, Ewan Harrison, David K. Jackson, Ian Johnston, Dominic Kwiatkowski, Cordelia Langford, John Sillitoe on behalf of the Wellcome Sanger Institute COVID-19 Surveillance Team |
| EPI_ISL_3109067 | Lighthouse Lab in Alderley Park | Wellcome Sanger Institute for the COVID-19 Genomics UK (COG-UK) Consortium | Jacquelyn Wynn, Mairead Hyland, The Lighthouse Lab in Alderley Park and Alex Alderton, Roberto Amato, Jeffrey Barrett, Sonia Goncalves, Ewan Harrison, David K. Jackson, Ian Johnston, Dominic Kwiatkowski, Cordelia Langford, John Sillitoe on behalf of the Wellcome Sanger Institute COVID-19 Surveillance Team |
| EPI_ISL_3109068, EPI_ISL_3109069, EPI_ISL_3109070, EPI_ISL_3109071, EPI_ISL_3109072, EPI_ISL_3109073, EPI_ISL_3109074 | Lighthouse Lab in Milton Keynes | Wellcome Sanger Institute for the COVID-19 Genomics UK (COG-UK) Consortium | The Lighthouse Lab in Milton Keynes and Alex Alderton, Roberto Amato, Jeffrey Barrett, Sonia Goncalves, Ewan Harrison, David K. Jackson, Ian Johnston, Dominic Kwiatkowski, Cordelia Langford, John Sillitoe on behalf of the Wellcome Sanger Institute COVID-19 Surveillance Team |
| EPI_ISL_3109075, EPI_ISL_3109076, EPI_ISL_3109077, EPI_ISL_3109078, EPI_ISL_3109079 | Lighthouse Lab in Alderley Park | Wellcome Sanger Institute for the COVID-19 Genomics UK (COG-UK) Consortium | Jacquelyn Wynn, Mairead Hyland, The Lighthouse Lab in Alderley Park and Alex Alderton, Roberto Amato, Jeffrey Barrett, Sonia Goncalves, Ewan Harrison, David K. Jackson, Ian Johnston, Dominic Kwiatkowski, Cordelia Langford, John Sillitoe on behalf of the Wellcome Sanger Institute COVID-19 Surveillance Team |
| EPI_ISL_3109080, EPI_ISL_3109081, EPI_ISL_3109082, EPI_ISL_3109083 | Lighthouse Lab in Milton Keynes | Wellcome Sanger Institute for the COVID-19 Genomics UK (COG-UK) Consortium | The Lighthouse Lab in Milton Keynes and Alex Alderton, Roberto Amato, Jeffrey Barrett, Sonia Goncalves, Ewan Harrison, David K. Jackson, Ian Johnston, Dominic Kwiatkowski, Cordelia Langford, John Sillitoe on behalf of the Wellcome Sanger Institute COVID-19 Surveillance Team |
| EPI_ISL_3109084 | Lighthouse Lab in Alderley Park | Wellcome Sanger Institute for the COVID-19 Genomics UK (COG-UK) Consortium | Jacquelyn Wynn, Mairead Hyland, The Lighthouse Lab in Alderley Park and Alex Alderton, Roberto Amato, Jeffrey Barrett, Sonia Goncalves, Ewan Harrison, David K. Jackson, Ian Johnston, Dominic Kwiatkowski, Cordelia Langford, John Sillitoe on behalf of the Wellcome Sanger Institute COVID-19 Surveillance Team |
| EPI_ISL_3109085 | Lighthouse Lab in Milton Keynes | Wellcome Sanger Institute for the COVID-19 Genomics UK (COG-UK) Consortium | The Lighthouse Lab in Milton Keynes and Alex Alderton, Roberto Amato, Jeffrey Barrett, Sonia Goncalves, Ewan Harrison, David K. Jackson, Ian Johnston, Dominic Kwiatkowski, Cordelia Langford, John Sillitoe on behalf of the Wellcome Sanger Institute COVID-19 Surveillance Team |
| EPI_ISL_3109086 | Lighthouse Lab in Alderley Park | Wellcome Sanger Institute for the COVID-19 Genomics UK (COG-UK) Consortium | Jacquelyn Wynn, Mairead Hyland, The Lighthouse Lab in Alderley Park and Alex Alderton, Roberto Amato, Jeffrey Barrett, Sonia Goncalves, Ewan Harrison, David K. Jackson, Ian Johnston, Dominic Kwiatkowski, Cordelia Langford, John Sillitoe on behalf of the Wellcome Sanger Institute COVID-19 Surveillance Team |
| EPI_ISL_3109087 | Lighthouse Lab in Milton Keynes | Wellcome Sanger Institute for the COVID-19 Genomics UK (COG-UK) Consortium | The Lighthouse Lab in Milton Keynes and Alex Alderton, Roberto Amato, Jeffrey Barrett, Sonia Goncalves, Ewan Harrison, David K. Jackson, Ian Johnston, Dominic Kwiatkowski, Cordelia Langford, John Sillitoe on behalf of the Wellcome Sanger Institute COVID-19 Surveillance Team |
| EPI_ISL_3109088 | Lighthouse Lab in Alderley Park | Wellcome Sanger Institute for the COVID-19 Genomics UK (COG-UK) Consortium | Jacquelyn Wynn, Mairead Hyland, The Lighthouse Lab in Alderley Park and Alex Alderton, Roberto Amato, Jeffrey Barrett, Sonia Goncalves, Ewan Harrison, David K. Jackson, Ian Johnston, Dominic Kwiatkowski, Cordelia Langford, John Sillitoe on behalf of the Wellcome Sanger Institute COVID-19 Surveillance Team |
| EPI_ISL_3109089, EPI_ISL_3109090, EPI_ISL_3109091 | Lighthouse Lab in Milton Keynes | Wellcome Sanger Institute for the COVID-19 Genomics UK (COG-UK) Consortium | The Lighthouse Lab in Milton Keynes and Alex Alderton, Roberto Amato, Jeffrey Barrett, Sonia Goncalves, Ewan Harrison, David K. Jackson, Ian Johnston, Dominic Kwiatkowski, Cordelia Langford, John Sillitoe on behalf of the Wellcome Sanger Institute COVID-19 Surveillance Team |
| EPI_ISL_3109092 | Lighthouse Lab in Alderley Park | Wellcome Sanger Institute for the COVID-19 Genomics UK (COG-UK) Consortium | Jacquelyn Wynn, Mairead Hyland, The Lighthouse Lab in Alderley Park and Alex Alderton, Roberto Amato, Jeffrey Barrett, Sonia Goncalves, Ewan Harrison, David K. Jackson, Ian Johnston, Dominic Kwiatkowski, Cordelia Langford, John Sillitoe on behalf of the Wellcome Sanger Institute COVID-19 Surveillance Team |
| EPI_ISL_3109093 | Lighthouse Lab in Milton Keynes | Wellcome Sanger Institute for the COVID-19 Genomics UK (COG-UK) Consortium | The Lighthouse Lab in Milton Keynes and Alex Alderton, Roberto Amato, Jeffrey Barrett, Sonia Goncalves, Ewan Harrison, David K. Jackson, Ian Johnston, Dominic Kwiatkowski, Cordelia Langford, John Sillitoe on behalf of the Wellcome Sanger Institute COVID-19 Surveillance Team |
| EPI_ISL_3109094 | Lighthouse Lab in Alderley Park | Wellcome Sanger Institute for the COVID-19 Genomics UK (COG-UK) Consortium | Jacquelyn Wynn, Mairead Hyland, The Lighthouse Lab in Alderley Park and Alex Alderton, Roberto Amato, Jeffrey Barrett, Sonia Goncalves, Ewan Harrison, David K. Jackson, Ian Johnston, Dominic Kwiatkowski, Cordelia Langford, John Sillitoe on behalf of the Wellcome Sanger Institute COVID-19 Surveillance Team |
| EPI_ISL_3109095 | Lighthouse Lab in Milton Keynes | Wellcome Sanger Institute for the COVID-19 Genomics UK (COG-UK) Consortium | The Lighthouse Lab in Milton Keynes and Alex Alderton, Roberto Amato, Jeffrey Barrett, Sonia Goncalves, Ewan Harrison, David K. Jackson, Ian Johnston, Dominic Kwiatkowski, Cordelia Langford, John Sillitoe on behalf of the Wellcome Sanger Institute COVID-19 Surveillance Team |
| EPI_ISL_3109096 | Lighthouse Lab in Glasgow | Wellcome Sanger Institute for the COVID-19 Genomics UK (COG-UK) Consortium | Harper VanSteenhouse, Yumi Kasai, David Gray, Carol Clugston, Anna Dominiczak and Alex Alderton, Roberto Amato, Jeffrey Barrett, Sonia Goncalves, Ewan Harrison, David K. Jackson, Ian Johnston, Dominic Kwiatkowski, Cordelia Langford, John Sillitoe on behalf of the Wellcome Sanger Institute COVID-19 Surveillance Team |
| EPI_ISL_3109097, EPI_ISL_3109098, EPI_ISL_3109099, EPI_ISL_3109100 | Lighthouse Lab in Milton Keynes | Wellcome Sanger Institute for the COVID-19 Genomics UK (COG-UK) Consortium | The Lighthouse Lab in Milton Keynes and Alex Alderton, Roberto Amato, Jeffrey Barrett, Sonia Goncalves, Ewan Harrison, David K. Jackson, Ian Johnston, Dominic Kwiatkowski, Cordelia Langford, John Sillitoe on behalf of the Wellcome Sanger Institute COVID-19 Surveillance Team |
| EPI_ISL_3109101 | Lighthouse Lab in Alderley Park | Wellcome Sanger Institute for the COVID-19 Genomics UK (COG-UK) Consortium | Jacquelyn Wynn, Mairead Hyland, The Lighthouse Lab in Alderley Park and Alex Alderton, Roberto Amato, Jeffrey Barrett, Sonia Goncalves, Ewan Harrison, David K. Jackson, Ian Johnston, Dominic Kwiatkowski, Cordelia Langford, John Sillitoe on behalf of the Wellcome Sanger Institute COVID-19 Surveillance Team |
| EPI_ISL_3109102 | Lighthouse Lab in Milton Keynes | Wellcome Sanger Institute for the COVID-19 Genomics UK (COG-UK) Consortium | The Lighthouse Lab in Milton Keynes and Alex Alderton, Roberto Amato, Jeffrey Barrett, Sonia Goncalves, Ewan Harrison, David K. Jackson, Ian Johnston, Dominic Kwiatkowski, Cordelia Langford, John Sillitoe on behalf of the Wellcome Sanger Institute COVID-19 Surveillance Team |
| EPI_ISL_3109103 | Lighthouse Lab in Alderley Park | Wellcome Sanger Institute for the COVID-19 Genomics UK (COG-UK) Consortium | Jacquelyn Wynn, Mairead Hyland, The Lighthouse Lab in Alderley Park and Alex Alderton, Roberto Amato, Jeffrey Barrett, Sonia Goncalves, Ewan Harrison, David K. Jackson, Ian Johnston, Dominic Kwiatkowski, Cordelia Langford, John Sillitoe on behalf of the Wellcome Sanger Institute COVID-19 Surveillance Team |
| EPI_ISL_3109104, EPI_ISL_3109105 | Lighthouse Lab in Milton Keynes | Wellcome Sanger Institute for the COVID-19 Genomics UK (COG-UK) Consortium | The Lighthouse Lab in Milton Keynes and Alex Alderton, Roberto Amato, Jeffrey Barrett, Sonia Goncalves, Ewan Harrison, David K. Jackson, Ian Johnston, Dominic Kwiatkowski, Cordelia Langford, John Sillitoe on behalf of the Wellcome Sanger Institute COVID-19 Surveillance Team |
| EPI_ISL_3109106 | Lighthouse Lab in Alderley Park | Wellcome Sanger Institute for the COVID-19 Genomics UK (COG-UK) Consortium | Jacquelyn Wynn, Mairead Hyland, The Lighthouse Lab in Alderley Park and Alex Alderton, Roberto Amato, Jeffrey Barrett, Sonia Goncalves, Ewan Harrison, David K. Jackson, Ian Johnston, Dominic Kwiatkowski, Cordelia Langford, John Sillitoe on behalf of the Wellcome Sanger Institute COVID-19 Surveillance Team |
| EPI_ISL_3109107 | Lighthouse Lab in Milton Keynes | Wellcome Sanger Institute for the COVID-19 Genomics UK (COG-UK) Consortium | The Lighthouse Lab in Milton Keynes and Alex Alderton, Roberto Amato, Jeffrey Barrett, Sonia Goncalves, Ewan Harrison, David K. Jackson, Ian Johnston, Dominic Kwiatkowski, Cordelia Langford, John Sillitoe on behalf of the Wellcome Sanger Institute COVID-19 Surveillance Team |
| EPI_ISL_3109108 | Lighthouse Lab in Alderley Park | Wellcome Sanger Institute for the COVID-19 Genomics UK (COG-UK) Consortium | Jacquelyn Wynn, Mairead Hyland, The Lighthouse Lab in Alderley Park and Alex Alderton, Roberto Amato, Jeffrey Barrett, Sonia Goncalves, Ewan Harrison, David K. Jackson, Ian Johnston, Dominic Kwiatkowski, Cordelia Langford, John Sillitoe on behalf of the Wellcome Sanger Institute COVID-19 Surveillance Team |

| | | | |
|---|---|---|---|
| EPI_ISL_3109109 | Lighthouse Lab in Milton Keynes | Wellcome Sanger Institute for the COVID-19 Genomics UK (COG-UK) Consortium | The Lighthouse Lab in Milton Keynes and Alex Alderton, Roberto Amato, Jeffrey Barrett, Sonia Goncalves, Ewan Harrison, David K. Jackson, Ian Johnston, Dominic Kwiatkowski, Cordelia Langford, John Sillitoe on behalf of the Wellcome Sanger Institute COVID-19 Surveillance Team |
| EPI_ISL_3109110, EPI_ISL_3109111 | Lighthouse Lab in Alderley Park | Wellcome Sanger Institute for the COVID-19 Genomics UK (COG-UK) Consortium | Jacquelyn Wynn, Mairead Hyland, The Lighthouse Lab in Alderley Park and Alex Alderton, Roberto Amato, Jeffrey Barrett, Sonia Goncalves, Ewan Harrison, David K. Jackson, Ian Johnston, Dominic Kwiatkowski, Cordelia Langford, John Sillitoe on behalf of the Wellcome Sanger Institute COVID-19 Surveillance Team |
| EPI_ISL_3109112, EPI_ISL_3109113 | Lighthouse Lab in Milton Keynes | Wellcome Sanger Institute for the COVID-19 Genomics UK (COG-UK) Consortium | The Lighthouse Lab in Milton Keynes and Alex Alderton, Roberto Amato, Jeffrey Barrett, Sonia Goncalves, Ewan Harrison, David K. Jackson, Ian Johnston, Dominic Kwiatkowski, Cordelia Langford, John Sillitoe on behalf of the Wellcome Sanger Institute COVID-19 Surveillance Team |
| EPI_ISL_3109114 | Lighthouse Lab in Alderley Park | Wellcome Sanger Institute for the COVID-19 Genomics UK (COG-UK) Consortium | Jacquelyn Wynn, Mairead Hyland, The Lighthouse Lab in Alderley Park and Alex Alderton, Roberto Amato, Jeffrey Barrett, Sonia Goncalves, Ewan Harrison, David K. Jackson, Ian Johnston, Dominic Kwiatkowski, Cordelia Langford, John Sillitoe on behalf of the Wellcome Sanger Institute COVID-19 Surveillance Team |
| EPI_ISL_3109115, EPI_ISL_3109116 | Lighthouse Lab in Milton Keynes | Wellcome Sanger Institute for the COVID-19 Genomics UK (COG-UK) Consortium | The Lighthouse Lab in Milton Keynes and Alex Alderton, Roberto Amato, Jeffrey Barrett, Sonia Goncalves, Ewan Harrison, David K. Jackson, Ian Johnston, Dominic Kwiatkowski, Cordelia Langford, John Sillitoe on behalf of the Wellcome Sanger Institute COVID-19 Surveillance Team |
| EPI_ISL_3109117, EPI_ISL_3109118 | Lighthouse Lab in Alderley Park | Wellcome Sanger Institute for the COVID-19 Genomics UK (COG-UK) Consortium | Jacquelyn Wynn, Mairead Hyland, The Lighthouse Lab in Alderley Park and Alex Alderton, Roberto Amato, Jeffrey Barrett, Sonia Goncalves, Ewan Harrison, David K. Jackson, Ian Johnston, Dominic Kwiatkowski, Cordelia Langford, John Sillitoe on behalf of the Wellcome Sanger Institute COVID-19 Surveillance Team |
| EPI_ISL_3109119 | Lighthouse Lab in Glasgow | Wellcome Sanger Institute for the COVID-19 Genomics UK (COG-UK) Consortium | Harper VanSteenhouse, Yumi Kasai, David Gray, Carol Clugston, Anna Dominiczak and Alex Alderton, Roberto Amato, Jeffrey Barrett, Sonia Goncalves, Ewan Harrison, David K. Jackson, Ian Johnston, Dominic Kwiatkowski, Cordelia Langford, John Sillitoe on behalf of the Wellcome Sanger Institute COVID-19 Surveillance Team |
| EPI_ISL_3109120, EPI_ISL_3109121, EPI_ISL_3109122, EPI_ISL_3109123, EPI_ISL_3109124 | Lighthouse Lab in Milton Keynes | Wellcome Sanger Institute for the COVID-19 Genomics UK (COG-UK) Consortium | The Lighthouse Lab in Milton Keynes and Alex Alderton, Roberto Amato, Jeffrey Barrett, Sonia Goncalves, Ewan Harrison, David K. Jackson, Ian Johnston, Dominic Kwiatkowski, Cordelia Langford, John Sillitoe on behalf of the Wellcome Sanger Institute COVID-19 Surveillance Team |
| EPI_ISL_3109125, EPI_ISL_3109126, EPI_ISL_3109127, EPI_ISL_3109128 | Lighthouse Lab in Alderley Park | Wellcome Sanger Institute for the COVID-19 Genomics UK (COG-UK) Consortium | Jacquelyn Wynn, Mairead Hyland, The Lighthouse Lab in Alderley Park and Alex Alderton, Roberto Amato, Jeffrey Barrett, Sonia Goncalves, Ewan Harrison, David K. Jackson, Ian Johnston, Dominic Kwiatkowski, Cordelia Langford, John Sillitoe on behalf of the Wellcome Sanger Institute COVID-19 Surveillance Team |
| EPI_ISL_3109129 | Berkshire and Surrey Pathology Services Lighthouse Laboratory | Wellcome Sanger Institute for the COVID-19 Genomics UK (COG-UK) Consortium | Berkshire and Surrey Pathology Services Lighthouse Laboratory and Alex Alderton, Roberto Amato, Jeffrey Barrett, Sonia Goncalves, Ewan Harrison, David K. Jackson, Ian Johnston, Dominic Kwiatkowski, Cordelia Langford, John Sillitoe on behalf of the Wellcome Sanger Institute COVID-19 Surveillance Team |
| EPI_ISL_3109130 | Lighthouse Lab in Glasgow | Wellcome Sanger Institute for the COVID-19 Genomics UK (COG-UK) Consortium | Harper VanSteenhouse, Yumi Kasai, David Gray, Carol Clugston, Anna Dominiczak and Alex Alderton, Roberto Amato, Jeffrey Barrett, Sonia Goncalves, Ewan Harrison, David K. Jackson, Ian Johnston, Dominic Kwiatkowski, Cordelia Langford, John Sillitoe on behalf of the Wellcome Sanger Institute COVID-19 Surveillance Team |
| EPI_ISL_3109131 | Berkshire and Surrey Pathology Services Lighthouse Laboratory | Wellcome Sanger Institute for the COVID-19 Genomics UK (COG-UK) Consortium | Berkshire and Surrey Pathology Services Lighthouse Laboratory and Alex Alderton, Roberto Amato, Jeffrey Barrett, Sonia Goncalves, Ewan Harrison, David K. Jackson, Ian Johnston, Dominic Kwiatkowski, Cordelia Langford, John Sillitoe on behalf of the Wellcome Sanger Institute COVID-19 Surveillance Team |
| EPI_ISL_3109132 | Lighthouse Lab in Glasgow | Wellcome Sanger Institute for the COVID-19 Genomics UK (COG-UK) Consortium | Harper VanSteenhouse, Yumi Kasai, David Gray, Carol Clugston, Anna Dominiczak and Alex Alderton, Roberto Amato, Jeffrey Barrett, Sonia Goncalves, Ewan Harrison, David K. Jackson, Ian Johnston, Dominic Kwiatkowski, Cordelia Langford, John Sillitoe on behalf of the Wellcome Sanger Institute COVID-19 Surveillance Team |
| EPI_ISL_3109133 | Berkshire and Surrey Pathology Services Lighthouse Laboratory | Wellcome Sanger Institute for the COVID-19 Genomics UK (COG-UK) Consortium | Berkshire and Surrey Pathology Services Lighthouse Laboratory and Alex Alderton, Roberto Amato, Jeffrey Barrett, Sonia Goncalves, Ewan Harrison, David K. Jackson, Ian Johnston, Dominic Kwiatkowski, Cordelia Langford, John Sillitoe on behalf of the Wellcome Sanger Institute COVID-19 Surveillance Team |
| EPI_ISL_3109134, EPI_ISL_3109135, EPI_ISL_3109136 | Lighthouse Lab in Alderley Park | Wellcome Sanger Institute for the COVID-19 Genomics UK (COG-UK) Consortium | Jacquelyn Wynn, Mairead Hyland, The Lighthouse Lab in Alderley Park and Alex Alderton, Roberto Amato, Jeffrey Barrett, Sonia Goncalves, Ewan Harrison, David K. Jackson, Ian Johnston, Dominic Kwiatkowski, Cordelia Langford, John Sillitoe on behalf of the Wellcome Sanger Institute COVID-19 Surveillance Team |
| EPI_ISL_3109137, EPI_ISL_3109138, EPI_ISL_3109139 | Berkshire and Surrey Pathology Services Lighthouse Laboratory | Wellcome Sanger Institute for the COVID-19 Genomics UK (COG-UK) Consortium | Berkshire and Surrey Pathology Services Lighthouse Laboratory and Alex Alderton, Roberto Amato, Jeffrey Barrett, Sonia Goncalves, Ewan Harrison, David K. Jackson, Ian Johnston, Dominic Kwiatkowski, Cordelia Langford, John Sillitoe on behalf of the Wellcome Sanger Institute COVID-19 Surveillance Team |
| EPI_ISL_3109140, EPI_ISL_3109141 | Lighthouse Lab in Alderley Park | Wellcome Sanger Institute for the COVID-19 Genomics UK (COG-UK) Consortium | Jacquelyn Wynn, Mairead Hyland, The Lighthouse Lab in Alderley Park and Alex Alderton, Roberto Amato, Jeffrey Barrett, Sonia Goncalves, Ewan Harrison, David K. Jackson, Ian Johnston, Dominic Kwiatkowski, Cordelia Langford, John Sillitoe on behalf of the Wellcome Sanger Institute COVID-19 Surveillance Team |
| EPI_ISL_3109142, EPI_ISL_3109143 | Berkshire and Surrey Pathology Services Lighthouse Laboratory | Wellcome Sanger Institute for the COVID-19 Genomics UK (COG-UK) Consortium | Berkshire and Surrey Pathology Services Lighthouse Laboratory and Alex Alderton, Roberto Amato, Jeffrey Barrett, Sonia Goncalves, Ewan Harrison, David K. Jackson, Ian Johnston, Dominic Kwiatkowski, Cordelia Langford, John Sillitoe on behalf of the Wellcome Sanger Institute COVID-19 Surveillance Team |
| EPI_ISL_3109144, EPI_ISL_3109145, EPI_ISL_3109146, EPI_ISL_3109147, EPI_ISL_3109148, EPI_ISL_3109149 | Lighthouse Lab in Alderley Park | Wellcome Sanger Institute for the COVID-19 Genomics UK (COG-UK) Consortium | Jacquelyn Wynn, Mairead Hyland, The Lighthouse Lab in Alderley Park and Alex Alderton, Roberto Amato, Jeffrey Barrett, Sonia Goncalves, Ewan Harrison, David K. Jackson, Ian Johnston, Dominic Kwiatkowski, Cordelia Langford, John Sillitoe on behalf of the Wellcome Sanger Institute COVID-19 Surveillance Team |
| EPI_ISL_3109150 | Berkshire and Surrey Pathology Services Lighthouse Laboratory | Wellcome Sanger Institute for the COVID-19 Genomics UK (COG-UK) Consortium | Berkshire and Surrey Pathology Services Lighthouse Laboratory and Alex Alderton, Roberto Amato, Jeffrey Barrett, Sonia Goncalves, Ewan Harrison, David K. Jackson, Ian Johnston, Dominic Kwiatkowski, Cordelia Langford, John Sillitoe on behalf of the Wellcome Sanger Institute COVID-19 Surveillance Team |
| EPI_ISL_3109151 | Lighthouse Lab in Alderley Park | Wellcome Sanger Institute for the COVID-19 Genomics UK (COG-UK) Consortium | Jacquelyn Wynn, Mairead Hyland, The Lighthouse Lab in Alderley Park and Alex Alderton, Roberto Amato, Jeffrey Barrett, Sonia Goncalves, Ewan Harrison, David K. Jackson, Ian Johnston, Dominic Kwiatkowski, Cordelia Langford, John Sillitoe on behalf of the Wellcome Sanger Institute COVID-19 Surveillance Team |
| EPI_ISL_3109152 | Berkshire and Surrey Pathology Services Lighthouse Laboratory | Wellcome Sanger Institute for the COVID-19 Genomics UK (COG-UK) Consortium | Berkshire and Surrey Pathology Services Lighthouse Laboratory and Alex Alderton, Roberto Amato, Jeffrey Barrett, Sonia Goncalves, Ewan Harrison, David K. Jackson, Ian Johnston, Dominic Kwiatkowski, Cordelia Langford, John Sillitoe on behalf of the Wellcome Sanger Institute COVID-19 Surveillance Team |
| EPI_ISL_3109153 | Lighthouse Lab in Alderley Park | Wellcome Sanger Institute for the COVID-19 Genomics UK (COG-UK) Consortium | Jacquelyn Wynn, Mairead Hyland, The Lighthouse Lab in Alderley Park and Alex Alderton, Roberto Amato, Jeffrey Barrett, Sonia Goncalves, Ewan Harrison, David K. Jackson, Ian Johnston, Dominic Kwiatkowski, Cordelia Langford, John Sillitoe on behalf of the Wellcome Sanger Institute COVID-19 Surveillance Team |
| EPI_ISL_3109154, EPI_ISL_3109155 | Berkshire and Surrey Pathology Services Lighthouse Laboratory | Wellcome Sanger Institute for the COVID-19 Genomics UK (COG-UK) Consortium | Berkshire and Surrey Pathology Services Lighthouse Laboratory and Alex Alderton, Roberto Amato, Jeffrey Barrett, Sonia Goncalves, Ewan Harrison, David K. Jackson, Ian Johnston, Dominic Kwiatkowski, Cordelia Langford, John Sillitoe on behalf of the Wellcome Sanger Institute COVID-19 Surveillance Team |
| EPI_ISL_3109156, EPI_ISL_3109157, EPI_ISL_3109158, EPI_ISL_3109159, EPI_ISL_3109160 | Lighthouse Lab in Alderley Park | Wellcome Sanger Institute for the COVID-19 Genomics UK (COG-UK) Consortium | Jacquelyn Wynn, Mairead Hyland, The Lighthouse Lab in Alderley Park and Alex Alderton, Roberto Amato, Jeffrey Barrett, Sonia Goncalves, Ewan Harrison, David K. Jackson, Ian Johnston, Dominic Kwiatkowski, Cordelia Langford, John Sillitoe on behalf of the Wellcome Sanger Institute COVID-19 Surveillance Team |
| EPI_ISL_3109161 | Berkshire and Surrey Pathology Services Lighthouse Laboratory | Wellcome Sanger Institute for the COVID-19 Genomics UK (COG-UK) Consortium | Berkshire and Surrey Pathology Services Lighthouse Laboratory and Alex Alderton, Roberto Amato, Jeffrey Barrett, Sonia Goncalves, Ewan Harrison, David K. Jackson, Ian Johnston, Dominic Kwiatkowski, Cordelia Langford, John Sillitoe on behalf of the Wellcome Sanger Institute COVID-19 Surveillance Team |
| EPI_ISL_3109162 | Lighthouse Lab in Alderley Park | Wellcome Sanger Institute for the COVID-19 Genomics UK (COG-UK) Consortium | Jacquelyn Wynn, Mairead Hyland, The Lighthouse Lab in Alderley Park and Alex Alderton, Roberto Amato, Jeffrey Barrett, Sonia Goncalves, Ewan Harrison, David K. Jackson, Ian Johnston, Dominic Kwiatkowski, Cordelia Langford, John Sillitoe on behalf of the Wellcome Sanger Institute COVID-19 Surveillance Team |
| EPI_ISL_3109163, EPI_ISL_3109164 | Berkshire and Surrey Pathology Services Lighthouse Laboratory | Wellcome Sanger Institute for the COVID-19 Genomics UK (COG-UK) Consortium | Berkshire and Surrey Pathology Services Lighthouse Laboratory and Alex Alderton, Roberto Amato, Jeffrey Barrett, Sonia Goncalves, Ewan Harrison, David K. Jackson, Ian Johnston, Dominic Kwiatkowski, Cordelia Langford, John Sillitoe on behalf of the Wellcome Sanger Institute COVID-19 Surveillance Team |
| EPI_ISL_3109165, EPI_ISL_3109166 | Lighthouse Lab in Alderley Park | Wellcome Sanger Institute for the COVID-19 Genomics UK (COG-UK) Consortium | Jacquelyn Wynn, Mairead Hyland, The Lighthouse Lab in Alderley Park and Alex Alderton, Roberto Amato, Jeffrey Barrett, Sonia Goncalves, Ewan Harrison, David K. Jackson, Ian Johnston, Dominic Kwiatkowski, Cordelia Langford, John Sillitoe on behalf of the Wellcome Sanger Institute COVID-19 Surveillance Team |
| EPI_ISL_3109167, EPI_ISL_3109168 | Berkshire and Surrey Pathology Services Lighthouse Laboratory | Wellcome Sanger Institute for the COVID-19 Genomics UK (COG-UK) Consortium | Berkshire and Surrey Pathology Services Lighthouse Laboratory and Alex Alderton, Roberto Amato, Jeffrey Barrett, Sonia Goncalves, Ewan Harrison, David K. Jackson, Ian Johnston, Dominic Kwiatkowski, Cordelia Langford, John Sillitoe on behalf of the Wellcome Sanger Institute COVID-19 Surveillance Team |
| EPI_ISL_3109169, EPI_ISL_3109170 | Lighthouse Lab in Alderley Park | Wellcome Sanger Institute for the COVID-19 Genomics UK (COG-UK) Consortium | Jacquelyn Wynn, Mairead Hyland, The Lighthouse Lab in Alderley Park and Alex Alderton, Roberto Amato, Jeffrey Barrett, Sonia Goncalves, Ewan Harrison, David K. Jackson, Ian Johnston, Dominic Kwiatkowski, Cordelia Langford, John Sillitoe on behalf of the Wellcome Sanger Institute COVID-19 Surveillance Team |
| EPI_ISL_3109171 | Berkshire and Surrey Pathology Services Lighthouse Laboratory | Wellcome Sanger Institute for the COVID-19 Genomics UK (COG-UK) Consortium | Berkshire and Surrey Pathology Services Lighthouse Laboratory and Alex Alderton, Roberto Amato, Jeffrey Barrett, Sonia Goncalves, Ewan Harrison, David K. Jackson, Ian Johnston, Dominic Kwiatkowski, Cordelia Langford, John Sillitoe on behalf of the Wellcome Sanger Institute COVID-19 Surveillance Team |
| EPI_ISL_3109172 | Lighthouse Lab in Glasgow | Wellcome Sanger Institute for the COVID-19 Genomics UK (COG-UK) Consortium | Harper VanSteenhouse, Yumi Kasai, David Gray, Carol Clugston, Anna Dominiczak and Alex Alderton, Roberto Amato, Jeffrey Barrett, Sonia Goncalves, Ewan Harrison, David K. Jackson, Ian Johnston, Dominic Kwiatkowski, Cordelia Langford, John Sillitoe on behalf of the Wellcome Sanger Institute COVID-19 Surveillance Team |
| EPI_ISL_3109173 | Lighthouse Lab in Alderley Park | Wellcome Sanger Institute for the COVID-19 Genomics UK (COG-UK) Consortium | Jacquelyn Wynn, Mairead Hyland, The Lighthouse Lab in Alderley Park and Alex Alderton, Roberto Amato, Jeffrey Barrett, Sonia Goncalves, Ewan Harrison, David K. Jackson, Ian Johnston, Dominic Kwiatkowski, Cordelia Langford, John Sillitoe on behalf of the Wellcome Sanger Institute COVID-19 Surveillance Team |
| EPI_ISL_3109174 | Lighthouse Lab in Glasgow | Wellcome Sanger Institute for the COVID-19 Genomics UK (COG-UK) Consortium | Harper VanSteenhouse, Yumi Kasai, David Gray, Carol Clugston, Anna Dominiczak and Alex Alderton, Roberto Amato, Jeffrey Barrett, Sonia Goncalves, Ewan Harrison, David K. Jackson, Ian Johnston, Dominic Kwiatkowski, Cordelia Langford, John Sillitoe on behalf of the Wellcome Sanger Institute COVID-19 Surveillance Team |
| EPI_ISL_3109175, EPI_ISL_3109176 | Lighthouse Lab in Alderley Park | Wellcome Sanger Institute for the COVID-19 Genomics UK (COG-UK) Consortium | Jacquelyn Wynn, Mairead Hyland, The Lighthouse Lab in Alderley Park and Alex Alderton, Roberto Amato, Jeffrey Barrett, Sonia Goncalves, Ewan Harrison, David K. Jackson, Ian Johnston, Dominic Kwiatkowski, Cordelia Langford, John Sillitoe on behalf of the Wellcome Sanger Institute COVID-19 Surveillance Team |
| EPI_ISL_3109177, EPI_ISL_3109178 | Lighthouse Lab in Glasgow | Wellcome Sanger Institute for the COVID-19 Genomics UK (COG-UK) Consortium | Harper VanSteenhouse, Yumi Kasai, David Gray, Carol Clugston, Anna Dominiczak and Alex Alderton, Roberto Amato, Jeffrey Barrett, Sonia Goncalves, Ewan Harrison, David K. Jackson, Ian Johnston, Dominic Kwiatkowski, Cordelia Langford, John Sillitoe on behalf of the Wellcome Sanger Institute COVID-19 Surveillance Team |
| EPI_ISL_3109179, EPI_ISL_3109180 | Lighthouse Lab in Alderley Park | Wellcome Sanger Institute for the COVID-19 Genomics UK (COG-UK) Consortium | Jacquelyn Wynn, Mairead Hyland, The Lighthouse Lab in Alderley Park and Alex Alderton, Roberto Amato, Jeffrey Barrett, Sonia Goncalves, Ewan Harrison, David K. Jackson, Ian Johnston, Dominic Kwiatkowski, Cordelia Langford, John Sillitoe on behalf of the Wellcome Sanger Institute COVID-19 Surveillance Team |
| EPI_ISL_3109181, EPI_ISL_3109182 | Berkshire and Surrey Pathology Services Lighthouse Laboratory | Wellcome Sanger Institute for the COVID-19 Genomics UK (COG-UK) Consortium | Berkshire and Surrey Pathology Services Lighthouse Laboratory and Alex Alderton, Roberto Amato, Jeffrey Barrett, Sonia Goncalves, Ewan Harrison, David K. Jackson, Ian Johnston, Dominic Kwiatkowski, Cordelia Langford, John Sillitoe on behalf of the Wellcome Sanger Institute COVID-19 Surveillance Team |

| | | | |
|---|---|---|---|
| EPI_ISL_3109183 | Lighthouse Lab in Glasgow | Wellcome Sanger Institute for the COVID-19 Genomics UK (COG-UK) Consortium | Harper VanSteenhouse, Yumi Kasai, David Gray, Carol Clugston, Anna Dominiczak and Alex Alderton, Roberto Amato, Jeffrey Barrett, Sonia Goncalves, Ewan Harrison, David K. Jackson, Ian Johnston, Dominic Kwiatkowski, Cordelia Langford, John Sillitoe on behalf of the Wellcome Sanger Institute COVID-19 Surveillance Team |
| EPI_ISL_3109184 | Lighthouse Lab in Alderley Park | Wellcome Sanger Institute for the COVID-19 Genomics UK (COG-UK) Consortium | Jacquelyn Wynn, Mairead Hyland, The Lighthouse Lab in Alderley Park and Alex Alderton, Roberto Amato, Jeffrey Barrett, Sonia Goncalves, Ewan Harrison, David K. Jackson, Ian Johnston, Dominic Kwiatkowski, Cordelia Langford, John Sillitoe on behalf of the Wellcome Sanger Institute COVID-19 Surveillance Team |
| EPI_ISL_3109185 | Lighthouse Lab in Glasgow | Wellcome Sanger Institute for the COVID-19 Genomics UK (COG-UK) Consortium | Harper VanSteenhouse, Yumi Kasai, David Gray, Carol Clugston, Anna Dominiczak and Alex Alderton, Roberto Amato, Jeffrey Barrett, Sonia Goncalves, Ewan Harrison, David K. Jackson, Ian Johnston, Dominic Kwiatkowski, Cordelia Langford, John Sillitoe on behalf of the Wellcome Sanger Institute COVID-19 Surveillance Team |
| EPI_ISL_3109186 | Lighthouse Lab in Alderley Park | Wellcome Sanger Institute for the COVID-19 Genomics UK (COG-UK) Consortium | Jacquelyn Wynn, Mairead Hyland, The Lighthouse Lab in Alderley Park and Alex Alderton, Roberto Amato, Jeffrey Barrett, Sonia Goncalves, Ewan Harrison, David K. Jackson, Ian Johnston, Dominic Kwiatkowski, Cordelia Langford, John Sillitoe on behalf of the Wellcome Sanger Institute COVID-19 Surveillance Team |
| EPI_ISL_3109187 | Lighthouse Lab in Glasgow | Wellcome Sanger Institute for the COVID-19 Genomics UK (COG-UK) Consortium | Harper VanSteenhouse, Yumi Kasai, David Gray, Carol Clugston, Anna Dominiczak and Alex Alderton, Roberto Amato, Jeffrey Barrett, Sonia Goncalves, Ewan Harrison, David K. Jackson, Ian Johnston, Dominic Kwiatkowski, Cordelia Langford, John Sillitoe on behalf of the Wellcome Sanger Institute COVID-19 Surveillance Team |
| EPI_ISL_3109188, EPI_ISL_3109189, EPI_ISL_3109190, EPI_ISL_3109191, EPI_ISL_3109192 | Lighthouse Lab in Alderley Park | Wellcome Sanger Institute for the COVID-19 Genomics UK (COG-UK) Consortium | Jacquelyn Wynn, Mairead Hyland, The Lighthouse Lab in Alderley Park and Alex Alderton, Roberto Amato, Jeffrey Barrett, Sonia Goncalves, Ewan Harrison, David K. Jackson, Ian Johnston, Dominic Kwiatkowski, Cordelia Langford, John Sillitoe on behalf of the Wellcome Sanger Institute COVID-19 Surveillance Team |
| EPI_ISL_3109193 | Berkshire and Surrey Pathology Services Lighthouse Laboratory | Wellcome Sanger Institute for the COVID-19 Genomics UK (COG-UK) Consortium | Berkshire and Surrey Pathology Services Lighthouse Laboratory and Alex Alderton, Roberto Amato, Jeffrey Barrett, Sonia Goncalves, Ewan Harrison, David K. Jackson, Ian Johnston, Dominic Kwiatkowski, Cordelia Langford, John Sillitoe on behalf of the Wellcome Sanger Institute COVID-19 Surveillance Team |
| EPI_ISL_3109194 | Lighthouse Lab in Alderley Park | Wellcome Sanger Institute for the COVID-19 Genomics UK (COG-UK) Consortium | Jacquelyn Wynn, Mairead Hyland, The Lighthouse Lab in Alderley Park and Alex Alderton, Roberto Amato, Jeffrey Barrett, Sonia Goncalves, Ewan Harrison, David K. Jackson, Ian Johnston, Dominic Kwiatkowski, Cordelia Langford, John Sillitoe on behalf of the Wellcome Sanger Institute COVID-19 Surveillance Team |
| EPI_ISL_3109195 | Berkshire and Surrey Pathology Services Lighthouse Laboratory | Wellcome Sanger Institute for the COVID-19 Genomics UK (COG-UK) Consortium | Berkshire and Surrey Pathology Services Lighthouse Laboratory and Alex Alderton, Roberto Amato, Jeffrey Barrett, Sonia Goncalves, Ewan Harrison, David K. Jackson, Ian Johnston, Dominic Kwiatkowski, Cordelia Langford, John Sillitoe on behalf of the Wellcome Sanger Institute COVID-19 Surveillance Team |
| EPI_ISL_3109196, EPI_ISL_3109197, EPI_ISL_3109198, EPI_ISL_3109199 | Lighthouse Lab in Alderley Park | Wellcome Sanger Institute for the COVID-19 Genomics UK (COG-UK) Consortium | Jacquelyn Wynn, Mairead Hyland, The Lighthouse Lab in Alderley Park and Alex Alderton, Roberto Amato, Jeffrey Barrett, Sonia Goncalves, Ewan Harrison, David K. Jackson, Ian Johnston, Dominic Kwiatkowski, Cordelia Langford, John Sillitoe on behalf of the Wellcome Sanger Institute COVID-19 Surveillance Team |
| EPI_ISL_3109200 | Lighthouse Lab in Glasgow | Wellcome Sanger Institute for the COVID-19 Genomics UK (COG-UK) Consortium | Harper VanSteenhouse, Yumi Kasai, David Gray, Carol Clugston, Anna Dominiczak and Alex Alderton, Roberto Amato, Jeffrey Barrett, Sonia Goncalves, Ewan Harrison, David K. Jackson, Ian Johnston, Dominic Kwiatkowski, Cordelia Langford, John Sillitoe on behalf of the Wellcome Sanger Institute COVID-19 Surveillance Team |
| EPI_ISL_3109201, EPI_ISL_3109202, EPI_ISL_3109203, EPI_ISL_3109204, EPI_ISL_3109205 | Lighthouse Lab in Alderley Park | Wellcome Sanger Institute for the COVID-19 Genomics UK (COG-UK) Consortium | Jacquelyn Wynn, Mairead Hyland, The Lighthouse Lab in Alderley Park and Alex Alderton, Roberto Amato, Jeffrey Barrett, Sonia Goncalves, Ewan Harrison, David K. Jackson, Ian Johnston, Dominic Kwiatkowski, Cordelia Langford, John Sillitoe on behalf of the Wellcome Sanger Institute COVID-19 Surveillance Team |
| EPI_ISL_3109206 | Lighthouse Lab in Glasgow | Wellcome Sanger Institute for the COVID-19 Genomics UK (COG-UK) Consortium | Harper VanSteenhouse, Yumi Kasai, David Gray, Carol Clugston, Anna Dominiczak and Alex Alderton, Roberto Amato, Jeffrey Barrett, Sonia Goncalves, Ewan Harrison, David K. Jackson, Ian Johnston, Dominic Kwiatkowski, Cordelia Langford, John Sillitoe on behalf of the Wellcome Sanger Institute COVID-19 Surveillance Team |
| EPI_ISL_3109207 | Lighthouse Lab in Alderley Park | Wellcome Sanger Institute for the COVID-19 Genomics UK (COG-UK) Consortium | Jacquelyn Wynn, Mairead Hyland, The Lighthouse Lab in Alderley Park and Alex Alderton, Roberto Amato, Jeffrey Barrett, Sonia Goncalves, Ewan Harrison, David K. Jackson, Ian Johnston, Dominic Kwiatkowski, Cordelia Langford, John Sillitoe on behalf of the Wellcome Sanger Institute COVID-19 Surveillance Team |
| EPI_ISL_3109208 | Berkshire and Surrey Pathology Services Lighthouse Laboratory | Wellcome Sanger Institute for the COVID-19 Genomics UK (COG-UK) Consortium | Berkshire and Surrey Pathology Services Lighthouse Laboratory and Alex Alderton, Roberto Amato, Jeffrey Barrett, Sonia Goncalves, Ewan Harrison, David K. Jackson, Ian Johnston, Dominic Kwiatkowski, Cordelia Langford, John Sillitoe on behalf of the Wellcome Sanger Institute COVID-19 Surveillance Team |
| EPI_ISL_3109209, EPI_ISL_3109210 | Lighthouse Lab in Alderley Park | Wellcome Sanger Institute for the COVID-19 Genomics UK (COG-UK) Consortium | Jacquelyn Wynn, Mairead Hyland, The Lighthouse Lab in Alderley Park and Alex Alderton, Roberto Amato, Jeffrey Barrett, Sonia Goncalves, Ewan Harrison, David K. Jackson, Ian Johnston, Dominic Kwiatkowski, Cordelia Langford, John Sillitoe on behalf of the Wellcome Sanger Institute COVID-19 Surveillance Team |
| EPI_ISL_3109211, EPI_ISL_3109212 | Berkshire and Surrey Pathology Services Lighthouse Laboratory | Wellcome Sanger Institute for the COVID-19 Genomics UK (COG-UK) Consortium | Berkshire and Surrey Pathology Services Lighthouse Laboratory and Alex Alderton, Roberto Amato, Jeffrey Barrett, Sonia Goncalves, Ewan Harrison, David K. Jackson, Ian Johnston, Dominic Kwiatkowski, Cordelia Langford, John Sillitoe on behalf of the Wellcome Sanger Institute COVID-19 Surveillance Team |
| EPI_ISL_3109213, EPI_ISL_3109214 | Lighthouse Lab in Alderley Park | Wellcome Sanger Institute for the COVID-19 Genomics UK (COG-UK) Consortium | Jacquelyn Wynn, Mairead Hyland, The Lighthouse Lab in Alderley Park and Alex Alderton, Roberto Amato, Jeffrey Barrett, Sonia Goncalves, Ewan Harrison, David K. Jackson, Ian Johnston, Dominic Kwiatkowski, Cordelia Langford, John Sillitoe on behalf of the Wellcome Sanger Institute COVID-19 Surveillance Team |
| EPI_ISL_3109215 | Lighthouse Lab in Glasgow | Wellcome Sanger Institute for the COVID-19 Genomics UK (COG-UK) Consortium | Harper VanSteenhouse, Yumi Kasai, David Gray, Carol Clugston, Anna Dominiczak and Alex Alderton, Roberto Amato, Jeffrey Barrett, Sonia Goncalves, Ewan Harrison, David K. Jackson, Ian Johnston, Dominic Kwiatkowski, Cordelia Langford, John Sillitoe on behalf of the Wellcome Sanger Institute COVID-19 Surveillance Team |
| EPI_ISL_3109216, EPI_ISL_3109217 | Lighthouse Lab in Alderley Park | Wellcome Sanger Institute for the COVID-19 Genomics UK (COG-UK) Consortium | Jacquelyn Wynn, Mairead Hyland, The Lighthouse Lab in Alderley Park and Alex Alderton, Roberto Amato, Jeffrey Barrett, Sonia Goncalves, Ewan Harrison, David K. Jackson, Ian Johnston, Dominic Kwiatkowski, Cordelia Langford, John Sillitoe on behalf of the Wellcome Sanger Institute COVID-19 Surveillance Team |
| EPI_ISL_3109218 | Berkshire and Surrey Pathology Services Lighthouse Laboratory | Wellcome Sanger Institute for the COVID-19 Genomics UK (COG-UK) Consortium | Berkshire and Surrey Pathology Services Lighthouse Laboratory and Alex Alderton, Roberto Amato, Jeffrey Barrett, Sonia Goncalves, Ewan Harrison, David K. Jackson, Ian Johnston, Dominic Kwiatkowski, Cordelia Langford, John Sillitoe on behalf of the Wellcome Sanger Institute COVID-19 Surveillance Team |
| EPI_ISL_3109219, EPI_ISL_3109220, EPI_ISL_3109221 | Lighthouse Lab in Alderley Park | Wellcome Sanger Institute for the COVID-19 Genomics UK (COG-UK) Consortium | Jacquelyn Wynn, Mairead Hyland, The Lighthouse Lab in Alderley Park and Alex Alderton, Roberto Amato, Jeffrey Barrett, Sonia Goncalves, Ewan Harrison, David K. Jackson, Ian Johnston, Dominic Kwiatkowski, Cordelia Langford, John Sillitoe on behalf of the Wellcome Sanger Institute COVID-19 Surveillance Team |
| EPI_ISL_3109222, EPI_ISL_3109223 | Berkshire and Surrey Pathology Services Lighthouse Laboratory | Wellcome Sanger Institute for the COVID-19 Genomics UK (COG-UK) Consortium | Berkshire and Surrey Pathology Services Lighthouse Laboratory and Alex Alderton, Roberto Amato, Jeffrey Barrett, Sonia Goncalves, Ewan Harrison, David K. Jackson, Ian Johnston, Dominic Kwiatkowski, Cordelia Langford, John Sillitoe on behalf of the Wellcome Sanger Institute COVID-19 Surveillance Team |
| EPI_ISL_3109224 | Lighthouse Lab in Alderley Park | Wellcome Sanger Institute for the COVID-19 Genomics UK (COG-UK) Consortium | Jacquelyn Wynn, Mairead Hyland, The Lighthouse Lab in Alderley Park and Alex Alderton, Roberto Amato, Jeffrey Barrett, Sonia Goncalves, Ewan Harrison, David K. Jackson, Ian Johnston, Dominic Kwiatkowski, Cordelia Langford, John Sillitoe on behalf of the Wellcome Sanger Institute COVID-19 Surveillance Team |
| EPI_ISL_3109225 | Lighthouse Lab in Glasgow | Wellcome Sanger Institute for the COVID-19 Genomics UK (COG-UK) Consortium | Harper VanSteenhouse, Yumi Kasai, David Gray, Carol Clugston, Anna Dominiczak and Alex Alderton, Roberto Amato, Jeffrey Barrett, Sonia Goncalves, Ewan Harrison, David K. Jackson, Ian Johnston, Dominic Kwiatkowski, Cordelia Langford, John Sillitoe on behalf of the Wellcome Sanger Institute COVID-19 Surveillance Team |
| EPI_ISL_3109226, EPI_ISL_3109227 | Lighthouse Lab in Alderley Park | Wellcome Sanger Institute for the COVID-19 Genomics UK (COG-UK) Consortium | Jacquelyn Wynn, Mairead Hyland, The Lighthouse Lab in Alderley Park and Alex Alderton, Roberto Amato, Jeffrey Barrett, Sonia Goncalves, Ewan Harrison, David K. Jackson, Ian Johnston, Dominic Kwiatkowski, Cordelia Langford, John Sillitoe on behalf of the Wellcome Sanger Institute COVID-19 Surveillance Team |
| EPI_ISL_3109228, EPI_ISL_3109229 | Berkshire and Surrey Pathology Services Lighthouse Laboratory | Wellcome Sanger Institute for the COVID-19 Genomics UK (COG-UK) Consortium | Berkshire and Surrey Pathology Services Lighthouse Laboratory and Alex Alderton, Roberto Amato, Jeffrey Barrett, Sonia Goncalves, Ewan Harrison, David K. Jackson, Ian Johnston, Dominic Kwiatkowski, Cordelia Langford, John Sillitoe on behalf of the Wellcome Sanger Institute COVID-19 Surveillance Team |
| EPI_ISL_3109230, EPI_ISL_3109231 | Lighthouse Lab in Alderley Park | Wellcome Sanger Institute for the COVID-19 Genomics UK (COG-UK) Consortium | Jacquelyn Wynn, Mairead Hyland, The Lighthouse Lab in Alderley Park and Alex Alderton, Roberto Amato, Jeffrey Barrett, Sonia Goncalves, Ewan Harrison, David K. Jackson, Ian Johnston, Dominic Kwiatkowski, Cordelia Langford, John Sillitoe on behalf of the Wellcome Sanger Institute COVID-19 Surveillance Team |
| EPI_ISL_3109232 | Berkshire and Surrey Pathology Services Lighthouse Laboratory | Wellcome Sanger Institute for the COVID-19 Genomics UK (COG-UK) Consortium | Berkshire and Surrey Pathology Services Lighthouse Laboratory and Alex Alderton, Roberto Amato, Jeffrey Barrett, Sonia Goncalves, Ewan Harrison, David K. Jackson, Ian Johnston, Dominic Kwiatkowski, Cordelia Langford, John Sillitoe on behalf of the Wellcome Sanger Institute COVID-19 Surveillance Team |
| EPI_ISL_3109233, EPI_ISL_3109234, EPI_ISL_3109235, EPI_ISL_3109236 | Lighthouse Lab in Alderley Park | Wellcome Sanger Institute for the COVID-19 Genomics UK (COG-UK) Consortium | Jacquelyn Wynn, Mairead Hyland, The Lighthouse Lab in Alderley Park and Alex Alderton, Roberto Amato, Jeffrey Barrett, Sonia Goncalves, Ewan Harrison, David K. Jackson, Ian Johnston, Dominic Kwiatkowski, Cordelia Langford, John Sillitoe on behalf of the Wellcome Sanger Institute COVID-19 Surveillance Team |
| EPI_ISL_3109237, EPI_ISL_3109238 | Berkshire and Surrey Pathology Services Lighthouse Laboratory | Wellcome Sanger Institute for the COVID-19 Genomics UK (COG-UK) Consortium | Berkshire and Surrey Pathology Services Lighthouse Laboratory and Alex Alderton, Roberto Amato, Jeffrey Barrett, Sonia Goncalves, Ewan Harrison, David K. Jackson, Ian Johnston, Dominic Kwiatkowski, Cordelia Langford, John Sillitoe on behalf of the Wellcome Sanger Institute COVID-19 Surveillance Team |
| EPI_ISL_3109239 | Lighthouse Lab in Glasgow | Wellcome Sanger Institute for the COVID-19 Genomics UK (COG-UK) Consortium | Harper VanSteenhouse, Yumi Kasai, David Gray, Carol Clugston, Anna Dominiczak and Alex Alderton, Roberto Amato, Jeffrey Barrett, Sonia Goncalves, Ewan Harrison, David K. Jackson, Ian Johnston, Dominic Kwiatkowski, Cordelia Langford, John Sillitoe on behalf of the Wellcome Sanger Institute COVID-19 Surveillance Team |
| EPI_ISL_3109240, EPI_ISL_3109241 | Lighthouse Lab in Alderley Park | Wellcome Sanger Institute for the COVID-19 Genomics UK (COG-UK) Consortium | Jacquelyn Wynn, Mairead Hyland, The Lighthouse Lab in Alderley Park and Alex Alderton, Roberto Amato, Jeffrey Barrett, Sonia Goncalves, Ewan Harrison, David K. Jackson, Ian Johnston, Dominic Kwiatkowski, Cordelia Langford, John Sillitoe on behalf of the Wellcome Sanger Institute COVID-19 Surveillance Team |
| EPI_ISL_3109242 | Lighthouse Lab in Glasgow | Wellcome Sanger Institute for the COVID-19 Genomics UK (COG-UK) Consortium | Harper VanSteenhouse, Yumi Kasai, David Gray, Carol Clugston, Anna Dominiczak and Alex Alderton, Roberto Amato, Jeffrey Barrett, Sonia Goncalves, Ewan Harrison, David K. Jackson, Ian Johnston, Dominic Kwiatkowski, Cordelia Langford, John Sillitoe on behalf of the Wellcome Sanger Institute COVID-19 Surveillance Team |
| EPI_ISL_3109243 | Berkshire and Surrey Pathology Services Lighthouse Laboratory | Wellcome Sanger Institute for the COVID-19 Genomics UK (COG-UK) Consortium | Berkshire and Surrey Pathology Services Lighthouse Laboratory and Alex Alderton, Roberto Amato, Jeffrey Barrett, Sonia Goncalves, Ewan Harrison, David K. Jackson, Ian Johnston, Dominic Kwiatkowski, Cordelia Langford, John Sillitoe on behalf of the Wellcome Sanger Institute COVID-19 Surveillance Team |
| EPI_ISL_3109244 | Lighthouse Lab in Alderley Park | Wellcome Sanger Institute for the COVID-19 Genomics UK (COG-UK) Consortium | Jacquelyn Wynn, Mairead Hyland, The Lighthouse Lab in Alderley Park and Alex Alderton, Roberto Amato, Jeffrey Barrett, Sonia Goncalves, Ewan Harrison, David K. Jackson, Ian Johnston, Dominic Kwiatkowski, Cordelia Langford, John Sillitoe on behalf of the Wellcome Sanger Institute COVID-19 Surveillance Team |
| EPI_ISL_3109245, EPI_ISL_3109246 | Berkshire and Surrey Pathology Services Lighthouse Laboratory | Wellcome Sanger Institute for the COVID-19 Genomics UK (COG-UK) Consortium | Berkshire and Surrey Pathology Services Lighthouse Laboratory and Alex Alderton, Roberto Amato, Jeffrey Barrett, Sonia Goncalves, Ewan Harrison, David K. Jackson, Ian Johnston, Dominic Kwiatkowski, Cordelia Langford, John Sillitoe on behalf of the Wellcome Sanger Institute COVID-19 Surveillance Team |
| EPI_ISL_3109247 | Lighthouse Lab in Alderley Park | Wellcome Sanger Institute for the COVID-19 Genomics UK (COG-UK) Consortium | Jacquelyn Wynn, Mairead Hyland, The Lighthouse Lab in Alderley Park and Alex Alderton, Roberto Amato, Jeffrey Barrett, Sonia Goncalves, Ewan Harrison, David K. Jackson, Ian Johnston, Dominic Kwiatkowski, Cordelia Langford, John Sillitoe on behalf of the Wellcome Sanger Institute COVID-19 Surveillance Team |
| EPI_ISL_3109248 | Berkshire and Surrey Pathology Services Lighthouse Laboratory | Wellcome Sanger Institute for the COVID-19 Genomics UK (COG-UK) Consortium | Berkshire and Surrey Pathology Services Lighthouse Laboratory and Alex Alderton, Roberto Amato, Jeffrey Barrett, Sonia Goncalves, Ewan Harrison, David K. Jackson, Ian Johnston, Dominic Kwiatkowski, Cordelia Langford, John Sillitoe on behalf of the Wellcome Sanger Institute COVID-19 Surveillance Team |
| EPI_ISL_3109249, EPI_ISL_3109250, EPI_ISL_3109251, EPI_ISL_3109252, | Lighthouse Lab in Alderley Park | Wellcome Sanger Institute for the COVID-19 Genomics UK (COG-UK) Consortium | Jacquelyn Wynn, Mairead Hyland, The Lighthouse Lab in Alderley Park and Alex Alderton, Roberto Amato, Jeffrey Barrett, Sonia Goncalves, Ewan Harrison, David K. Jackson, Ian Johnston, Dominic Kwiatkowski, Cordelia Langford, John Sillitoe on behalf of the Wellcome Sanger Institute COVID-19 Surveillance Team |

| | | | |
|---|---|---|---|
| EPI_ISL_3109253, EPI_ISL_3109254, EPI_ISL_3109255 | | | |
| EPI_ISL_3109256, EPI_ISL_3109257 | Berkshire and Surrey Pathology Services Lighthouse Laboratory | Wellcome Sanger Institute for the COVID-19 Genomics UK (COG-UK) Consortium | Berkshire and Surrey Pathology Services Lighthouse Laboratory and Alex Alderton, Roberto Amato, Jeffrey Barrett, Sonia Goncalves, Ewan Harrison, David K. Jackson, Ian Johnston, Dominic Kwiatkowski, Cordelia Langford, John Sillitoe on behalf of the Wellcome Sanger Institute COVID-19 Surveillance Team |
| EPI_ISL_3109258 | Lighthouse Lab in Glasgow | Wellcome Sanger Institute for the COVID-19 Genomics UK (COG-UK) Consortium | Harper VanSteenhouse, Yumi Kasai, David Gray, Carol Clugston, Anna Dominiczak and Alex Alderton, Roberto Amato, Jeffrey Barrett, Sonia Goncalves, Ewan Harrison, David K. Jackson, Ian Johnston, Dominic Kwiatkowski, Cordelia Langford, John Sillitoe on behalf of the Wellcome Sanger Institute COVID-19 Surveillance Team |
| EPI_ISL_3109259 | Lighthouse Lab in Alderley Park | Wellcome Sanger Institute for the COVID-19 Genomics UK (COG-UK) Consortium | Jacquelyn Wynn, Mairead Hyland, The Lighthouse Lab in Alderley Park and Alex Alderton, Roberto Amato, Jeffrey Barrett, Sonia Goncalves, Ewan Harrison, David K. Jackson, Ian Johnston, Dominic Kwiatkowski, Cordelia Langford, John Sillitoe on behalf of the Wellcome Sanger Institute COVID-19 Surveillance Team |
| EPI_ISL_3109260 | Berkshire and Surrey Pathology Services Lighthouse Laboratory | Wellcome Sanger Institute for the COVID-19 Genomics UK (COG-UK) Consortium | Berkshire and Surrey Pathology Services Lighthouse Laboratory and Alex Alderton, Roberto Amato, Jeffrey Barrett, Sonia Goncalves, Ewan Harrison, David K. Jackson, Ian Johnston, Dominic Kwiatkowski, Cordelia Langford, John Sillitoe on behalf of the Wellcome Sanger Institute COVID-19 Surveillance Team |
| EPI_ISL_3109261, EPI_ISL_3109262, EPI_ISL_3109263, EPI_ISL_3109264 | Lighthouse Lab in Alderley Park | Wellcome Sanger Institute for the COVID-19 Genomics UK (COG-UK) Consortium | Jacquelyn Wynn, Mairead Hyland, The Lighthouse Lab in Alderley Park and Alex Alderton, Roberto Amato, Jeffrey Barrett, Sonia Goncalves, Ewan Harrison, David K. Jackson, Ian Johnston, Dominic Kwiatkowski, Cordelia Langford, John Sillitoe on behalf of the Wellcome Sanger Institute COVID-19 Surveillance Team |
| EPI_ISL_3109265 | Berkshire and Surrey Pathology Services Lighthouse Laboratory | Wellcome Sanger Institute for the COVID-19 Genomics UK (COG-UK) Consortium | Berkshire and Surrey Pathology Services Lighthouse Laboratory and Alex Alderton, Roberto Amato, Jeffrey Barrett, Sonia Goncalves, Ewan Harrison, David K. Jackson, Ian Johnston, Dominic Kwiatkowski, Cordelia Langford, John Sillitoe on behalf of the Wellcome Sanger Institute COVID-19 Surveillance Team |
| EPI_ISL_3109266, EPI_ISL_3109267, EPI_ISL_3109268, EPI_ISL_3109269 | Lighthouse Lab in Alderley Park | Wellcome Sanger Institute for the COVID-19 Genomics UK (COG-UK) Consortium | Jacquelyn Wynn, Mairead Hyland, The Lighthouse Lab in Alderley Park and Alex Alderton, Roberto Amato, Jeffrey Barrett, Sonia Goncalves, Ewan Harrison, David K. Jackson, Ian Johnston, Dominic Kwiatkowski, Cordelia Langford, John Sillitoe on behalf of the Wellcome Sanger Institute COVID-19 Surveillance Team |
| EPI_ISL_3109270 | Lighthouse Lab in Glasgow | Wellcome Sanger Institute for the COVID-19 Genomics UK (COG-UK) Consortium | Harper VanSteenhouse, Yumi Kasai, David Gray, Carol Clugston, Anna Dominiczak and Alex Alderton, Roberto Amato, Jeffrey Barrett, Sonia Goncalves, Ewan Harrison, David K. Jackson, Ian Johnston, Dominic Kwiatkowski, Cordelia Langford, John Sillitoe on behalf of the Wellcome Sanger Institute COVID-19 Surveillance Team |
| EPI_ISL_3109271, EPI_ISL_3109272, EPI_ISL_3109273, EPI_ISL_3109274, EPI_ISL_3109275, EPI_ISL_3109276, EPI_ISL_3109277 | Lighthouse Lab in Alderley Park | Wellcome Sanger Institute for the COVID-19 Genomics UK (COG-UK) Consortium | Jacquelyn Wynn, Mairead Hyland, The Lighthouse Lab in Alderley Park and Alex Alderton, Roberto Amato, Jeffrey Barrett, Sonia Goncalves, Ewan Harrison, David K. Jackson, Ian Johnston, Dominic Kwiatkowski, Cordelia Langford, John Sillitoe on behalf of the Wellcome Sanger Institute COVID-19 Surveillance Team |
| EPI_ISL_3109278, EPI_ISL_3109279 | Berkshire and Surrey Pathology Services Lighthouse Laboratory | Wellcome Sanger Institute for the COVID-19 Genomics UK (COG-UK) Consortium | Berkshire and Surrey Pathology Services Lighthouse Laboratory and Alex Alderton, Roberto Amato, Jeffrey Barrett, Sonia Goncalves, Ewan Harrison, David K. Jackson, Ian Johnston, Dominic Kwiatkowski, Cordelia Langford, John Sillitoe on behalf of the Wellcome Sanger Institute COVID-19 Surveillance Team |
| EPI_ISL_3109280, EPI_ISL_3109281, EPI_ISL_3109282, EPI_ISL_3109283 | Lighthouse Lab in Alderley Park | Wellcome Sanger Institute for the COVID-19 Genomics UK (COG-UK) Consortium | Jacquelyn Wynn, Mairead Hyland, The Lighthouse Lab in Alderley Park and Alex Alderton, Roberto Amato, Jeffrey Barrett, Sonia Goncalves, Ewan Harrison, David K. Jackson, Ian Johnston, Dominic Kwiatkowski, Cordelia Langford, John Sillitoe on behalf of the Wellcome Sanger Institute COVID-19 Surveillance Team |
| EPI_ISL_3109284, EPI_ISL_3109285 | Berkshire and Surrey Pathology Services Lighthouse Laboratory | Wellcome Sanger Institute for the COVID-19 Genomics UK (COG-UK) Consortium | Berkshire and Surrey Pathology Services Lighthouse Laboratory and Alex Alderton, Roberto Amato, Jeffrey Barrett, Sonia Goncalves, Ewan Harrison, David K. Jackson, Ian Johnston, Dominic Kwiatkowski, Cordelia Langford, John Sillitoe on behalf of the Wellcome Sanger Institute COVID-19 Surveillance Team |
| EPI_ISL_3109286, EPI_ISL_3109287, EPI_ISL_3109288, EPI_ISL_3109289 | Lighthouse Lab in Alderley Park | Wellcome Sanger Institute for the COVID-19 Genomics UK (COG-UK) Consortium | Jacquelyn Wynn, Mairead Hyland, The Lighthouse Lab in Alderley Park and Alex Alderton, Roberto Amato, Jeffrey Barrett, Sonia Goncalves, Ewan Harrison, David K. Jackson, Ian Johnston, Dominic Kwiatkowski, Cordelia Langford, John Sillitoe on behalf of the Wellcome Sanger Institute COVID-19 Surveillance Team |
| EPI_ISL_3109290 | Lighthouse Lab in Glasgow | Wellcome Sanger Institute for the COVID-19 Genomics UK (COG-UK) Consortium | Harper VanSteenhouse, Yumi Kasai, David Gray, Carol Clugston, Anna Dominiczak and Alex Alderton, Roberto Amato, Jeffrey Barrett, Sonia Goncalves, Ewan Harrison, David K. Jackson, Ian Johnston, Dominic Kwiatkowski, Cordelia Langford, John Sillitoe on behalf of the Wellcome Sanger Institute COVID-19 Surveillance Team |
| EPI_ISL_3109291, EPI_ISL_3109292 | Lighthouse Lab in Alderley Park | Wellcome Sanger Institute for the COVID-19 Genomics UK (COG-UK) Consortium | Jacquelyn Wynn, Mairead Hyland, The Lighthouse Lab in Alderley Park and Alex Alderton, Roberto Amato, Jeffrey Barrett, Sonia Goncalves, Ewan Harrison, David K. Jackson, Ian Johnston, Dominic Kwiatkowski, Cordelia Langford, John Sillitoe on behalf of the Wellcome Sanger Institute COVID-19 Surveillance Team |
| EPI_ISL_3109293 | Berkshire and Surrey Pathology Services Lighthouse Laboratory | Wellcome Sanger Institute for the COVID-19 Genomics UK (COG-UK) Consortium | Berkshire and Surrey Pathology Services Lighthouse Laboratory and Alex Alderton, Roberto Amato, Jeffrey Barrett, Sonia Goncalves, Ewan Harrison, David K. Jackson, Ian Johnston, Dominic Kwiatkowski, Cordelia Langford, John Sillitoe on behalf of the Wellcome Sanger Institute COVID-19 Surveillance Team |
| EPI_ISL_3109294, EPI_ISL_3109295 | Lighthouse Lab in Alderley Park | Wellcome Sanger Institute for the COVID-19 Genomics UK (COG-UK) Consortium | Jacquelyn Wynn, Mairead Hyland, The Lighthouse Lab in Alderley Park and Alex Alderton, Roberto Amato, Jeffrey Barrett, Sonia Goncalves, Ewan Harrison, David K. Jackson, Ian Johnston, Dominic Kwiatkowski, Cordelia Langford, John Sillitoe on behalf of the Wellcome Sanger Institute COVID-19 Surveillance Team |
| EPI_ISL_3109296 | Berkshire and Surrey Pathology Services Lighthouse Laboratory | Wellcome Sanger Institute for the COVID-19 Genomics UK (COG-UK) Consortium | Berkshire and Surrey Pathology Services Lighthouse Laboratory and Alex Alderton, Roberto Amato, Jeffrey Barrett, Sonia Goncalves, Ewan Harrison, David K. Jackson, Ian Johnston, Dominic Kwiatkowski, Cordelia Langford, John Sillitoe on behalf of the Wellcome Sanger Institute COVID-19 Surveillance Team |
| EPI_ISL_3109297, EPI_ISL_3109298 | Lighthouse Lab in Alderley Park | Wellcome Sanger Institute for the COVID-19 Genomics UK (COG-UK) Consortium | Jacquelyn Wynn, Mairead Hyland, The Lighthouse Lab in Alderley Park and Alex Alderton, Roberto Amato, Jeffrey Barrett, Sonia Goncalves, Ewan Harrison, David K. Jackson, Ian Johnston, Dominic Kwiatkowski, Cordelia Langford, John Sillitoe on behalf of the Wellcome Sanger Institute COVID-19 Surveillance Team |
| EPI_ISL_3109299 | Berkshire and Surrey Pathology Services Lighthouse Laboratory | Wellcome Sanger Institute for the COVID-19 Genomics UK (COG-UK) Consortium | Berkshire and Surrey Pathology Services Lighthouse Laboratory and Alex Alderton, Roberto Amato, Jeffrey Barrett, Sonia Goncalves, Ewan Harrison, David K. Jackson, Ian Johnston, Dominic Kwiatkowski, Cordelia Langford, John Sillitoe on behalf of the Wellcome Sanger Institute COVID-19 Surveillance Team |
| EPI_ISL_3109300 | Lighthouse Lab in Glasgow | Wellcome Sanger Institute for the COVID-19 Genomics UK (COG-UK) Consortium | Harper VanSteenhouse, Yumi Kasai, David Gray, Carol Clugston, Anna Dominiczak and Alex Alderton, Roberto Amato, Jeffrey Barrett, Sonia Goncalves, Ewan Harrison, David K. Jackson, Ian Johnston, Dominic Kwiatkowski, Cordelia Langford, John Sillitoe on behalf of the Wellcome Sanger Institute COVID-19 Surveillance Team |
| EPI_ISL_3109301, EPI_ISL_3109302, EPI_ISL_3109303, EPI_ISL_3109304, EPI_ISL_3109305, EPI_ISL_3109306 | Lighthouse Lab in Alderley Park | Wellcome Sanger Institute for the COVID-19 Genomics UK (COG-UK) Consortium | Jacquelyn Wynn, Mairead Hyland, The Lighthouse Lab in Alderley Park and Alex Alderton, Roberto Amato, Jeffrey Barrett, Sonia Goncalves, Ewan Harrison, David K. Jackson, Ian Johnston, Dominic Kwiatkowski, Cordelia Langford, John Sillitoe on behalf of the Wellcome Sanger Institute COVID-19 Surveillance Team |
| EPI_ISL_3109307 | Lighthouse Lab in Glasgow | Wellcome Sanger Institute for the COVID-19 Genomics UK (COG-UK) Consortium | Harper VanSteenhouse, Yumi Kasai, David Gray, Carol Clugston, Anna Dominiczak and Alex Alderton, Roberto Amato, Jeffrey Barrett, Sonia Goncalves, Ewan Harrison, David K. Jackson, Ian Johnston, Dominic Kwiatkowski, Cordelia Langford, John Sillitoe on behalf of the Wellcome Sanger Institute COVID-19 Surveillance Team |
| EPI_ISL_3109308 | Berkshire and Surrey Pathology Services Lighthouse Laboratory | Wellcome Sanger Institute for the COVID-19 Genomics UK (COG-UK) Consortium | Berkshire and Surrey Pathology Services Lighthouse Laboratory and Alex Alderton, Roberto Amato, Jeffrey Barrett, Sonia Goncalves, Ewan Harrison, David K. Jackson, Ian Johnston, Dominic Kwiatkowski, Cordelia Langford, John Sillitoe on behalf of the Wellcome Sanger Institute COVID-19 Surveillance Team |
| EPI_ISL_3109309, EPI_ISL_3109310 | Lighthouse Lab in Alderley Park | Wellcome Sanger Institute for the COVID-19 Genomics UK (COG-UK) Consortium | Jacquelyn Wynn, Mairead Hyland, The Lighthouse Lab in Alderley Park and Alex Alderton, Roberto Amato, Jeffrey Barrett, Sonia Goncalves, Ewan Harrison, David K. Jackson, Ian Johnston, Dominic Kwiatkowski, Cordelia Langford, John Sillitoe on behalf of the Wellcome Sanger Institute COVID-19 Surveillance Team |
| EPI_ISL_3109311 | Lighthouse Lab in Glasgow | Wellcome Sanger Institute for the COVID-19 Genomics UK (COG-UK) Consortium | Harper VanSteenhouse, Yumi Kasai, David Gray, Carol Clugston, Anna Dominiczak and Alex Alderton, Roberto Amato, Jeffrey Barrett, Sonia Goncalves, Ewan Harrison, David K. Jackson, Ian Johnston, Dominic Kwiatkowski, Cordelia Langford, John Sillitoe on behalf of the Wellcome Sanger Institute COVID-19 Surveillance Team |
| EPI_ISL_3109312 | Lighthouse Lab in Alderley Park | Wellcome Sanger Institute for the COVID-19 Genomics UK (COG-UK) Consortium | Jacquelyn Wynn, Mairead Hyland, The Lighthouse Lab in Alderley Park and Alex Alderton, Roberto Amato, Jeffrey Barrett, Sonia Goncalves, Ewan Harrison, David K. Jackson, Ian Johnston, Dominic Kwiatkowski, Cordelia Langford, John Sillitoe on behalf of the Wellcome Sanger Institute COVID-19 Surveillance Team |
| EPI_ISL_3109313 | Berkshire and Surrey Pathology Services Lighthouse Laboratory | Wellcome Sanger Institute for the COVID-19 Genomics UK (COG-UK) Consortium | Berkshire and Surrey Pathology Services Lighthouse Laboratory and Alex Alderton, Roberto Amato, Jeffrey Barrett, Sonia Goncalves, Ewan Harrison, David K. Jackson, Ian Johnston, Dominic Kwiatkowski, Cordelia Langford, John Sillitoe on behalf of the Wellcome Sanger Institute COVID-19 Surveillance Team |
| EPI_ISL_3109314, EPI_ISL_3109315 | Lighthouse Lab in Alderley Park | Wellcome Sanger Institute for the COVID-19 Genomics UK (COG-UK) Consortium | Jacquelyn Wynn, Mairead Hyland, The Lighthouse Lab in Alderley Park and Alex Alderton, Roberto Amato, Jeffrey Barrett, Sonia Goncalves, Ewan Harrison, David K. Jackson, Ian Johnston, Dominic Kwiatkowski, Cordelia Langford, John Sillitoe on behalf of the Wellcome Sanger Institute COVID-19 Surveillance Team |
| EPI_ISL_3109316 | Berkshire and Surrey Pathology Services Lighthouse Laboratory | Wellcome Sanger Institute for the COVID-19 Genomics UK (COG-UK) Consortium | Berkshire and Surrey Pathology Services Lighthouse Laboratory and Alex Alderton, Roberto Amato, Jeffrey Barrett, Sonia Goncalves, Ewan Harrison, David K. Jackson, Ian Johnston, Dominic Kwiatkowski, Cordelia Langford, John Sillitoe on behalf of the Wellcome Sanger Institute COVID-19 Surveillance Team |
| EPI_ISL_3109317 | Lighthouse Lab in Alderley Park | Wellcome Sanger Institute for the COVID-19 Genomics UK (COG-UK) Consortium | Jacquelyn Wynn, Mairead Hyland, The Lighthouse Lab in Alderley Park and Alex Alderton, Roberto Amato, Jeffrey Barrett, Sonia Goncalves, Ewan Harrison, David K. Jackson, Ian Johnston, Dominic Kwiatkowski, Cordelia Langford, John Sillitoe on behalf of the Wellcome Sanger Institute COVID-19 Surveillance Team |
| EPI_ISL_3109318 | Berkshire and Surrey Pathology Services Lighthouse Laboratory | Wellcome Sanger Institute for the COVID-19 Genomics UK (COG-UK) Consortium | Berkshire and Surrey Pathology Services Lighthouse Laboratory and Alex Alderton, Roberto Amato, Jeffrey Barrett, Sonia Goncalves, Ewan Harrison, David K. Jackson, Ian Johnston, Dominic Kwiatkowski, Cordelia Langford, John Sillitoe on behalf of the Wellcome Sanger Institute COVID-19 Surveillance Team |
| EPI_ISL_3109319 | Lighthouse Lab in Alderley Park | Wellcome Sanger Institute for the COVID-19 Genomics UK (COG-UK) Consortium | Jacquelyn Wynn, Mairead Hyland, The Lighthouse Lab in Alderley Park and Alex Alderton, Roberto Amato, Jeffrey Barrett, Sonia Goncalves, Ewan Harrison, David K. Jackson, Ian Johnston, Dominic Kwiatkowski, Cordelia Langford, John Sillitoe on behalf of the Wellcome Sanger Institute COVID-19 Surveillance Team |
| EPI_ISL_3109320 | Berkshire and Surrey Pathology Services Lighthouse Laboratory | Wellcome Sanger Institute for the COVID-19 Genomics UK (COG-UK) Consortium | Berkshire and Surrey Pathology Services Lighthouse Laboratory and Alex Alderton, Roberto Amato, Jeffrey Barrett, Sonia Goncalves, Ewan Harrison, David K. Jackson, Ian Johnston, Dominic Kwiatkowski, Cordelia Langford, John Sillitoe on behalf of the Wellcome Sanger Institute COVID-19 Surveillance Team |
| EPI_ISL_3109321, EPI_ISL_3109322, EPI_ISL_3109323 | Lighthouse Lab in Alderley Park | Wellcome Sanger Institute for the COVID-19 Genomics UK (COG-UK) Consortium | Jacquelyn Wynn, Mairead Hyland, The Lighthouse Lab in Alderley Park and Alex Alderton, Roberto Amato, Jeffrey Barrett, Sonia Goncalves, Ewan Harrison, David K. Jackson, Ian Johnston, Dominic Kwiatkowski, Cordelia Langford, John Sillitoe on behalf of the Wellcome Sanger Institute COVID-19 Surveillance Team |
| EPI_ISL_3109324, EPI_ISL_3109325, EPI_ISL_3109326, | Lighthouse Lab in Glasgow | Wellcome Sanger Institute for the COVID-19 Genomics UK (COG-UK) Consortium | Harper VanSteenhouse, Yumi Kasai, David Gray, Carol Clugston, Anna Dominiczak and Alex Alderton, Roberto Amato, Jeffrey Barrett, Sonia Goncalves, Ewan Harrison, David K. Jackson, Ian Johnston, Dominic Kwiatkowski, Cordelia Langford, John Sillitoe on behalf of the Wellcome Sanger Institute COVID-19 Surveillance Team |

| | | | |
|---|---|---|---|
| EPI_ISL_3109327 | | | |
| EPI_ISL_3109328 | Lighthouse Lab in Alderley Park | Wellcome Sanger Institute for the COVID-19 Genomics UK (COG-UK) Consortium | Jacquelyn Wynn, Mairead Hyland, The Lighthouse Lab in Alderley Park and Alex Alderton, Roberto Amato, Jeffrey Barrett, Sonia Goncalves, Ewan Harrison, David K. Jackson, Ian Johnston, Dominic Kwiatkowski, Cordelia Langford, John Sillitoe on behalf of the Wellcome Sanger Institute COVID-19 Surveillance Team |
| EPI_ISL_3109329, EPI_ISL_3109330, EPI_ISL_3109331 | Berkshire and Surrey Pathology Services Lighthouse Laboratory | Wellcome Sanger Institute for the COVID-19 Genomics UK (COG-UK) Consortium | Berkshire and Surrey Pathology Services Lighthouse Laboratory and Alex Alderton, Roberto Amato, Jeffrey Barrett, Sonia Goncalves, Ewan Harrison, David K. Jackson, Ian Johnston, Dominic Kwiatkowski, Cordelia Langford, John Sillitoe on behalf of the Wellcome Sanger Institute COVID-19 Surveillance Team |
| EPI_ISL_3109332, EPI_ISL_3109333, EPI_ISL_3109334 | Lighthouse Lab in Alderley Park | Wellcome Sanger Institute for the COVID-19 Genomics UK (COG-UK) Consortium | Jacquelyn Wynn, Mairead Hyland, The Lighthouse Lab in Alderley Park and Alex Alderton, Roberto Amato, Jeffrey Barrett, Sonia Goncalves, Ewan Harrison, David K. Jackson, Ian Johnston, Dominic Kwiatkowski, Cordelia Langford, John Sillitoe on behalf of the Wellcome Sanger Institute COVID-19 Surveillance Team |
| EPI_ISL_3109335 | Lighthouse Lab in Glasgow | Wellcome Sanger Institute for the COVID-19 Genomics UK (COG-UK) Consortium | Harper VanSteenhouse, Yumi Kasai, David Gray, Carol Clugston, Anna Dominiczak and Alex Alderton, Roberto Amato, Jeffrey Barrett, Sonia Goncalves, Ewan Harrison, David K. Jackson, Ian Johnston, Dominic Kwiatkowski, Cordelia Langford, John Sillitoe on behalf of the Wellcome Sanger Institute COVID-19 Surveillance Team |
| EPI_ISL_3109336, EPI_ISL_3109337 | Berkshire and Surrey Pathology Services Lighthouse Laboratory | Wellcome Sanger Institute for the COVID-19 Genomics UK (COG-UK) Consortium | Berkshire and Surrey Pathology Services Lighthouse Laboratory and Alex Alderton, Roberto Amato, Jeffrey Barrett, Sonia Goncalves, Ewan Harrison, David K. Jackson, Ian Johnston, Dominic Kwiatkowski, Cordelia Langford, John Sillitoe on behalf of the Wellcome Sanger Institute COVID-19 Surveillance Team |
| EPI_ISL_3109338, EPI_ISL_3109339 | Lighthouse Lab in Alderley Park | Wellcome Sanger Institute for the COVID-19 Genomics UK (COG-UK) Consortium | Jacquelyn Wynn, Mairead Hyland, The Lighthouse Lab in Alderley Park and Alex Alderton, Roberto Amato, Jeffrey Barrett, Sonia Goncalves, Ewan Harrison, David K. Jackson, Ian Johnston, Dominic Kwiatkowski, Cordelia Langford, John Sillitoe on behalf of the Wellcome Sanger Institute COVID-19 Surveillance Team |
| EPI_ISL_3109340 | Lighthouse Lab in Glasgow | Wellcome Sanger Institute for the COVID-19 Genomics UK (COG-UK) Consortium | Harper VanSteenhouse, Yumi Kasai, David Gray, Carol Clugston, Anna Dominiczak and Alex Alderton, Roberto Amato, Jeffrey Barrett, Sonia Goncalves, Ewan Harrison, David K. Jackson, Ian Johnston, Dominic Kwiatkowski, Cordelia Langford, John Sillitoe on behalf of the Wellcome Sanger Institute COVID-19 Surveillance Team |
| EPI_ISL_3109341 | Berkshire and Surrey Pathology Services Lighthouse Laboratory | Wellcome Sanger Institute for the COVID-19 Genomics UK (COG-UK) Consortium | Berkshire and Surrey Pathology Services Lighthouse Laboratory and Alex Alderton, Roberto Amato, Jeffrey Barrett, Sonia Goncalves, Ewan Harrison, David K. Jackson, Ian Johnston, Dominic Kwiatkowski, Cordelia Langford, John Sillitoe on behalf of the Wellcome Sanger Institute COVID-19 Surveillance Team |
| EPI_ISL_3109342, EPI_ISL_3109343, EPI_ISL_3109344, EPI_ISL_3109345, EPI_ISL_3109346, EPI_ISL_3109347 | Lighthouse Lab in Alderley Park | Wellcome Sanger Institute for the COVID-19 Genomics UK (COG-UK) Consortium | Jacquelyn Wynn, Mairead Hyland, The Lighthouse Lab in Alderley Park and Alex Alderton, Roberto Amato, Jeffrey Barrett, Sonia Goncalves, Ewan Harrison, David K. Jackson, Ian Johnston, Dominic Kwiatkowski, Cordelia Langford, John Sillitoe on behalf of the Wellcome Sanger Institute COVID-19 Surveillance Team |
| EPI_ISL_3109348 | Berkshire and Surrey Pathology Services Lighthouse Laboratory | Wellcome Sanger Institute for the COVID-19 Genomics UK (COG-UK) Consortium | Berkshire and Surrey Pathology Services Lighthouse Laboratory and Alex Alderton, Roberto Amato, Jeffrey Barrett, Sonia Goncalves, Ewan Harrison, David K. Jackson, Ian Johnston, Dominic Kwiatkowski, Cordelia Langford, John Sillitoe on behalf of the Wellcome Sanger Institute COVID-19 Surveillance Team |
| EPI_ISL_3109349, EPI_ISL_3109350, EPI_ISL_3109351 | Lighthouse Lab in Alderley Park | Wellcome Sanger Institute for the COVID-19 Genomics UK (COG-UK) Consortium | Jacquelyn Wynn, Mairead Hyland, The Lighthouse Lab in Alderley Park and Alex Alderton, Roberto Amato, Jeffrey Barrett, Sonia Goncalves, Ewan Harrison, David K. Jackson, Ian Johnston, Dominic Kwiatkowski, Cordelia Langford, John Sillitoe on behalf of the Wellcome Sanger Institute COVID-19 Surveillance Team |
| EPI_ISL_3109352 | Lighthouse Lab in Glasgow | Wellcome Sanger Institute for the COVID-19 Genomics UK (COG-UK) Consortium | Harper VanSteenhouse, Yumi Kasai, David Gray, Carol Clugston, Anna Dominiczak and Alex Alderton, Roberto Amato, Jeffrey Barrett, Sonia Goncalves, Ewan Harrison, David K. Jackson, Ian Johnston, Dominic Kwiatkowski, Cordelia Langford, John Sillitoe on behalf of the Wellcome Sanger Institute COVID-19 Surveillance Team |
| EPI_ISL_3109353, EPI_ISL_3109354, EPI_ISL_3109355 | Lighthouse Lab in Alderley Park | Wellcome Sanger Institute for the COVID-19 Genomics UK (COG-UK) Consortium | Jacquelyn Wynn, Mairead Hyland, The Lighthouse Lab in Alderley Park and Alex Alderton, Roberto Amato, Jeffrey Barrett, Sonia Goncalves, Ewan Harrison, David K. Jackson, Ian Johnston, Dominic Kwiatkowski, Cordelia Langford, John Sillitoe on behalf of the Wellcome Sanger Institute COVID-19 Surveillance Team |
| EPI_ISL_3109356 | Berkshire and Surrey Pathology Services Lighthouse Laboratory | Wellcome Sanger Institute for the COVID-19 Genomics UK (COG-UK) Consortium | Berkshire and Surrey Pathology Services Lighthouse Laboratory and Alex Alderton, Roberto Amato, Jeffrey Barrett, Sonia Goncalves, Ewan Harrison, David K. Jackson, Ian Johnston, Dominic Kwiatkowski, Cordelia Langford, John Sillitoe on behalf of the Wellcome Sanger Institute COVID-19 Surveillance Team |
| EPI_ISL_3109357 | Lighthouse Lab in Glasgow | Wellcome Sanger Institute for the COVID-19 Genomics UK (COG-UK) Consortium | Harper VanSteenhouse, Yumi Kasai, David Gray, Carol Clugston, Anna Dominiczak and Alex Alderton, Roberto Amato, Jeffrey Barrett, Sonia Goncalves, Ewan Harrison, David K. Jackson, Ian Johnston, Dominic Kwiatkowski, Cordelia Langford, John Sillitoe on behalf of the Wellcome Sanger Institute COVID-19 Surveillance Team |
| EPI_ISL_3109358, EPI_ISL_3109359, EPI_ISL_3109360 | Lighthouse Lab in Alderley Park | Wellcome Sanger Institute for the COVID-19 Genomics UK (COG-UK) Consortium | Jacquelyn Wynn, Mairead Hyland, The Lighthouse Lab in Alderley Park and Alex Alderton, Roberto Amato, Jeffrey Barrett, Sonia Goncalves, Ewan Harrison, David K. Jackson, Ian Johnston, Dominic Kwiatkowski, Cordelia Langford, John Sillitoe on behalf of the Wellcome Sanger Institute COVID-19 Surveillance Team |
| EPI_ISL_3109361, EPI_ISL_3109362 | Berkshire and Surrey Pathology Services Lighthouse Laboratory | Wellcome Sanger Institute for the COVID-19 Genomics UK (COG-UK) Consortium | Berkshire and Surrey Pathology Services Lighthouse Laboratory and Alex Alderton, Roberto Amato, Jeffrey Barrett, Sonia Goncalves, Ewan Harrison, David K. Jackson, Ian Johnston, Dominic Kwiatkowski, Cordelia Langford, John Sillitoe on behalf of the Wellcome Sanger Institute COVID-19 Surveillance Team |
| EPI_ISL_3109363, EPI_ISL_3109364 | Lighthouse Lab in Alderley Park | Wellcome Sanger Institute for the COVID-19 Genomics UK (COG-UK) Consortium | Jacquelyn Wynn, Mairead Hyland, The Lighthouse Lab in Alderley Park and Alex Alderton, Roberto Amato, Jeffrey Barrett, Sonia Goncalves, Ewan Harrison, David K. Jackson, Ian Johnston, Dominic Kwiatkowski, Cordelia Langford, John Sillitoe on behalf of the Wellcome Sanger Institute COVID-19 Surveillance Team |
| EPI_ISL_3109365 | Berkshire and Surrey Pathology Services Lighthouse Laboratory | Wellcome Sanger Institute for the COVID-19 Genomics UK (COG-UK) Consortium | Berkshire and Surrey Pathology Services Lighthouse Laboratory and Alex Alderton, Roberto Amato, Jeffrey Barrett, Sonia Goncalves, Ewan Harrison, David K. Jackson, Ian Johnston, Dominic Kwiatkowski, Cordelia Langford, John Sillitoe on behalf of the Wellcome Sanger Institute COVID-19 Surveillance Team |
| EPI_ISL_3109366, EPI_ISL_3109367, EPI_ISL_3109368, EPI_ISL_3109369, EPI_ISL_3109370 | Lighthouse Lab in Alderley Park | Wellcome Sanger Institute for the COVID-19 Genomics UK (COG-UK) Consortium | Jacquelyn Wynn, Mairead Hyland, The Lighthouse Lab in Alderley Park and Alex Alderton, Roberto Amato, Jeffrey Barrett, Sonia Goncalves, Ewan Harrison, David K. Jackson, Ian Johnston, Dominic Kwiatkowski, Cordelia Langford, John Sillitoe on behalf of the Wellcome Sanger Institute COVID-19 Surveillance Team |
| EPI_ISL_3109371 | Berkshire and Surrey Pathology Services Lighthouse Laboratory | Wellcome Sanger Institute for the COVID-19 Genomics UK (COG-UK) Consortium | Berkshire and Surrey Pathology Services Lighthouse Laboratory and Alex Alderton, Roberto Amato, Jeffrey Barrett, Sonia Goncalves, Ewan Harrison, David K. Jackson, Ian Johnston, Dominic Kwiatkowski, Cordelia Langford, John Sillitoe on behalf of the Wellcome Sanger Institute COVID-19 Surveillance Team |
| EPI_ISL_3109372, EPI_ISL_3109373, EPI_ISL_3109374 | Lighthouse Lab in Alderley Park | Wellcome Sanger Institute for the COVID-19 Genomics UK (COG-UK) Consortium | Jacquelyn Wynn, Mairead Hyland, The Lighthouse Lab in Alderley Park and Alex Alderton, Roberto Amato, Jeffrey Barrett, Sonia Goncalves, Ewan Harrison, David K. Jackson, Ian Johnston, Dominic Kwiatkowski, Cordelia Langford, John Sillitoe on behalf of the Wellcome Sanger Institute COVID-19 Surveillance Team |
| EPI_ISL_3109375 | Berkshire and Surrey Pathology Services Lighthouse Laboratory | Wellcome Sanger Institute for the COVID-19 Genomics UK (COG-UK) Consortium | Berkshire and Surrey Pathology Services Lighthouse Laboratory and Alex Alderton, Roberto Amato, Jeffrey Barrett, Sonia Goncalves, Ewan Harrison, David K. Jackson, Ian Johnston, Dominic Kwiatkowski, Cordelia Langford, John Sillitoe on behalf of the Wellcome Sanger Institute COVID-19 Surveillance Team |
| EPI_ISL_3109376 | Lighthouse Lab in Alderley Park | Wellcome Sanger Institute for the COVID-19 Genomics UK (COG-UK) Consortium | Jacquelyn Wynn, Mairead Hyland, The Lighthouse Lab in Alderley Park and Alex Alderton, Roberto Amato, Jeffrey Barrett, Sonia Goncalves, Ewan Harrison, David K. Jackson, Ian Johnston, Dominic Kwiatkowski, Cordelia Langford, John Sillitoe on behalf of the Wellcome Sanger Institute COVID-19 Surveillance Team |
| EPI_ISL_3109377 | Lighthouse Lab in Glasgow | Wellcome Sanger Institute for the COVID-19 Genomics UK (COG-UK) Consortium | Harper VanSteenhouse, Yumi Kasai, David Gray, Carol Clugston, Anna Dominiczak and Alex Alderton, Roberto Amato, Jeffrey Barrett, Sonia Goncalves, Ewan Harrison, David K. Jackson, Ian Johnston, Dominic Kwiatkowski, Cordelia Langford, John Sillitoe on behalf of the Wellcome Sanger Institute COVID-19 Surveillance Team |
| EPI_ISL_3109378, EPI_ISL_3109379 | Lighthouse Lab in Alderley Park | Wellcome Sanger Institute for the COVID-19 Genomics UK (COG-UK) Consortium | Jacquelyn Wynn, Mairead Hyland, The Lighthouse Lab in Alderley Park and Alex Alderton, Roberto Amato, Jeffrey Barrett, Sonia Goncalves, Ewan Harrison, David K. Jackson, Ian Johnston, Dominic Kwiatkowski, Cordelia Langford, John Sillitoe on behalf of the Wellcome Sanger Institute COVID-19 Surveillance Team |
| EPI_ISL_3109380, EPI_ISL_3109381 | Berkshire and Surrey Pathology Services Lighthouse Laboratory | Wellcome Sanger Institute for the COVID-19 Genomics UK (COG-UK) Consortium | Berkshire and Surrey Pathology Services Lighthouse Laboratory and Alex Alderton, Roberto Amato, Jeffrey Barrett, Sonia Goncalves, Ewan Harrison, David K. Jackson, Ian Johnston, Dominic Kwiatkowski, Cordelia Langford, John Sillitoe on behalf of the Wellcome Sanger Institute COVID-19 Surveillance Team |
| EPI_ISL_3109382, EPI_ISL_3109383, EPI_ISL_3109384 | Lighthouse Lab in Alderley Park | Wellcome Sanger Institute for the COVID-19 Genomics UK (COG-UK) Consortium | Jacquelyn Wynn, Mairead Hyland, The Lighthouse Lab in Alderley Park and Alex Alderton, Roberto Amato, Jeffrey Barrett, Sonia Goncalves, Ewan Harrison, David K. Jackson, Ian Johnston, Dominic Kwiatkowski, Cordelia Langford, John Sillitoe on behalf of the Wellcome Sanger Institute COVID-19 Surveillance Team |
| EPI_ISL_3109385 | Berkshire and Surrey Pathology Services Lighthouse Laboratory | Wellcome Sanger Institute for the COVID-19 Genomics UK (COG-UK) Consortium | Berkshire and Surrey Pathology Services Lighthouse Laboratory and Alex Alderton, Roberto Amato, Jeffrey Barrett, Sonia Goncalves, Ewan Harrison, David K. Jackson, Ian Johnston, Dominic Kwiatkowski, Cordelia Langford, John Sillitoe on behalf of the Wellcome Sanger Institute COVID-19 Surveillance Team |
| EPI_ISL_3109386, EPI_ISL_3109387 | Lighthouse Lab in Alderley Park | Wellcome Sanger Institute for the COVID-19 Genomics UK (COG-UK) Consortium | Jacquelyn Wynn, Mairead Hyland, The Lighthouse Lab in Alderley Park and Alex Alderton, Roberto Amato, Jeffrey Barrett, Sonia Goncalves, Ewan Harrison, David K. Jackson, Ian Johnston, Dominic Kwiatkowski, Cordelia Langford, John Sillitoe on behalf of the Wellcome Sanger Institute COVID-19 Surveillance Team |
| EPI_ISL_3109388, EPI_ISL_3109389 | Berkshire and Surrey Pathology Services Lighthouse Laboratory | Wellcome Sanger Institute for the COVID-19 Genomics UK (COG-UK) Consortium | Berkshire and Surrey Pathology Services Lighthouse Laboratory and Alex Alderton, Roberto Amato, Jeffrey Barrett, Sonia Goncalves, Ewan Harrison, David K. Jackson, Ian Johnston, Dominic Kwiatkowski, Cordelia Langford, John Sillitoe on behalf of the Wellcome Sanger Institute COVID-19 Surveillance Team |
| EPI_ISL_3109390, EPI_ISL_3109391 | Lighthouse Lab in Alderley Park | Wellcome Sanger Institute for the COVID-19 Genomics UK (COG-UK) Consortium | Jacquelyn Wynn, Mairead Hyland, The Lighthouse Lab in Alderley Park and Alex Alderton, Roberto Amato, Jeffrey Barrett, Sonia Goncalves, Ewan Harrison, David K. Jackson, Ian Johnston, Dominic Kwiatkowski, Cordelia Langford, John Sillitoe on behalf of the Wellcome Sanger Institute COVID-19 Surveillance Team |
| EPI_ISL_3109392 | Berkshire and Surrey Pathology Services Lighthouse Laboratory | Wellcome Sanger Institute for the COVID-19 Genomics UK (COG-UK) Consortium | Berkshire and Surrey Pathology Services Lighthouse Laboratory and Alex Alderton, Roberto Amato, Jeffrey Barrett, Sonia Goncalves, Ewan Harrison, David K. Jackson, Ian Johnston, Dominic Kwiatkowski, Cordelia Langford, John Sillitoe on behalf of the Wellcome Sanger Institute COVID-19 Surveillance Team |
| EPI_ISL_3109393 | Lighthouse Lab in Glasgow | Wellcome Sanger Institute for the COVID-19 Genomics UK (COG-UK) Consortium | Harper VanSteenhouse, Yumi Kasai, David Gray, Carol Clugston, Anna Dominiczak and Alex Alderton, Roberto Amato, Jeffrey Barrett, Sonia Goncalves, Ewan Harrison, David K. Jackson, Ian Johnston, Dominic Kwiatkowski, Cordelia Langford, John Sillitoe on behalf of the Wellcome Sanger Institute COVID-19 Surveillance Team |
| EPI_ISL_3109394, EPI_ISL_3109395 | Lighthouse Lab in Alderley Park | Wellcome Sanger Institute for the COVID-19 Genomics UK (COG-UK) Consortium | Jacquelyn Wynn, Mairead Hyland, The Lighthouse Lab in Alderley Park and Alex Alderton, Roberto Amato, Jeffrey Barrett, Sonia Goncalves, Ewan Harrison, David K. Jackson, Ian Johnston, Dominic Kwiatkowski, Cordelia Langford, John Sillitoe on behalf of the Wellcome Sanger Institute COVID-19 Surveillance Team |
| EPI_ISL_3109396 | Berkshire and Surrey Pathology Services Lighthouse Laboratory | Wellcome Sanger Institute for the COVID-19 Genomics UK (COG-UK) Consortium | Berkshire and Surrey Pathology Services Lighthouse Laboratory and Alex Alderton, Roberto Amato, Jeffrey Barrett, Sonia Goncalves, Ewan Harrison, David K. Jackson, Ian Johnston, Dominic Kwiatkowski, Cordelia Langford, John Sillitoe on behalf of the Wellcome Sanger Institute COVID-19 Surveillance Team |
| EPI_ISL_3109397 | Lighthouse Lab in Alderley Park | Wellcome Sanger Institute for the COVID-19 Genomics UK (COG-UK) Consortium | Jacquelyn Wynn, Mairead Hyland, The Lighthouse Lab in Alderley Park and Alex Alderton, Roberto Amato, Jeffrey Barrett, Sonia Goncalves, Ewan Harrison, David K. Jackson, Ian Johnston, Dominic Kwiatkowski, Cordelia Langford, John Sillitoe on behalf of the Wellcome Sanger Institute COVID-19 Surveillance Team |
| EPI_ISL_3109398 | Lighthouse Lab in Glasgow | Wellcome Sanger Institute for the COVID-19 Genomics UK (COG-UK) Consortium | Harper VanSteenhouse, Yumi Kasai, David Gray, Carol Clugston, Anna Dominiczak and Alex Alderton, Roberto Amato, Jeffrey Barrett, Sonia Goncalves, Ewan Harrison, David K. Jackson, Ian Johnston, Dominic Kwiatkowski, Cordelia Langford, John Sillitoe on behalf of the Wellcome Sanger Institute COVID-19 Surveillance Team |
| EPI_ISL_3109399, EPI_ISL_3109400 | Lighthouse Lab in Alderley Park | Wellcome Sanger Institute for the COVID-19 Genomics UK (COG-UK) Consortium | Jacquelyn Wynn, Mairead Hyland, The Lighthouse Lab in Alderley Park and Alex Alderton, Roberto Amato, Jeffrey Barrett, Sonia Goncalves, Ewan Harrison, David K. Jackson, Ian Johnston, Dominic Kwiatkowski, Cordelia Langford, John Sillitoe on behalf of the Wellcome Sanger Institute COVID-19 Surveillance Team |

| | | | |
|---|---|---|---|
| EPI_ISL_3109401, EPI_ISL_3109402, EPI_ISL_3109403, EPI_ISL_3109404, EPI_ISL_3109405, EPI_ISL_3109406, EPI_ISL_3109407, EPI_ISL_3109408 | Lighthouse Lab in Glasgow | Wellcome Sanger Institute for the COVID-19 Genomics UK (COG-UK) Consortium | Harper VanSteenhouse, Yumi Kasai, David Gray, Carol Clugston, Anna Dominiczak and Alex Alderton, Roberto Amato, Jeffrey Barrett, Sonia Goncalves, Ewan Harrison, David K. Jackson, Ian Johnston, Dominic Kwiatkowski, Cordelia Langford, John Sillitoe on behalf of the Wellcome Sanger Institute COVID-19 Surveillance Team |
| EPI_ISL_3109409 | Lighthouse Laboratory Plymouth | Wellcome Sanger Institute for the COVID-19 Genomics UK (COG-UK) Consortium | Lighthouse Laboratory Plymouth and Alex Alderton, Roberto Amato, Jeffrey Barrett, Sonia Goncalves, Ewan Harrison, David K. Jackson, Ian Johnston, Dominic Kwiatkowski, Cordelia Langford, John Sillitoe on behalf of the Wellcome Sanger Institute COVID-19 Surveillance Team |
| EPI_ISL_3109410, EPI_ISL_3109411, EPI_ISL_3109412, EPI_ISL_3109413, EPI_ISL_3109414, EPI_ISL_3109415, EPI_ISL_3109416, EPI_ISL_3109417, EPI_ISL_3109418, EPI_ISL_3109419, EPI_ISL_3109420, EPI_ISL_3109421, EPI_ISL_3109422, EPI_ISL_3109423, EPI_ISL_3109424, EPI_ISL_3109425, EPI_ISL_3109426, EPI_ISL_3109427, EPI_ISL_3109428 | | | |
| see above | Lighthouse Lab in Glasgow | Wellcome Sanger Institute for the COVID-19 Genomics UK (COG-UK) Consortium | Harper VanSteenhouse, Yumi Kasai, David Gray, Carol Clugston, Anna Dominiczak and Alex Alderton, Roberto Amato, Jeffrey Barrett, Sonia Goncalves, Ewan Harrison, David K. Jackson, Ian Johnston, Dominic Kwiatkowski, Cordelia Langford, John Sillitoe on behalf of the Wellcome Sanger Institute COVID-19 Surveillance Team |
| EPI_ISL_3109429 | Lighthouse Lab in Milton Keynes | Wellcome Sanger Institute for the COVID-19 Genomics UK (COG-UK) Consortium | The Lighthouse Lab in Milton Keynes and Alex Alderton, Roberto Amato, Jeffrey Barrett, Sonia Goncalves, Ewan Harrison, David K. Jackson, Ian Johnston, Dominic Kwiatkowski, Cordelia Langford, John Sillitoe on behalf of the Wellcome Sanger Institute COVID-19 Surveillance Team |
| EPI_ISL_3109430, EPI_ISL_3109431 | Lighthouse Lab in Glasgow | Wellcome Sanger Institute for the COVID-19 Genomics UK (COG-UK) Consortium | Harper VanSteenhouse, Yumi Kasai, David Gray, Carol Clugston, Anna Dominiczak and Alex Alderton, Roberto Amato, Jeffrey Barrett, Sonia Goncalves, Ewan Harrison, David K. Jackson, Ian Johnston, Dominic Kwiatkowski, Cordelia Langford, John Sillitoe on behalf of the Wellcome Sanger Institute COVID-19 Surveillance Team |
| EPI_ISL_3109432 | Lighthouse Lab in Milton Keynes | Wellcome Sanger Institute for the COVID-19 Genomics UK (COG-UK) Consortium | The Lighthouse Lab in Milton Keynes and Alex Alderton, Roberto Amato, Jeffrey Barrett, Sonia Goncalves, Ewan Harrison, David K. Jackson, Ian Johnston, Dominic Kwiatkowski, Cordelia Langford, John Sillitoe on behalf of the Wellcome Sanger Institute COVID-19 Surveillance Team |
| EPI_ISL_3109433, EPI_ISL_3109434, EPI_ISL_3109435, EPI_ISL_3109436, EPI_ISL_3109437, EPI_ISL_3109438, EPI_ISL_3109439, EPI_ISL_3109440, EPI_ISL_3109441, EPI_ISL_3109442, EPI_ISL_3109443, EPI_ISL_3109444, EPI_ISL_3109445, EPI_ISL_3109446, EPI_ISL_3109447, EPI_ISL_3109448, EPI_ISL_3109449, EPI_ISL_3109450, EPI_ISL_3109451, EPI_ISL_3109452, EPI_ISL_3109453, EPI_ISL_3109454, EPI_ISL_3109455, EPI_ISL_3109456, EPI_ISL_3109457, EPI_ISL_3109458, EPI_ISL_3109459, EPI_ISL_3109460, EPI_ISL_3109461, EPI_ISL_3109462 | | | |
| see above | Lighthouse Lab in Glasgow | Wellcome Sanger Institute for the COVID-19 Genomics UK (COG-UK) Consortium | Harper VanSteenhouse, Yumi Kasai, David Gray, Carol Clugston, Anna Dominiczak and Alex Alderton, Roberto Amato, Jeffrey Barrett, Sonia Goncalves, Ewan Harrison, David K. Jackson, Ian Johnston, Dominic Kwiatkowski, Cordelia Langford, John Sillitoe on behalf of the Wellcome Sanger Institute COVID-19 Surveillance Team |
| EPI_ISL_3109463 | Lighthouse Laboratory Plymouth | Wellcome Sanger Institute for the COVID-19 Genomics UK (COG-UK) Consortium | Lighthouse Laboratory Plymouth and Alex Alderton, Roberto Amato, Jeffrey Barrett, Sonia Goncalves, Ewan Harrison, David K. Jackson, Ian Johnston, Dominic Kwiatkowski, Cordelia Langford, John Sillitoe on behalf of the Wellcome Sanger Institute COVID-19 Surveillance Team |
| EPI_ISL_3109464 | Lighthouse Lab in Glasgow | Wellcome Sanger Institute for the COVID-19 Genomics UK (COG-UK) Consortium | Harper VanSteenhouse, Yumi Kasai, David Gray, Carol Clugston, Anna Dominiczak and Alex Alderton, Roberto Amato, Jeffrey Barrett, Sonia Goncalves, Ewan Harrison, David K. Jackson, Ian Johnston, Dominic Kwiatkowski, Cordelia Langford, John Sillitoe on behalf of the Wellcome Sanger Institute COVID-19 Surveillance Team |
| EPI_ISL_3109465, EPI_ISL_3109466 | Lighthouse Lab in Milton Keynes | Wellcome Sanger Institute for the COVID-19 Genomics UK (COG-UK) Consortium | The Lighthouse Lab in Milton Keynes and Alex Alderton, Roberto Amato, Jeffrey Barrett, Sonia Goncalves, Ewan Harrison, David K. Jackson, Ian Johnston, Dominic Kwiatkowski, Cordelia Langford, John Sillitoe on behalf of the Wellcome Sanger Institute COVID-19 Surveillance Team |
| EPI_ISL_3109467, EPI_ISL_3109468 | Lighthouse Lab in Glasgow | Wellcome Sanger Institute for the COVID-19 Genomics UK (COG-UK) Consortium | Harper VanSteenhouse, Yumi Kasai, David Gray, Carol Clugston, Anna Dominiczak and Alex Alderton, Roberto Amato, Jeffrey Barrett, Sonia Goncalves, Ewan Harrison, David K. Jackson, Ian Johnston, Dominic Kwiatkowski, Cordelia Langford, John Sillitoe on behalf of the Wellcome Sanger Institute COVID-19 Surveillance Team |
| EPI_ISL_3109469 | Lighthouse Lab in Milton Keynes | Wellcome Sanger Institute for the COVID-19 Genomics UK (COG-UK) Consortium | The Lighthouse Lab in Milton Keynes and Alex Alderton, Roberto Amato, Jeffrey Barrett, Sonia Goncalves, Ewan Harrison, David K. Jackson, Ian Johnston, Dominic Kwiatkowski, Cordelia Langford, John Sillitoe on behalf of the Wellcome Sanger Institute COVID-19 Surveillance Team |
| EPI_ISL_3109470, EPI_ISL_3109471, EPI_ISL_3109472 | Lighthouse Lab in Glasgow | Wellcome Sanger Institute for the COVID-19 Genomics UK (COG-UK) Consortium | Harper VanSteenhouse, Yumi Kasai, David Gray, Carol Clugston, Anna Dominiczak and Alex Alderton, Roberto Amato, Jeffrey Barrett, Sonia Goncalves, Ewan Harrison, David K. Jackson, Ian Johnston, Dominic Kwiatkowski, Cordelia Langford, John Sillitoe on behalf of the Wellcome Sanger Institute COVID-19 Surveillance Team |
| EPI_ISL_3109473 | Lighthouse Lab in Milton Keynes | Wellcome Sanger Institute for the COVID-19 Genomics UK (COG-UK) Consortium | The Lighthouse Lab in Milton Keynes and Alex Alderton, Roberto Amato, Jeffrey Barrett, Sonia Goncalves, Ewan Harrison, David K. Jackson, Ian Johnston, Dominic Kwiatkowski, Cordelia Langford, John Sillitoe on behalf of the Wellcome Sanger Institute COVID-19 Surveillance Team |
| EPI_ISL_3109474, EPI_ISL_3109475 | Lighthouse Lab in Glasgow | Wellcome Sanger Institute for the COVID-19 Genomics UK (COG-UK) Consortium | Harper VanSteenhouse, Yumi Kasai, David Gray, Carol Clugston, Anna Dominiczak and Alex Alderton, Roberto Amato, Jeffrey Barrett, Sonia Goncalves, Ewan Harrison, David K. Jackson, Ian Johnston, Dominic Kwiatkowski, Cordelia Langford, John Sillitoe on behalf of the Wellcome Sanger Institute COVID-19 Surveillance Team |
| EPI_ISL_3109476 | Lighthouse Lab in Milton Keynes | Wellcome Sanger Institute for the COVID-19 Genomics UK (COG-UK) Consortium | The Lighthouse Lab in Milton Keynes and Alex Alderton, Roberto Amato, Jeffrey Barrett, Sonia Goncalves, Ewan Harrison, David K. Jackson, Ian Johnston, Dominic Kwiatkowski, Cordelia Langford, John Sillitoe on behalf of the Wellcome Sanger Institute COVID-19 Surveillance Team |
| EPI_ISL_3109477, EPI_ISL_3109478, EPI_ISL_3109479, EPI_ISL_3109480, EPI_ISL_3109481, EPI_ISL_3109482 | Lighthouse Lab in Glasgow | Wellcome Sanger Institute for the COVID-19 Genomics UK (COG-UK) Consortium | Harper VanSteenhouse, Yumi Kasai, David Gray, Carol Clugston, Anna Dominiczak and Alex Alderton, Roberto Amato, Jeffrey Barrett, Sonia Goncalves, Ewan Harrison, David K. Jackson, Ian Johnston, Dominic Kwiatkowski, Cordelia Langford, John Sillitoe on behalf of the Wellcome Sanger Institute COVID-19 Surveillance Team |
| EPI_ISL_3109483 | Lighthouse Lab in Milton Keynes | Wellcome Sanger Institute for the COVID-19 Genomics UK (COG-UK) Consortium | The Lighthouse Lab in Milton Keynes and Alex Alderton, Roberto Amato, Jeffrey Barrett, Sonia Goncalves, Ewan Harrison, David K. Jackson, Ian Johnston, Dominic Kwiatkowski, Cordelia Langford, John Sillitoe on behalf of the Wellcome Sanger Institute COVID-19 Surveillance Team |
| EPI_ISL_3109484, EPI_ISL_3109485, EPI_ISL_3109486 | Lighthouse Lab in Glasgow | Wellcome Sanger Institute for the COVID-19 Genomics UK (COG-UK) Consortium | Harper VanSteenhouse, Yumi Kasai, David Gray, Carol Clugston, Anna Dominiczak and Alex Alderton, Roberto Amato, Jeffrey Barrett, Sonia Goncalves, Ewan Harrison, David K. Jackson, Ian Johnston, Dominic Kwiatkowski, Cordelia Langford, John Sillitoe on behalf of the Wellcome Sanger Institute COVID-19 Surveillance Team |
| EPI_ISL_3109487, EPI_ISL_3109488, EPI_ISL_3109489 | Lighthouse Lab in Milton Keynes | Wellcome Sanger Institute for the COVID-19 Genomics UK (COG-UK) Consortium | The Lighthouse Lab in Milton Keynes and Alex Alderton, Roberto Amato, Jeffrey Barrett, Sonia Goncalves, Ewan Harrison, David K. Jackson, Ian Johnston, Dominic Kwiatkowski, Cordelia Langford, John Sillitoe on behalf of the Wellcome Sanger Institute COVID-19 Surveillance Team |
| EPI_ISL_3109490 | Lighthouse Lab in Glasgow | Wellcome Sanger Institute for the COVID-19 Genomics UK (COG-UK) Consortium | Harper VanSteenhouse, Yumi Kasai, David Gray, Carol Clugston, Anna Dominiczak and Alex Alderton, Roberto Amato, Jeffrey Barrett, Sonia Goncalves, Ewan Harrison, David K. Jackson, Ian Johnston, Dominic Kwiatkowski, Cordelia Langford, John Sillitoe on behalf of the Wellcome Sanger Institute COVID-19 Surveillance Team |
| EPI_ISL_3109491 | Lighthouse Lab in Milton Keynes | Wellcome Sanger Institute for the COVID-19 Genomics UK (COG-UK) Consortium | The Lighthouse Lab in Milton Keynes and Alex Alderton, Roberto Amato, Jeffrey Barrett, Sonia Goncalves, Ewan Harrison, David K. Jackson, Ian Johnston, Dominic Kwiatkowski, Cordelia Langford, John Sillitoe on behalf of the Wellcome Sanger Institute COVID-19 Surveillance Team |
| EPI_ISL_3109492 | Lighthouse Lab in Glasgow | Wellcome Sanger Institute for the COVID-19 Genomics UK (COG-UK) Consortium | Harper VanSteenhouse, Yumi Kasai, David Gray, Carol Clugston, Anna Dominiczak and Alex Alderton, Roberto Amato, Jeffrey Barrett, Sonia Goncalves, Ewan Harrison, David K. Jackson, Ian Johnston, Dominic Kwiatkowski, Cordelia Langford, John Sillitoe on behalf of the Wellcome Sanger Institute COVID-19 Surveillance Team |
| EPI_ISL_3109493 | Lighthouse Lab in Milton Keynes | Wellcome Sanger Institute for the COVID-19 Genomics UK (COG-UK) Consortium | The Lighthouse Lab in Milton Keynes and Alex Alderton, Roberto Amato, Jeffrey Barrett, Sonia Goncalves, Ewan Harrison, David K. Jackson, Ian Johnston, Dominic Kwiatkowski, Cordelia Langford, John Sillitoe on behalf of the Wellcome Sanger Institute COVID-19 Surveillance Team |
| EPI_ISL_3109494 | Lighthouse Lab in Glasgow | Wellcome Sanger Institute for the COVID-19 Genomics UK (COG-UK) Consortium | Harper VanSteenhouse, Yumi Kasai, David Gray, Carol Clugston, Anna Dominiczak and Alex Alderton, Roberto Amato, Jeffrey Barrett, Sonia Goncalves, Ewan Harrison, David K. Jackson, Ian Johnston, Dominic Kwiatkowski, Cordelia Langford, John Sillitoe on behalf of the Wellcome Sanger Institute COVID-19 Surveillance Team |
| EPI_ISL_3109495 | Lighthouse Lab in Milton Keynes | Wellcome Sanger Institute for the COVID-19 Genomics UK (COG-UK) Consortium | The Lighthouse Lab in Milton Keynes and Alex Alderton, Roberto Amato, Jeffrey Barrett, Sonia Goncalves, Ewan Harrison, David K. Jackson, Ian Johnston, Dominic Kwiatkowski, Cordelia Langford, John Sillitoe on behalf of the Wellcome Sanger Institute COVID-19 Surveillance Team |
| EPI_ISL_3109496, EPI_ISL_3109497 | Lighthouse Lab in Glasgow | Wellcome Sanger Institute for the COVID-19 Genomics UK (COG-UK) Consortium | Harper VanSteenhouse, Yumi Kasai, David Gray, Carol Clugston, Anna Dominiczak and Alex Alderton, Roberto Amato, Jeffrey Barrett, Sonia Goncalves, Ewan Harrison, David K. Jackson, Ian Johnston, Dominic Kwiatkowski, Cordelia Langford, John Sillitoe on behalf of the Wellcome Sanger Institute COVID-19 Surveillance Team |
| EPI_ISL_3109498, EPI_ISL_3109499 | Lighthouse Lab in Milton Keynes | Wellcome Sanger Institute for the COVID-19 Genomics UK (COG-UK) Consortium | The Lighthouse Lab in Milton Keynes and Alex Alderton, Roberto Amato, Jeffrey Barrett, Sonia Goncalves, Ewan Harrison, David K. Jackson, Ian Johnston, Dominic Kwiatkowski, Cordelia Langford, John Sillitoe on behalf of the Wellcome Sanger Institute COVID-19 Surveillance Team |
| EPI_ISL_3109500, EPI_ISL_3109501, EPI_ISL_3109502, EPI_ISL_3109503, EPI_ISL_3109504, EPI_ISL_3109505, EPI_ISL_3109506, EPI_ISL_3109507, EPI_ISL_3109508, EPI_ISL_3109509, EPI_ISL_3109510, EPI_ISL_3109511, EPI_ISL_3109512, EPI_ISL_3109513, EPI_ISL_3109514, EPI_ISL_3109515, EPI_ISL_3109516, EPI_ISL_3109517, EPI_ISL_3109518, EPI_ISL_3109519, EPI_ISL_3109520 | | | |
| see above | Lighthouse Lab in Glasgow | Wellcome Sanger Institute for the COVID-19 Genomics UK (COG-UK) Consortium | Harper VanSteenhouse, Yumi Kasai, David Gray, Carol Clugston, Anna Dominiczak and Alex Alderton, Roberto Amato, Jeffrey Barrett, Sonia Goncalves, Ewan Harrison, David K. Jackson, Ian Johnston, Dominic Kwiatkowski, Cordelia Langford, John Sillitoe on behalf of the Wellcome Sanger Institute COVID-19 Surveillance Team |
| EPI_ISL_3109521 | Berkshire and Surrey Pathology Services Lighthouse Laboratory | Wellcome Sanger Institute for the COVID-19 Genomics UK (COG-UK) Consortium | Berkshire and Surrey Pathology Services Lighthouse Laboratory and Alex Alderton, Roberto Amato, Jeffrey Barrett, Sonia Goncalves, Ewan Harrison, David K. Jackson, Ian Johnston, Dominic Kwiatkowski, Cordelia Langford, John Sillitoe on behalf of the Wellcome Sanger Institute COVID-19 Surveillance Team |
| EPI_ISL_3109522 | Lighthouse Lab in Alderley Park | Wellcome Sanger Institute for the COVID-19 Genomics UK (COG-UK) Consortium | Jacquelyn Wynn, Mairead Hyland, The Lighthouse Lab in Alderley Park and Alex Alderton, Roberto Amato, Jeffrey Barrett, Sonia Goncalves, Ewan Harrison, David K. Jackson, Ian Johnston, Dominic Kwiatkowski, Cordelia Langford, John Sillitoe on behalf of the Wellcome Sanger Institute COVID-19 Surveillance Team |
| EPI_ISL_3109523 | Lighthouse Laboratory Plymouth | Wellcome Sanger Institute for the COVID-19 Genomics UK (COG-UK) Consortium | Lighthouse Laboratory Plymouth and Alex Alderton, Roberto Amato, Jeffrey Barrett, Sonia Goncalves, Ewan Harrison, David K. Jackson, Ian Johnston, Dominic Kwiatkowski, Cordelia Langford, John Sillitoe on behalf of the Wellcome Sanger Institute COVID-19 Surveillance Team |
| EPI_ISL_3109524 | Berkshire and Surrey Pathology Services Lighthouse Laboratory | Wellcome Sanger Institute for the COVID-19 Genomics UK (COG-UK) Consortium | Berkshire and Surrey Pathology Services Lighthouse Laboratory and Alex Alderton, Roberto Amato, Jeffrey Barrett, Sonia Goncalves, Ewan Harrison, David K. Jackson, Ian Johnston, Dominic Kwiatkowski, Cordelia Langford, John Sillitoe on behalf of the Wellcome Sanger Institute COVID-19 Surveillance Team |
| EPI_ISL_3109525 | Lighthouse Lab in Alderley Park | Wellcome Sanger Institute for the COVID-19 Genomics UK (COG-UK) Consortium | Jacquelyn Wynn, Mairead Hyland, The Lighthouse Lab in Alderley Park and Alex Alderton, Roberto Amato, Jeffrey Barrett, Sonia Goncalves, Ewan Harrison, David K. Jackson, Ian Johnston, Dominic Kwiatkowski, Cordelia Langford, John Sillitoe on behalf of the Wellcome Sanger Institute COVID-19 Surveillance Team |
| EPI_ISL_3109526, EPI_ISL_3109527, EPI_ISL_3109528 | Lighthouse Lab in Milton Keynes | Wellcome Sanger Institute for the COVID-19 Genomics UK (COG-UK) Consortium | The Lighthouse Lab in Milton Keynes and Alex Alderton, Roberto Amato, Jeffrey Barrett, Sonia Goncalves, Ewan Harrison, David K. Jackson, Ian Johnston, Dominic Kwiatkowski, Cordelia Langford, John Sillitoe on behalf of the Wellcome Sanger Institute COVID-19 Surveillance Team |
| EPI_ISL_3109529 | Lighthouse Lab in Alderley Park | Wellcome Sanger Institute for the COVID-19 Genomics UK (COG-UK) Consortium | Jacquelyn Wynn, Mairead Hyland, The Lighthouse Lab in Alderley Park and Alex Alderton, Roberto Amato, Jeffrey Barrett, Sonia Goncalves, Ewan Harrison, David K. Jackson, Ian Johnston, Dominic Kwiatkowski, Cordelia Langford, John Sillitoe on behalf of the Wellcome Sanger Institute COVID-19 Surveillance Team |
| EPI_ISL_3109530 | Berkshire and Surrey Pathology Services Lighthouse Laboratory | Wellcome Sanger Institute for the COVID-19 Genomics UK (COG-UK) Consortium | Berkshire and Surrey Pathology Services Lighthouse Laboratory and Alex Alderton, Roberto Amato, Jeffrey Barrett, Sonia Goncalves, Ewan Harrison, David K. Jackson, Ian Johnston, Dominic Kwiatkowski, Cordelia Langford, John Sillitoe on behalf of the Wellcome Sanger Institute COVID-19 Surveillance Team |
| EPI_ISL_3109531, | Lighthouse Lab in Milton Keynes | Wellcome Sanger Institute for the COVID-19 Genomics UK (COG-UK) | The Lighthouse Lab in Milton Keynes and Alex Alderton, Roberto Amato, Jeffrey Barrett, Sonia Goncalves, Ewan Harrison, David K. Jackson, Ian Johnston, Dominic Kwiatkowski, Cordelia Langford, John Sillitoe on behalf of the Wellcome Sanger |

| | | | |
|---|---|---|---|
| EPI_ISL_3109532, EPI_ISL_3109533 | | Consortium | Institute COVID-19 Surveillance Team |
| EPI_ISL_3109534 | Lighthouse Lab in Alderley Park | Wellcome Sanger Institute for the COVID-19 Genomics UK (COG-UK) Consortium | Jacquelyn Wynn, Mairead Hyland, The Lighthouse Lab in Alderley Park and Alex Alderton, Roberto Amato, Jeffrey Barrett, Sonia Goncalves, Ewan Harrison, David K. Jackson, Ian Johnston, Dominic Kwiatkowski, Cordelia Langford, John Sillitoe on behalf of the Wellcome Sanger Institute COVID-19 Surveillance Team |
| EPI_ISL_3109535 | Lighthouse Lab in Milton Keynes | Wellcome Sanger Institute for the COVID-19 Genomics UK (COG-UK) Consortium | The Lighthouse Lab in Milton Keynes and Alex Alderton, Roberto Amato, Jeffrey Barrett, Sonia Goncalves, Ewan Harrison, David K. Jackson, Ian Johnston, Dominic Kwiatkowski, Cordelia Langford, John Sillitoe on behalf of the Wellcome Sanger Institute COVID-19 Surveillance Team |
| EPI_ISL_3109536 | Berkshire and Surrey Pathology Services Lighthouse Laboratory | Wellcome Sanger Institute for the COVID-19 Genomics UK (COG-UK) Consortium | Berkshire and Surrey Pathology Services Lighthouse Laboratory and Alex Alderton, Roberto Amato, Jeffrey Barrett, Sonia Goncalves, Ewan Harrison, David K. Jackson, Ian Johnston, Dominic Kwiatkowski, Cordelia Langford, John Sillitoe on behalf of the Wellcome Sanger Institute COVID-19 Surveillance Team |
| EPI_ISL_3109537, EPI_ISL_3109538 | Lighthouse Laboratory Plymouth | Wellcome Sanger Institute for the COVID-19 Genomics UK (COG-UK) Consortium | Lighthouse Laboratory Plymouth and Alex Alderton, Roberto Amato, Jeffrey Barrett, Sonia Goncalves, Ewan Harrison, David K. Jackson, Ian Johnston, Dominic Kwiatkowski, Cordelia Langford, John Sillitoe on behalf of the Wellcome Sanger Institute COVID-19 Surveillance Team |
| EPI_ISL_3109539 | Lighthouse Lab in Alderley Park | Wellcome Sanger Institute for the COVID-19 Genomics UK (COG-UK) Consortium | Jacquelyn Wynn, Mairead Hyland, The Lighthouse Lab in Alderley Park and Alex Alderton, Roberto Amato, Jeffrey Barrett, Sonia Goncalves, Ewan Harrison, David K. Jackson, Ian Johnston, Dominic Kwiatkowski, Cordelia Langford, John Sillitoe on behalf of the Wellcome Sanger Institute COVID-19 Surveillance Team |
| EPI_ISL_3109540 | Lighthouse Laboratory Plymouth | Wellcome Sanger Institute for the COVID-19 Genomics UK (COG-UK) Consortium | Lighthouse Laboratory Plymouth and Alex Alderton, Roberto Amato, Jeffrey Barrett, Sonia Goncalves, Ewan Harrison, David K. Jackson, Ian Johnston, Dominic Kwiatkowski, Cordelia Langford, John Sillitoe on behalf of the Wellcome Sanger Institute COVID-19 Surveillance Team |
| EPI_ISL_3109541 | Lighthouse Lab in Alderley Park | Wellcome Sanger Institute for the COVID-19 Genomics UK (COG-UK) Consortium | Jacquelyn Wynn, Mairead Hyland, The Lighthouse Lab in Alderley Park and Alex Alderton, Roberto Amato, Jeffrey Barrett, Sonia Goncalves, Ewan Harrison, David K. Jackson, Ian Johnston, Dominic Kwiatkowski, Cordelia Langford, John Sillitoe on behalf of the Wellcome Sanger Institute COVID-19 Surveillance Team |
| EPI_ISL_3109542 | Lighthouse Lab in Milton Keynes | Wellcome Sanger Institute for the COVID-19 Genomics UK (COG-UK) Consortium | The Lighthouse Lab in Milton Keynes and Alex Alderton, Roberto Amato, Jeffrey Barrett, Sonia Goncalves, Ewan Harrison, David K. Jackson, Ian Johnston, Dominic Kwiatkowski, Cordelia Langford, John Sillitoe on behalf of the Wellcome Sanger Institute COVID-19 Surveillance Team |
| EPI_ISL_3109543 | Berkshire and Surrey Pathology Services Lighthouse Laboratory | Wellcome Sanger Institute for the COVID-19 Genomics UK (COG-UK) Consortium | Berkshire and Surrey Pathology Services Lighthouse Laboratory and Alex Alderton, Roberto Amato, Jeffrey Barrett, Sonia Goncalves, Ewan Harrison, David K. Jackson, Ian Johnston, Dominic Kwiatkowski, Cordelia Langford, John Sillitoe on behalf of the Wellcome Sanger Institute COVID-19 Surveillance Team |
| EPI_ISL_3109544 | Lighthouse Lab in Alderley Park | Wellcome Sanger Institute for the COVID-19 Genomics UK (COG-UK) Consortium | Jacquelyn Wynn, Mairead Hyland, The Lighthouse Lab in Alderley Park and Alex Alderton, Roberto Amato, Jeffrey Barrett, Sonia Goncalves, Ewan Harrison, David K. Jackson, Ian Johnston, Dominic Kwiatkowski, Cordelia Langford, John Sillitoe on behalf of the Wellcome Sanger Institute COVID-19 Surveillance Team |
| EPI_ISL_3109545 | Lighthouse Lab in Milton Keynes | Wellcome Sanger Institute for the COVID-19 Genomics UK (COG-UK) Consortium | The Lighthouse Lab in Milton Keynes and Alex Alderton, Roberto Amato, Jeffrey Barrett, Sonia Goncalves, Ewan Harrison, David K. Jackson, Ian Johnston, Dominic Kwiatkowski, Cordelia Langford, John Sillitoe on behalf of the Wellcome Sanger Institute COVID-19 Surveillance Team |
| EPI_ISL_3109546 | Lighthouse Laboratory Plymouth | Wellcome Sanger Institute for the COVID-19 Genomics UK (COG-UK) Consortium | Lighthouse Laboratory Plymouth and Alex Alderton, Roberto Amato, Jeffrey Barrett, Sonia Goncalves, Ewan Harrison, David K. Jackson, Ian Johnston, Dominic Kwiatkowski, Cordelia Langford, John Sillitoe on behalf of the Wellcome Sanger Institute COVID-19 Surveillance Team |
| EPI_ISL_3109547 | Lighthouse Lab in Milton Keynes | Wellcome Sanger Institute for the COVID-19 Genomics UK (COG-UK) Consortium | The Lighthouse Lab in Milton Keynes and Alex Alderton, Roberto Amato, Jeffrey Barrett, Sonia Goncalves, Ewan Harrison, David K. Jackson, Ian Johnston, Dominic Kwiatkowski, Cordelia Langford, John Sillitoe on behalf of the Wellcome Sanger Institute COVID-19 Surveillance Team |
| EPI_ISL_3109548 | Lighthouse Lab in Alderley Park | Wellcome Sanger Institute for the COVID-19 Genomics UK (COG-UK) Consortium | Jacquelyn Wynn, Mairead Hyland, The Lighthouse Lab in Alderley Park and Alex Alderton, Roberto Amato, Jeffrey Barrett, Sonia Goncalves, Ewan Harrison, David K. Jackson, Ian Johnston, Dominic Kwiatkowski, Cordelia Langford, John Sillitoe on behalf of the Wellcome Sanger Institute COVID-19 Surveillance Team |
| EPI_ISL_3109549 | Berkshire and Surrey Pathology Services Lighthouse Laboratory | Wellcome Sanger Institute for the COVID-19 Genomics UK (COG-UK) Consortium | Berkshire and Surrey Pathology Services Lighthouse Laboratory and Alex Alderton, Roberto Amato, Jeffrey Barrett, Sonia Goncalves, Ewan Harrison, David K. Jackson, Ian Johnston, Dominic Kwiatkowski, Cordelia Langford, John Sillitoe on behalf of the Wellcome Sanger Institute COVID-19 Surveillance Team |
| EPI_ISL_3109550 | Lighthouse Lab in Milton Keynes | Wellcome Sanger Institute for the COVID-19 Genomics UK (COG-UK) Consortium | The Lighthouse Lab in Milton Keynes and Alex Alderton, Roberto Amato, Jeffrey Barrett, Sonia Goncalves, Ewan Harrison, David K. Jackson, Ian Johnston, Dominic Kwiatkowski, Cordelia Langford, John Sillitoe on behalf of the Wellcome Sanger Institute COVID-19 Surveillance Team |
| EPI_ISL_3109551 | Lighthouse Lab in Alderley Park | Wellcome Sanger Institute for the COVID-19 Genomics UK (COG-UK) Consortium | Jacquelyn Wynn, Mairead Hyland, The Lighthouse Lab in Alderley Park and Alex Alderton, Roberto Amato, Jeffrey Barrett, Sonia Goncalves, Ewan Harrison, David K. Jackson, Ian Johnston, Dominic Kwiatkowski, Cordelia Langford, John Sillitoe on behalf of the Wellcome Sanger Institute COVID-19 Surveillance Team |
| EPI_ISL_3109552, EPI_ISL_3109553, EPI_ISL_3109554, EPI_ISL_3109555 | Lighthouse Laboratory Plymouth | Wellcome Sanger Institute for the COVID-19 Genomics UK (COG-UK) Consortium | Lighthouse Laboratory Plymouth and Alex Alderton, Roberto Amato, Jeffrey Barrett, Sonia Goncalves, Ewan Harrison, David K. Jackson, Ian Johnston, Dominic Kwiatkowski, Cordelia Langford, John Sillitoe on behalf of the Wellcome Sanger Institute COVID-19 Surveillance Team |
| EPI_ISL_3109556, EPI_ISL_3109557 | Berkshire and Surrey Pathology Services Lighthouse Laboratory | Wellcome Sanger Institute for the COVID-19 Genomics UK (COG-UK) Consortium | Berkshire and Surrey Pathology Services Lighthouse Laboratory and Alex Alderton, Roberto Amato, Jeffrey Barrett, Sonia Goncalves, Ewan Harrison, David K. Jackson, Ian Johnston, Dominic Kwiatkowski, Cordelia Langford, John Sillitoe on behalf of the Wellcome Sanger Institute COVID-19 Surveillance Team |
| EPI_ISL_3109558, EPI_ISL_3109559 | Lighthouse Lab in Milton Keynes | Wellcome Sanger Institute for the COVID-19 Genomics UK (COG-UK) Consortium | The Lighthouse Lab in Milton Keynes and Alex Alderton, Roberto Amato, Jeffrey Barrett, Sonia Goncalves, Ewan Harrison, David K. Jackson, Ian Johnston, Dominic Kwiatkowski, Cordelia Langford, John Sillitoe on behalf of the Wellcome Sanger Institute COVID-19 Surveillance Team |
| EPI_ISL_3109560 | Lighthouse Laboratory Plymouth | Wellcome Sanger Institute for the COVID-19 Genomics UK (COG-UK) Consortium | Lighthouse Laboratory Plymouth and Alex Alderton, Roberto Amato, Jeffrey Barrett, Sonia Goncalves, Ewan Harrison, David K. Jackson, Ian Johnston, Dominic Kwiatkowski, Cordelia Langford, John Sillitoe on behalf of the Wellcome Sanger Institute COVID-19 Surveillance Team |
| EPI_ISL_3109561 | Lighthouse Lab in Alderley Park | Wellcome Sanger Institute for the COVID-19 Genomics UK (COG-UK) Consortium | Jacquelyn Wynn, Mairead Hyland, The Lighthouse Lab in Alderley Park and Alex Alderton, Roberto Amato, Jeffrey Barrett, Sonia Goncalves, Ewan Harrison, David K. Jackson, Ian Johnston, Dominic Kwiatkowski, Cordelia Langford, John Sillitoe on behalf of the Wellcome Sanger Institute COVID-19 Surveillance Team |
| EPI_ISL_3109562, EPI_ISL_3109563 | Berkshire and Surrey Pathology Services Lighthouse Laboratory | Wellcome Sanger Institute for the COVID-19 Genomics UK (COG-UK) Consortium | Berkshire and Surrey Pathology Services Lighthouse Laboratory and Alex Alderton, Roberto Amato, Jeffrey Barrett, Sonia Goncalves, Ewan Harrison, David K. Jackson, Ian Johnston, Dominic Kwiatkowski, Cordelia Langford, John Sillitoe on behalf of the Wellcome Sanger Institute COVID-19 Surveillance Team |
| EPI_ISL_3109564 | Lighthouse Lab in Alderley Park | Wellcome Sanger Institute for the COVID-19 Genomics UK (COG-UK) Consortium | Jacquelyn Wynn, Mairead Hyland, The Lighthouse Lab in Alderley Park and Alex Alderton, Roberto Amato, Jeffrey Barrett, Sonia Goncalves, Ewan Harrison, David K. Jackson, Ian Johnston, Dominic Kwiatkowski, Cordelia Langford, John Sillitoe on behalf of the Wellcome Sanger Institute COVID-19 Surveillance Team |
| EPI_ISL_3109565 | Lighthouse Lab in Milton Keynes | Wellcome Sanger Institute for the COVID-19 Genomics UK (COG-UK) Consortium | The Lighthouse Lab in Milton Keynes and Alex Alderton, Roberto Amato, Jeffrey Barrett, Sonia Goncalves, Ewan Harrison, David K. Jackson, Ian Johnston, Dominic Kwiatkowski, Cordelia Langford, John Sillitoe on behalf of the Wellcome Sanger Institute COVID-19 Surveillance Team |
| EPI_ISL_3109566 | Lighthouse Laboratory Plymouth | Wellcome Sanger Institute for the COVID-19 Genomics UK (COG-UK) Consortium | Lighthouse Laboratory Plymouth and Alex Alderton, Roberto Amato, Jeffrey Barrett, Sonia Goncalves, Ewan Harrison, David K. Jackson, Ian Johnston, Dominic Kwiatkowski, Cordelia Langford, John Sillitoe on behalf of the Wellcome Sanger Institute COVID-19 Surveillance Team |
| EPI_ISL_3109567 | Lighthouse Lab in Milton Keynes | Wellcome Sanger Institute for the COVID-19 Genomics UK (COG-UK) Consortium | The Lighthouse Lab in Milton Keynes and Alex Alderton, Roberto Amato, Jeffrey Barrett, Sonia Goncalves, Ewan Harrison, David K. Jackson, Ian Johnston, Dominic Kwiatkowski, Cordelia Langford, John Sillitoe on behalf of the Wellcome Sanger Institute COVID-19 Surveillance Team |
| EPI_ISL_3109568, EPI_ISL_3109569 | Lighthouse Laboratory Plymouth | Wellcome Sanger Institute for the COVID-19 Genomics UK (COG-UK) Consortium | Lighthouse Laboratory Plymouth and Alex Alderton, Roberto Amato, Jeffrey Barrett, Sonia Goncalves, Ewan Harrison, David K. Jackson, Ian Johnston, Dominic Kwiatkowski, Cordelia Langford, John Sillitoe on behalf of the Wellcome Sanger Institute COVID-19 Surveillance Team |
| EPI_ISL_3109570 | Lighthouse Lab in Alderley Park | Wellcome Sanger Institute for the COVID-19 Genomics UK (COG-UK) Consortium | Jacquelyn Wynn, Mairead Hyland, The Lighthouse Lab in Alderley Park and Alex Alderton, Roberto Amato, Jeffrey Barrett, Sonia Goncalves, Ewan Harrison, David K. Jackson, Ian Johnston, Dominic Kwiatkowski, Cordelia Langford, John Sillitoe on behalf of the Wellcome Sanger Institute COVID-19 Surveillance Team |
| EPI_ISL_3109571 | Lighthouse Laboratory Plymouth | Wellcome Sanger Institute for the COVID-19 Genomics UK (COG-UK) Consortium | Lighthouse Laboratory Plymouth and Alex Alderton, Roberto Amato, Jeffrey Barrett, Sonia Goncalves, Ewan Harrison, David K. Jackson, Ian Johnston, Dominic Kwiatkowski, Cordelia Langford, John Sillitoe on behalf of the Wellcome Sanger Institute COVID-19 Surveillance Team |
| EPI_ISL_3109572 | Lighthouse Lab in Alderley Park | Wellcome Sanger Institute for the COVID-19 Genomics UK (COG-UK) Consortium | Jacquelyn Wynn, Mairead Hyland, The Lighthouse Lab in Alderley Park and Alex Alderton, Roberto Amato, Jeffrey Barrett, Sonia Goncalves, Ewan Harrison, David K. Jackson, Ian Johnston, Dominic Kwiatkowski, Cordelia Langford, John Sillitoe on behalf of the Wellcome Sanger Institute COVID-19 Surveillance Team |
| EPI_ISL_3109573 | Lighthouse Laboratory Plymouth | Wellcome Sanger Institute for the COVID-19 Genomics UK (COG-UK) Consortium | Lighthouse Laboratory Plymouth and Alex Alderton, Roberto Amato, Jeffrey Barrett, Sonia Goncalves, Ewan Harrison, David K. Jackson, Ian Johnston, Dominic Kwiatkowski, Cordelia Langford, John Sillitoe on behalf of the Wellcome Sanger Institute COVID-19 Surveillance Team |
| EPI_ISL_3109574 | Berkshire and Surrey Pathology Services Lighthouse Laboratory | Wellcome Sanger Institute for the COVID-19 Genomics UK (COG-UK) Consortium | Berkshire and Surrey Pathology Services Lighthouse Laboratory and Alex Alderton, Roberto Amato, Jeffrey Barrett, Sonia Goncalves, Ewan Harrison, David K. Jackson, Ian Johnston, Dominic Kwiatkowski, Cordelia Langford, John Sillitoe on behalf of the Wellcome Sanger Institute COVID-19 Surveillance Team |
| EPI_ISL_3109575 | Lighthouse Lab in Alderley Park | Wellcome Sanger Institute for the COVID-19 Genomics UK (COG-UK) Consortium | Jacquelyn Wynn, Mairead Hyland, The Lighthouse Lab in Alderley Park and Alex Alderton, Roberto Amato, Jeffrey Barrett, Sonia Goncalves, Ewan Harrison, David K. Jackson, Ian Johnston, Dominic Kwiatkowski, Cordelia Langford, John Sillitoe on behalf of the Wellcome Sanger Institute COVID-19 Surveillance Team |
| EPI_ISL_3109576 | Lighthouse Lab in Milton Keynes | Wellcome Sanger Institute for the COVID-19 Genomics UK (COG-UK) Consortium | The Lighthouse Lab in Milton Keynes and Alex Alderton, Roberto Amato, Jeffrey Barrett, Sonia Goncalves, Ewan Harrison, David K. Jackson, Ian Johnston, Dominic Kwiatkowski, Cordelia Langford, John Sillitoe on behalf of the Wellcome Sanger Institute COVID-19 Surveillance Team |
| EPI_ISL_3109577 | Lighthouse Laboratory Plymouth | Wellcome Sanger Institute for the COVID-19 Genomics UK (COG-UK) Consortium | Lighthouse Laboratory Plymouth and Alex Alderton, Roberto Amato, Jeffrey Barrett, Sonia Goncalves, Ewan Harrison, David K. Jackson, Ian Johnston, Dominic Kwiatkowski, Cordelia Langford, John Sillitoe on behalf of the Wellcome Sanger Institute COVID-19 Surveillance Team |
| EPI_ISL_3109578 | Lighthouse Lab in Milton Keynes | Wellcome Sanger Institute for the COVID-19 Genomics UK (COG-UK) Consortium | The Lighthouse Lab in Milton Keynes and Alex Alderton, Roberto Amato, Jeffrey Barrett, Sonia Goncalves, Ewan Harrison, David K. Jackson, Ian Johnston, Dominic Kwiatkowski, Cordelia Langford, John Sillitoe on behalf of the Wellcome Sanger Institute COVID-19 Surveillance Team |
| EPI_ISL_3109579 | Lighthouse Laboratory Plymouth | Wellcome Sanger Institute for the COVID-19 Genomics UK (COG-UK) Consortium | Lighthouse Laboratory Plymouth and Alex Alderton, Roberto Amato, Jeffrey Barrett, Sonia Goncalves, Ewan Harrison, David K. Jackson, Ian Johnston, Dominic Kwiatkowski, Cordelia Langford, John Sillitoe on behalf of the Wellcome Sanger Institute COVID-19 Surveillance Team |
| EPI_ISL_3109580, EPI_ISL_3109581 | Lighthouse Lab in Milton Keynes | Wellcome Sanger Institute for the COVID-19 Genomics UK (COG-UK) Consortium | The Lighthouse Lab in Milton Keynes and Alex Alderton, Roberto Amato, Jeffrey Barrett, Sonia Goncalves, Ewan Harrison, David K. Jackson, Ian Johnston, Dominic Kwiatkowski, Cordelia Langford, John Sillitoe on behalf of the Wellcome Sanger Institute COVID-19 Surveillance Team |
| EPI_ISL_3109582 | Berkshire and Surrey Pathology Services Lighthouse Laboratory | Wellcome Sanger Institute for the COVID-19 Genomics UK (COG-UK) Consortium | Berkshire and Surrey Pathology Services Lighthouse Laboratory and Alex Alderton, Roberto Amato, Jeffrey Barrett, Sonia Goncalves, Ewan Harrison, David K. Jackson, Ian Johnston, Dominic Kwiatkowski, Cordelia Langford, John Sillitoe on behalf of the Wellcome Sanger Institute COVID-19 Surveillance Team |
| EPI_ISL_3109583, EPI_ISL_3109584 | Lighthouse Laboratory Plymouth | Wellcome Sanger Institute for the COVID-19 Genomics UK (COG-UK) Consortium | Lighthouse Laboratory Plymouth and Alex Alderton, Roberto Amato, Jeffrey Barrett, Sonia Goncalves, Ewan Harrison, David K. Jackson, Ian Johnston, Dominic Kwiatkowski, Cordelia Langford, John Sillitoe on behalf of the Wellcome Sanger Institute COVID-19 Surveillance Team |
| EPI_ISL_3109585 | Berkshire and Surrey Pathology Services Lighthouse Laboratory | Wellcome Sanger Institute for the COVID-19 Genomics UK (COG-UK) Consortium | Berkshire and Surrey Pathology Services Lighthouse Laboratory and Alex Alderton, Roberto Amato, Jeffrey Barrett, Sonia Goncalves, Ewan Harrison, David K. Jackson, Ian Johnston, Dominic Kwiatkowski, Cordelia Langford, John Sillitoe on behalf of the Wellcome Sanger Institute COVID-19 Surveillance Team |
| EPI_ISL_3109586 | Lighthouse Lab in Milton Keynes | Wellcome Sanger Institute for the COVID-19 Genomics UK (COG-UK) Consortium | The Lighthouse Lab in Milton Keynes and Alex Alderton, Roberto Amato, Jeffrey Barrett, Sonia Goncalves, Ewan Harrison, David K. Jackson, Ian Johnston, Dominic Kwiatkowski, Cordelia Langford, John Sillitoe on behalf of the Wellcome Sanger Institute COVID-19 Surveillance Team |

| | | | |
|---|---|---|---|
| EPI_ISL_3109587 | Lighthouse Laboratory Plymouth | Wellcome Sanger Institute for the COVID-19 Genomics UK (COG-UK) Consortium | Lighthouse Laboratory Plymouth and Alex Alderton, Roberto Amato, Jeffrey Barrett, Sonia Goncalves, Ewan Harrison, David K. Jackson, Ian Johnston, Dominic Kwiatkowski, Cordelia Langford, John Sillitoe on behalf of the Wellcome Sanger Institute COVID-19 Surveillance Team |
| EPI_ISL_3109588 | Berkshire and Surrey Pathology Services Lighthouse Laboratory | Wellcome Sanger Institute for the COVID-19 Genomics UK (COG-UK) Consortium | Berkshire and Surrey Pathology Services Lighthouse Laboratory and Alex Alderton, Roberto Amato, Jeffrey Barrett, Sonia Goncalves, Ewan Harrison, David K. Jackson, Ian Johnston, Dominic Kwiatkowski, Cordelia Langford, John Sillitoe on behalf of the Wellcome Sanger Institute COVID-19 Surveillance Team |
| EPI_ISL_3109589 | Lighthouse Lab in Alderley Park | Wellcome Sanger Institute for the COVID-19 Genomics UK (COG-UK) Consortium | Jacquelyn Wynn, Mairead Hyland, The Lighthouse Lab in Alderley Park and Alex Alderton, Roberto Amato, Jeffrey Barrett, Sonia Goncalves, Ewan Harrison, David K. Jackson, Ian Johnston, Dominic Kwiatkowski, Cordelia Langford, John Sillitoe on behalf of the Wellcome Sanger Institute COVID-19 Surveillance Team |
| EPI_ISL_3109590 | Lighthouse Lab in Milton Keynes | Wellcome Sanger Institute for the COVID-19 Genomics UK (COG-UK) Consortium | The Lighthouse Lab in Milton Keynes and Alex Alderton, Roberto Amato, Jeffrey Barrett, Sonia Goncalves, Ewan Harrison, David K. Jackson, Ian Johnston, Dominic Kwiatkowski, Cordelia Langford, John Sillitoe on behalf of the Wellcome Sanger Institute COVID-19 Surveillance Team |
| EPI_ISL_3109591 | Berkshire and Surrey Pathology Services Lighthouse Laboratory | Wellcome Sanger Institute for the COVID-19 Genomics UK (COG-UK) Consortium | Berkshire and Surrey Pathology Services Lighthouse Laboratory and Alex Alderton, Roberto Amato, Jeffrey Barrett, Sonia Goncalves, Ewan Harrison, David K. Jackson, Ian Johnston, Dominic Kwiatkowski, Cordelia Langford, John Sillitoe on behalf of the Wellcome Sanger Institute COVID-19 Surveillance Team |
| EPI_ISL_3109592 | Lighthouse Lab in Milton Keynes | Wellcome Sanger Institute for the COVID-19 Genomics UK (COG-UK) Consortium | The Lighthouse Lab in Milton Keynes and Alex Alderton, Roberto Amato, Jeffrey Barrett, Sonia Goncalves, Ewan Harrison, David K. Jackson, Ian Johnston, Dominic Kwiatkowski, Cordelia Langford, John Sillitoe on behalf of the Wellcome Sanger Institute COVID-19 Surveillance Team |
| EPI_ISL_3109593 | Lighthouse Lab in Alderley Park | Wellcome Sanger Institute for the COVID-19 Genomics UK (COG-UK) Consortium | Jacquelyn Wynn, Mairead Hyland, The Lighthouse Lab in Alderley Park and Alex Alderton, Roberto Amato, Jeffrey Barrett, Sonia Goncalves, Ewan Harrison, David K. Jackson, Ian Johnston, Dominic Kwiatkowski, Cordelia Langford, John Sillitoe on behalf of the Wellcome Sanger Institute COVID-19 Surveillance Team |
| EPI_ISL_3109594 | Berkshire and Surrey Pathology Services Lighthouse Laboratory | Wellcome Sanger Institute for the COVID-19 Genomics UK (COG-UK) Consortium | Berkshire and Surrey Pathology Services Lighthouse Laboratory and Alex Alderton, Roberto Amato, Jeffrey Barrett, Sonia Goncalves, Ewan Harrison, David K. Jackson, Ian Johnston, Dominic Kwiatkowski, Cordelia Langford, John Sillitoe on behalf of the Wellcome Sanger Institute COVID-19 Surveillance Team |
| EPI_ISL_3109595, EPI_ISL_3109596 | Lighthouse Lab in Milton Keynes | Wellcome Sanger Institute for the COVID-19 Genomics UK (COG-UK) Consortium | The Lighthouse Lab in Milton Keynes and Alex Alderton, Roberto Amato, Jeffrey Barrett, Sonia Goncalves, Ewan Harrison, David K. Jackson, Ian Johnston, Dominic Kwiatkowski, Cordelia Langford, John Sillitoe on behalf of the Wellcome Sanger Institute COVID-19 Surveillance Team |
| EPI_ISL_3109597, EPI_ISL_3109598 | Lighthouse Lab in Alderley Park | Wellcome Sanger Institute for the COVID-19 Genomics UK (COG-UK) Consortium | Jacquelyn Wynn, Mairead Hyland, The Lighthouse Lab in Alderley Park and Alex Alderton, Roberto Amato, Jeffrey Barrett, Sonia Goncalves, Ewan Harrison, David K. Jackson, Ian Johnston, Dominic Kwiatkowski, Cordelia Langford, John Sillitoe on behalf of the Wellcome Sanger Institute COVID-19 Surveillance Team |
| EPI_ISL_3109599 | Berkshire and Surrey Pathology Services Lighthouse Laboratory | Wellcome Sanger Institute for the COVID-19 Genomics UK (COG-UK) Consortium | Berkshire and Surrey Pathology Services Lighthouse Laboratory and Alex Alderton, Roberto Amato, Jeffrey Barrett, Sonia Goncalves, Ewan Harrison, David K. Jackson, Ian Johnston, Dominic Kwiatkowski, Cordelia Langford, John Sillitoe on behalf of the Wellcome Sanger Institute COVID-19 Surveillance Team |
| EPI_ISL_3109600 | Lighthouse Laboratory Plymouth | Wellcome Sanger Institute for the COVID-19 Genomics UK (COG-UK) Consortium | Lighthouse Laboratory Plymouth and Alex Alderton, Roberto Amato, Jeffrey Barrett, Sonia Goncalves, Ewan Harrison, David K. Jackson, Ian Johnston, Dominic Kwiatkowski, Cordelia Langford, John Sillitoe on behalf of the Wellcome Sanger Institute COVID-19 Surveillance Team |
| EPI_ISL_3109601 | Berkshire and Surrey Pathology Services Lighthouse Laboratory | Wellcome Sanger Institute for the COVID-19 Genomics UK (COG-UK) Consortium | Berkshire and Surrey Pathology Services Lighthouse Laboratory and Alex Alderton, Roberto Amato, Jeffrey Barrett, Sonia Goncalves, Ewan Harrison, David K. Jackson, Ian Johnston, Dominic Kwiatkowski, Cordelia Langford, John Sillitoe on behalf of the Wellcome Sanger Institute COVID-19 Surveillance Team |
| EPI_ISL_3109602 | Lighthouse Lab in Alderley Park | Wellcome Sanger Institute for the COVID-19 Genomics UK (COG-UK) Consortium | Jacquelyn Wynn, Mairead Hyland, The Lighthouse Lab in Alderley Park and Alex Alderton, Roberto Amato, Jeffrey Barrett, Sonia Goncalves, Ewan Harrison, David K. Jackson, Ian Johnston, Dominic Kwiatkowski, Cordelia Langford, John Sillitoe on behalf of the Wellcome Sanger Institute COVID-19 Surveillance Team |
| EPI_ISL_3109603 | Lighthouse Lab in Milton Keynes | Wellcome Sanger Institute for the COVID-19 Genomics UK (COG-UK) Consortium | The Lighthouse Lab in Milton Keynes and Alex Alderton, Roberto Amato, Jeffrey Barrett, Sonia Goncalves, Ewan Harrison, David K. Jackson, Ian Johnston, Dominic Kwiatkowski, Cordelia Langford, John Sillitoe on behalf of the Wellcome Sanger Institute COVID-19 Surveillance Team |
| EPI_ISL_3109604 | Berkshire and Surrey Pathology Services Lighthouse Laboratory | Wellcome Sanger Institute for the COVID-19 Genomics UK (COG-UK) Consortium | Berkshire and Surrey Pathology Services Lighthouse Laboratory and Alex Alderton, Roberto Amato, Jeffrey Barrett, Sonia Goncalves, Ewan Harrison, David K. Jackson, Ian Johnston, Dominic Kwiatkowski, Cordelia Langford, John Sillitoe on behalf of the Wellcome Sanger Institute COVID-19 Surveillance Team |
| EPI_ISL_3109605, EPI_ISL_3109606, EPI_ISL_3109607 | Lighthouse Lab in Milton Keynes | Wellcome Sanger Institute for the COVID-19 Genomics UK (COG-UK) Consortium | The Lighthouse Lab in Milton Keynes and Alex Alderton, Roberto Amato, Jeffrey Barrett, Sonia Goncalves, Ewan Harrison, David K. Jackson, Ian Johnston, Dominic Kwiatkowski, Cordelia Langford, John Sillitoe on behalf of the Wellcome Sanger Institute COVID-19 Surveillance Team |
| EPI_ISL_3109608, EPI_ISL_3109609, EPI_ISL_3109610 | Berkshire and Surrey Pathology Services Lighthouse Laboratory | Wellcome Sanger Institute for the COVID-19 Genomics UK (COG-UK) Consortium | Berkshire and Surrey Pathology Services Lighthouse Laboratory and Alex Alderton, Roberto Amato, Jeffrey Barrett, Sonia Goncalves, Ewan Harrison, David K. Jackson, Ian Johnston, Dominic Kwiatkowski, Cordelia Langford, John Sillitoe on behalf of the Wellcome Sanger Institute COVID-19 Surveillance Team |
| EPI_ISL_3109611, EPI_ISL_3109612 | Lighthouse Laboratory Plymouth | Wellcome Sanger Institute for the COVID-19 Genomics UK (COG-UK) Consortium | Lighthouse Laboratory Plymouth and Alex Alderton, Roberto Amato, Jeffrey Barrett, Sonia Goncalves, Ewan Harrison, David K. Jackson, Ian Johnston, Dominic Kwiatkowski, Cordelia Langford, John Sillitoe on behalf of the Wellcome Sanger Institute COVID-19 Surveillance Team |
| EPI_ISL_3109613 | Lighthouse Lab in Milton Keynes | Wellcome Sanger Institute for the COVID-19 Genomics UK (COG-UK) Consortium | The Lighthouse Lab in Milton Keynes and Alex Alderton, Roberto Amato, Jeffrey Barrett, Sonia Goncalves, Ewan Harrison, David K. Jackson, Ian Johnston, Dominic Kwiatkowski, Cordelia Langford, John Sillitoe on behalf of the Wellcome Sanger Institute COVID-19 Surveillance Team |
| EPI_ISL_3109614, EPI_ISL_3109615 | Lighthouse Lab in Alderley Park | Wellcome Sanger Institute for the COVID-19 Genomics UK (COG-UK) Consortium | Jacquelyn Wynn, Mairead Hyland, The Lighthouse Lab in Alderley Park and Alex Alderton, Roberto Amato, Jeffrey Barrett, Sonia Goncalves, Ewan Harrison, David K. Jackson, Ian Johnston, Dominic Kwiatkowski, Cordelia Langford, John Sillitoe on behalf of the Wellcome Sanger Institute COVID-19 Surveillance Team |
| EPI_ISL_3109616 | Lighthouse Laboratory Plymouth | Wellcome Sanger Institute for the COVID-19 Genomics UK (COG-UK) Consortium | Lighthouse Laboratory Plymouth and Alex Alderton, Roberto Amato, Jeffrey Barrett, Sonia Goncalves, Ewan Harrison, David K. Jackson, Ian Johnston, Dominic Kwiatkowski, Cordelia Langford, John Sillitoe on behalf of the Wellcome Sanger Institute COVID-19 Surveillance Team |
| EPI_ISL_3109617, EPI_ISL_3109618, EPI_ISL_3109619 | Berkshire and Surrey Pathology Services Lighthouse Laboratory | Wellcome Sanger Institute for the COVID-19 Genomics UK (COG-UK) Consortium | Berkshire and Surrey Pathology Services Lighthouse Laboratory and Alex Alderton, Roberto Amato, Jeffrey Barrett, Sonia Goncalves, Ewan Harrison, David K. Jackson, Ian Johnston, Dominic Kwiatkowski, Cordelia Langford, John Sillitoe on behalf of the Wellcome Sanger Institute COVID-19 Surveillance Team |
| EPI_ISL_3109620 | Lighthouse Laboratory Plymouth | Wellcome Sanger Institute for the COVID-19 Genomics UK (COG-UK) Consortium | Lighthouse Laboratory Plymouth and Alex Alderton, Roberto Amato, Jeffrey Barrett, Sonia Goncalves, Ewan Harrison, David K. Jackson, Ian Johnston, Dominic Kwiatkowski, Cordelia Langford, John Sillitoe on behalf of the Wellcome Sanger Institute COVID-19 Surveillance Team |
| EPI_ISL_3109621 | Lighthouse Lab in Milton Keynes | Wellcome Sanger Institute for the COVID-19 Genomics UK (COG-UK) Consortium | The Lighthouse Lab in Milton Keynes and Alex Alderton, Roberto Amato, Jeffrey Barrett, Sonia Goncalves, Ewan Harrison, David K. Jackson, Ian Johnston, Dominic Kwiatkowski, Cordelia Langford, John Sillitoe on behalf of the Wellcome Sanger Institute COVID-19 Surveillance Team |
| EPI_ISL_3109622 | Lighthouse Lab in Alderley Park | Wellcome Sanger Institute for the COVID-19 Genomics UK (COG-UK) Consortium | Jacquelyn Wynn, Mairead Hyland, The Lighthouse Lab in Alderley Park and Alex Alderton, Roberto Amato, Jeffrey Barrett, Sonia Goncalves, Ewan Harrison, David K. Jackson, Ian Johnston, Dominic Kwiatkowski, Cordelia Langford, John Sillitoe on behalf of the Wellcome Sanger Institute COVID-19 Surveillance Team |
| EPI_ISL_3109623 | Berkshire and Surrey Pathology Services Lighthouse Laboratory | Wellcome Sanger Institute for the COVID-19 Genomics UK (COG-UK) Consortium | Berkshire and Surrey Pathology Services Lighthouse Laboratory and Alex Alderton, Roberto Amato, Jeffrey Barrett, Sonia Goncalves, Ewan Harrison, David K. Jackson, Ian Johnston, Dominic Kwiatkowski, Cordelia Langford, John Sillitoe on behalf of the Wellcome Sanger Institute COVID-19 Surveillance Team |
| EPI_ISL_3109624 | Lighthouse Lab in Milton Keynes | Wellcome Sanger Institute for the COVID-19 Genomics UK (COG-UK) Consortium | The Lighthouse Lab in Milton Keynes and Alex Alderton, Roberto Amato, Jeffrey Barrett, Sonia Goncalves, Ewan Harrison, David K. Jackson, Ian Johnston, Dominic Kwiatkowski, Cordelia Langford, John Sillitoe on behalf of the Wellcome Sanger Institute COVID-19 Surveillance Team |
| EPI_ISL_3109625 | Lighthouse Lab in Alderley Park | Wellcome Sanger Institute for the COVID-19 Genomics UK (COG-UK) Consortium | Jacquelyn Wynn, Mairead Hyland, The Lighthouse Lab in Alderley Park and Alex Alderton, Roberto Amato, Jeffrey Barrett, Sonia Goncalves, Ewan Harrison, David K. Jackson, Ian Johnston, Dominic Kwiatkowski, Cordelia Langford, John Sillitoe on behalf of the Wellcome Sanger Institute COVID-19 Surveillance Team |
| EPI_ISL_3109626, EPI_ISL_3109627 | Berkshire and Surrey Pathology Services Lighthouse Laboratory | Wellcome Sanger Institute for the COVID-19 Genomics UK (COG-UK) Consortium | Berkshire and Surrey Pathology Services Lighthouse Laboratory and Alex Alderton, Roberto Amato, Jeffrey Barrett, Sonia Goncalves, Ewan Harrison, David K. Jackson, Ian Johnston, Dominic Kwiatkowski, Cordelia Langford, John Sillitoe on behalf of the Wellcome Sanger Institute COVID-19 Surveillance Team |
| EPI_ISL_3109628 | Lighthouse Laboratory Plymouth | Wellcome Sanger Institute for the COVID-19 Genomics UK (COG-UK) Consortium | Lighthouse Laboratory Plymouth and Alex Alderton, Roberto Amato, Jeffrey Barrett, Sonia Goncalves, Ewan Harrison, David K. Jackson, Ian Johnston, Dominic Kwiatkowski, Cordelia Langford, John Sillitoe on behalf of the Wellcome Sanger Institute COVID-19 Surveillance Team |
| EPI_ISL_3109629 | Lighthouse Lab in Milton Keynes | Wellcome Sanger Institute for the COVID-19 Genomics UK (COG-UK) Consortium | The Lighthouse Lab in Milton Keynes and Alex Alderton, Roberto Amato, Jeffrey Barrett, Sonia Goncalves, Ewan Harrison, David K. Jackson, Ian Johnston, Dominic Kwiatkowski, Cordelia Langford, John Sillitoe on behalf of the Wellcome Sanger Institute COVID-19 Surveillance Team |
| EPI_ISL_3109630, EPI_ISL_3109631 | Berkshire and Surrey Pathology Services Lighthouse Laboratory | Wellcome Sanger Institute for the COVID-19 Genomics UK (COG-UK) Consortium | Berkshire and Surrey Pathology Services Lighthouse Laboratory and Alex Alderton, Roberto Amato, Jeffrey Barrett, Sonia Goncalves, Ewan Harrison, David K. Jackson, Ian Johnston, Dominic Kwiatkowski, Cordelia Langford, John Sillitoe on behalf of the Wellcome Sanger Institute COVID-19 Surveillance Team |
| EPI_ISL_3109632 | Lighthouse Laboratory Plymouth | Wellcome Sanger Institute for the COVID-19 Genomics UK (COG-UK) Consortium | Lighthouse Laboratory Plymouth and Alex Alderton, Roberto Amato, Jeffrey Barrett, Sonia Goncalves, Ewan Harrison, David K. Jackson, Ian Johnston, Dominic Kwiatkowski, Cordelia Langford, John Sillitoe on behalf of the Wellcome Sanger Institute COVID-19 Surveillance Team |
| EPI_ISL_3109633 | Lighthouse Lab in Alderley Park | Wellcome Sanger Institute for the COVID-19 Genomics UK (COG-UK) Consortium | Jacquelyn Wynn, Mairead Hyland, The Lighthouse Lab in Alderley Park and Alex Alderton, Roberto Amato, Jeffrey Barrett, Sonia Goncalves, Ewan Harrison, David K. Jackson, Ian Johnston, Dominic Kwiatkowski, Cordelia Langford, John Sillitoe on behalf of the Wellcome Sanger Institute COVID-19 Surveillance Team |
| EPI_ISL_3109634 | Berkshire and Surrey Pathology Services Lighthouse Laboratory | Wellcome Sanger Institute for the COVID-19 Genomics UK (COG-UK) Consortium | Berkshire and Surrey Pathology Services Lighthouse Laboratory and Alex Alderton, Roberto Amato, Jeffrey Barrett, Sonia Goncalves, Ewan Harrison, David K. Jackson, Ian Johnston, Dominic Kwiatkowski, Cordelia Langford, John Sillitoe on behalf of the Wellcome Sanger Institute COVID-19 Surveillance Team |
| EPI_ISL_3109635 | Lighthouse Lab in Milton Keynes | Wellcome Sanger Institute for the COVID-19 Genomics UK (COG-UK) Consortium | The Lighthouse Lab in Milton Keynes and Alex Alderton, Roberto Amato, Jeffrey Barrett, Sonia Goncalves, Ewan Harrison, David K. Jackson, Ian Johnston, Dominic Kwiatkowski, Cordelia Langford, John Sillitoe on behalf of the Wellcome Sanger Institute COVID-19 Surveillance Team |
| EPI_ISL_3109636 | Berkshire and Surrey Pathology Services Lighthouse Laboratory | Wellcome Sanger Institute for the COVID-19 Genomics UK (COG-UK) Consortium | Berkshire and Surrey Pathology Services Lighthouse Laboratory and Alex Alderton, Roberto Amato, Jeffrey Barrett, Sonia Goncalves, Ewan Harrison, David K. Jackson, Ian Johnston, Dominic Kwiatkowski, Cordelia Langford, John Sillitoe on behalf of the Wellcome Sanger Institute COVID-19 Surveillance Team |
| EPI_ISL_3109637 | Lighthouse Lab in Alderley Park | Wellcome Sanger Institute for the COVID-19 Genomics UK (COG-UK) Consortium | Jacquelyn Wynn, Mairead Hyland, The Lighthouse Lab in Alderley Park and Alex Alderton, Roberto Amato, Jeffrey Barrett, Sonia Goncalves, Ewan Harrison, David K. Jackson, Ian Johnston, Dominic Kwiatkowski, Cordelia Langford, John Sillitoe on behalf of the Wellcome Sanger Institute COVID-19 Surveillance Team |
| EPI_ISL_3109638 | Berkshire and Surrey Pathology Services Lighthouse Laboratory | Wellcome Sanger Institute for the COVID-19 Genomics UK (COG-UK) Consortium | Berkshire and Surrey Pathology Services Lighthouse Laboratory and Alex Alderton, Roberto Amato, Jeffrey Barrett, Sonia Goncalves, Ewan Harrison, David K. Jackson, Ian Johnston, Dominic Kwiatkowski, Cordelia Langford, John Sillitoe on behalf of the Wellcome Sanger Institute COVID-19 Surveillance Team |
| EPI_ISL_3109640, EPI_ISL_3109641 | Lighthouse Lab in Milton Keynes | Wellcome Sanger Institute for the COVID-19 Genomics UK (COG-UK) Consortium | The Lighthouse Lab in Milton Keynes and Alex Alderton, Roberto Amato, Jeffrey Barrett, Sonia Goncalves, Ewan Harrison, David K. Jackson, Ian Johnston, Dominic Kwiatkowski, Cordelia Langford, John Sillitoe on behalf of the Wellcome Sanger Institute COVID-19 Surveillance Team |
| EPI_ISL_3109642 | Lighthouse Laboratory Plymouth | Wellcome Sanger Institute for the COVID-19 Genomics UK (COG-UK) Consortium | Lighthouse Laboratory Plymouth and Alex Alderton, Roberto Amato, Jeffrey Barrett, Sonia Goncalves, Ewan Harrison, David K. Jackson, Ian Johnston, Dominic Kwiatkowski, Cordelia Langford, John Sillitoe on behalf of the Wellcome Sanger Institute COVID-19 Surveillance Team |
| EPI_ISL_3109643, EPI_ISL_3109644 | Lighthouse Lab in Alderley Park | Wellcome Sanger Institute for the COVID-19 Genomics UK (COG-UK) Consortium | Jacquelyn Wynn, Mairead Hyland, The Lighthouse Lab in Alderley Park and Alex Alderton, Roberto Amato, Jeffrey Barrett, Sonia Goncalves, Ewan Harrison, David K. Jackson, Ian Johnston, Dominic Kwiatkowski, Cordelia Langford, John Sillitoe on behalf of the Wellcome Sanger Institute COVID-19 Surveillance Team |
| EPI_ISL_3109645, EPI_ISL_3109646 | Berkshire and Surrey Pathology Services Lighthouse Laboratory | Wellcome Sanger Institute for the COVID-19 Genomics UK (COG-UK) Consortium | Berkshire and Surrey Pathology Services Lighthouse Laboratory and Alex Alderton, Roberto Amato, Jeffrey Barrett, Sonia Goncalves, Ewan Harrison, David K. Jackson, Ian Johnston, Dominic Kwiatkowski, Cordelia Langford, John Sillitoe on behalf of the Wellcome Sanger Institute COVID-19 Surveillance Team |

| | | | |
|---|---|---|---|
| EPI_ISL_3109647 | Lighthouse Lab in Milton Keynes | Wellcome Sanger Institute for the COVID-19 Genomics UK (COG-UK) Consortium | The Lighthouse Lab in Milton Keynes and Alex Alderton, Roberto Amato, Jeffrey Barrett, Sonia Goncalves, Ewan Harrison, David K. Jackson, Ian Johnston, Dominic Kwiatkowski, Cordelia Langford, John Sillitoe on behalf of the Wellcome Sanger Institute COVID-19 Surveillance Team |
| EPI_ISL_3109648 | Lighthouse Lab in Alderley Park | Wellcome Sanger Institute for the COVID-19 Genomics UK (COG-UK) Consortium | Jacquelyn Wynn, Mairead Hyland, The Lighthouse Lab in Alderley Park and Alex Alderton, Roberto Amato, Jeffrey Barrett, Sonia Goncalves, Ewan Harrison, David K. Jackson, Ian Johnston, Dominic Kwiatkowski, Cordelia Langford, John Sillitoe on behalf of the Wellcome Sanger Institute COVID-19 Surveillance Team |
| EPI_ISL_3109649, EPI_ISL_3109650, EPI_ISL_3109651 | Lighthouse Lab in Milton Keynes | Wellcome Sanger Institute for the COVID-19 Genomics UK (COG-UK) Consortium | The Lighthouse Lab in Milton Keynes and Alex Alderton, Roberto Amato, Jeffrey Barrett, Sonia Goncalves, Ewan Harrison, David K. Jackson, Ian Johnston, Dominic Kwiatkowski, Cordelia Langford, John Sillitoe on behalf of the Wellcome Sanger Institute COVID-19 Surveillance Team |
| EPI_ISL_3109652, EPI_ISL_3109653 | Lighthouse Lab in Alderley Park | Wellcome Sanger Institute for the COVID-19 Genomics UK (COG-UK) Consortium | Jacquelyn Wynn, Mairead Hyland, The Lighthouse Lab in Alderley Park and Alex Alderton, Roberto Amato, Jeffrey Barrett, Sonia Goncalves, Ewan Harrison, David K. Jackson, Ian Johnston, Dominic Kwiatkowski, Cordelia Langford, John Sillitoe on behalf of the Wellcome Sanger Institute COVID-19 Surveillance Team |
| EPI_ISL_3109654 | Lighthouse Laboratory Plymouth | Wellcome Sanger Institute for the COVID-19 Genomics UK (COG-UK) Consortium | Lighthouse Laboratory Plymouth and Alex Alderton, Roberto Amato, Jeffrey Barrett, Sonia Goncalves, Ewan Harrison, David K. Jackson, Ian Johnston, Dominic Kwiatkowski, Cordelia Langford, John Sillitoe on behalf of the Wellcome Sanger Institute COVID-19 Surveillance Team |
| EPI_ISL_3109655, EPI_ISL_3109656 | Berkshire and Surrey Pathology Services Lighthouse Laboratory | Wellcome Sanger Institute for the COVID-19 Genomics UK (COG-UK) Consortium | Berkshire and Surrey Pathology Services Lighthouse Laboratory and Alex Alderton, Roberto Amato, Jeffrey Barrett, Sonia Goncalves, Ewan Harrison, David K. Jackson, Ian Johnston, Dominic Kwiatkowski, Cordelia Langford, John Sillitoe on behalf of the Wellcome Sanger Institute COVID-19 Surveillance Team |
| EPI_ISL_3109657 | Lighthouse Lab in Milton Keynes | Wellcome Sanger Institute for the COVID-19 Genomics UK (COG-UK) Consortium | The Lighthouse Lab in Milton Keynes and Alex Alderton, Roberto Amato, Jeffrey Barrett, Sonia Goncalves, Ewan Harrison, David K. Jackson, Ian Johnston, Dominic Kwiatkowski, Cordelia Langford, John Sillitoe on behalf of the Wellcome Sanger Institute COVID-19 Surveillance Team |
| EPI_ISL_3109658 | Lighthouse Laboratory Plymouth | Wellcome Sanger Institute for the COVID-19 Genomics UK (COG-UK) Consortium | Lighthouse Laboratory Plymouth and Alex Alderton, Roberto Amato, Jeffrey Barrett, Sonia Goncalves, Ewan Harrison, David K. Jackson, Ian Johnston, Dominic Kwiatkowski, Cordelia Langford, John Sillitoe on behalf of the Wellcome Sanger Institute COVID-19 Surveillance Team |
| EPI_ISL_3109659, EPI_ISL_3109660 | Berkshire and Surrey Pathology Services Lighthouse Laboratory | Wellcome Sanger Institute for the COVID-19 Genomics UK (COG-UK) Consortium | Berkshire and Surrey Pathology Services Lighthouse Laboratory and Alex Alderton, Roberto Amato, Jeffrey Barrett, Sonia Goncalves, Ewan Harrison, David K. Jackson, Ian Johnston, Dominic Kwiatkowski, Cordelia Langford, John Sillitoe on behalf of the Wellcome Sanger Institute COVID-19 Surveillance Team |
| EPI_ISL_3109661 | Lighthouse Lab in Alderley Park | Wellcome Sanger Institute for the COVID-19 Genomics UK (COG-UK) Consortium | Jacquelyn Wynn, Mairead Hyland, The Lighthouse Lab in Alderley Park and Alex Alderton, Roberto Amato, Jeffrey Barrett, Sonia Goncalves, Ewan Harrison, David K. Jackson, Ian Johnston, Dominic Kwiatkowski, Cordelia Langford, John Sillitoe on behalf of the Wellcome Sanger Institute COVID-19 Surveillance Team |
| EPI_ISL_3109662 | Lighthouse Lab in Milton Keynes | Wellcome Sanger Institute for the COVID-19 Genomics UK (COG-UK) Consortium | The Lighthouse Lab in Milton Keynes and Alex Alderton, Roberto Amato, Jeffrey Barrett, Sonia Goncalves, Ewan Harrison, David K. Jackson, Ian Johnston, Dominic Kwiatkowski, Cordelia Langford, John Sillitoe on behalf of the Wellcome Sanger Institute COVID-19 Surveillance Team |
| EPI_ISL_3109663, EPI_ISL_3109664 | Lighthouse Lab in Alderley Park | Wellcome Sanger Institute for the COVID-19 Genomics UK (COG-UK) Consortium | Jacquelyn Wynn, Mairead Hyland, The Lighthouse Lab in Alderley Park and Alex Alderton, Roberto Amato, Jeffrey Barrett, Sonia Goncalves, Ewan Harrison, David K. Jackson, Ian Johnston, Dominic Kwiatkowski, Cordelia Langford, John Sillitoe on behalf of the Wellcome Sanger Institute COVID-19 Surveillance Team |
| EPI_ISL_3109665, EPI_ISL_3109666 | Berkshire and Surrey Pathology Services Lighthouse Laboratory | Wellcome Sanger Institute for the COVID-19 Genomics UK (COG-UK) Consortium | Berkshire and Surrey Pathology Services Lighthouse Laboratory and Alex Alderton, Roberto Amato, Jeffrey Barrett, Sonia Goncalves, Ewan Harrison, David K. Jackson, Ian Johnston, Dominic Kwiatkowski, Cordelia Langford, John Sillitoe on behalf of the Wellcome Sanger Institute COVID-19 Surveillance Team |
| EPI_ISL_3109667, EPI_ISL_3109668, EPI_ISL_3109669 | Lighthouse Lab in Milton Keynes | Wellcome Sanger Institute for the COVID-19 Genomics UK (COG-UK) Consortium | The Lighthouse Lab in Milton Keynes and Alex Alderton, Roberto Amato, Jeffrey Barrett, Sonia Goncalves, Ewan Harrison, David K. Jackson, Ian Johnston, Dominic Kwiatkowski, Cordelia Langford, John Sillitoe on behalf of the Wellcome Sanger Institute COVID-19 Surveillance Team |
| EPI_ISL_3109670, EPI_ISL_3109671 | Lighthouse Lab in Alderley Park | Wellcome Sanger Institute for the COVID-19 Genomics UK (COG-UK) Consortium | Jacquelyn Wynn, Mairead Hyland, The Lighthouse Lab in Alderley Park and Alex Alderton, Roberto Amato, Jeffrey Barrett, Sonia Goncalves, Ewan Harrison, David K. Jackson, Ian Johnston, Dominic Kwiatkowski, Cordelia Langford, John Sillitoe on behalf of the Wellcome Sanger Institute COVID-19 Surveillance Team |
| EPI_ISL_3109672 | Berkshire and Surrey Pathology Services Lighthouse Laboratory | Wellcome Sanger Institute for the COVID-19 Genomics UK (COG-UK) Consortium | Berkshire and Surrey Pathology Services Lighthouse Laboratory and Alex Alderton, Roberto Amato, Jeffrey Barrett, Sonia Goncalves, Ewan Harrison, David K. Jackson, Ian Johnston, Dominic Kwiatkowski, Cordelia Langford, John Sillitoe on behalf of the Wellcome Sanger Institute COVID-19 Surveillance Team |
| EPI_ISL_3109673 | Lighthouse Lab in Alderley Park | Wellcome Sanger Institute for the COVID-19 Genomics UK (COG-UK) Consortium | Jacquelyn Wynn, Mairead Hyland, The Lighthouse Lab in Alderley Park and Alex Alderton, Roberto Amato, Jeffrey Barrett, Sonia Goncalves, Ewan Harrison, David K. Jackson, Ian Johnston, Dominic Kwiatkowski, Cordelia Langford, John Sillitoe on behalf of the Wellcome Sanger Institute COVID-19 Surveillance Team |
| EPI_ISL_3109674 | Berkshire and Surrey Pathology Services Lighthouse Laboratory | Wellcome Sanger Institute for the COVID-19 Genomics UK (COG-UK) Consortium | Berkshire and Surrey Pathology Services Lighthouse Laboratory and Alex Alderton, Roberto Amato, Jeffrey Barrett, Sonia Goncalves, Ewan Harrison, David K. Jackson, Ian Johnston, Dominic Kwiatkowski, Cordelia Langford, John Sillitoe on behalf of the Wellcome Sanger Institute COVID-19 Surveillance Team |
| EPI_ISL_3109675 | Lighthouse Laboratory Plymouth | Wellcome Sanger Institute for the COVID-19 Genomics UK (COG-UK) Consortium | Lighthouse Laboratory Plymouth and Alex Alderton, Roberto Amato, Jeffrey Barrett, Sonia Goncalves, Ewan Harrison, David K. Jackson, Ian Johnston, Dominic Kwiatkowski, Cordelia Langford, John Sillitoe on behalf of the Wellcome Sanger Institute COVID-19 Surveillance Team |
| EPI_ISL_3109676, EPI_ISL_3109677, EPI_ISL_3109678 | Lighthouse Lab in Alderley Park | Wellcome Sanger Institute for the COVID-19 Genomics UK (COG-UK) Consortium | Jacquelyn Wynn, Mairead Hyland, The Lighthouse Lab in Alderley Park and Alex Alderton, Roberto Amato, Jeffrey Barrett, Sonia Goncalves, Ewan Harrison, David K. Jackson, Ian Johnston, Dominic Kwiatkowski, Cordelia Langford, John Sillitoe on behalf of the Wellcome Sanger Institute COVID-19 Surveillance Team |
| EPI_ISL_3109679 | Berkshire and Surrey Pathology Services Lighthouse Laboratory | Wellcome Sanger Institute for the COVID-19 Genomics UK (COG-UK) Consortium | Berkshire and Surrey Pathology Services Lighthouse Laboratory and Alex Alderton, Roberto Amato, Jeffrey Barrett, Sonia Goncalves, Ewan Harrison, David K. Jackson, Ian Johnston, Dominic Kwiatkowski, Cordelia Langford, John Sillitoe on behalf of the Wellcome Sanger Institute COVID-19 Surveillance Team |
| EPI_ISL_3109680 | Lighthouse Lab in Alderley Park | Wellcome Sanger Institute for the COVID-19 Genomics UK (COG-UK) Consortium | Jacquelyn Wynn, Mairead Hyland, The Lighthouse Lab in Alderley Park and Alex Alderton, Roberto Amato, Jeffrey Barrett, Sonia Goncalves, Ewan Harrison, David K. Jackson, Ian Johnston, Dominic Kwiatkowski, Cordelia Langford, John Sillitoe on behalf of the Wellcome Sanger Institute COVID-19 Surveillance Team |
| EPI_ISL_3109681 | Berkshire and Surrey Pathology Services Lighthouse Laboratory | Wellcome Sanger Institute for the COVID-19 Genomics UK (COG-UK) Consortium | Berkshire and Surrey Pathology Services Lighthouse Laboratory and Alex Alderton, Roberto Amato, Jeffrey Barrett, Sonia Goncalves, Ewan Harrison, David K. Jackson, Ian Johnston, Dominic Kwiatkowski, Cordelia Langford, John Sillitoe on behalf of the Wellcome Sanger Institute COVID-19 Surveillance Team |
| EPI_ISL_3109682 | Lighthouse Laboratory Plymouth | Wellcome Sanger Institute for the COVID-19 Genomics UK (COG-UK) Consortium | Lighthouse Laboratory Plymouth and Alex Alderton, Roberto Amato, Jeffrey Barrett, Sonia Goncalves, Ewan Harrison, David K. Jackson, Ian Johnston, Dominic Kwiatkowski, Cordelia Langford, John Sillitoe on behalf of the Wellcome Sanger Institute COVID-19 Surveillance Team |
| EPI_ISL_3109683 | Berkshire and Surrey Pathology Services Lighthouse Laboratory | Wellcome Sanger Institute for the COVID-19 Genomics UK (COG-UK) Consortium | Berkshire and Surrey Pathology Services Lighthouse Laboratory and Alex Alderton, Roberto Amato, Jeffrey Barrett, Sonia Goncalves, Ewan Harrison, David K. Jackson, Ian Johnston, Dominic Kwiatkowski, Cordelia Langford, John Sillitoe on behalf of the Wellcome Sanger Institute COVID-19 Surveillance Team |
| EPI_ISL_3109684 | Lighthouse Lab in Alderley Park | Wellcome Sanger Institute for the COVID-19 Genomics UK (COG-UK) Consortium | Jacquelyn Wynn, Mairead Hyland, The Lighthouse Lab in Alderley Park and Alex Alderton, Roberto Amato, Jeffrey Barrett, Sonia Goncalves, Ewan Harrison, David K. Jackson, Ian Johnston, Dominic Kwiatkowski, Cordelia Langford, John Sillitoe on behalf of the Wellcome Sanger Institute COVID-19 Surveillance Team |
| EPI_ISL_3109685 | Lighthouse Laboratory Plymouth | Wellcome Sanger Institute for the COVID-19 Genomics UK (COG-UK) Consortium | Lighthouse Laboratory Plymouth and Alex Alderton, Roberto Amato, Jeffrey Barrett, Sonia Goncalves, Ewan Harrison, David K. Jackson, Ian Johnston, Dominic Kwiatkowski, Cordelia Langford, John Sillitoe on behalf of the Wellcome Sanger Institute COVID-19 Surveillance Team |
| EPI_ISL_3109686 | Lighthouse Lab in Milton Keynes | Wellcome Sanger Institute for the COVID-19 Genomics UK (COG-UK) Consortium | The Lighthouse Lab in Milton Keynes and Alex Alderton, Roberto Amato, Jeffrey Barrett, Sonia Goncalves, Ewan Harrison, David K. Jackson, Ian Johnston, Dominic Kwiatkowski, Cordelia Langford, John Sillitoe on behalf of the Wellcome Sanger Institute COVID-19 Surveillance Team |
| EPI_ISL_3109687 | Lighthouse Laboratory Plymouth | Wellcome Sanger Institute for the COVID-19 Genomics UK (COG-UK) Consortium | Lighthouse Laboratory Plymouth and Alex Alderton, Roberto Amato, Jeffrey Barrett, Sonia Goncalves, Ewan Harrison, David K. Jackson, Ian Johnston, Dominic Kwiatkowski, Cordelia Langford, John Sillitoe on behalf of the Wellcome Sanger Institute COVID-19 Surveillance Team |
| EPI_ISL_3109688, EPI_ISL_3109689 | Lighthouse Lab in Milton Keynes | Wellcome Sanger Institute for the COVID-19 Genomics UK (COG-UK) Consortium | The Lighthouse Lab in Milton Keynes and Alex Alderton, Roberto Amato, Jeffrey Barrett, Sonia Goncalves, Ewan Harrison, David K. Jackson, Ian Johnston, Dominic Kwiatkowski, Cordelia Langford, John Sillitoe on behalf of the Wellcome Sanger Institute COVID-19 Surveillance Team |
| EPI_ISL_3109690 | Berkshire and Surrey Pathology Services Lighthouse Laboratory | Wellcome Sanger Institute for the COVID-19 Genomics UK (COG-UK) Consortium | Berkshire and Surrey Pathology Services Lighthouse Laboratory and Alex Alderton, Roberto Amato, Jeffrey Barrett, Sonia Goncalves, Ewan Harrison, David K. Jackson, Ian Johnston, Dominic Kwiatkowski, Cordelia Langford, John Sillitoe on behalf of the Wellcome Sanger Institute COVID-19 Surveillance Team |
| EPI_ISL_3109691 | Lighthouse Lab in Alderley Park | Wellcome Sanger Institute for the COVID-19 Genomics UK (COG-UK) Consortium | Jacquelyn Wynn, Mairead Hyland, The Lighthouse Lab in Alderley Park and Alex Alderton, Roberto Amato, Jeffrey Barrett, Sonia Goncalves, Ewan Harrison, David K. Jackson, Ian Johnston, Dominic Kwiatkowski, Cordelia Langford, John Sillitoe on behalf of the Wellcome Sanger Institute COVID-19 Surveillance Team |
| EPI_ISL_3109692 | Lighthouse Laboratory Plymouth | Wellcome Sanger Institute for the COVID-19 Genomics UK (COG-UK) Consortium | Lighthouse Laboratory Plymouth and Alex Alderton, Roberto Amato, Jeffrey Barrett, Sonia Goncalves, Ewan Harrison, David K. Jackson, Ian Johnston, Dominic Kwiatkowski, Cordelia Langford, John Sillitoe on behalf of the Wellcome Sanger Institute COVID-19 Surveillance Team |
| EPI_ISL_3109693, EPI_ISL_3109694 | Lighthouse Lab in Alderley Park | Wellcome Sanger Institute for the COVID-19 Genomics UK (COG-UK) Consortium | Jacquelyn Wynn, Mairead Hyland, The Lighthouse Lab in Alderley Park and Alex Alderton, Roberto Amato, Jeffrey Barrett, Sonia Goncalves, Ewan Harrison, David K. Jackson, Ian Johnston, Dominic Kwiatkowski, Cordelia Langford, John Sillitoe on behalf of the Wellcome Sanger Institute COVID-19 Surveillance Team |
| EPI_ISL_3109695 | Lighthouse Laboratory Plymouth | Wellcome Sanger Institute for the COVID-19 Genomics UK (COG-UK) Consortium | Lighthouse Laboratory Plymouth and Alex Alderton, Roberto Amato, Jeffrey Barrett, Sonia Goncalves, Ewan Harrison, David K. Jackson, Ian Johnston, Dominic Kwiatkowski, Cordelia Langford, John Sillitoe on behalf of the Wellcome Sanger Institute COVID-19 Surveillance Team |
| EPI_ISL_3109696 | Berkshire and Surrey Pathology Services Lighthouse Laboratory | Wellcome Sanger Institute for the COVID-19 Genomics UK (COG-UK) Consortium | Berkshire and Surrey Pathology Services Lighthouse Laboratory and Alex Alderton, Roberto Amato, Jeffrey Barrett, Sonia Goncalves, Ewan Harrison, David K. Jackson, Ian Johnston, Dominic Kwiatkowski, Cordelia Langford, John Sillitoe on behalf of the Wellcome Sanger Institute COVID-19 Surveillance Team |
| EPI_ISL_3109697 | Lighthouse Lab in Alderley Park | Wellcome Sanger Institute for the COVID-19 Genomics UK (COG-UK) Consortium | Jacquelyn Wynn, Mairead Hyland, The Lighthouse Lab in Alderley Park and Alex Alderton, Roberto Amato, Jeffrey Barrett, Sonia Goncalves, Ewan Harrison, David K. Jackson, Ian Johnston, Dominic Kwiatkowski, Cordelia Langford, John Sillitoe on behalf of the Wellcome Sanger Institute COVID-19 Surveillance Team |
| EPI_ISL_3109698, EPI_ISL_3109699 | Berkshire and Surrey Pathology Services Lighthouse Laboratory | Wellcome Sanger Institute for the COVID-19 Genomics UK (COG-UK) Consortium | Berkshire and Surrey Pathology Services Lighthouse Laboratory and Alex Alderton, Roberto Amato, Jeffrey Barrett, Sonia Goncalves, Ewan Harrison, David K. Jackson, Ian Johnston, Dominic Kwiatkowski, Cordelia Langford, John Sillitoe on behalf of the Wellcome Sanger Institute COVID-19 Surveillance Team |
| EPI_ISL_3109700, EPI_ISL_3109701 | Lighthouse Lab in Alderley Park | Wellcome Sanger Institute for the COVID-19 Genomics UK (COG-UK) Consortium | Jacquelyn Wynn, Mairead Hyland, The Lighthouse Lab in Alderley Park and Alex Alderton, Roberto Amato, Jeffrey Barrett, Sonia Goncalves, Ewan Harrison, David K. Jackson, Ian Johnston, Dominic Kwiatkowski, Cordelia Langford, John Sillitoe on behalf of the Wellcome Sanger Institute COVID-19 Surveillance Team |
| EPI_ISL_3109702 | Lighthouse Lab in Milton Keynes | Wellcome Sanger Institute for the COVID-19 Genomics UK (COG-UK) Consortium | The Lighthouse Lab in Milton Keynes and Alex Alderton, Roberto Amato, Jeffrey Barrett, Sonia Goncalves, Ewan Harrison, David K. Jackson, Ian Johnston, Dominic Kwiatkowski, Cordelia Langford, John Sillitoe on behalf of the Wellcome Sanger Institute COVID-19 Surveillance Team |
| EPI_ISL_3109703, EPI_ISL_3109704 | Berkshire and Surrey Pathology Services Lighthouse Laboratory | Wellcome Sanger Institute for the COVID-19 Genomics UK (COG-UK) Consortium | Berkshire and Surrey Pathology Services Lighthouse Laboratory and Alex Alderton, Roberto Amato, Jeffrey Barrett, Sonia Goncalves, Ewan Harrison, David K. Jackson, Ian Johnston, Dominic Kwiatkowski, Cordelia Langford, John Sillitoe on behalf of the Wellcome Sanger Institute COVID-19 Surveillance Team |
| EPI_ISL_3109705 | Lighthouse Lab in Milton Keynes | Wellcome Sanger Institute for the COVID-19 Genomics UK (COG-UK) Consortium | The Lighthouse Lab in Milton Keynes and Alex Alderton, Roberto Amato, Jeffrey Barrett, Sonia Goncalves, Ewan Harrison, David K. Jackson, Ian Johnston, Dominic Kwiatkowski, Cordelia Langford, John Sillitoe on behalf of the Wellcome Sanger Institute COVID-19 Surveillance Team |
| EPI_ISL_3109706 | Lighthouse Laboratory Plymouth | Wellcome Sanger Institute for the COVID-19 Genomics UK (COG-UK) Consortium | Lighthouse Laboratory Plymouth and Alex Alderton, Roberto Amato, Jeffrey Barrett, Sonia Goncalves, Ewan Harrison, David K. Jackson, Ian Johnston, Dominic Kwiatkowski, Cordelia Langford, John Sillitoe on behalf of the Wellcome Sanger Institute COVID-19 Surveillance Team |
| EPI_ISL_3109707, EPI_ISL_3109708 | Lighthouse Lab in Alderley Park | Wellcome Sanger Institute for the COVID-19 Genomics UK (COG-UK) Consortium | Jacquelyn Wynn, Mairead Hyland, The Lighthouse Lab in Alderley Park and Alex Alderton, Roberto Amato, Jeffrey Barrett, Sonia Goncalves, Ewan Harrison, David K. Jackson, Ian Johnston, Dominic Kwiatkowski, Cordelia Langford, John Sillitoe on behalf of the Wellcome Sanger Institute COVID-19 Surveillance Team |

| | | | |
|---|---|---|---|
| EPI_ISL_3109709 | Lighthouse Lab in Milton Keynes | Wellcome Sanger Institute for the COVID-19 Genomics UK (COG-UK) Consortium | The Lighthouse Lab in Milton Keynes and Alex Alderton, Roberto Amato, Jeffrey Barrett, Sonia Goncalves, Ewan Harrison, David K. Jackson, Ian Johnston, Dominic Kwiatkowski, Cordelia Langford, John Sillitoe on behalf of the Wellcome Sanger Institute COVID-19 Surveillance Team |
| EPI_ISL_3109710 | Lighthouse Laboratory Plymouth | Wellcome Sanger Institute for the COVID-19 Genomics UK (COG-UK) Consortium | Lighthouse Laboratory Plymouth and Alex Alderton, Roberto Amato, Jeffrey Barrett, Sonia Goncalves, Ewan Harrison, David K. Jackson, Ian Johnston, Dominic Kwiatkowski, Cordelia Langford, John Sillitoe on behalf of the Wellcome Sanger Institute COVID-19 Surveillance Team |
| EPI_ISL_3109711 | Lighthouse Lab in Milton Keynes | Wellcome Sanger Institute for the COVID-19 Genomics UK (COG-UK) Consortium | The Lighthouse Lab in Milton Keynes and Alex Alderton, Roberto Amato, Jeffrey Barrett, Sonia Goncalves, Ewan Harrison, David K. Jackson, Ian Johnston, Dominic Kwiatkowski, Cordelia Langford, John Sillitoe on behalf of the Wellcome Sanger Institute COVID-19 Surveillance Team |
| EPI_ISL_3109712 | Berkshire and Surrey Pathology Services Lighthouse Laboratory | Wellcome Sanger Institute for the COVID-19 Genomics UK (COG-UK) Consortium | Berkshire and Surrey Pathology Services Lighthouse Laboratory and Alex Alderton, Roberto Amato, Jeffrey Barrett, Sonia Goncalves, Ewan Harrison, David K. Jackson, Ian Johnston, Dominic Kwiatkowski, Cordelia Langford, John Sillitoe on behalf of the Wellcome Sanger Institute COVID-19 Surveillance Team |
| EPI_ISL_3109713 | Lighthouse Laboratory Plymouth | Wellcome Sanger Institute for the COVID-19 Genomics UK (COG-UK) Consortium | Lighthouse Laboratory Plymouth and Alex Alderton, Roberto Amato, Jeffrey Barrett, Sonia Goncalves, Ewan Harrison, David K. Jackson, Ian Johnston, Dominic Kwiatkowski, Cordelia Langford, John Sillitoe on behalf of the Wellcome Sanger Institute COVID-19 Surveillance Team |
| EPI_ISL_3109714 | Lighthouse Lab in Alderley Park | Wellcome Sanger Institute for the COVID-19 Genomics UK (COG-UK) Consortium | Jacquelyn Wynn, Mairead Hyland, The Lighthouse Lab in Alderley Park and Alex Alderton, Roberto Amato, Jeffrey Barrett, Sonia Goncalves, Ewan Harrison, David K. Jackson, Ian Johnston, Dominic Kwiatkowski, Cordelia Langford, John Sillitoe on behalf of the Wellcome Sanger Institute COVID-19 Surveillance Team |
| EPI_ISL_3109715, EPI_ISL_3109716 | Lighthouse Lab in Milton Keynes | Wellcome Sanger Institute for the COVID-19 Genomics UK (COG-UK) Consortium | The Lighthouse Lab in Milton Keynes and Alex Alderton, Roberto Amato, Jeffrey Barrett, Sonia Goncalves, Ewan Harrison, David K. Jackson, Ian Johnston, Dominic Kwiatkowski, Cordelia Langford, John Sillitoe on behalf of the Wellcome Sanger Institute COVID-19 Surveillance Team |
| EPI_ISL_3109717 | Berkshire and Surrey Pathology Services Lighthouse Laboratory | Wellcome Sanger Institute for the COVID-19 Genomics UK (COG-UK) Consortium | Berkshire and Surrey Pathology Services Lighthouse Laboratory and Alex Alderton, Roberto Amato, Jeffrey Barrett, Sonia Goncalves, Ewan Harrison, David K. Jackson, Ian Johnston, Dominic Kwiatkowski, Cordelia Langford, John Sillitoe on behalf of the Wellcome Sanger Institute COVID-19 Surveillance Team |
| EPI_ISL_3109718 | Lighthouse Laboratory Plymouth | Wellcome Sanger Institute for the COVID-19 Genomics UK (COG-UK) Consortium | Lighthouse Laboratory Plymouth and Alex Alderton, Roberto Amato, Jeffrey Barrett, Sonia Goncalves, Ewan Harrison, David K. Jackson, Ian Johnston, Dominic Kwiatkowski, Cordelia Langford, John Sillitoe on behalf of the Wellcome Sanger Institute COVID-19 Surveillance Team |
| EPI_ISL_3109719 | Lighthouse Lab in Alderley Park | Wellcome Sanger Institute for the COVID-19 Genomics UK (COG-UK) Consortium | Jacquelyn Wynn, Mairead Hyland, The Lighthouse Lab in Alderley Park and Alex Alderton, Roberto Amato, Jeffrey Barrett, Sonia Goncalves, Ewan Harrison, David K. Jackson, Ian Johnston, Dominic Kwiatkowski, Cordelia Langford, John Sillitoe on behalf of the Wellcome Sanger Institute COVID-19 Surveillance Team |
| EPI_ISL_3109720 | Lighthouse Laboratory Plymouth | Wellcome Sanger Institute for the COVID-19 Genomics UK (COG-UK) Consortium | Lighthouse Laboratory Plymouth and Alex Alderton, Roberto Amato, Jeffrey Barrett, Sonia Goncalves, Ewan Harrison, David K. Jackson, Ian Johnston, Dominic Kwiatkowski, Cordelia Langford, John Sillitoe on behalf of the Wellcome Sanger Institute COVID-19 Surveillance Team |
| EPI_ISL_3109721, EPI_ISL_3109722 | Lighthouse Lab in Milton Keynes | Wellcome Sanger Institute for the COVID-19 Genomics UK (COG-UK) Consortium | The Lighthouse Lab in Milton Keynes and Alex Alderton, Roberto Amato, Jeffrey Barrett, Sonia Goncalves, Ewan Harrison, David K. Jackson, Ian Johnston, Dominic Kwiatkowski, Cordelia Langford, John Sillitoe on behalf of the Wellcome Sanger Institute COVID-19 Surveillance Team |
| EPI_ISL_3109723 | Lighthouse Laboratory Plymouth | Wellcome Sanger Institute for the COVID-19 Genomics UK (COG-UK) Consortium | Lighthouse Laboratory Plymouth and Alex Alderton, Roberto Amato, Jeffrey Barrett, Sonia Goncalves, Ewan Harrison, David K. Jackson, Ian Johnston, Dominic Kwiatkowski, Cordelia Langford, John Sillitoe on behalf of the Wellcome Sanger Institute COVID-19 Surveillance Team |
| EPI_ISL_3109724 | Lighthouse Lab in Alderley Park | Wellcome Sanger Institute for the COVID-19 Genomics UK (COG-UK) Consortium | Jacquelyn Wynn, Mairead Hyland, The Lighthouse Lab in Alderley Park and Alex Alderton, Roberto Amato, Jeffrey Barrett, Sonia Goncalves, Ewan Harrison, David K. Jackson, Ian Johnston, Dominic Kwiatkowski, Cordelia Langford, John Sillitoe on behalf of the Wellcome Sanger Institute COVID-19 Surveillance Team |
| EPI_ISL_3109725, EPI_ISL_3109726 | Lighthouse Laboratory Plymouth | Wellcome Sanger Institute for the COVID-19 Genomics UK (COG-UK) Consortium | Lighthouse Laboratory Plymouth and Alex Alderton, Roberto Amato, Jeffrey Barrett, Sonia Goncalves, Ewan Harrison, David K. Jackson, Ian Johnston, Dominic Kwiatkowski, Cordelia Langford, John Sillitoe on behalf of the Wellcome Sanger Institute COVID-19 Surveillance Team |
| EPI_ISL_3109727 | Lighthouse Lab in Alderley Park | Wellcome Sanger Institute for the COVID-19 Genomics UK (COG-UK) Consortium | Jacquelyn Wynn, Mairead Hyland, The Lighthouse Lab in Alderley Park and Alex Alderton, Roberto Amato, Jeffrey Barrett, Sonia Goncalves, Ewan Harrison, David K. Jackson, Ian Johnston, Dominic Kwiatkowski, Cordelia Langford, John Sillitoe on behalf of the Wellcome Sanger Institute COVID-19 Surveillance Team |
| EPI_ISL_3109728 | Lighthouse Laboratory Plymouth | Wellcome Sanger Institute for the COVID-19 Genomics UK (COG-UK) Consortium | Lighthouse Laboratory Plymouth and Alex Alderton, Roberto Amato, Jeffrey Barrett, Sonia Goncalves, Ewan Harrison, David K. Jackson, Ian Johnston, Dominic Kwiatkowski, Cordelia Langford, John Sillitoe on behalf of the Wellcome Sanger Institute COVID-19 Surveillance Team |
| EPI_ISL_3109729 | Lighthouse Lab in Alderley Park | Wellcome Sanger Institute for the COVID-19 Genomics UK (COG-UK) Consortium | Jacquelyn Wynn, Mairead Hyland, The Lighthouse Lab in Alderley Park and Alex Alderton, Roberto Amato, Jeffrey Barrett, Sonia Goncalves, Ewan Harrison, David K. Jackson, Ian Johnston, Dominic Kwiatkowski, Cordelia Langford, John Sillitoe on behalf of the Wellcome Sanger Institute COVID-19 Surveillance Team |
| EPI_ISL_3109730 | Lighthouse Laboratory Plymouth | Wellcome Sanger Institute for the COVID-19 Genomics UK (COG-UK) Consortium | Lighthouse Laboratory Plymouth and Alex Alderton, Roberto Amato, Jeffrey Barrett, Sonia Goncalves, Ewan Harrison, David K. Jackson, Ian Johnston, Dominic Kwiatkowski, Cordelia Langford, John Sillitoe on behalf of the Wellcome Sanger Institute COVID-19 Surveillance Team |
| EPI_ISL_3109731, EPI_ISL_3109732 | Berkshire and Surrey Pathology Services Lighthouse Laboratory | Wellcome Sanger Institute for the COVID-19 Genomics UK (COG-UK) Consortium | Berkshire and Surrey Pathology Services Lighthouse Laboratory and Alex Alderton, Roberto Amato, Jeffrey Barrett, Sonia Goncalves, Ewan Harrison, David K. Jackson, Ian Johnston, Dominic Kwiatkowski, Cordelia Langford, John Sillitoe on behalf of the Wellcome Sanger Institute COVID-19 Surveillance Team |
| EPI_ISL_3109733 | Lighthouse Lab in Alderley Park | Wellcome Sanger Institute for the COVID-19 Genomics UK (COG-UK) Consortium | Jacquelyn Wynn, Mairead Hyland, The Lighthouse Lab in Alderley Park and Alex Alderton, Roberto Amato, Jeffrey Barrett, Sonia Goncalves, Ewan Harrison, David K. Jackson, Ian Johnston, Dominic Kwiatkowski, Cordelia Langford, John Sillitoe on behalf of the Wellcome Sanger Institute COVID-19 Surveillance Team |
| EPI_ISL_3109734 | Lighthouse Lab in Milton Keynes | Wellcome Sanger Institute for the COVID-19 Genomics UK (COG-UK) Consortium | The Lighthouse Lab in Milton Keynes and Alex Alderton, Roberto Amato, Jeffrey Barrett, Sonia Goncalves, Ewan Harrison, David K. Jackson, Ian Johnston, Dominic Kwiatkowski, Cordelia Langford, John Sillitoe on behalf of the Wellcome Sanger Institute COVID-19 Surveillance Team |
| EPI_ISL_3109735 | Lighthouse Lab in Alderley Park | Wellcome Sanger Institute for the COVID-19 Genomics UK (COG-UK) Consortium | Jacquelyn Wynn, Mairead Hyland, The Lighthouse Lab in Alderley Park and Alex Alderton, Roberto Amato, Jeffrey Barrett, Sonia Goncalves, Ewan Harrison, David K. Jackson, Ian Johnston, Dominic Kwiatkowski, Cordelia Langford, John Sillitoe on behalf of the Wellcome Sanger Institute COVID-19 Surveillance Team |
| EPI_ISL_3109736, EPI_ISL_3109737 | Lighthouse Lab in Milton Keynes | Wellcome Sanger Institute for the COVID-19 Genomics UK (COG-UK) Consortium | The Lighthouse Lab in Milton Keynes and Alex Alderton, Roberto Amato, Jeffrey Barrett, Sonia Goncalves, Ewan Harrison, David K. Jackson, Ian Johnston, Dominic Kwiatkowski, Cordelia Langford, John Sillitoe on behalf of the Wellcome Sanger Institute COVID-19 Surveillance Team |
| EPI_ISL_3109738 | Berkshire and Surrey Pathology Services Lighthouse Laboratory | Wellcome Sanger Institute for the COVID-19 Genomics UK (COG-UK) Consortium | Berkshire and Surrey Pathology Services Lighthouse Laboratory and Alex Alderton, Roberto Amato, Jeffrey Barrett, Sonia Goncalves, Ewan Harrison, David K. Jackson, Ian Johnston, Dominic Kwiatkowski, Cordelia Langford, John Sillitoe on behalf of the Wellcome Sanger Institute COVID-19 Surveillance Team |
| EPI_ISL_3109739 | Lighthouse Laboratory Plymouth | Wellcome Sanger Institute for the COVID-19 Genomics UK (COG-UK) Consortium | Lighthouse Laboratory Plymouth and Alex Alderton, Roberto Amato, Jeffrey Barrett, Sonia Goncalves, Ewan Harrison, David K. Jackson, Ian Johnston, Dominic Kwiatkowski, Cordelia Langford, John Sillitoe on behalf of the Wellcome Sanger Institute COVID-19 Surveillance Team |
| EPI_ISL_3109740 | Lighthouse Lab in Milton Keynes | Wellcome Sanger Institute for the COVID-19 Genomics UK (COG-UK) Consortium | The Lighthouse Lab in Milton Keynes and Alex Alderton, Roberto Amato, Jeffrey Barrett, Sonia Goncalves, Ewan Harrison, David K. Jackson, Ian Johnston, Dominic Kwiatkowski, Cordelia Langford, John Sillitoe on behalf of the Wellcome Sanger Institute COVID-19 Surveillance Team |
| EPI_ISL_3109741 | Berkshire and Surrey Pathology Services Lighthouse Laboratory | Wellcome Sanger Institute for the COVID-19 Genomics UK (COG-UK) Consortium | Berkshire and Surrey Pathology Services Lighthouse Laboratory and Alex Alderton, Roberto Amato, Jeffrey Barrett, Sonia Goncalves, Ewan Harrison, David K. Jackson, Ian Johnston, Dominic Kwiatkowski, Cordelia Langford, John Sillitoe on behalf of the Wellcome Sanger Institute COVID-19 Surveillance Team |
| EPI_ISL_3109742 | Lighthouse Laboratory Plymouth | Wellcome Sanger Institute for the COVID-19 Genomics UK (COG-UK) Consortium | Lighthouse Laboratory Plymouth and Alex Alderton, Roberto Amato, Jeffrey Barrett, Sonia Goncalves, Ewan Harrison, David K. Jackson, Ian Johnston, Dominic Kwiatkowski, Cordelia Langford, John Sillitoe on behalf of the Wellcome Sanger Institute COVID-19 Surveillance Team |
| EPI_ISL_3109743 | Lighthouse Lab in Alderley Park | Wellcome Sanger Institute for the COVID-19 Genomics UK (COG-UK) Consortium | Jacquelyn Wynn, Mairead Hyland, The Lighthouse Lab in Alderley Park and Alex Alderton, Roberto Amato, Jeffrey Barrett, Sonia Goncalves, Ewan Harrison, David K. Jackson, Ian Johnston, Dominic Kwiatkowski, Cordelia Langford, John Sillitoe on behalf of the Wellcome Sanger Institute COVID-19 Surveillance Team |
| EPI_ISL_3109744, EPI_ISL_3109745, EPI_ISL_3109746 | Lighthouse Lab in Milton Keynes | Wellcome Sanger Institute for the COVID-19 Genomics UK (COG-UK) Consortium | The Lighthouse Lab in Milton Keynes and Alex Alderton, Roberto Amato, Jeffrey Barrett, Sonia Goncalves, Ewan Harrison, David K. Jackson, Ian Johnston, Dominic Kwiatkowski, Cordelia Langford, John Sillitoe on behalf of the Wellcome Sanger Institute COVID-19 Surveillance Team |
| EPI_ISL_3109747 | Berkshire and Surrey Pathology Services Lighthouse Laboratory | Wellcome Sanger Institute for the COVID-19 Genomics UK (COG-UK) Consortium | Berkshire and Surrey Pathology Services Lighthouse Laboratory and Alex Alderton, Roberto Amato, Jeffrey Barrett, Sonia Goncalves, Ewan Harrison, David K. Jackson, Ian Johnston, Dominic Kwiatkowski, Cordelia Langford, John Sillitoe on behalf of the Wellcome Sanger Institute COVID-19 Surveillance Team |
| EPI_ISL_3109748 | Lighthouse Laboratory Plymouth | Wellcome Sanger Institute for the COVID-19 Genomics UK (COG-UK) Consortium | Lighthouse Laboratory Plymouth and Alex Alderton, Roberto Amato, Jeffrey Barrett, Sonia Goncalves, Ewan Harrison, David K. Jackson, Ian Johnston, Dominic Kwiatkowski, Cordelia Langford, John Sillitoe on behalf of the Wellcome Sanger Institute COVID-19 Surveillance Team |
| EPI_ISL_3109749 | Lighthouse Lab in Milton Keynes | Wellcome Sanger Institute for the COVID-19 Genomics UK (COG-UK) Consortium | The Lighthouse Lab in Milton Keynes and Alex Alderton, Roberto Amato, Jeffrey Barrett, Sonia Goncalves, Ewan Harrison, David K. Jackson, Ian Johnston, Dominic Kwiatkowski, Cordelia Langford, John Sillitoe on behalf of the Wellcome Sanger Institute COVID-19 Surveillance Team |
| EPI_ISL_3109750 | Lighthouse Lab in Alderley Park | Wellcome Sanger Institute for the COVID-19 Genomics UK (COG-UK) Consortium | Jacquelyn Wynn, Mairead Hyland, The Lighthouse Lab in Alderley Park and Alex Alderton, Roberto Amato, Jeffrey Barrett, Sonia Goncalves, Ewan Harrison, David K. Jackson, Ian Johnston, Dominic Kwiatkowski, Cordelia Langford, John Sillitoe on behalf of the Wellcome Sanger Institute COVID-19 Surveillance Team |
| EPI_ISL_3109751 | Berkshire and Surrey Pathology Services Lighthouse Laboratory | Wellcome Sanger Institute for the COVID-19 Genomics UK (COG-UK) Consortium | Berkshire and Surrey Pathology Services Lighthouse Laboratory and Alex Alderton, Roberto Amato, Jeffrey Barrett, Sonia Goncalves, Ewan Harrison, David K. Jackson, Ian Johnston, Dominic Kwiatkowski, Cordelia Langford, John Sillitoe on behalf of the Wellcome Sanger Institute COVID-19 Surveillance Team |
| EPI_ISL_3109752 | Lighthouse Lab in Alderley Park | Wellcome Sanger Institute for the COVID-19 Genomics UK (COG-UK) Consortium | Jacquelyn Wynn, Mairead Hyland, The Lighthouse Lab in Alderley Park and Alex Alderton, Roberto Amato, Jeffrey Barrett, Sonia Goncalves, Ewan Harrison, David K. Jackson, Ian Johnston, Dominic Kwiatkowski, Cordelia Langford, John Sillitoe on behalf of the Wellcome Sanger Institute COVID-19 Surveillance Team |
| EPI_ISL_3109753, EPI_ISL_3109754 | Lighthouse Laboratory Plymouth | Wellcome Sanger Institute for the COVID-19 Genomics UK (COG-UK) Consortium | Lighthouse Laboratory Plymouth and Alex Alderton, Roberto Amato, Jeffrey Barrett, Sonia Goncalves, Ewan Harrison, David K. Jackson, Ian Johnston, Dominic Kwiatkowski, Cordelia Langford, John Sillitoe on behalf of the Wellcome Sanger Institute COVID-19 Surveillance Team |
| EPI_ISL_3109755 | Lighthouse Lab in Milton Keynes | Wellcome Sanger Institute for the COVID-19 Genomics UK (COG-UK) Consortium | The Lighthouse Lab in Milton Keynes and Alex Alderton, Roberto Amato, Jeffrey Barrett, Sonia Goncalves, Ewan Harrison, David K. Jackson, Ian Johnston, Dominic Kwiatkowski, Cordelia Langford, John Sillitoe on behalf of the Wellcome Sanger Institute COVID-19 Surveillance Team |
| EPI_ISL_3109756 | Lighthouse Laboratory Plymouth | Wellcome Sanger Institute for the COVID-19 Genomics UK (COG-UK) Consortium | Lighthouse Laboratory Plymouth and Alex Alderton, Roberto Amato, Jeffrey Barrett, Sonia Goncalves, Ewan Harrison, David K. Jackson, Ian Johnston, Dominic Kwiatkowski, Cordelia Langford, John Sillitoe on behalf of the Wellcome Sanger Institute COVID-19 Surveillance Team |
| EPI_ISL_3109757, EPI_ISL_3109758, EPI_ISL_3109759 | Lighthouse Lab in Alderley Park | Wellcome Sanger Institute for the COVID-19 Genomics UK (COG-UK) Consortium | Jacquelyn Wynn, Mairead Hyland, The Lighthouse Lab in Alderley Park and Alex Alderton, Roberto Amato, Jeffrey Barrett, Sonia Goncalves, Ewan Harrison, David K. Jackson, Ian Johnston, Dominic Kwiatkowski, Cordelia Langford, John Sillitoe on behalf of the Wellcome Sanger Institute COVID-19 Surveillance Team |
| EPI_ISL_3109760 | Lighthouse Lab in Milton Keynes | Wellcome Sanger Institute for the COVID-19 Genomics UK (COG-UK) Consortium | The Lighthouse Lab in Milton Keynes and Alex Alderton, Roberto Amato, Jeffrey Barrett, Sonia Goncalves, Ewan Harrison, David K. Jackson, Ian Johnston, Dominic Kwiatkowski, Cordelia Langford, John Sillitoe on behalf of the Wellcome Sanger Institute COVID-19 Surveillance Team |
| EPI_ISL_3109761, EPI_ISL_3109762, EPI_ISL_3109763 | Lighthouse Lab in Alderley Park | Wellcome Sanger Institute for the COVID-19 Genomics UK (COG-UK) Consortium | Jacquelyn Wynn, Mairead Hyland, The Lighthouse Lab in Alderley Park and Alex Alderton, Roberto Amato, Jeffrey Barrett, Sonia Goncalves, Ewan Harrison, David K. Jackson, Ian Johnston, Dominic Kwiatkowski, Cordelia Langford, John Sillitoe on behalf of the Wellcome Sanger Institute COVID-19 Surveillance Team |
| EPI_ISL_3109764 | Lighthouse Lab in Milton Keynes | Wellcome Sanger Institute for the COVID-19 Genomics UK (COG-UK) Consortium | The Lighthouse Lab in Milton Keynes and Alex Alderton, Roberto Amato, Jeffrey Barrett, Sonia Goncalves, Ewan Harrison, David K. Jackson, Ian Johnston, Dominic Kwiatkowski, Cordelia Langford, John Sillitoe on behalf of the Wellcome Sanger Institute COVID-19 Surveillance Team |

| | | | |
|---|---|---|---|
| EPI_ISL_3109765, EPI_ISL_3109766, EPI_ISL_3109767 | Lighthouse Laboratory Plymouth | Wellcome Sanger Institute for the COVID-19 Genomics UK (COG-UK) Consortium | Lighthouse Laboratory Plymouth and Alex Alderton, Roberto Amato, Jeffrey Barrett, Sonia Goncalves, Ewan Harrison, David K. Jackson, Ian Johnston, Dominic Kwiatkowski, Cordelia Langford, John Sillitoe on behalf of the Wellcome Sanger Institute COVID-19 Surveillance Team |
| EPI_ISL_3109768 | Berkshire and Surrey Pathology Services Lighthouse Laboratory | Wellcome Sanger Institute for the COVID-19 Genomics UK (COG-UK) Consortium | Berkshire and Surrey Pathology Services Lighthouse Laboratory and Alex Alderton, Roberto Amato, Jeffrey Barrett, Sonia Goncalves, Ewan Harrison, David K. Jackson, Ian Johnston, Dominic Kwiatkowski, Cordelia Langford, John Sillitoe on behalf of the Wellcome Sanger Institute COVID-19 Surveillance Team |
| EPI_ISL_3109769 | Lighthouse Lab in Alderley Park | Wellcome Sanger Institute for the COVID-19 Genomics UK (COG-UK) Consortium | Jacquelyn Wynn, Mairead Hyland, The Lighthouse Lab in Alderley Park and Alex Alderton, Roberto Amato, Jeffrey Barrett, Sonia Goncalves, Ewan Harrison, David K. Jackson, Ian Johnston, Dominic Kwiatkowski, Cordelia Langford, John Sillitoe on behalf of the Wellcome Sanger Institute COVID-19 Surveillance Team |
| EPI_ISL_3109770 | Lighthouse Lab in Milton Keynes | Wellcome Sanger Institute for the COVID-19 Genomics UK (COG-UK) Consortium | The Lighthouse Lab in Milton Keynes and Alex Alderton, Roberto Amato, Jeffrey Barrett, Sonia Goncalves, Ewan Harrison, David K. Jackson, Ian Johnston, Dominic Kwiatkowski, Cordelia Langford, John Sillitoe on behalf of the Wellcome Sanger Institute COVID-19 Surveillance Team |
| EPI_ISL_3109771 | Lighthouse Laboratory Plymouth | Wellcome Sanger Institute for the COVID-19 Genomics UK (COG-UK) Consortium | Lighthouse Laboratory Plymouth and Alex Alderton, Roberto Amato, Jeffrey Barrett, Sonia Goncalves, Ewan Harrison, David K. Jackson, Ian Johnston, Dominic Kwiatkowski, Cordelia Langford, John Sillitoe on behalf of the Wellcome Sanger Institute COVID-19 Surveillance Team |
| EPI_ISL_3109772 | Lighthouse Lab in Alderley Park | Wellcome Sanger Institute for the COVID-19 Genomics UK (COG-UK) Consortium | Jacquelyn Wynn, Mairead Hyland, The Lighthouse Lab in Alderley Park and Alex Alderton, Roberto Amato, Jeffrey Barrett, Sonia Goncalves, Ewan Harrison, David K. Jackson, Ian Johnston, Dominic Kwiatkowski, Cordelia Langford, John Sillitoe on behalf of the Wellcome Sanger Institute COVID-19 Surveillance Team |
| EPI_ISL_3109773 | Lighthouse Lab in Milton Keynes | Wellcome Sanger Institute for the COVID-19 Genomics UK (COG-UK) Consortium | The Lighthouse Lab in Milton Keynes and Alex Alderton, Roberto Amato, Jeffrey Barrett, Sonia Goncalves, Ewan Harrison, David K. Jackson, Ian Johnston, Dominic Kwiatkowski, Cordelia Langford, John Sillitoe on behalf of the Wellcome Sanger Institute COVID-19 Surveillance Team |
| EPI_ISL_3109774 | Lighthouse Lab in Alderley Park | Wellcome Sanger Institute for the COVID-19 Genomics UK (COG-UK) Consortium | Jacquelyn Wynn, Mairead Hyland, The Lighthouse Lab in Alderley Park and Alex Alderton, Roberto Amato, Jeffrey Barrett, Sonia Goncalves, Ewan Harrison, David K. Jackson, Ian Johnston, Dominic Kwiatkowski, Cordelia Langford, John Sillitoe on behalf of the Wellcome Sanger Institute COVID-19 Surveillance Team |
| EPI_ISL_3109775 | Lighthouse Laboratory Plymouth | Wellcome Sanger Institute for the COVID-19 Genomics UK (COG-UK) Consortium | Lighthouse Laboratory Plymouth and Alex Alderton, Roberto Amato, Jeffrey Barrett, Sonia Goncalves, Ewan Harrison, David K. Jackson, Ian Johnston, Dominic Kwiatkowski, Cordelia Langford, John Sillitoe on behalf of the Wellcome Sanger Institute COVID-19 Surveillance Team |
| EPI_ISL_3109776 | Berkshire and Surrey Pathology Services Lighthouse Laboratory | Wellcome Sanger Institute for the COVID-19 Genomics UK (COG-UK) Consortium | Berkshire and Surrey Pathology Services Lighthouse Laboratory and Alex Alderton, Roberto Amato, Jeffrey Barrett, Sonia Goncalves, Ewan Harrison, David K. Jackson, Ian Johnston, Dominic Kwiatkowski, Cordelia Langford, John Sillitoe on behalf of the Wellcome Sanger Institute COVID-19 Surveillance Team |
| EPI_ISL_3109777, EPI_ISL_3109778, EPI_ISL_3109779 | Lighthouse Lab in Alderley Park | Wellcome Sanger Institute for the COVID-19 Genomics UK (COG-UK) Consortium | Jacquelyn Wynn, Mairead Hyland, The Lighthouse Lab in Alderley Park and Alex Alderton, Roberto Amato, Jeffrey Barrett, Sonia Goncalves, Ewan Harrison, David K. Jackson, Ian Johnston, Dominic Kwiatkowski, Cordelia Langford, John Sillitoe on behalf of the Wellcome Sanger Institute COVID-19 Surveillance Team |
| EPI_ISL_3109780 | Lighthouse Lab in Milton Keynes | Wellcome Sanger Institute for the COVID-19 Genomics UK (COG-UK) Consortium | The Lighthouse Lab in Milton Keynes and Alex Alderton, Roberto Amato, Jeffrey Barrett, Sonia Goncalves, Ewan Harrison, David K. Jackson, Ian Johnston, Dominic Kwiatkowski, Cordelia Langford, John Sillitoe on behalf of the Wellcome Sanger Institute COVID-19 Surveillance Team |
| EPI_ISL_3109781 | Lighthouse Laboratory Plymouth | Wellcome Sanger Institute for the COVID-19 Genomics UK (COG-UK) Consortium | Lighthouse Laboratory Plymouth and Alex Alderton, Roberto Amato, Jeffrey Barrett, Sonia Goncalves, Ewan Harrison, David K. Jackson, Ian Johnston, Dominic Kwiatkowski, Cordelia Langford, John Sillitoe on behalf of the Wellcome Sanger Institute COVID-19 Surveillance Team |
| EPI_ISL_3109782 | Lighthouse Lab in Alderley Park | Wellcome Sanger Institute for the COVID-19 Genomics UK (COG-UK) Consortium | Jacquelyn Wynn, Mairead Hyland, The Lighthouse Lab in Alderley Park and Alex Alderton, Roberto Amato, Jeffrey Barrett, Sonia Goncalves, Ewan Harrison, David K. Jackson, Ian Johnston, Dominic Kwiatkowski, Cordelia Langford, John Sillitoe on behalf of the Wellcome Sanger Institute COVID-19 Surveillance Team |
| EPI_ISL_3109783, EPI_ISL_3109784 | Lighthouse Laboratory Plymouth | Wellcome Sanger Institute for the COVID-19 Genomics UK (COG-UK) Consortium | Lighthouse Laboratory Plymouth and Alex Alderton, Roberto Amato, Jeffrey Barrett, Sonia Goncalves, Ewan Harrison, David K. Jackson, Ian Johnston, Dominic Kwiatkowski, Cordelia Langford, John Sillitoe on behalf of the Wellcome Sanger Institute COVID-19 Surveillance Team |
| EPI_ISL_3109785, EPI_ISL_3109786 | Lighthouse Lab in Alderley Park | Wellcome Sanger Institute for the COVID-19 Genomics UK (COG-UK) Consortium | Jacquelyn Wynn, Mairead Hyland, The Lighthouse Lab in Alderley Park and Alex Alderton, Roberto Amato, Jeffrey Barrett, Sonia Goncalves, Ewan Harrison, David K. Jackson, Ian Johnston, Dominic Kwiatkowski, Cordelia Langford, John Sillitoe on behalf of the Wellcome Sanger Institute COVID-19 Surveillance Team |
| EPI_ISL_3109787 | Lighthouse Lab in Milton Keynes | Wellcome Sanger Institute for the COVID-19 Genomics UK (COG-UK) Consortium | The Lighthouse Lab in Milton Keynes and Alex Alderton, Roberto Amato, Jeffrey Barrett, Sonia Goncalves, Ewan Harrison, David K. Jackson, Ian Johnston, Dominic Kwiatkowski, Cordelia Langford, John Sillitoe on behalf of the Wellcome Sanger Institute COVID-19 Surveillance Team |
| EPI_ISL_3109788 | Lighthouse Laboratory Plymouth | Wellcome Sanger Institute for the COVID-19 Genomics UK (COG-UK) Consortium | Lighthouse Laboratory Plymouth and Alex Alderton, Roberto Amato, Jeffrey Barrett, Sonia Goncalves, Ewan Harrison, David K. Jackson, Ian Johnston, Dominic Kwiatkowski, Cordelia Langford, John Sillitoe on behalf of the Wellcome Sanger Institute COVID-19 Surveillance Team |
| EPI_ISL_3109789 | Lighthouse Lab in Alderley Park | Wellcome Sanger Institute for the COVID-19 Genomics UK (COG-UK) Consortium | Jacquelyn Wynn, Mairead Hyland, The Lighthouse Lab in Alderley Park and Alex Alderton, Roberto Amato, Jeffrey Barrett, Sonia Goncalves, Ewan Harrison, David K. Jackson, Ian Johnston, Dominic Kwiatkowski, Cordelia Langford, John Sillitoe on behalf of the Wellcome Sanger Institute COVID-19 Surveillance Team |
| EPI_ISL_3109790 | Lighthouse Lab in Milton Keynes | Wellcome Sanger Institute for the COVID-19 Genomics UK (COG-UK) Consortium | The Lighthouse Lab in Milton Keynes and Alex Alderton, Roberto Amato, Jeffrey Barrett, Sonia Goncalves, Ewan Harrison, David K. Jackson, Ian Johnston, Dominic Kwiatkowski, Cordelia Langford, John Sillitoe on behalf of the Wellcome Sanger Institute COVID-19 Surveillance Team |
| EPI_ISL_3109791 | Lighthouse Lab in Alderley Park | Wellcome Sanger Institute for the COVID-19 Genomics UK (COG-UK) Consortium | Jacquelyn Wynn, Mairead Hyland, The Lighthouse Lab in Alderley Park and Alex Alderton, Roberto Amato, Jeffrey Barrett, Sonia Goncalves, Ewan Harrison, David K. Jackson, Ian Johnston, Dominic Kwiatkowski, Cordelia Langford, John Sillitoe on behalf of the Wellcome Sanger Institute COVID-19 Surveillance Team |
| EPI_ISL_3109792 | Berkshire and Surrey Pathology Services Lighthouse Laboratory | Wellcome Sanger Institute for the COVID-19 Genomics UK (COG-UK) Consortium | Berkshire and Surrey Pathology Services Lighthouse Laboratory and Alex Alderton, Roberto Amato, Jeffrey Barrett, Sonia Goncalves, Ewan Harrison, David K. Jackson, Ian Johnston, Dominic Kwiatkowski, Cordelia Langford, John Sillitoe on behalf of the Wellcome Sanger Institute COVID-19 Surveillance Team |
| EPI_ISL_3109793 | Lighthouse Laboratory Plymouth | Wellcome Sanger Institute for the COVID-19 Genomics UK (COG-UK) Consortium | Lighthouse Laboratory Plymouth and Alex Alderton, Roberto Amato, Jeffrey Barrett, Sonia Goncalves, Ewan Harrison, David K. Jackson, Ian Johnston, Dominic Kwiatkowski, Cordelia Langford, John Sillitoe on behalf of the Wellcome Sanger Institute COVID-19 Surveillance Team |
| EPI_ISL_3109794 | Lighthouse Lab in Milton Keynes | Wellcome Sanger Institute for the COVID-19 Genomics UK (COG-UK) Consortium | The Lighthouse Lab in Milton Keynes and Alex Alderton, Roberto Amato, Jeffrey Barrett, Sonia Goncalves, Ewan Harrison, David K. Jackson, Ian Johnston, Dominic Kwiatkowski, Cordelia Langford, John Sillitoe on behalf of the Wellcome Sanger Institute COVID-19 Surveillance Team |
| EPI_ISL_3109795 | Berkshire and Surrey Pathology Services Lighthouse Laboratory | Wellcome Sanger Institute for the COVID-19 Genomics UK (COG-UK) Consortium | Berkshire and Surrey Pathology Services Lighthouse Laboratory and Alex Alderton, Roberto Amato, Jeffrey Barrett, Sonia Goncalves, Ewan Harrison, David K. Jackson, Ian Johnston, Dominic Kwiatkowski, Cordelia Langford, John Sillitoe on behalf of the Wellcome Sanger Institute COVID-19 Surveillance Team |
| EPI_ISL_3109796, EPI_ISL_3109797 | Lighthouse Laboratory Plymouth | Wellcome Sanger Institute for the COVID-19 Genomics UK (COG-UK) Consortium | Lighthouse Laboratory Plymouth and Alex Alderton, Roberto Amato, Jeffrey Barrett, Sonia Goncalves, Ewan Harrison, David K. Jackson, Ian Johnston, Dominic Kwiatkowski, Cordelia Langford, John Sillitoe on behalf of the Wellcome Sanger Institute COVID-19 Surveillance Team |
| EPI_ISL_3109798 | Berkshire and Surrey Pathology Services Lighthouse Laboratory | Wellcome Sanger Institute for the COVID-19 Genomics UK (COG-UK) Consortium | Berkshire and Surrey Pathology Services Lighthouse Laboratory and Alex Alderton, Roberto Amato, Jeffrey Barrett, Sonia Goncalves, Ewan Harrison, David K. Jackson, Ian Johnston, Dominic Kwiatkowski, Cordelia Langford, John Sillitoe on behalf of the Wellcome Sanger Institute COVID-19 Surveillance Team |
| EPI_ISL_3109799 | Lighthouse Lab in Milton Keynes | Wellcome Sanger Institute for the COVID-19 Genomics UK (COG-UK) Consortium | The Lighthouse Lab in Milton Keynes and Alex Alderton, Roberto Amato, Jeffrey Barrett, Sonia Goncalves, Ewan Harrison, David K. Jackson, Ian Johnston, Dominic Kwiatkowski, Cordelia Langford, John Sillitoe on behalf of the Wellcome Sanger Institute COVID-19 Surveillance Team |
| EPI_ISL_3109800 | Lighthouse Laboratory Plymouth | Wellcome Sanger Institute for the COVID-19 Genomics UK (COG-UK) Consortium | Lighthouse Laboratory Plymouth and Alex Alderton, Roberto Amato, Jeffrey Barrett, Sonia Goncalves, Ewan Harrison, David K. Jackson, Ian Johnston, Dominic Kwiatkowski, Cordelia Langford, John Sillitoe on behalf of the Wellcome Sanger Institute COVID-19 Surveillance Team |
| EPI_ISL_3109801 | Berkshire and Surrey Pathology Services Lighthouse Laboratory | Wellcome Sanger Institute for the COVID-19 Genomics UK (COG-UK) Consortium | Berkshire and Surrey Pathology Services Lighthouse Laboratory and Alex Alderton, Roberto Amato, Jeffrey Barrett, Sonia Goncalves, Ewan Harrison, David K. Jackson, Ian Johnston, Dominic Kwiatkowski, Cordelia Langford, John Sillitoe on behalf of the Wellcome Sanger Institute COVID-19 Surveillance Team |
| EPI_ISL_3109802 | Lighthouse Lab in Milton Keynes | Wellcome Sanger Institute for the COVID-19 Genomics UK (COG-UK) Consortium | The Lighthouse Lab in Milton Keynes and Alex Alderton, Roberto Amato, Jeffrey Barrett, Sonia Goncalves, Ewan Harrison, David K. Jackson, Ian Johnston, Dominic Kwiatkowski, Cordelia Langford, John Sillitoe on behalf of the Wellcome Sanger Institute COVID-19 Surveillance Team |
| EPI_ISL_3109803 | Lighthouse Lab in Alderley Park | Wellcome Sanger Institute for the COVID-19 Genomics UK (COG-UK) Consortium | Jacquelyn Wynn, Mairead Hyland, The Lighthouse Lab in Alderley Park and Alex Alderton, Roberto Amato, Jeffrey Barrett, Sonia Goncalves, Ewan Harrison, David K. Jackson, Ian Johnston, Dominic Kwiatkowski, Cordelia Langford, John Sillitoe on behalf of the Wellcome Sanger Institute COVID-19 Surveillance Team |
| EPI_ISL_3109804 | Lighthouse Laboratory Plymouth | Wellcome Sanger Institute for the COVID-19 Genomics UK (COG-UK) Consortium | Lighthouse Laboratory Plymouth and Alex Alderton, Roberto Amato, Jeffrey Barrett, Sonia Goncalves, Ewan Harrison, David K. Jackson, Ian Johnston, Dominic Kwiatkowski, Cordelia Langford, John Sillitoe on behalf of the Wellcome Sanger Institute COVID-19 Surveillance Team |
| EPI_ISL_3109805 | Lighthouse Lab in Milton Keynes | Wellcome Sanger Institute for the COVID-19 Genomics UK (COG-UK) Consortium | The Lighthouse Lab in Milton Keynes and Alex Alderton, Roberto Amato, Jeffrey Barrett, Sonia Goncalves, Ewan Harrison, David K. Jackson, Ian Johnston, Dominic Kwiatkowski, Cordelia Langford, John Sillitoe on behalf of the Wellcome Sanger Institute COVID-19 Surveillance Team |
| EPI_ISL_3109806 | Berkshire and Surrey Pathology Services Lighthouse Laboratory | Wellcome Sanger Institute for the COVID-19 Genomics UK (COG-UK) Consortium | Berkshire and Surrey Pathology Services Lighthouse Laboratory and Alex Alderton, Roberto Amato, Jeffrey Barrett, Sonia Goncalves, Ewan Harrison, David K. Jackson, Ian Johnston, Dominic Kwiatkowski, Cordelia Langford, John Sillitoe on behalf of the Wellcome Sanger Institute COVID-19 Surveillance Team |
| EPI_ISL_3109807 | Lighthouse Laboratory Plymouth | Wellcome Sanger Institute for the COVID-19 Genomics UK (COG-UK) Consortium | Lighthouse Laboratory Plymouth and Alex Alderton, Roberto Amato, Jeffrey Barrett, Sonia Goncalves, Ewan Harrison, David K. Jackson, Ian Johnston, Dominic Kwiatkowski, Cordelia Langford, John Sillitoe on behalf of the Wellcome Sanger Institute COVID-19 Surveillance Team |
| EPI_ISL_3109808 | Lighthouse Lab in Alderley Park | Wellcome Sanger Institute for the COVID-19 Genomics UK (COG-UK) Consortium | Jacquelyn Wynn, Mairead Hyland, The Lighthouse Lab in Alderley Park and Alex Alderton, Roberto Amato, Jeffrey Barrett, Sonia Goncalves, Ewan Harrison, David K. Jackson, Ian Johnston, Dominic Kwiatkowski, Cordelia Langford, John Sillitoe on behalf of the Wellcome Sanger Institute COVID-19 Surveillance Team |
| EPI_ISL_3109809 | Lighthouse Lab in Milton Keynes | Wellcome Sanger Institute for the COVID-19 Genomics UK (COG-UK) Consortium | The Lighthouse Lab in Milton Keynes and Alex Alderton, Roberto Amato, Jeffrey Barrett, Sonia Goncalves, Ewan Harrison, David K. Jackson, Ian Johnston, Dominic Kwiatkowski, Cordelia Langford, John Sillitoe on behalf of the Wellcome Sanger Institute COVID-19 Surveillance Team |
| EPI_ISL_3109810, EPI_ISL_3109811 | Berkshire and Surrey Pathology Services Lighthouse Laboratory | Wellcome Sanger Institute for the COVID-19 Genomics UK (COG-UK) Consortium | Berkshire and Surrey Pathology Services Lighthouse Laboratory and Alex Alderton, Roberto Amato, Jeffrey Barrett, Sonia Goncalves, Ewan Harrison, David K. Jackson, Ian Johnston, Dominic Kwiatkowski, Cordelia Langford, John Sillitoe on behalf of the Wellcome Sanger Institute COVID-19 Surveillance Team |
| EPI_ISL_3109812 | Lighthouse Lab in Alderley Park | Wellcome Sanger Institute for the COVID-19 Genomics UK (COG-UK) Consortium | Jacquelyn Wynn, Mairead Hyland, The Lighthouse Lab in Alderley Park and Alex Alderton, Roberto Amato, Jeffrey Barrett, Sonia Goncalves, Ewan Harrison, David K. Jackson, Ian Johnston, Dominic Kwiatkowski, Cordelia Langford, John Sillitoe on behalf of the Wellcome Sanger Institute COVID-19 Surveillance Team |
| EPI_ISL_3109813, EPI_ISL_3109814 | Berkshire and Surrey Pathology Services Lighthouse Laboratory | Wellcome Sanger Institute for the COVID-19 Genomics UK (COG-UK) Consortium | Berkshire and Surrey Pathology Services Lighthouse Laboratory and Alex Alderton, Roberto Amato, Jeffrey Barrett, Sonia Goncalves, Ewan Harrison, David K. Jackson, Ian Johnston, Dominic Kwiatkowski, Cordelia Langford, John Sillitoe on behalf of the Wellcome Sanger Institute COVID-19 Surveillance Team |
| EPI_ISL_3109815, EPI_ISL_3109816 | Lighthouse Laboratory Plymouth | Wellcome Sanger Institute for the COVID-19 Genomics UK (COG-UK) Consortium | Lighthouse Laboratory Plymouth and Alex Alderton, Roberto Amato, Jeffrey Barrett, Sonia Goncalves, Ewan Harrison, David K. Jackson, Ian Johnston, Dominic Kwiatkowski, Cordelia Langford, John Sillitoe on behalf of the Wellcome Sanger Institute COVID-19 Surveillance Team |
| EPI_ISL_3109817 | Lighthouse Lab in Alderley Park | Wellcome Sanger Institute for the COVID-19 Genomics UK (COG-UK) Consortium | Jacquelyn Wynn, Mairead Hyland, The Lighthouse Lab in Alderley Park and Alex Alderton, Roberto Amato, Jeffrey Barrett, Sonia Goncalves, Ewan Harrison, David K. Jackson, Ian Johnston, Dominic Kwiatkowski, Cordelia Langford, John Sillitoe on behalf of the Wellcome Sanger Institute COVID-19 Surveillance Team |
| EPI_ISL_3109818, EPI_ISL_3109819 | Lighthouse Laboratory Plymouth | Wellcome Sanger Institute for the COVID-19 Genomics UK (COG-UK) Consortium | Lighthouse Laboratory Plymouth and Alex Alderton, Roberto Amato, Jeffrey Barrett, Sonia Goncalves, Ewan Harrison, David K. Jackson, Ian Johnston, Dominic Kwiatkowski, Cordelia Langford, John Sillitoe on behalf of the Wellcome Sanger Institute COVID-19 Surveillance Team |
| EPI_ISL_3109820, | Lighthouse Lab in Milton Keynes | Wellcome Sanger Institute for the COVID-19 Genomics UK (COG-UK) | The Lighthouse Lab in Milton Keynes and Alex Alderton, Roberto Amato, Jeffrey Barrett, Sonia Goncalves, Ewan Harrison, David K. Jackson, Ian Johnston, Dominic Kwiatkowski, Cordelia Langford, John Sillitoe on behalf of the Wellcome Sanger |

| | | | |
|---|---|---|---|
| EPI_ISL_3109821, EPI_ISL_3109822, EPI_ISL_3109823 | | Consortium | Institute COVID-19 Surveillance Team |
| EPI_ISL_3109824 | Lighthouse Lab in Glasgow | Wellcome Sanger Institute for the COVID-19 Genomics UK (COG-UK) Consortium | Harper VanSteenhouse, Yumi Kasai, David Gray, Carol Clugston, Anna Dominiczak and Alex Alderton, Roberto Amato, Jeffrey Barrett, Sonia Goncalves, Ewan Harrison, David K. Jackson, Ian Johnston, Dominic Kwiatkowski, Cordelia Langford, John Sillitoe on behalf of the Wellcome Sanger Institute COVID-19 Surveillance Team |
| EPI_ISL_3109825 | Lighthouse Lab in Milton Keynes | Wellcome Sanger Institute for the COVID-19 Genomics UK (COG-UK) Consortium | The Lighthouse Lab in Milton Keynes and Alex Alderton, Roberto Amato, Jeffrey Barrett, Sonia Goncalves, Ewan Harrison, David K. Jackson, Ian Johnston, Dominic Kwiatkowski, Cordelia Langford, John Sillitoe on behalf of the Wellcome Sanger Institute COVID-19 Surveillance Team |
| EPI_ISL_3109826 | Lighthouse Lab in Alderley Park | Wellcome Sanger Institute for the COVID-19 Genomics UK (COG-UK) Consortium | Jacquelyn Wynn, Mairead Hyland, The Lighthouse Lab in Alderley Park and Alex Alderton, Roberto Amato, Jeffrey Barrett, Sonia Goncalves, Ewan Harrison, David K. Jackson, Ian Johnston, Dominic Kwiatkowski, Cordelia Langford, John Sillitoe on behalf of the Wellcome Sanger Institute COVID-19 Surveillance Team |
| EPI_ISL_3109827, EPI_ISL_3109828, EPI_ISL_3109829, EPI_ISL_3109830, EPI_ISL_3109831, EPI_ISL_3109832, EPI_ISL_3109833, EPI_ISL_3109834, EPI_ISL_3109835, EPI_ISL_3109836, EPI_ISL_3109837, EPI_ISL_3109838, EPI_ISL_3109839, EPI_ISL_3109840, EPI_ISL_3109841, EPI_ISL_3109842, EPI_ISL_3109843, EPI_ISL_3109844, EPI_ISL_3109845 | | | |
| see above | Lighthouse Lab in Glasgow | Wellcome Sanger Institute for the COVID-19 Genomics UK (COG-UK) Consortium | Harper VanSteenhouse, Yumi Kasai, David Gray, Carol Clugston, Anna Dominiczak and Alex Alderton, Roberto Amato, Jeffrey Barrett, Sonia Goncalves, Ewan Harrison, David K. Jackson, Ian Johnston, Dominic Kwiatkowski, Cordelia Langford, John Sillitoe on behalf of the Wellcome Sanger Institute COVID-19 Surveillance Team |
| EPI_ISL_3109846 | Lighthouse Lab in Alderley Park | Wellcome Sanger Institute for the COVID-19 Genomics UK (COG-UK) Consortium | Jacquelyn Wynn, Mairead Hyland, The Lighthouse Lab in Alderley Park and Alex Alderton, Roberto Amato, Jeffrey Barrett, Sonia Goncalves, Ewan Harrison, David K. Jackson, Ian Johnston, Dominic Kwiatkowski, Cordelia Langford, John Sillitoe on behalf of the Wellcome Sanger Institute COVID-19 Surveillance Team |
| EPI_ISL_3109847, EPI_ISL_3109848, EPI_ISL_3109849 | Lighthouse Lab in Glasgow | Wellcome Sanger Institute for the COVID-19 Genomics UK (COG-UK) Consortium | Harper VanSteenhouse, Yumi Kasai, David Gray, Carol Clugston, Anna Dominiczak and Alex Alderton, Roberto Amato, Jeffrey Barrett, Sonia Goncalves, Ewan Harrison, David K. Jackson, Ian Johnston, Dominic Kwiatkowski, Cordelia Langford, John Sillitoe on behalf of the Wellcome Sanger Institute COVID-19 Surveillance Team |
| EPI_ISL_3109850 | Lighthouse Lab in Alderley Park | Wellcome Sanger Institute for the COVID-19 Genomics UK (COG-UK) Consortium | Jacquelyn Wynn, Mairead Hyland, The Lighthouse Lab in Alderley Park and Alex Alderton, Roberto Amato, Jeffrey Barrett, Sonia Goncalves, Ewan Harrison, David K. Jackson, Ian Johnston, Dominic Kwiatkowski, Cordelia Langford, John Sillitoe on behalf of the Wellcome Sanger Institute COVID-19 Surveillance Team |
| EPI_ISL_3109851, EPI_ISL_3109852, EPI_ISL_3109853, EPI_ISL_3109854, EPI_ISL_3109855, EPI_ISL_3109856, EPI_ISL_3109857, EPI_ISL_3109858 | Lighthouse Lab in Glasgow | Wellcome Sanger Institute for the COVID-19 Genomics UK (COG-UK) Consortium | Harper VanSteenhouse, Yumi Kasai, David Gray, Carol Clugston, Anna Dominiczak and Alex Alderton, Roberto Amato, Jeffrey Barrett, Sonia Goncalves, Ewan Harrison, David K. Jackson, Ian Johnston, Dominic Kwiatkowski, Cordelia Langford, John Sillitoe on behalf of the Wellcome Sanger Institute COVID-19 Surveillance Team |
| EPI_ISL_3109859, EPI_ISL_3109860, EPI_ISL_3109861 | Lighthouse Lab in Milton Keynes | Wellcome Sanger Institute for the COVID-19 Genomics UK (COG-UK) Consortium | The Lighthouse Lab in Milton Keynes and Alex Alderton, Roberto Amato, Jeffrey Barrett, Sonia Goncalves, Ewan Harrison, David K. Jackson, Ian Johnston, Dominic Kwiatkowski, Cordelia Langford, John Sillitoe on behalf of the Wellcome Sanger Institute COVID-19 Surveillance Team |
| EPI_ISL_3109862, EPI_ISL_3109863, EPI_ISL_3109864, EPI_ISL_3109865, EPI_ISL_3109866 | Lighthouse Lab in Glasgow | Wellcome Sanger Institute for the COVID-19 Genomics UK (COG-UK) Consortium | Harper VanSteenhouse, Yumi Kasai, David Gray, Carol Clugston, Anna Dominiczak and Alex Alderton, Roberto Amato, Jeffrey Barrett, Sonia Goncalves, Ewan Harrison, David K. Jackson, Ian Johnston, Dominic Kwiatkowski, Cordelia Langford, John Sillitoe on behalf of the Wellcome Sanger Institute COVID-19 Surveillance Team |
| EPI_ISL_3109867 | Lighthouse Lab in Milton Keynes | Wellcome Sanger Institute for the COVID-19 Genomics UK (COG-UK) Consortium | The Lighthouse Lab in Milton Keynes and Alex Alderton, Roberto Amato, Jeffrey Barrett, Sonia Goncalves, Ewan Harrison, David K. Jackson, Ian Johnston, Dominic Kwiatkowski, Cordelia Langford, John Sillitoe on behalf of the Wellcome Sanger Institute COVID-19 Surveillance Team |
| EPI_ISL_3109868 | Lighthouse Lab in Glasgow | Wellcome Sanger Institute for the COVID-19 Genomics UK (COG-UK) Consortium | Harper VanSteenhouse, Yumi Kasai, David Gray, Carol Clugston, Anna Dominiczak and Alex Alderton, Roberto Amato, Jeffrey Barrett, Sonia Goncalves, Ewan Harrison, David K. Jackson, Ian Johnston, Dominic Kwiatkowski, Cordelia Langford, John Sillitoe on behalf of the Wellcome Sanger Institute COVID-19 Surveillance Team |
| EPI_ISL_3109869 | Lighthouse Lab in Milton Keynes | Wellcome Sanger Institute for the COVID-19 Genomics UK (COG-UK) Consortium | The Lighthouse Lab in Milton Keynes and Alex Alderton, Roberto Amato, Jeffrey Barrett, Sonia Goncalves, Ewan Harrison, David K. Jackson, Ian Johnston, Dominic Kwiatkowski, Cordelia Langford, John Sillitoe on behalf of the Wellcome Sanger Institute COVID-19 Surveillance Team |
| EPI_ISL_3109870, EPI_ISL_3109871 | Lighthouse Lab in Glasgow | Wellcome Sanger Institute for the COVID-19 Genomics UK (COG-UK) Consortium | Harper VanSteenhouse, Yumi Kasai, David Gray, Carol Clugston, Anna Dominiczak and Alex Alderton, Roberto Amato, Jeffrey Barrett, Sonia Goncalves, Ewan Harrison, David K. Jackson, Ian Johnston, Dominic Kwiatkowski, Cordelia Langford, John Sillitoe on behalf of the Wellcome Sanger Institute COVID-19 Surveillance Team |
| EPI_ISL_3109872, EPI_ISL_3109873 | Lighthouse Lab in Milton Keynes | Wellcome Sanger Institute for the COVID-19 Genomics UK (COG-UK) Consortium | The Lighthouse Lab in Milton Keynes and Alex Alderton, Roberto Amato, Jeffrey Barrett, Sonia Goncalves, Ewan Harrison, David K. Jackson, Ian Johnston, Dominic Kwiatkowski, Cordelia Langford, John Sillitoe on behalf of the Wellcome Sanger Institute COVID-19 Surveillance Team |
| EPI_ISL_3109874, EPI_ISL_3109875 | Lighthouse Lab in Glasgow | Wellcome Sanger Institute for the COVID-19 Genomics UK (COG-UK) Consortium | Harper VanSteenhouse, Yumi Kasai, David Gray, Carol Clugston, Anna Dominiczak and Alex Alderton, Roberto Amato, Jeffrey Barrett, Sonia Goncalves, Ewan Harrison, David K. Jackson, Ian Johnston, Dominic Kwiatkowski, Cordelia Langford, John Sillitoe on behalf of the Wellcome Sanger Institute COVID-19 Surveillance Team |
| EPI_ISL_3109876 | Lighthouse Lab in Milton Keynes | Wellcome Sanger Institute for the COVID-19 Genomics UK (COG-UK) Consortium | The Lighthouse Lab in Milton Keynes and Alex Alderton, Roberto Amato, Jeffrey Barrett, Sonia Goncalves, Ewan Harrison, David K. Jackson, Ian Johnston, Dominic Kwiatkowski, Cordelia Langford, John Sillitoe on behalf of the Wellcome Sanger Institute COVID-19 Surveillance Team |
| EPI_ISL_3109877 | Lighthouse Lab in Glasgow | Wellcome Sanger Institute for the COVID-19 Genomics UK (COG-UK) Consortium | Harper VanSteenhouse, Yumi Kasai, David Gray, Carol Clugston, Anna Dominiczak and Alex Alderton, Roberto Amato, Jeffrey Barrett, Sonia Goncalves, Ewan Harrison, David K. Jackson, Ian Johnston, Dominic Kwiatkowski, Cordelia Langford, John Sillitoe on behalf of the Wellcome Sanger Institute COVID-19 Surveillance Team |
| EPI_ISL_3109878 | Lighthouse Lab in Milton Keynes | Wellcome Sanger Institute for the COVID-19 Genomics UK (COG-UK) Consortium | The Lighthouse Lab in Milton Keynes and Alex Alderton, Roberto Amato, Jeffrey Barrett, Sonia Goncalves, Ewan Harrison, David K. Jackson, Ian Johnston, Dominic Kwiatkowski, Cordelia Langford, John Sillitoe on behalf of the Wellcome Sanger Institute COVID-19 Surveillance Team |
| EPI_ISL_3109879, EPI_ISL_3109880, EPI_ISL_3109881, EPI_ISL_3109882 | Lighthouse Lab in Glasgow | Wellcome Sanger Institute for the COVID-19 Genomics UK (COG-UK) Consortium | Harper VanSteenhouse, Yumi Kasai, David Gray, Carol Clugston, Anna Dominiczak and Alex Alderton, Roberto Amato, Jeffrey Barrett, Sonia Goncalves, Ewan Harrison, David K. Jackson, Ian Johnston, Dominic Kwiatkowski, Cordelia Langford, John Sillitoe on behalf of the Wellcome Sanger Institute COVID-19 Surveillance Team |
| EPI_ISL_3109883 | Lighthouse Lab in Milton Keynes | Wellcome Sanger Institute for the COVID-19 Genomics UK (COG-UK) Consortium | The Lighthouse Lab in Milton Keynes and Alex Alderton, Roberto Amato, Jeffrey Barrett, Sonia Goncalves, Ewan Harrison, David K. Jackson, Ian Johnston, Dominic Kwiatkowski, Cordelia Langford, John Sillitoe on behalf of the Wellcome Sanger Institute COVID-19 Surveillance Team |
| EPI_ISL_3109884, EPI_ISL_3109885, EPI_ISL_3109886, EPI_ISL_3109887 | Lighthouse Lab in Glasgow | Wellcome Sanger Institute for the COVID-19 Genomics UK (COG-UK) Consortium | Harper VanSteenhouse, Yumi Kasai, David Gray, Carol Clugston, Anna Dominiczak and Alex Alderton, Roberto Amato, Jeffrey Barrett, Sonia Goncalves, Ewan Harrison, David K. Jackson, Ian Johnston, Dominic Kwiatkowski, Cordelia Langford, John Sillitoe on behalf of the Wellcome Sanger Institute COVID-19 Surveillance Team |
| EPI_ISL_3109888 | Lighthouse Lab in Milton Keynes | Wellcome Sanger Institute for the COVID-19 Genomics UK (COG-UK) Consortium | The Lighthouse Lab in Milton Keynes and Alex Alderton, Roberto Amato, Jeffrey Barrett, Sonia Goncalves, Ewan Harrison, David K. Jackson, Ian Johnston, Dominic Kwiatkowski, Cordelia Langford, John Sillitoe on behalf of the Wellcome Sanger Institute COVID-19 Surveillance Team |
| EPI_ISL_3109889, EPI_ISL_3109890, EPI_ISL_3109891, EPI_ISL_3109892 | Lighthouse Lab in Glasgow | Wellcome Sanger Institute for the COVID-19 Genomics UK (COG-UK) Consortium | Harper VanSteenhouse, Yumi Kasai, David Gray, Carol Clugston, Anna Dominiczak and Alex Alderton, Roberto Amato, Jeffrey Barrett, Sonia Goncalves, Ewan Harrison, David K. Jackson, Ian Johnston, Dominic Kwiatkowski, Cordelia Langford, John Sillitoe on behalf of the Wellcome Sanger Institute COVID-19 Surveillance Team |
| EPI_ISL_3109893, EPI_ISL_3109894, EPI_ISL_3109895 | Lighthouse Lab in Milton Keynes | Wellcome Sanger Institute for the COVID-19 Genomics UK (COG-UK) Consortium | The Lighthouse Lab in Milton Keynes and Alex Alderton, Roberto Amato, Jeffrey Barrett, Sonia Goncalves, Ewan Harrison, David K. Jackson, Ian Johnston, Dominic Kwiatkowski, Cordelia Langford, John Sillitoe on behalf of the Wellcome Sanger Institute COVID-19 Surveillance Team |
| EPI_ISL_3109896, EPI_ISL_3109897, EPI_ISL_3109898, EPI_ISL_3109899, EPI_ISL_3109900, EPI_ISL_3109901, EPI_ISL_3109902, EPI_ISL_3109903 | Lighthouse Lab in Glasgow | Wellcome Sanger Institute for the COVID-19 Genomics UK (COG-UK) Consortium | Harper VanSteenhouse, Yumi Kasai, David Gray, Carol Clugston, Anna Dominiczak and Alex Alderton, Roberto Amato, Jeffrey Barrett, Sonia Goncalves, Ewan Harrison, David K. Jackson, Ian Johnston, Dominic Kwiatkowski, Cordelia Langford, John Sillitoe on behalf of the Wellcome Sanger Institute COVID-19 Surveillance Team |
| EPI_ISL_3109904 | Lighthouse Lab in Milton Keynes | Wellcome Sanger Institute for the COVID-19 Genomics UK (COG-UK) Consortium | The Lighthouse Lab in Milton Keynes and Alex Alderton, Roberto Amato, Jeffrey Barrett, Sonia Goncalves, Ewan Harrison, David K. Jackson, Ian Johnston, Dominic Kwiatkowski, Cordelia Langford, John Sillitoe on behalf of the Wellcome Sanger Institute COVID-19 Surveillance Team |
| EPI_ISL_3109905, EPI_ISL_3109906 | Lighthouse Lab in Alderley Park | Wellcome Sanger Institute for the COVID-19 Genomics UK (COG-UK) Consortium | Jacquelyn Wynn, Mairead Hyland, The Lighthouse Lab in Alderley Park and Alex Alderton, Roberto Amato, Jeffrey Barrett, Sonia Goncalves, Ewan Harrison, David K. Jackson, Ian Johnston, Dominic Kwiatkowski, Cordelia Langford, John Sillitoe on behalf of the Wellcome Sanger Institute COVID-19 Surveillance Team |
| EPI_ISL_3109907, EPI_ISL_3109908, EPI_ISL_3109909 | Lighthouse Lab in Glasgow | Wellcome Sanger Institute for the COVID-19 Genomics UK (COG-UK) Consortium | Harper VanSteenhouse, Yumi Kasai, David Gray, Carol Clugston, Anna Dominiczak and Alex Alderton, Roberto Amato, Jeffrey Barrett, Sonia Goncalves, Ewan Harrison, David K. Jackson, Ian Johnston, Dominic Kwiatkowski, Cordelia Langford, John Sillitoe on behalf of the Wellcome Sanger Institute COVID-19 Surveillance Team |
| EPI_ISL_3109910, EPI_ISL_3109911 | Lighthouse Lab in Milton Keynes | Wellcome Sanger Institute for the COVID-19 Genomics UK (COG-UK) Consortium | The Lighthouse Lab in Milton Keynes and Alex Alderton, Roberto Amato, Jeffrey Barrett, Sonia Goncalves, Ewan Harrison, David K. Jackson, Ian Johnston, Dominic Kwiatkowski, Cordelia Langford, John Sillitoe on behalf of the Wellcome Sanger Institute COVID-19 Surveillance Team |
| EPI_ISL_3109912, EPI_ISL_3109913, EPI_ISL_3109914, EPI_ISL_3109915, EPI_ISL_3109916, EPI_ISL_3109917, | Lighthouse Lab in Glasgow | Wellcome Sanger Institute for the COVID-19 Genomics UK (COG-UK) Consortium | Harper VanSteenhouse, Yumi Kasai, David Gray, Carol Clugston, Anna Dominiczak and Alex Alderton, Roberto Amato, Jeffrey Barrett, Sonia Goncalves, Ewan Harrison, David K. Jackson, Ian Johnston, Dominic Kwiatkowski, Cordelia Langford, John Sillitoe on behalf of the Wellcome Sanger Institute COVID-19 Surveillance Team |

| | | | |
|---|---|---|---|
| EPI_ISL_3109918, EPI_ISL_3109919 | | | |
| EPI_ISL_3109920 | Lighthouse Lab in Alderley Park | Wellcome Sanger Institute for the COVID-19 Genomics UK (COG-UK) Consortium | Jacquelyn Wynn, Mairead Hyland, The Lighthouse Lab in Alderley Park and Alex Alderton, Roberto Amato, Jeffrey Barrett, Sonia Goncalves, Ewan Harrison, David K. Jackson, Ian Johnston, Dominic Kwiatkowski, Cordelia Langford, John Sillitoe on behalf of the Wellcome Sanger Institute COVID-19 Surveillance Team |
| EPI_ISL_3109921 | Lighthouse Lab in Glasgow | Wellcome Sanger Institute for the COVID-19 Genomics UK (COG-UK) Consortium | Harper VanSteenhouse, Yumi Kasai, David Gray, Carol Clugston, Anna Dominiczak and Alex Alderton, Roberto Amato, Jeffrey Barrett, Sonia Goncalves, Ewan Harrison, David K. Jackson, Ian Johnston, Dominic Kwiatkowski, Cordelia Langford, John Sillitoe on behalf of the Wellcome Sanger Institute COVID-19 Surveillance Team |
| EPI_ISL_3109922, EPI_ISL_3109923 | Lighthouse Lab in Milton Keynes | Wellcome Sanger Institute for the COVID-19 Genomics UK (COG-UK) Consortium | The Lighthouse Lab in Milton Keynes and Alex Alderton, Roberto Amato, Jeffrey Barrett, Sonia Goncalves, Ewan Harrison, David K. Jackson, Ian Johnston, Dominic Kwiatkowski, Cordelia Langford, John Sillitoe on behalf of the Wellcome Sanger Institute COVID-19 Surveillance Team |
| EPI_ISL_3109924, EPI_ISL_3109925, EPI_ISL_3109926, EPI_ISL_3109927, EPI_ISL_3109928, EPI_ISL_3109929 | Lighthouse Lab in Glasgow | Wellcome Sanger Institute for the COVID-19 Genomics UK (COG-UK) Consortium | Harper VanSteenhouse, Yumi Kasai, David Gray, Carol Clugston, Anna Dominiczak and Alex Alderton, Roberto Amato, Jeffrey Barrett, Sonia Goncalves, Ewan Harrison, David K. Jackson, Ian Johnston, Dominic Kwiatkowski, Cordelia Langford, John Sillitoe on behalf of the Wellcome Sanger Institute COVID-19 Surveillance Team |
| EPI_ISL_3109930 | Lighthouse Lab in Milton Keynes | Wellcome Sanger Institute for the COVID-19 Genomics UK (COG-UK) Consortium | The Lighthouse Lab in Milton Keynes and Alex Alderton, Roberto Amato, Jeffrey Barrett, Sonia Goncalves, Ewan Harrison, David K. Jackson, Ian Johnston, Dominic Kwiatkowski, Cordelia Langford, John Sillitoe on behalf of the Wellcome Sanger Institute COVID-19 Surveillance Team |
| EPI_ISL_3109931, EPI_ISL_3109932 | Lighthouse Lab in Glasgow | Wellcome Sanger Institute for the COVID-19 Genomics UK (COG-UK) Consortium | Harper VanSteenhouse, Yumi Kasai, David Gray, Carol Clugston, Anna Dominiczak and Alex Alderton, Roberto Amato, Jeffrey Barrett, Sonia Goncalves, Ewan Harrison, David K. Jackson, Ian Johnston, Dominic Kwiatkowski, Cordelia Langford, John Sillitoe on behalf of the Wellcome Sanger Institute COVID-19 Surveillance Team |
| EPI_ISL_3109933 | Lighthouse Lab in Alderley Park | Wellcome Sanger Institute for the COVID-19 Genomics UK (COG-UK) Consortium | Jacquelyn Wynn, Mairead Hyland, The Lighthouse Lab in Alderley Park and Alex Alderton, Roberto Amato, Jeffrey Barrett, Sonia Goncalves, Ewan Harrison, David K. Jackson, Ian Johnston, Dominic Kwiatkowski, Cordelia Langford, John Sillitoe on behalf of the Wellcome Sanger Institute COVID-19 Surveillance Team |
| EPI_ISL_3109934 | Lighthouse Lab in Glasgow | Wellcome Sanger Institute for the COVID-19 Genomics UK (COG-UK) Consortium | Harper VanSteenhouse, Yumi Kasai, David Gray, Carol Clugston, Anna Dominiczak and Alex Alderton, Roberto Amato, Jeffrey Barrett, Sonia Goncalves, Ewan Harrison, David K. Jackson, Ian Johnston, Dominic Kwiatkowski, Cordelia Langford, John Sillitoe on behalf of the Wellcome Sanger Institute COVID-19 Surveillance Team |
| EPI_ISL_3109935, EPI_ISL_3109936 | Lighthouse Lab in Milton Keynes | Wellcome Sanger Institute for the COVID-19 Genomics UK (COG-UK) Consortium | The Lighthouse Lab in Milton Keynes and Alex Alderton, Roberto Amato, Jeffrey Barrett, Sonia Goncalves, Ewan Harrison, David K. Jackson, Ian Johnston, Dominic Kwiatkowski, Cordelia Langford, John Sillitoe on behalf of the Wellcome Sanger Institute COVID-19 Surveillance Team |
| EPI_ISL_3109937 | Lighthouse Lab in Alderley Park | Wellcome Sanger Institute for the COVID-19 Genomics UK (COG-UK) Consortium | Jacquelyn Wynn, Mairead Hyland, The Lighthouse Lab in Alderley Park and Alex Alderton, Roberto Amato, Jeffrey Barrett, Sonia Goncalves, Ewan Harrison, David K. Jackson, Ian Johnston, Dominic Kwiatkowski, Cordelia Langford, John Sillitoe on behalf of the Wellcome Sanger Institute COVID-19 Surveillance Team |
| EPI_ISL_3109938 | Lighthouse Lab in Milton Keynes | Wellcome Sanger Institute for the COVID-19 Genomics UK (COG-UK) Consortium | The Lighthouse Lab in Milton Keynes and Alex Alderton, Roberto Amato, Jeffrey Barrett, Sonia Goncalves, Ewan Harrison, David K. Jackson, Ian Johnston, Dominic Kwiatkowski, Cordelia Langford, John Sillitoe on behalf of the Wellcome Sanger Institute COVID-19 Surveillance Team |
| EPI_ISL_3109939 | Lighthouse Laboratory Plymouth | Wellcome Sanger Institute for the COVID-19 Genomics UK (COG-UK) Consortium | Lighthouse Laboratory Plymouth and Alex Alderton, Roberto Amato, Jeffrey Barrett, Sonia Goncalves, Ewan Harrison, David K. Jackson, Ian Johnston, Dominic Kwiatkowski, Cordelia Langford, John Sillitoe on behalf of the Wellcome Sanger Institute COVID-19 Surveillance Team |
| EPI_ISL_3109940 | Lighthouse Lab in Milton Keynes | Wellcome Sanger Institute for the COVID-19 Genomics UK (COG-UK) Consortium | The Lighthouse Lab in Milton Keynes and Alex Alderton, Roberto Amato, Jeffrey Barrett, Sonia Goncalves, Ewan Harrison, David K. Jackson, Ian Johnston, Dominic Kwiatkowski, Cordelia Langford, John Sillitoe on behalf of the Wellcome Sanger Institute COVID-19 Surveillance Team |
| EPI_ISL_3109941 | Lighthouse Laboratory Plymouth | Wellcome Sanger Institute for the COVID-19 Genomics UK (COG-UK) Consortium | Lighthouse Laboratory Plymouth and Alex Alderton, Roberto Amato, Jeffrey Barrett, Sonia Goncalves, Ewan Harrison, David K. Jackson, Ian Johnston, Dominic Kwiatkowski, Cordelia Langford, John Sillitoe on behalf of the Wellcome Sanger Institute COVID-19 Surveillance Team |
| EPI_ISL_3109942 | Lighthouse Lab in Milton Keynes | Wellcome Sanger Institute for the COVID-19 Genomics UK (COG-UK) Consortium | The Lighthouse Lab in Milton Keynes and Alex Alderton, Roberto Amato, Jeffrey Barrett, Sonia Goncalves, Ewan Harrison, David K. Jackson, Ian Johnston, Dominic Kwiatkowski, Cordelia Langford, John Sillitoe on behalf of the Wellcome Sanger Institute COVID-19 Surveillance Team |
| EPI_ISL_3109943 | Lighthouse Lab in Alderley Park | Wellcome Sanger Institute for the COVID-19 Genomics UK (COG-UK) Consortium | Jacquelyn Wynn, Mairead Hyland, The Lighthouse Lab in Alderley Park and Alex Alderton, Roberto Amato, Jeffrey Barrett, Sonia Goncalves, Ewan Harrison, David K. Jackson, Ian Johnston, Dominic Kwiatkowski, Cordelia Langford, John Sillitoe on behalf of the Wellcome Sanger Institute COVID-19 Surveillance Team |
| EPI_ISL_3109944 | Lighthouse Lab in Milton Keynes | Wellcome Sanger Institute for the COVID-19 Genomics UK (COG-UK) Consortium | The Lighthouse Lab in Milton Keynes and Alex Alderton, Roberto Amato, Jeffrey Barrett, Sonia Goncalves, Ewan Harrison, David K. Jackson, Ian Johnston, Dominic Kwiatkowski, Cordelia Langford, John Sillitoe on behalf of the Wellcome Sanger Institute COVID-19 Surveillance Team |
| EPI_ISL_3109945 | Lighthouse Laboratory Plymouth | Wellcome Sanger Institute for the COVID-19 Genomics UK (COG-UK) Consortium | Lighthouse Laboratory Plymouth and Alex Alderton, Roberto Amato, Jeffrey Barrett, Sonia Goncalves, Ewan Harrison, David K. Jackson, Ian Johnston, Dominic Kwiatkowski, Cordelia Langford, John Sillitoe on behalf of the Wellcome Sanger Institute COVID-19 Surveillance Team |
| EPI_ISL_3109946 | Lighthouse Lab in Alderley Park | Wellcome Sanger Institute for the COVID-19 Genomics UK (COG-UK) Consortium | Jacquelyn Wynn, Mairead Hyland, The Lighthouse Lab in Alderley Park and Alex Alderton, Roberto Amato, Jeffrey Barrett, Sonia Goncalves, Ewan Harrison, David K. Jackson, Ian Johnston, Dominic Kwiatkowski, Cordelia Langford, John Sillitoe on behalf of the Wellcome Sanger Institute COVID-19 Surveillance Team |
| EPI_ISL_3109947, EPI_ISL_3109948 | Lighthouse Lab in Glasgow | Wellcome Sanger Institute for the COVID-19 Genomics UK (COG-UK) Consortium | Harper VanSteenhouse, Yumi Kasai, David Gray, Carol Clugston, Anna Dominiczak and Alex Alderton, Roberto Amato, Jeffrey Barrett, Sonia Goncalves, Ewan Harrison, David K. Jackson, Ian Johnston, Dominic Kwiatkowski, Cordelia Langford, John Sillitoe on behalf of the Wellcome Sanger Institute COVID-19 Surveillance Team |
| EPI_ISL_3109949 | Lighthouse Laboratory Plymouth | Wellcome Sanger Institute for the COVID-19 Genomics UK (COG-UK) Consortium | Lighthouse Laboratory Plymouth and Alex Alderton, Roberto Amato, Jeffrey Barrett, Sonia Goncalves, Ewan Harrison, David K. Jackson, Ian Johnston, Dominic Kwiatkowski, Cordelia Langford, John Sillitoe on behalf of the Wellcome Sanger Institute COVID-19 Surveillance Team |
| EPI_ISL_3109950, EPI_ISL_3109951 | Lighthouse Lab in Alderley Park | Wellcome Sanger Institute for the COVID-19 Genomics UK (COG-UK) Consortium | Jacquelyn Wynn, Mairead Hyland, The Lighthouse Lab in Alderley Park and Alex Alderton, Roberto Amato, Jeffrey Barrett, Sonia Goncalves, Ewan Harrison, David K. Jackson, Ian Johnston, Dominic Kwiatkowski, Cordelia Langford, John Sillitoe on behalf of the Wellcome Sanger Institute COVID-19 Surveillance Team |
| EPI_ISL_3109952, EPI_ISL_3109953 | Lighthouse Lab in Milton Keynes | Wellcome Sanger Institute for the COVID-19 Genomics UK (COG-UK) Consortium | The Lighthouse Lab in Milton Keynes and Alex Alderton, Roberto Amato, Jeffrey Barrett, Sonia Goncalves, Ewan Harrison, David K. Jackson, Ian Johnston, Dominic Kwiatkowski, Cordelia Langford, John Sillitoe on behalf of the Wellcome Sanger Institute COVID-19 Surveillance Team |
| EPI_ISL_3109954 | Lighthouse Lab in Alderley Park | Wellcome Sanger Institute for the COVID-19 Genomics UK (COG-UK) Consortium | Jacquelyn Wynn, Mairead Hyland, The Lighthouse Lab in Alderley Park and Alex Alderton, Roberto Amato, Jeffrey Barrett, Sonia Goncalves, Ewan Harrison, David K. Jackson, Ian Johnston, Dominic Kwiatkowski, Cordelia Langford, John Sillitoe on behalf of the Wellcome Sanger Institute COVID-19 Surveillance Team |
| EPI_ISL_3109955 | Lighthouse Laboratory Plymouth | Wellcome Sanger Institute for the COVID-19 Genomics UK (COG-UK) Consortium | Lighthouse Laboratory Plymouth and Alex Alderton, Roberto Amato, Jeffrey Barrett, Sonia Goncalves, Ewan Harrison, David K. Jackson, Ian Johnston, Dominic Kwiatkowski, Cordelia Langford, John Sillitoe on behalf of the Wellcome Sanger Institute COVID-19 Surveillance Team |
| EPI_ISL_3109956 | Lighthouse Lab in Alderley Park | Wellcome Sanger Institute for the COVID-19 Genomics UK (COG-UK) Consortium | Jacquelyn Wynn, Mairead Hyland, The Lighthouse Lab in Alderley Park and Alex Alderton, Roberto Amato, Jeffrey Barrett, Sonia Goncalves, Ewan Harrison, David K. Jackson, Ian Johnston, Dominic Kwiatkowski, Cordelia Langford, John Sillitoe on behalf of the Wellcome Sanger Institute COVID-19 Surveillance Team |
| EPI_ISL_3109957 | Lighthouse Lab in Milton Keynes | Wellcome Sanger Institute for the COVID-19 Genomics UK (COG-UK) Consortium | The Lighthouse Lab in Milton Keynes and Alex Alderton, Roberto Amato, Jeffrey Barrett, Sonia Goncalves, Ewan Harrison, David K. Jackson, Ian Johnston, Dominic Kwiatkowski, Cordelia Langford, John Sillitoe on behalf of the Wellcome Sanger Institute COVID-19 Surveillance Team |
| EPI_ISL_3109958 | Lighthouse Lab in Alderley Park | Wellcome Sanger Institute for the COVID-19 Genomics UK (COG-UK) Consortium | Jacquelyn Wynn, Mairead Hyland, The Lighthouse Lab in Alderley Park and Alex Alderton, Roberto Amato, Jeffrey Barrett, Sonia Goncalves, Ewan Harrison, David K. Jackson, Ian Johnston, Dominic Kwiatkowski, Cordelia Langford, John Sillitoe on behalf of the Wellcome Sanger Institute COVID-19 Surveillance Team |
| EPI_ISL_3109959 | Lighthouse Laboratory Plymouth | Wellcome Sanger Institute for the COVID-19 Genomics UK (COG-UK) Consortium | Lighthouse Laboratory Plymouth and Alex Alderton, Roberto Amato, Jeffrey Barrett, Sonia Goncalves, Ewan Harrison, David K. Jackson, Ian Johnston, Dominic Kwiatkowski, Cordelia Langford, John Sillitoe on behalf of the Wellcome Sanger Institute COVID-19 Surveillance Team |
| EPI_ISL_3109960 | Lighthouse Lab in Glasgow | Wellcome Sanger Institute for the COVID-19 Genomics UK (COG-UK) Consortium | Harper VanSteenhouse, Yumi Kasai, David Gray, Carol Clugston, Anna Dominiczak and Alex Alderton, Roberto Amato, Jeffrey Barrett, Sonia Goncalves, Ewan Harrison, David K. Jackson, Ian Johnston, Dominic Kwiatkowski, Cordelia Langford, John Sillitoe on behalf of the Wellcome Sanger Institute COVID-19 Surveillance Team |
| EPI_ISL_3109961 | Lighthouse Laboratory Plymouth | Wellcome Sanger Institute for the COVID-19 Genomics UK (COG-UK) Consortium | Lighthouse Laboratory Plymouth and Alex Alderton, Roberto Amato, Jeffrey Barrett, Sonia Goncalves, Ewan Harrison, David K. Jackson, Ian Johnston, Dominic Kwiatkowski, Cordelia Langford, John Sillitoe on behalf of the Wellcome Sanger Institute COVID-19 Surveillance Team |
| EPI_ISL_3109962 | Lighthouse Lab in Alderley Park | Wellcome Sanger Institute for the COVID-19 Genomics UK (COG-UK) Consortium | Jacquelyn Wynn, Mairead Hyland, The Lighthouse Lab in Alderley Park and Alex Alderton, Roberto Amato, Jeffrey Barrett, Sonia Goncalves, Ewan Harrison, David K. Jackson, Ian Johnston, Dominic Kwiatkowski, Cordelia Langford, John Sillitoe on behalf of the Wellcome Sanger Institute COVID-19 Surveillance Team |
| EPI_ISL_3109963 | Lighthouse Laboratory Plymouth | Wellcome Sanger Institute for the COVID-19 Genomics UK (COG-UK) Consortium | Lighthouse Laboratory Plymouth and Alex Alderton, Roberto Amato, Jeffrey Barrett, Sonia Goncalves, Ewan Harrison, David K. Jackson, Ian Johnston, Dominic Kwiatkowski, Cordelia Langford, John Sillitoe on behalf of the Wellcome Sanger Institute COVID-19 Surveillance Team |
| EPI_ISL_3109964 | Lighthouse Lab in Milton Keynes | Wellcome Sanger Institute for the COVID-19 Genomics UK (COG-UK) Consortium | The Lighthouse Lab in Milton Keynes and Alex Alderton, Roberto Amato, Jeffrey Barrett, Sonia Goncalves, Ewan Harrison, David K. Jackson, Ian Johnston, Dominic Kwiatkowski, Cordelia Langford, John Sillitoe on behalf of the Wellcome Sanger Institute COVID-19 Surveillance Team |
| EPI_ISL_3109965 | Lighthouse Lab in Alderley Park | Wellcome Sanger Institute for the COVID-19 Genomics UK (COG-UK) Consortium | Jacquelyn Wynn, Mairead Hyland, The Lighthouse Lab in Alderley Park and Alex Alderton, Roberto Amato, Jeffrey Barrett, Sonia Goncalves, Ewan Harrison, David K. Jackson, Ian Johnston, Dominic Kwiatkowski, Cordelia Langford, John Sillitoe on behalf of the Wellcome Sanger Institute COVID-19 Surveillance Team |
| EPI_ISL_3109966 | Lighthouse Lab in Milton Keynes | Wellcome Sanger Institute for the COVID-19 Genomics UK (COG-UK) Consortium | The Lighthouse Lab in Milton Keynes and Alex Alderton, Roberto Amato, Jeffrey Barrett, Sonia Goncalves, Ewan Harrison, David K. Jackson, Ian Johnston, Dominic Kwiatkowski, Cordelia Langford, John Sillitoe on behalf of the Wellcome Sanger Institute COVID-19 Surveillance Team |
| EPI_ISL_3109967 | Lighthouse Lab in Alderley Park | Wellcome Sanger Institute for the COVID-19 Genomics UK (COG-UK) Consortium | Jacquelyn Wynn, Mairead Hyland, The Lighthouse Lab in Alderley Park and Alex Alderton, Roberto Amato, Jeffrey Barrett, Sonia Goncalves, Ewan Harrison, David K. Jackson, Ian Johnston, Dominic Kwiatkowski, Cordelia Langford, John Sillitoe on behalf of the Wellcome Sanger Institute COVID-19 Surveillance Team |
| EPI_ISL_3109968 | Lighthouse Lab in Milton Keynes | Wellcome Sanger Institute for the COVID-19 Genomics UK (COG-UK) Consortium | The Lighthouse Lab in Milton Keynes and Alex Alderton, Roberto Amato, Jeffrey Barrett, Sonia Goncalves, Ewan Harrison, David K. Jackson, Ian Johnston, Dominic Kwiatkowski, Cordelia Langford, John Sillitoe on behalf of the Wellcome Sanger Institute COVID-19 Surveillance Team |
| EPI_ISL_3109969 | Lighthouse Lab in Glasgow | Wellcome Sanger Institute for the COVID-19 Genomics UK (COG-UK) Consortium | Harper VanSteenhouse, Yumi Kasai, David Gray, Carol Clugston, Anna Dominiczak and Alex Alderton, Roberto Amato, Jeffrey Barrett, Sonia Goncalves, Ewan Harrison, David K. Jackson, Ian Johnston, Dominic Kwiatkowski, Cordelia Langford, John Sillitoe on behalf of the Wellcome Sanger Institute COVID-19 Surveillance Team |
| EPI_ISL_3109970 | Lighthouse Laboratory Plymouth | Wellcome Sanger Institute for the COVID-19 Genomics UK (COG-UK) Consortium | Lighthouse Laboratory Plymouth and Alex Alderton, Roberto Amato, Jeffrey Barrett, Sonia Goncalves, Ewan Harrison, David K. Jackson, Ian Johnston, Dominic Kwiatkowski, Cordelia Langford, John Sillitoe on behalf of the Wellcome Sanger Institute COVID-19 Surveillance Team |
| EPI_ISL_3109971 | Lighthouse Lab in Milton Keynes | Wellcome Sanger Institute for the COVID-19 Genomics UK (COG-UK) Consortium | The Lighthouse Lab in Milton Keynes and Alex Alderton, Roberto Amato, Jeffrey Barrett, Sonia Goncalves, Ewan Harrison, David K. Jackson, Ian Johnston, Dominic Kwiatkowski, Cordelia Langford, John Sillitoe on behalf of the Wellcome Sanger Institute COVID-19 Surveillance Team |
| EPI_ISL_3109972 | Lighthouse Lab in Alderley Park | Wellcome Sanger Institute for the COVID-19 Genomics UK (COG-UK) Consortium | Jacquelyn Wynn, Mairead Hyland, The Lighthouse Lab in Alderley Park and Alex Alderton, Roberto Amato, Jeffrey Barrett, Sonia Goncalves, Ewan Harrison, David K. Jackson, Ian Johnston, Dominic Kwiatkowski, Cordelia Langford, John Sillitoe on behalf of the Wellcome Sanger Institute COVID-19 Surveillance Team |
| EPI_ISL_3109973 | Lighthouse Lab in Glasgow | Wellcome Sanger Institute for the COVID-19 Genomics UK (COG-UK) | Harper VanSteenhouse, Yumi Kasai, David Gray, Carol Clugston, Anna Dominiczak and Alex Alderton, Roberto Amato, Jeffrey Barrett, Sonia Goncalves, Ewan Harrison, David K. Jackson, Ian Johnston, Dominic Kwiatkowski, Cordelia Langford, |

| | | | |
|---|---|---|---|
| | | Consortium | John Sillitoe on behalf of the Wellcome Sanger Institute COVID-19 Surveillance Team |
| EPI_ISL_3109974 | Lighthouse Laboratory Plymouth | Wellcome Sanger Institute for the COVID-19 Genomics UK (COG-UK) Consortium | Lighthouse Laboratory Plymouth and Alex Alderton, Roberto Amato, Jeffrey Barrett, Sonia Goncalves, Ewan Harrison, David K. Jackson, Ian Johnston, Dominic Kwiatkowski, Cordelia Langford, John Sillitoe on behalf of the Wellcome Sanger Institute COVID-19 Surveillance Team |
| EPI_ISL_3109975 | Lighthouse Lab in Glasgow | Wellcome Sanger Institute for the COVID-19 Genomics UK (COG-UK) Consortium | Harper VanSteenhouse, Yumi Kasai, David Gray, Carol Clugston, Anna Dominiczak and Alex Alderton, Roberto Amato, Jeffrey Barrett, Sonia Goncalves, Ewan Harrison, David K. Jackson, Ian Johnston, Dominic Kwiatkowski, Cordelia Langford, John Sillitoe on behalf of the Wellcome Sanger Institute COVID-19 Surveillance Team |
| EPI_ISL_3109976 | Lighthouse Laboratory Plymouth | Wellcome Sanger Institute for the COVID-19 Genomics UK (COG-UK) Consortium | Lighthouse Laboratory Plymouth and Alex Alderton, Roberto Amato, Jeffrey Barrett, Sonia Goncalves, Ewan Harrison, David K. Jackson, Ian Johnston, Dominic Kwiatkowski, Cordelia Langford, John Sillitoe on behalf of the Wellcome Sanger Institute COVID-19 Surveillance Team |
| EPI_ISL_3109977 | Lighthouse Lab in Alderley Park | Wellcome Sanger Institute for the COVID-19 Genomics UK (COG-UK) Consortium | Jacquelyn Wynn, Mairead Hyland, The Lighthouse Lab in Alderley Park and Alex Alderton, Roberto Amato, Jeffrey Barrett, Sonia Goncalves, Ewan Harrison, David K. Jackson, Ian Johnston, Dominic Kwiatkowski, Cordelia Langford, John Sillitoe on behalf of the Wellcome Sanger Institute COVID-19 Surveillance Team |
| EPI_ISL_3109978 | Lighthouse Lab in Milton Keynes | Wellcome Sanger Institute for the COVID-19 Genomics UK (COG-UK) Consortium | The Lighthouse Lab in Milton Keynes and Alex Alderton, Roberto Amato, Jeffrey Barrett, Sonia Goncalves, Ewan Harrison, David K. Jackson, Ian Johnston, Dominic Kwiatkowski, Cordelia Langford, John Sillitoe on behalf of the Wellcome Sanger Institute COVID-19 Surveillance Team |
| EPI_ISL_3109979 | Lighthouse Laboratory Plymouth | Wellcome Sanger Institute for the COVID-19 Genomics UK (COG-UK) Consortium | Lighthouse Laboratory Plymouth and Alex Alderton, Roberto Amato, Jeffrey Barrett, Sonia Goncalves, Ewan Harrison, David K. Jackson, Ian Johnston, Dominic Kwiatkowski, Cordelia Langford, John Sillitoe on behalf of the Wellcome Sanger Institute COVID-19 Surveillance Team |
| EPI_ISL_3109980 | Lighthouse Lab in Glasgow | Wellcome Sanger Institute for the COVID-19 Genomics UK (COG-UK) Consortium | Harper VanSteenhouse, Yumi Kasai, David Gray, Carol Clugston, Anna Dominiczak and Alex Alderton, Roberto Amato, Jeffrey Barrett, Sonia Goncalves, Ewan Harrison, David K. Jackson, Ian Johnston, Dominic Kwiatkowski, Cordelia Langford, John Sillitoe on behalf of the Wellcome Sanger Institute COVID-19 Surveillance Team |
| EPI_ISL_3109981 | Lighthouse Lab in Alderley Park | Wellcome Sanger Institute for the COVID-19 Genomics UK (COG-UK) Consortium | Jacquelyn Wynn, Mairead Hyland, The Lighthouse Lab in Alderley Park and Alex Alderton, Roberto Amato, Jeffrey Barrett, Sonia Goncalves, Ewan Harrison, David K. Jackson, Ian Johnston, Dominic Kwiatkowski, Cordelia Langford, John Sillitoe on behalf of the Wellcome Sanger Institute COVID-19 Surveillance Team |
| EPI_ISL_3109982, EPI_ISL_3109983 | Lighthouse Laboratory Plymouth | Wellcome Sanger Institute for the COVID-19 Genomics UK (COG-UK) Consortium | Lighthouse Laboratory Plymouth and Alex Alderton, Roberto Amato, Jeffrey Barrett, Sonia Goncalves, Ewan Harrison, David K. Jackson, Ian Johnston, Dominic Kwiatkowski, Cordelia Langford, John Sillitoe on behalf of the Wellcome Sanger Institute COVID-19 Surveillance Team |
| EPI_ISL_3109984, EPI_ISL_3109985 | Lighthouse Lab in Milton Keynes | Wellcome Sanger Institute for the COVID-19 Genomics UK (COG-UK) Consortium | The Lighthouse Lab in Milton Keynes and Alex Alderton, Roberto Amato, Jeffrey Barrett, Sonia Goncalves, Ewan Harrison, David K. Jackson, Ian Johnston, Dominic Kwiatkowski, Cordelia Langford, John Sillitoe on behalf of the Wellcome Sanger Institute COVID-19 Surveillance Team |
| EPI_ISL_3109986, EPI_ISL_3109987 | Lighthouse Lab in Alderley Park | Wellcome Sanger Institute for the COVID-19 Genomics UK (COG-UK) Consortium | Jacquelyn Wynn, Mairead Hyland, The Lighthouse Lab in Alderley Park and Alex Alderton, Roberto Amato, Jeffrey Barrett, Sonia Goncalves, Ewan Harrison, David K. Jackson, Ian Johnston, Dominic Kwiatkowski, Cordelia Langford, John Sillitoe on behalf of the Wellcome Sanger Institute COVID-19 Surveillance Team |
| EPI_ISL_3109988 | Lighthouse Laboratory Plymouth | Wellcome Sanger Institute for the COVID-19 Genomics UK (COG-UK) Consortium | Lighthouse Laboratory Plymouth and Alex Alderton, Roberto Amato, Jeffrey Barrett, Sonia Goncalves, Ewan Harrison, David K. Jackson, Ian Johnston, Dominic Kwiatkowski, Cordelia Langford, John Sillitoe on behalf of the Wellcome Sanger Institute COVID-19 Surveillance Team |
| EPI_ISL_3109989 | Lighthouse Lab in Milton Keynes | Wellcome Sanger Institute for the COVID-19 Genomics UK (COG-UK) Consortium | The Lighthouse Lab in Milton Keynes and Alex Alderton, Roberto Amato, Jeffrey Barrett, Sonia Goncalves, Ewan Harrison, David K. Jackson, Ian Johnston, Dominic Kwiatkowski, Cordelia Langford, John Sillitoe on behalf of the Wellcome Sanger Institute COVID-19 Surveillance Team |
| EPI_ISL_3109990 | Lighthouse Laboratory Plymouth | Wellcome Sanger Institute for the COVID-19 Genomics UK (COG-UK) Consortium | Lighthouse Laboratory Plymouth and Alex Alderton, Roberto Amato, Jeffrey Barrett, Sonia Goncalves, Ewan Harrison, David K. Jackson, Ian Johnston, Dominic Kwiatkowski, Cordelia Langford, John Sillitoe on behalf of the Wellcome Sanger Institute COVID-19 Surveillance Team |
| EPI_ISL_3109991 | Lighthouse Lab in Alderley Park | Wellcome Sanger Institute for the COVID-19 Genomics UK (COG-UK) Consortium | Jacquelyn Wynn, Mairead Hyland, The Lighthouse Lab in Alderley Park and Alex Alderton, Roberto Amato, Jeffrey Barrett, Sonia Goncalves, Ewan Harrison, David K. Jackson, Ian Johnston, Dominic Kwiatkowski, Cordelia Langford, John Sillitoe on behalf of the Wellcome Sanger Institute COVID-19 Surveillance Team |
| EPI_ISL_3109992, EPI_ISL_3109993 | Lighthouse Lab in Milton Keynes | Wellcome Sanger Institute for the COVID-19 Genomics UK (COG-UK) Consortium | The Lighthouse Lab in Milton Keynes and Alex Alderton, Roberto Amato, Jeffrey Barrett, Sonia Goncalves, Ewan Harrison, David K. Jackson, Ian Johnston, Dominic Kwiatkowski, Cordelia Langford, John Sillitoe on behalf of the Wellcome Sanger Institute COVID-19 Surveillance Team |
| EPI_ISL_3109994, EPI_ISL_3109995 | Lighthouse Laboratory Plymouth | Wellcome Sanger Institute for the COVID-19 Genomics UK (COG-UK) Consortium | Lighthouse Laboratory Plymouth and Alex Alderton, Roberto Amato, Jeffrey Barrett, Sonia Goncalves, Ewan Harrison, David K. Jackson, Ian Johnston, Dominic Kwiatkowski, Cordelia Langford, John Sillitoe on behalf of the Wellcome Sanger Institute COVID-19 Surveillance Team |
| EPI_ISL_3109996 | Lighthouse Lab in Glasgow | Wellcome Sanger Institute for the COVID-19 Genomics UK (COG-UK) Consortium | Harper VanSteenhouse, Yumi Kasai, David Gray, Carol Clugston, Anna Dominiczak and Alex Alderton, Roberto Amato, Jeffrey Barrett, Sonia Goncalves, Ewan Harrison, David K. Jackson, Ian Johnston, Dominic Kwiatkowski, Cordelia Langford, John Sillitoe on behalf of the Wellcome Sanger Institute COVID-19 Surveillance Team |
| EPI_ISL_3109997 | Lighthouse Lab in Alderley Park | Wellcome Sanger Institute for the COVID-19 Genomics UK (COG-UK) Consortium | Jacquelyn Wynn, Mairead Hyland, The Lighthouse Lab in Alderley Park and Alex Alderton, Roberto Amato, Jeffrey Barrett, Sonia Goncalves, Ewan Harrison, David K. Jackson, Ian Johnston, Dominic Kwiatkowski, Cordelia Langford, John Sillitoe on behalf of the Wellcome Sanger Institute COVID-19 Surveillance Team |
| EPI_ISL_3109998 | Lighthouse Lab in Glasgow | Wellcome Sanger Institute for the COVID-19 Genomics UK (COG-UK) Consortium | Harper VanSteenhouse, Yumi Kasai, David Gray, Carol Clugston, Anna Dominiczak and Alex Alderton, Roberto Amato, Jeffrey Barrett, Sonia Goncalves, Ewan Harrison, David K. Jackson, Ian Johnston, Dominic Kwiatkowski, Cordelia Langford, John Sillitoe on behalf of the Wellcome Sanger Institute COVID-19 Surveillance Team |
| EPI_ISL_3109999 | Lighthouse Lab in Milton Keynes | Wellcome Sanger Institute for the COVID-19 Genomics UK (COG-UK) Consortium | The Lighthouse Lab in Milton Keynes and Alex Alderton, Roberto Amato, Jeffrey Barrett, Sonia Goncalves, Ewan Harrison, David K. Jackson, Ian Johnston, Dominic Kwiatkowski, Cordelia Langford, John Sillitoe on behalf of the Wellcome Sanger Institute COVID-19 Surveillance Team |
| EPI_ISL_3110000 | Lighthouse Lab in Alderley Park | Wellcome Sanger Institute for the COVID-19 Genomics UK (COG-UK) Consortium | Jacquelyn Wynn, Mairead Hyland, The Lighthouse Lab in Alderley Park and Alex Alderton, Roberto Amato, Jeffrey Barrett, Sonia Goncalves, Ewan Harrison, David K. Jackson, Ian Johnston, Dominic Kwiatkowski, Cordelia Langford, John Sillitoe on behalf of the Wellcome Sanger Institute COVID-19 Surveillance Team |
| EPI_ISL_3110001, EPI_ISL_3110002 | Lighthouse Laboratory Plymouth | Wellcome Sanger Institute for the COVID-19 Genomics UK (COG-UK) Consortium | Lighthouse Laboratory Plymouth and Alex Alderton, Roberto Amato, Jeffrey Barrett, Sonia Goncalves, Ewan Harrison, David K. Jackson, Ian Johnston, Dominic Kwiatkowski, Cordelia Langford, John Sillitoe on behalf of the Wellcome Sanger Institute COVID-19 Surveillance Team |
| EPI_ISL_3110003 | Lighthouse Lab in Milton Keynes | Wellcome Sanger Institute for the COVID-19 Genomics UK (COG-UK) Consortium | The Lighthouse Lab in Milton Keynes and Alex Alderton, Roberto Amato, Jeffrey Barrett, Sonia Goncalves, Ewan Harrison, David K. Jackson, Ian Johnston, Dominic Kwiatkowski, Cordelia Langford, John Sillitoe on behalf of the Wellcome Sanger Institute COVID-19 Surveillance Team |
| EPI_ISL_3110004, EPI_ISL_3110005 | Lighthouse Laboratory Plymouth | Wellcome Sanger Institute for the COVID-19 Genomics UK (COG-UK) Consortium | Lighthouse Laboratory Plymouth and Alex Alderton, Roberto Amato, Jeffrey Barrett, Sonia Goncalves, Ewan Harrison, David K. Jackson, Ian Johnston, Dominic Kwiatkowski, Cordelia Langford, John Sillitoe on behalf of the Wellcome Sanger Institute COVID-19 Surveillance Team |
| EPI_ISL_3110006 | Lighthouse Lab in Glasgow | Wellcome Sanger Institute for the COVID-19 Genomics UK (COG-UK) Consortium | Harper VanSteenhouse, Yumi Kasai, David Gray, Carol Clugston, Anna Dominiczak and Alex Alderton, Roberto Amato, Jeffrey Barrett, Sonia Goncalves, Ewan Harrison, David K. Jackson, Ian Johnston, Dominic Kwiatkowski, Cordelia Langford, John Sillitoe on behalf of the Wellcome Sanger Institute COVID-19 Surveillance Team |
| EPI_ISL_3110007 | Lighthouse Laboratory Plymouth | Wellcome Sanger Institute for the COVID-19 Genomics UK (COG-UK) Consortium | Lighthouse Laboratory Plymouth and Alex Alderton, Roberto Amato, Jeffrey Barrett, Sonia Goncalves, Ewan Harrison, David K. Jackson, Ian Johnston, Dominic Kwiatkowski, Cordelia Langford, John Sillitoe on behalf of the Wellcome Sanger Institute COVID-19 Surveillance Team |
| EPI_ISL_3110008 | Lighthouse Lab in Alderley Park | Wellcome Sanger Institute for the COVID-19 Genomics UK (COG-UK) Consortium | Jacquelyn Wynn, Mairead Hyland, The Lighthouse Lab in Alderley Park and Alex Alderton, Roberto Amato, Jeffrey Barrett, Sonia Goncalves, Ewan Harrison, David K. Jackson, Ian Johnston, Dominic Kwiatkowski, Cordelia Langford, John Sillitoe on behalf of the Wellcome Sanger Institute COVID-19 Surveillance Team |
| EPI_ISL_3110009 | Lighthouse Laboratory Plymouth | Wellcome Sanger Institute for the COVID-19 Genomics UK (COG-UK) Consortium | Lighthouse Laboratory Plymouth and Alex Alderton, Roberto Amato, Jeffrey Barrett, Sonia Goncalves, Ewan Harrison, David K. Jackson, Ian Johnston, Dominic Kwiatkowski, Cordelia Langford, John Sillitoe on behalf of the Wellcome Sanger Institute COVID-19 Surveillance Team |
| EPI_ISL_3110010, EPI_ISL_3110011 | Lighthouse Lab in Alderley Park | Wellcome Sanger Institute for the COVID-19 Genomics UK (COG-UK) Consortium | Jacquelyn Wynn, Mairead Hyland, The Lighthouse Lab in Alderley Park and Alex Alderton, Roberto Amato, Jeffrey Barrett, Sonia Goncalves, Ewan Harrison, David K. Jackson, Ian Johnston, Dominic Kwiatkowski, Cordelia Langford, John Sillitoe on behalf of the Wellcome Sanger Institute COVID-19 Surveillance Team |
| EPI_ISL_3110012, EPI_ISL_3110013, EPI_ISL_3110014 | Lighthouse Laboratory Plymouth | Wellcome Sanger Institute for the COVID-19 Genomics UK (COG-UK) Consortium | Lighthouse Laboratory Plymouth and Alex Alderton, Roberto Amato, Jeffrey Barrett, Sonia Goncalves, Ewan Harrison, David K. Jackson, Ian Johnston, Dominic Kwiatkowski, Cordelia Langford, John Sillitoe on behalf of the Wellcome Sanger Institute COVID-19 Surveillance Team |
| EPI_ISL_3110015 | Lighthouse Lab in Alderley Park | Wellcome Sanger Institute for the COVID-19 Genomics UK (COG-UK) Consortium | Jacquelyn Wynn, Mairead Hyland, The Lighthouse Lab in Alderley Park and Alex Alderton, Roberto Amato, Jeffrey Barrett, Sonia Goncalves, Ewan Harrison, David K. Jackson, Ian Johnston, Dominic Kwiatkowski, Cordelia Langford, John Sillitoe on behalf of the Wellcome Sanger Institute COVID-19 Surveillance Team |
| EPI_ISL_3110016 | Lighthouse Laboratory Plymouth | Wellcome Sanger Institute for the COVID-19 Genomics UK (COG-UK) Consortium | Lighthouse Laboratory Plymouth and Alex Alderton, Roberto Amato, Jeffrey Barrett, Sonia Goncalves, Ewan Harrison, David K. Jackson, Ian Johnston, Dominic Kwiatkowski, Cordelia Langford, John Sillitoe on behalf of the Wellcome Sanger Institute COVID-19 Surveillance Team |
| EPI_ISL_3110017, EPI_ISL_3110018 | Lighthouse Lab in Milton Keynes | Wellcome Sanger Institute for the COVID-19 Genomics UK (COG-UK) Consortium | The Lighthouse Lab in Milton Keynes and Alex Alderton, Roberto Amato, Jeffrey Barrett, Sonia Goncalves, Ewan Harrison, David K. Jackson, Ian Johnston, Dominic Kwiatkowski, Cordelia Langford, John Sillitoe on behalf of the Wellcome Sanger Institute COVID-19 Surveillance Team |
| EPI_ISL_3110019 | Lighthouse Lab in Alderley Park | Wellcome Sanger Institute for the COVID-19 Genomics UK (COG-UK) Consortium | Jacquelyn Wynn, Mairead Hyland, The Lighthouse Lab in Alderley Park and Alex Alderton, Roberto Amato, Jeffrey Barrett, Sonia Goncalves, Ewan Harrison, David K. Jackson, Ian Johnston, Dominic Kwiatkowski, Cordelia Langford, John Sillitoe on behalf of the Wellcome Sanger Institute COVID-19 Surveillance Team |
| EPI_ISL_3110020 | Lighthouse Laboratory Plymouth | Wellcome Sanger Institute for the COVID-19 Genomics UK (COG-UK) Consortium | Lighthouse Laboratory Plymouth and Alex Alderton, Roberto Amato, Jeffrey Barrett, Sonia Goncalves, Ewan Harrison, David K. Jackson, Ian Johnston, Dominic Kwiatkowski, Cordelia Langford, John Sillitoe on behalf of the Wellcome Sanger Institute COVID-19 Surveillance Team |
| EPI_ISL_3110021 | Lighthouse Lab in Milton Keynes | Wellcome Sanger Institute for the COVID-19 Genomics UK (COG-UK) Consortium | The Lighthouse Lab in Milton Keynes and Alex Alderton, Roberto Amato, Jeffrey Barrett, Sonia Goncalves, Ewan Harrison, David K. Jackson, Ian Johnston, Dominic Kwiatkowski, Cordelia Langford, John Sillitoe on behalf of the Wellcome Sanger Institute COVID-19 Surveillance Team |
| EPI_ISL_3110022 | Lighthouse Laboratory Plymouth | Wellcome Sanger Institute for the COVID-19 Genomics UK (COG-UK) Consortium | Lighthouse Laboratory Plymouth and Alex Alderton, Roberto Amato, Jeffrey Barrett, Sonia Goncalves, Ewan Harrison, David K. Jackson, Ian Johnston, Dominic Kwiatkowski, Cordelia Langford, John Sillitoe on behalf of the Wellcome Sanger Institute COVID-19 Surveillance Team |
| EPI_ISL_3110023, EPI_ISL_3110024 | Lighthouse Lab in Alderley Park | Wellcome Sanger Institute for the COVID-19 Genomics UK (COG-UK) Consortium | Jacquelyn Wynn, Mairead Hyland, The Lighthouse Lab in Alderley Park and Alex Alderton, Roberto Amato, Jeffrey Barrett, Sonia Goncalves, Ewan Harrison, David K. Jackson, Ian Johnston, Dominic Kwiatkowski, Cordelia Langford, John Sillitoe on behalf of the Wellcome Sanger Institute COVID-19 Surveillance Team |
| EPI_ISL_3110025, EPI_ISL_3110026 | Lighthouse Laboratory Plymouth | Wellcome Sanger Institute for the COVID-19 Genomics UK (COG-UK) Consortium | Lighthouse Laboratory Plymouth and Alex Alderton, Roberto Amato, Jeffrey Barrett, Sonia Goncalves, Ewan Harrison, David K. Jackson, Ian Johnston, Dominic Kwiatkowski, Cordelia Langford, John Sillitoe on behalf of the Wellcome Sanger Institute COVID-19 Surveillance Team |
| EPI_ISL_3110027, EPI_ISL_3110028 | Lighthouse Lab in Milton Keynes | Wellcome Sanger Institute for the COVID-19 Genomics UK (COG-UK) Consortium | The Lighthouse Lab in Milton Keynes and Alex Alderton, Roberto Amato, Jeffrey Barrett, Sonia Goncalves, Ewan Harrison, David K. Jackson, Ian Johnston, Dominic Kwiatkowski, Cordelia Langford, John Sillitoe on behalf of the Wellcome Sanger Institute COVID-19 Surveillance Team |
| EPI_ISL_3110029, EPI_ISL_3110030 | Lighthouse Laboratory Plymouth | Wellcome Sanger Institute for the COVID-19 Genomics UK (COG-UK) Consortium | Lighthouse Laboratory Plymouth and Alex Alderton, Roberto Amato, Jeffrey Barrett, Sonia Goncalves, Ewan Harrison, David K. Jackson, Ian Johnston, Dominic Kwiatkowski, Cordelia Langford, John Sillitoe on behalf of the Wellcome Sanger Institute COVID-19 Surveillance Team |
| EPI_ISL_3110031, EPI_ISL_3110032 | Lighthouse Lab in Milton Keynes | Wellcome Sanger Institute for the COVID-19 Genomics UK (COG-UK) Consortium | The Lighthouse Lab in Milton Keynes and Alex Alderton, Roberto Amato, Jeffrey Barrett, Sonia Goncalves, Ewan Harrison, David K. Jackson, Ian Johnston, Dominic Kwiatkowski, Cordelia Langford, John Sillitoe on behalf of the Wellcome Sanger Institute COVID-19 Surveillance Team |
| EPI_ISL_3110033 | Lighthouse Lab in Alderley Park | Wellcome Sanger Institute for the COVID-19 Genomics UK (COG-UK) Consortium | Jacquelyn Wynn, Mairead Hyland, The Lighthouse Lab in Alderley Park and Alex Alderton, Roberto Amato, Jeffrey Barrett, Sonia Goncalves, Ewan Harrison, David K. Jackson, Ian Johnston, Dominic Kwiatkowski, Cordelia Langford, John Sillitoe on behalf of the Wellcome Sanger Institute COVID-19 Surveillance Team |

| | | | |
|---|---|---|---|
| EPI_ISL_3110034 | Lighthouse Lab in Milton Keynes | Wellcome Sanger Institute for the COVID-19 Genomics UK (COG-UK) Consortium | The Lighthouse Lab in Milton Keynes and Alex Alderton, Roberto Amato, Jeffrey Barrett, Sonia Goncalves, Ewan Harrison, David K. Jackson, Ian Johnston, Dominic Kwiatkowski, Cordelia Langford, John Sillitoe on behalf of the Wellcome Sanger Institute COVID-19 Surveillance Team |
| EPI_ISL_3110035 | Lighthouse Lab in Alderley Park | Wellcome Sanger Institute for the COVID-19 Genomics UK (COG-UK) Consortium | Jacquelyn Wynn, Mairead Hyland, The Lighthouse Lab in Alderley Park and Alex Alderton, Roberto Amato, Jeffrey Barrett, Sonia Goncalves, Ewan Harrison, David K. Jackson, Ian Johnston, Dominic Kwiatkowski, Cordelia Langford, John Sillitoe on behalf of the Wellcome Sanger Institute COVID-19 Surveillance Team |
| EPI_ISL_3110036 | Lighthouse Lab in Milton Keynes | Wellcome Sanger Institute for the COVID-19 Genomics UK (COG-UK) Consortium | The Lighthouse Lab in Milton Keynes and Alex Alderton, Roberto Amato, Jeffrey Barrett, Sonia Goncalves, Ewan Harrison, David K. Jackson, Ian Johnston, Dominic Kwiatkowski, Cordelia Langford, John Sillitoe on behalf of the Wellcome Sanger Institute COVID-19 Surveillance Team |
| EPI_ISL_3110037, EPI_ISL_3110038 | Lighthouse Lab in Alderley Park | Wellcome Sanger Institute for the COVID-19 Genomics UK (COG-UK) Consortium | Jacquelyn Wynn, Mairead Hyland, The Lighthouse Lab in Alderley Park and Alex Alderton, Roberto Amato, Jeffrey Barrett, Sonia Goncalves, Ewan Harrison, David K. Jackson, Ian Johnston, Dominic Kwiatkowski, Cordelia Langford, John Sillitoe on behalf of the Wellcome Sanger Institute COVID-19 Surveillance Team |
| EPI_ISL_3110039, EPI_ISL_3110040 | Lighthouse Laboratory Plymouth | Wellcome Sanger Institute for the COVID-19 Genomics UK (COG-UK) Consortium | Lighthouse Laboratory Plymouth and Alex Alderton, Roberto Amato, Jeffrey Barrett, Sonia Goncalves, Ewan Harrison, David K. Jackson, Ian Johnston, Dominic Kwiatkowski, Cordelia Langford, John Sillitoe on behalf of the Wellcome Sanger Institute COVID-19 Surveillance Team |
| EPI_ISL_3110041, EPI_ISL_3110042, EPI_ISL_3110043 | Lighthouse Lab in Alderley Park | Wellcome Sanger Institute for the COVID-19 Genomics UK (COG-UK) Consortium | Jacquelyn Wynn, Mairead Hyland, The Lighthouse Lab in Alderley Park and Alex Alderton, Roberto Amato, Jeffrey Barrett, Sonia Goncalves, Ewan Harrison, David K. Jackson, Ian Johnston, Dominic Kwiatkowski, Cordelia Langford, John Sillitoe on behalf of the Wellcome Sanger Institute COVID-19 Surveillance Team |
| EPI_ISL_3110044 | Lighthouse Lab in Glasgow | Wellcome Sanger Institute for the COVID-19 Genomics UK (COG-UK) Consortium | Harper VanSteenhouse, Yumi Kasai, David Gray, Carol Clugston, Anna Dominiczak and Alex Alderton, Roberto Amato, Jeffrey Barrett, Sonia Goncalves, Ewan Harrison, David K. Jackson, Ian Johnston, Dominic Kwiatkowski, Cordelia Langford, John Sillitoe on behalf of the Wellcome Sanger Institute COVID-19 Surveillance Team |
| EPI_ISL_3110045 | Lighthouse Lab in Milton Keynes | Wellcome Sanger Institute for the COVID-19 Genomics UK (COG-UK) Consortium | The Lighthouse Lab in Milton Keynes and Alex Alderton, Roberto Amato, Jeffrey Barrett, Sonia Goncalves, Ewan Harrison, David K. Jackson, Ian Johnston, Dominic Kwiatkowski, Cordelia Langford, John Sillitoe on behalf of the Wellcome Sanger Institute COVID-19 Surveillance Team |
| EPI_ISL_3110046 | Lighthouse Lab in Alderley Park | Wellcome Sanger Institute for the COVID-19 Genomics UK (COG-UK) Consortium | Jacquelyn Wynn, Mairead Hyland, The Lighthouse Lab in Alderley Park and Alex Alderton, Roberto Amato, Jeffrey Barrett, Sonia Goncalves, Ewan Harrison, David K. Jackson, Ian Johnston, Dominic Kwiatkowski, Cordelia Langford, John Sillitoe on behalf of the Wellcome Sanger Institute COVID-19 Surveillance Team |
| EPI_ISL_3110047 | Lighthouse Lab in Glasgow | Wellcome Sanger Institute for the COVID-19 Genomics UK (COG-UK) Consortium | Harper VanSteenhouse, Yumi Kasai, David Gray, Carol Clugston, Anna Dominiczak and Alex Alderton, Roberto Amato, Jeffrey Barrett, Sonia Goncalves, Ewan Harrison, David K. Jackson, Ian Johnston, Dominic Kwiatkowski, Cordelia Langford, John Sillitoe on behalf of the Wellcome Sanger Institute COVID-19 Surveillance Team |
| EPI_ISL_3110048, EPI_ISL_3110049 | Lighthouse Lab in Alderley Park | Wellcome Sanger Institute for the COVID-19 Genomics UK (COG-UK) Consortium | Jacquelyn Wynn, Mairead Hyland, The Lighthouse Lab in Alderley Park and Alex Alderton, Roberto Amato, Jeffrey Barrett, Sonia Goncalves, Ewan Harrison, David K. Jackson, Ian Johnston, Dominic Kwiatkowski, Cordelia Langford, John Sillitoe on behalf of the Wellcome Sanger Institute COVID-19 Surveillance Team |
| EPI_ISL_3110050 | Lighthouse Laboratory Plymouth | Wellcome Sanger Institute for the COVID-19 Genomics UK (COG-UK) Consortium | Lighthouse Laboratory Plymouth and Alex Alderton, Roberto Amato, Jeffrey Barrett, Sonia Goncalves, Ewan Harrison, David K. Jackson, Ian Johnston, Dominic Kwiatkowski, Cordelia Langford, John Sillitoe on behalf of the Wellcome Sanger Institute COVID-19 Surveillance Team |
| EPI_ISL_3110051 | Lighthouse Lab in Glasgow | Wellcome Sanger Institute for the COVID-19 Genomics UK (COG-UK) Consortium | Harper VanSteenhouse, Yumi Kasai, David Gray, Carol Clugston, Anna Dominiczak and Alex Alderton, Roberto Amato, Jeffrey Barrett, Sonia Goncalves, Ewan Harrison, David K. Jackson, Ian Johnston, Dominic Kwiatkowski, Cordelia Langford, John Sillitoe on behalf of the Wellcome Sanger Institute COVID-19 Surveillance Team |
| EPI_ISL_3110052 | Lighthouse Lab in Alderley Park | Wellcome Sanger Institute for the COVID-19 Genomics UK (COG-UK) Consortium | Jacquelyn Wynn, Mairead Hyland, The Lighthouse Lab in Alderley Park and Alex Alderton, Roberto Amato, Jeffrey Barrett, Sonia Goncalves, Ewan Harrison, David K. Jackson, Ian Johnston, Dominic Kwiatkowski, Cordelia Langford, John Sillitoe on behalf of the Wellcome Sanger Institute COVID-19 Surveillance Team |
| EPI_ISL_3110053 | Lighthouse Lab in Glasgow | Wellcome Sanger Institute for the COVID-19 Genomics UK (COG-UK) Consortium | Harper VanSteenhouse, Yumi Kasai, David Gray, Carol Clugston, Anna Dominiczak and Alex Alderton, Roberto Amato, Jeffrey Barrett, Sonia Goncalves, Ewan Harrison, David K. Jackson, Ian Johnston, Dominic Kwiatkowski, Cordelia Langford, John Sillitoe on behalf of the Wellcome Sanger Institute COVID-19 Surveillance Team |
| EPI_ISL_3110054, EPI_ISL_3110055 | Lighthouse Lab in Alderley Park | Wellcome Sanger Institute for the COVID-19 Genomics UK (COG-UK) Consortium | Jacquelyn Wynn, Mairead Hyland, The Lighthouse Lab in Alderley Park and Alex Alderton, Roberto Amato, Jeffrey Barrett, Sonia Goncalves, Ewan Harrison, David K. Jackson, Ian Johnston, Dominic Kwiatkowski, Cordelia Langford, John Sillitoe on behalf of the Wellcome Sanger Institute COVID-19 Surveillance Team |
| EPI_ISL_3110056 | Lighthouse Lab in Milton Keynes | Wellcome Sanger Institute for the COVID-19 Genomics UK (COG-UK) Consortium | The Lighthouse Lab in Milton Keynes and Alex Alderton, Roberto Amato, Jeffrey Barrett, Sonia Goncalves, Ewan Harrison, David K. Jackson, Ian Johnston, Dominic Kwiatkowski, Cordelia Langford, John Sillitoe on behalf of the Wellcome Sanger Institute COVID-19 Surveillance Team |
| EPI_ISL_3110057 | Lighthouse Lab in Alderley Park | Wellcome Sanger Institute for the COVID-19 Genomics UK (COG-UK) Consortium | Jacquelyn Wynn, Mairead Hyland, The Lighthouse Lab in Alderley Park and Alex Alderton, Roberto Amato, Jeffrey Barrett, Sonia Goncalves, Ewan Harrison, David K. Jackson, Ian Johnston, Dominic Kwiatkowski, Cordelia Langford, John Sillitoe on behalf of the Wellcome Sanger Institute COVID-19 Surveillance Team |
| EPI_ISL_3110058 | Lighthouse Lab in Glasgow | Wellcome Sanger Institute for the COVID-19 Genomics UK (COG-UK) Consortium | Harper VanSteenhouse, Yumi Kasai, David Gray, Carol Clugston, Anna Dominiczak and Alex Alderton, Roberto Amato, Jeffrey Barrett, Sonia Goncalves, Ewan Harrison, David K. Jackson, Ian Johnston, Dominic Kwiatkowski, Cordelia Langford, John Sillitoe on behalf of the Wellcome Sanger Institute COVID-19 Surveillance Team |
| EPI_ISL_3110059, EPI_ISL_3110060 | Lighthouse Lab in Alderley Park | Wellcome Sanger Institute for the COVID-19 Genomics UK (COG-UK) Consortium | Jacquelyn Wynn, Mairead Hyland, The Lighthouse Lab in Alderley Park and Alex Alderton, Roberto Amato, Jeffrey Barrett, Sonia Goncalves, Ewan Harrison, David K. Jackson, Ian Johnston, Dominic Kwiatkowski, Cordelia Langford, John Sillitoe on behalf of the Wellcome Sanger Institute COVID-19 Surveillance Team |
| EPI_ISL_3110061, EPI_ISL_3110062, EPI_ISL_3110063, EPI_ISL_3110064, EPI_ISL_3110065, EPI_ISL_3110066 | Lighthouse Lab in Milton Keynes | Wellcome Sanger Institute for the COVID-19 Genomics UK (COG-UK) Consortium | The Lighthouse Lab in Milton Keynes and Alex Alderton, Roberto Amato, Jeffrey Barrett, Sonia Goncalves, Ewan Harrison, David K. Jackson, Ian Johnston, Dominic Kwiatkowski, Cordelia Langford, John Sillitoe on behalf of the Wellcome Sanger Institute COVID-19 Surveillance Team |
| EPI_ISL_3110067 | Lighthouse Laboratory Plymouth | Wellcome Sanger Institute for the COVID-19 Genomics UK (COG-UK) Consortium | Lighthouse Laboratory Plymouth and Alex Alderton, Roberto Amato, Jeffrey Barrett, Sonia Goncalves, Ewan Harrison, David K. Jackson, Ian Johnston, Dominic Kwiatkowski, Cordelia Langford, John Sillitoe on behalf of the Wellcome Sanger Institute COVID-19 Surveillance Team |
| EPI_ISL_3110068, EPI_ISL_3110069 | Lighthouse Lab in Alderley Park | Wellcome Sanger Institute for the COVID-19 Genomics UK (COG-UK) Consortium | Jacquelyn Wynn, Mairead Hyland, The Lighthouse Lab in Alderley Park and Alex Alderton, Roberto Amato, Jeffrey Barrett, Sonia Goncalves, Ewan Harrison, David K. Jackson, Ian Johnston, Dominic Kwiatkowski, Cordelia Langford, John Sillitoe on behalf of the Wellcome Sanger Institute COVID-19 Surveillance Team |
| EPI_ISL_3110070 | Lighthouse Lab in Glasgow | Wellcome Sanger Institute for the COVID-19 Genomics UK (COG-UK) Consortium | Harper VanSteenhouse, Yumi Kasai, David Gray, Carol Clugston, Anna Dominiczak and Alex Alderton, Roberto Amato, Jeffrey Barrett, Sonia Goncalves, Ewan Harrison, David K. Jackson, Ian Johnston, Dominic Kwiatkowski, Cordelia Langford, John Sillitoe on behalf of the Wellcome Sanger Institute COVID-19 Surveillance Team |
| EPI_ISL_3110071 | Lighthouse Lab in Milton Keynes | Wellcome Sanger Institute for the COVID-19 Genomics UK (COG-UK) Consortium | The Lighthouse Lab in Milton Keynes and Alex Alderton, Roberto Amato, Jeffrey Barrett, Sonia Goncalves, Ewan Harrison, David K. Jackson, Ian Johnston, Dominic Kwiatkowski, Cordelia Langford, John Sillitoe on behalf of the Wellcome Sanger Institute COVID-19 Surveillance Team |
| EPI_ISL_3110072, EPI_ISL_3110073 | Lighthouse Lab in Alderley Park | Wellcome Sanger Institute for the COVID-19 Genomics UK (COG-UK) Consortium | Jacquelyn Wynn, Mairead Hyland, The Lighthouse Lab in Alderley Park and Alex Alderton, Roberto Amato, Jeffrey Barrett, Sonia Goncalves, Ewan Harrison, David K. Jackson, Ian Johnston, Dominic Kwiatkowski, Cordelia Langford, John Sillitoe on behalf of the Wellcome Sanger Institute COVID-19 Surveillance Team |
| EPI_ISL_3110074 | Lighthouse Lab in Milton Keynes | Wellcome Sanger Institute for the COVID-19 Genomics UK (COG-UK) Consortium | The Lighthouse Lab in Milton Keynes and Alex Alderton, Roberto Amato, Jeffrey Barrett, Sonia Goncalves, Ewan Harrison, David K. Jackson, Ian Johnston, Dominic Kwiatkowski, Cordelia Langford, John Sillitoe on behalf of the Wellcome Sanger Institute COVID-19 Surveillance Team |
| EPI_ISL_3110075 | Lighthouse Lab in Alderley Park | Wellcome Sanger Institute for the COVID-19 Genomics UK (COG-UK) Consortium | Jacquelyn Wynn, Mairead Hyland, The Lighthouse Lab in Alderley Park and Alex Alderton, Roberto Amato, Jeffrey Barrett, Sonia Goncalves, Ewan Harrison, David K. Jackson, Ian Johnston, Dominic Kwiatkowski, Cordelia Langford, John Sillitoe on behalf of the Wellcome Sanger Institute COVID-19 Surveillance Team |
| EPI_ISL_3110076 | Lighthouse Lab in Glasgow | Wellcome Sanger Institute for the COVID-19 Genomics UK (COG-UK) Consortium | Harper VanSteenhouse, Yumi Kasai, David Gray, Carol Clugston, Anna Dominiczak and Alex Alderton, Roberto Amato, Jeffrey Barrett, Sonia Goncalves, Ewan Harrison, David K. Jackson, Ian Johnston, Dominic Kwiatkowski, Cordelia Langford, John Sillitoe on behalf of the Wellcome Sanger Institute COVID-19 Surveillance Team |
| EPI_ISL_3110077, EPI_ISL_3110078, EPI_ISL_3110079, EPI_ISL_3110080 | Lighthouse Lab in Milton Keynes | Wellcome Sanger Institute for the COVID-19 Genomics UK (COG-UK) Consortium | The Lighthouse Lab in Milton Keynes and Alex Alderton, Roberto Amato, Jeffrey Barrett, Sonia Goncalves, Ewan Harrison, David K. Jackson, Ian Johnston, Dominic Kwiatkowski, Cordelia Langford, John Sillitoe on behalf of the Wellcome Sanger Institute COVID-19 Surveillance Team |
| EPI_ISL_3110081 | Lighthouse Lab in Alderley Park | Wellcome Sanger Institute for the COVID-19 Genomics UK (COG-UK) Consortium | Jacquelyn Wynn, Mairead Hyland, The Lighthouse Lab in Alderley Park and Alex Alderton, Roberto Amato, Jeffrey Barrett, Sonia Goncalves, Ewan Harrison, David K. Jackson, Ian Johnston, Dominic Kwiatkowski, Cordelia Langford, John Sillitoe on behalf of the Wellcome Sanger Institute COVID-19 Surveillance Team |
| EPI_ISL_3110082 | Lighthouse Lab in Glasgow | Wellcome Sanger Institute for the COVID-19 Genomics UK (COG-UK) Consortium | Harper VanSteenhouse, Yumi Kasai, David Gray, Carol Clugston, Anna Dominiczak and Alex Alderton, Roberto Amato, Jeffrey Barrett, Sonia Goncalves, Ewan Harrison, David K. Jackson, Ian Johnston, Dominic Kwiatkowski, Cordelia Langford, John Sillitoe on behalf of the Wellcome Sanger Institute COVID-19 Surveillance Team |
| EPI_ISL_3110083 | Lighthouse Lab in Milton Keynes | Wellcome Sanger Institute for the COVID-19 Genomics UK (COG-UK) Consortium | The Lighthouse Lab in Milton Keynes and Alex Alderton, Roberto Amato, Jeffrey Barrett, Sonia Goncalves, Ewan Harrison, David K. Jackson, Ian Johnston, Dominic Kwiatkowski, Cordelia Langford, John Sillitoe on behalf of the Wellcome Sanger Institute COVID-19 Surveillance Team |
| EPI_ISL_3110084 | Lighthouse Lab in Alderley Park | Wellcome Sanger Institute for the COVID-19 Genomics UK (COG-UK) Consortium | Jacquelyn Wynn, Mairead Hyland, The Lighthouse Lab in Alderley Park and Alex Alderton, Roberto Amato, Jeffrey Barrett, Sonia Goncalves, Ewan Harrison, David K. Jackson, Ian Johnston, Dominic Kwiatkowski, Cordelia Langford, John Sillitoe on behalf of the Wellcome Sanger Institute COVID-19 Surveillance Team |
| EPI_ISL_3110085 | Lighthouse Lab in Glasgow | Wellcome Sanger Institute for the COVID-19 Genomics UK (COG-UK) Consortium | Harper VanSteenhouse, Yumi Kasai, David Gray, Carol Clugston, Anna Dominiczak and Alex Alderton, Roberto Amato, Jeffrey Barrett, Sonia Goncalves, Ewan Harrison, David K. Jackson, Ian Johnston, Dominic Kwiatkowski, Cordelia Langford, John Sillitoe on behalf of the Wellcome Sanger Institute COVID-19 Surveillance Team |
| EPI_ISL_3110086 | Lighthouse Lab in Milton Keynes | Wellcome Sanger Institute for the COVID-19 Genomics UK (COG-UK) Consortium | The Lighthouse Lab in Milton Keynes and Alex Alderton, Roberto Amato, Jeffrey Barrett, Sonia Goncalves, Ewan Harrison, David K. Jackson, Ian Johnston, Dominic Kwiatkowski, Cordelia Langford, John Sillitoe on behalf of the Wellcome Sanger Institute COVID-19 Surveillance Team |
| EPI_ISL_3110087 | Lighthouse Lab in Alderley Park | Wellcome Sanger Institute for the COVID-19 Genomics UK (COG-UK) Consortium | Jacquelyn Wynn, Mairead Hyland, The Lighthouse Lab in Alderley Park and Alex Alderton, Roberto Amato, Jeffrey Barrett, Sonia Goncalves, Ewan Harrison, David K. Jackson, Ian Johnston, Dominic Kwiatkowski, Cordelia Langford, John Sillitoe on behalf of the Wellcome Sanger Institute COVID-19 Surveillance Team |
| EPI_ISL_3110088 | Lighthouse Lab in Milton Keynes | Wellcome Sanger Institute for the COVID-19 Genomics UK (COG-UK) Consortium | The Lighthouse Lab in Milton Keynes and Alex Alderton, Roberto Amato, Jeffrey Barrett, Sonia Goncalves, Ewan Harrison, David K. Jackson, Ian Johnston, Dominic Kwiatkowski, Cordelia Langford, John Sillitoe on behalf of the Wellcome Sanger Institute COVID-19 Surveillance Team |
| EPI_ISL_3110089 | Lighthouse Laboratory Plymouth | Wellcome Sanger Institute for the COVID-19 Genomics UK (COG-UK) Consortium | Lighthouse Laboratory Plymouth and Alex Alderton, Roberto Amato, Jeffrey Barrett, Sonia Goncalves, Ewan Harrison, David K. Jackson, Ian Johnston, Dominic Kwiatkowski, Cordelia Langford, John Sillitoe on behalf of the Wellcome Sanger Institute COVID-19 Surveillance Team |
| EPI_ISL_3110090 | Lighthouse Lab in Alderley Park | Wellcome Sanger Institute for the COVID-19 Genomics UK (COG-UK) Consortium | Jacquelyn Wynn, Mairead Hyland, The Lighthouse Lab in Alderley Park and Alex Alderton, Roberto Amato, Jeffrey Barrett, Sonia Goncalves, Ewan Harrison, David K. Jackson, Ian Johnston, Dominic Kwiatkowski, Cordelia Langford, John Sillitoe on behalf of the Wellcome Sanger Institute COVID-19 Surveillance Team |
| EPI_ISL_3110091 | Lighthouse Laboratory Plymouth | Wellcome Sanger Institute for the COVID-19 Genomics UK (COG-UK) Consortium | Lighthouse Laboratory Plymouth and Alex Alderton, Roberto Amato, Jeffrey Barrett, Sonia Goncalves, Ewan Harrison, David K. Jackson, Ian Johnston, Dominic Kwiatkowski, Cordelia Langford, John Sillitoe on behalf of the Wellcome Sanger Institute COVID-19 Surveillance Team |
| EPI_ISL_3110092 | Lighthouse Lab in Glasgow | Wellcome Sanger Institute for the COVID-19 Genomics UK (COG-UK) Consortium | Harper VanSteenhouse, Yumi Kasai, David Gray, Carol Clugston, Anna Dominiczak and Alex Alderton, Roberto Amato, Jeffrey Barrett, Sonia Goncalves, Ewan Harrison, David K. Jackson, Ian Johnston, Dominic Kwiatkowski, Cordelia Langford, John Sillitoe on behalf of the Wellcome Sanger Institute COVID-19 Surveillance Team |

| | | | |
|---|---|---|---|
| EPI_ISL_3110093 | Lighthouse Laboratory Plymouth | Wellcome Sanger Institute for the COVID-19 Genomics UK (COG-UK) Consortium | Lighthouse Laboratory Plymouth and Alex Alderton, Roberto Amato, Jeffrey Barrett, Sonia Goncalves, Ewan Harrison, David K. Jackson, Ian Johnston, Dominic Kwiatkowski, Cordelia Langford, John Sillitoe on behalf of the Wellcome Sanger Institute COVID-19 Surveillance Team |
| EPI_ISL_3110094 | Lighthouse Lab in Glasgow | Wellcome Sanger Institute for the COVID-19 Genomics UK (COG-UK) Consortium | Harper VanSteenhouse, Yumi Kasai, David Gray, Carol Clugston, Anna Dominiczak and Alex Alderton, Roberto Amato, Jeffrey Barrett, Sonia Goncalves, Ewan Harrison, David K. Jackson, Ian Johnston, Dominic Kwiatkowski, Cordelia Langford, John Sillitoe on behalf of the Wellcome Sanger Institute COVID-19 Surveillance Team |
| EPI_ISL_3110095, EPI_ISL_3110096 | Lighthouse Lab in Alderley Park | Wellcome Sanger Institute for the COVID-19 Genomics UK (COG-UK) Consortium | Jacquelyn Wynn, Mairead Hyland, The Lighthouse Lab in Alderley Park and Alex Alderton, Roberto Amato, Jeffrey Barrett, Sonia Goncalves, Ewan Harrison, David K. Jackson, Ian Johnston, Dominic Kwiatkowski, Cordelia Langford, John Sillitoe on behalf of the Wellcome Sanger Institute COVID-19 Surveillance Team |
| EPI_ISL_3110097 | Lighthouse Laboratory Plymouth | Wellcome Sanger Institute for the COVID-19 Genomics UK (COG-UK) Consortium | Lighthouse Laboratory Plymouth and Alex Alderton, Roberto Amato, Jeffrey Barrett, Sonia Goncalves, Ewan Harrison, David K. Jackson, Ian Johnston, Dominic Kwiatkowski, Cordelia Langford, John Sillitoe on behalf of the Wellcome Sanger Institute COVID-19 Surveillance Team |
| EPI_ISL_3110098, EPI_ISL_3110099 | Lighthouse Lab in Alderley Park | Wellcome Sanger Institute for the COVID-19 Genomics UK (COG-UK) Consortium | Jacquelyn Wynn, Mairead Hyland, The Lighthouse Lab in Alderley Park and Alex Alderton, Roberto Amato, Jeffrey Barrett, Sonia Goncalves, Ewan Harrison, David K. Jackson, Ian Johnston, Dominic Kwiatkowski, Cordelia Langford, John Sillitoe on behalf of the Wellcome Sanger Institute COVID-19 Surveillance Team |
| EPI_ISL_3110100 | Lighthouse Lab in Milton Keynes | Wellcome Sanger Institute for the COVID-19 Genomics UK (COG-UK) Consortium | The Lighthouse Lab in Milton Keynes and Alex Alderton, Roberto Amato, Jeffrey Barrett, Sonia Goncalves, Ewan Harrison, David K. Jackson, Ian Johnston, Dominic Kwiatkowski, Cordelia Langford, John Sillitoe on behalf of the Wellcome Sanger Institute COVID-19 Surveillance Team |
| EPI_ISL_3110101 | Lighthouse Laboratory Plymouth | Wellcome Sanger Institute for the COVID-19 Genomics UK (COG-UK) Consortium | Lighthouse Laboratory Plymouth and Alex Alderton, Roberto Amato, Jeffrey Barrett, Sonia Goncalves, Ewan Harrison, David K. Jackson, Ian Johnston, Dominic Kwiatkowski, Cordelia Langford, John Sillitoe on behalf of the Wellcome Sanger Institute COVID-19 Surveillance Team |
| EPI_ISL_3110102, EPI_ISL_3110103 | Lighthouse Lab in Milton Keynes | Wellcome Sanger Institute for the COVID-19 Genomics UK (COG-UK) Consortium | The Lighthouse Lab in Milton Keynes and Alex Alderton, Roberto Amato, Jeffrey Barrett, Sonia Goncalves, Ewan Harrison, David K. Jackson, Ian Johnston, Dominic Kwiatkowski, Cordelia Langford, John Sillitoe on behalf of the Wellcome Sanger Institute COVID-19 Surveillance Team |
| EPI_ISL_3110104 | Lighthouse Lab in Glasgow | Wellcome Sanger Institute for the COVID-19 Genomics UK (COG-UK) Consortium | Harper VanSteenhouse, Yumi Kasai, David Gray, Carol Clugston, Anna Dominiczak and Alex Alderton, Roberto Amato, Jeffrey Barrett, Sonia Goncalves, Ewan Harrison, David K. Jackson, Ian Johnston, Dominic Kwiatkowski, Cordelia Langford, John Sillitoe on behalf of the Wellcome Sanger Institute COVID-19 Surveillance Team |
| EPI_ISL_3110105, EPI_ISL_3110106 | Lighthouse Lab in Alderley Park | Wellcome Sanger Institute for the COVID-19 Genomics UK (COG-UK) Consortium | Jacquelyn Wynn, Mairead Hyland, The Lighthouse Lab in Alderley Park and Alex Alderton, Roberto Amato, Jeffrey Barrett, Sonia Goncalves, Ewan Harrison, David K. Jackson, Ian Johnston, Dominic Kwiatkowski, Cordelia Langford, John Sillitoe on behalf of the Wellcome Sanger Institute COVID-19 Surveillance Team |
| EPI_ISL_3110107 | Lighthouse Lab in Milton Keynes | Wellcome Sanger Institute for the COVID-19 Genomics UK (COG-UK) Consortium | The Lighthouse Lab in Milton Keynes and Alex Alderton, Roberto Amato, Jeffrey Barrett, Sonia Goncalves, Ewan Harrison, David K. Jackson, Ian Johnston, Dominic Kwiatkowski, Cordelia Langford, John Sillitoe on behalf of the Wellcome Sanger Institute COVID-19 Surveillance Team |
| EPI_ISL_3110108 | Lighthouse Lab in Glasgow | Wellcome Sanger Institute for the COVID-19 Genomics UK (COG-UK) Consortium | Harper VanSteenhouse, Yumi Kasai, David Gray, Carol Clugston, Anna Dominiczak and Alex Alderton, Roberto Amato, Jeffrey Barrett, Sonia Goncalves, Ewan Harrison, David K. Jackson, Ian Johnston, Dominic Kwiatkowski, Cordelia Langford, John Sillitoe on behalf of the Wellcome Sanger Institute COVID-19 Surveillance Team |
| EPI_ISL_3110109, EPI_ISL_3110110 | Lighthouse Lab in Milton Keynes | Wellcome Sanger Institute for the COVID-19 Genomics UK (COG-UK) Consortium | The Lighthouse Lab in Milton Keynes and Alex Alderton, Roberto Amato, Jeffrey Barrett, Sonia Goncalves, Ewan Harrison, David K. Jackson, Ian Johnston, Dominic Kwiatkowski, Cordelia Langford, John Sillitoe on behalf of the Wellcome Sanger Institute COVID-19 Surveillance Team |
| EPI_ISL_3110111, EPI_ISL_3110112, EPI_ISL_3110113 | Lighthouse Laboratory Plymouth | Wellcome Sanger Institute for the COVID-19 Genomics UK (COG-UK) Consortium | Lighthouse Laboratory Plymouth and Alex Alderton, Roberto Amato, Jeffrey Barrett, Sonia Goncalves, Ewan Harrison, David K. Jackson, Ian Johnston, Dominic Kwiatkowski, Cordelia Langford, John Sillitoe on behalf of the Wellcome Sanger Institute COVID-19 Surveillance Team |
| EPI_ISL_3110114, EPI_ISL_3110115 | Lighthouse Lab in Milton Keynes | Wellcome Sanger Institute for the COVID-19 Genomics UK (COG-UK) Consortium | The Lighthouse Lab in Milton Keynes and Alex Alderton, Roberto Amato, Jeffrey Barrett, Sonia Goncalves, Ewan Harrison, David K. Jackson, Ian Johnston, Dominic Kwiatkowski, Cordelia Langford, John Sillitoe on behalf of the Wellcome Sanger Institute COVID-19 Surveillance Team |
| EPI_ISL_3110116, EPI_ISL_3110117, EPI_ISL_3110118 | Lighthouse Lab in Alderley Park | Wellcome Sanger Institute for the COVID-19 Genomics UK (COG-UK) Consortium | Jacquelyn Wynn, Mairead Hyland, The Lighthouse Lab in Alderley Park and Alex Alderton, Roberto Amato, Jeffrey Barrett, Sonia Goncalves, Ewan Harrison, David K. Jackson, Ian Johnston, Dominic Kwiatkowski, Cordelia Langford, John Sillitoe on behalf of the Wellcome Sanger Institute COVID-19 Surveillance Team |
| EPI_ISL_3110119 | Lighthouse Laboratory Plymouth | Wellcome Sanger Institute for the COVID-19 Genomics UK (COG-UK) Consortium | Lighthouse Laboratory Plymouth and Alex Alderton, Roberto Amato, Jeffrey Barrett, Sonia Goncalves, Ewan Harrison, David K. Jackson, Ian Johnston, Dominic Kwiatkowski, Cordelia Langford, John Sillitoe on behalf of the Wellcome Sanger Institute COVID-19 Surveillance Team |
| EPI_ISL_3110120 | Lighthouse Lab in Alderley Park | Wellcome Sanger Institute for the COVID-19 Genomics UK (COG-UK) Consortium | Jacquelyn Wynn, Mairead Hyland, The Lighthouse Lab in Alderley Park and Alex Alderton, Roberto Amato, Jeffrey Barrett, Sonia Goncalves, Ewan Harrison, David K. Jackson, Ian Johnston, Dominic Kwiatkowski, Cordelia Langford, John Sillitoe on behalf of the Wellcome Sanger Institute COVID-19 Surveillance Team |
| EPI_ISL_3110121 | Lighthouse Laboratory Plymouth | Wellcome Sanger Institute for the COVID-19 Genomics UK (COG-UK) Consortium | Lighthouse Laboratory Plymouth and Alex Alderton, Roberto Amato, Jeffrey Barrett, Sonia Goncalves, Ewan Harrison, David K. Jackson, Ian Johnston, Dominic Kwiatkowski, Cordelia Langford, John Sillitoe on behalf of the Wellcome Sanger Institute COVID-19 Surveillance Team |
| EPI_ISL_3110122 | Lighthouse Lab in Milton Keynes | Wellcome Sanger Institute for the COVID-19 Genomics UK (COG-UK) Consortium | The Lighthouse Lab in Milton Keynes and Alex Alderton, Roberto Amato, Jeffrey Barrett, Sonia Goncalves, Ewan Harrison, David K. Jackson, Ian Johnston, Dominic Kwiatkowski, Cordelia Langford, John Sillitoe on behalf of the Wellcome Sanger Institute COVID-19 Surveillance Team |
| EPI_ISL_3110123 | Lighthouse Laboratory Plymouth | Wellcome Sanger Institute for the COVID-19 Genomics UK (COG-UK) Consortium | Lighthouse Laboratory Plymouth and Alex Alderton, Roberto Amato, Jeffrey Barrett, Sonia Goncalves, Ewan Harrison, David K. Jackson, Ian Johnston, Dominic Kwiatkowski, Cordelia Langford, John Sillitoe on behalf of the Wellcome Sanger Institute COVID-19 Surveillance Team |
| EPI_ISL_3110124 | Lighthouse Lab in Alderley Park | Wellcome Sanger Institute for the COVID-19 Genomics UK (COG-UK) Consortium | Jacquelyn Wynn, Mairead Hyland, The Lighthouse Lab in Alderley Park and Alex Alderton, Roberto Amato, Jeffrey Barrett, Sonia Goncalves, Ewan Harrison, David K. Jackson, Ian Johnston, Dominic Kwiatkowski, Cordelia Langford, John Sillitoe on behalf of the Wellcome Sanger Institute COVID-19 Surveillance Team |
| EPI_ISL_3110125 | Lighthouse Laboratory Plymouth | Wellcome Sanger Institute for the COVID-19 Genomics UK (COG-UK) Consortium | Lighthouse Laboratory Plymouth and Alex Alderton, Roberto Amato, Jeffrey Barrett, Sonia Goncalves, Ewan Harrison, David K. Jackson, Ian Johnston, Dominic Kwiatkowski, Cordelia Langford, John Sillitoe on behalf of the Wellcome Sanger Institute COVID-19 Surveillance Team |
| EPI_ISL_3110126 | Lighthouse Lab in Alderley Park | Wellcome Sanger Institute for the COVID-19 Genomics UK (COG-UK) Consortium | Jacquelyn Wynn, Mairead Hyland, The Lighthouse Lab in Alderley Park and Alex Alderton, Roberto Amato, Jeffrey Barrett, Sonia Goncalves, Ewan Harrison, David K. Jackson, Ian Johnston, Dominic Kwiatkowski, Cordelia Langford, John Sillitoe on behalf of the Wellcome Sanger Institute COVID-19 Surveillance Team |
| EPI_ISL_3110127, EPI_ISL_3110128, EPI_ISL_3110129 | Lighthouse Laboratory Plymouth | Wellcome Sanger Institute for the COVID-19 Genomics UK (COG-UK) Consortium | Lighthouse Laboratory Plymouth and Alex Alderton, Roberto Amato, Jeffrey Barrett, Sonia Goncalves, Ewan Harrison, David K. Jackson, Ian Johnston, Dominic Kwiatkowski, Cordelia Langford, John Sillitoe on behalf of the Wellcome Sanger Institute COVID-19 Surveillance Team |
| EPI_ISL_3110130 | Lighthouse Lab in Alderley Park | Wellcome Sanger Institute for the COVID-19 Genomics UK (COG-UK) Consortium | Jacquelyn Wynn, Mairead Hyland, The Lighthouse Lab in Alderley Park and Alex Alderton, Roberto Amato, Jeffrey Barrett, Sonia Goncalves, Ewan Harrison, David K. Jackson, Ian Johnston, Dominic Kwiatkowski, Cordelia Langford, John Sillitoe on behalf of the Wellcome Sanger Institute COVID-19 Surveillance Team |
| EPI_ISL_3110131 | Lighthouse Laboratory Plymouth | Wellcome Sanger Institute for the COVID-19 Genomics UK (COG-UK) Consortium | Lighthouse Laboratory Plymouth and Alex Alderton, Roberto Amato, Jeffrey Barrett, Sonia Goncalves, Ewan Harrison, David K. Jackson, Ian Johnston, Dominic Kwiatkowski, Cordelia Langford, John Sillitoe on behalf of the Wellcome Sanger Institute COVID-19 Surveillance Team |
| EPI_ISL_3110132 | Lighthouse Lab in Alderley Park | Wellcome Sanger Institute for the COVID-19 Genomics UK (COG-UK) Consortium | Jacquelyn Wynn, Mairead Hyland, The Lighthouse Lab in Alderley Park and Alex Alderton, Roberto Amato, Jeffrey Barrett, Sonia Goncalves, Ewan Harrison, David K. Jackson, Ian Johnston, Dominic Kwiatkowski, Cordelia Langford, John Sillitoe on behalf of the Wellcome Sanger Institute COVID-19 Surveillance Team |
| EPI_ISL_3110133 | Lighthouse Lab in Glasgow | Wellcome Sanger Institute for the COVID-19 Genomics UK (COG-UK) Consortium | Harper VanSteenhouse, Yumi Kasai, David Gray, Carol Clugston, Anna Dominiczak and Alex Alderton, Roberto Amato, Jeffrey Barrett, Sonia Goncalves, Ewan Harrison, David K. Jackson, Ian Johnston, Dominic Kwiatkowski, Cordelia Langford, John Sillitoe on behalf of the Wellcome Sanger Institute COVID-19 Surveillance Team |
| EPI_ISL_3110134, EPI_ISL_3110135 | Lighthouse Lab in Alderley Park | Wellcome Sanger Institute for the COVID-19 Genomics UK (COG-UK) Consortium | Jacquelyn Wynn, Mairead Hyland, The Lighthouse Lab in Alderley Park and Alex Alderton, Roberto Amato, Jeffrey Barrett, Sonia Goncalves, Ewan Harrison, David K. Jackson, Ian Johnston, Dominic Kwiatkowski, Cordelia Langford, John Sillitoe on behalf of the Wellcome Sanger Institute COVID-19 Surveillance Team |
| EPI_ISL_3110136, EPI_ISL_3110137 | Lighthouse Laboratory Plymouth | Wellcome Sanger Institute for the COVID-19 Genomics UK (COG-UK) Consortium | Lighthouse Laboratory Plymouth and Alex Alderton, Roberto Amato, Jeffrey Barrett, Sonia Goncalves, Ewan Harrison, David K. Jackson, Ian Johnston, Dominic Kwiatkowski, Cordelia Langford, John Sillitoe on behalf of the Wellcome Sanger Institute COVID-19 Surveillance Team |
| EPI_ISL_3110138 | Lighthouse Lab in Milton Keynes | Wellcome Sanger Institute for the COVID-19 Genomics UK (COG-UK) Consortium | The Lighthouse Lab in Milton Keynes and Alex Alderton, Roberto Amato, Jeffrey Barrett, Sonia Goncalves, Ewan Harrison, David K. Jackson, Ian Johnston, Dominic Kwiatkowski, Cordelia Langford, John Sillitoe on behalf of the Wellcome Sanger Institute COVID-19 Surveillance Team |
| EPI_ISL_3110139, EPI_ISL_3110140, EPI_ISL_3110141, EPI_ISL_3110142 | Lighthouse Lab in Alderley Park | Wellcome Sanger Institute for the COVID-19 Genomics UK (COG-UK) Consortium | Jacquelyn Wynn, Mairead Hyland, The Lighthouse Lab in Alderley Park and Alex Alderton, Roberto Amato, Jeffrey Barrett, Sonia Goncalves, Ewan Harrison, David K. Jackson, Ian Johnston, Dominic Kwiatkowski, Cordelia Langford, John Sillitoe on behalf of the Wellcome Sanger Institute COVID-19 Surveillance Team |
| EPI_ISL_3110143 | Lighthouse Laboratory Plymouth | Wellcome Sanger Institute for the COVID-19 Genomics UK (COG-UK) Consortium | Lighthouse Laboratory Plymouth and Alex Alderton, Roberto Amato, Jeffrey Barrett, Sonia Goncalves, Ewan Harrison, David K. Jackson, Ian Johnston, Dominic Kwiatkowski, Cordelia Langford, John Sillitoe on behalf of the Wellcome Sanger Institute COVID-19 Surveillance Team |
| EPI_ISL_3110144 | Lighthouse Lab in Alderley Park | Wellcome Sanger Institute for the COVID-19 Genomics UK (COG-UK) Consortium | Jacquelyn Wynn, Mairead Hyland, The Lighthouse Lab in Alderley Park and Alex Alderton, Roberto Amato, Jeffrey Barrett, Sonia Goncalves, Ewan Harrison, David K. Jackson, Ian Johnston, Dominic Kwiatkowski, Cordelia Langford, John Sillitoe on behalf of the Wellcome Sanger Institute COVID-19 Surveillance Team |
| EPI_ISL_3110145 | Lighthouse Lab in Milton Keynes | Wellcome Sanger Institute for the COVID-19 Genomics UK (COG-UK) Consortium | The Lighthouse Lab in Milton Keynes and Alex Alderton, Roberto Amato, Jeffrey Barrett, Sonia Goncalves, Ewan Harrison, David K. Jackson, Ian Johnston, Dominic Kwiatkowski, Cordelia Langford, John Sillitoe on behalf of the Wellcome Sanger Institute COVID-19 Surveillance Team |
| EPI_ISL_3110146 | Lighthouse Lab in Alderley Park | Wellcome Sanger Institute for the COVID-19 Genomics UK (COG-UK) Consortium | Jacquelyn Wynn, Mairead Hyland, The Lighthouse Lab in Alderley Park and Alex Alderton, Roberto Amato, Jeffrey Barrett, Sonia Goncalves, Ewan Harrison, David K. Jackson, Ian Johnston, Dominic Kwiatkowski, Cordelia Langford, John Sillitoe on behalf of the Wellcome Sanger Institute COVID-19 Surveillance Team |
| EPI_ISL_3110147 | Lighthouse Laboratory Plymouth | Wellcome Sanger Institute for the COVID-19 Genomics UK (COG-UK) Consortium | Lighthouse Laboratory Plymouth and Alex Alderton, Roberto Amato, Jeffrey Barrett, Sonia Goncalves, Ewan Harrison, David K. Jackson, Ian Johnston, Dominic Kwiatkowski, Cordelia Langford, John Sillitoe on behalf of the Wellcome Sanger Institute COVID-19 Surveillance Team |
| EPI_ISL_3110148, EPI_ISL_3110149 | Lighthouse Lab in Alderley Park | Wellcome Sanger Institute for the COVID-19 Genomics UK (COG-UK) Consortium | Jacquelyn Wynn, Mairead Hyland, The Lighthouse Lab in Alderley Park and Alex Alderton, Roberto Amato, Jeffrey Barrett, Sonia Goncalves, Ewan Harrison, David K. Jackson, Ian Johnston, Dominic Kwiatkowski, Cordelia Langford, John Sillitoe on behalf of the Wellcome Sanger Institute COVID-19 Surveillance Team |
| EPI_ISL_3110150 | Lighthouse Laboratory Plymouth | Wellcome Sanger Institute for the COVID-19 Genomics UK (COG-UK) Consortium | Lighthouse Laboratory Plymouth and Alex Alderton, Roberto Amato, Jeffrey Barrett, Sonia Goncalves, Ewan Harrison, David K. Jackson, Ian Johnston, Dominic Kwiatkowski, Cordelia Langford, John Sillitoe on behalf of the Wellcome Sanger Institute COVID-19 Surveillance Team |
| EPI_ISL_3110151, EPI_ISL_3110152 | Lighthouse Lab in Glasgow | Wellcome Sanger Institute for the COVID-19 Genomics UK (COG-UK) Consortium | Harper VanSteenhouse, Yumi Kasai, David Gray, Carol Clugston, Anna Dominiczak and Alex Alderton, Roberto Amato, Jeffrey Barrett, Sonia Goncalves, Ewan Harrison, David K. Jackson, Ian Johnston, Dominic Kwiatkowski, Cordelia Langford, John Sillitoe on behalf of the Wellcome Sanger Institute COVID-19 Surveillance Team |
| EPI_ISL_3110153 | Lighthouse Lab in Alderley Park | Wellcome Sanger Institute for the COVID-19 Genomics UK (COG-UK) Consortium | Jacquelyn Wynn, Mairead Hyland, The Lighthouse Lab in Alderley Park and Alex Alderton, Roberto Amato, Jeffrey Barrett, Sonia Goncalves, Ewan Harrison, David K. Jackson, Ian Johnston, Dominic Kwiatkowski, Cordelia Langford, John Sillitoe on behalf of the Wellcome Sanger Institute COVID-19 Surveillance Team |
| EPI_ISL_3110154 | Lighthouse Laboratory Plymouth | Wellcome Sanger Institute for the COVID-19 Genomics UK (COG-UK) Consortium | Lighthouse Laboratory Plymouth and Alex Alderton, Roberto Amato, Jeffrey Barrett, Sonia Goncalves, Ewan Harrison, David K. Jackson, Ian Johnston, Dominic Kwiatkowski, Cordelia Langford, John Sillitoe on behalf of the Wellcome Sanger Institute COVID-19 Surveillance Team |

| | | | |
|---|---|---|---|
| EPI_ISL_3110155, EPI_ISL_3110156 | Lighthouse Lab in Alderley Park | Wellcome Sanger Institute for the COVID-19 Genomics UK (COG-UK) Consortium | Jacquelyn Wynn, Mairead Hyland, The Lighthouse Lab in Alderley Park and Alex Alderton, Roberto Amato, Jeffrey Barrett, Sonia Goncalves, Ewan Harrison, David K. Jackson, Ian Johnston, Dominic Kwiatkowski, Cordelia Langford, John Sillitoe on behalf of the Wellcome Sanger Institute COVID-19 Surveillance Team |
| EPI_ISL_3110157 | Lighthouse Lab in Milton Keynes | Wellcome Sanger Institute for the COVID-19 Genomics UK (COG-UK) Consortium | The Lighthouse Lab in Milton Keynes and Alex Alderton, Roberto Amato, Jeffrey Barrett, Sonia Goncalves, Ewan Harrison, David K. Jackson, Ian Johnston, Dominic Kwiatkowski, Cordelia Langford, John Sillitoe on behalf of the Wellcome Sanger Institute COVID-19 Surveillance Team |
| EPI_ISL_3110158 | Lighthouse Lab in Glasgow | Wellcome Sanger Institute for the COVID-19 Genomics UK (COG-UK) Consortium | Harper VanSteenhouse, Yumi Kasai, David Gray, Carol Clugston, Anna Dominiczak and Alex Alderton, Roberto Amato, Jeffrey Barrett, Sonia Goncalves, Ewan Harrison, David K. Jackson, Ian Johnston, Dominic Kwiatkowski, Cordelia Langford, John Sillitoe on behalf of the Wellcome Sanger Institute COVID-19 Surveillance Team |
| EPI_ISL_3110159 | Lighthouse Lab in Milton Keynes | Wellcome Sanger Institute for the COVID-19 Genomics UK (COG-UK) Consortium | The Lighthouse Lab in Milton Keynes and Alex Alderton, Roberto Amato, Jeffrey Barrett, Sonia Goncalves, Ewan Harrison, David K. Jackson, Ian Johnston, Dominic Kwiatkowski, Cordelia Langford, John Sillitoe on behalf of the Wellcome Sanger Institute COVID-19 Surveillance Team |
| EPI_ISL_3110160 | Lighthouse Laboratory Plymouth | Wellcome Sanger Institute for the COVID-19 Genomics UK (COG-UK) Consortium | Lighthouse Laboratory Plymouth and Alex Alderton, Roberto Amato, Jeffrey Barrett, Sonia Goncalves, Ewan Harrison, David K. Jackson, Ian Johnston, Dominic Kwiatkowski, Cordelia Langford, John Sillitoe on behalf of the Wellcome Sanger Institute COVID-19 Surveillance Team |
| EPI_ISL_3110161 | Lighthouse Lab in Alderley Park | Wellcome Sanger Institute for the COVID-19 Genomics UK (COG-UK) Consortium | Jacquelyn Wynn, Mairead Hyland, The Lighthouse Lab in Alderley Park and Alex Alderton, Roberto Amato, Jeffrey Barrett, Sonia Goncalves, Ewan Harrison, David K. Jackson, Ian Johnston, Dominic Kwiatkowski, Cordelia Langford, John Sillitoe on behalf of the Wellcome Sanger Institute COVID-19 Surveillance Team |
| EPI_ISL_3110162, EPI_ISL_3110163 | Lighthouse Lab in Milton Keynes | Wellcome Sanger Institute for the COVID-19 Genomics UK (COG-UK) Consortium | The Lighthouse Lab in Milton Keynes and Alex Alderton, Roberto Amato, Jeffrey Barrett, Sonia Goncalves, Ewan Harrison, David K. Jackson, Ian Johnston, Dominic Kwiatkowski, Cordelia Langford, John Sillitoe on behalf of the Wellcome Sanger Institute COVID-19 Surveillance Team |
| EPI_ISL_3110164 | Lighthouse Laboratory Plymouth | Wellcome Sanger Institute for the COVID-19 Genomics UK (COG-UK) Consortium | Lighthouse Laboratory Plymouth and Alex Alderton, Roberto Amato, Jeffrey Barrett, Sonia Goncalves, Ewan Harrison, David K. Jackson, Ian Johnston, Dominic Kwiatkowski, Cordelia Langford, John Sillitoe on behalf of the Wellcome Sanger Institute COVID-19 Surveillance Team |
| EPI_ISL_3110165 | Lighthouse Lab in Milton Keynes | Wellcome Sanger Institute for the COVID-19 Genomics UK (COG-UK) Consortium | The Lighthouse Lab in Milton Keynes and Alex Alderton, Roberto Amato, Jeffrey Barrett, Sonia Goncalves, Ewan Harrison, David K. Jackson, Ian Johnston, Dominic Kwiatkowski, Cordelia Langford, John Sillitoe on behalf of the Wellcome Sanger Institute COVID-19 Surveillance Team |
| EPI_ISL_3110166 | Lighthouse Laboratory Plymouth | Wellcome Sanger Institute for the COVID-19 Genomics UK (COG-UK) Consortium | Lighthouse Laboratory Plymouth and Alex Alderton, Roberto Amato, Jeffrey Barrett, Sonia Goncalves, Ewan Harrison, David K. Jackson, Ian Johnston, Dominic Kwiatkowski, Cordelia Langford, John Sillitoe on behalf of the Wellcome Sanger Institute COVID-19 Surveillance Team |
| EPI_ISL_3110167 | Lighthouse Lab in Glasgow | Wellcome Sanger Institute for the COVID-19 Genomics UK (COG-UK) Consortium | Harper VanSteenhouse, Yumi Kasai, David Gray, Carol Clugston, Anna Dominiczak and Alex Alderton, Roberto Amato, Jeffrey Barrett, Sonia Goncalves, Ewan Harrison, David K. Jackson, Ian Johnston, Dominic Kwiatkowski, Cordelia Langford, John Sillitoe on behalf of the Wellcome Sanger Institute COVID-19 Surveillance Team |
| EPI_ISL_3110168 | Lighthouse Lab in Alderley Park | Wellcome Sanger Institute for the COVID-19 Genomics UK (COG-UK) Consortium | Jacquelyn Wynn, Mairead Hyland, The Lighthouse Lab in Alderley Park and Alex Alderton, Roberto Amato, Jeffrey Barrett, Sonia Goncalves, Ewan Harrison, David K. Jackson, Ian Johnston, Dominic Kwiatkowski, Cordelia Langford, John Sillitoe on behalf of the Wellcome Sanger Institute COVID-19 Surveillance Team |
| EPI_ISL_3110169 | Lighthouse Lab in Milton Keynes | Wellcome Sanger Institute for the COVID-19 Genomics UK (COG-UK) Consortium | The Lighthouse Lab in Milton Keynes and Alex Alderton, Roberto Amato, Jeffrey Barrett, Sonia Goncalves, Ewan Harrison, David K. Jackson, Ian Johnston, Dominic Kwiatkowski, Cordelia Langford, John Sillitoe on behalf of the Wellcome Sanger Institute COVID-19 Surveillance Team |
| EPI_ISL_3110170, EPI_ISL_3110171 | Lighthouse Lab in Glasgow | Wellcome Sanger Institute for the COVID-19 Genomics UK (COG-UK) Consortium | Harper VanSteenhouse, Yumi Kasai, David Gray, Carol Clugston, Anna Dominiczak and Alex Alderton, Roberto Amato, Jeffrey Barrett, Sonia Goncalves, Ewan Harrison, David K. Jackson, Ian Johnston, Dominic Kwiatkowski, Cordelia Langford, John Sillitoe on behalf of the Wellcome Sanger Institute COVID-19 Surveillance Team |
| EPI_ISL_3110172 | Lighthouse Lab in Alderley Park | Wellcome Sanger Institute for the COVID-19 Genomics UK (COG-UK) Consortium | Jacquelyn Wynn, Mairead Hyland, The Lighthouse Lab in Alderley Park and Alex Alderton, Roberto Amato, Jeffrey Barrett, Sonia Goncalves, Ewan Harrison, David K. Jackson, Ian Johnston, Dominic Kwiatkowski, Cordelia Langford, John Sillitoe on behalf of the Wellcome Sanger Institute COVID-19 Surveillance Team |
| EPI_ISL_3110173 | Lighthouse Laboratory Plymouth | Wellcome Sanger Institute for the COVID-19 Genomics UK (COG-UK) Consortium | Lighthouse Laboratory Plymouth and Alex Alderton, Roberto Amato, Jeffrey Barrett, Sonia Goncalves, Ewan Harrison, David K. Jackson, Ian Johnston, Dominic Kwiatkowski, Cordelia Langford, John Sillitoe on behalf of the Wellcome Sanger Institute COVID-19 Surveillance Team |
| EPI_ISL_3110174 | Lighthouse Lab in Alderley Park | Wellcome Sanger Institute for the COVID-19 Genomics UK (COG-UK) Consortium | Jacquelyn Wynn, Mairead Hyland, The Lighthouse Lab in Alderley Park and Alex Alderton, Roberto Amato, Jeffrey Barrett, Sonia Goncalves, Ewan Harrison, David K. Jackson, Ian Johnston, Dominic Kwiatkowski, Cordelia Langford, John Sillitoe on behalf of the Wellcome Sanger Institute COVID-19 Surveillance Team |
| EPI_ISL_3110175 | Lighthouse Laboratory Plymouth | Wellcome Sanger Institute for the COVID-19 Genomics UK (COG-UK) Consortium | Lighthouse Laboratory Plymouth and Alex Alderton, Roberto Amato, Jeffrey Barrett, Sonia Goncalves, Ewan Harrison, David K. Jackson, Ian Johnston, Dominic Kwiatkowski, Cordelia Langford, John Sillitoe on behalf of the Wellcome Sanger Institute COVID-19 Surveillance Team |
| EPI_ISL_3110176 | Lighthouse Lab in Alderley Park | Wellcome Sanger Institute for the COVID-19 Genomics UK (COG-UK) Consortium | Jacquelyn Wynn, Mairead Hyland, The Lighthouse Lab in Alderley Park and Alex Alderton, Roberto Amato, Jeffrey Barrett, Sonia Goncalves, Ewan Harrison, David K. Jackson, Ian Johnston, Dominic Kwiatkowski, Cordelia Langford, John Sillitoe on behalf of the Wellcome Sanger Institute COVID-19 Surveillance Team |
| EPI_ISL_3110177 | Lighthouse Laboratory Plymouth | Wellcome Sanger Institute for the COVID-19 Genomics UK (COG-UK) Consortium | Lighthouse Laboratory Plymouth and Alex Alderton, Roberto Amato, Jeffrey Barrett, Sonia Goncalves, Ewan Harrison, David K. Jackson, Ian Johnston, Dominic Kwiatkowski, Cordelia Langford, John Sillitoe on behalf of the Wellcome Sanger Institute COVID-19 Surveillance Team |
| EPI_ISL_3110178 | Lighthouse Lab in Alderley Park | Wellcome Sanger Institute for the COVID-19 Genomics UK (COG-UK) Consortium | Jacquelyn Wynn, Mairead Hyland, The Lighthouse Lab in Alderley Park and Alex Alderton, Roberto Amato, Jeffrey Barrett, Sonia Goncalves, Ewan Harrison, David K. Jackson, Ian Johnston, Dominic Kwiatkowski, Cordelia Langford, John Sillitoe on behalf of the Wellcome Sanger Institute COVID-19 Surveillance Team |
| EPI_ISL_3110179 | Lighthouse Lab in Milton Keynes | Wellcome Sanger Institute for the COVID-19 Genomics UK (COG-UK) Consortium | The Lighthouse Lab in Milton Keynes and Alex Alderton, Roberto Amato, Jeffrey Barrett, Sonia Goncalves, Ewan Harrison, David K. Jackson, Ian Johnston, Dominic Kwiatkowski, Cordelia Langford, John Sillitoe on behalf of the Wellcome Sanger Institute COVID-19 Surveillance Team |
| EPI_ISL_3110180, EPI_ISL_3110181 | Lighthouse Lab in Alderley Park | Wellcome Sanger Institute for the COVID-19 Genomics UK (COG-UK) Consortium | Jacquelyn Wynn, Mairead Hyland, The Lighthouse Lab in Alderley Park and Alex Alderton, Roberto Amato, Jeffrey Barrett, Sonia Goncalves, Ewan Harrison, David K. Jackson, Ian Johnston, Dominic Kwiatkowski, Cordelia Langford, John Sillitoe on behalf of the Wellcome Sanger Institute COVID-19 Surveillance Team |
| EPI_ISL_3110182, EPI_ISL_3110183 | Lighthouse Laboratory Plymouth | Wellcome Sanger Institute for the COVID-19 Genomics UK (COG-UK) Consortium | Lighthouse Laboratory Plymouth and Alex Alderton, Roberto Amato, Jeffrey Barrett, Sonia Goncalves, Ewan Harrison, David K. Jackson, Ian Johnston, Dominic Kwiatkowski, Cordelia Langford, John Sillitoe on behalf of the Wellcome Sanger Institute COVID-19 Surveillance Team |
| EPI_ISL_3110184, EPI_ISL_3110185 | Lighthouse Lab in Milton Keynes | Wellcome Sanger Institute for the COVID-19 Genomics UK (COG-UK) Consortium | The Lighthouse Lab in Milton Keynes and Alex Alderton, Roberto Amato, Jeffrey Barrett, Sonia Goncalves, Ewan Harrison, David K. Jackson, Ian Johnston, Dominic Kwiatkowski, Cordelia Langford, John Sillitoe on behalf of the Wellcome Sanger Institute COVID-19 Surveillance Team |
| EPI_ISL_3110186, EPI_ISL_3110187, EPI_ISL_3110188 | Lighthouse Lab in Alderley Park | Wellcome Sanger Institute for the COVID-19 Genomics UK (COG-UK) Consortium | Jacquelyn Wynn, Mairead Hyland, The Lighthouse Lab in Alderley Park and Alex Alderton, Roberto Amato, Jeffrey Barrett, Sonia Goncalves, Ewan Harrison, David K. Jackson, Ian Johnston, Dominic Kwiatkowski, Cordelia Langford, John Sillitoe on behalf of the Wellcome Sanger Institute COVID-19 Surveillance Team |
| EPI_ISL_3110189 | Lighthouse Laboratory Plymouth | Wellcome Sanger Institute for the COVID-19 Genomics UK (COG-UK) Consortium | Lighthouse Laboratory Plymouth and Alex Alderton, Roberto Amato, Jeffrey Barrett, Sonia Goncalves, Ewan Harrison, David K. Jackson, Ian Johnston, Dominic Kwiatkowski, Cordelia Langford, John Sillitoe on behalf of the Wellcome Sanger Institute COVID-19 Surveillance Team |
| EPI_ISL_3110190 | Lighthouse Lab in Alderley Park | Wellcome Sanger Institute for the COVID-19 Genomics UK (COG-UK) Consortium | Jacquelyn Wynn, Mairead Hyland, The Lighthouse Lab in Alderley Park and Alex Alderton, Roberto Amato, Jeffrey Barrett, Sonia Goncalves, Ewan Harrison, David K. Jackson, Ian Johnston, Dominic Kwiatkowski, Cordelia Langford, John Sillitoe on behalf of the Wellcome Sanger Institute COVID-19 Surveillance Team |
| EPI_ISL_3110191, EPI_ISL_3110192 | Lighthouse Laboratory Plymouth | Wellcome Sanger Institute for the COVID-19 Genomics UK (COG-UK) Consortium | Lighthouse Laboratory Plymouth and Alex Alderton, Roberto Amato, Jeffrey Barrett, Sonia Goncalves, Ewan Harrison, David K. Jackson, Ian Johnston, Dominic Kwiatkowski, Cordelia Langford, John Sillitoe on behalf of the Wellcome Sanger Institute COVID-19 Surveillance Team |
| EPI_ISL_3110193 | Lighthouse Lab in Alderley Park | Wellcome Sanger Institute for the COVID-19 Genomics UK (COG-UK) Consortium | Jacquelyn Wynn, Mairead Hyland, The Lighthouse Lab in Alderley Park and Alex Alderton, Roberto Amato, Jeffrey Barrett, Sonia Goncalves, Ewan Harrison, David K. Jackson, Ian Johnston, Dominic Kwiatkowski, Cordelia Langford, John Sillitoe on behalf of the Wellcome Sanger Institute COVID-19 Surveillance Team |
| EPI_ISL_3110194, EPI_ISL_3110195 | Lighthouse Lab in Milton Keynes | Wellcome Sanger Institute for the COVID-19 Genomics UK (COG-UK) Consortium | The Lighthouse Lab in Milton Keynes and Alex Alderton, Roberto Amato, Jeffrey Barrett, Sonia Goncalves, Ewan Harrison, David K. Jackson, Ian Johnston, Dominic Kwiatkowski, Cordelia Langford, John Sillitoe on behalf of the Wellcome Sanger Institute COVID-19 Surveillance Team |
| EPI_ISL_3110196 | Lighthouse Lab in Alderley Park | Wellcome Sanger Institute for the COVID-19 Genomics UK (COG-UK) Consortium | Jacquelyn Wynn, Mairead Hyland, The Lighthouse Lab in Alderley Park and Alex Alderton, Roberto Amato, Jeffrey Barrett, Sonia Goncalves, Ewan Harrison, David K. Jackson, Ian Johnston, Dominic Kwiatkowski, Cordelia Langford, John Sillitoe on behalf of the Wellcome Sanger Institute COVID-19 Surveillance Team |
| EPI_ISL_3110197, EPI_ISL_3110198 | Lighthouse Laboratory Plymouth | Wellcome Sanger Institute for the COVID-19 Genomics UK (COG-UK) Consortium | Lighthouse Laboratory Plymouth and Alex Alderton, Roberto Amato, Jeffrey Barrett, Sonia Goncalves, Ewan Harrison, David K. Jackson, Ian Johnston, Dominic Kwiatkowski, Cordelia Langford, John Sillitoe on behalf of the Wellcome Sanger Institute COVID-19 Surveillance Team |
| EPI_ISL_3110199 | Lighthouse Lab in Alderley Park | Wellcome Sanger Institute for the COVID-19 Genomics UK (COG-UK) Consortium | Jacquelyn Wynn, Mairead Hyland, The Lighthouse Lab in Alderley Park and Alex Alderton, Roberto Amato, Jeffrey Barrett, Sonia Goncalves, Ewan Harrison, David K. Jackson, Ian Johnston, Dominic Kwiatkowski, Cordelia Langford, John Sillitoe on behalf of the Wellcome Sanger Institute COVID-19 Surveillance Team |
| EPI_ISL_3110200 | Lighthouse Lab in Milton Keynes | Wellcome Sanger Institute for the COVID-19 Genomics UK (COG-UK) Consortium | The Lighthouse Lab in Milton Keynes and Alex Alderton, Roberto Amato, Jeffrey Barrett, Sonia Goncalves, Ewan Harrison, David K. Jackson, Ian Johnston, Dominic Kwiatkowski, Cordelia Langford, John Sillitoe on behalf of the Wellcome Sanger Institute COVID-19 Surveillance Team |
| EPI_ISL_3110201 | Lighthouse Lab in Alderley Park | Wellcome Sanger Institute for the COVID-19 Genomics UK (COG-UK) Consortium | Jacquelyn Wynn, Mairead Hyland, The Lighthouse Lab in Alderley Park and Alex Alderton, Roberto Amato, Jeffrey Barrett, Sonia Goncalves, Ewan Harrison, David K. Jackson, Ian Johnston, Dominic Kwiatkowski, Cordelia Langford, John Sillitoe on behalf of the Wellcome Sanger Institute COVID-19 Surveillance Team |
| EPI_ISL_3110202 | Lighthouse Laboratory Plymouth | Wellcome Sanger Institute for the COVID-19 Genomics UK (COG-UK) Consortium | Lighthouse Laboratory Plymouth and Alex Alderton, Roberto Amato, Jeffrey Barrett, Sonia Goncalves, Ewan Harrison, David K. Jackson, Ian Johnston, Dominic Kwiatkowski, Cordelia Langford, John Sillitoe on behalf of the Wellcome Sanger Institute COVID-19 Surveillance Team |
| EPI_ISL_3110203, EPI_ISL_3110204, EPI_ISL_3110205, EPI_ISL_3110206, EPI_ISL_3110207, EPI_ISL_3110208, EPI_ISL_3110209, EPI_ISL_3110210, EPI_ISL_3110211, EPI_ISL_3110212, EPI_ISL_3110213, EPI_ISL_3110214, EPI_ISL_3110215, EPI_ISL_3110216, EPI_ISL_3110217, EPI_ISL_3110218, EPI_ISL_3110219, EPI_ISL_3110220, EPI_ISL_3110221, EPI_ISL_3110222, EPI_ISL_3110223, EPI_ISL_3110224, EPI_ISL_3110225, EPI_ISL_3110226, EPI_ISL_3110227, EPI_ISL_3110228, EPI_ISL_3110229, EPI_ISL_3110230, EPI_ISL_3110231, EPI_ISL_3110232, EPI_ISL_3110233, EPI_ISL_3110234, EPI_ISL_3110235, EPI_ISL_3110236, EPI_ISL_3110237, EPI_ISL_3110238, EPI_ISL_3110239, EPI_ISL_3110240, EPI_ISL_3110241, EPI_ISL_3110242, EPI_ISL_3110243, EPI_ISL_3110244, EPI_ISL_3110245, EPI_ISL_3110246, EPI_ISL_3110247, EPI_ISL_3110248, EPI_ISL_3110249, EPI_ISL_3110250, EPI_ISL_3110251, EPI_ISL_3110252, EPI_ISL_3110253, EPI_ISL_3110254, EPI_ISL_3110255, EPI_ISL_3110256, EPI_ISL_3110257, EPI_ISL_3110258, EPI_ISL_3110259, EPI_ISL_3110260, EPI_ISL_3110261, EPI_ISL_3110262, EPI_ISL_3110263, EPI_ISL_3110264, EPI_ISL_3110265, EPI_ISL_3110266, EPI_ISL_3110267, EPI_ISL_3110268, EPI_ISL_3110269, EPI_ISL_3110270 | | | |
| see above | Lighthouse Lab in Glasgow | Wellcome Sanger Institute for the COVID-19 Genomics UK (COG-UK) Consortium | Harper VanSteenhouse, Yumi Kasai, David Gray, Carol Clugston, Anna Dominiczak and Alex Alderton, Roberto Amato, Jeffrey Barrett, Sonia Goncalves, Ewan Harrison, David K. Jackson, Ian Johnston, Dominic Kwiatkowski, Cordelia Langford, John Sillitoe on behalf of the Wellcome Sanger Institute COVID-19 Surveillance Team |
| EPI_ISL_3110272, EPI_ISL_3110273, EPI_ISL_3110274, EPI_ISL_3110275, EPI_ISL_3110276, EPI_ISL_3110277, EPI_ISL_3110278, EPI_ISL_3110279, EPI_ISL_3110280, EPI_ISL_3110281, EPI_ISL_3110282, EPI_ISL_3110283, EPI_ISL_3110284, EPI_ISL_3110285, EPI_ISL_3110286, EPI_ISL_3110288, EPI_ISL_3110289, EPI_ISL_3110291, EPI_ISL_3110292, EPI_ISL_3110293, EPI_ISL_3110294, EPI_ISL_3110295, EPI_ISL_3110296, EPI_ISL_3110297, EPI_ISL_3110298, EPI_ISL_3110299, EPI_ISL_3110300, EPI_ISL_3110301, EPI_ISL_3110302, EPI_ISL_3110303, EPI_ISL_3110304, EPI_ISL_3110305, EPI_ISL_3110306, EPI_ISL_3110307, EPI_ISL_3110308, EPI_ISL_3110309, EPI_ISL_3110310, EPI_ISL_3110311, EPI_ISL_3110312, EPI_ISL_3110313, EPI_ISL_3110314, EPI_ISL_3110316, EPI_ISL_3110317, EPI_ISL_3110318, EPI_ISL_3110319, EPI_ISL_3110320, EPI_ISL_3110321, EPI_ISL_3110322, EPI_ISL_3110323, EPI_ISL_3110324, EPI_ISL_3110325, EPI_ISL_3110326, EPI_ISL_3110327, EPI_ISL_3110328, EPI_ISL_3110329, EPI_ISL_3110330, EPI_ISL_3110331, EPI_ISL_3110332, EPI_ISL_3110333, EPI_ISL_3110334, EPI_ISL_3110335, EPI_ISL_3110336, EPI_ISL_3110337, EPI_ISL_3110338, EPI_ISL_3110339, EPI_ISL_3110340, EPI_ISL_3110341, EPI_ISL_3110342, EPI_ISL_3110343, EPI_ISL_3110344, EPI_ISL_3110346, EPI_ISL_3110347, EPI_ISL_3110348, EPI_ISL_3110349, EPI_ISL_3110350, EPI_ISL_3110351, EPI_ISL_3110352, EPI_ISL_3110353, EPI_ISL_3110354, EPI_ISL_3110355, EPI_ISL_3110356, EPI_ISL_3110357, EPI_ISL_3110358, EPI_ISL_3110359, EPI_ISL_3110360, EPI_ISL_3110361, EPI_ISL_3110362, EPI_ISL_3110363, EPI_ISL_3110364, EPI_ISL_3110365, EPI_ISL_3110366, EPI_ISL_3110367, EPI_ISL_3110368, EPI_ISL_3110369, EPI_ISL_3110370, EPI_ISL_3110371, EPI_ISL_3110372, EPI_ISL_3110373, EPI_ISL_3110374, EPI_ISL_3110375, EPI_ISL_3110376, EPI_ISL_3110377, EPI_ISL_3110378, EPI_ISL_3110379, EPI_ISL_3110380, EPI_ISL_3110381, EPI_ISL_3110382, EPI_ISL_3110383, EPI_ISL_3110384, EPI_ISL_3110385, EPI_ISL_3110386, EPI_ISL_3110387, EPI_ISL_3110389, EPI_ISL_3110390, EPI_ISL_3110391, EPI_ISL_3110392, EPI_ISL_3110393, EPI_ISL_3110394, EPI_ISL_3110395, EPI_ISL_3110396, EPI_ISL_3110397, EPI_ISL_3110398, EPI_ISL_3110399, EPI_ISL_3110400, EPI_ISL_3110401, EPI_ISL_3110402, EPI_ISL_3110403, EPI_ISL_3110404, EPI_ISL_3110405, EPI_ISL_3110406, EPI_ISL_3110407, EPI_ISL_3110408, EPI_ISL_3110409, EPI_ISL_3110410, EPI_ISL_3110411, EPI_ISL_3110412, EPI_ISL_3110413, EPI_ISL_3110414, EPI_ISL_3110415, EPI_ISL_3110416, EPI_ISL_3110417, EPI_ISL_3110418, EPI_ISL_3110419, EPI_ISL_3110420, EPI_ISL_3110421, EPI_ISL_3110422, EPI_ISL_3110423, EPI_ISL_3110424, EPI_ISL_3110425, EPI_ISL_3110426, EPI_ISL_3110427, EPI_ISL_3110428, EPI_ISL_3110429, EPI_ISL_3110430, EPI_ISL_3110431, EPI_ISL_3110432, EPI_ISL_3110433, EPI_ISL_3110434, EPI_ISL_3110435, EPI_ISL_3110436, EPI_ISL_3110438, EPI_ISL_3110439, EPI_ISL_3110440, EPI_ISL_3110441, EPI_ISL_3110442, EPI_ISL_3110443, EPI_ISL_3110444, EPI_ISL_3110445, EPI_ISL_3110446, EPI_ISL_3110447, EPI_ISL_3110448, EPI_ISL_3110449, EPI_ISL_3110450, EPI_ISL_3110451, EPI_ISL_3110453, EPI_ISL_3110454, EPI_ISL_3110455, EPI_ISL_3110456, EPI_ISL_3110457, EPI_ISL_3110458, EPI_ISL_3110459, EPI_ISL_3110460, EPI_ISL_3110461, EPI_ISL_3110462, EPI_ISL_3110463, EPI_ISL_3110464, EPI_ISL_3110465, EPI_ISL_3110466, EPI_ISL_3110467, EPI_ISL_3110468, EPI_ISL_3110470, EPI_ISL_3110471, EPI_ISL_3110472, EPI_ISL_3110473, EPI_ISL_3110474, EPI_ISL_3110475, EPI_ISL_3110476, EPI_ISL_3110477, EPI_ISL_3110478, EPI_ISL_3110479, EPI_ISL_3110480, EPI_ISL_3110481, EPI_ISL_3110482, EPI_ISL_3110484, EPI_ISL_3110486, EPI_ISL_3110487, EPI_ISL_3110488, EPI_ISL_3110489, EPI_ISL_3110490, EPI_ISL_3110491, EPI_ISL_3110492, EPI_ISL_3110493, EPI_ISL_3110494, EPI_ISL_3110495, EPI_ISL_3110496, EPI_ISL_3110497, EPI_ISL_3110498, EPI_ISL_3110499, EPI_ISL_3110500, EPI_ISL_3110502, EPI_ISL_3110503, EPI_ISL_3110504, EPI_ISL_3110505, EPI_ISL_3110506, EPI_ISL_3110507, EPI_ISL_3110508, EPI_ISL_3110509, EPI_ISL_3110510, EPI_ISL_3110511, EPI_ISL_3110512, EPI_ISL_3110513, EPI_ISL_3110514, EPI_ISL_3110515, EPI_ISL_3110516, EPI_ISL_3110517, EPI_ISL_3110518, EPI_ISL_3110519, EPI_ISL_3110520, EPI_ISL_3110521, EPI_ISL_3110522, EPI_ISL_3110523, EPI_ISL_3110524, EPI_ISL_3110525, EPI_ISL_3110526, EPI_ISL_3110527, EPI_ISL_3110528, EPI_ISL_3110529, EPI_ISL_3110530, EPI_ISL_3110531, EPI_ISL_3110532, EPI_ISL_3110533, EPI_ISL_3110534, EPI_ISL_3110535, EPI_ISL_3110536, EPI_ISL_3110537, EPI_ISL_3110538, EPI_ISL_3110539, EPI_ISL_3110540, EPI_ISL_3110541, EPI_ISL_3110542, EPI_ISL_3110543, EPI_ISL_3110544, EPI_ISL_3110545, EPI_ISL_3110546, | | | |

| | | | |
|---|---|---|---|
| EPI_ISL_3110547, EPI_ISL_3110548, EPI_ISL_3110549, EPI_ISL_3110550, EPI_ISL_3110551, EPI_ISL_3110552, EPI_ISL_3110553, EPI_ISL_3110555, EPI_ISL_3110556, EPI_ISL_3110557, EPI_ISL_3110558, EPI_ISL_3110560, EPI_ISL_3110561, EPI_ISL_3110562, EPI_ISL_3110563, EPI_ISL_3110565, EPI_ISL_3110567, EPI_ISL_3110568, EPI_ISL_3110569, EPI_ISL_3110570, EPI_ISL_3110571, EPI_ISL_3110572, EPI_ISL_3110573, EPI_ISL_3110574, EPI_ISL_3110575, EPI_ISL_3110576, EPI_ISL_3110577, EPI_ISL_3110578, EPI_ISL_3110579, EPI_ISL_3110580, EPI_ISL_3110581, EPI_ISL_3110582, EPI_ISL_3110583, EPI_ISL_3110584 | | | |
| see above | Fulgent Genetics | Centers for Disease Control and Prevention Division of Viral Diseases, Pathogen Discovery | Dakota Howard, Dhwani Batra, Peter W. Cook, Kara Moser, Adrian Paskey, Jason Caravas, Benjamin Rambo-Martin, Shatavia Morrison, Christopher Gulvick, Scott Sammons, Yvette Unoarumhi, Darlene Wagner, Matthew Schmerer, Harry Gao, Mickey Li, John Gao, Joseph Fierro, Benafsh Sapra, Becky Tsai, Yan Meng, Doreen Ng, James Xie, Clinton R. Paden, Duncan MacCannell |
| EPI_ISL_3110585, EPI_ISL_3110586, EPI_ISL_3110587, EPI_ISL_3110588, EPI_ISL_3110589, EPI_ISL_3110590, EPI_ISL_3110591, EPI_ISL_3110592, EPI_ISL_3110593, EPI_ISL_3110594, EPI_ISL_3110595, EPI_ISL_3110596, EPI_ISL_3110597, EPI_ISL_3110598, EPI_ISL_3110599, EPI_ISL_3110601, EPI_ISL_3110602, EPI_ISL_3110603, EPI_ISL_3110604, EPI_ISL_3110605, EPI_ISL_3110606, EPI_ISL_3110607, EPI_ISL_3110608, EPI_ISL_3110609, EPI_ISL_3110610, EPI_ISL_3110611, EPI_ISL_3110612, EPI_ISL_3110613, EPI_ISL_3110614, EPI_ISL_3110615, EPI_ISL_3110616, EPI_ISL_3110618, EPI_ISL_3110619, EPI_ISL_3110620, EPI_ISL_3110621, EPI_ISL_3110622, EPI_ISL_3110623, EPI_ISL_3110624, EPI_ISL_3110625, EPI_ISL_3110626, EPI_ISL_3110628, EPI_ISL_3110629, EPI_ISL_3110631, EPI_ISL_3110632, EPI_ISL_3110633, EPI_ISL_3110634, EPI_ISL_3110635, EPI_ISL_3110636, EPI_ISL_3110637, EPI_ISL_3110639, EPI_ISL_3110640, EPI_ISL_3110641, EPI_ISL_3110642, EPI_ISL_3110643, EPI_ISL_3110644, EPI_ISL_3110645, EPI_ISL_3110646, EPI_ISL_3110647, EPI_ISL_3110649, EPI_ISL_3110650, EPI_ISL_3110652, EPI_ISL_3110653, EPI_ISL_3110654, EPI_ISL_3110655, EPI_ISL_3110656, EPI_ISL_3110657, EPI_ISL_3110658, EPI_ISL_3110659, EPI_ISL_3110660, EPI_ISL_3110661, EPI_ISL_3110663, EPI_ISL_3110664, EPI_ISL_3110665, EPI_ISL_3110666, EPI_ISL_3110667, EPI_ISL_3110668, EPI_ISL_3110669, EPI_ISL_3110671, EPI_ISL_3110672, EPI_ISL_3110673, EPI_ISL_3110674, EPI_ISL_3110675, EPI_ISL_3110676, EPI_ISL_3110677, EPI_ISL_3110678, EPI_ISL_3110679, EPI_ISL_3110680, EPI_ISL_3110681, EPI_ISL_3110682, EPI_ISL_3110683, EPI_ISL_3110684, EPI_ISL_3110685, EPI_ISL_3110686, EPI_ISL_3110687, EPI_ISL_3110688, EPI_ISL_3110689, EPI_ISL_3110690, EPI_ISL_3110691, EPI_ISL_3110693, EPI_ISL_3110694, EPI_ISL_3110695, EPI_ISL_3110696, EPI_ISL_3110698, EPI_ISL_3110699, EPI_ISL_3110700, EPI_ISL_3110701, EPI_ISL_3110702, EPI_ISL_3110703, EPI_ISL_3110704, EPI_ISL_3110705, EPI_ISL_3110706, EPI_ISL_3110707, EPI_ISL_3110709, EPI_ISL_3110710, EPI_ISL_3110711, EPI_ISL_3110712, EPI_ISL_3110713, EPI_ISL_3110714, EPI_ISL_3110715, EPI_ISL_3110716, EPI_ISL_3110718, EPI_ISL_3110719, EPI_ISL_3110720, EPI_ISL_3110721, EPI_ISL_3110722, EPI_ISL_3110723, EPI_ISL_3110724, EPI_ISL_3110725, EPI_ISL_3110727, EPI_ISL_3110728, EPI_ISL_3110730, EPI_ISL_3110731, EPI_ISL_3110732, EPI_ISL_3110733, EPI_ISL_3110734, EPI_ISL_3110735, EPI_ISL_3110736, EPI_ISL_3110737, EPI_ISL_3110738, EPI_ISL_3110739, EPI_ISL_3110740, EPI_ISL_3110741, EPI_ISL_3110742, EPI_ISL_3110743, EPI_ISL_3110745, EPI_ISL_3110746, EPI_ISL_3110747, EPI_ISL_3110751, EPI_ISL_3110752, EPI_ISL_3110753, EPI_ISL_3110754, EPI_ISL_3110755, EPI_ISL_3110757, EPI_ISL_3110758, EPI_ISL_3110759, EPI_ISL_3110761, EPI_ISL_3110762, EPI_ISL_3110763, EPI_ISL_3110764, EPI_ISL_3110765, EPI_ISL_3110767, EPI_ISL_3110771, EPI_ISL_3110772, EPI_ISL_3110773, EPI_ISL_3110774, EPI_ISL_3110775, EPI_ISL_3110776, EPI_ISL_3110777, EPI_ISL_3110778, EPI_ISL_3110779, EPI_ISL_3110780, EPI_ISL_3110783, EPI_ISL_3110784, EPI_ISL_3110785, EPI_ISL_3110786, EPI_ISL_3110787, EPI_ISL_3110788, EPI_ISL_3110789, EPI_ISL_3110791, EPI_ISL_3110792, EPI_ISL_3110793, EPI_ISL_3110794, EPI_ISL_3110795, EPI_ISL_3110796, EPI_ISL_3110799, EPI_ISL_3110800, EPI_ISL_3110801, EPI_ISL_3110802, EPI_ISL_3110803, EPI_ISL_3110804, EPI_ISL_3110805, EPI_ISL_3110806, EPI_ISL_3110807, EPI_ISL_3110808, EPI_ISL_3110809, EPI_ISL_3110810, EPI_ISL_3110811, EPI_ISL_3110812, EPI_ISL_3110813, EPI_ISL_3110814, EPI_ISL_3110815, EPI_ISL_3110816, EPI_ISL_3110817, EPI_ISL_3110818, EPI_ISL_3110819, EPI_ISL_3110822, EPI_ISL_3110823, EPI_ISL_3110824, EPI_ISL_3110826, EPI_ISL_3110827, EPI_ISL_3110828, EPI_ISL_3110829, EPI_ISL_3110830, EPI_ISL_3110832, EPI_ISL_3110833, EPI_ISL_3110834, EPI_ISL_3110835, EPI_ISL_3110836, EPI_ISL_3110838, EPI_ISL_3110839, EPI_ISL_3110840, EPI_ISL_3110841, EPI_ISL_3110842, EPI_ISL_3110843, EPI_ISL_3110845, EPI_ISL_3110846, EPI_ISL_3110847, EPI_ISL_3110849, EPI_ISL_3110850, EPI_ISL_3110852, EPI_ISL_3110854, EPI_ISL_3110855, EPI_ISL_3110856, EPI_ISL_3110857, EPI_ISL_3110858, EPI_ISL_3110861, EPI_ISL_3110862, EPI_ISL_3110863, EPI_ISL_3110865, EPI_ISL_3110866, EPI_ISL_3110867, EPI_ISL_3110868, EPI_ISL_3110869, EPI_ISL_3110870, EPI_ISL_3110872, EPI_ISL_3110873, EPI_ISL_3110874, EPI_ISL_3110876, EPI_ISL_3110877, EPI_ISL_3110878, EPI_ISL_3110879, EPI_ISL_3110880, EPI_ISL_3110881, EPI_ISL_3110882, EPI_ISL_3110883, EPI_ISL_3110884, EPI_ISL_3110885, EPI_ISL_3110886, EPI_ISL_3110887, EPI_ISL_3110888, EPI_ISL_3110889, EPI_ISL_3110890, EPI_ISL_3110891, EPI_ISL_3110892, EPI_ISL_3110894, EPI_ISL_3110895, EPI_ISL_3110898, EPI_ISL_3110899, EPI_ISL_3110900, EPI_ISL_3110902, EPI_ISL_3110903, EPI_ISL_3110904, EPI_ISL_3110906, EPI_ISL_3110907, EPI_ISL_3110908, EPI_ISL_3110909, EPI_ISL_3110911, EPI_ISL_3110912, EPI_ISL_3110913, EPI_ISL_3110914, EPI_ISL_3110915, EPI_ISL_3110916, EPI_ISL_3110917, EPI_ISL_3110918, EPI_ISL_3110920, EPI_ISL_3110922, EPI_ISL_3110924, EPI_ISL_3110925, EPI_ISL_3110927, EPI_ISL_3110928, EPI_ISL_3110929, EPI_ISL_3110930, EPI_ISL_3110931, EPI_ISL_3110932, EPI_ISL_3110933, EPI_ISL_3110934, EPI_ISL_3110935, EPI_ISL_3110936, EPI_ISL_3110937, EPI_ISL_3110938, EPI_ISL_3110939, EPI_ISL_3110940, EPI_ISL_3110941, EPI_ISL_3110943, EPI_ISL_3110944, EPI_ISL_3110945, EPI_ISL_3110946, EPI_ISL_3110947, EPI_ISL_3110949, EPI_ISL_3110950, EPI_ISL_3110951, EPI_ISL_3110952, EPI_ISL_3110953, EPI_ISL_3110954, EPI_ISL_3110955, EPI_ISL_3110956, EPI_ISL_3110958, EPI_ISL_3110959, EPI_ISL_3110961, EPI_ISL_3110962, EPI_ISL_3110963, EPI_ISL_3110964, EPI_ISL_3110965, EPI_ISL_3110966, EPI_ISL_3110967, EPI_ISL_3110968, EPI_ISL_3110969, EPI_ISL_3110971, EPI_ISL_3110972, EPI_ISL_3110973, EPI_ISL_3110974, EPI_ISL_3110976, EPI_ISL_3110979, EPI_ISL_3110980 | | | |
| see above | Mako Medical | Centers for Disease Control and Prevention Division of Viral Diseases, Pathogen Discovery | Dakota Howard, Dhwani Batra, Peter W. Cook, Kara Moser, Adrian Paskey, Jason Caravas, Benjamin Rambo-Martin, Shatavia Morrison, Christopher Gulvick, Scott Sammons, Yvette Unoarumhi, Darlene Wagner, Matthew Schmerer,Matthew Tugwell, Lauren Moon, Clinton R. Paden, Duncan MacCannell |
| EPI_ISL_3110982, EPI_ISL_3110983, EPI_ISL_3110984, EPI_ISL_3110985, EPI_ISL_3110986, EPI_ISL_3110988, EPI_ISL_3110989, EPI_ISL_3110990, EPI_ISL_3110991, EPI_ISL_3110992, EPI_ISL_3110993, EPI_ISL_3110994, EPI_ISL_3110996, EPI_ISL_3110997, EPI_ISL_3110998, EPI_ISL_3110999, EPI_ISL_3111000, EPI_ISL_3111001, EPI_ISL_3111002, EPI_ISL_3111003, EPI_ISL_3111005, EPI_ISL_3111006, EPI_ISL_3111007, EPI_ISL_3111009, EPI_ISL_3111010, EPI_ISL_3111011, EPI_ISL_3111012, EPI_ISL_3111013, EPI_ISL_3111015, EPI_ISL_3111016, EPI_ISL_3111017, EPI_ISL_3111018, EPI_ISL_3111019, EPI_ISL_3111020, EPI_ISL_3111021, EPI_ISL_3111022, EPI_ISL_3111023, EPI_ISL_3111025, EPI_ISL_3111026, EPI_ISL_3111027, EPI_ISL_3111028, EPI_ISL_3111029, EPI_ISL_3111030, EPI_ISL_3111031, EPI_ISL_3111033, EPI_ISL_3111034, EPI_ISL_3111036, EPI_ISL_3111037, EPI_ISL_3111038, EPI_ISL_3111039, EPI_ISL_3111041, EPI_ISL_3111042, EPI_ISL_3111043, EPI_ISL_3111044, EPI_ISL_3111045, EPI_ISL_3111046, EPI_ISL_3111047, EPI_ISL_3111048, EPI_ISL_3111049, EPI_ISL_3111050, EPI_ISL_3111051, EPI_ISL_3111052, EPI_ISL_3111053, EPI_ISL_3111054, EPI_ISL_3111055, EPI_ISL_3111056, EPI_ISL_3111057, EPI_ISL_3111058, EPI_ISL_3111059, EPI_ISL_3111060, EPI_ISL_3111061, EPI_ISL_3111062, EPI_ISL_3111063, EPI_ISL_3111064, EPI_ISL_3111065, EPI_ISL_3111066, EPI_ISL_3111067, EPI_ISL_3111068, EPI_ISL_3111069, EPI_ISL_3111070, EPI_ISL_3111071, EPI_ISL_3111072, EPI_ISL_3111074, EPI_ISL_3111075, EPI_ISL_3111076, EPI_ISL_3111077, EPI_ISL_3111078, EPI_ISL_3111079, EPI_ISL_3111080, EPI_ISL_3111081, EPI_ISL_3111084, EPI_ISL_3111085, EPI_ISL_3111086, EPI_ISL_3111087, EPI_ISL_3111089, EPI_ISL_3111090, EPI_ISL_3111091, EPI_ISL_3111092, EPI_ISL_3111093 | | | |
| see above | Infinity Biologix | Centers for Disease Control and Prevention Division of Viral Diseases, Pathogen Discovery | Dakota Howard, Dhwani Batra, Peter W. Cook, Kara Moser, Adrian Paskey, Jason Caravas, Benjamin Rambo-Martin, Shatavia Morrison, Christopher Gulvick, Scott Sammons, Yvette Unoarumhi, Darlene Wagner, Matthew Schmerer, Christian Bixby, Yihe Wang, Jonathan Schultz, Chirayu Goswami, Russ Hager, Robin Grimwood, Clinton R. Paden, Duncan MacCannell |
| EPI_ISL_3111094, EPI_ISL_3111095, EPI_ISL_3111096, EPI_ISL_3111097, EPI_ISL_3111098, EPI_ISL_3111099, EPI_ISL_3111100, EPI_ISL_3111101, EPI_ISL_3111102, EPI_ISL_3111103, EPI_ISL_3111104, EPI_ISL_3111105, EPI_ISL_3111106, EPI_ISL_3111107, EPI_ISL_3111108, EPI_ISL_3111109, EPI_ISL_3111110, EPI_ISL_3111111, EPI_ISL_3111112, EPI_ISL_3111113, EPI_ISL_3111114, EPI_ISL_3111115, EPI_ISL_3111116, EPI_ISL_3111117, EPI_ISL_3111118, EPI_ISL_3111119, EPI_ISL_3111120, EPI_ISL_3111121, EPI_ISL_3111122, EPI_ISL_3111123, EPI_ISL_3111124, EPI_ISL_3111125, EPI_ISL_3111126, EPI_ISL_3111127, EPI_ISL_3111128, EPI_ISL_3111129, EPI_ISL_3111130, EPI_ISL_3111131, EPI_ISL_3111132, EPI_ISL_3111133, EPI_ISL_3111134, EPI_ISL_3111135, EPI_ISL_3111136, EPI_ISL_3111137, EPI_ISL_3111138, EPI_ISL_3111139, EPI_ISL_3111140, EPI_ISL_3111141, EPI_ISL_3111142, EPI_ISL_3111143, EPI_ISL_3111144, EPI_ISL_3111145, EPI_ISL_3111146, EPI_ISL_3111147, EPI_ISL_3111148, EPI_ISL_3111149, EPI_ISL_3111150, EPI_ISL_3111151, EPI_ISL_3111152, EPI_ISL_3111153, EPI_ISL_3111154, EPI_ISL_3111155, EPI_ISL_3111156, EPI_ISL_3111157, EPI_ISL_3111158, EPI_ISL_3111159, EPI_ISL_3111160, EPI_ISL_3111161, EPI_ISL_3111162, EPI_ISL_3111163, EPI_ISL_3111164, EPI_ISL_3111165, EPI_ISL_3111166, EPI_ISL_3111167, EPI_ISL_3111168, EPI_ISL_3111169, EPI_ISL_3111170, EPI_ISL_3111171, EPI_ISL_3111172, EPI_ISL_3111173, EPI_ISL_3111174, EPI_ISL_3111175, EPI_ISL_3111176, EPI_ISL_3111177, EPI_ISL_3111178, EPI_ISL_3111179, EPI_ISL_3111180, EPI_ISL_3111181, EPI_ISL_3111182, EPI_ISL_3111183, EPI_ISL_3111184, EPI_ISL_3111185, EPI_ISL_3111186, EPI_ISL_3111187, EPI_ISL_3111188, EPI_ISL_3111189, EPI_ISL_3111190, EPI_ISL_3111191, EPI_ISL_3111192, EPI_ISL_3111193, EPI_ISL_3111194, EPI_ISL_3111195, EPI_ISL_3111196, EPI_ISL_3111197, EPI_ISL_3111198, EPI_ISL_3111199, EPI_ISL_3111200, EPI_ISL_3111201, EPI_ISL_3111202, EPI_ISL_3111205, EPI_ISL_3111206, EPI_ISL_3111207, EPI_ISL_3111209, EPI_ISL_3111210, EPI_ISL_3111211, EPI_ISL_3111212, EPI_ISL_3111213, EPI_ISL_3111214, EPI_ISL_3111215, EPI_ISL_3111216, EPI_ISL_3111217, EPI_ISL_3111218, EPI_ISL_3111219, EPI_ISL_3111220, EPI_ISL_3111221, EPI_ISL_3111223, EPI_ISL_3111224, EPI_ISL_3111225, EPI_ISL_3111226, EPI_ISL_3111227, EPI_ISL_3111228, EPI_ISL_3111229, EPI_ISL_3111230, EPI_ISL_3111231, EPI_ISL_3111232, EPI_ISL_3111233, EPI_ISL_3111234, EPI_ISL_3111235, EPI_ISL_3111236, EPI_ISL_3111237, EPI_ISL_3111238, EPI_ISL_3111239, EPI_ISL_3111240, EPI_ISL_3111241, EPI_ISL_3111242, EPI_ISL_3111243, EPI_ISL_3111244, EPI_ISL_3111245, EPI_ISL_3111246, EPI_ISL_3111247, EPI_ISL_3111248, EPI_ISL_3111249, EPI_ISL_3111250, EPI_ISL_3111251, EPI_ISL_3111252, EPI_ISL_3111253, EPI_ISL_3111254, EPI_ISL_3111255, EPI_ISL_3111256, EPI_ISL_3111257, EPI_ISL_3111259, EPI_ISL_3111260, EPI_ISL_3111261, EPI_ISL_3111262, EPI_ISL_3111263, EPI_ISL_3111264, EPI_ISL_3111265, EPI_ISL_3111266, EPI_ISL_3111267, EPI_ISL_3111268, EPI_ISL_3111269, EPI_ISL_3111270, EPI_ISL_3111271, EPI_ISL_3111272, EPI_ISL_3111273, EPI_ISL_3111274, EPI_ISL_3111275, EPI_ISL_3111276, EPI_ISL_3111277, EPI_ISL_3111280, EPI_ISL_3111281, EPI_ISL_3111283, EPI_ISL_3111284, EPI_ISL_3111285, EPI_ISL_3111287, EPI_ISL_3111288, EPI_ISL_3111290, EPI_ISL_3111291, EPI_ISL_3111292, EPI_ISL_3111293, EPI_ISL_3111294, EPI_ISL_3111295, EPI_ISL_3111296, EPI_ISL_3111297, EPI_ISL_3111298, EPI_ISL_3111299, EPI_ISL_3111300, EPI_ISL_3111301, EPI_ISL_3111302, EPI_ISL_3111303, EPI_ISL_3111304, EPI_ISL_3111305, EPI_ISL_3111306, EPI_ISL_3111307, EPI_ISL_3111308, EPI_ISL_3111309, EPI_ISL_3111310, EPI_ISL_3111311, EPI_ISL_3111312, EPI_ISL_3111313, EPI_ISL_3111314, EPI_ISL_3111315, EPI_ISL_3111316, EPI_ISL_3111317, EPI_ISL_3111318, EPI_ISL_3111320, EPI_ISL_3111321, EPI_ISL_3111322, EPI_ISL_3111323, EPI_ISL_3111324, EPI_ISL_3111325, EPI_ISL_3111326, EPI_ISL_3111327, EPI_ISL_3111328, EPI_ISL_3111329, EPI_ISL_3111332, EPI_ISL_3111333, EPI_ISL_3111334, EPI_ISL_3111337, EPI_ISL_3111340, EPI_ISL_3111341, EPI_ISL_3111342, EPI_ISL_3111343, EPI_ISL_3111344, EPI_ISL_3111345, EPI_ISL_3111346, EPI_ISL_3111347, EPI_ISL_3111348, EPI_ISL_3111349, EPI_ISL_3111350, EPI_ISL_3111351, EPI_ISL_3111352, EPI_ISL_3111353, EPI_ISL_3111354, EPI_ISL_3111355, EPI_ISL_3111356, EPI_ISL_3111357, EPI_ISL_3111359, EPI_ISL_3111360, EPI_ISL_3111361, EPI_ISL_3111362, EPI_ISL_3111363, EPI_ISL_3111364, EPI_ISL_3111365, EPI_ISL_3111366, EPI_ISL_3111368, EPI_ISL_3111369, EPI_ISL_3111370, EPI_ISL_3111371, EPI_ISL_3111372, EPI_ISL_3111373, EPI_ISL_3111374, EPI_ISL_3111375, EPI_ISL_3111376, EPI_ISL_3111377, EPI_ISL_3111378, EPI_ISL_3111379, EPI_ISL_3111380, EPI_ISL_3111381, EPI_ISL_3111382, EPI_ISL_3111383, EPI_ISL_3111384, EPI_ISL_3111385, EPI_ISL_3111386, EPI_ISL_3111387, EPI_ISL_3111388, EPI_ISL_3111389, EPI_ISL_3111390, EPI_ISL_3111391, EPI_ISL_3111392, EPI_ISL_3111393, EPI_ISL_3111394, EPI_ISL_3111395, EPI_ISL_3111396, EPI_ISL_3111397, EPI_ISL_3111398, EPI_ISL_3111399, EPI_ISL_3111400, EPI_ISL_3111401, EPI_ISL_3111402, EPI_ISL_3111403, EPI_ISL_3111404, EPI_ISL_3111405, EPI_ISL_3111406, EPI_ISL_3111407, EPI_ISL_3111408, EPI_ISL_3111409, EPI_ISL_3111410, EPI_ISL_3111411, EPI_ISL_3111412, EPI_ISL_3111413, EPI_ISL_3111414, EPI_ISL_3111415, EPI_ISL_3111416, EPI_ISL_3111417, EPI_ISL_3111418, EPI_ISL_3111420, EPI_ISL_3111421, EPI_ISL_3111423, EPI_ISL_3111424, EPI_ISL_3111426, EPI_ISL_3111427, EPI_ISL_3111428, EPI_ISL_3111429, EPI_ISL_3111430, EPI_ISL_3111431, EPI_ISL_3111432, EPI_ISL_3111434, EPI_ISL_3111435, EPI_ISL_3111437, EPI_ISL_3111438, EPI_ISL_3111439, EPI_ISL_3111440, EPI_ISL_3111443, EPI_ISL_3111444, EPI_ISL_3111445, EPI_ISL_3111446, EPI_ISL_3111447, EPI_ISL_3111448, EPI_ISL_3111449, EPI_ISL_3111450, EPI_ISL_3111451, EPI_ISL_3111452, EPI_ISL_3111453, EPI_ISL_3111454, EPI_ISL_3111456, EPI_ISL_3111458, EPI_ISL_3111460, EPI_ISL_3111461, EPI_ISL_3111463, EPI_ISL_3111465, EPI_ISL_3111466, EPI_ISL_3111467, EPI_ISL_3111469, EPI_ISL_3111470, EPI_ISL_3111472, EPI_ISL_3111473, EPI_ISL_3111474, EPI_ISL_3111475, EPI_ISL_3111476, EPI_ISL_3111477, EPI_ISL_3111478, EPI_ISL_3111479, EPI_ISL_3111481, EPI_ISL_3111482, EPI_ISL_3111483, EPI_ISL_3111484, EPI_ISL_3111486, EPI_ISL_3111487, EPI_ISL_3111488, EPI_ISL_3111490, EPI_ISL_3111493, EPI_ISL_3111494, EPI_ISL_3111495, EPI_ISL_3111496, EPI_ISL_3111497, EPI_ISL_3111498, EPI_ISL_3111500, EPI_ISL_3111501, EPI_ISL_3111502, EPI_ISL_3111503, EPI_ISL_3111504, EPI_ISL_3111505, EPI_ISL_3111506, EPI_ISL_3111507, EPI_ISL_3111510, EPI_ISL_3111511, EPI_ISL_3111512, EPI_ISL_3111513, EPI_ISL_3111514, EPI_ISL_3111515, EPI_ISL_3111516, EPI_ISL_3111517, EPI_ISL_3111518, EPI_ISL_3111519, EPI_ISL_3111520, EPI_ISL_3111521, EPI_ISL_3111523, EPI_ISL_3111524, EPI_ISL_3111525, EPI_ISL_3111526, EPI_ISL_3111528, EPI_ISL_3111529, EPI_ISL_3111530, EPI_ISL_3111531, EPI_ISL_3111532, EPI_ISL_3111535, EPI_ISL_3111538, EPI_ISL_3111539, EPI_ISL_3111540, EPI_ISL_3111541, EPI_ISL_3111544, EPI_ISL_3111545, EPI_ISL_3111546, EPI_ISL_3111547, EPI_ISL_3111548, EPI_ISL_3111549, EPI_ISL_3111550, EPI_ISL_3111551, EPI_ISL_3111552, EPI_ISL_3111553, EPI_ISL_3111554, EPI_ISL_3111555, EPI_ISL_3111557, EPI_ISL_3111558, EPI_ISL_3111559, EPI_ISL_3111561, EPI_ISL_3111562, EPI_ISL_3111563, EPI_ISL_3111564, EPI_ISL_3111565, EPI_ISL_3111566, EPI_ISL_3111567, EPI_ISL_3111568, EPI_ISL_3111569, EPI_ISL_3111571, EPI_ISL_3111575, EPI_ISL_3111577, EPI_ISL_3111578, EPI_ISL_3111579, EPI_ISL_3111580, EPI_ISL_3111581, EPI_ISL_3111584, EPI_ISL_3111585, EPI_ISL_3111587, EPI_ISL_3111588, EPI_ISL_3111589, EPI_ISL_3111590, EPI_ISL_3111591, EPI_ISL_3111592, EPI_ISL_3111593, EPI_ISL_3111594, EPI_ISL_3111595, EPI_ISL_3111597, EPI_ISL_3111599, EPI_ISL_3111603, EPI_ISL_3111604, EPI_ISL_3111607, EPI_ISL_3111608, EPI_ISL_3111610, EPI_ISL_3111611, EPI_ISL_3111612, EPI_ISL_3111613, EPI_ISL_3111614, EPI_ISL_3111615, EPI_ISL_3111616, EPI_ISL_3111617, EPI_ISL_3111618, EPI_ISL_3111619, EPI_ISL_3111621, EPI_ISL_3111622, EPI_ISL_3111623, EPI_ISL_3111624, EPI_ISL_3111626, EPI_ISL_3111627, EPI_ISL_3111629, EPI_ISL_3111630, EPI_ISL_3111631, EPI_ISL_3111632, EPI_ISL_3111633, EPI_ISL_3111634, EPI_ISL_3111635, EPI_ISL_3111636, EPI_ISL_3111637, EPI_ISL_3111638, EPI_ISL_3111639, EPI_ISL_3111640, EPI_ISL_3111641, EPI_ISL_3111642, EPI_ISL_3111643, EPI_ISL_3111644, EPI_ISL_3111645, EPI_ISL_3111646, EPI_ISL_3111647, EPI_ISL_3111648, EPI_ISL_3111649, EPI_ISL_3111650, EPI_ISL_3111651, EPI_ISL_3111652, EPI_ISL_3111653, EPI_ISL_3111654, EPI_ISL_3111655, EPI_ISL_3111656, EPI_ISL_3111657, EPI_ISL_3111658, EPI_ISL_3111659, EPI_ISL_3111660, EPI_ISL_3111661, EPI_ISL_3111662, EPI_ISL_3111663, EPI_ISL_3111664, EPI_ISL_3111665, EPI_ISL_3111666, EPI_ISL_3111667, EPI_ISL_3111668, EPI_ISL_3111669, EPI_ISL_3111670, EPI_ISL_3111671, EPI_ISL_3111672, EPI_ISL_3111673, EPI_ISL_3111674, EPI_ISL_3111675, EPI_ISL_3111676, EPI_ISL_3111678, EPI_ISL_3111679, EPI_ISL_3111680, EPI_ISL_3111681, EPI_ISL_3111682, EPI_ISL_3111683, EPI_ISL_3111684, EPI_ISL_3111685, EPI_ISL_3111687, EPI_ISL_3111688, EPI_ISL_3111689, EPI_ISL_3111690, EPI_ISL_3111691, EPI_ISL_3111692, EPI_ISL_3111693, EPI_ISL_3111694, EPI_ISL_3111696, EPI_ISL_3111697, EPI_ISL_3111698, EPI_ISL_3111700, EPI_ISL_3111701, EPI_ISL_3111702, EPI_ISL_3111705, EPI_ISL_3111706, EPI_ISL_3111708, EPI_ISL_3111710, EPI_ISL_3111711, EPI_ISL_3111712, EPI_ISL_3111713, EPI_ISL_3111714, EPI_ISL_3111715, EPI_ISL_3111717, EPI_ISL_3111718, EPI_ISL_3111719, EPI_ISL_3111720, EPI_ISL_3111721, EPI_ISL_3111722, EPI_ISL_3111723, EPI_ISL_3111724, EPI_ISL_3111725, EPI_ISL_3111726, EPI_ISL_3111727, EPI_ISL_3111728, EPI_ISL_3111730, EPI_ISL_3111731, EPI_ISL_3111732, EPI_ISL_3111733, EPI_ISL_3111734, EPI_ISL_3111735, EPI_ISL_3111736, EPI_ISL_3111737, EPI_ISL_3111738, EPI_ISL_3111739, EPI_ISL_3111740, EPI_ISL_3111741, EPI_ISL_3111742, EPI_ISL_3111743, EPI_ISL_3111744, EPI_ISL_3111745, EPI_ISL_3111746, EPI_ISL_3111747, EPI_ISL_3111748, EPI_ISL_3111749, EPI_ISL_3111750, EPI_ISL_3111751, EPI_ISL_3111752, EPI_ISL_3111753, EPI_ISL_3111754, EPI_ISL_3111755, EPI_ISL_3111756, EPI_ISL_3111757, EPI_ISL_3111758, EPI_ISL_3111759, EPI_ISL_3111760, EPI_ISL_3111763, EPI_ISL_3111764, EPI_ISL_3111765, EPI_ISL_3111766, EPI_ISL_3111767, EPI_ISL_3111768, EPI_ISL_3111769, EPI_ISL_3111770, EPI_ISL_3111771, EPI_ISL_3111772, EPI_ISL_3111773, EPI_ISL_3111774, EPI_ISL_3111775, EPI_ISL_3111776, EPI_ISL_3111777, EPI_ISL_3111778, EPI_ISL_3111780, EPI_ISL_3111781, EPI_ISL_3111782, EPI_ISL_3111783, EPI_ISL_3111784, EPI_ISL_3111785, EPI_ISL_3111786, EPI_ISL_3111787, EPI_ISL_3111788, EPI_ISL_3111789, EPI_ISL_3111790, EPI_ISL_3111791, EPI_ISL_3111792, EPI_ISL_3111793, EPI_ISL_3111794, EPI_ISL_3111795, EPI_ISL_3111796, EPI_ISL_3111797, EPI_ISL_3111798, EPI_ISL_3111799, EPI_ISL_3111800, EPI_ISL_3111801, EPI_ISL_3111802, EPI_ISL_3111803, EPI_ISL_3111806, EPI_ISL_3111807, EPI_ISL_3111808, EPI_ISL_3111809, EPI_ISL_3111810, EPI_ISL_3111812, EPI_ISL_3111813, EPI_ISL_3111815, EPI_ISL_3111816, EPI_ISL_3111817, EPI_ISL_3111818, EPI_ISL_3111819, EPI_ISL_3111820, EPI_ISL_3111821, EPI_ISL_3111822, EPI_ISL_3111823, EPI_ISL_3111825, EPI_ISL_3111826, EPI_ISL_3111827, EPI_ISL_3111828, EPI_ISL_3111829, EPI_ISL_3111830, EPI_ISL_3111831, EPI_ISL_3111832, EPI_ISL_3111833, EPI_ISL_3111834, EPI_ISL_3111835, EPI_ISL_3111836, EPI_ISL_3111837, EPI_ISL_3111838, EPI_ISL_3111839, EPI_ISL_3111840, EPI_ISL_3111841, EPI_ISL_3111842, EPI_ISL_3111843, EPI_ISL_3111844, EPI_ISL_3111845, EPI_ISL_3111846, EPI_ISL_3111849, EPI_ISL_3111850, EPI_ISL_3111851, EPI_ISL_3111852, EPI_ISL_3111853, EPI_ISL_3111854, EPI_ISL_3111855, EPI_ISL_3111856, EPI_ISL_3111858, EPI_ISL_3111860, EPI_ISL_3111861, EPI_ISL_3111862, EPI_ISL_3111863, EPI_ISL_3111864, EPI_ISL_3111865, EPI_ISL_3111866, EPI_ISL_3111867, EPI_ISL_3111868, EPI_ISL_3111869, EPI_ISL_3111871, EPI_ISL_3111872, EPI_ISL_3111873, EPI_ISL_3111874, EPI_ISL_3111875, EPI_ISL_3111877, EPI_ISL_3111878, EPI_ISL_3111879, EPI_ISL_3111880, EPI_ISL_3111881, EPI_ISL_3111882, EPI_ISL_3111883, EPI_ISL_3111884, EPI_ISL_3111885, EPI_ISL_3111886, EPI_ISL_3111887, EPI_ISL_3111888, EPI_ISL_3111889, EPI_ISL_3111890, EPI_ISL_3111891, EPI_ISL_3111892, EPI_ISL_3111893, EPI_ISL_3111894, EPI_ISL_3111895, EPI_ISL_3111896, EPI_ISL_3111897, EPI_ISL_3111898, EPI_ISL_3111899, EPI_ISL_3111900, EPI_ISL_3111901, EPI_ISL_3111903, EPI_ISL_3111904, EPI_ISL_3111905, EPI_ISL_3111906, EPI_ISL_3111907, EPI_ISL_3111908, EPI_ISL_3111909, EPI_ISL_3111910, EPI_ISL_3111911, EPI_ISL_3111912, EPI_ISL_3111913, EPI_ISL_3111914, EPI_ISL_3111915, EPI_ISL_3111916, EPI_ISL_3111917, EPI_ISL_3111918, EPI_ISL_3111919, EPI_ISL_3111920, EPI_ISL_3111921, EPI_ISL_3111922, EPI_ISL_3111923, EPI_ISL_3111924, EPI_ISL_3111925, EPI_ISL_3111926, EPI_ISL_3111927, EPI_ISL_3111928, EPI_ISL_3111929, EPI_ISL_3111930, EPI_ISL_3111931, EPI_ISL_3111932, EPI_ISL_3111933, EPI_ISL_3111934, EPI_ISL_3111935, EPI_ISL_3111936, EPI_ISL_3111939, EPI_ISL_3111940, EPI_ISL_3111942, EPI_ISL_3111943, EPI_ISL_3111944, EPI_ISL_3111946, EPI_ISL_3111947, EPI_ISL_3111948, EPI_ISL_3111949, EPI_ISL_3111950, EPI_ISL_3111951, EPI_ISL_3111952, EPI_ISL_3111954, EPI_ISL_3111956, EPI_ISL_3111957, EPI_ISL_3111958, EPI_ISL_3111959, EPI_ISL_3111960, EPI_ISL_3111961, EPI_ISL_3111962, EPI_ISL_3111963, EPI_ISL_3111964, EPI_ISL_3111965, EPI_ISL_3111966, EPI_ISL_3111967, EPI_ISL_3111968, EPI_ISL_3111970, EPI_ISL_3111971, EPI_ISL_3111972, EPI_ISL_3111973, EPI_ISL_3111974, EPI_ISL_3111975, EPI_ISL_3111976, EPI_ISL_3111977, EPI_ISL_3111978, EPI_ISL_3111979, EPI_ISL_3111980, EPI_ISL_3111981, EPI_ISL_3111982, EPI_ISL_3111983, EPI_ISL_3111984, EPI_ISL_3111985, EPI_ISL_3111986, EPI_ISL_3111987, EPI_ISL_3111988, EPI_ISL_3111989, EPI_ISL_3111990, EPI_ISL_3111991, EPI_ISL_3111992, EPI_ISL_3111993, EPI_ISL_3111994, EPI_ISL_3111995, EPI_ISL_3111996, EPI_ISL_3111997, EPI_ISL_3111998, EPI_ISL_3111999, EPI_ISL_3112000, EPI_ISL_3112001, EPI_ISL_3112002, EPI_ISL_3112003, EPI_ISL_3112004, EPI_ISL_3112005, EPI_ISL_3112006, EPI_ISL_3112007, EPI_ISL_3112008, EPI_ISL_3112009, EPI_ISL_3112010, EPI_ISL_3112011, EPI_ISL_3112012, EPI_ISL_3112013, EPI_ISL_3112014, EPI_ISL_3112015, EPI_ISL_3112016, EPI_ISL_3112017, EPI_ISL_3112018, EPI_ISL_3112019, EPI_ISL_3112020, EPI_ISL_3112021, EPI_ISL_3112022, EPI_ISL_3112023, EPI_ISL_3112024, EPI_ISL_3112025, EPI_ISL_3112026, EPI_ISL_3112027, EPI_ISL_3112028, EPI_ISL_3112029, EPI_ISL_3112030, EPI_ISL_3112031, EPI_ISL_3112032, EPI_ISL_3112033, EPI_ISL_3112034, EPI_ISL_3112035, EPI_ISL_3112036, EPI_ISL_3112037, EPI_ISL_3112038, EPI_ISL_3112039, EPI_ISL_3112040, EPI_ISL_3112041, EPI_ISL_3112042, EPI_ISL_3112043, EPI_ISL_3112044, EPI_ISL_3112045, EPI_ISL_3112046, EPI_ISL_3112047, EPI_ISL_3112048, EPI_ISL_3112049, EPI_ISL_3112050, EPI_ISL_3112051, EPI_ISL_3112052, EPI_ISL_3112053, EPI_ISL_3112054, EPI_ISL_3112055, EPI_ISL_3112057, EPI_ISL_3112058, EPI_ISL_3112059, EPI_ISL_3112060, EPI_ISL_3112061, EPI_ISL_3112062, EPI_ISL_3112063, EPI_ISL_3112064, EPI_ISL_3112065, EPI_ISL_3112066, EPI_ISL_3112067, EPI_ISL_3112068, EPI_ISL_3112069, EPI_ISL_3112070, EPI_ISL_3112071, EPI_ISL_3112072, EPI_ISL_3112073, EPI_ISL_3112074, EPI_ISL_3112075, EPI_ISL_3112076, EPI_ISL_3112077, EPI_ISL_3112078, EPI_ISL_3112079, EPI_ISL_3112080, EPI_ISL_3112081, EPI_ISL_3112082, EPI_ISL_3112083, EPI_ISL_3112084, EPI_ISL_3112085, EPI_ISL_3112086, EPI_ISL_3112087, EPI_ISL_3112088, EPI_ISL_3112089, EPI_ISL_3112090, EPI_ISL_3112091, EPI_ISL_3112092, EPI_ISL_3112093, EPI_ISL_3112094, EPI_ISL_3112095, EPI_ISL_3112096, EPI_ISL_3112097, EPI_ISL_3112098, EPI_ISL_3112099, EPI_ISL_3112100, EPI_ISL_3112101, EPI_ISL_3112102, EPI_ISL_3112103, EPI_ISL_3112104, EPI_ISL_3112105, EPI_ISL_3112106, EPI_ISL_3112107, EPI_ISL_3112108, EPI_ISL_3112109, EPI_ISL_3112110, EPI_ISL_3112111, EPI_ISL_3112112, EPI_ISL_3112113, EPI_ISL_3112114, EPI_ISL_3112115, EPI_ISL_3112116, EPI_ISL_3112117, EPI_ISL_3112118, EPI_ISL_3112119, EPI_ISL_3112120, EPI_ISL_3112121, EPI_ISL_3112122, EPI_ISL_3112123, EPI_ISL_3112124, EPI_ISL_3112125, EPI_ISL_3112126, EPI_ISL_3112127, EPI_ISL_3112128, EPI_ISL_3112129, EPI_ISL_3112130, EPI_ISL_3112131, EPI_ISL_3112132, EPI_ISL_3112133, EPI_ISL_3112134, EPI_ISL_3112135, EPI_ISL_3112136, EPI_ISL_3112137, EPI_ISL_3112138, EPI_ISL_3112139, EPI_ISL_3112140, EPI_ISL_3112141, EPI_ISL_3112142, EPI_ISL_3112143, EPI_ISL_3112144, EPI_ISL_3112145, EPI_ISL_3112146, EPI_ISL_3112147, EPI_ISL_3112148, EPI_ISL_3112149, EPI_ISL_3112150, EPI_ISL_3112151, EPI_ISL_3112152, EPI_ISL_3112153, EPI_ISL_3112154, EPI_ISL_3112155, EPI_ISL_3112156, EPI_ISL_3112157, EPI_ISL_3112158, EPI_ISL_3112159, EPI_ISL_3112160, EPI_ISL_3112161, EPI_ISL_3112162, EPI_ISL_3112163, EPI_ISL_3112164, EPI_ISL_3112165, EPI_ISL_3112166, EPI_ISL_3112167, EPI_ISL_3112168, EPI_ISL_3112169, EPI_ISL_3112170, EPI_ISL_3112174, EPI_ISL_3112175, EPI_ISL_3112176, EPI_ISL_3112177, EPI_ISL_3112178, EPI_ISL_3112179, EPI_ISL_3112181, EPI_ISL_3112182, EPI_ISL_3112184, EPI_ISL_3112185, EPI_ISL_3112187, EPI_ISL_3112188, EPI_ISL_3112189, EPI_ISL_3112190, EPI_ISL_3112191, EPI_ISL_3112193, EPI_ISL_3112196, EPI_ISL_3112198, EPI_ISL_3112201, EPI_ISL_3112202, EPI_ISL_3112203, EPI_ISL_3112204, EPI_ISL_3112205, EPI_ISL_3112206, EPI_ISL_3112207, EPI_ISL_3112208, EPI_ISL_3112209, EPI_ISL_3112210, EPI_ISL_3112211, EPI_ISL_3112212, EPI_ISL_3112213, EPI_ISL_3112214, EPI_ISL_3112215, EPI_ISL_3112216, EPI_ISL_3112217, EPI_ISL_3112218, EPI_ISL_3112219, EPI_ISL_3112220, EPI_ISL_3112221, EPI_ISL_3112222, EPI_ISL_3112224, EPI_ISL_3112225, EPI_ISL_3112226, EPI_ISL_3112227, EPI_ISL_3112228, EPI_ISL_3112229, EPI_ISL_3112231, EPI_ISL_3112233, EPI_ISL_3112234, EPI_ISL_3112235, EPI_ISL_3112236, EPI_ISL_3112237, EPI_ISL_3112238, EPI_ISL_3112239, EPI_ISL_3112240, EPI_ISL_3112241, EPI_ISL_3112242, EPI_ISL_3112243, EPI_ISL_3112245, EPI_ISL_3112246, EPI_ISL_3112247, EPI_ISL_3112248, EPI_ISL_3112249, EPI_ISL_3112251, EPI_ISL_3112252, EPI_ISL_3112253, EPI_ISL_3112254, EPI_ISL_3112255, EPI_ISL_3112256, EPI_ISL_3112257, EPI_ISL_3112258, EPI_ISL_3112260, EPI_ISL_3112261, EPI_ISL_3112262, EPI_ISL_3112263, EPI_ISL_3112264, EPI_ISL_3112265, EPI_ISL_3112266, EPI_ISL_3112267, EPI_ISL_3112268, EPI_ISL_3112269, EPI_ISL_3112270, EPI_ISL_3112271, EPI_ISL_3112272, EPI_ISL_3112273, EPI_ISL_3112274, EPI_ISL_3112275, EPI_ISL_3112277, EPI_ISL_3112278, EPI_ISL_3112279, EPI_ISL_3112280, EPI_ISL_3112281, EPI_ISL_3112283, EPI_ISL_3112284, EPI_ISL_3112285, EPI_ISL_3112286, EPI_ISL_3112287, EPI_ISL_3112288, EPI_ISL_3112291, EPI_ISL_3112293, EPI_ISL_3112295, EPI_ISL_3112296, EPI_ISL_3112297, EPI_ISL_3112298, EPI_ISL_3112299, EPI_ISL_3112301, EPI_ISL_3112303, EPI_ISL_3112306, EPI_ISL_3112307, EPI_ISL_3112308, EPI_ISL_3112310, EPI_ISL_3112311, EPI_ISL_3112312, EPI_ISL_3112313, EPI_ISL_3112316, EPI_ISL_3112317, EPI_ISL_3112318, EPI_ISL_3112319, EPI_ISL_3112320, EPI_ISL_3112321, EPI_ISL_3112322, EPI_ISL_3112324, EPI_ISL_3112325, EPI_ISL_3112326, EPI_ISL_3112327, EPI_ISL_3112328, EPI_ISL_3112329, EPI_ISL_3112330, EPI_ISL_3112331, EPI_ISL_3112332, EPI_ISL_3112333, EPI_ISL_3112334, EPI_ISL_3112337, EPI_ISL_3112338, EPI_ISL_3112339 | | | |
| see above | Laboratory Corporation of America | Centers for Disease Control and Prevention Division of Viral Diseases, Pathogen Discovery | Dakota Howard, Dhwani Batra, Peter W. Cook, Kara Moser, Adrian Paskey, Jason Caravas, Benjamin Rambo-Martin, Shatavia Morrison, Christopher Gulvick, Scott Sammons, Yvette Unoarumhi, Darlene Wagner, Matthew Schmerer, Minoo Agarwal, Eyad Almasri, Debbie Boles, Ayla Burns, Nuthawin Charoensri, Oren Cohen, Susan Countryman, Mary Ann Cristobal, Bobbi Croy, Suzanne Dale, Hrushikesh Deshmukh, Amanda Douglas, Vincent Drouillon, Marcia Eisenberg, Howard Engler, Rama Ghatti, Prashant Gupta, Susan Hicks, Jake Humphrey, Lax Iyer, Lisa Pfefferle, Manoj Jain, Matthew Robinson, Mohan Kolli, Brian Krueger, Tim Kuphal, Stanley Letovsky, Michael Levandoski, Craig Lukasik, Jonathan Meltzer, Brian Norvell, Mindy Nye, Scott Parker, Christos Petropoulos, John Pruitt, Steven Ragan, Scott Ryan, Mike Sapeta, Jana Schroth, Suresh Babu Selvaraju, Goran Stevovic, Amanda Suchanek, Andrea Throop, Lyndon Tilson, Thomas Urban, Joe Voshell, Kimberly Wagner, Jonathan Williams, Mary Williamson, Qian Zeng, Tricia Zwiefelhofer, Clinton R. Paden, Duncan MacCannell |
| EPI_ISL_3112340 | Institute for Infectious Diseases | Institute for Infectious Diseases | Stefan Neuenschwander, Christian Baumann, Miguel A Terrazos Miani, Cora SÃ¤gesser, Pascal Bittel, Peter Keller, Franziska Suter-Riniker, Stephen L Leib, Alban Ramette |
| EPI_ISL_3112341 | Laboratory Corporation of America | Centers for Disease Control and Prevention Division of Viral Diseases, Pathogen Discovery | Dakota Howard, Dhwani Batra, Peter W. Cook, Kara Moser, Adrian Paskey, Jason Caravas, Benjamin Rambo-Martin, Shatavia Morrison, Christopher Gulvick, Scott Sammons, Yvette Unoarumhi, Darlene Wagner, Matthew Schmerer, Minoo Agarwal, Eyad Almasri, Debbie Boles, Ayla Burns, Nuthawin Charoensri, Oren Cohen, Susan Countryman, Mary Ann Cristobal, Bobbi Croy, Suzanne Dale, Hrushikesh Deshmukh, Amanda Douglas, Vincent Drouillon, Marcia Eisenberg, Howard Engler, Rama Ghatti, Prashant Gupta, Susan Hicks, Jake Humphrey, Lax Iyer, Lisa Pfefferle, Manoj Jain, Matthew Robinson, Mohan Kolli, Brian Krueger, Tim Kuphal, Stanley Letovsky, Michael Levandoski, Craig Lukasik, Jonathan Meltzer, Brian Norvell, Mindy Nye, Scott Parker, Christos Petropoulos, John Pruitt, Steven Ragan, Scott Ryan, Mike Sapeta, Jana Schroth, Suresh Babu Selvaraju, Goran Stevovic, Amanda Suchanek, Andrea Throop, Lyndon Tilson, Thomas Urban, Joe Voshell, Kimberly Wagner, Jonathan Williams, Mary Williamson, Qian Zeng, Tricia Zwiefelhofer, Clinton R. Paden, Duncan MacCannell |
| EPI_ISL_3112342, EPI_ISL_3112343, EPI_ISL_3112345, EPI_ISL_3112346 | Institute for Infectious Diseases | Institute for Infectious Diseases | Stefan Neuenschwander, Christian Baumann, Miguel A Terrazos Miani, Cora SÃ¤gesser, Pascal Bittel, Peter Keller, Franziska Suter-Riniker, Stephen L Leib, Alban Ramette |
| EPI_ISL_3112347 | Laboratory Corporation of America | Centers for Disease Control and Prevention Division of Viral Diseases, Pathogen Discovery | Dakota Howard, Dhwani Batra, Peter W. Cook, Kara Moser, Adrian Paskey, Jason Caravas, Benjamin Rambo-Martin, Shatavia Morrison, Christopher Gulvick, Scott Sammons, Yvette Unoarumhi, Darlene Wagner, Matthew Schmerer, Minoo Agarwal, Eyad Almasri, Debbie Boles, Ayla Burns, Nuthawin Charoensri, Oren Cohen, Susan Countryman, Mary Ann Cristobal, Bobbi Croy, Suzanne Dale, Hrushikesh Deshmukh, Amanda Douglas, Vincent Drouillon, Marcia Eisenberg, Howard Engler, Rama Ghatti, Prashant Gupta, Susan Hicks, Jake Humphrey, Lax Iyer, Lisa Pfefferle, Manoj Jain, Matthew Robinson, Mohan Kolli, Brian Krueger, Tim Kuphal, Stanley Letovsky, Michael Levandoski, Craig Lukasik, Jonathan Meltzer, Brian Norvell, Mindy Nye, Scott Parker, Christos Petropoulos, John Pruitt, Steven Ragan, Scott Ryan, Mike Sapeta, Jana Schroth, Suresh Babu Selvaraju, Goran Stevovic, Amanda Suchanek, Andrea Throop, Lyndon Tilson, Thomas Urban, Joe Voshell, Kimberly Wagner, Jonathan Williams, Mary Williamson, Qian Zeng, Tricia Zwiefelhofer, Clinton R. Paden, Duncan MacCannell |
| EPI_ISL_3112348, EPI_ISL_3112349 | Institute for Infectious Diseases | Institute for Infectious Diseases | Stefan Neuenschwander, Christian Baumann, Miguel A Terrazos Miani, Cora SÃ¤gesser, Pascal Bittel, Peter Keller, Franziska Suter-Riniker, Stephen L Leib, Alban Ramette |
| EPI_ISL_3112350 | Laboratory Corporation of America | Centers for Disease Control and Prevention Division of Viral Diseases, Pathogen Discovery | Dakota Howard, Dhwani Batra, Peter W. Cook, Kara Moser, Adrian Paskey, Jason Caravas, Benjamin Rambo-Martin, Shatavia Morrison, Christopher Gulvick, Scott Sammons, Yvette Unoarumhi, Darlene Wagner, Matthew Schmerer, Minoo Agarwal, Eyad Almasri, Debbie Boles, Ayla Burns, Nuthawin Charoensri, Oren Cohen, Susan Countryman, Mary Ann Cristobal, Bobbi Croy, Suzanne Dale, Hrushikesh Deshmukh, Amanda Douglas, Vincent Drouillon, Marcia Eisenberg, Howard |

| | | | |
|---|---|---|---|
| | | | Engler, Rama Ghatti, Prashant Gupta, Susan Hicks, Jake Humphrey, Lax Iyer, Lisa Pfefferle, Manoj Jain, Matthew Robinson, Mohan Kolli, Brian Krueger, Tim Kuphal, Stanley Letovsky, Michael Levandoski, Craig Lukasik, Jonathan Meltzer, Brian Norvell, Mindy Nye, Scott Parker, Christos Petropoulos, John Pruitt, Steven Ragan, Scott Ryan, Mike Sapeta, Jana Schroth, Suresh Babu Selvaraju, Goran Stevovic, Amanda Suchanek, Andrea Throop, Lyndon Tilson, Thomas Urban, Joe Voshell, Kimberly Wagner, Jonathan Williams, Mary Williamson, Qian Zeng, Tricia Zwiefelhofer, Clinton R. Paden, Duncan MacCannell |
| EPI_ISL_3112351 | Institute for Infectious Diseases | Institute for Infectious Diseases | Stefan Neuenschwander, Christian Baumann, Miguel A Terrazos Miani, Cora SÃ¤gesser, Pascal Bittel, Peter Keller, Franziska Suter-Riniker, Stephen L Leib, Alban Ramette |
| EPI_ISL_3112352 | Laboratory Corporation of America | Centers for Disease Control and Prevention Division of Viral Diseases, Pathogen Discovery | Dakota Howard, Dhwani Batra, Peter W. Cook, Kara Moser, Adrian Paskey, Jason Caravas, Benjamin Rambo-Martin, Shatavia Morrison, Christopher Gulvick, Scott Sammons, Yvette Unoarumhi, Darlene Wagner, Matthew Schmerer, Minoo Agarwal, Eyad Almasri, Debbie Boles, Ayla Burns, Nuthawin Charoensri, Oren Cohen, Susan Countryman, Mary Ann Cristobal, Bobbi Croy, Suzanne Dale, Hrushikesh Deshmukh, Amanda Douglas, Vincent Drouillon, Marcia Eisenberg, Howard Engler, Rama Ghatti, Prashant Gupta, Susan Hicks, Jake Humphrey, Lax Iyer, Lisa Pfefferle, Manoj Jain, Matthew Robinson, Mohan Kolli, Brian Krueger, Tim Kuphal, Stanley Letovsky, Michael Levandoski, Craig Lukasik, Jonathan Meltzer, Brian Norvell, Mindy Nye, Scott Parker, Christos Petropoulos, John Pruitt, Steven Ragan, Scott Ryan, Mike Sapeta, Jana Schroth, Suresh Babu Selvaraju, Goran Stevovic, Amanda Suchanek, Andrea Throop, Lyndon Tilson, Thomas Urban, Joe Voshell, Kimberly Wagner, Jonathan Williams, Mary Williamson, Qian Zeng, Tricia Zwiefelhofer, Clinton R. Paden, Duncan MacCannell |
| EPI_ISL_3112353, EPI_ISL_3112354 | Institute for Infectious Diseases | Institute for Infectious Diseases | Stefan Neuenschwander, Christian Baumann, Miguel A Terrazos Miani, Cora SÃ¤gesser, Pascal Bittel, Peter Keller, Franziska Suter-Riniker, Stephen L Leib, Alban Ramette |
| EPI_ISL_3112355 | Laboratory Corporation of America | Centers for Disease Control and Prevention Division of Viral Diseases, Pathogen Discovery | Dakota Howard, Dhwani Batra, Peter W. Cook, Kara Moser, Adrian Paskey, Jason Caravas, Benjamin Rambo-Martin, Shatavia Morrison, Christopher Gulvick, Scott Sammons, Yvette Unoarumhi, Darlene Wagner, Matthew Schmerer, Minoo Agarwal, Eyad Almasri, Debbie Boles, Ayla Burns, Nuthawin Charoensri, Oren Cohen, Susan Countryman, Mary Ann Cristobal, Bobbi Croy, Suzanne Dale, Hrushikesh Deshmukh, Amanda Douglas, Vincent Drouillon, Marcia Eisenberg, Howard Engler, Rama Ghatti, Prashant Gupta, Susan Hicks, Jake Humphrey, Lax Iyer, Lisa Pfefferle, Manoj Jain, Matthew Robinson, Mohan Kolli, Brian Krueger, Tim Kuphal, Stanley Letovsky, Michael Levandoski, Craig Lukasik, Jonathan Meltzer, Brian Norvell, Mindy Nye, Scott Parker, Christos Petropoulos, John Pruitt, Steven Ragan, Scott Ryan, Mike Sapeta, Jana Schroth, Suresh Babu Selvaraju, Goran Stevovic, Amanda Suchanek, Andrea Throop, Lyndon Tilson, Thomas Urban, Joe Voshell, Kimberly Wagner, Jonathan Williams, Mary Williamson, Qian Zeng, Tricia Zwiefelhofer, Clinton R. Paden, Duncan MacCannell |
| EPI_ISL_3112356, EPI_ISL_3112357 | Institute for Infectious Diseases | Institute for Infectious Diseases | Stefan Neuenschwander, Christian Baumann, Miguel A Terrazos Miani, Cora SÃ¤gesser, Pascal Bittel, Peter Keller, Franziska Suter-Riniker, Stephen L Leib, Alban Ramette |
| EPI_ISL_3112358 | Laboratory Corporation of America | Centers for Disease Control and Prevention Division of Viral Diseases, Pathogen Discovery | Dakota Howard, Dhwani Batra, Peter W. Cook, Kara Moser, Adrian Paskey, Jason Caravas, Benjamin Rambo-Martin, Shatavia Morrison, Christopher Gulvick, Scott Sammons, Yvette Unoarumhi, Darlene Wagner, Matthew Schmerer, Minoo Agarwal, Eyad Almasri, Debbie Boles, Ayla Burns, Nuthawin Charoensri, Oren Cohen, Susan Countryman, Mary Ann Cristobal, Bobbi Croy, Suzanne Dale, Hrushikesh Deshmukh, Amanda Douglas, Vincent Drouillon, Marcia Eisenberg, Howard Engler, Rama Ghatti, Prashant Gupta, Susan Hicks, Jake Humphrey, Lax Iyer, Lisa Pfefferle, Manoj Jain, Matthew Robinson, Mohan Kolli, Brian Krueger, Tim Kuphal, Stanley Letovsky, Michael Levandoski, Craig Lukasik, Jonathan Meltzer, Brian Norvell, Mindy Nye, Scott Parker, Christos Petropoulos, John Pruitt, Steven Ragan, Scott Ryan, Mike Sapeta, Jana Schroth, Suresh Babu Selvaraju, Goran Stevovic, Amanda Suchanek, Andrea Throop, Lyndon Tilson, Thomas Urban, Joe Voshell, Kimberly Wagner, Jonathan Williams, Mary Williamson, Qian Zeng, Tricia Zwiefelhofer, Clinton R. Paden, Duncan MacCannell |
| EPI_ISL_3112359, EPI_ISL_3112360 | Institute for Infectious Diseases | Institute for Infectious Diseases | Stefan Neuenschwander, Christian Baumann, Miguel A Terrazos Miani, Cora SÃ¤gesser, Pascal Bittel, Peter Keller, Franziska Suter-Riniker, Stephen L Leib, Alban Ramette |
| EPI_ISL_3112361 | Laboratory Corporation of America | Centers for Disease Control and Prevention Division of Viral Diseases, Pathogen Discovery | Dakota Howard, Dhwani Batra, Peter W. Cook, Kara Moser, Adrian Paskey, Jason Caravas, Benjamin Rambo-Martin, Shatavia Morrison, Christopher Gulvick, Scott Sammons, Yvette Unoarumhi, Darlene Wagner, Matthew Schmerer, Minoo Agarwal, Eyad Almasri, Debbie Boles, Ayla Burns, Nuthawin Charoensri, Oren Cohen, Susan Countryman, Mary Ann Cristobal, Bobbi Croy, Suzanne Dale, Hrushikesh Deshmukh, Amanda Douglas, Vincent Drouillon, Marcia Eisenberg, Howard Engler, Rama Ghatti, Prashant Gupta, Susan Hicks, Jake Humphrey, Lax Iyer, Lisa Pfefferle, Manoj Jain, Matthew Robinson, Mohan Kolli, Brian Krueger, Tim Kuphal, Stanley Letovsky, Michael Levandoski, Craig Lukasik, Jonathan Meltzer, Brian Norvell, Mindy Nye, Scott Parker, Christos Petropoulos, John Pruitt, Steven Ragan, Scott Ryan, Mike Sapeta, Jana Schroth, Suresh Babu Selvaraju, Goran Stevovic, Amanda Suchanek, Andrea Throop, Lyndon Tilson, Thomas Urban, Joe Voshell, Kimberly Wagner, Jonathan Williams, Mary Williamson, Qian Zeng, Tricia Zwiefelhofer, Clinton R. Paden, Duncan MacCannell |
| EPI_ISL_3112362 | Institute for Infectious Diseases | Institute for Infectious Diseases | Stefan Neuenschwander, Christian Baumann, Miguel A Terrazos Miani, Cora SÃ¤gesser, Pascal Bittel, Peter Keller, Franziska Suter-Riniker, Stephen L Leib, Alban Ramette |
| EPI_ISL_3112363 | Laboratory Corporation of America | Centers for Disease Control and Prevention Division of Viral Diseases, Pathogen Discovery | Dakota Howard, Dhwani Batra, Peter W. Cook, Kara Moser, Adrian Paskey, Jason Caravas, Benjamin Rambo-Martin, Shatavia Morrison, Christopher Gulvick, Scott Sammons, Yvette Unoarumhi, Darlene Wagner, Matthew Schmerer, Minoo Agarwal, Eyad Almasri, Debbie Boles, Ayla Burns, Nuthawin Charoensri, Oren Cohen, Susan Countryman, Mary Ann Cristobal, Bobbi Croy, Suzanne Dale, Hrushikesh Deshmukh, Amanda Douglas, Vincent Drouillon, Marcia Eisenberg, Howard Engler, Rama Ghatti, Prashant Gupta, Susan Hicks, Jake Humphrey, Lax Iyer, Lisa Pfefferle, Manoj Jain, Matthew Robinson, Mohan Kolli, Brian Krueger, Tim Kuphal, Stanley Letovsky, Michael Levandoski, Craig Lukasik, Jonathan Meltzer, Brian Norvell, Mindy Nye, Scott Parker, Christos Petropoulos, John Pruitt, Steven Ragan, Scott Ryan, Mike Sapeta, Jana Schroth, Suresh Babu Selvaraju, Goran Stevovic, Amanda Suchanek, Andrea Throop, Lyndon Tilson, Thomas Urban, Joe Voshell, Kimberly Wagner, Jonathan Williams, Mary Williamson, Qian Zeng, Tricia Zwiefelhofer, Clinton R. Paden, Duncan MacCannell |
| EPI_ISL_3112364, EPI_ISL_3112365 | Institute for Infectious Diseases | Institute for Infectious Diseases | Stefan Neuenschwander, Christian Baumann, Miguel A Terrazos Miani, Cora SÃ¤gesser, Pascal Bittel, Peter Keller, Franziska Suter-Riniker, Stephen L Leib, Alban Ramette |
| EPI_ISL_3112366 | Laboratory Corporation of America | Centers for Disease Control and Prevention Division of Viral Diseases, Pathogen Discovery | Dakota Howard, Dhwani Batra, Peter W. Cook, Kara Moser, Adrian Paskey, Jason Caravas, Benjamin Rambo-Martin, Shatavia Morrison, Christopher Gulvick, Scott Sammons, Yvette Unoarumhi, Darlene Wagner, Matthew Schmerer, Minoo Agarwal, Eyad Almasri, Debbie Boles, Ayla Burns, Nuthawin Charoensri, Oren Cohen, Susan Countryman, Mary Ann Cristobal, Bobbi Croy, Suzanne Dale, Hrushikesh Deshmukh, Amanda Douglas, Vincent Drouillon, Marcia Eisenberg, Howard Engler, Rama Ghatti, Prashant Gupta, Susan Hicks, Jake Humphrey, Lax Iyer, Lisa Pfefferle, Manoj Jain, Matthew Robinson, Mohan Kolli, Brian Krueger, Tim Kuphal, Stanley Letovsky, Michael Levandoski, Craig Lukasik, Jonathan Meltzer, Brian Norvell, Mindy Nye, Scott Parker, Christos Petropoulos, John Pruitt, Steven Ragan, Scott Ryan, Mike Sapeta, Jana Schroth, Suresh Babu Selvaraju, Goran Stevovic, Amanda Suchanek, Andrea Throop, Lyndon Tilson, Thomas Urban, Joe Voshell, Kimberly Wagner, Jonathan Williams, Mary Williamson, Qian Zeng, Tricia Zwiefelhofer, Clinton R. Paden, Duncan MacCannell |
| EPI_ISL_3112368, EPI_ISL_3112369, EPI_ISL_3112370, EPI_ISL_3112371 | Institute for Infectious Diseases | Institute for Infectious Diseases | Stefan Neuenschwander, Christian Baumann, Miguel A Terrazos Miani, Cora SÃ¤gesser, Pascal Bittel, Peter Keller, Franziska Suter-Riniker, Stephen L Leib, Alban Ramette |
| EPI_ISL_3112372 | Laboratory Corporation of America | Centers for Disease Control and Prevention Division of Viral Diseases, Pathogen Discovery | Dakota Howard, Dhwani Batra, Peter W. Cook, Kara Moser, Adrian Paskey, Jason Caravas, Benjamin Rambo-Martin, Shatavia Morrison, Christopher Gulvick, Scott Sammons, Yvette Unoarumhi, Darlene Wagner, Matthew Schmerer, Minoo Agarwal, Eyad Almasri, Debbie Boles, Ayla Burns, Nuthawin Charoensri, Oren Cohen, Susan Countryman, Mary Ann Cristobal, Bobbi Croy, Suzanne Dale, Hrushikesh Deshmukh, Amanda Douglas, Vincent Drouillon, Marcia Eisenberg, Howard Engler, Rama Ghatti, Prashant Gupta, Susan Hicks, Jake Humphrey, Lax Iyer, Lisa Pfefferle, Manoj Jain, Matthew Robinson, Mohan Kolli, Brian Krueger, Tim Kuphal, Stanley Letovsky, Michael Levandoski, Craig Lukasik, Jonathan Meltzer, Brian Norvell, Mindy Nye, Scott Parker, Christos Petropoulos, John Pruitt, Steven Ragan, Scott Ryan, Mike Sapeta, Jana Schroth, Suresh Babu Selvaraju, Goran Stevovic, Amanda Suchanek, Andrea Throop, Lyndon Tilson, Thomas Urban, Joe Voshell, Kimberly Wagner, Jonathan Williams, Mary Williamson, Qian Zeng, Tricia Zwiefelhofer, Clinton R. Paden, Duncan MacCannell |
| EPI_ISL_3112373, EPI_ISL_3112374 | Institute for Infectious Diseases | Institute for Infectious Diseases | Stefan Neuenschwander, Christian Baumann, Miguel A Terrazos Miani, Cora SÃ¤gesser, Pascal Bittel, Peter Keller, Franziska Suter-Riniker, Stephen L Leib, Alban Ramette |
| EPI_ISL_3112375 | Laboratory Corporation of America | Centers for Disease Control and Prevention Division of Viral Diseases, Pathogen Discovery | Dakota Howard, Dhwani Batra, Peter W. Cook, Kara Moser, Adrian Paskey, Jason Caravas, Benjamin Rambo-Martin, Shatavia Morrison, Christopher Gulvick, Scott Sammons, Yvette Unoarumhi, Darlene Wagner, Matthew Schmerer, Minoo Agarwal, Eyad Almasri, Debbie Boles, Ayla Burns, Nuthawin Charoensri, Oren Cohen, Susan Countryman, Mary Ann Cristobal, Bobbi Croy, Suzanne Dale, Hrushikesh Deshmukh, Amanda Douglas, Vincent Drouillon, Marcia Eisenberg, Howard Engler, Rama Ghatti, Prashant Gupta, Susan Hicks, Jake Humphrey, Lax Iyer, Lisa Pfefferle, Manoj Jain, Matthew Robinson, Mohan Kolli, Brian Krueger, Tim Kuphal, Stanley Letovsky, Michael Levandoski, Craig Lukasik, Jonathan Meltzer, Brian Norvell, Mindy Nye, Scott Parker, Christos Petropoulos, John Pruitt, Steven Ragan, Scott Ryan, Mike Sapeta, Jana Schroth, Suresh Babu Selvaraju, Goran Stevovic, Amanda Suchanek, Andrea Throop, Lyndon Tilson, Thomas Urban, Joe Voshell, Kimberly Wagner, Jonathan Williams, Mary Williamson, Qian Zeng, Tricia Zwiefelhofer, Clinton R. Paden, Duncan MacCannell |
| EPI_ISL_3112376, EPI_ISL_3112377 | Institute for Infectious Diseases | Institute for Infectious Diseases | Stefan Neuenschwander, Christian Baumann, Miguel A Terrazos Miani, Cora SÃ¤gesser, Pascal Bittel, Peter Keller, Franziska Suter-Riniker, Stephen L Leib, Alban Ramette |
| EPI_ISL_3112378 | Laboratory Corporation of America | Centers for Disease Control and Prevention Division of Viral Diseases, Pathogen Discovery | Dakota Howard, Dhwani Batra, Peter W. Cook, Kara Moser, Adrian Paskey, Jason Caravas, Benjamin Rambo-Martin, Shatavia Morrison, Christopher Gulvick, Scott Sammons, Yvette Unoarumhi, Darlene Wagner, Matthew Schmerer, Minoo Agarwal, Eyad Almasri, Debbie Boles, Ayla Burns, Nuthawin Charoensri, Oren Cohen, Susan Countryman, Mary Ann Cristobal, Bobbi Croy, Suzanne Dale, Hrushikesh Deshmukh, Amanda Douglas, Vincent Drouillon, Marcia Eisenberg, Howard Engler, Rama Ghatti, Prashant Gupta, Susan Hicks, Jake Humphrey, Lax Iyer, Lisa Pfefferle, Manoj Jain, Matthew Robinson, Mohan Kolli, Brian Krueger, Tim Kuphal, Stanley Letovsky, Michael Levandoski, Craig Lukasik, Jonathan Meltzer, Brian Norvell, Mindy Nye, Scott Parker, Christos Petropoulos, John Pruitt, Steven Ragan, Scott Ryan, Mike Sapeta, Jana Schroth, Suresh Babu Selvaraju, Goran Stevovic, Amanda Suchanek, Andrea Throop, Lyndon Tilson, Thomas Urban, Joe Voshell, Kimberly Wagner, Jonathan Williams, Mary Williamson, Qian Zeng, Tricia Zwiefelhofer, Clinton R. Paden, Duncan MacCannell |
| EPI_ISL_3112379 | Institute for Infectious Diseases | Institute for Infectious Diseases | Stefan Neuenschwander, Christian Baumann, Miguel A Terrazos Miani, Cora SÃ¤gesser, Pascal Bittel, Peter Keller, Franziska Suter-Riniker, Stephen L Leib, Alban Ramette |
| EPI_ISL_3112380 | Laboratory Corporation of America | Centers for Disease Control and Prevention Division of Viral Diseases, Pathogen Discovery | Dakota Howard, Dhwani Batra, Peter W. Cook, Kara Moser, Adrian Paskey, Jason Caravas, Benjamin Rambo-Martin, Shatavia Morrison, Christopher Gulvick, Scott Sammons, Yvette Unoarumhi, Darlene Wagner, Matthew Schmerer, Minoo Agarwal, Eyad Almasri, Debbie Boles, Ayla Burns, Nuthawin Charoensri, Oren Cohen, Susan Countryman, Mary Ann Cristobal, Bobbi Croy, Suzanne Dale, Hrushikesh Deshmukh, Amanda Douglas, Vincent Drouillon, Marcia Eisenberg, Howard Engler, Rama Ghatti, Prashant Gupta, Susan Hicks, Jake Humphrey, Lax Iyer, Lisa Pfefferle, Manoj Jain, Matthew Robinson, Mohan Kolli, Brian Krueger, Tim Kuphal, Stanley Letovsky, Michael Levandoski, Craig Lukasik, Jonathan Meltzer, Brian Norvell, Mindy Nye, Scott Parker, Christos Petropoulos, John Pruitt, Steven Ragan, Scott Ryan, Mike Sapeta, Jana Schroth, Suresh Babu Selvaraju, Goran Stevovic, Amanda Suchanek, Andrea Throop, Lyndon Tilson, Thomas Urban, Joe Voshell, Kimberly Wagner, Jonathan Williams, Mary Williamson, Qian Zeng, Tricia Zwiefelhofer, Clinton R. Paden, Duncan MacCannell |
| EPI_ISL_3112381, EPI_ISL_3112382 | Institute for Infectious Diseases | Institute for Infectious Diseases | Stefan Neuenschwander, Christian Baumann, Miguel A Terrazos Miani, Cora SÃ¤gesser, Pascal Bittel, Peter Keller, Franziska Suter-Riniker, Stephen L Leib, Alban Ramette |
| EPI_ISL_3112383 | Laboratory Corporation of America | Centers for Disease Control and Prevention Division of Viral Diseases, Pathogen Discovery | Dakota Howard, Dhwani Batra, Peter W. Cook, Kara Moser, Adrian Paskey, Jason Caravas, Benjamin Rambo-Martin, Shatavia Morrison, Christopher Gulvick, Scott Sammons, Yvette Unoarumhi, Darlene Wagner, Matthew Schmerer, Minoo Agarwal, Eyad Almasri, Debbie Boles, Ayla Burns, Nuthawin Charoensri, Oren Cohen, Susan Countryman, Mary Ann Cristobal, Bobbi Croy, Suzanne Dale, Hrushikesh Deshmukh, Amanda Douglas, Vincent Drouillon, Marcia Eisenberg, Howard Engler, Rama Ghatti, Prashant Gupta, Susan Hicks, Jake Humphrey, Lax Iyer, Lisa Pfefferle, Manoj Jain, Matthew Robinson, Mohan Kolli, Brian Krueger, Tim Kuphal, Stanley Letovsky, Michael Levandoski, Craig Lukasik, Jonathan Meltzer, Brian Norvell, Mindy Nye, Scott Parker, Christos Petropoulos, John Pruitt, Steven Ragan, Scott Ryan, Mike Sapeta, Jana Schroth, Suresh Babu Selvaraju, Goran Stevovic, Amanda Suchanek, Andrea Throop, Lyndon Tilson, Thomas Urban, Joe Voshell, Kimberly Wagner, Jonathan Williams, Mary Williamson, Qian Zeng, Tricia Zwiefelhofer, Clinton R. Paden, Duncan MacCannell |
| EPI_ISL_3112384, EPI_ISL_3112385 | Institute for Infectious Diseases | Institute for Infectious Diseases | Stefan Neuenschwander, Christian Baumann, Miguel A Terrazos Miani, Cora SÃ¤gesser, Pascal Bittel, Peter Keller, Franziska Suter-Riniker, Stephen L Leib, Alban Ramette |
| EPI_ISL_3112386 | Laboratory Corporation of America | Centers for Disease Control and Prevention Division of Viral Diseases, Pathogen Discovery | Dakota Howard, Dhwani Batra, Peter W. Cook, Kara Moser, Adrian Paskey, Jason Caravas, Benjamin Rambo-Martin, Shatavia Morrison, Christopher Gulvick, Scott Sammons, Yvette Unoarumhi, Darlene Wagner, Matthew Schmerer, Minoo Agarwal, Eyad Almasri, Debbie Boles, Ayla Burns, Nuthawin Charoensri, Oren Cohen, Susan Countryman, Mary Ann Cristobal, Bobbi Croy, Suzanne Dale, Hrushikesh Deshmukh, Amanda Douglas, Vincent Drouillon, Marcia Eisenberg, Howard Engler, Rama Ghatti, Prashant Gupta, Susan Hicks, Jake Humphrey, Lax Iyer, Lisa Pfefferle, Manoj Jain, Matthew Robinson, Mohan Kolli, Brian Krueger, Tim Kuphal, Stanley Letovsky, Michael Levandoski, Craig Lukasik, Jonathan Meltzer, Brian Norvell, Mindy Nye, Scott Parker, Christos Petropoulos, John Pruitt, Steven Ragan, Scott Ryan, Mike Sapeta, Jana Schroth, Suresh Babu Selvaraju, Goran Stevovic, Amanda Suchanek, Andrea Throop, Lyndon Tilson, Thomas Urban, Joe Voshell, Kimberly Wagner, Jonathan Williams, Mary Williamson, Qian Zeng, Tricia Zwiefelhofer, Clinton R. Paden, Duncan MacCannell |
| EPI_ISL_3112387, EPI_ISL_3112388 | Institute for Infectious Diseases | Institute for Infectious Diseases | Stefan Neuenschwander, Christian Baumann, Miguel A Terrazos Miani, Cora SÃ¤gesser, Pascal Bittel, Peter Keller, Franziska Suter-Riniker, Stephen L Leib, Alban Ramette |
| EPI_ISL_3112389 | Laboratory Corporation of America | Centers for Disease Control and Prevention Division of Viral Diseases, Pathogen Discovery | Dakota Howard, Dhwani Batra, Peter W. Cook, Kara Moser, Adrian Paskey, Jason Caravas, Benjamin Rambo-Martin, Shatavia Morrison, Christopher Gulvick, Scott Sammons, Yvette Unoarumhi, Darlene Wagner, Matthew Schmerer, Minoo Agarwal, Eyad Almasri, Debbie Boles, Ayla Burns, Nuthawin Charoensri, Oren Cohen, Susan Countryman, Mary Ann Cristobal, Bobbi Croy, Suzanne Dale, Hrushikesh Deshmukh, Amanda Douglas, Vincent Drouillon, Marcia Eisenberg, Howard Engler, Rama Ghatti, Prashant Gupta, Susan Hicks, Jake Humphrey, Lax Iyer, Lisa Pfefferle, Manoj Jain, Matthew Robinson, Mohan Kolli, Brian Krueger, Tim Kuphal, Stanley Letovsky, Michael Levandoski, Craig Lukasik, Jonathan Meltzer, Brian Norvell, Mindy Nye, Scott Parker, Christos Petropoulos, John Pruitt, Steven Ragan, Scott Ryan, Mike Sapeta, Jana Schroth, Suresh Babu Selvaraju, Goran Stevovic, Amanda Suchanek, Andrea Throop, Lyndon Tilson, Thomas Urban, Joe Voshell, Kimberly Wagner, Jonathan Williams, Mary Williamson, Qian Zeng, Tricia Zwiefelhofer, Clinton R. Paden, Duncan MacCannell |
| EPI_ISL_3112390 | Institute for Infectious Diseases | Institute for Infectious Diseases | Stefan Neuenschwander, Christian Baumann, Miguel A Terrazos Miani, Cora SÃ¤gesser, Pascal Bittel, Peter Keller, Franziska Suter-Riniker, Stephen L Leib, Alban Ramette |
| EPI_ISL_3112391 | Laboratory Corporation of America | Centers for Disease Control and Prevention Division of Viral Diseases, Pathogen Discovery | Dakota Howard, Dhwani Batra, Peter W. Cook, Kara Moser, Adrian Paskey, Jason Caravas, Benjamin Rambo-Martin, Shatavia Morrison, Christopher Gulvick, Scott Sammons, Yvette Unoarumhi, Darlene Wagner, Matthew Schmerer, Minoo Agarwal, Eyad Almasri, Debbie Boles, Ayla Burns, Nuthawin Charoensri, Oren Cohen, Susan Countryman, Mary Ann Cristobal, Bobbi Croy, Suzanne Dale, Hrushikesh Deshmukh, Amanda Douglas, Vincent Drouillon, Marcia Eisenberg, Howard Engler, Rama Ghatti, Prashant Gupta, Susan Hicks, Jake Humphrey, Lax Iyer, Lisa Pfefferle, Manoj Jain, Matthew Robinson, Mohan Kolli, Brian Krueger, Tim Kuphal, Stanley Letovsky, Michael Levandoski, Craig Lukasik, Jonathan Meltzer, Brian |

| | | | |
|---|---|---|---|
| | | | Norvell, Mindy Nye, Scott Parker, Christos Petropoulos, John Pruitt, Steven Ragan, Scott Ryan, Mike Sapeta, Jana Schroth, Suresh Babu Selvaraju, Goran Stevovic, Amanda Suchanek, Andrea Throop, Lyndon Tilson, Thomas Urban, Joe Voshell, Kimberly Wagner, Jonathan Williams, Mary Williamson, Qian Zeng, Tricia Zwiefelhofer, Clinton R. Paden, Duncan MacCannell |
| EPI_ISL_3112392, EPI_ISL_3112393 | Institute for Infectious Diseases | Institute for Infectious Diseases | Stefan Neuenschwander, Christian Baumann, Miguel A Terrazos Miani, Cora SÃ¤gesser, Pascal Bittel, Peter Keller, Franziska Suter-Riniker, Stephen L Leib, Alban Ramette |
| EPI_ISL_3112394 | Laboratory Corporation of America | Centers for Disease Control and Prevention Division of Viral Diseases, Pathogen Discovery | Dakota Howard, Dhwani Batra, Peter W. Cook, Kara Moser, Adrian Paskey, Jason Caravas, Benjamin Rambo-Martin, Shatavia Morrison, Christopher Gulvick, Scott Sammons, Yvette Unoarumhi, Darlene Wagner, Matthew Schmerer, Minoo Agarwal, Eyad Almasri, Debbie Boles, Ayla Burns, Nuthawin Charoensri, Oren Cohen, Susan Countryman, Mary Ann Cristobal, Bobbi Croy, Suzanne Dale, Hrushikesh Deshmukh, Amanda Douglas, Vincent Drouillon, Marcia Eisenberg, Howard Engler, Rama Ghatti, Prashant Gupta, Susan Hicks, Jake Humphrey, Lax Iyer, Lisa Pfefferle, Manoj Jain, Matthew Robinson, Mohan Kolli, Brian Krueger, Tim Kuphal, Stanley Letovsky, Michael Levandoski, Craig Lukasik, Jonathan Meltzer, Brian Norvell, Mindy Nye, Scott Parker, Christos Petropoulos, John Pruitt, Steven Ragan, Scott Ryan, Mike Sapeta, Jana Schroth, Suresh Babu Selvaraju, Goran Stevovic, Amanda Suchanek, Andrea Throop, Lyndon Tilson, Thomas Urban, Joe Voshell, Kimberly Wagner, Jonathan Williams, Mary Williamson, Qian Zeng, Tricia Zwiefelhofer, Clinton R. Paden, Duncan MacCannell |
| EPI_ISL_3112395, EPI_ISL_3112396 | Institute for Infectious Diseases | Institute for Infectious Diseases | Stefan Neuenschwander, Christian Baumann, Miguel A Terrazos Miani, Cora SÃ¤gesser, Pascal Bittel, Peter Keller, Franziska Suter-Riniker, Stephen L Leib, Alban Ramette |
| EPI_ISL_3112397 | Laboratory Corporation of America | Centers for Disease Control and Prevention Division of Viral Diseases, Pathogen Discovery | Dakota Howard, Dhwani Batra, Peter W. Cook, Kara Moser, Adrian Paskey, Jason Caravas, Benjamin Rambo-Martin, Shatavia Morrison, Christopher Gulvick, Scott Sammons, Yvette Unoarumhi, Darlene Wagner, Matthew Schmerer, Minoo Agarwal, Eyad Almasri, Debbie Boles, Ayla Burns, Nuthawin Charoensri, Oren Cohen, Susan Countryman, Mary Ann Cristobal, Bobbi Croy, Suzanne Dale, Hrushikesh Deshmukh, Amanda Douglas, Vincent Drouillon, Marcia Eisenberg, Howard Engler, Rama Ghatti, Prashant Gupta, Susan Hicks, Jake Humphrey, Lax Iyer, Lisa Pfefferle, Manoj Jain, Matthew Robinson, Mohan Kolli, Brian Krueger, Tim Kuphal, Stanley Letovsky, Michael Levandoski, Craig Lukasik, Jonathan Meltzer, Brian Norvell, Mindy Nye, Scott Parker, Christos Petropoulos, John Pruitt, Steven Ragan, Scott Ryan, Mike Sapeta, Jana Schroth, Suresh Babu Selvaraju, Goran Stevovic, Amanda Suchanek, Andrea Throop, Lyndon Tilson, Thomas Urban, Joe Voshell, Kimberly Wagner, Jonathan Williams, Mary Williamson, Qian Zeng, Tricia Zwiefelhofer, Clinton R. Paden, Duncan MacCannell |
| EPI_ISL_3112398, EPI_ISL_3112399 | Institute for Infectious Diseases | Institute for Infectious Diseases | Stefan Neuenschwander, Christian Baumann, Miguel A Terrazos Miani, Cora SÃ¤gesser, Pascal Bittel, Peter Keller, Franziska Suter-Riniker, Stephen L Leib, Alban Ramette |
| EPI_ISL_3112400 | Laboratory Corporation of America | Centers for Disease Control and Prevention Division of Viral Diseases, Pathogen Discovery | Dakota Howard, Dhwani Batra, Peter W. Cook, Kara Moser, Adrian Paskey, Jason Caravas, Benjamin Rambo-Martin, Shatavia Morrison, Christopher Gulvick, Scott Sammons, Yvette Unoarumhi, Darlene Wagner, Matthew Schmerer, Minoo Agarwal, Eyad Almasri, Debbie Boles, Ayla Burns, Nuthawin Charoensri, Oren Cohen, Susan Countryman, Mary Ann Cristobal, Bobbi Croy, Suzanne Dale, Hrushikesh Deshmukh, Amanda Douglas, Vincent Drouillon, Marcia Eisenberg, Howard Engler, Rama Ghatti, Prashant Gupta, Susan Hicks, Jake Humphrey, Lax Iyer, Lisa Pfefferle, Manoj Jain, Matthew Robinson, Mohan Kolli, Brian Krueger, Tim Kuphal, Stanley Letovsky, Michael Levandoski, Craig Lukasik, Jonathan Meltzer, Brian Norvell, Mindy Nye, Scott Parker, Christos Petropoulos, John Pruitt, Steven Ragan, Scott Ryan, Mike Sapeta, Jana Schroth, Suresh Babu Selvaraju, Goran Stevovic, Amanda Suchanek, Andrea Throop, Lyndon Tilson, Thomas Urban, Joe Voshell, Kimberly Wagner, Jonathan Williams, Mary Williamson, Qian Zeng, Tricia Zwiefelhofer, Clinton R. Paden, Duncan MacCannell |
| EPI_ISL_3112401, EPI_ISL_3112403, EPI_ISL_3112404 | Institute for Infectious Diseases | Institute for Infectious Diseases | Stefan Neuenschwander, Christian Baumann, Miguel A Terrazos Miani, Cora SÃ¤gesser, Pascal Bittel, Peter Keller, Franziska Suter-Riniker, Stephen L Leib, Alban Ramette |
| EPI_ISL_3112405 | Laboratory Corporation of America | Centers for Disease Control and Prevention Division of Viral Diseases, Pathogen Discovery | Dakota Howard, Dhwani Batra, Peter W. Cook, Kara Moser, Adrian Paskey, Jason Caravas, Benjamin Rambo-Martin, Shatavia Morrison, Christopher Gulvick, Scott Sammons, Yvette Unoarumhi, Darlene Wagner, Matthew Schmerer, Minoo Agarwal, Eyad Almasri, Debbie Boles, Ayla Burns, Nuthawin Charoensri, Oren Cohen, Susan Countryman, Mary Ann Cristobal, Bobbi Croy, Suzanne Dale, Hrushikesh Deshmukh, Amanda Douglas, Vincent Drouillon, Marcia Eisenberg, Howard Engler, Rama Ghatti, Prashant Gupta, Susan Hicks, Jake Humphrey, Lax Iyer, Lisa Pfefferle, Manoj Jain, Matthew Robinson, Mohan Kolli, Brian Krueger, Tim Kuphal, Stanley Letovsky, Michael Levandoski, Craig Lukasik, Jonathan Meltzer, Brian Norvell, Mindy Nye, Scott Parker, Christos Petropoulos, John Pruitt, Steven Ragan, Scott Ryan, Mike Sapeta, Jana Schroth, Suresh Babu Selvaraju, Goran Stevovic, Amanda Suchanek, Andrea Throop, Lyndon Tilson, Thomas Urban, Joe Voshell, Kimberly Wagner, Jonathan Williams, Mary Williamson, Qian Zeng, Tricia Zwiefelhofer, Clinton R. Paden, Duncan MacCannell |
| EPI_ISL_3112406 | Institute for Infectious Diseases | Institute for Infectious Diseases | Stefan Neuenschwander, Christian Baumann, Miguel A Terrazos Miani, Cora SÃ¤gesser, Pascal Bittel, Peter Keller, Franziska Suter-Riniker, Stephen L Leib, Alban Ramette |
| EPI_ISL_3112408 | Laboratory Corporation of America | Centers for Disease Control and Prevention Division of Viral Diseases, Pathogen Discovery | Dakota Howard, Dhwani Batra, Peter W. Cook, Kara Moser, Adrian Paskey, Jason Caravas, Benjamin Rambo-Martin, Shatavia Morrison, Christopher Gulvick, Scott Sammons, Yvette Unoarumhi, Darlene Wagner, Matthew Schmerer, Minoo Agarwal, Eyad Almasri, Debbie Boles, Ayla Burns, Nuthawin Charoensri, Oren Cohen, Susan Countryman, Mary Ann Cristobal, Bobbi Croy, Suzanne Dale, Hrushikesh Deshmukh, Amanda Douglas, Vincent Drouillon, Marcia Eisenberg, Howard Engler, Rama Ghatti, Prashant Gupta, Susan Hicks, Jake Humphrey, Lax Iyer, Lisa Pfefferle, Manoj Jain, Matthew Robinson, Mohan Kolli, Brian Krueger, Tim Kuphal, Stanley Letovsky, Michael Levandoski, Craig Lukasik, Jonathan Meltzer, Brian Norvell, Mindy Nye, Scott Parker, Christos Petropoulos, John Pruitt, Steven Ragan, Scott Ryan, Mike Sapeta, Jana Schroth, Suresh Babu Selvaraju, Goran Stevovic, Amanda Suchanek, Andrea Throop, Lyndon Tilson, Thomas Urban, Joe Voshell, Kimberly Wagner, Jonathan Williams, Mary Williamson, Qian Zeng, Tricia Zwiefelhofer, Clinton R. Paden, Duncan MacCannell |
| EPI_ISL_3112409 | Institute for Infectious Diseases | Institute for Infectious Diseases | Stefan Neuenschwander, Christian Baumann, Miguel A Terrazos Miani, Cora SÃ¤gesser, Pascal Bittel, Peter Keller, Franziska Suter-Riniker, Stephen L Leib, Alban Ramette |
| EPI_ISL_3112410, EPI_ISL_3112411, EPI_ISL_3112412, EPI_ISL_3112415, EPI_ISL_3112417, EPI_ISL_3112418, EPI_ISL_3112420, EPI_ISL_3112421, EPI_ISL_3112422, EPI_ISL_3112424, EPI_ISL_3112425, EPI_ISL_3112426, EPI_ISL_3112427, EPI_ISL_3112428, EPI_ISL_3112429, EPI_ISL_3112431, EPI_ISL_3112432, EPI_ISL_3112433, EPI_ISL_3112434, EPI_ISL_3112435, EPI_ISL_3112436, EPI_ISL_3112438, EPI_ISL_3112439, EPI_ISL_3112440, EPI_ISL_3112441, EPI_ISL_3112442, EPI_ISL_3112443, EPI_ISL_3112444, EPI_ISL_3112445, EPI_ISL_3112446, EPI_ISL_3112447, EPI_ISL_3112448, EPI_ISL_3112449, EPI_ISL_3112450, EPI_ISL_3112451, EPI_ISL_3112452, EPI_ISL_3112453, EPI_ISL_3112454, EPI_ISL_3112456, EPI_ISL_3112457, EPI_ISL_3112458, EPI_ISL_3112460, EPI_ISL_3112461, EPI_ISL_3112462, EPI_ISL_3112463, EPI_ISL_3112464, EPI_ISL_3112465, EPI_ISL_3112466, EPI_ISL_3112467, EPI_ISL_3112468, EPI_ISL_3112469, EPI_ISL_3112470, EPI_ISL_3112471, EPI_ISL_3112473, EPI_ISL_3112475, EPI_ISL_3112476, EPI_ISL_3112477, EPI_ISL_3112480, EPI_ISL_3112481, EPI_ISL_3112482, EPI_ISL_3112483, EPI_ISL_3112484, EPI_ISL_3112486, EPI_ISL_3112487, EPI_ISL_3112488, EPI_ISL_3112489, EPI_ISL_3112490, EPI_ISL_3112491, EPI_ISL_3112492, EPI_ISL_3112493, EPI_ISL_3112494, EPI_ISL_3112495, EPI_ISL_3112496, EPI_ISL_3112497, EPI_ISL_3112498, EPI_ISL_3112499, EPI_ISL_3112500, EPI_ISL_3112501, EPI_ISL_3112502, EPI_ISL_3112504, EPI_ISL_3112505, EPI_ISL_3112506, EPI_ISL_3112507, EPI_ISL_3112508, EPI_ISL_3112509, EPI_ISL_3112510, EPI_ISL_3112511, EPI_ISL_3112512, EPI_ISL_3112513, EPI_ISL_3112514, EPI_ISL_3112516, EPI_ISL_3112517, EPI_ISL_3112519, EPI_ISL_3112520, EPI_ISL_3112521, EPI_ISL_3112524, EPI_ISL_3112525, EPI_ISL_3112526, EPI_ISL_3112527, EPI_ISL_3112528, EPI_ISL_3112529, EPI_ISL_3112530, EPI_ISL_3112531, EPI_ISL_3112532, EPI_ISL_3112533, EPI_ISL_3112535, EPI_ISL_3112536, EPI_ISL_3112537, EPI_ISL_3112538, EPI_ISL_3112539, EPI_ISL_3112541, EPI_ISL_3112542, EPI_ISL_3112543, EPI_ISL_3112544, EPI_ISL_3112545, EPI_ISL_3112547, EPI_ISL_3112548, EPI_ISL_3112549, EPI_ISL_3112550, EPI_ISL_3112551, EPI_ISL_3112553, EPI_ISL_3112554, EPI_ISL_3112555, EPI_ISL_3112556, EPI_ISL_3112557, EPI_ISL_3112558, EPI_ISL_3112560, EPI_ISL_3112561, EPI_ISL_3112562, EPI_ISL_3112564, EPI_ISL_3112565, EPI_ISL_3112566, EPI_ISL_3112570, EPI_ISL_3112571, EPI_ISL_3112572, EPI_ISL_3112573, EPI_ISL_3112574, EPI_ISL_3112575, EPI_ISL_3112576, EPI_ISL_3112577, EPI_ISL_3112578, EPI_ISL_3112579, EPI_ISL_3112580, EPI_ISL_3112581, EPI_ISL_3112582, EPI_ISL_3112583, EPI_ISL_3112584, EPI_ISL_3112585, EPI_ISL_3112586, EPI_ISL_3112587, EPI_ISL_3112588, EPI_ISL_3112589, EPI_ISL_3112590, EPI_ISL_3112591, EPI_ISL_3112592, EPI_ISL_3112593, EPI_ISL_3112594, EPI_ISL_3112595, EPI_ISL_3112596, EPI_ISL_3112597, EPI_ISL_3112598, EPI_ISL_3112599, EPI_ISL_3112600, EPI_ISL_3112601, EPI_ISL_3112602, EPI_ISL_3112603, EPI_ISL_3112605, EPI_ISL_3112606, EPI_ISL_3112607, EPI_ISL_3112608, EPI_ISL_3112609, EPI_ISL_3112611, EPI_ISL_3112612, EPI_ISL_3112613, EPI_ISL_3112614, EPI_ISL_3112615, EPI_ISL_3112616, EPI_ISL_3112617, EPI_ISL_3112618, EPI_ISL_3112619, EPI_ISL_3112621, EPI_ISL_3112622, EPI_ISL_3112623, EPI_ISL_3112624, EPI_ISL_3112625, EPI_ISL_3112626, EPI_ISL_3112627, EPI_ISL_3112628, EPI_ISL_3112629, EPI_ISL_3112630, EPI_ISL_3112631, EPI_ISL_3112632, EPI_ISL_3112633, EPI_ISL_3112634, EPI_ISL_3112635, EPI_ISL_3112636, EPI_ISL_3112637, EPI_ISL_3112638, EPI_ISL_3112639, EPI_ISL_3112640, EPI_ISL_3112641, EPI_ISL_3112642, EPI_ISL_3112643, EPI_ISL_3112644, EPI_ISL_3112645, EPI_ISL_3112646, EPI_ISL_3112647, EPI_ISL_3112648, EPI_ISL_3112649, EPI_ISL_3112650, EPI_ISL_3112651, EPI_ISL_3112653, EPI_ISL_3112654, EPI_ISL_3112655, EPI_ISL_3112657, EPI_ISL_3112658, EPI_ISL_3112659, EPI_ISL_3112660, EPI_ISL_3112661, EPI_ISL_3112662, EPI_ISL_3112663, EPI_ISL_3112664, EPI_ISL_3112665, EPI_ISL_3112666, EPI_ISL_3112667, EPI_ISL_3112668, EPI_ISL_3112670, EPI_ISL_3112671, EPI_ISL_3112672, EPI_ISL_3112673, EPI_ISL_3112674, EPI_ISL_3112675, EPI_ISL_3112676, EPI_ISL_3112677, EPI_ISL_3112678, EPI_ISL_3112679, EPI_ISL_3112680, EPI_ISL_3112681, EPI_ISL_3112682, EPI_ISL_3112683, EPI_ISL_3112684, EPI_ISL_3112685, EPI_ISL_3112686, EPI_ISL_3112687, EPI_ISL_3112688, EPI_ISL_3112690, EPI_ISL_3112691, EPI_ISL_3112692, EPI_ISL_3112693, EPI_ISL_3112694, EPI_ISL_3112695, EPI_ISL_3112696, EPI_ISL_3112697, EPI_ISL_3112698, EPI_ISL_3112699, EPI_ISL_3112700, EPI_ISL_3112701, EPI_ISL_3112702, EPI_ISL_3112703, EPI_ISL_3112704, EPI_ISL_3112707, EPI_ISL_3112708, EPI_ISL_3112709, EPI_ISL_3112710, EPI_ISL_3112711, EPI_ISL_3112712, EPI_ISL_3112713, EPI_ISL_3112714, EPI_ISL_3112715, EPI_ISL_3112716, EPI_ISL_3112717, EPI_ISL_3112718, EPI_ISL_3112719, EPI_ISL_3112720, EPI_ISL_3112721, EPI_ISL_3112722, EPI_ISL_3112723, EPI_ISL_3112724, EPI_ISL_3112725, EPI_ISL_3112726, EPI_ISL_3112728, EPI_ISL_3112729, EPI_ISL_3112730, EPI_ISL_3112731, EPI_ISL_3112732, EPI_ISL_3112733, EPI_ISL_3112734, EPI_ISL_3112735, EPI_ISL_3112736, EPI_ISL_3112737, EPI_ISL_3112739, EPI_ISL_3112741, EPI_ISL_3112742, EPI_ISL_3112743, EPI_ISL_3112744, EPI_ISL_3112745, EPI_ISL_3112746, EPI_ISL_3112747, EPI_ISL_3112748, EPI_ISL_3112751, EPI_ISL_3112752, EPI_ISL_3112753, EPI_ISL_3112754, EPI_ISL_3112755, EPI_ISL_3112756, EPI_ISL_3112757, EPI_ISL_3112758, EPI_ISL_3112759, EPI_ISL_3112760, EPI_ISL_3112762, EPI_ISL_3112763, EPI_ISL_3112765, EPI_ISL_3112766, EPI_ISL_3112769, EPI_ISL_3112770, EPI_ISL_3112771, EPI_ISL_3112772, EPI_ISL_3112773, EPI_ISL_3112774, EPI_ISL_3112775, EPI_ISL_3112776, EPI_ISL_3112777, EPI_ISL_3112778, EPI_ISL_3112779, EPI_ISL_3112780, EPI_ISL_3112781, EPI_ISL_3112783, EPI_ISL_3112784, EPI_ISL_3112785, EPI_ISL_3112786, EPI_ISL_3112787, EPI_ISL_3112788, EPI_ISL_3112789, EPI_ISL_3112791, EPI_ISL_3112792, EPI_ISL_3112793, EPI_ISL_3112794, EPI_ISL_3112796, EPI_ISL_3112798, EPI_ISL_3112799, EPI_ISL_3112800, EPI_ISL_3112803, EPI_ISL_3112804, EPI_ISL_3112805, EPI_ISL_3112806, EPI_ISL_3112807, EPI_ISL_3112808, EPI_ISL_3112809, EPI_ISL_3112810, EPI_ISL_3112811, EPI_ISL_3112812, EPI_ISL_3112813, EPI_ISL_3112814, EPI_ISL_3112815, EPI_ISL_3112816, EPI_ISL_3112817, EPI_ISL_3112818, EPI_ISL_3112821, EPI_ISL_3112822, EPI_ISL_3112823, EPI_ISL_3112824, EPI_ISL_3112825, EPI_ISL_3112826, EPI_ISL_3112827, EPI_ISL_3112828, EPI_ISL_3112829, EPI_ISL_3112830, EPI_ISL_3112833, EPI_ISL_3112834, EPI_ISL_3112835, EPI_ISL_3112836, EPI_ISL_3112837, EPI_ISL_3112838, EPI_ISL_3112839, EPI_ISL_3112840, EPI_ISL_3112841, EPI_ISL_3112842, EPI_ISL_3112843, EPI_ISL_3112844, EPI_ISL_3112845, EPI_ISL_3112846, EPI_ISL_3112847, EPI_ISL_3112848, EPI_ISL_3112849, EPI_ISL_3112850, EPI_ISL_3112851, EPI_ISL_3112852, EPI_ISL_3112853, EPI_ISL_3112854, EPI_ISL_3112855, EPI_ISL_3112856, EPI_ISL_3112857, EPI_ISL_3112858, EPI_ISL_3112859, EPI_ISL_3112860, EPI_ISL_3112861, EPI_ISL_3112862, EPI_ISL_3112863, EPI_ISL_3112864, EPI_ISL_3112865, EPI_ISL_3112867, EPI_ISL_3112868, EPI_ISL_3112869, EPI_ISL_3112870, EPI_ISL_3112871, EPI_ISL_3112872, EPI_ISL_3112873, EPI_ISL_3112874, EPI_ISL_3112875, EPI_ISL_3112876, EPI_ISL_3112877, EPI_ISL_3112878, EPI_ISL_3112879, EPI_ISL_3112880, EPI_ISL_3112881, EPI_ISL_3112882, EPI_ISL_3112883, EPI_ISL_3112885, EPI_ISL_3112886, EPI_ISL_3112887, EPI_ISL_3112888, EPI_ISL_3112890, EPI_ISL_3112891, EPI_ISL_3112893, EPI_ISL_3112894, EPI_ISL_3112895, EPI_ISL_3112896, EPI_ISL_3112897, EPI_ISL_3112898, EPI_ISL_3112899, EPI_ISL_3112900, EPI_ISL_3112901, EPI_ISL_3112902, EPI_ISL_3112903, EPI_ISL_3112904, EPI_ISL_3112905, EPI_ISL_3112906, EPI_ISL_3112907, EPI_ISL_3112908, EPI_ISL_3112909, EPI_ISL_3112910, EPI_ISL_3112911, EPI_ISL_3112913, EPI_ISL_3112914, EPI_ISL_3112916, EPI_ISL_3112917, EPI_ISL_3112918, EPI_ISL_3112920, EPI_ISL_3112921, EPI_ISL_3112923, EPI_ISL_3112924, EPI_ISL_3112925, EPI_ISL_3112927, EPI_ISL_3112928, EPI_ISL_3112929, EPI_ISL_3112930, EPI_ISL_3112931, EPI_ISL_3112932, EPI_ISL_3112933, EPI_ISL_3112934, EPI_ISL_3112935, EPI_ISL_3112936, EPI_ISL_3112938, EPI_ISL_3112940, EPI_ISL_3112941, EPI_ISL_3112942, EPI_ISL_3112943, EPI_ISL_3112944, EPI_ISL_3112945, EPI_ISL_3112946, EPI_ISL_3112947, EPI_ISL_3112948, EPI_ISL_3112949, EPI_ISL_3112950, EPI_ISL_3112951, EPI_ISL_3112952, EPI_ISL_3112953, EPI_ISL_3112954, EPI_ISL_3112955, EPI_ISL_3112959, EPI_ISL_3112961, EPI_ISL_3112962, EPI_ISL_3112964, EPI_ISL_3112965, EPI_ISL_3112967, EPI_ISL_3112968, EPI_ISL_3112969, EPI_ISL_3112970, EPI_ISL_3112971, EPI_ISL_3112972, EPI_ISL_3112973, EPI_ISL_3112974, EPI_ISL_3112975, EPI_ISL_3112977, EPI_ISL_3112978, EPI_ISL_3112979, EPI_ISL_3112980, EPI_ISL_3112981, EPI_ISL_3112982, EPI_ISL_3112985, EPI_ISL_3112986, EPI_ISL_3112987, EPI_ISL_3112988, EPI_ISL_3112992, EPI_ISL_3112994, EPI_ISL_3112996, EPI_ISL_3112999, EPI_ISL_3113001, EPI_ISL_3113003, EPI_ISL_3113004, EPI_ISL_3113005, EPI_ISL_3113006, EPI_ISL_3113008, EPI_ISL_3113009, EPI_ISL_3113010, EPI_ISL_3113011, EPI_ISL_3113012, EPI_ISL_3113013, EPI_ISL_3113015, EPI_ISL_3113016, EPI_ISL_3113017, EPI_ISL_3113018, EPI_ISL_3113019, EPI_ISL_3113020, EPI_ISL_3113021, EPI_ISL_3113022, EPI_ISL_3113023, EPI_ISL_3113024, EPI_ISL_3113025, EPI_ISL_3113026, EPI_ISL_3113028, EPI_ISL_3113029, EPI_ISL_3113030, EPI_ISL_3113031, EPI_ISL_3113032, EPI_ISL_3113037, EPI_ISL_3113039, EPI_ISL_3113040, EPI_ISL_3113041, EPI_ISL_3113044, EPI_ISL_3113045, EPI_ISL_3113046, EPI_ISL_3113047, EPI_ISL_3113048, EPI_ISL_3113049, EPI_ISL_3113050, EPI_ISL_3113051, EPI_ISL_3113053, EPI_ISL_3113055, EPI_ISL_3113056, EPI_ISL_3113057, EPI_ISL_3113058, EPI_ISL_3113059, EPI_ISL_3113062, EPI_ISL_3113064, EPI_ISL_3113065, EPI_ISL_3113066, EPI_ISL_3113067, EPI_ISL_3113069, EPI_ISL_3113070, EPI_ISL_3113072, EPI_ISL_3113073, EPI_ISL_3113074, EPI_ISL_3113075, EPI_ISL_3113076, EPI_ISL_3113077, EPI_ISL_3113078, EPI_ISL_3113079, EPI_ISL_3113080, EPI_ISL_3113083, EPI_ISL_3113084, EPI_ISL_3113085, EPI_ISL_3113086, EPI_ISL_3113087, EPI_ISL_3113088, EPI_ISL_3113089, EPI_ISL_3113090, EPI_ISL_3113091, EPI_ISL_3113092, EPI_ISL_3113093, EPI_ISL_3113094, EPI_ISL_3113095, EPI_ISL_3113097, EPI_ISL_3113098, EPI_ISL_3113099, EPI_ISL_3113100, EPI_ISL_3113104, EPI_ISL_3113105, EPI_ISL_3113106, EPI_ISL_3113107, EPI_ISL_3113108, EPI_ISL_3113109, EPI_ISL_3113110, EPI_ISL_3113111, EPI_ISL_3113112, EPI_ISL_3113113, EPI_ISL_3113114, EPI_ISL_3113115, EPI_ISL_3113116, EPI_ISL_3113117, EPI_ISL_3113119, EPI_ISL_3113121, EPI_ISL_3113122, EPI_ISL_3113123, EPI_ISL_3113124, EPI_ISL_3113125, EPI_ISL_3113126, EPI_ISL_3113127, EPI_ISL_3113128, EPI_ISL_3113129, EPI_ISL_3113130, EPI_ISL_3113131, EPI_ISL_3113132, EPI_ISL_3113133, EPI_ISL_3113135, EPI_ISL_3113136, EPI_ISL_3113138 | | | |
| see above | Laboratory Corporation of America | Centers for Disease Control and Prevention Division of Viral Diseases, Pathogen Discovery | Dakota Howard, Dhwani Batra, Peter W. Cook, Kara Moser, Adrian Paskey, Jason Caravas, Benjamin Rambo-Martin, Shatavia Morrison, Christopher Gulvick, Scott Sammons, Yvette Unoarumhi, Darlene Wagner, Matthew Schmerer, Minoo Agarwal, Eyad Almasri, Debbie Boles, Ayla Burns, Nuthawin Charoensri, Oren Cohen, Susan Countryman, Mary Ann Cristobal, Bobbi Croy, Suzanne Dale, Hrushikesh Deshmukh, Amanda Douglas, Vincent Drouillon, Marcia Eisenberg, Howard Engler, Rama Ghatti, Prashant Gupta, Susan Hicks, Jake Humphrey, Lax Iyer, Lisa Pfefferle, Manoj Jain, Matthew Robinson, Mohan Kolli, Brian Krueger, Tim Kuphal, Stanley Letovsky, Michael Levandoski, Craig Lukasik, Jonathan Meltzer, Brian Norvell, Mindy Nye, Scott Parker, Christos Petropoulos, John Pruitt, Steven Ragan, Scott Ryan, Mike Sapeta, Jana Schroth, Suresh Babu Selvaraju, Goran Stevovic, Amanda Suchanek, Andrea Throop, Lyndon Tilson, Thomas Urban, Joe Voshell, Kimberly Wagner, Jonathan Williams, Mary Williamson, Qian Zeng, Tricia Zwiefelhofer, Clinton R. Paden, Duncan MacCannell |
| EPI_ISL_3113139 | Labor Berlin Charité Vivantes GmbH / Institut für Virologie | Charité Universitätsmedizin Berlin, Institut für Virologie/Labor Berlin | Peter Menzel, Christine Stephan, Rolf Schwarzer, Victor M Corman, Barbara Mühlemann, Terry Jones, Christian Drosten |
| EPI_ISL_3113140 | Laboratory Corporation of America | Centers for Disease Control and Prevention Division of Viral Diseases, Pathogen Discovery | Dakota Howard, Dhwani Batra, Peter W. Cook, Kara Moser, Adrian Paskey, Jason Caravas, Benjamin Rambo-Martin, Shatavia Morrison, Christopher Gulvick, Scott Sammons, Yvette Unoarumhi, Darlene Wagner, Matthew Schmerer, Minoo Agarwal, Eyad Almasri, Debbie Boles, Ayla Burns, Nuthawin Charoensri, Oren Cohen, Susan Countryman, Mary Ann Cristobal, Bobbi Croy, Suzanne Dale, Hrushikesh Deshmukh, Amanda Douglas, Vincent Drouillon, Marcia Eisenberg, Howard Engler, Rama Ghatti, Prashant Gupta, Susan Hicks, Jake Humphrey, Lax Iyer, Lisa Pfefferle, Manoj Jain, Matthew Robinson, Mohan Kolli, Brian Krueger, Tim Kuphal, Stanley Letovsky, Michael Levandoski, Craig Lukasik, Jonathan Meltzer, Brian Norvell, Mindy Nye, Scott Parker, Christos Petropoulos, John Pruitt, Steven Ragan, Scott Ryan, Mike Sapeta, Jana Schroth, Suresh Babu Selvaraju, Goran Stevovic, Amanda Suchanek, Andrea Throop, Lyndon Tilson, Thomas Urban, Joe Voshell, Kimberly Wagner, Jonathan Williams, Mary Williamson, Qian Zeng, Tricia Zwiefelhofer, Clinton R. Paden, Duncan MacCannell |
| EPI_ISL_3113141 | Labor Berlin Charité Vivantes GmbH / Institut für Virologie | Charité Universitätsmedizin Berlin, Institut für Virologie/Labor Berlin | Peter Menzel, Christine Stephan, Rolf Schwarzer, Victor M Corman, Barbara Mühlemann, Terry Jones, Christian Drosten |
| EPI_ISL_3113143 | Laboratory Corporation of America | Centers for Disease Control and Prevention Division of Viral Diseases, Pathogen Discovery | Dakota Howard, Dhwani Batra, Peter W. Cook, Kara Moser, Adrian Paskey, Jason Caravas, Benjamin Rambo-Martin, Shatavia Morrison, Christopher Gulvick, Scott Sammons, Yvette Unoarumhi, Darlene Wagner, Matthew Schmerer, Minoo Agarwal, Eyad Almasri, Debbie Boles, Ayla Burns, Nuthawin Charoensri, Oren Cohen, Susan Countryman, Mary Ann Cristobal, Bobbi Croy, Suzanne Dale, Hrushikesh Deshmukh, Amanda Douglas, Vincent Drouillon, Marcia Eisenberg, Howard Engler, Rama Ghatti, Prashant Gupta, Susan Hicks, Jake Humphrey, Lax Iyer, Lisa Pfefferle, Manoj Jain, Matthew Robinson, Mohan Kolli, Brian Krueger, Tim Kuphal, Stanley Letovsky, Michael Levandoski, Craig Lukasik, Jonathan Meltzer, Brian Norvell, Mindy Nye, Scott Parker, Christos Petropoulos, John Pruitt, Steven Ragan, Scott Ryan, Mike Sapeta, Jana Schroth, Suresh Babu Selvaraju, Goran Stevovic, Amanda Suchanek, Andrea Throop, Lyndon Tilson, Thomas Urban, Joe Voshell, Kimberly Wagner, Jonathan Williams, Mary Williamson, Qian Zeng, Tricia Zwiefelhofer, Clinton R. Paden, Duncan MacCannell |
| EPI_ISL_3113145, EPI_ISL_3113146, EPI_ISL_3113148 | Labor Berlin Charité Vivantes GmbH / Institut für Virologie | Charité Universitätsmedizin Berlin, Institut für Virologie/Labor Berlin | Peter Menzel, Christine Stephan, Rolf Schwarzer, Victor M Corman, Barbara Mühlemann, Terry Jones, Christian Drosten |
| EPI_ISL_3113149 | Laboratory Corporation of America | Centers for Disease Control and Prevention Division of Viral Diseases, Pathogen Discovery | Dakota Howard, Dhwani Batra, Peter W. Cook, Kara Moser, Adrian Paskey, Jason Caravas, Benjamin Rambo-Martin, Shatavia Morrison, Christopher Gulvick, Scott Sammons, Yvette Unoarumhi, Darlene Wagner, Matthew Schmerer, Minoo Agarwal, Eyad Almasri, Debbie Boles, Ayla Burns, Nuthawin Charoensri, Oren Cohen, Susan Countryman, Mary Ann Cristobal, Bobbi Croy, Suzanne Dale, Hrushikesh Deshmukh, Amanda Douglas, Vincent Drouillon, Marcia Eisenberg, Howard Engler, Rama Ghatti, Prashant Gupta, Susan Hicks, Jake Humphrey, Lax Iyer, Lisa Pfefferle, Manoj Jain, Matthew Robinson, Mohan Kolli, Brian Krueger, Tim Kuphal, Stanley Letovsky, Michael Levandoski, Craig Lukasik, Jonathan Meltzer, Brian Norvell, Mindy Nye, Scott Parker, Christos Petropoulos, John Pruitt, Steven Ragan, Scott Ryan, Mike Sapeta, Jana Schroth, Suresh Babu Selvaraju, Goran Stevovic, Amanda Suchanek, Andrea Throop, Lyndon Tilson, Thomas Urban, Joe Voshell, Kimberly Wagner, Jonathan Williams, Mary Williamson, Qian Zeng, Tricia Zwiefelhofer, Clinton R. Paden, Duncan MacCannell |
| EPI_ISL_3113151 | Labor Berlin Charité Vivantes GmbH / Institut für Virologie | Charité Universitätsmedizin Berlin, Institut für Virologie/Labor Berlin | Peter Menzel, Christine Stephan, Rolf Schwarzer, Victor M Corman, Barbara Mühlemann, Terry Jones, Christian Drosten |
| EPI_ISL_3113152 | Laboratory Corporation of America | Centers for Disease Control and Prevention Division of Viral Diseases, Pathogen Discovery | Dakota Howard, Dhwani Batra, Peter W. Cook, Kara Moser, Adrian Paskey, Jason Caravas, Benjamin Rambo-Martin, Shatavia Morrison, Christopher Gulvick, Scott Sammons, Yvette Unoarumhi, Darlene Wagner, Matthew Schmerer, Minoo Agarwal, Eyad Almasri, Debbie Boles, Ayla Burns, Nuthawin Charoensri, Oren Cohen, Susan Countryman, Mary Ann Cristobal, Bobbi Croy, Suzanne Dale, Hrushikesh Deshmukh, Amanda Douglas, Vincent Drouillon, Marcia Eisenberg, Howard Engler, Rama Ghatti, Prashant Gupta, Susan Hicks, Jake Humphrey, Lax Iyer, Lisa Pfefferle, Manoj Jain, Matthew Robinson, Mohan Kolli, Brian Krueger, Tim Kuphal, Stanley Letovsky, Michael Levandoski, Craig Lukasik, Jonathan Meltzer, Brian Norvell, Mindy Nye, Scott Parker, Christos Petropoulos, John Pruitt, Steven Ragan, Scott Ryan, Mike Sapeta, Jana Schroth, Suresh Babu Selvaraju, Goran Stevovic, Amanda Suchanek, Andrea Throop, Lyndon Tilson, Thomas Urban, Joe Voshell, Kimberly Wagner, Jonathan Williams, Mary Williamson, Qian Zeng, Tricia Zwiefelhofer, Clinton R. Paden, Duncan MacCannell |
| EPI_ISL_3113153 | Labor Berlin Charité Vivantes GmbH / Institut für Virologie | Charité Universitätsmedizin Berlin, Institut für Virologie/Labor Berlin | Peter Menzel, Christine Stephan, Rolf Schwarzer, Victor M Corman, Barbara Mühlemann, Terry Jones, Christian Drosten |
| EPI_ISL_3113154 | Laboratory Corporation of America | Centers for Disease Control and Prevention Division of Viral | Dakota Howard, Dhwani Batra, Peter W. Cook, Kara Moser, Adrian Paskey, Jason Caravas, Benjamin Rambo-Martin, Shatavia Morrison, Christopher Gulvick, Scott Sammons, Yvette Unoarumhi, Darlene Wagner, Matthew Schmerer, Minoo |

| | | | |
|---|---|---|---|
| | | Diseases, Pathogen Discovery | Agarwal, Eyad Almasri, Debbie Boles, Ayla Burns, Nuthawin Charoensri, Oren Cohen, Susan Countryman, Mary Ann Cristobal, Bobbi Croy, Suzanne Dale, Hrushikesh Deshmukh, Amanda Douglas, Vincent Drouillon, Marcia Eisenberg, Howard Engler, Rama Ghatti, Prashant Gupta, Susan Hicks, Jake Humphrey, Lax Iyer, Lisa Pfefferle, Manoj Jain, Matthew Robinson, Mohan Kolli, Brian Krueger, Tim Kuphal, Stanley Letovsky, Michael Levandoski, Craig Lukasik, Jonathan Meltzer, Brian Norvell, Mindy Nye, Scott Parker, Christos Petropoulos, John Pruitt, Steven Ragan, Scott Ryan, Mike Sapeta, Jana Schroth, Suresh Babu Selvaraju, Goran Stevovic, Amanda Suchanek, Andrea Throop, Lyndon Tilson, Thomas Urban, Joe Voshell, Kimberly Wagner, Jonathan Williams, Mary Williamson, Qian Zeng, Tricia Zwiefelhofer, Clinton R. Paden, Duncan MacCannell |
| EPI_ISL_3113155 | Labor Berlin Charité Vivantes GmbH / Institut für Virologie | Charité Universitätsmedizin Berlin, Institut für Virologie/Labor Berlin | Peter Menzel, Christine Stephan, Rolf Schwarzer, Victor M Corman, Barbara Mühlemann, Terry Jones, Christian Drosten |
| EPI_ISL_3113157 | Laboratory Corporation of America | Centers for Disease Control and Prevention Division of Viral Diseases, Pathogen Discovery | Dakota Howard, Dhwani Batra, Peter W. Cook, Kara Moser, Adrian Paskey, Jason Caravas, Benjamin Rambo-Martin, Shatavia Morrison, Christopher Gulvick, Scott Sammons, Yvette Unoarumhi, Darlene Wagner, Matthew Schmerer, Minoo Agarwal, Eyad Almasri, Debbie Boles, Ayla Burns, Nuthawin Charoensri, Oren Cohen, Susan Countryman, Mary Ann Cristobal, Bobbi Croy, Suzanne Dale, Hrushikesh Deshmukh, Amanda Douglas, Vincent Drouillon, Marcia Eisenberg, Howard Engler, Rama Ghatti, Prashant Gupta, Susan Hicks, Jake Humphrey, Lax Iyer, Lisa Pfefferle, Manoj Jain, Matthew Robinson, Mohan Kolli, Brian Krueger, Tim Kuphal, Stanley Letovsky, Michael Levandoski, Craig Lukasik, Jonathan Meltzer, Brian Norvell, Mindy Nye, Scott Parker, Christos Petropoulos, John Pruitt, Steven Ragan, Scott Ryan, Mike Sapeta, Jana Schroth, Suresh Babu Selvaraju, Goran Stevovic, Amanda Suchanek, Andrea Throop, Lyndon Tilson, Thomas Urban, Joe Voshell, Kimberly Wagner, Jonathan Williams, Mary Williamson, Qian Zeng, Tricia Zwiefelhofer, Clinton R. Paden, Duncan MacCannell |
| EPI_ISL_3113158 | Labor Berlin Charité Vivantes GmbH / Institut für Virologie | Charité Universitätsmedizin Berlin, Institut für Virologie/Labor Berlin | Peter Menzel, Christine Stephan, Rolf Schwarzer, Victor M Corman, Barbara Mühlemann, Terry Jones, Christian Drosten |
| EPI_ISL_3113159 | Laboratory Corporation of America | Centers for Disease Control and Prevention Division of Viral Diseases, Pathogen Discovery | Dakota Howard, Dhwani Batra, Peter W. Cook, Kara Moser, Adrian Paskey, Jason Caravas, Benjamin Rambo-Martin, Shatavia Morrison, Christopher Gulvick, Scott Sammons, Yvette Unoarumhi, Darlene Wagner, Matthew Schmerer, Minoo Agarwal, Eyad Almasri, Debbie Boles, Ayla Burns, Nuthawin Charoensri, Oren Cohen, Susan Countryman, Mary Ann Cristobal, Bobbi Croy, Suzanne Dale, Hrushikesh Deshmukh, Amanda Douglas, Vincent Drouillon, Marcia Eisenberg, Howard Engler, Rama Ghatti, Prashant Gupta, Susan Hicks, Jake Humphrey, Lax Iyer, Lisa Pfefferle, Manoj Jain, Matthew Robinson, Mohan Kolli, Brian Krueger, Tim Kuphal, Stanley Letovsky, Michael Levandoski, Craig Lukasik, Jonathan Meltzer, Brian Norvell, Mindy Nye, Scott Parker, Christos Petropoulos, John Pruitt, Steven Ragan, Scott Ryan, Mike Sapeta, Jana Schroth, Suresh Babu Selvaraju, Goran Stevovic, Amanda Suchanek, Andrea Throop, Lyndon Tilson, Thomas Urban, Joe Voshell, Kimberly Wagner, Jonathan Williams, Mary Williamson, Qian Zeng, Tricia Zwiefelhofer, Clinton R. Paden, Duncan MacCannell |
| EPI_ISL_3113160 | Labor Berlin Charité Vivantes GmbH / Institut für Virologie | Charité Universitätsmedizin Berlin, Institut für Virologie/Labor Berlin | Peter Menzel, Christine Stephan, Rolf Schwarzer, Victor M Corman, Barbara Mühlemann, Terry Jones, Christian Drosten |
| EPI_ISL_3113162, EPI_ISL_3113164 | Laboratory Corporation of America | Centers for Disease Control and Prevention Division of Viral Diseases, Pathogen Discovery | Dakota Howard, Dhwani Batra, Peter W. Cook, Kara Moser, Adrian Paskey, Jason Caravas, Benjamin Rambo-Martin, Shatavia Morrison, Christopher Gulvick, Scott Sammons, Yvette Unoarumhi, Darlene Wagner, Matthew Schmerer, Minoo Agarwal, Eyad Almasri, Debbie Boles, Ayla Burns, Nuthawin Charoensri, Oren Cohen, Susan Countryman, Mary Ann Cristobal, Bobbi Croy, Suzanne Dale, Hrushikesh Deshmukh, Amanda Douglas, Vincent Drouillon, Marcia Eisenberg, Howard Engler, Rama Ghatti, Prashant Gupta, Susan Hicks, Jake Humphrey, Lax Iyer, Lisa Pfefferle, Manoj Jain, Matthew Robinson, Mohan Kolli, Brian Krueger, Tim Kuphal, Stanley Letovsky, Michael Levandoski, Craig Lukasik, Jonathan Meltzer, Brian Norvell, Mindy Nye, Scott Parker, Christos Petropoulos, John Pruitt, Steven Ragan, Scott Ryan, Mike Sapeta, Jana Schroth, Suresh Babu Selvaraju, Goran Stevovic, Amanda Suchanek, Andrea Throop, Lyndon Tilson, Thomas Urban, Joe Voshell, Kimberly Wagner, Jonathan Williams, Mary Williamson, Qian Zeng, Tricia Zwiefelhofer, Clinton R. Paden, Duncan MacCannell |
| EPI_ISL_3113165 | Labor Berlin Charité Vivantes GmbH / Institut für Virologie | Charité Universitätsmedizin Berlin, Institut für Virologie/Labor Berlin | Peter Menzel, Christine Stephan, Rolf Schwarzer, Victor M Corman, Barbara Mühlemann, Terry Jones, Christian Drosten |
| EPI_ISL_3113166 | Laboratory Corporation of America | Centers for Disease Control and Prevention Division of Viral Diseases, Pathogen Discovery | Dakota Howard, Dhwani Batra, Peter W. Cook, Kara Moser, Adrian Paskey, Jason Caravas, Benjamin Rambo-Martin, Shatavia Morrison, Christopher Gulvick, Scott Sammons, Yvette Unoarumhi, Darlene Wagner, Matthew Schmerer, Minoo Agarwal, Eyad Almasri, Debbie Boles, Ayla Burns, Nuthawin Charoensri, Oren Cohen, Susan Countryman, Mary Ann Cristobal, Bobbi Croy, Suzanne Dale, Hrushikesh Deshmukh, Amanda Douglas, Vincent Drouillon, Marcia Eisenberg, Howard Engler, Rama Ghatti, Prashant Gupta, Susan Hicks, Jake Humphrey, Lax Iyer, Lisa Pfefferle, Manoj Jain, Matthew Robinson, Mohan Kolli, Brian Krueger, Tim Kuphal, Stanley Letovsky, Michael Levandoski, Craig Lukasik, Jonathan Meltzer, Brian Norvell, Mindy Nye, Scott Parker, Christos Petropoulos, John Pruitt, Steven Ragan, Scott Ryan, Mike Sapeta, Jana Schroth, Suresh Babu Selvaraju, Goran Stevovic, Amanda Suchanek, Andrea Throop, Lyndon Tilson, Thomas Urban, Joe Voshell, Kimberly Wagner, Jonathan Williams, Mary Williamson, Qian Zeng, Tricia Zwiefelhofer, Clinton R. Paden, Duncan MacCannell |
| EPI_ISL_3113167, EPI_ISL_3113168 | Labor Berlin Charité Vivantes GmbH / Institut für Virologie | Charité Universitätsmedizin Berlin, Institut für Virologie/Labor Berlin | Peter Menzel, Christine Stephan, Rolf Schwarzer, Victor M Corman, Barbara Mühlemann, Terry Jones, Christian Drosten |
| EPI_ISL_3113169 | Laboratory Corporation of America | Centers for Disease Control and Prevention Division of Viral Diseases, Pathogen Discovery | Dakota Howard, Dhwani Batra, Peter W. Cook, Kara Moser, Adrian Paskey, Jason Caravas, Benjamin Rambo-Martin, Shatavia Morrison, Christopher Gulvick, Scott Sammons, Yvette Unoarumhi, Darlene Wagner, Matthew Schmerer, Minoo Agarwal, Eyad Almasri, Debbie Boles, Ayla Burns, Nuthawin Charoensri, Oren Cohen, Susan Countryman, Mary Ann Cristobal, Bobbi Croy, Suzanne Dale, Hrushikesh Deshmukh, Amanda Douglas, Vincent Drouillon, Marcia Eisenberg, Howard Engler, Rama Ghatti, Prashant Gupta, Susan Hicks, Jake Humphrey, Lax Iyer, Lisa Pfefferle, Manoj Jain, Matthew Robinson, Mohan Kolli, Brian Krueger, Tim Kuphal, Stanley Letovsky, Michael Levandoski, Craig Lukasik, Jonathan Meltzer, Brian Norvell, Mindy Nye, Scott Parker, Christos Petropoulos, John Pruitt, Steven Ragan, Scott Ryan, Mike Sapeta, Jana Schroth, Suresh Babu Selvaraju, Goran Stevovic, Amanda Suchanek, Andrea Throop, Lyndon Tilson, Thomas Urban, Joe Voshell, Kimberly Wagner, Jonathan Williams, Mary Williamson, Qian Zeng, Tricia Zwiefelhofer, Clinton R. Paden, Duncan MacCannell |
| EPI_ISL_3113170 | Labor Berlin Charité Vivantes GmbH / Institut für Virologie | Charité Universitätsmedizin Berlin, Institut für Virologie/Labor Berlin | Peter Menzel, Christine Stephan, Rolf Schwarzer, Victor M Corman, Barbara Mühlemann, Terry Jones, Christian Drosten |
| EPI_ISL_3113171 | Laboratory Corporation of America | Centers for Disease Control and Prevention Division of Viral Diseases, Pathogen Discovery | Dakota Howard, Dhwani Batra, Peter W. Cook, Kara Moser, Adrian Paskey, Jason Caravas, Benjamin Rambo-Martin, Shatavia Morrison, Christopher Gulvick, Scott Sammons, Yvette Unoarumhi, Darlene Wagner, Matthew Schmerer, Minoo Agarwal, Eyad Almasri, Debbie Boles, Ayla Burns, Nuthawin Charoensri, Oren Cohen, Susan Countryman, Mary Ann Cristobal, Bobbi Croy, Suzanne Dale, Hrushikesh Deshmukh, Amanda Douglas, Vincent Drouillon, Marcia Eisenberg, Howard Engler, Rama Ghatti, Prashant Gupta, Susan Hicks, Jake Humphrey, Lax Iyer, Lisa Pfefferle, Manoj Jain, Matthew Robinson, Mohan Kolli, Brian Krueger, Tim Kuphal, Stanley Letovsky, Michael Levandoski, Craig Lukasik, Jonathan Meltzer, Brian Norvell, Mindy Nye, Scott Parker, Christos Petropoulos, John Pruitt, Steven Ragan, Scott Ryan, Mike Sapeta, Jana Schroth, Suresh Babu Selvaraju, Goran Stevovic, Amanda Suchanek, Andrea Throop, Lyndon Tilson, Thomas Urban, Joe Voshell, Kimberly Wagner, Jonathan Williams, Mary Williamson, Qian Zeng, Tricia Zwiefelhofer, Clinton R. Paden, Duncan MacCannell |
| EPI_ISL_3113172 | Labor Berlin Charité Vivantes GmbH / Institut für Virologie | Charité Universitätsmedizin Berlin, Institut für Virologie/Labor Berlin | Peter Menzel, Christine Stephan, Rolf Schwarzer, Victor M Corman, Barbara Mühlemann, Terry Jones, Christian Drosten |
| EPI_ISL_3113173 | Laboratory Corporation of America | Centers for Disease Control and Prevention Division of Viral Diseases, Pathogen Discovery | Dakota Howard, Dhwani Batra, Peter W. Cook, Kara Moser, Adrian Paskey, Jason Caravas, Benjamin Rambo-Martin, Shatavia Morrison, Christopher Gulvick, Scott Sammons, Yvette Unoarumhi, Darlene Wagner, Matthew Schmerer, Minoo Agarwal, Eyad Almasri, Debbie Boles, Ayla Burns, Nuthawin Charoensri, Oren Cohen, Susan Countryman, Mary Ann Cristobal, Bobbi Croy, Suzanne Dale, Hrushikesh Deshmukh, Amanda Douglas, Vincent Drouillon, Marcia Eisenberg, Howard Engler, Rama Ghatti, Prashant Gupta, Susan Hicks, Jake Humphrey, Lax Iyer, Lisa Pfefferle, Manoj Jain, Matthew Robinson, Mohan Kolli, Brian Krueger, Tim Kuphal, Stanley Letovsky, Michael Levandoski, Craig Lukasik, Jonathan Meltzer, Brian Norvell, Mindy Nye, Scott Parker, Christos Petropoulos, John Pruitt, Steven Ragan, Scott Ryan, Mike Sapeta, Jana Schroth, Suresh Babu Selvaraju, Goran Stevovic, Amanda Suchanek, Andrea Throop, Lyndon Tilson, Thomas Urban, Joe Voshell, Kimberly Wagner, Jonathan Williams, Mary Williamson, Qian Zeng, Tricia Zwiefelhofer, Clinton R. Paden, Duncan MacCannell |
| EPI_ISL_3113174, EPI_ISL_3113175 | Labor Berlin Charité Vivantes GmbH / Institut für Virologie | Charité Universitätsmedizin Berlin, Institut für Virologie/Labor Berlin | Peter Menzel, Christine Stephan, Rolf Schwarzer, Victor M Corman, Barbara Mühlemann, Terry Jones, Christian Drosten |
| EPI_ISL_3113176 | Laboratory Corporation of America | Centers for Disease Control and Prevention Division of Viral Diseases, Pathogen Discovery | Dakota Howard, Dhwani Batra, Peter W. Cook, Kara Moser, Adrian Paskey, Jason Caravas, Benjamin Rambo-Martin, Shatavia Morrison, Christopher Gulvick, Scott Sammons, Yvette Unoarumhi, Darlene Wagner, Matthew Schmerer, Minoo Agarwal, Eyad Almasri, Debbie Boles, Ayla Burns, Nuthawin Charoensri, Oren Cohen, Susan Countryman, Mary Ann Cristobal, Bobbi Croy, Suzanne Dale, Hrushikesh Deshmukh, Amanda Douglas, Vincent Drouillon, Marcia Eisenberg, Howard Engler, Rama Ghatti, Prashant Gupta, Susan Hicks, Jake Humphrey, Lax Iyer, Lisa Pfefferle, Manoj Jain, Matthew Robinson, Mohan Kolli, Brian Krueger, Tim Kuphal, Stanley Letovsky, Michael Levandoski, Craig Lukasik, Jonathan Meltzer, Brian Norvell, Mindy Nye, Scott Parker, Christos Petropoulos, John Pruitt, Steven Ragan, Scott Ryan, Mike Sapeta, Jana Schroth, Suresh Babu Selvaraju, Goran Stevovic, Amanda Suchanek, Andrea Throop, Lyndon Tilson, Thomas Urban, Joe Voshell, Kimberly Wagner, Jonathan Williams, Mary Williamson, Qian Zeng, Tricia Zwiefelhofer, Clinton R. Paden, Duncan MacCannell |
| EPI_ISL_3113177, EPI_ISL_3113178 | Labor Berlin Charité Vivantes GmbH / Institut für Virologie | Charité Universitätsmedizin Berlin, Institut für Virologie/Labor Berlin | Peter Menzel, Christine Stephan, Rolf Schwarzer, Victor M Corman, Barbara Mühlemann, Terry Jones, Christian Drosten |
| EPI_ISL_3113179, EPI_ISL_3113181 | Laboratory Corporation of America | Centers for Disease Control and Prevention Division of Viral Diseases, Pathogen Discovery | Dakota Howard, Dhwani Batra, Peter W. Cook, Kara Moser, Adrian Paskey, Jason Caravas, Benjamin Rambo-Martin, Shatavia Morrison, Christopher Gulvick, Scott Sammons, Yvette Unoarumhi, Darlene Wagner, Matthew Schmerer, Minoo Agarwal, Eyad Almasri, Debbie Boles, Ayla Burns, Nuthawin Charoensri, Oren Cohen, Susan Countryman, Mary Ann Cristobal, Bobbi Croy, Suzanne Dale, Hrushikesh Deshmukh, Amanda Douglas, Vincent Drouillon, Marcia Eisenberg, Howard Engler, Rama Ghatti, Prashant Gupta, Susan Hicks, Jake Humphrey, Lax Iyer, Lisa Pfefferle, Manoj Jain, Matthew Robinson, Mohan Kolli, Brian Krueger, Tim Kuphal, Stanley Letovsky, Michael Levandoski, Craig Lukasik, Jonathan Meltzer, Brian Norvell, Mindy Nye, Scott Parker, Christos Petropoulos, John Pruitt, Steven Ragan, Scott Ryan, Mike Sapeta, Jana Schroth, Suresh Babu Selvaraju, Goran Stevovic, Amanda Suchanek, Andrea Throop, Lyndon Tilson, Thomas Urban, Joe Voshell, Kimberly Wagner, Jonathan Williams, Mary Williamson, Qian Zeng, Tricia Zwiefelhofer, Clinton R. Paden, Duncan MacCannell |
| EPI_ISL_3113182 | Labor Berlin Charité Vivantes GmbH / Institut für Virologie | Charité Universitätsmedizin Berlin, Institut für Virologie/Labor Berlin | Peter Menzel, Christine Stephan, Rolf Schwarzer, Victor M Corman, Barbara Mühlemann, Terry Jones, Christian Drosten |
| EPI_ISL_3113184, EPI_ISL_3113186 | Laboratory Corporation of America | Centers for Disease Control and Prevention Division of Viral Diseases, Pathogen Discovery | Dakota Howard, Dhwani Batra, Peter W. Cook, Kara Moser, Adrian Paskey, Jason Caravas, Benjamin Rambo-Martin, Shatavia Morrison, Christopher Gulvick, Scott Sammons, Yvette Unoarumhi, Darlene Wagner, Matthew Schmerer, Minoo Agarwal, Eyad Almasri, Debbie Boles, Ayla Burns, Nuthawin Charoensri, Oren Cohen, Susan Countryman, Mary Ann Cristobal, Bobbi Croy, Suzanne Dale, Hrushikesh Deshmukh, Amanda Douglas, Vincent Drouillon, Marcia Eisenberg, Howard Engler, Rama Ghatti, Prashant Gupta, Susan Hicks, Jake Humphrey, Lax Iyer, Lisa Pfefferle, Manoj Jain, Matthew Robinson, Mohan Kolli, Brian Krueger, Tim Kuphal, Stanley Letovsky, Michael Levandoski, Craig Lukasik, Jonathan Meltzer, Brian Norvell, Mindy Nye, Scott Parker, Christos Petropoulos, John Pruitt, Steven Ragan, Scott Ryan, Mike Sapeta, Jana Schroth, Suresh Babu Selvaraju, Goran Stevovic, Amanda Suchanek, Andrea Throop, Lyndon Tilson, Thomas Urban, Joe Voshell, Kimberly Wagner, Jonathan Williams, Mary Williamson, Qian Zeng, Tricia Zwiefelhofer, Clinton R. Paden, Duncan MacCannell |
| EPI_ISL_3113187 | Labor Berlin Charité Vivantes GmbH / Institut für Virologie | Charité Universitätsmedizin Berlin, Institut für Virologie/Labor Berlin | Peter Menzel, Christine Stephan, Rolf Schwarzer, Victor M Corman, Barbara Mühlemann, Terry Jones, Christian Drosten |
| EPI_ISL_3113189 | Laboratory Corporation of America | Centers for Disease Control and Prevention Division of Viral Diseases, Pathogen Discovery | Dakota Howard, Dhwani Batra, Peter W. Cook, Kara Moser, Adrian Paskey, Jason Caravas, Benjamin Rambo-Martin, Shatavia Morrison, Christopher Gulvick, Scott Sammons, Yvette Unoarumhi, Darlene Wagner, Matthew Schmerer, Minoo Agarwal, Eyad Almasri, Debbie Boles, Ayla Burns, Nuthawin Charoensri, Oren Cohen, Susan Countryman, Mary Ann Cristobal, Bobbi Croy, Suzanne Dale, Hrushikesh Deshmukh, Amanda Douglas, Vincent Drouillon, Marcia Eisenberg, Howard Engler, Rama Ghatti, Prashant Gupta, Susan Hicks, Jake Humphrey, Lax Iyer, Lisa Pfefferle, Manoj Jain, Matthew Robinson, Mohan Kolli, Brian Krueger, Tim Kuphal, Stanley Letovsky, Michael Levandoski, Craig Lukasik, Jonathan Meltzer, Brian Norvell, Mindy Nye, Scott Parker, Christos Petropoulos, John Pruitt, Steven Ragan, Scott Ryan, Mike Sapeta, Jana Schroth, Suresh Babu Selvaraju, Goran Stevovic, Amanda Suchanek, Andrea Throop, Lyndon Tilson, Thomas Urban, Joe Voshell, Kimberly Wagner, Jonathan Williams, Mary Williamson, Qian Zeng, Tricia Zwiefelhofer, Clinton R. Paden, Duncan MacCannell |
| EPI_ISL_3113191 | Labor Berlin Charité Vivantes GmbH / Institut für Virologie | Charité Universitätsmedizin Berlin, Institut für Virologie/Labor Berlin | Peter Menzel, Christine Stephan, Rolf Schwarzer, Victor M Corman, Barbara Mühlemann, Terry Jones, Christian Drosten |
| EPI_ISL_3113192 | Laboratory Corporation of America | Centers for Disease Control and Prevention Division of Viral Diseases, Pathogen Discovery | Dakota Howard, Dhwani Batra, Peter W. Cook, Kara Moser, Adrian Paskey, Jason Caravas, Benjamin Rambo-Martin, Shatavia Morrison, Christopher Gulvick, Scott Sammons, Yvette Unoarumhi, Darlene Wagner, Matthew Schmerer, Minoo Agarwal, Eyad Almasri, Debbie Boles, Ayla Burns, Nuthawin Charoensri, Oren Cohen, Susan Countryman, Mary Ann Cristobal, Bobbi Croy, Suzanne Dale, Hrushikesh Deshmukh, Amanda Douglas, Vincent Drouillon, Marcia Eisenberg, Howard Engler, Rama Ghatti, Prashant Gupta, Susan Hicks, Jake Humphrey, Lax Iyer, Lisa Pfefferle, Manoj Jain, Matthew Robinson, Mohan Kolli, Brian Krueger, Tim Kuphal, Stanley Letovsky, Michael Levandoski, Craig Lukasik, Jonathan Meltzer, Brian Norvell, Mindy Nye, Scott Parker, Christos Petropoulos, John Pruitt, Steven Ragan, Scott Ryan, Mike Sapeta, Jana Schroth, Suresh Babu Selvaraju, Goran Stevovic, Amanda Suchanek, Andrea Throop, Lyndon Tilson, Thomas Urban, Joe Voshell, Kimberly Wagner, Jonathan Williams, Mary Williamson, Qian Zeng, Tricia Zwiefelhofer, Clinton R. Paden, Duncan MacCannell |
| EPI_ISL_3113193 | Labor Berlin Charité Vivantes GmbH / Institut für Virologie | Charité Universitätsmedizin Berlin, Institut für Virologie/Labor Berlin | Peter Menzel, Christine Stephan, Rolf Schwarzer, Victor M Corman, Barbara Mühlemann, Terry Jones, Christian Drosten |
| EPI_ISL_3113194 | Laboratory Corporation of America | Centers for Disease Control and Prevention Division of Viral Diseases, Pathogen Discovery | Dakota Howard, Dhwani Batra, Peter W. Cook, Kara Moser, Adrian Paskey, Jason Caravas, Benjamin Rambo-Martin, Shatavia Morrison, Christopher Gulvick, Scott Sammons, Yvette Unoarumhi, Darlene Wagner, Matthew Schmerer, Minoo Agarwal, Eyad Almasri, Debbie Boles, Ayla Burns, Nuthawin Charoensri, Oren Cohen, Susan Countryman, Mary Ann Cristobal, Bobbi Croy, Suzanne Dale, Hrushikesh Deshmukh, Amanda Douglas, Vincent Drouillon, Marcia Eisenberg, Howard Engler, Rama Ghatti, Prashant Gupta, Susan Hicks, Jake Humphrey, Lax Iyer, Lisa Pfefferle, Manoj Jain, Matthew Robinson, Mohan Kolli, Brian Krueger, Tim Kuphal, Stanley Letovsky, Michael Levandoski, Craig Lukasik, Jonathan Meltzer, Brian Norvell, Mindy Nye, Scott Parker, Christos Petropoulos, John Pruitt, Steven Ragan, Scott Ryan, Mike Sapeta, Jana Schroth, Suresh Babu Selvaraju, Goran Stevovic, Amanda Suchanek, Andrea Throop, Lyndon Tilson, Thomas Urban, Joe Voshell, Kimberly Wagner, Jonathan Williams, Mary Williamson, Qian Zeng, Tricia Zwiefelhofer, Clinton R. Paden, Duncan MacCannell |
| EPI_ISL_3113195 | Labor Berlin Charité Vivantes GmbH / Institut für Virologie | Charité Universitätsmedizin Berlin, Institut für Virologie/Labor Berlin | Peter Menzel, Christine Stephan, Rolf Schwarzer, Victor M Corman, Barbara Mühlemann, Terry Jones, Christian Drosten |
| EPI_ISL_3113196 | Laboratory Corporation of America | Centers for Disease Control and Prevention Division of Viral Diseases, Pathogen Discovery | Dakota Howard, Dhwani Batra, Peter W. Cook, Kara Moser, Adrian Paskey, Jason Caravas, Benjamin Rambo-Martin, Shatavia Morrison, Christopher Gulvick, Scott Sammons, Yvette Unoarumhi, Darlene Wagner, Matthew Schmerer, Minoo Agarwal, Eyad Almasri, Debbie Boles, Ayla Burns, Nuthawin Charoensri, Oren Cohen, Susan Countryman, Mary Ann Cristobal, Bobbi Croy, Suzanne Dale, Hrushikesh Deshmukh, Amanda Douglas, Vincent Drouillon, Marcia Eisenberg, Howard Engler, Rama Ghatti, Prashant Gupta, Susan Hicks, Jake Humphrey, Lax Iyer, Lisa Pfefferle, Manoj Jain, Matthew Robinson, Mohan Kolli, Brian Krueger, Tim Kuphal, Stanley Letovsky, Michael Levandoski, Craig Lukasik, Jonathan Meltzer, Brian Norvell, Mindy Nye, Scott Parker, Christos Petropoulos, John Pruitt, Steven Ragan, Scott Ryan, Mike Sapeta, Jana Schroth, Suresh Babu Selvaraju, Goran Stevovic, Amanda Suchanek, Andrea Throop, Lyndon Tilson, Thomas Urban, Joe Voshell, Kimberly Wagner, Jonathan Williams, Mary Williamson, Qian Zeng, Tricia Zwiefelhofer, Clinton R. Paden, Duncan MacCannell |
| EPI_ISL_3113197 | Labor Berlin Charité Vivantes GmbH / Institut für Virologie | Charité Universitätsmedizin Berlin, Institut für Virologie/Labor Berlin | Peter Menzel, Christine Stephan, Rolf Schwarzer, Victor M Corman, Barbara Mühlemann, Terry Jones, Christian Drosten |
| EPI_ISL_3113198 | Laboratory Corporation of America | Centers for Disease Control and Prevention Division of Viral Diseases, Pathogen Discovery | Dakota Howard, Dhwani Batra, Peter W. Cook, Kara Moser, Adrian Paskey, Jason Caravas, Benjamin Rambo-Martin, Shatavia Morrison, Christopher Gulvick, Scott Sammons, Yvette Unoarumhi, Darlene Wagner, Matthew Schmerer, Minoo Agarwal, Eyad Almasri, Debbie Boles, Ayla Burns, Nuthawin Charoensri, Oren Cohen, Susan Countryman, Mary Ann Cristobal, Bobbi Croy, Suzanne Dale, Hrushikesh Deshmukh, Amanda Douglas, Vincent Drouillon, Marcia Eisenberg, Howard Engler, Rama Ghatti, Prashant Gupta, Susan Hicks, Jake Humphrey, Lax Iyer, Lisa Pfefferle, Manoj Jain, Matthew Robinson, Mohan Kolli, Brian Krueger, Tim Kuphal, Stanley Letovsky, Michael Levandoski, Craig Lukasik, Jonathan Meltzer, Brian Norvell, Mindy Nye, Scott Parker, Christos Petropoulos, John Pruitt, Steven Ragan, Scott Ryan, Mike Sapeta, Jana Schroth, Suresh Babu Selvaraju, Goran Stevovic, Amanda Suchanek, Andrea Throop, Lyndon Tilson, Thomas Urban, Joe Voshell, |

| | | | |
|---|---|---|---|
| | | | Kimberly Wagner, Jonathan Williams, Mary Williamson, Qian Zeng, Tricia Zwiefelhofer, Clinton R. Paden, Duncan MacCannell |
| EPI_ISL_3113199 | Labor Berlin Charité Vivantes GmbH / Institut für Virologie | Charité Universitätsmedizin Berlin, Institut für Virologie/Labor Berlin | Peter Menzel, Christine Stephan, Rolf Schwarzer, Victor M Corman, Barbara Mühlemann, Terry Jones, Christian Drosten |
| EPI_ISL_3113201 | Laboratory Corporation of America | Centers for Disease Control and Prevention Division of Viral Diseases, Pathogen Discovery | Dakota Howard, Dhwani Batra, Peter W. Cook, Kara Moser, Adrian Paskey, Jason Caravas, Benjamin Rambo-Martin, Shatavia Morrison, Christopher Gulvick, Scott Sammons, Yvette Unoarumhi, Darlene Wagner, Matthew Schmerer, Minoo Agarwal, Eyad Almasri, Debbie Boles, Ayla Burns, Nuthawin Charoensri, Oren Cohen, Susan Countryman, Mary Ann Cristobal, Bobbi Croy, Suzanne Dale, Hrushikesh Deshmukh, Amanda Douglas, Vincent Drouillon, Marcia Eisenberg, Howard Engler, Rama Ghatti, Prashant Gupta, Susan Hicks, Jake Humphrey, Lax Iyer, Lisa Pfefferle, Manoj Jain, Matthew Robinson, Mohan Kolli, Brian Krueger, Tim Kuphal, Stanley Letovsky, Michael Levandoski, Craig Lukasik, Jonathan Meltzer, Brian Norvell, Mindy Nye, Scott Parker, Christos Petropoulos, John Pruitt, Steven Ragan, Scott Ryan, Mike Sapeta, Jana Schroth, Suresh Babu Selvaraju, Goran Stevovic, Amanda Suchanek, Andrea Throop, Lyndon Tilson, Thomas Urban, Joe Voshell, Kimberly Wagner, Jonathan Williams, Mary Williamson, Qian Zeng, Tricia Zwiefelhofer, Clinton R. Paden, Duncan MacCannell |
| EPI_ISL_3113204, EPI_ISL_3113205, EPI_ISL_3113207, EPI_ISL_3113208 | Labor Berlin Charité Vivantes GmbH / Institut für Virologie | Charité Universitätsmedizin Berlin, Institut für Virologie/Labor Berlin | Peter Menzel, Christine Stephan, Rolf Schwarzer, Victor M Corman, Barbara Mühlemann, Terry Jones, Christian Drosten |
| EPI_ISL_3113209 | Laboratory Corporation of America | Centers for Disease Control and Prevention Division of Viral Diseases, Pathogen Discovery | Dakota Howard, Dhwani Batra, Peter W. Cook, Kara Moser, Adrian Paskey, Jason Caravas, Benjamin Rambo-Martin, Shatavia Morrison, Christopher Gulvick, Scott Sammons, Yvette Unoarumhi, Darlene Wagner, Matthew Schmerer, Minoo Agarwal, Eyad Almasri, Debbie Boles, Ayla Burns, Nuthawin Charoensri, Oren Cohen, Susan Countryman, Mary Ann Cristobal, Bobbi Croy, Suzanne Dale, Hrushikesh Deshmukh, Amanda Douglas, Vincent Drouillon, Marcia Eisenberg, Howard Engler, Rama Ghatti, Prashant Gupta, Susan Hicks, Jake Humphrey, Lax Iyer, Lisa Pfefferle, Manoj Jain, Matthew Robinson, Mohan Kolli, Brian Krueger, Tim Kuphal, Stanley Letovsky, Michael Levandoski, Craig Lukasik, Jonathan Meltzer, Brian Norvell, Mindy Nye, Scott Parker, Christos Petropoulos, John Pruitt, Steven Ragan, Scott Ryan, Mike Sapeta, Jana Schroth, Suresh Babu Selvaraju, Goran Stevovic, Amanda Suchanek, Andrea Throop, Lyndon Tilson, Thomas Urban, Joe Voshell, Kimberly Wagner, Jonathan Williams, Mary Williamson, Qian Zeng, Tricia Zwiefelhofer, Clinton R. Paden, Duncan MacCannell |
| EPI_ISL_3113210 | Labor Berlin Charité Vivantes GmbH / Institut für Virologie | Charité Universitätsmedizin Berlin, Institut für Virologie/Labor Berlin | Peter Menzel, Christine Stephan, Rolf Schwarzer, Victor M Corman, Barbara Mühlemann, Terry Jones, Christian Drosten |
| EPI_ISL_3113211 | Laboratory Corporation of America | Centers for Disease Control and Prevention Division of Viral Diseases, Pathogen Discovery | Dakota Howard, Dhwani Batra, Peter W. Cook, Kara Moser, Adrian Paskey, Jason Caravas, Benjamin Rambo-Martin, Shatavia Morrison, Christopher Gulvick, Scott Sammons, Yvette Unoarumhi, Darlene Wagner, Matthew Schmerer, Minoo Agarwal, Eyad Almasri, Debbie Boles, Ayla Burns, Nuthawin Charoensri, Oren Cohen, Susan Countryman, Mary Ann Cristobal, Bobbi Croy, Suzanne Dale, Hrushikesh Deshmukh, Amanda Douglas, Vincent Drouillon, Marcia Eisenberg, Howard Engler, Rama Ghatti, Prashant Gupta, Susan Hicks, Jake Humphrey, Lax Iyer, Lisa Pfefferle, Manoj Jain, Matthew Robinson, Mohan Kolli, Brian Krueger, Tim Kuphal, Stanley Letovsky, Michael Levandoski, Craig Lukasik, Jonathan Meltzer, Brian Norvell, Mindy Nye, Scott Parker, Christos Petropoulos, John Pruitt, Steven Ragan, Scott Ryan, Mike Sapeta, Jana Schroth, Suresh Babu Selvaraju, Goran Stevovic, Amanda Suchanek, Andrea Throop, Lyndon Tilson, Thomas Urban, Joe Voshell, Kimberly Wagner, Jonathan Williams, Mary Williamson, Qian Zeng, Tricia Zwiefelhofer, Clinton R. Paden, Duncan MacCannell |
| EPI_ISL_3113212 | Labor Berlin Charité Vivantes GmbH / Institut für Virologie | Charité Universitätsmedizin Berlin, Institut für Virologie/Labor Berlin | Peter Menzel, Christine Stephan, Rolf Schwarzer, Victor M Corman, Barbara Mühlemann, Terry Jones, Christian Drosten |
| EPI_ISL_3113214 | Laboratory Corporation of America | Centers for Disease Control and Prevention Division of Viral Diseases, Pathogen Discovery | Dakota Howard, Dhwani Batra, Peter W. Cook, Kara Moser, Adrian Paskey, Jason Caravas, Benjamin Rambo-Martin, Shatavia Morrison, Christopher Gulvick, Scott Sammons, Yvette Unoarumhi, Darlene Wagner, Matthew Schmerer, Minoo Agarwal, Eyad Almasri, Debbie Boles, Ayla Burns, Nuthawin Charoensri, Oren Cohen, Susan Countryman, Mary Ann Cristobal, Bobbi Croy, Suzanne Dale, Hrushikesh Deshmukh, Amanda Douglas, Vincent Drouillon, Marcia Eisenberg, Howard Engler, Rama Ghatti, Prashant Gupta, Susan Hicks, Jake Humphrey, Lax Iyer, Lisa Pfefferle, Manoj Jain, Matthew Robinson, Mohan Kolli, Brian Krueger, Tim Kuphal, Stanley Letovsky, Michael Levandoski, Craig Lukasik, Jonathan Meltzer, Brian Norvell, Mindy Nye, Scott Parker, Christos Petropoulos, John Pruitt, Steven Ragan, Scott Ryan, Mike Sapeta, Jana Schroth, Suresh Babu Selvaraju, Goran Stevovic, Amanda Suchanek, Andrea Throop, Lyndon Tilson, Thomas Urban, Joe Voshell, Kimberly Wagner, Jonathan Williams, Mary Williamson, Qian Zeng, Tricia Zwiefelhofer, Clinton R. Paden, Duncan MacCannell |
| EPI_ISL_3113215 | Labor Berlin Charité Vivantes GmbH / Institut für Virologie | Charité Universitätsmedizin Berlin, Institut für Virologie/Labor Berlin | Peter Menzel, Christine Stephan, Rolf Schwarzer, Victor M Corman, Barbara Mühlemann, Terry Jones, Christian Drosten |
| EPI_ISL_3113218, EPI_ISL_3113220, EPI_ISL_3113221, EPI_ISL_3113222, EPI_ISL_3113223, EPI_ISL_3113224, EPI_ISL_3113225, EPI_ISL_3113226, EPI_ISL_3113227, EPI_ISL_3113228, EPI_ISL_3113229, EPI_ISL_3113230, EPI_ISL_3113231, EPI_ISL_3113232, EPI_ISL_3113233, EPI_ISL_3113234 | | | |
| see above | Laboratory Corporation of America | Centers for Disease Control and Prevention Division of Viral Diseases, Pathogen Discovery | Dakota Howard, Dhwani Batra, Peter W. Cook, Kara Moser, Adrian Paskey, Jason Caravas, Benjamin Rambo-Martin, Shatavia Morrison, Christopher Gulvick, Scott Sammons, Yvette Unoarumhi, Darlene Wagner, Matthew Schmerer, Minoo Agarwal, Eyad Almasri, Debbie Boles, Ayla Burns, Nuthawin Charoensri, Oren Cohen, Susan Countryman, Mary Ann Cristobal, Bobbi Croy, Suzanne Dale, Hrushikesh Deshmukh, Amanda Douglas, Vincent Drouillon, Marcia Eisenberg, Howard Engler, Rama Ghatti, Prashant Gupta, Susan Hicks, Jake Humphrey, Lax Iyer, Lisa Pfefferle, Manoj Jain, Matthew Robinson, Mohan Kolli, Brian Krueger, Tim Kuphal, Stanley Letovsky, Michael Levandoski, Craig Lukasik, Jonathan Meltzer, Brian Norvell, Mindy Nye, Scott Parker, Christos Petropoulos, John Pruitt, Steven Ragan, Scott Ryan, Mike Sapeta, Jana Schroth, Suresh Babu Selvaraju, Goran Stevovic, Amanda Suchanek, Andrea Throop, Lyndon Tilson, Thomas Urban, Joe Voshell, Kimberly Wagner, Jonathan Williams, Mary Williamson, Qian Zeng, Tricia Zwiefelhofer, Clinton R. Paden, Duncan MacCannell |
| EPI_ISL_3113235 | Labor Berlin Charité Vivantes GmbH / Institut für Virologie | Charité Universitätsmedizin Berlin, Institut für Virologie/Labor Berlin | Peter Menzel, Christine Stephan, Rolf Schwarzer, Victor M Corman, Barbara Mühlemann, Terry Jones, Christian Drosten |
| EPI_ISL_3113236 | Laboratory Corporation of America | Centers for Disease Control and Prevention Division of Viral Diseases, Pathogen Discovery | Dakota Howard, Dhwani Batra, Peter W. Cook, Kara Moser, Adrian Paskey, Jason Caravas, Benjamin Rambo-Martin, Shatavia Morrison, Christopher Gulvick, Scott Sammons, Yvette Unoarumhi, Darlene Wagner, Matthew Schmerer, Minoo Agarwal, Eyad Almasri, Debbie Boles, Ayla Burns, Nuthawin Charoensri, Oren Cohen, Susan Countryman, Mary Ann Cristobal, Bobbi Croy, Suzanne Dale, Hrushikesh Deshmukh, Amanda Douglas, Vincent Drouillon, Marcia Eisenberg, Howard Engler, Rama Ghatti, Prashant Gupta, Susan Hicks, Jake Humphrey, Lax Iyer, Lisa Pfefferle, Manoj Jain, Matthew Robinson, Mohan Kolli, Brian Krueger, Tim Kuphal, Stanley Letovsky, Michael Levandoski, Craig Lukasik, Jonathan Meltzer, Brian Norvell, Mindy Nye, Scott Parker, Christos Petropoulos, John Pruitt, Steven Ragan, Scott Ryan, Mike Sapeta, Jana Schroth, Suresh Babu Selvaraju, Goran Stevovic, Amanda Suchanek, Andrea Throop, Lyndon Tilson, Thomas Urban, Joe Voshell, Kimberly Wagner, Jonathan Williams, Mary Williamson, Qian Zeng, Tricia Zwiefelhofer, Clinton R. Paden, Duncan MacCannell |
| EPI_ISL_3113237, EPI_ISL_3113238 | Labor Berlin Charité Vivantes GmbH / Institut für Virologie | Charité Universitätsmedizin Berlin, Institut für Virologie/Labor Berlin | Peter Menzel, Christine Stephan, Rolf Schwarzer, Victor M Corman, Barbara Mühlemann, Terry Jones, Christian Drosten |
| EPI_ISL_3113243 | Laboratory Corporation of America | Centers for Disease Control and Prevention Division of Viral Diseases, Pathogen Discovery | Dakota Howard, Dhwani Batra, Peter W. Cook, Kara Moser, Adrian Paskey, Jason Caravas, Benjamin Rambo-Martin, Shatavia Morrison, Christopher Gulvick, Scott Sammons, Yvette Unoarumhi, Darlene Wagner, Matthew Schmerer, Minoo Agarwal, Eyad Almasri, Debbie Boles, Ayla Burns, Nuthawin Charoensri, Oren Cohen, Susan Countryman, Mary Ann Cristobal, Bobbi Croy, Suzanne Dale, Hrushikesh Deshmukh, Amanda Douglas, Vincent Drouillon, Marcia Eisenberg, Howard Engler, Rama Ghatti, Prashant Gupta, Susan Hicks, Jake Humphrey, Lax Iyer, Lisa Pfefferle, Manoj Jain, Matthew Robinson, Mohan Kolli, Brian Krueger, Tim Kuphal, Stanley Letovsky, Michael Levandoski, Craig Lukasik, Jonathan Meltzer, Brian Norvell, Mindy Nye, Scott Parker, Christos Petropoulos, John Pruitt, Steven Ragan, Scott Ryan, Mike Sapeta, Jana Schroth, Suresh Babu Selvaraju, Goran Stevovic, Amanda Suchanek, Andrea Throop, Lyndon Tilson, Thomas Urban, Joe Voshell, Kimberly Wagner, Jonathan Williams, Mary Williamson, Qian Zeng, Tricia Zwiefelhofer, Clinton R. Paden, Duncan MacCannell |
| EPI_ISL_3113244, EPI_ISL_3113245 | Labor Berlin Charité Vivantes GmbH / Institut für Virologie | Charité Universitätsmedizin Berlin, Institut für Virologie/Labor Berlin | Peter Menzel, Christine Stephan, Rolf Schwarzer, Victor M Corman, Barbara Mühlemann, Terry Jones, Christian Drosten |
| EPI_ISL_3113246 | Laboratory Corporation of America | Centers for Disease Control and Prevention Division of Viral Diseases, Pathogen Discovery | Dakota Howard, Dhwani Batra, Peter W. Cook, Kara Moser, Adrian Paskey, Jason Caravas, Benjamin Rambo-Martin, Shatavia Morrison, Christopher Gulvick, Scott Sammons, Yvette Unoarumhi, Darlene Wagner, Matthew Schmerer, Minoo Agarwal, Eyad Almasri, Debbie Boles, Ayla Burns, Nuthawin Charoensri, Oren Cohen, Susan Countryman, Mary Ann Cristobal, Bobbi Croy, Suzanne Dale, Hrushikesh Deshmukh, Amanda Douglas, Vincent Drouillon, Marcia Eisenberg, Howard Engler, Rama Ghatti, Prashant Gupta, Susan Hicks, Jake Humphrey, Lax Iyer, Lisa Pfefferle, Manoj Jain, Matthew Robinson, Mohan Kolli, Brian Krueger, Tim Kuphal, Stanley Letovsky, Michael Levandoski, Craig Lukasik, Jonathan Meltzer, Brian Norvell, Mindy Nye, Scott Parker, Christos Petropoulos, John Pruitt, Steven Ragan, Scott Ryan, Mike Sapeta, Jana Schroth, Suresh Babu Selvaraju, Goran Stevovic, Amanda Suchanek, Andrea Throop, Lyndon Tilson, Thomas Urban, Joe Voshell, Kimberly Wagner, Jonathan Williams, Mary Williamson, Qian Zeng, Tricia Zwiefelhofer, Clinton R. Paden, Duncan MacCannell |
| EPI_ISL_3113247, EPI_ISL_3113248 | Labor Berlin Charité Vivantes GmbH / Institut für Virologie | Charité Universitätsmedizin Berlin, Institut für Virologie/Labor Berlin | Peter Menzel, Christine Stephan, Rolf Schwarzer, Victor M Corman, Barbara Mühlemann, Terry Jones, Christian Drosten |
| EPI_ISL_3113249 | Laboratory Corporation of America | Centers for Disease Control and Prevention Division of Viral Diseases, Pathogen Discovery | Dakota Howard, Dhwani Batra, Peter W. Cook, Kara Moser, Adrian Paskey, Jason Caravas, Benjamin Rambo-Martin, Shatavia Morrison, Christopher Gulvick, Scott Sammons, Yvette Unoarumhi, Darlene Wagner, Matthew Schmerer, Minoo Agarwal, Eyad Almasri, Debbie Boles, Ayla Burns, Nuthawin Charoensri, Oren Cohen, Susan Countryman, Mary Ann Cristobal, Bobbi Croy, Suzanne Dale, Hrushikesh Deshmukh, Amanda Douglas, Vincent Drouillon, Marcia Eisenberg, Howard Engler, Rama Ghatti, Prashant Gupta, Susan Hicks, Jake Humphrey, Lax Iyer, Lisa Pfefferle, Manoj Jain, Matthew Robinson, Mohan Kolli, Brian Krueger, Tim Kuphal, Stanley Letovsky, Michael Levandoski, Craig Lukasik, Jonathan Meltzer, Brian Norvell, Mindy Nye, Scott Parker, Christos Petropoulos, John Pruitt, Steven Ragan, Scott Ryan, Mike Sapeta, Jana Schroth, Suresh Babu Selvaraju, Goran Stevovic, Amanda Suchanek, Andrea Throop, Lyndon Tilson, Thomas Urban, Joe Voshell, Kimberly Wagner, Jonathan Williams, Mary Williamson, Qian Zeng, Tricia Zwiefelhofer, Clinton R. Paden, Duncan MacCannell |
| EPI_ISL_3113250, EPI_ISL_3113252, EPI_ISL_3113253 | Labor Berlin Charité Vivantes GmbH / Institut für Virologie | Charité Universitätsmedizin Berlin, Institut für Virologie/Labor Berlin | Peter Menzel, Christine Stephan, Rolf Schwarzer, Victor M Corman, Barbara Mühlemann, Terry Jones, Christian Drosten |
| EPI_ISL_3113254 | Laboratory Corporation of America | Centers for Disease Control and Prevention Division of Viral Diseases, Pathogen Discovery | Dakota Howard, Dhwani Batra, Peter W. Cook, Kara Moser, Adrian Paskey, Jason Caravas, Benjamin Rambo-Martin, Shatavia Morrison, Christopher Gulvick, Scott Sammons, Yvette Unoarumhi, Darlene Wagner, Matthew Schmerer, Minoo Agarwal, Eyad Almasri, Debbie Boles, Ayla Burns, Nuthawin Charoensri, Oren Cohen, Susan Countryman, Mary Ann Cristobal, Bobbi Croy, Suzanne Dale, Hrushikesh Deshmukh, Amanda Douglas, Vincent Drouillon, Marcia Eisenberg, Howard Engler, Rama Ghatti, Prashant Gupta, Susan Hicks, Jake Humphrey, Lax Iyer, Lisa Pfefferle, Manoj Jain, Matthew Robinson, Mohan Kolli, Brian Krueger, Tim Kuphal, Stanley Letovsky, Michael Levandoski, Craig Lukasik, Jonathan Meltzer, Brian Norvell, Mindy Nye, Scott Parker, Christos Petropoulos, John Pruitt, Steven Ragan, Scott Ryan, Mike Sapeta, Jana Schroth, Suresh Babu Selvaraju, Goran Stevovic, Amanda Suchanek, Andrea Throop, Lyndon Tilson, Thomas Urban, Joe Voshell, Kimberly Wagner, Jonathan Williams, Mary Williamson, Qian Zeng, Tricia Zwiefelhofer, Clinton R. Paden, Duncan MacCannell |
| EPI_ISL_3113255 | Labor Berlin Charité Vivantes GmbH / Institut für Virologie | Charité Universitätsmedizin Berlin, Institut für Virologie/Labor Berlin | Peter Menzel, Christine Stephan, Rolf Schwarzer, Victor M Corman, Barbara Mühlemann, Terry Jones, Christian Drosten |
| EPI_ISL_3113256 | Laboratory Corporation of America | Centers for Disease Control and Prevention Division of Viral Diseases, Pathogen Discovery | Dakota Howard, Dhwani Batra, Peter W. Cook, Kara Moser, Adrian Paskey, Jason Caravas, Benjamin Rambo-Martin, Shatavia Morrison, Christopher Gulvick, Scott Sammons, Yvette Unoarumhi, Darlene Wagner, Matthew Schmerer, Minoo Agarwal, Eyad Almasri, Debbie Boles, Ayla Burns, Nuthawin Charoensri, Oren Cohen, Susan Countryman, Mary Ann Cristobal, Bobbi Croy, Suzanne Dale, Hrushikesh Deshmukh, Amanda Douglas, Vincent Drouillon, Marcia Eisenberg, Howard Engler, Rama Ghatti, Prashant Gupta, Susan Hicks, Jake Humphrey, Lax Iyer, Lisa Pfefferle, Manoj Jain, Matthew Robinson, Mohan Kolli, Brian Krueger, Tim Kuphal, Stanley Letovsky, Michael Levandoski, Craig Lukasik, Jonathan Meltzer, Brian Norvell, Mindy Nye, Scott Parker, Christos Petropoulos, John Pruitt, Steven Ragan, Scott Ryan, Mike Sapeta, Jana Schroth, Suresh Babu Selvaraju, Goran Stevovic, Amanda Suchanek, Andrea Throop, Lyndon Tilson, Thomas Urban, Joe Voshell, Kimberly Wagner, Jonathan Williams, Mary Williamson, Qian Zeng, Tricia Zwiefelhofer, Clinton R. Paden, Duncan MacCannell |
| EPI_ISL_3113257 | Labor Berlin Charité Vivantes GmbH / Institut für Virologie | Charité Universitätsmedizin Berlin, Institut für Virologie/Labor Berlin | Peter Menzel, Christine Stephan, Rolf Schwarzer, Victor M Corman, Barbara Mühlemann, Terry Jones, Christian Drosten |
| EPI_ISL_3113258 | Laboratory Corporation of America | Centers for Disease Control and Prevention Division of Viral Diseases, Pathogen Discovery | Dakota Howard, Dhwani Batra, Peter W. Cook, Kara Moser, Adrian Paskey, Jason Caravas, Benjamin Rambo-Martin, Shatavia Morrison, Christopher Gulvick, Scott Sammons, Yvette Unoarumhi, Darlene Wagner, Matthew Schmerer, Minoo Agarwal, Eyad Almasri, Debbie Boles, Ayla Burns, Nuthawin Charoensri, Oren Cohen, Susan Countryman, Mary Ann Cristobal, Bobbi Croy, Suzanne Dale, Hrushikesh Deshmukh, Amanda Douglas, Vincent Drouillon, Marcia Eisenberg, Howard Engler, Rama Ghatti, Prashant Gupta, Susan Hicks, Jake Humphrey, Lax Iyer, Lisa Pfefferle, Manoj Jain, Matthew Robinson, Mohan Kolli, Brian Krueger, Tim Kuphal, Stanley Letovsky, Michael Levandoski, Craig Lukasik, Jonathan Meltzer, Brian Norvell, Mindy Nye, Scott Parker, Christos Petropoulos, John Pruitt, Steven Ragan, Scott Ryan, Mike Sapeta, Jana Schroth, Suresh Babu Selvaraju, Goran Stevovic, Amanda Suchanek, Andrea Throop, Lyndon Tilson, Thomas Urban, Joe Voshell, Kimberly Wagner, Jonathan Williams, Mary Williamson, Qian Zeng, Tricia Zwiefelhofer, Clinton R. Paden, Duncan MacCannell |
| EPI_ISL_3113259, EPI_ISL_3113260 | Labor Berlin Charité Vivantes GmbH / Institut für Virologie | Charité Universitätsmedizin Berlin, Institut für Virologie/Labor Berlin | Peter Menzel, Christine Stephan, Rolf Schwarzer, Victor M Corman, Barbara Mühlemann, Terry Jones, Christian Drosten |
| EPI_ISL_3113261 | Laboratory Corporation of America | Centers for Disease Control and Prevention Division of Viral Diseases, Pathogen Discovery | Dakota Howard, Dhwani Batra, Peter W. Cook, Kara Moser, Adrian Paskey, Jason Caravas, Benjamin Rambo-Martin, Shatavia Morrison, Christopher Gulvick, Scott Sammons, Yvette Unoarumhi, Darlene Wagner, Matthew Schmerer, Minoo Agarwal, Eyad Almasri, Debbie Boles, Ayla Burns, Nuthawin Charoensri, Oren Cohen, Susan Countryman, Mary Ann Cristobal, Bobbi Croy, Suzanne Dale, Hrushikesh Deshmukh, Amanda Douglas, Vincent Drouillon, Marcia Eisenberg, Howard Engler, Rama Ghatti, Prashant Gupta, Susan Hicks, Jake Humphrey, Lax Iyer, Lisa Pfefferle, Manoj Jain, Matthew Robinson, Mohan Kolli, Brian Krueger, Tim Kuphal, Stanley Letovsky, Michael Levandoski, Craig Lukasik, Jonathan Meltzer, Brian Norvell, Mindy Nye, Scott Parker, Christos Petropoulos, John Pruitt, Steven Ragan, Scott Ryan, Mike Sapeta, Jana Schroth, Suresh Babu Selvaraju, Goran Stevovic, Amanda Suchanek, Andrea Throop, Lyndon Tilson, Thomas Urban, Joe Voshell, Kimberly Wagner, Jonathan Williams, Mary Williamson, Qian Zeng, Tricia Zwiefelhofer, Clinton R. Paden, Duncan MacCannell |
| EPI_ISL_3113262, EPI_ISL_3113263 | Labor Berlin Charité Vivantes GmbH / Institut für Virologie | Charité Universitätsmedizin Berlin, Institut für Virologie/Labor Berlin | Peter Menzel, Christine Stephan, Rolf Schwarzer, Victor M Corman, Barbara Mühlemann, Terry Jones, Christian Drosten |
| EPI_ISL_3113264 | Laboratory Corporation of America | Centers for Disease Control and Prevention Division of Viral Diseases, Pathogen Discovery | Dakota Howard, Dhwani Batra, Peter W. Cook, Kara Moser, Adrian Paskey, Jason Caravas, Benjamin Rambo-Martin, Shatavia Morrison, Christopher Gulvick, Scott Sammons, Yvette Unoarumhi, Darlene Wagner, Matthew Schmerer, Minoo Agarwal, Eyad Almasri, Debbie Boles, Ayla Burns, Nuthawin Charoensri, Oren Cohen, Susan Countryman, Mary Ann Cristobal, Bobbi Croy, Suzanne Dale, Hrushikesh Deshmukh, Amanda Douglas, Vincent Drouillon, Marcia Eisenberg, Howard Engler, Rama Ghatti, Prashant Gupta, Susan Hicks, Jake Humphrey, Lax Iyer, Lisa Pfefferle, Manoj Jain, Matthew Robinson, Mohan Kolli, Brian Krueger, Tim Kuphal, Stanley Letovsky, Michael Levandoski, Craig Lukasik, Jonathan Meltzer, Brian Norvell, Mindy Nye, Scott Parker, Christos Petropoulos, John Pruitt, Steven Ragan, Scott Ryan, Mike Sapeta, Jana Schroth, Suresh Babu Selvaraju, Goran Stevovic, Amanda Suchanek, Andrea Throop, Lyndon Tilson, Thomas Urban, Joe Voshell, Kimberly Wagner, Jonathan Williams, Mary Williamson, Qian Zeng, Tricia Zwiefelhofer, Clinton R. Paden, Duncan MacCannell |
| EPI_ISL_3113265 | Labor Berlin Charité Vivantes GmbH / Institut für Virologie | Charité Universitätsmedizin Berlin, Institut für Virologie/Labor Berlin | Peter Menzel, Christine Stephan, Rolf Schwarzer, Victor M Corman, Barbara Mühlemann, Terry Jones, Christian Drosten |

| | | | |
|---|---|---|---|
| EPI_ISL_3113266 | Laboratory Corporation of America | Centers for Disease Control and Prevention Division of Viral Diseases, Pathogen Discovery | Dakota Howard, Dhwani Batra, Peter W. Cook, Kara Moser, Adrian Paskey, Jason Caravas, Benjamin Rambo-Martin, Shatavia Morrison, Christopher Gulvick, Scott Sammons, Yvette Unoarumhi, Darlene Wagner, Matthew Schmerer, Minoo Agarwal, Eyad Almasri, Debbie Boles, Ayla Burns, Nuthawin Charoensri, Oren Cohen, Susan Countryman, Mary Ann Cristobal, Bobbi Croy, Suzanne Dale, Hrushikesh Deshmukh, Amanda Douglas, Vincent Drouillon, Marcia Eisenberg, Howard Engler, Rama Ghatti, Prashant Gupta, Susan Hicks, Jake Humphrey, Lax Iyer, Lisa Pfefferle, Manoj Jain, Matthew Robinson, Mohan Kolli, Brian Krueger, Tim Kuphal, Stanley Letovsky, Michael Levandoski, Craig Lukasik, Jonathan Meltzer, Brian Norvell, Mindy Nye, Scott Parker, Christos Petropoulos, John Pruitt, Steven Ragan, Scott Ryan, Mike Sapeta, Jana Schroth, Suresh Babu Selvaraju, Goran Stevovic, Amanda Suchanek, Andrea Throop, Lyndon Tilson, Thomas Urban, Joe Voshell, Kimberly Wagner, Jonathan Williams, Mary Williamson, Qian Zeng, Tricia Zwiefelhofer, Clinton R. Paden, Duncan MacCannell |
| EPI_ISL_3113267 | Labor Berlin Charité Vivantes GmbH / Institut für Virologie | Charité Universitätsmedizin Berlin, Institut für Virologie/Labor Berlin | Peter Menzel, Christine Stephan, Rolf Schwarzer, Victor M Corman, Barbara Mühlemann, Terry Jones, Christian Drosten |
| EPI_ISL_3113268 | Laboratory Corporation of America | Centers for Disease Control and Prevention Division of Viral Diseases, Pathogen Discovery | Dakota Howard, Dhwani Batra, Peter W. Cook, Kara Moser, Adrian Paskey, Jason Caravas, Benjamin Rambo-Martin, Shatavia Morrison, Christopher Gulvick, Scott Sammons, Yvette Unoarumhi, Darlene Wagner, Matthew Schmerer, Minoo Agarwal, Eyad Almasri, Debbie Boles, Ayla Burns, Nuthawin Charoensri, Oren Cohen, Susan Countryman, Mary Ann Cristobal, Bobbi Croy, Suzanne Dale, Hrushikesh Deshmukh, Amanda Douglas, Vincent Drouillon, Marcia Eisenberg, Howard Engler, Rama Ghatti, Prashant Gupta, Susan Hicks, Jake Humphrey, Lax Iyer, Lisa Pfefferle, Manoj Jain, Matthew Robinson, Mohan Kolli, Brian Krueger, Tim Kuphal, Stanley Letovsky, Michael Levandoski, Craig Lukasik, Jonathan Meltzer, Brian Norvell, Mindy Nye, Scott Parker, Christos Petropoulos, John Pruitt, Steven Ragan, Scott Ryan, Mike Sapeta, Jana Schroth, Suresh Babu Selvaraju, Goran Stevovic, Amanda Suchanek, Andrea Throop, Lyndon Tilson, Thomas Urban, Joe Voshell, Kimberly Wagner, Jonathan Williams, Mary Williamson, Qian Zeng, Tricia Zwiefelhofer, Clinton R. Paden, Duncan MacCannell |
| EPI_ISL_3113269 | Labor Berlin Charité Vivantes GmbH / Institut für Virologie | Charité Universitätsmedizin Berlin, Institut für Virologie/Labor Berlin | Peter Menzel, Christine Stephan, Rolf Schwarzer, Victor M Corman, Barbara Mühlemann, Terry Jones, Christian Drosten |
| EPI_ISL_3113270 | Laboratory Corporation of America | Centers for Disease Control and Prevention Division of Viral Diseases, Pathogen Discovery | Dakota Howard, Dhwani Batra, Peter W. Cook, Kara Moser, Adrian Paskey, Jason Caravas, Benjamin Rambo-Martin, Shatavia Morrison, Christopher Gulvick, Scott Sammons, Yvette Unoarumhi, Darlene Wagner, Matthew Schmerer, Minoo Agarwal, Eyad Almasri, Debbie Boles, Ayla Burns, Nuthawin Charoensri, Oren Cohen, Susan Countryman, Mary Ann Cristobal, Bobbi Croy, Suzanne Dale, Hrushikesh Deshmukh, Amanda Douglas, Vincent Drouillon, Marcia Eisenberg, Howard Engler, Rama Ghatti, Prashant Gupta, Susan Hicks, Jake Humphrey, Lax Iyer, Lisa Pfefferle, Manoj Jain, Matthew Robinson, Mohan Kolli, Brian Krueger, Tim Kuphal, Stanley Letovsky, Michael Levandoski, Craig Lukasik, Jonathan Meltzer, Brian Norvell, Mindy Nye, Scott Parker, Christos Petropoulos, John Pruitt, Steven Ragan, Scott Ryan, Mike Sapeta, Jana Schroth, Suresh Babu Selvaraju, Goran Stevovic, Amanda Suchanek, Andrea Throop, Lyndon Tilson, Thomas Urban, Joe Voshell, Kimberly Wagner, Jonathan Williams, Mary Williamson, Qian Zeng, Tricia Zwiefelhofer, Clinton R. Paden, Duncan MacCannell |
| EPI_ISL_3113271, EPI_ISL_3113272 | Labor Berlin Charité Vivantes GmbH / Institut für Virologie | Charité Universitätsmedizin Berlin, Institut für Virologie/Labor Berlin | Peter Menzel, Christine Stephan, Rolf Schwarzer, Victor M Corman, Barbara Mühlemann, Terry Jones, Christian Drosten |
| EPI_ISL_3113275 | Laboratory Corporation of America | Centers for Disease Control and Prevention Division of Viral Diseases, Pathogen Discovery | Dakota Howard, Dhwani Batra, Peter W. Cook, Kara Moser, Adrian Paskey, Jason Caravas, Benjamin Rambo-Martin, Shatavia Morrison, Christopher Gulvick, Scott Sammons, Yvette Unoarumhi, Darlene Wagner, Matthew Schmerer, Minoo Agarwal, Eyad Almasri, Debbie Boles, Ayla Burns, Nuthawin Charoensri, Oren Cohen, Susan Countryman, Mary Ann Cristobal, Bobbi Croy, Suzanne Dale, Hrushikesh Deshmukh, Amanda Douglas, Vincent Drouillon, Marcia Eisenberg, Howard Engler, Rama Ghatti, Prashant Gupta, Susan Hicks, Jake Humphrey, Lax Iyer, Lisa Pfefferle, Manoj Jain, Matthew Robinson, Mohan Kolli, Brian Krueger, Tim Kuphal, Stanley Letovsky, Michael Levandoski, Craig Lukasik, Jonathan Meltzer, Brian Norvell, Mindy Nye, Scott Parker, Christos Petropoulos, John Pruitt, Steven Ragan, Scott Ryan, Mike Sapeta, Jana Schroth, Suresh Babu Selvaraju, Goran Stevovic, Amanda Suchanek, Andrea Throop, Lyndon Tilson, Thomas Urban, Joe Voshell, Kimberly Wagner, Jonathan Williams, Mary Williamson, Qian Zeng, Tricia Zwiefelhofer, Clinton R. Paden, Duncan MacCannell |
| EPI_ISL_3113276 | Labor Berlin Charité Vivantes GmbH / Institut für Virologie | Charité Universitätsmedizin Berlin, Institut für Virologie/Labor Berlin | Peter Menzel, Christine Stephan, Rolf Schwarzer, Victor M Corman, Barbara Mühlemann, Terry Jones, Christian Drosten |
| EPI_ISL_3113277 | Laboratory Corporation of America | Centers for Disease Control and Prevention Division of Viral Diseases, Pathogen Discovery | Dakota Howard, Dhwani Batra, Peter W. Cook, Kara Moser, Adrian Paskey, Jason Caravas, Benjamin Rambo-Martin, Shatavia Morrison, Christopher Gulvick, Scott Sammons, Yvette Unoarumhi, Darlene Wagner, Matthew Schmerer, Minoo Agarwal, Eyad Almasri, Debbie Boles, Ayla Burns, Nuthawin Charoensri, Oren Cohen, Susan Countryman, Mary Ann Cristobal, Bobbi Croy, Suzanne Dale, Hrushikesh Deshmukh, Amanda Douglas, Vincent Drouillon, Marcia Eisenberg, Howard Engler, Rama Ghatti, Prashant Gupta, Susan Hicks, Jake Humphrey, Lax Iyer, Lisa Pfefferle, Manoj Jain, Matthew Robinson, Mohan Kolli, Brian Krueger, Tim Kuphal, Stanley Letovsky, Michael Levandoski, Craig Lukasik, Jonathan Meltzer, Brian Norvell, Mindy Nye, Scott Parker, Christos Petropoulos, John Pruitt, Steven Ragan, Scott Ryan, Mike Sapeta, Jana Schroth, Suresh Babu Selvaraju, Goran Stevovic, Amanda Suchanek, Andrea Throop, Lyndon Tilson, Thomas Urban, Joe Voshell, Kimberly Wagner, Jonathan Williams, Mary Williamson, Qian Zeng, Tricia Zwiefelhofer, Clinton R. Paden, Duncan MacCannell |
| EPI_ISL_3113278, EPI_ISL_3113279, EPI_ISL_3113281 | Labor Berlin Charité Vivantes GmbH / Institut für Virologie | Charité Universitätsmedizin Berlin, Institut für Virologie/Labor Berlin | Peter Menzel, Christine Stephan, Rolf Schwarzer, Victor M Corman, Barbara Mühlemann, Terry Jones, Christian Drosten |
| EPI_ISL_3113282 | Laboratory Corporation of America | Centers for Disease Control and Prevention Division of Viral Diseases, Pathogen Discovery | Dakota Howard, Dhwani Batra, Peter W. Cook, Kara Moser, Adrian Paskey, Jason Caravas, Benjamin Rambo-Martin, Shatavia Morrison, Christopher Gulvick, Scott Sammons, Yvette Unoarumhi, Darlene Wagner, Matthew Schmerer, Minoo Agarwal, Eyad Almasri, Debbie Boles, Ayla Burns, Nuthawin Charoensri, Oren Cohen, Susan Countryman, Mary Ann Cristobal, Bobbi Croy, Suzanne Dale, Hrushikesh Deshmukh, Amanda Douglas, Vincent Drouillon, Marcia Eisenberg, Howard Engler, Rama Ghatti, Prashant Gupta, Susan Hicks, Jake Humphrey, Lax Iyer, Lisa Pfefferle, Manoj Jain, Matthew Robinson, Mohan Kolli, Brian Krueger, Tim Kuphal, Stanley Letovsky, Michael Levandoski, Craig Lukasik, Jonathan Meltzer, Brian Norvell, Mindy Nye, Scott Parker, Christos Petropoulos, John Pruitt, Steven Ragan, Scott Ryan, Mike Sapeta, Jana Schroth, Suresh Babu Selvaraju, Goran Stevovic, Amanda Suchanek, Andrea Throop, Lyndon Tilson, Thomas Urban, Joe Voshell, Kimberly Wagner, Jonathan Williams, Mary Williamson, Qian Zeng, Tricia Zwiefelhofer, Clinton R. Paden, Duncan MacCannell |
| EPI_ISL_3113283 | Labor Berlin Charité Vivantes GmbH / Institut für Virologie | Charité Universitätsmedizin Berlin, Institut für Virologie/Labor Berlin | Peter Menzel, Christine Stephan, Rolf Schwarzer, Victor M Corman, Barbara Mühlemann, Terry Jones, Christian Drosten |
| EPI_ISL_3113284 | Laboratory Corporation of America | Centers for Disease Control and Prevention Division of Viral Diseases, Pathogen Discovery | Dakota Howard, Dhwani Batra, Peter W. Cook, Kara Moser, Adrian Paskey, Jason Caravas, Benjamin Rambo-Martin, Shatavia Morrison, Christopher Gulvick, Scott Sammons, Yvette Unoarumhi, Darlene Wagner, Matthew Schmerer, Minoo Agarwal, Eyad Almasri, Debbie Boles, Ayla Burns, Nuthawin Charoensri, Oren Cohen, Susan Countryman, Mary Ann Cristobal, Bobbi Croy, Suzanne Dale, Hrushikesh Deshmukh, Amanda Douglas, Vincent Drouillon, Marcia Eisenberg, Howard Engler, Rama Ghatti, Prashant Gupta, Susan Hicks, Jake Humphrey, Lax Iyer, Lisa Pfefferle, Manoj Jain, Matthew Robinson, Mohan Kolli, Brian Krueger, Tim Kuphal, Stanley Letovsky, Michael Levandoski, Craig Lukasik, Jonathan Meltzer, Brian Norvell, Mindy Nye, Scott Parker, Christos Petropoulos, John Pruitt, Steven Ragan, Scott Ryan, Mike Sapeta, Jana Schroth, Suresh Babu Selvaraju, Goran Stevovic, Amanda Suchanek, Andrea Throop, Lyndon Tilson, Thomas Urban, Joe Voshell, Kimberly Wagner, Jonathan Williams, Mary Williamson, Qian Zeng, Tricia Zwiefelhofer, Clinton R. Paden, Duncan MacCannell |
| EPI_ISL_3113285, EPI_ISL_3113286, EPI_ISL_3113288 | Labor Berlin Charité Vivantes GmbH / Institut für Virologie | Charité Universitätsmedizin Berlin, Institut für Virologie/Labor Berlin | Peter Menzel, Christine Stephan, Rolf Schwarzer, Victor M Corman, Barbara Mühlemann, Terry Jones, Christian Drosten |
| EPI_ISL_3113289 | Laboratory Corporation of America | Centers for Disease Control and Prevention Division of Viral Diseases, Pathogen Discovery | Dakota Howard, Dhwani Batra, Peter W. Cook, Kara Moser, Adrian Paskey, Jason Caravas, Benjamin Rambo-Martin, Shatavia Morrison, Christopher Gulvick, Scott Sammons, Yvette Unoarumhi, Darlene Wagner, Matthew Schmerer, Minoo Agarwal, Eyad Almasri, Debbie Boles, Ayla Burns, Nuthawin Charoensri, Oren Cohen, Susan Countryman, Mary Ann Cristobal, Bobbi Croy, Suzanne Dale, Hrushikesh Deshmukh, Amanda Douglas, Vincent Drouillon, Marcia Eisenberg, Howard Engler, Rama Ghatti, Prashant Gupta, Susan Hicks, Jake Humphrey, Lax Iyer, Lisa Pfefferle, Manoj Jain, Matthew Robinson, Mohan Kolli, Brian Krueger, Tim Kuphal, Stanley Letovsky, Michael Levandoski, Craig Lukasik, Jonathan Meltzer, Brian Norvell, Mindy Nye, Scott Parker, Christos Petropoulos, John Pruitt, Steven Ragan, Scott Ryan, Mike Sapeta, Jana Schroth, Suresh Babu Selvaraju, Goran Stevovic, Amanda Suchanek, Andrea Throop, Lyndon Tilson, Thomas Urban, Joe Voshell, Kimberly Wagner, Jonathan Williams, Mary Williamson, Qian Zeng, Tricia Zwiefelhofer, Clinton R. Paden, Duncan MacCannell |
| EPI_ISL_3113290, EPI_ISL_3113291 | Labor Berlin Charité Vivantes GmbH / Institut für Virologie | Charité Universitätsmedizin Berlin, Institut für Virologie/Labor Berlin | Peter Menzel, Christine Stephan, Rolf Schwarzer, Victor M Corman, Barbara Mühlemann, Terry Jones, Christian Drosten |
| EPI_ISL_3113292 | Laboratory Corporation of America | Centers for Disease Control and Prevention Division of Viral Diseases, Pathogen Discovery | Dakota Howard, Dhwani Batra, Peter W. Cook, Kara Moser, Adrian Paskey, Jason Caravas, Benjamin Rambo-Martin, Shatavia Morrison, Christopher Gulvick, Scott Sammons, Yvette Unoarumhi, Darlene Wagner, Matthew Schmerer, Minoo Agarwal, Eyad Almasri, Debbie Boles, Ayla Burns, Nuthawin Charoensri, Oren Cohen, Susan Countryman, Mary Ann Cristobal, Bobbi Croy, Suzanne Dale, Hrushikesh Deshmukh, Amanda Douglas, Vincent Drouillon, Marcia Eisenberg, Howard Engler, Rama Ghatti, Prashant Gupta, Susan Hicks, Jake Humphrey, Lax Iyer, Lisa Pfefferle, Manoj Jain, Matthew Robinson, Mohan Kolli, Brian Krueger, Tim Kuphal, Stanley Letovsky, Michael Levandoski, Craig Lukasik, Jonathan Meltzer, Brian Norvell, Mindy Nye, Scott Parker, Christos Petropoulos, John Pruitt, Steven Ragan, Scott Ryan, Mike Sapeta, Jana Schroth, Suresh Babu Selvaraju, Goran Stevovic, Amanda Suchanek, Andrea Throop, Lyndon Tilson, Thomas Urban, Joe Voshell, Kimberly Wagner, Jonathan Williams, Mary Williamson, Qian Zeng, Tricia Zwiefelhofer, Clinton R. Paden, Duncan MacCannell |
| EPI_ISL_3113293, EPI_ISL_3113296 | Labor Berlin Charité Vivantes GmbH / Institut für Virologie | Charité Universitätsmedizin Berlin, Institut für Virologie/Labor Berlin | Peter Menzel, Christine Stephan, Rolf Schwarzer, Victor M Corman, Barbara Mühlemann, Terry Jones, Christian Drosten |
| EPI_ISL_3113297 | Laboratory Corporation of America | Centers for Disease Control and Prevention Division of Viral Diseases, Pathogen Discovery | Dakota Howard, Dhwani Batra, Peter W. Cook, Kara Moser, Adrian Paskey, Jason Caravas, Benjamin Rambo-Martin, Shatavia Morrison, Christopher Gulvick, Scott Sammons, Yvette Unoarumhi, Darlene Wagner, Matthew Schmerer, Minoo Agarwal, Eyad Almasri, Debbie Boles, Ayla Burns, Nuthawin Charoensri, Oren Cohen, Susan Countryman, Mary Ann Cristobal, Bobbi Croy, Suzanne Dale, Hrushikesh Deshmukh, Amanda Douglas, Vincent Drouillon, Marcia Eisenberg, Howard Engler, Rama Ghatti, Prashant Gupta, Susan Hicks, Jake Humphrey, Lax Iyer, Lisa Pfefferle, Manoj Jain, Matthew Robinson, Mohan Kolli, Brian Krueger, Tim Kuphal, Stanley Letovsky, Michael Levandoski, Craig Lukasik, Jonathan Meltzer, Brian Norvell, Mindy Nye, Scott Parker, Christos Petropoulos, John Pruitt, Steven Ragan, Scott Ryan, Mike Sapeta, Jana Schroth, Suresh Babu Selvaraju, Goran Stevovic, Amanda Suchanek, Andrea Throop, Lyndon Tilson, Thomas Urban, Joe Voshell, Kimberly Wagner, Jonathan Williams, Mary Williamson, Qian Zeng, Tricia Zwiefelhofer, Clinton R. Paden, Duncan MacCannell |
| EPI_ISL_3113298 | Labor Berlin Charité Vivantes GmbH / Institut für Virologie | Charité Universitätsmedizin Berlin, Institut für Virologie/Labor Berlin | Peter Menzel, Christine Stephan, Rolf Schwarzer, Victor M Corman, Barbara Mühlemann, Terry Jones, Christian Drosten |
| EPI_ISL_3113299 | Laboratory Corporation of America | Centers for Disease Control and Prevention Division of Viral Diseases, Pathogen Discovery | Dakota Howard, Dhwani Batra, Peter W. Cook, Kara Moser, Adrian Paskey, Jason Caravas, Benjamin Rambo-Martin, Shatavia Morrison, Christopher Gulvick, Scott Sammons, Yvette Unoarumhi, Darlene Wagner, Matthew Schmerer, Minoo Agarwal, Eyad Almasri, Debbie Boles, Ayla Burns, Nuthawin Charoensri, Oren Cohen, Susan Countryman, Mary Ann Cristobal, Bobbi Croy, Suzanne Dale, Hrushikesh Deshmukh, Amanda Douglas, Vincent Drouillon, Marcia Eisenberg, Howard Engler, Rama Ghatti, Prashant Gupta, Susan Hicks, Jake Humphrey, Lax Iyer, Lisa Pfefferle, Manoj Jain, Matthew Robinson, Mohan Kolli, Brian Krueger, Tim Kuphal, Stanley Letovsky, Michael Levandoski, Craig Lukasik, Jonathan Meltzer, Brian Norvell, Mindy Nye, Scott Parker, Christos Petropoulos, John Pruitt, Steven Ragan, Scott Ryan, Mike Sapeta, Jana Schroth, Suresh Babu Selvaraju, Goran Stevovic, Amanda Suchanek, Andrea Throop, Lyndon Tilson, Thomas Urban, Joe Voshell, Kimberly Wagner, Jonathan Williams, Mary Williamson, Qian Zeng, Tricia Zwiefelhofer, Clinton R. Paden, Duncan MacCannell |
| EPI_ISL_3113300 | Labor Berlin Charité Vivantes GmbH / Institut für Virologie | Charité Universitätsmedizin Berlin, Institut für Virologie/Labor Berlin | Peter Menzel, Christine Stephan, Rolf Schwarzer, Victor M Corman, Barbara Mühlemann, Terry Jones, Christian Drosten |
| EPI_ISL_3113301 | Laboratory Corporation of America | Centers for Disease Control and Prevention Division of Viral Diseases, Pathogen Discovery | Dakota Howard, Dhwani Batra, Peter W. Cook, Kara Moser, Adrian Paskey, Jason Caravas, Benjamin Rambo-Martin, Shatavia Morrison, Christopher Gulvick, Scott Sammons, Yvette Unoarumhi, Darlene Wagner, Matthew Schmerer, Minoo Agarwal, Eyad Almasri, Debbie Boles, Ayla Burns, Nuthawin Charoensri, Oren Cohen, Susan Countryman, Mary Ann Cristobal, Bobbi Croy, Suzanne Dale, Hrushikesh Deshmukh, Amanda Douglas, Vincent Drouillon, Marcia Eisenberg, Howard Engler, Rama Ghatti, Prashant Gupta, Susan Hicks, Jake Humphrey, Lax Iyer, Lisa Pfefferle, Manoj Jain, Matthew Robinson, Mohan Kolli, Brian Krueger, Tim Kuphal, Stanley Letovsky, Michael Levandoski, Craig Lukasik, Jonathan Meltzer, Brian Norvell, Mindy Nye, Scott Parker, Christos Petropoulos, John Pruitt, Steven Ragan, Scott Ryan, Mike Sapeta, Jana Schroth, Suresh Babu Selvaraju, Goran Stevovic, Amanda Suchanek, Andrea Throop, Lyndon Tilson, Thomas Urban, Joe Voshell, Kimberly Wagner, Jonathan Williams, Mary Williamson, Qian Zeng, Tricia Zwiefelhofer, Clinton R. Paden, Duncan MacCannell |
| EPI_ISL_3113302 | Labor Berlin Charité Vivantes GmbH / Institut für Virologie | Charité Universitätsmedizin Berlin, Institut für Virologie/Labor Berlin | Peter Menzel, Christine Stephan, Rolf Schwarzer, Victor M Corman, Barbara Mühlemann, Terry Jones, Christian Drosten |
| EPI_ISL_3113303 | Laboratory Corporation of America | Centers for Disease Control and Prevention Division of Viral Diseases, Pathogen Discovery | Dakota Howard, Dhwani Batra, Peter W. Cook, Kara Moser, Adrian Paskey, Jason Caravas, Benjamin Rambo-Martin, Shatavia Morrison, Christopher Gulvick, Scott Sammons, Yvette Unoarumhi, Darlene Wagner, Matthew Schmerer, Minoo Agarwal, Eyad Almasri, Debbie Boles, Ayla Burns, Nuthawin Charoensri, Oren Cohen, Susan Countryman, Mary Ann Cristobal, Bobbi Croy, Suzanne Dale, Hrushikesh Deshmukh, Amanda Douglas, Vincent Drouillon, Marcia Eisenberg, Howard Engler, Rama Ghatti, Prashant Gupta, Susan Hicks, Jake Humphrey, Lax Iyer, Lisa Pfefferle, Manoj Jain, Matthew Robinson, Mohan Kolli, Brian Krueger, Tim Kuphal, Stanley Letovsky, Michael Levandoski, Craig Lukasik, Jonathan Meltzer, Brian Norvell, Mindy Nye, Scott Parker, Christos Petropoulos, John Pruitt, Steven Ragan, Scott Ryan, Mike Sapeta, Jana Schroth, Suresh Babu Selvaraju, Goran Stevovic, Amanda Suchanek, Andrea Throop, Lyndon Tilson, Thomas Urban, Joe Voshell, Kimberly Wagner, Jonathan Williams, Mary Williamson, Qian Zeng, Tricia Zwiefelhofer, Clinton R. Paden, Duncan MacCannell |
| EPI_ISL_3113304, EPI_ISL_3113305 | Labor Berlin Charité Vivantes GmbH / Institut für Virologie | Charité Universitätsmedizin Berlin, Institut für Virologie/Labor Berlin | Peter Menzel, Christine Stephan, Rolf Schwarzer, Victor M Corman, Barbara Mühlemann, Terry Jones, Christian Drosten |
| EPI_ISL_3113306, EPI_ISL_3113308 | Laboratory Corporation of America | Centers for Disease Control and Prevention Division of Viral Diseases, Pathogen Discovery | Dakota Howard, Dhwani Batra, Peter W. Cook, Kara Moser, Adrian Paskey, Jason Caravas, Benjamin Rambo-Martin, Shatavia Morrison, Christopher Gulvick, Scott Sammons, Yvette Unoarumhi, Darlene Wagner, Matthew Schmerer, Minoo Agarwal, Eyad Almasri, Debbie Boles, Ayla Burns, Nuthawin Charoensri, Oren Cohen, Susan Countryman, Mary Ann Cristobal, Bobbi Croy, Suzanne Dale, Hrushikesh Deshmukh, Amanda Douglas, Vincent Drouillon, Marcia Eisenberg, Howard Engler, Rama Ghatti, Prashant Gupta, Susan Hicks, Jake Humphrey, Lax Iyer, Lisa Pfefferle, Manoj Jain, Matthew Robinson, Mohan Kolli, Brian Krueger, Tim Kuphal, Stanley Letovsky, Michael Levandoski, Craig Lukasik, Jonathan Meltzer, Brian Norvell, Mindy Nye, Scott Parker, Christos Petropoulos, John Pruitt, Steven Ragan, Scott Ryan, Mike Sapeta, Jana Schroth, Suresh Babu Selvaraju, Goran Stevovic, Amanda Suchanek, Andrea Throop, Lyndon Tilson, Thomas Urban, Joe Voshell, Kimberly Wagner, Jonathan Williams, Mary Williamson, Qian Zeng, Tricia Zwiefelhofer, Clinton R. Paden, Duncan MacCannell |
| EPI_ISL_3113309, EPI_ISL_3113310 | Labor Berlin Charité Vivantes GmbH / Institut für Virologie | Charité Universitätsmedizin Berlin, Institut für Virologie/Labor Berlin | Peter Menzel, Christine Stephan, Rolf Schwarzer, Victor M Corman, Barbara Mühlemann, Terry Jones, Christian Drosten |
| EPI_ISL_3113311 | Laboratory Corporation of America | Centers for Disease Control and Prevention Division of Viral Diseases, Pathogen Discovery | Dakota Howard, Dhwani Batra, Peter W. Cook, Kara Moser, Adrian Paskey, Jason Caravas, Benjamin Rambo-Martin, Shatavia Morrison, Christopher Gulvick, Scott Sammons, Yvette Unoarumhi, Darlene Wagner, Matthew Schmerer, Minoo Agarwal, Eyad Almasri, Debbie Boles, Ayla Burns, Nuthawin Charoensri, Oren Cohen, Susan Countryman, Mary Ann Cristobal, Bobbi Croy, Suzanne Dale, Hrushikesh Deshmukh, Amanda Douglas, Vincent Drouillon, Marcia Eisenberg, Howard Engler, Rama Ghatti, Prashant Gupta, Susan Hicks, Jake Humphrey, Lax Iyer, Lisa Pfefferle, Manoj Jain, Matthew Robinson, Mohan Kolli, Brian Krueger, Tim Kuphal, Stanley Letovsky, Michael Levandoski, Craig Lukasik, Jonathan Meltzer, Brian Norvell, Mindy Nye, Scott Parker, Christos Petropoulos, John Pruitt, Steven Ragan, Scott Ryan, Mike Sapeta, Jana Schroth, Suresh Babu Selvaraju, Goran Stevovic, Amanda Suchanek, Andrea Throop, Lyndon Tilson, Thomas Urban, Joe Voshell, Kimberly Wagner, Jonathan Williams, Mary Williamson, Qian Zeng, Tricia Zwiefelhofer, Clinton R. Paden, Duncan MacCannell |

| | | | |
|---|---|---|---|
| EPI_ISL_3113312 | Labor Berlin Charité Vivantes GmbH / Institut für Virologie | Charité Universitätsmedizin Berlin, Institut für Virologie/Labor Berlin | Peter Menzel, Christine Stephan, Rolf Schwarzer, Victor M Corman, Barbara Mühlemann, Terry Jones, Christian Drosten |
| EPI_ISL_3113314 | Laboratory Corporation of America | Centers for Disease Control and Prevention Division of Viral Diseases, Pathogen Discovery | Dakota Howard, Dhwani Batra, Peter W. Cook, Kara Moser, Adrian Paskey, Jason Caravas, Benjamin Rambo-Martin, Shatavia Morrison, Christopher Gulvick, Scott Sammons, Yvette Unoarumhi, Darlene Wagner, Matthew Schmerer, Minoo Agarwal, Eyad Almasri, Debbie Boles, Ayla Burns, Nuthawin Charoensri, Oren Cohen, Susan Countryman, Mary Ann Cristobal, Bobbi Croy, Suzanne Dale, Hrushikesh Deshmukh, Amanda Douglas, Vincent Drouillon, Marcia Eisenberg, Howard Engler, Rama Ghatti, Prashant Gupta, Susan Hicks, Jake Humphrey, Lax Iyer, Lisa Pfefferle, Manoj Jain, Matthew Robinson, Mohan Kolli, Brian Krueger, Tim Kuphal, Stanley Letovsky, Michael Levandoski, Craig Lukasik, Jonathan Meltzer, Brian Norvell, Mindy Nye, Scott Parker, Christos Petropoulos, John Pruitt, Steven Ragan, Scott Ryan, Mike Sapeta, Jana Schroth, Suresh Babu Selvaraju, Goran Stevovic, Amanda Suchanek, Andrea Throop, Lyndon Tilson, Thomas Urban, Joe Voshell, Kimberly Wagner, Jonathan Williams, Mary Williamson, Qian Zeng, Tricia Zwiefelhofer, Clinton R. Paden, Duncan MacCannell |
| EPI_ISL_3113315 | Labor Berlin Charité Vivantes GmbH / Institut für Virologie | Charité Universitätsmedizin Berlin, Institut für Virologie/Labor Berlin | Peter Menzel, Christine Stephan, Rolf Schwarzer, Victor M Corman, Barbara Mühlemann, Terry Jones, Christian Drosten |
| EPI_ISL_3113316 | Laboratory Corporation of America | Centers for Disease Control and Prevention Division of Viral Diseases, Pathogen Discovery | Dakota Howard, Dhwani Batra, Peter W. Cook, Kara Moser, Adrian Paskey, Jason Caravas, Benjamin Rambo-Martin, Shatavia Morrison, Christopher Gulvick, Scott Sammons, Yvette Unoarumhi, Darlene Wagner, Matthew Schmerer, Minoo Agarwal, Eyad Almasri, Debbie Boles, Ayla Burns, Nuthawin Charoensri, Oren Cohen, Susan Countryman, Mary Ann Cristobal, Bobbi Croy, Suzanne Dale, Hrushikesh Deshmukh, Amanda Douglas, Vincent Drouillon, Marcia Eisenberg, Howard Engler, Rama Ghatti, Prashant Gupta, Susan Hicks, Jake Humphrey, Lax Iyer, Lisa Pfefferle, Manoj Jain, Matthew Robinson, Mohan Kolli, Brian Krueger, Tim Kuphal, Stanley Letovsky, Michael Levandoski, Craig Lukasik, Jonathan Meltzer, Brian Norvell, Mindy Nye, Scott Parker, Christos Petropoulos, John Pruitt, Steven Ragan, Scott Ryan, Mike Sapeta, Jana Schroth, Suresh Babu Selvaraju, Goran Stevovic, Amanda Suchanek, Andrea Throop, Lyndon Tilson, Thomas Urban, Joe Voshell, Kimberly Wagner, Jonathan Williams, Mary Williamson, Qian Zeng, Tricia Zwiefelhofer, Clinton R. Paden, Duncan MacCannell |
| EPI_ISL_3113317, EPI_ISL_3113318 | Labor Berlin Charité Vivantes GmbH / Institut für Virologie | Charité Universitätsmedizin Berlin, Institut für Virologie/Labor Berlin | Peter Menzel, Christine Stephan, Rolf Schwarzer, Victor M Corman, Barbara Mühlemann, Terry Jones, Christian Drosten |
| EPI_ISL_3113319 | Laboratory Corporation of America | Centers for Disease Control and Prevention Division of Viral Diseases, Pathogen Discovery | Dakota Howard, Dhwani Batra, Peter W. Cook, Kara Moser, Adrian Paskey, Jason Caravas, Benjamin Rambo-Martin, Shatavia Morrison, Christopher Gulvick, Scott Sammons, Yvette Unoarumhi, Darlene Wagner, Matthew Schmerer, Minoo Agarwal, Eyad Almasri, Debbie Boles, Ayla Burns, Nuthawin Charoensri, Oren Cohen, Susan Countryman, Mary Ann Cristobal, Bobbi Croy, Suzanne Dale, Hrushikesh Deshmukh, Amanda Douglas, Vincent Drouillon, Marcia Eisenberg, Howard Engler, Rama Ghatti, Prashant Gupta, Susan Hicks, Jake Humphrey, Lax Iyer, Lisa Pfefferle, Manoj Jain, Matthew Robinson, Mohan Kolli, Brian Krueger, Tim Kuphal, Stanley Letovsky, Michael Levandoski, Craig Lukasik, Jonathan Meltzer, Brian Norvell, Mindy Nye, Scott Parker, Christos Petropoulos, John Pruitt, Steven Ragan, Scott Ryan, Mike Sapeta, Jana Schroth, Suresh Babu Selvaraju, Goran Stevovic, Amanda Suchanek, Andrea Throop, Lyndon Tilson, Thomas Urban, Joe Voshell, Kimberly Wagner, Jonathan Williams, Mary Williamson, Qian Zeng, Tricia Zwiefelhofer, Clinton R. Paden, Duncan MacCannell |
| EPI_ISL_3113320, EPI_ISL_3113322 | Labor Berlin Charité Vivantes GmbH / Institut für Virologie | Charité Universitätsmedizin Berlin, Institut für Virologie/Labor Berlin | Peter Menzel, Christine Stephan, Rolf Schwarzer, Victor M Corman, Barbara Mühlemann, Terry Jones, Christian Drosten |
| EPI_ISL_3113324 | Laboratory Corporation of America | Centers for Disease Control and Prevention Division of Viral Diseases, Pathogen Discovery | Dakota Howard, Dhwani Batra, Peter W. Cook, Kara Moser, Adrian Paskey, Jason Caravas, Benjamin Rambo-Martin, Shatavia Morrison, Christopher Gulvick, Scott Sammons, Yvette Unoarumhi, Darlene Wagner, Matthew Schmerer, Minoo Agarwal, Eyad Almasri, Debbie Boles, Ayla Burns, Nuthawin Charoensri, Oren Cohen, Susan Countryman, Mary Ann Cristobal, Bobbi Croy, Suzanne Dale, Hrushikesh Deshmukh, Amanda Douglas, Vincent Drouillon, Marcia Eisenberg, Howard Engler, Rama Ghatti, Prashant Gupta, Susan Hicks, Jake Humphrey, Lax Iyer, Lisa Pfefferle, Manoj Jain, Matthew Robinson, Mohan Kolli, Brian Krueger, Tim Kuphal, Stanley Letovsky, Michael Levandoski, Craig Lukasik, Jonathan Meltzer, Brian Norvell, Mindy Nye, Scott Parker, Christos Petropoulos, John Pruitt, Steven Ragan, Scott Ryan, Mike Sapeta, Jana Schroth, Suresh Babu Selvaraju, Goran Stevovic, Amanda Suchanek, Andrea Throop, Lyndon Tilson, Thomas Urban, Joe Voshell, Kimberly Wagner, Jonathan Williams, Mary Williamson, Qian Zeng, Tricia Zwiefelhofer, Clinton R. Paden, Duncan MacCannell |
| EPI_ISL_3113325 | Labor Berlin Charité Vivantes GmbH / Institut für Virologie | Charité Universitätsmedizin Berlin, Institut für Virologie/Labor Berlin | Peter Menzel, Christine Stephan, Rolf Schwarzer, Victor M Corman, Barbara Mühlemann, Terry Jones, Christian Drosten |
| EPI_ISL_3113326 | Laboratory Corporation of America | Centers for Disease Control and Prevention Division of Viral Diseases, Pathogen Discovery | Dakota Howard, Dhwani Batra, Peter W. Cook, Kara Moser, Adrian Paskey, Jason Caravas, Benjamin Rambo-Martin, Shatavia Morrison, Christopher Gulvick, Scott Sammons, Yvette Unoarumhi, Darlene Wagner, Matthew Schmerer, Minoo Agarwal, Eyad Almasri, Debbie Boles, Ayla Burns, Nuthawin Charoensri, Oren Cohen, Susan Countryman, Mary Ann Cristobal, Bobbi Croy, Suzanne Dale, Hrushikesh Deshmukh, Amanda Douglas, Vincent Drouillon, Marcia Eisenberg, Howard Engler, Rama Ghatti, Prashant Gupta, Susan Hicks, Jake Humphrey, Lax Iyer, Lisa Pfefferle, Manoj Jain, Matthew Robinson, Mohan Kolli, Brian Krueger, Tim Kuphal, Stanley Letovsky, Michael Levandoski, Craig Lukasik, Jonathan Meltzer, Brian Norvell, Mindy Nye, Scott Parker, Christos Petropoulos, John Pruitt, Steven Ragan, Scott Ryan, Mike Sapeta, Jana Schroth, Suresh Babu Selvaraju, Goran Stevovic, Amanda Suchanek, Andrea Throop, Lyndon Tilson, Thomas Urban, Joe Voshell, Kimberly Wagner, Jonathan Williams, Mary Williamson, Qian Zeng, Tricia Zwiefelhofer, Clinton R. Paden, Duncan MacCannell |
| EPI_ISL_3113327, EPI_ISL_3113328 | Labor Berlin Charité Vivantes GmbH / Institut für Virologie | Charité Universitätsmedizin Berlin, Institut für Virologie/Labor Berlin | Peter Menzel, Christine Stephan, Rolf Schwarzer, Victor M Corman, Barbara Mühlemann, Terry Jones, Christian Drosten |

EPI_ISL_3113329, EPI_ISL_3113330, EPI_ISL_3113331, EPI_ISL_3113332, EPI_ISL_3113333, EPI_ISL_3113334, EPI_ISL_3113336, EPI_ISL_3113337, EPI_ISL_3113338, EPI_ISL_3113339, EPI_ISL_3113340, EPI_ISL_3113341, EPI_ISL_3113342, EPI_ISL_3113343, EPI_ISL_3113344, EPI_ISL_3113345, EPI_ISL_3113346, EPI_ISL_3113347, EPI_ISL_3113348, EPI_ISL_3113349, EPI_ISL_3113350, EPI_ISL_3113351, EPI_ISL_3113352, EPI_ISL_3113353, EPI_ISL_3113354, EPI_ISL_3113355, EPI_ISL_3113356, EPI_ISL_3113361, EPI_ISL_3113363, EPI_ISL_3113364, EPI_ISL_3113365, EPI_ISL_3113367, EPI_ISL_3113368, EPI_ISL_3113369, EPI_ISL_3113370, EPI_ISL_3113371, EPI_ISL_3113373, EPI_ISL_3113374, EPI_ISL_3113375, EPI_ISL_3113376, EPI_ISL_3113377, EPI_ISL_3113378, EPI_ISL_3113379, EPI_ISL_3113380, EPI_ISL_3113381, EPI_ISL_3113382, EPI_ISL_3113383, EPI_ISL_3113384, EPI_ISL_3113385, EPI_ISL_3113390, EPI_ISL_3113391, EPI_ISL_3113392, EPI_ISL_3113393, EPI_ISL_3113394, EPI_ISL_3113395, EPI_ISL_3113396, EPI_ISL_3113398, EPI_ISL_3113399, EPI_ISL_3113401, EPI_ISL_3113402, EPI_ISL_3113403, EPI_ISL_3113404, EPI_ISL_3113405, EPI_ISL_3113407, EPI_ISL_3113409, EPI_ISL_3113410, EPI_ISL_3113411, EPI_ISL_3113412, EPI_ISL_3113415, EPI_ISL_3113416, EPI_ISL_3113417, EPI_ISL_3113420, EPI_ISL_3113421, EPI_ISL_3113423, EPI_ISL_3113424, EPI_ISL_3113425, EPI_ISL_3113426, EPI_ISL_3113427, EPI_ISL_3113431, EPI_ISL_3113432, EPI_ISL_3113434, EPI_ISL_3113435, EPI_ISL_3113437, EPI_ISL_3113438, EPI_ISL_3113439, EPI_ISL_3113441, EPI_ISL_3113442, EPI_ISL_3113444, EPI_ISL_3113445, EPI_ISL_3113446, EPI_ISL_3113447, EPI_ISL_3113448, EPI_ISL_3113449, EPI_ISL_3113451, EPI_ISL_3113452, EPI_ISL_3113454, EPI_ISL_3113455, EPI_ISL_3113458, EPI_ISL_3113461, EPI_ISL_3113462, EPI_ISL_3113465, EPI_ISL_3113469, EPI_ISL_3113470, EPI_ISL_3113471, EPI_ISL_3113472, EPI_ISL_3113476, EPI_ISL_3113477, EPI_ISL_3113478, EPI_ISL_3113479, EPI_ISL_3113480, EPI_ISL_3113481, EPI_ISL_3113482, EPI_ISL_3113484, EPI_ISL_3113485, EPI_ISL_3113486, EPI_ISL_3113487, EPI_ISL_3113488, EPI_ISL_3113489, EPI_ISL_3113490, EPI_ISL_3113491, EPI_ISL_3113493, EPI_ISL_3113494, EPI_ISL_3113496, EPI_ISL_3113498, EPI_ISL_3113499, EPI_ISL_3113500, EPI_ISL_3113501, EPI_ISL_3113503, EPI_ISL_3113504, EPI_ISL_3113505, EPI_ISL_3113506, EPI_ISL_3113508, EPI_ISL_3113509, EPI_ISL_3113510, EPI_ISL_3113511, EPI_ISL_3113512, EPI_ISL_3113513, EPI_ISL_3113514, EPI_ISL_3113515, EPI_ISL_3113516, EPI_ISL_3113517, EPI_ISL_3113518, EPI_ISL_3113519, EPI_ISL_3113520, EPI_ISL_3113521, EPI_ISL_3113523, EPI_ISL_3113524, EPI_ISL_3113525, EPI_ISL_3113526, EPI_ISL_3113527, EPI_ISL_3113528, EPI_ISL_3113530, EPI_ISL_3113531, EPI_ISL_3113532, EPI_ISL_3113533, EPI_ISL_3113535, EPI_ISL_3113537, EPI_ISL_3113538, EPI_ISL_3113539, EPI_ISL_3113540, EPI_ISL_3113541, EPI_ISL_3113542, EPI_ISL_3113543, EPI_ISL_3113544, EPI_ISL_3113545, EPI_ISL_3113546, EPI_ISL_3113547, EPI_ISL_3113548, EPI_ISL_3113549, EPI_ISL_3113551, EPI_ISL_3113552, EPI_ISL_3113553, EPI_ISL_3113554, EPI_ISL_3113555, EPI_ISL_3113556, EPI_ISL_3113557, EPI_ISL_3113558, EPI_ISL_3113559, EPI_ISL_3113560, EPI_ISL_3113561, EPI_ISL_3113562, EPI_ISL_3113563, EPI_ISL_3113564, EPI_ISL_3113565, EPI_ISL_3113566, EPI_ISL_3113567, EPI_ISL_3113569, EPI_ISL_3113570, EPI_ISL_3113571, EPI_ISL_3113572, EPI_ISL_3113573, EPI_ISL_3113574, EPI_ISL_3113576, EPI_ISL_3113577, EPI_ISL_3113580, EPI_ISL_3113581, EPI_ISL_3113582, EPI_ISL_3113583, EPI_ISL_3113585, EPI_ISL_3113586, EPI_ISL_3113587, EPI_ISL_3113588, EPI_ISL_3113589, EPI_ISL_3113590, EPI_ISL_3113591, EPI_ISL_3113592, EPI_ISL_3113593, EPI_ISL_3113594, EPI_ISL_3113595, EPI_ISL_3113596, EPI_ISL_3113598, EPI_ISL_3113599, EPI_ISL_3113600, EPI_ISL_3113601, EPI_ISL_3113602, EPI_ISL_3113603, EPI_ISL_3113605, EPI_ISL_3113606, EPI_ISL_3113607, EPI_ISL_3113609, EPI_ISL_3113610, EPI_ISL_3113611, EPI_ISL_3113612, EPI_ISL_3113613, EPI_ISL_3113614, EPI_ISL_3113615, EPI_ISL_3113616, EPI_ISL_3113618, EPI_ISL_3113619, EPI_ISL_3113620, EPI_ISL_3113621, EPI_ISL_3113623, EPI_ISL_3113624, EPI_ISL_3113625, EPI_ISL_3113626, EPI_ISL_3113627, EPI_ISL_3113630, EPI_ISL_3113631, EPI_ISL_3113632, EPI_ISL_3113633, EPI_ISL_3113634, EPI_ISL_3113635, EPI_ISL_3113636, EPI_ISL_3113637, EPI_ISL_3113638, EPI_ISL_3113639, EPI_ISL_3113640, EPI_ISL_3113642, EPI_ISL_3113643, EPI_ISL_3113644, EPI_ISL_3113645, EPI_ISL_3113646, EPI_ISL_3113647, EPI_ISL_3113648, EPI_ISL_3113649, EPI_ISL_3113650, EPI_ISL_3113651, EPI_ISL_3113652, EPI_ISL_3113653, EPI_ISL_3113654, EPI_ISL_3113655, EPI_ISL_3113656, EPI_ISL_3113657, EPI_ISL_3113658, EPI_ISL_3113661, EPI_ISL_3113662, EPI_ISL_3113663, EPI_ISL_3113664, EPI_ISL_3113665, EPI_ISL_3113666, EPI_ISL_3113667, EPI_ISL_3113668, EPI_ISL_3113669, EPI_ISL_3113670, EPI_ISL_3113671, EPI_ISL_3113672, EPI_ISL_3113673, EPI_ISL_3113674, EPI_ISL_3113675, EPI_ISL_3113676, EPI_ISL_3113677, EPI_ISL_3113678, EPI_ISL_3113679, EPI_ISL_3113680, EPI_ISL_3113682, EPI_ISL_3113683, EPI_ISL_3113684, EPI_ISL_3113685, EPI_ISL_3113687, EPI_ISL_3113688, EPI_ISL_3113689, EPI_ISL_3113690, EPI_ISL_3113691, EPI_ISL_3113692, EPI_ISL_3113693, EPI_ISL_3113694, EPI_ISL_3113695, EPI_ISL_3113696, EPI_ISL_3113697, EPI_ISL_3113698, EPI_ISL_3113699, EPI_ISL_3113700, EPI_ISL_3113701, EPI_ISL_3113702, EPI_ISL_3113704, EPI_ISL_3113705, EPI_ISL_3113706, EPI_ISL_3113707, EPI_ISL_3113709, EPI_ISL_3113711, EPI_ISL_3113712, EPI_ISL_3113714, EPI_ISL_3113715, EPI_ISL_3113716, EPI_ISL_3113717, EPI_ISL_3113718, EPI_ISL_3113719, EPI_ISL_3113721, EPI_ISL_3113722, EPI_ISL_3113723, EPI_ISL_3113724, EPI_ISL_3113725, EPI_ISL_3113727, EPI_ISL_3113728, EPI_ISL_3113729, EPI_ISL_3113730, EPI_ISL_3113731, EPI_ISL_3113732, EPI_ISL_3113733, EPI_ISL_3113734, EPI_ISL_3113736, EPI_ISL_3113737, EPI_ISL_3113738, EPI_ISL_3113739, EPI_ISL_3113740, EPI_ISL_3113743, EPI_ISL_3113744, EPI_ISL_3113745, EPI_ISL_3113746, EPI_ISL_3113747, EPI_ISL_3113748, EPI_ISL_3113749, EPI_ISL_3113750, EPI_ISL_3113752, EPI_ISL_3113753, EPI_ISL_3113754, EPI_ISL_3113755, EPI_ISL_3113756, EPI_ISL_3113757, EPI_ISL_3113758, EPI_ISL_3113759, EPI_ISL_3113760

| | | | |
|---|---|---|---|
| see above | Laboratory Corporation of America | Centers for Disease Control and Prevention Division of Viral Diseases, Pathogen Discovery | Dakota Howard, Dhwani Batra, Peter W. Cook, Kara Moser, Adrian Paskey, Jason Caravas, Benjamin Rambo-Martin, Shatavia Morrison, Christopher Gulvick, Scott Sammons, Yvette Unoarumhi, Darlene Wagner, Matthew Schmerer, Minoo Agarwal, Eyad Almasri, Debbie Boles, Ayla Burns, Nuthawin Charoensri, Oren Cohen, Susan Countryman, Mary Ann Cristobal, Bobbi Croy, Suzanne Dale, Hrushikesh Deshmukh, Amanda Douglas, Vincent Drouillon, Marcia Eisenberg, Howard Engler, Rama Ghatti, Prashant Gupta, Susan Hicks, Jake Humphrey, Lax Iyer, Lisa Pfefferle, Manoj Jain, Matthew Robinson, Mohan Kolli, Brian Krueger, Tim Kuphal, Stanley Letovsky, Michael Levandoski, Craig Lukasik, Jonathan Meltzer, Brian Norvell, Mindy Nye, Scott Parker, Christos Petropoulos, John Pruitt, Steven Ragan, Scott Ryan, Mike Sapeta, Jana Schroth, Suresh Babu Selvaraju, Goran Stevovic, Amanda Suchanek, Andrea Throop, Lyndon Tilson, Thomas Urban, Joe Voshell, Kimberly Wagner, Jonathan Williams, Mary Williamson, Qian Zeng, Tricia Zwiefelhofer, Clinton R. Paden, Duncan MacCannell |

EPI_ISL_3113761, EPI_ISL_3113762, EPI_ISL_3113763, EPI_ISL_3113764, EPI_ISL_3113765, EPI_ISL_3113766, EPI_ISL_3113767, EPI_ISL_3113768, EPI_ISL_3113769, EPI_ISL_3113770, EPI_ISL_3113771, EPI_ISL_3113772, EPI_ISL_3113773, EPI_ISL_3113774, EPI_ISL_3113775, EPI_ISL_3113776, EPI_ISL_3113777, EPI_ISL_3113778, EPI_ISL_3113779, EPI_ISL_3113780, EPI_ISL_3113781, EPI_ISL_3113782, EPI_ISL_3113783, EPI_ISL_3113784, EPI_ISL_3113785, EPI_ISL_3113786, EPI_ISL_3113787, EPI_ISL_3113788, EPI_ISL_3113789, EPI_ISL_3113791, EPI_ISL_3113792, EPI_ISL_3113793, EPI_ISL_3113794, EPI_ISL_3113795, EPI_ISL_3113796, EPI_ISL_3113797, EPI_ISL_3113798, EPI_ISL_3113799, EPI_ISL_3113800, EPI_ISL_3113801, EPI_ISL_3113802, EPI_ISL_3113803, EPI_ISL_3113804, EPI_ISL_3113805, EPI_ISL_3113806, EPI_ISL_3113807, EPI_ISL_3113808, EPI_ISL_3113810, EPI_ISL_3113811, EPI_ISL_3113812, EPI_ISL_3113813, EPI_ISL_3113814, EPI_ISL_3113815, EPI_ISL_3113816, EPI_ISL_3113817, EPI_ISL_3113819, EPI_ISL_3113820, EPI_ISL_3113822, EPI_ISL_3113823, EPI_ISL_3113825, EPI_ISL_3113826, EPI_ISL_3113828, EPI_ISL_3113829, EPI_ISL_3113830, EPI_ISL_3113831, EPI_ISL_3113832, EPI_ISL_3113833, EPI_ISL_3113834, EPI_ISL_3113835, EPI_ISL_3113836, EPI_ISL_3113837, EPI_ISL_3113838, EPI_ISL_3113839, EPI_ISL_3113840, EPI_ISL_3113841, EPI_ISL_3113842, EPI_ISL_3113843, EPI_ISL_3113844, EPI_ISL_3113846, EPI_ISL_3113847, EPI_ISL_3113848, EPI_ISL_3113849, EPI_ISL_3113850, EPI_ISL_3113852, EPI_ISL_3113853, EPI_ISL_3113854, EPI_ISL_3113855, EPI_ISL_3113856, EPI_ISL_3113857, EPI_ISL_3113858, EPI_ISL_3113859, EPI_ISL_3113860, EPI_ISL_3113861, EPI_ISL_3113863, EPI_ISL_3113864, EPI_ISL_3113865, EPI_ISL_3113866, EPI_ISL_3113868, EPI_ISL_3113869, EPI_ISL_3113870, EPI_ISL_3113871, EPI_ISL_3113872, EPI_ISL_3113873, EPI_ISL_3113875, EPI_ISL_3113876, EPI_ISL_3113877, EPI_ISL_3113878, EPI_ISL_3113879, EPI_ISL_3113880, EPI_ISL_3113881, EPI_ISL_3113882, EPI_ISL_3113883, EPI_ISL_3113884, EPI_ISL_3113885, EPI_ISL_3113887, EPI_ISL_3113888, EPI_ISL_3113889, EPI_ISL_3113890, EPI_ISL_3113891, EPI_ISL_3113892, EPI_ISL_3113893, EPI_ISL_3113894, EPI_ISL_3113895, EPI_ISL_3113896, EPI_ISL_3113897, EPI_ISL_3113898, EPI_ISL_3113899, EPI_ISL_3113900, EPI_ISL_3113901, EPI_ISL_3113902, EPI_ISL_3113903, EPI_ISL_3113906, EPI_ISL_3113907, EPI_ISL_3113908, EPI_ISL_3113909, EPI_ISL_3113910, EPI_ISL_3113911, EPI_ISL_3113912, EPI_ISL_3113913, EPI_ISL_3113915, EPI_ISL_3113916, EPI_ISL_3113917, EPI_ISL_3113918, EPI_ISL_3113919, EPI_ISL_3113920, EPI_ISL_3113921, EPI_ISL_3113922, EPI_ISL_3113923, EPI_ISL_3113924, EPI_ISL_3113925, EPI_ISL_3113926, EPI_ISL_3113927, EPI_ISL_3113928, EPI_ISL_3113929, EPI_ISL_3113930, EPI_ISL_3113931, EPI_ISL_3113932, EPI_ISL_3113933, EPI_ISL_3113934, EPI_ISL_3113935, EPI_ISL_3113936, EPI_ISL_3113937, EPI_ISL_3113938, EPI_ISL_3113939, EPI_ISL_3113940, EPI_ISL_3113943, EPI_ISL_3113944, EPI_ISL_3113945, EPI_ISL_3113946, EPI_ISL_3113947, EPI_ISL_3113948, EPI_ISL_3113949, EPI_ISL_3113950, EPI_ISL_3113951, EPI_ISL_3113952, EPI_ISL_3113954, EPI_ISL_3113955, EPI_ISL_3113957, EPI_ISL_3113958, EPI_ISL_3113959, EPI_ISL_3113961, EPI_ISL_3113962, EPI_ISL_3113964, EPI_ISL_3113965, EPI_ISL_3113966, EPI_ISL_3113967, EPI_ISL_3113968, EPI_ISL_3113969, EPI_ISL_3113970, EPI_ISL_3113971, EPI_ISL_3113972, EPI_ISL_3113973, EPI_ISL_3113974, EPI_ISL_3113975, EPI_ISL_3113976, EPI_ISL_3113977, EPI_ISL_3113978, EPI_ISL_3113979, EPI_ISL_3113980, EPI_ISL_3113981, EPI_ISL_3113982, EPI_ISL_3113983, EPI_ISL_3113984, EPI_ISL_3113985, EPI_ISL_3113987, EPI_ISL_3113988, EPI_ISL_3113989, EPI_ISL_3113990, EPI_ISL_3113991, EPI_ISL_3113992, EPI_ISL_3113993, EPI_ISL_3113994, EPI_ISL_3113995, EPI_ISL_3113996, EPI_ISL_3113997, EPI_ISL_3113998, EPI_ISL_3114002, EPI_ISL_3114003, EPI_ISL_3114004, EPI_ISL_3114006, EPI_ISL_3114010, EPI_ISL_3114011, EPI_ISL_3114012, EPI_ISL_3114013, EPI_ISL_3114014, EPI_ISL_3114016, EPI_ISL_3114017, EPI_ISL_3114018, EPI_ISL_3114019, EPI_ISL_3114021, EPI_ISL_3114022, EPI_ISL_3114023, EPI_ISL_3114024, EPI_ISL_3114025, EPI_ISL_3114026, EPI_ISL_3114027, EPI_ISL_3114028, EPI_ISL_3114029, EPI_ISL_3114030, EPI_ISL_3114031, EPI_ISL_3114032, EPI_ISL_3114033, EPI_ISL_3114034, EPI_ISL_3114035, EPI_ISL_3114036, EPI_ISL_3114037, EPI_ISL_3114038, EPI_ISL_3114039, EPI_ISL_3114040, EPI_ISL_3114041, EPI_ISL_3114042, EPI_ISL_3114043, EPI_ISL_3114044, EPI_ISL_3114045, EPI_ISL_3114047, EPI_ISL_3114048, EPI_ISL_3114049, EPI_ISL_3114050, EPI_ISL_3114052, EPI_ISL_3114053, EPI_ISL_3114054, EPI_ISL_3114055, EPI_ISL_3114056, EPI_ISL_3114057, EPI_ISL_3114058, EPI_ISL_3114059, EPI_ISL_3114060, EPI_ISL_3114061, EPI_ISL_3114062, EPI_ISL_3114064, EPI_ISL_3114065, EPI_ISL_3114066, EPI_ISL_3114067, EPI_ISL_3114068, EPI_ISL_3114070, EPI_ISL_3114071, EPI_ISL_3114072, EPI_ISL_3114073, EPI_ISL_3114074, EPI_ISL_3114075, EPI_ISL_3114076, EPI_ISL_3114077, EPI_ISL_3114078, EPI_ISL_3114080, EPI_ISL_3114081, EPI_ISL_3114082, EPI_ISL_3114083, EPI_ISL_3114084, EPI_ISL_3114086, EPI_ISL_3114088, EPI_ISL_3114089, EPI_ISL_3114090, EPI_ISL_3114091, EPI_ISL_3114092, EPI_ISL_3114093, EPI_ISL_3114094, EPI_ISL_3114095, EPI_ISL_3114096, EPI_ISL_3114097, EPI_ISL_3114098, EPI_ISL_3114100, EPI_ISL_3114101, EPI_ISL_3114102, EPI_ISL_3114103, EPI_ISL_3114104, EPI_ISL_3114105, EPI_ISL_3114106, EPI_ISL_3114107, EPI_ISL_3114108, EPI_ISL_3114109, EPI_ISL_3114110, EPI_ISL_3114111, EPI_ISL_3114113, EPI_ISL_3114114, EPI_ISL_3114115, EPI_ISL_3114116, EPI_ISL_3114118, EPI_ISL_3114119, EPI_ISL_3114120, EPI_ISL_3114121, EPI_ISL_3114122, EPI_ISL_3114124, EPI_ISL_3114125, EPI_ISL_3114126, EPI_ISL_3114127, EPI_ISL_3114128, EPI_ISL_3114129, EPI_ISL_3114131, EPI_ISL_3114132, EPI_ISL_3114133, EPI_ISL_3114134, EPI_ISL_3114135, EPI_ISL_3114136, EPI_ISL_3114137, EPI_ISL_3114138, EPI_ISL_3114139, EPI_ISL_3114140, EPI_ISL_3114141, EPI_ISL_3114142, EPI_ISL_3114143, EPI_ISL_3114144, EPI_ISL_3114145, EPI_ISL_3114147, EPI_ISL_3114148, EPI_ISL_3114149, EPI_ISL_3114150, EPI_ISL_3114151, EPI_ISL_3114156, EPI_ISL_3114157, EPI_ISL_3114158, EPI_ISL_3114160, EPI_ISL_3114161, EPI_ISL_3114163, EPI_ISL_3114166, EPI_ISL_3114168, EPI_ISL_3114169, EPI_ISL_3114170, EPI_ISL_3114171, EPI_ISL_3114172, EPI_ISL_3114173, EPI_ISL_3114174, EPI_ISL_3114175, EPI_ISL_3114176, EPI_ISL_3114177, EPI_ISL_3114178, EPI_ISL_3114179, EPI_ISL_3114180, EPI_ISL_3114183, EPI_ISL_3114184, EPI_ISL_3114185, EPI_ISL_3114188, EPI_ISL_3114189, EPI_ISL_3114190, EPI_ISL_3114191, EPI_ISL_3114192, EPI_ISL_3114193, EPI_ISL_3114194, EPI_ISL_3114195, EPI_ISL_3114197, EPI_ISL_3114198, EPI_ISL_3114202, EPI_ISL_3114203, EPI_ISL_3114204, EPI_ISL_3114205, EPI_ISL_3114206, EPI_ISL_3114208, EPI_ISL_3114209, EPI_ISL_3114210, EPI_ISL_3114211, EPI_ISL_3114212, EPI_ISL_3114213, EPI_ISL_3114214, EPI_ISL_3114215, EPI_ISL_3114216, EPI_ISL_3114217, EPI_ISL_3114218, EPI_ISL_3114219, EPI_ISL_3114220, EPI_ISL_3114222, EPI_ISL_3114223, EPI_ISL_3114224, EPI_ISL_3114225, EPI_ISL_3114227, EPI_ISL_3114228, EPI_ISL_3114230, EPI_ISL_3114231, EPI_ISL_3114232, EPI_ISL_3114233, EPI_ISL_3114234, EPI_ISL_3114235, EPI_ISL_3114236, EPI_ISL_3114237, EPI_ISL_3114239, EPI_ISL_3114241, EPI_ISL_3114242, EPI_ISL_3114243, EPI_ISL_3114244, EPI_ISL_3114245, EPI_ISL_3114246, EPI_ISL_3114247, EPI_ISL_3114248, EPI_ISL_3114249, EPI_ISL_3114250, EPI_ISL_3114251, EPI_ISL_3114252, EPI_ISL_3114253, EPI_ISL_3114254, EPI_ISL_3114256, EPI_ISL_3114257, EPI_ISL_3114258, EPI_ISL_3114259, EPI_ISL_3114260, EPI_ISL_3114261, EPI_ISL_3114262, EPI_ISL_3114263, EPI_ISL_3114264, EPI_ISL_3114265, EPI_ISL_3114266, EPI_ISL_3114267, EPI_ISL_3114268, EPI_ISL_3114269, EPI_ISL_3114272, EPI_ISL_3114273, EPI_ISL_3114274, EPI_ISL_3114275, EPI_ISL_3114277, EPI_ISL_3114279, EPI_ISL_3114280, EPI_ISL_3114281, EPI_ISL_3114283, EPI_ISL_3114284, EPI_ISL_3114286, EPI_ISL_3114287, EPI_ISL_3114288, EPI_ISL_3114289, EPI_ISL_3114290, EPI_ISL_3114291, EPI_ISL_3114292, EPI_ISL_3114293, EPI_ISL_3114294, EPI_ISL_3114295, EPI_ISL_3114296, EPI_ISL_3114297, EPI_ISL_3114298, EPI_ISL_3114300, EPI_ISL_3114301, EPI_ISL_3114302, EPI_ISL_3114304, EPI_ISL_3114305, EPI_ISL_3114306, EPI_ISL_3114310, EPI_ISL_3114311, EPI_ISL_3114312, EPI_ISL_3114313, EPI_ISL_3114314, EPI_ISL_3114315, EPI_ISL_3114316, EPI_ISL_3114318, EPI_ISL_3114319, EPI_ISL_3114320, EPI_ISL_3114321, EPI_ISL_3114322, EPI_ISL_3114323, EPI_ISL_3114324, EPI_ISL_3114325, EPI_ISL_3114326, EPI_ISL_3114327, EPI_ISL_3114328, EPI_ISL_3114329, EPI_ISL_3114330, EPI_ISL_3114331, EPI_ISL_3114333, EPI_ISL_3114335, EPI_ISL_3114336, EPI_ISL_3114337, EPI_ISL_3114338, EPI_ISL_3114339, EPI_ISL_3114340, EPI_ISL_3114341, EPI_ISL_3114342, EPI_ISL_3114343, EPI_ISL_3114344, EPI_ISL_3114345, EPI_ISL_3114346, EPI_ISL_3114347, EPI_ISL_3114348, EPI_ISL_3114349, EPI_ISL_3114350, EPI_ISL_3114352, EPI_ISL_3114353, EPI_ISL_3114354, EPI_ISL_3114355, EPI_ISL_3114356, EPI_ISL_3114357, EPI_ISL_3114359, EPI_ISL_3114360, EPI_ISL_3114361, EPI_ISL_3114362, EPI_ISL_3114363, EPI_ISL_3114364, EPI_ISL_3114365, EPI_ISL_3114366, EPI_ISL_3114367, EPI_ISL_3114369, EPI_ISL_3114370, EPI_ISL_3114371, EPI_ISL_3114372, EPI_ISL_3114373, EPI_ISL_3114375, EPI_ISL_3114376, EPI_ISL_3114377, EPI_ISL_3114378, EPI_ISL_3114379, EPI_ISL_3114381, EPI_ISL_3114382, EPI_ISL_3114383, EPI_ISL_3114385, EPI_ISL_3114386, EPI_ISL_3114387, EPI_ISL_3114388, EPI_ISL_3114389, EPI_ISL_3114390, EPI_ISL_3114391, EPI_ISL_3114392, EPI_ISL_3114393, EPI_ISL_3114395, EPI_ISL_3114397, EPI_ISL_3114399, EPI_ISL_3114401, EPI_ISL_3114402, EPI_ISL_3114404, EPI_ISL_3114406, EPI_ISL_3114407, EPI_ISL_3114408, EPI_ISL_3114409, EPI_ISL_3114410, EPI_ISL_3114411, EPI_ISL_3114412, EPI_ISL_3114413, EPI_ISL_3114414, EPI_ISL_3114415, EPI_ISL_3114416, EPI_ISL_3114417, EPI_ISL_3114418, EPI_ISL_3114419, EPI_ISL_3114420, EPI_ISL_3114421, EPI_ISL_3114422, EPI_ISL_3114424, EPI_ISL_3114425, EPI_ISL_3114426, EPI_ISL_3114427, EPI_ISL_3114428, EPI_ISL_3114429, EPI_ISL_3114430, EPI_ISL_3114431, EPI_ISL_3114432, EPI_ISL_3114433, EPI_ISL_3114434, EPI_ISL_3114435, EPI_ISL_3114436, EPI_ISL_3114437, EPI_ISL_3114439, EPI_ISL_3114440, EPI_ISL_3114441, EPI_ISL_3114442, EPI_ISL_3114445, EPI_ISL_3114446, EPI_ISL_3114447, EPI_ISL_3114448, EPI_ISL_3114449, EPI_ISL_3114451, EPI_ISL_3114452, EPI_ISL_3114453, EPI_ISL_3114454, EPI_ISL_3114455, EPI_ISL_3114456, EPI_ISL_3114457, EPI_ISL_3114458, EPI_ISL_3114459, EPI_ISL_3114460, EPI_ISL_3114461, EPI_ISL_3114462, EPI_ISL_3114463, EPI_ISL_3114464, EPI_ISL_3114465, EPI_ISL_3114466, EPI_ISL_3114467, EPI_ISL_3114468, EPI_ISL_3114469, EPI_ISL_3114470, EPI_ISL_3114471, EPI_ISL_3114472, EPI_ISL_3114473, EPI_ISL_3114474, EPI_ISL_3114475, EPI_ISL_3114476, EPI_ISL_3114477, EPI_ISL_3114478, EPI_ISL_3114479, EPI_ISL_3114481, EPI_ISL_3114483, EPI_ISL_3114484, EPI_ISL_3114485, EPI_ISL_3114486, EPI_ISL_3114487, EPI_ISL_3114488, EPI_ISL_3114489, EPI_ISL_3114490, EPI_ISL_3114491, EPI_ISL_3114492, EPI_ISL_3114493, EPI_ISL_3114494, EPI_ISL_3114495, EPI_ISL_3114496, EPI_ISL_3114498, EPI_ISL_3114499, EPI_ISL_3114500, EPI_ISL_3114501, EPI_ISL_3114502, EPI_ISL_3114503, EPI_ISL_3114504, EPI_ISL_3114505, EPI_ISL_3114507, EPI_ISL_3114508, EPI_ISL_3114509, EPI_ISL_3114510, EPI_ISL_3114511, EPI_ISL_3114512, EPI_ISL_3114515, EPI_ISL_3114516, EPI_ISL_3114518, EPI_ISL_3114519, EPI_ISL_3114520, EPI_ISL_3114521, EPI_ISL_3114522, EPI_ISL_3114523, EPI_ISL_3114527, EPI_ISL_3114528, EPI_ISL_3114529, EPI_ISL_3114530, EPI_ISL_3114531, EPI_ISL_3114534, EPI_ISL_3114535, EPI_ISL_3114536, EPI_ISL_3114540, EPI_ISL_3114541, EPI_ISL_3114542, EPI_ISL_3114543, EPI_ISL_3114544, EPI_ISL_3114545, EPI_ISL_3114546, EPI_ISL_3114547, EPI_ISL_3114548, EPI_ISL_3114550, EPI_ISL_3114552, EPI_ISL_3114553, EPI_ISL_3114554, EPI_ISL_3114555, EPI_ISL_3114557, EPI_ISL_3114558, EPI_ISL_3114559, EPI_ISL_3114560, EPI_ISL_3114561, EPI_ISL_3114563, EPI_ISL_3114564, EPI_ISL_3114567, EPI_ISL_3114568, EPI_ISL_3114570, EPI_ISL_3114571, EPI_ISL_3114572, EPI_ISL_3114573, EPI_ISL_3114574, EPI_ISL_3114576, EPI_ISL_3114578, EPI_ISL_3114579, EPI_ISL_3114580, EPI_ISL_3114581, EPI_ISL_3114582, EPI_ISL_3114583, EPI_ISL_3114584, EPI_ISL_3114586, EPI_ISL_3114587, EPI_ISL_3114588, EPI_ISL_3114589, EPI_ISL_3114590, EPI_ISL_3114591, EPI_ISL_3114592, EPI_ISL_3114593, EPI_ISL_3114594, EPI_ISL_3114595, EPI_ISL_3114596, EPI_ISL_3114597, EPI_ISL_3114598, EPI_ISL_3114599, EPI_ISL_3114601, EPI_ISL_3114603, EPI_ISL_3114604, EPI_ISL_3114605, EPI_ISL_3114606, EPI_ISL_3114607, EPI_ISL_3114608, EPI_ISL_3114609, EPI_ISL_3114610, EPI_ISL_3114611, EPI_ISL_3114612, EPI_ISL_3114613, EPI_ISL_3114614, EPI_ISL_3114615, EPI_ISL_3114616, EPI_ISL_3114617, EPI_ISL_3114618, EPI_ISL_3114619, EPI_ISL_3114620, EPI_ISL_3114621, EPI_ISL_3114622, EPI_ISL_3114623, EPI_ISL_3114624, EPI_ISL_3114625, EPI_ISL_3114626, EPI_ISL_3114627, EPI_ISL_3114628, EPI_ISL_3114629, EPI_ISL_3114632, EPI_ISL_3114633, EPI_ISL_3114634, EPI_ISL_3114635, EPI_ISL_3114636, EPI_ISL_3114638, EPI_ISL_3114639, EPI_ISL_3114640, EPI_ISL_3114641, EPI_ISL_3114642, EPI_ISL_3114643, EPI_ISL_3114644, EPI_ISL_3114645, EPI_ISL_3114646, EPI_ISL_3114649, EPI_ISL_3114652, EPI_ISL_3114653, EPI_ISL_3114654, EPI_ISL_3114655, EPI_ISL_3114656, EPI_ISL_3114657, EPI_ISL_3114658, EPI_ISL_3114659, EPI_ISL_3114660, EPI_ISL_3114661, EPI_ISL_3114662, EPI_ISL_3114664, EPI_ISL_3114665, EPI_ISL_3114668, EPI_ISL_3114669, EPI_ISL_3114670, EPI_ISL_3114672, EPI_ISL_3114673, EPI_ISL_3114674, EPI_ISL_3114675, EPI_ISL_3114676, EPI_ISL_3114677, EPI_ISL_3114679, EPI_ISL_3114680, EPI_ISL_3114681, EPI_ISL_3114682, EPI_ISL_3114683, EPI_ISL_3114684, EPI_ISL_3114685, EPI_ISL_3114686, EPI_ISL_3114688, EPI_ISL_3114689, EPI_ISL_3114690, EPI_ISL_3114691, EPI_ISL_3114692, EPI_ISL_3114693, EPI_ISL_3114694, EPI_ISL_3114695, EPI_ISL_3114696, EPI_ISL_3114697, EPI_ISL_3114698, EPI_ISL_3114699, EPI_ISL_3114700, EPI_ISL_3114701, EPI_ISL_3114702, EPI_ISL_3114703, EPI_ISL_3114704, EPI_ISL_3114705, EPI_ISL_3114706, EPI_ISL_3114707, EPI_ISL_3114708, EPI_ISL_3114709, EPI_ISL_3114710, EPI_ISL_3114711, EPI_ISL_3114712, EPI_ISL_3114713, EPI_ISL_3114714, EPI_ISL_3114715, EPI_ISL_3114716, EPI_ISL_3114717, EPI_ISL_3114718, EPI_ISL_3114719, EPI_ISL_3114720, EPI_ISL_3114722, EPI_ISL_3114725, EPI_ISL_3114727, EPI_ISL_3114728, EPI_ISL_3114729, EPI_ISL_3114730, EPI_ISL_3114731, EPI_ISL_3114732, EPI_ISL_3114733, EPI_ISL_3114734, EPI_ISL_3114735, EPI_ISL_3114736, EPI_ISL_3114737, EPI_ISL_3114738, EPI_ISL_3114740, EPI_ISL_3114741, EPI_ISL_3114742, EPI_ISL_3114743, EPI_ISL_3114744, EPI_ISL_3114745, EPI_ISL_3114747, EPI_ISL_3114748, EPI_ISL_3114749, EPI_ISL_3114750, EPI_ISL_3114751, EPI_ISL_3114752, EPI_ISL_3114753, EPI_ISL_3114754, EPI_ISL_3114755, EPI_ISL_3114756, EPI_ISL_3114757, EPI_ISL_3114760, EPI_ISL_3114761, EPI_ISL_3114762, EPI_ISL_3114763, EPI_ISL_3114764, EPI_ISL_3114765, EPI_ISL_3114766, EPI_ISL_3114767, EPI_ISL_3114768, EPI_ISL_3114770, EPI_ISL_3114771, EPI_ISL_3114772, EPI_ISL_3114773, EPI_ISL_3114774, EPI_ISL_3114775, EPI_ISL_3114776, EPI_ISL_3114777, EPI_ISL_3114778, EPI_ISL_3114779, EPI_ISL_3114780, EPI_ISL_3114782, EPI_ISL_3114783, EPI_ISL_3114784, EPI_ISL_3114785, EPI_ISL_3114786, EPI_ISL_3114787, EPI_ISL_3114788, EPI_ISL_3114790, EPI_ISL_3114792, EPI_ISL_3114793, EPI_ISL_3114794, EPI_ISL_3114797, EPI_ISL_3114798, EPI_ISL_3114799, EPI_ISL_3114801, EPI_ISL_3114802, EPI_ISL_3114803, EPI_ISL_3114804, EPI_ISL_3114805, EPI_ISL_3114806, EPI_ISL_3114807, EPI_ISL_3114808, EPI_ISL_3114809, EPI_ISL_3114810, EPI_ISL_3114812, EPI_ISL_3114814, EPI_ISL_3114815, EPI_ISL_3114816, EPI_ISL_3114817, EPI_ISL_3114818, EPI_ISL_3114819, EPI_ISL_3114820, EPI_ISL_3114821, EPI_ISL_3114822, EPI_ISL_3114823, EPI_ISL_3114824, EPI_ISL_3114825, EPI_ISL_3114826, EPI_ISL_3114827, EPI_ISL_3114828, EPI_ISL_3114829, EPI_ISL_3114830, EPI_ISL_3114831, EPI_ISL_3114832, EPI_ISL_3114833, EPI_ISL_3114834, EPI_ISL_3114835, EPI_ISL_3114836, EPI_ISL_3114837, EPI_ISL_3114838, EPI_ISL_3114839, EPI_ISL_3114840, EPI_ISL_3114841, EPI_ISL_3114842, EPI_ISL_3114843, EPI_ISL_3114844, EPI_ISL_3114846, EPI_ISL_3114847, EPI_ISL_3114848, EPI_ISL_3114849, EPI_ISL_3114850, EPI_ISL_3114851, EPI_ISL_3114852, EPI_ISL_3114853, EPI_ISL_3114854, EPI_ISL_3114855, EPI_ISL_3114856, EPI_ISL_3114857, EPI_ISL_3114858, EPI_ISL_3114859, EPI_ISL_3114860, EPI_ISL_3114861, EPI_ISL_3114862, EPI_ISL_3114863, EPI_ISL_3114864, EPI_ISL_3114865, EPI_ISL_3114866, EPI_ISL_3114867, EPI_ISL_3114868, EPI_ISL_3114870, EPI_ISL_3114871, EPI_ISL_3114872, EPI_ISL_3114873, EPI_ISL_3114874, EPI_ISL_3114875, EPI_ISL_3114876, EPI_ISL_3114878, EPI_ISL_3114879, EPI_ISL_3114880, EPI_ISL_3114881, EPI_ISL_3114882, EPI_ISL_3114883, EPI_ISL_3114884, EPI_ISL_3114885, EPI_ISL_3114886, EPI_ISL_3114887, EPI_ISL_3114888, EPI_ISL_3114889, EPI_ISL_3114890, EPI_ISL_3114891, EPI_ISL_3114892, EPI_ISL_3114893, EPI_ISL_3114894, EPI_ISL_3114895, EPI_ISL_3114896, EPI_ISL_3114897, EPI_ISL_3114898, EPI_ISL_3114899, EPI_ISL_3114900, EPI_ISL_3114901, EPI_ISL_3114902, EPI_ISL_3114903, EPI_ISL_3114905, EPI_ISL_3114907, EPI_ISL_3114908, EPI_ISL_3114909, EPI_ISL_3114910, EPI_ISL_3114911, EPI_ISL_3114912, EPI_ISL_3114913, EPI_ISL_3114914, EPI_ISL_3114915, EPI_ISL_3114916, EPI_ISL_3114917, EPI_ISL_3114918, EPI_ISL_3114919, EPI_ISL_3114920, EPI_ISL_3114921, EPI_ISL_3114922, EPI_ISL_3114923, EPI_ISL_3114925, EPI_ISL_3114926, EPI_ISL_3114927, EPI_ISL_3114928, EPI_ISL_3114929, EPI_ISL_3114930, EPI_ISL_3114931, EPI_ISL_3114932, EPI_ISL_3114933, EPI_ISL_3114934, EPI_ISL_3114935, EPI_ISL_3114936, EPI_ISL_3114937, EPI_ISL_3114938, EPI_ISL_3114939, EPI_ISL_3114940, EPI_ISL_3114941, EPI_ISL_3114942, EPI_ISL_3114943, EPI_ISL_3114944, EPI_ISL_3114945, EPI_ISL_3114946, EPI_ISL_3114948, EPI_ISL_3114949, EPI_ISL_3114950, EPI_ISL_3114952, EPI_ISL_3114953, EPI_ISL_3114954, EPI_ISL_3114955, EPI_ISL_3114956, EPI_ISL_3114957, EPI_ISL_3114958, EPI_ISL_3114959, EPI_ISL_3114960, EPI_ISL_3114961, EPI_ISL_3114962, EPI_ISL_3114963, EPI_ISL_3114964, EPI_ISL_3114965, EPI_ISL_3114966, EPI_ISL_3114967, EPI_ISL_3114968, EPI_ISL_3114969, EPI_ISL_3114970, EPI_ISL_3114971, EPI_ISL_3114972, EPI_ISL_3114974, EPI_ISL_3114975, EPI_ISL_3114976, EPI_ISL_3114977, EPI_ISL_3114978, EPI_ISL_3114979, EPI_ISL_3114980, EPI_ISL_3114982,

| | | | |
|---|---|---|---|
| EPI_ISL_3114983, EPI_ISL_3114984, EPI_ISL_3114985, EPI_ISL_3114986, EPI_ISL_3114987, EPI_ISL_3114988, EPI_ISL_3114989, EPI_ISL_3114990, EPI_ISL_3114991, EPI_ISL_3114993, EPI_ISL_3114994, EPI_ISL_3114995, EPI_ISL_3114996, EPI_ISL_3114997, EPI_ISL_3114998, EPI_ISL_3114999, EPI_ISL_3115000, EPI_ISL_3115001, EPI_ISL_3115002, EPI_ISL_3115003, EPI_ISL_3115004, EPI_ISL_3115005, EPI_ISL_3115006, EPI_ISL_3115007, EPI_ISL_3115008, EPI_ISL_3115009, EPI_ISL_3115010, EPI_ISL_3115011, EPI_ISL_3115012, EPI_ISL_3115013, EPI_ISL_3115014, EPI_ISL_3115015, EPI_ISL_3115017, EPI_ISL_3115018 | | | |
| see above | Quest Diagnostics Incorporated | Centers for Disease Control and Prevention Division of Viral Diseases, Pathogen Discovery | Dakota Howard, Dhwani Batra, Peter W. Cook, Kara Moser, Adrian Paskey, Jason Caravas, Benjamin Rambo-Martin, Shatavia Morrison, Christopher Gulvick, Scott Sammons, Yvette Unoarumhi, Darlene Wagner, Matthew Schmerer, S. H. Rosenthal, A. Gerasimova, R. M. Kagan, B. Anderson, M. Hua, Y. Liu, L.E. Bernstein, K.E. Livingston, A. Perez, I. A. Shlyakhter, R. V. Rolando, R. Owen, P. Tanpaiboon, F. Lacbawan, Clinton R. Paden, Duncan MacCannell |
| EPI_ISL_3115019, EPI_ISL_3115020 | Algemeen Klinisch Labo | Labo Klinische Biologie, UZA | Marie Le Mercier, Jasmine Coppens, Kathleen Holemans, Ines Verbesselt, Basil Britto Xavier, Christine Lammens, Veerle Matheeussen, Herman Goossens |
| EPI_ISL_3115021, EPI_ISL_3115022, EPI_ISL_3115024, EPI_ISL_3115025, EPI_ISL_3115026, EPI_ISL_3115027 | Centrum voor Medische Analyse | Labo Klinische Biologie, UZA | Marie Le Mercier, Jasmine Coppens, Kathleen Holemans, Ines Verbesselt, Basil Britto Xavier, Christine Lammens, Veerle Matheeussen, Herman Goossens |
| EPI_ISL_3115028, EPI_ISL_3115029, EPI_ISL_3115030, EPI_ISL_3115031, EPI_ISL_3115032, EPI_ISL_3115033, EPI_ISL_3115034, EPI_ISL_3115035, EPI_ISL_3115036, EPI_ISL_3115037, EPI_ISL_3115038, EPI_ISL_3115039, EPI_ISL_3115040, EPI_ISL_3115041, EPI_ISL_3115042, EPI_ISL_3115043, EPI_ISL_3115045, EPI_ISL_3115046, EPI_ISL_3115047, EPI_ISL_3115048, EPI_ISL_3115049, EPI_ISL_3115050, EPI_ISL_3115051, EPI_ISL_3115052, EPI_ISL_3115053, EPI_ISL_3115055, EPI_ISL_3115056, EPI_ISL_3115057, EPI_ISL_3115058, EPI_ISL_3115059, EPI_ISL_3115060 | | | |
| see above | LHUB-ULB | Labo Klinische Biologie, UZA | Marie Le Mercier, Jasmine Coppens, Kathleen Holemans, Ines Verbesselt, Basil Britto Xavier, Christine Lammens, Veerle Matheeussen, Herman Goossens |
| EPI_ISL_3115061, EPI_ISL_3115062, EPI_ISL_3115063, EPI_ISL_3115064, EPI_ISL_3115065, EPI_ISL_3115066, EPI_ISL_3115067, EPI_ISL_3115068, EPI_ISL_3115069, EPI_ISL_3115070, EPI_ISL_3115071, EPI_ISL_3115072, EPI_ISL_3115073, EPI_ISL_3115074, EPI_ISL_3115075, EPI_ISL_3115076, EPI_ISL_3115077, EPI_ISL_3115078, EPI_ISL_3115079, EPI_ISL_3115080, EPI_ISL_3115081, EPI_ISL_3115082, EPI_ISL_3115083, EPI_ISL_3115084, EPI_ISL_3115085, EPI_ISL_3115086, EPI_ISL_3115087, EPI_ISL_3115088, EPI_ISL_3115089, EPI_ISL_3115090, EPI_ISL_3115091, EPI_ISL_3115092, EPI_ISL_3115093, EPI_ISL_3115094, EPI_ISL_3115095, EPI_ISL_3115096, EPI_ISL_3115097, EPI_ISL_3115098, EPI_ISL_3115099, EPI_ISL_3115100, EPI_ISL_3115101, EPI_ISL_3115102, EPI_ISL_3115104, EPI_ISL_3115105, EPI_ISL_3115106, EPI_ISL_3115107, EPI_ISL_3115108, EPI_ISL_3115109, EPI_ISL_3115110, EPI_ISL_3115111, EPI_ISL_3115112, EPI_ISL_3115113, EPI_ISL_3115114, EPI_ISL_3115115, EPI_ISL_3115116, EPI_ISL_3115117, EPI_ISL_3115118, EPI_ISL_3115119, EPI_ISL_3115120, EPI_ISL_3115121, EPI_ISL_3115122, EPI_ISL_3115123, EPI_ISL_3115124, EPI_ISL_3115125, EPI_ISL_3115126, EPI_ISL_3115127, EPI_ISL_3115128, EPI_ISL_3115129, EPI_ISL_3115130, EPI_ISL_3115131, EPI_ISL_3115132, EPI_ISL_3115133, EPI_ISL_3115134, EPI_ISL_3115135, EPI_ISL_3115136, EPI_ISL_3115137, EPI_ISL_3115138, EPI_ISL_3115139, EPI_ISL_3115140, EPI_ISL_3115141, EPI_ISL_3115142, EPI_ISL_3115143, EPI_ISL_3115144, EPI_ISL_3115145, EPI_ISL_3115146, EPI_ISL_3115147, EPI_ISL_3115148, EPI_ISL_3115149, EPI_ISL_3115150, EPI_ISL_3115151, EPI_ISL_3115152, EPI_ISL_3115153, EPI_ISL_3115154, EPI_ISL_3115155, EPI_ISL_3115156, EPI_ISL_3115158, EPI_ISL_3115159, EPI_ISL_3115160, EPI_ISL_3115161, EPI_ISL_3115162, EPI_ISL_3115163, EPI_ISL_3115164, EPI_ISL_3115165, EPI_ISL_3115166, EPI_ISL_3115167, EPI_ISL_3115168, EPI_ISL_3115169, EPI_ISL_3115170, EPI_ISL_3115171, EPI_ISL_3115172, EPI_ISL_3115173, EPI_ISL_3115174, EPI_ISL_3115175, EPI_ISL_3115176, EPI_ISL_3115177, EPI_ISL_3115178, EPI_ISL_3115179, EPI_ISL_3115180, EPI_ISL_3115181, EPI_ISL_3115182, EPI_ISL_3115183, EPI_ISL_3115184, EPI_ISL_3115185, EPI_ISL_3115186, EPI_ISL_3115187 | | | |
| see above | Platform BIS UZA/UAntwerpen | Labo Klinische Biologie, UZA | Marie Le Mercier, Jasmine Coppens, Kathleen Holemans, Ines Verbesselt, Basil Britto Xavier, Christine Lammens, Veerle Matheeussen, Herman Goossens |
| EPI_ISL_3115189, EPI_ISL_3115197 | Willis-Knighton Medical Center Hospital Laboratory | LSUHS Emerging Viral Threat Laboratory | Gregory L. Ware, Alexander Mijalis, Jeremy P. Kamil, Jennifer L. Carroll, Maarten Van Diest, Rona S. Scott, Andrew D. Yurochko, Christopher G. Kevil, John A. Vanchiere |
| EPI_ISL_3115201 | Louisiana Office of Public Heath Laboratory | LSUHS Emerging Viral Threat Laboratory | Gregory L. Ware, Alexander Mijalis, Jeremy P. Kamil, Jennifer L. Carroll, Maarten Van Diest, Rona S. Scott, Andrew D. Yurochko, Christopher G. Kevil, John A. Vanchiere |
| EPI_ISL_3115204 | Willis-Knighton Medical Center Hospital Laboratory | LSUHS Emerging Viral Threat Laboratory | Gregory L. Ware, Alexander Mijalis, Jeremy P. Kamil, Jennifer L. Carroll, Maarten Van Diest, Rona S. Scott, Andrew D. Yurochko, Christopher G. Kevil, John A. Vanchiere |
| EPI_ISL_3115208 | Louisiana Office of Public Heath Laboratory | LSUHS Emerging Viral Threat Laboratory | Gregory L. Ware, Alexander Mijalis, Jeremy P. Kamil, Jennifer L. Carroll, Maarten Van Diest, Rona S. Scott, Andrew D. Yurochko, Christopher G. Kevil, John A. Vanchiere |
| EPI_ISL_3115210 | Willis-Knighton Medical Center Hospital Laboratory | LSUHS Emerging Viral Threat Laboratory | Gregory L. Ware, Alexander Mijalis, Jeremy P. Kamil, Jennifer L. Carroll, Maarten Van Diest, Rona S. Scott, Andrew D. Yurochko, Christopher G. Kevil, John A. Vanchiere |
| EPI_ISL_3115213 | Louisiana Office of Public Heath Laboratory | LSUHS Emerging Viral Threat Laboratory | Gregory L. Ware, Alexander Mijalis, Jeremy P. Kamil, Jennifer L. Carroll, Maarten Van Diest, Rona S. Scott, Andrew D. Yurochko, Christopher G. Kevil, John A. Vanchiere |
| EPI_ISL_3115215, EPI_ISL_3115218 | Willis-Knighton Medical Center Hospital Laboratory | LSUHS Emerging Viral Threat Laboratory | Gregory L. Ware, Alexander Mijalis, Jeremy P. Kamil, Jennifer L. Carroll, Maarten Van Diest, Rona S. Scott, Andrew D. Yurochko, Christopher G. Kevil, John A. Vanchiere |
| EPI_ISL_3115220 | Louisiana Office of Public Heath Laboratory | LSUHS Emerging Viral Threat Laboratory | Gregory L. Ware, Alexander Mijalis, Jeremy P. Kamil, Jennifer L. Carroll, Maarten Van Diest, Rona S. Scott, Andrew D. Yurochko, Christopher G. Kevil, John A. Vanchiere |
| EPI_ISL_3115221, EPI_ISL_3115222, EPI_ISL_3115223 | Willis-Knighton Medical Center Hospital Laboratory | LSUHS Emerging Viral Threat Laboratory | Gregory L. Ware, Alexander Mijalis, Jeremy P. Kamil, Jennifer L. Carroll, Maarten Van Diest, Rona S. Scott, Andrew D. Yurochko, Christopher G. Kevil, John A. Vanchiere |
| EPI_ISL_3115224 | Louisiana Office of Public Heath Laboratory | LSUHS Emerging Viral Threat Laboratory | Gregory L. Ware, Alexander Mijalis, Jeremy P. Kamil, Jennifer L. Carroll, Maarten Van Diest, Rona S. Scott, Andrew D. Yurochko, Christopher G. Kevil, John A. Vanchiere |
| EPI_ISL_3115225, EPI_ISL_3115226, EPI_ISL_3115227, EPI_ISL_3115228, EPI_ISL_3115229, EPI_ISL_3115230 | Willis-Knighton Medical Center Hospital Laboratory | LSUHS Emerging Viral Threat Laboratory | Gregory L. Ware, Alexander Mijalis, Jeremy P. Kamil, Jennifer L. Carroll, Maarten Van Diest, Rona S. Scott, Andrew D. Yurochko, Christopher G. Kevil, John A. Vanchiere |
| EPI_ISL_3115232 | Louisiana Office of Public Heath Laboratory | LSUHS Emerging Viral Threat Laboratory | Gregory L. Ware, Alexander Mijalis, Jeremy P. Kamil, Jennifer L. Carroll, Maarten Van Diest, Rona S. Scott, Andrew D. Yurochko, Christopher G. Kevil, John A. Vanchiere |
| EPI_ISL_3115233, EPI_ISL_3115234, EPI_ISL_3115235 | Willis-Knighton Medical Center Hospital Laboratory | LSUHS Emerging Viral Threat Laboratory | Gregory L. Ware, Alexander Mijalis, Jeremy P. Kamil, Jennifer L. Carroll, Maarten Van Diest, Rona S. Scott, Andrew D. Yurochko, Christopher G. Kevil, John A. Vanchiere |
| EPI_ISL_3115236, EPI_ISL_3115238 | LSUHS Emerging Viral Threat Laboratory | LSUHS Emerging Viral Threat Laboratory | Gregory L. Ware, Alexander Mijalis, Jeremy P. Kamil, Jennifer L. Carroll, Maarten Van Diest, Rona S. Scott, Andrew D. Yurochko, Christopher G. Kevil, John A. Vanchiere |
| EPI_ISL_3115239, EPI_ISL_3115240 | Willis-Knighton Medical Center Hospital Laboratory | LSUHS Emerging Viral Threat Laboratory | Gregory L. Ware, Alexander Mijalis, Jeremy P. Kamil, Jennifer L. Carroll, Maarten Van Diest, Rona S. Scott, Andrew D. Yurochko, Christopher G. Kevil, John A. Vanchiere |
| EPI_ISL_3115241 | Louisiana Office of Public Heath Laboratory | LSUHS Emerging Viral Threat Laboratory | Gregory L. Ware, Alexander Mijalis, Jeremy P. Kamil, Jennifer L. Carroll, Maarten Van Diest, Rona S. Scott, Andrew D. Yurochko, Christopher G. Kevil, John A. Vanchiere |
| EPI_ISL_3115242 | Willis-Knighton Medical Center Hospital Laboratory | LSUHS Emerging Viral Threat Laboratory | Gregory L. Ware, Alexander Mijalis, Jeremy P. Kamil, Jennifer L. Carroll, Maarten Van Diest, Rona S. Scott, Andrew D. Yurochko, Christopher G. Kevil, John A. Vanchiere |
| EPI_ISL_3115243 | Louisiana Office of Public Heath Laboratory | LSUHS Emerging Viral Threat Laboratory | Gregory L. Ware, Alexander Mijalis, Jeremy P. Kamil, Jennifer L. Carroll, Maarten Van Diest, Rona S. Scott, Andrew D. Yurochko, Christopher G. Kevil, John A. Vanchiere |
| EPI_ISL_3115244 | LSUHS Emerging Viral Threat Laboratory | LSUHS Emerging Viral Threat Laboratory | Gregory L. Ware, Alexander Mijalis, Jeremy P. Kamil, Jennifer L. Carroll, Maarten Van Diest, Rona S. Scott, Andrew D. Yurochko, Christopher G. Kevil, John A. Vanchiere |
| EPI_ISL_3115245, EPI_ISL_3115246, EPI_ISL_3115247 | Louisiana Office of Public Heath Laboratory | LSUHS Emerging Viral Threat Laboratory | Gregory L. Ware, Alexander Mijalis, Jeremy P. Kamil, Jennifer L. Carroll, Maarten Van Diest, Rona S. Scott, Andrew D. Yurochko, Christopher G. Kevil, John A. Vanchiere |
| EPI_ISL_3115248 | LSUHS Emerging Viral Threat Laboratory | LSUHS Emerging Viral Threat Laboratory | Gregory L. Ware, Alexander Mijalis, Jeremy P. Kamil, Jennifer L. Carroll, Maarten Van Diest, Rona S. Scott, Andrew D. Yurochko, Christopher G. Kevil, John A. Vanchiere |
| EPI_ISL_3115249 | Louisiana Office of Public Heath Laboratory | LSUHS Emerging Viral Threat Laboratory | Gregory L. Ware, Alexander Mijalis, Jeremy P. Kamil, Jennifer L. Carroll, Maarten Van Diest, Rona S. Scott, Andrew D. Yurochko, Christopher G. Kevil, John A. Vanchiere |
| EPI_ISL_3115250 | LSUHS Emerging Viral Threat Laboratory | LSUHS Emerging Viral Threat Laboratory | Gregory L. Ware, Alexander Mijalis, Jeremy P. Kamil, Jennifer L. Carroll, Maarten Van Diest, Rona S. Scott, Andrew D. Yurochko, Christopher G. Kevil, John A. Vanchiere |
| EPI_ISL_3115251 | Willis-Knighton Medical Center Hospital Laboratory | LSUHS Emerging Viral Threat Laboratory | Gregory L. Ware, Alexander Mijalis, Jeremy P. Kamil, Jennifer L. Carroll, Maarten Van Diest, Rona S. Scott, Andrew D. Yurochko, Christopher G. Kevil, John A. Vanchiere |
| EPI_ISL_3115252, EPI_ISL_3115253 | Louisiana Office of Public Heath Laboratory | LSUHS Emerging Viral Threat Laboratory | Gregory L. Ware, Alexander Mijalis, Jeremy P. Kamil, Jennifer L. Carroll, Maarten Van Diest, Rona S. Scott, Andrew D. Yurochko, Christopher G. Kevil, John A. Vanchiere |
| EPI_ISL_3115254 | LSUHS Emerging Viral Threat Laboratory | LSUHS Emerging Viral Threat Laboratory | Gregory L. Ware, Alexander Mijalis, Jeremy P. Kamil, Jennifer L. Carroll, Maarten Van Diest, Rona S. Scott, Andrew D. Yurochko, Christopher G. Kevil, John A. Vanchiere |
| EPI_ISL_3115255, EPI_ISL_3115256 | Willis-Knighton Medical Center Hospital Laboratory | LSUHS Emerging Viral Threat Laboratory | Gregory L. Ware, Alexander Mijalis, Jeremy P. Kamil, Jennifer L. Carroll, Maarten Van Diest, Rona S. Scott, Andrew D. Yurochko, Christopher G. Kevil, John A. Vanchiere |
| EPI_ISL_3115258 | Louisiana Office of Public Heath Laboratory | LSUHS Emerging Viral Threat Laboratory | Gregory L. Ware, Alexander Mijalis, Jeremy P. Kamil, Jennifer L. Carroll, Maarten Van Diest, Rona S. Scott, Andrew D. Yurochko, Christopher G. Kevil, John A. Vanchiere |
| EPI_ISL_3115259 | Willis-Knighton Medical Center Hospital Laboratory | LSUHS Emerging Viral Threat Laboratory | Gregory L. Ware, Alexander Mijalis, Jeremy P. Kamil, Jennifer L. Carroll, Maarten Van Diest, Rona S. Scott, Andrew D. Yurochko, Christopher G. Kevil, John A. Vanchiere |
| EPI_ISL_3115260 | Louisiana Office of Public Heath Laboratory | LSUHS Emerging Viral Threat Laboratory | Gregory L. Ware, Alexander Mijalis, Jeremy P. Kamil, Jennifer L. Carroll, Maarten Van Diest, Rona S. Scott, Andrew D. Yurochko, Christopher G. Kevil, John A. Vanchiere |
| EPI_ISL_3115261, EPI_ISL_3115262, EPI_ISL_3115264, EPI_ISL_3115266, EPI_ISL_3115267 | Willis-Knighton Medical Center Hospital Laboratory | LSUHS Emerging Viral Threat Laboratory | Gregory L. Ware, Alexander Mijalis, Jeremy P. Kamil, Jennifer L. Carroll, Maarten Van Diest, Rona S. Scott, Andrew D. Yurochko, Christopher G. Kevil, John A. Vanchiere |
| EPI_ISL_3115268 | Louisiana Office of Public Heath Laboratory | LSUHS Emerging Viral Threat Laboratory | Gregory L. Ware, Alexander Mijalis, Jeremy P. Kamil, Jennifer L. Carroll, Maarten Van Diest, Rona S. Scott, Andrew D. Yurochko, Christopher G. Kevil, John A. Vanchiere |
| EPI_ISL_3115269 | Willis-Knighton Medical Center Hospital Laboratory | LSUHS Emerging Viral Threat Laboratory | Gregory L. Ware, Alexander Mijalis, Jeremy P. Kamil, Jennifer L. Carroll, Maarten Van Diest, Rona S. Scott, Andrew D. Yurochko, Christopher G. Kevil, John A. Vanchiere |
| EPI_ISL_3115270, EPI_ISL_3115271 | Louisiana Office of Public Heath Laboratory | LSUHS Emerging Viral Threat Laboratory | Gregory L. Ware, Alexander Mijalis, Jeremy P. Kamil, Jennifer L. Carroll, Maarten Van Diest, Rona S. Scott, Andrew D. Yurochko, Christopher G. Kevil, John A. Vanchiere |
| EPI_ISL_3115272 | LSUHS Emerging Viral Threat Laboratory | LSUHS Emerging Viral Threat Laboratory | Gregory L. Ware, Alexander Mijalis, Jeremy P. Kamil, Jennifer L. Carroll, Maarten Van Diest, Rona S. Scott, Andrew D. Yurochko, Christopher G. Kevil, John A. Vanchiere |
| EPI_ISL_3115273, EPI_ISL_3115274 | Willis-Knighton Medical Center Hospital Laboratory | LSUHS Emerging Viral Threat Laboratory | Gregory L. Ware, Alexander Mijalis, Jeremy P. Kamil, Jennifer L. Carroll, Maarten Van Diest, Rona S. Scott, Andrew D. Yurochko, Christopher G. Kevil, John A. Vanchiere |
| EPI_ISL_3115275 | LSUHS Emerging Viral Threat Laboratory | LSUHS Emerging Viral Threat Laboratory | Gregory L. Ware, Alexander Mijalis, Jeremy P. Kamil, Jennifer L. Carroll, Maarten Van Diest, Rona S. Scott, Andrew D. Yurochko, Christopher G. Kevil, John A. Vanchiere |
| EPI_ISL_3115276, EPI_ISL_3115277 | Louisiana Office of Public Heath Laboratory | LSUHS Emerging Viral Threat Laboratory | Gregory L. Ware, Alexander Mijalis, Jeremy P. Kamil, Jennifer L. Carroll, Maarten Van Diest, Rona S. Scott, Andrew D. Yurochko, Christopher G. Kevil, John A. Vanchiere |
| EPI_ISL_3115278 | Willis-Knighton Medical Center Hospital Laboratory | LSUHS Emerging Viral Threat Laboratory | Gregory L. Ware, Alexander Mijalis, Jeremy P. Kamil, Jennifer L. Carroll, Maarten Van Diest, Rona S. Scott, Andrew D. Yurochko, Christopher G. Kevil, John A. Vanchiere |
| EPI_ISL_3115279 | Louisiana Office of Public Heath Laboratory | LSUHS Emerging Viral Threat Laboratory | Gregory L. Ware, Alexander Mijalis, Jeremy P. Kamil, Jennifer L. Carroll, Maarten Van Diest, Rona S. Scott, Andrew D. Yurochko, Christopher G. Kevil, John A. Vanchiere |
| EPI_ISL_3115280 | LSUHS Emerging Viral Threat Laboratory | LSUHS Emerging Viral Threat Laboratory | Gregory L. Ware, Alexander Mijalis, Jeremy P. Kamil, Jennifer L. Carroll, Maarten Van Diest, Rona S. Scott, Andrew D. Yurochko, Christopher G. Kevil, John A. Vanchiere |
| EPI_ISL_3115281 | Willis-Knighton Medical Center Hospital Laboratory | LSUHS Emerging Viral Threat Laboratory | Gregory L. Ware, Alexander Mijalis, Jeremy P. Kamil, Jennifer L. Carroll, Maarten Van Diest, Rona S. Scott, Andrew D. Yurochko, Christopher G. Kevil, John A. Vanchiere |

| | | | |
|---|---|---|---|
| EPI_ISL_3115282 | Louisiana Office of Public Heath Laboratory | LSUHS Emerging Viral Threat Laboratory | Gregory L. Ware, Alexander Mijalis, Jeremy P. Kamil, Jennifer L. Carroll, Maarten Van Diest, Rona S. Scott, Andrew D. Yurochko, Christopher G. Kevil, John A. Vanchiere |
| EPI_ISL_3115283 | Willis-Knighton Medical Center Hospital Laboratory | LSUHS Emerging Viral Threat Laboratory | Gregory L. Ware, Alexander Mijalis, Jeremy P. Kamil, Jennifer L. Carroll, Maarten Van Diest, Rona S. Scott, Andrew D. Yurochko, Christopher G. Kevil, John A. Vanchiere |
| EPI_ISL_3115284, EPI_ISL_3115285, EPI_ISL_3115286 | Louisiana Office of Public Heath Laboratory | LSUHS Emerging Viral Threat Laboratory | Gregory L. Ware, Alexander Mijalis, Jeremy P. Kamil, Jennifer L. Carroll, Maarten Van Diest, Rona S. Scott, Andrew D. Yurochko, Christopher G. Kevil, John A. Vanchiere |
| EPI_ISL_3115287, EPI_ISL_3115288 | Willis-Knighton Medical Center Hospital Laboratory | LSUHS Emerging Viral Threat Laboratory | Gregory L. Ware, Alexander Mijalis, Jeremy P. Kamil, Jennifer L. Carroll, Maarten Van Diest, Rona S. Scott, Andrew D. Yurochko, Christopher G. Kevil, John A. Vanchiere |
| EPI_ISL_3115289, EPI_ISL_3115290, EPI_ISL_3115291 | Louisiana Office of Public Heath Laboratory | LSUHS Emerging Viral Threat Laboratory | Gregory L. Ware, Alexander Mijalis, Jeremy P. Kamil, Jennifer L. Carroll, Maarten Van Diest, Rona S. Scott, Andrew D. Yurochko, Christopher G. Kevil, John A. Vanchiere |
| EPI_ISL_3115292, EPI_ISL_3115293, EPI_ISL_3115294, EPI_ISL_3115295 | Willis-Knighton Medical Center Hospital Laboratory | LSUHS Emerging Viral Threat Laboratory | Gregory L. Ware, Alexander Mijalis, Jeremy P. Kamil, Jennifer L. Carroll, Maarten Van Diest, Rona S. Scott, Andrew D. Yurochko, Christopher G. Kevil, John A. Vanchiere |
| EPI_ISL_3115296 | Louisiana Office of Public Heath Laboratory | LSUHS Emerging Viral Threat Laboratory | Gregory L. Ware, Alexander Mijalis, Jeremy P. Kamil, Jennifer L. Carroll, Maarten Van Diest, Rona S. Scott, Andrew D. Yurochko, Christopher G. Kevil, John A. Vanchiere |
| EPI_ISL_3115297 | Willis-Knighton Medical Center Hospital Laboratory | LSUHS Emerging Viral Threat Laboratory | Gregory L. Ware, Alexander Mijalis, Jeremy P. Kamil, Jennifer L. Carroll, Maarten Van Diest, Rona S. Scott, Andrew D. Yurochko, Christopher G. Kevil, John A. Vanchiere |
| EPI_ISL_3115298, EPI_ISL_3115299 | Louisiana Office of Public Heath Laboratory | LSUHS Emerging Viral Threat Laboratory | Gregory L. Ware, Alexander Mijalis, Jeremy P. Kamil, Jennifer L. Carroll, Maarten Van Diest, Rona S. Scott, Andrew D. Yurochko, Christopher G. Kevil, John A. Vanchiere |
| EPI_ISL_3115300 | Willis-Knighton Medical Center Hospital Laboratory | LSUHS Emerging Viral Threat Laboratory | Gregory L. Ware, Alexander Mijalis, Jeremy P. Kamil, Jennifer L. Carroll, Maarten Van Diest, Rona S. Scott, Andrew D. Yurochko, Christopher G. Kevil, John A. Vanchiere |
| EPI_ISL_3115301 | Louisiana Office of Public Heath Laboratory | LSUHS Emerging Viral Threat Laboratory | Gregory L. Ware, Alexander Mijalis, Jeremy P. Kamil, Jennifer L. Carroll, Maarten Van Diest, Rona S. Scott, Andrew D. Yurochko, Christopher G. Kevil, John A. Vanchiere |
| EPI_ISL_3115302, EPI_ISL_3115303 | Willis-Knighton Medical Center Hospital Laboratory | LSUHS Emerging Viral Threat Laboratory | Gregory L. Ware, Alexander Mijalis, Jeremy P. Kamil, Jennifer L. Carroll, Maarten Van Diest, Rona S. Scott, Andrew D. Yurochko, Christopher G. Kevil, John A. Vanchiere |
| EPI_ISL_3115304, EPI_ISL_3115305 | Louisiana Office of Public Heath Laboratory | LSUHS Emerging Viral Threat Laboratory | Gregory L. Ware, Alexander Mijalis, Jeremy P. Kamil, Jennifer L. Carroll, Maarten Van Diest, Rona S. Scott, Andrew D. Yurochko, Christopher G. Kevil, John A. Vanchiere |
| EPI_ISL_3115306 | LSUHS Emerging Viral Threat Laboratory | LSUHS Emerging Viral Threat Laboratory | Gregory L. Ware, Alexander Mijalis, Jeremy P. Kamil, Jennifer L. Carroll, Maarten Van Diest, Rona S. Scott, Andrew D. Yurochko, Christopher G. Kevil, John A. Vanchiere |
| EPI_ISL_3115307, EPI_ISL_3115308, EPI_ISL_3115309 | Willis-Knighton Medical Center Hospital Laboratory | LSUHS Emerging Viral Threat Laboratory | Gregory L. Ware, Alexander Mijalis, Jeremy P. Kamil, Jennifer L. Carroll, Maarten Van Diest, Rona S. Scott, Andrew D. Yurochko, Christopher G. Kevil, John A. Vanchiere |
| EPI_ISL_3115310 | LSUHS Emerging Viral Threat Laboratory | LSUHS Emerging Viral Threat Laboratory | Gregory L. Ware, Alexander Mijalis, Jeremy P. Kamil, Jennifer L. Carroll, Maarten Van Diest, Rona S. Scott, Andrew D. Yurochko, Christopher G. Kevil, John A. Vanchiere |
| EPI_ISL_3115311, EPI_ISL_3115312, EPI_ISL_3115313, EPI_ISL_3115314, EPI_ISL_3115315, EPI_ISL_3115316 | Willis-Knighton Medical Center Hospital Laboratory | LSUHS Emerging Viral Threat Laboratory | Gregory L. Ware, Alexander Mijalis, Jeremy P. Kamil, Jennifer L. Carroll, Maarten Van Diest, Rona S. Scott, Andrew D. Yurochko, Christopher G. Kevil, John A. Vanchiere |
| EPI_ISL_3115317 | Louisiana Office of Public Heath Laboratory | LSUHS Emerging Viral Threat Laboratory | Gregory L. Ware, Alexander Mijalis, Jeremy P. Kamil, Jennifer L. Carroll, Maarten Van Diest, Rona S. Scott, Andrew D. Yurochko, Christopher G. Kevil, John A. Vanchiere |
| EPI_ISL_3115318, EPI_ISL_3115319 | Willis-Knighton Medical Center Hospital Laboratory | LSUHS Emerging Viral Threat Laboratory | Gregory L. Ware, Alexander Mijalis, Jeremy P. Kamil, Jennifer L. Carroll, Maarten Van Diest, Rona S. Scott, Andrew D. Yurochko, Christopher G. Kevil, John A. Vanchiere |
| EPI_ISL_3115320 | LSUHS Emerging Viral Threat Laboratory | LSUHS Emerging Viral Threat Laboratory | Gregory L. Ware, Alexander Mijalis, Jeremy P. Kamil, Jennifer L. Carroll, Maarten Van Diest, Rona S. Scott, Andrew D. Yurochko, Christopher G. Kevil, John A. Vanchiere |
| EPI_ISL_3115321 | Willis-Knighton Medical Center Hospital Laboratory | LSUHS Emerging Viral Threat Laboratory | Gregory L. Ware, Alexander Mijalis, Jeremy P. Kamil, Jennifer L. Carroll, Maarten Van Diest, Rona S. Scott, Andrew D. Yurochko, Christopher G. Kevil, John A. Vanchiere |
| EPI_ISL_3115322, EPI_ISL_3115323 | Louisiana Office of Public Heath Laboratory | LSUHS Emerging Viral Threat Laboratory | Gregory L. Ware, Alexander Mijalis, Jeremy P. Kamil, Jennifer L. Carroll, Maarten Van Diest, Rona S. Scott, Andrew D. Yurochko, Christopher G. Kevil, John A. Vanchiere |
| EPI_ISL_3115324 | LSUHS Emerging Viral Threat Laboratory | LSUHS Emerging Viral Threat Laboratory | Gregory L. Ware, Alexander Mijalis, Jeremy P. Kamil, Jennifer L. Carroll, Maarten Van Diest, Rona S. Scott, Andrew D. Yurochko, Christopher G. Kevil, John A. Vanchiere |
| EPI_ISL_3115325, EPI_ISL_3115326, EPI_ISL_3115327 | Willis-Knighton Medical Center Hospital Laboratory | LSUHS Emerging Viral Threat Laboratory | Gregory L. Ware, Alexander Mijalis, Jeremy P. Kamil, Jennifer L. Carroll, Maarten Van Diest, Rona S. Scott, Andrew D. Yurochko, Christopher G. Kevil, John A. Vanchiere |
| EPI_ISL_3115328, EPI_ISL_3115329 | Louisiana Office of Public Heath Laboratory | LSUHS Emerging Viral Threat Laboratory | Gregory L. Ware, Alexander Mijalis, Jeremy P. Kamil, Jennifer L. Carroll, Maarten Van Diest, Rona S. Scott, Andrew D. Yurochko, Christopher G. Kevil, John A. Vanchiere |
| EPI_ISL_3115330 | Willis-Knighton Medical Center Hospital Laboratory | LSUHS Emerging Viral Threat Laboratory | Gregory L. Ware, Alexander Mijalis, Jeremy P. Kamil, Jennifer L. Carroll, Maarten Van Diest, Rona S. Scott, Andrew D. Yurochko, Christopher G. Kevil, John A. Vanchiere |
| EPI_ISL_3115332 | LSUHS Emerging Viral Threat Laboratory | LSUHS Emerging Viral Threat Laboratory | Gregory L. Ware, Alexander Mijalis, Jeremy P. Kamil, Jennifer L. Carroll, Maarten Van Diest, Rona S. Scott, Andrew D. Yurochko, Christopher G. Kevil, John A. Vanchiere |
| EPI_ISL_3115333, EPI_ISL_3115334 | Louisiana Office of Public Heath Laboratory | LSUHS Emerging Viral Threat Laboratory | Gregory L. Ware, Alexander Mijalis, Jeremy P. Kamil, Jennifer L. Carroll, Maarten Van Diest, Rona S. Scott, Andrew D. Yurochko, Christopher G. Kevil, John A. Vanchiere |
| EPI_ISL_3115335, EPI_ISL_3115336 | Willis-Knighton Medical Center Hospital Laboratory | LSUHS Emerging Viral Threat Laboratory | Gregory L. Ware, Alexander Mijalis, Jeremy P. Kamil, Jennifer L. Carroll, Maarten Van Diest, Rona S. Scott, Andrew D. Yurochko, Christopher G. Kevil, John A. Vanchiere |
| EPI_ISL_3115337, EPI_ISL_3115338 | LSUHS Emerging Viral Threat Laboratory | LSUHS Emerging Viral Threat Laboratory | Gregory L. Ware, Alexander Mijalis, Jeremy P. Kamil, Jennifer L. Carroll, Maarten Van Diest, Rona S. Scott, Andrew D. Yurochko, Christopher G. Kevil, John A. Vanchiere |
| EPI_ISL_3115339 | Louisiana Office of Public Heath Laboratory | LSUHS Emerging Viral Threat Laboratory | Gregory L. Ware, Alexander Mijalis, Jeremy P. Kamil, Jennifer L. Carroll, Maarten Van Diest, Rona S. Scott, Andrew D. Yurochko, Christopher G. Kevil, John A. Vanchiere |
| EPI_ISL_3115340 | LSUHS Emerging Viral Threat Laboratory | LSUHS Emerging Viral Threat Laboratory | Gregory L. Ware, Alexander Mijalis, Jeremy P. Kamil, Jennifer L. Carroll, Maarten Van Diest, Rona S. Scott, Andrew D. Yurochko, Christopher G. Kevil, John A. Vanchiere |
| EPI_ISL_3115341 | Louisiana Office of Public Heath Laboratory | LSUHS Emerging Viral Threat Laboratory | Gregory L. Ware, Alexander Mijalis, Jeremy P. Kamil, Jennifer L. Carroll, Maarten Van Diest, Rona S. Scott, Andrew D. Yurochko, Christopher G. Kevil, John A. Vanchiere |
| EPI_ISL_3115342 | LSUHS Emerging Viral Threat Laboratory | LSUHS Emerging Viral Threat Laboratory | Gregory L. Ware, Alexander Mijalis, Jeremy P. Kamil, Jennifer L. Carroll, Maarten Van Diest, Rona S. Scott, Andrew D. Yurochko, Christopher G. Kevil, John A. Vanchiere |
| EPI_ISL_3115343 | Willis-Knighton Medical Center Hospital Laboratory | LSUHS Emerging Viral Threat Laboratory | Gregory L. Ware, Alexander Mijalis, Jeremy P. Kamil, Jennifer L. Carroll, Maarten Van Diest, Rona S. Scott, Andrew D. Yurochko, Christopher G. Kevil, John A. Vanchiere |
| EPI_ISL_3115344 | LSUHS Emerging Viral Threat Laboratory | LSUHS Emerging Viral Threat Laboratory | Gregory L. Ware, Alexander Mijalis, Jeremy P. Kamil, Jennifer L. Carroll, Maarten Van Diest, Rona S. Scott, Andrew D. Yurochko, Christopher G. Kevil, John A. Vanchiere |
| EPI_ISL_3115345, EPI_ISL_3115346, EPI_ISL_3115347 | Clínica Rotger Laboratorio de Análisis Clínicos Dpto. Biología Molecular | Clínica Rotger Laboratorio de Análisis Clínicos Dpto. Biología Molecular | Francisco Aliaga-Lozano, Chaymaá Gammani-Chairi, Laura Gómez, Juana González-Calderón, Marina Barceló-Vives, Catalina Vidal-Gómez, Marina Kleber-Pol, Esperança Frau-Fuster, Carmen Morales-González, Esther Nieto-Molina |
| EPI_ISL_3115361, EPI_ISL_3115362, EPI_ISL_3115363 | Biorepository Clinical Virology Laboratory, Ibadan | Africa Centre for Excellence for Genomics of Infectious Diseases (ACEGID), Redeemer's University | Olawoye, I.B., Oluniyi, P.E., Eromon, P.E., Oguzie, J.U., Kayode, A.T., Uwanibe, J.N., Ugwu, C.A., Akano, K.O., Ajogbasile, F.V., Abechi, P.S., Olumade, T.J., Nosamiefan, I., Folarin, O., Happi, C.T. |
| EPI_ISL_3115515, EPI_ISL_3115560, EPI_ISL_3115561, EPI_ISL_3115562, EPI_ISL_3115563, EPI_ISL_3115564, EPI_ISL_3115565 | Clínica Rotger Laboratorio de Análisis Clínicos Dpto. Biología Molecular | Clínica Rotger Laboratorio de Análisis Clínicos Dpto. Biología Molecular | Francisco Aliaga-Lozano, Chaymaá Gammani-Chairi, Laura Gómez, Juana González-Calderón, Marina Barceló-Vives, Catalina Vidal-Gómez, Marina Kleber-Pol, Esperança Frau-Fuster, Carmen Morales-González, Esther Nieto-Molina |
| EPI_ISL_3115566, EPI_ISL_3115567, EPI_ISL_3115568, EPI_ISL_3115569 | Lafene Health Center | Kansas State Veterinary Diagnostic Laboratory | Tyler Doerksen, Andreea Lu, Lance Noll, Elizabeth Porter, Emily Cox, Jianfa Bai, Jamie Henningson, Rachel Palinski |
| EPI_ISL_3115586, EPI_ISL_3115593, EPI_ISL_3115594, EPI_ISL_3115648, EPI_ISL_3115689, EPI_ISL_3115692, EPI_ISL_3115697 | ZOTZ KLIMAS MVZ Düsseldorf-Centrum GbR ÜBAG für Labormedizin, Genetik, Zytologie, Pathologie | Center of Medical Microbiology, Virology, and Hospital Hygiene, University of Duesseldorf | Maximilian Damagnez;Alexander Dilthey;Patrick Finzer;Katrin Hoffmann;Torsten Houwaart;Lisanna Hülse;Malte Kohns Vasconcelos;Nadine Lübke;Jessica Nicolai;Klaus Pfeffer;Daniel Strelow;Jörg Timm;Andreas Walker;Tobias Wienemann;Rainer Zotz |
| EPI_ISL_3115699, EPI_ISL_3115700, EPI_ISL_3115701, EPI_ISL_3115702, EPI_ISL_3115703, EPI_ISL_3115704, EPI_ISL_3115705, EPI_ISL_3115706, EPI_ISL_3115707, EPI_ISL_3115708, EPI_ISL_3115709, EPI_ISL_3115710, EPI_ISL_3115711, EPI_ISL_3115712, EPI_ISL_3115713, EPI_ISL_3115714, EPI_ISL_3115716, EPI_ISL_3115718, EPI_ISL_3115719, EPI_ISL_3115720, EPI_ISL_3115721, EPI_ISL_3115722, EPI_ISL_3115723, EPI_ISL_3115724, EPI_ISL_3115725, EPI_ISL_3115726, EPI_ISL_3115727, EPI_ISL_3115728, EPI_ISL_3115729, EPI_ISL_3115730 | | | |
| see above | IN State Department of Health Laboratory Services | IN State Department of Health Laboratory Services | Cassandra Campion, Jamie Yeadon, Brian Pope, Lixia Liu, Kyle Brownlee, Melissa Hindenlang, Mark Glazier |
| EPI_ISL_3115750, EPI_ISL_3115765 | ZOTZ KLIMAS MVZ Düsseldorf-Centrum GbR ÜBAG für Labormedizin, Genetik, Zytologie, Pathologie | Center of Medical Microbiology, Virology, and Hospital Hygiene, University of Duesseldorf | Maximilian Damagnez;Alexander Dilthey;Patrick Finzer;Katrin Hoffmann;Torsten Houwaart;Lisanna Hülse;Malte Kohns Vasconcelos;Nadine Lübke;Jessica Nicolai;Klaus Pfeffer;Daniel Strelow;Jörg Timm;Andreas Walker;Tobias Wienemann;Rainer Zotz |

| | | | |
|---|---|---|---|
| EPI_ISL_3115770 | Center of Medical Microbiology, Virology, and Hospital Hygiene, University of Duesseldorf | Center of Medical Microbiology, Virology, and Hospital Hygiene, University of Duesseldorf | Maximilian Damagnez;Alexander Dilthey;Torsten Houwaart;Lisanna Hülse;Malte Kohns Vasconcelos;Nadine Lübke;Jessica Nicolai;Klaus Pfeffer;Daniel Strelow;Jörg Timm;Andreas Walker;Tobias Wienemann |
| EPI_ISL_3115775, EPI_ISL_3115781, EPI_ISL_3115800, EPI_ISL_3115801, EPI_ISL_3115802, EPI_ISL_3115803, EPI_ISL_3115809, EPI_ISL_3115810, EPI_ISL_3115813, EPI_ISL_3115814, EPI_ISL_3115815 | | | |
| see above | ZOTZ KLIMAS MVZ Düsseldorf-Centrum GbR ÜBAG für Labormedizin, Genetik, Zytologie, Pathologie | Center of Medical Microbiology, Virology, and Hospital Hygiene, University of Duesseldorf | Maximilian Damagnez;Alexander Dilthey;Patrick Finzer;Katrin Hoffmann;Torsten Houwaart;Lisanna Hülse;Malte Kohns Vasconcelos;Nadine Lübke;Jessica Nicolai;Klaus Pfeffer;Daniel Strelow;Jörg Timm;Andreas Walker;Tobias Wienemann;Rainer Zotz |
| EPI_ISL_3116024, EPI_ISL_3116025, EPI_ISL_3116026, EPI_ISL_3116027, EPI_ISL_3116028, EPI_ISL_3116029, EPI_ISL_3116030, EPI_ISL_3116031, EPI_ISL_3116032, EPI_ISL_3116033, EPI_ISL_3116034, EPI_ISL_3116035, EPI_ISL_3116036, EPI_ISL_3116037, EPI_ISL_3116038, EPI_ISL_3116039, EPI_ISL_3116040, EPI_ISL_3116041, EPI_ISL_3116042, EPI_ISL_3116043, EPI_ISL_3116044, EPI_ISL_3116045, EPI_ISL_3116046, EPI_ISL_3116047, EPI_ISL_3116048, EPI_ISL_3116049, EPI_ISL_3116050, EPI_ISL_3116051, EPI_ISL_3116052, EPI_ISL_3116053, EPI_ISL_3116054, EPI_ISL_3116055, EPI_ISL_3116056, EPI_ISL_3116057, EPI_ISL_3116058, EPI_ISL_3116059, EPI_ISL_3116060, EPI_ISL_3116061, EPI_ISL_3116062, EPI_ISL_3116063, EPI_ISL_3116064, EPI_ISL_3116065, EPI_ISL_3116066, EPI_ISL_3116067, EPI_ISL_3116068, EPI_ISL_3116069, EPI_ISL_3116070, EPI_ISL_3116071, EPI_ISL_3116072, EPI_ISL_3116073, EPI_ISL_3116074, EPI_ISL_3116075, EPI_ISL_3116076, EPI_ISL_3116077, EPI_ISL_3116078, EPI_ISL_3116079, EPI_ISL_3116080, EPI_ISL_3116081, EPI_ISL_3116082, EPI_ISL_3116083, EPI_ISL_3116084, EPI_ISL_3116085, EPI_ISL_3116086, EPI_ISL_3116087, EPI_ISL_3116088, EPI_ISL_3116089, EPI_ISL_3116090, EPI_ISL_3116091, EPI_ISL_3116092, EPI_ISL_3116093, EPI_ISL_3116094, EPI_ISL_3116095, EPI_ISL_3116096, EPI_ISL_3116097, EPI_ISL_3116098, EPI_ISL_3116099, EPI_ISL_3116100, EPI_ISL_3116101, EPI_ISL_3116102, EPI_ISL_3116103, EPI_ISL_3116104, EPI_ISL_3116105, EPI_ISL_3116106, EPI_ISL_3116107, EPI_ISL_3116108, EPI_ISL_3116109, EPI_ISL_3116110, EPI_ISL_3116111, EPI_ISL_3116112, EPI_ISL_3116113, EPI_ISL_3116114, EPI_ISL_3116115, EPI_ISL_3116116, EPI_ISL_3116117, EPI_ISL_3116118, EPI_ISL_3116119, EPI_ISL_3116120, EPI_ISL_3116121, EPI_ISL_3116122, EPI_ISL_3116123, EPI_ISL_3116124, EPI_ISL_3116125, EPI_ISL_3116126, EPI_ISL_3116127, EPI_ISL_3116128, EPI_ISL_3116129, EPI_ISL_3116130, EPI_ISL_3116131, EPI_ISL_3116132, EPI_ISL_3116133, EPI_ISL_3116134, EPI_ISL_3116135, EPI_ISL_3116136, EPI_ISL_3116137, EPI_ISL_3116138, EPI_ISL_3116139, EPI_ISL_3116140, EPI_ISL_3116141, EPI_ISL_3116142, EPI_ISL_3116143, EPI_ISL_3116144, EPI_ISL_3116145, EPI_ISL_3116146, EPI_ISL_3116147, EPI_ISL_3116148, EPI_ISL_3116149, EPI_ISL_3116150, EPI_ISL_3116151, EPI_ISL_3116152, EPI_ISL_3116153, EPI_ISL_3116154, EPI_ISL_3116155, EPI_ISL_3116156, EPI_ISL_3116157, EPI_ISL_3116158, EPI_ISL_3116159, EPI_ISL_3116160, EPI_ISL_3116161, EPI_ISL_3116162, EPI_ISL_3116163, EPI_ISL_3116164, EPI_ISL_3116165, EPI_ISL_3116166, EPI_ISL_3116167, EPI_ISL_3116168, EPI_ISL_3116169, EPI_ISL_3116170, EPI_ISL_3116171, EPI_ISL_3116172, EPI_ISL_3116173, EPI_ISL_3116174, EPI_ISL_3116175, EPI_ISL_3116176, EPI_ISL_3116177, EPI_ISL_3116178, EPI_ISL_3116179, EPI_ISL_3116180, EPI_ISL_3116181, EPI_ISL_3116182, EPI_ISL_3116183, EPI_ISL_3116184, EPI_ISL_3116185, EPI_ISL_3116186, EPI_ISL_3116187, EPI_ISL_3116188, EPI_ISL_3116189, EPI_ISL_3116190, EPI_ISL_3116191, EPI_ISL_3116192, EPI_ISL_3116193, EPI_ISL_3116194, EPI_ISL_3116195, EPI_ISL_3116196, EPI_ISL_3116197, EPI_ISL_3116198, EPI_ISL_3116199, EPI_ISL_3116200, EPI_ISL_3116201, EPI_ISL_3116202, EPI_ISL_3116203, EPI_ISL_3116204, EPI_ISL_3116205, EPI_ISL_3116206, EPI_ISL_3116207, EPI_ISL_3116208, EPI_ISL_3116209, EPI_ISL_3116210, EPI_ISL_3116211, EPI_ISL_3116212, EPI_ISL_3116213, EPI_ISL_3116214, EPI_ISL_3116215, EPI_ISL_3116216, EPI_ISL_3116217, EPI_ISL_3116218, EPI_ISL_3116219, EPI_ISL_3116220, EPI_ISL_3116221, EPI_ISL_3116222, EPI_ISL_3116223, EPI_ISL_3116224, EPI_ISL_3116225, EPI_ISL_3116226, EPI_ISL_3116227, EPI_ISL_3116228, EPI_ISL_3116229, EPI_ISL_3116230, EPI_ISL_3116231, EPI_ISL_3116232, EPI_ISL_3116233, EPI_ISL_3116234, EPI_ISL_3116235, EPI_ISL_3116236, EPI_ISL_3116237, EPI_ISL_3116238, EPI_ISL_3116239, EPI_ISL_3116240, EPI_ISL_3116241, EPI_ISL_3116242, EPI_ISL_3116243, EPI_ISL_3116244, EPI_ISL_3116245, EPI_ISL_3116246, EPI_ISL_3116247, EPI_ISL_3116248, EPI_ISL_3116249, EPI_ISL_3116250, EPI_ISL_3116251, EPI_ISL_3116252, EPI_ISL_3116253, EPI_ISL_3116254, EPI_ISL_3116255, EPI_ISL_3116256, EPI_ISL_3116257, EPI_ISL_3116258, EPI_ISL_3116259, EPI_ISL_3116260, EPI_ISL_3116261, EPI_ISL_3116262, EPI_ISL_3116263, EPI_ISL_3116264, EPI_ISL_3116265, EPI_ISL_3116266, EPI_ISL_3116267, EPI_ISL_3116268, EPI_ISL_3116269, EPI_ISL_3116270, EPI_ISL_3116271, EPI_ISL_3116272, EPI_ISL_3116273, EPI_ISL_3116274, EPI_ISL_3116275, EPI_ISL_3116276, EPI_ISL_3116277, EPI_ISL_3116278, EPI_ISL_3116279, EPI_ISL_3116280, EPI_ISL_3116281, EPI_ISL_3116282, EPI_ISL_3116283, EPI_ISL_3116284, EPI_ISL_3116285, EPI_ISL_3116286, EPI_ISL_3116287, EPI_ISL_3116288, EPI_ISL_3116289, EPI_ISL_3116290, EPI_ISL_3116291, EPI_ISL_3116292, EPI_ISL_3116293, EPI_ISL_3116294, EPI_ISL_3116295, EPI_ISL_3116296, EPI_ISL_3116297, EPI_ISL_3116298, EPI_ISL_3116299, EPI_ISL_3116300, EPI_ISL_3116301, EPI_ISL_3116302, EPI_ISL_3116303, EPI_ISL_3116304, EPI_ISL_3116305, EPI_ISL_3116306, EPI_ISL_3116307, EPI_ISL_3116308, EPI_ISL_3116309, EPI_ISL_3116310, EPI_ISL_3116311, EPI_ISL_3116312, EPI_ISL_3116313, EPI_ISL_3116314, EPI_ISL_3116315, EPI_ISL_3116316, EPI_ISL_3116317, EPI_ISL_3116318, EPI_ISL_3116319, EPI_ISL_3116320, EPI_ISL_3116321, EPI_ISL_3116322, EPI_ISL_3116323, EPI_ISL_3116324, EPI_ISL_3116325, EPI_ISL_3116326, EPI_ISL_3116327, EPI_ISL_3116328, EPI_ISL_3116329, EPI_ISL_3116330, EPI_ISL_3116331, EPI_ISL_3116332, EPI_ISL_3116333, EPI_ISL_3116334, EPI_ISL_3116335, EPI_ISL_3116336, EPI_ISL_3116337, EPI_ISL_3116338, EPI_ISL_3116339, EPI_ISL_3116340, EPI_ISL_3116341, EPI_ISL_3116342, EPI_ISL_3116343, EPI_ISL_3116344, EPI_ISL_3116345, EPI_ISL_3116346, EPI_ISL_3116347, EPI_ISL_3116348, EPI_ISL_3116349, EPI_ISL_3116350, EPI_ISL_3116351, EPI_ISL_3116352, EPI_ISL_3116353, EPI_ISL_3116354, EPI_ISL_3116355, EPI_ISL_3116356, EPI_ISL_3116357, EPI_ISL_3116358, EPI_ISL_3116359, EPI_ISL_3116360, EPI_ISL_3116361, EPI_ISL_3116362, EPI_ISL_3116363, EPI_ISL_3116364, EPI_ISL_3116365, EPI_ISL_3116366, EPI_ISL_3116367, EPI_ISL_3116368, EPI_ISL_3116369, EPI_ISL_3116370, EPI_ISL_3116371, EPI_ISL_3116372, EPI_ISL_3116373, EPI_ISL_3116374, EPI_ISL_3116375, EPI_ISL_3116376, EPI_ISL_3116377, EPI_ISL_3116378, EPI_ISL_3116379, EPI_ISL_3116380, EPI_ISL_3116381, EPI_ISL_3116382, EPI_ISL_3116383, EPI_ISL_3116384, EPI_ISL_3116385, EPI_ISL_3116386, EPI_ISL_3116387, EPI_ISL_3116388, EPI_ISL_3116389, EPI_ISL_3116390, EPI_ISL_3116391, EPI_ISL_3116392, EPI_ISL_3116393, EPI_ISL_3116394, EPI_ISL_3116395, EPI_ISL_3116396, EPI_ISL_3116397, EPI_ISL_3116398, EPI_ISL_3116399, EPI_ISL_3116400, EPI_ISL_3116401, EPI_ISL_3116402, EPI_ISL_3116403, EPI_ISL_3116404, EPI_ISL_3116405, EPI_ISL_3116406, EPI_ISL_3116407, EPI_ISL_3116408, EPI_ISL_3116409, EPI_ISL_3116410, EPI_ISL_3116411, EPI_ISL_3116412, EPI_ISL_3116413, EPI_ISL_3116414, EPI_ISL_3116415, EPI_ISL_3116416, EPI_ISL_3116417, EPI_ISL_3116418, EPI_ISL_3116419, EPI_ISL_3116420, EPI_ISL_3116421, EPI_ISL_3116422, EPI_ISL_3116423, EPI_ISL_3116424, EPI_ISL_3116425, EPI_ISL_3116426, EPI_ISL_3116427, EPI_ISL_3116428, EPI_ISL_3116429, EPI_ISL_3116430, EPI_ISL_3116431, EPI_ISL_3116432, EPI_ISL_3116433, EPI_ISL_3116434, EPI_ISL_3116435, EPI_ISL_3116436, EPI_ISL_3116437, EPI_ISL_3116438, EPI_ISL_3116439, EPI_ISL_3116440, EPI_ISL_3116441, EPI_ISL_3116442, EPI_ISL_3116443, EPI_ISL_3116444, EPI_ISL_3116445, EPI_ISL_3116446, EPI_ISL_3116447, EPI_ISL_3116448, EPI_ISL_3116449, EPI_ISL_3116450, EPI_ISL_3116451, EPI_ISL_3116452, EPI_ISL_3116453, EPI_ISL_3116454, EPI_ISL_3116455, EPI_ISL_3116456, EPI_ISL_3116457, EPI_ISL_3116458, EPI_ISL_3116459, EPI_ISL_3116460, EPI_ISL_3116461, EPI_ISL_3116462, EPI_ISL_3116463, EPI_ISL_3116464, EPI_ISL_3116465, EPI_ISL_3116466, EPI_ISL_3116467, EPI_ISL_3116468, EPI_ISL_3116469, EPI_ISL_3116470, EPI_ISL_3116471, EPI_ISL_3116472, EPI_ISL_3116473, EPI_ISL_3116474, EPI_ISL_3116475, EPI_ISL_3116476, EPI_ISL_3116477, EPI_ISL_3116478, EPI_ISL_3116479, EPI_ISL_3116480, EPI_ISL_3116481, EPI_ISL_3116482, EPI_ISL_3116483, EPI_ISL_3116484, EPI_ISL_3116485 | | | |
| see above | Edmonton Provincial Lab | Public Health Agency of Canada (PHAC) National Microbiology Laboratory | Buss, E, Croxen M, Deo A, Dieu P, Gill K, Ferrato C, Khan F, Koleva P, Li V, Lloyd C, Lynch T, Ma R, Murphy S, Pabbaraju K, Shokoples S, Tipples G, Thayer J, Whitehouse M, Wong A, Yu C, Zelyas N |
| EPI_ISL_3116597, EPI_ISL_3116600, EPI_ISL_3116602, EPI_ISL_3116603, EPI_ISL_3116606, EPI_ISL_3116608, EPI_ISL_3116610, EPI_ISL_3116619, EPI_ISL_3116636, EPI_ISL_3116638, EPI_ISL_3116639, EPI_ISL_3116653, EPI_ISL_3116654, EPI_ISL_3116656, EPI_ISL_3116657, EPI_ISL_3116659, EPI_ISL_3116660 | | | |
| see above | ZOTZ KLIMAS MVZ Düsseldorf-Centrum GbR ÜBAG für Labormedizin, Genetik, Zytologie, Pathologie | Center of Medical Microbiology, Virology, and Hospital Hygiene, University of Duesseldorf | Maximilian Damagnez;Alexander Dilthey;Patrick Finzer;Katrin Hoffmann;Torsten Houwaart;Lisanna Hülse;Malte Kohns Vasconcelos;Nadine Lübke;Jessica Nicolai;Klaus Pfeffer;Daniel Strelow;Jörg Timm;Andreas Walker;Tobias Wienemann;Rainer Zotz |
| EPI_ISL_3116788, EPI_ISL_3116789, EPI_ISL_3116790, EPI_ISL_3116791, EPI_ISL_3116792, EPI_ISL_3116793, EPI_ISL_3116794, EPI_ISL_3116795, EPI_ISL_3116796, EPI_ISL_3116797, EPI_ISL_3116798, EPI_ISL_3116799, EPI_ISL_3116800, EPI_ISL_3116801, EPI_ISL_3116802, EPI_ISL_3116803, EPI_ISL_3116804, EPI_ISL_3116805, EPI_ISL_3116806, EPI_ISL_3116807, EPI_ISL_3116808, EPI_ISL_3116809, EPI_ISL_3116810, EPI_ISL_3116811, EPI_ISL_3116812, EPI_ISL_3116813, EPI_ISL_3116814, EPI_ISL_3116815, EPI_ISL_3116816, EPI_ISL_3116817, EPI_ISL_3116818, EPI_ISL_3116890, EPI_ISL_3116893, EPI_ISL_3116903, EPI_ISL_3116904, EPI_ISL_3116905, EPI_ISL_3116906, EPI_ISL_3116907, EPI_ISL_3117238, EPI_ISL_3117239, EPI_ISL_3117240, EPI_ISL_3117241, EPI_ISL_3117242, EPI_ISL_3117243, EPI_ISL_3117244, EPI_ISL_3117245, EPI_ISL_3117246, EPI_ISL_3117247, EPI_ISL_3117248, EPI_ISL_3117249, EPI_ISL_3117250, EPI_ISL_3117251, EPI_ISL_3117252, EPI_ISL_3117253, EPI_ISL_3117254, EPI_ISL_3117255, EPI_ISL_3117274, EPI_ISL_3117275, EPI_ISL_3117276, EPI_ISL_3117277, EPI_ISL_3117278, EPI_ISL_3117279, EPI_ISL_3117280, EPI_ISL_3117281, EPI_ISL_3117282, EPI_ISL_3117283, EPI_ISL_3117284, EPI_ISL_3117285, EPI_ISL_3117286, EPI_ISL_3117287, EPI_ISL_3117288, EPI_ISL_3117289, EPI_ISL_3117296, EPI_ISL_3117311, EPI_ISL_3117312, EPI_ISL_3117313, EPI_ISL_3117314, EPI_ISL_3117315, EPI_ISL_3117316, EPI_ISL_3117317, EPI_ISL_3117318, EPI_ISL_3117319, EPI_ISL_3117320, EPI_ISL_3117321, EPI_ISL_3117322, EPI_ISL_3117323, EPI_ISL_3117324, EPI_ISL_3117325, EPI_ISL_3117326, EPI_ISL_3117327, EPI_ISL_3117328, EPI_ISL_3117329, EPI_ISL_3117330, EPI_ISL_3117331, EPI_ISL_3117332, EPI_ISL_3117333, EPI_ISL_3117334, EPI_ISL_3117335, EPI_ISL_3117336, EPI_ISL_3117337, EPI_ISL_3117338, EPI_ISL_3117339, EPI_ISL_3117340, EPI_ISL_3117341, EPI_ISL_3117342, EPI_ISL_3117343, EPI_ISL_3117344, EPI_ISL_3117345, EPI_ISL_3117346, EPI_ISL_3117347, EPI_ISL_3117348, EPI_ISL_3117349, EPI_ISL_3117350, EPI_ISL_3117351, EPI_ISL_3117352, EPI_ISL_3117353, EPI_ISL_3117354, EPI_ISL_3117355, EPI_ISL_3117356, EPI_ISL_3117357, EPI_ISL_3117358, EPI_ISL_3117359, EPI_ISL_3117360, EPI_ISL_3117361, EPI_ISL_3117362, EPI_ISL_3117363, EPI_ISL_3117364, EPI_ISL_3117365, EPI_ISL_3117366, EPI_ISL_3117367, EPI_ISL_3117368, EPI_ISL_3117369, EPI_ISL_3117370, EPI_ISL_3117371, EPI_ISL_3117372, EPI_ISL_3117381, EPI_ISL_3117384, EPI_ISL_3117385, EPI_ISL_3117386, EPI_ISL_3117387, EPI_ISL_3117392, EPI_ISL_3117393, EPI_ISL_3117394, EPI_ISL_3117395, EPI_ISL_3117410, EPI_ISL_3117411, EPI_ISL_3117412, EPI_ISL_3117413, EPI_ISL_3117419, EPI_ISL_3117423 | | | |
| see above | Edmonton Provincial Lab | Public Health Agency of Canada (PHAC) National Microbiology Laboratory | Buss, E, Croxen M, Deo A, Dieu P, Gill K, Ferrato C, Khan F, Koleva P, Li V, Lloyd C, Lynch T, Ma R, Murphy S, Pabbaraju K, Shokoples S, Tipples G, Thayer J, Whitehouse M, Wong A, Yu C, Zelyas N |
| EPI_ISL_3117447, EPI_ISL_3117458, EPI_ISL_3117459, EPI_ISL_3117460, EPI_ISL_3117462, EPI_ISL_3117463, EPI_ISL_3117465, EPI_ISL_3117466, EPI_ISL_3117467, EPI_ISL_3117468, EPI_ISL_3117469, EPI_ISL_3117470, EPI_ISL_3117471, EPI_ISL_3117472, EPI_ISL_3117473, EPI_ISL_3117474, EPI_ISL_3117475, EPI_ISL_3117476, EPI_ISL_3117477, EPI_ISL_3117478, EPI_ISL_3117479, EPI_ISL_3117480, EPI_ISL_3117481, EPI_ISL_3117482, EPI_ISL_3117483, EPI_ISL_3117484, EPI_ISL_3117485, EPI_ISL_3117486, EPI_ISL_3117487, EPI_ISL_3117488, EPI_ISL_3117489, EPI_ISL_3117490, EPI_ISL_3117491, EPI_ISL_3117492, EPI_ISL_3117493, EPI_ISL_3117494, EPI_ISL_3117495, EPI_ISL_3117496, EPI_ISL_3117497, EPI_ISL_3117499, EPI_ISL_3117500, EPI_ISL_3117501, EPI_ISL_3117502, EPI_ISL_3117503, EPI_ISL_3117504, EPI_ISL_3117505, EPI_ISL_3117506, EPI_ISL_3117507, EPI_ISL_3117508, EPI_ISL_3117509, EPI_ISL_3117510, EPI_ISL_3117511, EPI_ISL_3117512, EPI_ISL_3117513, EPI_ISL_3117514, EPI_ISL_3117515, EPI_ISL_3117516, EPI_ISL_3117517 | | | |
| see above | ZOTZ KLIMAS MVZ Düsseldorf-Centrum GbR ÜBAG für Labormedizin, Genetik, Zytologie, Pathologie | Center of Medical Microbiology, Virology, and Hospital Hygiene, University of Duesseldorf | Maximilian Damagnez;Alexander Dilthey;Patrick Finzer;Katrin Hoffmann;Torsten Houwaart;Lisanna Hülse;Malte Kohns Vasconcelos;Nadine Lübke;Jessica Nicolai;Klaus Pfeffer;Daniel Strelow;Jörg Timm;Andreas Walker;Tobias Wienemann;Rainer Zotz |
| EPI_ISL_3117519, EPI_ISL_3117520, EPI_ISL_3117521, EPI_ISL_3117522, EPI_ISL_3117523, EPI_ISL_3117524, EPI_ISL_3117525, EPI_ISL_3117526, EPI_ISL_3117527, EPI_ISL_3117528, EPI_ISL_3117529, EPI_ISL_3117530, EPI_ISL_3117531, EPI_ISL_3117532, EPI_ISL_3117533, EPI_ISL_3117534, EPI_ISL_3117535, EPI_ISL_3117536, EPI_ISL_3117537, EPI_ISL_3117539, EPI_ISL_3117540, EPI_ISL_3117541, EPI_ISL_3117542 | | | |
| see above | CHU Pontchaillou | CHU Pontchaillou | GROLHIER Claire, DENOUAL Florent, ETCHEVERRY Amandine, SASSI Mohamed, JAGLINE Steven, FEBREAU Christine, PRONIER Charlotte, GALIBERT Marie Dominique, DE TAYRAC Marie, THIBAULT Vincent |
| EPI_ISL_3117545, EPI_ISL_3117554, EPI_ISL_3117555, EPI_ISL_3117556 | University of Wisconsin-Madison AIDS Vaccine Research Laboratories | University of Wisconsin-Madison AIDS Vaccine Research Laboratories | Gage Moreno, Katarina Braun, et al. AIDS Vaccine Research Laboratories |
| EPI_ISL_3117561, EPI_ISL_3117562, EPI_ISL_3117563, EPI_ISL_3117565, EPI_ISL_3117566, EPI_ISL_3117567, EPI_ISL_3117568, EPI_ISL_3117569, EPI_ISL_3117570, EPI_ISL_3117571, EPI_ISL_3117572, EPI_ISL_3117574, EPI_ISL_3117575, EPI_ISL_3117576, EPI_ISL_3117577, EPI_ISL_3117578, EPI_ISL_3117579, EPI_ISL_3117580, EPI_ISL_3117581, EPI_ISL_3117582, EPI_ISL_3117584, EPI_ISL_3117585, EPI_ISL_3117586, EPI_ISL_3117588, EPI_ISL_3117590, EPI_ISL_3117591, EPI_ISL_3117592, EPI_ISL_3117593, EPI_ISL_3117594, EPI_ISL_3117595, EPI_ISL_3117596, EPI_ISL_3117597, EPI_ISL_3117599, EPI_ISL_3117600, EPI_ISL_3117601, EPI_ISL_3117602 | | | |
| see above | Virginia Division of Consolidated Laboratory Services | Virginia Division of Consolidated Laboratory Services | Virginia Division of Consolidated Laboratory Services |
| EPI_ISL_3117603, EPI_ISL_3117604, EPI_ISL_3117605, EPI_ISL_3117606, EPI_ISL_3117607, EPI_ISL_3117608, EPI_ISL_3117609, EPI_ISL_3117610, EPI_ISL_3117611, EPI_ISL_3117612, EPI_ISL_3117613, EPI_ISL_3117614, EPI_ISL_3117615, EPI_ISL_3117616, EPI_ISL_3117617, EPI_ISL_3117618, EPI_ISL_3117619, EPI_ISL_3117620, EPI_ISL_3117621, EPI_ISL_3117622, EPI_ISL_3117623, EPI_ISL_3117624 | | | |
| see above | Molecular Diagnostic and Research Laboratory, GIMS, Greater Noida | Virology Laboratory, Department of Microbiology, King George's Medical University | Vivek Gupta, Shantanu Prakash, Suruchi Shukla, Amita Jain |
| EPI_ISL_3117625, EPI_ISL_3117626, EPI_ISL_3117627 | Department of Microbiology, Command Hospital Central Command | Virology Laboratory, Department of Microbiology, King George's Medical University | Mahesh Reddy, Shantanu Prakash, Suruchi Shukla, Amita Jain |
| EPI_ISL_3117628, EPI_ISL_3117629, EPI_ISL_3117630, EPI_ISL_3117631, EPI_ISL_3117632, EPI_ISL_3117633, EPI_ISL_3117634, EPI_ISL_3117635, EPI_ISL_3117636, EPI_ISL_3117637, EPI_ISL_3117638, EPI_ISL_3117639, EPI_ISL_3117640, EPI_ISL_3117641, EPI_ISL_3117642, EPI_ISL_3117643, EPI_ISL_3117644 | | | |
| see above | Virology Laboratory, Department of Microbiology, King George's Medical University | Virology Laboratory, Department of Microbiology, King George's Medical University | Shantanu Prakash, Hricha Mishra, Suruchi Shukla, Amita Jain |
| EPI_ISL_3117645, EPI_ISL_3117646 | Department of Microbiology, Dr. Ram Manohar Lohia Institute of Medical Sciences | Virology Laboratory, Department of Microbiology, King George's Medical University | Jyotsna Agarwal, Shantanu Prakash, Suruchi Shukla, Amita Jain |
| EPI_ISL_3117647, EPI_ISL_3117648, EPI_ISL_3117649, EPI_ISL_3117650, EPI_ISL_3117651, EPI_ISL_3117652, EPI_ISL_3117653, EPI_ISL_3117654, EPI_ISL_3117655 | Department of Microbiology, AIIMS, Raebareli | Virology Laboratory, Department of Microbiology, King George's Medical University | Shefali Gupta, Shantanu Prakash, Suruchi Shukla, Amita Jain |
| EPI_ISL_3117656, EPI_ISL_3117657, EPI_ISL_3117658, EPI_ISL_3117659, EPI_ISL_3117660, EPI_ISL_3117661, EPI_ISL_3117662, EPI_ISL_3117663, EPI_ISL_3117664, EPI_ISL_3117665, EPI_ISL_3117666, EPI_ISL_3117667, EPI_ISL_3117668, EPI_ISL_3117669, EPI_ISL_3117670, EPI_ISL_3117671, EPI_ISL_3117672, EPI_ISL_3117673, EPI_ISL_3117674, EPI_ISL_3117675, EPI_ISL_3117676, EPI_ISL_3117677, EPI_ISL_3117678, EPI_ISL_3117679 | | | |
| see above | CSIR-Central Drug Research Institute | Virology Laboratory, Department of Microbiology, King George's Medical University | Raj Kamal Tripathi, Prem Narayan Yadav, Shantanu Prakash, Suruchi Shukla, Amita Jain |
| EPI_ISL_3117680, EPI_ISL_3117681, EPI_ISL_3117682 | Molecular Diagnostic and Research Laboratory, GIMS, Greater Noida | Virology Laboratory, Department of Microbiology, King George's Medical University | Vivek Gupta, Shantanu Prakash, Suruchi Shukla, Amita Jain |
| EPI_ISL_3117683, EPI_ISL_3117684, EPI_ISL_3117685, EPI_ISL_3117686, EPI_ISL_3117687, EPI_ISL_3117688, EPI_ISL_3117689, EPI_ISL_3117690, EPI_ISL_3117691, EPI_ISL_3117692 | National Institute of Cancer Prevention and Research | Virology Laboratory, Department of Microbiology, King George's Medical University | Shalini Singh, Shantanu Prakash, Suruchi Shukla, Amita Jain |
| EPI_ISL_3117693, EPI_ISL_3117694, EPI_ISL_3117695, EPI_ISL_3117696, EPI_ISL_3117697, EPI_ISL_3117698 | Department of Microbiology, LLRM Medical College | Virology Laboratory, Department of Microbiology, King George's Medical University | Amit Garg, Shantanu Prakash, Suruchi Shukla, Amita Jain |

| | | | |
|---|---|---|---|
| EPI_ISL_3117699, EPI_ISL_3117700, EPI_ISL_3117701, EPI_ISL_3117702, EPI_ISL_3117703, EPI_ISL_3117704 | National Institute of Biologicals | Virology Laboratory, Department of Microbiology, King George's Medical University | Harish Chander, Shantanu Prakash, Suruchi Shukla, Amita Jain |
| EPI_ISL_3117705, EPI_ISL_3117706, EPI_ISL_3117707, EPI_ISL_3117708, EPI_ISL_3117709, EPI_ISL_3117710, EPI_ISL_3117711, EPI_ISL_3117712, EPI_ISL_3117713, EPI_ISL_3117714 | Department of Microbiology, SSPHPGTI | Virology Laboratory, Department of Microbiology, King George's Medical University | Sumi Nandwani, Shantanu Prakash, Suruchi Shukla, Amita Jain |
| EPI_ISL_3117715, EPI_ISL_3117716, EPI_ISL_3117717, EPI_ISL_3117718, EPI_ISL_3117719, EPI_ISL_3117720, EPI_ISL_3117721, EPI_ISL_3117722, EPI_ISL_3117723, EPI_ISL_3117724, EPI_ISL_3117725, EPI_ISL_3117726, EPI_ISL_3117727, EPI_ISL_3117728, EPI_ISL_3117729, EPI_ISL_3117730 | | | |
| see above | Department of Microbiology, Veterinary College Mathura, DUVASU | Virology Laboratory, Department of Microbiology, King George's Medical University | Rashmi Singh, Shantanu Prakash, Suruchi Shukla, Amita Jain |
| EPI_ISL_3117731, EPI_ISL_3117732, EPI_ISL_3117733, EPI_ISL_3117734 | Department of Microbiology, Autonomous State Medical College Firozabad | Virology Laboratory, Department of Microbiology, King George's Medical University | Hariom Sharan, Lekha Tuli, Shantanu Prakash, Suruchi Shukla, Amita Jain |
| EPI_ISL_3117742, EPI_ISL_3117743, EPI_ISL_3117744, EPI_ISL_3117746, EPI_ISL_3117747, EPI_ISL_3117748, EPI_ISL_3117749, EPI_ISL_3117750, EPI_ISL_3117751, EPI_ISL_3117752, EPI_ISL_3117753 | | | |
| see above | Department of Microbiology, S.N. Medical College | Virology Laboratory, Department of Microbiology, King George's Medical University | Arti Agrawal, Shantanu Prakash, Suruchi Shukla, Amita Jain |
| EPI_ISL_3117754 | Department of Microbiology, LLRM Medical College | Virology Laboratory, Department of Microbiology, King George's Medical University | Amit Garg, Shantanu Prakash, Suruchi Shukla, Amita Jain |
| EPI_ISL_3117755, EPI_ISL_3117756, EPI_ISL_3117757, EPI_ISL_3117758 | CSIR-Indian Institute of Toxicology Research | Virology Laboratory, Department of Microbiology, King George's Medical University | Aditya Bhushan Pant, Shantanu Prakash, Suruchi Shukla, Amita Jain |
| EPI_ISL_3117759, EPI_ISL_3117760, EPI_ISL_3117761, EPI_ISL_3117763, EPI_ISL_3117764 | Virology Laboratory, Department of Microbiology, King George's Medical University | Virology Laboratory, Department of Microbiology, King George's Medical University | Shantanu Prakash, Hricha Mishra, Suruchi Shukla, Amita Jain |
| EPI_ISL_3117765 | Department of Microbiology, Government Medical College, Budaun | Virology Laboratory, Department of Microbiology, King George's Medical University | Vashishth Mishra, Shantanu Prakash, Suruchi Shukla, Amita Jain |
| EPI_ISL_3117825, EPI_ISL_3117827, EPI_ISL_3117841, EPI_ISL_3117842, EPI_ISL_3117843, EPI_ISL_3117844, EPI_ISL_3117845, EPI_ISL_3117848, EPI_ISL_3117849, EPI_ISL_3117850, EPI_ISL_3117851, EPI_ISL_3117852, EPI_ISL_3117857, EPI_ISL_3117859, EPI_ISL_3117864, EPI_ISL_3117867, EPI_ISL_3117869, EPI_ISL_3117870, EPI_ISL_3117871, EPI_ISL_3117872, EPI_ISL_3117873, EPI_ISL_3117874, EPI_ISL_3117875, EPI_ISL_3117876, EPI_ISL_3117877, EPI_ISL_3117878, EPI_ISL_3117879, EPI_ISL_3117882, EPI_ISL_3117884, EPI_ISL_3117885, EPI_ISL_3117886, EPI_ISL_3117887, EPI_ISL_3117888, EPI_ISL_3117889, EPI_ISL_3117891, EPI_ISL_3117893, EPI_ISL_3117894, EPI_ISL_3117895, EPI_ISL_3117896, EPI_ISL_3117897, EPI_ISL_3117900, EPI_ISL_3117901, EPI_ISL_3117903, EPI_ISL_3117904, EPI_ISL_3117909, EPI_ISL_3117911, EPI_ISL_3117912, EPI_ISL_3117913, EPI_ISL_3117914, EPI_ISL_3117916, EPI_ISL_3117917, EPI_ISL_3117918, EPI_ISL_3117919, EPI_ISL_3117922, EPI_ISL_3117923, EPI_ISL_3117924, EPI_ISL_3117925, EPI_ISL_3117926, EPI_ISL_3117927, EPI_ISL_3117928, EPI_ISL_3117929, EPI_ISL_3117930, EPI_ISL_3117931, EPI_ISL_3117932, EPI_ISL_3117933, EPI_ISL_3117934, EPI_ISL_3117935, EPI_ISL_3117937, EPI_ISL_3117938, EPI_ISL_3117939, EPI_ISL_3117940, EPI_ISL_3117941, EPI_ISL_3117942, EPI_ISL_3117943, EPI_ISL_3117944, EPI_ISL_3117945, EPI_ISL_3117946, EPI_ISL_3117947, EPI_ISL_3117948, EPI_ISL_3117953, EPI_ISL_3117954, EPI_ISL_3117956, EPI_ISL_3117957, EPI_ISL_3117958, EPI_ISL_3117960, EPI_ISL_3117961, EPI_ISL_3117962, EPI_ISL_3117963, EPI_ISL_3117964, EPI_ISL_3117965, EPI_ISL_3117966, EPI_ISL_3117967, EPI_ISL_3117968, EPI_ISL_3117969, EPI_ISL_3117970, EPI_ISL_3117971, EPI_ISL_3117972, EPI_ISL_3117973 | | | |
| see above | Public Health Authority of the Slovak Republic | Laboratory of Genomics and Bioinformatics, Comenius University Science Park | Tatiana Sedláčková, Diana Rusňáková, Miroslav Böhmer, Anna Gičová, Jaroslav Budiš, Tomáš Szemes, Jakub Styk |
| EPI_ISL_3117977, EPI_ISL_3117978, EPI_ISL_3117979, EPI_ISL_3117980, EPI_ISL_3117981, EPI_ISL_3117982, EPI_ISL_3117983, EPI_ISL_3117984, EPI_ISL_3117985, EPI_ISL_3117986, EPI_ISL_3117987, EPI_ISL_3117988, EPI_ISL_3117989, EPI_ISL_3117990, EPI_ISL_3117991, EPI_ISL_3117992, EPI_ISL_3117993, EPI_ISL_3117994, EPI_ISL_3117995, EPI_ISL_3117996, EPI_ISL_3117997, EPI_ISL_3117998, EPI_ISL_3117999, EPI_ISL_3118000, EPI_ISL_3118001, EPI_ISL_3118002, EPI_ISL_3118003, EPI_ISL_3118004, EPI_ISL_3118005, EPI_ISL_3118006, EPI_ISL_3118007, EPI_ISL_3118008, EPI_ISL_3118009, EPI_ISL_3118010, EPI_ISL_3118011, EPI_ISL_3118012, EPI_ISL_3118013, EPI_ISL_3118014, EPI_ISL_3118015, EPI_ISL_3118016, EPI_ISL_3118017, EPI_ISL_3118019, EPI_ISL_3118020, EPI_ISL_3118021, EPI_ISL_3118022, EPI_ISL_3118023, EPI_ISL_3118024, EPI_ISL_3118025, EPI_ISL_3118026, EPI_ISL_3118027, EPI_ISL_3118028, EPI_ISL_3118029, EPI_ISL_3118030, EPI_ISL_3118031, EPI_ISL_3118032, EPI_ISL_3118033, EPI_ISL_3118034, EPI_ISL_3118035, EPI_ISL_3118036, EPI_ISL_3118037, EPI_ISL_3118038, EPI_ISL_3118039, EPI_ISL_3118040, EPI_ISL_3118042, EPI_ISL_3118043, EPI_ISL_3118044, EPI_ISL_3118045, EPI_ISL_3118046, EPI_ISL_3118047, EPI_ISL_3118048, EPI_ISL_3118049, EPI_ISL_3118050, EPI_ISL_3118051, EPI_ISL_3118052, EPI_ISL_3118053, EPI_ISL_3118054, EPI_ISL_3118055, EPI_ISL_3118056, EPI_ISL_3118057, EPI_ISL_3118058, EPI_ISL_3118059, EPI_ISL_3118060, EPI_ISL_3118062, EPI_ISL_3118063, EPI_ISL_3118064, EPI_ISL_3118065, EPI_ISL_3118066, EPI_ISL_3118067, EPI_ISL_3118068, EPI_ISL_3118069, EPI_ISL_3118070, EPI_ISL_3118071, EPI_ISL_3118072, EPI_ISL_3118073, EPI_ISL_3118075, EPI_ISL_3118076, EPI_ISL_3118077, EPI_ISL_3118078, EPI_ISL_3118079, EPI_ISL_3118080, EPI_ISL_3118081, EPI_ISL_3118082, EPI_ISL_3118083, EPI_ISL_3118084, EPI_ISL_3118085, EPI_ISL_3118086, EPI_ISL_3118087, EPI_ISL_3118088, EPI_ISL_3118089, EPI_ISL_3118090, EPI_ISL_3118091, EPI_ISL_3118092, EPI_ISL_3118093, EPI_ISL_3118094, EPI_ISL_3118095, EPI_ISL_3118096, EPI_ISL_3118097, EPI_ISL_3118098, EPI_ISL_3118099, EPI_ISL_3118100, EPI_ISL_3118101, EPI_ISL_3118102, EPI_ISL_3118103, EPI_ISL_3118104, EPI_ISL_3118105, EPI_ISL_3118106, EPI_ISL_3118107, EPI_ISL_3118108, EPI_ISL_3118109, EPI_ISL_3118110, EPI_ISL_3118111, EPI_ISL_3118112, EPI_ISL_3118113, EPI_ISL_3118114, EPI_ISL_3118115, EPI_ISL_3118116, EPI_ISL_3118117, EPI_ISL_3118118, EPI_ISL_3118119, EPI_ISL_3118120, EPI_ISL_3118121, EPI_ISL_3118122, EPI_ISL_3118123, EPI_ISL_3118124, EPI_ISL_3118125, EPI_ISL_3118126, EPI_ISL_3118127, EPI_ISL_3118128, EPI_ISL_3118129, EPI_ISL_3118130, EPI_ISL_3118131, EPI_ISL_3118132, EPI_ISL_3118133, EPI_ISL_3118134, EPI_ISL_3118135, EPI_ISL_3118136, EPI_ISL_3118137, EPI_ISL_3118138, EPI_ISL_3118139, EPI_ISL_3118140, EPI_ISL_3118142, EPI_ISL_3118143, EPI_ISL_3118144, EPI_ISL_3118145, EPI_ISL_3118146, EPI_ISL_3118147, EPI_ISL_3118148, EPI_ISL_3118149, EPI_ISL_3118151, EPI_ISL_3118152, EPI_ISL_3118153, EPI_ISL_3118154, EPI_ISL_3118155, EPI_ISL_3118156, EPI_ISL_3118157, EPI_ISL_3118158, EPI_ISL_3118160, EPI_ISL_3118161, EPI_ISL_3118162, EPI_ISL_3118163, EPI_ISL_3118164, EPI_ISL_3118165, EPI_ISL_3118166, EPI_ISL_3118167, EPI_ISL_3118168, EPI_ISL_3118169, EPI_ISL_3118170, EPI_ISL_3118171, EPI_ISL_3118172, EPI_ISL_3118173, EPI_ISL_3118174, EPI_ISL_3118175, EPI_ISL_3118176, EPI_ISL_3118177, EPI_ISL_3118178, EPI_ISL_3118179, EPI_ISL_3118180, EPI_ISL_3118181, EPI_ISL_3118182, EPI_ISL_3118183, EPI_ISL_3118184, EPI_ISL_3118186, EPI_ISL_3118187, EPI_ISL_3118188, EPI_ISL_3118189, EPI_ISL_3118190, EPI_ISL_3118191, EPI_ISL_3118192, EPI_ISL_3118193, EPI_ISL_3118194, EPI_ISL_3118195, EPI_ISL_3118196, EPI_ISL_3118197, EPI_ISL_3118199, EPI_ISL_3118200, EPI_ISL_3118201, EPI_ISL_3118202, EPI_ISL_3118203, EPI_ISL_3118204, EPI_ISL_3118205, EPI_ISL_3118206, EPI_ISL_3118207, EPI_ISL_3118208, EPI_ISL_3118209, EPI_ISL_3118210, EPI_ISL_3118211, EPI_ISL_3118212, EPI_ISL_3118213, EPI_ISL_3118214, EPI_ISL_3118215 | | | |
| see above | National Virus Reference Laboratory | National Virus Reference Laboratory | Zoe Yandle, Charlene Bennett, Gabriel Gonzalez, Michael Carr, Jonathan Dean, Cillian F De Gascun |
| EPI_ISL_3118216, EPI_ISL_3118217, EPI_ISL_3118218, EPI_ISL_3118219, EPI_ISL_3118220, EPI_ISL_3118221 | Oddělení klinické mikrobiologie | Institute of Medical Microbiology and Virology, University Hospital Carl Gustav Carus, TU Dresden | Marlena Stadtmüller, Alexa Laubner, Johanna Beil, Fabian Rost, Sylvia Klemroth, Grit Mehnert, Montserrat Palau de Miquel, Susanne Reinhardt, Leo Büttner, Anett Zabzinski, Sylke Winkler, Alexander Dalpke, Eva Patrasová, Lenka Šimůnková, Lenka Dvořáková |
| EPI_ISL_3118222, EPI_ISL_3118223, EPI_ISL_3118224 | Zdravotní ústav Ústí nad Labem | Institute of Medical Microbiology and Virology, University Hospital Carl Gustav Carus, TU Dresden | Marlena Stadtmüller, Alexa Laubner, Johanna Beil, Fabian Rost, Sylvia Klemroth, Grit Mehnert, Montserrat Palau de Miquel, Susanne Reinhardt, Leo Büttner, Anett Zabzinski, Sylke Winkler, Alexander Dalpke, Romana Mikešová, Jitka Pohořská, Ivana Stiborová, Eva Patrasová, Lenka Šimůnková |
| EPI_ISL_3118274, EPI_ISL_3118275, EPI_ISL_3118276, EPI_ISL_3118277, EPI_ISL_3118278, EPI_ISL_3118279, EPI_ISL_3118280, EPI_ISL_3118281, EPI_ISL_3118282, EPI_ISL_3118283 | CLILAB | Microbiology Department | Sara Marti, Aida Gonzalez-Diaz, Yolanda Hernandez, Laura Calatayud, Jordi Niubó, Carmen Ardanuy, Jordi Camara, M Angeles Domínguez |
| EPI_ISL_3118285, EPI_ISL_3118286, EPI_ISL_3118287, EPI_ISL_3118288, EPI_ISL_3118289, EPI_ISL_3118290, EPI_ISL_3118291, EPI_ISL_3118292, EPI_ISL_3118293, EPI_ISL_3118294, EPI_ISL_3118295, EPI_ISL_3118296, EPI_ISL_3118297, EPI_ISL_3118298, EPI_ISL_3118299, EPI_ISL_3118300, EPI_ISL_3118301, EPI_ISL_3118302, EPI_ISL_3118303, EPI_ISL_3118304, EPI_ISL_3118305, EPI_ISL_3118306, EPI_ISL_3118307, EPI_ISL_3118308, EPI_ISL_3118309, EPI_ISL_3118310, EPI_ISL_3118311, EPI_ISL_3118312, EPI_ISL_3118313, EPI_ISL_3118314, EPI_ISL_3118315, EPI_ISL_3118316, EPI_ISL_3118317, EPI_ISL_3118319, EPI_ISL_3118320, EPI_ISL_3118321, EPI_ISL_3118322, EPI_ISL_3118323, EPI_ISL_3118324, EPI_ISL_3118325, EPI_ISL_3118326, EPI_ISL_3118327, EPI_ISL_3118328, EPI_ISL_3118329, EPI_ISL_3118330, EPI_ISL_3118331, EPI_ISL_3118332, EPI_ISL_3118333, EPI_ISL_3118334, EPI_ISL_3118336, EPI_ISL_3118337, EPI_ISL_3118338, EPI_ISL_3118339, EPI_ISL_3118340, EPI_ISL_3118341, EPI_ISL_3118342, EPI_ISL_3118343, EPI_ISL_3118344, EPI_ISL_3118345, EPI_ISL_3118346, EPI_ISL_3118347, EPI_ISL_3118348, EPI_ISL_3118350, EPI_ISL_3118351, EPI_ISL_3118352, EPI_ISL_3118353, EPI_ISL_3118354, EPI_ISL_3118355, EPI_ISL_3118357, EPI_ISL_3118359 | | | |
| see above | Hospital Universitat de Bellvitge | Microbiology Department | Sara Marti, Aida Gonzalez-Diaz, Yolanda Hernandez, Laura Calatayud, Jordi Niubó, Carmen Ardanuy, Jordi Camara, M Angeles Domínguez |
| EPI_ISL_3118365 | Center of Advanced Studies and Technology, Molecular Genetics Laboratory | Center of Advanced Studies and Technology, Molecular Genetics Laboratory | Anaclerio Federico, De Fabritiis Simone |
| EPI_ISL_3118366, EPI_ISL_3118367, EPI_ISL_3118368, EPI_ISL_3118369 | Banteay Meanchey Regional Laboratory | Virology Unit, Institut Pasteur du Cambodge | Leakhena Pum, Sophoannadedh Rath, Cecile Troupin, Ly Sovann, Kraing Sidonn, Yi Sengdoeurn, Chin Savuth, Chau Darapheak, Veasna Duong, Erik A Karlsson |
| EPI_ISL_3118370, EPI_ISL_3118371, EPI_ISL_3118372 | Battambang Repid Response Team | Virology Unit, Institut Pasteur du Cambodge | Leakhena Pum, Sophoannadedh Rath, Cecile Troupin, Ly Sovann, Kraing Sidonn, Yi Sengdoeurn, Chin Savuth, Chau Darapheak, Veasna Duong, Erik A Karlsson |
| EPI_ISL_3118373 | Kampong Cham Rapid Response Team | Virology Unit, Institut Pasteur du Cambodge | Leakhena Pum, Sophoannadedh Rath, Cecile Troupin, Ly Sovann, Kraing Sidonn, Yi Sengdoeurn, Chin Savuth, Chau Darapheak, Veasna Duong, Erik A Karlsson |
| EPI_ISL_3118374 | Kampomg Thom Rapid Response Team | Virology Unit, Institut Pasteur du Cambodge | Leakhena Pum, Sophoannadedh Rath, Cecile Troupin, Ly Sovann, Kraing Sidonn, Yi Sengdoeurn, Chin Savuth, Chau Darapheak, Veasna Duong, Erik A Karlsson |
| EPI_ISL_3118375, EPI_ISL_3118376 | Pailin Rapid Response Team | Virology Unit, Institut Pasteur du Cambodge | Leakhena Pum, Sophoannadedh Rath, Cecile Troupin, Ly Sovann, Kraing Sidonn, Yi Sengdoeurn, Chin Savuth, Chau Darapheak, Veasna Duong, Erik A Karlsson |
| EPI_ISL_3118378, EPI_ISL_3118379, EPI_ISL_3118380, | Oddat Meanchey Rapid Response Team | Virology Unit, Institut Pasteur du Cambodge | Leakhena Pum, Sophoannadedh Rath, Cecile Troupin, Ly Sovann, Kraing Sidonn, Yi Sengdoeurn, Chin Savuth, Chau Darapheak, Veasna Duong, Erik A Karlsson |

| | | | |
|---|---|---|---|
| EPI_ISL_3118381, EPI_ISL_3118382, EPI_ISL_3118383 | | | |
| EPI_ISL_3118385, EPI_ISL_3118396, EPI_ISL_3118397, EPI_ISL_3118398, EPI_ISL_3118402, EPI_ISL_3118403, EPI_ISL_3118405 | Virology Unit, Institut Pasteur du Cambodge | Virology Unit, Institut Pasteur du Cambodge | Leakhena Pum, Sophoannadedh Rath, Cecile Troupin, Ly Sovann, Kraing Sidonn, Yi Sengdoeurn, Chin Savuth, Chau Darapheak, Veasna Duong, Erik A Karlsson |
| EPI_ISL_3118406, EPI_ISL_3118407 | Prey Veng Rapid Response Team | Virology Unit, Institut Pasteur du Cambodge | Leakhena Pum, Sophoannadedh Rath, Cecile Troupin, Ly Sovann, Kraing Sidonn, Yi Sengdoeurn, Chin Savuth, Chau Darapheak, Veasna Duong, Erik A Karlsson |
| EPI_ISL_3118408, EPI_ISL_3118409, EPI_ISL_3118410, EPI_ISL_3118411 | Pailin Rapid Response Team | Virology Unit, Institut Pasteur du Cambodge | Leakhena Pum, Sophoannadedh Rath, Cecile Troupin, Ly Sovann, Kraing Sidonn, Yi Sengdoeurn, Chin Savuth, Chau Darapheak, Veasna Duong, Erik A Karlsson |
| EPI_ISL_3118460, EPI_ISL_3118461 | Lafene Health Center | Kansas State Veterinary Diagnostic Laboratory | Tyler Doerksen, Andreea Lu, Lance Noll, Elizabeth Porter, Emily Cox, Jianfa Bai, Jamie Henningson, Rachel Palinski |
| EPI_ISL_3118495, EPI_ISL_3118496 | Mayo Clinic & Mayo Clinic Laboratories | Minnesota Department of Health, Public Health Laboratory | Alexandra Lorentz, Jacob Garfin, Matt Plumb, and Xiong Wang |
| EPI_ISL_3118497, EPI_ISL_3118498, EPI_ISL_3118499, EPI_ISL_3118500, EPI_ISL_3118501, EPI_ISL_3118502 | Allina Health Laboratory | Minnesota Department of Health, Public Health Laboratory | Alexandra Lorentz, Jacob Garfin, Matt Plumb, and Xiong Wang |
| EPI_ISL_3118503, EPI_ISL_3118504, EPI_ISL_3118505 | Minnesota Department of Health, Public Health Laboratory | Minnesota Department of Health, Public Health Laboratory | Alexandra Lorentz, Jacob Garfin, Matt Plumb, and Xiong Wang |
| EPI_ISL_3118507, EPI_ISL_3118508, EPI_ISL_3118509, EPI_ISL_3118510 | Regions Hospital | Minnesota Department of Health, Public Health Laboratory | Alexandra Lorentz, Jacob Garfin, Matt Plumb, and Xiong Wang |
| EPI_ISL_3118511, EPI_ISL_3118512 | Hennepin County Medical Center | Minnesota Department of Health, Public Health Laboratory | Alexandra Lorentz, Jacob Garfin, Matt Plumb, and Xiong Wang |
| EPI_ISL_3118513 | Centracare Laboratory Services | Minnesota Department of Health, Public Health Laboratory | Alexandra Lorentz, Jacob Garfin, Matt Plumb, and Xiong Wang |
| EPI_ISL_3118514, EPI_ISL_3118515 | Minnesota Department of Health, Public Health Laboratory | Minnesota Department of Health, Public Health Laboratory | Alexandra Lorentz, Jacob Garfin, Matt Plumb, and Xiong Wang |
| EPI_ISL_3118516, EPI_ISL_3118517 | Essentia Health-St. Mary's Medical Center | Minnesota Department of Health, Public Health Laboratory | Alexandra Lorentz, Jacob Garfin, Matt Plumb, and Xiong Wang |
| EPI_ISL_3118518, EPI_ISL_3118519 | Park Nicollet Methodist Hospital | Minnesota Department of Health, Public Health Laboratory | Alexandra Lorentz, Jacob Garfin, Matt Plumb, and Xiong Wang |
| EPI_ISL_3118522 | Allina Health Laboratory | Minnesota Department of Health, Public Health Laboratory | Alexandra Lorentz, Jacob Garfin, Matt Plumb, and Xiong Wang |
| EPI_ISL_3118524 | Minnesota Department of Health, Public Health Laboratory | Minnesota Department of Health, Public Health Laboratory | Alexandra Lorentz, Jacob Garfin, Matt Plumb, and Xiong Wang |
| EPI_ISL_3118525 | Mayo Clinic & Mayo Clinic Laboratories | Minnesota Department of Health, Public Health Laboratory | Alexandra Lorentz, Jacob Garfin, Matt Plumb, and Xiong Wang |
| EPI_ISL_3118526, EPI_ISL_3118527, EPI_ISL_3118529 | HealthPartners Central Lab | Minnesota Department of Health, Public Health Laboratory | Alexandra Lorentz, Jacob Garfin, Matt Plumb, and Xiong Wang |
| EPI_ISL_3118530, EPI_ISL_3118531, EPI_ISL_3118532, EPI_ISL_3118533, EPI_ISL_3118534, EPI_ISL_3118536, EPI_ISL_3118541, EPI_ISL_3118542, EPI_ISL_3118543, EPI_ISL_3118544, EPI_ISL_3118545, EPI_ISL_3118546, EPI_ISL_3118547, EPI_ISL_3118548, EPI_ISL_3118549, EPI_ISL_3118550, EPI_ISL_3118551, EPI_ISL_3118552, EPI_ISL_3118553, EPI_ISL_3118554, EPI_ISL_3118555, EPI_ISL_3118556, EPI_ISL_3118557, EPI_ISL_3118558, EPI_ISL_3118559, EPI_ISL_3118560, EPI_ISL_3118561, EPI_ISL_3118562, EPI_ISL_3118563, EPI_ISL_3118564, EPI_ISL_3118565, EPI_ISL_3118567, EPI_ISL_3118568 | | | |
| see above | M Health Fairview | Minnesota Department of Health, Public Health Laboratory | Alexandra Lorentz, Jacob Garfin, Matt Plumb, and Xiong Wang |
| EPI_ISL_3118569, EPI_ISL_3118570, EPI_ISL_3118571, EPI_ISL_3118575, EPI_ISL_3118577, EPI_ISL_3118579, EPI_ISL_3118581, EPI_ISL_3118582, EPI_ISL_3118583, EPI_ISL_3118584, EPI_ISL_3118585, EPI_ISL_3118586, EPI_ISL_3118587, EPI_ISL_3118588, EPI_ISL_3118589, EPI_ISL_3118613, EPI_ISL_3118618, EPI_ISL_3118629, EPI_ISL_3118630, EPI_ISL_3118638, EPI_ISL_3118639, EPI_ISL_3118640, EPI_ISL_3118641, EPI_ISL_3118642, EPI_ISL_3118643, EPI_ISL_3118644, EPI_ISL_3118645, EPI_ISL_3118646, EPI_ISL_3118647, EPI_ISL_3118648 | | | |
| see above | Alaska State Virology Laboratory | Alaska State Virology Laboratory | Stephanie DeRonde, Elva House, Jacob Zidek, Lisa Smith, Ph.D., Jack Chen, Ph.D. |
| EPI_ISL_3118687, EPI_ISL_3118688 | AHRI- Alex Sigal lab (Sandile) | KRISP, KZn Research Innovation and Sequencing Platform | Sigal A, Sandile C, Farina K, Khadija K, Mallory B, Giandhari J, Pillay S, Yajna R, Lessells R, Mdlalose K, York D, Khan S, Emmanuel SJ, Tegally H, Wilkinson E, de Oliveira T |
| EPI_ISL_3118697 | Netcare | KRISP, KZn Research Innovation and Sequencing Platform | Omar S, Brown MJ, Wadula J, Giandhari J, Pillay S, Yajna R, Lessells R, Emmanuel SJ, Tegally H, Wilkinson E, de Oliveira T |
| EPI_ISL_3118698, EPI_ISL_3118699, EPI_ISL_3118700, EPI_ISL_3118701, EPI_ISL_3118702, EPI_ISL_3118703, EPI_ISL_3118704, EPI_ISL_3118705, EPI_ISL_3118706, EPI_ISL_3118707, EPI_ISL_3118708, EPI_ISL_3118709, EPI_ISL_3118710, EPI_ISL_3118711, EPI_ISL_3118712, EPI_ISL_3118713, EPI_ISL_3118714, EPI_ISL_3118715, EPI_ISL_3118716, EPI_ISL_3118717, EPI_ISL_3118718, EPI_ISL_3118721, EPI_ISL_3118722, EPI_ISL_3118723, EPI_ISL_3118724, EPI_ISL_3118725, EPI_ISL_3118726, EPI_ISL_3118727, EPI_ISL_3118728, EPI_ISL_3118729, EPI_ISL_3118730, EPI_ISL_3118731, EPI_ISL_3118732, EPI_ISL_3118733, EPI_ISL_3118734, EPI_ISL_3118735, EPI_ISL_3118736, EPI_ISL_3118737, EPI_ISL_3118738, EPI_ISL_3118739, EPI_ISL_3118740, EPI_ISL_3118741, EPI_ISL_3118742, EPI_ISL_3118743, EPI_ISL_3118744, EPI_ISL_3118745, EPI_ISL_3118746, EPI_ISL_3118747, EPI_ISL_3118748, EPI_ISL_3118749, EPI_ISL_3118750, EPI_ISL_3118751, EPI_ISL_3118753, EPI_ISL_3118754, EPI_ISL_3118755, EPI_ISL_3118756, EPI_ISL_3118757, EPI_ISL_3118758, EPI_ISL_3118759, EPI_ISL_3118760, EPI_ISL_3118761, EPI_ISL_3118762, EPI_ISL_3118763, EPI_ISL_3118764, EPI_ISL_3118765, EPI_ISL_3118766, EPI_ISL_3118767, EPI_ISL_3118768, EPI_ISL_3118769, EPI_ISL_3118770, EPI_ISL_3118771, EPI_ISL_3118772, EPI_ISL_3118773, EPI_ISL_3118774, EPI_ISL_3118775, EPI_ISL_3118776, EPI_ISL_3118777, EPI_ISL_3118778, EPI_ISL_3118779, EPI_ISL_3118780, EPI_ISL_3118781, EPI_ISL_3118782, EPI_ISL_3118783 | | | |
| see above | Division of Medical Virology, National Health Laboratory Service (NHLS), Tygerberg Hospital / Stellenbosch University | Division of Medical Virology, National Health Laboratory Service (NHLS), Tygerberg Hospital / Stellenbosch University | Susan Engelbrecht, Tongai Maponga, Tania Stander, Kamela Mahlakwane, Gert van Zyl, Wolfgang Preiser |
| EPI_ISL_3119268, EPI_ISL_3119269, EPI_ISL_3119270, EPI_ISL_3119271, EPI_ISL_3119272, EPI_ISL_3119273, EPI_ISL_3119274, EPI_ISL_3119275, EPI_ISL_3119277, EPI_ISL_3119278, EPI_ISL_3119279, EPI_ISL_3119280, EPI_ISL_3119281, EPI_ISL_3119282, EPI_ISL_3119283, EPI_ISL_3119284, EPI_ISL_3119285, EPI_ISL_3119286, EPI_ISL_3119287, EPI_ISL_3119288, EPI_ISL_3119289, EPI_ISL_3119290, EPI_ISL_3119291, EPI_ISL_3119292, EPI_ISL_3119293, EPI_ISL_3119294, EPI_ISL_3119295, EPI_ISL_3119296, EPI_ISL_3119297, EPI_ISL_3119298, EPI_ISL_3119299, EPI_ISL_3119300, EPI_ISL_3119301, EPI_ISL_3119302, EPI_ISL_3119303, EPI_ISL_3119304, EPI_ISL_3119305, EPI_ISL_3119306, EPI_ISL_3119307, EPI_ISL_3119308, EPI_ISL_3119309, EPI_ISL_3119310, EPI_ISL_3119311, EPI_ISL_3119312, EPI_ISL_3119313, EPI_ISL_3119314, EPI_ISL_3119315, EPI_ISL_3119317, EPI_ISL_3119318, EPI_ISL_3119319, EPI_ISL_3119320, EPI_ISL_3119321, EPI_ISL_3119322, EPI_ISL_3119323, EPI_ISL_3119324, EPI_ISL_3119325, EPI_ISL_3119326, EPI_ISL_3119327, EPI_ISL_3119328, EPI_ISL_3119329, EPI_ISL_3119330, EPI_ISL_3119331, EPI_ISL_3119332, EPI_ISL_3119333, EPI_ISL_3119334, EPI_ISL_3119335, EPI_ISL_3119336, EPI_ISL_3119337, EPI_ISL_3119338, EPI_ISL_3119339, EPI_ISL_3119340, EPI_ISL_3119341, EPI_ISL_3119342, EPI_ISL_3119343, EPI_ISL_3119344, EPI_ISL_3119345, EPI_ISL_3119346, EPI_ISL_3119347, EPI_ISL_3119348, EPI_ISL_3119349, EPI_ISL_3119350, EPI_ISL_3119351, EPI_ISL_3119352, EPI_ISL_3119353, EPI_ISL_3119354, EPI_ISL_3119355, EPI_ISL_3119356, EPI_ISL_3119357 | | | |
| see above | Lifebrain Covid Labor GmbH | Dept. of Laboratory Medicine | Harald Esterbauer, Anna Gschaider, Petra Jurkowitsch, Fabian Konig, Sabina Plumer, Robert Strassl, Claudia Weber, Oswald Wagner |
| EPI_ISL_3119424, EPI_ISL_3119425, EPI_ISL_3119426, EPI_ISL_3119427, EPI_ISL_3119428, EPI_ISL_3119430, EPI_ISL_3119431, EPI_ISL_3119432, EPI_ISL_3119433, EPI_ISL_3119434, EPI_ISL_3119435, EPI_ISL_3119436, EPI_ISL_3119437, EPI_ISL_3119438, EPI_ISL_3119439, EPI_ISL_3119440, EPI_ISL_3119441, EPI_ISL_3119442, EPI_ISL_3119443, EPI_ISL_3119444, EPI_ISL_3119445, EPI_ISL_3119446, EPI_ISL_3119447, EPI_ISL_3119448, EPI_ISL_3119449, EPI_ISL_3119450, EPI_ISL_3119451, EPI_ISL_3119452, EPI_ISL_3119453, EPI_ISL_3119454, EPI_ISL_3119455, EPI_ISL_3119456, EPI_ISL_3119457, EPI_ISL_3119458, EPI_ISL_3119459, EPI_ISL_3119460 | | | |
| see above | Dept. of Microbiology and Infection Control, Akershus University Hospital HF | Dept. of Microbiology and Infection Control, Akershus University Hospital HF | Hege Vangstein Aamot, Alexander Hesselberg Løvestad |
| EPI_ISL_3119461, EPI_ISL_3119462, EPI_ISL_3119463 | Laverty Pathology | NSW Health Pathology - Institute of Clinical Pathology and Medical Research; Westmead Hospital; University of Sydney | CIDM-PH et al. |
| EPI_ISL_3119464 | Sydney South West Pathology Service (SSWPS) - Liverpool Hospital - NSW Health Pathology | NSW Health Pathology - Institute of Clinical Pathology and Medical Research; Westmead Hospital; University of Sydney | CIDM-PH et al. |
| EPI_ISL_3119465 | St Vincent's Pathology (SydPath) | NSW Health Pathology - Institute of Clinical Pathology and Medical Research; Westmead Hospital; University of Sydney | CIDM-PH et al. |
| EPI_ISL_3119466 | 4Cyte Pathology | NSW Health Pathology - Institute of Clinical Pathology and Medical Research; Westmead Hospital; University of Sydney | CIDM-PH et al. |
| EPI_ISL_3119467, EPI_ISL_3119469, EPI_ISL_3119470, EPI_ISL_3119472, EPI_ISL_3119473, EPI_ISL_3119474, EPI_ISL_3119475, EPI_ISL_3119476, EPI_ISL_3119477, EPI_ISL_3119478, EPI_ISL_3119480, EPI_ISL_3119481, EPI_ISL_3119482, EPI_ISL_3119483, EPI_ISL_3119484, EPI_ISL_3119485, EPI_ISL_3119486, EPI_ISL_3119487, EPI_ISL_3119488, EPI_ISL_3119489, EPI_ISL_3119490, EPI_ISL_3119491, EPI_ISL_3119492, EPI_ISL_3119493, EPI_ISL_3119494, EPI_ISL_3119495, EPI_ISL_3119496, EPI_ISL_3119497, EPI_ISL_3119498, EPI_ISL_3119499, EPI_ISL_3119500, EPI_ISL_3119501, EPI_ISL_3119502, EPI_ISL_3119503, EPI_ISL_3119505, EPI_ISL_3119508, EPI_ISL_3119509, EPI_ISL_3119510, EPI_ISL_3119511, EPI_ISL_3119512, EPI_ISL_3119513, EPI_ISL_3119514, EPI_ISL_3119515, EPI_ISL_3119516, EPI_ISL_3119517, EPI_ISL_3119518, EPI_ISL_3119519, EPI_ISL_3119520, EPI_ISL_3119521, EPI_ISL_3119522, EPI_ISL_3119524, EPI_ISL_3119525, EPI_ISL_3119526, EPI_ISL_3119527, EPI_ISL_3119530, EPI_ISL_3119531, EPI_ISL_3119533, EPI_ISL_3119535, EPI_ISL_3119539, EPI_ISL_3119540, EPI_ISL_3119541, EPI_ISL_3119542, EPI_ISL_3119543, EPI_ISL_3119545, EPI_ISL_3119547, EPI_ISL_3119567, EPI_ISL_3119568, EPI_ISL_3119569, EPI_ISL_3119570, EPI_ISL_3119573, EPI_ISL_3119574, EPI_ISL_3119575, EPI_ISL_3119576, EPI_ISL_3119577, EPI_ISL_3119578, EPI_ISL_3119579, EPI_ISL_3119582, EPI_ISL_3119586, EPI_ISL_3119587, EPI_ISL_3119589, EPI_ISL_3119590, EPI_ISL_3119591, EPI_ISL_3119592, EPI_ISL_3119593, EPI_ISL_3119594, EPI_ISL_3119595, EPI_ISL_3119596, EPI_ISL_3119597, EPI_ISL_3119598, EPI_ISL_3119599, EPI_ISL_3119600, EPI_ISL_3119601, EPI_ISL_3119602, EPI_ISL_3119603, EPI_ISL_3119604, EPI_ISL_3119605, EPI_ISL_3119606, EPI_ISL_3119607, EPI_ISL_3119608, EPI_ISL_3119609, EPI_ISL_3119610, EPI_ISL_3119611, EPI_ISL_3119612, EPI_ISL_3119613, EPI_ISL_3119614, EPI_ISL_3119615, EPI_ISL_3119616, EPI_ISL_3119617, EPI_ISL_3119618, EPI_ISL_3119619, EPI_ISL_3119620, EPI_ISL_3119621, EPI_ISL_3119622, EPI_ISL_3119623, EPI_ISL_3119624, EPI_ISL_3119625, EPI_ISL_3119626, EPI_ISL_3119627, EPI_ISL_3119628, EPI_ISL_3119629, EPI_ISL_3119630, EPI_ISL_3119631, EPI_ISL_3119632, EPI_ISL_3119634, EPI_ISL_3119635, EPI_ISL_3119636, EPI_ISL_3119637, EPI_ISL_3119638, EPI_ISL_3119639, EPI_ISL_3119640, EPI_ISL_3119641, EPI_ISL_3119642, EPI_ISL_3119643, EPI_ISL_3119644, EPI_ISL_3119645, EPI_ISL_3119646, EPI_ISL_3119647, EPI_ISL_3119648, EPI_ISL_3119649, EPI_ISL_3119650, EPI_ISL_3119651, EPI_ISL_3119652, EPI_ISL_3119653, EPI_ISL_3119654, EPI_ISL_3119655, EPI_ISL_3119656, EPI_ISL_3119657, EPI_ISL_3119658, EPI_ISL_3119659, EPI_ISL_3119660, EPI_ISL_3119661, EPI_ISL_3119662, EPI_ISL_3119663, EPI_ISL_3119664, EPI_ISL_3119665, EPI_ISL_3119666, EPI_ISL_3119667, EPI_ISL_3119668, EPI_ISL_3119669, EPI_ISL_3119670, EPI_ISL_3119671, EPI_ISL_3119672, EPI_ISL_3119673, EPI_ISL_3119674, EPI_ISL_3119675, EPI_ISL_3119676, EPI_ISL_3119677, EPI_ISL_3119678, EPI_ISL_3119679, EPI_ISL_3119680, EPI_ISL_3119681, EPI_ISL_3119682, EPI_ISL_3119683, EPI_ISL_3119684, EPI_ISL_3119685, EPI_ISL_3119686, EPI_ISL_3119687, EPI_ISL_3119688, EPI_ISL_3119689, EPI_ISL_3119691, EPI_ISL_3119692, EPI_ISL_3119693, EPI_ISL_3119694 | | | |
| see above | Indian Council of Medical Research-National Institute of Virology, Microbial Containment Complex | Indian Council of Medical Research-National Institute of Virology, Microbial Containment Complex | Pragya D Yadav |
| EPI_ISL_3119695, EPI_ISL_3119696, EPI_ISL_3119697, EPI_ISL_3119698, EPI_ISL_3119699, EPI_ISL_3119700, EPI_ISL_3119701, EPI_ISL_3119702, EPI_ISL_3119703, EPI_ISL_3119704, EPI_ISL_3119705, EPI_ISL_3119706, EPI_ISL_3119707 | | | |
| see above | AULSS 2 Marca Trevigiana | Istituto Zooprofilattico Sperimentale delle Venezie | Adelaide Milani, Alessia Schivo, Annalisa Salviato, Elisa Palumbo, Erika Giorgia Quaranta, Luca Tassoni, Ambra Pastori, Edoardo Giussani, Alice Fusaro, Isabella Monne, Calogero Terregino, Antonia Ricci |

| | | | |
|---|---|---|---|
| EPI_ISL_3119708, EPI_ISL_3119709, EPI_ISL_3119710, EPI_ISL_3119711, EPI_ISL_3119712, EPI_ISL_3119713, EPI_ISL_3119714, EPI_ISL_3119715 | AULSS 1 Dolomiti | Istituto Zooprofilattico Sperimentale delle Venezie | Adelaide Milani, Alessia Schivo, Annalisa Salviato, Elisa Palumbo, Erika Giorgia Quaranta, Luca Tassoni, Ambra Pastori, Edoardo Giussani, Alice Fusaro, Isabella Monne, Calogero Terregino, Antonia Ricci |
| EPI_ISL_3119716, EPI_ISL_3119717, EPI_ISL_3119718, EPI_ISL_3119720, EPI_ISL_3119721, EPI_ISL_3119722, EPI_ISL_3119723, EPI_ISL_3119724, EPI_ISL_3119725, EPI_ISL_3119726, EPI_ISL_3119727, EPI_ISL_3119728, EPI_ISL_3119729 | | | |
| see above | AULSS 3 Venezia | Istituto Zooprofilattico Sperimentale delle Venezie | Adelaide Milani, Alessia Schivo, Annalisa Salviato, Elisa Palumbo, Erika Giorgia Quaranta, Luca Tassoni, Ambra Pastori, Edoardo Giussani, Alice Fusaro, Isabella Monne, Calogero Terregino, Antonia Ricci |
| EPI_ISL_3119730, EPI_ISL_3119731, EPI_ISL_3119732, EPI_ISL_3119733, EPI_ISL_3119734, EPI_ISL_3119735, EPI_ISL_3119736, EPI_ISL_3119737, EPI_ISL_3119738 | AULSS 9 Scaligera - San Bonifacio | Istituto Zooprofilattico Sperimentale delle Venezie | Adelaide Milani, Alessia Schivo, Annalisa Salviato, Elisa Palumbo, Erika Giorgia Quaranta, Luca Tassoni, Ambra Pastori, Edoardo Giussani, Alice Fusaro, Isabella Monne, Calogero Terregino, Antonia Ricci |
| EPI_ISL_3119739, EPI_ISL_3119740, EPI_ISL_3119741, EPI_ISL_3119742, EPI_ISL_3119743, EPI_ISL_3119744, EPI_ISL_3119745, EPI_ISL_3119746, EPI_ISL_3119747, EPI_ISL_3119748, EPI_ISL_3119749, EPI_ISL_3119750, EPI_ISL_3119751, EPI_ISL_3119752 | | | |
| see above | AULSS 8 Berica | Istituto Zooprofilattico Sperimentale delle Venezie | Adelaide Milani, Alessia Schivo, Annalisa Salviato, Elisa Palumbo, Erika Giorgia Quaranta, Luca Tassoni, Ambra Pastori, Edoardo Giussani, Alice Fusaro, Isabella Monne, Calogero Terregino, Antonia Ricci |
| EPI_ISL_3119753, EPI_ISL_3119754, EPI_ISL_3119755, EPI_ISL_3119756, EPI_ISL_3119757, EPI_ISL_3119758, EPI_ISL_3119759, EPI_ISL_3119760, EPI_ISL_3119761, EPI_ISL_3119762, EPI_ISL_3119763, EPI_ISL_3119764 | | | |
| see above | AULSS 9 Scaligera - Legnago | Istituto Zooprofilattico Sperimentale delle Venezie | Adelaide Milani, Alessia Schivo, Annalisa Salviato, Elisa Palumbo, Erika Giorgia Quaranta, Luca Tassoni, Ambra Pastori, Edoardo Giussani, Alice Fusaro, Isabella Monne, Calogero Terregino, Antonia Ricci |
| EPI_ISL_3119765, EPI_ISL_3119766 | AULSS 6 Euganea | Istituto Zooprofilattico Sperimentale delle Venezie | Adelaide Milani, Alessia Schivo, Annalisa Salviato, Elisa Palumbo, Erika Giorgia Quaranta, Luca Tassoni, Ambra Pastori, Edoardo Giussani, Alice Fusaro, Isabella Monne, Calogero Terregino, Antonia Ricci |
| EPI_ISL_3119767, EPI_ISL_3119768 | AULSS 9 Scaligera - Legnago | Istituto Zooprofilattico Sperimentale delle Venezie | Adelaide Milani, Alessia Schivo, Annalisa Salviato, Elisa Palumbo, Erika Giorgia Quaranta, Luca Tassoni, Ambra Pastori, Edoardo Giussani, Alice Fusaro, Isabella Monne, Calogero Terregino, Antonia Ricci |
| EPI_ISL_3119769, EPI_ISL_3119770, EPI_ISL_3119771, EPI_ISL_3119772, EPI_ISL_3119773, EPI_ISL_3119774, EPI_ISL_3119775, EPI_ISL_3119776, EPI_ISL_3119777, EPI_ISL_3119778, EPI_ISL_3119779, EPI_ISL_3119780 | | | |
| see above | AULSS 6 Euganea | Istituto Zooprofilattico Sperimentale delle Venezie | Adelaide Milani, Alessia Schivo, Annalisa Salviato, Elisa Palumbo, Erika Giorgia Quaranta, Luca Tassoni, Ambra Pastori, Edoardo Giussani, Alice Fusaro, Isabella Monne, Calogero Terregino, Antonia Ricci |
| EPI_ISL_3119781, EPI_ISL_3119782, EPI_ISL_3119783, EPI_ISL_3119784, EPI_ISL_3119785, EPI_ISL_3119786, EPI_ISL_3119787, EPI_ISL_3119788, EPI_ISL_3119789, EPI_ISL_3119790, EPI_ISL_3119791, EPI_ISL_3119792, EPI_ISL_3119793, EPI_ISL_3119794 | | | |
| see above | AULSS 9 Scaligera - Villafranca | Istituto Zooprofilattico Sperimentale delle Venezie | Adelaide Milani, Alessia Schivo, Annalisa Salviato, Elisa Palumbo, Erika Giorgia Quaranta, Luca Tassoni, Ambra Pastori, Edoardo Giussani, Alice Fusaro, Isabella Monne, Calogero Terregino, Antonia Ricci |
| EPI_ISL_3119795, EPI_ISL_3119796, EPI_ISL_3119797, EPI_ISL_3119798, EPI_ISL_3119799 | AULSS 7 Pedemontana - Santorso | Istituto Zooprofilattico Sperimentale delle Venezie | Adelaide Milani, Alessia Schivo, Annalisa Salviato, Elisa Palumbo, Erika Giorgia Quaranta, Luca Tassoni, Ambra Pastori, Edoardo Giussani, Alice Fusaro, Isabella Monne, Calogero Terregino, Antonia Ricci |
| EPI_ISL_3119800 | AULSS 7 Pedemontana - Bassano | Istituto Zooprofilattico Sperimentale delle Venezie | Adelaide Milani, Alessia Schivo, Annalisa Salviato, Elisa Palumbo, Erika Giorgia Quaranta, Luca Tassoni, Ambra Pastori, Edoardo Giussani, Alice Fusaro, Isabella Monne, Calogero Terregino, Antonia Ricci |
| EPI_ISL_3119801, EPI_ISL_3119802, EPI_ISL_3119803, EPI_ISL_3119804, EPI_ISL_3119805 | AOUIVR | Istituto Zooprofilattico Sperimentale delle Venezie | Adelaide Milani, Alessia Schivo, Annalisa Salviato, Elisa Palumbo, Erika Giorgia Quaranta, Luca Tassoni, Ambra Pastori, Edoardo Giussani, Alice Fusaro, Isabella Monne, Calogero Terregino, Antonia Ricci |
| EPI_ISL_3119806, EPI_ISL_3119807, EPI_ISL_3119808, EPI_ISL_3119809, EPI_ISL_3119810, EPI_ISL_3119812, EPI_ISL_3119813, EPI_ISL_3119815, EPI_ISL_3119816, EPI_ISL_3119817, EPI_ISL_3119818, EPI_ISL_3119819, EPI_ISL_3119820, EPI_ISL_3119821, EPI_ISL_3119822, EPI_ISL_3119823 | | | |
| see above | AULSS 5 Polesana | Istituto Zooprofilattico Sperimentale delle Venezie | Adelaide Milani, Alessia Schivo, Annalisa Salviato, Elisa Palumbo, Erika Giorgia Quaranta, Luca Tassoni, Ambra Pastori, Edoardo Giussani, Alice Fusaro, Isabella Monne, Calogero Terregino, Antonia Ricci |
| EPI_ISL_3119824, EPI_ISL_3119825, EPI_ISL_3119826, EPI_ISL_3119827, EPI_ISL_3119828, EPI_ISL_3119829, EPI_ISL_3119830, EPI_ISL_3119831, EPI_ISL_3119832, EPI_ISL_3119833, EPI_ISL_3119834 | | | |
| see above | AOUIVR | Istituto Zooprofilattico Sperimentale delle Venezie | Adelaide Milani, Alessia Schivo, Annalisa Salviato, Elisa Palumbo, Erika Giorgia Quaranta, Luca Tassoni, Ambra Pastori, Edoardo Giussani, Alice Fusaro, Isabella Monne, Calogero Terregino, Antonia Ricci |
| EPI_ISL_3119835, EPI_ISL_3119836, EPI_ISL_3119837, EPI_ISL_3119838, EPI_ISL_3119839, EPI_ISL_3119840, EPI_ISL_3119841, EPI_ISL_3119843, EPI_ISL_3119844 | AULSS 9 Scaligera - San Bonifacio | Istituto Zooprofilattico Sperimentale delle Venezie | Adelaide Milani, Alessia Schivo, Annalisa Salviato, Elisa Palumbo, Erika Giorgia Quaranta, Luca Tassoni, Ambra Pastori, Edoardo Giussani, Alice Fusaro, Isabella Monne, Calogero Terregino, Antonia Ricci |
| EPI_ISL_3119845, EPI_ISL_3119846, EPI_ISL_3119847, EPI_ISL_3119848, EPI_ISL_3119849, EPI_ISL_3119850 | AULSS 7 Pedemontana - Santorso | Istituto Zooprofilattico Sperimentale delle Venezie | Adelaide Milani, Alessia Schivo, Annalisa Salviato, Elisa Palumbo, Erika Giorgia Quaranta, Luca Tassoni, Ambra Pastori, Edoardo Giussani, Alice Fusaro, Isabella Monne, Calogero Terregino, Antonia Ricci |
| EPI_ISL_3119851 | AULSS 6 Euganea | Istituto Zooprofilattico Sperimentale delle Venezie | Adelaide Milani, Alessia Schivo, Annalisa Salviato, Elisa Palumbo, Erika Giorgia Quaranta, Luca Tassoni, Ambra Pastori, Edoardo Giussani, Alice Fusaro, Isabella Monne, Calogero Terregino, Antonia Ricci |
| EPI_ISL_3119852, EPI_ISL_3119853 | AULSS 7 Pedemontana - Santorso | Istituto Zooprofilattico Sperimentale delle Venezie | Adelaide Milani, Alessia Schivo, Annalisa Salviato, Elisa Palumbo, Erika Giorgia Quaranta, Luca Tassoni, Ambra Pastori, Edoardo Giussani, Alice Fusaro, Isabella Monne, Calogero Terregino, Antonia Ricci |
| EPI_ISL_3119854, EPI_ISL_3119855 | AULSS 8 Berica | Istituto Zooprofilattico Sperimentale delle Venezie | Adelaide Milani, Alessia Schivo, Annalisa Salviato, Elisa Palumbo, Erika Giorgia Quaranta, Luca Tassoni, Ambra Pastori, Edoardo Giussani, Alice Fusaro, Isabella Monne, Calogero Terregino, Antonia Ricci |
| EPI_ISL_3119856 | AULSS 6 Euganea | Istituto Zooprofilattico Sperimentale delle Venezie | Adelaide Milani, Alessia Schivo, Annalisa Salviato, Elisa Palumbo, Erika Giorgia Quaranta, Luca Tassoni, Ambra Pastori, Edoardo Giussani, Alice Fusaro, Isabella Monne, Calogero Terregino, Antonia Ricci |
| EPI_ISL_3119858 | AULSS 8 Berica | Istituto Zooprofilattico Sperimentale delle Venezie | Adelaide Milani, Alessia Schivo, Annalisa Salviato, Elisa Palumbo, Erika Giorgia Quaranta, Luca Tassoni, Ambra Pastori, Edoardo Giussani, Alice Fusaro, Isabella Monne, Calogero Terregino, Antonia Ricci |
| EPI_ISL_3119859 | AULSS 7 Pedemontana - Bassano | Istituto Zooprofilattico Sperimentale delle Venezie | Adelaide Milani, Alessia Schivo, Annalisa Salviato, Elisa Palumbo, Erika Giorgia Quaranta, Luca Tassoni, Ambra Pastori, Edoardo Giussani, Alice Fusaro, Isabella Monne, Calogero Terregino, Antonia Ricci |
| EPI_ISL_3119861 | AULSS 8 Berica | Istituto Zooprofilattico Sperimentale delle Venezie | Adelaide Milani, Alessia Schivo, Annalisa Salviato, Elisa Palumbo, Erika Giorgia Quaranta, Luca Tassoni, Ambra Pastori, Edoardo Giussani, Alice Fusaro, Isabella Monne, Calogero Terregino, Antonia Ricci |
| EPI_ISL_3119862 | AULSS 6 Euganea | Istituto Zooprofilattico Sperimentale delle Venezie | Adelaide Milani, Alessia Schivo, Annalisa Salviato, Elisa Palumbo, Erika Giorgia Quaranta, Luca Tassoni, Ambra Pastori, Edoardo Giussani, Alice Fusaro, Isabella Monne, Calogero Terregino, Antonia Ricci |
| EPI_ISL_3119863 | AULSS 7 Pedemontana - Bassano | Istituto Zooprofilattico Sperimentale delle Venezie | Adelaide Milani, Alessia Schivo, Annalisa Salviato, Elisa Palumbo, Erika Giorgia Quaranta, Luca Tassoni, Ambra Pastori, Edoardo Giussani, Alice Fusaro, Isabella Monne, Calogero Terregino, Antonia Ricci |
| EPI_ISL_3119864, EPI_ISL_3119865, EPI_ISL_3119866, EPI_ISL_3119867, EPI_ISL_3119868 | AULSS 8 Berica | Istituto Zooprofilattico Sperimentale delle Venezie | Adelaide Milani, Alessia Schivo, Annalisa Salviato, Elisa Palumbo, Erika Giorgia Quaranta, Luca Tassoni, Ambra Pastori, Edoardo Giussani, Alice Fusaro, Isabella Monne, Calogero Terregino, Antonia Ricci |
| EPI_ISL_3119869 | AULSS 6 Euganea | Istituto Zooprofilattico Sperimentale delle Venezie | Adelaide Milani, Alessia Schivo, Annalisa Salviato, Elisa Palumbo, Erika Giorgia Quaranta, Luca Tassoni, Ambra Pastori, Edoardo Giussani, Alice Fusaro, Isabella Monne, Calogero Terregino, Antonia Ricci |
| EPI_ISL_3119871, EPI_ISL_3119872, EPI_ISL_3119873, EPI_ISL_3119874, EPI_ISL_3119875, EPI_ISL_3119876, EPI_ISL_3119877 | AULSS 8 Berica | Istituto Zooprofilattico Sperimentale delle Venezie | Adelaide Milani, Alessia Schivo, Annalisa Salviato, Elisa Palumbo, Erika Giorgia Quaranta, Luca Tassoni, Ambra Pastori, Edoardo Giussani, Alice Fusaro, Isabella Monne, Calogero Terregino, Antonia Ricci |
| EPI_ISL_3119879 | AULSS 6 Euganea | Istituto Zooprofilattico Sperimentale delle Venezie | Adelaide Milani, Alessia Schivo, Annalisa Salviato, Elisa Palumbo, Erika Giorgia Quaranta, Luca Tassoni, Ambra Pastori, Edoardo Giussani, Alice Fusaro, Isabella Monne, Calogero Terregino, Antonia Ricci |
| EPI_ISL_3119880, EPI_ISL_3119881, EPI_ISL_3119882, EPI_ISL_3119883, EPI_ISL_3119884, EPI_ISL_3119885 | AULSS 8 Berica | Istituto Zooprofilattico Sperimentale delle Venezie | Adelaide Milani, Alessia Schivo, Annalisa Salviato, Elisa Palumbo, Erika Giorgia Quaranta, Luca Tassoni, Ambra Pastori, Edoardo Giussani, Alice Fusaro, Isabella Monne, Calogero Terregino, Antonia Ricci |
| EPI_ISL_3119886, | AULSS 6 Euganea | Istituto Zooprofilattico Sperimentale delle Venezie | Adelaide Milani, Alessia Schivo, Annalisa Salviato, Elisa Palumbo, Erika Giorgia Quaranta, Luca Tassoni, Ambra Pastori, Edoardo Giussani, Alice Fusaro, Isabella Monne, Calogero Terregino, Antonia Ricci |

| | | | |
|---|---|---|---|
| EPI_ISL_3119887 | | | |
| EPI_ISL_3119888 | AULSS 7 Pedemontana - Bassano | Istituto Zooprofilattico Sperimentale delle Venezie | Adelaide Milani, Alessia Schivo, Annalisa Salviato, Elisa Palumbo, Erika Giorgia Quaranta, Luca Tassoni, Ambra Pastori, Edoardo Giussani, Alice Fusaro, Isabella Monne, Calogero Terregino, Antonia Ricci |
| EPI_ISL_3119889 | AULSS 8 Berica | Istituto Zooprofilattico Sperimentale delle Venezie | Adelaide Milani, Alessia Schivo, Annalisa Salviato, Elisa Palumbo, Erika Giorgia Quaranta, Luca Tassoni, Ambra Pastori, Edoardo Giussani, Alice Fusaro, Isabella Monne, Calogero Terregino, Antonia Ricci |
| EPI_ISL_3119890 | AULSS 7 Pedemontana - Bassano | Istituto Zooprofilattico Sperimentale delle Venezie | Adelaide Milani, Alessia Schivo, Annalisa Salviato, Elisa Palumbo, Erika Giorgia Quaranta, Luca Tassoni, Ambra Pastori, Edoardo Giussani, Alice Fusaro, Isabella Monne, Calogero Terregino, Antonia Ricci |
| EPI_ISL_3119891, EPI_ISL_3119892, EPI_ISL_3119893, EPI_ISL_3119894 | AULSS 8 Berica | Istituto Zooprofilattico Sperimentale delle Venezie | Adelaide Milani, Alessia Schivo, Annalisa Salviato, Elisa Palumbo, Erika Giorgia Quaranta, Luca Tassoni, Ambra Pastori, Edoardo Giussani, Alice Fusaro, Isabella Monne, Calogero Terregino, Antonia Ricci |
| EPI_ISL_3119895 | AULSS 6 Euganea | Istituto Zooprofilattico Sperimentale delle Venezie | Adelaide Milani, Alessia Schivo, Annalisa Salviato, Elisa Palumbo, Erika Giorgia Quaranta, Luca Tassoni, Ambra Pastori, Edoardo Giussani, Alice Fusaro, Isabella Monne, Calogero Terregino, Antonia Ricci |
| EPI_ISL_3119896 | AULSS 8 Berica | Istituto Zooprofilattico Sperimentale delle Venezie | Adelaide Milani, Alessia Schivo, Annalisa Salviato, Elisa Palumbo, Erika Giorgia Quaranta, Luca Tassoni, Ambra Pastori, Edoardo Giussani, Alice Fusaro, Isabella Monne, Calogero Terregino, Antonia Ricci |
| EPI_ISL_3119897, EPI_ISL_3119898 | AULSS 7 Pedemontana - Bassano | Istituto Zooprofilattico Sperimentale delle Venezie | Adelaide Milani, Alessia Schivo, Annalisa Salviato, Elisa Palumbo, Erika Giorgia Quaranta, Luca Tassoni, Ambra Pastori, Edoardo Giussani, Alice Fusaro, Isabella Monne, Calogero Terregino, Antonia Ricci |
| EPI_ISL_3119899 | AULSS 6 Euganea | Istituto Zooprofilattico Sperimentale delle Venezie | Adelaide Milani, Alessia Schivo, Annalisa Salviato, Elisa Palumbo, Erika Giorgia Quaranta, Luca Tassoni, Ambra Pastori, Edoardo Giussani, Alice Fusaro, Isabella Monne, Calogero Terregino, Antonia Ricci |
| EPI_ISL_3119900, EPI_ISL_3119901, EPI_ISL_3119902 | AULSS 8 Berica | Istituto Zooprofilattico Sperimentale delle Venezie | Adelaide Milani, Alessia Schivo, Annalisa Salviato, Elisa Palumbo, Erika Giorgia Quaranta, Luca Tassoni, Ambra Pastori, Edoardo Giussani, Alice Fusaro, Isabella Monne, Calogero Terregino, Antonia Ricci |
| EPI_ISL_3119903, EPI_ISL_3119904 | AULSS 6 Euganea | Istituto Zooprofilattico Sperimentale delle Venezie | Adelaide Milani, Alessia Schivo, Annalisa Salviato, Elisa Palumbo, Erika Giorgia Quaranta, Luca Tassoni, Ambra Pastori, Edoardo Giussani, Alice Fusaro, Isabella Monne, Calogero Terregino, Antonia Ricci |
| EPI_ISL_3119905, EPI_ISL_3119906, EPI_ISL_3119907, EPI_ISL_3119908, EPI_ISL_3119909 | AULSS 8 Berica | Istituto Zooprofilattico Sperimentale delle Venezie | Adelaide Milani, Alessia Schivo, Annalisa Salviato, Elisa Palumbo, Erika Giorgia Quaranta, Luca Tassoni, Ambra Pastori, Edoardo Giussani, Alice Fusaro, Isabella Monne, Calogero Terregino, Antonia Ricci |
| EPI_ISL_3119911, EPI_ISL_3119912, EPI_ISL_3119913, EPI_ISL_3119914, EPI_ISL_3119915, EPI_ISL_3119916, EPI_ISL_3119917 | AULSS 5 Polesana | Istituto Zooprofilattico Sperimentale delle Venezie | Adelaide Milani, Alessia Schivo, Annalisa Salviato, Elisa Palumbo, Erika Giorgia Quaranta, Luca Tassoni, Ambra Pastori, Edoardo Giussani, Alice Fusaro, Isabella Monne, Calogero Terregino, Antonia Ricci |
| EPI_ISL_3119918 | RM2/A.U. Tor Vergata | National Institute for Infectious Diseases (INMI) L. Spallanzani I.R.C.C.S | CEM Gruber, F Messina, F Santini, M Rueca, G Bonfiglio, O Butera, B Bartolini, E Giombini, MR Capobianchi, A Di Caro |
| EPI_ISL_3119919 | RM5/Osp. S.Giovanni Evangelista | National Institute for Infectious Diseases (INMI) L. Spallanzani I.R.C.C.S | CEM Gruber, F Messina, F Santini, M Rueca, G Bonfiglio, O Butera, B Bartolini, E Giombini, MR Capobianchi, A Di Caro |
| EPI_ISL_3119920, EPI_ISL_3119921, EPI_ISL_3119922, EPI_ISL_3119923, EPI_ISL_3119924 | RM2/Pol.U. Campus Biomedico | National Institute for Infectious Diseases (INMI) L. Spallanzani I.R.C.C.S | CEM Gruber, F Messina, F Santini, M Rueca, G Bonfiglio, O Butera, B Bartolini, E Giombini, MR Capobianchi, A Di Caro |
| EPI_ISL_3119925 | RM5/Osp. S.Giovanni Evangelista | National Institute for Infectious Diseases (INMI) L. Spallanzani I.R.C.C.S | CEM Gruber, F Messina, F Santini, M Rueca, G Bonfiglio, O Butera, B Bartolini, E Giombini, MR Capobianchi, A Di Caro |
| EPI_ISL_3119926 | RM2/Osp. S.Eugenio | National Institute for Infectious Diseases (INMI) L. Spallanzani I.R.C.C.S | CEM Gruber, F Messina, F Santini, M Rueca, G Bonfiglio, O Butera, B Bartolini, E Giombini, MR Capobianchi, A Di Caro |
| EPI_ISL_3119927 | RM5/Osp. S.Giovanni Evangelista | National Institute for Infectious Diseases (INMI) L. Spallanzani I.R.C.C.S | CEM Gruber, F Messina, F Santini, M Rueca, G Bonfiglio, O Butera, B Bartolini, E Giombini, MR Capobianchi, A Di Caro |
| EPI_ISL_3119928 | RM2/Pol.U. Campus Biomedico | National Institute for Infectious Diseases (INMI) L. Spallanzani I.R.C.C.S | CEM Gruber, F Messina, F Santini, M Rueca, G Bonfiglio, O Butera, B Bartolini, E Giombini, MR Capobianchi, A Di Caro |
| EPI_ISL_3119929 | RM5/Osp. S.Giovanni Evangelista | National Institute for Infectious Diseases (INMI) L. Spallanzani I.R.C.C.S | CEM Gruber, F Messina, F Santini, M Rueca, G Bonfiglio, O Butera, B Bartolini, E Giombini, MR Capobianchi, A Di Caro |
| EPI_ISL_3119930 | RM2/A.U. Tor Vergata | National Institute for Infectious Diseases (INMI) L. Spallanzani I.R.C.C.S | CEM Gruber, F Messina, F Santini, M Rueca, G Bonfiglio, O Butera, B Bartolini, E Giombini, MR Capobianchi, A Di Caro |
| EPI_ISL_3119931 | RM2/Osp. S.Eugenio | National Institute for Infectious Diseases (INMI) L. Spallanzani I.R.C.C.S | CEM Gruber, F Messina, F Santini, M Rueca, G Bonfiglio, O Butera, B Bartolini, E Giombini, MR Capobianchi, A Di Caro |
| EPI_ISL_3119932, EPI_ISL_3119933 | RM5/Osp. S.Giovanni Evangelista | National Institute for Infectious Diseases (INMI) L. Spallanzani I.R.C.C.S | CEM Gruber, F Messina, F Santini, M Rueca, G Bonfiglio, O Butera, B Bartolini, E Giombini, MR Capobianchi, A Di Caro |
| EPI_ISL_3119934, EPI_ISL_3119935, EPI_ISL_3119936, EPI_ISL_3119937, EPI_ISL_3119938, EPI_ISL_3119939, EPI_ISL_3119940, EPI_ISL_3119941, EPI_ISL_3119942, EPI_ISL_3119943, EPI_ISL_3119944, EPI_ISL_3119945, EPI_ISL_3119947 | | | |
| see above | RM2/A.U. Tor Vergata | National Institute for Infectious Diseases (INMI) L. Spallanzani I.R.C.C.S | CEM Gruber, F Messina, F Santini, M Rueca, G Bonfiglio, O Butera, B Bartolini, E Giombini, MR Capobianchi, A Di Caro |
| EPI_ISL_3119948 | RM6/Genzano | National Institute for Infectious Diseases (INMI) L. Spallanzani I.R.C.C.S | CEM Gruber, F Messina, F Santini, M Rueca, G Bonfiglio, O Butera, B Bartolini, E Giombini, MR Capobianchi, A Di Caro |
| EPI_ISL_3119949 | RM1/Sant'Andrea | National Institute for Infectious Diseases (INMI) L. Spallanzani I.R.C.C.S | CEM Gruber, F Messina, F Santini, M Rueca, G Bonfiglio, O Butera, B Bartolini, E Giombini, MR Capobianchi, A Di Caro |
| EPI_ISL_3119950, EPI_ISL_3119951, EPI_ISL_3119952 | FR/Osp. F.Spaziani | National Institute for Infectious Diseases (INMI) L. Spallanzani I.R.C.C.S | CEM Gruber, F Messina, F Santini, M Rueca, G Bonfiglio, O Butera, B Bartolini, E Giombini, MR Capobianchi, A Di Caro |
| EPI_ISL_3119953, EPI_ISL_3119954, EPI_ISL_3119955, EPI_ISL_3119956, EPI_ISL_3119957, EPI_ISL_3119959, EPI_ISL_3119960, EPI_ISL_3119961, EPI_ISL_3119962, EPI_ISL_3119963, EPI_ISL_3119964, EPI_ISL_3119965, EPI_ISL_3119966 | | | |
| see above | RM2/Osp. S.Pertini | National Institute for Infectious Diseases (INMI) L. Spallanzani I.R.C.C.S | CEM Gruber, F Messina, F Santini, M Rueca, G Bonfiglio, O Butera, B Bartolini, E Giombini, MR Capobianchi, A Di Caro |
| EPI_ISL_3119967, EPI_ISL_3119968, EPI_ISL_3119969, EPI_ISL_3119970, EPI_ISL_3119971, EPI_ISL_3119972, EPI_ISL_3119973, EPI_ISL_3119974, EPI_ISL_3119975 | VT/Osp. Di Belcolle | National Institute for Infectious Diseases (INMI) L. Spallanzani I.R.C.C.S | CEM Gruber, F Messina, F Santini, M Rueca, G Bonfiglio, O Butera, B Bartolini, E Giombini, MR Capobianchi, A Di Caro |
| EPI_ISL_3119976 | RM2/Osp. S.Pertini | National Institute for Infectious Diseases (INMI) L. Spallanzani I.R.C.C.S | CEM Gruber, F Messina, F Santini, M Rueca, G Bonfiglio, O Butera, B Bartolini, E Giombini, MR Capobianchi, A Di Caro |
| EPI_ISL_3119977, EPI_ISL_3119978, EPI_ISL_3119980 | RM3/IRCCS INMI Lazzaro Spallanzani | National Institute for Infectious Diseases (INMI) L. Spallanzani I.R.C.C.S | CEM Gruber, F Messina, F Santini, M Rueca, G Bonfiglio, O Butera, B Bartolini, E Giombini, MR Capobianchi, A Di Caro |
| EPI_ISL_3119981 | RM1/IRCCS IDI | National Institute for Infectious Diseases (INMI) L. Spallanzani I.R.C.C.S | CEM Gruber, F Messina, F Santini, M Rueca, G Bonfiglio, O Butera, B Bartolini, E Giombini, MR Capobianchi, A Di Caro |
| EPI_ISL_3119982, EPI_ISL_3119983, EPI_ISL_3119984, EPI_ISL_3119985, EPI_ISL_3119986, EPI_ISL_3119987, EPI_ISL_3119988, EPI_ISL_3119989, EPI_ISL_3119991, EPI_ISL_3119992, EPI_ISL_3119993, EPI_ISL_3119994, EPI_ISL_3119995, EPI_ISL_3119996, EPI_ISL_3119997 | | | |
| see above | RM3/IFO | National Institute for Infectious Diseases (INMI) L. Spallanzani I.R.C.C.S | CEM Gruber, F Messina, F Santini, M Rueca, G Bonfiglio, O Butera, B Bartolini, E Giombini, MR Capobianchi, A Di Caro |
| EPI_ISL_3119999 | A.S.L. CITTA' DI TORINO-ASO MAURIZIANO | Fondazione del Piemonte per l'Oncologia IRCCS | Antonino Sottile, Silvia Brossa, Paola Marino, Giorgio Giardina |
| EPI_ISL_3120000 | A.S.L. ALESSANDRIA | Fondazione del Piemonte per l'Oncologia IRCCS | Antonino Sottile, Silvia Brossa, Paola Marino, Giorgio Giardina |
| EPI_ISL_3120001, EPI_ISL_3120002 | A.S.L. CITTA' DI TORINO-ASO MAURIZIANO | Fondazione del Piemonte per l'Oncologia IRCCS | Antonino Sottile, Silvia Brossa, Paola Marino, Giorgio Giardina |
| EPI_ISL_3120003, | A.S.L. ALESSANDRIA | Fondazione del Piemonte per l'Oncologia IRCCS | Antonino Sottile, Silvia Brossa, Paola Marino, Giorgio Giardina |

| | | | |
|---|---|---|---|
| EPI_ISL_3120004, EPI_ISL_3120005, EPI_ISL_3120006, EPI_ISL_3120007 | | | |
| EPI_ISL_3120008, EPI_ISL_3120009 | A.S.L. ASTI | Fondazione del Piemonte per l'Oncologia IRCCS | Antonino Sottile, Silvia Brossa, Paola Marino, Giorgio Giardina |
| EPI_ISL_3120011 | A.S.L. TO4 | Fondazione del Piemonte per l'Oncologia IRCCS | Antonino Sottile, Silvia Brossa, Paola Marino, Giorgio Giardina |
| EPI_ISL_3120012 | A.S.L. ALESSANDRIA | Fondazione del Piemonte per l'Oncologia IRCCS | Antonino Sottile, Silvia Brossa, Paola Marino, Giorgio Giardina |
| EPI_ISL_3120013, EPI_ISL_3120014, EPI_ISL_3120015, EPI_ISL_3120016, EPI_ISL_3120017 | A.S.L. TO4 | Fondazione del Piemonte per l'Oncologia IRCCS | Antonino Sottile, Silvia Brossa, Paola Marino, Giorgio Giardina |
| EPI_ISL_3120018, EPI_ISL_3120020, EPI_ISL_3120021 | A.S.L. TORINO 5 | Fondazione del Piemonte per l'Oncologia IRCCS | Antonino Sottile, Silvia Brossa, Paola Marino, Giorgio Giardina |
| EPI_ISL_3120022 | A.S.L. BIELLA - OSPEDALE DEGLI INFERMI - SC LABORATORIO ANALISI - | Fondazione del Piemonte per l'Oncologia IRCCS | Antonino Sottile, Silvia Brossa, Paola Marino, Giorgio Giardina |
| EPI_ISL_3120023 | A.S.L. CUNEO 1 - - S.S. LABORATORIO ANALISI - | Fondazione del Piemonte per l'Oncologia IRCCS | Antonino Sottile, Silvia Brossa, Paola Marino, Giorgio Giardina |
| EPI_ISL_3120024, EPI_ISL_3120025 | AZIENDA OSP. S.CROCE E CARLE - AZ. OSPEDAL. S. CROCE E CARLE - S.C. LABORATORIO ANALISI CHIMICO CLINICHE E MICROBIOLOGIA - | Fondazione del Piemonte per l'Oncologia IRCCS | Antonino Sottile, Silvia Brossa, Paola Marino, Giorgio Giardina |
| EPI_ISL_3120026, EPI_ISL_3120027 | A.S.L. CUNEO 1 - - S.S. LABORATORIO ANALISI - | Fondazione del Piemonte per l'Oncologia IRCCS | Antonino Sottile, Silvia Brossa, Paola Marino, Giorgio Giardina |
| EPI_ISL_3120028, EPI_ISL_3120029, EPI_ISL_3120030, EPI_ISL_3120031, EPI_ISL_3120032 | A.S.L. CITTA' DI TORINO - OSPEDALE AMEDEO DI SAVOIA - SC MICROBIOLOGIA E VIROLOGIA - | Fondazione del Piemonte per l'Oncologia IRCCS | Antonino Sottile, Silvia Brossa, Paola Marino, Giorgio Giardina |
| EPI_ISL_3120033 | A.S.L. TO3 | Fondazione del Piemonte per l'Oncologia IRCCS | Antonino Sottile, Silvia Brossa, Paola Marino, Giorgio Giardina |
| EPI_ISL_3120034 | A.S.L. CITTA' DI TORINO - OSPEDALE AMEDEO DI SAVOIA - SC MICROBIOLOGIA E VIROLOGIA - | Fondazione del Piemonte per l'Oncologia IRCCS | Antonino Sottile, Silvia Brossa, Paola Marino, Giorgio Giardina |
| EPI_ISL_3120035, EPI_ISL_3120036 | A.S.L. TO3 | Fondazione del Piemonte per l'Oncologia IRCCS | Antonino Sottile, Silvia Brossa, Paola Marino, Giorgio Giardina |
| EPI_ISL_3120037 | A.S.L. TORINO 5 | Fondazione del Piemonte per l'Oncologia IRCCS | Antonino Sottile, Silvia Brossa, Paola Marino, Giorgio Giardina |
| EPI_ISL_3120038 | A.S.L. TO4 | Fondazione del Piemonte per l'Oncologia IRCCS | Antonino Sottile, Silvia Brossa, Paola Marino, Giorgio Giardina |
| EPI_ISL_3120039 | A.S.L. CUNEO 1 | Fondazione del Piemonte per l'Oncologia IRCCS | Antonino Sottile, Silvia Brossa, Paola Marino, Giorgio Giardina |
| EPI_ISL_3120040, EPI_ISL_3120041 | A.S.L. TO4 | Fondazione del Piemonte per l'Oncologia IRCCS | Antonino Sottile, Silvia Brossa, Paola Marino, Giorgio Giardina |
| EPI_ISL_3120042, EPI_ISL_3120043, EPI_ISL_3120044, EPI_ISL_3120047 | A.S.L. CUNEO 1 | Fondazione del Piemonte per l'Oncologia IRCCS | Antonino Sottile, Silvia Brossa, Paola Marino, Giorgio Giardina |
| EPI_ISL_3120048, EPI_ISL_3120049, EPI_ISL_3120050, EPI_ISL_3120051, EPI_ISL_3120052, EPI_ISL_3120053, EPI_ISL_3120054, EPI_ISL_3120057, EPI_ISL_3120059, EPI_ISL_3120060 | A.S.L. TO4 | Fondazione del Piemonte per l'Oncologia IRCCS | Antonino Sottile, Silvia Brossa, Paola Marino, Giorgio Giardina |
| EPI_ISL_3120061 | A.S.L. TO3 | Fondazione del Piemonte per l'Oncologia IRCCS | Antonino Sottile, Silvia Brossa, Paola Marino, Giorgio Giardina |
| EPI_ISL_3120062, EPI_ISL_3120063 | A.S.L. TO4 | Fondazione del Piemonte per l'Oncologia IRCCS | Antonino Sottile, Silvia Brossa, Paola Marino, Giorgio Giardina |
| EPI_ISL_3120064, EPI_ISL_3120065, EPI_ISL_3120066 | A.S.L. TO3 | Fondazione del Piemonte per l'Oncologia IRCCS | Antonino Sottile, Silvia Brossa, Paola Marino, Giorgio Giardina |
| EPI_ISL_3120067, EPI_ISL_3120069, EPI_ISL_3120070, EPI_ISL_3120071 | A.S.L. ALESSANDRIA | Fondazione del Piemonte per l'Oncologia IRCCS | Antonino Sottile, Silvia Brossa, Paola Marino, Giorgio Giardina |
| EPI_ISL_3120072, EPI_ISL_3120073, EPI_ISL_3120074, EPI_ISL_3120075, EPI_ISL_3120076, EPI_ISL_3120077, EPI_ISL_3120078, EPI_ISL_3120079, EPI_ISL_3120080 | A.S.L. CUNEO 1 | Fondazione del Piemonte per l'Oncologia IRCCS | Antonino Sottile, Silvia Brossa, Paola Marino, Giorgio Giardina |
| EPI_ISL_3120081 | A.S.L. NOVARA - ISTITUTO S.S. TRINITA' - BORGOMANERO - S.S. LABORATORIO ANALISI CHIMICO CLINICHE - | Fondazione del Piemonte per l'Oncologia IRCCS | Antonino Sottile, Silvia Brossa, Paola Marino, Giorgio Giardina |
| EPI_ISL_3120082, EPI_ISL_3120084 | A.S.L. CUNEO 1 | Fondazione del Piemonte per l'Oncologia IRCCS | Antonino Sottile, Silvia Brossa, Paola Marino, Giorgio Giardina |
| EPI_ISL_3120085 | A.S.L. NOVARA - ISTITUTO S.S. TRINITA' - BORGOMANERO - S.S. LABORATORIO ANALISI CHIMICO CLINICHE - | Fondazione del Piemonte per l'Oncologia IRCCS | Antonino Sottile, Silvia Brossa, Paola Marino, Giorgio Giardina |
| EPI_ISL_3120086 | A.S.L. CUNEO 1 | Fondazione del Piemonte per l'Oncologia IRCCS | Antonino Sottile, Silvia Brossa, Paola Marino, Giorgio Giardina |
| EPI_ISL_3120087, EPI_ISL_3120088, EPI_ISL_3120089, EPI_ISL_3120091, EPI_ISL_3120092, EPI_ISL_3120093, EPI_ISL_3120094, EPI_ISL_3120095, EPI_ISL_3120096, EPI_ISL_3120097 | A.S.L. NOVARA - ISTITUTO S.S. TRINITA' - BORGOMANERO - S.S. LABORATORIO ANALISI CHIMICO CLINICHE - | Fondazione del Piemonte per l'Oncologia IRCCS | Antonino Sottile, Silvia Brossa, Paola Marino, Giorgio Giardina |
| EPI_ISL_3120105 | Policlinico Ponte S. Pietro | Laboratory of Clinical Microbiology, Virology and Bioemergencies, ASST Fatebenefratelli Sacco - Sacco Hospital | Valeria Micheli |
| EPI_ISL_3120106 | Ospedale Galeazzi | Laboratory of Clinical Microbiology, Virology and Bioemergencies, ASST Fatebenefratelli Sacco - Sacco Hospital | Valeria Micheli |
| EPI_ISL_3120107 | Policlinico San Donato | Laboratory of Clinical Microbiology, Virology and Bioemergencies, ASST Fatebenefratelli Sacco - Sacco Hospital | Valeria Micheli |
| EPI_ISL_3120109 | Ospedale di Vimercate | Laboratory of Clinical Microbiology, Virology and Bioemergencies, | Valeria Micheli |

| | | | |
|---|---|---|---|
| | | ASST Fatebenefratelli Sacco - Sacco Hospital | |
| EPI_ISL_3120110 | Raja Perempuan Zainab II Hospital | Institute for Medical Research, Infectious Disease Research Centre, National Institutes of Health, Ministry of Health Malaysia | Suppiah J, Kamel K, Mohd Zawawi Z, Azizan MA, Ramly N, Robert F, Anasir MI, Thayan R |
| EPI_ISL_3120111 | Selayang Hospital | Institute for Medical Research, Infectious Disease Research Centre, National Institutes of Health, Ministry of Health Malaysia | Suppiah J, Kamel K, Mohd Zawawi Z, Azizan MA, Ramly N, Robert F, Anasir MI, Thayan R |
| EPI_ISL_3120112, EPI_ISL_3120113 | Lembah Pantai Health Office | Institute for Medical Research, Infectious Disease Research Centre, National Institutes of Health, Ministry of Health Malaysia | Suppiah J, Kamel K, Mohd Zawawi Z, Azizan MA, Ramly N, Robert F, Anasir MI, Thayan R |
| EPI_ISL_3120114, EPI_ISL_3120115 | Tengku Ampuan Rahimah Hospital | Institute for Medical Research, Infectious Disease Research Centre, National Institutes of Health, Ministry of Health Malaysia | Suppiah J, Kamel K, Mohd Zawawi Z, Azizan MA, Ramly N, Robert F, Anasir MI, Thayan R |
| EPI_ISL_3120116 | Sultan Abdul Halim Hospital | Institute for Medical Research, Infectious Disease Research Centre, National Institutes of Health, Ministry of Health Malaysia | Suppiah J, Kamel K, Mohd Zawawi Z, Azizan MA, Ramly N, Robert F, Anasir MI, Thayan R |
| EPI_ISL_3120117, EPI_ISL_3120118, EPI_ISL_3120119, EPI_ISL_3120120, EPI_ISL_3120121, EPI_ISL_3120122, EPI_ISL_3120123, EPI_ISL_3120124, EPI_ISL_3120125, EPI_ISL_3120126, EPI_ISL_3120127, EPI_ISL_3120128, EPI_ISL_3120129, EPI_ISL_3120130, EPI_ISL_3120131, EPI_ISL_3120132, EPI_ISL_3120133, EPI_ISL_3120134, EPI_ISL_3120135, EPI_ISL_3120136, EPI_ISL_3120137 | | | |
| see above | Dipartimento di Medicina di Laboratorio, Azienda sanitaria universitaria Friuli Centrale (ASU FC) | Dipartimento di Medicina di Laboratorio, Azienda sanitaria universitaria Friuli Centrale (ASU FC) | Corrado Pipan, Catia Mio, Chiara Dal Secco, Stefania Marzinotto, Francesco Curcio |
| EPI_ISL_3120139, EPI_ISL_3120140, EPI_ISL_3120141, EPI_ISL_3120142 | ASST FBF Sacco | Laboratory of Clinical Microbiology, Virology and Bioemergencies, ASST Fatebenefratelli Sacco - Sacco Hospital | Valeria Micheli |
| EPI_ISL_3120143 | Sultanah Bahiyah Hospital, Alor Setar | Institute for Medical Research, Infectious Disease Research Centre, National Institutes of Health, Ministry of Health Malaysia | Suppiah J, Kamel K, Mohd Zawawi Z, Azizan MA, Ramly N, Robert F, Anasir MI, Thayan R |
| EPI_ISL_3120144, EPI_ISL_3120145, EPI_ISL_3120146, EPI_ISL_3120147, EPI_ISL_3120148, EPI_ISL_3120149, EPI_ISL_3120150, EPI_ISL_3120151, EPI_ISL_3120152, EPI_ISL_3120153, EPI_ISL_3120154, EPI_ISL_3120155, EPI_ISL_3120156, EPI_ISL_3120157, EPI_ISL_3120158, EPI_ISL_3120159, EPI_ISL_3120160, EPI_ISL_3120161, EPI_ISL_3120162, EPI_ISL_3120163, EPI_ISL_3120164, EPI_ISL_3120165, EPI_ISL_3120166, EPI_ISL_3120167, EPI_ISL_3120168, EPI_ISL_3120169, EPI_ISL_3120171, EPI_ISL_3120172, EPI_ISL_3120173, EPI_ISL_3120174, EPI_ISL_3120175, EPI_ISL_3120176, EPI_ISL_3120177, EPI_ISL_3120178, EPI_ISL_3120179, EPI_ISL_3120180, EPI_ISL_3120181, EPI_ISL_3120182, EPI_ISL_3120183, EPI_ISL_3120184, EPI_ISL_3120185, EPI_ISL_3120186, EPI_ISL_3120187, EPI_ISL_3120188, EPI_ISL_3120189, EPI_ISL_3120190, EPI_ISL_3120191, EPI_ISL_3120192, EPI_ISL_3120193, EPI_ISL_3120194, EPI_ISL_3120195, EPI_ISL_3120196, EPI_ISL_3120197, EPI_ISL_3120198, EPI_ISL_3120199, EPI_ISL_3120200, EPI_ISL_3120201, EPI_ISL_3120202, EPI_ISL_3120203, EPI_ISL_3120204, EPI_ISL_3120205, EPI_ISL_3120206, EPI_ISL_3120207 | | | |
| see above | University of Liège COVID-19 testing center | GIGA Medical Genomics | Keith Durkin, Maria Artesi, Bouchra Boujemla, Nathalie Renotte, Nadine Cambisano, Sébastien Bontems, Cécile Meex, Claire Gourzonès, Olivier Ek, Laurent Gillet, Marie-Pierre Hayette, Vincent Bours |
| EPI_ISL_3120208, EPI_ISL_3120209 | Basurto University Hospital: Clinical Microbiology Laboratory | Biocruces Bizkaia | Mikel Urrutikoetxea-Gutiérrez, Mª Carmen Nieto Toboso, Estibaliz Ugalde Zarraga, Mikel Gallego Rodrigo, Ana de la Hoz, José Luis Díaz de Tuesta del Arco |
| EPI_ISL_3120210, EPI_ISL_3120212, EPI_ISL_3120213, EPI_ISL_3120214, EPI_ISL_3120215 | Plateforme de testing Namuroise | Plateforme de testing Namuroise | Lesly Nyinkeu Kemamen; Demars Aurore ; Maschietto Céline ; Nobis Chloé ; Otto Gaetan ; Denis Olivier ; Degosserie Jonathan ; Mullier François |
| EPI_ISL_3120216 | Marche en Famenne | Plateforme de testing Namuroise | Lesly Nyinkeu Kemamen; Demars Aurore ; Maschietto Céline ; Nobis Chloé ; Otto Gaetan ; Denis Olivier ; Degosserie Jonathan ; Mullier François |
| EPI_ISL_3120217, EPI_ISL_3120219, EPI_ISL_3120220 | Plateforme de testing Namuroise | Plateforme de testing Namuroise | Lesly Nyinkeu Kemamen; Demars Aurore ; Maschietto Céline ; Nobis Chloé ; Otto Gaetan ; Denis Olivier ; Degosserie Jonathan ; Mullier François |
| EPI_ISL_3120221 | Marche en Famenne | Plateforme de testing Namuroise | Lesly Nyinkeu Kemamen; Demars Aurore ; Maschietto Céline ; Nobis Chloé ; Otto Gaetan ; Denis Olivier ; Degosserie Jonathan ; Mullier François |
| EPI_ISL_3120222, EPI_ISL_3120223, EPI_ISL_3120226, EPI_ISL_3120227, EPI_ISL_3120228, EPI_ISL_3120229 | Plateforme de testing Namuroise | Plateforme de testing Namuroise | Lesly Nyinkeu Kemamen; Demars Aurore ; Maschietto Céline ; Nobis Chloé ; Otto Gaetan ; Denis Olivier ; Degosserie Jonathan ; Mullier François |
| EPI_ISL_3120230 | Marche en Famenne | Plateforme de testing Namuroise | Lesly Nyinkeu Kemamen; Demars Aurore ; Maschietto Céline ; Nobis Chloé ; Otto Gaetan ; Denis Olivier ; Degosserie Jonathan ; Mullier François |
| EPI_ISL_3120231, EPI_ISL_3120232, EPI_ISL_3120233 | Plateforme de testing Namuroise | Plateforme de testing Namuroise | Lesly Nyinkeu Kemamen; Demars Aurore ; Maschietto Céline ; Nobis Chloé ; Otto Gaetan ; Denis Olivier ; Degosserie Jonathan ; Mullier François |
| EPI_ISL_3120234 | Marche en Famenne | Plateforme de testing Namuroise | Lesly Nyinkeu Kemamen; Demars Aurore ; Maschietto Céline ; Nobis Chloé ; Otto Gaetan ; Denis Olivier ; Degosserie Jonathan ; Mullier François |
| EPI_ISL_3120235 | Plateforme de testing Namuroise | Plateforme de testing Namuroise | Lesly Nyinkeu Kemamen; Demars Aurore ; Maschietto Céline ; Nobis Chloé ; Otto Gaetan ; Denis Olivier ; Degosserie Jonathan ; Mullier François |
| EPI_ISL_3120237 | Sungai Buloh Hospital | Institute for Medical Research, Infectious Disease Research Centre, National Institutes of Health, Ministry of Health Malaysia | Suppiah J, Kamel K, Mohd Zawawi Z, Azizan MA, Ramly N, Robert F, Anasir MI, Thayan R |
| EPI_ISL_3120238, EPI_ISL_3120239, EPI_ISL_3120240, EPI_ISL_3120241, EPI_ISL_3120242, EPI_ISL_3120243, EPI_ISL_3120244, EPI_ISL_3120245, EPI_ISL_3120246, EPI_ISL_3120247, EPI_ISL_3120248, EPI_ISL_3120249, EPI_ISL_3120250, EPI_ISL_3120251, EPI_ISL_3120252, EPI_ISL_3120253, EPI_ISL_3120254, EPI_ISL_3120255, EPI_ISL_3120256, EPI_ISL_3120257, EPI_ISL_3120258, EPI_ISL_3120259, EPI_ISL_3120260, EPI_ISL_3120261, EPI_ISL_3120262, EPI_ISL_3120263, EPI_ISL_3120264, EPI_ISL_3120266, EPI_ISL_3120267, EPI_ISL_3120268, EPI_ISL_3120269, EPI_ISL_3120270, EPI_ISL_3120271, EPI_ISL_3120272, EPI_ISL_3120273, EPI_ISL_3120274, EPI_ISL_3120275, EPI_ISL_3120276, EPI_ISL_3120277, EPI_ISL_3120278, EPI_ISL_3120280, EPI_ISL_3120281, EPI_ISL_3120283, EPI_ISL_3120284, EPI_ISL_3120286, EPI_ISL_3120287, EPI_ISL_3120288, EPI_ISL_3120289, EPI_ISL_3120290, EPI_ISL_3120291, EPI_ISL_3120292, EPI_ISL_3120293, EPI_ISL_3120294, EPI_ISL_3120295, EPI_ISL_3120296, EPI_ISL_3120297, EPI_ISL_3120298, EPI_ISL_3120299, EPI_ISL_3120300, EPI_ISL_3120301, EPI_ISL_3120302, EPI_ISL_3120303, EPI_ISL_3120304, EPI_ISL_3120305, EPI_ISL_3120306, EPI_ISL_3120307, EPI_ISL_3120308, EPI_ISL_3120309, EPI_ISL_3120310, EPI_ISL_3120311, EPI_ISL_3120312 | | | |
| see above | AZDelta | AZ Delta Medical Laboratories in Roeselare, Belgium | Geert Martens, Dieter De Smet, Frederik Van Hoecke, on behalf of AZ Delta COVID-19 Genomics core (member of Genomic surveillance of SARS-CoV-2 in Belgium network) |
| EPI_ISL_3120313 | Basurto University Hospital: Clinical Microbiology Laboratory | Biocruces Bizkaia | Mikel Urrutikoetxea-Gutiérrez, Mª Carmen Nieto Toboso, Estibaliz Ugalde Zarraga, Mikel Gallego Rodrigo, Ana de la Hoz, José Luis Díaz de Tuesta del Arco |
| EPI_ISL_3120753 | Synlab MVZ Augsburg | Robert Koch Institute | unknown |
| EPI_ISL_3120795, EPI_ISL_3120800, EPI_ISL_3120805 | SYNLAB MVZ Weiden | Robert Koch Institute | unknown |
| EPI_ISL_3120815, EPI_ISL_3120816, EPI_ISL_3120817 | Institut für Labormedizin Mikrobiologie und Hygiene | Robert Koch Institute | unknown |
| EPI_ISL_3120818, EPI_ISL_3120819, EPI_ISL_3120820 | Landesamt für Verbraucherschutz des Landes Sachsen-Anhalt | Robert Koch Institute | unknown |
| EPI_ISL_3120821 | MVZ für Laboratoriumsmedizin und Mikrobiologie Würzburg | Robert Koch Institute | unknown |
| EPI_ISL_3120822, EPI_ISL_3120823, EPI_ISL_3120824, EPI_ISL_3120825 | MVZ Dr. Eberhard & Partner Dortmund | Robert Koch Institute | unknown |
| EPI_ISL_3120826, EPI_ISL_3120827 | MVZ für Laboratoriumsmedizin und Mikrobiologie Würzburg | Robert Koch Institute | unknown |
| EPI_ISL_3120828, EPI_ISL_3120829, EPI_ISL_3120830, EPI_ISL_3120831, EPI_ISL_3120832 | MVZ Dr. Eberhard & Partner Dortmund | Robert Koch Institute | unknown |
| EPI_ISL_3120833, EPI_ISL_3120834, EPI_ISL_3120835, EPI_ISL_3120836, EPI_ISL_3120837, EPI_ISL_3120838, EPI_ISL_3120839, EPI_ISL_3120840 | Klinikum Ernst von Bergmann gemeinnützige GmbH - stationärer Bereich | Robert Koch Institute | unknown |
| EPI_ISL_3120841, EPI_ISL_3120842, EPI_ISL_3120843, EPI_ISL_3120844, EPI_ISL_3120846 | Bioscientia Labor Wermsdorf | Robert Koch Institute | unknown |
| EPI_ISL_3120847 | Sonic - MVZ Medizinisches Labor Bremen GmbH | Robert Koch Institute | unknown |
| EPI_ISL_3120848 | Sonic - Dr. Staber & Kollegen GmbH München | Robert Koch Institute | unknown |

| | | | |
|---|---|---|---|
| EPI_ISL_3120849 | Sonic - MVZ Medizinisches Labor Bremen GmbH | Robert Koch Institute | unknown |
| EPI_ISL_3120850 | Bioscientia Labor Wermsdorf | Robert Koch Institute | unknown |
| EPI_ISL_3120851 | Bioscientia MVZ Labor Karlsruhe GmbH | Robert Koch Institute | unknown |
| EPI_ISL_3120852 | Sonic - MVZ Medizinisches Labor Bremen GmbH | Robert Koch Institute | unknown |
| EPI_ISL_3120853 | Bioscientia Labor Wermsdorf | Robert Koch Institute | unknown |
| EPI_ISL_3120854, EPI_ISL_3120855 | Sonic - MVZ Medizinisches Labor Bremen GmbH | Robert Koch Institute | unknown |
| EPI_ISL_3120856, EPI_ISL_3120858, EPI_ISL_3120859, EPI_ISL_3120860, EPI_ISL_3120861 | Bioscientia Labor Wermsdorf | Robert Koch Institute | unknown |
| EPI_ISL_3120862, EPI_ISL_3120863 | Sonic - MVZ Medizinisches Labor Bremen GmbH | Robert Koch Institute | unknown |
| EPI_ISL_3120865 | Sonic - Dr. Staber & Kollegen GmbH München | Robert Koch Institute | unknown |
| EPI_ISL_3120866, EPI_ISL_3120867 | Sonic - MVZ Medizinisches Labor Bremen GmbH | Robert Koch Institute | unknown |
| EPI_ISL_3120868 | Sonic - Labor Staber Nürnberg | Robert Koch Institute | unknown |
| EPI_ISL_3120870, EPI_ISL_3120871, EPI_ISL_3120872, EPI_ISL_3120873 | Bioscientia Labor Wermsdorf | Robert Koch Institute | unknown |
| EPI_ISL_3120875, EPI_ISL_3120876 | Sonic - MVZ Medizinisches Labor Bremen GmbH | Robert Koch Institute | unknown |
| EPI_ISL_3120877, EPI_ISL_3120878 | Bioscientia Labor Wermsdorf | Robert Koch Institute | unknown |
| EPI_ISL_3120879, EPI_ISL_3120880, EPI_ISL_3120881, EPI_ISL_3120882 | Sonic - MVZ Medizinisches Labor Bremen GmbH | Robert Koch Institute | unknown |
| EPI_ISL_3120883, EPI_ISL_3120885 | Bioscientia Labor Wermsdorf | Robert Koch Institute | unknown |
| EPI_ISL_3120886 | Klinikum Fulda gAG | Robert Koch Institute | unknown |
| EPI_ISL_3120887 | Bioscientia Labor Wermsdorf | Robert Koch Institute | unknown |
| EPI_ISL_3120888 | Sonic - Labor Staber Dresden (Klipphausen) | Robert Koch Institute | unknown |
| EPI_ISL_3120889 | Bioscientia Labor Wermsdorf | Robert Koch Institute | unknown |
| EPI_ISL_3120890, EPI_ISL_3120891 | Sonic - MVZ Medizinisches Labor Bremen GmbH | Robert Koch Institute | unknown |
| EPI_ISL_3120892, EPI_ISL_3120893 | Bioscientia Labor Wermsdorf | Robert Koch Institute | unknown |
| EPI_ISL_3120894, EPI_ISL_3120895 | Sonic - MVZ Medizinisches Labor Bremen GmbH | Robert Koch Institute | unknown |
| EPI_ISL_3120896, EPI_ISL_3120897, EPI_ISL_3120898 | Bioscientia Labor Wermsdorf | Robert Koch Institute | unknown |
| EPI_ISL_3120899 | Sonic - MVZ Medizinisches Labor Bremen GmbH | Robert Koch Institute | unknown |
| EPI_ISL_3120900, EPI_ISL_3120902 | Bioscientia Labor Wermsdorf | Robert Koch Institute | unknown |
| EPI_ISL_3120903 | Sonic - MVZ Medizinisches Labor Bremen GmbH | Robert Koch Institute | unknown |
| EPI_ISL_3120904 | Bioscientia Labor Wermsdorf | Robert Koch Institute | unknown |
| EPI_ISL_3120905 | Sonic - MVZ Medizinisches Labor Bremen GmbH | Robert Koch Institute | unknown |
| EPI_ISL_3120907 | Sonic - Labor Staber Dresden (Klipphausen) | Robert Koch Institute | unknown |
| EPI_ISL_3120908 | Sonic - MVZ Medizinisches Labor Bremen GmbH | Robert Koch Institute | unknown |
| EPI_ISL_3120909 | SYNLAB MVZ Weiden | Robert Koch Institute | unknown |
| EPI_ISL_3120910 | SYNLAB MVZ Leinfelden-Echterdingen | Robert Koch Institute | unknown |
| EPI_ISL_3120911, EPI_ISL_3120912 | SYNLAB MVZ Leverkusen | Robert Koch Institute | unknown |
| EPI_ISL_3120913 | SYNLAB MVZ Weiden | Robert Koch Institute | unknown |
| EPI_ISL_3120914 | SYNLAB MVZ Leinfelden-Echterdingen | Robert Koch Institute | unknown |
| EPI_ISL_3120915 | Synlab MVZ Augsburg | Robert Koch Institute | unknown |
| EPI_ISL_3120916 | SYNLAB MVZ Weiden | Robert Koch Institute | unknown |
| EPI_ISL_3120917 | Synlab MVZ Augsburg | Robert Koch Institute | unknown |
| EPI_ISL_3120918 | SYNLAB MVZ Weiden | Robert Koch Institute | unknown |
| EPI_ISL_3120919 | SYNLAB MVZ Leverkusen | Robert Koch Institute | unknown |
| EPI_ISL_3120920 | SYNLAB MVZ Leinfelden-Echterdingen | Robert Koch Institute | unknown |
| EPI_ISL_3120921 | Synlab MVZ Augsburg | Robert Koch Institute | unknown |
| EPI_ISL_3120922 | SYNLAB MVZ Leinfelden-Echterdingen | Robert Koch Institute | unknown |
| EPI_ISL_3120924 | SYNLAB MVZ Weiden | Robert Koch Institute | unknown |
| EPI_ISL_3120925 | SYNLAB MVZ Leverkusen | Robert Koch Institute | unknown |
| EPI_ISL_3120926 | SYNLAB MVZ Weiden | Robert Koch Institute | unknown |
| EPI_ISL_3120927 | SYNLAB MVZ Leinfelden-Echterdingen | Robert Koch Institute | unknown |
| EPI_ISL_3120928 | SYNLAB MVZ Berlin | Robert Koch Institute | unknown |
| EPI_ISL_3120929 | SYNLAB MVZ Weiden | Robert Koch Institute | unknown |
| EPI_ISL_3120930 | SYNLAB MVZ Leverkusen | Robert Koch Institute | unknown |
| EPI_ISL_3120931 | SYNLAB MVZ Weiden | Robert Koch Institute | unknown |
| EPI_ISL_3120932, EPI_ISL_3120933 | SYNLAB MVZ Leverkusen | Robert Koch Institute | unknown |
| EPI_ISL_3120934, EPI_ISL_3120935 | SYNLAB MVZ Weiden | Robert Koch Institute | unknown |
| EPI_ISL_3120936 | SYNLAB MVZ Trier | Robert Koch Institute | unknown |
| EPI_ISL_3120937 | SYNLAB MVZ Leinfelden-Echterdingen | Robert Koch Institute | unknown |
| EPI_ISL_3120938, EPI_ISL_3120939 | SYNLAB MVZ Leverkusen | Robert Koch Institute | unknown |
| EPI_ISL_3120940 | Synlab MVZ Augsburg | Robert Koch Institute | unknown |
| EPI_ISL_3120941 | SYNLAB MVZ Weiden | Robert Koch Institute | unknown |

| | | | |
|---|---|---|---|
| EPI_ISL_3120942 | SYNLAB MVZ Berlin | Robert Koch Institute | unknown |
| EPI_ISL_3120943, EPI_ISL_3120944 | SYNLAB MVZ Leverkusen | Robert Koch Institute | unknown |
| EPI_ISL_3120945 | Synlab MVZ Augsburg | Robert Koch Institute | unknown |
| EPI_ISL_3120946 | SYNLAB MVZ Leverkusen | Robert Koch Institute | unknown |
| EPI_ISL_3120947 | Synlab MVZ Augsburg | Robert Koch Institute | unknown |
| EPI_ISL_3120948 | SYNLAB MVZ Leverkusen | Robert Koch Institute | unknown |
| EPI_ISL_3120949 | Synlab MVZ Augsburg | Robert Koch Institute | unknown |
| EPI_ISL_3120951, EPI_ISL_3120952, EPI_ISL_3120953 | SYNLAB MVZ Leverkusen | Robert Koch Institute | unknown |
| EPI_ISL_3120954 | SYNLAB MVZ Leinfelden-Echterdingen | Robert Koch Institute | unknown |
| EPI_ISL_3120955, EPI_ISL_3120956 | SYNLAB MVZ Leverkusen | Robert Koch Institute | unknown |
| EPI_ISL_3120957 | SYNLAB MVZ Trier | Robert Koch Institute | unknown |
| EPI_ISL_3120958, EPI_ISL_3120959 | SYNLAB MVZ Leverkusen | Robert Koch Institute | unknown |
| EPI_ISL_3120960 | SYNLAB MVZ Dachau | Robert Koch Institute | unknown |
| EPI_ISL_3120961, EPI_ISL_3120962, EPI_ISL_3120964, EPI_ISL_3120965 | SYNLAB MVZ Leverkusen | Robert Koch Institute | unknown |
| EPI_ISL_3120966, EPI_ISL_3120967 | SYNLAB MVZ Weiden | Robert Koch Institute | unknown |
| EPI_ISL_3120968 | SYNLAB MVZ Leverkusen | Robert Koch Institute | unknown |
| EPI_ISL_3120969 | SYNLAB MVZ Hamburg | Robert Koch Institute | unknown |
| EPI_ISL_3120970 | SYNLAB MVZ Leinfelden-Echterdingen | Robert Koch Institute | unknown |
| EPI_ISL_3120971 | SYNLAB MVZ Leverkusen | Robert Koch Institute | unknown |
| EPI_ISL_3120972 | SYNLAB MVZ Weiden | Robert Koch Institute | unknown |
| EPI_ISL_3120973, EPI_ISL_3120974 | SYNLAB MVZ Leverkusen | Robert Koch Institute | unknown |
| EPI_ISL_3120975 | SYNLAB MVZ Weiden | Robert Koch Institute | unknown |
| EPI_ISL_3120976 | SYNLAB MVZ Trier | Robert Koch Institute | unknown |
| EPI_ISL_3120977, EPI_ISL_3120978 | SYNLAB MVZ Weiden | Robert Koch Institute | unknown |
| EPI_ISL_3120979, EPI_ISL_3120980, EPI_ISL_3120981 | SYNLAB MVZ Leinfelden-Echterdingen | Robert Koch Institute | unknown |
| EPI_ISL_3120982 | SYNLAB MVZ Weiden | Robert Koch Institute | unknown |
| EPI_ISL_3120983 | SYNLAB MVZ Leverkusen | Robert Koch Institute | unknown |
| EPI_ISL_3120984 | SYNLAB MVZ Berlin | Robert Koch Institute | unknown |
| EPI_ISL_3120985, EPI_ISL_3120986 | SYNLAB MVZ Leverkusen | Robert Koch Institute | unknown |
| EPI_ISL_3120987 | SYNLAB MVZ Weiden | Robert Koch Institute | unknown |
| EPI_ISL_3120989, EPI_ISL_3120990 | SYNLAB MVZ Leverkusen | Robert Koch Institute | unknown |
| EPI_ISL_3120991 | Synlab MVZ Augsburg | Robert Koch Institute | unknown |
| EPI_ISL_3120993, EPI_ISL_3120994, EPI_ISL_3120995 | SYNLAB MVZ Weiden | Robert Koch Institute | unknown |
| EPI_ISL_3120996 | Synlab MVZ Augsburg | Robert Koch Institute | unknown |
| EPI_ISL_3120997 | SYNLAB MVZ Weiden | Robert Koch Institute | unknown |
| EPI_ISL_3120999 | SYNLAB MVZ Trier | Robert Koch Institute | unknown |
| EPI_ISL_3121000 | SYNLAB MVZ Hamburg | Robert Koch Institute | unknown |
| EPI_ISL_3121001 | SYNLAB MVZ Leverkusen | Robert Koch Institute | unknown |
| EPI_ISL_3121002 | SYNLAB MVZ Trier | Robert Koch Institute | unknown |
| EPI_ISL_3121004 | SYNLAB MVZ Leverkusen | Robert Koch Institute | unknown |
| EPI_ISL_3121005 | SYNLAB MVZ Berlin | Robert Koch Institute | unknown |
| EPI_ISL_3121006 | SYNLAB MVZ Weiden | Robert Koch Institute | unknown |
| EPI_ISL_3121007 | SYNLAB MVZ Leverkusen | Robert Koch Institute | unknown |
| EPI_ISL_3121008 | SYNLAB MVZ Trier | Robert Koch Institute | unknown |
| EPI_ISL_3121009 | SYNLAB MVZ Hamburg | Robert Koch Institute | unknown |
| EPI_ISL_3121010, EPI_ISL_3121011, EPI_ISL_3121012, EPI_ISL_3121013 | SYNLAB MVZ Leverkusen | Robert Koch Institute | unknown |
| EPI_ISL_3121014 | Synlab MVZ Augsburg | Robert Koch Institute | unknown |
| EPI_ISL_3121015 | SYNLAB MVZ Leinfelden-Echterdingen | Robert Koch Institute | unknown |
| EPI_ISL_3121016, EPI_ISL_3121017 | SYNLAB MVZ Weiden | Robert Koch Institute | unknown |
| EPI_ISL_3121018, EPI_ISL_3121019, EPI_ISL_3121020, EPI_ISL_3121021, EPI_ISL_3121022 | SYNLAB MVZ Leverkusen | Robert Koch Institute | unknown |
| EPI_ISL_3121023 | SYNLAB MVZ Hamburg | Robert Koch Institute | unknown |
| EPI_ISL_3121024 | SYNLAB MVZ Leinfelden-Echterdingen | Robert Koch Institute | unknown |
| EPI_ISL_3121025 | SYNLAB Jena Oncoscreen | Robert Koch Institute | unknown |
| EPI_ISL_3121026 | SYNLAB MVZ Trier | Robert Koch Institute | unknown |
| EPI_ISL_3121027 | SYNLAB MVZ Leverkusen | Robert Koch Institute | unknown |
| EPI_ISL_3121029, EPI_ISL_3121030 | SYNLAB MVZ Weiden | Robert Koch Institute | unknown |
| EPI_ISL_3121031, EPI_ISL_3121032, EPI_ISL_3121033, | SYNLAB MVZ Leverkusen | Robert Koch Institute | unknown |

| | | | |
|---|---|---|---|
| EPI_ISL_3121034, EPI_ISL_3121035, EPI_ISL_3121036 | | | |
| EPI_ISL_3121037 | SYNLAB MVZ Weiden | Robert Koch Institute | unknown |
| EPI_ISL_3121038, EPI_ISL_3121039, EPI_ISL_3121040 | SYNLAB MVZ Leverkusen | Robert Koch Institute | unknown |
| EPI_ISL_3121041 | SYNLAB MVZ Trier | Robert Koch Institute | unknown |
| EPI_ISL_3121042, EPI_ISL_3121043 | SYNLAB MVZ Leverkusen | Robert Koch Institute | unknown |
| EPI_ISL_3121044 | SYNLAB MVZ Weiden | Robert Koch Institute | unknown |
| EPI_ISL_3121045 | SYNLAB MVZ Leverkusen | Robert Koch Institute | unknown |
| EPI_ISL_3121046 | SYNLAB MVZ Trier | Robert Koch Institute | unknown |
| EPI_ISL_3121047 | SYNLAB MVZ Leverkusen | Robert Koch Institute | unknown |
| EPI_ISL_3121048 | Synlab MVZ Augsburg | Robert Koch Institute | unknown |
| EPI_ISL_3121049 | SYNLAB MVZ Leverkusen | Robert Koch Institute | unknown |
| EPI_ISL_3121050 | SYNLAB MVZ Weiden | Robert Koch Institute | unknown |
| EPI_ISL_3121051 | SYNLAB MVZ Leverkusen | Robert Koch Institute | unknown |
| EPI_ISL_3121052 | SYNLAB MVZ Weiden | Robert Koch Institute | unknown |
| EPI_ISL_3121053, EPI_ISL_3121054 | SYNLAB MVZ Leverkusen | Robert Koch Institute | unknown |
| EPI_ISL_3121055 | SYNLAB MVZ Weiden | Robert Koch Institute | unknown |
| EPI_ISL_3121056 | SYNLAB MVZ Leverkusen | Robert Koch Institute | unknown |
| EPI_ISL_3121057 | SYNLAB MVZ Weiden | Robert Koch Institute | unknown |
| EPI_ISL_3121058, EPI_ISL_3121059 | SYNLAB MVZ Trier | Robert Koch Institute | unknown |
| EPI_ISL_3121060, EPI_ISL_3121061 | SYNLAB MVZ Leverkusen | Robert Koch Institute | unknown |
| EPI_ISL_3121062, EPI_ISL_3121063 | SYNLAB MVZ Trier | Robert Koch Institute | unknown |
| EPI_ISL_3121064, EPI_ISL_3121065, EPI_ISL_3121066, EPI_ISL_3121067, EPI_ISL_3121068, EPI_ISL_3121069 | SYNLAB MVZ Leverkusen | Robert Koch Institute | unknown |
| EPI_ISL_3121070 | SYNLAB MVZ Weiden | Robert Koch Institute | unknown |
| EPI_ISL_3121071, EPI_ISL_3121072, EPI_ISL_3121073 | SYNLAB MVZ Leverkusen | Robert Koch Institute | unknown |
| EPI_ISL_3121074 | SYNLAB MVZ Leinfelden-Echterdingen | Robert Koch Institute | unknown |
| EPI_ISL_3121075 | SYNLAB MVZ Weiden | Robert Koch Institute | unknown |
| EPI_ISL_3121076 | SYNLAB MVZ Trier | Robert Koch Institute | unknown |
| EPI_ISL_3121077 | SYNLAB MVZ Leverkusen | Robert Koch Institute | unknown |
| EPI_ISL_3121078, EPI_ISL_3121079, EPI_ISL_3121080 | SYNLAB MVZ Trier | Robert Koch Institute | unknown |
| EPI_ISL_3121081 | SYNLAB MVZ Weiden | Robert Koch Institute | unknown |
| EPI_ISL_3121082, EPI_ISL_3121083, EPI_ISL_3121084 | SYNLAB MVZ Trier | Robert Koch Institute | unknown |
| EPI_ISL_3121085 | SYNLAB MVZ Weiden | Robert Koch Institute | unknown |
| EPI_ISL_3121086, EPI_ISL_3121087, EPI_ISL_3121088 | SYNLAB MVZ Leverkusen | Robert Koch Institute | unknown |
| EPI_ISL_3121089 | Synlab MVZ Augsburg | Robert Koch Institute | unknown |
| EPI_ISL_3121091 | SYNLAB MVZ Leverkusen | Robert Koch Institute | unknown |
| EPI_ISL_3121092 | SYNLAB MVZ Weiden | Robert Koch Institute | unknown |
| EPI_ISL_3121093, EPI_ISL_3121094 | SYNLAB MVZ Leverkusen | Robert Koch Institute | unknown |
| EPI_ISL_3121095, EPI_ISL_3121096, EPI_ISL_3121097 | SYNLAB MVZ Weiden | Robert Koch Institute | unknown |
| EPI_ISL_3121098 | SYNLAB MVZ Leverkusen | Robert Koch Institute | unknown |
| EPI_ISL_3121100 | SYNLAB MVZ Trier | Robert Koch Institute | unknown |
| EPI_ISL_3121101 | SYNLAB MVZ Leverkusen | Robert Koch Institute | unknown |
| EPI_ISL_3121102 | SYNLAB MVZ Weiden | Robert Koch Institute | unknown |
| EPI_ISL_3121103 | SYNLAB Jena Oncoscreen | Robert Koch Institute | unknown |
| EPI_ISL_3121105 | SYNLAB MVZ Weiden | Robert Koch Institute | unknown |
| EPI_ISL_3121106 | SYNLAB MVZ Berlin | Robert Koch Institute | unknown |
| EPI_ISL_3121107 | SYNLAB MVZ Weiden | Robert Koch Institute | unknown |
| EPI_ISL_3121108 | SYNLAB MVZ Leverkusen | Robert Koch Institute | unknown |
| EPI_ISL_3121109 | SYNLAB MVZ Weiden | Robert Koch Institute | unknown |
| EPI_ISL_3121110 | Synlab MVZ Augsburg | Robert Koch Institute | unknown |
| EPI_ISL_3121111, EPI_ISL_3121112 | SYNLAB MVZ Weiden | Robert Koch Institute | unknown |
| EPI_ISL_3121113 | SYNLAB MVZ Leinfelden-Echterdingen | Robert Koch Institute | unknown |
| EPI_ISL_3121114, EPI_ISL_3121116 | SYNLAB MVZ Leverkusen | Robert Koch Institute | unknown |
| EPI_ISL_3121117 | SYNLAB MVZ Hamburg | Robert Koch Institute | unknown |
| EPI_ISL_3121118, EPI_ISL_3121119 | SYNLAB MVZ Leverkusen | Robert Koch Institute | unknown |
| EPI_ISL_3121120 | SYNLAB MVZ Weiden | Robert Koch Institute | unknown |
| EPI_ISL_3121121, EPI_ISL_3121122 | SYNLAB MVZ Leverkusen | Robert Koch Institute | unknown |

| | | | |
|---|---|---|---|
| EPI_ISL_3121123 | SYNLAB MVZ Weiden | Robert Koch Institute | unknown |
| EPI_ISL_3121124 | SYNLAB MVZ Trier | Robert Koch Institute | unknown |
| EPI_ISL_3121125 | SYNLAB MVZ Weiden | Robert Koch Institute | unknown |
| EPI_ISL_3121126 | Synlab MVZ Augsburg | Robert Koch Institute | unknown |
| EPI_ISL_3121127 | SYNLAB MVZ Weiden | Robert Koch Institute | unknown |
| EPI_ISL_3121128 | Universitätsklinikum des Saarlandes - Institut für Virologie | Robert Koch Institute | unknown |
| EPI_ISL_3121129 | Labormedizin Darmstadt | Robert Koch Institute | unknown |
| EPI_ISL_3121130 | IMD Labor Frankfurt | Robert Koch Institute | unknown |
| EPI_ISL_3121132 | Labormedizin Darmstadt | Robert Koch Institute | unknown |
| EPI_ISL_3121133 | Universitätsklinikum des Saarlandes - Institut für Virologie | Robert Koch Institute | unknown |
| EPI_ISL_3121134 | Labormedizin Darmstadt | Robert Koch Institute | unknown |
| EPI_ISL_3121136 | IMD Labor Frankfurt | Robert Koch Institute | unknown |
| EPI_ISL_3121137 | Labormedizin Darmstadt | Robert Koch Institute | unknown |
| EPI_ISL_3121138, EPI_ISL_3121140 | IMD Labor Frankfurt | Robert Koch Institute | unknown |
| EPI_ISL_3121141 | Labormedizin Darmstadt | Robert Koch Institute | unknown |
| EPI_ISL_3121142, EPI_ISL_3121143 | IMD Labor Frankfurt | Robert Koch Institute | unknown |
| EPI_ISL_3121144, EPI_ISL_3121145, EPI_ISL_3121146 | Labormedizin Darmstadt | Robert Koch Institute | unknown |
| EPI_ISL_3121147, EPI_ISL_3121148, EPI_ISL_3121149, EPI_ISL_3121150, EPI_ISL_3121151, EPI_ISL_3121152, EPI_ISL_3121153, EPI_ISL_3121154, EPI_ISL_3121155, EPI_ISL_3121156, EPI_ISL_3121157 | | | |
| see above | Chemische und Veterinäruntersuchungsamt Münsterland-Emscher-Lippe (CVUA-MEL) | Robert Koch Institute | unknown |
| EPI_ISL_3121158, EPI_ISL_3121159, EPI_ISL_3121160, EPI_ISL_3121161, EPI_ISL_3121162, EPI_ISL_3121163, EPI_ISL_3121164, EPI_ISL_3121165, EPI_ISL_3121166, EPI_ISL_3121167, EPI_ISL_3121168, EPI_ISL_3121169, EPI_ISL_3121170, EPI_ISL_3121171, EPI_ISL_3121172, EPI_ISL_3121173, EPI_ISL_3121175, EPI_ISL_3121176 | | | |
| see above | Labor Dr. Heidrich & Kollegen MVZ GmbH Hamburg | Robert Koch Institute | unknown |
| EPI_ISL_3121177, EPI_ISL_3121178 | Laborarztpraxis Dres. med. Walther Weindel & Kollegen | Robert Koch Institute | unknown |
| EPI_ISL_3121179 | alphaomega Labor GbR (Medizinische Kooperationsgemeinschaft Dres. Ackermann/Tennert/Zaiß) | Robert Koch Institute | unknown |
| EPI_ISL_3121180 | Labor für medizinische Mikrobiologie Bakteriologie Mykologie Virologie & Infektionsserologie - Labor Mölbis | Robert Koch Institute | unknown |
| EPI_ISL_3121181, EPI_ISL_3121182 | alphaomega Labor GbR (Medizinische Kooperationsgemeinschaft Dres. Ackermann/Tennert/Zaiß) | Robert Koch Institute | unknown |
| EPI_ISL_3121183 | amedes MVZ für Labordiagnostik Rhein-Main | Robert Koch Institute | unknown |
| EPI_ISL_3121184, EPI_ISL_3121185, EPI_ISL_3121186, EPI_ISL_3121187 | amedes MVZ DIAMEDIS Sennestadt | Robert Koch Institute | unknown |
| EPI_ISL_3121188 | amedes MVZ für Laboratoriumsmedizin Rhein/Ruhr | Robert Koch Institute | unknown |
| EPI_ISL_3121189 | amedes MVZ für Labordiagnostik Rhein-Main | Robert Koch Institute | unknown |
| EPI_ISL_3121190, EPI_ISL_3121191, EPI_ISL_3121192, EPI_ISL_3121193, EPI_ISL_3121194 | amedes MVZ DIAMEDIS Sennestadt | Robert Koch Institute | unknown |
| EPI_ISL_3121195 | amedes MVZ Halle/Leipzig | Robert Koch Institute | unknown |
| EPI_ISL_3121196 | amedes MVZ DIAMEDIS Sennestadt | Robert Koch Institute | unknown |
| EPI_ISL_3121197 | amedes MVZ für Labordiagnostik Rhein-Main | Robert Koch Institute | unknown |
| EPI_ISL_3121198 | amedes MVZ Hannover | Robert Koch Institute | unknown |
| EPI_ISL_3121199 | amedes MVZ DIAMEDIS Sennestadt | Robert Koch Institute | unknown |
| EPI_ISL_3121200 | amedes Göttingen | Robert Koch Institute | unknown |
| EPI_ISL_3121201 | amedes MVZ Hannover | Robert Koch Institute | unknown |
| EPI_ISL_3121202 | amedes Göttingen | Robert Koch Institute | unknown |
| EPI_ISL_3121203 | amedes MVZ DIAMEDIS Sennestadt | Robert Koch Institute | unknown |
| EPI_ISL_3121204 | amedes Medlab Arnold Analytik MVZ GmbH | Robert Koch Institute | unknown |
| EPI_ISL_3121205, EPI_ISL_3121206 | amedes Göttingen | Robert Koch Institute | unknown |
| EPI_ISL_3121207 | amedes MVZ DIAMEDIS Sennestadt | Robert Koch Institute | unknown |
| EPI_ISL_3121208, EPI_ISL_3121209 | amedes Göttingen | Robert Koch Institute | unknown |
| EPI_ISL_3121210 | amedes MVZ für Laboratoriumsdiagnostik Raubling GmbH | Robert Koch Institute | unknown |
| EPI_ISL_3121211 | amedes MVZ DIAMEDIS Sennestadt | Robert Koch Institute | unknown |
| EPI_ISL_3121212 | amedes MVZ Jena | Robert Koch Institute | unknown |
| EPI_ISL_3121213 | amedes MVZ für Labordiagnostik Rhein-Main | Robert Koch Institute | unknown |
| EPI_ISL_3121214 | amedes MVZ Halle/Leipzig | Robert Koch Institute | unknown |
| EPI_ISL_3121215, EPI_ISL_3121216 | amedes MVZ DIAMEDIS Sennestadt | Robert Koch Institute | unknown |
| EPI_ISL_3121217 | amedes MVZ Jena | Robert Koch Institute | unknown |
| EPI_ISL_3121218 | nordlab - Partnerschaftspraxis für Laboratoriumsmedizin | Robert Koch Institute | unknown |
| EPI_ISL_3121220 | amedes MVZ DIAMEDIS Sennestadt | Robert Koch Institute | unknown |
| EPI_ISL_3121221 | nordlab - Partnerschaftspraxis für Laboratoriumsmedizin | Robert Koch Institute | unknown |
| EPI_ISL_3121222 | amedes MVZ für Laboratoriumsdiagnostik Raubling GmbH | Robert Koch Institute | unknown |
| EPI_ISL_3121224 | amedes MVZ für Labordiagnostik Rhein-Main | Robert Koch Institute | unknown |
| EPI_ISL_3121225 | nordlab - Partnerschaftspraxis für Laboratoriumsmedizin | Robert Koch Institute | unknown |
| EPI_ISL_3121226 | amedes MVZ Jena | Robert Koch Institute | unknown |
| EPI_ISL_3121227 | amedes MVZ DIAMEDIS Sennestadt | Robert Koch Institute | unknown |
| EPI_ISL_3121228 | nordlab - Partnerschaftspraxis für Laboratoriumsmedizin | Robert Koch Institute | unknown |
| EPI_ISL_3121229 | amedes MVZ DIAMEDIS Sennestadt | Robert Koch Institute | unknown |
| EPI_ISL_3121230, EPI_ISL_3121231 | nordlab - Partnerschaftspraxis für Laboratoriumsmedizin | Robert Koch Institute | unknown |
| EPI_ISL_3121232 | amedes MVZ DIAMEDIS Sennestadt | Robert Koch Institute | unknown |

| | | | |
|---|---|---|---|
| EPI_ISL_3121233 | amedes MVZ für Labordiagnostik Rhein-Main | Robert Koch Institute | unknown |
| EPI_ISL_3121234 | amedes MVZ DIAMEDIS Sennestadt | Robert Koch Institute | unknown |
| EPI_ISL_3121235 | amedes MVZ Zentrum für Pathologie und Zytodiagnostik GmbH (Köln) | Robert Koch Institute | unknown |
| EPI_ISL_3121236 | amedes MVZ DIAMEDIS Sennestadt | Robert Koch Institute | unknown |
| EPI_ISL_3121237 | amedes MVZ Zentrum für Pathologie und Zytodiagnostik GmbH (Köln) | Robert Koch Institute | unknown |
| EPI_ISL_3121238 | amedes MVZ DIAMEDIS Sennestadt | Robert Koch Institute | unknown |
| EPI_ISL_3121239 | amedes MVZ Zentrum für Pathologie und Zytodiagnostik GmbH (Köln) | Robert Koch Institute | unknown |
| EPI_ISL_3121240 | amedes MVZ DIAMEDIS Sennestadt | Robert Koch Institute | unknown |
| EPI_ISL_3121241 | amedes MVZ Hannover | Robert Koch Institute | unknown |
| EPI_ISL_3121242 | amedes MVZ für Laboratoriumsdiagnostik Raubling GmbH | Robert Koch Institute | unknown |
| EPI_ISL_3121243 | amedes MVZ Zentrum für Pathologie und Zytodiagnostik GmbH (Köln) | Robert Koch Institute | unknown |
| EPI_ISL_3121246 | amedes MVZ für Laboratoriumsmedizin Fürstenfeldbruck | Robert Koch Institute | unknown |
| EPI_ISL_3121247 | amedes MVZ Jena | Robert Koch Institute | unknown |
| EPI_ISL_3121248 | amedes MVZ DIAMEDIS Sennestadt | Robert Koch Institute | unknown |
| EPI_ISL_3121249 | amedes Medlab Arnold Analytik MVZ GmbH | Robert Koch Institute | unknown |
| EPI_ISL_3121250 | amedes MVZ DIAMEDIS Sennestadt | Robert Koch Institute | unknown |
| EPI_ISL_3121251 | amedes MVZ für Laboratoriumsmedizin Rhein/Ruhr | Robert Koch Institute | unknown |
| EPI_ISL_3121252 | amedes MVZ DIAMEDIS Sennestadt | Robert Koch Institute | unknown |
| EPI_ISL_3121253 | nordlab - Partnerschaftspraxis für Laboratoriumsmedizin | Robert Koch Institute | unknown |
| EPI_ISL_3121254 | amedes MVZ DIAMEDIS Sennestadt | Robert Koch Institute | unknown |
| EPI_ISL_3121255 | amedes MVZ Hannover | Robert Koch Institute | unknown |
| EPI_ISL_3121256 | amedes MVZ DIAMEDIS Sennestadt | Robert Koch Institute | unknown |
| EPI_ISL_3121257, EPI_ISL_3121258, EPI_ISL_3121259 | nordlab - Partnerschaftspraxis für Laboratoriumsmedizin | Robert Koch Institute | unknown |
| EPI_ISL_3121260 | amedes MVZ DIAMEDIS Sennestadt | Robert Koch Institute | unknown |
| EPI_ISL_3121261 | amedes MVZ Jena | Robert Koch Institute | unknown |
| EPI_ISL_3121262 | amedes MVZ DIAMEDIS Sennestadt | Robert Koch Institute | unknown |
| EPI_ISL_3121263 | nordlab - Partnerschaftspraxis für Laboratoriumsmedizin | Robert Koch Institute | unknown |
| EPI_ISL_3121264 | amedes Medlab Arnold Analytik MVZ GmbH | Robert Koch Institute | unknown |
| EPI_ISL_3121265 | nordlab - Partnerschaftspraxis für Laboratoriumsmedizin | Robert Koch Institute | unknown |
| EPI_ISL_3121266, EPI_ISL_3121267, EPI_ISL_3121268, EPI_ISL_3121269, EPI_ISL_3121270 | Niedersächsisches Landesgesundheitsamt (NLGA) | Robert Koch Institute | unknown |
| EPI_ISL_3121271, EPI_ISL_3121272 | Universitätsklinikum Schleswig-Holstein (standortübergreifend inkl. Kiel) - Institut für Klinische Chemie und Laboratoriumsdiagnostik | Robert Koch Institute | unknown |
| EPI_ISL_3121274, EPI_ISL_3121275, EPI_ISL_3121276, EPI_ISL_3121277, EPI_ISL_3121278, EPI_ISL_3121279, EPI_ISL_3121280, EPI_ISL_3121281, EPI_ISL_3121282, EPI_ISL_3121283, EPI_ISL_3121285, EPI_ISL_3121286, EPI_ISL_3121287, EPI_ISL_3121288, EPI_ISL_3121289, EPI_ISL_3121290, EPI_ISL_3121291, EPI_ISL_3121292, EPI_ISL_3121293, EPI_ISL_3121294, EPI_ISL_3121295, EPI_ISL_3121296, EPI_ISL_3121297, EPI_ISL_3121298, EPI_ISL_3121299, EPI_ISL_3121300, EPI_ISL_3121301, EPI_ISL_3121302, EPI_ISL_3121303, EPI_ISL_3121304, EPI_ISL_3121305, EPI_ISL_3121306, EPI_ISL_3121307, EPI_ISL_3121308, EPI_ISL_3121309, EPI_ISL_3121310, EPI_ISL_3121311, EPI_ISL_3121312, EPI_ISL_3121313, EPI_ISL_3121314, EPI_ISL_3121315, EPI_ISL_3121316 | | | |
| see above | Medizinische Laboratorien Düsseldorf | Robert Koch Institute | unknown |
| EPI_ISL_3121317 | Laborarztpraxis Dres. med. Walther Weindel & Kollegen | Robert Koch Institute | unknown |
| EPI_ISL_3121318, EPI_ISL_3121319, EPI_ISL_3121320, EPI_ISL_3121321, EPI_ISL_3121322, EPI_ISL_3121323, EPI_ISL_3121324, EPI_ISL_3121325, EPI_ISL_3121326, EPI_ISL_3121327, EPI_ISL_3121328, EPI_ISL_3121329 | | | |
| see above | Helios Universitätsklinikum Wuppertal: Universität Witten/Herdecke: Institut für Medizinische Labordiagnostik | Robert Koch Institute | unknown |
| EPI_ISL_3121330 | Thüringer Landesamtes für Verbraucherschutz | Robert Koch Institute | unknown |
| EPI_ISL_3121331, EPI_ISL_3121332, EPI_ISL_3121333, EPI_ISL_3121335, EPI_ISL_3121336, EPI_ISL_3121337, EPI_ISL_3121338, EPI_ISL_3121339, EPI_ISL_3121340, EPI_ISL_3121341, EPI_ISL_3121342, EPI_ISL_3121343, EPI_ISL_3121345, EPI_ISL_3121346, EPI_ISL_3121347, EPI_ISL_3121348, EPI_ISL_3121349, EPI_ISL_3121350, EPI_ISL_3121351, EPI_ISL_3121352, EPI_ISL_3121353, EPI_ISL_3121355, EPI_ISL_3121356, EPI_ISL_3121357, EPI_ISL_3121358, EPI_ISL_3121359, EPI_ISL_3121360, EPI_ISL_3121361, EPI_ISL_3121362, EPI_ISL_3121364, EPI_ISL_3121365, EPI_ISL_3121366, EPI_ISL_3121367, EPI_ISL_3121368, EPI_ISL_3121369, EPI_ISL_3121370, EPI_ISL_3121371, EPI_ISL_3121372, EPI_ISL_3121373, EPI_ISL_3121374, EPI_ISL_3121375, EPI_ISL_3121376, EPI_ISL_3121377, EPI_ISL_3121378, EPI_ISL_3121379, EPI_ISL_3121380, EPI_ISL_3121381, EPI_ISL_3121382, EPI_ISL_3121383, EPI_ISL_3121384, EPI_ISL_3121385, EPI_ISL_3121386, EPI_ISL_3121387, EPI_ISL_3121388, EPI_ISL_3121389, EPI_ISL_3121390, EPI_ISL_3121392, EPI_ISL_3121393, EPI_ISL_3121394, EPI_ISL_3121395, EPI_ISL_3121396, EPI_ISL_3121397, EPI_ISL_3121398, EPI_ISL_3121399, EPI_ISL_3121400, EPI_ISL_3121401, EPI_ISL_3121402, EPI_ISL_3121403, EPI_ISL_3121404, EPI_ISL_3121405, EPI_ISL_3121406, EPI_ISL_3121407, EPI_ISL_3121408, EPI_ISL_3121409, EPI_ISL_3121410, EPI_ISL_3121411, EPI_ISL_3121412, EPI_ISL_3121413, EPI_ISL_3121414, EPI_ISL_3121415, EPI_ISL_3121417, EPI_ISL_3121418, EPI_ISL_3121419, EPI_ISL_3121420, EPI_ISL_3121421, EPI_ISL_3121422, EPI_ISL_3121423, EPI_ISL_3121424, EPI_ISL_3121425, EPI_ISL_3121426, EPI_ISL_3121427, EPI_ISL_3121428, EPI_ISL_3121429, EPI_ISL_3121430, EPI_ISL_3121431, EPI_ISL_3121432, EPI_ISL_3121433, EPI_ISL_3121434, EPI_ISL_3121435, EPI_ISL_3121436, EPI_ISL_3121437, EPI_ISL_3121438, EPI_ISL_3121439, EPI_ISL_3121440, EPI_ISL_3121441, EPI_ISL_3121442, EPI_ISL_3121443, EPI_ISL_3121444, EPI_ISL_3121445, EPI_ISL_3121446, EPI_ISL_3121447, EPI_ISL_3121448, EPI_ISL_3121450, EPI_ISL_3121451, EPI_ISL_3121452, EPI_ISL_3121453, EPI_ISL_3121454, EPI_ISL_3121455, EPI_ISL_3121456, EPI_ISL_3121457, EPI_ISL_3121458, EPI_ISL_3121459, EPI_ISL_3121460, EPI_ISL_3121461, EPI_ISL_3121462, EPI_ISL_3121464, EPI_ISL_3121465, EPI_ISL_3121466, EPI_ISL_3121467, EPI_ISL_3121468, EPI_ISL_3121469, EPI_ISL_3121470, EPI_ISL_3121471, EPI_ISL_3121472, EPI_ISL_3121474, EPI_ISL_3121475, EPI_ISL_3121476, EPI_ISL_3121477 | | | |
| see above | CENTOGENE Frankfurt Laboratory: Niederlassung Industriepark Höchst | Robert Koch Institute | unknown |
| EPI_ISL_3121478 | MVZ Dr. Eberhard & Partner Dortmund | Robert Koch Institute | unknown |
| EPI_ISL_3121479, EPI_ISL_3121480 | Laborarztpraxis Osnabrück | Robert Koch Institute | unknown |
| EPI_ISL_3121481 | MVZ Dr. Eberhard & Partner Dortmund | Robert Koch Institute | unknown |
| EPI_ISL_3121482, EPI_ISL_3121483, EPI_ISL_3121484, EPI_ISL_3121485, EPI_ISL_3121486, EPI_ISL_3121487 | Laborarztpraxis Osnabrück | Robert Koch Institute | unknown |
| EPI_ISL_3121488 | MVZ Dr. Eberhard & Partner Dortmund | Robert Koch Institute | unknown |
| EPI_ISL_3121489, EPI_ISL_3121490, EPI_ISL_3121491, EPI_ISL_3121492, EPI_ISL_3121493, EPI_ISL_3121494, EPI_ISL_3121495, EPI_ISL_3121496, EPI_ISL_3121497, EPI_ISL_3121498 | Laborarztpraxis Osnabrück | Robert Koch Institute | unknown |
| EPI_ISL_3121499 | labopart - Medizinische Laboratorien Dresden | Robert Koch Institute | unknown |
| EPI_ISL_3121501, EPI_ISL_3121502, EPI_ISL_3121503 | Laborarztpraxis Osnabrück | Robert Koch Institute | unknown |
| EPI_ISL_3121504 | MVZ Dr. Eberhard & Partner Dortmund | Robert Koch Institute | unknown |
| EPI_ISL_3121505 | Labor Lübeck bzw. Laborärztliche Gemeinschaftspraxis Lübeck | Robert Koch Institute | unknown |
| EPI_ISL_3121506 | Laborarztpraxis Osnabrück | Robert Koch Institute | unknown |

| | | | |
|---|---|---|---|
| EPI_ISL_3121507 | labopart - Medizinische Laboratorien Dresden | Robert Koch Institute | unknown |
| EPI_ISL_3121508 | Labor Lübeck bzw. Laborärztliche Gemeinschaftspraxis Lübeck | Robert Koch Institute | unknown |
| EPI_ISL_3121509 | Laborarztpraxis Osnabrück | Robert Koch Institute | unknown |
| EPI_ISL_3121510 | MVZ Dr. Eberhard & Partner Dortmund | Robert Koch Institute | unknown |
| EPI_ISL_3121511, EPI_ISL_3121512, EPI_ISL_3121514, EPI_ISL_3121515, EPI_ISL_3121516, EPI_ISL_3121517 | Laborarztpraxis Osnabrück | Robert Koch Institute | unknown |
| EPI_ISL_3121518 | labopart - Medizinische Laboratorien Dresden | Robert Koch Institute | unknown |
| EPI_ISL_3121519 | Laborarztpraxis Osnabrück | Robert Koch Institute | unknown |
| EPI_ISL_3121520, EPI_ISL_3121521, EPI_ISL_3121522, EPI_ISL_3121523, EPI_ISL_3121524, EPI_ISL_3121525, EPI_ISL_3121526, EPI_ISL_3121527, EPI_ISL_3121528, EPI_ISL_3121529, EPI_ISL_3121530, EPI_ISL_3121531, EPI_ISL_3121532 | | | |
| see above | MVZ Labor Dr. Limbach & Kollegen GbR | Robert Koch Institute | unknown |
| EPI_ISL_3121533 | Bioscientia Labor Wermsdorf | Robert Koch Institute | unknown |
| EPI_ISL_3121534 | LabKom - Labor Augsburg MVZ GmbH | Robert Koch Institute | unknown |
| EPI_ISL_3121535, EPI_ISL_3121536 | Sonic - MVZ Medizinisches Labor Bremen GmbH | Robert Koch Institute | unknown |
| EPI_ISL_3121537 | Bioscientia Labor Wermsdorf | Robert Koch Institute | unknown |
| EPI_ISL_3121538 | LabKom - Labor Augsburg MVZ GmbH | Robert Koch Institute | unknown |
| EPI_ISL_3121540 | Bioscientia Labor Wermsdorf | Robert Koch Institute | unknown |
| EPI_ISL_3121541 | Sonic - Bioscientia - MVZ Labor Saar GmbH | Robert Koch Institute | unknown |
| EPI_ISL_3121542, EPI_ISL_3121543, EPI_ISL_3121544 | LabKom - Labor Augsburg MVZ GmbH | Robert Koch Institute | unknown |
| EPI_ISL_3121545 | Bioscientia MVZ Labor Karlsruhe GmbH | Robert Koch Institute | unknown |
| EPI_ISL_3121546 | Bioscientia Labor Wermsdorf | Robert Koch Institute | unknown |
| EPI_ISL_3121547 | LabKom - Labor Augsburg MVZ GmbH | Robert Koch Institute | unknown |
| EPI_ISL_3121548 | Bioscientia MVZ Labor Karlsruhe GmbH | Robert Koch Institute | unknown |
| EPI_ISL_3121549 | Bioscientia Labor Wermsdorf | Robert Koch Institute | unknown |
| EPI_ISL_3121550 | Universitätsklinikum Frankfurt - Institut für Medizinische Virologie | Robert Koch Institute | unknown |
| EPI_ISL_3121551 | LabKom - Labor Augsburg MVZ GmbH | Robert Koch Institute | unknown |
| EPI_ISL_3121552 | Labor 28 MVZ GmbH | Robert Koch Institute | unknown |
| EPI_ISL_3121553 | Sonic - Labor Staber Nürnberg | Robert Koch Institute | unknown |
| EPI_ISL_3121554 | LabKom - Labor Augsburg MVZ GmbH | Robert Koch Institute | unknown |
| EPI_ISL_3121555 | Sonic - Labor Staber Dresden (Klipphausen) | Robert Koch Institute | unknown |
| EPI_ISL_3121556, EPI_ISL_3121557 | LabKom - Labor Augsburg MVZ GmbH | Robert Koch Institute | unknown |
| EPI_ISL_3121559 | Labor 28 MVZ GmbH | Robert Koch Institute | unknown |
| EPI_ISL_3121560 | LabKom - Labor Augsburg MVZ GmbH | Robert Koch Institute | unknown |
| EPI_ISL_3121561 | Bioscientia MVZ Labor Karlsruhe GmbH | Robert Koch Institute | unknown |
| EPI_ISL_3121562 | Labor 28 MVZ GmbH | Robert Koch Institute | unknown |
| EPI_ISL_3121563 | Bioscientia MVZ Labor Karlsruhe GmbH | Robert Koch Institute | unknown |
| EPI_ISL_3121564 | Sonic - MVZ Medizinisches Labor Bremen GmbH | Robert Koch Institute | unknown |
| EPI_ISL_3121565, EPI_ISL_3121567 | Bioscientia Labor Wermsdorf | Robert Koch Institute | unknown |
| EPI_ISL_3121568 | LabKom - Labor Augsburg MVZ GmbH | Robert Koch Institute | unknown |
| EPI_ISL_3121569, EPI_ISL_3121570 | Bioscientia Labor Wermsdorf | Robert Koch Institute | unknown |
| EPI_ISL_3121571, EPI_ISL_3121572 | Labor 28 MVZ GmbH | Robert Koch Institute | unknown |
| EPI_ISL_3121573 | LabKom - Labor Augsburg MVZ GmbH | Robert Koch Institute | unknown |
| EPI_ISL_3121574, EPI_ISL_3121575, EPI_ISL_3121576 | Bioscientia Labor Wermsdorf | Robert Koch Institute | unknown |
| EPI_ISL_3121578 | Bioscientia MVZ Labor Karlsruhe GmbH | Robert Koch Institute | unknown |
| EPI_ISL_3121579 | LabKom - Labor Augsburg MVZ GmbH | Robert Koch Institute | unknown |
| EPI_ISL_3121580 | Bioscientia Labor Wermsdorf | Robert Koch Institute | unknown |
| EPI_ISL_3121581 | Labor 28 MVZ GmbH | Robert Koch Institute | unknown |
| EPI_ISL_3121582 | Sonic - MVZ Medizinisches Labor Bremen GmbH | Robert Koch Institute | unknown |
| EPI_ISL_3121583 | LabKom - Labor Augsburg MVZ GmbH | Robert Koch Institute | unknown |
| EPI_ISL_3121584 | Sonic - MVZ Medizinisches Labor Bremen GmbH | Robert Koch Institute | unknown |
| EPI_ISL_3121585 | Labor 28 MVZ GmbH | Robert Koch Institute | unknown |
| EPI_ISL_3121586 | LabKom - Labor Augsburg MVZ GmbH | Robert Koch Institute | unknown |
| EPI_ISL_3121587, EPI_ISL_3121588 | Sonic - MVZ Medizinisches Labor Bremen GmbH | Robert Koch Institute | unknown |
| EPI_ISL_3121589 | Sonic - Labor Staber Bayreuth | Robert Koch Institute | unknown |
| EPI_ISL_3121590 | Bioscientia Labor Wermsdorf | Robert Koch Institute | unknown |
| EPI_ISL_3121591, EPI_ISL_3121592 | Bioscientia MVZ Labor Karlsruhe GmbH | Robert Koch Institute | unknown |
| EPI_ISL_3121593 | LabKom - Labor Mainz MVZ GmbH | Robert Koch Institute | unknown |
| EPI_ISL_3121594 | LabKom - Labor Augsburg MVZ GmbH | Robert Koch Institute | unknown |
| EPI_ISL_3121595 | Sonic - Bioscientia - MVZ Labor Saar GmbH | Robert Koch Institute | unknown |
| EPI_ISL_3121596 | Bioscientia MVZ Labor Karlsruhe GmbH | Robert Koch Institute | unknown |
| EPI_ISL_3121597 | LabKom - Labor Augsburg MVZ GmbH | Robert Koch Institute | unknown |
| EPI_ISL_3121598, EPI_ISL_3121599, EPI_ISL_3121600, EPI_ISL_3121601, EPI_ISL_3121602, EPI_ISL_3121603 | Bioscientia Labor Wermsdorf | Robert Koch Institute | unknown |
| EPI_ISL_3121604 | LabKom - Labor Augsburg MVZ GmbH | Robert Koch Institute | unknown |

| | | | |
|---|---|---|---|
| EPI_ISL_3121605, EPI_ISL_3121606 | Bioscientia Labor Wermsdorf | Robert Koch Institute | unknown |
| EPI_ISL_3121607 | Labor 28 MVZ GmbH | Robert Koch Institute | unknown |
| EPI_ISL_3121608 | Sonic - Dr. Staber & Kollegen GmbH München | Robert Koch Institute | unknown |
| EPI_ISL_3121609 | Sonic - MVZ Medizinisches Labor Bremen GmbH | Robert Koch Institute | unknown |
| EPI_ISL_3121610 | Bioscientia Labor Wermsdorf | Robert Koch Institute | unknown |
| EPI_ISL_3121611 | Sonic - MVZ Medizinisches Labor Bremen GmbH | Robert Koch Institute | unknown |
| EPI_ISL_3121612 | Sonic - Bioscientia - MVZ Labor Saar GmbH | Robert Koch Institute | unknown |
| EPI_ISL_3121613, EPI_ISL_3121615 | Bioscientia Labor Wermsdorf | Robert Koch Institute | unknown |
| EPI_ISL_3121616 | Bioscientia MVZ Labor Karlsruhe GmbH | Robert Koch Institute | unknown |
| EPI_ISL_3121617, EPI_ISL_3121618, EPI_ISL_3121619 | Bioscientia Labor Wermsdorf | Robert Koch Institute | unknown |
| EPI_ISL_3121620, EPI_ISL_3121621 | LabKom - Labor Augsburg MVZ GmbH | Robert Koch Institute | unknown |
| EPI_ISL_3121622 | Sonic - Bioscientia - MVZ Labor Saar GmbH | Robert Koch Institute | unknown |
| EPI_ISL_3121623 | Bioscientia MVZ Labor Karlsruhe GmbH | Robert Koch Institute | unknown |
| EPI_ISL_3121624, EPI_ISL_3121625, EPI_ISL_3121626 | Bioscientia Labor Wermsdorf | Robert Koch Institute | unknown |
| EPI_ISL_3121627 | Labor 28 MVZ GmbH | Robert Koch Institute | unknown |
| EPI_ISL_3121628 | Bioscientia Labor Wermsdorf | Robert Koch Institute | unknown |
| EPI_ISL_3121629 | Sonic - MVZ Medizinisches Labor Bremen GmbH | Robert Koch Institute | unknown |
| EPI_ISL_3121630, EPI_ISL_3121631 | Bioscientia MVZ Labor Karlsruhe GmbH | Robert Koch Institute | unknown |
| EPI_ISL_3121632, EPI_ISL_3121633, EPI_ISL_3121634, EPI_ISL_3121636 | Bioscientia Labor Wermsdorf | Robert Koch Institute | unknown |
| EPI_ISL_3121637 | Sonic - Dr. Staber & Kollegen GmbH München | Robert Koch Institute | unknown |
| EPI_ISL_3121638 | Sonic - MVZ Medizinisches Labor Bremen GmbH | Robert Koch Institute | unknown |
| EPI_ISL_3121639 | Labor 28 MVZ GmbH | Robert Koch Institute | unknown |
| EPI_ISL_3121640, EPI_ISL_3121641 | Bioscientia Labor Wermsdorf | Robert Koch Institute | unknown |
| EPI_ISL_3121642 | LabKom - Labor Augsburg MVZ GmbH | Robert Koch Institute | unknown |
| EPI_ISL_3121643 | Bioscientia Labor Wermsdorf | Robert Koch Institute | unknown |
| EPI_ISL_3121644 | Sonic - Labor Staber Nürnberg | Robert Koch Institute | unknown |
| EPI_ISL_3121645, EPI_ISL_3121646 | Bioscientia Labor Wermsdorf | Robert Koch Institute | unknown |
| EPI_ISL_3121648 | Bioscientia MVZ Labor Karlsruhe GmbH | Robert Koch Institute | unknown |
| EPI_ISL_3121649 | Sonic - Bioscientia - MVZ Labor Saar GmbH | Robert Koch Institute | unknown |
| EPI_ISL_3121650 | Labor 28 MVZ GmbH | Robert Koch Institute | unknown |
| EPI_ISL_3121651 | Bioscientia MVZ Labor Karlsruhe GmbH | Robert Koch Institute | unknown |
| EPI_ISL_3121652 | Bioscientia Labor Wermsdorf | Robert Koch Institute | unknown |
| EPI_ISL_3121653 | Sonic - MVZ Medizinisches Labor Bremen GmbH | Robert Koch Institute | unknown |
| EPI_ISL_3121654 | Bioscientia Labor Wermsdorf | Robert Koch Institute | unknown |
| EPI_ISL_3121655 | LabKom - Labor Augsburg MVZ GmbH | Robert Koch Institute | unknown |
| EPI_ISL_3121656 | Labor 28 MVZ GmbH | Robert Koch Institute | unknown |
| EPI_ISL_3121657 | Bioscientia MVZ Labor Karlsruhe GmbH | Robert Koch Institute | unknown |
| EPI_ISL_3121658 | LabKom - Labor Augsburg MVZ GmbH | Robert Koch Institute | unknown |
| EPI_ISL_3121659 | Labor 28 MVZ GmbH | Robert Koch Institute | unknown |
| EPI_ISL_3121660, EPI_ISL_3121661, EPI_ISL_3121662 | Bioscientia Labor Wermsdorf | Robert Koch Institute | unknown |
| EPI_ISL_3121663 | Labor 28 MVZ GmbH | Robert Koch Institute | unknown |
| EPI_ISL_3121664 | Bioscientia MVZ Labor Karlsruhe GmbH | Robert Koch Institute | unknown |
| EPI_ISL_3121665 | Bioscientia Labor Wermsdorf | Robert Koch Institute | unknown |
| EPI_ISL_3121666, EPI_ISL_3121667 | Labor 28 MVZ GmbH | Robert Koch Institute | unknown |
| EPI_ISL_3121668 | Bioscientia MVZ Labor Karlsruhe GmbH | Robert Koch Institute | unknown |
| EPI_ISL_3121669 | Sonic - Dr. Staber & Kollegen GmbH München | Robert Koch Institute | unknown |
| EPI_ISL_3121670, EPI_ISL_3121671 | Bioscientia MVZ Labor Karlsruhe GmbH | Robert Koch Institute | unknown |
| EPI_ISL_3121672 | Sonic - Labor Staber Dresden (Klipphausen) | Robert Koch Institute | unknown |
| EPI_ISL_3121673, EPI_ISL_3121674 | Bioscientia Labor Wermsdorf | Robert Koch Institute | unknown |
| EPI_ISL_3121675 | Sonic - Bioscientia - MVZ Labor Saar GmbH | Robert Koch Institute | unknown |
| EPI_ISL_3121676 | Sonic - Labor Staber Nürnberg | Robert Koch Institute | unknown |
| EPI_ISL_3121677 | Bioscientia MVZ Labor Karlsruhe GmbH | Robert Koch Institute | unknown |
| EPI_ISL_3121678 | Sonic - Bioscientia - MVZ Labor Saar GmbH | Robert Koch Institute | unknown |
| EPI_ISL_3121679 | Sonic - MVZ Medizinisches Labor Bremen GmbH | Robert Koch Institute | unknown |
| EPI_ISL_3121680, EPI_ISL_3121681 | Bioscientia Labor Wermsdorf | Robert Koch Institute | unknown |
| EPI_ISL_3121682, EPI_ISL_3121683 | LabKom - Labor Augsburg MVZ GmbH | Robert Koch Institute | unknown |
| EPI_ISL_3121684 | Sonic - Dr. Staber & Kollegen GmbH München | Robert Koch Institute | unknown |
| EPI_ISL_3121685, EPI_ISL_3121686 | Labor 28 MVZ GmbH | Robert Koch Institute | unknown |
| EPI_ISL_3121687, EPI_ISL_3121688, EPI_ISL_3121689, EPI_ISL_3121690, EPI_ISL_3121692 | Bioscientia Labor Wermsdorf | Robert Koch Institute | unknown |

| | | | |
|---|---|---|---|
| EPI_ISL_3121693 | LabKom - Labor Augsburg MVZ GmbH | Robert Koch Institute | unknown |
| EPI_ISL_3121694, EPI_ISL_3121695, EPI_ISL_3121696, EPI_ISL_3121697 | Bioscientia Labor Wermsdorf | Robert Koch Institute | unknown |
| EPI_ISL_3121698 | Labor 28 MVZ GmbH | Robert Koch Institute | unknown |
| EPI_ISL_3121699 | Sonic - Labor Staber Nürnberg | Robert Koch Institute | unknown |
| EPI_ISL_3121700 | Bioscientia MVZ Labor Karlsruhe GmbH | Robert Koch Institute | unknown |
| EPI_ISL_3121701, EPI_ISL_3121702, EPI_ISL_3121703 | Bioscientia Labor Wermsdorf | Robert Koch Institute | unknown |
| EPI_ISL_3121704 | Sonic - Bioscientia - MVZ Labor Saar GmbH | Robert Koch Institute | unknown |
| EPI_ISL_3121705 | Labor 28 MVZ GmbH | Robert Koch Institute | unknown |
| EPI_ISL_3121706 | Bioscientia MVZ Labor Karlsruhe GmbH | Robert Koch Institute | unknown |
| EPI_ISL_3121707 | Labor 28 MVZ GmbH | Robert Koch Institute | unknown |
| EPI_ISL_3121708 | Bioscientia Labor Wermsdorf | Robert Koch Institute | unknown |
| EPI_ISL_3121709 | Labor 28 MVZ GmbH | Robert Koch Institute | unknown |
| EPI_ISL_3121710 | LabKom - Labor Hannover MVZ GmbH | Robert Koch Institute | unknown |
| EPI_ISL_3121711 | Bioscientia Labor Wermsdorf | Robert Koch Institute | unknown |
| EPI_ISL_3121712 | Sonic - Labor Staber MVZ Kassel | Robert Koch Institute | unknown |
| EPI_ISL_3121713 | Bioscientia MVZ Labor Karlsruhe GmbH | Robert Koch Institute | unknown |
| EPI_ISL_3121714 | Sonic - Bioscientia - MVZ Labor Saar GmbH | Robert Koch Institute | unknown |
| EPI_ISL_3121715 | Bioscientia MVZ Labor Karlsruhe GmbH | Robert Koch Institute | unknown |
| EPI_ISL_3121716 | Sonic - MVZ Medizinisches Labor Bremen GmbH | Robert Koch Institute | unknown |
| EPI_ISL_3121717 | Bioscientia MVZ Labor Karlsruhe GmbH | Robert Koch Institute | unknown |
| EPI_ISL_3121718 | Bioscientia Labor Wermsdorf | Robert Koch Institute | unknown |
| EPI_ISL_3121719 | Sonic - Labor Staber Dresden (Klipphausen) | Robert Koch Institute | unknown |
| EPI_ISL_3121720 | LabKom - Labor Augsburg MVZ GmbH | Robert Koch Institute | unknown |
| EPI_ISL_3121721 | Bioscientia MVZ Labor Karlsruhe GmbH | Robert Koch Institute | unknown |
| EPI_ISL_3121722 | Sonic - Bioscientia - MVZ Labor Saar GmbH | Robert Koch Institute | unknown |
| EPI_ISL_3121723, EPI_ISL_3121724 | Bioscientia Labor Wermsdorf | Robert Koch Institute | unknown |
| EPI_ISL_3121725, EPI_ISL_3121726, EPI_ISL_3121727 | LabKom - Labor Augsburg MVZ GmbH | Robert Koch Institute | unknown |
| EPI_ISL_3121728, EPI_ISL_3121730 | Bioscientia MVZ Labor Karlsruhe GmbH | Robert Koch Institute | unknown |
| EPI_ISL_3121731, EPI_ISL_3121732 | Bioscientia Labor Wermsdorf | Robert Koch Institute | unknown |
| EPI_ISL_3121733 | Sonic - MVZ Medizinisches Labor Bremen GmbH | Robert Koch Institute | unknown |
| EPI_ISL_3121734, EPI_ISL_3121735 | Labor 28 MVZ GmbH | Robert Koch Institute | unknown |
| EPI_ISL_3121736 | Bioscientia Labor Wermsdorf | Robert Koch Institute | unknown |
| EPI_ISL_3121737 | Bioscientia MVZ Labor Karlsruhe GmbH | Robert Koch Institute | unknown |
| EPI_ISL_3121738 | Sonic - Bioscientia - MVZ Labor Saar GmbH | Robert Koch Institute | unknown |
| EPI_ISL_3121739 | Labor 28 MVZ GmbH | Robert Koch Institute | unknown |
| EPI_ISL_3121740, EPI_ISL_3121741 | Bioscientia Labor Wermsdorf | Robert Koch Institute | unknown |
| EPI_ISL_3121742 | Labor 28 MVZ GmbH | Robert Koch Institute | unknown |
| EPI_ISL_3121743 | Sonic - Bioscientia - MVZ Labor Saar GmbH | Robert Koch Institute | unknown |
| EPI_ISL_3121744 | Bioscientia Labor Wermsdorf | Robert Koch Institute | unknown |
| EPI_ISL_3121745 | LabKom - Labor Mainz MVZ GmbH | Robert Koch Institute | unknown |
| EPI_ISL_3121746 | Bioscientia MVZ Labor Karlsruhe GmbH | Robert Koch Institute | unknown |
| EPI_ISL_3121747 | Sonic - MVZ Medizinisches Labor Bremen GmbH | Robert Koch Institute | unknown |
| EPI_ISL_3121748 | Bioscientia MVZ Labor Karlsruhe GmbH | Robert Koch Institute | unknown |
| EPI_ISL_3121749, EPI_ISL_3121750 | LabKom - Labor Augsburg MVZ GmbH | Robert Koch Institute | unknown |
| EPI_ISL_3121751 | Sonic - Bioscientia - MVZ Labor Saar GmbH | Robert Koch Institute | unknown |
| EPI_ISL_3121752 | Bioscientia Labor Wermsdorf | Robert Koch Institute | unknown |
| EPI_ISL_3121753 | Universitätsklinikum Frankfurt - Institut für Medizinische Virologie | Robert Koch Institute | unknown |
| EPI_ISL_3121754, EPI_ISL_3121755 | Bioscientia MVZ Labor Karlsruhe GmbH | Robert Koch Institute | unknown |
| EPI_ISL_3121756 | LabKom - Labor Augsburg MVZ GmbH | Robert Koch Institute | unknown |
| EPI_ISL_3121757 | Labor 28 MVZ GmbH | Robert Koch Institute | unknown |
| EPI_ISL_3121758, EPI_ISL_3121759, EPI_ISL_3121760 | Bioscientia Labor Wermsdorf | Robert Koch Institute | unknown |
| EPI_ISL_3121761 | Labor 28 MVZ GmbH | Robert Koch Institute | unknown |
| EPI_ISL_3121762, EPI_ISL_3121763, EPI_ISL_3121764 | Bioscientia MVZ Labor Karlsruhe GmbH | Robert Koch Institute | unknown |
| EPI_ISL_3121765, EPI_ISL_3121766, EPI_ISL_3121767 | Bioscientia Labor Wermsdorf | Robert Koch Institute | unknown |
| EPI_ISL_3121768 | Robert-Bosch-Krankenhaus | Robert Koch Institute | unknown |
| EPI_ISL_3121769 | Bioscientia Labor Wermsdorf | Robert Koch Institute | unknown |
| EPI_ISL_3121770 | LabKom - Labor Augsburg MVZ GmbH | Robert Koch Institute | unknown |
| EPI_ISL_3121771 | Bioscientia MVZ Labor Karlsruhe GmbH | Robert Koch Institute | unknown |
| EPI_ISL_3121772 | LabKom - Labor Augsburg MVZ GmbH | Robert Koch Institute | unknown |
| EPI_ISL_3121773 | Sonic - MVZ Medizinisches Labor Bremen GmbH | Robert Koch Institute | unknown |
| EPI_ISL_3121774 | Labor 28 MVZ GmbH | Robert Koch Institute | unknown |
| EPI_ISL_3121775 | Bioscientia Labor Wermsdorf | Robert Koch Institute | unknown |

| | | | |
|---|---|---|---|
| EPI_ISL_3121776 | Labor 28 MVZ GmbH | Robert Koch Institute | unknown |
| EPI_ISL_3121777, EPI_ISL_3121778 | Bioscientia MVZ Labor Karlsruhe GmbH | Robert Koch Institute | unknown |
| EPI_ISL_3121779, EPI_ISL_3121780 | Bioscientia Labor Wermsdorf | Robert Koch Institute | unknown |
| EPI_ISL_3121781 | Labor 28 MVZ GmbH | Robert Koch Institute | unknown |
| EPI_ISL_3121782 | Bioscientia MVZ Labor Karlsruhe GmbH | Robert Koch Institute | unknown |
| EPI_ISL_3121783, EPI_ISL_3121784, EPI_ISL_3121785 | alphaomega Labor GbR (Medizinische Kooperationsgemeinschaft Dres. Ackermann/Tennert/Zaiß) | Robert Koch Institute | unknown |
| EPI_ISL_3121786 | MVZ Labor Dr. Limbach & Kollegen GbR | Robert Koch Institute | unknown |
| EPI_ISL_3121787 | Limbach - MVZ Labor Eveld & Kollegen Essen | Robert Koch Institute | unknown |
| EPI_ISL_3121789, EPI_ISL_3121790, EPI_ISL_3121791, EPI_ISL_3121793, EPI_ISL_3121794, EPI_ISL_3121795, EPI_ISL_3121796 | MVZ Labor Dr. Limbach & Kollegen GbR | Robert Koch Institute | unknown |
| EPI_ISL_3121797 | Limbach - MVZ Labor Eveld & Kollegen Essen | Robert Koch Institute | unknown |
| EPI_ISL_3121798 | MVZ Labor Dr. Limbach & Kollegen GbR | Robert Koch Institute | unknown |
| EPI_ISL_3121799 | MVZ Medizinisches Labor Hannover GmbH | Robert Koch Institute | unknown |
| EPI_ISL_3121800, EPI_ISL_3121801, EPI_ISL_3121802 | MVZ für Laboratoriumsmedizin und Mikrobiologie Koblenz-Mittelrhein (Labor Koblenz) | Robert Koch Institute | unknown |
| EPI_ISL_3121803 | MVZ Labor Dr. Limbach & Kollegen GbR | Robert Koch Institute | unknown |
| EPI_ISL_3121804, EPI_ISL_3121806 | MVZ für Laboratoriumsmedizin und Mikrobiologie Koblenz-Mittelrhein (Labor Koblenz) | Robert Koch Institute | unknown |
| EPI_ISL_3121807, EPI_ISL_3121808 | MVZ Labor Dr. Limbach & Kollegen GbR | Robert Koch Institute | unknown |
| EPI_ISL_3121809, EPI_ISL_3121811 | MDI Limbach Berlin GmbH; MVZ Labor Berlin | Robert Koch Institute | unknown |
| EPI_ISL_3121812 | Labor Mönchengladbach MVZ Dr. Stein + Kollegen GbR | Robert Koch Institute | unknown |
| EPI_ISL_3121813 | MVZ Labor Dr. Limbach & Kollegen GbR | Robert Koch Institute | unknown |
| EPI_ISL_3121814 | MVZ Medizinisches Labor Hannover GmbH | Robert Koch Institute | unknown |
| EPI_ISL_3121815, EPI_ISL_3121816 | MVZ Labor Dr. Limbach & Kollegen GbR | Robert Koch Institute | unknown |
| EPI_ISL_3121817, EPI_ISL_3121818, EPI_ISL_3121819 | MVZ Medizinisches Labor Hannover GmbH | Robert Koch Institute | unknown |
| EPI_ISL_3121820 | MVZ Labor Dr. Limbach & Kollegen GbR | Robert Koch Institute | unknown |
| EPI_ISL_3121821 | MVZ Medizinisches Labor Hannover GmbH | Robert Koch Institute | unknown |
| EPI_ISL_3121822 | MVZ Labor Dr. Limbach & Kollegen GbR | Robert Koch Institute | unknown |
| EPI_ISL_3121823, EPI_ISL_3121824 | MVZ Medizinisches Labor Hannover GmbH | Robert Koch Institute | unknown |
| EPI_ISL_3121825 | MVZ Labor Dr. Limbach & Kollegen GbR | Robert Koch Institute | unknown |
| EPI_ISL_3121826 | MVZ Medizinisches Labor Hannover GmbH | Robert Koch Institute | unknown |
| EPI_ISL_3121827 | Limbach - MVZ Labor Eveld & Kollegen Essen | Robert Koch Institute | unknown |
| EPI_ISL_3121828 | Kerckhoff-Klinik GmbH | Robert Koch Institute | unknown |
| EPI_ISL_3121829, EPI_ISL_3121830 | MVZ Medizinisches Labor Hannover GmbH | Robert Koch Institute | unknown |
| EPI_ISL_3121831 | Limbach - MVZ Labor Eveld & Kollegen Essen | Robert Koch Institute | unknown |
| EPI_ISL_3121832, EPI_ISL_3121833, EPI_ISL_3121834, EPI_ISL_3121835, EPI_ISL_3121836, EPI_ISL_3121837, EPI_ISL_3121838 | Labor Mönchengladbach MVZ Dr. Stein + Kollegen GbR | Robert Koch Institute | unknown |
| EPI_ISL_3121839 | MVZ Labor Dr. Limbach & Kollegen GbR | Robert Koch Institute | unknown |
| EPI_ISL_3121840, EPI_ISL_3121841, EPI_ISL_3121842, EPI_ISL_3121843 | Labor Mönchengladbach MVZ Dr. Stein + Kollegen GbR | Robert Koch Institute | unknown |
| EPI_ISL_3121844 | Limbach - MVZ Labor Eveld & Kollegen Essen | Robert Koch Institute | unknown |
| EPI_ISL_3121845, EPI_ISL_3121846, EPI_ISL_3121847, EPI_ISL_3121849, EPI_ISL_3121850, EPI_ISL_3121851, EPI_ISL_3121853, EPI_ISL_3121854, EPI_ISL_3121855, EPI_ISL_3121856, EPI_ISL_3121857, EPI_ISL_3121858, EPI_ISL_3121859, EPI_ISL_3121860, EPI_ISL_3121861, EPI_ISL_3121862, EPI_ISL_3121863, EPI_ISL_3121864, EPI_ISL_3121865, EPI_ISL_3121866, EPI_ISL_3121867, EPI_ISL_3121868, EPI_ISL_3121869, EPI_ISL_3121870, EPI_ISL_3121871, EPI_ISL_3121872, EPI_ISL_3121873, EPI_ISL_3121874, EPI_ISL_3121875, EPI_ISL_3121876, EPI_ISL_3121877 | | | |
| see above | Labor Mönchengladbach MVZ Dr. Stein + Kollegen GbR | Robert Koch Institute | unknown |
| EPI_ISL_3121878 | MVZ Labor Dr. Limbach & Kollegen GbR | Robert Koch Institute | unknown |
| EPI_ISL_3121879, EPI_ISL_3121881, EPI_ISL_3121882, EPI_ISL_3121884, EPI_ISL_3121885, EPI_ISL_3121886, EPI_ISL_3121887, EPI_ISL_3121888 | Klinikum Stuttgart | Robert Koch Institute | unknown |
| EPI_ISL_3121889, EPI_ISL_3121890, EPI_ISL_3121891, EPI_ISL_3121892, EPI_ISL_3121893, EPI_ISL_3121894, EPI_ISL_3121895, EPI_ISL_3121896, EPI_ISL_3121897, EPI_ISL_3121898, EPI_ISL_3121899, EPI_ISL_3121900, EPI_ISL_3121901, EPI_ISL_3121902, EPI_ISL_3121903, EPI_ISL_3121904, EPI_ISL_3121905, EPI_ISL_3121906, EPI_ISL_3121907, EPI_ISL_3121908, EPI_ISL_3121909, EPI_ISL_3121910, EPI_ISL_3121911, EPI_ISL_3121912, EPI_ISL_3121913, EPI_ISL_3121914, EPI_ISL_3121915, EPI_ISL_3121916, EPI_ISL_3121917, EPI_ISL_3121918, EPI_ISL_3121920, EPI_ISL_3121921, EPI_ISL_3121922, EPI_ISL_3121923, EPI_ISL_3121924, EPI_ISL_3121925, EPI_ISL_3121926, EPI_ISL_3121927, EPI_ISL_3121928, EPI_ISL_3121929, EPI_ISL_3121930, EPI_ISL_3121931 | | | |
| see above | Eurofins MVZ Labor Gelsenkirchen | Robert Koch Institute | unknown |
| EPI_ISL_3121932, EPI_ISL_3121933, EPI_ISL_3121934, EPI_ISL_3121935, EPI_ISL_3121936, EPI_ISL_3121937, EPI_ISL_3121938, EPI_ISL_3121939, EPI_ISL_3121940, EPI_ISL_3121941, EPI_ISL_3121942, EPI_ISL_3121943, EPI_ISL_3121944, EPI_ISL_3121945, EPI_ISL_3121946, EPI_ISL_3121947, EPI_ISL_3121948, EPI_ISL_3121949, EPI_ISL_3121950, EPI_ISL_3121951, EPI_ISL_3121952, EPI_ISL_3121953, EPI_ISL_3121954, EPI_ISL_3121955, EPI_ISL_3121956, EPI_ISL_3121957, EPI_ISL_3121958, EPI_ISL_3121959, EPI_ISL_3121960, EPI_ISL_3121961, EPI_ISL_3121962, EPI_ISL_3121963, EPI_ISL_3121964, EPI_ISL_3121965, EPI_ISL_3121966, EPI_ISL_3121968, EPI_ISL_3121969, EPI_ISL_3121970, EPI_ISL_3121971, EPI_ISL_3121972, EPI_ISL_3121973, EPI_ISL_3121974 | | | |
| see above | MVZ Labor Krone GbR | Robert Koch Institute | unknown |
| EPI_ISL_3121975 | MVZ Medizinisches Labor Hannover GmbH | Robert Koch Institute | unknown |
| EPI_ISL_3121976, EPI_ISL_3121977, EPI_ISL_3121978, EPI_ISL_3121979, EPI_ISL_3121980, EPI_ISL_3121981, EPI_ISL_3121982, EPI_ISL_3121983, EPI_ISL_3121984, EPI_ISL_3121985, EPI_ISL_3121986, EPI_ISL_3121987, EPI_ISL_3121988, EPI_ISL_3121989 | | | |
| see above | MDI Limbach Berlin GmbH; MVZ Labor Berlin | Robert Koch Institute | unknown |
| EPI_ISL_3121990 | Wyoming Public Health Laboratory | Wyoming Public Health Laboratory | Jim Mildenberger, Taylor Fearing, Channing Weber, Ashley Norberg, Chayse Rowley, Marley Goetz, Brian Dominguez, Elliot Thomasson, Sam Britz, Cari Sloma, Robert Petit, and Rob Christensen |
| EPI_ISL_3121991, EPI_ISL_3121992, | MDI Limbach Berlin GmbH; MVZ Labor Berlin | Robert Koch Institute | unknown |

| | | | |
|---|---|---|---|
| EPI_ISL_3121993 | | | |
| EPI_ISL_3121994, EPI_ISL_3121995 | Wyoming Public Health Laboratory | Wyoming Public Health Laboratory | Jim Mildenberger, Taylor Fearing, Channing Weber, Ashley Norberg, Chayse Rowley, Marley Goetz, Brian Dominguez, Elliot Thomasson, Sam Britz, Cari Sloma, Robert Petit, and Rob Christensen |
| EPI_ISL_3121996, EPI_ISL_3121997 | MDI Limbach Berlin GmbH; MVZ Labor Berlin | Robert Koch Institute | unknown |
| EPI_ISL_3121998 | Wyoming Public Health Laboratory | Wyoming Public Health Laboratory | Jim Mildenberger, Taylor Fearing, Channing Weber, Ashley Norberg, Chayse Rowley, Marley Goetz, Brian Dominguez, Elliot Thomasson, Sam Britz, Cari Sloma, Robert Petit, and Rob Christensen |
| EPI_ISL_3121999 | MDI Limbach Berlin GmbH; MVZ Labor Berlin | Robert Koch Institute | unknown |
| EPI_ISL_3122000 | Wyoming Public Health Laboratory | Wyoming Public Health Laboratory | Jim Mildenberger, Taylor Fearing, Channing Weber, Ashley Norberg, Chayse Rowley, Marley Goetz, Brian Dominguez, Elliot Thomasson, Sam Britz, Cari Sloma, Robert Petit, and Rob Christensen |
| EPI_ISL_3122001 | MDI Limbach Berlin GmbH; MVZ Labor Berlin | Robert Koch Institute | unknown |
| EPI_ISL_3122002 | Wyoming Public Health Laboratory | Wyoming Public Health Laboratory | Jim Mildenberger, Taylor Fearing, Channing Weber, Ashley Norberg, Chayse Rowley, Marley Goetz, Brian Dominguez, Elliot Thomasson, Sam Britz, Cari Sloma, Robert Petit, and Rob Christensen |
| EPI_ISL_3122003 | MDI Limbach Berlin GmbH; MVZ Labor Berlin | Robert Koch Institute | unknown |
| EPI_ISL_3122004 | Wyoming Public Health Laboratory | Wyoming Public Health Laboratory | Jim Mildenberger, Taylor Fearing, Channing Weber, Ashley Norberg, Chayse Rowley, Marley Goetz, Brian Dominguez, Elliot Thomasson, Sam Britz, Cari Sloma, Robert Petit, and Rob Christensen |
| EPI_ISL_3122005 | MDI Limbach Berlin GmbH; MVZ Labor Berlin | Robert Koch Institute | unknown |
| EPI_ISL_3122006 | Wyoming Public Health Laboratory | Wyoming Public Health Laboratory | Jim Mildenberger, Taylor Fearing, Channing Weber, Ashley Norberg, Chayse Rowley, Marley Goetz, Brian Dominguez, Elliot Thomasson, Sam Britz, Cari Sloma, Robert Petit, and Rob Christensen |
| EPI_ISL_3122007 | MDI Limbach Berlin GmbH; MVZ Labor Berlin | Robert Koch Institute | unknown |
| EPI_ISL_3122008 | Wyoming Public Health Laboratory | Wyoming Public Health Laboratory | Jim Mildenberger, Taylor Fearing, Channing Weber, Ashley Norberg, Chayse Rowley, Marley Goetz, Brian Dominguez, Elliot Thomasson, Sam Britz, Cari Sloma, Robert Petit, and Rob Christensen |
| EPI_ISL_3122009 | MDI Limbach Berlin GmbH; MVZ Labor Berlin | Robert Koch Institute | unknown |
| EPI_ISL_3122010 | Wyoming Public Health Laboratory | Wyoming Public Health Laboratory | Jim Mildenberger, Taylor Fearing, Channing Weber, Ashley Norberg, Chayse Rowley, Marley Goetz, Brian Dominguez, Elliot Thomasson, Sam Britz, Cari Sloma, Robert Petit, and Rob Christensen |
| EPI_ISL_3122011 | MDI Limbach Berlin GmbH; MVZ Labor Berlin | Robert Koch Institute | unknown |
| EPI_ISL_3122012 | Wyoming Public Health Laboratory | Wyoming Public Health Laboratory | Jim Mildenberger, Taylor Fearing, Channing Weber, Ashley Norberg, Chayse Rowley, Marley Goetz, Brian Dominguez, Elliot Thomasson, Sam Britz, Cari Sloma, Robert Petit, and Rob Christensen |
| EPI_ISL_3122013 | MDI Limbach Berlin GmbH; MVZ Labor Berlin | Robert Koch Institute | unknown |
| EPI_ISL_3122014 | Wyoming Public Health Laboratory | Wyoming Public Health Laboratory | Jim Mildenberger, Taylor Fearing, Channing Weber, Ashley Norberg, Chayse Rowley, Marley Goetz, Brian Dominguez, Elliot Thomasson, Sam Britz, Cari Sloma, Robert Petit, and Rob Christensen |
| EPI_ISL_3122015 | MVZ Medizinisches Labor Hannover GmbH | Robert Koch Institute | unknown |
| EPI_ISL_3122016 | Wyoming Public Health Laboratory | Wyoming Public Health Laboratory | Jim Mildenberger, Taylor Fearing, Channing Weber, Ashley Norberg, Chayse Rowley, Marley Goetz, Brian Dominguez, Elliot Thomasson, Sam Britz, Cari Sloma, Robert Petit, and Rob Christensen |
| EPI_ISL_3122017 | MDI Limbach Berlin GmbH; MVZ Labor Berlin | Robert Koch Institute | unknown |
| EPI_ISL_3122018 | Wyoming Public Health Laboratory | Wyoming Public Health Laboratory | Jim Mildenberger, Taylor Fearing, Channing Weber, Ashley Norberg, Chayse Rowley, Marley Goetz, Brian Dominguez, Elliot Thomasson, Sam Britz, Cari Sloma, Robert Petit, and Rob Christensen |
| EPI_ISL_3122019, EPI_ISL_3122020 | MDI Limbach Berlin GmbH; MVZ Labor Berlin | Robert Koch Institute | unknown |
| EPI_ISL_3122021 | Wyoming Public Health Laboratory | Wyoming Public Health Laboratory | Jim Mildenberger, Taylor Fearing, Channing Weber, Ashley Norberg, Chayse Rowley, Marley Goetz, Brian Dominguez, Elliot Thomasson, Sam Britz, Cari Sloma, Robert Petit, and Rob Christensen |
| EPI_ISL_3122022 | MDI Limbach Berlin GmbH; MVZ Labor Berlin | Robert Koch Institute | unknown |
| EPI_ISL_3122023 | Wyoming Public Health Laboratory | Wyoming Public Health Laboratory | Jim Mildenberger, Taylor Fearing, Channing Weber, Ashley Norberg, Chayse Rowley, Marley Goetz, Brian Dominguez, Elliot Thomasson, Sam Britz, Cari Sloma, Robert Petit, and Rob Christensen |
| EPI_ISL_3122024 | MDI Limbach Berlin GmbH; MVZ Labor Berlin | Robert Koch Institute | unknown |
| EPI_ISL_3122025, EPI_ISL_3122026 | Wyoming Public Health Laboratory | Wyoming Public Health Laboratory | Jim Mildenberger, Taylor Fearing, Channing Weber, Ashley Norberg, Chayse Rowley, Marley Goetz, Brian Dominguez, Elliot Thomasson, Sam Britz, Cari Sloma, Robert Petit, and Rob Christensen |
| EPI_ISL_3122027, EPI_ISL_3122028 | MDI Limbach Berlin GmbH; MVZ Labor Berlin | Robert Koch Institute | unknown |
| EPI_ISL_3122029 | Wyoming Public Health Laboratory | Wyoming Public Health Laboratory | Jim Mildenberger, Taylor Fearing, Channing Weber, Ashley Norberg, Chayse Rowley, Marley Goetz, Brian Dominguez, Elliot Thomasson, Sam Britz, Cari Sloma, Robert Petit, and Rob Christensen |
| EPI_ISL_3122030 | MDI Limbach Berlin GmbH; MVZ Labor Berlin | Robert Koch Institute | unknown |
| EPI_ISL_3122031 | Wyoming Public Health Laboratory | Wyoming Public Health Laboratory | Jim Mildenberger, Taylor Fearing, Channing Weber, Ashley Norberg, Chayse Rowley, Marley Goetz, Brian Dominguez, Elliot Thomasson, Sam Britz, Cari Sloma, Robert Petit, and Rob Christensen |
| EPI_ISL_3122032 | MDI Limbach Berlin GmbH; MVZ Labor Berlin | Robert Koch Institute | unknown |
| EPI_ISL_3122033 | Wyoming Public Health Laboratory | Wyoming Public Health Laboratory | Jim Mildenberger, Taylor Fearing, Channing Weber, Ashley Norberg, Chayse Rowley, Marley Goetz, Brian Dominguez, Elliot Thomasson, Sam Britz, Cari Sloma, Robert Petit, and Rob Christensen |
| EPI_ISL_3122034, EPI_ISL_3122036, EPI_ISL_3122038, EPI_ISL_3122039 | MDI Limbach Berlin GmbH; MVZ Labor Berlin | Robert Koch Institute | unknown |
| EPI_ISL_3122040, EPI_ISL_3122041 | Wyoming Public Health Laboratory | Wyoming Public Health Laboratory | Jim Mildenberger, Taylor Fearing, Channing Weber, Ashley Norberg, Chayse Rowley, Marley Goetz, Brian Dominguez, Elliot Thomasson, Sam Britz, Cari Sloma, Robert Petit, and Rob Christensen |
| EPI_ISL_3122042, EPI_ISL_3122044 | MDI Limbach Berlin GmbH; MVZ Labor Berlin | Robert Koch Institute | unknown |
| EPI_ISL_3122046 | MVZ Medizinisches Labor Hannover GmbH | Robert Koch Institute | unknown |
| EPI_ISL_3122048 | MDI Limbach Berlin GmbH; MVZ Labor Berlin | Robert Koch Institute | unknown |
| EPI_ISL_3122050 | MVZ Medizinisches Labor Hannover GmbH | Robert Koch Institute | unknown |
| EPI_ISL_3122051 | MDI Limbach Berlin GmbH; MVZ Labor Berlin | Robert Koch Institute | unknown |
| EPI_ISL_3122053, EPI_ISL_3122055, EPI_ISL_3122056, EPI_ISL_3122057, EPI_ISL_3122058, EPI_ISL_3122059 | MVZ Medizinisches Labor Hannover GmbH | Robert Koch Institute | unknown |
| EPI_ISL_3122060 | Limbach - MVZ Gemeinschaftslabor Suhl | Robert Koch Institute | unknown |
| EPI_ISL_3122061 | MVZ Medizinisches Labor Hannover GmbH | Robert Koch Institute | unknown |
| EPI_ISL_3122062 | Limbach - MVZ Gemeinschaftslabor Suhl | Robert Koch Institute | unknown |
| EPI_ISL_3122063, EPI_ISL_3122064 | MVZ Medizinisches Labor Hannover GmbH | Robert Koch Institute | unknown |
| EPI_ISL_3122065 | MDI Limbach Berlin GmbH; MVZ Labor Berlin | Robert Koch Institute | unknown |
| EPI_ISL_3122066, EPI_ISL_3122067, EPI_ISL_3122068, EPI_ISL_3122069, EPI_ISL_3122070 | MVZ Medizinisches Labor Hannover GmbH | Robert Koch Institute | unknown |
| EPI_ISL_3122071 | MDI Limbach Berlin GmbH; MVZ Labor Berlin | Robert Koch Institute | unknown |
| EPI_ISL_3122072 | MVZ Medizinisches Labor Hannover GmbH | Robert Koch Institute | unknown |
| EPI_ISL_3122073 | MDI Limbach Berlin GmbH; MVZ Labor Berlin | Robert Koch Institute | unknown |
| EPI_ISL_3122074, EPI_ISL_3122075, EPI_ISL_3122076 | MVZ Medizinisches Labor Hannover GmbH | Robert Koch Institute | unknown |
| EPI_ISL_3122077 | MDI Limbach Berlin GmbH; MVZ Labor Berlin | Robert Koch Institute | unknown |
| EPI_ISL_3122078 | MVZ Medizinisches Labor Hannover GmbH | Robert Koch Institute | unknown |
| EPI_ISL_3122079 | MDI Limbach Berlin GmbH; MVZ Labor Berlin | Robert Koch Institute | unknown |
| EPI_ISL_3122080, EPI_ISL_3122081 | MVZ Medizinisches Labor Hannover GmbH | Robert Koch Institute | unknown |
| EPI_ISL_3122082 | MDI Limbach Berlin GmbH; MVZ Labor Berlin | Robert Koch Institute | unknown |
| EPI_ISL_3122083, | MVZ Medizinisches Labor Hannover GmbH | Robert Koch Institute | unknown |

| | | | |
|---|---|---|---|
| EPI_ISL_3122084 | | | |
| EPI_ISL_3122085 | MDI Limbach Berlin GmbH; MVZ Labor Berlin | Robert Koch Institute | unknown |
| EPI_ISL_3122086, EPI_ISL_3122087, EPI_ISL_3122088 | MVZ Medizinisches Labor Hannover GmbH | Robert Koch Institute | unknown |
| EPI_ISL_3122089 | MDI Limbach Berlin GmbH; MVZ Labor Berlin | Robert Koch Institute | unknown |
| EPI_ISL_3122090, EPI_ISL_3122091, EPI_ISL_3122092, EPI_ISL_3122093 | MVZ Medizinisches Labor Hannover GmbH | Robert Koch Institute | unknown |
| EPI_ISL_3122094 | Klinikum Darmstadt GmbH | Robert Koch Institute | unknown |
| EPI_ISL_3122095, EPI_ISL_3122096, EPI_ISL_3122097, EPI_ISL_3122098, EPI_ISL_3122099, EPI_ISL_3122100, EPI_ISL_3122101, EPI_ISL_3122102, EPI_ISL_3122103, EPI_ISL_3122104, EPI_ISL_3122105, EPI_ISL_3122106, EPI_ISL_3122107, EPI_ISL_3122108, EPI_ISL_3122109, EPI_ISL_3122110, EPI_ISL_3122111, EPI_ISL_3122112, EPI_ISL_3122113, EPI_ISL_3122114, EPI_ISL_3122115, EPI_ISL_3122116, EPI_ISL_3122117 | | | |
| see above | MDI Limbach Berlin GmbH; MVZ Labor Berlin | Robert Koch Institute | unknown |
| EPI_ISL_3122119, EPI_ISL_3122120, EPI_ISL_3122121, EPI_ISL_3122122, EPI_ISL_3122123 | IMD - MVZ Labor Greifswald | Robert Koch Institute | unknown |
| EPI_ISL_3122124, EPI_ISL_3122125 | IMD - MVZ Labor Martinsried | Robert Koch Institute | unknown |
| EPI_ISL_3122126, EPI_ISL_3122127, EPI_ISL_3122128, EPI_ISL_3122129 | IMD - MVZ Labor Greifswald | Robert Koch Institute | unknown |
| EPI_ISL_3122130 | IMD - Labor Oderland | Robert Koch Institute | unknown |
| EPI_ISL_3122131, EPI_ISL_3122132, EPI_ISL_3122133 | IMD - MVZ Labor Greifswald | Robert Koch Institute | unknown |
| EPI_ISL_3122134 | MVZ Labor Dr. Limbach & Kollegen GbR | Robert Koch Institute | unknown |
| EPI_ISL_3122135, EPI_ISL_3122136, EPI_ISL_3122137, EPI_ISL_3122138, EPI_ISL_3122139, EPI_ISL_3122140, EPI_ISL_3122141, EPI_ISL_3122142, EPI_ISL_3122143, EPI_ISL_3122144, EPI_ISL_3122145, EPI_ISL_3122146, EPI_ISL_3122147, EPI_ISL_3122148, EPI_ISL_3122150, EPI_ISL_3122151, EPI_ISL_3122152, EPI_ISL_3122153, EPI_ISL_3122154, EPI_ISL_3122155, EPI_ISL_3122156, EPI_ISL_3122157, EPI_ISL_3122158, EPI_ISL_3122159, EPI_ISL_3122160, EPI_ISL_3122161, EPI_ISL_3122162, EPI_ISL_3122163, EPI_ISL_3122164, EPI_ISL_3122165, EPI_ISL_3122166, EPI_ISL_3122167, EPI_ISL_3122168, EPI_ISL_3122169, EPI_ISL_3122170, EPI_ISL_3122171, EPI_ISL_3122172, EPI_ISL_3122173, EPI_ISL_3122174, EPI_ISL_3122175, EPI_ISL_3122176, EPI_ISL_3122177, EPI_ISL_3122178, EPI_ISL_3122179, EPI_ISL_3122180, EPI_ISL_3122181, EPI_ISL_3122182, EPI_ISL_3122183, EPI_ISL_3122184, EPI_ISL_3122186, EPI_ISL_3122187, EPI_ISL_3122188, EPI_ISL_3122189, EPI_ISL_3122190, EPI_ISL_3122191, EPI_ISL_3122192, EPI_ISL_3122193, EPI_ISL_3122194, EPI_ISL_3122196, EPI_ISL_3122197, EPI_ISL_3122198, EPI_ISL_3122199, EPI_ISL_3122200, EPI_ISL_3122201, EPI_ISL_3122202, EPI_ISL_3122203, EPI_ISL_3122204, EPI_ISL_3122205, EPI_ISL_3122206, EPI_ISL_3122207, EPI_ISL_3122208, EPI_ISL_3122209, EPI_ISL_3122210, EPI_ISL_3122211, EPI_ISL_3122212, EPI_ISL_3122213, EPI_ISL_3122214, EPI_ISL_3122215, EPI_ISL_3122216, EPI_ISL_3122217, EPI_ISL_3122218, EPI_ISL_3122219, EPI_ISL_3122220, EPI_ISL_3122221, EPI_ISL_3122222, EPI_ISL_3122223, EPI_ISL_3122224, EPI_ISL_3122225, EPI_ISL_3122226, EPI_ISL_3122227, EPI_ISL_3122228, EPI_ISL_3122229, EPI_ISL_3122230, EPI_ISL_3122231 | | | |
| see above | Eurofins LifeCodexx GmbH | Robert Koch Institute | unknown |
| EPI_ISL_3122232, EPI_ISL_3122233, EPI_ISL_3122235, EPI_ISL_3122236 | Bayerisches Landesamt für Gesundheit und Lebensmittelsicherheit (LGL) | Robert Koch Institute | unknown |
| EPI_ISL_3122240, EPI_ISL_3122241, EPI_ISL_3122243, EPI_ISL_3122244, EPI_ISL_3122245, EPI_ISL_3122246, EPI_ISL_3122247, EPI_ISL_3122248, EPI_ISL_3122249, EPI_ISL_3122250, EPI_ISL_3122251, EPI_ISL_3122252, EPI_ISL_3122253, EPI_ISL_3122254, EPI_ISL_3122255, EPI_ISL_3122256, EPI_ISL_3122257, EPI_ISL_3122258, EPI_ISL_3122259, EPI_ISL_3122260, EPI_ISL_3122261, EPI_ISL_3122262, EPI_ISL_3122263, EPI_ISL_3122264, EPI_ISL_3122265, EPI_ISL_3122266, EPI_ISL_3122267 | | | |
| see above | MVZ Labor Dr. Fenner und Kollegen (Standort Hamburg) | Robert Koch Institute | unknown |
| EPI_ISL_3122268 | SYNLAB MVZ Weiden | Robert Koch Institute | unknown |
| EPI_ISL_3122269 | SYNLAB Labor München Zentrum LMZ | Robert Koch Institute | unknown |
| EPI_ISL_3122270, EPI_ISL_3122271, EPI_ISL_3122272, EPI_ISL_3122273 | SYNLAB MVZ Leverkusen | Robert Koch Institute | unknown |
| EPI_ISL_3122275 | SYNLAB MVZ Weiden | Robert Koch Institute | unknown |
| EPI_ISL_3122276 | SYNLAB MVZ Trier | Robert Koch Institute | unknown |
| EPI_ISL_3122277, EPI_ISL_3122278 | SYNLAB MVZ Leverkusen | Robert Koch Institute | unknown |
| EPI_ISL_3122279 | SYNLAB Labor München Zentrum LMZ | Robert Koch Institute | unknown |
| EPI_ISL_3122280 | SYNLAB MVZ Weiden | Robert Koch Institute | unknown |
| EPI_ISL_3122281 | SYNLAB MVZ Leverkusen | Robert Koch Institute | unknown |
| EPI_ISL_3122283, EPI_ISL_3122284 | SYNLAB Labor München Zentrum LMZ | Robert Koch Institute | unknown |
| EPI_ISL_3122285 | SYNLAB MVZ Weiden | Robert Koch Institute | unknown |
| EPI_ISL_3122286 | SYNLAB MVZ Trier | Robert Koch Institute | unknown |
| EPI_ISL_3122287, EPI_ISL_3122288 | SYNLAB MVZ Weiden | Robert Koch Institute | unknown |
| EPI_ISL_3122289, EPI_ISL_3122290, EPI_ISL_3122291, EPI_ISL_3122293, EPI_ISL_3122294 | SYNLAB MVZ Leverkusen | Robert Koch Institute | unknown |
| EPI_ISL_3122295 | SYNLAB MVZ Trier | Robert Koch Institute | unknown |
| EPI_ISL_3122296 | SYNLAB MVZ Leverkusen | Robert Koch Institute | unknown |
| EPI_ISL_3122297 | SYNLAB MVZ Weiden | Robert Koch Institute | unknown |
| EPI_ISL_3122298 | SYNLAB MVZ Dachau | Robert Koch Institute | unknown |
| EPI_ISL_3122299 | SYNLAB MVZ Leinfelden-Echterdingen | Robert Koch Institute | unknown |
| EPI_ISL_3122300, EPI_ISL_3122301 | SYNLAB MVZ Leverkusen | Robert Koch Institute | unknown |
| EPI_ISL_3122302 | Synlab MVZ Augsburg | Robert Koch Institute | unknown |
| EPI_ISL_3122303, EPI_ISL_3122304 | SYNLAB MVZ Leverkusen | Robert Koch Institute | unknown |
| EPI_ISL_3122305 | SYNLAB MVZ Weiden | Robert Koch Institute | unknown |
| EPI_ISL_3122306, EPI_ISL_3122307 | SYNLAB MVZ Leverkusen | Robert Koch Institute | unknown |
| EPI_ISL_3122308 | SYNLAB MVZ Trier | Robert Koch Institute | unknown |
| EPI_ISL_3122309 | SYNLAB Jena Oncoscreen | Robert Koch Institute | unknown |
| EPI_ISL_3122310, EPI_ISL_3122311, EPI_ISL_3122312, EPI_ISL_3122313, EPI_ISL_3122314, EPI_ISL_3122315, EPI_ISL_3122316 | SYNLAB MVZ Leverkusen | Robert Koch Institute | unknown |
| EPI_ISL_3122317 | Synlab MVZ Augsburg | Robert Koch Institute | unknown |

| | | | |
|---|---|---|---|
| EPI_ISL_3122318 | SYNLAB MVZ Leinfelden-Echterdingen | Robert Koch Institute | unknown |
| EPI_ISL_3122319 | SYNLAB MVZ Leverkusen | Robert Koch Institute | unknown |
| EPI_ISL_3122320, EPI_ISL_3122321 | SYNLAB MVZ Weiden | Robert Koch Institute | unknown |
| EPI_ISL_3122322, EPI_ISL_3122323, EPI_ISL_3122324, EPI_ISL_3122325 | SYNLAB MVZ Leverkusen | Robert Koch Institute | unknown |
| EPI_ISL_3122326 | SYNLAB MVZ Weiden | Robert Koch Institute | unknown |
| EPI_ISL_3122327 | Synlab MVZ Augsburg | Robert Koch Institute | unknown |
| EPI_ISL_3122328 | SYNLAB MVZ Leverkusen | Robert Koch Institute | unknown |
| EPI_ISL_3122329, EPI_ISL_3122330 | SYNLAB MVZ Weiden | Robert Koch Institute | unknown |
| EPI_ISL_3122331, EPI_ISL_3122332, EPI_ISL_3122333, EPI_ISL_3122334 | SYNLAB MVZ Leverkusen | Robert Koch Institute | unknown |
| EPI_ISL_3122335 | SYNLAB Labor München Zentrum LMZ | Robert Koch Institute | unknown |
| EPI_ISL_3122336 | SYNLAB MVZ Weiden | Robert Koch Institute | unknown |
| EPI_ISL_3122337 | SYNLAB MVZ Leverkusen | Robert Koch Institute | unknown |
| EPI_ISL_3122338 | SYNLAB MVZ Weiden | Robert Koch Institute | unknown |
| EPI_ISL_3122339 | SYNLAB MVZ Leverkusen | Robert Koch Institute | unknown |
| EPI_ISL_3122340 | SYNLAB Labor München Zentrum LMZ | Robert Koch Institute | unknown |
| EPI_ISL_3122341 | SYNLAB MVZ Weiden | Robert Koch Institute | unknown |
| EPI_ISL_3122342 | SYNLAB MVZ Trier | Robert Koch Institute | unknown |
| EPI_ISL_3122343 | SYNLAB MVZ Weiden | Robert Koch Institute | unknown |
| EPI_ISL_3122344 | SYNLAB Jena Oncoscreen | Robert Koch Institute | unknown |
| EPI_ISL_3122345 | SYNLAB Labor München Zentrum LMZ | Robert Koch Institute | unknown |
| EPI_ISL_3122346 | SYNLAB MVZ Leverkusen | Robert Koch Institute | unknown |
| EPI_ISL_3122347 | SYNLAB MVZ Weiden | Robert Koch Institute | unknown |
| EPI_ISL_3122348 | SYNLAB MVZ Leverkusen | Robert Koch Institute | unknown |
| EPI_ISL_3122349 | SYNLAB MVZ Hamburg | Robert Koch Institute | unknown |
| EPI_ISL_3122350 | SYNLAB MVZ Weiden | Robert Koch Institute | unknown |
| EPI_ISL_3122351 | SYNLAB MVZ Trier | Robert Koch Institute | unknown |
| EPI_ISL_3122352 | Synlab MVZ Augsburg | Robert Koch Institute | unknown |
| EPI_ISL_3122353 | SYNLAB MVZ Weiden | Robert Koch Institute | unknown |
| EPI_ISL_3122354 | SYNLAB MVZ Leverkusen | Robert Koch Institute | unknown |
| EPI_ISL_3122355 | SYNLAB MVZ Leinfelden-Echterdingen | Robert Koch Institute | unknown |
| EPI_ISL_3122356 | SYNLAB MVZ Weiden | Robert Koch Institute | unknown |
| EPI_ISL_3122357 | SYNLAB MVZ Leverkusen | Robert Koch Institute | unknown |
| EPI_ISL_3122358 | SYNLAB MVZ Hamburg | Robert Koch Institute | unknown |
| EPI_ISL_3122359, EPI_ISL_3122360 | Synlab MVZ Augsburg | Robert Koch Institute | unknown |
| EPI_ISL_3122361 | SYNLAB MVZ Leverkusen | Robert Koch Institute | unknown |
| EPI_ISL_3122362 | SYNLAB MVZ Weiden | Robert Koch Institute | unknown |
| EPI_ISL_3122363 | Synlab MVZ Augsburg | Robert Koch Institute | unknown |
| EPI_ISL_3122364, EPI_ISL_3122365, EPI_ISL_3122366, EPI_ISL_3122367, EPI_ISL_3122368 | SYNLAB MVZ Leverkusen | Robert Koch Institute | unknown |
| EPI_ISL_3122369 | Synlab MVZ Augsburg | Robert Koch Institute | unknown |
| EPI_ISL_3122370, EPI_ISL_3122371, EPI_ISL_3122372 | SYNLAB MVZ Leverkusen | Robert Koch Institute | unknown |
| EPI_ISL_3122373 | SYNLAB MVZ Hamburg | Robert Koch Institute | unknown |
| EPI_ISL_3122375 | SYNLAB MVZ Leverkusen | Robert Koch Institute | unknown |
| EPI_ISL_3122376 | SYNLAB Labor München Zentrum LMZ | Robert Koch Institute | unknown |
| EPI_ISL_3122377 | SYNLAB MVZ Leverkusen | Robert Koch Institute | unknown |
| EPI_ISL_3122378, EPI_ISL_3122379, EPI_ISL_3122380 | SYNLAB Labor München Zentrum LMZ | Robert Koch Institute | unknown |
| EPI_ISL_3122381, EPI_ISL_3122382, EPI_ISL_3122383, EPI_ISL_3122384 | SYNLAB MVZ Leverkusen | Robert Koch Institute | unknown |
| EPI_ISL_3122385 | SYNLAB MVZ Trier | Robert Koch Institute | unknown |
| EPI_ISL_3122386 | SYNLAB MVZ Leverkusen | Robert Koch Institute | unknown |
| EPI_ISL_3122387 | SYNLAB MVZ Trier | Robert Koch Institute | unknown |
| EPI_ISL_3122388, EPI_ISL_3122389 | SYNLAB MVZ Leinfelden-Echterdingen | Robert Koch Institute | unknown |
| EPI_ISL_3122390 | SYNLAB MVZ Leverkusen | Robert Koch Institute | unknown |
| EPI_ISL_3122391 | SYNLAB Labor München Zentrum LMZ | Robert Koch Institute | unknown |
| EPI_ISL_3122392 | SYNLAB MVZ Leverkusen | Robert Koch Institute | unknown |
| EPI_ISL_3122393 | SYNLAB Labor München Zentrum LMZ | Robert Koch Institute | unknown |
| EPI_ISL_3122394 | SYNLAB MVZ Leverkusen | Robert Koch Institute | unknown |
| EPI_ISL_3122395 | SYNLAB Labor München Zentrum LMZ | Robert Koch Institute | unknown |
| EPI_ISL_3122396, EPI_ISL_3122397 | SYNLAB MVZ Weiden | Robert Koch Institute | unknown |
| EPI_ISL_3122398, EPI_ISL_3122399, EPI_ISL_3122400 | SYNLAB MVZ Trier | Robert Koch Institute | unknown |
| EPI_ISL_3122401 | SYNLAB MVZ Weiden | Robert Koch Institute | unknown |
| EPI_ISL_3122402 | SYNLAB MVZ Trier | Robert Koch Institute | unknown |

| | | | |
|---|---|---|---|
| EPI_ISL_3122403, EPI_ISL_3122404, EPI_ISL_3122405, EPI_ISL_3122406 | SYNLAB MVZ Leverkusen | Robert Koch Institute | unknown |
| EPI_ISL_3122407, EPI_ISL_3122408 | SYNLAB MVZ Berlin | Robert Koch Institute | unknown |
| EPI_ISL_3122410, EPI_ISL_3122411, EPI_ISL_3122412 | SYNLAB MVZ Leverkusen | Robert Koch Institute | unknown |
| EPI_ISL_3122413 | SYNLAB Labor München Zentrum LMZ | Robert Koch Institute | unknown |
| EPI_ISL_3122414, EPI_ISL_3122415, EPI_ISL_3122416, EPI_ISL_3122417 | SYNLAB MVZ Leverkusen | Robert Koch Institute | unknown |
| EPI_ISL_3122418 | SYNLAB Labor München Zentrum LMZ | Robert Koch Institute | unknown |
| EPI_ISL_3122419 | SYNLAB MVZ Weiden | Robert Koch Institute | unknown |
| EPI_ISL_3122420 | SYNLAB MVZ Leverkusen | Robert Koch Institute | unknown |
| EPI_ISL_3122421 | SYNLAB MVZ Trier | Robert Koch Institute | unknown |
| EPI_ISL_3122422 | SYNLAB MVZ Weiden | Robert Koch Institute | unknown |
| EPI_ISL_3122423, EPI_ISL_3122424 | SYNLAB MVZ Leverkusen | Robert Koch Institute | unknown |
| EPI_ISL_3122425 | Praxisgemeinschaft für Laboratoriumsmedizin Labor Blumenstraße; Praxis Dr. Sel und Dr. Späte | Robert Koch Institute | unknown |
| EPI_ISL_3122426 | SYNLAB MVZ Leinfelden-Echterdingen | Robert Koch Institute | unknown |
| EPI_ISL_3122427, EPI_ISL_3122428 | SYNLAB MVZ Leverkusen | Robert Koch Institute | unknown |
| EPI_ISL_3122429 | SYNLAB MVZ Trier | Robert Koch Institute | unknown |
| EPI_ISL_3122430 | SYNLAB MVZ Leverkusen | Robert Koch Institute | unknown |
| EPI_ISL_3122431 | SYNLAB MVZ Leinfelden-Echterdingen | Robert Koch Institute | unknown |
| EPI_ISL_3122432, EPI_ISL_3122433 | SYNLAB MVZ Leverkusen | Robert Koch Institute | unknown |
| EPI_ISL_3122434 | SYNLAB MVZ Weiden | Robert Koch Institute | unknown |
| EPI_ISL_3122435 | SYNLAB MVZ Leverkusen | Robert Koch Institute | unknown |
| EPI_ISL_3122436 | SYNLAB MVZ Trier | Robert Koch Institute | unknown |
| EPI_ISL_3122437, EPI_ISL_3122438 | SYNLAB MVZ Leinfelden-Echterdingen | Robert Koch Institute | unknown |
| EPI_ISL_3122439 | SYNLAB MVZ Leverkusen | Robert Koch Institute | unknown |
| EPI_ISL_3122440 | SYNLAB MVZ Leinfelden-Echterdingen | Robert Koch Institute | unknown |
| EPI_ISL_3122441 | SYNLAB MVZ Leverkusen | Robert Koch Institute | unknown |
| EPI_ISL_3122442 | Synlab MVZ Augsburg | Robert Koch Institute | unknown |
| EPI_ISL_3122443 | SYNLAB MVZ Weiden | Robert Koch Institute | unknown |
| EPI_ISL_3122444 | SYNLAB MVZ Trier | Robert Koch Institute | unknown |
| EPI_ISL_3122445 | SYNLAB MVZ Weiden | Robert Koch Institute | unknown |
| EPI_ISL_3122446, EPI_ISL_3122447, EPI_ISL_3122448, EPI_ISL_3122449 | SYNLAB MVZ Leverkusen | Robert Koch Institute | unknown |
| EPI_ISL_3122450 | SYNLAB MVZ Leinfelden-Echterdingen | Robert Koch Institute | unknown |
| EPI_ISL_3122451 | SYNLAB MVZ Leverkusen | Robert Koch Institute | unknown |
| EPI_ISL_3122452 | SYNLAB MVZ Berlin | Robert Koch Institute | unknown |
| EPI_ISL_3122453, EPI_ISL_3122454 | SYNLAB MVZ Hamburg | Robert Koch Institute | unknown |
| EPI_ISL_3122455 | SYNLAB MVZ Leverkusen | Robert Koch Institute | unknown |
| EPI_ISL_3122456 | SYNLAB Labor München Zentrum LMZ | Robert Koch Institute | unknown |
| EPI_ISL_3122457, EPI_ISL_3122458, EPI_ISL_3122459 | SYNLAB MVZ Leinfelden-Echterdingen | Robert Koch Institute | unknown |
| EPI_ISL_3122460 | SYNLAB MVZ Leverkusen | Robert Koch Institute | unknown |
| EPI_ISL_3122461 | SYNLAB MVZ Weiden | Robert Koch Institute | unknown |
| EPI_ISL_3122462 | SYNLAB MVZ Trier | Robert Koch Institute | unknown |
| EPI_ISL_3122463, EPI_ISL_3122464 | SYNLAB Labor München Zentrum LMZ | Robert Koch Institute | unknown |
| EPI_ISL_3122465 | SYNLAB MVZ Berlin | Robert Koch Institute | unknown |
| EPI_ISL_3122466 | SYNLAB Labor München Zentrum LMZ | Robert Koch Institute | unknown |
| EPI_ISL_3122467 | SYNLAB MVZ Leverkusen | Robert Koch Institute | unknown |
| EPI_ISL_3122468 | SYNLAB MVZ Trier | Robert Koch Institute | unknown |
| EPI_ISL_3122470, EPI_ISL_3122471 | Synlab MVZ Augsburg | Robert Koch Institute | unknown |
| EPI_ISL_3122472, EPI_ISL_3122473 | SYNLAB MVZ Leverkusen | Robert Koch Institute | unknown |
| EPI_ISL_3122474 | SYNLAB MVZ Leinfelden-Echterdingen | Robert Koch Institute | unknown |
| EPI_ISL_3122475 | SYNLAB MVZ Leverkusen | Robert Koch Institute | unknown |
| EPI_ISL_3122476, EPI_ISL_3122477, EPI_ISL_3122478 | SYNLAB MVZ Leinfelden-Echterdingen | Robert Koch Institute | unknown |
| EPI_ISL_3122479 | SYNLAB MVZ Leverkusen | Robert Koch Institute | unknown |
| EPI_ISL_3122481 | Synlab MVZ Augsburg | Robert Koch Institute | unknown |
| EPI_ISL_3122482 | SYNLAB MVZ Leverkusen | Robert Koch Institute | unknown |
| EPI_ISL_3122483 | SYNLAB MVZ Weiden | Robert Koch Institute | unknown |
| EPI_ISL_3122485 | Synlab MVZ Augsburg | Robert Koch Institute | unknown |
| EPI_ISL_3122486 | SYNLAB MVZ Weiden | Robert Koch Institute | unknown |
| EPI_ISL_3122487 | Synlab MVZ Augsburg | Robert Koch Institute | unknown |
| EPI_ISL_3122488 | SYNLAB MVZ Weiden | Robert Koch Institute | unknown |
| EPI_ISL_3122489, | SYNLAB Labor München Zentrum LMZ | Robert Koch Institute | unknown |

| | | | |
|---|---|---|---|
| EPI_ISL_3122490 | | | |
| EPI_ISL_3122491 | SYNLAB MVZ Berlin | Robert Koch Institute | unknown |
| EPI_ISL_3122493 | SYNLAB MVZ Leinfelden-Echterdingen | Robert Koch Institute | unknown |
| EPI_ISL_3122494, EPI_ISL_3122495 | SYNLAB MVZ Leverkusen | Robert Koch Institute | unknown |
| EPI_ISL_3122496 | SYNLAB MVZ Weiden | Robert Koch Institute | unknown |
| EPI_ISL_3122497, EPI_ISL_3122498 | SYNLAB MVZ Leverkusen | Robert Koch Institute | unknown |
| EPI_ISL_3122499, EPI_ISL_3122500, EPI_ISL_3122501, EPI_ISL_3122502 | Labor Dr. Wisplinghoff - Köln | Robert Koch Institute | unknown |
| EPI_ISL_3122503 | Labor Mönchengladbach MVZ Dr. Stein + Kollegen GbR | Robert Koch Institute | unknown |
| EPI_ISL_3122504, EPI_ISL_3122505, EPI_ISL_3122506, EPI_ISL_3122507, EPI_ISL_3122508 | MVZ Labor Dr. Limbach & Kollegen GbR | Robert Koch Institute | unknown |
| EPI_ISL_3122509 | Labor Mönchengladbach MVZ Dr. Stein + Kollegen GbR | Robert Koch Institute | unknown |
| EPI_ISL_3122510, EPI_ISL_3122513, EPI_ISL_3122514 | MVZ Labor Dr. Limbach & Kollegen GbR | Robert Koch Institute | unknown |
| EPI_ISL_3122515 | Labor Mönchengladbach MVZ Dr. Stein + Kollegen GbR | Robert Koch Institute | unknown |
| EPI_ISL_3122516, EPI_ISL_3122517 | MVZ Labor Dr. Limbach & Kollegen GbR | Robert Koch Institute | unknown |
| EPI_ISL_3122518 | Limbach - MVZ Labor Eveld & Kollegen Essen | Robert Koch Institute | unknown |
| EPI_ISL_3122519, EPI_ISL_3122520 | MVZ Labor Dr. Limbach & Kollegen GbR | Robert Koch Institute | unknown |
| EPI_ISL_3122521, EPI_ISL_3122522 | Labor Mönchengladbach MVZ Dr. Stein + Kollegen GbR | Robert Koch Institute | unknown |
| EPI_ISL_3122524 | MVZ Labor Dr. Limbach & Kollegen GbR | Robert Koch Institute | unknown |
| EPI_ISL_3122525, EPI_ISL_3122526, EPI_ISL_3122527, EPI_ISL_3122528, EPI_ISL_3122529, EPI_ISL_3122530, EPI_ISL_3122531, EPI_ISL_3122532, EPI_ISL_3122533, EPI_ISL_3122534, EPI_ISL_3122535, EPI_ISL_3122536, EPI_ISL_3122537, EPI_ISL_3122540, EPI_ISL_3122541, EPI_ISL_3122542, EPI_ISL_3122543, EPI_ISL_3122544, EPI_ISL_3122545, EPI_ISL_3122546 | | | |
| see above | Labor Mönchengladbach MVZ Dr. Stein + Kollegen GbR | Robert Koch Institute | unknown |
| EPI_ISL_3122547 | MDI Limbach Berlin GmbH; MVZ Labor Berlin | Robert Koch Institute | unknown |
| EPI_ISL_3122548, EPI_ISL_3122549 | MVZ für Laboratoriumsmedizin und Mikrobiologie Koblenz-Mittelrhein (Labor Koblenz) | Robert Koch Institute | unknown |
| EPI_ISL_3122550 | MDI Limbach Berlin GmbH; MVZ Labor Berlin | Robert Koch Institute | unknown |
| EPI_ISL_3122551, EPI_ISL_3122552 | MVZ für Laboratoriumsmedizin und Mikrobiologie Koblenz-Mittelrhein (Labor Koblenz) | Robert Koch Institute | unknown |
| EPI_ISL_3122553 | Labor Prof. Dr. G. Enders MVZ GbR | Robert Koch Institute | unknown |
| EPI_ISL_3122554 | MVZ für Laboratoriumsmedizin und Mikrobiologie Koblenz-Mittelrhein (Labor Koblenz) | Robert Koch Institute | unknown |
| EPI_ISL_3122555 | MDI Limbach Berlin GmbH; MVZ Labor Berlin | Robert Koch Institute | unknown |
| EPI_ISL_3122557, EPI_ISL_3122558 | Labor Prof. Dr. G. Enders MVZ GbR | Robert Koch Institute | unknown |
| EPI_ISL_3122559, EPI_ISL_3122560 | MDI Limbach Berlin GmbH; MVZ Labor Berlin | Robert Koch Institute | unknown |
| EPI_ISL_3122561, EPI_ISL_3122562, EPI_ISL_3122563 | Labor Prof. Dr. G. Enders MVZ GbR | Robert Koch Institute | unknown |
| EPI_ISL_3122564 | MVZ für Laboratoriumsmedizin und Mikrobiologie Koblenz-Mittelrhein (Labor Koblenz) | Robert Koch Institute | unknown |
| EPI_ISL_3122565 | Labor Prof. Dr. G. Enders MVZ GbR | Robert Koch Institute | unknown |
| EPI_ISL_3122566 | MVZ für Laboratoriumsmedizin und Mikrobiologie Koblenz-Mittelrhein (Labor Koblenz) | Robert Koch Institute | unknown |
| EPI_ISL_3122567 | Labor Prof. Dr. G. Enders MVZ GbR | Robert Koch Institute | unknown |
| EPI_ISL_3122568, EPI_ISL_3122569, EPI_ISL_3122571, EPI_ISL_3122572, EPI_ISL_3122573, EPI_ISL_3122574, EPI_ISL_3122575, EPI_ISL_3122576, EPI_ISL_3122581, EPI_ISL_3122582, EPI_ISL_3122583, EPI_ISL_3122584, EPI_ISL_3122585, EPI_ISL_3122586, EPI_ISL_3122587, EPI_ISL_3122588, EPI_ISL_3122589 | | | |
| see above | MDI Limbach Berlin GmbH; MVZ Labor Berlin | Robert Koch Institute | unknown |
| EPI_ISL_3122590 | MVZ Medizinisches Labor Hannover GmbH | Robert Koch Institute | unknown |
| EPI_ISL_3122591, EPI_ISL_3122592 | MDI Limbach Berlin GmbH; MVZ Labor Berlin | Robert Koch Institute | unknown |
| EPI_ISL_3122593 | MVZ Medizinisches Labor Hannover GmbH | Robert Koch Institute | unknown |
| EPI_ISL_3122594, EPI_ISL_3122595, EPI_ISL_3122596, EPI_ISL_3122597 | MDI Limbach Berlin GmbH; MVZ Labor Berlin | Robert Koch Institute | unknown |
| EPI_ISL_3122598 | MVZ Medizinisches Labor Hannover GmbH | Robert Koch Institute | unknown |
| EPI_ISL_3122599, EPI_ISL_3122600, EPI_ISL_3122601, EPI_ISL_3122602, EPI_ISL_3122603, EPI_ISL_3122604, EPI_ISL_3122605, EPI_ISL_3122606 | MDI Limbach Berlin GmbH; MVZ Labor Berlin | Robert Koch Institute | unknown |
| EPI_ISL_3122607 | MVZ Medizinisches Labor Hannover GmbH | Robert Koch Institute | unknown |
| EPI_ISL_3122608, EPI_ISL_3122609, EPI_ISL_3122610, EPI_ISL_3122611, EPI_ISL_3122612, EPI_ISL_3122613 | MDI Limbach Berlin GmbH; MVZ Labor Berlin | Robert Koch Institute | unknown |
| EPI_ISL_3122614 | MVZ Medizinisches Labor Hannover GmbH | Robert Koch Institute | unknown |
| EPI_ISL_3122615, EPI_ISL_3122616, EPI_ISL_3122617, EPI_ISL_3122618, EPI_ISL_3122619, EPI_ISL_3122620, EPI_ISL_3122621, | MDI Limbach Berlin GmbH; MVZ Labor Berlin | Robert Koch Institute | unknown |

| | | | |
|---|---|---|---|
| EPI_ISL_3122622 | | | |
| EPI_ISL_3122623 | MVZ Medizinisches Labor Hannover GmbH | Robert Koch Institute | unknown |
| EPI_ISL_3122624, EPI_ISL_3122625, EPI_ISL_3122626 | MDI Limbach Berlin GmbH; MVZ Labor Berlin | Robert Koch Institute | unknown |
| EPI_ISL_3122627, EPI_ISL_3122628 | MVZ Labor Dr. Limbach & Kollegen GbR | Robert Koch Institute | unknown |
| EPI_ISL_3122629 | MVZ Medizinisches Labor Hannover GmbH | Robert Koch Institute | unknown |
| EPI_ISL_3122630 | MVZ Labor Dr. Limbach & Kollegen GbR | Robert Koch Institute | unknown |
| EPI_ISL_3122632 | MVZ Medizinisches Labor Hannover GmbH | Robert Koch Institute | unknown |
| EPI_ISL_3122633 | MVZ Labor Dr. Limbach & Kollegen GbR | Robert Koch Institute | unknown |
| EPI_ISL_3122634, EPI_ISL_3122635 | MVZ Medizinisches Labor Hannover GmbH | Robert Koch Institute | unknown |
| EPI_ISL_3122636 | MVZ Labor Dr. Limbach & Kollegen GbR | Robert Koch Institute | unknown |
| EPI_ISL_3122637 | MVZ Medizinisches Labor Hannover GmbH | Robert Koch Institute | unknown |
| EPI_ISL_3122638 | Klinikum Darmstadt GmbH | Robert Koch Institute | unknown |
| EPI_ISL_3122639 | MVZ Labor Dr. Limbach & Kollegen GbR | Robert Koch Institute | unknown |
| EPI_ISL_3122640 | MVZ Medizinisches Labor Hannover GmbH | Robert Koch Institute | unknown |
| EPI_ISL_3122641, EPI_ISL_3122642 | MVZ Labor Dr. Limbach & Kollegen GbR | Robert Koch Institute | unknown |
| EPI_ISL_3122643, EPI_ISL_3122644 | MVZ Medizinisches Labor Hannover GmbH | Robert Koch Institute | unknown |
| EPI_ISL_3122645, EPI_ISL_3122646 | MVZ Labor Dr. Limbach & Kollegen GbR | Robert Koch Institute | unknown |
| EPI_ISL_3122647 | MVZ Medizinisches Labor Hannover GmbH | Robert Koch Institute | unknown |
| EPI_ISL_3122648, EPI_ISL_3122649, EPI_ISL_3122650 | MVZ Labor Dr. Limbach & Kollegen GbR | Robert Koch Institute | unknown |
| EPI_ISL_3122651 | MVZ für Laboratoriumsmedizin und Mikrobiologie Koblenz-Mittelrhein (Labor Koblenz) | Robert Koch Institute | unknown |
| EPI_ISL_3122652, EPI_ISL_3122653 | MVZ Labor Dr. Limbach & Kollegen GbR | Robert Koch Institute | unknown |
| EPI_ISL_3122654, EPI_ISL_3122655, EPI_ISL_3122656, EPI_ISL_3122657, EPI_ISL_3122658, EPI_ISL_3122659, EPI_ISL_3122660 | MVZ für Laboratoriumsmedizin und Mikrobiologie Koblenz-Mittelrhein (Labor Koblenz) | Robert Koch Institute | unknown |
| EPI_ISL_3122661 | MVZ Labor Dr. Limbach & Kollegen GbR | Robert Koch Institute | unknown |
| EPI_ISL_3122662 | MVZ für Laboratoriumsmedizin und Mikrobiologie Koblenz-Mittelrhein (Labor Koblenz) | Robert Koch Institute | unknown |
| EPI_ISL_3122663, EPI_ISL_3122664 | MVZ Labor Dr. Limbach & Kollegen GbR | Robert Koch Institute | unknown |
| EPI_ISL_3122665, EPI_ISL_3122666 | Labor Mönchengladbach MVZ Dr. Stein + Kollegen GbR | Robert Koch Institute | unknown |
| EPI_ISL_3122667 | MVZ Labor Dr. Limbach & Kollegen GbR | Robert Koch Institute | unknown |
| EPI_ISL_3122668, EPI_ISL_3122669, EPI_ISL_3122670, EPI_ISL_3122671 | Labor Mönchengladbach MVZ Dr. Stein + Kollegen GbR | Robert Koch Institute | unknown |
| EPI_ISL_3122672 | MVZ Labor Dr. Limbach & Kollegen GbR | Robert Koch Institute | unknown |
| EPI_ISL_3122673 | Labor Mönchengladbach MVZ Dr. Stein + Kollegen GbR | Robert Koch Institute | unknown |
| EPI_ISL_3122675, EPI_ISL_3122676, EPI_ISL_3122677, EPI_ISL_3122678, EPI_ISL_3122680, EPI_ISL_3122681, EPI_ISL_3122682, EPI_ISL_3122683, EPI_ISL_3122684 | MVZ für Laboratoriumsmedizin und Mikrobiologie Koblenz-Mittelrhein (Labor Koblenz) | Robert Koch Institute | unknown |
| EPI_ISL_3122725, EPI_ISL_3122726, EPI_ISL_3122727, EPI_ISL_3122728, EPI_ISL_3122802, EPI_ISL_3122803, EPI_ISL_3122804, EPI_ISL_3122805, EPI_ISL_3122806, EPI_ISL_3122809, EPI_ISL_3122810, EPI_ISL_3122811, EPI_ISL_3122812, EPI_ISL_3122813, EPI_ISL_3122814, EPI_ISL_3122815 | | | |
| see above | ASST GOM NIGUARDA | ASST Grande ospedale Metropolitano Niguarda | Alice Nava |
| EPI_ISL_3122852, EPI_ISL_3122853, EPI_ISL_3122854, EPI_ISL_3122856, EPI_ISL_3122857, EPI_ISL_3122858 | Infectious Diseases Hospital | WHO National Influenza Centre Russian Federation | Andrey Komissarov, Artem Fadeev, Kseniya Komissarova, Oula Mansour, Kirill Varchenko, Mikhail Bakaev, Tamila Musaeva, Veronika Eder, Maria Pisareva, Nikita Yolshin, Daria Danilenko, Maria Baturova, Alexey Masharsky, Ksenia Safina, Elena Nabieva, Georgii Bazykin, Dmitry Lioznov |
| EPI_ISL_3122859, EPI_ISL_3122860 | Kursk Dermatovenerologic Dispensary | WHO National Influenza Centre Russian Federation | Andrey Komissarov, Artem Fadeev, Kseniya Komissarova, Oula Mansour, Kirill Varchenko, Mikhail Bakaev, Tamila Musaeva, Veronika Eder, Maria Pisareva, Nikita Yolshin, Daria Danilenko, Maria Baturova, Alexey Masharsky, Ksenia Safina, Elena Nabieva, Georgii Bazykin, Dmitry Lioznov |
| EPI_ISL_3122861, EPI_ISL_3122863, EPI_ISL_3122864, EPI_ISL_3122865, EPI_ISL_3122866, EPI_ISL_3122867, EPI_ISL_3122868, EPI_ISL_3122869, EPI_ISL_3122870 | Krai Clinical Hospital No 2 | WHO National Influenza Centre Russian Federation | Andrey Komissarov, Artem Fadeev, Kseniya Komissarova, Oula Mansour, Kirill Varchenko, Mikhail Bakaev, Tamila Musaeva, Veronika Eder, Maria Pisareva, Nikita Yolshin, Daria Danilenko, Maria Baturova, Alexey Masharsky, Ksenia Safina, Elena Nabieva, Georgii Bazykin, Dmitry Lioznov |
| EPI_ISL_3122871, EPI_ISL_3122872, EPI_ISL_3122873, EPI_ISL_3122874, EPI_ISL_3122875, EPI_ISL_3122876, EPI_ISL_3122877, EPI_ISL_3122878, EPI_ISL_3122879, EPI_ISL_3122880, EPI_ISL_3122881 | | | |
| see above | Ulyanovsk Regional Clinical Hospital | WHO National Influenza Centre Russian Federation | Andrey Komissarov, Artem Fadeev, Kseniya Komissarova, Oula Mansour, Kirill Varchenko, Mikhail Bakaev, Tamila Musaeva, Veronika Eder, Maria Pisareva, Nikita Yolshin, Daria Danilenko, Maria Baturova, Alexey Masharsky, Ksenia Safina, Elena Nabieva, Georgii Bazykin, Dmitry Lioznov |
| EPI_ISL_3122882, EPI_ISL_3122883, EPI_ISL_3122884, EPI_ISL_3122885, EPI_ISL_3122886, EPI_ISL_3122887, EPI_ISL_3122888, EPI_ISL_3122889, EPI_ISL_3122890, EPI_ISL_3122891, EPI_ISL_3122892, EPI_ISL_3122893, EPI_ISL_3122894, EPI_ISL_3122895, EPI_ISL_3122896, EPI_ISL_3122898 | | | |
| see above | Perinatal Center | WHO National Influenza Centre Russian Federation | Andrey Komissarov, Artem Fadeev, Kseniya Komissarova, Oula Mansour, Kirill Varchenko, Mikhail Bakaev, Tamila Musaeva, Veronika Eder, Maria Pisareva, Nikita Yolshin, Daria Danilenko, Maria Baturova, Alexey Masharsky, Ksenia Safina, Elena Nabieva, Georgii Bazykin, Dmitry Lioznov |
| EPI_ISL_3122899, EPI_ISL_3122900 | Engels City Hospital No 2 | WHO National Influenza Centre Russian Federation | Andrey Komissarov, Artem Fadeev, Kseniya Komissarova, Oula Mansour, Kirill Varchenko, Mikhail Bakaev, Tamila Musaeva, Veronika Eder, Maria Pisareva, Nikita Yolshin, Daria Danilenko, Maria Baturova, Alexey Masharsky, Ksenia Safina, Elena Nabieva, Georgii Bazykin, Dmitry Lioznov |
| EPI_ISL_3122901, EPI_ISL_3122902, EPI_ISL_3122904, | Voronezh Regional Clinical Consultative and Diagnostic Center | WHO National Influenza Centre Russian Federation | Andrey Komissarov, Artem Fadeev, Kseniya Komissarova, Oula Mansour, Kirill Varchenko, Mikhail Bakaev, Tamila Musaeva, Veronika Eder, Maria Pisareva, Nikita Yolshin, Daria Danilenko, Maria Baturova, Alexey Masharsky, Ksenia Safina, Elena Nabieva, Georgii Bazykin, Dmitry Lioznov |

| | | | |
|---|---|---|---|
| EPI_ISL_3122905, EPI_ISL_3122906, EPI_ISL_3122907, EPI_ISL_3122908 | | | |
| EPI_ISL_3122909, EPI_ISL_3122911, EPI_ISL_3122912 | Kaliningrad Region Infectious Diseases Hospital | WHO National Influenza Centre Russian Federation | Andrey Komissarov, Artem Fadeev, Kseniya Komissarova, Oula Mansour, Kirill Varchenko, Mikhail Bakaev, Tamila Musaeva, Veronika Eder, Maria Pisareva, Nikita Yolshin, Daria Danilenko, Maria Baturova, Alexey Masharsky, Ksenia Safina, Elena Nabieva, Georgii Bazykin, Dmitry Lioznov |
| EPI_ISL_3122913, EPI_ISL_3122914, EPI_ISL_3122915, EPI_ISL_3122916, EPI_ISL_3122917, EPI_ISL_3122918, EPI_ISL_3122919, EPI_ISL_3122920, EPI_ISL_3122921, EPI_ISL_3122922, EPI_ISL_3122923, EPI_ISL_3122924, EPI_ISL_3122925, EPI_ISL_3122926, EPI_ISL_3122927 | | | |
| see above | Vologda City Polyclinic No 1 | WHO National Influenza Centre Russian Federation | Andrey Komissarov, Artem Fadeev, Kseniya Komissarova, Oula Mansour, Kirill Varchenko, Mikhail Bakaev, Tamila Musaeva, Veronika Eder, Maria Pisareva, Nikita Yolshin, Daria Danilenko, Maria Baturova, Alexey Masharsky, Ksenia Safina, Elena Nabieva, Georgii Bazykin, Dmitry Lioznov |
| EPI_ISL_3122928, EPI_ISL_3122929, EPI_ISL_3122931, EPI_ISL_3122932, EPI_ISL_3122933 | Novgorod Regional Clinical Hospital | WHO National Influenza Centre Russian Federation | Andrey Komissarov, Artem Fadeev, Kseniya Komissarova, Oula Mansour, Kirill Varchenko, Mikhail Bakaev, Tamila Musaeva, Veronika Eder, Maria Pisareva, Nikita Yolshin, Daria Danilenko, Maria Baturova, Alexey Masharsky, Ksenia Safina, Elena Nabieva, Georgii Bazykin, Dmitry Lioznov |
| EPI_ISL_3122934, EPI_ISL_3122935, EPI_ISL_3122936, EPI_ISL_3122937, EPI_ISL_3122938, EPI_ISL_3122939, EPI_ISL_3122940, EPI_ISL_3122941, EPI_ISL_3122942, EPI_ISL_3122943 | Tumen Regional Clinical Hospital No 1 | WHO National Influenza Centre Russian Federation | Andrey Komissarov, Artem Fadeev, Kseniya Komissarova, Oula Mansour, Kirill Varchenko, Mikhail Bakaev, Tamila Musaeva, Veronika Eder, Maria Pisareva, Nikita Yolshin, Daria Danilenko, Maria Baturova, Alexey Masharsky, Ksenia Safina, Elena Nabieva, Georgii Bazykin, Dmitry Lioznov |
| EPI_ISL_3122944, EPI_ISL_3122945, EPI_ISL_3122946, EPI_ISL_3122947, EPI_ISL_3122948, EPI_ISL_3122949, EPI_ISL_3122950, EPI_ISL_3122951, EPI_ISL_3122952, EPI_ISL_3122953, EPI_ISL_3122954, EPI_ISL_3122955 | | | |
| see above | Surgut Country Clinical Hospital | WHO National Influenza Centre Russian Federation | Andrey Komissarov, Artem Fadeev, Kseniya Komissarova, Oula Mansour, Kirill Varchenko, Mikhail Bakaev, Tamila Musaeva, Veronika Eder, Maria Pisareva, Nikita Yolshin, Daria Danilenko, Maria Baturova, Alexey Masharsky, Ksenia Safina, Elena Nabieva, Georgii Bazykin, Dmitry Lioznov |
| EPI_ISL_3122956 | Republican Medicogenetic Center | WHO National Influenza Centre Russian Federation | Andrey Komissarov, Artem Fadeev, Kseniya Komissarova, Oula Mansour, Kirill Varchenko, Mikhail Bakaev, Tamila Musaeva, Veronika Eder, Maria Pisareva, Nikita Yolshin, Daria Danilenko, Maria Baturova, Alexey Masharsky, Ksenia Safina, Elena Nabieva, Georgii Bazykin, Dmitry Lioznov |
| EPI_ISL_3122957, EPI_ISL_3122958, EPI_ISL_3122959, EPI_ISL_3122960, EPI_ISL_3122961, EPI_ISL_3122962 | Ufa City Clinical Hospital No 13 | WHO National Influenza Centre Russian Federation | Andrey Komissarov, Artem Fadeev, Kseniya Komissarova, Oula Mansour, Kirill Varchenko, Mikhail Bakaev, Tamila Musaeva, Veronika Eder, Maria Pisareva, Nikita Yolshin, Daria Danilenko, Maria Baturova, Alexey Masharsky, Ksenia Safina, Elena Nabieva, Georgii Bazykin, Dmitry Lioznov |
| EPI_ISL_3122963, EPI_ISL_3122964 | Republican Clinical Infectious Diseases Hospital | WHO National Influenza Centre Russian Federation | Andrey Komissarov, Artem Fadeev, Kseniya Komissarova, Oula Mansour, Kirill Varchenko, Mikhail Bakaev, Tamila Musaeva, Veronika Eder, Maria Pisareva, Nikita Yolshin, Daria Danilenko, Maria Baturova, Alexey Masharsky, Ksenia Safina, Elena Nabieva, Georgii Bazykin, Dmitry Lioznov |
| EPI_ISL_3122965, EPI_ISL_3122966 | Magadan Regional Dispensary of Phthisiology and Infectious Diseases | WHO National Influenza Centre Russian Federation | Andrey Komissarov, Artem Fadeev, Kseniya Komissarova, Oula Mansour, Kirill Varchenko, Mikhail Bakaev, Tamila Musaeva, Veronika Eder, Maria Pisareva, Nikita Yolshin, Daria Danilenko, Maria Baturova, Alexey Masharsky, Ksenia Safina, Elena Nabieva, Georgii Bazykin, Dmitry Lioznov |
| EPI_ISL_3122967, EPI_ISL_3122968, EPI_ISL_3122969, EPI_ISL_3122970 | Chelyabinsk Regional Dermatovenerologic Dispensary | WHO National Influenza Centre Russian Federation | Andrey Komissarov, Artem Fadeev, Kseniya Komissarova, Oula Mansour, Kirill Varchenko, Mikhail Bakaev, Tamila Musaeva, Veronika Eder, Maria Pisareva, Nikita Yolshin, Daria Danilenko, Maria Baturova, Alexey Masharsky, Ksenia Safina, Elena Nabieva, Georgii Bazykin, Dmitry Lioznov |
| EPI_ISL_3122971, EPI_ISL_3122972, EPI_ISL_3122973, EPI_ISL_3122974, EPI_ISL_3122975, EPI_ISL_3122976, EPI_ISL_3122978, EPI_ISL_3122979, EPI_ISL_3122980, EPI_ISL_3122981, EPI_ISL_3122982, EPI_ISL_3122983, EPI_ISL_3122984 | | | |
| see above | City Clinical Hospital No 1 | WHO National Influenza Centre Russian Federation | Andrey Komissarov, Artem Fadeev, Kseniya Komissarova, Oula Mansour, Kirill Varchenko, Mikhail Bakaev, Tamila Musaeva, Veronika Eder, Maria Pisareva, Nikita Yolshin, Daria Danilenko, Maria Baturova, Alexey Masharsky, Ksenia Safina, Elena Nabieva, Georgii Bazykin, Dmitry Lioznov |
| EPI_ISL_3122985, EPI_ISL_3122986, EPI_ISL_3122987 | N. A. Semashko Republican Clinical Hospital | WHO National Influenza Centre Russian Federation | Andrey Komissarov, Artem Fadeev, Kseniya Komissarova, Oula Mansour, Kirill Varchenko, Mikhail Bakaev, Tamila Musaeva, Veronika Eder, Maria Pisareva, Nikita Yolshin, Daria Danilenko, Maria Baturova, Alexey Masharsky, Ksenia Safina, Elena Nabieva, Georgii Bazykin, Dmitry Lioznov |
| EPI_ISL_3122988, EPI_ISL_3122989, EPI_ISL_3122990, EPI_ISL_3122991, EPI_ISL_3122992, EPI_ISL_3122993, EPI_ISL_3122994, EPI_ISL_3122995, EPI_ISL_3122996, EPI_ISL_3122997, EPI_ISL_3122998, EPI_ISL_3122999, EPI_ISL_3123000, EPI_ISL_3123001, EPI_ISL_3123002, EPI_ISL_3123003, EPI_ISL_3123005, EPI_ISL_3123006, EPI_ISL_3123007, EPI_ISL_3123008 | | | |
| see above | Republican Children's Clinical Infectious Diseases Hospital | WHO National Influenza Centre Russian Federation | Andrey Komissarov, Artem Fadeev, Kseniya Komissarova, Oula Mansour, Kirill Varchenko, Mikhail Bakaev, Tamila Musaeva, Veronika Eder, Maria Pisareva, Nikita Yolshin, Daria Danilenko, Maria Baturova, Alexey Masharsky, Ksenia Safina, Elena Nabieva, Georgii Bazykin, Dmitry Lioznov |
| EPI_ISL_3123009, EPI_ISL_3123010, EPI_ISL_3123011, EPI_ISL_3123012, EPI_ISL_3123013, EPI_ISL_3123014 | Rostov-on-Don Regional Clinical Consultative and Diagnostic Center | WHO National Influenza Centre Russian Federation | Andrey Komissarov, Artem Fadeev, Kseniya Komissarova, Oula Mansour, Kirill Varchenko, Mikhail Bakaev, Tamila Musaeva, Veronika Eder, Maria Pisareva, Nikita Yolshin, Daria Danilenko, Maria Baturova, Alexey Masharsky, Ksenia Safina, Elena Nabieva, Georgii Bazykin, Dmitry Lioznov |
| EPI_ISL_3123017, EPI_ISL_3123018 | Petropavlosk-Kamchatsky City Polyclinic No 3 | WHO National Influenza Centre Russian Federation | Andrey Komissarov, Artem Fadeev, Kseniya Komissarova, Oula Mansour, Kirill Varchenko, Mikhail Bakaev, Tamila Musaeva, Veronika Eder, Maria Pisareva, Nikita Yolshin, Daria Danilenko, Maria Baturova, Alexey Masharsky, Ksenia Safina, Elena Nabieva, Georgii Bazykin, Dmitry Lioznov |
| EPI_ISL_3123019, EPI_ISL_3123020, EPI_ISL_3123021, EPI_ISL_3123022 | Petropavlosk-Kamchatsky City Polyclinic No 1 | WHO National Influenza Centre Russian Federation | Andrey Komissarov, Artem Fadeev, Kseniya Komissarova, Oula Mansour, Kirill Varchenko, Mikhail Bakaev, Tamila Musaeva, Veronika Eder, Maria Pisareva, Nikita Yolshin, Daria Danilenko, Maria Baturova, Alexey Masharsky, Ksenia Safina, Elena Nabieva, Georgii Bazykin, Dmitry Lioznov |
| EPI_ISL_3123023, EPI_ISL_3123024 | Kamchatka Krai Birth Center | WHO National Influenza Centre Russian Federation | Andrey Komissarov, Artem Fadeev, Kseniya Komissarova, Oula Mansour, Kirill Varchenko, Mikhail Bakaev, Tamila Musaeva, Veronika Eder, Maria Pisareva, Nikita Yolshin, Daria Danilenko, Maria Baturova, Alexey Masharsky, Ksenia Safina, Elena Nabieva, Georgii Bazykin, Dmitry Lioznov |
| EPI_ISL_3123025 | Petropavlosk-Kamchatsky City Polyclinic No 2 | WHO National Influenza Centre Russian Federation | Andrey Komissarov, Artem Fadeev, Kseniya Komissarova, Oula Mansour, Kirill Varchenko, Mikhail Bakaev, Tamila Musaeva, Veronika Eder, Maria Pisareva, Nikita Yolshin, Daria Danilenko, Maria Baturova, Alexey Masharsky, Ksenia Safina, Elena Nabieva, Georgii Bazykin, Dmitry Lioznov |
| EPI_ISL_3123026 | Kamchatka Krai Children's Infectious Diseases Hospital | WHO National Influenza Centre Russian Federation | Andrey Komissarov, Artem Fadeev, Kseniya Komissarova, Oula Mansour, Kirill Varchenko, Mikhail Bakaev, Tamila Musaeva, Veronika Eder, Maria Pisareva, Nikita Yolshin, Daria Danilenko, Maria Baturova, Alexey Masharsky, Ksenia Safina, Elena Nabieva, Georgii Bazykin, Dmitry Lioznov |
| EPI_ISL_3123027, EPI_ISL_3123028, EPI_ISL_3123029, EPI_ISL_3123030, EPI_ISL_3123031, EPI_ISL_3123032 | Petropavlosk-Kamchatsky City Polyclinic No 3 | WHO National Influenza Centre Russian Federation | Andrey Komissarov, Artem Fadeev, Kseniya Komissarova, Oula Mansour, Kirill Varchenko, Mikhail Bakaev, Tamila Musaeva, Veronika Eder, Maria Pisareva, Nikita Yolshin, Daria Danilenko, Maria Baturova, Alexey Masharsky, Ksenia Safina, Elena Nabieva, Georgii Bazykin, Dmitry Lioznov |
| EPI_ISL_3123033, EPI_ISL_3123034, EPI_ISL_3123035, EPI_ISL_3123036, EPI_ISL_3123037, EPI_ISL_3123038, EPI_ISL_3123039, EPI_ISL_3123040, EPI_ISL_3123041, EPI_ISL_3123042, EPI_ISL_3123043, EPI_ISL_3123044, EPI_ISL_3123045, EPI_ISL_3123046, EPI_ISL_3123047, EPI_ISL_3123048, EPI_ISL_3123049, EPI_ISL_3123050, EPI_ISL_3123051, EPI_ISL_3123052, EPI_ISL_3123053, EPI_ISL_3123054, EPI_ISL_3123055, EPI_ISL_3123056, EPI_ISL_3123057, EPI_ISL_3123058, EPI_ISL_3123059, EPI_ISL_3123060, EPI_ISL_3123061, EPI_ISL_3123062, EPI_ISL_3123063, EPI_ISL_3123064, EPI_ISL_3123065, EPI_ISL_3123066, EPI_ISL_3123067, EPI_ISL_3123068, EPI_ISL_3123069, EPI_ISL_3123070, EPI_ISL_3123071, EPI_ISL_3123072, EPI_ISL_3123073, EPI_ISL_3123074, EPI_ISL_3123075, EPI_ISL_3123076, EPI_ISL_3123077, EPI_ISL_3123079, EPI_ISL_3123080, EPI_ISL_3123081, EPI_ISL_3123082, EPI_ISL_3123083, EPI_ISL_3123084, EPI_ISL_3123085, EPI_ISL_3123086, EPI_ISL_3123087, EPI_ISL_3123088, EPI_ISL_3123089, EPI_ISL_3123090, EPI_ISL_3123091, EPI_ISL_3123092, EPI_ISL_3123093, EPI_ISL_3123094, EPI_ISL_3123095, EPI_ISL_3123096 | | | |
| see above | N.F. Gamaleya Research Center for Epidemiology and Microbiology | WHO National Influenza Centre Russian Federation | Andrey Komissarov, Artem Fadeev, Kseniya Komissarova, Oula Mansour, Kirill Varchenko, Mikhail Bakaev, Tamila Musaeva, Veronika Eder, Maria Pisareva, Nikita Yolshin, Daria Danilenko, Maria Baturova, Alexey Masharsky, Elizaveta Divisenko, Alina Odintsova, Elena Shidlovskaya, Ludmila Vasilchenko, Nadezhda Kuznetsova, Maria Nikiforova, Alexey Shchetinin, Artem Tkachuk, Andrei Pochtovyi, Andrei Siniavin, Denis Kleymenov, Evgeniia Bykonia, Liubov Popova, Elena Mazunina, Inna Dolzhikova, Anna Kovyrshina, Andrei Botikov, Dmitry Shcheblyakov, Vladimir Gushchin, Denis Logunov, Alexander Gintsburg, Ksenia Safina, Elena Nabieva, Georgii Bazykin, Dmitry Lioznov |
| EPI_ISL_3123097 | Apollo Institute of Medical Sciences and Research | CDFD | Murali Bashyam, Asmita Gupta, Reelina Basu, Vinay Donipadi, Nagamani Kammili, Divya Vashisht, Ashwin Dalal |
| EPI_ISL_3123098, EPI_ISL_3123101, EPI_ISL_3123102, EPI_ISL_3123104, EPI_ISL_3123105, EPI_ISL_3123106, EPI_ISL_3123107, EPI_ISL_3123108, EPI_ISL_3123112, EPI_ISL_3123114, EPI_ISL_3123115, EPI_ISL_3123116, EPI_ISL_3123117, EPI_ISL_3123118, EPI_ISL_3123122, EPI_ISL_3123123, EPI_ISL_3123124, EPI_ISL_3123125, EPI_ISL_3123126, EPI_ISL_3123127, EPI_ISL_3123128, EPI_ISL_3123129, EPI_ISL_3123130, EPI_ISL_3123131, EPI_ISL_3123132, EPI_ISL_3123133, EPI_ISL_3123134, EPI_ISL_3123135, EPI_ISL_3123136, EPI_ISL_3123137, EPI_ISL_3123138, EPI_ISL_3123139, EPI_ISL_3123140, EPI_ISL_3123142, EPI_ISL_3123143, EPI_ISL_3123144, EPI_ISL_3123145, EPI_ISL_3123146, EPI_ISL_3123147, EPI_ISL_3123148, EPI_ISL_3123149, EPI_ISL_3123150, EPI_ISL_3123151, EPI_ISL_3123152, EPI_ISL_3123156, EPI_ISL_3123157, EPI_ISL_3123158, EPI_ISL_3123159, EPI_ISL_3123160, EPI_ISL_3123161, EPI_ISL_3123162, EPI_ISL_3123166, EPI_ISL_3123167, EPI_ISL_3123168, EPI_ISL_3123170, EPI_ISL_3123171, EPI_ISL_3123174, EPI_ISL_3123175, EPI_ISL_3123176, EPI_ISL_3123177, EPI_ISL_3123178, EPI_ISL_3123179, EPI_ISL_3123180, EPI_ISL_3123181, EPI_ISL_3123182, EPI_ISL_3123183, EPI_ISL_3123184, EPI_ISL_3123185, EPI_ISL_3123186, EPI_ISL_3123187, EPI_ISL_3123188, EPI_ISL_3123189, EPI_ISL_3123191, EPI_ISL_3123192, EPI_ISL_3123193, EPI_ISL_3123194, EPI_ISL_3123195, EPI_ISL_3123196, EPI_ISL_3123197, EPI_ISL_3123198, EPI_ISL_3123199, EPI_ISL_3123200, EPI_ISL_3123201, EPI_ISL_3123202, EPI_ISL_3123203, EPI_ISL_3123204, EPI_ISL_3123206, EPI_ISL_3123207, EPI_ISL_3123208, EPI_ISL_3123209, EPI_ISL_3123210, EPI_ISL_3123211, EPI_ISL_3123212, EPI_ISL_3123213, EPI_ISL_3123214, EPI_ISL_3123215, EPI_ISL_3123216, EPI_ISL_3123217, EPI_ISL_3123218, EPI_ISL_3123219, EPI_ISL_3123220, EPI_ISL_3123221, EPI_ISL_3123222, EPI_ISL_3123223, EPI_ISL_3123224, EPI_ISL_3123226, EPI_ISL_3123227, EPI_ISL_3123228, EPI_ISL_3123229, EPI_ISL_3123230, EPI_ISL_3123231, EPI_ISL_3123233, EPI_ISL_3123234, EPI_ISL_3123235, EPI_ISL_3123236, EPI_ISL_3123237, EPI_ISL_3123238, EPI_ISL_3123239, EPI_ISL_3123240, EPI_ISL_3123241, EPI_ISL_3123242, EPI_ISL_3123243, EPI_ISL_3123244, EPI_ISL_3123245, EPI_ISL_3123246, EPI_ISL_3123247, EPI_ISL_3123248, EPI_ISL_3123249, EPI_ISL_3123250, EPI_ISL_3123251, EPI_ISL_3123252, EPI_ISL_3123253, EPI_ISL_3123254, EPI_ISL_3123255, EPI_ISL_3123256, EPI_ISL_3123257, EPI_ISL_3123258, EPI_ISL_3123259, EPI_ISL_3123260, EPI_ISL_3123261, EPI_ISL_3123262, EPI_ISL_3123263, EPI_ISL_3123264, EPI_ISL_3123265, EPI_ISL_3123266, EPI_ISL_3123267, EPI_ISL_3123269, EPI_ISL_3123270, EPI_ISL_3123271, EPI_ISL_3123272, EPI_ISL_3123273, EPI_ISL_3123274, EPI_ISL_3123276, EPI_ISL_3123277, EPI_ISL_3123278, EPI_ISL_3123279, EPI_ISL_3123280, EPI_ISL_3123281, EPI_ISL_3123282, EPI_ISL_3123283, EPI_ISL_3123284, EPI_ISL_3123285, EPI_ISL_3123286, EPI_ISL_3123287, EPI_ISL_3123288, EPI_ISL_3123289, EPI_ISL_3123290, EPI_ISL_3123291, EPI_ISL_3123292, EPI_ISL_3123293, EPI_ISL_3123294, EPI_ISL_3123295, EPI_ISL_3123296, EPI_ISL_3123297, EPI_ISL_3123298, EPI_ISL_3123299, EPI_ISL_3123300, EPI_ISL_3123301, EPI_ISL_3123302, EPI_ISL_3123303, EPI_ISL_3123304, EPI_ISL_3123305, EPI_ISL_3123306, EPI_ISL_3123307, EPI_ISL_3123308, EPI_ISL_3123309, EPI_ISL_3123310, EPI_ISL_3123311, EPI_ISL_3123312, EPI_ISL_3123313, EPI_ISL_3123314, EPI_ISL_3123315, EPI_ISL_3123316, EPI_ISL_3123317, EPI_ISL_3123318, EPI_ISL_3123319, EPI_ISL_3123320, EPI_ISL_3123321, EPI_ISL_3123322, EPI_ISL_3123323 | | | |
| see above | Austrian Agency for Health and Food Safety (AGES) | Bergthaler laboratory, CeMM Research Center for Molecular Medicine of the Austrian Academy of Sciences | Lukas Endler, Anna Schedl, Fabian Amman, Petr Triska, Matthew Thornton, Thomas Penz, Benedikt Agerer, Maelle Le Moing, Michael Schuster, Bekir Erguner, Jan Laine, Martin Senekowitsch, Christoph Bock, Andreas Bergthaler |

| | | | |
|---|---|---|---|
| EPI_ISL_3123332, EPI_ISL_3123333, EPI_ISL_3123334, EPI_ISL_3123336, EPI_ISL_3123337, EPI_ISL_3123338, EPI_ISL_3123339, EPI_ISL_3123340, EPI_ISL_3123341, EPI_ISL_3123342 | Universität Innsbruck, Institut für Mikrobiologie | Bergthaler laboratory, CeMM Research Center for Molecular Medicine of the Austrian Academy of Sciences | Lukas Endler, Anna Schedl, Fabian Amman, Petr Triska, Matthew Thornton, Thomas Penz, Benedikt Agerer, Maelle Le Moing, Michael Schuster, Bekir Erguner, Jan Laine, Martin Senekowitsch, Christoph Bock, Andreas Bergthaler |
| EPI_ISL_3123344 | Institute of Legal Medicine, Medical University of Innsbruck | Bergthaler laboratory, CeMM Research Center for Molecular Medicine of the Austrian Academy of Sciences | Lukas Endler, Anna Schedl, Fabian Amman, Petr Triska, Matthew Thornton, Thomas Penz, Benedikt Agerer, Maelle Le Moing, Michael Schuster, Bekir Erguner, Jan Laine, Martin Senekowitsch, Christoph Bock, Andreas Bergthaler |
| EPI_ISL_3123356 | Institute for Water Quality and Resource Management, Technical University Vienna | Bergthaler laboratory, CeMM Research Center for Molecular Medicine of the Austrian Academy of Sciences | Lukas Endler, Anna Schedl, Fabian Amman, Petr Triska, Matthew Thornton, Thomas Penz, Benedikt Agerer, Maelle Le Moing, Michael Schuster, Bekir Erguner, Jan Laine, Martin Senekowitsch, Christoph Bock, Andreas Bergthaler |
| EPI_ISL_3123357, EPI_ISL_3123358, EPI_ISL_3123359, EPI_ISL_3123360 | WSSE w Bialymstoku | Wojewodzka Stacja Sanitarno-Epidemiologiczna w Olsztynie, Laboratorium Badan Epidemiologiczno-Klinicznych | Sylwia Krzetowska, Monika Czerminska, Ewa Liszewska, Tomasz Jakubczak, Marta Lukian, Patryk Bielecki, Aleksandra Kobiatko, Emilia Tarabasz, Paulina Rozycka, Barbara Dolinska |
| EPI_ISL_3123361 | WSSE w Bydgoszczy | Wojewodzka Stacja Sanitarno-Epidemiologiczna w Olsztynie, Laboratorium Badan Epidemiologiczno-Klinicznych | Sylwia Krzetowska, Monika Czerminska, Ewa Liszewska, Tomasz Jakubczak, Marta Lukian, Patryk Bielecki, Aleksandra Kobiatko, Emilia Tarabasz, Paulina Rozycka, Barbara Dolinska |
| EPI_ISL_3123362, EPI_ISL_3123363, EPI_ISL_3123364, EPI_ISL_3123365, EPI_ISL_3123366 | WSSE w Gdansku | Wojewodzka Stacja Sanitarno-Epidemiologiczna w Olsztynie, Laboratorium Badan Epidemiologiczno-Klinicznych | Sylwia Krzetowska, Monika Czerminska, Ewa Liszewska, Tomasz Jakubczak, Marta Lukian, Patryk Bielecki, Aleksandra Kobiatko, Emilia Tarabasz, Paulina Rozycka, Barbara Dolinska |
| EPI_ISL_3123369, EPI_ISL_3123370, EPI_ISL_3123371 | WSSE w Gorzowie Wielkopolskim | Wojewodzka Stacja Sanitarno-Epidemiologiczna w Olsztynie, Laboratorium Badan Epidemiologiczno-Klinicznych | Sylwia Krzetowska, Monika Czerminska, Ewa Liszewska, Tomasz Jakubczak, Marta Lukian, Patryk Bielecki, Aleksandra Kobiatko, Emilia Tarabasz, Paulina Rozycka, Barbara Dolinska |
| EPI_ISL_3123373, EPI_ISL_3123374, EPI_ISL_3123375, EPI_ISL_3123376, EPI_ISL_3123377, EPI_ISL_3123378, EPI_ISL_3123379, EPI_ISL_3123382 | WSSE w Lublinie | Wojewodzka Stacja Sanitarno-Epidemiologiczna w Olsztynie, Laboratorium Badan Epidemiologiczno-Klinicznych | Sylwia Krzetowska, Monika Czerminska, Ewa Liszewska, Tomasz Jakubczak, Marta Lukian, Patryk Bielecki, Aleksandra Kobiatko, Emilia Tarabasz, Paulina Rozycka, Barbara Dolinska |
| EPI_ISL_3123384, EPI_ISL_3123385, EPI_ISL_3123386, EPI_ISL_3123388, EPI_ISL_3123389 | WSSE w Poznaniu | Wojewodzka Stacja Sanitarno-Epidemiologiczna w Olsztynie, Laboratorium Badan Epidemiologiczno-Klinicznych | Sylwia Krzetowska, Monika Czerminska, Ewa Liszewska, Tomasz Jakubczak, Marta Lukian, Patryk Bielecki, Aleksandra Kobiatko, Emilia Tarabasz, Paulina Rozycka, Barbara Dolinska |
| EPI_ISL_3123390, EPI_ISL_3123391, EPI_ISL_3123392, EPI_ISL_3123393, EPI_ISL_3123394, EPI_ISL_3123395, EPI_ISL_3123396, EPI_ISL_3123397 | WSSE w Rzeszowie | Wojewodzka Stacja Sanitarno-Epidemiologiczna w Olsztynie, Laboratorium Badan Epidemiologiczno-Klinicznych | Sylwia Krzetowska, Monika Czerminska, Ewa Liszewska, Tomasz Jakubczak, Marta Lukian, Patryk Bielecki, Aleksandra Kobiatko, Emilia Tarabasz, Paulina Rozycka, Barbara Dolinska |
| EPI_ISL_3123398, EPI_ISL_3123400, EPI_ISL_3123401 | WSSE w Szczecinie | Wojewodzka Stacja Sanitarno-Epidemiologiczna w Olsztynie, Laboratorium Badan Epidemiologiczno-Klinicznych | Sylwia Krzetowska, Monika Czerminska, Ewa Liszewska, Tomasz Jakubczak, Marta Lukian, Patryk Bielecki, Aleksandra Kobiatko, Emilia Tarabasz, Paulina Rozycka, Barbara Dolinska |
| EPI_ISL_3123402, EPI_ISL_3123403, EPI_ISL_3123404, EPI_ISL_3123405, EPI_ISL_3123406 | WSSE w Warszawie | Wojewodzka Stacja Sanitarno-Epidemiologiczna w Olsztynie, Laboratorium Badan Epidemiologiczno-Klinicznych | Sylwia Krzetowska, Monika Czerminska, Ewa Liszewska, Tomasz Jakubczak, Marta Lukian, Patryk Bielecki, Aleksandra Kobiatko, Emilia Tarabasz, Paulina Rozycka, Barbara Dolinska |
| EPI_ISL_3123407, EPI_ISL_3123408, EPI_ISL_3123411, EPI_ISL_3123412, EPI_ISL_3123413, EPI_ISL_3123414, EPI_ISL_3123415, EPI_ISL_3123416, EPI_ISL_3123417, EPI_ISL_3123418 | WSSE we Wroclawiu | Wojewodzka Stacja Sanitarno-Epidemiologiczna w Olsztynie, Laboratorium Badan Epidemiologiczno-Klinicznych | Sylwia Krzetowska, Monika Czerminska, Ewa Liszewska, Tomasz Jakubczak, Marta Lukian, Patryk Bielecki, Aleksandra Kobiatko, Emilia Tarabasz, Paulina Rozycka, Barbara Dolinska |
| EPI_ISL_3123419, EPI_ISL_3123421, EPI_ISL_3123422, EPI_ISL_3123423, EPI_ISL_3123424 | WSSE w Lodzi | Wojewodzka Stacja Sanitarno-Epidemiologiczna w Olsztynie, Laboratorium Badan Epidemiologiczno-Klinicznych | Sylwia Krzetowska, Monika Czerminska, Ewa Liszewska, Tomasz Jakubczak, Marta Lukian, Patryk Bielecki, Aleksandra Kobiatko, Emilia Tarabasz, Paulina Rozycka, Barbara Dolinska |
| EPI_ISL_3123425 | WSSE w Opolu | Wojewodzka Stacja Sanitarno-Epidemiologiczna w Olsztynie, Laboratorium Badan Epidemiologiczno-Klinicznych | Sylwia Krzetowska, Monika Czerminska, Ewa Liszewska, Tomasz Jakubczak, Marta Lukian, Patryk Bielecki, Aleksandra Kobiatko, Emilia Tarabasz, Paulina Rozycka, Barbara Dolinska |
| EPI_ISL_3123426, EPI_ISL_3123428, EPI_ISL_3123429, EPI_ISL_3123431, EPI_ISL_3123432, EPI_ISL_3123433, EPI_ISL_3123434, EPI_ISL_3123436, EPI_ISL_3123437, EPI_ISL_3123438, EPI_ISL_3123439, EPI_ISL_3123440, EPI_ISL_3123441, EPI_ISL_3123442, EPI_ISL_3123443, EPI_ISL_3123444, EPI_ISL_3123445, EPI_ISL_3123446, EPI_ISL_3123447, EPI_ISL_3123448, EPI_ISL_3123449, EPI_ISL_3123450, EPI_ISL_3123451, EPI_ISL_3123452, EPI_ISL_3123453, EPI_ISL_3123454, EPI_ISL_3123455, EPI_ISL_3123457, EPI_ISL_3123458, EPI_ISL_3123459, EPI_ISL_3123460, EPI_ISL_3123461, EPI_ISL_3123462, EPI_ISL_3123463, EPI_ISL_3123467 | | | |
| see above | Virginia Division of Consolidated Laboratory Services | Virginia Division of Consolidated Laboratory Services | Virginia Division of Consolidated Laboratory Services |
| EPI_ISL_3123468, EPI_ISL_3123469, EPI_ISL_3123471, EPI_ISL_3123472, EPI_ISL_3123473, EPI_ISL_3123474, EPI_ISL_3123475, EPI_ISL_3123476, EPI_ISL_3123477, EPI_ISL_3123478, EPI_ISL_3123479, EPI_ISL_3123480 | | | |
| see above | AULSS 1 Dolomiti | Istituto Zooprofilattico Sperimentale delle Venezie | Adelaide Milani, Alessia Schivo, Annalisa Salviato, Elisa Palumbo, Erika Giorgia Quaranta, Luca Tassoni, Ambra Pastori, Edoardo Giussani, Alice Fusaro, Isabella Monne, Calogero Terregino, Antonia Ricci |
| EPI_ISL_3123481 | AULSS 5 Polesana | Istituto Zooprofilattico Sperimentale delle Venezie | Adelaide Milani, Alessia Schivo, Annalisa Salviato, Elisa Palumbo, Erika Giorgia Quaranta, Luca Tassoni, Ambra Pastori, Edoardo Giussani, Alice Fusaro, Isabella Monne, Calogero Terregino, Antonia Ricci |
| EPI_ISL_3123482, EPI_ISL_3123483 | AULSS 1 Dolomiti | Istituto Zooprofilattico Sperimentale delle Venezie | Adelaide Milani, Alessia Schivo, Annalisa Salviato, Elisa Palumbo, Erika Giorgia Quaranta, Luca Tassoni, Ambra Pastori, Edoardo Giussani, Alice Fusaro, Isabella Monne, Calogero Terregino, Antonia Ricci |
| EPI_ISL_3123485, EPI_ISL_3123487, EPI_ISL_3123488, EPI_ISL_3123489, EPI_ISL_3123490 | AULSS 5 Polesana | Istituto Zooprofilattico Sperimentale delle Venezie | Adelaide Milani, Alessia Schivo, Annalisa Salviato, Elisa Palumbo, Erika Giorgia Quaranta, Luca Tassoni, Ambra Pastori, Edoardo Giussani, Alice Fusaro, Isabella Monne, Calogero Terregino, Antonia Ricci |
| EPI_ISL_3123491 | AULSS 9 Scaligera - San Bonifacio | Istituto Zooprofilattico Sperimentale delle Venezie | Adelaide Milani, Alessia Schivo, Annalisa Salviato, Elisa Palumbo, Erika Giorgia Quaranta, Luca Tassoni, Ambra Pastori, Edoardo Giussani, Alice Fusaro, Isabella Monne, Calogero Terregino, Antonia Ricci |
| EPI_ISL_3123492 | AULSS 5 Polesana | Istituto Zooprofilattico Sperimentale delle Venezie | Adelaide Milani, Alessia Schivo, Annalisa Salviato, Elisa Palumbo, Erika Giorgia Quaranta, Luca Tassoni, Ambra Pastori, Edoardo Giussani, Alice Fusaro, Isabella Monne, Calogero Terregino, Antonia Ricci |
| EPI_ISL_3123493, EPI_ISL_3123494, EPI_ISL_3123495, EPI_ISL_3123496, EPI_ISL_3123497, EPI_ISL_3123498, EPI_ISL_3123499, EPI_ISL_3123500, EPI_ISL_3123501 | AULSS 9 Scaligera - San Bonifacio | Istituto Zooprofilattico Sperimentale delle Venezie | Adelaide Milani, Alessia Schivo, Annalisa Salviato, Elisa Palumbo, Erika Giorgia Quaranta, Luca Tassoni, Ambra Pastori, Edoardo Giussani, Alice Fusaro, Isabella Monne, Calogero Terregino, Antonia Ricci |
| EPI_ISL_3123502, EPI_ISL_3123503, EPI_ISL_3123504, EPI_ISL_3123505, EPI_ISL_3123506, EPI_ISL_3123507, EPI_ISL_3123508, EPI_ISL_3123509, EPI_ISL_3123510, EPI_ISL_3123511, EPI_ISL_3123512, EPI_ISL_3123513, EPI_ISL_3123514, EPI_ISL_3123515 | | | |

| | | | |
|---|---|---|---|
| see above | AULSS 7 Pedemontana - Santorso | Istituto Zooprofilattico Sperimentale delle Venezie | Adelaide Milani, Alessia Schivo, Annalisa Salviato, Elisa Palumbo, Erika Giorgia Quaranta, Luca Tassoni, Ambra Pastori, Edoardo Giussani, Alice Fusaro, Isabella Monne, Calogero Terregino, Antonia Ricci |
| EPI_ISL_3123516 | AULSS 8 Berica | Istituto Zooprofilattico Sperimentale delle Venezie | Adelaide Milani, Alessia Schivo, Annalisa Salviato, Elisa Palumbo, Erika Giorgia Quaranta, Luca Tassoni, Ambra Pastori, Edoardo Giussani, Alice Fusaro, Isabella Monne, Calogero Terregino, Antonia Ricci |
| EPI_ISL_3123517 | AULSS 7 Pedemontana - Santorso | Istituto Zooprofilattico Sperimentale delle Venezie | Adelaide Milani, Alessia Schivo, Annalisa Salviato, Elisa Palumbo, Erika Giorgia Quaranta, Luca Tassoni, Ambra Pastori, Edoardo Giussani, Alice Fusaro, Isabella Monne, Calogero Terregino, Antonia Ricci |
| EPI_ISL_3123518, EPI_ISL_3123519, EPI_ISL_3123520, EPI_ISL_3123522, EPI_ISL_3123523 | AULSS 8 Berica | Istituto Zooprofilattico Sperimentale delle Venezie | Adelaide Milani, Alessia Schivo, Annalisa Salviato, Elisa Palumbo, Erika Giorgia Quaranta, Luca Tassoni, Ambra Pastori, Edoardo Giussani, Alice Fusaro, Isabella Monne, Calogero Terregino, Antonia Ricci |
| EPI_ISL_3123524 | AULSS 1 Dolomiti | Istituto Zooprofilattico Sperimentale delle Venezie | Adelaide Milani, Alessia Schivo, Annalisa Salviato, Elisa Palumbo, Erika Giorgia Quaranta, Luca Tassoni, Ambra Pastori, Edoardo Giussani, Alice Fusaro, Isabella Monne, Calogero Terregino, Antonia Ricci |
| EPI_ISL_3123525 | AULSS 8 Berica | Istituto Zooprofilattico Sperimentale delle Venezie | Adelaide Milani, Alessia Schivo, Annalisa Salviato, Elisa Palumbo, Erika Giorgia Quaranta, Luca Tassoni, Ambra Pastori, Edoardo Giussani, Alice Fusaro, Isabella Monne, Calogero Terregino, Antonia Ricci |
| EPI_ISL_3123526 | AULSS 1 Dolomiti | Istituto Zooprofilattico Sperimentale delle Venezie | Adelaide Milani, Alessia Schivo, Annalisa Salviato, Elisa Palumbo, Erika Giorgia Quaranta, Luca Tassoni, Ambra Pastori, Edoardo Giussani, Alice Fusaro, Isabella Monne, Calogero Terregino, Antonia Ricci |
| EPI_ISL_3123527, EPI_ISL_3123528, EPI_ISL_3123529, EPI_ISL_3123530, EPI_ISL_3123531, EPI_ISL_3123532, EPI_ISL_3123533, EPI_ISL_3123534, EPI_ISL_3123535, EPI_ISL_3123536, EPI_ISL_3123537, EPI_ISL_3123538, EPI_ISL_3123539, EPI_ISL_3123540, EPI_ISL_3123541, EPI_ISL_3123542, EPI_ISL_3123543, EPI_ISL_3123544, EPI_ISL_3123545, EPI_ISL_3123546, EPI_ISL_3123547, EPI_ISL_3123548, EPI_ISL_3123549, EPI_ISL_3123550, EPI_ISL_3123551, EPI_ISL_3123552, EPI_ISL_3123553, EPI_ISL_3123554, EPI_ISL_3123555, EPI_ISL_3123556, EPI_ISL_3123557, EPI_ISL_3123558 | | | |
| see above | AULSS 2 Marca Trevigiana | Istituto Zooprofilattico Sperimentale delle Venezie | Adelaide Milani, Alessia Schivo, Annalisa Salviato, Elisa Palumbo, Erika Giorgia Quaranta, Luca Tassoni, Ambra Pastori, Edoardo Giussani, Alice Fusaro, Isabella Monne, Calogero Terregino, Antonia Ricci |
| EPI_ISL_3123559, EPI_ISL_3123560, EPI_ISL_3123561, EPI_ISL_3123562, EPI_ISL_3123563, EPI_ISL_3123564, EPI_ISL_3123565, EPI_ISL_3123566 | Unit of laboratory surveillance of viral emergent disease Control of Communicable Diseases Departament National Institute of Public Health Albania | Charité Universitätsmedizin Berlin, Institut für Virologie | Victor M Corman, Barbara Mühlemann, Jörn Beheim-Schwarzbach, Talitha Veith, Julia Schneider, Terry Jones, Iris Hasibra, Christian Drosten |
| EPI_ISL_3123567 | Lighthouse Lab in Glasgow | Wellcome Sanger Institute for the COVID-19 Genomics UK (COG-UK) Consortium | Harper VanSteenhouse, Yumi Kasai, David Gray, Carol Clugston, Anna Dominiczak and Alex Alderton, Roberto Amato, Jeffrey Barrett, Sonia Goncalves, Ewan Harrison, David K. Jackson, Ian Johnston, Dominic Kwiatkowski, Cordelia Langford, John Sillitoe on behalf of the Wellcome Sanger Institute COVID-19 Surveillance Team |
| EPI_ISL_3123568 | Lighthouse Lab in Milton Keynes | Wellcome Sanger Institute for the COVID-19 Genomics UK (COG-UK) Consortium | The Lighthouse Lab in Milton Keynes and Alex Alderton, Roberto Amato, Jeffrey Barrett, Sonia Goncalves, Ewan Harrison, David K. Jackson, Ian Johnston, Dominic Kwiatkowski, Cordelia Langford, John Sillitoe on behalf of the Wellcome Sanger Institute COVID-19 Surveillance Team |
| EPI_ISL_3123569, EPI_ISL_3123570, EPI_ISL_3123571, EPI_ISL_3123572 | Lighthouse Lab in Glasgow | Wellcome Sanger Institute for the COVID-19 Genomics UK (COG-UK) Consortium | Harper VanSteenhouse, Yumi Kasai, David Gray, Carol Clugston, Anna Dominiczak and Alex Alderton, Roberto Amato, Jeffrey Barrett, Sonia Goncalves, Ewan Harrison, David K. Jackson, Ian Johnston, Dominic Kwiatkowski, Cordelia Langford, John Sillitoe on behalf of the Wellcome Sanger Institute COVID-19 Surveillance Team |
| EPI_ISL_3123573 | Lighthouse Lab in Milton Keynes | Wellcome Sanger Institute for the COVID-19 Genomics UK (COG-UK) Consortium | The Lighthouse Lab in Milton Keynes and Alex Alderton, Roberto Amato, Jeffrey Barrett, Sonia Goncalves, Ewan Harrison, David K. Jackson, Ian Johnston, Dominic Kwiatkowski, Cordelia Langford, John Sillitoe on behalf of the Wellcome Sanger Institute COVID-19 Surveillance Team |
| EPI_ISL_3123574 | Lighthouse Lab in Glasgow | Wellcome Sanger Institute for the COVID-19 Genomics UK (COG-UK) Consortium | Harper VanSteenhouse, Yumi Kasai, David Gray, Carol Clugston, Anna Dominiczak and Alex Alderton, Roberto Amato, Jeffrey Barrett, Sonia Goncalves, Ewan Harrison, David K. Jackson, Ian Johnston, Dominic Kwiatkowski, Cordelia Langford, John Sillitoe on behalf of the Wellcome Sanger Institute COVID-19 Surveillance Team |
| EPI_ISL_3123575, EPI_ISL_3123576 | Lighthouse Lab in Milton Keynes | Wellcome Sanger Institute for the COVID-19 Genomics UK (COG-UK) Consortium | The Lighthouse Lab in Milton Keynes and Alex Alderton, Roberto Amato, Jeffrey Barrett, Sonia Goncalves, Ewan Harrison, David K. Jackson, Ian Johnston, Dominic Kwiatkowski, Cordelia Langford, John Sillitoe on behalf of the Wellcome Sanger Institute COVID-19 Surveillance Team |
| EPI_ISL_3123577 | Berkshire and Surrey Pathology Services Lighthouse Laboratory | Wellcome Sanger Institute for the COVID-19 Genomics UK (COG-UK) Consortium | Berkshire and Surrey Pathology Services Lighthouse Laboratory and Alex Alderton, Roberto Amato, Jeffrey Barrett, Sonia Goncalves, Ewan Harrison, David K. Jackson, Ian Johnston, Dominic Kwiatkowski, Cordelia Langford, John Sillitoe on behalf of the Wellcome Sanger Institute COVID-19 Surveillance Team |
| EPI_ISL_3123578, EPI_ISL_3123579 | Lighthouse Lab in Milton Keynes | Wellcome Sanger Institute for the COVID-19 Genomics UK (COG-UK) Consortium | The Lighthouse Lab in Milton Keynes and Alex Alderton, Roberto Amato, Jeffrey Barrett, Sonia Goncalves, Ewan Harrison, David K. Jackson, Ian Johnston, Dominic Kwiatkowski, Cordelia Langford, John Sillitoe on behalf of the Wellcome Sanger Institute COVID-19 Surveillance Team |
| EPI_ISL_3123580, EPI_ISL_3123581 | Lighthouse Lab in Glasgow | Wellcome Sanger Institute for the COVID-19 Genomics UK (COG-UK) Consortium | Harper VanSteenhouse, Yumi Kasai, David Gray, Carol Clugston, Anna Dominiczak and Alex Alderton, Roberto Amato, Jeffrey Barrett, Sonia Goncalves, Ewan Harrison, David K. Jackson, Ian Johnston, Dominic Kwiatkowski, Cordelia Langford, John Sillitoe on behalf of the Wellcome Sanger Institute COVID-19 Surveillance Team |
| EPI_ISL_3123582 | Berkshire and Surrey Pathology Services Lighthouse Laboratory | Wellcome Sanger Institute for the COVID-19 Genomics UK (COG-UK) Consortium | Berkshire and Surrey Pathology Services Lighthouse Laboratory and Alex Alderton, Roberto Amato, Jeffrey Barrett, Sonia Goncalves, Ewan Harrison, David K. Jackson, Ian Johnston, Dominic Kwiatkowski, Cordelia Langford, John Sillitoe on behalf of the Wellcome Sanger Institute COVID-19 Surveillance Team |
| EPI_ISL_3123583 | Lighthouse Lab in Milton Keynes | Wellcome Sanger Institute for the COVID-19 Genomics UK (COG-UK) Consortium | The Lighthouse Lab in Milton Keynes and Alex Alderton, Roberto Amato, Jeffrey Barrett, Sonia Goncalves, Ewan Harrison, David K. Jackson, Ian Johnston, Dominic Kwiatkowski, Cordelia Langford, John Sillitoe on behalf of the Wellcome Sanger Institute COVID-19 Surveillance Team |
| EPI_ISL_3123584 | Lighthouse Lab in Glasgow | Wellcome Sanger Institute for the COVID-19 Genomics UK (COG-UK) Consortium | Harper VanSteenhouse, Yumi Kasai, David Gray, Carol Clugston, Anna Dominiczak and Alex Alderton, Roberto Amato, Jeffrey Barrett, Sonia Goncalves, Ewan Harrison, David K. Jackson, Ian Johnston, Dominic Kwiatkowski, Cordelia Langford, John Sillitoe on behalf of the Wellcome Sanger Institute COVID-19 Surveillance Team |
| EPI_ISL_3123585, EPI_ISL_3123586 | Lighthouse Lab in Milton Keynes | Wellcome Sanger Institute for the COVID-19 Genomics UK (COG-UK) Consortium | The Lighthouse Lab in Milton Keynes and Alex Alderton, Roberto Amato, Jeffrey Barrett, Sonia Goncalves, Ewan Harrison, David K. Jackson, Ian Johnston, Dominic Kwiatkowski, Cordelia Langford, John Sillitoe on behalf of the Wellcome Sanger Institute COVID-19 Surveillance Team |
| EPI_ISL_3123587 | Berkshire and Surrey Pathology Services Lighthouse Laboratory | Wellcome Sanger Institute for the COVID-19 Genomics UK (COG-UK) Consortium | Berkshire and Surrey Pathology Services Lighthouse Laboratory and Alex Alderton, Roberto Amato, Jeffrey Barrett, Sonia Goncalves, Ewan Harrison, David K. Jackson, Ian Johnston, Dominic Kwiatkowski, Cordelia Langford, John Sillitoe on behalf of the Wellcome Sanger Institute COVID-19 Surveillance Team |
| EPI_ISL_3123588 | Lighthouse Lab in Glasgow | Wellcome Sanger Institute for the COVID-19 Genomics UK (COG-UK) Consortium | Harper VanSteenhouse, Yumi Kasai, David Gray, Carol Clugston, Anna Dominiczak and Alex Alderton, Roberto Amato, Jeffrey Barrett, Sonia Goncalves, Ewan Harrison, David K. Jackson, Ian Johnston, Dominic Kwiatkowski, Cordelia Langford, John Sillitoe on behalf of the Wellcome Sanger Institute COVID-19 Surveillance Team |
| EPI_ISL_3123589 | Berkshire and Surrey Pathology Services Lighthouse Laboratory | Wellcome Sanger Institute for the COVID-19 Genomics UK (COG-UK) Consortium | Berkshire and Surrey Pathology Services Lighthouse Laboratory and Alex Alderton, Roberto Amato, Jeffrey Barrett, Sonia Goncalves, Ewan Harrison, David K. Jackson, Ian Johnston, Dominic Kwiatkowski, Cordelia Langford, John Sillitoe on behalf of the Wellcome Sanger Institute COVID-19 Surveillance Team |
| EPI_ISL_3123590 | Lighthouse Lab in Milton Keynes | Wellcome Sanger Institute for the COVID-19 Genomics UK (COG-UK) Consortium | The Lighthouse Lab in Milton Keynes and Alex Alderton, Roberto Amato, Jeffrey Barrett, Sonia Goncalves, Ewan Harrison, David K. Jackson, Ian Johnston, Dominic Kwiatkowski, Cordelia Langford, John Sillitoe on behalf of the Wellcome Sanger Institute COVID-19 Surveillance Team |
| EPI_ISL_3123591 | Berkshire and Surrey Pathology Services Lighthouse Laboratory | Wellcome Sanger Institute for the COVID-19 Genomics UK (COG-UK) Consortium | Berkshire and Surrey Pathology Services Lighthouse Laboratory and Alex Alderton, Roberto Amato, Jeffrey Barrett, Sonia Goncalves, Ewan Harrison, David K. Jackson, Ian Johnston, Dominic Kwiatkowski, Cordelia Langford, John Sillitoe on behalf of the Wellcome Sanger Institute COVID-19 Surveillance Team |
| EPI_ISL_3123592 | Lighthouse Lab in Milton Keynes | Wellcome Sanger Institute for the COVID-19 Genomics UK (COG-UK) Consortium | The Lighthouse Lab in Milton Keynes and Alex Alderton, Roberto Amato, Jeffrey Barrett, Sonia Goncalves, Ewan Harrison, David K. Jackson, Ian Johnston, Dominic Kwiatkowski, Cordelia Langford, John Sillitoe on behalf of the Wellcome Sanger Institute COVID-19 Surveillance Team |
| EPI_ISL_3123593, EPI_ISL_3123594 | Berkshire and Surrey Pathology Services Lighthouse Laboratory | Wellcome Sanger Institute for the COVID-19 Genomics UK (COG-UK) Consortium | Berkshire and Surrey Pathology Services Lighthouse Laboratory and Alex Alderton, Roberto Amato, Jeffrey Barrett, Sonia Goncalves, Ewan Harrison, David K. Jackson, Ian Johnston, Dominic Kwiatkowski, Cordelia Langford, John Sillitoe on behalf of the Wellcome Sanger Institute COVID-19 Surveillance Team |
| EPI_ISL_3123595 | Lighthouse Lab in Glasgow | Wellcome Sanger Institute for the COVID-19 Genomics UK (COG-UK) Consortium | Harper VanSteenhouse, Yumi Kasai, David Gray, Carol Clugston, Anna Dominiczak and Alex Alderton, Roberto Amato, Jeffrey Barrett, Sonia Goncalves, Ewan Harrison, David K. Jackson, Ian Johnston, Dominic Kwiatkowski, Cordelia Langford, John Sillitoe on behalf of the Wellcome Sanger Institute COVID-19 Surveillance Team |
| EPI_ISL_3123596, EPI_ISL_3123597 | Lighthouse Lab in Milton Keynes | Wellcome Sanger Institute for the COVID-19 Genomics UK (COG-UK) Consortium | The Lighthouse Lab in Milton Keynes and Alex Alderton, Roberto Amato, Jeffrey Barrett, Sonia Goncalves, Ewan Harrison, David K. Jackson, Ian Johnston, Dominic Kwiatkowski, Cordelia Langford, John Sillitoe on behalf of the Wellcome Sanger Institute COVID-19 Surveillance Team |
| EPI_ISL_3123598 | Berkshire and Surrey Pathology Services Lighthouse Laboratory | Wellcome Sanger Institute for the COVID-19 Genomics UK (COG-UK) Consortium | Berkshire and Surrey Pathology Services Lighthouse Laboratory and Alex Alderton, Roberto Amato, Jeffrey Barrett, Sonia Goncalves, Ewan Harrison, David K. Jackson, Ian Johnston, Dominic Kwiatkowski, Cordelia Langford, John Sillitoe on behalf of the Wellcome Sanger Institute COVID-19 Surveillance Team |
| EPI_ISL_3123599 | Lighthouse Lab in Milton Keynes | Wellcome Sanger Institute for the COVID-19 Genomics UK (COG-UK) Consortium | The Lighthouse Lab in Milton Keynes and Alex Alderton, Roberto Amato, Jeffrey Barrett, Sonia Goncalves, Ewan Harrison, David K. Jackson, Ian Johnston, Dominic Kwiatkowski, Cordelia Langford, John Sillitoe on behalf of the Wellcome Sanger Institute COVID-19 Surveillance Team |
| EPI_ISL_3123600 | Berkshire and Surrey Pathology Services Lighthouse Laboratory | Wellcome Sanger Institute for the COVID-19 Genomics UK (COG-UK) Consortium | Berkshire and Surrey Pathology Services Lighthouse Laboratory and Alex Alderton, Roberto Amato, Jeffrey Barrett, Sonia Goncalves, Ewan Harrison, David K. Jackson, Ian Johnston, Dominic Kwiatkowski, Cordelia Langford, John Sillitoe on behalf of the Wellcome Sanger Institute COVID-19 Surveillance Team |
| EPI_ISL_3123601 | Lighthouse Lab in Milton Keynes | Wellcome Sanger Institute for the COVID-19 Genomics UK (COG-UK) Consortium | The Lighthouse Lab in Milton Keynes and Alex Alderton, Roberto Amato, Jeffrey Barrett, Sonia Goncalves, Ewan Harrison, David K. Jackson, Ian Johnston, Dominic Kwiatkowski, Cordelia Langford, John Sillitoe on behalf of the Wellcome Sanger Institute COVID-19 Surveillance Team |
| EPI_ISL_3123602 | Berkshire and Surrey Pathology Services Lighthouse Laboratory | Wellcome Sanger Institute for the COVID-19 Genomics UK (COG-UK) Consortium | Berkshire and Surrey Pathology Services Lighthouse Laboratory and Alex Alderton, Roberto Amato, Jeffrey Barrett, Sonia Goncalves, Ewan Harrison, David K. Jackson, Ian Johnston, Dominic Kwiatkowski, Cordelia Langford, John Sillitoe on behalf of the Wellcome Sanger Institute COVID-19 Surveillance Team |
| EPI_ISL_3123603 | Lighthouse Lab in Glasgow | Wellcome Sanger Institute for the COVID-19 Genomics UK (COG-UK) Consortium | Harper VanSteenhouse, Yumi Kasai, David Gray, Carol Clugston, Anna Dominiczak and Alex Alderton, Roberto Amato, Jeffrey Barrett, Sonia Goncalves, Ewan Harrison, David K. Jackson, Ian Johnston, Dominic Kwiatkowski, Cordelia Langford, John Sillitoe on behalf of the Wellcome Sanger Institute COVID-19 Surveillance Team |
| EPI_ISL_3123604 | Lighthouse Lab in Milton Keynes | Wellcome Sanger Institute for the COVID-19 Genomics UK (COG-UK) Consortium | The Lighthouse Lab in Milton Keynes and Alex Alderton, Roberto Amato, Jeffrey Barrett, Sonia Goncalves, Ewan Harrison, David K. Jackson, Ian Johnston, Dominic Kwiatkowski, Cordelia Langford, John Sillitoe on behalf of the Wellcome Sanger Institute COVID-19 Surveillance Team |
| EPI_ISL_3123605, EPI_ISL_3123606 | Lighthouse Lab in Glasgow | Wellcome Sanger Institute for the COVID-19 Genomics UK (COG-UK) Consortium | Harper VanSteenhouse, Yumi Kasai, David Gray, Carol Clugston, Anna Dominiczak and Alex Alderton, Roberto Amato, Jeffrey Barrett, Sonia Goncalves, Ewan Harrison, David K. Jackson, Ian Johnston, Dominic Kwiatkowski, Cordelia Langford, John Sillitoe on behalf of the Wellcome Sanger Institute COVID-19 Surveillance Team |
| EPI_ISL_3123607, EPI_ISL_3123608 | Lighthouse Lab in Milton Keynes | Wellcome Sanger Institute for the COVID-19 Genomics UK (COG-UK) Consortium | The Lighthouse Lab in Milton Keynes and Alex Alderton, Roberto Amato, Jeffrey Barrett, Sonia Goncalves, Ewan Harrison, David K. Jackson, Ian Johnston, Dominic Kwiatkowski, Cordelia Langford, John Sillitoe on behalf of the Wellcome Sanger Institute COVID-19 Surveillance Team |
| EPI_ISL_3123609 | Berkshire and Surrey Pathology Services Lighthouse Laboratory | Wellcome Sanger Institute for the COVID-19 Genomics UK (COG-UK) Consortium | Berkshire and Surrey Pathology Services Lighthouse Laboratory and Alex Alderton, Roberto Amato, Jeffrey Barrett, Sonia Goncalves, Ewan Harrison, David K. Jackson, Ian Johnston, Dominic Kwiatkowski, Cordelia Langford, John Sillitoe on behalf of the Wellcome Sanger Institute COVID-19 Surveillance Team |
| EPI_ISL_3123610, EPI_ISL_3123611, EPI_ISL_3123612 | Lighthouse Lab in Glasgow | Wellcome Sanger Institute for the COVID-19 Genomics UK (COG-UK) Consortium | Harper VanSteenhouse, Yumi Kasai, David Gray, Carol Clugston, Anna Dominiczak and Alex Alderton, Roberto Amato, Jeffrey Barrett, Sonia Goncalves, Ewan Harrison, David K. Jackson, Ian Johnston, Dominic Kwiatkowski, Cordelia Langford, John Sillitoe on behalf of the Wellcome Sanger Institute COVID-19 Surveillance Team |

| | | | |
|---|---|---|---|
| EPI_ISL_3123613 | Lighthouse Lab in Milton Keynes | Wellcome Sanger Institute for the COVID-19 Genomics UK (COG-UK) Consortium | The Lighthouse Lab in Milton Keynes and Alex Alderton, Roberto Amato, Jeffrey Barrett, Sonia Goncalves, Ewan Harrison, David K. Jackson, Ian Johnston, Dominic Kwiatkowski, Cordelia Langford, John Sillitoe on behalf of the Wellcome Sanger Institute COVID-19 Surveillance Team |
| EPI_ISL_3123614 | Lighthouse Lab in Glasgow | Wellcome Sanger Institute for the COVID-19 Genomics UK (COG-UK) Consortium | Harper VanSteenhouse, Yumi Kasai, David Gray, Carol Clugston, Anna Dominiczak and Alex Alderton, Roberto Amato, Jeffrey Barrett, Sonia Goncalves, Ewan Harrison, David K. Jackson, Ian Johnston, Dominic Kwiatkowski, Cordelia Langford, John Sillitoe on behalf of the Wellcome Sanger Institute COVID-19 Surveillance Team |
| EPI_ISL_3123615 | Berkshire and Surrey Pathology Services Lighthouse Laboratory | Wellcome Sanger Institute for the COVID-19 Genomics UK (COG-UK) Consortium | Berkshire and Surrey Pathology Services Lighthouse Laboratory and Alex Alderton, Roberto Amato, Jeffrey Barrett, Sonia Goncalves, Ewan Harrison, David K. Jackson, Ian Johnston, Dominic Kwiatkowski, Cordelia Langford, John Sillitoe on behalf of the Wellcome Sanger Institute COVID-19 Surveillance Team |
| EPI_ISL_3123616, EPI_ISL_3123617 | Lighthouse Lab in Milton Keynes | Wellcome Sanger Institute for the COVID-19 Genomics UK (COG-UK) Consortium | The Lighthouse Lab in Milton Keynes and Alex Alderton, Roberto Amato, Jeffrey Barrett, Sonia Goncalves, Ewan Harrison, David K. Jackson, Ian Johnston, Dominic Kwiatkowski, Cordelia Langford, John Sillitoe on behalf of the Wellcome Sanger Institute COVID-19 Surveillance Team |
| EPI_ISL_3123618, EPI_ISL_3123619, EPI_ISL_3123620, EPI_ISL_3123621 | Lighthouse Lab in Glasgow | Wellcome Sanger Institute for the COVID-19 Genomics UK (COG-UK) Consortium | Harper VanSteenhouse, Yumi Kasai, David Gray, Carol Clugston, Anna Dominiczak and Alex Alderton, Roberto Amato, Jeffrey Barrett, Sonia Goncalves, Ewan Harrison, David K. Jackson, Ian Johnston, Dominic Kwiatkowski, Cordelia Langford, John Sillitoe on behalf of the Wellcome Sanger Institute COVID-19 Surveillance Team |
| EPI_ISL_3123622 | Lighthouse Lab in Milton Keynes | Wellcome Sanger Institute for the COVID-19 Genomics UK (COG-UK) Consortium | The Lighthouse Lab in Milton Keynes and Alex Alderton, Roberto Amato, Jeffrey Barrett, Sonia Goncalves, Ewan Harrison, David K. Jackson, Ian Johnston, Dominic Kwiatkowski, Cordelia Langford, John Sillitoe on behalf of the Wellcome Sanger Institute COVID-19 Surveillance Team |
| EPI_ISL_3123623, EPI_ISL_3123624, EPI_ISL_3123625 | Berkshire and Surrey Pathology Services Lighthouse Laboratory | Wellcome Sanger Institute for the COVID-19 Genomics UK (COG-UK) Consortium | Berkshire and Surrey Pathology Services Lighthouse Laboratory and Alex Alderton, Roberto Amato, Jeffrey Barrett, Sonia Goncalves, Ewan Harrison, David K. Jackson, Ian Johnston, Dominic Kwiatkowski, Cordelia Langford, John Sillitoe on behalf of the Wellcome Sanger Institute COVID-19 Surveillance Team |
| EPI_ISL_3123626 | Lighthouse Lab in Milton Keynes | Wellcome Sanger Institute for the COVID-19 Genomics UK (COG-UK) Consortium | The Lighthouse Lab in Milton Keynes and Alex Alderton, Roberto Amato, Jeffrey Barrett, Sonia Goncalves, Ewan Harrison, David K. Jackson, Ian Johnston, Dominic Kwiatkowski, Cordelia Langford, John Sillitoe on behalf of the Wellcome Sanger Institute COVID-19 Surveillance Team |
| EPI_ISL_3123627 | Berkshire and Surrey Pathology Services Lighthouse Laboratory | Wellcome Sanger Institute for the COVID-19 Genomics UK (COG-UK) Consortium | Berkshire and Surrey Pathology Services Lighthouse Laboratory and Alex Alderton, Roberto Amato, Jeffrey Barrett, Sonia Goncalves, Ewan Harrison, David K. Jackson, Ian Johnston, Dominic Kwiatkowski, Cordelia Langford, John Sillitoe on behalf of the Wellcome Sanger Institute COVID-19 Surveillance Team |
| EPI_ISL_3123628 | Lighthouse Lab in Glasgow | Wellcome Sanger Institute for the COVID-19 Genomics UK (COG-UK) Consortium | Harper VanSteenhouse, Yumi Kasai, David Gray, Carol Clugston, Anna Dominiczak and Alex Alderton, Roberto Amato, Jeffrey Barrett, Sonia Goncalves, Ewan Harrison, David K. Jackson, Ian Johnston, Dominic Kwiatkowski, Cordelia Langford, John Sillitoe on behalf of the Wellcome Sanger Institute COVID-19 Surveillance Team |
| EPI_ISL_3123629, EPI_ISL_3123630 | Lighthouse Lab in Milton Keynes | Wellcome Sanger Institute for the COVID-19 Genomics UK (COG-UK) Consortium | The Lighthouse Lab in Milton Keynes and Alex Alderton, Roberto Amato, Jeffrey Barrett, Sonia Goncalves, Ewan Harrison, David K. Jackson, Ian Johnston, Dominic Kwiatkowski, Cordelia Langford, John Sillitoe on behalf of the Wellcome Sanger Institute COVID-19 Surveillance Team |
| EPI_ISL_3123631, EPI_ISL_3123632 | Berkshire and Surrey Pathology Services Lighthouse Laboratory | Wellcome Sanger Institute for the COVID-19 Genomics UK (COG-UK) Consortium | Berkshire and Surrey Pathology Services Lighthouse Laboratory and Alex Alderton, Roberto Amato, Jeffrey Barrett, Sonia Goncalves, Ewan Harrison, David K. Jackson, Ian Johnston, Dominic Kwiatkowski, Cordelia Langford, John Sillitoe on behalf of the Wellcome Sanger Institute COVID-19 Surveillance Team |
| EPI_ISL_3123633 | Lighthouse Lab in Milton Keynes | Wellcome Sanger Institute for the COVID-19 Genomics UK (COG-UK) Consortium | The Lighthouse Lab in Milton Keynes and Alex Alderton, Roberto Amato, Jeffrey Barrett, Sonia Goncalves, Ewan Harrison, David K. Jackson, Ian Johnston, Dominic Kwiatkowski, Cordelia Langford, John Sillitoe on behalf of the Wellcome Sanger Institute COVID-19 Surveillance Team |
| EPI_ISL_3123634, EPI_ISL_3123635, EPI_ISL_3123636 | Berkshire and Surrey Pathology Services Lighthouse Laboratory | Wellcome Sanger Institute for the COVID-19 Genomics UK (COG-UK) Consortium | Berkshire and Surrey Pathology Services Lighthouse Laboratory and Alex Alderton, Roberto Amato, Jeffrey Barrett, Sonia Goncalves, Ewan Harrison, David K. Jackson, Ian Johnston, Dominic Kwiatkowski, Cordelia Langford, John Sillitoe on behalf of the Wellcome Sanger Institute COVID-19 Surveillance Team |
| EPI_ISL_3123637 | Lighthouse Lab in Milton Keynes | Wellcome Sanger Institute for the COVID-19 Genomics UK (COG-UK) Consortium | The Lighthouse Lab in Milton Keynes and Alex Alderton, Roberto Amato, Jeffrey Barrett, Sonia Goncalves, Ewan Harrison, David K. Jackson, Ian Johnston, Dominic Kwiatkowski, Cordelia Langford, John Sillitoe on behalf of the Wellcome Sanger Institute COVID-19 Surveillance Team |
| EPI_ISL_3123638, EPI_ISL_3123639, EPI_ISL_3123640 | Lighthouse Lab in Glasgow | Wellcome Sanger Institute for the COVID-19 Genomics UK (COG-UK) Consortium | Harper VanSteenhouse, Yumi Kasai, David Gray, Carol Clugston, Anna Dominiczak and Alex Alderton, Roberto Amato, Jeffrey Barrett, Sonia Goncalves, Ewan Harrison, David K. Jackson, Ian Johnston, Dominic Kwiatkowski, Cordelia Langford, John Sillitoe on behalf of the Wellcome Sanger Institute COVID-19 Surveillance Team |
| EPI_ISL_3123641, EPI_ISL_3123642 | Lighthouse Lab in Milton Keynes | Wellcome Sanger Institute for the COVID-19 Genomics UK (COG-UK) Consortium | The Lighthouse Lab in Milton Keynes and Alex Alderton, Roberto Amato, Jeffrey Barrett, Sonia Goncalves, Ewan Harrison, David K. Jackson, Ian Johnston, Dominic Kwiatkowski, Cordelia Langford, John Sillitoe on behalf of the Wellcome Sanger Institute COVID-19 Surveillance Team |
| EPI_ISL_3123643 | Lighthouse Lab in Glasgow | Wellcome Sanger Institute for the COVID-19 Genomics UK (COG-UK) Consortium | Harper VanSteenhouse, Yumi Kasai, David Gray, Carol Clugston, Anna Dominiczak and Alex Alderton, Roberto Amato, Jeffrey Barrett, Sonia Goncalves, Ewan Harrison, David K. Jackson, Ian Johnston, Dominic Kwiatkowski, Cordelia Langford, John Sillitoe on behalf of the Wellcome Sanger Institute COVID-19 Surveillance Team |
| EPI_ISL_3123644, EPI_ISL_3123645 | Lighthouse Lab in Milton Keynes | Wellcome Sanger Institute for the COVID-19 Genomics UK (COG-UK) Consortium | The Lighthouse Lab in Milton Keynes and Alex Alderton, Roberto Amato, Jeffrey Barrett, Sonia Goncalves, Ewan Harrison, David K. Jackson, Ian Johnston, Dominic Kwiatkowski, Cordelia Langford, John Sillitoe on behalf of the Wellcome Sanger Institute COVID-19 Surveillance Team |
| EPI_ISL_3123646 | Berkshire and Surrey Pathology Services Lighthouse Laboratory | Wellcome Sanger Institute for the COVID-19 Genomics UK (COG-UK) Consortium | Berkshire and Surrey Pathology Services Lighthouse Laboratory and Alex Alderton, Roberto Amato, Jeffrey Barrett, Sonia Goncalves, Ewan Harrison, David K. Jackson, Ian Johnston, Dominic Kwiatkowski, Cordelia Langford, John Sillitoe on behalf of the Wellcome Sanger Institute COVID-19 Surveillance Team |
| EPI_ISL_3123647 | Lighthouse Lab in Milton Keynes | Wellcome Sanger Institute for the COVID-19 Genomics UK (COG-UK) Consortium | The Lighthouse Lab in Milton Keynes and Alex Alderton, Roberto Amato, Jeffrey Barrett, Sonia Goncalves, Ewan Harrison, David K. Jackson, Ian Johnston, Dominic Kwiatkowski, Cordelia Langford, John Sillitoe on behalf of the Wellcome Sanger Institute COVID-19 Surveillance Team |
| EPI_ISL_3123648, EPI_ISL_3123649, EPI_ISL_3123650 | Berkshire and Surrey Pathology Services Lighthouse Laboratory | Wellcome Sanger Institute for the COVID-19 Genomics UK (COG-UK) Consortium | Berkshire and Surrey Pathology Services Lighthouse Laboratory and Alex Alderton, Roberto Amato, Jeffrey Barrett, Sonia Goncalves, Ewan Harrison, David K. Jackson, Ian Johnston, Dominic Kwiatkowski, Cordelia Langford, John Sillitoe on behalf of the Wellcome Sanger Institute COVID-19 Surveillance Team |
| EPI_ISL_3123651 | Lighthouse Lab in Glasgow | Wellcome Sanger Institute for the COVID-19 Genomics UK (COG-UK) Consortium | Harper VanSteenhouse, Yumi Kasai, David Gray, Carol Clugston, Anna Dominiczak and Alex Alderton, Roberto Amato, Jeffrey Barrett, Sonia Goncalves, Ewan Harrison, David K. Jackson, Ian Johnston, Dominic Kwiatkowski, Cordelia Langford, John Sillitoe on behalf of the Wellcome Sanger Institute COVID-19 Surveillance Team |
| EPI_ISL_3123652 | Berkshire and Surrey Pathology Services Lighthouse Laboratory | Wellcome Sanger Institute for the COVID-19 Genomics UK (COG-UK) Consortium | Berkshire and Surrey Pathology Services Lighthouse Laboratory and Alex Alderton, Roberto Amato, Jeffrey Barrett, Sonia Goncalves, Ewan Harrison, David K. Jackson, Ian Johnston, Dominic Kwiatkowski, Cordelia Langford, John Sillitoe on behalf of the Wellcome Sanger Institute COVID-19 Surveillance Team |
| EPI_ISL_3123653 | Lighthouse Lab in Milton Keynes | Wellcome Sanger Institute for the COVID-19 Genomics UK (COG-UK) Consortium | The Lighthouse Lab in Milton Keynes and Alex Alderton, Roberto Amato, Jeffrey Barrett, Sonia Goncalves, Ewan Harrison, David K. Jackson, Ian Johnston, Dominic Kwiatkowski, Cordelia Langford, John Sillitoe on behalf of the Wellcome Sanger Institute COVID-19 Surveillance Team |
| EPI_ISL_3123654, EPI_ISL_3123655, EPI_ISL_3123656 | Lighthouse Lab in Glasgow | Wellcome Sanger Institute for the COVID-19 Genomics UK (COG-UK) Consortium | Harper VanSteenhouse, Yumi Kasai, David Gray, Carol Clugston, Anna Dominiczak and Alex Alderton, Roberto Amato, Jeffrey Barrett, Sonia Goncalves, Ewan Harrison, David K. Jackson, Ian Johnston, Dominic Kwiatkowski, Cordelia Langford, John Sillitoe on behalf of the Wellcome Sanger Institute COVID-19 Surveillance Team |
| EPI_ISL_3123657 | Berkshire and Surrey Pathology Services Lighthouse Laboratory | Wellcome Sanger Institute for the COVID-19 Genomics UK (COG-UK) Consortium | Berkshire and Surrey Pathology Services Lighthouse Laboratory and Alex Alderton, Roberto Amato, Jeffrey Barrett, Sonia Goncalves, Ewan Harrison, David K. Jackson, Ian Johnston, Dominic Kwiatkowski, Cordelia Langford, John Sillitoe on behalf of the Wellcome Sanger Institute COVID-19 Surveillance Team |
| EPI_ISL_3123658 | Lighthouse Lab in Milton Keynes | Wellcome Sanger Institute for the COVID-19 Genomics UK (COG-UK) Consortium | The Lighthouse Lab in Milton Keynes and Alex Alderton, Roberto Amato, Jeffrey Barrett, Sonia Goncalves, Ewan Harrison, David K. Jackson, Ian Johnston, Dominic Kwiatkowski, Cordelia Langford, John Sillitoe on behalf of the Wellcome Sanger Institute COVID-19 Surveillance Team |
| EPI_ISL_3123659 | Lighthouse Lab in Glasgow | Wellcome Sanger Institute for the COVID-19 Genomics UK (COG-UK) Consortium | Harper VanSteenhouse, Yumi Kasai, David Gray, Carol Clugston, Anna Dominiczak and Alex Alderton, Roberto Amato, Jeffrey Barrett, Sonia Goncalves, Ewan Harrison, David K. Jackson, Ian Johnston, Dominic Kwiatkowski, Cordelia Langford, John Sillitoe on behalf of the Wellcome Sanger Institute COVID-19 Surveillance Team |
| EPI_ISL_3123660 | Berkshire and Surrey Pathology Services Lighthouse Laboratory | Wellcome Sanger Institute for the COVID-19 Genomics UK (COG-UK) Consortium | Berkshire and Surrey Pathology Services Lighthouse Laboratory and Alex Alderton, Roberto Amato, Jeffrey Barrett, Sonia Goncalves, Ewan Harrison, David K. Jackson, Ian Johnston, Dominic Kwiatkowski, Cordelia Langford, John Sillitoe on behalf of the Wellcome Sanger Institute COVID-19 Surveillance Team |
| EPI_ISL_3123661, EPI_ISL_3123662, EPI_ISL_3123663 | Lighthouse Lab in Milton Keynes | Wellcome Sanger Institute for the COVID-19 Genomics UK (COG-UK) Consortium | The Lighthouse Lab in Milton Keynes and Alex Alderton, Roberto Amato, Jeffrey Barrett, Sonia Goncalves, Ewan Harrison, David K. Jackson, Ian Johnston, Dominic Kwiatkowski, Cordelia Langford, John Sillitoe on behalf of the Wellcome Sanger Institute COVID-19 Surveillance Team |
| EPI_ISL_3123664, EPI_ISL_3123665 | Lighthouse Lab in Glasgow | Wellcome Sanger Institute for the COVID-19 Genomics UK (COG-UK) Consortium | Harper VanSteenhouse, Yumi Kasai, David Gray, Carol Clugston, Anna Dominiczak and Alex Alderton, Roberto Amato, Jeffrey Barrett, Sonia Goncalves, Ewan Harrison, David K. Jackson, Ian Johnston, Dominic Kwiatkowski, Cordelia Langford, John Sillitoe on behalf of the Wellcome Sanger Institute COVID-19 Surveillance Team |
| EPI_ISL_3123666 | Berkshire and Surrey Pathology Services Lighthouse Laboratory | Wellcome Sanger Institute for the COVID-19 Genomics UK (COG-UK) Consortium | Berkshire and Surrey Pathology Services Lighthouse Laboratory and Alex Alderton, Roberto Amato, Jeffrey Barrett, Sonia Goncalves, Ewan Harrison, David K. Jackson, Ian Johnston, Dominic Kwiatkowski, Cordelia Langford, John Sillitoe on behalf of the Wellcome Sanger Institute COVID-19 Surveillance Team |
| EPI_ISL_3123667 | Lighthouse Lab in Milton Keynes | Wellcome Sanger Institute for the COVID-19 Genomics UK (COG-UK) Consortium | The Lighthouse Lab in Milton Keynes and Alex Alderton, Roberto Amato, Jeffrey Barrett, Sonia Goncalves, Ewan Harrison, David K. Jackson, Ian Johnston, Dominic Kwiatkowski, Cordelia Langford, John Sillitoe on behalf of the Wellcome Sanger Institute COVID-19 Surveillance Team |
| EPI_ISL_3123668, EPI_ISL_3123669, EPI_ISL_3123670, EPI_ISL_3123671, EPI_ISL_3123672 | Lighthouse Lab in Glasgow | Wellcome Sanger Institute for the COVID-19 Genomics UK (COG-UK) Consortium | Harper VanSteenhouse, Yumi Kasai, David Gray, Carol Clugston, Anna Dominiczak and Alex Alderton, Roberto Amato, Jeffrey Barrett, Sonia Goncalves, Ewan Harrison, David K. Jackson, Ian Johnston, Dominic Kwiatkowski, Cordelia Langford, John Sillitoe on behalf of the Wellcome Sanger Institute COVID-19 Surveillance Team |
| EPI_ISL_3123673 | Lighthouse Lab in Milton Keynes | Wellcome Sanger Institute for the COVID-19 Genomics UK (COG-UK) Consortium | The Lighthouse Lab in Milton Keynes and Alex Alderton, Roberto Amato, Jeffrey Barrett, Sonia Goncalves, Ewan Harrison, David K. Jackson, Ian Johnston, Dominic Kwiatkowski, Cordelia Langford, John Sillitoe on behalf of the Wellcome Sanger Institute COVID-19 Surveillance Team |
| EPI_ISL_3123674 | Lighthouse Lab in Glasgow | Wellcome Sanger Institute for the COVID-19 Genomics UK (COG-UK) Consortium | Harper VanSteenhouse, Yumi Kasai, David Gray, Carol Clugston, Anna Dominiczak and Alex Alderton, Roberto Amato, Jeffrey Barrett, Sonia Goncalves, Ewan Harrison, David K. Jackson, Ian Johnston, Dominic Kwiatkowski, Cordelia Langford, John Sillitoe on behalf of the Wellcome Sanger Institute COVID-19 Surveillance Team |
| EPI_ISL_3123675 | Lighthouse Lab in Milton Keynes | Wellcome Sanger Institute for the COVID-19 Genomics UK (COG-UK) Consortium | The Lighthouse Lab in Milton Keynes and Alex Alderton, Roberto Amato, Jeffrey Barrett, Sonia Goncalves, Ewan Harrison, David K. Jackson, Ian Johnston, Dominic Kwiatkowski, Cordelia Langford, John Sillitoe on behalf of the Wellcome Sanger Institute COVID-19 Surveillance Team |
| EPI_ISL_3123676 | Berkshire and Surrey Pathology Services Lighthouse Laboratory | Wellcome Sanger Institute for the COVID-19 Genomics UK (COG-UK) Consortium | Berkshire and Surrey Pathology Services Lighthouse Laboratory and Alex Alderton, Roberto Amato, Jeffrey Barrett, Sonia Goncalves, Ewan Harrison, David K. Jackson, Ian Johnston, Dominic Kwiatkowski, Cordelia Langford, John Sillitoe on behalf of the Wellcome Sanger Institute COVID-19 Surveillance Team |
| EPI_ISL_3123677 | Lighthouse Lab in Glasgow | Wellcome Sanger Institute for the COVID-19 Genomics UK (COG-UK) Consortium | Harper VanSteenhouse, Yumi Kasai, David Gray, Carol Clugston, Anna Dominiczak and Alex Alderton, Roberto Amato, Jeffrey Barrett, Sonia Goncalves, Ewan Harrison, David K. Jackson, Ian Johnston, Dominic Kwiatkowski, Cordelia Langford, John Sillitoe on behalf of the Wellcome Sanger Institute COVID-19 Surveillance Team |
| EPI_ISL_3123678 | Lighthouse Lab in Milton Keynes | Wellcome Sanger Institute for the COVID-19 Genomics UK (COG-UK) | The Lighthouse Lab in Milton Keynes and Alex Alderton, Roberto Amato, Jeffrey Barrett, Sonia Goncalves, Ewan Harrison, David K. Jackson, Ian Johnston, Dominic Kwiatkowski, Cordelia Langford, John Sillitoe on behalf of the Wellcome Sanger |

| | | | |
|---|---|---|---|
| | | Consortium | Institute COVID-19 Surveillance Team |
| EPI_ISL_3123679, EPI_ISL_3123680, EPI_ISL_3123681, EPI_ISL_3123682 | Berkshire and Surrey Pathology Services Lighthouse Laboratory | Wellcome Sanger Institute for the COVID-19 Genomics UK (COG-UK) Consortium | Berkshire and Surrey Pathology Services Lighthouse Laboratory and Alex Alderton, Roberto Amato, Jeffrey Barrett, Sonia Goncalves, Ewan Harrison, David K. Jackson, Ian Johnston, Dominic Kwiatkowski, Cordelia Langford, John Sillitoe on behalf of the Wellcome Sanger Institute COVID-19 Surveillance Team |
| EPI_ISL_3123683 | Lighthouse Lab in Glasgow | Wellcome Sanger Institute for the COVID-19 Genomics UK (COG-UK) Consortium | Harper VanSteenhouse, Yumi Kasai, David Gray, Carol Clugston, Anna Dominiczak and Alex Alderton, Roberto Amato, Jeffrey Barrett, Sonia Goncalves, Ewan Harrison, David K. Jackson, Ian Johnston, Dominic Kwiatkowski, Cordelia Langford, John Sillitoe on behalf of the Wellcome Sanger Institute COVID-19 Surveillance Team |
| EPI_ISL_3123684, EPI_ISL_3123685 | Lighthouse Lab in Milton Keynes | Wellcome Sanger Institute for the COVID-19 Genomics UK (COG-UK) Consortium | The Lighthouse Lab in Milton Keynes and Alex Alderton, Roberto Amato, Jeffrey Barrett, Sonia Goncalves, Ewan Harrison, David K. Jackson, Ian Johnston, Dominic Kwiatkowski, Cordelia Langford, John Sillitoe on behalf of the Wellcome Sanger Institute COVID-19 Surveillance Team |
| EPI_ISL_3123686 | Lighthouse Lab in Glasgow | Wellcome Sanger Institute for the COVID-19 Genomics UK (COG-UK) Consortium | Harper VanSteenhouse, Yumi Kasai, David Gray, Carol Clugston, Anna Dominiczak and Alex Alderton, Roberto Amato, Jeffrey Barrett, Sonia Goncalves, Ewan Harrison, David K. Jackson, Ian Johnston, Dominic Kwiatkowski, Cordelia Langford, John Sillitoe on behalf of the Wellcome Sanger Institute COVID-19 Surveillance Team |
| EPI_ISL_3123687 | Lighthouse Lab in Milton Keynes | Wellcome Sanger Institute for the COVID-19 Genomics UK (COG-UK) Consortium | The Lighthouse Lab in Milton Keynes and Alex Alderton, Roberto Amato, Jeffrey Barrett, Sonia Goncalves, Ewan Harrison, David K. Jackson, Ian Johnston, Dominic Kwiatkowski, Cordelia Langford, John Sillitoe on behalf of the Wellcome Sanger Institute COVID-19 Surveillance Team |
| EPI_ISL_3123688, EPI_ISL_3123689 | Lighthouse Lab in Glasgow | Wellcome Sanger Institute for the COVID-19 Genomics UK (COG-UK) Consortium | Harper VanSteenhouse, Yumi Kasai, David Gray, Carol Clugston, Anna Dominiczak and Alex Alderton, Roberto Amato, Jeffrey Barrett, Sonia Goncalves, Ewan Harrison, David K. Jackson, Ian Johnston, Dominic Kwiatkowski, Cordelia Langford, John Sillitoe on behalf of the Wellcome Sanger Institute COVID-19 Surveillance Team |
| EPI_ISL_3123690 | Berkshire and Surrey Pathology Services Lighthouse Laboratory | Wellcome Sanger Institute for the COVID-19 Genomics UK (COG-UK) Consortium | Berkshire and Surrey Pathology Services Lighthouse Laboratory and Alex Alderton, Roberto Amato, Jeffrey Barrett, Sonia Goncalves, Ewan Harrison, David K. Jackson, Ian Johnston, Dominic Kwiatkowski, Cordelia Langford, John Sillitoe on behalf of the Wellcome Sanger Institute COVID-19 Surveillance Team |
| EPI_ISL_3123691 | Lighthouse Lab in Glasgow | Wellcome Sanger Institute for the COVID-19 Genomics UK (COG-UK) Consortium | Harper VanSteenhouse, Yumi Kasai, David Gray, Carol Clugston, Anna Dominiczak and Alex Alderton, Roberto Amato, Jeffrey Barrett, Sonia Goncalves, Ewan Harrison, David K. Jackson, Ian Johnston, Dominic Kwiatkowski, Cordelia Langford, John Sillitoe on behalf of the Wellcome Sanger Institute COVID-19 Surveillance Team |
| EPI_ISL_3123692 | Berkshire and Surrey Pathology Services Lighthouse Laboratory | Wellcome Sanger Institute for the COVID-19 Genomics UK (COG-UK) Consortium | Berkshire and Surrey Pathology Services Lighthouse Laboratory and Alex Alderton, Roberto Amato, Jeffrey Barrett, Sonia Goncalves, Ewan Harrison, David K. Jackson, Ian Johnston, Dominic Kwiatkowski, Cordelia Langford, John Sillitoe on behalf of the Wellcome Sanger Institute COVID-19 Surveillance Team |
| EPI_ISL_3123693 | Lighthouse Lab in Milton Keynes | Wellcome Sanger Institute for the COVID-19 Genomics UK (COG-UK) Consortium | The Lighthouse Lab in Milton Keynes and Alex Alderton, Roberto Amato, Jeffrey Barrett, Sonia Goncalves, Ewan Harrison, David K. Jackson, Ian Johnston, Dominic Kwiatkowski, Cordelia Langford, John Sillitoe on behalf of the Wellcome Sanger Institute COVID-19 Surveillance Team |
| EPI_ISL_3123694, EPI_ISL_3123695 | Lighthouse Lab in Glasgow | Wellcome Sanger Institute for the COVID-19 Genomics UK (COG-UK) Consortium | Harper VanSteenhouse, Yumi Kasai, David Gray, Carol Clugston, Anna Dominiczak and Alex Alderton, Roberto Amato, Jeffrey Barrett, Sonia Goncalves, Ewan Harrison, David K. Jackson, Ian Johnston, Dominic Kwiatkowski, Cordelia Langford, John Sillitoe on behalf of the Wellcome Sanger Institute COVID-19 Surveillance Team |
| EPI_ISL_3123696 | Lighthouse Lab in Milton Keynes | Wellcome Sanger Institute for the COVID-19 Genomics UK (COG-UK) Consortium | The Lighthouse Lab in Milton Keynes and Alex Alderton, Roberto Amato, Jeffrey Barrett, Sonia Goncalves, Ewan Harrison, David K. Jackson, Ian Johnston, Dominic Kwiatkowski, Cordelia Langford, John Sillitoe on behalf of the Wellcome Sanger Institute COVID-19 Surveillance Team |
| EPI_ISL_3123697, EPI_ISL_3123698, EPI_ISL_3123699, EPI_ISL_3123700 | Lighthouse Lab in Glasgow | Wellcome Sanger Institute for the COVID-19 Genomics UK (COG-UK) Consortium | Harper VanSteenhouse, Yumi Kasai, David Gray, Carol Clugston, Anna Dominiczak and Alex Alderton, Roberto Amato, Jeffrey Barrett, Sonia Goncalves, Ewan Harrison, David K. Jackson, Ian Johnston, Dominic Kwiatkowski, Cordelia Langford, John Sillitoe on behalf of the Wellcome Sanger Institute COVID-19 Surveillance Team |
| EPI_ISL_3123701 | Lighthouse Lab in Milton Keynes | Wellcome Sanger Institute for the COVID-19 Genomics UK (COG-UK) Consortium | The Lighthouse Lab in Milton Keynes and Alex Alderton, Roberto Amato, Jeffrey Barrett, Sonia Goncalves, Ewan Harrison, David K. Jackson, Ian Johnston, Dominic Kwiatkowski, Cordelia Langford, John Sillitoe on behalf of the Wellcome Sanger Institute COVID-19 Surveillance Team |
| EPI_ISL_3123702 | Berkshire and Surrey Pathology Services Lighthouse Laboratory | Wellcome Sanger Institute for the COVID-19 Genomics UK (COG-UK) Consortium | Berkshire and Surrey Pathology Services Lighthouse Laboratory and Alex Alderton, Roberto Amato, Jeffrey Barrett, Sonia Goncalves, Ewan Harrison, David K. Jackson, Ian Johnston, Dominic Kwiatkowski, Cordelia Langford, John Sillitoe on behalf of the Wellcome Sanger Institute COVID-19 Surveillance Team |
| EPI_ISL_3123703 | Lighthouse Lab in Milton Keynes | Wellcome Sanger Institute for the COVID-19 Genomics UK (COG-UK) Consortium | The Lighthouse Lab in Milton Keynes and Alex Alderton, Roberto Amato, Jeffrey Barrett, Sonia Goncalves, Ewan Harrison, David K. Jackson, Ian Johnston, Dominic Kwiatkowski, Cordelia Langford, John Sillitoe on behalf of the Wellcome Sanger Institute COVID-19 Surveillance Team |
| EPI_ISL_3123704, EPI_ISL_3123705, EPI_ISL_3123706, EPI_ISL_3123707 | Lighthouse Lab in Glasgow | Wellcome Sanger Institute for the COVID-19 Genomics UK (COG-UK) Consortium | Harper VanSteenhouse, Yumi Kasai, David Gray, Carol Clugston, Anna Dominiczak and Alex Alderton, Roberto Amato, Jeffrey Barrett, Sonia Goncalves, Ewan Harrison, David K. Jackson, Ian Johnston, Dominic Kwiatkowski, Cordelia Langford, John Sillitoe on behalf of the Wellcome Sanger Institute COVID-19 Surveillance Team |
| EPI_ISL_3123708 | Berkshire and Surrey Pathology Services Lighthouse Laboratory | Wellcome Sanger Institute for the COVID-19 Genomics UK (COG-UK) Consortium | Berkshire and Surrey Pathology Services Lighthouse Laboratory and Alex Alderton, Roberto Amato, Jeffrey Barrett, Sonia Goncalves, Ewan Harrison, David K. Jackson, Ian Johnston, Dominic Kwiatkowski, Cordelia Langford, John Sillitoe on behalf of the Wellcome Sanger Institute COVID-19 Surveillance Team |
| EPI_ISL_3123709 | Lighthouse Lab in Milton Keynes | Wellcome Sanger Institute for the COVID-19 Genomics UK (COG-UK) Consortium | The Lighthouse Lab in Milton Keynes and Alex Alderton, Roberto Amato, Jeffrey Barrett, Sonia Goncalves, Ewan Harrison, David K. Jackson, Ian Johnston, Dominic Kwiatkowski, Cordelia Langford, John Sillitoe on behalf of the Wellcome Sanger Institute COVID-19 Surveillance Team |
| EPI_ISL_3123710 | Lighthouse Lab in Glasgow | Wellcome Sanger Institute for the COVID-19 Genomics UK (COG-UK) Consortium | Harper VanSteenhouse, Yumi Kasai, David Gray, Carol Clugston, Anna Dominiczak and Alex Alderton, Roberto Amato, Jeffrey Barrett, Sonia Goncalves, Ewan Harrison, David K. Jackson, Ian Johnston, Dominic Kwiatkowski, Cordelia Langford, John Sillitoe on behalf of the Wellcome Sanger Institute COVID-19 Surveillance Team |
| EPI_ISL_3123711 | Berkshire and Surrey Pathology Services Lighthouse Laboratory | Wellcome Sanger Institute for the COVID-19 Genomics UK (COG-UK) Consortium | Berkshire and Surrey Pathology Services Lighthouse Laboratory and Alex Alderton, Roberto Amato, Jeffrey Barrett, Sonia Goncalves, Ewan Harrison, David K. Jackson, Ian Johnston, Dominic Kwiatkowski, Cordelia Langford, John Sillitoe on behalf of the Wellcome Sanger Institute COVID-19 Surveillance Team |
| EPI_ISL_3123712 | Lighthouse Lab in Milton Keynes | Wellcome Sanger Institute for the COVID-19 Genomics UK (COG-UK) Consortium | The Lighthouse Lab in Milton Keynes and Alex Alderton, Roberto Amato, Jeffrey Barrett, Sonia Goncalves, Ewan Harrison, David K. Jackson, Ian Johnston, Dominic Kwiatkowski, Cordelia Langford, John Sillitoe on behalf of the Wellcome Sanger Institute COVID-19 Surveillance Team |
| EPI_ISL_3123713 | Lighthouse Lab in Glasgow | Wellcome Sanger Institute for the COVID-19 Genomics UK (COG-UK) Consortium | Harper VanSteenhouse, Yumi Kasai, David Gray, Carol Clugston, Anna Dominiczak and Alex Alderton, Roberto Amato, Jeffrey Barrett, Sonia Goncalves, Ewan Harrison, David K. Jackson, Ian Johnston, Dominic Kwiatkowski, Cordelia Langford, John Sillitoe on behalf of the Wellcome Sanger Institute COVID-19 Surveillance Team |
| EPI_ISL_3123714 | Lighthouse Lab in Milton Keynes | Wellcome Sanger Institute for the COVID-19 Genomics UK (COG-UK) Consortium | The Lighthouse Lab in Milton Keynes and Alex Alderton, Roberto Amato, Jeffrey Barrett, Sonia Goncalves, Ewan Harrison, David K. Jackson, Ian Johnston, Dominic Kwiatkowski, Cordelia Langford, John Sillitoe on behalf of the Wellcome Sanger Institute COVID-19 Surveillance Team |
| EPI_ISL_3123715 | Lighthouse Lab in Glasgow | Wellcome Sanger Institute for the COVID-19 Genomics UK (COG-UK) Consortium | Harper VanSteenhouse, Yumi Kasai, David Gray, Carol Clugston, Anna Dominiczak and Alex Alderton, Roberto Amato, Jeffrey Barrett, Sonia Goncalves, Ewan Harrison, David K. Jackson, Ian Johnston, Dominic Kwiatkowski, Cordelia Langford, John Sillitoe on behalf of the Wellcome Sanger Institute COVID-19 Surveillance Team |
| EPI_ISL_3123716, EPI_ISL_3123717, EPI_ISL_3123718, EPI_ISL_3123719 | Lighthouse Lab in Milton Keynes | Wellcome Sanger Institute for the COVID-19 Genomics UK (COG-UK) Consortium | The Lighthouse Lab in Milton Keynes and Alex Alderton, Roberto Amato, Jeffrey Barrett, Sonia Goncalves, Ewan Harrison, David K. Jackson, Ian Johnston, Dominic Kwiatkowski, Cordelia Langford, John Sillitoe on behalf of the Wellcome Sanger Institute COVID-19 Surveillance Team |
| EPI_ISL_3123720, EPI_ISL_3123721, EPI_ISL_3123722, EPI_ISL_3123723 | Lighthouse Lab in Glasgow | Wellcome Sanger Institute for the COVID-19 Genomics UK (COG-UK) Consortium | Harper VanSteenhouse, Yumi Kasai, David Gray, Carol Clugston, Anna Dominiczak and Alex Alderton, Roberto Amato, Jeffrey Barrett, Sonia Goncalves, Ewan Harrison, David K. Jackson, Ian Johnston, Dominic Kwiatkowski, Cordelia Langford, John Sillitoe on behalf of the Wellcome Sanger Institute COVID-19 Surveillance Team |
| EPI_ISL_3123724 | Berkshire and Surrey Pathology Services Lighthouse Laboratory | Wellcome Sanger Institute for the COVID-19 Genomics UK (COG-UK) Consortium | Berkshire and Surrey Pathology Services Lighthouse Laboratory and Alex Alderton, Roberto Amato, Jeffrey Barrett, Sonia Goncalves, Ewan Harrison, David K. Jackson, Ian Johnston, Dominic Kwiatkowski, Cordelia Langford, John Sillitoe on behalf of the Wellcome Sanger Institute COVID-19 Surveillance Team |
| EPI_ISL_3123725 | Lighthouse Lab in Glasgow | Wellcome Sanger Institute for the COVID-19 Genomics UK (COG-UK) Consortium | Harper VanSteenhouse, Yumi Kasai, David Gray, Carol Clugston, Anna Dominiczak and Alex Alderton, Roberto Amato, Jeffrey Barrett, Sonia Goncalves, Ewan Harrison, David K. Jackson, Ian Johnston, Dominic Kwiatkowski, Cordelia Langford, John Sillitoe on behalf of the Wellcome Sanger Institute COVID-19 Surveillance Team |
| EPI_ISL_3123726 | Lighthouse Lab in Milton Keynes | Wellcome Sanger Institute for the COVID-19 Genomics UK (COG-UK) Consortium | The Lighthouse Lab in Milton Keynes and Alex Alderton, Roberto Amato, Jeffrey Barrett, Sonia Goncalves, Ewan Harrison, David K. Jackson, Ian Johnston, Dominic Kwiatkowski, Cordelia Langford, John Sillitoe on behalf of the Wellcome Sanger Institute COVID-19 Surveillance Team |
| EPI_ISL_3123727 | Lighthouse Lab in Glasgow | Wellcome Sanger Institute for the COVID-19 Genomics UK (COG-UK) Consortium | Harper VanSteenhouse, Yumi Kasai, David Gray, Carol Clugston, Anna Dominiczak and Alex Alderton, Roberto Amato, Jeffrey Barrett, Sonia Goncalves, Ewan Harrison, David K. Jackson, Ian Johnston, Dominic Kwiatkowski, Cordelia Langford, John Sillitoe on behalf of the Wellcome Sanger Institute COVID-19 Surveillance Team |
| EPI_ISL_3123728 | Lighthouse Lab in Milton Keynes | Wellcome Sanger Institute for the COVID-19 Genomics UK (COG-UK) Consortium | The Lighthouse Lab in Milton Keynes and Alex Alderton, Roberto Amato, Jeffrey Barrett, Sonia Goncalves, Ewan Harrison, David K. Jackson, Ian Johnston, Dominic Kwiatkowski, Cordelia Langford, John Sillitoe on behalf of the Wellcome Sanger Institute COVID-19 Surveillance Team |
| EPI_ISL_3123729 | Berkshire and Surrey Pathology Services Lighthouse Laboratory | Wellcome Sanger Institute for the COVID-19 Genomics UK (COG-UK) Consortium | Berkshire and Surrey Pathology Services Lighthouse Laboratory and Alex Alderton, Roberto Amato, Jeffrey Barrett, Sonia Goncalves, Ewan Harrison, David K. Jackson, Ian Johnston, Dominic Kwiatkowski, Cordelia Langford, John Sillitoe on behalf of the Wellcome Sanger Institute COVID-19 Surveillance Team |
| EPI_ISL_3123730 | Lighthouse Lab in Milton Keynes | Wellcome Sanger Institute for the COVID-19 Genomics UK (COG-UK) Consortium | The Lighthouse Lab in Milton Keynes and Alex Alderton, Roberto Amato, Jeffrey Barrett, Sonia Goncalves, Ewan Harrison, David K. Jackson, Ian Johnston, Dominic Kwiatkowski, Cordelia Langford, John Sillitoe on behalf of the Wellcome Sanger Institute COVID-19 Surveillance Team |
| EPI_ISL_3123731, EPI_ISL_3123732, EPI_ISL_3123733 | Lighthouse Lab in Glasgow | Wellcome Sanger Institute for the COVID-19 Genomics UK (COG-UK) Consortium | Harper VanSteenhouse, Yumi Kasai, David Gray, Carol Clugston, Anna Dominiczak and Alex Alderton, Roberto Amato, Jeffrey Barrett, Sonia Goncalves, Ewan Harrison, David K. Jackson, Ian Johnston, Dominic Kwiatkowski, Cordelia Langford, John Sillitoe on behalf of the Wellcome Sanger Institute COVID-19 Surveillance Team |
| EPI_ISL_3123734 | Lighthouse Lab in Milton Keynes | Wellcome Sanger Institute for the COVID-19 Genomics UK (COG-UK) Consortium | The Lighthouse Lab in Milton Keynes and Alex Alderton, Roberto Amato, Jeffrey Barrett, Sonia Goncalves, Ewan Harrison, David K. Jackson, Ian Johnston, Dominic Kwiatkowski, Cordelia Langford, John Sillitoe on behalf of the Wellcome Sanger Institute COVID-19 Surveillance Team |
| EPI_ISL_3123735 | Lighthouse Lab in Glasgow | Wellcome Sanger Institute for the COVID-19 Genomics UK (COG-UK) Consortium | Harper VanSteenhouse, Yumi Kasai, David Gray, Carol Clugston, Anna Dominiczak and Alex Alderton, Roberto Amato, Jeffrey Barrett, Sonia Goncalves, Ewan Harrison, David K. Jackson, Ian Johnston, Dominic Kwiatkowski, Cordelia Langford, John Sillitoe on behalf of the Wellcome Sanger Institute COVID-19 Surveillance Team |
| EPI_ISL_3123736, EPI_ISL_3123737 | Berkshire and Surrey Pathology Services Lighthouse Laboratory | Wellcome Sanger Institute for the COVID-19 Genomics UK (COG-UK) Consortium | Berkshire and Surrey Pathology Services Lighthouse Laboratory and Alex Alderton, Roberto Amato, Jeffrey Barrett, Sonia Goncalves, Ewan Harrison, David K. Jackson, Ian Johnston, Dominic Kwiatkowski, Cordelia Langford, John Sillitoe on behalf of the Wellcome Sanger Institute COVID-19 Surveillance Team |
| EPI_ISL_3123738 | Lighthouse Lab in Milton Keynes | Wellcome Sanger Institute for the COVID-19 Genomics UK (COG-UK) Consortium | The Lighthouse Lab in Milton Keynes and Alex Alderton, Roberto Amato, Jeffrey Barrett, Sonia Goncalves, Ewan Harrison, David K. Jackson, Ian Johnston, Dominic Kwiatkowski, Cordelia Langford, John Sillitoe on behalf of the Wellcome Sanger Institute COVID-19 Surveillance Team |
| EPI_ISL_3123739 | Lighthouse Lab in Glasgow | Wellcome Sanger Institute for the COVID-19 Genomics UK (COG-UK) Consortium | Harper VanSteenhouse, Yumi Kasai, David Gray, Carol Clugston, Anna Dominiczak and Alex Alderton, Roberto Amato, Jeffrey Barrett, Sonia Goncalves, Ewan Harrison, David K. Jackson, Ian Johnston, Dominic Kwiatkowski, Cordelia Langford, John Sillitoe on behalf of the Wellcome Sanger Institute COVID-19 Surveillance Team |

| | | | |
|---|---|---|---|
| EPI_ISL_3123740 | Berkshire and Surrey Pathology Services Lighthouse Laboratory | Wellcome Sanger Institute for the COVID-19 Genomics UK (COG-UK) Consortium | Berkshire and Surrey Pathology Services Lighthouse Laboratory and Alex Alderton, Roberto Amato, Jeffrey Barrett, Sonia Goncalves, Ewan Harrison, David K. Jackson, Ian Johnston, Dominic Kwiatkowski, Cordelia Langford, John Sillitoe on behalf of the Wellcome Sanger Institute COVID-19 Surveillance Team |
| EPI_ISL_3123741, EPI_ISL_3123742 | Lighthouse Lab in Glasgow | Wellcome Sanger Institute for the COVID-19 Genomics UK (COG-UK) Consortium | Harper VanSteenhouse, Yumi Kasai, David Gray, Carol Clugston, Anna Dominiczak and Alex Alderton, Roberto Amato, Jeffrey Barrett, Sonia Goncalves, Ewan Harrison, David K. Jackson, Ian Johnston, Dominic Kwiatkowski, Cordelia Langford, John Sillitoe on behalf of the Wellcome Sanger Institute COVID-19 Surveillance Team |
| EPI_ISL_3123743 | Lighthouse Lab in Milton Keynes | Wellcome Sanger Institute for the COVID-19 Genomics UK (COG-UK) Consortium | The Lighthouse Lab in Milton Keynes and Alex Alderton, Roberto Amato, Jeffrey Barrett, Sonia Goncalves, Ewan Harrison, David K. Jackson, Ian Johnston, Dominic Kwiatkowski, Cordelia Langford, John Sillitoe on behalf of the Wellcome Sanger Institute COVID-19 Surveillance Team |
| EPI_ISL_3123744, EPI_ISL_3123745, EPI_ISL_3123746, EPI_ISL_3123747 | Lighthouse Lab in Glasgow | Wellcome Sanger Institute for the COVID-19 Genomics UK (COG-UK) Consortium | Harper VanSteenhouse, Yumi Kasai, David Gray, Carol Clugston, Anna Dominiczak and Alex Alderton, Roberto Amato, Jeffrey Barrett, Sonia Goncalves, Ewan Harrison, David K. Jackson, Ian Johnston, Dominic Kwiatkowski, Cordelia Langford, John Sillitoe on behalf of the Wellcome Sanger Institute COVID-19 Surveillance Team |
| EPI_ISL_3123748 | Berkshire and Surrey Pathology Services Lighthouse Laboratory | Wellcome Sanger Institute for the COVID-19 Genomics UK (COG-UK) Consortium | Berkshire and Surrey Pathology Services Lighthouse Laboratory and Alex Alderton, Roberto Amato, Jeffrey Barrett, Sonia Goncalves, Ewan Harrison, David K. Jackson, Ian Johnston, Dominic Kwiatkowski, Cordelia Langford, John Sillitoe on behalf of the Wellcome Sanger Institute COVID-19 Surveillance Team |
| EPI_ISL_3123749, EPI_ISL_3123750 | Lighthouse Lab in Glasgow | Wellcome Sanger Institute for the COVID-19 Genomics UK (COG-UK) Consortium | Harper VanSteenhouse, Yumi Kasai, David Gray, Carol Clugston, Anna Dominiczak and Alex Alderton, Roberto Amato, Jeffrey Barrett, Sonia Goncalves, Ewan Harrison, David K. Jackson, Ian Johnston, Dominic Kwiatkowski, Cordelia Langford, John Sillitoe on behalf of the Wellcome Sanger Institute COVID-19 Surveillance Team |
| EPI_ISL_3123751 | Lighthouse Lab in Milton Keynes | Wellcome Sanger Institute for the COVID-19 Genomics UK (COG-UK) Consortium | The Lighthouse Lab in Milton Keynes and Alex Alderton, Roberto Amato, Jeffrey Barrett, Sonia Goncalves, Ewan Harrison, David K. Jackson, Ian Johnston, Dominic Kwiatkowski, Cordelia Langford, John Sillitoe on behalf of the Wellcome Sanger Institute COVID-19 Surveillance Team |
| EPI_ISL_3123752, EPI_ISL_3123753 | Lighthouse Lab in Glasgow | Wellcome Sanger Institute for the COVID-19 Genomics UK (COG-UK) Consortium | Harper VanSteenhouse, Yumi Kasai, David Gray, Carol Clugston, Anna Dominiczak and Alex Alderton, Roberto Amato, Jeffrey Barrett, Sonia Goncalves, Ewan Harrison, David K. Jackson, Ian Johnston, Dominic Kwiatkowski, Cordelia Langford, John Sillitoe on behalf of the Wellcome Sanger Institute COVID-19 Surveillance Team |
| EPI_ISL_3123754 | Berkshire and Surrey Pathology Services Lighthouse Laboratory | Wellcome Sanger Institute for the COVID-19 Genomics UK (COG-UK) Consortium | Berkshire and Surrey Pathology Services Lighthouse Laboratory and Alex Alderton, Roberto Amato, Jeffrey Barrett, Sonia Goncalves, Ewan Harrison, David K. Jackson, Ian Johnston, Dominic Kwiatkowski, Cordelia Langford, John Sillitoe on behalf of the Wellcome Sanger Institute COVID-19 Surveillance Team |
| EPI_ISL_3123755 | Lighthouse Lab in Milton Keynes | Wellcome Sanger Institute for the COVID-19 Genomics UK (COG-UK) Consortium | The Lighthouse Lab in Milton Keynes and Alex Alderton, Roberto Amato, Jeffrey Barrett, Sonia Goncalves, Ewan Harrison, David K. Jackson, Ian Johnston, Dominic Kwiatkowski, Cordelia Langford, John Sillitoe on behalf of the Wellcome Sanger Institute COVID-19 Surveillance Team |
| EPI_ISL_3123756 | Lighthouse Lab in Glasgow | Wellcome Sanger Institute for the COVID-19 Genomics UK (COG-UK) Consortium | Harper VanSteenhouse, Yumi Kasai, David Gray, Carol Clugston, Anna Dominiczak and Alex Alderton, Roberto Amato, Jeffrey Barrett, Sonia Goncalves, Ewan Harrison, David K. Jackson, Ian Johnston, Dominic Kwiatkowski, Cordelia Langford, John Sillitoe on behalf of the Wellcome Sanger Institute COVID-19 Surveillance Team |
| EPI_ISL_3123757 | Lighthouse Lab in Milton Keynes | Wellcome Sanger Institute for the COVID-19 Genomics UK (COG-UK) Consortium | The Lighthouse Lab in Milton Keynes and Alex Alderton, Roberto Amato, Jeffrey Barrett, Sonia Goncalves, Ewan Harrison, David K. Jackson, Ian Johnston, Dominic Kwiatkowski, Cordelia Langford, John Sillitoe on behalf of the Wellcome Sanger Institute COVID-19 Surveillance Team |
| EPI_ISL_3123758, EPI_ISL_3123759 | Lighthouse Lab in Glasgow | Wellcome Sanger Institute for the COVID-19 Genomics UK (COG-UK) Consortium | Harper VanSteenhouse, Yumi Kasai, David Gray, Carol Clugston, Anna Dominiczak and Alex Alderton, Roberto Amato, Jeffrey Barrett, Sonia Goncalves, Ewan Harrison, David K. Jackson, Ian Johnston, Dominic Kwiatkowski, Cordelia Langford, John Sillitoe on behalf of the Wellcome Sanger Institute COVID-19 Surveillance Team |
| EPI_ISL_3123760 | Berkshire and Surrey Pathology Services Lighthouse Laboratory | Wellcome Sanger Institute for the COVID-19 Genomics UK (COG-UK) Consortium | Berkshire and Surrey Pathology Services Lighthouse Laboratory and Alex Alderton, Roberto Amato, Jeffrey Barrett, Sonia Goncalves, Ewan Harrison, David K. Jackson, Ian Johnston, Dominic Kwiatkowski, Cordelia Langford, John Sillitoe on behalf of the Wellcome Sanger Institute COVID-19 Surveillance Team |
| EPI_ISL_3123761 | Lighthouse Lab in Milton Keynes | Wellcome Sanger Institute for the COVID-19 Genomics UK (COG-UK) Consortium | The Lighthouse Lab in Milton Keynes and Alex Alderton, Roberto Amato, Jeffrey Barrett, Sonia Goncalves, Ewan Harrison, David K. Jackson, Ian Johnston, Dominic Kwiatkowski, Cordelia Langford, John Sillitoe on behalf of the Wellcome Sanger Institute COVID-19 Surveillance Team |
| EPI_ISL_3123762 | Berkshire and Surrey Pathology Services Lighthouse Laboratory | Wellcome Sanger Institute for the COVID-19 Genomics UK (COG-UK) Consortium | Berkshire and Surrey Pathology Services Lighthouse Laboratory and Alex Alderton, Roberto Amato, Jeffrey Barrett, Sonia Goncalves, Ewan Harrison, David K. Jackson, Ian Johnston, Dominic Kwiatkowski, Cordelia Langford, John Sillitoe on behalf of the Wellcome Sanger Institute COVID-19 Surveillance Team |
| EPI_ISL_3123763 | Lighthouse Lab in Milton Keynes | Wellcome Sanger Institute for the COVID-19 Genomics UK (COG-UK) Consortium | The Lighthouse Lab in Milton Keynes and Alex Alderton, Roberto Amato, Jeffrey Barrett, Sonia Goncalves, Ewan Harrison, David K. Jackson, Ian Johnston, Dominic Kwiatkowski, Cordelia Langford, John Sillitoe on behalf of the Wellcome Sanger Institute COVID-19 Surveillance Team |
| EPI_ISL_3123764, EPI_ISL_3123765 | Berkshire and Surrey Pathology Services Lighthouse Laboratory | Wellcome Sanger Institute for the COVID-19 Genomics UK (COG-UK) Consortium | Berkshire and Surrey Pathology Services Lighthouse Laboratory and Alex Alderton, Roberto Amato, Jeffrey Barrett, Sonia Goncalves, Ewan Harrison, David K. Jackson, Ian Johnston, Dominic Kwiatkowski, Cordelia Langford, John Sillitoe on behalf of the Wellcome Sanger Institute COVID-19 Surveillance Team |
| EPI_ISL_3123766 | Lighthouse Lab in Glasgow | Wellcome Sanger Institute for the COVID-19 Genomics UK (COG-UK) Consortium | Harper VanSteenhouse, Yumi Kasai, David Gray, Carol Clugston, Anna Dominiczak and Alex Alderton, Roberto Amato, Jeffrey Barrett, Sonia Goncalves, Ewan Harrison, David K. Jackson, Ian Johnston, Dominic Kwiatkowski, Cordelia Langford, John Sillitoe on behalf of the Wellcome Sanger Institute COVID-19 Surveillance Team |
| EPI_ISL_3123767 | Lighthouse Lab in Milton Keynes | Wellcome Sanger Institute for the COVID-19 Genomics UK (COG-UK) Consortium | The Lighthouse Lab in Milton Keynes and Alex Alderton, Roberto Amato, Jeffrey Barrett, Sonia Goncalves, Ewan Harrison, David K. Jackson, Ian Johnston, Dominic Kwiatkowski, Cordelia Langford, John Sillitoe on behalf of the Wellcome Sanger Institute COVID-19 Surveillance Team |
| EPI_ISL_3123768 | Lighthouse Lab in Glasgow | Wellcome Sanger Institute for the COVID-19 Genomics UK (COG-UK) Consortium | Harper VanSteenhouse, Yumi Kasai, David Gray, Carol Clugston, Anna Dominiczak and Alex Alderton, Roberto Amato, Jeffrey Barrett, Sonia Goncalves, Ewan Harrison, David K. Jackson, Ian Johnston, Dominic Kwiatkowski, Cordelia Langford, John Sillitoe on behalf of the Wellcome Sanger Institute COVID-19 Surveillance Team |
| EPI_ISL_3123769 | Berkshire and Surrey Pathology Services Lighthouse Laboratory | Wellcome Sanger Institute for the COVID-19 Genomics UK (COG-UK) Consortium | Berkshire and Surrey Pathology Services Lighthouse Laboratory and Alex Alderton, Roberto Amato, Jeffrey Barrett, Sonia Goncalves, Ewan Harrison, David K. Jackson, Ian Johnston, Dominic Kwiatkowski, Cordelia Langford, John Sillitoe on behalf of the Wellcome Sanger Institute COVID-19 Surveillance Team |
| EPI_ISL_3123770 | Lighthouse Lab in Glasgow | Wellcome Sanger Institute for the COVID-19 Genomics UK (COG-UK) Consortium | Harper VanSteenhouse, Yumi Kasai, David Gray, Carol Clugston, Anna Dominiczak and Alex Alderton, Roberto Amato, Jeffrey Barrett, Sonia Goncalves, Ewan Harrison, David K. Jackson, Ian Johnston, Dominic Kwiatkowski, Cordelia Langford, John Sillitoe on behalf of the Wellcome Sanger Institute COVID-19 Surveillance Team |
| EPI_ISL_3123771 | Berkshire and Surrey Pathology Services Lighthouse Laboratory | Wellcome Sanger Institute for the COVID-19 Genomics UK (COG-UK) Consortium | Berkshire and Surrey Pathology Services Lighthouse Laboratory and Alex Alderton, Roberto Amato, Jeffrey Barrett, Sonia Goncalves, Ewan Harrison, David K. Jackson, Ian Johnston, Dominic Kwiatkowski, Cordelia Langford, John Sillitoe on behalf of the Wellcome Sanger Institute COVID-19 Surveillance Team |
| EPI_ISL_3123772 | Lighthouse Lab in Milton Keynes | Wellcome Sanger Institute for the COVID-19 Genomics UK (COG-UK) Consortium | The Lighthouse Lab in Milton Keynes and Alex Alderton, Roberto Amato, Jeffrey Barrett, Sonia Goncalves, Ewan Harrison, David K. Jackson, Ian Johnston, Dominic Kwiatkowski, Cordelia Langford, John Sillitoe on behalf of the Wellcome Sanger Institute COVID-19 Surveillance Team |
| EPI_ISL_3123773, EPI_ISL_3123774 | Berkshire and Surrey Pathology Services Lighthouse Laboratory | Wellcome Sanger Institute for the COVID-19 Genomics UK (COG-UK) Consortium | Berkshire and Surrey Pathology Services Lighthouse Laboratory and Alex Alderton, Roberto Amato, Jeffrey Barrett, Sonia Goncalves, Ewan Harrison, David K. Jackson, Ian Johnston, Dominic Kwiatkowski, Cordelia Langford, John Sillitoe on behalf of the Wellcome Sanger Institute COVID-19 Surveillance Team |
| EPI_ISL_3123775, EPI_ISL_3123776 | Lighthouse Lab in Glasgow | Wellcome Sanger Institute for the COVID-19 Genomics UK (COG-UK) Consortium | Harper VanSteenhouse, Yumi Kasai, David Gray, Carol Clugston, Anna Dominiczak and Alex Alderton, Roberto Amato, Jeffrey Barrett, Sonia Goncalves, Ewan Harrison, David K. Jackson, Ian Johnston, Dominic Kwiatkowski, Cordelia Langford, John Sillitoe on behalf of the Wellcome Sanger Institute COVID-19 Surveillance Team |
| EPI_ISL_3123777 | Berkshire and Surrey Pathology Services Lighthouse Laboratory | Wellcome Sanger Institute for the COVID-19 Genomics UK (COG-UK) Consortium | Berkshire and Surrey Pathology Services Lighthouse Laboratory and Alex Alderton, Roberto Amato, Jeffrey Barrett, Sonia Goncalves, Ewan Harrison, David K. Jackson, Ian Johnston, Dominic Kwiatkowski, Cordelia Langford, John Sillitoe on behalf of the Wellcome Sanger Institute COVID-19 Surveillance Team |
| EPI_ISL_3123778 | Lighthouse Lab in Glasgow | Wellcome Sanger Institute for the COVID-19 Genomics UK (COG-UK) Consortium | Harper VanSteenhouse, Yumi Kasai, David Gray, Carol Clugston, Anna Dominiczak and Alex Alderton, Roberto Amato, Jeffrey Barrett, Sonia Goncalves, Ewan Harrison, David K. Jackson, Ian Johnston, Dominic Kwiatkowski, Cordelia Langford, John Sillitoe on behalf of the Wellcome Sanger Institute COVID-19 Surveillance Team |
| EPI_ISL_3123779 | Lighthouse Lab in Milton Keynes | Wellcome Sanger Institute for the COVID-19 Genomics UK (COG-UK) Consortium | The Lighthouse Lab in Milton Keynes and Alex Alderton, Roberto Amato, Jeffrey Barrett, Sonia Goncalves, Ewan Harrison, David K. Jackson, Ian Johnston, Dominic Kwiatkowski, Cordelia Langford, John Sillitoe on behalf of the Wellcome Sanger Institute COVID-19 Surveillance Team |
| EPI_ISL_3123780 | Berkshire and Surrey Pathology Services Lighthouse Laboratory | Wellcome Sanger Institute for the COVID-19 Genomics UK (COG-UK) Consortium | Berkshire and Surrey Pathology Services Lighthouse Laboratory and Alex Alderton, Roberto Amato, Jeffrey Barrett, Sonia Goncalves, Ewan Harrison, David K. Jackson, Ian Johnston, Dominic Kwiatkowski, Cordelia Langford, John Sillitoe on behalf of the Wellcome Sanger Institute COVID-19 Surveillance Team |
| EPI_ISL_3123781, EPI_ISL_3123782 | Lighthouse Lab in Glasgow | Wellcome Sanger Institute for the COVID-19 Genomics UK (COG-UK) Consortium | Harper VanSteenhouse, Yumi Kasai, David Gray, Carol Clugston, Anna Dominiczak and Alex Alderton, Roberto Amato, Jeffrey Barrett, Sonia Goncalves, Ewan Harrison, David K. Jackson, Ian Johnston, Dominic Kwiatkowski, Cordelia Langford, John Sillitoe on behalf of the Wellcome Sanger Institute COVID-19 Surveillance Team |
| EPI_ISL_3123783 | Berkshire and Surrey Pathology Services Lighthouse Laboratory | Wellcome Sanger Institute for the COVID-19 Genomics UK (COG-UK) Consortium | Berkshire and Surrey Pathology Services Lighthouse Laboratory and Alex Alderton, Roberto Amato, Jeffrey Barrett, Sonia Goncalves, Ewan Harrison, David K. Jackson, Ian Johnston, Dominic Kwiatkowski, Cordelia Langford, John Sillitoe on behalf of the Wellcome Sanger Institute COVID-19 Surveillance Team |
| EPI_ISL_3123784, EPI_ISL_3123785 | Lighthouse Lab in Milton Keynes | Wellcome Sanger Institute for the COVID-19 Genomics UK (COG-UK) Consortium | The Lighthouse Lab in Milton Keynes and Alex Alderton, Roberto Amato, Jeffrey Barrett, Sonia Goncalves, Ewan Harrison, David K. Jackson, Ian Johnston, Dominic Kwiatkowski, Cordelia Langford, John Sillitoe on behalf of the Wellcome Sanger Institute COVID-19 Surveillance Team |
| EPI_ISL_3123786 | Lighthouse Lab in Glasgow | Wellcome Sanger Institute for the COVID-19 Genomics UK (COG-UK) Consortium | Harper VanSteenhouse, Yumi Kasai, David Gray, Carol Clugston, Anna Dominiczak and Alex Alderton, Roberto Amato, Jeffrey Barrett, Sonia Goncalves, Ewan Harrison, David K. Jackson, Ian Johnston, Dominic Kwiatkowski, Cordelia Langford, John Sillitoe on behalf of the Wellcome Sanger Institute COVID-19 Surveillance Team |
| EPI_ISL_3123787 | Berkshire and Surrey Pathology Services Lighthouse Laboratory | Wellcome Sanger Institute for the COVID-19 Genomics UK (COG-UK) Consortium | Berkshire and Surrey Pathology Services Lighthouse Laboratory and Alex Alderton, Roberto Amato, Jeffrey Barrett, Sonia Goncalves, Ewan Harrison, David K. Jackson, Ian Johnston, Dominic Kwiatkowski, Cordelia Langford, John Sillitoe on behalf of the Wellcome Sanger Institute COVID-19 Surveillance Team |
| EPI_ISL_3123788 | Lighthouse Lab in Glasgow | Wellcome Sanger Institute for the COVID-19 Genomics UK (COG-UK) Consortium | Harper VanSteenhouse, Yumi Kasai, David Gray, Carol Clugston, Anna Dominiczak and Alex Alderton, Roberto Amato, Jeffrey Barrett, Sonia Goncalves, Ewan Harrison, David K. Jackson, Ian Johnston, Dominic Kwiatkowski, Cordelia Langford, John Sillitoe on behalf of the Wellcome Sanger Institute COVID-19 Surveillance Team |
| EPI_ISL_3123789, EPI_ISL_3123790 | Lighthouse Lab in Milton Keynes | Wellcome Sanger Institute for the COVID-19 Genomics UK (COG-UK) Consortium | The Lighthouse Lab in Milton Keynes and Alex Alderton, Roberto Amato, Jeffrey Barrett, Sonia Goncalves, Ewan Harrison, David K. Jackson, Ian Johnston, Dominic Kwiatkowski, Cordelia Langford, John Sillitoe on behalf of the Wellcome Sanger Institute COVID-19 Surveillance Team |
| EPI_ISL_3123791 | Lighthouse Lab in Glasgow | Wellcome Sanger Institute for the COVID-19 Genomics UK (COG-UK) Consortium | Harper VanSteenhouse, Yumi Kasai, David Gray, Carol Clugston, Anna Dominiczak and Alex Alderton, Roberto Amato, Jeffrey Barrett, Sonia Goncalves, Ewan Harrison, David K. Jackson, Ian Johnston, Dominic Kwiatkowski, Cordelia Langford, John Sillitoe on behalf of the Wellcome Sanger Institute COVID-19 Surveillance Team |
| EPI_ISL_3123792 | Lighthouse Lab in Milton Keynes | Wellcome Sanger Institute for the COVID-19 Genomics UK (COG-UK) Consortium | The Lighthouse Lab in Milton Keynes and Alex Alderton, Roberto Amato, Jeffrey Barrett, Sonia Goncalves, Ewan Harrison, David K. Jackson, Ian Johnston, Dominic Kwiatkowski, Cordelia Langford, John Sillitoe on behalf of the Wellcome Sanger Institute COVID-19 Surveillance Team |
| EPI_ISL_3123793 | Berkshire and Surrey Pathology Services Lighthouse Laboratory | Wellcome Sanger Institute for the COVID-19 Genomics UK (COG-UK) Consortium | Berkshire and Surrey Pathology Services Lighthouse Laboratory and Alex Alderton, Roberto Amato, Jeffrey Barrett, Sonia Goncalves, Ewan Harrison, David K. Jackson, Ian Johnston, Dominic Kwiatkowski, Cordelia Langford, John Sillitoe on behalf of the Wellcome Sanger Institute COVID-19 Surveillance Team |
| EPI_ISL_3123794, EPI_ISL_3123795 | Lighthouse Lab in Glasgow | Wellcome Sanger Institute for the COVID-19 Genomics UK (COG-UK) Consortium | Harper VanSteenhouse, Yumi Kasai, David Gray, Carol Clugston, Anna Dominiczak and Alex Alderton, Roberto Amato, Jeffrey Barrett, Sonia Goncalves, Ewan Harrison, David K. Jackson, Ian Johnston, Dominic Kwiatkowski, Cordelia Langford, John Sillitoe on behalf of the Wellcome Sanger Institute COVID-19 Surveillance Team |
| EPI_ISL_3123796 | Berkshire and Surrey Pathology Services Lighthouse Laboratory | Wellcome Sanger Institute for the COVID-19 Genomics UK (COG-UK) Consortium | Berkshire and Surrey Pathology Services Lighthouse Laboratory and Alex Alderton, Roberto Amato, Jeffrey Barrett, Sonia Goncalves, Ewan Harrison, David K. Jackson, Ian Johnston, Dominic Kwiatkowski, Cordelia Langford, John Sillitoe on behalf of the Wellcome Sanger Institute COVID-19 Surveillance Team |
| EPI_ISL_3123797 | Lighthouse Lab in Milton Keynes | Wellcome Sanger Institute for the COVID-19 Genomics UK (COG-UK) Consortium | The Lighthouse Lab in Milton Keynes and Alex Alderton, Roberto Amato, Jeffrey Barrett, Sonia Goncalves, Ewan Harrison, David K. Jackson, Ian Johnston, Dominic Kwiatkowski, Cordelia Langford, John Sillitoe on behalf of the Wellcome Sanger Institute COVID-19 Surveillance Team |
| EPI_ISL_3123798 | Lighthouse Lab in Glasgow | Wellcome Sanger Institute for the COVID-19 Genomics UK (COG-UK) | Harper VanSteenhouse, Yumi Kasai, David Gray, Carol Clugston, Anna Dominiczak and Alex Alderton, Roberto Amato, Jeffrey Barrett, Sonia Goncalves, Ewan Harrison, David K. Jackson, Ian Johnston, Dominic Kwiatkowski, Cordelia Langford, |

| | | | |
|---|---|---|---|
| | | Consortium | John Sillitoe on behalf of the Wellcome Sanger Institute COVID-19 Surveillance Team |
| EPI_ISL_3123799 | Lighthouse Lab in Milton Keynes | Wellcome Sanger Institute for the COVID-19 Genomics UK (COG-UK) Consortium | The Lighthouse Lab in Milton Keynes and Alex Alderton, Roberto Amato, Jeffrey Barrett, Sonia Goncalves, Ewan Harrison, David K. Jackson, Ian Johnston, Dominic Kwiatkowski, Cordelia Langford, John Sillitoe on behalf of the Wellcome Sanger Institute COVID-19 Surveillance Team |
| EPI_ISL_3123800 | Lighthouse Lab in Glasgow | Wellcome Sanger Institute for the COVID-19 Genomics UK (COG-UK) Consortium | Harper VanSteenhouse, Yumi Kasai, David Gray, Carol Clugston, Anna Dominiczak and Alex Alderton, Roberto Amato, Jeffrey Barrett, Sonia Goncalves, Ewan Harrison, David K. Jackson, Ian Johnston, Dominic Kwiatkowski, Cordelia Langford, John Sillitoe on behalf of the Wellcome Sanger Institute COVID-19 Surveillance Team |
| EPI_ISL_3123801 | Berkshire and Surrey Pathology Services Lighthouse Laboratory | Wellcome Sanger Institute for the COVID-19 Genomics UK (COG-UK) Consortium | Berkshire and Surrey Pathology Services Lighthouse Laboratory and Alex Alderton, Roberto Amato, Jeffrey Barrett, Sonia Goncalves, Ewan Harrison, David K. Jackson, Ian Johnston, Dominic Kwiatkowski, Cordelia Langford, John Sillitoe on behalf of the Wellcome Sanger Institute COVID-19 Surveillance Team |
| EPI_ISL_3123802 | Lighthouse Lab in Milton Keynes | Wellcome Sanger Institute for the COVID-19 Genomics UK (COG-UK) Consortium | The Lighthouse Lab in Milton Keynes and Alex Alderton, Roberto Amato, Jeffrey Barrett, Sonia Goncalves, Ewan Harrison, David K. Jackson, Ian Johnston, Dominic Kwiatkowski, Cordelia Langford, John Sillitoe on behalf of the Wellcome Sanger Institute COVID-19 Surveillance Team |
| EPI_ISL_3123803 | Lighthouse Lab in Glasgow | Wellcome Sanger Institute for the COVID-19 Genomics UK (COG-UK) Consortium | Harper VanSteenhouse, Yumi Kasai, David Gray, Carol Clugston, Anna Dominiczak and Alex Alderton, Roberto Amato, Jeffrey Barrett, Sonia Goncalves, Ewan Harrison, David K. Jackson, Ian Johnston, Dominic Kwiatkowski, Cordelia Langford, John Sillitoe on behalf of the Wellcome Sanger Institute COVID-19 Surveillance Team |
| EPI_ISL_3123804 | Lighthouse Lab in Milton Keynes | Wellcome Sanger Institute for the COVID-19 Genomics UK (COG-UK) Consortium | The Lighthouse Lab in Milton Keynes and Alex Alderton, Roberto Amato, Jeffrey Barrett, Sonia Goncalves, Ewan Harrison, David K. Jackson, Ian Johnston, Dominic Kwiatkowski, Cordelia Langford, John Sillitoe on behalf of the Wellcome Sanger Institute COVID-19 Surveillance Team |
| EPI_ISL_3123805 | Lighthouse Lab in Glasgow | Wellcome Sanger Institute for the COVID-19 Genomics UK (COG-UK) Consortium | Harper VanSteenhouse, Yumi Kasai, David Gray, Carol Clugston, Anna Dominiczak and Alex Alderton, Roberto Amato, Jeffrey Barrett, Sonia Goncalves, Ewan Harrison, David K. Jackson, Ian Johnston, Dominic Kwiatkowski, Cordelia Langford, John Sillitoe on behalf of the Wellcome Sanger Institute COVID-19 Surveillance Team |
| EPI_ISL_3123806, EPI_ISL_3123807, EPI_ISL_3123808 | Berkshire and Surrey Pathology Services Lighthouse Laboratory | Wellcome Sanger Institute for the COVID-19 Genomics UK (COG-UK) Consortium | Berkshire and Surrey Pathology Services Lighthouse Laboratory and Alex Alderton, Roberto Amato, Jeffrey Barrett, Sonia Goncalves, Ewan Harrison, David K. Jackson, Ian Johnston, Dominic Kwiatkowski, Cordelia Langford, John Sillitoe on behalf of the Wellcome Sanger Institute COVID-19 Surveillance Team |
| EPI_ISL_3123809, EPI_ISL_3123810, EPI_ISL_3123811, EPI_ISL_3123812, EPI_ISL_3123813 | Lighthouse Lab in Glasgow | Wellcome Sanger Institute for the COVID-19 Genomics UK (COG-UK) Consortium | Harper VanSteenhouse, Yumi Kasai, David Gray, Carol Clugston, Anna Dominiczak and Alex Alderton, Roberto Amato, Jeffrey Barrett, Sonia Goncalves, Ewan Harrison, David K. Jackson, Ian Johnston, Dominic Kwiatkowski, Cordelia Langford, John Sillitoe on behalf of the Wellcome Sanger Institute COVID-19 Surveillance Team |
| EPI_ISL_3123814, EPI_ISL_3123815 | Berkshire and Surrey Pathology Services Lighthouse Laboratory | Wellcome Sanger Institute for the COVID-19 Genomics UK (COG-UK) Consortium | Berkshire and Surrey Pathology Services Lighthouse Laboratory and Alex Alderton, Roberto Amato, Jeffrey Barrett, Sonia Goncalves, Ewan Harrison, David K. Jackson, Ian Johnston, Dominic Kwiatkowski, Cordelia Langford, John Sillitoe on behalf of the Wellcome Sanger Institute COVID-19 Surveillance Team |
| EPI_ISL_3123816 | Lighthouse Lab in Glasgow | Wellcome Sanger Institute for the COVID-19 Genomics UK (COG-UK) Consortium | Harper VanSteenhouse, Yumi Kasai, David Gray, Carol Clugston, Anna Dominiczak and Alex Alderton, Roberto Amato, Jeffrey Barrett, Sonia Goncalves, Ewan Harrison, David K. Jackson, Ian Johnston, Dominic Kwiatkowski, Cordelia Langford, John Sillitoe on behalf of the Wellcome Sanger Institute COVID-19 Surveillance Team |
| EPI_ISL_3123817 | Lighthouse Lab in Milton Keynes | Wellcome Sanger Institute for the COVID-19 Genomics UK (COG-UK) Consortium | The Lighthouse Lab in Milton Keynes and Alex Alderton, Roberto Amato, Jeffrey Barrett, Sonia Goncalves, Ewan Harrison, David K. Jackson, Ian Johnston, Dominic Kwiatkowski, Cordelia Langford, John Sillitoe on behalf of the Wellcome Sanger Institute COVID-19 Surveillance Team |
| EPI_ISL_3123818, EPI_ISL_3123819 | Lighthouse Lab in Glasgow | Wellcome Sanger Institute for the COVID-19 Genomics UK (COG-UK) Consortium | Harper VanSteenhouse, Yumi Kasai, David Gray, Carol Clugston, Anna Dominiczak and Alex Alderton, Roberto Amato, Jeffrey Barrett, Sonia Goncalves, Ewan Harrison, David K. Jackson, Ian Johnston, Dominic Kwiatkowski, Cordelia Langford, John Sillitoe on behalf of the Wellcome Sanger Institute COVID-19 Surveillance Team |
| EPI_ISL_3123820, EPI_ISL_3123821, EPI_ISL_3123822 | Berkshire and Surrey Pathology Services Lighthouse Laboratory | Wellcome Sanger Institute for the COVID-19 Genomics UK (COG-UK) Consortium | Berkshire and Surrey Pathology Services Lighthouse Laboratory and Alex Alderton, Roberto Amato, Jeffrey Barrett, Sonia Goncalves, Ewan Harrison, David K. Jackson, Ian Johnston, Dominic Kwiatkowski, Cordelia Langford, John Sillitoe on behalf of the Wellcome Sanger Institute COVID-19 Surveillance Team |
| EPI_ISL_3123823, EPI_ISL_3123824 | Lighthouse Lab in Glasgow | Wellcome Sanger Institute for the COVID-19 Genomics UK (COG-UK) Consortium | Harper VanSteenhouse, Yumi Kasai, David Gray, Carol Clugston, Anna Dominiczak and Alex Alderton, Roberto Amato, Jeffrey Barrett, Sonia Goncalves, Ewan Harrison, David K. Jackson, Ian Johnston, Dominic Kwiatkowski, Cordelia Langford, John Sillitoe on behalf of the Wellcome Sanger Institute COVID-19 Surveillance Team |
| EPI_ISL_3123825 | Lighthouse Lab in Milton Keynes | Wellcome Sanger Institute for the COVID-19 Genomics UK (COG-UK) Consortium | The Lighthouse Lab in Milton Keynes and Alex Alderton, Roberto Amato, Jeffrey Barrett, Sonia Goncalves, Ewan Harrison, David K. Jackson, Ian Johnston, Dominic Kwiatkowski, Cordelia Langford, John Sillitoe on behalf of the Wellcome Sanger Institute COVID-19 Surveillance Team |
| EPI_ISL_3123826 | Berkshire and Surrey Pathology Services Lighthouse Laboratory | Wellcome Sanger Institute for the COVID-19 Genomics UK (COG-UK) Consortium | Berkshire and Surrey Pathology Services Lighthouse Laboratory and Alex Alderton, Roberto Amato, Jeffrey Barrett, Sonia Goncalves, Ewan Harrison, David K. Jackson, Ian Johnston, Dominic Kwiatkowski, Cordelia Langford, John Sillitoe on behalf of the Wellcome Sanger Institute COVID-19 Surveillance Team |
| EPI_ISL_3123827, EPI_ISL_3123828 | Lighthouse Lab in Milton Keynes | Wellcome Sanger Institute for the COVID-19 Genomics UK (COG-UK) Consortium | The Lighthouse Lab in Milton Keynes and Alex Alderton, Roberto Amato, Jeffrey Barrett, Sonia Goncalves, Ewan Harrison, David K. Jackson, Ian Johnston, Dominic Kwiatkowski, Cordelia Langford, John Sillitoe on behalf of the Wellcome Sanger Institute COVID-19 Surveillance Team |
| EPI_ISL_3123829, EPI_ISL_3123830 | Berkshire and Surrey Pathology Services Lighthouse Laboratory | Wellcome Sanger Institute for the COVID-19 Genomics UK (COG-UK) Consortium | Berkshire and Surrey Pathology Services Lighthouse Laboratory and Alex Alderton, Roberto Amato, Jeffrey Barrett, Sonia Goncalves, Ewan Harrison, David K. Jackson, Ian Johnston, Dominic Kwiatkowski, Cordelia Langford, John Sillitoe on behalf of the Wellcome Sanger Institute COVID-19 Surveillance Team |
| EPI_ISL_3123831, EPI_ISL_3123832 | Lighthouse Lab in Milton Keynes | Wellcome Sanger Institute for the COVID-19 Genomics UK (COG-UK) Consortium | The Lighthouse Lab in Milton Keynes and Alex Alderton, Roberto Amato, Jeffrey Barrett, Sonia Goncalves, Ewan Harrison, David K. Jackson, Ian Johnston, Dominic Kwiatkowski, Cordelia Langford, John Sillitoe on behalf of the Wellcome Sanger Institute COVID-19 Surveillance Team |
| EPI_ISL_3123833 | Lighthouse Lab in Glasgow | Wellcome Sanger Institute for the COVID-19 Genomics UK (COG-UK) Consortium | Harper VanSteenhouse, Yumi Kasai, David Gray, Carol Clugston, Anna Dominiczak and Alex Alderton, Roberto Amato, Jeffrey Barrett, Sonia Goncalves, Ewan Harrison, David K. Jackson, Ian Johnston, Dominic Kwiatkowski, Cordelia Langford, John Sillitoe on behalf of the Wellcome Sanger Institute COVID-19 Surveillance Team |
| EPI_ISL_3123834 | Lighthouse Lab in Milton Keynes | Wellcome Sanger Institute for the COVID-19 Genomics UK (COG-UK) Consortium | The Lighthouse Lab in Milton Keynes and Alex Alderton, Roberto Amato, Jeffrey Barrett, Sonia Goncalves, Ewan Harrison, David K. Jackson, Ian Johnston, Dominic Kwiatkowski, Cordelia Langford, John Sillitoe on behalf of the Wellcome Sanger Institute COVID-19 Surveillance Team |
| EPI_ISL_3123835, EPI_ISL_3123836 | Berkshire and Surrey Pathology Services Lighthouse Laboratory | Wellcome Sanger Institute for the COVID-19 Genomics UK (COG-UK) Consortium | Berkshire and Surrey Pathology Services Lighthouse Laboratory and Alex Alderton, Roberto Amato, Jeffrey Barrett, Sonia Goncalves, Ewan Harrison, David K. Jackson, Ian Johnston, Dominic Kwiatkowski, Cordelia Langford, John Sillitoe on behalf of the Wellcome Sanger Institute COVID-19 Surveillance Team |
| EPI_ISL_3123837, EPI_ISL_3123838, EPI_ISL_3123839, EPI_ISL_3123840 | Lighthouse Lab in Glasgow | Wellcome Sanger Institute for the COVID-19 Genomics UK (COG-UK) Consortium | Harper VanSteenhouse, Yumi Kasai, David Gray, Carol Clugston, Anna Dominiczak and Alex Alderton, Roberto Amato, Jeffrey Barrett, Sonia Goncalves, Ewan Harrison, David K. Jackson, Ian Johnston, Dominic Kwiatkowski, Cordelia Langford, John Sillitoe on behalf of the Wellcome Sanger Institute COVID-19 Surveillance Team |
| EPI_ISL_3123841 | Berkshire and Surrey Pathology Services Lighthouse Laboratory | Wellcome Sanger Institute for the COVID-19 Genomics UK (COG-UK) Consortium | Berkshire and Surrey Pathology Services Lighthouse Laboratory and Alex Alderton, Roberto Amato, Jeffrey Barrett, Sonia Goncalves, Ewan Harrison, David K. Jackson, Ian Johnston, Dominic Kwiatkowski, Cordelia Langford, John Sillitoe on behalf of the Wellcome Sanger Institute COVID-19 Surveillance Team |
| EPI_ISL_3123842, EPI_ISL_3123843, EPI_ISL_3123844, EPI_ISL_3123845, EPI_ISL_3123846 | Lighthouse Lab in Glasgow | Wellcome Sanger Institute for the COVID-19 Genomics UK (COG-UK) Consortium | Harper VanSteenhouse, Yumi Kasai, David Gray, Carol Clugston, Anna Dominiczak and Alex Alderton, Roberto Amato, Jeffrey Barrett, Sonia Goncalves, Ewan Harrison, David K. Jackson, Ian Johnston, Dominic Kwiatkowski, Cordelia Langford, John Sillitoe on behalf of the Wellcome Sanger Institute COVID-19 Surveillance Team |
| EPI_ISL_3123847 | Berkshire and Surrey Pathology Services Lighthouse Laboratory | Wellcome Sanger Institute for the COVID-19 Genomics UK (COG-UK) Consortium | Berkshire and Surrey Pathology Services Lighthouse Laboratory and Alex Alderton, Roberto Amato, Jeffrey Barrett, Sonia Goncalves, Ewan Harrison, David K. Jackson, Ian Johnston, Dominic Kwiatkowski, Cordelia Langford, John Sillitoe on behalf of the Wellcome Sanger Institute COVID-19 Surveillance Team |
| EPI_ISL_3123848, EPI_ISL_3123849, EPI_ISL_3123850, EPI_ISL_3123851, EPI_ISL_3123852 | Lighthouse Lab in Glasgow | Wellcome Sanger Institute for the COVID-19 Genomics UK (COG-UK) Consortium | Harper VanSteenhouse, Yumi Kasai, David Gray, Carol Clugston, Anna Dominiczak and Alex Alderton, Roberto Amato, Jeffrey Barrett, Sonia Goncalves, Ewan Harrison, David K. Jackson, Ian Johnston, Dominic Kwiatkowski, Cordelia Langford, John Sillitoe on behalf of the Wellcome Sanger Institute COVID-19 Surveillance Team |
| EPI_ISL_3123853 | Berkshire and Surrey Pathology Services Lighthouse Laboratory | Wellcome Sanger Institute for the COVID-19 Genomics UK (COG-UK) Consortium | Berkshire and Surrey Pathology Services Lighthouse Laboratory and Alex Alderton, Roberto Amato, Jeffrey Barrett, Sonia Goncalves, Ewan Harrison, David K. Jackson, Ian Johnston, Dominic Kwiatkowski, Cordelia Langford, John Sillitoe on behalf of the Wellcome Sanger Institute COVID-19 Surveillance Team |
| EPI_ISL_3123854, EPI_ISL_3123855, EPI_ISL_3123856, EPI_ISL_3123857 | Lighthouse Lab in Glasgow | Wellcome Sanger Institute for the COVID-19 Genomics UK (COG-UK) Consortium | Harper VanSteenhouse, Yumi Kasai, David Gray, Carol Clugston, Anna Dominiczak and Alex Alderton, Roberto Amato, Jeffrey Barrett, Sonia Goncalves, Ewan Harrison, David K. Jackson, Ian Johnston, Dominic Kwiatkowski, Cordelia Langford, John Sillitoe on behalf of the Wellcome Sanger Institute COVID-19 Surveillance Team |
| EPI_ISL_3123858 | Berkshire and Surrey Pathology Services Lighthouse Laboratory | Wellcome Sanger Institute for the COVID-19 Genomics UK (COG-UK) Consortium | Berkshire and Surrey Pathology Services Lighthouse Laboratory and Alex Alderton, Roberto Amato, Jeffrey Barrett, Sonia Goncalves, Ewan Harrison, David K. Jackson, Ian Johnston, Dominic Kwiatkowski, Cordelia Langford, John Sillitoe on behalf of the Wellcome Sanger Institute COVID-19 Surveillance Team |
| EPI_ISL_3123859, EPI_ISL_3123860 | Lighthouse Lab in Glasgow | Wellcome Sanger Institute for the COVID-19 Genomics UK (COG-UK) Consortium | Harper VanSteenhouse, Yumi Kasai, David Gray, Carol Clugston, Anna Dominiczak and Alex Alderton, Roberto Amato, Jeffrey Barrett, Sonia Goncalves, Ewan Harrison, David K. Jackson, Ian Johnston, Dominic Kwiatkowski, Cordelia Langford, John Sillitoe on behalf of the Wellcome Sanger Institute COVID-19 Surveillance Team |
| EPI_ISL_3123861, EPI_ISL_3123862, EPI_ISL_3123863, EPI_ISL_3123864 | Berkshire and Surrey Pathology Services Lighthouse Laboratory | Wellcome Sanger Institute for the COVID-19 Genomics UK (COG-UK) Consortium | Berkshire and Surrey Pathology Services Lighthouse Laboratory and Alex Alderton, Roberto Amato, Jeffrey Barrett, Sonia Goncalves, Ewan Harrison, David K. Jackson, Ian Johnston, Dominic Kwiatkowski, Cordelia Langford, John Sillitoe on behalf of the Wellcome Sanger Institute COVID-19 Surveillance Team |
| EPI_ISL_3123865, EPI_ISL_3123866, EPI_ISL_3123867 | Lighthouse Lab in Glasgow | Wellcome Sanger Institute for the COVID-19 Genomics UK (COG-UK) Consortium | Harper VanSteenhouse, Yumi Kasai, David Gray, Carol Clugston, Anna Dominiczak and Alex Alderton, Roberto Amato, Jeffrey Barrett, Sonia Goncalves, Ewan Harrison, David K. Jackson, Ian Johnston, Dominic Kwiatkowski, Cordelia Langford, John Sillitoe on behalf of the Wellcome Sanger Institute COVID-19 Surveillance Team |
| EPI_ISL_3123868 | Berkshire and Surrey Pathology Services Lighthouse Laboratory | Wellcome Sanger Institute for the COVID-19 Genomics UK (COG-UK) Consortium | Berkshire and Surrey Pathology Services Lighthouse Laboratory and Alex Alderton, Roberto Amato, Jeffrey Barrett, Sonia Goncalves, Ewan Harrison, David K. Jackson, Ian Johnston, Dominic Kwiatkowski, Cordelia Langford, John Sillitoe on behalf of the Wellcome Sanger Institute COVID-19 Surveillance Team |
| EPI_ISL_3123869, EPI_ISL_3123870, EPI_ISL_3123871, EPI_ISL_3123872, EPI_ISL_3123873, | Lighthouse Lab in Glasgow | Wellcome Sanger Institute for the COVID-19 Genomics UK (COG-UK) Consortium | Harper VanSteenhouse, Yumi Kasai, David Gray, Carol Clugston, Anna Dominiczak and Alex Alderton, Roberto Amato, Jeffrey Barrett, Sonia Goncalves, Ewan Harrison, David K. Jackson, Ian Johnston, Dominic Kwiatkowski, Cordelia Langford, John Sillitoe on behalf of the Wellcome Sanger Institute COVID-19 Surveillance Team |

| | | | |
|---|---|---|---|
| EPI_ISL_3123874, EPI_ISL_3123875 | | | |
| EPI_ISL_3123876, EPI_ISL_3123877 | Berkshire and Surrey Pathology Services Lighthouse Laboratory | Wellcome Sanger Institute for the COVID-19 Genomics UK (COG-UK) Consortium | Berkshire and Surrey Pathology Services Lighthouse Laboratory and Alex Alderton, Roberto Amato, Jeffrey Barrett, Sonia Goncalves, Ewan Harrison, David K. Jackson, Ian Johnston, Dominic Kwiatkowski, Cordelia Langford, John Sillitoe on behalf of the Wellcome Sanger Institute COVID-19 Surveillance Team |
| EPI_ISL_3123878, EPI_ISL_3123879 | Lighthouse Lab in Glasgow | Wellcome Sanger Institute for the COVID-19 Genomics UK (COG-UK) Consortium | Harper VanSteenhouse, Yumi Kasai, David Gray, Carol Clugston, Anna Dominiczak and Alex Alderton, Roberto Amato, Jeffrey Barrett, Sonia Goncalves, Ewan Harrison, David K. Jackson, Ian Johnston, Dominic Kwiatkowski, Cordelia Langford, John Sillitoe on behalf of the Wellcome Sanger Institute COVID-19 Surveillance Team |
| EPI_ISL_3123880, EPI_ISL_3123881 | Berkshire and Surrey Pathology Services Lighthouse Laboratory | Wellcome Sanger Institute for the COVID-19 Genomics UK (COG-UK) Consortium | Berkshire and Surrey Pathology Services Lighthouse Laboratory and Alex Alderton, Roberto Amato, Jeffrey Barrett, Sonia Goncalves, Ewan Harrison, David K. Jackson, Ian Johnston, Dominic Kwiatkowski, Cordelia Langford, John Sillitoe on behalf of the Wellcome Sanger Institute COVID-19 Surveillance Team |
| EPI_ISL_3123882, EPI_ISL_3123883, EPI_ISL_3123884, EPI_ISL_3123885, EPI_ISL_3123886, EPI_ISL_3123887, EPI_ISL_3123888, EPI_ISL_3123889, EPI_ISL_3123890, EPI_ISL_3123891, EPI_ISL_3123892, EPI_ISL_3123893, EPI_ISL_3123894, EPI_ISL_3123895, EPI_ISL_3123896, EPI_ISL_3123897, EPI_ISL_3123898 | | | |
| see above | Lighthouse Lab in Glasgow | Wellcome Sanger Institute for the COVID-19 Genomics UK (COG-UK) Consortium | Harper VanSteenhouse, Yumi Kasai, David Gray, Carol Clugston, Anna Dominiczak and Alex Alderton, Roberto Amato, Jeffrey Barrett, Sonia Goncalves, Ewan Harrison, David K. Jackson, Ian Johnston, Dominic Kwiatkowski, Cordelia Langford, John Sillitoe on behalf of the Wellcome Sanger Institute COVID-19 Surveillance Team |
| EPI_ISL_3123899, EPI_ISL_3123900 | Berkshire and Surrey Pathology Services Lighthouse Laboratory | Wellcome Sanger Institute for the COVID-19 Genomics UK (COG-UK) Consortium | Berkshire and Surrey Pathology Services Lighthouse Laboratory and Alex Alderton, Roberto Amato, Jeffrey Barrett, Sonia Goncalves, Ewan Harrison, David K. Jackson, Ian Johnston, Dominic Kwiatkowski, Cordelia Langford, John Sillitoe on behalf of the Wellcome Sanger Institute COVID-19 Surveillance Team |
| EPI_ISL_3123901, EPI_ISL_3123902, EPI_ISL_3123903 | Lighthouse Lab in Glasgow | Wellcome Sanger Institute for the COVID-19 Genomics UK (COG-UK) Consortium | Harper VanSteenhouse, Yumi Kasai, David Gray, Carol Clugston, Anna Dominiczak and Alex Alderton, Roberto Amato, Jeffrey Barrett, Sonia Goncalves, Ewan Harrison, David K. Jackson, Ian Johnston, Dominic Kwiatkowski, Cordelia Langford, John Sillitoe on behalf of the Wellcome Sanger Institute COVID-19 Surveillance Team |
| EPI_ISL_3123904 | Berkshire and Surrey Pathology Services Lighthouse Laboratory | Wellcome Sanger Institute for the COVID-19 Genomics UK (COG-UK) Consortium | Berkshire and Surrey Pathology Services Lighthouse Laboratory and Alex Alderton, Roberto Amato, Jeffrey Barrett, Sonia Goncalves, Ewan Harrison, David K. Jackson, Ian Johnston, Dominic Kwiatkowski, Cordelia Langford, John Sillitoe on behalf of the Wellcome Sanger Institute COVID-19 Surveillance Team |
| EPI_ISL_3123905 | Lighthouse Lab in Glasgow | Wellcome Sanger Institute for the COVID-19 Genomics UK (COG-UK) Consortium | Harper VanSteenhouse, Yumi Kasai, David Gray, Carol Clugston, Anna Dominiczak and Alex Alderton, Roberto Amato, Jeffrey Barrett, Sonia Goncalves, Ewan Harrison, David K. Jackson, Ian Johnston, Dominic Kwiatkowski, Cordelia Langford, John Sillitoe on behalf of the Wellcome Sanger Institute COVID-19 Surveillance Team |
| EPI_ISL_3123906 | Berkshire and Surrey Pathology Services Lighthouse Laboratory | Wellcome Sanger Institute for the COVID-19 Genomics UK (COG-UK) Consortium | Berkshire and Surrey Pathology Services Lighthouse Laboratory and Alex Alderton, Roberto Amato, Jeffrey Barrett, Sonia Goncalves, Ewan Harrison, David K. Jackson, Ian Johnston, Dominic Kwiatkowski, Cordelia Langford, John Sillitoe on behalf of the Wellcome Sanger Institute COVID-19 Surveillance Team |
| EPI_ISL_3123907 | Lighthouse Lab in Glasgow | Wellcome Sanger Institute for the COVID-19 Genomics UK (COG-UK) Consortium | Harper VanSteenhouse, Yumi Kasai, David Gray, Carol Clugston, Anna Dominiczak and Alex Alderton, Roberto Amato, Jeffrey Barrett, Sonia Goncalves, Ewan Harrison, David K. Jackson, Ian Johnston, Dominic Kwiatkowski, Cordelia Langford, John Sillitoe on behalf of the Wellcome Sanger Institute COVID-19 Surveillance Team |
| EPI_ISL_3123908 | Berkshire and Surrey Pathology Services Lighthouse Laboratory | Wellcome Sanger Institute for the COVID-19 Genomics UK (COG-UK) Consortium | Berkshire and Surrey Pathology Services Lighthouse Laboratory and Alex Alderton, Roberto Amato, Jeffrey Barrett, Sonia Goncalves, Ewan Harrison, David K. Jackson, Ian Johnston, Dominic Kwiatkowski, Cordelia Langford, John Sillitoe on behalf of the Wellcome Sanger Institute COVID-19 Surveillance Team |
| EPI_ISL_3123909, EPI_ISL_3123910 | Lighthouse Lab in Glasgow | Wellcome Sanger Institute for the COVID-19 Genomics UK (COG-UK) Consortium | Harper VanSteenhouse, Yumi Kasai, David Gray, Carol Clugston, Anna Dominiczak and Alex Alderton, Roberto Amato, Jeffrey Barrett, Sonia Goncalves, Ewan Harrison, David K. Jackson, Ian Johnston, Dominic Kwiatkowski, Cordelia Langford, John Sillitoe on behalf of the Wellcome Sanger Institute COVID-19 Surveillance Team |
| EPI_ISL_3123911 | Berkshire and Surrey Pathology Services Lighthouse Laboratory | Wellcome Sanger Institute for the COVID-19 Genomics UK (COG-UK) Consortium | Berkshire and Surrey Pathology Services Lighthouse Laboratory and Alex Alderton, Roberto Amato, Jeffrey Barrett, Sonia Goncalves, Ewan Harrison, David K. Jackson, Ian Johnston, Dominic Kwiatkowski, Cordelia Langford, John Sillitoe on behalf of the Wellcome Sanger Institute COVID-19 Surveillance Team |
| EPI_ISL_3123912, EPI_ISL_3123913, EPI_ISL_3123914 | Lighthouse Lab in Glasgow | Wellcome Sanger Institute for the COVID-19 Genomics UK (COG-UK) Consortium | Harper VanSteenhouse, Yumi Kasai, David Gray, Carol Clugston, Anna Dominiczak and Alex Alderton, Roberto Amato, Jeffrey Barrett, Sonia Goncalves, Ewan Harrison, David K. Jackson, Ian Johnston, Dominic Kwiatkowski, Cordelia Langford, John Sillitoe on behalf of the Wellcome Sanger Institute COVID-19 Surveillance Team |
| EPI_ISL_3123915, EPI_ISL_3123916 | Berkshire and Surrey Pathology Services Lighthouse Laboratory | Wellcome Sanger Institute for the COVID-19 Genomics UK (COG-UK) Consortium | Berkshire and Surrey Pathology Services Lighthouse Laboratory and Alex Alderton, Roberto Amato, Jeffrey Barrett, Sonia Goncalves, Ewan Harrison, David K. Jackson, Ian Johnston, Dominic Kwiatkowski, Cordelia Langford, John Sillitoe on behalf of the Wellcome Sanger Institute COVID-19 Surveillance Team |
| EPI_ISL_3123917 | Lighthouse Lab in Glasgow | Wellcome Sanger Institute for the COVID-19 Genomics UK (COG-UK) Consortium | Harper VanSteenhouse, Yumi Kasai, David Gray, Carol Clugston, Anna Dominiczak and Alex Alderton, Roberto Amato, Jeffrey Barrett, Sonia Goncalves, Ewan Harrison, David K. Jackson, Ian Johnston, Dominic Kwiatkowski, Cordelia Langford, John Sillitoe on behalf of the Wellcome Sanger Institute COVID-19 Surveillance Team |
| EPI_ISL_3123918, EPI_ISL_3123919 | Berkshire and Surrey Pathology Services Lighthouse Laboratory | Wellcome Sanger Institute for the COVID-19 Genomics UK (COG-UK) Consortium | Berkshire and Surrey Pathology Services Lighthouse Laboratory and Alex Alderton, Roberto Amato, Jeffrey Barrett, Sonia Goncalves, Ewan Harrison, David K. Jackson, Ian Johnston, Dominic Kwiatkowski, Cordelia Langford, John Sillitoe on behalf of the Wellcome Sanger Institute COVID-19 Surveillance Team |
| EPI_ISL_3123920 | Lighthouse Lab in Glasgow | Wellcome Sanger Institute for the COVID-19 Genomics UK (COG-UK) Consortium | Harper VanSteenhouse, Yumi Kasai, David Gray, Carol Clugston, Anna Dominiczak and Alex Alderton, Roberto Amato, Jeffrey Barrett, Sonia Goncalves, Ewan Harrison, David K. Jackson, Ian Johnston, Dominic Kwiatkowski, Cordelia Langford, John Sillitoe on behalf of the Wellcome Sanger Institute COVID-19 Surveillance Team |
| EPI_ISL_3123921 | Berkshire and Surrey Pathology Services Lighthouse Laboratory | Wellcome Sanger Institute for the COVID-19 Genomics UK (COG-UK) Consortium | Berkshire and Surrey Pathology Services Lighthouse Laboratory and Alex Alderton, Roberto Amato, Jeffrey Barrett, Sonia Goncalves, Ewan Harrison, David K. Jackson, Ian Johnston, Dominic Kwiatkowski, Cordelia Langford, John Sillitoe on behalf of the Wellcome Sanger Institute COVID-19 Surveillance Team |
| EPI_ISL_3123922, EPI_ISL_3123923 | Lighthouse Lab in Glasgow | Wellcome Sanger Institute for the COVID-19 Genomics UK (COG-UK) Consortium | Harper VanSteenhouse, Yumi Kasai, David Gray, Carol Clugston, Anna Dominiczak and Alex Alderton, Roberto Amato, Jeffrey Barrett, Sonia Goncalves, Ewan Harrison, David K. Jackson, Ian Johnston, Dominic Kwiatkowski, Cordelia Langford, John Sillitoe on behalf of the Wellcome Sanger Institute COVID-19 Surveillance Team |
| EPI_ISL_3123924 | Berkshire and Surrey Pathology Services Lighthouse Laboratory | Wellcome Sanger Institute for the COVID-19 Genomics UK (COG-UK) Consortium | Berkshire and Surrey Pathology Services Lighthouse Laboratory and Alex Alderton, Roberto Amato, Jeffrey Barrett, Sonia Goncalves, Ewan Harrison, David K. Jackson, Ian Johnston, Dominic Kwiatkowski, Cordelia Langford, John Sillitoe on behalf of the Wellcome Sanger Institute COVID-19 Surveillance Team |
| EPI_ISL_3123925, EPI_ISL_3123926 | Lighthouse Lab in Glasgow | Wellcome Sanger Institute for the COVID-19 Genomics UK (COG-UK) Consortium | Harper VanSteenhouse, Yumi Kasai, David Gray, Carol Clugston, Anna Dominiczak and Alex Alderton, Roberto Amato, Jeffrey Barrett, Sonia Goncalves, Ewan Harrison, David K. Jackson, Ian Johnston, Dominic Kwiatkowski, Cordelia Langford, John Sillitoe on behalf of the Wellcome Sanger Institute COVID-19 Surveillance Team |
| EPI_ISL_3123927 | Berkshire and Surrey Pathology Services Lighthouse Laboratory | Wellcome Sanger Institute for the COVID-19 Genomics UK (COG-UK) Consortium | Berkshire and Surrey Pathology Services Lighthouse Laboratory and Alex Alderton, Roberto Amato, Jeffrey Barrett, Sonia Goncalves, Ewan Harrison, David K. Jackson, Ian Johnston, Dominic Kwiatkowski, Cordelia Langford, John Sillitoe on behalf of the Wellcome Sanger Institute COVID-19 Surveillance Team |
| EPI_ISL_3123928, EPI_ISL_3123929, EPI_ISL_3123930, EPI_ISL_3123931 | Lighthouse Lab in Glasgow | Wellcome Sanger Institute for the COVID-19 Genomics UK (COG-UK) Consortium | Harper VanSteenhouse, Yumi Kasai, David Gray, Carol Clugston, Anna Dominiczak and Alex Alderton, Roberto Amato, Jeffrey Barrett, Sonia Goncalves, Ewan Harrison, David K. Jackson, Ian Johnston, Dominic Kwiatkowski, Cordelia Langford, John Sillitoe on behalf of the Wellcome Sanger Institute COVID-19 Surveillance Team |
| EPI_ISL_3123932 | Berkshire and Surrey Pathology Services Lighthouse Laboratory | Wellcome Sanger Institute for the COVID-19 Genomics UK (COG-UK) Consortium | Berkshire and Surrey Pathology Services Lighthouse Laboratory and Alex Alderton, Roberto Amato, Jeffrey Barrett, Sonia Goncalves, Ewan Harrison, David K. Jackson, Ian Johnston, Dominic Kwiatkowski, Cordelia Langford, John Sillitoe on behalf of the Wellcome Sanger Institute COVID-19 Surveillance Team |
| EPI_ISL_3123933, EPI_ISL_3123934 | Lighthouse Lab in Glasgow | Wellcome Sanger Institute for the COVID-19 Genomics UK (COG-UK) Consortium | Harper VanSteenhouse, Yumi Kasai, David Gray, Carol Clugston, Anna Dominiczak and Alex Alderton, Roberto Amato, Jeffrey Barrett, Sonia Goncalves, Ewan Harrison, David K. Jackson, Ian Johnston, Dominic Kwiatkowski, Cordelia Langford, John Sillitoe on behalf of the Wellcome Sanger Institute COVID-19 Surveillance Team |
| EPI_ISL_3123935 | Berkshire and Surrey Pathology Services Lighthouse Laboratory | Wellcome Sanger Institute for the COVID-19 Genomics UK (COG-UK) Consortium | Berkshire and Surrey Pathology Services Lighthouse Laboratory and Alex Alderton, Roberto Amato, Jeffrey Barrett, Sonia Goncalves, Ewan Harrison, David K. Jackson, Ian Johnston, Dominic Kwiatkowski, Cordelia Langford, John Sillitoe on behalf of the Wellcome Sanger Institute COVID-19 Surveillance Team |
| EPI_ISL_3123937, EPI_ISL_3123938, EPI_ISL_3123939 | Lighthouse Lab in Glasgow | Wellcome Sanger Institute for the COVID-19 Genomics UK (COG-UK) Consortium | Harper VanSteenhouse, Yumi Kasai, David Gray, Carol Clugston, Anna Dominiczak and Alex Alderton, Roberto Amato, Jeffrey Barrett, Sonia Goncalves, Ewan Harrison, David K. Jackson, Ian Johnston, Dominic Kwiatkowski, Cordelia Langford, John Sillitoe on behalf of the Wellcome Sanger Institute COVID-19 Surveillance Team |
| EPI_ISL_3123940 | Berkshire and Surrey Pathology Services Lighthouse Laboratory | Wellcome Sanger Institute for the COVID-19 Genomics UK (COG-UK) Consortium | Berkshire and Surrey Pathology Services Lighthouse Laboratory and Alex Alderton, Roberto Amato, Jeffrey Barrett, Sonia Goncalves, Ewan Harrison, David K. Jackson, Ian Johnston, Dominic Kwiatkowski, Cordelia Langford, John Sillitoe on behalf of the Wellcome Sanger Institute COVID-19 Surveillance Team |
| EPI_ISL_3123941, EPI_ISL_3123942, EPI_ISL_3123943, EPI_ISL_3123944, EPI_ISL_3123945, EPI_ISL_3123946, EPI_ISL_3123947, EPI_ISL_3123948, EPI_ISL_3123949 | Lighthouse Lab in Glasgow | Wellcome Sanger Institute for the COVID-19 Genomics UK (COG-UK) Consortium | Harper VanSteenhouse, Yumi Kasai, David Gray, Carol Clugston, Anna Dominiczak and Alex Alderton, Roberto Amato, Jeffrey Barrett, Sonia Goncalves, Ewan Harrison, David K. Jackson, Ian Johnston, Dominic Kwiatkowski, Cordelia Langford, John Sillitoe on behalf of the Wellcome Sanger Institute COVID-19 Surveillance Team |
| EPI_ISL_3123950 | Berkshire and Surrey Pathology Services Lighthouse Laboratory | Wellcome Sanger Institute for the COVID-19 Genomics UK (COG-UK) Consortium | Berkshire and Surrey Pathology Services Lighthouse Laboratory and Alex Alderton, Roberto Amato, Jeffrey Barrett, Sonia Goncalves, Ewan Harrison, David K. Jackson, Ian Johnston, Dominic Kwiatkowski, Cordelia Langford, John Sillitoe on behalf of the Wellcome Sanger Institute COVID-19 Surveillance Team |
| EPI_ISL_3123951, EPI_ISL_3123952, EPI_ISL_3123953, EPI_ISL_3123954, EPI_ISL_3123955, EPI_ISL_3123956 | Lighthouse Lab in Glasgow | Wellcome Sanger Institute for the COVID-19 Genomics UK (COG-UK) Consortium | Harper VanSteenhouse, Yumi Kasai, David Gray, Carol Clugston, Anna Dominiczak and Alex Alderton, Roberto Amato, Jeffrey Barrett, Sonia Goncalves, Ewan Harrison, David K. Jackson, Ian Johnston, Dominic Kwiatkowski, Cordelia Langford, John Sillitoe on behalf of the Wellcome Sanger Institute COVID-19 Surveillance Team |
| EPI_ISL_3123957, EPI_ISL_3123958, EPI_ISL_3123959 | Berkshire and Surrey Pathology Services Lighthouse Laboratory | Wellcome Sanger Institute for the COVID-19 Genomics UK (COG-UK) Consortium | Berkshire and Surrey Pathology Services Lighthouse Laboratory and Alex Alderton, Roberto Amato, Jeffrey Barrett, Sonia Goncalves, Ewan Harrison, David K. Jackson, Ian Johnston, Dominic Kwiatkowski, Cordelia Langford, John Sillitoe on behalf of the Wellcome Sanger Institute COVID-19 Surveillance Team |
| EPI_ISL_3123960, EPI_ISL_3123961 | Lighthouse Lab in Glasgow | Wellcome Sanger Institute for the COVID-19 Genomics UK (COG-UK) Consortium | Harper VanSteenhouse, Yumi Kasai, David Gray, Carol Clugston, Anna Dominiczak and Alex Alderton, Roberto Amato, Jeffrey Barrett, Sonia Goncalves, Ewan Harrison, David K. Jackson, Ian Johnston, Dominic Kwiatkowski, Cordelia Langford, John Sillitoe on behalf of the Wellcome Sanger Institute COVID-19 Surveillance Team |
| EPI_ISL_3123962, EPI_ISL_3123963 | Berkshire and Surrey Pathology Services Lighthouse Laboratory | Wellcome Sanger Institute for the COVID-19 Genomics UK (COG-UK) Consortium | Berkshire and Surrey Pathology Services Lighthouse Laboratory and Alex Alderton, Roberto Amato, Jeffrey Barrett, Sonia Goncalves, Ewan Harrison, David K. Jackson, Ian Johnston, Dominic Kwiatkowski, Cordelia Langford, John Sillitoe on behalf of the Wellcome Sanger Institute COVID-19 Surveillance Team |
| EPI_ISL_3123964, EPI_ISL_3123965 | Lighthouse Lab in Glasgow | Wellcome Sanger Institute for the COVID-19 Genomics UK (COG-UK) Consortium | Harper VanSteenhouse, Yumi Kasai, David Gray, Carol Clugston, Anna Dominiczak and Alex Alderton, Roberto Amato, Jeffrey Barrett, Sonia Goncalves, Ewan Harrison, David K. Jackson, Ian Johnston, Dominic Kwiatkowski, Cordelia Langford, John Sillitoe on behalf of the Wellcome Sanger Institute COVID-19 Surveillance Team |
| EPI_ISL_3123966 | Berkshire and Surrey Pathology Services Lighthouse Laboratory | Wellcome Sanger Institute for the COVID-19 Genomics UK (COG-UK) | Berkshire and Surrey Pathology Services Lighthouse Laboratory and Alex Alderton, Roberto Amato, Jeffrey Barrett, Sonia Goncalves, Ewan Harrison, David K. Jackson, Ian Johnston, Dominic Kwiatkowski, Cordelia Langford, John Sillitoe on behalf |

| | | | |
|---|---|---|---|
| | | Consortium | of the Wellcome Sanger Institute COVID-19 Surveillance Team |
| EPI_ISL_3123967, EPI_ISL_3123968, EPI_ISL_3123969, EPI_ISL_3123970, EPI_ISL_3123971, EPI_ISL_3123972 | Lighthouse Lab in Glasgow | Wellcome Sanger Institute for the COVID-19 Genomics UK (COG-UK) Consortium | Harper VanSteenhouse, Yumi Kasai, David Gray, Carol Clugston, Anna Dominiczak and Alex Alderton, Roberto Amato, Jeffrey Barrett, Sonia Goncalves, Ewan Harrison, David K. Jackson, Ian Johnston, Dominic Kwiatkowski, Cordelia Langford, John Sillitoe on behalf of the Wellcome Sanger Institute COVID-19 Surveillance Team |
| EPI_ISL_3123973, EPI_ISL_3123974 | Berkshire and Surrey Pathology Services Lighthouse Laboratory | Wellcome Sanger Institute for the COVID-19 Genomics UK (COG-UK) Consortium | Berkshire and Surrey Pathology Services Lighthouse Laboratory and Alex Alderton, Roberto Amato, Jeffrey Barrett, Sonia Goncalves, Ewan Harrison, David K. Jackson, Ian Johnston, Dominic Kwiatkowski, Cordelia Langford, John Sillitoe on behalf of the Wellcome Sanger Institute COVID-19 Surveillance Team |
| EPI_ISL_3123975, EPI_ISL_3123976 | Lighthouse Lab in Glasgow | Wellcome Sanger Institute for the COVID-19 Genomics UK (COG-UK) Consortium | Harper VanSteenhouse, Yumi Kasai, David Gray, Carol Clugston, Anna Dominiczak and Alex Alderton, Roberto Amato, Jeffrey Barrett, Sonia Goncalves, Ewan Harrison, David K. Jackson, Ian Johnston, Dominic Kwiatkowski, Cordelia Langford, John Sillitoe on behalf of the Wellcome Sanger Institute COVID-19 Surveillance Team |
| EPI_ISL_3123977, EPI_ISL_3123978 | Berkshire and Surrey Pathology Services Lighthouse Laboratory | Wellcome Sanger Institute for the COVID-19 Genomics UK (COG-UK) Consortium | Berkshire and Surrey Pathology Services Lighthouse Laboratory and Alex Alderton, Roberto Amato, Jeffrey Barrett, Sonia Goncalves, Ewan Harrison, David K. Jackson, Ian Johnston, Dominic Kwiatkowski, Cordelia Langford, John Sillitoe on behalf of the Wellcome Sanger Institute COVID-19 Surveillance Team |
| EPI_ISL_3123979, EPI_ISL_3123980 | Lighthouse Lab in Glasgow | Wellcome Sanger Institute for the COVID-19 Genomics UK (COG-UK) Consortium | Harper VanSteenhouse, Yumi Kasai, David Gray, Carol Clugston, Anna Dominiczak and Alex Alderton, Roberto Amato, Jeffrey Barrett, Sonia Goncalves, Ewan Harrison, David K. Jackson, Ian Johnston, Dominic Kwiatkowski, Cordelia Langford, John Sillitoe on behalf of the Wellcome Sanger Institute COVID-19 Surveillance Team |
| EPI_ISL_3123981 | Berkshire and Surrey Pathology Services Lighthouse Laboratory | Wellcome Sanger Institute for the COVID-19 Genomics UK (COG-UK) Consortium | Berkshire and Surrey Pathology Services Lighthouse Laboratory and Alex Alderton, Roberto Amato, Jeffrey Barrett, Sonia Goncalves, Ewan Harrison, David K. Jackson, Ian Johnston, Dominic Kwiatkowski, Cordelia Langford, John Sillitoe on behalf of the Wellcome Sanger Institute COVID-19 Surveillance Team |
| EPI_ISL_3123982, EPI_ISL_3123983 | Lighthouse Lab in Glasgow | Wellcome Sanger Institute for the COVID-19 Genomics UK (COG-UK) Consortium | Harper VanSteenhouse, Yumi Kasai, David Gray, Carol Clugston, Anna Dominiczak and Alex Alderton, Roberto Amato, Jeffrey Barrett, Sonia Goncalves, Ewan Harrison, David K. Jackson, Ian Johnston, Dominic Kwiatkowski, Cordelia Langford, John Sillitoe on behalf of the Wellcome Sanger Institute COVID-19 Surveillance Team |
| EPI_ISL_3123984, EPI_ISL_3123985 | Berkshire and Surrey Pathology Services Lighthouse Laboratory | Wellcome Sanger Institute for the COVID-19 Genomics UK (COG-UK) Consortium | Berkshire and Surrey Pathology Services Lighthouse Laboratory and Alex Alderton, Roberto Amato, Jeffrey Barrett, Sonia Goncalves, Ewan Harrison, David K. Jackson, Ian Johnston, Dominic Kwiatkowski, Cordelia Langford, John Sillitoe on behalf of the Wellcome Sanger Institute COVID-19 Surveillance Team |
| EPI_ISL_3123986, EPI_ISL_3123987 | Lighthouse Lab in Glasgow | Wellcome Sanger Institute for the COVID-19 Genomics UK (COG-UK) Consortium | Harper VanSteenhouse, Yumi Kasai, David Gray, Carol Clugston, Anna Dominiczak and Alex Alderton, Roberto Amato, Jeffrey Barrett, Sonia Goncalves, Ewan Harrison, David K. Jackson, Ian Johnston, Dominic Kwiatkowski, Cordelia Langford, John Sillitoe on behalf of the Wellcome Sanger Institute COVID-19 Surveillance Team |
| EPI_ISL_3123988, EPI_ISL_3123989 | Berkshire and Surrey Pathology Services Lighthouse Laboratory | Wellcome Sanger Institute for the COVID-19 Genomics UK (COG-UK) Consortium | Berkshire and Surrey Pathology Services Lighthouse Laboratory and Alex Alderton, Roberto Amato, Jeffrey Barrett, Sonia Goncalves, Ewan Harrison, David K. Jackson, Ian Johnston, Dominic Kwiatkowski, Cordelia Langford, John Sillitoe on behalf of the Wellcome Sanger Institute COVID-19 Surveillance Team |
| EPI_ISL_3123990 | Lighthouse Lab in Glasgow | Wellcome Sanger Institute for the COVID-19 Genomics UK (COG-UK) Consortium | Harper VanSteenhouse, Yumi Kasai, David Gray, Carol Clugston, Anna Dominiczak and Alex Alderton, Roberto Amato, Jeffrey Barrett, Sonia Goncalves, Ewan Harrison, David K. Jackson, Ian Johnston, Dominic Kwiatkowski, Cordelia Langford, John Sillitoe on behalf of the Wellcome Sanger Institute COVID-19 Surveillance Team |
| EPI_ISL_3123991 | Berkshire and Surrey Pathology Services Lighthouse Laboratory | Wellcome Sanger Institute for the COVID-19 Genomics UK (COG-UK) Consortium | Berkshire and Surrey Pathology Services Lighthouse Laboratory and Alex Alderton, Roberto Amato, Jeffrey Barrett, Sonia Goncalves, Ewan Harrison, David K. Jackson, Ian Johnston, Dominic Kwiatkowski, Cordelia Langford, John Sillitoe on behalf of the Wellcome Sanger Institute COVID-19 Surveillance Team |
| EPI_ISL_3123992 | Lighthouse Lab in Glasgow | Wellcome Sanger Institute for the COVID-19 Genomics UK (COG-UK) Consortium | Harper VanSteenhouse, Yumi Kasai, David Gray, Carol Clugston, Anna Dominiczak and Alex Alderton, Roberto Amato, Jeffrey Barrett, Sonia Goncalves, Ewan Harrison, David K. Jackson, Ian Johnston, Dominic Kwiatkowski, Cordelia Langford, John Sillitoe on behalf of the Wellcome Sanger Institute COVID-19 Surveillance Team |
| EPI_ISL_3123993, EPI_ISL_3123994, EPI_ISL_3123995 | Berkshire and Surrey Pathology Services Lighthouse Laboratory | Wellcome Sanger Institute for the COVID-19 Genomics UK (COG-UK) Consortium | Berkshire and Surrey Pathology Services Lighthouse Laboratory and Alex Alderton, Roberto Amato, Jeffrey Barrett, Sonia Goncalves, Ewan Harrison, David K. Jackson, Ian Johnston, Dominic Kwiatkowski, Cordelia Langford, John Sillitoe on behalf of the Wellcome Sanger Institute COVID-19 Surveillance Team |
| EPI_ISL_3123996 | Lighthouse Lab in Glasgow | Wellcome Sanger Institute for the COVID-19 Genomics UK (COG-UK) Consortium | Harper VanSteenhouse, Yumi Kasai, David Gray, Carol Clugston, Anna Dominiczak and Alex Alderton, Roberto Amato, Jeffrey Barrett, Sonia Goncalves, Ewan Harrison, David K. Jackson, Ian Johnston, Dominic Kwiatkowski, Cordelia Langford, John Sillitoe on behalf of the Wellcome Sanger Institute COVID-19 Surveillance Team |
| EPI_ISL_3123997, EPI_ISL_3123998 | Berkshire and Surrey Pathology Services Lighthouse Laboratory | Wellcome Sanger Institute for the COVID-19 Genomics UK (COG-UK) Consortium | Berkshire and Surrey Pathology Services Lighthouse Laboratory and Alex Alderton, Roberto Amato, Jeffrey Barrett, Sonia Goncalves, Ewan Harrison, David K. Jackson, Ian Johnston, Dominic Kwiatkowski, Cordelia Langford, John Sillitoe on behalf of the Wellcome Sanger Institute COVID-19 Surveillance Team |
| EPI_ISL_3123999, EPI_ISL_3124000, EPI_ISL_3124001, EPI_ISL_3124002, EPI_ISL_3124003, EPI_ISL_3124004, EPI_ISL_3124005, EPI_ISL_3124006, EPI_ISL_3124007, EPI_ISL_3124008 | Lighthouse Lab in Glasgow | Wellcome Sanger Institute for the COVID-19 Genomics UK (COG-UK) Consortium | Harper VanSteenhouse, Yumi Kasai, David Gray, Carol Clugston, Anna Dominiczak and Alex Alderton, Roberto Amato, Jeffrey Barrett, Sonia Goncalves, Ewan Harrison, David K. Jackson, Ian Johnston, Dominic Kwiatkowski, Cordelia Langford, John Sillitoe on behalf of the Wellcome Sanger Institute COVID-19 Surveillance Team |
| EPI_ISL_3124009, EPI_ISL_3124010 | Lighthouse Laboratory Plymouth | Wellcome Sanger Institute for the COVID-19 Genomics UK (COG-UK) Consortium | Lighthouse Laboratory Plymouth and Alex Alderton, Roberto Amato, Jeffrey Barrett, Sonia Goncalves, Ewan Harrison, David K. Jackson, Ian Johnston, Dominic Kwiatkowski, Cordelia Langford, John Sillitoe on behalf of the Wellcome Sanger Institute COVID-19 Surveillance Team |
| EPI_ISL_3124011 | Berkshire and Surrey Pathology Services Lighthouse Laboratory | Wellcome Sanger Institute for the COVID-19 Genomics UK (COG-UK) Consortium | Berkshire and Surrey Pathology Services Lighthouse Laboratory and Alex Alderton, Roberto Amato, Jeffrey Barrett, Sonia Goncalves, Ewan Harrison, David K. Jackson, Ian Johnston, Dominic Kwiatkowski, Cordelia Langford, John Sillitoe on behalf of the Wellcome Sanger Institute COVID-19 Surveillance Team |
| EPI_ISL_3124012, EPI_ISL_3124013, EPI_ISL_3124014, EPI_ISL_3124015 | Lighthouse Laboratory Plymouth | Wellcome Sanger Institute for the COVID-19 Genomics UK (COG-UK) Consortium | Lighthouse Laboratory Plymouth and Alex Alderton, Roberto Amato, Jeffrey Barrett, Sonia Goncalves, Ewan Harrison, David K. Jackson, Ian Johnston, Dominic Kwiatkowski, Cordelia Langford, John Sillitoe on behalf of the Wellcome Sanger Institute COVID-19 Surveillance Team |
| EPI_ISL_3124016, EPI_ISL_3124017, EPI_ISL_3124018 | Berkshire and Surrey Pathology Services Lighthouse Laboratory | Wellcome Sanger Institute for the COVID-19 Genomics UK (COG-UK) Consortium | Berkshire and Surrey Pathology Services Lighthouse Laboratory and Alex Alderton, Roberto Amato, Jeffrey Barrett, Sonia Goncalves, Ewan Harrison, David K. Jackson, Ian Johnston, Dominic Kwiatkowski, Cordelia Langford, John Sillitoe on behalf of the Wellcome Sanger Institute COVID-19 Surveillance Team |
| EPI_ISL_3124019, EPI_ISL_3124020, EPI_ISL_3124021 | Lighthouse Laboratory Plymouth | Wellcome Sanger Institute for the COVID-19 Genomics UK (COG-UK) Consortium | Lighthouse Laboratory Plymouth and Alex Alderton, Roberto Amato, Jeffrey Barrett, Sonia Goncalves, Ewan Harrison, David K. Jackson, Ian Johnston, Dominic Kwiatkowski, Cordelia Langford, John Sillitoe on behalf of the Wellcome Sanger Institute COVID-19 Surveillance Team |
| EPI_ISL_3124022 | Berkshire and Surrey Pathology Services Lighthouse Laboratory | Wellcome Sanger Institute for the COVID-19 Genomics UK (COG-UK) Consortium | Berkshire and Surrey Pathology Services Lighthouse Laboratory and Alex Alderton, Roberto Amato, Jeffrey Barrett, Sonia Goncalves, Ewan Harrison, David K. Jackson, Ian Johnston, Dominic Kwiatkowski, Cordelia Langford, John Sillitoe on behalf of the Wellcome Sanger Institute COVID-19 Surveillance Team |
| EPI_ISL_3124023 | Lighthouse Laboratory Plymouth | Wellcome Sanger Institute for the COVID-19 Genomics UK (COG-UK) Consortium | Lighthouse Laboratory Plymouth and Alex Alderton, Roberto Amato, Jeffrey Barrett, Sonia Goncalves, Ewan Harrison, David K. Jackson, Ian Johnston, Dominic Kwiatkowski, Cordelia Langford, John Sillitoe on behalf of the Wellcome Sanger Institute COVID-19 Surveillance Team |
| EPI_ISL_3124024 | Berkshire and Surrey Pathology Services Lighthouse Laboratory | Wellcome Sanger Institute for the COVID-19 Genomics UK (COG-UK) Consortium | Berkshire and Surrey Pathology Services Lighthouse Laboratory and Alex Alderton, Roberto Amato, Jeffrey Barrett, Sonia Goncalves, Ewan Harrison, David K. Jackson, Ian Johnston, Dominic Kwiatkowski, Cordelia Langford, John Sillitoe on behalf of the Wellcome Sanger Institute COVID-19 Surveillance Team |
| EPI_ISL_3124025, EPI_ISL_3124026 | Lighthouse Laboratory Plymouth | Wellcome Sanger Institute for the COVID-19 Genomics UK (COG-UK) Consortium | Lighthouse Laboratory Plymouth and Alex Alderton, Roberto Amato, Jeffrey Barrett, Sonia Goncalves, Ewan Harrison, David K. Jackson, Ian Johnston, Dominic Kwiatkowski, Cordelia Langford, John Sillitoe on behalf of the Wellcome Sanger Institute COVID-19 Surveillance Team |
| EPI_ISL_3124027 | Berkshire and Surrey Pathology Services Lighthouse Laboratory | Wellcome Sanger Institute for the COVID-19 Genomics UK (COG-UK) Consortium | Berkshire and Surrey Pathology Services Lighthouse Laboratory and Alex Alderton, Roberto Amato, Jeffrey Barrett, Sonia Goncalves, Ewan Harrison, David K. Jackson, Ian Johnston, Dominic Kwiatkowski, Cordelia Langford, John Sillitoe on behalf of the Wellcome Sanger Institute COVID-19 Surveillance Team |
| EPI_ISL_3124028, EPI_ISL_3124029 | Lighthouse Laboratory Plymouth | Wellcome Sanger Institute for the COVID-19 Genomics UK (COG-UK) Consortium | Lighthouse Laboratory Plymouth and Alex Alderton, Roberto Amato, Jeffrey Barrett, Sonia Goncalves, Ewan Harrison, David K. Jackson, Ian Johnston, Dominic Kwiatkowski, Cordelia Langford, John Sillitoe on behalf of the Wellcome Sanger Institute COVID-19 Surveillance Team |
| EPI_ISL_3124030, EPI_ISL_3124031, EPI_ISL_3124032, EPI_ISL_3124033, EPI_ISL_3124034 | Berkshire and Surrey Pathology Services Lighthouse Laboratory | Wellcome Sanger Institute for the COVID-19 Genomics UK (COG-UK) Consortium | Berkshire and Surrey Pathology Services Lighthouse Laboratory and Alex Alderton, Roberto Amato, Jeffrey Barrett, Sonia Goncalves, Ewan Harrison, David K. Jackson, Ian Johnston, Dominic Kwiatkowski, Cordelia Langford, John Sillitoe on behalf of the Wellcome Sanger Institute COVID-19 Surveillance Team |
| EPI_ISL_3124035, EPI_ISL_3124036, EPI_ISL_3124037, EPI_ISL_3124038, EPI_ISL_3124039 | Lighthouse Laboratory Plymouth | Wellcome Sanger Institute for the COVID-19 Genomics UK (COG-UK) Consortium | Lighthouse Laboratory Plymouth and Alex Alderton, Roberto Amato, Jeffrey Barrett, Sonia Goncalves, Ewan Harrison, David K. Jackson, Ian Johnston, Dominic Kwiatkowski, Cordelia Langford, John Sillitoe on behalf of the Wellcome Sanger Institute COVID-19 Surveillance Team |
| EPI_ISL_3124040 | Berkshire and Surrey Pathology Services Lighthouse Laboratory | Wellcome Sanger Institute for the COVID-19 Genomics UK (COG-UK) Consortium | Berkshire and Surrey Pathology Services Lighthouse Laboratory and Alex Alderton, Roberto Amato, Jeffrey Barrett, Sonia Goncalves, Ewan Harrison, David K. Jackson, Ian Johnston, Dominic Kwiatkowski, Cordelia Langford, John Sillitoe on behalf of the Wellcome Sanger Institute COVID-19 Surveillance Team |
| EPI_ISL_3124041 | Unit of laboratory surveillance of viral emergent disease Control of Communicable Diseases Departament National Institute of Public Health Albania | Charité Universitätsmedizin Berlin, Institut für Virologie | Victor M Corman, Barbara Mühlemann, Jörn Beheim-Schwarzbach, Talitha Veith, Julia Schneider, Terry Jones, Iris Hasibra, Christian Drosten |
| EPI_ISL_3124042, EPI_ISL_3124043, EPI_ISL_3124044, EPI_ISL_3124045, EPI_ISL_3124046, EPI_ISL_3124047 | Lighthouse Laboratory Plymouth | Wellcome Sanger Institute for the COVID-19 Genomics UK (COG-UK) Consortium | Lighthouse Laboratory Plymouth and Alex Alderton, Roberto Amato, Jeffrey Barrett, Sonia Goncalves, Ewan Harrison, David K. Jackson, Ian Johnston, Dominic Kwiatkowski, Cordelia Langford, John Sillitoe on behalf of the Wellcome Sanger Institute COVID-19 Surveillance Team |

| | | | |
|---|---|---|---|
| EPI_ISL_3124048 | Berkshire and Surrey Pathology Services Lighthouse Laboratory | Wellcome Sanger Institute for the COVID-19 Genomics UK (COG-UK) Consortium | Berkshire and Surrey Pathology Services Lighthouse Laboratory and Alex Alderton, Roberto Amato, Jeffrey Barrett, Sonia Goncalves, Ewan Harrison, David K. Jackson, Ian Johnston, Dominic Kwiatkowski, Cordelia Langford, John Sillitoe on behalf of the Wellcome Sanger Institute COVID-19 Surveillance Team |
| EPI_ISL_3124049, EPI_ISL_3124050, EPI_ISL_3124051, EPI_ISL_3124052, EPI_ISL_3124053 | Lighthouse Laboratory Plymouth | Wellcome Sanger Institute for the COVID-19 Genomics UK (COG-UK) Consortium | Lighthouse Laboratory Plymouth and Alex Alderton, Roberto Amato, Jeffrey Barrett, Sonia Goncalves, Ewan Harrison, David K. Jackson, Ian Johnston, Dominic Kwiatkowski, Cordelia Langford, John Sillitoe on behalf of the Wellcome Sanger Institute COVID-19 Surveillance Team |
| EPI_ISL_3124054 | Berkshire and Surrey Pathology Services Lighthouse Laboratory | Wellcome Sanger Institute for the COVID-19 Genomics UK (COG-UK) Consortium | Berkshire and Surrey Pathology Services Lighthouse Laboratory and Alex Alderton, Roberto Amato, Jeffrey Barrett, Sonia Goncalves, Ewan Harrison, David K. Jackson, Ian Johnston, Dominic Kwiatkowski, Cordelia Langford, John Sillitoe on behalf of the Wellcome Sanger Institute COVID-19 Surveillance Team |
| EPI_ISL_3124055 | Lighthouse Laboratory Plymouth | Wellcome Sanger Institute for the COVID-19 Genomics UK (COG-UK) Consortium | Lighthouse Laboratory Plymouth and Alex Alderton, Roberto Amato, Jeffrey Barrett, Sonia Goncalves, Ewan Harrison, David K. Jackson, Ian Johnston, Dominic Kwiatkowski, Cordelia Langford, John Sillitoe on behalf of the Wellcome Sanger Institute COVID-19 Surveillance Team |
| EPI_ISL_3124056 | Berkshire and Surrey Pathology Services Lighthouse Laboratory | Wellcome Sanger Institute for the COVID-19 Genomics UK (COG-UK) Consortium | Berkshire and Surrey Pathology Services Lighthouse Laboratory and Alex Alderton, Roberto Amato, Jeffrey Barrett, Sonia Goncalves, Ewan Harrison, David K. Jackson, Ian Johnston, Dominic Kwiatkowski, Cordelia Langford, John Sillitoe on behalf of the Wellcome Sanger Institute COVID-19 Surveillance Team |
| EPI_ISL_3124057, EPI_ISL_3124058, EPI_ISL_3124059, EPI_ISL_3124060, EPI_ISL_3124061, EPI_ISL_3124062, EPI_ISL_3124063, EPI_ISL_3124064, EPI_ISL_3124065, EPI_ISL_3124066, EPI_ISL_3124067, EPI_ISL_3124068 | | | |
| see above | Lighthouse Laboratory Plymouth | Wellcome Sanger Institute for the COVID-19 Genomics UK (COG-UK) Consortium | Lighthouse Laboratory Plymouth and Alex Alderton, Roberto Amato, Jeffrey Barrett, Sonia Goncalves, Ewan Harrison, David K. Jackson, Ian Johnston, Dominic Kwiatkowski, Cordelia Langford, John Sillitoe on behalf of the Wellcome Sanger Institute COVID-19 Surveillance Team |
| EPI_ISL_3124069, EPI_ISL_3124070 | Berkshire and Surrey Pathology Services Lighthouse Laboratory | Wellcome Sanger Institute for the COVID-19 Genomics UK (COG-UK) Consortium | Berkshire and Surrey Pathology Services Lighthouse Laboratory and Alex Alderton, Roberto Amato, Jeffrey Barrett, Sonia Goncalves, Ewan Harrison, David K. Jackson, Ian Johnston, Dominic Kwiatkowski, Cordelia Langford, John Sillitoe on behalf of the Wellcome Sanger Institute COVID-19 Surveillance Team |
| EPI_ISL_3124071, EPI_ISL_3124072, EPI_ISL_3124073, EPI_ISL_3124074, EPI_ISL_3124075, EPI_ISL_3124076, EPI_ISL_3124077, EPI_ISL_3124078, EPI_ISL_3124079, EPI_ISL_3124080, EPI_ISL_3124081, EPI_ISL_3124082, EPI_ISL_3124083, EPI_ISL_3124084, EPI_ISL_3124085, EPI_ISL_3124086 | | | |
| see above | Lighthouse Laboratory Plymouth | Wellcome Sanger Institute for the COVID-19 Genomics UK (COG-UK) Consortium | Lighthouse Laboratory Plymouth and Alex Alderton, Roberto Amato, Jeffrey Barrett, Sonia Goncalves, Ewan Harrison, David K. Jackson, Ian Johnston, Dominic Kwiatkowski, Cordelia Langford, John Sillitoe on behalf of the Wellcome Sanger Institute COVID-19 Surveillance Team |
| EPI_ISL_3124087 | Berkshire and Surrey Pathology Services Lighthouse Laboratory | Wellcome Sanger Institute for the COVID-19 Genomics UK (COG-UK) Consortium | Berkshire and Surrey Pathology Services Lighthouse Laboratory and Alex Alderton, Roberto Amato, Jeffrey Barrett, Sonia Goncalves, Ewan Harrison, David K. Jackson, Ian Johnston, Dominic Kwiatkowski, Cordelia Langford, John Sillitoe on behalf of the Wellcome Sanger Institute COVID-19 Surveillance Team |
| EPI_ISL_3124088, EPI_ISL_3124089, EPI_ISL_3124090, EPI_ISL_3124091 | Lighthouse Laboratory Plymouth | Wellcome Sanger Institute for the COVID-19 Genomics UK (COG-UK) Consortium | Lighthouse Laboratory Plymouth and Alex Alderton, Roberto Amato, Jeffrey Barrett, Sonia Goncalves, Ewan Harrison, David K. Jackson, Ian Johnston, Dominic Kwiatkowski, Cordelia Langford, John Sillitoe on behalf of the Wellcome Sanger Institute COVID-19 Surveillance Team |
| EPI_ISL_3124092, EPI_ISL_3124093 | Berkshire and Surrey Pathology Services Lighthouse Laboratory | Wellcome Sanger Institute for the COVID-19 Genomics UK (COG-UK) Consortium | Berkshire and Surrey Pathology Services Lighthouse Laboratory and Alex Alderton, Roberto Amato, Jeffrey Barrett, Sonia Goncalves, Ewan Harrison, David K. Jackson, Ian Johnston, Dominic Kwiatkowski, Cordelia Langford, John Sillitoe on behalf of the Wellcome Sanger Institute COVID-19 Surveillance Team |
| EPI_ISL_3124094, EPI_ISL_3124095, EPI_ISL_3124096, EPI_ISL_3124097, EPI_ISL_3124098, EPI_ISL_3124099, EPI_ISL_3124100, EPI_ISL_3124101, EPI_ISL_3124102, EPI_ISL_3124103, EPI_ISL_3124104, EPI_ISL_3124105, EPI_ISL_3124106, EPI_ISL_3124107, EPI_ISL_3124108, EPI_ISL_3124109, EPI_ISL_3124110, EPI_ISL_3124111, EPI_ISL_3124112, EPI_ISL_3124113, EPI_ISL_3124114, EPI_ISL_3124115 | | | |
| see above | Lighthouse Laboratory Plymouth | Wellcome Sanger Institute for the COVID-19 Genomics UK (COG-UK) Consortium | Lighthouse Laboratory Plymouth and Alex Alderton, Roberto Amato, Jeffrey Barrett, Sonia Goncalves, Ewan Harrison, David K. Jackson, Ian Johnston, Dominic Kwiatkowski, Cordelia Langford, John Sillitoe on behalf of the Wellcome Sanger Institute COVID-19 Surveillance Team |
| EPI_ISL_3124116 | Berkshire and Surrey Pathology Services Lighthouse Laboratory | Wellcome Sanger Institute for the COVID-19 Genomics UK (COG-UK) Consortium | Berkshire and Surrey Pathology Services Lighthouse Laboratory and Alex Alderton, Roberto Amato, Jeffrey Barrett, Sonia Goncalves, Ewan Harrison, David K. Jackson, Ian Johnston, Dominic Kwiatkowski, Cordelia Langford, John Sillitoe on behalf of the Wellcome Sanger Institute COVID-19 Surveillance Team |
| EPI_ISL_3124117, EPI_ISL_3124118, EPI_ISL_3124119, EPI_ISL_3124120 | Lighthouse Laboratory Plymouth | Wellcome Sanger Institute for the COVID-19 Genomics UK (COG-UK) Consortium | Lighthouse Laboratory Plymouth and Alex Alderton, Roberto Amato, Jeffrey Barrett, Sonia Goncalves, Ewan Harrison, David K. Jackson, Ian Johnston, Dominic Kwiatkowski, Cordelia Langford, John Sillitoe on behalf of the Wellcome Sanger Institute COVID-19 Surveillance Team |
| EPI_ISL_3124121, EPI_ISL_3124122 | Berkshire and Surrey Pathology Services Lighthouse Laboratory | Wellcome Sanger Institute for the COVID-19 Genomics UK (COG-UK) Consortium | Berkshire and Surrey Pathology Services Lighthouse Laboratory and Alex Alderton, Roberto Amato, Jeffrey Barrett, Sonia Goncalves, Ewan Harrison, David K. Jackson, Ian Johnston, Dominic Kwiatkowski, Cordelia Langford, John Sillitoe on behalf of the Wellcome Sanger Institute COVID-19 Surveillance Team |
| EPI_ISL_3124123, EPI_ISL_3124124, EPI_ISL_3124125, EPI_ISL_3124126, EPI_ISL_3124127, EPI_ISL_3124128, EPI_ISL_3124129, EPI_ISL_3124130, EPI_ISL_3124131, EPI_ISL_3124132, EPI_ISL_3124133, EPI_ISL_3124134, EPI_ISL_3124135 | | | |
| see above | Lighthouse Laboratory Plymouth | Wellcome Sanger Institute for the COVID-19 Genomics UK (COG-UK) Consortium | Lighthouse Laboratory Plymouth and Alex Alderton, Roberto Amato, Jeffrey Barrett, Sonia Goncalves, Ewan Harrison, David K. Jackson, Ian Johnston, Dominic Kwiatkowski, Cordelia Langford, John Sillitoe on behalf of the Wellcome Sanger Institute COVID-19 Surveillance Team |
| EPI_ISL_3124136 | Berkshire and Surrey Pathology Services Lighthouse Laboratory | Wellcome Sanger Institute for the COVID-19 Genomics UK (COG-UK) Consortium | Berkshire and Surrey Pathology Services Lighthouse Laboratory and Alex Alderton, Roberto Amato, Jeffrey Barrett, Sonia Goncalves, Ewan Harrison, David K. Jackson, Ian Johnston, Dominic Kwiatkowski, Cordelia Langford, John Sillitoe on behalf of the Wellcome Sanger Institute COVID-19 Surveillance Team |
| EPI_ISL_3124137 | Lighthouse Laboratory Plymouth | Wellcome Sanger Institute for the COVID-19 Genomics UK (COG-UK) Consortium | Lighthouse Laboratory Plymouth and Alex Alderton, Roberto Amato, Jeffrey Barrett, Sonia Goncalves, Ewan Harrison, David K. Jackson, Ian Johnston, Dominic Kwiatkowski, Cordelia Langford, John Sillitoe on behalf of the Wellcome Sanger Institute COVID-19 Surveillance Team |
| EPI_ISL_3124138 | Berkshire and Surrey Pathology Services Lighthouse Laboratory | Wellcome Sanger Institute for the COVID-19 Genomics UK (COG-UK) Consortium | Berkshire and Surrey Pathology Services Lighthouse Laboratory and Alex Alderton, Roberto Amato, Jeffrey Barrett, Sonia Goncalves, Ewan Harrison, David K. Jackson, Ian Johnston, Dominic Kwiatkowski, Cordelia Langford, John Sillitoe on behalf of the Wellcome Sanger Institute COVID-19 Surveillance Team |
| EPI_ISL_3124139, EPI_ISL_3124140 | Lighthouse Laboratory Plymouth | Wellcome Sanger Institute for the COVID-19 Genomics UK (COG-UK) Consortium | Lighthouse Laboratory Plymouth and Alex Alderton, Roberto Amato, Jeffrey Barrett, Sonia Goncalves, Ewan Harrison, David K. Jackson, Ian Johnston, Dominic Kwiatkowski, Cordelia Langford, John Sillitoe on behalf of the Wellcome Sanger Institute COVID-19 Surveillance Team |
| EPI_ISL_3124141 | Berkshire and Surrey Pathology Services Lighthouse Laboratory | Wellcome Sanger Institute for the COVID-19 Genomics UK (COG-UK) Consortium | Berkshire and Surrey Pathology Services Lighthouse Laboratory and Alex Alderton, Roberto Amato, Jeffrey Barrett, Sonia Goncalves, Ewan Harrison, David K. Jackson, Ian Johnston, Dominic Kwiatkowski, Cordelia Langford, John Sillitoe on behalf of the Wellcome Sanger Institute COVID-19 Surveillance Team |
| EPI_ISL_3124142, EPI_ISL_3124143, EPI_ISL_3124144, EPI_ISL_3124145 | Lighthouse Laboratory Plymouth | Wellcome Sanger Institute for the COVID-19 Genomics UK (COG-UK) Consortium | Lighthouse Laboratory Plymouth and Alex Alderton, Roberto Amato, Jeffrey Barrett, Sonia Goncalves, Ewan Harrison, David K. Jackson, Ian Johnston, Dominic Kwiatkowski, Cordelia Langford, John Sillitoe on behalf of the Wellcome Sanger Institute COVID-19 Surveillance Team |
| EPI_ISL_3124146 | Berkshire and Surrey Pathology Services Lighthouse Laboratory | Wellcome Sanger Institute for the COVID-19 Genomics UK (COG-UK) Consortium | Berkshire and Surrey Pathology Services Lighthouse Laboratory and Alex Alderton, Roberto Amato, Jeffrey Barrett, Sonia Goncalves, Ewan Harrison, David K. Jackson, Ian Johnston, Dominic Kwiatkowski, Cordelia Langford, John Sillitoe on behalf of the Wellcome Sanger Institute COVID-19 Surveillance Team |
| EPI_ISL_3124147, EPI_ISL_3124148, EPI_ISL_3124149, EPI_ISL_3124150, EPI_ISL_3124151, EPI_ISL_3124152, EPI_ISL_3124153, EPI_ISL_3124154 | Lighthouse Laboratory Plymouth | Wellcome Sanger Institute for the COVID-19 Genomics UK (COG-UK) Consortium | Lighthouse Laboratory Plymouth and Alex Alderton, Roberto Amato, Jeffrey Barrett, Sonia Goncalves, Ewan Harrison, David K. Jackson, Ian Johnston, Dominic Kwiatkowski, Cordelia Langford, John Sillitoe on behalf of the Wellcome Sanger Institute COVID-19 Surveillance Team |
| EPI_ISL_3124155 | Berkshire and Surrey Pathology Services Lighthouse Laboratory | Wellcome Sanger Institute for the COVID-19 Genomics UK (COG-UK) Consortium | Berkshire and Surrey Pathology Services Lighthouse Laboratory and Alex Alderton, Roberto Amato, Jeffrey Barrett, Sonia Goncalves, Ewan Harrison, David K. Jackson, Ian Johnston, Dominic Kwiatkowski, Cordelia Langford, John Sillitoe on behalf of the Wellcome Sanger Institute COVID-19 Surveillance Team |
| EPI_ISL_3124156, EPI_ISL_3124157, EPI_ISL_3124158, EPI_ISL_3124159, EPI_ISL_3124160, EPI_ISL_3124161, EPI_ISL_3124162, EPI_ISL_3124163, EPI_ISL_3124164, EPI_ISL_3124165 | Lighthouse Laboratory Plymouth | Wellcome Sanger Institute for the COVID-19 Genomics UK (COG-UK) Consortium | Lighthouse Laboratory Plymouth and Alex Alderton, Roberto Amato, Jeffrey Barrett, Sonia Goncalves, Ewan Harrison, David K. Jackson, Ian Johnston, Dominic Kwiatkowski, Cordelia Langford, John Sillitoe on behalf of the Wellcome Sanger Institute COVID-19 Surveillance Team |
| EPI_ISL_3124166 | Berkshire and Surrey Pathology Services Lighthouse Laboratory | Wellcome Sanger Institute for the COVID-19 Genomics UK (COG-UK) Consortium | Berkshire and Surrey Pathology Services Lighthouse Laboratory and Alex Alderton, Roberto Amato, Jeffrey Barrett, Sonia Goncalves, Ewan Harrison, David K. Jackson, Ian Johnston, Dominic Kwiatkowski, Cordelia Langford, John Sillitoe on behalf of the Wellcome Sanger Institute COVID-19 Surveillance Team |
| EPI_ISL_3124167, EPI_ISL_3124168, EPI_ISL_3124169, EPI_ISL_3124170, EPI_ISL_3124171, EPI_ISL_3124172, EPI_ISL_3124173, EPI_ISL_3124174 | Lighthouse Laboratory Plymouth | Wellcome Sanger Institute for the COVID-19 Genomics UK (COG-UK) Consortium | Lighthouse Laboratory Plymouth and Alex Alderton, Roberto Amato, Jeffrey Barrett, Sonia Goncalves, Ewan Harrison, David K. Jackson, Ian Johnston, Dominic Kwiatkowski, Cordelia Langford, John Sillitoe on behalf of the Wellcome Sanger Institute COVID-19 Surveillance Team |
| EPI_ISL_3124175 | Berkshire and Surrey Pathology Services Lighthouse Laboratory | Wellcome Sanger Institute for the COVID-19 Genomics UK (COG-UK) Consortium | Berkshire and Surrey Pathology Services Lighthouse Laboratory and Alex Alderton, Roberto Amato, Jeffrey Barrett, Sonia Goncalves, Ewan Harrison, David K. Jackson, Ian Johnston, Dominic Kwiatkowski, Cordelia Langford, John Sillitoe on behalf of the Wellcome Sanger Institute COVID-19 Surveillance Team |
| EPI_ISL_3124176, EPI_ISL_3124177, EPI_ISL_3124178, EPI_ISL_3124179 | Lighthouse Laboratory Plymouth | Wellcome Sanger Institute for the COVID-19 Genomics UK (COG-UK) Consortium | Lighthouse Laboratory Plymouth and Alex Alderton, Roberto Amato, Jeffrey Barrett, Sonia Goncalves, Ewan Harrison, David K. Jackson, Ian Johnston, Dominic Kwiatkowski, Cordelia Langford, John Sillitoe on behalf of the Wellcome Sanger Institute COVID-19 Surveillance Team |

| | | | |
|---|---|---|---|
| EPI_ISL_3124180, EPI_ISL_3124181 | Berkshire and Surrey Pathology Services Lighthouse Laboratory | Wellcome Sanger Institute for the COVID-19 Genomics UK (COG-UK) Consortium | Berkshire and Surrey Pathology Services Lighthouse Laboratory and Alex Alderton, Roberto Amato, Jeffrey Barrett, Sonia Goncalves, Ewan Harrison, David K. Jackson, Ian Johnston, Dominic Kwiatkowski, Cordelia Langford, John Sillitoe on behalf of the Wellcome Sanger Institute COVID-19 Surveillance Team |
| EPI_ISL_3124182 | Lighthouse Laboratory Plymouth | Wellcome Sanger Institute for the COVID-19 Genomics UK (COG-UK) Consortium | Lighthouse Laboratory Plymouth and Alex Alderton, Roberto Amato, Jeffrey Barrett, Sonia Goncalves, Ewan Harrison, David K. Jackson, Ian Johnston, Dominic Kwiatkowski, Cordelia Langford, John Sillitoe on behalf of the Wellcome Sanger Institute COVID-19 Surveillance Team |
| EPI_ISL_3124183, EPI_ISL_3124184 | Berkshire and Surrey Pathology Services Lighthouse Laboratory | Wellcome Sanger Institute for the COVID-19 Genomics UK (COG-UK) Consortium | Berkshire and Surrey Pathology Services Lighthouse Laboratory and Alex Alderton, Roberto Amato, Jeffrey Barrett, Sonia Goncalves, Ewan Harrison, David K. Jackson, Ian Johnston, Dominic Kwiatkowski, Cordelia Langford, John Sillitoe on behalf of the Wellcome Sanger Institute COVID-19 Surveillance Team |
| EPI_ISL_3124185, EPI_ISL_3124186, EPI_ISL_3124187, EPI_ISL_3124188, EPI_ISL_3124189, EPI_ISL_3124190, EPI_ISL_3124191, EPI_ISL_3124192, EPI_ISL_3124193, EPI_ISL_3124194, EPI_ISL_3124195, EPI_ISL_3124196, EPI_ISL_3124197, EPI_ISL_3124198, EPI_ISL_3124199, EPI_ISL_3124200, EPI_ISL_3124201, EPI_ISL_3124202 | | | |
| see above | Lighthouse Laboratory Plymouth | Wellcome Sanger Institute for the COVID-19 Genomics UK (COG-UK) Consortium | Lighthouse Laboratory Plymouth and Alex Alderton, Roberto Amato, Jeffrey Barrett, Sonia Goncalves, Ewan Harrison, David K. Jackson, Ian Johnston, Dominic Kwiatkowski, Cordelia Langford, John Sillitoe on behalf of the Wellcome Sanger Institute COVID-19 Surveillance Team |
| EPI_ISL_3124203, EPI_ISL_3124204 | Berkshire and Surrey Pathology Services Lighthouse Laboratory | Wellcome Sanger Institute for the COVID-19 Genomics UK (COG-UK) Consortium | Berkshire and Surrey Pathology Services Lighthouse Laboratory and Alex Alderton, Roberto Amato, Jeffrey Barrett, Sonia Goncalves, Ewan Harrison, David K. Jackson, Ian Johnston, Dominic Kwiatkowski, Cordelia Langford, John Sillitoe on behalf of the Wellcome Sanger Institute COVID-19 Surveillance Team |
| EPI_ISL_3124205, EPI_ISL_3124206, EPI_ISL_3124207, EPI_ISL_3124208, EPI_ISL_3124209, EPI_ISL_3124210, EPI_ISL_3124211 | Lighthouse Laboratory Plymouth | Wellcome Sanger Institute for the COVID-19 Genomics UK (COG-UK) Consortium | Lighthouse Laboratory Plymouth and Alex Alderton, Roberto Amato, Jeffrey Barrett, Sonia Goncalves, Ewan Harrison, David K. Jackson, Ian Johnston, Dominic Kwiatkowski, Cordelia Langford, John Sillitoe on behalf of the Wellcome Sanger Institute COVID-19 Surveillance Team |
| EPI_ISL_3124212, EPI_ISL_3124213 | Berkshire and Surrey Pathology Services Lighthouse Laboratory | Wellcome Sanger Institute for the COVID-19 Genomics UK (COG-UK) Consortium | Berkshire and Surrey Pathology Services Lighthouse Laboratory and Alex Alderton, Roberto Amato, Jeffrey Barrett, Sonia Goncalves, Ewan Harrison, David K. Jackson, Ian Johnston, Dominic Kwiatkowski, Cordelia Langford, John Sillitoe on behalf of the Wellcome Sanger Institute COVID-19 Surveillance Team |
| EPI_ISL_3124214, EPI_ISL_3124215, EPI_ISL_3124216, EPI_ISL_3124217, EPI_ISL_3124218, EPI_ISL_3124219, EPI_ISL_3124220, EPI_ISL_3124221, EPI_ISL_3124222, EPI_ISL_3124223, EPI_ISL_3124224, EPI_ISL_3124225, EPI_ISL_3124226 | | | |
| see above | Lighthouse Laboratory Plymouth | Wellcome Sanger Institute for the COVID-19 Genomics UK (COG-UK) Consortium | Lighthouse Laboratory Plymouth and Alex Alderton, Roberto Amato, Jeffrey Barrett, Sonia Goncalves, Ewan Harrison, David K. Jackson, Ian Johnston, Dominic Kwiatkowski, Cordelia Langford, John Sillitoe on behalf of the Wellcome Sanger Institute COVID-19 Surveillance Team |
| EPI_ISL_3124227 | Berkshire and Surrey Pathology Services Lighthouse Laboratory | Wellcome Sanger Institute for the COVID-19 Genomics UK (COG-UK) Consortium | Berkshire and Surrey Pathology Services Lighthouse Laboratory and Alex Alderton, Roberto Amato, Jeffrey Barrett, Sonia Goncalves, Ewan Harrison, David K. Jackson, Ian Johnston, Dominic Kwiatkowski, Cordelia Langford, John Sillitoe on behalf of the Wellcome Sanger Institute COVID-19 Surveillance Team |
| EPI_ISL_3124228, EPI_ISL_3124229 | Lighthouse Laboratory Plymouth | Wellcome Sanger Institute for the COVID-19 Genomics UK (COG-UK) Consortium | Lighthouse Laboratory Plymouth and Alex Alderton, Roberto Amato, Jeffrey Barrett, Sonia Goncalves, Ewan Harrison, David K. Jackson, Ian Johnston, Dominic Kwiatkowski, Cordelia Langford, John Sillitoe on behalf of the Wellcome Sanger Institute COVID-19 Surveillance Team |
| EPI_ISL_3124230 | Berkshire and Surrey Pathology Services Lighthouse Laboratory | Wellcome Sanger Institute for the COVID-19 Genomics UK (COG-UK) Consortium | Berkshire and Surrey Pathology Services Lighthouse Laboratory and Alex Alderton, Roberto Amato, Jeffrey Barrett, Sonia Goncalves, Ewan Harrison, David K. Jackson, Ian Johnston, Dominic Kwiatkowski, Cordelia Langford, John Sillitoe on behalf of the Wellcome Sanger Institute COVID-19 Surveillance Team |
| EPI_ISL_3124231, EPI_ISL_3124232 | Lighthouse Laboratory Plymouth | Wellcome Sanger Institute for the COVID-19 Genomics UK (COG-UK) Consortium | Lighthouse Laboratory Plymouth and Alex Alderton, Roberto Amato, Jeffrey Barrett, Sonia Goncalves, Ewan Harrison, David K. Jackson, Ian Johnston, Dominic Kwiatkowski, Cordelia Langford, John Sillitoe on behalf of the Wellcome Sanger Institute COVID-19 Surveillance Team |
| EPI_ISL_3124233 | Berkshire and Surrey Pathology Services Lighthouse Laboratory | Wellcome Sanger Institute for the COVID-19 Genomics UK (COG-UK) Consortium | Berkshire and Surrey Pathology Services Lighthouse Laboratory and Alex Alderton, Roberto Amato, Jeffrey Barrett, Sonia Goncalves, Ewan Harrison, David K. Jackson, Ian Johnston, Dominic Kwiatkowski, Cordelia Langford, John Sillitoe on behalf of the Wellcome Sanger Institute COVID-19 Surveillance Team |
| EPI_ISL_3124234, EPI_ISL_3124235, EPI_ISL_3124236, EPI_ISL_3124237 | Lighthouse Laboratory Plymouth | Wellcome Sanger Institute for the COVID-19 Genomics UK (COG-UK) Consortium | Lighthouse Laboratory Plymouth and Alex Alderton, Roberto Amato, Jeffrey Barrett, Sonia Goncalves, Ewan Harrison, David K. Jackson, Ian Johnston, Dominic Kwiatkowski, Cordelia Langford, John Sillitoe on behalf of the Wellcome Sanger Institute COVID-19 Surveillance Team |
| EPI_ISL_3124238, EPI_ISL_3124239 | Berkshire and Surrey Pathology Services Lighthouse Laboratory | Wellcome Sanger Institute for the COVID-19 Genomics UK (COG-UK) Consortium | Berkshire and Surrey Pathology Services Lighthouse Laboratory and Alex Alderton, Roberto Amato, Jeffrey Barrett, Sonia Goncalves, Ewan Harrison, David K. Jackson, Ian Johnston, Dominic Kwiatkowski, Cordelia Langford, John Sillitoe on behalf of the Wellcome Sanger Institute COVID-19 Surveillance Team |
| EPI_ISL_3124240, EPI_ISL_3124241 | Lighthouse Laboratory Plymouth | Wellcome Sanger Institute for the COVID-19 Genomics UK (COG-UK) Consortium | Lighthouse Laboratory Plymouth and Alex Alderton, Roberto Amato, Jeffrey Barrett, Sonia Goncalves, Ewan Harrison, David K. Jackson, Ian Johnston, Dominic Kwiatkowski, Cordelia Langford, John Sillitoe on behalf of the Wellcome Sanger Institute COVID-19 Surveillance Team |
| EPI_ISL_3124242 | Berkshire and Surrey Pathology Services Lighthouse Laboratory | Wellcome Sanger Institute for the COVID-19 Genomics UK (COG-UK) Consortium | Berkshire and Surrey Pathology Services Lighthouse Laboratory and Alex Alderton, Roberto Amato, Jeffrey Barrett, Sonia Goncalves, Ewan Harrison, David K. Jackson, Ian Johnston, Dominic Kwiatkowski, Cordelia Langford, John Sillitoe on behalf of the Wellcome Sanger Institute COVID-19 Surveillance Team |
| EPI_ISL_3124243, EPI_ISL_3124244, EPI_ISL_3124245, EPI_ISL_3124246, EPI_ISL_3124247 | Lighthouse Laboratory Plymouth | Wellcome Sanger Institute for the COVID-19 Genomics UK (COG-UK) Consortium | Lighthouse Laboratory Plymouth and Alex Alderton, Roberto Amato, Jeffrey Barrett, Sonia Goncalves, Ewan Harrison, David K. Jackson, Ian Johnston, Dominic Kwiatkowski, Cordelia Langford, John Sillitoe on behalf of the Wellcome Sanger Institute COVID-19 Surveillance Team |
| EPI_ISL_3124248 | Berkshire and Surrey Pathology Services Lighthouse Laboratory | Wellcome Sanger Institute for the COVID-19 Genomics UK (COG-UK) Consortium | Berkshire and Surrey Pathology Services Lighthouse Laboratory and Alex Alderton, Roberto Amato, Jeffrey Barrett, Sonia Goncalves, Ewan Harrison, David K. Jackson, Ian Johnston, Dominic Kwiatkowski, Cordelia Langford, John Sillitoe on behalf of the Wellcome Sanger Institute COVID-19 Surveillance Team |
| EPI_ISL_3124249 | Lighthouse Laboratory Plymouth | Wellcome Sanger Institute for the COVID-19 Genomics UK (COG-UK) Consortium | Lighthouse Laboratory Plymouth and Alex Alderton, Roberto Amato, Jeffrey Barrett, Sonia Goncalves, Ewan Harrison, David K. Jackson, Ian Johnston, Dominic Kwiatkowski, Cordelia Langford, John Sillitoe on behalf of the Wellcome Sanger Institute COVID-19 Surveillance Team |
| EPI_ISL_3124250, EPI_ISL_3124251 | Berkshire and Surrey Pathology Services Lighthouse Laboratory | Wellcome Sanger Institute for the COVID-19 Genomics UK (COG-UK) Consortium | Berkshire and Surrey Pathology Services Lighthouse Laboratory and Alex Alderton, Roberto Amato, Jeffrey Barrett, Sonia Goncalves, Ewan Harrison, David K. Jackson, Ian Johnston, Dominic Kwiatkowski, Cordelia Langford, John Sillitoe on behalf of the Wellcome Sanger Institute COVID-19 Surveillance Team |
| EPI_ISL_3124252, EPI_ISL_3124253, EPI_ISL_3124254, EPI_ISL_3124255, EPI_ISL_3124256, EPI_ISL_3124257, EPI_ISL_3124258, EPI_ISL_3124259, EPI_ISL_3124260 | Lighthouse Laboratory Plymouth | Wellcome Sanger Institute for the COVID-19 Genomics UK (COG-UK) Consortium | Lighthouse Laboratory Plymouth and Alex Alderton, Roberto Amato, Jeffrey Barrett, Sonia Goncalves, Ewan Harrison, David K. Jackson, Ian Johnston, Dominic Kwiatkowski, Cordelia Langford, John Sillitoe on behalf of the Wellcome Sanger Institute COVID-19 Surveillance Team |
| EPI_ISL_3124261 | Berkshire and Surrey Pathology Services Lighthouse Laboratory | Wellcome Sanger Institute for the COVID-19 Genomics UK (COG-UK) Consortium | Berkshire and Surrey Pathology Services Lighthouse Laboratory and Alex Alderton, Roberto Amato, Jeffrey Barrett, Sonia Goncalves, Ewan Harrison, David K. Jackson, Ian Johnston, Dominic Kwiatkowski, Cordelia Langford, John Sillitoe on behalf of the Wellcome Sanger Institute COVID-19 Surveillance Team |
| EPI_ISL_3124262 | Lighthouse Laboratory Plymouth | Wellcome Sanger Institute for the COVID-19 Genomics UK (COG-UK) Consortium | Lighthouse Laboratory Plymouth and Alex Alderton, Roberto Amato, Jeffrey Barrett, Sonia Goncalves, Ewan Harrison, David K. Jackson, Ian Johnston, Dominic Kwiatkowski, Cordelia Langford, John Sillitoe on behalf of the Wellcome Sanger Institute COVID-19 Surveillance Team |
| EPI_ISL_3124263 | Berkshire and Surrey Pathology Services Lighthouse Laboratory | Wellcome Sanger Institute for the COVID-19 Genomics UK (COG-UK) Consortium | Berkshire and Surrey Pathology Services Lighthouse Laboratory and Alex Alderton, Roberto Amato, Jeffrey Barrett, Sonia Goncalves, Ewan Harrison, David K. Jackson, Ian Johnston, Dominic Kwiatkowski, Cordelia Langford, John Sillitoe on behalf of the Wellcome Sanger Institute COVID-19 Surveillance Team |
| EPI_ISL_3124264, EPI_ISL_3124265, EPI_ISL_3124266, EPI_ISL_3124267, EPI_ISL_3124268, EPI_ISL_3124269, EPI_ISL_3124270, EPI_ISL_3124271 | Lighthouse Laboratory Plymouth | Wellcome Sanger Institute for the COVID-19 Genomics UK (COG-UK) Consortium | Lighthouse Laboratory Plymouth and Alex Alderton, Roberto Amato, Jeffrey Barrett, Sonia Goncalves, Ewan Harrison, David K. Jackson, Ian Johnston, Dominic Kwiatkowski, Cordelia Langford, John Sillitoe on behalf of the Wellcome Sanger Institute COVID-19 Surveillance Team |
| EPI_ISL_3124272 | Berkshire and Surrey Pathology Services Lighthouse Laboratory | Wellcome Sanger Institute for the COVID-19 Genomics UK (COG-UK) Consortium | Berkshire and Surrey Pathology Services Lighthouse Laboratory and Alex Alderton, Roberto Amato, Jeffrey Barrett, Sonia Goncalves, Ewan Harrison, David K. Jackson, Ian Johnston, Dominic Kwiatkowski, Cordelia Langford, John Sillitoe on behalf of the Wellcome Sanger Institute COVID-19 Surveillance Team |
| EPI_ISL_3124273, EPI_ISL_3124274, EPI_ISL_3124275, EPI_ISL_3124276, EPI_ISL_3124277, EPI_ISL_3124278, EPI_ISL_3124279, EPI_ISL_3124280, EPI_ISL_3124281, EPI_ISL_3124282, EPI_ISL_3124283, EPI_ISL_3124284, EPI_ISL_3124285 | | | |
| see above | Lighthouse Laboratory Plymouth | Wellcome Sanger Institute for the COVID-19 Genomics UK (COG-UK) Consortium | Lighthouse Laboratory Plymouth and Alex Alderton, Roberto Amato, Jeffrey Barrett, Sonia Goncalves, Ewan Harrison, David K. Jackson, Ian Johnston, Dominic Kwiatkowski, Cordelia Langford, John Sillitoe on behalf of the Wellcome Sanger Institute COVID-19 Surveillance Team |
| EPI_ISL_3124286 | Berkshire and Surrey Pathology Services Lighthouse Laboratory | Wellcome Sanger Institute for the COVID-19 Genomics UK (COG-UK) Consortium | Berkshire and Surrey Pathology Services Lighthouse Laboratory and Alex Alderton, Roberto Amato, Jeffrey Barrett, Sonia Goncalves, Ewan Harrison, David K. Jackson, Ian Johnston, Dominic Kwiatkowski, Cordelia Langford, John Sillitoe on behalf of the Wellcome Sanger Institute COVID-19 Surveillance Team |
| EPI_ISL_3124287, EPI_ISL_3124288, EPI_ISL_3124289, EPI_ISL_3124290, EPI_ISL_3124291, EPI_ISL_3124292, EPI_ISL_3124293, EPI_ISL_3124294, EPI_ISL_3124295 | Lighthouse Laboratory Plymouth | Wellcome Sanger Institute for the COVID-19 Genomics UK (COG-UK) Consortium | Lighthouse Laboratory Plymouth and Alex Alderton, Roberto Amato, Jeffrey Barrett, Sonia Goncalves, Ewan Harrison, David K. Jackson, Ian Johnston, Dominic Kwiatkowski, Cordelia Langford, John Sillitoe on behalf of the Wellcome Sanger Institute COVID-19 Surveillance Team |
| EPI_ISL_3124296 | Berkshire and Surrey Pathology Services Lighthouse Laboratory | Wellcome Sanger Institute for the COVID-19 Genomics UK (COG-UK) Consortium | Berkshire and Surrey Pathology Services Lighthouse Laboratory and Alex Alderton, Roberto Amato, Jeffrey Barrett, Sonia Goncalves, Ewan Harrison, David K. Jackson, Ian Johnston, Dominic Kwiatkowski, Cordelia Langford, John Sillitoe on behalf of the Wellcome Sanger Institute COVID-19 Surveillance Team |

| | | | |
|---|---|---|---|
| EPI_ISL_3124297 | Lighthouse Lab in Glasgow | Wellcome Sanger Institute for the COVID-19 Genomics UK (COG-UK) Consortium | Harper VanSteenhouse, Yumi Kasai, David Gray, Carol Clugston, Anna Dominiczak and Alex Alderton, Roberto Amato, Jeffrey Barrett, Sonia Goncalves, Ewan Harrison, David K. Jackson, Ian Johnston, Dominic Kwiatkowski, Cordelia Langford, John Sillitoe on behalf of the Wellcome Sanger Institute COVID-19 Surveillance Team |
| EPI_ISL_3124298, EPI_ISL_3124299 | Berkshire and Surrey Pathology Services Lighthouse Laboratory | Wellcome Sanger Institute for the COVID-19 Genomics UK (COG-UK) Consortium | Berkshire and Surrey Pathology Services Lighthouse Laboratory and Alex Alderton, Roberto Amato, Jeffrey Barrett, Sonia Goncalves, Ewan Harrison, David K. Jackson, Ian Johnston, Dominic Kwiatkowski, Cordelia Langford, John Sillitoe on behalf of the Wellcome Sanger Institute COVID-19 Surveillance Team |
| EPI_ISL_3124300 | Lighthouse Laboratory Plymouth | Wellcome Sanger Institute for the COVID-19 Genomics UK (COG-UK) Consortium | Lighthouse Laboratory Plymouth and Alex Alderton, Roberto Amato, Jeffrey Barrett, Sonia Goncalves, Ewan Harrison, David K. Jackson, Ian Johnston, Dominic Kwiatkowski, Cordelia Langford, John Sillitoe on behalf of the Wellcome Sanger Institute COVID-19 Surveillance Team |
| EPI_ISL_3124301 | Berkshire and Surrey Pathology Services Lighthouse Laboratory | Wellcome Sanger Institute for the COVID-19 Genomics UK (COG-UK) Consortium | Berkshire and Surrey Pathology Services Lighthouse Laboratory and Alex Alderton, Roberto Amato, Jeffrey Barrett, Sonia Goncalves, Ewan Harrison, David K. Jackson, Ian Johnston, Dominic Kwiatkowski, Cordelia Langford, John Sillitoe on behalf of the Wellcome Sanger Institute COVID-19 Surveillance Team |
| EPI_ISL_3124302 | Lighthouse Laboratory Plymouth | Wellcome Sanger Institute for the COVID-19 Genomics UK (COG-UK) Consortium | Lighthouse Laboratory Plymouth and Alex Alderton, Roberto Amato, Jeffrey Barrett, Sonia Goncalves, Ewan Harrison, David K. Jackson, Ian Johnston, Dominic Kwiatkowski, Cordelia Langford, John Sillitoe on behalf of the Wellcome Sanger Institute COVID-19 Surveillance Team |
| EPI_ISL_3124303 | Lighthouse Lab in Milton Keynes | Wellcome Sanger Institute for the COVID-19 Genomics UK (COG-UK) Consortium | The Lighthouse Lab in Milton Keynes and Alex Alderton, Roberto Amato, Jeffrey Barrett, Sonia Goncalves, Ewan Harrison, David K. Jackson, Ian Johnston, Dominic Kwiatkowski, Cordelia Langford, John Sillitoe on behalf of the Wellcome Sanger Institute COVID-19 Surveillance Team |
| EPI_ISL_3124304 | Berkshire and Surrey Pathology Services Lighthouse Laboratory | Wellcome Sanger Institute for the COVID-19 Genomics UK (COG-UK) Consortium | Berkshire and Surrey Pathology Services Lighthouse Laboratory and Alex Alderton, Roberto Amato, Jeffrey Barrett, Sonia Goncalves, Ewan Harrison, David K. Jackson, Ian Johnston, Dominic Kwiatkowski, Cordelia Langford, John Sillitoe on behalf of the Wellcome Sanger Institute COVID-19 Surveillance Team |
| EPI_ISL_3124305 | Lighthouse Lab in Glasgow | Wellcome Sanger Institute for the COVID-19 Genomics UK (COG-UK) Consortium | Harper VanSteenhouse, Yumi Kasai, David Gray, Carol Clugston, Anna Dominiczak and Alex Alderton, Roberto Amato, Jeffrey Barrett, Sonia Goncalves, Ewan Harrison, David K. Jackson, Ian Johnston, Dominic Kwiatkowski, Cordelia Langford, John Sillitoe on behalf of the Wellcome Sanger Institute COVID-19 Surveillance Team |
| EPI_ISL_3124306 | Lighthouse Lab in Milton Keynes | Wellcome Sanger Institute for the COVID-19 Genomics UK (COG-UK) Consortium | The Lighthouse Lab in Milton Keynes and Alex Alderton, Roberto Amato, Jeffrey Barrett, Sonia Goncalves, Ewan Harrison, David K. Jackson, Ian Johnston, Dominic Kwiatkowski, Cordelia Langford, John Sillitoe on behalf of the Wellcome Sanger Institute COVID-19 Surveillance Team |
| EPI_ISL_3124307 | Berkshire and Surrey Pathology Services Lighthouse Laboratory | Wellcome Sanger Institute for the COVID-19 Genomics UK (COG-UK) Consortium | Berkshire and Surrey Pathology Services Lighthouse Laboratory and Alex Alderton, Roberto Amato, Jeffrey Barrett, Sonia Goncalves, Ewan Harrison, David K. Jackson, Ian Johnston, Dominic Kwiatkowski, Cordelia Langford, John Sillitoe on behalf of the Wellcome Sanger Institute COVID-19 Surveillance Team |
| EPI_ISL_3124308 | Lighthouse Laboratory Plymouth | Wellcome Sanger Institute for the COVID-19 Genomics UK (COG-UK) Consortium | Lighthouse Laboratory Plymouth and Alex Alderton, Roberto Amato, Jeffrey Barrett, Sonia Goncalves, Ewan Harrison, David K. Jackson, Ian Johnston, Dominic Kwiatkowski, Cordelia Langford, John Sillitoe on behalf of the Wellcome Sanger Institute COVID-19 Surveillance Team |
| EPI_ISL_3124309, EPI_ISL_3124310 | Lighthouse Lab in Milton Keynes | Wellcome Sanger Institute for the COVID-19 Genomics UK (COG-UK) Consortium | The Lighthouse Lab in Milton Keynes and Alex Alderton, Roberto Amato, Jeffrey Barrett, Sonia Goncalves, Ewan Harrison, David K. Jackson, Ian Johnston, Dominic Kwiatkowski, Cordelia Langford, John Sillitoe on behalf of the Wellcome Sanger Institute COVID-19 Surveillance Team |
| EPI_ISL_3124311, EPI_ISL_3124312 | Berkshire and Surrey Pathology Services Lighthouse Laboratory | Wellcome Sanger Institute for the COVID-19 Genomics UK (COG-UK) Consortium | Berkshire and Surrey Pathology Services Lighthouse Laboratory and Alex Alderton, Roberto Amato, Jeffrey Barrett, Sonia Goncalves, Ewan Harrison, David K. Jackson, Ian Johnston, Dominic Kwiatkowski, Cordelia Langford, John Sillitoe on behalf of the Wellcome Sanger Institute COVID-19 Surveillance Team |
| EPI_ISL_3124313, EPI_ISL_3124314 | Lighthouse Lab in Milton Keynes | Wellcome Sanger Institute for the COVID-19 Genomics UK (COG-UK) Consortium | The Lighthouse Lab in Milton Keynes and Alex Alderton, Roberto Amato, Jeffrey Barrett, Sonia Goncalves, Ewan Harrison, David K. Jackson, Ian Johnston, Dominic Kwiatkowski, Cordelia Langford, John Sillitoe on behalf of the Wellcome Sanger Institute COVID-19 Surveillance Team |
| EPI_ISL_3124315 | Berkshire and Surrey Pathology Services Lighthouse Laboratory | Wellcome Sanger Institute for the COVID-19 Genomics UK (COG-UK) Consortium | Berkshire and Surrey Pathology Services Lighthouse Laboratory and Alex Alderton, Roberto Amato, Jeffrey Barrett, Sonia Goncalves, Ewan Harrison, David K. Jackson, Ian Johnston, Dominic Kwiatkowski, Cordelia Langford, John Sillitoe on behalf of the Wellcome Sanger Institute COVID-19 Surveillance Team |
| EPI_ISL_3124316, EPI_ISL_3124317 | Lighthouse Lab in Milton Keynes | Wellcome Sanger Institute for the COVID-19 Genomics UK (COG-UK) Consortium | The Lighthouse Lab in Milton Keynes and Alex Alderton, Roberto Amato, Jeffrey Barrett, Sonia Goncalves, Ewan Harrison, David K. Jackson, Ian Johnston, Dominic Kwiatkowski, Cordelia Langford, John Sillitoe on behalf of the Wellcome Sanger Institute COVID-19 Surveillance Team |
| EPI_ISL_3124318 | Lighthouse Lab in Glasgow | Wellcome Sanger Institute for the COVID-19 Genomics UK (COG-UK) Consortium | Harper VanSteenhouse, Yumi Kasai, David Gray, Carol Clugston, Anna Dominiczak and Alex Alderton, Roberto Amato, Jeffrey Barrett, Sonia Goncalves, Ewan Harrison, David K. Jackson, Ian Johnston, Dominic Kwiatkowski, Cordelia Langford, John Sillitoe on behalf of the Wellcome Sanger Institute COVID-19 Surveillance Team |
| EPI_ISL_3124319, EPI_ISL_3124320, EPI_ISL_3124321, EPI_ISL_3124322 | Lighthouse Lab in Milton Keynes | Wellcome Sanger Institute for the COVID-19 Genomics UK (COG-UK) Consortium | The Lighthouse Lab in Milton Keynes and Alex Alderton, Roberto Amato, Jeffrey Barrett, Sonia Goncalves, Ewan Harrison, David K. Jackson, Ian Johnston, Dominic Kwiatkowski, Cordelia Langford, John Sillitoe on behalf of the Wellcome Sanger Institute COVID-19 Surveillance Team |
| EPI_ISL_3124323 | Lighthouse Lab in Glasgow | Wellcome Sanger Institute for the COVID-19 Genomics UK (COG-UK) Consortium | Harper VanSteenhouse, Yumi Kasai, David Gray, Carol Clugston, Anna Dominiczak and Alex Alderton, Roberto Amato, Jeffrey Barrett, Sonia Goncalves, Ewan Harrison, David K. Jackson, Ian Johnston, Dominic Kwiatkowski, Cordelia Langford, John Sillitoe on behalf of the Wellcome Sanger Institute COVID-19 Surveillance Team |
| EPI_ISL_3124324 | Lighthouse Lab in Milton Keynes | Wellcome Sanger Institute for the COVID-19 Genomics UK (COG-UK) Consortium | The Lighthouse Lab in Milton Keynes and Alex Alderton, Roberto Amato, Jeffrey Barrett, Sonia Goncalves, Ewan Harrison, David K. Jackson, Ian Johnston, Dominic Kwiatkowski, Cordelia Langford, John Sillitoe on behalf of the Wellcome Sanger Institute COVID-19 Surveillance Team |
| EPI_ISL_3124325 | Lighthouse Lab in Glasgow | Wellcome Sanger Institute for the COVID-19 Genomics UK (COG-UK) Consortium | Harper VanSteenhouse, Yumi Kasai, David Gray, Carol Clugston, Anna Dominiczak and Alex Alderton, Roberto Amato, Jeffrey Barrett, Sonia Goncalves, Ewan Harrison, David K. Jackson, Ian Johnston, Dominic Kwiatkowski, Cordelia Langford, John Sillitoe on behalf of the Wellcome Sanger Institute COVID-19 Surveillance Team |
| EPI_ISL_3124326 | Lighthouse Lab in Milton Keynes | Wellcome Sanger Institute for the COVID-19 Genomics UK (COG-UK) Consortium | The Lighthouse Lab in Milton Keynes and Alex Alderton, Roberto Amato, Jeffrey Barrett, Sonia Goncalves, Ewan Harrison, David K. Jackson, Ian Johnston, Dominic Kwiatkowski, Cordelia Langford, John Sillitoe on behalf of the Wellcome Sanger Institute COVID-19 Surveillance Team |
| EPI_ISL_3124327 | Lighthouse Lab in Glasgow | Wellcome Sanger Institute for the COVID-19 Genomics UK (COG-UK) Consortium | Harper VanSteenhouse, Yumi Kasai, David Gray, Carol Clugston, Anna Dominiczak and Alex Alderton, Roberto Amato, Jeffrey Barrett, Sonia Goncalves, Ewan Harrison, David K. Jackson, Ian Johnston, Dominic Kwiatkowski, Cordelia Langford, John Sillitoe on behalf of the Wellcome Sanger Institute COVID-19 Surveillance Team |
| EPI_ISL_3124328, EPI_ISL_3124329, EPI_ISL_3124330 | Berkshire and Surrey Pathology Services Lighthouse Laboratory | Wellcome Sanger Institute for the COVID-19 Genomics UK (COG-UK) Consortium | Berkshire and Surrey Pathology Services Lighthouse Laboratory and Alex Alderton, Roberto Amato, Jeffrey Barrett, Sonia Goncalves, Ewan Harrison, David K. Jackson, Ian Johnston, Dominic Kwiatkowski, Cordelia Langford, John Sillitoe on behalf of the Wellcome Sanger Institute COVID-19 Surveillance Team |
| EPI_ISL_3124331 | Lighthouse Lab in Milton Keynes | Wellcome Sanger Institute for the COVID-19 Genomics UK (COG-UK) Consortium | The Lighthouse Lab in Milton Keynes and Alex Alderton, Roberto Amato, Jeffrey Barrett, Sonia Goncalves, Ewan Harrison, David K. Jackson, Ian Johnston, Dominic Kwiatkowski, Cordelia Langford, John Sillitoe on behalf of the Wellcome Sanger Institute COVID-19 Surveillance Team |
| EPI_ISL_3124332, EPI_ISL_3124333 | Lighthouse Laboratory Plymouth | Wellcome Sanger Institute for the COVID-19 Genomics UK (COG-UK) Consortium | Lighthouse Laboratory Plymouth and Alex Alderton, Roberto Amato, Jeffrey Barrett, Sonia Goncalves, Ewan Harrison, David K. Jackson, Ian Johnston, Dominic Kwiatkowski, Cordelia Langford, John Sillitoe on behalf of the Wellcome Sanger Institute COVID-19 Surveillance Team |
| EPI_ISL_3124334 | Berkshire and Surrey Pathology Services Lighthouse Laboratory | Wellcome Sanger Institute for the COVID-19 Genomics UK (COG-UK) Consortium | Berkshire and Surrey Pathology Services Lighthouse Laboratory and Alex Alderton, Roberto Amato, Jeffrey Barrett, Sonia Goncalves, Ewan Harrison, David K. Jackson, Ian Johnston, Dominic Kwiatkowski, Cordelia Langford, John Sillitoe on behalf of the Wellcome Sanger Institute COVID-19 Surveillance Team |
| EPI_ISL_3124335 | Lighthouse Lab in Glasgow | Wellcome Sanger Institute for the COVID-19 Genomics UK (COG-UK) Consortium | Harper VanSteenhouse, Yumi Kasai, David Gray, Carol Clugston, Anna Dominiczak and Alex Alderton, Roberto Amato, Jeffrey Barrett, Sonia Goncalves, Ewan Harrison, David K. Jackson, Ian Johnston, Dominic Kwiatkowski, Cordelia Langford, John Sillitoe on behalf of the Wellcome Sanger Institute COVID-19 Surveillance Team |
| EPI_ISL_3124336, EPI_ISL_3124337, EPI_ISL_3124338 | Lighthouse Laboratory Plymouth | Wellcome Sanger Institute for the COVID-19 Genomics UK (COG-UK) Consortium | Lighthouse Laboratory Plymouth and Alex Alderton, Roberto Amato, Jeffrey Barrett, Sonia Goncalves, Ewan Harrison, David K. Jackson, Ian Johnston, Dominic Kwiatkowski, Cordelia Langford, John Sillitoe on behalf of the Wellcome Sanger Institute COVID-19 Surveillance Team |
| EPI_ISL_3124339 | Berkshire and Surrey Pathology Services Lighthouse Laboratory | Wellcome Sanger Institute for the COVID-19 Genomics UK (COG-UK) Consortium | Berkshire and Surrey Pathology Services Lighthouse Laboratory and Alex Alderton, Roberto Amato, Jeffrey Barrett, Sonia Goncalves, Ewan Harrison, David K. Jackson, Ian Johnston, Dominic Kwiatkowski, Cordelia Langford, John Sillitoe on behalf of the Wellcome Sanger Institute COVID-19 Surveillance Team |
| EPI_ISL_3124340, EPI_ISL_3124341 | Lighthouse Lab in Glasgow | Wellcome Sanger Institute for the COVID-19 Genomics UK (COG-UK) Consortium | Harper VanSteenhouse, Yumi Kasai, David Gray, Carol Clugston, Anna Dominiczak and Alex Alderton, Roberto Amato, Jeffrey Barrett, Sonia Goncalves, Ewan Harrison, David K. Jackson, Ian Johnston, Dominic Kwiatkowski, Cordelia Langford, John Sillitoe on behalf of the Wellcome Sanger Institute COVID-19 Surveillance Team |
| EPI_ISL_3124342, EPI_ISL_3124343 | Lighthouse Lab in Milton Keynes | Wellcome Sanger Institute for the COVID-19 Genomics UK (COG-UK) Consortium | The Lighthouse Lab in Milton Keynes and Alex Alderton, Roberto Amato, Jeffrey Barrett, Sonia Goncalves, Ewan Harrison, David K. Jackson, Ian Johnston, Dominic Kwiatkowski, Cordelia Langford, John Sillitoe on behalf of the Wellcome Sanger Institute COVID-19 Surveillance Team |
| EPI_ISL_3124344 | Lighthouse Laboratory Plymouth | Wellcome Sanger Institute for the COVID-19 Genomics UK (COG-UK) Consortium | Lighthouse Laboratory Plymouth and Alex Alderton, Roberto Amato, Jeffrey Barrett, Sonia Goncalves, Ewan Harrison, David K. Jackson, Ian Johnston, Dominic Kwiatkowski, Cordelia Langford, John Sillitoe on behalf of the Wellcome Sanger Institute COVID-19 Surveillance Team |
| EPI_ISL_3124345, EPI_ISL_3124346 | Lighthouse Lab in Glasgow | Wellcome Sanger Institute for the COVID-19 Genomics UK (COG-UK) Consortium | Harper VanSteenhouse, Yumi Kasai, David Gray, Carol Clugston, Anna Dominiczak and Alex Alderton, Roberto Amato, Jeffrey Barrett, Sonia Goncalves, Ewan Harrison, David K. Jackson, Ian Johnston, Dominic Kwiatkowski, Cordelia Langford, John Sillitoe on behalf of the Wellcome Sanger Institute COVID-19 Surveillance Team |
| EPI_ISL_3124347 | Lighthouse Lab in Milton Keynes | Wellcome Sanger Institute for the COVID-19 Genomics UK (COG-UK) Consortium | The Lighthouse Lab in Milton Keynes and Alex Alderton, Roberto Amato, Jeffrey Barrett, Sonia Goncalves, Ewan Harrison, David K. Jackson, Ian Johnston, Dominic Kwiatkowski, Cordelia Langford, John Sillitoe on behalf of the Wellcome Sanger Institute COVID-19 Surveillance Team |
| EPI_ISL_3124348 | Lighthouse Lab in Glasgow | Wellcome Sanger Institute for the COVID-19 Genomics UK (COG-UK) Consortium | Harper VanSteenhouse, Yumi Kasai, David Gray, Carol Clugston, Anna Dominiczak and Alex Alderton, Roberto Amato, Jeffrey Barrett, Sonia Goncalves, Ewan Harrison, David K. Jackson, Ian Johnston, Dominic Kwiatkowski, Cordelia Langford, John Sillitoe on behalf of the Wellcome Sanger Institute COVID-19 Surveillance Team |
| EPI_ISL_3124349 | Lighthouse Lab in Milton Keynes | Wellcome Sanger Institute for the COVID-19 Genomics UK (COG-UK) Consortium | The Lighthouse Lab in Milton Keynes and Alex Alderton, Roberto Amato, Jeffrey Barrett, Sonia Goncalves, Ewan Harrison, David K. Jackson, Ian Johnston, Dominic Kwiatkowski, Cordelia Langford, John Sillitoe on behalf of the Wellcome Sanger Institute COVID-19 Surveillance Team |
| EPI_ISL_3124350 | Lighthouse Laboratory Plymouth | Wellcome Sanger Institute for the COVID-19 Genomics UK (COG-UK) Consortium | Lighthouse Laboratory Plymouth and Alex Alderton, Roberto Amato, Jeffrey Barrett, Sonia Goncalves, Ewan Harrison, David K. Jackson, Ian Johnston, Dominic Kwiatkowski, Cordelia Langford, John Sillitoe on behalf of the Wellcome Sanger Institute COVID-19 Surveillance Team |
| EPI_ISL_3124351, EPI_ISL_3124352, EPI_ISL_3124353 | Berkshire and Surrey Pathology Services Lighthouse Laboratory | Wellcome Sanger Institute for the COVID-19 Genomics UK (COG-UK) Consortium | Berkshire and Surrey Pathology Services Lighthouse Laboratory and Alex Alderton, Roberto Amato, Jeffrey Barrett, Sonia Goncalves, Ewan Harrison, David K. Jackson, Ian Johnston, Dominic Kwiatkowski, Cordelia Langford, John Sillitoe on behalf of the Wellcome Sanger Institute COVID-19 Surveillance Team |
| EPI_ISL_3124354, EPI_ISL_3124355 | Lighthouse Lab in Milton Keynes | Wellcome Sanger Institute for the COVID-19 Genomics UK (COG-UK) Consortium | The Lighthouse Lab in Milton Keynes and Alex Alderton, Roberto Amato, Jeffrey Barrett, Sonia Goncalves, Ewan Harrison, David K. Jackson, Ian Johnston, Dominic Kwiatkowski, Cordelia Langford, John Sillitoe on behalf of the Wellcome Sanger Institute COVID-19 Surveillance Team |
| EPI_ISL_3124356 | Lighthouse Laboratory Plymouth | Wellcome Sanger Institute for the COVID-19 Genomics UK (COG-UK) Consortium | Lighthouse Laboratory Plymouth and Alex Alderton, Roberto Amato, Jeffrey Barrett, Sonia Goncalves, Ewan Harrison, David K. Jackson, Ian Johnston, Dominic Kwiatkowski, Cordelia Langford, John Sillitoe on behalf of the Wellcome Sanger Institute COVID-19 Surveillance Team |
| EPI_ISL_3124357 | Berkshire and Surrey Pathology Services Lighthouse Laboratory | Wellcome Sanger Institute for the COVID-19 Genomics UK (COG-UK) Consortium | Berkshire and Surrey Pathology Services Lighthouse Laboratory and Alex Alderton, Roberto Amato, Jeffrey Barrett, Sonia Goncalves, Ewan Harrison, David K. Jackson, Ian Johnston, Dominic Kwiatkowski, Cordelia Langford, John Sillitoe on behalf of the Wellcome Sanger Institute COVID-19 Surveillance Team |
| EPI_ISL_3124358 | Lighthouse Lab in Milton Keynes | Wellcome Sanger Institute for the COVID-19 Genomics UK (COG-UK) Consortium | The Lighthouse Lab in Milton Keynes and Alex Alderton, Roberto Amato, Jeffrey Barrett, Sonia Goncalves, Ewan Harrison, David K. Jackson, Ian Johnston, Dominic Kwiatkowski, Cordelia Langford, John Sillitoe on behalf of the Wellcome Sanger Institute COVID-19 Surveillance Team |

| | | | |
|---|---|---|---|
| EPI_ISL_3124359, EPI_ISL_3124360 | Lighthouse Laboratory Plymouth | Wellcome Sanger Institute for the COVID-19 Genomics UK (COG-UK) Consortium | Lighthouse Laboratory Plymouth and Alex Alderton, Roberto Amato, Jeffrey Barrett, Sonia Goncalves, Ewan Harrison, David K. Jackson, Ian Johnston, Dominic Kwiatkowski, Cordelia Langford, John Sillitoe on behalf of the Wellcome Sanger Institute COVID-19 Surveillance Team |
| EPI_ISL_3124361 | Lighthouse Lab in Glasgow | Wellcome Sanger Institute for the COVID-19 Genomics UK (COG-UK) Consortium | Harper VanSteenhouse, Yumi Kasai, David Gray, Carol Clugston, Anna Dominiczak and Alex Alderton, Roberto Amato, Jeffrey Barrett, Sonia Goncalves, Ewan Harrison, David K. Jackson, Ian Johnston, Dominic Kwiatkowski, Cordelia Langford, John Sillitoe on behalf of the Wellcome Sanger Institute COVID-19 Surveillance Team |
| EPI_ISL_3124362 | Lighthouse Laboratory Plymouth | Wellcome Sanger Institute for the COVID-19 Genomics UK (COG-UK) Consortium | Lighthouse Laboratory Plymouth and Alex Alderton, Roberto Amato, Jeffrey Barrett, Sonia Goncalves, Ewan Harrison, David K. Jackson, Ian Johnston, Dominic Kwiatkowski, Cordelia Langford, John Sillitoe on behalf of the Wellcome Sanger Institute COVID-19 Surveillance Team |
| EPI_ISL_3124363 | Lighthouse Lab in Milton Keynes | Wellcome Sanger Institute for the COVID-19 Genomics UK (COG-UK) Consortium | The Lighthouse Lab in Milton Keynes and Alex Alderton, Roberto Amato, Jeffrey Barrett, Sonia Goncalves, Ewan Harrison, David K. Jackson, Ian Johnston, Dominic Kwiatkowski, Cordelia Langford, John Sillitoe on behalf of the Wellcome Sanger Institute COVID-19 Surveillance Team |
| EPI_ISL_3124364 | Lighthouse Lab in Glasgow | Wellcome Sanger Institute for the COVID-19 Genomics UK (COG-UK) Consortium | Harper VanSteenhouse, Yumi Kasai, David Gray, Carol Clugston, Anna Dominiczak and Alex Alderton, Roberto Amato, Jeffrey Barrett, Sonia Goncalves, Ewan Harrison, David K. Jackson, Ian Johnston, Dominic Kwiatkowski, Cordelia Langford, John Sillitoe on behalf of the Wellcome Sanger Institute COVID-19 Surveillance Team |
| EPI_ISL_3124365 | Lighthouse Lab in Milton Keynes | Wellcome Sanger Institute for the COVID-19 Genomics UK (COG-UK) Consortium | The Lighthouse Lab in Milton Keynes and Alex Alderton, Roberto Amato, Jeffrey Barrett, Sonia Goncalves, Ewan Harrison, David K. Jackson, Ian Johnston, Dominic Kwiatkowski, Cordelia Langford, John Sillitoe on behalf of the Wellcome Sanger Institute COVID-19 Surveillance Team |
| EPI_ISL_3124366 | Lighthouse Lab in Glasgow | Wellcome Sanger Institute for the COVID-19 Genomics UK (COG-UK) Consortium | Harper VanSteenhouse, Yumi Kasai, David Gray, Carol Clugston, Anna Dominiczak and Alex Alderton, Roberto Amato, Jeffrey Barrett, Sonia Goncalves, Ewan Harrison, David K. Jackson, Ian Johnston, Dominic Kwiatkowski, Cordelia Langford, John Sillitoe on behalf of the Wellcome Sanger Institute COVID-19 Surveillance Team |
| EPI_ISL_3124367 | Berkshire and Surrey Pathology Services Lighthouse Laboratory | Wellcome Sanger Institute for the COVID-19 Genomics UK (COG-UK) Consortium | Berkshire and Surrey Pathology Services Lighthouse Laboratory and Alex Alderton, Roberto Amato, Jeffrey Barrett, Sonia Goncalves, Ewan Harrison, David K. Jackson, Ian Johnston, Dominic Kwiatkowski, Cordelia Langford, John Sillitoe on behalf of the Wellcome Sanger Institute COVID-19 Surveillance Team |
| EPI_ISL_3124368, EPI_ISL_3124369 | Lighthouse Laboratory Plymouth | Wellcome Sanger Institute for the COVID-19 Genomics UK (COG-UK) Consortium | Lighthouse Laboratory Plymouth and Alex Alderton, Roberto Amato, Jeffrey Barrett, Sonia Goncalves, Ewan Harrison, David K. Jackson, Ian Johnston, Dominic Kwiatkowski, Cordelia Langford, John Sillitoe on behalf of the Wellcome Sanger Institute COVID-19 Surveillance Team |
| EPI_ISL_3124370, EPI_ISL_3124371 | Berkshire and Surrey Pathology Services Lighthouse Laboratory | Wellcome Sanger Institute for the COVID-19 Genomics UK (COG-UK) Consortium | Berkshire and Surrey Pathology Services Lighthouse Laboratory and Alex Alderton, Roberto Amato, Jeffrey Barrett, Sonia Goncalves, Ewan Harrison, David K. Jackson, Ian Johnston, Dominic Kwiatkowski, Cordelia Langford, John Sillitoe on behalf of the Wellcome Sanger Institute COVID-19 Surveillance Team |
| EPI_ISL_3124372 | Lighthouse Lab in Glasgow | Wellcome Sanger Institute for the COVID-19 Genomics UK (COG-UK) Consortium | Harper VanSteenhouse, Yumi Kasai, David Gray, Carol Clugston, Anna Dominiczak and Alex Alderton, Roberto Amato, Jeffrey Barrett, Sonia Goncalves, Ewan Harrison, David K. Jackson, Ian Johnston, Dominic Kwiatkowski, Cordelia Langford, John Sillitoe on behalf of the Wellcome Sanger Institute COVID-19 Surveillance Team |
| EPI_ISL_3124373 | Berkshire and Surrey Pathology Services Lighthouse Laboratory | Wellcome Sanger Institute for the COVID-19 Genomics UK (COG-UK) Consortium | Berkshire and Surrey Pathology Services Lighthouse Laboratory and Alex Alderton, Roberto Amato, Jeffrey Barrett, Sonia Goncalves, Ewan Harrison, David K. Jackson, Ian Johnston, Dominic Kwiatkowski, Cordelia Langford, John Sillitoe on behalf of the Wellcome Sanger Institute COVID-19 Surveillance Team |
| EPI_ISL_3124374 | Lighthouse Lab in Glasgow | Wellcome Sanger Institute for the COVID-19 Genomics UK (COG-UK) Consortium | Harper VanSteenhouse, Yumi Kasai, David Gray, Carol Clugston, Anna Dominiczak and Alex Alderton, Roberto Amato, Jeffrey Barrett, Sonia Goncalves, Ewan Harrison, David K. Jackson, Ian Johnston, Dominic Kwiatkowski, Cordelia Langford, John Sillitoe on behalf of the Wellcome Sanger Institute COVID-19 Surveillance Team |
| EPI_ISL_3124375, EPI_ISL_3124376, EPI_ISL_3124377 | Lighthouse Laboratory Plymouth | Wellcome Sanger Institute for the COVID-19 Genomics UK (COG-UK) Consortium | Lighthouse Laboratory Plymouth and Alex Alderton, Roberto Amato, Jeffrey Barrett, Sonia Goncalves, Ewan Harrison, David K. Jackson, Ian Johnston, Dominic Kwiatkowski, Cordelia Langford, John Sillitoe on behalf of the Wellcome Sanger Institute COVID-19 Surveillance Team |
| EPI_ISL_3124378 | Lighthouse Lab in Milton Keynes | Wellcome Sanger Institute for the COVID-19 Genomics UK (COG-UK) Consortium | The Lighthouse Lab in Milton Keynes and Alex Alderton, Roberto Amato, Jeffrey Barrett, Sonia Goncalves, Ewan Harrison, David K. Jackson, Ian Johnston, Dominic Kwiatkowski, Cordelia Langford, John Sillitoe on behalf of the Wellcome Sanger Institute COVID-19 Surveillance Team |
| EPI_ISL_3124379 | Lighthouse Lab in Glasgow | Wellcome Sanger Institute for the COVID-19 Genomics UK (COG-UK) Consortium | Harper VanSteenhouse, Yumi Kasai, David Gray, Carol Clugston, Anna Dominiczak and Alex Alderton, Roberto Amato, Jeffrey Barrett, Sonia Goncalves, Ewan Harrison, David K. Jackson, Ian Johnston, Dominic Kwiatkowski, Cordelia Langford, John Sillitoe on behalf of the Wellcome Sanger Institute COVID-19 Surveillance Team |
| EPI_ISL_3124380 | Berkshire and Surrey Pathology Services Lighthouse Laboratory | Wellcome Sanger Institute for the COVID-19 Genomics UK (COG-UK) Consortium | Berkshire and Surrey Pathology Services Lighthouse Laboratory and Alex Alderton, Roberto Amato, Jeffrey Barrett, Sonia Goncalves, Ewan Harrison, David K. Jackson, Ian Johnston, Dominic Kwiatkowski, Cordelia Langford, John Sillitoe on behalf of the Wellcome Sanger Institute COVID-19 Surveillance Team |
| EPI_ISL_3124381, EPI_ISL_3124382, EPI_ISL_3124383, EPI_ISL_3124384 | Lighthouse Laboratory Plymouth | Wellcome Sanger Institute for the COVID-19 Genomics UK (COG-UK) Consortium | Lighthouse Laboratory Plymouth and Alex Alderton, Roberto Amato, Jeffrey Barrett, Sonia Goncalves, Ewan Harrison, David K. Jackson, Ian Johnston, Dominic Kwiatkowski, Cordelia Langford, John Sillitoe on behalf of the Wellcome Sanger Institute COVID-19 Surveillance Team |
| EPI_ISL_3124385 | Berkshire and Surrey Pathology Services Lighthouse Laboratory | Wellcome Sanger Institute for the COVID-19 Genomics UK (COG-UK) Consortium | Berkshire and Surrey Pathology Services Lighthouse Laboratory and Alex Alderton, Roberto Amato, Jeffrey Barrett, Sonia Goncalves, Ewan Harrison, David K. Jackson, Ian Johnston, Dominic Kwiatkowski, Cordelia Langford, John Sillitoe on behalf of the Wellcome Sanger Institute COVID-19 Surveillance Team |
| EPI_ISL_3124386, EPI_ISL_3124387, EPI_ISL_3124388 | Lighthouse Laboratory Plymouth | Wellcome Sanger Institute for the COVID-19 Genomics UK (COG-UK) Consortium | Lighthouse Laboratory Plymouth and Alex Alderton, Roberto Amato, Jeffrey Barrett, Sonia Goncalves, Ewan Harrison, David K. Jackson, Ian Johnston, Dominic Kwiatkowski, Cordelia Langford, John Sillitoe on behalf of the Wellcome Sanger Institute COVID-19 Surveillance Team |
| EPI_ISL_3124389 | Lighthouse Lab in Glasgow | Wellcome Sanger Institute for the COVID-19 Genomics UK (COG-UK) Consortium | Harper VanSteenhouse, Yumi Kasai, David Gray, Carol Clugston, Anna Dominiczak and Alex Alderton, Roberto Amato, Jeffrey Barrett, Sonia Goncalves, Ewan Harrison, David K. Jackson, Ian Johnston, Dominic Kwiatkowski, Cordelia Langford, John Sillitoe on behalf of the Wellcome Sanger Institute COVID-19 Surveillance Team |
| EPI_ISL_3124390 | Lighthouse Laboratory Plymouth | Wellcome Sanger Institute for the COVID-19 Genomics UK (COG-UK) Consortium | Lighthouse Laboratory Plymouth and Alex Alderton, Roberto Amato, Jeffrey Barrett, Sonia Goncalves, Ewan Harrison, David K. Jackson, Ian Johnston, Dominic Kwiatkowski, Cordelia Langford, John Sillitoe on behalf of the Wellcome Sanger Institute COVID-19 Surveillance Team |
| EPI_ISL_3124391, EPI_ISL_3124392 | Lighthouse Lab in Glasgow | Wellcome Sanger Institute for the COVID-19 Genomics UK (COG-UK) Consortium | Harper VanSteenhouse, Yumi Kasai, David Gray, Carol Clugston, Anna Dominiczak and Alex Alderton, Roberto Amato, Jeffrey Barrett, Sonia Goncalves, Ewan Harrison, David K. Jackson, Ian Johnston, Dominic Kwiatkowski, Cordelia Langford, John Sillitoe on behalf of the Wellcome Sanger Institute COVID-19 Surveillance Team |
| EPI_ISL_3124393, EPI_ISL_3124394 | Lighthouse Lab in Milton Keynes | Wellcome Sanger Institute for the COVID-19 Genomics UK (COG-UK) Consortium | The Lighthouse Lab in Milton Keynes and Alex Alderton, Roberto Amato, Jeffrey Barrett, Sonia Goncalves, Ewan Harrison, David K. Jackson, Ian Johnston, Dominic Kwiatkowski, Cordelia Langford, John Sillitoe on behalf of the Wellcome Sanger Institute COVID-19 Surveillance Team |
| EPI_ISL_3124395, EPI_ISL_3124396 | Berkshire and Surrey Pathology Services Lighthouse Laboratory | Wellcome Sanger Institute for the COVID-19 Genomics UK (COG-UK) Consortium | Berkshire and Surrey Pathology Services Lighthouse Laboratory and Alex Alderton, Roberto Amato, Jeffrey Barrett, Sonia Goncalves, Ewan Harrison, David K. Jackson, Ian Johnston, Dominic Kwiatkowski, Cordelia Langford, John Sillitoe on behalf of the Wellcome Sanger Institute COVID-19 Surveillance Team |
| EPI_ISL_3124397 | Lighthouse Laboratory Plymouth | Wellcome Sanger Institute for the COVID-19 Genomics UK (COG-UK) Consortium | Lighthouse Laboratory Plymouth and Alex Alderton, Roberto Amato, Jeffrey Barrett, Sonia Goncalves, Ewan Harrison, David K. Jackson, Ian Johnston, Dominic Kwiatkowski, Cordelia Langford, John Sillitoe on behalf of the Wellcome Sanger Institute COVID-19 Surveillance Team |
| EPI_ISL_3124398 | Lighthouse Lab in Milton Keynes | Wellcome Sanger Institute for the COVID-19 Genomics UK (COG-UK) Consortium | The Lighthouse Lab in Milton Keynes and Alex Alderton, Roberto Amato, Jeffrey Barrett, Sonia Goncalves, Ewan Harrison, David K. Jackson, Ian Johnston, Dominic Kwiatkowski, Cordelia Langford, John Sillitoe on behalf of the Wellcome Sanger Institute COVID-19 Surveillance Team |
| EPI_ISL_3124399 | Lighthouse Lab in Glasgow | Wellcome Sanger Institute for the COVID-19 Genomics UK (COG-UK) Consortium | Harper VanSteenhouse, Yumi Kasai, David Gray, Carol Clugston, Anna Dominiczak and Alex Alderton, Roberto Amato, Jeffrey Barrett, Sonia Goncalves, Ewan Harrison, David K. Jackson, Ian Johnston, Dominic Kwiatkowski, Cordelia Langford, John Sillitoe on behalf of the Wellcome Sanger Institute COVID-19 Surveillance Team |
| EPI_ISL_3124400 | Lighthouse Lab in Milton Keynes | Wellcome Sanger Institute for the COVID-19 Genomics UK (COG-UK) Consortium | The Lighthouse Lab in Milton Keynes and Alex Alderton, Roberto Amato, Jeffrey Barrett, Sonia Goncalves, Ewan Harrison, David K. Jackson, Ian Johnston, Dominic Kwiatkowski, Cordelia Langford, John Sillitoe on behalf of the Wellcome Sanger Institute COVID-19 Surveillance Team |
| EPI_ISL_3124401, EPI_ISL_3124402 | Lighthouse Laboratory Plymouth | Wellcome Sanger Institute for the COVID-19 Genomics UK (COG-UK) Consortium | Lighthouse Laboratory Plymouth and Alex Alderton, Roberto Amato, Jeffrey Barrett, Sonia Goncalves, Ewan Harrison, David K. Jackson, Ian Johnston, Dominic Kwiatkowski, Cordelia Langford, John Sillitoe on behalf of the Wellcome Sanger Institute COVID-19 Surveillance Team |
| EPI_ISL_3124403 | Berkshire and Surrey Pathology Services Lighthouse Laboratory | Wellcome Sanger Institute for the COVID-19 Genomics UK (COG-UK) Consortium | Berkshire and Surrey Pathology Services Lighthouse Laboratory and Alex Alderton, Roberto Amato, Jeffrey Barrett, Sonia Goncalves, Ewan Harrison, David K. Jackson, Ian Johnston, Dominic Kwiatkowski, Cordelia Langford, John Sillitoe on behalf of the Wellcome Sanger Institute COVID-19 Surveillance Team |
| EPI_ISL_3124404, EPI_ISL_3124405 | Lighthouse Laboratory Plymouth | Wellcome Sanger Institute for the COVID-19 Genomics UK (COG-UK) Consortium | Lighthouse Laboratory Plymouth and Alex Alderton, Roberto Amato, Jeffrey Barrett, Sonia Goncalves, Ewan Harrison, David K. Jackson, Ian Johnston, Dominic Kwiatkowski, Cordelia Langford, John Sillitoe on behalf of the Wellcome Sanger Institute COVID-19 Surveillance Team |
| EPI_ISL_3124406, EPI_ISL_3124407 | Lighthouse Lab in Glasgow | Wellcome Sanger Institute for the COVID-19 Genomics UK (COG-UK) Consortium | Harper VanSteenhouse, Yumi Kasai, David Gray, Carol Clugston, Anna Dominiczak and Alex Alderton, Roberto Amato, Jeffrey Barrett, Sonia Goncalves, Ewan Harrison, David K. Jackson, Ian Johnston, Dominic Kwiatkowski, Cordelia Langford, John Sillitoe on behalf of the Wellcome Sanger Institute COVID-19 Surveillance Team |
| EPI_ISL_3124408 | Lighthouse Lab in Milton Keynes | Wellcome Sanger Institute for the COVID-19 Genomics UK (COG-UK) Consortium | The Lighthouse Lab in Milton Keynes and Alex Alderton, Roberto Amato, Jeffrey Barrett, Sonia Goncalves, Ewan Harrison, David K. Jackson, Ian Johnston, Dominic Kwiatkowski, Cordelia Langford, John Sillitoe on behalf of the Wellcome Sanger Institute COVID-19 Surveillance Team |
| EPI_ISL_3124409 | Lighthouse Lab in Glasgow | Wellcome Sanger Institute for the COVID-19 Genomics UK (COG-UK) Consortium | Harper VanSteenhouse, Yumi Kasai, David Gray, Carol Clugston, Anna Dominiczak and Alex Alderton, Roberto Amato, Jeffrey Barrett, Sonia Goncalves, Ewan Harrison, David K. Jackson, Ian Johnston, Dominic Kwiatkowski, Cordelia Langford, John Sillitoe on behalf of the Wellcome Sanger Institute COVID-19 Surveillance Team |
| EPI_ISL_3124410 | Lighthouse Laboratory Plymouth | Wellcome Sanger Institute for the COVID-19 Genomics UK (COG-UK) Consortium | Lighthouse Laboratory Plymouth and Alex Alderton, Roberto Amato, Jeffrey Barrett, Sonia Goncalves, Ewan Harrison, David K. Jackson, Ian Johnston, Dominic Kwiatkowski, Cordelia Langford, John Sillitoe on behalf of the Wellcome Sanger Institute COVID-19 Surveillance Team |
| EPI_ISL_3124411 | Lighthouse Lab in Glasgow | Wellcome Sanger Institute for the COVID-19 Genomics UK (COG-UK) Consortium | Harper VanSteenhouse, Yumi Kasai, David Gray, Carol Clugston, Anna Dominiczak and Alex Alderton, Roberto Amato, Jeffrey Barrett, Sonia Goncalves, Ewan Harrison, David K. Jackson, Ian Johnston, Dominic Kwiatkowski, Cordelia Langford, John Sillitoe on behalf of the Wellcome Sanger Institute COVID-19 Surveillance Team |
| EPI_ISL_3124412 | Lighthouse Laboratory Plymouth | Wellcome Sanger Institute for the COVID-19 Genomics UK (COG-UK) Consortium | Lighthouse Laboratory Plymouth and Alex Alderton, Roberto Amato, Jeffrey Barrett, Sonia Goncalves, Ewan Harrison, David K. Jackson, Ian Johnston, Dominic Kwiatkowski, Cordelia Langford, John Sillitoe on behalf of the Wellcome Sanger Institute COVID-19 Surveillance Team |
| EPI_ISL_3124413, EPI_ISL_3124414 | Berkshire and Surrey Pathology Services Lighthouse Laboratory | Wellcome Sanger Institute for the COVID-19 Genomics UK (COG-UK) Consortium | Berkshire and Surrey Pathology Services Lighthouse Laboratory and Alex Alderton, Roberto Amato, Jeffrey Barrett, Sonia Goncalves, Ewan Harrison, David K. Jackson, Ian Johnston, Dominic Kwiatkowski, Cordelia Langford, John Sillitoe on behalf of the Wellcome Sanger Institute COVID-19 Surveillance Team |
| EPI_ISL_3124415, EPI_ISL_3124416 | Lighthouse Lab in Milton Keynes | Wellcome Sanger Institute for the COVID-19 Genomics UK (COG-UK) Consortium | The Lighthouse Lab in Milton Keynes and Alex Alderton, Roberto Amato, Jeffrey Barrett, Sonia Goncalves, Ewan Harrison, David K. Jackson, Ian Johnston, Dominic Kwiatkowski, Cordelia Langford, John Sillitoe on behalf of the Wellcome Sanger Institute COVID-19 Surveillance Team |
| EPI_ISL_3124417 | Berkshire and Surrey Pathology Services Lighthouse Laboratory | Wellcome Sanger Institute for the COVID-19 Genomics UK (COG-UK) Consortium | Berkshire and Surrey Pathology Services Lighthouse Laboratory and Alex Alderton, Roberto Amato, Jeffrey Barrett, Sonia Goncalves, Ewan Harrison, David K. Jackson, Ian Johnston, Dominic Kwiatkowski, Cordelia Langford, John Sillitoe on behalf of the Wellcome Sanger Institute COVID-19 Surveillance Team |
| EPI_ISL_3124418 | Lighthouse Lab in Glasgow | Wellcome Sanger Institute for the COVID-19 Genomics UK (COG-UK) Consortium | Harper VanSteenhouse, Yumi Kasai, David Gray, Carol Clugston, Anna Dominiczak and Alex Alderton, Roberto Amato, Jeffrey Barrett, Sonia Goncalves, Ewan Harrison, David K. Jackson, Ian Johnston, Dominic Kwiatkowski, Cordelia Langford, John Sillitoe on behalf of the Wellcome Sanger Institute COVID-19 Surveillance Team |
| EPI_ISL_3124419 | Lighthouse Laboratory Plymouth | Wellcome Sanger Institute for the COVID-19 Genomics UK (COG-UK) Consortium | Lighthouse Laboratory Plymouth and Alex Alderton, Roberto Amato, Jeffrey Barrett, Sonia Goncalves, Ewan Harrison, David K. Jackson, Ian Johnston, Dominic Kwiatkowski, Cordelia Langford, John Sillitoe on behalf of the Wellcome Sanger Institute COVID-19 Surveillance Team |
| EPI_ISL_3124420 | Lighthouse Lab in Milton Keynes | Wellcome Sanger Institute for the COVID-19 Genomics UK (COG-UK) | The Lighthouse Lab in Milton Keynes and Alex Alderton, Roberto Amato, Jeffrey Barrett, Sonia Goncalves, Ewan Harrison, David K. Jackson, Ian Johnston, Dominic Kwiatkowski, Cordelia Langford, John Sillitoe on behalf of the Wellcome Sanger |

| | | | |
|---|---|---|---|
| | | Consortium | Institute COVID-19 Surveillance Team |
| EPI_ISL_3124421 | Berkshire and Surrey Pathology Services Lighthouse Laboratory | Wellcome Sanger Institute for the COVID-19 Genomics UK (COG-UK) Consortium | Berkshire and Surrey Pathology Services Lighthouse Laboratory and Alex Alderton, Roberto Amato, Jeffrey Barrett, Sonia Goncalves, Ewan Harrison, David K. Jackson, Ian Johnston, Dominic Kwiatkowski, Cordelia Langford, John Sillitoe on behalf of the Wellcome Sanger Institute COVID-19 Surveillance Team |
| EPI_ISL_3124422, EPI_ISL_3124423 | Lighthouse Lab in Milton Keynes | Wellcome Sanger Institute for the COVID-19 Genomics UK (COG-UK) Consortium | The Lighthouse Lab in Milton Keynes and Alex Alderton, Roberto Amato, Jeffrey Barrett, Sonia Goncalves, Ewan Harrison, David K. Jackson, Ian Johnston, Dominic Kwiatkowski, Cordelia Langford, John Sillitoe on behalf of the Wellcome Sanger Institute COVID-19 Surveillance Team |
| EPI_ISL_3124424 | Berkshire and Surrey Pathology Services Lighthouse Laboratory | Wellcome Sanger Institute for the COVID-19 Genomics UK (COG-UK) Consortium | Berkshire and Surrey Pathology Services Lighthouse Laboratory and Alex Alderton, Roberto Amato, Jeffrey Barrett, Sonia Goncalves, Ewan Harrison, David K. Jackson, Ian Johnston, Dominic Kwiatkowski, Cordelia Langford, John Sillitoe on behalf of the Wellcome Sanger Institute COVID-19 Surveillance Team |
| EPI_ISL_3124425 | Lighthouse Lab in Milton Keynes | Wellcome Sanger Institute for the COVID-19 Genomics UK (COG-UK) Consortium | The Lighthouse Lab in Milton Keynes and Alex Alderton, Roberto Amato, Jeffrey Barrett, Sonia Goncalves, Ewan Harrison, David K. Jackson, Ian Johnston, Dominic Kwiatkowski, Cordelia Langford, John Sillitoe on behalf of the Wellcome Sanger Institute COVID-19 Surveillance Team |
| EPI_ISL_3124426 | Lighthouse Lab in Glasgow | Wellcome Sanger Institute for the COVID-19 Genomics UK (COG-UK) Consortium | Harper VanSteenhouse, Yumi Kasai, David Gray, Carol Clugston, Anna Dominiczak and Alex Alderton, Roberto Amato, Jeffrey Barrett, Sonia Goncalves, Ewan Harrison, David K. Jackson, Ian Johnston, Dominic Kwiatkowski, Cordelia Langford, John Sillitoe on behalf of the Wellcome Sanger Institute COVID-19 Surveillance Team |
| EPI_ISL_3124427, EPI_ISL_3124428 | Lighthouse Lab in Milton Keynes | Wellcome Sanger Institute for the COVID-19 Genomics UK (COG-UK) Consortium | The Lighthouse Lab in Milton Keynes and Alex Alderton, Roberto Amato, Jeffrey Barrett, Sonia Goncalves, Ewan Harrison, David K. Jackson, Ian Johnston, Dominic Kwiatkowski, Cordelia Langford, John Sillitoe on behalf of the Wellcome Sanger Institute COVID-19 Surveillance Team |
| EPI_ISL_3124429 | Lighthouse Lab in Glasgow | Wellcome Sanger Institute for the COVID-19 Genomics UK (COG-UK) Consortium | Harper VanSteenhouse, Yumi Kasai, David Gray, Carol Clugston, Anna Dominiczak and Alex Alderton, Roberto Amato, Jeffrey Barrett, Sonia Goncalves, Ewan Harrison, David K. Jackson, Ian Johnston, Dominic Kwiatkowski, Cordelia Langford, John Sillitoe on behalf of the Wellcome Sanger Institute COVID-19 Surveillance Team |
| EPI_ISL_3124430 | Lighthouse Laboratory Plymouth | Wellcome Sanger Institute for the COVID-19 Genomics UK (COG-UK) Consortium | Lighthouse Laboratory Plymouth and Alex Alderton, Roberto Amato, Jeffrey Barrett, Sonia Goncalves, Ewan Harrison, David K. Jackson, Ian Johnston, Dominic Kwiatkowski, Cordelia Langford, John Sillitoe on behalf of the Wellcome Sanger Institute COVID-19 Surveillance Team |
| EPI_ISL_3124431 | Lighthouse Lab in Glasgow | Wellcome Sanger Institute for the COVID-19 Genomics UK (COG-UK) Consortium | Harper VanSteenhouse, Yumi Kasai, David Gray, Carol Clugston, Anna Dominiczak and Alex Alderton, Roberto Amato, Jeffrey Barrett, Sonia Goncalves, Ewan Harrison, David K. Jackson, Ian Johnston, Dominic Kwiatkowski, Cordelia Langford, John Sillitoe on behalf of the Wellcome Sanger Institute COVID-19 Surveillance Team |
| EPI_ISL_3124432, EPI_ISL_3124433, EPI_ISL_3124434 | Lighthouse Laboratory Plymouth | Wellcome Sanger Institute for the COVID-19 Genomics UK (COG-UK) Consortium | Lighthouse Laboratory Plymouth and Alex Alderton, Roberto Amato, Jeffrey Barrett, Sonia Goncalves, Ewan Harrison, David K. Jackson, Ian Johnston, Dominic Kwiatkowski, Cordelia Langford, John Sillitoe on behalf of the Wellcome Sanger Institute COVID-19 Surveillance Team |
| EPI_ISL_3124435 | Lighthouse Lab in Glasgow | Wellcome Sanger Institute for the COVID-19 Genomics UK (COG-UK) Consortium | Harper VanSteenhouse, Yumi Kasai, David Gray, Carol Clugston, Anna Dominiczak and Alex Alderton, Roberto Amato, Jeffrey Barrett, Sonia Goncalves, Ewan Harrison, David K. Jackson, Ian Johnston, Dominic Kwiatkowski, Cordelia Langford, John Sillitoe on behalf of the Wellcome Sanger Institute COVID-19 Surveillance Team |
| EPI_ISL_3124436 | Lighthouse Laboratory Plymouth | Wellcome Sanger Institute for the COVID-19 Genomics UK (COG-UK) Consortium | Lighthouse Laboratory Plymouth and Alex Alderton, Roberto Amato, Jeffrey Barrett, Sonia Goncalves, Ewan Harrison, David K. Jackson, Ian Johnston, Dominic Kwiatkowski, Cordelia Langford, John Sillitoe on behalf of the Wellcome Sanger Institute COVID-19 Surveillance Team |
| EPI_ISL_3124437 | Berkshire and Surrey Pathology Services Lighthouse Laboratory | Wellcome Sanger Institute for the COVID-19 Genomics UK (COG-UK) Consortium | Berkshire and Surrey Pathology Services Lighthouse Laboratory and Alex Alderton, Roberto Amato, Jeffrey Barrett, Sonia Goncalves, Ewan Harrison, David K. Jackson, Ian Johnston, Dominic Kwiatkowski, Cordelia Langford, John Sillitoe on behalf of the Wellcome Sanger Institute COVID-19 Surveillance Team |
| EPI_ISL_3124438 | Lighthouse Lab in Milton Keynes | Wellcome Sanger Institute for the COVID-19 Genomics UK (COG-UK) Consortium | The Lighthouse Lab in Milton Keynes and Alex Alderton, Roberto Amato, Jeffrey Barrett, Sonia Goncalves, Ewan Harrison, David K. Jackson, Ian Johnston, Dominic Kwiatkowski, Cordelia Langford, John Sillitoe on behalf of the Wellcome Sanger Institute COVID-19 Surveillance Team |
| EPI_ISL_3124439 | Lighthouse Lab in Glasgow | Wellcome Sanger Institute for the COVID-19 Genomics UK (COG-UK) Consortium | Harper VanSteenhouse, Yumi Kasai, David Gray, Carol Clugston, Anna Dominiczak and Alex Alderton, Roberto Amato, Jeffrey Barrett, Sonia Goncalves, Ewan Harrison, David K. Jackson, Ian Johnston, Dominic Kwiatkowski, Cordelia Langford, John Sillitoe on behalf of the Wellcome Sanger Institute COVID-19 Surveillance Team |
| EPI_ISL_3124440 | Lighthouse Lab in Milton Keynes | Wellcome Sanger Institute for the COVID-19 Genomics UK (COG-UK) Consortium | The Lighthouse Lab in Milton Keynes and Alex Alderton, Roberto Amato, Jeffrey Barrett, Sonia Goncalves, Ewan Harrison, David K. Jackson, Ian Johnston, Dominic Kwiatkowski, Cordelia Langford, John Sillitoe on behalf of the Wellcome Sanger Institute COVID-19 Surveillance Team |
| EPI_ISL_3124441 | Berkshire and Surrey Pathology Services Lighthouse Laboratory | Wellcome Sanger Institute for the COVID-19 Genomics UK (COG-UK) Consortium | Berkshire and Surrey Pathology Services Lighthouse Laboratory and Alex Alderton, Roberto Amato, Jeffrey Barrett, Sonia Goncalves, Ewan Harrison, David K. Jackson, Ian Johnston, Dominic Kwiatkowski, Cordelia Langford, John Sillitoe on behalf of the Wellcome Sanger Institute COVID-19 Surveillance Team |
| EPI_ISL_3124442, EPI_ISL_3124443 | Lighthouse Lab in Milton Keynes | Wellcome Sanger Institute for the COVID-19 Genomics UK (COG-UK) Consortium | The Lighthouse Lab in Milton Keynes and Alex Alderton, Roberto Amato, Jeffrey Barrett, Sonia Goncalves, Ewan Harrison, David K. Jackson, Ian Johnston, Dominic Kwiatkowski, Cordelia Langford, John Sillitoe on behalf of the Wellcome Sanger Institute COVID-19 Surveillance Team |
| EPI_ISL_3124444 | Lighthouse Lab in Glasgow | Wellcome Sanger Institute for the COVID-19 Genomics UK (COG-UK) Consortium | Harper VanSteenhouse, Yumi Kasai, David Gray, Carol Clugston, Anna Dominiczak and Alex Alderton, Roberto Amato, Jeffrey Barrett, Sonia Goncalves, Ewan Harrison, David K. Jackson, Ian Johnston, Dominic Kwiatkowski, Cordelia Langford, John Sillitoe on behalf of the Wellcome Sanger Institute COVID-19 Surveillance Team |
| EPI_ISL_3124445 | Lighthouse Laboratory Plymouth | Wellcome Sanger Institute for the COVID-19 Genomics UK (COG-UK) Consortium | Lighthouse Laboratory Plymouth and Alex Alderton, Roberto Amato, Jeffrey Barrett, Sonia Goncalves, Ewan Harrison, David K. Jackson, Ian Johnston, Dominic Kwiatkowski, Cordelia Langford, John Sillitoe on behalf of the Wellcome Sanger Institute COVID-19 Surveillance Team |
| EPI_ISL_3124446, EPI_ISL_3124447 | Lighthouse Lab in Glasgow | Wellcome Sanger Institute for the COVID-19 Genomics UK (COG-UK) Consortium | Harper VanSteenhouse, Yumi Kasai, David Gray, Carol Clugston, Anna Dominiczak and Alex Alderton, Roberto Amato, Jeffrey Barrett, Sonia Goncalves, Ewan Harrison, David K. Jackson, Ian Johnston, Dominic Kwiatkowski, Cordelia Langford, John Sillitoe on behalf of the Wellcome Sanger Institute COVID-19 Surveillance Team |
| EPI_ISL_3124448 | Berkshire and Surrey Pathology Services Lighthouse Laboratory | Wellcome Sanger Institute for the COVID-19 Genomics UK (COG-UK) Consortium | Berkshire and Surrey Pathology Services Lighthouse Laboratory and Alex Alderton, Roberto Amato, Jeffrey Barrett, Sonia Goncalves, Ewan Harrison, David K. Jackson, Ian Johnston, Dominic Kwiatkowski, Cordelia Langford, John Sillitoe on behalf of the Wellcome Sanger Institute COVID-19 Surveillance Team |
| EPI_ISL_3124449 | Lighthouse Laboratory Plymouth | Wellcome Sanger Institute for the COVID-19 Genomics UK (COG-UK) Consortium | Lighthouse Laboratory Plymouth and Alex Alderton, Roberto Amato, Jeffrey Barrett, Sonia Goncalves, Ewan Harrison, David K. Jackson, Ian Johnston, Dominic Kwiatkowski, Cordelia Langford, John Sillitoe on behalf of the Wellcome Sanger Institute COVID-19 Surveillance Team |
| EPI_ISL_3124450 | Lighthouse Lab in Glasgow | Wellcome Sanger Institute for the COVID-19 Genomics UK (COG-UK) Consortium | Harper VanSteenhouse, Yumi Kasai, David Gray, Carol Clugston, Anna Dominiczak and Alex Alderton, Roberto Amato, Jeffrey Barrett, Sonia Goncalves, Ewan Harrison, David K. Jackson, Ian Johnston, Dominic Kwiatkowski, Cordelia Langford, John Sillitoe on behalf of the Wellcome Sanger Institute COVID-19 Surveillance Team |
| EPI_ISL_3124451 | Lighthouse Lab in Milton Keynes | Wellcome Sanger Institute for the COVID-19 Genomics UK (COG-UK) Consortium | The Lighthouse Lab in Milton Keynes and Alex Alderton, Roberto Amato, Jeffrey Barrett, Sonia Goncalves, Ewan Harrison, David K. Jackson, Ian Johnston, Dominic Kwiatkowski, Cordelia Langford, John Sillitoe on behalf of the Wellcome Sanger Institute COVID-19 Surveillance Team |
| EPI_ISL_3124452 | Lighthouse Lab in Glasgow | Wellcome Sanger Institute for the COVID-19 Genomics UK (COG-UK) Consortium | Harper VanSteenhouse, Yumi Kasai, David Gray, Carol Clugston, Anna Dominiczak and Alex Alderton, Roberto Amato, Jeffrey Barrett, Sonia Goncalves, Ewan Harrison, David K. Jackson, Ian Johnston, Dominic Kwiatkowski, Cordelia Langford, John Sillitoe on behalf of the Wellcome Sanger Institute COVID-19 Surveillance Team |
| EPI_ISL_3124453 | Lighthouse Lab in Milton Keynes | Wellcome Sanger Institute for the COVID-19 Genomics UK (COG-UK) Consortium | The Lighthouse Lab in Milton Keynes and Alex Alderton, Roberto Amato, Jeffrey Barrett, Sonia Goncalves, Ewan Harrison, David K. Jackson, Ian Johnston, Dominic Kwiatkowski, Cordelia Langford, John Sillitoe on behalf of the Wellcome Sanger Institute COVID-19 Surveillance Team |
| EPI_ISL_3124454 | Lighthouse Lab in Glasgow | Wellcome Sanger Institute for the COVID-19 Genomics UK (COG-UK) Consortium | Harper VanSteenhouse, Yumi Kasai, David Gray, Carol Clugston, Anna Dominiczak and Alex Alderton, Roberto Amato, Jeffrey Barrett, Sonia Goncalves, Ewan Harrison, David K. Jackson, Ian Johnston, Dominic Kwiatkowski, Cordelia Langford, John Sillitoe on behalf of the Wellcome Sanger Institute COVID-19 Surveillance Team |
| EPI_ISL_3124455, EPI_ISL_3124456 | Berkshire and Surrey Pathology Services Lighthouse Laboratory | Wellcome Sanger Institute for the COVID-19 Genomics UK (COG-UK) Consortium | Berkshire and Surrey Pathology Services Lighthouse Laboratory and Alex Alderton, Roberto Amato, Jeffrey Barrett, Sonia Goncalves, Ewan Harrison, David K. Jackson, Ian Johnston, Dominic Kwiatkowski, Cordelia Langford, John Sillitoe on behalf of the Wellcome Sanger Institute COVID-19 Surveillance Team |
| EPI_ISL_3124457 | Lighthouse Laboratory Plymouth | Wellcome Sanger Institute for the COVID-19 Genomics UK (COG-UK) Consortium | Lighthouse Laboratory Plymouth and Alex Alderton, Roberto Amato, Jeffrey Barrett, Sonia Goncalves, Ewan Harrison, David K. Jackson, Ian Johnston, Dominic Kwiatkowski, Cordelia Langford, John Sillitoe on behalf of the Wellcome Sanger Institute COVID-19 Surveillance Team |
| EPI_ISL_3124458, EPI_ISL_3124459 | Lighthouse Lab in Milton Keynes | Wellcome Sanger Institute for the COVID-19 Genomics UK (COG-UK) Consortium | The Lighthouse Lab in Milton Keynes and Alex Alderton, Roberto Amato, Jeffrey Barrett, Sonia Goncalves, Ewan Harrison, David K. Jackson, Ian Johnston, Dominic Kwiatkowski, Cordelia Langford, John Sillitoe on behalf of the Wellcome Sanger Institute COVID-19 Surveillance Team |
| EPI_ISL_3124460, EPI_ISL_3124461, EPI_ISL_3124462 | Lighthouse Laboratory Plymouth | Wellcome Sanger Institute for the COVID-19 Genomics UK (COG-UK) Consortium | Lighthouse Laboratory Plymouth and Alex Alderton, Roberto Amato, Jeffrey Barrett, Sonia Goncalves, Ewan Harrison, David K. Jackson, Ian Johnston, Dominic Kwiatkowski, Cordelia Langford, John Sillitoe on behalf of the Wellcome Sanger Institute COVID-19 Surveillance Team |
| EPI_ISL_3124463 | Lighthouse Lab in Glasgow | Wellcome Sanger Institute for the COVID-19 Genomics UK (COG-UK) Consortium | Harper VanSteenhouse, Yumi Kasai, David Gray, Carol Clugston, Anna Dominiczak and Alex Alderton, Roberto Amato, Jeffrey Barrett, Sonia Goncalves, Ewan Harrison, David K. Jackson, Ian Johnston, Dominic Kwiatkowski, Cordelia Langford, John Sillitoe on behalf of the Wellcome Sanger Institute COVID-19 Surveillance Team |
| EPI_ISL_3124464 | Lighthouse Laboratory Plymouth | Wellcome Sanger Institute for the COVID-19 Genomics UK (COG-UK) Consortium | Lighthouse Laboratory Plymouth and Alex Alderton, Roberto Amato, Jeffrey Barrett, Sonia Goncalves, Ewan Harrison, David K. Jackson, Ian Johnston, Dominic Kwiatkowski, Cordelia Langford, John Sillitoe on behalf of the Wellcome Sanger Institute COVID-19 Surveillance Team |
| EPI_ISL_3124465 | Lighthouse Lab in Glasgow | Wellcome Sanger Institute for the COVID-19 Genomics UK (COG-UK) Consortium | Harper VanSteenhouse, Yumi Kasai, David Gray, Carol Clugston, Anna Dominiczak and Alex Alderton, Roberto Amato, Jeffrey Barrett, Sonia Goncalves, Ewan Harrison, David K. Jackson, Ian Johnston, Dominic Kwiatkowski, Cordelia Langford, John Sillitoe on behalf of the Wellcome Sanger Institute COVID-19 Surveillance Team |
| EPI_ISL_3124466 | Berkshire and Surrey Pathology Services Lighthouse Laboratory | Wellcome Sanger Institute for the COVID-19 Genomics UK (COG-UK) Consortium | Berkshire and Surrey Pathology Services Lighthouse Laboratory and Alex Alderton, Roberto Amato, Jeffrey Barrett, Sonia Goncalves, Ewan Harrison, David K. Jackson, Ian Johnston, Dominic Kwiatkowski, Cordelia Langford, John Sillitoe on behalf of the Wellcome Sanger Institute COVID-19 Surveillance Team |
| EPI_ISL_3124467 | Lighthouse Lab in Glasgow | Wellcome Sanger Institute for the COVID-19 Genomics UK (COG-UK) Consortium | Harper VanSteenhouse, Yumi Kasai, David Gray, Carol Clugston, Anna Dominiczak and Alex Alderton, Roberto Amato, Jeffrey Barrett, Sonia Goncalves, Ewan Harrison, David K. Jackson, Ian Johnston, Dominic Kwiatkowski, Cordelia Langford, John Sillitoe on behalf of the Wellcome Sanger Institute COVID-19 Surveillance Team |
| EPI_ISL_3124468 | Lighthouse Laboratory Plymouth | Wellcome Sanger Institute for the COVID-19 Genomics UK (COG-UK) Consortium | Lighthouse Laboratory Plymouth and Alex Alderton, Roberto Amato, Jeffrey Barrett, Sonia Goncalves, Ewan Harrison, David K. Jackson, Ian Johnston, Dominic Kwiatkowski, Cordelia Langford, John Sillitoe on behalf of the Wellcome Sanger Institute COVID-19 Surveillance Team |
| EPI_ISL_3124469 | Berkshire and Surrey Pathology Services Lighthouse Laboratory | Wellcome Sanger Institute for the COVID-19 Genomics UK (COG-UK) Consortium | Berkshire and Surrey Pathology Services Lighthouse Laboratory and Alex Alderton, Roberto Amato, Jeffrey Barrett, Sonia Goncalves, Ewan Harrison, David K. Jackson, Ian Johnston, Dominic Kwiatkowski, Cordelia Langford, John Sillitoe on behalf of the Wellcome Sanger Institute COVID-19 Surveillance Team |
| EPI_ISL_3124470, EPI_ISL_3124471, EPI_ISL_3124472 | Lighthouse Lab in Glasgow | Wellcome Sanger Institute for the COVID-19 Genomics UK (COG-UK) Consortium | Harper VanSteenhouse, Yumi Kasai, David Gray, Carol Clugston, Anna Dominiczak and Alex Alderton, Roberto Amato, Jeffrey Barrett, Sonia Goncalves, Ewan Harrison, David K. Jackson, Ian Johnston, Dominic Kwiatkowski, Cordelia Langford, John Sillitoe on behalf of the Wellcome Sanger Institute COVID-19 Surveillance Team |
| EPI_ISL_3124473 | Lighthouse Laboratory Plymouth | Wellcome Sanger Institute for the COVID-19 Genomics UK (COG-UK) Consortium | Lighthouse Laboratory Plymouth and Alex Alderton, Roberto Amato, Jeffrey Barrett, Sonia Goncalves, Ewan Harrison, David K. Jackson, Ian Johnston, Dominic Kwiatkowski, Cordelia Langford, John Sillitoe on behalf of the Wellcome Sanger Institute COVID-19 Surveillance Team |
| EPI_ISL_3124474 | Berkshire and Surrey Pathology Services Lighthouse Laboratory | Wellcome Sanger Institute for the COVID-19 Genomics UK (COG-UK) Consortium | Berkshire and Surrey Pathology Services Lighthouse Laboratory and Alex Alderton, Roberto Amato, Jeffrey Barrett, Sonia Goncalves, Ewan Harrison, David K. Jackson, Ian Johnston, Dominic Kwiatkowski, Cordelia Langford, John Sillitoe on behalf of the Wellcome Sanger Institute COVID-19 Surveillance Team |
| EPI_ISL_3124475 | Lighthouse Lab in Glasgow | Wellcome Sanger Institute for the COVID-19 Genomics UK (COG-UK) Consortium | Harper VanSteenhouse, Yumi Kasai, David Gray, Carol Clugston, Anna Dominiczak and Alex Alderton, Roberto Amato, Jeffrey Barrett, Sonia Goncalves, Ewan Harrison, David K. Jackson, Ian Johnston, Dominic Kwiatkowski, Cordelia Langford, John Sillitoe on behalf of the Wellcome Sanger Institute COVID-19 Surveillance Team |
| EPI_ISL_3124476 | Berkshire and Surrey Pathology Services Lighthouse Laboratory | Wellcome Sanger Institute for the COVID-19 Genomics UK (COG-UK) | Berkshire and Surrey Pathology Services Lighthouse Laboratory and Alex Alderton, Roberto Amato, Jeffrey Barrett, Sonia Goncalves, Ewan Harrison, David K. Jackson, Ian Johnston, Dominic Kwiatkowski, Cordelia Langford, John Sillitoe on behalf |

| | | | |
|---|---|---|---|
| | | Consortium | of the Wellcome Sanger Institute COVID-19 Surveillance Team |
| EPI_ISL_3124477 | Lighthouse Laboratory Plymouth | Wellcome Sanger Institute for the COVID-19 Genomics UK (COG-UK) Consortium | Lighthouse Laboratory Plymouth and Alex Alderton, Roberto Amato, Jeffrey Barrett, Sonia Goncalves, Ewan Harrison, David K. Jackson, Ian Johnston, Dominic Kwiatkowski, Cordelia Langford, John Sillitoe on behalf of the Wellcome Sanger Institute COVID-19 Surveillance Team |
| EPI_ISL_3124478 | Lighthouse Lab in Milton Keynes | Wellcome Sanger Institute for the COVID-19 Genomics UK (COG-UK) Consortium | The Lighthouse Lab in Milton Keynes and Alex Alderton, Roberto Amato, Jeffrey Barrett, Sonia Goncalves, Ewan Harrison, David K. Jackson, Ian Johnston, Dominic Kwiatkowski, Cordelia Langford, John Sillitoe on behalf of the Wellcome Sanger Institute COVID-19 Surveillance Team |
| EPI_ISL_3124479, EPI_ISL_3124480 | Lighthouse Lab in Glasgow | Wellcome Sanger Institute for the COVID-19 Genomics UK (COG-UK) Consortium | Harper VanSteenhouse, Yumi Kasai, David Gray, Carol Clugston, Anna Dominiczak and Alex Alderton, Roberto Amato, Jeffrey Barrett, Sonia Goncalves, Ewan Harrison, David K. Jackson, Ian Johnston, Dominic Kwiatkowski, Cordelia Langford, John Sillitoe on behalf of the Wellcome Sanger Institute COVID-19 Surveillance Team |
| EPI_ISL_3124481, EPI_ISL_3124482 | Lighthouse Laboratory Plymouth | Wellcome Sanger Institute for the COVID-19 Genomics UK (COG-UK) Consortium | Lighthouse Laboratory Plymouth and Alex Alderton, Roberto Amato, Jeffrey Barrett, Sonia Goncalves, Ewan Harrison, David K. Jackson, Ian Johnston, Dominic Kwiatkowski, Cordelia Langford, John Sillitoe on behalf of the Wellcome Sanger Institute COVID-19 Surveillance Team |
| EPI_ISL_3124483, EPI_ISL_3124484 | Lighthouse Lab in Milton Keynes | Wellcome Sanger Institute for the COVID-19 Genomics UK (COG-UK) Consortium | The Lighthouse Lab in Milton Keynes and Alex Alderton, Roberto Amato, Jeffrey Barrett, Sonia Goncalves, Ewan Harrison, David K. Jackson, Ian Johnston, Dominic Kwiatkowski, Cordelia Langford, John Sillitoe on behalf of the Wellcome Sanger Institute COVID-19 Surveillance Team |
| EPI_ISL_3124485 | Lighthouse Laboratory Plymouth | Wellcome Sanger Institute for the COVID-19 Genomics UK (COG-UK) Consortium | Lighthouse Laboratory Plymouth and Alex Alderton, Roberto Amato, Jeffrey Barrett, Sonia Goncalves, Ewan Harrison, David K. Jackson, Ian Johnston, Dominic Kwiatkowski, Cordelia Langford, John Sillitoe on behalf of the Wellcome Sanger Institute COVID-19 Surveillance Team |
| EPI_ISL_3124486 | Lighthouse Lab in Glasgow | Wellcome Sanger Institute for the COVID-19 Genomics UK (COG-UK) Consortium | Harper VanSteenhouse, Yumi Kasai, David Gray, Carol Clugston, Anna Dominiczak and Alex Alderton, Roberto Amato, Jeffrey Barrett, Sonia Goncalves, Ewan Harrison, David K. Jackson, Ian Johnston, Dominic Kwiatkowski, Cordelia Langford, John Sillitoe on behalf of the Wellcome Sanger Institute COVID-19 Surveillance Team |
| EPI_ISL_3124487, EPI_ISL_3124488 | Lighthouse Laboratory Plymouth | Wellcome Sanger Institute for the COVID-19 Genomics UK (COG-UK) Consortium | Lighthouse Laboratory Plymouth and Alex Alderton, Roberto Amato, Jeffrey Barrett, Sonia Goncalves, Ewan Harrison, David K. Jackson, Ian Johnston, Dominic Kwiatkowski, Cordelia Langford, John Sillitoe on behalf of the Wellcome Sanger Institute COVID-19 Surveillance Team |
| EPI_ISL_3124489 | Berkshire and Surrey Pathology Services Lighthouse Laboratory | Wellcome Sanger Institute for the COVID-19 Genomics UK (COG-UK) Consortium | Berkshire and Surrey Pathology Services Lighthouse Laboratory and Alex Alderton, Roberto Amato, Jeffrey Barrett, Sonia Goncalves, Ewan Harrison, David K. Jackson, Ian Johnston, Dominic Kwiatkowski, Cordelia Langford, John Sillitoe on behalf of the Wellcome Sanger Institute COVID-19 Surveillance Team |
| EPI_ISL_3124490 | Lighthouse Lab in Milton Keynes | Wellcome Sanger Institute for the COVID-19 Genomics UK (COG-UK) Consortium | The Lighthouse Lab in Milton Keynes and Alex Alderton, Roberto Amato, Jeffrey Barrett, Sonia Goncalves, Ewan Harrison, David K. Jackson, Ian Johnston, Dominic Kwiatkowski, Cordelia Langford, John Sillitoe on behalf of the Wellcome Sanger Institute COVID-19 Surveillance Team |
| EPI_ISL_3124491, EPI_ISL_3124492, EPI_ISL_3124493, EPI_ISL_3124494, EPI_ISL_3124495 | Lighthouse Laboratory Plymouth | Wellcome Sanger Institute for the COVID-19 Genomics UK (COG-UK) Consortium | Lighthouse Laboratory Plymouth and Alex Alderton, Roberto Amato, Jeffrey Barrett, Sonia Goncalves, Ewan Harrison, David K. Jackson, Ian Johnston, Dominic Kwiatkowski, Cordelia Langford, John Sillitoe on behalf of the Wellcome Sanger Institute COVID-19 Surveillance Team |
| EPI_ISL_3124496 | Berkshire and Surrey Pathology Services Lighthouse Laboratory | Wellcome Sanger Institute for the COVID-19 Genomics UK (COG-UK) Consortium | Berkshire and Surrey Pathology Services Lighthouse Laboratory and Alex Alderton, Roberto Amato, Jeffrey Barrett, Sonia Goncalves, Ewan Harrison, David K. Jackson, Ian Johnston, Dominic Kwiatkowski, Cordelia Langford, John Sillitoe on behalf of the Wellcome Sanger Institute COVID-19 Surveillance Team |
| EPI_ISL_3124497 | Lighthouse Lab in Milton Keynes | Wellcome Sanger Institute for the COVID-19 Genomics UK (COG-UK) Consortium | The Lighthouse Lab in Milton Keynes and Alex Alderton, Roberto Amato, Jeffrey Barrett, Sonia Goncalves, Ewan Harrison, David K. Jackson, Ian Johnston, Dominic Kwiatkowski, Cordelia Langford, John Sillitoe on behalf of the Wellcome Sanger Institute COVID-19 Surveillance Team |
| EPI_ISL_3124498, EPI_ISL_3124499 | Lighthouse Laboratory Plymouth | Wellcome Sanger Institute for the COVID-19 Genomics UK (COG-UK) Consortium | Lighthouse Laboratory Plymouth and Alex Alderton, Roberto Amato, Jeffrey Barrett, Sonia Goncalves, Ewan Harrison, David K. Jackson, Ian Johnston, Dominic Kwiatkowski, Cordelia Langford, John Sillitoe on behalf of the Wellcome Sanger Institute COVID-19 Surveillance Team |
| EPI_ISL_3124500 | Berkshire and Surrey Pathology Services Lighthouse Laboratory | Wellcome Sanger Institute for the COVID-19 Genomics UK (COG-UK) Consortium | Berkshire and Surrey Pathology Services Lighthouse Laboratory and Alex Alderton, Roberto Amato, Jeffrey Barrett, Sonia Goncalves, Ewan Harrison, David K. Jackson, Ian Johnston, Dominic Kwiatkowski, Cordelia Langford, John Sillitoe on behalf of the Wellcome Sanger Institute COVID-19 Surveillance Team |
| EPI_ISL_3124501, EPI_ISL_3124502 | Lighthouse Lab in Milton Keynes | Wellcome Sanger Institute for the COVID-19 Genomics UK (COG-UK) Consortium | The Lighthouse Lab in Milton Keynes and Alex Alderton, Roberto Amato, Jeffrey Barrett, Sonia Goncalves, Ewan Harrison, David K. Jackson, Ian Johnston, Dominic Kwiatkowski, Cordelia Langford, John Sillitoe on behalf of the Wellcome Sanger Institute COVID-19 Surveillance Team |
| EPI_ISL_3124503 | Berkshire and Surrey Pathology Services Lighthouse Laboratory | Wellcome Sanger Institute for the COVID-19 Genomics UK (COG-UK) Consortium | Berkshire and Surrey Pathology Services Lighthouse Laboratory and Alex Alderton, Roberto Amato, Jeffrey Barrett, Sonia Goncalves, Ewan Harrison, David K. Jackson, Ian Johnston, Dominic Kwiatkowski, Cordelia Langford, John Sillitoe on behalf of the Wellcome Sanger Institute COVID-19 Surveillance Team |
| EPI_ISL_3124504, EPI_ISL_3124505, EPI_ISL_3124506 | Lighthouse Lab in Milton Keynes | Wellcome Sanger Institute for the COVID-19 Genomics UK (COG-UK) Consortium | The Lighthouse Lab in Milton Keynes and Alex Alderton, Roberto Amato, Jeffrey Barrett, Sonia Goncalves, Ewan Harrison, David K. Jackson, Ian Johnston, Dominic Kwiatkowski, Cordelia Langford, John Sillitoe on behalf of the Wellcome Sanger Institute COVID-19 Surveillance Team |
| EPI_ISL_3124507 | Lighthouse Laboratory Plymouth | Wellcome Sanger Institute for the COVID-19 Genomics UK (COG-UK) Consortium | Lighthouse Laboratory Plymouth and Alex Alderton, Roberto Amato, Jeffrey Barrett, Sonia Goncalves, Ewan Harrison, David K. Jackson, Ian Johnston, Dominic Kwiatkowski, Cordelia Langford, John Sillitoe on behalf of the Wellcome Sanger Institute COVID-19 Surveillance Team |
| EPI_ISL_3124508, EPI_ISL_3124509 | Lighthouse Lab in Milton Keynes | Wellcome Sanger Institute for the COVID-19 Genomics UK (COG-UK) Consortium | The Lighthouse Lab in Milton Keynes and Alex Alderton, Roberto Amato, Jeffrey Barrett, Sonia Goncalves, Ewan Harrison, David K. Jackson, Ian Johnston, Dominic Kwiatkowski, Cordelia Langford, John Sillitoe on behalf of the Wellcome Sanger Institute COVID-19 Surveillance Team |
| EPI_ISL_3124510 | Berkshire and Surrey Pathology Services Lighthouse Laboratory | Wellcome Sanger Institute for the COVID-19 Genomics UK (COG-UK) Consortium | Berkshire and Surrey Pathology Services Lighthouse Laboratory and Alex Alderton, Roberto Amato, Jeffrey Barrett, Sonia Goncalves, Ewan Harrison, David K. Jackson, Ian Johnston, Dominic Kwiatkowski, Cordelia Langford, John Sillitoe on behalf of the Wellcome Sanger Institute COVID-19 Surveillance Team |
| EPI_ISL_3124511 | Lighthouse Lab in Glasgow | Wellcome Sanger Institute for the COVID-19 Genomics UK (COG-UK) Consortium | Harper VanSteenhouse, Yumi Kasai, David Gray, Carol Clugston, Anna Dominiczak and Alex Alderton, Roberto Amato, Jeffrey Barrett, Sonia Goncalves, Ewan Harrison, David K. Jackson, Ian Johnston, Dominic Kwiatkowski, Cordelia Langford, John Sillitoe on behalf of the Wellcome Sanger Institute COVID-19 Surveillance Team |
| EPI_ISL_3124512, EPI_ISL_3124513 | Lighthouse Lab in Milton Keynes | Wellcome Sanger Institute for the COVID-19 Genomics UK (COG-UK) Consortium | The Lighthouse Lab in Milton Keynes and Alex Alderton, Roberto Amato, Jeffrey Barrett, Sonia Goncalves, Ewan Harrison, David K. Jackson, Ian Johnston, Dominic Kwiatkowski, Cordelia Langford, John Sillitoe on behalf of the Wellcome Sanger Institute COVID-19 Surveillance Team |
| EPI_ISL_3124514, EPI_ISL_3124515 | Lighthouse Lab in Glasgow | Wellcome Sanger Institute for the COVID-19 Genomics UK (COG-UK) Consortium | Harper VanSteenhouse, Yumi Kasai, David Gray, Carol Clugston, Anna Dominiczak and Alex Alderton, Roberto Amato, Jeffrey Barrett, Sonia Goncalves, Ewan Harrison, David K. Jackson, Ian Johnston, Dominic Kwiatkowski, Cordelia Langford, John Sillitoe on behalf of the Wellcome Sanger Institute COVID-19 Surveillance Team |
| EPI_ISL_3124516 | Lighthouse Lab in Milton Keynes | Wellcome Sanger Institute for the COVID-19 Genomics UK (COG-UK) Consortium | The Lighthouse Lab in Milton Keynes and Alex Alderton, Roberto Amato, Jeffrey Barrett, Sonia Goncalves, Ewan Harrison, David K. Jackson, Ian Johnston, Dominic Kwiatkowski, Cordelia Langford, John Sillitoe on behalf of the Wellcome Sanger Institute COVID-19 Surveillance Team |
| EPI_ISL_3124517 | Lighthouse Laboratory Plymouth | Wellcome Sanger Institute for the COVID-19 Genomics UK (COG-UK) Consortium | Lighthouse Laboratory Plymouth and Alex Alderton, Roberto Amato, Jeffrey Barrett, Sonia Goncalves, Ewan Harrison, David K. Jackson, Ian Johnston, Dominic Kwiatkowski, Cordelia Langford, John Sillitoe on behalf of the Wellcome Sanger Institute COVID-19 Surveillance Team |
| EPI_ISL_3124518, EPI_ISL_3124519 | Lighthouse Lab in Glasgow | Wellcome Sanger Institute for the COVID-19 Genomics UK (COG-UK) Consortium | Harper VanSteenhouse, Yumi Kasai, David Gray, Carol Clugston, Anna Dominiczak and Alex Alderton, Roberto Amato, Jeffrey Barrett, Sonia Goncalves, Ewan Harrison, David K. Jackson, Ian Johnston, Dominic Kwiatkowski, Cordelia Langford, John Sillitoe on behalf of the Wellcome Sanger Institute COVID-19 Surveillance Team |
| EPI_ISL_3124520 | Lighthouse Lab in Milton Keynes | Wellcome Sanger Institute for the COVID-19 Genomics UK (COG-UK) Consortium | The Lighthouse Lab in Milton Keynes and Alex Alderton, Roberto Amato, Jeffrey Barrett, Sonia Goncalves, Ewan Harrison, David K. Jackson, Ian Johnston, Dominic Kwiatkowski, Cordelia Langford, John Sillitoe on behalf of the Wellcome Sanger Institute COVID-19 Surveillance Team |
| EPI_ISL_3124521, EPI_ISL_3124522 | Lighthouse Laboratory Plymouth | Wellcome Sanger Institute for the COVID-19 Genomics UK (COG-UK) Consortium | Lighthouse Laboratory Plymouth and Alex Alderton, Roberto Amato, Jeffrey Barrett, Sonia Goncalves, Ewan Harrison, David K. Jackson, Ian Johnston, Dominic Kwiatkowski, Cordelia Langford, John Sillitoe on behalf of the Wellcome Sanger Institute COVID-19 Surveillance Team |
| EPI_ISL_3124523 | Lighthouse Lab in Glasgow | Wellcome Sanger Institute for the COVID-19 Genomics UK (COG-UK) Consortium | Harper VanSteenhouse, Yumi Kasai, David Gray, Carol Clugston, Anna Dominiczak and Alex Alderton, Roberto Amato, Jeffrey Barrett, Sonia Goncalves, Ewan Harrison, David K. Jackson, Ian Johnston, Dominic Kwiatkowski, Cordelia Langford, John Sillitoe on behalf of the Wellcome Sanger Institute COVID-19 Surveillance Team |
| EPI_ISL_3124524 | Berkshire and Surrey Pathology Services Lighthouse Laboratory | Wellcome Sanger Institute for the COVID-19 Genomics UK (COG-UK) Consortium | Berkshire and Surrey Pathology Services Lighthouse Laboratory and Alex Alderton, Roberto Amato, Jeffrey Barrett, Sonia Goncalves, Ewan Harrison, David K. Jackson, Ian Johnston, Dominic Kwiatkowski, Cordelia Langford, John Sillitoe on behalf of the Wellcome Sanger Institute COVID-19 Surveillance Team |
| EPI_ISL_3124525 | Lighthouse Laboratory Plymouth | Wellcome Sanger Institute for the COVID-19 Genomics UK (COG-UK) Consortium | Lighthouse Laboratory Plymouth and Alex Alderton, Roberto Amato, Jeffrey Barrett, Sonia Goncalves, Ewan Harrison, David K. Jackson, Ian Johnston, Dominic Kwiatkowski, Cordelia Langford, John Sillitoe on behalf of the Wellcome Sanger Institute COVID-19 Surveillance Team |
| EPI_ISL_3124526 | Lighthouse Lab in Milton Keynes | Wellcome Sanger Institute for the COVID-19 Genomics UK (COG-UK) Consortium | The Lighthouse Lab in Milton Keynes and Alex Alderton, Roberto Amato, Jeffrey Barrett, Sonia Goncalves, Ewan Harrison, David K. Jackson, Ian Johnston, Dominic Kwiatkowski, Cordelia Langford, John Sillitoe on behalf of the Wellcome Sanger Institute COVID-19 Surveillance Team |
| EPI_ISL_3124527 | Berkshire and Surrey Pathology Services Lighthouse Laboratory | Wellcome Sanger Institute for the COVID-19 Genomics UK (COG-UK) Consortium | Berkshire and Surrey Pathology Services Lighthouse Laboratory and Alex Alderton, Roberto Amato, Jeffrey Barrett, Sonia Goncalves, Ewan Harrison, David K. Jackson, Ian Johnston, Dominic Kwiatkowski, Cordelia Langford, John Sillitoe on behalf of the Wellcome Sanger Institute COVID-19 Surveillance Team |
| EPI_ISL_3124528, EPI_ISL_3124529 | Lighthouse Lab in Milton Keynes | Wellcome Sanger Institute for the COVID-19 Genomics UK (COG-UK) Consortium | The Lighthouse Lab in Milton Keynes and Alex Alderton, Roberto Amato, Jeffrey Barrett, Sonia Goncalves, Ewan Harrison, David K. Jackson, Ian Johnston, Dominic Kwiatkowski, Cordelia Langford, John Sillitoe on behalf of the Wellcome Sanger Institute COVID-19 Surveillance Team |
| EPI_ISL_3124530 | Berkshire and Surrey Pathology Services Lighthouse Laboratory | Wellcome Sanger Institute for the COVID-19 Genomics UK (COG-UK) Consortium | Berkshire and Surrey Pathology Services Lighthouse Laboratory and Alex Alderton, Roberto Amato, Jeffrey Barrett, Sonia Goncalves, Ewan Harrison, David K. Jackson, Ian Johnston, Dominic Kwiatkowski, Cordelia Langford, John Sillitoe on behalf of the Wellcome Sanger Institute COVID-19 Surveillance Team |
| EPI_ISL_3124531 | Lighthouse Lab in Glasgow | Wellcome Sanger Institute for the COVID-19 Genomics UK (COG-UK) Consortium | Harper VanSteenhouse, Yumi Kasai, David Gray, Carol Clugston, Anna Dominiczak and Alex Alderton, Roberto Amato, Jeffrey Barrett, Sonia Goncalves, Ewan Harrison, David K. Jackson, Ian Johnston, Dominic Kwiatkowski, Cordelia Langford, John Sillitoe on behalf of the Wellcome Sanger Institute COVID-19 Surveillance Team |
| EPI_ISL_3124532 | Lighthouse Lab in Milton Keynes | Wellcome Sanger Institute for the COVID-19 Genomics UK (COG-UK) Consortium | The Lighthouse Lab in Milton Keynes and Alex Alderton, Roberto Amato, Jeffrey Barrett, Sonia Goncalves, Ewan Harrison, David K. Jackson, Ian Johnston, Dominic Kwiatkowski, Cordelia Langford, John Sillitoe on behalf of the Wellcome Sanger Institute COVID-19 Surveillance Team |
| EPI_ISL_3124533, EPI_ISL_3124534 | Lighthouse Laboratory Plymouth | Wellcome Sanger Institute for the COVID-19 Genomics UK (COG-UK) Consortium | Lighthouse Laboratory Plymouth and Alex Alderton, Roberto Amato, Jeffrey Barrett, Sonia Goncalves, Ewan Harrison, David K. Jackson, Ian Johnston, Dominic Kwiatkowski, Cordelia Langford, John Sillitoe on behalf of the Wellcome Sanger Institute COVID-19 Surveillance Team |
| EPI_ISL_3124535 | Berkshire and Surrey Pathology Services Lighthouse Laboratory | Wellcome Sanger Institute for the COVID-19 Genomics UK (COG-UK) Consortium | Berkshire and Surrey Pathology Services Lighthouse Laboratory and Alex Alderton, Roberto Amato, Jeffrey Barrett, Sonia Goncalves, Ewan Harrison, David K. Jackson, Ian Johnston, Dominic Kwiatkowski, Cordelia Langford, John Sillitoe on behalf of the Wellcome Sanger Institute COVID-19 Surveillance Team |
| EPI_ISL_3124536 | Lighthouse Laboratory Plymouth | Wellcome Sanger Institute for the COVID-19 Genomics UK (COG-UK) Consortium | Lighthouse Laboratory Plymouth and Alex Alderton, Roberto Amato, Jeffrey Barrett, Sonia Goncalves, Ewan Harrison, David K. Jackson, Ian Johnston, Dominic Kwiatkowski, Cordelia Langford, John Sillitoe on behalf of the Wellcome Sanger Institute COVID-19 Surveillance Team |
| EPI_ISL_3124537 | Lighthouse Lab in Milton Keynes | Wellcome Sanger Institute for the COVID-19 Genomics UK (COG-UK) Consortium | The Lighthouse Lab in Milton Keynes and Alex Alderton, Roberto Amato, Jeffrey Barrett, Sonia Goncalves, Ewan Harrison, David K. Jackson, Ian Johnston, Dominic Kwiatkowski, Cordelia Langford, John Sillitoe on behalf of the Wellcome Sanger Institute COVID-19 Surveillance Team |
| EPI_ISL_3124538 | Lighthouse Lab in Glasgow | Wellcome Sanger Institute for the COVID-19 Genomics UK (COG-UK) Consortium | Harper VanSteenhouse, Yumi Kasai, David Gray, Carol Clugston, Anna Dominiczak and Alex Alderton, Roberto Amato, Jeffrey Barrett, Sonia Goncalves, Ewan Harrison, David K. Jackson, Ian Johnston, Dominic Kwiatkowski, Cordelia Langford, John Sillitoe on behalf of the Wellcome Sanger Institute COVID-19 Surveillance Team |

| | | | |
|---|---|---|---|
| EPI_ISL_3124539 | Lighthouse Lab in Milton Keynes | Wellcome Sanger Institute for the COVID-19 Genomics UK (COG-UK) Consortium | The Lighthouse Lab in Milton Keynes and Alex Alderton, Roberto Amato, Jeffrey Barrett, Sonia Goncalves, Ewan Harrison, David K. Jackson, Ian Johnston, Dominic Kwiatkowski, Cordelia Langford, John Sillitoe on behalf of the Wellcome Sanger Institute COVID-19 Surveillance Team |
| EPI_ISL_3124540 | Lighthouse Laboratory Plymouth | Wellcome Sanger Institute for the COVID-19 Genomics UK (COG-UK) Consortium | Lighthouse Laboratory Plymouth and Alex Alderton, Roberto Amato, Jeffrey Barrett, Sonia Goncalves, Ewan Harrison, David K. Jackson, Ian Johnston, Dominic Kwiatkowski, Cordelia Langford, John Sillitoe on behalf of the Wellcome Sanger Institute COVID-19 Surveillance Team |
| EPI_ISL_3124541 | Lighthouse Lab in Milton Keynes | Wellcome Sanger Institute for the COVID-19 Genomics UK (COG-UK) Consortium | The Lighthouse Lab in Milton Keynes and Alex Alderton, Roberto Amato, Jeffrey Barrett, Sonia Goncalves, Ewan Harrison, David K. Jackson, Ian Johnston, Dominic Kwiatkowski, Cordelia Langford, John Sillitoe on behalf of the Wellcome Sanger Institute COVID-19 Surveillance Team |
| EPI_ISL_3124542 | Lighthouse Laboratory Plymouth | Wellcome Sanger Institute for the COVID-19 Genomics UK (COG-UK) Consortium | Lighthouse Laboratory Plymouth and Alex Alderton, Roberto Amato, Jeffrey Barrett, Sonia Goncalves, Ewan Harrison, David K. Jackson, Ian Johnston, Dominic Kwiatkowski, Cordelia Langford, John Sillitoe on behalf of the Wellcome Sanger Institute COVID-19 Surveillance Team |
| EPI_ISL_3124543, EPI_ISL_3124544, EPI_ISL_3124545 | Lighthouse Lab in Milton Keynes | Wellcome Sanger Institute for the COVID-19 Genomics UK (COG-UK) Consortium | The Lighthouse Lab in Milton Keynes and Alex Alderton, Roberto Amato, Jeffrey Barrett, Sonia Goncalves, Ewan Harrison, David K. Jackson, Ian Johnston, Dominic Kwiatkowski, Cordelia Langford, John Sillitoe on behalf of the Wellcome Sanger Institute COVID-19 Surveillance Team |
| EPI_ISL_3124546 | Lighthouse Laboratory Plymouth | Wellcome Sanger Institute for the COVID-19 Genomics UK (COG-UK) Consortium | Lighthouse Laboratory Plymouth and Alex Alderton, Roberto Amato, Jeffrey Barrett, Sonia Goncalves, Ewan Harrison, David K. Jackson, Ian Johnston, Dominic Kwiatkowski, Cordelia Langford, John Sillitoe on behalf of the Wellcome Sanger Institute COVID-19 Surveillance Team |
| EPI_ISL_3124547 | Lighthouse Lab in Glasgow | Wellcome Sanger Institute for the COVID-19 Genomics UK (COG-UK) Consortium | Harper VanSteenhouse, Yumi Kasai, David Gray, Carol Clugston, Anna Dominiczak and Alex Alderton, Roberto Amato, Jeffrey Barrett, Sonia Goncalves, Ewan Harrison, David K. Jackson, Ian Johnston, Dominic Kwiatkowski, Cordelia Langford, John Sillitoe on behalf of the Wellcome Sanger Institute COVID-19 Surveillance Team |
| EPI_ISL_3124548, EPI_ISL_3124549 | Lighthouse Laboratory Plymouth | Wellcome Sanger Institute for the COVID-19 Genomics UK (COG-UK) Consortium | Lighthouse Laboratory Plymouth and Alex Alderton, Roberto Amato, Jeffrey Barrett, Sonia Goncalves, Ewan Harrison, David K. Jackson, Ian Johnston, Dominic Kwiatkowski, Cordelia Langford, John Sillitoe on behalf of the Wellcome Sanger Institute COVID-19 Surveillance Team |
| EPI_ISL_3124550 | Lighthouse Lab in Milton Keynes | Wellcome Sanger Institute for the COVID-19 Genomics UK (COG-UK) Consortium | The Lighthouse Lab in Milton Keynes and Alex Alderton, Roberto Amato, Jeffrey Barrett, Sonia Goncalves, Ewan Harrison, David K. Jackson, Ian Johnston, Dominic Kwiatkowski, Cordelia Langford, John Sillitoe on behalf of the Wellcome Sanger Institute COVID-19 Surveillance Team |
| EPI_ISL_3124551 | Lighthouse Lab in Glasgow | Wellcome Sanger Institute for the COVID-19 Genomics UK (COG-UK) Consortium | Harper VanSteenhouse, Yumi Kasai, David Gray, Carol Clugston, Anna Dominiczak and Alex Alderton, Roberto Amato, Jeffrey Barrett, Sonia Goncalves, Ewan Harrison, David K. Jackson, Ian Johnston, Dominic Kwiatkowski, Cordelia Langford, John Sillitoe on behalf of the Wellcome Sanger Institute COVID-19 Surveillance Team |
| EPI_ISL_3124552, EPI_ISL_3124553 | Lighthouse Laboratory Plymouth | Wellcome Sanger Institute for the COVID-19 Genomics UK (COG-UK) Consortium | Lighthouse Laboratory Plymouth and Alex Alderton, Roberto Amato, Jeffrey Barrett, Sonia Goncalves, Ewan Harrison, David K. Jackson, Ian Johnston, Dominic Kwiatkowski, Cordelia Langford, John Sillitoe on behalf of the Wellcome Sanger Institute COVID-19 Surveillance Team |
| EPI_ISL_3124554 | Berkshire and Surrey Pathology Services Lighthouse Laboratory | Wellcome Sanger Institute for the COVID-19 Genomics UK (COG-UK) Consortium | Berkshire and Surrey Pathology Services Lighthouse Laboratory and Alex Alderton, Roberto Amato, Jeffrey Barrett, Sonia Goncalves, Ewan Harrison, David K. Jackson, Ian Johnston, Dominic Kwiatkowski, Cordelia Langford, John Sillitoe on behalf of the Wellcome Sanger Institute COVID-19 Surveillance Team |
| EPI_ISL_3124555 | Lighthouse Lab in Milton Keynes | Wellcome Sanger Institute for the COVID-19 Genomics UK (COG-UK) Consortium | The Lighthouse Lab in Milton Keynes and Alex Alderton, Roberto Amato, Jeffrey Barrett, Sonia Goncalves, Ewan Harrison, David K. Jackson, Ian Johnston, Dominic Kwiatkowski, Cordelia Langford, John Sillitoe on behalf of the Wellcome Sanger Institute COVID-19 Surveillance Team |
| EPI_ISL_3124556, EPI_ISL_3124557, EPI_ISL_3124558 | Lighthouse Lab in Glasgow | Wellcome Sanger Institute for the COVID-19 Genomics UK (COG-UK) Consortium | Harper VanSteenhouse, Yumi Kasai, David Gray, Carol Clugston, Anna Dominiczak and Alex Alderton, Roberto Amato, Jeffrey Barrett, Sonia Goncalves, Ewan Harrison, David K. Jackson, Ian Johnston, Dominic Kwiatkowski, Cordelia Langford, John Sillitoe on behalf of the Wellcome Sanger Institute COVID-19 Surveillance Team |
| EPI_ISL_3124559 | Berkshire and Surrey Pathology Services Lighthouse Laboratory | Wellcome Sanger Institute for the COVID-19 Genomics UK (COG-UK) Consortium | Berkshire and Surrey Pathology Services Lighthouse Laboratory and Alex Alderton, Roberto Amato, Jeffrey Barrett, Sonia Goncalves, Ewan Harrison, David K. Jackson, Ian Johnston, Dominic Kwiatkowski, Cordelia Langford, John Sillitoe on behalf of the Wellcome Sanger Institute COVID-19 Surveillance Team |
| EPI_ISL_3124560, EPI_ISL_3124561 | Lighthouse Lab in Glasgow | Wellcome Sanger Institute for the COVID-19 Genomics UK (COG-UK) Consortium | Harper VanSteenhouse, Yumi Kasai, David Gray, Carol Clugston, Anna Dominiczak and Alex Alderton, Roberto Amato, Jeffrey Barrett, Sonia Goncalves, Ewan Harrison, David K. Jackson, Ian Johnston, Dominic Kwiatkowski, Cordelia Langford, John Sillitoe on behalf of the Wellcome Sanger Institute COVID-19 Surveillance Team |
| EPI_ISL_3124562 | Lighthouse Lab in Milton Keynes | Wellcome Sanger Institute for the COVID-19 Genomics UK (COG-UK) Consortium | The Lighthouse Lab in Milton Keynes and Alex Alderton, Roberto Amato, Jeffrey Barrett, Sonia Goncalves, Ewan Harrison, David K. Jackson, Ian Johnston, Dominic Kwiatkowski, Cordelia Langford, John Sillitoe on behalf of the Wellcome Sanger Institute COVID-19 Surveillance Team |
| EPI_ISL_3124563 | Lighthouse Lab in Glasgow | Wellcome Sanger Institute for the COVID-19 Genomics UK (COG-UK) Consortium | Harper VanSteenhouse, Yumi Kasai, David Gray, Carol Clugston, Anna Dominiczak and Alex Alderton, Roberto Amato, Jeffrey Barrett, Sonia Goncalves, Ewan Harrison, David K. Jackson, Ian Johnston, Dominic Kwiatkowski, Cordelia Langford, John Sillitoe on behalf of the Wellcome Sanger Institute COVID-19 Surveillance Team |
| EPI_ISL_3124564 | Berkshire and Surrey Pathology Services Lighthouse Laboratory | Wellcome Sanger Institute for the COVID-19 Genomics UK (COG-UK) Consortium | Berkshire and Surrey Pathology Services Lighthouse Laboratory and Alex Alderton, Roberto Amato, Jeffrey Barrett, Sonia Goncalves, Ewan Harrison, David K. Jackson, Ian Johnston, Dominic Kwiatkowski, Cordelia Langford, John Sillitoe on behalf of the Wellcome Sanger Institute COVID-19 Surveillance Team |
| EPI_ISL_3124565 | Lighthouse Laboratory Plymouth | Wellcome Sanger Institute for the COVID-19 Genomics UK (COG-UK) Consortium | Lighthouse Laboratory Plymouth and Alex Alderton, Roberto Amato, Jeffrey Barrett, Sonia Goncalves, Ewan Harrison, David K. Jackson, Ian Johnston, Dominic Kwiatkowski, Cordelia Langford, John Sillitoe on behalf of the Wellcome Sanger Institute COVID-19 Surveillance Team |
| EPI_ISL_3124566, EPI_ISL_3124567, EPI_ISL_3124568 | Lighthouse Lab in Milton Keynes | Wellcome Sanger Institute for the COVID-19 Genomics UK (COG-UK) Consortium | The Lighthouse Lab in Milton Keynes and Alex Alderton, Roberto Amato, Jeffrey Barrett, Sonia Goncalves, Ewan Harrison, David K. Jackson, Ian Johnston, Dominic Kwiatkowski, Cordelia Langford, John Sillitoe on behalf of the Wellcome Sanger Institute COVID-19 Surveillance Team |
| EPI_ISL_3124569 | Berkshire and Surrey Pathology Services Lighthouse Laboratory | Wellcome Sanger Institute for the COVID-19 Genomics UK (COG-UK) Consortium | Berkshire and Surrey Pathology Services Lighthouse Laboratory and Alex Alderton, Roberto Amato, Jeffrey Barrett, Sonia Goncalves, Ewan Harrison, David K. Jackson, Ian Johnston, Dominic Kwiatkowski, Cordelia Langford, John Sillitoe on behalf of the Wellcome Sanger Institute COVID-19 Surveillance Team |
| EPI_ISL_3124570 | Lighthouse Lab in Milton Keynes | Wellcome Sanger Institute for the COVID-19 Genomics UK (COG-UK) Consortium | The Lighthouse Lab in Milton Keynes and Alex Alderton, Roberto Amato, Jeffrey Barrett, Sonia Goncalves, Ewan Harrison, David K. Jackson, Ian Johnston, Dominic Kwiatkowski, Cordelia Langford, John Sillitoe on behalf of the Wellcome Sanger Institute COVID-19 Surveillance Team |
| EPI_ISL_3124571 | Lighthouse Laboratory Plymouth | Wellcome Sanger Institute for the COVID-19 Genomics UK (COG-UK) Consortium | Lighthouse Laboratory Plymouth and Alex Alderton, Roberto Amato, Jeffrey Barrett, Sonia Goncalves, Ewan Harrison, David K. Jackson, Ian Johnston, Dominic Kwiatkowski, Cordelia Langford, John Sillitoe on behalf of the Wellcome Sanger Institute COVID-19 Surveillance Team |
| EPI_ISL_3124572 | Lighthouse Lab in Glasgow | Wellcome Sanger Institute for the COVID-19 Genomics UK (COG-UK) Consortium | Harper VanSteenhouse, Yumi Kasai, David Gray, Carol Clugston, Anna Dominiczak and Alex Alderton, Roberto Amato, Jeffrey Barrett, Sonia Goncalves, Ewan Harrison, David K. Jackson, Ian Johnston, Dominic Kwiatkowski, Cordelia Langford, John Sillitoe on behalf of the Wellcome Sanger Institute COVID-19 Surveillance Team |
| EPI_ISL_3124573, EPI_ISL_3124574 | Lighthouse Laboratory Plymouth | Wellcome Sanger Institute for the COVID-19 Genomics UK (COG-UK) Consortium | Lighthouse Laboratory Plymouth and Alex Alderton, Roberto Amato, Jeffrey Barrett, Sonia Goncalves, Ewan Harrison, David K. Jackson, Ian Johnston, Dominic Kwiatkowski, Cordelia Langford, John Sillitoe on behalf of the Wellcome Sanger Institute COVID-19 Surveillance Team |
| EPI_ISL_3124575, EPI_ISL_3124576, EPI_ISL_3124577 | Lighthouse Lab in Milton Keynes | Wellcome Sanger Institute for the COVID-19 Genomics UK (COG-UK) Consortium | The Lighthouse Lab in Milton Keynes and Alex Alderton, Roberto Amato, Jeffrey Barrett, Sonia Goncalves, Ewan Harrison, David K. Jackson, Ian Johnston, Dominic Kwiatkowski, Cordelia Langford, John Sillitoe on behalf of the Wellcome Sanger Institute COVID-19 Surveillance Team |
| EPI_ISL_3124578 | Lighthouse Lab in Glasgow | Wellcome Sanger Institute for the COVID-19 Genomics UK (COG-UK) Consortium | Harper VanSteenhouse, Yumi Kasai, David Gray, Carol Clugston, Anna Dominiczak and Alex Alderton, Roberto Amato, Jeffrey Barrett, Sonia Goncalves, Ewan Harrison, David K. Jackson, Ian Johnston, Dominic Kwiatkowski, Cordelia Langford, John Sillitoe on behalf of the Wellcome Sanger Institute COVID-19 Surveillance Team |
| EPI_ISL_3124579 | Lighthouse Laboratory Plymouth | Wellcome Sanger Institute for the COVID-19 Genomics UK (COG-UK) Consortium | Lighthouse Laboratory Plymouth and Alex Alderton, Roberto Amato, Jeffrey Barrett, Sonia Goncalves, Ewan Harrison, David K. Jackson, Ian Johnston, Dominic Kwiatkowski, Cordelia Langford, John Sillitoe on behalf of the Wellcome Sanger Institute COVID-19 Surveillance Team |
| EPI_ISL_3124580, EPI_ISL_3124581 | Berkshire and Surrey Pathology Services Lighthouse Laboratory | Wellcome Sanger Institute for the COVID-19 Genomics UK (COG-UK) Consortium | Berkshire and Surrey Pathology Services Lighthouse Laboratory and Alex Alderton, Roberto Amato, Jeffrey Barrett, Sonia Goncalves, Ewan Harrison, David K. Jackson, Ian Johnston, Dominic Kwiatkowski, Cordelia Langford, John Sillitoe on behalf of the Wellcome Sanger Institute COVID-19 Surveillance Team |
| EPI_ISL_3124582 | Lighthouse Laboratory Plymouth | Wellcome Sanger Institute for the COVID-19 Genomics UK (COG-UK) Consortium | Lighthouse Laboratory Plymouth and Alex Alderton, Roberto Amato, Jeffrey Barrett, Sonia Goncalves, Ewan Harrison, David K. Jackson, Ian Johnston, Dominic Kwiatkowski, Cordelia Langford, John Sillitoe on behalf of the Wellcome Sanger Institute COVID-19 Surveillance Team |
| EPI_ISL_3124583 | Lighthouse Lab in Milton Keynes | Wellcome Sanger Institute for the COVID-19 Genomics UK (COG-UK) Consortium | The Lighthouse Lab in Milton Keynes and Alex Alderton, Roberto Amato, Jeffrey Barrett, Sonia Goncalves, Ewan Harrison, David K. Jackson, Ian Johnston, Dominic Kwiatkowski, Cordelia Langford, John Sillitoe on behalf of the Wellcome Sanger Institute COVID-19 Surveillance Team |
| EPI_ISL_3124584, EPI_ISL_3124585, EPI_ISL_3124586 | Lighthouse Laboratory Plymouth | Wellcome Sanger Institute for the COVID-19 Genomics UK (COG-UK) Consortium | Lighthouse Laboratory Plymouth and Alex Alderton, Roberto Amato, Jeffrey Barrett, Sonia Goncalves, Ewan Harrison, David K. Jackson, Ian Johnston, Dominic Kwiatkowski, Cordelia Langford, John Sillitoe on behalf of the Wellcome Sanger Institute COVID-19 Surveillance Team |
| EPI_ISL_3124587 | Lighthouse Lab in Milton Keynes | Wellcome Sanger Institute for the COVID-19 Genomics UK (COG-UK) Consortium | The Lighthouse Lab in Milton Keynes and Alex Alderton, Roberto Amato, Jeffrey Barrett, Sonia Goncalves, Ewan Harrison, David K. Jackson, Ian Johnston, Dominic Kwiatkowski, Cordelia Langford, John Sillitoe on behalf of the Wellcome Sanger Institute COVID-19 Surveillance Team |
| EPI_ISL_3124588 | Lighthouse Lab in Glasgow | Wellcome Sanger Institute for the COVID-19 Genomics UK (COG-UK) Consortium | Harper VanSteenhouse, Yumi Kasai, David Gray, Carol Clugston, Anna Dominiczak and Alex Alderton, Roberto Amato, Jeffrey Barrett, Sonia Goncalves, Ewan Harrison, David K. Jackson, Ian Johnston, Dominic Kwiatkowski, Cordelia Langford, John Sillitoe on behalf of the Wellcome Sanger Institute COVID-19 Surveillance Team |
| EPI_ISL_3124589 | Lighthouse Lab in Milton Keynes | Wellcome Sanger Institute for the COVID-19 Genomics UK (COG-UK) Consortium | The Lighthouse Lab in Milton Keynes and Alex Alderton, Roberto Amato, Jeffrey Barrett, Sonia Goncalves, Ewan Harrison, David K. Jackson, Ian Johnston, Dominic Kwiatkowski, Cordelia Langford, John Sillitoe on behalf of the Wellcome Sanger Institute COVID-19 Surveillance Team |
| EPI_ISL_3124590 | Lighthouse Laboratory Plymouth | Wellcome Sanger Institute for the COVID-19 Genomics UK (COG-UK) Consortium | Lighthouse Laboratory Plymouth and Alex Alderton, Roberto Amato, Jeffrey Barrett, Sonia Goncalves, Ewan Harrison, David K. Jackson, Ian Johnston, Dominic Kwiatkowski, Cordelia Langford, John Sillitoe on behalf of the Wellcome Sanger Institute COVID-19 Surveillance Team |
| EPI_ISL_3124591, EPI_ISL_3124592 | Berkshire and Surrey Pathology Services Lighthouse Laboratory | Wellcome Sanger Institute for the COVID-19 Genomics UK (COG-UK) Consortium | Berkshire and Surrey Pathology Services Lighthouse Laboratory and Alex Alderton, Roberto Amato, Jeffrey Barrett, Sonia Goncalves, Ewan Harrison, David K. Jackson, Ian Johnston, Dominic Kwiatkowski, Cordelia Langford, John Sillitoe on behalf of the Wellcome Sanger Institute COVID-19 Surveillance Team |
| EPI_ISL_3124593, EPI_ISL_3124594 | Lighthouse Laboratory Plymouth | Wellcome Sanger Institute for the COVID-19 Genomics UK (COG-UK) Consortium | Lighthouse Laboratory Plymouth and Alex Alderton, Roberto Amato, Jeffrey Barrett, Sonia Goncalves, Ewan Harrison, David K. Jackson, Ian Johnston, Dominic Kwiatkowski, Cordelia Langford, John Sillitoe on behalf of the Wellcome Sanger Institute COVID-19 Surveillance Team |
| EPI_ISL_3124595, EPI_ISL_3124596 | Lighthouse Lab in Milton Keynes | Wellcome Sanger Institute for the COVID-19 Genomics UK (COG-UK) Consortium | The Lighthouse Lab in Milton Keynes and Alex Alderton, Roberto Amato, Jeffrey Barrett, Sonia Goncalves, Ewan Harrison, David K. Jackson, Ian Johnston, Dominic Kwiatkowski, Cordelia Langford, John Sillitoe on behalf of the Wellcome Sanger Institute COVID-19 Surveillance Team |
| EPI_ISL_3124597 | Lighthouse Laboratory Plymouth | Wellcome Sanger Institute for the COVID-19 Genomics UK (COG-UK) Consortium | Lighthouse Laboratory Plymouth and Alex Alderton, Roberto Amato, Jeffrey Barrett, Sonia Goncalves, Ewan Harrison, David K. Jackson, Ian Johnston, Dominic Kwiatkowski, Cordelia Langford, John Sillitoe on behalf of the Wellcome Sanger Institute COVID-19 Surveillance Team |
| EPI_ISL_3124598 | Lighthouse Lab in Milton Keynes | Wellcome Sanger Institute for the COVID-19 Genomics UK (COG-UK) Consortium | The Lighthouse Lab in Milton Keynes and Alex Alderton, Roberto Amato, Jeffrey Barrett, Sonia Goncalves, Ewan Harrison, David K. Jackson, Ian Johnston, Dominic Kwiatkowski, Cordelia Langford, John Sillitoe on behalf of the Wellcome Sanger Institute COVID-19 Surveillance Team |
| EPI_ISL_3124599 | Lighthouse Lab in Glasgow | Wellcome Sanger Institute for the COVID-19 Genomics UK (COG-UK) Consortium | Harper VanSteenhouse, Yumi Kasai, David Gray, Carol Clugston, Anna Dominiczak and Alex Alderton, Roberto Amato, Jeffrey Barrett, Sonia Goncalves, Ewan Harrison, David K. Jackson, Ian Johnston, Dominic Kwiatkowski, Cordelia Langford, John Sillitoe on behalf of the Wellcome Sanger Institute COVID-19 Surveillance Team |

| | | | |
|---|---|---|---|
| EPI_ISL_3124600 | Berkshire and Surrey Pathology Services Lighthouse Laboratory | Wellcome Sanger Institute for the COVID-19 Genomics UK (COG-UK) Consortium | Berkshire and Surrey Pathology Services Lighthouse Laboratory and Alex Alderton, Roberto Amato, Jeffrey Barrett, Sonia Goncalves, Ewan Harrison, David K. Jackson, Ian Johnston, Dominic Kwiatkowski, Cordelia Langford, John Sillitoe on behalf of the Wellcome Sanger Institute COVID-19 Surveillance Team |
| EPI_ISL_3124601, EPI_ISL_3124602 | Lighthouse Laboratory Plymouth | Wellcome Sanger Institute for the COVID-19 Genomics UK (COG-UK) Consortium | Lighthouse Laboratory Plymouth and Alex Alderton, Roberto Amato, Jeffrey Barrett, Sonia Goncalves, Ewan Harrison, David K. Jackson, Ian Johnston, Dominic Kwiatkowski, Cordelia Langford, John Sillitoe on behalf of the Wellcome Sanger Institute COVID-19 Surveillance Team |
| EPI_ISL_3124603 | Lighthouse Lab in Glasgow | Wellcome Sanger Institute for the COVID-19 Genomics UK (COG-UK) Consortium | Harper VanSteenhouse, Yumi Kasai, David Gray, Carol Clugston, Anna Dominiczak and Alex Alderton, Roberto Amato, Jeffrey Barrett, Sonia Goncalves, Ewan Harrison, David K. Jackson, Ian Johnston, Dominic Kwiatkowski, Cordelia Langford, John Sillitoe on behalf of the Wellcome Sanger Institute COVID-19 Surveillance Team |
| EPI_ISL_3124604, EPI_ISL_3124605, EPI_ISL_3124606 | Lighthouse Lab in Milton Keynes | Wellcome Sanger Institute for the COVID-19 Genomics UK (COG-UK) Consortium | The Lighthouse Lab in Milton Keynes and Alex Alderton, Roberto Amato, Jeffrey Barrett, Sonia Goncalves, Ewan Harrison, David K. Jackson, Ian Johnston, Dominic Kwiatkowski, Cordelia Langford, John Sillitoe on behalf of the Wellcome Sanger Institute COVID-19 Surveillance Team |
| EPI_ISL_3124607 | Lighthouse Lab in Alderley Park | Wellcome Sanger Institute for the COVID-19 Genomics UK (COG-UK) Consortium | Jacquelyn Wynn, Mairead Hyland, The Lighthouse Lab in Alderley Park and Alex Alderton, Roberto Amato, Jeffrey Barrett, Sonia Goncalves, Ewan Harrison, David K. Jackson, Ian Johnston, Dominic Kwiatkowski, Cordelia Langford, John Sillitoe on behalf of the Wellcome Sanger Institute COVID-19 Surveillance Team |
| EPI_ISL_3124608, EPI_ISL_3124609, EPI_ISL_3124610, EPI_ISL_3124611, EPI_ISL_3124612, EPI_ISL_3124613 | Lighthouse Lab in Milton Keynes | Wellcome Sanger Institute for the COVID-19 Genomics UK (COG-UK) Consortium | The Lighthouse Lab in Milton Keynes and Alex Alderton, Roberto Amato, Jeffrey Barrett, Sonia Goncalves, Ewan Harrison, David K. Jackson, Ian Johnston, Dominic Kwiatkowski, Cordelia Langford, John Sillitoe on behalf of the Wellcome Sanger Institute COVID-19 Surveillance Team |
| EPI_ISL_3124614 | Lighthouse Lab in Alderley Park | Wellcome Sanger Institute for the COVID-19 Genomics UK (COG-UK) Consortium | Jacquelyn Wynn, Mairead Hyland, The Lighthouse Lab in Alderley Park and Alex Alderton, Roberto Amato, Jeffrey Barrett, Sonia Goncalves, Ewan Harrison, David K. Jackson, Ian Johnston, Dominic Kwiatkowski, Cordelia Langford, John Sillitoe on behalf of the Wellcome Sanger Institute COVID-19 Surveillance Team |
| EPI_ISL_3124615 | Lighthouse Lab in Milton Keynes | Wellcome Sanger Institute for the COVID-19 Genomics UK (COG-UK) Consortium | The Lighthouse Lab in Milton Keynes and Alex Alderton, Roberto Amato, Jeffrey Barrett, Sonia Goncalves, Ewan Harrison, David K. Jackson, Ian Johnston, Dominic Kwiatkowski, Cordelia Langford, John Sillitoe on behalf of the Wellcome Sanger Institute COVID-19 Surveillance Team |
| EPI_ISL_3124616, EPI_ISL_3124617 | Lighthouse Lab in Alderley Park | Wellcome Sanger Institute for the COVID-19 Genomics UK (COG-UK) Consortium | Jacquelyn Wynn, Mairead Hyland, The Lighthouse Lab in Alderley Park and Alex Alderton, Roberto Amato, Jeffrey Barrett, Sonia Goncalves, Ewan Harrison, David K. Jackson, Ian Johnston, Dominic Kwiatkowski, Cordelia Langford, John Sillitoe on behalf of the Wellcome Sanger Institute COVID-19 Surveillance Team |
| EPI_ISL_3124618, EPI_ISL_3124619 | Lighthouse Lab in Glasgow | Wellcome Sanger Institute for the COVID-19 Genomics UK (COG-UK) Consortium | Harper VanSteenhouse, Yumi Kasai, David Gray, Carol Clugston, Anna Dominiczak and Alex Alderton, Roberto Amato, Jeffrey Barrett, Sonia Goncalves, Ewan Harrison, David K. Jackson, Ian Johnston, Dominic Kwiatkowski, Cordelia Langford, John Sillitoe on behalf of the Wellcome Sanger Institute COVID-19 Surveillance Team |
| EPI_ISL_3124620, EPI_ISL_3124621, EPI_ISL_3124622 | Lighthouse Lab in Alderley Park | Wellcome Sanger Institute for the COVID-19 Genomics UK (COG-UK) Consortium | Jacquelyn Wynn, Mairead Hyland, The Lighthouse Lab in Alderley Park and Alex Alderton, Roberto Amato, Jeffrey Barrett, Sonia Goncalves, Ewan Harrison, David K. Jackson, Ian Johnston, Dominic Kwiatkowski, Cordelia Langford, John Sillitoe on behalf of the Wellcome Sanger Institute COVID-19 Surveillance Team |
| EPI_ISL_3124623 | Lighthouse Lab in Glasgow | Wellcome Sanger Institute for the COVID-19 Genomics UK (COG-UK) Consortium | Harper VanSteenhouse, Yumi Kasai, David Gray, Carol Clugston, Anna Dominiczak and Alex Alderton, Roberto Amato, Jeffrey Barrett, Sonia Goncalves, Ewan Harrison, David K. Jackson, Ian Johnston, Dominic Kwiatkowski, Cordelia Langford, John Sillitoe on behalf of the Wellcome Sanger Institute COVID-19 Surveillance Team |
| EPI_ISL_3124624, EPI_ISL_3124625, EPI_ISL_3124626, EPI_ISL_3124627 | Lighthouse Lab in Alderley Park | Wellcome Sanger Institute for the COVID-19 Genomics UK (COG-UK) Consortium | Jacquelyn Wynn, Mairead Hyland, The Lighthouse Lab in Alderley Park and Alex Alderton, Roberto Amato, Jeffrey Barrett, Sonia Goncalves, Ewan Harrison, David K. Jackson, Ian Johnston, Dominic Kwiatkowski, Cordelia Langford, John Sillitoe on behalf of the Wellcome Sanger Institute COVID-19 Surveillance Team |
| EPI_ISL_3124628 | Lighthouse Lab in Glasgow | Wellcome Sanger Institute for the COVID-19 Genomics UK (COG-UK) Consortium | Harper VanSteenhouse, Yumi Kasai, David Gray, Carol Clugston, Anna Dominiczak and Alex Alderton, Roberto Amato, Jeffrey Barrett, Sonia Goncalves, Ewan Harrison, David K. Jackson, Ian Johnston, Dominic Kwiatkowski, Cordelia Langford, John Sillitoe on behalf of the Wellcome Sanger Institute COVID-19 Surveillance Team |
| EPI_ISL_3124629 | Lighthouse Lab in Milton Keynes | Wellcome Sanger Institute for the COVID-19 Genomics UK (COG-UK) Consortium | The Lighthouse Lab in Milton Keynes and Alex Alderton, Roberto Amato, Jeffrey Barrett, Sonia Goncalves, Ewan Harrison, David K. Jackson, Ian Johnston, Dominic Kwiatkowski, Cordelia Langford, John Sillitoe on behalf of the Wellcome Sanger Institute COVID-19 Surveillance Team |
| EPI_ISL_3124630 | Lighthouse Lab in Glasgow | Wellcome Sanger Institute for the COVID-19 Genomics UK (COG-UK) Consortium | Harper VanSteenhouse, Yumi Kasai, David Gray, Carol Clugston, Anna Dominiczak and Alex Alderton, Roberto Amato, Jeffrey Barrett, Sonia Goncalves, Ewan Harrison, David K. Jackson, Ian Johnston, Dominic Kwiatkowski, Cordelia Langford, John Sillitoe on behalf of the Wellcome Sanger Institute COVID-19 Surveillance Team |
| EPI_ISL_3124631 | Lighthouse Lab in Milton Keynes | Wellcome Sanger Institute for the COVID-19 Genomics UK (COG-UK) Consortium | The Lighthouse Lab in Milton Keynes and Alex Alderton, Roberto Amato, Jeffrey Barrett, Sonia Goncalves, Ewan Harrison, David K. Jackson, Ian Johnston, Dominic Kwiatkowski, Cordelia Langford, John Sillitoe on behalf of the Wellcome Sanger Institute COVID-19 Surveillance Team |
| EPI_ISL_3124632 | Lighthouse Lab in Alderley Park | Wellcome Sanger Institute for the COVID-19 Genomics UK (COG-UK) Consortium | Jacquelyn Wynn, Mairead Hyland, The Lighthouse Lab in Alderley Park and Alex Alderton, Roberto Amato, Jeffrey Barrett, Sonia Goncalves, Ewan Harrison, David K. Jackson, Ian Johnston, Dominic Kwiatkowski, Cordelia Langford, John Sillitoe on behalf of the Wellcome Sanger Institute COVID-19 Surveillance Team |
| EPI_ISL_3124633, EPI_ISL_3124634 | Lighthouse Lab in Milton Keynes | Wellcome Sanger Institute for the COVID-19 Genomics UK (COG-UK) Consortium | The Lighthouse Lab in Milton Keynes and Alex Alderton, Roberto Amato, Jeffrey Barrett, Sonia Goncalves, Ewan Harrison, David K. Jackson, Ian Johnston, Dominic Kwiatkowski, Cordelia Langford, John Sillitoe on behalf of the Wellcome Sanger Institute COVID-19 Surveillance Team |
| EPI_ISL_3124635 | Lighthouse Lab in Glasgow | Wellcome Sanger Institute for the COVID-19 Genomics UK (COG-UK) Consortium | Harper VanSteenhouse, Yumi Kasai, David Gray, Carol Clugston, Anna Dominiczak and Alex Alderton, Roberto Amato, Jeffrey Barrett, Sonia Goncalves, Ewan Harrison, David K. Jackson, Ian Johnston, Dominic Kwiatkowski, Cordelia Langford, John Sillitoe on behalf of the Wellcome Sanger Institute COVID-19 Surveillance Team |
| EPI_ISL_3124636 | Lighthouse Lab in Alderley Park | Wellcome Sanger Institute for the COVID-19 Genomics UK (COG-UK) Consortium | Jacquelyn Wynn, Mairead Hyland, The Lighthouse Lab in Alderley Park and Alex Alderton, Roberto Amato, Jeffrey Barrett, Sonia Goncalves, Ewan Harrison, David K. Jackson, Ian Johnston, Dominic Kwiatkowski, Cordelia Langford, John Sillitoe on behalf of the Wellcome Sanger Institute COVID-19 Surveillance Team |
| EPI_ISL_3124637 | Lighthouse Lab in Milton Keynes | Wellcome Sanger Institute for the COVID-19 Genomics UK (COG-UK) Consortium | The Lighthouse Lab in Milton Keynes and Alex Alderton, Roberto Amato, Jeffrey Barrett, Sonia Goncalves, Ewan Harrison, David K. Jackson, Ian Johnston, Dominic Kwiatkowski, Cordelia Langford, John Sillitoe on behalf of the Wellcome Sanger Institute COVID-19 Surveillance Team |
| EPI_ISL_3124638 | Lighthouse Lab in Alderley Park | Wellcome Sanger Institute for the COVID-19 Genomics UK (COG-UK) Consortium | Jacquelyn Wynn, Mairead Hyland, The Lighthouse Lab in Alderley Park and Alex Alderton, Roberto Amato, Jeffrey Barrett, Sonia Goncalves, Ewan Harrison, David K. Jackson, Ian Johnston, Dominic Kwiatkowski, Cordelia Langford, John Sillitoe on behalf of the Wellcome Sanger Institute COVID-19 Surveillance Team |
| EPI_ISL_3124639 | Lighthouse Lab in Milton Keynes | Wellcome Sanger Institute for the COVID-19 Genomics UK (COG-UK) Consortium | The Lighthouse Lab in Milton Keynes and Alex Alderton, Roberto Amato, Jeffrey Barrett, Sonia Goncalves, Ewan Harrison, David K. Jackson, Ian Johnston, Dominic Kwiatkowski, Cordelia Langford, John Sillitoe on behalf of the Wellcome Sanger Institute COVID-19 Surveillance Team |
| EPI_ISL_3124640, EPI_ISL_3124641 | Lighthouse Lab in Glasgow | Wellcome Sanger Institute for the COVID-19 Genomics UK (COG-UK) Consortium | Harper VanSteenhouse, Yumi Kasai, David Gray, Carol Clugston, Anna Dominiczak and Alex Alderton, Roberto Amato, Jeffrey Barrett, Sonia Goncalves, Ewan Harrison, David K. Jackson, Ian Johnston, Dominic Kwiatkowski, Cordelia Langford, John Sillitoe on behalf of the Wellcome Sanger Institute COVID-19 Surveillance Team |
| EPI_ISL_3124642 | Lighthouse Lab in Alderley Park | Wellcome Sanger Institute for the COVID-19 Genomics UK (COG-UK) Consortium | Jacquelyn Wynn, Mairead Hyland, The Lighthouse Lab in Alderley Park and Alex Alderton, Roberto Amato, Jeffrey Barrett, Sonia Goncalves, Ewan Harrison, David K. Jackson, Ian Johnston, Dominic Kwiatkowski, Cordelia Langford, John Sillitoe on behalf of the Wellcome Sanger Institute COVID-19 Surveillance Team |
| EPI_ISL_3124643 | Lighthouse Lab in Glasgow | Wellcome Sanger Institute for the COVID-19 Genomics UK (COG-UK) Consortium | Harper VanSteenhouse, Yumi Kasai, David Gray, Carol Clugston, Anna Dominiczak and Alex Alderton, Roberto Amato, Jeffrey Barrett, Sonia Goncalves, Ewan Harrison, David K. Jackson, Ian Johnston, Dominic Kwiatkowski, Cordelia Langford, John Sillitoe on behalf of the Wellcome Sanger Institute COVID-19 Surveillance Team |
| EPI_ISL_3124644, EPI_ISL_3124645 | Lighthouse Lab in Milton Keynes | Wellcome Sanger Institute for the COVID-19 Genomics UK (COG-UK) Consortium | The Lighthouse Lab in Milton Keynes and Alex Alderton, Roberto Amato, Jeffrey Barrett, Sonia Goncalves, Ewan Harrison, David K. Jackson, Ian Johnston, Dominic Kwiatkowski, Cordelia Langford, John Sillitoe on behalf of the Wellcome Sanger Institute COVID-19 Surveillance Team |
| EPI_ISL_3124646 | Lighthouse Lab in Glasgow | Wellcome Sanger Institute for the COVID-19 Genomics UK (COG-UK) Consortium | Harper VanSteenhouse, Yumi Kasai, David Gray, Carol Clugston, Anna Dominiczak and Alex Alderton, Roberto Amato, Jeffrey Barrett, Sonia Goncalves, Ewan Harrison, David K. Jackson, Ian Johnston, Dominic Kwiatkowski, Cordelia Langford, John Sillitoe on behalf of the Wellcome Sanger Institute COVID-19 Surveillance Team |
| EPI_ISL_3124647 | Lighthouse Lab in Alderley Park | Wellcome Sanger Institute for the COVID-19 Genomics UK (COG-UK) Consortium | Jacquelyn Wynn, Mairead Hyland, The Lighthouse Lab in Alderley Park and Alex Alderton, Roberto Amato, Jeffrey Barrett, Sonia Goncalves, Ewan Harrison, David K. Jackson, Ian Johnston, Dominic Kwiatkowski, Cordelia Langford, John Sillitoe on behalf of the Wellcome Sanger Institute COVID-19 Surveillance Team |
| EPI_ISL_3124648 | Lighthouse Lab in Glasgow | Wellcome Sanger Institute for the COVID-19 Genomics UK (COG-UK) Consortium | Harper VanSteenhouse, Yumi Kasai, David Gray, Carol Clugston, Anna Dominiczak and Alex Alderton, Roberto Amato, Jeffrey Barrett, Sonia Goncalves, Ewan Harrison, David K. Jackson, Ian Johnston, Dominic Kwiatkowski, Cordelia Langford, John Sillitoe on behalf of the Wellcome Sanger Institute COVID-19 Surveillance Team |
| EPI_ISL_3124649 | Lighthouse Lab in Alderley Park | Wellcome Sanger Institute for the COVID-19 Genomics UK (COG-UK) Consortium | Jacquelyn Wynn, Mairead Hyland, The Lighthouse Lab in Alderley Park and Alex Alderton, Roberto Amato, Jeffrey Barrett, Sonia Goncalves, Ewan Harrison, David K. Jackson, Ian Johnston, Dominic Kwiatkowski, Cordelia Langford, John Sillitoe on behalf of the Wellcome Sanger Institute COVID-19 Surveillance Team |
| EPI_ISL_3124650, EPI_ISL_3124651 | Lighthouse Lab in Milton Keynes | Wellcome Sanger Institute for the COVID-19 Genomics UK (COG-UK) Consortium | The Lighthouse Lab in Milton Keynes and Alex Alderton, Roberto Amato, Jeffrey Barrett, Sonia Goncalves, Ewan Harrison, David K. Jackson, Ian Johnston, Dominic Kwiatkowski, Cordelia Langford, John Sillitoe on behalf of the Wellcome Sanger Institute COVID-19 Surveillance Team |
| EPI_ISL_3124652, EPI_ISL_3124653 | Lighthouse Lab in Alderley Park | Wellcome Sanger Institute for the COVID-19 Genomics UK (COG-UK) Consortium | Jacquelyn Wynn, Mairead Hyland, The Lighthouse Lab in Alderley Park and Alex Alderton, Roberto Amato, Jeffrey Barrett, Sonia Goncalves, Ewan Harrison, David K. Jackson, Ian Johnston, Dominic Kwiatkowski, Cordelia Langford, John Sillitoe on behalf of the Wellcome Sanger Institute COVID-19 Surveillance Team |
| EPI_ISL_3124654 | Lighthouse Lab in Milton Keynes | Wellcome Sanger Institute for the COVID-19 Genomics UK (COG-UK) Consortium | The Lighthouse Lab in Milton Keynes and Alex Alderton, Roberto Amato, Jeffrey Barrett, Sonia Goncalves, Ewan Harrison, David K. Jackson, Ian Johnston, Dominic Kwiatkowski, Cordelia Langford, John Sillitoe on behalf of the Wellcome Sanger Institute COVID-19 Surveillance Team |
| EPI_ISL_3124655 | Lighthouse Lab in Glasgow | Wellcome Sanger Institute for the COVID-19 Genomics UK (COG-UK) Consortium | Harper VanSteenhouse, Yumi Kasai, David Gray, Carol Clugston, Anna Dominiczak and Alex Alderton, Roberto Amato, Jeffrey Barrett, Sonia Goncalves, Ewan Harrison, David K. Jackson, Ian Johnston, Dominic Kwiatkowski, Cordelia Langford, John Sillitoe on behalf of the Wellcome Sanger Institute COVID-19 Surveillance Team |
| EPI_ISL_3124656 | Lighthouse Lab in Milton Keynes | Wellcome Sanger Institute for the COVID-19 Genomics UK (COG-UK) Consortium | The Lighthouse Lab in Milton Keynes and Alex Alderton, Roberto Amato, Jeffrey Barrett, Sonia Goncalves, Ewan Harrison, David K. Jackson, Ian Johnston, Dominic Kwiatkowski, Cordelia Langford, John Sillitoe on behalf of the Wellcome Sanger Institute COVID-19 Surveillance Team |
| EPI_ISL_3124657 | Lighthouse Lab in Glasgow | Wellcome Sanger Institute for the COVID-19 Genomics UK (COG-UK) Consortium | Harper VanSteenhouse, Yumi Kasai, David Gray, Carol Clugston, Anna Dominiczak and Alex Alderton, Roberto Amato, Jeffrey Barrett, Sonia Goncalves, Ewan Harrison, David K. Jackson, Ian Johnston, Dominic Kwiatkowski, Cordelia Langford, John Sillitoe on behalf of the Wellcome Sanger Institute COVID-19 Surveillance Team |
| EPI_ISL_3124658, EPI_ISL_3124659 | Lighthouse Lab in Alderley Park | Wellcome Sanger Institute for the COVID-19 Genomics UK (COG-UK) Consortium | Jacquelyn Wynn, Mairead Hyland, The Lighthouse Lab in Alderley Park and Alex Alderton, Roberto Amato, Jeffrey Barrett, Sonia Goncalves, Ewan Harrison, David K. Jackson, Ian Johnston, Dominic Kwiatkowski, Cordelia Langford, John Sillitoe on behalf of the Wellcome Sanger Institute COVID-19 Surveillance Team |
| EPI_ISL_3124660, EPI_ISL_3124661 | Lighthouse Lab in Milton Keynes | Wellcome Sanger Institute for the COVID-19 Genomics UK (COG-UK) Consortium | The Lighthouse Lab in Milton Keynes and Alex Alderton, Roberto Amato, Jeffrey Barrett, Sonia Goncalves, Ewan Harrison, David K. Jackson, Ian Johnston, Dominic Kwiatkowski, Cordelia Langford, John Sillitoe on behalf of the Wellcome Sanger Institute COVID-19 Surveillance Team |
| EPI_ISL_3124662 | Lighthouse Lab in Glasgow | Wellcome Sanger Institute for the COVID-19 Genomics UK (COG-UK) Consortium | Harper VanSteenhouse, Yumi Kasai, David Gray, Carol Clugston, Anna Dominiczak and Alex Alderton, Roberto Amato, Jeffrey Barrett, Sonia Goncalves, Ewan Harrison, David K. Jackson, Ian Johnston, Dominic Kwiatkowski, Cordelia Langford, John Sillitoe on behalf of the Wellcome Sanger Institute COVID-19 Surveillance Team |
| EPI_ISL_3124663 | Lighthouse Lab in Alderley Park | Wellcome Sanger Institute for the COVID-19 Genomics UK (COG-UK) | Jacquelyn Wynn, Mairead Hyland, The Lighthouse Lab in Alderley Park and Alex Alderton, Roberto Amato, Jeffrey Barrett, Sonia Goncalves, Ewan Harrison, David K. Jackson, Ian Johnston, Dominic Kwiatkowski, Cordelia Langford, John Sillitoe on |

| | | | |
|---|---|---|---|
| | | Consortium | behalf of the Wellcome Sanger Institute COVID-19 Surveillance Team |
| EPI_ISL_3124664, EPI_ISL_3124665 | Lighthouse Lab in Glasgow | Wellcome Sanger Institute for the COVID-19 Genomics UK (COG-UK) Consortium | Harper VanSteenhouse, Yumi Kasai, David Gray, Carol Clugston, Anna Dominiczak and Alex Alderton, Roberto Amato, Jeffrey Barrett, Sonia Goncalves, Ewan Harrison, David K. Jackson, Ian Johnston, Dominic Kwiatkowski, Cordelia Langford, John Sillitoe on behalf of the Wellcome Sanger Institute COVID-19 Surveillance Team |
| EPI_ISL_3124666, EPI_ISL_3124667, EPI_ISL_3124668 | Lighthouse Lab in Milton Keynes | Wellcome Sanger Institute for the COVID-19 Genomics UK (COG-UK) Consortium | The Lighthouse Lab in Milton Keynes and Alex Alderton, Roberto Amato, Jeffrey Barrett, Sonia Goncalves, Ewan Harrison, David K. Jackson, Ian Johnston, Dominic Kwiatkowski, Cordelia Langford, John Sillitoe on behalf of the Wellcome Sanger Institute COVID-19 Surveillance Team |
| EPI_ISL_3124669 | Lighthouse Lab in Alderley Park | Wellcome Sanger Institute for the COVID-19 Genomics UK (COG-UK) Consortium | Jacquelyn Wynn, Mairead Hyland, The Lighthouse Lab in Alderley Park and Alex Alderton, Roberto Amato, Jeffrey Barrett, Sonia Goncalves, Ewan Harrison, David K. Jackson, Ian Johnston, Dominic Kwiatkowski, Cordelia Langford, John Sillitoe on behalf of the Wellcome Sanger Institute COVID-19 Surveillance Team |
| EPI_ISL_3124670 | Lighthouse Lab in Glasgow | Wellcome Sanger Institute for the COVID-19 Genomics UK (COG-UK) Consortium | Harper VanSteenhouse, Yumi Kasai, David Gray, Carol Clugston, Anna Dominiczak and Alex Alderton, Roberto Amato, Jeffrey Barrett, Sonia Goncalves, Ewan Harrison, David K. Jackson, Ian Johnston, Dominic Kwiatkowski, Cordelia Langford, John Sillitoe on behalf of the Wellcome Sanger Institute COVID-19 Surveillance Team |
| EPI_ISL_3124671 | Lighthouse Lab in Alderley Park | Wellcome Sanger Institute for the COVID-19 Genomics UK (COG-UK) Consortium | Jacquelyn Wynn, Mairead Hyland, The Lighthouse Lab in Alderley Park and Alex Alderton, Roberto Amato, Jeffrey Barrett, Sonia Goncalves, Ewan Harrison, David K. Jackson, Ian Johnston, Dominic Kwiatkowski, Cordelia Langford, John Sillitoe on behalf of the Wellcome Sanger Institute COVID-19 Surveillance Team |
| EPI_ISL_3124672 | Lighthouse Lab in Glasgow | Wellcome Sanger Institute for the COVID-19 Genomics UK (COG-UK) Consortium | Harper VanSteenhouse, Yumi Kasai, David Gray, Carol Clugston, Anna Dominiczak and Alex Alderton, Roberto Amato, Jeffrey Barrett, Sonia Goncalves, Ewan Harrison, David K. Jackson, Ian Johnston, Dominic Kwiatkowski, Cordelia Langford, John Sillitoe on behalf of the Wellcome Sanger Institute COVID-19 Surveillance Team |
| EPI_ISL_3124673 | Lighthouse Lab in Milton Keynes | Wellcome Sanger Institute for the COVID-19 Genomics UK (COG-UK) Consortium | The Lighthouse Lab in Milton Keynes and Alex Alderton, Roberto Amato, Jeffrey Barrett, Sonia Goncalves, Ewan Harrison, David K. Jackson, Ian Johnston, Dominic Kwiatkowski, Cordelia Langford, John Sillitoe on behalf of the Wellcome Sanger Institute COVID-19 Surveillance Team |
| EPI_ISL_3124674 | Lighthouse Lab in Glasgow | Wellcome Sanger Institute for the COVID-19 Genomics UK (COG-UK) Consortium | Harper VanSteenhouse, Yumi Kasai, David Gray, Carol Clugston, Anna Dominiczak and Alex Alderton, Roberto Amato, Jeffrey Barrett, Sonia Goncalves, Ewan Harrison, David K. Jackson, Ian Johnston, Dominic Kwiatkowski, Cordelia Langford, John Sillitoe on behalf of the Wellcome Sanger Institute COVID-19 Surveillance Team |
| EPI_ISL_3124675 | Lighthouse Lab in Alderley Park | Wellcome Sanger Institute for the COVID-19 Genomics UK (COG-UK) Consortium | Jacquelyn Wynn, Mairead Hyland, The Lighthouse Lab in Alderley Park and Alex Alderton, Roberto Amato, Jeffrey Barrett, Sonia Goncalves, Ewan Harrison, David K. Jackson, Ian Johnston, Dominic Kwiatkowski, Cordelia Langford, John Sillitoe on behalf of the Wellcome Sanger Institute COVID-19 Surveillance Team |
| EPI_ISL_3124676 | Lighthouse Lab in Glasgow | Wellcome Sanger Institute for the COVID-19 Genomics UK (COG-UK) Consortium | Harper VanSteenhouse, Yumi Kasai, David Gray, Carol Clugston, Anna Dominiczak and Alex Alderton, Roberto Amato, Jeffrey Barrett, Sonia Goncalves, Ewan Harrison, David K. Jackson, Ian Johnston, Dominic Kwiatkowski, Cordelia Langford, John Sillitoe on behalf of the Wellcome Sanger Institute COVID-19 Surveillance Team |
| EPI_ISL_3124677 | Lighthouse Lab in Alderley Park | Wellcome Sanger Institute for the COVID-19 Genomics UK (COG-UK) Consortium | Jacquelyn Wynn, Mairead Hyland, The Lighthouse Lab in Alderley Park and Alex Alderton, Roberto Amato, Jeffrey Barrett, Sonia Goncalves, Ewan Harrison, David K. Jackson, Ian Johnston, Dominic Kwiatkowski, Cordelia Langford, John Sillitoe on behalf of the Wellcome Sanger Institute COVID-19 Surveillance Team |
| EPI_ISL_3124678, EPI_ISL_3124679, EPI_ISL_3124680, EPI_ISL_3124681 | Lighthouse Lab in Glasgow | Wellcome Sanger Institute for the COVID-19 Genomics UK (COG-UK) Consortium | Harper VanSteenhouse, Yumi Kasai, David Gray, Carol Clugston, Anna Dominiczak and Alex Alderton, Roberto Amato, Jeffrey Barrett, Sonia Goncalves, Ewan Harrison, David K. Jackson, Ian Johnston, Dominic Kwiatkowski, Cordelia Langford, John Sillitoe on behalf of the Wellcome Sanger Institute COVID-19 Surveillance Team |
| EPI_ISL_3124682 | Lighthouse Lab in Milton Keynes | Wellcome Sanger Institute for the COVID-19 Genomics UK (COG-UK) Consortium | The Lighthouse Lab in Milton Keynes and Alex Alderton, Roberto Amato, Jeffrey Barrett, Sonia Goncalves, Ewan Harrison, David K. Jackson, Ian Johnston, Dominic Kwiatkowski, Cordelia Langford, John Sillitoe on behalf of the Wellcome Sanger Institute COVID-19 Surveillance Team |
| EPI_ISL_3124683 | Lighthouse Lab in Alderley Park | Wellcome Sanger Institute for the COVID-19 Genomics UK (COG-UK) Consortium | Jacquelyn Wynn, Mairead Hyland, The Lighthouse Lab in Alderley Park and Alex Alderton, Roberto Amato, Jeffrey Barrett, Sonia Goncalves, Ewan Harrison, David K. Jackson, Ian Johnston, Dominic Kwiatkowski, Cordelia Langford, John Sillitoe on behalf of the Wellcome Sanger Institute COVID-19 Surveillance Team |
| EPI_ISL_3124684 | Lighthouse Lab in Glasgow | Wellcome Sanger Institute for the COVID-19 Genomics UK (COG-UK) Consortium | Harper VanSteenhouse, Yumi Kasai, David Gray, Carol Clugston, Anna Dominiczak and Alex Alderton, Roberto Amato, Jeffrey Barrett, Sonia Goncalves, Ewan Harrison, David K. Jackson, Ian Johnston, Dominic Kwiatkowski, Cordelia Langford, John Sillitoe on behalf of the Wellcome Sanger Institute COVID-19 Surveillance Team |
| EPI_ISL_3124685, EPI_ISL_3124686, EPI_ISL_3124687, EPI_ISL_3124688 | Lighthouse Lab in Alderley Park | Wellcome Sanger Institute for the COVID-19 Genomics UK (COG-UK) Consortium | Jacquelyn Wynn, Mairead Hyland, The Lighthouse Lab in Alderley Park and Alex Alderton, Roberto Amato, Jeffrey Barrett, Sonia Goncalves, Ewan Harrison, David K. Jackson, Ian Johnston, Dominic Kwiatkowski, Cordelia Langford, John Sillitoe on behalf of the Wellcome Sanger Institute COVID-19 Surveillance Team |
| EPI_ISL_3124689 | Lighthouse Lab in Glasgow | Wellcome Sanger Institute for the COVID-19 Genomics UK (COG-UK) Consortium | Harper VanSteenhouse, Yumi Kasai, David Gray, Carol Clugston, Anna Dominiczak and Alex Alderton, Roberto Amato, Jeffrey Barrett, Sonia Goncalves, Ewan Harrison, David K. Jackson, Ian Johnston, Dominic Kwiatkowski, Cordelia Langford, John Sillitoe on behalf of the Wellcome Sanger Institute COVID-19 Surveillance Team |
| EPI_ISL_3124690 | Lighthouse Lab in Alderley Park | Wellcome Sanger Institute for the COVID-19 Genomics UK (COG-UK) Consortium | Jacquelyn Wynn, Mairead Hyland, The Lighthouse Lab in Alderley Park and Alex Alderton, Roberto Amato, Jeffrey Barrett, Sonia Goncalves, Ewan Harrison, David K. Jackson, Ian Johnston, Dominic Kwiatkowski, Cordelia Langford, John Sillitoe on behalf of the Wellcome Sanger Institute COVID-19 Surveillance Team |
| EPI_ISL_3124691 | Lighthouse Lab in Milton Keynes | Wellcome Sanger Institute for the COVID-19 Genomics UK (COG-UK) Consortium | The Lighthouse Lab in Milton Keynes and Alex Alderton, Roberto Amato, Jeffrey Barrett, Sonia Goncalves, Ewan Harrison, David K. Jackson, Ian Johnston, Dominic Kwiatkowski, Cordelia Langford, John Sillitoe on behalf of the Wellcome Sanger Institute COVID-19 Surveillance Team |
| EPI_ISL_3124692, EPI_ISL_3124693, EPI_ISL_3124694, EPI_ISL_3124695, EPI_ISL_3124696 | Lighthouse Lab in Alderley Park | Wellcome Sanger Institute for the COVID-19 Genomics UK (COG-UK) Consortium | Jacquelyn Wynn, Mairead Hyland, The Lighthouse Lab in Alderley Park and Alex Alderton, Roberto Amato, Jeffrey Barrett, Sonia Goncalves, Ewan Harrison, David K. Jackson, Ian Johnston, Dominic Kwiatkowski, Cordelia Langford, John Sillitoe on behalf of the Wellcome Sanger Institute COVID-19 Surveillance Team |
| EPI_ISL_3124697 | Lighthouse Lab in Milton Keynes | Wellcome Sanger Institute for the COVID-19 Genomics UK (COG-UK) Consortium | The Lighthouse Lab in Milton Keynes and Alex Alderton, Roberto Amato, Jeffrey Barrett, Sonia Goncalves, Ewan Harrison, David K. Jackson, Ian Johnston, Dominic Kwiatkowski, Cordelia Langford, John Sillitoe on behalf of the Wellcome Sanger Institute COVID-19 Surveillance Team |
| EPI_ISL_3124698 | Lighthouse Lab in Glasgow | Wellcome Sanger Institute for the COVID-19 Genomics UK (COG-UK) Consortium | Harper VanSteenhouse, Yumi Kasai, David Gray, Carol Clugston, Anna Dominiczak and Alex Alderton, Roberto Amato, Jeffrey Barrett, Sonia Goncalves, Ewan Harrison, David K. Jackson, Ian Johnston, Dominic Kwiatkowski, Cordelia Langford, John Sillitoe on behalf of the Wellcome Sanger Institute COVID-19 Surveillance Team |
| EPI_ISL_3124699 | Lighthouse Lab in Alderley Park | Wellcome Sanger Institute for the COVID-19 Genomics UK (COG-UK) Consortium | Jacquelyn Wynn, Mairead Hyland, The Lighthouse Lab in Alderley Park and Alex Alderton, Roberto Amato, Jeffrey Barrett, Sonia Goncalves, Ewan Harrison, David K. Jackson, Ian Johnston, Dominic Kwiatkowski, Cordelia Langford, John Sillitoe on behalf of the Wellcome Sanger Institute COVID-19 Surveillance Team |
| EPI_ISL_3124700 | Lighthouse Lab in Glasgow | Wellcome Sanger Institute for the COVID-19 Genomics UK (COG-UK) Consortium | Harper VanSteenhouse, Yumi Kasai, David Gray, Carol Clugston, Anna Dominiczak and Alex Alderton, Roberto Amato, Jeffrey Barrett, Sonia Goncalves, Ewan Harrison, David K. Jackson, Ian Johnston, Dominic Kwiatkowski, Cordelia Langford, John Sillitoe on behalf of the Wellcome Sanger Institute COVID-19 Surveillance Team |
| EPI_ISL_3124701 | Lighthouse Lab in Milton Keynes | Wellcome Sanger Institute for the COVID-19 Genomics UK (COG-UK) Consortium | The Lighthouse Lab in Milton Keynes and Alex Alderton, Roberto Amato, Jeffrey Barrett, Sonia Goncalves, Ewan Harrison, David K. Jackson, Ian Johnston, Dominic Kwiatkowski, Cordelia Langford, John Sillitoe on behalf of the Wellcome Sanger Institute COVID-19 Surveillance Team |
| EPI_ISL_3124702, EPI_ISL_3124703, EPI_ISL_3124704 | Lighthouse Lab in Glasgow | Wellcome Sanger Institute for the COVID-19 Genomics UK (COG-UK) Consortium | Harper VanSteenhouse, Yumi Kasai, David Gray, Carol Clugston, Anna Dominiczak and Alex Alderton, Roberto Amato, Jeffrey Barrett, Sonia Goncalves, Ewan Harrison, David K. Jackson, Ian Johnston, Dominic Kwiatkowski, Cordelia Langford, John Sillitoe on behalf of the Wellcome Sanger Institute COVID-19 Surveillance Team |
| EPI_ISL_3124705 | Lighthouse Lab in Alderley Park | Wellcome Sanger Institute for the COVID-19 Genomics UK (COG-UK) Consortium | Jacquelyn Wynn, Mairead Hyland, The Lighthouse Lab in Alderley Park and Alex Alderton, Roberto Amato, Jeffrey Barrett, Sonia Goncalves, Ewan Harrison, David K. Jackson, Ian Johnston, Dominic Kwiatkowski, Cordelia Langford, John Sillitoe on behalf of the Wellcome Sanger Institute COVID-19 Surveillance Team |
| EPI_ISL_3124706, EPI_ISL_3124707 | Lighthouse Lab in Glasgow | Wellcome Sanger Institute for the COVID-19 Genomics UK (COG-UK) Consortium | Harper VanSteenhouse, Yumi Kasai, David Gray, Carol Clugston, Anna Dominiczak and Alex Alderton, Roberto Amato, Jeffrey Barrett, Sonia Goncalves, Ewan Harrison, David K. Jackson, Ian Johnston, Dominic Kwiatkowski, Cordelia Langford, John Sillitoe on behalf of the Wellcome Sanger Institute COVID-19 Surveillance Team |
| EPI_ISL_3124708 | Lighthouse Lab in Alderley Park | Wellcome Sanger Institute for the COVID-19 Genomics UK (COG-UK) Consortium | Jacquelyn Wynn, Mairead Hyland, The Lighthouse Lab in Alderley Park and Alex Alderton, Roberto Amato, Jeffrey Barrett, Sonia Goncalves, Ewan Harrison, David K. Jackson, Ian Johnston, Dominic Kwiatkowski, Cordelia Langford, John Sillitoe on behalf of the Wellcome Sanger Institute COVID-19 Surveillance Team |
| EPI_ISL_3124709, EPI_ISL_3124710 | Lighthouse Lab in Glasgow | Wellcome Sanger Institute for the COVID-19 Genomics UK (COG-UK) Consortium | Harper VanSteenhouse, Yumi Kasai, David Gray, Carol Clugston, Anna Dominiczak and Alex Alderton, Roberto Amato, Jeffrey Barrett, Sonia Goncalves, Ewan Harrison, David K. Jackson, Ian Johnston, Dominic Kwiatkowski, Cordelia Langford, John Sillitoe on behalf of the Wellcome Sanger Institute COVID-19 Surveillance Team |
| EPI_ISL_3124711 | Lighthouse Lab in Alderley Park | Wellcome Sanger Institute for the COVID-19 Genomics UK (COG-UK) Consortium | Jacquelyn Wynn, Mairead Hyland, The Lighthouse Lab in Alderley Park and Alex Alderton, Roberto Amato, Jeffrey Barrett, Sonia Goncalves, Ewan Harrison, David K. Jackson, Ian Johnston, Dominic Kwiatkowski, Cordelia Langford, John Sillitoe on behalf of the Wellcome Sanger Institute COVID-19 Surveillance Team |
| EPI_ISL_3124712 | Lighthouse Lab in Milton Keynes | Wellcome Sanger Institute for the COVID-19 Genomics UK (COG-UK) Consortium | The Lighthouse Lab in Milton Keynes and Alex Alderton, Roberto Amato, Jeffrey Barrett, Sonia Goncalves, Ewan Harrison, David K. Jackson, Ian Johnston, Dominic Kwiatkowski, Cordelia Langford, John Sillitoe on behalf of the Wellcome Sanger Institute COVID-19 Surveillance Team |
| EPI_ISL_3124713, EPI_ISL_3124714 | Lighthouse Lab in Alderley Park | Wellcome Sanger Institute for the COVID-19 Genomics UK (COG-UK) Consortium | Jacquelyn Wynn, Mairead Hyland, The Lighthouse Lab in Alderley Park and Alex Alderton, Roberto Amato, Jeffrey Barrett, Sonia Goncalves, Ewan Harrison, David K. Jackson, Ian Johnston, Dominic Kwiatkowski, Cordelia Langford, John Sillitoe on behalf of the Wellcome Sanger Institute COVID-19 Surveillance Team |
| EPI_ISL_3124715 | Lighthouse Lab in Milton Keynes | Wellcome Sanger Institute for the COVID-19 Genomics UK (COG-UK) Consortium | The Lighthouse Lab in Milton Keynes and Alex Alderton, Roberto Amato, Jeffrey Barrett, Sonia Goncalves, Ewan Harrison, David K. Jackson, Ian Johnston, Dominic Kwiatkowski, Cordelia Langford, John Sillitoe on behalf of the Wellcome Sanger Institute COVID-19 Surveillance Team |
| EPI_ISL_3124716, EPI_ISL_3124717 | Lighthouse Lab in Alderley Park | Wellcome Sanger Institute for the COVID-19 Genomics UK (COG-UK) Consortium | Jacquelyn Wynn, Mairead Hyland, The Lighthouse Lab in Alderley Park and Alex Alderton, Roberto Amato, Jeffrey Barrett, Sonia Goncalves, Ewan Harrison, David K. Jackson, Ian Johnston, Dominic Kwiatkowski, Cordelia Langford, John Sillitoe on behalf of the Wellcome Sanger Institute COVID-19 Surveillance Team |
| EPI_ISL_3124718 | Lighthouse Lab in Milton Keynes | Wellcome Sanger Institute for the COVID-19 Genomics UK (COG-UK) Consortium | The Lighthouse Lab in Milton Keynes and Alex Alderton, Roberto Amato, Jeffrey Barrett, Sonia Goncalves, Ewan Harrison, David K. Jackson, Ian Johnston, Dominic Kwiatkowski, Cordelia Langford, John Sillitoe on behalf of the Wellcome Sanger Institute COVID-19 Surveillance Team |
| EPI_ISL_3124719, EPI_ISL_3124720, EPI_ISL_3124721 | Lighthouse Lab in Alderley Park | Wellcome Sanger Institute for the COVID-19 Genomics UK (COG-UK) Consortium | Jacquelyn Wynn, Mairead Hyland, The Lighthouse Lab in Alderley Park and Alex Alderton, Roberto Amato, Jeffrey Barrett, Sonia Goncalves, Ewan Harrison, David K. Jackson, Ian Johnston, Dominic Kwiatkowski, Cordelia Langford, John Sillitoe on behalf of the Wellcome Sanger Institute COVID-19 Surveillance Team |
| EPI_ISL_3124722 | Lighthouse Lab in Milton Keynes | Wellcome Sanger Institute for the COVID-19 Genomics UK (COG-UK) Consortium | The Lighthouse Lab in Milton Keynes and Alex Alderton, Roberto Amato, Jeffrey Barrett, Sonia Goncalves, Ewan Harrison, David K. Jackson, Ian Johnston, Dominic Kwiatkowski, Cordelia Langford, John Sillitoe on behalf of the Wellcome Sanger Institute COVID-19 Surveillance Team |
| EPI_ISL_3124723 | Lighthouse Lab in Glasgow | Wellcome Sanger Institute for the COVID-19 Genomics UK (COG-UK) Consortium | Harper VanSteenhouse, Yumi Kasai, David Gray, Carol Clugston, Anna Dominiczak and Alex Alderton, Roberto Amato, Jeffrey Barrett, Sonia Goncalves, Ewan Harrison, David K. Jackson, Ian Johnston, Dominic Kwiatkowski, Cordelia Langford, John Sillitoe on behalf of the Wellcome Sanger Institute COVID-19 Surveillance Team |
| EPI_ISL_3124724 | Lighthouse Lab in Alderley Park | Wellcome Sanger Institute for the COVID-19 Genomics UK (COG-UK) Consortium | Jacquelyn Wynn, Mairead Hyland, The Lighthouse Lab in Alderley Park and Alex Alderton, Roberto Amato, Jeffrey Barrett, Sonia Goncalves, Ewan Harrison, David K. Jackson, Ian Johnston, Dominic Kwiatkowski, Cordelia Langford, John Sillitoe on behalf of the Wellcome Sanger Institute COVID-19 Surveillance Team |
| EPI_ISL_3124725 | Lighthouse Lab in Glasgow | Wellcome Sanger Institute for the COVID-19 Genomics UK (COG-UK) | Harper VanSteenhouse, Yumi Kasai, David Gray, Carol Clugston, Anna Dominiczak and Alex Alderton, Roberto Amato, Jeffrey Barrett, Sonia Goncalves, Ewan Harrison, David K. Jackson, Ian Johnston, Dominic Kwiatkowski, Cordelia Langford, |

| | | | |
|---|---|---|---|
| | | Consortium | John Sillitoe on behalf of the Wellcome Sanger Institute COVID-19 Surveillance Team |
| EPI_ISL_3124726 | Lighthouse Lab in Milton Keynes | Wellcome Sanger Institute for the COVID-19 Genomics UK (COG-UK) Consortium | The Lighthouse Lab in Milton Keynes and Alex Alderton, Roberto Amato, Jeffrey Barrett, Sonia Goncalves, Ewan Harrison, David K. Jackson, Ian Johnston, Dominic Kwiatkowski, Cordelia Langford, John Sillitoe on behalf of the Wellcome Sanger Institute COVID-19 Surveillance Team |
| EPI_ISL_3124727 | Lighthouse Lab in Alderley Park | Wellcome Sanger Institute for the COVID-19 Genomics UK (COG-UK) Consortium | Jacquelyn Wynn, Mairead Hyland, The Lighthouse Lab in Alderley Park and Alex Alderton, Roberto Amato, Jeffrey Barrett, Sonia Goncalves, Ewan Harrison, David K. Jackson, Ian Johnston, Dominic Kwiatkowski, Cordelia Langford, John Sillitoe on behalf of the Wellcome Sanger Institute COVID-19 Surveillance Team |
| EPI_ISL_3124728 | Lighthouse Lab in Milton Keynes | Wellcome Sanger Institute for the COVID-19 Genomics UK (COG-UK) Consortium | The Lighthouse Lab in Milton Keynes and Alex Alderton, Roberto Amato, Jeffrey Barrett, Sonia Goncalves, Ewan Harrison, David K. Jackson, Ian Johnston, Dominic Kwiatkowski, Cordelia Langford, John Sillitoe on behalf of the Wellcome Sanger Institute COVID-19 Surveillance Team |
| EPI_ISL_3124729 | Lighthouse Lab in Glasgow | Wellcome Sanger Institute for the COVID-19 Genomics UK (COG-UK) Consortium | Harper VanSteenhouse, Yumi Kasai, David Gray, Carol Clugston, Anna Dominiczak and Alex Alderton, Roberto Amato, Jeffrey Barrett, Sonia Goncalves, Ewan Harrison, David K. Jackson, Ian Johnston, Dominic Kwiatkowski, Cordelia Langford, John Sillitoe on behalf of the Wellcome Sanger Institute COVID-19 Surveillance Team |
| EPI_ISL_3124730, EPI_ISL_3124731, EPI_ISL_3124732 | Lighthouse Lab in Alderley Park | Wellcome Sanger Institute for the COVID-19 Genomics UK (COG-UK) Consortium | Jacquelyn Wynn, Mairead Hyland, The Lighthouse Lab in Alderley Park and Alex Alderton, Roberto Amato, Jeffrey Barrett, Sonia Goncalves, Ewan Harrison, David K. Jackson, Ian Johnston, Dominic Kwiatkowski, Cordelia Langford, John Sillitoe on behalf of the Wellcome Sanger Institute COVID-19 Surveillance Team |
| EPI_ISL_3124733, EPI_ISL_3124734 | Lighthouse Lab in Glasgow | Wellcome Sanger Institute for the COVID-19 Genomics UK (COG-UK) Consortium | Harper VanSteenhouse, Yumi Kasai, David Gray, Carol Clugston, Anna Dominiczak and Alex Alderton, Roberto Amato, Jeffrey Barrett, Sonia Goncalves, Ewan Harrison, David K. Jackson, Ian Johnston, Dominic Kwiatkowski, Cordelia Langford, John Sillitoe on behalf of the Wellcome Sanger Institute COVID-19 Surveillance Team |
| EPI_ISL_3124735 | Lighthouse Lab in Milton Keynes | Wellcome Sanger Institute for the COVID-19 Genomics UK (COG-UK) Consortium | The Lighthouse Lab in Milton Keynes and Alex Alderton, Roberto Amato, Jeffrey Barrett, Sonia Goncalves, Ewan Harrison, David K. Jackson, Ian Johnston, Dominic Kwiatkowski, Cordelia Langford, John Sillitoe on behalf of the Wellcome Sanger Institute COVID-19 Surveillance Team |
| EPI_ISL_3124736 | Lighthouse Lab in Glasgow | Wellcome Sanger Institute for the COVID-19 Genomics UK (COG-UK) Consortium | Harper VanSteenhouse, Yumi Kasai, David Gray, Carol Clugston, Anna Dominiczak and Alex Alderton, Roberto Amato, Jeffrey Barrett, Sonia Goncalves, Ewan Harrison, David K. Jackson, Ian Johnston, Dominic Kwiatkowski, Cordelia Langford, John Sillitoe on behalf of the Wellcome Sanger Institute COVID-19 Surveillance Team |
| EPI_ISL_3124737 | Lighthouse Lab in Milton Keynes | Wellcome Sanger Institute for the COVID-19 Genomics UK (COG-UK) Consortium | The Lighthouse Lab in Milton Keynes and Alex Alderton, Roberto Amato, Jeffrey Barrett, Sonia Goncalves, Ewan Harrison, David K. Jackson, Ian Johnston, Dominic Kwiatkowski, Cordelia Langford, John Sillitoe on behalf of the Wellcome Sanger Institute COVID-19 Surveillance Team |
| EPI_ISL_3124738 | Lighthouse Lab in Glasgow | Wellcome Sanger Institute for the COVID-19 Genomics UK (COG-UK) Consortium | Harper VanSteenhouse, Yumi Kasai, David Gray, Carol Clugston, Anna Dominiczak and Alex Alderton, Roberto Amato, Jeffrey Barrett, Sonia Goncalves, Ewan Harrison, David K. Jackson, Ian Johnston, Dominic Kwiatkowski, Cordelia Langford, John Sillitoe on behalf of the Wellcome Sanger Institute COVID-19 Surveillance Team |
| EPI_ISL_3124739 | Lighthouse Lab in Alderley Park | Wellcome Sanger Institute for the COVID-19 Genomics UK (COG-UK) Consortium | Jacquelyn Wynn, Mairead Hyland, The Lighthouse Lab in Alderley Park and Alex Alderton, Roberto Amato, Jeffrey Barrett, Sonia Goncalves, Ewan Harrison, David K. Jackson, Ian Johnston, Dominic Kwiatkowski, Cordelia Langford, John Sillitoe on behalf of the Wellcome Sanger Institute COVID-19 Surveillance Team |
| EPI_ISL_3124740 | Lighthouse Lab in Glasgow | Wellcome Sanger Institute for the COVID-19 Genomics UK (COG-UK) Consortium | Harper VanSteenhouse, Yumi Kasai, David Gray, Carol Clugston, Anna Dominiczak and Alex Alderton, Roberto Amato, Jeffrey Barrett, Sonia Goncalves, Ewan Harrison, David K. Jackson, Ian Johnston, Dominic Kwiatkowski, Cordelia Langford, John Sillitoe on behalf of the Wellcome Sanger Institute COVID-19 Surveillance Team |
| EPI_ISL_3124741 | Lighthouse Lab in Milton Keynes | Wellcome Sanger Institute for the COVID-19 Genomics UK (COG-UK) Consortium | The Lighthouse Lab in Milton Keynes and Alex Alderton, Roberto Amato, Jeffrey Barrett, Sonia Goncalves, Ewan Harrison, David K. Jackson, Ian Johnston, Dominic Kwiatkowski, Cordelia Langford, John Sillitoe on behalf of the Wellcome Sanger Institute COVID-19 Surveillance Team |
| EPI_ISL_3124742 | Lighthouse Lab in Glasgow | Wellcome Sanger Institute for the COVID-19 Genomics UK (COG-UK) Consortium | Harper VanSteenhouse, Yumi Kasai, David Gray, Carol Clugston, Anna Dominiczak and Alex Alderton, Roberto Amato, Jeffrey Barrett, Sonia Goncalves, Ewan Harrison, David K. Jackson, Ian Johnston, Dominic Kwiatkowski, Cordelia Langford, John Sillitoe on behalf of the Wellcome Sanger Institute COVID-19 Surveillance Team |
| EPI_ISL_3124743, EPI_ISL_3124744 | Lighthouse Lab in Milton Keynes | Wellcome Sanger Institute for the COVID-19 Genomics UK (COG-UK) Consortium | The Lighthouse Lab in Milton Keynes and Alex Alderton, Roberto Amato, Jeffrey Barrett, Sonia Goncalves, Ewan Harrison, David K. Jackson, Ian Johnston, Dominic Kwiatkowski, Cordelia Langford, John Sillitoe on behalf of the Wellcome Sanger Institute COVID-19 Surveillance Team |
| EPI_ISL_3124745, EPI_ISL_3124746 | Lighthouse Lab in Alderley Park | Wellcome Sanger Institute for the COVID-19 Genomics UK (COG-UK) Consortium | Jacquelyn Wynn, Mairead Hyland, The Lighthouse Lab in Alderley Park and Alex Alderton, Roberto Amato, Jeffrey Barrett, Sonia Goncalves, Ewan Harrison, David K. Jackson, Ian Johnston, Dominic Kwiatkowski, Cordelia Langford, John Sillitoe on behalf of the Wellcome Sanger Institute COVID-19 Surveillance Team |
| EPI_ISL_3124747 | Lighthouse Lab in Milton Keynes | Wellcome Sanger Institute for the COVID-19 Genomics UK (COG-UK) Consortium | The Lighthouse Lab in Milton Keynes and Alex Alderton, Roberto Amato, Jeffrey Barrett, Sonia Goncalves, Ewan Harrison, David K. Jackson, Ian Johnston, Dominic Kwiatkowski, Cordelia Langford, John Sillitoe on behalf of the Wellcome Sanger Institute COVID-19 Surveillance Team |
| EPI_ISL_3124748 | Lighthouse Lab in Glasgow | Wellcome Sanger Institute for the COVID-19 Genomics UK (COG-UK) Consortium | Harper VanSteenhouse, Yumi Kasai, David Gray, Carol Clugston, Anna Dominiczak and Alex Alderton, Roberto Amato, Jeffrey Barrett, Sonia Goncalves, Ewan Harrison, David K. Jackson, Ian Johnston, Dominic Kwiatkowski, Cordelia Langford, John Sillitoe on behalf of the Wellcome Sanger Institute COVID-19 Surveillance Team |
| EPI_ISL_3124749 | Lighthouse Lab in Alderley Park | Wellcome Sanger Institute for the COVID-19 Genomics UK (COG-UK) Consortium | Jacquelyn Wynn, Mairead Hyland, The Lighthouse Lab in Alderley Park and Alex Alderton, Roberto Amato, Jeffrey Barrett, Sonia Goncalves, Ewan Harrison, David K. Jackson, Ian Johnston, Dominic Kwiatkowski, Cordelia Langford, John Sillitoe on behalf of the Wellcome Sanger Institute COVID-19 Surveillance Team |
| EPI_ISL_3124750 | Lighthouse Lab in Glasgow | Wellcome Sanger Institute for the COVID-19 Genomics UK (COG-UK) Consortium | Harper VanSteenhouse, Yumi Kasai, David Gray, Carol Clugston, Anna Dominiczak and Alex Alderton, Roberto Amato, Jeffrey Barrett, Sonia Goncalves, Ewan Harrison, David K. Jackson, Ian Johnston, Dominic Kwiatkowski, Cordelia Langford, John Sillitoe on behalf of the Wellcome Sanger Institute COVID-19 Surveillance Team |
| EPI_ISL_3124751 | Lighthouse Lab in Milton Keynes | Wellcome Sanger Institute for the COVID-19 Genomics UK (COG-UK) Consortium | The Lighthouse Lab in Milton Keynes and Alex Alderton, Roberto Amato, Jeffrey Barrett, Sonia Goncalves, Ewan Harrison, David K. Jackson, Ian Johnston, Dominic Kwiatkowski, Cordelia Langford, John Sillitoe on behalf of the Wellcome Sanger Institute COVID-19 Surveillance Team |
| EPI_ISL_3124752 | Lighthouse Lab in Glasgow | Wellcome Sanger Institute for the COVID-19 Genomics UK (COG-UK) Consortium | Harper VanSteenhouse, Yumi Kasai, David Gray, Carol Clugston, Anna Dominiczak and Alex Alderton, Roberto Amato, Jeffrey Barrett, Sonia Goncalves, Ewan Harrison, David K. Jackson, Ian Johnston, Dominic Kwiatkowski, Cordelia Langford, John Sillitoe on behalf of the Wellcome Sanger Institute COVID-19 Surveillance Team |
| EPI_ISL_3124753, EPI_ISL_3124754 | Lighthouse Lab in Milton Keynes | Wellcome Sanger Institute for the COVID-19 Genomics UK (COG-UK) Consortium | The Lighthouse Lab in Milton Keynes and Alex Alderton, Roberto Amato, Jeffrey Barrett, Sonia Goncalves, Ewan Harrison, David K. Jackson, Ian Johnston, Dominic Kwiatkowski, Cordelia Langford, John Sillitoe on behalf of the Wellcome Sanger Institute COVID-19 Surveillance Team |
| EPI_ISL_3124755 | Lighthouse Lab in Alderley Park | Wellcome Sanger Institute for the COVID-19 Genomics UK (COG-UK) Consortium | Jacquelyn Wynn, Mairead Hyland, The Lighthouse Lab in Alderley Park and Alex Alderton, Roberto Amato, Jeffrey Barrett, Sonia Goncalves, Ewan Harrison, David K. Jackson, Ian Johnston, Dominic Kwiatkowski, Cordelia Langford, John Sillitoe on behalf of the Wellcome Sanger Institute COVID-19 Surveillance Team |
| EPI_ISL_3124756, EPI_ISL_3124757 | Lighthouse Lab in Glasgow | Wellcome Sanger Institute for the COVID-19 Genomics UK (COG-UK) Consortium | Harper VanSteenhouse, Yumi Kasai, David Gray, Carol Clugston, Anna Dominiczak and Alex Alderton, Roberto Amato, Jeffrey Barrett, Sonia Goncalves, Ewan Harrison, David K. Jackson, Ian Johnston, Dominic Kwiatkowski, Cordelia Langford, John Sillitoe on behalf of the Wellcome Sanger Institute COVID-19 Surveillance Team |
| EPI_ISL_3124758 | Lighthouse Lab in Alderley Park | Wellcome Sanger Institute for the COVID-19 Genomics UK (COG-UK) Consortium | Jacquelyn Wynn, Mairead Hyland, The Lighthouse Lab in Alderley Park and Alex Alderton, Roberto Amato, Jeffrey Barrett, Sonia Goncalves, Ewan Harrison, David K. Jackson, Ian Johnston, Dominic Kwiatkowski, Cordelia Langford, John Sillitoe on behalf of the Wellcome Sanger Institute COVID-19 Surveillance Team |
| EPI_ISL_3124759, EPI_ISL_3124760 | Lighthouse Lab in Glasgow | Wellcome Sanger Institute for the COVID-19 Genomics UK (COG-UK) Consortium | Harper VanSteenhouse, Yumi Kasai, David Gray, Carol Clugston, Anna Dominiczak and Alex Alderton, Roberto Amato, Jeffrey Barrett, Sonia Goncalves, Ewan Harrison, David K. Jackson, Ian Johnston, Dominic Kwiatkowski, Cordelia Langford, John Sillitoe on behalf of the Wellcome Sanger Institute COVID-19 Surveillance Team |
| EPI_ISL_3124761, EPI_ISL_3124762, EPI_ISL_3124763 | Lighthouse Lab in Milton Keynes | Wellcome Sanger Institute for the COVID-19 Genomics UK (COG-UK) Consortium | The Lighthouse Lab in Milton Keynes and Alex Alderton, Roberto Amato, Jeffrey Barrett, Sonia Goncalves, Ewan Harrison, David K. Jackson, Ian Johnston, Dominic Kwiatkowski, Cordelia Langford, John Sillitoe on behalf of the Wellcome Sanger Institute COVID-19 Surveillance Team |
| EPI_ISL_3124764, EPI_ISL_3124765 | Lighthouse Lab in Glasgow | Wellcome Sanger Institute for the COVID-19 Genomics UK (COG-UK) Consortium | Harper VanSteenhouse, Yumi Kasai, David Gray, Carol Clugston, Anna Dominiczak and Alex Alderton, Roberto Amato, Jeffrey Barrett, Sonia Goncalves, Ewan Harrison, David K. Jackson, Ian Johnston, Dominic Kwiatkowski, Cordelia Langford, John Sillitoe on behalf of the Wellcome Sanger Institute COVID-19 Surveillance Team |
| EPI_ISL_3124766, EPI_ISL_3124767 | Lighthouse Lab in Alderley Park | Wellcome Sanger Institute for the COVID-19 Genomics UK (COG-UK) Consortium | Jacquelyn Wynn, Mairead Hyland, The Lighthouse Lab in Alderley Park and Alex Alderton, Roberto Amato, Jeffrey Barrett, Sonia Goncalves, Ewan Harrison, David K. Jackson, Ian Johnston, Dominic Kwiatkowski, Cordelia Langford, John Sillitoe on behalf of the Wellcome Sanger Institute COVID-19 Surveillance Team |
| EPI_ISL_3124768 | Lighthouse Lab in Glasgow | Wellcome Sanger Institute for the COVID-19 Genomics UK (COG-UK) Consortium | Harper VanSteenhouse, Yumi Kasai, David Gray, Carol Clugston, Anna Dominiczak and Alex Alderton, Roberto Amato, Jeffrey Barrett, Sonia Goncalves, Ewan Harrison, David K. Jackson, Ian Johnston, Dominic Kwiatkowski, Cordelia Langford, John Sillitoe on behalf of the Wellcome Sanger Institute COVID-19 Surveillance Team |
| EPI_ISL_3124769 | Lighthouse Lab in Milton Keynes | Wellcome Sanger Institute for the COVID-19 Genomics UK (COG-UK) Consortium | The Lighthouse Lab in Milton Keynes and Alex Alderton, Roberto Amato, Jeffrey Barrett, Sonia Goncalves, Ewan Harrison, David K. Jackson, Ian Johnston, Dominic Kwiatkowski, Cordelia Langford, John Sillitoe on behalf of the Wellcome Sanger Institute COVID-19 Surveillance Team |
| EPI_ISL_3124770 | Lighthouse Lab in Glasgow | Wellcome Sanger Institute for the COVID-19 Genomics UK (COG-UK) Consortium | Harper VanSteenhouse, Yumi Kasai, David Gray, Carol Clugston, Anna Dominiczak and Alex Alderton, Roberto Amato, Jeffrey Barrett, Sonia Goncalves, Ewan Harrison, David K. Jackson, Ian Johnston, Dominic Kwiatkowski, Cordelia Langford, John Sillitoe on behalf of the Wellcome Sanger Institute COVID-19 Surveillance Team |
| EPI_ISL_3124771 | Lighthouse Lab in Milton Keynes | Wellcome Sanger Institute for the COVID-19 Genomics UK (COG-UK) Consortium | The Lighthouse Lab in Milton Keynes and Alex Alderton, Roberto Amato, Jeffrey Barrett, Sonia Goncalves, Ewan Harrison, David K. Jackson, Ian Johnston, Dominic Kwiatkowski, Cordelia Langford, John Sillitoe on behalf of the Wellcome Sanger Institute COVID-19 Surveillance Team |
| EPI_ISL_3124772, EPI_ISL_3124773 | Lighthouse Lab in Glasgow | Wellcome Sanger Institute for the COVID-19 Genomics UK (COG-UK) Consortium | Harper VanSteenhouse, Yumi Kasai, David Gray, Carol Clugston, Anna Dominiczak and Alex Alderton, Roberto Amato, Jeffrey Barrett, Sonia Goncalves, Ewan Harrison, David K. Jackson, Ian Johnston, Dominic Kwiatkowski, Cordelia Langford, John Sillitoe on behalf of the Wellcome Sanger Institute COVID-19 Surveillance Team |
| EPI_ISL_3124774, EPI_ISL_3124775 | Lighthouse Lab in Alderley Park | Wellcome Sanger Institute for the COVID-19 Genomics UK (COG-UK) Consortium | Jacquelyn Wynn, Mairead Hyland, The Lighthouse Lab in Alderley Park and Alex Alderton, Roberto Amato, Jeffrey Barrett, Sonia Goncalves, Ewan Harrison, David K. Jackson, Ian Johnston, Dominic Kwiatkowski, Cordelia Langford, John Sillitoe on behalf of the Wellcome Sanger Institute COVID-19 Surveillance Team |
| EPI_ISL_3124776 | Lighthouse Lab in Glasgow | Wellcome Sanger Institute for the COVID-19 Genomics UK (COG-UK) Consortium | Harper VanSteenhouse, Yumi Kasai, David Gray, Carol Clugston, Anna Dominiczak and Alex Alderton, Roberto Amato, Jeffrey Barrett, Sonia Goncalves, Ewan Harrison, David K. Jackson, Ian Johnston, Dominic Kwiatkowski, Cordelia Langford, John Sillitoe on behalf of the Wellcome Sanger Institute COVID-19 Surveillance Team |
| EPI_ISL_3124777 | Lighthouse Lab in Milton Keynes | Wellcome Sanger Institute for the COVID-19 Genomics UK (COG-UK) Consortium | The Lighthouse Lab in Milton Keynes and Alex Alderton, Roberto Amato, Jeffrey Barrett, Sonia Goncalves, Ewan Harrison, David K. Jackson, Ian Johnston, Dominic Kwiatkowski, Cordelia Langford, John Sillitoe on behalf of the Wellcome Sanger Institute COVID-19 Surveillance Team |
| EPI_ISL_3124778, EPI_ISL_3124779 | Lighthouse Lab in Glasgow | Wellcome Sanger Institute for the COVID-19 Genomics UK (COG-UK) Consortium | Harper VanSteenhouse, Yumi Kasai, David Gray, Carol Clugston, Anna Dominiczak and Alex Alderton, Roberto Amato, Jeffrey Barrett, Sonia Goncalves, Ewan Harrison, David K. Jackson, Ian Johnston, Dominic Kwiatkowski, Cordelia Langford, John Sillitoe on behalf of the Wellcome Sanger Institute COVID-19 Surveillance Team |
| EPI_ISL_3124780 | Lighthouse Lab in Milton Keynes | Wellcome Sanger Institute for the COVID-19 Genomics UK (COG-UK) Consortium | The Lighthouse Lab in Milton Keynes and Alex Alderton, Roberto Amato, Jeffrey Barrett, Sonia Goncalves, Ewan Harrison, David K. Jackson, Ian Johnston, Dominic Kwiatkowski, Cordelia Langford, John Sillitoe on behalf of the Wellcome Sanger Institute COVID-19 Surveillance Team |
| EPI_ISL_3124781, EPI_ISL_3124782 | Lighthouse Lab in Alderley Park | Wellcome Sanger Institute for the COVID-19 Genomics UK (COG-UK) Consortium | Jacquelyn Wynn, Mairead Hyland, The Lighthouse Lab in Alderley Park and Alex Alderton, Roberto Amato, Jeffrey Barrett, Sonia Goncalves, Ewan Harrison, David K. Jackson, Ian Johnston, Dominic Kwiatkowski, Cordelia Langford, John Sillitoe on behalf of the Wellcome Sanger Institute COVID-19 Surveillance Team |
| EPI_ISL_3124783 | Lighthouse Lab in Glasgow | Wellcome Sanger Institute for the COVID-19 Genomics UK (COG-UK) Consortium | Harper VanSteenhouse, Yumi Kasai, David Gray, Carol Clugston, Anna Dominiczak and Alex Alderton, Roberto Amato, Jeffrey Barrett, Sonia Goncalves, Ewan Harrison, David K. Jackson, Ian Johnston, Dominic Kwiatkowski, Cordelia Langford, John Sillitoe on behalf of the Wellcome Sanger Institute COVID-19 Surveillance Team |
| EPI_ISL_3124784 | Lighthouse Lab in Milton Keynes | Wellcome Sanger Institute for the COVID-19 Genomics UK (COG-UK) Consortium | The Lighthouse Lab in Milton Keynes and Alex Alderton, Roberto Amato, Jeffrey Barrett, Sonia Goncalves, Ewan Harrison, David K. Jackson, Ian Johnston, Dominic Kwiatkowski, Cordelia Langford, John Sillitoe on behalf of the Wellcome Sanger Institute COVID-19 Surveillance Team |
| EPI_ISL_3124785 | Lighthouse Lab in Glasgow | Wellcome Sanger Institute for the COVID-19 Genomics UK (COG-UK) Consortium | Harper VanSteenhouse, Yumi Kasai, David Gray, Carol Clugston, Anna Dominiczak and Alex Alderton, Roberto Amato, Jeffrey Barrett, Sonia Goncalves, Ewan Harrison, David K. Jackson, Ian Johnston, Dominic Kwiatkowski, Cordelia Langford, John Sillitoe on behalf of the Wellcome Sanger Institute COVID-19 Surveillance Team |

| | | | |
|---|---|---|---|
| EPI_ISL_3124786 | Lighthouse Lab in Alderley Park | Wellcome Sanger Institute for the COVID-19 Genomics UK (COG-UK) Consortium | Jacquelyn Wynn, Mairead Hyland, The Lighthouse Lab in Alderley Park and Alex Alderton, Roberto Amato, Jeffrey Barrett, Sonia Goncalves, Ewan Harrison, David K. Jackson, Ian Johnston, Dominic Kwiatkowski, Cordelia Langford, John Sillitoe on behalf of the Wellcome Sanger Institute COVID-19 Surveillance Team |
| EPI_ISL_3124787, EPI_ISL_3124788 | Lighthouse Lab in Glasgow | Wellcome Sanger Institute for the COVID-19 Genomics UK (COG-UK) Consortium | Harper VanSteenhouse, Yumi Kasai, David Gray, Carol Clugston, Anna Dominiczak and Alex Alderton, Roberto Amato, Jeffrey Barrett, Sonia Goncalves, Ewan Harrison, David K. Jackson, Ian Johnston, Dominic Kwiatkowski, Cordelia Langford, John Sillitoe on behalf of the Wellcome Sanger Institute COVID-19 Surveillance Team |
| EPI_ISL_3124789 | Lighthouse Lab in Alderley Park | Wellcome Sanger Institute for the COVID-19 Genomics UK (COG-UK) Consortium | Jacquelyn Wynn, Mairead Hyland, The Lighthouse Lab in Alderley Park and Alex Alderton, Roberto Amato, Jeffrey Barrett, Sonia Goncalves, Ewan Harrison, David K. Jackson, Ian Johnston, Dominic Kwiatkowski, Cordelia Langford, John Sillitoe on behalf of the Wellcome Sanger Institute COVID-19 Surveillance Team |
| EPI_ISL_3124790 | Lighthouse Lab in Milton Keynes | Wellcome Sanger Institute for the COVID-19 Genomics UK (COG-UK) Consortium | The Lighthouse Lab in Milton Keynes and Alex Alderton, Roberto Amato, Jeffrey Barrett, Sonia Goncalves, Ewan Harrison, David K. Jackson, Ian Johnston, Dominic Kwiatkowski, Cordelia Langford, John Sillitoe on behalf of the Wellcome Sanger Institute COVID-19 Surveillance Team |
| EPI_ISL_3124791 | Lighthouse Lab in Glasgow | Wellcome Sanger Institute for the COVID-19 Genomics UK (COG-UK) Consortium | Harper VanSteenhouse, Yumi Kasai, David Gray, Carol Clugston, Anna Dominiczak and Alex Alderton, Roberto Amato, Jeffrey Barrett, Sonia Goncalves, Ewan Harrison, David K. Jackson, Ian Johnston, Dominic Kwiatkowski, Cordelia Langford, John Sillitoe on behalf of the Wellcome Sanger Institute COVID-19 Surveillance Team |
| EPI_ISL_3124792 | Lighthouse Lab in Milton Keynes | Wellcome Sanger Institute for the COVID-19 Genomics UK (COG-UK) Consortium | The Lighthouse Lab in Milton Keynes and Alex Alderton, Roberto Amato, Jeffrey Barrett, Sonia Goncalves, Ewan Harrison, David K. Jackson, Ian Johnston, Dominic Kwiatkowski, Cordelia Langford, John Sillitoe on behalf of the Wellcome Sanger Institute COVID-19 Surveillance Team |
| EPI_ISL_3124793, EPI_ISL_3124794, EPI_ISL_3124795, EPI_ISL_3124796, EPI_ISL_3124797, EPI_ISL_3124798, EPI_ISL_3124799, EPI_ISL_3124800 | Lighthouse Lab in Glasgow | Wellcome Sanger Institute for the COVID-19 Genomics UK (COG-UK) Consortium | Harper VanSteenhouse, Yumi Kasai, David Gray, Carol Clugston, Anna Dominiczak and Alex Alderton, Roberto Amato, Jeffrey Barrett, Sonia Goncalves, Ewan Harrison, David K. Jackson, Ian Johnston, Dominic Kwiatkowski, Cordelia Langford, John Sillitoe on behalf of the Wellcome Sanger Institute COVID-19 Surveillance Team |
| EPI_ISL_3124801 | Lighthouse Lab in Alderley Park | Wellcome Sanger Institute for the COVID-19 Genomics UK (COG-UK) Consortium | Jacquelyn Wynn, Mairead Hyland, The Lighthouse Lab in Alderley Park and Alex Alderton, Roberto Amato, Jeffrey Barrett, Sonia Goncalves, Ewan Harrison, David K. Jackson, Ian Johnston, Dominic Kwiatkowski, Cordelia Langford, John Sillitoe on behalf of the Wellcome Sanger Institute COVID-19 Surveillance Team |
| EPI_ISL_3124802, EPI_ISL_3124803 | Lighthouse Lab in Glasgow | Wellcome Sanger Institute for the COVID-19 Genomics UK (COG-UK) Consortium | Harper VanSteenhouse, Yumi Kasai, David Gray, Carol Clugston, Anna Dominiczak and Alex Alderton, Roberto Amato, Jeffrey Barrett, Sonia Goncalves, Ewan Harrison, David K. Jackson, Ian Johnston, Dominic Kwiatkowski, Cordelia Langford, John Sillitoe on behalf of the Wellcome Sanger Institute COVID-19 Surveillance Team |
| EPI_ISL_3124804 | Lighthouse Lab in Milton Keynes | Wellcome Sanger Institute for the COVID-19 Genomics UK (COG-UK) Consortium | The Lighthouse Lab in Milton Keynes and Alex Alderton, Roberto Amato, Jeffrey Barrett, Sonia Goncalves, Ewan Harrison, David K. Jackson, Ian Johnston, Dominic Kwiatkowski, Cordelia Langford, John Sillitoe on behalf of the Wellcome Sanger Institute COVID-19 Surveillance Team |
| EPI_ISL_3124805, EPI_ISL_3124806 | Lighthouse Lab in Glasgow | Wellcome Sanger Institute for the COVID-19 Genomics UK (COG-UK) Consortium | Harper VanSteenhouse, Yumi Kasai, David Gray, Carol Clugston, Anna Dominiczak and Alex Alderton, Roberto Amato, Jeffrey Barrett, Sonia Goncalves, Ewan Harrison, David K. Jackson, Ian Johnston, Dominic Kwiatkowski, Cordelia Langford, John Sillitoe on behalf of the Wellcome Sanger Institute COVID-19 Surveillance Team |
| EPI_ISL_3124807, EPI_ISL_3124808 | Lighthouse Lab in Milton Keynes | Wellcome Sanger Institute for the COVID-19 Genomics UK (COG-UK) Consortium | The Lighthouse Lab in Milton Keynes and Alex Alderton, Roberto Amato, Jeffrey Barrett, Sonia Goncalves, Ewan Harrison, David K. Jackson, Ian Johnston, Dominic Kwiatkowski, Cordelia Langford, John Sillitoe on behalf of the Wellcome Sanger Institute COVID-19 Surveillance Team |
| EPI_ISL_3124809 | Lighthouse Lab in Glasgow | Wellcome Sanger Institute for the COVID-19 Genomics UK (COG-UK) Consortium | Harper VanSteenhouse, Yumi Kasai, David Gray, Carol Clugston, Anna Dominiczak and Alex Alderton, Roberto Amato, Jeffrey Barrett, Sonia Goncalves, Ewan Harrison, David K. Jackson, Ian Johnston, Dominic Kwiatkowski, Cordelia Langford, John Sillitoe on behalf of the Wellcome Sanger Institute COVID-19 Surveillance Team |
| EPI_ISL_3124810 | Lighthouse Lab in Milton Keynes | Wellcome Sanger Institute for the COVID-19 Genomics UK (COG-UK) Consortium | The Lighthouse Lab in Milton Keynes and Alex Alderton, Roberto Amato, Jeffrey Barrett, Sonia Goncalves, Ewan Harrison, David K. Jackson, Ian Johnston, Dominic Kwiatkowski, Cordelia Langford, John Sillitoe on behalf of the Wellcome Sanger Institute COVID-19 Surveillance Team |
| EPI_ISL_3124811 | Lighthouse Lab in Alderley Park | Wellcome Sanger Institute for the COVID-19 Genomics UK (COG-UK) Consortium | Jacquelyn Wynn, Mairead Hyland, The Lighthouse Lab in Alderley Park and Alex Alderton, Roberto Amato, Jeffrey Barrett, Sonia Goncalves, Ewan Harrison, David K. Jackson, Ian Johnston, Dominic Kwiatkowski, Cordelia Langford, John Sillitoe on behalf of the Wellcome Sanger Institute COVID-19 Surveillance Team |
| EPI_ISL_3124812 | Lighthouse Lab in Glasgow | Wellcome Sanger Institute for the COVID-19 Genomics UK (COG-UK) Consortium | Harper VanSteenhouse, Yumi Kasai, David Gray, Carol Clugston, Anna Dominiczak and Alex Alderton, Roberto Amato, Jeffrey Barrett, Sonia Goncalves, Ewan Harrison, David K. Jackson, Ian Johnston, Dominic Kwiatkowski, Cordelia Langford, John Sillitoe on behalf of the Wellcome Sanger Institute COVID-19 Surveillance Team |
| EPI_ISL_3124813, EPI_ISL_3124814 | Lighthouse Lab in Alderley Park | Wellcome Sanger Institute for the COVID-19 Genomics UK (COG-UK) Consortium | Jacquelyn Wynn, Mairead Hyland, The Lighthouse Lab in Alderley Park and Alex Alderton, Roberto Amato, Jeffrey Barrett, Sonia Goncalves, Ewan Harrison, David K. Jackson, Ian Johnston, Dominic Kwiatkowski, Cordelia Langford, John Sillitoe on behalf of the Wellcome Sanger Institute COVID-19 Surveillance Team |
| EPI_ISL_3124815 | Lighthouse Lab in Milton Keynes | Wellcome Sanger Institute for the COVID-19 Genomics UK (COG-UK) Consortium | The Lighthouse Lab in Milton Keynes and Alex Alderton, Roberto Amato, Jeffrey Barrett, Sonia Goncalves, Ewan Harrison, David K. Jackson, Ian Johnston, Dominic Kwiatkowski, Cordelia Langford, John Sillitoe on behalf of the Wellcome Sanger Institute COVID-19 Surveillance Team |
| EPI_ISL_3124816, EPI_ISL_3124817 | Lighthouse Lab in Alderley Park | Wellcome Sanger Institute for the COVID-19 Genomics UK (COG-UK) Consortium | Jacquelyn Wynn, Mairead Hyland, The Lighthouse Lab in Alderley Park and Alex Alderton, Roberto Amato, Jeffrey Barrett, Sonia Goncalves, Ewan Harrison, David K. Jackson, Ian Johnston, Dominic Kwiatkowski, Cordelia Langford, John Sillitoe on behalf of the Wellcome Sanger Institute COVID-19 Surveillance Team |
| EPI_ISL_3124818 | Lighthouse Lab in Glasgow | Wellcome Sanger Institute for the COVID-19 Genomics UK (COG-UK) Consortium | Harper VanSteenhouse, Yumi Kasai, David Gray, Carol Clugston, Anna Dominiczak and Alex Alderton, Roberto Amato, Jeffrey Barrett, Sonia Goncalves, Ewan Harrison, David K. Jackson, Ian Johnston, Dominic Kwiatkowski, Cordelia Langford, John Sillitoe on behalf of the Wellcome Sanger Institute COVID-19 Surveillance Team |
| EPI_ISL_3124819 | Lighthouse Lab in Milton Keynes | Wellcome Sanger Institute for the COVID-19 Genomics UK (COG-UK) Consortium | The Lighthouse Lab in Milton Keynes and Alex Alderton, Roberto Amato, Jeffrey Barrett, Sonia Goncalves, Ewan Harrison, David K. Jackson, Ian Johnston, Dominic Kwiatkowski, Cordelia Langford, John Sillitoe on behalf of the Wellcome Sanger Institute COVID-19 Surveillance Team |
| EPI_ISL_3124820 | Lighthouse Lab in Glasgow | Wellcome Sanger Institute for the COVID-19 Genomics UK (COG-UK) Consortium | Harper VanSteenhouse, Yumi Kasai, David Gray, Carol Clugston, Anna Dominiczak and Alex Alderton, Roberto Amato, Jeffrey Barrett, Sonia Goncalves, Ewan Harrison, David K. Jackson, Ian Johnston, Dominic Kwiatkowski, Cordelia Langford, John Sillitoe on behalf of the Wellcome Sanger Institute COVID-19 Surveillance Team |
| EPI_ISL_3124821 | Lighthouse Lab in Alderley Park | Wellcome Sanger Institute for the COVID-19 Genomics UK (COG-UK) Consortium | Jacquelyn Wynn, Mairead Hyland, The Lighthouse Lab in Alderley Park and Alex Alderton, Roberto Amato, Jeffrey Barrett, Sonia Goncalves, Ewan Harrison, David K. Jackson, Ian Johnston, Dominic Kwiatkowski, Cordelia Langford, John Sillitoe on behalf of the Wellcome Sanger Institute COVID-19 Surveillance Team |
| EPI_ISL_3124822, EPI_ISL_3124823, EPI_ISL_3124824, EPI_ISL_3124825, EPI_ISL_3124826 | Lighthouse Lab in Glasgow | Wellcome Sanger Institute for the COVID-19 Genomics UK (COG-UK) Consortium | Harper VanSteenhouse, Yumi Kasai, David Gray, Carol Clugston, Anna Dominiczak and Alex Alderton, Roberto Amato, Jeffrey Barrett, Sonia Goncalves, Ewan Harrison, David K. Jackson, Ian Johnston, Dominic Kwiatkowski, Cordelia Langford, John Sillitoe on behalf of the Wellcome Sanger Institute COVID-19 Surveillance Team |
| EPI_ISL_3124827, EPI_ISL_3124828 | Lighthouse Lab in Milton Keynes | Wellcome Sanger Institute for the COVID-19 Genomics UK (COG-UK) Consortium | The Lighthouse Lab in Milton Keynes and Alex Alderton, Roberto Amato, Jeffrey Barrett, Sonia Goncalves, Ewan Harrison, David K. Jackson, Ian Johnston, Dominic Kwiatkowski, Cordelia Langford, John Sillitoe on behalf of the Wellcome Sanger Institute COVID-19 Surveillance Team |
| EPI_ISL_3124829, EPI_ISL_3124830 | Lighthouse Lab in Alderley Park | Wellcome Sanger Institute for the COVID-19 Genomics UK (COG-UK) Consortium | Jacquelyn Wynn, Mairead Hyland, The Lighthouse Lab in Alderley Park and Alex Alderton, Roberto Amato, Jeffrey Barrett, Sonia Goncalves, Ewan Harrison, David K. Jackson, Ian Johnston, Dominic Kwiatkowski, Cordelia Langford, John Sillitoe on behalf of the Wellcome Sanger Institute COVID-19 Surveillance Team |
| EPI_ISL_3124831 | Lighthouse Lab in Glasgow | Wellcome Sanger Institute for the COVID-19 Genomics UK (COG-UK) Consortium | Harper VanSteenhouse, Yumi Kasai, David Gray, Carol Clugston, Anna Dominiczak and Alex Alderton, Roberto Amato, Jeffrey Barrett, Sonia Goncalves, Ewan Harrison, David K. Jackson, Ian Johnston, Dominic Kwiatkowski, Cordelia Langford, John Sillitoe on behalf of the Wellcome Sanger Institute COVID-19 Surveillance Team |
| EPI_ISL_3124832 | Lighthouse Lab in Milton Keynes | Wellcome Sanger Institute for the COVID-19 Genomics UK (COG-UK) Consortium | The Lighthouse Lab in Milton Keynes and Alex Alderton, Roberto Amato, Jeffrey Barrett, Sonia Goncalves, Ewan Harrison, David K. Jackson, Ian Johnston, Dominic Kwiatkowski, Cordelia Langford, John Sillitoe on behalf of the Wellcome Sanger Institute COVID-19 Surveillance Team |
| EPI_ISL_3124833 | Lighthouse Lab in Alderley Park | Wellcome Sanger Institute for the COVID-19 Genomics UK (COG-UK) Consortium | Jacquelyn Wynn, Mairead Hyland, The Lighthouse Lab in Alderley Park and Alex Alderton, Roberto Amato, Jeffrey Barrett, Sonia Goncalves, Ewan Harrison, David K. Jackson, Ian Johnston, Dominic Kwiatkowski, Cordelia Langford, John Sillitoe on behalf of the Wellcome Sanger Institute COVID-19 Surveillance Team |
| EPI_ISL_3124834 | Lighthouse Lab in Milton Keynes | Wellcome Sanger Institute for the COVID-19 Genomics UK (COG-UK) Consortium | The Lighthouse Lab in Milton Keynes and Alex Alderton, Roberto Amato, Jeffrey Barrett, Sonia Goncalves, Ewan Harrison, David K. Jackson, Ian Johnston, Dominic Kwiatkowski, Cordelia Langford, John Sillitoe on behalf of the Wellcome Sanger Institute COVID-19 Surveillance Team |
| EPI_ISL_3124835, EPI_ISL_3124836, EPI_ISL_3124837, EPI_ISL_3124838, EPI_ISL_3124839 | Lighthouse Lab in Glasgow | Wellcome Sanger Institute for the COVID-19 Genomics UK (COG-UK) Consortium | Harper VanSteenhouse, Yumi Kasai, David Gray, Carol Clugston, Anna Dominiczak and Alex Alderton, Roberto Amato, Jeffrey Barrett, Sonia Goncalves, Ewan Harrison, David K. Jackson, Ian Johnston, Dominic Kwiatkowski, Cordelia Langford, John Sillitoe on behalf of the Wellcome Sanger Institute COVID-19 Surveillance Team |
| EPI_ISL_3124840 | Lighthouse Lab in Alderley Park | Wellcome Sanger Institute for the COVID-19 Genomics UK (COG-UK) Consortium | Jacquelyn Wynn, Mairead Hyland, The Lighthouse Lab in Alderley Park and Alex Alderton, Roberto Amato, Jeffrey Barrett, Sonia Goncalves, Ewan Harrison, David K. Jackson, Ian Johnston, Dominic Kwiatkowski, Cordelia Langford, John Sillitoe on behalf of the Wellcome Sanger Institute COVID-19 Surveillance Team |
| EPI_ISL_3124841 | Lighthouse Lab in Glasgow | Wellcome Sanger Institute for the COVID-19 Genomics UK (COG-UK) Consortium | Harper VanSteenhouse, Yumi Kasai, David Gray, Carol Clugston, Anna Dominiczak and Alex Alderton, Roberto Amato, Jeffrey Barrett, Sonia Goncalves, Ewan Harrison, David K. Jackson, Ian Johnston, Dominic Kwiatkowski, Cordelia Langford, John Sillitoe on behalf of the Wellcome Sanger Institute COVID-19 Surveillance Team |
| EPI_ISL_3124842 | Lighthouse Lab in Alderley Park | Wellcome Sanger Institute for the COVID-19 Genomics UK (COG-UK) Consortium | Jacquelyn Wynn, Mairead Hyland, The Lighthouse Lab in Alderley Park and Alex Alderton, Roberto Amato, Jeffrey Barrett, Sonia Goncalves, Ewan Harrison, David K. Jackson, Ian Johnston, Dominic Kwiatkowski, Cordelia Langford, John Sillitoe on behalf of the Wellcome Sanger Institute COVID-19 Surveillance Team |
| EPI_ISL_3124843 | Lighthouse Lab in Glasgow | Wellcome Sanger Institute for the COVID-19 Genomics UK (COG-UK) Consortium | Harper VanSteenhouse, Yumi Kasai, David Gray, Carol Clugston, Anna Dominiczak and Alex Alderton, Roberto Amato, Jeffrey Barrett, Sonia Goncalves, Ewan Harrison, David K. Jackson, Ian Johnston, Dominic Kwiatkowski, Cordelia Langford, John Sillitoe on behalf of the Wellcome Sanger Institute COVID-19 Surveillance Team |
| EPI_ISL_3124844, EPI_ISL_3124845 | Lighthouse Lab in Milton Keynes | Wellcome Sanger Institute for the COVID-19 Genomics UK (COG-UK) Consortium | The Lighthouse Lab in Milton Keynes and Alex Alderton, Roberto Amato, Jeffrey Barrett, Sonia Goncalves, Ewan Harrison, David K. Jackson, Ian Johnston, Dominic Kwiatkowski, Cordelia Langford, John Sillitoe on behalf of the Wellcome Sanger Institute COVID-19 Surveillance Team |
| EPI_ISL_3124846, EPI_ISL_3124847 | Lighthouse Lab in Glasgow | Wellcome Sanger Institute for the COVID-19 Genomics UK (COG-UK) Consortium | Harper VanSteenhouse, Yumi Kasai, David Gray, Carol Clugston, Anna Dominiczak and Alex Alderton, Roberto Amato, Jeffrey Barrett, Sonia Goncalves, Ewan Harrison, David K. Jackson, Ian Johnston, Dominic Kwiatkowski, Cordelia Langford, John Sillitoe on behalf of the Wellcome Sanger Institute COVID-19 Surveillance Team |
| EPI_ISL_3124848 | Lighthouse Lab in Milton Keynes | Wellcome Sanger Institute for the COVID-19 Genomics UK (COG-UK) Consortium | The Lighthouse Lab in Milton Keynes and Alex Alderton, Roberto Amato, Jeffrey Barrett, Sonia Goncalves, Ewan Harrison, David K. Jackson, Ian Johnston, Dominic Kwiatkowski, Cordelia Langford, John Sillitoe on behalf of the Wellcome Sanger Institute COVID-19 Surveillance Team |
| EPI_ISL_3124849, EPI_ISL_3124850, EPI_ISL_3124851, EPI_ISL_3124852, EPI_ISL_3124853 | Lighthouse Lab in Glasgow | Wellcome Sanger Institute for the COVID-19 Genomics UK (COG-UK) Consortium | Harper VanSteenhouse, Yumi Kasai, David Gray, Carol Clugston, Anna Dominiczak and Alex Alderton, Roberto Amato, Jeffrey Barrett, Sonia Goncalves, Ewan Harrison, David K. Jackson, Ian Johnston, Dominic Kwiatkowski, Cordelia Langford, John Sillitoe on behalf of the Wellcome Sanger Institute COVID-19 Surveillance Team |

| | | | |
|---|---|---|---|
| EPI_ISL_3124854 | Lighthouse Lab in Milton Keynes | Wellcome Sanger Institute for the COVID-19 Genomics UK (COG-UK) Consortium | The Lighthouse Lab in Milton Keynes and Alex Alderton, Roberto Amato, Jeffrey Barrett, Sonia Goncalves, Ewan Harrison, David K. Jackson, Ian Johnston, Dominic Kwiatkowski, Cordelia Langford, John Sillitoe on behalf of the Wellcome Sanger Institute COVID-19 Surveillance Team |
| EPI_ISL_3124855, EPI_ISL_3124856 | Lighthouse Lab in Glasgow | Wellcome Sanger Institute for the COVID-19 Genomics UK (COG-UK) Consortium | Harper VanSteenhouse, Yumi Kasai, David Gray, Carol Clugston, Anna Dominiczak and Alex Alderton, Roberto Amato, Jeffrey Barrett, Sonia Goncalves, Ewan Harrison, David K. Jackson, Ian Johnston, Dominic Kwiatkowski, Cordelia Langford, John Sillitoe on behalf of the Wellcome Sanger Institute COVID-19 Surveillance Team |
| EPI_ISL_3124857 | Lighthouse Lab in Alderley Park | Wellcome Sanger Institute for the COVID-19 Genomics UK (COG-UK) Consortium | Jacquelyn Wynn, Mairead Hyland, The Lighthouse Lab in Alderley Park and Alex Alderton, Roberto Amato, Jeffrey Barrett, Sonia Goncalves, Ewan Harrison, David K. Jackson, Ian Johnston, Dominic Kwiatkowski, Cordelia Langford, John Sillitoe on behalf of the Wellcome Sanger Institute COVID-19 Surveillance Team |
| EPI_ISL_3124858, EPI_ISL_3124859, EPI_ISL_3124860, EPI_ISL_3124861, EPI_ISL_3124862, EPI_ISL_3124863, EPI_ISL_3124864 | Lighthouse Lab in Glasgow | Wellcome Sanger Institute for the COVID-19 Genomics UK (COG-UK) Consortium | Harper VanSteenhouse, Yumi Kasai, David Gray, Carol Clugston, Anna Dominiczak and Alex Alderton, Roberto Amato, Jeffrey Barrett, Sonia Goncalves, Ewan Harrison, David K. Jackson, Ian Johnston, Dominic Kwiatkowski, Cordelia Langford, John Sillitoe on behalf of the Wellcome Sanger Institute COVID-19 Surveillance Team |
| EPI_ISL_3124865, EPI_ISL_3124866 | Lighthouse Lab in Alderley Park | Wellcome Sanger Institute for the COVID-19 Genomics UK (COG-UK) Consortium | Jacquelyn Wynn, Mairead Hyland, The Lighthouse Lab in Alderley Park and Alex Alderton, Roberto Amato, Jeffrey Barrett, Sonia Goncalves, Ewan Harrison, David K. Jackson, Ian Johnston, Dominic Kwiatkowski, Cordelia Langford, John Sillitoe on behalf of the Wellcome Sanger Institute COVID-19 Surveillance Team |
| EPI_ISL_3124867 | Lighthouse Lab in Milton Keynes | Wellcome Sanger Institute for the COVID-19 Genomics UK (COG-UK) Consortium | The Lighthouse Lab in Milton Keynes and Alex Alderton, Roberto Amato, Jeffrey Barrett, Sonia Goncalves, Ewan Harrison, David K. Jackson, Ian Johnston, Dominic Kwiatkowski, Cordelia Langford, John Sillitoe on behalf of the Wellcome Sanger Institute COVID-19 Surveillance Team |
| EPI_ISL_3124868, EPI_ISL_3124869 | Lighthouse Lab in Alderley Park | Wellcome Sanger Institute for the COVID-19 Genomics UK (COG-UK) Consortium | Jacquelyn Wynn, Mairead Hyland, The Lighthouse Lab in Alderley Park and Alex Alderton, Roberto Amato, Jeffrey Barrett, Sonia Goncalves, Ewan Harrison, David K. Jackson, Ian Johnston, Dominic Kwiatkowski, Cordelia Langford, John Sillitoe on behalf of the Wellcome Sanger Institute COVID-19 Surveillance Team |
| EPI_ISL_3124870, EPI_ISL_3124871 | Lighthouse Lab in Glasgow | Wellcome Sanger Institute for the COVID-19 Genomics UK (COG-UK) Consortium | Harper VanSteenhouse, Yumi Kasai, David Gray, Carol Clugston, Anna Dominiczak and Alex Alderton, Roberto Amato, Jeffrey Barrett, Sonia Goncalves, Ewan Harrison, David K. Jackson, Ian Johnston, Dominic Kwiatkowski, Cordelia Langford, John Sillitoe on behalf of the Wellcome Sanger Institute COVID-19 Surveillance Team |
| EPI_ISL_3124872 | Lighthouse Lab in Milton Keynes | Wellcome Sanger Institute for the COVID-19 Genomics UK (COG-UK) Consortium | The Lighthouse Lab in Milton Keynes and Alex Alderton, Roberto Amato, Jeffrey Barrett, Sonia Goncalves, Ewan Harrison, David K. Jackson, Ian Johnston, Dominic Kwiatkowski, Cordelia Langford, John Sillitoe on behalf of the Wellcome Sanger Institute COVID-19 Surveillance Team |
| EPI_ISL_3124873 | Lighthouse Lab in Alderley Park | Wellcome Sanger Institute for the COVID-19 Genomics UK (COG-UK) Consortium | Jacquelyn Wynn, Mairead Hyland, The Lighthouse Lab in Alderley Park and Alex Alderton, Roberto Amato, Jeffrey Barrett, Sonia Goncalves, Ewan Harrison, David K. Jackson, Ian Johnston, Dominic Kwiatkowski, Cordelia Langford, John Sillitoe on behalf of the Wellcome Sanger Institute COVID-19 Surveillance Team |
| EPI_ISL_3124874 | Lighthouse Lab in Glasgow | Wellcome Sanger Institute for the COVID-19 Genomics UK (COG-UK) Consortium | Harper VanSteenhouse, Yumi Kasai, David Gray, Carol Clugston, Anna Dominiczak and Alex Alderton, Roberto Amato, Jeffrey Barrett, Sonia Goncalves, Ewan Harrison, David K. Jackson, Ian Johnston, Dominic Kwiatkowski, Cordelia Langford, John Sillitoe on behalf of the Wellcome Sanger Institute COVID-19 Surveillance Team |
| EPI_ISL_3124875 | Lighthouse Lab in Alderley Park | Wellcome Sanger Institute for the COVID-19 Genomics UK (COG-UK) Consortium | Jacquelyn Wynn, Mairead Hyland, The Lighthouse Lab in Alderley Park and Alex Alderton, Roberto Amato, Jeffrey Barrett, Sonia Goncalves, Ewan Harrison, David K. Jackson, Ian Johnston, Dominic Kwiatkowski, Cordelia Langford, John Sillitoe on behalf of the Wellcome Sanger Institute COVID-19 Surveillance Team |
| EPI_ISL_3124876, EPI_ISL_3124877 | Lighthouse Lab in Glasgow | Wellcome Sanger Institute for the COVID-19 Genomics UK (COG-UK) Consortium | Harper VanSteenhouse, Yumi Kasai, David Gray, Carol Clugston, Anna Dominiczak and Alex Alderton, Roberto Amato, Jeffrey Barrett, Sonia Goncalves, Ewan Harrison, David K. Jackson, Ian Johnston, Dominic Kwiatkowski, Cordelia Langford, John Sillitoe on behalf of the Wellcome Sanger Institute COVID-19 Surveillance Team |
| EPI_ISL_3124878 | Lighthouse Lab in Alderley Park | Wellcome Sanger Institute for the COVID-19 Genomics UK (COG-UK) Consortium | Jacquelyn Wynn, Mairead Hyland, The Lighthouse Lab in Alderley Park and Alex Alderton, Roberto Amato, Jeffrey Barrett, Sonia Goncalves, Ewan Harrison, David K. Jackson, Ian Johnston, Dominic Kwiatkowski, Cordelia Langford, John Sillitoe on behalf of the Wellcome Sanger Institute COVID-19 Surveillance Team |
| EPI_ISL_3124879, EPI_ISL_3124880 | Lighthouse Lab in Milton Keynes | Wellcome Sanger Institute for the COVID-19 Genomics UK (COG-UK) Consortium | The Lighthouse Lab in Milton Keynes and Alex Alderton, Roberto Amato, Jeffrey Barrett, Sonia Goncalves, Ewan Harrison, David K. Jackson, Ian Johnston, Dominic Kwiatkowski, Cordelia Langford, John Sillitoe on behalf of the Wellcome Sanger Institute COVID-19 Surveillance Team |
| EPI_ISL_3124881, EPI_ISL_3124882, EPI_ISL_3124883 | Lighthouse Lab in Glasgow | Wellcome Sanger Institute for the COVID-19 Genomics UK (COG-UK) Consortium | Harper VanSteenhouse, Yumi Kasai, David Gray, Carol Clugston, Anna Dominiczak and Alex Alderton, Roberto Amato, Jeffrey Barrett, Sonia Goncalves, Ewan Harrison, David K. Jackson, Ian Johnston, Dominic Kwiatkowski, Cordelia Langford, John Sillitoe on behalf of the Wellcome Sanger Institute COVID-19 Surveillance Team |
| EPI_ISL_3124884 | Lighthouse Lab in Alderley Park | Wellcome Sanger Institute for the COVID-19 Genomics UK (COG-UK) Consortium | Jacquelyn Wynn, Mairead Hyland, The Lighthouse Lab in Alderley Park and Alex Alderton, Roberto Amato, Jeffrey Barrett, Sonia Goncalves, Ewan Harrison, David K. Jackson, Ian Johnston, Dominic Kwiatkowski, Cordelia Langford, John Sillitoe on behalf of the Wellcome Sanger Institute COVID-19 Surveillance Team |
| EPI_ISL_3124885, EPI_ISL_3124886 | Lighthouse Lab in Glasgow | Wellcome Sanger Institute for the COVID-19 Genomics UK (COG-UK) Consortium | Harper VanSteenhouse, Yumi Kasai, David Gray, Carol Clugston, Anna Dominiczak and Alex Alderton, Roberto Amato, Jeffrey Barrett, Sonia Goncalves, Ewan Harrison, David K. Jackson, Ian Johnston, Dominic Kwiatkowski, Cordelia Langford, John Sillitoe on behalf of the Wellcome Sanger Institute COVID-19 Surveillance Team |
| EPI_ISL_3124887, EPI_ISL_3124888, EPI_ISL_3124889 | Lighthouse Lab in Alderley Park | Wellcome Sanger Institute for the COVID-19 Genomics UK (COG-UK) Consortium | Jacquelyn Wynn, Mairead Hyland, The Lighthouse Lab in Alderley Park and Alex Alderton, Roberto Amato, Jeffrey Barrett, Sonia Goncalves, Ewan Harrison, David K. Jackson, Ian Johnston, Dominic Kwiatkowski, Cordelia Langford, John Sillitoe on behalf of the Wellcome Sanger Institute COVID-19 Surveillance Team |
| EPI_ISL_3124890, EPI_ISL_3124891, EPI_ISL_3124892 | Lighthouse Lab in Milton Keynes | Wellcome Sanger Institute for the COVID-19 Genomics UK (COG-UK) Consortium | The Lighthouse Lab in Milton Keynes and Alex Alderton, Roberto Amato, Jeffrey Barrett, Sonia Goncalves, Ewan Harrison, David K. Jackson, Ian Johnston, Dominic Kwiatkowski, Cordelia Langford, John Sillitoe on behalf of the Wellcome Sanger Institute COVID-19 Surveillance Team |
| EPI_ISL_3124893 | Lighthouse Lab in Glasgow | Wellcome Sanger Institute for the COVID-19 Genomics UK (COG-UK) Consortium | Harper VanSteenhouse, Yumi Kasai, David Gray, Carol Clugston, Anna Dominiczak and Alex Alderton, Roberto Amato, Jeffrey Barrett, Sonia Goncalves, Ewan Harrison, David K. Jackson, Ian Johnston, Dominic Kwiatkowski, Cordelia Langford, John Sillitoe on behalf of the Wellcome Sanger Institute COVID-19 Surveillance Team |
| EPI_ISL_3124894 | Lighthouse Lab in Alderley Park | Wellcome Sanger Institute for the COVID-19 Genomics UK (COG-UK) Consortium | Jacquelyn Wynn, Mairead Hyland, The Lighthouse Lab in Alderley Park and Alex Alderton, Roberto Amato, Jeffrey Barrett, Sonia Goncalves, Ewan Harrison, David K. Jackson, Ian Johnston, Dominic Kwiatkowski, Cordelia Langford, John Sillitoe on behalf of the Wellcome Sanger Institute COVID-19 Surveillance Team |
| EPI_ISL_3124895, EPI_ISL_3124896 | Lighthouse Lab in Glasgow | Wellcome Sanger Institute for the COVID-19 Genomics UK (COG-UK) Consortium | Harper VanSteenhouse, Yumi Kasai, David Gray, Carol Clugston, Anna Dominiczak and Alex Alderton, Roberto Amato, Jeffrey Barrett, Sonia Goncalves, Ewan Harrison, David K. Jackson, Ian Johnston, Dominic Kwiatkowski, Cordelia Langford, John Sillitoe on behalf of the Wellcome Sanger Institute COVID-19 Surveillance Team |
| EPI_ISL_3124897, EPI_ISL_3124898 | Lighthouse Lab in Milton Keynes | Wellcome Sanger Institute for the COVID-19 Genomics UK (COG-UK) Consortium | The Lighthouse Lab in Milton Keynes and Alex Alderton, Roberto Amato, Jeffrey Barrett, Sonia Goncalves, Ewan Harrison, David K. Jackson, Ian Johnston, Dominic Kwiatkowski, Cordelia Langford, John Sillitoe on behalf of the Wellcome Sanger Institute COVID-19 Surveillance Team |
| EPI_ISL_3124899 | Lighthouse Lab in Glasgow | Wellcome Sanger Institute for the COVID-19 Genomics UK (COG-UK) Consortium | Harper VanSteenhouse, Yumi Kasai, David Gray, Carol Clugston, Anna Dominiczak and Alex Alderton, Roberto Amato, Jeffrey Barrett, Sonia Goncalves, Ewan Harrison, David K. Jackson, Ian Johnston, Dominic Kwiatkowski, Cordelia Langford, John Sillitoe on behalf of the Wellcome Sanger Institute COVID-19 Surveillance Team |
| EPI_ISL_3124900 | Lighthouse Lab in Milton Keynes | Wellcome Sanger Institute for the COVID-19 Genomics UK (COG-UK) Consortium | The Lighthouse Lab in Milton Keynes and Alex Alderton, Roberto Amato, Jeffrey Barrett, Sonia Goncalves, Ewan Harrison, David K. Jackson, Ian Johnston, Dominic Kwiatkowski, Cordelia Langford, John Sillitoe on behalf of the Wellcome Sanger Institute COVID-19 Surveillance Team |
| EPI_ISL_3124901 | Lighthouse Lab in Glasgow | Wellcome Sanger Institute for the COVID-19 Genomics UK (COG-UK) Consortium | Harper VanSteenhouse, Yumi Kasai, David Gray, Carol Clugston, Anna Dominiczak and Alex Alderton, Roberto Amato, Jeffrey Barrett, Sonia Goncalves, Ewan Harrison, David K. Jackson, Ian Johnston, Dominic Kwiatkowski, Cordelia Langford, John Sillitoe on behalf of the Wellcome Sanger Institute COVID-19 Surveillance Team |
| EPI_ISL_3124902 | Lighthouse Lab in Milton Keynes | Wellcome Sanger Institute for the COVID-19 Genomics UK (COG-UK) Consortium | The Lighthouse Lab in Milton Keynes and Alex Alderton, Roberto Amato, Jeffrey Barrett, Sonia Goncalves, Ewan Harrison, David K. Jackson, Ian Johnston, Dominic Kwiatkowski, Cordelia Langford, John Sillitoe on behalf of the Wellcome Sanger Institute COVID-19 Surveillance Team |
| EPI_ISL_3124903, EPI_ISL_3124904 | Lighthouse Lab in Glasgow | Wellcome Sanger Institute for the COVID-19 Genomics UK (COG-UK) Consortium | Harper VanSteenhouse, Yumi Kasai, David Gray, Carol Clugston, Anna Dominiczak and Alex Alderton, Roberto Amato, Jeffrey Barrett, Sonia Goncalves, Ewan Harrison, David K. Jackson, Ian Johnston, Dominic Kwiatkowski, Cordelia Langford, John Sillitoe on behalf of the Wellcome Sanger Institute COVID-19 Surveillance Team |
| EPI_ISL_3124905 | Lighthouse Lab in Alderley Park | Wellcome Sanger Institute for the COVID-19 Genomics UK (COG-UK) Consortium | Jacquelyn Wynn, Mairead Hyland, The Lighthouse Lab in Alderley Park and Alex Alderton, Roberto Amato, Jeffrey Barrett, Sonia Goncalves, Ewan Harrison, David K. Jackson, Ian Johnston, Dominic Kwiatkowski, Cordelia Langford, John Sillitoe on behalf of the Wellcome Sanger Institute COVID-19 Surveillance Team |
| EPI_ISL_3124906 | Lighthouse Lab in Glasgow | Wellcome Sanger Institute for the COVID-19 Genomics UK (COG-UK) Consortium | Harper VanSteenhouse, Yumi Kasai, David Gray, Carol Clugston, Anna Dominiczak and Alex Alderton, Roberto Amato, Jeffrey Barrett, Sonia Goncalves, Ewan Harrison, David K. Jackson, Ian Johnston, Dominic Kwiatkowski, Cordelia Langford, John Sillitoe on behalf of the Wellcome Sanger Institute COVID-19 Surveillance Team |
| EPI_ISL_3124907, EPI_ISL_3124908, EPI_ISL_3124909 | Lighthouse Lab in Alderley Park | Wellcome Sanger Institute for the COVID-19 Genomics UK (COG-UK) Consortium | Jacquelyn Wynn, Mairead Hyland, The Lighthouse Lab in Alderley Park and Alex Alderton, Roberto Amato, Jeffrey Barrett, Sonia Goncalves, Ewan Harrison, David K. Jackson, Ian Johnston, Dominic Kwiatkowski, Cordelia Langford, John Sillitoe on behalf of the Wellcome Sanger Institute COVID-19 Surveillance Team |
| EPI_ISL_3124910, EPI_ISL_3124911 | Lighthouse Lab in Glasgow | Wellcome Sanger Institute for the COVID-19 Genomics UK (COG-UK) Consortium | Harper VanSteenhouse, Yumi Kasai, David Gray, Carol Clugston, Anna Dominiczak and Alex Alderton, Roberto Amato, Jeffrey Barrett, Sonia Goncalves, Ewan Harrison, David K. Jackson, Ian Johnston, Dominic Kwiatkowski, Cordelia Langford, John Sillitoe on behalf of the Wellcome Sanger Institute COVID-19 Surveillance Team |
| EPI_ISL_3124912, EPI_ISL_3124913 | Lighthouse Lab in Alderley Park | Wellcome Sanger Institute for the COVID-19 Genomics UK (COG-UK) Consortium | Jacquelyn Wynn, Mairead Hyland, The Lighthouse Lab in Alderley Park and Alex Alderton, Roberto Amato, Jeffrey Barrett, Sonia Goncalves, Ewan Harrison, David K. Jackson, Ian Johnston, Dominic Kwiatkowski, Cordelia Langford, John Sillitoe on behalf of the Wellcome Sanger Institute COVID-19 Surveillance Team |
| EPI_ISL_3124914, EPI_ISL_3124915 | Lighthouse Lab in Milton Keynes | Wellcome Sanger Institute for the COVID-19 Genomics UK (COG-UK) Consortium | The Lighthouse Lab in Milton Keynes and Alex Alderton, Roberto Amato, Jeffrey Barrett, Sonia Goncalves, Ewan Harrison, David K. Jackson, Ian Johnston, Dominic Kwiatkowski, Cordelia Langford, John Sillitoe on behalf of the Wellcome Sanger Institute COVID-19 Surveillance Team |
| EPI_ISL_3124916 | Lighthouse Lab in Glasgow | Wellcome Sanger Institute for the COVID-19 Genomics UK (COG-UK) Consortium | Harper VanSteenhouse, Yumi Kasai, David Gray, Carol Clugston, Anna Dominiczak and Alex Alderton, Roberto Amato, Jeffrey Barrett, Sonia Goncalves, Ewan Harrison, David K. Jackson, Ian Johnston, Dominic Kwiatkowski, Cordelia Langford, John Sillitoe on behalf of the Wellcome Sanger Institute COVID-19 Surveillance Team |
| EPI_ISL_3124917, EPI_ISL_3124918 | Lighthouse Lab in Alderley Park | Wellcome Sanger Institute for the COVID-19 Genomics UK (COG-UK) Consortium | Jacquelyn Wynn, Mairead Hyland, The Lighthouse Lab in Alderley Park and Alex Alderton, Roberto Amato, Jeffrey Barrett, Sonia Goncalves, Ewan Harrison, David K. Jackson, Ian Johnston, Dominic Kwiatkowski, Cordelia Langford, John Sillitoe on behalf of the Wellcome Sanger Institute COVID-19 Surveillance Team |
| EPI_ISL_3124919, EPI_ISL_3124920 | Lighthouse Lab in Milton Keynes | Wellcome Sanger Institute for the COVID-19 Genomics UK (COG-UK) Consortium | The Lighthouse Lab in Milton Keynes and Alex Alderton, Roberto Amato, Jeffrey Barrett, Sonia Goncalves, Ewan Harrison, David K. Jackson, Ian Johnston, Dominic Kwiatkowski, Cordelia Langford, John Sillitoe on behalf of the Wellcome Sanger Institute COVID-19 Surveillance Team |
| EPI_ISL_3124921, EPI_ISL_3124922 | Lighthouse Lab in Glasgow | Wellcome Sanger Institute for the COVID-19 Genomics UK (COG-UK) Consortium | Harper VanSteenhouse, Yumi Kasai, David Gray, Carol Clugston, Anna Dominiczak and Alex Alderton, Roberto Amato, Jeffrey Barrett, Sonia Goncalves, Ewan Harrison, David K. Jackson, Ian Johnston, Dominic Kwiatkowski, Cordelia Langford, John Sillitoe on behalf of the Wellcome Sanger Institute COVID-19 Surveillance Team |
| EPI_ISL_3124923, EPI_ISL_3124924, EPI_ISL_3124926 | Lighthouse Lab in Milton Keynes | Wellcome Sanger Institute for the COVID-19 Genomics UK (COG-UK) Consortium | The Lighthouse Lab in Milton Keynes and Alex Alderton, Roberto Amato, Jeffrey Barrett, Sonia Goncalves, Ewan Harrison, David K. Jackson, Ian Johnston, Dominic Kwiatkowski, Cordelia Langford, John Sillitoe on behalf of the Wellcome Sanger Institute COVID-19 Surveillance Team |
| EPI_ISL_3124927 | Lighthouse Lab in Alderley Park | Wellcome Sanger Institute for the COVID-19 Genomics UK (COG-UK) Consortium | Jacquelyn Wynn, Mairead Hyland, The Lighthouse Lab in Alderley Park and Alex Alderton, Roberto Amato, Jeffrey Barrett, Sonia Goncalves, Ewan Harrison, David K. Jackson, Ian Johnston, Dominic Kwiatkowski, Cordelia Langford, John Sillitoe on behalf of the Wellcome Sanger Institute COVID-19 Surveillance Team |

| | | | |
|---|---|---|---|
| EPI_ISL_3124928 | Lighthouse Lab in Milton Keynes | Wellcome Sanger Institute for the COVID-19 Genomics UK (COG-UK) Consortium | The Lighthouse Lab in Milton Keynes and Alex Alderton, Roberto Amato, Jeffrey Barrett, Sonia Goncalves, Ewan Harrison, David K. Jackson, Ian Johnston, Dominic Kwiatkowski, Cordelia Langford, John Sillitoe on behalf of the Wellcome Sanger Institute COVID-19 Surveillance Team |
| EPI_ISL_3124929 | Lighthouse Lab in Glasgow | Wellcome Sanger Institute for the COVID-19 Genomics UK (COG-UK) Consortium | Harper VanSteenhouse, Yumi Kasai, David Gray, Carol Clugston, Anna Dominiczak and Alex Alderton, Roberto Amato, Jeffrey Barrett, Sonia Goncalves, Ewan Harrison, David K. Jackson, Ian Johnston, Dominic Kwiatkowski, Cordelia Langford, John Sillitoe on behalf of the Wellcome Sanger Institute COVID-19 Surveillance Team |
| EPI_ISL_3124930 | Lighthouse Lab in Alderley Park | Wellcome Sanger Institute for the COVID-19 Genomics UK (COG-UK) Consortium | Jacquelyn Wynn, Mairead Hyland, The Lighthouse Lab in Alderley Park and Alex Alderton, Roberto Amato, Jeffrey Barrett, Sonia Goncalves, Ewan Harrison, David K. Jackson, Ian Johnston, Dominic Kwiatkowski, Cordelia Langford, John Sillitoe on behalf of the Wellcome Sanger Institute COVID-19 Surveillance Team |
| EPI_ISL_3124931 | Lighthouse Lab in Milton Keynes | Wellcome Sanger Institute for the COVID-19 Genomics UK (COG-UK) Consortium | The Lighthouse Lab in Milton Keynes and Alex Alderton, Roberto Amato, Jeffrey Barrett, Sonia Goncalves, Ewan Harrison, David K. Jackson, Ian Johnston, Dominic Kwiatkowski, Cordelia Langford, John Sillitoe on behalf of the Wellcome Sanger Institute COVID-19 Surveillance Team |
| EPI_ISL_3124932 | Lighthouse Lab in Glasgow | Wellcome Sanger Institute for the COVID-19 Genomics UK (COG-UK) Consortium | Harper VanSteenhouse, Yumi Kasai, David Gray, Carol Clugston, Anna Dominiczak and Alex Alderton, Roberto Amato, Jeffrey Barrett, Sonia Goncalves, Ewan Harrison, David K. Jackson, Ian Johnston, Dominic Kwiatkowski, Cordelia Langford, John Sillitoe on behalf of the Wellcome Sanger Institute COVID-19 Surveillance Team |
| EPI_ISL_3124933 | Lighthouse Lab in Milton Keynes | Wellcome Sanger Institute for the COVID-19 Genomics UK (COG-UK) Consortium | The Lighthouse Lab in Milton Keynes and Alex Alderton, Roberto Amato, Jeffrey Barrett, Sonia Goncalves, Ewan Harrison, David K. Jackson, Ian Johnston, Dominic Kwiatkowski, Cordelia Langford, John Sillitoe on behalf of the Wellcome Sanger Institute COVID-19 Surveillance Team |
| EPI_ISL_3124934 | Lighthouse Lab in Glasgow | Wellcome Sanger Institute for the COVID-19 Genomics UK (COG-UK) Consortium | Harper VanSteenhouse, Yumi Kasai, David Gray, Carol Clugston, Anna Dominiczak and Alex Alderton, Roberto Amato, Jeffrey Barrett, Sonia Goncalves, Ewan Harrison, David K. Jackson, Ian Johnston, Dominic Kwiatkowski, Cordelia Langford, John Sillitoe on behalf of the Wellcome Sanger Institute COVID-19 Surveillance Team |
| EPI_ISL_3124935 | Lighthouse Lab in Milton Keynes | Wellcome Sanger Institute for the COVID-19 Genomics UK (COG-UK) Consortium | The Lighthouse Lab in Milton Keynes and Alex Alderton, Roberto Amato, Jeffrey Barrett, Sonia Goncalves, Ewan Harrison, David K. Jackson, Ian Johnston, Dominic Kwiatkowski, Cordelia Langford, John Sillitoe on behalf of the Wellcome Sanger Institute COVID-19 Surveillance Team |
| EPI_ISL_3124936 | Lighthouse Lab in Alderley Park | Wellcome Sanger Institute for the COVID-19 Genomics UK (COG-UK) Consortium | Jacquelyn Wynn, Mairead Hyland, The Lighthouse Lab in Alderley Park and Alex Alderton, Roberto Amato, Jeffrey Barrett, Sonia Goncalves, Ewan Harrison, David K. Jackson, Ian Johnston, Dominic Kwiatkowski, Cordelia Langford, John Sillitoe on behalf of the Wellcome Sanger Institute COVID-19 Surveillance Team |
| EPI_ISL_3124937, EPI_ISL_3124938 | Lighthouse Lab in Glasgow | Wellcome Sanger Institute for the COVID-19 Genomics UK (COG-UK) Consortium | Harper VanSteenhouse, Yumi Kasai, David Gray, Carol Clugston, Anna Dominiczak and Alex Alderton, Roberto Amato, Jeffrey Barrett, Sonia Goncalves, Ewan Harrison, David K. Jackson, Ian Johnston, Dominic Kwiatkowski, Cordelia Langford, John Sillitoe on behalf of the Wellcome Sanger Institute COVID-19 Surveillance Team |
| EPI_ISL_3124939, EPI_ISL_3124940 | Lighthouse Lab in Alderley Park | Wellcome Sanger Institute for the COVID-19 Genomics UK (COG-UK) Consortium | Jacquelyn Wynn, Mairead Hyland, The Lighthouse Lab in Alderley Park and Alex Alderton, Roberto Amato, Jeffrey Barrett, Sonia Goncalves, Ewan Harrison, David K. Jackson, Ian Johnston, Dominic Kwiatkowski, Cordelia Langford, John Sillitoe on behalf of the Wellcome Sanger Institute COVID-19 Surveillance Team |
| EPI_ISL_3124941 | Lighthouse Lab in Milton Keynes | Wellcome Sanger Institute for the COVID-19 Genomics UK (COG-UK) Consortium | The Lighthouse Lab in Milton Keynes and Alex Alderton, Roberto Amato, Jeffrey Barrett, Sonia Goncalves, Ewan Harrison, David K. Jackson, Ian Johnston, Dominic Kwiatkowski, Cordelia Langford, John Sillitoe on behalf of the Wellcome Sanger Institute COVID-19 Surveillance Team |
| EPI_ISL_3124942 | Lighthouse Lab in Alderley Park | Wellcome Sanger Institute for the COVID-19 Genomics UK (COG-UK) Consortium | Jacquelyn Wynn, Mairead Hyland, The Lighthouse Lab in Alderley Park and Alex Alderton, Roberto Amato, Jeffrey Barrett, Sonia Goncalves, Ewan Harrison, David K. Jackson, Ian Johnston, Dominic Kwiatkowski, Cordelia Langford, John Sillitoe on behalf of the Wellcome Sanger Institute COVID-19 Surveillance Team |
| EPI_ISL_3124943 | Lighthouse Lab in Milton Keynes | Wellcome Sanger Institute for the COVID-19 Genomics UK (COG-UK) Consortium | The Lighthouse Lab in Milton Keynes and Alex Alderton, Roberto Amato, Jeffrey Barrett, Sonia Goncalves, Ewan Harrison, David K. Jackson, Ian Johnston, Dominic Kwiatkowski, Cordelia Langford, John Sillitoe on behalf of the Wellcome Sanger Institute COVID-19 Surveillance Team |
| EPI_ISL_3124944, EPI_ISL_3124945, EPI_ISL_3124946, EPI_ISL_3124947 | Lighthouse Lab in Alderley Park | Wellcome Sanger Institute for the COVID-19 Genomics UK (COG-UK) Consortium | Jacquelyn Wynn, Mairead Hyland, The Lighthouse Lab in Alderley Park and Alex Alderton, Roberto Amato, Jeffrey Barrett, Sonia Goncalves, Ewan Harrison, David K. Jackson, Ian Johnston, Dominic Kwiatkowski, Cordelia Langford, John Sillitoe on behalf of the Wellcome Sanger Institute COVID-19 Surveillance Team |
| EPI_ISL_3124948, EPI_ISL_3124949 | Lighthouse Lab in Glasgow | Wellcome Sanger Institute for the COVID-19 Genomics UK (COG-UK) Consortium | Harper VanSteenhouse, Yumi Kasai, David Gray, Carol Clugston, Anna Dominiczak and Alex Alderton, Roberto Amato, Jeffrey Barrett, Sonia Goncalves, Ewan Harrison, David K. Jackson, Ian Johnston, Dominic Kwiatkowski, Cordelia Langford, John Sillitoe on behalf of the Wellcome Sanger Institute COVID-19 Surveillance Team |
| EPI_ISL_3124950 | Lighthouse Lab in Alderley Park | Wellcome Sanger Institute for the COVID-19 Genomics UK (COG-UK) Consortium | Jacquelyn Wynn, Mairead Hyland, The Lighthouse Lab in Alderley Park and Alex Alderton, Roberto Amato, Jeffrey Barrett, Sonia Goncalves, Ewan Harrison, David K. Jackson, Ian Johnston, Dominic Kwiatkowski, Cordelia Langford, John Sillitoe on behalf of the Wellcome Sanger Institute COVID-19 Surveillance Team |
| EPI_ISL_3124951 | Lighthouse Lab in Milton Keynes | Wellcome Sanger Institute for the COVID-19 Genomics UK (COG-UK) Consortium | The Lighthouse Lab in Milton Keynes and Alex Alderton, Roberto Amato, Jeffrey Barrett, Sonia Goncalves, Ewan Harrison, David K. Jackson, Ian Johnston, Dominic Kwiatkowski, Cordelia Langford, John Sillitoe on behalf of the Wellcome Sanger Institute COVID-19 Surveillance Team |
| EPI_ISL_3124952 | Lighthouse Lab in Glasgow | Wellcome Sanger Institute for the COVID-19 Genomics UK (COG-UK) Consortium | Harper VanSteenhouse, Yumi Kasai, David Gray, Carol Clugston, Anna Dominiczak and Alex Alderton, Roberto Amato, Jeffrey Barrett, Sonia Goncalves, Ewan Harrison, David K. Jackson, Ian Johnston, Dominic Kwiatkowski, Cordelia Langford, John Sillitoe on behalf of the Wellcome Sanger Institute COVID-19 Surveillance Team |
| EPI_ISL_3124953 | Lighthouse Lab in Milton Keynes | Wellcome Sanger Institute for the COVID-19 Genomics UK (COG-UK) Consortium | The Lighthouse Lab in Milton Keynes and Alex Alderton, Roberto Amato, Jeffrey Barrett, Sonia Goncalves, Ewan Harrison, David K. Jackson, Ian Johnston, Dominic Kwiatkowski, Cordelia Langford, John Sillitoe on behalf of the Wellcome Sanger Institute COVID-19 Surveillance Team |
| EPI_ISL_3124954 | Lighthouse Lab in Glasgow | Wellcome Sanger Institute for the COVID-19 Genomics UK (COG-UK) Consortium | Harper VanSteenhouse, Yumi Kasai, David Gray, Carol Clugston, Anna Dominiczak and Alex Alderton, Roberto Amato, Jeffrey Barrett, Sonia Goncalves, Ewan Harrison, David K. Jackson, Ian Johnston, Dominic Kwiatkowski, Cordelia Langford, John Sillitoe on behalf of the Wellcome Sanger Institute COVID-19 Surveillance Team |
| EPI_ISL_3124955 | Lighthouse Lab in Alderley Park | Wellcome Sanger Institute for the COVID-19 Genomics UK (COG-UK) Consortium | Jacquelyn Wynn, Mairead Hyland, The Lighthouse Lab in Alderley Park and Alex Alderton, Roberto Amato, Jeffrey Barrett, Sonia Goncalves, Ewan Harrison, David K. Jackson, Ian Johnston, Dominic Kwiatkowski, Cordelia Langford, John Sillitoe on behalf of the Wellcome Sanger Institute COVID-19 Surveillance Team |
| EPI_ISL_3124956 | Lighthouse Lab in Milton Keynes | Wellcome Sanger Institute for the COVID-19 Genomics UK (COG-UK) Consortium | The Lighthouse Lab in Milton Keynes and Alex Alderton, Roberto Amato, Jeffrey Barrett, Sonia Goncalves, Ewan Harrison, David K. Jackson, Ian Johnston, Dominic Kwiatkowski, Cordelia Langford, John Sillitoe on behalf of the Wellcome Sanger Institute COVID-19 Surveillance Team |
| EPI_ISL_3124957, EPI_ISL_3124958, EPI_ISL_3124959 | Lighthouse Lab in Glasgow | Wellcome Sanger Institute for the COVID-19 Genomics UK (COG-UK) Consortium | Harper VanSteenhouse, Yumi Kasai, David Gray, Carol Clugston, Anna Dominiczak and Alex Alderton, Roberto Amato, Jeffrey Barrett, Sonia Goncalves, Ewan Harrison, David K. Jackson, Ian Johnston, Dominic Kwiatkowski, Cordelia Langford, John Sillitoe on behalf of the Wellcome Sanger Institute COVID-19 Surveillance Team |
| EPI_ISL_3124960 | Lighthouse Lab in Milton Keynes | Wellcome Sanger Institute for the COVID-19 Genomics UK (COG-UK) Consortium | The Lighthouse Lab in Milton Keynes and Alex Alderton, Roberto Amato, Jeffrey Barrett, Sonia Goncalves, Ewan Harrison, David K. Jackson, Ian Johnston, Dominic Kwiatkowski, Cordelia Langford, John Sillitoe on behalf of the Wellcome Sanger Institute COVID-19 Surveillance Team |
| EPI_ISL_3124961 | Lighthouse Lab in Glasgow | Wellcome Sanger Institute for the COVID-19 Genomics UK (COG-UK) Consortium | Harper VanSteenhouse, Yumi Kasai, David Gray, Carol Clugston, Anna Dominiczak and Alex Alderton, Roberto Amato, Jeffrey Barrett, Sonia Goncalves, Ewan Harrison, David K. Jackson, Ian Johnston, Dominic Kwiatkowski, Cordelia Langford, John Sillitoe on behalf of the Wellcome Sanger Institute COVID-19 Surveillance Team |
| EPI_ISL_3124962, EPI_ISL_3124963 | Lighthouse Lab in Alderley Park | Wellcome Sanger Institute for the COVID-19 Genomics UK (COG-UK) Consortium | Jacquelyn Wynn, Mairead Hyland, The Lighthouse Lab in Alderley Park and Alex Alderton, Roberto Amato, Jeffrey Barrett, Sonia Goncalves, Ewan Harrison, David K. Jackson, Ian Johnston, Dominic Kwiatkowski, Cordelia Langford, John Sillitoe on behalf of the Wellcome Sanger Institute COVID-19 Surveillance Team |
| EPI_ISL_3124964 | Lighthouse Lab in Glasgow | Wellcome Sanger Institute for the COVID-19 Genomics UK (COG-UK) Consortium | Harper VanSteenhouse, Yumi Kasai, David Gray, Carol Clugston, Anna Dominiczak and Alex Alderton, Roberto Amato, Jeffrey Barrett, Sonia Goncalves, Ewan Harrison, David K. Jackson, Ian Johnston, Dominic Kwiatkowski, Cordelia Langford, John Sillitoe on behalf of the Wellcome Sanger Institute COVID-19 Surveillance Team |
| EPI_ISL_3124965 | Lighthouse Lab in Milton Keynes | Wellcome Sanger Institute for the COVID-19 Genomics UK (COG-UK) Consortium | The Lighthouse Lab in Milton Keynes and Alex Alderton, Roberto Amato, Jeffrey Barrett, Sonia Goncalves, Ewan Harrison, David K. Jackson, Ian Johnston, Dominic Kwiatkowski, Cordelia Langford, John Sillitoe on behalf of the Wellcome Sanger Institute COVID-19 Surveillance Team |
| EPI_ISL_3124966 | Lighthouse Lab in Glasgow | Wellcome Sanger Institute for the COVID-19 Genomics UK (COG-UK) Consortium | Harper VanSteenhouse, Yumi Kasai, David Gray, Carol Clugston, Anna Dominiczak and Alex Alderton, Roberto Amato, Jeffrey Barrett, Sonia Goncalves, Ewan Harrison, David K. Jackson, Ian Johnston, Dominic Kwiatkowski, Cordelia Langford, John Sillitoe on behalf of the Wellcome Sanger Institute COVID-19 Surveillance Team |
| EPI_ISL_3124967 | Lighthouse Lab in Alderley Park | Wellcome Sanger Institute for the COVID-19 Genomics UK (COG-UK) Consortium | Jacquelyn Wynn, Mairead Hyland, The Lighthouse Lab in Alderley Park and Alex Alderton, Roberto Amato, Jeffrey Barrett, Sonia Goncalves, Ewan Harrison, David K. Jackson, Ian Johnston, Dominic Kwiatkowski, Cordelia Langford, John Sillitoe on behalf of the Wellcome Sanger Institute COVID-19 Surveillance Team |
| EPI_ISL_3124968, EPI_ISL_3124969, EPI_ISL_3124970 | Lighthouse Lab in Glasgow | Wellcome Sanger Institute for the COVID-19 Genomics UK (COG-UK) Consortium | Harper VanSteenhouse, Yumi Kasai, David Gray, Carol Clugston, Anna Dominiczak and Alex Alderton, Roberto Amato, Jeffrey Barrett, Sonia Goncalves, Ewan Harrison, David K. Jackson, Ian Johnston, Dominic Kwiatkowski, Cordelia Langford, John Sillitoe on behalf of the Wellcome Sanger Institute COVID-19 Surveillance Team |
| EPI_ISL_3124971, EPI_ISL_3124972 | Lighthouse Lab in Alderley Park | Wellcome Sanger Institute for the COVID-19 Genomics UK (COG-UK) Consortium | Jacquelyn Wynn, Mairead Hyland, The Lighthouse Lab in Alderley Park and Alex Alderton, Roberto Amato, Jeffrey Barrett, Sonia Goncalves, Ewan Harrison, David K. Jackson, Ian Johnston, Dominic Kwiatkowski, Cordelia Langford, John Sillitoe on behalf of the Wellcome Sanger Institute COVID-19 Surveillance Team |
| EPI_ISL_3124973 | Lighthouse Lab in Glasgow | Wellcome Sanger Institute for the COVID-19 Genomics UK (COG-UK) Consortium | Harper VanSteenhouse, Yumi Kasai, David Gray, Carol Clugston, Anna Dominiczak and Alex Alderton, Roberto Amato, Jeffrey Barrett, Sonia Goncalves, Ewan Harrison, David K. Jackson, Ian Johnston, Dominic Kwiatkowski, Cordelia Langford, John Sillitoe on behalf of the Wellcome Sanger Institute COVID-19 Surveillance Team |
| EPI_ISL_3124974, EPI_ISL_3124975, EPI_ISL_3124976, EPI_ISL_3124977 | Lighthouse Lab in Milton Keynes | Wellcome Sanger Institute for the COVID-19 Genomics UK (COG-UK) Consortium | The Lighthouse Lab in Milton Keynes and Alex Alderton, Roberto Amato, Jeffrey Barrett, Sonia Goncalves, Ewan Harrison, David K. Jackson, Ian Johnston, Dominic Kwiatkowski, Cordelia Langford, John Sillitoe on behalf of the Wellcome Sanger Institute COVID-19 Surveillance Team |
| EPI_ISL_3124978, EPI_ISL_3124979, EPI_ISL_3124980 | Lighthouse Lab in Alderley Park | Wellcome Sanger Institute for the COVID-19 Genomics UK (COG-UK) Consortium | Jacquelyn Wynn, Mairead Hyland, The Lighthouse Lab in Alderley Park and Alex Alderton, Roberto Amato, Jeffrey Barrett, Sonia Goncalves, Ewan Harrison, David K. Jackson, Ian Johnston, Dominic Kwiatkowski, Cordelia Langford, John Sillitoe on behalf of the Wellcome Sanger Institute COVID-19 Surveillance Team |
| EPI_ISL_3124981, EPI_ISL_3124982, EPI_ISL_3124983 | Lighthouse Lab in Glasgow | Wellcome Sanger Institute for the COVID-19 Genomics UK (COG-UK) Consortium | Harper VanSteenhouse, Yumi Kasai, David Gray, Carol Clugston, Anna Dominiczak and Alex Alderton, Roberto Amato, Jeffrey Barrett, Sonia Goncalves, Ewan Harrison, David K. Jackson, Ian Johnston, Dominic Kwiatkowski, Cordelia Langford, John Sillitoe on behalf of the Wellcome Sanger Institute COVID-19 Surveillance Team |
| EPI_ISL_3124984 | Lighthouse Lab in Milton Keynes | Wellcome Sanger Institute for the COVID-19 Genomics UK (COG-UK) Consortium | The Lighthouse Lab in Milton Keynes and Alex Alderton, Roberto Amato, Jeffrey Barrett, Sonia Goncalves, Ewan Harrison, David K. Jackson, Ian Johnston, Dominic Kwiatkowski, Cordelia Langford, John Sillitoe on behalf of the Wellcome Sanger Institute COVID-19 Surveillance Team |
| EPI_ISL_3124985 | Lighthouse Lab in Alderley Park | Wellcome Sanger Institute for the COVID-19 Genomics UK (COG-UK) Consortium | Jacquelyn Wynn, Mairead Hyland, The Lighthouse Lab in Alderley Park and Alex Alderton, Roberto Amato, Jeffrey Barrett, Sonia Goncalves, Ewan Harrison, David K. Jackson, Ian Johnston, Dominic Kwiatkowski, Cordelia Langford, John Sillitoe on behalf of the Wellcome Sanger Institute COVID-19 Surveillance Team |
| EPI_ISL_3124986, EPI_ISL_3124987, EPI_ISL_3124988, EPI_ISL_3124989, EPI_ISL_3124990 | Lighthouse Lab in Glasgow | Wellcome Sanger Institute for the COVID-19 Genomics UK (COG-UK) Consortium | Harper VanSteenhouse, Yumi Kasai, David Gray, Carol Clugston, Anna Dominiczak and Alex Alderton, Roberto Amato, Jeffrey Barrett, Sonia Goncalves, Ewan Harrison, David K. Jackson, Ian Johnston, Dominic Kwiatkowski, Cordelia Langford, John Sillitoe on behalf of the Wellcome Sanger Institute COVID-19 Surveillance Team |
| EPI_ISL_3124991 | Lighthouse Lab in Alderley Park | Wellcome Sanger Institute for the COVID-19 Genomics UK (COG-UK) | Jacquelyn Wynn, Mairead Hyland, The Lighthouse Lab in Alderley Park and Alex Alderton, Roberto Amato, Jeffrey Barrett, Sonia Goncalves, Ewan Harrison, David K. Jackson, Ian Johnston, Dominic Kwiatkowski, Cordelia Langford, John Sillitoe on |

| | | | |
|---|---|---|---|
| | | Consortium | behalf of the Wellcome Sanger Institute COVID-19 Surveillance Team |
| EPI_ISL_3124992, EPI_ISL_3124993 | Lighthouse Lab in Glasgow | Wellcome Sanger Institute for the COVID-19 Genomics UK (COG-UK) Consortium | Harper VanSteenhouse, Yumi Kasai, David Gray, Carol Clugston, Anna Dominiczak and Alex Alderton, Roberto Amato, Jeffrey Barrett, Sonia Goncalves, Ewan Harrison, David K. Jackson, Ian Johnston, Dominic Kwiatkowski, Cordelia Langford, John Sillitoe on behalf of the Wellcome Sanger Institute COVID-19 Surveillance Team |
| EPI_ISL_3124994 | Lighthouse Lab in Milton Keynes | Wellcome Sanger Institute for the COVID-19 Genomics UK (COG-UK) Consortium | The Lighthouse Lab in Milton Keynes and Alex Alderton, Roberto Amato, Jeffrey Barrett, Sonia Goncalves, Ewan Harrison, David K. Jackson, Ian Johnston, Dominic Kwiatkowski, Cordelia Langford, John Sillitoe on behalf of the Wellcome Sanger Institute COVID-19 Surveillance Team |
| EPI_ISL_3124995 | Lighthouse Lab in Alderley Park | Wellcome Sanger Institute for the COVID-19 Genomics UK (COG-UK) Consortium | Jacquelyn Wynn, Mairead Hyland, The Lighthouse Lab in Alderley Park and Alex Alderton, Roberto Amato, Jeffrey Barrett, Sonia Goncalves, Ewan Harrison, David K. Jackson, Ian Johnston, Dominic Kwiatkowski, Cordelia Langford, John Sillitoe on behalf of the Wellcome Sanger Institute COVID-19 Surveillance Team |
| EPI_ISL_3124996 | Lighthouse Lab in Glasgow | Wellcome Sanger Institute for the COVID-19 Genomics UK (COG-UK) Consortium | Harper VanSteenhouse, Yumi Kasai, David Gray, Carol Clugston, Anna Dominiczak and Alex Alderton, Roberto Amato, Jeffrey Barrett, Sonia Goncalves, Ewan Harrison, David K. Jackson, Ian Johnston, Dominic Kwiatkowski, Cordelia Langford, John Sillitoe on behalf of the Wellcome Sanger Institute COVID-19 Surveillance Team |
| EPI_ISL_3124997 | Lighthouse Lab in Milton Keynes | Wellcome Sanger Institute for the COVID-19 Genomics UK (COG-UK) Consortium | The Lighthouse Lab in Milton Keynes and Alex Alderton, Roberto Amato, Jeffrey Barrett, Sonia Goncalves, Ewan Harrison, David K. Jackson, Ian Johnston, Dominic Kwiatkowski, Cordelia Langford, John Sillitoe on behalf of the Wellcome Sanger Institute COVID-19 Surveillance Team |
| EPI_ISL_3124998 | Lighthouse Lab in Glasgow | Wellcome Sanger Institute for the COVID-19 Genomics UK (COG-UK) Consortium | Harper VanSteenhouse, Yumi Kasai, David Gray, Carol Clugston, Anna Dominiczak and Alex Alderton, Roberto Amato, Jeffrey Barrett, Sonia Goncalves, Ewan Harrison, David K. Jackson, Ian Johnston, Dominic Kwiatkowski, Cordelia Langford, John Sillitoe on behalf of the Wellcome Sanger Institute COVID-19 Surveillance Team |
| EPI_ISL_3124999 | Lighthouse Lab in Milton Keynes | Wellcome Sanger Institute for the COVID-19 Genomics UK (COG-UK) Consortium | The Lighthouse Lab in Milton Keynes and Alex Alderton, Roberto Amato, Jeffrey Barrett, Sonia Goncalves, Ewan Harrison, David K. Jackson, Ian Johnston, Dominic Kwiatkowski, Cordelia Langford, John Sillitoe on behalf of the Wellcome Sanger Institute COVID-19 Surveillance Team |
| EPI_ISL_3125000, EPI_ISL_3125001 | Lighthouse Lab in Glasgow | Wellcome Sanger Institute for the COVID-19 Genomics UK (COG-UK) Consortium | Harper VanSteenhouse, Yumi Kasai, David Gray, Carol Clugston, Anna Dominiczak and Alex Alderton, Roberto Amato, Jeffrey Barrett, Sonia Goncalves, Ewan Harrison, David K. Jackson, Ian Johnston, Dominic Kwiatkowski, Cordelia Langford, John Sillitoe on behalf of the Wellcome Sanger Institute COVID-19 Surveillance Team |
| EPI_ISL_3125002, EPI_ISL_3125003 | Lighthouse Lab in Milton Keynes | Wellcome Sanger Institute for the COVID-19 Genomics UK (COG-UK) Consortium | The Lighthouse Lab in Milton Keynes and Alex Alderton, Roberto Amato, Jeffrey Barrett, Sonia Goncalves, Ewan Harrison, David K. Jackson, Ian Johnston, Dominic Kwiatkowski, Cordelia Langford, John Sillitoe on behalf of the Wellcome Sanger Institute COVID-19 Surveillance Team |
| EPI_ISL_3125004 | Lighthouse Lab in Glasgow | Wellcome Sanger Institute for the COVID-19 Genomics UK (COG-UK) Consortium | Harper VanSteenhouse, Yumi Kasai, David Gray, Carol Clugston, Anna Dominiczak and Alex Alderton, Roberto Amato, Jeffrey Barrett, Sonia Goncalves, Ewan Harrison, David K. Jackson, Ian Johnston, Dominic Kwiatkowski, Cordelia Langford, John Sillitoe on behalf of the Wellcome Sanger Institute COVID-19 Surveillance Team |
| EPI_ISL_3125005 | Lighthouse Lab in Milton Keynes | Wellcome Sanger Institute for the COVID-19 Genomics UK (COG-UK) Consortium | The Lighthouse Lab in Milton Keynes and Alex Alderton, Roberto Amato, Jeffrey Barrett, Sonia Goncalves, Ewan Harrison, David K. Jackson, Ian Johnston, Dominic Kwiatkowski, Cordelia Langford, John Sillitoe on behalf of the Wellcome Sanger Institute COVID-19 Surveillance Team |
| EPI_ISL_3125006 | Lighthouse Lab in Glasgow | Wellcome Sanger Institute for the COVID-19 Genomics UK (COG-UK) Consortium | Harper VanSteenhouse, Yumi Kasai, David Gray, Carol Clugston, Anna Dominiczak and Alex Alderton, Roberto Amato, Jeffrey Barrett, Sonia Goncalves, Ewan Harrison, David K. Jackson, Ian Johnston, Dominic Kwiatkowski, Cordelia Langford, John Sillitoe on behalf of the Wellcome Sanger Institute COVID-19 Surveillance Team |
| EPI_ISL_3125007 | Lighthouse Lab in Milton Keynes | Wellcome Sanger Institute for the COVID-19 Genomics UK (COG-UK) Consortium | The Lighthouse Lab in Milton Keynes and Alex Alderton, Roberto Amato, Jeffrey Barrett, Sonia Goncalves, Ewan Harrison, David K. Jackson, Ian Johnston, Dominic Kwiatkowski, Cordelia Langford, John Sillitoe on behalf of the Wellcome Sanger Institute COVID-19 Surveillance Team |
| EPI_ISL_3125008 | Lighthouse Lab in Glasgow | Wellcome Sanger Institute for the COVID-19 Genomics UK (COG-UK) Consortium | Harper VanSteenhouse, Yumi Kasai, David Gray, Carol Clugston, Anna Dominiczak and Alex Alderton, Roberto Amato, Jeffrey Barrett, Sonia Goncalves, Ewan Harrison, David K. Jackson, Ian Johnston, Dominic Kwiatkowski, Cordelia Langford, John Sillitoe on behalf of the Wellcome Sanger Institute COVID-19 Surveillance Team |
| EPI_ISL_3125009, EPI_ISL_3125010 | Lighthouse Lab in Milton Keynes | Wellcome Sanger Institute for the COVID-19 Genomics UK (COG-UK) Consortium | The Lighthouse Lab in Milton Keynes and Alex Alderton, Roberto Amato, Jeffrey Barrett, Sonia Goncalves, Ewan Harrison, David K. Jackson, Ian Johnston, Dominic Kwiatkowski, Cordelia Langford, John Sillitoe on behalf of the Wellcome Sanger Institute COVID-19 Surveillance Team |
| EPI_ISL_3125011, EPI_ISL_3125012, EPI_ISL_3125013 | Lighthouse Lab in Glasgow | Wellcome Sanger Institute for the COVID-19 Genomics UK (COG-UK) Consortium | Harper VanSteenhouse, Yumi Kasai, David Gray, Carol Clugston, Anna Dominiczak and Alex Alderton, Roberto Amato, Jeffrey Barrett, Sonia Goncalves, Ewan Harrison, David K. Jackson, Ian Johnston, Dominic Kwiatkowski, Cordelia Langford, John Sillitoe on behalf of the Wellcome Sanger Institute COVID-19 Surveillance Team |
| EPI_ISL_3125014 | Lighthouse Lab in Alderley Park | Wellcome Sanger Institute for the COVID-19 Genomics UK (COG-UK) Consortium | Jacquelyn Wynn, Mairead Hyland, The Lighthouse Lab in Alderley Park and Alex Alderton, Roberto Amato, Jeffrey Barrett, Sonia Goncalves, Ewan Harrison, David K. Jackson, Ian Johnston, Dominic Kwiatkowski, Cordelia Langford, John Sillitoe on behalf of the Wellcome Sanger Institute COVID-19 Surveillance Team |
| EPI_ISL_3125015 | Lighthouse Lab in Glasgow | Wellcome Sanger Institute for the COVID-19 Genomics UK (COG-UK) Consortium | Harper VanSteenhouse, Yumi Kasai, David Gray, Carol Clugston, Anna Dominiczak and Alex Alderton, Roberto Amato, Jeffrey Barrett, Sonia Goncalves, Ewan Harrison, David K. Jackson, Ian Johnston, Dominic Kwiatkowski, Cordelia Langford, John Sillitoe on behalf of the Wellcome Sanger Institute COVID-19 Surveillance Team |
| EPI_ISL_3125016 | Lighthouse Lab in Milton Keynes | Wellcome Sanger Institute for the COVID-19 Genomics UK (COG-UK) Consortium | The Lighthouse Lab in Milton Keynes and Alex Alderton, Roberto Amato, Jeffrey Barrett, Sonia Goncalves, Ewan Harrison, David K. Jackson, Ian Johnston, Dominic Kwiatkowski, Cordelia Langford, John Sillitoe on behalf of the Wellcome Sanger Institute COVID-19 Surveillance Team |
| EPI_ISL_3125017, EPI_ISL_3125018 | Lighthouse Lab in Glasgow | Wellcome Sanger Institute for the COVID-19 Genomics UK (COG-UK) Consortium | Harper VanSteenhouse, Yumi Kasai, David Gray, Carol Clugston, Anna Dominiczak and Alex Alderton, Roberto Amato, Jeffrey Barrett, Sonia Goncalves, Ewan Harrison, David K. Jackson, Ian Johnston, Dominic Kwiatkowski, Cordelia Langford, John Sillitoe on behalf of the Wellcome Sanger Institute COVID-19 Surveillance Team |
| EPI_ISL_3125019, EPI_ISL_3125020 | Lighthouse Lab in Milton Keynes | Wellcome Sanger Institute for the COVID-19 Genomics UK (COG-UK) Consortium | The Lighthouse Lab in Milton Keynes and Alex Alderton, Roberto Amato, Jeffrey Barrett, Sonia Goncalves, Ewan Harrison, David K. Jackson, Ian Johnston, Dominic Kwiatkowski, Cordelia Langford, John Sillitoe on behalf of the Wellcome Sanger Institute COVID-19 Surveillance Team |
| EPI_ISL_3125021 | Lighthouse Lab in Glasgow | Wellcome Sanger Institute for the COVID-19 Genomics UK (COG-UK) Consortium | Harper VanSteenhouse, Yumi Kasai, David Gray, Carol Clugston, Anna Dominiczak and Alex Alderton, Roberto Amato, Jeffrey Barrett, Sonia Goncalves, Ewan Harrison, David K. Jackson, Ian Johnston, Dominic Kwiatkowski, Cordelia Langford, John Sillitoe on behalf of the Wellcome Sanger Institute COVID-19 Surveillance Team |
| EPI_ISL_3125022 | Lighthouse Lab in Milton Keynes | Wellcome Sanger Institute for the COVID-19 Genomics UK (COG-UK) Consortium | The Lighthouse Lab in Milton Keynes and Alex Alderton, Roberto Amato, Jeffrey Barrett, Sonia Goncalves, Ewan Harrison, David K. Jackson, Ian Johnston, Dominic Kwiatkowski, Cordelia Langford, John Sillitoe on behalf of the Wellcome Sanger Institute COVID-19 Surveillance Team |
| EPI_ISL_3125023 | Lighthouse Lab in Glasgow | Wellcome Sanger Institute for the COVID-19 Genomics UK (COG-UK) Consortium | Harper VanSteenhouse, Yumi Kasai, David Gray, Carol Clugston, Anna Dominiczak and Alex Alderton, Roberto Amato, Jeffrey Barrett, Sonia Goncalves, Ewan Harrison, David K. Jackson, Ian Johnston, Dominic Kwiatkowski, Cordelia Langford, John Sillitoe on behalf of the Wellcome Sanger Institute COVID-19 Surveillance Team |
| EPI_ISL_3125024 | Lighthouse Lab in Milton Keynes | Wellcome Sanger Institute for the COVID-19 Genomics UK (COG-UK) Consortium | The Lighthouse Lab in Milton Keynes and Alex Alderton, Roberto Amato, Jeffrey Barrett, Sonia Goncalves, Ewan Harrison, David K. Jackson, Ian Johnston, Dominic Kwiatkowski, Cordelia Langford, John Sillitoe on behalf of the Wellcome Sanger Institute COVID-19 Surveillance Team |
| EPI_ISL_3125025, EPI_ISL_3125026 | Lighthouse Lab in Alderley Park | Wellcome Sanger Institute for the COVID-19 Genomics UK (COG-UK) Consortium | Jacquelyn Wynn, Mairead Hyland, The Lighthouse Lab in Alderley Park and Alex Alderton, Roberto Amato, Jeffrey Barrett, Sonia Goncalves, Ewan Harrison, David K. Jackson, Ian Johnston, Dominic Kwiatkowski, Cordelia Langford, John Sillitoe on behalf of the Wellcome Sanger Institute COVID-19 Surveillance Team |
| EPI_ISL_3125027 | Lighthouse Lab in Glasgow | Wellcome Sanger Institute for the COVID-19 Genomics UK (COG-UK) Consortium | Harper VanSteenhouse, Yumi Kasai, David Gray, Carol Clugston, Anna Dominiczak and Alex Alderton, Roberto Amato, Jeffrey Barrett, Sonia Goncalves, Ewan Harrison, David K. Jackson, Ian Johnston, Dominic Kwiatkowski, Cordelia Langford, John Sillitoe on behalf of the Wellcome Sanger Institute COVID-19 Surveillance Team |
| EPI_ISL_3125028 | Lighthouse Lab in Alderley Park | Wellcome Sanger Institute for the COVID-19 Genomics UK (COG-UK) Consortium | Jacquelyn Wynn, Mairead Hyland, The Lighthouse Lab in Alderley Park and Alex Alderton, Roberto Amato, Jeffrey Barrett, Sonia Goncalves, Ewan Harrison, David K. Jackson, Ian Johnston, Dominic Kwiatkowski, Cordelia Langford, John Sillitoe on behalf of the Wellcome Sanger Institute COVID-19 Surveillance Team |
| EPI_ISL_3125029 | Lighthouse Lab in Glasgow | Wellcome Sanger Institute for the COVID-19 Genomics UK (COG-UK) Consortium | Harper VanSteenhouse, Yumi Kasai, David Gray, Carol Clugston, Anna Dominiczak and Alex Alderton, Roberto Amato, Jeffrey Barrett, Sonia Goncalves, Ewan Harrison, David K. Jackson, Ian Johnston, Dominic Kwiatkowski, Cordelia Langford, John Sillitoe on behalf of the Wellcome Sanger Institute COVID-19 Surveillance Team |
| EPI_ISL_3125030 | Lighthouse Lab in Milton Keynes | Wellcome Sanger Institute for the COVID-19 Genomics UK (COG-UK) Consortium | The Lighthouse Lab in Milton Keynes and Alex Alderton, Roberto Amato, Jeffrey Barrett, Sonia Goncalves, Ewan Harrison, David K. Jackson, Ian Johnston, Dominic Kwiatkowski, Cordelia Langford, John Sillitoe on behalf of the Wellcome Sanger Institute COVID-19 Surveillance Team |
| EPI_ISL_3125031 | Lighthouse Lab in Glasgow | Wellcome Sanger Institute for the COVID-19 Genomics UK (COG-UK) Consortium | Harper VanSteenhouse, Yumi Kasai, David Gray, Carol Clugston, Anna Dominiczak and Alex Alderton, Roberto Amato, Jeffrey Barrett, Sonia Goncalves, Ewan Harrison, David K. Jackson, Ian Johnston, Dominic Kwiatkowski, Cordelia Langford, John Sillitoe on behalf of the Wellcome Sanger Institute COVID-19 Surveillance Team |
| EPI_ISL_3125032, EPI_ISL_3125033 | Lighthouse Lab in Milton Keynes | Wellcome Sanger Institute for the COVID-19 Genomics UK (COG-UK) Consortium | The Lighthouse Lab in Milton Keynes and Alex Alderton, Roberto Amato, Jeffrey Barrett, Sonia Goncalves, Ewan Harrison, David K. Jackson, Ian Johnston, Dominic Kwiatkowski, Cordelia Langford, John Sillitoe on behalf of the Wellcome Sanger Institute COVID-19 Surveillance Team |
| EPI_ISL_3125034, EPI_ISL_3125035, EPI_ISL_3125036 | Lighthouse Lab in Glasgow | Wellcome Sanger Institute for the COVID-19 Genomics UK (COG-UK) Consortium | Harper VanSteenhouse, Yumi Kasai, David Gray, Carol Clugston, Anna Dominiczak and Alex Alderton, Roberto Amato, Jeffrey Barrett, Sonia Goncalves, Ewan Harrison, David K. Jackson, Ian Johnston, Dominic Kwiatkowski, Cordelia Langford, John Sillitoe on behalf of the Wellcome Sanger Institute COVID-19 Surveillance Team |
| EPI_ISL_3125037 | Lighthouse Lab in Alderley Park | Wellcome Sanger Institute for the COVID-19 Genomics UK (COG-UK) Consortium | Jacquelyn Wynn, Mairead Hyland, The Lighthouse Lab in Alderley Park and Alex Alderton, Roberto Amato, Jeffrey Barrett, Sonia Goncalves, Ewan Harrison, David K. Jackson, Ian Johnston, Dominic Kwiatkowski, Cordelia Langford, John Sillitoe on behalf of the Wellcome Sanger Institute COVID-19 Surveillance Team |
| EPI_ISL_3125038 | Lighthouse Lab in Milton Keynes | Wellcome Sanger Institute for the COVID-19 Genomics UK (COG-UK) Consortium | The Lighthouse Lab in Milton Keynes and Alex Alderton, Roberto Amato, Jeffrey Barrett, Sonia Goncalves, Ewan Harrison, David K. Jackson, Ian Johnston, Dominic Kwiatkowski, Cordelia Langford, John Sillitoe on behalf of the Wellcome Sanger Institute COVID-19 Surveillance Team |
| EPI_ISL_3125039 | Lighthouse Lab in Glasgow | Wellcome Sanger Institute for the COVID-19 Genomics UK (COG-UK) Consortium | Harper VanSteenhouse, Yumi Kasai, David Gray, Carol Clugston, Anna Dominiczak and Alex Alderton, Roberto Amato, Jeffrey Barrett, Sonia Goncalves, Ewan Harrison, David K. Jackson, Ian Johnston, Dominic Kwiatkowski, Cordelia Langford, John Sillitoe on behalf of the Wellcome Sanger Institute COVID-19 Surveillance Team |
| EPI_ISL_3125040 | Lighthouse Lab in Alderley Park | Wellcome Sanger Institute for the COVID-19 Genomics UK (COG-UK) Consortium | Jacquelyn Wynn, Mairead Hyland, The Lighthouse Lab in Alderley Park and Alex Alderton, Roberto Amato, Jeffrey Barrett, Sonia Goncalves, Ewan Harrison, David K. Jackson, Ian Johnston, Dominic Kwiatkowski, Cordelia Langford, John Sillitoe on behalf of the Wellcome Sanger Institute COVID-19 Surveillance Team |
| EPI_ISL_3125041, EPI_ISL_3125042 | Lighthouse Lab in Glasgow | Wellcome Sanger Institute for the COVID-19 Genomics UK (COG-UK) Consortium | Harper VanSteenhouse, Yumi Kasai, David Gray, Carol Clugston, Anna Dominiczak and Alex Alderton, Roberto Amato, Jeffrey Barrett, Sonia Goncalves, Ewan Harrison, David K. Jackson, Ian Johnston, Dominic Kwiatkowski, Cordelia Langford, John Sillitoe on behalf of the Wellcome Sanger Institute COVID-19 Surveillance Team |
| EPI_ISL_3125043, EPI_ISL_3125044 | Lighthouse Lab in Alderley Park | Wellcome Sanger Institute for the COVID-19 Genomics UK (COG-UK) Consortium | Jacquelyn Wynn, Mairead Hyland, The Lighthouse Lab in Alderley Park and Alex Alderton, Roberto Amato, Jeffrey Barrett, Sonia Goncalves, Ewan Harrison, David K. Jackson, Ian Johnston, Dominic Kwiatkowski, Cordelia Langford, John Sillitoe on behalf of the Wellcome Sanger Institute COVID-19 Surveillance Team |
| EPI_ISL_3125045 | Lighthouse Lab in Glasgow | Wellcome Sanger Institute for the COVID-19 Genomics UK (COG-UK) Consortium | Harper VanSteenhouse, Yumi Kasai, David Gray, Carol Clugston, Anna Dominiczak and Alex Alderton, Roberto Amato, Jeffrey Barrett, Sonia Goncalves, Ewan Harrison, David K. Jackson, Ian Johnston, Dominic Kwiatkowski, Cordelia Langford, John Sillitoe on behalf of the Wellcome Sanger Institute COVID-19 Surveillance Team |
| EPI_ISL_3125046 | Lighthouse Lab in Milton Keynes | Wellcome Sanger Institute for the COVID-19 Genomics UK (COG-UK) Consortium | The Lighthouse Lab in Milton Keynes and Alex Alderton, Roberto Amato, Jeffrey Barrett, Sonia Goncalves, Ewan Harrison, David K. Jackson, Ian Johnston, Dominic Kwiatkowski, Cordelia Langford, John Sillitoe on behalf of the Wellcome Sanger Institute COVID-19 Surveillance Team |
| EPI_ISL_3125047 | Lighthouse Lab in Alderley Park | Wellcome Sanger Institute for the COVID-19 Genomics UK (COG-UK) Consortium | Jacquelyn Wynn, Mairead Hyland, The Lighthouse Lab in Alderley Park and Alex Alderton, Roberto Amato, Jeffrey Barrett, Sonia Goncalves, Ewan Harrison, David K. Jackson, Ian Johnston, Dominic Kwiatkowski, Cordelia Langford, John Sillitoe on behalf of the Wellcome Sanger Institute COVID-19 Surveillance Team |
| EPI_ISL_3125048 | Lighthouse Lab in Milton Keynes | Wellcome Sanger Institute for the COVID-19 Genomics UK (COG-UK) Consortium | The Lighthouse Lab in Milton Keynes and Alex Alderton, Roberto Amato, Jeffrey Barrett, Sonia Goncalves, Ewan Harrison, David K. Jackson, Ian Johnston, Dominic Kwiatkowski, Cordelia Langford, John Sillitoe on behalf of the Wellcome Sanger Institute COVID-19 Surveillance Team |
| EPI_ISL_3125049 | Lighthouse Lab in Alderley Park | Wellcome Sanger Institute for the COVID-19 Genomics UK (COG-UK) Consortium | Jacquelyn Wynn, Mairead Hyland, The Lighthouse Lab in Alderley Park and Alex Alderton, Roberto Amato, Jeffrey Barrett, Sonia Goncalves, Ewan Harrison, David K. Jackson, Ian Johnston, Dominic Kwiatkowski, Cordelia Langford, John Sillitoe on behalf of the Wellcome Sanger Institute COVID-19 Surveillance Team |

| | | | |
|---|---|---|---|
| EPI_ISL_3125050, EPI_ISL_3125051 | Lighthouse Lab in Glasgow | Wellcome Sanger Institute for the COVID-19 Genomics UK (COG-UK) Consortium | Harper VanSteenhouse, Yumi Kasai, David Gray, Carol Clugston, Anna Dominiczak and Alex Alderton, Roberto Amato, Jeffrey Barrett, Sonia Goncalves, Ewan Harrison, David K. Jackson, Ian Johnston, Dominic Kwiatkowski, Cordelia Langford, John Sillitoe on behalf of the Wellcome Sanger Institute COVID-19 Surveillance Team |
| EPI_ISL_3125052 | Lighthouse Lab in Alderley Park | Wellcome Sanger Institute for the COVID-19 Genomics UK (COG-UK) Consortium | Jacquelyn Wynn, Mairead Hyland, The Lighthouse Lab in Alderley Park and Alex Alderton, Roberto Amato, Jeffrey Barrett, Sonia Goncalves, Ewan Harrison, David K. Jackson, Ian Johnston, Dominic Kwiatkowski, Cordelia Langford, John Sillitoe on behalf of the Wellcome Sanger Institute COVID-19 Surveillance Team |
| EPI_ISL_3125053 | Lighthouse Lab in Milton Keynes | Wellcome Sanger Institute for the COVID-19 Genomics UK (COG-UK) Consortium | The Lighthouse Lab in Milton Keynes and Alex Alderton, Roberto Amato, Jeffrey Barrett, Sonia Goncalves, Ewan Harrison, David K. Jackson, Ian Johnston, Dominic Kwiatkowski, Cordelia Langford, John Sillitoe on behalf of the Wellcome Sanger Institute COVID-19 Surveillance Team |
| EPI_ISL_3125054, EPI_ISL_3125055 | Lighthouse Lab in Glasgow | Wellcome Sanger Institute for the COVID-19 Genomics UK (COG-UK) Consortium | Harper VanSteenhouse, Yumi Kasai, David Gray, Carol Clugston, Anna Dominiczak and Alex Alderton, Roberto Amato, Jeffrey Barrett, Sonia Goncalves, Ewan Harrison, David K. Jackson, Ian Johnston, Dominic Kwiatkowski, Cordelia Langford, John Sillitoe on behalf of the Wellcome Sanger Institute COVID-19 Surveillance Team |
| EPI_ISL_3125056 | Lighthouse Lab in Alderley Park | Wellcome Sanger Institute for the COVID-19 Genomics UK (COG-UK) Consortium | Jacquelyn Wynn, Mairead Hyland, The Lighthouse Lab in Alderley Park and Alex Alderton, Roberto Amato, Jeffrey Barrett, Sonia Goncalves, Ewan Harrison, David K. Jackson, Ian Johnston, Dominic Kwiatkowski, Cordelia Langford, John Sillitoe on behalf of the Wellcome Sanger Institute COVID-19 Surveillance Team |
| EPI_ISL_3125057 | Lighthouse Lab in Glasgow | Wellcome Sanger Institute for the COVID-19 Genomics UK (COG-UK) Consortium | Harper VanSteenhouse, Yumi Kasai, David Gray, Carol Clugston, Anna Dominiczak and Alex Alderton, Roberto Amato, Jeffrey Barrett, Sonia Goncalves, Ewan Harrison, David K. Jackson, Ian Johnston, Dominic Kwiatkowski, Cordelia Langford, John Sillitoe on behalf of the Wellcome Sanger Institute COVID-19 Surveillance Team |
| EPI_ISL_3125058 | Lighthouse Lab in Milton Keynes | Wellcome Sanger Institute for the COVID-19 Genomics UK (COG-UK) Consortium | The Lighthouse Lab in Milton Keynes and Alex Alderton, Roberto Amato, Jeffrey Barrett, Sonia Goncalves, Ewan Harrison, David K. Jackson, Ian Johnston, Dominic Kwiatkowski, Cordelia Langford, John Sillitoe on behalf of the Wellcome Sanger Institute COVID-19 Surveillance Team |
| EPI_ISL_3125059 | Lighthouse Lab in Glasgow | Wellcome Sanger Institute for the COVID-19 Genomics UK (COG-UK) Consortium | Harper VanSteenhouse, Yumi Kasai, David Gray, Carol Clugston, Anna Dominiczak and Alex Alderton, Roberto Amato, Jeffrey Barrett, Sonia Goncalves, Ewan Harrison, David K. Jackson, Ian Johnston, Dominic Kwiatkowski, Cordelia Langford, John Sillitoe on behalf of the Wellcome Sanger Institute COVID-19 Surveillance Team |
| EPI_ISL_3125060 | Lighthouse Lab in Milton Keynes | Wellcome Sanger Institute for the COVID-19 Genomics UK (COG-UK) Consortium | The Lighthouse Lab in Milton Keynes and Alex Alderton, Roberto Amato, Jeffrey Barrett, Sonia Goncalves, Ewan Harrison, David K. Jackson, Ian Johnston, Dominic Kwiatkowski, Cordelia Langford, John Sillitoe on behalf of the Wellcome Sanger Institute COVID-19 Surveillance Team |
| EPI_ISL_3125061 | Lighthouse Lab in Glasgow | Wellcome Sanger Institute for the COVID-19 Genomics UK (COG-UK) Consortium | Harper VanSteenhouse, Yumi Kasai, David Gray, Carol Clugston, Anna Dominiczak and Alex Alderton, Roberto Amato, Jeffrey Barrett, Sonia Goncalves, Ewan Harrison, David K. Jackson, Ian Johnston, Dominic Kwiatkowski, Cordelia Langford, John Sillitoe on behalf of the Wellcome Sanger Institute COVID-19 Surveillance Team |
| EPI_ISL_3125062, EPI_ISL_3125063, EPI_ISL_3125064, EPI_ISL_3125065 | Lighthouse Lab in Alderley Park | Wellcome Sanger Institute for the COVID-19 Genomics UK (COG-UK) Consortium | Jacquelyn Wynn, Mairead Hyland, The Lighthouse Lab in Alderley Park and Alex Alderton, Roberto Amato, Jeffrey Barrett, Sonia Goncalves, Ewan Harrison, David K. Jackson, Ian Johnston, Dominic Kwiatkowski, Cordelia Langford, John Sillitoe on behalf of the Wellcome Sanger Institute COVID-19 Surveillance Team |
| EPI_ISL_3125066, EPI_ISL_3125067 | Lighthouse Lab in Milton Keynes | Wellcome Sanger Institute for the COVID-19 Genomics UK (COG-UK) Consortium | The Lighthouse Lab in Milton Keynes and Alex Alderton, Roberto Amato, Jeffrey Barrett, Sonia Goncalves, Ewan Harrison, David K. Jackson, Ian Johnston, Dominic Kwiatkowski, Cordelia Langford, John Sillitoe on behalf of the Wellcome Sanger Institute COVID-19 Surveillance Team |
| EPI_ISL_3125068, EPI_ISL_3125069 | Lighthouse Lab in Alderley Park | Wellcome Sanger Institute for the COVID-19 Genomics UK (COG-UK) Consortium | Jacquelyn Wynn, Mairead Hyland, The Lighthouse Lab in Alderley Park and Alex Alderton, Roberto Amato, Jeffrey Barrett, Sonia Goncalves, Ewan Harrison, David K. Jackson, Ian Johnston, Dominic Kwiatkowski, Cordelia Langford, John Sillitoe on behalf of the Wellcome Sanger Institute COVID-19 Surveillance Team |
| EPI_ISL_3125070 | Lighthouse Lab in Milton Keynes | Wellcome Sanger Institute for the COVID-19 Genomics UK (COG-UK) Consortium | The Lighthouse Lab in Milton Keynes and Alex Alderton, Roberto Amato, Jeffrey Barrett, Sonia Goncalves, Ewan Harrison, David K. Jackson, Ian Johnston, Dominic Kwiatkowski, Cordelia Langford, John Sillitoe on behalf of the Wellcome Sanger Institute COVID-19 Surveillance Team |
| EPI_ISL_3125071 | Lighthouse Lab in Alderley Park | Wellcome Sanger Institute for the COVID-19 Genomics UK (COG-UK) Consortium | Jacquelyn Wynn, Mairead Hyland, The Lighthouse Lab in Alderley Park and Alex Alderton, Roberto Amato, Jeffrey Barrett, Sonia Goncalves, Ewan Harrison, David K. Jackson, Ian Johnston, Dominic Kwiatkowski, Cordelia Langford, John Sillitoe on behalf of the Wellcome Sanger Institute COVID-19 Surveillance Team |
| EPI_ISL_3125072, EPI_ISL_3125073 | Lighthouse Lab in Glasgow | Wellcome Sanger Institute for the COVID-19 Genomics UK (COG-UK) Consortium | Harper VanSteenhouse, Yumi Kasai, David Gray, Carol Clugston, Anna Dominiczak and Alex Alderton, Roberto Amato, Jeffrey Barrett, Sonia Goncalves, Ewan Harrison, David K. Jackson, Ian Johnston, Dominic Kwiatkowski, Cordelia Langford, John Sillitoe on behalf of the Wellcome Sanger Institute COVID-19 Surveillance Team |
| EPI_ISL_3125074 | Lighthouse Lab in Milton Keynes | Wellcome Sanger Institute for the COVID-19 Genomics UK (COG-UK) Consortium | The Lighthouse Lab in Milton Keynes and Alex Alderton, Roberto Amato, Jeffrey Barrett, Sonia Goncalves, Ewan Harrison, David K. Jackson, Ian Johnston, Dominic Kwiatkowski, Cordelia Langford, John Sillitoe on behalf of the Wellcome Sanger Institute COVID-19 Surveillance Team |
| EPI_ISL_3125075, EPI_ISL_3125076 | Lighthouse Lab in Glasgow | Wellcome Sanger Institute for the COVID-19 Genomics UK (COG-UK) Consortium | Harper VanSteenhouse, Yumi Kasai, David Gray, Carol Clugston, Anna Dominiczak and Alex Alderton, Roberto Amato, Jeffrey Barrett, Sonia Goncalves, Ewan Harrison, David K. Jackson, Ian Johnston, Dominic Kwiatkowski, Cordelia Langford, John Sillitoe on behalf of the Wellcome Sanger Institute COVID-19 Surveillance Team |
| EPI_ISL_3125077 | Lighthouse Lab in Milton Keynes | Wellcome Sanger Institute for the COVID-19 Genomics UK (COG-UK) Consortium | The Lighthouse Lab in Milton Keynes and Alex Alderton, Roberto Amato, Jeffrey Barrett, Sonia Goncalves, Ewan Harrison, David K. Jackson, Ian Johnston, Dominic Kwiatkowski, Cordelia Langford, John Sillitoe on behalf of the Wellcome Sanger Institute COVID-19 Surveillance Team |
| EPI_ISL_3125078 | Lighthouse Lab in Alderley Park | Wellcome Sanger Institute for the COVID-19 Genomics UK (COG-UK) Consortium | Jacquelyn Wynn, Mairead Hyland, The Lighthouse Lab in Alderley Park and Alex Alderton, Roberto Amato, Jeffrey Barrett, Sonia Goncalves, Ewan Harrison, David K. Jackson, Ian Johnston, Dominic Kwiatkowski, Cordelia Langford, John Sillitoe on behalf of the Wellcome Sanger Institute COVID-19 Surveillance Team |
| EPI_ISL_3125079 | Lighthouse Lab in Glasgow | Wellcome Sanger Institute for the COVID-19 Genomics UK (COG-UK) Consortium | Harper VanSteenhouse, Yumi Kasai, David Gray, Carol Clugston, Anna Dominiczak and Alex Alderton, Roberto Amato, Jeffrey Barrett, Sonia Goncalves, Ewan Harrison, David K. Jackson, Ian Johnston, Dominic Kwiatkowski, Cordelia Langford, John Sillitoe on behalf of the Wellcome Sanger Institute COVID-19 Surveillance Team |
| EPI_ISL_3125080 | Lighthouse Lab in Milton Keynes | Wellcome Sanger Institute for the COVID-19 Genomics UK (COG-UK) Consortium | The Lighthouse Lab in Milton Keynes and Alex Alderton, Roberto Amato, Jeffrey Barrett, Sonia Goncalves, Ewan Harrison, David K. Jackson, Ian Johnston, Dominic Kwiatkowski, Cordelia Langford, John Sillitoe on behalf of the Wellcome Sanger Institute COVID-19 Surveillance Team |
| EPI_ISL_3125081, EPI_ISL_3125082, EPI_ISL_3125083, EPI_ISL_3125084 | Lighthouse Lab in Glasgow | Wellcome Sanger Institute for the COVID-19 Genomics UK (COG-UK) Consortium | Harper VanSteenhouse, Yumi Kasai, David Gray, Carol Clugston, Anna Dominiczak and Alex Alderton, Roberto Amato, Jeffrey Barrett, Sonia Goncalves, Ewan Harrison, David K. Jackson, Ian Johnston, Dominic Kwiatkowski, Cordelia Langford, John Sillitoe on behalf of the Wellcome Sanger Institute COVID-19 Surveillance Team |
| EPI_ISL_3125085 | Lighthouse Lab in Milton Keynes | Wellcome Sanger Institute for the COVID-19 Genomics UK (COG-UK) Consortium | The Lighthouse Lab in Milton Keynes and Alex Alderton, Roberto Amato, Jeffrey Barrett, Sonia Goncalves, Ewan Harrison, David K. Jackson, Ian Johnston, Dominic Kwiatkowski, Cordelia Langford, John Sillitoe on behalf of the Wellcome Sanger Institute COVID-19 Surveillance Team |
| EPI_ISL_3125086 | Lighthouse Lab in Glasgow | Wellcome Sanger Institute for the COVID-19 Genomics UK (COG-UK) Consortium | Harper VanSteenhouse, Yumi Kasai, David Gray, Carol Clugston, Anna Dominiczak and Alex Alderton, Roberto Amato, Jeffrey Barrett, Sonia Goncalves, Ewan Harrison, David K. Jackson, Ian Johnston, Dominic Kwiatkowski, Cordelia Langford, John Sillitoe on behalf of the Wellcome Sanger Institute COVID-19 Surveillance Team |
| EPI_ISL_3125087, EPI_ISL_3125088, EPI_ISL_3125089 | Lighthouse Lab in Milton Keynes | Wellcome Sanger Institute for the COVID-19 Genomics UK (COG-UK) Consortium | The Lighthouse Lab in Milton Keynes and Alex Alderton, Roberto Amato, Jeffrey Barrett, Sonia Goncalves, Ewan Harrison, David K. Jackson, Ian Johnston, Dominic Kwiatkowski, Cordelia Langford, John Sillitoe on behalf of the Wellcome Sanger Institute COVID-19 Surveillance Team |
| EPI_ISL_3125090, EPI_ISL_3125091 | Lighthouse Lab in Alderley Park | Wellcome Sanger Institute for the COVID-19 Genomics UK (COG-UK) Consortium | Jacquelyn Wynn, Mairead Hyland, The Lighthouse Lab in Alderley Park and Alex Alderton, Roberto Amato, Jeffrey Barrett, Sonia Goncalves, Ewan Harrison, David K. Jackson, Ian Johnston, Dominic Kwiatkowski, Cordelia Langford, John Sillitoe on behalf of the Wellcome Sanger Institute COVID-19 Surveillance Team |
| EPI_ISL_3125092 | Lighthouse Lab in Glasgow | Wellcome Sanger Institute for the COVID-19 Genomics UK (COG-UK) Consortium | Harper VanSteenhouse, Yumi Kasai, David Gray, Carol Clugston, Anna Dominiczak and Alex Alderton, Roberto Amato, Jeffrey Barrett, Sonia Goncalves, Ewan Harrison, David K. Jackson, Ian Johnston, Dominic Kwiatkowski, Cordelia Langford, John Sillitoe on behalf of the Wellcome Sanger Institute COVID-19 Surveillance Team |
| EPI_ISL_3125093 | Lighthouse Lab in Alderley Park | Wellcome Sanger Institute for the COVID-19 Genomics UK (COG-UK) Consortium | Jacquelyn Wynn, Mairead Hyland, The Lighthouse Lab in Alderley Park and Alex Alderton, Roberto Amato, Jeffrey Barrett, Sonia Goncalves, Ewan Harrison, David K. Jackson, Ian Johnston, Dominic Kwiatkowski, Cordelia Langford, John Sillitoe on behalf of the Wellcome Sanger Institute COVID-19 Surveillance Team |
| EPI_ISL_3125094, EPI_ISL_3125095 | Lighthouse Lab in Milton Keynes | Wellcome Sanger Institute for the COVID-19 Genomics UK (COG-UK) Consortium | The Lighthouse Lab in Milton Keynes and Alex Alderton, Roberto Amato, Jeffrey Barrett, Sonia Goncalves, Ewan Harrison, David K. Jackson, Ian Johnston, Dominic Kwiatkowski, Cordelia Langford, John Sillitoe on behalf of the Wellcome Sanger Institute COVID-19 Surveillance Team |
| EPI_ISL_3125096 | Lighthouse Lab in Glasgow | Wellcome Sanger Institute for the COVID-19 Genomics UK (COG-UK) Consortium | Harper VanSteenhouse, Yumi Kasai, David Gray, Carol Clugston, Anna Dominiczak and Alex Alderton, Roberto Amato, Jeffrey Barrett, Sonia Goncalves, Ewan Harrison, David K. Jackson, Ian Johnston, Dominic Kwiatkowski, Cordelia Langford, John Sillitoe on behalf of the Wellcome Sanger Institute COVID-19 Surveillance Team |
| EPI_ISL_3125097 | Lighthouse Lab in Alderley Park | Wellcome Sanger Institute for the COVID-19 Genomics UK (COG-UK) Consortium | Jacquelyn Wynn, Mairead Hyland, The Lighthouse Lab in Alderley Park and Alex Alderton, Roberto Amato, Jeffrey Barrett, Sonia Goncalves, Ewan Harrison, David K. Jackson, Ian Johnston, Dominic Kwiatkowski, Cordelia Langford, John Sillitoe on behalf of the Wellcome Sanger Institute COVID-19 Surveillance Team |
| EPI_ISL_3125098 | Lighthouse Lab in Glasgow | Wellcome Sanger Institute for the COVID-19 Genomics UK (COG-UK) Consortium | Harper VanSteenhouse, Yumi Kasai, David Gray, Carol Clugston, Anna Dominiczak and Alex Alderton, Roberto Amato, Jeffrey Barrett, Sonia Goncalves, Ewan Harrison, David K. Jackson, Ian Johnston, Dominic Kwiatkowski, Cordelia Langford, John Sillitoe on behalf of the Wellcome Sanger Institute COVID-19 Surveillance Team |
| EPI_ISL_3125099, EPI_ISL_3125100 | Lighthouse Lab in Alderley Park | Wellcome Sanger Institute for the COVID-19 Genomics UK (COG-UK) Consortium | Jacquelyn Wynn, Mairead Hyland, The Lighthouse Lab in Alderley Park and Alex Alderton, Roberto Amato, Jeffrey Barrett, Sonia Goncalves, Ewan Harrison, David K. Jackson, Ian Johnston, Dominic Kwiatkowski, Cordelia Langford, John Sillitoe on behalf of the Wellcome Sanger Institute COVID-19 Surveillance Team |
| EPI_ISL_3125101 | Lighthouse Lab in Glasgow | Wellcome Sanger Institute for the COVID-19 Genomics UK (COG-UK) Consortium | Harper VanSteenhouse, Yumi Kasai, David Gray, Carol Clugston, Anna Dominiczak and Alex Alderton, Roberto Amato, Jeffrey Barrett, Sonia Goncalves, Ewan Harrison, David K. Jackson, Ian Johnston, Dominic Kwiatkowski, Cordelia Langford, John Sillitoe on behalf of the Wellcome Sanger Institute COVID-19 Surveillance Team |
| EPI_ISL_3125102 | Lighthouse Lab in Milton Keynes | Wellcome Sanger Institute for the COVID-19 Genomics UK (COG-UK) Consortium | The Lighthouse Lab in Milton Keynes and Alex Alderton, Roberto Amato, Jeffrey Barrett, Sonia Goncalves, Ewan Harrison, David K. Jackson, Ian Johnston, Dominic Kwiatkowski, Cordelia Langford, John Sillitoe on behalf of the Wellcome Sanger Institute COVID-19 Surveillance Team |
| EPI_ISL_3125103 | Lighthouse Lab in Alderley Park | Wellcome Sanger Institute for the COVID-19 Genomics UK (COG-UK) Consortium | Jacquelyn Wynn, Mairead Hyland, The Lighthouse Lab in Alderley Park and Alex Alderton, Roberto Amato, Jeffrey Barrett, Sonia Goncalves, Ewan Harrison, David K. Jackson, Ian Johnston, Dominic Kwiatkowski, Cordelia Langford, John Sillitoe on behalf of the Wellcome Sanger Institute COVID-19 Surveillance Team |
| EPI_ISL_3125104 | Lighthouse Lab in Glasgow | Wellcome Sanger Institute for the COVID-19 Genomics UK (COG-UK) Consortium | Harper VanSteenhouse, Yumi Kasai, David Gray, Carol Clugston, Anna Dominiczak and Alex Alderton, Roberto Amato, Jeffrey Barrett, Sonia Goncalves, Ewan Harrison, David K. Jackson, Ian Johnston, Dominic Kwiatkowski, Cordelia Langford, John Sillitoe on behalf of the Wellcome Sanger Institute COVID-19 Surveillance Team |
| EPI_ISL_3125105 | Lighthouse Lab in Alderley Park | Wellcome Sanger Institute for the COVID-19 Genomics UK (COG-UK) Consortium | Jacquelyn Wynn, Mairead Hyland, The Lighthouse Lab in Alderley Park and Alex Alderton, Roberto Amato, Jeffrey Barrett, Sonia Goncalves, Ewan Harrison, David K. Jackson, Ian Johnston, Dominic Kwiatkowski, Cordelia Langford, John Sillitoe on behalf of the Wellcome Sanger Institute COVID-19 Surveillance Team |
| EPI_ISL_3125106 | Lighthouse Lab in Glasgow | Wellcome Sanger Institute for the COVID-19 Genomics UK (COG-UK) Consortium | Harper VanSteenhouse, Yumi Kasai, David Gray, Carol Clugston, Anna Dominiczak and Alex Alderton, Roberto Amato, Jeffrey Barrett, Sonia Goncalves, Ewan Harrison, David K. Jackson, Ian Johnston, Dominic Kwiatkowski, Cordelia Langford, John Sillitoe on behalf of the Wellcome Sanger Institute COVID-19 Surveillance Team |
| EPI_ISL_3125107 | Lighthouse Lab in Alderley Park | Wellcome Sanger Institute for the COVID-19 Genomics UK (COG-UK) Consortium | Jacquelyn Wynn, Mairead Hyland, The Lighthouse Lab in Alderley Park and Alex Alderton, Roberto Amato, Jeffrey Barrett, Sonia Goncalves, Ewan Harrison, David K. Jackson, Ian Johnston, Dominic Kwiatkowski, Cordelia Langford, John Sillitoe on behalf of the Wellcome Sanger Institute COVID-19 Surveillance Team |
| EPI_ISL_3125108 | Lighthouse Lab in Milton Keynes | Wellcome Sanger Institute for the COVID-19 Genomics UK (COG-UK) Consortium | The Lighthouse Lab in Milton Keynes and Alex Alderton, Roberto Amato, Jeffrey Barrett, Sonia Goncalves, Ewan Harrison, David K. Jackson, Ian Johnston, Dominic Kwiatkowski, Cordelia Langford, John Sillitoe on behalf of the Wellcome Sanger Institute COVID-19 Surveillance Team |
| EPI_ISL_3125109, EPI_ISL_3125110 | Lighthouse Lab in Glasgow | Wellcome Sanger Institute for the COVID-19 Genomics UK (COG-UK) Consortium | Harper VanSteenhouse, Yumi Kasai, David Gray, Carol Clugston, Anna Dominiczak and Alex Alderton, Roberto Amato, Jeffrey Barrett, Sonia Goncalves, Ewan Harrison, David K. Jackson, Ian Johnston, Dominic Kwiatkowski, Cordelia Langford, John Sillitoe on behalf of the Wellcome Sanger Institute COVID-19 Surveillance Team |

| | | | |
|---|---|---|---|
| EPI_ISL_3125111, EPI_ISL_3125112 | Lighthouse Lab in Alderley Park | Wellcome Sanger Institute for the COVID-19 Genomics UK (COG-UK) Consortium | Jacquelyn Wynn, Mairead Hyland, The Lighthouse Lab in Alderley Park and Alex Alderton, Roberto Amato, Jeffrey Barrett, Sonia Goncalves, Ewan Harrison, David K. Jackson, Ian Johnston, Dominic Kwiatkowski, Cordelia Langford, John Sillitoe on behalf of the Wellcome Sanger Institute COVID-19 Surveillance Team |
| EPI_ISL_3125113 | Lighthouse Lab in Glasgow | Wellcome Sanger Institute for the COVID-19 Genomics UK (COG-UK) Consortium | Harper VanSteenhouse, Yumi Kasai, David Gray, Carol Clugston, Anna Dominiczak and Alex Alderton, Roberto Amato, Jeffrey Barrett, Sonia Goncalves, Ewan Harrison, David K. Jackson, Ian Johnston, Dominic Kwiatkowski, Cordelia Langford, John Sillitoe on behalf of the Wellcome Sanger Institute COVID-19 Surveillance Team |
| EPI_ISL_3125114 | Lighthouse Lab in Milton Keynes | Wellcome Sanger Institute for the COVID-19 Genomics UK (COG-UK) Consortium | The Lighthouse Lab in Milton Keynes and Alex Alderton, Roberto Amato, Jeffrey Barrett, Sonia Goncalves, Ewan Harrison, David K. Jackson, Ian Johnston, Dominic Kwiatkowski, Cordelia Langford, John Sillitoe on behalf of the Wellcome Sanger Institute COVID-19 Surveillance Team |
| EPI_ISL_3125115 | Lighthouse Lab in Alderley Park | Wellcome Sanger Institute for the COVID-19 Genomics UK (COG-UK) Consortium | Jacquelyn Wynn, Mairead Hyland, The Lighthouse Lab in Alderley Park and Alex Alderton, Roberto Amato, Jeffrey Barrett, Sonia Goncalves, Ewan Harrison, David K. Jackson, Ian Johnston, Dominic Kwiatkowski, Cordelia Langford, John Sillitoe on behalf of the Wellcome Sanger Institute COVID-19 Surveillance Team |
| EPI_ISL_3125116 | Lighthouse Lab in Glasgow | Wellcome Sanger Institute for the COVID-19 Genomics UK (COG-UK) Consortium | Harper VanSteenhouse, Yumi Kasai, David Gray, Carol Clugston, Anna Dominiczak and Alex Alderton, Roberto Amato, Jeffrey Barrett, Sonia Goncalves, Ewan Harrison, David K. Jackson, Ian Johnston, Dominic Kwiatkowski, Cordelia Langford, John Sillitoe on behalf of the Wellcome Sanger Institute COVID-19 Surveillance Team |
| EPI_ISL_3125117 | Lighthouse Lab in Milton Keynes | Wellcome Sanger Institute for the COVID-19 Genomics UK (COG-UK) Consortium | The Lighthouse Lab in Milton Keynes and Alex Alderton, Roberto Amato, Jeffrey Barrett, Sonia Goncalves, Ewan Harrison, David K. Jackson, Ian Johnston, Dominic Kwiatkowski, Cordelia Langford, John Sillitoe on behalf of the Wellcome Sanger Institute COVID-19 Surveillance Team |
| EPI_ISL_3125118 | Lighthouse Lab in Alderley Park | Wellcome Sanger Institute for the COVID-19 Genomics UK (COG-UK) Consortium | Jacquelyn Wynn, Mairead Hyland, The Lighthouse Lab in Alderley Park and Alex Alderton, Roberto Amato, Jeffrey Barrett, Sonia Goncalves, Ewan Harrison, David K. Jackson, Ian Johnston, Dominic Kwiatkowski, Cordelia Langford, John Sillitoe on behalf of the Wellcome Sanger Institute COVID-19 Surveillance Team |
| EPI_ISL_3125119 | Lighthouse Lab in Milton Keynes | Wellcome Sanger Institute for the COVID-19 Genomics UK (COG-UK) Consortium | The Lighthouse Lab in Milton Keynes and Alex Alderton, Roberto Amato, Jeffrey Barrett, Sonia Goncalves, Ewan Harrison, David K. Jackson, Ian Johnston, Dominic Kwiatkowski, Cordelia Langford, John Sillitoe on behalf of the Wellcome Sanger Institute COVID-19 Surveillance Team |
| EPI_ISL_3125120, EPI_ISL_3125121, EPI_ISL_3125122 | Lighthouse Lab in Glasgow | Wellcome Sanger Institute for the COVID-19 Genomics UK (COG-UK) Consortium | Harper VanSteenhouse, Yumi Kasai, David Gray, Carol Clugston, Anna Dominiczak and Alex Alderton, Roberto Amato, Jeffrey Barrett, Sonia Goncalves, Ewan Harrison, David K. Jackson, Ian Johnston, Dominic Kwiatkowski, Cordelia Langford, John Sillitoe on behalf of the Wellcome Sanger Institute COVID-19 Surveillance Team |
| EPI_ISL_3125123, EPI_ISL_3125124 | Lighthouse Lab in Alderley Park | Wellcome Sanger Institute for the COVID-19 Genomics UK (COG-UK) Consortium | Jacquelyn Wynn, Mairead Hyland, The Lighthouse Lab in Alderley Park and Alex Alderton, Roberto Amato, Jeffrey Barrett, Sonia Goncalves, Ewan Harrison, David K. Jackson, Ian Johnston, Dominic Kwiatkowski, Cordelia Langford, John Sillitoe on behalf of the Wellcome Sanger Institute COVID-19 Surveillance Team |
| EPI_ISL_3125125, EPI_ISL_3125126 | Lighthouse Lab in Milton Keynes | Wellcome Sanger Institute for the COVID-19 Genomics UK (COG-UK) Consortium | The Lighthouse Lab in Milton Keynes and Alex Alderton, Roberto Amato, Jeffrey Barrett, Sonia Goncalves, Ewan Harrison, David K. Jackson, Ian Johnston, Dominic Kwiatkowski, Cordelia Langford, John Sillitoe on behalf of the Wellcome Sanger Institute COVID-19 Surveillance Team |
| EPI_ISL_3125127 | Lighthouse Lab in Glasgow | Wellcome Sanger Institute for the COVID-19 Genomics UK (COG-UK) Consortium | Harper VanSteenhouse, Yumi Kasai, David Gray, Carol Clugston, Anna Dominiczak and Alex Alderton, Roberto Amato, Jeffrey Barrett, Sonia Goncalves, Ewan Harrison, David K. Jackson, Ian Johnston, Dominic Kwiatkowski, Cordelia Langford, John Sillitoe on behalf of the Wellcome Sanger Institute COVID-19 Surveillance Team |
| EPI_ISL_3125128, EPI_ISL_3125129, EPI_ISL_3125130 | Lighthouse Lab in Alderley Park | Wellcome Sanger Institute for the COVID-19 Genomics UK (COG-UK) Consortium | Jacquelyn Wynn, Mairead Hyland, The Lighthouse Lab in Alderley Park and Alex Alderton, Roberto Amato, Jeffrey Barrett, Sonia Goncalves, Ewan Harrison, David K. Jackson, Ian Johnston, Dominic Kwiatkowski, Cordelia Langford, John Sillitoe on behalf of the Wellcome Sanger Institute COVID-19 Surveillance Team |
| EPI_ISL_3125131 | Lighthouse Lab in Milton Keynes | Wellcome Sanger Institute for the COVID-19 Genomics UK (COG-UK) Consortium | The Lighthouse Lab in Milton Keynes and Alex Alderton, Roberto Amato, Jeffrey Barrett, Sonia Goncalves, Ewan Harrison, David K. Jackson, Ian Johnston, Dominic Kwiatkowski, Cordelia Langford, John Sillitoe on behalf of the Wellcome Sanger Institute COVID-19 Surveillance Team |
| EPI_ISL_3125132 | Lighthouse Lab in Glasgow | Wellcome Sanger Institute for the COVID-19 Genomics UK (COG-UK) Consortium | Harper VanSteenhouse, Yumi Kasai, David Gray, Carol Clugston, Anna Dominiczak and Alex Alderton, Roberto Amato, Jeffrey Barrett, Sonia Goncalves, Ewan Harrison, David K. Jackson, Ian Johnston, Dominic Kwiatkowski, Cordelia Langford, John Sillitoe on behalf of the Wellcome Sanger Institute COVID-19 Surveillance Team |
| EPI_ISL_3125133 | Lighthouse Lab in Alderley Park | Wellcome Sanger Institute for the COVID-19 Genomics UK (COG-UK) Consortium | Jacquelyn Wynn, Mairead Hyland, The Lighthouse Lab in Alderley Park and Alex Alderton, Roberto Amato, Jeffrey Barrett, Sonia Goncalves, Ewan Harrison, David K. Jackson, Ian Johnston, Dominic Kwiatkowski, Cordelia Langford, John Sillitoe on behalf of the Wellcome Sanger Institute COVID-19 Surveillance Team |
| EPI_ISL_3125134 | Lighthouse Lab in Glasgow | Wellcome Sanger Institute for the COVID-19 Genomics UK (COG-UK) Consortium | Harper VanSteenhouse, Yumi Kasai, David Gray, Carol Clugston, Anna Dominiczak and Alex Alderton, Roberto Amato, Jeffrey Barrett, Sonia Goncalves, Ewan Harrison, David K. Jackson, Ian Johnston, Dominic Kwiatkowski, Cordelia Langford, John Sillitoe on behalf of the Wellcome Sanger Institute COVID-19 Surveillance Team |
| EPI_ISL_3125135 | Lighthouse Lab in Milton Keynes | Wellcome Sanger Institute for the COVID-19 Genomics UK (COG-UK) Consortium | The Lighthouse Lab in Milton Keynes and Alex Alderton, Roberto Amato, Jeffrey Barrett, Sonia Goncalves, Ewan Harrison, David K. Jackson, Ian Johnston, Dominic Kwiatkowski, Cordelia Langford, John Sillitoe on behalf of the Wellcome Sanger Institute COVID-19 Surveillance Team |
| EPI_ISL_3125136 | Lighthouse Lab in Glasgow | Wellcome Sanger Institute for the COVID-19 Genomics UK (COG-UK) Consortium | Harper VanSteenhouse, Yumi Kasai, David Gray, Carol Clugston, Anna Dominiczak and Alex Alderton, Roberto Amato, Jeffrey Barrett, Sonia Goncalves, Ewan Harrison, David K. Jackson, Ian Johnston, Dominic Kwiatkowski, Cordelia Langford, John Sillitoe on behalf of the Wellcome Sanger Institute COVID-19 Surveillance Team |
| EPI_ISL_3125137 | Lighthouse Lab in Alderley Park | Wellcome Sanger Institute for the COVID-19 Genomics UK (COG-UK) Consortium | Jacquelyn Wynn, Mairead Hyland, The Lighthouse Lab in Alderley Park and Alex Alderton, Roberto Amato, Jeffrey Barrett, Sonia Goncalves, Ewan Harrison, David K. Jackson, Ian Johnston, Dominic Kwiatkowski, Cordelia Langford, John Sillitoe on behalf of the Wellcome Sanger Institute COVID-19 Surveillance Team |
| EPI_ISL_3125138, EPI_ISL_3125139 | Lighthouse Lab in Glasgow | Wellcome Sanger Institute for the COVID-19 Genomics UK (COG-UK) Consortium | Harper VanSteenhouse, Yumi Kasai, David Gray, Carol Clugston, Anna Dominiczak and Alex Alderton, Roberto Amato, Jeffrey Barrett, Sonia Goncalves, Ewan Harrison, David K. Jackson, Ian Johnston, Dominic Kwiatkowski, Cordelia Langford, John Sillitoe on behalf of the Wellcome Sanger Institute COVID-19 Surveillance Team |
| EPI_ISL_3125140 | Lighthouse Lab in Alderley Park | Wellcome Sanger Institute for the COVID-19 Genomics UK (COG-UK) Consortium | Jacquelyn Wynn, Mairead Hyland, The Lighthouse Lab in Alderley Park and Alex Alderton, Roberto Amato, Jeffrey Barrett, Sonia Goncalves, Ewan Harrison, David K. Jackson, Ian Johnston, Dominic Kwiatkowski, Cordelia Langford, John Sillitoe on behalf of the Wellcome Sanger Institute COVID-19 Surveillance Team |
| EPI_ISL_3125141 | Lighthouse Lab in Milton Keynes | Wellcome Sanger Institute for the COVID-19 Genomics UK (COG-UK) Consortium | The Lighthouse Lab in Milton Keynes and Alex Alderton, Roberto Amato, Jeffrey Barrett, Sonia Goncalves, Ewan Harrison, David K. Jackson, Ian Johnston, Dominic Kwiatkowski, Cordelia Langford, John Sillitoe on behalf of the Wellcome Sanger Institute COVID-19 Surveillance Team |
| EPI_ISL_3125142 | Lighthouse Lab in Glasgow | Wellcome Sanger Institute for the COVID-19 Genomics UK (COG-UK) Consortium | Harper VanSteenhouse, Yumi Kasai, David Gray, Carol Clugston, Anna Dominiczak and Alex Alderton, Roberto Amato, Jeffrey Barrett, Sonia Goncalves, Ewan Harrison, David K. Jackson, Ian Johnston, Dominic Kwiatkowski, Cordelia Langford, John Sillitoe on behalf of the Wellcome Sanger Institute COVID-19 Surveillance Team |
| EPI_ISL_3125143 | Lighthouse Lab in Milton Keynes | Wellcome Sanger Institute for the COVID-19 Genomics UK (COG-UK) Consortium | The Lighthouse Lab in Milton Keynes and Alex Alderton, Roberto Amato, Jeffrey Barrett, Sonia Goncalves, Ewan Harrison, David K. Jackson, Ian Johnston, Dominic Kwiatkowski, Cordelia Langford, John Sillitoe on behalf of the Wellcome Sanger Institute COVID-19 Surveillance Team |
| EPI_ISL_3125144, EPI_ISL_3125145, EPI_ISL_3125146 | Lighthouse Lab in Glasgow | Wellcome Sanger Institute for the COVID-19 Genomics UK (COG-UK) Consortium | Harper VanSteenhouse, Yumi Kasai, David Gray, Carol Clugston, Anna Dominiczak and Alex Alderton, Roberto Amato, Jeffrey Barrett, Sonia Goncalves, Ewan Harrison, David K. Jackson, Ian Johnston, Dominic Kwiatkowski, Cordelia Langford, John Sillitoe on behalf of the Wellcome Sanger Institute COVID-19 Surveillance Team |
| EPI_ISL_3125147 | Lighthouse Lab in Alderley Park | Wellcome Sanger Institute for the COVID-19 Genomics UK (COG-UK) Consortium | Jacquelyn Wynn, Mairead Hyland, The Lighthouse Lab in Alderley Park and Alex Alderton, Roberto Amato, Jeffrey Barrett, Sonia Goncalves, Ewan Harrison, David K. Jackson, Ian Johnston, Dominic Kwiatkowski, Cordelia Langford, John Sillitoe on behalf of the Wellcome Sanger Institute COVID-19 Surveillance Team |
| EPI_ISL_3125148 | Lighthouse Lab in Glasgow | Wellcome Sanger Institute for the COVID-19 Genomics UK (COG-UK) Consortium | Harper VanSteenhouse, Yumi Kasai, David Gray, Carol Clugston, Anna Dominiczak and Alex Alderton, Roberto Amato, Jeffrey Barrett, Sonia Goncalves, Ewan Harrison, David K. Jackson, Ian Johnston, Dominic Kwiatkowski, Cordelia Langford, John Sillitoe on behalf of the Wellcome Sanger Institute COVID-19 Surveillance Team |
| EPI_ISL_3125149 | Lighthouse Lab in Milton Keynes | Wellcome Sanger Institute for the COVID-19 Genomics UK (COG-UK) Consortium | The Lighthouse Lab in Milton Keynes and Alex Alderton, Roberto Amato, Jeffrey Barrett, Sonia Goncalves, Ewan Harrison, David K. Jackson, Ian Johnston, Dominic Kwiatkowski, Cordelia Langford, John Sillitoe on behalf of the Wellcome Sanger Institute COVID-19 Surveillance Team |
| EPI_ISL_3125150 | Lighthouse Lab in Alderley Park | Wellcome Sanger Institute for the COVID-19 Genomics UK (COG-UK) Consortium | Jacquelyn Wynn, Mairead Hyland, The Lighthouse Lab in Alderley Park and Alex Alderton, Roberto Amato, Jeffrey Barrett, Sonia Goncalves, Ewan Harrison, David K. Jackson, Ian Johnston, Dominic Kwiatkowski, Cordelia Langford, John Sillitoe on behalf of the Wellcome Sanger Institute COVID-19 Surveillance Team |
| EPI_ISL_3125151 | Lighthouse Lab in Glasgow | Wellcome Sanger Institute for the COVID-19 Genomics UK (COG-UK) Consortium | Harper VanSteenhouse, Yumi Kasai, David Gray, Carol Clugston, Anna Dominiczak and Alex Alderton, Roberto Amato, Jeffrey Barrett, Sonia Goncalves, Ewan Harrison, David K. Jackson, Ian Johnston, Dominic Kwiatkowski, Cordelia Langford, John Sillitoe on behalf of the Wellcome Sanger Institute COVID-19 Surveillance Team |
| EPI_ISL_3125152 | Lighthouse Lab in Alderley Park | Wellcome Sanger Institute for the COVID-19 Genomics UK (COG-UK) Consortium | Jacquelyn Wynn, Mairead Hyland, The Lighthouse Lab in Alderley Park and Alex Alderton, Roberto Amato, Jeffrey Barrett, Sonia Goncalves, Ewan Harrison, David K. Jackson, Ian Johnston, Dominic Kwiatkowski, Cordelia Langford, John Sillitoe on behalf of the Wellcome Sanger Institute COVID-19 Surveillance Team |
| EPI_ISL_3125153 | Lighthouse Lab in Glasgow | Wellcome Sanger Institute for the COVID-19 Genomics UK (COG-UK) Consortium | Harper VanSteenhouse, Yumi Kasai, David Gray, Carol Clugston, Anna Dominiczak and Alex Alderton, Roberto Amato, Jeffrey Barrett, Sonia Goncalves, Ewan Harrison, David K. Jackson, Ian Johnston, Dominic Kwiatkowski, Cordelia Langford, John Sillitoe on behalf of the Wellcome Sanger Institute COVID-19 Surveillance Team |
| EPI_ISL_3125154 | Lighthouse Lab in Milton Keynes | Wellcome Sanger Institute for the COVID-19 Genomics UK (COG-UK) Consortium | The Lighthouse Lab in Milton Keynes and Alex Alderton, Roberto Amato, Jeffrey Barrett, Sonia Goncalves, Ewan Harrison, David K. Jackson, Ian Johnston, Dominic Kwiatkowski, Cordelia Langford, John Sillitoe on behalf of the Wellcome Sanger Institute COVID-19 Surveillance Team |
| EPI_ISL_3125155 | Lighthouse Lab in Alderley Park | Wellcome Sanger Institute for the COVID-19 Genomics UK (COG-UK) Consortium | Jacquelyn Wynn, Mairead Hyland, The Lighthouse Lab in Alderley Park and Alex Alderton, Roberto Amato, Jeffrey Barrett, Sonia Goncalves, Ewan Harrison, David K. Jackson, Ian Johnston, Dominic Kwiatkowski, Cordelia Langford, John Sillitoe on behalf of the Wellcome Sanger Institute COVID-19 Surveillance Team |
| EPI_ISL_3125156 | Lighthouse Lab in Glasgow | Wellcome Sanger Institute for the COVID-19 Genomics UK (COG-UK) Consortium | Harper VanSteenhouse, Yumi Kasai, David Gray, Carol Clugston, Anna Dominiczak and Alex Alderton, Roberto Amato, Jeffrey Barrett, Sonia Goncalves, Ewan Harrison, David K. Jackson, Ian Johnston, Dominic Kwiatkowski, Cordelia Langford, John Sillitoe on behalf of the Wellcome Sanger Institute COVID-19 Surveillance Team |
| EPI_ISL_3125157 | Lighthouse Lab in Milton Keynes | Wellcome Sanger Institute for the COVID-19 Genomics UK (COG-UK) Consortium | The Lighthouse Lab in Milton Keynes and Alex Alderton, Roberto Amato, Jeffrey Barrett, Sonia Goncalves, Ewan Harrison, David K. Jackson, Ian Johnston, Dominic Kwiatkowski, Cordelia Langford, John Sillitoe on behalf of the Wellcome Sanger Institute COVID-19 Surveillance Team |
| EPI_ISL_3125158, EPI_ISL_3125159 | Lighthouse Lab in Glasgow | Wellcome Sanger Institute for the COVID-19 Genomics UK (COG-UK) Consortium | Harper VanSteenhouse, Yumi Kasai, David Gray, Carol Clugston, Anna Dominiczak and Alex Alderton, Roberto Amato, Jeffrey Barrett, Sonia Goncalves, Ewan Harrison, David K. Jackson, Ian Johnston, Dominic Kwiatkowski, Cordelia Langford, John Sillitoe on behalf of the Wellcome Sanger Institute COVID-19 Surveillance Team |
| EPI_ISL_3125160, EPI_ISL_3125161, EPI_ISL_3125162, EPI_ISL_3125163 | Lighthouse Lab in Alderley Park | Wellcome Sanger Institute for the COVID-19 Genomics UK (COG-UK) Consortium | Jacquelyn Wynn, Mairead Hyland, The Lighthouse Lab in Alderley Park and Alex Alderton, Roberto Amato, Jeffrey Barrett, Sonia Goncalves, Ewan Harrison, David K. Jackson, Ian Johnston, Dominic Kwiatkowski, Cordelia Langford, John Sillitoe on behalf of the Wellcome Sanger Institute COVID-19 Surveillance Team |
| EPI_ISL_3125164, EPI_ISL_3125165 | Lighthouse Lab in Milton Keynes | Wellcome Sanger Institute for the COVID-19 Genomics UK (COG-UK) Consortium | The Lighthouse Lab in Milton Keynes and Alex Alderton, Roberto Amato, Jeffrey Barrett, Sonia Goncalves, Ewan Harrison, David K. Jackson, Ian Johnston, Dominic Kwiatkowski, Cordelia Langford, John Sillitoe on behalf of the Wellcome Sanger Institute COVID-19 Surveillance Team |
| EPI_ISL_3125166 | Lighthouse Lab in Glasgow | Wellcome Sanger Institute for the COVID-19 Genomics UK (COG-UK) Consortium | Harper VanSteenhouse, Yumi Kasai, David Gray, Carol Clugston, Anna Dominiczak and Alex Alderton, Roberto Amato, Jeffrey Barrett, Sonia Goncalves, Ewan Harrison, David K. Jackson, Ian Johnston, Dominic Kwiatkowski, Cordelia Langford, John Sillitoe on behalf of the Wellcome Sanger Institute COVID-19 Surveillance Team |
| EPI_ISL_3125167 | Lighthouse Lab in Milton Keynes | Wellcome Sanger Institute for the COVID-19 Genomics UK (COG-UK) Consortium | The Lighthouse Lab in Milton Keynes and Alex Alderton, Roberto Amato, Jeffrey Barrett, Sonia Goncalves, Ewan Harrison, David K. Jackson, Ian Johnston, Dominic Kwiatkowski, Cordelia Langford, John Sillitoe on behalf of the Wellcome Sanger Institute COVID-19 Surveillance Team |
| EPI_ISL_3125168, EPI_ISL_3125169 | Lighthouse Lab in Glasgow | Wellcome Sanger Institute for the COVID-19 Genomics UK (COG-UK) Consortium | Harper VanSteenhouse, Yumi Kasai, David Gray, Carol Clugston, Anna Dominiczak and Alex Alderton, Roberto Amato, Jeffrey Barrett, Sonia Goncalves, Ewan Harrison, David K. Jackson, Ian Johnston, Dominic Kwiatkowski, Cordelia Langford, John Sillitoe on behalf of the Wellcome Sanger Institute COVID-19 Surveillance Team |

| | | | |
|---|---|---|---|
| EPI_ISL_3125170, EPI_ISL_3125171 | Lighthouse Lab in Alderley Park | Wellcome Sanger Institute for the COVID-19 Genomics UK (COG-UK) Consortium | Jacquelyn Wynn, Mairead Hyland, The Lighthouse Lab in Alderley Park and Alex Alderton, Roberto Amato, Jeffrey Barrett, Sonia Goncalves, Ewan Harrison, David K. Jackson, Ian Johnston, Dominic Kwiatkowski, Cordelia Langford, John Sillitoe on behalf of the Wellcome Sanger Institute COVID-19 Surveillance Team |
| EPI_ISL_3125172 | Lighthouse Lab in Milton Keynes | Wellcome Sanger Institute for the COVID-19 Genomics UK (COG-UK) Consortium | The Lighthouse Lab in Milton Keynes and Alex Alderton, Roberto Amato, Jeffrey Barrett, Sonia Goncalves, Ewan Harrison, David K. Jackson, Ian Johnston, Dominic Kwiatkowski, Cordelia Langford, John Sillitoe on behalf of the Wellcome Sanger Institute COVID-19 Surveillance Team |
| EPI_ISL_3125173, EPI_ISL_3125174 | Lighthouse Lab in Glasgow | Wellcome Sanger Institute for the COVID-19 Genomics UK (COG-UK) Consortium | Harper VanSteenhouse, Yumi Kasai, David Gray, Carol Clugston, Anna Dominiczak and Alex Alderton, Roberto Amato, Jeffrey Barrett, Sonia Goncalves, Ewan Harrison, David K. Jackson, Ian Johnston, Dominic Kwiatkowski, Cordelia Langford, John Sillitoe on behalf of the Wellcome Sanger Institute COVID-19 Surveillance Team |
| EPI_ISL_3125175 | Lighthouse Lab in Milton Keynes | Wellcome Sanger Institute for the COVID-19 Genomics UK (COG-UK) Consortium | The Lighthouse Lab in Milton Keynes and Alex Alderton, Roberto Amato, Jeffrey Barrett, Sonia Goncalves, Ewan Harrison, David K. Jackson, Ian Johnston, Dominic Kwiatkowski, Cordelia Langford, John Sillitoe on behalf of the Wellcome Sanger Institute COVID-19 Surveillance Team |
| EPI_ISL_3125176, EPI_ISL_3125177 | Lighthouse Lab in Glasgow | Wellcome Sanger Institute for the COVID-19 Genomics UK (COG-UK) Consortium | Harper VanSteenhouse, Yumi Kasai, David Gray, Carol Clugston, Anna Dominiczak and Alex Alderton, Roberto Amato, Jeffrey Barrett, Sonia Goncalves, Ewan Harrison, David K. Jackson, Ian Johnston, Dominic Kwiatkowski, Cordelia Langford, John Sillitoe on behalf of the Wellcome Sanger Institute COVID-19 Surveillance Team |
| EPI_ISL_3125178 | Lighthouse Lab in Milton Keynes | Wellcome Sanger Institute for the COVID-19 Genomics UK (COG-UK) Consortium | The Lighthouse Lab in Milton Keynes and Alex Alderton, Roberto Amato, Jeffrey Barrett, Sonia Goncalves, Ewan Harrison, David K. Jackson, Ian Johnston, Dominic Kwiatkowski, Cordelia Langford, John Sillitoe on behalf of the Wellcome Sanger Institute COVID-19 Surveillance Team |
| EPI_ISL_3125179 | Lighthouse Lab in Alderley Park | Wellcome Sanger Institute for the COVID-19 Genomics UK (COG-UK) Consortium | Jacquelyn Wynn, Mairead Hyland, The Lighthouse Lab in Alderley Park and Alex Alderton, Roberto Amato, Jeffrey Barrett, Sonia Goncalves, Ewan Harrison, David K. Jackson, Ian Johnston, Dominic Kwiatkowski, Cordelia Langford, John Sillitoe on behalf of the Wellcome Sanger Institute COVID-19 Surveillance Team |
| EPI_ISL_3125180 | Lighthouse Lab in Milton Keynes | Wellcome Sanger Institute for the COVID-19 Genomics UK (COG-UK) Consortium | The Lighthouse Lab in Milton Keynes and Alex Alderton, Roberto Amato, Jeffrey Barrett, Sonia Goncalves, Ewan Harrison, David K. Jackson, Ian Johnston, Dominic Kwiatkowski, Cordelia Langford, John Sillitoe on behalf of the Wellcome Sanger Institute COVID-19 Surveillance Team |
| EPI_ISL_3125181, EPI_ISL_3125182 | Lighthouse Lab in Alderley Park | Wellcome Sanger Institute for the COVID-19 Genomics UK (COG-UK) Consortium | Jacquelyn Wynn, Mairead Hyland, The Lighthouse Lab in Alderley Park and Alex Alderton, Roberto Amato, Jeffrey Barrett, Sonia Goncalves, Ewan Harrison, David K. Jackson, Ian Johnston, Dominic Kwiatkowski, Cordelia Langford, John Sillitoe on behalf of the Wellcome Sanger Institute COVID-19 Surveillance Team |
| EPI_ISL_3125183 | Lighthouse Lab in Milton Keynes | Wellcome Sanger Institute for the COVID-19 Genomics UK (COG-UK) Consortium | The Lighthouse Lab in Milton Keynes and Alex Alderton, Roberto Amato, Jeffrey Barrett, Sonia Goncalves, Ewan Harrison, David K. Jackson, Ian Johnston, Dominic Kwiatkowski, Cordelia Langford, John Sillitoe on behalf of the Wellcome Sanger Institute COVID-19 Surveillance Team |
| EPI_ISL_3125184 | Lighthouse Lab in Alderley Park | Wellcome Sanger Institute for the COVID-19 Genomics UK (COG-UK) Consortium | Jacquelyn Wynn, Mairead Hyland, The Lighthouse Lab in Alderley Park and Alex Alderton, Roberto Amato, Jeffrey Barrett, Sonia Goncalves, Ewan Harrison, David K. Jackson, Ian Johnston, Dominic Kwiatkowski, Cordelia Langford, John Sillitoe on behalf of the Wellcome Sanger Institute COVID-19 Surveillance Team |
| EPI_ISL_3125185 | Lighthouse Lab in Glasgow | Wellcome Sanger Institute for the COVID-19 Genomics UK (COG-UK) Consortium | Harper VanSteenhouse, Yumi Kasai, David Gray, Carol Clugston, Anna Dominiczak and Alex Alderton, Roberto Amato, Jeffrey Barrett, Sonia Goncalves, Ewan Harrison, David K. Jackson, Ian Johnston, Dominic Kwiatkowski, Cordelia Langford, John Sillitoe on behalf of the Wellcome Sanger Institute COVID-19 Surveillance Team |
| EPI_ISL_3125186, EPI_ISL_3125187, EPI_ISL_3125188 | Lighthouse Lab in Milton Keynes | Wellcome Sanger Institute for the COVID-19 Genomics UK (COG-UK) Consortium | The Lighthouse Lab in Milton Keynes and Alex Alderton, Roberto Amato, Jeffrey Barrett, Sonia Goncalves, Ewan Harrison, David K. Jackson, Ian Johnston, Dominic Kwiatkowski, Cordelia Langford, John Sillitoe on behalf of the Wellcome Sanger Institute COVID-19 Surveillance Team |
| EPI_ISL_3125189, EPI_ISL_3125190 | Lighthouse Lab in Glasgow | Wellcome Sanger Institute for the COVID-19 Genomics UK (COG-UK) Consortium | Harper VanSteenhouse, Yumi Kasai, David Gray, Carol Clugston, Anna Dominiczak and Alex Alderton, Roberto Amato, Jeffrey Barrett, Sonia Goncalves, Ewan Harrison, David K. Jackson, Ian Johnston, Dominic Kwiatkowski, Cordelia Langford, John Sillitoe on behalf of the Wellcome Sanger Institute COVID-19 Surveillance Team |
| EPI_ISL_3125191 | Lighthouse Lab in Milton Keynes | Wellcome Sanger Institute for the COVID-19 Genomics UK (COG-UK) Consortium | The Lighthouse Lab in Milton Keynes and Alex Alderton, Roberto Amato, Jeffrey Barrett, Sonia Goncalves, Ewan Harrison, David K. Jackson, Ian Johnston, Dominic Kwiatkowski, Cordelia Langford, John Sillitoe on behalf of the Wellcome Sanger Institute COVID-19 Surveillance Team |
| EPI_ISL_3125192 | Lighthouse Lab in Alderley Park | Wellcome Sanger Institute for the COVID-19 Genomics UK (COG-UK) Consortium | Jacquelyn Wynn, Mairead Hyland, The Lighthouse Lab in Alderley Park and Alex Alderton, Roberto Amato, Jeffrey Barrett, Sonia Goncalves, Ewan Harrison, David K. Jackson, Ian Johnston, Dominic Kwiatkowski, Cordelia Langford, John Sillitoe on behalf of the Wellcome Sanger Institute COVID-19 Surveillance Team |
| EPI_ISL_3125193 | Lighthouse Lab in Glasgow | Wellcome Sanger Institute for the COVID-19 Genomics UK (COG-UK) Consortium | Harper VanSteenhouse, Yumi Kasai, David Gray, Carol Clugston, Anna Dominiczak and Alex Alderton, Roberto Amato, Jeffrey Barrett, Sonia Goncalves, Ewan Harrison, David K. Jackson, Ian Johnston, Dominic Kwiatkowski, Cordelia Langford, John Sillitoe on behalf of the Wellcome Sanger Institute COVID-19 Surveillance Team |
| EPI_ISL_3125194, EPI_ISL_3125195 | Lighthouse Lab in Alderley Park | Wellcome Sanger Institute for the COVID-19 Genomics UK (COG-UK) Consortium | Jacquelyn Wynn, Mairead Hyland, The Lighthouse Lab in Alderley Park and Alex Alderton, Roberto Amato, Jeffrey Barrett, Sonia Goncalves, Ewan Harrison, David K. Jackson, Ian Johnston, Dominic Kwiatkowski, Cordelia Langford, John Sillitoe on behalf of the Wellcome Sanger Institute COVID-19 Surveillance Team |
| EPI_ISL_3125196 | Lighthouse Lab in Glasgow | Wellcome Sanger Institute for the COVID-19 Genomics UK (COG-UK) Consortium | Harper VanSteenhouse, Yumi Kasai, David Gray, Carol Clugston, Anna Dominiczak and Alex Alderton, Roberto Amato, Jeffrey Barrett, Sonia Goncalves, Ewan Harrison, David K. Jackson, Ian Johnston, Dominic Kwiatkowski, Cordelia Langford, John Sillitoe on behalf of the Wellcome Sanger Institute COVID-19 Surveillance Team |
| EPI_ISL_3125197 | Lighthouse Lab in Milton Keynes | Wellcome Sanger Institute for the COVID-19 Genomics UK (COG-UK) Consortium | The Lighthouse Lab in Milton Keynes and Alex Alderton, Roberto Amato, Jeffrey Barrett, Sonia Goncalves, Ewan Harrison, David K. Jackson, Ian Johnston, Dominic Kwiatkowski, Cordelia Langford, John Sillitoe on behalf of the Wellcome Sanger Institute COVID-19 Surveillance Team |
| EPI_ISL_3125198, EPI_ISL_3125199 | Lighthouse Lab in Glasgow | Wellcome Sanger Institute for the COVID-19 Genomics UK (COG-UK) Consortium | Harper VanSteenhouse, Yumi Kasai, David Gray, Carol Clugston, Anna Dominiczak and Alex Alderton, Roberto Amato, Jeffrey Barrett, Sonia Goncalves, Ewan Harrison, David K. Jackson, Ian Johnston, Dominic Kwiatkowski, Cordelia Langford, John Sillitoe on behalf of the Wellcome Sanger Institute COVID-19 Surveillance Team |
| EPI_ISL_3125200 | Lighthouse Lab in Milton Keynes | Wellcome Sanger Institute for the COVID-19 Genomics UK (COG-UK) Consortium | The Lighthouse Lab in Milton Keynes and Alex Alderton, Roberto Amato, Jeffrey Barrett, Sonia Goncalves, Ewan Harrison, David K. Jackson, Ian Johnston, Dominic Kwiatkowski, Cordelia Langford, John Sillitoe on behalf of the Wellcome Sanger Institute COVID-19 Surveillance Team |
| EPI_ISL_3125201, EPI_ISL_3125202, EPI_ISL_3125203 | Lighthouse Lab in Glasgow | Wellcome Sanger Institute for the COVID-19 Genomics UK (COG-UK) Consortium | Harper VanSteenhouse, Yumi Kasai, David Gray, Carol Clugston, Anna Dominiczak and Alex Alderton, Roberto Amato, Jeffrey Barrett, Sonia Goncalves, Ewan Harrison, David K. Jackson, Ian Johnston, Dominic Kwiatkowski, Cordelia Langford, John Sillitoe on behalf of the Wellcome Sanger Institute COVID-19 Surveillance Team |
| EPI_ISL_3125204, EPI_ISL_3125205 | Lighthouse Lab in Milton Keynes | Wellcome Sanger Institute for the COVID-19 Genomics UK (COG-UK) Consortium | The Lighthouse Lab in Milton Keynes and Alex Alderton, Roberto Amato, Jeffrey Barrett, Sonia Goncalves, Ewan Harrison, David K. Jackson, Ian Johnston, Dominic Kwiatkowski, Cordelia Langford, John Sillitoe on behalf of the Wellcome Sanger Institute COVID-19 Surveillance Team |
| EPI_ISL_3125206, EPI_ISL_3125207, EPI_ISL_3125208 | Lighthouse Lab in Glasgow | Wellcome Sanger Institute for the COVID-19 Genomics UK (COG-UK) Consortium | Harper VanSteenhouse, Yumi Kasai, David Gray, Carol Clugston, Anna Dominiczak and Alex Alderton, Roberto Amato, Jeffrey Barrett, Sonia Goncalves, Ewan Harrison, David K. Jackson, Ian Johnston, Dominic Kwiatkowski, Cordelia Langford, John Sillitoe on behalf of the Wellcome Sanger Institute COVID-19 Surveillance Team |
| EPI_ISL_3125209 | Lighthouse Lab in Alderley Park | Wellcome Sanger Institute for the COVID-19 Genomics UK (COG-UK) Consortium | Jacquelyn Wynn, Mairead Hyland, The Lighthouse Lab in Alderley Park and Alex Alderton, Roberto Amato, Jeffrey Barrett, Sonia Goncalves, Ewan Harrison, David K. Jackson, Ian Johnston, Dominic Kwiatkowski, Cordelia Langford, John Sillitoe on behalf of the Wellcome Sanger Institute COVID-19 Surveillance Team |
| EPI_ISL_3125210, EPI_ISL_3125211 | Lighthouse Lab in Milton Keynes | Wellcome Sanger Institute for the COVID-19 Genomics UK (COG-UK) Consortium | The Lighthouse Lab in Milton Keynes and Alex Alderton, Roberto Amato, Jeffrey Barrett, Sonia Goncalves, Ewan Harrison, David K. Jackson, Ian Johnston, Dominic Kwiatkowski, Cordelia Langford, John Sillitoe on behalf of the Wellcome Sanger Institute COVID-19 Surveillance Team |
| EPI_ISL_3125212, EPI_ISL_3125213 | Lighthouse Lab in Glasgow | Wellcome Sanger Institute for the COVID-19 Genomics UK (COG-UK) Consortium | Harper VanSteenhouse, Yumi Kasai, David Gray, Carol Clugston, Anna Dominiczak and Alex Alderton, Roberto Amato, Jeffrey Barrett, Sonia Goncalves, Ewan Harrison, David K. Jackson, Ian Johnston, Dominic Kwiatkowski, Cordelia Langford, John Sillitoe on behalf of the Wellcome Sanger Institute COVID-19 Surveillance Team |
| EPI_ISL_3125214 | Lighthouse Lab in Alderley Park | Wellcome Sanger Institute for the COVID-19 Genomics UK (COG-UK) Consortium | Jacquelyn Wynn, Mairead Hyland, The Lighthouse Lab in Alderley Park and Alex Alderton, Roberto Amato, Jeffrey Barrett, Sonia Goncalves, Ewan Harrison, David K. Jackson, Ian Johnston, Dominic Kwiatkowski, Cordelia Langford, John Sillitoe on behalf of the Wellcome Sanger Institute COVID-19 Surveillance Team |
| EPI_ISL_3125215 | Lighthouse Lab in Milton Keynes | Wellcome Sanger Institute for the COVID-19 Genomics UK (COG-UK) Consortium | The Lighthouse Lab in Milton Keynes and Alex Alderton, Roberto Amato, Jeffrey Barrett, Sonia Goncalves, Ewan Harrison, David K. Jackson, Ian Johnston, Dominic Kwiatkowski, Cordelia Langford, John Sillitoe on behalf of the Wellcome Sanger Institute COVID-19 Surveillance Team |
| EPI_ISL_3125216, EPI_ISL_3125217 | Lighthouse Lab in Glasgow | Wellcome Sanger Institute for the COVID-19 Genomics UK (COG-UK) Consortium | Harper VanSteenhouse, Yumi Kasai, David Gray, Carol Clugston, Anna Dominiczak and Alex Alderton, Roberto Amato, Jeffrey Barrett, Sonia Goncalves, Ewan Harrison, David K. Jackson, Ian Johnston, Dominic Kwiatkowski, Cordelia Langford, John Sillitoe on behalf of the Wellcome Sanger Institute COVID-19 Surveillance Team |
| EPI_ISL_3125218 | Lighthouse Lab in Alderley Park | Wellcome Sanger Institute for the COVID-19 Genomics UK (COG-UK) Consortium | Jacquelyn Wynn, Mairead Hyland, The Lighthouse Lab in Alderley Park and Alex Alderton, Roberto Amato, Jeffrey Barrett, Sonia Goncalves, Ewan Harrison, David K. Jackson, Ian Johnston, Dominic Kwiatkowski, Cordelia Langford, John Sillitoe on behalf of the Wellcome Sanger Institute COVID-19 Surveillance Team |
| EPI_ISL_3125219 | Lighthouse Lab in Milton Keynes | Wellcome Sanger Institute for the COVID-19 Genomics UK (COG-UK) Consortium | The Lighthouse Lab in Milton Keynes and Alex Alderton, Roberto Amato, Jeffrey Barrett, Sonia Goncalves, Ewan Harrison, David K. Jackson, Ian Johnston, Dominic Kwiatkowski, Cordelia Langford, John Sillitoe on behalf of the Wellcome Sanger Institute COVID-19 Surveillance Team |
| EPI_ISL_3125220, EPI_ISL_3125221 | Lighthouse Lab in Alderley Park | Wellcome Sanger Institute for the COVID-19 Genomics UK (COG-UK) Consortium | Jacquelyn Wynn, Mairead Hyland, The Lighthouse Lab in Alderley Park and Alex Alderton, Roberto Amato, Jeffrey Barrett, Sonia Goncalves, Ewan Harrison, David K. Jackson, Ian Johnston, Dominic Kwiatkowski, Cordelia Langford, John Sillitoe on behalf of the Wellcome Sanger Institute COVID-19 Surveillance Team |
| EPI_ISL_3125222, EPI_ISL_3125223, EPI_ISL_3125224, EPI_ISL_3125225, EPI_ISL_3125226 | Lighthouse Lab in Glasgow | Wellcome Sanger Institute for the COVID-19 Genomics UK (COG-UK) Consortium | Harper VanSteenhouse, Yumi Kasai, David Gray, Carol Clugston, Anna Dominiczak and Alex Alderton, Roberto Amato, Jeffrey Barrett, Sonia Goncalves, Ewan Harrison, David K. Jackson, Ian Johnston, Dominic Kwiatkowski, Cordelia Langford, John Sillitoe on behalf of the Wellcome Sanger Institute COVID-19 Surveillance Team |
| EPI_ISL_3125227, EPI_ISL_3125228, EPI_ISL_3125229, EPI_ISL_3125230, EPI_ISL_3125231, EPI_ISL_3125232 | Lighthouse Lab in Alderley Park | Wellcome Sanger Institute for the COVID-19 Genomics UK (COG-UK) Consortium | Jacquelyn Wynn, Mairead Hyland, The Lighthouse Lab in Alderley Park and Alex Alderton, Roberto Amato, Jeffrey Barrett, Sonia Goncalves, Ewan Harrison, David K. Jackson, Ian Johnston, Dominic Kwiatkowski, Cordelia Langford, John Sillitoe on behalf of the Wellcome Sanger Institute COVID-19 Surveillance Team |
| EPI_ISL_3125233 | Lighthouse Lab in Milton Keynes | Wellcome Sanger Institute for the COVID-19 Genomics UK (COG-UK) Consortium | The Lighthouse Lab in Milton Keynes and Alex Alderton, Roberto Amato, Jeffrey Barrett, Sonia Goncalves, Ewan Harrison, David K. Jackson, Ian Johnston, Dominic Kwiatkowski, Cordelia Langford, John Sillitoe on behalf of the Wellcome Sanger Institute COVID-19 Surveillance Team |
| EPI_ISL_3125234, EPI_ISL_3125235, EPI_ISL_3125236 | Lighthouse Lab in Alderley Park | Wellcome Sanger Institute for the COVID-19 Genomics UK (COG-UK) Consortium | Jacquelyn Wynn, Mairead Hyland, The Lighthouse Lab in Alderley Park and Alex Alderton, Roberto Amato, Jeffrey Barrett, Sonia Goncalves, Ewan Harrison, David K. Jackson, Ian Johnston, Dominic Kwiatkowski, Cordelia Langford, John Sillitoe on behalf of the Wellcome Sanger Institute COVID-19 Surveillance Team |
| EPI_ISL_3125237, EPI_ISL_3125238 | Berkshire and Surrey Pathology Services Lighthouse Laboratory | Wellcome Sanger Institute for the COVID-19 Genomics UK (COG-UK) Consortium | Berkshire and Surrey Pathology Services Lighthouse Laboratory and Alex Alderton, Roberto Amato, Jeffrey Barrett, Sonia Goncalves, Ewan Harrison, David K. Jackson, Ian Johnston, Dominic Kwiatkowski, Cordelia Langford, John Sillitoe on behalf of the Wellcome Sanger Institute COVID-19 Surveillance Team |
| EPI_ISL_3125239, EPI_ISL_3125240, EPI_ISL_3125241, EPI_ISL_3125242 | Lighthouse Lab in Alderley Park | Wellcome Sanger Institute for the COVID-19 Genomics UK (COG-UK) Consortium | Jacquelyn Wynn, Mairead Hyland, The Lighthouse Lab in Alderley Park and Alex Alderton, Roberto Amato, Jeffrey Barrett, Sonia Goncalves, Ewan Harrison, David K. Jackson, Ian Johnston, Dominic Kwiatkowski, Cordelia Langford, John Sillitoe on behalf of the Wellcome Sanger Institute COVID-19 Surveillance Team |

| | | | |
|---|---|---|---|
| EPI_ISL_3125243, EPI_ISL_3125244, EPI_ISL_3125245 | Berkshire and Surrey Pathology Services Lighthouse Laboratory | Wellcome Sanger Institute for the COVID-19 Genomics UK (COG-UK) Consortium | Berkshire and Surrey Pathology Services Lighthouse Laboratory and Alex Alderton, Roberto Amato, Jeffrey Barrett, Sonia Goncalves, Ewan Harrison, David K. Jackson, Ian Johnston, Dominic Kwiatkowski, Cordelia Langford, John Sillitoe on behalf of the Wellcome Sanger Institute COVID-19 Surveillance Team |
| EPI_ISL_3125246, EPI_ISL_3125247, EPI_ISL_3125248 | Lighthouse Lab in Alderley Park | Wellcome Sanger Institute for the COVID-19 Genomics UK (COG-UK) Consortium | Jacquelyn Wynn, Mairead Hyland, The Lighthouse Lab in Alderley Park and Alex Alderton, Roberto Amato, Jeffrey Barrett, Sonia Goncalves, Ewan Harrison, David K. Jackson, Ian Johnston, Dominic Kwiatkowski, Cordelia Langford, John Sillitoe on behalf of the Wellcome Sanger Institute COVID-19 Surveillance Team |
| EPI_ISL_3125249 | Lighthouse Lab in Milton Keynes | Wellcome Sanger Institute for the COVID-19 Genomics UK (COG-UK) Consortium | The Lighthouse Lab in Milton Keynes and Alex Alderton, Roberto Amato, Jeffrey Barrett, Sonia Goncalves, Ewan Harrison, David K. Jackson, Ian Johnston, Dominic Kwiatkowski, Cordelia Langford, John Sillitoe on behalf of the Wellcome Sanger Institute COVID-19 Surveillance Team |
| EPI_ISL_3125250, EPI_ISL_3125251 | Lighthouse Lab in Alderley Park | Wellcome Sanger Institute for the COVID-19 Genomics UK (COG-UK) Consortium | Jacquelyn Wynn, Mairead Hyland, The Lighthouse Lab in Alderley Park and Alex Alderton, Roberto Amato, Jeffrey Barrett, Sonia Goncalves, Ewan Harrison, David K. Jackson, Ian Johnston, Dominic Kwiatkowski, Cordelia Langford, John Sillitoe on behalf of the Wellcome Sanger Institute COVID-19 Surveillance Team |
| EPI_ISL_3125252 | Lighthouse Lab in Milton Keynes | Wellcome Sanger Institute for the COVID-19 Genomics UK (COG-UK) Consortium | The Lighthouse Lab in Milton Keynes and Alex Alderton, Roberto Amato, Jeffrey Barrett, Sonia Goncalves, Ewan Harrison, David K. Jackson, Ian Johnston, Dominic Kwiatkowski, Cordelia Langford, John Sillitoe on behalf of the Wellcome Sanger Institute COVID-19 Surveillance Team |
| EPI_ISL_3125253 | Berkshire and Surrey Pathology Services Lighthouse Laboratory | Wellcome Sanger Institute for the COVID-19 Genomics UK (COG-UK) Consortium | Berkshire and Surrey Pathology Services Lighthouse Laboratory and Alex Alderton, Roberto Amato, Jeffrey Barrett, Sonia Goncalves, Ewan Harrison, David K. Jackson, Ian Johnston, Dominic Kwiatkowski, Cordelia Langford, John Sillitoe on behalf of the Wellcome Sanger Institute COVID-19 Surveillance Team |
| EPI_ISL_3125254, EPI_ISL_3125255 | Lighthouse Lab in Milton Keynes | Wellcome Sanger Institute for the COVID-19 Genomics UK (COG-UK) Consortium | The Lighthouse Lab in Milton Keynes and Alex Alderton, Roberto Amato, Jeffrey Barrett, Sonia Goncalves, Ewan Harrison, David K. Jackson, Ian Johnston, Dominic Kwiatkowski, Cordelia Langford, John Sillitoe on behalf of the Wellcome Sanger Institute COVID-19 Surveillance Team |
| EPI_ISL_3125256, EPI_ISL_3125257 | Lighthouse Lab in Alderley Park | Wellcome Sanger Institute for the COVID-19 Genomics UK (COG-UK) Consortium | Jacquelyn Wynn, Mairead Hyland, The Lighthouse Lab in Alderley Park and Alex Alderton, Roberto Amato, Jeffrey Barrett, Sonia Goncalves, Ewan Harrison, David K. Jackson, Ian Johnston, Dominic Kwiatkowski, Cordelia Langford, John Sillitoe on behalf of the Wellcome Sanger Institute COVID-19 Surveillance Team |
| EPI_ISL_3125258 | Berkshire and Surrey Pathology Services Lighthouse Laboratory | Wellcome Sanger Institute for the COVID-19 Genomics UK (COG-UK) Consortium | Berkshire and Surrey Pathology Services Lighthouse Laboratory and Alex Alderton, Roberto Amato, Jeffrey Barrett, Sonia Goncalves, Ewan Harrison, David K. Jackson, Ian Johnston, Dominic Kwiatkowski, Cordelia Langford, John Sillitoe on behalf of the Wellcome Sanger Institute COVID-19 Surveillance Team |
| EPI_ISL_3125259 | Lighthouse Lab in Milton Keynes | Wellcome Sanger Institute for the COVID-19 Genomics UK (COG-UK) Consortium | The Lighthouse Lab in Milton Keynes and Alex Alderton, Roberto Amato, Jeffrey Barrett, Sonia Goncalves, Ewan Harrison, David K. Jackson, Ian Johnston, Dominic Kwiatkowski, Cordelia Langford, John Sillitoe on behalf of the Wellcome Sanger Institute COVID-19 Surveillance Team |
| EPI_ISL_3125260, EPI_ISL_3125261, EPI_ISL_3125262 | Lighthouse Lab in Alderley Park | Wellcome Sanger Institute for the COVID-19 Genomics UK (COG-UK) Consortium | Jacquelyn Wynn, Mairead Hyland, The Lighthouse Lab in Alderley Park and Alex Alderton, Roberto Amato, Jeffrey Barrett, Sonia Goncalves, Ewan Harrison, David K. Jackson, Ian Johnston, Dominic Kwiatkowski, Cordelia Langford, John Sillitoe on behalf of the Wellcome Sanger Institute COVID-19 Surveillance Team |
| EPI_ISL_3125263 | Berkshire and Surrey Pathology Services Lighthouse Laboratory | Wellcome Sanger Institute for the COVID-19 Genomics UK (COG-UK) Consortium | Berkshire and Surrey Pathology Services Lighthouse Laboratory and Alex Alderton, Roberto Amato, Jeffrey Barrett, Sonia Goncalves, Ewan Harrison, David K. Jackson, Ian Johnston, Dominic Kwiatkowski, Cordelia Langford, John Sillitoe on behalf of the Wellcome Sanger Institute COVID-19 Surveillance Team |
| EPI_ISL_3125264 | Lighthouse Lab in Alderley Park | Wellcome Sanger Institute for the COVID-19 Genomics UK (COG-UK) Consortium | Jacquelyn Wynn, Mairead Hyland, The Lighthouse Lab in Alderley Park and Alex Alderton, Roberto Amato, Jeffrey Barrett, Sonia Goncalves, Ewan Harrison, David K. Jackson, Ian Johnston, Dominic Kwiatkowski, Cordelia Langford, John Sillitoe on behalf of the Wellcome Sanger Institute COVID-19 Surveillance Team |
| EPI_ISL_3125265 | Lighthouse Lab in Milton Keynes | Wellcome Sanger Institute for the COVID-19 Genomics UK (COG-UK) Consortium | The Lighthouse Lab in Milton Keynes and Alex Alderton, Roberto Amato, Jeffrey Barrett, Sonia Goncalves, Ewan Harrison, David K. Jackson, Ian Johnston, Dominic Kwiatkowski, Cordelia Langford, John Sillitoe on behalf of the Wellcome Sanger Institute COVID-19 Surveillance Team |
| EPI_ISL_3125266 | Lighthouse Lab in Alderley Park | Wellcome Sanger Institute for the COVID-19 Genomics UK (COG-UK) Consortium | Jacquelyn Wynn, Mairead Hyland, The Lighthouse Lab in Alderley Park and Alex Alderton, Roberto Amato, Jeffrey Barrett, Sonia Goncalves, Ewan Harrison, David K. Jackson, Ian Johnston, Dominic Kwiatkowski, Cordelia Langford, John Sillitoe on behalf of the Wellcome Sanger Institute COVID-19 Surveillance Team |
| EPI_ISL_3125267 | Lighthouse Lab in Milton Keynes | Wellcome Sanger Institute for the COVID-19 Genomics UK (COG-UK) Consortium | The Lighthouse Lab in Milton Keynes and Alex Alderton, Roberto Amato, Jeffrey Barrett, Sonia Goncalves, Ewan Harrison, David K. Jackson, Ian Johnston, Dominic Kwiatkowski, Cordelia Langford, John Sillitoe on behalf of the Wellcome Sanger Institute COVID-19 Surveillance Team |
| EPI_ISL_3125268, EPI_ISL_3125269, EPI_ISL_3125270 | Lighthouse Lab in Alderley Park | Wellcome Sanger Institute for the COVID-19 Genomics UK (COG-UK) Consortium | Jacquelyn Wynn, Mairead Hyland, The Lighthouse Lab in Alderley Park and Alex Alderton, Roberto Amato, Jeffrey Barrett, Sonia Goncalves, Ewan Harrison, David K. Jackson, Ian Johnston, Dominic Kwiatkowski, Cordelia Langford, John Sillitoe on behalf of the Wellcome Sanger Institute COVID-19 Surveillance Team |
| EPI_ISL_3125271 | Berkshire and Surrey Pathology Services Lighthouse Laboratory | Wellcome Sanger Institute for the COVID-19 Genomics UK (COG-UK) Consortium | Berkshire and Surrey Pathology Services Lighthouse Laboratory and Alex Alderton, Roberto Amato, Jeffrey Barrett, Sonia Goncalves, Ewan Harrison, David K. Jackson, Ian Johnston, Dominic Kwiatkowski, Cordelia Langford, John Sillitoe on behalf of the Wellcome Sanger Institute COVID-19 Surveillance Team |
| EPI_ISL_3125272 | Lighthouse Lab in Alderley Park | Wellcome Sanger Institute for the COVID-19 Genomics UK (COG-UK) Consortium | Jacquelyn Wynn, Mairead Hyland, The Lighthouse Lab in Alderley Park and Alex Alderton, Roberto Amato, Jeffrey Barrett, Sonia Goncalves, Ewan Harrison, David K. Jackson, Ian Johnston, Dominic Kwiatkowski, Cordelia Langford, John Sillitoe on behalf of the Wellcome Sanger Institute COVID-19 Surveillance Team |
| EPI_ISL_3125273 | Lighthouse Lab in Milton Keynes | Wellcome Sanger Institute for the COVID-19 Genomics UK (COG-UK) Consortium | The Lighthouse Lab in Milton Keynes and Alex Alderton, Roberto Amato, Jeffrey Barrett, Sonia Goncalves, Ewan Harrison, David K. Jackson, Ian Johnston, Dominic Kwiatkowski, Cordelia Langford, John Sillitoe on behalf of the Wellcome Sanger Institute COVID-19 Surveillance Team |
| EPI_ISL_3125274 | Berkshire and Surrey Pathology Services Lighthouse Laboratory | Wellcome Sanger Institute for the COVID-19 Genomics UK (COG-UK) Consortium | Berkshire and Surrey Pathology Services Lighthouse Laboratory and Alex Alderton, Roberto Amato, Jeffrey Barrett, Sonia Goncalves, Ewan Harrison, David K. Jackson, Ian Johnston, Dominic Kwiatkowski, Cordelia Langford, John Sillitoe on behalf of the Wellcome Sanger Institute COVID-19 Surveillance Team |
| EPI_ISL_3125275, EPI_ISL_3125276 | Lighthouse Lab in Alderley Park | Wellcome Sanger Institute for the COVID-19 Genomics UK (COG-UK) Consortium | Jacquelyn Wynn, Mairead Hyland, The Lighthouse Lab in Alderley Park and Alex Alderton, Roberto Amato, Jeffrey Barrett, Sonia Goncalves, Ewan Harrison, David K. Jackson, Ian Johnston, Dominic Kwiatkowski, Cordelia Langford, John Sillitoe on behalf of the Wellcome Sanger Institute COVID-19 Surveillance Team |
| EPI_ISL_3125277 | Berkshire and Surrey Pathology Services Lighthouse Laboratory | Wellcome Sanger Institute for the COVID-19 Genomics UK (COG-UK) Consortium | Berkshire and Surrey Pathology Services Lighthouse Laboratory and Alex Alderton, Roberto Amato, Jeffrey Barrett, Sonia Goncalves, Ewan Harrison, David K. Jackson, Ian Johnston, Dominic Kwiatkowski, Cordelia Langford, John Sillitoe on behalf of the Wellcome Sanger Institute COVID-19 Surveillance Team |
| EPI_ISL_3125278, EPI_ISL_3125279 | Lighthouse Lab in Milton Keynes | Wellcome Sanger Institute for the COVID-19 Genomics UK (COG-UK) Consortium | The Lighthouse Lab in Milton Keynes and Alex Alderton, Roberto Amato, Jeffrey Barrett, Sonia Goncalves, Ewan Harrison, David K. Jackson, Ian Johnston, Dominic Kwiatkowski, Cordelia Langford, John Sillitoe on behalf of the Wellcome Sanger Institute COVID-19 Surveillance Team |
| EPI_ISL_3125280, EPI_ISL_3125281 | Lighthouse Lab in Alderley Park | Wellcome Sanger Institute for the COVID-19 Genomics UK (COG-UK) Consortium | Jacquelyn Wynn, Mairead Hyland, The Lighthouse Lab in Alderley Park and Alex Alderton, Roberto Amato, Jeffrey Barrett, Sonia Goncalves, Ewan Harrison, David K. Jackson, Ian Johnston, Dominic Kwiatkowski, Cordelia Langford, John Sillitoe on behalf of the Wellcome Sanger Institute COVID-19 Surveillance Team |
| EPI_ISL_3125282, EPI_ISL_3125283, EPI_ISL_3125284 | Lighthouse Lab in Milton Keynes | Wellcome Sanger Institute for the COVID-19 Genomics UK (COG-UK) Consortium | The Lighthouse Lab in Milton Keynes and Alex Alderton, Roberto Amato, Jeffrey Barrett, Sonia Goncalves, Ewan Harrison, David K. Jackson, Ian Johnston, Dominic Kwiatkowski, Cordelia Langford, John Sillitoe on behalf of the Wellcome Sanger Institute COVID-19 Surveillance Team |
| EPI_ISL_3125285 | Berkshire and Surrey Pathology Services Lighthouse Laboratory | Wellcome Sanger Institute for the COVID-19 Genomics UK (COG-UK) Consortium | Berkshire and Surrey Pathology Services Lighthouse Laboratory and Alex Alderton, Roberto Amato, Jeffrey Barrett, Sonia Goncalves, Ewan Harrison, David K. Jackson, Ian Johnston, Dominic Kwiatkowski, Cordelia Langford, John Sillitoe on behalf of the Wellcome Sanger Institute COVID-19 Surveillance Team |
| EPI_ISL_3125286 | Lighthouse Lab in Milton Keynes | Wellcome Sanger Institute for the COVID-19 Genomics UK (COG-UK) Consortium | The Lighthouse Lab in Milton Keynes and Alex Alderton, Roberto Amato, Jeffrey Barrett, Sonia Goncalves, Ewan Harrison, David K. Jackson, Ian Johnston, Dominic Kwiatkowski, Cordelia Langford, John Sillitoe on behalf of the Wellcome Sanger Institute COVID-19 Surveillance Team |
| EPI_ISL_3125287, EPI_ISL_3125288, EPI_ISL_3125289, EPI_ISL_3125290 | Lighthouse Lab in Alderley Park | Wellcome Sanger Institute for the COVID-19 Genomics UK (COG-UK) Consortium | Jacquelyn Wynn, Mairead Hyland, The Lighthouse Lab in Alderley Park and Alex Alderton, Roberto Amato, Jeffrey Barrett, Sonia Goncalves, Ewan Harrison, David K. Jackson, Ian Johnston, Dominic Kwiatkowski, Cordelia Langford, John Sillitoe on behalf of the Wellcome Sanger Institute COVID-19 Surveillance Team |
| EPI_ISL_3125291 | Berkshire and Surrey Pathology Services Lighthouse Laboratory | Wellcome Sanger Institute for the COVID-19 Genomics UK (COG-UK) Consortium | Berkshire and Surrey Pathology Services Lighthouse Laboratory and Alex Alderton, Roberto Amato, Jeffrey Barrett, Sonia Goncalves, Ewan Harrison, David K. Jackson, Ian Johnston, Dominic Kwiatkowski, Cordelia Langford, John Sillitoe on behalf of the Wellcome Sanger Institute COVID-19 Surveillance Team |
| EPI_ISL_3125292 | Lighthouse Lab in Alderley Park | Wellcome Sanger Institute for the COVID-19 Genomics UK (COG-UK) Consortium | Jacquelyn Wynn, Mairead Hyland, The Lighthouse Lab in Alderley Park and Alex Alderton, Roberto Amato, Jeffrey Barrett, Sonia Goncalves, Ewan Harrison, David K. Jackson, Ian Johnston, Dominic Kwiatkowski, Cordelia Langford, John Sillitoe on behalf of the Wellcome Sanger Institute COVID-19 Surveillance Team |
| EPI_ISL_3125293 | Lighthouse Lab in Milton Keynes | Wellcome Sanger Institute for the COVID-19 Genomics UK (COG-UK) Consortium | The Lighthouse Lab in Milton Keynes and Alex Alderton, Roberto Amato, Jeffrey Barrett, Sonia Goncalves, Ewan Harrison, David K. Jackson, Ian Johnston, Dominic Kwiatkowski, Cordelia Langford, John Sillitoe on behalf of the Wellcome Sanger Institute COVID-19 Surveillance Team |
| EPI_ISL_3125294, EPI_ISL_3125295, EPI_ISL_3125296, EPI_ISL_3125297, EPI_ISL_3125298 | Lighthouse Lab in Alderley Park | Wellcome Sanger Institute for the COVID-19 Genomics UK (COG-UK) Consortium | Jacquelyn Wynn, Mairead Hyland, The Lighthouse Lab in Alderley Park and Alex Alderton, Roberto Amato, Jeffrey Barrett, Sonia Goncalves, Ewan Harrison, David K. Jackson, Ian Johnston, Dominic Kwiatkowski, Cordelia Langford, John Sillitoe on behalf of the Wellcome Sanger Institute COVID-19 Surveillance Team |
| EPI_ISL_3125299 | Lighthouse Lab in Milton Keynes | Wellcome Sanger Institute for the COVID-19 Genomics UK (COG-UK) Consortium | The Lighthouse Lab in Milton Keynes and Alex Alderton, Roberto Amato, Jeffrey Barrett, Sonia Goncalves, Ewan Harrison, David K. Jackson, Ian Johnston, Dominic Kwiatkowski, Cordelia Langford, John Sillitoe on behalf of the Wellcome Sanger Institute COVID-19 Surveillance Team |
| EPI_ISL_3125300, EPI_ISL_3125301 | Lighthouse Lab in Alderley Park | Wellcome Sanger Institute for the COVID-19 Genomics UK (COG-UK) Consortium | Jacquelyn Wynn, Mairead Hyland, The Lighthouse Lab in Alderley Park and Alex Alderton, Roberto Amato, Jeffrey Barrett, Sonia Goncalves, Ewan Harrison, David K. Jackson, Ian Johnston, Dominic Kwiatkowski, Cordelia Langford, John Sillitoe on behalf of the Wellcome Sanger Institute COVID-19 Surveillance Team |
| EPI_ISL_3125302 | Berkshire and Surrey Pathology Services Lighthouse Laboratory | Wellcome Sanger Institute for the COVID-19 Genomics UK (COG-UK) Consortium | Berkshire and Surrey Pathology Services Lighthouse Laboratory and Alex Alderton, Roberto Amato, Jeffrey Barrett, Sonia Goncalves, Ewan Harrison, David K. Jackson, Ian Johnston, Dominic Kwiatkowski, Cordelia Langford, John Sillitoe on behalf of the Wellcome Sanger Institute COVID-19 Surveillance Team |
| EPI_ISL_3125303, EPI_ISL_3125304 | Lighthouse Lab in Alderley Park | Wellcome Sanger Institute for the COVID-19 Genomics UK (COG-UK) Consortium | Jacquelyn Wynn, Mairead Hyland, The Lighthouse Lab in Alderley Park and Alex Alderton, Roberto Amato, Jeffrey Barrett, Sonia Goncalves, Ewan Harrison, David K. Jackson, Ian Johnston, Dominic Kwiatkowski, Cordelia Langford, John Sillitoe on behalf of the Wellcome Sanger Institute COVID-19 Surveillance Team |
| EPI_ISL_3125305, EPI_ISL_3125306 | Berkshire and Surrey Pathology Services Lighthouse Laboratory | Wellcome Sanger Institute for the COVID-19 Genomics UK (COG-UK) Consortium | Berkshire and Surrey Pathology Services Lighthouse Laboratory and Alex Alderton, Roberto Amato, Jeffrey Barrett, Sonia Goncalves, Ewan Harrison, David K. Jackson, Ian Johnston, Dominic Kwiatkowski, Cordelia Langford, John Sillitoe on behalf of the Wellcome Sanger Institute COVID-19 Surveillance Team |
| EPI_ISL_3125307 | Lighthouse Lab in Milton Keynes | Wellcome Sanger Institute for the COVID-19 Genomics UK (COG-UK) Consortium | The Lighthouse Lab in Milton Keynes and Alex Alderton, Roberto Amato, Jeffrey Barrett, Sonia Goncalves, Ewan Harrison, David K. Jackson, Ian Johnston, Dominic Kwiatkowski, Cordelia Langford, John Sillitoe on behalf of the Wellcome Sanger Institute COVID-19 Surveillance Team |
| EPI_ISL_3125308 | Lighthouse Lab in Alderley Park | Wellcome Sanger Institute for the COVID-19 Genomics UK (COG-UK) Consortium | Jacquelyn Wynn, Mairead Hyland, The Lighthouse Lab in Alderley Park and Alex Alderton, Roberto Amato, Jeffrey Barrett, Sonia Goncalves, Ewan Harrison, David K. Jackson, Ian Johnston, Dominic Kwiatkowski, Cordelia Langford, John Sillitoe on behalf of the Wellcome Sanger Institute COVID-19 Surveillance Team |
| EPI_ISL_3125309 | Berkshire and Surrey Pathology Services Lighthouse Laboratory | Wellcome Sanger Institute for the COVID-19 Genomics UK (COG-UK) | Berkshire and Surrey Pathology Services Lighthouse Laboratory and Alex Alderton, Roberto Amato, Jeffrey Barrett, Sonia Goncalves, Ewan Harrison, David K. Jackson, Ian Johnston, Dominic Kwiatkowski, Cordelia Langford, John Sillitoe on behalf |

| | | | |
|---|---|---|---|
| | | Consortium | of the Wellcome Sanger Institute COVID-19 Surveillance Team |
| EPI_ISL_3125310 | Lighthouse Lab in Milton Keynes | Wellcome Sanger Institute for the COVID-19 Genomics UK (COG-UK) Consortium | The Lighthouse Lab in Milton Keynes and Alex Alderton, Roberto Amato, Jeffrey Barrett, Sonia Goncalves, Ewan Harrison, David K. Jackson, Ian Johnston, Dominic Kwiatkowski, Cordelia Langford, John Sillitoe on behalf of the Wellcome Sanger Institute COVID-19 Surveillance Team |
| EPI_ISL_3125311 | Berkshire and Surrey Pathology Services Lighthouse Laboratory | Wellcome Sanger Institute for the COVID-19 Genomics UK (COG-UK) Consortium | Berkshire and Surrey Pathology Services Lighthouse Laboratory and Alex Alderton, Roberto Amato, Jeffrey Barrett, Sonia Goncalves, Ewan Harrison, David K. Jackson, Ian Johnston, Dominic Kwiatkowski, Cordelia Langford, John Sillitoe on behalf of the Wellcome Sanger Institute COVID-19 Surveillance Team |
| EPI_ISL_3125312, EPI_ISL_3125313, EPI_ISL_3125314 | Lighthouse Lab in Alderley Park | Wellcome Sanger Institute for the COVID-19 Genomics UK (COG-UK) Consortium | Jacquelyn Wynn, Mairead Hyland, The Lighthouse Lab in Alderley Park and Alex Alderton, Roberto Amato, Jeffrey Barrett, Sonia Goncalves, Ewan Harrison, David K. Jackson, Ian Johnston, Dominic Kwiatkowski, Cordelia Langford, John Sillitoe on behalf of the Wellcome Sanger Institute COVID-19 Surveillance Team |
| EPI_ISL_3125315 | Lighthouse Lab in Milton Keynes | Wellcome Sanger Institute for the COVID-19 Genomics UK (COG-UK) Consortium | The Lighthouse Lab in Milton Keynes and Alex Alderton, Roberto Amato, Jeffrey Barrett, Sonia Goncalves, Ewan Harrison, David K. Jackson, Ian Johnston, Dominic Kwiatkowski, Cordelia Langford, John Sillitoe on behalf of the Wellcome Sanger Institute COVID-19 Surveillance Team |
| EPI_ISL_3125316 | Lighthouse Lab in Alderley Park | Wellcome Sanger Institute for the COVID-19 Genomics UK (COG-UK) Consortium | Jacquelyn Wynn, Mairead Hyland, The Lighthouse Lab in Alderley Park and Alex Alderton, Roberto Amato, Jeffrey Barrett, Sonia Goncalves, Ewan Harrison, David K. Jackson, Ian Johnston, Dominic Kwiatkowski, Cordelia Langford, John Sillitoe on behalf of the Wellcome Sanger Institute COVID-19 Surveillance Team |
| EPI_ISL_3125317 | Berkshire and Surrey Pathology Services Lighthouse Laboratory | Wellcome Sanger Institute for the COVID-19 Genomics UK (COG-UK) Consortium | Berkshire and Surrey Pathology Services Lighthouse Laboratory and Alex Alderton, Roberto Amato, Jeffrey Barrett, Sonia Goncalves, Ewan Harrison, David K. Jackson, Ian Johnston, Dominic Kwiatkowski, Cordelia Langford, John Sillitoe on behalf of the Wellcome Sanger Institute COVID-19 Surveillance Team |
| EPI_ISL_3125318 | Lighthouse Lab in Alderley Park | Wellcome Sanger Institute for the COVID-19 Genomics UK (COG-UK) Consortium | Jacquelyn Wynn, Mairead Hyland, The Lighthouse Lab in Alderley Park and Alex Alderton, Roberto Amato, Jeffrey Barrett, Sonia Goncalves, Ewan Harrison, David K. Jackson, Ian Johnston, Dominic Kwiatkowski, Cordelia Langford, John Sillitoe on behalf of the Wellcome Sanger Institute COVID-19 Surveillance Team |
| EPI_ISL_3125319 | Lighthouse Lab in Milton Keynes | Wellcome Sanger Institute for the COVID-19 Genomics UK (COG-UK) Consortium | The Lighthouse Lab in Milton Keynes and Alex Alderton, Roberto Amato, Jeffrey Barrett, Sonia Goncalves, Ewan Harrison, David K. Jackson, Ian Johnston, Dominic Kwiatkowski, Cordelia Langford, John Sillitoe on behalf of the Wellcome Sanger Institute COVID-19 Surveillance Team |
| EPI_ISL_3125320 | Lighthouse Lab in Alderley Park | Wellcome Sanger Institute for the COVID-19 Genomics UK (COG-UK) Consortium | Jacquelyn Wynn, Mairead Hyland, The Lighthouse Lab in Alderley Park and Alex Alderton, Roberto Amato, Jeffrey Barrett, Sonia Goncalves, Ewan Harrison, David K. Jackson, Ian Johnston, Dominic Kwiatkowski, Cordelia Langford, John Sillitoe on behalf of the Wellcome Sanger Institute COVID-19 Surveillance Team |
| EPI_ISL_3125321 | Berkshire and Surrey Pathology Services Lighthouse Laboratory | Wellcome Sanger Institute for the COVID-19 Genomics UK (COG-UK) Consortium | Berkshire and Surrey Pathology Services Lighthouse Laboratory and Alex Alderton, Roberto Amato, Jeffrey Barrett, Sonia Goncalves, Ewan Harrison, David K. Jackson, Ian Johnston, Dominic Kwiatkowski, Cordelia Langford, John Sillitoe on behalf of the Wellcome Sanger Institute COVID-19 Surveillance Team |
| EPI_ISL_3125322, EPI_ISL_3125323 | Lighthouse Lab in Alderley Park | Wellcome Sanger Institute for the COVID-19 Genomics UK (COG-UK) Consortium | Jacquelyn Wynn, Mairead Hyland, The Lighthouse Lab in Alderley Park and Alex Alderton, Roberto Amato, Jeffrey Barrett, Sonia Goncalves, Ewan Harrison, David K. Jackson, Ian Johnston, Dominic Kwiatkowski, Cordelia Langford, John Sillitoe on behalf of the Wellcome Sanger Institute COVID-19 Surveillance Team |
| EPI_ISL_3125324 | Berkshire and Surrey Pathology Services Lighthouse Laboratory | Wellcome Sanger Institute for the COVID-19 Genomics UK (COG-UK) Consortium | Berkshire and Surrey Pathology Services Lighthouse Laboratory and Alex Alderton, Roberto Amato, Jeffrey Barrett, Sonia Goncalves, Ewan Harrison, David K. Jackson, Ian Johnston, Dominic Kwiatkowski, Cordelia Langford, John Sillitoe on behalf of the Wellcome Sanger Institute COVID-19 Surveillance Team |
| EPI_ISL_3125325 | Lighthouse Lab in Milton Keynes | Wellcome Sanger Institute for the COVID-19 Genomics UK (COG-UK) Consortium | The Lighthouse Lab in Milton Keynes and Alex Alderton, Roberto Amato, Jeffrey Barrett, Sonia Goncalves, Ewan Harrison, David K. Jackson, Ian Johnston, Dominic Kwiatkowski, Cordelia Langford, John Sillitoe on behalf of the Wellcome Sanger Institute COVID-19 Surveillance Team |
| EPI_ISL_3125326 | Berkshire and Surrey Pathology Services Lighthouse Laboratory | Wellcome Sanger Institute for the COVID-19 Genomics UK (COG-UK) Consortium | Berkshire and Surrey Pathology Services Lighthouse Laboratory and Alex Alderton, Roberto Amato, Jeffrey Barrett, Sonia Goncalves, Ewan Harrison, David K. Jackson, Ian Johnston, Dominic Kwiatkowski, Cordelia Langford, John Sillitoe on behalf of the Wellcome Sanger Institute COVID-19 Surveillance Team |
| EPI_ISL_3125327 | Lighthouse Lab in Alderley Park | Wellcome Sanger Institute for the COVID-19 Genomics UK (COG-UK) Consortium | Jacquelyn Wynn, Mairead Hyland, The Lighthouse Lab in Alderley Park and Alex Alderton, Roberto Amato, Jeffrey Barrett, Sonia Goncalves, Ewan Harrison, David K. Jackson, Ian Johnston, Dominic Kwiatkowski, Cordelia Langford, John Sillitoe on behalf of the Wellcome Sanger Institute COVID-19 Surveillance Team |
| EPI_ISL_3125328 | Berkshire and Surrey Pathology Services Lighthouse Laboratory | Wellcome Sanger Institute for the COVID-19 Genomics UK (COG-UK) Consortium | Berkshire and Surrey Pathology Services Lighthouse Laboratory and Alex Alderton, Roberto Amato, Jeffrey Barrett, Sonia Goncalves, Ewan Harrison, David K. Jackson, Ian Johnston, Dominic Kwiatkowski, Cordelia Langford, John Sillitoe on behalf of the Wellcome Sanger Institute COVID-19 Surveillance Team |
| EPI_ISL_3125329, EPI_ISL_3125330 | Lighthouse Lab in Milton Keynes | Wellcome Sanger Institute for the COVID-19 Genomics UK (COG-UK) Consortium | The Lighthouse Lab in Milton Keynes and Alex Alderton, Roberto Amato, Jeffrey Barrett, Sonia Goncalves, Ewan Harrison, David K. Jackson, Ian Johnston, Dominic Kwiatkowski, Cordelia Langford, John Sillitoe on behalf of the Wellcome Sanger Institute COVID-19 Surveillance Team |
| EPI_ISL_3125331 | Berkshire and Surrey Pathology Services Lighthouse Laboratory | Wellcome Sanger Institute for the COVID-19 Genomics UK (COG-UK) Consortium | Berkshire and Surrey Pathology Services Lighthouse Laboratory and Alex Alderton, Roberto Amato, Jeffrey Barrett, Sonia Goncalves, Ewan Harrison, David K. Jackson, Ian Johnston, Dominic Kwiatkowski, Cordelia Langford, John Sillitoe on behalf of the Wellcome Sanger Institute COVID-19 Surveillance Team |
| EPI_ISL_3125332 | Lighthouse Lab in Alderley Park | Wellcome Sanger Institute for the COVID-19 Genomics UK (COG-UK) Consortium | Jacquelyn Wynn, Mairead Hyland, The Lighthouse Lab in Alderley Park and Alex Alderton, Roberto Amato, Jeffrey Barrett, Sonia Goncalves, Ewan Harrison, David K. Jackson, Ian Johnston, Dominic Kwiatkowski, Cordelia Langford, John Sillitoe on behalf of the Wellcome Sanger Institute COVID-19 Surveillance Team |
| EPI_ISL_3125333, EPI_ISL_3125334 | Berkshire and Surrey Pathology Services Lighthouse Laboratory | Wellcome Sanger Institute for the COVID-19 Genomics UK (COG-UK) Consortium | Berkshire and Surrey Pathology Services Lighthouse Laboratory and Alex Alderton, Roberto Amato, Jeffrey Barrett, Sonia Goncalves, Ewan Harrison, David K. Jackson, Ian Johnston, Dominic Kwiatkowski, Cordelia Langford, John Sillitoe on behalf of the Wellcome Sanger Institute COVID-19 Surveillance Team |
| EPI_ISL_3125335 | Lighthouse Lab in Alderley Park | Wellcome Sanger Institute for the COVID-19 Genomics UK (COG-UK) Consortium | Jacquelyn Wynn, Mairead Hyland, The Lighthouse Lab in Alderley Park and Alex Alderton, Roberto Amato, Jeffrey Barrett, Sonia Goncalves, Ewan Harrison, David K. Jackson, Ian Johnston, Dominic Kwiatkowski, Cordelia Langford, John Sillitoe on behalf of the Wellcome Sanger Institute COVID-19 Surveillance Team |
| EPI_ISL_3125336 | Lighthouse Lab in Milton Keynes | Wellcome Sanger Institute for the COVID-19 Genomics UK (COG-UK) Consortium | The Lighthouse Lab in Milton Keynes and Alex Alderton, Roberto Amato, Jeffrey Barrett, Sonia Goncalves, Ewan Harrison, David K. Jackson, Ian Johnston, Dominic Kwiatkowski, Cordelia Langford, John Sillitoe on behalf of the Wellcome Sanger Institute COVID-19 Surveillance Team |
| EPI_ISL_3125337, EPI_ISL_3125338 | Lighthouse Lab in Alderley Park | Wellcome Sanger Institute for the COVID-19 Genomics UK (COG-UK) Consortium | Jacquelyn Wynn, Mairead Hyland, The Lighthouse Lab in Alderley Park and Alex Alderton, Roberto Amato, Jeffrey Barrett, Sonia Goncalves, Ewan Harrison, David K. Jackson, Ian Johnston, Dominic Kwiatkowski, Cordelia Langford, John Sillitoe on behalf of the Wellcome Sanger Institute COVID-19 Surveillance Team |
| EPI_ISL_3125339 | Berkshire and Surrey Pathology Services Lighthouse Laboratory | Wellcome Sanger Institute for the COVID-19 Genomics UK (COG-UK) Consortium | Berkshire and Surrey Pathology Services Lighthouse Laboratory and Alex Alderton, Roberto Amato, Jeffrey Barrett, Sonia Goncalves, Ewan Harrison, David K. Jackson, Ian Johnston, Dominic Kwiatkowski, Cordelia Langford, John Sillitoe on behalf of the Wellcome Sanger Institute COVID-19 Surveillance Team |
| EPI_ISL_3125340, EPI_ISL_3125341, EPI_ISL_3125342, EPI_ISL_3125343, EPI_ISL_3125344, EPI_ISL_3125345 | Lighthouse Lab in Alderley Park | Wellcome Sanger Institute for the COVID-19 Genomics UK (COG-UK) Consortium | Jacquelyn Wynn, Mairead Hyland, The Lighthouse Lab in Alderley Park and Alex Alderton, Roberto Amato, Jeffrey Barrett, Sonia Goncalves, Ewan Harrison, David K. Jackson, Ian Johnston, Dominic Kwiatkowski, Cordelia Langford, John Sillitoe on behalf of the Wellcome Sanger Institute COVID-19 Surveillance Team |
| EPI_ISL_3125346, EPI_ISL_3125347 | Lighthouse Lab in Milton Keynes | Wellcome Sanger Institute for the COVID-19 Genomics UK (COG-UK) Consortium | The Lighthouse Lab in Milton Keynes and Alex Alderton, Roberto Amato, Jeffrey Barrett, Sonia Goncalves, Ewan Harrison, David K. Jackson, Ian Johnston, Dominic Kwiatkowski, Cordelia Langford, John Sillitoe on behalf of the Wellcome Sanger Institute COVID-19 Surveillance Team |
| EPI_ISL_3125348 | Berkshire and Surrey Pathology Services Lighthouse Laboratory | Wellcome Sanger Institute for the COVID-19 Genomics UK (COG-UK) Consortium | Berkshire and Surrey Pathology Services Lighthouse Laboratory and Alex Alderton, Roberto Amato, Jeffrey Barrett, Sonia Goncalves, Ewan Harrison, David K. Jackson, Ian Johnston, Dominic Kwiatkowski, Cordelia Langford, John Sillitoe on behalf of the Wellcome Sanger Institute COVID-19 Surveillance Team |
| EPI_ISL_3125349 | Lighthouse Lab in Alderley Park | Wellcome Sanger Institute for the COVID-19 Genomics UK (COG-UK) Consortium | Jacquelyn Wynn, Mairead Hyland, The Lighthouse Lab in Alderley Park and Alex Alderton, Roberto Amato, Jeffrey Barrett, Sonia Goncalves, Ewan Harrison, David K. Jackson, Ian Johnston, Dominic Kwiatkowski, Cordelia Langford, John Sillitoe on behalf of the Wellcome Sanger Institute COVID-19 Surveillance Team |
| EPI_ISL_3125350, EPI_ISL_3125351 | Lighthouse Lab in Milton Keynes | Wellcome Sanger Institute for the COVID-19 Genomics UK (COG-UK) Consortium | The Lighthouse Lab in Milton Keynes and Alex Alderton, Roberto Amato, Jeffrey Barrett, Sonia Goncalves, Ewan Harrison, David K. Jackson, Ian Johnston, Dominic Kwiatkowski, Cordelia Langford, John Sillitoe on behalf of the Wellcome Sanger Institute COVID-19 Surveillance Team |
| EPI_ISL_3125352 | Lighthouse Lab in Alderley Park | Wellcome Sanger Institute for the COVID-19 Genomics UK (COG-UK) Consortium | Jacquelyn Wynn, Mairead Hyland, The Lighthouse Lab in Alderley Park and Alex Alderton, Roberto Amato, Jeffrey Barrett, Sonia Goncalves, Ewan Harrison, David K. Jackson, Ian Johnston, Dominic Kwiatkowski, Cordelia Langford, John Sillitoe on behalf of the Wellcome Sanger Institute COVID-19 Surveillance Team |
| EPI_ISL_3125353, EPI_ISL_3125354 | Berkshire and Surrey Pathology Services Lighthouse Laboratory | Wellcome Sanger Institute for the COVID-19 Genomics UK (COG-UK) Consortium | Berkshire and Surrey Pathology Services Lighthouse Laboratory and Alex Alderton, Roberto Amato, Jeffrey Barrett, Sonia Goncalves, Ewan Harrison, David K. Jackson, Ian Johnston, Dominic Kwiatkowski, Cordelia Langford, John Sillitoe on behalf of the Wellcome Sanger Institute COVID-19 Surveillance Team |
| EPI_ISL_3125355, EPI_ISL_3125356 | Lighthouse Lab in Alderley Park | Wellcome Sanger Institute for the COVID-19 Genomics UK (COG-UK) Consortium | Jacquelyn Wynn, Mairead Hyland, The Lighthouse Lab in Alderley Park and Alex Alderton, Roberto Amato, Jeffrey Barrett, Sonia Goncalves, Ewan Harrison, David K. Jackson, Ian Johnston, Dominic Kwiatkowski, Cordelia Langford, John Sillitoe on behalf of the Wellcome Sanger Institute COVID-19 Surveillance Team |
| EPI_ISL_3125357 | Berkshire and Surrey Pathology Services Lighthouse Laboratory | Wellcome Sanger Institute for the COVID-19 Genomics UK (COG-UK) Consortium | Berkshire and Surrey Pathology Services Lighthouse Laboratory and Alex Alderton, Roberto Amato, Jeffrey Barrett, Sonia Goncalves, Ewan Harrison, David K. Jackson, Ian Johnston, Dominic Kwiatkowski, Cordelia Langford, John Sillitoe on behalf of the Wellcome Sanger Institute COVID-19 Surveillance Team |
| EPI_ISL_3125358, EPI_ISL_3125359 | Lighthouse Lab in Alderley Park | Wellcome Sanger Institute for the COVID-19 Genomics UK (COG-UK) Consortium | Jacquelyn Wynn, Mairead Hyland, The Lighthouse Lab in Alderley Park and Alex Alderton, Roberto Amato, Jeffrey Barrett, Sonia Goncalves, Ewan Harrison, David K. Jackson, Ian Johnston, Dominic Kwiatkowski, Cordelia Langford, John Sillitoe on behalf of the Wellcome Sanger Institute COVID-19 Surveillance Team |
| EPI_ISL_3125360 | Lighthouse Lab in Milton Keynes | Wellcome Sanger Institute for the COVID-19 Genomics UK (COG-UK) Consortium | The Lighthouse Lab in Milton Keynes and Alex Alderton, Roberto Amato, Jeffrey Barrett, Sonia Goncalves, Ewan Harrison, David K. Jackson, Ian Johnston, Dominic Kwiatkowski, Cordelia Langford, John Sillitoe on behalf of the Wellcome Sanger Institute COVID-19 Surveillance Team |
| EPI_ISL_3125361 | Berkshire and Surrey Pathology Services Lighthouse Laboratory | Wellcome Sanger Institute for the COVID-19 Genomics UK (COG-UK) Consortium | Berkshire and Surrey Pathology Services Lighthouse Laboratory and Alex Alderton, Roberto Amato, Jeffrey Barrett, Sonia Goncalves, Ewan Harrison, David K. Jackson, Ian Johnston, Dominic Kwiatkowski, Cordelia Langford, John Sillitoe on behalf of the Wellcome Sanger Institute COVID-19 Surveillance Team |
| EPI_ISL_3125362, EPI_ISL_3125363 | Lighthouse Lab in Alderley Park | Wellcome Sanger Institute for the COVID-19 Genomics UK (COG-UK) Consortium | Jacquelyn Wynn, Mairead Hyland, The Lighthouse Lab in Alderley Park and Alex Alderton, Roberto Amato, Jeffrey Barrett, Sonia Goncalves, Ewan Harrison, David K. Jackson, Ian Johnston, Dominic Kwiatkowski, Cordelia Langford, John Sillitoe on behalf of the Wellcome Sanger Institute COVID-19 Surveillance Team |
| EPI_ISL_3125364 | Berkshire and Surrey Pathology Services Lighthouse Laboratory | Wellcome Sanger Institute for the COVID-19 Genomics UK (COG-UK) Consortium | Berkshire and Surrey Pathology Services Lighthouse Laboratory and Alex Alderton, Roberto Amato, Jeffrey Barrett, Sonia Goncalves, Ewan Harrison, David K. Jackson, Ian Johnston, Dominic Kwiatkowski, Cordelia Langford, John Sillitoe on behalf of the Wellcome Sanger Institute COVID-19 Surveillance Team |
| EPI_ISL_3125365, EPI_ISL_3125366, EPI_ISL_3125367, EPI_ISL_3125368, EPI_ISL_3125369, EPI_ISL_3125370 | Lighthouse Lab in Alderley Park | Wellcome Sanger Institute for the COVID-19 Genomics UK (COG-UK) Consortium | Jacquelyn Wynn, Mairead Hyland, The Lighthouse Lab in Alderley Park and Alex Alderton, Roberto Amato, Jeffrey Barrett, Sonia Goncalves, Ewan Harrison, David K. Jackson, Ian Johnston, Dominic Kwiatkowski, Cordelia Langford, John Sillitoe on behalf of the Wellcome Sanger Institute COVID-19 Surveillance Team |
| EPI_ISL_3125371 | Berkshire and Surrey Pathology Services Lighthouse Laboratory | Wellcome Sanger Institute for the COVID-19 Genomics UK (COG-UK) Consortium | Berkshire and Surrey Pathology Services Lighthouse Laboratory and Alex Alderton, Roberto Amato, Jeffrey Barrett, Sonia Goncalves, Ewan Harrison, David K. Jackson, Ian Johnston, Dominic Kwiatkowski, Cordelia Langford, John Sillitoe on behalf of the Wellcome Sanger Institute COVID-19 Surveillance Team |
| EPI_ISL_3125372 | Lighthouse Lab in Alderley Park | Wellcome Sanger Institute for the COVID-19 Genomics UK (COG-UK) Consortium | Jacquelyn Wynn, Mairead Hyland, The Lighthouse Lab in Alderley Park and Alex Alderton, Roberto Amato, Jeffrey Barrett, Sonia Goncalves, Ewan Harrison, David K. Jackson, Ian Johnston, Dominic Kwiatkowski, Cordelia Langford, John Sillitoe on behalf of the Wellcome Sanger Institute COVID-19 Surveillance Team |

| | | | |
|---|---|---|---|
| EPI_ISL_3125373, EPI_ISL_3125374, EPI_ISL_3125375 | Berkshire and Surrey Pathology Services Lighthouse Laboratory | Wellcome Sanger Institute for the COVID-19 Genomics UK (COG-UK) Consortium | Berkshire and Surrey Pathology Services Lighthouse Laboratory and Alex Alderton, Roberto Amato, Jeffrey Barrett, Sonia Goncalves, Ewan Harrison, David K. Jackson, Ian Johnston, Dominic Kwiatkowski, Cordelia Langford, John Sillitoe on behalf of the Wellcome Sanger Institute COVID-19 Surveillance Team |
| EPI_ISL_3125376 | Lighthouse Lab in Milton Keynes | Wellcome Sanger Institute for the COVID-19 Genomics UK (COG-UK) Consortium | The Lighthouse Lab in Milton Keynes and Alex Alderton, Roberto Amato, Jeffrey Barrett, Sonia Goncalves, Ewan Harrison, David K. Jackson, Ian Johnston, Dominic Kwiatkowski, Cordelia Langford, John Sillitoe on behalf of the Wellcome Sanger Institute COVID-19 Surveillance Team |
| EPI_ISL_3125377 | Lighthouse Lab in Alderley Park | Wellcome Sanger Institute for the COVID-19 Genomics UK (COG-UK) Consortium | Jacquelyn Wynn, Mairead Hyland, The Lighthouse Lab in Alderley Park and Alex Alderton, Roberto Amato, Jeffrey Barrett, Sonia Goncalves, Ewan Harrison, David K. Jackson, Ian Johnston, Dominic Kwiatkowski, Cordelia Langford, John Sillitoe on behalf of the Wellcome Sanger Institute COVID-19 Surveillance Team |
| EPI_ISL_3125378, EPI_ISL_3125379 | Lighthouse Lab in Milton Keynes | Wellcome Sanger Institute for the COVID-19 Genomics UK (COG-UK) Consortium | The Lighthouse Lab in Milton Keynes and Alex Alderton, Roberto Amato, Jeffrey Barrett, Sonia Goncalves, Ewan Harrison, David K. Jackson, Ian Johnston, Dominic Kwiatkowski, Cordelia Langford, John Sillitoe on behalf of the Wellcome Sanger Institute COVID-19 Surveillance Team |
| EPI_ISL_3125380 | Lighthouse Lab in Alderley Park | Wellcome Sanger Institute for the COVID-19 Genomics UK (COG-UK) Consortium | Jacquelyn Wynn, Mairead Hyland, The Lighthouse Lab in Alderley Park and Alex Alderton, Roberto Amato, Jeffrey Barrett, Sonia Goncalves, Ewan Harrison, David K. Jackson, Ian Johnston, Dominic Kwiatkowski, Cordelia Langford, John Sillitoe on behalf of the Wellcome Sanger Institute COVID-19 Surveillance Team |
| EPI_ISL_3125381, EPI_ISL_3125382 | Berkshire and Surrey Pathology Services Lighthouse Laboratory | Wellcome Sanger Institute for the COVID-19 Genomics UK (COG-UK) Consortium | Berkshire and Surrey Pathology Services Lighthouse Laboratory and Alex Alderton, Roberto Amato, Jeffrey Barrett, Sonia Goncalves, Ewan Harrison, David K. Jackson, Ian Johnston, Dominic Kwiatkowski, Cordelia Langford, John Sillitoe on behalf of the Wellcome Sanger Institute COVID-19 Surveillance Team |
| EPI_ISL_3125383 | Lighthouse Lab in Milton Keynes | Wellcome Sanger Institute for the COVID-19 Genomics UK (COG-UK) Consortium | The Lighthouse Lab in Milton Keynes and Alex Alderton, Roberto Amato, Jeffrey Barrett, Sonia Goncalves, Ewan Harrison, David K. Jackson, Ian Johnston, Dominic Kwiatkowski, Cordelia Langford, John Sillitoe on behalf of the Wellcome Sanger Institute COVID-19 Surveillance Team |
| EPI_ISL_3125384 | Berkshire and Surrey Pathology Services Lighthouse Laboratory | Wellcome Sanger Institute for the COVID-19 Genomics UK (COG-UK) Consortium | Berkshire and Surrey Pathology Services Lighthouse Laboratory and Alex Alderton, Roberto Amato, Jeffrey Barrett, Sonia Goncalves, Ewan Harrison, David K. Jackson, Ian Johnston, Dominic Kwiatkowski, Cordelia Langford, John Sillitoe on behalf of the Wellcome Sanger Institute COVID-19 Surveillance Team |
| EPI_ISL_3125385 | Lighthouse Lab in Alderley Park | Wellcome Sanger Institute for the COVID-19 Genomics UK (COG-UK) Consortium | Jacquelyn Wynn, Mairead Hyland, The Lighthouse Lab in Alderley Park and Alex Alderton, Roberto Amato, Jeffrey Barrett, Sonia Goncalves, Ewan Harrison, David K. Jackson, Ian Johnston, Dominic Kwiatkowski, Cordelia Langford, John Sillitoe on behalf of the Wellcome Sanger Institute COVID-19 Surveillance Team |
| EPI_ISL_3125386 | Lighthouse Lab in Milton Keynes | Wellcome Sanger Institute for the COVID-19 Genomics UK (COG-UK) Consortium | The Lighthouse Lab in Milton Keynes and Alex Alderton, Roberto Amato, Jeffrey Barrett, Sonia Goncalves, Ewan Harrison, David K. Jackson, Ian Johnston, Dominic Kwiatkowski, Cordelia Langford, John Sillitoe on behalf of the Wellcome Sanger Institute COVID-19 Surveillance Team |
| EPI_ISL_3125387, EPI_ISL_3125388, EPI_ISL_3125389 | Lighthouse Lab in Alderley Park | Wellcome Sanger Institute for the COVID-19 Genomics UK (COG-UK) Consortium | Jacquelyn Wynn, Mairead Hyland, The Lighthouse Lab in Alderley Park and Alex Alderton, Roberto Amato, Jeffrey Barrett, Sonia Goncalves, Ewan Harrison, David K. Jackson, Ian Johnston, Dominic Kwiatkowski, Cordelia Langford, John Sillitoe on behalf of the Wellcome Sanger Institute COVID-19 Surveillance Team |
| EPI_ISL_3125390 | Lighthouse Lab in Milton Keynes | Wellcome Sanger Institute for the COVID-19 Genomics UK (COG-UK) Consortium | The Lighthouse Lab in Milton Keynes and Alex Alderton, Roberto Amato, Jeffrey Barrett, Sonia Goncalves, Ewan Harrison, David K. Jackson, Ian Johnston, Dominic Kwiatkowski, Cordelia Langford, John Sillitoe on behalf of the Wellcome Sanger Institute COVID-19 Surveillance Team |
| EPI_ISL_3125391, EPI_ISL_3125392 | Lighthouse Lab in Alderley Park | Wellcome Sanger Institute for the COVID-19 Genomics UK (COG-UK) Consortium | Jacquelyn Wynn, Mairead Hyland, The Lighthouse Lab in Alderley Park and Alex Alderton, Roberto Amato, Jeffrey Barrett, Sonia Goncalves, Ewan Harrison, David K. Jackson, Ian Johnston, Dominic Kwiatkowski, Cordelia Langford, John Sillitoe on behalf of the Wellcome Sanger Institute COVID-19 Surveillance Team |
| EPI_ISL_3125393 | Lighthouse Lab in Milton Keynes | Wellcome Sanger Institute for the COVID-19 Genomics UK (COG-UK) Consortium | The Lighthouse Lab in Milton Keynes and Alex Alderton, Roberto Amato, Jeffrey Barrett, Sonia Goncalves, Ewan Harrison, David K. Jackson, Ian Johnston, Dominic Kwiatkowski, Cordelia Langford, John Sillitoe on behalf of the Wellcome Sanger Institute COVID-19 Surveillance Team |
| EPI_ISL_3125394, EPI_ISL_3125395 | Lighthouse Lab in Alderley Park | Wellcome Sanger Institute for the COVID-19 Genomics UK (COG-UK) Consortium | Jacquelyn Wynn, Mairead Hyland, The Lighthouse Lab in Alderley Park and Alex Alderton, Roberto Amato, Jeffrey Barrett, Sonia Goncalves, Ewan Harrison, David K. Jackson, Ian Johnston, Dominic Kwiatkowski, Cordelia Langford, John Sillitoe on behalf of the Wellcome Sanger Institute COVID-19 Surveillance Team |
| EPI_ISL_3125396, EPI_ISL_3125397 | Lighthouse Lab in Milton Keynes | Wellcome Sanger Institute for the COVID-19 Genomics UK (COG-UK) Consortium | The Lighthouse Lab in Milton Keynes and Alex Alderton, Roberto Amato, Jeffrey Barrett, Sonia Goncalves, Ewan Harrison, David K. Jackson, Ian Johnston, Dominic Kwiatkowski, Cordelia Langford, John Sillitoe on behalf of the Wellcome Sanger Institute COVID-19 Surveillance Team |
| EPI_ISL_3125398 | Lighthouse Lab in Alderley Park | Wellcome Sanger Institute for the COVID-19 Genomics UK (COG-UK) Consortium | Jacquelyn Wynn, Mairead Hyland, The Lighthouse Lab in Alderley Park and Alex Alderton, Roberto Amato, Jeffrey Barrett, Sonia Goncalves, Ewan Harrison, David K. Jackson, Ian Johnston, Dominic Kwiatkowski, Cordelia Langford, John Sillitoe on behalf of the Wellcome Sanger Institute COVID-19 Surveillance Team |
| EPI_ISL_3125399, EPI_ISL_3125400, EPI_ISL_3125401 | Lighthouse Lab in Milton Keynes | Wellcome Sanger Institute for the COVID-19 Genomics UK (COG-UK) Consortium | The Lighthouse Lab in Milton Keynes and Alex Alderton, Roberto Amato, Jeffrey Barrett, Sonia Goncalves, Ewan Harrison, David K. Jackson, Ian Johnston, Dominic Kwiatkowski, Cordelia Langford, John Sillitoe on behalf of the Wellcome Sanger Institute COVID-19 Surveillance Team |
| EPI_ISL_3125402, EPI_ISL_3125403, EPI_ISL_3125404, EPI_ISL_3125405, EPI_ISL_3125406, EPI_ISL_3125407 | Berkshire and Surrey Pathology Services Lighthouse Laboratory | Wellcome Sanger Institute for the COVID-19 Genomics UK (COG-UK) Consortium | Berkshire and Surrey Pathology Services Lighthouse Laboratory and Alex Alderton, Roberto Amato, Jeffrey Barrett, Sonia Goncalves, Ewan Harrison, David K. Jackson, Ian Johnston, Dominic Kwiatkowski, Cordelia Langford, John Sillitoe on behalf of the Wellcome Sanger Institute COVID-19 Surveillance Team |
| EPI_ISL_3125408 | Lighthouse Lab in Milton Keynes | Wellcome Sanger Institute for the COVID-19 Genomics UK (COG-UK) Consortium | The Lighthouse Lab in Milton Keynes and Alex Alderton, Roberto Amato, Jeffrey Barrett, Sonia Goncalves, Ewan Harrison, David K. Jackson, Ian Johnston, Dominic Kwiatkowski, Cordelia Langford, John Sillitoe on behalf of the Wellcome Sanger Institute COVID-19 Surveillance Team |
| EPI_ISL_3125409 | Lighthouse Lab in Alderley Park | Wellcome Sanger Institute for the COVID-19 Genomics UK (COG-UK) Consortium | Jacquelyn Wynn, Mairead Hyland, The Lighthouse Lab in Alderley Park and Alex Alderton, Roberto Amato, Jeffrey Barrett, Sonia Goncalves, Ewan Harrison, David K. Jackson, Ian Johnston, Dominic Kwiatkowski, Cordelia Langford, John Sillitoe on behalf of the Wellcome Sanger Institute COVID-19 Surveillance Team |
| EPI_ISL_3125410, EPI_ISL_3125411 | Berkshire and Surrey Pathology Services Lighthouse Laboratory | Wellcome Sanger Institute for the COVID-19 Genomics UK (COG-UK) Consortium | Berkshire and Surrey Pathology Services Lighthouse Laboratory and Alex Alderton, Roberto Amato, Jeffrey Barrett, Sonia Goncalves, Ewan Harrison, David K. Jackson, Ian Johnston, Dominic Kwiatkowski, Cordelia Langford, John Sillitoe on behalf of the Wellcome Sanger Institute COVID-19 Surveillance Team |
| EPI_ISL_3125412, EPI_ISL_3125413 | Lighthouse Lab in Alderley Park | Wellcome Sanger Institute for the COVID-19 Genomics UK (COG-UK) Consortium | Jacquelyn Wynn, Mairead Hyland, The Lighthouse Lab in Alderley Park and Alex Alderton, Roberto Amato, Jeffrey Barrett, Sonia Goncalves, Ewan Harrison, David K. Jackson, Ian Johnston, Dominic Kwiatkowski, Cordelia Langford, John Sillitoe on behalf of the Wellcome Sanger Institute COVID-19 Surveillance Team |
| EPI_ISL_3125414, EPI_ISL_3125415 | Lighthouse Lab in Milton Keynes | Wellcome Sanger Institute for the COVID-19 Genomics UK (COG-UK) Consortium | The Lighthouse Lab in Milton Keynes and Alex Alderton, Roberto Amato, Jeffrey Barrett, Sonia Goncalves, Ewan Harrison, David K. Jackson, Ian Johnston, Dominic Kwiatkowski, Cordelia Langford, John Sillitoe on behalf of the Wellcome Sanger Institute COVID-19 Surveillance Team |
| EPI_ISL_3125416 | Lighthouse Lab in Alderley Park | Wellcome Sanger Institute for the COVID-19 Genomics UK (COG-UK) Consortium | Jacquelyn Wynn, Mairead Hyland, The Lighthouse Lab in Alderley Park and Alex Alderton, Roberto Amato, Jeffrey Barrett, Sonia Goncalves, Ewan Harrison, David K. Jackson, Ian Johnston, Dominic Kwiatkowski, Cordelia Langford, John Sillitoe on behalf of the Wellcome Sanger Institute COVID-19 Surveillance Team |
| EPI_ISL_3125417, EPI_ISL_3125418, EPI_ISL_3125419 | Lighthouse Lab in Milton Keynes | Wellcome Sanger Institute for the COVID-19 Genomics UK (COG-UK) Consortium | The Lighthouse Lab in Milton Keynes and Alex Alderton, Roberto Amato, Jeffrey Barrett, Sonia Goncalves, Ewan Harrison, David K. Jackson, Ian Johnston, Dominic Kwiatkowski, Cordelia Langford, John Sillitoe on behalf of the Wellcome Sanger Institute COVID-19 Surveillance Team |
| EPI_ISL_3125420 | Berkshire and Surrey Pathology Services Lighthouse Laboratory | Wellcome Sanger Institute for the COVID-19 Genomics UK (COG-UK) Consortium | Berkshire and Surrey Pathology Services Lighthouse Laboratory and Alex Alderton, Roberto Amato, Jeffrey Barrett, Sonia Goncalves, Ewan Harrison, David K. Jackson, Ian Johnston, Dominic Kwiatkowski, Cordelia Langford, John Sillitoe on behalf of the Wellcome Sanger Institute COVID-19 Surveillance Team |
| EPI_ISL_3125421, EPI_ISL_3125422, EPI_ISL_3125423 | Lighthouse Lab in Alderley Park | Wellcome Sanger Institute for the COVID-19 Genomics UK (COG-UK) Consortium | Jacquelyn Wynn, Mairead Hyland, The Lighthouse Lab in Alderley Park and Alex Alderton, Roberto Amato, Jeffrey Barrett, Sonia Goncalves, Ewan Harrison, David K. Jackson, Ian Johnston, Dominic Kwiatkowski, Cordelia Langford, John Sillitoe on behalf of the Wellcome Sanger Institute COVID-19 Surveillance Team |
| EPI_ISL_3125424 | Lighthouse Lab in Milton Keynes | Wellcome Sanger Institute for the COVID-19 Genomics UK (COG-UK) Consortium | The Lighthouse Lab in Milton Keynes and Alex Alderton, Roberto Amato, Jeffrey Barrett, Sonia Goncalves, Ewan Harrison, David K. Jackson, Ian Johnston, Dominic Kwiatkowski, Cordelia Langford, John Sillitoe on behalf of the Wellcome Sanger Institute COVID-19 Surveillance Team |
| EPI_ISL_3125425 | Lighthouse Lab in Alderley Park | Wellcome Sanger Institute for the COVID-19 Genomics UK (COG-UK) Consortium | Jacquelyn Wynn, Mairead Hyland, The Lighthouse Lab in Alderley Park and Alex Alderton, Roberto Amato, Jeffrey Barrett, Sonia Goncalves, Ewan Harrison, David K. Jackson, Ian Johnston, Dominic Kwiatkowski, Cordelia Langford, John Sillitoe on behalf of the Wellcome Sanger Institute COVID-19 Surveillance Team |
| EPI_ISL_3125426 | Lighthouse Lab in Milton Keynes | Wellcome Sanger Institute for the COVID-19 Genomics UK (COG-UK) Consortium | The Lighthouse Lab in Milton Keynes and Alex Alderton, Roberto Amato, Jeffrey Barrett, Sonia Goncalves, Ewan Harrison, David K. Jackson, Ian Johnston, Dominic Kwiatkowski, Cordelia Langford, John Sillitoe on behalf of the Wellcome Sanger Institute COVID-19 Surveillance Team |
| EPI_ISL_3125427, EPI_ISL_3125428 | Lighthouse Lab in Alderley Park | Wellcome Sanger Institute for the COVID-19 Genomics UK (COG-UK) Consortium | Jacquelyn Wynn, Mairead Hyland, The Lighthouse Lab in Alderley Park and Alex Alderton, Roberto Amato, Jeffrey Barrett, Sonia Goncalves, Ewan Harrison, David K. Jackson, Ian Johnston, Dominic Kwiatkowski, Cordelia Langford, John Sillitoe on behalf of the Wellcome Sanger Institute COVID-19 Surveillance Team |
| EPI_ISL_3125429 | Berkshire and Surrey Pathology Services Lighthouse Laboratory | Wellcome Sanger Institute for the COVID-19 Genomics UK (COG-UK) Consortium | Berkshire and Surrey Pathology Services Lighthouse Laboratory and Alex Alderton, Roberto Amato, Jeffrey Barrett, Sonia Goncalves, Ewan Harrison, David K. Jackson, Ian Johnston, Dominic Kwiatkowski, Cordelia Langford, John Sillitoe on behalf of the Wellcome Sanger Institute COVID-19 Surveillance Team |
| EPI_ISL_3125430, EPI_ISL_3125431, EPI_ISL_3125432 | Lighthouse Lab in Alderley Park | Wellcome Sanger Institute for the COVID-19 Genomics UK (COG-UK) Consortium | Jacquelyn Wynn, Mairead Hyland, The Lighthouse Lab in Alderley Park and Alex Alderton, Roberto Amato, Jeffrey Barrett, Sonia Goncalves, Ewan Harrison, David K. Jackson, Ian Johnston, Dominic Kwiatkowski, Cordelia Langford, John Sillitoe on behalf of the Wellcome Sanger Institute COVID-19 Surveillance Team |
| EPI_ISL_3125433 | Berkshire and Surrey Pathology Services Lighthouse Laboratory | Wellcome Sanger Institute for the COVID-19 Genomics UK (COG-UK) Consortium | Berkshire and Surrey Pathology Services Lighthouse Laboratory and Alex Alderton, Roberto Amato, Jeffrey Barrett, Sonia Goncalves, Ewan Harrison, David K. Jackson, Ian Johnston, Dominic Kwiatkowski, Cordelia Langford, John Sillitoe on behalf of the Wellcome Sanger Institute COVID-19 Surveillance Team |
| EPI_ISL_3125434, EPI_ISL_3125435, EPI_ISL_3125436, EPI_ISL_3125437 | Lighthouse Lab in Alderley Park | Wellcome Sanger Institute for the COVID-19 Genomics UK (COG-UK) Consortium | Jacquelyn Wynn, Mairead Hyland, The Lighthouse Lab in Alderley Park and Alex Alderton, Roberto Amato, Jeffrey Barrett, Sonia Goncalves, Ewan Harrison, David K. Jackson, Ian Johnston, Dominic Kwiatkowski, Cordelia Langford, John Sillitoe on behalf of the Wellcome Sanger Institute COVID-19 Surveillance Team |
| EPI_ISL_3125438 | Berkshire and Surrey Pathology Services Lighthouse Laboratory | Wellcome Sanger Institute for the COVID-19 Genomics UK (COG-UK) Consortium | Berkshire and Surrey Pathology Services Lighthouse Laboratory and Alex Alderton, Roberto Amato, Jeffrey Barrett, Sonia Goncalves, Ewan Harrison, David K. Jackson, Ian Johnston, Dominic Kwiatkowski, Cordelia Langford, John Sillitoe on behalf of the Wellcome Sanger Institute COVID-19 Surveillance Team |
| EPI_ISL_3125439 | Lighthouse Lab in Milton Keynes | Wellcome Sanger Institute for the COVID-19 Genomics UK (COG-UK) Consortium | The Lighthouse Lab in Milton Keynes and Alex Alderton, Roberto Amato, Jeffrey Barrett, Sonia Goncalves, Ewan Harrison, David K. Jackson, Ian Johnston, Dominic Kwiatkowski, Cordelia Langford, John Sillitoe on behalf of the Wellcome Sanger Institute COVID-19 Surveillance Team |
| EPI_ISL_3125440, EPI_ISL_3125441, EPI_ISL_3125442, EPI_ISL_3125443 | Lighthouse Lab in Alderley Park | Wellcome Sanger Institute for the COVID-19 Genomics UK (COG-UK) Consortium | Jacquelyn Wynn, Mairead Hyland, The Lighthouse Lab in Alderley Park and Alex Alderton, Roberto Amato, Jeffrey Barrett, Sonia Goncalves, Ewan Harrison, David K. Jackson, Ian Johnston, Dominic Kwiatkowski, Cordelia Langford, John Sillitoe on behalf of the Wellcome Sanger Institute COVID-19 Surveillance Team |
| EPI_ISL_3125444 | Lighthouse Lab in Milton Keynes | Wellcome Sanger Institute for the COVID-19 Genomics UK (COG-UK) | The Lighthouse Lab in Milton Keynes and Alex Alderton, Roberto Amato, Jeffrey Barrett, Sonia Goncalves, Ewan Harrison, David K. Jackson, Ian Johnston, Dominic Kwiatkowski, Cordelia Langford, John Sillitoe on behalf of the Wellcome Sanger |

| | | | |
|---|---|---|---|
| | | Consortium | Institute COVID-19 Surveillance Team |
| EPI_ISL_3125445 | Lighthouse Lab in Alderley Park | Wellcome Sanger Institute for the COVID-19 Genomics UK (COG-UK) Consortium | Jacquelyn Wynn, Mairead Hyland, The Lighthouse Lab in Alderley Park and Alex Alderton, Roberto Amato, Jeffrey Barrett, Sonia Goncalves, Ewan Harrison, David K. Jackson, Ian Johnston, Dominic Kwiatkowski, Cordelia Langford, John Sillitoe on behalf of the Wellcome Sanger Institute COVID-19 Surveillance Team |
| EPI_ISL_3125446 | Lighthouse Lab in Milton Keynes | Wellcome Sanger Institute for the COVID-19 Genomics UK (COG-UK) Consortium | The Lighthouse Lab in Milton Keynes and Alex Alderton, Roberto Amato, Jeffrey Barrett, Sonia Goncalves, Ewan Harrison, David K. Jackson, Ian Johnston, Dominic Kwiatkowski, Cordelia Langford, John Sillitoe on behalf of the Wellcome Sanger Institute COVID-19 Surveillance Team |
| EPI_ISL_3125447, EPI_ISL_3125448 | Berkshire and Surrey Pathology Services Lighthouse Laboratory | Wellcome Sanger Institute for the COVID-19 Genomics UK (COG-UK) Consortium | Berkshire and Surrey Pathology Services Lighthouse Laboratory and Alex Alderton, Roberto Amato, Jeffrey Barrett, Sonia Goncalves, Ewan Harrison, David K. Jackson, Ian Johnston, Dominic Kwiatkowski, Cordelia Langford, John Sillitoe on behalf of the Wellcome Sanger Institute COVID-19 Surveillance Team |
| EPI_ISL_3125449 | Lighthouse Lab in Alderley Park | Wellcome Sanger Institute for the COVID-19 Genomics UK (COG-UK) Consortium | Jacquelyn Wynn, Mairead Hyland, The Lighthouse Lab in Alderley Park and Alex Alderton, Roberto Amato, Jeffrey Barrett, Sonia Goncalves, Ewan Harrison, David K. Jackson, Ian Johnston, Dominic Kwiatkowski, Cordelia Langford, John Sillitoe on behalf of the Wellcome Sanger Institute COVID-19 Surveillance Team |
| EPI_ISL_3125450 | Lighthouse Lab in Milton Keynes | Wellcome Sanger Institute for the COVID-19 Genomics UK (COG-UK) Consortium | The Lighthouse Lab in Milton Keynes and Alex Alderton, Roberto Amato, Jeffrey Barrett, Sonia Goncalves, Ewan Harrison, David K. Jackson, Ian Johnston, Dominic Kwiatkowski, Cordelia Langford, John Sillitoe on behalf of the Wellcome Sanger Institute COVID-19 Surveillance Team |
| EPI_ISL_3125451 | Lighthouse Lab in Alderley Park | Wellcome Sanger Institute for the COVID-19 Genomics UK (COG-UK) Consortium | Jacquelyn Wynn, Mairead Hyland, The Lighthouse Lab in Alderley Park and Alex Alderton, Roberto Amato, Jeffrey Barrett, Sonia Goncalves, Ewan Harrison, David K. Jackson, Ian Johnston, Dominic Kwiatkowski, Cordelia Langford, John Sillitoe on behalf of the Wellcome Sanger Institute COVID-19 Surveillance Team |
| EPI_ISL_3125452 | Berkshire and Surrey Pathology Services Lighthouse Laboratory | Wellcome Sanger Institute for the COVID-19 Genomics UK (COG-UK) Consortium | Berkshire and Surrey Pathology Services Lighthouse Laboratory and Alex Alderton, Roberto Amato, Jeffrey Barrett, Sonia Goncalves, Ewan Harrison, David K. Jackson, Ian Johnston, Dominic Kwiatkowski, Cordelia Langford, John Sillitoe on behalf of the Wellcome Sanger Institute COVID-19 Surveillance Team |
| EPI_ISL_3125453, EPI_ISL_3125454 | Lighthouse Lab in Alderley Park | Wellcome Sanger Institute for the COVID-19 Genomics UK (COG-UK) Consortium | Jacquelyn Wynn, Mairead Hyland, The Lighthouse Lab in Alderley Park and Alex Alderton, Roberto Amato, Jeffrey Barrett, Sonia Goncalves, Ewan Harrison, David K. Jackson, Ian Johnston, Dominic Kwiatkowski, Cordelia Langford, John Sillitoe on behalf of the Wellcome Sanger Institute COVID-19 Surveillance Team |
| EPI_ISL_3125455 | Lighthouse Lab in Milton Keynes | Wellcome Sanger Institute for the COVID-19 Genomics UK (COG-UK) Consortium | The Lighthouse Lab in Milton Keynes and Alex Alderton, Roberto Amato, Jeffrey Barrett, Sonia Goncalves, Ewan Harrison, David K. Jackson, Ian Johnston, Dominic Kwiatkowski, Cordelia Langford, John Sillitoe on behalf of the Wellcome Sanger Institute COVID-19 Surveillance Team |
| EPI_ISL_3125456 | Berkshire and Surrey Pathology Services Lighthouse Laboratory | Wellcome Sanger Institute for the COVID-19 Genomics UK (COG-UK) Consortium | Berkshire and Surrey Pathology Services Lighthouse Laboratory and Alex Alderton, Roberto Amato, Jeffrey Barrett, Sonia Goncalves, Ewan Harrison, David K. Jackson, Ian Johnston, Dominic Kwiatkowski, Cordelia Langford, John Sillitoe on behalf of the Wellcome Sanger Institute COVID-19 Surveillance Team |
| EPI_ISL_3125457 | Lighthouse Lab in Alderley Park | Wellcome Sanger Institute for the COVID-19 Genomics UK (COG-UK) Consortium | Jacquelyn Wynn, Mairead Hyland, The Lighthouse Lab in Alderley Park and Alex Alderton, Roberto Amato, Jeffrey Barrett, Sonia Goncalves, Ewan Harrison, David K. Jackson, Ian Johnston, Dominic Kwiatkowski, Cordelia Langford, John Sillitoe on behalf of the Wellcome Sanger Institute COVID-19 Surveillance Team |
| EPI_ISL_3125458 | Lighthouse Lab in Milton Keynes | Wellcome Sanger Institute for the COVID-19 Genomics UK (COG-UK) Consortium | The Lighthouse Lab in Milton Keynes and Alex Alderton, Roberto Amato, Jeffrey Barrett, Sonia Goncalves, Ewan Harrison, David K. Jackson, Ian Johnston, Dominic Kwiatkowski, Cordelia Langford, John Sillitoe on behalf of the Wellcome Sanger Institute COVID-19 Surveillance Team |
| EPI_ISL_3125459 | Lighthouse Lab in Alderley Park | Wellcome Sanger Institute for the COVID-19 Genomics UK (COG-UK) Consortium | Jacquelyn Wynn, Mairead Hyland, The Lighthouse Lab in Alderley Park and Alex Alderton, Roberto Amato, Jeffrey Barrett, Sonia Goncalves, Ewan Harrison, David K. Jackson, Ian Johnston, Dominic Kwiatkowski, Cordelia Langford, John Sillitoe on behalf of the Wellcome Sanger Institute COVID-19 Surveillance Team |
| EPI_ISL_3125460 | Berkshire and Surrey Pathology Services Lighthouse Laboratory | Wellcome Sanger Institute for the COVID-19 Genomics UK (COG-UK) Consortium | Berkshire and Surrey Pathology Services Lighthouse Laboratory and Alex Alderton, Roberto Amato, Jeffrey Barrett, Sonia Goncalves, Ewan Harrison, David K. Jackson, Ian Johnston, Dominic Kwiatkowski, Cordelia Langford, John Sillitoe on behalf of the Wellcome Sanger Institute COVID-19 Surveillance Team |
| EPI_ISL_3125461 | Lighthouse Lab in Alderley Park | Wellcome Sanger Institute for the COVID-19 Genomics UK (COG-UK) Consortium | Jacquelyn Wynn, Mairead Hyland, The Lighthouse Lab in Alderley Park and Alex Alderton, Roberto Amato, Jeffrey Barrett, Sonia Goncalves, Ewan Harrison, David K. Jackson, Ian Johnston, Dominic Kwiatkowski, Cordelia Langford, John Sillitoe on behalf of the Wellcome Sanger Institute COVID-19 Surveillance Team |
| EPI_ISL_3125462 | Berkshire and Surrey Pathology Services Lighthouse Laboratory | Wellcome Sanger Institute for the COVID-19 Genomics UK (COG-UK) Consortium | Berkshire and Surrey Pathology Services Lighthouse Laboratory and Alex Alderton, Roberto Amato, Jeffrey Barrett, Sonia Goncalves, Ewan Harrison, David K. Jackson, Ian Johnston, Dominic Kwiatkowski, Cordelia Langford, John Sillitoe on behalf of the Wellcome Sanger Institute COVID-19 Surveillance Team |
| EPI_ISL_3125463 | Lighthouse Lab in Alderley Park | Wellcome Sanger Institute for the COVID-19 Genomics UK (COG-UK) Consortium | Jacquelyn Wynn, Mairead Hyland, The Lighthouse Lab in Alderley Park and Alex Alderton, Roberto Amato, Jeffrey Barrett, Sonia Goncalves, Ewan Harrison, David K. Jackson, Ian Johnston, Dominic Kwiatkowski, Cordelia Langford, John Sillitoe on behalf of the Wellcome Sanger Institute COVID-19 Surveillance Team |
| EPI_ISL_3125464 | Lighthouse Lab in Milton Keynes | Wellcome Sanger Institute for the COVID-19 Genomics UK (COG-UK) Consortium | The Lighthouse Lab in Milton Keynes and Alex Alderton, Roberto Amato, Jeffrey Barrett, Sonia Goncalves, Ewan Harrison, David K. Jackson, Ian Johnston, Dominic Kwiatkowski, Cordelia Langford, John Sillitoe on behalf of the Wellcome Sanger Institute COVID-19 Surveillance Team |
| EPI_ISL_3125465, EPI_ISL_3125466 | Lighthouse Lab in Alderley Park | Wellcome Sanger Institute for the COVID-19 Genomics UK (COG-UK) Consortium | Jacquelyn Wynn, Mairead Hyland, The Lighthouse Lab in Alderley Park and Alex Alderton, Roberto Amato, Jeffrey Barrett, Sonia Goncalves, Ewan Harrison, David K. Jackson, Ian Johnston, Dominic Kwiatkowski, Cordelia Langford, John Sillitoe on behalf of the Wellcome Sanger Institute COVID-19 Surveillance Team |
| EPI_ISL_3125467 | Lighthouse Lab in Milton Keynes | Wellcome Sanger Institute for the COVID-19 Genomics UK (COG-UK) Consortium | The Lighthouse Lab in Milton Keynes and Alex Alderton, Roberto Amato, Jeffrey Barrett, Sonia Goncalves, Ewan Harrison, David K. Jackson, Ian Johnston, Dominic Kwiatkowski, Cordelia Langford, John Sillitoe on behalf of the Wellcome Sanger Institute COVID-19 Surveillance Team |
| EPI_ISL_3125468 | Lighthouse Lab in Alderley Park | Wellcome Sanger Institute for the COVID-19 Genomics UK (COG-UK) Consortium | Jacquelyn Wynn, Mairead Hyland, The Lighthouse Lab in Alderley Park and Alex Alderton, Roberto Amato, Jeffrey Barrett, Sonia Goncalves, Ewan Harrison, David K. Jackson, Ian Johnston, Dominic Kwiatkowski, Cordelia Langford, John Sillitoe on behalf of the Wellcome Sanger Institute COVID-19 Surveillance Team |
| EPI_ISL_3125469 | Lighthouse Lab in Milton Keynes | Wellcome Sanger Institute for the COVID-19 Genomics UK (COG-UK) Consortium | The Lighthouse Lab in Milton Keynes and Alex Alderton, Roberto Amato, Jeffrey Barrett, Sonia Goncalves, Ewan Harrison, David K. Jackson, Ian Johnston, Dominic Kwiatkowski, Cordelia Langford, John Sillitoe on behalf of the Wellcome Sanger Institute COVID-19 Surveillance Team |
| EPI_ISL_3125470 | Lighthouse Lab in Alderley Park | Wellcome Sanger Institute for the COVID-19 Genomics UK (COG-UK) Consortium | Jacquelyn Wynn, Mairead Hyland, The Lighthouse Lab in Alderley Park and Alex Alderton, Roberto Amato, Jeffrey Barrett, Sonia Goncalves, Ewan Harrison, David K. Jackson, Ian Johnston, Dominic Kwiatkowski, Cordelia Langford, John Sillitoe on behalf of the Wellcome Sanger Institute COVID-19 Surveillance Team |
| EPI_ISL_3125471 | Berkshire and Surrey Pathology Services Lighthouse Laboratory | Wellcome Sanger Institute for the COVID-19 Genomics UK (COG-UK) Consortium | Berkshire and Surrey Pathology Services Lighthouse Laboratory and Alex Alderton, Roberto Amato, Jeffrey Barrett, Sonia Goncalves, Ewan Harrison, David K. Jackson, Ian Johnston, Dominic Kwiatkowski, Cordelia Langford, John Sillitoe on behalf of the Wellcome Sanger Institute COVID-19 Surveillance Team |
| EPI_ISL_3125472 | Lighthouse Lab in Alderley Park | Wellcome Sanger Institute for the COVID-19 Genomics UK (COG-UK) Consortium | Jacquelyn Wynn, Mairead Hyland, The Lighthouse Lab in Alderley Park and Alex Alderton, Roberto Amato, Jeffrey Barrett, Sonia Goncalves, Ewan Harrison, David K. Jackson, Ian Johnston, Dominic Kwiatkowski, Cordelia Langford, John Sillitoe on behalf of the Wellcome Sanger Institute COVID-19 Surveillance Team |
| EPI_ISL_3125473 | Berkshire and Surrey Pathology Services Lighthouse Laboratory | Wellcome Sanger Institute for the COVID-19 Genomics UK (COG-UK) Consortium | Berkshire and Surrey Pathology Services Lighthouse Laboratory and Alex Alderton, Roberto Amato, Jeffrey Barrett, Sonia Goncalves, Ewan Harrison, David K. Jackson, Ian Johnston, Dominic Kwiatkowski, Cordelia Langford, John Sillitoe on behalf of the Wellcome Sanger Institute COVID-19 Surveillance Team |
| EPI_ISL_3125474 | Lighthouse Lab in Milton Keynes | Wellcome Sanger Institute for the COVID-19 Genomics UK (COG-UK) Consortium | The Lighthouse Lab in Milton Keynes and Alex Alderton, Roberto Amato, Jeffrey Barrett, Sonia Goncalves, Ewan Harrison, David K. Jackson, Ian Johnston, Dominic Kwiatkowski, Cordelia Langford, John Sillitoe on behalf of the Wellcome Sanger Institute COVID-19 Surveillance Team |
| EPI_ISL_3125475, EPI_ISL_3125476 | Lighthouse Lab in Alderley Park | Wellcome Sanger Institute for the COVID-19 Genomics UK (COG-UK) Consortium | Jacquelyn Wynn, Mairead Hyland, The Lighthouse Lab in Alderley Park and Alex Alderton, Roberto Amato, Jeffrey Barrett, Sonia Goncalves, Ewan Harrison, David K. Jackson, Ian Johnston, Dominic Kwiatkowski, Cordelia Langford, John Sillitoe on behalf of the Wellcome Sanger Institute COVID-19 Surveillance Team |
| EPI_ISL_3125477, EPI_ISL_3125478 | Lighthouse Lab in Milton Keynes | Wellcome Sanger Institute for the COVID-19 Genomics UK (COG-UK) Consortium | The Lighthouse Lab in Milton Keynes and Alex Alderton, Roberto Amato, Jeffrey Barrett, Sonia Goncalves, Ewan Harrison, David K. Jackson, Ian Johnston, Dominic Kwiatkowski, Cordelia Langford, John Sillitoe on behalf of the Wellcome Sanger Institute COVID-19 Surveillance Team |
| EPI_ISL_3125479 | Lighthouse Lab in Alderley Park | Wellcome Sanger Institute for the COVID-19 Genomics UK (COG-UK) Consortium | Jacquelyn Wynn, Mairead Hyland, The Lighthouse Lab in Alderley Park and Alex Alderton, Roberto Amato, Jeffrey Barrett, Sonia Goncalves, Ewan Harrison, David K. Jackson, Ian Johnston, Dominic Kwiatkowski, Cordelia Langford, John Sillitoe on behalf of the Wellcome Sanger Institute COVID-19 Surveillance Team |
| EPI_ISL_3125480, EPI_ISL_3125481 | Berkshire and Surrey Pathology Services Lighthouse Laboratory | Wellcome Sanger Institute for the COVID-19 Genomics UK (COG-UK) Consortium | Berkshire and Surrey Pathology Services Lighthouse Laboratory and Alex Alderton, Roberto Amato, Jeffrey Barrett, Sonia Goncalves, Ewan Harrison, David K. Jackson, Ian Johnston, Dominic Kwiatkowski, Cordelia Langford, John Sillitoe on behalf of the Wellcome Sanger Institute COVID-19 Surveillance Team |
| EPI_ISL_3125482 | Lighthouse Lab in Alderley Park | Wellcome Sanger Institute for the COVID-19 Genomics UK (COG-UK) Consortium | Jacquelyn Wynn, Mairead Hyland, The Lighthouse Lab in Alderley Park and Alex Alderton, Roberto Amato, Jeffrey Barrett, Sonia Goncalves, Ewan Harrison, David K. Jackson, Ian Johnston, Dominic Kwiatkowski, Cordelia Langford, John Sillitoe on behalf of the Wellcome Sanger Institute COVID-19 Surveillance Team |
| EPI_ISL_3125483, EPI_ISL_3125484 | Berkshire and Surrey Pathology Services Lighthouse Laboratory | Wellcome Sanger Institute for the COVID-19 Genomics UK (COG-UK) Consortium | Berkshire and Surrey Pathology Services Lighthouse Laboratory and Alex Alderton, Roberto Amato, Jeffrey Barrett, Sonia Goncalves, Ewan Harrison, David K. Jackson, Ian Johnston, Dominic Kwiatkowski, Cordelia Langford, John Sillitoe on behalf of the Wellcome Sanger Institute COVID-19 Surveillance Team |
| EPI_ISL_3125485, EPI_ISL_3125486, EPI_ISL_3125487 | Lighthouse Lab in Alderley Park | Wellcome Sanger Institute for the COVID-19 Genomics UK (COG-UK) Consortium | Jacquelyn Wynn, Mairead Hyland, The Lighthouse Lab in Alderley Park and Alex Alderton, Roberto Amato, Jeffrey Barrett, Sonia Goncalves, Ewan Harrison, David K. Jackson, Ian Johnston, Dominic Kwiatkowski, Cordelia Langford, John Sillitoe on behalf of the Wellcome Sanger Institute COVID-19 Surveillance Team |
| EPI_ISL_3125488 | Lighthouse Lab in Milton Keynes | Wellcome Sanger Institute for the COVID-19 Genomics UK (COG-UK) Consortium | The Lighthouse Lab in Milton Keynes and Alex Alderton, Roberto Amato, Jeffrey Barrett, Sonia Goncalves, Ewan Harrison, David K. Jackson, Ian Johnston, Dominic Kwiatkowski, Cordelia Langford, John Sillitoe on behalf of the Wellcome Sanger Institute COVID-19 Surveillance Team |
| EPI_ISL_3125489, EPI_ISL_3125490 | Lighthouse Lab in Alderley Park | Wellcome Sanger Institute for the COVID-19 Genomics UK (COG-UK) Consortium | Jacquelyn Wynn, Mairead Hyland, The Lighthouse Lab in Alderley Park and Alex Alderton, Roberto Amato, Jeffrey Barrett, Sonia Goncalves, Ewan Harrison, David K. Jackson, Ian Johnston, Dominic Kwiatkowski, Cordelia Langford, John Sillitoe on behalf of the Wellcome Sanger Institute COVID-19 Surveillance Team |
| EPI_ISL_3125491, EPI_ISL_3125492, EPI_ISL_3125493 | Lighthouse Lab in Milton Keynes | Wellcome Sanger Institute for the COVID-19 Genomics UK (COG-UK) Consortium | The Lighthouse Lab in Milton Keynes and Alex Alderton, Roberto Amato, Jeffrey Barrett, Sonia Goncalves, Ewan Harrison, David K. Jackson, Ian Johnston, Dominic Kwiatkowski, Cordelia Langford, John Sillitoe on behalf of the Wellcome Sanger Institute COVID-19 Surveillance Team |
| EPI_ISL_3125494 | Lighthouse Lab in Alderley Park | Wellcome Sanger Institute for the COVID-19 Genomics UK (COG-UK) Consortium | Jacquelyn Wynn, Mairead Hyland, The Lighthouse Lab in Alderley Park and Alex Alderton, Roberto Amato, Jeffrey Barrett, Sonia Goncalves, Ewan Harrison, David K. Jackson, Ian Johnston, Dominic Kwiatkowski, Cordelia Langford, John Sillitoe on behalf of the Wellcome Sanger Institute COVID-19 Surveillance Team |
| EPI_ISL_3125495, EPI_ISL_3125496 | Berkshire and Surrey Pathology Services Lighthouse Laboratory | Wellcome Sanger Institute for the COVID-19 Genomics UK (COG-UK) Consortium | Berkshire and Surrey Pathology Services Lighthouse Laboratory and Alex Alderton, Roberto Amato, Jeffrey Barrett, Sonia Goncalves, Ewan Harrison, David K. Jackson, Ian Johnston, Dominic Kwiatkowski, Cordelia Langford, John Sillitoe on behalf of the Wellcome Sanger Institute COVID-19 Surveillance Team |
| EPI_ISL_3125497 | Lighthouse Lab in Alderley Park | Wellcome Sanger Institute for the COVID-19 Genomics UK (COG-UK) Consortium | Jacquelyn Wynn, Mairead Hyland, The Lighthouse Lab in Alderley Park and Alex Alderton, Roberto Amato, Jeffrey Barrett, Sonia Goncalves, Ewan Harrison, David K. Jackson, Ian Johnston, Dominic Kwiatkowski, Cordelia Langford, John Sillitoe on behalf of the Wellcome Sanger Institute COVID-19 Surveillance Team |
| EPI_ISL_3125498 | Lighthouse Lab in Milton Keynes | Wellcome Sanger Institute for the COVID-19 Genomics UK (COG-UK) Consortium | The Lighthouse Lab in Milton Keynes and Alex Alderton, Roberto Amato, Jeffrey Barrett, Sonia Goncalves, Ewan Harrison, David K. Jackson, Ian Johnston, Dominic Kwiatkowski, Cordelia Langford, John Sillitoe on behalf of the Wellcome Sanger Institute COVID-19 Surveillance Team |
| EPI_ISL_3125499 | Lighthouse Lab in Alderley Park | Wellcome Sanger Institute for the COVID-19 Genomics UK (COG-UK) Consortium | Jacquelyn Wynn, Mairead Hyland, The Lighthouse Lab in Alderley Park and Alex Alderton, Roberto Amato, Jeffrey Barrett, Sonia Goncalves, Ewan Harrison, David K. Jackson, Ian Johnston, Dominic Kwiatkowski, Cordelia Langford, John Sillitoe on behalf of the Wellcome Sanger Institute COVID-19 Surveillance Team |
| EPI_ISL_3125500 | Lighthouse Lab in Milton Keynes | Wellcome Sanger Institute for the COVID-19 Genomics UK (COG-UK) Consortium | The Lighthouse Lab in Milton Keynes and Alex Alderton, Roberto Amato, Jeffrey Barrett, Sonia Goncalves, Ewan Harrison, David K. Jackson, Ian Johnston, Dominic Kwiatkowski, Cordelia Langford, John Sillitoe on behalf of the Wellcome Sanger Institute COVID-19 Surveillance Team |
| EPI_ISL_3125501, | Lighthouse Lab in Alderley Park | Wellcome Sanger Institute for the COVID-19 Genomics UK (COG-UK) | Jacquelyn Wynn, Mairead Hyland, The Lighthouse Lab in Alderley Park and Alex Alderton, Roberto Amato, Jeffrey Barrett, Sonia Goncalves, Ewan Harrison, David K. Jackson, Ian Johnston, Dominic Kwiatkowski, Cordelia Langford, John Sillitoe on |

| | | | |
|---|---|---|---|
| EPI_ISL_3125502, EPI_ISL_3125503 | | Consortium | behalf of the Wellcome Sanger Institute COVID-19 Surveillance Team |
| EPI_ISL_3125504, EPI_ISL_3125505 | Lighthouse Lab in Milton Keynes | Wellcome Sanger Institute for the COVID-19 Genomics UK (COG-UK) Consortium | The Lighthouse Lab in Milton Keynes and Alex Alderton, Roberto Amato, Jeffrey Barrett, Sonia Goncalves, Ewan Harrison, David K. Jackson, Ian Johnston, Dominic Kwiatkowski, Cordelia Langford, John Sillitoe on behalf of the Wellcome Sanger Institute COVID-19 Surveillance Team |
| EPI_ISL_3125506, EPI_ISL_3125507 | Lighthouse Lab in Alderley Park | Wellcome Sanger Institute for the COVID-19 Genomics UK (COG-UK) Consortium | Jacquelyn Wynn, Mairead Hyland, The Lighthouse Lab in Alderley Park and Alex Alderton, Roberto Amato, Jeffrey Barrett, Sonia Goncalves, Ewan Harrison, David K. Jackson, Ian Johnston, Dominic Kwiatkowski, Cordelia Langford, John Sillitoe on behalf of the Wellcome Sanger Institute COVID-19 Surveillance Team |
| EPI_ISL_3125508 | Berkshire and Surrey Pathology Services Lighthouse Laboratory | Wellcome Sanger Institute for the COVID-19 Genomics UK (COG-UK) Consortium | Berkshire and Surrey Pathology Services Lighthouse Laboratory and Alex Alderton, Roberto Amato, Jeffrey Barrett, Sonia Goncalves, Ewan Harrison, David K. Jackson, Ian Johnston, Dominic Kwiatkowski, Cordelia Langford, John Sillitoe on behalf of the Wellcome Sanger Institute COVID-19 Surveillance Team |
| EPI_ISL_3125509 | Lighthouse Lab in Milton Keynes | Wellcome Sanger Institute for the COVID-19 Genomics UK (COG-UK) Consortium | The Lighthouse Lab in Milton Keynes and Alex Alderton, Roberto Amato, Jeffrey Barrett, Sonia Goncalves, Ewan Harrison, David K. Jackson, Ian Johnston, Dominic Kwiatkowski, Cordelia Langford, John Sillitoe on behalf of the Wellcome Sanger Institute COVID-19 Surveillance Team |
| EPI_ISL_3125510, EPI_ISL_3125511 | Lighthouse Lab in Alderley Park | Wellcome Sanger Institute for the COVID-19 Genomics UK (COG-UK) Consortium | Jacquelyn Wynn, Mairead Hyland, The Lighthouse Lab in Alderley Park and Alex Alderton, Roberto Amato, Jeffrey Barrett, Sonia Goncalves, Ewan Harrison, David K. Jackson, Ian Johnston, Dominic Kwiatkowski, Cordelia Langford, John Sillitoe on behalf of the Wellcome Sanger Institute COVID-19 Surveillance Team |
| EPI_ISL_3125512 | Lighthouse Lab in Milton Keynes | Wellcome Sanger Institute for the COVID-19 Genomics UK (COG-UK) Consortium | The Lighthouse Lab in Milton Keynes and Alex Alderton, Roberto Amato, Jeffrey Barrett, Sonia Goncalves, Ewan Harrison, David K. Jackson, Ian Johnston, Dominic Kwiatkowski, Cordelia Langford, John Sillitoe on behalf of the Wellcome Sanger Institute COVID-19 Surveillance Team |
| EPI_ISL_3125513, EPI_ISL_3125514 | Berkshire and Surrey Pathology Services Lighthouse Laboratory | Wellcome Sanger Institute for the COVID-19 Genomics UK (COG-UK) Consortium | Berkshire and Surrey Pathology Services Lighthouse Laboratory and Alex Alderton, Roberto Amato, Jeffrey Barrett, Sonia Goncalves, Ewan Harrison, David K. Jackson, Ian Johnston, Dominic Kwiatkowski, Cordelia Langford, John Sillitoe on behalf of the Wellcome Sanger Institute COVID-19 Surveillance Team |
| EPI_ISL_3125515, EPI_ISL_3125516 | Lighthouse Lab in Milton Keynes | Wellcome Sanger Institute for the COVID-19 Genomics UK (COG-UK) Consortium | The Lighthouse Lab in Milton Keynes and Alex Alderton, Roberto Amato, Jeffrey Barrett, Sonia Goncalves, Ewan Harrison, David K. Jackson, Ian Johnston, Dominic Kwiatkowski, Cordelia Langford, John Sillitoe on behalf of the Wellcome Sanger Institute COVID-19 Surveillance Team |
| EPI_ISL_3125517 | Lighthouse Lab in Alderley Park | Wellcome Sanger Institute for the COVID-19 Genomics UK (COG-UK) Consortium | Jacquelyn Wynn, Mairead Hyland, The Lighthouse Lab in Alderley Park and Alex Alderton, Roberto Amato, Jeffrey Barrett, Sonia Goncalves, Ewan Harrison, David K. Jackson, Ian Johnston, Dominic Kwiatkowski, Cordelia Langford, John Sillitoe on behalf of the Wellcome Sanger Institute COVID-19 Surveillance Team |
| EPI_ISL_3125518 | Berkshire and Surrey Pathology Services Lighthouse Laboratory | Wellcome Sanger Institute for the COVID-19 Genomics UK (COG-UK) Consortium | Berkshire and Surrey Pathology Services Lighthouse Laboratory and Alex Alderton, Roberto Amato, Jeffrey Barrett, Sonia Goncalves, Ewan Harrison, David K. Jackson, Ian Johnston, Dominic Kwiatkowski, Cordelia Langford, John Sillitoe on behalf of the Wellcome Sanger Institute COVID-19 Surveillance Team |
| EPI_ISL_3125519 | Lighthouse Lab in Alderley Park | Wellcome Sanger Institute for the COVID-19 Genomics UK (COG-UK) Consortium | Jacquelyn Wynn, Mairead Hyland, The Lighthouse Lab in Alderley Park and Alex Alderton, Roberto Amato, Jeffrey Barrett, Sonia Goncalves, Ewan Harrison, David K. Jackson, Ian Johnston, Dominic Kwiatkowski, Cordelia Langford, John Sillitoe on behalf of the Wellcome Sanger Institute COVID-19 Surveillance Team |
| EPI_ISL_3125520 | Lighthouse Lab in Milton Keynes | Wellcome Sanger Institute for the COVID-19 Genomics UK (COG-UK) Consortium | The Lighthouse Lab in Milton Keynes and Alex Alderton, Roberto Amato, Jeffrey Barrett, Sonia Goncalves, Ewan Harrison, David K. Jackson, Ian Johnston, Dominic Kwiatkowski, Cordelia Langford, John Sillitoe on behalf of the Wellcome Sanger Institute COVID-19 Surveillance Team |
| EPI_ISL_3125521, EPI_ISL_3125522, EPI_ISL_3125523, EPI_ISL_3125524 | Lighthouse Lab in Alderley Park | Wellcome Sanger Institute for the COVID-19 Genomics UK (COG-UK) Consortium | Jacquelyn Wynn, Mairead Hyland, The Lighthouse Lab in Alderley Park and Alex Alderton, Roberto Amato, Jeffrey Barrett, Sonia Goncalves, Ewan Harrison, David K. Jackson, Ian Johnston, Dominic Kwiatkowski, Cordelia Langford, John Sillitoe on behalf of the Wellcome Sanger Institute COVID-19 Surveillance Team |
| EPI_ISL_3125525 | Berkshire and Surrey Pathology Services Lighthouse Laboratory | Wellcome Sanger Institute for the COVID-19 Genomics UK (COG-UK) Consortium | Berkshire and Surrey Pathology Services Lighthouse Laboratory and Alex Alderton, Roberto Amato, Jeffrey Barrett, Sonia Goncalves, Ewan Harrison, David K. Jackson, Ian Johnston, Dominic Kwiatkowski, Cordelia Langford, John Sillitoe on behalf of the Wellcome Sanger Institute COVID-19 Surveillance Team |
| EPI_ISL_3125526 | Lighthouse Lab in Alderley Park | Wellcome Sanger Institute for the COVID-19 Genomics UK (COG-UK) Consortium | Jacquelyn Wynn, Mairead Hyland, The Lighthouse Lab in Alderley Park and Alex Alderton, Roberto Amato, Jeffrey Barrett, Sonia Goncalves, Ewan Harrison, David K. Jackson, Ian Johnston, Dominic Kwiatkowski, Cordelia Langford, John Sillitoe on behalf of the Wellcome Sanger Institute COVID-19 Surveillance Team |
| EPI_ISL_3125527 | Lighthouse Lab in Milton Keynes | Wellcome Sanger Institute for the COVID-19 Genomics UK (COG-UK) Consortium | The Lighthouse Lab in Milton Keynes and Alex Alderton, Roberto Amato, Jeffrey Barrett, Sonia Goncalves, Ewan Harrison, David K. Jackson, Ian Johnston, Dominic Kwiatkowski, Cordelia Langford, John Sillitoe on behalf of the Wellcome Sanger Institute COVID-19 Surveillance Team |
| EPI_ISL_3125528 | Berkshire and Surrey Pathology Services Lighthouse Laboratory | Wellcome Sanger Institute for the COVID-19 Genomics UK (COG-UK) Consortium | Berkshire and Surrey Pathology Services Lighthouse Laboratory and Alex Alderton, Roberto Amato, Jeffrey Barrett, Sonia Goncalves, Ewan Harrison, David K. Jackson, Ian Johnston, Dominic Kwiatkowski, Cordelia Langford, John Sillitoe on behalf of the Wellcome Sanger Institute COVID-19 Surveillance Team |
| EPI_ISL_3125529 | Lighthouse Lab in Alderley Park | Wellcome Sanger Institute for the COVID-19 Genomics UK (COG-UK) Consortium | Jacquelyn Wynn, Mairead Hyland, The Lighthouse Lab in Alderley Park and Alex Alderton, Roberto Amato, Jeffrey Barrett, Sonia Goncalves, Ewan Harrison, David K. Jackson, Ian Johnston, Dominic Kwiatkowski, Cordelia Langford, John Sillitoe on behalf of the Wellcome Sanger Institute COVID-19 Surveillance Team |
| EPI_ISL_3125530 | Berkshire and Surrey Pathology Services Lighthouse Laboratory | Wellcome Sanger Institute for the COVID-19 Genomics UK (COG-UK) Consortium | Berkshire and Surrey Pathology Services Lighthouse Laboratory and Alex Alderton, Roberto Amato, Jeffrey Barrett, Sonia Goncalves, Ewan Harrison, David K. Jackson, Ian Johnston, Dominic Kwiatkowski, Cordelia Langford, John Sillitoe on behalf of the Wellcome Sanger Institute COVID-19 Surveillance Team |
| EPI_ISL_3125531, EPI_ISL_3125532, EPI_ISL_3125533, EPI_ISL_3125534, EPI_ISL_3125535 | Lighthouse Lab in Alderley Park | Wellcome Sanger Institute for the COVID-19 Genomics UK (COG-UK) Consortium | Jacquelyn Wynn, Mairead Hyland, The Lighthouse Lab in Alderley Park and Alex Alderton, Roberto Amato, Jeffrey Barrett, Sonia Goncalves, Ewan Harrison, David K. Jackson, Ian Johnston, Dominic Kwiatkowski, Cordelia Langford, John Sillitoe on behalf of the Wellcome Sanger Institute COVID-19 Surveillance Team |
| EPI_ISL_3125536 | Lighthouse Lab in Milton Keynes | Wellcome Sanger Institute for the COVID-19 Genomics UK (COG-UK) Consortium | The Lighthouse Lab in Milton Keynes and Alex Alderton, Roberto Amato, Jeffrey Barrett, Sonia Goncalves, Ewan Harrison, David K. Jackson, Ian Johnston, Dominic Kwiatkowski, Cordelia Langford, John Sillitoe on behalf of the Wellcome Sanger Institute COVID-19 Surveillance Team |
| EPI_ISL_3125537 | Berkshire and Surrey Pathology Services Lighthouse Laboratory | Wellcome Sanger Institute for the COVID-19 Genomics UK (COG-UK) Consortium | Berkshire and Surrey Pathology Services Lighthouse Laboratory and Alex Alderton, Roberto Amato, Jeffrey Barrett, Sonia Goncalves, Ewan Harrison, David K. Jackson, Ian Johnston, Dominic Kwiatkowski, Cordelia Langford, John Sillitoe on behalf of the Wellcome Sanger Institute COVID-19 Surveillance Team |
| EPI_ISL_3125538 | Lighthouse Lab in Milton Keynes | Wellcome Sanger Institute for the COVID-19 Genomics UK (COG-UK) Consortium | The Lighthouse Lab in Milton Keynes and Alex Alderton, Roberto Amato, Jeffrey Barrett, Sonia Goncalves, Ewan Harrison, David K. Jackson, Ian Johnston, Dominic Kwiatkowski, Cordelia Langford, John Sillitoe on behalf of the Wellcome Sanger Institute COVID-19 Surveillance Team |
| EPI_ISL_3125539, EPI_ISL_3125540, EPI_ISL_3125541 | Berkshire and Surrey Pathology Services Lighthouse Laboratory | Wellcome Sanger Institute for the COVID-19 Genomics UK (COG-UK) Consortium | Berkshire and Surrey Pathology Services Lighthouse Laboratory and Alex Alderton, Roberto Amato, Jeffrey Barrett, Sonia Goncalves, Ewan Harrison, David K. Jackson, Ian Johnston, Dominic Kwiatkowski, Cordelia Langford, John Sillitoe on behalf of the Wellcome Sanger Institute COVID-19 Surveillance Team |
| EPI_ISL_3125542 | Lighthouse Lab in Alderley Park | Wellcome Sanger Institute for the COVID-19 Genomics UK (COG-UK) Consortium | Jacquelyn Wynn, Mairead Hyland, The Lighthouse Lab in Alderley Park and Alex Alderton, Roberto Amato, Jeffrey Barrett, Sonia Goncalves, Ewan Harrison, David K. Jackson, Ian Johnston, Dominic Kwiatkowski, Cordelia Langford, John Sillitoe on behalf of the Wellcome Sanger Institute COVID-19 Surveillance Team |
| EPI_ISL_3125543 | Lighthouse Lab in Milton Keynes | Wellcome Sanger Institute for the COVID-19 Genomics UK (COG-UK) Consortium | The Lighthouse Lab in Milton Keynes and Alex Alderton, Roberto Amato, Jeffrey Barrett, Sonia Goncalves, Ewan Harrison, David K. Jackson, Ian Johnston, Dominic Kwiatkowski, Cordelia Langford, John Sillitoe on behalf of the Wellcome Sanger Institute COVID-19 Surveillance Team |
| EPI_ISL_3125544, EPI_ISL_3125545, EPI_ISL_3125546, EPI_ISL_3125547, EPI_ISL_3125548 | Lighthouse Lab in Alderley Park | Wellcome Sanger Institute for the COVID-19 Genomics UK (COG-UK) Consortium | Jacquelyn Wynn, Mairead Hyland, The Lighthouse Lab in Alderley Park and Alex Alderton, Roberto Amato, Jeffrey Barrett, Sonia Goncalves, Ewan Harrison, David K. Jackson, Ian Johnston, Dominic Kwiatkowski, Cordelia Langford, John Sillitoe on behalf of the Wellcome Sanger Institute COVID-19 Surveillance Team |
| EPI_ISL_3125549 | Lighthouse Lab in Milton Keynes | Wellcome Sanger Institute for the COVID-19 Genomics UK (COG-UK) Consortium | The Lighthouse Lab in Milton Keynes and Alex Alderton, Roberto Amato, Jeffrey Barrett, Sonia Goncalves, Ewan Harrison, David K. Jackson, Ian Johnston, Dominic Kwiatkowski, Cordelia Langford, John Sillitoe on behalf of the Wellcome Sanger Institute COVID-19 Surveillance Team |
| EPI_ISL_3125550 | Berkshire and Surrey Pathology Services Lighthouse Laboratory | Wellcome Sanger Institute for the COVID-19 Genomics UK (COG-UK) Consortium | Berkshire and Surrey Pathology Services Lighthouse Laboratory and Alex Alderton, Roberto Amato, Jeffrey Barrett, Sonia Goncalves, Ewan Harrison, David K. Jackson, Ian Johnston, Dominic Kwiatkowski, Cordelia Langford, John Sillitoe on behalf of the Wellcome Sanger Institute COVID-19 Surveillance Team |
| EPI_ISL_3125551 | Lighthouse Lab in Milton Keynes | Wellcome Sanger Institute for the COVID-19 Genomics UK (COG-UK) Consortium | The Lighthouse Lab in Milton Keynes and Alex Alderton, Roberto Amato, Jeffrey Barrett, Sonia Goncalves, Ewan Harrison, David K. Jackson, Ian Johnston, Dominic Kwiatkowski, Cordelia Langford, John Sillitoe on behalf of the Wellcome Sanger Institute COVID-19 Surveillance Team |
| EPI_ISL_3125552 | Berkshire and Surrey Pathology Services Lighthouse Laboratory | Wellcome Sanger Institute for the COVID-19 Genomics UK (COG-UK) Consortium | Berkshire and Surrey Pathology Services Lighthouse Laboratory and Alex Alderton, Roberto Amato, Jeffrey Barrett, Sonia Goncalves, Ewan Harrison, David K. Jackson, Ian Johnston, Dominic Kwiatkowski, Cordelia Langford, John Sillitoe on behalf of the Wellcome Sanger Institute COVID-19 Surveillance Team |
| EPI_ISL_3125553, EPI_ISL_3125554, EPI_ISL_3125555 | Lighthouse Lab in Milton Keynes | Wellcome Sanger Institute for the COVID-19 Genomics UK (COG-UK) Consortium | The Lighthouse Lab in Milton Keynes and Alex Alderton, Roberto Amato, Jeffrey Barrett, Sonia Goncalves, Ewan Harrison, David K. Jackson, Ian Johnston, Dominic Kwiatkowski, Cordelia Langford, John Sillitoe on behalf of the Wellcome Sanger Institute COVID-19 Surveillance Team |
| EPI_ISL_3125556, EPI_ISL_3125557 | Lighthouse Lab in Alderley Park | Wellcome Sanger Institute for the COVID-19 Genomics UK (COG-UK) Consortium | Jacquelyn Wynn, Mairead Hyland, The Lighthouse Lab in Alderley Park and Alex Alderton, Roberto Amato, Jeffrey Barrett, Sonia Goncalves, Ewan Harrison, David K. Jackson, Ian Johnston, Dominic Kwiatkowski, Cordelia Langford, John Sillitoe on behalf of the Wellcome Sanger Institute COVID-19 Surveillance Team |
| EPI_ISL_3125558 | Lighthouse Lab in Milton Keynes | Wellcome Sanger Institute for the COVID-19 Genomics UK (COG-UK) Consortium | The Lighthouse Lab in Milton Keynes and Alex Alderton, Roberto Amato, Jeffrey Barrett, Sonia Goncalves, Ewan Harrison, David K. Jackson, Ian Johnston, Dominic Kwiatkowski, Cordelia Langford, John Sillitoe on behalf of the Wellcome Sanger Institute COVID-19 Surveillance Team |
| EPI_ISL_3125559, EPI_ISL_3125560, EPI_ISL_3125561, EPI_ISL_3125562 | Lighthouse Lab in Alderley Park | Wellcome Sanger Institute for the COVID-19 Genomics UK (COG-UK) Consortium | Jacquelyn Wynn, Mairead Hyland, The Lighthouse Lab in Alderley Park and Alex Alderton, Roberto Amato, Jeffrey Barrett, Sonia Goncalves, Ewan Harrison, David K. Jackson, Ian Johnston, Dominic Kwiatkowski, Cordelia Langford, John Sillitoe on behalf of the Wellcome Sanger Institute COVID-19 Surveillance Team |
| EPI_ISL_3125563 | Lighthouse Lab in Milton Keynes | Wellcome Sanger Institute for the COVID-19 Genomics UK (COG-UK) Consortium | The Lighthouse Lab in Milton Keynes and Alex Alderton, Roberto Amato, Jeffrey Barrett, Sonia Goncalves, Ewan Harrison, David K. Jackson, Ian Johnston, Dominic Kwiatkowski, Cordelia Langford, John Sillitoe on behalf of the Wellcome Sanger Institute COVID-19 Surveillance Team |
| EPI_ISL_3125564 | Lighthouse Lab in Alderley Park | Wellcome Sanger Institute for the COVID-19 Genomics UK (COG-UK) Consortium | Jacquelyn Wynn, Mairead Hyland, The Lighthouse Lab in Alderley Park and Alex Alderton, Roberto Amato, Jeffrey Barrett, Sonia Goncalves, Ewan Harrison, David K. Jackson, Ian Johnston, Dominic Kwiatkowski, Cordelia Langford, John Sillitoe on behalf of the Wellcome Sanger Institute COVID-19 Surveillance Team |
| EPI_ISL_3125565 | Lighthouse Lab in Milton Keynes | Wellcome Sanger Institute for the COVID-19 Genomics UK (COG-UK) Consortium | The Lighthouse Lab in Milton Keynes and Alex Alderton, Roberto Amato, Jeffrey Barrett, Sonia Goncalves, Ewan Harrison, David K. Jackson, Ian Johnston, Dominic Kwiatkowski, Cordelia Langford, John Sillitoe on behalf of the Wellcome Sanger Institute COVID-19 Surveillance Team |
| EPI_ISL_3125566 | Lighthouse Lab in Glasgow | Wellcome Sanger Institute for the COVID-19 Genomics UK (COG-UK) Consortium | Harper VanSteenhouse, Yumi Kasai, David Gray, Carol Clugston, Anna Dominiczak and Alex Alderton, Roberto Amato, Jeffrey Barrett, Sonia Goncalves, Ewan Harrison, David K. Jackson, Ian Johnston, Dominic Kwiatkowski, Cordelia Langford, John Sillitoe on behalf of the Wellcome Sanger Institute COVID-19 Surveillance Team |

| | | | |
|---|---|---|---|
| EPI_ISL_3125567, EPI_ISL_3125568, EPI_ISL_3125569, EPI_ISL_3125570, EPI_ISL_3125571 | Lighthouse Lab in Milton Keynes | Wellcome Sanger Institute for the COVID-19 Genomics UK (COG-UK) Consortium | The Lighthouse Lab in Milton Keynes and Alex Alderton, Roberto Amato, Jeffrey Barrett, Sonia Goncalves, Ewan Harrison, David K. Jackson, Ian Johnston, Dominic Kwiatkowski, Cordelia Langford, John Sillitoe on behalf of the Wellcome Sanger Institute COVID-19 Surveillance Team |
| EPI_ISL_3125572 | Berkshire and Surrey Pathology Services Lighthouse Laboratory | Wellcome Sanger Institute for the COVID-19 Genomics UK (COG-UK) Consortium | Berkshire and Surrey Pathology Services Lighthouse Laboratory and Alex Alderton, Roberto Amato, Jeffrey Barrett, Sonia Goncalves, Ewan Harrison, David K. Jackson, Ian Johnston, Dominic Kwiatkowski, Cordelia Langford, John Sillitoe on behalf of the Wellcome Sanger Institute COVID-19 Surveillance Team |
| EPI_ISL_3125573, EPI_ISL_3125574 | Lighthouse Lab in Milton Keynes | Wellcome Sanger Institute for the COVID-19 Genomics UK (COG-UK) Consortium | The Lighthouse Lab in Milton Keynes and Alex Alderton, Roberto Amato, Jeffrey Barrett, Sonia Goncalves, Ewan Harrison, David K. Jackson, Ian Johnston, Dominic Kwiatkowski, Cordelia Langford, John Sillitoe on behalf of the Wellcome Sanger Institute COVID-19 Surveillance Team |
| EPI_ISL_3125575 | Lighthouse Lab in Glasgow | Wellcome Sanger Institute for the COVID-19 Genomics UK (COG-UK) Consortium | Harper VanSteenhouse, Yumi Kasai, David Gray, Carol Clugston, Anna Dominiczak and Alex Alderton, Roberto Amato, Jeffrey Barrett, Sonia Goncalves, Ewan Harrison, David K. Jackson, Ian Johnston, Dominic Kwiatkowski, Cordelia Langford, John Sillitoe on behalf of the Wellcome Sanger Institute COVID-19 Surveillance Team |
| EPI_ISL_3125576, EPI_ISL_3125577 | Lighthouse Lab in Milton Keynes | Wellcome Sanger Institute for the COVID-19 Genomics UK (COG-UK) Consortium | The Lighthouse Lab in Milton Keynes and Alex Alderton, Roberto Amato, Jeffrey Barrett, Sonia Goncalves, Ewan Harrison, David K. Jackson, Ian Johnston, Dominic Kwiatkowski, Cordelia Langford, John Sillitoe on behalf of the Wellcome Sanger Institute COVID-19 Surveillance Team |
| EPI_ISL_3125578 | Berkshire and Surrey Pathology Services Lighthouse Laboratory | Wellcome Sanger Institute for the COVID-19 Genomics UK (COG-UK) Consortium | Berkshire and Surrey Pathology Services Lighthouse Laboratory and Alex Alderton, Roberto Amato, Jeffrey Barrett, Sonia Goncalves, Ewan Harrison, David K. Jackson, Ian Johnston, Dominic Kwiatkowski, Cordelia Langford, John Sillitoe on behalf of the Wellcome Sanger Institute COVID-19 Surveillance Team |
| EPI_ISL_3125579 | Lighthouse Lab in Milton Keynes | Wellcome Sanger Institute for the COVID-19 Genomics UK (COG-UK) Consortium | The Lighthouse Lab in Milton Keynes and Alex Alderton, Roberto Amato, Jeffrey Barrett, Sonia Goncalves, Ewan Harrison, David K. Jackson, Ian Johnston, Dominic Kwiatkowski, Cordelia Langford, John Sillitoe on behalf of the Wellcome Sanger Institute COVID-19 Surveillance Team |
| EPI_ISL_3125580 | Berkshire and Surrey Pathology Services Lighthouse Laboratory | Wellcome Sanger Institute for the COVID-19 Genomics UK (COG-UK) Consortium | Berkshire and Surrey Pathology Services Lighthouse Laboratory and Alex Alderton, Roberto Amato, Jeffrey Barrett, Sonia Goncalves, Ewan Harrison, David K. Jackson, Ian Johnston, Dominic Kwiatkowski, Cordelia Langford, John Sillitoe on behalf of the Wellcome Sanger Institute COVID-19 Surveillance Team |
| EPI_ISL_3125581, EPI_ISL_3125582, EPI_ISL_3125583, EPI_ISL_3125584 | Lighthouse Lab in Milton Keynes | Wellcome Sanger Institute for the COVID-19 Genomics UK (COG-UK) Consortium | The Lighthouse Lab in Milton Keynes and Alex Alderton, Roberto Amato, Jeffrey Barrett, Sonia Goncalves, Ewan Harrison, David K. Jackson, Ian Johnston, Dominic Kwiatkowski, Cordelia Langford, John Sillitoe on behalf of the Wellcome Sanger Institute COVID-19 Surveillance Team |
| EPI_ISL_3125585 | Lighthouse Lab in Glasgow | Wellcome Sanger Institute for the COVID-19 Genomics UK (COG-UK) Consortium | Harper VanSteenhouse, Yumi Kasai, David Gray, Carol Clugston, Anna Dominiczak and Alex Alderton, Roberto Amato, Jeffrey Barrett, Sonia Goncalves, Ewan Harrison, David K. Jackson, Ian Johnston, Dominic Kwiatkowski, Cordelia Langford, John Sillitoe on behalf of the Wellcome Sanger Institute COVID-19 Surveillance Team |
| EPI_ISL_3125586 | Lighthouse Lab in Milton Keynes | Wellcome Sanger Institute for the COVID-19 Genomics UK (COG-UK) Consortium | The Lighthouse Lab in Milton Keynes and Alex Alderton, Roberto Amato, Jeffrey Barrett, Sonia Goncalves, Ewan Harrison, David K. Jackson, Ian Johnston, Dominic Kwiatkowski, Cordelia Langford, John Sillitoe on behalf of the Wellcome Sanger Institute COVID-19 Surveillance Team |
| EPI_ISL_3125587 | Berkshire and Surrey Pathology Services Lighthouse Laboratory | Wellcome Sanger Institute for the COVID-19 Genomics UK (COG-UK) Consortium | Berkshire and Surrey Pathology Services Lighthouse Laboratory and Alex Alderton, Roberto Amato, Jeffrey Barrett, Sonia Goncalves, Ewan Harrison, David K. Jackson, Ian Johnston, Dominic Kwiatkowski, Cordelia Langford, John Sillitoe on behalf of the Wellcome Sanger Institute COVID-19 Surveillance Team |
| EPI_ISL_3125588, EPI_ISL_3125589 | Lighthouse Lab in Milton Keynes | Wellcome Sanger Institute for the COVID-19 Genomics UK (COG-UK) Consortium | The Lighthouse Lab in Milton Keynes and Alex Alderton, Roberto Amato, Jeffrey Barrett, Sonia Goncalves, Ewan Harrison, David K. Jackson, Ian Johnston, Dominic Kwiatkowski, Cordelia Langford, John Sillitoe on behalf of the Wellcome Sanger Institute COVID-19 Surveillance Team |
| EPI_ISL_3125590, EPI_ISL_3125591, EPI_ISL_3125592 | Berkshire and Surrey Pathology Services Lighthouse Laboratory | Wellcome Sanger Institute for the COVID-19 Genomics UK (COG-UK) Consortium | Berkshire and Surrey Pathology Services Lighthouse Laboratory and Alex Alderton, Roberto Amato, Jeffrey Barrett, Sonia Goncalves, Ewan Harrison, David K. Jackson, Ian Johnston, Dominic Kwiatkowski, Cordelia Langford, John Sillitoe on behalf of the Wellcome Sanger Institute COVID-19 Surveillance Team |
| EPI_ISL_3125593, EPI_ISL_3125594 | Lighthouse Lab in Milton Keynes | Wellcome Sanger Institute for the COVID-19 Genomics UK (COG-UK) Consortium | The Lighthouse Lab in Milton Keynes and Alex Alderton, Roberto Amato, Jeffrey Barrett, Sonia Goncalves, Ewan Harrison, David K. Jackson, Ian Johnston, Dominic Kwiatkowski, Cordelia Langford, John Sillitoe on behalf of the Wellcome Sanger Institute COVID-19 Surveillance Team |
| EPI_ISL_3125595, EPI_ISL_3125596, EPI_ISL_3125597 | Berkshire and Surrey Pathology Services Lighthouse Laboratory | Wellcome Sanger Institute for the COVID-19 Genomics UK (COG-UK) Consortium | Berkshire and Surrey Pathology Services Lighthouse Laboratory and Alex Alderton, Roberto Amato, Jeffrey Barrett, Sonia Goncalves, Ewan Harrison, David K. Jackson, Ian Johnston, Dominic Kwiatkowski, Cordelia Langford, John Sillitoe on behalf of the Wellcome Sanger Institute COVID-19 Surveillance Team |
| EPI_ISL_3125598 | Lighthouse Lab in Milton Keynes | Wellcome Sanger Institute for the COVID-19 Genomics UK (COG-UK) Consortium | The Lighthouse Lab in Milton Keynes and Alex Alderton, Roberto Amato, Jeffrey Barrett, Sonia Goncalves, Ewan Harrison, David K. Jackson, Ian Johnston, Dominic Kwiatkowski, Cordelia Langford, John Sillitoe on behalf of the Wellcome Sanger Institute COVID-19 Surveillance Team |
| EPI_ISL_3125599, EPI_ISL_3125600 | Berkshire and Surrey Pathology Services Lighthouse Laboratory | Wellcome Sanger Institute for the COVID-19 Genomics UK (COG-UK) Consortium | Berkshire and Surrey Pathology Services Lighthouse Laboratory and Alex Alderton, Roberto Amato, Jeffrey Barrett, Sonia Goncalves, Ewan Harrison, David K. Jackson, Ian Johnston, Dominic Kwiatkowski, Cordelia Langford, John Sillitoe on behalf of the Wellcome Sanger Institute COVID-19 Surveillance Team |
| EPI_ISL_3125601, EPI_ISL_3125602, EPI_ISL_3125603, EPI_ISL_3125604, EPI_ISL_3125605 | Lighthouse Lab in Milton Keynes | Wellcome Sanger Institute for the COVID-19 Genomics UK (COG-UK) Consortium | The Lighthouse Lab in Milton Keynes and Alex Alderton, Roberto Amato, Jeffrey Barrett, Sonia Goncalves, Ewan Harrison, David K. Jackson, Ian Johnston, Dominic Kwiatkowski, Cordelia Langford, John Sillitoe on behalf of the Wellcome Sanger Institute COVID-19 Surveillance Team |
| EPI_ISL_3125606, EPI_ISL_3125607 | Lighthouse Lab in Glasgow | Wellcome Sanger Institute for the COVID-19 Genomics UK (COG-UK) Consortium | Harper VanSteenhouse, Yumi Kasai, David Gray, Carol Clugston, Anna Dominiczak and Alex Alderton, Roberto Amato, Jeffrey Barrett, Sonia Goncalves, Ewan Harrison, David K. Jackson, Ian Johnston, Dominic Kwiatkowski, Cordelia Langford, John Sillitoe on behalf of the Wellcome Sanger Institute COVID-19 Surveillance Team |
| EPI_ISL_3125608, EPI_ISL_3125609 | Lighthouse Lab in Milton Keynes | Wellcome Sanger Institute for the COVID-19 Genomics UK (COG-UK) Consortium | The Lighthouse Lab in Milton Keynes and Alex Alderton, Roberto Amato, Jeffrey Barrett, Sonia Goncalves, Ewan Harrison, David K. Jackson, Ian Johnston, Dominic Kwiatkowski, Cordelia Langford, John Sillitoe on behalf of the Wellcome Sanger Institute COVID-19 Surveillance Team |
| EPI_ISL_3125610 | Berkshire and Surrey Pathology Services Lighthouse Laboratory | Wellcome Sanger Institute for the COVID-19 Genomics UK (COG-UK) Consortium | Berkshire and Surrey Pathology Services Lighthouse Laboratory and Alex Alderton, Roberto Amato, Jeffrey Barrett, Sonia Goncalves, Ewan Harrison, David K. Jackson, Ian Johnston, Dominic Kwiatkowski, Cordelia Langford, John Sillitoe on behalf of the Wellcome Sanger Institute COVID-19 Surveillance Team |
| EPI_ISL_3125611 | Lighthouse Lab in Glasgow | Wellcome Sanger Institute for the COVID-19 Genomics UK (COG-UK) Consortium | Harper VanSteenhouse, Yumi Kasai, David Gray, Carol Clugston, Anna Dominiczak and Alex Alderton, Roberto Amato, Jeffrey Barrett, Sonia Goncalves, Ewan Harrison, David K. Jackson, Ian Johnston, Dominic Kwiatkowski, Cordelia Langford, John Sillitoe on behalf of the Wellcome Sanger Institute COVID-19 Surveillance Team |
| EPI_ISL_3125612 | Berkshire and Surrey Pathology Services Lighthouse Laboratory | Wellcome Sanger Institute for the COVID-19 Genomics UK (COG-UK) Consortium | Berkshire and Surrey Pathology Services Lighthouse Laboratory and Alex Alderton, Roberto Amato, Jeffrey Barrett, Sonia Goncalves, Ewan Harrison, David K. Jackson, Ian Johnston, Dominic Kwiatkowski, Cordelia Langford, John Sillitoe on behalf of the Wellcome Sanger Institute COVID-19 Surveillance Team |
| EPI_ISL_3125613, EPI_ISL_3125614, EPI_ISL_3125615 | Lighthouse Lab in Milton Keynes | Wellcome Sanger Institute for the COVID-19 Genomics UK (COG-UK) Consortium | The Lighthouse Lab in Milton Keynes and Alex Alderton, Roberto Amato, Jeffrey Barrett, Sonia Goncalves, Ewan Harrison, David K. Jackson, Ian Johnston, Dominic Kwiatkowski, Cordelia Langford, John Sillitoe on behalf of the Wellcome Sanger Institute COVID-19 Surveillance Team |
| EPI_ISL_3125616 | Berkshire and Surrey Pathology Services Lighthouse Laboratory | Wellcome Sanger Institute for the COVID-19 Genomics UK (COG-UK) Consortium | Berkshire and Surrey Pathology Services Lighthouse Laboratory and Alex Alderton, Roberto Amato, Jeffrey Barrett, Sonia Goncalves, Ewan Harrison, David K. Jackson, Ian Johnston, Dominic Kwiatkowski, Cordelia Langford, John Sillitoe on behalf of the Wellcome Sanger Institute COVID-19 Surveillance Team |
| EPI_ISL_3125617 | Lighthouse Lab in Milton Keynes | Wellcome Sanger Institute for the COVID-19 Genomics UK (COG-UK) Consortium | The Lighthouse Lab in Milton Keynes and Alex Alderton, Roberto Amato, Jeffrey Barrett, Sonia Goncalves, Ewan Harrison, David K. Jackson, Ian Johnston, Dominic Kwiatkowski, Cordelia Langford, John Sillitoe on behalf of the Wellcome Sanger Institute COVID-19 Surveillance Team |
| EPI_ISL_3125618 | Lighthouse Lab in Glasgow | Wellcome Sanger Institute for the COVID-19 Genomics UK (COG-UK) Consortium | Harper VanSteenhouse, Yumi Kasai, David Gray, Carol Clugston, Anna Dominiczak and Alex Alderton, Roberto Amato, Jeffrey Barrett, Sonia Goncalves, Ewan Harrison, David K. Jackson, Ian Johnston, Dominic Kwiatkowski, Cordelia Langford, John Sillitoe on behalf of the Wellcome Sanger Institute COVID-19 Surveillance Team |
| EPI_ISL_3125619, EPI_ISL_3125620 | Berkshire and Surrey Pathology Services Lighthouse Laboratory | Wellcome Sanger Institute for the COVID-19 Genomics UK (COG-UK) Consortium | Berkshire and Surrey Pathology Services Lighthouse Laboratory and Alex Alderton, Roberto Amato, Jeffrey Barrett, Sonia Goncalves, Ewan Harrison, David K. Jackson, Ian Johnston, Dominic Kwiatkowski, Cordelia Langford, John Sillitoe on behalf of the Wellcome Sanger Institute COVID-19 Surveillance Team |
| EPI_ISL_3125621, EPI_ISL_3125622 | Lighthouse Lab in Glasgow | Wellcome Sanger Institute for the COVID-19 Genomics UK (COG-UK) Consortium | Harper VanSteenhouse, Yumi Kasai, David Gray, Carol Clugston, Anna Dominiczak and Alex Alderton, Roberto Amato, Jeffrey Barrett, Sonia Goncalves, Ewan Harrison, David K. Jackson, Ian Johnston, Dominic Kwiatkowski, Cordelia Langford, John Sillitoe on behalf of the Wellcome Sanger Institute COVID-19 Surveillance Team |
| EPI_ISL_3125623 | Berkshire and Surrey Pathology Services Lighthouse Laboratory | Wellcome Sanger Institute for the COVID-19 Genomics UK (COG-UK) Consortium | Berkshire and Surrey Pathology Services Lighthouse Laboratory and Alex Alderton, Roberto Amato, Jeffrey Barrett, Sonia Goncalves, Ewan Harrison, David K. Jackson, Ian Johnston, Dominic Kwiatkowski, Cordelia Langford, John Sillitoe on behalf of the Wellcome Sanger Institute COVID-19 Surveillance Team |
| EPI_ISL_3125624 | Lighthouse Lab in Milton Keynes | Wellcome Sanger Institute for the COVID-19 Genomics UK (COG-UK) Consortium | The Lighthouse Lab in Milton Keynes and Alex Alderton, Roberto Amato, Jeffrey Barrett, Sonia Goncalves, Ewan Harrison, David K. Jackson, Ian Johnston, Dominic Kwiatkowski, Cordelia Langford, John Sillitoe on behalf of the Wellcome Sanger Institute COVID-19 Surveillance Team |
| EPI_ISL_3125625 | Berkshire and Surrey Pathology Services Lighthouse Laboratory | Wellcome Sanger Institute for the COVID-19 Genomics UK (COG-UK) Consortium | Berkshire and Surrey Pathology Services Lighthouse Laboratory and Alex Alderton, Roberto Amato, Jeffrey Barrett, Sonia Goncalves, Ewan Harrison, David K. Jackson, Ian Johnston, Dominic Kwiatkowski, Cordelia Langford, John Sillitoe on behalf of the Wellcome Sanger Institute COVID-19 Surveillance Team |
| EPI_ISL_3125626, EPI_ISL_3125627 | Lighthouse Lab in Milton Keynes | Wellcome Sanger Institute for the COVID-19 Genomics UK (COG-UK) Consortium | The Lighthouse Lab in Milton Keynes and Alex Alderton, Roberto Amato, Jeffrey Barrett, Sonia Goncalves, Ewan Harrison, David K. Jackson, Ian Johnston, Dominic Kwiatkowski, Cordelia Langford, John Sillitoe on behalf of the Wellcome Sanger Institute COVID-19 Surveillance Team |
| EPI_ISL_3125628 | Berkshire and Surrey Pathology Services Lighthouse Laboratory | Wellcome Sanger Institute for the COVID-19 Genomics UK (COG-UK) Consortium | Berkshire and Surrey Pathology Services Lighthouse Laboratory and Alex Alderton, Roberto Amato, Jeffrey Barrett, Sonia Goncalves, Ewan Harrison, David K. Jackson, Ian Johnston, Dominic Kwiatkowski, Cordelia Langford, John Sillitoe on behalf of the Wellcome Sanger Institute COVID-19 Surveillance Team |
| EPI_ISL_3125629, EPI_ISL_3125630, EPI_ISL_3125631, EPI_ISL_3125632, EPI_ISL_3125633 | Lighthouse Lab in Milton Keynes | Wellcome Sanger Institute for the COVID-19 Genomics UK (COG-UK) Consortium | The Lighthouse Lab in Milton Keynes and Alex Alderton, Roberto Amato, Jeffrey Barrett, Sonia Goncalves, Ewan Harrison, David K. Jackson, Ian Johnston, Dominic Kwiatkowski, Cordelia Langford, John Sillitoe on behalf of the Wellcome Sanger Institute COVID-19 Surveillance Team |
| EPI_ISL_3125634 | Lighthouse Lab in Glasgow | Wellcome Sanger Institute for the COVID-19 Genomics UK (COG-UK) Consortium | Harper VanSteenhouse, Yumi Kasai, David Gray, Carol Clugston, Anna Dominiczak and Alex Alderton, Roberto Amato, Jeffrey Barrett, Sonia Goncalves, Ewan Harrison, David K. Jackson, Ian Johnston, Dominic Kwiatkowski, Cordelia Langford, John Sillitoe on behalf of the Wellcome Sanger Institute COVID-19 Surveillance Team |
| EPI_ISL_3125635 | Lighthouse Lab in Milton Keynes | Wellcome Sanger Institute for the COVID-19 Genomics UK (COG-UK) Consortium | The Lighthouse Lab in Milton Keynes and Alex Alderton, Roberto Amato, Jeffrey Barrett, Sonia Goncalves, Ewan Harrison, David K. Jackson, Ian Johnston, Dominic Kwiatkowski, Cordelia Langford, John Sillitoe on behalf of the Wellcome Sanger Institute COVID-19 Surveillance Team |
| EPI_ISL_3125636 | Lighthouse Lab in Glasgow | Wellcome Sanger Institute for the COVID-19 Genomics UK (COG-UK) Consortium | Harper VanSteenhouse, Yumi Kasai, David Gray, Carol Clugston, Anna Dominiczak and Alex Alderton, Roberto Amato, Jeffrey Barrett, Sonia Goncalves, Ewan Harrison, David K. Jackson, Ian Johnston, Dominic Kwiatkowski, Cordelia Langford, John Sillitoe on behalf of the Wellcome Sanger Institute COVID-19 Surveillance Team |
| EPI_ISL_3125637 | Lighthouse Lab in Milton Keynes | Wellcome Sanger Institute for the COVID-19 Genomics UK (COG-UK) | The Lighthouse Lab in Milton Keynes and Alex Alderton, Roberto Amato, Jeffrey Barrett, Sonia Goncalves, Ewan Harrison, David K. Jackson, Ian Johnston, Dominic Kwiatkowski, Cordelia Langford, John Sillitoe on behalf of the Wellcome Sanger |

| | | | |
|---|---|---|---|
| | | Consortium | Institute COVID-19 Surveillance Team |
| EPI_ISL_3125638, EPI_ISL_3125639 | Lighthouse Lab in Glasgow | Wellcome Sanger Institute for the COVID-19 Genomics UK (COG-UK) Consortium | Harper VanSteenhouse, Yumi Kasai, David Gray, Carol Clugston, Anna Dominiczak and Alex Alderton, Roberto Amato, Jeffrey Barrett, Sonia Goncalves, Ewan Harrison, David K. Jackson, Ian Johnston, Dominic Kwiatkowski, Cordelia Langford, John Sillitoe on behalf of the Wellcome Sanger Institute COVID-19 Surveillance Team |
| EPI_ISL_3125640, EPI_ISL_3125641 | Lighthouse Lab in Milton Keynes | Wellcome Sanger Institute for the COVID-19 Genomics UK (COG-UK) Consortium | The Lighthouse Lab in Milton Keynes and Alex Alderton, Roberto Amato, Jeffrey Barrett, Sonia Goncalves, Ewan Harrison, David K. Jackson, Ian Johnston, Dominic Kwiatkowski, Cordelia Langford, John Sillitoe on behalf of the Wellcome Sanger Institute COVID-19 Surveillance Team |
| EPI_ISL_3125642 | Lighthouse Lab in Glasgow | Wellcome Sanger Institute for the COVID-19 Genomics UK (COG-UK) Consortium | Harper VanSteenhouse, Yumi Kasai, David Gray, Carol Clugston, Anna Dominiczak and Alex Alderton, Roberto Amato, Jeffrey Barrett, Sonia Goncalves, Ewan Harrison, David K. Jackson, Ian Johnston, Dominic Kwiatkowski, Cordelia Langford, John Sillitoe on behalf of the Wellcome Sanger Institute COVID-19 Surveillance Team |
| EPI_ISL_3125643, EPI_ISL_3125644, EPI_ISL_3125645, EPI_ISL_3125646 | Lighthouse Lab in Milton Keynes | Wellcome Sanger Institute for the COVID-19 Genomics UK (COG-UK) Consortium | The Lighthouse Lab in Milton Keynes and Alex Alderton, Roberto Amato, Jeffrey Barrett, Sonia Goncalves, Ewan Harrison, David K. Jackson, Ian Johnston, Dominic Kwiatkowski, Cordelia Langford, John Sillitoe on behalf of the Wellcome Sanger Institute COVID-19 Surveillance Team |
| EPI_ISL_3125647 | Lighthouse Lab in Glasgow | Wellcome Sanger Institute for the COVID-19 Genomics UK (COG-UK) Consortium | Harper VanSteenhouse, Yumi Kasai, David Gray, Carol Clugston, Anna Dominiczak and Alex Alderton, Roberto Amato, Jeffrey Barrett, Sonia Goncalves, Ewan Harrison, David K. Jackson, Ian Johnston, Dominic Kwiatkowski, Cordelia Langford, John Sillitoe on behalf of the Wellcome Sanger Institute COVID-19 Surveillance Team |
| EPI_ISL_3125648, EPI_ISL_3125649 | Lighthouse Lab in Milton Keynes | Wellcome Sanger Institute for the COVID-19 Genomics UK (COG-UK) Consortium | The Lighthouse Lab in Milton Keynes and Alex Alderton, Roberto Amato, Jeffrey Barrett, Sonia Goncalves, Ewan Harrison, David K. Jackson, Ian Johnston, Dominic Kwiatkowski, Cordelia Langford, John Sillitoe on behalf of the Wellcome Sanger Institute COVID-19 Surveillance Team |
| EPI_ISL_3125650 | Berkshire and Surrey Pathology Services Lighthouse Laboratory | Wellcome Sanger Institute for the COVID-19 Genomics UK (COG-UK) Consortium | Berkshire and Surrey Pathology Services Lighthouse Laboratory and Alex Alderton, Roberto Amato, Jeffrey Barrett, Sonia Goncalves, Ewan Harrison, David K. Jackson, Ian Johnston, Dominic Kwiatkowski, Cordelia Langford, John Sillitoe on behalf of the Wellcome Sanger Institute COVID-19 Surveillance Team |
| EPI_ISL_3125651, EPI_ISL_3125652 | Lighthouse Lab in Milton Keynes | Wellcome Sanger Institute for the COVID-19 Genomics UK (COG-UK) Consortium | The Lighthouse Lab in Milton Keynes and Alex Alderton, Roberto Amato, Jeffrey Barrett, Sonia Goncalves, Ewan Harrison, David K. Jackson, Ian Johnston, Dominic Kwiatkowski, Cordelia Langford, John Sillitoe on behalf of the Wellcome Sanger Institute COVID-19 Surveillance Team |
| EPI_ISL_3125653, EPI_ISL_3125654 | Berkshire and Surrey Pathology Services Lighthouse Laboratory | Wellcome Sanger Institute for the COVID-19 Genomics UK (COG-UK) Consortium | Berkshire and Surrey Pathology Services Lighthouse Laboratory and Alex Alderton, Roberto Amato, Jeffrey Barrett, Sonia Goncalves, Ewan Harrison, David K. Jackson, Ian Johnston, Dominic Kwiatkowski, Cordelia Langford, John Sillitoe on behalf of the Wellcome Sanger Institute COVID-19 Surveillance Team |
| EPI_ISL_3125655, EPI_ISL_3125656 | Lighthouse Lab in Milton Keynes | Wellcome Sanger Institute for the COVID-19 Genomics UK (COG-UK) Consortium | The Lighthouse Lab in Milton Keynes and Alex Alderton, Roberto Amato, Jeffrey Barrett, Sonia Goncalves, Ewan Harrison, David K. Jackson, Ian Johnston, Dominic Kwiatkowski, Cordelia Langford, John Sillitoe on behalf of the Wellcome Sanger Institute COVID-19 Surveillance Team |
| EPI_ISL_3125657 | Berkshire and Surrey Pathology Services Lighthouse Laboratory | Wellcome Sanger Institute for the COVID-19 Genomics UK (COG-UK) Consortium | Berkshire and Surrey Pathology Services Lighthouse Laboratory and Alex Alderton, Roberto Amato, Jeffrey Barrett, Sonia Goncalves, Ewan Harrison, David K. Jackson, Ian Johnston, Dominic Kwiatkowski, Cordelia Langford, John Sillitoe on behalf of the Wellcome Sanger Institute COVID-19 Surveillance Team |
| EPI_ISL_3125658, EPI_ISL_3125659 | Lighthouse Lab in Milton Keynes | Wellcome Sanger Institute for the COVID-19 Genomics UK (COG-UK) Consortium | The Lighthouse Lab in Milton Keynes and Alex Alderton, Roberto Amato, Jeffrey Barrett, Sonia Goncalves, Ewan Harrison, David K. Jackson, Ian Johnston, Dominic Kwiatkowski, Cordelia Langford, John Sillitoe on behalf of the Wellcome Sanger Institute COVID-19 Surveillance Team |
| EPI_ISL_3125660 | Berkshire and Surrey Pathology Services Lighthouse Laboratory | Wellcome Sanger Institute for the COVID-19 Genomics UK (COG-UK) Consortium | Berkshire and Surrey Pathology Services Lighthouse Laboratory and Alex Alderton, Roberto Amato, Jeffrey Barrett, Sonia Goncalves, Ewan Harrison, David K. Jackson, Ian Johnston, Dominic Kwiatkowski, Cordelia Langford, John Sillitoe on behalf of the Wellcome Sanger Institute COVID-19 Surveillance Team |
| EPI_ISL_3125661 | Lighthouse Lab in Milton Keynes | Wellcome Sanger Institute for the COVID-19 Genomics UK (COG-UK) Consortium | The Lighthouse Lab in Milton Keynes and Alex Alderton, Roberto Amato, Jeffrey Barrett, Sonia Goncalves, Ewan Harrison, David K. Jackson, Ian Johnston, Dominic Kwiatkowski, Cordelia Langford, John Sillitoe on behalf of the Wellcome Sanger Institute COVID-19 Surveillance Team |
| EPI_ISL_3125662 | Berkshire and Surrey Pathology Services Lighthouse Laboratory | Wellcome Sanger Institute for the COVID-19 Genomics UK (COG-UK) Consortium | Berkshire and Surrey Pathology Services Lighthouse Laboratory and Alex Alderton, Roberto Amato, Jeffrey Barrett, Sonia Goncalves, Ewan Harrison, David K. Jackson, Ian Johnston, Dominic Kwiatkowski, Cordelia Langford, John Sillitoe on behalf of the Wellcome Sanger Institute COVID-19 Surveillance Team |
| EPI_ISL_3125663 | Lighthouse Lab in Milton Keynes | Wellcome Sanger Institute for the COVID-19 Genomics UK (COG-UK) Consortium | The Lighthouse Lab in Milton Keynes and Alex Alderton, Roberto Amato, Jeffrey Barrett, Sonia Goncalves, Ewan Harrison, David K. Jackson, Ian Johnston, Dominic Kwiatkowski, Cordelia Langford, John Sillitoe on behalf of the Wellcome Sanger Institute COVID-19 Surveillance Team |
| EPI_ISL_3125664 | Berkshire and Surrey Pathology Services Lighthouse Laboratory | Wellcome Sanger Institute for the COVID-19 Genomics UK (COG-UK) Consortium | Berkshire and Surrey Pathology Services Lighthouse Laboratory and Alex Alderton, Roberto Amato, Jeffrey Barrett, Sonia Goncalves, Ewan Harrison, David K. Jackson, Ian Johnston, Dominic Kwiatkowski, Cordelia Langford, John Sillitoe on behalf of the Wellcome Sanger Institute COVID-19 Surveillance Team |
| EPI_ISL_3125665, EPI_ISL_3125666 | Lighthouse Lab in Milton Keynes | Wellcome Sanger Institute for the COVID-19 Genomics UK (COG-UK) Consortium | The Lighthouse Lab in Milton Keynes and Alex Alderton, Roberto Amato, Jeffrey Barrett, Sonia Goncalves, Ewan Harrison, David K. Jackson, Ian Johnston, Dominic Kwiatkowski, Cordelia Langford, John Sillitoe on behalf of the Wellcome Sanger Institute COVID-19 Surveillance Team |
| EPI_ISL_3125667 | Lighthouse Lab in Glasgow | Wellcome Sanger Institute for the COVID-19 Genomics UK (COG-UK) Consortium | Harper VanSteenhouse, Yumi Kasai, David Gray, Carol Clugston, Anna Dominiczak and Alex Alderton, Roberto Amato, Jeffrey Barrett, Sonia Goncalves, Ewan Harrison, David K. Jackson, Ian Johnston, Dominic Kwiatkowski, Cordelia Langford, John Sillitoe on behalf of the Wellcome Sanger Institute COVID-19 Surveillance Team |
| EPI_ISL_3125668, EPI_ISL_3125669, EPI_ISL_3125670, EPI_ISL_3125671 | Lighthouse Lab in Milton Keynes | Wellcome Sanger Institute for the COVID-19 Genomics UK (COG-UK) Consortium | The Lighthouse Lab in Milton Keynes and Alex Alderton, Roberto Amato, Jeffrey Barrett, Sonia Goncalves, Ewan Harrison, David K. Jackson, Ian Johnston, Dominic Kwiatkowski, Cordelia Langford, John Sillitoe on behalf of the Wellcome Sanger Institute COVID-19 Surveillance Team |
| EPI_ISL_3125672 | Berkshire and Surrey Pathology Services Lighthouse Laboratory | Wellcome Sanger Institute for the COVID-19 Genomics UK (COG-UK) Consortium | Berkshire and Surrey Pathology Services Lighthouse Laboratory and Alex Alderton, Roberto Amato, Jeffrey Barrett, Sonia Goncalves, Ewan Harrison, David K. Jackson, Ian Johnston, Dominic Kwiatkowski, Cordelia Langford, John Sillitoe on behalf of the Wellcome Sanger Institute COVID-19 Surveillance Team |
| EPI_ISL_3125673, EPI_ISL_3125674, EPI_ISL_3125675, EPI_ISL_3125676, EPI_ISL_3125677, EPI_ISL_3125678 | Lighthouse Lab in Milton Keynes | Wellcome Sanger Institute for the COVID-19 Genomics UK (COG-UK) Consortium | The Lighthouse Lab in Milton Keynes and Alex Alderton, Roberto Amato, Jeffrey Barrett, Sonia Goncalves, Ewan Harrison, David K. Jackson, Ian Johnston, Dominic Kwiatkowski, Cordelia Langford, John Sillitoe on behalf of the Wellcome Sanger Institute COVID-19 Surveillance Team |
| EPI_ISL_3125679 | Berkshire and Surrey Pathology Services Lighthouse Laboratory | Wellcome Sanger Institute for the COVID-19 Genomics UK (COG-UK) Consortium | Berkshire and Surrey Pathology Services Lighthouse Laboratory and Alex Alderton, Roberto Amato, Jeffrey Barrett, Sonia Goncalves, Ewan Harrison, David K. Jackson, Ian Johnston, Dominic Kwiatkowski, Cordelia Langford, John Sillitoe on behalf of the Wellcome Sanger Institute COVID-19 Surveillance Team |
| EPI_ISL_3125680 | Lighthouse Lab in Glasgow | Wellcome Sanger Institute for the COVID-19 Genomics UK (COG-UK) Consortium | Harper VanSteenhouse, Yumi Kasai, David Gray, Carol Clugston, Anna Dominiczak and Alex Alderton, Roberto Amato, Jeffrey Barrett, Sonia Goncalves, Ewan Harrison, David K. Jackson, Ian Johnston, Dominic Kwiatkowski, Cordelia Langford, John Sillitoe on behalf of the Wellcome Sanger Institute COVID-19 Surveillance Team |
| EPI_ISL_3125681, EPI_ISL_3125682 | Lighthouse Lab in Milton Keynes | Wellcome Sanger Institute for the COVID-19 Genomics UK (COG-UK) Consortium | The Lighthouse Lab in Milton Keynes and Alex Alderton, Roberto Amato, Jeffrey Barrett, Sonia Goncalves, Ewan Harrison, David K. Jackson, Ian Johnston, Dominic Kwiatkowski, Cordelia Langford, John Sillitoe on behalf of the Wellcome Sanger Institute COVID-19 Surveillance Team |
| EPI_ISL_3125683, EPI_ISL_3125684 | Berkshire and Surrey Pathology Services Lighthouse Laboratory | Wellcome Sanger Institute for the COVID-19 Genomics UK (COG-UK) Consortium | Berkshire and Surrey Pathology Services Lighthouse Laboratory and Alex Alderton, Roberto Amato, Jeffrey Barrett, Sonia Goncalves, Ewan Harrison, David K. Jackson, Ian Johnston, Dominic Kwiatkowski, Cordelia Langford, John Sillitoe on behalf of the Wellcome Sanger Institute COVID-19 Surveillance Team |
| EPI_ISL_3125685 | Lighthouse Lab in Milton Keynes | Wellcome Sanger Institute for the COVID-19 Genomics UK (COG-UK) Consortium | The Lighthouse Lab in Milton Keynes and Alex Alderton, Roberto Amato, Jeffrey Barrett, Sonia Goncalves, Ewan Harrison, David K. Jackson, Ian Johnston, Dominic Kwiatkowski, Cordelia Langford, John Sillitoe on behalf of the Wellcome Sanger Institute COVID-19 Surveillance Team |
| EPI_ISL_3125686 | Berkshire and Surrey Pathology Services Lighthouse Laboratory | Wellcome Sanger Institute for the COVID-19 Genomics UK (COG-UK) Consortium | Berkshire and Surrey Pathology Services Lighthouse Laboratory and Alex Alderton, Roberto Amato, Jeffrey Barrett, Sonia Goncalves, Ewan Harrison, David K. Jackson, Ian Johnston, Dominic Kwiatkowski, Cordelia Langford, John Sillitoe on behalf of the Wellcome Sanger Institute COVID-19 Surveillance Team |
| EPI_ISL_3125687, EPI_ISL_3125688, EPI_ISL_3125689 | Lighthouse Lab in Milton Keynes | Wellcome Sanger Institute for the COVID-19 Genomics UK (COG-UK) Consortium | The Lighthouse Lab in Milton Keynes and Alex Alderton, Roberto Amato, Jeffrey Barrett, Sonia Goncalves, Ewan Harrison, David K. Jackson, Ian Johnston, Dominic Kwiatkowski, Cordelia Langford, John Sillitoe on behalf of the Wellcome Sanger Institute COVID-19 Surveillance Team |
| EPI_ISL_3125690, EPI_ISL_3125691 | Berkshire and Surrey Pathology Services Lighthouse Laboratory | Wellcome Sanger Institute for the COVID-19 Genomics UK (COG-UK) Consortium | Berkshire and Surrey Pathology Services Lighthouse Laboratory and Alex Alderton, Roberto Amato, Jeffrey Barrett, Sonia Goncalves, Ewan Harrison, David K. Jackson, Ian Johnston, Dominic Kwiatkowski, Cordelia Langford, John Sillitoe on behalf of the Wellcome Sanger Institute COVID-19 Surveillance Team |
| EPI_ISL_3125692 | Lighthouse Lab in Milton Keynes | Wellcome Sanger Institute for the COVID-19 Genomics UK (COG-UK) Consortium | The Lighthouse Lab in Milton Keynes and Alex Alderton, Roberto Amato, Jeffrey Barrett, Sonia Goncalves, Ewan Harrison, David K. Jackson, Ian Johnston, Dominic Kwiatkowski, Cordelia Langford, John Sillitoe on behalf of the Wellcome Sanger Institute COVID-19 Surveillance Team |
| EPI_ISL_3125693 | Berkshire and Surrey Pathology Services Lighthouse Laboratory | Wellcome Sanger Institute for the COVID-19 Genomics UK (COG-UK) Consortium | Berkshire and Surrey Pathology Services Lighthouse Laboratory and Alex Alderton, Roberto Amato, Jeffrey Barrett, Sonia Goncalves, Ewan Harrison, David K. Jackson, Ian Johnston, Dominic Kwiatkowski, Cordelia Langford, John Sillitoe on behalf of the Wellcome Sanger Institute COVID-19 Surveillance Team |
| EPI_ISL_3125694 | Lighthouse Lab in Milton Keynes | Wellcome Sanger Institute for the COVID-19 Genomics UK (COG-UK) Consortium | The Lighthouse Lab in Milton Keynes and Alex Alderton, Roberto Amato, Jeffrey Barrett, Sonia Goncalves, Ewan Harrison, David K. Jackson, Ian Johnston, Dominic Kwiatkowski, Cordelia Langford, John Sillitoe on behalf of the Wellcome Sanger Institute COVID-19 Surveillance Team |
| EPI_ISL_3125695, EPI_ISL_3125696, EPI_ISL_3125697 | Berkshire and Surrey Pathology Services Lighthouse Laboratory | Wellcome Sanger Institute for the COVID-19 Genomics UK (COG-UK) Consortium | Berkshire and Surrey Pathology Services Lighthouse Laboratory and Alex Alderton, Roberto Amato, Jeffrey Barrett, Sonia Goncalves, Ewan Harrison, David K. Jackson, Ian Johnston, Dominic Kwiatkowski, Cordelia Langford, John Sillitoe on behalf of the Wellcome Sanger Institute COVID-19 Surveillance Team |
| EPI_ISL_3125698, EPI_ISL_3125699 | Lighthouse Lab in Milton Keynes | Wellcome Sanger Institute for the COVID-19 Genomics UK (COG-UK) Consortium | The Lighthouse Lab in Milton Keynes and Alex Alderton, Roberto Amato, Jeffrey Barrett, Sonia Goncalves, Ewan Harrison, David K. Jackson, Ian Johnston, Dominic Kwiatkowski, Cordelia Langford, John Sillitoe on behalf of the Wellcome Sanger Institute COVID-19 Surveillance Team |
| EPI_ISL_3125700, EPI_ISL_3125701 | Berkshire and Surrey Pathology Services Lighthouse Laboratory | Wellcome Sanger Institute for the COVID-19 Genomics UK (COG-UK) Consortium | Berkshire and Surrey Pathology Services Lighthouse Laboratory and Alex Alderton, Roberto Amato, Jeffrey Barrett, Sonia Goncalves, Ewan Harrison, David K. Jackson, Ian Johnston, Dominic Kwiatkowski, Cordelia Langford, John Sillitoe on behalf of the Wellcome Sanger Institute COVID-19 Surveillance Team |
| EPI_ISL_3125702, EPI_ISL_3125703 | Lighthouse Lab in Milton Keynes | Wellcome Sanger Institute for the COVID-19 Genomics UK (COG-UK) Consortium | The Lighthouse Lab in Milton Keynes and Alex Alderton, Roberto Amato, Jeffrey Barrett, Sonia Goncalves, Ewan Harrison, David K. Jackson, Ian Johnston, Dominic Kwiatkowski, Cordelia Langford, John Sillitoe on behalf of the Wellcome Sanger Institute COVID-19 Surveillance Team |
| EPI_ISL_3125704, EPI_ISL_3125705 | Berkshire and Surrey Pathology Services Lighthouse Laboratory | Wellcome Sanger Institute for the COVID-19 Genomics UK (COG-UK) Consortium | Berkshire and Surrey Pathology Services Lighthouse Laboratory and Alex Alderton, Roberto Amato, Jeffrey Barrett, Sonia Goncalves, Ewan Harrison, David K. Jackson, Ian Johnston, Dominic Kwiatkowski, Cordelia Langford, John Sillitoe on behalf of the Wellcome Sanger Institute COVID-19 Surveillance Team |
| EPI_ISL_3125706 | Lighthouse Lab in Milton Keynes | Wellcome Sanger Institute for the COVID-19 Genomics UK (COG-UK) Consortium | The Lighthouse Lab in Milton Keynes and Alex Alderton, Roberto Amato, Jeffrey Barrett, Sonia Goncalves, Ewan Harrison, David K. Jackson, Ian Johnston, Dominic Kwiatkowski, Cordelia Langford, John Sillitoe on behalf of the Wellcome Sanger Institute COVID-19 Surveillance Team |
| EPI_ISL_3125707 | Lighthouse Lab in Glasgow | Wellcome Sanger Institute for the COVID-19 Genomics UK (COG-UK) Consortium | Harper VanSteenhouse, Yumi Kasai, David Gray, Carol Clugston, Anna Dominiczak and Alex Alderton, Roberto Amato, Jeffrey Barrett, Sonia Goncalves, Ewan Harrison, David K. Jackson, Ian Johnston, Dominic Kwiatkowski, Cordelia Langford, John Sillitoe on behalf of the Wellcome Sanger Institute COVID-19 Surveillance Team |
| EPI_ISL_3125708 | Berkshire and Surrey Pathology Services Lighthouse Laboratory | Wellcome Sanger Institute for the COVID-19 Genomics UK (COG-UK) | Berkshire and Surrey Pathology Services Lighthouse Laboratory and Alex Alderton, Roberto Amato, Jeffrey Barrett, Sonia Goncalves, Ewan Harrison, David K. Jackson, Ian Johnston, Dominic Kwiatkowski, Cordelia Langford, John Sillitoe on behalf |

| | | | |
|---|---|---|---|
| | | Consortium | of the Wellcome Sanger Institute COVID-19 Surveillance Team |
| EPI_ISL_3125709, EPI_ISL_3125710, EPI_ISL_3125711 | Lighthouse Lab in Milton Keynes | Wellcome Sanger Institute for the COVID-19 Genomics UK (COG-UK) Consortium | The Lighthouse Lab in Milton Keynes and Alex Alderton, Roberto Amato, Jeffrey Barrett, Sonia Goncalves, Ewan Harrison, David K. Jackson, Ian Johnston, Dominic Kwiatkowski, Cordelia Langford, John Sillitoe on behalf of the Wellcome Sanger Institute COVID-19 Surveillance Team |
| EPI_ISL_3125712 | Lighthouse Lab in Glasgow | Wellcome Sanger Institute for the COVID-19 Genomics UK (COG-UK) Consortium | Harper VanSteenhouse, Yumi Kasai, David Gray, Carol Clugston, Anna Dominiczak and Alex Alderton, Roberto Amato, Jeffrey Barrett, Sonia Goncalves, Ewan Harrison, David K. Jackson, Ian Johnston, Dominic Kwiatkowski, Cordelia Langford, John Sillitoe on behalf of the Wellcome Sanger Institute COVID-19 Surveillance Team |
| EPI_ISL_3125713 | Lighthouse Lab in Milton Keynes | Wellcome Sanger Institute for the COVID-19 Genomics UK (COG-UK) Consortium | The Lighthouse Lab in Milton Keynes and Alex Alderton, Roberto Amato, Jeffrey Barrett, Sonia Goncalves, Ewan Harrison, David K. Jackson, Ian Johnston, Dominic Kwiatkowski, Cordelia Langford, John Sillitoe on behalf of the Wellcome Sanger Institute COVID-19 Surveillance Team |
| EPI_ISL_3125714 | Lighthouse Lab in Glasgow | Wellcome Sanger Institute for the COVID-19 Genomics UK (COG-UK) Consortium | Harper VanSteenhouse, Yumi Kasai, David Gray, Carol Clugston, Anna Dominiczak and Alex Alderton, Roberto Amato, Jeffrey Barrett, Sonia Goncalves, Ewan Harrison, David K. Jackson, Ian Johnston, Dominic Kwiatkowski, Cordelia Langford, John Sillitoe on behalf of the Wellcome Sanger Institute COVID-19 Surveillance Team |
| EPI_ISL_3125715 | Lighthouse Lab in Milton Keynes | Wellcome Sanger Institute for the COVID-19 Genomics UK (COG-UK) Consortium | The Lighthouse Lab in Milton Keynes and Alex Alderton, Roberto Amato, Jeffrey Barrett, Sonia Goncalves, Ewan Harrison, David K. Jackson, Ian Johnston, Dominic Kwiatkowski, Cordelia Langford, John Sillitoe on behalf of the Wellcome Sanger Institute COVID-19 Surveillance Team |
| EPI_ISL_3125716 | Lighthouse Lab in Glasgow | Wellcome Sanger Institute for the COVID-19 Genomics UK (COG-UK) Consortium | Harper VanSteenhouse, Yumi Kasai, David Gray, Carol Clugston, Anna Dominiczak and Alex Alderton, Roberto Amato, Jeffrey Barrett, Sonia Goncalves, Ewan Harrison, David K. Jackson, Ian Johnston, Dominic Kwiatkowski, Cordelia Langford, John Sillitoe on behalf of the Wellcome Sanger Institute COVID-19 Surveillance Team |
| EPI_ISL_3125717, EPI_ISL_3125718, EPI_ISL_3125719, EPI_ISL_3125720, EPI_ISL_3125721, EPI_ISL_3125722 | Lighthouse Lab in Milton Keynes | Wellcome Sanger Institute for the COVID-19 Genomics UK (COG-UK) Consortium | The Lighthouse Lab in Milton Keynes and Alex Alderton, Roberto Amato, Jeffrey Barrett, Sonia Goncalves, Ewan Harrison, David K. Jackson, Ian Johnston, Dominic Kwiatkowski, Cordelia Langford, John Sillitoe on behalf of the Wellcome Sanger Institute COVID-19 Surveillance Team |
| EPI_ISL_3125723, EPI_ISL_3125724 | Berkshire and Surrey Pathology Services Lighthouse Laboratory | Wellcome Sanger Institute for the COVID-19 Genomics UK (COG-UK) Consortium | Berkshire and Surrey Pathology Services Lighthouse Laboratory and Alex Alderton, Roberto Amato, Jeffrey Barrett, Sonia Goncalves, Ewan Harrison, David K. Jackson, Ian Johnston, Dominic Kwiatkowski, Cordelia Langford, John Sillitoe on behalf of the Wellcome Sanger Institute COVID-19 Surveillance Team |
| EPI_ISL_3125725 | Lighthouse Lab in Glasgow | Wellcome Sanger Institute for the COVID-19 Genomics UK (COG-UK) Consortium | Harper VanSteenhouse, Yumi Kasai, David Gray, Carol Clugston, Anna Dominiczak and Alex Alderton, Roberto Amato, Jeffrey Barrett, Sonia Goncalves, Ewan Harrison, David K. Jackson, Ian Johnston, Dominic Kwiatkowski, Cordelia Langford, John Sillitoe on behalf of the Wellcome Sanger Institute COVID-19 Surveillance Team |
| EPI_ISL_3125726, EPI_ISL_3125727 | Lighthouse Lab in Milton Keynes | Wellcome Sanger Institute for the COVID-19 Genomics UK (COG-UK) Consortium | The Lighthouse Lab in Milton Keynes and Alex Alderton, Roberto Amato, Jeffrey Barrett, Sonia Goncalves, Ewan Harrison, David K. Jackson, Ian Johnston, Dominic Kwiatkowski, Cordelia Langford, John Sillitoe on behalf of the Wellcome Sanger Institute COVID-19 Surveillance Team |
| EPI_ISL_3125728 | Berkshire and Surrey Pathology Services Lighthouse Laboratory | Wellcome Sanger Institute for the COVID-19 Genomics UK (COG-UK) Consortium | Berkshire and Surrey Pathology Services Lighthouse Laboratory and Alex Alderton, Roberto Amato, Jeffrey Barrett, Sonia Goncalves, Ewan Harrison, David K. Jackson, Ian Johnston, Dominic Kwiatkowski, Cordelia Langford, John Sillitoe on behalf of the Wellcome Sanger Institute COVID-19 Surveillance Team |
| EPI_ISL_3125729, EPI_ISL_3125730, EPI_ISL_3125731 | Lighthouse Lab in Milton Keynes | Wellcome Sanger Institute for the COVID-19 Genomics UK (COG-UK) Consortium | The Lighthouse Lab in Milton Keynes and Alex Alderton, Roberto Amato, Jeffrey Barrett, Sonia Goncalves, Ewan Harrison, David K. Jackson, Ian Johnston, Dominic Kwiatkowski, Cordelia Langford, John Sillitoe on behalf of the Wellcome Sanger Institute COVID-19 Surveillance Team |
| EPI_ISL_3125732, EPI_ISL_3125733, EPI_ISL_3125734 | Lighthouse Lab in Glasgow | Wellcome Sanger Institute for the COVID-19 Genomics UK (COG-UK) Consortium | Harper VanSteenhouse, Yumi Kasai, David Gray, Carol Clugston, Anna Dominiczak and Alex Alderton, Roberto Amato, Jeffrey Barrett, Sonia Goncalves, Ewan Harrison, David K. Jackson, Ian Johnston, Dominic Kwiatkowski, Cordelia Langford, John Sillitoe on behalf of the Wellcome Sanger Institute COVID-19 Surveillance Team |
| EPI_ISL_3125735 | Lighthouse Lab in Milton Keynes | Wellcome Sanger Institute for the COVID-19 Genomics UK (COG-UK) Consortium | The Lighthouse Lab in Milton Keynes and Alex Alderton, Roberto Amato, Jeffrey Barrett, Sonia Goncalves, Ewan Harrison, David K. Jackson, Ian Johnston, Dominic Kwiatkowski, Cordelia Langford, John Sillitoe on behalf of the Wellcome Sanger Institute COVID-19 Surveillance Team |
| EPI_ISL_3125736 | Berkshire and Surrey Pathology Services Lighthouse Laboratory | Wellcome Sanger Institute for the COVID-19 Genomics UK (COG-UK) Consortium | Berkshire and Surrey Pathology Services Lighthouse Laboratory and Alex Alderton, Roberto Amato, Jeffrey Barrett, Sonia Goncalves, Ewan Harrison, David K. Jackson, Ian Johnston, Dominic Kwiatkowski, Cordelia Langford, John Sillitoe on behalf of the Wellcome Sanger Institute COVID-19 Surveillance Team |
| EPI_ISL_3125737, EPI_ISL_3125738, EPI_ISL_3125739, EPI_ISL_3125740 | Lighthouse Lab in Milton Keynes | Wellcome Sanger Institute for the COVID-19 Genomics UK (COG-UK) Consortium | The Lighthouse Lab in Milton Keynes and Alex Alderton, Roberto Amato, Jeffrey Barrett, Sonia Goncalves, Ewan Harrison, David K. Jackson, Ian Johnston, Dominic Kwiatkowski, Cordelia Langford, John Sillitoe on behalf of the Wellcome Sanger Institute COVID-19 Surveillance Team |
| EPI_ISL_3125741 | Berkshire and Surrey Pathology Services Lighthouse Laboratory | Wellcome Sanger Institute for the COVID-19 Genomics UK (COG-UK) Consortium | Berkshire and Surrey Pathology Services Lighthouse Laboratory and Alex Alderton, Roberto Amato, Jeffrey Barrett, Sonia Goncalves, Ewan Harrison, David K. Jackson, Ian Johnston, Dominic Kwiatkowski, Cordelia Langford, John Sillitoe on behalf of the Wellcome Sanger Institute COVID-19 Surveillance Team |
| EPI_ISL_3125742 | Lighthouse Lab in Milton Keynes | Wellcome Sanger Institute for the COVID-19 Genomics UK (COG-UK) Consortium | The Lighthouse Lab in Milton Keynes and Alex Alderton, Roberto Amato, Jeffrey Barrett, Sonia Goncalves, Ewan Harrison, David K. Jackson, Ian Johnston, Dominic Kwiatkowski, Cordelia Langford, John Sillitoe on behalf of the Wellcome Sanger Institute COVID-19 Surveillance Team |
| EPI_ISL_3125743, EPI_ISL_3125744 | Berkshire and Surrey Pathology Services Lighthouse Laboratory | Wellcome Sanger Institute for the COVID-19 Genomics UK (COG-UK) Consortium | Berkshire and Surrey Pathology Services Lighthouse Laboratory and Alex Alderton, Roberto Amato, Jeffrey Barrett, Sonia Goncalves, Ewan Harrison, David K. Jackson, Ian Johnston, Dominic Kwiatkowski, Cordelia Langford, John Sillitoe on behalf of the Wellcome Sanger Institute COVID-19 Surveillance Team |
| EPI_ISL_3125745 | Lighthouse Lab in Glasgow | Wellcome Sanger Institute for the COVID-19 Genomics UK (COG-UK) Consortium | Harper VanSteenhouse, Yumi Kasai, David Gray, Carol Clugston, Anna Dominiczak and Alex Alderton, Roberto Amato, Jeffrey Barrett, Sonia Goncalves, Ewan Harrison, David K. Jackson, Ian Johnston, Dominic Kwiatkowski, Cordelia Langford, John Sillitoe on behalf of the Wellcome Sanger Institute COVID-19 Surveillance Team |
| EPI_ISL_3125746, EPI_ISL_3125747 | Lighthouse Lab in Milton Keynes | Wellcome Sanger Institute for the COVID-19 Genomics UK (COG-UK) Consortium | The Lighthouse Lab in Milton Keynes and Alex Alderton, Roberto Amato, Jeffrey Barrett, Sonia Goncalves, Ewan Harrison, David K. Jackson, Ian Johnston, Dominic Kwiatkowski, Cordelia Langford, John Sillitoe on behalf of the Wellcome Sanger Institute COVID-19 Surveillance Team |
| EPI_ISL_3125748 | Lighthouse Lab in Glasgow | Wellcome Sanger Institute for the COVID-19 Genomics UK (COG-UK) Consortium | Harper VanSteenhouse, Yumi Kasai, David Gray, Carol Clugston, Anna Dominiczak and Alex Alderton, Roberto Amato, Jeffrey Barrett, Sonia Goncalves, Ewan Harrison, David K. Jackson, Ian Johnston, Dominic Kwiatkowski, Cordelia Langford, John Sillitoe on behalf of the Wellcome Sanger Institute COVID-19 Surveillance Team |
| EPI_ISL_3125749 | Lighthouse Lab in Milton Keynes | Wellcome Sanger Institute for the COVID-19 Genomics UK (COG-UK) Consortium | The Lighthouse Lab in Milton Keynes and Alex Alderton, Roberto Amato, Jeffrey Barrett, Sonia Goncalves, Ewan Harrison, David K. Jackson, Ian Johnston, Dominic Kwiatkowski, Cordelia Langford, John Sillitoe on behalf of the Wellcome Sanger Institute COVID-19 Surveillance Team |
| EPI_ISL_3125750 | Berkshire and Surrey Pathology Services Lighthouse Laboratory | Wellcome Sanger Institute for the COVID-19 Genomics UK (COG-UK) Consortium | Berkshire and Surrey Pathology Services Lighthouse Laboratory and Alex Alderton, Roberto Amato, Jeffrey Barrett, Sonia Goncalves, Ewan Harrison, David K. Jackson, Ian Johnston, Dominic Kwiatkowski, Cordelia Langford, John Sillitoe on behalf of the Wellcome Sanger Institute COVID-19 Surveillance Team |
| EPI_ISL_3125751 | Lighthouse Lab in Milton Keynes | Wellcome Sanger Institute for the COVID-19 Genomics UK (COG-UK) Consortium | The Lighthouse Lab in Milton Keynes and Alex Alderton, Roberto Amato, Jeffrey Barrett, Sonia Goncalves, Ewan Harrison, David K. Jackson, Ian Johnston, Dominic Kwiatkowski, Cordelia Langford, John Sillitoe on behalf of the Wellcome Sanger Institute COVID-19 Surveillance Team |
| EPI_ISL_3125752, EPI_ISL_3125753 | Berkshire and Surrey Pathology Services Lighthouse Laboratory | Wellcome Sanger Institute for the COVID-19 Genomics UK (COG-UK) Consortium | Berkshire and Surrey Pathology Services Lighthouse Laboratory and Alex Alderton, Roberto Amato, Jeffrey Barrett, Sonia Goncalves, Ewan Harrison, David K. Jackson, Ian Johnston, Dominic Kwiatkowski, Cordelia Langford, John Sillitoe on behalf of the Wellcome Sanger Institute COVID-19 Surveillance Team |
| EPI_ISL_3125754 | Lighthouse Lab in Glasgow | Wellcome Sanger Institute for the COVID-19 Genomics UK (COG-UK) Consortium | Harper VanSteenhouse, Yumi Kasai, David Gray, Carol Clugston, Anna Dominiczak and Alex Alderton, Roberto Amato, Jeffrey Barrett, Sonia Goncalves, Ewan Harrison, David K. Jackson, Ian Johnston, Dominic Kwiatkowski, Cordelia Langford, John Sillitoe on behalf of the Wellcome Sanger Institute COVID-19 Surveillance Team |
| EPI_ISL_3125755, EPI_ISL_3125756, EPI_ISL_3125757, EPI_ISL_3125758, EPI_ISL_3125759, EPI_ISL_3125760, EPI_ISL_3125761, EPI_ISL_3125762 | Lighthouse Lab in Milton Keynes | Wellcome Sanger Institute for the COVID-19 Genomics UK (COG-UK) Consortium | The Lighthouse Lab in Milton Keynes and Alex Alderton, Roberto Amato, Jeffrey Barrett, Sonia Goncalves, Ewan Harrison, David K. Jackson, Ian Johnston, Dominic Kwiatkowski, Cordelia Langford, John Sillitoe on behalf of the Wellcome Sanger Institute COVID-19 Surveillance Team |
| EPI_ISL_3125763 | Berkshire and Surrey Pathology Services Lighthouse Laboratory | Wellcome Sanger Institute for the COVID-19 Genomics UK (COG-UK) Consortium | Berkshire and Surrey Pathology Services Lighthouse Laboratory and Alex Alderton, Roberto Amato, Jeffrey Barrett, Sonia Goncalves, Ewan Harrison, David K. Jackson, Ian Johnston, Dominic Kwiatkowski, Cordelia Langford, John Sillitoe on behalf of the Wellcome Sanger Institute COVID-19 Surveillance Team |
| EPI_ISL_3125764, EPI_ISL_3125765 | Lighthouse Lab in Milton Keynes | Wellcome Sanger Institute for the COVID-19 Genomics UK (COG-UK) Consortium | The Lighthouse Lab in Milton Keynes and Alex Alderton, Roberto Amato, Jeffrey Barrett, Sonia Goncalves, Ewan Harrison, David K. Jackson, Ian Johnston, Dominic Kwiatkowski, Cordelia Langford, John Sillitoe on behalf of the Wellcome Sanger Institute COVID-19 Surveillance Team |
| EPI_ISL_3125766 | Berkshire and Surrey Pathology Services Lighthouse Laboratory | Wellcome Sanger Institute for the COVID-19 Genomics UK (COG-UK) Consortium | Berkshire and Surrey Pathology Services Lighthouse Laboratory and Alex Alderton, Roberto Amato, Jeffrey Barrett, Sonia Goncalves, Ewan Harrison, David K. Jackson, Ian Johnston, Dominic Kwiatkowski, Cordelia Langford, John Sillitoe on behalf of the Wellcome Sanger Institute COVID-19 Surveillance Team |
| EPI_ISL_3125767 | Lighthouse Lab in Milton Keynes | Wellcome Sanger Institute for the COVID-19 Genomics UK (COG-UK) Consortium | The Lighthouse Lab in Milton Keynes and Alex Alderton, Roberto Amato, Jeffrey Barrett, Sonia Goncalves, Ewan Harrison, David K. Jackson, Ian Johnston, Dominic Kwiatkowski, Cordelia Langford, John Sillitoe on behalf of the Wellcome Sanger Institute COVID-19 Surveillance Team |
| EPI_ISL_3125768, EPI_ISL_3125769, EPI_ISL_3125770 | Lighthouse Lab in Glasgow | Wellcome Sanger Institute for the COVID-19 Genomics UK (COG-UK) Consortium | Harper VanSteenhouse, Yumi Kasai, David Gray, Carol Clugston, Anna Dominiczak and Alex Alderton, Roberto Amato, Jeffrey Barrett, Sonia Goncalves, Ewan Harrison, David K. Jackson, Ian Johnston, Dominic Kwiatkowski, Cordelia Langford, John Sillitoe on behalf of the Wellcome Sanger Institute COVID-19 Surveillance Team |
| EPI_ISL_3125771, EPI_ISL_3125772, EPI_ISL_3125773, EPI_ISL_3125774, EPI_ISL_3125775, EPI_ISL_3125776 | Lighthouse Lab in Milton Keynes | Wellcome Sanger Institute for the COVID-19 Genomics UK (COG-UK) Consortium | The Lighthouse Lab in Milton Keynes and Alex Alderton, Roberto Amato, Jeffrey Barrett, Sonia Goncalves, Ewan Harrison, David K. Jackson, Ian Johnston, Dominic Kwiatkowski, Cordelia Langford, John Sillitoe on behalf of the Wellcome Sanger Institute COVID-19 Surveillance Team |
| EPI_ISL_3125777 | Berkshire and Surrey Pathology Services Lighthouse Laboratory | Wellcome Sanger Institute for the COVID-19 Genomics UK (COG-UK) Consortium | Berkshire and Surrey Pathology Services Lighthouse Laboratory and Alex Alderton, Roberto Amato, Jeffrey Barrett, Sonia Goncalves, Ewan Harrison, David K. Jackson, Ian Johnston, Dominic Kwiatkowski, Cordelia Langford, John Sillitoe on behalf of the Wellcome Sanger Institute COVID-19 Surveillance Team |
| EPI_ISL_3125778 | Lighthouse Lab in Milton Keynes | Wellcome Sanger Institute for the COVID-19 Genomics UK (COG-UK) Consortium | The Lighthouse Lab in Milton Keynes and Alex Alderton, Roberto Amato, Jeffrey Barrett, Sonia Goncalves, Ewan Harrison, David K. Jackson, Ian Johnston, Dominic Kwiatkowski, Cordelia Langford, John Sillitoe on behalf of the Wellcome Sanger Institute COVID-19 Surveillance Team |
| EPI_ISL_3125779 | Berkshire and Surrey Pathology Services Lighthouse Laboratory | Wellcome Sanger Institute for the COVID-19 Genomics UK (COG-UK) | Berkshire and Surrey Pathology Services Lighthouse Laboratory and Alex Alderton, Roberto Amato, Jeffrey Barrett, Sonia Goncalves, Ewan Harrison, David K. Jackson, Ian Johnston, Dominic Kwiatkowski, Cordelia Langford, John Sillitoe on behalf |

| | | | |
|---|---|---|---|
| | | Consortium | of the Wellcome Sanger Institute COVID-19 Surveillance Team |
| EPI_ISL_3125780 | Lighthouse Lab in Milton Keynes | Wellcome Sanger Institute for the COVID-19 Genomics UK (COG-UK) Consortium | The Lighthouse Lab in Milton Keynes and Alex Alderton, Roberto Amato, Jeffrey Barrett, Sonia Goncalves, Ewan Harrison, David K. Jackson, Ian Johnston, Dominic Kwiatkowski, Cordelia Langford, John Sillitoe on behalf of the Wellcome Sanger Institute COVID-19 Surveillance Team |
| EPI_ISL_3125781 | Lighthouse Lab in Glasgow | Wellcome Sanger Institute for the COVID-19 Genomics UK (COG-UK) Consortium | Harper VanSteenhouse, Yumi Kasai, David Gray, Carol Clugston, Anna Dominiczak and Alex Alderton, Roberto Amato, Jeffrey Barrett, Sonia Goncalves, Ewan Harrison, David K. Jackson, Ian Johnston, Dominic Kwiatkowski, Cordelia Langford, John Sillitoe on behalf of the Wellcome Sanger Institute COVID-19 Surveillance Team |
| EPI_ISL_3125782 | Berkshire and Surrey Pathology Services Lighthouse Laboratory | Wellcome Sanger Institute for the COVID-19 Genomics UK (COG-UK) Consortium | Berkshire and Surrey Pathology Services Lighthouse Laboratory and Alex Alderton, Roberto Amato, Jeffrey Barrett, Sonia Goncalves, Ewan Harrison, David K. Jackson, Ian Johnston, Dominic Kwiatkowski, Cordelia Langford, John Sillitoe on behalf of the Wellcome Sanger Institute COVID-19 Surveillance Team |
| EPI_ISL_3125783 | Lighthouse Lab in Milton Keynes | Wellcome Sanger Institute for the COVID-19 Genomics UK (COG-UK) Consortium | The Lighthouse Lab in Milton Keynes and Alex Alderton, Roberto Amato, Jeffrey Barrett, Sonia Goncalves, Ewan Harrison, David K. Jackson, Ian Johnston, Dominic Kwiatkowski, Cordelia Langford, John Sillitoe on behalf of the Wellcome Sanger Institute COVID-19 Surveillance Team |
| EPI_ISL_3125784 | Lighthouse Lab in Glasgow | Wellcome Sanger Institute for the COVID-19 Genomics UK (COG-UK) Consortium | Harper VanSteenhouse, Yumi Kasai, David Gray, Carol Clugston, Anna Dominiczak and Alex Alderton, Roberto Amato, Jeffrey Barrett, Sonia Goncalves, Ewan Harrison, David K. Jackson, Ian Johnston, Dominic Kwiatkowski, Cordelia Langford, John Sillitoe on behalf of the Wellcome Sanger Institute COVID-19 Surveillance Team |
| EPI_ISL_3125785, EPI_ISL_3125786 | Lighthouse Lab in Milton Keynes | Wellcome Sanger Institute for the COVID-19 Genomics UK (COG-UK) Consortium | The Lighthouse Lab in Milton Keynes and Alex Alderton, Roberto Amato, Jeffrey Barrett, Sonia Goncalves, Ewan Harrison, David K. Jackson, Ian Johnston, Dominic Kwiatkowski, Cordelia Langford, John Sillitoe on behalf of the Wellcome Sanger Institute COVID-19 Surveillance Team |
| EPI_ISL_3125787 | Lighthouse Lab in Glasgow | Wellcome Sanger Institute for the COVID-19 Genomics UK (COG-UK) Consortium | Harper VanSteenhouse, Yumi Kasai, David Gray, Carol Clugston, Anna Dominiczak and Alex Alderton, Roberto Amato, Jeffrey Barrett, Sonia Goncalves, Ewan Harrison, David K. Jackson, Ian Johnston, Dominic Kwiatkowski, Cordelia Langford, John Sillitoe on behalf of the Wellcome Sanger Institute COVID-19 Surveillance Team |
| EPI_ISL_3125788 | Lighthouse Lab in Milton Keynes | Wellcome Sanger Institute for the COVID-19 Genomics UK (COG-UK) Consortium | The Lighthouse Lab in Milton Keynes and Alex Alderton, Roberto Amato, Jeffrey Barrett, Sonia Goncalves, Ewan Harrison, David K. Jackson, Ian Johnston, Dominic Kwiatkowski, Cordelia Langford, John Sillitoe on behalf of the Wellcome Sanger Institute COVID-19 Surveillance Team |
| EPI_ISL_3125789 | Lighthouse Lab in Glasgow | Wellcome Sanger Institute for the COVID-19 Genomics UK (COG-UK) Consortium | Harper VanSteenhouse, Yumi Kasai, David Gray, Carol Clugston, Anna Dominiczak and Alex Alderton, Roberto Amato, Jeffrey Barrett, Sonia Goncalves, Ewan Harrison, David K. Jackson, Ian Johnston, Dominic Kwiatkowski, Cordelia Langford, John Sillitoe on behalf of the Wellcome Sanger Institute COVID-19 Surveillance Team |
| EPI_ISL_3125790 | Berkshire and Surrey Pathology Services Lighthouse Laboratory | Wellcome Sanger Institute for the COVID-19 Genomics UK (COG-UK) Consortium | Berkshire and Surrey Pathology Services Lighthouse Laboratory and Alex Alderton, Roberto Amato, Jeffrey Barrett, Sonia Goncalves, Ewan Harrison, David K. Jackson, Ian Johnston, Dominic Kwiatkowski, Cordelia Langford, John Sillitoe on behalf of the Wellcome Sanger Institute COVID-19 Surveillance Team |
| EPI_ISL_3125791, EPI_ISL_3125792, EPI_ISL_3125793, EPI_ISL_3125794 | Lighthouse Lab in Milton Keynes | Wellcome Sanger Institute for the COVID-19 Genomics UK (COG-UK) Consortium | The Lighthouse Lab in Milton Keynes and Alex Alderton, Roberto Amato, Jeffrey Barrett, Sonia Goncalves, Ewan Harrison, David K. Jackson, Ian Johnston, Dominic Kwiatkowski, Cordelia Langford, John Sillitoe on behalf of the Wellcome Sanger Institute COVID-19 Surveillance Team |
| EPI_ISL_3125795, EPI_ISL_3125796 | Lighthouse Lab in Glasgow | Wellcome Sanger Institute for the COVID-19 Genomics UK (COG-UK) Consortium | Harper VanSteenhouse, Yumi Kasai, David Gray, Carol Clugston, Anna Dominiczak and Alex Alderton, Roberto Amato, Jeffrey Barrett, Sonia Goncalves, Ewan Harrison, David K. Jackson, Ian Johnston, Dominic Kwiatkowski, Cordelia Langford, John Sillitoe on behalf of the Wellcome Sanger Institute COVID-19 Surveillance Team |
| EPI_ISL_3125797 | Lighthouse Lab in Milton Keynes | Wellcome Sanger Institute for the COVID-19 Genomics UK (COG-UK) Consortium | The Lighthouse Lab in Milton Keynes and Alex Alderton, Roberto Amato, Jeffrey Barrett, Sonia Goncalves, Ewan Harrison, David K. Jackson, Ian Johnston, Dominic Kwiatkowski, Cordelia Langford, John Sillitoe on behalf of the Wellcome Sanger Institute COVID-19 Surveillance Team |
| EPI_ISL_3125798 | Berkshire and Surrey Pathology Services Lighthouse Laboratory | Wellcome Sanger Institute for the COVID-19 Genomics UK (COG-UK) Consortium | Berkshire and Surrey Pathology Services Lighthouse Laboratory and Alex Alderton, Roberto Amato, Jeffrey Barrett, Sonia Goncalves, Ewan Harrison, David K. Jackson, Ian Johnston, Dominic Kwiatkowski, Cordelia Langford, John Sillitoe on behalf of the Wellcome Sanger Institute COVID-19 Surveillance Team |
| EPI_ISL_3125799 | Lighthouse Lab in Glasgow | Wellcome Sanger Institute for the COVID-19 Genomics UK (COG-UK) Consortium | Harper VanSteenhouse, Yumi Kasai, David Gray, Carol Clugston, Anna Dominiczak and Alex Alderton, Roberto Amato, Jeffrey Barrett, Sonia Goncalves, Ewan Harrison, David K. Jackson, Ian Johnston, Dominic Kwiatkowski, Cordelia Langford, John Sillitoe on behalf of the Wellcome Sanger Institute COVID-19 Surveillance Team |
| EPI_ISL_3125800 | Lighthouse Lab in Milton Keynes | Wellcome Sanger Institute for the COVID-19 Genomics UK (COG-UK) Consortium | The Lighthouse Lab in Milton Keynes and Alex Alderton, Roberto Amato, Jeffrey Barrett, Sonia Goncalves, Ewan Harrison, David K. Jackson, Ian Johnston, Dominic Kwiatkowski, Cordelia Langford, John Sillitoe on behalf of the Wellcome Sanger Institute COVID-19 Surveillance Team |
| EPI_ISL_3125801 | Berkshire and Surrey Pathology Services Lighthouse Laboratory | Wellcome Sanger Institute for the COVID-19 Genomics UK (COG-UK) Consortium | Berkshire and Surrey Pathology Services Lighthouse Laboratory and Alex Alderton, Roberto Amato, Jeffrey Barrett, Sonia Goncalves, Ewan Harrison, David K. Jackson, Ian Johnston, Dominic Kwiatkowski, Cordelia Langford, John Sillitoe on behalf of the Wellcome Sanger Institute COVID-19 Surveillance Team |
| EPI_ISL_3125802, EPI_ISL_3125803, EPI_ISL_3125804, EPI_ISL_3125805, EPI_ISL_3125806 | Lighthouse Lab in Milton Keynes | Wellcome Sanger Institute for the COVID-19 Genomics UK (COG-UK) Consortium | The Lighthouse Lab in Milton Keynes and Alex Alderton, Roberto Amato, Jeffrey Barrett, Sonia Goncalves, Ewan Harrison, David K. Jackson, Ian Johnston, Dominic Kwiatkowski, Cordelia Langford, John Sillitoe on behalf of the Wellcome Sanger Institute COVID-19 Surveillance Team |
| EPI_ISL_3125807 | Lighthouse Lab in Glasgow | Wellcome Sanger Institute for the COVID-19 Genomics UK (COG-UK) Consortium | Harper VanSteenhouse, Yumi Kasai, David Gray, Carol Clugston, Anna Dominiczak and Alex Alderton, Roberto Amato, Jeffrey Barrett, Sonia Goncalves, Ewan Harrison, David K. Jackson, Ian Johnston, Dominic Kwiatkowski, Cordelia Langford, John Sillitoe on behalf of the Wellcome Sanger Institute COVID-19 Surveillance Team |
| EPI_ISL_3125808, EPI_ISL_3125809, EPI_ISL_3125810 | Lighthouse Lab in Milton Keynes | Wellcome Sanger Institute for the COVID-19 Genomics UK (COG-UK) Consortium | The Lighthouse Lab in Milton Keynes and Alex Alderton, Roberto Amato, Jeffrey Barrett, Sonia Goncalves, Ewan Harrison, David K. Jackson, Ian Johnston, Dominic Kwiatkowski, Cordelia Langford, John Sillitoe on behalf of the Wellcome Sanger Institute COVID-19 Surveillance Team |
| EPI_ISL_3125811, EPI_ISL_3125812 | Berkshire and Surrey Pathology Services Lighthouse Laboratory | Wellcome Sanger Institute for the COVID-19 Genomics UK (COG-UK) Consortium | Berkshire and Surrey Pathology Services Lighthouse Laboratory and Alex Alderton, Roberto Amato, Jeffrey Barrett, Sonia Goncalves, Ewan Harrison, David K. Jackson, Ian Johnston, Dominic Kwiatkowski, Cordelia Langford, John Sillitoe on behalf of the Wellcome Sanger Institute COVID-19 Surveillance Team |
| EPI_ISL_3125813, EPI_ISL_3125814, EPI_ISL_3125815 | Lighthouse Lab in Milton Keynes | Wellcome Sanger Institute for the COVID-19 Genomics UK (COG-UK) Consortium | The Lighthouse Lab in Milton Keynes and Alex Alderton, Roberto Amato, Jeffrey Barrett, Sonia Goncalves, Ewan Harrison, David K. Jackson, Ian Johnston, Dominic Kwiatkowski, Cordelia Langford, John Sillitoe on behalf of the Wellcome Sanger Institute COVID-19 Surveillance Team |
| EPI_ISL_3125816 | Unit of laboratory surveillance of viral emergent disease Control of Communicable Diseases Departament National Institute of Public Health Albania | Charité Universitätsmedizin Berlin, Institut für Virologie | Victor M Corman, Barbara Mühlemann, Jörn Beheim-Schwarzbach, Talitha Veith, Julia Schneider, Terry Jones, Iris Hasibra, Christian Drosten |
| EPI_ISL_3125817, EPI_ISL_3125818 | Lighthouse Lab in Milton Keynes | Wellcome Sanger Institute for the COVID-19 Genomics UK (COG-UK) Consortium | The Lighthouse Lab in Milton Keynes and Alex Alderton, Roberto Amato, Jeffrey Barrett, Sonia Goncalves, Ewan Harrison, David K. Jackson, Ian Johnston, Dominic Kwiatkowski, Cordelia Langford, John Sillitoe on behalf of the Wellcome Sanger Institute COVID-19 Surveillance Team |
| EPI_ISL_3125819 | Lighthouse Lab in Glasgow | Wellcome Sanger Institute for the COVID-19 Genomics UK (COG-UK) Consortium | Harper VanSteenhouse, Yumi Kasai, David Gray, Carol Clugston, Anna Dominiczak and Alex Alderton, Roberto Amato, Jeffrey Barrett, Sonia Goncalves, Ewan Harrison, David K. Jackson, Ian Johnston, Dominic Kwiatkowski, Cordelia Langford, John Sillitoe on behalf of the Wellcome Sanger Institute COVID-19 Surveillance Team |
| EPI_ISL_3125820, EPI_ISL_3125821, EPI_ISL_3125822 | Lighthouse Lab in Milton Keynes | Wellcome Sanger Institute for the COVID-19 Genomics UK (COG-UK) Consortium | The Lighthouse Lab in Milton Keynes and Alex Alderton, Roberto Amato, Jeffrey Barrett, Sonia Goncalves, Ewan Harrison, David K. Jackson, Ian Johnston, Dominic Kwiatkowski, Cordelia Langford, John Sillitoe on behalf of the Wellcome Sanger Institute COVID-19 Surveillance Team |
| EPI_ISL_3125823 | Berkshire and Surrey Pathology Services Lighthouse Laboratory | Wellcome Sanger Institute for the COVID-19 Genomics UK (COG-UK) Consortium | Berkshire and Surrey Pathology Services Lighthouse Laboratory and Alex Alderton, Roberto Amato, Jeffrey Barrett, Sonia Goncalves, Ewan Harrison, David K. Jackson, Ian Johnston, Dominic Kwiatkowski, Cordelia Langford, John Sillitoe on behalf of the Wellcome Sanger Institute COVID-19 Surveillance Team |
| EPI_ISL_3125824 | Lighthouse Lab in Milton Keynes | Wellcome Sanger Institute for the COVID-19 Genomics UK (COG-UK) Consortium | The Lighthouse Lab in Milton Keynes and Alex Alderton, Roberto Amato, Jeffrey Barrett, Sonia Goncalves, Ewan Harrison, David K. Jackson, Ian Johnston, Dominic Kwiatkowski, Cordelia Langford, John Sillitoe on behalf of the Wellcome Sanger Institute COVID-19 Surveillance Team |
| EPI_ISL_3125825, EPI_ISL_3125826 | Berkshire and Surrey Pathology Services Lighthouse Laboratory | Wellcome Sanger Institute for the COVID-19 Genomics UK (COG-UK) Consortium | Berkshire and Surrey Pathology Services Lighthouse Laboratory and Alex Alderton, Roberto Amato, Jeffrey Barrett, Sonia Goncalves, Ewan Harrison, David K. Jackson, Ian Johnston, Dominic Kwiatkowski, Cordelia Langford, John Sillitoe on behalf of the Wellcome Sanger Institute COVID-19 Surveillance Team |
| EPI_ISL_3125827, EPI_ISL_3125828 | Lighthouse Lab in Milton Keynes | Wellcome Sanger Institute for the COVID-19 Genomics UK (COG-UK) Consortium | The Lighthouse Lab in Milton Keynes and Alex Alderton, Roberto Amato, Jeffrey Barrett, Sonia Goncalves, Ewan Harrison, David K. Jackson, Ian Johnston, Dominic Kwiatkowski, Cordelia Langford, John Sillitoe on behalf of the Wellcome Sanger Institute COVID-19 Surveillance Team |
| EPI_ISL_3125829 | Berkshire and Surrey Pathology Services Lighthouse Laboratory | Wellcome Sanger Institute for the COVID-19 Genomics UK (COG-UK) Consortium | Berkshire and Surrey Pathology Services Lighthouse Laboratory and Alex Alderton, Roberto Amato, Jeffrey Barrett, Sonia Goncalves, Ewan Harrison, David K. Jackson, Ian Johnston, Dominic Kwiatkowski, Cordelia Langford, John Sillitoe on behalf of the Wellcome Sanger Institute COVID-19 Surveillance Team |
| EPI_ISL_3125830 | Lighthouse Lab in Milton Keynes | Wellcome Sanger Institute for the COVID-19 Genomics UK (COG-UK) Consortium | The Lighthouse Lab in Milton Keynes and Alex Alderton, Roberto Amato, Jeffrey Barrett, Sonia Goncalves, Ewan Harrison, David K. Jackson, Ian Johnston, Dominic Kwiatkowski, Cordelia Langford, John Sillitoe on behalf of the Wellcome Sanger Institute COVID-19 Surveillance Team |
| EPI_ISL_3125831 | Lighthouse Lab in Alderley Park | Wellcome Sanger Institute for the COVID-19 Genomics UK (COG-UK) Consortium | Jacquelyn Wynn, Mairead Hyland, The Lighthouse Lab in Alderley Park and Alex Alderton, Roberto Amato, Jeffrey Barrett, Sonia Goncalves, Ewan Harrison, David K. Jackson, Ian Johnston, Dominic Kwiatkowski, Cordelia Langford, John Sillitoe on behalf of the Wellcome Sanger Institute COVID-19 Surveillance Team |
| EPI_ISL_3125832, EPI_ISL_3125833, EPI_ISL_3125834, EPI_ISL_3125835, EPI_ISL_3125836, EPI_ISL_3125837, EPI_ISL_3125838 | Lighthouse Lab in Milton Keynes | Wellcome Sanger Institute for the COVID-19 Genomics UK (COG-UK) Consortium | The Lighthouse Lab in Milton Keynes and Alex Alderton, Roberto Amato, Jeffrey Barrett, Sonia Goncalves, Ewan Harrison, David K. Jackson, Ian Johnston, Dominic Kwiatkowski, Cordelia Langford, John Sillitoe on behalf of the Wellcome Sanger Institute COVID-19 Surveillance Team |
| EPI_ISL_3125839 | Lighthouse Lab in Glasgow | Wellcome Sanger Institute for the COVID-19 Genomics UK (COG-UK) Consortium | Harper VanSteenhouse, Yumi Kasai, David Gray, Carol Clugston, Anna Dominiczak and Alex Alderton, Roberto Amato, Jeffrey Barrett, Sonia Goncalves, Ewan Harrison, David K. Jackson, Ian Johnston, Dominic Kwiatkowski, Cordelia Langford, John Sillitoe on behalf of the Wellcome Sanger Institute COVID-19 Surveillance Team |
| EPI_ISL_3125840, EPI_ISL_3125841 | Lighthouse Lab in Milton Keynes | Wellcome Sanger Institute for the COVID-19 Genomics UK (COG-UK) Consortium | The Lighthouse Lab in Milton Keynes and Alex Alderton, Roberto Amato, Jeffrey Barrett, Sonia Goncalves, Ewan Harrison, David K. Jackson, Ian Johnston, Dominic Kwiatkowski, Cordelia Langford, John Sillitoe on behalf of the Wellcome Sanger Institute COVID-19 Surveillance Team |
| EPI_ISL_3125842 | Lighthouse Lab in Alderley Park | Wellcome Sanger Institute for the COVID-19 Genomics UK (COG-UK) Consortium | Jacquelyn Wynn, Mairead Hyland, The Lighthouse Lab in Alderley Park and Alex Alderton, Roberto Amato, Jeffrey Barrett, Sonia Goncalves, Ewan Harrison, David K. Jackson, Ian Johnston, Dominic Kwiatkowski, Cordelia Langford, John Sillitoe on behalf of the Wellcome Sanger Institute COVID-19 Surveillance Team |
| EPI_ISL_3125843 | Lighthouse Lab in Milton Keynes | Wellcome Sanger Institute for the COVID-19 Genomics UK (COG-UK) Consortium | The Lighthouse Lab in Milton Keynes and Alex Alderton, Roberto Amato, Jeffrey Barrett, Sonia Goncalves, Ewan Harrison, David K. Jackson, Ian Johnston, Dominic Kwiatkowski, Cordelia Langford, John Sillitoe on behalf of the Wellcome Sanger Institute COVID-19 Surveillance Team |
| EPI_ISL_3125844, EPI_ISL_3125845, | Lighthouse Lab in Glasgow | Wellcome Sanger Institute for the COVID-19 Genomics UK (COG-UK) Consortium | Harper VanSteenhouse, Yumi Kasai, David Gray, Carol Clugston, Anna Dominiczak and Alex Alderton, Roberto Amato, Jeffrey Barrett, Sonia Goncalves, Ewan Harrison, David K. Jackson, Ian Johnston, Dominic Kwiatkowski, Cordelia Langford, John Sillitoe on behalf of the Wellcome Sanger Institute COVID-19 Surveillance Team |

| | | | |
|---|---|---|---|
| EPI_ISL_3125846, EPI_ISL_3125847, EPI_ISL_3125848, EPI_ISL_3125849, EPI_ISL_3125850, EPI_ISL_3125851 | | | |
| EPI_ISL_3125852 | Lighthouse Lab in Alderley Park | Wellcome Sanger Institute for the COVID-19 Genomics UK (COG-UK) Consortium | Jacquelyn Wynn, Mairead Hyland, The Lighthouse Lab in Alderley Park and Alex Alderton, Roberto Amato, Jeffrey Barrett, Sonia Goncalves, Ewan Harrison, David K. Jackson, Ian Johnston, Dominic Kwiatkowski, Cordelia Langford, John Sillitoe on behalf of the Wellcome Sanger Institute COVID-19 Surveillance Team |
| EPI_ISL_3125853 | Lighthouse Lab in Glasgow | Wellcome Sanger Institute for the COVID-19 Genomics UK (COG-UK) Consortium | Harper VanSteenhouse, Yumi Kasai, David Gray, Carol Clugston, Anna Dominiczak and Alex Alderton, Roberto Amato, Jeffrey Barrett, Sonia Goncalves, Ewan Harrison, David K. Jackson, Ian Johnston, Dominic Kwiatkowski, Cordelia Langford, John Sillitoe on behalf of the Wellcome Sanger Institute COVID-19 Surveillance Team |
| EPI_ISL_3125854, EPI_ISL_3125855 | Lighthouse Lab in Alderley Park | Wellcome Sanger Institute for the COVID-19 Genomics UK (COG-UK) Consortium | Jacquelyn Wynn, Mairead Hyland, The Lighthouse Lab in Alderley Park and Alex Alderton, Roberto Amato, Jeffrey Barrett, Sonia Goncalves, Ewan Harrison, David K. Jackson, Ian Johnston, Dominic Kwiatkowski, Cordelia Langford, John Sillitoe on behalf of the Wellcome Sanger Institute COVID-19 Surveillance Team |
| EPI_ISL_3125856 | Lighthouse Lab in Glasgow | Wellcome Sanger Institute for the COVID-19 Genomics UK (COG-UK) Consortium | Harper VanSteenhouse, Yumi Kasai, David Gray, Carol Clugston, Anna Dominiczak and Alex Alderton, Roberto Amato, Jeffrey Barrett, Sonia Goncalves, Ewan Harrison, David K. Jackson, Ian Johnston, Dominic Kwiatkowski, Cordelia Langford, John Sillitoe on behalf of the Wellcome Sanger Institute COVID-19 Surveillance Team |
| EPI_ISL_3125857, EPI_ISL_3125858 | Lighthouse Lab in Alderley Park | Wellcome Sanger Institute for the COVID-19 Genomics UK (COG-UK) Consortium | Jacquelyn Wynn, Mairead Hyland, The Lighthouse Lab in Alderley Park and Alex Alderton, Roberto Amato, Jeffrey Barrett, Sonia Goncalves, Ewan Harrison, David K. Jackson, Ian Johnston, Dominic Kwiatkowski, Cordelia Langford, John Sillitoe on behalf of the Wellcome Sanger Institute COVID-19 Surveillance Team |
| EPI_ISL_3125859, EPI_ISL_3125860 | Lighthouse Lab in Glasgow | Wellcome Sanger Institute for the COVID-19 Genomics UK (COG-UK) Consortium | Harper VanSteenhouse, Yumi Kasai, David Gray, Carol Clugston, Anna Dominiczak and Alex Alderton, Roberto Amato, Jeffrey Barrett, Sonia Goncalves, Ewan Harrison, David K. Jackson, Ian Johnston, Dominic Kwiatkowski, Cordelia Langford, John Sillitoe on behalf of the Wellcome Sanger Institute COVID-19 Surveillance Team |
| EPI_ISL_3125861 | Lighthouse Lab in Alderley Park | Wellcome Sanger Institute for the COVID-19 Genomics UK (COG-UK) Consortium | Jacquelyn Wynn, Mairead Hyland, The Lighthouse Lab in Alderley Park and Alex Alderton, Roberto Amato, Jeffrey Barrett, Sonia Goncalves, Ewan Harrison, David K. Jackson, Ian Johnston, Dominic Kwiatkowski, Cordelia Langford, John Sillitoe on behalf of the Wellcome Sanger Institute COVID-19 Surveillance Team |
| EPI_ISL_3125862, EPI_ISL_3125863 | Lighthouse Lab in Glasgow | Wellcome Sanger Institute for the COVID-19 Genomics UK (COG-UK) Consortium | Harper VanSteenhouse, Yumi Kasai, David Gray, Carol Clugston, Anna Dominiczak and Alex Alderton, Roberto Amato, Jeffrey Barrett, Sonia Goncalves, Ewan Harrison, David K. Jackson, Ian Johnston, Dominic Kwiatkowski, Cordelia Langford, John Sillitoe on behalf of the Wellcome Sanger Institute COVID-19 Surveillance Team |
| EPI_ISL_3125864, EPI_ISL_3125865, EPI_ISL_3125866, EPI_ISL_3125867 | Lighthouse Lab in Alderley Park | Wellcome Sanger Institute for the COVID-19 Genomics UK (COG-UK) Consortium | Jacquelyn Wynn, Mairead Hyland, The Lighthouse Lab in Alderley Park and Alex Alderton, Roberto Amato, Jeffrey Barrett, Sonia Goncalves, Ewan Harrison, David K. Jackson, Ian Johnston, Dominic Kwiatkowski, Cordelia Langford, John Sillitoe on behalf of the Wellcome Sanger Institute COVID-19 Surveillance Team |
| EPI_ISL_3125868 | Lighthouse Lab in Glasgow | Wellcome Sanger Institute for the COVID-19 Genomics UK (COG-UK) Consortium | Harper VanSteenhouse, Yumi Kasai, David Gray, Carol Clugston, Anna Dominiczak and Alex Alderton, Roberto Amato, Jeffrey Barrett, Sonia Goncalves, Ewan Harrison, David K. Jackson, Ian Johnston, Dominic Kwiatkowski, Cordelia Langford, John Sillitoe on behalf of the Wellcome Sanger Institute COVID-19 Surveillance Team |
| EPI_ISL_3125869, EPI_ISL_3125870 | Lighthouse Lab in Alderley Park | Wellcome Sanger Institute for the COVID-19 Genomics UK (COG-UK) Consortium | Jacquelyn Wynn, Mairead Hyland, The Lighthouse Lab in Alderley Park and Alex Alderton, Roberto Amato, Jeffrey Barrett, Sonia Goncalves, Ewan Harrison, David K. Jackson, Ian Johnston, Dominic Kwiatkowski, Cordelia Langford, John Sillitoe on behalf of the Wellcome Sanger Institute COVID-19 Surveillance Team |
| EPI_ISL_3125871 | Lighthouse Lab in Glasgow | Wellcome Sanger Institute for the COVID-19 Genomics UK (COG-UK) Consortium | Harper VanSteenhouse, Yumi Kasai, David Gray, Carol Clugston, Anna Dominiczak and Alex Alderton, Roberto Amato, Jeffrey Barrett, Sonia Goncalves, Ewan Harrison, David K. Jackson, Ian Johnston, Dominic Kwiatkowski, Cordelia Langford, John Sillitoe on behalf of the Wellcome Sanger Institute COVID-19 Surveillance Team |
| EPI_ISL_3125872 | Lighthouse Lab in Alderley Park | Wellcome Sanger Institute for the COVID-19 Genomics UK (COG-UK) Consortium | Jacquelyn Wynn, Mairead Hyland, The Lighthouse Lab in Alderley Park and Alex Alderton, Roberto Amato, Jeffrey Barrett, Sonia Goncalves, Ewan Harrison, David K. Jackson, Ian Johnston, Dominic Kwiatkowski, Cordelia Langford, John Sillitoe on behalf of the Wellcome Sanger Institute COVID-19 Surveillance Team |
| EPI_ISL_3125873 | Lighthouse Lab in Glasgow | Wellcome Sanger Institute for the COVID-19 Genomics UK (COG-UK) Consortium | Harper VanSteenhouse, Yumi Kasai, David Gray, Carol Clugston, Anna Dominiczak and Alex Alderton, Roberto Amato, Jeffrey Barrett, Sonia Goncalves, Ewan Harrison, David K. Jackson, Ian Johnston, Dominic Kwiatkowski, Cordelia Langford, John Sillitoe on behalf of the Wellcome Sanger Institute COVID-19 Surveillance Team |
| EPI_ISL_3125874, EPI_ISL_3125875, EPI_ISL_3125876 | Lighthouse Lab in Alderley Park | Wellcome Sanger Institute for the COVID-19 Genomics UK (COG-UK) Consortium | Jacquelyn Wynn, Mairead Hyland, The Lighthouse Lab in Alderley Park and Alex Alderton, Roberto Amato, Jeffrey Barrett, Sonia Goncalves, Ewan Harrison, David K. Jackson, Ian Johnston, Dominic Kwiatkowski, Cordelia Langford, John Sillitoe on behalf of the Wellcome Sanger Institute COVID-19 Surveillance Team |
| EPI_ISL_3125877 | Lighthouse Lab in Glasgow | Wellcome Sanger Institute for the COVID-19 Genomics UK (COG-UK) Consortium | Harper VanSteenhouse, Yumi Kasai, David Gray, Carol Clugston, Anna Dominiczak and Alex Alderton, Roberto Amato, Jeffrey Barrett, Sonia Goncalves, Ewan Harrison, David K. Jackson, Ian Johnston, Dominic Kwiatkowski, Cordelia Langford, John Sillitoe on behalf of the Wellcome Sanger Institute COVID-19 Surveillance Team |
| EPI_ISL_3125878, EPI_ISL_3125879 | Lighthouse Lab in Alderley Park | Wellcome Sanger Institute for the COVID-19 Genomics UK (COG-UK) Consortium | Jacquelyn Wynn, Mairead Hyland, The Lighthouse Lab in Alderley Park and Alex Alderton, Roberto Amato, Jeffrey Barrett, Sonia Goncalves, Ewan Harrison, David K. Jackson, Ian Johnston, Dominic Kwiatkowski, Cordelia Langford, John Sillitoe on behalf of the Wellcome Sanger Institute COVID-19 Surveillance Team |
| EPI_ISL_3125880, EPI_ISL_3125881, EPI_ISL_3125882 | Lighthouse Lab in Glasgow | Wellcome Sanger Institute for the COVID-19 Genomics UK (COG-UK) Consortium | Harper VanSteenhouse, Yumi Kasai, David Gray, Carol Clugston, Anna Dominiczak and Alex Alderton, Roberto Amato, Jeffrey Barrett, Sonia Goncalves, Ewan Harrison, David K. Jackson, Ian Johnston, Dominic Kwiatkowski, Cordelia Langford, John Sillitoe on behalf of the Wellcome Sanger Institute COVID-19 Surveillance Team |
| EPI_ISL_3125883, EPI_ISL_3125884 | Lighthouse Lab in Alderley Park | Wellcome Sanger Institute for the COVID-19 Genomics UK (COG-UK) Consortium | Jacquelyn Wynn, Mairead Hyland, The Lighthouse Lab in Alderley Park and Alex Alderton, Roberto Amato, Jeffrey Barrett, Sonia Goncalves, Ewan Harrison, David K. Jackson, Ian Johnston, Dominic Kwiatkowski, Cordelia Langford, John Sillitoe on behalf of the Wellcome Sanger Institute COVID-19 Surveillance Team |
| EPI_ISL_3125885, EPI_ISL_3125886 | Lighthouse Lab in Glasgow | Wellcome Sanger Institute for the COVID-19 Genomics UK (COG-UK) Consortium | Harper VanSteenhouse, Yumi Kasai, David Gray, Carol Clugston, Anna Dominiczak and Alex Alderton, Roberto Amato, Jeffrey Barrett, Sonia Goncalves, Ewan Harrison, David K. Jackson, Ian Johnston, Dominic Kwiatkowski, Cordelia Langford, John Sillitoe on behalf of the Wellcome Sanger Institute COVID-19 Surveillance Team |
| EPI_ISL_3125887, EPI_ISL_3125888 | Lighthouse Lab in Alderley Park | Wellcome Sanger Institute for the COVID-19 Genomics UK (COG-UK) Consortium | Jacquelyn Wynn, Mairead Hyland, The Lighthouse Lab in Alderley Park and Alex Alderton, Roberto Amato, Jeffrey Barrett, Sonia Goncalves, Ewan Harrison, David K. Jackson, Ian Johnston, Dominic Kwiatkowski, Cordelia Langford, John Sillitoe on behalf of the Wellcome Sanger Institute COVID-19 Surveillance Team |
| EPI_ISL_3125889 | Lighthouse Lab in Glasgow | Wellcome Sanger Institute for the COVID-19 Genomics UK (COG-UK) Consortium | Harper VanSteenhouse, Yumi Kasai, David Gray, Carol Clugston, Anna Dominiczak and Alex Alderton, Roberto Amato, Jeffrey Barrett, Sonia Goncalves, Ewan Harrison, David K. Jackson, Ian Johnston, Dominic Kwiatkowski, Cordelia Langford, John Sillitoe on behalf of the Wellcome Sanger Institute COVID-19 Surveillance Team |
| EPI_ISL_3125890, EPI_ISL_3125891, EPI_ISL_3125892, EPI_ISL_3125893, EPI_ISL_3125894, EPI_ISL_3125895 | Lighthouse Lab in Alderley Park | Wellcome Sanger Institute for the COVID-19 Genomics UK (COG-UK) Consortium | Jacquelyn Wynn, Mairead Hyland, The Lighthouse Lab in Alderley Park and Alex Alderton, Roberto Amato, Jeffrey Barrett, Sonia Goncalves, Ewan Harrison, David K. Jackson, Ian Johnston, Dominic Kwiatkowski, Cordelia Langford, John Sillitoe on behalf of the Wellcome Sanger Institute COVID-19 Surveillance Team |
| EPI_ISL_3125896, EPI_ISL_3125897 | Lighthouse Lab in Glasgow | Wellcome Sanger Institute for the COVID-19 Genomics UK (COG-UK) Consortium | Harper VanSteenhouse, Yumi Kasai, David Gray, Carol Clugston, Anna Dominiczak and Alex Alderton, Roberto Amato, Jeffrey Barrett, Sonia Goncalves, Ewan Harrison, David K. Jackson, Ian Johnston, Dominic Kwiatkowski, Cordelia Langford, John Sillitoe on behalf of the Wellcome Sanger Institute COVID-19 Surveillance Team |
| EPI_ISL_3125898, EPI_ISL_3125899 | Lighthouse Lab in Alderley Park | Wellcome Sanger Institute for the COVID-19 Genomics UK (COG-UK) Consortium | Jacquelyn Wynn, Mairead Hyland, The Lighthouse Lab in Alderley Park and Alex Alderton, Roberto Amato, Jeffrey Barrett, Sonia Goncalves, Ewan Harrison, David K. Jackson, Ian Johnston, Dominic Kwiatkowski, Cordelia Langford, John Sillitoe on behalf of the Wellcome Sanger Institute COVID-19 Surveillance Team |
| EPI_ISL_3125900 | Lighthouse Lab in Glasgow | Wellcome Sanger Institute for the COVID-19 Genomics UK (COG-UK) Consortium | Harper VanSteenhouse, Yumi Kasai, David Gray, Carol Clugston, Anna Dominiczak and Alex Alderton, Roberto Amato, Jeffrey Barrett, Sonia Goncalves, Ewan Harrison, David K. Jackson, Ian Johnston, Dominic Kwiatkowski, Cordelia Langford, John Sillitoe on behalf of the Wellcome Sanger Institute COVID-19 Surveillance Team |
| EPI_ISL_3125901, EPI_ISL_3125902, EPI_ISL_3125903 | Lighthouse Lab in Alderley Park | Wellcome Sanger Institute for the COVID-19 Genomics UK (COG-UK) Consortium | Jacquelyn Wynn, Mairead Hyland, The Lighthouse Lab in Alderley Park and Alex Alderton, Roberto Amato, Jeffrey Barrett, Sonia Goncalves, Ewan Harrison, David K. Jackson, Ian Johnston, Dominic Kwiatkowski, Cordelia Langford, John Sillitoe on behalf of the Wellcome Sanger Institute COVID-19 Surveillance Team |
| EPI_ISL_3125904 | Lighthouse Lab in Glasgow | Wellcome Sanger Institute for the COVID-19 Genomics UK (COG-UK) Consortium | Harper VanSteenhouse, Yumi Kasai, David Gray, Carol Clugston, Anna Dominiczak and Alex Alderton, Roberto Amato, Jeffrey Barrett, Sonia Goncalves, Ewan Harrison, David K. Jackson, Ian Johnston, Dominic Kwiatkowski, Cordelia Langford, John Sillitoe on behalf of the Wellcome Sanger Institute COVID-19 Surveillance Team |
| EPI_ISL_3125905 | Lighthouse Lab in Alderley Park | Wellcome Sanger Institute for the COVID-19 Genomics UK (COG-UK) Consortium | Jacquelyn Wynn, Mairead Hyland, The Lighthouse Lab in Alderley Park and Alex Alderton, Roberto Amato, Jeffrey Barrett, Sonia Goncalves, Ewan Harrison, David K. Jackson, Ian Johnston, Dominic Kwiatkowski, Cordelia Langford, John Sillitoe on behalf of the Wellcome Sanger Institute COVID-19 Surveillance Team |
| EPI_ISL_3125906, EPI_ISL_3125907 | Lighthouse Lab in Glasgow | Wellcome Sanger Institute for the COVID-19 Genomics UK (COG-UK) Consortium | Harper VanSteenhouse, Yumi Kasai, David Gray, Carol Clugston, Anna Dominiczak and Alex Alderton, Roberto Amato, Jeffrey Barrett, Sonia Goncalves, Ewan Harrison, David K. Jackson, Ian Johnston, Dominic Kwiatkowski, Cordelia Langford, John Sillitoe on behalf of the Wellcome Sanger Institute COVID-19 Surveillance Team |
| EPI_ISL_3125908 | Lighthouse Lab in Alderley Park | Wellcome Sanger Institute for the COVID-19 Genomics UK (COG-UK) Consortium | Jacquelyn Wynn, Mairead Hyland, The Lighthouse Lab in Alderley Park and Alex Alderton, Roberto Amato, Jeffrey Barrett, Sonia Goncalves, Ewan Harrison, David K. Jackson, Ian Johnston, Dominic Kwiatkowski, Cordelia Langford, John Sillitoe on behalf of the Wellcome Sanger Institute COVID-19 Surveillance Team |
| EPI_ISL_3125909, EPI_ISL_3125910, EPI_ISL_3125911, EPI_ISL_3125912, EPI_ISL_3125913, EPI_ISL_3125914 | Lighthouse Lab in Glasgow | Wellcome Sanger Institute for the COVID-19 Genomics UK (COG-UK) Consortium | Harper VanSteenhouse, Yumi Kasai, David Gray, Carol Clugston, Anna Dominiczak and Alex Alderton, Roberto Amato, Jeffrey Barrett, Sonia Goncalves, Ewan Harrison, David K. Jackson, Ian Johnston, Dominic Kwiatkowski, Cordelia Langford, John Sillitoe on behalf of the Wellcome Sanger Institute COVID-19 Surveillance Team |
| EPI_ISL_3125915 | Lighthouse Lab in Alderley Park | Wellcome Sanger Institute for the COVID-19 Genomics UK (COG-UK) Consortium | Jacquelyn Wynn, Mairead Hyland, The Lighthouse Lab in Alderley Park and Alex Alderton, Roberto Amato, Jeffrey Barrett, Sonia Goncalves, Ewan Harrison, David K. Jackson, Ian Johnston, Dominic Kwiatkowski, Cordelia Langford, John Sillitoe on behalf of the Wellcome Sanger Institute COVID-19 Surveillance Team |
| EPI_ISL_3125916 | Lighthouse Lab in Glasgow | Wellcome Sanger Institute for the COVID-19 Genomics UK (COG-UK) Consortium | Harper VanSteenhouse, Yumi Kasai, David Gray, Carol Clugston, Anna Dominiczak and Alex Alderton, Roberto Amato, Jeffrey Barrett, Sonia Goncalves, Ewan Harrison, David K. Jackson, Ian Johnston, Dominic Kwiatkowski, Cordelia Langford, John Sillitoe on behalf of the Wellcome Sanger Institute COVID-19 Surveillance Team |
| EPI_ISL_3125917, EPI_ISL_3125918 | Lighthouse Lab in Alderley Park | Wellcome Sanger Institute for the COVID-19 Genomics UK (COG-UK) Consortium | Jacquelyn Wynn, Mairead Hyland, The Lighthouse Lab in Alderley Park and Alex Alderton, Roberto Amato, Jeffrey Barrett, Sonia Goncalves, Ewan Harrison, David K. Jackson, Ian Johnston, Dominic Kwiatkowski, Cordelia Langford, John Sillitoe on behalf of the Wellcome Sanger Institute COVID-19 Surveillance Team |
| EPI_ISL_3125919 | Lighthouse Lab in Glasgow | Wellcome Sanger Institute for the COVID-19 Genomics UK (COG-UK) Consortium | Harper VanSteenhouse, Yumi Kasai, David Gray, Carol Clugston, Anna Dominiczak and Alex Alderton, Roberto Amato, Jeffrey Barrett, Sonia Goncalves, Ewan Harrison, David K. Jackson, Ian Johnston, Dominic Kwiatkowski, Cordelia Langford, John Sillitoe on behalf of the Wellcome Sanger Institute COVID-19 Surveillance Team |
| EPI_ISL_3125920 | Lighthouse Lab in Alderley Park | Wellcome Sanger Institute for the COVID-19 Genomics UK (COG-UK) Consortium | Jacquelyn Wynn, Mairead Hyland, The Lighthouse Lab in Alderley Park and Alex Alderton, Roberto Amato, Jeffrey Barrett, Sonia Goncalves, Ewan Harrison, David K. Jackson, Ian Johnston, Dominic Kwiatkowski, Cordelia Langford, John Sillitoe on behalf of the Wellcome Sanger Institute COVID-19 Surveillance Team |

| | | | |
|---|---|---|---|
| EPI_ISL_3125921, EPI_ISL_3125922 | Lighthouse Lab in Glasgow | Wellcome Sanger Institute for the COVID-19 Genomics UK (COG-UK) Consortium | Harper VanSteenhouse, Yumi Kasai, David Gray, Carol Clugston, Anna Dominiczak and Alex Alderton, Roberto Amato, Jeffrey Barrett, Sonia Goncalves, Ewan Harrison, David K. Jackson, Ian Johnston, Dominic Kwiatkowski, Cordelia Langford, John Sillitoe on behalf of the Wellcome Sanger Institute COVID-19 Surveillance Team |
| EPI_ISL_3125923 | Lighthouse Lab in Alderley Park | Wellcome Sanger Institute for the COVID-19 Genomics UK (COG-UK) Consortium | Jacquelyn Wynn, Mairead Hyland, The Lighthouse Lab in Alderley Park and Alex Alderton, Roberto Amato, Jeffrey Barrett, Sonia Goncalves, Ewan Harrison, David K. Jackson, Ian Johnston, Dominic Kwiatkowski, Cordelia Langford, John Sillitoe on behalf of the Wellcome Sanger Institute COVID-19 Surveillance Team |
| EPI_ISL_3125924, EPI_ISL_3125925 | Lighthouse Lab in Glasgow | Wellcome Sanger Institute for the COVID-19 Genomics UK (COG-UK) Consortium | Harper VanSteenhouse, Yumi Kasai, David Gray, Carol Clugston, Anna Dominiczak and Alex Alderton, Roberto Amato, Jeffrey Barrett, Sonia Goncalves, Ewan Harrison, David K. Jackson, Ian Johnston, Dominic Kwiatkowski, Cordelia Langford, John Sillitoe on behalf of the Wellcome Sanger Institute COVID-19 Surveillance Team |
| EPI_ISL_3125926 | Lighthouse Lab in Alderley Park | Wellcome Sanger Institute for the COVID-19 Genomics UK (COG-UK) Consortium | Jacquelyn Wynn, Mairead Hyland, The Lighthouse Lab in Alderley Park and Alex Alderton, Roberto Amato, Jeffrey Barrett, Sonia Goncalves, Ewan Harrison, David K. Jackson, Ian Johnston, Dominic Kwiatkowski, Cordelia Langford, John Sillitoe on behalf of the Wellcome Sanger Institute COVID-19 Surveillance Team |
| EPI_ISL_3125927, EPI_ISL_3125928 | Lighthouse Lab in Glasgow | Wellcome Sanger Institute for the COVID-19 Genomics UK (COG-UK) Consortium | Harper VanSteenhouse, Yumi Kasai, David Gray, Carol Clugston, Anna Dominiczak and Alex Alderton, Roberto Amato, Jeffrey Barrett, Sonia Goncalves, Ewan Harrison, David K. Jackson, Ian Johnston, Dominic Kwiatkowski, Cordelia Langford, John Sillitoe on behalf of the Wellcome Sanger Institute COVID-19 Surveillance Team |
| EPI_ISL_3125929, EPI_ISL_3125930 | Lighthouse Lab in Alderley Park | Wellcome Sanger Institute for the COVID-19 Genomics UK (COG-UK) Consortium | Jacquelyn Wynn, Mairead Hyland, The Lighthouse Lab in Alderley Park and Alex Alderton, Roberto Amato, Jeffrey Barrett, Sonia Goncalves, Ewan Harrison, David K. Jackson, Ian Johnston, Dominic Kwiatkowski, Cordelia Langford, John Sillitoe on behalf of the Wellcome Sanger Institute COVID-19 Surveillance Team |
| EPI_ISL_3125931, EPI_ISL_3125932 | Lighthouse Lab in Glasgow | Wellcome Sanger Institute for the COVID-19 Genomics UK (COG-UK) Consortium | Harper VanSteenhouse, Yumi Kasai, David Gray, Carol Clugston, Anna Dominiczak and Alex Alderton, Roberto Amato, Jeffrey Barrett, Sonia Goncalves, Ewan Harrison, David K. Jackson, Ian Johnston, Dominic Kwiatkowski, Cordelia Langford, John Sillitoe on behalf of the Wellcome Sanger Institute COVID-19 Surveillance Team |
| EPI_ISL_3125933, EPI_ISL_3125934, EPI_ISL_3125935, EPI_ISL_3125936, EPI_ISL_3125937, EPI_ISL_3125938, EPI_ISL_3125939 | Lighthouse Lab in Alderley Park | Wellcome Sanger Institute for the COVID-19 Genomics UK (COG-UK) Consortium | Jacquelyn Wynn, Mairead Hyland, The Lighthouse Lab in Alderley Park and Alex Alderton, Roberto Amato, Jeffrey Barrett, Sonia Goncalves, Ewan Harrison, David K. Jackson, Ian Johnston, Dominic Kwiatkowski, Cordelia Langford, John Sillitoe on behalf of the Wellcome Sanger Institute COVID-19 Surveillance Team |
| EPI_ISL_3125940 | Lighthouse Lab in Glasgow | Wellcome Sanger Institute for the COVID-19 Genomics UK (COG-UK) Consortium | Harper VanSteenhouse, Yumi Kasai, David Gray, Carol Clugston, Anna Dominiczak and Alex Alderton, Roberto Amato, Jeffrey Barrett, Sonia Goncalves, Ewan Harrison, David K. Jackson, Ian Johnston, Dominic Kwiatkowski, Cordelia Langford, John Sillitoe on behalf of the Wellcome Sanger Institute COVID-19 Surveillance Team |
| EPI_ISL_3125941, EPI_ISL_3125942, EPI_ISL_3125943, EPI_ISL_3125944 | Lighthouse Lab in Alderley Park | Wellcome Sanger Institute for the COVID-19 Genomics UK (COG-UK) Consortium | Jacquelyn Wynn, Mairead Hyland, The Lighthouse Lab in Alderley Park and Alex Alderton, Roberto Amato, Jeffrey Barrett, Sonia Goncalves, Ewan Harrison, David K. Jackson, Ian Johnston, Dominic Kwiatkowski, Cordelia Langford, John Sillitoe on behalf of the Wellcome Sanger Institute COVID-19 Surveillance Team |
| EPI_ISL_3125945, EPI_ISL_3125946 | Lighthouse Lab in Glasgow | Wellcome Sanger Institute for the COVID-19 Genomics UK (COG-UK) Consortium | Harper VanSteenhouse, Yumi Kasai, David Gray, Carol Clugston, Anna Dominiczak and Alex Alderton, Roberto Amato, Jeffrey Barrett, Sonia Goncalves, Ewan Harrison, David K. Jackson, Ian Johnston, Dominic Kwiatkowski, Cordelia Langford, John Sillitoe on behalf of the Wellcome Sanger Institute COVID-19 Surveillance Team |
| EPI_ISL_3125947, EPI_ISL_3125948, EPI_ISL_3125949, EPI_ISL_3125950, EPI_ISL_3125951 | Lighthouse Lab in Alderley Park | Wellcome Sanger Institute for the COVID-19 Genomics UK (COG-UK) Consortium | Jacquelyn Wynn, Mairead Hyland, The Lighthouse Lab in Alderley Park and Alex Alderton, Roberto Amato, Jeffrey Barrett, Sonia Goncalves, Ewan Harrison, David K. Jackson, Ian Johnston, Dominic Kwiatkowski, Cordelia Langford, John Sillitoe on behalf of the Wellcome Sanger Institute COVID-19 Surveillance Team |
| EPI_ISL_3125952 | Lighthouse Lab in Glasgow | Wellcome Sanger Institute for the COVID-19 Genomics UK (COG-UK) Consortium | Harper VanSteenhouse, Yumi Kasai, David Gray, Carol Clugston, Anna Dominiczak and Alex Alderton, Roberto Amato, Jeffrey Barrett, Sonia Goncalves, Ewan Harrison, David K. Jackson, Ian Johnston, Dominic Kwiatkowski, Cordelia Langford, John Sillitoe on behalf of the Wellcome Sanger Institute COVID-19 Surveillance Team |
| EPI_ISL_3125953 | Lighthouse Lab in Alderley Park | Wellcome Sanger Institute for the COVID-19 Genomics UK (COG-UK) Consortium | Jacquelyn Wynn, Mairead Hyland, The Lighthouse Lab in Alderley Park and Alex Alderton, Roberto Amato, Jeffrey Barrett, Sonia Goncalves, Ewan Harrison, David K. Jackson, Ian Johnston, Dominic Kwiatkowski, Cordelia Langford, John Sillitoe on behalf of the Wellcome Sanger Institute COVID-19 Surveillance Team |
| EPI_ISL_3125954 | Lighthouse Lab in Glasgow | Wellcome Sanger Institute for the COVID-19 Genomics UK (COG-UK) Consortium | Harper VanSteenhouse, Yumi Kasai, David Gray, Carol Clugston, Anna Dominiczak and Alex Alderton, Roberto Amato, Jeffrey Barrett, Sonia Goncalves, Ewan Harrison, David K. Jackson, Ian Johnston, Dominic Kwiatkowski, Cordelia Langford, John Sillitoe on behalf of the Wellcome Sanger Institute COVID-19 Surveillance Team |
| EPI_ISL_3125955, EPI_ISL_3125956 | Lighthouse Lab in Alderley Park | Wellcome Sanger Institute for the COVID-19 Genomics UK (COG-UK) Consortium | Jacquelyn Wynn, Mairead Hyland, The Lighthouse Lab in Alderley Park and Alex Alderton, Roberto Amato, Jeffrey Barrett, Sonia Goncalves, Ewan Harrison, David K. Jackson, Ian Johnston, Dominic Kwiatkowski, Cordelia Langford, John Sillitoe on behalf of the Wellcome Sanger Institute COVID-19 Surveillance Team |
| EPI_ISL_3125957, EPI_ISL_3125958 | Lighthouse Lab in Glasgow | Wellcome Sanger Institute for the COVID-19 Genomics UK (COG-UK) Consortium | Harper VanSteenhouse, Yumi Kasai, David Gray, Carol Clugston, Anna Dominiczak and Alex Alderton, Roberto Amato, Jeffrey Barrett, Sonia Goncalves, Ewan Harrison, David K. Jackson, Ian Johnston, Dominic Kwiatkowski, Cordelia Langford, John Sillitoe on behalf of the Wellcome Sanger Institute COVID-19 Surveillance Team |
| EPI_ISL_3125959 | Lighthouse Lab in Alderley Park | Wellcome Sanger Institute for the COVID-19 Genomics UK (COG-UK) Consortium | Jacquelyn Wynn, Mairead Hyland, The Lighthouse Lab in Alderley Park and Alex Alderton, Roberto Amato, Jeffrey Barrett, Sonia Goncalves, Ewan Harrison, David K. Jackson, Ian Johnston, Dominic Kwiatkowski, Cordelia Langford, John Sillitoe on behalf of the Wellcome Sanger Institute COVID-19 Surveillance Team |
| EPI_ISL_3125960 | Lighthouse Lab in Glasgow | Wellcome Sanger Institute for the COVID-19 Genomics UK (COG-UK) Consortium | Harper VanSteenhouse, Yumi Kasai, David Gray, Carol Clugston, Anna Dominiczak and Alex Alderton, Roberto Amato, Jeffrey Barrett, Sonia Goncalves, Ewan Harrison, David K. Jackson, Ian Johnston, Dominic Kwiatkowski, Cordelia Langford, John Sillitoe on behalf of the Wellcome Sanger Institute COVID-19 Surveillance Team |
| EPI_ISL_3125961, EPI_ISL_3125962, EPI_ISL_3125963, EPI_ISL_3125964, EPI_ISL_3125965 | Lighthouse Lab in Alderley Park | Wellcome Sanger Institute for the COVID-19 Genomics UK (COG-UK) Consortium | Jacquelyn Wynn, Mairead Hyland, The Lighthouse Lab in Alderley Park and Alex Alderton, Roberto Amato, Jeffrey Barrett, Sonia Goncalves, Ewan Harrison, David K. Jackson, Ian Johnston, Dominic Kwiatkowski, Cordelia Langford, John Sillitoe on behalf of the Wellcome Sanger Institute COVID-19 Surveillance Team |
| EPI_ISL_3125966 | Lighthouse Lab in Glasgow | Wellcome Sanger Institute for the COVID-19 Genomics UK (COG-UK) Consortium | Harper VanSteenhouse, Yumi Kasai, David Gray, Carol Clugston, Anna Dominiczak and Alex Alderton, Roberto Amato, Jeffrey Barrett, Sonia Goncalves, Ewan Harrison, David K. Jackson, Ian Johnston, Dominic Kwiatkowski, Cordelia Langford, John Sillitoe on behalf of the Wellcome Sanger Institute COVID-19 Surveillance Team |
| EPI_ISL_3125967 | Lighthouse Lab in Alderley Park | Wellcome Sanger Institute for the COVID-19 Genomics UK (COG-UK) Consortium | Jacquelyn Wynn, Mairead Hyland, The Lighthouse Lab in Alderley Park and Alex Alderton, Roberto Amato, Jeffrey Barrett, Sonia Goncalves, Ewan Harrison, David K. Jackson, Ian Johnston, Dominic Kwiatkowski, Cordelia Langford, John Sillitoe on behalf of the Wellcome Sanger Institute COVID-19 Surveillance Team |
| EPI_ISL_3125968 | Lighthouse Lab in Glasgow | Wellcome Sanger Institute for the COVID-19 Genomics UK (COG-UK) Consortium | Harper VanSteenhouse, Yumi Kasai, David Gray, Carol Clugston, Anna Dominiczak and Alex Alderton, Roberto Amato, Jeffrey Barrett, Sonia Goncalves, Ewan Harrison, David K. Jackson, Ian Johnston, Dominic Kwiatkowski, Cordelia Langford, John Sillitoe on behalf of the Wellcome Sanger Institute COVID-19 Surveillance Team |
| EPI_ISL_3125969, EPI_ISL_3125970 | Lighthouse Lab in Alderley Park | Wellcome Sanger Institute for the COVID-19 Genomics UK (COG-UK) Consortium | Jacquelyn Wynn, Mairead Hyland, The Lighthouse Lab in Alderley Park and Alex Alderton, Roberto Amato, Jeffrey Barrett, Sonia Goncalves, Ewan Harrison, David K. Jackson, Ian Johnston, Dominic Kwiatkowski, Cordelia Langford, John Sillitoe on behalf of the Wellcome Sanger Institute COVID-19 Surveillance Team |
| EPI_ISL_3125971 | Lighthouse Lab in Glasgow | Wellcome Sanger Institute for the COVID-19 Genomics UK (COG-UK) Consortium | Harper VanSteenhouse, Yumi Kasai, David Gray, Carol Clugston, Anna Dominiczak and Alex Alderton, Roberto Amato, Jeffrey Barrett, Sonia Goncalves, Ewan Harrison, David K. Jackson, Ian Johnston, Dominic Kwiatkowski, Cordelia Langford, John Sillitoe on behalf of the Wellcome Sanger Institute COVID-19 Surveillance Team |
| EPI_ISL_3125972 | Lighthouse Lab in Alderley Park | Wellcome Sanger Institute for the COVID-19 Genomics UK (COG-UK) Consortium | Jacquelyn Wynn, Mairead Hyland, The Lighthouse Lab in Alderley Park and Alex Alderton, Roberto Amato, Jeffrey Barrett, Sonia Goncalves, Ewan Harrison, David K. Jackson, Ian Johnston, Dominic Kwiatkowski, Cordelia Langford, John Sillitoe on behalf of the Wellcome Sanger Institute COVID-19 Surveillance Team |
| EPI_ISL_3125973 | Lighthouse Lab in Glasgow | Wellcome Sanger Institute for the COVID-19 Genomics UK (COG-UK) Consortium | Harper VanSteenhouse, Yumi Kasai, David Gray, Carol Clugston, Anna Dominiczak and Alex Alderton, Roberto Amato, Jeffrey Barrett, Sonia Goncalves, Ewan Harrison, David K. Jackson, Ian Johnston, Dominic Kwiatkowski, Cordelia Langford, John Sillitoe on behalf of the Wellcome Sanger Institute COVID-19 Surveillance Team |
| EPI_ISL_3125974 | Lighthouse Lab in Alderley Park | Wellcome Sanger Institute for the COVID-19 Genomics UK (COG-UK) Consortium | Jacquelyn Wynn, Mairead Hyland, The Lighthouse Lab in Alderley Park and Alex Alderton, Roberto Amato, Jeffrey Barrett, Sonia Goncalves, Ewan Harrison, David K. Jackson, Ian Johnston, Dominic Kwiatkowski, Cordelia Langford, John Sillitoe on behalf of the Wellcome Sanger Institute COVID-19 Surveillance Team |
| EPI_ISL_3125975, EPI_ISL_3125976, EPI_ISL_3125977, EPI_ISL_3125978 | Lighthouse Lab in Glasgow | Wellcome Sanger Institute for the COVID-19 Genomics UK (COG-UK) Consortium | Harper VanSteenhouse, Yumi Kasai, David Gray, Carol Clugston, Anna Dominiczak and Alex Alderton, Roberto Amato, Jeffrey Barrett, Sonia Goncalves, Ewan Harrison, David K. Jackson, Ian Johnston, Dominic Kwiatkowski, Cordelia Langford, John Sillitoe on behalf of the Wellcome Sanger Institute COVID-19 Surveillance Team |
| EPI_ISL_3125979, EPI_ISL_3125980, EPI_ISL_3125981, EPI_ISL_3125982, EPI_ISL_3125983, EPI_ISL_3125984 | Lighthouse Lab in Alderley Park | Wellcome Sanger Institute for the COVID-19 Genomics UK (COG-UK) Consortium | Jacquelyn Wynn, Mairead Hyland, The Lighthouse Lab in Alderley Park and Alex Alderton, Roberto Amato, Jeffrey Barrett, Sonia Goncalves, Ewan Harrison, David K. Jackson, Ian Johnston, Dominic Kwiatkowski, Cordelia Langford, John Sillitoe on behalf of the Wellcome Sanger Institute COVID-19 Surveillance Team |
| EPI_ISL_3125985, EPI_ISL_3125986 | Lighthouse Lab in Glasgow | Wellcome Sanger Institute for the COVID-19 Genomics UK (COG-UK) Consortium | Harper VanSteenhouse, Yumi Kasai, David Gray, Carol Clugston, Anna Dominiczak and Alex Alderton, Roberto Amato, Jeffrey Barrett, Sonia Goncalves, Ewan Harrison, David K. Jackson, Ian Johnston, Dominic Kwiatkowski, Cordelia Langford, John Sillitoe on behalf of the Wellcome Sanger Institute COVID-19 Surveillance Team |
| EPI_ISL_3125987 | Lighthouse Lab in Alderley Park | Wellcome Sanger Institute for the COVID-19 Genomics UK (COG-UK) Consortium | Jacquelyn Wynn, Mairead Hyland, The Lighthouse Lab in Alderley Park and Alex Alderton, Roberto Amato, Jeffrey Barrett, Sonia Goncalves, Ewan Harrison, David K. Jackson, Ian Johnston, Dominic Kwiatkowski, Cordelia Langford, John Sillitoe on behalf of the Wellcome Sanger Institute COVID-19 Surveillance Team |
| EPI_ISL_3125988 | Lighthouse Lab in Glasgow | Wellcome Sanger Institute for the COVID-19 Genomics UK (COG-UK) Consortium | Harper VanSteenhouse, Yumi Kasai, David Gray, Carol Clugston, Anna Dominiczak and Alex Alderton, Roberto Amato, Jeffrey Barrett, Sonia Goncalves, Ewan Harrison, David K. Jackson, Ian Johnston, Dominic Kwiatkowski, Cordelia Langford, John Sillitoe on behalf of the Wellcome Sanger Institute COVID-19 Surveillance Team |
| EPI_ISL_3125989, EPI_ISL_3125990, EPI_ISL_3125991, EPI_ISL_3125992 | Lighthouse Lab in Alderley Park | Wellcome Sanger Institute for the COVID-19 Genomics UK (COG-UK) Consortium | Jacquelyn Wynn, Mairead Hyland, The Lighthouse Lab in Alderley Park and Alex Alderton, Roberto Amato, Jeffrey Barrett, Sonia Goncalves, Ewan Harrison, David K. Jackson, Ian Johnston, Dominic Kwiatkowski, Cordelia Langford, John Sillitoe on behalf of the Wellcome Sanger Institute COVID-19 Surveillance Team |
| EPI_ISL_3125993, EPI_ISL_3125994 | Lighthouse Lab in Glasgow | Wellcome Sanger Institute for the COVID-19 Genomics UK (COG-UK) Consortium | Harper VanSteenhouse, Yumi Kasai, David Gray, Carol Clugston, Anna Dominiczak and Alex Alderton, Roberto Amato, Jeffrey Barrett, Sonia Goncalves, Ewan Harrison, David K. Jackson, Ian Johnston, Dominic Kwiatkowski, Cordelia Langford, John Sillitoe on behalf of the Wellcome Sanger Institute COVID-19 Surveillance Team |
| EPI_ISL_3125995, EPI_ISL_3125996, EPI_ISL_3125997, EPI_ISL_3125998 | Lighthouse Lab in Alderley Park | Wellcome Sanger Institute for the COVID-19 Genomics UK (COG-UK) Consortium | Jacquelyn Wynn, Mairead Hyland, The Lighthouse Lab in Alderley Park and Alex Alderton, Roberto Amato, Jeffrey Barrett, Sonia Goncalves, Ewan Harrison, David K. Jackson, Ian Johnston, Dominic Kwiatkowski, Cordelia Langford, John Sillitoe on behalf of the Wellcome Sanger Institute COVID-19 Surveillance Team |

| | | | |
|---|---|---|---|
| EPI_ISL_3125999, EPI_ISL_3126000 | Lighthouse Lab in Glasgow | Wellcome Sanger Institute for the COVID-19 Genomics UK (COG-UK) Consortium | Harper VanSteenhouse, Yumi Kasai, David Gray, Carol Clugston, Anna Dominiczak and Alex Alderton, Roberto Amato, Jeffrey Barrett, Sonia Goncalves, Ewan Harrison, David K. Jackson, Ian Johnston, Dominic Kwiatkowski, Cordelia Langford, John Sillitoe on behalf of the Wellcome Sanger Institute COVID-19 Surveillance Team |
| EPI_ISL_3126001, EPI_ISL_3126002 | Lighthouse Lab in Alderley Park | Wellcome Sanger Institute for the COVID-19 Genomics UK (COG-UK) Consortium | Jacquelyn Wynn, Mairead Hyland, The Lighthouse Lab in Alderley Park and Alex Alderton, Roberto Amato, Jeffrey Barrett, Sonia Goncalves, Ewan Harrison, David K. Jackson, Ian Johnston, Dominic Kwiatkowski, Cordelia Langford, John Sillitoe on behalf of the Wellcome Sanger Institute COVID-19 Surveillance Team |
| EPI_ISL_3126003 | Lighthouse Lab in Glasgow | Wellcome Sanger Institute for the COVID-19 Genomics UK (COG-UK) Consortium | Harper VanSteenhouse, Yumi Kasai, David Gray, Carol Clugston, Anna Dominiczak and Alex Alderton, Roberto Amato, Jeffrey Barrett, Sonia Goncalves, Ewan Harrison, David K. Jackson, Ian Johnston, Dominic Kwiatkowski, Cordelia Langford, John Sillitoe on behalf of the Wellcome Sanger Institute COVID-19 Surveillance Team |
| EPI_ISL_3126004, EPI_ISL_3126005 | Lighthouse Lab in Alderley Park | Wellcome Sanger Institute for the COVID-19 Genomics UK (COG-UK) Consortium | Jacquelyn Wynn, Mairead Hyland, The Lighthouse Lab in Alderley Park and Alex Alderton, Roberto Amato, Jeffrey Barrett, Sonia Goncalves, Ewan Harrison, David K. Jackson, Ian Johnston, Dominic Kwiatkowski, Cordelia Langford, John Sillitoe on behalf of the Wellcome Sanger Institute COVID-19 Surveillance Team |
| EPI_ISL_3126006, EPI_ISL_3126007, EPI_ISL_3126008, EPI_ISL_3126009 | Lighthouse Lab in Glasgow | Wellcome Sanger Institute for the COVID-19 Genomics UK (COG-UK) Consortium | Harper VanSteenhouse, Yumi Kasai, David Gray, Carol Clugston, Anna Dominiczak and Alex Alderton, Roberto Amato, Jeffrey Barrett, Sonia Goncalves, Ewan Harrison, David K. Jackson, Ian Johnston, Dominic Kwiatkowski, Cordelia Langford, John Sillitoe on behalf of the Wellcome Sanger Institute COVID-19 Surveillance Team |
| EPI_ISL_3126010, EPI_ISL_3126011 | Lighthouse Lab in Alderley Park | Wellcome Sanger Institute for the COVID-19 Genomics UK (COG-UK) Consortium | Jacquelyn Wynn, Mairead Hyland, The Lighthouse Lab in Alderley Park and Alex Alderton, Roberto Amato, Jeffrey Barrett, Sonia Goncalves, Ewan Harrison, David K. Jackson, Ian Johnston, Dominic Kwiatkowski, Cordelia Langford, John Sillitoe on behalf of the Wellcome Sanger Institute COVID-19 Surveillance Team |
| EPI_ISL_3126012, EPI_ISL_3126013, EPI_ISL_3126014 | Lighthouse Lab in Glasgow | Wellcome Sanger Institute for the COVID-19 Genomics UK (COG-UK) Consortium | Harper VanSteenhouse, Yumi Kasai, David Gray, Carol Clugston, Anna Dominiczak and Alex Alderton, Roberto Amato, Jeffrey Barrett, Sonia Goncalves, Ewan Harrison, David K. Jackson, Ian Johnston, Dominic Kwiatkowski, Cordelia Langford, John Sillitoe on behalf of the Wellcome Sanger Institute COVID-19 Surveillance Team |
| EPI_ISL_3126015, EPI_ISL_3126016 | Lighthouse Lab in Alderley Park | Wellcome Sanger Institute for the COVID-19 Genomics UK (COG-UK) Consortium | Jacquelyn Wynn, Mairead Hyland, The Lighthouse Lab in Alderley Park and Alex Alderton, Roberto Amato, Jeffrey Barrett, Sonia Goncalves, Ewan Harrison, David K. Jackson, Ian Johnston, Dominic Kwiatkowski, Cordelia Langford, John Sillitoe on behalf of the Wellcome Sanger Institute COVID-19 Surveillance Team |
| EPI_ISL_3126017 | Lighthouse Lab in Glasgow | Wellcome Sanger Institute for the COVID-19 Genomics UK (COG-UK) Consortium | Harper VanSteenhouse, Yumi Kasai, David Gray, Carol Clugston, Anna Dominiczak and Alex Alderton, Roberto Amato, Jeffrey Barrett, Sonia Goncalves, Ewan Harrison, David K. Jackson, Ian Johnston, Dominic Kwiatkowski, Cordelia Langford, John Sillitoe on behalf of the Wellcome Sanger Institute COVID-19 Surveillance Team |
| EPI_ISL_3126018 | Lighthouse Lab in Alderley Park | Wellcome Sanger Institute for the COVID-19 Genomics UK (COG-UK) Consortium | Jacquelyn Wynn, Mairead Hyland, The Lighthouse Lab in Alderley Park and Alex Alderton, Roberto Amato, Jeffrey Barrett, Sonia Goncalves, Ewan Harrison, David K. Jackson, Ian Johnston, Dominic Kwiatkowski, Cordelia Langford, John Sillitoe on behalf of the Wellcome Sanger Institute COVID-19 Surveillance Team |
| EPI_ISL_3126019 | Lighthouse Lab in Glasgow | Wellcome Sanger Institute for the COVID-19 Genomics UK (COG-UK) Consortium | Harper VanSteenhouse, Yumi Kasai, David Gray, Carol Clugston, Anna Dominiczak and Alex Alderton, Roberto Amato, Jeffrey Barrett, Sonia Goncalves, Ewan Harrison, David K. Jackson, Ian Johnston, Dominic Kwiatkowski, Cordelia Langford, John Sillitoe on behalf of the Wellcome Sanger Institute COVID-19 Surveillance Team |
| EPI_ISL_3126020, EPI_ISL_3126021 | Lighthouse Lab in Alderley Park | Wellcome Sanger Institute for the COVID-19 Genomics UK (COG-UK) Consortium | Jacquelyn Wynn, Mairead Hyland, The Lighthouse Lab in Alderley Park and Alex Alderton, Roberto Amato, Jeffrey Barrett, Sonia Goncalves, Ewan Harrison, David K. Jackson, Ian Johnston, Dominic Kwiatkowski, Cordelia Langford, John Sillitoe on behalf of the Wellcome Sanger Institute COVID-19 Surveillance Team |
| EPI_ISL_3126022 | Lighthouse Lab in Glasgow | Wellcome Sanger Institute for the COVID-19 Genomics UK (COG-UK) Consortium | Harper VanSteenhouse, Yumi Kasai, David Gray, Carol Clugston, Anna Dominiczak and Alex Alderton, Roberto Amato, Jeffrey Barrett, Sonia Goncalves, Ewan Harrison, David K. Jackson, Ian Johnston, Dominic Kwiatkowski, Cordelia Langford, John Sillitoe on behalf of the Wellcome Sanger Institute COVID-19 Surveillance Team |
| EPI_ISL_3126023 | Lighthouse Lab in Alderley Park | Wellcome Sanger Institute for the COVID-19 Genomics UK (COG-UK) Consortium | Jacquelyn Wynn, Mairead Hyland, The Lighthouse Lab in Alderley Park and Alex Alderton, Roberto Amato, Jeffrey Barrett, Sonia Goncalves, Ewan Harrison, David K. Jackson, Ian Johnston, Dominic Kwiatkowski, Cordelia Langford, John Sillitoe on behalf of the Wellcome Sanger Institute COVID-19 Surveillance Team |
| EPI_ISL_3126024 | Lighthouse Lab in Glasgow | Wellcome Sanger Institute for the COVID-19 Genomics UK (COG-UK) Consortium | Harper VanSteenhouse, Yumi Kasai, David Gray, Carol Clugston, Anna Dominiczak and Alex Alderton, Roberto Amato, Jeffrey Barrett, Sonia Goncalves, Ewan Harrison, David K. Jackson, Ian Johnston, Dominic Kwiatkowski, Cordelia Langford, John Sillitoe on behalf of the Wellcome Sanger Institute COVID-19 Surveillance Team |
| EPI_ISL_3126025, EPI_ISL_3126026 | Lighthouse Lab in Alderley Park | Wellcome Sanger Institute for the COVID-19 Genomics UK (COG-UK) Consortium | Jacquelyn Wynn, Mairead Hyland, The Lighthouse Lab in Alderley Park and Alex Alderton, Roberto Amato, Jeffrey Barrett, Sonia Goncalves, Ewan Harrison, David K. Jackson, Ian Johnston, Dominic Kwiatkowski, Cordelia Langford, John Sillitoe on behalf of the Wellcome Sanger Institute COVID-19 Surveillance Team |
| EPI_ISL_3126027 | Lighthouse Lab in Glasgow | Wellcome Sanger Institute for the COVID-19 Genomics UK (COG-UK) Consortium | Harper VanSteenhouse, Yumi Kasai, David Gray, Carol Clugston, Anna Dominiczak and Alex Alderton, Roberto Amato, Jeffrey Barrett, Sonia Goncalves, Ewan Harrison, David K. Jackson, Ian Johnston, Dominic Kwiatkowski, Cordelia Langford, John Sillitoe on behalf of the Wellcome Sanger Institute COVID-19 Surveillance Team |
| EPI_ISL_3126028 | Lighthouse Lab in Alderley Park | Wellcome Sanger Institute for the COVID-19 Genomics UK (COG-UK) Consortium | Jacquelyn Wynn, Mairead Hyland, The Lighthouse Lab in Alderley Park and Alex Alderton, Roberto Amato, Jeffrey Barrett, Sonia Goncalves, Ewan Harrison, David K. Jackson, Ian Johnston, Dominic Kwiatkowski, Cordelia Langford, John Sillitoe on behalf of the Wellcome Sanger Institute COVID-19 Surveillance Team |
| EPI_ISL_3126029 | Lighthouse Lab in Glasgow | Wellcome Sanger Institute for the COVID-19 Genomics UK (COG-UK) Consortium | Harper VanSteenhouse, Yumi Kasai, David Gray, Carol Clugston, Anna Dominiczak and Alex Alderton, Roberto Amato, Jeffrey Barrett, Sonia Goncalves, Ewan Harrison, David K. Jackson, Ian Johnston, Dominic Kwiatkowski, Cordelia Langford, John Sillitoe on behalf of the Wellcome Sanger Institute COVID-19 Surveillance Team |
| EPI_ISL_3126030, EPI_ISL_3126031 | Lighthouse Lab in Alderley Park | Wellcome Sanger Institute for the COVID-19 Genomics UK (COG-UK) Consortium | Jacquelyn Wynn, Mairead Hyland, The Lighthouse Lab in Alderley Park and Alex Alderton, Roberto Amato, Jeffrey Barrett, Sonia Goncalves, Ewan Harrison, David K. Jackson, Ian Johnston, Dominic Kwiatkowski, Cordelia Langford, John Sillitoe on behalf of the Wellcome Sanger Institute COVID-19 Surveillance Team |
| EPI_ISL_3126032 | Lighthouse Lab in Glasgow | Wellcome Sanger Institute for the COVID-19 Genomics UK (COG-UK) Consortium | Harper VanSteenhouse, Yumi Kasai, David Gray, Carol Clugston, Anna Dominiczak and Alex Alderton, Roberto Amato, Jeffrey Barrett, Sonia Goncalves, Ewan Harrison, David K. Jackson, Ian Johnston, Dominic Kwiatkowski, Cordelia Langford, John Sillitoe on behalf of the Wellcome Sanger Institute COVID-19 Surveillance Team |
| EPI_ISL_3126033 | Lighthouse Lab in Alderley Park | Wellcome Sanger Institute for the COVID-19 Genomics UK (COG-UK) Consortium | Jacquelyn Wynn, Mairead Hyland, The Lighthouse Lab in Alderley Park and Alex Alderton, Roberto Amato, Jeffrey Barrett, Sonia Goncalves, Ewan Harrison, David K. Jackson, Ian Johnston, Dominic Kwiatkowski, Cordelia Langford, John Sillitoe on behalf of the Wellcome Sanger Institute COVID-19 Surveillance Team |
| EPI_ISL_3126034 | Lighthouse Lab in Glasgow | Wellcome Sanger Institute for the COVID-19 Genomics UK (COG-UK) Consortium | Harper VanSteenhouse, Yumi Kasai, David Gray, Carol Clugston, Anna Dominiczak and Alex Alderton, Roberto Amato, Jeffrey Barrett, Sonia Goncalves, Ewan Harrison, David K. Jackson, Ian Johnston, Dominic Kwiatkowski, Cordelia Langford, John Sillitoe on behalf of the Wellcome Sanger Institute COVID-19 Surveillance Team |
| EPI_ISL_3126035, EPI_ISL_3126036, EPI_ISL_3126037 | Lighthouse Lab in Alderley Park | Wellcome Sanger Institute for the COVID-19 Genomics UK (COG-UK) Consortium | Jacquelyn Wynn, Mairead Hyland, The Lighthouse Lab in Alderley Park and Alex Alderton, Roberto Amato, Jeffrey Barrett, Sonia Goncalves, Ewan Harrison, David K. Jackson, Ian Johnston, Dominic Kwiatkowski, Cordelia Langford, John Sillitoe on behalf of the Wellcome Sanger Institute COVID-19 Surveillance Team |
| EPI_ISL_3126038 | Lighthouse Lab in Glasgow | Wellcome Sanger Institute for the COVID-19 Genomics UK (COG-UK) Consortium | Harper VanSteenhouse, Yumi Kasai, David Gray, Carol Clugston, Anna Dominiczak and Alex Alderton, Roberto Amato, Jeffrey Barrett, Sonia Goncalves, Ewan Harrison, David K. Jackson, Ian Johnston, Dominic Kwiatkowski, Cordelia Langford, John Sillitoe on behalf of the Wellcome Sanger Institute COVID-19 Surveillance Team |
| EPI_ISL_3126039, EPI_ISL_3126040, EPI_ISL_3126041 | Lighthouse Lab in Alderley Park | Wellcome Sanger Institute for the COVID-19 Genomics UK (COG-UK) Consortium | Jacquelyn Wynn, Mairead Hyland, The Lighthouse Lab in Alderley Park and Alex Alderton, Roberto Amato, Jeffrey Barrett, Sonia Goncalves, Ewan Harrison, David K. Jackson, Ian Johnston, Dominic Kwiatkowski, Cordelia Langford, John Sillitoe on behalf of the Wellcome Sanger Institute COVID-19 Surveillance Team |
| EPI_ISL_3126042 | Lighthouse Lab in Glasgow | Wellcome Sanger Institute for the COVID-19 Genomics UK (COG-UK) Consortium | Harper VanSteenhouse, Yumi Kasai, David Gray, Carol Clugston, Anna Dominiczak and Alex Alderton, Roberto Amato, Jeffrey Barrett, Sonia Goncalves, Ewan Harrison, David K. Jackson, Ian Johnston, Dominic Kwiatkowski, Cordelia Langford, John Sillitoe on behalf of the Wellcome Sanger Institute COVID-19 Surveillance Team |
| EPI_ISL_3126043, EPI_ISL_3126044, EPI_ISL_3126045, EPI_ISL_3126046 | Lighthouse Lab in Alderley Park | Wellcome Sanger Institute for the COVID-19 Genomics UK (COG-UK) Consortium | Jacquelyn Wynn, Mairead Hyland, The Lighthouse Lab in Alderley Park and Alex Alderton, Roberto Amato, Jeffrey Barrett, Sonia Goncalves, Ewan Harrison, David K. Jackson, Ian Johnston, Dominic Kwiatkowski, Cordelia Langford, John Sillitoe on behalf of the Wellcome Sanger Institute COVID-19 Surveillance Team |
| EPI_ISL_3126047 | Lighthouse Lab in Glasgow | Wellcome Sanger Institute for the COVID-19 Genomics UK (COG-UK) Consortium | Harper VanSteenhouse, Yumi Kasai, David Gray, Carol Clugston, Anna Dominiczak and Alex Alderton, Roberto Amato, Jeffrey Barrett, Sonia Goncalves, Ewan Harrison, David K. Jackson, Ian Johnston, Dominic Kwiatkowski, Cordelia Langford, John Sillitoe on behalf of the Wellcome Sanger Institute COVID-19 Surveillance Team |
| EPI_ISL_3126048 | Lighthouse Lab in Alderley Park | Wellcome Sanger Institute for the COVID-19 Genomics UK (COG-UK) Consortium | Jacquelyn Wynn, Mairead Hyland, The Lighthouse Lab in Alderley Park and Alex Alderton, Roberto Amato, Jeffrey Barrett, Sonia Goncalves, Ewan Harrison, David K. Jackson, Ian Johnston, Dominic Kwiatkowski, Cordelia Langford, John Sillitoe on behalf of the Wellcome Sanger Institute COVID-19 Surveillance Team |
| EPI_ISL_3126049 | Lighthouse Lab in Glasgow | Wellcome Sanger Institute for the COVID-19 Genomics UK (COG-UK) Consortium | Harper VanSteenhouse, Yumi Kasai, David Gray, Carol Clugston, Anna Dominiczak and Alex Alderton, Roberto Amato, Jeffrey Barrett, Sonia Goncalves, Ewan Harrison, David K. Jackson, Ian Johnston, Dominic Kwiatkowski, Cordelia Langford, John Sillitoe on behalf of the Wellcome Sanger Institute COVID-19 Surveillance Team |
| EPI_ISL_3126050, EPI_ISL_3126051, EPI_ISL_3126052 | Lighthouse Lab in Alderley Park | Wellcome Sanger Institute for the COVID-19 Genomics UK (COG-UK) Consortium | Jacquelyn Wynn, Mairead Hyland, The Lighthouse Lab in Alderley Park and Alex Alderton, Roberto Amato, Jeffrey Barrett, Sonia Goncalves, Ewan Harrison, David K. Jackson, Ian Johnston, Dominic Kwiatkowski, Cordelia Langford, John Sillitoe on behalf of the Wellcome Sanger Institute COVID-19 Surveillance Team |
| EPI_ISL_3126053 | Lighthouse Lab in Glasgow | Wellcome Sanger Institute for the COVID-19 Genomics UK (COG-UK) Consortium | Harper VanSteenhouse, Yumi Kasai, David Gray, Carol Clugston, Anna Dominiczak and Alex Alderton, Roberto Amato, Jeffrey Barrett, Sonia Goncalves, Ewan Harrison, David K. Jackson, Ian Johnston, Dominic Kwiatkowski, Cordelia Langford, John Sillitoe on behalf of the Wellcome Sanger Institute COVID-19 Surveillance Team |
| EPI_ISL_3126054, EPI_ISL_3126055 | Lighthouse Lab in Alderley Park | Wellcome Sanger Institute for the COVID-19 Genomics UK (COG-UK) Consortium | Jacquelyn Wynn, Mairead Hyland, The Lighthouse Lab in Alderley Park and Alex Alderton, Roberto Amato, Jeffrey Barrett, Sonia Goncalves, Ewan Harrison, David K. Jackson, Ian Johnston, Dominic Kwiatkowski, Cordelia Langford, John Sillitoe on behalf of the Wellcome Sanger Institute COVID-19 Surveillance Team |
| EPI_ISL_3126056 | Lighthouse Lab in Glasgow | Wellcome Sanger Institute for the COVID-19 Genomics UK (COG-UK) Consortium | Harper VanSteenhouse, Yumi Kasai, David Gray, Carol Clugston, Anna Dominiczak and Alex Alderton, Roberto Amato, Jeffrey Barrett, Sonia Goncalves, Ewan Harrison, David K. Jackson, Ian Johnston, Dominic Kwiatkowski, Cordelia Langford, John Sillitoe on behalf of the Wellcome Sanger Institute COVID-19 Surveillance Team |
| EPI_ISL_3126057, EPI_ISL_3126058, EPI_ISL_3126059 | Lighthouse Lab in Alderley Park | Wellcome Sanger Institute for the COVID-19 Genomics UK (COG-UK) Consortium | Jacquelyn Wynn, Mairead Hyland, The Lighthouse Lab in Alderley Park and Alex Alderton, Roberto Amato, Jeffrey Barrett, Sonia Goncalves, Ewan Harrison, David K. Jackson, Ian Johnston, Dominic Kwiatkowski, Cordelia Langford, John Sillitoe on behalf of the Wellcome Sanger Institute COVID-19 Surveillance Team |
| EPI_ISL_3126060, EPI_ISL_3126061, EPI_ISL_3126062 | Lighthouse Lab in Glasgow | Wellcome Sanger Institute for the COVID-19 Genomics UK (COG-UK) Consortium | Harper VanSteenhouse, Yumi Kasai, David Gray, Carol Clugston, Anna Dominiczak and Alex Alderton, Roberto Amato, Jeffrey Barrett, Sonia Goncalves, Ewan Harrison, David K. Jackson, Ian Johnston, Dominic Kwiatkowski, Cordelia Langford, John Sillitoe on behalf of the Wellcome Sanger Institute COVID-19 Surveillance Team |
| EPI_ISL_3126063, EPI_ISL_3126064, EPI_ISL_3126065, EPI_ISL_3126066 | Lighthouse Lab in Alderley Park | Wellcome Sanger Institute for the COVID-19 Genomics UK (COG-UK) Consortium | Jacquelyn Wynn, Mairead Hyland, The Lighthouse Lab in Alderley Park and Alex Alderton, Roberto Amato, Jeffrey Barrett, Sonia Goncalves, Ewan Harrison, David K. Jackson, Ian Johnston, Dominic Kwiatkowski, Cordelia Langford, John Sillitoe on behalf of the Wellcome Sanger Institute COVID-19 Surveillance Team |
| EPI_ISL_3126067 | Lighthouse Lab in Glasgow | Wellcome Sanger Institute for the COVID-19 Genomics UK (COG-UK) Consortium | Harper VanSteenhouse, Yumi Kasai, David Gray, Carol Clugston, Anna Dominiczak and Alex Alderton, Roberto Amato, Jeffrey Barrett, Sonia Goncalves, Ewan Harrison, David K. Jackson, Ian Johnston, Dominic Kwiatkowski, Cordelia Langford, John Sillitoe on behalf of the Wellcome Sanger Institute COVID-19 Surveillance Team |
| EPI_ISL_3126068 | Lighthouse Lab in Alderley Park | Wellcome Sanger Institute for the COVID-19 Genomics UK (COG-UK) Consortium | Jacquelyn Wynn, Mairead Hyland, The Lighthouse Lab in Alderley Park and Alex Alderton, Roberto Amato, Jeffrey Barrett, Sonia Goncalves, Ewan Harrison, David K. Jackson, Ian Johnston, Dominic Kwiatkowski, Cordelia Langford, John Sillitoe on behalf of the Wellcome Sanger Institute COVID-19 Surveillance Team |

| | | | |
|---|---|---|---|
| EPI_ISL_3126069 | Lighthouse Lab in Glasgow | Wellcome Sanger Institute for the COVID-19 Genomics UK (COG-UK) Consortium | Harper VanSteenhouse, Yumi Kasai, David Gray, Carol Clugston, Anna Dominiczak and Alex Alderton, Roberto Amato, Jeffrey Barrett, Sonia Goncalves, Ewan Harrison, David K. Jackson, Ian Johnston, Dominic Kwiatkowski, Cordelia Langford, John Sillitoe on behalf of the Wellcome Sanger Institute COVID-19 Surveillance Team |
| EPI_ISL_3126070 | Lighthouse Lab in Alderley Park | Wellcome Sanger Institute for the COVID-19 Genomics UK (COG-UK) Consortium | Jacquelyn Wynn, Mairead Hyland, The Lighthouse Lab in Alderley Park and Alex Alderton, Roberto Amato, Jeffrey Barrett, Sonia Goncalves, Ewan Harrison, David K. Jackson, Ian Johnston, Dominic Kwiatkowski, Cordelia Langford, John Sillitoe on behalf of the Wellcome Sanger Institute COVID-19 Surveillance Team |
| EPI_ISL_3126071 | Lighthouse Lab in Glasgow | Wellcome Sanger Institute for the COVID-19 Genomics UK (COG-UK) Consortium | Harper VanSteenhouse, Yumi Kasai, David Gray, Carol Clugston, Anna Dominiczak and Alex Alderton, Roberto Amato, Jeffrey Barrett, Sonia Goncalves, Ewan Harrison, David K. Jackson, Ian Johnston, Dominic Kwiatkowski, Cordelia Langford, John Sillitoe on behalf of the Wellcome Sanger Institute COVID-19 Surveillance Team |
| EPI_ISL_3126072 | Lighthouse Lab in Alderley Park | Wellcome Sanger Institute for the COVID-19 Genomics UK (COG-UK) Consortium | Jacquelyn Wynn, Mairead Hyland, The Lighthouse Lab in Alderley Park and Alex Alderton, Roberto Amato, Jeffrey Barrett, Sonia Goncalves, Ewan Harrison, David K. Jackson, Ian Johnston, Dominic Kwiatkowski, Cordelia Langford, John Sillitoe on behalf of the Wellcome Sanger Institute COVID-19 Surveillance Team |
| EPI_ISL_3126073, EPI_ISL_3126074, EPI_ISL_3126075, EPI_ISL_3126076, EPI_ISL_3126077 | Lighthouse Lab in Glasgow | Wellcome Sanger Institute for the COVID-19 Genomics UK (COG-UK) Consortium | Harper VanSteenhouse, Yumi Kasai, David Gray, Carol Clugston, Anna Dominiczak and Alex Alderton, Roberto Amato, Jeffrey Barrett, Sonia Goncalves, Ewan Harrison, David K. Jackson, Ian Johnston, Dominic Kwiatkowski, Cordelia Langford, John Sillitoe on behalf of the Wellcome Sanger Institute COVID-19 Surveillance Team |
| EPI_ISL_3126078 | Lighthouse Lab in Alderley Park | Wellcome Sanger Institute for the COVID-19 Genomics UK (COG-UK) Consortium | Jacquelyn Wynn, Mairead Hyland, The Lighthouse Lab in Alderley Park and Alex Alderton, Roberto Amato, Jeffrey Barrett, Sonia Goncalves, Ewan Harrison, David K. Jackson, Ian Johnston, Dominic Kwiatkowski, Cordelia Langford, John Sillitoe on behalf of the Wellcome Sanger Institute COVID-19 Surveillance Team |
| EPI_ISL_3126079, EPI_ISL_3126080, EPI_ISL_3126081 | Lighthouse Lab in Glasgow | Wellcome Sanger Institute for the COVID-19 Genomics UK (COG-UK) Consortium | Harper VanSteenhouse, Yumi Kasai, David Gray, Carol Clugston, Anna Dominiczak and Alex Alderton, Roberto Amato, Jeffrey Barrett, Sonia Goncalves, Ewan Harrison, David K. Jackson, Ian Johnston, Dominic Kwiatkowski, Cordelia Langford, John Sillitoe on behalf of the Wellcome Sanger Institute COVID-19 Surveillance Team |
| EPI_ISL_3126082, EPI_ISL_3126083 | Lighthouse Lab in Alderley Park | Wellcome Sanger Institute for the COVID-19 Genomics UK (COG-UK) Consortium | Jacquelyn Wynn, Mairead Hyland, The Lighthouse Lab in Alderley Park and Alex Alderton, Roberto Amato, Jeffrey Barrett, Sonia Goncalves, Ewan Harrison, David K. Jackson, Ian Johnston, Dominic Kwiatkowski, Cordelia Langford, John Sillitoe on behalf of the Wellcome Sanger Institute COVID-19 Surveillance Team |
| EPI_ISL_3126084 | Lighthouse Lab in Glasgow | Wellcome Sanger Institute for the COVID-19 Genomics UK (COG-UK) Consortium | Harper VanSteenhouse, Yumi Kasai, David Gray, Carol Clugston, Anna Dominiczak and Alex Alderton, Roberto Amato, Jeffrey Barrett, Sonia Goncalves, Ewan Harrison, David K. Jackson, Ian Johnston, Dominic Kwiatkowski, Cordelia Langford, John Sillitoe on behalf of the Wellcome Sanger Institute COVID-19 Surveillance Team |
| EPI_ISL_3126085, EPI_ISL_3126086 | Lighthouse Lab in Alderley Park | Wellcome Sanger Institute for the COVID-19 Genomics UK (COG-UK) Consortium | Jacquelyn Wynn, Mairead Hyland, The Lighthouse Lab in Alderley Park and Alex Alderton, Roberto Amato, Jeffrey Barrett, Sonia Goncalves, Ewan Harrison, David K. Jackson, Ian Johnston, Dominic Kwiatkowski, Cordelia Langford, John Sillitoe on behalf of the Wellcome Sanger Institute COVID-19 Surveillance Team |
| EPI_ISL_3126087, EPI_ISL_3126088 | Lighthouse Lab in Glasgow | Wellcome Sanger Institute for the COVID-19 Genomics UK (COG-UK) Consortium | Harper VanSteenhouse, Yumi Kasai, David Gray, Carol Clugston, Anna Dominiczak and Alex Alderton, Roberto Amato, Jeffrey Barrett, Sonia Goncalves, Ewan Harrison, David K. Jackson, Ian Johnston, Dominic Kwiatkowski, Cordelia Langford, John Sillitoe on behalf of the Wellcome Sanger Institute COVID-19 Surveillance Team |
| EPI_ISL_3126089 | Lighthouse Lab in Alderley Park | Wellcome Sanger Institute for the COVID-19 Genomics UK (COG-UK) Consortium | Jacquelyn Wynn, Mairead Hyland, The Lighthouse Lab in Alderley Park and Alex Alderton, Roberto Amato, Jeffrey Barrett, Sonia Goncalves, Ewan Harrison, David K. Jackson, Ian Johnston, Dominic Kwiatkowski, Cordelia Langford, John Sillitoe on behalf of the Wellcome Sanger Institute COVID-19 Surveillance Team |
| EPI_ISL_3126090, EPI_ISL_3126091, EPI_ISL_3126092 | Lighthouse Lab in Glasgow | Wellcome Sanger Institute for the COVID-19 Genomics UK (COG-UK) Consortium | Harper VanSteenhouse, Yumi Kasai, David Gray, Carol Clugston, Anna Dominiczak and Alex Alderton, Roberto Amato, Jeffrey Barrett, Sonia Goncalves, Ewan Harrison, David K. Jackson, Ian Johnston, Dominic Kwiatkowski, Cordelia Langford, John Sillitoe on behalf of the Wellcome Sanger Institute COVID-19 Surveillance Team |
| EPI_ISL_3126093 | Lighthouse Lab in Alderley Park | Wellcome Sanger Institute for the COVID-19 Genomics UK (COG-UK) Consortium | Jacquelyn Wynn, Mairead Hyland, The Lighthouse Lab in Alderley Park and Alex Alderton, Roberto Amato, Jeffrey Barrett, Sonia Goncalves, Ewan Harrison, David K. Jackson, Ian Johnston, Dominic Kwiatkowski, Cordelia Langford, John Sillitoe on behalf of the Wellcome Sanger Institute COVID-19 Surveillance Team |
| EPI_ISL_3126094, EPI_ISL_3126095 | Lighthouse Lab in Glasgow | Wellcome Sanger Institute for the COVID-19 Genomics UK (COG-UK) Consortium | Harper VanSteenhouse, Yumi Kasai, David Gray, Carol Clugston, Anna Dominiczak and Alex Alderton, Roberto Amato, Jeffrey Barrett, Sonia Goncalves, Ewan Harrison, David K. Jackson, Ian Johnston, Dominic Kwiatkowski, Cordelia Langford, John Sillitoe on behalf of the Wellcome Sanger Institute COVID-19 Surveillance Team |
| EPI_ISL_3126096, EPI_ISL_3126097, EPI_ISL_3126098 | Lighthouse Lab in Alderley Park | Wellcome Sanger Institute for the COVID-19 Genomics UK (COG-UK) Consortium | Jacquelyn Wynn, Mairead Hyland, The Lighthouse Lab in Alderley Park and Alex Alderton, Roberto Amato, Jeffrey Barrett, Sonia Goncalves, Ewan Harrison, David K. Jackson, Ian Johnston, Dominic Kwiatkowski, Cordelia Langford, John Sillitoe on behalf of the Wellcome Sanger Institute COVID-19 Surveillance Team |
| EPI_ISL_3126099, EPI_ISL_3126100 | Lighthouse Lab in Glasgow | Wellcome Sanger Institute for the COVID-19 Genomics UK (COG-UK) Consortium | Harper VanSteenhouse, Yumi Kasai, David Gray, Carol Clugston, Anna Dominiczak and Alex Alderton, Roberto Amato, Jeffrey Barrett, Sonia Goncalves, Ewan Harrison, David K. Jackson, Ian Johnston, Dominic Kwiatkowski, Cordelia Langford, John Sillitoe on behalf of the Wellcome Sanger Institute COVID-19 Surveillance Team |
| EPI_ISL_3126101, EPI_ISL_3126102, EPI_ISL_3126103, EPI_ISL_3126104, EPI_ISL_3126105, EPI_ISL_3126106, EPI_ISL_3126107, EPI_ISL_3126108, EPI_ISL_3126109, EPI_ISL_3126110, EPI_ISL_3126111, EPI_ISL_3126112, EPI_ISL_3126113, EPI_ISL_3126114, EPI_ISL_3126115, EPI_ISL_3126116, EPI_ISL_3126117 | | | |
| see above | Lighthouse Lab in Alderley Park | Wellcome Sanger Institute for the COVID-19 Genomics UK (COG-UK) Consortium | Jacquelyn Wynn, Mairead Hyland, The Lighthouse Lab in Alderley Park and Alex Alderton, Roberto Amato, Jeffrey Barrett, Sonia Goncalves, Ewan Harrison, David K. Jackson, Ian Johnston, Dominic Kwiatkowski, Cordelia Langford, John Sillitoe on behalf of the Wellcome Sanger Institute COVID-19 Surveillance Team |
| EPI_ISL_3126118 | Lighthouse Lab in Glasgow | Wellcome Sanger Institute for the COVID-19 Genomics UK (COG-UK) Consortium | Harper VanSteenhouse, Yumi Kasai, David Gray, Carol Clugston, Anna Dominiczak and Alex Alderton, Roberto Amato, Jeffrey Barrett, Sonia Goncalves, Ewan Harrison, David K. Jackson, Ian Johnston, Dominic Kwiatkowski, Cordelia Langford, John Sillitoe on behalf of the Wellcome Sanger Institute COVID-19 Surveillance Team |
| EPI_ISL_3126119 | Lighthouse Lab in Alderley Park | Wellcome Sanger Institute for the COVID-19 Genomics UK (COG-UK) Consortium | Jacquelyn Wynn, Mairead Hyland, The Lighthouse Lab in Alderley Park and Alex Alderton, Roberto Amato, Jeffrey Barrett, Sonia Goncalves, Ewan Harrison, David K. Jackson, Ian Johnston, Dominic Kwiatkowski, Cordelia Langford, John Sillitoe on behalf of the Wellcome Sanger Institute COVID-19 Surveillance Team |
| EPI_ISL_3126120 | Lighthouse Lab in Glasgow | Wellcome Sanger Institute for the COVID-19 Genomics UK (COG-UK) Consortium | Harper VanSteenhouse, Yumi Kasai, David Gray, Carol Clugston, Anna Dominiczak and Alex Alderton, Roberto Amato, Jeffrey Barrett, Sonia Goncalves, Ewan Harrison, David K. Jackson, Ian Johnston, Dominic Kwiatkowski, Cordelia Langford, John Sillitoe on behalf of the Wellcome Sanger Institute COVID-19 Surveillance Team |
| EPI_ISL_3126121 | Lighthouse Lab in Alderley Park | Wellcome Sanger Institute for the COVID-19 Genomics UK (COG-UK) Consortium | Jacquelyn Wynn, Mairead Hyland, The Lighthouse Lab in Alderley Park and Alex Alderton, Roberto Amato, Jeffrey Barrett, Sonia Goncalves, Ewan Harrison, David K. Jackson, Ian Johnston, Dominic Kwiatkowski, Cordelia Langford, John Sillitoe on behalf of the Wellcome Sanger Institute COVID-19 Surveillance Team |
| EPI_ISL_3126122 | Lighthouse Lab in Glasgow | Wellcome Sanger Institute for the COVID-19 Genomics UK (COG-UK) Consortium | Harper VanSteenhouse, Yumi Kasai, David Gray, Carol Clugston, Anna Dominiczak and Alex Alderton, Roberto Amato, Jeffrey Barrett, Sonia Goncalves, Ewan Harrison, David K. Jackson, Ian Johnston, Dominic Kwiatkowski, Cordelia Langford, John Sillitoe on behalf of the Wellcome Sanger Institute COVID-19 Surveillance Team |
| EPI_ISL_3126123 | Lighthouse Lab in Alderley Park | Wellcome Sanger Institute for the COVID-19 Genomics UK (COG-UK) Consortium | Jacquelyn Wynn, Mairead Hyland, The Lighthouse Lab in Alderley Park and Alex Alderton, Roberto Amato, Jeffrey Barrett, Sonia Goncalves, Ewan Harrison, David K. Jackson, Ian Johnston, Dominic Kwiatkowski, Cordelia Langford, John Sillitoe on behalf of the Wellcome Sanger Institute COVID-19 Surveillance Team |
| EPI_ISL_3126124, EPI_ISL_3126125 | Lighthouse Lab in Glasgow | Wellcome Sanger Institute for the COVID-19 Genomics UK (COG-UK) Consortium | Harper VanSteenhouse, Yumi Kasai, David Gray, Carol Clugston, Anna Dominiczak and Alex Alderton, Roberto Amato, Jeffrey Barrett, Sonia Goncalves, Ewan Harrison, David K. Jackson, Ian Johnston, Dominic Kwiatkowski, Cordelia Langford, John Sillitoe on behalf of the Wellcome Sanger Institute COVID-19 Surveillance Team |
| EPI_ISL_3126126 | Lighthouse Lab in Alderley Park | Wellcome Sanger Institute for the COVID-19 Genomics UK (COG-UK) Consortium | Jacquelyn Wynn, Mairead Hyland, The Lighthouse Lab in Alderley Park and Alex Alderton, Roberto Amato, Jeffrey Barrett, Sonia Goncalves, Ewan Harrison, David K. Jackson, Ian Johnston, Dominic Kwiatkowski, Cordelia Langford, John Sillitoe on behalf of the Wellcome Sanger Institute COVID-19 Surveillance Team |
| EPI_ISL_3126127 | Lighthouse Lab in Glasgow | Wellcome Sanger Institute for the COVID-19 Genomics UK (COG-UK) Consortium | Harper VanSteenhouse, Yumi Kasai, David Gray, Carol Clugston, Anna Dominiczak and Alex Alderton, Roberto Amato, Jeffrey Barrett, Sonia Goncalves, Ewan Harrison, David K. Jackson, Ian Johnston, Dominic Kwiatkowski, Cordelia Langford, John Sillitoe on behalf of the Wellcome Sanger Institute COVID-19 Surveillance Team |
| EPI_ISL_3126128, EPI_ISL_3126129, EPI_ISL_3126130 | Lighthouse Lab in Alderley Park | Wellcome Sanger Institute for the COVID-19 Genomics UK (COG-UK) Consortium | Jacquelyn Wynn, Mairead Hyland, The Lighthouse Lab in Alderley Park and Alex Alderton, Roberto Amato, Jeffrey Barrett, Sonia Goncalves, Ewan Harrison, David K. Jackson, Ian Johnston, Dominic Kwiatkowski, Cordelia Langford, John Sillitoe on behalf of the Wellcome Sanger Institute COVID-19 Surveillance Team |
| EPI_ISL_3126131 | Lighthouse Lab in Glasgow | Wellcome Sanger Institute for the COVID-19 Genomics UK (COG-UK) Consortium | Harper VanSteenhouse, Yumi Kasai, David Gray, Carol Clugston, Anna Dominiczak and Alex Alderton, Roberto Amato, Jeffrey Barrett, Sonia Goncalves, Ewan Harrison, David K. Jackson, Ian Johnston, Dominic Kwiatkowski, Cordelia Langford, John Sillitoe on behalf of the Wellcome Sanger Institute COVID-19 Surveillance Team |
| EPI_ISL_3126132, EPI_ISL_3126133, EPI_ISL_3126134 | Lighthouse Lab in Alderley Park | Wellcome Sanger Institute for the COVID-19 Genomics UK (COG-UK) Consortium | Jacquelyn Wynn, Mairead Hyland, The Lighthouse Lab in Alderley Park and Alex Alderton, Roberto Amato, Jeffrey Barrett, Sonia Goncalves, Ewan Harrison, David K. Jackson, Ian Johnston, Dominic Kwiatkowski, Cordelia Langford, John Sillitoe on behalf of the Wellcome Sanger Institute COVID-19 Surveillance Team |
| EPI_ISL_3126135 | Lighthouse Lab in Glasgow | Wellcome Sanger Institute for the COVID-19 Genomics UK (COG-UK) Consortium | Harper VanSteenhouse, Yumi Kasai, David Gray, Carol Clugston, Anna Dominiczak and Alex Alderton, Roberto Amato, Jeffrey Barrett, Sonia Goncalves, Ewan Harrison, David K. Jackson, Ian Johnston, Dominic Kwiatkowski, Cordelia Langford, John Sillitoe on behalf of the Wellcome Sanger Institute COVID-19 Surveillance Team |
| EPI_ISL_3126136 | Lighthouse Lab in Alderley Park | Wellcome Sanger Institute for the COVID-19 Genomics UK (COG-UK) Consortium | Jacquelyn Wynn, Mairead Hyland, The Lighthouse Lab in Alderley Park and Alex Alderton, Roberto Amato, Jeffrey Barrett, Sonia Goncalves, Ewan Harrison, David K. Jackson, Ian Johnston, Dominic Kwiatkowski, Cordelia Langford, John Sillitoe on behalf of the Wellcome Sanger Institute COVID-19 Surveillance Team |
| EPI_ISL_3126137 | Lighthouse Lab in Glasgow | Wellcome Sanger Institute for the COVID-19 Genomics UK (COG-UK) Consortium | Harper VanSteenhouse, Yumi Kasai, David Gray, Carol Clugston, Anna Dominiczak and Alex Alderton, Roberto Amato, Jeffrey Barrett, Sonia Goncalves, Ewan Harrison, David K. Jackson, Ian Johnston, Dominic Kwiatkowski, Cordelia Langford, John Sillitoe on behalf of the Wellcome Sanger Institute COVID-19 Surveillance Team |
| EPI_ISL_3126138 | Lighthouse Lab in Alderley Park | Wellcome Sanger Institute for the COVID-19 Genomics UK (COG-UK) Consortium | Jacquelyn Wynn, Mairead Hyland, The Lighthouse Lab in Alderley Park and Alex Alderton, Roberto Amato, Jeffrey Barrett, Sonia Goncalves, Ewan Harrison, David K. Jackson, Ian Johnston, Dominic Kwiatkowski, Cordelia Langford, John Sillitoe on behalf of the Wellcome Sanger Institute COVID-19 Surveillance Team |
| EPI_ISL_3126139, EPI_ISL_3126140, EPI_ISL_3126141, EPI_ISL_3126142, EPI_ISL_3126143 | Lighthouse Lab in Glasgow | Wellcome Sanger Institute for the COVID-19 Genomics UK (COG-UK) Consortium | Harper VanSteenhouse, Yumi Kasai, David Gray, Carol Clugston, Anna Dominiczak and Alex Alderton, Roberto Amato, Jeffrey Barrett, Sonia Goncalves, Ewan Harrison, David K. Jackson, Ian Johnston, Dominic Kwiatkowski, Cordelia Langford, John Sillitoe on behalf of the Wellcome Sanger Institute COVID-19 Surveillance Team |
| EPI_ISL_3126144 | Lighthouse Lab in Milton Keynes | Wellcome Sanger Institute for the COVID-19 Genomics UK (COG-UK) Consortium | The Lighthouse Lab in Milton Keynes and Alex Alderton, Roberto Amato, Jeffrey Barrett, Sonia Goncalves, Ewan Harrison, David K. Jackson, Ian Johnston, Dominic Kwiatkowski, Cordelia Langford, John Sillitoe on behalf of the Wellcome Sanger Institute COVID-19 Surveillance Team |
| EPI_ISL_3126145, EPI_ISL_3126146 | Lighthouse Lab in Alderley Park | Wellcome Sanger Institute for the COVID-19 Genomics UK (COG-UK) Consortium | Jacquelyn Wynn, Mairead Hyland, The Lighthouse Lab in Alderley Park and Alex Alderton, Roberto Amato, Jeffrey Barrett, Sonia Goncalves, Ewan Harrison, David K. Jackson, Ian Johnston, Dominic Kwiatkowski, Cordelia Langford, John Sillitoe on behalf of the Wellcome Sanger Institute COVID-19 Surveillance Team |
| EPI_ISL_3126147 | Berkshire and Surrey Pathology Services Lighthouse Laboratory | Wellcome Sanger Institute for the COVID-19 Genomics UK (COG-UK) Consortium | Berkshire and Surrey Pathology Services Lighthouse Laboratory and Alex Alderton, Roberto Amato, Jeffrey Barrett, Sonia Goncalves, Ewan Harrison, David K. Jackson, Ian Johnston, Dominic Kwiatkowski, Cordelia Langford, John Sillitoe on behalf of the Wellcome Sanger Institute COVID-19 Surveillance Team |
| EPI_ISL_3126148, EPI_ISL_3126149, EPI_ISL_3126150 | Lighthouse Lab in Alderley Park | Wellcome Sanger Institute for the COVID-19 Genomics UK (COG-UK) Consortium | Jacquelyn Wynn, Mairead Hyland, The Lighthouse Lab in Alderley Park and Alex Alderton, Roberto Amato, Jeffrey Barrett, Sonia Goncalves, Ewan Harrison, David K. Jackson, Ian Johnston, Dominic Kwiatkowski, Cordelia Langford, John Sillitoe on behalf of the Wellcome Sanger Institute COVID-19 Surveillance Team |
| EPI_ISL_3126151 | Lighthouse Lab in Milton Keynes | Wellcome Sanger Institute for the COVID-19 Genomics UK (COG-UK) | The Lighthouse Lab in Milton Keynes and Alex Alderton, Roberto Amato, Jeffrey Barrett, Sonia Goncalves, Ewan Harrison, David K. Jackson, Ian Johnston, Dominic Kwiatkowski, Cordelia Langford, John Sillitoe on behalf of the Wellcome Sanger |

| | | | |
|---|---|---|---|
| | | Consortium | Institute COVID-19 Surveillance Team |
| EPI_ISL_3126152, EPI_ISL_3126153, EPI_ISL_3126154, EPI_ISL_3126155 | Lighthouse Lab in Alderley Park | Wellcome Sanger Institute for the COVID-19 Genomics UK (COG-UK) Consortium | Jacquelyn Wynn, Mairead Hyland, The Lighthouse Lab in Alderley Park and Alex Alderton, Roberto Amato, Jeffrey Barrett, Sonia Goncalves, Ewan Harrison, David K. Jackson, Ian Johnston, Dominic Kwiatkowski, Cordelia Langford, John Sillitoe on behalf of the Wellcome Sanger Institute COVID-19 Surveillance Team |
| EPI_ISL_3126156 | Berkshire and Surrey Pathology Services Lighthouse Laboratory | Wellcome Sanger Institute for the COVID-19 Genomics UK (COG-UK) Consortium | Berkshire and Surrey Pathology Services Lighthouse Laboratory and Alex Alderton, Roberto Amato, Jeffrey Barrett, Sonia Goncalves, Ewan Harrison, David K. Jackson, Ian Johnston, Dominic Kwiatkowski, Cordelia Langford, John Sillitoe on behalf of the Wellcome Sanger Institute COVID-19 Surveillance Team |
| EPI_ISL_3126157, EPI_ISL_3126158 | Lighthouse Lab in Alderley Park | Wellcome Sanger Institute for the COVID-19 Genomics UK (COG-UK) Consortium | Jacquelyn Wynn, Mairead Hyland, The Lighthouse Lab in Alderley Park and Alex Alderton, Roberto Amato, Jeffrey Barrett, Sonia Goncalves, Ewan Harrison, David K. Jackson, Ian Johnston, Dominic Kwiatkowski, Cordelia Langford, John Sillitoe on behalf of the Wellcome Sanger Institute COVID-19 Surveillance Team |
| EPI_ISL_3126159, EPI_ISL_3126160 | Berkshire and Surrey Pathology Services Lighthouse Laboratory | Wellcome Sanger Institute for the COVID-19 Genomics UK (COG-UK) Consortium | Berkshire and Surrey Pathology Services Lighthouse Laboratory and Alex Alderton, Roberto Amato, Jeffrey Barrett, Sonia Goncalves, Ewan Harrison, David K. Jackson, Ian Johnston, Dominic Kwiatkowski, Cordelia Langford, John Sillitoe on behalf of the Wellcome Sanger Institute COVID-19 Surveillance Team |
| EPI_ISL_3126162, EPI_ISL_3126163, EPI_ISL_3126164 | Lighthouse Lab in Alderley Park | Wellcome Sanger Institute for the COVID-19 Genomics UK (COG-UK) Consortium | Jacquelyn Wynn, Mairead Hyland, The Lighthouse Lab in Alderley Park and Alex Alderton, Roberto Amato, Jeffrey Barrett, Sonia Goncalves, Ewan Harrison, David K. Jackson, Ian Johnston, Dominic Kwiatkowski, Cordelia Langford, John Sillitoe on behalf of the Wellcome Sanger Institute COVID-19 Surveillance Team |
| EPI_ISL_3126165 | Berkshire and Surrey Pathology Services Lighthouse Laboratory | Wellcome Sanger Institute for the COVID-19 Genomics UK (COG-UK) Consortium | Berkshire and Surrey Pathology Services Lighthouse Laboratory and Alex Alderton, Roberto Amato, Jeffrey Barrett, Sonia Goncalves, Ewan Harrison, David K. Jackson, Ian Johnston, Dominic Kwiatkowski, Cordelia Langford, John Sillitoe on behalf of the Wellcome Sanger Institute COVID-19 Surveillance Team |
| EPI_ISL_3126166, EPI_ISL_3126167 | Lighthouse Lab in Alderley Park | Wellcome Sanger Institute for the COVID-19 Genomics UK (COG-UK) Consortium | Jacquelyn Wynn, Mairead Hyland, The Lighthouse Lab in Alderley Park and Alex Alderton, Roberto Amato, Jeffrey Barrett, Sonia Goncalves, Ewan Harrison, David K. Jackson, Ian Johnston, Dominic Kwiatkowski, Cordelia Langford, John Sillitoe on behalf of the Wellcome Sanger Institute COVID-19 Surveillance Team |
| EPI_ISL_3126168 | Lighthouse Lab in Milton Keynes | Wellcome Sanger Institute for the COVID-19 Genomics UK (COG-UK) Consortium | The Lighthouse Lab in Milton Keynes and Alex Alderton, Roberto Amato, Jeffrey Barrett, Sonia Goncalves, Ewan Harrison, David K. Jackson, Ian Johnston, Dominic Kwiatkowski, Cordelia Langford, John Sillitoe on behalf of the Wellcome Sanger Institute COVID-19 Surveillance Team |
| EPI_ISL_3126169, EPI_ISL_3126170, EPI_ISL_3126171, EPI_ISL_3126172 | Lighthouse Lab in Alderley Park | Wellcome Sanger Institute for the COVID-19 Genomics UK (COG-UK) Consortium | Jacquelyn Wynn, Mairead Hyland, The Lighthouse Lab in Alderley Park and Alex Alderton, Roberto Amato, Jeffrey Barrett, Sonia Goncalves, Ewan Harrison, David K. Jackson, Ian Johnston, Dominic Kwiatkowski, Cordelia Langford, John Sillitoe on behalf of the Wellcome Sanger Institute COVID-19 Surveillance Team |
| EPI_ISL_3126173 | Lighthouse Lab in Milton Keynes | Wellcome Sanger Institute for the COVID-19 Genomics UK (COG-UK) Consortium | The Lighthouse Lab in Milton Keynes and Alex Alderton, Roberto Amato, Jeffrey Barrett, Sonia Goncalves, Ewan Harrison, David K. Jackson, Ian Johnston, Dominic Kwiatkowski, Cordelia Langford, John Sillitoe on behalf of the Wellcome Sanger Institute COVID-19 Surveillance Team |
| EPI_ISL_3126174, EPI_ISL_3126175, EPI_ISL_3126176, EPI_ISL_3126177, EPI_ISL_3126178 | Lighthouse Lab in Alderley Park | Wellcome Sanger Institute for the COVID-19 Genomics UK (COG-UK) Consortium | Jacquelyn Wynn, Mairead Hyland, The Lighthouse Lab in Alderley Park and Alex Alderton, Roberto Amato, Jeffrey Barrett, Sonia Goncalves, Ewan Harrison, David K. Jackson, Ian Johnston, Dominic Kwiatkowski, Cordelia Langford, John Sillitoe on behalf of the Wellcome Sanger Institute COVID-19 Surveillance Team |
| EPI_ISL_3126179 | Berkshire and Surrey Pathology Services Lighthouse Laboratory | Wellcome Sanger Institute for the COVID-19 Genomics UK (COG-UK) Consortium | Berkshire and Surrey Pathology Services Lighthouse Laboratory and Alex Alderton, Roberto Amato, Jeffrey Barrett, Sonia Goncalves, Ewan Harrison, David K. Jackson, Ian Johnston, Dominic Kwiatkowski, Cordelia Langford, John Sillitoe on behalf of the Wellcome Sanger Institute COVID-19 Surveillance Team |
| EPI_ISL_3126180 | Lighthouse Lab in Milton Keynes | Wellcome Sanger Institute for the COVID-19 Genomics UK (COG-UK) Consortium | The Lighthouse Lab in Milton Keynes and Alex Alderton, Roberto Amato, Jeffrey Barrett, Sonia Goncalves, Ewan Harrison, David K. Jackson, Ian Johnston, Dominic Kwiatkowski, Cordelia Langford, John Sillitoe on behalf of the Wellcome Sanger Institute COVID-19 Surveillance Team |
| EPI_ISL_3126181 | Berkshire and Surrey Pathology Services Lighthouse Laboratory | Wellcome Sanger Institute for the COVID-19 Genomics UK (COG-UK) Consortium | Berkshire and Surrey Pathology Services Lighthouse Laboratory and Alex Alderton, Roberto Amato, Jeffrey Barrett, Sonia Goncalves, Ewan Harrison, David K. Jackson, Ian Johnston, Dominic Kwiatkowski, Cordelia Langford, John Sillitoe on behalf of the Wellcome Sanger Institute COVID-19 Surveillance Team |
| EPI_ISL_3126182 | Lighthouse Lab in Milton Keynes | Wellcome Sanger Institute for the COVID-19 Genomics UK (COG-UK) Consortium | The Lighthouse Lab in Milton Keynes and Alex Alderton, Roberto Amato, Jeffrey Barrett, Sonia Goncalves, Ewan Harrison, David K. Jackson, Ian Johnston, Dominic Kwiatkowski, Cordelia Langford, John Sillitoe on behalf of the Wellcome Sanger Institute COVID-19 Surveillance Team |
| EPI_ISL_3126183 | Lighthouse Lab in Alderley Park | Wellcome Sanger Institute for the COVID-19 Genomics UK (COG-UK) Consortium | Jacquelyn Wynn, Mairead Hyland, The Lighthouse Lab in Alderley Park and Alex Alderton, Roberto Amato, Jeffrey Barrett, Sonia Goncalves, Ewan Harrison, David K. Jackson, Ian Johnston, Dominic Kwiatkowski, Cordelia Langford, John Sillitoe on behalf of the Wellcome Sanger Institute COVID-19 Surveillance Team |
| EPI_ISL_3126184 | Lighthouse Lab in Milton Keynes | Wellcome Sanger Institute for the COVID-19 Genomics UK (COG-UK) Consortium | The Lighthouse Lab in Milton Keynes and Alex Alderton, Roberto Amato, Jeffrey Barrett, Sonia Goncalves, Ewan Harrison, David K. Jackson, Ian Johnston, Dominic Kwiatkowski, Cordelia Langford, John Sillitoe on behalf of the Wellcome Sanger Institute COVID-19 Surveillance Team |
| EPI_ISL_3126185, EPI_ISL_3126186 | Lighthouse Lab in Alderley Park | Wellcome Sanger Institute for the COVID-19 Genomics UK (COG-UK) Consortium | Jacquelyn Wynn, Mairead Hyland, The Lighthouse Lab in Alderley Park and Alex Alderton, Roberto Amato, Jeffrey Barrett, Sonia Goncalves, Ewan Harrison, David K. Jackson, Ian Johnston, Dominic Kwiatkowski, Cordelia Langford, John Sillitoe on behalf of the Wellcome Sanger Institute COVID-19 Surveillance Team |
| EPI_ISL_3126187, EPI_ISL_3126188 | Lighthouse Lab in Milton Keynes | Wellcome Sanger Institute for the COVID-19 Genomics UK (COG-UK) Consortium | The Lighthouse Lab in Milton Keynes and Alex Alderton, Roberto Amato, Jeffrey Barrett, Sonia Goncalves, Ewan Harrison, David K. Jackson, Ian Johnston, Dominic Kwiatkowski, Cordelia Langford, John Sillitoe on behalf of the Wellcome Sanger Institute COVID-19 Surveillance Team |
| EPI_ISL_3126189, EPI_ISL_3126190 | Lighthouse Lab in Alderley Park | Wellcome Sanger Institute for the COVID-19 Genomics UK (COG-UK) Consortium | Jacquelyn Wynn, Mairead Hyland, The Lighthouse Lab in Alderley Park and Alex Alderton, Roberto Amato, Jeffrey Barrett, Sonia Goncalves, Ewan Harrison, David K. Jackson, Ian Johnston, Dominic Kwiatkowski, Cordelia Langford, John Sillitoe on behalf of the Wellcome Sanger Institute COVID-19 Surveillance Team |
| EPI_ISL_3126191 | Lighthouse Lab in Milton Keynes | Wellcome Sanger Institute for the COVID-19 Genomics UK (COG-UK) Consortium | The Lighthouse Lab in Milton Keynes and Alex Alderton, Roberto Amato, Jeffrey Barrett, Sonia Goncalves, Ewan Harrison, David K. Jackson, Ian Johnston, Dominic Kwiatkowski, Cordelia Langford, John Sillitoe on behalf of the Wellcome Sanger Institute COVID-19 Surveillance Team |
| EPI_ISL_3126192 | Lighthouse Lab in Alderley Park | Wellcome Sanger Institute for the COVID-19 Genomics UK (COG-UK) Consortium | Jacquelyn Wynn, Mairead Hyland, The Lighthouse Lab in Alderley Park and Alex Alderton, Roberto Amato, Jeffrey Barrett, Sonia Goncalves, Ewan Harrison, David K. Jackson, Ian Johnston, Dominic Kwiatkowski, Cordelia Langford, John Sillitoe on behalf of the Wellcome Sanger Institute COVID-19 Surveillance Team |
| EPI_ISL_3126193 | Lighthouse Lab in Milton Keynes | Wellcome Sanger Institute for the COVID-19 Genomics UK (COG-UK) Consortium | The Lighthouse Lab in Milton Keynes and Alex Alderton, Roberto Amato, Jeffrey Barrett, Sonia Goncalves, Ewan Harrison, David K. Jackson, Ian Johnston, Dominic Kwiatkowski, Cordelia Langford, John Sillitoe on behalf of the Wellcome Sanger Institute COVID-19 Surveillance Team |
| EPI_ISL_3126194 | Berkshire and Surrey Pathology Services Lighthouse Laboratory | Wellcome Sanger Institute for the COVID-19 Genomics UK (COG-UK) Consortium | Berkshire and Surrey Pathology Services Lighthouse Laboratory and Alex Alderton, Roberto Amato, Jeffrey Barrett, Sonia Goncalves, Ewan Harrison, David K. Jackson, Ian Johnston, Dominic Kwiatkowski, Cordelia Langford, John Sillitoe on behalf of the Wellcome Sanger Institute COVID-19 Surveillance Team |
| EPI_ISL_3126195 | Lighthouse Lab in Milton Keynes | Wellcome Sanger Institute for the COVID-19 Genomics UK (COG-UK) Consortium | The Lighthouse Lab in Milton Keynes and Alex Alderton, Roberto Amato, Jeffrey Barrett, Sonia Goncalves, Ewan Harrison, David K. Jackson, Ian Johnston, Dominic Kwiatkowski, Cordelia Langford, John Sillitoe on behalf of the Wellcome Sanger Institute COVID-19 Surveillance Team |
| EPI_ISL_3126196, EPI_ISL_3126197, EPI_ISL_3126198 | Lighthouse Lab in Alderley Park | Wellcome Sanger Institute for the COVID-19 Genomics UK (COG-UK) Consortium | Jacquelyn Wynn, Mairead Hyland, The Lighthouse Lab in Alderley Park and Alex Alderton, Roberto Amato, Jeffrey Barrett, Sonia Goncalves, Ewan Harrison, David K. Jackson, Ian Johnston, Dominic Kwiatkowski, Cordelia Langford, John Sillitoe on behalf of the Wellcome Sanger Institute COVID-19 Surveillance Team |
| EPI_ISL_3126199, EPI_ISL_3126200 | Lighthouse Lab in Milton Keynes | Wellcome Sanger Institute for the COVID-19 Genomics UK (COG-UK) Consortium | The Lighthouse Lab in Milton Keynes and Alex Alderton, Roberto Amato, Jeffrey Barrett, Sonia Goncalves, Ewan Harrison, David K. Jackson, Ian Johnston, Dominic Kwiatkowski, Cordelia Langford, John Sillitoe on behalf of the Wellcome Sanger Institute COVID-19 Surveillance Team |
| EPI_ISL_3126201, EPI_ISL_3126202 | Lighthouse Lab in Alderley Park | Wellcome Sanger Institute for the COVID-19 Genomics UK (COG-UK) Consortium | Jacquelyn Wynn, Mairead Hyland, The Lighthouse Lab in Alderley Park and Alex Alderton, Roberto Amato, Jeffrey Barrett, Sonia Goncalves, Ewan Harrison, David K. Jackson, Ian Johnston, Dominic Kwiatkowski, Cordelia Langford, John Sillitoe on behalf of the Wellcome Sanger Institute COVID-19 Surveillance Team |
| EPI_ISL_3126203, EPI_ISL_3126204 | Berkshire and Surrey Pathology Services Lighthouse Laboratory | Wellcome Sanger Institute for the COVID-19 Genomics UK (COG-UK) Consortium | Berkshire and Surrey Pathology Services Lighthouse Laboratory and Alex Alderton, Roberto Amato, Jeffrey Barrett, Sonia Goncalves, Ewan Harrison, David K. Jackson, Ian Johnston, Dominic Kwiatkowski, Cordelia Langford, John Sillitoe on behalf of the Wellcome Sanger Institute COVID-19 Surveillance Team |
| EPI_ISL_3126205, EPI_ISL_3126206, EPI_ISL_3126207, EPI_ISL_3126208 | Lighthouse Lab in Alderley Park | Wellcome Sanger Institute for the COVID-19 Genomics UK (COG-UK) Consortium | Jacquelyn Wynn, Mairead Hyland, The Lighthouse Lab in Alderley Park and Alex Alderton, Roberto Amato, Jeffrey Barrett, Sonia Goncalves, Ewan Harrison, David K. Jackson, Ian Johnston, Dominic Kwiatkowski, Cordelia Langford, John Sillitoe on behalf of the Wellcome Sanger Institute COVID-19 Surveillance Team |
| EPI_ISL_3126209, EPI_ISL_3126210 | Lighthouse Lab in Milton Keynes | Wellcome Sanger Institute for the COVID-19 Genomics UK (COG-UK) Consortium | The Lighthouse Lab in Milton Keynes and Alex Alderton, Roberto Amato, Jeffrey Barrett, Sonia Goncalves, Ewan Harrison, David K. Jackson, Ian Johnston, Dominic Kwiatkowski, Cordelia Langford, John Sillitoe on behalf of the Wellcome Sanger Institute COVID-19 Surveillance Team |
| EPI_ISL_3126211 | Lighthouse Lab in Alderley Park | Wellcome Sanger Institute for the COVID-19 Genomics UK (COG-UK) Consortium | Jacquelyn Wynn, Mairead Hyland, The Lighthouse Lab in Alderley Park and Alex Alderton, Roberto Amato, Jeffrey Barrett, Sonia Goncalves, Ewan Harrison, David K. Jackson, Ian Johnston, Dominic Kwiatkowski, Cordelia Langford, John Sillitoe on behalf of the Wellcome Sanger Institute COVID-19 Surveillance Team |
| EPI_ISL_3126212 | Berkshire and Surrey Pathology Services Lighthouse Laboratory | Wellcome Sanger Institute for the COVID-19 Genomics UK (COG-UK) Consortium | Berkshire and Surrey Pathology Services Lighthouse Laboratory and Alex Alderton, Roberto Amato, Jeffrey Barrett, Sonia Goncalves, Ewan Harrison, David K. Jackson, Ian Johnston, Dominic Kwiatkowski, Cordelia Langford, John Sillitoe on behalf of the Wellcome Sanger Institute COVID-19 Surveillance Team |
| EPI_ISL_3126213 | Lighthouse Lab in Alderley Park | Wellcome Sanger Institute for the COVID-19 Genomics UK (COG-UK) Consortium | Jacquelyn Wynn, Mairead Hyland, The Lighthouse Lab in Alderley Park and Alex Alderton, Roberto Amato, Jeffrey Barrett, Sonia Goncalves, Ewan Harrison, David K. Jackson, Ian Johnston, Dominic Kwiatkowski, Cordelia Langford, John Sillitoe on behalf of the Wellcome Sanger Institute COVID-19 Surveillance Team |
| EPI_ISL_3126214 | Lighthouse Lab in Milton Keynes | Wellcome Sanger Institute for the COVID-19 Genomics UK (COG-UK) Consortium | The Lighthouse Lab in Milton Keynes and Alex Alderton, Roberto Amato, Jeffrey Barrett, Sonia Goncalves, Ewan Harrison, David K. Jackson, Ian Johnston, Dominic Kwiatkowski, Cordelia Langford, John Sillitoe on behalf of the Wellcome Sanger Institute COVID-19 Surveillance Team |
| EPI_ISL_3126215 | Lighthouse Lab in Alderley Park | Wellcome Sanger Institute for the COVID-19 Genomics UK (COG-UK) Consortium | Jacquelyn Wynn, Mairead Hyland, The Lighthouse Lab in Alderley Park and Alex Alderton, Roberto Amato, Jeffrey Barrett, Sonia Goncalves, Ewan Harrison, David K. Jackson, Ian Johnston, Dominic Kwiatkowski, Cordelia Langford, John Sillitoe on behalf of the Wellcome Sanger Institute COVID-19 Surveillance Team |
| EPI_ISL_3126216 | Lighthouse Lab in Milton Keynes | Wellcome Sanger Institute for the COVID-19 Genomics UK (COG-UK) Consortium | The Lighthouse Lab in Milton Keynes and Alex Alderton, Roberto Amato, Jeffrey Barrett, Sonia Goncalves, Ewan Harrison, David K. Jackson, Ian Johnston, Dominic Kwiatkowski, Cordelia Langford, John Sillitoe on behalf of the Wellcome Sanger Institute COVID-19 Surveillance Team |
| EPI_ISL_3126217 | Lighthouse Lab in Alderley Park | Wellcome Sanger Institute for the COVID-19 Genomics UK (COG-UK) Consortium | Jacquelyn Wynn, Mairead Hyland, The Lighthouse Lab in Alderley Park and Alex Alderton, Roberto Amato, Jeffrey Barrett, Sonia Goncalves, Ewan Harrison, David K. Jackson, Ian Johnston, Dominic Kwiatkowski, Cordelia Langford, John Sillitoe on behalf of the Wellcome Sanger Institute COVID-19 Surveillance Team |
| EPI_ISL_3126218 | Lighthouse Lab in Milton Keynes | Wellcome Sanger Institute for the COVID-19 Genomics UK (COG-UK) Consortium | The Lighthouse Lab in Milton Keynes and Alex Alderton, Roberto Amato, Jeffrey Barrett, Sonia Goncalves, Ewan Harrison, David K. Jackson, Ian Johnston, Dominic Kwiatkowski, Cordelia Langford, John Sillitoe on behalf of the Wellcome Sanger Institute COVID-19 Surveillance Team |
| EPI_ISL_3126219, EPI_ISL_3126220 | Lighthouse Lab in Alderley Park | Wellcome Sanger Institute for the COVID-19 Genomics UK (COG-UK) Consortium | Jacquelyn Wynn, Mairead Hyland, The Lighthouse Lab in Alderley Park and Alex Alderton, Roberto Amato, Jeffrey Barrett, Sonia Goncalves, Ewan Harrison, David K. Jackson, Ian Johnston, Dominic Kwiatkowski, Cordelia Langford, John Sillitoe on behalf of the Wellcome Sanger Institute COVID-19 Surveillance Team |

| | | | |
|---|---|---|---|
| EPI_ISL_3126221 | Berkshire and Surrey Pathology Services Lighthouse Laboratory | Wellcome Sanger Institute for the COVID-19 Genomics UK (COG-UK) Consortium | Berkshire and Surrey Pathology Services Lighthouse Laboratory and Alex Alderton, Roberto Amato, Jeffrey Barrett, Sonia Goncalves, Ewan Harrison, David K. Jackson, Ian Johnston, Dominic Kwiatkowski, Cordelia Langford, John Sillitoe on behalf of the Wellcome Sanger Institute COVID-19 Surveillance Team |
| EPI_ISL_3126222, EPI_ISL_3126223 | Lighthouse Lab in Milton Keynes | Wellcome Sanger Institute for the COVID-19 Genomics UK (COG-UK) Consortium | The Lighthouse Lab in Milton Keynes and Alex Alderton, Roberto Amato, Jeffrey Barrett, Sonia Goncalves, Ewan Harrison, David K. Jackson, Ian Johnston, Dominic Kwiatkowski, Cordelia Langford, John Sillitoe on behalf of the Wellcome Sanger Institute COVID-19 Surveillance Team |
| EPI_ISL_3126224, EPI_ISL_3126225, EPI_ISL_3126226 | Lighthouse Lab in Alderley Park | Wellcome Sanger Institute for the COVID-19 Genomics UK (COG-UK) Consortium | Jacquelyn Wynn, Mairead Hyland, The Lighthouse Lab in Alderley Park and Alex Alderton, Roberto Amato, Jeffrey Barrett, Sonia Goncalves, Ewan Harrison, David K. Jackson, Ian Johnston, Dominic Kwiatkowski, Cordelia Langford, John Sillitoe on behalf of the Wellcome Sanger Institute COVID-19 Surveillance Team |
| EPI_ISL_3126227 | Berkshire and Surrey Pathology Services Lighthouse Laboratory | Wellcome Sanger Institute for the COVID-19 Genomics UK (COG-UK) Consortium | Berkshire and Surrey Pathology Services Lighthouse Laboratory and Alex Alderton, Roberto Amato, Jeffrey Barrett, Sonia Goncalves, Ewan Harrison, David K. Jackson, Ian Johnston, Dominic Kwiatkowski, Cordelia Langford, John Sillitoe on behalf of the Wellcome Sanger Institute COVID-19 Surveillance Team |
| EPI_ISL_3126228 | Lighthouse Lab in Milton Keynes | Wellcome Sanger Institute for the COVID-19 Genomics UK (COG-UK) Consortium | The Lighthouse Lab in Milton Keynes and Alex Alderton, Roberto Amato, Jeffrey Barrett, Sonia Goncalves, Ewan Harrison, David K. Jackson, Ian Johnston, Dominic Kwiatkowski, Cordelia Langford, John Sillitoe on behalf of the Wellcome Sanger Institute COVID-19 Surveillance Team |
| EPI_ISL_3126229, EPI_ISL_3126230, EPI_ISL_3126231 | Lighthouse Lab in Alderley Park | Wellcome Sanger Institute for the COVID-19 Genomics UK (COG-UK) Consortium | Jacquelyn Wynn, Mairead Hyland, The Lighthouse Lab in Alderley Park and Alex Alderton, Roberto Amato, Jeffrey Barrett, Sonia Goncalves, Ewan Harrison, David K. Jackson, Ian Johnston, Dominic Kwiatkowski, Cordelia Langford, John Sillitoe on behalf of the Wellcome Sanger Institute COVID-19 Surveillance Team |
| EPI_ISL_3126232 | Berkshire and Surrey Pathology Services Lighthouse Laboratory | Wellcome Sanger Institute for the COVID-19 Genomics UK (COG-UK) Consortium | Berkshire and Surrey Pathology Services Lighthouse Laboratory and Alex Alderton, Roberto Amato, Jeffrey Barrett, Sonia Goncalves, Ewan Harrison, David K. Jackson, Ian Johnston, Dominic Kwiatkowski, Cordelia Langford, John Sillitoe on behalf of the Wellcome Sanger Institute COVID-19 Surveillance Team |
| EPI_ISL_3126233, EPI_ISL_3126234 | Lighthouse Lab in Milton Keynes | Wellcome Sanger Institute for the COVID-19 Genomics UK (COG-UK) Consortium | The Lighthouse Lab in Milton Keynes and Alex Alderton, Roberto Amato, Jeffrey Barrett, Sonia Goncalves, Ewan Harrison, David K. Jackson, Ian Johnston, Dominic Kwiatkowski, Cordelia Langford, John Sillitoe on behalf of the Wellcome Sanger Institute COVID-19 Surveillance Team |
| EPI_ISL_3126235 | Lighthouse Lab in Alderley Park | Wellcome Sanger Institute for the COVID-19 Genomics UK (COG-UK) Consortium | Jacquelyn Wynn, Mairead Hyland, The Lighthouse Lab in Alderley Park and Alex Alderton, Roberto Amato, Jeffrey Barrett, Sonia Goncalves, Ewan Harrison, David K. Jackson, Ian Johnston, Dominic Kwiatkowski, Cordelia Langford, John Sillitoe on behalf of the Wellcome Sanger Institute COVID-19 Surveillance Team |
| EPI_ISL_3126236 | Berkshire and Surrey Pathology Services Lighthouse Laboratory | Wellcome Sanger Institute for the COVID-19 Genomics UK (COG-UK) Consortium | Berkshire and Surrey Pathology Services Lighthouse Laboratory and Alex Alderton, Roberto Amato, Jeffrey Barrett, Sonia Goncalves, Ewan Harrison, David K. Jackson, Ian Johnston, Dominic Kwiatkowski, Cordelia Langford, John Sillitoe on behalf of the Wellcome Sanger Institute COVID-19 Surveillance Team |
| EPI_ISL_3126237, EPI_ISL_3126238 | Lighthouse Lab in Alderley Park | Wellcome Sanger Institute for the COVID-19 Genomics UK (COG-UK) Consortium | Jacquelyn Wynn, Mairead Hyland, The Lighthouse Lab in Alderley Park and Alex Alderton, Roberto Amato, Jeffrey Barrett, Sonia Goncalves, Ewan Harrison, David K. Jackson, Ian Johnston, Dominic Kwiatkowski, Cordelia Langford, John Sillitoe on behalf of the Wellcome Sanger Institute COVID-19 Surveillance Team |
| EPI_ISL_3126239, EPI_ISL_3126240 | Lighthouse Lab in Milton Keynes | Wellcome Sanger Institute for the COVID-19 Genomics UK (COG-UK) Consortium | The Lighthouse Lab in Milton Keynes and Alex Alderton, Roberto Amato, Jeffrey Barrett, Sonia Goncalves, Ewan Harrison, David K. Jackson, Ian Johnston, Dominic Kwiatkowski, Cordelia Langford, John Sillitoe on behalf of the Wellcome Sanger Institute COVID-19 Surveillance Team |
| EPI_ISL_3126241, EPI_ISL_3126242 | Lighthouse Lab in Alderley Park | Wellcome Sanger Institute for the COVID-19 Genomics UK (COG-UK) Consortium | Jacquelyn Wynn, Mairead Hyland, The Lighthouse Lab in Alderley Park and Alex Alderton, Roberto Amato, Jeffrey Barrett, Sonia Goncalves, Ewan Harrison, David K. Jackson, Ian Johnston, Dominic Kwiatkowski, Cordelia Langford, John Sillitoe on behalf of the Wellcome Sanger Institute COVID-19 Surveillance Team |
| EPI_ISL_3126243 | Berkshire and Surrey Pathology Services Lighthouse Laboratory | Wellcome Sanger Institute for the COVID-19 Genomics UK (COG-UK) Consortium | Berkshire and Surrey Pathology Services Lighthouse Laboratory and Alex Alderton, Roberto Amato, Jeffrey Barrett, Sonia Goncalves, Ewan Harrison, David K. Jackson, Ian Johnston, Dominic Kwiatkowski, Cordelia Langford, John Sillitoe on behalf of the Wellcome Sanger Institute COVID-19 Surveillance Team |
| EPI_ISL_3126244, EPI_ISL_3126245 | Lighthouse Lab in Alderley Park | Wellcome Sanger Institute for the COVID-19 Genomics UK (COG-UK) Consortium | Jacquelyn Wynn, Mairead Hyland, The Lighthouse Lab in Alderley Park and Alex Alderton, Roberto Amato, Jeffrey Barrett, Sonia Goncalves, Ewan Harrison, David K. Jackson, Ian Johnston, Dominic Kwiatkowski, Cordelia Langford, John Sillitoe on behalf of the Wellcome Sanger Institute COVID-19 Surveillance Team |
| EPI_ISL_3126246, EPI_ISL_3126247 | Lighthouse Lab in Milton Keynes | Wellcome Sanger Institute for the COVID-19 Genomics UK (COG-UK) Consortium | The Lighthouse Lab in Milton Keynes and Alex Alderton, Roberto Amato, Jeffrey Barrett, Sonia Goncalves, Ewan Harrison, David K. Jackson, Ian Johnston, Dominic Kwiatkowski, Cordelia Langford, John Sillitoe on behalf of the Wellcome Sanger Institute COVID-19 Surveillance Team |
| EPI_ISL_3126248 | Berkshire and Surrey Pathology Services Lighthouse Laboratory | Wellcome Sanger Institute for the COVID-19 Genomics UK (COG-UK) Consortium | Berkshire and Surrey Pathology Services Lighthouse Laboratory and Alex Alderton, Roberto Amato, Jeffrey Barrett, Sonia Goncalves, Ewan Harrison, David K. Jackson, Ian Johnston, Dominic Kwiatkowski, Cordelia Langford, John Sillitoe on behalf of the Wellcome Sanger Institute COVID-19 Surveillance Team |
| EPI_ISL_3126249, EPI_ISL_3126250 | Lighthouse Lab in Milton Keynes | Wellcome Sanger Institute for the COVID-19 Genomics UK (COG-UK) Consortium | The Lighthouse Lab in Milton Keynes and Alex Alderton, Roberto Amato, Jeffrey Barrett, Sonia Goncalves, Ewan Harrison, David K. Jackson, Ian Johnston, Dominic Kwiatkowski, Cordelia Langford, John Sillitoe on behalf of the Wellcome Sanger Institute COVID-19 Surveillance Team |
| EPI_ISL_3126251, EPI_ISL_3126252, EPI_ISL_3126253, EPI_ISL_3126254 | Lighthouse Lab in Alderley Park | Wellcome Sanger Institute for the COVID-19 Genomics UK (COG-UK) Consortium | Jacquelyn Wynn, Mairead Hyland, The Lighthouse Lab in Alderley Park and Alex Alderton, Roberto Amato, Jeffrey Barrett, Sonia Goncalves, Ewan Harrison, David K. Jackson, Ian Johnston, Dominic Kwiatkowski, Cordelia Langford, John Sillitoe on behalf of the Wellcome Sanger Institute COVID-19 Surveillance Team |
| EPI_ISL_3126255 | Berkshire and Surrey Pathology Services Lighthouse Laboratory | Wellcome Sanger Institute for the COVID-19 Genomics UK (COG-UK) Consortium | Berkshire and Surrey Pathology Services Lighthouse Laboratory and Alex Alderton, Roberto Amato, Jeffrey Barrett, Sonia Goncalves, Ewan Harrison, David K. Jackson, Ian Johnston, Dominic Kwiatkowski, Cordelia Langford, John Sillitoe on behalf of the Wellcome Sanger Institute COVID-19 Surveillance Team |
| EPI_ISL_3126256, EPI_ISL_3126257 | Lighthouse Lab in Milton Keynes | Wellcome Sanger Institute for the COVID-19 Genomics UK (COG-UK) Consortium | The Lighthouse Lab in Milton Keynes and Alex Alderton, Roberto Amato, Jeffrey Barrett, Sonia Goncalves, Ewan Harrison, David K. Jackson, Ian Johnston, Dominic Kwiatkowski, Cordelia Langford, John Sillitoe on behalf of the Wellcome Sanger Institute COVID-19 Surveillance Team |
| EPI_ISL_3126258 | Berkshire and Surrey Pathology Services Lighthouse Laboratory | Wellcome Sanger Institute for the COVID-19 Genomics UK (COG-UK) Consortium | Berkshire and Surrey Pathology Services Lighthouse Laboratory and Alex Alderton, Roberto Amato, Jeffrey Barrett, Sonia Goncalves, Ewan Harrison, David K. Jackson, Ian Johnston, Dominic Kwiatkowski, Cordelia Langford, John Sillitoe on behalf of the Wellcome Sanger Institute COVID-19 Surveillance Team |
| EPI_ISL_3126259 | Lighthouse Lab in Milton Keynes | Wellcome Sanger Institute for the COVID-19 Genomics UK (COG-UK) Consortium | The Lighthouse Lab in Milton Keynes and Alex Alderton, Roberto Amato, Jeffrey Barrett, Sonia Goncalves, Ewan Harrison, David K. Jackson, Ian Johnston, Dominic Kwiatkowski, Cordelia Langford, John Sillitoe on behalf of the Wellcome Sanger Institute COVID-19 Surveillance Team |
| EPI_ISL_3126260, EPI_ISL_3126261, EPI_ISL_3126262 | Lighthouse Lab in Alderley Park | Wellcome Sanger Institute for the COVID-19 Genomics UK (COG-UK) Consortium | Jacquelyn Wynn, Mairead Hyland, The Lighthouse Lab in Alderley Park and Alex Alderton, Roberto Amato, Jeffrey Barrett, Sonia Goncalves, Ewan Harrison, David K. Jackson, Ian Johnston, Dominic Kwiatkowski, Cordelia Langford, John Sillitoe on behalf of the Wellcome Sanger Institute COVID-19 Surveillance Team |
| EPI_ISL_3126263 | Lighthouse Lab in Milton Keynes | Wellcome Sanger Institute for the COVID-19 Genomics UK (COG-UK) Consortium | The Lighthouse Lab in Milton Keynes and Alex Alderton, Roberto Amato, Jeffrey Barrett, Sonia Goncalves, Ewan Harrison, David K. Jackson, Ian Johnston, Dominic Kwiatkowski, Cordelia Langford, John Sillitoe on behalf of the Wellcome Sanger Institute COVID-19 Surveillance Team |
| EPI_ISL_3126264 | Berkshire and Surrey Pathology Services Lighthouse Laboratory | Wellcome Sanger Institute for the COVID-19 Genomics UK (COG-UK) Consortium | Berkshire and Surrey Pathology Services Lighthouse Laboratory and Alex Alderton, Roberto Amato, Jeffrey Barrett, Sonia Goncalves, Ewan Harrison, David K. Jackson, Ian Johnston, Dominic Kwiatkowski, Cordelia Langford, John Sillitoe on behalf of the Wellcome Sanger Institute COVID-19 Surveillance Team |
| EPI_ISL_3126265 | Lighthouse Lab in Milton Keynes | Wellcome Sanger Institute for the COVID-19 Genomics UK (COG-UK) Consortium | The Lighthouse Lab in Milton Keynes and Alex Alderton, Roberto Amato, Jeffrey Barrett, Sonia Goncalves, Ewan Harrison, David K. Jackson, Ian Johnston, Dominic Kwiatkowski, Cordelia Langford, John Sillitoe on behalf of the Wellcome Sanger Institute COVID-19 Surveillance Team |
| EPI_ISL_3126266 | Lighthouse Lab in Alderley Park | Wellcome Sanger Institute for the COVID-19 Genomics UK (COG-UK) Consortium | Jacquelyn Wynn, Mairead Hyland, The Lighthouse Lab in Alderley Park and Alex Alderton, Roberto Amato, Jeffrey Barrett, Sonia Goncalves, Ewan Harrison, David K. Jackson, Ian Johnston, Dominic Kwiatkowski, Cordelia Langford, John Sillitoe on behalf of the Wellcome Sanger Institute COVID-19 Surveillance Team |
| EPI_ISL_3126267, EPI_ISL_3126268 | Lighthouse Lab in Milton Keynes | Wellcome Sanger Institute for the COVID-19 Genomics UK (COG-UK) Consortium | The Lighthouse Lab in Milton Keynes and Alex Alderton, Roberto Amato, Jeffrey Barrett, Sonia Goncalves, Ewan Harrison, David K. Jackson, Ian Johnston, Dominic Kwiatkowski, Cordelia Langford, John Sillitoe on behalf of the Wellcome Sanger Institute COVID-19 Surveillance Team |
| EPI_ISL_3126269, EPI_ISL_3126270, EPI_ISL_3126271, EPI_ISL_3126272, EPI_ISL_3126273 | Lighthouse Lab in Alderley Park | Wellcome Sanger Institute for the COVID-19 Genomics UK (COG-UK) Consortium | Jacquelyn Wynn, Mairead Hyland, The Lighthouse Lab in Alderley Park and Alex Alderton, Roberto Amato, Jeffrey Barrett, Sonia Goncalves, Ewan Harrison, David K. Jackson, Ian Johnston, Dominic Kwiatkowski, Cordelia Langford, John Sillitoe on behalf of the Wellcome Sanger Institute COVID-19 Surveillance Team |
| EPI_ISL_3126274 | Lighthouse Lab in Milton Keynes | Wellcome Sanger Institute for the COVID-19 Genomics UK (COG-UK) Consortium | The Lighthouse Lab in Milton Keynes and Alex Alderton, Roberto Amato, Jeffrey Barrett, Sonia Goncalves, Ewan Harrison, David K. Jackson, Ian Johnston, Dominic Kwiatkowski, Cordelia Langford, John Sillitoe on behalf of the Wellcome Sanger Institute COVID-19 Surveillance Team |
| EPI_ISL_3126275 | Berkshire and Surrey Pathology Services Lighthouse Laboratory | Wellcome Sanger Institute for the COVID-19 Genomics UK (COG-UK) Consortium | Berkshire and Surrey Pathology Services Lighthouse Laboratory and Alex Alderton, Roberto Amato, Jeffrey Barrett, Sonia Goncalves, Ewan Harrison, David K. Jackson, Ian Johnston, Dominic Kwiatkowski, Cordelia Langford, John Sillitoe on behalf of the Wellcome Sanger Institute COVID-19 Surveillance Team |
| EPI_ISL_3126276, EPI_ISL_3126277 | Lighthouse Lab in Alderley Park | Wellcome Sanger Institute for the COVID-19 Genomics UK (COG-UK) Consortium | Jacquelyn Wynn, Mairead Hyland, The Lighthouse Lab in Alderley Park and Alex Alderton, Roberto Amato, Jeffrey Barrett, Sonia Goncalves, Ewan Harrison, David K. Jackson, Ian Johnston, Dominic Kwiatkowski, Cordelia Langford, John Sillitoe on behalf of the Wellcome Sanger Institute COVID-19 Surveillance Team |
| EPI_ISL_3126278, EPI_ISL_3126279 | Lighthouse Lab in Milton Keynes | Wellcome Sanger Institute for the COVID-19 Genomics UK (COG-UK) Consortium | The Lighthouse Lab in Milton Keynes and Alex Alderton, Roberto Amato, Jeffrey Barrett, Sonia Goncalves, Ewan Harrison, David K. Jackson, Ian Johnston, Dominic Kwiatkowski, Cordelia Langford, John Sillitoe on behalf of the Wellcome Sanger Institute COVID-19 Surveillance Team |
| EPI_ISL_3126280 | Berkshire and Surrey Pathology Services Lighthouse Laboratory | Wellcome Sanger Institute for the COVID-19 Genomics UK (COG-UK) Consortium | Berkshire and Surrey Pathology Services Lighthouse Laboratory and Alex Alderton, Roberto Amato, Jeffrey Barrett, Sonia Goncalves, Ewan Harrison, David K. Jackson, Ian Johnston, Dominic Kwiatkowski, Cordelia Langford, John Sillitoe on behalf of the Wellcome Sanger Institute COVID-19 Surveillance Team |
| EPI_ISL_3126281 | Lighthouse Lab in Milton Keynes | Wellcome Sanger Institute for the COVID-19 Genomics UK (COG-UK) Consortium | The Lighthouse Lab in Milton Keynes and Alex Alderton, Roberto Amato, Jeffrey Barrett, Sonia Goncalves, Ewan Harrison, David K. Jackson, Ian Johnston, Dominic Kwiatkowski, Cordelia Langford, John Sillitoe on behalf of the Wellcome Sanger Institute COVID-19 Surveillance Team |
| EPI_ISL_3126282 | Lighthouse Lab in Alderley Park | Wellcome Sanger Institute for the COVID-19 Genomics UK (COG-UK) Consortium | Jacquelyn Wynn, Mairead Hyland, The Lighthouse Lab in Alderley Park and Alex Alderton, Roberto Amato, Jeffrey Barrett, Sonia Goncalves, Ewan Harrison, David K. Jackson, Ian Johnston, Dominic Kwiatkowski, Cordelia Langford, John Sillitoe on behalf of the Wellcome Sanger Institute COVID-19 Surveillance Team |
| EPI_ISL_3126283, EPI_ISL_3126284, EPI_ISL_3126285 | Lighthouse Lab in Milton Keynes | Wellcome Sanger Institute for the COVID-19 Genomics UK (COG-UK) Consortium | The Lighthouse Lab in Milton Keynes and Alex Alderton, Roberto Amato, Jeffrey Barrett, Sonia Goncalves, Ewan Harrison, David K. Jackson, Ian Johnston, Dominic Kwiatkowski, Cordelia Langford, John Sillitoe on behalf of the Wellcome Sanger Institute COVID-19 Surveillance Team |
| EPI_ISL_3126286 | Berkshire and Surrey Pathology Services Lighthouse Laboratory | Wellcome Sanger Institute for the COVID-19 Genomics UK (COG-UK) Consortium | Berkshire and Surrey Pathology Services Lighthouse Laboratory and Alex Alderton, Roberto Amato, Jeffrey Barrett, Sonia Goncalves, Ewan Harrison, David K. Jackson, Ian Johnston, Dominic Kwiatkowski, Cordelia Langford, John Sillitoe on behalf of the Wellcome Sanger Institute COVID-19 Surveillance Team |
| EPI_ISL_3126287, EPI_ISL_3126288 | Lighthouse Lab in Alderley Park | Wellcome Sanger Institute for the COVID-19 Genomics UK (COG-UK) Consortium | Jacquelyn Wynn, Mairead Hyland, The Lighthouse Lab in Alderley Park and Alex Alderton, Roberto Amato, Jeffrey Barrett, Sonia Goncalves, Ewan Harrison, David K. Jackson, Ian Johnston, Dominic Kwiatkowski, Cordelia Langford, John Sillitoe on behalf of the Wellcome Sanger Institute COVID-19 Surveillance Team |
| EPI_ISL_3126289 | Lighthouse Lab in Milton Keynes | Wellcome Sanger Institute for the COVID-19 Genomics UK (COG-UK) Consortium | The Lighthouse Lab in Milton Keynes and Alex Alderton, Roberto Amato, Jeffrey Barrett, Sonia Goncalves, Ewan Harrison, David K. Jackson, Ian Johnston, Dominic Kwiatkowski, Cordelia Langford, John Sillitoe on behalf of the Wellcome Sanger Institute COVID-19 Surveillance Team |
| EPI_ISL_3126290 | Lighthouse Lab in Alderley Park | Wellcome Sanger Institute for the COVID-19 Genomics UK (COG-UK) | Jacquelyn Wynn, Mairead Hyland, The Lighthouse Lab in Alderley Park and Alex Alderton, Roberto Amato, Jeffrey Barrett, Sonia Goncalves, Ewan Harrison, David K. Jackson, Ian Johnston, Dominic Kwiatkowski, Cordelia Langford, John Sillitoe on |

| | | | |
|---|---|---|---|
| | | Consortium | behalf of the Wellcome Sanger Institute COVID-19 Surveillance Team |
| EPI_ISL_3126291, EPI_ISL_3126292 | Berkshire and Surrey Pathology Services Lighthouse Laboratory | Wellcome Sanger Institute for the COVID-19 Genomics UK (COG-UK) Consortium | Berkshire and Surrey Pathology Services Lighthouse Laboratory and Alex Alderton, Roberto Amato, Jeffrey Barrett, Sonia Goncalves, Ewan Harrison, David K. Jackson, Ian Johnston, Dominic Kwiatkowski, Cordelia Langford, John Sillitoe on behalf of the Wellcome Sanger Institute COVID-19 Surveillance Team |
| EPI_ISL_3126293, EPI_ISL_3126294 | Lighthouse Lab in Milton Keynes | Wellcome Sanger Institute for the COVID-19 Genomics UK (COG-UK) Consortium | The Lighthouse Lab in Milton Keynes and Alex Alderton, Roberto Amato, Jeffrey Barrett, Sonia Goncalves, Ewan Harrison, David K. Jackson, Ian Johnston, Dominic Kwiatkowski, Cordelia Langford, John Sillitoe on behalf of the Wellcome Sanger Institute COVID-19 Surveillance Team |
| EPI_ISL_3126295 | Lighthouse Lab in Alderley Park | Wellcome Sanger Institute for the COVID-19 Genomics UK (COG-UK) Consortium | Jacquelyn Wynn, Mairead Hyland, The Lighthouse Lab in Alderley Park and Alex Alderton, Roberto Amato, Jeffrey Barrett, Sonia Goncalves, Ewan Harrison, David K. Jackson, Ian Johnston, Dominic Kwiatkowski, Cordelia Langford, John Sillitoe on behalf of the Wellcome Sanger Institute COVID-19 Surveillance Team |
| EPI_ISL_3126296 | Lighthouse Lab in Milton Keynes | Wellcome Sanger Institute for the COVID-19 Genomics UK (COG-UK) Consortium | The Lighthouse Lab in Milton Keynes and Alex Alderton, Roberto Amato, Jeffrey Barrett, Sonia Goncalves, Ewan Harrison, David K. Jackson, Ian Johnston, Dominic Kwiatkowski, Cordelia Langford, John Sillitoe on behalf of the Wellcome Sanger Institute COVID-19 Surveillance Team |
| EPI_ISL_3126297 | Berkshire and Surrey Pathology Services Lighthouse Laboratory | Wellcome Sanger Institute for the COVID-19 Genomics UK (COG-UK) Consortium | Berkshire and Surrey Pathology Services Lighthouse Laboratory and Alex Alderton, Roberto Amato, Jeffrey Barrett, Sonia Goncalves, Ewan Harrison, David K. Jackson, Ian Johnston, Dominic Kwiatkowski, Cordelia Langford, John Sillitoe on behalf of the Wellcome Sanger Institute COVID-19 Surveillance Team |
| EPI_ISL_3126298, EPI_ISL_3126299 | Lighthouse Lab in Alderley Park | Wellcome Sanger Institute for the COVID-19 Genomics UK (COG-UK) Consortium | Jacquelyn Wynn, Mairead Hyland, The Lighthouse Lab in Alderley Park and Alex Alderton, Roberto Amato, Jeffrey Barrett, Sonia Goncalves, Ewan Harrison, David K. Jackson, Ian Johnston, Dominic Kwiatkowski, Cordelia Langford, John Sillitoe on behalf of the Wellcome Sanger Institute COVID-19 Surveillance Team |
| EPI_ISL_3126300 | Lighthouse Lab in Milton Keynes | Wellcome Sanger Institute for the COVID-19 Genomics UK (COG-UK) Consortium | The Lighthouse Lab in Milton Keynes and Alex Alderton, Roberto Amato, Jeffrey Barrett, Sonia Goncalves, Ewan Harrison, David K. Jackson, Ian Johnston, Dominic Kwiatkowski, Cordelia Langford, John Sillitoe on behalf of the Wellcome Sanger Institute COVID-19 Surveillance Team |
| EPI_ISL_3126301, EPI_ISL_3126302 | Lighthouse Lab in Alderley Park | Wellcome Sanger Institute for the COVID-19 Genomics UK (COG-UK) Consortium | Jacquelyn Wynn, Mairead Hyland, The Lighthouse Lab in Alderley Park and Alex Alderton, Roberto Amato, Jeffrey Barrett, Sonia Goncalves, Ewan Harrison, David K. Jackson, Ian Johnston, Dominic Kwiatkowski, Cordelia Langford, John Sillitoe on behalf of the Wellcome Sanger Institute COVID-19 Surveillance Team |
| EPI_ISL_3126303 | Berkshire and Surrey Pathology Services Lighthouse Laboratory | Wellcome Sanger Institute for the COVID-19 Genomics UK (COG-UK) Consortium | Berkshire and Surrey Pathology Services Lighthouse Laboratory and Alex Alderton, Roberto Amato, Jeffrey Barrett, Sonia Goncalves, Ewan Harrison, David K. Jackson, Ian Johnston, Dominic Kwiatkowski, Cordelia Langford, John Sillitoe on behalf of the Wellcome Sanger Institute COVID-19 Surveillance Team |
| EPI_ISL_3126304 | Lighthouse Lab in Milton Keynes | Wellcome Sanger Institute for the COVID-19 Genomics UK (COG-UK) Consortium | The Lighthouse Lab in Milton Keynes and Alex Alderton, Roberto Amato, Jeffrey Barrett, Sonia Goncalves, Ewan Harrison, David K. Jackson, Ian Johnston, Dominic Kwiatkowski, Cordelia Langford, John Sillitoe on behalf of the Wellcome Sanger Institute COVID-19 Surveillance Team |
| EPI_ISL_3126305 | Lighthouse Lab in Alderley Park | Wellcome Sanger Institute for the COVID-19 Genomics UK (COG-UK) Consortium | Jacquelyn Wynn, Mairead Hyland, The Lighthouse Lab in Alderley Park and Alex Alderton, Roberto Amato, Jeffrey Barrett, Sonia Goncalves, Ewan Harrison, David K. Jackson, Ian Johnston, Dominic Kwiatkowski, Cordelia Langford, John Sillitoe on behalf of the Wellcome Sanger Institute COVID-19 Surveillance Team |
| EPI_ISL_3126306 | Berkshire and Surrey Pathology Services Lighthouse Laboratory | Wellcome Sanger Institute for the COVID-19 Genomics UK (COG-UK) Consortium | Berkshire and Surrey Pathology Services Lighthouse Laboratory and Alex Alderton, Roberto Amato, Jeffrey Barrett, Sonia Goncalves, Ewan Harrison, David K. Jackson, Ian Johnston, Dominic Kwiatkowski, Cordelia Langford, John Sillitoe on behalf of the Wellcome Sanger Institute COVID-19 Surveillance Team |
| EPI_ISL_3126307, EPI_ISL_3126308 | Lighthouse Lab in Alderley Park | Wellcome Sanger Institute for the COVID-19 Genomics UK (COG-UK) Consortium | Jacquelyn Wynn, Mairead Hyland, The Lighthouse Lab in Alderley Park and Alex Alderton, Roberto Amato, Jeffrey Barrett, Sonia Goncalves, Ewan Harrison, David K. Jackson, Ian Johnston, Dominic Kwiatkowski, Cordelia Langford, John Sillitoe on behalf of the Wellcome Sanger Institute COVID-19 Surveillance Team |
| EPI_ISL_3126309 | Berkshire and Surrey Pathology Services Lighthouse Laboratory | Wellcome Sanger Institute for the COVID-19 Genomics UK (COG-UK) Consortium | Berkshire and Surrey Pathology Services Lighthouse Laboratory and Alex Alderton, Roberto Amato, Jeffrey Barrett, Sonia Goncalves, Ewan Harrison, David K. Jackson, Ian Johnston, Dominic Kwiatkowski, Cordelia Langford, John Sillitoe on behalf of the Wellcome Sanger Institute COVID-19 Surveillance Team |
| EPI_ISL_3126310, EPI_ISL_3126311, EPI_ISL_3126312, EPI_ISL_3126313 | Lighthouse Lab in Alderley Park | Wellcome Sanger Institute for the COVID-19 Genomics UK (COG-UK) Consortium | Jacquelyn Wynn, Mairead Hyland, The Lighthouse Lab in Alderley Park and Alex Alderton, Roberto Amato, Jeffrey Barrett, Sonia Goncalves, Ewan Harrison, David K. Jackson, Ian Johnston, Dominic Kwiatkowski, Cordelia Langford, John Sillitoe on behalf of the Wellcome Sanger Institute COVID-19 Surveillance Team |
| EPI_ISL_3126314, EPI_ISL_3126315 | Lighthouse Lab in Milton Keynes | Wellcome Sanger Institute for the COVID-19 Genomics UK (COG-UK) Consortium | The Lighthouse Lab in Milton Keynes and Alex Alderton, Roberto Amato, Jeffrey Barrett, Sonia Goncalves, Ewan Harrison, David K. Jackson, Ian Johnston, Dominic Kwiatkowski, Cordelia Langford, John Sillitoe on behalf of the Wellcome Sanger Institute COVID-19 Surveillance Team |
| EPI_ISL_3126316 | Lighthouse Lab in Alderley Park | Wellcome Sanger Institute for the COVID-19 Genomics UK (COG-UK) Consortium | Jacquelyn Wynn, Mairead Hyland, The Lighthouse Lab in Alderley Park and Alex Alderton, Roberto Amato, Jeffrey Barrett, Sonia Goncalves, Ewan Harrison, David K. Jackson, Ian Johnston, Dominic Kwiatkowski, Cordelia Langford, John Sillitoe on behalf of the Wellcome Sanger Institute COVID-19 Surveillance Team |
| EPI_ISL_3126317 | Lighthouse Lab in Milton Keynes | Wellcome Sanger Institute for the COVID-19 Genomics UK (COG-UK) Consortium | The Lighthouse Lab in Milton Keynes and Alex Alderton, Roberto Amato, Jeffrey Barrett, Sonia Goncalves, Ewan Harrison, David K. Jackson, Ian Johnston, Dominic Kwiatkowski, Cordelia Langford, John Sillitoe on behalf of the Wellcome Sanger Institute COVID-19 Surveillance Team |
| EPI_ISL_3126318 | Lighthouse Lab in Alderley Park | Wellcome Sanger Institute for the COVID-19 Genomics UK (COG-UK) Consortium | Jacquelyn Wynn, Mairead Hyland, The Lighthouse Lab in Alderley Park and Alex Alderton, Roberto Amato, Jeffrey Barrett, Sonia Goncalves, Ewan Harrison, David K. Jackson, Ian Johnston, Dominic Kwiatkowski, Cordelia Langford, John Sillitoe on behalf of the Wellcome Sanger Institute COVID-19 Surveillance Team |
| EPI_ISL_3126319 | Berkshire and Surrey Pathology Services Lighthouse Laboratory | Wellcome Sanger Institute for the COVID-19 Genomics UK (COG-UK) Consortium | Berkshire and Surrey Pathology Services Lighthouse Laboratory and Alex Alderton, Roberto Amato, Jeffrey Barrett, Sonia Goncalves, Ewan Harrison, David K. Jackson, Ian Johnston, Dominic Kwiatkowski, Cordelia Langford, John Sillitoe on behalf of the Wellcome Sanger Institute COVID-19 Surveillance Team |
| EPI_ISL_3126320, EPI_ISL_3126321, EPI_ISL_3126322, EPI_ISL_3126323, EPI_ISL_3126324, EPI_ISL_3126325 | Lighthouse Lab in Alderley Park | Wellcome Sanger Institute for the COVID-19 Genomics UK (COG-UK) Consortium | Jacquelyn Wynn, Mairead Hyland, The Lighthouse Lab in Alderley Park and Alex Alderton, Roberto Amato, Jeffrey Barrett, Sonia Goncalves, Ewan Harrison, David K. Jackson, Ian Johnston, Dominic Kwiatkowski, Cordelia Langford, John Sillitoe on behalf of the Wellcome Sanger Institute COVID-19 Surveillance Team |
| EPI_ISL_3126326, EPI_ISL_3126327 | Berkshire and Surrey Pathology Services Lighthouse Laboratory | Wellcome Sanger Institute for the COVID-19 Genomics UK (COG-UK) Consortium | Berkshire and Surrey Pathology Services Lighthouse Laboratory and Alex Alderton, Roberto Amato, Jeffrey Barrett, Sonia Goncalves, Ewan Harrison, David K. Jackson, Ian Johnston, Dominic Kwiatkowski, Cordelia Langford, John Sillitoe on behalf of the Wellcome Sanger Institute COVID-19 Surveillance Team |
| EPI_ISL_3126328 | Lighthouse Lab in Alderley Park | Wellcome Sanger Institute for the COVID-19 Genomics UK (COG-UK) Consortium | Jacquelyn Wynn, Mairead Hyland, The Lighthouse Lab in Alderley Park and Alex Alderton, Roberto Amato, Jeffrey Barrett, Sonia Goncalves, Ewan Harrison, David K. Jackson, Ian Johnston, Dominic Kwiatkowski, Cordelia Langford, John Sillitoe on behalf of the Wellcome Sanger Institute COVID-19 Surveillance Team |
| EPI_ISL_3126329 | Berkshire and Surrey Pathology Services Lighthouse Laboratory | Wellcome Sanger Institute for the COVID-19 Genomics UK (COG-UK) Consortium | Berkshire and Surrey Pathology Services Lighthouse Laboratory and Alex Alderton, Roberto Amato, Jeffrey Barrett, Sonia Goncalves, Ewan Harrison, David K. Jackson, Ian Johnston, Dominic Kwiatkowski, Cordelia Langford, John Sillitoe on behalf of the Wellcome Sanger Institute COVID-19 Surveillance Team |
| EPI_ISL_3126330, EPI_ISL_3126331 | Lighthouse Lab in Alderley Park | Wellcome Sanger Institute for the COVID-19 Genomics UK (COG-UK) Consortium | Jacquelyn Wynn, Mairead Hyland, The Lighthouse Lab in Alderley Park and Alex Alderton, Roberto Amato, Jeffrey Barrett, Sonia Goncalves, Ewan Harrison, David K. Jackson, Ian Johnston, Dominic Kwiatkowski, Cordelia Langford, John Sillitoe on behalf of the Wellcome Sanger Institute COVID-19 Surveillance Team |
| EPI_ISL_3126332, EPI_ISL_3126333 | Lighthouse Lab in Milton Keynes | Wellcome Sanger Institute for the COVID-19 Genomics UK (COG-UK) Consortium | The Lighthouse Lab in Milton Keynes and Alex Alderton, Roberto Amato, Jeffrey Barrett, Sonia Goncalves, Ewan Harrison, David K. Jackson, Ian Johnston, Dominic Kwiatkowski, Cordelia Langford, John Sillitoe on behalf of the Wellcome Sanger Institute COVID-19 Surveillance Team |
| EPI_ISL_3126334 | Berkshire and Surrey Pathology Services Lighthouse Laboratory | Wellcome Sanger Institute for the COVID-19 Genomics UK (COG-UK) Consortium | Berkshire and Surrey Pathology Services Lighthouse Laboratory and Alex Alderton, Roberto Amato, Jeffrey Barrett, Sonia Goncalves, Ewan Harrison, David K. Jackson, Ian Johnston, Dominic Kwiatkowski, Cordelia Langford, John Sillitoe on behalf of the Wellcome Sanger Institute COVID-19 Surveillance Team |
| EPI_ISL_3126335 | Lighthouse Lab in Alderley Park | Wellcome Sanger Institute for the COVID-19 Genomics UK (COG-UK) Consortium | Jacquelyn Wynn, Mairead Hyland, The Lighthouse Lab in Alderley Park and Alex Alderton, Roberto Amato, Jeffrey Barrett, Sonia Goncalves, Ewan Harrison, David K. Jackson, Ian Johnston, Dominic Kwiatkowski, Cordelia Langford, John Sillitoe on behalf of the Wellcome Sanger Institute COVID-19 Surveillance Team |
| EPI_ISL_3126336 | Berkshire and Surrey Pathology Services Lighthouse Laboratory | Wellcome Sanger Institute for the COVID-19 Genomics UK (COG-UK) Consortium | Berkshire and Surrey Pathology Services Lighthouse Laboratory and Alex Alderton, Roberto Amato, Jeffrey Barrett, Sonia Goncalves, Ewan Harrison, David K. Jackson, Ian Johnston, Dominic Kwiatkowski, Cordelia Langford, John Sillitoe on behalf of the Wellcome Sanger Institute COVID-19 Surveillance Team |
| EPI_ISL_3126337 | Lighthouse Lab in Milton Keynes | Wellcome Sanger Institute for the COVID-19 Genomics UK (COG-UK) Consortium | The Lighthouse Lab in Milton Keynes and Alex Alderton, Roberto Amato, Jeffrey Barrett, Sonia Goncalves, Ewan Harrison, David K. Jackson, Ian Johnston, Dominic Kwiatkowski, Cordelia Langford, John Sillitoe on behalf of the Wellcome Sanger Institute COVID-19 Surveillance Team |
| EPI_ISL_3126338 | Lighthouse Lab in Alderley Park | Wellcome Sanger Institute for the COVID-19 Genomics UK (COG-UK) Consortium | Jacquelyn Wynn, Mairead Hyland, The Lighthouse Lab in Alderley Park and Alex Alderton, Roberto Amato, Jeffrey Barrett, Sonia Goncalves, Ewan Harrison, David K. Jackson, Ian Johnston, Dominic Kwiatkowski, Cordelia Langford, John Sillitoe on behalf of the Wellcome Sanger Institute COVID-19 Surveillance Team |
| EPI_ISL_3126339, EPI_ISL_3126340, EPI_ISL_3126341 | Lighthouse Lab in Milton Keynes | Wellcome Sanger Institute for the COVID-19 Genomics UK (COG-UK) Consortium | The Lighthouse Lab in Milton Keynes and Alex Alderton, Roberto Amato, Jeffrey Barrett, Sonia Goncalves, Ewan Harrison, David K. Jackson, Ian Johnston, Dominic Kwiatkowski, Cordelia Langford, John Sillitoe on behalf of the Wellcome Sanger Institute COVID-19 Surveillance Team |
| EPI_ISL_3126342 | Lighthouse Lab in Alderley Park | Wellcome Sanger Institute for the COVID-19 Genomics UK (COG-UK) Consortium | Jacquelyn Wynn, Mairead Hyland, The Lighthouse Lab in Alderley Park and Alex Alderton, Roberto Amato, Jeffrey Barrett, Sonia Goncalves, Ewan Harrison, David K. Jackson, Ian Johnston, Dominic Kwiatkowski, Cordelia Langford, John Sillitoe on behalf of the Wellcome Sanger Institute COVID-19 Surveillance Team |
| EPI_ISL_3126343 | Berkshire and Surrey Pathology Services Lighthouse Laboratory | Wellcome Sanger Institute for the COVID-19 Genomics UK (COG-UK) Consortium | Berkshire and Surrey Pathology Services Lighthouse Laboratory and Alex Alderton, Roberto Amato, Jeffrey Barrett, Sonia Goncalves, Ewan Harrison, David K. Jackson, Ian Johnston, Dominic Kwiatkowski, Cordelia Langford, John Sillitoe on behalf of the Wellcome Sanger Institute COVID-19 Surveillance Team |
| EPI_ISL_3126344 | Lighthouse Lab in Alderley Park | Wellcome Sanger Institute for the COVID-19 Genomics UK (COG-UK) Consortium | Jacquelyn Wynn, Mairead Hyland, The Lighthouse Lab in Alderley Park and Alex Alderton, Roberto Amato, Jeffrey Barrett, Sonia Goncalves, Ewan Harrison, David K. Jackson, Ian Johnston, Dominic Kwiatkowski, Cordelia Langford, John Sillitoe on behalf of the Wellcome Sanger Institute COVID-19 Surveillance Team |
| EPI_ISL_3126345, EPI_ISL_3126346 | Berkshire and Surrey Pathology Services Lighthouse Laboratory | Wellcome Sanger Institute for the COVID-19 Genomics UK (COG-UK) Consortium | Berkshire and Surrey Pathology Services Lighthouse Laboratory and Alex Alderton, Roberto Amato, Jeffrey Barrett, Sonia Goncalves, Ewan Harrison, David K. Jackson, Ian Johnston, Dominic Kwiatkowski, Cordelia Langford, John Sillitoe on behalf of the Wellcome Sanger Institute COVID-19 Surveillance Team |
| EPI_ISL_3126347, EPI_ISL_3126348, EPI_ISL_3126349 | Lighthouse Lab in Alderley Park | Wellcome Sanger Institute for the COVID-19 Genomics UK (COG-UK) Consortium | Jacquelyn Wynn, Mairead Hyland, The Lighthouse Lab in Alderley Park and Alex Alderton, Roberto Amato, Jeffrey Barrett, Sonia Goncalves, Ewan Harrison, David K. Jackson, Ian Johnston, Dominic Kwiatkowski, Cordelia Langford, John Sillitoe on behalf of the Wellcome Sanger Institute COVID-19 Surveillance Team |
| EPI_ISL_3126350 | Lighthouse Lab in Milton Keynes | Wellcome Sanger Institute for the COVID-19 Genomics UK (COG-UK) Consortium | The Lighthouse Lab in Milton Keynes and Alex Alderton, Roberto Amato, Jeffrey Barrett, Sonia Goncalves, Ewan Harrison, David K. Jackson, Ian Johnston, Dominic Kwiatkowski, Cordelia Langford, John Sillitoe on behalf of the Wellcome Sanger Institute COVID-19 Surveillance Team |
| EPI_ISL_3126351, EPI_ISL_3126352, EPI_ISL_3126353, EPI_ISL_3126354, EPI_ISL_3126355, EPI_ISL_3126356 | Lighthouse Lab in Alderley Park | Wellcome Sanger Institute for the COVID-19 Genomics UK (COG-UK) Consortium | Jacquelyn Wynn, Mairead Hyland, The Lighthouse Lab in Alderley Park and Alex Alderton, Roberto Amato, Jeffrey Barrett, Sonia Goncalves, Ewan Harrison, David K. Jackson, Ian Johnston, Dominic Kwiatkowski, Cordelia Langford, John Sillitoe on behalf of the Wellcome Sanger Institute COVID-19 Surveillance Team |
| EPI_ISL_3126357 | Berkshire and Surrey Pathology Services Lighthouse Laboratory | Wellcome Sanger Institute for the COVID-19 Genomics UK (COG-UK) | Berkshire and Surrey Pathology Services Lighthouse Laboratory and Alex Alderton, Roberto Amato, Jeffrey Barrett, Sonia Goncalves, Ewan Harrison, David K. Jackson, Ian Johnston, Dominic Kwiatkowski, Cordelia Langford, John Sillitoe on behalf |

| | | | |
|---|---|---|---|
| | | Consortium | of the Wellcome Sanger Institute COVID-19 Surveillance Team |
| EPI_ISL_3126358, EPI_ISL_3126359 | Lighthouse Lab in Alderley Park | Wellcome Sanger Institute for the COVID-19 Genomics UK (COG-UK) Consortium | Jacquelyn Wynn, Mairead Hyland, The Lighthouse Lab in Alderley Park and Alex Alderton, Roberto Amato, Jeffrey Barrett, Sonia Goncalves, Ewan Harrison, David K. Jackson, Ian Johnston, Dominic Kwiatkowski, Cordelia Langford, John Sillitoe on behalf of the Wellcome Sanger Institute COVID-19 Surveillance Team |
| EPI_ISL_3126360, EPI_ISL_3126361 | Lighthouse Lab in Milton Keynes | Wellcome Sanger Institute for the COVID-19 Genomics UK (COG-UK) Consortium | The Lighthouse Lab in Milton Keynes and Alex Alderton, Roberto Amato, Jeffrey Barrett, Sonia Goncalves, Ewan Harrison, David K. Jackson, Ian Johnston, Dominic Kwiatkowski, Cordelia Langford, John Sillitoe on behalf of the Wellcome Sanger Institute COVID-19 Surveillance Team |
| EPI_ISL_3126362, EPI_ISL_3126363 | Lighthouse Lab in Alderley Park | Wellcome Sanger Institute for the COVID-19 Genomics UK (COG-UK) Consortium | Jacquelyn Wynn, Mairead Hyland, The Lighthouse Lab in Alderley Park and Alex Alderton, Roberto Amato, Jeffrey Barrett, Sonia Goncalves, Ewan Harrison, David K. Jackson, Ian Johnston, Dominic Kwiatkowski, Cordelia Langford, John Sillitoe on behalf of the Wellcome Sanger Institute COVID-19 Surveillance Team |
| EPI_ISL_3126364 | Lighthouse Lab in Milton Keynes | Wellcome Sanger Institute for the COVID-19 Genomics UK (COG-UK) Consortium | The Lighthouse Lab in Milton Keynes and Alex Alderton, Roberto Amato, Jeffrey Barrett, Sonia Goncalves, Ewan Harrison, David K. Jackson, Ian Johnston, Dominic Kwiatkowski, Cordelia Langford, John Sillitoe on behalf of the Wellcome Sanger Institute COVID-19 Surveillance Team |
| EPI_ISL_3126365 | Berkshire and Surrey Pathology Services Lighthouse Laboratory | Wellcome Sanger Institute for the COVID-19 Genomics UK (COG-UK) Consortium | Berkshire and Surrey Pathology Services Lighthouse Laboratory and Alex Alderton, Roberto Amato, Jeffrey Barrett, Sonia Goncalves, Ewan Harrison, David K. Jackson, Ian Johnston, Dominic Kwiatkowski, Cordelia Langford, John Sillitoe on behalf of the Wellcome Sanger Institute COVID-19 Surveillance Team |
| EPI_ISL_3126366 | Lighthouse Lab in Alderley Park | Wellcome Sanger Institute for the COVID-19 Genomics UK (COG-UK) Consortium | Jacquelyn Wynn, Mairead Hyland, The Lighthouse Lab in Alderley Park and Alex Alderton, Roberto Amato, Jeffrey Barrett, Sonia Goncalves, Ewan Harrison, David K. Jackson, Ian Johnston, Dominic Kwiatkowski, Cordelia Langford, John Sillitoe on behalf of the Wellcome Sanger Institute COVID-19 Surveillance Team |
| EPI_ISL_3126367 | Lighthouse Lab in Milton Keynes | Wellcome Sanger Institute for the COVID-19 Genomics UK (COG-UK) Consortium | The Lighthouse Lab in Milton Keynes and Alex Alderton, Roberto Amato, Jeffrey Barrett, Sonia Goncalves, Ewan Harrison, David K. Jackson, Ian Johnston, Dominic Kwiatkowski, Cordelia Langford, John Sillitoe on behalf of the Wellcome Sanger Institute COVID-19 Surveillance Team |
| EPI_ISL_3126368 | Lighthouse Lab in Alderley Park | Wellcome Sanger Institute for the COVID-19 Genomics UK (COG-UK) Consortium | Jacquelyn Wynn, Mairead Hyland, The Lighthouse Lab in Alderley Park and Alex Alderton, Roberto Amato, Jeffrey Barrett, Sonia Goncalves, Ewan Harrison, David K. Jackson, Ian Johnston, Dominic Kwiatkowski, Cordelia Langford, John Sillitoe on behalf of the Wellcome Sanger Institute COVID-19 Surveillance Team |
| EPI_ISL_3126369, EPI_ISL_3126370 | Berkshire and Surrey Pathology Services Lighthouse Laboratory | Wellcome Sanger Institute for the COVID-19 Genomics UK (COG-UK) Consortium | Berkshire and Surrey Pathology Services Lighthouse Laboratory and Alex Alderton, Roberto Amato, Jeffrey Barrett, Sonia Goncalves, Ewan Harrison, David K. Jackson, Ian Johnston, Dominic Kwiatkowski, Cordelia Langford, John Sillitoe on behalf of the Wellcome Sanger Institute COVID-19 Surveillance Team |
| EPI_ISL_3126371, EPI_ISL_3126372 | Lighthouse Lab in Alderley Park | Wellcome Sanger Institute for the COVID-19 Genomics UK (COG-UK) Consortium | Jacquelyn Wynn, Mairead Hyland, The Lighthouse Lab in Alderley Park and Alex Alderton, Roberto Amato, Jeffrey Barrett, Sonia Goncalves, Ewan Harrison, David K. Jackson, Ian Johnston, Dominic Kwiatkowski, Cordelia Langford, John Sillitoe on behalf of the Wellcome Sanger Institute COVID-19 Surveillance Team |
| EPI_ISL_3126373, EPI_ISL_3126374, EPI_ISL_3126375 | Lighthouse Lab in Milton Keynes | Wellcome Sanger Institute for the COVID-19 Genomics UK (COG-UK) Consortium | The Lighthouse Lab in Milton Keynes and Alex Alderton, Roberto Amato, Jeffrey Barrett, Sonia Goncalves, Ewan Harrison, David K. Jackson, Ian Johnston, Dominic Kwiatkowski, Cordelia Langford, John Sillitoe on behalf of the Wellcome Sanger Institute COVID-19 Surveillance Team |
| EPI_ISL_3126376, EPI_ISL_3126377 | Berkshire and Surrey Pathology Services Lighthouse Laboratory | Wellcome Sanger Institute for the COVID-19 Genomics UK (COG-UK) Consortium | Berkshire and Surrey Pathology Services Lighthouse Laboratory and Alex Alderton, Roberto Amato, Jeffrey Barrett, Sonia Goncalves, Ewan Harrison, David K. Jackson, Ian Johnston, Dominic Kwiatkowski, Cordelia Langford, John Sillitoe on behalf of the Wellcome Sanger Institute COVID-19 Surveillance Team |
| EPI_ISL_3126378 | Lighthouse Lab in Milton Keynes | Wellcome Sanger Institute for the COVID-19 Genomics UK (COG-UK) Consortium | The Lighthouse Lab in Milton Keynes and Alex Alderton, Roberto Amato, Jeffrey Barrett, Sonia Goncalves, Ewan Harrison, David K. Jackson, Ian Johnston, Dominic Kwiatkowski, Cordelia Langford, John Sillitoe on behalf of the Wellcome Sanger Institute COVID-19 Surveillance Team |
| EPI_ISL_3126379, EPI_ISL_3126380, EPI_ISL_3126381, EPI_ISL_3126382 | Lighthouse Lab in Alderley Park | Wellcome Sanger Institute for the COVID-19 Genomics UK (COG-UK) Consortium | Jacquelyn Wynn, Mairead Hyland, The Lighthouse Lab in Alderley Park and Alex Alderton, Roberto Amato, Jeffrey Barrett, Sonia Goncalves, Ewan Harrison, David K. Jackson, Ian Johnston, Dominic Kwiatkowski, Cordelia Langford, John Sillitoe on behalf of the Wellcome Sanger Institute COVID-19 Surveillance Team |
| EPI_ISL_3126383 | Lighthouse Lab in Milton Keynes | Wellcome Sanger Institute for the COVID-19 Genomics UK (COG-UK) Consortium | The Lighthouse Lab in Milton Keynes and Alex Alderton, Roberto Amato, Jeffrey Barrett, Sonia Goncalves, Ewan Harrison, David K. Jackson, Ian Johnston, Dominic Kwiatkowski, Cordelia Langford, John Sillitoe on behalf of the Wellcome Sanger Institute COVID-19 Surveillance Team |
| EPI_ISL_3126384, EPI_ISL_3126385 | Berkshire and Surrey Pathology Services Lighthouse Laboratory | Wellcome Sanger Institute for the COVID-19 Genomics UK (COG-UK) Consortium | Berkshire and Surrey Pathology Services Lighthouse Laboratory and Alex Alderton, Roberto Amato, Jeffrey Barrett, Sonia Goncalves, Ewan Harrison, David K. Jackson, Ian Johnston, Dominic Kwiatkowski, Cordelia Langford, John Sillitoe on behalf of the Wellcome Sanger Institute COVID-19 Surveillance Team |
| EPI_ISL_3126386 | Lighthouse Lab in Milton Keynes | Wellcome Sanger Institute for the COVID-19 Genomics UK (COG-UK) Consortium | The Lighthouse Lab in Milton Keynes and Alex Alderton, Roberto Amato, Jeffrey Barrett, Sonia Goncalves, Ewan Harrison, David K. Jackson, Ian Johnston, Dominic Kwiatkowski, Cordelia Langford, John Sillitoe on behalf of the Wellcome Sanger Institute COVID-19 Surveillance Team |
| EPI_ISL_3126387 | Lighthouse Lab in Alderley Park | Wellcome Sanger Institute for the COVID-19 Genomics UK (COG-UK) Consortium | Jacquelyn Wynn, Mairead Hyland, The Lighthouse Lab in Alderley Park and Alex Alderton, Roberto Amato, Jeffrey Barrett, Sonia Goncalves, Ewan Harrison, David K. Jackson, Ian Johnston, Dominic Kwiatkowski, Cordelia Langford, John Sillitoe on behalf of the Wellcome Sanger Institute COVID-19 Surveillance Team |
| EPI_ISL_3126388 | Berkshire and Surrey Pathology Services Lighthouse Laboratory | Wellcome Sanger Institute for the COVID-19 Genomics UK (COG-UK) Consortium | Berkshire and Surrey Pathology Services Lighthouse Laboratory and Alex Alderton, Roberto Amato, Jeffrey Barrett, Sonia Goncalves, Ewan Harrison, David K. Jackson, Ian Johnston, Dominic Kwiatkowski, Cordelia Langford, John Sillitoe on behalf of the Wellcome Sanger Institute COVID-19 Surveillance Team |
| EPI_ISL_3126389 | Lighthouse Lab in Alderley Park | Wellcome Sanger Institute for the COVID-19 Genomics UK (COG-UK) Consortium | Jacquelyn Wynn, Mairead Hyland, The Lighthouse Lab in Alderley Park and Alex Alderton, Roberto Amato, Jeffrey Barrett, Sonia Goncalves, Ewan Harrison, David K. Jackson, Ian Johnston, Dominic Kwiatkowski, Cordelia Langford, John Sillitoe on behalf of the Wellcome Sanger Institute COVID-19 Surveillance Team |
| EPI_ISL_3126390, EPI_ISL_3126391, EPI_ISL_3126392 | Lighthouse Lab in Milton Keynes | Wellcome Sanger Institute for the COVID-19 Genomics UK (COG-UK) Consortium | The Lighthouse Lab in Milton Keynes and Alex Alderton, Roberto Amato, Jeffrey Barrett, Sonia Goncalves, Ewan Harrison, David K. Jackson, Ian Johnston, Dominic Kwiatkowski, Cordelia Langford, John Sillitoe on behalf of the Wellcome Sanger Institute COVID-19 Surveillance Team |
| EPI_ISL_3126393 | Berkshire and Surrey Pathology Services Lighthouse Laboratory | Wellcome Sanger Institute for the COVID-19 Genomics UK (COG-UK) Consortium | Berkshire and Surrey Pathology Services Lighthouse Laboratory and Alex Alderton, Roberto Amato, Jeffrey Barrett, Sonia Goncalves, Ewan Harrison, David K. Jackson, Ian Johnston, Dominic Kwiatkowski, Cordelia Langford, John Sillitoe on behalf of the Wellcome Sanger Institute COVID-19 Surveillance Team |
| EPI_ISL_3126394 | Lighthouse Lab in Milton Keynes | Wellcome Sanger Institute for the COVID-19 Genomics UK (COG-UK) Consortium | The Lighthouse Lab in Milton Keynes and Alex Alderton, Roberto Amato, Jeffrey Barrett, Sonia Goncalves, Ewan Harrison, David K. Jackson, Ian Johnston, Dominic Kwiatkowski, Cordelia Langford, John Sillitoe on behalf of the Wellcome Sanger Institute COVID-19 Surveillance Team |
| EPI_ISL_3126395 | Lighthouse Lab in Alderley Park | Wellcome Sanger Institute for the COVID-19 Genomics UK (COG-UK) Consortium | Jacquelyn Wynn, Mairead Hyland, The Lighthouse Lab in Alderley Park and Alex Alderton, Roberto Amato, Jeffrey Barrett, Sonia Goncalves, Ewan Harrison, David K. Jackson, Ian Johnston, Dominic Kwiatkowski, Cordelia Langford, John Sillitoe on behalf of the Wellcome Sanger Institute COVID-19 Surveillance Team |
| EPI_ISL_3126396, EPI_ISL_3126397 | Berkshire and Surrey Pathology Services Lighthouse Laboratory | Wellcome Sanger Institute for the COVID-19 Genomics UK (COG-UK) Consortium | Berkshire and Surrey Pathology Services Lighthouse Laboratory and Alex Alderton, Roberto Amato, Jeffrey Barrett, Sonia Goncalves, Ewan Harrison, David K. Jackson, Ian Johnston, Dominic Kwiatkowski, Cordelia Langford, John Sillitoe on behalf of the Wellcome Sanger Institute COVID-19 Surveillance Team |
| EPI_ISL_3126398 | Lighthouse Lab in Alderley Park | Wellcome Sanger Institute for the COVID-19 Genomics UK (COG-UK) Consortium | Jacquelyn Wynn, Mairead Hyland, The Lighthouse Lab in Alderley Park and Alex Alderton, Roberto Amato, Jeffrey Barrett, Sonia Goncalves, Ewan Harrison, David K. Jackson, Ian Johnston, Dominic Kwiatkowski, Cordelia Langford, John Sillitoe on behalf of the Wellcome Sanger Institute COVID-19 Surveillance Team |
| EPI_ISL_3126399 | Berkshire and Surrey Pathology Services Lighthouse Laboratory | Wellcome Sanger Institute for the COVID-19 Genomics UK (COG-UK) Consortium | Berkshire and Surrey Pathology Services Lighthouse Laboratory and Alex Alderton, Roberto Amato, Jeffrey Barrett, Sonia Goncalves, Ewan Harrison, David K. Jackson, Ian Johnston, Dominic Kwiatkowski, Cordelia Langford, John Sillitoe on behalf of the Wellcome Sanger Institute COVID-19 Surveillance Team |
| EPI_ISL_3126400 | Lighthouse Lab in Alderley Park | Wellcome Sanger Institute for the COVID-19 Genomics UK (COG-UK) Consortium | Jacquelyn Wynn, Mairead Hyland, The Lighthouse Lab in Alderley Park and Alex Alderton, Roberto Amato, Jeffrey Barrett, Sonia Goncalves, Ewan Harrison, David K. Jackson, Ian Johnston, Dominic Kwiatkowski, Cordelia Langford, John Sillitoe on behalf of the Wellcome Sanger Institute COVID-19 Surveillance Team |
| EPI_ISL_3126401, EPI_ISL_3126402 | Berkshire and Surrey Pathology Services Lighthouse Laboratory | Wellcome Sanger Institute for the COVID-19 Genomics UK (COG-UK) Consortium | Berkshire and Surrey Pathology Services Lighthouse Laboratory and Alex Alderton, Roberto Amato, Jeffrey Barrett, Sonia Goncalves, Ewan Harrison, David K. Jackson, Ian Johnston, Dominic Kwiatkowski, Cordelia Langford, John Sillitoe on behalf of the Wellcome Sanger Institute COVID-19 Surveillance Team |
| EPI_ISL_3126403, EPI_ISL_3126404, EPI_ISL_3126405 | Lighthouse Lab in Milton Keynes | Wellcome Sanger Institute for the COVID-19 Genomics UK (COG-UK) Consortium | The Lighthouse Lab in Milton Keynes and Alex Alderton, Roberto Amato, Jeffrey Barrett, Sonia Goncalves, Ewan Harrison, David K. Jackson, Ian Johnston, Dominic Kwiatkowski, Cordelia Langford, John Sillitoe on behalf of the Wellcome Sanger Institute COVID-19 Surveillance Team |
| EPI_ISL_3126406 | Lighthouse Lab in Alderley Park | Wellcome Sanger Institute for the COVID-19 Genomics UK (COG-UK) Consortium | Jacquelyn Wynn, Mairead Hyland, The Lighthouse Lab in Alderley Park and Alex Alderton, Roberto Amato, Jeffrey Barrett, Sonia Goncalves, Ewan Harrison, David K. Jackson, Ian Johnston, Dominic Kwiatkowski, Cordelia Langford, John Sillitoe on behalf of the Wellcome Sanger Institute COVID-19 Surveillance Team |
| EPI_ISL_3126407 | Lighthouse Lab in Milton Keynes | Wellcome Sanger Institute for the COVID-19 Genomics UK (COG-UK) Consortium | The Lighthouse Lab in Milton Keynes and Alex Alderton, Roberto Amato, Jeffrey Barrett, Sonia Goncalves, Ewan Harrison, David K. Jackson, Ian Johnston, Dominic Kwiatkowski, Cordelia Langford, John Sillitoe on behalf of the Wellcome Sanger Institute COVID-19 Surveillance Team |
| EPI_ISL_3126408 | Berkshire and Surrey Pathology Services Lighthouse Laboratory | Wellcome Sanger Institute for the COVID-19 Genomics UK (COG-UK) Consortium | Berkshire and Surrey Pathology Services Lighthouse Laboratory and Alex Alderton, Roberto Amato, Jeffrey Barrett, Sonia Goncalves, Ewan Harrison, David K. Jackson, Ian Johnston, Dominic Kwiatkowski, Cordelia Langford, John Sillitoe on behalf of the Wellcome Sanger Institute COVID-19 Surveillance Team |
| EPI_ISL_3126409, EPI_ISL_3126410 | Lighthouse Lab in Alderley Park | Wellcome Sanger Institute for the COVID-19 Genomics UK (COG-UK) Consortium | Jacquelyn Wynn, Mairead Hyland, The Lighthouse Lab in Alderley Park and Alex Alderton, Roberto Amato, Jeffrey Barrett, Sonia Goncalves, Ewan Harrison, David K. Jackson, Ian Johnston, Dominic Kwiatkowski, Cordelia Langford, John Sillitoe on behalf of the Wellcome Sanger Institute COVID-19 Surveillance Team |
| EPI_ISL_3126411, EPI_ISL_3126412 | Berkshire and Surrey Pathology Services Lighthouse Laboratory | Wellcome Sanger Institute for the COVID-19 Genomics UK (COG-UK) Consortium | Berkshire and Surrey Pathology Services Lighthouse Laboratory and Alex Alderton, Roberto Amato, Jeffrey Barrett, Sonia Goncalves, Ewan Harrison, David K. Jackson, Ian Johnston, Dominic Kwiatkowski, Cordelia Langford, John Sillitoe on behalf of the Wellcome Sanger Institute COVID-19 Surveillance Team |
| EPI_ISL_3126413 | Lighthouse Lab in Milton Keynes | Wellcome Sanger Institute for the COVID-19 Genomics UK (COG-UK) Consortium | The Lighthouse Lab in Milton Keynes and Alex Alderton, Roberto Amato, Jeffrey Barrett, Sonia Goncalves, Ewan Harrison, David K. Jackson, Ian Johnston, Dominic Kwiatkowski, Cordelia Langford, John Sillitoe on behalf of the Wellcome Sanger Institute COVID-19 Surveillance Team |
| EPI_ISL_3126414, EPI_ISL_3126415, EPI_ISL_3126416, EPI_ISL_3126417, EPI_ISL_3126418 | Lighthouse Lab in Alderley Park | Wellcome Sanger Institute for the COVID-19 Genomics UK (COG-UK) Consortium | Jacquelyn Wynn, Mairead Hyland, The Lighthouse Lab in Alderley Park and Alex Alderton, Roberto Amato, Jeffrey Barrett, Sonia Goncalves, Ewan Harrison, David K. Jackson, Ian Johnston, Dominic Kwiatkowski, Cordelia Langford, John Sillitoe on behalf of the Wellcome Sanger Institute COVID-19 Surveillance Team |
| EPI_ISL_3126419 | Lighthouse Lab in Milton Keynes | Wellcome Sanger Institute for the COVID-19 Genomics UK (COG-UK) Consortium | The Lighthouse Lab in Milton Keynes and Alex Alderton, Roberto Amato, Jeffrey Barrett, Sonia Goncalves, Ewan Harrison, David K. Jackson, Ian Johnston, Dominic Kwiatkowski, Cordelia Langford, John Sillitoe on behalf of the Wellcome Sanger Institute COVID-19 Surveillance Team |
| EPI_ISL_3126420, EPI_ISL_3126421, EPI_ISL_3126422, EPI_ISL_3126423 | Lighthouse Lab in Alderley Park | Wellcome Sanger Institute for the COVID-19 Genomics UK (COG-UK) Consortium | Jacquelyn Wynn, Mairead Hyland, The Lighthouse Lab in Alderley Park and Alex Alderton, Roberto Amato, Jeffrey Barrett, Sonia Goncalves, Ewan Harrison, David K. Jackson, Ian Johnston, Dominic Kwiatkowski, Cordelia Langford, John Sillitoe on behalf of the Wellcome Sanger Institute COVID-19 Surveillance Team |
| EPI_ISL_3126424 | Lighthouse Lab in Milton Keynes | Wellcome Sanger Institute for the COVID-19 Genomics UK (COG-UK) Consortium | The Lighthouse Lab in Milton Keynes and Alex Alderton, Roberto Amato, Jeffrey Barrett, Sonia Goncalves, Ewan Harrison, David K. Jackson, Ian Johnston, Dominic Kwiatkowski, Cordelia Langford, John Sillitoe on behalf of the Wellcome Sanger Institute COVID-19 Surveillance Team |

| | | | |
|---|---|---|---|
| EPI_ISL_3126425 | Lighthouse Lab in Alderley Park | Wellcome Sanger Institute for the COVID-19 Genomics UK (COG-UK) Consortium | Jacquelyn Wynn, Mairead Hyland, The Lighthouse Lab in Alderley Park and Alex Alderton, Roberto Amato, Jeffrey Barrett, Sonia Goncalves, Ewan Harrison, David K. Jackson, Ian Johnston, Dominic Kwiatkowski, Cordelia Langford, John Sillitoe on behalf of the Wellcome Sanger Institute COVID-19 Surveillance Team |
| EPI_ISL_3126426 | Berkshire and Surrey Pathology Services Lighthouse Laboratory | Wellcome Sanger Institute for the COVID-19 Genomics UK (COG-UK) Consortium | Berkshire and Surrey Pathology Services Lighthouse Laboratory and Alex Alderton, Roberto Amato, Jeffrey Barrett, Sonia Goncalves, Ewan Harrison, David K. Jackson, Ian Johnston, Dominic Kwiatkowski, Cordelia Langford, John Sillitoe on behalf of the Wellcome Sanger Institute COVID-19 Surveillance Team |
| EPI_ISL_3126427 | Lighthouse Lab in Alderley Park | Wellcome Sanger Institute for the COVID-19 Genomics UK (COG-UK) Consortium | Jacquelyn Wynn, Mairead Hyland, The Lighthouse Lab in Alderley Park and Alex Alderton, Roberto Amato, Jeffrey Barrett, Sonia Goncalves, Ewan Harrison, David K. Jackson, Ian Johnston, Dominic Kwiatkowski, Cordelia Langford, John Sillitoe on behalf of the Wellcome Sanger Institute COVID-19 Surveillance Team |
| EPI_ISL_3126428 | Berkshire and Surrey Pathology Services Lighthouse Laboratory | Wellcome Sanger Institute for the COVID-19 Genomics UK (COG-UK) Consortium | Berkshire and Surrey Pathology Services Lighthouse Laboratory and Alex Alderton, Roberto Amato, Jeffrey Barrett, Sonia Goncalves, Ewan Harrison, David K. Jackson, Ian Johnston, Dominic Kwiatkowski, Cordelia Langford, John Sillitoe on behalf of the Wellcome Sanger Institute COVID-19 Surveillance Team |
| EPI_ISL_3126429, EPI_ISL_3126430, EPI_ISL_3126431, EPI_ISL_3126432 | Lighthouse Lab in Alderley Park | Wellcome Sanger Institute for the COVID-19 Genomics UK (COG-UK) Consortium | Jacquelyn Wynn, Mairead Hyland, The Lighthouse Lab in Alderley Park and Alex Alderton, Roberto Amato, Jeffrey Barrett, Sonia Goncalves, Ewan Harrison, David K. Jackson, Ian Johnston, Dominic Kwiatkowski, Cordelia Langford, John Sillitoe on behalf of the Wellcome Sanger Institute COVID-19 Surveillance Team |
| EPI_ISL_3126433 | Berkshire and Surrey Pathology Services Lighthouse Laboratory | Wellcome Sanger Institute for the COVID-19 Genomics UK (COG-UK) Consortium | Berkshire and Surrey Pathology Services Lighthouse Laboratory and Alex Alderton, Roberto Amato, Jeffrey Barrett, Sonia Goncalves, Ewan Harrison, David K. Jackson, Ian Johnston, Dominic Kwiatkowski, Cordelia Langford, John Sillitoe on behalf of the Wellcome Sanger Institute COVID-19 Surveillance Team |
| EPI_ISL_3126434 | Lighthouse Lab in Alderley Park | Wellcome Sanger Institute for the COVID-19 Genomics UK (COG-UK) Consortium | Jacquelyn Wynn, Mairead Hyland, The Lighthouse Lab in Alderley Park and Alex Alderton, Roberto Amato, Jeffrey Barrett, Sonia Goncalves, Ewan Harrison, David K. Jackson, Ian Johnston, Dominic Kwiatkowski, Cordelia Langford, John Sillitoe on behalf of the Wellcome Sanger Institute COVID-19 Surveillance Team |
| EPI_ISL_3126435 | Lighthouse Lab in Glasgow | Wellcome Sanger Institute for the COVID-19 Genomics UK (COG-UK) Consortium | Harper VanSteenhouse, Yumi Kasai, David Gray, Carol Clugston, Anna Dominiczak and Alex Alderton, Roberto Amato, Jeffrey Barrett, Sonia Goncalves, Ewan Harrison, David K. Jackson, Ian Johnston, Dominic Kwiatkowski, Cordelia Langford, John Sillitoe on behalf of the Wellcome Sanger Institute COVID-19 Surveillance Team |
| EPI_ISL_3126436 | Lighthouse Lab in Alderley Park | Wellcome Sanger Institute for the COVID-19 Genomics UK (COG-UK) Consortium | Jacquelyn Wynn, Mairead Hyland, The Lighthouse Lab in Alderley Park and Alex Alderton, Roberto Amato, Jeffrey Barrett, Sonia Goncalves, Ewan Harrison, David K. Jackson, Ian Johnston, Dominic Kwiatkowski, Cordelia Langford, John Sillitoe on behalf of the Wellcome Sanger Institute COVID-19 Surveillance Team |
| EPI_ISL_3126437 | Berkshire and Surrey Pathology Services Lighthouse Laboratory | Wellcome Sanger Institute for the COVID-19 Genomics UK (COG-UK) Consortium | Berkshire and Surrey Pathology Services Lighthouse Laboratory and Alex Alderton, Roberto Amato, Jeffrey Barrett, Sonia Goncalves, Ewan Harrison, David K. Jackson, Ian Johnston, Dominic Kwiatkowski, Cordelia Langford, John Sillitoe on behalf of the Wellcome Sanger Institute COVID-19 Surveillance Team |
| EPI_ISL_3126438 | Lighthouse Lab in Alderley Park | Wellcome Sanger Institute for the COVID-19 Genomics UK (COG-UK) Consortium | Jacquelyn Wynn, Mairead Hyland, The Lighthouse Lab in Alderley Park and Alex Alderton, Roberto Amato, Jeffrey Barrett, Sonia Goncalves, Ewan Harrison, David K. Jackson, Ian Johnston, Dominic Kwiatkowski, Cordelia Langford, John Sillitoe on behalf of the Wellcome Sanger Institute COVID-19 Surveillance Team |
| EPI_ISL_3126439 | Berkshire and Surrey Pathology Services Lighthouse Laboratory | Wellcome Sanger Institute for the COVID-19 Genomics UK (COG-UK) Consortium | Berkshire and Surrey Pathology Services Lighthouse Laboratory and Alex Alderton, Roberto Amato, Jeffrey Barrett, Sonia Goncalves, Ewan Harrison, David K. Jackson, Ian Johnston, Dominic Kwiatkowski, Cordelia Langford, John Sillitoe on behalf of the Wellcome Sanger Institute COVID-19 Surveillance Team |
| EPI_ISL_3126440 | Lighthouse Lab in Alderley Park | Wellcome Sanger Institute for the COVID-19 Genomics UK (COG-UK) Consortium | Jacquelyn Wynn, Mairead Hyland, The Lighthouse Lab in Alderley Park and Alex Alderton, Roberto Amato, Jeffrey Barrett, Sonia Goncalves, Ewan Harrison, David K. Jackson, Ian Johnston, Dominic Kwiatkowski, Cordelia Langford, John Sillitoe on behalf of the Wellcome Sanger Institute COVID-19 Surveillance Team |
| EPI_ISL_3126441 | Berkshire and Surrey Pathology Services Lighthouse Laboratory | Wellcome Sanger Institute for the COVID-19 Genomics UK (COG-UK) Consortium | Berkshire and Surrey Pathology Services Lighthouse Laboratory and Alex Alderton, Roberto Amato, Jeffrey Barrett, Sonia Goncalves, Ewan Harrison, David K. Jackson, Ian Johnston, Dominic Kwiatkowski, Cordelia Langford, John Sillitoe on behalf of the Wellcome Sanger Institute COVID-19 Surveillance Team |
| EPI_ISL_3126442 | Lighthouse Lab in Alderley Park | Wellcome Sanger Institute for the COVID-19 Genomics UK (COG-UK) Consortium | Jacquelyn Wynn, Mairead Hyland, The Lighthouse Lab in Alderley Park and Alex Alderton, Roberto Amato, Jeffrey Barrett, Sonia Goncalves, Ewan Harrison, David K. Jackson, Ian Johnston, Dominic Kwiatkowski, Cordelia Langford, John Sillitoe on behalf of the Wellcome Sanger Institute COVID-19 Surveillance Team |
| EPI_ISL_3126443 | Lighthouse Lab in Glasgow | Wellcome Sanger Institute for the COVID-19 Genomics UK (COG-UK) Consortium | Harper VanSteenhouse, Yumi Kasai, David Gray, Carol Clugston, Anna Dominiczak and Alex Alderton, Roberto Amato, Jeffrey Barrett, Sonia Goncalves, Ewan Harrison, David K. Jackson, Ian Johnston, Dominic Kwiatkowski, Cordelia Langford, John Sillitoe on behalf of the Wellcome Sanger Institute COVID-19 Surveillance Team |
| EPI_ISL_3126444 | Lighthouse Lab in Alderley Park | Wellcome Sanger Institute for the COVID-19 Genomics UK (COG-UK) Consortium | Jacquelyn Wynn, Mairead Hyland, The Lighthouse Lab in Alderley Park and Alex Alderton, Roberto Amato, Jeffrey Barrett, Sonia Goncalves, Ewan Harrison, David K. Jackson, Ian Johnston, Dominic Kwiatkowski, Cordelia Langford, John Sillitoe on behalf of the Wellcome Sanger Institute COVID-19 Surveillance Team |
| EPI_ISL_3126445, EPI_ISL_3126446, EPI_ISL_3126447, EPI_ISL_3126448 | Lighthouse Lab in Glasgow | Wellcome Sanger Institute for the COVID-19 Genomics UK (COG-UK) Consortium | Harper VanSteenhouse, Yumi Kasai, David Gray, Carol Clugston, Anna Dominiczak and Alex Alderton, Roberto Amato, Jeffrey Barrett, Sonia Goncalves, Ewan Harrison, David K. Jackson, Ian Johnston, Dominic Kwiatkowski, Cordelia Langford, John Sillitoe on behalf of the Wellcome Sanger Institute COVID-19 Surveillance Team |
| EPI_ISL_3126449, EPI_ISL_3126450 | Berkshire and Surrey Pathology Services Lighthouse Laboratory | Wellcome Sanger Institute for the COVID-19 Genomics UK (COG-UK) Consortium | Berkshire and Surrey Pathology Services Lighthouse Laboratory and Alex Alderton, Roberto Amato, Jeffrey Barrett, Sonia Goncalves, Ewan Harrison, David K. Jackson, Ian Johnston, Dominic Kwiatkowski, Cordelia Langford, John Sillitoe on behalf of the Wellcome Sanger Institute COVID-19 Surveillance Team |
| EPI_ISL_3126451 | Lighthouse Lab in Glasgow | Wellcome Sanger Institute for the COVID-19 Genomics UK (COG-UK) Consortium | Harper VanSteenhouse, Yumi Kasai, David Gray, Carol Clugston, Anna Dominiczak and Alex Alderton, Roberto Amato, Jeffrey Barrett, Sonia Goncalves, Ewan Harrison, David K. Jackson, Ian Johnston, Dominic Kwiatkowski, Cordelia Langford, John Sillitoe on behalf of the Wellcome Sanger Institute COVID-19 Surveillance Team |
| EPI_ISL_3126452 | Berkshire and Surrey Pathology Services Lighthouse Laboratory | Wellcome Sanger Institute for the COVID-19 Genomics UK (COG-UK) Consortium | Berkshire and Surrey Pathology Services Lighthouse Laboratory and Alex Alderton, Roberto Amato, Jeffrey Barrett, Sonia Goncalves, Ewan Harrison, David K. Jackson, Ian Johnston, Dominic Kwiatkowski, Cordelia Langford, John Sillitoe on behalf of the Wellcome Sanger Institute COVID-19 Surveillance Team |
| EPI_ISL_3126453 | Lighthouse Lab in Glasgow | Wellcome Sanger Institute for the COVID-19 Genomics UK (COG-UK) Consortium | Harper VanSteenhouse, Yumi Kasai, David Gray, Carol Clugston, Anna Dominiczak and Alex Alderton, Roberto Amato, Jeffrey Barrett, Sonia Goncalves, Ewan Harrison, David K. Jackson, Ian Johnston, Dominic Kwiatkowski, Cordelia Langford, John Sillitoe on behalf of the Wellcome Sanger Institute COVID-19 Surveillance Team |
| EPI_ISL_3126454, EPI_ISL_3126455, EPI_ISL_3126456 | Lighthouse Lab in Alderley Park | Wellcome Sanger Institute for the COVID-19 Genomics UK (COG-UK) Consortium | Jacquelyn Wynn, Mairead Hyland, The Lighthouse Lab in Alderley Park and Alex Alderton, Roberto Amato, Jeffrey Barrett, Sonia Goncalves, Ewan Harrison, David K. Jackson, Ian Johnston, Dominic Kwiatkowski, Cordelia Langford, John Sillitoe on behalf of the Wellcome Sanger Institute COVID-19 Surveillance Team |
| EPI_ISL_3126457 | Berkshire and Surrey Pathology Services Lighthouse Laboratory | Wellcome Sanger Institute for the COVID-19 Genomics UK (COG-UK) Consortium | Berkshire and Surrey Pathology Services Lighthouse Laboratory and Alex Alderton, Roberto Amato, Jeffrey Barrett, Sonia Goncalves, Ewan Harrison, David K. Jackson, Ian Johnston, Dominic Kwiatkowski, Cordelia Langford, John Sillitoe on behalf of the Wellcome Sanger Institute COVID-19 Surveillance Team |
| EPI_ISL_3126458 | Lighthouse Lab in Glasgow | Wellcome Sanger Institute for the COVID-19 Genomics UK (COG-UK) Consortium | Harper VanSteenhouse, Yumi Kasai, David Gray, Carol Clugston, Anna Dominiczak and Alex Alderton, Roberto Amato, Jeffrey Barrett, Sonia Goncalves, Ewan Harrison, David K. Jackson, Ian Johnston, Dominic Kwiatkowski, Cordelia Langford, John Sillitoe on behalf of the Wellcome Sanger Institute COVID-19 Surveillance Team |
| EPI_ISL_3126459 | Berkshire and Surrey Pathology Services Lighthouse Laboratory | Wellcome Sanger Institute for the COVID-19 Genomics UK (COG-UK) Consortium | Berkshire and Surrey Pathology Services Lighthouse Laboratory and Alex Alderton, Roberto Amato, Jeffrey Barrett, Sonia Goncalves, Ewan Harrison, David K. Jackson, Ian Johnston, Dominic Kwiatkowski, Cordelia Langford, John Sillitoe on behalf of the Wellcome Sanger Institute COVID-19 Surveillance Team |
| EPI_ISL_3126460 | Lighthouse Lab in Alderley Park | Wellcome Sanger Institute for the COVID-19 Genomics UK (COG-UK) Consortium | Jacquelyn Wynn, Mairead Hyland, The Lighthouse Lab in Alderley Park and Alex Alderton, Roberto Amato, Jeffrey Barrett, Sonia Goncalves, Ewan Harrison, David K. Jackson, Ian Johnston, Dominic Kwiatkowski, Cordelia Langford, John Sillitoe on behalf of the Wellcome Sanger Institute COVID-19 Surveillance Team |
| EPI_ISL_3126461, EPI_ISL_3126462, EPI_ISL_3126463 | Lighthouse Lab in Glasgow | Wellcome Sanger Institute for the COVID-19 Genomics UK (COG-UK) Consortium | Harper VanSteenhouse, Yumi Kasai, David Gray, Carol Clugston, Anna Dominiczak and Alex Alderton, Roberto Amato, Jeffrey Barrett, Sonia Goncalves, Ewan Harrison, David K. Jackson, Ian Johnston, Dominic Kwiatkowski, Cordelia Langford, John Sillitoe on behalf of the Wellcome Sanger Institute COVID-19 Surveillance Team |
| EPI_ISL_3126464, EPI_ISL_3126465 | Lighthouse Lab in Alderley Park | Wellcome Sanger Institute for the COVID-19 Genomics UK (COG-UK) Consortium | Jacquelyn Wynn, Mairead Hyland, The Lighthouse Lab in Alderley Park and Alex Alderton, Roberto Amato, Jeffrey Barrett, Sonia Goncalves, Ewan Harrison, David K. Jackson, Ian Johnston, Dominic Kwiatkowski, Cordelia Langford, John Sillitoe on behalf of the Wellcome Sanger Institute COVID-19 Surveillance Team |
| EPI_ISL_3126466 | Lighthouse Lab in Glasgow | Wellcome Sanger Institute for the COVID-19 Genomics UK (COG-UK) Consortium | Harper VanSteenhouse, Yumi Kasai, David Gray, Carol Clugston, Anna Dominiczak and Alex Alderton, Roberto Amato, Jeffrey Barrett, Sonia Goncalves, Ewan Harrison, David K. Jackson, Ian Johnston, Dominic Kwiatkowski, Cordelia Langford, John Sillitoe on behalf of the Wellcome Sanger Institute COVID-19 Surveillance Team |
| EPI_ISL_3126467 | Lighthouse Lab in Alderley Park | Wellcome Sanger Institute for the COVID-19 Genomics UK (COG-UK) Consortium | Jacquelyn Wynn, Mairead Hyland, The Lighthouse Lab in Alderley Park and Alex Alderton, Roberto Amato, Jeffrey Barrett, Sonia Goncalves, Ewan Harrison, David K. Jackson, Ian Johnston, Dominic Kwiatkowski, Cordelia Langford, John Sillitoe on behalf of the Wellcome Sanger Institute COVID-19 Surveillance Team |
| EPI_ISL_3126468 | Berkshire and Surrey Pathology Services Lighthouse Laboratory | Wellcome Sanger Institute for the COVID-19 Genomics UK (COG-UK) Consortium | Berkshire and Surrey Pathology Services Lighthouse Laboratory and Alex Alderton, Roberto Amato, Jeffrey Barrett, Sonia Goncalves, Ewan Harrison, David K. Jackson, Ian Johnston, Dominic Kwiatkowski, Cordelia Langford, John Sillitoe on behalf of the Wellcome Sanger Institute COVID-19 Surveillance Team |
| EPI_ISL_3126469 | Lighthouse Lab in Glasgow | Wellcome Sanger Institute for the COVID-19 Genomics UK (COG-UK) Consortium | Harper VanSteenhouse, Yumi Kasai, David Gray, Carol Clugston, Anna Dominiczak and Alex Alderton, Roberto Amato, Jeffrey Barrett, Sonia Goncalves, Ewan Harrison, David K. Jackson, Ian Johnston, Dominic Kwiatkowski, Cordelia Langford, John Sillitoe on behalf of the Wellcome Sanger Institute COVID-19 Surveillance Team |
| EPI_ISL_3126470 | Lighthouse Lab in Alderley Park | Wellcome Sanger Institute for the COVID-19 Genomics UK (COG-UK) Consortium | Jacquelyn Wynn, Mairead Hyland, The Lighthouse Lab in Alderley Park and Alex Alderton, Roberto Amato, Jeffrey Barrett, Sonia Goncalves, Ewan Harrison, David K. Jackson, Ian Johnston, Dominic Kwiatkowski, Cordelia Langford, John Sillitoe on behalf of the Wellcome Sanger Institute COVID-19 Surveillance Team |
| EPI_ISL_3126471 | Berkshire and Surrey Pathology Services Lighthouse Laboratory | Wellcome Sanger Institute for the COVID-19 Genomics UK (COG-UK) Consortium | Berkshire and Surrey Pathology Services Lighthouse Laboratory and Alex Alderton, Roberto Amato, Jeffrey Barrett, Sonia Goncalves, Ewan Harrison, David K. Jackson, Ian Johnston, Dominic Kwiatkowski, Cordelia Langford, John Sillitoe on behalf of the Wellcome Sanger Institute COVID-19 Surveillance Team |
| EPI_ISL_3126472 | Lighthouse Lab in Glasgow | Wellcome Sanger Institute for the COVID-19 Genomics UK (COG-UK) Consortium | Harper VanSteenhouse, Yumi Kasai, David Gray, Carol Clugston, Anna Dominiczak and Alex Alderton, Roberto Amato, Jeffrey Barrett, Sonia Goncalves, Ewan Harrison, David K. Jackson, Ian Johnston, Dominic Kwiatkowski, Cordelia Langford, John Sillitoe on behalf of the Wellcome Sanger Institute COVID-19 Surveillance Team |
| EPI_ISL_3126473 | Berkshire and Surrey Pathology Services Lighthouse Laboratory | Wellcome Sanger Institute for the COVID-19 Genomics UK (COG-UK) Consortium | Berkshire and Surrey Pathology Services Lighthouse Laboratory and Alex Alderton, Roberto Amato, Jeffrey Barrett, Sonia Goncalves, Ewan Harrison, David K. Jackson, Ian Johnston, Dominic Kwiatkowski, Cordelia Langford, John Sillitoe on behalf of the Wellcome Sanger Institute COVID-19 Surveillance Team |
| EPI_ISL_3126474 | Lighthouse Lab in Glasgow | Wellcome Sanger Institute for the COVID-19 Genomics UK (COG-UK) Consortium | Harper VanSteenhouse, Yumi Kasai, David Gray, Carol Clugston, Anna Dominiczak and Alex Alderton, Roberto Amato, Jeffrey Barrett, Sonia Goncalves, Ewan Harrison, David K. Jackson, Ian Johnston, Dominic Kwiatkowski, Cordelia Langford, John Sillitoe on behalf of the Wellcome Sanger Institute COVID-19 Surveillance Team |
| EPI_ISL_3126475 | Berkshire and Surrey Pathology Services Lighthouse Laboratory | Wellcome Sanger Institute for the COVID-19 Genomics UK (COG-UK) Consortium | Berkshire and Surrey Pathology Services Lighthouse Laboratory and Alex Alderton, Roberto Amato, Jeffrey Barrett, Sonia Goncalves, Ewan Harrison, David K. Jackson, Ian Johnston, Dominic Kwiatkowski, Cordelia Langford, John Sillitoe on behalf of the Wellcome Sanger Institute COVID-19 Surveillance Team |
| EPI_ISL_3126476, EPI_ISL_3126477, EPI_ISL_3126478, EPI_ISL_3126479 | Lighthouse Lab in Glasgow | Wellcome Sanger Institute for the COVID-19 Genomics UK (COG-UK) Consortium | Harper VanSteenhouse, Yumi Kasai, David Gray, Carol Clugston, Anna Dominiczak and Alex Alderton, Roberto Amato, Jeffrey Barrett, Sonia Goncalves, Ewan Harrison, David K. Jackson, Ian Johnston, Dominic Kwiatkowski, Cordelia Langford, John Sillitoe on behalf of the Wellcome Sanger Institute COVID-19 Surveillance Team |
| EPI_ISL_3126480 | Lighthouse Lab in Alderley Park | Wellcome Sanger Institute for the COVID-19 Genomics UK (COG-UK) Consortium | Jacquelyn Wynn, Mairead Hyland, The Lighthouse Lab in Alderley Park and Alex Alderton, Roberto Amato, Jeffrey Barrett, Sonia Goncalves, Ewan Harrison, David K. Jackson, Ian Johnston, Dominic Kwiatkowski, Cordelia Langford, John Sillitoe on behalf of the Wellcome Sanger Institute COVID-19 Surveillance Team |
| EPI_ISL_3126481, | Lighthouse Lab in Glasgow | Wellcome Sanger Institute for the COVID-19 Genomics UK (COG-UK) | Harper VanSteenhouse, Yumi Kasai, David Gray, Carol Clugston, Anna Dominiczak and Alex Alderton, Roberto Amato, Jeffrey Barrett, Sonia Goncalves, Ewan Harrison, David K. Jackson, Ian Johnston, Dominic Kwiatkowski, Cordelia Langford, |

| | | | |
|---|---|---|---|
| EPI_ISL_3126482 | | Consortium | John Sillitoe on behalf of the Wellcome Sanger Institute COVID-19 Surveillance Team |
| EPI_ISL_3126483 | Lighthouse Lab in Alderley Park | Wellcome Sanger Institute for the COVID-19 Genomics UK (COG-UK) Consortium | Jacquelyn Wynn, Mairead Hyland, The Lighthouse Lab in Alderley Park and Alex Alderton, Roberto Amato, Jeffrey Barrett, Sonia Goncalves, Ewan Harrison, David K. Jackson, Ian Johnston, Dominic Kwiatkowski, Cordelia Langford, John Sillitoe on behalf of the Wellcome Sanger Institute COVID-19 Surveillance Team |
| EPI_ISL_3126484 | Lighthouse Lab in Glasgow | Wellcome Sanger Institute for the COVID-19 Genomics UK (COG-UK) Consortium | Harper VanSteenhouse, Yumi Kasai, David Gray, Carol Clugston, Anna Dominiczak and Alex Alderton, Roberto Amato, Jeffrey Barrett, Sonia Goncalves, Ewan Harrison, David K. Jackson, Ian Johnston, Dominic Kwiatkowski, Cordelia Langford, John Sillitoe on behalf of the Wellcome Sanger Institute COVID-19 Surveillance Team |
| EPI_ISL_3126485 | Berkshire and Surrey Pathology Services Lighthouse Laboratory | Wellcome Sanger Institute for the COVID-19 Genomics UK (COG-UK) Consortium | Berkshire and Surrey Pathology Services Lighthouse Laboratory and Alex Alderton, Roberto Amato, Jeffrey Barrett, Sonia Goncalves, Ewan Harrison, David K. Jackson, Ian Johnston, Dominic Kwiatkowski, Cordelia Langford, John Sillitoe on behalf of the Wellcome Sanger Institute COVID-19 Surveillance Team |
| EPI_ISL_3126486 | Lighthouse Lab in Glasgow | Wellcome Sanger Institute for the COVID-19 Genomics UK (COG-UK) Consortium | Harper VanSteenhouse, Yumi Kasai, David Gray, Carol Clugston, Anna Dominiczak and Alex Alderton, Roberto Amato, Jeffrey Barrett, Sonia Goncalves, Ewan Harrison, David K. Jackson, Ian Johnston, Dominic Kwiatkowski, Cordelia Langford, John Sillitoe on behalf of the Wellcome Sanger Institute COVID-19 Surveillance Team |
| EPI_ISL_3126487 | Lighthouse Lab in Alderley Park | Wellcome Sanger Institute for the COVID-19 Genomics UK (COG-UK) Consortium | Jacquelyn Wynn, Mairead Hyland, The Lighthouse Lab in Alderley Park and Alex Alderton, Roberto Amato, Jeffrey Barrett, Sonia Goncalves, Ewan Harrison, David K. Jackson, Ian Johnston, Dominic Kwiatkowski, Cordelia Langford, John Sillitoe on behalf of the Wellcome Sanger Institute COVID-19 Surveillance Team |
| EPI_ISL_3126488 | Lighthouse Lab in Glasgow | Wellcome Sanger Institute for the COVID-19 Genomics UK (COG-UK) Consortium | Harper VanSteenhouse, Yumi Kasai, David Gray, Carol Clugston, Anna Dominiczak and Alex Alderton, Roberto Amato, Jeffrey Barrett, Sonia Goncalves, Ewan Harrison, David K. Jackson, Ian Johnston, Dominic Kwiatkowski, Cordelia Langford, John Sillitoe on behalf of the Wellcome Sanger Institute COVID-19 Surveillance Team |
| EPI_ISL_3126489 | Lighthouse Lab in Alderley Park | Wellcome Sanger Institute for the COVID-19 Genomics UK (COG-UK) Consortium | Jacquelyn Wynn, Mairead Hyland, The Lighthouse Lab in Alderley Park and Alex Alderton, Roberto Amato, Jeffrey Barrett, Sonia Goncalves, Ewan Harrison, David K. Jackson, Ian Johnston, Dominic Kwiatkowski, Cordelia Langford, John Sillitoe on behalf of the Wellcome Sanger Institute COVID-19 Surveillance Team |
| EPI_ISL_3126490, EPI_ISL_3126491, EPI_ISL_3126492, EPI_ISL_3126493 | Lighthouse Lab in Glasgow | Wellcome Sanger Institute for the COVID-19 Genomics UK (COG-UK) Consortium | Harper VanSteenhouse, Yumi Kasai, David Gray, Carol Clugston, Anna Dominiczak and Alex Alderton, Roberto Amato, Jeffrey Barrett, Sonia Goncalves, Ewan Harrison, David K. Jackson, Ian Johnston, Dominic Kwiatkowski, Cordelia Langford, John Sillitoe on behalf of the Wellcome Sanger Institute COVID-19 Surveillance Team |
| EPI_ISL_3126494 | Berkshire and Surrey Pathology Services Lighthouse Laboratory | Wellcome Sanger Institute for the COVID-19 Genomics UK (COG-UK) Consortium | Berkshire and Surrey Pathology Services Lighthouse Laboratory and Alex Alderton, Roberto Amato, Jeffrey Barrett, Sonia Goncalves, Ewan Harrison, David K. Jackson, Ian Johnston, Dominic Kwiatkowski, Cordelia Langford, John Sillitoe on behalf of the Wellcome Sanger Institute COVID-19 Surveillance Team |
| EPI_ISL_3126495 | Lighthouse Lab in Alderley Park | Wellcome Sanger Institute for the COVID-19 Genomics UK (COG-UK) Consortium | Jacquelyn Wynn, Mairead Hyland, The Lighthouse Lab in Alderley Park and Alex Alderton, Roberto Amato, Jeffrey Barrett, Sonia Goncalves, Ewan Harrison, David K. Jackson, Ian Johnston, Dominic Kwiatkowski, Cordelia Langford, John Sillitoe on behalf of the Wellcome Sanger Institute COVID-19 Surveillance Team |
| EPI_ISL_3126496, EPI_ISL_3126497 | Lighthouse Lab in Glasgow | Wellcome Sanger Institute for the COVID-19 Genomics UK (COG-UK) Consortium | Harper VanSteenhouse, Yumi Kasai, David Gray, Carol Clugston, Anna Dominiczak and Alex Alderton, Roberto Amato, Jeffrey Barrett, Sonia Goncalves, Ewan Harrison, David K. Jackson, Ian Johnston, Dominic Kwiatkowski, Cordelia Langford, John Sillitoe on behalf of the Wellcome Sanger Institute COVID-19 Surveillance Team |
| EPI_ISL_3126498 | Berkshire and Surrey Pathology Services Lighthouse Laboratory | Wellcome Sanger Institute for the COVID-19 Genomics UK (COG-UK) Consortium | Berkshire and Surrey Pathology Services Lighthouse Laboratory and Alex Alderton, Roberto Amato, Jeffrey Barrett, Sonia Goncalves, Ewan Harrison, David K. Jackson, Ian Johnston, Dominic Kwiatkowski, Cordelia Langford, John Sillitoe on behalf of the Wellcome Sanger Institute COVID-19 Surveillance Team |
| EPI_ISL_3126499 | Lighthouse Lab in Glasgow | Wellcome Sanger Institute for the COVID-19 Genomics UK (COG-UK) Consortium | Harper VanSteenhouse, Yumi Kasai, David Gray, Carol Clugston, Anna Dominiczak and Alex Alderton, Roberto Amato, Jeffrey Barrett, Sonia Goncalves, Ewan Harrison, David K. Jackson, Ian Johnston, Dominic Kwiatkowski, Cordelia Langford, John Sillitoe on behalf of the Wellcome Sanger Institute COVID-19 Surveillance Team |
| EPI_ISL_3126500 | Lighthouse Lab in Alderley Park | Wellcome Sanger Institute for the COVID-19 Genomics UK (COG-UK) Consortium | Jacquelyn Wynn, Mairead Hyland, The Lighthouse Lab in Alderley Park and Alex Alderton, Roberto Amato, Jeffrey Barrett, Sonia Goncalves, Ewan Harrison, David K. Jackson, Ian Johnston, Dominic Kwiatkowski, Cordelia Langford, John Sillitoe on behalf of the Wellcome Sanger Institute COVID-19 Surveillance Team |
| EPI_ISL_3126501 | Lighthouse Lab in Glasgow | Wellcome Sanger Institute for the COVID-19 Genomics UK (COG-UK) Consortium | Harper VanSteenhouse, Yumi Kasai, David Gray, Carol Clugston, Anna Dominiczak and Alex Alderton, Roberto Amato, Jeffrey Barrett, Sonia Goncalves, Ewan Harrison, David K. Jackson, Ian Johnston, Dominic Kwiatkowski, Cordelia Langford, John Sillitoe on behalf of the Wellcome Sanger Institute COVID-19 Surveillance Team |
| EPI_ISL_3126502 | Lighthouse Lab in Alderley Park | Wellcome Sanger Institute for the COVID-19 Genomics UK (COG-UK) Consortium | Jacquelyn Wynn, Mairead Hyland, The Lighthouse Lab in Alderley Park and Alex Alderton, Roberto Amato, Jeffrey Barrett, Sonia Goncalves, Ewan Harrison, David K. Jackson, Ian Johnston, Dominic Kwiatkowski, Cordelia Langford, John Sillitoe on behalf of the Wellcome Sanger Institute COVID-19 Surveillance Team |
| EPI_ISL_3126503, EPI_ISL_3126504 | Lighthouse Lab in Glasgow | Wellcome Sanger Institute for the COVID-19 Genomics UK (COG-UK) Consortium | Harper VanSteenhouse, Yumi Kasai, David Gray, Carol Clugston, Anna Dominiczak and Alex Alderton, Roberto Amato, Jeffrey Barrett, Sonia Goncalves, Ewan Harrison, David K. Jackson, Ian Johnston, Dominic Kwiatkowski, Cordelia Langford, John Sillitoe on behalf of the Wellcome Sanger Institute COVID-19 Surveillance Team |
| EPI_ISL_3126505 | Berkshire and Surrey Pathology Services Lighthouse Laboratory | Wellcome Sanger Institute for the COVID-19 Genomics UK (COG-UK) Consortium | Berkshire and Surrey Pathology Services Lighthouse Laboratory and Alex Alderton, Roberto Amato, Jeffrey Barrett, Sonia Goncalves, Ewan Harrison, David K. Jackson, Ian Johnston, Dominic Kwiatkowski, Cordelia Langford, John Sillitoe on behalf of the Wellcome Sanger Institute COVID-19 Surveillance Team |
| EPI_ISL_3126506 | Lighthouse Lab in Glasgow | Wellcome Sanger Institute for the COVID-19 Genomics UK (COG-UK) Consortium | Harper VanSteenhouse, Yumi Kasai, David Gray, Carol Clugston, Anna Dominiczak and Alex Alderton, Roberto Amato, Jeffrey Barrett, Sonia Goncalves, Ewan Harrison, David K. Jackson, Ian Johnston, Dominic Kwiatkowski, Cordelia Langford, John Sillitoe on behalf of the Wellcome Sanger Institute COVID-19 Surveillance Team |
| EPI_ISL_3126507 | Berkshire and Surrey Pathology Services Lighthouse Laboratory | Wellcome Sanger Institute for the COVID-19 Genomics UK (COG-UK) Consortium | Berkshire and Surrey Pathology Services Lighthouse Laboratory and Alex Alderton, Roberto Amato, Jeffrey Barrett, Sonia Goncalves, Ewan Harrison, David K. Jackson, Ian Johnston, Dominic Kwiatkowski, Cordelia Langford, John Sillitoe on behalf of the Wellcome Sanger Institute COVID-19 Surveillance Team |
| EPI_ISL_3126508 | Lighthouse Lab in Alderley Park | Wellcome Sanger Institute for the COVID-19 Genomics UK (COG-UK) Consortium | Jacquelyn Wynn, Mairead Hyland, The Lighthouse Lab in Alderley Park and Alex Alderton, Roberto Amato, Jeffrey Barrett, Sonia Goncalves, Ewan Harrison, David K. Jackson, Ian Johnston, Dominic Kwiatkowski, Cordelia Langford, John Sillitoe on behalf of the Wellcome Sanger Institute COVID-19 Surveillance Team |
| EPI_ISL_3126509 | Lighthouse Lab in Glasgow | Wellcome Sanger Institute for the COVID-19 Genomics UK (COG-UK) Consortium | Harper VanSteenhouse, Yumi Kasai, David Gray, Carol Clugston, Anna Dominiczak and Alex Alderton, Roberto Amato, Jeffrey Barrett, Sonia Goncalves, Ewan Harrison, David K. Jackson, Ian Johnston, Dominic Kwiatkowski, Cordelia Langford, John Sillitoe on behalf of the Wellcome Sanger Institute COVID-19 Surveillance Team |
| EPI_ISL_3126510 | Lighthouse Lab in Alderley Park | Wellcome Sanger Institute for the COVID-19 Genomics UK (COG-UK) Consortium | Jacquelyn Wynn, Mairead Hyland, The Lighthouse Lab in Alderley Park and Alex Alderton, Roberto Amato, Jeffrey Barrett, Sonia Goncalves, Ewan Harrison, David K. Jackson, Ian Johnston, Dominic Kwiatkowski, Cordelia Langford, John Sillitoe on behalf of the Wellcome Sanger Institute COVID-19 Surveillance Team |
| EPI_ISL_3126511 | Berkshire and Surrey Pathology Services Lighthouse Laboratory | Wellcome Sanger Institute for the COVID-19 Genomics UK (COG-UK) Consortium | Berkshire and Surrey Pathology Services Lighthouse Laboratory and Alex Alderton, Roberto Amato, Jeffrey Barrett, Sonia Goncalves, Ewan Harrison, David K. Jackson, Ian Johnston, Dominic Kwiatkowski, Cordelia Langford, John Sillitoe on behalf of the Wellcome Sanger Institute COVID-19 Surveillance Team |
| EPI_ISL_3126512, EPI_ISL_3126513 | Lighthouse Lab in Alderley Park | Wellcome Sanger Institute for the COVID-19 Genomics UK (COG-UK) Consortium | Jacquelyn Wynn, Mairead Hyland, The Lighthouse Lab in Alderley Park and Alex Alderton, Roberto Amato, Jeffrey Barrett, Sonia Goncalves, Ewan Harrison, David K. Jackson, Ian Johnston, Dominic Kwiatkowski, Cordelia Langford, John Sillitoe on behalf of the Wellcome Sanger Institute COVID-19 Surveillance Team |
| EPI_ISL_3126514 | Lighthouse Lab in Glasgow | Wellcome Sanger Institute for the COVID-19 Genomics UK (COG-UK) Consortium | Harper VanSteenhouse, Yumi Kasai, David Gray, Carol Clugston, Anna Dominiczak and Alex Alderton, Roberto Amato, Jeffrey Barrett, Sonia Goncalves, Ewan Harrison, David K. Jackson, Ian Johnston, Dominic Kwiatkowski, Cordelia Langford, John Sillitoe on behalf of the Wellcome Sanger Institute COVID-19 Surveillance Team |
| EPI_ISL_3126515 | Lighthouse Lab in Alderley Park | Wellcome Sanger Institute for the COVID-19 Genomics UK (COG-UK) Consortium | Jacquelyn Wynn, Mairead Hyland, The Lighthouse Lab in Alderley Park and Alex Alderton, Roberto Amato, Jeffrey Barrett, Sonia Goncalves, Ewan Harrison, David K. Jackson, Ian Johnston, Dominic Kwiatkowski, Cordelia Langford, John Sillitoe on behalf of the Wellcome Sanger Institute COVID-19 Surveillance Team |
| EPI_ISL_3126516, EPI_ISL_3126517 | Lighthouse Lab in Glasgow | Wellcome Sanger Institute for the COVID-19 Genomics UK (COG-UK) Consortium | Harper VanSteenhouse, Yumi Kasai, David Gray, Carol Clugston, Anna Dominiczak and Alex Alderton, Roberto Amato, Jeffrey Barrett, Sonia Goncalves, Ewan Harrison, David K. Jackson, Ian Johnston, Dominic Kwiatkowski, Cordelia Langford, John Sillitoe on behalf of the Wellcome Sanger Institute COVID-19 Surveillance Team |
| EPI_ISL_3126518, EPI_ISL_3126519 | Berkshire and Surrey Pathology Services Lighthouse Laboratory | Wellcome Sanger Institute for the COVID-19 Genomics UK (COG-UK) Consortium | Berkshire and Surrey Pathology Services Lighthouse Laboratory and Alex Alderton, Roberto Amato, Jeffrey Barrett, Sonia Goncalves, Ewan Harrison, David K. Jackson, Ian Johnston, Dominic Kwiatkowski, Cordelia Langford, John Sillitoe on behalf of the Wellcome Sanger Institute COVID-19 Surveillance Team |
| EPI_ISL_3126520 | Lighthouse Lab in Glasgow | Wellcome Sanger Institute for the COVID-19 Genomics UK (COG-UK) Consortium | Harper VanSteenhouse, Yumi Kasai, David Gray, Carol Clugston, Anna Dominiczak and Alex Alderton, Roberto Amato, Jeffrey Barrett, Sonia Goncalves, Ewan Harrison, David K. Jackson, Ian Johnston, Dominic Kwiatkowski, Cordelia Langford, John Sillitoe on behalf of the Wellcome Sanger Institute COVID-19 Surveillance Team |
| EPI_ISL_3126521 | Berkshire and Surrey Pathology Services Lighthouse Laboratory | Wellcome Sanger Institute for the COVID-19 Genomics UK (COG-UK) Consortium | Berkshire and Surrey Pathology Services Lighthouse Laboratory and Alex Alderton, Roberto Amato, Jeffrey Barrett, Sonia Goncalves, Ewan Harrison, David K. Jackson, Ian Johnston, Dominic Kwiatkowski, Cordelia Langford, John Sillitoe on behalf of the Wellcome Sanger Institute COVID-19 Surveillance Team |
| EPI_ISL_3126522, EPI_ISL_3126523 | Lighthouse Lab in Glasgow | Wellcome Sanger Institute for the COVID-19 Genomics UK (COG-UK) Consortium | Harper VanSteenhouse, Yumi Kasai, David Gray, Carol Clugston, Anna Dominiczak and Alex Alderton, Roberto Amato, Jeffrey Barrett, Sonia Goncalves, Ewan Harrison, David K. Jackson, Ian Johnston, Dominic Kwiatkowski, Cordelia Langford, John Sillitoe on behalf of the Wellcome Sanger Institute COVID-19 Surveillance Team |
| EPI_ISL_3126524 | Lighthouse Lab in Alderley Park | Wellcome Sanger Institute for the COVID-19 Genomics UK (COG-UK) Consortium | Jacquelyn Wynn, Mairead Hyland, The Lighthouse Lab in Alderley Park and Alex Alderton, Roberto Amato, Jeffrey Barrett, Sonia Goncalves, Ewan Harrison, David K. Jackson, Ian Johnston, Dominic Kwiatkowski, Cordelia Langford, John Sillitoe on behalf of the Wellcome Sanger Institute COVID-19 Surveillance Team |
| EPI_ISL_3126525 | Berkshire and Surrey Pathology Services Lighthouse Laboratory | Wellcome Sanger Institute for the COVID-19 Genomics UK (COG-UK) Consortium | Berkshire and Surrey Pathology Services Lighthouse Laboratory and Alex Alderton, Roberto Amato, Jeffrey Barrett, Sonia Goncalves, Ewan Harrison, David K. Jackson, Ian Johnston, Dominic Kwiatkowski, Cordelia Langford, John Sillitoe on behalf of the Wellcome Sanger Institute COVID-19 Surveillance Team |
| EPI_ISL_3126526 | Lighthouse Lab in Glasgow | Wellcome Sanger Institute for the COVID-19 Genomics UK (COG-UK) Consortium | Harper VanSteenhouse, Yumi Kasai, David Gray, Carol Clugston, Anna Dominiczak and Alex Alderton, Roberto Amato, Jeffrey Barrett, Sonia Goncalves, Ewan Harrison, David K. Jackson, Ian Johnston, Dominic Kwiatkowski, Cordelia Langford, John Sillitoe on behalf of the Wellcome Sanger Institute COVID-19 Surveillance Team |
| EPI_ISL_3126527, EPI_ISL_3126528, EPI_ISL_3126529 | Lighthouse Lab in Alderley Park | Wellcome Sanger Institute for the COVID-19 Genomics UK (COG-UK) Consortium | Jacquelyn Wynn, Mairead Hyland, The Lighthouse Lab in Alderley Park and Alex Alderton, Roberto Amato, Jeffrey Barrett, Sonia Goncalves, Ewan Harrison, David K. Jackson, Ian Johnston, Dominic Kwiatkowski, Cordelia Langford, John Sillitoe on behalf of the Wellcome Sanger Institute COVID-19 Surveillance Team |
| EPI_ISL_3126530, EPI_ISL_3126531, EPI_ISL_3126532, EPI_ISL_3126533 | Lighthouse Lab in Glasgow | Wellcome Sanger Institute for the COVID-19 Genomics UK (COG-UK) Consortium | Harper VanSteenhouse, Yumi Kasai, David Gray, Carol Clugston, Anna Dominiczak and Alex Alderton, Roberto Amato, Jeffrey Barrett, Sonia Goncalves, Ewan Harrison, David K. Jackson, Ian Johnston, Dominic Kwiatkowski, Cordelia Langford, John Sillitoe on behalf of the Wellcome Sanger Institute COVID-19 Surveillance Team |
| EPI_ISL_3126534 | Berkshire and Surrey Pathology Services Lighthouse Laboratory | Wellcome Sanger Institute for the COVID-19 Genomics UK (COG-UK) Consortium | Berkshire and Surrey Pathology Services Lighthouse Laboratory and Alex Alderton, Roberto Amato, Jeffrey Barrett, Sonia Goncalves, Ewan Harrison, David K. Jackson, Ian Johnston, Dominic Kwiatkowski, Cordelia Langford, John Sillitoe on behalf of the Wellcome Sanger Institute COVID-19 Surveillance Team |
| EPI_ISL_3126535 | Lighthouse Lab in Glasgow | Wellcome Sanger Institute for the COVID-19 Genomics UK (COG-UK) Consortium | Harper VanSteenhouse, Yumi Kasai, David Gray, Carol Clugston, Anna Dominiczak and Alex Alderton, Roberto Amato, Jeffrey Barrett, Sonia Goncalves, Ewan Harrison, David K. Jackson, Ian Johnston, Dominic Kwiatkowski, Cordelia Langford, John Sillitoe on behalf of the Wellcome Sanger Institute COVID-19 Surveillance Team |
| EPI_ISL_3126536, EPI_ISL_3126537 | Berkshire and Surrey Pathology Services Lighthouse Laboratory | Wellcome Sanger Institute for the COVID-19 Genomics UK (COG-UK) Consortium | Berkshire and Surrey Pathology Services Lighthouse Laboratory and Alex Alderton, Roberto Amato, Jeffrey Barrett, Sonia Goncalves, Ewan Harrison, David K. Jackson, Ian Johnston, Dominic Kwiatkowski, Cordelia Langford, John Sillitoe on behalf of the Wellcome Sanger Institute COVID-19 Surveillance Team |
| EPI_ISL_3126538 | Lighthouse Lab in Glasgow | Wellcome Sanger Institute for the COVID-19 Genomics UK (COG-UK) Consortium | Harper VanSteenhouse, Yumi Kasai, David Gray, Carol Clugston, Anna Dominiczak and Alex Alderton, Roberto Amato, Jeffrey Barrett, Sonia Goncalves, Ewan Harrison, David K. Jackson, Ian Johnston, Dominic Kwiatkowski, Cordelia Langford, John Sillitoe on behalf of the Wellcome Sanger Institute COVID-19 Surveillance Team |
| EPI_ISL_3126539 | Lighthouse Lab in Alderley Park | Wellcome Sanger Institute for the COVID-19 Genomics UK (COG-UK) Consortium | Jacquelyn Wynn, Mairead Hyland, The Lighthouse Lab in Alderley Park and Alex Alderton, Roberto Amato, Jeffrey Barrett, Sonia Goncalves, Ewan Harrison, David K. Jackson, Ian Johnston, Dominic Kwiatkowski, Cordelia Langford, John Sillitoe on behalf of the Wellcome Sanger Institute COVID-19 Surveillance Team |

| | | | |
|---|---|---|---|
| EPI_ISL_3126540 | Berkshire and Surrey Pathology Services Lighthouse Laboratory | Wellcome Sanger Institute for the COVID-19 Genomics UK (COG-UK) Consortium | Berkshire and Surrey Pathology Services Lighthouse Laboratory and Alex Alderton, Roberto Amato, Jeffrey Barrett, Sonia Goncalves, Ewan Harrison, David K. Jackson, Ian Johnston, Dominic Kwiatkowski, Cordelia Langford, John Sillitoe on behalf of the Wellcome Sanger Institute COVID-19 Surveillance Team |
| EPI_ISL_3126541 | Lighthouse Lab in Glasgow | Wellcome Sanger Institute for the COVID-19 Genomics UK (COG-UK) Consortium | Harper VanSteenhouse, Yumi Kasai, David Gray, Carol Clugston, Anna Dominiczak and Alex Alderton, Roberto Amato, Jeffrey Barrett, Sonia Goncalves, Ewan Harrison, David K. Jackson, Ian Johnston, Dominic Kwiatkowski, Cordelia Langford, John Sillitoe on behalf of the Wellcome Sanger Institute COVID-19 Surveillance Team |
| EPI_ISL_3126542 | Lighthouse Lab in Alderley Park | Wellcome Sanger Institute for the COVID-19 Genomics UK (COG-UK) Consortium | Jacquelyn Wynn, Mairead Hyland, The Lighthouse Lab in Alderley Park and Alex Alderton, Roberto Amato, Jeffrey Barrett, Sonia Goncalves, Ewan Harrison, David K. Jackson, Ian Johnston, Dominic Kwiatkowski, Cordelia Langford, John Sillitoe on behalf of the Wellcome Sanger Institute COVID-19 Surveillance Team |
| EPI_ISL_3126543, EPI_ISL_3126544, EPI_ISL_3126545 | Lighthouse Lab in Glasgow | Wellcome Sanger Institute for the COVID-19 Genomics UK (COG-UK) Consortium | Harper VanSteenhouse, Yumi Kasai, David Gray, Carol Clugston, Anna Dominiczak and Alex Alderton, Roberto Amato, Jeffrey Barrett, Sonia Goncalves, Ewan Harrison, David K. Jackson, Ian Johnston, Dominic Kwiatkowski, Cordelia Langford, John Sillitoe on behalf of the Wellcome Sanger Institute COVID-19 Surveillance Team |
| EPI_ISL_3126546 | Berkshire and Surrey Pathology Services Lighthouse Laboratory | Wellcome Sanger Institute for the COVID-19 Genomics UK (COG-UK) Consortium | Berkshire and Surrey Pathology Services Lighthouse Laboratory and Alex Alderton, Roberto Amato, Jeffrey Barrett, Sonia Goncalves, Ewan Harrison, David K. Jackson, Ian Johnston, Dominic Kwiatkowski, Cordelia Langford, John Sillitoe on behalf of the Wellcome Sanger Institute COVID-19 Surveillance Team |
| EPI_ISL_3126547, EPI_ISL_3126548, EPI_ISL_3126549, EPI_ISL_3126550, EPI_ISL_3126551, EPI_ISL_3126552 | Lighthouse Lab in Glasgow | Wellcome Sanger Institute for the COVID-19 Genomics UK (COG-UK) Consortium | Harper VanSteenhouse, Yumi Kasai, David Gray, Carol Clugston, Anna Dominiczak and Alex Alderton, Roberto Amato, Jeffrey Barrett, Sonia Goncalves, Ewan Harrison, David K. Jackson, Ian Johnston, Dominic Kwiatkowski, Cordelia Langford, John Sillitoe on behalf of the Wellcome Sanger Institute COVID-19 Surveillance Team |
| EPI_ISL_3126553 | Lighthouse Lab in Alderley Park | Wellcome Sanger Institute for the COVID-19 Genomics UK (COG-UK) Consortium | Jacquelyn Wynn, Mairead Hyland, The Lighthouse Lab in Alderley Park and Alex Alderton, Roberto Amato, Jeffrey Barrett, Sonia Goncalves, Ewan Harrison, David K. Jackson, Ian Johnston, Dominic Kwiatkowski, Cordelia Langford, John Sillitoe on behalf of the Wellcome Sanger Institute COVID-19 Surveillance Team |
| EPI_ISL_3126554 | Lighthouse Lab in Glasgow | Wellcome Sanger Institute for the COVID-19 Genomics UK (COG-UK) Consortium | Harper VanSteenhouse, Yumi Kasai, David Gray, Carol Clugston, Anna Dominiczak and Alex Alderton, Roberto Amato, Jeffrey Barrett, Sonia Goncalves, Ewan Harrison, David K. Jackson, Ian Johnston, Dominic Kwiatkowski, Cordelia Langford, John Sillitoe on behalf of the Wellcome Sanger Institute COVID-19 Surveillance Team |
| EPI_ISL_3126555, EPI_ISL_3126556 | Lighthouse Lab in Alderley Park | Wellcome Sanger Institute for the COVID-19 Genomics UK (COG-UK) Consortium | Jacquelyn Wynn, Mairead Hyland, The Lighthouse Lab in Alderley Park and Alex Alderton, Roberto Amato, Jeffrey Barrett, Sonia Goncalves, Ewan Harrison, David K. Jackson, Ian Johnston, Dominic Kwiatkowski, Cordelia Langford, John Sillitoe on behalf of the Wellcome Sanger Institute COVID-19 Surveillance Team |
| EPI_ISL_3126557 | Lighthouse Lab in Glasgow | Wellcome Sanger Institute for the COVID-19 Genomics UK (COG-UK) Consortium | Harper VanSteenhouse, Yumi Kasai, David Gray, Carol Clugston, Anna Dominiczak and Alex Alderton, Roberto Amato, Jeffrey Barrett, Sonia Goncalves, Ewan Harrison, David K. Jackson, Ian Johnston, Dominic Kwiatkowski, Cordelia Langford, John Sillitoe on behalf of the Wellcome Sanger Institute COVID-19 Surveillance Team |
| EPI_ISL_3126558, EPI_ISL_3126559 | Lighthouse Lab in Alderley Park | Wellcome Sanger Institute for the COVID-19 Genomics UK (COG-UK) Consortium | Jacquelyn Wynn, Mairead Hyland, The Lighthouse Lab in Alderley Park and Alex Alderton, Roberto Amato, Jeffrey Barrett, Sonia Goncalves, Ewan Harrison, David K. Jackson, Ian Johnston, Dominic Kwiatkowski, Cordelia Langford, John Sillitoe on behalf of the Wellcome Sanger Institute COVID-19 Surveillance Team |
| EPI_ISL_3126560 | Lighthouse Lab in Glasgow | Wellcome Sanger Institute for the COVID-19 Genomics UK (COG-UK) Consortium | Harper VanSteenhouse, Yumi Kasai, David Gray, Carol Clugston, Anna Dominiczak and Alex Alderton, Roberto Amato, Jeffrey Barrett, Sonia Goncalves, Ewan Harrison, David K. Jackson, Ian Johnston, Dominic Kwiatkowski, Cordelia Langford, John Sillitoe on behalf of the Wellcome Sanger Institute COVID-19 Surveillance Team |
| EPI_ISL_3126561 | Berkshire and Surrey Pathology Services Lighthouse Laboratory | Wellcome Sanger Institute for the COVID-19 Genomics UK (COG-UK) Consortium | Berkshire and Surrey Pathology Services Lighthouse Laboratory and Alex Alderton, Roberto Amato, Jeffrey Barrett, Sonia Goncalves, Ewan Harrison, David K. Jackson, Ian Johnston, Dominic Kwiatkowski, Cordelia Langford, John Sillitoe on behalf of the Wellcome Sanger Institute COVID-19 Surveillance Team |
| EPI_ISL_3126562, EPI_ISL_3126563 | Lighthouse Lab in Glasgow | Wellcome Sanger Institute for the COVID-19 Genomics UK (COG-UK) Consortium | Harper VanSteenhouse, Yumi Kasai, David Gray, Carol Clugston, Anna Dominiczak and Alex Alderton, Roberto Amato, Jeffrey Barrett, Sonia Goncalves, Ewan Harrison, David K. Jackson, Ian Johnston, Dominic Kwiatkowski, Cordelia Langford, John Sillitoe on behalf of the Wellcome Sanger Institute COVID-19 Surveillance Team |
| EPI_ISL_3126564, EPI_ISL_3126565, EPI_ISL_3126566, EPI_ISL_3126567, EPI_ISL_3126568 | Lighthouse Lab in Alderley Park | Wellcome Sanger Institute for the COVID-19 Genomics UK (COG-UK) Consortium | Jacquelyn Wynn, Mairead Hyland, The Lighthouse Lab in Alderley Park and Alex Alderton, Roberto Amato, Jeffrey Barrett, Sonia Goncalves, Ewan Harrison, David K. Jackson, Ian Johnston, Dominic Kwiatkowski, Cordelia Langford, John Sillitoe on behalf of the Wellcome Sanger Institute COVID-19 Surveillance Team |
| EPI_ISL_3126569, EPI_ISL_3126570 | Berkshire and Surrey Pathology Services Lighthouse Laboratory | Wellcome Sanger Institute for the COVID-19 Genomics UK (COG-UK) Consortium | Berkshire and Surrey Pathology Services Lighthouse Laboratory and Alex Alderton, Roberto Amato, Jeffrey Barrett, Sonia Goncalves, Ewan Harrison, David K. Jackson, Ian Johnston, Dominic Kwiatkowski, Cordelia Langford, John Sillitoe on behalf of the Wellcome Sanger Institute COVID-19 Surveillance Team |
| EPI_ISL_3126571, EPI_ISL_3126572 | Lighthouse Lab in Glasgow | Wellcome Sanger Institute for the COVID-19 Genomics UK (COG-UK) Consortium | Harper VanSteenhouse, Yumi Kasai, David Gray, Carol Clugston, Anna Dominiczak and Alex Alderton, Roberto Amato, Jeffrey Barrett, Sonia Goncalves, Ewan Harrison, David K. Jackson, Ian Johnston, Dominic Kwiatkowski, Cordelia Langford, John Sillitoe on behalf of the Wellcome Sanger Institute COVID-19 Surveillance Team |
| EPI_ISL_3126573 | Berkshire and Surrey Pathology Services Lighthouse Laboratory | Wellcome Sanger Institute for the COVID-19 Genomics UK (COG-UK) Consortium | Berkshire and Surrey Pathology Services Lighthouse Laboratory and Alex Alderton, Roberto Amato, Jeffrey Barrett, Sonia Goncalves, Ewan Harrison, David K. Jackson, Ian Johnston, Dominic Kwiatkowski, Cordelia Langford, John Sillitoe on behalf of the Wellcome Sanger Institute COVID-19 Surveillance Team |
| EPI_ISL_3126574 | Lighthouse Lab in Glasgow | Wellcome Sanger Institute for the COVID-19 Genomics UK (COG-UK) Consortium | Harper VanSteenhouse, Yumi Kasai, David Gray, Carol Clugston, Anna Dominiczak and Alex Alderton, Roberto Amato, Jeffrey Barrett, Sonia Goncalves, Ewan Harrison, David K. Jackson, Ian Johnston, Dominic Kwiatkowski, Cordelia Langford, John Sillitoe on behalf of the Wellcome Sanger Institute COVID-19 Surveillance Team |
| EPI_ISL_3126575, EPI_ISL_3126576 | Berkshire and Surrey Pathology Services Lighthouse Laboratory | Wellcome Sanger Institute for the COVID-19 Genomics UK (COG-UK) Consortium | Berkshire and Surrey Pathology Services Lighthouse Laboratory and Alex Alderton, Roberto Amato, Jeffrey Barrett, Sonia Goncalves, Ewan Harrison, David K. Jackson, Ian Johnston, Dominic Kwiatkowski, Cordelia Langford, John Sillitoe on behalf of the Wellcome Sanger Institute COVID-19 Surveillance Team |
| EPI_ISL_3126577, EPI_ISL_3126578, EPI_ISL_3126579, EPI_ISL_3126580, EPI_ISL_3126581, EPI_ISL_3126582 | Lighthouse Lab in Glasgow | Wellcome Sanger Institute for the COVID-19 Genomics UK (COG-UK) Consortium | Harper VanSteenhouse, Yumi Kasai, David Gray, Carol Clugston, Anna Dominiczak and Alex Alderton, Roberto Amato, Jeffrey Barrett, Sonia Goncalves, Ewan Harrison, David K. Jackson, Ian Johnston, Dominic Kwiatkowski, Cordelia Langford, John Sillitoe on behalf of the Wellcome Sanger Institute COVID-19 Surveillance Team |
| EPI_ISL_3126583, EPI_ISL_3126584 | Berkshire and Surrey Pathology Services Lighthouse Laboratory | Wellcome Sanger Institute for the COVID-19 Genomics UK (COG-UK) Consortium | Berkshire and Surrey Pathology Services Lighthouse Laboratory and Alex Alderton, Roberto Amato, Jeffrey Barrett, Sonia Goncalves, Ewan Harrison, David K. Jackson, Ian Johnston, Dominic Kwiatkowski, Cordelia Langford, John Sillitoe on behalf of the Wellcome Sanger Institute COVID-19 Surveillance Team |
| EPI_ISL_3126585, EPI_ISL_3126586, EPI_ISL_3126587, EPI_ISL_3126588 | Lighthouse Lab in Glasgow | Wellcome Sanger Institute for the COVID-19 Genomics UK (COG-UK) Consortium | Harper VanSteenhouse, Yumi Kasai, David Gray, Carol Clugston, Anna Dominiczak and Alex Alderton, Roberto Amato, Jeffrey Barrett, Sonia Goncalves, Ewan Harrison, David K. Jackson, Ian Johnston, Dominic Kwiatkowski, Cordelia Langford, John Sillitoe on behalf of the Wellcome Sanger Institute COVID-19 Surveillance Team |
| EPI_ISL_3126589 | Berkshire and Surrey Pathology Services Lighthouse Laboratory | Wellcome Sanger Institute for the COVID-19 Genomics UK (COG-UK) Consortium | Berkshire and Surrey Pathology Services Lighthouse Laboratory and Alex Alderton, Roberto Amato, Jeffrey Barrett, Sonia Goncalves, Ewan Harrison, David K. Jackson, Ian Johnston, Dominic Kwiatkowski, Cordelia Langford, John Sillitoe on behalf of the Wellcome Sanger Institute COVID-19 Surveillance Team |
| EPI_ISL_3126590 | Lighthouse Lab in Glasgow | Wellcome Sanger Institute for the COVID-19 Genomics UK (COG-UK) Consortium | Harper VanSteenhouse, Yumi Kasai, David Gray, Carol Clugston, Anna Dominiczak and Alex Alderton, Roberto Amato, Jeffrey Barrett, Sonia Goncalves, Ewan Harrison, David K. Jackson, Ian Johnston, Dominic Kwiatkowski, Cordelia Langford, John Sillitoe on behalf of the Wellcome Sanger Institute COVID-19 Surveillance Team |
| EPI_ISL_3126591, EPI_ISL_3126592 | Lighthouse Lab in Alderley Park | Wellcome Sanger Institute for the COVID-19 Genomics UK (COG-UK) Consortium | Jacquelyn Wynn, Mairead Hyland, The Lighthouse Lab in Alderley Park and Alex Alderton, Roberto Amato, Jeffrey Barrett, Sonia Goncalves, Ewan Harrison, David K. Jackson, Ian Johnston, Dominic Kwiatkowski, Cordelia Langford, John Sillitoe on behalf of the Wellcome Sanger Institute COVID-19 Surveillance Team |
| EPI_ISL_3126593, EPI_ISL_3126594 | Lighthouse Lab in Glasgow | Wellcome Sanger Institute for the COVID-19 Genomics UK (COG-UK) Consortium | Harper VanSteenhouse, Yumi Kasai, David Gray, Carol Clugston, Anna Dominiczak and Alex Alderton, Roberto Amato, Jeffrey Barrett, Sonia Goncalves, Ewan Harrison, David K. Jackson, Ian Johnston, Dominic Kwiatkowski, Cordelia Langford, John Sillitoe on behalf of the Wellcome Sanger Institute COVID-19 Surveillance Team |
| EPI_ISL_3126595 | Lighthouse Lab in Alderley Park | Wellcome Sanger Institute for the COVID-19 Genomics UK (COG-UK) Consortium | Jacquelyn Wynn, Mairead Hyland, The Lighthouse Lab in Alderley Park and Alex Alderton, Roberto Amato, Jeffrey Barrett, Sonia Goncalves, Ewan Harrison, David K. Jackson, Ian Johnston, Dominic Kwiatkowski, Cordelia Langford, John Sillitoe on behalf of the Wellcome Sanger Institute COVID-19 Surveillance Team |
| EPI_ISL_3126596, EPI_ISL_3126597, EPI_ISL_3126598, EPI_ISL_3126599 | Lighthouse Lab in Glasgow | Wellcome Sanger Institute for the COVID-19 Genomics UK (COG-UK) Consortium | Harper VanSteenhouse, Yumi Kasai, David Gray, Carol Clugston, Anna Dominiczak and Alex Alderton, Roberto Amato, Jeffrey Barrett, Sonia Goncalves, Ewan Harrison, David K. Jackson, Ian Johnston, Dominic Kwiatkowski, Cordelia Langford, John Sillitoe on behalf of the Wellcome Sanger Institute COVID-19 Surveillance Team |
| EPI_ISL_3126600, EPI_ISL_3126601 | Lighthouse Lab in Alderley Park | Wellcome Sanger Institute for the COVID-19 Genomics UK (COG-UK) Consortium | Jacquelyn Wynn, Mairead Hyland, The Lighthouse Lab in Alderley Park and Alex Alderton, Roberto Amato, Jeffrey Barrett, Sonia Goncalves, Ewan Harrison, David K. Jackson, Ian Johnston, Dominic Kwiatkowski, Cordelia Langford, John Sillitoe on behalf of the Wellcome Sanger Institute COVID-19 Surveillance Team |
| EPI_ISL_3126602 | Berkshire and Surrey Pathology Services Lighthouse Laboratory | Wellcome Sanger Institute for the COVID-19 Genomics UK (COG-UK) Consortium | Berkshire and Surrey Pathology Services Lighthouse Laboratory and Alex Alderton, Roberto Amato, Jeffrey Barrett, Sonia Goncalves, Ewan Harrison, David K. Jackson, Ian Johnston, Dominic Kwiatkowski, Cordelia Langford, John Sillitoe on behalf of the Wellcome Sanger Institute COVID-19 Surveillance Team |
| EPI_ISL_3126603 | Lighthouse Lab in Glasgow | Wellcome Sanger Institute for the COVID-19 Genomics UK (COG-UK) Consortium | Harper VanSteenhouse, Yumi Kasai, David Gray, Carol Clugston, Anna Dominiczak and Alex Alderton, Roberto Amato, Jeffrey Barrett, Sonia Goncalves, Ewan Harrison, David K. Jackson, Ian Johnston, Dominic Kwiatkowski, Cordelia Langford, John Sillitoe on behalf of the Wellcome Sanger Institute COVID-19 Surveillance Team |
| EPI_ISL_3126604 | Berkshire and Surrey Pathology Services Lighthouse Laboratory | Wellcome Sanger Institute for the COVID-19 Genomics UK (COG-UK) Consortium | Berkshire and Surrey Pathology Services Lighthouse Laboratory and Alex Alderton, Roberto Amato, Jeffrey Barrett, Sonia Goncalves, Ewan Harrison, David K. Jackson, Ian Johnston, Dominic Kwiatkowski, Cordelia Langford, John Sillitoe on behalf of the Wellcome Sanger Institute COVID-19 Surveillance Team |
| EPI_ISL_3126605 | Lighthouse Lab in Glasgow | Wellcome Sanger Institute for the COVID-19 Genomics UK (COG-UK) Consortium | Harper VanSteenhouse, Yumi Kasai, David Gray, Carol Clugston, Anna Dominiczak and Alex Alderton, Roberto Amato, Jeffrey Barrett, Sonia Goncalves, Ewan Harrison, David K. Jackson, Ian Johnston, Dominic Kwiatkowski, Cordelia Langford, John Sillitoe on behalf of the Wellcome Sanger Institute COVID-19 Surveillance Team |
| EPI_ISL_3126606 | Lighthouse Lab in Alderley Park | Wellcome Sanger Institute for the COVID-19 Genomics UK (COG-UK) Consortium | Jacquelyn Wynn, Mairead Hyland, The Lighthouse Lab in Alderley Park and Alex Alderton, Roberto Amato, Jeffrey Barrett, Sonia Goncalves, Ewan Harrison, David K. Jackson, Ian Johnston, Dominic Kwiatkowski, Cordelia Langford, John Sillitoe on behalf of the Wellcome Sanger Institute COVID-19 Surveillance Team |
| EPI_ISL_3126607 | Lighthouse Lab in Glasgow | Wellcome Sanger Institute for the COVID-19 Genomics UK (COG-UK) Consortium | Harper VanSteenhouse, Yumi Kasai, David Gray, Carol Clugston, Anna Dominiczak and Alex Alderton, Roberto Amato, Jeffrey Barrett, Sonia Goncalves, Ewan Harrison, David K. Jackson, Ian Johnston, Dominic Kwiatkowski, Cordelia Langford, John Sillitoe on behalf of the Wellcome Sanger Institute COVID-19 Surveillance Team |
| EPI_ISL_3126608 | Lighthouse Lab in Alderley Park | Wellcome Sanger Institute for the COVID-19 Genomics UK (COG-UK) Consortium | Jacquelyn Wynn, Mairead Hyland, The Lighthouse Lab in Alderley Park and Alex Alderton, Roberto Amato, Jeffrey Barrett, Sonia Goncalves, Ewan Harrison, David K. Jackson, Ian Johnston, Dominic Kwiatkowski, Cordelia Langford, John Sillitoe on behalf of the Wellcome Sanger Institute COVID-19 Surveillance Team |
| EPI_ISL_3126609 | Lighthouse Lab in Glasgow | Wellcome Sanger Institute for the COVID-19 Genomics UK (COG-UK) Consortium | Harper VanSteenhouse, Yumi Kasai, David Gray, Carol Clugston, Anna Dominiczak and Alex Alderton, Roberto Amato, Jeffrey Barrett, Sonia Goncalves, Ewan Harrison, David K. Jackson, Ian Johnston, Dominic Kwiatkowski, Cordelia Langford, John Sillitoe on behalf of the Wellcome Sanger Institute COVID-19 Surveillance Team |
| EPI_ISL_3126610 | Lighthouse Lab in Alderley Park | Wellcome Sanger Institute for the COVID-19 Genomics UK (COG-UK) | Jacquelyn Wynn, Mairead Hyland, The Lighthouse Lab in Alderley Park and Alex Alderton, Roberto Amato, Jeffrey Barrett, Sonia Goncalves, Ewan Harrison, David K. Jackson, Ian Johnston, Dominic Kwiatkowski, Cordelia Langford, John Sillitoe on |

| | | | |
|---|---|---|---|
| | | Consortium | behalf of the Wellcome Sanger Institute COVID-19 Surveillance Team |
| EPI_ISL_3126611, EPI_ISL_3126612, EPI_ISL_3126613 | Berkshire and Surrey Pathology Services Lighthouse Laboratory | Wellcome Sanger Institute for the COVID-19 Genomics UK (COG-UK) Consortium | Berkshire and Surrey Pathology Services Lighthouse Laboratory and Alex Alderton, Roberto Amato, Jeffrey Barrett, Sonia Goncalves, Ewan Harrison, David K. Jackson, Ian Johnston, Dominic Kwiatkowski, Cordelia Langford, John Sillitoe on behalf of the Wellcome Sanger Institute COVID-19 Surveillance Team |
| EPI_ISL_3126614 | Lighthouse Lab in Glasgow | Wellcome Sanger Institute for the COVID-19 Genomics UK (COG-UK) Consortium | Harper VanSteenhouse, Yumi Kasai, David Gray, Carol Clugston, Anna Dominiczak and Alex Alderton, Roberto Amato, Jeffrey Barrett, Sonia Goncalves, Ewan Harrison, David K. Jackson, Ian Johnston, Dominic Kwiatkowski, Cordelia Langford, John Sillitoe on behalf of the Wellcome Sanger Institute COVID-19 Surveillance Team |
| EPI_ISL_3126615 | Berkshire and Surrey Pathology Services Lighthouse Laboratory | Wellcome Sanger Institute for the COVID-19 Genomics UK (COG-UK) Consortium | Berkshire and Surrey Pathology Services Lighthouse Laboratory and Alex Alderton, Roberto Amato, Jeffrey Barrett, Sonia Goncalves, Ewan Harrison, David K. Jackson, Ian Johnston, Dominic Kwiatkowski, Cordelia Langford, John Sillitoe on behalf of the Wellcome Sanger Institute COVID-19 Surveillance Team |
| EPI_ISL_3126616, EPI_ISL_3126617 | Lighthouse Lab in Alderley Park | Wellcome Sanger Institute for the COVID-19 Genomics UK (COG-UK) Consortium | Jacquelyn Wynn, Mairead Hyland, The Lighthouse Lab in Alderley Park and Alex Alderton, Roberto Amato, Jeffrey Barrett, Sonia Goncalves, Ewan Harrison, David K. Jackson, Ian Johnston, Dominic Kwiatkowski, Cordelia Langford, John Sillitoe on behalf of the Wellcome Sanger Institute COVID-19 Surveillance Team |
| EPI_ISL_3126618 | Lighthouse Lab in Glasgow | Wellcome Sanger Institute for the COVID-19 Genomics UK (COG-UK) Consortium | Harper VanSteenhouse, Yumi Kasai, David Gray, Carol Clugston, Anna Dominiczak and Alex Alderton, Roberto Amato, Jeffrey Barrett, Sonia Goncalves, Ewan Harrison, David K. Jackson, Ian Johnston, Dominic Kwiatkowski, Cordelia Langford, John Sillitoe on behalf of the Wellcome Sanger Institute COVID-19 Surveillance Team |
| EPI_ISL_3126619 | Berkshire and Surrey Pathology Services Lighthouse Laboratory | Wellcome Sanger Institute for the COVID-19 Genomics UK (COG-UK) Consortium | Berkshire and Surrey Pathology Services Lighthouse Laboratory and Alex Alderton, Roberto Amato, Jeffrey Barrett, Sonia Goncalves, Ewan Harrison, David K. Jackson, Ian Johnston, Dominic Kwiatkowski, Cordelia Langford, John Sillitoe on behalf of the Wellcome Sanger Institute COVID-19 Surveillance Team |
| EPI_ISL_3126620, EPI_ISL_3126621 | Lighthouse Lab in Glasgow | Wellcome Sanger Institute for the COVID-19 Genomics UK (COG-UK) Consortium | Harper VanSteenhouse, Yumi Kasai, David Gray, Carol Clugston, Anna Dominiczak and Alex Alderton, Roberto Amato, Jeffrey Barrett, Sonia Goncalves, Ewan Harrison, David K. Jackson, Ian Johnston, Dominic Kwiatkowski, Cordelia Langford, John Sillitoe on behalf of the Wellcome Sanger Institute COVID-19 Surveillance Team |
| EPI_ISL_3126622 | Lighthouse Lab in Alderley Park | Wellcome Sanger Institute for the COVID-19 Genomics UK (COG-UK) Consortium | Jacquelyn Wynn, Mairead Hyland, The Lighthouse Lab in Alderley Park and Alex Alderton, Roberto Amato, Jeffrey Barrett, Sonia Goncalves, Ewan Harrison, David K. Jackson, Ian Johnston, Dominic Kwiatkowski, Cordelia Langford, John Sillitoe on behalf of the Wellcome Sanger Institute COVID-19 Surveillance Team |
| EPI_ISL_3126623 | Lighthouse Lab in Glasgow | Wellcome Sanger Institute for the COVID-19 Genomics UK (COG-UK) Consortium | Harper VanSteenhouse, Yumi Kasai, David Gray, Carol Clugston, Anna Dominiczak and Alex Alderton, Roberto Amato, Jeffrey Barrett, Sonia Goncalves, Ewan Harrison, David K. Jackson, Ian Johnston, Dominic Kwiatkowski, Cordelia Langford, John Sillitoe on behalf of the Wellcome Sanger Institute COVID-19 Surveillance Team |
| EPI_ISL_3126624 | Lighthouse Lab in Alderley Park | Wellcome Sanger Institute for the COVID-19 Genomics UK (COG-UK) Consortium | Jacquelyn Wynn, Mairead Hyland, The Lighthouse Lab in Alderley Park and Alex Alderton, Roberto Amato, Jeffrey Barrett, Sonia Goncalves, Ewan Harrison, David K. Jackson, Ian Johnston, Dominic Kwiatkowski, Cordelia Langford, John Sillitoe on behalf of the Wellcome Sanger Institute COVID-19 Surveillance Team |
| EPI_ISL_3126625, EPI_ISL_3126626 | Lighthouse Lab in Glasgow | Wellcome Sanger Institute for the COVID-19 Genomics UK (COG-UK) Consortium | Harper VanSteenhouse, Yumi Kasai, David Gray, Carol Clugston, Anna Dominiczak and Alex Alderton, Roberto Amato, Jeffrey Barrett, Sonia Goncalves, Ewan Harrison, David K. Jackson, Ian Johnston, Dominic Kwiatkowski, Cordelia Langford, John Sillitoe on behalf of the Wellcome Sanger Institute COVID-19 Surveillance Team |
| EPI_ISL_3126627 | Berkshire and Surrey Pathology Services Lighthouse Laboratory | Wellcome Sanger Institute for the COVID-19 Genomics UK (COG-UK) Consortium | Berkshire and Surrey Pathology Services Lighthouse Laboratory and Alex Alderton, Roberto Amato, Jeffrey Barrett, Sonia Goncalves, Ewan Harrison, David K. Jackson, Ian Johnston, Dominic Kwiatkowski, Cordelia Langford, John Sillitoe on behalf of the Wellcome Sanger Institute COVID-19 Surveillance Team |
| EPI_ISL_3126628, EPI_ISL_3126629 | Lighthouse Lab in Glasgow | Wellcome Sanger Institute for the COVID-19 Genomics UK (COG-UK) Consortium | Harper VanSteenhouse, Yumi Kasai, David Gray, Carol Clugston, Anna Dominiczak and Alex Alderton, Roberto Amato, Jeffrey Barrett, Sonia Goncalves, Ewan Harrison, David K. Jackson, Ian Johnston, Dominic Kwiatkowski, Cordelia Langford, John Sillitoe on behalf of the Wellcome Sanger Institute COVID-19 Surveillance Team |
| EPI_ISL_3126630 | Lighthouse Lab in Alderley Park | Wellcome Sanger Institute for the COVID-19 Genomics UK (COG-UK) Consortium | Jacquelyn Wynn, Mairead Hyland, The Lighthouse Lab in Alderley Park and Alex Alderton, Roberto Amato, Jeffrey Barrett, Sonia Goncalves, Ewan Harrison, David K. Jackson, Ian Johnston, Dominic Kwiatkowski, Cordelia Langford, John Sillitoe on behalf of the Wellcome Sanger Institute COVID-19 Surveillance Team |
| EPI_ISL_3126631 | Berkshire and Surrey Pathology Services Lighthouse Laboratory | Wellcome Sanger Institute for the COVID-19 Genomics UK (COG-UK) Consortium | Berkshire and Surrey Pathology Services Lighthouse Laboratory and Alex Alderton, Roberto Amato, Jeffrey Barrett, Sonia Goncalves, Ewan Harrison, David K. Jackson, Ian Johnston, Dominic Kwiatkowski, Cordelia Langford, John Sillitoe on behalf of the Wellcome Sanger Institute COVID-19 Surveillance Team |
| EPI_ISL_3126632 | Lighthouse Lab in Glasgow | Wellcome Sanger Institute for the COVID-19 Genomics UK (COG-UK) Consortium | Harper VanSteenhouse, Yumi Kasai, David Gray, Carol Clugston, Anna Dominiczak and Alex Alderton, Roberto Amato, Jeffrey Barrett, Sonia Goncalves, Ewan Harrison, David K. Jackson, Ian Johnston, Dominic Kwiatkowski, Cordelia Langford, John Sillitoe on behalf of the Wellcome Sanger Institute COVID-19 Surveillance Team |
| EPI_ISL_3126633 | Lighthouse Lab in Alderley Park | Wellcome Sanger Institute for the COVID-19 Genomics UK (COG-UK) Consortium | Jacquelyn Wynn, Mairead Hyland, The Lighthouse Lab in Alderley Park and Alex Alderton, Roberto Amato, Jeffrey Barrett, Sonia Goncalves, Ewan Harrison, David K. Jackson, Ian Johnston, Dominic Kwiatkowski, Cordelia Langford, John Sillitoe on behalf of the Wellcome Sanger Institute COVID-19 Surveillance Team |
| EPI_ISL_3126634 | Lighthouse Lab in Glasgow | Wellcome Sanger Institute for the COVID-19 Genomics UK (COG-UK) Consortium | Harper VanSteenhouse, Yumi Kasai, David Gray, Carol Clugston, Anna Dominiczak and Alex Alderton, Roberto Amato, Jeffrey Barrett, Sonia Goncalves, Ewan Harrison, David K. Jackson, Ian Johnston, Dominic Kwiatkowski, Cordelia Langford, John Sillitoe on behalf of the Wellcome Sanger Institute COVID-19 Surveillance Team |
| EPI_ISL_3126635 | Berkshire and Surrey Pathology Services Lighthouse Laboratory | Wellcome Sanger Institute for the COVID-19 Genomics UK (COG-UK) Consortium | Berkshire and Surrey Pathology Services Lighthouse Laboratory and Alex Alderton, Roberto Amato, Jeffrey Barrett, Sonia Goncalves, Ewan Harrison, David K. Jackson, Ian Johnston, Dominic Kwiatkowski, Cordelia Langford, John Sillitoe on behalf of the Wellcome Sanger Institute COVID-19 Surveillance Team |
| EPI_ISL_3126636 | Lighthouse Lab in Glasgow | Wellcome Sanger Institute for the COVID-19 Genomics UK (COG-UK) Consortium | Harper VanSteenhouse, Yumi Kasai, David Gray, Carol Clugston, Anna Dominiczak and Alex Alderton, Roberto Amato, Jeffrey Barrett, Sonia Goncalves, Ewan Harrison, David K. Jackson, Ian Johnston, Dominic Kwiatkowski, Cordelia Langford, John Sillitoe on behalf of the Wellcome Sanger Institute COVID-19 Surveillance Team |
| EPI_ISL_3126637 | Lighthouse Lab in Alderley Park | Wellcome Sanger Institute for the COVID-19 Genomics UK (COG-UK) Consortium | Jacquelyn Wynn, Mairead Hyland, The Lighthouse Lab in Alderley Park and Alex Alderton, Roberto Amato, Jeffrey Barrett, Sonia Goncalves, Ewan Harrison, David K. Jackson, Ian Johnston, Dominic Kwiatkowski, Cordelia Langford, John Sillitoe on behalf of the Wellcome Sanger Institute COVID-19 Surveillance Team |
| EPI_ISL_3126638 | Lighthouse Lab in Glasgow | Wellcome Sanger Institute for the COVID-19 Genomics UK (COG-UK) Consortium | Harper VanSteenhouse, Yumi Kasai, David Gray, Carol Clugston, Anna Dominiczak and Alex Alderton, Roberto Amato, Jeffrey Barrett, Sonia Goncalves, Ewan Harrison, David K. Jackson, Ian Johnston, Dominic Kwiatkowski, Cordelia Langford, John Sillitoe on behalf of the Wellcome Sanger Institute COVID-19 Surveillance Team |
| EPI_ISL_3126639, EPI_ISL_3126640 | Berkshire and Surrey Pathology Services Lighthouse Laboratory | Wellcome Sanger Institute for the COVID-19 Genomics UK (COG-UK) Consortium | Berkshire and Surrey Pathology Services Lighthouse Laboratory and Alex Alderton, Roberto Amato, Jeffrey Barrett, Sonia Goncalves, Ewan Harrison, David K. Jackson, Ian Johnston, Dominic Kwiatkowski, Cordelia Langford, John Sillitoe on behalf of the Wellcome Sanger Institute COVID-19 Surveillance Team |
| EPI_ISL_3126641 | Lighthouse Lab in Alderley Park | Wellcome Sanger Institute for the COVID-19 Genomics UK (COG-UK) Consortium | Jacquelyn Wynn, Mairead Hyland, The Lighthouse Lab in Alderley Park and Alex Alderton, Roberto Amato, Jeffrey Barrett, Sonia Goncalves, Ewan Harrison, David K. Jackson, Ian Johnston, Dominic Kwiatkowski, Cordelia Langford, John Sillitoe on behalf of the Wellcome Sanger Institute COVID-19 Surveillance Team |
| EPI_ISL_3126642 | Lighthouse Lab in Glasgow | Wellcome Sanger Institute for the COVID-19 Genomics UK (COG-UK) Consortium | Harper VanSteenhouse, Yumi Kasai, David Gray, Carol Clugston, Anna Dominiczak and Alex Alderton, Roberto Amato, Jeffrey Barrett, Sonia Goncalves, Ewan Harrison, David K. Jackson, Ian Johnston, Dominic Kwiatkowski, Cordelia Langford, John Sillitoe on behalf of the Wellcome Sanger Institute COVID-19 Surveillance Team |
| EPI_ISL_3126643 | Unit of laboratory surveillance of viral emergent disease Control of Communicable Diseases Departament National Institute of Public Health Albania | Charité Universitätsmedizin Berlin, Institut für Virologie | Victor M Corman, Barbara Mühlemann, Jörn Beheim-Schwarzbach, Talitha Veith, Julia Schneider, Terry Jones, Iris Hasibra, Christian Drosten |
| EPI_ISL_3126644 | Lighthouse Lab in Alderley Park | Wellcome Sanger Institute for the COVID-19 Genomics UK (COG-UK) Consortium | Jacquelyn Wynn, Mairead Hyland, The Lighthouse Lab in Alderley Park and Alex Alderton, Roberto Amato, Jeffrey Barrett, Sonia Goncalves, Ewan Harrison, David K. Jackson, Ian Johnston, Dominic Kwiatkowski, Cordelia Langford, John Sillitoe on behalf of the Wellcome Sanger Institute COVID-19 Surveillance Team |
| EPI_ISL_3126645 | Lighthouse Lab in Glasgow | Wellcome Sanger Institute for the COVID-19 Genomics UK (COG-UK) Consortium | Harper VanSteenhouse, Yumi Kasai, David Gray, Carol Clugston, Anna Dominiczak and Alex Alderton, Roberto Amato, Jeffrey Barrett, Sonia Goncalves, Ewan Harrison, David K. Jackson, Ian Johnston, Dominic Kwiatkowski, Cordelia Langford, John Sillitoe on behalf of the Wellcome Sanger Institute COVID-19 Surveillance Team |
| EPI_ISL_3126646 | Lighthouse Lab in Alderley Park | Wellcome Sanger Institute for the COVID-19 Genomics UK (COG-UK) Consortium | Jacquelyn Wynn, Mairead Hyland, The Lighthouse Lab in Alderley Park and Alex Alderton, Roberto Amato, Jeffrey Barrett, Sonia Goncalves, Ewan Harrison, David K. Jackson, Ian Johnston, Dominic Kwiatkowski, Cordelia Langford, John Sillitoe on behalf of the Wellcome Sanger Institute COVID-19 Surveillance Team |
| EPI_ISL_3126647 | Berkshire and Surrey Pathology Services Lighthouse Laboratory | Wellcome Sanger Institute for the COVID-19 Genomics UK (COG-UK) Consortium | Berkshire and Surrey Pathology Services Lighthouse Laboratory and Alex Alderton, Roberto Amato, Jeffrey Barrett, Sonia Goncalves, Ewan Harrison, David K. Jackson, Ian Johnston, Dominic Kwiatkowski, Cordelia Langford, John Sillitoe on behalf of the Wellcome Sanger Institute COVID-19 Surveillance Team |
| EPI_ISL_3126648 | Lighthouse Lab in Alderley Park | Wellcome Sanger Institute for the COVID-19 Genomics UK (COG-UK) Consortium | Jacquelyn Wynn, Mairead Hyland, The Lighthouse Lab in Alderley Park and Alex Alderton, Roberto Amato, Jeffrey Barrett, Sonia Goncalves, Ewan Harrison, David K. Jackson, Ian Johnston, Dominic Kwiatkowski, Cordelia Langford, John Sillitoe on behalf of the Wellcome Sanger Institute COVID-19 Surveillance Team |
| EPI_ISL_3126649, EPI_ISL_3126650, EPI_ISL_3126651 | Lighthouse Lab in Glasgow | Wellcome Sanger Institute for the COVID-19 Genomics UK (COG-UK) Consortium | Harper VanSteenhouse, Yumi Kasai, David Gray, Carol Clugston, Anna Dominiczak and Alex Alderton, Roberto Amato, Jeffrey Barrett, Sonia Goncalves, Ewan Harrison, David K. Jackson, Ian Johnston, Dominic Kwiatkowski, Cordelia Langford, John Sillitoe on behalf of the Wellcome Sanger Institute COVID-19 Surveillance Team |
| EPI_ISL_3126652 | Lighthouse Lab in Alderley Park | Wellcome Sanger Institute for the COVID-19 Genomics UK (COG-UK) Consortium | Jacquelyn Wynn, Mairead Hyland, The Lighthouse Lab in Alderley Park and Alex Alderton, Roberto Amato, Jeffrey Barrett, Sonia Goncalves, Ewan Harrison, David K. Jackson, Ian Johnston, Dominic Kwiatkowski, Cordelia Langford, John Sillitoe on behalf of the Wellcome Sanger Institute COVID-19 Surveillance Team |
| EPI_ISL_3126653 | Lighthouse Lab in Glasgow | Wellcome Sanger Institute for the COVID-19 Genomics UK (COG-UK) Consortium | Harper VanSteenhouse, Yumi Kasai, David Gray, Carol Clugston, Anna Dominiczak and Alex Alderton, Roberto Amato, Jeffrey Barrett, Sonia Goncalves, Ewan Harrison, David K. Jackson, Ian Johnston, Dominic Kwiatkowski, Cordelia Langford, John Sillitoe on behalf of the Wellcome Sanger Institute COVID-19 Surveillance Team |
| EPI_ISL_3126654 | Lighthouse Lab in Alderley Park | Wellcome Sanger Institute for the COVID-19 Genomics UK (COG-UK) Consortium | Jacquelyn Wynn, Mairead Hyland, The Lighthouse Lab in Alderley Park and Alex Alderton, Roberto Amato, Jeffrey Barrett, Sonia Goncalves, Ewan Harrison, David K. Jackson, Ian Johnston, Dominic Kwiatkowski, Cordelia Langford, John Sillitoe on behalf of the Wellcome Sanger Institute COVID-19 Surveillance Team |
| EPI_ISL_3126655, EPI_ISL_3126656 | Lighthouse Lab in Glasgow | Wellcome Sanger Institute for the COVID-19 Genomics UK (COG-UK) Consortium | Harper VanSteenhouse, Yumi Kasai, David Gray, Carol Clugston, Anna Dominiczak and Alex Alderton, Roberto Amato, Jeffrey Barrett, Sonia Goncalves, Ewan Harrison, David K. Jackson, Ian Johnston, Dominic Kwiatkowski, Cordelia Langford, John Sillitoe on behalf of the Wellcome Sanger Institute COVID-19 Surveillance Team |
| EPI_ISL_3126657, EPI_ISL_3126658 | Berkshire and Surrey Pathology Services Lighthouse Laboratory | Wellcome Sanger Institute for the COVID-19 Genomics UK (COG-UK) Consortium | Berkshire and Surrey Pathology Services Lighthouse Laboratory and Alex Alderton, Roberto Amato, Jeffrey Barrett, Sonia Goncalves, Ewan Harrison, David K. Jackson, Ian Johnston, Dominic Kwiatkowski, Cordelia Langford, John Sillitoe on behalf of the Wellcome Sanger Institute COVID-19 Surveillance Team |
| EPI_ISL_3126659 | Lighthouse Lab in Glasgow | Wellcome Sanger Institute for the COVID-19 Genomics UK (COG-UK) Consortium | Harper VanSteenhouse, Yumi Kasai, David Gray, Carol Clugston, Anna Dominiczak and Alex Alderton, Roberto Amato, Jeffrey Barrett, Sonia Goncalves, Ewan Harrison, David K. Jackson, Ian Johnston, Dominic Kwiatkowski, Cordelia Langford, John Sillitoe on behalf of the Wellcome Sanger Institute COVID-19 Surveillance Team |
| EPI_ISL_3126660 | Lighthouse Lab in Alderley Park | Wellcome Sanger Institute for the COVID-19 Genomics UK (COG-UK) Consortium | Jacquelyn Wynn, Mairead Hyland, The Lighthouse Lab in Alderley Park and Alex Alderton, Roberto Amato, Jeffrey Barrett, Sonia Goncalves, Ewan Harrison, David K. Jackson, Ian Johnston, Dominic Kwiatkowski, Cordelia Langford, John Sillitoe on behalf of the Wellcome Sanger Institute COVID-19 Surveillance Team |
| EPI_ISL_3126661 | Lighthouse Lab in Glasgow | Wellcome Sanger Institute for the COVID-19 Genomics UK (COG-UK) Consortium | Harper VanSteenhouse, Yumi Kasai, David Gray, Carol Clugston, Anna Dominiczak and Alex Alderton, Roberto Amato, Jeffrey Barrett, Sonia Goncalves, Ewan Harrison, David K. Jackson, Ian Johnston, Dominic Kwiatkowski, Cordelia Langford, John Sillitoe on behalf of the Wellcome Sanger Institute COVID-19 Surveillance Team |
| EPI_ISL_3126662 | Lighthouse Lab in Alderley Park | Wellcome Sanger Institute for the COVID-19 Genomics UK (COG-UK) Consortium | Jacquelyn Wynn, Mairead Hyland, The Lighthouse Lab in Alderley Park and Alex Alderton, Roberto Amato, Jeffrey Barrett, Sonia Goncalves, Ewan Harrison, David K. Jackson, Ian Johnston, Dominic Kwiatkowski, Cordelia Langford, John Sillitoe on behalf of the Wellcome Sanger Institute COVID-19 Surveillance Team |
| EPI_ISL_3126663 | Lighthouse Lab in Glasgow | Wellcome Sanger Institute for the COVID-19 Genomics UK (COG-UK) Consortium | Harper VanSteenhouse, Yumi Kasai, David Gray, Carol Clugston, Anna Dominiczak and Alex Alderton, Roberto Amato, Jeffrey Barrett, Sonia Goncalves, Ewan Harrison, David K. Jackson, Ian Johnston, Dominic Kwiatkowski, Cordelia Langford, John Sillitoe on behalf of the Wellcome Sanger Institute COVID-19 Surveillance Team |
| EPI_ISL_3126664 | Berkshire and Surrey Pathology Services Lighthouse Laboratory | Wellcome Sanger Institute for the COVID-19 Genomics UK (COG-UK) Consortium | Berkshire and Surrey Pathology Services Lighthouse Laboratory and Alex Alderton, Roberto Amato, Jeffrey Barrett, Sonia Goncalves, Ewan Harrison, David K. Jackson, Ian Johnston, Dominic Kwiatkowski, Cordelia Langford, John Sillitoe on behalf of the Wellcome Sanger Institute COVID-19 Surveillance Team |

| | | | |
|---|---|---|---|
| EPI_ISL_3126665 | Lighthouse Lab in Alderley Park | Wellcome Sanger Institute for the COVID-19 Genomics UK (COG-UK) Consortium | Jacquelyn Wynn, Mairead Hyland, The Lighthouse Lab in Alderley Park and Alex Alderton, Roberto Amato, Jeffrey Barrett, Sonia Goncalves, Ewan Harrison, David K. Jackson, Ian Johnston, Dominic Kwiatkowski, Cordelia Langford, John Sillitoe on behalf of the Wellcome Sanger Institute COVID-19 Surveillance Team |
| EPI_ISL_3126666, EPI_ISL_3126667, EPI_ISL_3126668, EPI_ISL_3126669, EPI_ISL_3126670, EPI_ISL_3126671, EPI_ISL_3126672 | Lighthouse Lab in Glasgow | Wellcome Sanger Institute for the COVID-19 Genomics UK (COG-UK) Consortium | Harper VanSteenhouse, Yumi Kasai, David Gray, Carol Clugston, Anna Dominiczak and Alex Alderton, Roberto Amato, Jeffrey Barrett, Sonia Goncalves, Ewan Harrison, David K. Jackson, Ian Johnston, Dominic Kwiatkowski, Cordelia Langford, John Sillitoe on behalf of the Wellcome Sanger Institute COVID-19 Surveillance Team |
| EPI_ISL_3126673 | Lighthouse Lab in Alderley Park | Wellcome Sanger Institute for the COVID-19 Genomics UK (COG-UK) Consortium | Jacquelyn Wynn, Mairead Hyland, The Lighthouse Lab in Alderley Park and Alex Alderton, Roberto Amato, Jeffrey Barrett, Sonia Goncalves, Ewan Harrison, David K. Jackson, Ian Johnston, Dominic Kwiatkowski, Cordelia Langford, John Sillitoe on behalf of the Wellcome Sanger Institute COVID-19 Surveillance Team |
| EPI_ISL_3126674 | Lighthouse Lab in Glasgow | Wellcome Sanger Institute for the COVID-19 Genomics UK (COG-UK) Consortium | Harper VanSteenhouse, Yumi Kasai, David Gray, Carol Clugston, Anna Dominiczak and Alex Alderton, Roberto Amato, Jeffrey Barrett, Sonia Goncalves, Ewan Harrison, David K. Jackson, Ian Johnston, Dominic Kwiatkowski, Cordelia Langford, John Sillitoe on behalf of the Wellcome Sanger Institute COVID-19 Surveillance Team |
| EPI_ISL_3126675 | Berkshire and Surrey Pathology Services Lighthouse Laboratory | Wellcome Sanger Institute for the COVID-19 Genomics UK (COG-UK) Consortium | Berkshire and Surrey Pathology Services Lighthouse Laboratory and Alex Alderton, Roberto Amato, Jeffrey Barrett, Sonia Goncalves, Ewan Harrison, David K. Jackson, Ian Johnston, Dominic Kwiatkowski, Cordelia Langford, John Sillitoe on behalf of the Wellcome Sanger Institute COVID-19 Surveillance Team |
| EPI_ISL_3126676 | Lighthouse Lab in Glasgow | Wellcome Sanger Institute for the COVID-19 Genomics UK (COG-UK) Consortium | Harper VanSteenhouse, Yumi Kasai, David Gray, Carol Clugston, Anna Dominiczak and Alex Alderton, Roberto Amato, Jeffrey Barrett, Sonia Goncalves, Ewan Harrison, David K. Jackson, Ian Johnston, Dominic Kwiatkowski, Cordelia Langford, John Sillitoe on behalf of the Wellcome Sanger Institute COVID-19 Surveillance Team |
| EPI_ISL_3126677 | Lighthouse Lab in Alderley Park | Wellcome Sanger Institute for the COVID-19 Genomics UK (COG-UK) Consortium | Jacquelyn Wynn, Mairead Hyland, The Lighthouse Lab in Alderley Park and Alex Alderton, Roberto Amato, Jeffrey Barrett, Sonia Goncalves, Ewan Harrison, David K. Jackson, Ian Johnston, Dominic Kwiatkowski, Cordelia Langford, John Sillitoe on behalf of the Wellcome Sanger Institute COVID-19 Surveillance Team |
| EPI_ISL_3126678, EPI_ISL_3126679, EPI_ISL_3126680, EPI_ISL_3126681, EPI_ISL_3126682 | Lighthouse Lab in Glasgow | Wellcome Sanger Institute for the COVID-19 Genomics UK (COG-UK) Consortium | Harper VanSteenhouse, Yumi Kasai, David Gray, Carol Clugston, Anna Dominiczak and Alex Alderton, Roberto Amato, Jeffrey Barrett, Sonia Goncalves, Ewan Harrison, David K. Jackson, Ian Johnston, Dominic Kwiatkowski, Cordelia Langford, John Sillitoe on behalf of the Wellcome Sanger Institute COVID-19 Surveillance Team |
| EPI_ISL_3126683 | Lighthouse Lab in Alderley Park | Wellcome Sanger Institute for the COVID-19 Genomics UK (COG-UK) Consortium | Jacquelyn Wynn, Mairead Hyland, The Lighthouse Lab in Alderley Park and Alex Alderton, Roberto Amato, Jeffrey Barrett, Sonia Goncalves, Ewan Harrison, David K. Jackson, Ian Johnston, Dominic Kwiatkowski, Cordelia Langford, John Sillitoe on behalf of the Wellcome Sanger Institute COVID-19 Surveillance Team |
| EPI_ISL_3126684, EPI_ISL_3126685, EPI_ISL_3126686 | Berkshire and Surrey Pathology Services Lighthouse Laboratory | Wellcome Sanger Institute for the COVID-19 Genomics UK (COG-UK) Consortium | Berkshire and Surrey Pathology Services Lighthouse Laboratory and Alex Alderton, Roberto Amato, Jeffrey Barrett, Sonia Goncalves, Ewan Harrison, David K. Jackson, Ian Johnston, Dominic Kwiatkowski, Cordelia Langford, John Sillitoe on behalf of the Wellcome Sanger Institute COVID-19 Surveillance Team |
| EPI_ISL_3126687 | Lighthouse Lab in Alderley Park | Wellcome Sanger Institute for the COVID-19 Genomics UK (COG-UK) Consortium | Jacquelyn Wynn, Mairead Hyland, The Lighthouse Lab in Alderley Park and Alex Alderton, Roberto Amato, Jeffrey Barrett, Sonia Goncalves, Ewan Harrison, David K. Jackson, Ian Johnston, Dominic Kwiatkowski, Cordelia Langford, John Sillitoe on behalf of the Wellcome Sanger Institute COVID-19 Surveillance Team |
| EPI_ISL_3126688 | Berkshire and Surrey Pathology Services Lighthouse Laboratory | Wellcome Sanger Institute for the COVID-19 Genomics UK (COG-UK) Consortium | Berkshire and Surrey Pathology Services Lighthouse Laboratory and Alex Alderton, Roberto Amato, Jeffrey Barrett, Sonia Goncalves, Ewan Harrison, David K. Jackson, Ian Johnston, Dominic Kwiatkowski, Cordelia Langford, John Sillitoe on behalf of the Wellcome Sanger Institute COVID-19 Surveillance Team |
| EPI_ISL_3126689 | Lighthouse Lab in Glasgow | Wellcome Sanger Institute for the COVID-19 Genomics UK (COG-UK) Consortium | Harper VanSteenhouse, Yumi Kasai, David Gray, Carol Clugston, Anna Dominiczak and Alex Alderton, Roberto Amato, Jeffrey Barrett, Sonia Goncalves, Ewan Harrison, David K. Jackson, Ian Johnston, Dominic Kwiatkowski, Cordelia Langford, John Sillitoe on behalf of the Wellcome Sanger Institute COVID-19 Surveillance Team |
| EPI_ISL_3126690 | Lighthouse Lab in Alderley Park | Wellcome Sanger Institute for the COVID-19 Genomics UK (COG-UK) Consortium | Jacquelyn Wynn, Mairead Hyland, The Lighthouse Lab in Alderley Park and Alex Alderton, Roberto Amato, Jeffrey Barrett, Sonia Goncalves, Ewan Harrison, David K. Jackson, Ian Johnston, Dominic Kwiatkowski, Cordelia Langford, John Sillitoe on behalf of the Wellcome Sanger Institute COVID-19 Surveillance Team |
| EPI_ISL_3126691, EPI_ISL_3126692, EPI_ISL_3126693 | Lighthouse Lab in Glasgow | Wellcome Sanger Institute for the COVID-19 Genomics UK (COG-UK) Consortium | Harper VanSteenhouse, Yumi Kasai, David Gray, Carol Clugston, Anna Dominiczak and Alex Alderton, Roberto Amato, Jeffrey Barrett, Sonia Goncalves, Ewan Harrison, David K. Jackson, Ian Johnston, Dominic Kwiatkowski, Cordelia Langford, John Sillitoe on behalf of the Wellcome Sanger Institute COVID-19 Surveillance Team |
| EPI_ISL_3126694 | Berkshire and Surrey Pathology Services Lighthouse Laboratory | Wellcome Sanger Institute for the COVID-19 Genomics UK (COG-UK) Consortium | Berkshire and Surrey Pathology Services Lighthouse Laboratory and Alex Alderton, Roberto Amato, Jeffrey Barrett, Sonia Goncalves, Ewan Harrison, David K. Jackson, Ian Johnston, Dominic Kwiatkowski, Cordelia Langford, John Sillitoe on behalf of the Wellcome Sanger Institute COVID-19 Surveillance Team |
| EPI_ISL_3126695, EPI_ISL_3126696 | Lighthouse Lab in Glasgow | Wellcome Sanger Institute for the COVID-19 Genomics UK (COG-UK) Consortium | Harper VanSteenhouse, Yumi Kasai, David Gray, Carol Clugston, Anna Dominiczak and Alex Alderton, Roberto Amato, Jeffrey Barrett, Sonia Goncalves, Ewan Harrison, David K. Jackson, Ian Johnston, Dominic Kwiatkowski, Cordelia Langford, John Sillitoe on behalf of the Wellcome Sanger Institute COVID-19 Surveillance Team |
| EPI_ISL_3126697 | Lighthouse Lab in Alderley Park | Wellcome Sanger Institute for the COVID-19 Genomics UK (COG-UK) Consortium | Jacquelyn Wynn, Mairead Hyland, The Lighthouse Lab in Alderley Park and Alex Alderton, Roberto Amato, Jeffrey Barrett, Sonia Goncalves, Ewan Harrison, David K. Jackson, Ian Johnston, Dominic Kwiatkowski, Cordelia Langford, John Sillitoe on behalf of the Wellcome Sanger Institute COVID-19 Surveillance Team |
| EPI_ISL_3126698, EPI_ISL_3126699 | Lighthouse Lab in Glasgow | Wellcome Sanger Institute for the COVID-19 Genomics UK (COG-UK) Consortium | Harper VanSteenhouse, Yumi Kasai, David Gray, Carol Clugston, Anna Dominiczak and Alex Alderton, Roberto Amato, Jeffrey Barrett, Sonia Goncalves, Ewan Harrison, David K. Jackson, Ian Johnston, Dominic Kwiatkowski, Cordelia Langford, John Sillitoe on behalf of the Wellcome Sanger Institute COVID-19 Surveillance Team |
| EPI_ISL_3126700 | Lighthouse Lab in Alderley Park | Wellcome Sanger Institute for the COVID-19 Genomics UK (COG-UK) Consortium | Jacquelyn Wynn, Mairead Hyland, The Lighthouse Lab in Alderley Park and Alex Alderton, Roberto Amato, Jeffrey Barrett, Sonia Goncalves, Ewan Harrison, David K. Jackson, Ian Johnston, Dominic Kwiatkowski, Cordelia Langford, John Sillitoe on behalf of the Wellcome Sanger Institute COVID-19 Surveillance Team |
| EPI_ISL_3126701, EPI_ISL_3126702 | Lighthouse Lab in Glasgow | Wellcome Sanger Institute for the COVID-19 Genomics UK (COG-UK) Consortium | Harper VanSteenhouse, Yumi Kasai, David Gray, Carol Clugston, Anna Dominiczak and Alex Alderton, Roberto Amato, Jeffrey Barrett, Sonia Goncalves, Ewan Harrison, David K. Jackson, Ian Johnston, Dominic Kwiatkowski, Cordelia Langford, John Sillitoe on behalf of the Wellcome Sanger Institute COVID-19 Surveillance Team |
| EPI_ISL_3126703 | Lighthouse Lab in Alderley Park | Wellcome Sanger Institute for the COVID-19 Genomics UK (COG-UK) Consortium | Jacquelyn Wynn, Mairead Hyland, The Lighthouse Lab in Alderley Park and Alex Alderton, Roberto Amato, Jeffrey Barrett, Sonia Goncalves, Ewan Harrison, David K. Jackson, Ian Johnston, Dominic Kwiatkowski, Cordelia Langford, John Sillitoe on behalf of the Wellcome Sanger Institute COVID-19 Surveillance Team |
| EPI_ISL_3126704 | Lighthouse Lab in Glasgow | Wellcome Sanger Institute for the COVID-19 Genomics UK (COG-UK) Consortium | Harper VanSteenhouse, Yumi Kasai, David Gray, Carol Clugston, Anna Dominiczak and Alex Alderton, Roberto Amato, Jeffrey Barrett, Sonia Goncalves, Ewan Harrison, David K. Jackson, Ian Johnston, Dominic Kwiatkowski, Cordelia Langford, John Sillitoe on behalf of the Wellcome Sanger Institute COVID-19 Surveillance Team |
| EPI_ISL_3126705, EPI_ISL_3126706, EPI_ISL_3126707, EPI_ISL_3126708, EPI_ISL_3126709 | Lighthouse Lab in Alderley Park | Wellcome Sanger Institute for the COVID-19 Genomics UK (COG-UK) Consortium | Jacquelyn Wynn, Mairead Hyland, The Lighthouse Lab in Alderley Park and Alex Alderton, Roberto Amato, Jeffrey Barrett, Sonia Goncalves, Ewan Harrison, David K. Jackson, Ian Johnston, Dominic Kwiatkowski, Cordelia Langford, John Sillitoe on behalf of the Wellcome Sanger Institute COVID-19 Surveillance Team |
| EPI_ISL_3126710 | Lighthouse Lab in Milton Keynes | Wellcome Sanger Institute for the COVID-19 Genomics UK (COG-UK) Consortium | The Lighthouse Lab in Milton Keynes and Alex Alderton, Roberto Amato, Jeffrey Barrett, Sonia Goncalves, Ewan Harrison, David K. Jackson, Ian Johnston, Dominic Kwiatkowski, Cordelia Langford, John Sillitoe on behalf of the Wellcome Sanger Institute COVID-19 Surveillance Team |
| EPI_ISL_3126711 | Lighthouse Lab in Glasgow | Wellcome Sanger Institute for the COVID-19 Genomics UK (COG-UK) Consortium | Harper VanSteenhouse, Yumi Kasai, David Gray, Carol Clugston, Anna Dominiczak and Alex Alderton, Roberto Amato, Jeffrey Barrett, Sonia Goncalves, Ewan Harrison, David K. Jackson, Ian Johnston, Dominic Kwiatkowski, Cordelia Langford, John Sillitoe on behalf of the Wellcome Sanger Institute COVID-19 Surveillance Team |
| EPI_ISL_3126712 | Lighthouse Lab in Alderley Park | Wellcome Sanger Institute for the COVID-19 Genomics UK (COG-UK) Consortium | Jacquelyn Wynn, Mairead Hyland, The Lighthouse Lab in Alderley Park and Alex Alderton, Roberto Amato, Jeffrey Barrett, Sonia Goncalves, Ewan Harrison, David K. Jackson, Ian Johnston, Dominic Kwiatkowski, Cordelia Langford, John Sillitoe on behalf of the Wellcome Sanger Institute COVID-19 Surveillance Team |
| EPI_ISL_3126713 | Lighthouse Lab in Milton Keynes | Wellcome Sanger Institute for the COVID-19 Genomics UK (COG-UK) Consortium | The Lighthouse Lab in Milton Keynes and Alex Alderton, Roberto Amato, Jeffrey Barrett, Sonia Goncalves, Ewan Harrison, David K. Jackson, Ian Johnston, Dominic Kwiatkowski, Cordelia Langford, John Sillitoe on behalf of the Wellcome Sanger Institute COVID-19 Surveillance Team |
| EPI_ISL_3126714, EPI_ISL_3126715, EPI_ISL_3126716 | Lighthouse Lab in Alderley Park | Wellcome Sanger Institute for the COVID-19 Genomics UK (COG-UK) Consortium | Jacquelyn Wynn, Mairead Hyland, The Lighthouse Lab in Alderley Park and Alex Alderton, Roberto Amato, Jeffrey Barrett, Sonia Goncalves, Ewan Harrison, David K. Jackson, Ian Johnston, Dominic Kwiatkowski, Cordelia Langford, John Sillitoe on behalf of the Wellcome Sanger Institute COVID-19 Surveillance Team |
| EPI_ISL_3126717 | Lighthouse Lab in Glasgow | Wellcome Sanger Institute for the COVID-19 Genomics UK (COG-UK) Consortium | Harper VanSteenhouse, Yumi Kasai, David Gray, Carol Clugston, Anna Dominiczak and Alex Alderton, Roberto Amato, Jeffrey Barrett, Sonia Goncalves, Ewan Harrison, David K. Jackson, Ian Johnston, Dominic Kwiatkowski, Cordelia Langford, John Sillitoe on behalf of the Wellcome Sanger Institute COVID-19 Surveillance Team |
| EPI_ISL_3126718 | Lighthouse Lab in Alderley Park | Wellcome Sanger Institute for the COVID-19 Genomics UK (COG-UK) Consortium | Jacquelyn Wynn, Mairead Hyland, The Lighthouse Lab in Alderley Park and Alex Alderton, Roberto Amato, Jeffrey Barrett, Sonia Goncalves, Ewan Harrison, David K. Jackson, Ian Johnston, Dominic Kwiatkowski, Cordelia Langford, John Sillitoe on behalf of the Wellcome Sanger Institute COVID-19 Surveillance Team |
| EPI_ISL_3126719, EPI_ISL_3126720 | Lighthouse Lab in Milton Keynes | Wellcome Sanger Institute for the COVID-19 Genomics UK (COG-UK) Consortium | The Lighthouse Lab in Milton Keynes and Alex Alderton, Roberto Amato, Jeffrey Barrett, Sonia Goncalves, Ewan Harrison, David K. Jackson, Ian Johnston, Dominic Kwiatkowski, Cordelia Langford, John Sillitoe on behalf of the Wellcome Sanger Institute COVID-19 Surveillance Team |
| EPI_ISL_3126721, EPI_ISL_3126722 | Lighthouse Lab in Alderley Park | Wellcome Sanger Institute for the COVID-19 Genomics UK (COG-UK) Consortium | Jacquelyn Wynn, Mairead Hyland, The Lighthouse Lab in Alderley Park and Alex Alderton, Roberto Amato, Jeffrey Barrett, Sonia Goncalves, Ewan Harrison, David K. Jackson, Ian Johnston, Dominic Kwiatkowski, Cordelia Langford, John Sillitoe on behalf of the Wellcome Sanger Institute COVID-19 Surveillance Team |
| EPI_ISL_3126723 | Berkshire and Surrey Pathology Services Lighthouse Laboratory | Wellcome Sanger Institute for the COVID-19 Genomics UK (COG-UK) Consortium | Berkshire and Surrey Pathology Services Lighthouse Laboratory and Alex Alderton, Roberto Amato, Jeffrey Barrett, Sonia Goncalves, Ewan Harrison, David K. Jackson, Ian Johnston, Dominic Kwiatkowski, Cordelia Langford, John Sillitoe on behalf of the Wellcome Sanger Institute COVID-19 Surveillance Team |
| EPI_ISL_3126724, EPI_ISL_3126725 | Lighthouse Lab in Alderley Park | Wellcome Sanger Institute for the COVID-19 Genomics UK (COG-UK) Consortium | Jacquelyn Wynn, Mairead Hyland, The Lighthouse Lab in Alderley Park and Alex Alderton, Roberto Amato, Jeffrey Barrett, Sonia Goncalves, Ewan Harrison, David K. Jackson, Ian Johnston, Dominic Kwiatkowski, Cordelia Langford, John Sillitoe on behalf of the Wellcome Sanger Institute COVID-19 Surveillance Team |
| EPI_ISL_3126726 | Berkshire and Surrey Pathology Services Lighthouse Laboratory | Wellcome Sanger Institute for the COVID-19 Genomics UK (COG-UK) Consortium | Berkshire and Surrey Pathology Services Lighthouse Laboratory and Alex Alderton, Roberto Amato, Jeffrey Barrett, Sonia Goncalves, Ewan Harrison, David K. Jackson, Ian Johnston, Dominic Kwiatkowski, Cordelia Langford, John Sillitoe on behalf of the Wellcome Sanger Institute COVID-19 Surveillance Team |
| EPI_ISL_3126727 | Lighthouse Lab in Alderley Park | Wellcome Sanger Institute for the COVID-19 Genomics UK (COG-UK) Consortium | Jacquelyn Wynn, Mairead Hyland, The Lighthouse Lab in Alderley Park and Alex Alderton, Roberto Amato, Jeffrey Barrett, Sonia Goncalves, Ewan Harrison, David K. Jackson, Ian Johnston, Dominic Kwiatkowski, Cordelia Langford, John Sillitoe on behalf of the Wellcome Sanger Institute COVID-19 Surveillance Team |
| EPI_ISL_3126728 | Berkshire and Surrey Pathology Services Lighthouse Laboratory | Wellcome Sanger Institute for the COVID-19 Genomics UK (COG-UK) Consortium | Berkshire and Surrey Pathology Services Lighthouse Laboratory and Alex Alderton, Roberto Amato, Jeffrey Barrett, Sonia Goncalves, Ewan Harrison, David K. Jackson, Ian Johnston, Dominic Kwiatkowski, Cordelia Langford, John Sillitoe on behalf of the Wellcome Sanger Institute COVID-19 Surveillance Team |
| EPI_ISL_3126729 | Lighthouse Lab in Alderley Park | Wellcome Sanger Institute for the COVID-19 Genomics UK (COG-UK) Consortium | Jacquelyn Wynn, Mairead Hyland, The Lighthouse Lab in Alderley Park and Alex Alderton, Roberto Amato, Jeffrey Barrett, Sonia Goncalves, Ewan Harrison, David K. Jackson, Ian Johnston, Dominic Kwiatkowski, Cordelia Langford, John Sillitoe on behalf of the Wellcome Sanger Institute COVID-19 Surveillance Team |
| EPI_ISL_3126730 | Lighthouse Lab in Milton Keynes | Wellcome Sanger Institute for the COVID-19 Genomics UK (COG-UK) | The Lighthouse Lab in Milton Keynes and Alex Alderton, Roberto Amato, Jeffrey Barrett, Sonia Goncalves, Ewan Harrison, David K. Jackson, Ian Johnston, Dominic Kwiatkowski, Cordelia Langford, John Sillitoe on behalf of the Wellcome Sanger |

| | | | |
|---|---|---|---|
| | | Consortium | Institute COVID-19 Surveillance Team |
| EPI_ISL_3126731, EPI_ISL_3126732 | Lighthouse Lab in Alderley Park | Wellcome Sanger Institute for the COVID-19 Genomics UK (COG-UK) Consortium | Jacquelyn Wynn, Mairead Hyland, The Lighthouse Lab in Alderley Park and Alex Alderton, Roberto Amato, Jeffrey Barrett, Sonia Goncalves, Ewan Harrison, David K. Jackson, Ian Johnston, Dominic Kwiatkowski, Cordelia Langford, John Sillitoe on behalf of the Wellcome Sanger Institute COVID-19 Surveillance Team |
| EPI_ISL_3126733 | Berkshire and Surrey Pathology Services Lighthouse Laboratory | Wellcome Sanger Institute for the COVID-19 Genomics UK (COG-UK) Consortium | Berkshire and Surrey Pathology Services Lighthouse Laboratory and Alex Alderton, Roberto Amato, Jeffrey Barrett, Sonia Goncalves, Ewan Harrison, David K. Jackson, Ian Johnston, Dominic Kwiatkowski, Cordelia Langford, John Sillitoe on behalf of the Wellcome Sanger Institute COVID-19 Surveillance Team |
| EPI_ISL_3126734, EPI_ISL_3126735 | Lighthouse Lab in Alderley Park | Wellcome Sanger Institute for the COVID-19 Genomics UK (COG-UK) Consortium | Jacquelyn Wynn, Mairead Hyland, The Lighthouse Lab in Alderley Park and Alex Alderton, Roberto Amato, Jeffrey Barrett, Sonia Goncalves, Ewan Harrison, David K. Jackson, Ian Johnston, Dominic Kwiatkowski, Cordelia Langford, John Sillitoe on behalf of the Wellcome Sanger Institute COVID-19 Surveillance Team |
| EPI_ISL_3126736 | Lighthouse Lab in Milton Keynes | Wellcome Sanger Institute for the COVID-19 Genomics UK (COG-UK) Consortium | The Lighthouse Lab in Milton Keynes and Alex Alderton, Roberto Amato, Jeffrey Barrett, Sonia Goncalves, Ewan Harrison, David K. Jackson, Ian Johnston, Dominic Kwiatkowski, Cordelia Langford, John Sillitoe on behalf of the Wellcome Sanger Institute COVID-19 Surveillance Team |
| EPI_ISL_3126737 | Lighthouse Lab in Alderley Park | Wellcome Sanger Institute for the COVID-19 Genomics UK (COG-UK) Consortium | Jacquelyn Wynn, Mairead Hyland, The Lighthouse Lab in Alderley Park and Alex Alderton, Roberto Amato, Jeffrey Barrett, Sonia Goncalves, Ewan Harrison, David K. Jackson, Ian Johnston, Dominic Kwiatkowski, Cordelia Langford, John Sillitoe on behalf of the Wellcome Sanger Institute COVID-19 Surveillance Team |
| EPI_ISL_3126738 | Lighthouse Lab in Milton Keynes | Wellcome Sanger Institute for the COVID-19 Genomics UK (COG-UK) Consortium | The Lighthouse Lab in Milton Keynes and Alex Alderton, Roberto Amato, Jeffrey Barrett, Sonia Goncalves, Ewan Harrison, David K. Jackson, Ian Johnston, Dominic Kwiatkowski, Cordelia Langford, John Sillitoe on behalf of the Wellcome Sanger Institute COVID-19 Surveillance Team |
| EPI_ISL_3126739 | Lighthouse Lab in Alderley Park | Wellcome Sanger Institute for the COVID-19 Genomics UK (COG-UK) Consortium | Jacquelyn Wynn, Mairead Hyland, The Lighthouse Lab in Alderley Park and Alex Alderton, Roberto Amato, Jeffrey Barrett, Sonia Goncalves, Ewan Harrison, David K. Jackson, Ian Johnston, Dominic Kwiatkowski, Cordelia Langford, John Sillitoe on behalf of the Wellcome Sanger Institute COVID-19 Surveillance Team |
| EPI_ISL_3126740 | Berkshire and Surrey Pathology Services Lighthouse Laboratory | Wellcome Sanger Institute for the COVID-19 Genomics UK (COG-UK) Consortium | Berkshire and Surrey Pathology Services Lighthouse Laboratory and Alex Alderton, Roberto Amato, Jeffrey Barrett, Sonia Goncalves, Ewan Harrison, David K. Jackson, Ian Johnston, Dominic Kwiatkowski, Cordelia Langford, John Sillitoe on behalf of the Wellcome Sanger Institute COVID-19 Surveillance Team |
| EPI_ISL_3126741, EPI_ISL_3126742, EPI_ISL_3126743 | Lighthouse Lab in Milton Keynes | Wellcome Sanger Institute for the COVID-19 Genomics UK (COG-UK) Consortium | The Lighthouse Lab in Milton Keynes and Alex Alderton, Roberto Amato, Jeffrey Barrett, Sonia Goncalves, Ewan Harrison, David K. Jackson, Ian Johnston, Dominic Kwiatkowski, Cordelia Langford, John Sillitoe on behalf of the Wellcome Sanger Institute COVID-19 Surveillance Team |
| EPI_ISL_3126744, EPI_ISL_3126745, EPI_ISL_3126746 | Lighthouse Lab in Glasgow | Wellcome Sanger Institute for the COVID-19 Genomics UK (COG-UK) Consortium | Harper VanSteenhouse, Yumi Kasai, David Gray, Carol Clugston, Anna Dominiczak and Alex Alderton, Roberto Amato, Jeffrey Barrett, Sonia Goncalves, Ewan Harrison, David K. Jackson, Ian Johnston, Dominic Kwiatkowski, Cordelia Langford, John Sillitoe on behalf of the Wellcome Sanger Institute COVID-19 Surveillance Team |
| EPI_ISL_3126747, EPI_ISL_3126748, EPI_ISL_3126749, EPI_ISL_3126750 | Lighthouse Lab in Milton Keynes | Wellcome Sanger Institute for the COVID-19 Genomics UK (COG-UK) Consortium | The Lighthouse Lab in Milton Keynes and Alex Alderton, Roberto Amato, Jeffrey Barrett, Sonia Goncalves, Ewan Harrison, David K. Jackson, Ian Johnston, Dominic Kwiatkowski, Cordelia Langford, John Sillitoe on behalf of the Wellcome Sanger Institute COVID-19 Surveillance Team |
| EPI_ISL_3126751 | Lighthouse Lab in Alderley Park | Wellcome Sanger Institute for the COVID-19 Genomics UK (COG-UK) Consortium | Jacquelyn Wynn, Mairead Hyland, The Lighthouse Lab in Alderley Park and Alex Alderton, Roberto Amato, Jeffrey Barrett, Sonia Goncalves, Ewan Harrison, David K. Jackson, Ian Johnston, Dominic Kwiatkowski, Cordelia Langford, John Sillitoe on behalf of the Wellcome Sanger Institute COVID-19 Surveillance Team |
| EPI_ISL_3126752, EPI_ISL_3126753, EPI_ISL_3126754, EPI_ISL_3126755 | Lighthouse Lab in Milton Keynes | Wellcome Sanger Institute for the COVID-19 Genomics UK (COG-UK) Consortium | The Lighthouse Lab in Milton Keynes and Alex Alderton, Roberto Amato, Jeffrey Barrett, Sonia Goncalves, Ewan Harrison, David K. Jackson, Ian Johnston, Dominic Kwiatkowski, Cordelia Langford, John Sillitoe on behalf of the Wellcome Sanger Institute COVID-19 Surveillance Team |
| EPI_ISL_3126756, EPI_ISL_3126757, EPI_ISL_3126758, EPI_ISL_3126759, EPI_ISL_3126760 | Berkshire and Surrey Pathology Services Lighthouse Laboratory | Wellcome Sanger Institute for the COVID-19 Genomics UK (COG-UK) Consortium | Berkshire and Surrey Pathology Services Lighthouse Laboratory and Alex Alderton, Roberto Amato, Jeffrey Barrett, Sonia Goncalves, Ewan Harrison, David K. Jackson, Ian Johnston, Dominic Kwiatkowski, Cordelia Langford, John Sillitoe on behalf of the Wellcome Sanger Institute COVID-19 Surveillance Team |
| EPI_ISL_3126761, EPI_ISL_3126762, EPI_ISL_3126763 | Lighthouse Lab in Alderley Park | Wellcome Sanger Institute for the COVID-19 Genomics UK (COG-UK) Consortium | Jacquelyn Wynn, Mairead Hyland, The Lighthouse Lab in Alderley Park and Alex Alderton, Roberto Amato, Jeffrey Barrett, Sonia Goncalves, Ewan Harrison, David K. Jackson, Ian Johnston, Dominic Kwiatkowski, Cordelia Langford, John Sillitoe on behalf of the Wellcome Sanger Institute COVID-19 Surveillance Team |
| EPI_ISL_3126764, EPI_ISL_3126765, EPI_ISL_3126766 | Lighthouse Lab in Milton Keynes | Wellcome Sanger Institute for the COVID-19 Genomics UK (COG-UK) Consortium | The Lighthouse Lab in Milton Keynes and Alex Alderton, Roberto Amato, Jeffrey Barrett, Sonia Goncalves, Ewan Harrison, David K. Jackson, Ian Johnston, Dominic Kwiatkowski, Cordelia Langford, John Sillitoe on behalf of the Wellcome Sanger Institute COVID-19 Surveillance Team |
| EPI_ISL_3126767 | Lighthouse Lab in Alderley Park | Wellcome Sanger Institute for the COVID-19 Genomics UK (COG-UK) Consortium | Jacquelyn Wynn, Mairead Hyland, The Lighthouse Lab in Alderley Park and Alex Alderton, Roberto Amato, Jeffrey Barrett, Sonia Goncalves, Ewan Harrison, David K. Jackson, Ian Johnston, Dominic Kwiatkowski, Cordelia Langford, John Sillitoe on behalf of the Wellcome Sanger Institute COVID-19 Surveillance Team |
| EPI_ISL_3126768, EPI_ISL_3126769, EPI_ISL_3126770 | Berkshire and Surrey Pathology Services Lighthouse Laboratory | Wellcome Sanger Institute for the COVID-19 Genomics UK (COG-UK) Consortium | Berkshire and Surrey Pathology Services Lighthouse Laboratory and Alex Alderton, Roberto Amato, Jeffrey Barrett, Sonia Goncalves, Ewan Harrison, David K. Jackson, Ian Johnston, Dominic Kwiatkowski, Cordelia Langford, John Sillitoe on behalf of the Wellcome Sanger Institute COVID-19 Surveillance Team |
| EPI_ISL_3126771, EPI_ISL_3126772, EPI_ISL_3126773, EPI_ISL_3126774, EPI_ISL_3126775, EPI_ISL_3126776 | Lighthouse Lab in Alderley Park | Wellcome Sanger Institute for the COVID-19 Genomics UK (COG-UK) Consortium | Jacquelyn Wynn, Mairead Hyland, The Lighthouse Lab in Alderley Park and Alex Alderton, Roberto Amato, Jeffrey Barrett, Sonia Goncalves, Ewan Harrison, David K. Jackson, Ian Johnston, Dominic Kwiatkowski, Cordelia Langford, John Sillitoe on behalf of the Wellcome Sanger Institute COVID-19 Surveillance Team |
| EPI_ISL_3126777, EPI_ISL_3126778, EPI_ISL_3126779 | Lighthouse Lab in Milton Keynes | Wellcome Sanger Institute for the COVID-19 Genomics UK (COG-UK) Consortium | The Lighthouse Lab in Milton Keynes and Alex Alderton, Roberto Amato, Jeffrey Barrett, Sonia Goncalves, Ewan Harrison, David K. Jackson, Ian Johnston, Dominic Kwiatkowski, Cordelia Langford, John Sillitoe on behalf of the Wellcome Sanger Institute COVID-19 Surveillance Team |
| EPI_ISL_3126780 | Lighthouse Lab in Glasgow | Wellcome Sanger Institute for the COVID-19 Genomics UK (COG-UK) Consortium | Harper VanSteenhouse, Yumi Kasai, David Gray, Carol Clugston, Anna Dominiczak and Alex Alderton, Roberto Amato, Jeffrey Barrett, Sonia Goncalves, Ewan Harrison, David K. Jackson, Ian Johnston, Dominic Kwiatkowski, Cordelia Langford, John Sillitoe on behalf of the Wellcome Sanger Institute COVID-19 Surveillance Team |
| EPI_ISL_3126781 | Lighthouse Lab in Milton Keynes | Wellcome Sanger Institute for the COVID-19 Genomics UK (COG-UK) Consortium | The Lighthouse Lab in Milton Keynes and Alex Alderton, Roberto Amato, Jeffrey Barrett, Sonia Goncalves, Ewan Harrison, David K. Jackson, Ian Johnston, Dominic Kwiatkowski, Cordelia Langford, John Sillitoe on behalf of the Wellcome Sanger Institute COVID-19 Surveillance Team |
| EPI_ISL_3126782 | Lighthouse Lab in Glasgow | Wellcome Sanger Institute for the COVID-19 Genomics UK (COG-UK) Consortium | Harper VanSteenhouse, Yumi Kasai, David Gray, Carol Clugston, Anna Dominiczak and Alex Alderton, Roberto Amato, Jeffrey Barrett, Sonia Goncalves, Ewan Harrison, David K. Jackson, Ian Johnston, Dominic Kwiatkowski, Cordelia Langford, John Sillitoe on behalf of the Wellcome Sanger Institute COVID-19 Surveillance Team |
| EPI_ISL_3126783, EPI_ISL_3126784, EPI_ISL_3126785, EPI_ISL_3126786, EPI_ISL_3126787, EPI_ISL_3126788, EPI_ISL_3126789, EPI_ISL_3126790 | Lighthouse Lab in Milton Keynes | Wellcome Sanger Institute for the COVID-19 Genomics UK (COG-UK) Consortium | The Lighthouse Lab in Milton Keynes and Alex Alderton, Roberto Amato, Jeffrey Barrett, Sonia Goncalves, Ewan Harrison, David K. Jackson, Ian Johnston, Dominic Kwiatkowski, Cordelia Langford, John Sillitoe on behalf of the Wellcome Sanger Institute COVID-19 Surveillance Team |
| EPI_ISL_3126791, EPI_ISL_3126792, EPI_ISL_3126793, EPI_ISL_3126794, EPI_ISL_3126795, EPI_ISL_3126796, EPI_ISL_3126797, EPI_ISL_3126798, EPI_ISL_3126799, EPI_ISL_3126800, EPI_ISL_3126801, EPI_ISL_3126802, EPI_ISL_3126803, EPI_ISL_3126804, EPI_ISL_3126805, EPI_ISL_3126806, EPI_ISL_3126807, EPI_ISL_3126808, EPI_ISL_3126809, EPI_ISL_3126810, EPI_ISL_3126811, EPI_ISL_3126812, EPI_ISL_3126813, EPI_ISL_3126814, EPI_ISL_3126815, EPI_ISL_3126816 | | | |
| see above | Lighthouse Lab in Alderley Park | Wellcome Sanger Institute for the COVID-19 Genomics UK (COG-UK) Consortium | Jacquelyn Wynn, Mairead Hyland, The Lighthouse Lab in Alderley Park and Alex Alderton, Roberto Amato, Jeffrey Barrett, Sonia Goncalves, Ewan Harrison, David K. Jackson, Ian Johnston, Dominic Kwiatkowski, Cordelia Langford, John Sillitoe on behalf of the Wellcome Sanger Institute COVID-19 Surveillance Team |
| EPI_ISL_3126817 | Lighthouse Lab in Glasgow | Wellcome Sanger Institute for the COVID-19 Genomics UK (COG-UK) Consortium | Harper VanSteenhouse, Yumi Kasai, David Gray, Carol Clugston, Anna Dominiczak and Alex Alderton, Roberto Amato, Jeffrey Barrett, Sonia Goncalves, Ewan Harrison, David K. Jackson, Ian Johnston, Dominic Kwiatkowski, Cordelia Langford, John Sillitoe on behalf of the Wellcome Sanger Institute COVID-19 Surveillance Team |
| EPI_ISL_3126818 | Lighthouse Lab in Alderley Park | Wellcome Sanger Institute for the COVID-19 Genomics UK (COG-UK) Consortium | Jacquelyn Wynn, Mairead Hyland, The Lighthouse Lab in Alderley Park and Alex Alderton, Roberto Amato, Jeffrey Barrett, Sonia Goncalves, Ewan Harrison, David K. Jackson, Ian Johnston, Dominic Kwiatkowski, Cordelia Langford, John Sillitoe on behalf of the Wellcome Sanger Institute COVID-19 Surveillance Team |
| EPI_ISL_3126819, EPI_ISL_3126820 | Lighthouse Lab in Milton Keynes | Wellcome Sanger Institute for the COVID-19 Genomics UK (COG-UK) Consortium | The Lighthouse Lab in Milton Keynes and Alex Alderton, Roberto Amato, Jeffrey Barrett, Sonia Goncalves, Ewan Harrison, David K. Jackson, Ian Johnston, Dominic Kwiatkowski, Cordelia Langford, John Sillitoe on behalf of the Wellcome Sanger Institute COVID-19 Surveillance Team |
| EPI_ISL_3126821 | Berkshire and Surrey Pathology Services Lighthouse Laboratory | Wellcome Sanger Institute for the COVID-19 Genomics UK (COG-UK) Consortium | Berkshire and Surrey Pathology Services Lighthouse Laboratory and Alex Alderton, Roberto Amato, Jeffrey Barrett, Sonia Goncalves, Ewan Harrison, David K. Jackson, Ian Johnston, Dominic Kwiatkowski, Cordelia Langford, John Sillitoe on behalf of the Wellcome Sanger Institute COVID-19 Surveillance Team |
| EPI_ISL_3126822 | Lighthouse Lab in Milton Keynes | Wellcome Sanger Institute for the COVID-19 Genomics UK (COG-UK) Consortium | The Lighthouse Lab in Milton Keynes and Alex Alderton, Roberto Amato, Jeffrey Barrett, Sonia Goncalves, Ewan Harrison, David K. Jackson, Ian Johnston, Dominic Kwiatkowski, Cordelia Langford, John Sillitoe on behalf of the Wellcome Sanger Institute COVID-19 Surveillance Team |
| EPI_ISL_3126823 | Berkshire and Surrey Pathology Services Lighthouse Laboratory | Wellcome Sanger Institute for the COVID-19 Genomics UK (COG-UK) Consortium | Berkshire and Surrey Pathology Services Lighthouse Laboratory and Alex Alderton, Roberto Amato, Jeffrey Barrett, Sonia Goncalves, Ewan Harrison, David K. Jackson, Ian Johnston, Dominic Kwiatkowski, Cordelia Langford, John Sillitoe on behalf of the Wellcome Sanger Institute COVID-19 Surveillance Team |
| EPI_ISL_3126824, EPI_ISL_3126825, EPI_ISL_3126826 | Lighthouse Lab in Alderley Park | Wellcome Sanger Institute for the COVID-19 Genomics UK (COG-UK) Consortium | Jacquelyn Wynn, Mairead Hyland, The Lighthouse Lab in Alderley Park and Alex Alderton, Roberto Amato, Jeffrey Barrett, Sonia Goncalves, Ewan Harrison, David K. Jackson, Ian Johnston, Dominic Kwiatkowski, Cordelia Langford, John Sillitoe on behalf of the Wellcome Sanger Institute COVID-19 Surveillance Team |
| EPI_ISL_3126827, EPI_ISL_3126828, EPI_ISL_3126829, EPI_ISL_3126830, | Lighthouse Lab in Glasgow | Wellcome Sanger Institute for the COVID-19 Genomics UK (COG-UK) Consortium | Harper VanSteenhouse, Yumi Kasai, David Gray, Carol Clugston, Anna Dominiczak and Alex Alderton, Roberto Amato, Jeffrey Barrett, Sonia Goncalves, Ewan Harrison, David K. Jackson, Ian Johnston, Dominic Kwiatkowski, Cordelia Langford, John Sillitoe on behalf of the Wellcome Sanger Institute COVID-19 Surveillance Team |

| | | | |
|---|---|---|---|
| EPI_ISL_3126831, EPI_ISL_3126832, EPI_ISL_3126833, EPI_ISL_3126834 | | | |
| EPI_ISL_3126835, EPI_ISL_3126836, EPI_ISL_3126837 | Lighthouse Laboratory Plymouth | Wellcome Sanger Institute for the COVID-19 Genomics UK (COG-UK) Consortium | Lighthouse Laboratory Plymouth and Alex Alderton, Roberto Amato, Jeffrey Barrett, Sonia Goncalves, Ewan Harrison, David K. Jackson, Ian Johnston, Dominic Kwiatkowski, Cordelia Langford, John Sillitoe on behalf of the Wellcome Sanger Institute COVID-19 Surveillance Team |
| EPI_ISL_3126838 | Lighthouse Lab in Milton Keynes | Wellcome Sanger Institute for the COVID-19 Genomics UK (COG-UK) Consortium | The Lighthouse Lab in Milton Keynes and Alex Alderton, Roberto Amato, Jeffrey Barrett, Sonia Goncalves, Ewan Harrison, David K. Jackson, Ian Johnston, Dominic Kwiatkowski, Cordelia Langford, John Sillitoe on behalf of the Wellcome Sanger Institute COVID-19 Surveillance Team |
| EPI_ISL_3126839, EPI_ISL_3126840 | Lighthouse Lab in Glasgow | Wellcome Sanger Institute for the COVID-19 Genomics UK (COG-UK) Consortium | Harper VanSteenhouse, Yumi Kasai, David Gray, Carol Clugston, Anna Dominiczak and Alex Alderton, Roberto Amato, Jeffrey Barrett, Sonia Goncalves, Ewan Harrison, David K. Jackson, Ian Johnston, Dominic Kwiatkowski, Cordelia Langford, John Sillitoe on behalf of the Wellcome Sanger Institute COVID-19 Surveillance Team |
| EPI_ISL_3126841, EPI_ISL_3126842 | Lighthouse Laboratory Plymouth | Wellcome Sanger Institute for the COVID-19 Genomics UK (COG-UK) Consortium | Lighthouse Laboratory Plymouth and Alex Alderton, Roberto Amato, Jeffrey Barrett, Sonia Goncalves, Ewan Harrison, David K. Jackson, Ian Johnston, Dominic Kwiatkowski, Cordelia Langford, John Sillitoe on behalf of the Wellcome Sanger Institute COVID-19 Surveillance Team |
| EPI_ISL_3126843 | Lighthouse Lab in Milton Keynes | Wellcome Sanger Institute for the COVID-19 Genomics UK (COG-UK) Consortium | The Lighthouse Lab in Milton Keynes and Alex Alderton, Roberto Amato, Jeffrey Barrett, Sonia Goncalves, Ewan Harrison, David K. Jackson, Ian Johnston, Dominic Kwiatkowski, Cordelia Langford, John Sillitoe on behalf of the Wellcome Sanger Institute COVID-19 Surveillance Team |
| EPI_ISL_3126844, EPI_ISL_3126845 | Lighthouse Lab in Glasgow | Wellcome Sanger Institute for the COVID-19 Genomics UK (COG-UK) Consortium | Harper VanSteenhouse, Yumi Kasai, David Gray, Carol Clugston, Anna Dominiczak and Alex Alderton, Roberto Amato, Jeffrey Barrett, Sonia Goncalves, Ewan Harrison, David K. Jackson, Ian Johnston, Dominic Kwiatkowski, Cordelia Langford, John Sillitoe on behalf of the Wellcome Sanger Institute COVID-19 Surveillance Team |
| EPI_ISL_3126846 | Lighthouse Lab in Milton Keynes | Wellcome Sanger Institute for the COVID-19 Genomics UK (COG-UK) Consortium | The Lighthouse Lab in Milton Keynes and Alex Alderton, Roberto Amato, Jeffrey Barrett, Sonia Goncalves, Ewan Harrison, David K. Jackson, Ian Johnston, Dominic Kwiatkowski, Cordelia Langford, John Sillitoe on behalf of the Wellcome Sanger Institute COVID-19 Surveillance Team |
| EPI_ISL_3126847 | ASST FBF Sacco | Laboratory of Clinical Microbiology, Virology and Bioemergencies, ASST Fatebenefratelli Sacco - Sacco Hospital | Valeria Micheli |
| EPI_ISL_3126849, EPI_ISL_3126850, EPI_ISL_3126851, EPI_ISL_3126852, EPI_ISL_3126853, EPI_ISL_3126854, EPI_ISL_3126855, EPI_ISL_3126856, EPI_ISL_3126857, EPI_ISL_3126858, EPI_ISL_3126859, EPI_ISL_3126860, EPI_ISL_3126861, EPI_ISL_3126862, EPI_ISL_3126863, EPI_ISL_3126864, EPI_ISL_3126865, EPI_ISL_3126866, EPI_ISL_3126867, EPI_ISL_3126868, EPI_ISL_3126869, EPI_ISL_3126870, EPI_ISL_3126871, EPI_ISL_3126872, EPI_ISL_3126874, EPI_ISL_3126875, EPI_ISL_3126876, EPI_ISL_3126877, EPI_ISL_3126878, EPI_ISL_3126879, EPI_ISL_3126880, EPI_ISL_3126881, EPI_ISL_3126882, EPI_ISL_3126883, EPI_ISL_3126884, EPI_ISL_3126885, EPI_ISL_3126886, EPI_ISL_3126887, EPI_ISL_3126888, EPI_ISL_3126889, EPI_ISL_3126890, EPI_ISL_3126891, EPI_ISL_3126892, EPI_ISL_3126893, EPI_ISL_3126894, EPI_ISL_3126895, EPI_ISL_3126896, EPI_ISL_3126897, EPI_ISL_3126898, EPI_ISL_3126899, EPI_ISL_3126900, EPI_ISL_3126901, EPI_ISL_3126902, EPI_ISL_3126903, EPI_ISL_3126904, EPI_ISL_3126905, EPI_ISL_3126906, EPI_ISL_3126907, EPI_ISL_3126908, EPI_ISL_3126909, EPI_ISL_3126910, EPI_ISL_3126911, EPI_ISL_3126912, EPI_ISL_3126913, EPI_ISL_3126914, EPI_ISL_3126915, EPI_ISL_3126916, EPI_ISL_3126917, EPI_ISL_3126918, EPI_ISL_3126919, EPI_ISL_3126920 | | | |
| see above | Department of Bacteria, Parasites and Fungi, Statens Serum Institut, Copenhagen, Denmark | Statens Serum Institut Bioinformatics and Microbial Genomics | Danish Covid-19 Genome Consortium |
| EPI_ISL_3126921 | Molekylær Medicinsk Afdeling, Aarhus University Hospital, Aarhus, Denmark | Statens Serum Institut Bioinformatics and Microbial Genomics | Danish Covid-19 Genome Consortium |
| EPI_ISL_3126922, EPI_ISL_3126923, EPI_ISL_3126924, EPI_ISL_3126925, EPI_ISL_3126926, EPI_ISL_3126927, EPI_ISL_3126928, EPI_ISL_3126929, EPI_ISL_3126930, EPI_ISL_3126931, EPI_ISL_3126932, EPI_ISL_3126933, EPI_ISL_3126934, EPI_ISL_3126935, EPI_ISL_3126936, EPI_ISL_3126937, EPI_ISL_3126938, EPI_ISL_3126939, EPI_ISL_3126940, EPI_ISL_3126941, EPI_ISL_3126942, EPI_ISL_3126943, EPI_ISL_3126944, EPI_ISL_3126945, EPI_ISL_3126946, EPI_ISL_3126947, EPI_ISL_3126948 | | | |
| see above | Department of Bacteria, Parasites and Fungi, Statens Serum Institut, Copenhagen, Denmark | Statens Serum Institut Bioinformatics and Microbial Genomics | Danish Covid-19 Genome Consortium |
| EPI_ISL_3126949 | Molekylær Medicinsk Afdeling, Aarhus University Hospital, Aarhus, Denmark | Statens Serum Institut Bioinformatics and Microbial Genomics | Danish Covid-19 Genome Consortium |
| EPI_ISL_3126950, EPI_ISL_3126951, EPI_ISL_3126952, EPI_ISL_3126953, EPI_ISL_3126954, EPI_ISL_3126955, EPI_ISL_3126956, EPI_ISL_3126957, EPI_ISL_3126958, EPI_ISL_3126959, EPI_ISL_3126960, EPI_ISL_3126961, EPI_ISL_3126962, EPI_ISL_3126963, EPI_ISL_3126964, EPI_ISL_3126965, EPI_ISL_3126966, EPI_ISL_3126967, EPI_ISL_3126968, EPI_ISL_3126969, EPI_ISL_3126970, EPI_ISL_3126971, EPI_ISL_3126972, EPI_ISL_3126973, EPI_ISL_3126974, EPI_ISL_3126975, EPI_ISL_3126976, EPI_ISL_3126977, EPI_ISL_3126978, EPI_ISL_3126979, EPI_ISL_3126980, EPI_ISL_3126981, EPI_ISL_3126982, EPI_ISL_3126983 | | | |
| see above | Department of Bacteria, Parasites and Fungi, Statens Serum Institut, Copenhagen, Denmark | Statens Serum Institut Bioinformatics and Microbial Genomics | Danish Covid-19 Genome Consortium |
| EPI_ISL_3126984 | Molekylær Medicinsk Afdeling, Aarhus University Hospital, Aarhus, Denmark | Statens Serum Institut Bioinformatics and Microbial Genomics | Danish Covid-19 Genome Consortium |
| EPI_ISL_3126985, EPI_ISL_3126986, EPI_ISL_3126987, EPI_ISL_3126988, EPI_ISL_3126989, EPI_ISL_3126990, EPI_ISL_3126991, EPI_ISL_3126992, EPI_ISL_3126993, EPI_ISL_3126994, EPI_ISL_3126995, EPI_ISL_3126996, EPI_ISL_3126997, EPI_ISL_3126998, EPI_ISL_3126999, EPI_ISL_3127000, EPI_ISL_3127001, EPI_ISL_3127002, EPI_ISL_3127003, EPI_ISL_3127004, EPI_ISL_3127005, EPI_ISL_3127006, EPI_ISL_3127007, EPI_ISL_3127008, EPI_ISL_3127009, EPI_ISL_3127010, EPI_ISL_3127011, EPI_ISL_3127012, EPI_ISL_3127013, EPI_ISL_3127014, EPI_ISL_3127015, EPI_ISL_3127017, EPI_ISL_3127018, EPI_ISL_3127019, EPI_ISL_3127020, EPI_ISL_3127021, EPI_ISL_3127022, EPI_ISL_3127023, EPI_ISL_3127024, EPI_ISL_3127025, EPI_ISL_3127026, EPI_ISL_3127027, EPI_ISL_3127028, EPI_ISL_3127029, EPI_ISL_3127030, EPI_ISL_3127031, EPI_ISL_3127032, EPI_ISL_3127033, EPI_ISL_3127034, EPI_ISL_3127035, EPI_ISL_3127036 | | | |
| see above | Department of Bacteria, Parasites and Fungi, Statens Serum Institut, Copenhagen, Denmark | Statens Serum Institut Bioinformatics and Microbial Genomics | Danish Covid-19 Genome Consortium |
| EPI_ISL_3127037 | Molekylær Medicinsk Afdeling, Aarhus University Hospital, Aarhus, Denmark | Statens Serum Institut Bioinformatics and Microbial Genomics | Danish Covid-19 Genome Consortium |
| EPI_ISL_3127038, EPI_ISL_3127039, EPI_ISL_3127040, EPI_ISL_3127041, EPI_ISL_3127042, EPI_ISL_3127043, EPI_ISL_3127044, EPI_ISL_3127045, EPI_ISL_3127046, EPI_ISL_3127047, EPI_ISL_3127048, EPI_ISL_3127049, EPI_ISL_3127050, EPI_ISL_3127051, EPI_ISL_3127052, EPI_ISL_3127053, EPI_ISL_3127054, EPI_ISL_3127055, EPI_ISL_3127056, EPI_ISL_3127057, EPI_ISL_3127058, EPI_ISL_3127059, EPI_ISL_3127060, EPI_ISL_3127061, EPI_ISL_3127062, EPI_ISL_3127063, EPI_ISL_3127064, EPI_ISL_3127065, EPI_ISL_3127066, EPI_ISL_3127067, EPI_ISL_3127068, EPI_ISL_3127070, EPI_ISL_3127071, EPI_ISL_3127072, EPI_ISL_3127073, EPI_ISL_3127074, EPI_ISL_3127075, EPI_ISL_3127076, EPI_ISL_3127077, EPI_ISL_3127078, EPI_ISL_3127079, EPI_ISL_3127080, EPI_ISL_3127081, EPI_ISL_3127082, EPI_ISL_3127083, EPI_ISL_3127084, EPI_ISL_3127085, EPI_ISL_3127086, EPI_ISL_3127087, EPI_ISL_3127088, EPI_ISL_3127089, EPI_ISL_3127090, EPI_ISL_3127091, EPI_ISL_3127093, EPI_ISL_3127094, EPI_ISL_3127095, EPI_ISL_3127096, EPI_ISL_3127097, EPI_ISL_3127098, EPI_ISL_3127099, EPI_ISL_3127100, EPI_ISL_3127101, EPI_ISL_3127102, EPI_ISL_3127103, EPI_ISL_3127104, EPI_ISL_3127105, EPI_ISL_3127106, EPI_ISL_3127107, EPI_ISL_3127108, EPI_ISL_3127109, EPI_ISL_3127110, EPI_ISL_3127111, EPI_ISL_3127112, EPI_ISL_3127113, EPI_ISL_3127114, EPI_ISL_3127115, EPI_ISL_3127116, EPI_ISL_3127117 | | | |
| see above | Department of Bacteria, Parasites and Fungi, Statens Serum Institut, Copenhagen, Denmark | Statens Serum Institut Bioinformatics and Microbial Genomics | Danish Covid-19 Genome Consortium |
| EPI_ISL_3127119, EPI_ISL_3127121, EPI_ISL_3127122, EPI_ISL_3127123, EPI_ISL_3127125, EPI_ISL_3127126, EPI_ISL_3127127, EPI_ISL_3127129, EPI_ISL_3127130, EPI_ISL_3127131, EPI_ISL_3127132, EPI_ISL_3127133, EPI_ISL_3127134, EPI_ISL_3127137, EPI_ISL_3127140, EPI_ISL_3127142, EPI_ISL_3127143, EPI_ISL_3127144, EPI_ISL_3127145, EPI_ISL_3127146, EPI_ISL_3127147, EPI_ISL_3127149, EPI_ISL_3127150, EPI_ISL_3127151, EPI_ISL_3127154, EPI_ISL_3127155, EPI_ISL_3127157, EPI_ISL_3127158, EPI_ISL_3127159, EPI_ISL_3127161, EPI_ISL_3127162, EPI_ISL_3127165, EPI_ISL_3127166, EPI_ISL_3127167, EPI_ISL_3127168, EPI_ISL_3127169, EPI_ISL_3127170, EPI_ISL_3127171, EPI_ISL_3127172, EPI_ISL_3127173, EPI_ISL_3127174, EPI_ISL_3127175, EPI_ISL_3127176, EPI_ISL_3127177, EPI_ISL_3127179, EPI_ISL_3127180, EPI_ISL_3127181, EPI_ISL_3127182, EPI_ISL_3127183, EPI_ISL_3127184, EPI_ISL_3127185, EPI_ISL_3127186, EPI_ISL_3127187, EPI_ISL_3127188, EPI_ISL_3127189, EPI_ISL_3127190, EPI_ISL_3127191, EPI_ISL_3127192, EPI_ISL_3127193, EPI_ISL_3127194, EPI_ISL_3127196, EPI_ISL_3127197, EPI_ISL_3127198, EPI_ISL_3127199, EPI_ISL_3127200 | | | |
| see above | New Mexico Department of Health Scientific Laboratory | New Mexico Department of Health Scientific Laboratory | Ellie Johnson, D'eldra Malone, Jennifer Benoit, Ratheesh Rajan, Linda Salazar, T. Mark Willmon, Anastacia Griego-Fisher |
| EPI_ISL_3127201, EPI_ISL_3127202, EPI_ISL_3127203, EPI_ISL_3127204, EPI_ISL_3127205, EPI_ISL_3127206, EPI_ISL_3127207, EPI_ISL_3127208, EPI_ISL_3127209, EPI_ISL_3127210, EPI_ISL_3127211, EPI_ISL_3127212, EPI_ISL_3127213, EPI_ISL_3127214, EPI_ISL_3127215, EPI_ISL_3127216, EPI_ISL_3127217, EPI_ISL_3127218, EPI_ISL_3127219, EPI_ISL_3127220, EPI_ISL_3127221, EPI_ISL_3127222, EPI_ISL_3127223, EPI_ISL_3127224, EPI_ISL_3127225, EPI_ISL_3127226, EPI_ISL_3127227, EPI_ISL_3127228, EPI_ISL_3127229, EPI_ISL_3127230, EPI_ISL_3127231, EPI_ISL_3127232, EPI_ISL_3127233, EPI_ISL_3127234, EPI_ISL_3127235, EPI_ISL_3127236, EPI_ISL_3127237, EPI_ISL_3127238, EPI_ISL_3127239, EPI_ISL_3127240, EPI_ISL_3127242, EPI_ISL_3127243, EPI_ISL_3127244, EPI_ISL_3127245, EPI_ISL_3127246, EPI_ISL_3127247, EPI_ISL_3127248, EPI_ISL_3127249, EPI_ISL_3127250, EPI_ISL_3127251, EPI_ISL_3127252, EPI_ISL_3127253, EPI_ISL_3127254, EPI_ISL_3127255, EPI_ISL_3127256, EPI_ISL_3127257, EPI_ISL_3127258, EPI_ISL_3127259, EPI_ISL_3127260, EPI_ISL_3127261, EPI_ISL_3127262, EPI_ISL_3127263, EPI_ISL_3127264, EPI_ISL_3127265, EPI_ISL_3127266, EPI_ISL_3127267, EPI_ISL_3127268, EPI_ISL_3127269, EPI_ISL_3127270, EPI_ISL_3127271, EPI_ISL_3127272, EPI_ISL_3127273, EPI_ISL_3127274, EPI_ISL_3127275, EPI_ISL_3127276, EPI_ISL_3127277, EPI_ISL_3127278, EPI_ISL_3127279, EPI_ISL_3127280, EPI_ISL_3127281, EPI_ISL_3127282, EPI_ISL_3127283, EPI_ISL_3127284, EPI_ISL_3127285, EPI_ISL_3127286, EPI_ISL_3127287, EPI_ISL_3127288, EPI_ISL_3127289, EPI_ISL_3127290, EPI_ISL_3127291, EPI_ISL_3127292, EPI_ISL_3127294, EPI_ISL_3127295, EPI_ISL_3127296, EPI_ISL_3127297, EPI_ISL_3127298, EPI_ISL_3127299, EPI_ISL_3127300, EPI_ISL_3127301, EPI_ISL_3127302, EPI_ISL_3127303, EPI_ISL_3127304, EPI_ISL_3127305, EPI_ISL_3127306, EPI_ISL_3127307, EPI_ISL_3127308, EPI_ISL_3127309, EPI_ISL_3127310, EPI_ISL_3127311, EPI_ISL_3127312, EPI_ISL_3127313, EPI_ISL_3127314, EPI_ISL_3127315, EPI_ISL_3127316, EPI_ISL_3127317, EPI_ISL_3127318, EPI_ISL_3127319, EPI_ISL_3127320, EPI_ISL_3127321, EPI_ISL_3127322, EPI_ISL_3127323, EPI_ISL_3127324, EPI_ISL_3127325, EPI_ISL_3127326, EPI_ISL_3127327, EPI_ISL_3127328, EPI_ISL_3127329, EPI_ISL_3127330, EPI_ISL_3127331, EPI_ISL_3127332, EPI_ISL_3127333, EPI_ISL_3127334, EPI_ISL_3127335, EPI_ISL_3127336, EPI_ISL_3127337, EPI_ISL_3127338, EPI_ISL_3127339, EPI_ISL_3127340, EPI_ISL_3127341, EPI_ISL_3127342, EPI_ISL_3127343, EPI_ISL_3127344, EPI_ISL_3127345, EPI_ISL_3127346, EPI_ISL_3127347, EPI_ISL_3127348, EPI_ISL_3127349, EPI_ISL_3127350, EPI_ISL_3127351, EPI_ISL_3127352, EPI_ISL_3127353, EPI_ISL_3127354, EPI_ISL_3127355, EPI_ISL_3127356, EPI_ISL_3127357, EPI_ISL_3127358, EPI_ISL_3127359, EPI_ISL_3127360, EPI_ISL_3127361, EPI_ISL_3127362, EPI_ISL_3127363, EPI_ISL_3127364, EPI_ISL_3127365, EPI_ISL_3127366, EPI_ISL_3127367, EPI_ISL_3127368, EPI_ISL_3127369, EPI_ISL_3127370, EPI_ISL_3127371, EPI_ISL_3127372, EPI_ISL_3127373, EPI_ISL_3127374, EPI_ISL_3127375, EPI_ISL_3127376, EPI_ISL_3127377, EPI_ISL_3127378, EPI_ISL_3127379, EPI_ISL_3127380, EPI_ISL_3127381, EPI_ISL_3127382, EPI_ISL_3127383, EPI_ISL_3127384, EPI_ISL_3127385, EPI_ISL_3127386, EPI_ISL_3127387, EPI_ISL_3127388, EPI_ISL_3127390, EPI_ISL_3127391, EPI_ISL_3127392, EPI_ISL_3127393, EPI_ISL_3127394, EPI_ISL_3127395, EPI_ISL_3127396, EPI_ISL_3127397, EPI_ISL_3127398, EPI_ISL_3127399, EPI_ISL_3127400, EPI_ISL_3127401 | | | |
| see above | Broad Institute Clinical Research Sequencing Platform | Infectious Disease Program, Broad Institute of Harvard and MIT | Siddle,K.J., Adams,G., Pearlman,L., Gladden-Young,A., Vicente,G., Blumenstiel,B., DeFelice,M., Lee,M., McGovern,S., Lagerborg,K., Rudy,M., DeRuff,K., Carter,A., Normandin,E., Bauer,M., Reilly,S., Tomkins-Tinch,C., Loreth,C., Chaluvadi,S., Meldrim,J., Granger,B., Lemieux,J.E., Birren,B.W., Sabeti,P.C., Larkin,K., Dodge,S., Lennon,N., Madoff,L., Brown,C., Gallagher,G., Smole,S., Park,D.J., Gabriel,S., and MacInnis,B.L. |
| EPI_ISL_3127402, EPI_ISL_3127403, EPI_ISL_3127404, EPI_ISL_3127405, EPI_ISL_3127406, EPI_ISL_3127407, EPI_ISL_3127409, EPI_ISL_3127410, EPI_ISL_3127411, EPI_ISL_3127412, EPI_ISL_3127413, EPI_ISL_3127415 | | | |
| see above | A.S.L. NOVARA - ISTITUTO S.S. TRINITA' - BORGOMANERO - S.S. LABORATORIO ANALISI CHIMICO CLINICHE - | Fondazione del Piemonte per l'Oncologia IRCCS | Antonino Sottile, Silvia Brossa, Paola Marino, Giorgio Giardina |
| EPI_ISL_3127417, EPI_ISL_3127418 | A.S.L. TO4 | Fondazione del Piemonte per l'Oncologia IRCCS | Antonino Sottile, Silvia Brossa, Paola Marino, Giorgio Giardina |
| EPI_ISL_3127422, EPI_ISL_3127429, EPI_ISL_3127442 | Presidio Ospedaliero "Madonna delle Grazie" di Matera | ASM - P.O. Madonna delle Grazie - Matera | Massimo Dell'Edera |
| EPI_ISL_3127444 | State Key Laboratory for Diagnosis and Treatment of Infectious Diseases, National Clinical Research Center for Infectious Diseases, First Affiliated Hospital, Zhejiang University School of Medicine, Hangzhou, China. 310003 | State Key Laboratory for Diagnosis and Treatment of Infectious Diseases, National Clinical Research Center for Infectious Diseases, First Affiliated Hospital, Zhejiang University School of Medicine, Hangzhou, China. 310003 | Hangping Yao, Nanping Wu, Shibo Li, Xiangyun Lu, Keda Chen，Zhigang Wu, Tianhao Weng，Danrong Shi，Linfang Cheng, Fumin Liu, Haibo Wu,Changzhong Jin, Min Zheng, Lanjuan Li |
| EPI_ISL_3127445 | State Key Laboratory for Diagnosis and Treatment of Infectious Diseases, National Clinical Research Center for Infectious Diseases, First Affiliated Hospital, Zhejiang University School of Medicine, Hangzhou, China. 310003 | State Key Laboratory for Diagnosis and Treatment of Infectious Diseases, National Clinical Research Center for Infectious Diseases, First Affiliated Hospital, Zhejiang University School of Medicine, Hangzhou, China. 310003 | Hangping Yao, Nanping Wu, Shibo Li, Xiangyun Lu, Keda Chen, Zhigang Wu, Tianhao Weng, Danrong Shi, Linfang Cheng, Fumin Liu, Haibo Wu, Changzhong Jin, Min Zheng, Lanjuan Li |
| EPI_ISL_3127446, EPI_ISL_3127447, EPI_ISL_3127449 | NCSLPH | NCSLPH | Chase K, Miller MC, Greene S, Glover W |
| EPI_ISL_3127451 | WSSE w Poznaniu | Wojewodzka Stacja Sanitarno-Epidemiologiczna w Olsztynie, Laboratorium Badan Epidemiologiczno-Klinicznych | Sylwia Krzetowska, Monika Czerminska, Ewa Liszewska, Tomasz Jakubczak, Marta Lukian, Patryk Bielecki, Aleksandra Kobiatko, Emilia Tarabasz, Paulina Rozycka, Barbara Dolinska |
| EPI_ISL_3127453 | WSSE we Wroclawiu | Wojewodzka Stacja Sanitarno-Epidemiologiczna w Olsztynie, Laboratorium Badan Epidemiologiczno-Klinicznych | Sylwia Krzetowska, Monika Czerminska, Ewa Liszewska, Tomasz Jakubczak, Marta Lukian, Patryk Bielecki, Aleksandra Kobiatko, Emilia Tarabasz, Paulina Rozycka, Barbara Dolinska |
| EPI_ISL_3127454 | WSSE w Opolu | Wojewodzka Stacja Sanitarno-Epidemiologiczna w Olsztynie, | Sylwia Krzetowska, Monika Czerminska, Ewa Liszewska, Tomasz Jakubczak, Marta Lukian, Patryk Bielecki, Aleksandra Kobiatko, Emilia Tarabasz, Paulina Rozycka, Barbara Dolinska |

| | | | |
|---|---|---|---|
| | | Laboratorium Badan Epidemiologiczno-Klinicznych | |
| EPI_ISL_3127455, EPI_ISL_3127759, EPI_ISL_3127760, EPI_ISL_3127761 | NCSLPH | NCSLPH | Chase K, Miller MC, Greene S, Glover W |
| EPI_ISL_3127762, EPI_ISL_3127763, EPI_ISL_3127766 | Ospedale Regionale Umberto Parini | U.O. Genomica - Istituto Zooprofilattico Sperimentale del Piemonte, Liguria e Valle d'Aosta | Simone Peletto |
| EPI_ISL_3127767, EPI_ISL_3127768, EPI_ISL_3127769, EPI_ISL_3127770, EPI_ISL_3127771, EPI_ISL_3127772, EPI_ISL_3127773, EPI_ISL_3127774, EPI_ISL_3127775, EPI_ISL_3127777, EPI_ISL_3127778, EPI_ISL_3127779, EPI_ISL_3127780, EPI_ISL_3127781 | | | |
| see above | AULSS 9 Scaligera - Villafranca | UOSD Genetica e Citogenetica - Azienda ULSS 3 Serenissima; Istituto Zooprofilattico Sperimentale delle Venezie | Mosé Favarato, Elisa Squarcina, Laura Squarzon, Claudia Perini, Laura Bevilacqua, Adelaide Milani, Alessia Schivo, Annalisa Salviato, Elisa Palumbo, Erika Giorgia Quaranta, Luca Tassoni, Ambra Pastori, Edoardo Giussani, Alice Fusaro, Isabella Monne, Calogero Terregino, Antonia Ricci |
| EPI_ISL_3127782, EPI_ISL_3127783, EPI_ISL_3127784, EPI_ISL_3127785 | AULSS 3 Venezia | UOSD Genetica e Citogenetica - Azienda ULSS 3 Serenissima; Istituto Zooprofilattico Sperimentale delle Venezie | Mosé Favarato, Elisa Squarcina, Laura Squarzon, Claudia Perini, Laura Bevilacqua, Adelaide Milani, Alessia Schivo, Annalisa Salviato, Elisa Palumbo, Erika Giorgia Quaranta, Luca Tassoni, Ambra Pastori, Edoardo Giussani, Alice Fusaro, Isabella Monne, Calogero Terregino, Antonia Ricci |
| EPI_ISL_3127786 | AULSS 9 Scaligera - Villafranca | UOSD Genetica e Citogenetica - Azienda ULSS 3 Serenissima; Istituto Zooprofilattico Sperimentale delle Venezie | Mosé Favarato, Elisa Squarcina, Laura Squarzon, Claudia Perini, Laura Bevilacqua, Adelaide Milani, Alessia Schivo, Annalisa Salviato, Elisa Palumbo, Erika Giorgia Quaranta, Luca Tassoni, Ambra Pastori, Edoardo Giussani, Alice Fusaro, Isabella Monne, Calogero Terregino, Antonia Ricci |
| EPI_ISL_3127788, EPI_ISL_3127789, EPI_ISL_3127790, EPI_ISL_3127791, EPI_ISL_3127792, EPI_ISL_3127793, EPI_ISL_3127794, EPI_ISL_3127795, EPI_ISL_3127796, EPI_ISL_3127797, EPI_ISL_3127798 | | | |
| see above | AULSS 3 Venezia | UOSD Genetica e Citogenetica - Azienda ULSS 3 Serenissima; Istituto Zooprofilattico Sperimentale delle Venezie | Mosé Favarato, Elisa Squarcina, Laura Squarzon, Claudia Perini, Laura Bevilacqua, Adelaide Milani, Alessia Schivo, Annalisa Salviato, Elisa Palumbo, Erika Giorgia Quaranta, Luca Tassoni, Ambra Pastori, Edoardo Giussani, Alice Fusaro, Isabella Monne, Calogero Terregino, Antonia Ricci |
| EPI_ISL_3127799 | AULSS 4 Veneto Orientale | UOSD Genetica e Citogenetica - Azienda ULSS 3 Serenissima; Istituto Zooprofilattico Sperimentale delle Venezie | Mosé Favarato, Elisa Squarcina, Laura Squarzon, Claudia Perini, Laura Bevilacqua, Adelaide Milani, Alessia Schivo, Annalisa Salviato, Elisa Palumbo, Erika Giorgia Quaranta, Luca Tassoni, Ambra Pastori, Edoardo Giussani, Alice Fusaro, Isabella Monne, Calogero Terregino, Antonia Ricci |
| EPI_ISL_3127800, EPI_ISL_3127801, EPI_ISL_3127803 | AULSS 3 Venezia | UOSD Genetica e Citogenetica - Azienda ULSS 3 Serenissima; Istituto Zooprofilattico Sperimentale delle Venezie | Mosé Favarato, Elisa Squarcina, Laura Squarzon, Claudia Perini, Laura Bevilacqua, Adelaide Milani, Alessia Schivo, Annalisa Salviato, Elisa Palumbo, Erika Giorgia Quaranta, Luca Tassoni, Ambra Pastori, Edoardo Giussani, Alice Fusaro, Isabella Monne, Calogero Terregino, Antonia Ricci |
| EPI_ISL_3127804, EPI_ISL_3127805, EPI_ISL_3127806 | Center of Medical Microbiology, Virology, and Hospital Hygiene, University of Duesseldorf | Center of Medical Microbiology, Virology, and Hospital Hygiene, University of Duesseldorf | Maximilian Damagnez;Alexander Dilthey;Torsten Houwaart;Lisanna Hülse;Malte Kohns Vasconcelos;Nadine Lübke;Jessica Nicolai;Klaus Pfeffer;Daniel Strelow;Jörg Timm;Andreas Walker;Tobias Wienemann |
| EPI_ISL_3127814, EPI_ISL_3127815, EPI_ISL_3127816, EPI_ISL_3127817, EPI_ISL_3127818, EPI_ISL_3127819, EPI_ISL_3127820, EPI_ISL_3127821, EPI_ISL_3127822, EPI_ISL_3127823, EPI_ISL_3127824, EPI_ISL_3127825, EPI_ISL_3127826, EPI_ISL_3127827, EPI_ISL_3127828, EPI_ISL_3127829, EPI_ISL_3127830, EPI_ISL_3127831, EPI_ISL_3127832, EPI_ISL_3127833, EPI_ISL_3127839, EPI_ISL_3127840, EPI_ISL_3127841, EPI_ISL_3127843, EPI_ISL_3127844, EPI_ISL_3127845, EPI_ISL_3127846 | | | |
| see above | ZOTZ KLIMAS MVZ Düsseldorf-Centrum GbR ÜBAG für Labormedizin, Genetik, Zytologie, Pathologie | Center of Medical Microbiology, Virology, and Hospital Hygiene, University of Duesseldorf | Maximilian Damagnez;Alexander Dilthey;Patrick Finzer;Katrin Hoffmann;Torsten Houwaart;Lisanna Hülse;Malte Kohns Vasconcelos;Nadine Lübke;Jessica Nicolai;Klaus Pfeffer;Daniel Strelow;Jörg Timm;Andreas Walker;Tobias Wienemann;Rainer Zotz |
| EPI_ISL_3127847, EPI_ISL_3127848 | NCSLPH | NCSLPH | Chase K, Miller MC, Greene S, Glover W |
| EPI_ISL_3127853 | SD Public Health Laboratory | Centers for Disease Control and Prevention Division of Viral Diseases, Pathogen Discovery | Mili Sheth, Sarah Nobles, Jasmine Padilla, Alex Burgin, Meghan Bentz, Mark Burroughs, Peter Cook, Clinton R. Paden, Dhwani Batra, Krista Queen, Kristen Knipe, Dakota Howard, Yvette Unoarumhi, Matthew Schmerer, Ben L. Rambo-Martin, Kristine Lacek, Sam Shepard, Dave Wentworth, Vivien Dugan, Suxiang Tong, Justin Lee |
| EPI_ISL_3127854, EPI_ISL_3127855 | GA Department of Public Health Laboratory | Centers for Disease Control and Prevention Division of Viral Diseases, Pathogen Discovery | Mili Sheth, Sarah Nobles, Jasmine Padilla, Alex Burgin, Meghan Bentz, Mark Burroughs, Peter Cook, Clinton R. Paden, Dhwani Batra, Krista Queen, Kristen Knipe, Dakota Howard, Yvette Unoarumhi, Matthew Schmerer, Ben L. Rambo-Martin, Kristine Lacek, Sam Shepard, Dave Wentworth, Vivien Dugan, Suxiang Tong, Justin Lee |
| EPI_ISL_3127856 | ND Dept. of Health Laboratory Services-Microbiology | Centers for Disease Control and Prevention Division of Viral Diseases, Pathogen Discovery | Mili Sheth, Sarah Nobles, Jasmine Padilla, Alex Burgin, Meghan Bentz, Mark Burroughs, Peter Cook, Clinton R. Paden, Dhwani Batra, Krista Queen, Kristen Knipe, Dakota Howard, Yvette Unoarumhi, Matthew Schmerer, Ben L. Rambo-Martin, Kristine Lacek, Sam Shepard, Dave Wentworth, Vivien Dugan, Suxiang Tong, Justin Lee |
| EPI_ISL_3127858, EPI_ISL_3127859 | KS Health and Environmental Laboratories | Centers for Disease Control and Prevention Division of Viral Diseases, Pathogen Discovery | Mili Sheth, Sarah Nobles, Jasmine Padilla, Alex Burgin, Meghan Bentz, Mark Burroughs, Peter Cook, Clinton R. Paden, Dhwani Batra, Krista Queen, Kristen Knipe, Dakota Howard, Yvette Unoarumhi, Matthew Schmerer, Ben L. Rambo-Martin, Kristine Lacek, Sam Shepard, Dave Wentworth, Vivien Dugan, Suxiang Tong, Justin Lee |
| EPI_ISL_3127860, EPI_ISL_3127861, EPI_ISL_3127862, EPI_ISL_3127863, EPI_ISL_3127864 | RI State Health Laboratories | Centers for Disease Control and Prevention Division of Viral Diseases, Pathogen Discovery | Mili Sheth, Sarah Nobles, Jasmine Padilla, Alex Burgin, Meghan Bentz, Mark Burroughs, Peter Cook, Clinton R. Paden, Dhwani Batra, Krista Queen, Kristen Knipe, Dakota Howard, Yvette Unoarumhi, Matthew Schmerer, Ben L. Rambo-Martin, Kristine Lacek, Sam Shepard, Dave Wentworth, Vivien Dugan, Suxiang Tong, Justin Lee |
| EPI_ISL_3127867, EPI_ISL_3127871, EPI_ISL_3127874, EPI_ISL_3127877, EPI_ISL_3127880, EPI_ISL_3127881, EPI_ISL_3127883, EPI_ISL_3127886 | NV-Southern Nevada Public Health Laboratory | Centers for Disease Control and Prevention Division of Viral Diseases, Pathogen Discovery | Mili Sheth, Sarah Nobles, Jasmine Padilla, Alex Burgin, Meghan Bentz, Mark Burroughs, Peter Cook, Clinton R. Paden, Dhwani Batra, Krista Queen, Kristen Knipe, Dakota Howard, Yvette Unoarumhi, Matthew Schmerer, Ben L. Rambo-Martin, Kristine Lacek, Sam Shepard, Dave Wentworth, Vivien Dugan, Suxiang Tong, Justin Lee |
| EPI_ISL_3127892, EPI_ISL_3127893, EPI_ISL_3127894, EPI_ISL_3127895, EPI_ISL_3127896, EPI_ISL_3127897 | AL Dept. of Public Health Bureau of Clinical Laboratories | Centers for Disease Control and Prevention Division of Viral Diseases, Pathogen Discovery | Mili Sheth, Sarah Nobles, Jasmine Padilla, Alex Burgin, Meghan Bentz, Mark Burroughs, Peter Cook, Clinton R. Paden, Dhwani Batra, Krista Queen, Kristen Knipe, Dakota Howard, Yvette Unoarumhi, Matthew Schmerer, Ben L. Rambo-Martin, Kristine Lacek, Sam Shepard, Dave Wentworth, Vivien Dugan, Suxiang Tong, Justin Lee |
| EPI_ISL_3127898 | Commonwealth Healthcare Center | Centers for Disease Control and Prevention Division of Viral Diseases, Pathogen Discovery | Mili Sheth, Sarah Nobles, Jasmine Padilla, Alex Burgin, Meghan Bentz, Mark Burroughs, Peter Cook, Clinton R. Paden, Dhwani Batra, Krista Queen, Kristen Knipe, Dakota Howard, Yvette Unoarumhi, Matthew Schmerer, Ben L. Rambo-Martin, Kristine Lacek, Sam Shepard, Dave Wentworth, Vivien Dugan, Suxiang Tong, Justin Lee |
| EPI_ISL_3127899, EPI_ISL_3127900, EPI_ISL_3127901, EPI_ISL_3127902, EPI_ISL_3127903, EPI_ISL_3127904 | KY State Public Health Lab | Centers for Disease Control and Prevention Division of Viral Diseases, Pathogen Discovery | Mili Sheth, Sarah Nobles, Jasmine Padilla, Alex Burgin, Meghan Bentz, Mark Burroughs, Peter Cook, Clinton R. Paden, Dhwani Batra, Krista Queen, Kristen Knipe, Dakota Howard, Yvette Unoarumhi, Matthew Schmerer, Ben L. Rambo-Martin, Kristine Lacek, Sam Shepard, Dave Wentworth, Vivien Dugan, Suxiang Tong, Justin Lee |
| EPI_ISL_3127914, EPI_ISL_3127916, EPI_ISL_3127917, EPI_ISL_3127919, EPI_ISL_3127921, EPI_ISL_3127922, EPI_ISL_3127923, EPI_ISL_3127924, EPI_ISL_3127925, EPI_ISL_3127926 | PR Public Health Lab | Centers for Disease Control and Prevention Division of Viral Diseases, Pathogen Discovery | Mili Sheth, Sarah Nobles, Jasmine Padilla, Alex Burgin, Meghan Bentz, Mark Burroughs, Peter Cook, Clinton R. Paden, Dhwani Batra, Krista Queen, Kristen Knipe, Dakota Howard, Yvette Unoarumhi, Matthew Schmerer, Ben L. Rambo-Martin, Kristine Lacek, Sam Shepard, Dave Wentworth, Vivien Dugan, Suxiang Tong, Justin Lee |
| EPI_ISL_3127930, EPI_ISL_3127931, EPI_ISL_3127932, EPI_ISL_3127933, EPI_ISL_3127934, EPI_ISL_3127935, EPI_ISL_3127936, EPI_ISL_3127937, EPI_ISL_3127938, EPI_ISL_3127939, EPI_ISL_3127940, EPI_ISL_3127941, EPI_ISL_3127942, EPI_ISL_3127943, EPI_ISL_3127944, EPI_ISL_3127945, EPI_ISL_3127946, EPI_ISL_3127947, EPI_ISL_3127948, EPI_ISL_3127949, EPI_ISL_3127950 | | | |
| see above | "CO Dept. of Public Health and Environment, Lab Services Division" | Centers for Disease Control and Prevention Division of Viral Diseases, Pathogen Discovery | Mili Sheth, Sarah Nobles, Jasmine Padilla, Alex Burgin, Meghan Bentz, Mark Burroughs, Peter Cook, Clinton R. Paden, Dhwani Batra, Krista Queen, Kristen Knipe, Dakota Howard, Yvette Unoarumhi, Matthew Schmerer, Ben L. Rambo-Martin, Kristine Lacek, Sam Shepard, Dave Wentworth, Vivien Dugan, Suxiang Tong, Justin Lee |
| EPI_ISL_3127951, EPI_ISL_3127952, EPI_ISL_3127953, EPI_ISL_3127954, EPI_ISL_3127955, EPI_ISL_3127956 | "NM Dept. Health, Scientific Laboratory Division " | Centers for Disease Control and Prevention Division of Viral Diseases, Pathogen Discovery | Mili Sheth, Sarah Nobles, Jasmine Padilla, Alex Burgin, Meghan Bentz, Mark Burroughs, Peter Cook, Clinton R. Paden, Dhwani Batra, Krista Queen, Kristen Knipe, Dakota Howard, Yvette Unoarumhi, Matthew Schmerer, Ben L. Rambo-Martin, Kristine Lacek, Sam Shepard, Dave Wentworth, Vivien Dugan, Suxiang Tong, Justin Lee |
| EPI_ISL_3127957 | WA State Department of Health | Centers for Disease Control and Prevention Division of Viral Diseases, Pathogen Discovery | Mili Sheth, Sarah Nobles, Jasmine Padilla, Alex Burgin, Meghan Bentz, Mark Burroughs, Peter Cook, Clinton R. Paden, Dhwani Batra, Krista Queen, Kristen Knipe, Dakota Howard, Yvette Unoarumhi, Matthew Schmerer, Ben L. Rambo-Martin, Kristine Lacek, Sam Shepard, Dave Wentworth, Vivien Dugan, Suxiang Tong, Justin Lee |
| EPI_ISL_3127958 | UW Virology Lab | UW Virology Lab | Pavitra Roychoudhury, Shah Mohamed Bakhash, Ricardo Perez, Maria Lukes, Hong Xie, Tien V. Nguyen, Noah R. Baker, Sean Ellis, Meei-Li Huang, Keith R Jerome, Alexander Greninger |
| EPI_ISL_3127960 | WA State Department of Health | Centers for Disease Control and Prevention Division of Viral Diseases, Pathogen Discovery | Mili Sheth, Sarah Nobles, Jasmine Padilla, Alex Burgin, Meghan Bentz, Mark Burroughs, Peter Cook, Clinton R. Paden, Dhwani Batra, Krista Queen, Kristen Knipe, Dakota Howard, Yvette Unoarumhi, Matthew Schmerer, Ben L. Rambo-Martin, Kristine Lacek, Sam Shepard, Dave Wentworth, Vivien Dugan, Suxiang Tong, Justin Lee |
| EPI_ISL_3127962 | UW Virology Lab | UW Virology Lab | Pavitra Roychoudhury, Shah Mohamed Bakhash, Ricardo Perez, Maria Lukes, Hong Xie, Tien V. Nguyen, Noah R. Baker, Sean Ellis, Meei-Li Huang, Keith R Jerome, Alexander Greninger |
| EPI_ISL_3127963, EPI_ISL_3127966, | WA State Department of Health | Centers for Disease Control and Prevention Division of Viral Diseases, Pathogen Discovery | Mili Sheth, Sarah Nobles, Jasmine Padilla, Alex Burgin, Meghan Bentz, Mark Burroughs, Peter Cook, Clinton R. Paden, Dhwani Batra, Krista Queen, Kristen Knipe, Dakota Howard, Yvette Unoarumhi, Matthew Schmerer, Ben L. Rambo-Martin, Kristine Lacek, Sam Shepard, Dave Wentworth, Vivien Dugan, Suxiang Tong, Justin Lee |

| | | | |
|---|---|---|---|
| EPI_ISL_3127969, EPI_ISL_3127970 | | | |
| EPI_ISL_3127974, EPI_ISL_3127975, EPI_ISL_3127976, EPI_ISL_3127977, EPI_ISL_3127978, EPI_ISL_3127979, EPI_ISL_3127980, EPI_ISL_3127981, EPI_ISL_3127982 | SD Public Health Laboratory | Centers for Disease Control and Prevention Division of Viral Diseases, Pathogen Discovery | Mili Sheth, Sarah Nobles, Jasmine Padilla, Alex Burgin, Meghan Bentz, Mark Burroughs, Peter Cook, Clinton R. Paden, Dhwani Batra, Krista Queen, Kristen Knipe, Dakota Howard, Yvette Unoarumhi, Matthew Schmerer, Ben L. Rambo-Martin, Kristine Lacek, Sam Shepard, Dave Wentworth, Vivien Dugan, Suxiang Tong, Justin Lee |
| EPI_ISL_3127983, EPI_ISL_3127985, EPI_ISL_3127986, EPI_ISL_3127990, EPI_ISL_3127992, EPI_ISL_3127993 | IA State Hygienic Laboratory | Centers for Disease Control and Prevention Division of Viral Diseases, Pathogen Discovery | Mili Sheth, Sarah Nobles, Jasmine Padilla, Alex Burgin, Meghan Bentz, Mark Burroughs, Peter Cook, Clinton R. Paden, Dhwani Batra, Krista Queen, Kristen Knipe, Dakota Howard, Yvette Unoarumhi, Matthew Schmerer, Ben L. Rambo-Martin, Kristine Lacek, Sam Shepard, Dave Wentworth, Vivien Dugan, Suxiang Tong, Justin Lee |
| EPI_ISL_3128000, EPI_ISL_3128001, EPI_ISL_3128002 | CT-Dr. Katherine A. Kelley State Public Health Lab | Centers for Disease Control and Prevention Division of Viral Diseases, Pathogen Discovery | Mili Sheth, Sarah Nobles, Jasmine Padilla, Alex Burgin, Meghan Bentz, Mark Burroughs, Peter Cook, Clinton R. Paden, Dhwani Batra, Krista Queen, Kristen Knipe, Dakota Howard, Yvette Unoarumhi, Matthew Schmerer, Ben L. Rambo-Martin, Kristine Lacek, Sam Shepard, Dave Wentworth, Vivien Dugan, Suxiang Tong, Justin Lee |
| EPI_ISL_3128003, EPI_ISL_3128004, EPI_ISL_3128005, EPI_ISL_3128006, EPI_ISL_3128007, EPI_ISL_3128008, EPI_ISL_3128009, EPI_ISL_3128010, EPI_ISL_3128011, EPI_ISL_3128012, EPI_ISL_3128013, EPI_ISL_3128014, EPI_ISL_3128015, EPI_ISL_3128016 | | | |
| see above | UT-Unified State Labs: Public Health Utah DOH | Centers for Disease Control and Prevention Division of Viral Diseases, Pathogen Discovery | Mili Sheth, Sarah Nobles, Jasmine Padilla, Alex Burgin, Meghan Bentz, Mark Burroughs, Peter Cook, Clinton R. Paden, Dhwani Batra, Krista Queen, Kristen Knipe, Dakota Howard, Yvette Unoarumhi, Matthew Schmerer, Ben L. Rambo-Martin, Kristine Lacek, Sam Shepard, Dave Wentworth, Vivien Dugan, Suxiang Tong, Justin Lee |
| EPI_ISL_3128017 | NCSLPH | NCSLPH | Chase K, Miller MC, Greene S, Glover W |
| EPI_ISL_3128018, EPI_ISL_3128019, EPI_ISL_3128020, EPI_ISL_3128021, EPI_ISL_3128022 | UT-Unified State Labs: Public Health Utah DOH | Centers for Disease Control and Prevention Division of Viral Diseases, Pathogen Discovery | Mili Sheth, Sarah Nobles, Jasmine Padilla, Alex Burgin, Meghan Bentz, Mark Burroughs, Peter Cook, Clinton R. Paden, Dhwani Batra, Krista Queen, Kristen Knipe, Dakota Howard, Yvette Unoarumhi, Matthew Schmerer, Ben L. Rambo-Martin, Kristine Lacek, Sam Shepard, Dave Wentworth, Vivien Dugan, Suxiang Tong, Justin Lee |
| EPI_ISL_3128025, EPI_ISL_3128027, EPI_ISL_3128037 | ID Bureau of Laboratories | Centers for Disease Control and Prevention Division of Viral Diseases, Pathogen Discovery | Mili Sheth, Sarah Nobles, Jasmine Padilla, Alex Burgin, Meghan Bentz, Mark Burroughs, Peter Cook, Clinton R. Paden, Dhwani Batra, Krista Queen, Kristen Knipe, Dakota Howard, Yvette Unoarumhi, Matthew Schmerer, Ben L. Rambo-Martin, Kristine Lacek, Sam Shepard, Dave Wentworth, Vivien Dugan, Suxiang Tong, Justin Lee |
| EPI_ISL_3128038, EPI_ISL_3128039, EPI_ISL_3128040, EPI_ISL_3128041, EPI_ISL_3128042, EPI_ISL_3128043, EPI_ISL_3128044 | WY Public Health Laboratory | Centers for Disease Control and Prevention Division of Viral Diseases, Pathogen Discovery | Mili Sheth, Sarah Nobles, Jasmine Padilla, Alex Burgin, Meghan Bentz, Mark Burroughs, Peter Cook, Clinton R. Paden, Dhwani Batra, Krista Queen, Kristen Knipe, Dakota Howard, Yvette Unoarumhi, Matthew Schmerer, Ben L. Rambo-Martin, Kristine Lacek, Sam Shepard, Dave Wentworth, Vivien Dugan, Suxiang Tong, Justin Lee |
| EPI_ISL_3128057, EPI_ISL_3128058, EPI_ISL_3128059, EPI_ISL_3128060, EPI_ISL_3128061, EPI_ISL_3128062, EPI_ISL_3128063 | "AR Dept. of Health-PHL, Molecular Diagnostics" | Centers for Disease Control and Prevention Division of Viral Diseases, Pathogen Discovery | Mili Sheth, Sarah Nobles, Jasmine Padilla, Alex Burgin, Meghan Bentz, Mark Burroughs, Peter Cook, Clinton R. Paden, Dhwani Batra, Krista Queen, Kristen Knipe, Dakota Howard, Yvette Unoarumhi, Matthew Schmerer, Ben L. Rambo-Martin, Kristine Lacek, Sam Shepard, Dave Wentworth, Vivien Dugan, Suxiang Tong, Justin Lee |
| EPI_ISL_3128064, EPI_ISL_3128065, EPI_ISL_3128066, EPI_ISL_3128068, EPI_ISL_3128070, EPI_ISL_3128071 | NE Public Health Laboratory | Centers for Disease Control and Prevention Division of Viral Diseases, Pathogen Discovery | Mili Sheth, Sarah Nobles, Jasmine Padilla, Alex Burgin, Meghan Bentz, Mark Burroughs, Peter Cook, Clinton R. Paden, Dhwani Batra, Krista Queen, Kristen Knipe, Dakota Howard, Yvette Unoarumhi, Matthew Schmerer, Ben L. Rambo-Martin, Kristine Lacek, Sam Shepard, Dave Wentworth, Vivien Dugan, Suxiang Tong, Justin Lee |
| EPI_ISL_3128072, EPI_ISL_3128074, EPI_ISL_3128076 | MI - Michigan Department of Health and Human Services - Bureau of Laboratories | Centers for Disease Control and Prevention Division of Viral Diseases, Pathogen Discovery | Mili Sheth, Sarah Nobles, Jasmine Padilla, Alex Burgin, Meghan Bentz, Mark Burroughs, Peter Cook, Clinton R. Paden, Dhwani Batra, Krista Queen, Kristen Knipe, Dakota Howard, Yvette Unoarumhi, Matthew Schmerer, Ben L. Rambo-Martin, Kristine Lacek, Sam Shepard, Dave Wentworth, Vivien Dugan, Suxiang Tong, Justin Lee |
| EPI_ISL_3128083, EPI_ISL_3128084, EPI_ISL_3128089, EPI_ISL_3128090, EPI_ISL_3128093, EPI_ISL_3128095, EPI_ISL_3128098, EPI_ISL_3128099, EPI_ISL_3128104, EPI_ISL_3128105 | GA Department of Public Health Laboratory | Centers for Disease Control and Prevention Division of Viral Diseases, Pathogen Discovery | Mili Sheth, Sarah Nobles, Jasmine Padilla, Alex Burgin, Meghan Bentz, Mark Burroughs, Peter Cook, Clinton R. Paden, Dhwani Batra, Krista Queen, Kristen Knipe, Dakota Howard, Yvette Unoarumhi, Matthew Schmerer, Ben L. Rambo-Martin, Kristine Lacek, Sam Shepard, Dave Wentworth, Vivien Dugan, Suxiang Tong, Justin Lee |
| EPI_ISL_3128111, EPI_ISL_3128112, EPI_ISL_3128113, EPI_ISL_3128114 | "OK Public Health Laboratory, Oklahoma State DOH" | Centers for Disease Control and Prevention Division of Viral Diseases, Pathogen Discovery | Mili Sheth, Sarah Nobles, Jasmine Padilla, Alex Burgin, Meghan Bentz, Mark Burroughs, Peter Cook, Clinton R. Paden, Dhwani Batra, Krista Queen, Kristen Knipe, Dakota Howard, Yvette Unoarumhi, Matthew Schmerer, Ben L. Rambo-Martin, Kristine Lacek, Sam Shepard, Dave Wentworth, Vivien Dugan, Suxiang Tong, Justin Lee |
| EPI_ISL_3128115, EPI_ISL_3128116, EPI_ISL_3128117, EPI_ISL_3128118, EPI_ISL_3128121, EPI_ISL_3128123 | LA Office of Public Health Laboratories | Centers for Disease Control and Prevention Division of Viral Diseases, Pathogen Discovery | Mili Sheth, Sarah Nobles, Jasmine Padilla, Alex Burgin, Meghan Bentz, Mark Burroughs, Peter Cook, Clinton R. Paden, Dhwani Batra, Krista Queen, Kristen Knipe, Dakota Howard, Yvette Unoarumhi, Matthew Schmerer, Ben L. Rambo-Martin, Kristine Lacek, Sam Shepard, Dave Wentworth, Vivien Dugan, Suxiang Tong, Justin Lee |
| EPI_ISL_3128125, EPI_ISL_3128126, EPI_ISL_3128127, EPI_ISL_3128128, EPI_ISL_3128129 | ND Dept. of Health Laboratory Services-Microbiology | Centers for Disease Control and Prevention Division of Viral Diseases, Pathogen Discovery | Mili Sheth, Sarah Nobles, Jasmine Padilla, Alex Burgin, Meghan Bentz, Mark Burroughs, Peter Cook, Clinton R. Paden, Dhwani Batra, Krista Queen, Kristen Knipe, Dakota Howard, Yvette Unoarumhi, Matthew Schmerer, Ben L. Rambo-Martin, Kristine Lacek, Sam Shepard, Dave Wentworth, Vivien Dugan, Suxiang Tong, Justin Lee |
| EPI_ISL_3128130, EPI_ISL_3128133, EPI_ISL_3128134, EPI_ISL_3128135, EPI_ISL_3128137 | "AZ SPHL, Arizona Department of Health Services" | Centers for Disease Control and Prevention Division of Viral Diseases, Pathogen Discovery | Mili Sheth, Sarah Nobles, Jasmine Padilla, Alex Burgin, Meghan Bentz, Mark Burroughs, Peter Cook, Clinton R. Paden, Dhwani Batra, Krista Queen, Kristen Knipe, Dakota Howard, Yvette Unoarumhi, Matthew Schmerer, Ben L. Rambo-Martin, Kristine Lacek, Sam Shepard, Dave Wentworth, Vivien Dugan, Suxiang Tong, Justin Lee |
| EPI_ISL_3128138 | OH Department of Health Laboratory | Centers for Disease Control and Prevention Division of Viral Diseases, Pathogen Discovery | Mili Sheth, Sarah Nobles, Jasmine Padilla, Alex Burgin, Meghan Bentz, Mark Burroughs, Peter Cook, Clinton R. Paden, Dhwani Batra, Krista Queen, Kristen Knipe, Dakota Howard, Yvette Unoarumhi, Matthew Schmerer, Ben L. Rambo-Martin, Kristine Lacek, Sam Shepard, Dave Wentworth, Vivien Dugan, Suxiang Tong, Justin Lee |
| EPI_ISL_3128143 | KS Health and Environmental Laboratories | Centers for Disease Control and Prevention Division of Viral Diseases, Pathogen Discovery | Mili Sheth, Sarah Nobles, Jasmine Padilla, Alex Burgin, Meghan Bentz, Mark Burroughs, Peter Cook, Clinton R. Paden, Dhwani Batra, Krista Queen, Kristen Knipe, Dakota Howard, Yvette Unoarumhi, Matthew Schmerer, Ben L. Rambo-Martin, Kristine Lacek, Sam Shepard, Dave Wentworth, Vivien Dugan, Suxiang Tong, Justin Lee |
| EPI_ISL_3128144 | NCSLPH | NCSLPH | Chase K, Miller MC, Greene S, Glover W |
| EPI_ISL_3128145, EPI_ISL_3128146 | ZOTZ KLIMAS MVZ Düsseldorf-Centrum GbR ÜBAG für Labormedizin, Genetik, Zytologie, Pathologie | Center of Medical Microbiology, Virology, and Hospital Hygiene, University of Duesseldorf | Maximilian Damagnez;Alexander Dilthey;Patrick Finzer;Katrin Hoffmann;Torsten Houwaart;Lisanna Hülse;Malte Kohns Vasconcelos;Nadine Lübke;Jessica Nicolai;Klaus Pfeffer;Daniel Strelow;Jörg Timm;Andreas Walker;Tobias Wienemann;Rainer Zotz |
| EPI_ISL_3128147, EPI_ISL_3128149, EPI_ISL_3128150, EPI_ISL_3128151, EPI_ISL_3128153 | NCSLPH | NCSLPH | Chase K, Miller MC, Greene S, Glover W |
| EPI_ISL_3128459 | ASL GENZANO | University Campus Bio-Medico of Rome (UCBM) | Riva E., De Florio L., Fogolari M., Francesconi M., Lintas C., Veralli R., Angeletti S. |
| EPI_ISL_3128461, EPI_ISL_3128462, EPI_ISL_3128463, EPI_ISL_3128464, EPI_ISL_3128465, EPI_ISL_3128466, EPI_ISL_3128467, EPI_ISL_3128468, EPI_ISL_3128469, EPI_ISL_3128470, EPI_ISL_3128472, EPI_ISL_3128475, EPI_ISL_3128477, EPI_ISL_3128478, EPI_ISL_3128479, EPI_ISL_3128481, EPI_ISL_3128482, EPI_ISL_3128493, EPI_ISL_3128494, EPI_ISL_3128496, EPI_ISL_3128499, EPI_ISL_3128500, EPI_ISL_3128502, EPI_ISL_3128503, EPI_ISL_3128504, EPI_ISL_3128505, EPI_ISL_3128506, EPI_ISL_3128507, EPI_ISL_3128508, EPI_ISL_3128509, EPI_ISL_3128510, EPI_ISL_3128511, EPI_ISL_3128513, EPI_ISL_3128515, EPI_ISL_3128516, EPI_ISL_3128517, EPI_ISL_3128518, EPI_ISL_3128519, EPI_ISL_3128520 | | | |

| | | | |
|---|---|---|---|
| see above | Laboratorio di Riferimento Regionale della Sicilia Occidentale per l'Emergenza COVID-19 | Laboratorio di Riferimento Regionale della Sicilia Occidentale per l'Emergenza COVID-19 | Fabio Tramuto, Carmelo Massimo Maida, Daniela Di Naro, Giulia Randazzo, Walter Mazzucco, Giorgio Graziano, Vincenzo Restivo, Claudio Costantino, Francesco Vitale |
| EPI_ISL_3128536 | Sciensano - Animal diseases | Sciensano - Animal diseases | Elisabeth Mathijs, Frank Vandenbussche, Nadège Balmelle, Anne Declercq, Ann Brigitte Cay |
| EPI_ISL_3128540 | Centre Hospitalier Universitaire Clermont-Ferrand | CHU Clermont-Ferrand, service de virologie | Bisseux Maxime, Mirand Audrey, Combes Patricia, Henquell Cécile |
| EPI_ISL_3128541 | Hospital de Vichy | CHU Clermont-Ferrand, service de virologie | Bisseux Maxime, Mirand Audrey, Combes Patricia, Henquell Cécile |
| EPI_ISL_3128542 | Centre Hospitalier Universitaire Clermont-Ferrand | CHU Clermont-Ferrand, service de virologie | Bisseux Maxime, Mirand Audrey, Combes Patricia, Henquell Cécile |
| EPI_ISL_3128543 | Hospital de Vichy | CHU Clermont-Ferrand, service de virologie | Bisseux Maxime, Mirand Audrey, Combes Patricia, Henquell Cécile |
| EPI_ISL_3128544, EPI_ISL_3128545, EPI_ISL_3128546, EPI_ISL_3128547, EPI_ISL_3128548, EPI_ISL_3128549, EPI_ISL_3128550 | Centre Hospitalier Universitaire Clermont-Ferrand | CHU Clermont-Ferrand, service de virologie | Bisseux Maxime, Mirand Audrey, Combes Patricia, Henquell Cécile |
| EPI_ISL_3128560 | Center of Medical Microbiology, Virology, and Hospital Hygiene, University of Duesseldorf | Center of Medical Microbiology, Virology, and Hospital Hygiene, University of Duesseldorf | Maximilian Damagnez;Alexander Dilthey;Torsten Houwaart;Lisanna Hülse;Malte Kohns Vasconcelos;Nadine Lübke;Jessica Nicolai;Klaus Pfeffer;Daniel Strelow;Jörg Timm;Andreas Walker;Tobias Wienemann |
| EPI_ISL_3128588, EPI_ISL_3128589, EPI_ISL_3128590, EPI_ISL_3128591, EPI_ISL_3128592, EPI_ISL_3128593, EPI_ISL_3128594, EPI_ISL_3128595, EPI_ISL_3128596, EPI_ISL_3128597, EPI_ISL_3128599, EPI_ISL_3128600, EPI_ISL_3128601, EPI_ISL_3128602, EPI_ISL_3128606, EPI_ISL_3128607, EPI_ISL_3128608, EPI_ISL_3128611, EPI_ISL_3128612, EPI_ISL_3128613, EPI_ISL_3128614, EPI_ISL_3128615, EPI_ISL_3128617, EPI_ISL_3128618, EPI_ISL_3128619, EPI_ISL_3128622, EPI_ISL_3128623, EPI_ISL_3128624, EPI_ISL_3128625, EPI_ISL_3128626 | | | |
| see above | ZOTZ KLIMAS MVZ Düsseldorf-Centrum GbR ÜBAG für Labormedizin, Genetik, Zytologie, Pathologie | Center of Medical Microbiology, Virology, and Hospital Hygiene, University of Duesseldorf | Maximilian Damagnez;Alexander Dilthey;Patrick Finzer;Katrin Hoffmann;Torsten Houwaart;Lisanna Hülse;Malte Kohns Vasconcelos;Nadine Lübke;Jessica Nicolai;Klaus Pfeffer;Daniel Strelow;Jörg Timm;Andreas Walker;Tobias Wienemann;Rainer Zotz |
| EPI_ISL_3128722, EPI_ISL_3128723, EPI_ISL_3128724 | Stadtspital Triemli | Institute of Medical Virology | Daniel Ehrsam, Isabel Stürmer, Catharine Aquino, Joel Wirz, Weihong Qi, Hubert Rehrauer, Verena Kufner, Gabriela Ziltener, Maryam Zaheri, Stefan Schmutz, Annette Audigé, Maria Grünberg, Kevin Steiner, Jon Huder, Cyril Shah, Riccarda Capaul, Guido Bloemberg, Jürg Böni, Michael Huber, Alexandra Trkola |
| EPI_ISL_3128725 | UniversitätsSpital Zürich | Institute of Medical Virology | Daniel Ehrsam, Isabel Stürmer, Catharine Aquino, Joel Wirz, Weihong Qi, Hubert Rehrauer, Verena Kufner, Gabriela Ziltener, Maryam Zaheri, Stefan Schmutz, Annette Audigé, Maria Grünberg, Kevin Steiner, Jon Huder, Cyril Shah, Riccarda Capaul, Guido Bloemberg, Jürg Böni, Michael Huber, Alexandra Trkola |
| EPI_ISL_3128726, EPI_ISL_3128727 | Spital Limmattal | Institute of Medical Virology | Daniel Ehrsam, Isabel Stürmer, Catharine Aquino, Joel Wirz, Weihong Qi, Hubert Rehrauer, Verena Kufner, Gabriela Ziltener, Maryam Zaheri, Stefan Schmutz, Annette Audigé, Maria Grünberg, Kevin Steiner, Jon Huder, Cyril Shah, Riccarda Capaul, Guido Bloemberg, Jürg Böni, Michael Huber, Alexandra Trkola |
| EPI_ISL_3128728, EPI_ISL_3128729 | UniversitätsSpital Zürich | Institute of Medical Virology | Daniel Ehrsam, Isabel Stürmer, Catharine Aquino, Joel Wirz, Weihong Qi, Hubert Rehrauer, Verena Kufner, Gabriela Ziltener, Maryam Zaheri, Stefan Schmutz, Annette Audigé, Maria Grünberg, Kevin Steiner, Jon Huder, Cyril Shah, Riccarda Capaul, Guido Bloemberg, Jürg Böni, Michael Huber, Alexandra Trkola |
| EPI_ISL_3128730 | Zentrallabor Zürich | Institute of Medical Virology | Daniel Ehrsam, Isabel Stürmer, Catharine Aquino, Joel Wirz, Weihong Qi, Hubert Rehrauer, Verena Kufner, Gabriela Ziltener, Maryam Zaheri, Stefan Schmutz, Annette Audigé, Maria Grünberg, Kevin Steiner, Jon Huder, Cyril Shah, Riccarda Capaul, Guido Bloemberg, Jürg Böni, Michael Huber, Alexandra Trkola |
| EPI_ISL_3128731, EPI_ISL_3128732, EPI_ISL_3128733, EPI_ISL_3128734, EPI_ISL_3128735 | Spital Limmattal | Institute of Medical Virology | Daniel Ehrsam, Isabel Stürmer, Catharine Aquino, Joel Wirz, Weihong Qi, Hubert Rehrauer, Verena Kufner, Gabriela Ziltener, Maryam Zaheri, Stefan Schmutz, Annette Audigé, Maria Grünberg, Kevin Steiner, Jon Huder, Cyril Shah, Riccarda Capaul, Guido Bloemberg, Jürg Böni, Michael Huber, Alexandra Trkola |
| EPI_ISL_3128736, EPI_ISL_3128737, EPI_ISL_3128738, EPI_ISL_3128739, EPI_ISL_3128740, EPI_ISL_3128741, EPI_ISL_3128742, EPI_ISL_3128743, EPI_ISL_3128744, EPI_ISL_3128745, EPI_ISL_3128746, EPI_ISL_3128747 | | | |
| see above | Stadtspital Triemli | Institute of Medical Virology | Daniel Ehrsam, Isabel Stürmer, Catharine Aquino, Joel Wirz, Weihong Qi, Hubert Rehrauer, Verena Kufner, Gabriela Ziltener, Maryam Zaheri, Stefan Schmutz, Annette Audigé, Maria Grünberg, Kevin Steiner, Jon Huder, Cyril Shah, Riccarda Capaul, Guido Bloemberg, Jürg Böni, Michael Huber, Alexandra Trkola |
| EPI_ISL_3128748, EPI_ISL_3128749 | UniversitätsSpital Zürich | Institute of Medical Virology | Daniel Ehrsam, Isabel Stürmer, Catharine Aquino, Joel Wirz, Weihong Qi, Hubert Rehrauer, Verena Kufner, Gabriela Ziltener, Maryam Zaheri, Stefan Schmutz, Annette Audigé, Maria Grünberg, Kevin Steiner, Jon Huder, Cyril Shah, Riccarda Capaul, Guido Bloemberg, Jürg Böni, Michael Huber, Alexandra Trkola |
| EPI_ISL_3128750 | Spital Männedorf AG | Institute of Medical Virology | Daniel Ehrsam, Isabel Stürmer, Catharine Aquino, Joel Wirz, Weihong Qi, Hubert Rehrauer, Verena Kufner, Gabriela Ziltener, Maryam Zaheri, Stefan Schmutz, Annette Audigé, Maria Grünberg, Kevin Steiner, Jon Huder, Cyril Shah, Riccarda Capaul, Guido Bloemberg, Jürg Böni, Michael Huber, Alexandra Trkola |
| EPI_ISL_3128751 | Arzthaus Zürich Stadelhofen | Institute of Medical Virology | Daniel Ehrsam, Isabel Stürmer, Catharine Aquino, Joel Wirz, Weihong Qi, Hubert Rehrauer, Verena Kufner, Gabriela Ziltener, Maryam Zaheri, Stefan Schmutz, Annette Audigé, Maria Grünberg, Kevin Steiner, Jon Huder, Cyril Shah, Riccarda Capaul, Guido Bloemberg, Jürg Böni, Michael Huber, Alexandra Trkola |
| EPI_ISL_3128752, EPI_ISL_3128753, EPI_ISL_3128754 | Spital Limmattal | Institute of Medical Virology | Daniel Ehrsam, Isabel Stürmer, Catharine Aquino, Joel Wirz, Weihong Qi, Hubert Rehrauer, Verena Kufner, Gabriela Ziltener, Maryam Zaheri, Stefan Schmutz, Annette Audigé, Maria Grünberg, Kevin Steiner, Jon Huder, Cyril Shah, Riccarda Capaul, Guido Bloemberg, Jürg Böni, Michael Huber, Alexandra Trkola |
| EPI_ISL_3128755, EPI_ISL_3128756 | UniversitätsSpital Zürich | Institute of Medical Virology | Daniel Ehrsam, Isabel Stürmer, Catharine Aquino, Joel Wirz, Weihong Qi, Hubert Rehrauer, Verena Kufner, Gabriela Ziltener, Maryam Zaheri, Stefan Schmutz, Annette Audigé, Maria Grünberg, Kevin Steiner, Jon Huder, Cyril Shah, Riccarda Capaul, Guido Bloemberg, Jürg Böni, Michael Huber, Alexandra Trkola |
| EPI_ISL_3128757 | Arzthaus Zürich Stadelhofen | Institute of Medical Virology | Daniel Ehrsam, Isabel Stürmer, Catharine Aquino, Joel Wirz, Weihong Qi, Hubert Rehrauer, Verena Kufner, Gabriela Ziltener, Maryam Zaheri, Stefan Schmutz, Annette Audigé, Maria Grünberg, Kevin Steiner, Jon Huder, Cyril Shah, Riccarda Capaul, Guido Bloemberg, Jürg Böni, Michael Huber, Alexandra Trkola |
| EPI_ISL_3128758, EPI_ISL_3128759, EPI_ISL_3128760, EPI_ISL_3128761, EPI_ISL_3128762 | Universitäts-Kinderspital Zürich | Institute of Medical Virology | Daniel Ehrsam, Isabel Stürmer, Catharine Aquino, Joel Wirz, Weihong Qi, Hubert Rehrauer, Verena Kufner, Gabriela Ziltener, Maryam Zaheri, Stefan Schmutz, Annette Audigé, Maria Grünberg, Kevin Steiner, Jon Huder, Cyril Shah, Riccarda Capaul, Guido Bloemberg, Jürg Böni, Michael Huber, Alexandra Trkola |
| EPI_ISL_3128763 | Spital Limmattal | Institute of Medical Virology | Daniel Ehrsam, Isabel Stürmer, Catharine Aquino, Joel Wirz, Weihong Qi, Hubert Rehrauer, Verena Kufner, Gabriela Ziltener, Maryam Zaheri, Stefan Schmutz, Annette Audigé, Maria Grünberg, Kevin Steiner, Jon Huder, Cyril Shah, Riccarda Capaul, Guido Bloemberg, Jürg Böni, Michael Huber, Alexandra Trkola |
| EPI_ISL_3128764 | Sanatorium Kilchberg | Institute of Medical Virology | Daniel Ehrsam, Isabel Stürmer, Catharine Aquino, Joel Wirz, Weihong Qi, Hubert Rehrauer, Verena Kufner, Gabriela Ziltener, Maryam Zaheri, Stefan Schmutz, Annette Audigé, Maria Grünberg, Kevin Steiner, Jon Huder, Cyril Shah, Riccarda Capaul, Guido Bloemberg, Jürg Böni, Michael Huber, Alexandra Trkola |
| EPI_ISL_3128765 | Spital Limmattal | Institute of Medical Virology | Daniel Ehrsam, Isabel Stürmer, Catharine Aquino, Joel Wirz, Weihong Qi, Hubert Rehrauer, Verena Kufner, Gabriela Ziltener, Maryam Zaheri, Stefan Schmutz, Annette Audigé, Maria Grünberg, Kevin Steiner, Jon Huder, Cyril Shah, Riccarda Capaul, Guido Bloemberg, Jürg Böni, Michael Huber, Alexandra Trkola |
| EPI_ISL_3128766, EPI_ISL_3128767, EPI_ISL_3128768, EPI_ISL_3128769 | UniversitätsSpital Zürich | Institute of Medical Virology | Daniel Ehrsam, Isabel Stürmer, Catharine Aquino, Joel Wirz, Weihong Qi, Hubert Rehrauer, Verena Kufner, Gabriela Ziltener, Maryam Zaheri, Stefan Schmutz, Annette Audigé, Maria Grünberg, Kevin Steiner, Jon Huder, Cyril Shah, Riccarda Capaul, Guido Bloemberg, Jürg Böni, Michael Huber, Alexandra Trkola |
| EPI_ISL_3128770, EPI_ISL_3128771, EPI_ISL_3128772, EPI_ISL_3128773, EPI_ISL_3128774, EPI_ISL_3128776, EPI_ISL_3128777, EPI_ISL_3128778, EPI_ISL_3128779, EPI_ISL_3128780, EPI_ISL_3128781, EPI_ISL_3128782, EPI_ISL_3128783, EPI_ISL_3128784, EPI_ISL_3128785, EPI_ISL_3128786, EPI_ISL_3128787, EPI_ISL_3128788, EPI_ISL_3128789, EPI_ISL_3128790, EPI_ISL_3128791, EPI_ISL_3128792 | | | |
| see above | Stadtspital Triemli | Institute of Medical Virology | Daniel Ehrsam, Isabel Stürmer, Catharine Aquino, Joel Wirz, Weihong Qi, Hubert Rehrauer, Verena Kufner, Gabriela Ziltener, Maryam Zaheri, Stefan Schmutz, Annette Audigé, Maria Grünberg, Kevin Steiner, Jon Huder, Cyril Shah, Riccarda Capaul, Guido Bloemberg, Jürg Böni, Michael Huber, Alexandra Trkola |
| EPI_ISL_3128794 | Kantonsspital Baden AG | Institute of Medical Virology | Verena Kufner, Gabriela Ziltener, Maryam Zaheri, Stefan Schmutz, Annette Audigé, Maria Grünberg, Kevin Steiner, Jon Huder, Cyril Shah, Riccarda Capaul, Guido Bloemberg, Jürg Böni, Michael Huber, Alexandra Trkola |
| EPI_ISL_3128795 | Invenimus AG | Institute of Medical Virology | Verena Kufner, Gabriela Ziltener, Maryam Zaheri, Stefan Schmutz, Annette Audigé, Maria Grünberg, Kevin Steiner, Jon Huder, Cyril Shah, Riccarda Capaul, Guido Bloemberg, Jürg Böni, Michael Huber, Alexandra Trkola |
| EPI_ISL_3128796 | Spital Männedorf AG | Institute of Medical Virology | Verena Kufner, Gabriela Ziltener, Maryam Zaheri, Stefan Schmutz, Annette Audigé, Maria Grünberg, Kevin Steiner, Jon Huder, Cyril Shah, Riccarda Capaul, Guido Bloemberg, Jürg Böni, Michael Huber, Alexandra Trkola |
| EPI_ISL_3128797, EPI_ISL_3128798 | Analytica | Institute of Medical Virology | Verena Kufner, Gabriela Ziltener, Maryam Zaheri, Stefan Schmutz, Annette Audigé, Maria Grünberg, Kevin Steiner, Jon Huder, Cyril Shah, Riccarda Capaul, Guido Bloemberg, Jürg Böni, Michael Huber, Alexandra Trkola |
| EPI_ISL_3128799 | UniversitätsSpital Zürich | Institute of Medical Virology | Verena Kufner, Gabriela Ziltener, Maryam Zaheri, Stefan Schmutz, Annette Audigé, Maria Grünberg, Kevin Steiner, Jon Huder, Cyril Shah, Riccarda Capaul, Guido Bloemberg, Jürg Böni, Michael Huber, Alexandra Trkola |
| EPI_ISL_3128800 | Kantonsspital Aarau | Institute of Medical Virology | Verena Kufner, Gabriela Ziltener, Maryam Zaheri, Stefan Schmutz, Annette Audigé, Maria Grünberg, Kevin Steiner, Jon Huder, Cyril Shah, Riccarda Capaul, Guido Bloemberg, Jürg Böni, Michael Huber, Alexandra Trkola |
| EPI_ISL_3128801 | Medica | Institute of Medical Virology | Verena Kufner, Gabriela Ziltener, Maryam Zaheri, Stefan Schmutz, Annette Audigé, Maria Grünberg, Kevin Steiner, Jon Huder, Cyril Shah, Riccarda Capaul, Guido Bloemberg, Jürg Böni, Michael Huber, Alexandra Trkola |
| EPI_ISL_3128803 | Zentrallabor Zürich | Institute of Medical Virology | Verena Kufner, Gabriela Ziltener, Maryam Zaheri, Stefan Schmutz, Annette Audigé, Maria Grünberg, Kevin Steiner, Jon Huder, Cyril Shah, Riccarda Capaul, Guido Bloemberg, Jürg Böni, Michael Huber, Alexandra Trkola |
| EPI_ISL_3128804 | Analytica | Institute of Medical Virology | Verena Kufner, Gabriela Ziltener, Maryam Zaheri, Stefan Schmutz, Annette Audigé, Maria Grünberg, Kevin Steiner, Jon Huder, Cyril Shah, Riccarda Capaul, Guido Bloemberg, Jürg Böni, Michael Huber, Alexandra Trkola |
| EPI_ISL_3128805 | UniversitätsSpital Zürich | Institute of Medical Virology | Verena Kufner, Gabriela Ziltener, Maryam Zaheri, Stefan Schmutz, Annette Audigé, Maria Grünberg, Kevin Steiner, Jon Huder, Cyril Shah, Riccarda Capaul, Guido Bloemberg, Jürg Böni, Michael Huber, Alexandra Trkola |
| EPI_ISL_3128806 | Universitätsspital Zürich 046 | Institute of Medical Virology | Verena Kufner, Gabriela Ziltener, Maryam Zaheri, Stefan Schmutz, Annette Audigé, Maria Grünberg, Kevin Steiner, Jon Huder, Cyril Shah, Riccarda Capaul, Guido Bloemberg, Jürg Böni, Michael Huber, Alexandra Trkola |
| EPI_ISL_3128807 | Gesundheitsdirektion Zürich | Institute of Medical Virology | Verena Kufner, Gabriela Ziltener, Maryam Zaheri, Stefan Schmutz, Annette Audigé, Maria Grünberg, Kevin Steiner, Jon Huder, Cyril Shah, Riccarda Capaul, Guido Bloemberg, Jürg Böni, Michael Huber, Alexandra Trkola |
| EPI_ISL_3128808 | Spital Langenthal | Institute of Medical Virology | Verena Kufner, Gabriela Ziltener, Maryam Zaheri, Stefan Schmutz, Annette Audigé, Maria Grünberg, Kevin Steiner, Jon Huder, Cyril Shah, Riccarda Capaul, Guido Bloemberg, Jürg Böni, Michael Huber, Alexandra Trkola |
| EPI_ISL_3128809, EPI_ISL_3128810 | UniversitätsSpital Zürich | Institute of Medical Virology | Verena Kufner, Gabriela Ziltener, Maryam Zaheri, Stefan Schmutz, Annette Audigé, Maria Grünberg, Kevin Steiner, Jon Huder, Cyril Shah, Riccarda Capaul, Guido Bloemberg, Jürg Böni, Michael Huber, Alexandra Trkola |
| EPI_ISL_3128811 | Analytica | Institute of Medical Virology | Verena Kufner, Gabriela Ziltener, Maryam Zaheri, Stefan Schmutz, Annette Audigé, Maria Grünberg, Kevin Steiner, Jon Huder, Cyril Shah, Riccarda Capaul, Guido Bloemberg, Jürg Böni, Michael Huber, Alexandra Trkola |
| EPI_ISL_3128812 | Unilabs | Institute of Medical Virology | Verena Kufner, Gabriela Ziltener, Maryam Zaheri, Stefan Schmutz, Annette Audigé, Maria Grünberg, Kevin Steiner, Jon Huder, Cyril Shah, Riccarda Capaul, Guido Bloemberg, Jürg Böni, Michael Huber, Alexandra Trkola |
| EPI_ISL_3128813 | Kantonsspital Aarau | Institute of Medical Virology | Verena Kufner, Gabriela Ziltener, Maryam Zaheri, Stefan Schmutz, Annette Audigé, Maria Grünberg, Kevin Steiner, Jon Huder, Cyril Shah, Riccarda Capaul, Guido Bloemberg, Jürg Böni, Michael Huber, Alexandra Trkola |

| | | | |
|---|---|---|---|
| EPI_ISL_3128814 | UniversitätsSpital Zürich | Institute of Medical Virology | Verena Kufner, Gabriela Ziltener, Maryam Zaheri, Stefan Schmutz, Annette Audigé, Maria Grünberg, Kevin Steiner, Jon Huder, Cyril Shah, Riccarda Capaul, Guido Bloemberg, Jürg Böni, Michael Huber, Alexandra Trkola |
| EPI_ISL_3128815 | Medica | Institute of Medical Virology | Verena Kufner, Gabriela Ziltener, Maryam Zaheri, Stefan Schmutz, Annette Audigé, Maria Grünberg, Kevin Steiner, Jon Huder, Cyril Shah, Riccarda Capaul, Guido Bloemberg, Jürg Böni, Michael Huber, Alexandra Trkola |
| EPI_ISL_3128816 | Spital Region Oberaargau | Institute of Medical Virology | Verena Kufner, Gabriela Ziltener, Maryam Zaheri, Stefan Schmutz, Annette Audigé, Maria Grünberg, Kevin Steiner, Jon Huder, Cyril Shah, Riccarda Capaul, Guido Bloemberg, Jürg Böni, Michael Huber, Alexandra Trkola |
| EPI_ISL_3128817 | Unilabs | Institute of Medical Virology | Verena Kufner, Gabriela Ziltener, Maryam Zaheri, Stefan Schmutz, Annette Audigé, Maria Grünberg, Kevin Steiner, Jon Huder, Cyril Shah, Riccarda Capaul, Guido Bloemberg, Jürg Böni, Michael Huber, Alexandra Trkola |
| EPI_ISL_3128818 | Kantonsspital Baden AG | Institute of Medical Virology | Verena Kufner, Gabriela Ziltener, Maryam Zaheri, Stefan Schmutz, Annette Audigé, Maria Grünberg, Kevin Steiner, Jon Huder, Cyril Shah, Riccarda Capaul, Guido Bloemberg, Jürg Böni, Michael Huber, Alexandra Trkola |
| EPI_ISL_3128819 | Laborgemeinschaft 1 | Institute of Medical Virology | Verena Kufner, Gabriela Ziltener, Maryam Zaheri, Stefan Schmutz, Annette Audigé, Maria Grünberg, Kevin Steiner, Jon Huder, Cyril Shah, Riccarda Capaul, Guido Bloemberg, Jürg Böni, Michael Huber, Alexandra Trkola |
| EPI_ISL_3128820, EPI_ISL_3128821 | Zentrallabor Zürich | Institute of Medical Virology | Verena Kufner, Gabriela Ziltener, Maryam Zaheri, Stefan Schmutz, Annette Audigé, Maria Grünberg, Kevin Steiner, Jon Huder, Cyril Shah, Riccarda Capaul, Guido Bloemberg, Jürg Böni, Michael Huber, Alexandra Trkola |
| EPI_ISL_3128822, EPI_ISL_3128823, EPI_ISL_3128824 | Spital Limmattal | Institute of Medical Virology | Verena Kufner, Gabriela Ziltener, Maryam Zaheri, Stefan Schmutz, Annette Audigé, Maria Grünberg, Kevin Steiner, Jon Huder, Cyril Shah, Riccarda Capaul, Guido Bloemberg, Jürg Böni, Michael Huber, Alexandra Trkola |
| EPI_ISL_3128825 | Analytica | Institute of Medical Virology | Verena Kufner, Gabriela Ziltener, Maryam Zaheri, Stefan Schmutz, Annette Audigé, Maria Grünberg, Kevin Steiner, Jon Huder, Cyril Shah, Riccarda Capaul, Guido Bloemberg, Jürg Böni, Michael Huber, Alexandra Trkola |
| EPI_ISL_3128826 | UniversitätsSpital Zürich | Institute of Medical Virology | Verena Kufner, Gabriela Ziltener, Maryam Zaheri, Stefan Schmutz, Annette Audigé, Maria Grünberg, Kevin Steiner, Jon Huder, Cyril Shah, Riccarda Capaul, Guido Bloemberg, Jürg Böni, Michael Huber, Alexandra Trkola |
| EPI_ISL_3128827, EPI_ISL_3128828 | Medica | Institute of Medical Virology | Verena Kufner, Gabriela Ziltener, Maryam Zaheri, Stefan Schmutz, Annette Audigé, Maria Grünberg, Kevin Steiner, Jon Huder, Cyril Shah, Riccarda Capaul, Guido Bloemberg, Jürg Böni, Michael Huber, Alexandra Trkola |
| EPI_ISL_3128829, EPI_ISL_3128830 | UniversitätsSpital Zürich | Institute of Medical Virology | Verena Kufner, Gabriela Ziltener, Maryam Zaheri, Stefan Schmutz, Annette Audigé, Maria Grünberg, Kevin Steiner, Jon Huder, Cyril Shah, Riccarda Capaul, Guido Bloemberg, Jürg Böni, Michael Huber, Alexandra Trkola |
| EPI_ISL_3128831 | UniversitätsSpital Zürich 064.3 | Institute of Medical Virology | Verena Kufner, Gabriela Ziltener, Maryam Zaheri, Stefan Schmutz, Annette Audigé, Maria Grünberg, Kevin Steiner, Jon Huder, Cyril Shah, Riccarda Capaul, Guido Bloemberg, Jürg Böni, Michael Huber, Alexandra Trkola |
| EPI_ISL_3128832 | UniversitätsSpital Zürich 225 | Institute of Medical Virology | Verena Kufner, Gabriela Ziltener, Maryam Zaheri, Stefan Schmutz, Annette Audigé, Maria Grünberg, Kevin Steiner, Jon Huder, Cyril Shah, Riccarda Capaul, Guido Bloemberg, Jürg Böni, Michael Huber, Alexandra Trkola |
| EPI_ISL_3129549, EPI_ISL_3129635, EPI_ISL_3129751, EPI_ISL_3129797, EPI_ISL_3129798, EPI_ISL_3129799, EPI_ISL_3129800, EPI_ISL_3129801, EPI_ISL_3129802, EPI_ISL_3129804, EPI_ISL_3129806, EPI_ISL_3129809, EPI_ISL_3129812, EPI_ISL_3129813, EPI_ISL_3129814, EPI_ISL_3129815, EPI_ISL_3129816, EPI_ISL_3129817, EPI_ISL_3129818, EPI_ISL_3129820, EPI_ISL_3129821, EPI_ISL_3129822, EPI_ISL_3129823, EPI_ISL_3129824, EPI_ISL_3129825, EPI_ISL_3129826, EPI_ISL_3129827, EPI_ISL_3129828, EPI_ISL_3129830, EPI_ISL_3129831, EPI_ISL_3129834, EPI_ISL_3129836, EPI_ISL_3129837, EPI_ISL_3129838, EPI_ISL_3129839, EPI_ISL_3129840, EPI_ISL_3129841, EPI_ISL_3129842, EPI_ISL_3129843, EPI_ISL_3129847, EPI_ISL_3129848, EPI_ISL_3129849, EPI_ISL_3129850, EPI_ISL_3129851, EPI_ISL_3129852, EPI_ISL_3129853, EPI_ISL_3129854, EPI_ISL_3129855 | | | |
| see above | Michigan Department of Health and Human Services, Bureau of Laboratories | Michigan Department of Health and Human Services, Bureau of Laboratories | Blankenship HM, Riner D, Soehnlen MK |
| EPI_ISL_3129863, EPI_ISL_3129864, EPI_ISL_3129865, EPI_ISL_3129867, EPI_ISL_3129868, EPI_ISL_3129893, EPI_ISL_3129895, EPI_ISL_3129896, EPI_ISL_3129911, EPI_ISL_3129912, EPI_ISL_3129914, EPI_ISL_3129915, EPI_ISL_3129916, EPI_ISL_3129918, EPI_ISL_3129920, EPI_ISL_3129925, EPI_ISL_3129926, EPI_ISL_3129928 | | | |
| see above | Centro de Investigación Biomédica de La Rioja - Hospital San Pedro Logroño | SeqCOVID-SPAIN consortium/IBV(CSIC) | María de Toro, José Manuel Azcona Gutiérrez, María Pilar Bea Escudero, Miriam Blasco Alberdi and SeqCOVID-SPAIN consortium |
| EPI_ISL_3129955 | CNR Institut Pasteur de la Guyane | Institut Pasteur de la Guyane | Anne Lavergne, Dominique Rousset, Antoine Enfissi, Arielle Salmier |
| EPI_ISL_3129971 | Laboratoire Carage | Institut Pasteur de la Guyane | Anne Lavergne, Dominique Rousset, Antoine Enfissi, Arielle Salmier |
| EPI_ISL_3129973, EPI_ISL_3129974, EPI_ISL_3129975, EPI_ISL_3129976, EPI_ISL_3129977, EPI_ISL_3129979, EPI_ISL_3129980, EPI_ISL_3129981, EPI_ISL_3129983, EPI_ISL_3129984, EPI_ISL_3129985, EPI_ISL_3129987, EPI_ISL_3129989, EPI_ISL_3129990, EPI_ISL_3129991, EPI_ISL_3129993, EPI_ISL_3129994, EPI_ISL_3129995, EPI_ISL_3129996, EPI_ISL_3129997, EPI_ISL_3129998, EPI_ISL_3129999, EPI_ISL_3130002, EPI_ISL_3130004, EPI_ISL_3130005, EPI_ISL_3130006, EPI_ISL_3130007, EPI_ISL_3130008, EPI_ISL_3130009, EPI_ISL_3130010, EPI_ISL_3130011, EPI_ISL_3130012, EPI_ISL_3130013, EPI_ISL_3130014, EPI_ISL_3130015, EPI_ISL_3130016, EPI_ISL_3130017, EPI_ISL_3130018, EPI_ISL_3130019, EPI_ISL_3130020, EPI_ISL_3130021, EPI_ISL_3130022, EPI_ISL_3130023, EPI_ISL_3130024, EPI_ISL_3130025, EPI_ISL_3130026, EPI_ISL_3130027, EPI_ISL_3130028, EPI_ISL_3130029, EPI_ISL_3130030, EPI_ISL_3130031, EPI_ISL_3130032, EPI_ISL_3130033, EPI_ISL_3130037, EPI_ISL_3130038, EPI_ISL_3130039, EPI_ISL_3130040, EPI_ISL_3130041, EPI_ISL_3130042, EPI_ISL_3130044 | | | |
| see above | NJDOH, Public Health and Environmental Laboratories | NJ_PHEL | Lindsey Bodnar, Shiv K. Verma, Ryan Pachucki, Chelsea San Filippo, Jacquelyn Deverell, Dana Woell, Allison Roder, Byeong Jeong |
| EPI_ISL_3130045, EPI_ISL_3130046, EPI_ISL_3130047, EPI_ISL_3130048, EPI_ISL_3130049, EPI_ISL_3130050, EPI_ISL_3130051, EPI_ISL_3130052, EPI_ISL_3130053, EPI_ISL_3130054, EPI_ISL_3130055, EPI_ISL_3130056, EPI_ISL_3130057, EPI_ISL_3130058, EPI_ISL_3130059, EPI_ISL_3130060, EPI_ISL_3130061, EPI_ISL_3130062, EPI_ISL_3130063, EPI_ISL_3130064, EPI_ISL_3130065, EPI_ISL_3130066, EPI_ISL_3130067, EPI_ISL_3130068, EPI_ISL_3130069, EPI_ISL_3130070, EPI_ISL_3130071, EPI_ISL_3130072, EPI_ISL_3130073, EPI_ISL_3130074, EPI_ISL_3130075, EPI_ISL_3130076 | | | |
| see above | IN State Department of Health Laboratory Services | IN State Department of Health Laboratory Services | Cassandra Campion, Jamie Yeadon, Brian Pope, Lixia Liu, Kyle Brownlee, Melissa Hindenlang, Mark Glazier |
| EPI_ISL_3130078 | AX BIO OCEAN | CNR Virus des Infections Respiratoires - France SUD | Antonin Bal, Gregory Destras, Gwendolyne Burfin, Hadrien Regue, Quentin Semanas, Martine Valette, Bruno Lina, Laurence Josset |
| EPI_ISL_3130079 | LABORATOIRE FORTE BIO ST PAUL LES DAX | CNR Virus des Infections Respiratoires - France SUD | Antonin Bal, Gregory Destras, Gwendolyne Burfin, Hadrien Regue, Quentin Semanas, Martine Valette, Bruno Lina, Laurence Josset |
| EPI_ISL_3130082 | CENTRE HOSPITALIER PIERRE OUDOT | CNR Virus des Infections Respiratoires - France SUD | Antonin Bal, Gregory Destras, Gwendolyne Burfin, Hadrien Regue, Quentin Semanas, Martine Valette, Bruno Lina, Laurence Josset |
| EPI_ISL_3130084 | ASTRALAB TASSIGNY | CNR Virus des Infections Respiratoires - France SUD | Antonin Bal, Gregory Destras, Gwendolyne Burfin, Hadrien Regue, Quentin Semanas, Martine Valette, Bruno Lina, Laurence Josset |
| EPI_ISL_3130089, EPI_ISL_3130090 | BIOMNIS LYON | CNR Virus des Infections Respiratoires - France SUD | Antonin Bal, Gregory Destras, Gwendolyne Burfin, Hadrien Regue, Quentin Semanas, Martine Valette, Bruno Lina, Laurence Josset |
| EPI_ISL_3130091 | LABORATOIRE UNILIANS DECINES | CNR Virus des Infections Respiratoires - France SUD | Antonin Bal, Gregory Destras, Gwendolyne Burfin, Hadrien Regue, Quentin Semanas, Martine Valette, Bruno Lina, Laurence Josset |
| EPI_ISL_3130092 | MIRIALIS CLUSES BECHET | CNR Virus des Infections Respiratoires - France SUD | Antonin Bal, Gregory Destras, Gwendolyne Burfin, Hadrien Regue, Quentin Semanas, Martine Valette, Bruno Lina, Laurence Josset |
| EPI_ISL_3130094, EPI_ISL_3130095, EPI_ISL_3130097 | BIOMNIS LYON | CNR Virus des Infections Respiratoires - France SUD | Antonin Bal, Gregory Destras, Gwendolyne Burfin, Hadrien Regue, Quentin Semanas, Martine Valette, Bruno Lina, Laurence Josset |
| EPI_ISL_3130101 | ORIAPOLE | CNR Virus des Infections Respiratoires - France SUD | Antonin Bal, Gregory Destras, Gwendolyne Burfin, Hadrien Regue, Quentin Semanas, Martine Valette, Bruno Lina, Laurence Josset |
| EPI_ISL_3130102 | LABORATOIRE DES CEDRES | CNR Virus des Infections Respiratoires - France SUD | Antonin Bal, Gregory Destras, Gwendolyne Burfin, Hadrien Regue, Quentin Semanas, Martine Valette, Bruno Lina, Laurence Josset |
| EPI_ISL_3130104 | CNR Virus des Infections Respiratoires - France SUD | CNR Virus des Infections Respiratoires - France SUD | Antonin Bal, Gregory Destras, Gwendolyne Burfin, Hadrien Regue, Quentin Semanas, Martine Valette, Bruno Lina, Laurence Josset |
| EPI_ISL_3130105 | BIOLITTORAL BIOGROUP PLATEAU TECH | CNR Virus des Infections Respiratoires - France SUD | Antonin Bal, Gregory Destras, Gwendolyne Burfin, Hadrien Regue, Quentin Semanas, Martine Valette, Bruno Lina, Laurence Josset |
| EPI_ISL_3130106 | CERBALLIANCE CHARENTES SAINTES | CNR Virus des Infections Respiratoires - France SUD | Antonin Bal, Gregory Destras, Gwendolyne Burfin, Hadrien Regue, Quentin Semanas, Martine Valette, Bruno Lina, Laurence Josset |
| EPI_ISL_3130107 | BIOMNIS LYON | CNR Virus des Infections Respiratoires - France SUD | Antonin Bal, Gregory Destras, Gwendolyne Burfin, Hadrien Regue, Quentin Semanas, Martine Valette, Bruno Lina, Laurence Josset |
| EPI_ISL_3130108 | LABORATOIRE MAYMAT | CNR Virus des Infections Respiratoires - France SUD | Antonin Bal, Gregory Destras, Gwendolyne Burfin, Hadrien Regue, Quentin Semanas, Martine Valette, Bruno Lina, Laurence Josset |
| EPI_ISL_3130109 | CHU ST ETIENNE HOPITAL NORD | CNR Virus des Infections Respiratoires - France SUD | Antonin Bal, Gregory Destras, Gwendolyne Burfin, Hadrien Regue, Quentin Semanas, Martine Valette, Bruno Lina, Laurence Josset |
| EPI_ISL_3130111, EPI_ISL_3130112, EPI_ISL_3130113, EPI_ISL_3130114, EPI_ISL_3130115, EPI_ISL_3130116, EPI_ISL_3130117, EPI_ISL_3130118, EPI_ISL_3130119, EPI_ISL_3130120, EPI_ISL_3130121, EPI_ISL_3130122, EPI_ISL_3130123 | | | |
| see above | AX BIO OCEAN | CNR Virus des Infections Respiratoires - France SUD | Antonin Bal, Gregory Destras, Gwendolyne Burfin, Hadrien Regue, Quentin Semanas, Martine Valette, Bruno Lina, Laurence Josset |
| EPI_ISL_3130126 | LABORATOIRE BIOESTEREL | CNR Virus des Infections Respiratoires - France SUD | Antonin Bal, Gregory Destras, Gwendolyne Burfin, Hadrien Regue, Quentin Semanas, Martine Valette, Bruno Lina, Laurence Josset |
| EPI_ISL_3130127, EPI_ISL_3130128, EPI_ISL_3130129, EPI_ISL_3130130, EPI_ISL_3130131, EPI_ISL_3130132, EPI_ISL_3130133, EPI_ISL_3130134, EPI_ISL_3130135 | LABORATOIRE CBM DE MURET | CNR Virus des Infections Respiratoires - France SUD | Antonin Bal, Gregory Destras, Gwendolyne Burfin, Hadrien Regue, Quentin Semanas, Martine Valette, Bruno Lina, Laurence Josset |
| EPI_ISL_3130136 | CH ANTIBES - JUAN LES PINS | CNR Virus des Infections Respiratoires - France SUD | Antonin Bal, Gregory Destras, Gwendolyne Burfin, Hadrien Regue, Quentin Semanas, Martine Valette, Bruno Lina, Laurence Josset |
| EPI_ISL_3130142 | LABORATOIRE BOUVIER | CNR Virus des Infections Respiratoires - France SUD | Antonin Bal, Gregory Destras, Gwendolyne Burfin, Hadrien Regue, Quentin Semanas, Martine Valette, Bruno Lina, Laurence Josset |
| EPI_ISL_3130143 | GPT DE LABORATOIRES DE BIOLOGIE MED | CNR Virus des Infections Respiratoires - France SUD | Antonin Bal, Gregory Destras, Gwendolyne Burfin, Hadrien Regue, Quentin Semanas, Martine Valette, Bruno Lina, Laurence Josset |
| EPI_ISL_3130144, EPI_ISL_3130145, EPI_ISL_3130146, EPI_ISL_3130148, EPI_ISL_3130150, EPI_ISL_3130151, EPI_ISL_3130152, EPI_ISL_3130153, EPI_ISL_3130154, EPI_ISL_3130155, EPI_ISL_3130156, EPI_ISL_3130157, EPI_ISL_3130158, EPI_ISL_3130159, EPI_ISL_3130160, EPI_ISL_3130161, EPI_ISL_3130162, EPI_ISL_3130163 | | | |
| see above | MIRIALIS CLUSES BECHET | CNR Virus des Infections Respiratoires - France SUD | Antonin Bal, Gregory Destras, Gwendolyne Burfin, Hadrien Regue, Quentin Semanas, Martine Valette, Bruno Lina, Laurence Josset |
| EPI_ISL_3130165, EPI_ISL_3130166 | LABORATOIRE MAYMAT | CNR Virus des Infections Respiratoires - France SUD | Antonin Bal, Gregory Destras, Gwendolyne Burfin, Hadrien Regue, Quentin Semanas, Martine Valette, Bruno Lina, Laurence Josset |
| EPI_ISL_3130168 | ASTRALAB TASSIGNY | CNR Virus des Infections Respiratoires - France SUD | Antonin Bal, Gregory Destras, Gwendolyne Burfin, Hadrien Regue, Quentin Semanas, Martine Valette, Bruno Lina, Laurence Josset |
| EPI_ISL_3130169, EPI_ISL_3130171 | LABORATOIRE CERBALLIANCE PLT VILLON | CNR Virus des Infections Respiratoires - France SUD | Antonin Bal, Gregory Destras, Gwendolyne Burfin, Hadrien Regue, Quentin Semanas, Martine Valette, Bruno Lina, Laurence Josset |
| EPI_ISL_3130172 | LABORATOIRE DU C.H.A.R.M.E. | CNR Virus des Infections Respiratoires - France SUD | Antonin Bal, Gregory Destras, Gwendolyne Burfin, Hadrien Regue, Quentin Semanas, Martine Valette, Bruno Lina, Laurence Josset |
| EPI_ISL_3130174, EPI_ISL_3130175, EPI_ISL_3130176 | LAM BAYARD-BIOPYRENEES | CNR Virus des Infections Respiratoires - France SUD | Antonin Bal, Gregory Destras, Gwendolyne Burfin, Hadrien Regue, Quentin Semanas, Martine Valette, Bruno Lina, Laurence Josset |
| EPI_ISL_3130178 | CH VALENCE | CNR Virus des Infections Respiratoires - France SUD | Antonin Bal, Gregory Destras, Gwendolyne Burfin, Hadrien Regue, Quentin Semanas, Martine Valette, Bruno Lina, Laurence Josset |
| EPI_ISL_3130179 | LABORATOIRE ACCOLAB SUD OUEST | CNR Virus des Infections Respiratoires - France SUD | Antonin Bal, Gregory Destras, Gwendolyne Burfin, Hadrien Regue, Quentin Semanas, Martine Valette, Bruno Lina, Laurence Josset |
| EPI_ISL_3130181 | LAM BAYARD-BIOPYRENEES | CNR Virus des Infections Respiratoires - France SUD | Antonin Bal, Gregory Destras, Gwendolyne Burfin, Hadrien Regue, Quentin Semanas, Martine Valette, Bruno Lina, Laurence Josset |
| EPI_ISL_3130190, EPI_ISL_3130192, EPI_ISL_3130193, EPI_ISL_3130196, EPI_ISL_3130198, EPI_ISL_3130200, EPI_ISL_3130205, EPI_ISL_3130208, EPI_ISL_3130209, EPI_ISL_3130212, EPI_ISL_3130214, EPI_ISL_3130230, EPI_ISL_3130233, EPI_ISL_3130235, EPI_ISL_3130236, EPI_ISL_3130237, EPI_ISL_3130238, EPI_ISL_3130239, EPI_ISL_3130241, EPI_ISL_3130243, EPI_ISL_3130244, EPI_ISL_3130245, | | | |

| | | | |
|---|---|---|---|
| EPI_ISL_3130246, EPI_ISL_3130247 | | | |
| see above | LABORATOIRE CERBALLIANCE PLT VILLON | CNR Virus des Infections Respiratoires - France SUD | Antonin Bal, Gregory Destras, Gwendolyne Burfin, Hadrien Regue, Quentin Semanas, Martine Valette, Bruno Lina, Laurence Josset |
| EPI_ISL_3130248, EPI_ISL_3130249, EPI_ISL_3130250 | LAM BIORYLIS | CNR Virus des Infections Respiratoires - France SUD | Antonin Bal, Gregory Destras, Gwendolyne Burfin, Hadrien Regue, Quentin Semanas, Martine Valette, Bruno Lina, Laurence Josset |
| EPI_ISL_3130262, EPI_ISL_3130264, EPI_ISL_3130265, EPI_ISL_3130267, EPI_ISL_3130269, EPI_ISL_3130270, EPI_ISL_3130271, EPI_ISL_3130273, EPI_ISL_3130274, EPI_ISL_3130275, EPI_ISL_3130276 | | | |
| see above | LABORATOIRE CERBALLIANCE PLT VILLON | CNR Virus des Infections Respiratoires - France SUD | Antonin Bal, Gregory Destras, Gwendolyne Burfin, Hadrien Regue, Quentin Semanas, Martine Valette, Bruno Lina, Laurence Josset |
| EPI_ISL_3130278, EPI_ISL_3130279, EPI_ISL_3130281, EPI_ISL_3130282, EPI_ISL_3130283, EPI_ISL_3130291, EPI_ISL_3130292 | LABORATOIRE UNILIANS DECINES | CNR Virus des Infections Respiratoires - France SUD | Antonin Bal, Gregory Destras, Gwendolyne Burfin, Hadrien Regue, Quentin Semanas, Martine Valette, Bruno Lina, Laurence Josset |
| EPI_ISL_3130293, EPI_ISL_3130295 | AX BIO OCEAN | CNR Virus des Infections Respiratoires - France SUD | Antonin Bal, Gregory Destras, Gwendolyne Burfin, Hadrien Regue, Quentin Semanas, Martine Valette, Bruno Lina, Laurence Josset |
| EPI_ISL_3130296 | LABORATOIRE UNILIANS DECINES | CNR Virus des Infections Respiratoires - France SUD | Antonin Bal, Gregory Destras, Gwendolyne Burfin, Hadrien Regue, Quentin Semanas, Martine Valette, Bruno Lina, Laurence Josset |
| EPI_ISL_3130298, EPI_ISL_3130300, EPI_ISL_3130301, EPI_ISL_3130303 | AX BIO OCEAN | CNR Virus des Infections Respiratoires - France SUD | Antonin Bal, Gregory Destras, Gwendolyne Burfin, Hadrien Regue, Quentin Semanas, Martine Valette, Bruno Lina, Laurence Josset |
| EPI_ISL_3130304, EPI_ISL_3130305 | LABORATOIRE BIO-VAL | CNR Virus des Infections Respiratoires - France SUD | Antonin Bal, Gregory Destras, Gwendolyne Burfin, Hadrien Regue, Quentin Semanas, Martine Valette, Bruno Lina, Laurence Josset |
| EPI_ISL_3130306, EPI_ISL_3130307 | AX BIO OCEAN | CNR Virus des Infections Respiratoires - France SUD | Antonin Bal, Gregory Destras, Gwendolyne Burfin, Hadrien Regue, Quentin Semanas, Martine Valette, Bruno Lina, Laurence Josset |
| EPI_ISL_3130308, EPI_ISL_3130309 | LABORATOIRE BIO-VAL | CNR Virus des Infections Respiratoires - France SUD | Antonin Bal, Gregory Destras, Gwendolyne Burfin, Hadrien Regue, Quentin Semanas, Martine Valette, Bruno Lina, Laurence Josset |
| EPI_ISL_3130310 | AX BIO OCEAN | CNR Virus des Infections Respiratoires - France SUD | Antonin Bal, Gregory Destras, Gwendolyne Burfin, Hadrien Regue, Quentin Semanas, Martine Valette, Bruno Lina, Laurence Josset |
| EPI_ISL_3130311, EPI_ISL_3130312 | LABORATOIRE BIO-VAL | CNR Virus des Infections Respiratoires - France SUD | Antonin Bal, Gregory Destras, Gwendolyne Burfin, Hadrien Regue, Quentin Semanas, Martine Valette, Bruno Lina, Laurence Josset |
| EPI_ISL_3130313, EPI_ISL_3130314, EPI_ISL_3130315, EPI_ISL_3130316, EPI_ISL_3130317 | LABORATOIRE d'ANALYSES de BIOLOGIE MEDICALES | CNR Virus des Infections Respiratoires - France SUD | Antonin Bal, Gregory Destras, Gwendolyne Burfin, Hadrien Regue, Quentin Semanas, Martine Valette, Bruno Lina, Laurence Josset |
| EPI_ISL_3130318, EPI_ISL_3130319, EPI_ISL_3130320, EPI_ISL_3130321 | LABORATOIRE BOUVIER | CNR Virus des Infections Respiratoires - France SUD | Antonin Bal, Gregory Destras, Gwendolyne Burfin, Hadrien Regue, Quentin Semanas, Martine Valette, Bruno Lina, Laurence Josset |
| EPI_ISL_3130322 | ASTRALAB TASSIGNY | CNR Virus des Infections Respiratoires - France SUD | Antonin Bal, Gregory Destras, Gwendolyne Burfin, Hadrien Regue, Quentin Semanas, Martine Valette, Bruno Lina, Laurence Josset |
| EPI_ISL_3130323 | LAM DE BIO HD | CNR Virus des Infections Respiratoires - France SUD | Antonin Bal, Gregory Destras, Gwendolyne Burfin, Hadrien Regue, Quentin Semanas, Martine Valette, Bruno Lina, Laurence Josset |
| EPI_ISL_3130325 | LABORATOIRE MAYMAT | CNR Virus des Infections Respiratoires - France SUD | Antonin Bal, Gregory Destras, Gwendolyne Burfin, Hadrien Regue, Quentin Semanas, Martine Valette, Bruno Lina, Laurence Josset |
| EPI_ISL_3130335, EPI_ISL_3130336, EPI_ISL_3130337, EPI_ISL_3130338, EPI_ISL_3130339, EPI_ISL_3130340, EPI_ISL_3130341, EPI_ISL_3130342 | LABORATOIRE d'ANALYSES de BIOLOGIE MEDICALES | CNR Virus des Infections Respiratoires - France SUD | Antonin Bal, Gregory Destras, Gwendolyne Burfin, Hadrien Regue, Quentin Semanas, Martine Valette, Bruno Lina, Laurence Josset |
| EPI_ISL_3130343, EPI_ISL_3130344, EPI_ISL_3130345, EPI_ISL_3130346, EPI_ISL_3130347, EPI_ISL_3130349 | LABORATOIRE CBM DE MURET | CNR Virus des Infections Respiratoires - France SUD | Antonin Bal, Gregory Destras, Gwendolyne Burfin, Hadrien Regue, Quentin Semanas, Martine Valette, Bruno Lina, Laurence Josset |
| EPI_ISL_3130352, EPI_ISL_3130354, EPI_ISL_3130356, EPI_ISL_3130362 | LABORATOIRE DES CEDRES | CNR Virus des Infections Respiratoires - France SUD | Antonin Bal, Gregory Destras, Gwendolyne Burfin, Hadrien Regue, Quentin Semanas, Martine Valette, Bruno Lina, Laurence Josset |
| EPI_ISL_3130377 | CNR Virus des Infections Respiratoires - France SUD | CNR Virus des Infections Respiratoires - France SUD | Antonin Bal, Gregory Destras, Gwendolyne Burfin, Hadrien Regue, Quentin Semanas, Martine Valette, Bruno Lina, Laurence Josset |
| EPI_ISL_3130379, EPI_ISL_3130380, EPI_ISL_3130381, EPI_ISL_3130382 | BIOLITTORAL BIOGROUP PLATEAU TECH | CNR Virus des Infections Respiratoires - France SUD | Antonin Bal, Gregory Destras, Gwendolyne Burfin, Hadrien Regue, Quentin Semanas, Martine Valette, Bruno Lina, Laurence Josset |
| EPI_ISL_3130383, EPI_ISL_3130384, EPI_ISL_3130385, EPI_ISL_3130386, EPI_ISL_3130391 | LAM GEN-BIO REPUBLIQUE | CNR Virus des Infections Respiratoires - France SUD | Antonin Bal, Gregory Destras, Gwendolyne Burfin, Hadrien Regue, Quentin Semanas, Martine Valette, Bruno Lina, Laurence Josset |
| EPI_ISL_3130392, EPI_ISL_3130394, EPI_ISL_3130395 | BIOMNIS LYON | CNR Virus des Infections Respiratoires - France SUD | Antonin Bal, Gregory Destras, Gwendolyne Burfin, Hadrien Regue, Quentin Semanas, Martine Valette, Bruno Lina, Laurence Josset |
| EPI_ISL_3130397 | LABORATOIRE CBM DE MURET | CNR Virus des Infections Respiratoires - France SUD | Antonin Bal, Gregory Destras, Gwendolyne Burfin, Hadrien Regue, Quentin Semanas, Martine Valette, Bruno Lina, Laurence Josset |
| EPI_ISL_3130398 | BIOMNIS LYON | CNR Virus des Infections Respiratoires - France SUD | Antonin Bal, Gregory Destras, Gwendolyne Burfin, Hadrien Regue, Quentin Semanas, Martine Valette, Bruno Lina, Laurence Josset |
| EPI_ISL_3130399 | LABORATOIRE CBM DE MURET | CNR Virus des Infections Respiratoires - France SUD | Antonin Bal, Gregory Destras, Gwendolyne Burfin, Hadrien Regue, Quentin Semanas, Martine Valette, Bruno Lina, Laurence Josset |
| EPI_ISL_3130400 | BIOMNIS LYON | CNR Virus des Infections Respiratoires - France SUD | Antonin Bal, Gregory Destras, Gwendolyne Burfin, Hadrien Regue, Quentin Semanas, Martine Valette, Bruno Lina, Laurence Josset |
| EPI_ISL_3130401 | LABORATOIRE CBM DE MURET | CNR Virus des Infections Respiratoires - France SUD | Antonin Bal, Gregory Destras, Gwendolyne Burfin, Hadrien Regue, Quentin Semanas, Martine Valette, Bruno Lina, Laurence Josset |
| EPI_ISL_3130402 | BIOMNIS LYON | CNR Virus des Infections Respiratoires - France SUD | Antonin Bal, Gregory Destras, Gwendolyne Burfin, Hadrien Regue, Quentin Semanas, Martine Valette, Bruno Lina, Laurence Josset |
| EPI_ISL_3130404 | LABORATOIRE CBM DE MURET | CNR Virus des Infections Respiratoires - France SUD | Antonin Bal, Gregory Destras, Gwendolyne Burfin, Hadrien Regue, Quentin Semanas, Martine Valette, Bruno Lina, Laurence Josset |
| EPI_ISL_3130408 | BIOLITTORAL BIOGROUP PLATEAU TECH | CNR Virus des Infections Respiratoires - France SUD | Antonin Bal, Gregory Destras, Gwendolyne Burfin, Hadrien Regue, Quentin Semanas, Martine Valette, Bruno Lina, Laurence Josset |
| EPI_ISL_3130410, EPI_ISL_3130411 | LABORATOIRE BIOESTEREL | CNR Virus des Infections Respiratoires - France SUD | Antonin Bal, Gregory Destras, Gwendolyne Burfin, Hadrien Regue, Quentin Semanas, Martine Valette, Bruno Lina, Laurence Josset |
| EPI_ISL_3130412 | CHU TOULOUSE | CNR Virus des Infections Respiratoires - France SUD | Antonin Bal, Gregory Destras, Gwendolyne Burfin, Hadrien Regue, Quentin Semanas, Martine Valette, Bruno Lina, Laurence Josset |
| EPI_ISL_3130413 | LABORATOIRE BIOESTEREL | CNR Virus des Infections Respiratoires - France SUD | Antonin Bal, Gregory Destras, Gwendolyne Burfin, Hadrien Regue, Quentin Semanas, Martine Valette, Bruno Lina, Laurence Josset |
| EPI_ISL_3130415 | CHU TOULOUSE | CNR Virus des Infections Respiratoires - France SUD | Antonin Bal, Gregory Destras, Gwendolyne Burfin, Hadrien Regue, Quentin Semanas, Martine Valette, Bruno Lina, Laurence Josset |
| EPI_ISL_3130416, EPI_ISL_3130418 | BIOLITTORAL BIOGROUP PLATEAU TECH | CNR Virus des Infections Respiratoires - France SUD | Antonin Bal, Gregory Destras, Gwendolyne Burfin, Hadrien Regue, Quentin Semanas, Martine Valette, Bruno Lina, Laurence Josset |
| EPI_ISL_3130419, EPI_ISL_3130422, EPI_ISL_3130424 | CHU TOULOUSE | CNR Virus des Infections Respiratoires - France SUD | Antonin Bal, Gregory Destras, Gwendolyne Burfin, Hadrien Regue, Quentin Semanas, Martine Valette, Bruno Lina, Laurence Josset |
| EPI_ISL_3130425 | BIOLITTORAL BIOGROUP PLATEAU TECH | CNR Virus des Infections Respiratoires - France SUD | Antonin Bal, Gregory Destras, Gwendolyne Burfin, Hadrien Regue, Quentin Semanas, Martine Valette, Bruno Lina, Laurence Josset |

| | | | |
|---|---|---|---|
| EPI_ISL_3130428, EPI_ISL_3130429, EPI_ISL_3130431 | LABORATOIRE BIOESTEREL | CNR Virus des Infections Respiratoires - France SUD | Antonin Bal, Gregory Destras, Gwendolyne Burfin, Hadrien Regue, Quentin Semanas, Martine Valette, Bruno Lina, Laurence Josset |
| EPI_ISL_3130432 | LABORATOIRE DES CEDRES | CNR Virus des Infections Respiratoires - France SUD | Antonin Bal, Gregory Destras, Gwendolyne Burfin, Hadrien Regue, Quentin Semanas, Martine Valette, Bruno Lina, Laurence Josset |
| EPI_ISL_3130433 | LABORATOIRE BIOESTEREL | CNR Virus des Infections Respiratoires - France SUD | Antonin Bal, Gregory Destras, Gwendolyne Burfin, Hadrien Regue, Quentin Semanas, Martine Valette, Bruno Lina, Laurence Josset |
| EPI_ISL_3130434, EPI_ISL_3130436 | LABORATOIRE DES CEDRES | CNR Virus des Infections Respiratoires - France SUD | Antonin Bal, Gregory Destras, Gwendolyne Burfin, Hadrien Regue, Quentin Semanas, Martine Valette, Bruno Lina, Laurence Josset |
| EPI_ISL_3130437 | LABORATOIRE BIOESTEREL | CNR Virus des Infections Respiratoires - France SUD | Antonin Bal, Gregory Destras, Gwendolyne Burfin, Hadrien Regue, Quentin Semanas, Martine Valette, Bruno Lina, Laurence Josset |
| EPI_ISL_3130439 | LABORATOIRE DES CEDRES | CNR Virus des Infections Respiratoires - France SUD | Antonin Bal, Gregory Destras, Gwendolyne Burfin, Hadrien Regue, Quentin Semanas, Martine Valette, Bruno Lina, Laurence Josset |
| EPI_ISL_3130440, EPI_ISL_3130441 | LABORATOIRE BIOESTEREL | CNR Virus des Infections Respiratoires - France SUD | Antonin Bal, Gregory Destras, Gwendolyne Burfin, Hadrien Regue, Quentin Semanas, Martine Valette, Bruno Lina, Laurence Josset |
| EPI_ISL_3130442, EPI_ISL_3130443 | LABORATOIRE DES CEDRES | CNR Virus des Infections Respiratoires - France SUD | Antonin Bal, Gregory Destras, Gwendolyne Burfin, Hadrien Regue, Quentin Semanas, Martine Valette, Bruno Lina, Laurence Josset |
| EPI_ISL_3130444 | LABORATOIRE FORTE BIO DAX | CNR Virus des Infections Respiratoires - France SUD | Antonin Bal, Gregory Destras, Gwendolyne Burfin, Hadrien Regue, Quentin Semanas, Martine Valette, Bruno Lina, Laurence Josset |
| EPI_ISL_3130445 | LABORATOIRE BIOESTEREL | CNR Virus des Infections Respiratoires - France SUD | Antonin Bal, Gregory Destras, Gwendolyne Burfin, Hadrien Regue, Quentin Semanas, Martine Valette, Bruno Lina, Laurence Josset |
| EPI_ISL_3130446 | LABORATOIRE FORTE BIO DAX | CNR Virus des Infections Respiratoires - France SUD | Antonin Bal, Gregory Destras, Gwendolyne Burfin, Hadrien Regue, Quentin Semanas, Martine Valette, Bruno Lina, Laurence Josset |
| EPI_ISL_3130447 | LABORATOIRE BIOESTEREL | CNR Virus des Infections Respiratoires - France SUD | Antonin Bal, Gregory Destras, Gwendolyne Burfin, Hadrien Regue, Quentin Semanas, Martine Valette, Bruno Lina, Laurence Josset |
| EPI_ISL_3130448 | LABORATOIRE FORTE BIO DAX | CNR Virus des Infections Respiratoires - France SUD | Antonin Bal, Gregory Destras, Gwendolyne Burfin, Hadrien Regue, Quentin Semanas, Martine Valette, Bruno Lina, Laurence Josset |
| EPI_ISL_3130449, EPI_ISL_3130450 | LABORATOIRE BIOESTEREL | CNR Virus des Infections Respiratoires - France SUD | Antonin Bal, Gregory Destras, Gwendolyne Burfin, Hadrien Regue, Quentin Semanas, Martine Valette, Bruno Lina, Laurence Josset |
| EPI_ISL_3130451 | LABORATOIRE FORTE BIO DAX | CNR Virus des Infections Respiratoires - France SUD | Antonin Bal, Gregory Destras, Gwendolyne Burfin, Hadrien Regue, Quentin Semanas, Martine Valette, Bruno Lina, Laurence Josset |
| EPI_ISL_3130453 | LABORATOIRE BIOESTEREL | CNR Virus des Infections Respiratoires - France SUD | Antonin Bal, Gregory Destras, Gwendolyne Burfin, Hadrien Regue, Quentin Semanas, Martine Valette, Bruno Lina, Laurence Josset |
| EPI_ISL_3130454 | LABORATOIRE FORTE BIO DAX | CNR Virus des Infections Respiratoires - France SUD | Antonin Bal, Gregory Destras, Gwendolyne Burfin, Hadrien Regue, Quentin Semanas, Martine Valette, Bruno Lina, Laurence Josset |
| EPI_ISL_3130455, EPI_ISL_3130456 | LABORATOIRE BIOESTEREL | CNR Virus des Infections Respiratoires - France SUD | Antonin Bal, Gregory Destras, Gwendolyne Burfin, Hadrien Regue, Quentin Semanas, Martine Valette, Bruno Lina, Laurence Josset |
| EPI_ISL_3130457 | LABORATOIRE FORTE BIO DAX | CNR Virus des Infections Respiratoires - France SUD | Antonin Bal, Gregory Destras, Gwendolyne Burfin, Hadrien Regue, Quentin Semanas, Martine Valette, Bruno Lina, Laurence Josset |
| EPI_ISL_3130458, EPI_ISL_3130460 | LABORATOIRE BIOESTEREL | CNR Virus des Infections Respiratoires - France SUD | Antonin Bal, Gregory Destras, Gwendolyne Burfin, Hadrien Regue, Quentin Semanas, Martine Valette, Bruno Lina, Laurence Josset |
| EPI_ISL_3130463 | LABORATOIRE FORTE BIO DAX | CNR Virus des Infections Respiratoires - France SUD | Antonin Bal, Gregory Destras, Gwendolyne Burfin, Hadrien Regue, Quentin Semanas, Martine Valette, Bruno Lina, Laurence Josset |
| EPI_ISL_3130464, EPI_ISL_3130465 | LABORATOIRE BIOESTEREL | CNR Virus des Infections Respiratoires - France SUD | Antonin Bal, Gregory Destras, Gwendolyne Burfin, Hadrien Regue, Quentin Semanas, Martine Valette, Bruno Lina, Laurence Josset |
| EPI_ISL_3130466 | LABORATOIRE FORTE BIO DAX | CNR Virus des Infections Respiratoires - France SUD | Antonin Bal, Gregory Destras, Gwendolyne Burfin, Hadrien Regue, Quentin Semanas, Martine Valette, Bruno Lina, Laurence Josset |
| EPI_ISL_3130467, EPI_ISL_3130468, EPI_ISL_3130469 | LABORATOIRE BIOESTEREL | CNR Virus des Infections Respiratoires - France SUD | Antonin Bal, Gregory Destras, Gwendolyne Burfin, Hadrien Regue, Quentin Semanas, Martine Valette, Bruno Lina, Laurence Josset |
| EPI_ISL_3130470 | CERBALLIANCE MIDI-PYRENEES | CNR Virus des Infections Respiratoires - France SUD | Antonin Bal, Gregory Destras, Gwendolyne Burfin, Hadrien Regue, Quentin Semanas, Martine Valette, Bruno Lina, Laurence Josset |
| EPI_ISL_3130471 | LABORATOIRE FORTE BIO DAX | CNR Virus des Infections Respiratoires - France SUD | Antonin Bal, Gregory Destras, Gwendolyne Burfin, Hadrien Regue, Quentin Semanas, Martine Valette, Bruno Lina, Laurence Josset |
| EPI_ISL_3130472, EPI_ISL_3130473, EPI_ISL_3130474 | CERBALLIANCE MIDI-PYRENEES | CNR Virus des Infections Respiratoires - France SUD | Antonin Bal, Gregory Destras, Gwendolyne Burfin, Hadrien Regue, Quentin Semanas, Martine Valette, Bruno Lina, Laurence Josset |
| EPI_ISL_3130475 | LABORATOIRE BIOESTEREL | CNR Virus des Infections Respiratoires - France SUD | Antonin Bal, Gregory Destras, Gwendolyne Burfin, Hadrien Regue, Quentin Semanas, Martine Valette, Bruno Lina, Laurence Josset |
| EPI_ISL_3130476 | LABORATOIRE FORTE BIO DAX | CNR Virus des Infections Respiratoires - France SUD | Antonin Bal, Gregory Destras, Gwendolyne Burfin, Hadrien Regue, Quentin Semanas, Martine Valette, Bruno Lina, Laurence Josset |
| EPI_ISL_3130477, EPI_ISL_3130479 | LABORATOIRE BIOESTEREL | CNR Virus des Infections Respiratoires - France SUD | Antonin Bal, Gregory Destras, Gwendolyne Burfin, Hadrien Regue, Quentin Semanas, Martine Valette, Bruno Lina, Laurence Josset |
| EPI_ISL_3130480 | CERBALLIANCE MIDI-PYRENEES | CNR Virus des Infections Respiratoires - France SUD | Antonin Bal, Gregory Destras, Gwendolyne Burfin, Hadrien Regue, Quentin Semanas, Martine Valette, Bruno Lina, Laurence Josset |
| EPI_ISL_3130482 | LABORATOIRE BIOESTEREL | CNR Virus des Infections Respiratoires - France SUD | Antonin Bal, Gregory Destras, Gwendolyne Burfin, Hadrien Regue, Quentin Semanas, Martine Valette, Bruno Lina, Laurence Josset |
| EPI_ISL_3130483 | CERBALLIANCE MIDI-PYRENEES | CNR Virus des Infections Respiratoires - France SUD | Antonin Bal, Gregory Destras, Gwendolyne Burfin, Hadrien Regue, Quentin Semanas, Martine Valette, Bruno Lina, Laurence Josset |
| EPI_ISL_3130484 | LABORATOIRE BIOESTEREL | CNR Virus des Infections Respiratoires - France SUD | Antonin Bal, Gregory Destras, Gwendolyne Burfin, Hadrien Regue, Quentin Semanas, Martine Valette, Bruno Lina, Laurence Josset |
| EPI_ISL_3130485, EPI_ISL_3130486 | CERBALLIANCE MIDI-PYRENEES | CNR Virus des Infections Respiratoires - France SUD | Antonin Bal, Gregory Destras, Gwendolyne Burfin, Hadrien Regue, Quentin Semanas, Martine Valette, Bruno Lina, Laurence Josset |
| EPI_ISL_3130488 | LABORATOIRE CERBALLIANCE PLT VILLON | CNR Virus des Infections Respiratoires - France SUD | Antonin Bal, Gregory Destras, Gwendolyne Burfin, Hadrien Regue, Quentin Semanas, Martine Valette, Bruno Lina, Laurence Josset |
| EPI_ISL_3130489 | CERBALLIANCE MIDI-PYRENEES | CNR Virus des Infections Respiratoires - France SUD | Antonin Bal, Gregory Destras, Gwendolyne Burfin, Hadrien Regue, Quentin Semanas, Martine Valette, Bruno Lina, Laurence Josset |
| EPI_ISL_3130490, EPI_ISL_3130491, EPI_ISL_3130492, EPI_ISL_3130493, EPI_ISL_3130494 | LABORATOIRE CERBALLIANCE PLT VILLON | CNR Virus des Infections Respiratoires - France SUD | Antonin Bal, Gregory Destras, Gwendolyne Burfin, Hadrien Regue, Quentin Semanas, Martine Valette, Bruno Lina, Laurence Josset |
| EPI_ISL_3130495 | CERBALLIANCE MIDI-PYRENEES | CNR Virus des Infections Respiratoires - France SUD | Antonin Bal, Gregory Destras, Gwendolyne Burfin, Hadrien Regue, Quentin Semanas, Martine Valette, Bruno Lina, Laurence Josset |
| EPI_ISL_3130496, EPI_ISL_3130497, EPI_ISL_3130498 | LABORATOIRE CERBALLIANCE PLT VILLON | CNR Virus des Infections Respiratoires - France SUD | Antonin Bal, Gregory Destras, Gwendolyne Burfin, Hadrien Regue, Quentin Semanas, Martine Valette, Bruno Lina, Laurence Josset |
| EPI_ISL_3130499 | CERBALLIANCE MIDI-PYRENEES | CNR Virus des Infections Respiratoires - France SUD | Antonin Bal, Gregory Destras, Gwendolyne Burfin, Hadrien Regue, Quentin Semanas, Martine Valette, Bruno Lina, Laurence Josset |
| EPI_ISL_3130500, EPI_ISL_3130502 | LABORATOIRE CERBALLIANCE PLT VILLON | CNR Virus des Infections Respiratoires - France SUD | Antonin Bal, Gregory Destras, Gwendolyne Burfin, Hadrien Regue, Quentin Semanas, Martine Valette, Bruno Lina, Laurence Josset |
| EPI_ISL_3130503 | CERBALLIANCE MIDI-PYRENEES | CNR Virus des Infections Respiratoires - France SUD | Antonin Bal, Gregory Destras, Gwendolyne Burfin, Hadrien Regue, Quentin Semanas, Martine Valette, Bruno Lina, Laurence Josset |
| EPI_ISL_3130504, EPI_ISL_3130505 | LABORATOIRE CERBALLIANCE PLT VILLON | CNR Virus des Infections Respiratoires - France SUD | Antonin Bal, Gregory Destras, Gwendolyne Burfin, Hadrien Regue, Quentin Semanas, Martine Valette, Bruno Lina, Laurence Josset |
| EPI_ISL_3130506, EPI_ISL_3130507 | CERBALLIANCE MIDI-PYRENEES | CNR Virus des Infections Respiratoires - France SUD | Antonin Bal, Gregory Destras, Gwendolyne Burfin, Hadrien Regue, Quentin Semanas, Martine Valette, Bruno Lina, Laurence Josset |
| EPI_ISL_3130508, EPI_ISL_3130509 | LABORATOIRE CERBALLIANCE PLT VILLON | CNR Virus des Infections Respiratoires - France SUD | Antonin Bal, Gregory Destras, Gwendolyne Burfin, Hadrien Regue, Quentin Semanas, Martine Valette, Bruno Lina, Laurence Josset |
| EPI_ISL_3130510 | CERBALLIANCE MIDI-PYRENEES | CNR Virus des Infections Respiratoires - France SUD | Antonin Bal, Gregory Destras, Gwendolyne Burfin, Hadrien Regue, Quentin Semanas, Martine Valette, Bruno Lina, Laurence Josset |
| EPI_ISL_3130511, EPI_ISL_3130512, EPI_ISL_3130513, EPI_ISL_3130515, EPI_ISL_3130516, EPI_ISL_3130519 | LABORATOIRE CERBALLIANCE PLT VILLON | CNR Virus des Infections Respiratoires - France SUD | Antonin Bal, Gregory Destras, Gwendolyne Burfin, Hadrien Regue, Quentin Semanas, Martine Valette, Bruno Lina, Laurence Josset |
| EPI_ISL_3130520 | CERBALLIANCE MIDI-PYRENEES | CNR Virus des Infections Respiratoires - France SUD | Antonin Bal, Gregory Destras, Gwendolyne Burfin, Hadrien Regue, Quentin Semanas, Martine Valette, Bruno Lina, Laurence Josset |
| EPI_ISL_3130523 | LABORATOIRE CERBALLIANCE PLT VILLON | CNR Virus des Infections Respiratoires - France SUD | Antonin Bal, Gregory Destras, Gwendolyne Burfin, Hadrien Regue, Quentin Semanas, Martine Valette, Bruno Lina, Laurence Josset |
| EPI_ISL_3130526 | CENTRE HOSPITALIER DE LA COTE BASQUE | CNR Virus des Infections Respiratoires - France SUD | Antonin Bal, Gregory Destras, Gwendolyne Burfin, Hadrien Regue, Quentin Semanas, Martine Valette, Bruno Lina, Laurence Josset |
| EPI_ISL_3130527 | LABORATOIRE CERBALLIANCE PLT VILLON | CNR Virus des Infections Respiratoires - France SUD | Antonin Bal, Gregory Destras, Gwendolyne Burfin, Hadrien Regue, Quentin Semanas, Martine Valette, Bruno Lina, Laurence Josset |
| EPI_ISL_3130528 | CERBALLIANCE CHARENTES SAINTES | CNR Virus des Infections Respiratoires - France SUD | Antonin Bal, Gregory Destras, Gwendolyne Burfin, Hadrien Regue, Quentin Semanas, Martine Valette, Bruno Lina, Laurence Josset |
| EPI_ISL_3130530 | LABORATOIRE CERBALLIANCE PLT VILLON | CNR Virus des Infections Respiratoires - France SUD | Antonin Bal, Gregory Destras, Gwendolyne Burfin, Hadrien Regue, Quentin Semanas, Martine Valette, Bruno Lina, Laurence Josset |
| EPI_ISL_3130532 | CENTRE HOSPITALIER DE LA COTE BASQUE | CNR Virus des Infections Respiratoires - France SUD | Antonin Bal, Gregory Destras, Gwendolyne Burfin, Hadrien Regue, Quentin Semanas, Martine Valette, Bruno Lina, Laurence Josset |
| EPI_ISL_3130533, | LABORATOIRE CERBALLIANCE PLT VILLON | CNR Virus des Infections Respiratoires - France SUD | Antonin Bal, Gregory Destras, Gwendolyne Burfin, Hadrien Regue, Quentin Semanas, Martine Valette, Bruno Lina, Laurence Josset |

| | | | |
|---|---|---|---|
| EPI_ISL_3130535, EPI_ISL_3130536, EPI_ISL_3130537, EPI_ISL_3130539, EPI_ISL_3130540, EPI_ISL_3130543 | | | |
| EPI_ISL_3130544 | CEDIBIO-UNILABS-TOUNIS | CNR Virus des Infections Respiratoires - France SUD | Antonin Bal, Gregory Destras, Gwendolyne Burfin, Hadrien Regue, Quentin Semanas, Martine Valette, Bruno Lina, Laurence Josset |
| EPI_ISL_3130545, EPI_ISL_3130546 | LABORATOIRE CERBALLIANCE PLT VILLON | CNR Virus des Infections Respiratoires - France SUD | Antonin Bal, Gregory Destras, Gwendolyne Burfin, Hadrien Regue, Quentin Semanas, Martine Valette, Bruno Lina, Laurence Josset |
| EPI_ISL_3130547 | CEDIBIO-UNILABS-TOUNIS | CNR Virus des Infections Respiratoires - France SUD | Antonin Bal, Gregory Destras, Gwendolyne Burfin, Hadrien Regue, Quentin Semanas, Martine Valette, Bruno Lina, Laurence Josset |
| EPI_ISL_3130548 | LABORATOIRE CERBALLIANCE PLT VILLON | CNR Virus des Infections Respiratoires - France SUD | Antonin Bal, Gregory Destras, Gwendolyne Burfin, Hadrien Regue, Quentin Semanas, Martine Valette, Bruno Lina, Laurence Josset |
| EPI_ISL_3130549 | CEDIBIO-UNILABS-TOUNIS | CNR Virus des Infections Respiratoires - France SUD | Antonin Bal, Gregory Destras, Gwendolyne Burfin, Hadrien Regue, Quentin Semanas, Martine Valette, Bruno Lina, Laurence Josset |
| EPI_ISL_3130550, EPI_ISL_3130551 | LABORATOIRE CERBALLIANCE PLT VILLON | CNR Virus des Infections Respiratoires - France SUD | Antonin Bal, Gregory Destras, Gwendolyne Burfin, Hadrien Regue, Quentin Semanas, Martine Valette, Bruno Lina, Laurence Josset |
| EPI_ISL_3130552 | BIOLITTORAL BIOGROUP PLATEAU TECH | CNR Virus des Infections Respiratoires - France SUD | Antonin Bal, Gregory Destras, Gwendolyne Burfin, Hadrien Regue, Quentin Semanas, Martine Valette, Bruno Lina, Laurence Josset |
| EPI_ISL_3130553, EPI_ISL_3130554, EPI_ISL_3130555 | LABORATOIRE BOUVIER | CNR Virus des Infections Respiratoires - France SUD | Antonin Bal, Gregory Destras, Gwendolyne Burfin, Hadrien Regue, Quentin Semanas, Martine Valette, Bruno Lina, Laurence Josset |
| EPI_ISL_3130558 | LABORATOIRE BIOESTEREL | CNR Virus des Infections Respiratoires - France SUD | Antonin Bal, Gregory Destras, Gwendolyne Burfin, Hadrien Regue, Quentin Semanas, Martine Valette, Bruno Lina, Laurence Josset |
| EPI_ISL_3130560 | LABORATOIRE CBM DE MURET | CNR Virus des Infections Respiratoires - France SUD | Antonin Bal, Gregory Destras, Gwendolyne Burfin, Hadrien Regue, Quentin Semanas, Martine Valette, Bruno Lina, Laurence Josset |
| EPI_ISL_3130561 | LABORATOIRE BIOESTEREL | CNR Virus des Infections Respiratoires - France SUD | Antonin Bal, Gregory Destras, Gwendolyne Burfin, Hadrien Regue, Quentin Semanas, Martine Valette, Bruno Lina, Laurence Josset |
| EPI_ISL_3130562 | LABORATOIRE CBM DE MURET | CNR Virus des Infections Respiratoires - France SUD | Antonin Bal, Gregory Destras, Gwendolyne Burfin, Hadrien Regue, Quentin Semanas, Martine Valette, Bruno Lina, Laurence Josset |
| EPI_ISL_3130563, EPI_ISL_3130564, EPI_ISL_3130566, EPI_ISL_3130567, EPI_ISL_3130568 | LABORATOIRE BIOESTEREL | CNR Virus des Infections Respiratoires - France SUD | Antonin Bal, Gregory Destras, Gwendolyne Burfin, Hadrien Regue, Quentin Semanas, Martine Valette, Bruno Lina, Laurence Josset |
| EPI_ISL_3130570 | CNR Virus des Infections Respiratoires - France SUD | CNR Virus des Infections Respiratoires - France SUD | Antonin Bal, Gregory Destras, Gwendolyne Burfin, Hadrien Regue, Quentin Semanas, Martine Valette, Bruno Lina, Laurence Josset |
| EPI_ISL_3130571 | LABORATOIRE NOVELAB | CNR Virus des Infections Respiratoires - France SUD | Antonin Bal, Gregory Destras, Gwendolyne Burfin, Hadrien Regue, Quentin Semanas, Martine Valette, Bruno Lina, Laurence Josset |
| EPI_ISL_3130573 | C.H.R.U. MONTPIED | CNR Virus des Infections Respiratoires - France SUD | Antonin Bal, Gregory Destras, Gwendolyne Burfin, Hadrien Regue, Quentin Semanas, Martine Valette, Bruno Lina, Laurence Josset |
| EPI_ISL_3130574 | LAM CERBALLIANCE | CNR Virus des Infections Respiratoires - France SUD | Antonin Bal, Gregory Destras, Gwendolyne Burfin, Hadrien Regue, Quentin Semanas, Martine Valette, Bruno Lina, Laurence Josset |
| EPI_ISL_3130576 | C.H.R.U. MONTPIED | CNR Virus des Infections Respiratoires - France SUD | Antonin Bal, Gregory Destras, Gwendolyne Burfin, Hadrien Regue, Quentin Semanas, Martine Valette, Bruno Lina, Laurence Josset |
| EPI_ISL_3130577 | LAM CERBALLIANCE | CNR Virus des Infections Respiratoires - France SUD | Antonin Bal, Gregory Destras, Gwendolyne Burfin, Hadrien Regue, Quentin Semanas, Martine Valette, Bruno Lina, Laurence Josset |
| EPI_ISL_3130578 | LABORATOIRE UNILIANS DECINES | CNR Virus des Infections Respiratoires - France SUD | Antonin Bal, Gregory Destras, Gwendolyne Burfin, Hadrien Regue, Quentin Semanas, Martine Valette, Bruno Lina, Laurence Josset |
| EPI_ISL_3130579 | SYNLAB GASCOGNE | CNR Virus des Infections Respiratoires - France SUD | Antonin Bal, Gregory Destras, Gwendolyne Burfin, Hadrien Regue, Quentin Semanas, Martine Valette, Bruno Lina, Laurence Josset |
| EPI_ISL_3130580, EPI_ISL_3130582 | LABORATOIRE UNILIANS DECINES | CNR Virus des Infections Respiratoires - France SUD | Antonin Bal, Gregory Destras, Gwendolyne Burfin, Hadrien Regue, Quentin Semanas, Martine Valette, Bruno Lina, Laurence Josset |
| EPI_ISL_3130583 | AX BIO OCEAN | CNR Virus des Infections Respiratoires - France SUD | Antonin Bal, Gregory Destras, Gwendolyne Burfin, Hadrien Regue, Quentin Semanas, Martine Valette, Bruno Lina, Laurence Josset |
| EPI_ISL_3130584 | SYNLAB GASCOGNE | CNR Virus des Infections Respiratoires - France SUD | Antonin Bal, Gregory Destras, Gwendolyne Burfin, Hadrien Regue, Quentin Semanas, Martine Valette, Bruno Lina, Laurence Josset |
| EPI_ISL_3130585 | LAM CERBALLIANCE | CNR Virus des Infections Respiratoires - France SUD | Antonin Bal, Gregory Destras, Gwendolyne Burfin, Hadrien Regue, Quentin Semanas, Martine Valette, Bruno Lina, Laurence Josset |
| EPI_ISL_3130587 | AX BIO OCEAN | CNR Virus des Infections Respiratoires - France SUD | Antonin Bal, Gregory Destras, Gwendolyne Burfin, Hadrien Regue, Quentin Semanas, Martine Valette, Bruno Lina, Laurence Josset |
| EPI_ISL_3130588 | SYNLAB GASCOGNE | CNR Virus des Infections Respiratoires - France SUD | Antonin Bal, Gregory Destras, Gwendolyne Burfin, Hadrien Regue, Quentin Semanas, Martine Valette, Bruno Lina, Laurence Josset |
| EPI_ISL_3130590, EPI_ISL_3130591 | AX BIO OCEAN | CNR Virus des Infections Respiratoires - France SUD | Antonin Bal, Gregory Destras, Gwendolyne Burfin, Hadrien Regue, Quentin Semanas, Martine Valette, Bruno Lina, Laurence Josset |
| EPI_ISL_3130592, EPI_ISL_3130594 | LABORATOIRE UNILIANS DECINES | CNR Virus des Infections Respiratoires - France SUD | Antonin Bal, Gregory Destras, Gwendolyne Burfin, Hadrien Regue, Quentin Semanas, Martine Valette, Bruno Lina, Laurence Josset |
| EPI_ISL_3130595 | AX BIO OCEAN | CNR Virus des Infections Respiratoires - France SUD | Antonin Bal, Gregory Destras, Gwendolyne Burfin, Hadrien Regue, Quentin Semanas, Martine Valette, Bruno Lina, Laurence Josset |
| EPI_ISL_3130596 | SYNLAB GASCOGNE | CNR Virus des Infections Respiratoires - France SUD | Antonin Bal, Gregory Destras, Gwendolyne Burfin, Hadrien Regue, Quentin Semanas, Martine Valette, Bruno Lina, Laurence Josset |
| EPI_ISL_3130597, EPI_ISL_3130599 | LABORATOIRE UNILIANS DECINES | CNR Virus des Infections Respiratoires - France SUD | Antonin Bal, Gregory Destras, Gwendolyne Burfin, Hadrien Regue, Quentin Semanas, Martine Valette, Bruno Lina, Laurence Josset |
| EPI_ISL_3130601 | SYNLAB GASCOGNE | CNR Virus des Infections Respiratoires - France SUD | Antonin Bal, Gregory Destras, Gwendolyne Burfin, Hadrien Regue, Quentin Semanas, Martine Valette, Bruno Lina, Laurence Josset |
| EPI_ISL_3130607 | LABORATOIRE UNILIANS DECINES | CNR Virus des Infections Respiratoires - France SUD | Antonin Bal, Gregory Destras, Gwendolyne Burfin, Hadrien Regue, Quentin Semanas, Martine Valette, Bruno Lina, Laurence Josset |
| EPI_ISL_3130609 | SYNLAB GASCOGNE | CNR Virus des Infections Respiratoires - France SUD | Antonin Bal, Gregory Destras, Gwendolyne Burfin, Hadrien Regue, Quentin Semanas, Martine Valette, Bruno Lina, Laurence Josset |
| EPI_ISL_3130612 | LABORATOIRE UNILIANS DECINES | CNR Virus des Infections Respiratoires - France SUD | Antonin Bal, Gregory Destras, Gwendolyne Burfin, Hadrien Regue, Quentin Semanas, Martine Valette, Bruno Lina, Laurence Josset |
| EPI_ISL_3130614, EPI_ISL_3130623 | AX BIO OCEAN | CNR Virus des Infections Respiratoires - France SUD | Antonin Bal, Gregory Destras, Gwendolyne Burfin, Hadrien Regue, Quentin Semanas, Martine Valette, Bruno Lina, Laurence Josset |
| EPI_ISL_3130625 | LAM BIORYLIS | CNR Virus des Infections Respiratoires - France SUD | Antonin Bal, Gregory Destras, Gwendolyne Burfin, Hadrien Regue, Quentin Semanas, Martine Valette, Bruno Lina, Laurence Josset |
| EPI_ISL_3130627 | AX BIO OCEAN | CNR Virus des Infections Respiratoires - France SUD | Antonin Bal, Gregory Destras, Gwendolyne Burfin, Hadrien Regue, Quentin Semanas, Martine Valette, Bruno Lina, Laurence Josset |
| EPI_ISL_3130629 | LAM CERBALLIANCE | CNR Virus des Infections Respiratoires - France SUD | Antonin Bal, Gregory Destras, Gwendolyne Burfin, Hadrien Regue, Quentin Semanas, Martine Valette, Bruno Lina, Laurence Josset |
| EPI_ISL_3130630, EPI_ISL_3130632 | AX BIO OCEAN | CNR Virus des Infections Respiratoires - France SUD | Antonin Bal, Gregory Destras, Gwendolyne Burfin, Hadrien Regue, Quentin Semanas, Martine Valette, Bruno Lina, Laurence Josset |
| EPI_ISL_3130633 | LAM CERBALLIANCE | CNR Virus des Infections Respiratoires - France SUD | Antonin Bal, Gregory Destras, Gwendolyne Burfin, Hadrien Regue, Quentin Semanas, Martine Valette, Bruno Lina, Laurence Josset |
| EPI_ISL_3130634, EPI_ISL_3130636 | AX BIO OCEAN | CNR Virus des Infections Respiratoires - France SUD | Antonin Bal, Gregory Destras, Gwendolyne Burfin, Hadrien Regue, Quentin Semanas, Martine Valette, Bruno Lina, Laurence Josset |
| EPI_ISL_3130640, EPI_ISL_3130643 | LAM CERBALLIANCE | CNR Virus des Infections Respiratoires - France SUD | Antonin Bal, Gregory Destras, Gwendolyne Burfin, Hadrien Regue, Quentin Semanas, Martine Valette, Bruno Lina, Laurence Josset |
| EPI_ISL_3130646 | ASTRALAB TASSIGNY | CNR Virus des Infections Respiratoires - France SUD | Antonin Bal, Gregory Destras, Gwendolyne Burfin, Hadrien Regue, Quentin Semanas, Martine Valette, Bruno Lina, Laurence Josset |
| EPI_ISL_3130648 | LAM CERBALLIANCE | CNR Virus des Infections Respiratoires - France SUD | Antonin Bal, Gregory Destras, Gwendolyne Burfin, Hadrien Regue, Quentin Semanas, Martine Valette, Bruno Lina, Laurence Josset |
| EPI_ISL_3130649 | ASTRALAB TASSIGNY | CNR Virus des Infections Respiratoires - France SUD | Antonin Bal, Gregory Destras, Gwendolyne Burfin, Hadrien Regue, Quentin Semanas, Martine Valette, Bruno Lina, Laurence Josset |
| EPI_ISL_3130650 | LAM CERBALLIANCE | CNR Virus des Infections Respiratoires - France SUD | Antonin Bal, Gregory Destras, Gwendolyne Burfin, Hadrien Regue, Quentin Semanas, Martine Valette, Bruno Lina, Laurence Josset |
| EPI_ISL_3130651, EPI_ISL_3130652, EPI_ISL_3130653 | ASTRALAB TASSIGNY | CNR Virus des Infections Respiratoires - France SUD | Antonin Bal, Gregory Destras, Gwendolyne Burfin, Hadrien Regue, Quentin Semanas, Martine Valette, Bruno Lina, Laurence Josset |
| EPI_ISL_3130654 | LAM CERBALLIANCE | CNR Virus des Infections Respiratoires - France SUD | Antonin Bal, Gregory Destras, Gwendolyne Burfin, Hadrien Regue, Quentin Semanas, Martine Valette, Bruno Lina, Laurence Josset |
| EPI_ISL_3130655 | LAM EIMER LENYS | CNR Virus des Infections Respiratoires - France SUD | Antonin Bal, Gregory Destras, Gwendolyne Burfin, Hadrien Regue, Quentin Semanas, Martine Valette, Bruno Lina, Laurence Josset |
| EPI_ISL_3130656, EPI_ISL_3130657 | LAM CERBALLIANCE | CNR Virus des Infections Respiratoires - France SUD | Antonin Bal, Gregory Destras, Gwendolyne Burfin, Hadrien Regue, Quentin Semanas, Martine Valette, Bruno Lina, Laurence Josset |
| EPI_ISL_3130658, EPI_ISL_3130659 | LAM EIMER LENYS | CNR Virus des Infections Respiratoires - France SUD | Antonin Bal, Gregory Destras, Gwendolyne Burfin, Hadrien Regue, Quentin Semanas, Martine Valette, Bruno Lina, Laurence Josset |
| EPI_ISL_3130660, EPI_ISL_3130662 | LAM CERBALLIANCE | CNR Virus des Infections Respiratoires - France SUD | Antonin Bal, Gregory Destras, Gwendolyne Burfin, Hadrien Regue, Quentin Semanas, Martine Valette, Bruno Lina, Laurence Josset |
| EPI_ISL_3130663, EPI_ISL_3130664 | LAM EIMER LENYS | CNR Virus des Infections Respiratoires - France SUD | Antonin Bal, Gregory Destras, Gwendolyne Burfin, Hadrien Regue, Quentin Semanas, Martine Valette, Bruno Lina, Laurence Josset |
| EPI_ISL_3130665 | LAM CERBALLIANCE | CNR Virus des Infections Respiratoires - France SUD | Antonin Bal, Gregory Destras, Gwendolyne Burfin, Hadrien Regue, Quentin Semanas, Martine Valette, Bruno Lina, Laurence Josset |
| EPI_ISL_3130666 | LAM EIMER LENYS | CNR Virus des Infections Respiratoires - France SUD | Antonin Bal, Gregory Destras, Gwendolyne Burfin, Hadrien Regue, Quentin Semanas, Martine Valette, Bruno Lina, Laurence Josset |

| | | | |
|---|---|---|---|
| EPI_ISL_3130667 | LAM CERBALLIANCE | CNR Virus des Infections Respiratoires - France SUD | Antonin Bal, Gregory Destras, Gwendolyne Burfin, Hadrien Regue, Quentin Semanas, Martine Valette, Bruno Lina, Laurence Josset |
| EPI_ISL_3130668 | LAM EIMER LENYS | CNR Virus des Infections Respiratoires - France SUD | Antonin Bal, Gregory Destras, Gwendolyne Burfin, Hadrien Regue, Quentin Semanas, Martine Valette, Bruno Lina, Laurence Josset |
| EPI_ISL_3130669, EPI_ISL_3130670 | LAM CERBALLIANCE | CNR Virus des Infections Respiratoires - France SUD | Antonin Bal, Gregory Destras, Gwendolyne Burfin, Hadrien Regue, Quentin Semanas, Martine Valette, Bruno Lina, Laurence Josset |
| EPI_ISL_3130675 | CENTRE HOSPITALIER ST JOSEPH ST LUC | CNR Virus des Infections Respiratoires - France SUD | Antonin Bal, Gregory Destras, Gwendolyne Burfin, Hadrien Regue, Quentin Semanas, Martine Valette, Bruno Lina, Laurence Josset |
| EPI_ISL_3130676, EPI_ISL_3130677, EPI_ISL_3130679, EPI_ISL_3130680, EPI_ISL_3130681 | LABORATOIRE d'ANALYSES de BIOLOGIE MEDICALES | CNR Virus des Infections Respiratoires - France SUD | Antonin Bal, Gregory Destras, Gwendolyne Burfin, Hadrien Regue, Quentin Semanas, Martine Valette, Bruno Lina, Laurence Josset |
| EPI_ISL_3130693, EPI_ISL_3130694, EPI_ISL_3130695, EPI_ISL_3130696, EPI_ISL_3130700, EPI_ISL_3130701 | DYOMEDEA-LABORATOIRE DE LA SAUVEGARDE | CNR Virus des Infections Respiratoires - France SUD | Antonin Bal, Gregory Destras, Gwendolyne Burfin, Hadrien Regue, Quentin Semanas, Martine Valette, Bruno Lina, Laurence Josset |
| EPI_ISL_3130702, EPI_ISL_3130703 | HOPITAL SAINT ANDRE | CNR Virus des Infections Respiratoires - France SUD | Antonin Bal, Gregory Destras, Gwendolyne Burfin, Hadrien Regue, Quentin Semanas, Martine Valette, Bruno Lina, Laurence Josset |
| EPI_ISL_3130704 | DYOMEDEA-LABORATOIRE DE LA SAUVEGARDE | CNR Virus des Infections Respiratoires - France SUD | Antonin Bal, Gregory Destras, Gwendolyne Burfin, Hadrien Regue, Quentin Semanas, Martine Valette, Bruno Lina, Laurence Josset |
| EPI_ISL_3130705 | LABORATOIRE BIOMEDICA | CNR Virus des Infections Respiratoires - France SUD | Antonin Bal, Gregory Destras, Gwendolyne Burfin, Hadrien Regue, Quentin Semanas, Martine Valette, Bruno Lina, Laurence Josset |
| EPI_ISL_3130706 | LABORATOIRE MAYMAT | CNR Virus des Infections Respiratoires - France SUD | Antonin Bal, Gregory Destras, Gwendolyne Burfin, Hadrien Regue, Quentin Semanas, Martine Valette, Bruno Lina, Laurence Josset |
| EPI_ISL_3130707, EPI_ISL_3130709 | LABORATOIRE BIOMEDICA | CNR Virus des Infections Respiratoires - France SUD | Antonin Bal, Gregory Destras, Gwendolyne Burfin, Hadrien Regue, Quentin Semanas, Martine Valette, Bruno Lina, Laurence Josset |
| EPI_ISL_3130712 | MIRIALIS CLUSES BECHET | CNR Virus des Infections Respiratoires - France SUD | Antonin Bal, Gregory Destras, Gwendolyne Burfin, Hadrien Regue, Quentin Semanas, Martine Valette, Bruno Lina, Laurence Josset |
| EPI_ISL_3130713, EPI_ISL_3130714 | LX BIO | CNR Virus des Infections Respiratoires - France SUD | Antonin Bal, Gregory Destras, Gwendolyne Burfin, Hadrien Regue, Quentin Semanas, Martine Valette, Bruno Lina, Laurence Josset |
| EPI_ISL_3130715 | MIRIALIS CLUSES BECHET | CNR Virus des Infections Respiratoires - France SUD | Antonin Bal, Gregory Destras, Gwendolyne Burfin, Hadrien Regue, Quentin Semanas, Martine Valette, Bruno Lina, Laurence Josset |
| EPI_ISL_3130716 | LX BIO | CNR Virus des Infections Respiratoires - France SUD | Antonin Bal, Gregory Destras, Gwendolyne Burfin, Hadrien Regue, Quentin Semanas, Martine Valette, Bruno Lina, Laurence Josset |
| EPI_ISL_3130717 | MIRIALIS CLUSES BECHET | CNR Virus des Infections Respiratoires - France SUD | Antonin Bal, Gregory Destras, Gwendolyne Burfin, Hadrien Regue, Quentin Semanas, Martine Valette, Bruno Lina, Laurence Josset |
| EPI_ISL_3130718 | LX BIO | CNR Virus des Infections Respiratoires - France SUD | Antonin Bal, Gregory Destras, Gwendolyne Burfin, Hadrien Regue, Quentin Semanas, Martine Valette, Bruno Lina, Laurence Josset |
| EPI_ISL_3130719 | MIRIALIS CLUSES BECHET | CNR Virus des Infections Respiratoires - France SUD | Antonin Bal, Gregory Destras, Gwendolyne Burfin, Hadrien Regue, Quentin Semanas, Martine Valette, Bruno Lina, Laurence Josset |
| EPI_ISL_3130721 | SCM B12 | CNR Virus des Infections Respiratoires - France SUD | Antonin Bal, Gregory Destras, Gwendolyne Burfin, Hadrien Regue, Quentin Semanas, Martine Valette, Bruno Lina, Laurence Josset |
| EPI_ISL_3130722 | BIOMNIS LYON | CNR Virus des Infections Respiratoires - France SUD | Antonin Bal, Gregory Destras, Gwendolyne Burfin, Hadrien Regue, Quentin Semanas, Martine Valette, Bruno Lina, Laurence Josset |
| EPI_ISL_3130723 | SCM B12 | CNR Virus des Infections Respiratoires - France SUD | Antonin Bal, Gregory Destras, Gwendolyne Burfin, Hadrien Regue, Quentin Semanas, Martine Valette, Bruno Lina, Laurence Josset |
| EPI_ISL_3130724 | BIOMNIS LYON | CNR Virus des Infections Respiratoires - France SUD | Antonin Bal, Gregory Destras, Gwendolyne Burfin, Hadrien Regue, Quentin Semanas, Martine Valette, Bruno Lina, Laurence Josset |
| EPI_ISL_3130725 | MIRIALIS CLUSES BECHET | CNR Virus des Infections Respiratoires - France SUD | Antonin Bal, Gregory Destras, Gwendolyne Burfin, Hadrien Regue, Quentin Semanas, Martine Valette, Bruno Lina, Laurence Josset |
| EPI_ISL_3130726 | SCM B12 | CNR Virus des Infections Respiratoires - France SUD | Antonin Bal, Gregory Destras, Gwendolyne Burfin, Hadrien Regue, Quentin Semanas, Martine Valette, Bruno Lina, Laurence Josset |
| EPI_ISL_3130727 | MIRIALIS CLUSES BECHET | CNR Virus des Infections Respiratoires - France SUD | Antonin Bal, Gregory Destras, Gwendolyne Burfin, Hadrien Regue, Quentin Semanas, Martine Valette, Bruno Lina, Laurence Josset |
| EPI_ISL_3130728 | SCM B12 | CNR Virus des Infections Respiratoires - France SUD | Antonin Bal, Gregory Destras, Gwendolyne Burfin, Hadrien Regue, Quentin Semanas, Martine Valette, Bruno Lina, Laurence Josset |
| EPI_ISL_3130729, EPI_ISL_3130730 | MIRIALIS CLUSES BECHET | CNR Virus des Infections Respiratoires - France SUD | Antonin Bal, Gregory Destras, Gwendolyne Burfin, Hadrien Regue, Quentin Semanas, Martine Valette, Bruno Lina, Laurence Josset |
| EPI_ISL_3130731 | SCM B12 | CNR Virus des Infections Respiratoires - France SUD | Antonin Bal, Gregory Destras, Gwendolyne Burfin, Hadrien Regue, Quentin Semanas, Martine Valette, Bruno Lina, Laurence Josset |
| EPI_ISL_3130732 | MIRIALIS CLUSES BECHET | CNR Virus des Infections Respiratoires - France SUD | Antonin Bal, Gregory Destras, Gwendolyne Burfin, Hadrien Regue, Quentin Semanas, Martine Valette, Bruno Lina, Laurence Josset |
| EPI_ISL_3130733 | SCM B12 | CNR Virus des Infections Respiratoires - France SUD | Antonin Bal, Gregory Destras, Gwendolyne Burfin, Hadrien Regue, Quentin Semanas, Martine Valette, Bruno Lina, Laurence Josset |
| EPI_ISL_3130734 | MIRIALIS CLUSES BECHET | CNR Virus des Infections Respiratoires - France SUD | Antonin Bal, Gregory Destras, Gwendolyne Burfin, Hadrien Regue, Quentin Semanas, Martine Valette, Bruno Lina, Laurence Josset |
| EPI_ISL_3130735, EPI_ISL_3130736 | LABORATOIRE ANALYSEO | CNR Virus des Infections Respiratoires - France SUD | Antonin Bal, Gregory Destras, Gwendolyne Burfin, Hadrien Regue, Quentin Semanas, Martine Valette, Bruno Lina, Laurence Josset |
| EPI_ISL_3130737 | SCM B12 | CNR Virus des Infections Respiratoires - France SUD | Antonin Bal, Gregory Destras, Gwendolyne Burfin, Hadrien Regue, Quentin Semanas, Martine Valette, Bruno Lina, Laurence Josset |
| EPI_ISL_3130738 | LABORATOIRE ANALYSEO | CNR Virus des Infections Respiratoires - France SUD | Antonin Bal, Gregory Destras, Gwendolyne Burfin, Hadrien Regue, Quentin Semanas, Martine Valette, Bruno Lina, Laurence Josset |
| EPI_ISL_3130739 | SCM B12 | CNR Virus des Infections Respiratoires - France SUD | Antonin Bal, Gregory Destras, Gwendolyne Burfin, Hadrien Regue, Quentin Semanas, Martine Valette, Bruno Lina, Laurence Josset |
| EPI_ISL_3130740 | MIRIALIS CLUSES BECHET | CNR Virus des Infections Respiratoires - France SUD | Antonin Bal, Gregory Destras, Gwendolyne Burfin, Hadrien Regue, Quentin Semanas, Martine Valette, Bruno Lina, Laurence Josset |
| EPI_ISL_3130741 | LABORATOIRE ANALYSEO | CNR Virus des Infections Respiratoires - France SUD | Antonin Bal, Gregory Destras, Gwendolyne Burfin, Hadrien Regue, Quentin Semanas, Martine Valette, Bruno Lina, Laurence Josset |
| EPI_ISL_3130742 | MIRIALIS CLUSES BECHET | CNR Virus des Infections Respiratoires - France SUD | Antonin Bal, Gregory Destras, Gwendolyne Burfin, Hadrien Regue, Quentin Semanas, Martine Valette, Bruno Lina, Laurence Josset |
| EPI_ISL_3130743 | SCM B12 | CNR Virus des Infections Respiratoires - France SUD | Antonin Bal, Gregory Destras, Gwendolyne Burfin, Hadrien Regue, Quentin Semanas, Martine Valette, Bruno Lina, Laurence Josset |
| EPI_ISL_3130744 | LABORATOIRE ANALYSEO | CNR Virus des Infections Respiratoires - France SUD | Antonin Bal, Gregory Destras, Gwendolyne Burfin, Hadrien Regue, Quentin Semanas, Martine Valette, Bruno Lina, Laurence Josset |
| EPI_ISL_3130746, EPI_ISL_3130747, EPI_ISL_3130748, EPI_ISL_3130751, EPI_ISL_3130752, EPI_ISL_3130754, EPI_ISL_3130755, EPI_ISL_3130756, EPI_ISL_3130757, EPI_ISL_3130758, EPI_ISL_3130759, EPI_ISL_3130760, EPI_ISL_3130761, EPI_ISL_3130762, EPI_ISL_3130763, EPI_ISL_3130764, EPI_ISL_3130766, EPI_ISL_3130767, EPI_ISL_3130768 | | | |
| see above | MIRIALIS CLUSES BECHET | CNR Virus des Infections Respiratoires - France SUD | Antonin Bal, Gregory Destras, Gwendolyne Burfin, Hadrien Regue, Quentin Semanas, Martine Valette, Bruno Lina, Laurence Josset |
| EPI_ISL_3130769, EPI_ISL_3130770, EPI_ISL_3130773, EPI_ISL_3130774, EPI_ISL_3130775, EPI_ISL_3130776 | DYOMEDEA-LABORATOIRE DE LA SAUVEGARDE | CNR Virus des Infections Respiratoires - France SUD | Antonin Bal, Gregory Destras, Gwendolyne Burfin, Hadrien Regue, Quentin Semanas, Martine Valette, Bruno Lina, Laurence Josset |
| EPI_ISL_3130777 | ASTRALAB TASSIGNY | CNR Virus des Infections Respiratoires - France SUD | Antonin Bal, Gregory Destras, Gwendolyne Burfin, Hadrien Regue, Quentin Semanas, Martine Valette, Bruno Lina, Laurence Josset |
| EPI_ISL_3130778, EPI_ISL_3130779, EPI_ISL_3130780, EPI_ISL_3130781, EPI_ISL_3130782 | DYOMEDEA-LABORATOIRE DE LA SAUVEGARDE | CNR Virus des Infections Respiratoires - France SUD | Antonin Bal, Gregory Destras, Gwendolyne Burfin, Hadrien Regue, Quentin Semanas, Martine Valette, Bruno Lina, Laurence Josset |
| EPI_ISL_3130783 | ASTRALAB TASSIGNY | CNR Virus des Infections Respiratoires - France SUD | Antonin Bal, Gregory Destras, Gwendolyne Burfin, Hadrien Regue, Quentin Semanas, Martine Valette, Bruno Lina, Laurence Josset |
| EPI_ISL_3130784, EPI_ISL_3130785, EPI_ISL_3130786, EPI_ISL_3130787 | DYOMEDEA-LABORATOIRE DE LA SAUVEGARDE | CNR Virus des Infections Respiratoires - France SUD | Antonin Bal, Gregory Destras, Gwendolyne Burfin, Hadrien Regue, Quentin Semanas, Martine Valette, Bruno Lina, Laurence Josset |
| EPI_ISL_3130789, EPI_ISL_3130790 | CNR Virus des Infections Respiratoires - France SUD | CNR Virus des Infections Respiratoires - France SUD | Antonin Bal, Gregory Destras, Gwendolyne Burfin, Hadrien Regue, Quentin Semanas, Martine Valette, Bruno Lina, Laurence Josset |
| EPI_ISL_3130791, EPI_ISL_3130792 | BIOLITTORAL BIOGROUP PLATEAU TECH | CNR Virus des Infections Respiratoires - France SUD | Antonin Bal, Gregory Destras, Gwendolyne Burfin, Hadrien Regue, Quentin Semanas, Martine Valette, Bruno Lina, Laurence Josset |
| EPI_ISL_3130793 | CNR Virus des Infections Respiratoires - France SUD | CNR Virus des Infections Respiratoires - France SUD | Antonin Bal, Gregory Destras, Gwendolyne Burfin, Hadrien Regue, Quentin Semanas, Martine Valette, Bruno Lina, Laurence Josset |
| EPI_ISL_3130794 | BIOLITTORAL BIOGROUP PLATEAU TECH | CNR Virus des Infections Respiratoires - France SUD | Antonin Bal, Gregory Destras, Gwendolyne Burfin, Hadrien Regue, Quentin Semanas, Martine Valette, Bruno Lina, Laurence Josset |
| EPI_ISL_3130795, EPI_ISL_3130796, EPI_ISL_3130798, EPI_ISL_3130799 | ASTRALAB TASSIGNY | CNR Virus des Infections Respiratoires - France SUD | Antonin Bal, Gregory Destras, Gwendolyne Burfin, Hadrien Regue, Quentin Semanas, Martine Valette, Bruno Lina, Laurence Josset |
| EPI_ISL_3130801, EPI_ISL_3130802, | SCM B12 | CNR Virus des Infections Respiratoires - France SUD | Antonin Bal, Gregory Destras, Gwendolyne Burfin, Hadrien Regue, Quentin Semanas, Martine Valette, Bruno Lina, Laurence Josset |

| | | | |
|---|---|---|---|
| EPI_ISL_3130803, EPI_ISL_3130804, EPI_ISL_3130805, EPI_ISL_3130806, EPI_ISL_3130807 | | | |
| EPI_ISL_3130808 | CNR Virus des Infections Respiratoires - France SUD | CNR Virus des Infections Respiratoires - France SUD | Antonin Bal, Gregory Destras, Gwendolyne Burfin, Hadrien Regue, Quentin Semanas, Martine Valette, Bruno Lina, Laurence Josset |
| EPI_ISL_3130810, EPI_ISL_3130811 | SCM B12 | CNR Virus des Infections Respiratoires - France SUD | Antonin Bal, Gregory Destras, Gwendolyne Burfin, Hadrien Regue, Quentin Semanas, Martine Valette, Bruno Lina, Laurence Josset |
| EPI_ISL_3130812 | CH VALENCE | CNR Virus des Infections Respiratoires - France SUD | Antonin Bal, Gregory Destras, Gwendolyne Burfin, Hadrien Regue, Quentin Semanas, Martine Valette, Bruno Lina, Laurence Josset |
| EPI_ISL_3130813, EPI_ISL_3130814, EPI_ISL_3130815, EPI_ISL_3130816, EPI_ISL_3130817, EPI_ISL_3130818, EPI_ISL_3130819, EPI_ISL_3130820, EPI_ISL_3130821, EPI_ISL_3130822, EPI_ISL_3130823, EPI_ISL_3130824, EPI_ISL_3130825, EPI_ISL_3130826, EPI_ISL_3130827, EPI_ISL_3130828, EPI_ISL_3130829 | | | |
| see above | SCM B12 | CNR Virus des Infections Respiratoires - France SUD | Antonin Bal, Gregory Destras, Gwendolyne Burfin, Hadrien Regue, Quentin Semanas, Martine Valette, Bruno Lina, Laurence Josset |
| EPI_ISL_3130830 | CENTRE HOSPITALIER PIERRE OUDOT | CNR Virus des Infections Respiratoires - France SUD | Antonin Bal, Gregory Destras, Gwendolyne Burfin, Hadrien Regue, Quentin Semanas, Martine Valette, Bruno Lina, Laurence Josset |
| EPI_ISL_3130831 | CH DE ROANNE | CNR Virus des Infections Respiratoires - France SUD | Antonin Bal, Gregory Destras, Gwendolyne Burfin, Hadrien Regue, Quentin Semanas, Martine Valette, Bruno Lina, Laurence Josset |
| EPI_ISL_3130832 | SYNLAB GASCOGNE | CNR Virus des Infections Respiratoires - France SUD | Antonin Bal, Gregory Destras, Gwendolyne Burfin, Hadrien Regue, Quentin Semanas, Martine Valette, Bruno Lina, Laurence Josset |
| EPI_ISL_3130833 | CHU ST ETIENNE HOPITAL NORD | CNR Virus des Infections Respiratoires - France SUD | Antonin Bal, Gregory Destras, Gwendolyne Burfin, Hadrien Regue, Quentin Semanas, Martine Valette, Bruno Lina, Laurence Josset |
| EPI_ISL_3130834, EPI_ISL_3130835, EPI_ISL_3130836, EPI_ISL_3130837, EPI_ISL_3130838 | SYNLAB GASCOGNE | CNR Virus des Infections Respiratoires - France SUD | Antonin Bal, Gregory Destras, Gwendolyne Burfin, Hadrien Regue, Quentin Semanas, Martine Valette, Bruno Lina, Laurence Josset |
| EPI_ISL_3130839 | DYOMEDEA-LABORATOIRE DE LA SAUVEGARDE | CNR Virus des Infections Respiratoires - France SUD | Antonin Bal, Gregory Destras, Gwendolyne Burfin, Hadrien Regue, Quentin Semanas, Martine Valette, Bruno Lina, Laurence Josset |
| EPI_ISL_3130840, EPI_ISL_3130841, EPI_ISL_3130842 | ASTRALAB TASSIGNY | CNR Virus des Infections Respiratoires - France SUD | Antonin Bal, Gregory Destras, Gwendolyne Burfin, Hadrien Regue, Quentin Semanas, Martine Valette, Bruno Lina, Laurence Josset |
| EPI_ISL_3130844 | LABORATOIRE CERBALLIANCE PLT VILLON | CNR Virus des Infections Respiratoires - France SUD | Antonin Bal, Gregory Destras, Gwendolyne Burfin, Hadrien Regue, Quentin Semanas, Martine Valette, Bruno Lina, Laurence Josset |
| EPI_ISL_3130845 | LAM BIOPOLE ORTHEZ | CNR Virus des Infections Respiratoires - France SUD | Antonin Bal, Gregory Destras, Gwendolyne Burfin, Hadrien Regue, Quentin Semanas, Martine Valette, Bruno Lina, Laurence Josset |
| EPI_ISL_3130846 | LABORATOIRE CERBALLIANCE PLT VILLON | CNR Virus des Infections Respiratoires - France SUD | Antonin Bal, Gregory Destras, Gwendolyne Burfin, Hadrien Regue, Quentin Semanas, Martine Valette, Bruno Lina, Laurence Josset |
| EPI_ISL_3130847 | LAM BIOPOLE ORTHEZ | CNR Virus des Infections Respiratoires - France SUD | Antonin Bal, Gregory Destras, Gwendolyne Burfin, Hadrien Regue, Quentin Semanas, Martine Valette, Bruno Lina, Laurence Josset |
| EPI_ISL_3130848 | LABORATOIRE CERBALLIANCE PLT VILLON | CNR Virus des Infections Respiratoires - France SUD | Antonin Bal, Gregory Destras, Gwendolyne Burfin, Hadrien Regue, Quentin Semanas, Martine Valette, Bruno Lina, Laurence Josset |
| EPI_ISL_3130849 | LAM BIOPOLE ORTHEZ | CNR Virus des Infections Respiratoires - France SUD | Antonin Bal, Gregory Destras, Gwendolyne Burfin, Hadrien Regue, Quentin Semanas, Martine Valette, Bruno Lina, Laurence Josset |
| EPI_ISL_3130850 | LABORATOIRE CERBALLIANCE PLT VILLON | CNR Virus des Infections Respiratoires - France SUD | Antonin Bal, Gregory Destras, Gwendolyne Burfin, Hadrien Regue, Quentin Semanas, Martine Valette, Bruno Lina, Laurence Josset |
| EPI_ISL_3130851 | LAM BIOPOLE ORTHEZ | CNR Virus des Infections Respiratoires - France SUD | Antonin Bal, Gregory Destras, Gwendolyne Burfin, Hadrien Regue, Quentin Semanas, Martine Valette, Bruno Lina, Laurence Josset |
| EPI_ISL_3130852 | LABORATOIRE CERBALLIANCE PLT VILLON | CNR Virus des Infections Respiratoires - France SUD | Antonin Bal, Gregory Destras, Gwendolyne Burfin, Hadrien Regue, Quentin Semanas, Martine Valette, Bruno Lina, Laurence Josset |
| EPI_ISL_3130853 | CENTRE HOSPITALIER DE BASTIA | CNR Virus des Infections Respiratoires - France SUD | Antonin Bal, Gregory Destras, Gwendolyne Burfin, Hadrien Regue, Quentin Semanas, Martine Valette, Bruno Lina, Laurence Josset |
| EPI_ISL_3130854 | LABORATOIRE CERBALLIANCE PLT VILLON | CNR Virus des Infections Respiratoires - France SUD | Antonin Bal, Gregory Destras, Gwendolyne Burfin, Hadrien Regue, Quentin Semanas, Martine Valette, Bruno Lina, Laurence Josset |
| EPI_ISL_3130855 | MIRIALIS CLUSES BECHET | CNR Virus des Infections Respiratoires - France SUD | Antonin Bal, Gregory Destras, Gwendolyne Burfin, Hadrien Regue, Quentin Semanas, Martine Valette, Bruno Lina, Laurence Josset |
| EPI_ISL_3130856 | LABORATOIRE CERBALLIANCE PLT VILLON | CNR Virus des Infections Respiratoires - France SUD | Antonin Bal, Gregory Destras, Gwendolyne Burfin, Hadrien Regue, Quentin Semanas, Martine Valette, Bruno Lina, Laurence Josset |
| EPI_ISL_3130857 | MIRIALIS CLUSES BECHET | CNR Virus des Infections Respiratoires - France SUD | Antonin Bal, Gregory Destras, Gwendolyne Burfin, Hadrien Regue, Quentin Semanas, Martine Valette, Bruno Lina, Laurence Josset |
| EPI_ISL_3130858 | LABORATOIRE CERBALLIANCE PLT VILLON | CNR Virus des Infections Respiratoires - France SUD | Antonin Bal, Gregory Destras, Gwendolyne Burfin, Hadrien Regue, Quentin Semanas, Martine Valette, Bruno Lina, Laurence Josset |
| EPI_ISL_3130859 | MIRIALIS CLUSES BECHET | CNR Virus des Infections Respiratoires - France SUD | Antonin Bal, Gregory Destras, Gwendolyne Burfin, Hadrien Regue, Quentin Semanas, Martine Valette, Bruno Lina, Laurence Josset |
| EPI_ISL_3130860 | LABORATOIRE CERBALLIANCE PLT VILLON | CNR Virus des Infections Respiratoires - France SUD | Antonin Bal, Gregory Destras, Gwendolyne Burfin, Hadrien Regue, Quentin Semanas, Martine Valette, Bruno Lina, Laurence Josset |
| EPI_ISL_3130861 | MIRIALIS CLUSES BECHET | CNR Virus des Infections Respiratoires - France SUD | Antonin Bal, Gregory Destras, Gwendolyne Burfin, Hadrien Regue, Quentin Semanas, Martine Valette, Bruno Lina, Laurence Josset |
| EPI_ISL_3130862, EPI_ISL_3130863 | LABORATOIRE CERBALLIANCE PLT VILLON | CNR Virus des Infections Respiratoires - France SUD | Antonin Bal, Gregory Destras, Gwendolyne Burfin, Hadrien Regue, Quentin Semanas, Martine Valette, Bruno Lina, Laurence Josset |
| EPI_ISL_3130864 | MIRIALIS CLUSES BECHET | CNR Virus des Infections Respiratoires - France SUD | Antonin Bal, Gregory Destras, Gwendolyne Burfin, Hadrien Regue, Quentin Semanas, Martine Valette, Bruno Lina, Laurence Josset |
| EPI_ISL_3130866, EPI_ISL_3130867 | LABORATOIRE CERBALLIANCE PLT VILLON | CNR Virus des Infections Respiratoires - France SUD | Antonin Bal, Gregory Destras, Gwendolyne Burfin, Hadrien Regue, Quentin Semanas, Martine Valette, Bruno Lina, Laurence Josset |
| EPI_ISL_3130868 | MIRIALIS CLUSES BECHET | CNR Virus des Infections Respiratoires - France SUD | Antonin Bal, Gregory Destras, Gwendolyne Burfin, Hadrien Regue, Quentin Semanas, Martine Valette, Bruno Lina, Laurence Josset |
| EPI_ISL_3130869 | LABORATOIRE CERBALLIANCE PLT VILLON | CNR Virus des Infections Respiratoires - France SUD | Antonin Bal, Gregory Destras, Gwendolyne Burfin, Hadrien Regue, Quentin Semanas, Martine Valette, Bruno Lina, Laurence Josset |
| EPI_ISL_3130870 | MIRIALIS CLUSES BECHET | CNR Virus des Infections Respiratoires - France SUD | Antonin Bal, Gregory Destras, Gwendolyne Burfin, Hadrien Regue, Quentin Semanas, Martine Valette, Bruno Lina, Laurence Josset |
| EPI_ISL_3130871 | CENTRE HOSPITALIER PIERRE OUDOT | CNR Virus des Infections Respiratoires - France SUD | Antonin Bal, Gregory Destras, Gwendolyne Burfin, Hadrien Regue, Quentin Semanas, Martine Valette, Bruno Lina, Laurence Josset |
| EPI_ISL_3130872 | MIRIALIS CLUSES BECHET | CNR Virus des Infections Respiratoires - France SUD | Antonin Bal, Gregory Destras, Gwendolyne Burfin, Hadrien Regue, Quentin Semanas, Martine Valette, Bruno Lina, Laurence Josset |
| EPI_ISL_3130873 | LABORATOIRE CERBALLIANCE PLT VILLON | CNR Virus des Infections Respiratoires - France SUD | Antonin Bal, Gregory Destras, Gwendolyne Burfin, Hadrien Regue, Quentin Semanas, Martine Valette, Bruno Lina, Laurence Josset |
| EPI_ISL_3130874, EPI_ISL_3130876 | MIRIALIS CLUSES BECHET | CNR Virus des Infections Respiratoires - France SUD | Antonin Bal, Gregory Destras, Gwendolyne Burfin, Hadrien Regue, Quentin Semanas, Martine Valette, Bruno Lina, Laurence Josset |
| EPI_ISL_3130877 | LABORATOIRE CERBALLIANCE PLT VILLON | CNR Virus des Infections Respiratoires - France SUD | Antonin Bal, Gregory Destras, Gwendolyne Burfin, Hadrien Regue, Quentin Semanas, Martine Valette, Bruno Lina, Laurence Josset |
| EPI_ISL_3130878 | MIRIALIS CLUSES BECHET | CNR Virus des Infections Respiratoires - France SUD | Antonin Bal, Gregory Destras, Gwendolyne Burfin, Hadrien Regue, Quentin Semanas, Martine Valette, Bruno Lina, Laurence Josset |
| EPI_ISL_3130879 | LABORATOIRE CERBALLIANCE PLT VILLON | CNR Virus des Infections Respiratoires - France SUD | Antonin Bal, Gregory Destras, Gwendolyne Burfin, Hadrien Regue, Quentin Semanas, Martine Valette, Bruno Lina, Laurence Josset |
| EPI_ISL_3130880 | MIRIALIS CLUSES BECHET | CNR Virus des Infections Respiratoires - France SUD | Antonin Bal, Gregory Destras, Gwendolyne Burfin, Hadrien Regue, Quentin Semanas, Martine Valette, Bruno Lina, Laurence Josset |
| EPI_ISL_3130881 | LABORATOIRE CERBALLIANCE PLT VILLON | CNR Virus des Infections Respiratoires - France SUD | Antonin Bal, Gregory Destras, Gwendolyne Burfin, Hadrien Regue, Quentin Semanas, Martine Valette, Bruno Lina, Laurence Josset |
| EPI_ISL_3130882 | MIRIALIS CLUSES BECHET | CNR Virus des Infections Respiratoires - France SUD | Antonin Bal, Gregory Destras, Gwendolyne Burfin, Hadrien Regue, Quentin Semanas, Martine Valette, Bruno Lina, Laurence Josset |
| EPI_ISL_3130883 | LABORATOIRE CERBALLIANCE PLT VILLON | CNR Virus des Infections Respiratoires - France SUD | Antonin Bal, Gregory Destras, Gwendolyne Burfin, Hadrien Regue, Quentin Semanas, Martine Valette, Bruno Lina, Laurence Josset |
| EPI_ISL_3130884 | LABORATOIRE BOUVIER | CNR Virus des Infections Respiratoires - France SUD | Antonin Bal, Gregory Destras, Gwendolyne Burfin, Hadrien Regue, Quentin Semanas, Martine Valette, Bruno Lina, Laurence Josset |
| EPI_ISL_3130885, EPI_ISL_3130886, EPI_ISL_3130888 | LABORATOIRE CERBALLIANCE PLT VILLON | CNR Virus des Infections Respiratoires - France SUD | Antonin Bal, Gregory Destras, Gwendolyne Burfin, Hadrien Regue, Quentin Semanas, Martine Valette, Bruno Lina, Laurence Josset |
| EPI_ISL_3130890 | LABORATOIRE BOUVIER | CNR Virus des Infections Respiratoires - France SUD | Antonin Bal, Gregory Destras, Gwendolyne Burfin, Hadrien Regue, Quentin Semanas, Martine Valette, Bruno Lina, Laurence Josset |
| EPI_ISL_3130891 | LABORATOIRE CERBALLIANCE PLT VILLON | CNR Virus des Infections Respiratoires - France SUD | Antonin Bal, Gregory Destras, Gwendolyne Burfin, Hadrien Regue, Quentin Semanas, Martine Valette, Bruno Lina, Laurence Josset |
| EPI_ISL_3130892 | MIRIALIS CLUSES BECHET | CNR Virus des Infections Respiratoires - France SUD | Antonin Bal, Gregory Destras, Gwendolyne Burfin, Hadrien Regue, Quentin Semanas, Martine Valette, Bruno Lina, Laurence Josset |
| EPI_ISL_3130893 | LABORATOIRE CERBALLIANCE PLT VILLON | CNR Virus des Infections Respiratoires - France SUD | Antonin Bal, Gregory Destras, Gwendolyne Burfin, Hadrien Regue, Quentin Semanas, Martine Valette, Bruno Lina, Laurence Josset |
| EPI_ISL_3130895 | LBM SEALAB DARRASSE ET ASSOCIES | CNR Virus des Infections Respiratoires - France SUD | Antonin Bal, Gregory Destras, Gwendolyne Burfin, Hadrien Regue, Quentin Semanas, Martine Valette, Bruno Lina, Laurence Josset |
| EPI_ISL_3130896, EPI_ISL_3130897 | LABORATOIRE CERBALLIANCE PLT VILLON | CNR Virus des Infections Respiratoires - France SUD | Antonin Bal, Gregory Destras, Gwendolyne Burfin, Hadrien Regue, Quentin Semanas, Martine Valette, Bruno Lina, Laurence Josset |
| EPI_ISL_3130898 | MIRIALIS CLUSES BECHET | CNR Virus des Infections Respiratoires - France SUD | Antonin Bal, Gregory Destras, Gwendolyne Burfin, Hadrien Regue, Quentin Semanas, Martine Valette, Bruno Lina, Laurence Josset |
| EPI_ISL_3130899 | LBM SEALAB DARRASSE ET ASSOCIES | CNR Virus des Infections Respiratoires - France SUD | Antonin Bal, Gregory Destras, Gwendolyne Burfin, Hadrien Regue, Quentin Semanas, Martine Valette, Bruno Lina, Laurence Josset |
| EPI_ISL_3130900 | LABORATOIRE BOUVIER | CNR Virus des Infections Respiratoires - France SUD | Antonin Bal, Gregory Destras, Gwendolyne Burfin, Hadrien Regue, Quentin Semanas, Martine Valette, Bruno Lina, Laurence Josset |
| EPI_ISL_3130901 | CH ANTIBES - JUAN LES PINS | CNR Virus des Infections Respiratoires - France SUD | Antonin Bal, Gregory Destras, Gwendolyne Burfin, Hadrien Regue, Quentin Semanas, Martine Valette, Bruno Lina, Laurence Josset |
| EPI_ISL_3130902 | MIRIALIS CLUSES BECHET | CNR Virus des Infections Respiratoires - France SUD | Antonin Bal, Gregory Destras, Gwendolyne Burfin, Hadrien Regue, Quentin Semanas, Martine Valette, Bruno Lina, Laurence Josset |
| EPI_ISL_3130903 | LABORATOIRE BOUVIER | CNR Virus des Infections Respiratoires - France SUD | Antonin Bal, Gregory Destras, Gwendolyne Burfin, Hadrien Regue, Quentin Semanas, Martine Valette, Bruno Lina, Laurence Josset |
| EPI_ISL_3130904 | CENTRE HOSPITALIER BRIVE | CNR Virus des Infections Respiratoires - France SUD | Antonin Bal, Gregory Destras, Gwendolyne Burfin, Hadrien Regue, Quentin Semanas, Martine Valette, Bruno Lina, Laurence Josset |
| EPI_ISL_3130906 | MIRIALIS CLUSES BECHET | CNR Virus des Infections Respiratoires - France SUD | Antonin Bal, Gregory Destras, Gwendolyne Burfin, Hadrien Regue, Quentin Semanas, Martine Valette, Bruno Lina, Laurence Josset |
| EPI_ISL_3130907 | CENTRE HOSPITALIER DE ROCHEFORT | CNR Virus des Infections Respiratoires - France SUD | Antonin Bal, Gregory Destras, Gwendolyne Burfin, Hadrien Regue, Quentin Semanas, Martine Valette, Bruno Lina, Laurence Josset |

| | | | |
|---|---|---|---|
| EPI_ISL_3130909, EPI_ISL_3130911, EPI_ISL_3130912, EPI_ISL_3130913, EPI_ISL_3130915, EPI_ISL_3130916, EPI_ISL_3130917, EPI_ISL_3130918 | MIRIALIS CLUSES BECHET | CNR Virus des Infections Respiratoires - France SUD | Antonin Bal, Gregory Destras, Gwendolyne Burfin, Hadrien Regue, Quentin Semanas, Martine Valette, Bruno Lina, Laurence Josset |
| EPI_ISL_3130919 | CENTRE HOSPITALIER DE BASTIA | CNR Virus des Infections Respiratoires - France SUD | Antonin Bal, Gregory Destras, Gwendolyne Burfin, Hadrien Regue, Quentin Semanas, Martine Valette, Bruno Lina, Laurence Josset |
| EPI_ISL_3130920, EPI_ISL_3130921, EPI_ISL_3130922, EPI_ISL_3130923, EPI_ISL_3130924, EPI_ISL_3130925, EPI_ISL_3130926 | MIRIALIS CLUSES BECHET | CNR Virus des Infections Respiratoires - France SUD | Antonin Bal, Gregory Destras, Gwendolyne Burfin, Hadrien Regue, Quentin Semanas, Martine Valette, Bruno Lina, Laurence Josset |
| EPI_ISL_3130927 | CENTRE HOSPITALIER LUCIEN HUSSEL | CNR Virus des Infections Respiratoires - France SUD | Antonin Bal, Gregory Destras, Gwendolyne Burfin, Hadrien Regue, Quentin Semanas, Martine Valette, Bruno Lina, Laurence Josset |
| EPI_ISL_3130928, EPI_ISL_3130929, EPI_ISL_3130931, EPI_ISL_3130933, EPI_ISL_3130934, EPI_ISL_3130935, EPI_ISL_3130936 | MIRIALIS CLUSES BECHET | CNR Virus des Infections Respiratoires - France SUD | Antonin Bal, Gregory Destras, Gwendolyne Burfin, Hadrien Regue, Quentin Semanas, Martine Valette, Bruno Lina, Laurence Josset |
| EPI_ISL_3130938 | LBM SEALAB DARRASSE ET ASSOCIES | CNR Virus des Infections Respiratoires - France SUD | Antonin Bal, Gregory Destras, Gwendolyne Burfin, Hadrien Regue, Quentin Semanas, Martine Valette, Bruno Lina, Laurence Josset |
| EPI_ISL_3130939, EPI_ISL_3130940, EPI_ISL_3130941, EPI_ISL_3130942, EPI_ISL_3130945 | MIRIALIS CLUSES BECHET | CNR Virus des Infections Respiratoires - France SUD | Antonin Bal, Gregory Destras, Gwendolyne Burfin, Hadrien Regue, Quentin Semanas, Martine Valette, Bruno Lina, Laurence Josset |
| EPI_ISL_3130946 | CH ANTIBES - JUAN LES PINS | CNR Virus des Infections Respiratoires - France SUD | Antonin Bal, Gregory Destras, Gwendolyne Burfin, Hadrien Regue, Quentin Semanas, Martine Valette, Bruno Lina, Laurence Josset |
| EPI_ISL_3130947, EPI_ISL_3130948, EPI_ISL_3130949, EPI_ISL_3130950, EPI_ISL_3130951, EPI_ISL_3130952, EPI_ISL_3130953, EPI_ISL_3130955, EPI_ISL_3130956, EPI_ISL_3130958, EPI_ISL_3130960, EPI_ISL_3130961, EPI_ISL_3130962, EPI_ISL_3130963, EPI_ISL_3130964 | | | |
| see above | MIRIALIS CLUSES BECHET | CNR Virus des Infections Respiratoires - France SUD | Antonin Bal, Gregory Destras, Gwendolyne Burfin, Hadrien Regue, Quentin Semanas, Martine Valette, Bruno Lina, Laurence Josset |
| EPI_ISL_3130975 | CH GIVORS | CNR Virus des Infections Respiratoires - France SUD | Antonin Bal, Gregory Destras, Gwendolyne Burfin, Hadrien Regue, Quentin Semanas, Martine Valette, Bruno Lina, Laurence Josset |
| EPI_ISL_3130994 | CNR Virus des Infections Respiratoires - France SUD | CNR Virus des Infections Respiratoires - France SUD | Antonin Bal, Gregory Destras, Gwendolyne Burfin, Hadrien Regue, Quentin Semanas, Martine Valette, Bruno Lina, Laurence Josset |
| EPI_ISL_3130995 | CH GIVORS | CNR Virus des Infections Respiratoires - France SUD | Antonin Bal, Gregory Destras, Gwendolyne Burfin, Hadrien Regue, Quentin Semanas, Martine Valette, Bruno Lina, Laurence Josset |
| EPI_ISL_3130997, EPI_ISL_3130998, EPI_ISL_3130999 | LABORATOIRE MAYMAT | CNR Virus des Infections Respiratoires - France SUD | Antonin Bal, Gregory Destras, Gwendolyne Burfin, Hadrien Regue, Quentin Semanas, Martine Valette, Bruno Lina, Laurence Josset |
| EPI_ISL_3131000, EPI_ISL_3131001, EPI_ISL_3131003 | LBM SEALAB DARRASSE ET ASSOCIES | CNR Virus des Infections Respiratoires - France SUD | Antonin Bal, Gregory Destras, Gwendolyne Burfin, Hadrien Regue, Quentin Semanas, Martine Valette, Bruno Lina, Laurence Josset |
| EPI_ISL_3131004 | LABORATOIRE MAYMAT | CNR Virus des Infections Respiratoires - France SUD | Antonin Bal, Gregory Destras, Gwendolyne Burfin, Hadrien Regue, Quentin Semanas, Martine Valette, Bruno Lina, Laurence Josset |
| EPI_ISL_3131005 | CH VALENCE | CNR Virus des Infections Respiratoires - France SUD | Antonin Bal, Gregory Destras, Gwendolyne Burfin, Hadrien Regue, Quentin Semanas, Martine Valette, Bruno Lina, Laurence Josset |
| EPI_ISL_3131006, EPI_ISL_3131008, EPI_ISL_3131009, EPI_ISL_3131010, EPI_ISL_3131011, EPI_ISL_3131012, EPI_ISL_3131013, EPI_ISL_3131015, EPI_ISL_3131016, EPI_ISL_3131018, EPI_ISL_3131021, EPI_ISL_3131022 | | | |
| see above | LABORATOIRE ACCOLAB SUD OUEST | CNR Virus des Infections Respiratoires - France SUD | Antonin Bal, Gregory Destras, Gwendolyne Burfin, Hadrien Regue, Quentin Semanas, Martine Valette, Bruno Lina, Laurence Josset |
| EPI_ISL_3131024 | LBM SEALAB DARRASSE ET ASSOCIES | CNR Virus des Infections Respiratoires - France SUD | Antonin Bal, Gregory Destras, Gwendolyne Burfin, Hadrien Regue, Quentin Semanas, Martine Valette, Bruno Lina, Laurence Josset |
| EPI_ISL_3131025, EPI_ISL_3131026, EPI_ISL_3131027 | LABORATOIRE CERBALLIANCE PLT VILLON | CNR Virus des Infections Respiratoires - France SUD | Antonin Bal, Gregory Destras, Gwendolyne Burfin, Hadrien Regue, Quentin Semanas, Martine Valette, Bruno Lina, Laurence Josset |
| EPI_ISL_3131028 | LBM SEALAB DARRASSE ET ASSOCIES | CNR Virus des Infections Respiratoires - France SUD | Antonin Bal, Gregory Destras, Gwendolyne Burfin, Hadrien Regue, Quentin Semanas, Martine Valette, Bruno Lina, Laurence Josset |
| EPI_ISL_3131029, EPI_ISL_3131030, EPI_ISL_3131031, EPI_ISL_3131032 | LABORATOIRE CERBALLIANCE PLT VILLON | CNR Virus des Infections Respiratoires - France SUD | Antonin Bal, Gregory Destras, Gwendolyne Burfin, Hadrien Regue, Quentin Semanas, Martine Valette, Bruno Lina, Laurence Josset |
| EPI_ISL_3131033 | LBM SEALAB DARRASSE ET ASSOCIES | CNR Virus des Infections Respiratoires - France SUD | Antonin Bal, Gregory Destras, Gwendolyne Burfin, Hadrien Regue, Quentin Semanas, Martine Valette, Bruno Lina, Laurence Josset |
| EPI_ISL_3131034, EPI_ISL_3131035 | LABORATOIRE CERBALLIANCE PLT VILLON | CNR Virus des Infections Respiratoires - France SUD | Antonin Bal, Gregory Destras, Gwendolyne Burfin, Hadrien Regue, Quentin Semanas, Martine Valette, Bruno Lina, Laurence Josset |
| EPI_ISL_3131036 | LBM SEALAB DARRASSE ET ASSOCIES | CNR Virus des Infections Respiratoires - France SUD | Antonin Bal, Gregory Destras, Gwendolyne Burfin, Hadrien Regue, Quentin Semanas, Martine Valette, Bruno Lina, Laurence Josset |
| EPI_ISL_3131037 | LABORATOIRE CERBALLIANCE PLT VILLON | CNR Virus des Infections Respiratoires - France SUD | Antonin Bal, Gregory Destras, Gwendolyne Burfin, Hadrien Regue, Quentin Semanas, Martine Valette, Bruno Lina, Laurence Josset |
| EPI_ISL_3131039 | LBM SEALAB DARRASSE ET ASSOCIES | CNR Virus des Infections Respiratoires - France SUD | Antonin Bal, Gregory Destras, Gwendolyne Burfin, Hadrien Regue, Quentin Semanas, Martine Valette, Bruno Lina, Laurence Josset |
| EPI_ISL_3131040, EPI_ISL_3131041, EPI_ISL_3131043, EPI_ISL_3131044, EPI_ISL_3131045, EPI_ISL_3131046, EPI_ISL_3131047, EPI_ISL_3131048, EPI_ISL_3131049, EPI_ISL_3131050, EPI_ISL_3131051, EPI_ISL_3131052, EPI_ISL_3131053, EPI_ISL_3131054 | | | |
| see above | LABORATOIRE CERBALLIANCE PLT VILLON | CNR Virus des Infections Respiratoires - France SUD | Antonin Bal, Gregory Destras, Gwendolyne Burfin, Hadrien Regue, Quentin Semanas, Martine Valette, Bruno Lina, Laurence Josset |
| EPI_ISL_3131055 | CNR Virus des Infections Respiratoires - France SUD | CNR Virus des Infections Respiratoires - France SUD | Antonin Bal, Gregory Destras, Gwendolyne Burfin, Hadrien Regue, Quentin Semanas, Martine Valette, Bruno Lina, Laurence Josset |
| EPI_ISL_3131059, EPI_ISL_3131063 | LBM SEALAB DARRASSE ET ASSOCIES | CNR Virus des Infections Respiratoires - France SUD | Antonin Bal, Gregory Destras, Gwendolyne Burfin, Hadrien Regue, Quentin Semanas, Martine Valette, Bruno Lina, Laurence Josset |
| EPI_ISL_3131064 | NEOLAB JASSANS | CNR Virus des Infections Respiratoires - France SUD | Antonin Bal, Gregory Destras, Gwendolyne Burfin, Hadrien Regue, Quentin Semanas, Martine Valette, Bruno Lina, Laurence Josset |
| EPI_ISL_3131065, EPI_ISL_3131066 | CNR Virus des Infections Respiratoires - France SUD | CNR Virus des Infections Respiratoires - France SUD | Antonin Bal, Gregory Destras, Gwendolyne Burfin, Hadrien Regue, Quentin Semanas, Martine Valette, Bruno Lina, Laurence Josset |
| EPI_ISL_3131067, EPI_ISL_3131068 | LBM BIOLYSS | CNR Virus des Infections Respiratoires - France SUD | Antonin Bal, Gregory Destras, Gwendolyne Burfin, Hadrien Regue, Quentin Semanas, Martine Valette, Bruno Lina, Laurence Josset |
| EPI_ISL_3131069 | CHU GRENOBLE | CNR Virus des Infections Respiratoires - France SUD | Antonin Bal, Gregory Destras, Gwendolyne Burfin, Hadrien Regue, Quentin Semanas, Martine Valette, Bruno Lina, Laurence Josset |
| EPI_ISL_3131072 | CHU ST ETIENNE HOPITAL NORD | CNR Virus des Infections Respiratoires - France SUD | Antonin Bal, Gregory Destras, Gwendolyne Burfin, Hadrien Regue, Quentin Semanas, Martine Valette, Bruno Lina, Laurence Josset |
| EPI_ISL_3131073, EPI_ISL_3131074, EPI_ISL_3131075 | LAM BIORYLIS | CNR Virus des Infections Respiratoires - France SUD | Antonin Bal, Gregory Destras, Gwendolyne Burfin, Hadrien Regue, Quentin Semanas, Martine Valette, Bruno Lina, Laurence Josset |
| EPI_ISL_3131076, EPI_ISL_3131077, EPI_ISL_3131078, EPI_ISL_3131079, EPI_ISL_3131080, EPI_ISL_3131081 | LABM BIO 17 | CNR Virus des Infections Respiratoires - France SUD | Antonin Bal, Gregory Destras, Gwendolyne Burfin, Hadrien Regue, Quentin Semanas, Martine Valette, Bruno Lina, Laurence Josset |
| EPI_ISL_3131082 | DYOMEDEA-LABORATOIRE DE LA SAUVEGARDE | CNR Virus des Infections Respiratoires - France SUD | Antonin Bal, Gregory Destras, Gwendolyne Burfin, Hadrien Regue, Quentin Semanas, Martine Valette, Bruno Lina, Laurence Josset |
| EPI_ISL_3131083, EPI_ISL_3131084 | LABM BIO 17 | CNR Virus des Infections Respiratoires - France SUD | Antonin Bal, Gregory Destras, Gwendolyne Burfin, Hadrien Regue, Quentin Semanas, Martine Valette, Bruno Lina, Laurence Josset |
| EPI_ISL_3131085 | DYOMEDEA-LABORATOIRE DE LA SAUVEGARDE | CNR Virus des Infections Respiratoires - France SUD | Antonin Bal, Gregory Destras, Gwendolyne Burfin, Hadrien Regue, Quentin Semanas, Martine Valette, Bruno Lina, Laurence Josset |
| EPI_ISL_3131086 | LABM BIO 17 | CNR Virus des Infections Respiratoires - France SUD | Antonin Bal, Gregory Destras, Gwendolyne Burfin, Hadrien Regue, Quentin Semanas, Martine Valette, Bruno Lina, Laurence Josset |
| EPI_ISL_3131087 | DYOMEDEA-LABORATOIRE DE LA SAUVEGARDE | CNR Virus des Infections Respiratoires - France SUD | Antonin Bal, Gregory Destras, Gwendolyne Burfin, Hadrien Regue, Quentin Semanas, Martine Valette, Bruno Lina, Laurence Josset |
| EPI_ISL_3131088, EPI_ISL_3131089 | LABM BIO 17 | CNR Virus des Infections Respiratoires - France SUD | Antonin Bal, Gregory Destras, Gwendolyne Burfin, Hadrien Regue, Quentin Semanas, Martine Valette, Bruno Lina, Laurence Josset |

| | | | |
|---|---|---|---|
| EPI_ISL_3131090 | DYOMEDEA-LABORATOIRE DE LA SAUVEGARDE | CNR Virus des Infections Respiratoires - France SUD | Antonin Bal, Gregory Destras, Gwendolyne Burfin, Hadrien Regue, Quentin Semanas, Martine Valette, Bruno Lina, Laurence Josset |
| EPI_ISL_3131091, EPI_ISL_3131092 | LABM BIO 17 | CNR Virus des Infections Respiratoires - France SUD | Antonin Bal, Gregory Destras, Gwendolyne Burfin, Hadrien Regue, Quentin Semanas, Martine Valette, Bruno Lina, Laurence Josset |
| EPI_ISL_3131093 | DYOMEDEA-LABORATOIRE DE LA SAUVEGARDE | CNR Virus des Infections Respiratoires - France SUD | Antonin Bal, Gregory Destras, Gwendolyne Burfin, Hadrien Regue, Quentin Semanas, Martine Valette, Bruno Lina, Laurence Josset |
| EPI_ISL_3131094, EPI_ISL_3131095 | LABM BIO 17 | CNR Virus des Infections Respiratoires - France SUD | Antonin Bal, Gregory Destras, Gwendolyne Burfin, Hadrien Regue, Quentin Semanas, Martine Valette, Bruno Lina, Laurence Josset |
| EPI_ISL_3131096 | DYOMEDEA-LABORATOIRE DE LA SAUVEGARDE | CNR Virus des Infections Respiratoires - France SUD | Antonin Bal, Gregory Destras, Gwendolyne Burfin, Hadrien Regue, Quentin Semanas, Martine Valette, Bruno Lina, Laurence Josset |
| EPI_ISL_3131097, EPI_ISL_3131098 | LABM BIO 17 | CNR Virus des Infections Respiratoires - France SUD | Antonin Bal, Gregory Destras, Gwendolyne Burfin, Hadrien Regue, Quentin Semanas, Martine Valette, Bruno Lina, Laurence Josset |
| EPI_ISL_3131099 | DYOMEDEA-LABORATOIRE DE LA SAUVEGARDE | CNR Virus des Infections Respiratoires - France SUD | Antonin Bal, Gregory Destras, Gwendolyne Burfin, Hadrien Regue, Quentin Semanas, Martine Valette, Bruno Lina, Laurence Josset |
| EPI_ISL_3131100 | LABM BIO 17 | CNR Virus des Infections Respiratoires - France SUD | Antonin Bal, Gregory Destras, Gwendolyne Burfin, Hadrien Regue, Quentin Semanas, Martine Valette, Bruno Lina, Laurence Josset |
| EPI_ISL_3131101, EPI_ISL_3131102 | DYOMEDEA-LABORATOIRE DE LA SAUVEGARDE | CNR Virus des Infections Respiratoires - France SUD | Antonin Bal, Gregory Destras, Gwendolyne Burfin, Hadrien Regue, Quentin Semanas, Martine Valette, Bruno Lina, Laurence Josset |
| EPI_ISL_3131103, EPI_ISL_3131104 | LABM BIO 17 | CNR Virus des Infections Respiratoires - France SUD | Antonin Bal, Gregory Destras, Gwendolyne Burfin, Hadrien Regue, Quentin Semanas, Martine Valette, Bruno Lina, Laurence Josset |
| EPI_ISL_3131105 | DYOMEDEA-LABORATOIRE DE LA SAUVEGARDE | CNR Virus des Infections Respiratoires - France SUD | Antonin Bal, Gregory Destras, Gwendolyne Burfin, Hadrien Regue, Quentin Semanas, Martine Valette, Bruno Lina, Laurence Josset |
| EPI_ISL_3131106 | LABM BIO 17 | CNR Virus des Infections Respiratoires - France SUD | Antonin Bal, Gregory Destras, Gwendolyne Burfin, Hadrien Regue, Quentin Semanas, Martine Valette, Bruno Lina, Laurence Josset |
| EPI_ISL_3131107 | DYOMEDEA-LABORATOIRE DE LA SAUVEGARDE | CNR Virus des Infections Respiratoires - France SUD | Antonin Bal, Gregory Destras, Gwendolyne Burfin, Hadrien Regue, Quentin Semanas, Martine Valette, Bruno Lina, Laurence Josset |
| EPI_ISL_3131108, EPI_ISL_3131109 | LABM BIO 17 | CNR Virus des Infections Respiratoires - France SUD | Antonin Bal, Gregory Destras, Gwendolyne Burfin, Hadrien Regue, Quentin Semanas, Martine Valette, Bruno Lina, Laurence Josset |
| EPI_ISL_3131110 | DYOMEDEA-LABORATOIRE DE LA SAUVEGARDE | CNR Virus des Infections Respiratoires - France SUD | Antonin Bal, Gregory Destras, Gwendolyne Burfin, Hadrien Regue, Quentin Semanas, Martine Valette, Bruno Lina, Laurence Josset |
| EPI_ISL_3131111 | LABM BIO 17 | CNR Virus des Infections Respiratoires - France SUD | Antonin Bal, Gregory Destras, Gwendolyne Burfin, Hadrien Regue, Quentin Semanas, Martine Valette, Bruno Lina, Laurence Josset |
| EPI_ISL_3131112 | BIOMER LABORATOIRE ST DIE | CNR Virus des Infections Respiratoires - France SUD | Antonin Bal, Gregory Destras, Gwendolyne Burfin, Hadrien Regue, Quentin Semanas, Martine Valette, Bruno Lina, Laurence Josset |
| EPI_ISL_3131113, EPI_ISL_3131114, EPI_ISL_3131115, EPI_ISL_3131116, EPI_ISL_3131117, EPI_ISL_3131118, EPI_ISL_3131119, EPI_ISL_3131120 | LABM BIO 17 | CNR Virus des Infections Respiratoires - France SUD | Antonin Bal, Gregory Destras, Gwendolyne Burfin, Hadrien Regue, Quentin Semanas, Martine Valette, Bruno Lina, Laurence Josset |
| EPI_ISL_3131121, EPI_ISL_3131122, EPI_ISL_3131123, EPI_ISL_3131124 | LABORATOIRE UNILIANS DECINES | CNR Virus des Infections Respiratoires - France SUD | Antonin Bal, Gregory Destras, Gwendolyne Burfin, Hadrien Regue, Quentin Semanas, Martine Valette, Bruno Lina, Laurence Josset |
| EPI_ISL_3131125, EPI_ISL_3131127, EPI_ISL_3131128, EPI_ISL_3131129, EPI_ISL_3131130, EPI_ISL_3131131, EPI_ISL_3131132, EPI_ISL_3131133 | ASTRALAB TASSIGNY | CNR Virus des Infections Respiratoires - France SUD | Antonin Bal, Gregory Destras, Gwendolyne Burfin, Hadrien Regue, Quentin Semanas, Martine Valette, Bruno Lina, Laurence Josset |
| EPI_ISL_3131134 | LABORATOIRE BOUVIER | CNR Virus des Infections Respiratoires - France SUD | Antonin Bal, Gregory Destras, Gwendolyne Burfin, Hadrien Regue, Quentin Semanas, Martine Valette, Bruno Lina, Laurence Josset |
| EPI_ISL_3131135, EPI_ISL_3131136 | ASTRALAB TASSIGNY | CNR Virus des Infections Respiratoires - France SUD | Antonin Bal, Gregory Destras, Gwendolyne Burfin, Hadrien Regue, Quentin Semanas, Martine Valette, Bruno Lina, Laurence Josset |
| EPI_ISL_3131137 | LABORATOIRE BOUVIER | CNR Virus des Infections Respiratoires - France SUD | Antonin Bal, Gregory Destras, Gwendolyne Burfin, Hadrien Regue, Quentin Semanas, Martine Valette, Bruno Lina, Laurence Josset |
| EPI_ISL_3131138, EPI_ISL_3131139, EPI_ISL_3131140 | ASTRALAB TASSIGNY | CNR Virus des Infections Respiratoires - France SUD | Antonin Bal, Gregory Destras, Gwendolyne Burfin, Hadrien Regue, Quentin Semanas, Martine Valette, Bruno Lina, Laurence Josset |
| EPI_ISL_3131141, EPI_ISL_3131143, EPI_ISL_3131144 | LABORATOIRE BOUVIER | CNR Virus des Infections Respiratoires - France SUD | Antonin Bal, Gregory Destras, Gwendolyne Burfin, Hadrien Regue, Quentin Semanas, Martine Valette, Bruno Lina, Laurence Josset |
| EPI_ISL_3131145 | LAM EIMER LENYS | CNR Virus des Infections Respiratoires - France SUD | Antonin Bal, Gregory Destras, Gwendolyne Burfin, Hadrien Regue, Quentin Semanas, Martine Valette, Bruno Lina, Laurence Josset |
| EPI_ISL_3131146 | LABORATOIRE BOUVIER | CNR Virus des Infections Respiratoires - France SUD | Antonin Bal, Gregory Destras, Gwendolyne Burfin, Hadrien Regue, Quentin Semanas, Martine Valette, Bruno Lina, Laurence Josset |
| EPI_ISL_3131147, EPI_ISL_3131148 | LAM EIMER LENYS | CNR Virus des Infections Respiratoires - France SUD | Antonin Bal, Gregory Destras, Gwendolyne Burfin, Hadrien Regue, Quentin Semanas, Martine Valette, Bruno Lina, Laurence Josset |
| EPI_ISL_3131149 | LABORATOIRE BOUVIER | CNR Virus des Infections Respiratoires - France SUD | Antonin Bal, Gregory Destras, Gwendolyne Burfin, Hadrien Regue, Quentin Semanas, Martine Valette, Bruno Lina, Laurence Josset |
| EPI_ISL_3131150, EPI_ISL_3131151 | LAM EIMER LENYS | CNR Virus des Infections Respiratoires - France SUD | Antonin Bal, Gregory Destras, Gwendolyne Burfin, Hadrien Regue, Quentin Semanas, Martine Valette, Bruno Lina, Laurence Josset |
| EPI_ISL_3131152 | LABORATOIRE CERDIBIO-CHARENTES | CNR Virus des Infections Respiratoires - France SUD | Antonin Bal, Gregory Destras, Gwendolyne Burfin, Hadrien Regue, Quentin Semanas, Martine Valette, Bruno Lina, Laurence Josset |
| EPI_ISL_3131153 | LABORATOIRE LABAZUR | CNR Virus des Infections Respiratoires - France SUD | Antonin Bal, Gregory Destras, Gwendolyne Burfin, Hadrien Regue, Quentin Semanas, Martine Valette, Bruno Lina, Laurence Josset |
| EPI_ISL_3131154 | LABORATOIRE CERDIBIO-CHARENTES | CNR Virus des Infections Respiratoires - France SUD | Antonin Bal, Gregory Destras, Gwendolyne Burfin, Hadrien Regue, Quentin Semanas, Martine Valette, Bruno Lina, Laurence Josset |
| EPI_ISL_3131155 | LABORATOIRE LABAZUR | CNR Virus des Infections Respiratoires - France SUD | Antonin Bal, Gregory Destras, Gwendolyne Burfin, Hadrien Regue, Quentin Semanas, Martine Valette, Bruno Lina, Laurence Josset |
| EPI_ISL_3131156 | LABORATOIRE CERDIBIO-CHARENTES | CNR Virus des Infections Respiratoires - France SUD | Antonin Bal, Gregory Destras, Gwendolyne Burfin, Hadrien Regue, Quentin Semanas, Martine Valette, Bruno Lina, Laurence Josset |
| EPI_ISL_3131157 | LABORATOIRE LABAZUR | CNR Virus des Infections Respiratoires - France SUD | Antonin Bal, Gregory Destras, Gwendolyne Burfin, Hadrien Regue, Quentin Semanas, Martine Valette, Bruno Lina, Laurence Josset |
| EPI_ISL_3131158 | LABORATOIRE CERDIBIO-CHARENTES | CNR Virus des Infections Respiratoires - France SUD | Antonin Bal, Gregory Destras, Gwendolyne Burfin, Hadrien Regue, Quentin Semanas, Martine Valette, Bruno Lina, Laurence Josset |
| EPI_ISL_3131159 | LABORATOIRE LABAZUR | CNR Virus des Infections Respiratoires - France SUD | Antonin Bal, Gregory Destras, Gwendolyne Burfin, Hadrien Regue, Quentin Semanas, Martine Valette, Bruno Lina, Laurence Josset |
| EPI_ISL_3131160, EPI_ISL_3131161 | LABORATOIRE CERDIBIO-CHARENTES | CNR Virus des Infections Respiratoires - France SUD | Antonin Bal, Gregory Destras, Gwendolyne Burfin, Hadrien Regue, Quentin Semanas, Martine Valette, Bruno Lina, Laurence Josset |
| EPI_ISL_3131163 | LABORATOIRE LABAZUR | CNR Virus des Infections Respiratoires - France SUD | Antonin Bal, Gregory Destras, Gwendolyne Burfin, Hadrien Regue, Quentin Semanas, Martine Valette, Bruno Lina, Laurence Josset |
| EPI_ISL_3131164 | CERBALLIANCE CHARENTES SAINTES | CNR Virus des Infections Respiratoires - France SUD | Antonin Bal, Gregory Destras, Gwendolyne Burfin, Hadrien Regue, Quentin Semanas, Martine Valette, Bruno Lina, Laurence Josset |
| EPI_ISL_3131165 | LABORATOIRE LABAZUR | CNR Virus des Infections Respiratoires - France SUD | Antonin Bal, Gregory Destras, Gwendolyne Burfin, Hadrien Regue, Quentin Semanas, Martine Valette, Bruno Lina, Laurence Josset |
| EPI_ISL_3131166 | LABORATOIRE CERDIBIO-CHARENTES | CNR Virus des Infections Respiratoires - France SUD | Antonin Bal, Gregory Destras, Gwendolyne Burfin, Hadrien Regue, Quentin Semanas, Martine Valette, Bruno Lina, Laurence Josset |
| EPI_ISL_3131167 | LABORATOIRE LABAZUR | CNR Virus des Infections Respiratoires - France SUD | Antonin Bal, Gregory Destras, Gwendolyne Burfin, Hadrien Regue, Quentin Semanas, Martine Valette, Bruno Lina, Laurence Josset |
| EPI_ISL_3131168, EPI_ISL_3131169, EPI_ISL_3131170, EPI_ISL_3131172, EPI_ISL_3131173, EPI_ISL_3131174 | LABORATOIRE CERDIBIO-CHARENTES | CNR Virus des Infections Respiratoires - France SUD | Antonin Bal, Gregory Destras, Gwendolyne Burfin, Hadrien Regue, Quentin Semanas, Martine Valette, Bruno Lina, Laurence Josset |
| EPI_ISL_3131175 | CNR Virus des Infections Respiratoires - France SUD | CNR Virus des Infections Respiratoires - France SUD | Antonin Bal, Gregory Destras, Gwendolyne Burfin, Hadrien Regue, Quentin Semanas, Martine Valette, Bruno Lina, Laurence Josset |
| EPI_ISL_3131176, EPI_ISL_3131177 | LABORATOIRE CERDIBIO-CHARENTES | CNR Virus des Infections Respiratoires - France SUD | Antonin Bal, Gregory Destras, Gwendolyne Burfin, Hadrien Regue, Quentin Semanas, Martine Valette, Bruno Lina, Laurence Josset |
| EPI_ISL_3131178 | HOPITAL SAINT ANDRE | CNR Virus des Infections Respiratoires - France SUD | Antonin Bal, Gregory Destras, Gwendolyne Burfin, Hadrien Regue, Quentin Semanas, Martine Valette, Bruno Lina, Laurence Josset |
| EPI_ISL_3131179, EPI_ISL_3131180, EPI_ISL_3131181, | LABORATOIRE CERDIBIO-CHARENTES | CNR Virus des Infections Respiratoires - France SUD | Antonin Bal, Gregory Destras, Gwendolyne Burfin, Hadrien Regue, Quentin Semanas, Martine Valette, Bruno Lina, Laurence Josset |

| | | | |
|---|---|---|---|
| EPI_ISL_3131183 | | | |
| EPI_ISL_3131185 | HOPITAL SAINT ANDRE | CNR Virus des Infections Respiratoires - France SUD | Antonin Bal, Gregory Destras, Gwendolyne Burfin, Hadrien Regue, Quentin Semanas, Martine Valette, Bruno Lina, Laurence Josset |
| EPI_ISL_3131186, EPI_ISL_3131187 | SYNLAB GASCOGNE | CNR Virus des Infections Respiratoires - France SUD | Antonin Bal, Gregory Destras, Gwendolyne Burfin, Hadrien Regue, Quentin Semanas, Martine Valette, Bruno Lina, Laurence Josset |
| EPI_ISL_3131188, EPI_ISL_3131189, EPI_ISL_3131190, EPI_ISL_3131191 | HOPITAL SAINT ANDRE | CNR Virus des Infections Respiratoires - France SUD | Antonin Bal, Gregory Destras, Gwendolyne Burfin, Hadrien Regue, Quentin Semanas, Martine Valette, Bruno Lina, Laurence Josset |
| EPI_ISL_3131192 | SYNLAB GASCOGNE | CNR Virus des Infections Respiratoires - France SUD | Antonin Bal, Gregory Destras, Gwendolyne Burfin, Hadrien Regue, Quentin Semanas, Martine Valette, Bruno Lina, Laurence Josset |
| EPI_ISL_3131193 | HOPITAL SAINT ANDRE | CNR Virus des Infections Respiratoires - France SUD | Antonin Bal, Gregory Destras, Gwendolyne Burfin, Hadrien Regue, Quentin Semanas, Martine Valette, Bruno Lina, Laurence Josset |
| EPI_ISL_3131194 | LABORATOIRE UNILIANS DECINES | CNR Virus des Infections Respiratoires - France SUD | Antonin Bal, Gregory Destras, Gwendolyne Burfin, Hadrien Regue, Quentin Semanas, Martine Valette, Bruno Lina, Laurence Josset |
| EPI_ISL_3131195 | HOPITAL SAINT ANDRE | CNR Virus des Infections Respiratoires - France SUD | Antonin Bal, Gregory Destras, Gwendolyne Burfin, Hadrien Regue, Quentin Semanas, Martine Valette, Bruno Lina, Laurence Josset |
| EPI_ISL_3131196 | LABORATOIRE UNILIANS DECINES | CNR Virus des Infections Respiratoires - France SUD | Antonin Bal, Gregory Destras, Gwendolyne Burfin, Hadrien Regue, Quentin Semanas, Martine Valette, Bruno Lina, Laurence Josset |
| EPI_ISL_3131197 | HOPITAL SAINT ANDRE | CNR Virus des Infections Respiratoires - France SUD | Antonin Bal, Gregory Destras, Gwendolyne Burfin, Hadrien Regue, Quentin Semanas, Martine Valette, Bruno Lina, Laurence Josset |
| EPI_ISL_3131198, EPI_ISL_3131199 | LABORATOIRE UNILIANS DECINES | CNR Virus des Infections Respiratoires - France SUD | Antonin Bal, Gregory Destras, Gwendolyne Burfin, Hadrien Regue, Quentin Semanas, Martine Valette, Bruno Lina, Laurence Josset |
| EPI_ISL_3131200, EPI_ISL_3131201, EPI_ISL_3131203 | HOPITAL SAINT ANDRE | CNR Virus des Infections Respiratoires - France SUD | Antonin Bal, Gregory Destras, Gwendolyne Burfin, Hadrien Regue, Quentin Semanas, Martine Valette, Bruno Lina, Laurence Josset |
| EPI_ISL_3131204 | LAM BAYARD-BIOPYRENEES | CNR Virus des Infections Respiratoires - France SUD | Antonin Bal, Gregory Destras, Gwendolyne Burfin, Hadrien Regue, Quentin Semanas, Martine Valette, Bruno Lina, Laurence Josset |
| EPI_ISL_3131205 | HOPITAL SAINT ANDRE | CNR Virus des Infections Respiratoires - France SUD | Antonin Bal, Gregory Destras, Gwendolyne Burfin, Hadrien Regue, Quentin Semanas, Martine Valette, Bruno Lina, Laurence Josset |
| EPI_ISL_3131206 | LAM BAYARD-BIOPYRENEES | CNR Virus des Infections Respiratoires - France SUD | Antonin Bal, Gregory Destras, Gwendolyne Burfin, Hadrien Regue, Quentin Semanas, Martine Valette, Bruno Lina, Laurence Josset |
| EPI_ISL_3131207 | HOPITAL SAINT ANDRE | CNR Virus des Infections Respiratoires - France SUD | Antonin Bal, Gregory Destras, Gwendolyne Burfin, Hadrien Regue, Quentin Semanas, Martine Valette, Bruno Lina, Laurence Josset |
| EPI_ISL_3131208, EPI_ISL_3131209, EPI_ISL_3131211, EPI_ISL_3131213 | LAM BAYARD-BIOPYRENEES | CNR Virus des Infections Respiratoires - France SUD | Antonin Bal, Gregory Destras, Gwendolyne Burfin, Hadrien Regue, Quentin Semanas, Martine Valette, Bruno Lina, Laurence Josset |
| EPI_ISL_3131214 | HOPITAL SAINT ANDRE | CNR Virus des Infections Respiratoires - France SUD | Antonin Bal, Gregory Destras, Gwendolyne Burfin, Hadrien Regue, Quentin Semanas, Martine Valette, Bruno Lina, Laurence Josset |
| EPI_ISL_3131215, EPI_ISL_3131216, EPI_ISL_3131217, EPI_ISL_3131218 | LAM BAYARD-BIOPYRENEES | CNR Virus des Infections Respiratoires - France SUD | Antonin Bal, Gregory Destras, Gwendolyne Burfin, Hadrien Regue, Quentin Semanas, Martine Valette, Bruno Lina, Laurence Josset |
| EPI_ISL_3131219 | HOPITAL SAINT ANDRE | CNR Virus des Infections Respiratoires - France SUD | Antonin Bal, Gregory Destras, Gwendolyne Burfin, Hadrien Regue, Quentin Semanas, Martine Valette, Bruno Lina, Laurence Josset |
| EPI_ISL_3131220, EPI_ISL_3131221, EPI_ISL_3131222, EPI_ISL_3131223, EPI_ISL_3131224, EPI_ISL_3131225, EPI_ISL_3131226, EPI_ISL_3131227, EPI_ISL_3131228 | LAM BAYARD-BIOPYRENEES | CNR Virus des Infections Respiratoires - France SUD | Antonin Bal, Gregory Destras, Gwendolyne Burfin, Hadrien Regue, Quentin Semanas, Martine Valette, Bruno Lina, Laurence Josset |
| EPI_ISL_3131229, EPI_ISL_3131230 | LABORATOIRE CBM DE MURET | CNR Virus des Infections Respiratoires - France SUD | Antonin Bal, Gregory Destras, Gwendolyne Burfin, Hadrien Regue, Quentin Semanas, Martine Valette, Bruno Lina, Laurence Josset |
| EPI_ISL_3131231 | CNR Virus des Infections Respiratoires - France SUD | CNR Virus des Infections Respiratoires - France SUD | Antonin Bal, Gregory Destras, Gwendolyne Burfin, Hadrien Regue, Quentin Semanas, Martine Valette, Bruno Lina, Laurence Josset |
| EPI_ISL_3131232, EPI_ISL_3131235, EPI_ISL_3131236, EPI_ISL_3131237, EPI_ISL_3131238, EPI_ISL_3131239, EPI_ISL_3131240, EPI_ISL_3131241, EPI_ISL_3131242, EPI_ISL_3131243, EPI_ISL_3131244, EPI_ISL_3131245, EPI_ISL_3131250, EPI_ISL_3131251, EPI_ISL_3131252, EPI_ISL_3131253, EPI_ISL_3131254, EPI_ISL_3131255 | | | |
| see above | LABORATOIRE CBM DE MURET | CNR Virus des Infections Respiratoires - France SUD | Antonin Bal, Gregory Destras, Gwendolyne Burfin, Hadrien Regue, Quentin Semanas, Martine Valette, Bruno Lina, Laurence Josset |
| EPI_ISL_3131256 | LBM CHARENTE-MARITIME NORD | CNR Virus des Infections Respiratoires - France SUD | Antonin Bal, Gregory Destras, Gwendolyne Burfin, Hadrien Regue, Quentin Semanas, Martine Valette, Bruno Lina, Laurence Josset |
| EPI_ISL_3131257, EPI_ISL_3131259, EPI_ISL_3131260, EPI_ISL_3131261 | LABORATOIRE CBM DE MURET | CNR Virus des Infections Respiratoires - France SUD | Antonin Bal, Gregory Destras, Gwendolyne Burfin, Hadrien Regue, Quentin Semanas, Martine Valette, Bruno Lina, Laurence Josset |
| EPI_ISL_3131262, EPI_ISL_3131263, EPI_ISL_3131264, EPI_ISL_3131265 | CNR Virus des Infections Respiratoires - France SUD | CNR Virus des Infections Respiratoires - France SUD | Antonin Bal, Gregory Destras, Gwendolyne Burfin, Hadrien Regue, Quentin Semanas, Martine Valette, Bruno Lina, Laurence Josset |
| EPI_ISL_3131266, EPI_ISL_3131267, EPI_ISL_3131268, EPI_ISL_3131269, EPI_ISL_3131270, EPI_ISL_3131271, EPI_ISL_3131272, EPI_ISL_3131273, EPI_ISL_3131274, EPI_ISL_3131275, EPI_ISL_3131276, EPI_ISL_3131277, EPI_ISL_3131278 | | | |
| see above | LABORATOIRE CBM DE MURET | CNR Virus des Infections Respiratoires - France SUD | Antonin Bal, Gregory Destras, Gwendolyne Burfin, Hadrien Regue, Quentin Semanas, Martine Valette, Bruno Lina, Laurence Josset |
| EPI_ISL_3131279 | BIOLITTORAL BIOGROUP PLATEAU TECH | CNR Virus des Infections Respiratoires - France SUD | Antonin Bal, Gregory Destras, Gwendolyne Burfin, Hadrien Regue, Quentin Semanas, Martine Valette, Bruno Lina, Laurence Josset |
| EPI_ISL_3131280 | LABORATOIRE CBM DE MURET | CNR Virus des Infections Respiratoires - France SUD | Antonin Bal, Gregory Destras, Gwendolyne Burfin, Hadrien Regue, Quentin Semanas, Martine Valette, Bruno Lina, Laurence Josset |
| EPI_ISL_3131282, EPI_ISL_3131283, EPI_ISL_3131284, EPI_ISL_3131285, EPI_ISL_3131286, EPI_ISL_3131287 | BIOLITTORAL BIOGROUP PLATEAU TECH | CNR Virus des Infections Respiratoires - France SUD | Antonin Bal, Gregory Destras, Gwendolyne Burfin, Hadrien Regue, Quentin Semanas, Martine Valette, Bruno Lina, Laurence Josset |
| EPI_ISL_3131289, EPI_ISL_3131290, EPI_ISL_3131291, EPI_ISL_3131292, EPI_ISL_3131293, EPI_ISL_3131294, EPI_ISL_3131295, EPI_ISL_3131296, EPI_ISL_3131297 | BIOLITTORAL SANARY ROSE | CNR Virus des Infections Respiratoires - France SUD | Antonin Bal, Gregory Destras, Gwendolyne Burfin, Hadrien Regue, Quentin Semanas, Martine Valette, Bruno Lina, Laurence Josset |
| EPI_ISL_3131298 | CNR Virus des Infections Respiratoires - France SUD | CNR Virus des Infections Respiratoires - France SUD | Antonin Bal, Gregory Destras, Gwendolyne Burfin, Hadrien Regue, Quentin Semanas, Martine Valette, Bruno Lina, Laurence Josset |
| EPI_ISL_3131299, EPI_ISL_3131300, EPI_ISL_3131301 | BIOLITTORAL SANARY ROSE | CNR Virus des Infections Respiratoires - France SUD | Antonin Bal, Gregory Destras, Gwendolyne Burfin, Hadrien Regue, Quentin Semanas, Martine Valette, Bruno Lina, Laurence Josset |
| EPI_ISL_3131302 | CNR Virus des Infections Respiratoires - France SUD | CNR Virus des Infections Respiratoires - France SUD | Antonin Bal, Gregory Destras, Gwendolyne Burfin, Hadrien Regue, Quentin Semanas, Martine Valette, Bruno Lina, Laurence Josset |
| EPI_ISL_3131303, EPI_ISL_3131304 | CH GIVORS | CNR Virus des Infections Respiratoires - France SUD | Antonin Bal, Gregory Destras, Gwendolyne Burfin, Hadrien Regue, Quentin Semanas, Martine Valette, Bruno Lina, Laurence Josset |
| EPI_ISL_3131305 | CNR Virus des Infections Respiratoires - France SUD | CNR Virus des Infections Respiratoires - France SUD | Antonin Bal, Gregory Destras, Gwendolyne Burfin, Hadrien Regue, Quentin Semanas, Martine Valette, Bruno Lina, Laurence Josset |
| EPI_ISL_3131306, EPI_ISL_3131307, EPI_ISL_3131308, EPI_ISL_3131309, EPI_ISL_3131310, EPI_ISL_3131312, EPI_ISL_3131313, EPI_ISL_3131314, EPI_ISL_3131315, EPI_ISL_3131316, EPI_ISL_3131317 | | | |
| see above | BIOLITTORAL BIOGROUP PLATEAU TECH | CNR Virus des Infections Respiratoires - France SUD | Antonin Bal, Gregory Destras, Gwendolyne Burfin, Hadrien Regue, Quentin Semanas, Martine Valette, Bruno Lina, Laurence Josset |
| EPI_ISL_3131318 | LABORATOIRE ACCOLAB SUD OUEST | CNR Virus des Infections Respiratoires - France SUD | Antonin Bal, Gregory Destras, Gwendolyne Burfin, Hadrien Regue, Quentin Semanas, Martine Valette, Bruno Lina, Laurence Josset |
| EPI_ISL_3131319, EPI_ISL_3131320, EPI_ISL_3131321, EPI_ISL_3131322, EPI_ISL_3131323 | BIOLITTORAL BIOGROUP PLATEAU TECH | CNR Virus des Infections Respiratoires - France SUD | Antonin Bal, Gregory Destras, Gwendolyne Burfin, Hadrien Regue, Quentin Semanas, Martine Valette, Bruno Lina, Laurence Josset |

| | | | |
|---|---|---|---|
| EPI_ISL_3131324, EPI_ISL_3131325, EPI_ISL_3131326, EPI_ISL_3131327 | LABORATOIRE ACCOLAB SUD OUEST | CNR Virus des Infections Respiratoires - France SUD | Antonin Bal, Gregory Destras, Gwendolyne Burfin, Hadrien Regue, Quentin Semanas, Martine Valette, Bruno Lina, Laurence Josset |
| EPI_ISL_3131328 | CHU TOULOUSE | CNR Virus des Infections Respiratoires - France SUD | Antonin Bal, Gregory Destras, Gwendolyne Burfin, Hadrien Regue, Quentin Semanas, Martine Valette, Bruno Lina, Laurence Josset |
| EPI_ISL_3131330 | LABORATOIRE ACCOLAB SUD OUEST | CNR Virus des Infections Respiratoires - France SUD | Antonin Bal, Gregory Destras, Gwendolyne Burfin, Hadrien Regue, Quentin Semanas, Martine Valette, Bruno Lina, Laurence Josset |
| EPI_ISL_3131331, EPI_ISL_3131332 | CHU TOULOUSE | CNR Virus des Infections Respiratoires - France SUD | Antonin Bal, Gregory Destras, Gwendolyne Burfin, Hadrien Regue, Quentin Semanas, Martine Valette, Bruno Lina, Laurence Josset |
| EPI_ISL_3131333 | LABORATOIRE ACCOLAB SUD OUEST | CNR Virus des Infections Respiratoires - France SUD | Antonin Bal, Gregory Destras, Gwendolyne Burfin, Hadrien Regue, Quentin Semanas, Martine Valette, Bruno Lina, Laurence Josset |
| EPI_ISL_3131334, EPI_ISL_3131335 | CHU TOULOUSE | CNR Virus des Infections Respiratoires - France SUD | Antonin Bal, Gregory Destras, Gwendolyne Burfin, Hadrien Regue, Quentin Semanas, Martine Valette, Bruno Lina, Laurence Josset |
| EPI_ISL_3131336 | LABORATOIRE ACCOLAB SUD OUEST | CNR Virus des Infections Respiratoires - France SUD | Antonin Bal, Gregory Destras, Gwendolyne Burfin, Hadrien Regue, Quentin Semanas, Martine Valette, Bruno Lina, Laurence Josset |
| EPI_ISL_3131337, EPI_ISL_3131338, EPI_ISL_3131340 | CHU TOULOUSE | CNR Virus des Infections Respiratoires - France SUD | Antonin Bal, Gregory Destras, Gwendolyne Burfin, Hadrien Regue, Quentin Semanas, Martine Valette, Bruno Lina, Laurence Josset |
| EPI_ISL_3131341 | LABORATOIRE ACCOLAB SUD OUEST | CNR Virus des Infections Respiratoires - France SUD | Antonin Bal, Gregory Destras, Gwendolyne Burfin, Hadrien Regue, Quentin Semanas, Martine Valette, Bruno Lina, Laurence Josset |
| EPI_ISL_3131343, EPI_ISL_3131344 | CHU TOULOUSE | CNR Virus des Infections Respiratoires - France SUD | Antonin Bal, Gregory Destras, Gwendolyne Burfin, Hadrien Regue, Quentin Semanas, Martine Valette, Bruno Lina, Laurence Josset |
| EPI_ISL_3131345 | LABORATOIRE ACCOLAB SUD OUEST | CNR Virus des Infections Respiratoires - France SUD | Antonin Bal, Gregory Destras, Gwendolyne Burfin, Hadrien Regue, Quentin Semanas, Martine Valette, Bruno Lina, Laurence Josset |
| EPI_ISL_3131346, EPI_ISL_3131347 | CHU TOULOUSE | CNR Virus des Infections Respiratoires - France SUD | Antonin Bal, Gregory Destras, Gwendolyne Burfin, Hadrien Regue, Quentin Semanas, Martine Valette, Bruno Lina, Laurence Josset |
| EPI_ISL_3131348 | LABORATOIRE ACCOLAB SUD OUEST | CNR Virus des Infections Respiratoires - France SUD | Antonin Bal, Gregory Destras, Gwendolyne Burfin, Hadrien Regue, Quentin Semanas, Martine Valette, Bruno Lina, Laurence Josset |
| EPI_ISL_3131349, EPI_ISL_3131350 | CHU TOULOUSE | CNR Virus des Infections Respiratoires - France SUD | Antonin Bal, Gregory Destras, Gwendolyne Burfin, Hadrien Regue, Quentin Semanas, Martine Valette, Bruno Lina, Laurence Josset |
| EPI_ISL_3131353 | LABORATOIRE ACCOLAB SUD OUEST | CNR Virus des Infections Respiratoires - France SUD | Antonin Bal, Gregory Destras, Gwendolyne Burfin, Hadrien Regue, Quentin Semanas, Martine Valette, Bruno Lina, Laurence Josset |
| EPI_ISL_3131356, EPI_ISL_3131357 | CERBALLIANCE MIDI-PYRENEES | CNR Virus des Infections Respiratoires - France SUD | Antonin Bal, Gregory Destras, Gwendolyne Burfin, Hadrien Regue, Quentin Semanas, Martine Valette, Bruno Lina, Laurence Josset |
| EPI_ISL_3131358, EPI_ISL_3131360 | LABORATOIRE ACCOLAB SUD OUEST | CNR Virus des Infections Respiratoires - France SUD | Antonin Bal, Gregory Destras, Gwendolyne Burfin, Hadrien Regue, Quentin Semanas, Martine Valette, Bruno Lina, Laurence Josset |
| EPI_ISL_3131361, EPI_ISL_3131362, EPI_ISL_3131364, EPI_ISL_3131365, EPI_ISL_3131366, EPI_ISL_3131367 | CERBALLIANCE MIDI-PYRENEES | CNR Virus des Infections Respiratoires - France SUD | Antonin Bal, Gregory Destras, Gwendolyne Burfin, Hadrien Regue, Quentin Semanas, Martine Valette, Bruno Lina, Laurence Josset |
| EPI_ISL_3131369, EPI_ISL_3131370, EPI_ISL_3131371, EPI_ISL_3131372, EPI_ISL_3131374, EPI_ISL_3131375, EPI_ISL_3131376, EPI_ISL_3131378 | LAM CBM | CNR Virus des Infections Respiratoires - France SUD | Antonin Bal, Gregory Destras, Gwendolyne Burfin, Hadrien Regue, Quentin Semanas, Martine Valette, Bruno Lina, Laurence Josset |
| EPI_ISL_3131379, EPI_ISL_3131380, EPI_ISL_3131381, EPI_ISL_3131382, EPI_ISL_3131383, EPI_ISL_3131384, EPI_ISL_3131385 | LABORATOIRE BOUVIER | CNR Virus des Infections Respiratoires - France SUD | Antonin Bal, Gregory Destras, Gwendolyne Burfin, Hadrien Regue, Quentin Semanas, Martine Valette, Bruno Lina, Laurence Josset |
| EPI_ISL_3131386 | LABORATOIRE d'ANALYSES de BIOLOGIE MEDICALES | CNR Virus des Infections Respiratoires - France SUD | Antonin Bal, Gregory Destras, Gwendolyne Burfin, Hadrien Regue, Quentin Semanas, Martine Valette, Bruno Lina, Laurence Josset |
| EPI_ISL_3131387 | LABORATOIRE DES CARMES | CNR Virus des Infections Respiratoires - France SUD | Antonin Bal, Gregory Destras, Gwendolyne Burfin, Hadrien Regue, Quentin Semanas, Martine Valette, Bruno Lina, Laurence Josset |
| EPI_ISL_3131388 | LABORATOIRE d'ANALYSES de BIOLOGIE MEDICALES | CNR Virus des Infections Respiratoires - France SUD | Antonin Bal, Gregory Destras, Gwendolyne Burfin, Hadrien Regue, Quentin Semanas, Martine Valette, Bruno Lina, Laurence Josset |
| EPI_ISL_3131389 | LABORATOIRE DES CARMES | CNR Virus des Infections Respiratoires - France SUD | Antonin Bal, Gregory Destras, Gwendolyne Burfin, Hadrien Regue, Quentin Semanas, Martine Valette, Bruno Lina, Laurence Josset |
| EPI_ISL_3131390 | LABORATOIRE d'ANALYSES de BIOLOGIE MEDICALES | CNR Virus des Infections Respiratoires - France SUD | Antonin Bal, Gregory Destras, Gwendolyne Burfin, Hadrien Regue, Quentin Semanas, Martine Valette, Bruno Lina, Laurence Josset |
| EPI_ISL_3131391 | CENTRE JEAN PERRIN | CNR Virus des Infections Respiratoires - France SUD | Antonin Bal, Gregory Destras, Gwendolyne Burfin, Hadrien Regue, Quentin Semanas, Martine Valette, Bruno Lina, Laurence Josset |
| EPI_ISL_3131392, EPI_ISL_3131394 | LABORATOIRE d'ANALYSES de BIOLOGIE MEDICALES | CNR Virus des Infections Respiratoires - France SUD | Antonin Bal, Gregory Destras, Gwendolyne Burfin, Hadrien Regue, Quentin Semanas, Martine Valette, Bruno Lina, Laurence Josset |
| EPI_ISL_3131395 | LABORATOIRE DES CARMES | CNR Virus des Infections Respiratoires - France SUD | Antonin Bal, Gregory Destras, Gwendolyne Burfin, Hadrien Regue, Quentin Semanas, Martine Valette, Bruno Lina, Laurence Josset |
| EPI_ISL_3131396 | LABORATOIRE d'ANALYSES de BIOLOGIE MEDICALES | CNR Virus des Infections Respiratoires - France SUD | Antonin Bal, Gregory Destras, Gwendolyne Burfin, Hadrien Regue, Quentin Semanas, Martine Valette, Bruno Lina, Laurence Josset |
| EPI_ISL_3131397 | LABORATOIRE DES CARMES | CNR Virus des Infections Respiratoires - France SUD | Antonin Bal, Gregory Destras, Gwendolyne Burfin, Hadrien Regue, Quentin Semanas, Martine Valette, Bruno Lina, Laurence Josset |
| EPI_ISL_3131398 | LABORATOIRE d'ANALYSES de BIOLOGIE MEDICALES | CNR Virus des Infections Respiratoires - France SUD | Antonin Bal, Gregory Destras, Gwendolyne Burfin, Hadrien Regue, Quentin Semanas, Martine Valette, Bruno Lina, Laurence Josset |
| EPI_ISL_3131399 | LABORATOIRE DES CARMES | CNR Virus des Infections Respiratoires - France SUD | Antonin Bal, Gregory Destras, Gwendolyne Burfin, Hadrien Regue, Quentin Semanas, Martine Valette, Bruno Lina, Laurence Josset |
| EPI_ISL_3131400 | LABORATOIRE d'ANALYSES de BIOLOGIE MEDICALES | CNR Virus des Infections Respiratoires - France SUD | Antonin Bal, Gregory Destras, Gwendolyne Burfin, Hadrien Regue, Quentin Semanas, Martine Valette, Bruno Lina, Laurence Josset |
| EPI_ISL_3131401 | LABORATOIRE FORTE BIO DAX | CNR Virus des Infections Respiratoires - France SUD | Antonin Bal, Gregory Destras, Gwendolyne Burfin, Hadrien Regue, Quentin Semanas, Martine Valette, Bruno Lina, Laurence Josset |
| EPI_ISL_3131403, EPI_ISL_3131404 | LABORATOIRE DES CARMES | CNR Virus des Infections Respiratoires - France SUD | Antonin Bal, Gregory Destras, Gwendolyne Burfin, Hadrien Regue, Quentin Semanas, Martine Valette, Bruno Lina, Laurence Josset |
| EPI_ISL_3131405 | LABORATOIRE d'ANALYSES de BIOLOGIE MEDICALES | CNR Virus des Infections Respiratoires - France SUD | Antonin Bal, Gregory Destras, Gwendolyne Burfin, Hadrien Regue, Quentin Semanas, Martine Valette, Bruno Lina, Laurence Josset |
| EPI_ISL_3131406 | LABORATOIRE DES CARMES | CNR Virus des Infections Respiratoires - France SUD | Antonin Bal, Gregory Destras, Gwendolyne Burfin, Hadrien Regue, Quentin Semanas, Martine Valette, Bruno Lina, Laurence Josset |
| EPI_ISL_3131407 | LABORATOIRE d'ANALYSES de BIOLOGIE MEDICALES | CNR Virus des Infections Respiratoires - France SUD | Antonin Bal, Gregory Destras, Gwendolyne Burfin, Hadrien Regue, Quentin Semanas, Martine Valette, Bruno Lina, Laurence Josset |
| EPI_ISL_3131408, EPI_ISL_3131409, EPI_ISL_3131410 | LABORATOIRE DES CARMES | CNR Virus des Infections Respiratoires - France SUD | Antonin Bal, Gregory Destras, Gwendolyne Burfin, Hadrien Regue, Quentin Semanas, Martine Valette, Bruno Lina, Laurence Josset |
| EPI_ISL_3131411, EPI_ISL_3131412, EPI_ISL_3131413 | LABORATOIRE d'ANALYSES de BIOLOGIE MEDICALES | CNR Virus des Infections Respiratoires - France SUD | Antonin Bal, Gregory Destras, Gwendolyne Burfin, Hadrien Regue, Quentin Semanas, Martine Valette, Bruno Lina, Laurence Josset |
| EPI_ISL_3131414 | LABORATOIRE FORTE BIO DAX | CNR Virus des Infections Respiratoires - France SUD | Antonin Bal, Gregory Destras, Gwendolyne Burfin, Hadrien Regue, Quentin Semanas, Martine Valette, Bruno Lina, Laurence Josset |
| EPI_ISL_3131415 | CENTRE HOSPITALIER DE SAINTONGE | CNR Virus des Infections Respiratoires - France SUD | Antonin Bal, Gregory Destras, Gwendolyne Burfin, Hadrien Regue, Quentin Semanas, Martine Valette, Bruno Lina, Laurence Josset |
| EPI_ISL_3131416 | LABORATOIRE FORTE BIO DAX | CNR Virus des Infections Respiratoires - France SUD | Antonin Bal, Gregory Destras, Gwendolyne Burfin, Hadrien Regue, Quentin Semanas, Martine Valette, Bruno Lina, Laurence Josset |
| EPI_ISL_3131417 | LABORATOIRE d'ANALYSES de BIOLOGIE MEDICALES | CNR Virus des Infections Respiratoires - France SUD | Antonin Bal, Gregory Destras, Gwendolyne Burfin, Hadrien Regue, Quentin Semanas, Martine Valette, Bruno Lina, Laurence Josset |
| EPI_ISL_3131418, EPI_ISL_3131419, EPI_ISL_3131420, EPI_ISL_3131421, EPI_ISL_3131422 | LABORATOIRE FORTE BIO DAX | CNR Virus des Infections Respiratoires - France SUD | Antonin Bal, Gregory Destras, Gwendolyne Burfin, Hadrien Regue, Quentin Semanas, Martine Valette, Bruno Lina, Laurence Josset |
| EPI_ISL_3131423 | LABORATOIRE d'ANALYSES de BIOLOGIE MEDICALES | CNR Virus des Infections Respiratoires - France SUD | Antonin Bal, Gregory Destras, Gwendolyne Burfin, Hadrien Regue, Quentin Semanas, Martine Valette, Bruno Lina, Laurence Josset |
| EPI_ISL_3131424 | LABORATOIRE FORTE BIO DAX | CNR Virus des Infections Respiratoires - France SUD | Antonin Bal, Gregory Destras, Gwendolyne Burfin, Hadrien Regue, Quentin Semanas, Martine Valette, Bruno Lina, Laurence Josset |
| EPI_ISL_3131425 | LABORATOIRE BIOESTEREL | CNR Virus des Infections Respiratoires - France SUD | Antonin Bal, Gregory Destras, Gwendolyne Burfin, Hadrien Regue, Quentin Semanas, Martine Valette, Bruno Lina, Laurence Josset |

| | | | |
|---|---|---|---|
| EPI_ISL_3131426 | LABORATOIRE FORTE BIO DAX | CNR Virus des Infections Respiratoires - France SUD | Antonin Bal, Gregory Destras, Gwendolyne Burfin, Hadrien Regue, Quentin Semanas, Martine Valette, Bruno Lina, Laurence Josset |
| EPI_ISL_3131427 | LABORATOIRE d'ANALYSES de BIOLOGIE MEDICALES | CNR Virus des Infections Respiratoires - France SUD | Antonin Bal, Gregory Destras, Gwendolyne Burfin, Hadrien Regue, Quentin Semanas, Martine Valette, Bruno Lina, Laurence Josset |
| EPI_ISL_3131428, EPI_ISL_3131429, EPI_ISL_3131430 | LABORATOIRE FORTE BIO DAX | CNR Virus des Infections Respiratoires - France SUD | Antonin Bal, Gregory Destras, Gwendolyne Burfin, Hadrien Regue, Quentin Semanas, Martine Valette, Bruno Lina, Laurence Josset |
| EPI_ISL_3131431 | LABORATOIRE d'ANALYSES de BIOLOGIE MEDICALES | CNR Virus des Infections Respiratoires - France SUD | Antonin Bal, Gregory Destras, Gwendolyne Burfin, Hadrien Regue, Quentin Semanas, Martine Valette, Bruno Lina, Laurence Josset |
| EPI_ISL_3131432, EPI_ISL_3131433, EPI_ISL_3131434 | LABORATOIRE FORTE BIO DAX | CNR Virus des Infections Respiratoires - France SUD | Antonin Bal, Gregory Destras, Gwendolyne Burfin, Hadrien Regue, Quentin Semanas, Martine Valette, Bruno Lina, Laurence Josset |
| EPI_ISL_3131435 | LABORATOIRE d'ANALYSES de BIOLOGIE MEDICALES | CNR Virus des Infections Respiratoires - France SUD | Antonin Bal, Gregory Destras, Gwendolyne Burfin, Hadrien Regue, Quentin Semanas, Martine Valette, Bruno Lina, Laurence Josset |
| EPI_ISL_3131436 | LABORATOIRE BIOESTEREL | CNR Virus des Infections Respiratoires - France SUD | Antonin Bal, Gregory Destras, Gwendolyne Burfin, Hadrien Regue, Quentin Semanas, Martine Valette, Bruno Lina, Laurence Josset |
| EPI_ISL_3131438 | LABORATOIRE FORTE BIO DAX | CNR Virus des Infections Respiratoires - France SUD | Antonin Bal, Gregory Destras, Gwendolyne Burfin, Hadrien Regue, Quentin Semanas, Martine Valette, Bruno Lina, Laurence Josset |
| EPI_ISL_3131443 | LABORATOIRE d'ANALYSES de BIOLOGIE MEDICALES | CNR Virus des Infections Respiratoires - France SUD | Antonin Bal, Gregory Destras, Gwendolyne Burfin, Hadrien Regue, Quentin Semanas, Martine Valette, Bruno Lina, Laurence Josset |
| EPI_ISL_3131446 | LABORATOIRE FORTE BIO DAX | CNR Virus des Infections Respiratoires - France SUD | Antonin Bal, Gregory Destras, Gwendolyne Burfin, Hadrien Regue, Quentin Semanas, Martine Valette, Bruno Lina, Laurence Josset |
| EPI_ISL_3131451 | LABORATOIRE BIOESTEREL | CNR Virus des Infections Respiratoires - France SUD | Antonin Bal, Gregory Destras, Gwendolyne Burfin, Hadrien Regue, Quentin Semanas, Martine Valette, Bruno Lina, Laurence Josset |
| EPI_ISL_3131453 | LABORATOIRE d'ANALYSES de BIOLOGIE MEDICALES | CNR Virus des Infections Respiratoires - France SUD | Antonin Bal, Gregory Destras, Gwendolyne Burfin, Hadrien Regue, Quentin Semanas, Martine Valette, Bruno Lina, Laurence Josset |
| EPI_ISL_3131455 | LABORATOIRE FORTE BIO DAX | CNR Virus des Infections Respiratoires - France SUD | Antonin Bal, Gregory Destras, Gwendolyne Burfin, Hadrien Regue, Quentin Semanas, Martine Valette, Bruno Lina, Laurence Josset |
| EPI_ISL_3131457 | LABORATOIRE BIOESTEREL | CNR Virus des Infections Respiratoires - France SUD | Antonin Bal, Gregory Destras, Gwendolyne Burfin, Hadrien Regue, Quentin Semanas, Martine Valette, Bruno Lina, Laurence Josset |
| EPI_ISL_3131459 | CERBALLIANCE RHONE ALPES | CNR Virus des Infections Respiratoires - France SUD | Antonin Bal, Gregory Destras, Gwendolyne Burfin, Hadrien Regue, Quentin Semanas, Martine Valette, Bruno Lina, Laurence Josset |
| EPI_ISL_3131464 | LABORATOIRE FORTE BIO DAX | CNR Virus des Infections Respiratoires - France SUD | Antonin Bal, Gregory Destras, Gwendolyne Burfin, Hadrien Regue, Quentin Semanas, Martine Valette, Bruno Lina, Laurence Josset |
| EPI_ISL_3131467 | LABORATOIRE d'ANALYSES de BIOLOGIE MEDICALES | CNR Virus des Infections Respiratoires - France SUD | Antonin Bal, Gregory Destras, Gwendolyne Burfin, Hadrien Regue, Quentin Semanas, Martine Valette, Bruno Lina, Laurence Josset |
| EPI_ISL_3131469 | LABORATOIRE FORTE BIO DAX | CNR Virus des Infections Respiratoires - France SUD | Antonin Bal, Gregory Destras, Gwendolyne Burfin, Hadrien Regue, Quentin Semanas, Martine Valette, Bruno Lina, Laurence Josset |
| EPI_ISL_3131471 | CERBALLIANCE RHONE ALPES | CNR Virus des Infections Respiratoires - France SUD | Antonin Bal, Gregory Destras, Gwendolyne Burfin, Hadrien Regue, Quentin Semanas, Martine Valette, Bruno Lina, Laurence Josset |
| EPI_ISL_3131472, EPI_ISL_3131473, EPI_ISL_3131475 | ZOTZ KLIMAS MVZ Düsseldorf-Centrum GbR ÜBAG für Labormedizin, Genetik, Zytologie, Pathologie | Center of Medical Microbiology, Virology, and Hospital Hygiene, University of Duesseldorf | Maximilian Damagnez;Alexander Dilthey;Patrick Finzer;Katrin Hoffmann;Torsten Houwaart;Lisanna Hülse;Malte Kohns Vasconcelos;Nadine Lübke;Jessica Nicolai;Klaus Pfeffer;Daniel Strelow;Jörg Timm;Andreas Walker;Tobias Wienemann;Rainer Zotz |
| EPI_ISL_3131479 | LABORATOIRE d'ANALYSES de BIOLOGIE MEDICALES | CNR Virus des Infections Respiratoires - France SUD | Antonin Bal, Gregory Destras, Gwendolyne Burfin, Hadrien Regue, Quentin Semanas, Martine Valette, Bruno Lina, Laurence Josset |
| EPI_ISL_3131481 | CERBALLIANCE RHONE ALPES | CNR Virus des Infections Respiratoires - France SUD | Antonin Bal, Gregory Destras, Gwendolyne Burfin, Hadrien Regue, Quentin Semanas, Martine Valette, Bruno Lina, Laurence Josset |
| EPI_ISL_3131483 | LABORATOIRE UNILIANS DECINES | CNR Virus des Infections Respiratoires - France SUD | Antonin Bal, Gregory Destras, Gwendolyne Burfin, Hadrien Regue, Quentin Semanas, Martine Valette, Bruno Lina, Laurence Josset |
| EPI_ISL_3131484 | ZOTZ KLIMAS MVZ Düsseldorf-Centrum GbR ÜBAG für Labormedizin, Genetik, Zytologie, Pathologie | Center of Medical Microbiology, Virology, and Hospital Hygiene, University of Duesseldorf | Maximilian Damagnez;Alexander Dilthey;Patrick Finzer;Katrin Hoffmann;Torsten Houwaart;Lisanna Hülse;Malte Kohns Vasconcelos;Nadine Lübke;Jessica Nicolai;Klaus Pfeffer;Daniel Strelow;Jörg Timm;Andreas Walker;Tobias Wienemann;Rainer Zotz |
| EPI_ISL_3131485 | CERBALLIANCE RHONE ALPES | CNR Virus des Infections Respiratoires - France SUD | Antonin Bal, Gregory Destras, Gwendolyne Burfin, Hadrien Regue, Quentin Semanas, Martine Valette, Bruno Lina, Laurence Josset |
| EPI_ISL_3131486 | ZOTZ KLIMAS MVZ Düsseldorf-Centrum GbR ÜBAG für Labormedizin, Genetik, Zytologie, Pathologie | Center of Medical Microbiology, Virology, and Hospital Hygiene, University of Duesseldorf | Maximilian Damagnez;Alexander Dilthey;Patrick Finzer;Katrin Hoffmann;Torsten Houwaart;Lisanna Hülse;Malte Kohns Vasconcelos;Nadine Lübke;Jessica Nicolai;Klaus Pfeffer;Daniel Strelow;Jörg Timm;Andreas Walker;Tobias Wienemann;Rainer Zotz |
| EPI_ISL_3131488 | LABORATOIRE d'ANALYSES de BIOLOGIE MEDICALES | CNR Virus des Infections Respiratoires - France SUD | Antonin Bal, Gregory Destras, Gwendolyne Burfin, Hadrien Regue, Quentin Semanas, Martine Valette, Bruno Lina, Laurence Josset |
| EPI_ISL_3131489, EPI_ISL_3131491 | ZOTZ KLIMAS MVZ Düsseldorf-Centrum GbR ÜBAG für Labormedizin, Genetik, Zytologie, Pathologie | Center of Medical Microbiology, Virology, and Hospital Hygiene, University of Duesseldorf | Maximilian Damagnez;Alexander Dilthey;Patrick Finzer;Katrin Hoffmann;Torsten Houwaart;Lisanna Hülse;Malte Kohns Vasconcelos;Nadine Lübke;Jessica Nicolai;Klaus Pfeffer;Daniel Strelow;Jörg Timm;Andreas Walker;Tobias Wienemann;Rainer Zotz |
| EPI_ISL_3131492 | CERBALLIANCE RHONE ALPES | CNR Virus des Infections Respiratoires - France SUD | Antonin Bal, Gregory Destras, Gwendolyne Burfin, Hadrien Regue, Quentin Semanas, Martine Valette, Bruno Lina, Laurence Josset |
| EPI_ISL_3131493, EPI_ISL_3131494 | ZOTZ KLIMAS MVZ Düsseldorf-Centrum GbR ÜBAG für Labormedizin, Genetik, Zytologie, Pathologie | Center of Medical Microbiology, Virology, and Hospital Hygiene, University of Duesseldorf | Maximilian Damagnez;Alexander Dilthey;Patrick Finzer;Katrin Hoffmann;Torsten Houwaart;Lisanna Hülse;Malte Kohns Vasconcelos;Nadine Lübke;Jessica Nicolai;Klaus Pfeffer;Daniel Strelow;Jörg Timm;Andreas Walker;Tobias Wienemann;Rainer Zotz |
| EPI_ISL_3131495 | LABORATOIRE UNILIANS DECINES | CNR Virus des Infections Respiratoires - France SUD | Antonin Bal, Gregory Destras, Gwendolyne Burfin, Hadrien Regue, Quentin Semanas, Martine Valette, Bruno Lina, Laurence Josset |
| EPI_ISL_3131497 | CERBALLIANCE RHONE ALPES | CNR Virus des Infections Respiratoires - France SUD | Antonin Bal, Gregory Destras, Gwendolyne Burfin, Hadrien Regue, Quentin Semanas, Martine Valette, Bruno Lina, Laurence Josset |
| EPI_ISL_3131498, EPI_ISL_3131499, EPI_ISL_3131500 | ZOTZ KLIMAS MVZ Düsseldorf-Centrum GbR ÜBAG für Labormedizin, Genetik, Zytologie, Pathologie | Center of Medical Microbiology, Virology, and Hospital Hygiene, University of Duesseldorf | Maximilian Damagnez;Alexander Dilthey;Patrick Finzer;Katrin Hoffmann;Torsten Houwaart;Lisanna Hülse;Malte Kohns Vasconcelos;Nadine Lübke;Jessica Nicolai;Klaus Pfeffer;Daniel Strelow;Jörg Timm;Andreas Walker;Tobias Wienemann;Rainer Zotz |
| EPI_ISL_3131501 | LABORATOIRE UNILIANS DECINES | CNR Virus des Infections Respiratoires - France SUD | Antonin Bal, Gregory Destras, Gwendolyne Burfin, Hadrien Regue, Quentin Semanas, Martine Valette, Bruno Lina, Laurence Josset |
| EPI_ISL_3131505 | ZOTZ KLIMAS MVZ Düsseldorf-Centrum GbR ÜBAG für Labormedizin, Genetik, Zytologie, Pathologie | Center of Medical Microbiology, Virology, and Hospital Hygiene, University of Duesseldorf | Maximilian Damagnez;Alexander Dilthey;Patrick Finzer;Katrin Hoffmann;Torsten Houwaart;Lisanna Hülse;Malte Kohns Vasconcelos;Nadine Lübke;Jessica Nicolai;Klaus Pfeffer;Daniel Strelow;Jörg Timm;Andreas Walker;Tobias Wienemann;Rainer Zotz |
| EPI_ISL_3131506 | CERBALLIANCE RHONE ALPES | CNR Virus des Infections Respiratoires - France SUD | Antonin Bal, Gregory Destras, Gwendolyne Burfin, Hadrien Regue, Quentin Semanas, Martine Valette, Bruno Lina, Laurence Josset |
| EPI_ISL_3131507 | ZOTZ KLIMAS MVZ Düsseldorf-Centrum GbR ÜBAG für Labormedizin, Genetik, Zytologie, Pathologie | Center of Medical Microbiology, Virology, and Hospital Hygiene, University of Duesseldorf | Maximilian Damagnez;Alexander Dilthey;Patrick Finzer;Katrin Hoffmann;Torsten Houwaart;Lisanna Hülse;Malte Kohns Vasconcelos;Nadine Lübke;Jessica Nicolai;Klaus Pfeffer;Daniel Strelow;Jörg Timm;Andreas Walker;Tobias Wienemann;Rainer Zotz |
| EPI_ISL_3131508 | LABORATOIRE FORTE BIO DAX | CNR Virus des Infections Respiratoires - France SUD | Antonin Bal, Gregory Destras, Gwendolyne Burfin, Hadrien Regue, Quentin Semanas, Martine Valette, Bruno Lina, Laurence Josset |
| EPI_ISL_3131509 | CERBALLIANCE RHONE ALPES | CNR Virus des Infections Respiratoires - France SUD | Antonin Bal, Gregory Destras, Gwendolyne Burfin, Hadrien Regue, Quentin Semanas, Martine Valette, Bruno Lina, Laurence Josset |
| EPI_ISL_3131510 | LABORATOIRE BIOMEDICA | CNR Virus des Infections Respiratoires - France SUD | Antonin Bal, Gregory Destras, Gwendolyne Burfin, Hadrien Regue, Quentin Semanas, Martine Valette, Bruno Lina, Laurence Josset |
| EPI_ISL_3131511 | CERBALLIANCE RHONE ALPES | CNR Virus des Infections Respiratoires - France SUD | Antonin Bal, Gregory Destras, Gwendolyne Burfin, Hadrien Regue, Quentin Semanas, Martine Valette, Bruno Lina, Laurence Josset |
| EPI_ISL_3131512, EPI_ISL_3131513 | LABORATOIRE FORTE BIO DAX | CNR Virus des Infections Respiratoires - France SUD | Antonin Bal, Gregory Destras, Gwendolyne Burfin, Hadrien Regue, Quentin Semanas, Martine Valette, Bruno Lina, Laurence Josset |
| EPI_ISL_3131515 | LABORATOIRE BIOMEDICA | CNR Virus des Infections Respiratoires - France SUD | Antonin Bal, Gregory Destras, Gwendolyne Burfin, Hadrien Regue, Quentin Semanas, Martine Valette, Bruno Lina, Laurence Josset |
| EPI_ISL_3131516 | LABORATOIRE FORTE BIO DAX | CNR Virus des Infections Respiratoires - France SUD | Antonin Bal, Gregory Destras, Gwendolyne Burfin, Hadrien Regue, Quentin Semanas, Martine Valette, Bruno Lina, Laurence Josset |
| EPI_ISL_3131517 | CERBALLIANCE RHONE ALPES | CNR Virus des Infections Respiratoires - France SUD | Antonin Bal, Gregory Destras, Gwendolyne Burfin, Hadrien Regue, Quentin Semanas, Martine Valette, Bruno Lina, Laurence Josset |
| EPI_ISL_3131518 | LABORATOIRE BIOMEDICA | CNR Virus des Infections Respiratoires - France SUD | Antonin Bal, Gregory Destras, Gwendolyne Burfin, Hadrien Regue, Quentin Semanas, Martine Valette, Bruno Lina, Laurence Josset |
| EPI_ISL_3131519 | LABORATOIRE UNILIANS DECINES | CNR Virus des Infections Respiratoires - France SUD | Antonin Bal, Gregory Destras, Gwendolyne Burfin, Hadrien Regue, Quentin Semanas, Martine Valette, Bruno Lina, Laurence Josset |
| EPI_ISL_3131520, EPI_ISL_3131521 | LABORATOIRE FORTE BIO DAX | CNR Virus des Infections Respiratoires - France SUD | Antonin Bal, Gregory Destras, Gwendolyne Burfin, Hadrien Regue, Quentin Semanas, Martine Valette, Bruno Lina, Laurence Josset |
| EPI_ISL_3131522 | LABORATOIRE BIOMEDICA | CNR Virus des Infections Respiratoires - France SUD | Antonin Bal, Gregory Destras, Gwendolyne Burfin, Hadrien Regue, Quentin Semanas, Martine Valette, Bruno Lina, Laurence Josset |
| EPI_ISL_3131523 | LABORATOIRE FORTE BIO DAX | CNR Virus des Infections Respiratoires - France SUD | Antonin Bal, Gregory Destras, Gwendolyne Burfin, Hadrien Regue, Quentin Semanas, Martine Valette, Bruno Lina, Laurence Josset |
| EPI_ISL_3131524 | CERBALLIANCE RHONE ALPES | CNR Virus des Infections Respiratoires - France SUD | Antonin Bal, Gregory Destras, Gwendolyne Burfin, Hadrien Regue, Quentin Semanas, Martine Valette, Bruno Lina, Laurence Josset |
| EPI_ISL_3131525 | LABORATOIRE UNILIANS DECINES | CNR Virus des Infections Respiratoires - France SUD | Antonin Bal, Gregory Destras, Gwendolyne Burfin, Hadrien Regue, Quentin Semanas, Martine Valette, Bruno Lina, Laurence Josset |
| EPI_ISL_3131526 | LABORATOIRE FORTE BIO DAX | CNR Virus des Infections Respiratoires - France SUD | Antonin Bal, Gregory Destras, Gwendolyne Burfin, Hadrien Regue, Quentin Semanas, Martine Valette, Bruno Lina, Laurence Josset |
| EPI_ISL_3131527 | LABORATOIRE BIOMEDICA | CNR Virus des Infections Respiratoires - France SUD | Antonin Bal, Gregory Destras, Gwendolyne Burfin, Hadrien Regue, Quentin Semanas, Martine Valette, Bruno Lina, Laurence Josset |
| EPI_ISL_3131528 | CERBALLIANCE RHONE ALPES | CNR Virus des Infections Respiratoires - France SUD | Antonin Bal, Gregory Destras, Gwendolyne Burfin, Hadrien Regue, Quentin Semanas, Martine Valette, Bruno Lina, Laurence Josset |
| EPI_ISL_3131529 | LABORATOIRE FORTE BIO DAX | CNR Virus des Infections Respiratoires - France SUD | Antonin Bal, Gregory Destras, Gwendolyne Burfin, Hadrien Regue, Quentin Semanas, Martine Valette, Bruno Lina, Laurence Josset |
| EPI_ISL_3131530 | LABORATOIRE BIOMEDICA | CNR Virus des Infections Respiratoires - France SUD | Antonin Bal, Gregory Destras, Gwendolyne Burfin, Hadrien Regue, Quentin Semanas, Martine Valette, Bruno Lina, Laurence Josset |
| EPI_ISL_3131531 | LABORATOIRE UNILIANS DECINES | CNR Virus des Infections Respiratoires - France SUD | Antonin Bal, Gregory Destras, Gwendolyne Burfin, Hadrien Regue, Quentin Semanas, Martine Valette, Bruno Lina, Laurence Josset |
| EPI_ISL_3131532 | CERBALLIANCE RHONE ALPES | CNR Virus des Infections Respiratoires - France SUD | Antonin Bal, Gregory Destras, Gwendolyne Burfin, Hadrien Regue, Quentin Semanas, Martine Valette, Bruno Lina, Laurence Josset |
| EPI_ISL_3131533 | LABORATOIRE UNILIANS DECINES | CNR Virus des Infections Respiratoires - France SUD | Antonin Bal, Gregory Destras, Gwendolyne Burfin, Hadrien Regue, Quentin Semanas, Martine Valette, Bruno Lina, Laurence Josset |
| EPI_ISL_3131534, EPI_ISL_3131535 | CERBALLIANCE RHONE ALPES | CNR Virus des Infections Respiratoires - France SUD | Antonin Bal, Gregory Destras, Gwendolyne Burfin, Hadrien Regue, Quentin Semanas, Martine Valette, Bruno Lina, Laurence Josset |
| EPI_ISL_3131536, EPI_ISL_3131537 | LABORATOIRE FORTE BIO DAX | CNR Virus des Infections Respiratoires - France SUD | Antonin Bal, Gregory Destras, Gwendolyne Burfin, Hadrien Regue, Quentin Semanas, Martine Valette, Bruno Lina, Laurence Josset |
| EPI_ISL_3131538 | LABORATOIRE UNILIANS DECINES | CNR Virus des Infections Respiratoires - France SUD | Antonin Bal, Gregory Destras, Gwendolyne Burfin, Hadrien Regue, Quentin Semanas, Martine Valette, Bruno Lina, Laurence Josset |
| EPI_ISL_3131541 | LABORATOIRE FORTE BIO DAX | CNR Virus des Infections Respiratoires - France SUD | Antonin Bal, Gregory Destras, Gwendolyne Burfin, Hadrien Regue, Quentin Semanas, Martine Valette, Bruno Lina, Laurence Josset |

| | | | |
|---|---|---|---|
| EPI_ISL_3131542 | LABORATOIRE UNILIANS DECINES | CNR Virus des Infections Respiratoires - France SUD | Antonin Bal, Gregory Destras, Gwendolyne Burfin, Hadrien Regue, Quentin Semanas, Martine Valette, Bruno Lina, Laurence Josset |
| EPI_ISL_3131544 | LABORATOIRE FORTE BIO DAX | CNR Virus des Infections Respiratoires - France SUD | Antonin Bal, Gregory Destras, Gwendolyne Burfin, Hadrien Regue, Quentin Semanas, Martine Valette, Bruno Lina, Laurence Josset |
| EPI_ISL_3131545 | LABM BIO 17 | CNR Virus des Infections Respiratoires - France SUD | Antonin Bal, Gregory Destras, Gwendolyne Burfin, Hadrien Regue, Quentin Semanas, Martine Valette, Bruno Lina, Laurence Josset |
| EPI_ISL_3131546 | CERBALLIANCE RHONE ALPES | CNR Virus des Infections Respiratoires - France SUD | Antonin Bal, Gregory Destras, Gwendolyne Burfin, Hadrien Regue, Quentin Semanas, Martine Valette, Bruno Lina, Laurence Josset |
| EPI_ISL_3131547 | LABORATOIRE FORTE BIO DAX | CNR Virus des Infections Respiratoires - France SUD | Antonin Bal, Gregory Destras, Gwendolyne Burfin, Hadrien Regue, Quentin Semanas, Martine Valette, Bruno Lina, Laurence Josset |
| EPI_ISL_3131548 | LABM BIO 17 | CNR Virus des Infections Respiratoires - France SUD | Antonin Bal, Gregory Destras, Gwendolyne Burfin, Hadrien Regue, Quentin Semanas, Martine Valette, Bruno Lina, Laurence Josset |
| EPI_ISL_3131549 | CERBALLIANCE RHONE ALPES | CNR Virus des Infections Respiratoires - France SUD | Antonin Bal, Gregory Destras, Gwendolyne Burfin, Hadrien Regue, Quentin Semanas, Martine Valette, Bruno Lina, Laurence Josset |
| EPI_ISL_3131550 | LABORATOIRE FORTE BIO DAX | CNR Virus des Infections Respiratoires - France SUD | Antonin Bal, Gregory Destras, Gwendolyne Burfin, Hadrien Regue, Quentin Semanas, Martine Valette, Bruno Lina, Laurence Josset |
| EPI_ISL_3131551 | LABORATOIRE UNILIANS DECINES | CNR Virus des Infections Respiratoires - France SUD | Antonin Bal, Gregory Destras, Gwendolyne Burfin, Hadrien Regue, Quentin Semanas, Martine Valette, Bruno Lina, Laurence Josset |
| EPI_ISL_3131552 | LABM BIO 17 | CNR Virus des Infections Respiratoires - France SUD | Antonin Bal, Gregory Destras, Gwendolyne Burfin, Hadrien Regue, Quentin Semanas, Martine Valette, Bruno Lina, Laurence Josset |
| EPI_ISL_3131553 | CERBALLIANCE RHONE ALPES | CNR Virus des Infections Respiratoires - France SUD | Antonin Bal, Gregory Destras, Gwendolyne Burfin, Hadrien Regue, Quentin Semanas, Martine Valette, Bruno Lina, Laurence Josset |
| EPI_ISL_3131554 | LABORATOIRE FORTE BIO DAX | CNR Virus des Infections Respiratoires - France SUD | Antonin Bal, Gregory Destras, Gwendolyne Burfin, Hadrien Regue, Quentin Semanas, Martine Valette, Bruno Lina, Laurence Josset |
| EPI_ISL_3131555 | LABM BIO 17 | CNR Virus des Infections Respiratoires - France SUD | Antonin Bal, Gregory Destras, Gwendolyne Burfin, Hadrien Regue, Quentin Semanas, Martine Valette, Bruno Lina, Laurence Josset |
| EPI_ISL_3131556 | CERBALLIANCE RHONE ALPES | CNR Virus des Infections Respiratoires - France SUD | Antonin Bal, Gregory Destras, Gwendolyne Burfin, Hadrien Regue, Quentin Semanas, Martine Valette, Bruno Lina, Laurence Josset |
| EPI_ISL_3131558 | LABM BIO 17 | CNR Virus des Infections Respiratoires - France SUD | Antonin Bal, Gregory Destras, Gwendolyne Burfin, Hadrien Regue, Quentin Semanas, Martine Valette, Bruno Lina, Laurence Josset |
| EPI_ISL_3131559 | MIRIALIS CLUSES BECHET | CNR Virus des Infections Respiratoires - France SUD | Antonin Bal, Gregory Destras, Gwendolyne Burfin, Hadrien Regue, Quentin Semanas, Martine Valette, Bruno Lina, Laurence Josset |
| EPI_ISL_3131560 | LABM BIO 17 | CNR Virus des Infections Respiratoires - France SUD | Antonin Bal, Gregory Destras, Gwendolyne Burfin, Hadrien Regue, Quentin Semanas, Martine Valette, Bruno Lina, Laurence Josset |
| EPI_ISL_3131561 | LABORATOIRE FORTE BIO DAX | CNR Virus des Infections Respiratoires - France SUD | Antonin Bal, Gregory Destras, Gwendolyne Burfin, Hadrien Regue, Quentin Semanas, Martine Valette, Bruno Lina, Laurence Josset |
| EPI_ISL_3131563 | LABM BIO 17 | CNR Virus des Infections Respiratoires - France SUD | Antonin Bal, Gregory Destras, Gwendolyne Burfin, Hadrien Regue, Quentin Semanas, Martine Valette, Bruno Lina, Laurence Josset |
| EPI_ISL_3131564 | LABORATOIRE FORTE BIO DAX | CNR Virus des Infections Respiratoires - France SUD | Antonin Bal, Gregory Destras, Gwendolyne Burfin, Hadrien Regue, Quentin Semanas, Martine Valette, Bruno Lina, Laurence Josset |
| EPI_ISL_3131565 | LABM BIO 17 | CNR Virus des Infections Respiratoires - France SUD | Antonin Bal, Gregory Destras, Gwendolyne Burfin, Hadrien Regue, Quentin Semanas, Martine Valette, Bruno Lina, Laurence Josset |
| EPI_ISL_3131566 | LABORATOIRE UNILIANS DECINES | CNR Virus des Infections Respiratoires - France SUD | Antonin Bal, Gregory Destras, Gwendolyne Burfin, Hadrien Regue, Quentin Semanas, Martine Valette, Bruno Lina, Laurence Josset |
| EPI_ISL_3131567 | CERBALLIANCE RHONE ALPES | CNR Virus des Infections Respiratoires - France SUD | Antonin Bal, Gregory Destras, Gwendolyne Burfin, Hadrien Regue, Quentin Semanas, Martine Valette, Bruno Lina, Laurence Josset |
| EPI_ISL_3131568 | LABM BIO 17 | CNR Virus des Infections Respiratoires - France SUD | Antonin Bal, Gregory Destras, Gwendolyne Burfin, Hadrien Regue, Quentin Semanas, Martine Valette, Bruno Lina, Laurence Josset |
| EPI_ISL_3131569 | LABORATOIRE FORTE BIO DAX | CNR Virus des Infections Respiratoires - France SUD | Antonin Bal, Gregory Destras, Gwendolyne Burfin, Hadrien Regue, Quentin Semanas, Martine Valette, Bruno Lina, Laurence Josset |
| EPI_ISL_3131570 | LABM BIO 17 | CNR Virus des Infections Respiratoires - France SUD | Antonin Bal, Gregory Destras, Gwendolyne Burfin, Hadrien Regue, Quentin Semanas, Martine Valette, Bruno Lina, Laurence Josset |
| EPI_ISL_3131571 | LABORATOIRE FORTE BIO DAX | CNR Virus des Infections Respiratoires - France SUD | Antonin Bal, Gregory Destras, Gwendolyne Burfin, Hadrien Regue, Quentin Semanas, Martine Valette, Bruno Lina, Laurence Josset |
| EPI_ISL_3131572 | CERBALLIANCE RHONE ALPES | CNR Virus des Infections Respiratoires - France SUD | Antonin Bal, Gregory Destras, Gwendolyne Burfin, Hadrien Regue, Quentin Semanas, Martine Valette, Bruno Lina, Laurence Josset |
| EPI_ISL_3131573 | LABM BIO 17 | CNR Virus des Infections Respiratoires - France SUD | Antonin Bal, Gregory Destras, Gwendolyne Burfin, Hadrien Regue, Quentin Semanas, Martine Valette, Bruno Lina, Laurence Josset |
| EPI_ISL_3131574 | LABORATOIRE UNILIANS DECINES | CNR Virus des Infections Respiratoires - France SUD | Antonin Bal, Gregory Destras, Gwendolyne Burfin, Hadrien Regue, Quentin Semanas, Martine Valette, Bruno Lina, Laurence Josset |
| EPI_ISL_3131575 | CERBALLIANCE RHONE ALPES | CNR Virus des Infections Respiratoires - France SUD | Antonin Bal, Gregory Destras, Gwendolyne Burfin, Hadrien Regue, Quentin Semanas, Martine Valette, Bruno Lina, Laurence Josset |
| EPI_ISL_3131576 | LABORATOIRE FORTE BIO DAX | CNR Virus des Infections Respiratoires - France SUD | Antonin Bal, Gregory Destras, Gwendolyne Burfin, Hadrien Regue, Quentin Semanas, Martine Valette, Bruno Lina, Laurence Josset |
| EPI_ISL_3131577 | LABM BIO 17 | CNR Virus des Infections Respiratoires - France SUD | Antonin Bal, Gregory Destras, Gwendolyne Burfin, Hadrien Regue, Quentin Semanas, Martine Valette, Bruno Lina, Laurence Josset |
| EPI_ISL_3131578 | LABORATOIRE FORTE BIO DAX | CNR Virus des Infections Respiratoires - France SUD | Antonin Bal, Gregory Destras, Gwendolyne Burfin, Hadrien Regue, Quentin Semanas, Martine Valette, Bruno Lina, Laurence Josset |
| EPI_ISL_3131579 | CERBALLIANCE RHONE ALPES | CNR Virus des Infections Respiratoires - France SUD | Antonin Bal, Gregory Destras, Gwendolyne Burfin, Hadrien Regue, Quentin Semanas, Martine Valette, Bruno Lina, Laurence Josset |
| EPI_ISL_3131580 | LABORATOIRE UNILIANS DECINES | CNR Virus des Infections Respiratoires - France SUD | Antonin Bal, Gregory Destras, Gwendolyne Burfin, Hadrien Regue, Quentin Semanas, Martine Valette, Bruno Lina, Laurence Josset |
| EPI_ISL_3131581 | LABORATOIRE FORTE BIO DAX | CNR Virus des Infections Respiratoires - France SUD | Antonin Bal, Gregory Destras, Gwendolyne Burfin, Hadrien Regue, Quentin Semanas, Martine Valette, Bruno Lina, Laurence Josset |
| EPI_ISL_3131582 | LABORATOIRE UNILIANS DECINES | CNR Virus des Infections Respiratoires - France SUD | Antonin Bal, Gregory Destras, Gwendolyne Burfin, Hadrien Regue, Quentin Semanas, Martine Valette, Bruno Lina, Laurence Josset |
| EPI_ISL_3131583 | CERBALLIANCE RHONE ALPES | CNR Virus des Infections Respiratoires - France SUD | Antonin Bal, Gregory Destras, Gwendolyne Burfin, Hadrien Regue, Quentin Semanas, Martine Valette, Bruno Lina, Laurence Josset |
| EPI_ISL_3131584 | LABORATOIRE FORTE BIO DAX | CNR Virus des Infections Respiratoires - France SUD | Antonin Bal, Gregory Destras, Gwendolyne Burfin, Hadrien Regue, Quentin Semanas, Martine Valette, Bruno Lina, Laurence Josset |
| EPI_ISL_3131585 | LABORATOIRE UNILIANS DECINES | CNR Virus des Infections Respiratoires - France SUD | Antonin Bal, Gregory Destras, Gwendolyne Burfin, Hadrien Regue, Quentin Semanas, Martine Valette, Bruno Lina, Laurence Josset |
| EPI_ISL_3131586 | LABORATOIRE FORTE BIO DAX | CNR Virus des Infections Respiratoires - France SUD | Antonin Bal, Gregory Destras, Gwendolyne Burfin, Hadrien Regue, Quentin Semanas, Martine Valette, Bruno Lina, Laurence Josset |
| EPI_ISL_3131587 | CERBALLIANCE RHONE ALPES | CNR Virus des Infections Respiratoires - France SUD | Antonin Bal, Gregory Destras, Gwendolyne Burfin, Hadrien Regue, Quentin Semanas, Martine Valette, Bruno Lina, Laurence Josset |
| EPI_ISL_3131588 | MIRIALIS CLUSES BECHET | CNR Virus des Infections Respiratoires - France SUD | Antonin Bal, Gregory Destras, Gwendolyne Burfin, Hadrien Regue, Quentin Semanas, Martine Valette, Bruno Lina, Laurence Josset |
| EPI_ISL_3131590 | CERBALLIANCE RHONE ALPES | CNR Virus des Infections Respiratoires - France SUD | Antonin Bal, Gregory Destras, Gwendolyne Burfin, Hadrien Regue, Quentin Semanas, Martine Valette, Bruno Lina, Laurence Josset |
| EPI_ISL_3131591 | MIRIALIS CLUSES BECHET | CNR Virus des Infections Respiratoires - France SUD | Antonin Bal, Gregory Destras, Gwendolyne Burfin, Hadrien Regue, Quentin Semanas, Martine Valette, Bruno Lina, Laurence Josset |
| EPI_ISL_3131592 | LABORATOIRE FORTE BIO DAX | CNR Virus des Infections Respiratoires - France SUD | Antonin Bal, Gregory Destras, Gwendolyne Burfin, Hadrien Regue, Quentin Semanas, Martine Valette, Bruno Lina, Laurence Josset |
| EPI_ISL_3131593 | MIRIALIS CLUSES BECHET | CNR Virus des Infections Respiratoires - France SUD | Antonin Bal, Gregory Destras, Gwendolyne Burfin, Hadrien Regue, Quentin Semanas, Martine Valette, Bruno Lina, Laurence Josset |
| EPI_ISL_3131594 | CERBALLIANCE RHONE ALPES | CNR Virus des Infections Respiratoires - France SUD | Antonin Bal, Gregory Destras, Gwendolyne Burfin, Hadrien Regue, Quentin Semanas, Martine Valette, Bruno Lina, Laurence Josset |
| EPI_ISL_3131595 | LABORATOIRE FORTE BIO DAX | CNR Virus des Infections Respiratoires - France SUD | Antonin Bal, Gregory Destras, Gwendolyne Burfin, Hadrien Regue, Quentin Semanas, Martine Valette, Bruno Lina, Laurence Josset |
| EPI_ISL_3131596 | CERBALLIANCE RHONE ALPES | CNR Virus des Infections Respiratoires - France SUD | Antonin Bal, Gregory Destras, Gwendolyne Burfin, Hadrien Regue, Quentin Semanas, Martine Valette, Bruno Lina, Laurence Josset |
| EPI_ISL_3131597 | LABORATOIRE FORTE BIO DAX | CNR Virus des Infections Respiratoires - France SUD | Antonin Bal, Gregory Destras, Gwendolyne Burfin, Hadrien Regue, Quentin Semanas, Martine Valette, Bruno Lina, Laurence Josset |
| EPI_ISL_3131598 | MIRIALIS CLUSES BECHET | CNR Virus des Infections Respiratoires - France SUD | Antonin Bal, Gregory Destras, Gwendolyne Burfin, Hadrien Regue, Quentin Semanas, Martine Valette, Bruno Lina, Laurence Josset |
| EPI_ISL_3131599 | CERBALLIANCE RHONE ALPES | CNR Virus des Infections Respiratoires - France SUD | Antonin Bal, Gregory Destras, Gwendolyne Burfin, Hadrien Regue, Quentin Semanas, Martine Valette, Bruno Lina, Laurence Josset |
| EPI_ISL_3131600 | LABORATOIRE FORTE BIO DAX | CNR Virus des Infections Respiratoires - France SUD | Antonin Bal, Gregory Destras, Gwendolyne Burfin, Hadrien Regue, Quentin Semanas, Martine Valette, Bruno Lina, Laurence Josset |
| EPI_ISL_3131601 | MIRIALIS CLUSES BECHET | CNR Virus des Infections Respiratoires - France SUD | Antonin Bal, Gregory Destras, Gwendolyne Burfin, Hadrien Regue, Quentin Semanas, Martine Valette, Bruno Lina, Laurence Josset |
| EPI_ISL_3131602 | LABORATOIRE FORTE BIO DAX | CNR Virus des Infections Respiratoires - France SUD | Antonin Bal, Gregory Destras, Gwendolyne Burfin, Hadrien Regue, Quentin Semanas, Martine Valette, Bruno Lina, Laurence Josset |
| EPI_ISL_3131603 | CERBALLIANCE RHONE ALPES | CNR Virus des Infections Respiratoires - France SUD | Antonin Bal, Gregory Destras, Gwendolyne Burfin, Hadrien Regue, Quentin Semanas, Martine Valette, Bruno Lina, Laurence Josset |
| EPI_ISL_3131604 | LABORATOIRE FORTE BIO DAX | CNR Virus des Infections Respiratoires - France SUD | Antonin Bal, Gregory Destras, Gwendolyne Burfin, Hadrien Regue, Quentin Semanas, Martine Valette, Bruno Lina, Laurence Josset |
| EPI_ISL_3131605 | CENTRE HOSPITALIER DE BASTIA | CNR Virus des Infections Respiratoires - France SUD | Antonin Bal, Gregory Destras, Gwendolyne Burfin, Hadrien Regue, Quentin Semanas, Martine Valette, Bruno Lina, Laurence Josset |
| EPI_ISL_3131606 | CERBALLIANCE RHONE ALPES | CNR Virus des Infections Respiratoires - France SUD | Antonin Bal, Gregory Destras, Gwendolyne Burfin, Hadrien Regue, Quentin Semanas, Martine Valette, Bruno Lina, Laurence Josset |
| EPI_ISL_3131607 | LABORATOIRE FORTE BIO DAX | CNR Virus des Infections Respiratoires - France SUD | Antonin Bal, Gregory Destras, Gwendolyne Burfin, Hadrien Regue, Quentin Semanas, Martine Valette, Bruno Lina, Laurence Josset |
| EPI_ISL_3131608 | CENTRE HOSPITALIER DE BASTIA | CNR Virus des Infections Respiratoires - France SUD | Antonin Bal, Gregory Destras, Gwendolyne Burfin, Hadrien Regue, Quentin Semanas, Martine Valette, Bruno Lina, Laurence Josset |
| EPI_ISL_3131609 | CERBALLIANCE RHONE ALPES | CNR Virus des Infections Respiratoires - France SUD | Antonin Bal, Gregory Destras, Gwendolyne Burfin, Hadrien Regue, Quentin Semanas, Martine Valette, Bruno Lina, Laurence Josset |
| EPI_ISL_3131610 | LABORATOIRE FORTE BIO DAX | CNR Virus des Infections Respiratoires - France SUD | Antonin Bal, Gregory Destras, Gwendolyne Burfin, Hadrien Regue, Quentin Semanas, Martine Valette, Bruno Lina, Laurence Josset |
| EPI_ISL_3131611 | CENTRE HOSPITALIER DE BASTIA | CNR Virus des Infections Respiratoires - France SUD | Antonin Bal, Gregory Destras, Gwendolyne Burfin, Hadrien Regue, Quentin Semanas, Martine Valette, Bruno Lina, Laurence Josset |
| EPI_ISL_3131612 | CERBALLIANCE RHONE ALPES | CNR Virus des Infections Respiratoires - France SUD | Antonin Bal, Gregory Destras, Gwendolyne Burfin, Hadrien Regue, Quentin Semanas, Martine Valette, Bruno Lina, Laurence Josset |
| EPI_ISL_3131613 | CENTRE HOSPITALIER DE BASTIA | CNR Virus des Infections Respiratoires - France SUD | Antonin Bal, Gregory Destras, Gwendolyne Burfin, Hadrien Regue, Quentin Semanas, Martine Valette, Bruno Lina, Laurence Josset |
| EPI_ISL_3131614 | CERBALLIANCE RHONE ALPES | CNR Virus des Infections Respiratoires - France SUD | Antonin Bal, Gregory Destras, Gwendolyne Burfin, Hadrien Regue, Quentin Semanas, Martine Valette, Bruno Lina, Laurence Josset |
| EPI_ISL_3131615, EPI_ISL_3131616, EPI_ISL_3131617, EPI_ISL_3131618, EPI_ISL_3131619, EPI_ISL_3131620 | AX BIO OCEAN | CNR Virus des Infections Respiratoires - France SUD | Antonin Bal, Gregory Destras, Gwendolyne Burfin, Hadrien Regue, Quentin Semanas, Martine Valette, Bruno Lina, Laurence Josset |
| EPI_ISL_3131621 | CERBALLIANCE RHONE ALPES | CNR Virus des Infections Respiratoires - France SUD | Antonin Bal, Gregory Destras, Gwendolyne Burfin, Hadrien Regue, Quentin Semanas, Martine Valette, Bruno Lina, Laurence Josset |
| EPI_ISL_3131622, EPI_ISL_3131623 | AX BIO OCEAN | CNR Virus des Infections Respiratoires - France SUD | Antonin Bal, Gregory Destras, Gwendolyne Burfin, Hadrien Regue, Quentin Semanas, Martine Valette, Bruno Lina, Laurence Josset |

| | | | |
|---|---|---|---|
| EPI_ISL_3131624 | CERBALLIANCE RHONE ALPES | CNR Virus des Infections Respiratoires - France SUD | Antonin Bal, Gregory Destras, Gwendolyne Burfin, Hadrien Regue, Quentin Semanas, Martine Valette, Bruno Lina, Laurence Josset |
| EPI_ISL_3131625, EPI_ISL_3131626, EPI_ISL_3131628 | AX BIO OCEAN | CNR Virus des Infections Respiratoires - France SUD | Antonin Bal, Gregory Destras, Gwendolyne Burfin, Hadrien Regue, Quentin Semanas, Martine Valette, Bruno Lina, Laurence Josset |
| EPI_ISL_3131629 | LABORATOIRE UNILIANS DECINES | CNR Virus des Infections Respiratoires - France SUD | Antonin Bal, Gregory Destras, Gwendolyne Burfin, Hadrien Regue, Quentin Semanas, Martine Valette, Bruno Lina, Laurence Josset |
| EPI_ISL_3131630, EPI_ISL_3131631, EPI_ISL_3131632 | AX BIO OCEAN | CNR Virus des Infections Respiratoires - France SUD | Antonin Bal, Gregory Destras, Gwendolyne Burfin, Hadrien Regue, Quentin Semanas, Martine Valette, Bruno Lina, Laurence Josset |
| EPI_ISL_3131633, EPI_ISL_3131634, EPI_ISL_3131635, EPI_ISL_3131636, EPI_ISL_3131637, EPI_ISL_3131638, EPI_ISL_3131639, EPI_ISL_3131640 | MIRIALIS CLUSES BECHET | CNR Virus des Infections Respiratoires - France SUD | Antonin Bal, Gregory Destras, Gwendolyne Burfin, Hadrien Regue, Quentin Semanas, Martine Valette, Bruno Lina, Laurence Josset |
| EPI_ISL_3131641, EPI_ISL_3131643 | LABORATOIRE UNILIANS DECINES | CNR Virus des Infections Respiratoires - France SUD | Antonin Bal, Gregory Destras, Gwendolyne Burfin, Hadrien Regue, Quentin Semanas, Martine Valette, Bruno Lina, Laurence Josset |
| EPI_ISL_3131644, EPI_ISL_3131645 | MIRIALIS CLUSES BECHET | CNR Virus des Infections Respiratoires - France SUD | Antonin Bal, Gregory Destras, Gwendolyne Burfin, Hadrien Regue, Quentin Semanas, Martine Valette, Bruno Lina, Laurence Josset |
| EPI_ISL_3131646 | LABORATOIRE UNILIANS DECINES | CNR Virus des Infections Respiratoires - France SUD | Antonin Bal, Gregory Destras, Gwendolyne Burfin, Hadrien Regue, Quentin Semanas, Martine Valette, Bruno Lina, Laurence Josset |
| EPI_ISL_3131647, EPI_ISL_3131648 | MIRIALIS CLUSES BECHET | CNR Virus des Infections Respiratoires - France SUD | Antonin Bal, Gregory Destras, Gwendolyne Burfin, Hadrien Regue, Quentin Semanas, Martine Valette, Bruno Lina, Laurence Josset |
| EPI_ISL_3131649 | LABORATOIRE UNILIANS DECINES | CNR Virus des Infections Respiratoires - France SUD | Antonin Bal, Gregory Destras, Gwendolyne Burfin, Hadrien Regue, Quentin Semanas, Martine Valette, Bruno Lina, Laurence Josset |
| EPI_ISL_3131650 | MIRIALIS CLUSES BECHET | CNR Virus des Infections Respiratoires - France SUD | Antonin Bal, Gregory Destras, Gwendolyne Burfin, Hadrien Regue, Quentin Semanas, Martine Valette, Bruno Lina, Laurence Josset |
| EPI_ISL_3131651, EPI_ISL_3131652, EPI_ISL_3131653, EPI_ISL_3131654, EPI_ISL_3131655, EPI_ISL_3131656, EPI_ISL_3131657, EPI_ISL_3131658, EPI_ISL_3131659, EPI_ISL_3131660, EPI_ISL_3131661, EPI_ISL_3131662, EPI_ISL_3131663, EPI_ISL_3131664, EPI_ISL_3131665 | | | |
| see above | LABORATOIRE UNILIANS DECINES | CNR Virus des Infections Respiratoires - France SUD | Antonin Bal, Gregory Destras, Gwendolyne Burfin, Hadrien Regue, Quentin Semanas, Martine Valette, Bruno Lina, Laurence Josset |
| EPI_ISL_3131666, EPI_ISL_3131667, EPI_ISL_3131668, EPI_ISL_3131669, EPI_ISL_3131671, EPI_ISL_3131672, EPI_ISL_3131674, EPI_ISL_3131675, EPI_ISL_3131677, EPI_ISL_3131678, EPI_ISL_3131679, EPI_ISL_3131681, EPI_ISL_3131682, EPI_ISL_3131683 | | | |
| see above | LABORATOIRE BIOFUSION | CNR Virus des Infections Respiratoires - France SUD | Antonin Bal, Gregory Destras, Gwendolyne Burfin, Hadrien Regue, Quentin Semanas, Martine Valette, Bruno Lina, Laurence Josset |
| EPI_ISL_3131684, EPI_ISL_3131685, EPI_ISL_3131686 | DYOMEDEA-LABORATOIRE DE LA SAUVEGARDE | CNR Virus des Infections Respiratoires - France SUD | Antonin Bal, Gregory Destras, Gwendolyne Burfin, Hadrien Regue, Quentin Semanas, Martine Valette, Bruno Lina, Laurence Josset |
| EPI_ISL_3131687, EPI_ISL_3131688, EPI_ISL_3131689 | LABORATOIRE UNILIANS DECINES | CNR Virus des Infections Respiratoires - France SUD | Antonin Bal, Gregory Destras, Gwendolyne Burfin, Hadrien Regue, Quentin Semanas, Martine Valette, Bruno Lina, Laurence Josset |
| EPI_ISL_3131690 | DYOMEDEA-LABORATOIRE DE LA SAUVEGARDE | CNR Virus des Infections Respiratoires - France SUD | Antonin Bal, Gregory Destras, Gwendolyne Burfin, Hadrien Regue, Quentin Semanas, Martine Valette, Bruno Lina, Laurence Josset |
| EPI_ISL_3131691, EPI_ISL_3131692, EPI_ISL_3131693, EPI_ISL_3131694, EPI_ISL_3131695, EPI_ISL_3131696, EPI_ISL_3131697, EPI_ISL_3131698 | LABORATOIRE UNILIANS DECINES | CNR Virus des Infections Respiratoires - France SUD | Antonin Bal, Gregory Destras, Gwendolyne Burfin, Hadrien Regue, Quentin Semanas, Martine Valette, Bruno Lina, Laurence Josset |
| EPI_ISL_3131699 | LABORATOIRE d'ANALYSES de BIOLOGIE MEDICALES | CNR Virus des Infections Respiratoires - France SUD | Antonin Bal, Gregory Destras, Gwendolyne Burfin, Hadrien Regue, Quentin Semanas, Martine Valette, Bruno Lina, Laurence Josset |
| EPI_ISL_3131700 | LABORATOIRE UNILIANS DECINES | CNR Virus des Infections Respiratoires - France SUD | Antonin Bal, Gregory Destras, Gwendolyne Burfin, Hadrien Regue, Quentin Semanas, Martine Valette, Bruno Lina, Laurence Josset |
| EPI_ISL_3131701 | LABORATOIRE d'ANALYSES de BIOLOGIE MEDICALES | CNR Virus des Infections Respiratoires - France SUD | Antonin Bal, Gregory Destras, Gwendolyne Burfin, Hadrien Regue, Quentin Semanas, Martine Valette, Bruno Lina, Laurence Josset |
| EPI_ISL_3131702, EPI_ISL_3131703, EPI_ISL_3131706, EPI_ISL_3131707, EPI_ISL_3131709, EPI_ISL_3131710, EPI_ISL_3131711, EPI_ISL_3131712, EPI_ISL_3131713, EPI_ISL_3131714, EPI_ISL_3131715, EPI_ISL_3131716, EPI_ISL_3131717, EPI_ISL_3131718, EPI_ISL_3131719, EPI_ISL_3131721, EPI_ISL_3131722, EPI_ISL_3131723, EPI_ISL_3131724, EPI_ISL_3131725, EPI_ISL_3131726, EPI_ISL_3131727, EPI_ISL_3131728, EPI_ISL_3131729, EPI_ISL_3131730, EPI_ISL_3131731, EPI_ISL_3131732, EPI_ISL_3131733, EPI_ISL_3131734, EPI_ISL_3131735 | | | |
| see above | CNR Virus des Infections Respiratoires - France SUD | CNR Virus des Infections Respiratoires - France SUD | Antonin Bal, Gregory Destras, Gwendolyne Burfin, Hadrien Regue, Quentin Semanas, Martine Valette, Bruno Lina, Laurence Josset |
| EPI_ISL_3131736, EPI_ISL_3131737, EPI_ISL_3131738, EPI_ISL_3131739, EPI_ISL_3131740, EPI_ISL_3131741, EPI_ISL_3131742, EPI_ISL_3131743, EPI_ISL_3131744, EPI_ISL_3131745, EPI_ISL_3131746, EPI_ISL_3131747, EPI_ISL_3131748, EPI_ISL_3131749, EPI_ISL_3131750, EPI_ISL_3131751, EPI_ISL_3131752, EPI_ISL_3131753, EPI_ISL_3131754, EPI_ISL_3131755, EPI_ISL_3131756, EPI_ISL_3131757, EPI_ISL_3131758, EPI_ISL_3131759, EPI_ISL_3131760, EPI_ISL_3131761, EPI_ISL_3131762, EPI_ISL_3131763, EPI_ISL_3131764 | | | |
| see above | Nebraska Public Health Laboratory | NPHL COVID-19 Response Team | NPHL COVID-19 Response Team |
| EPI_ISL_3131765, EPI_ISL_3131766, EPI_ISL_3131767, EPI_ISL_3131768, EPI_ISL_3131769, EPI_ISL_3131770, EPI_ISL_3131772, EPI_ISL_3131773 | ZOTZ KLIMAS MVZ Düsseldorf-Centrum GbR ÜBAG für Labormedizin, Genetik, Zytologie, Pathologie | Center of Medical Microbiology, Virology, and Hospital Hygiene, University of Duesseldorf | Maximilian Damagnez;Alexander Dilthey;Patrick Finzer;Katrin Hoffmann;Torsten Houwaart;Lisanna Hülse;Malte Kohns Vasconcelos;Nadine Lübke;Jessica Nicolai;Klaus Pfeffer;Daniel Strelow;Jörg Timm;Andreas Walker;Tobias Wienemann;Rainer Zotz |
| EPI_ISL_3131776 | MANKWENG LABORATORY | National Institute for Communicable Diseases of the National Health Laboratory Service | Amoako DG, Everatt J, Scheepers C, Mohale T, Ntuli N, Mahlangu B, Mnguni A, Ismail A, Bhiman JN |
| EPI_ISL_3131908, EPI_ISL_3131909, EPI_ISL_3131911, EPI_ISL_3131917, EPI_ISL_3131919, EPI_ISL_3131922, EPI_ISL_3131926, EPI_ISL_3131927, EPI_ISL_3131931, EPI_ISL_3131932, EPI_ISL_3131933, EPI_ISL_3131937, EPI_ISL_3131940, EPI_ISL_3131941, EPI_ISL_3131942, EPI_ISL_3131943, EPI_ISL_3131944, EPI_ISL_3131946, EPI_ISL_3131947, EPI_ISL_3131948, EPI_ISL_3131950, EPI_ISL_3131952, EPI_ISL_3131953, EPI_ISL_3131954, EPI_ISL_3131955, EPI_ISL_3131956, EPI_ISL_3131957, EPI_ISL_3131958, EPI_ISL_3131959, EPI_ISL_3131961, EPI_ISL_3131962, EPI_ISL_3131963, EPI_ISL_3131964, EPI_ISL_3131965, EPI_ISL_3131966, EPI_ISL_3131969, EPI_ISL_3131971, EPI_ISL_3131972, EPI_ISL_3131973, EPI_ISL_3131974, EPI_ISL_3131976, EPI_ISL_3131977, EPI_ISL_3131978, EPI_ISL_3131979, EPI_ISL_3131980, EPI_ISL_3131981, EPI_ISL_3131983, EPI_ISL_3131984, EPI_ISL_3131985, EPI_ISL_3131986, EPI_ISL_3131987, EPI_ISL_3131988, EPI_ISL_3131989, EPI_ISL_3131990, EPI_ISL_3131991, EPI_ISL_3131992, EPI_ISL_3131993, EPI_ISL_3131997, EPI_ISL_3131998, EPI_ISL_3131999, EPI_ISL_3132000, EPI_ISL_3132001, EPI_ISL_3132002, EPI_ISL_3132003, EPI_ISL_3132004, EPI_ISL_3132005, EPI_ISL_3132006, EPI_ISL_3132007, EPI_ISL_3132008, EPI_ISL_3132009, EPI_ISL_3132010, EPI_ISL_3132011, EPI_ISL_3132012, EPI_ISL_3132014, EPI_ISL_3132015, EPI_ISL_3132016, EPI_ISL_3132017, EPI_ISL_3132018, EPI_ISL_3132019, EPI_ISL_3132020, EPI_ISL_3132021, EPI_ISL_3132022, EPI_ISL_3132023, EPI_ISL_3132024, EPI_ISL_3132025, EPI_ISL_3132026, EPI_ISL_3132027, EPI_ISL_3132028, EPI_ISL_3132029, EPI_ISL_3132030, EPI_ISL_3132031, EPI_ISL_3132032, EPI_ISL_3132033, EPI_ISL_3132034, EPI_ISL_3132035, EPI_ISL_3132036, EPI_ISL_3132037, EPI_ISL_3132038, EPI_ISL_3132039, EPI_ISL_3132040, EPI_ISL_3132041, EPI_ISL_3132042, EPI_ISL_3132044, EPI_ISL_3132047, EPI_ISL_3132048, EPI_ISL_3132049, EPI_ISL_3132050, EPI_ISL_3132051, EPI_ISL_3132052, EPI_ISL_3132053, EPI_ISL_3132054, EPI_ISL_3132055, EPI_ISL_3132056, EPI_ISL_3132057, EPI_ISL_3132058, EPI_ISL_3132059, EPI_ISL_3132060, EPI_ISL_3132061, EPI_ISL_3132062, EPI_ISL_3132063, EPI_ISL_3132064, EPI_ISL_3132065, EPI_ISL_3132066, EPI_ISL_3132067, EPI_ISL_3132068, EPI_ISL_3132069, EPI_ISL_3132070, EPI_ISL_3132071, EPI_ISL_3132072, EPI_ISL_3132073, EPI_ISL_3132074, EPI_ISL_3132075, EPI_ISL_3132076, EPI_ISL_3132077, EPI_ISL_3132078, EPI_ISL_3132079, EPI_ISL_3132080, EPI_ISL_3132081, EPI_ISL_3132082, EPI_ISL_3132083, EPI_ISL_3132084, EPI_ISL_3132085, EPI_ISL_3132086, EPI_ISL_3132087, EPI_ISL_3132088, EPI_ISL_3132089, EPI_ISL_3132090, EPI_ISL_3132092, EPI_ISL_3132093, EPI_ISL_3132094, EPI_ISL_3132095, EPI_ISL_3132096, EPI_ISL_3132097, EPI_ISL_3132098, EPI_ISL_3132099, EPI_ISL_3132100, EPI_ISL_3132101, EPI_ISL_3132102, EPI_ISL_3132103, EPI_ISL_3132104, EPI_ISL_3132105, EPI_ISL_3132106, EPI_ISL_3132107, EPI_ISL_3132108 | | | |
| see above | Wisconsin State Laboratory of Hygiene Communicable Disease Division | Wisconsin State Laboratory of Hygiene Communicable Disease Division | Abigail C. Shockey, Alicia J. Mooney, Erika M. Hanson, Tonya Danz, Richard Griesser, Sara Wagner, Kelsey R. Florek |
| EPI_ISL_3132109, EPI_ISL_3132110, EPI_ISL_3132111, EPI_ISL_3132112, EPI_ISL_3132113, EPI_ISL_3132116, EPI_ISL_3132117, EPI_ISL_3132118, EPI_ISL_3132119, EPI_ISL_3132120, EPI_ISL_3132122, EPI_ISL_3132123, EPI_ISL_3132124, EPI_ISL_3132125, EPI_ISL_3132127, EPI_ISL_3132128, EPI_ISL_3132129, EPI_ISL_3132132, EPI_ISL_3132133 | | | |
| see above | TXDSHS | TXDSHS | Rashmi Tuladhar, Bonnie Oh, Jenny Zhang, Maliha Rahman, Mayela Pedrueza, Anita Pokharel, Karen Bobier, Lorraine Rodriguez, Myong Koag, Chun Wang, Rachel Lee, Grace Kubin |
| EPI_ISL_3132134, EPI_ISL_3132135, EPI_ISL_3132136, EPI_ISL_3132137, EPI_ISL_3132138, EPI_ISL_3132139, EPI_ISL_3132140, EPI_ISL_3132141, EPI_ISL_3132142, EPI_ISL_3132143, EPI_ISL_3132144, EPI_ISL_3132145, EPI_ISL_3132146, EPI_ISL_3132147, EPI_ISL_3132148, EPI_ISL_3132149, EPI_ISL_3132150, EPI_ISL_3132151, EPI_ISL_3132152, EPI_ISL_3132153, EPI_ISL_3132154, EPI_ISL_3132155, EPI_ISL_3132156, EPI_ISL_3132157, EPI_ISL_3132158, EPI_ISL_3132159, EPI_ISL_3132160, EPI_ISL_3132161, EPI_ISL_3132162, EPI_ISL_3132163, EPI_ISL_3132164, EPI_ISL_3132165, EPI_ISL_3132166 | | | |
| see above | Wyoming Public Health Laboratory | Wyoming Public Health Laboratory | Jim Mildenberger, Taylor Fearing, Channing Weber, Ashley Norberg, Chayse Rowley, Marley Goetz, Brian Dominguez, Elliot Thomasson, Sam Britz, Cari Sloma, Robert Petit, and Rob Christensen |
| EPI_ISL_3132167, EPI_ISL_3132168, EPI_ISL_3132169, EPI_ISL_3132170, EPI_ISL_3132171 | LCSP | IICS-UNA | Magaly Martinez, Adriana Valenzuela, Alejandra Rojas, Chyntia Diaz, Eva Nara, Fatima Cardozo, Florencia del Puerto, Joel Ortiz, Jonas Fernandez, Laura Franco, Leticia Rojas, Maria Eugenia Galeano, Cynthia Vazquez, Andrea Gomez de la Fuente, Shirley Villalba, Juan Torales, Maria Liz Gamarra, Andrea Ojeda, Guillermo Sequera. |
| EPI_ISL_3132172 | MeyerLab | IICS-UNA | Magaly Martinez, Adriana Valenzuela, Alejandra Rojas, Chyntia Diaz, Eva Nara, Fatima Cardozo, Florencia del Puerto, Joel Ortiz, Jonas Fernandez, Laura Franco, Leticia Rojas, Maria Eugenia Galeano, Cynthia Vazquez, Andrea Gomez de la Fuente, Shirley Villalba, Juan Torales, Maria Liz Gamarra, Andrea Ojeda, Guillermo Sequera. |
| EPI_ISL_3132191, EPI_ISL_3132192, EPI_ISL_3132193, EPI_ISL_3132194, EPI_ISL_3132195 | Puerto Rico Department of Health | Centers for Disease Control and Prevention, Dengue Branch | Gilberto A. Santiago, Glenda Gonzalez, Betzabel Flores, Keyla Charriez, Gabriela Paz-Bailey, Jorge L. Munoz-Jordan |
| EPI_ISL_3132196 | Centers for Disease Control and Prevention, Dengue Branch | Centers for Disease Control and Prevention, Dengue Branch | Gilberto A. Santiago, Glenda Gonzalez, Betzabel Flores, Keyla Charriez, Gabriela Paz-Bailey, Jorge L. Munoz-Jordan |
| EPI_ISL_3132252, EPI_ISL_3132253, | Genetica Molecular and Subdepartamento de Virologia ISP Chile | Instituto de Salud Publica de Chile | Karen Orostica, Constanza Campano, Barbara Parra, Loredana Arata, Gisselle Barra, Patricia Bustos, Rodrigo Fasce, Javier Tognarelli, Andres Castillo, Soledad Ulloa, Jorge Fernandez |

| | | | |
|---|---|---|---|
| EPI_ISL_3132255, EPI_ISL_3132256, EPI_ISL_3132258, EPI_ISL_3132259, EPI_ISL_3132260, EPI_ISL_3132261, EPI_ISL_3132262 | | | |
| EPI_ISL_3132295, EPI_ISL_3132296, EPI_ISL_3132297, EPI_ISL_3132298, EPI_ISL_3132299, EPI_ISL_3132300, EPI_ISL_3132301, EPI_ISL_3132302, EPI_ISL_3132303, EPI_ISL_3132304, EPI_ISL_3132305, EPI_ISL_3132306, EPI_ISL_3132307, EPI_ISL_3132308, EPI_ISL_3132309, EPI_ISL_3132310, EPI_ISL_3132311, EPI_ISL_3132312, EPI_ISL_3132313, EPI_ISL_3132314, EPI_ISL_3132315, EPI_ISL_3132316, EPI_ISL_3132317, EPI_ISL_3132318, EPI_ISL_3132319, EPI_ISL_3132320, EPI_ISL_3132321, EPI_ISL_3132322, EPI_ISL_3132323, EPI_ISL_3132324, EPI_ISL_3132325, EPI_ISL_3132326, EPI_ISL_3132327, EPI_ISL_3132328, EPI_ISL_3132329, EPI_ISL_3132330, EPI_ISL_3132331, EPI_ISL_3132332, EPI_ISL_3132334, EPI_ISL_3132335, EPI_ISL_3132336, EPI_ISL_3132337, EPI_ISL_3132338, EPI_ISL_3132339, EPI_ISL_3132340, EPI_ISL_3132341 | | | |
| see above | MRCG at LSHTM Genomics lab | MRCG at LSHTM Genomics lab | Abdul Karim sesay, Abdoulie Kanteh, Jarra Manneh, Mariama Kujabi, Bakary Sanyang,Dabiri Damilari |
| EPI_ISL_3132368 | Institute of Microbiology, Universidad San Francisco de Quito | Institute of Microbiology, Universidad San Francisco de Quito | Erika Muñoz, Juan José Guadalupe, Sully Márquez, Belén Prado-Vivar, Monica Becerra-Wong, Fernanda Zurita, Bernardo Gutiérrez, Andrea Pinos, Verónica Barragán, Patricio Rojas-Silva, Gabriel Trueba, Michelle Grunauer, Paúl Cárdenas |
| EPI_ISL_3132374, EPI_ISL_3132375, EPI_ISL_3132376, EPI_ISL_3132432, EPI_ISL_3132434, EPI_ISL_3132435, EPI_ISL_3132436, EPI_ISL_3132437 | North Dakota Department of Health, Public Health Laboratory | North Dakota Department of Health, Public Health Laboratory | Lisa Wingerter |
| EPI_ISL_3132495, EPI_ISL_3132496, EPI_ISL_3132497, EPI_ISL_3132498, EPI_ISL_3132500, EPI_ISL_3132501, EPI_ISL_3132502, EPI_ISL_3132503, EPI_ISL_3132504, EPI_ISL_3132505, EPI_ISL_3132506, EPI_ISL_3132507, EPI_ISL_3132508, EPI_ISL_3132509, EPI_ISL_3132510, EPI_ISL_3132511, EPI_ISL_3132512, EPI_ISL_3132513, EPI_ISL_3132514, EPI_ISL_3132515, EPI_ISL_3132516, EPI_ISL_3132517, EPI_ISL_3132518, EPI_ISL_3132519, EPI_ISL_3132520, EPI_ISL_3132521, EPI_ISL_3132523, EPI_ISL_3132524, EPI_ISL_3132525, EPI_ISL_3132526, EPI_ISL_3132527, EPI_ISL_3132528, EPI_ISL_3132530, EPI_ISL_3132531, EPI_ISL_3132532, EPI_ISL_3132533, EPI_ISL_3132534, EPI_ISL_3132535, EPI_ISL_3132536, EPI_ISL_3132537, EPI_ISL_3132538, EPI_ISL_3132539, EPI_ISL_3132541, EPI_ISL_3132542, EPI_ISL_3132543, EPI_ISL_3132544, EPI_ISL_3132546, EPI_ISL_3132547, EPI_ISL_3132548, EPI_ISL_3132549, EPI_ISL_3132550, EPI_ISL_3132551, EPI_ISL_3132553, EPI_ISL_3132554, EPI_ISL_3132555, EPI_ISL_3132556, EPI_ISL_3132557, EPI_ISL_3132558, EPI_ISL_3132559, EPI_ISL_3132560, EPI_ISL_3132561, EPI_ISL_3132562, EPI_ISL_3132563, EPI_ISL_3132565, EPI_ISL_3132567, EPI_ISL_3132568, EPI_ISL_3132569, EPI_ISL_3132570, EPI_ISL_3132571, EPI_ISL_3132572, EPI_ISL_3132573 | | | |
| see above | NHLS Charlotte Maxeke Johannesburg Academic Hospital and the University of the Witwatersrand | KRISP, KZn Research Innovation and Sequencing Platform | Florette Treurnicht, Bulelani Manene, Kathleen Subramoney, Giandhari Jennifer, Pillay Sureshnee, Yajna Ramphal, Naidoo Yeshnee, Tshabuila Derek,Tegally Houriiyah, San James, Wilkinson Eduan, de Oliveira Tulio |
| EPI_ISL_3132574, EPI_ISL_3132575, EPI_ISL_3132576, EPI_ISL_3132577, EPI_ISL_3132578, EPI_ISL_3132579, EPI_ISL_3132580, EPI_ISL_3132581, EPI_ISL_3132582, EPI_ISL_3132583, EPI_ISL_3132584, EPI_ISL_3132585, EPI_ISL_3132587, EPI_ISL_3132588, EPI_ISL_3132589, EPI_ISL_3132590, EPI_ISL_3132591, EPI_ISL_3132592, EPI_ISL_3132593, EPI_ISL_3132594, EPI_ISL_3132595, EPI_ISL_3132596, EPI_ISL_3132597, EPI_ISL_3132598, EPI_ISL_3132599, EPI_ISL_3132600, EPI_ISL_3132601, EPI_ISL_3132603, EPI_ISL_3132604, EPI_ISL_3132605, EPI_ISL_3132606, EPI_ISL_3132607, EPI_ISL_3132609, EPI_ISL_3132610, EPI_ISL_3132611, EPI_ISL_3132612, EPI_ISL_3132613, EPI_ISL_3132614, EPI_ISL_3132615, EPI_ISL_3132616, EPI_ISL_3132617, EPI_ISL_3132618, EPI_ISL_3132619, EPI_ISL_3132620, EPI_ISL_3132621, EPI_ISL_3132622 | | | |
| see above | National Health Laboratory Service, South Africa | KRISP, KZn Research Innovation and Sequencing Platform | Giandhari Jennifer, Pillay Sureshnee, Yajna Ramphal, Naidoo Yeshnee, Tshabuila Derek,Tegally Houriiyah, San James, Wilkinson Eduan, de Oliveira Tulio |
| EPI_ISL_3132624, EPI_ISL_3132625, EPI_ISL_3132626, EPI_ISL_3132627 | NHLS Charlotte Maxeke Johannesburg Academic Hospital and the University of the Witwatersrand | KRISP, KZn Research Innovation and Sequencing Platform | Florette Treurnicht, Bulelani Manene, Kathleen Subramoney, Giandhari Jennifer, Pillay Sureshnee, Yajna Ramphal, Naidoo Yeshnee, Tshabuila Derek,Tegally Houriiyah, San James, Wilkinson Eduan, de Oliveira Tulio |
| EPI_ISL_3132628 | Netcare Kingsway Hospital | KRISP, KZn Research Innovation and Sequencing Platform | Giandhari Jennifer, Pillay Sureshnee, Yajna Ramphal, Naidoo Yeshnee, Tshabuila Derek,Tegally Houriiyah, San James, Wilkinson Eduan, de Oliveira Tulio |
| EPI_ISL_3132630 | Netcare Parklands Hospital | KRISP, KZn Research Innovation and Sequencing Platform | Giandhari Jennifer, Pillay Sureshnee, Yajna Ramphal, Naidoo Yeshnee, Tshabuila Derek,Tegally Houriiyah, San James, Wilkinson Eduan, de Oliveira Tulio |
| EPI_ISL_3132632, EPI_ISL_3132633, EPI_ISL_3132634 | Netcare St Augustines | KRISP, KZn Research Innovation and Sequencing Platform | Giandhari Jennifer, Pillay Sureshnee, Yajna Ramphal, Naidoo Yeshnee, Tshabuila Derek,Tegally Houriiyah, San James, Wilkinson Eduan, de Oliveira Tulio |
| EPI_ISL_3132699, EPI_ISL_3132700, EPI_ISL_3132701, EPI_ISL_3132702, EPI_ISL_3132703, EPI_ISL_3132704, EPI_ISL_3132705, EPI_ISL_3132706, EPI_ISL_3132707, EPI_ISL_3132708, EPI_ISL_3132709, EPI_ISL_3132710, EPI_ISL_3132713, EPI_ISL_3132714, EPI_ISL_3132715, EPI_ISL_3132716, EPI_ISL_3132717, EPI_ISL_3132718, EPI_ISL_3132719, EPI_ISL_3132720, EPI_ISL_3132721, EPI_ISL_3132723, EPI_ISL_3132724, EPI_ISL_3132725, EPI_ISL_3132726, EPI_ISL_3132727, EPI_ISL_3132729, EPI_ISL_3132730, EPI_ISL_3132733, EPI_ISL_3132735, EPI_ISL_3132738, EPI_ISL_3132742, EPI_ISL_3132744, EPI_ISL_3132745, EPI_ISL_3132747, EPI_ISL_3132748, EPI_ISL_3132749, EPI_ISL_3132751, EPI_ISL_3132753, EPI_ISL_3132754, EPI_ISL_3132755, EPI_ISL_3132756, EPI_ISL_3132758, EPI_ISL_3132759, EPI_ISL_3132760, EPI_ISL_3132762, EPI_ISL_3132763, EPI_ISL_3132764, EPI_ISL_3132766, EPI_ISL_3132767, EPI_ISL_3132768, EPI_ISL_3132769, EPI_ISL_3132770, EPI_ISL_3132771, EPI_ISL_3132772, EPI_ISL_3132773, EPI_ISL_3132774, EPI_ISL_3132775, EPI_ISL_3132779, EPI_ISL_3132780, EPI_ISL_3132781, EPI_ISL_3132783, EPI_ISL_3132784, EPI_ISL_3132788, EPI_ISL_3132789, EPI_ISL_3132790, EPI_ISL_3132792, EPI_ISL_3132793, EPI_ISL_3132794, EPI_ISL_3132795, EPI_ISL_3132797, EPI_ISL_3132798, EPI_ISL_3132799, EPI_ISL_3132800, EPI_ISL_3132801, EPI_ISL_3132802, EPI_ISL_3132803, EPI_ISL_3132804, EPI_ISL_3132809, EPI_ISL_3132810, EPI_ISL_3132811, EPI_ISL_3132813, EPI_ISL_3132814, EPI_ISL_3132815, EPI_ISL_3132816, EPI_ISL_3132817, EPI_ISL_3132818, EPI_ISL_3132820, EPI_ISL_3132821, EPI_ISL_3132822, EPI_ISL_3132824, EPI_ISL_3132825, EPI_ISL_3132826, EPI_ISL_3132827, EPI_ISL_3132830, EPI_ISL_3132831, EPI_ISL_3132832, EPI_ISL_3132833, EPI_ISL_3132834, EPI_ISL_3132835, EPI_ISL_3132836, EPI_ISL_3132838, EPI_ISL_3132839, EPI_ISL_3132841, EPI_ISL_3132842, EPI_ISL_3132843, EPI_ISL_3132844, EPI_ISL_3132846, EPI_ISL_3132847, EPI_ISL_3132848, EPI_ISL_3132849, EPI_ISL_3132850, EPI_ISL_3132851, EPI_ISL_3132853, EPI_ISL_3132854, EPI_ISL_3132856, EPI_ISL_3132857, EPI_ISL_3132858, EPI_ISL_3132859, EPI_ISL_3132860, EPI_ISL_3132861, EPI_ISL_3132862, EPI_ISL_3132863, EPI_ISL_3132865, EPI_ISL_3132867, EPI_ISL_3132868, EPI_ISL_3132870, EPI_ISL_3132872, EPI_ISL_3132873, EPI_ISL_3132874, EPI_ISL_3132876, EPI_ISL_3132878, EPI_ISL_3132879, EPI_ISL_3132880, EPI_ISL_3132881, EPI_ISL_3132882, EPI_ISL_3132884, EPI_ISL_3132885, EPI_ISL_3132886, EPI_ISL_3132887, EPI_ISL_3132888, EPI_ISL_3132889, EPI_ISL_3132891, EPI_ISL_3132892, EPI_ISL_3132893, EPI_ISL_3132894, EPI_ISL_3132895, EPI_ISL_3132896, EPI_ISL_3132897, EPI_ISL_3132898, EPI_ISL_3132899, EPI_ISL_3132901, EPI_ISL_3132902, EPI_ISL_3132903, EPI_ISL_3132905, EPI_ISL_3132906, EPI_ISL_3132908, EPI_ISL_3132910, EPI_ISL_3132911, EPI_ISL_3132912, EPI_ISL_3132913, EPI_ISL_3132915, EPI_ISL_3132916, EPI_ISL_3132917, EPI_ISL_3132919, EPI_ISL_3132920, EPI_ISL_3132921, EPI_ISL_3132922, EPI_ISL_3132924, EPI_ISL_3132925, EPI_ISL_3132927, EPI_ISL_3132928, EPI_ISL_3132929, EPI_ISL_3132930, EPI_ISL_3132931, EPI_ISL_3132933, EPI_ISL_3132934, EPI_ISL_3132936, EPI_ISL_3132938, EPI_ISL_3132939, EPI_ISL_3132940, EPI_ISL_3132941, EPI_ISL_3132942, EPI_ISL_3132944, EPI_ISL_3132945, EPI_ISL_3132946, EPI_ISL_3132947, EPI_ISL_3132948, EPI_ISL_3132949, EPI_ISL_3132950, EPI_ISL_3132951, EPI_ISL_3132952, EPI_ISL_3132953, EPI_ISL_3132954, EPI_ISL_3132957, EPI_ISL_3132959, EPI_ISL_3132960, EPI_ISL_3132961, EPI_ISL_3132962, EPI_ISL_3132966, EPI_ISL_3132967, EPI_ISL_3132971, EPI_ISL_3132972, EPI_ISL_3132974, EPI_ISL_3132975, EPI_ISL_3132976, EPI_ISL_3132978, EPI_ISL_3132979, EPI_ISL_3132980, EPI_ISL_3132981, EPI_ISL_3132986, EPI_ISL_3132987, EPI_ISL_3132988, EPI_ISL_3132989, EPI_ISL_3132990, EPI_ISL_3132991, EPI_ISL_3132992, EPI_ISL_3132993, EPI_ISL_3132994, EPI_ISL_3132995, EPI_ISL_3132996, EPI_ISL_3132997, EPI_ISL_3132998, EPI_ISL_3132999, EPI_ISL_3133001, EPI_ISL_3133003, EPI_ISL_3133005, EPI_ISL_3133008, EPI_ISL_3133012, EPI_ISL_3133020, EPI_ISL_3133021 | | | |
| see above | Utah Public Health Laboratory | Utah Public Health Laboratory | Olinto Linares-Perdomo, Erin L. Young, Kelly F. Oakeson |
| EPI_ISL_3133022 | Yale Clinical Virology Lab | Grubaugh Lab - Yale School of Public Health | Joseph Fauver, Mallery Breban, Isabel Ott, Tara Alpert, Mary Petrone, Anderson Brito, Chantal Vogels, Annie Watkins, Chaney Kalinich, Jessica Rothman, Kendall Billig, Tobias Koch, Marie L. Landry, Nathan Grubaugh |
| EPI_ISL_3133026, EPI_ISL_3133027, EPI_ISL_3133028, EPI_ISL_3133029, EPI_ISL_3133030, EPI_ISL_3133032, EPI_ISL_3133033, EPI_ISL_3133034, EPI_ISL_3133036, EPI_ISL_3133038, EPI_ISL_3133040, EPI_ISL_3133041, EPI_ISL_3133042, EPI_ISL_3133043, EPI_ISL_3133045, EPI_ISL_3133046, EPI_ISL_3133047, EPI_ISL_3133048, EPI_ISL_3133049, EPI_ISL_3133050, EPI_ISL_3133051, EPI_ISL_3133052, EPI_ISL_3133053, EPI_ISL_3133054, EPI_ISL_3133056, EPI_ISL_3133057, EPI_ISL_3133058, EPI_ISL_3133059 | | | |
| see above | National Platform bis UMONS/Jolimont | National Platform bis UMONS/Jolimont | François Dufrasne, Guillaume Bayon-Vicente, Florian Juszczak, Laetitia Gheysen, Eric Tarantino, Gautier Detry, Ruddy Wattiez |
| EPI_ISL_3133060, EPI_ISL_3133061, EPI_ISL_3133062, EPI_ISL_3133063, EPI_ISL_3133064, EPI_ISL_3133065, EPI_ISL_3133066, EPI_ISL_3133067, EPI_ISL_3133069, EPI_ISL_3133070, EPI_ISL_3133071, EPI_ISL_3133072, EPI_ISL_3133073, EPI_ISL_3133074, EPI_ISL_3133075, EPI_ISL_3133076, EPI_ISL_3133077, EPI_ISL_3133078, EPI_ISL_3133079, EPI_ISL_3133080, EPI_ISL_3133081, EPI_ISL_3133082, EPI_ISL_3133083, EPI_ISL_3133084, EPI_ISL_3133085, EPI_ISL_3133086, EPI_ISL_3133087, EPI_ISL_3133088, EPI_ISL_3133089, EPI_ISL_3133090, EPI_ISL_3133091, EPI_ISL_3133092, EPI_ISL_3133093, EPI_ISL_3133094, EPI_ISL_3133095, EPI_ISL_3133096, EPI_ISL_3133097, EPI_ISL_3133098, EPI_ISL_3133099, EPI_ISL_3133100, EPI_ISL_3133101, EPI_ISL_3133102, EPI_ISL_3133103, EPI_ISL_3133104, EPI_ISL_3133105, EPI_ISL_3133106, EPI_ISL_3133107, EPI_ISL_3133108, EPI_ISL_3133109, EPI_ISL_3133110, EPI_ISL_3133111, EPI_ISL_3133112, EPI_ISL_3133113, EPI_ISL_3133114, EPI_ISL_3133115, EPI_ISL_3133116, EPI_ISL_3133117, EPI_ISL_3133118, EPI_ISL_3133119, EPI_ISL_3133120, EPI_ISL_3133121, EPI_ISL_3133122, EPI_ISL_3133123, EPI_ISL_3133124, EPI_ISL_3133125, EPI_ISL_3133126, EPI_ISL_3133128, EPI_ISL_3133130 | | | |
| see above | Cliniques universitaires Saint-Luc | UCLouvain/IREC/MBLG | Jean Ruelle, Bertrand Bearzatto, Benoit Kabamba Mukadi |
| EPI_ISL_3133131, EPI_ISL_3133132, EPI_ISL_3133133, EPI_ISL_3133134, EPI_ISL_3133135, EPI_ISL_3133136, EPI_ISL_3133137, EPI_ISL_3133138, EPI_ISL_3133139, EPI_ISL_3133140, EPI_ISL_3133141, EPI_ISL_3133142, EPI_ISL_3133143, EPI_ISL_3133144, EPI_ISL_3133145, EPI_ISL_3133146, EPI_ISL_3133147, EPI_ISL_3133148, EPI_ISL_3133149, EPI_ISL_3133150, EPI_ISL_3133151 | | | |
| see above | Clinique Saint-Pierre Ottignies | UCLouvain/IREC/MBLG | Jean Ruelle, Bertrand Bearzatto, Benoit Kabamba Mukadi |
| EPI_ISL_3133152, EPI_ISL_3133153, EPI_ISL_3133155, EPI_ISL_3133157, EPI_ISL_3133158, EPI_ISL_3133160, EPI_ISL_3133161, EPI_ISL_3133162, EPI_ISL_3133165, EPI_ISL_3133166, EPI_ISL_3133167, EPI_ISL_3133168, EPI_ISL_3133169, EPI_ISL_3133171, EPI_ISL_3133172, EPI_ISL_3133173, EPI_ISL_3133174, EPI_ISL_3133175, EPI_ISL_3133176, EPI_ISL_3133177, EPI_ISL_3133178, EPI_ISL_3133179, EPI_ISL_3133180, EPI_ISL_3133181, EPI_ISL_3133182 | | | |
| see above | Montana Public Health Laboratory | Montana Public Health Laboratory | Joy Ritter, Michelle Mozer, Carrie Biskupiak, Deborah Gibson, Michael Dills |
| EPI_ISL_3133228, EPI_ISL_3133229, EPI_ISL_3133230, EPI_ISL_3133231, EPI_ISL_3133232, EPI_ISL_3133233, EPI_ISL_3133234, EPI_ISL_3133235, EPI_ISL_3133236, EPI_ISL_3133237, EPI_ISL_3133238, EPI_ISL_3133239, EPI_ISL_3133240, EPI_ISL_3133241, EPI_ISL_3133242, EPI_ISL_3133243, EPI_ISL_3133244, EPI_ISL_3133245, EPI_ISL_3133246, EPI_ISL_3133247, EPI_ISL_3133248, EPI_ISL_3133249, EPI_ISL_3133250, EPI_ISL_3133251, EPI_ISL_3133252, EPI_ISL_3133253, EPI_ISL_3133254, EPI_ISL_3133255, EPI_ISL_3133256, EPI_ISL_3133257, EPI_ISL_3133258, EPI_ISL_3133259, EPI_ISL_3133260 | | | |
| see above | Kansas Health and Environmental Lab | Kansas Health and Environmental Lab | Mike Grose, Katherine Wiggins, Jonathan Barnell, Ben Olsen, and Phil Adam |
| EPI_ISL_3133261, EPI_ISL_3133262, EPI_ISL_3133263, EPI_ISL_3133264, EPI_ISL_3133265, EPI_ISL_3133266, EPI_ISL_3133267, EPI_ISL_3133268, EPI_ISL_3133269, EPI_ISL_3133270, EPI_ISL_3133271, EPI_ISL_3133272, EPI_ISL_3133274, EPI_ISL_3133275, EPI_ISL_3133276, EPI_ISL_3133277, EPI_ISL_3133278, EPI_ISL_3133279, EPI_ISL_3133280, EPI_ISL_3133281, EPI_ISL_3133282, EPI_ISL_3133283, EPI_ISL_3133284, EPI_ISL_3133285, EPI_ISL_3133286, EPI_ISL_3133287, EPI_ISL_3133288, EPI_ISL_3133289, EPI_ISL_3133290, EPI_ISL_3133291, EPI_ISL_3133292, EPI_ISL_3133293, EPI_ISL_3133294, EPI_ISL_3133295, EPI_ISL_3133296, EPI_ISL_3133297, EPI_ISL_3133299, EPI_ISL_3133300, EPI_ISL_3133301, EPI_ISL_3133302, EPI_ISL_3133303, EPI_ISL_3133304, EPI_ISL_3133306, EPI_ISL_3133308, EPI_ISL_3133309, EPI_ISL_3133310, EPI_ISL_3133311, EPI_ISL_3133312, EPI_ISL_3133313, EPI_ISL_3133314, EPI_ISL_3133315, EPI_ISL_3133316, EPI_ISL_3133317, EPI_ISL_3133318, EPI_ISL_3133320, EPI_ISL_3133321, EPI_ISL_3133322, EPI_ISL_3133323, EPI_ISL_3133324, EPI_ISL_3133325, EPI_ISL_3133326, EPI_ISL_3133327, EPI_ISL_3133328, EPI_ISL_3133330, EPI_ISL_3133331, EPI_ISL_3133332, EPI_ISL_3133333, EPI_ISL_3133334, EPI_ISL_3133335, EPI_ISL_3133336, EPI_ISL_3133337, EPI_ISL_3133338 | | | |
| see above | UW Virology Lab | UW Virology Lab | Pavitra Roychoudhury, Shah Mohamed Bakhash, Ricardo Perez, Maria Lukes, Hong Xie, Tien V. Nguyen, Noah R. Baker, Sean Ellis, Meei-Li Huang, Keith R Jerome, Alexander Greninger |
| EPI_ISL_3133339, EPI_ISL_3133340, EPI_ISL_3133342, EPI_ISL_3133343, EPI_ISL_3133344, EPI_ISL_3133345, EPI_ISL_3133346, EPI_ISL_3133347, EPI_ISL_3133348, EPI_ISL_3133349, EPI_ISL_3133350, EPI_ISL_3133354, EPI_ISL_3133356, EPI_ISL_3133357, EPI_ISL_3133358, EPI_ISL_3133359, EPI_ISL_3133360, EPI_ISL_3133361, EPI_ISL_3133362, EPI_ISL_3133363, EPI_ISL_3133364, EPI_ISL_3133365, EPI_ISL_3133366, EPI_ISL_3133367, EPI_ISL_3133368, EPI_ISL_3133369 | | | |
| see above | Oregon State Public Health Laboratory | Oregon State Public Health Laboratory | Rafia Razzaque, Eugene Yeboah, Vanda Makris, Laura Tsaknaridis, John Fontana and Shane Sevey |
| EPI_ISL_3133413, EPI_ISL_3133414, EPI_ISL_3133415, EPI_ISL_3133416, EPI_ISL_3133417, EPI_ISL_3133418, EPI_ISL_3133419, EPI_ISL_3133420, EPI_ISL_3133421, EPI_ISL_3133422, EPI_ISL_3133423, EPI_ISL_3133424, EPI_ISL_3133425, EPI_ISL_3133426, EPI_ISL_3133427, EPI_ISL_3133428, EPI_ISL_3133429, EPI_ISL_3133430, EPI_ISL_3133431, EPI_ISL_3133432, EPI_ISL_3133434, EPI_ISL_3133435, EPI_ISL_3133437, EPI_ISL_3133438, EPI_ISL_3133439, EPI_ISL_3133441, EPI_ISL_3133442, EPI_ISL_3133444, EPI_ISL_3133445, EPI_ISL_3133446, EPI_ISL_3133447, EPI_ISL_3133448, EPI_ISL_3133449, EPI_ISL_3133450, EPI_ISL_3133451 | | | |
| see above | Tampa General Hospital Esoteric Lab | Tampa General Hospital Esoteric Research & Development Lab | Jon Faughn, Marci O'Driscoll, Deanna Becker, Dominic Uy, Jorge Mecias-Frias, Elaine Vendrone, Vicki Healer, Amorce Lima, Suzane Silbert |
| EPI_ISL_3133452, EPI_ISL_3133453, EPI_ISL_3133454, EPI_ISL_3133455, EPI_ISL_3133456, EPI_ISL_3133457, EPI_ISL_3133458, EPI_ISL_3133459, EPI_ISL_3133460, EPI_ISL_3133461, EPI_ISL_3133462, EPI_ISL_3133463, EPI_ISL_3133464, EPI_ISL_3133465, EPI_ISL_3133466, EPI_ISL_3133467, EPI_ISL_3133468, EPI_ISL_3133469, EPI_ISL_3133470, EPI_ISL_3133471, EPI_ISL_3133472, EPI_ISL_3133473, EPI_ISL_3133474, EPI_ISL_3133475, EPI_ISL_3133476, EPI_ISL_3133477, EPI_ISL_3133478, EPI_ISL_3133479, EPI_ISL_3133480, EPI_ISL_3133481, EPI_ISL_3133482, EPI_ISL_3133483, EPI_ISL_3133484, EPI_ISL_3133485, EPI_ISL_3133486, EPI_ISL_3133487, EPI_ISL_3133488, EPI_ISL_3133489, EPI_ISL_3133490, EPI_ISL_3133491, EPI_ISL_3133492, EPI_ISL_3133493, EPI_ISL_3133494, EPI_ISL_3133495, EPI_ISL_3133496, EPI_ISL_3133497, EPI_ISL_3133498, EPI_ISL_3133499, EPI_ISL_3133500, EPI_ISL_3133501, EPI_ISL_3133502, EPI_ISL_3133503, EPI_ISL_3133504, EPI_ISL_3133505, EPI_ISL_3133506, EPI_ISL_3133507, EPI_ISL_3133508, EPI_ISL_3133509, EPI_ISL_3133510, EPI_ISL_3133511, EPI_ISL_3133512, EPI_ISL_3133513, EPI_ISL_3133514, EPI_ISL_3133515, EPI_ISL_3133516, EPI_ISL_3133517, EPI_ISL_3133518, EPI_ISL_3133519, EPI_ISL_3133520, EPI_ISL_3133521, EPI_ISL_3133522, EPI_ISL_3133525, EPI_ISL_3133526, EPI_ISL_3133527, EPI_ISL_3133528, EPI_ISL_3133529, EPI_ISL_3133530, EPI_ISL_3133531, EPI_ISL_3133532, EPI_ISL_3133533, EPI_ISL_3133534, EPI_ISL_3133535, EPI_ISL_3133536, EPI_ISL_3133537 | | | |
| see above | Illinois Department of Public Health - Springfield Lab | Illinois Department of Public Health - Springfield Lab | Bryan Sim, Gordon McCall |
| EPI_ISL_3133538, EPI_ISL_3133539, EPI_ISL_3133540 | Furst Medical Laboratory | Norwegian Institute of Public Health, Department of Virology | Kathrine Stene-Johansen, Kamilla Heddeland Instefjord, Hilde Elshaug, Garcia Llorente Ignacio, Jon Bråte, Engebretsen Serina Beate,Pedersen Benedikte Nevjen, Line Victoria Moen, Debech Nadia, Atiya R Ali,Marie Paulsen Madsen, Rasmus Riis Kopperud, Hilde Vollan, Karoline Bragstad, Olav Hungnes |
| EPI_ISL_3133541, EPI_ISL_3133542, EPI_ISL_3133543 | Haukeland University Hospital, Dept. of Microbiology | Norwegian Institute of Public Health, Department of Virology | Kathrine Stene-Johansen, Kamilla Heddeland Instefjord, Hilde Elshaug, Garcia Llorente Ignacio, Jon Bråte, Engebretsen Serina Beate,Pedersen Benedikte Nevjen, Line Victoria Moen, Debech Nadia, Atiya R Ali,Marie Paulsen Madsen, Rasmus Riis Kopperud, Hilde Vollan, Karoline Bragstad, Olav Hungnes |
| EPI_ISL_3133546, EPI_ISL_3133548, EPI_ISL_3133549, EPI_ISL_3133550 | Ostfold Hospital Trust - Kalnes, Centre for Laboratory Medicine, Section for gene technology and infection serology | Norwegian Institute of Public Health, Department of Virology | Kathrine Stene-Johansen, Kamilla Heddeland Instefjord, Hilde Elshaug, Garcia Llorente Ignacio, Jon Bråte, Engebretsen Serina Beate,Pedersen Benedikte Nevjen, Line Victoria Moen, Debech Nadia, Atiya R Ali,Marie Paulsen Madsen, Rasmus Riis Kopperud, Hilde Vollan, Karoline Bragstad, Olav Hungnes |

| | | | |
|---|---|---|---|
| EPI_ISL_3133553, EPI_ISL_3133554, EPI_ISL_3133555, EPI_ISL_3133556 | Furst Medical Laboratory | Norwegian Institute of Public Health, Department of Virology | Kathrine Stene-Johansen, Kamilla Heddeland Instefjord, Hilde Elshaug, Garcia Llorente Ignacio, Jon Bråte, Engebretsen Serina Beate,Pedersen Benedikte Nevjen, Line Victoria Moen, Debech Nadia, Atiya R Ali,Marie Paulsen Madsen, Rasmus Riis Kopperud, Hilde Vollan, Karoline Bragstad, Olav Hungnes |
| EPI_ISL_3133557 | Ostfold Hospital Trust - Kalnes, Centre for Laboratory Medicine, Section for gene technology and infection serology | Norwegian Institute of Public Health, Department of Virology | Kathrine Stene-Johansen, Kamilla Heddeland Instefjord, Hilde Elshaug, Garcia Llorente Ignacio, Jon Bråte, Engebretsen Serina Beate,Pedersen Benedikte Nevjen, Line Victoria Moen, Debech Nadia, Atiya R Ali,Marie Paulsen Madsen, Rasmus Riis Kopperud, Hilde Vollan, Karoline Bragstad, Olav Hungnes |
| EPI_ISL_3133558 | Furst Medical Laboratory | Norwegian Institute of Public Health, Department of Virology | Kathrine Stene-Johansen, Kamilla Heddeland Instefjord, Hilde Elshaug, Garcia Llorente Ignacio, Jon Bråte, Engebretsen Serina Beate,Pedersen Benedikte Nevjen, Line Victoria Moen, Debech Nadia, Atiya R Ali,Marie Paulsen Madsen, Rasmus Riis Kopperud, Hilde Vollan, Karoline Bragstad, Olav Hungnes |
| EPI_ISL_3133559, EPI_ISL_3133560, EPI_ISL_3133561 | Ostfold Hospital Trust - Kalnes, Centre for Laboratory Medicine, Section for gene technology and infection serology | Norwegian Institute of Public Health, Department of Virology | Kathrine Stene-Johansen, Kamilla Heddeland Instefjord, Hilde Elshaug, Garcia Llorente Ignacio, Jon Bråte, Engebretsen Serina Beate,Pedersen Benedikte Nevjen, Line Victoria Moen, Debech Nadia, Atiya R Ali,Marie Paulsen Madsen, Rasmus Riis Kopperud, Hilde Vollan, Karoline Bragstad, Olav Hungnes |
| EPI_ISL_3133562 | Furst Medical Laboratory | Norwegian Institute of Public Health, Department of Virology | Kathrine Stene-Johansen, Kamilla Heddeland Instefjord, Hilde Elshaug, Garcia Llorente Ignacio, Jon Bråte, Engebretsen Serina Beate,Pedersen Benedikte Nevjen, Line Victoria Moen, Debech Nadia, Atiya R Ali,Marie Paulsen Madsen, Rasmus Riis Kopperud, Hilde Vollan, Karoline Bragstad, Olav Hungnes |
| EPI_ISL_3133563, EPI_ISL_3133564 | Ostfold Hospital Trust - Kalnes, Centre for Laboratory Medicine, Section for gene technology and infection serology | Norwegian Institute of Public Health, Department of Virology | Kathrine Stene-Johansen, Kamilla Heddeland Instefjord, Hilde Elshaug, Garcia Llorente Ignacio, Jon Bråte, Engebretsen Serina Beate,Pedersen Benedikte Nevjen, Line Victoria Moen, Debech Nadia, Atiya R Ali,Marie Paulsen Madsen, Rasmus Riis Kopperud, Hilde Vollan, Karoline Bragstad, Olav Hungnes |
| EPI_ISL_3133565 | Furst Medical Laboratory | Norwegian Institute of Public Health, Department of Virology | Kathrine Stene-Johansen, Kamilla Heddeland Instefjord, Hilde Elshaug, Garcia Llorente Ignacio, Jon Bråte, Engebretsen Serina Beate,Pedersen Benedikte Nevjen, Line Victoria Moen, Debech Nadia, Atiya R Ali,Marie Paulsen Madsen, Rasmus Riis Kopperud, Hilde Vollan, Karoline Bragstad, Olav Hungnes |
| EPI_ISL_3133566, EPI_ISL_3133567, EPI_ISL_3133571, EPI_ISL_3133572, EPI_ISL_3133573, EPI_ISL_3133574, EPI_ISL_3133575, EPI_ISL_3133576, EPI_ISL_3133577, EPI_ISL_3133578, EPI_ISL_3133580, EPI_ISL_3133581, EPI_ISL_3133582, EPI_ISL_3133583, EPI_ISL_3133584, EPI_ISL_3133585, EPI_ISL_3133586, EPI_ISL_3133587, EPI_ISL_3133588, EPI_ISL_3133589, EPI_ISL_3133590, EPI_ISL_3133591, EPI_ISL_3133593, EPI_ISL_3133594, EPI_ISL_3133595, EPI_ISL_3133596, EPI_ISL_3133597, EPI_ISL_3133598, EPI_ISL_3133599, EPI_ISL_3133601, EPI_ISL_3133602, EPI_ISL_3133603, EPI_ISL_3133604, EPI_ISL_3133605, EPI_ISL_3133606, EPI_ISL_3133607, EPI_ISL_3133608, EPI_ISL_3133609 | | | |
| see above | Microvida | Microvida | Suzan D. Pas, Jaco J. Verweij, Joep J. J. M. Stohr |
| EPI_ISL_3133712, EPI_ISL_3133718 | Servizo de Microbioloxía. Complexo Hospitalario de Santiago de Compostela | Servizo de Microbioloxía. Complexo Hospitalario de Santiago de Compostela | Xosé Costa, Antonio Aguilera, Gema Barbeito, Amparo Coira, Carlos García_Riestra, Manuela Hernández, José Llovo, Teresa Lopez_Valiño, Laura Millán, Daniel Navarro, Rocío trastoy, Mercedes Treviño, María Luisa Pérez_del_Molino |
| EPI_ISL_3133775, EPI_ISL_3133778 | Laboratorio Central de Epidemiología (LCE) | Instituto de Biotecnología de la UNAM | Consorcio Mexicano de Vigilancia Genómica (CoViGen-Mex). Authors (in alphabetical order): Julio Elias Alvarado-Yaah, Carlos F. Arias, Santiago Ávila-Ríos, Eduardo Becerril-Vargas, Víctor Hugo Borja-Aburto, Celia Boukadida, Cristóbal Cháidez-Quiróz, Juan Bautista Chale-Dzul, Ricardo Ciria Merce, Andreu Comas-García, Célida Duque Molina, Julissa Enciso-Ibarra, José Antonio Enciso-Moreno, Gloria Elena Espinosa-Ayala, Fernando Fontove-Herrera, Daniel Fregoso-Rueda, Victor Eduardo García-Arias, Alejandra García-Gasca, Bruno Gómez-Gil, Jean Pierre González, Irvin González-López, Concepción Grajales-Muñiz, Ricardo Grande, Rosa María Gutiérrez Rios, Jesús Hernández, Alejandra Hernández-Terán, Alfredo Herrera-Estrella, Carla Ivón Herrera-Najera, Pavel Isa, Verónica Mata-Haro, Daniel Lira Morales, Susana Lopez, Antonio Loza Román, Brenda Irasema Maldonado-Meza, Bernardo Martínez-Miguel, José Arturo Martínez-Orozco, Célida Martínez-Rodríguez, Margarita Matías-Florentino, Fidencio Mejía-Nepomuceno, María Guadalupe de Jesús Mireles-Rivera, Gloria María Molina-Salinas, Hector Montoya-Fuentes, , Mario Mújica-Sánchez, José Esteban Muñoz-Medina, José de Jesús Nuñez-Contreras, Alicia Ocaña-Mondragón, Luis Alberto Ochoa-Carrera, Hector Esteban Paz-Juárez, Marissa Perez-Garcia, Francisco Pulido, Helen Haydee Fernanda Ramírez-Plascencia, Jorge Salas-Hernández, Angel Gustavo Salas-Lais, Alejandro Sánchez-Flores, Clara Esperanza Santacruz-Tinoco, María Guadalupe Santiago-Mauricio, Selene Zárate, Nelly Sélem-Mojica, Blanca Taboada, Gloria Vazquez, Joel Armando Vázquez-Pérez. |
| EPI_ISL_3133795 | Furst Medical Laboratory | Norwegian Institute of Public Health, Department of Virology | Kathrine Stene-Johansen, Kamilla Heddeland Instefjord, Hilde Elshaug, Garcia Llorente Ignacio, Jon Bråte, Engebretsen Serina Beate,Pedersen Benedikte Nevjen, Line Victoria Moen, Debech Nadia, Atiya R Ali,Marie Paulsen Madsen, Rasmus Riis Kopperud, Hilde Vollan, Karoline Bragstad, Olav Hungnes |
| EPI_ISL_3133796 | Haukeland University Hospital, Dept. of Microbiology | Norwegian Institute of Public Health, Department of Virology | Kathrine Stene-Johansen, Kamilla Heddeland Instefjord, Hilde Elshaug, Garcia Llorente Ignacio, Jon Bråte, Engebretsen Serina Beate,Pedersen Benedikte Nevjen, Line Victoria Moen, Debech Nadia, Atiya R Ali,Marie Paulsen Madsen, Rasmus Riis Kopperud, Hilde Vollan, Karoline Bragstad, Olav Hungnes |
| EPI_ISL_3133797 | Furst Medical Laboratory | Norwegian Institute of Public Health, Department of Virology | Kathrine Stene-Johansen, Kamilla Heddeland Instefjord, Hilde Elshaug, Garcia Llorente Ignacio, Jon Bråte, Engebretsen Serina Beate,Pedersen Benedikte Nevjen, Line Victoria Moen, Debech Nadia, Atiya R Ali,Marie Paulsen Madsen, Rasmus Riis Kopperud, Hilde Vollan, Karoline Bragstad, Olav Hungnes |
| EPI_ISL_3133798 | Haukeland University Hospital, Dept. of Microbiology | Norwegian Institute of Public Health, Department of Virology | Kathrine Stene-Johansen, Kamilla Heddeland Instefjord, Hilde Elshaug, Garcia Llorente Ignacio, Jon Bråte, Engebretsen Serina Beate,Pedersen Benedikte Nevjen, Line Victoria Moen, Debech Nadia, Atiya R Ali,Marie Paulsen Madsen, Rasmus Riis Kopperud, Hilde Vollan, Karoline Bragstad, Olav Hungnes |
| EPI_ISL_3133799, EPI_ISL_3133802, EPI_ISL_3133804 | Furst Medical Laboratory | Norwegian Institute of Public Health, Department of Virology | Kathrine Stene-Johansen, Kamilla Heddeland Instefjord, Hilde Elshaug, Garcia Llorente Ignacio, Jon Bråte, Engebretsen Serina Beate,Pedersen Benedikte Nevjen, Line Victoria Moen, Debech Nadia, Atiya R Ali,Marie Paulsen Madsen, Rasmus Riis Kopperud, Hilde Vollan, Karoline Bragstad, Olav Hungnes |
| EPI_ISL_3133806 | Ostfold Hospital Trust - Kalnes, Centre for Laboratory Medicine, Section for gene technology and infection serology | Norwegian Institute of Public Health, Department of Virology | Kathrine Stene-Johansen, Kamilla Heddeland Instefjord, Hilde Elshaug, Garcia Llorente Ignacio, Jon Bråte, Engebretsen Serina Beate,Pedersen Benedikte Nevjen, Line Victoria Moen, Debech Nadia, Atiya R Ali,Marie Paulsen Madsen, Rasmus Riis Kopperud, Hilde Vollan, Karoline Bragstad, Olav Hungnes |
| EPI_ISL_3133807, EPI_ISL_3133811 | Furst Medical Laboratory | Norwegian Institute of Public Health, Department of Virology | Kathrine Stene-Johansen, Kamilla Heddeland Instefjord, Hilde Elshaug, Garcia Llorente Ignacio, Jon Bråte, Engebretsen Serina Beate,Pedersen Benedikte Nevjen, Line Victoria Moen, Debech Nadia, Atiya R Ali,Marie Paulsen Madsen, Rasmus Riis Kopperud, Hilde Vollan, Karoline Bragstad, Olav Hungnes |
| EPI_ISL_3133813 | Medical Microbiology Unit, Department for Laboratory Medicine, Drammen Hospital, Vestre Viken Health Trust, | Norwegian Institute of Public Health, Department of Virology | Kathrine Stene-Johansen, Kamilla Heddeland Instefjord, Hilde Elshaug, Garcia Llorente Ignacio, Jon Bråte, Engebretsen Serina Beate,Pedersen Benedikte Nevjen, Line Victoria Moen, Debech Nadia, Atiya R Ali,Marie Paulsen Madsen, Rasmus Riis Kopperud, Hilde Vollan, Karoline Bragstad, Olav Hungnes |
| EPI_ISL_3133814 | Ostfold Hospital Trust - Kalnes, Centre for Laboratory Medicine, Section for gene technology and infection serology | Norwegian Institute of Public Health, Department of Virology | Kathrine Stene-Johansen, Kamilla Heddeland Instefjord, Hilde Elshaug, Garcia Llorente Ignacio, Jon Bråte, Engebretsen Serina Beate,Pedersen Benedikte Nevjen, Line Victoria Moen, Debech Nadia, Atiya R Ali,Marie Paulsen Madsen, Rasmus Riis Kopperud, Hilde Vollan, Karoline Bragstad, Olav Hungnes |
| EPI_ISL_3133815, EPI_ISL_3133817, EPI_ISL_3133818 | Furst Medical Laboratory | Norwegian Institute of Public Health, Department of Virology | Kathrine Stene-Johansen, Kamilla Heddeland Instefjord, Hilde Elshaug, Garcia Llorente Ignacio, Jon Bråte, Engebretsen Serina Beate,Pedersen Benedikte Nevjen, Line Victoria Moen, Debech Nadia, Atiya R Ali,Marie Paulsen Madsen, Rasmus Riis Kopperud, Hilde Vollan, Karoline Bragstad, Olav Hungnes |
| EPI_ISL_3133819 | Medical Microbiology Unit, Department for Laboratory Medicine, Drammen Hospital, Vestre Viken Health Trust, | Norwegian Institute of Public Health, Department of Virology | Kathrine Stene-Johansen, Kamilla Heddeland Instefjord, Hilde Elshaug, Garcia Llorente Ignacio, Jon Bråte, Engebretsen Serina Beate,Pedersen Benedikte Nevjen, Line Victoria Moen, Debech Nadia, Atiya R Ali,Marie Paulsen Madsen, Rasmus Riis Kopperud, Hilde Vollan, Karoline Bragstad, Olav Hungnes |
| EPI_ISL_3133820, EPI_ISL_3133821 | Furst Medical Laboratory | Norwegian Institute of Public Health, Department of Virology | Kathrine Stene-Johansen, Kamilla Heddeland Instefjord, Hilde Elshaug, Garcia Llorente Ignacio, Jon Bråte, Engebretsen Serina Beate,Pedersen Benedikte Nevjen, Line Victoria Moen, Debech Nadia, Atiya R Ali,Marie Paulsen Madsen, Rasmus Riis Kopperud, Hilde Vollan, Karoline Bragstad, Olav Hungnes |
| EPI_ISL_3133823 | Ostfold Hospital Trust - Kalnes, Centre for Laboratory Medicine, Section for gene technology and infection serology | Norwegian Institute of Public Health, Department of Virology | Kathrine Stene-Johansen, Kamilla Heddeland Instefjord, Hilde Elshaug, Garcia Llorente Ignacio, Jon Bråte, Engebretsen Serina Beate,Pedersen Benedikte Nevjen, Line Victoria Moen, Debech Nadia, Atiya R Ali,Marie Paulsen Madsen, Rasmus Riis Kopperud, Hilde Vollan, Karoline Bragstad, Olav Hungnes |
| EPI_ISL_3133824 | Furst Medical Laboratory | Norwegian Institute of Public Health, Department of Virology | Kathrine Stene-Johansen, Kamilla Heddeland Instefjord, Hilde Elshaug, Garcia Llorente Ignacio, Jon Bråte, Engebretsen Serina Beate,Pedersen Benedikte Nevjen, Line Victoria Moen, Debech Nadia, Atiya R Ali,Marie Paulsen Madsen, Rasmus Riis Kopperud, Hilde Vollan, Karoline Bragstad, Olav Hungnes |
| EPI_ISL_3133826 | Medical Microbiology Unit, Department for Laboratory Medicine, Drammen Hospital, Vestre Viken Health Trust, | Norwegian Institute of Public Health, Department of Virology | Kathrine Stene-Johansen, Kamilla Heddeland Instefjord, Hilde Elshaug, Garcia Llorente Ignacio, Jon Bråte, Engebretsen Serina Beate,Pedersen Benedikte Nevjen, Line Victoria Moen, Debech Nadia, Atiya R Ali,Marie Paulsen Madsen, Rasmus Riis Kopperud, Hilde Vollan, Karoline Bragstad, Olav Hungnes |
| EPI_ISL_3133827 | Furst Medical Laboratory | Norwegian Institute of Public Health, Department of Virology | Kathrine Stene-Johansen, Kamilla Heddeland Instefjord, Hilde Elshaug, Garcia Llorente Ignacio, Jon Bråte, Engebretsen Serina Beate,Pedersen Benedikte Nevjen, Line Victoria Moen, Debech Nadia, Atiya R Ali,Marie Paulsen Madsen, Rasmus Riis Kopperud, Hilde Vollan, Karoline Bragstad, Olav Hungnes |
| EPI_ISL_3133830 | Dept. of Medical Microbiology, Stavanger University Hospital, Helse Stavanger HF | Norwegian Institute of Public Health, Department of Virology | Kathrine Stene-Johansen, Kamilla Heddeland Instefjord, Hilde Elshaug, Garcia Llorente Ignacio, Jon Bråte, Engebretsen Serina Beate,Pedersen Benedikte Nevjen, Line Victoria Moen, Debech Nadia, Atiya R Ali,Marie Paulsen Madsen, Rasmus Riis Kopperud, Hilde Vollan, Karoline Bragstad, Olav Hungnes |
| EPI_ISL_3133832 | Furst Medical Laboratory | Norwegian Institute of Public Health, Department of Virology | Kathrine Stene-Johansen, Kamilla Heddeland Instefjord, Hilde Elshaug, Garcia Llorente Ignacio, Jon Bråte, Engebretsen Serina Beate,Pedersen Benedikte Nevjen, Line Victoria Moen, Debech Nadia, Atiya R Ali,Marie Paulsen Madsen, Rasmus Riis Kopperud, Hilde Vollan, Karoline Bragstad, Olav Hungnes |
| EPI_ISL_3133833 | Medical Microbiology Unit, Department for Laboratory Medicine, Drammen Hospital, Vestre Viken Health Trust, | Norwegian Institute of Public Health, Department of Virology | Kathrine Stene-Johansen, Kamilla Heddeland Instefjord, Hilde Elshaug, Garcia Llorente Ignacio, Jon Bråte, Engebretsen Serina Beate,Pedersen Benedikte Nevjen, Line Victoria Moen, Debech Nadia, Atiya R Ali,Marie Paulsen Madsen, Rasmus Riis Kopperud, Hilde Vollan, Karoline Bragstad, Olav Hungnes |
| EPI_ISL_3133834 | Dept. of Medical Microbiology, Stavanger University Hospital, Helse Stavanger HF | Norwegian Institute of Public Health, Department of Virology | Kathrine Stene-Johansen, Kamilla Heddeland Instefjord, Hilde Elshaug, Garcia Llorente Ignacio, Jon Bråte, Engebretsen Serina Beate,Pedersen Benedikte Nevjen, Line Victoria Moen, Debech Nadia, Atiya R Ali,Marie Paulsen Madsen, Rasmus Riis Kopperud, Hilde Vollan, Karoline Bragstad, Olav Hungnes |
| EPI_ISL_3133835, EPI_ISL_3133836, EPI_ISL_3133837 | Furst Medical Laboratory | Norwegian Institute of Public Health, Department of Virology | Kathrine Stene-Johansen, Kamilla Heddeland Instefjord, Hilde Elshaug, Garcia Llorente Ignacio, Jon Bråte, Engebretsen Serina Beate,Pedersen Benedikte Nevjen, Line Victoria Moen, Debech Nadia, Atiya R Ali,Marie Paulsen Madsen, Rasmus Riis Kopperud, Hilde Vollan, Karoline Bragstad, Olav Hungnes |
| EPI_ISL_3133838 | Medical Microbiology Unit, Department for Laboratory Medicine, Drammen Hospital, Vestre Viken Health Trust, | Norwegian Institute of Public Health, Department of Virology | Kathrine Stene-Johansen, Kamilla Heddeland Instefjord, Hilde Elshaug, Garcia Llorente Ignacio, Jon Bråte, Engebretsen Serina Beate,Pedersen Benedikte Nevjen, Line Victoria Moen, Debech Nadia, Atiya R Ali,Marie Paulsen Madsen, Rasmus Riis Kopperud, Hilde Vollan, Karoline Bragstad, Olav Hungnes |
| EPI_ISL_3133840, EPI_ISL_3133841 | Furst Medical Laboratory | Norwegian Institute of Public Health, Department of Virology | Kathrine Stene-Johansen, Kamilla Heddeland Instefjord, Hilde Elshaug, Garcia Llorente Ignacio, Jon Bråte, Engebretsen Serina Beate,Pedersen Benedikte Nevjen, Line Victoria Moen, Debech Nadia, Atiya R Ali,Marie Paulsen Madsen, Rasmus Riis Kopperud, Hilde Vollan, Karoline Bragstad, Olav Hungnes |
| EPI_ISL_3133842, EPI_ISL_3133843 | Medical Microbiology Unit, Department for Laboratory Medicine, Drammen Hospital, Vestre Viken Health Trust, | Norwegian Institute of Public Health, Department of Virology | Kathrine Stene-Johansen, Kamilla Heddeland Instefjord, Hilde Elshaug, Garcia Llorente Ignacio, Jon Bråte, Engebretsen Serina Beate,Pedersen Benedikte Nevjen, Line Victoria Moen, Debech Nadia, Atiya R Ali,Marie Paulsen Madsen, Rasmus Riis Kopperud, Hilde Vollan, Karoline Bragstad, Olav Hungnes |
| EPI_ISL_3133845 | Furst Medical Laboratory | Norwegian Institute of Public Health, Department of Virology | Kathrine Stene-Johansen, Kamilla Heddeland Instefjord, Hilde Elshaug, Garcia Llorente Ignacio, Jon Bråte, Engebretsen Serina Beate,Pedersen Benedikte Nevjen, Line Victoria Moen, Debech Nadia, Atiya R Ali,Marie Paulsen Madsen, Rasmus Riis Kopperud, Hilde Vollan, Karoline Bragstad, Olav Hungnes |
| EPI_ISL_3133846, EPI_ISL_3133847, EPI_ISL_3133848, EPI_ISL_3133849 | Medical Microbiology Unit, Department for Laboratory Medicine, Drammen Hospital, Vestre Viken Health Trust, | Norwegian Institute of Public Health, Department of Virology | Kathrine Stene-Johansen, Kamilla Heddeland Instefjord, Hilde Elshaug, Garcia Llorente Ignacio, Jon Bråte, Engebretsen Serina Beate,Pedersen Benedikte Nevjen, Line Victoria Moen, Debech Nadia, Atiya R Ali,Marie Paulsen Madsen, Rasmus Riis Kopperud, Hilde Vollan, Karoline Bragstad, Olav Hungnes |
| EPI_ISL_3133855 | Vestfold Hospital, Toensberg Department of Microbiology | Norwegian Institute of Public Health, Department of Virology | Kathrine Stene-Johansen, Kamilla Heddeland Instefjord, Hilde Elshaug, Garcia Llorente Ignacio, Jon Bråte, Engebretsen Serina Beate,Pedersen Benedikte Nevjen, Line Victoria Moen, Debech Nadia, Atiya R Ali,Marie Paulsen Madsen, Rasmus Riis Kopperud, Hilde Vollan, Karoline Bragstad, Olav Hungnes |
| EPI_ISL_3133860, EPI_ISL_3133862, EPI_ISL_3133864, EPI_ISL_3133865, EPI_ISL_3133866, | Furst Medical Laboratory | Norwegian Institute of Public Health, Department of Virology | Kathrine Stene-Johansen, Kamilla Heddeland Instefjord, Hilde Elshaug, Garcia Llorente Ignacio, Jon Bråte, Engebretsen Serina Beate,Pedersen Benedikte Nevjen, Line Victoria Moen, Debech Nadia, Atiya R Ali,Marie Paulsen Madsen, Rasmus Riis Kopperud, Hilde Vollan, Karoline Bragstad, Olav Hungnes |

| | | | |
|---|---|---|---|
| EPI_ISL_3133868 | | | |
| EPI_ISL_3133869 | Ostfold Hospital Trust - Kalnes, Centre for Laboratory Medicine, Section for gene technology and infection serology | Norwegian Institute of Public Health, Department of Virology | Kathrine Stene-Johansen, Kamilla Heddeland Instefjord, Hilde Elshaug, Garcia Llorente Ignacio, Jon Bråte, Engebretsen Serina Beate,Pedersen Benedikte Nevjen, Line Victoria Moen, Debech Nadia, Atiya R Ali,Marie Paulsen Madsen, Rasmus Riis Kopperud, Hilde Vollan, Karoline Bragstad, Olav Hungnes |
| EPI_ISL_3133871 | Furst Medical Laboratory | Norwegian Institute of Public Health, Department of Virology | Kathrine Stene-Johansen, Kamilla Heddeland Instefjord, Hilde Elshaug, Garcia Llorente Ignacio, Jon Bråte, Engebretsen Serina Beate,Pedersen Benedikte Nevjen, Line Victoria Moen, Debech Nadia, Atiya R Ali,Marie Paulsen Madsen, Rasmus Riis Kopperud, Hilde Vollan, Karoline Bragstad, Olav Hungnes |
| EPI_ISL_3133872 | Ostfold Hospital Trust - Kalnes, Centre for Laboratory Medicine, Section for gene technology and infection serology | Norwegian Institute of Public Health, Department of Virology | Kathrine Stene-Johansen, Kamilla Heddeland Instefjord, Hilde Elshaug, Garcia Llorente Ignacio, Jon Bråte, Engebretsen Serina Beate,Pedersen Benedikte Nevjen, Line Victoria Moen, Debech Nadia, Atiya R Ali,Marie Paulsen Madsen, Rasmus Riis Kopperud, Hilde Vollan, Karoline Bragstad, Olav Hungnes |
| EPI_ISL_3133873 | Unilabs Laboratory Medicine | Norwegian Institute of Public Health, Department of Virology | Kathrine Stene-Johansen, Kamilla Heddeland Instefjord, Hilde Elshaug, Garcia Llorente Ignacio, Jon Bråte, Engebretsen Serina Beate,Pedersen Benedikte Nevjen, Line Victoria Moen, Debech Nadia, Atiya R Ali,Marie Paulsen Madsen, Rasmus Riis Kopperud, Hilde Vollan, Karoline Bragstad, Olav Hungnes |
| EPI_ISL_3133874, EPI_ISL_3133876 | Furst Medical Laboratory | Norwegian Institute of Public Health, Department of Virology | Kathrine Stene-Johansen, Kamilla Heddeland Instefjord, Hilde Elshaug, Garcia Llorente Ignacio, Jon Bråte, Engebretsen Serina Beate,Pedersen Benedikte Nevjen, Line Victoria Moen, Debech Nadia, Atiya R Ali,Marie Paulsen Madsen, Rasmus Riis Kopperud, Hilde Vollan, Karoline Bragstad, Olav Hungnes |
| EPI_ISL_3133885, EPI_ISL_3133886, EPI_ISL_3133887, EPI_ISL_3133888, EPI_ISL_3133889, EPI_ISL_3133890, EPI_ISL_3133891, EPI_ISL_3133892, EPI_ISL_3133893, EPI_ISL_3133894, EPI_ISL_3133895, EPI_ISL_3133896, EPI_ISL_3133897, EPI_ISL_3133898, EPI_ISL_3133899, EPI_ISL_3133900, EPI_ISL_3133901, EPI_ISL_3133902, EPI_ISL_3133903, EPI_ISL_3133904, EPI_ISL_3133905, EPI_ISL_3133906, EPI_ISL_3133907, EPI_ISL_3133910, EPI_ISL_3133911, EPI_ISL_3133912, EPI_ISL_3133913, EPI_ISL_3133914, EPI_ISL_3133915, EPI_ISL_3133916, EPI_ISL_3133917, EPI_ISL_3133918, EPI_ISL_3133919, EPI_ISL_3133920, EPI_ISL_3133921, EPI_ISL_3133922, EPI_ISL_3133923, EPI_ISL_3133924, EPI_ISL_3133925, EPI_ISL_3133930, EPI_ISL_3133931, EPI_ISL_3133932, EPI_ISL_3133933, EPI_ISL_3133934, EPI_ISL_3133935, EPI_ISL_3133936, EPI_ISL_3133937, EPI_ISL_3133938, EPI_ISL_3133939, EPI_ISL_3133940, EPI_ISL_3133941, EPI_ISL_3133942, EPI_ISL_3133943, EPI_ISL_3133944, EPI_ISL_3133946, EPI_ISL_3133947 | | | |
| see above | Servicio Microbiología Hospital La Paz | Servicio Microbiología Hospital La Paz | Fernando Lázaro, Rubén Cáceres, Iván Bloise, Jesús Mingorance Cruz, Elie Dahdouh |
| EPI_ISL_3134062, EPI_ISL_3134078, EPI_ISL_3134081, EPI_ISL_3134085, EPI_ISL_3134086, EPI_ISL_3134088, EPI_ISL_3134089, EPI_ISL_3134098, EPI_ISL_3134103, EPI_ISL_3134112, EPI_ISL_3134115, EPI_ISL_3134122, EPI_ISL_3134132, EPI_ISL_3134142, EPI_ISL_3134143, EPI_ISL_3134144, EPI_ISL_3134146, EPI_ISL_3134150, EPI_ISL_3134153, EPI_ISL_3134159 | | | |
| see above | California Department of Public Health Valencia Branch Laboratory | California Department of Public Health | CDPH-COVIDNet,UCSF Center for Advanced Technology |
| EPI_ISL_3134163, EPI_ISL_3134164, EPI_ISL_3134165, EPI_ISL_3134166, EPI_ISL_3134167, EPI_ISL_3134168, EPI_ISL_3134169, EPI_ISL_3134170, EPI_ISL_3134171, EPI_ISL_3134172, EPI_ISL_3134174, EPI_ISL_3134175, EPI_ISL_3134176 | | | |
| see above | Ayass Bioscience LLC | Ayass Bioscience LLC | Mohamad Ammar Ayass, Natalya Griko, Kevin Zhu, Nazanin Taheri, Lina Abi Mosleh |
| EPI_ISL_3134177, EPI_ISL_3134178, EPI_ISL_3134179, EPI_ISL_3134180, EPI_ISL_3134181, EPI_ISL_3134182, EPI_ISL_3134183, EPI_ISL_3134184, EPI_ISL_3134185, EPI_ISL_3134187, EPI_ISL_3134188, EPI_ISL_3134189, EPI_ISL_3134190, EPI_ISL_3134191, EPI_ISL_3134192, EPI_ISL_3134193, EPI_ISL_3134194, EPI_ISL_3134196, EPI_ISL_3134197, EPI_ISL_3134198, EPI_ISL_3134199, EPI_ISL_3134200, EPI_ISL_3134201, EPI_ISL_3134202, EPI_ISL_3134203, EPI_ISL_3134206, EPI_ISL_3134207, EPI_ISL_3134208, EPI_ISL_3134209, EPI_ISL_3134210, EPI_ISL_3134211, EPI_ISL_3134212, EPI_ISL_3134213, EPI_ISL_3134214, EPI_ISL_3134215, EPI_ISL_3134216, EPI_ISL_3134217, EPI_ISL_3134218, EPI_ISL_3134219, EPI_ISL_3134220, EPI_ISL_3134221, EPI_ISL_3134222, EPI_ISL_3134223, EPI_ISL_3134224, EPI_ISL_3134225, EPI_ISL_3134226, EPI_ISL_3134227, EPI_ISL_3134228, EPI_ISL_3134229, EPI_ISL_3134230, EPI_ISL_3134231, EPI_ISL_3134232, EPI_ISL_3134233, EPI_ISL_3134234, EPI_ISL_3134235, EPI_ISL_3134237, EPI_ISL_3134238, EPI_ISL_3134241, EPI_ISL_3134242, EPI_ISL_3134243, EPI_ISL_3134244, EPI_ISL_3134247 | | | |
| see above | GA Department of Public Health | GA Department of Public Health | Stacy Reeves, Sharmila Talekar, Jonathan Edwards, Cynthia Dixey, Tonia Parrott, Aliyah Fields, Taylor Smith |
| EPI_ISL_3134282, EPI_ISL_3134283, EPI_ISL_3134284, EPI_ISL_3134285, EPI_ISL_3134286, EPI_ISL_3134287, EPI_ISL_3134288, EPI_ISL_3134289, EPI_ISL_3134290, EPI_ISL_3134291, EPI_ISL_3134292, EPI_ISL_3134293, EPI_ISL_3134294, EPI_ISL_3134295, EPI_ISL_3134296, EPI_ISL_3134297, EPI_ISL_3134298, EPI_ISL_3134299, EPI_ISL_3134300, EPI_ISL_3134301, EPI_ISL_3134302, EPI_ISL_3134303, EPI_ISL_3134304, EPI_ISL_3134305, EPI_ISL_3134306, EPI_ISL_3134307, EPI_ISL_3134308, EPI_ISL_3134309, EPI_ISL_3134310, EPI_ISL_3134311, EPI_ISL_3134312, EPI_ISL_3134313, EPI_ISL_3134314, EPI_ISL_3134315, EPI_ISL_3134316, EPI_ISL_3134317, EPI_ISL_3134318, EPI_ISL_3134319, EPI_ISL_3134320, EPI_ISL_3134321, EPI_ISL_3134322, EPI_ISL_3134323, EPI_ISL_3134324, EPI_ISL_3134325, EPI_ISL_3134326, EPI_ISL_3134327, EPI_ISL_3134328, EPI_ISL_3134329, EPI_ISL_3134330, EPI_ISL_3134331, EPI_ISL_3134332, EPI_ISL_3134334, EPI_ISL_3134335, EPI_ISL_3134336, EPI_ISL_3134337, EPI_ISL_3134338, EPI_ISL_3134339, EPI_ISL_3134340, EPI_ISL_3134341, EPI_ISL_3134342, EPI_ISL_3134343, EPI_ISL_3134344, EPI_ISL_3134345, EPI_ISL_3134346, EPI_ISL_3134347, EPI_ISL_3134348, EPI_ISL_3134349, EPI_ISL_3134350, EPI_ISL_3134351, EPI_ISL_3134352, EPI_ISL_3134353, EPI_ISL_3134354, EPI_ISL_3134355, EPI_ISL_3134356, EPI_ISL_3134357, EPI_ISL_3134358, EPI_ISL_3134359, EPI_ISL_3134361, EPI_ISL_3134362, EPI_ISL_3134363, EPI_ISL_3134364, EPI_ISL_3134365, EPI_ISL_3134366, EPI_ISL_3134367, EPI_ISL_3134368, EPI_ISL_3134369, EPI_ISL_3134370, EPI_ISL_3134371, EPI_ISL_3134372, EPI_ISL_3134373, EPI_ISL_3134374, EPI_ISL_3134375, EPI_ISL_3134376, EPI_ISL_3134377, EPI_ISL_3134378, EPI_ISL_3134379, EPI_ISL_3134380, EPI_ISL_3134381, EPI_ISL_3134382, EPI_ISL_3134383, EPI_ISL_3134384, EPI_ISL_3134385, EPI_ISL_3134386, EPI_ISL_3134387, EPI_ISL_3134388, EPI_ISL_3134389, EPI_ISL_3134390, EPI_ISL_3134391, EPI_ISL_3134392, EPI_ISL_3134393, EPI_ISL_3134394, EPI_ISL_3134395, EPI_ISL_3134396, EPI_ISL_3134398, EPI_ISL_3134399, EPI_ISL_3134400, EPI_ISL_3134401, EPI_ISL_3134402, EPI_ISL_3134403, EPI_ISL_3134404, EPI_ISL_3134405, EPI_ISL_3134406, EPI_ISL_3134407, EPI_ISL_3134408, EPI_ISL_3134409, EPI_ISL_3134410, EPI_ISL_3134411, EPI_ISL_3134412, EPI_ISL_3134413, EPI_ISL_3134414, EPI_ISL_3134415, EPI_ISL_3134416, EPI_ISL_3134417, EPI_ISL_3134418, EPI_ISL_3134419, EPI_ISL_3134420, EPI_ISL_3134421, EPI_ISL_3134422, EPI_ISL_3134423, EPI_ISL_3134424, EPI_ISL_3134425, EPI_ISL_3134426, EPI_ISL_3134427, EPI_ISL_3134428, EPI_ISL_3134429, EPI_ISL_3134430, EPI_ISL_3134431, EPI_ISL_3134432, EPI_ISL_3134433, EPI_ISL_3134434, EPI_ISL_3134435, EPI_ISL_3134436, EPI_ISL_3134437, EPI_ISL_3134438, EPI_ISL_3134439, EPI_ISL_3134440, EPI_ISL_3134441, EPI_ISL_3134442, EPI_ISL_3134443, EPI_ISL_3134444, EPI_ISL_3134445, EPI_ISL_3134446, EPI_ISL_3134447, EPI_ISL_3134449, EPI_ISL_3134450, EPI_ISL_3134451, EPI_ISL_3134452, EPI_ISL_3134453, EPI_ISL_3134454, EPI_ISL_3134455, EPI_ISL_3134456, EPI_ISL_3134457, EPI_ISL_3134458, EPI_ISL_3134459, EPI_ISL_3134460, EPI_ISL_3134461, EPI_ISL_3134462, EPI_ISL_3134463, EPI_ISL_3134464, EPI_ISL_3134465, EPI_ISL_3134466, EPI_ISL_3134467, EPI_ISL_3134468, EPI_ISL_3134469, EPI_ISL_3134470, EPI_ISL_3134471, EPI_ISL_3134472, EPI_ISL_3134473, EPI_ISL_3134474, EPI_ISL_3134475, EPI_ISL_3134476, EPI_ISL_3134477, EPI_ISL_3134478, EPI_ISL_3134479, EPI_ISL_3134480, EPI_ISL_3134481, EPI_ISL_3134482, EPI_ISL_3134483, EPI_ISL_3134484, EPI_ISL_3134485, EPI_ISL_3134486, EPI_ISL_3134488, EPI_ISL_3134489, EPI_ISL_3134490, EPI_ISL_3134491, EPI_ISL_3134492, EPI_ISL_3134493, EPI_ISL_3134494, EPI_ISL_3134495, EPI_ISL_3134496, EPI_ISL_3134497, EPI_ISL_3134498, EPI_ISL_3134499, EPI_ISL_3134500, EPI_ISL_3134501, EPI_ISL_3134502, EPI_ISL_3134503, EPI_ISL_3134504, EPI_ISL_3134505, EPI_ISL_3134506, EPI_ISL_3134507, EPI_ISL_3134508, EPI_ISL_3134509, EPI_ISL_3134510, EPI_ISL_3134511, EPI_ISL_3134512, EPI_ISL_3134513, EPI_ISL_3134514, EPI_ISL_3134515, EPI_ISL_3134516, EPI_ISL_3134517, EPI_ISL_3134518, EPI_ISL_3134519, EPI_ISL_3134520, EPI_ISL_3134521, EPI_ISL_3134522, EPI_ISL_3134523, EPI_ISL_3134524, EPI_ISL_3134525, EPI_ISL_3134526, EPI_ISL_3134527, EPI_ISL_3134528, EPI_ISL_3134529, EPI_ISL_3134530, EPI_ISL_3134531, EPI_ISL_3134532, EPI_ISL_3134533, EPI_ISL_3134534, EPI_ISL_3134535, EPI_ISL_3134536, EPI_ISL_3134537, EPI_ISL_3134538, EPI_ISL_3134539, EPI_ISL_3134540, EPI_ISL_3134541, EPI_ISL_3134542, EPI_ISL_3134544, EPI_ISL_3134545, EPI_ISL_3134546, EPI_ISL_3134547, EPI_ISL_3134548, EPI_ISL_3134549 | | | |
| see above | Broad Institute Clinical Research Sequencing Platform | Infectious Disease Program, Broad Institute of Harvard and MIT | Siddle,K.J., Adams,G., Pearlman,L., Gladden-Young,A., Vicente,G., Blumenstiel,B., DeFelice,M., Lee,M., McGovern,S., Lagerborg,K., Rudy,M., DeRuff,K., Carter,A., Normandin,E., Bauer,M., Reilly,S., Tomkins-Tinch,C., Loreth,C., Chaluvadi,S., Meldrim,J., Granger,B., Lemieux,J.E., Birren,B.W., Sabeti,P.C., Larkin,K., Dodge,S., Lennon,N., Madoff,L., Brown,C., Gallagher,G., Smole,S., Park,D.J., Gabriel,S., and MacInnis,B.L. |
| EPI_ISL_3134853, EPI_ISL_3134854, EPI_ISL_3134855 | LACEN/PE | WallauLab on behalf of Fiocruz COVID-19 Genomic Surveillance Network | Marcelo Henrique dos Santos Paiva, Cassia Docena, Matheus Filgueira Bezerra, Filipe Zimmer Dezordi, Lais Ceschini Machado, Lilian Caroliny Amorim Silva, Gustavo Barbosa de Lima, Alexandre Freitas da Silva, Sinval Pinto Brandão Filho, Constância Flávia Junqueira Ayres, Gabriel Luz Wallau |
| EPI_ISL_3134913, EPI_ISL_3134917 | Microbiological Diagnostic Unit - Public Health Laboratory (MDU-PHL) | MDU-PHL | Seemann T., Sait, M.L., Sherry, N.L. |
| EPI_ISL_3134918 | Royal Darwin Hospital Pathology | MDU-PHL | Meumann, E., Caly L., Seemann T., Sait, M.L., Druce J., Sherry, N.L. |
| EPI_ISL_3134919, EPI_ISL_3134920 | Victorian Infectious Diseases Reference Laboratory (VIDRL) | VIDRL and MDU-PHL | Caly L., Seemann T., Sait, M.L., Druce J., Sherry, N.L. |
| EPI_ISL_3134947, EPI_ISL_3134951, EPI_ISL_3134959, EPI_ISL_3134964 | LATE - Laboratório de Técnicas Especiais - Hospital Israelita Albert Einstein | LATE - Laboratório de Técnicas Especiais - Hospital Israelita Albert Einstein | Deyvid Amgarten, Fernanda de Mello Malta, Raquel Riyuzo, Ana Paula Moreira Salles, Pedro Henrique Sebe Rodrigues, Alexandre Hideaki Takara, Anelisie da Silva Santos, Erick Gustavo Dorlass, Marcio Anunciacao Menezes, João Renato Rebello Pinho |
| EPI_ISL_3134981, EPI_ISL_3134982, EPI_ISL_3134984, EPI_ISL_3134985, EPI_ISL_3134987, EPI_ISL_3134988, EPI_ISL_3134989, EPI_ISL_3134991, EPI_ISL_3134992, EPI_ISL_3134993, EPI_ISL_3134994, EPI_ISL_3134995, EPI_ISL_3134996, EPI_ISL_3134997, EPI_ISL_3134998, EPI_ISL_3135000, EPI_ISL_3135002, EPI_ISL_3135003, EPI_ISL_3135004, EPI_ISL_3135005, EPI_ISL_3135007, EPI_ISL_3135008, EPI_ISL_3135009, EPI_ISL_3135010, EPI_ISL_3135012, EPI_ISL_3135013, EPI_ISL_3135015, EPI_ISL_3135016, EPI_ISL_3135017, EPI_ISL_3135018, EPI_ISL_3135019, EPI_ISL_3135020, EPI_ISL_3135021, EPI_ISL_3135022, EPI_ISL_3135026, EPI_ISL_3135028, EPI_ISL_3135029, EPI_ISL_3135030, EPI_ISL_3135031, EPI_ISL_3135032, EPI_ISL_3135033, EPI_ISL_3135034, EPI_ISL_3135035, EPI_ISL_3135036, EPI_ISL_3135038, EPI_ISL_3135040, EPI_ISL_3135041, EPI_ISL_3135042, EPI_ISL_3135043, EPI_ISL_3135044, EPI_ISL_3135045, EPI_ISL_3135049, EPI_ISL_3135050, EPI_ISL_3135051, EPI_ISL_3135052, EPI_ISL_3135053, EPI_ISL_3135054, EPI_ISL_3135055, EPI_ISL_3135056, EPI_ISL_3135057, EPI_ISL_3135059, EPI_ISL_3135060, EPI_ISL_3135061, EPI_ISL_3135062, EPI_ISL_3135064, EPI_ISL_3135065, EPI_ISL_3135067, EPI_ISL_3135068, EPI_ISL_3135069, EPI_ISL_3135070, EPI_ISL_3135071, EPI_ISL_3135072, EPI_ISL_3135074, EPI_ISL_3135075, EPI_ISL_3135076, EPI_ISL_3135077, EPI_ISL_3135078, EPI_ISL_3135079, EPI_ISL_3135080, EPI_ISL_3135081, EPI_ISL_3135083, EPI_ISL_3135084, EPI_ISL_3135085, EPI_ISL_3135086, EPI_ISL_3135087, EPI_ISL_3135088, EPI_ISL_3135093, EPI_ISL_3135094, EPI_ISL_3135095, EPI_ISL_3135096, EPI_ISL_3135097, EPI_ISL_3135098, EPI_ISL_3135099, EPI_ISL_3135100, EPI_ISL_3135101, EPI_ISL_3135102, EPI_ISL_3135103, EPI_ISL_3135105, EPI_ISL_3135106, EPI_ISL_3135107, EPI_ISL_3135108, EPI_ISL_3135109, EPI_ISL_3135111, EPI_ISL_3135112, EPI_ISL_3135113, EPI_ISL_3135114, EPI_ISL_3135115, EPI_ISL_3135117, EPI_ISL_3135118, EPI_ISL_3135119, EPI_ISL_3135120, EPI_ISL_3135121, EPI_ISL_3135122, EPI_ISL_3135124, EPI_ISL_3135125, EPI_ISL_3135126, EPI_ISL_3135127, EPI_ISL_3135128, EPI_ISL_3135129, EPI_ISL_3135130, EPI_ISL_3135131, EPI_ISL_3135132, EPI_ISL_3135133, EPI_ISL_3135135, EPI_ISL_3135136, EPI_ISL_3135137, EPI_ISL_3135139, EPI_ISL_3135140, EPI_ISL_3135141, EPI_ISL_3135142, EPI_ISL_3135144, EPI_ISL_3135145, EPI_ISL_3135146, EPI_ISL_3135148, EPI_ISL_3135149, EPI_ISL_3135151, EPI_ISL_3135152, EPI_ISL_3135154, EPI_ISL_3135155, EPI_ISL_3135156, EPI_ISL_3135157, EPI_ISL_3135158, EPI_ISL_3135161, EPI_ISL_3135162, EPI_ISL_3135164, EPI_ISL_3135165, EPI_ISL_3135168, EPI_ISL_3135169, EPI_ISL_3135170, EPI_ISL_3135171, EPI_ISL_3135173, EPI_ISL_3135174, EPI_ISL_3135175, EPI_ISL_3135176, EPI_ISL_3135177, EPI_ISL_3135178, EPI_ISL_3135179, EPI_ISL_3135180, EPI_ISL_3135181, EPI_ISL_3135182, EPI_ISL_3135183, EPI_ISL_3135184, EPI_ISL_3135185, EPI_ISL_3135186, EPI_ISL_3135187, EPI_ISL_3135188, EPI_ISL_3135189, EPI_ISL_3135190, EPI_ISL_3135192, EPI_ISL_3135193, EPI_ISL_3135195, EPI_ISL_3135197, EPI_ISL_3135198, EPI_ISL_3135199, EPI_ISL_3135200, EPI_ISL_3135201, EPI_ISL_3135202, EPI_ISL_3135203, EPI_ISL_3135204, EPI_ISL_3135205, EPI_ISL_3135207, EPI_ISL_3135208, EPI_ISL_3135210, EPI_ISL_3135211, EPI_ISL_3135212, EPI_ISL_3135213, EPI_ISL_3135215, EPI_ISL_3135216, EPI_ISL_3135218, EPI_ISL_3135219, EPI_ISL_3135220, EPI_ISL_3135221, EPI_ISL_3135222, EPI_ISL_3135223, EPI_ISL_3135224, EPI_ISL_3135225, EPI_ISL_3135226, EPI_ISL_3135227, EPI_ISL_3135228, EPI_ISL_3135229, EPI_ISL_3135230, EPI_ISL_3135231, EPI_ISL_3135232, EPI_ISL_3135233, EPI_ISL_3135234, EPI_ISL_3135235, EPI_ISL_3135236, EPI_ISL_3135237, EPI_ISL_3135238, EPI_ISL_3135239, EPI_ISL_3135240, EPI_ISL_3135241, EPI_ISL_3135242, EPI_ISL_3135243, EPI_ISL_3135244, EPI_ISL_3135245, EPI_ISL_3135246, EPI_ISL_3135247, EPI_ISL_3135248, EPI_ISL_3135249, EPI_ISL_3135250, EPI_ISL_3135251, EPI_ISL_3135252, EPI_ISL_3135253, EPI_ISL_3135254, EPI_ISL_3135255, EPI_ISL_3135256, EPI_ISL_3135257, EPI_ISL_3135258, EPI_ISL_3135259, EPI_ISL_3135260, EPI_ISL_3135261, EPI_ISL_3135262, EPI_ISL_3135263, EPI_ISL_3135264, EPI_ISL_3135265, EPI_ISL_3135266, EPI_ISL_3135268, EPI_ISL_3135269, EPI_ISL_3135270, EPI_ISL_3135271, EPI_ISL_3135272, EPI_ISL_3135273, EPI_ISL_3135274, EPI_ISL_3135275, EPI_ISL_3135276, EPI_ISL_3135277, EPI_ISL_3135278, EPI_ISL_3135279, EPI_ISL_3135280, EPI_ISL_3135281, EPI_ISL_3135282, EPI_ISL_3135283, EPI_ISL_3135284, EPI_ISL_3135285, EPI_ISL_3135286, EPI_ISL_3135287, EPI_ISL_3135288, EPI_ISL_3135289, EPI_ISL_3135290, EPI_ISL_3135291, EPI_ISL_3135292, EPI_ISL_3135293, EPI_ISL_3135294, EPI_ISL_3135295, EPI_ISL_3135296, EPI_ISL_3135297, EPI_ISL_3135298, EPI_ISL_3135299, EPI_ISL_3135300, EPI_ISL_3135301, EPI_ISL_3135302, EPI_ISL_3135303, EPI_ISL_3135304, EPI_ISL_3135305, EPI_ISL_3135306, EPI_ISL_3135307, EPI_ISL_3135308, EPI_ISL_3135309, EPI_ISL_3135310, EPI_ISL_3135311, EPI_ISL_3135312, EPI_ISL_3135313, EPI_ISL_3135314, EPI_ISL_3135315, EPI_ISL_3135316, EPI_ISL_3135317, EPI_ISL_3135318, EPI_ISL_3135320, EPI_ISL_3135322, EPI_ISL_3135323, EPI_ISL_3135324, EPI_ISL_3135325, EPI_ISL_3135326, EPI_ISL_3135327, EPI_ISL_3135329, EPI_ISL_3135330, EPI_ISL_3135331, EPI_ISL_3135332, EPI_ISL_3135333, EPI_ISL_3135335, EPI_ISL_3135336, EPI_ISL_3135337, EPI_ISL_3135338, EPI_ISL_3135339, EPI_ISL_3135340, EPI_ISL_3135341, EPI_ISL_3135343, EPI_ISL_3135344, EPI_ISL_3135346, EPI_ISL_3135347, EPI_ISL_3135348, EPI_ISL_3135349, EPI_ISL_3135350, EPI_ISL_3135353, EPI_ISL_3135354, EPI_ISL_3135355, EPI_ISL_3135356, EPI_ISL_3135357, EPI_ISL_3135359, EPI_ISL_3135360, EPI_ISL_3135361, EPI_ISL_3135362, EPI_ISL_3135363, EPI_ISL_3135364, EPI_ISL_3135367, EPI_ISL_3135368, EPI_ISL_3135369, EPI_ISL_3135370, EPI_ISL_3135371, EPI_ISL_3135372, EPI_ISL_3135373, EPI_ISL_3135374, EPI_ISL_3135375, EPI_ISL_3135376, EPI_ISL_3135377, EPI_ISL_3135378, EPI_ISL_3135379, EPI_ISL_3135380, EPI_ISL_3135381, EPI_ISL_3135382, EPI_ISL_3135383, EPI_ISL_3135384, EPI_ISL_3135385, EPI_ISL_3135386, EPI_ISL_3135387, EPI_ISL_3135388, EPI_ISL_3135389, EPI_ISL_3135390, EPI_ISL_3135391, EPI_ISL_3135392, EPI_ISL_3135393, EPI_ISL_3135394, EPI_ISL_3135395, EPI_ISL_3135396, EPI_ISL_3135397, EPI_ISL_3135398, EPI_ISL_3135399, EPI_ISL_3135400, EPI_ISL_3135401, EPI_ISL_3135402, EPI_ISL_3135403, EPI_ISL_3135404, EPI_ISL_3135405, EPI_ISL_3135406, EPI_ISL_3135407, EPI_ISL_3135408, EPI_ISL_3135412, EPI_ISL_3135413, EPI_ISL_3135414, EPI_ISL_3135415, EPI_ISL_3135416, EPI_ISL_3135417, EPI_ISL_3135418, EPI_ISL_3135419, EPI_ISL_3135420, EPI_ISL_3135421, EPI_ISL_3135422, EPI_ISL_3135423, EPI_ISL_3135424, EPI_ISL_3135425, EPI_ISL_3135426, EPI_ISL_3135427, EPI_ISL_3135428, EPI_ISL_3135429, EPI_ISL_3135430, EPI_ISL_3135431, EPI_ISL_3135432, EPI_ISL_3135433, EPI_ISL_3135434, EPI_ISL_3135435, EPI_ISL_3135436, EPI_ISL_3135437, EPI_ISL_3135438, EPI_ISL_3135439, EPI_ISL_3135440, EPI_ISL_3135441, EPI_ISL_3135442, EPI_ISL_3135443, EPI_ISL_3135444, EPI_ISL_3135445, EPI_ISL_3135446, EPI_ISL_3135447, EPI_ISL_3135448, EPI_ISL_3135449, EPI_ISL_3135450, EPI_ISL_3135451, EPI_ISL_3135452, EPI_ISL_3135453, EPI_ISL_3135454, EPI_ISL_3135455, EPI_ISL_3135456, EPI_ISL_3135457, EPI_ISL_3135458, EPI_ISL_3135459, EPI_ISL_3135460, EPI_ISL_3135461, EPI_ISL_3135462, EPI_ISL_3135463, EPI_ISL_3135464, EPI_ISL_3135465, EPI_ISL_3135466, EPI_ISL_3135467, EPI_ISL_3135468, EPI_ISL_3135469, EPI_ISL_3135470, EPI_ISL_3135471, EPI_ISL_3135472, EPI_ISL_3135473, EPI_ISL_3135474, EPI_ISL_3135475, EPI_ISL_3135476, EPI_ISL_3135477, EPI_ISL_3135478, EPI_ISL_3135479, EPI_ISL_3135480, EPI_ISL_3135481, EPI_ISL_3135482, EPI_ISL_3135483, EPI_ISL_3135484, EPI_ISL_3135485, EPI_ISL_3135486, EPI_ISL_3135487, EPI_ISL_3135488, EPI_ISL_3135489, EPI_ISL_3135490, EPI_ISL_3135491, EPI_ISL_3135492, EPI_ISL_3135493, EPI_ISL_3135494, EPI_ISL_3135495, EPI_ISL_3135496, EPI_ISL_3135497, EPI_ISL_3135498, EPI_ISL_3135499, EPI_ISL_3135500, EPI_ISL_3135501, EPI_ISL_3135502, EPI_ISL_3135503, EPI_ISL_3135504, EPI_ISL_3135505, EPI_ISL_3135506, EPI_ISL_3135507, EPI_ISL_3135508, EPI_ISL_3135509, EPI_ISL_3135510, EPI_ISL_3135511, EPI_ISL_3135512, EPI_ISL_3135513, EPI_ISL_3135514, EPI_ISL_3135515, EPI_ISL_3135516, EPI_ISL_3135517, EPI_ISL_3135518, EPI_ISL_3135519, EPI_ISL_3135520, EPI_ISL_3135521, EPI_ISL_3135522, EPI_ISL_3135523, EPI_ISL_3135524, EPI_ISL_3135525, EPI_ISL_3135526, EPI_ISL_3135527, EPI_ISL_3135528, EPI_ISL_3135529, EPI_ISL_3135530, EPI_ISL_3135531, EPI_ISL_3135532, EPI_ISL_3135533, EPI_ISL_3135534, EPI_ISL_3135535, EPI_ISL_3135536, EPI_ISL_3135537, EPI_ISL_3135538, EPI_ISL_3135539, EPI_ISL_3135540, EPI_ISL_3135541, EPI_ISL_3135542, EPI_ISL_3135543, EPI_ISL_3135544, EPI_ISL_3135545, EPI_ISL_3135546, EPI_ISL_3135547, EPI_ISL_3135548, EPI_ISL_3135549, EPI_ISL_3135550, EPI_ISL_3135551, EPI_ISL_3135552, EPI_ISL_3135553, EPI_ISL_3135554, EPI_ISL_3135555, EPI_ISL_3135556, EPI_ISL_3135557, EPI_ISL_3135558, EPI_ISL_3135559, EPI_ISL_3135560, EPI_ISL_3135561, EPI_ISL_3135562, EPI_ISL_3135563, EPI_ISL_3135564, EPI_ISL_3135565, EPI_ISL_3135566, EPI_ISL_3135567, EPI_ISL_3135568, EPI_ISL_3135569, EPI_ISL_3135570, EPI_ISL_3135571, EPI_ISL_3135572, EPI_ISL_3135573, EPI_ISL_3135574, EPI_ISL_3135575, EPI_ISL_3135576, EPI_ISL_3135577, EPI_ISL_3135578, EPI_ISL_3135579, EPI_ISL_3135580, EPI_ISL_3135581, EPI_ISL_3135582, EPI_ISL_3135583, EPI_ISL_3135584, EPI_ISL_3135585, EPI_ISL_3135586, EPI_ISL_3135587, EPI_ISL_3135588, EPI_ISL_3135589, EPI_ISL_3135590, EPI_ISL_3135591, EPI_ISL_3135592, EPI_ISL_3135593, EPI_ISL_3135594, EPI_ISL_3135595, EPI_ISL_3135596, EPI_ISL_3135597, EPI_ISL_3135598, EPI_ISL_3135599, EPI_ISL_3135600, EPI_ISL_3135601, EPI_ISL_3135602, EPI_ISL_3135603, EPI_ISL_3135604, EPI_ISL_3135605, EPI_ISL_3135606, EPI_ISL_3135607, EPI_ISL_3135608, EPI_ISL_3135609, EPI_ISL_3135610, EPI_ISL_3135611, EPI_ISL_3135612, EPI_ISL_3135613, EPI_ISL_3135614, EPI_ISL_3135615, EPI_ISL_3135616, EPI_ISL_3135617, EPI_ISL_3135618, EPI_ISL_3135619, EPI_ISL_3135620, EPI_ISL_3135621, EPI_ISL_3135622, EPI_ISL_3135623, EPI_ISL_3135624, EPI_ISL_3135625, EPI_ISL_3135626, EPI_ISL_3135627, EPI_ISL_3135628, EPI_ISL_3135629, EPI_ISL_3135630, EPI_ISL_3135631, EPI_ISL_3135632, EPI_ISL_3135633, EPI_ISL_3135634, EPI_ISL_3135635, EPI_ISL_3135637, EPI_ISL_3135639, EPI_ISL_3135640, EPI_ISL_3135641, EPI_ISL_3135643, EPI_ISL_3135644, EPI_ISL_3135645, EPI_ISL_3135646, EPI_ISL_3135647, EPI_ISL_3135648, EPI_ISL_3135649, EPI_ISL_3135650, EPI_ISL_3135651, EPI_ISL_3135652, EPI_ISL_3135653, EPI_ISL_3135654, EPI_ISL_3135655, EPI_ISL_3135656, EPI_ISL_3135657, EPI_ISL_3135658, EPI_ISL_3135661, EPI_ISL_3135662, EPI_ISL_3135663, EPI_ISL_3135664, EPI_ISL_3135665, EPI_ISL_3135666, EPI_ISL_3135667, EPI_ISL_3135669, EPI_ISL_3135670, EPI_ISL_3135671, EPI_ISL_3135672, EPI_ISL_3135674, EPI_ISL_3135675, EPI_ISL_3135677, EPI_ISL_3135678, EPI_ISL_3135679, EPI_ISL_3135680, EPI_ISL_3135681, EPI_ISL_3135682, EPI_ISL_3135684, EPI_ISL_3135685, EPI_ISL_3135687, EPI_ISL_3135688, EPI_ISL_3135689, EPI_ISL_3135690, EPI_ISL_3135691, EPI_ISL_3135692, EPI_ISL_3135694, EPI_ISL_3135695, EPI_ISL_3135697, EPI_ISL_3135698, EPI_ISL_3135699, EPI_ISL_3135700, EPI_ISL_3135701, EPI_ISL_3135703, EPI_ISL_3135704, EPI_ISL_3135705, EPI_ISL_3135706, EPI_ISL_3135707, EPI_ISL_3135708, EPI_ISL_3135710, EPI_ISL_3135713, EPI_ISL_3135714, EPI_ISL_3135715, EPI_ISL_3135716, EPI_ISL_3135717, EPI_ISL_3135718, EPI_ISL_3135719, EPI_ISL_3135720, EPI_ISL_3135721, EPI_ISL_3135722, EPI_ISL_3135723, EPI_ISL_3135724, EPI_ISL_3135725, EPI_ISL_3135726, EPI_ISL_3135727, EPI_ISL_3135728, EPI_ISL_3135729, EPI_ISL_3135731, EPI_ISL_3135732, EPI_ISL_3135733, EPI_ISL_3135734, EPI_ISL_3135735, EPI_ISL_3135736, EPI_ISL_3135737, EPI_ISL_3135738, EPI_ISL_3135739, EPI_ISL_3135740, EPI_ISL_3135741, EPI_ISL_3135742, EPI_ISL_3135743, EPI_ISL_3135745, EPI_ISL_3135746, EPI_ISL_3135749, EPI_ISL_3135750, EPI_ISL_3135752, EPI_ISL_3135753, EPI_ISL_3135754, EPI_ISL_3135755, EPI_ISL_3135756, EPI_ISL_3135757, EPI_ISL_3135758, EPI_ISL_3135759, EPI_ISL_3135760, EPI_ISL_3135761, EPI_ISL_3135762, EPI_ISL_3135763, EPI_ISL_3135764, EPI_ISL_3135765, EPI_ISL_3135766, EPI_ISL_3135767, EPI_ISL_3135768, EPI_ISL_3135769, EPI_ISL_3135770, EPI_ISL_3135771, EPI_ISL_3135772, EPI_ISL_3135773, EPI_ISL_3135774, EPI_ISL_3135775, EPI_ISL_3135776, EPI_ISL_3135777, EPI_ISL_3135778, EPI_ISL_3135779, EPI_ISL_3135780, EPI_ISL_3135781, EPI_ISL_3135782, EPI_ISL_3135783, EPI_ISL_3135784, EPI_ISL_3135786, EPI_ISL_3135787, EPI_ISL_3135789, EPI_ISL_3135790, EPI_ISL_3135791, EPI_ISL_3135792, EPI_ISL_3135793, EPI_ISL_3135794, EPI_ISL_3135795, EPI_ISL_3135796, EPI_ISL_3135797, EPI_ISL_3135798, EPI_ISL_3135799, EPI_ISL_3135800, EPI_ISL_3135803, EPI_ISL_3135805, EPI_ISL_3135806, EPI_ISL_3135807, EPI_ISL_3135808, EPI_ISL_3135809, EPI_ISL_3135811, EPI_ISL_3135812, EPI_ISL_3135813, EPI_ISL_3135814, EPI_ISL_3135815, EPI_ISL_3135817, EPI_ISL_3135818, EPI_ISL_3135819, EPI_ISL_3135820, EPI_ISL_3135821, EPI_ISL_3135823, EPI_ISL_3135824, EPI_ISL_3135825, EPI_ISL_3135826, EPI_ISL_3135827, EPI_ISL_3135828, EPI_ISL_3135829, EPI_ISL_3135830, EPI_ISL_3135831, EPI_ISL_3135835, EPI_ISL_3135836, EPI_ISL_3135837, EPI_ISL_3135838, EPI_ISL_3135842, EPI_ISL_3135843, EPI_ISL_3135845, EPI_ISL_3135846, EPI_ISL_3135847, EPI_ISL_3135848, EPI_ISL_3135849, EPI_ISL_3135850, EPI_ISL_3135851, EPI_ISL_3135852, EPI_ISL_3135853, EPI_ISL_3135854, EPI_ISL_3135855, EPI_ISL_3135856, EPI_ISL_3135857, EPI_ISL_3135858, EPI_ISL_3135859, EPI_ISL_3135860, EPI_ISL_3135861, EPI_ISL_3135862, EPI_ISL_3135864, EPI_ISL_3135865, EPI_ISL_3135866, EPI_ISL_3135868, EPI_ISL_3135869, EPI_ISL_3135870, EPI_ISL_3135871, EPI_ISL_3135872, EPI_ISL_3135873, EPI_ISL_3135874, EPI_ISL_3135876, EPI_ISL_3135877, EPI_ISL_3135878, EPI_ISL_3135880, EPI_ISL_3135881, EPI_ISL_3135882, EPI_ISL_3135883, EPI_ISL_3135884, EPI_ISL_3135885, EPI_ISL_3135887, EPI_ISL_3135888, EPI_ISL_3135889, EPI_ISL_3135890, EPI_ISL_3135892, EPI_ISL_3135893, EPI_ISL_3135895, EPI_ISL_3135896 | | | |
| see above | Department of Virology and Immunology, University of Helsinki and Helsinki University Hospital, Huslab Finland | Department of Virology, Faculty of Medicine, University of Helsinki, Helsinki, Finland | Teemu Smura, Ravi Kant, Phuoc Truong, Hussein Alburkat, Hannimari Kallio-Kokko, Jenni Virtanen, Maija Suvanto, Essi Korhonen, Sari Hannula, Harri Kangas, Hanna Liimatainen, Satu Kurkela, Hanna Jarva, Maija Lappalainen, Pekka Ellonen, Olli Vapalahti |
| EPI_ISL_3135898, EPI_ISL_3135899 | Laboratory Medicine and Molecular Diagnostics | Laboratory Medicine and Molecular Diagnostics | Dayakar Seetha, Heera R Pillai, Sanughosh Kalpothodi, Radhakrishna R Nair |
| EPI_ISL_3135900 | Laboratory Medicine and Molecular Diagnostics | Laboratory Medicine and Molecular Diagnostics | Dayakar Seetha, Heera R Pillai, Sanughosh Kalpothodi, Radhakrishnan R Nair |
| EPI_ISL_3135901, EPI_ISL_3135902 | Laboratory Medicine and Molecular Diagnostics | Laboratory Medicine and Molecular Diagnostics | Dayakar Seetha, Heera R Pillai, Sanughosh Kalpothodi, Radhakrishna R Nair |
| EPI_ISL_3135903, EPI_ISL_3135904 | Laboratory Medicine and Molecular Diagnostics | Laboratory Medicine and Molecular Diagnostics | Dayakar Seetha, Heera R Pillai, Sanughosh Kalpothodi, Radhakrishnan R Nair |
| EPI_ISL_3135905, EPI_ISL_3135906, EPI_ISL_3135907, EPI_ISL_3135908, EPI_ISL_3135909, EPI_ISL_3135910, EPI_ISL_3135911, EPI_ISL_3135912, EPI_ISL_3135913, EPI_ISL_3135914, EPI_ISL_3135915, EPI_ISL_3135916, EPI_ISL_3135917, EPI_ISL_3135918, EPI_ISL_3135919, EPI_ISL_3135920, EPI_ISL_3135921, EPI_ISL_3135922, EPI_ISL_3135923, EPI_ISL_3135924 | | | |

| | | | |
|---|---|---|---|
| see above | Wyoming Public Health Laboratory | Wyoming Public Health Laboratory | Jim Mildenberger, Taylor Fearing, Channing Weber, Ashley Norberg, Chayse Rowley, Marley Goetz, Brian Dominguez, Elliot Thomasson, Sam Britz, Cari Sloma, Robert Petit, and Rob Christensen |
| EPI_ISL_3135925, EPI_ISL_3135927, EPI_ISL_3135928, EPI_ISL_3135929, EPI_ISL_3135932 | Laboratory Medicine and Molecular Diagnostics | Laboratory Medicine and Molecular Diagnostics | Dayakar Seetha, Heera R Pillai, Sanughosh Kalpothodi, Radhakrishnan R Nair |
| EPI_ISL_3135933, EPI_ISL_3135934, EPI_ISL_3135935, EPI_ISL_3135936, EPI_ISL_3135937, EPI_ISL_3135939, EPI_ISL_3135940, EPI_ISL_3135941, EPI_ISL_3135944, EPI_ISL_3135945, EPI_ISL_3135946, EPI_ISL_3135950, EPI_ISL_3135954, EPI_ISL_3135955, EPI_ISL_3135956, EPI_ISL_3135957, EPI_ISL_3135959, EPI_ISL_3135960, EPI_ISL_3135961, EPI_ISL_3135962, EPI_ISL_3135963, EPI_ISL_3135966, EPI_ISL_3135967, EPI_ISL_3135968, EPI_ISL_3135969, EPI_ISL_3135970, EPI_ISL_3135971, EPI_ISL_3135972, EPI_ISL_3135973, EPI_ISL_3135974, EPI_ISL_3135975, EPI_ISL_3135976, EPI_ISL_3135977, EPI_ISL_3135978, EPI_ISL_3135979, EPI_ISL_3135981, EPI_ISL_3135982, EPI_ISL_3135985, EPI_ISL_3135986, EPI_ISL_3135987, EPI_ISL_3135988, EPI_ISL_3135989, EPI_ISL_3135990, EPI_ISL_3135991, EPI_ISL_3135992, EPI_ISL_3135993, EPI_ISL_3135995, EPI_ISL_3135996, EPI_ISL_3135997, EPI_ISL_3135998, EPI_ISL_3135999, EPI_ISL_3136000, EPI_ISL_3136002, EPI_ISL_3136004, EPI_ISL_3136005, EPI_ISL_3136006, EPI_ISL_3136007, EPI_ISL_3136008, EPI_ISL_3136010, EPI_ISL_3136011, EPI_ISL_3136012, EPI_ISL_3136014, EPI_ISL_3136015, EPI_ISL_3136016, EPI_ISL_3136017, EPI_ISL_3136018, EPI_ISL_3136019, EPI_ISL_3136020, EPI_ISL_3136021, EPI_ISL_3136022, EPI_ISL_3136023, EPI_ISL_3136024, EPI_ISL_3136025, EPI_ISL_3136026, EPI_ISL_3136028, EPI_ISL_3136029, EPI_ISL_3136030, EPI_ISL_3136031, EPI_ISL_3136032, EPI_ISL_3136033, EPI_ISL_3136034, EPI_ISL_3136035, EPI_ISL_3136037, EPI_ISL_3136038, EPI_ISL_3136039, EPI_ISL_3136040, EPI_ISL_3136041, EPI_ISL_3136042, EPI_ISL_3136043, EPI_ISL_3136044, EPI_ISL_3136047, EPI_ISL_3136048, EPI_ISL_3136049, EPI_ISL_3136050, EPI_ISL_3136051, EPI_ISL_3136052, EPI_ISL_3136053, EPI_ISL_3136054, EPI_ISL_3136055, EPI_ISL_3136056, EPI_ISL_3136057, EPI_ISL_3136058, EPI_ISL_3136060, EPI_ISL_3136061, EPI_ISL_3136062, EPI_ISL_3136063, EPI_ISL_3136064, EPI_ISL_3136065, EPI_ISL_3136066, EPI_ISL_3136067, EPI_ISL_3136068, EPI_ISL_3136069, EPI_ISL_3136070, EPI_ISL_3136071, EPI_ISL_3136072, EPI_ISL_3136073, EPI_ISL_3136074, EPI_ISL_3136075, EPI_ISL_3136076, EPI_ISL_3136077, EPI_ISL_3136078, EPI_ISL_3136079, EPI_ISL_3136080, EPI_ISL_3136081, EPI_ISL_3136082, EPI_ISL_3136083, EPI_ISL_3136084, EPI_ISL_3136085, EPI_ISL_3136086, EPI_ISL_3136087, EPI_ISL_3136089, EPI_ISL_3136090, EPI_ISL_3136091, EPI_ISL_3136092, EPI_ISL_3136093, EPI_ISL_3136094, EPI_ISL_3136095, EPI_ISL_3136096, EPI_ISL_3136097, EPI_ISL_3136098, EPI_ISL_3136099, EPI_ISL_3136100, EPI_ISL_3136101, EPI_ISL_3136103, EPI_ISL_3136104, EPI_ISL_3136105, EPI_ISL_3136106, EPI_ISL_3136107, EPI_ISL_3136108, EPI_ISL_3136109, EPI_ISL_3136110, EPI_ISL_3136111, EPI_ISL_3136112, EPI_ISL_3136113, EPI_ISL_3136114, EPI_ISL_3136115, EPI_ISL_3136116, EPI_ISL_3136117, EPI_ISL_3136118, EPI_ISL_3136119, EPI_ISL_3136120, EPI_ISL_3136121, EPI_ISL_3136122, EPI_ISL_3136123, EPI_ISL_3136124, EPI_ISL_3136125 | | | |
| see above | Dutch COVID-19 response team | Erasmus Medical Center | Bas Oude Munnink, Reina Sikkema, David Nieuwenhuijse, Irina Chestakova, Anne van der Linden, Marjan Boter, Emmanuelle Munger, Corine GeurtsvanKessel, Annemiek van der Eijk, Richard Molenkamp, Marion Koopmans, on behalf of the Dutch national COVID-19 response team. |
| EPI_ISL_3136174, EPI_ISL_3136175, EPI_ISL_3136176, EPI_ISL_3136178 | Fakultní nemocnice Hradec Králové | University Hospital Hradec Kralove | Marketa Gancarcikova, Helena Kovarikova |
| EPI_ISL_3136179, EPI_ISL_3136180, EPI_ISL_3136181, EPI_ISL_3136182, EPI_ISL_3136183, EPI_ISL_3136184, EPI_ISL_3136185, EPI_ISL_3136186, EPI_ISL_3136187, EPI_ISL_3136188, EPI_ISL_3136189, EPI_ISL_3136190, EPI_ISL_3136191, EPI_ISL_3136192, EPI_ISL_3136193 | | | |
| see above | Pardubická nemocnice | University Hospital Hradec Kralove | Marketa Gancarcikova, Helena Kovarikova |
| EPI_ISL_3136194, EPI_ISL_3136195 | Oblastní nemocnice Trutnov a.s. | University Hospital Hradec Kralove | Marketa Gancarcikova, Helena Kovarikova |
| EPI_ISL_3136196, EPI_ISL_3136197, EPI_ISL_3136198 | Pardubická nemocnice | University Hospital Hradec Kralove | Marketa Gancarcikova, Helena Kovarikova |
| EPI_ISL_3136199 | Orlickoústecká nemocnice | University Hospital Hradec Kralove | Marketa Gancarcikova, Helena Kovarikova |
| EPI_ISL_3136200 | Oblastní nemocnice Náchod a.s. | University Hospital Hradec Kralove | Marketa Gancarcikova, Helena Kovarikova |
| EPI_ISL_3136201, EPI_ISL_3136202 | Fakultní nemocnice Hradec Králové | University Hospital Hradec Kralove | Marketa Gancarcikova, Helena Kovarikova |
| EPI_ISL_3136203 | Laboratory Medicine and Molecular Diagnostics | Laboratory Medicine and Molecular Diagnostics | Dayakar Seetha, Heera R Pillai, Sanughosh Kalpothodi, Radhakrishnan R Nair |
| EPI_ISL_3136208 | National Virus Reference Laboratory | National Virus Reference Laboratory | Zoe Yandle, Charlene Bennett, Gabriel Gonzalez, Michael Carr, Jonathan Dean, Cillian F De Gascun |
| EPI_ISL_3136215, EPI_ISL_3136216, EPI_ISL_3136217, EPI_ISL_3136218, EPI_ISL_3136234, EPI_ISL_3136235, EPI_ISL_3136236, EPI_ISL_3136237, EPI_ISL_3136238, EPI_ISL_3136241 | National Virus Reference Laboratory | National Virus Reference Laboratory | Guerrino Macori, Gabriel Gonzalez, Michael Carr, Zoe Yandle, Charlene Bennett, Jonathan Dean, Seamus Fanning, Cillian F De Gascun |
| EPI_ISL_3136245, EPI_ISL_3136246, EPI_ISL_3136247, EPI_ISL_3136248, EPI_ISL_3136249, EPI_ISL_3136250, EPI_ISL_3136251, EPI_ISL_3136252, EPI_ISL_3136253, EPI_ISL_3136254, EPI_ISL_3136255, EPI_ISL_3136256, EPI_ISL_3136257, EPI_ISL_3136258, EPI_ISL_3136259, EPI_ISL_3136260, EPI_ISL_3136261, EPI_ISL_3136262, EPI_ISL_3136263, EPI_ISL_3136264, EPI_ISL_3136265, EPI_ISL_3136266, EPI_ISL_3136267, EPI_ISL_3136268, EPI_ISL_3136269, EPI_ISL_3136270, EPI_ISL_3136271, EPI_ISL_3136272, EPI_ISL_3136273, EPI_ISL_3136274, EPI_ISL_3136275, EPI_ISL_3136276, EPI_ISL_3136277, EPI_ISL_3136278, EPI_ISL_3136279, EPI_ISL_3136280, EPI_ISL_3136281, EPI_ISL_3136282, EPI_ISL_3136283, EPI_ISL_3136284, EPI_ISL_3136285, EPI_ISL_3136286, EPI_ISL_3136287, EPI_ISL_3136288, EPI_ISL_3136289, EPI_ISL_3136290, EPI_ISL_3136291, EPI_ISL_3136292, EPI_ISL_3136293, EPI_ISL_3136294, EPI_ISL_3136295, EPI_ISL_3136296, EPI_ISL_3136297, EPI_ISL_3136298, EPI_ISL_3136299, EPI_ISL_3136300, EPI_ISL_3136301, EPI_ISL_3136302, EPI_ISL_3136303, EPI_ISL_3136304, EPI_ISL_3136305, EPI_ISL_3136306, EPI_ISL_3136307, EPI_ISL_3136308, EPI_ISL_3136309, EPI_ISL_3136310, EPI_ISL_3136311, EPI_ISL_3136312, EPI_ISL_3136314, EPI_ISL_3136315, EPI_ISL_3136316, EPI_ISL_3136317, EPI_ISL_3136318, EPI_ISL_3136319, EPI_ISL_3136320, EPI_ISL_3136321, EPI_ISL_3136322, EPI_ISL_3136325, EPI_ISL_3136326, EPI_ISL_3136327, EPI_ISL_3136328, EPI_ISL_3136329, EPI_ISL_3136330, EPI_ISL_3136331, EPI_ISL_3136332, EPI_ISL_3136333, EPI_ISL_3136334, EPI_ISL_3136335, EPI_ISL_3136336, EPI_ISL_3136337, EPI_ISL_3136338, EPI_ISL_3136339, EPI_ISL_3136340, EPI_ISL_3136341, EPI_ISL_3136342, EPI_ISL_3136344, EPI_ISL_3136346, EPI_ISL_3136347, EPI_ISL_3136348, EPI_ISL_3136349, EPI_ISL_3136350, EPI_ISL_3136352, EPI_ISL_3136353, EPI_ISL_3136354, EPI_ISL_3136355, EPI_ISL_3136356, EPI_ISL_3136357, EPI_ISL_3136358, EPI_ISL_3136359, EPI_ISL_3136360, EPI_ISL_3136361, EPI_ISL_3136362, EPI_ISL_3136363, EPI_ISL_3136364, EPI_ISL_3136365, EPI_ISL_3136366, EPI_ISL_3136367, EPI_ISL_3136368, EPI_ISL_3136369, EPI_ISL_3136370, EPI_ISL_3136371, EPI_ISL_3136372, EPI_ISL_3136373, EPI_ISL_3136374, EPI_ISL_3136375, EPI_ISL_3136376, EPI_ISL_3136377, EPI_ISL_3136378, EPI_ISL_3136379, EPI_ISL_3136380, EPI_ISL_3136381, EPI_ISL_3136382, EPI_ISL_3136383, EPI_ISL_3136384, EPI_ISL_3136385, EPI_ISL_3136386, EPI_ISL_3136387, EPI_ISL_3136388, EPI_ISL_3136389, EPI_ISL_3136390, EPI_ISL_3136391, EPI_ISL_3136392, EPI_ISL_3136393, EPI_ISL_3136394, EPI_ISL_3136395, EPI_ISL_3136396, EPI_ISL_3136397, EPI_ISL_3136398, EPI_ISL_3136399, EPI_ISL_3136400, EPI_ISL_3136401, EPI_ISL_3136402, EPI_ISL_3136403, EPI_ISL_3136404, EPI_ISL_3136405, EPI_ISL_3136406, EPI_ISL_3136407, EPI_ISL_3136408, EPI_ISL_3136409, EPI_ISL_3136410, EPI_ISL_3136411, EPI_ISL_3136412, EPI_ISL_3136413, EPI_ISL_3136414, EPI_ISL_3136415, EPI_ISL_3136416, EPI_ISL_3136417, EPI_ISL_3136418, EPI_ISL_3136419, EPI_ISL_3136420, EPI_ISL_3136421, EPI_ISL_3136422, EPI_ISL_3136423, EPI_ISL_3136424, EPI_ISL_3136425, EPI_ISL_3136426, EPI_ISL_3136431, EPI_ISL_3136432, EPI_ISL_3136434, EPI_ISL_3136435, EPI_ISL_3136437, EPI_ISL_3136442, EPI_ISL_3136443, EPI_ISL_3136444, EPI_ISL_3136445, EPI_ISL_3136446, EPI_ISL_3136447, EPI_ISL_3136449, EPI_ISL_3136450, EPI_ISL_3136451, EPI_ISL_3136452, EPI_ISL_3136453, EPI_ISL_3136454, EPI_ISL_3136455, EPI_ISL_3136456, EPI_ISL_3136458, EPI_ISL_3136460, EPI_ISL_3136461, EPI_ISL_3136462, EPI_ISL_3136463, EPI_ISL_3136464, EPI_ISL_3136465, EPI_ISL_3136466, EPI_ISL_3136467, EPI_ISL_3136468, EPI_ISL_3136469, EPI_ISL_3136470, EPI_ISL_3136471, EPI_ISL_3136472 | | | |
| see above | National Virus Reference Laboratory | National Virus Reference Laboratory | Zoe Yandle, Charlene Bennett, Gabriel Gonzalez, Michael Carr, Jonathan Dean, Cillian F De Gascun |
| EPI_ISL_3136475, EPI_ISL_3136476, EPI_ISL_3136477, EPI_ISL_3136478, EPI_ISL_3136479, EPI_ISL_3136480, EPI_ISL_3136481, EPI_ISL_3136482, EPI_ISL_3136483, EPI_ISL_3136484, EPI_ISL_3136486, EPI_ISL_3136488, EPI_ISL_3136489, EPI_ISL_3136490, EPI_ISL_3136492, EPI_ISL_3136493, EPI_ISL_3136494, EPI_ISL_3136495, EPI_ISL_3136496, EPI_ISL_3136497, EPI_ISL_3136498, EPI_ISL_3136499, EPI_ISL_3136500, EPI_ISL_3136501, EPI_ISL_3136502, EPI_ISL_3136503, EPI_ISL_3136504, EPI_ISL_3136506, EPI_ISL_3136507, EPI_ISL_3136508, EPI_ISL_3136510, EPI_ISL_3136511, EPI_ISL_3136512, EPI_ISL_3136513, EPI_ISL_3136514, EPI_ISL_3136515, EPI_ISL_3136516, EPI_ISL_3136517, EPI_ISL_3136518, EPI_ISL_3136519, EPI_ISL_3136520, EPI_ISL_3136521, EPI_ISL_3136522, EPI_ISL_3136523, EPI_ISL_3136524, EPI_ISL_3136525, EPI_ISL_3136526, EPI_ISL_3136527, EPI_ISL_3136528, EPI_ISL_3136529, EPI_ISL_3136530, EPI_ISL_3136531, EPI_ISL_3136532, EPI_ISL_3136534, EPI_ISL_3136535, EPI_ISL_3136536, EPI_ISL_3136537, EPI_ISL_3136538, EPI_ISL_3136539, EPI_ISL_3136540, EPI_ISL_3136541, EPI_ISL_3136542, EPI_ISL_3136543, EPI_ISL_3136544, EPI_ISL_3136546, EPI_ISL_3136547, EPI_ISL_3136548, EPI_ISL_3136549, EPI_ISL_3136550 | | | |
| see above | Microvida | Microvida | Suzan D. Pas, Jaco J. Verweij, Joep J. J. M. Stohr |
| EPI_ISL_3136552, EPI_ISL_3136553, EPI_ISL_3136555 | National Platform bis UMONS/Jolimont | National Platform bis UMONS/Jolimont | François Dufrasne, Guillaume Bayon-Vicente, Florian Juszczak, Laetitia Gheysen, Eric Tarantino, Gautier Detry, Ruddy Wattiez |
| EPI_ISL_3136557, EPI_ISL_3136558, EPI_ISL_3136559, EPI_ISL_3136561, EPI_ISL_3136563, EPI_ISL_3136564, EPI_ISL_3136565, EPI_ISL_3136566, EPI_ISL_3136567, EPI_ISL_3136568, EPI_ISL_3136570, EPI_ISL_3136571, EPI_ISL_3136572, EPI_ISL_3136573, EPI_ISL_3136574, EPI_ISL_3136575, EPI_ISL_3136576, EPI_ISL_3136577, EPI_ISL_3136579, EPI_ISL_3136580, EPI_ISL_3136581, EPI_ISL_3136582, EPI_ISL_3136583, EPI_ISL_3136584, EPI_ISL_3136586, EPI_ISL_3136587, EPI_ISL_3136588, EPI_ISL_3136589, EPI_ISL_3136590, EPI_ISL_3136591, EPI_ISL_3136592, EPI_ISL_3136593, EPI_ISL_3136594, EPI_ISL_3136595, EPI_ISL_3136596, EPI_ISL_3136598, EPI_ISL_3136599, EPI_ISL_3136600, EPI_ISL_3136601, EPI_ISL_3136602, EPI_ISL_3136603, EPI_ISL_3136604, EPI_ISL_3136606, EPI_ISL_3136607, EPI_ISL_3136609, EPI_ISL_3136611, EPI_ISL_3136612, EPI_ISL_3136613, EPI_ISL_3136614, EPI_ISL_3136616, EPI_ISL_3136618, EPI_ISL_3136619, EPI_ISL_3136621, EPI_ISL_3136622, EPI_ISL_3136623, EPI_ISL_3136625, EPI_ISL_3136626, EPI_ISL_3136627, EPI_ISL_3136628, EPI_ISL_3136629, EPI_ISL_3136630, EPI_ISL_3136631, EPI_ISL_3136632, EPI_ISL_3136633, EPI_ISL_3136634, EPI_ISL_3136635, EPI_ISL_3136636, EPI_ISL_3136637, EPI_ISL_3136638, EPI_ISL_3136639, EPI_ISL_3136640, EPI_ISL_3136641, EPI_ISL_3136644, EPI_ISL_3136645, EPI_ISL_3136646, EPI_ISL_3136647, EPI_ISL_3136648, EPI_ISL_3136649, EPI_ISL_3136650, EPI_ISL_3136651, EPI_ISL_3136652, EPI_ISL_3136653, EPI_ISL_3136654, EPI_ISL_3136655, EPI_ISL_3136656, EPI_ISL_3136657, EPI_ISL_3136658, EPI_ISL_3136659, EPI_ISL_3136660, EPI_ISL_3136661, EPI_ISL_3136662, EPI_ISL_3136663, EPI_ISL_3136664, EPI_ISL_3136666, EPI_ISL_3136667, EPI_ISL_3136668, EPI_ISL_3136669, EPI_ISL_3136670, EPI_ISL_3136671, EPI_ISL_3136672, EPI_ISL_3136673, EPI_ISL_3136674, EPI_ISL_3136675, EPI_ISL_3136676, EPI_ISL_3136677, EPI_ISL_3136678, EPI_ISL_3136679, EPI_ISL_3136680, EPI_ISL_3136681, EPI_ISL_3136682, EPI_ISL_3136683, EPI_ISL_3136684, EPI_ISL_3136685, EPI_ISL_3136686, EPI_ISL_3136687, EPI_ISL_3136688, EPI_ISL_3136689, EPI_ISL_3136690, EPI_ISL_3136691, EPI_ISL_3136692, EPI_ISL_3136693, EPI_ISL_3136694, EPI_ISL_3136695, EPI_ISL_3136696, EPI_ISL_3136697, EPI_ISL_3136698, EPI_ISL_3136699, EPI_ISL_3136700, EPI_ISL_3136702, EPI_ISL_3136703, EPI_ISL_3136704, EPI_ISL_3136705, EPI_ISL_3136706, EPI_ISL_3136707, EPI_ISL_3136708, EPI_ISL_3136709, EPI_ISL_3136710, EPI_ISL_3136711, EPI_ISL_3136712, EPI_ISL_3136713, EPI_ISL_3136714, EPI_ISL_3136715, EPI_ISL_3136716, EPI_ISL_3136717, EPI_ISL_3136719, EPI_ISL_3136720, EPI_ISL_3136721, EPI_ISL_3136722, EPI_ISL_3136723, EPI_ISL_3136724, EPI_ISL_3136725, EPI_ISL_3136726, EPI_ISL_3136727, EPI_ISL_3136729, EPI_ISL_3136730, EPI_ISL_3136731, EPI_ISL_3136732, EPI_ISL_3136734, EPI_ISL_3136735, EPI_ISL_3136736, EPI_ISL_3136737, EPI_ISL_3136738, EPI_ISL_3136739, EPI_ISL_3136740, EPI_ISL_3136741, EPI_ISL_3136742, EPI_ISL_3136743, EPI_ISL_3136744, EPI_ISL_3136745, EPI_ISL_3136746, EPI_ISL_3136747, EPI_ISL_3136748, EPI_ISL_3136749, EPI_ISL_3136750, EPI_ISL_3136751, EPI_ISL_3136752, EPI_ISL_3136753, EPI_ISL_3136754, EPI_ISL_3136755, EPI_ISL_3136756, EPI_ISL_3136757, EPI_ISL_3136758, EPI_ISL_3136759, EPI_ISL_3136760, EPI_ISL_3136762, EPI_ISL_3136763, EPI_ISL_3136764, EPI_ISL_3136765, EPI_ISL_3136766, EPI_ISL_3136767, EPI_ISL_3136768, EPI_ISL_3136769, EPI_ISL_3136770, EPI_ISL_3136771, EPI_ISL_3136772, EPI_ISL_3136773, EPI_ISL_3136774, EPI_ISL_3136775, EPI_ISL_3136776, EPI_ISL_3136778, EPI_ISL_3136779, EPI_ISL_3136780, EPI_ISL_3136781, EPI_ISL_3136782, EPI_ISL_3136783, EPI_ISL_3136784, EPI_ISL_3136785, EPI_ISL_3136786, EPI_ISL_3136787, EPI_ISL_3136788, EPI_ISL_3136789, EPI_ISL_3136790, EPI_ISL_3136791, EPI_ISL_3136792, EPI_ISL_3136793, EPI_ISL_3136795, EPI_ISL_3136796, EPI_ISL_3136797, EPI_ISL_3136798, EPI_ISL_3136799, EPI_ISL_3136800, EPI_ISL_3136801, EPI_ISL_3136803, EPI_ISL_3136804, EPI_ISL_3136805, EPI_ISL_3136806, EPI_ISL_3136807, EPI_ISL_3136808, EPI_ISL_3136809, EPI_ISL_3136810, EPI_ISL_3136811, EPI_ISL_3136812, EPI_ISL_3136813, EPI_ISL_3136814, EPI_ISL_3136815, EPI_ISL_3136816, EPI_ISL_3136817, EPI_ISL_3136818, EPI_ISL_3136819, EPI_ISL_3136820, EPI_ISL_3136821, EPI_ISL_3136822, EPI_ISL_3136823, EPI_ISL_3136824, EPI_ISL_3136825, EPI_ISL_3136826, EPI_ISL_3136827, EPI_ISL_3136828, EPI_ISL_3136829, EPI_ISL_3136830, EPI_ISL_3136831, EPI_ISL_3136832, EPI_ISL_3136833, EPI_ISL_3136834, EPI_ISL_3136835, EPI_ISL_3136837, EPI_ISL_3136838, EPI_ISL_3136839, EPI_ISL_3136840, EPI_ISL_3136841, EPI_ISL_3136842, EPI_ISL_3136843, EPI_ISL_3136844, EPI_ISL_3136845, EPI_ISL_3136846, EPI_ISL_3136847, EPI_ISL_3136848, EPI_ISL_3136849, EPI_ISL_3136850, EPI_ISL_3136851, EPI_ISL_3136852, EPI_ISL_3136853, EPI_ISL_3136854, EPI_ISL_3136855, EPI_ISL_3136856, EPI_ISL_3136857, EPI_ISL_3136858, EPI_ISL_3136859, EPI_ISL_3136860, EPI_ISL_3136861, EPI_ISL_3136862, EPI_ISL_3136863, EPI_ISL_3136864, EPI_ISL_3136865, EPI_ISL_3136866, EPI_ISL_3136867, EPI_ISL_3136868, EPI_ISL_3136869, EPI_ISL_3136870, EPI_ISL_3136871, EPI_ISL_3136872, EPI_ISL_3136873, EPI_ISL_3136874, EPI_ISL_3136875, EPI_ISL_3136877, EPI_ISL_3136878, EPI_ISL_3136879, EPI_ISL_3136880, EPI_ISL_3136881, EPI_ISL_3136882, EPI_ISL_3136883, EPI_ISL_3136884, EPI_ISL_3136885, EPI_ISL_3136886, EPI_ISL_3136887, EPI_ISL_3136888, EPI_ISL_3136889, EPI_ISL_3136890, EPI_ISL_3136891, EPI_ISL_3136892, EPI_ISL_3136893, EPI_ISL_3136894, EPI_ISL_3136895, EPI_ISL_3136896, EPI_ISL_3136897, EPI_ISL_3136898, EPI_ISL_3136899, EPI_ISL_3136900, EPI_ISL_3136901, EPI_ISL_3136902, EPI_ISL_3136903, EPI_ISL_3136904, EPI_ISL_3136905, EPI_ISL_3136906, EPI_ISL_3136907, EPI_ISL_3136908, EPI_ISL_3136909, EPI_ISL_3136910, EPI_ISL_3136911, EPI_ISL_3136912, EPI_ISL_3136913, EPI_ISL_3136914, EPI_ISL_3136915, EPI_ISL_3136916, EPI_ISL_3136917, EPI_ISL_3136918, EPI_ISL_3136919, EPI_ISL_3136921, EPI_ISL_3136922, EPI_ISL_3136923, EPI_ISL_3136924, EPI_ISL_3136925, EPI_ISL_3136926, EPI_ISL_3136927, EPI_ISL_3136928, EPI_ISL_3136930, EPI_ISL_3136931, EPI_ISL_3136932, EPI_ISL_3136933, EPI_ISL_3136934, EPI_ISL_3136935, EPI_ISL_3136936, EPI_ISL_3136937, EPI_ISL_3136938, EPI_ISL_3136939, EPI_ISL_3136940, EPI_ISL_3136941, EPI_ISL_3136942, EPI_ISL_3136943, EPI_ISL_3136944, EPI_ISL_3136945, EPI_ISL_3136946, EPI_ISL_3136947, EPI_ISL_3136948, EPI_ISL_3136949, EPI_ISL_3136950, EPI_ISL_3136951, EPI_ISL_3136952, EPI_ISL_3136953, EPI_ISL_3136954, EPI_ISL_3136955, EPI_ISL_3136956, EPI_ISL_3136957, EPI_ISL_3136958, EPI_ISL_3136959, EPI_ISL_3136960, EPI_ISL_3136961, EPI_ISL_3136962, EPI_ISL_3136963, EPI_ISL_3136964, EPI_ISL_3136965, EPI_ISL_3136966, EPI_ISL_3136967, EPI_ISL_3136968, EPI_ISL_3136969, EPI_ISL_3136970, EPI_ISL_3136971, EPI_ISL_3136972, EPI_ISL_3136974, EPI_ISL_3136975, EPI_ISL_3136976, EPI_ISL_3136977, EPI_ISL_3136978, EPI_ISL_3136979, EPI_ISL_3136980, EPI_ISL_3136981, EPI_ISL_3136982, EPI_ISL_3136984, EPI_ISL_3136985, EPI_ISL_3136986, EPI_ISL_3136987, EPI_ISL_3136988, EPI_ISL_3136989, EPI_ISL_3136991, EPI_ISL_3136992, EPI_ISL_3136993, EPI_ISL_3136994, EPI_ISL_3136995, EPI_ISL_3136996, EPI_ISL_3136997, EPI_ISL_3136998, EPI_ISL_3136999, EPI_ISL_3137000, EPI_ISL_3137001, EPI_ISL_3137002, EPI_ISL_3137003, EPI_ISL_3137004, EPI_ISL_3137005, EPI_ISL_3137006, EPI_ISL_3137007, EPI_ISL_3137008, EPI_ISL_3137010, EPI_ISL_3137011, EPI_ISL_3137012, EPI_ISL_3137013, EPI_ISL_3137015, EPI_ISL_3137016, EPI_ISL_3137017, EPI_ISL_3137018, EPI_ISL_3137019, EPI_ISL_3137020, EPI_ISL_3137021, EPI_ISL_3137022, EPI_ISL_3137023, EPI_ISL_3137024, EPI_ISL_3137026, EPI_ISL_3137027, EPI_ISL_3137028, EPI_ISL_3137029, EPI_ISL_3137030, EPI_ISL_3137031, EPI_ISL_3137032, EPI_ISL_3137033, EPI_ISL_3137034, EPI_ISL_3137035, EPI_ISL_3137036, EPI_ISL_3137038, EPI_ISL_3137039, EPI_ISL_3137040, EPI_ISL_3137041, EPI_ISL_3137042, EPI_ISL_3137043, EPI_ISL_3137044, EPI_ISL_3137045, EPI_ISL_3137046, EPI_ISL_3137047, EPI_ISL_3137049, EPI_ISL_3137050, EPI_ISL_3137051, EPI_ISL_3137052, EPI_ISL_3137053, EPI_ISL_3137054, EPI_ISL_3137055, EPI_ISL_3137056, EPI_ISL_3137057, EPI_ISL_3137059, EPI_ISL_3137060, EPI_ISL_3137061, EPI_ISL_3137062, EPI_ISL_3137063, EPI_ISL_3137064, EPI_ISL_3137065, EPI_ISL_3137066, EPI_ISL_3137067, EPI_ISL_3137068, EPI_ISL_3137071, EPI_ISL_3137072, EPI_ISL_3137073, EPI_ISL_3137074, EPI_ISL_3137075, EPI_ISL_3137076, EPI_ISL_3137077, EPI_ISL_3137078, EPI_ISL_3137079, EPI_ISL_3137080, EPI_ISL_3137081, EPI_ISL_3137083, EPI_ISL_3137084, EPI_ISL_3137085, EPI_ISL_3137086, EPI_ISL_3137088, EPI_ISL_3137089, EPI_ISL_3137091, EPI_ISL_3137092, EPI_ISL_3137093, EPI_ISL_3137095, EPI_ISL_3137096, EPI_ISL_3137097, EPI_ISL_3137098, EPI_ISL_3137099, EPI_ISL_3137100, EPI_ISL_3137101, EPI_ISL_3137102, EPI_ISL_3137103, EPI_ISL_3137104, EPI_ISL_3137105, EPI_ISL_3137107, EPI_ISL_3137109, EPI_ISL_3137110, EPI_ISL_3137111, EPI_ISL_3137113, EPI_ISL_3137114, EPI_ISL_3137115, EPI_ISL_3137116, EPI_ISL_3137117, EPI_ISL_3137118, EPI_ISL_3137119, EPI_ISL_3137120, EPI_ISL_3137121, EPI_ISL_3137122, EPI_ISL_3137123, EPI_ISL_3137124, EPI_ISL_3137126, EPI_ISL_3137127, EPI_ISL_3137128, EPI_ISL_3137129, EPI_ISL_3137130, EPI_ISL_3137131, EPI_ISL_3137132, EPI_ISL_3137133, EPI_ISL_3137134, EPI_ISL_3137135, EPI_ISL_3137136, EPI_ISL_3137137, EPI_ISL_3137138, EPI_ISL_3137139, EPI_ISL_3137140, EPI_ISL_3137142, EPI_ISL_3137143, EPI_ISL_3137145, EPI_ISL_3137146, EPI_ISL_3137147, EPI_ISL_3137148, EPI_ISL_3137149, EPI_ISL_3137150, EPI_ISL_3137151, EPI_ISL_3137152, EPI_ISL_3137154, EPI_ISL_3137155, EPI_ISL_3137156, EPI_ISL_3137157, EPI_ISL_3137161, EPI_ISL_3137162, EPI_ISL_3137163, EPI_ISL_3137164, EPI_ISL_3137165, EPI_ISL_3137166, EPI_ISL_3137167, EPI_ISL_3137168, EPI_ISL_3137169, EPI_ISL_3137170, EPI_ISL_3137171, EPI_ISL_3137173, EPI_ISL_3137174, EPI_ISL_3137177, EPI_ISL_3137178, EPI_ISL_3137179, EPI_ISL_3137180, EPI_ISL_3137181, EPI_ISL_3137182, EPI_ISL_3137183, EPI_ISL_3137184, EPI_ISL_3137185, EPI_ISL_3137186, EPI_ISL_3137189, EPI_ISL_3137190, EPI_ISL_3137191, EPI_ISL_3137192, EPI_ISL_3137193, EPI_ISL_3137194, EPI_ISL_3137195, EPI_ISL_3137196, EPI_ISL_3137197, EPI_ISL_3137201, EPI_ISL_3137202, EPI_ISL_3137203, EPI_ISL_3137205, EPI_ISL_3137206, EPI_ISL_3137207, EPI_ISL_3137208, EPI_ISL_3137211, EPI_ISL_3137212, EPI_ISL_3137213, EPI_ISL_3137214, EPI_ISL_3137215, EPI_ISL_3137216, EPI_ISL_3137217, EPI_ISL_3137218, EPI_ISL_3137219, EPI_ISL_3137222, EPI_ISL_3137223, EPI_ISL_3137224, EPI_ISL_3137225, EPI_ISL_3137226, EPI_ISL_3137227, EPI_ISL_3137228, EPI_ISL_3137229, EPI_ISL_3137230, EPI_ISL_3137231, EPI_ISL_3137233, EPI_ISL_3137235, EPI_ISL_3137236, EPI_ISL_3137237, EPI_ISL_3137238, EPI_ISL_3137239, EPI_ISL_3137240, EPI_ISL_3137241, EPI_ISL_3137242, EPI_ISL_3137243, EPI_ISL_3137247, EPI_ISL_3137248, EPI_ISL_3137249, EPI_ISL_3137250, EPI_ISL_3137251, EPI_ISL_3137253, EPI_ISL_3137254, EPI_ISL_3137255, EPI_ISL_3137256, EPI_ISL_3137259, EPI_ISL_3137260, EPI_ISL_3137261, EPI_ISL_3137262, EPI_ISL_3137263, EPI_ISL_3137264, EPI_ISL_3137266, EPI_ISL_3137268, EPI_ISL_3137269, EPI_ISL_3137270, EPI_ISL_3137271, EPI_ISL_3137272, EPI_ISL_3137273, EPI_ISL_3137274, EPI_ISL_3137275, EPI_ISL_3137276, EPI_ISL_3137277, EPI_ISL_3137278, EPI_ISL_3137279, EPI_ISL_3137280, EPI_ISL_3137281, EPI_ISL_3137283, EPI_ISL_3137284, EPI_ISL_3137285, EPI_ISL_3137286, EPI_ISL_3137287, EPI_ISL_3137289, EPI_ISL_3137290, EPI_ISL_3137291, EPI_ISL_3137292, EPI_ISL_3137293, EPI_ISL_3137294, EPI_ISL_3137295, EPI_ISL_3137296, EPI_ISL_3137297, EPI_ISL_3137298, EPI_ISL_3137299, EPI_ISL_3137300, EPI_ISL_3137301, EPI_ISL_3137302, EPI_ISL_3137304, EPI_ISL_3137305, EPI_ISL_3137306, EPI_ISL_3137307, EPI_ISL_3137309, EPI_ISL_3137310, EPI_ISL_3137311, EPI_ISL_3137312, EPI_ISL_3137313, EPI_ISL_3137314, EPI_ISL_3137316, EPI_ISL_3137317, EPI_ISL_3137318, EPI_ISL_3137319, EPI_ISL_3137321, EPI_ISL_3137322, EPI_ISL_3137323, EPI_ISL_3137325, EPI_ISL_3137326, EPI_ISL_3137327, EPI_ISL_3137328, EPI_ISL_3137330, EPI_ISL_3137331, EPI_ISL_3137333, EPI_ISL_3137334, EPI_ISL_3137335, EPI_ISL_3137336, EPI_ISL_3137337, EPI_ISL_3137338, EPI_ISL_3137340, EPI_ISL_3137341, EPI_ISL_3137342, EPI_ISL_3137343, EPI_ISL_3137344, EPI_ISL_3137345, EPI_ISL_3137346, EPI_ISL_3137347, EPI_ISL_3137349, EPI_ISL_3137350, EPI_ISL_3137351, EPI_ISL_3137353, EPI_ISL_3137354, EPI_ISL_3137356, EPI_ISL_3137357, EPI_ISL_3137358, EPI_ISL_3137359, EPI_ISL_3137360, EPI_ISL_3137361, EPI_ISL_3137362, EPI_ISL_3137365, EPI_ISL_3137366, EPI_ISL_3137367, EPI_ISL_3137368, EPI_ISL_3137369, EPI_ISL_3137370, EPI_ISL_3137371, EPI_ISL_3137372, EPI_ISL_3137373, EPI_ISL_3137374, EPI_ISL_3137375, EPI_ISL_3137376, EPI_ISL_3137377, EPI_ISL_3137378, EPI_ISL_3137379, EPI_ISL_3137380, EPI_ISL_3137381, EPI_ISL_3137382, EPI_ISL_3137383, EPI_ISL_3137384, EPI_ISL_3137385, EPI_ISL_3137386, EPI_ISL_3137387, EPI_ISL_3137388, EPI_ISL_3137389, EPI_ISL_3137390, EPI_ISL_3137391, EPI_ISL_3137392, EPI_ISL_3137393, EPI_ISL_3137394, EPI_ISL_3137395, EPI_ISL_3137396, EPI_ISL_3137397, EPI_ISL_3137398, EPI_ISL_3137399, EPI_ISL_3137400, EPI_ISL_3137401, EPI_ISL_3137402, EPI_ISL_3137403, EPI_ISL_3137404, EPI_ISL_3137405, EPI_ISL_3137406, EPI_ISL_3137407, EPI_ISL_3137408, EPI_ISL_3137409, EPI_ISL_3137410, EPI_ISL_3137411, EPI_ISL_3137412, EPI_ISL_3137413, EPI_ISL_3137414, EPI_ISL_3137415, EPI_ISL_3137416, EPI_ISL_3137417, EPI_ISL_3137418, EPI_ISL_3137419, EPI_ISL_3137420, EPI_ISL_3137421, EPI_ISL_3137422, EPI_ISL_3137423, EPI_ISL_3137424, EPI_ISL_3137425, EPI_ISL_3137426, EPI_ISL_3137427, EPI_ISL_3137428, EPI_ISL_3137429, EPI_ISL_3137430, EPI_ISL_3137431, EPI_ISL_3137432, EPI_ISL_3137433, EPI_ISL_3137434, EPI_ISL_3137435, EPI_ISL_3137437, EPI_ISL_3137438, EPI_ISL_3137440, EPI_ISL_3137441, EPI_ISL_3137442, EPI_ISL_3137443, EPI_ISL_3137444, EPI_ISL_3137445, EPI_ISL_3137446, EPI_ISL_3137447, EPI_ISL_3137448, EPI_ISL_3137449, EPI_ISL_3137450, EPI_ISL_3137451, EPI_ISL_3137452, EPI_ISL_3137453, EPI_ISL_3137454, EPI_ISL_3137455, EPI_ISL_3137457, EPI_ISL_3137458, EPI_ISL_3137459, EPI_ISL_3137460, EPI_ISL_3137461, EPI_ISL_3137463, EPI_ISL_3137464, EPI_ISL_3137465, EPI_ISL_3137466, EPI_ISL_3137467, EPI_ISL_3137468, EPI_ISL_3137469, EPI_ISL_3137471, EPI_ISL_3137472, EPI_ISL_3137474, EPI_ISL_3137475, EPI_ISL_3137476, EPI_ISL_3137477, EPI_ISL_3137478, EPI_ISL_3137479, EPI_ISL_3137480, EPI_ISL_3137481, EPI_ISL_3137482, EPI_ISL_3137483, EPI_ISL_3137484, EPI_ISL_3137485, EPI_ISL_3137486, EPI_ISL_3137487, EPI_ISL_3137489, EPI_ISL_3137490, EPI_ISL_3137491, EPI_ISL_3137492, EPI_ISL_3137493, EPI_ISL_3137494, EPI_ISL_3137495, EPI_ISL_3137496, EPI_ISL_3137497, EPI_ISL_3137498, EPI_ISL_3137499, EPI_ISL_3137500, EPI_ISL_3137501, EPI_ISL_3137502, EPI_ISL_3137503, EPI_ISL_3137504, EPI_ISL_3137505, EPI_ISL_3137506, EPI_ISL_3137507, EPI_ISL_3137508, EPI_ISL_3137510, EPI_ISL_3137511, EPI_ISL_3137512, EPI_ISL_3137513, | | | |

| | | | |
|---|---|---|---|
| EPI_ISL_3137514, EPI_ISL_3137515, EPI_ISL_3137516, EPI_ISL_3137517, EPI_ISL_3137518, EPI_ISL_3137519, EPI_ISL_3137520, EPI_ISL_3137521, EPI_ISL_3137522, EPI_ISL_3137523, EPI_ISL_3137524, EPI_ISL_3137525, EPI_ISL_3137526, EPI_ISL_3137527, EPI_ISL_3137528, EPI_ISL_3137529, EPI_ISL_3137530, EPI_ISL_3137531, EPI_ISL_3137532, EPI_ISL_3137533, EPI_ISL_3137534, EPI_ISL_3137535, EPI_ISL_3137536, EPI_ISL_3137537, EPI_ISL_3137538, EPI_ISL_3137539, EPI_ISL_3137540, EPI_ISL_3137541, EPI_ISL_3137543, EPI_ISL_3137544, EPI_ISL_3137545, EPI_ISL_3137547, EPI_ISL_3137548, EPI_ISL_3137549, EPI_ISL_3137550, EPI_ISL_3137551, EPI_ISL_3137552, EPI_ISL_3137553, EPI_ISL_3137554, EPI_ISL_3137555, EPI_ISL_3137557, EPI_ISL_3137558, EPI_ISL_3137559, EPI_ISL_3137560, EPI_ISL_3137561, EPI_ISL_3137562, EPI_ISL_3137563, EPI_ISL_3137564 | | | |
| see above | Dutch COVID-19 response team | National Institute for Public Health and the Environment (RIVM) | Adam Meijer, Harry Vennema, Dirk Eggink, Jeroen Cremer, Sharon van den Brink, Bas van der Veer, AnneMarie van den Brandt, Lisa Wijsman, Kim Freriks, Ryanne Jaarsma, Euníce Then, Lynn Aarts, Sanne Bos, Melissa van Tuil, Florian Zwagemaker, Dennis Schmitz, Annelies Kroneman, Karim Hajji, Chantal Reusken, on behalf of the national COVID-19 response team |
| EPI_ISL_3137565 | Ospedale San Filippo Neri Microvir | University Campus Bio-Medico of Rome (UCBM) | Riva E., De Florio L., Fogolari M., Francesconi M., Lintas C., Veralli R., Angeletti S. |
| EPI_ISL_3137566, EPI_ISL_3137567, EPI_ISL_3137568, EPI_ISL_3137570, EPI_ISL_3137571, EPI_ISL_3137572, EPI_ISL_3137573, EPI_ISL_3137574, EPI_ISL_3137575, EPI_ISL_3137576, EPI_ISL_3137577, EPI_ISL_3137578, EPI_ISL_3137579, EPI_ISL_3137580, EPI_ISL_3137581, EPI_ISL_3137582, EPI_ISL_3137583, EPI_ISL_3137584, EPI_ISL_3137585, EPI_ISL_3137586, EPI_ISL_3137587, EPI_ISL_3137588, EPI_ISL_3137589, EPI_ISL_3137590, EPI_ISL_3137591, EPI_ISL_3137592, EPI_ISL_3137593, EPI_ISL_3137595, EPI_ISL_3137596, EPI_ISL_3137597, EPI_ISL_3137598, EPI_ISL_3137599, EPI_ISL_3137600, EPI_ISL_3137601, EPI_ISL_3137602, EPI_ISL_3137603, EPI_ISL_3137604, EPI_ISL_3137605, EPI_ISL_3137607, EPI_ISL_3137608, EPI_ISL_3137609, EPI_ISL_3137610, EPI_ISL_3137611, EPI_ISL_3137612, EPI_ISL_3137613, EPI_ISL_3137614, EPI_ISL_3137615, EPI_ISL_3137616, EPI_ISL_3137617, EPI_ISL_3137618, EPI_ISL_3137619, EPI_ISL_3137620, EPI_ISL_3137622, EPI_ISL_3137623, EPI_ISL_3137624, EPI_ISL_3137625, EPI_ISL_3137626, EPI_ISL_3137627, EPI_ISL_3137628, EPI_ISL_3137629, EPI_ISL_3137630, EPI_ISL_3137632, EPI_ISL_3137633, EPI_ISL_3137634, EPI_ISL_3137635, EPI_ISL_3137636, EPI_ISL_3137637, EPI_ISL_3137638, EPI_ISL_3137639, EPI_ISL_3137640, EPI_ISL_3137641, EPI_ISL_3137643, EPI_ISL_3137645, EPI_ISL_3137646, EPI_ISL_3137647, EPI_ISL_3137649, EPI_ISL_3137650, EPI_ISL_3137651, EPI_ISL_3137652, EPI_ISL_3137653, EPI_ISL_3137655, EPI_ISL_3137657, EPI_ISL_3137658, EPI_ISL_3137659, EPI_ISL_3137660, EPI_ISL_3137661, EPI_ISL_3137662, EPI_ISL_3137663, EPI_ISL_3137665, EPI_ISL_3137666, EPI_ISL_3137667, EPI_ISL_3137668, EPI_ISL_3137669, EPI_ISL_3137670, EPI_ISL_3137671, EPI_ISL_3137672, EPI_ISL_3137673, EPI_ISL_3137674, EPI_ISL_3137676, EPI_ISL_3137678, EPI_ISL_3137680, EPI_ISL_3137682, EPI_ISL_3137684, EPI_ISL_3137685, EPI_ISL_3137686, EPI_ISL_3137687, EPI_ISL_3137688, EPI_ISL_3137689, EPI_ISL_3137690, EPI_ISL_3137691, EPI_ISL_3137692, EPI_ISL_3137693, EPI_ISL_3137694, EPI_ISL_3137695, EPI_ISL_3137696, EPI_ISL_3137698, EPI_ISL_3137699, EPI_ISL_3137700, EPI_ISL_3137701, EPI_ISL_3137702, EPI_ISL_3137703, EPI_ISL_3137704, EPI_ISL_3137705, EPI_ISL_3137706, EPI_ISL_3137707, EPI_ISL_3137708, EPI_ISL_3137710, EPI_ISL_3137711, EPI_ISL_3137712, EPI_ISL_3137713, EPI_ISL_3137714, EPI_ISL_3137715, EPI_ISL_3137716, EPI_ISL_3137717, EPI_ISL_3137718, EPI_ISL_3137720, EPI_ISL_3137721, EPI_ISL_3137722, EPI_ISL_3137723, EPI_ISL_3137725, EPI_ISL_3137726, EPI_ISL_3137727, EPI_ISL_3137728, EPI_ISL_3137729, EPI_ISL_3137731, EPI_ISL_3137732, EPI_ISL_3137733, EPI_ISL_3137734, EPI_ISL_3137735, EPI_ISL_3137736, EPI_ISL_3137737, EPI_ISL_3137738, EPI_ISL_3137739, EPI_ISL_3137741, EPI_ISL_3137742, EPI_ISL_3137743, EPI_ISL_3137744, EPI_ISL_3137745, EPI_ISL_3137746, EPI_ISL_3137747, EPI_ISL_3137748, EPI_ISL_3137749, EPI_ISL_3137750, EPI_ISL_3137751, EPI_ISL_3137755, EPI_ISL_3137756, EPI_ISL_3137757, EPI_ISL_3137758, EPI_ISL_3137759, EPI_ISL_3137760, EPI_ISL_3137761, EPI_ISL_3137762, EPI_ISL_3137764, EPI_ISL_3137766, EPI_ISL_3137767, EPI_ISL_3137768, EPI_ISL_3137770, EPI_ISL_3137772, EPI_ISL_3137773, EPI_ISL_3137774, EPI_ISL_3137775, EPI_ISL_3137776, EPI_ISL_3137777, EPI_ISL_3137778, EPI_ISL_3137779, EPI_ISL_3137781, EPI_ISL_3137783, EPI_ISL_3137784, EPI_ISL_3137785, EPI_ISL_3137786, EPI_ISL_3137787, EPI_ISL_3137788, EPI_ISL_3137789, EPI_ISL_3137791, EPI_ISL_3137792, EPI_ISL_3137793, EPI_ISL_3137794, EPI_ISL_3137795, EPI_ISL_3137796, EPI_ISL_3137797, EPI_ISL_3137798, EPI_ISL_3137799, EPI_ISL_3137800, EPI_ISL_3137801, EPI_ISL_3137802, EPI_ISL_3137803, EPI_ISL_3137804, EPI_ISL_3137805, EPI_ISL_3137806, EPI_ISL_3137807, EPI_ISL_3137808, EPI_ISL_3137809, EPI_ISL_3137810, EPI_ISL_3137811, EPI_ISL_3137812, EPI_ISL_3137813, EPI_ISL_3137814, EPI_ISL_3137815, EPI_ISL_3137816, EPI_ISL_3137817, EPI_ISL_3137818, EPI_ISL_3137819, EPI_ISL_3137820, EPI_ISL_3137821, EPI_ISL_3137822, EPI_ISL_3137823, EPI_ISL_3137824, EPI_ISL_3137825, EPI_ISL_3137826, EPI_ISL_3137827, EPI_ISL_3137828, EPI_ISL_3137829, EPI_ISL_3137830, EPI_ISL_3137831, EPI_ISL_3137832, EPI_ISL_3137833, EPI_ISL_3137834, EPI_ISL_3137835, EPI_ISL_3137836, EPI_ISL_3137837, EPI_ISL_3137838, EPI_ISL_3137839, EPI_ISL_3137840, EPI_ISL_3137841, EPI_ISL_3137842, EPI_ISL_3137843, EPI_ISL_3137844, EPI_ISL_3137845, EPI_ISL_3137846, EPI_ISL_3137847, EPI_ISL_3137848, EPI_ISL_3137849, EPI_ISL_3137850, EPI_ISL_3137851, EPI_ISL_3137853, EPI_ISL_3137854, EPI_ISL_3137855, EPI_ISL_3137856, EPI_ISL_3137857, EPI_ISL_3137858, EPI_ISL_3137859, EPI_ISL_3137860, EPI_ISL_3137861, EPI_ISL_3137862, EPI_ISL_3137863, EPI_ISL_3137864, EPI_ISL_3137865, EPI_ISL_3137866, EPI_ISL_3137867, EPI_ISL_3137868, EPI_ISL_3137869, EPI_ISL_3137870, EPI_ISL_3137871, EPI_ISL_3137872, EPI_ISL_3137873, EPI_ISL_3137875, EPI_ISL_3137876, EPI_ISL_3137877, EPI_ISL_3137878, EPI_ISL_3137879, EPI_ISL_3137880, EPI_ISL_3137881, EPI_ISL_3137882, EPI_ISL_3137883, EPI_ISL_3137884, EPI_ISL_3137885, EPI_ISL_3137886, EPI_ISL_3137887, EPI_ISL_3137888, EPI_ISL_3137889, EPI_ISL_3137891, EPI_ISL_3137892, EPI_ISL_3137893, EPI_ISL_3137895, EPI_ISL_3137896, EPI_ISL_3137897, EPI_ISL_3137898, EPI_ISL_3137899, EPI_ISL_3137900, EPI_ISL_3137901, EPI_ISL_3137902, EPI_ISL_3137903, EPI_ISL_3137904, EPI_ISL_3137905, EPI_ISL_3137906, EPI_ISL_3137907, EPI_ISL_3137908, EPI_ISL_3137909, EPI_ISL_3137910, EPI_ISL_3137911, EPI_ISL_3137912, EPI_ISL_3137913, EPI_ISL_3137914, EPI_ISL_3137915, EPI_ISL_3137916, EPI_ISL_3137917, EPI_ISL_3137918, EPI_ISL_3137919, EPI_ISL_3137920, EPI_ISL_3137922, EPI_ISL_3137923, EPI_ISL_3137924, EPI_ISL_3137925, EPI_ISL_3137927, EPI_ISL_3137928, EPI_ISL_3137929, EPI_ISL_3137930, EPI_ISL_3137931, EPI_ISL_3137932, EPI_ISL_3137933, EPI_ISL_3137934, EPI_ISL_3137935, EPI_ISL_3137936, EPI_ISL_3137937, EPI_ISL_3137938, EPI_ISL_3137939, EPI_ISL_3137940, EPI_ISL_3137941, EPI_ISL_3137942, EPI_ISL_3137944, EPI_ISL_3137945, EPI_ISL_3137946, EPI_ISL_3137947, EPI_ISL_3137948 | | | |
| see above | Dutch COVID-19 response team | National Institute for Public Health and the Environment (RIVM) | Adam Meijer, Harry Vennema, Dirk Eggink, Jeroen Cremer, Sharon van den Brink, Bas van der Veer, AnneMarie van den Brandt, Lisa Wijsman, Kim Freriks, Ryanne Jaarsma, Euníce Then, Lynn Aarts, Sanne Bos, Melissa van Tuil, Florian Zwagemaker, Dennis Schmitz, Annelies Kroneman, Karim Hajji, Chantal Reusken, on behalf of the national COVID-19 response team |
| EPI_ISL_3137949 | Ospedale San Filippo Neri Microvir | University Campus Bio-Medico of Rome (UCBM) | Riva E., De Florio L., Fogolari M., Francesconi M., Lintas C., Veralli R., Angeletti S. |
| EPI_ISL_3137950, EPI_ISL_3137951, EPI_ISL_3137952, EPI_ISL_3137953, EPI_ISL_3137954, EPI_ISL_3137955, EPI_ISL_3137956, EPI_ISL_3137958, EPI_ISL_3137959, EPI_ISL_3137960, EPI_ISL_3137961, EPI_ISL_3137962, EPI_ISL_3137963, EPI_ISL_3137964, EPI_ISL_3137965, EPI_ISL_3137966, EPI_ISL_3137967, EPI_ISL_3137968, EPI_ISL_3137969, EPI_ISL_3137971, EPI_ISL_3137972, EPI_ISL_3137973, EPI_ISL_3137974, EPI_ISL_3137975, EPI_ISL_3137976, EPI_ISL_3137977, EPI_ISL_3137978, EPI_ISL_3137979, EPI_ISL_3137980, EPI_ISL_3137981, EPI_ISL_3137982, EPI_ISL_3137983, EPI_ISL_3137984, EPI_ISL_3137985, EPI_ISL_3137986, EPI_ISL_3137987, EPI_ISL_3137988, EPI_ISL_3137989, EPI_ISL_3137990, EPI_ISL_3137991, EPI_ISL_3137992, EPI_ISL_3137993, EPI_ISL_3137994, EPI_ISL_3137995, EPI_ISL_3137996, EPI_ISL_3137998, EPI_ISL_3137999, EPI_ISL_3138000, EPI_ISL_3138001, EPI_ISL_3138002, EPI_ISL_3138003, EPI_ISL_3138004, EPI_ISL_3138005, EPI_ISL_3138006, EPI_ISL_3138007, EPI_ISL_3138008, EPI_ISL_3138009, EPI_ISL_3138010, EPI_ISL_3138011, EPI_ISL_3138012, EPI_ISL_3138013, EPI_ISL_3138014, EPI_ISL_3138015, EPI_ISL_3138016, EPI_ISL_3138017, EPI_ISL_3138018, EPI_ISL_3138019, EPI_ISL_3138021, EPI_ISL_3138022, EPI_ISL_3138023, EPI_ISL_3138024, EPI_ISL_3138025, EPI_ISL_3138026, EPI_ISL_3138027, EPI_ISL_3138028, EPI_ISL_3138029, EPI_ISL_3138030, EPI_ISL_3138031, EPI_ISL_3138032, EPI_ISL_3138033, EPI_ISL_3138034, EPI_ISL_3138035, EPI_ISL_3138036, EPI_ISL_3138037, EPI_ISL_3138038, EPI_ISL_3138039, EPI_ISL_3138040, EPI_ISL_3138041, EPI_ISL_3138042, EPI_ISL_3138043, EPI_ISL_3138044, EPI_ISL_3138045, EPI_ISL_3138046, EPI_ISL_3138047, EPI_ISL_3138048, EPI_ISL_3138049, EPI_ISL_3138050, EPI_ISL_3138052, EPI_ISL_3138053, EPI_ISL_3138054, EPI_ISL_3138055, EPI_ISL_3138056, EPI_ISL_3138057, EPI_ISL_3138058, EPI_ISL_3138059, EPI_ISL_3138060, EPI_ISL_3138061, EPI_ISL_3138062, EPI_ISL_3138063, EPI_ISL_3138064, EPI_ISL_3138065, EPI_ISL_3138066, EPI_ISL_3138067, EPI_ISL_3138068, EPI_ISL_3138069, EPI_ISL_3138070, EPI_ISL_3138071, EPI_ISL_3138072, EPI_ISL_3138073, EPI_ISL_3138074, EPI_ISL_3138075, EPI_ISL_3138076, EPI_ISL_3138077, EPI_ISL_3138078, EPI_ISL_3138079, EPI_ISL_3138080, EPI_ISL_3138081, EPI_ISL_3138082, EPI_ISL_3138083, EPI_ISL_3138084, EPI_ISL_3138085, EPI_ISL_3138086, EPI_ISL_3138087, EPI_ISL_3138088, EPI_ISL_3138089, EPI_ISL_3138090, EPI_ISL_3138091, EPI_ISL_3138092, EPI_ISL_3138093, EPI_ISL_3138094, EPI_ISL_3138095, EPI_ISL_3138096, EPI_ISL_3138097, EPI_ISL_3138101, EPI_ISL_3138102, EPI_ISL_3138104, EPI_ISL_3138105, EPI_ISL_3138106, EPI_ISL_3138107, EPI_ISL_3138108, EPI_ISL_3138109, EPI_ISL_3138112, EPI_ISL_3138113, EPI_ISL_3138115, EPI_ISL_3138116, EPI_ISL_3138117, EPI_ISL_3138118, EPI_ISL_3138119, EPI_ISL_3138120, EPI_ISL_3138121, EPI_ISL_3138122, EPI_ISL_3138123, EPI_ISL_3138124, EPI_ISL_3138125, EPI_ISL_3138126, EPI_ISL_3138128, EPI_ISL_3138132, EPI_ISL_3138133, EPI_ISL_3138134, EPI_ISL_3138135, EPI_ISL_3138136, EPI_ISL_3138137, EPI_ISL_3138139, EPI_ISL_3138141, EPI_ISL_3138142, EPI_ISL_3138143, EPI_ISL_3138144, EPI_ISL_3138145, EPI_ISL_3138146, EPI_ISL_3138148, EPI_ISL_3138149, EPI_ISL_3138150, EPI_ISL_3138152, EPI_ISL_3138153, EPI_ISL_3138154, EPI_ISL_3138155, EPI_ISL_3138156, EPI_ISL_3138157, EPI_ISL_3138158, EPI_ISL_3138160, EPI_ISL_3138161, EPI_ISL_3138162, EPI_ISL_3138163, EPI_ISL_3138164, EPI_ISL_3138165, EPI_ISL_3138166, EPI_ISL_3138167, EPI_ISL_3138168, EPI_ISL_3138171, EPI_ISL_3138172, EPI_ISL_3138173 | | | |
| see above | Dutch COVID-19 response team | National Institute for Public Health and the Environment (RIVM) | Adam Meijer, Harry Vennema, Dirk Eggink, Jeroen Cremer, Sharon van den Brink, Bas van der Veer, AnneMarie van den Brandt, Lisa Wijsman, Kim Freriks, Ryanne Jaarsma, Euníce Then, Lynn Aarts, Sanne Bos, Melissa van Tuil, Florian Zwagemaker, Dennis Schmitz, Annelies Kroneman, Karim Hajji, Chantal Reusken, on behalf of the national COVID-19 response team |
| EPI_ISL_3138174 | Ospedale San Filippo Neri Microvir | University Campus Bio-Medico of Rome (UCBM) | Riva E., De Florio L., Fogolari M., Francesconi M., Lintas C., Veralli R., Angeletti S. |
| EPI_ISL_3138175, EPI_ISL_3138176, EPI_ISL_3138177, EPI_ISL_3138178, EPI_ISL_3138179, EPI_ISL_3138180, EPI_ISL_3138181, EPI_ISL_3138182, EPI_ISL_3138183, EPI_ISL_3138184, EPI_ISL_3138185, EPI_ISL_3138186, EPI_ISL_3138187, EPI_ISL_3138188, EPI_ISL_3138189, EPI_ISL_3138190, EPI_ISL_3138191, EPI_ISL_3138192, EPI_ISL_3138193, EPI_ISL_3138194, EPI_ISL_3138195, EPI_ISL_3138196, EPI_ISL_3138197, EPI_ISL_3138198, EPI_ISL_3138199, EPI_ISL_3138200, EPI_ISL_3138201, EPI_ISL_3138202, EPI_ISL_3138204, EPI_ISL_3138205, EPI_ISL_3138206, EPI_ISL_3138207, EPI_ISL_3138208, EPI_ISL_3138209, EPI_ISL_3138210, EPI_ISL_3138211, EPI_ISL_3138212, EPI_ISL_3138213, EPI_ISL_3138214, EPI_ISL_3138216, EPI_ISL_3138217, EPI_ISL_3138218, EPI_ISL_3138219, EPI_ISL_3138220, EPI_ISL_3138221, EPI_ISL_3138222, EPI_ISL_3138223, EPI_ISL_3138224, EPI_ISL_3138225, EPI_ISL_3138227, EPI_ISL_3138228, EPI_ISL_3138229, EPI_ISL_3138230, EPI_ISL_3138231, EPI_ISL_3138232, EPI_ISL_3138233, EPI_ISL_3138234, EPI_ISL_3138235, EPI_ISL_3138236, EPI_ISL_3138238, EPI_ISL_3138239, EPI_ISL_3138240, EPI_ISL_3138241, EPI_ISL_3138242, EPI_ISL_3138243, EPI_ISL_3138244, EPI_ISL_3138245, EPI_ISL_3138246, EPI_ISL_3138248, EPI_ISL_3138249, EPI_ISL_3138250, EPI_ISL_3138251, EPI_ISL_3138252, EPI_ISL_3138253, EPI_ISL_3138254, EPI_ISL_3138255, EPI_ISL_3138256, EPI_ISL_3138257, EPI_ISL_3138261, EPI_ISL_3138262, EPI_ISL_3138263, EPI_ISL_3138265, EPI_ISL_3138266, EPI_ISL_3138267, EPI_ISL_3138268, EPI_ISL_3138271, EPI_ISL_3138272, EPI_ISL_3138273, EPI_ISL_3138274, EPI_ISL_3138275, EPI_ISL_3138276, EPI_ISL_3138277, EPI_ISL_3138278, EPI_ISL_3138281, EPI_ISL_3138282, EPI_ISL_3138283, EPI_ISL_3138284, EPI_ISL_3138285, EPI_ISL_3138286, EPI_ISL_3138288, EPI_ISL_3138291, EPI_ISL_3138292, EPI_ISL_3138293, EPI_ISL_3138294, EPI_ISL_3138296, EPI_ISL_3138297, EPI_ISL_3138303, EPI_ISL_3138304, EPI_ISL_3138305, EPI_ISL_3138306, EPI_ISL_3138307, EPI_ISL_3138308, EPI_ISL_3138309, EPI_ISL_3138313, EPI_ISL_3138314, EPI_ISL_3138315, EPI_ISL_3138316, EPI_ISL_3138317, EPI_ISL_3138318, EPI_ISL_3138319, EPI_ISL_3138321, EPI_ISL_3138324, EPI_ISL_3138326, EPI_ISL_3138327, EPI_ISL_3138328, EPI_ISL_3138329, EPI_ISL_3138330, EPI_ISL_3138331, EPI_ISL_3138332, EPI_ISL_3138333, EPI_ISL_3138335, EPI_ISL_3138336, EPI_ISL_3138338, EPI_ISL_3138339, EPI_ISL_3138340, EPI_ISL_3138341, EPI_ISL_3138346, EPI_ISL_3138347, EPI_ISL_3138348, EPI_ISL_3138349, EPI_ISL_3138352, EPI_ISL_3138353, EPI_ISL_3138354, EPI_ISL_3138355, EPI_ISL_3138356, EPI_ISL_3138357, EPI_ISL_3138358, EPI_ISL_3138359, EPI_ISL_3138361, EPI_ISL_3138364, EPI_ISL_3138365, EPI_ISL_3138369, EPI_ISL_3138371, EPI_ISL_3138372, EPI_ISL_3138373, EPI_ISL_3138375, EPI_ISL_3138376, EPI_ISL_3138377, EPI_ISL_3138379, EPI_ISL_3138380, EPI_ISL_3138381, EPI_ISL_3138382, EPI_ISL_3138383, EPI_ISL_3138384, EPI_ISL_3138385, EPI_ISL_3138387, EPI_ISL_3138388, EPI_ISL_3138389, EPI_ISL_3138390, EPI_ISL_3138391, EPI_ISL_3138392, EPI_ISL_3138393, EPI_ISL_3138394, EPI_ISL_3138395, EPI_ISL_3138396, EPI_ISL_3138397, EPI_ISL_3138399, EPI_ISL_3138400, EPI_ISL_3138401, EPI_ISL_3138403, EPI_ISL_3138404, EPI_ISL_3138407, EPI_ISL_3138408, EPI_ISL_3138411, EPI_ISL_3138412, EPI_ISL_3138413, EPI_ISL_3138414, EPI_ISL_3138415, EPI_ISL_3138416, EPI_ISL_3138417, EPI_ISL_3138418, EPI_ISL_3138421, EPI_ISL_3138422, EPI_ISL_3138423, EPI_ISL_3138425, EPI_ISL_3138426, EPI_ISL_3138427, EPI_ISL_3138428, EPI_ISL_3138431, EPI_ISL_3138433, EPI_ISL_3138434, EPI_ISL_3138435, EPI_ISL_3138436, EPI_ISL_3138437, EPI_ISL_3138438, EPI_ISL_3138439, EPI_ISL_3138440, EPI_ISL_3138441, EPI_ISL_3138442, EPI_ISL_3138443, EPI_ISL_3138444, EPI_ISL_3138445, EPI_ISL_3138446, EPI_ISL_3138447, EPI_ISL_3138448, EPI_ISL_3138450, EPI_ISL_3138451, EPI_ISL_3138452, EPI_ISL_3138453, EPI_ISL_3138454, EPI_ISL_3138455, EPI_ISL_3138456, EPI_ISL_3138457, EPI_ISL_3138458, EPI_ISL_3138459, EPI_ISL_3138460, EPI_ISL_3138461, EPI_ISL_3138462, EPI_ISL_3138463, EPI_ISL_3138464, EPI_ISL_3138465, EPI_ISL_3138466, EPI_ISL_3138467, EPI_ISL_3138468, EPI_ISL_3138469, EPI_ISL_3138470, EPI_ISL_3138471, EPI_ISL_3138472, EPI_ISL_3138473, EPI_ISL_3138474, EPI_ISL_3138475, EPI_ISL_3138476, EPI_ISL_3138477, EPI_ISL_3138478, EPI_ISL_3138479, EPI_ISL_3138480, EPI_ISL_3138481, EPI_ISL_3138482, EPI_ISL_3138483, EPI_ISL_3138484, EPI_ISL_3138485, EPI_ISL_3138486, EPI_ISL_3138487, EPI_ISL_3138488, EPI_ISL_3138489, EPI_ISL_3138490, EPI_ISL_3138491, EPI_ISL_3138492, EPI_ISL_3138493, EPI_ISL_3138494 | | | |
| see above | Dutch COVID-19 response team | National Institute for Public Health and the Environment (RIVM) | Adam Meijer, Harry Vennema, Dirk Eggink, Jeroen Cremer, Sharon van den Brink, Bas van der Veer, AnneMarie van den Brandt, Lisa Wijsman, Kim Freriks, Ryanne Jaarsma, Euníce Then, Lynn Aarts, Sanne Bos, Melissa van Tuil, Florian Zwagemaker, Dennis Schmitz, Annelies Kroneman, Karim Hajji, Chantal Reusken, on behalf of the national COVID-19 response team |
| EPI_ISL_3138495, EPI_ISL_3138496, EPI_ISL_3138497, EPI_ISL_3138498, EPI_ISL_3138499, EPI_ISL_3138500 | ZOL | Jessa | Severine Berden et al. on behalf of the Jessa_cmdLab |
| EPI_ISL_3138501, EPI_ISL_3138502 | LKO | Jessa | Severine Berden et al. on behalf of the Jessa_cmdLab |
| EPI_ISL_3138503, EPI_ISL_3138504, EPI_ISL_3138505, EPI_ISL_3138506, EPI_ISL_3138507, EPI_ISL_3138508, EPI_ISL_3138509, EPI_ISL_3138510, EPI_ISL_3138511, EPI_ISL_3138512 | ZOL | Jessa | Severine Berden et al. on behalf of the Jessa_cmdLab |
| EPI_ISL_3138515, EPI_ISL_3138516, EPI_ISL_3138517, EPI_ISL_3138518, EPI_ISL_3138519, EPI_ISL_3138520, EPI_ISL_3138521, EPI_ISL_3138522, EPI_ISL_3138523, EPI_ISL_3138524, EPI_ISL_3138525, EPI_ISL_3138526, EPI_ISL_3138527, EPI_ISL_3138528, EPI_ISL_3138529, EPI_ISL_3138530, EPI_ISL_3138531, EPI_ISL_3138532, EPI_ISL_3138533, EPI_ISL_3138534, EPI_ISL_3138535, EPI_ISL_3138536, EPI_ISL_3138537, EPI_ISL_3138538, EPI_ISL_3138539 | | | |
| see above | Jessa | Jessa | Severine Berden et al. on behalf of the Jessa_cmdLab |
| EPI_ISL_3138540, EPI_ISL_3138541, EPI_ISL_3138542, EPI_ISL_3138543, EPI_ISL_3138544, EPI_ISL_3138545, EPI_ISL_3138546 | Ospedale San Filippo Neri Microvir | University Campus Bio-Medico of Rome (UCBM) | Riva E., De Florio L., Fogolari M., Francesconi M., Lintas C., Veralli R., Angeletti S. |
| EPI_ISL_3138547 | AULSS 7 Pedemontana - Bassano | Istituto Zooprofilattico Sperimentale delle Venezie | Adelaide Milani, Alessia Schivo, Annalisa Salviato, Elisa Palumbo, Erika Giorgia Quaranta, Luca Tassoni, Ambra Pastori, Edoardo Giussani, Alice Fusaro, Isabella Monne, Calogero Terregino, Antonia Ricci |
| EPI_ISL_3138548, EPI_ISL_3138549, EPI_ISL_3138550, EPI_ISL_3138551, EPI_ISL_3138552 | AOUIVR | Istituto Zooprofilattico Sperimentale delle Venezie | Adelaide Milani, Alessia Schivo, Annalisa Salviato, Elisa Palumbo, Erika Giorgia Quaranta, Luca Tassoni, Ambra Pastori, Edoardo Giussani, Alice Fusaro, Isabella Monne, Calogero Terregino, Antonia Ricci |
| EPI_ISL_3138553, EPI_ISL_3138554, EPI_ISL_3138556 | AULSS 7 Pedemontana - Bassano | Istituto Zooprofilattico Sperimentale delle Venezie | Adelaide Milani, Alessia Schivo, Annalisa Salviato, Elisa Palumbo, Erika Giorgia Quaranta, Luca Tassoni, Ambra Pastori, Edoardo Giussani, Alice Fusaro, Isabella Monne, Calogero Terregino, Antonia Ricci |
| EPI_ISL_3138557, EPI_ISL_3138558, EPI_ISL_3138559 | AULSS 7 Pedemontana - Santorso | Istituto Zooprofilattico Sperimentale delle Venezie | Adelaide Milani, Alessia Schivo, Annalisa Salviato, Elisa Palumbo, Erika Giorgia Quaranta, Luca Tassoni, Ambra Pastori, Edoardo Giussani, Alice Fusaro, Isabella Monne, Calogero Terregino, Antonia Ricci |
| EPI_ISL_3138560 | AULSS 9 Scaligera - San Bonifacio | Istituto Zooprofilattico Sperimentale delle Venezie | Adelaide Milani, Alessia Schivo, Annalisa Salviato, Elisa Palumbo, Erika Giorgia Quaranta, Luca Tassoni, Ambra Pastori, Edoardo Giussani, Alice Fusaro, Isabella Monne, Calogero Terregino, Antonia Ricci |
| EPI_ISL_3138561, EPI_ISL_3138562 | AULSS 1 Dolomiti | Istituto Zooprofilattico Sperimentale delle Venezie | Adelaide Milani, Alessia Schivo, Annalisa Salviato, Elisa Palumbo, Erika Giorgia Quaranta, Luca Tassoni, Ambra Pastori, Edoardo Giussani, Alice Fusaro, Isabella Monne, Calogero Terregino, Antonia Ricci |
| EPI_ISL_3138563 | Ospedale San Filippo Neri Microvir | University Campus Bio-Medico of Rome (UCBM) | Riva E., De Florio L., Fogolari M., Francesconi M., Lintas C., Veralli R., Angeletti S. |
| EPI_ISL_3138564 | AULSS 1 Dolomiti | Istituto Zooprofilattico Sperimentale delle Venezie | Adelaide Milani, Alessia Schivo, Annalisa Salviato, Elisa Palumbo, Erika Giorgia Quaranta, Luca Tassoni, Ambra Pastori, Edoardo Giussani, Alice Fusaro, Isabella Monne, Calogero Terregino, Antonia Ricci |
| EPI_ISL_3138565 | Ospedale San Filippo Neri Microvir | University Campus Bio-Medico of Rome (UCBM) | Riva E., De Florio L., Fogolari M., Francesconi M., Lintas C., Veralli R., Angeletti S. |
| EPI_ISL_3138568, EPI_ISL_3138569, EPI_ISL_3138570, EPI_ISL_3138571, EPI_ISL_3138572, EPI_ISL_3138573, EPI_ISL_3138574, EPI_ISL_3138575, EPI_ISL_3138576, EPI_ISL_3138577, EPI_ISL_3138578, EPI_ISL_3138579, EPI_ISL_3138580, EPI_ISL_3138581, EPI_ISL_3138582, EPI_ISL_3138583, EPI_ISL_3138584, EPI_ISL_3138585, EPI_ISL_3138586, EPI_ISL_3138587, EPI_ISL_3138588, EPI_ISL_3138590, | | | |

| | | | |
|---|---|---|---|
| EPI_ISL_3138591, EPI_ISL_3138592, EPI_ISL_3138593, EPI_ISL_3138594, EPI_ISL_3138595, EPI_ISL_3138596, EPI_ISL_3138597, EPI_ISL_3138598, EPI_ISL_3138599, EPI_ISL_3138600, EPI_ISL_3138601, EPI_ISL_3138602, EPI_ISL_3138603, EPI_ISL_3138604, EPI_ISL_3138605, EPI_ISL_3138606, EPI_ISL_3138607 | | | |
| see above | Dipartimento di Medicina di Laboratorio, Azienda sanitaria universitaria Friuli Centrale (ASU FC) | Dipartimento di Medicina di Laboratorio, Azienda sanitaria universitaria Friuli Centrale (ASU FC) | Corrado Pipan, Catia Mio, Chiara Dal Secco, Stefania Marzinotto, Francesco Curcio |
| EPI_ISL_3138611 | AULSS 3 Venezia | UOSD Genetica e Citogenetica - Azienda ULSS 3 Serenissima; Istituto Zooprofilattico Sperimentale delle Venezie | Mosé Favarato, Elisa Squarcina, Laura Squarzon, Claudia Perini, Laura Bevilacqua, Adelaide Milani, Alessia Schivo, Annalisa Salviato, Elisa Palumbo, Erika Giorgia Quaranta, Luca Tassoni, Ambra Pastori, Edoardo Giussani, Alice Fusaro, Isabella Monne, Calogero Terregino, Antonia Ricci |
| EPI_ISL_3138612 | AULSS 9 Scaligera - Villafranca | UOSD Genetica e Citogenetica - Azienda ULSS 3 Serenissima; Istituto Zooprofilattico Sperimentale delle Venezie | Mosé Favarato, Elisa Squarcina, Laura Squarzon, Claudia Perini, Laura Bevilacqua, Adelaide Milani, Alessia Schivo, Annalisa Salviato, Elisa Palumbo, Erika Giorgia Quaranta, Luca Tassoni, Ambra Pastori, Edoardo Giussani, Alice Fusaro, Isabella Monne, Calogero Terregino, Antonia Ricci |
| EPI_ISL_3138613, EPI_ISL_3138614 | AULSS 4 Veneto Orientale | UOSD Genetica e Citogenetica - Azienda ULSS 3 Serenissima; Istituto Zooprofilattico Sperimentale delle Venezie | Mosé Favarato, Elisa Squarcina, Laura Squarzon, Claudia Perini, Laura Bevilacqua, Adelaide Milani, Alessia Schivo, Annalisa Salviato, Elisa Palumbo, Erika Giorgia Quaranta, Luca Tassoni, Ambra Pastori, Edoardo Giussani, Alice Fusaro, Isabella Monne, Calogero Terregino, Antonia Ricci |
| EPI_ISL_3138616, EPI_ISL_3138617, EPI_ISL_3138619, EPI_ISL_3138621, EPI_ISL_3138622, EPI_ISL_3138623, EPI_ISL_3138624 | University Campus Bio-Medico of Rome (UCBM) | University Campus Bio-Medico of Rome (UCBM) | Riva E., De Florio L., Fogolari M., Francesconi M., Lintas C., Veralli R., Angeletti S. |
| EPI_ISL_3138632, EPI_ISL_3138634, EPI_ISL_3138636, EPI_ISL_3138638, EPI_ISL_3138639 | Servizo de Microbioloxía. Complexo Hospitalario de Santiago de Compostela | Servizo de Microbioloxía. Complexo Hospitalario de Santiago de Compostela | Xosé Costa, Antonio Aguilera, Gema Barbeito, Amparo Coira, Carlos García_Riestra, Manuela Hernández, José Llovo, Teresa Lopez_Valiño, Laura Millán, Daniel Navarro, Rocío trastoy, Mercedes Treviño, María Luisa Pérez_del_Molino |
| EPI_ISL_3138642, EPI_ISL_3138643, EPI_ISL_3138644, EPI_ISL_3138645, EPI_ISL_3138646, EPI_ISL_3138647, EPI_ISL_3138648, EPI_ISL_3138649, EPI_ISL_3138650, EPI_ISL_3138651, EPI_ISL_3138652, EPI_ISL_3138653, EPI_ISL_3138654 | | | |
| see above | University Campus Bio-Medico of Rome (UCBM) | University Campus Bio-Medico of Rome (UCBM) | Riva E., De Florio L., Fogolari M., Francesconi M., Lintas C., Veralli R., Angeletti S. |
| EPI_ISL_3138741, EPI_ISL_3138742, EPI_ISL_3138743, EPI_ISL_3138744, EPI_ISL_3138745, EPI_ISL_3138746, EPI_ISL_3138747, EPI_ISL_3138748 | University Hospital Brno, OKMI | University Hospital Brno, CMBG | Matej Bezdicek, Jan Svaton, Martina Lengerova, Kristyna Dufkova |
| EPI_ISL_3138749 | Nemocnice TGM Hodonin | University Hospital Brno, CMBG | Matej Bezdicek, Jan Svaton, Martina Lengerova, Kristyna Dufkova |
| EPI_ISL_3138750, EPI_ISL_3138751 | Nemocnice Hodonin | University Hospital Brno, CMBG | Matej Bezdicek, Jan Svaton, Martina Lengerova, Kristyna Dufkova |
| EPI_ISL_3138752, EPI_ISL_3138753 | Cytogeneticka laborator Brno | University Hospital Brno, CMBG | Matej Bezdicek, Jan Svaton, Martina Lengerova, Kristyna Dufkova |
| EPI_ISL_3138754, EPI_ISL_3138755, EPI_ISL_3138756, EPI_ISL_3138757, EPI_ISL_3138758, EPI_ISL_3138759, EPI_ISL_3138760, EPI_ISL_3138761, EPI_ISL_3138762, EPI_ISL_3138766 | Laboratory Medicine and Molecular Diagnostics | Laboratory Medicine and Molecular Diagnostics | Dayakar Seetha, Heera R Pillai, Sanughosh Kalpothodi, Radhakrishnan R Nair |
| EPI_ISL_3138805 | RSPI Sulianti Saroso | National Institute of Health Research and Development | Arie Ardiansyah Nugraha, Hana Apsari Pawestri, Subangkit, Kartika Dewi Puspa, Hartanti Dian Ikawati, Krisna Pangesti, Triyani Soekarso, Fajar Nur Sulistiyohadi, Nelly Puspandari, Vivi Setiawaty |
| EPI_ISL_3138823, EPI_ISL_3138824, EPI_ISL_3138825, EPI_ISL_3138826, EPI_ISL_3138828, EPI_ISL_3138831 | GSI LAB | National Institute of Health Research and Development | Arie Ardiansyah Nugraha, Hana Apsari Pawestri, Subangkit, Kartika Dewi Puspa, Hartanti Dian Ikawati, Krisna Pangesti, Triyani Soekarso, Fajar Nur Sulistiyohadi, Nelly Puspandari, Vivi Setiawaty |
| EPI_ISL_3138843, EPI_ISL_3138844, EPI_ISL_3138845, EPI_ISL_3138846, EPI_ISL_3138847, EPI_ISL_3138848, EPI_ISL_3138849, EPI_ISL_3138850 | Labkes Provinsi Bali | National Institute of Health Research and Development | Arie Ardiansyah Nugraha, Hana Apsari Pawestri, Subangkit, Kartika Dewi Puspa, Hartanti Dian Ikawati, Krisna Pangesti, Triyani Soekarso, Fajar Nur Sulistiyohadi, Nelly Puspandari, Vivi Setiawaty |
| EPI_ISL_3138851 | RS Bali Jimbaran | National Institute of Health Research and Development | Arie Ardiansyah Nugraha, Hana Apsari Pawestri, Subangkit, Kartika Dewi Puspa, Hartanti Dian Ikawati, Krisna Pangesti, Triyani Soekarso, Fajar Nur Sulistiyohadi, Nelly Puspandari, Vivi Setiawaty |
| EPI_ISL_3138852 | Labkes Provinsi Bali | National Institute of Health Research and Development | Arie Ardiansyah Nugraha, Hana Apsari Pawestri, Subangkit, Kartika Dewi Puspa, Hartanti Dian Ikawati, Krisna Pangesti, Triyani Soekarso, Fajar Nur Sulistiyohadi, Nelly Puspandari, Vivi Setiawaty |
| EPI_ISL_3138853 | RS Balimed Karangasem | National Institute of Health Research and Development | Arie Ardiansyah Nugraha, Hana Apsari Pawestri, Subangkit, Kartika Dewi Puspa, Hartanti Dian Ikawati, Krisna Pangesti, Triyani Soekarso, Fajar Nur Sulistiyohadi, Nelly Puspandari, Vivi Setiawaty |
| EPI_ISL_3138854 | RS Balimed Karangasem | National Institute of Health Research and Development | Subangkit, Hana Apsari Pawestri, , Kartika Dewi Puspa, Arie Ardiansyah Nugraha, Hartanti Dian Ikawati, Fajar Nur Sulistiyohadi, Krisna Pangesti, Triyani Soekarso, Nelly Puspandari, Vivi Setiawaty |
| EPI_ISL_3138855, EPI_ISL_3138856, EPI_ISL_3138857, EPI_ISL_3138858, EPI_ISL_3138859 | Dinas Kesehatan Klungkung | National Institute of Health Research and Development | Subangkit, Hana Apsari Pawestri, , Kartika Dewi Puspa, Arie Ardiansyah Nugraha, Hartanti Dian Ikawati, Fajar Nur Sulistiyohadi, Krisna Pangesti, Triyani Soekarso, Nelly Puspandari, Vivi Setiawaty |
| EPI_ISL_3138860, EPI_ISL_3138861, EPI_ISL_3138862 | RS Jantung dan Pembulu Darah Harapan Kita | National Institute of Health Research and Development | Subangkit, Hana Apsari Pawestri, , Kartika Dewi Puspa, Arie Ardiansyah Nugraha, Hartanti Dian Ikawati, Fajar Nur Sulistiyohadi, Krisna Pangesti, Triyani Soekarso, Nelly Puspandari, Vivi Setiawaty |
| EPI_ISL_3138863, EPI_ISL_3138864, EPI_ISL_3138865 | RS Universitas Brawijaya | National Institute of Health Research and Development | Subangkit, Hana Apsari Pawestri, , Kartika Dewi Puspa, Arie Ardiansyah Nugraha, Hartanti Dian Ikawati, Fajar Nur Sulistiyohadi, Krisna Pangesti, Triyani Soekarso, Nelly Puspandari, Vivi Setiawaty |
| EPI_ISL_3138866 | BBTKL PP Banjarbaru | National Institute of Health Research and Development | Subangkit, Hana Apsari Pawestri, , Kartika Dewi Puspa, Arie Ardiansyah Nugraha, Hartanti Dian Ikawati, Fajar Nur Sulistiyohadi, Krisna Pangesti, Triyani Soekarso, Nelly Puspandari, Vivi Setiawaty |
| EPI_ISL_3138875, EPI_ISL_3138876, EPI_ISL_3138877, EPI_ISL_3138879, EPI_ISL_3138880, EPI_ISL_3138881, EPI_ISL_3138882, EPI_ISL_3138883, EPI_ISL_3138884 | BTKL PP Ambon | National Institute of Health Research and Development | Kartika Dewi Puspa, Hana Apsari Pawestri,Subangkit, Arie Ardiansyah Nugraha, Hartanti Dian Ikawati, Fajar Nur Sulistiyohadi, Krisna Pangesti, Triyani Soekarso, Nelly Puspandari, Vivi Setiawaty |
| EPI_ISL_3138885 | RSUD Mangusada Badung | National Institute of Health Research and Development | Kartika Dewi Puspa, Hana Apsari Pawestri,Subangkit, Arie Ardiansyah Nugraha, Hartanti Dian Ikawati, Fajar Nur Sulistiyohadi, Krisna Pangesti, Triyani Soekarso, Nelly Puspandari, Vivi Setiawaty |
| EPI_ISL_3138886 | BTKL PP Kelas 1 Manado | National Institute of Health Research and Development | Kartika Dewi Puspa, Hana Apsari Pawestri,Subangkit, Arie Ardiansyah Nugraha, Hartanti Dian Ikawati, Fajar Nur Sulistiyohadi, Krisna Pangesti, Triyani Soekarso, Nelly Puspandari, Vivi Setiawaty |
| EPI_ISL_3138888, EPI_ISL_3138889, EPI_ISL_3138890, EPI_ISL_3138891, EPI_ISL_3138892, EPI_ISL_3138893 | BTKL PP Kelas 1 Manado | National Institute of Health Research and Development | Vivi Setiawaty, Hana Apsari Pawestri, Subangkit, Kartika Dewi Puspa, Arie Ardiansyah Nugraha, Hartanti Dian Ikawati, Krisna Pangesti, Triyani Soekarso, Fajar Nur Sulistiyohadi, Nelly Puspandari |
| EPI_ISL_3138894, EPI_ISL_3138895, EPI_ISL_3138896 | Labkesda DKI Jakarta | National Institute of Health Research and Development | Vivi Setiawaty, Hana Apsari Pawestri, Subangkit, Kartika Dewi Puspa, Arie Ardiansyah Nugraha, Hartanti Dian Ikawati, Krisna Pangesti, Triyani Soekarso, Fajar Nur Sulistiyohadi, Nelly Puspandari |

| | | | |
|---|---|---|---|
| EPI_ISL_3138898, EPI_ISL_3138899, EPI_ISL_3138901, EPI_ISL_3138902, EPI_ISL_3138903, EPI_ISL_3138904, EPI_ISL_3138905, EPI_ISL_3138906 | Dinkes Provinsi Kalimantan Utara | National Institute of Health Research and Development | Vivi Setiawaty, Hana Apsari Pawestri, Subangkit, Kartika Dewi Puspa, Arie Ardiansyah Nugraha, Hartanti Dian Ikawati, Krisna Pangesti, Triyani Soekarso, Fajar Nur Sulistiyohadi, Nelly Puspandari |
| EPI_ISL_3138907, EPI_ISL_3138908 | RSUD Tarutung | National Institute of Health Research and Development | Vivi Setiawaty, Hana Apsari Pawestri, Subangkit, Kartika Dewi Puspa, Arie Ardiansyah Nugraha, Hartanti Dian Ikawati, Krisna Pangesti, Triyani Soekarso, Fajar Nur Sulistiyohadi, Nelly Puspandari |
| EPI_ISL_3138918, EPI_ISL_3138919, EPI_ISL_3138920, EPI_ISL_3138922, EPI_ISL_3138923, EPI_ISL_3138924 | RSUP Dr Kariadi Semarang | National Institute of Health Research and Development | Vivi Setiawaty, Hana Apsari Pawestri, Subangkit, Kartika Dewi Puspa, Arie Ardiansyah Nugraha, Hartanti Dian Ikawati, Krisna Pangesti, Triyani Soekarso, Fajar Nur Sulistiyohadi, Nelly Puspandari |
| EPI_ISL_3138942, EPI_ISL_3138943 | RSUD Tarakan | National Institute of Health Research and Development | Hana Apsari Pawestri, Subangkit, Arie Ardiansyah Nugraha, Kartika Dewi Puspa, Hartanti Dian Ikawati, Krisna Pangesti, Triyani Soekarso, Fajar Nur Sulistiyohadi, Nelly Puspandari, Vivi Setiawaty |
| EPI_ISL_3138944, EPI_ISL_3138945, EPI_ISL_3138946, EPI_ISL_3138948 | RSUP Nasional Dr. Cipto Mangunkusumo | National Institute of Health Research and Development | Hana Apsari Pawestri, Subangkit, Arie Ardiansyah Nugraha, Kartika Dewi Puspa, Hartanti Dian Ikawati, Krisna Pangesti, Triyani Soekarso, Fajar Nur Sulistiyohadi, Nelly Puspandari, Vivi Setiawaty |
| EPI_ISL_3138950 | Department of Virology and Immunology, University of Helsinki and Helsinki University Hospital, Huslab Finland | Department of Virology, Faculty of Medicine, University of Helsinki, Helsinki, Finland | Teemu Smura, Ravi Kant, Phuoc Truong, Hussein Alburkat, Hannimari Kallio-Kokko, Jenni Virtanen, Maija Suvanto, Essi Korhonen, Sari Hannula, Harri Kangas, Hanna Liimatainen, Satu Kurkela, Hanna Jarva, Maija Lappalainen, Pekka Ellonen, Olli Vapalahti |
| EPI_ISL_3138972 | Tivoli Fondazione Policlinico Universitario Agostino Gemelli IRCCS | Fondazione Policlinico Universitario Agostino Gemelli IRCCS | Istituto di Microbiologia e Virologia |
| EPI_ISL_3138973, EPI_ISL_3138974 | Lighthouse Lab in Glasgow | Wellcome Sanger Institute for the COVID-19 Genomics UK (COG-UK) Consortium | Harper VanSteenhouse, Yumi Kasai, David Gray, Carol Clugston, Anna Dominiczak and Alex Alderton, Roberto Amato, Jeffrey Barrett, Sonia Goncalves, Ewan Harrison, David K. Jackson, Ian Johnston, Dominic Kwiatkowski, Cordelia Langford, John Sillitoe on behalf of the Wellcome Sanger Institute COVID-19 Surveillance Team |
| EPI_ISL_3138975 | Berkshire and Surrey Pathology Services Lighthouse Laboratory | Wellcome Sanger Institute for the COVID-19 Genomics UK (COG-UK) Consortium | Berkshire and Surrey Pathology Services Lighthouse Laboratory and Alex Alderton, Roberto Amato, Jeffrey Barrett, Sonia Goncalves, Ewan Harrison, David K. Jackson, Ian Johnston, Dominic Kwiatkowski, Cordelia Langford, John Sillitoe on behalf of the Wellcome Sanger Institute COVID-19 Surveillance Team |
| EPI_ISL_3138976, EPI_ISL_3138977, EPI_ISL_3138978 | Lighthouse Lab in Glasgow | Wellcome Sanger Institute for the COVID-19 Genomics UK (COG-UK) Consortium | Harper VanSteenhouse, Yumi Kasai, David Gray, Carol Clugston, Anna Dominiczak and Alex Alderton, Roberto Amato, Jeffrey Barrett, Sonia Goncalves, Ewan Harrison, David K. Jackson, Ian Johnston, Dominic Kwiatkowski, Cordelia Langford, John Sillitoe on behalf of the Wellcome Sanger Institute COVID-19 Surveillance Team |
| EPI_ISL_3138979, EPI_ISL_3138980, EPI_ISL_3138981 | Berkshire and Surrey Pathology Services Lighthouse Laboratory | Wellcome Sanger Institute for the COVID-19 Genomics UK (COG-UK) Consortium | Berkshire and Surrey Pathology Services Lighthouse Laboratory and Alex Alderton, Roberto Amato, Jeffrey Barrett, Sonia Goncalves, Ewan Harrison, David K. Jackson, Ian Johnston, Dominic Kwiatkowski, Cordelia Langford, John Sillitoe on behalf of the Wellcome Sanger Institute COVID-19 Surveillance Team |
| EPI_ISL_3138982, EPI_ISL_3138983 | Lighthouse Lab in Glasgow | Wellcome Sanger Institute for the COVID-19 Genomics UK (COG-UK) Consortium | Harper VanSteenhouse, Yumi Kasai, David Gray, Carol Clugston, Anna Dominiczak and Alex Alderton, Roberto Amato, Jeffrey Barrett, Sonia Goncalves, Ewan Harrison, David K. Jackson, Ian Johnston, Dominic Kwiatkowski, Cordelia Langford, John Sillitoe on behalf of the Wellcome Sanger Institute COVID-19 Surveillance Team |
| EPI_ISL_3138984, EPI_ISL_3138985 | Berkshire and Surrey Pathology Services Lighthouse Laboratory | Wellcome Sanger Institute for the COVID-19 Genomics UK (COG-UK) Consortium | Berkshire and Surrey Pathology Services Lighthouse Laboratory and Alex Alderton, Roberto Amato, Jeffrey Barrett, Sonia Goncalves, Ewan Harrison, David K. Jackson, Ian Johnston, Dominic Kwiatkowski, Cordelia Langford, John Sillitoe on behalf of the Wellcome Sanger Institute COVID-19 Surveillance Team |
| EPI_ISL_3138986 | Lighthouse Lab in Glasgow | Wellcome Sanger Institute for the COVID-19 Genomics UK (COG-UK) Consortium | Harper VanSteenhouse, Yumi Kasai, David Gray, Carol Clugston, Anna Dominiczak and Alex Alderton, Roberto Amato, Jeffrey Barrett, Sonia Goncalves, Ewan Harrison, David K. Jackson, Ian Johnston, Dominic Kwiatkowski, Cordelia Langford, John Sillitoe on behalf of the Wellcome Sanger Institute COVID-19 Surveillance Team |
| EPI_ISL_3138987 | Berkshire and Surrey Pathology Services Lighthouse Laboratory | Wellcome Sanger Institute for the COVID-19 Genomics UK (COG-UK) Consortium | Berkshire and Surrey Pathology Services Lighthouse Laboratory and Alex Alderton, Roberto Amato, Jeffrey Barrett, Sonia Goncalves, Ewan Harrison, David K. Jackson, Ian Johnston, Dominic Kwiatkowski, Cordelia Langford, John Sillitoe on behalf of the Wellcome Sanger Institute COVID-19 Surveillance Team |
| EPI_ISL_3138988 | Lighthouse Lab in Glasgow | Wellcome Sanger Institute for the COVID-19 Genomics UK (COG-UK) Consortium | Harper VanSteenhouse, Yumi Kasai, David Gray, Carol Clugston, Anna Dominiczak and Alex Alderton, Roberto Amato, Jeffrey Barrett, Sonia Goncalves, Ewan Harrison, David K. Jackson, Ian Johnston, Dominic Kwiatkowski, Cordelia Langford, John Sillitoe on behalf of the Wellcome Sanger Institute COVID-19 Surveillance Team |
| EPI_ISL_3138989, EPI_ISL_3138990 | Berkshire and Surrey Pathology Services Lighthouse Laboratory | Wellcome Sanger Institute for the COVID-19 Genomics UK (COG-UK) Consortium | Berkshire and Surrey Pathology Services Lighthouse Laboratory and Alex Alderton, Roberto Amato, Jeffrey Barrett, Sonia Goncalves, Ewan Harrison, David K. Jackson, Ian Johnston, Dominic Kwiatkowski, Cordelia Langford, John Sillitoe on behalf of the Wellcome Sanger Institute COVID-19 Surveillance Team |
| EPI_ISL_3138991 | Lighthouse Lab in Glasgow | Wellcome Sanger Institute for the COVID-19 Genomics UK (COG-UK) Consortium | Harper VanSteenhouse, Yumi Kasai, David Gray, Carol Clugston, Anna Dominiczak and Alex Alderton, Roberto Amato, Jeffrey Barrett, Sonia Goncalves, Ewan Harrison, David K. Jackson, Ian Johnston, Dominic Kwiatkowski, Cordelia Langford, John Sillitoe on behalf of the Wellcome Sanger Institute COVID-19 Surveillance Team |
| EPI_ISL_3138992, EPI_ISL_3138993, EPI_ISL_3138994 | Berkshire and Surrey Pathology Services Lighthouse Laboratory | Wellcome Sanger Institute for the COVID-19 Genomics UK (COG-UK) Consortium | Berkshire and Surrey Pathology Services Lighthouse Laboratory and Alex Alderton, Roberto Amato, Jeffrey Barrett, Sonia Goncalves, Ewan Harrison, David K. Jackson, Ian Johnston, Dominic Kwiatkowski, Cordelia Langford, John Sillitoe on behalf of the Wellcome Sanger Institute COVID-19 Surveillance Team |
| EPI_ISL_3138995 | Lighthouse Lab in Glasgow | Wellcome Sanger Institute for the COVID-19 Genomics UK (COG-UK) Consortium | Harper VanSteenhouse, Yumi Kasai, David Gray, Carol Clugston, Anna Dominiczak and Alex Alderton, Roberto Amato, Jeffrey Barrett, Sonia Goncalves, Ewan Harrison, David K. Jackson, Ian Johnston, Dominic Kwiatkowski, Cordelia Langford, John Sillitoe on behalf of the Wellcome Sanger Institute COVID-19 Surveillance Team |
| EPI_ISL_3138996 | Berkshire and Surrey Pathology Services Lighthouse Laboratory | Wellcome Sanger Institute for the COVID-19 Genomics UK (COG-UK) Consortium | Berkshire and Surrey Pathology Services Lighthouse Laboratory and Alex Alderton, Roberto Amato, Jeffrey Barrett, Sonia Goncalves, Ewan Harrison, David K. Jackson, Ian Johnston, Dominic Kwiatkowski, Cordelia Langford, John Sillitoe on behalf of the Wellcome Sanger Institute COVID-19 Surveillance Team |
| EPI_ISL_3138997, EPI_ISL_3138998 | Lighthouse Lab in Glasgow | Wellcome Sanger Institute for the COVID-19 Genomics UK (COG-UK) Consortium | Harper VanSteenhouse, Yumi Kasai, David Gray, Carol Clugston, Anna Dominiczak and Alex Alderton, Roberto Amato, Jeffrey Barrett, Sonia Goncalves, Ewan Harrison, David K. Jackson, Ian Johnston, Dominic Kwiatkowski, Cordelia Langford, John Sillitoe on behalf of the Wellcome Sanger Institute COVID-19 Surveillance Team |
| EPI_ISL_3138999, EPI_ISL_3139000, EPI_ISL_3139001, EPI_ISL_3139002 | Berkshire and Surrey Pathology Services Lighthouse Laboratory | Wellcome Sanger Institute for the COVID-19 Genomics UK (COG-UK) Consortium | Berkshire and Surrey Pathology Services Lighthouse Laboratory and Alex Alderton, Roberto Amato, Jeffrey Barrett, Sonia Goncalves, Ewan Harrison, David K. Jackson, Ian Johnston, Dominic Kwiatkowski, Cordelia Langford, John Sillitoe on behalf of the Wellcome Sanger Institute COVID-19 Surveillance Team |
| EPI_ISL_3139003, EPI_ISL_3139004, EPI_ISL_3139005, EPI_ISL_3139006, EPI_ISL_3139007 | Lighthouse Lab in Glasgow | Wellcome Sanger Institute for the COVID-19 Genomics UK (COG-UK) Consortium | Harper VanSteenhouse, Yumi Kasai, David Gray, Carol Clugston, Anna Dominiczak and Alex Alderton, Roberto Amato, Jeffrey Barrett, Sonia Goncalves, Ewan Harrison, David K. Jackson, Ian Johnston, Dominic Kwiatkowski, Cordelia Langford, John Sillitoe on behalf of the Wellcome Sanger Institute COVID-19 Surveillance Team |
| EPI_ISL_3139008 | Berkshire and Surrey Pathology Services Lighthouse Laboratory | Wellcome Sanger Institute for the COVID-19 Genomics UK (COG-UK) Consortium | Berkshire and Surrey Pathology Services Lighthouse Laboratory and Alex Alderton, Roberto Amato, Jeffrey Barrett, Sonia Goncalves, Ewan Harrison, David K. Jackson, Ian Johnston, Dominic Kwiatkowski, Cordelia Langford, John Sillitoe on behalf of the Wellcome Sanger Institute COVID-19 Surveillance Team |
| EPI_ISL_3139009 | Lighthouse Lab in Glasgow | Wellcome Sanger Institute for the COVID-19 Genomics UK (COG-UK) Consortium | Harper VanSteenhouse, Yumi Kasai, David Gray, Carol Clugston, Anna Dominiczak and Alex Alderton, Roberto Amato, Jeffrey Barrett, Sonia Goncalves, Ewan Harrison, David K. Jackson, Ian Johnston, Dominic Kwiatkowski, Cordelia Langford, John Sillitoe on behalf of the Wellcome Sanger Institute COVID-19 Surveillance Team |
| EPI_ISL_3139010 | Berkshire and Surrey Pathology Services Lighthouse Laboratory | Wellcome Sanger Institute for the COVID-19 Genomics UK (COG-UK) Consortium | Berkshire and Surrey Pathology Services Lighthouse Laboratory and Alex Alderton, Roberto Amato, Jeffrey Barrett, Sonia Goncalves, Ewan Harrison, David K. Jackson, Ian Johnston, Dominic Kwiatkowski, Cordelia Langford, John Sillitoe on behalf of the Wellcome Sanger Institute COVID-19 Surveillance Team |
| EPI_ISL_3139011 | Lighthouse Lab in Glasgow | Wellcome Sanger Institute for the COVID-19 Genomics UK (COG-UK) Consortium | Harper VanSteenhouse, Yumi Kasai, David Gray, Carol Clugston, Anna Dominiczak and Alex Alderton, Roberto Amato, Jeffrey Barrett, Sonia Goncalves, Ewan Harrison, David K. Jackson, Ian Johnston, Dominic Kwiatkowski, Cordelia Langford, John Sillitoe on behalf of the Wellcome Sanger Institute COVID-19 Surveillance Team |
| EPI_ISL_3139012, EPI_ISL_3139013 | Berkshire and Surrey Pathology Services Lighthouse Laboratory | Wellcome Sanger Institute for the COVID-19 Genomics UK (COG-UK) Consortium | Berkshire and Surrey Pathology Services Lighthouse Laboratory and Alex Alderton, Roberto Amato, Jeffrey Barrett, Sonia Goncalves, Ewan Harrison, David K. Jackson, Ian Johnston, Dominic Kwiatkowski, Cordelia Langford, John Sillitoe on behalf of the Wellcome Sanger Institute COVID-19 Surveillance Team |
| EPI_ISL_3139014, EPI_ISL_3139015 | Lighthouse Lab in Glasgow | Wellcome Sanger Institute for the COVID-19 Genomics UK (COG-UK) Consortium | Harper VanSteenhouse, Yumi Kasai, David Gray, Carol Clugston, Anna Dominiczak and Alex Alderton, Roberto Amato, Jeffrey Barrett, Sonia Goncalves, Ewan Harrison, David K. Jackson, Ian Johnston, Dominic Kwiatkowski, Cordelia Langford, John Sillitoe on behalf of the Wellcome Sanger Institute COVID-19 Surveillance Team |
| EPI_ISL_3139016, EPI_ISL_3139017, EPI_ISL_3139018 | Berkshire and Surrey Pathology Services Lighthouse Laboratory | Wellcome Sanger Institute for the COVID-19 Genomics UK (COG-UK) Consortium | Berkshire and Surrey Pathology Services Lighthouse Laboratory and Alex Alderton, Roberto Amato, Jeffrey Barrett, Sonia Goncalves, Ewan Harrison, David K. Jackson, Ian Johnston, Dominic Kwiatkowski, Cordelia Langford, John Sillitoe on behalf of the Wellcome Sanger Institute COVID-19 Surveillance Team |
| EPI_ISL_3139019 | Lighthouse Lab in Glasgow | Wellcome Sanger Institute for the COVID-19 Genomics UK (COG-UK) Consortium | Harper VanSteenhouse, Yumi Kasai, David Gray, Carol Clugston, Anna Dominiczak and Alex Alderton, Roberto Amato, Jeffrey Barrett, Sonia Goncalves, Ewan Harrison, David K. Jackson, Ian Johnston, Dominic Kwiatkowski, Cordelia Langford, John Sillitoe on behalf of the Wellcome Sanger Institute COVID-19 Surveillance Team |
| EPI_ISL_3139020 | Berkshire and Surrey Pathology Services Lighthouse Laboratory | Wellcome Sanger Institute for the COVID-19 Genomics UK (COG-UK) Consortium | Berkshire and Surrey Pathology Services Lighthouse Laboratory and Alex Alderton, Roberto Amato, Jeffrey Barrett, Sonia Goncalves, Ewan Harrison, David K. Jackson, Ian Johnston, Dominic Kwiatkowski, Cordelia Langford, John Sillitoe on behalf of the Wellcome Sanger Institute COVID-19 Surveillance Team |
| EPI_ISL_3139021, EPI_ISL_3139022 | Lighthouse Lab in Glasgow | Wellcome Sanger Institute for the COVID-19 Genomics UK (COG-UK) Consortium | Harper VanSteenhouse, Yumi Kasai, David Gray, Carol Clugston, Anna Dominiczak and Alex Alderton, Roberto Amato, Jeffrey Barrett, Sonia Goncalves, Ewan Harrison, David K. Jackson, Ian Johnston, Dominic Kwiatkowski, Cordelia Langford, John Sillitoe on behalf of the Wellcome Sanger Institute COVID-19 Surveillance Team |
| EPI_ISL_3139023 | Berkshire and Surrey Pathology Services Lighthouse Laboratory | Wellcome Sanger Institute for the COVID-19 Genomics UK (COG-UK) Consortium | Berkshire and Surrey Pathology Services Lighthouse Laboratory and Alex Alderton, Roberto Amato, Jeffrey Barrett, Sonia Goncalves, Ewan Harrison, David K. Jackson, Ian Johnston, Dominic Kwiatkowski, Cordelia Langford, John Sillitoe on behalf of the Wellcome Sanger Institute COVID-19 Surveillance Team |
| EPI_ISL_3139024, EPI_ISL_3139025, EPI_ISL_3139026 | Lighthouse Lab in Glasgow | Wellcome Sanger Institute for the COVID-19 Genomics UK (COG-UK) Consortium | Harper VanSteenhouse, Yumi Kasai, David Gray, Carol Clugston, Anna Dominiczak and Alex Alderton, Roberto Amato, Jeffrey Barrett, Sonia Goncalves, Ewan Harrison, David K. Jackson, Ian Johnston, Dominic Kwiatkowski, Cordelia Langford, John Sillitoe on behalf of the Wellcome Sanger Institute COVID-19 Surveillance Team |

| | | | |
|---|---|---|---|
| EPI_ISL_3139027 | Berkshire and Surrey Pathology Services Lighthouse Laboratory | Wellcome Sanger Institute for the COVID-19 Genomics UK (COG-UK) Consortium | Berkshire and Surrey Pathology Services Lighthouse Laboratory and Alex Alderton, Roberto Amato, Jeffrey Barrett, Sonia Goncalves, Ewan Harrison, David K. Jackson, Ian Johnston, Dominic Kwiatkowski, Cordelia Langford, John Sillitoe on behalf of the Wellcome Sanger Institute COVID-19 Surveillance Team |
| EPI_ISL_3139028 | Lighthouse Lab in Glasgow | Wellcome Sanger Institute for the COVID-19 Genomics UK (COG-UK) Consortium | Harper VanSteenhouse, Yumi Kasai, David Gray, Carol Clugston, Anna Dominiczak and Alex Alderton, Roberto Amato, Jeffrey Barrett, Sonia Goncalves, Ewan Harrison, David K. Jackson, Ian Johnston, Dominic Kwiatkowski, Cordelia Langford, John Sillitoe on behalf of the Wellcome Sanger Institute COVID-19 Surveillance Team |
| EPI_ISL_3139029 | Berkshire and Surrey Pathology Services Lighthouse Laboratory | Wellcome Sanger Institute for the COVID-19 Genomics UK (COG-UK) Consortium | Berkshire and Surrey Pathology Services Lighthouse Laboratory and Alex Alderton, Roberto Amato, Jeffrey Barrett, Sonia Goncalves, Ewan Harrison, David K. Jackson, Ian Johnston, Dominic Kwiatkowski, Cordelia Langford, John Sillitoe on behalf of the Wellcome Sanger Institute COVID-19 Surveillance Team |
| EPI_ISL_3139030 | Lighthouse Lab in Glasgow | Wellcome Sanger Institute for the COVID-19 Genomics UK (COG-UK) Consortium | Harper VanSteenhouse, Yumi Kasai, David Gray, Carol Clugston, Anna Dominiczak and Alex Alderton, Roberto Amato, Jeffrey Barrett, Sonia Goncalves, Ewan Harrison, David K. Jackson, Ian Johnston, Dominic Kwiatkowski, Cordelia Langford, John Sillitoe on behalf of the Wellcome Sanger Institute COVID-19 Surveillance Team |
| EPI_ISL_3139031 | Berkshire and Surrey Pathology Services Lighthouse Laboratory | Wellcome Sanger Institute for the COVID-19 Genomics UK (COG-UK) Consortium | Berkshire and Surrey Pathology Services Lighthouse Laboratory and Alex Alderton, Roberto Amato, Jeffrey Barrett, Sonia Goncalves, Ewan Harrison, David K. Jackson, Ian Johnston, Dominic Kwiatkowski, Cordelia Langford, John Sillitoe on behalf of the Wellcome Sanger Institute COVID-19 Surveillance Team |
| EPI_ISL_3139032 | Lighthouse Lab in Glasgow | Wellcome Sanger Institute for the COVID-19 Genomics UK (COG-UK) Consortium | Harper VanSteenhouse, Yumi Kasai, David Gray, Carol Clugston, Anna Dominiczak and Alex Alderton, Roberto Amato, Jeffrey Barrett, Sonia Goncalves, Ewan Harrison, David K. Jackson, Ian Johnston, Dominic Kwiatkowski, Cordelia Langford, John Sillitoe on behalf of the Wellcome Sanger Institute COVID-19 Surveillance Team |
| EPI_ISL_3139033, EPI_ISL_3139034, EPI_ISL_3139035, EPI_ISL_3139036 | Berkshire and Surrey Pathology Services Lighthouse Laboratory | Wellcome Sanger Institute for the COVID-19 Genomics UK (COG-UK) Consortium | Berkshire and Surrey Pathology Services Lighthouse Laboratory and Alex Alderton, Roberto Amato, Jeffrey Barrett, Sonia Goncalves, Ewan Harrison, David K. Jackson, Ian Johnston, Dominic Kwiatkowski, Cordelia Langford, John Sillitoe on behalf of the Wellcome Sanger Institute COVID-19 Surveillance Team |
| EPI_ISL_3139037, EPI_ISL_3139038 | Lighthouse Lab in Glasgow | Wellcome Sanger Institute for the COVID-19 Genomics UK (COG-UK) Consortium | Harper VanSteenhouse, Yumi Kasai, David Gray, Carol Clugston, Anna Dominiczak and Alex Alderton, Roberto Amato, Jeffrey Barrett, Sonia Goncalves, Ewan Harrison, David K. Jackson, Ian Johnston, Dominic Kwiatkowski, Cordelia Langford, John Sillitoe on behalf of the Wellcome Sanger Institute COVID-19 Surveillance Team |
| EPI_ISL_3139039, EPI_ISL_3139040 | Berkshire and Surrey Pathology Services Lighthouse Laboratory | Wellcome Sanger Institute for the COVID-19 Genomics UK (COG-UK) Consortium | Berkshire and Surrey Pathology Services Lighthouse Laboratory and Alex Alderton, Roberto Amato, Jeffrey Barrett, Sonia Goncalves, Ewan Harrison, David K. Jackson, Ian Johnston, Dominic Kwiatkowski, Cordelia Langford, John Sillitoe on behalf of the Wellcome Sanger Institute COVID-19 Surveillance Team |
| EPI_ISL_3139041, EPI_ISL_3139042, EPI_ISL_3139043, EPI_ISL_3139044 | Lighthouse Lab in Glasgow | Wellcome Sanger Institute for the COVID-19 Genomics UK (COG-UK) Consortium | Harper VanSteenhouse, Yumi Kasai, David Gray, Carol Clugston, Anna Dominiczak and Alex Alderton, Roberto Amato, Jeffrey Barrett, Sonia Goncalves, Ewan Harrison, David K. Jackson, Ian Johnston, Dominic Kwiatkowski, Cordelia Langford, John Sillitoe on behalf of the Wellcome Sanger Institute COVID-19 Surveillance Team |
| EPI_ISL_3139045 | Berkshire and Surrey Pathology Services Lighthouse Laboratory | Wellcome Sanger Institute for the COVID-19 Genomics UK (COG-UK) Consortium | Berkshire and Surrey Pathology Services Lighthouse Laboratory and Alex Alderton, Roberto Amato, Jeffrey Barrett, Sonia Goncalves, Ewan Harrison, David K. Jackson, Ian Johnston, Dominic Kwiatkowski, Cordelia Langford, John Sillitoe on behalf of the Wellcome Sanger Institute COVID-19 Surveillance Team |
| EPI_ISL_3139046, EPI_ISL_3139047 | Lighthouse Lab in Glasgow | Wellcome Sanger Institute for the COVID-19 Genomics UK (COG-UK) Consortium | Harper VanSteenhouse, Yumi Kasai, David Gray, Carol Clugston, Anna Dominiczak and Alex Alderton, Roberto Amato, Jeffrey Barrett, Sonia Goncalves, Ewan Harrison, David K. Jackson, Ian Johnston, Dominic Kwiatkowski, Cordelia Langford, John Sillitoe on behalf of the Wellcome Sanger Institute COVID-19 Surveillance Team |
| EPI_ISL_3139048, EPI_ISL_3139049 | Berkshire and Surrey Pathology Services Lighthouse Laboratory | Wellcome Sanger Institute for the COVID-19 Genomics UK (COG-UK) Consortium | Berkshire and Surrey Pathology Services Lighthouse Laboratory and Alex Alderton, Roberto Amato, Jeffrey Barrett, Sonia Goncalves, Ewan Harrison, David K. Jackson, Ian Johnston, Dominic Kwiatkowski, Cordelia Langford, John Sillitoe on behalf of the Wellcome Sanger Institute COVID-19 Surveillance Team |
| EPI_ISL_3139050 | Lighthouse Lab in Glasgow | Wellcome Sanger Institute for the COVID-19 Genomics UK (COG-UK) Consortium | Harper VanSteenhouse, Yumi Kasai, David Gray, Carol Clugston, Anna Dominiczak and Alex Alderton, Roberto Amato, Jeffrey Barrett, Sonia Goncalves, Ewan Harrison, David K. Jackson, Ian Johnston, Dominic Kwiatkowski, Cordelia Langford, John Sillitoe on behalf of the Wellcome Sanger Institute COVID-19 Surveillance Team |
| EPI_ISL_3139051 | Berkshire and Surrey Pathology Services Lighthouse Laboratory | Wellcome Sanger Institute for the COVID-19 Genomics UK (COG-UK) Consortium | Berkshire and Surrey Pathology Services Lighthouse Laboratory and Alex Alderton, Roberto Amato, Jeffrey Barrett, Sonia Goncalves, Ewan Harrison, David K. Jackson, Ian Johnston, Dominic Kwiatkowski, Cordelia Langford, John Sillitoe on behalf of the Wellcome Sanger Institute COVID-19 Surveillance Team |
| EPI_ISL_3139052, EPI_ISL_3139053 | Lighthouse Lab in Glasgow | Wellcome Sanger Institute for the COVID-19 Genomics UK (COG-UK) Consortium | Harper VanSteenhouse, Yumi Kasai, David Gray, Carol Clugston, Anna Dominiczak and Alex Alderton, Roberto Amato, Jeffrey Barrett, Sonia Goncalves, Ewan Harrison, David K. Jackson, Ian Johnston, Dominic Kwiatkowski, Cordelia Langford, John Sillitoe on behalf of the Wellcome Sanger Institute COVID-19 Surveillance Team |
| EPI_ISL_3139054, EPI_ISL_3139055 | Berkshire and Surrey Pathology Services Lighthouse Laboratory | Wellcome Sanger Institute for the COVID-19 Genomics UK (COG-UK) Consortium | Berkshire and Surrey Pathology Services Lighthouse Laboratory and Alex Alderton, Roberto Amato, Jeffrey Barrett, Sonia Goncalves, Ewan Harrison, David K. Jackson, Ian Johnston, Dominic Kwiatkowski, Cordelia Langford, John Sillitoe on behalf of the Wellcome Sanger Institute COVID-19 Surveillance Team |
| EPI_ISL_3139056 | Lighthouse Lab in Glasgow | Wellcome Sanger Institute for the COVID-19 Genomics UK (COG-UK) Consortium | Harper VanSteenhouse, Yumi Kasai, David Gray, Carol Clugston, Anna Dominiczak and Alex Alderton, Roberto Amato, Jeffrey Barrett, Sonia Goncalves, Ewan Harrison, David K. Jackson, Ian Johnston, Dominic Kwiatkowski, Cordelia Langford, John Sillitoe on behalf of the Wellcome Sanger Institute COVID-19 Surveillance Team |
| EPI_ISL_3139057, EPI_ISL_3139058 | Berkshire and Surrey Pathology Services Lighthouse Laboratory | Wellcome Sanger Institute for the COVID-19 Genomics UK (COG-UK) Consortium | Berkshire and Surrey Pathology Services Lighthouse Laboratory and Alex Alderton, Roberto Amato, Jeffrey Barrett, Sonia Goncalves, Ewan Harrison, David K. Jackson, Ian Johnston, Dominic Kwiatkowski, Cordelia Langford, John Sillitoe on behalf of the Wellcome Sanger Institute COVID-19 Surveillance Team |
| EPI_ISL_3139059, EPI_ISL_3139060, EPI_ISL_3139061, EPI_ISL_3139062, EPI_ISL_3139063, EPI_ISL_3139064, EPI_ISL_3139065 | Lighthouse Lab in Glasgow | Wellcome Sanger Institute for the COVID-19 Genomics UK (COG-UK) Consortium | Harper VanSteenhouse, Yumi Kasai, David Gray, Carol Clugston, Anna Dominiczak and Alex Alderton, Roberto Amato, Jeffrey Barrett, Sonia Goncalves, Ewan Harrison, David K. Jackson, Ian Johnston, Dominic Kwiatkowski, Cordelia Langford, John Sillitoe on behalf of the Wellcome Sanger Institute COVID-19 Surveillance Team |
| EPI_ISL_3139066, EPI_ISL_3139067, EPI_ISL_3139068, EPI_ISL_3139069, EPI_ISL_3139070 | Berkshire and Surrey Pathology Services Lighthouse Laboratory | Wellcome Sanger Institute for the COVID-19 Genomics UK (COG-UK) Consortium | Berkshire and Surrey Pathology Services Lighthouse Laboratory and Alex Alderton, Roberto Amato, Jeffrey Barrett, Sonia Goncalves, Ewan Harrison, David K. Jackson, Ian Johnston, Dominic Kwiatkowski, Cordelia Langford, John Sillitoe on behalf of the Wellcome Sanger Institute COVID-19 Surveillance Team |
| EPI_ISL_3139071, EPI_ISL_3139072, EPI_ISL_3139073 | Lighthouse Lab in Glasgow | Wellcome Sanger Institute for the COVID-19 Genomics UK (COG-UK) Consortium | Harper VanSteenhouse, Yumi Kasai, David Gray, Carol Clugston, Anna Dominiczak and Alex Alderton, Roberto Amato, Jeffrey Barrett, Sonia Goncalves, Ewan Harrison, David K. Jackson, Ian Johnston, Dominic Kwiatkowski, Cordelia Langford, John Sillitoe on behalf of the Wellcome Sanger Institute COVID-19 Surveillance Team |
| EPI_ISL_3139074 | Berkshire and Surrey Pathology Services Lighthouse Laboratory | Wellcome Sanger Institute for the COVID-19 Genomics UK (COG-UK) Consortium | Berkshire and Surrey Pathology Services Lighthouse Laboratory and Alex Alderton, Roberto Amato, Jeffrey Barrett, Sonia Goncalves, Ewan Harrison, David K. Jackson, Ian Johnston, Dominic Kwiatkowski, Cordelia Langford, John Sillitoe on behalf of the Wellcome Sanger Institute COVID-19 Surveillance Team |
| EPI_ISL_3139075 | Lighthouse Lab in Glasgow | Wellcome Sanger Institute for the COVID-19 Genomics UK (COG-UK) Consortium | Harper VanSteenhouse, Yumi Kasai, David Gray, Carol Clugston, Anna Dominiczak and Alex Alderton, Roberto Amato, Jeffrey Barrett, Sonia Goncalves, Ewan Harrison, David K. Jackson, Ian Johnston, Dominic Kwiatkowski, Cordelia Langford, John Sillitoe on behalf of the Wellcome Sanger Institute COVID-19 Surveillance Team |
| EPI_ISL_3139076 | Berkshire and Surrey Pathology Services Lighthouse Laboratory | Wellcome Sanger Institute for the COVID-19 Genomics UK (COG-UK) Consortium | Berkshire and Surrey Pathology Services Lighthouse Laboratory and Alex Alderton, Roberto Amato, Jeffrey Barrett, Sonia Goncalves, Ewan Harrison, David K. Jackson, Ian Johnston, Dominic Kwiatkowski, Cordelia Langford, John Sillitoe on behalf of the Wellcome Sanger Institute COVID-19 Surveillance Team |
| EPI_ISL_3139077, EPI_ISL_3139078, EPI_ISL_3139079, EPI_ISL_3139080, EPI_ISL_3139081, EPI_ISL_3139082, EPI_ISL_3139083, EPI_ISL_3139084 | Lighthouse Lab in Glasgow | Wellcome Sanger Institute for the COVID-19 Genomics UK (COG-UK) Consortium | Harper VanSteenhouse, Yumi Kasai, David Gray, Carol Clugston, Anna Dominiczak and Alex Alderton, Roberto Amato, Jeffrey Barrett, Sonia Goncalves, Ewan Harrison, David K. Jackson, Ian Johnston, Dominic Kwiatkowski, Cordelia Langford, John Sillitoe on behalf of the Wellcome Sanger Institute COVID-19 Surveillance Team |
| EPI_ISL_3139085 | Berkshire and Surrey Pathology Services Lighthouse Laboratory | Wellcome Sanger Institute for the COVID-19 Genomics UK (COG-UK) Consortium | Berkshire and Surrey Pathology Services Lighthouse Laboratory and Alex Alderton, Roberto Amato, Jeffrey Barrett, Sonia Goncalves, Ewan Harrison, David K. Jackson, Ian Johnston, Dominic Kwiatkowski, Cordelia Langford, John Sillitoe on behalf of the Wellcome Sanger Institute COVID-19 Surveillance Team |
| EPI_ISL_3139086, EPI_ISL_3139087, EPI_ISL_3139088, EPI_ISL_3139089 | Lighthouse Lab in Glasgow | Wellcome Sanger Institute for the COVID-19 Genomics UK (COG-UK) Consortium | Harper VanSteenhouse, Yumi Kasai, David Gray, Carol Clugston, Anna Dominiczak and Alex Alderton, Roberto Amato, Jeffrey Barrett, Sonia Goncalves, Ewan Harrison, David K. Jackson, Ian Johnston, Dominic Kwiatkowski, Cordelia Langford, John Sillitoe on behalf of the Wellcome Sanger Institute COVID-19 Surveillance Team |
| EPI_ISL_3139090 | Berkshire and Surrey Pathology Services Lighthouse Laboratory | Wellcome Sanger Institute for the COVID-19 Genomics UK (COG-UK) Consortium | Berkshire and Surrey Pathology Services Lighthouse Laboratory and Alex Alderton, Roberto Amato, Jeffrey Barrett, Sonia Goncalves, Ewan Harrison, David K. Jackson, Ian Johnston, Dominic Kwiatkowski, Cordelia Langford, John Sillitoe on behalf of the Wellcome Sanger Institute COVID-19 Surveillance Team |
| EPI_ISL_3139091, EPI_ISL_3139092 | Lighthouse Lab in Glasgow | Wellcome Sanger Institute for the COVID-19 Genomics UK (COG-UK) Consortium | Harper VanSteenhouse, Yumi Kasai, David Gray, Carol Clugston, Anna Dominiczak and Alex Alderton, Roberto Amato, Jeffrey Barrett, Sonia Goncalves, Ewan Harrison, David K. Jackson, Ian Johnston, Dominic Kwiatkowski, Cordelia Langford, John Sillitoe on behalf of the Wellcome Sanger Institute COVID-19 Surveillance Team |
| EPI_ISL_3139093 | Berkshire and Surrey Pathology Services Lighthouse Laboratory | Wellcome Sanger Institute for the COVID-19 Genomics UK (COG-UK) Consortium | Berkshire and Surrey Pathology Services Lighthouse Laboratory and Alex Alderton, Roberto Amato, Jeffrey Barrett, Sonia Goncalves, Ewan Harrison, David K. Jackson, Ian Johnston, Dominic Kwiatkowski, Cordelia Langford, John Sillitoe on behalf of the Wellcome Sanger Institute COVID-19 Surveillance Team |
| EPI_ISL_3139094 | Lighthouse Lab in Glasgow | Wellcome Sanger Institute for the COVID-19 Genomics UK (COG-UK) Consortium | Harper VanSteenhouse, Yumi Kasai, David Gray, Carol Clugston, Anna Dominiczak and Alex Alderton, Roberto Amato, Jeffrey Barrett, Sonia Goncalves, Ewan Harrison, David K. Jackson, Ian Johnston, Dominic Kwiatkowski, Cordelia Langford, John Sillitoe on behalf of the Wellcome Sanger Institute COVID-19 Surveillance Team |
| EPI_ISL_3139095 | Berkshire and Surrey Pathology Services Lighthouse Laboratory | Wellcome Sanger Institute for the COVID-19 Genomics UK (COG-UK) Consortium | Berkshire and Surrey Pathology Services Lighthouse Laboratory and Alex Alderton, Roberto Amato, Jeffrey Barrett, Sonia Goncalves, Ewan Harrison, David K. Jackson, Ian Johnston, Dominic Kwiatkowski, Cordelia Langford, John Sillitoe on behalf of the Wellcome Sanger Institute COVID-19 Surveillance Team |
| EPI_ISL_3139096 | Lighthouse Lab in Glasgow | Wellcome Sanger Institute for the COVID-19 Genomics UK (COG-UK) Consortium | Harper VanSteenhouse, Yumi Kasai, David Gray, Carol Clugston, Anna Dominiczak and Alex Alderton, Roberto Amato, Jeffrey Barrett, Sonia Goncalves, Ewan Harrison, David K. Jackson, Ian Johnston, Dominic Kwiatkowski, Cordelia Langford, John Sillitoe on behalf of the Wellcome Sanger Institute COVID-19 Surveillance Team |
| EPI_ISL_3139097 | Berkshire and Surrey Pathology Services Lighthouse Laboratory | Wellcome Sanger Institute for the COVID-19 Genomics UK (COG-UK) Consortium | Berkshire and Surrey Pathology Services Lighthouse Laboratory and Alex Alderton, Roberto Amato, Jeffrey Barrett, Sonia Goncalves, Ewan Harrison, David K. Jackson, Ian Johnston, Dominic Kwiatkowski, Cordelia Langford, John Sillitoe on behalf of the Wellcome Sanger Institute COVID-19 Surveillance Team |
| EPI_ISL_3139098, EPI_ISL_3139099 | Lighthouse Lab in Glasgow | Wellcome Sanger Institute for the COVID-19 Genomics UK (COG-UK) Consortium | Harper VanSteenhouse, Yumi Kasai, David Gray, Carol Clugston, Anna Dominiczak and Alex Alderton, Roberto Amato, Jeffrey Barrett, Sonia Goncalves, Ewan Harrison, David K. Jackson, Ian Johnston, Dominic Kwiatkowski, Cordelia Langford, John Sillitoe on behalf of the Wellcome Sanger Institute COVID-19 Surveillance Team |
| EPI_ISL_3139100, | Berkshire and Surrey Pathology Services Lighthouse Laboratory | Wellcome Sanger Institute for the COVID-19 Genomics UK (COG-UK) | Berkshire and Surrey Pathology Services Lighthouse Laboratory and Alex Alderton, Roberto Amato, Jeffrey Barrett, Sonia Goncalves, Ewan Harrison, David K. Jackson, Ian Johnston, Dominic Kwiatkowski, Cordelia Langford, John Sillitoe on behalf |

| | | | |
|---|---|---|---|
| EPI_ISL_3139101, EPI_ISL_3139102 | | Consortium | of the Wellcome Sanger Institute COVID-19 Surveillance Team |
| EPI_ISL_3139103 | Lighthouse Lab in Glasgow | Wellcome Sanger Institute for the COVID-19 Genomics UK (COG-UK) Consortium | Harper VanSteenhouse, Yumi Kasai, David Gray, Carol Clugston, Anna Dominiczak and Alex Alderton, Roberto Amato, Jeffrey Barrett, Sonia Goncalves, Ewan Harrison, David K. Jackson, Ian Johnston, Dominic Kwiatkowski, Cordelia Langford, John Sillitoe on behalf of the Wellcome Sanger Institute COVID-19 Surveillance Team |
| EPI_ISL_3139104, EPI_ISL_3139105, EPI_ISL_3139106 | Berkshire and Surrey Pathology Services Lighthouse Laboratory | Wellcome Sanger Institute for the COVID-19 Genomics UK (COG-UK) Consortium | Berkshire and Surrey Pathology Services Lighthouse Laboratory and Alex Alderton, Roberto Amato, Jeffrey Barrett, Sonia Goncalves, Ewan Harrison, David K. Jackson, Ian Johnston, Dominic Kwiatkowski, Cordelia Langford, John Sillitoe on behalf of the Wellcome Sanger Institute COVID-19 Surveillance Team |
| EPI_ISL_3139107, EPI_ISL_3139108, EPI_ISL_3139109 | Lighthouse Lab in Glasgow | Wellcome Sanger Institute for the COVID-19 Genomics UK (COG-UK) Consortium | Harper VanSteenhouse, Yumi Kasai, David Gray, Carol Clugston, Anna Dominiczak and Alex Alderton, Roberto Amato, Jeffrey Barrett, Sonia Goncalves, Ewan Harrison, David K. Jackson, Ian Johnston, Dominic Kwiatkowski, Cordelia Langford, John Sillitoe on behalf of the Wellcome Sanger Institute COVID-19 Surveillance Team |
| EPI_ISL_3139110 | Berkshire and Surrey Pathology Services Lighthouse Laboratory | Wellcome Sanger Institute for the COVID-19 Genomics UK (COG-UK) Consortium | Berkshire and Surrey Pathology Services Lighthouse Laboratory and Alex Alderton, Roberto Amato, Jeffrey Barrett, Sonia Goncalves, Ewan Harrison, David K. Jackson, Ian Johnston, Dominic Kwiatkowski, Cordelia Langford, John Sillitoe on behalf of the Wellcome Sanger Institute COVID-19 Surveillance Team |
| EPI_ISL_3139111, EPI_ISL_3139112 | Lighthouse Lab in Glasgow | Wellcome Sanger Institute for the COVID-19 Genomics UK (COG-UK) Consortium | Harper VanSteenhouse, Yumi Kasai, David Gray, Carol Clugston, Anna Dominiczak and Alex Alderton, Roberto Amato, Jeffrey Barrett, Sonia Goncalves, Ewan Harrison, David K. Jackson, Ian Johnston, Dominic Kwiatkowski, Cordelia Langford, John Sillitoe on behalf of the Wellcome Sanger Institute COVID-19 Surveillance Team |
| EPI_ISL_3139113, EPI_ISL_3139114, EPI_ISL_3139115 | Berkshire and Surrey Pathology Services Lighthouse Laboratory | Wellcome Sanger Institute for the COVID-19 Genomics UK (COG-UK) Consortium | Berkshire and Surrey Pathology Services Lighthouse Laboratory and Alex Alderton, Roberto Amato, Jeffrey Barrett, Sonia Goncalves, Ewan Harrison, David K. Jackson, Ian Johnston, Dominic Kwiatkowski, Cordelia Langford, John Sillitoe on behalf of the Wellcome Sanger Institute COVID-19 Surveillance Team |
| EPI_ISL_3139116, EPI_ISL_3139117, EPI_ISL_3139118 | Lighthouse Lab in Glasgow | Wellcome Sanger Institute for the COVID-19 Genomics UK (COG-UK) Consortium | Harper VanSteenhouse, Yumi Kasai, David Gray, Carol Clugston, Anna Dominiczak and Alex Alderton, Roberto Amato, Jeffrey Barrett, Sonia Goncalves, Ewan Harrison, David K. Jackson, Ian Johnston, Dominic Kwiatkowski, Cordelia Langford, John Sillitoe on behalf of the Wellcome Sanger Institute COVID-19 Surveillance Team |
| EPI_ISL_3139119, EPI_ISL_3139120, EPI_ISL_3139121, EPI_ISL_3139122 | Berkshire and Surrey Pathology Services Lighthouse Laboratory | Wellcome Sanger Institute for the COVID-19 Genomics UK (COG-UK) Consortium | Berkshire and Surrey Pathology Services Lighthouse Laboratory and Alex Alderton, Roberto Amato, Jeffrey Barrett, Sonia Goncalves, Ewan Harrison, David K. Jackson, Ian Johnston, Dominic Kwiatkowski, Cordelia Langford, John Sillitoe on behalf of the Wellcome Sanger Institute COVID-19 Surveillance Team |
| EPI_ISL_3139123 | Lighthouse Lab in Glasgow | Wellcome Sanger Institute for the COVID-19 Genomics UK (COG-UK) Consortium | Harper VanSteenhouse, Yumi Kasai, David Gray, Carol Clugston, Anna Dominiczak and Alex Alderton, Roberto Amato, Jeffrey Barrett, Sonia Goncalves, Ewan Harrison, David K. Jackson, Ian Johnston, Dominic Kwiatkowski, Cordelia Langford, John Sillitoe on behalf of the Wellcome Sanger Institute COVID-19 Surveillance Team |
| EPI_ISL_3139124 | Berkshire and Surrey Pathology Services Lighthouse Laboratory | Wellcome Sanger Institute for the COVID-19 Genomics UK (COG-UK) Consortium | Berkshire and Surrey Pathology Services Lighthouse Laboratory and Alex Alderton, Roberto Amato, Jeffrey Barrett, Sonia Goncalves, Ewan Harrison, David K. Jackson, Ian Johnston, Dominic Kwiatkowski, Cordelia Langford, John Sillitoe on behalf of the Wellcome Sanger Institute COVID-19 Surveillance Team |
| EPI_ISL_3139125, EPI_ISL_3139126, EPI_ISL_3139127 | Lighthouse Lab in Glasgow | Wellcome Sanger Institute for the COVID-19 Genomics UK (COG-UK) Consortium | Harper VanSteenhouse, Yumi Kasai, David Gray, Carol Clugston, Anna Dominiczak and Alex Alderton, Roberto Amato, Jeffrey Barrett, Sonia Goncalves, Ewan Harrison, David K. Jackson, Ian Johnston, Dominic Kwiatkowski, Cordelia Langford, John Sillitoe on behalf of the Wellcome Sanger Institute COVID-19 Surveillance Team |
| EPI_ISL_3139128 | Berkshire and Surrey Pathology Services Lighthouse Laboratory | Wellcome Sanger Institute for the COVID-19 Genomics UK (COG-UK) Consortium | Berkshire and Surrey Pathology Services Lighthouse Laboratory and Alex Alderton, Roberto Amato, Jeffrey Barrett, Sonia Goncalves, Ewan Harrison, David K. Jackson, Ian Johnston, Dominic Kwiatkowski, Cordelia Langford, John Sillitoe on behalf of the Wellcome Sanger Institute COVID-19 Surveillance Team |
| EPI_ISL_3139129 | Lighthouse Lab in Glasgow | Wellcome Sanger Institute for the COVID-19 Genomics UK (COG-UK) Consortium | Harper VanSteenhouse, Yumi Kasai, David Gray, Carol Clugston, Anna Dominiczak and Alex Alderton, Roberto Amato, Jeffrey Barrett, Sonia Goncalves, Ewan Harrison, David K. Jackson, Ian Johnston, Dominic Kwiatkowski, Cordelia Langford, John Sillitoe on behalf of the Wellcome Sanger Institute COVID-19 Surveillance Team |
| EPI_ISL_3139130, EPI_ISL_3139131, EPI_ISL_3139132 | Berkshire and Surrey Pathology Services Lighthouse Laboratory | Wellcome Sanger Institute for the COVID-19 Genomics UK (COG-UK) Consortium | Berkshire and Surrey Pathology Services Lighthouse Laboratory and Alex Alderton, Roberto Amato, Jeffrey Barrett, Sonia Goncalves, Ewan Harrison, David K. Jackson, Ian Johnston, Dominic Kwiatkowski, Cordelia Langford, John Sillitoe on behalf of the Wellcome Sanger Institute COVID-19 Surveillance Team |
| EPI_ISL_3139133, EPI_ISL_3139134, EPI_ISL_3139135 | Lighthouse Lab in Glasgow | Wellcome Sanger Institute for the COVID-19 Genomics UK (COG-UK) Consortium | Harper VanSteenhouse, Yumi Kasai, David Gray, Carol Clugston, Anna Dominiczak and Alex Alderton, Roberto Amato, Jeffrey Barrett, Sonia Goncalves, Ewan Harrison, David K. Jackson, Ian Johnston, Dominic Kwiatkowski, Cordelia Langford, John Sillitoe on behalf of the Wellcome Sanger Institute COVID-19 Surveillance Team |
| EPI_ISL_3139136, EPI_ISL_3139137 | Berkshire and Surrey Pathology Services Lighthouse Laboratory | Wellcome Sanger Institute for the COVID-19 Genomics UK (COG-UK) Consortium | Berkshire and Surrey Pathology Services Lighthouse Laboratory and Alex Alderton, Roberto Amato, Jeffrey Barrett, Sonia Goncalves, Ewan Harrison, David K. Jackson, Ian Johnston, Dominic Kwiatkowski, Cordelia Langford, John Sillitoe on behalf of the Wellcome Sanger Institute COVID-19 Surveillance Team |
| EPI_ISL_3139138, EPI_ISL_3139139, EPI_ISL_3139140, EPI_ISL_3139141, EPI_ISL_3139142, EPI_ISL_3139143 | Lighthouse Lab in Glasgow | Wellcome Sanger Institute for the COVID-19 Genomics UK (COG-UK) Consortium | Harper VanSteenhouse, Yumi Kasai, David Gray, Carol Clugston, Anna Dominiczak and Alex Alderton, Roberto Amato, Jeffrey Barrett, Sonia Goncalves, Ewan Harrison, David K. Jackson, Ian Johnston, Dominic Kwiatkowski, Cordelia Langford, John Sillitoe on behalf of the Wellcome Sanger Institute COVID-19 Surveillance Team |
| EPI_ISL_3139144, EPI_ISL_3139145, EPI_ISL_3139146 | Berkshire and Surrey Pathology Services Lighthouse Laboratory | Wellcome Sanger Institute for the COVID-19 Genomics UK (COG-UK) Consortium | Berkshire and Surrey Pathology Services Lighthouse Laboratory and Alex Alderton, Roberto Amato, Jeffrey Barrett, Sonia Goncalves, Ewan Harrison, David K. Jackson, Ian Johnston, Dominic Kwiatkowski, Cordelia Langford, John Sillitoe on behalf of the Wellcome Sanger Institute COVID-19 Surveillance Team |
| EPI_ISL_3139147, EPI_ISL_3139148 | Lighthouse Lab in Glasgow | Wellcome Sanger Institute for the COVID-19 Genomics UK (COG-UK) Consortium | Harper VanSteenhouse, Yumi Kasai, David Gray, Carol Clugston, Anna Dominiczak and Alex Alderton, Roberto Amato, Jeffrey Barrett, Sonia Goncalves, Ewan Harrison, David K. Jackson, Ian Johnston, Dominic Kwiatkowski, Cordelia Langford, John Sillitoe on behalf of the Wellcome Sanger Institute COVID-19 Surveillance Team |
| EPI_ISL_3139149, EPI_ISL_3139150 | Berkshire and Surrey Pathology Services Lighthouse Laboratory | Wellcome Sanger Institute for the COVID-19 Genomics UK (COG-UK) Consortium | Berkshire and Surrey Pathology Services Lighthouse Laboratory and Alex Alderton, Roberto Amato, Jeffrey Barrett, Sonia Goncalves, Ewan Harrison, David K. Jackson, Ian Johnston, Dominic Kwiatkowski, Cordelia Langford, John Sillitoe on behalf of the Wellcome Sanger Institute COVID-19 Surveillance Team |
| EPI_ISL_3139151 | Lighthouse Lab in Glasgow | Wellcome Sanger Institute for the COVID-19 Genomics UK (COG-UK) Consortium | Harper VanSteenhouse, Yumi Kasai, David Gray, Carol Clugston, Anna Dominiczak and Alex Alderton, Roberto Amato, Jeffrey Barrett, Sonia Goncalves, Ewan Harrison, David K. Jackson, Ian Johnston, Dominic Kwiatkowski, Cordelia Langford, John Sillitoe on behalf of the Wellcome Sanger Institute COVID-19 Surveillance Team |
| EPI_ISL_3139152 | Berkshire and Surrey Pathology Services Lighthouse Laboratory | Wellcome Sanger Institute for the COVID-19 Genomics UK (COG-UK) Consortium | Berkshire and Surrey Pathology Services Lighthouse Laboratory and Alex Alderton, Roberto Amato, Jeffrey Barrett, Sonia Goncalves, Ewan Harrison, David K. Jackson, Ian Johnston, Dominic Kwiatkowski, Cordelia Langford, John Sillitoe on behalf of the Wellcome Sanger Institute COVID-19 Surveillance Team |
| EPI_ISL_3139153 | Lighthouse Lab in Glasgow | Wellcome Sanger Institute for the COVID-19 Genomics UK (COG-UK) Consortium | Harper VanSteenhouse, Yumi Kasai, David Gray, Carol Clugston, Anna Dominiczak and Alex Alderton, Roberto Amato, Jeffrey Barrett, Sonia Goncalves, Ewan Harrison, David K. Jackson, Ian Johnston, Dominic Kwiatkowski, Cordelia Langford, John Sillitoe on behalf of the Wellcome Sanger Institute COVID-19 Surveillance Team |
| EPI_ISL_3139154, EPI_ISL_3139155, EPI_ISL_3139156, EPI_ISL_3139157, EPI_ISL_3139158 | Berkshire and Surrey Pathology Services Lighthouse Laboratory | Wellcome Sanger Institute for the COVID-19 Genomics UK (COG-UK) Consortium | Berkshire and Surrey Pathology Services Lighthouse Laboratory and Alex Alderton, Roberto Amato, Jeffrey Barrett, Sonia Goncalves, Ewan Harrison, David K. Jackson, Ian Johnston, Dominic Kwiatkowski, Cordelia Langford, John Sillitoe on behalf of the Wellcome Sanger Institute COVID-19 Surveillance Team |
| EPI_ISL_3139159 | Lighthouse Lab in Glasgow | Wellcome Sanger Institute for the COVID-19 Genomics UK (COG-UK) Consortium | Harper VanSteenhouse, Yumi Kasai, David Gray, Carol Clugston, Anna Dominiczak and Alex Alderton, Roberto Amato, Jeffrey Barrett, Sonia Goncalves, Ewan Harrison, David K. Jackson, Ian Johnston, Dominic Kwiatkowski, Cordelia Langford, John Sillitoe on behalf of the Wellcome Sanger Institute COVID-19 Surveillance Team |
| EPI_ISL_3139160 | Berkshire and Surrey Pathology Services Lighthouse Laboratory | Wellcome Sanger Institute for the COVID-19 Genomics UK (COG-UK) Consortium | Berkshire and Surrey Pathology Services Lighthouse Laboratory and Alex Alderton, Roberto Amato, Jeffrey Barrett, Sonia Goncalves, Ewan Harrison, David K. Jackson, Ian Johnston, Dominic Kwiatkowski, Cordelia Langford, John Sillitoe on behalf of the Wellcome Sanger Institute COVID-19 Surveillance Team |
| EPI_ISL_3139161, EPI_ISL_3139162 | Lighthouse Lab in Glasgow | Wellcome Sanger Institute for the COVID-19 Genomics UK (COG-UK) Consortium | Harper VanSteenhouse, Yumi Kasai, David Gray, Carol Clugston, Anna Dominiczak and Alex Alderton, Roberto Amato, Jeffrey Barrett, Sonia Goncalves, Ewan Harrison, David K. Jackson, Ian Johnston, Dominic Kwiatkowski, Cordelia Langford, John Sillitoe on behalf of the Wellcome Sanger Institute COVID-19 Surveillance Team |
| EPI_ISL_3139163, EPI_ISL_3139164, EPI_ISL_3139165, EPI_ISL_3139166 | Berkshire and Surrey Pathology Services Lighthouse Laboratory | Wellcome Sanger Institute for the COVID-19 Genomics UK (COG-UK) Consortium | Berkshire and Surrey Pathology Services Lighthouse Laboratory and Alex Alderton, Roberto Amato, Jeffrey Barrett, Sonia Goncalves, Ewan Harrison, David K. Jackson, Ian Johnston, Dominic Kwiatkowski, Cordelia Langford, John Sillitoe on behalf of the Wellcome Sanger Institute COVID-19 Surveillance Team |
| EPI_ISL_3139167, EPI_ISL_3139168 | Lighthouse Lab in Glasgow | Wellcome Sanger Institute for the COVID-19 Genomics UK (COG-UK) Consortium | Harper VanSteenhouse, Yumi Kasai, David Gray, Carol Clugston, Anna Dominiczak and Alex Alderton, Roberto Amato, Jeffrey Barrett, Sonia Goncalves, Ewan Harrison, David K. Jackson, Ian Johnston, Dominic Kwiatkowski, Cordelia Langford, John Sillitoe on behalf of the Wellcome Sanger Institute COVID-19 Surveillance Team |
| EPI_ISL_3139169, EPI_ISL_3139170, EPI_ISL_3139171 | Berkshire and Surrey Pathology Services Lighthouse Laboratory | Wellcome Sanger Institute for the COVID-19 Genomics UK (COG-UK) Consortium | Berkshire and Surrey Pathology Services Lighthouse Laboratory and Alex Alderton, Roberto Amato, Jeffrey Barrett, Sonia Goncalves, Ewan Harrison, David K. Jackson, Ian Johnston, Dominic Kwiatkowski, Cordelia Langford, John Sillitoe on behalf of the Wellcome Sanger Institute COVID-19 Surveillance Team |
| EPI_ISL_3139172, EPI_ISL_3139173, EPI_ISL_3139174 | Lighthouse Lab in Glasgow | Wellcome Sanger Institute for the COVID-19 Genomics UK (COG-UK) Consortium | Harper VanSteenhouse, Yumi Kasai, David Gray, Carol Clugston, Anna Dominiczak and Alex Alderton, Roberto Amato, Jeffrey Barrett, Sonia Goncalves, Ewan Harrison, David K. Jackson, Ian Johnston, Dominic Kwiatkowski, Cordelia Langford, John Sillitoe on behalf of the Wellcome Sanger Institute COVID-19 Surveillance Team |
| EPI_ISL_3139175 | Berkshire and Surrey Pathology Services Lighthouse Laboratory | Wellcome Sanger Institute for the COVID-19 Genomics UK (COG-UK) Consortium | Berkshire and Surrey Pathology Services Lighthouse Laboratory and Alex Alderton, Roberto Amato, Jeffrey Barrett, Sonia Goncalves, Ewan Harrison, David K. Jackson, Ian Johnston, Dominic Kwiatkowski, Cordelia Langford, John Sillitoe on behalf of the Wellcome Sanger Institute COVID-19 Surveillance Team |
| EPI_ISL_3139176 | Lighthouse Lab in Glasgow | Wellcome Sanger Institute for the COVID-19 Genomics UK (COG-UK) Consortium | Harper VanSteenhouse, Yumi Kasai, David Gray, Carol Clugston, Anna Dominiczak and Alex Alderton, Roberto Amato, Jeffrey Barrett, Sonia Goncalves, Ewan Harrison, David K. Jackson, Ian Johnston, Dominic Kwiatkowski, Cordelia Langford, John Sillitoe on behalf of the Wellcome Sanger Institute COVID-19 Surveillance Team |
| EPI_ISL_3139177 | Berkshire and Surrey Pathology Services Lighthouse Laboratory | Wellcome Sanger Institute for the COVID-19 Genomics UK (COG-UK) Consortium | Berkshire and Surrey Pathology Services Lighthouse Laboratory and Alex Alderton, Roberto Amato, Jeffrey Barrett, Sonia Goncalves, Ewan Harrison, David K. Jackson, Ian Johnston, Dominic Kwiatkowski, Cordelia Langford, John Sillitoe on behalf of the Wellcome Sanger Institute COVID-19 Surveillance Team |
| EPI_ISL_3139178, EPI_ISL_3139179 | Lighthouse Lab in Glasgow | Wellcome Sanger Institute for the COVID-19 Genomics UK (COG-UK) Consortium | Harper VanSteenhouse, Yumi Kasai, David Gray, Carol Clugston, Anna Dominiczak and Alex Alderton, Roberto Amato, Jeffrey Barrett, Sonia Goncalves, Ewan Harrison, David K. Jackson, Ian Johnston, Dominic Kwiatkowski, Cordelia Langford, John Sillitoe on behalf of the Wellcome Sanger Institute COVID-19 Surveillance Team |
| EPI_ISL_3139180, | Berkshire and Surrey Pathology Services Lighthouse Laboratory | Wellcome Sanger Institute for the COVID-19 Genomics UK (COG-UK) | Berkshire and Surrey Pathology Services Lighthouse Laboratory and Alex Alderton, Roberto Amato, Jeffrey Barrett, Sonia Goncalves, Ewan Harrison, David K. Jackson, Ian Johnston, Dominic Kwiatkowski, Cordelia Langford, John Sillitoe on behalf |

| | | | |
|---|---|---|---|
| EPI_ISL_3139181 | | Consortium | of the Wellcome Sanger Institute COVID-19 Surveillance Team |
| EPI_ISL_3139182, EPI_ISL_3139183 | Lighthouse Lab in Glasgow | Wellcome Sanger Institute for the COVID-19 Genomics UK (COG-UK) Consortium | Harper VanSteenhouse, Yumi Kasai, David Gray, Carol Clugston, Anna Dominiczak and Alex Alderton, Roberto Amato, Jeffrey Barrett, Sonia Goncalves, Ewan Harrison, David K. Jackson, Ian Johnston, Dominic Kwiatkowski, Cordelia Langford, John Sillitoe on behalf of the Wellcome Sanger Institute COVID-19 Surveillance Team |
| EPI_ISL_3139184, EPI_ISL_3139185 | Berkshire and Surrey Pathology Services Lighthouse Laboratory | Wellcome Sanger Institute for the COVID-19 Genomics UK (COG-UK) Consortium | Berkshire and Surrey Pathology Services Lighthouse Laboratory and Alex Alderton, Roberto Amato, Jeffrey Barrett, Sonia Goncalves, Ewan Harrison, David K. Jackson, Ian Johnston, Dominic Kwiatkowski, Cordelia Langford, John Sillitoe on behalf of the Wellcome Sanger Institute COVID-19 Surveillance Team |
| EPI_ISL_3139186, EPI_ISL_3139187, EPI_ISL_3139188, EPI_ISL_3139189, EPI_ISL_3139190, EPI_ISL_3139191 | Lighthouse Lab in Glasgow | Wellcome Sanger Institute for the COVID-19 Genomics UK (COG-UK) Consortium | Harper VanSteenhouse, Yumi Kasai, David Gray, Carol Clugston, Anna Dominiczak and Alex Alderton, Roberto Amato, Jeffrey Barrett, Sonia Goncalves, Ewan Harrison, David K. Jackson, Ian Johnston, Dominic Kwiatkowski, Cordelia Langford, John Sillitoe on behalf of the Wellcome Sanger Institute COVID-19 Surveillance Team |
| EPI_ISL_3139192, EPI_ISL_3139193, EPI_ISL_3139194 | Berkshire and Surrey Pathology Services Lighthouse Laboratory | Wellcome Sanger Institute for the COVID-19 Genomics UK (COG-UK) Consortium | Berkshire and Surrey Pathology Services Lighthouse Laboratory and Alex Alderton, Roberto Amato, Jeffrey Barrett, Sonia Goncalves, Ewan Harrison, David K. Jackson, Ian Johnston, Dominic Kwiatkowski, Cordelia Langford, John Sillitoe on behalf of the Wellcome Sanger Institute COVID-19 Surveillance Team |
| EPI_ISL_3139195, EPI_ISL_3139196 | Lighthouse Lab in Glasgow | Wellcome Sanger Institute for the COVID-19 Genomics UK (COG-UK) Consortium | Harper VanSteenhouse, Yumi Kasai, David Gray, Carol Clugston, Anna Dominiczak and Alex Alderton, Roberto Amato, Jeffrey Barrett, Sonia Goncalves, Ewan Harrison, David K. Jackson, Ian Johnston, Dominic Kwiatkowski, Cordelia Langford, John Sillitoe on behalf of the Wellcome Sanger Institute COVID-19 Surveillance Team |
| EPI_ISL_3139197, EPI_ISL_3139198, EPI_ISL_3139199 | Berkshire and Surrey Pathology Services Lighthouse Laboratory | Wellcome Sanger Institute for the COVID-19 Genomics UK (COG-UK) Consortium | Berkshire and Surrey Pathology Services Lighthouse Laboratory and Alex Alderton, Roberto Amato, Jeffrey Barrett, Sonia Goncalves, Ewan Harrison, David K. Jackson, Ian Johnston, Dominic Kwiatkowski, Cordelia Langford, John Sillitoe on behalf of the Wellcome Sanger Institute COVID-19 Surveillance Team |
| EPI_ISL_3139200, EPI_ISL_3139201 | Lighthouse Lab in Glasgow | Wellcome Sanger Institute for the COVID-19 Genomics UK (COG-UK) Consortium | Harper VanSteenhouse, Yumi Kasai, David Gray, Carol Clugston, Anna Dominiczak and Alex Alderton, Roberto Amato, Jeffrey Barrett, Sonia Goncalves, Ewan Harrison, David K. Jackson, Ian Johnston, Dominic Kwiatkowski, Cordelia Langford, John Sillitoe on behalf of the Wellcome Sanger Institute COVID-19 Surveillance Team |
| EPI_ISL_3139202 | Berkshire and Surrey Pathology Services Lighthouse Laboratory | Wellcome Sanger Institute for the COVID-19 Genomics UK (COG-UK) Consortium | Berkshire and Surrey Pathology Services Lighthouse Laboratory and Alex Alderton, Roberto Amato, Jeffrey Barrett, Sonia Goncalves, Ewan Harrison, David K. Jackson, Ian Johnston, Dominic Kwiatkowski, Cordelia Langford, John Sillitoe on behalf of the Wellcome Sanger Institute COVID-19 Surveillance Team |
| EPI_ISL_3139203, EPI_ISL_3139204 | Lighthouse Lab in Glasgow | Wellcome Sanger Institute for the COVID-19 Genomics UK (COG-UK) Consortium | Harper VanSteenhouse, Yumi Kasai, David Gray, Carol Clugston, Anna Dominiczak and Alex Alderton, Roberto Amato, Jeffrey Barrett, Sonia Goncalves, Ewan Harrison, David K. Jackson, Ian Johnston, Dominic Kwiatkowski, Cordelia Langford, John Sillitoe on behalf of the Wellcome Sanger Institute COVID-19 Surveillance Team |
| EPI_ISL_3139205, EPI_ISL_3139206, EPI_ISL_3139207, EPI_ISL_3139208 | Berkshire and Surrey Pathology Services Lighthouse Laboratory | Wellcome Sanger Institute for the COVID-19 Genomics UK (COG-UK) Consortium | Berkshire and Surrey Pathology Services Lighthouse Laboratory and Alex Alderton, Roberto Amato, Jeffrey Barrett, Sonia Goncalves, Ewan Harrison, David K. Jackson, Ian Johnston, Dominic Kwiatkowski, Cordelia Langford, John Sillitoe on behalf of the Wellcome Sanger Institute COVID-19 Surveillance Team |
| EPI_ISL_3139209 | Lighthouse Lab in Glasgow | Wellcome Sanger Institute for the COVID-19 Genomics UK (COG-UK) Consortium | Harper VanSteenhouse, Yumi Kasai, David Gray, Carol Clugston, Anna Dominiczak and Alex Alderton, Roberto Amato, Jeffrey Barrett, Sonia Goncalves, Ewan Harrison, David K. Jackson, Ian Johnston, Dominic Kwiatkowski, Cordelia Langford, John Sillitoe on behalf of the Wellcome Sanger Institute COVID-19 Surveillance Team |
| EPI_ISL_3139210 | Berkshire and Surrey Pathology Services Lighthouse Laboratory | Wellcome Sanger Institute for the COVID-19 Genomics UK (COG-UK) Consortium | Berkshire and Surrey Pathology Services Lighthouse Laboratory and Alex Alderton, Roberto Amato, Jeffrey Barrett, Sonia Goncalves, Ewan Harrison, David K. Jackson, Ian Johnston, Dominic Kwiatkowski, Cordelia Langford, John Sillitoe on behalf of the Wellcome Sanger Institute COVID-19 Surveillance Team |
| EPI_ISL_3139211, EPI_ISL_3139212, EPI_ISL_3139213, EPI_ISL_3139214 | Lighthouse Lab in Glasgow | Wellcome Sanger Institute for the COVID-19 Genomics UK (COG-UK) Consortium | Harper VanSteenhouse, Yumi Kasai, David Gray, Carol Clugston, Anna Dominiczak and Alex Alderton, Roberto Amato, Jeffrey Barrett, Sonia Goncalves, Ewan Harrison, David K. Jackson, Ian Johnston, Dominic Kwiatkowski, Cordelia Langford, John Sillitoe on behalf of the Wellcome Sanger Institute COVID-19 Surveillance Team |
| EPI_ISL_3139215 | Berkshire and Surrey Pathology Services Lighthouse Laboratory | Wellcome Sanger Institute for the COVID-19 Genomics UK (COG-UK) Consortium | Berkshire and Surrey Pathology Services Lighthouse Laboratory and Alex Alderton, Roberto Amato, Jeffrey Barrett, Sonia Goncalves, Ewan Harrison, David K. Jackson, Ian Johnston, Dominic Kwiatkowski, Cordelia Langford, John Sillitoe on behalf of the Wellcome Sanger Institute COVID-19 Surveillance Team |
| EPI_ISL_3139216, EPI_ISL_3139217, EPI_ISL_3139218, EPI_ISL_3139219, EPI_ISL_3139220 | Lighthouse Lab in Glasgow | Wellcome Sanger Institute for the COVID-19 Genomics UK (COG-UK) Consortium | Harper VanSteenhouse, Yumi Kasai, David Gray, Carol Clugston, Anna Dominiczak and Alex Alderton, Roberto Amato, Jeffrey Barrett, Sonia Goncalves, Ewan Harrison, David K. Jackson, Ian Johnston, Dominic Kwiatkowski, Cordelia Langford, John Sillitoe on behalf of the Wellcome Sanger Institute COVID-19 Surveillance Team |
| EPI_ISL_3139221 | Berkshire and Surrey Pathology Services Lighthouse Laboratory | Wellcome Sanger Institute for the COVID-19 Genomics UK (COG-UK) Consortium | Berkshire and Surrey Pathology Services Lighthouse Laboratory and Alex Alderton, Roberto Amato, Jeffrey Barrett, Sonia Goncalves, Ewan Harrison, David K. Jackson, Ian Johnston, Dominic Kwiatkowski, Cordelia Langford, John Sillitoe on behalf of the Wellcome Sanger Institute COVID-19 Surveillance Team |
| EPI_ISL_3139222, EPI_ISL_3139223, EPI_ISL_3139224, EPI_ISL_3139225, EPI_ISL_3139226, EPI_ISL_3139227, EPI_ISL_3139228, EPI_ISL_3139229 | Lighthouse Lab in Glasgow | Wellcome Sanger Institute for the COVID-19 Genomics UK (COG-UK) Consortium | Harper VanSteenhouse, Yumi Kasai, David Gray, Carol Clugston, Anna Dominiczak and Alex Alderton, Roberto Amato, Jeffrey Barrett, Sonia Goncalves, Ewan Harrison, David K. Jackson, Ian Johnston, Dominic Kwiatkowski, Cordelia Langford, John Sillitoe on behalf of the Wellcome Sanger Institute COVID-19 Surveillance Team |
| EPI_ISL_3139230 | Berkshire and Surrey Pathology Services Lighthouse Laboratory | Wellcome Sanger Institute for the COVID-19 Genomics UK (COG-UK) Consortium | Berkshire and Surrey Pathology Services Lighthouse Laboratory and Alex Alderton, Roberto Amato, Jeffrey Barrett, Sonia Goncalves, Ewan Harrison, David K. Jackson, Ian Johnston, Dominic Kwiatkowski, Cordelia Langford, John Sillitoe on behalf of the Wellcome Sanger Institute COVID-19 Surveillance Team |
| EPI_ISL_3139231 | Lighthouse Lab in Glasgow | Wellcome Sanger Institute for the COVID-19 Genomics UK (COG-UK) Consortium | Harper VanSteenhouse, Yumi Kasai, David Gray, Carol Clugston, Anna Dominiczak and Alex Alderton, Roberto Amato, Jeffrey Barrett, Sonia Goncalves, Ewan Harrison, David K. Jackson, Ian Johnston, Dominic Kwiatkowski, Cordelia Langford, John Sillitoe on behalf of the Wellcome Sanger Institute COVID-19 Surveillance Team |
| EPI_ISL_3139232, EPI_ISL_3139233 | Berkshire and Surrey Pathology Services Lighthouse Laboratory | Wellcome Sanger Institute for the COVID-19 Genomics UK (COG-UK) Consortium | Berkshire and Surrey Pathology Services Lighthouse Laboratory and Alex Alderton, Roberto Amato, Jeffrey Barrett, Sonia Goncalves, Ewan Harrison, David K. Jackson, Ian Johnston, Dominic Kwiatkowski, Cordelia Langford, John Sillitoe on behalf of the Wellcome Sanger Institute COVID-19 Surveillance Team |
| EPI_ISL_3139234 | Lighthouse Lab in Glasgow | Wellcome Sanger Institute for the COVID-19 Genomics UK (COG-UK) Consortium | Harper VanSteenhouse, Yumi Kasai, David Gray, Carol Clugston, Anna Dominiczak and Alex Alderton, Roberto Amato, Jeffrey Barrett, Sonia Goncalves, Ewan Harrison, David K. Jackson, Ian Johnston, Dominic Kwiatkowski, Cordelia Langford, John Sillitoe on behalf of the Wellcome Sanger Institute COVID-19 Surveillance Team |
| EPI_ISL_3139235 | Berkshire and Surrey Pathology Services Lighthouse Laboratory | Wellcome Sanger Institute for the COVID-19 Genomics UK (COG-UK) Consortium | Berkshire and Surrey Pathology Services Lighthouse Laboratory and Alex Alderton, Roberto Amato, Jeffrey Barrett, Sonia Goncalves, Ewan Harrison, David K. Jackson, Ian Johnston, Dominic Kwiatkowski, Cordelia Langford, John Sillitoe on behalf of the Wellcome Sanger Institute COVID-19 Surveillance Team |
| EPI_ISL_3139236 | Lighthouse Lab in Glasgow | Wellcome Sanger Institute for the COVID-19 Genomics UK (COG-UK) Consortium | Harper VanSteenhouse, Yumi Kasai, David Gray, Carol Clugston, Anna Dominiczak and Alex Alderton, Roberto Amato, Jeffrey Barrett, Sonia Goncalves, Ewan Harrison, David K. Jackson, Ian Johnston, Dominic Kwiatkowski, Cordelia Langford, John Sillitoe on behalf of the Wellcome Sanger Institute COVID-19 Surveillance Team |
| EPI_ISL_3139237, EPI_ISL_3139238 | Berkshire and Surrey Pathology Services Lighthouse Laboratory | Wellcome Sanger Institute for the COVID-19 Genomics UK (COG-UK) Consortium | Berkshire and Surrey Pathology Services Lighthouse Laboratory and Alex Alderton, Roberto Amato, Jeffrey Barrett, Sonia Goncalves, Ewan Harrison, David K. Jackson, Ian Johnston, Dominic Kwiatkowski, Cordelia Langford, John Sillitoe on behalf of the Wellcome Sanger Institute COVID-19 Surveillance Team |
| EPI_ISL_3139239, EPI_ISL_3139240, EPI_ISL_3139241, EPI_ISL_3139242, EPI_ISL_3139243 | Lighthouse Lab in Glasgow | Wellcome Sanger Institute for the COVID-19 Genomics UK (COG-UK) Consortium | Harper VanSteenhouse, Yumi Kasai, David Gray, Carol Clugston, Anna Dominiczak and Alex Alderton, Roberto Amato, Jeffrey Barrett, Sonia Goncalves, Ewan Harrison, David K. Jackson, Ian Johnston, Dominic Kwiatkowski, Cordelia Langford, John Sillitoe on behalf of the Wellcome Sanger Institute COVID-19 Surveillance Team |
| EPI_ISL_3139244 | Berkshire and Surrey Pathology Services Lighthouse Laboratory | Wellcome Sanger Institute for the COVID-19 Genomics UK (COG-UK) Consortium | Berkshire and Surrey Pathology Services Lighthouse Laboratory and Alex Alderton, Roberto Amato, Jeffrey Barrett, Sonia Goncalves, Ewan Harrison, David K. Jackson, Ian Johnston, Dominic Kwiatkowski, Cordelia Langford, John Sillitoe on behalf of the Wellcome Sanger Institute COVID-19 Surveillance Team |
| EPI_ISL_3139245, EPI_ISL_3139246, EPI_ISL_3139247 | Lighthouse Lab in Glasgow | Wellcome Sanger Institute for the COVID-19 Genomics UK (COG-UK) Consortium | Harper VanSteenhouse, Yumi Kasai, David Gray, Carol Clugston, Anna Dominiczak and Alex Alderton, Roberto Amato, Jeffrey Barrett, Sonia Goncalves, Ewan Harrison, David K. Jackson, Ian Johnston, Dominic Kwiatkowski, Cordelia Langford, John Sillitoe on behalf of the Wellcome Sanger Institute COVID-19 Surveillance Team |
| EPI_ISL_3139248, EPI_ISL_3139249, EPI_ISL_3139250 | Berkshire and Surrey Pathology Services Lighthouse Laboratory | Wellcome Sanger Institute for the COVID-19 Genomics UK (COG-UK) Consortium | Berkshire and Surrey Pathology Services Lighthouse Laboratory and Alex Alderton, Roberto Amato, Jeffrey Barrett, Sonia Goncalves, Ewan Harrison, David K. Jackson, Ian Johnston, Dominic Kwiatkowski, Cordelia Langford, John Sillitoe on behalf of the Wellcome Sanger Institute COVID-19 Surveillance Team |
| EPI_ISL_3139251 | Lighthouse Lab in Glasgow | Wellcome Sanger Institute for the COVID-19 Genomics UK (COG-UK) Consortium | Harper VanSteenhouse, Yumi Kasai, David Gray, Carol Clugston, Anna Dominiczak and Alex Alderton, Roberto Amato, Jeffrey Barrett, Sonia Goncalves, Ewan Harrison, David K. Jackson, Ian Johnston, Dominic Kwiatkowski, Cordelia Langford, John Sillitoe on behalf of the Wellcome Sanger Institute COVID-19 Surveillance Team |
| EPI_ISL_3139252, EPI_ISL_3139253, EPI_ISL_3139254, EPI_ISL_3139255 | Berkshire and Surrey Pathology Services Lighthouse Laboratory | Wellcome Sanger Institute for the COVID-19 Genomics UK (COG-UK) Consortium | Berkshire and Surrey Pathology Services Lighthouse Laboratory and Alex Alderton, Roberto Amato, Jeffrey Barrett, Sonia Goncalves, Ewan Harrison, David K. Jackson, Ian Johnston, Dominic Kwiatkowski, Cordelia Langford, John Sillitoe on behalf of the Wellcome Sanger Institute COVID-19 Surveillance Team |
| EPI_ISL_3139256, EPI_ISL_3139257, EPI_ISL_3139258, EPI_ISL_3139259, EPI_ISL_3139260, EPI_ISL_3139261 | Lighthouse Lab in Glasgow | Wellcome Sanger Institute for the COVID-19 Genomics UK (COG-UK) Consortium | Harper VanSteenhouse, Yumi Kasai, David Gray, Carol Clugston, Anna Dominiczak and Alex Alderton, Roberto Amato, Jeffrey Barrett, Sonia Goncalves, Ewan Harrison, David K. Jackson, Ian Johnston, Dominic Kwiatkowski, Cordelia Langford, John Sillitoe on behalf of the Wellcome Sanger Institute COVID-19 Surveillance Team |
| EPI_ISL_3139262 | Berkshire and Surrey Pathology Services Lighthouse Laboratory | Wellcome Sanger Institute for the COVID-19 Genomics UK (COG-UK) Consortium | Berkshire and Surrey Pathology Services Lighthouse Laboratory and Alex Alderton, Roberto Amato, Jeffrey Barrett, Sonia Goncalves, Ewan Harrison, David K. Jackson, Ian Johnston, Dominic Kwiatkowski, Cordelia Langford, John Sillitoe on behalf of the Wellcome Sanger Institute COVID-19 Surveillance Team |

| | | | |
|---|---|---|---|
| EPI_ISL_3139263 | Lighthouse Lab in Glasgow | Wellcome Sanger Institute for the COVID-19 Genomics UK (COG-UK) Consortium | Harper VanSteenhouse, Yumi Kasai, David Gray, Carol Clugston, Anna Dominiczak and Alex Alderton, Roberto Amato, Jeffrey Barrett, Sonia Goncalves, Ewan Harrison, David K. Jackson, Ian Johnston, Dominic Kwiatkowski, Cordelia Langford, John Sillitoe on behalf of the Wellcome Sanger Institute COVID-19 Surveillance Team |
| EPI_ISL_3139264 | Lighthouse Laboratory Plymouth | Wellcome Sanger Institute for the COVID-19 Genomics UK (COG-UK) Consortium | Lighthouse Laboratory Plymouth and Alex Alderton, Roberto Amato, Jeffrey Barrett, Sonia Goncalves, Ewan Harrison, David K. Jackson, Ian Johnston, Dominic Kwiatkowski, Cordelia Langford, John Sillitoe on behalf of the Wellcome Sanger Institute COVID-19 Surveillance Team |
| EPI_ISL_3139265 | Lighthouse Lab in Milton Keynes | Wellcome Sanger Institute for the COVID-19 Genomics UK (COG-UK) Consortium | The Lighthouse Lab in Milton Keynes and Alex Alderton, Roberto Amato, Jeffrey Barrett, Sonia Goncalves, Ewan Harrison, David K. Jackson, Ian Johnston, Dominic Kwiatkowski, Cordelia Langford, John Sillitoe on behalf of the Wellcome Sanger Institute COVID-19 Surveillance Team |
| EPI_ISL_3139266 | Lighthouse Laboratory Plymouth | Wellcome Sanger Institute for the COVID-19 Genomics UK (COG-UK) Consortium | Lighthouse Laboratory Plymouth and Alex Alderton, Roberto Amato, Jeffrey Barrett, Sonia Goncalves, Ewan Harrison, David K. Jackson, Ian Johnston, Dominic Kwiatkowski, Cordelia Langford, John Sillitoe on behalf of the Wellcome Sanger Institute COVID-19 Surveillance Team |
| EPI_ISL_3139267 | Lighthouse Lab in Milton Keynes | Wellcome Sanger Institute for the COVID-19 Genomics UK (COG-UK) Consortium | The Lighthouse Lab in Milton Keynes and Alex Alderton, Roberto Amato, Jeffrey Barrett, Sonia Goncalves, Ewan Harrison, David K. Jackson, Ian Johnston, Dominic Kwiatkowski, Cordelia Langford, John Sillitoe on behalf of the Wellcome Sanger Institute COVID-19 Surveillance Team |
| EPI_ISL_3139268, EPI_ISL_3139269, EPI_ISL_3139270 | Lighthouse Laboratory Plymouth | Wellcome Sanger Institute for the COVID-19 Genomics UK (COG-UK) Consortium | Lighthouse Laboratory Plymouth and Alex Alderton, Roberto Amato, Jeffrey Barrett, Sonia Goncalves, Ewan Harrison, David K. Jackson, Ian Johnston, Dominic Kwiatkowski, Cordelia Langford, John Sillitoe on behalf of the Wellcome Sanger Institute COVID-19 Surveillance Team |
| EPI_ISL_3139271 | Lighthouse Lab in Glasgow | Wellcome Sanger Institute for the COVID-19 Genomics UK (COG-UK) Consortium | Harper VanSteenhouse, Yumi Kasai, David Gray, Carol Clugston, Anna Dominiczak and Alex Alderton, Roberto Amato, Jeffrey Barrett, Sonia Goncalves, Ewan Harrison, David K. Jackson, Ian Johnston, Dominic Kwiatkowski, Cordelia Langford, John Sillitoe on behalf of the Wellcome Sanger Institute COVID-19 Surveillance Team |
| EPI_ISL_3139272, EPI_ISL_3139273, EPI_ISL_3139274 | Lighthouse Laboratory Plymouth | Wellcome Sanger Institute for the COVID-19 Genomics UK (COG-UK) Consortium | Lighthouse Laboratory Plymouth and Alex Alderton, Roberto Amato, Jeffrey Barrett, Sonia Goncalves, Ewan Harrison, David K. Jackson, Ian Johnston, Dominic Kwiatkowski, Cordelia Langford, John Sillitoe on behalf of the Wellcome Sanger Institute COVID-19 Surveillance Team |
| EPI_ISL_3139275 | Lighthouse Lab in Glasgow | Wellcome Sanger Institute for the COVID-19 Genomics UK (COG-UK) Consortium | Harper VanSteenhouse, Yumi Kasai, David Gray, Carol Clugston, Anna Dominiczak and Alex Alderton, Roberto Amato, Jeffrey Barrett, Sonia Goncalves, Ewan Harrison, David K. Jackson, Ian Johnston, Dominic Kwiatkowski, Cordelia Langford, John Sillitoe on behalf of the Wellcome Sanger Institute COVID-19 Surveillance Team |
| EPI_ISL_3139276 | Lighthouse Laboratory Plymouth | Wellcome Sanger Institute for the COVID-19 Genomics UK (COG-UK) Consortium | Lighthouse Laboratory Plymouth and Alex Alderton, Roberto Amato, Jeffrey Barrett, Sonia Goncalves, Ewan Harrison, David K. Jackson, Ian Johnston, Dominic Kwiatkowski, Cordelia Langford, John Sillitoe on behalf of the Wellcome Sanger Institute COVID-19 Surveillance Team |
| EPI_ISL_3139277, EPI_ISL_3139278 | Lighthouse Lab in Milton Keynes | Wellcome Sanger Institute for the COVID-19 Genomics UK (COG-UK) Consortium | The Lighthouse Lab in Milton Keynes and Alex Alderton, Roberto Amato, Jeffrey Barrett, Sonia Goncalves, Ewan Harrison, David K. Jackson, Ian Johnston, Dominic Kwiatkowski, Cordelia Langford, John Sillitoe on behalf of the Wellcome Sanger Institute COVID-19 Surveillance Team |
| EPI_ISL_3139279, EPI_ISL_3139280, EPI_ISL_3139281, EPI_ISL_3139282 | Lighthouse Laboratory Plymouth | Wellcome Sanger Institute for the COVID-19 Genomics UK (COG-UK) Consortium | Lighthouse Laboratory Plymouth and Alex Alderton, Roberto Amato, Jeffrey Barrett, Sonia Goncalves, Ewan Harrison, David K. Jackson, Ian Johnston, Dominic Kwiatkowski, Cordelia Langford, John Sillitoe on behalf of the Wellcome Sanger Institute COVID-19 Surveillance Team |
| EPI_ISL_3139283 | Lighthouse Lab in Glasgow | Wellcome Sanger Institute for the COVID-19 Genomics UK (COG-UK) Consortium | Harper VanSteenhouse, Yumi Kasai, David Gray, Carol Clugston, Anna Dominiczak and Alex Alderton, Roberto Amato, Jeffrey Barrett, Sonia Goncalves, Ewan Harrison, David K. Jackson, Ian Johnston, Dominic Kwiatkowski, Cordelia Langford, John Sillitoe on behalf of the Wellcome Sanger Institute COVID-19 Surveillance Team |
| EPI_ISL_3139284, EPI_ISL_3139285, EPI_ISL_3139286 | Lighthouse Laboratory Plymouth | Wellcome Sanger Institute for the COVID-19 Genomics UK (COG-UK) Consortium | Lighthouse Laboratory Plymouth and Alex Alderton, Roberto Amato, Jeffrey Barrett, Sonia Goncalves, Ewan Harrison, David K. Jackson, Ian Johnston, Dominic Kwiatkowski, Cordelia Langford, John Sillitoe on behalf of the Wellcome Sanger Institute COVID-19 Surveillance Team |
| EPI_ISL_3139287 | Lighthouse Lab in Milton Keynes | Wellcome Sanger Institute for the COVID-19 Genomics UK (COG-UK) Consortium | The Lighthouse Lab in Milton Keynes and Alex Alderton, Roberto Amato, Jeffrey Barrett, Sonia Goncalves, Ewan Harrison, David K. Jackson, Ian Johnston, Dominic Kwiatkowski, Cordelia Langford, John Sillitoe on behalf of the Wellcome Sanger Institute COVID-19 Surveillance Team |
| EPI_ISL_3139288 | Lighthouse Lab in Glasgow | Wellcome Sanger Institute for the COVID-19 Genomics UK (COG-UK) Consortium | Harper VanSteenhouse, Yumi Kasai, David Gray, Carol Clugston, Anna Dominiczak and Alex Alderton, Roberto Amato, Jeffrey Barrett, Sonia Goncalves, Ewan Harrison, David K. Jackson, Ian Johnston, Dominic Kwiatkowski, Cordelia Langford, John Sillitoe on behalf of the Wellcome Sanger Institute COVID-19 Surveillance Team |
| EPI_ISL_3139289 | Lighthouse Laboratory Plymouth | Wellcome Sanger Institute for the COVID-19 Genomics UK (COG-UK) Consortium | Lighthouse Laboratory Plymouth and Alex Alderton, Roberto Amato, Jeffrey Barrett, Sonia Goncalves, Ewan Harrison, David K. Jackson, Ian Johnston, Dominic Kwiatkowski, Cordelia Langford, John Sillitoe on behalf of the Wellcome Sanger Institute COVID-19 Surveillance Team |
| EPI_ISL_3139290 | Lighthouse Lab in Milton Keynes | Wellcome Sanger Institute for the COVID-19 Genomics UK (COG-UK) Consortium | The Lighthouse Lab in Milton Keynes and Alex Alderton, Roberto Amato, Jeffrey Barrett, Sonia Goncalves, Ewan Harrison, David K. Jackson, Ian Johnston, Dominic Kwiatkowski, Cordelia Langford, John Sillitoe on behalf of the Wellcome Sanger Institute COVID-19 Surveillance Team |
| EPI_ISL_3139291, EPI_ISL_3139292 | Lighthouse Laboratory Plymouth | Wellcome Sanger Institute for the COVID-19 Genomics UK (COG-UK) Consortium | Lighthouse Laboratory Plymouth and Alex Alderton, Roberto Amato, Jeffrey Barrett, Sonia Goncalves, Ewan Harrison, David K. Jackson, Ian Johnston, Dominic Kwiatkowski, Cordelia Langford, John Sillitoe on behalf of the Wellcome Sanger Institute COVID-19 Surveillance Team |
| EPI_ISL_3139293 | Lighthouse Lab in Glasgow | Wellcome Sanger Institute for the COVID-19 Genomics UK (COG-UK) Consortium | Harper VanSteenhouse, Yumi Kasai, David Gray, Carol Clugston, Anna Dominiczak and Alex Alderton, Roberto Amato, Jeffrey Barrett, Sonia Goncalves, Ewan Harrison, David K. Jackson, Ian Johnston, Dominic Kwiatkowski, Cordelia Langford, John Sillitoe on behalf of the Wellcome Sanger Institute COVID-19 Surveillance Team |
| EPI_ISL_3139294, EPI_ISL_3139295 | Lighthouse Laboratory Plymouth | Wellcome Sanger Institute for the COVID-19 Genomics UK (COG-UK) Consortium | Lighthouse Laboratory Plymouth and Alex Alderton, Roberto Amato, Jeffrey Barrett, Sonia Goncalves, Ewan Harrison, David K. Jackson, Ian Johnston, Dominic Kwiatkowski, Cordelia Langford, John Sillitoe on behalf of the Wellcome Sanger Institute COVID-19 Surveillance Team |
| EPI_ISL_3139296 | Lighthouse Lab in Glasgow | Wellcome Sanger Institute for the COVID-19 Genomics UK (COG-UK) Consortium | Harper VanSteenhouse, Yumi Kasai, David Gray, Carol Clugston, Anna Dominiczak and Alex Alderton, Roberto Amato, Jeffrey Barrett, Sonia Goncalves, Ewan Harrison, David K. Jackson, Ian Johnston, Dominic Kwiatkowski, Cordelia Langford, John Sillitoe on behalf of the Wellcome Sanger Institute COVID-19 Surveillance Team |
| EPI_ISL_3139297 | Lighthouse Laboratory Plymouth | Wellcome Sanger Institute for the COVID-19 Genomics UK (COG-UK) Consortium | Lighthouse Laboratory Plymouth and Alex Alderton, Roberto Amato, Jeffrey Barrett, Sonia Goncalves, Ewan Harrison, David K. Jackson, Ian Johnston, Dominic Kwiatkowski, Cordelia Langford, John Sillitoe on behalf of the Wellcome Sanger Institute COVID-19 Surveillance Team |
| EPI_ISL_3139298 | Lighthouse Lab in Milton Keynes | Wellcome Sanger Institute for the COVID-19 Genomics UK (COG-UK) Consortium | The Lighthouse Lab in Milton Keynes and Alex Alderton, Roberto Amato, Jeffrey Barrett, Sonia Goncalves, Ewan Harrison, David K. Jackson, Ian Johnston, Dominic Kwiatkowski, Cordelia Langford, John Sillitoe on behalf of the Wellcome Sanger Institute COVID-19 Surveillance Team |
| EPI_ISL_3139299 | Lighthouse Laboratory Plymouth | Wellcome Sanger Institute for the COVID-19 Genomics UK (COG-UK) Consortium | Lighthouse Laboratory Plymouth and Alex Alderton, Roberto Amato, Jeffrey Barrett, Sonia Goncalves, Ewan Harrison, David K. Jackson, Ian Johnston, Dominic Kwiatkowski, Cordelia Langford, John Sillitoe on behalf of the Wellcome Sanger Institute COVID-19 Surveillance Team |
| EPI_ISL_3139300 | Lighthouse Lab in Milton Keynes | Wellcome Sanger Institute for the COVID-19 Genomics UK (COG-UK) Consortium | The Lighthouse Lab in Milton Keynes and Alex Alderton, Roberto Amato, Jeffrey Barrett, Sonia Goncalves, Ewan Harrison, David K. Jackson, Ian Johnston, Dominic Kwiatkowski, Cordelia Langford, John Sillitoe on behalf of the Wellcome Sanger Institute COVID-19 Surveillance Team |
| EPI_ISL_3139301, EPI_ISL_3139302 | Lighthouse Laboratory Plymouth | Wellcome Sanger Institute for the COVID-19 Genomics UK (COG-UK) Consortium | Lighthouse Laboratory Plymouth and Alex Alderton, Roberto Amato, Jeffrey Barrett, Sonia Goncalves, Ewan Harrison, David K. Jackson, Ian Johnston, Dominic Kwiatkowski, Cordelia Langford, John Sillitoe on behalf of the Wellcome Sanger Institute COVID-19 Surveillance Team |
| EPI_ISL_3139303 | Lighthouse Lab in Glasgow | Wellcome Sanger Institute for the COVID-19 Genomics UK (COG-UK) Consortium | Harper VanSteenhouse, Yumi Kasai, David Gray, Carol Clugston, Anna Dominiczak and Alex Alderton, Roberto Amato, Jeffrey Barrett, Sonia Goncalves, Ewan Harrison, David K. Jackson, Ian Johnston, Dominic Kwiatkowski, Cordelia Langford, John Sillitoe on behalf of the Wellcome Sanger Institute COVID-19 Surveillance Team |
| EPI_ISL_3139304, EPI_ISL_3139305 | Lighthouse Laboratory Plymouth | Wellcome Sanger Institute for the COVID-19 Genomics UK (COG-UK) Consortium | Lighthouse Laboratory Plymouth and Alex Alderton, Roberto Amato, Jeffrey Barrett, Sonia Goncalves, Ewan Harrison, David K. Jackson, Ian Johnston, Dominic Kwiatkowski, Cordelia Langford, John Sillitoe on behalf of the Wellcome Sanger Institute COVID-19 Surveillance Team |
| EPI_ISL_3139306 | Lighthouse Lab in Milton Keynes | Wellcome Sanger Institute for the COVID-19 Genomics UK (COG-UK) Consortium | The Lighthouse Lab in Milton Keynes and Alex Alderton, Roberto Amato, Jeffrey Barrett, Sonia Goncalves, Ewan Harrison, David K. Jackson, Ian Johnston, Dominic Kwiatkowski, Cordelia Langford, John Sillitoe on behalf of the Wellcome Sanger Institute COVID-19 Surveillance Team |
| EPI_ISL_3139307 | Lighthouse Laboratory Plymouth | Wellcome Sanger Institute for the COVID-19 Genomics UK (COG-UK) Consortium | Lighthouse Laboratory Plymouth and Alex Alderton, Roberto Amato, Jeffrey Barrett, Sonia Goncalves, Ewan Harrison, David K. Jackson, Ian Johnston, Dominic Kwiatkowski, Cordelia Langford, John Sillitoe on behalf of the Wellcome Sanger Institute COVID-19 Surveillance Team |
| EPI_ISL_3139308 | Lighthouse Lab in Milton Keynes | Wellcome Sanger Institute for the COVID-19 Genomics UK (COG-UK) Consortium | The Lighthouse Lab in Milton Keynes and Alex Alderton, Roberto Amato, Jeffrey Barrett, Sonia Goncalves, Ewan Harrison, David K. Jackson, Ian Johnston, Dominic Kwiatkowski, Cordelia Langford, John Sillitoe on behalf of the Wellcome Sanger Institute COVID-19 Surveillance Team |
| EPI_ISL_3139309, EPI_ISL_3139310 | Lighthouse Laboratory Plymouth | Wellcome Sanger Institute for the COVID-19 Genomics UK (COG-UK) Consortium | Lighthouse Laboratory Plymouth and Alex Alderton, Roberto Amato, Jeffrey Barrett, Sonia Goncalves, Ewan Harrison, David K. Jackson, Ian Johnston, Dominic Kwiatkowski, Cordelia Langford, John Sillitoe on behalf of the Wellcome Sanger Institute COVID-19 Surveillance Team |
| EPI_ISL_3139311, EPI_ISL_3139312, EPI_ISL_3139313 | Lighthouse Lab in Glasgow | Wellcome Sanger Institute for the COVID-19 Genomics UK (COG-UK) Consortium | Harper VanSteenhouse, Yumi Kasai, David Gray, Carol Clugston, Anna Dominiczak and Alex Alderton, Roberto Amato, Jeffrey Barrett, Sonia Goncalves, Ewan Harrison, David K. Jackson, Ian Johnston, Dominic Kwiatkowski, Cordelia Langford, John Sillitoe on behalf of the Wellcome Sanger Institute COVID-19 Surveillance Team |
| EPI_ISL_3139314, EPI_ISL_3139315, EPI_ISL_3139316 | Lighthouse Laboratory Plymouth | Wellcome Sanger Institute for the COVID-19 Genomics UK (COG-UK) Consortium | Lighthouse Laboratory Plymouth and Alex Alderton, Roberto Amato, Jeffrey Barrett, Sonia Goncalves, Ewan Harrison, David K. Jackson, Ian Johnston, Dominic Kwiatkowski, Cordelia Langford, John Sillitoe on behalf of the Wellcome Sanger Institute COVID-19 Surveillance Team |
| EPI_ISL_3139317 | Lighthouse Lab in Milton Keynes | Wellcome Sanger Institute for the COVID-19 Genomics UK (COG-UK) Consortium | The Lighthouse Lab in Milton Keynes and Alex Alderton, Roberto Amato, Jeffrey Barrett, Sonia Goncalves, Ewan Harrison, David K. Jackson, Ian Johnston, Dominic Kwiatkowski, Cordelia Langford, John Sillitoe on behalf of the Wellcome Sanger Institute COVID-19 Surveillance Team |
| EPI_ISL_3139318 | Lighthouse Laboratory Plymouth | Wellcome Sanger Institute for the COVID-19 Genomics UK (COG-UK) Consortium | Lighthouse Laboratory Plymouth and Alex Alderton, Roberto Amato, Jeffrey Barrett, Sonia Goncalves, Ewan Harrison, David K. Jackson, Ian Johnston, Dominic Kwiatkowski, Cordelia Langford, John Sillitoe on behalf of the Wellcome Sanger Institute COVID-19 Surveillance Team |
| EPI_ISL_3139319, EPI_ISL_3139320, EPI_ISL_3139321, EPI_ISL_3139322 | Lighthouse Lab in Glasgow | Wellcome Sanger Institute for the COVID-19 Genomics UK (COG-UK) Consortium | Harper VanSteenhouse, Yumi Kasai, David Gray, Carol Clugston, Anna Dominiczak and Alex Alderton, Roberto Amato, Jeffrey Barrett, Sonia Goncalves, Ewan Harrison, David K. Jackson, Ian Johnston, Dominic Kwiatkowski, Cordelia Langford, John Sillitoe on behalf of the Wellcome Sanger Institute COVID-19 Surveillance Team |
| EPI_ISL_3139323 | Lighthouse Laboratory Plymouth | Wellcome Sanger Institute for the COVID-19 Genomics UK (COG-UK) Consortium | Lighthouse Laboratory Plymouth and Alex Alderton, Roberto Amato, Jeffrey Barrett, Sonia Goncalves, Ewan Harrison, David K. Jackson, Ian Johnston, Dominic Kwiatkowski, Cordelia Langford, John Sillitoe on behalf of the Wellcome Sanger Institute COVID-19 Surveillance Team |
| EPI_ISL_3139324 | Lighthouse Lab in Milton Keynes | Wellcome Sanger Institute for the COVID-19 Genomics UK (COG-UK) Consortium | The Lighthouse Lab in Milton Keynes and Alex Alderton, Roberto Amato, Jeffrey Barrett, Sonia Goncalves, Ewan Harrison, David K. Jackson, Ian Johnston, Dominic Kwiatkowski, Cordelia Langford, John Sillitoe on behalf of the Wellcome Sanger Institute COVID-19 Surveillance Team |
| EPI_ISL_3139325, EPI_ISL_3139326, EPI_ISL_3139327, | Lighthouse Laboratory Plymouth | Wellcome Sanger Institute for the COVID-19 Genomics UK (COG-UK) Consortium | Lighthouse Laboratory Plymouth and Alex Alderton, Roberto Amato, Jeffrey Barrett, Sonia Goncalves, Ewan Harrison, David K. Jackson, Ian Johnston, Dominic Kwiatkowski, Cordelia Langford, John Sillitoe on behalf of the Wellcome Sanger Institute COVID-19 Surveillance Team |

| | | | |
|---|---|---|---|
| EPI_ISL_3139328, EPI_ISL_3139329 | | | |
| EPI_ISL_3139330 | Lighthouse Lab in Glasgow | Wellcome Sanger Institute for the COVID-19 Genomics UK (COG-UK) Consortium | Harper VanSteenhouse, Yumi Kasai, David Gray, Carol Clugston, Anna Dominiczak and Alex Alderton, Roberto Amato, Jeffrey Barrett, Sonia Goncalves, Ewan Harrison, David K. Jackson, Ian Johnston, Dominic Kwiatkowski, Cordelia Langford, John Sillitoe on behalf of the Wellcome Sanger Institute COVID-19 Surveillance Team |
| EPI_ISL_3139331, EPI_ISL_3139332 | Lighthouse Laboratory Plymouth | Wellcome Sanger Institute for the COVID-19 Genomics UK (COG-UK) Consortium | Lighthouse Laboratory Plymouth and Alex Alderton, Roberto Amato, Jeffrey Barrett, Sonia Goncalves, Ewan Harrison, David K. Jackson, Ian Johnston, Dominic Kwiatkowski, Cordelia Langford, John Sillitoe on behalf of the Wellcome Sanger Institute COVID-19 Surveillance Team |
| EPI_ISL_3139333 | Lighthouse Lab in Milton Keynes | Wellcome Sanger Institute for the COVID-19 Genomics UK (COG-UK) Consortium | The Lighthouse Lab in Milton Keynes and Alex Alderton, Roberto Amato, Jeffrey Barrett, Sonia Goncalves, Ewan Harrison, David K. Jackson, Ian Johnston, Dominic Kwiatkowski, Cordelia Langford, John Sillitoe on behalf of the Wellcome Sanger Institute COVID-19 Surveillance Team |
| EPI_ISL_3139334 | Lighthouse Laboratory Plymouth | Wellcome Sanger Institute for the COVID-19 Genomics UK (COG-UK) Consortium | Lighthouse Laboratory Plymouth and Alex Alderton, Roberto Amato, Jeffrey Barrett, Sonia Goncalves, Ewan Harrison, David K. Jackson, Ian Johnston, Dominic Kwiatkowski, Cordelia Langford, John Sillitoe on behalf of the Wellcome Sanger Institute COVID-19 Surveillance Team |
| EPI_ISL_3139335, EPI_ISL_3139336, EPI_ISL_3139337 | Lighthouse Lab in Glasgow | Wellcome Sanger Institute for the COVID-19 Genomics UK (COG-UK) Consortium | Harper VanSteenhouse, Yumi Kasai, David Gray, Carol Clugston, Anna Dominiczak and Alex Alderton, Roberto Amato, Jeffrey Barrett, Sonia Goncalves, Ewan Harrison, David K. Jackson, Ian Johnston, Dominic Kwiatkowski, Cordelia Langford, John Sillitoe on behalf of the Wellcome Sanger Institute COVID-19 Surveillance Team |
| EPI_ISL_3139338, EPI_ISL_3139339 | Lighthouse Laboratory Plymouth | Wellcome Sanger Institute for the COVID-19 Genomics UK (COG-UK) Consortium | Lighthouse Laboratory Plymouth and Alex Alderton, Roberto Amato, Jeffrey Barrett, Sonia Goncalves, Ewan Harrison, David K. Jackson, Ian Johnston, Dominic Kwiatkowski, Cordelia Langford, John Sillitoe on behalf of the Wellcome Sanger Institute COVID-19 Surveillance Team |
| EPI_ISL_3139340 | Lighthouse Lab in Glasgow | Wellcome Sanger Institute for the COVID-19 Genomics UK (COG-UK) Consortium | Harper VanSteenhouse, Yumi Kasai, David Gray, Carol Clugston, Anna Dominiczak and Alex Alderton, Roberto Amato, Jeffrey Barrett, Sonia Goncalves, Ewan Harrison, David K. Jackson, Ian Johnston, Dominic Kwiatkowski, Cordelia Langford, John Sillitoe on behalf of the Wellcome Sanger Institute COVID-19 Surveillance Team |
| EPI_ISL_3139341, EPI_ISL_3139342, EPI_ISL_3139343, EPI_ISL_3139344, EPI_ISL_3139345, EPI_ISL_3139346, EPI_ISL_3139347, EPI_ISL_3139348, EPI_ISL_3139349 | Lighthouse Laboratory Plymouth | Wellcome Sanger Institute for the COVID-19 Genomics UK (COG-UK) Consortium | Lighthouse Laboratory Plymouth and Alex Alderton, Roberto Amato, Jeffrey Barrett, Sonia Goncalves, Ewan Harrison, David K. Jackson, Ian Johnston, Dominic Kwiatkowski, Cordelia Langford, John Sillitoe on behalf of the Wellcome Sanger Institute COVID-19 Surveillance Team |
| EPI_ISL_3139350, EPI_ISL_3139351, EPI_ISL_3139352 | Lighthouse Lab in Glasgow | Wellcome Sanger Institute for the COVID-19 Genomics UK (COG-UK) Consortium | Harper VanSteenhouse, Yumi Kasai, David Gray, Carol Clugston, Anna Dominiczak and Alex Alderton, Roberto Amato, Jeffrey Barrett, Sonia Goncalves, Ewan Harrison, David K. Jackson, Ian Johnston, Dominic Kwiatkowski, Cordelia Langford, John Sillitoe on behalf of the Wellcome Sanger Institute COVID-19 Surveillance Team |
| EPI_ISL_3139353 | Lighthouse Lab in Milton Keynes | Wellcome Sanger Institute for the COVID-19 Genomics UK (COG-UK) Consortium | The Lighthouse Lab in Milton Keynes and Alex Alderton, Roberto Amato, Jeffrey Barrett, Sonia Goncalves, Ewan Harrison, David K. Jackson, Ian Johnston, Dominic Kwiatkowski, Cordelia Langford, John Sillitoe on behalf of the Wellcome Sanger Institute COVID-19 Surveillance Team |
| EPI_ISL_3139354 | Lighthouse Lab in Glasgow | Wellcome Sanger Institute for the COVID-19 Genomics UK (COG-UK) Consortium | Harper VanSteenhouse, Yumi Kasai, David Gray, Carol Clugston, Anna Dominiczak and Alex Alderton, Roberto Amato, Jeffrey Barrett, Sonia Goncalves, Ewan Harrison, David K. Jackson, Ian Johnston, Dominic Kwiatkowski, Cordelia Langford, John Sillitoe on behalf of the Wellcome Sanger Institute COVID-19 Surveillance Team |
| EPI_ISL_3139355 | Lighthouse Laboratory Plymouth | Wellcome Sanger Institute for the COVID-19 Genomics UK (COG-UK) Consortium | Lighthouse Laboratory Plymouth and Alex Alderton, Roberto Amato, Jeffrey Barrett, Sonia Goncalves, Ewan Harrison, David K. Jackson, Ian Johnston, Dominic Kwiatkowski, Cordelia Langford, John Sillitoe on behalf of the Wellcome Sanger Institute COVID-19 Surveillance Team |
| EPI_ISL_3139356, EPI_ISL_3139357, EPI_ISL_3139358 | Lighthouse Lab in Glasgow | Wellcome Sanger Institute for the COVID-19 Genomics UK (COG-UK) Consortium | Harper VanSteenhouse, Yumi Kasai, David Gray, Carol Clugston, Anna Dominiczak and Alex Alderton, Roberto Amato, Jeffrey Barrett, Sonia Goncalves, Ewan Harrison, David K. Jackson, Ian Johnston, Dominic Kwiatkowski, Cordelia Langford, John Sillitoe on behalf of the Wellcome Sanger Institute COVID-19 Surveillance Team |
| EPI_ISL_3139359, EPI_ISL_3139360 | Lighthouse Lab in Milton Keynes | Wellcome Sanger Institute for the COVID-19 Genomics UK (COG-UK) Consortium | The Lighthouse Lab in Milton Keynes and Alex Alderton, Roberto Amato, Jeffrey Barrett, Sonia Goncalves, Ewan Harrison, David K. Jackson, Ian Johnston, Dominic Kwiatkowski, Cordelia Langford, John Sillitoe on behalf of the Wellcome Sanger Institute COVID-19 Surveillance Team |
| EPI_ISL_3139361 | Lighthouse Laboratory Plymouth | Wellcome Sanger Institute for the COVID-19 Genomics UK (COG-UK) Consortium | Lighthouse Laboratory Plymouth and Alex Alderton, Roberto Amato, Jeffrey Barrett, Sonia Goncalves, Ewan Harrison, David K. Jackson, Ian Johnston, Dominic Kwiatkowski, Cordelia Langford, John Sillitoe on behalf of the Wellcome Sanger Institute COVID-19 Surveillance Team |
| EPI_ISL_3139362 | Lighthouse Lab in Glasgow | Wellcome Sanger Institute for the COVID-19 Genomics UK (COG-UK) Consortium | Harper VanSteenhouse, Yumi Kasai, David Gray, Carol Clugston, Anna Dominiczak and Alex Alderton, Roberto Amato, Jeffrey Barrett, Sonia Goncalves, Ewan Harrison, David K. Jackson, Ian Johnston, Dominic Kwiatkowski, Cordelia Langford, John Sillitoe on behalf of the Wellcome Sanger Institute COVID-19 Surveillance Team |
| EPI_ISL_3139363 | Lighthouse Lab in Milton Keynes | Wellcome Sanger Institute for the COVID-19 Genomics UK (COG-UK) Consortium | The Lighthouse Lab in Milton Keynes and Alex Alderton, Roberto Amato, Jeffrey Barrett, Sonia Goncalves, Ewan Harrison, David K. Jackson, Ian Johnston, Dominic Kwiatkowski, Cordelia Langford, John Sillitoe on behalf of the Wellcome Sanger Institute COVID-19 Surveillance Team |
| EPI_ISL_3139364, EPI_ISL_3139365 | Lighthouse Laboratory Plymouth | Wellcome Sanger Institute for the COVID-19 Genomics UK (COG-UK) Consortium | Lighthouse Laboratory Plymouth and Alex Alderton, Roberto Amato, Jeffrey Barrett, Sonia Goncalves, Ewan Harrison, David K. Jackson, Ian Johnston, Dominic Kwiatkowski, Cordelia Langford, John Sillitoe on behalf of the Wellcome Sanger Institute COVID-19 Surveillance Team |
| EPI_ISL_3139366, EPI_ISL_3139367 | Lighthouse Lab in Glasgow | Wellcome Sanger Institute for the COVID-19 Genomics UK (COG-UK) Consortium | Harper VanSteenhouse, Yumi Kasai, David Gray, Carol Clugston, Anna Dominiczak and Alex Alderton, Roberto Amato, Jeffrey Barrett, Sonia Goncalves, Ewan Harrison, David K. Jackson, Ian Johnston, Dominic Kwiatkowski, Cordelia Langford, John Sillitoe on behalf of the Wellcome Sanger Institute COVID-19 Surveillance Team |
| EPI_ISL_3139368, EPI_ISL_3139369 | Lighthouse Laboratory Plymouth | Wellcome Sanger Institute for the COVID-19 Genomics UK (COG-UK) Consortium | Lighthouse Laboratory Plymouth and Alex Alderton, Roberto Amato, Jeffrey Barrett, Sonia Goncalves, Ewan Harrison, David K. Jackson, Ian Johnston, Dominic Kwiatkowski, Cordelia Langford, John Sillitoe on behalf of the Wellcome Sanger Institute COVID-19 Surveillance Team |
| EPI_ISL_3139370 | Lighthouse Lab in Glasgow | Wellcome Sanger Institute for the COVID-19 Genomics UK (COG-UK) Consortium | Harper VanSteenhouse, Yumi Kasai, David Gray, Carol Clugston, Anna Dominiczak and Alex Alderton, Roberto Amato, Jeffrey Barrett, Sonia Goncalves, Ewan Harrison, David K. Jackson, Ian Johnston, Dominic Kwiatkowski, Cordelia Langford, John Sillitoe on behalf of the Wellcome Sanger Institute COVID-19 Surveillance Team |
| EPI_ISL_3139371, EPI_ISL_3139372 | Lighthouse Lab in Milton Keynes | Wellcome Sanger Institute for the COVID-19 Genomics UK (COG-UK) Consortium | The Lighthouse Lab in Milton Keynes and Alex Alderton, Roberto Amato, Jeffrey Barrett, Sonia Goncalves, Ewan Harrison, David K. Jackson, Ian Johnston, Dominic Kwiatkowski, Cordelia Langford, John Sillitoe on behalf of the Wellcome Sanger Institute COVID-19 Surveillance Team |
| EPI_ISL_3139373 | Lighthouse Laboratory Plymouth | Wellcome Sanger Institute for the COVID-19 Genomics UK (COG-UK) Consortium | Lighthouse Laboratory Plymouth and Alex Alderton, Roberto Amato, Jeffrey Barrett, Sonia Goncalves, Ewan Harrison, David K. Jackson, Ian Johnston, Dominic Kwiatkowski, Cordelia Langford, John Sillitoe on behalf of the Wellcome Sanger Institute COVID-19 Surveillance Team |
| EPI_ISL_3139374, EPI_ISL_3139375 | Lighthouse Lab in Glasgow | Wellcome Sanger Institute for the COVID-19 Genomics UK (COG-UK) Consortium | Harper VanSteenhouse, Yumi Kasai, David Gray, Carol Clugston, Anna Dominiczak and Alex Alderton, Roberto Amato, Jeffrey Barrett, Sonia Goncalves, Ewan Harrison, David K. Jackson, Ian Johnston, Dominic Kwiatkowski, Cordelia Langford, John Sillitoe on behalf of the Wellcome Sanger Institute COVID-19 Surveillance Team |
| EPI_ISL_3139376, EPI_ISL_3139377, EPI_ISL_3139378 | Lighthouse Laboratory Plymouth | Wellcome Sanger Institute for the COVID-19 Genomics UK (COG-UK) Consortium | Lighthouse Laboratory Plymouth and Alex Alderton, Roberto Amato, Jeffrey Barrett, Sonia Goncalves, Ewan Harrison, David K. Jackson, Ian Johnston, Dominic Kwiatkowski, Cordelia Langford, John Sillitoe on behalf of the Wellcome Sanger Institute COVID-19 Surveillance Team |
| EPI_ISL_3139379 | Lighthouse Lab in Milton Keynes | Wellcome Sanger Institute for the COVID-19 Genomics UK (COG-UK) Consortium | The Lighthouse Lab in Milton Keynes and Alex Alderton, Roberto Amato, Jeffrey Barrett, Sonia Goncalves, Ewan Harrison, David K. Jackson, Ian Johnston, Dominic Kwiatkowski, Cordelia Langford, John Sillitoe on behalf of the Wellcome Sanger Institute COVID-19 Surveillance Team |
| EPI_ISL_3139380, EPI_ISL_3139381, EPI_ISL_3139382 | Lighthouse Laboratory Plymouth | Wellcome Sanger Institute for the COVID-19 Genomics UK (COG-UK) Consortium | Lighthouse Laboratory Plymouth and Alex Alderton, Roberto Amato, Jeffrey Barrett, Sonia Goncalves, Ewan Harrison, David K. Jackson, Ian Johnston, Dominic Kwiatkowski, Cordelia Langford, John Sillitoe on behalf of the Wellcome Sanger Institute COVID-19 Surveillance Team |
| EPI_ISL_3139383 | Lighthouse Lab in Glasgow | Wellcome Sanger Institute for the COVID-19 Genomics UK (COG-UK) Consortium | Harper VanSteenhouse, Yumi Kasai, David Gray, Carol Clugston, Anna Dominiczak and Alex Alderton, Roberto Amato, Jeffrey Barrett, Sonia Goncalves, Ewan Harrison, David K. Jackson, Ian Johnston, Dominic Kwiatkowski, Cordelia Langford, John Sillitoe on behalf of the Wellcome Sanger Institute COVID-19 Surveillance Team |
| EPI_ISL_3139384 | Lighthouse Laboratory Plymouth | Wellcome Sanger Institute for the COVID-19 Genomics UK (COG-UK) Consortium | Lighthouse Laboratory Plymouth and Alex Alderton, Roberto Amato, Jeffrey Barrett, Sonia Goncalves, Ewan Harrison, David K. Jackson, Ian Johnston, Dominic Kwiatkowski, Cordelia Langford, John Sillitoe on behalf of the Wellcome Sanger Institute COVID-19 Surveillance Team |
| EPI_ISL_3139385, EPI_ISL_3139386 | Lighthouse Lab in Milton Keynes | Wellcome Sanger Institute for the COVID-19 Genomics UK (COG-UK) Consortium | The Lighthouse Lab in Milton Keynes and Alex Alderton, Roberto Amato, Jeffrey Barrett, Sonia Goncalves, Ewan Harrison, David K. Jackson, Ian Johnston, Dominic Kwiatkowski, Cordelia Langford, John Sillitoe on behalf of the Wellcome Sanger Institute COVID-19 Surveillance Team |
| EPI_ISL_3139387 | Lighthouse Laboratory Plymouth | Wellcome Sanger Institute for the COVID-19 Genomics UK (COG-UK) Consortium | Lighthouse Laboratory Plymouth and Alex Alderton, Roberto Amato, Jeffrey Barrett, Sonia Goncalves, Ewan Harrison, David K. Jackson, Ian Johnston, Dominic Kwiatkowski, Cordelia Langford, John Sillitoe on behalf of the Wellcome Sanger Institute COVID-19 Surveillance Team |
| EPI_ISL_3139388, EPI_ISL_3139389 | Lighthouse Lab in Glasgow | Wellcome Sanger Institute for the COVID-19 Genomics UK (COG-UK) Consortium | Harper VanSteenhouse, Yumi Kasai, David Gray, Carol Clugston, Anna Dominiczak and Alex Alderton, Roberto Amato, Jeffrey Barrett, Sonia Goncalves, Ewan Harrison, David K. Jackson, Ian Johnston, Dominic Kwiatkowski, Cordelia Langford, John Sillitoe on behalf of the Wellcome Sanger Institute COVID-19 Surveillance Team |
| EPI_ISL_3139390 | Lighthouse Lab in Milton Keynes | Wellcome Sanger Institute for the COVID-19 Genomics UK (COG-UK) Consortium | The Lighthouse Lab in Milton Keynes and Alex Alderton, Roberto Amato, Jeffrey Barrett, Sonia Goncalves, Ewan Harrison, David K. Jackson, Ian Johnston, Dominic Kwiatkowski, Cordelia Langford, John Sillitoe on behalf of the Wellcome Sanger Institute COVID-19 Surveillance Team |
| EPI_ISL_3139391, EPI_ISL_3139392, EPI_ISL_3139393, EPI_ISL_3139394 | Lighthouse Laboratory Plymouth | Wellcome Sanger Institute for the COVID-19 Genomics UK (COG-UK) Consortium | Lighthouse Laboratory Plymouth and Alex Alderton, Roberto Amato, Jeffrey Barrett, Sonia Goncalves, Ewan Harrison, David K. Jackson, Ian Johnston, Dominic Kwiatkowski, Cordelia Langford, John Sillitoe on behalf of the Wellcome Sanger Institute COVID-19 Surveillance Team |
| EPI_ISL_3139395, EPI_ISL_3139396 | Lighthouse Lab in Milton Keynes | Wellcome Sanger Institute for the COVID-19 Genomics UK (COG-UK) Consortium | The Lighthouse Lab in Milton Keynes and Alex Alderton, Roberto Amato, Jeffrey Barrett, Sonia Goncalves, Ewan Harrison, David K. Jackson, Ian Johnston, Dominic Kwiatkowski, Cordelia Langford, John Sillitoe on behalf of the Wellcome Sanger Institute COVID-19 Surveillance Team |
| EPI_ISL_3139397 | Lighthouse Lab in Glasgow | Wellcome Sanger Institute for the COVID-19 Genomics UK (COG-UK) Consortium | Harper VanSteenhouse, Yumi Kasai, David Gray, Carol Clugston, Anna Dominiczak and Alex Alderton, Roberto Amato, Jeffrey Barrett, Sonia Goncalves, Ewan Harrison, David K. Jackson, Ian Johnston, Dominic Kwiatkowski, Cordelia Langford, John Sillitoe on behalf of the Wellcome Sanger Institute COVID-19 Surveillance Team |
| EPI_ISL_3139398, EPI_ISL_3139399 | Lighthouse Lab in Milton Keynes | Wellcome Sanger Institute for the COVID-19 Genomics UK (COG-UK) Consortium | The Lighthouse Lab in Milton Keynes and Alex Alderton, Roberto Amato, Jeffrey Barrett, Sonia Goncalves, Ewan Harrison, David K. Jackson, Ian Johnston, Dominic Kwiatkowski, Cordelia Langford, John Sillitoe on behalf of the Wellcome Sanger Institute COVID-19 Surveillance Team |
| EPI_ISL_3139400 | Lighthouse Lab in Glasgow | Wellcome Sanger Institute for the COVID-19 Genomics UK (COG-UK) Consortium | Harper VanSteenhouse, Yumi Kasai, David Gray, Carol Clugston, Anna Dominiczak and Alex Alderton, Roberto Amato, Jeffrey Barrett, Sonia Goncalves, Ewan Harrison, David K. Jackson, Ian Johnston, Dominic Kwiatkowski, Cordelia Langford, John Sillitoe on behalf of the Wellcome Sanger Institute COVID-19 Surveillance Team |
| EPI_ISL_3139401, | Lighthouse Laboratory Plymouth | Wellcome Sanger Institute for the COVID-19 Genomics UK (COG-UK) | Lighthouse Laboratory Plymouth and Alex Alderton, Roberto Amato, Jeffrey Barrett, Sonia Goncalves, Ewan Harrison, David K. Jackson, Ian Johnston, Dominic Kwiatkowski, Cordelia Langford, John Sillitoe on behalf of the Wellcome Sanger |

| | | | |
|---|---|---|---|
| EPI_ISL_3139402, EPI_ISL_3139403 | | Consortium | Institute COVID-19 Surveillance Team |
| EPI_ISL_3139404 | Lighthouse Lab in Glasgow | Wellcome Sanger Institute for the COVID-19 Genomics UK (COG-UK) Consortium | Harper VanSteenhouse, Yumi Kasai, David Gray, Carol Clugston, Anna Dominiczak and Alex Alderton, Roberto Amato, Jeffrey Barrett, Sonia Goncalves, Ewan Harrison, David K. Jackson, Ian Johnston, Dominic Kwiatkowski, Cordelia Langford, John Sillitoe on behalf of the Wellcome Sanger Institute COVID-19 Surveillance Team |
| EPI_ISL_3139405 | Lighthouse Laboratory Plymouth | Wellcome Sanger Institute for the COVID-19 Genomics UK (COG-UK) Consortium | Lighthouse Laboratory Plymouth and Alex Alderton, Roberto Amato, Jeffrey Barrett, Sonia Goncalves, Ewan Harrison, David K. Jackson, Ian Johnston, Dominic Kwiatkowski, Cordelia Langford, John Sillitoe on behalf of the Wellcome Sanger Institute COVID-19 Surveillance Team |
| EPI_ISL_3139406 | Lighthouse Lab in Glasgow | Wellcome Sanger Institute for the COVID-19 Genomics UK (COG-UK) Consortium | Harper VanSteenhouse, Yumi Kasai, David Gray, Carol Clugston, Anna Dominiczak and Alex Alderton, Roberto Amato, Jeffrey Barrett, Sonia Goncalves, Ewan Harrison, David K. Jackson, Ian Johnston, Dominic Kwiatkowski, Cordelia Langford, John Sillitoe on behalf of the Wellcome Sanger Institute COVID-19 Surveillance Team |
| EPI_ISL_3139407 | Lighthouse Lab in Milton Keynes | Wellcome Sanger Institute for the COVID-19 Genomics UK (COG-UK) Consortium | The Lighthouse Lab in Milton Keynes and Alex Alderton, Roberto Amato, Jeffrey Barrett, Sonia Goncalves, Ewan Harrison, David K. Jackson, Ian Johnston, Dominic Kwiatkowski, Cordelia Langford, John Sillitoe on behalf of the Wellcome Sanger Institute COVID-19 Surveillance Team |
| EPI_ISL_3139408, EPI_ISL_3139409 | Lighthouse Laboratory Plymouth | Wellcome Sanger Institute for the COVID-19 Genomics UK (COG-UK) Consortium | Lighthouse Laboratory Plymouth and Alex Alderton, Roberto Amato, Jeffrey Barrett, Sonia Goncalves, Ewan Harrison, David K. Jackson, Ian Johnston, Dominic Kwiatkowski, Cordelia Langford, John Sillitoe on behalf of the Wellcome Sanger Institute COVID-19 Surveillance Team |
| EPI_ISL_3139410 | Lighthouse Lab in Glasgow | Wellcome Sanger Institute for the COVID-19 Genomics UK (COG-UK) Consortium | Harper VanSteenhouse, Yumi Kasai, David Gray, Carol Clugston, Anna Dominiczak and Alex Alderton, Roberto Amato, Jeffrey Barrett, Sonia Goncalves, Ewan Harrison, David K. Jackson, Ian Johnston, Dominic Kwiatkowski, Cordelia Langford, John Sillitoe on behalf of the Wellcome Sanger Institute COVID-19 Surveillance Team |
| EPI_ISL_3139411 | Lighthouse Lab in Milton Keynes | Wellcome Sanger Institute for the COVID-19 Genomics UK (COG-UK) Consortium | The Lighthouse Lab in Milton Keynes and Alex Alderton, Roberto Amato, Jeffrey Barrett, Sonia Goncalves, Ewan Harrison, David K. Jackson, Ian Johnston, Dominic Kwiatkowski, Cordelia Langford, John Sillitoe on behalf of the Wellcome Sanger Institute COVID-19 Surveillance Team |
| EPI_ISL_3139412 | Lighthouse Lab in Glasgow | Wellcome Sanger Institute for the COVID-19 Genomics UK (COG-UK) Consortium | Harper VanSteenhouse, Yumi Kasai, David Gray, Carol Clugston, Anna Dominiczak and Alex Alderton, Roberto Amato, Jeffrey Barrett, Sonia Goncalves, Ewan Harrison, David K. Jackson, Ian Johnston, Dominic Kwiatkowski, Cordelia Langford, John Sillitoe on behalf of the Wellcome Sanger Institute COVID-19 Surveillance Team |
| EPI_ISL_3139413, EPI_ISL_3139414 | Lighthouse Lab in Milton Keynes | Wellcome Sanger Institute for the COVID-19 Genomics UK (COG-UK) Consortium | The Lighthouse Lab in Milton Keynes and Alex Alderton, Roberto Amato, Jeffrey Barrett, Sonia Goncalves, Ewan Harrison, David K. Jackson, Ian Johnston, Dominic Kwiatkowski, Cordelia Langford, John Sillitoe on behalf of the Wellcome Sanger Institute COVID-19 Surveillance Team |
| EPI_ISL_3139415 | Lighthouse Lab in Glasgow | Wellcome Sanger Institute for the COVID-19 Genomics UK (COG-UK) Consortium | Harper VanSteenhouse, Yumi Kasai, David Gray, Carol Clugston, Anna Dominiczak and Alex Alderton, Roberto Amato, Jeffrey Barrett, Sonia Goncalves, Ewan Harrison, David K. Jackson, Ian Johnston, Dominic Kwiatkowski, Cordelia Langford, John Sillitoe on behalf of the Wellcome Sanger Institute COVID-19 Surveillance Team |
| EPI_ISL_3139416 | Lighthouse Laboratory Plymouth | Wellcome Sanger Institute for the COVID-19 Genomics UK (COG-UK) Consortium | Lighthouse Laboratory Plymouth and Alex Alderton, Roberto Amato, Jeffrey Barrett, Sonia Goncalves, Ewan Harrison, David K. Jackson, Ian Johnston, Dominic Kwiatkowski, Cordelia Langford, John Sillitoe on behalf of the Wellcome Sanger Institute COVID-19 Surveillance Team |
| EPI_ISL_3139417 | Lighthouse Lab in Milton Keynes | Wellcome Sanger Institute for the COVID-19 Genomics UK (COG-UK) Consortium | The Lighthouse Lab in Milton Keynes and Alex Alderton, Roberto Amato, Jeffrey Barrett, Sonia Goncalves, Ewan Harrison, David K. Jackson, Ian Johnston, Dominic Kwiatkowski, Cordelia Langford, John Sillitoe on behalf of the Wellcome Sanger Institute COVID-19 Surveillance Team |
| EPI_ISL_3139418, EPI_ISL_3139419 | Lighthouse Laboratory Plymouth | Wellcome Sanger Institute for the COVID-19 Genomics UK (COG-UK) Consortium | Lighthouse Laboratory Plymouth and Alex Alderton, Roberto Amato, Jeffrey Barrett, Sonia Goncalves, Ewan Harrison, David K. Jackson, Ian Johnston, Dominic Kwiatkowski, Cordelia Langford, John Sillitoe on behalf of the Wellcome Sanger Institute COVID-19 Surveillance Team |
| EPI_ISL_3139420, EPI_ISL_3139421, EPI_ISL_3139422, EPI_ISL_3139423, EPI_ISL_3139424 | Lighthouse Lab in Glasgow | Wellcome Sanger Institute for the COVID-19 Genomics UK (COG-UK) Consortium | Harper VanSteenhouse, Yumi Kasai, David Gray, Carol Clugston, Anna Dominiczak and Alex Alderton, Roberto Amato, Jeffrey Barrett, Sonia Goncalves, Ewan Harrison, David K. Jackson, Ian Johnston, Dominic Kwiatkowski, Cordelia Langford, John Sillitoe on behalf of the Wellcome Sanger Institute COVID-19 Surveillance Team |
| EPI_ISL_3139425, EPI_ISL_3139426 | Lighthouse Laboratory Plymouth | Wellcome Sanger Institute for the COVID-19 Genomics UK (COG-UK) Consortium | Lighthouse Laboratory Plymouth and Alex Alderton, Roberto Amato, Jeffrey Barrett, Sonia Goncalves, Ewan Harrison, David K. Jackson, Ian Johnston, Dominic Kwiatkowski, Cordelia Langford, John Sillitoe on behalf of the Wellcome Sanger Institute COVID-19 Surveillance Team |
| EPI_ISL_3139427, EPI_ISL_3139428 | Lighthouse Lab in Glasgow | Wellcome Sanger Institute for the COVID-19 Genomics UK (COG-UK) Consortium | Harper VanSteenhouse, Yumi Kasai, David Gray, Carol Clugston, Anna Dominiczak and Alex Alderton, Roberto Amato, Jeffrey Barrett, Sonia Goncalves, Ewan Harrison, David K. Jackson, Ian Johnston, Dominic Kwiatkowski, Cordelia Langford, John Sillitoe on behalf of the Wellcome Sanger Institute COVID-19 Surveillance Team |
| EPI_ISL_3139429 | Lighthouse Laboratory Plymouth | Wellcome Sanger Institute for the COVID-19 Genomics UK (COG-UK) Consortium | Lighthouse Laboratory Plymouth and Alex Alderton, Roberto Amato, Jeffrey Barrett, Sonia Goncalves, Ewan Harrison, David K. Jackson, Ian Johnston, Dominic Kwiatkowski, Cordelia Langford, John Sillitoe on behalf of the Wellcome Sanger Institute COVID-19 Surveillance Team |
| EPI_ISL_3139430 | Lighthouse Lab in Glasgow | Wellcome Sanger Institute for the COVID-19 Genomics UK (COG-UK) Consortium | Harper VanSteenhouse, Yumi Kasai, David Gray, Carol Clugston, Anna Dominiczak and Alex Alderton, Roberto Amato, Jeffrey Barrett, Sonia Goncalves, Ewan Harrison, David K. Jackson, Ian Johnston, Dominic Kwiatkowski, Cordelia Langford, John Sillitoe on behalf of the Wellcome Sanger Institute COVID-19 Surveillance Team |
| EPI_ISL_3139431, EPI_ISL_3139432 | Lighthouse Laboratory Plymouth | Wellcome Sanger Institute for the COVID-19 Genomics UK (COG-UK) Consortium | Lighthouse Laboratory Plymouth and Alex Alderton, Roberto Amato, Jeffrey Barrett, Sonia Goncalves, Ewan Harrison, David K. Jackson, Ian Johnston, Dominic Kwiatkowski, Cordelia Langford, John Sillitoe on behalf of the Wellcome Sanger Institute COVID-19 Surveillance Team |
| EPI_ISL_3139433, EPI_ISL_3139434, EPI_ISL_3139435, EPI_ISL_3139436 | Lighthouse Lab in Glasgow | Wellcome Sanger Institute for the COVID-19 Genomics UK (COG-UK) Consortium | Harper VanSteenhouse, Yumi Kasai, David Gray, Carol Clugston, Anna Dominiczak and Alex Alderton, Roberto Amato, Jeffrey Barrett, Sonia Goncalves, Ewan Harrison, David K. Jackson, Ian Johnston, Dominic Kwiatkowski, Cordelia Langford, John Sillitoe on behalf of the Wellcome Sanger Institute COVID-19 Surveillance Team |
| EPI_ISL_3139437, EPI_ISL_3139438, EPI_ISL_3139439 | Lighthouse Lab in Milton Keynes | Wellcome Sanger Institute for the COVID-19 Genomics UK (COG-UK) Consortium | The Lighthouse Lab in Milton Keynes and Alex Alderton, Roberto Amato, Jeffrey Barrett, Sonia Goncalves, Ewan Harrison, David K. Jackson, Ian Johnston, Dominic Kwiatkowski, Cordelia Langford, John Sillitoe on behalf of the Wellcome Sanger Institute COVID-19 Surveillance Team |
| EPI_ISL_3139440, EPI_ISL_3139441 | Lighthouse Laboratory Plymouth | Wellcome Sanger Institute for the COVID-19 Genomics UK (COG-UK) Consortium | Lighthouse Laboratory Plymouth and Alex Alderton, Roberto Amato, Jeffrey Barrett, Sonia Goncalves, Ewan Harrison, David K. Jackson, Ian Johnston, Dominic Kwiatkowski, Cordelia Langford, John Sillitoe on behalf of the Wellcome Sanger Institute COVID-19 Surveillance Team |
| EPI_ISL_3139442 | Lighthouse Lab in Milton Keynes | Wellcome Sanger Institute for the COVID-19 Genomics UK (COG-UK) Consortium | The Lighthouse Lab in Milton Keynes and Alex Alderton, Roberto Amato, Jeffrey Barrett, Sonia Goncalves, Ewan Harrison, David K. Jackson, Ian Johnston, Dominic Kwiatkowski, Cordelia Langford, John Sillitoe on behalf of the Wellcome Sanger Institute COVID-19 Surveillance Team |
| EPI_ISL_3139443 | Lighthouse Lab in Glasgow | Wellcome Sanger Institute for the COVID-19 Genomics UK (COG-UK) Consortium | Harper VanSteenhouse, Yumi Kasai, David Gray, Carol Clugston, Anna Dominiczak and Alex Alderton, Roberto Amato, Jeffrey Barrett, Sonia Goncalves, Ewan Harrison, David K. Jackson, Ian Johnston, Dominic Kwiatkowski, Cordelia Langford, John Sillitoe on behalf of the Wellcome Sanger Institute COVID-19 Surveillance Team |
| EPI_ISL_3139444 | Lighthouse Laboratory Plymouth | Wellcome Sanger Institute for the COVID-19 Genomics UK (COG-UK) Consortium | Lighthouse Laboratory Plymouth and Alex Alderton, Roberto Amato, Jeffrey Barrett, Sonia Goncalves, Ewan Harrison, David K. Jackson, Ian Johnston, Dominic Kwiatkowski, Cordelia Langford, John Sillitoe on behalf of the Wellcome Sanger Institute COVID-19 Surveillance Team |
| EPI_ISL_3139445 | Lighthouse Lab in Milton Keynes | Wellcome Sanger Institute for the COVID-19 Genomics UK (COG-UK) Consortium | The Lighthouse Lab in Milton Keynes and Alex Alderton, Roberto Amato, Jeffrey Barrett, Sonia Goncalves, Ewan Harrison, David K. Jackson, Ian Johnston, Dominic Kwiatkowski, Cordelia Langford, John Sillitoe on behalf of the Wellcome Sanger Institute COVID-19 Surveillance Team |
| EPI_ISL_3139446 | Lighthouse Laboratory Plymouth | Wellcome Sanger Institute for the COVID-19 Genomics UK (COG-UK) Consortium | Lighthouse Laboratory Plymouth and Alex Alderton, Roberto Amato, Jeffrey Barrett, Sonia Goncalves, Ewan Harrison, David K. Jackson, Ian Johnston, Dominic Kwiatkowski, Cordelia Langford, John Sillitoe on behalf of the Wellcome Sanger Institute COVID-19 Surveillance Team |
| EPI_ISL_3139447 | Lighthouse Lab in Glasgow | Wellcome Sanger Institute for the COVID-19 Genomics UK (COG-UK) Consortium | Harper VanSteenhouse, Yumi Kasai, David Gray, Carol Clugston, Anna Dominiczak and Alex Alderton, Roberto Amato, Jeffrey Barrett, Sonia Goncalves, Ewan Harrison, David K. Jackson, Ian Johnston, Dominic Kwiatkowski, Cordelia Langford, John Sillitoe on behalf of the Wellcome Sanger Institute COVID-19 Surveillance Team |
| EPI_ISL_3139448, EPI_ISL_3139449 | Lighthouse Laboratory Plymouth | Wellcome Sanger Institute for the COVID-19 Genomics UK (COG-UK) Consortium | Lighthouse Laboratory Plymouth and Alex Alderton, Roberto Amato, Jeffrey Barrett, Sonia Goncalves, Ewan Harrison, David K. Jackson, Ian Johnston, Dominic Kwiatkowski, Cordelia Langford, John Sillitoe on behalf of the Wellcome Sanger Institute COVID-19 Surveillance Team |
| EPI_ISL_3139450 | Lighthouse Lab in Milton Keynes | Wellcome Sanger Institute for the COVID-19 Genomics UK (COG-UK) Consortium | The Lighthouse Lab in Milton Keynes and Alex Alderton, Roberto Amato, Jeffrey Barrett, Sonia Goncalves, Ewan Harrison, David K. Jackson, Ian Johnston, Dominic Kwiatkowski, Cordelia Langford, John Sillitoe on behalf of the Wellcome Sanger Institute COVID-19 Surveillance Team |
| EPI_ISL_3139451 | Lighthouse Lab in Glasgow | Wellcome Sanger Institute for the COVID-19 Genomics UK (COG-UK) Consortium | Harper VanSteenhouse, Yumi Kasai, David Gray, Carol Clugston, Anna Dominiczak and Alex Alderton, Roberto Amato, Jeffrey Barrett, Sonia Goncalves, Ewan Harrison, David K. Jackson, Ian Johnston, Dominic Kwiatkowski, Cordelia Langford, John Sillitoe on behalf of the Wellcome Sanger Institute COVID-19 Surveillance Team |
| EPI_ISL_3139452 | Lighthouse Lab in Milton Keynes | Wellcome Sanger Institute for the COVID-19 Genomics UK (COG-UK) Consortium | The Lighthouse Lab in Milton Keynes and Alex Alderton, Roberto Amato, Jeffrey Barrett, Sonia Goncalves, Ewan Harrison, David K. Jackson, Ian Johnston, Dominic Kwiatkowski, Cordelia Langford, John Sillitoe on behalf of the Wellcome Sanger Institute COVID-19 Surveillance Team |
| EPI_ISL_3139453 | Lighthouse Laboratory Plymouth | Wellcome Sanger Institute for the COVID-19 Genomics UK (COG-UK) Consortium | Lighthouse Laboratory Plymouth and Alex Alderton, Roberto Amato, Jeffrey Barrett, Sonia Goncalves, Ewan Harrison, David K. Jackson, Ian Johnston, Dominic Kwiatkowski, Cordelia Langford, John Sillitoe on behalf of the Wellcome Sanger Institute COVID-19 Surveillance Team |
| EPI_ISL_3139454 | Lighthouse Lab in Milton Keynes | Wellcome Sanger Institute for the COVID-19 Genomics UK (COG-UK) Consortium | The Lighthouse Lab in Milton Keynes and Alex Alderton, Roberto Amato, Jeffrey Barrett, Sonia Goncalves, Ewan Harrison, David K. Jackson, Ian Johnston, Dominic Kwiatkowski, Cordelia Langford, John Sillitoe on behalf of the Wellcome Sanger Institute COVID-19 Surveillance Team |
| EPI_ISL_3139455, EPI_ISL_3139456, EPI_ISL_3139457 | Lighthouse Laboratory Plymouth | Wellcome Sanger Institute for the COVID-19 Genomics UK (COG-UK) Consortium | Lighthouse Laboratory Plymouth and Alex Alderton, Roberto Amato, Jeffrey Barrett, Sonia Goncalves, Ewan Harrison, David K. Jackson, Ian Johnston, Dominic Kwiatkowski, Cordelia Langford, John Sillitoe on behalf of the Wellcome Sanger Institute COVID-19 Surveillance Team |
| EPI_ISL_3139458 | Lighthouse Lab in Milton Keynes | Wellcome Sanger Institute for the COVID-19 Genomics UK (COG-UK) Consortium | The Lighthouse Lab in Milton Keynes and Alex Alderton, Roberto Amato, Jeffrey Barrett, Sonia Goncalves, Ewan Harrison, David K. Jackson, Ian Johnston, Dominic Kwiatkowski, Cordelia Langford, John Sillitoe on behalf of the Wellcome Sanger Institute COVID-19 Surveillance Team |
| EPI_ISL_3139459 | Lighthouse Lab in Glasgow | Wellcome Sanger Institute for the COVID-19 Genomics UK (COG-UK) Consortium | Harper VanSteenhouse, Yumi Kasai, David Gray, Carol Clugston, Anna Dominiczak and Alex Alderton, Roberto Amato, Jeffrey Barrett, Sonia Goncalves, Ewan Harrison, David K. Jackson, Ian Johnston, Dominic Kwiatkowski, Cordelia Langford, John Sillitoe on behalf of the Wellcome Sanger Institute COVID-19 Surveillance Team |
| EPI_ISL_3139460, EPI_ISL_3139461, EPI_ISL_3139462 | Lighthouse Laboratory Plymouth | Wellcome Sanger Institute for the COVID-19 Genomics UK (COG-UK) Consortium | Lighthouse Laboratory Plymouth and Alex Alderton, Roberto Amato, Jeffrey Barrett, Sonia Goncalves, Ewan Harrison, David K. Jackson, Ian Johnston, Dominic Kwiatkowski, Cordelia Langford, John Sillitoe on behalf of the Wellcome Sanger Institute COVID-19 Surveillance Team |
| EPI_ISL_3139463 | Lighthouse Lab in Glasgow | Wellcome Sanger Institute for the COVID-19 Genomics UK (COG-UK) Consortium | Harper VanSteenhouse, Yumi Kasai, David Gray, Carol Clugston, Anna Dominiczak and Alex Alderton, Roberto Amato, Jeffrey Barrett, Sonia Goncalves, Ewan Harrison, David K. Jackson, Ian Johnston, Dominic Kwiatkowski, Cordelia Langford, John Sillitoe on behalf of the Wellcome Sanger Institute COVID-19 Surveillance Team |
| EPI_ISL_3139464 | Lighthouse Laboratory Plymouth | Wellcome Sanger Institute for the COVID-19 Genomics UK (COG-UK) Consortium | Lighthouse Laboratory Plymouth and Alex Alderton, Roberto Amato, Jeffrey Barrett, Sonia Goncalves, Ewan Harrison, David K. Jackson, Ian Johnston, Dominic Kwiatkowski, Cordelia Langford, John Sillitoe on behalf of the Wellcome Sanger Institute COVID-19 Surveillance Team |
| EPI_ISL_3139465 | Lighthouse Lab in Milton Keynes | Wellcome Sanger Institute for the COVID-19 Genomics UK (COG-UK) Consortium | The Lighthouse Lab in Milton Keynes and Alex Alderton, Roberto Amato, Jeffrey Barrett, Sonia Goncalves, Ewan Harrison, David K. Jackson, Ian Johnston, Dominic Kwiatkowski, Cordelia Langford, John Sillitoe on behalf of the Wellcome Sanger Institute COVID-19 Surveillance Team |

| | | | |
|---|---|---|---|
| EPI_ISL_3139466 | Lighthouse Laboratory Plymouth | Wellcome Sanger Institute for the COVID-19 Genomics UK (COG-UK) Consortium | Lighthouse Laboratory Plymouth and Alex Alderton, Roberto Amato, Jeffrey Barrett, Sonia Goncalves, Ewan Harrison, David K. Jackson, Ian Johnston, Dominic Kwiatkowski, Cordelia Langford, John Sillitoe on behalf of the Wellcome Sanger Institute COVID-19 Surveillance Team |
| EPI_ISL_3139467 | Lighthouse Lab in Milton Keynes | Wellcome Sanger Institute for the COVID-19 Genomics UK (COG-UK) Consortium | The Lighthouse Lab in Milton Keynes and Alex Alderton, Roberto Amato, Jeffrey Barrett, Sonia Goncalves, Ewan Harrison, David K. Jackson, Ian Johnston, Dominic Kwiatkowski, Cordelia Langford, John Sillitoe on behalf of the Wellcome Sanger Institute COVID-19 Surveillance Team |
| EPI_ISL_3139468, EPI_ISL_3139469 | Lighthouse Laboratory Plymouth | Wellcome Sanger Institute for the COVID-19 Genomics UK (COG-UK) Consortium | Lighthouse Laboratory Plymouth and Alex Alderton, Roberto Amato, Jeffrey Barrett, Sonia Goncalves, Ewan Harrison, David K. Jackson, Ian Johnston, Dominic Kwiatkowski, Cordelia Langford, John Sillitoe on behalf of the Wellcome Sanger Institute COVID-19 Surveillance Team |
| EPI_ISL_3139470 | Lighthouse Lab in Milton Keynes | Wellcome Sanger Institute for the COVID-19 Genomics UK (COG-UK) Consortium | The Lighthouse Lab in Milton Keynes and Alex Alderton, Roberto Amato, Jeffrey Barrett, Sonia Goncalves, Ewan Harrison, David K. Jackson, Ian Johnston, Dominic Kwiatkowski, Cordelia Langford, John Sillitoe on behalf of the Wellcome Sanger Institute COVID-19 Surveillance Team |
| EPI_ISL_3139471, EPI_ISL_3139472 | Lighthouse Laboratory Plymouth | Wellcome Sanger Institute for the COVID-19 Genomics UK (COG-UK) Consortium | Lighthouse Laboratory Plymouth and Alex Alderton, Roberto Amato, Jeffrey Barrett, Sonia Goncalves, Ewan Harrison, David K. Jackson, Ian Johnston, Dominic Kwiatkowski, Cordelia Langford, John Sillitoe on behalf of the Wellcome Sanger Institute COVID-19 Surveillance Team |
| EPI_ISL_3139473, EPI_ISL_3139474, EPI_ISL_3139475 | Lighthouse Lab in Milton Keynes | Wellcome Sanger Institute for the COVID-19 Genomics UK (COG-UK) Consortium | The Lighthouse Lab in Milton Keynes and Alex Alderton, Roberto Amato, Jeffrey Barrett, Sonia Goncalves, Ewan Harrison, David K. Jackson, Ian Johnston, Dominic Kwiatkowski, Cordelia Langford, John Sillitoe on behalf of the Wellcome Sanger Institute COVID-19 Surveillance Team |
| EPI_ISL_3139476 | Lighthouse Laboratory Plymouth | Wellcome Sanger Institute for the COVID-19 Genomics UK (COG-UK) Consortium | Lighthouse Laboratory Plymouth and Alex Alderton, Roberto Amato, Jeffrey Barrett, Sonia Goncalves, Ewan Harrison, David K. Jackson, Ian Johnston, Dominic Kwiatkowski, Cordelia Langford, John Sillitoe on behalf of the Wellcome Sanger Institute COVID-19 Surveillance Team |
| EPI_ISL_3139477 | Lighthouse Lab in Glasgow | Wellcome Sanger Institute for the COVID-19 Genomics UK (COG-UK) Consortium | Harper VanSteenhouse, Yumi Kasai, David Gray, Carol Clugston, Anna Dominiczak and Alex Alderton, Roberto Amato, Jeffrey Barrett, Sonia Goncalves, Ewan Harrison, David K. Jackson, Ian Johnston, Dominic Kwiatkowski, Cordelia Langford, John Sillitoe on behalf of the Wellcome Sanger Institute COVID-19 Surveillance Team |
| EPI_ISL_3139478 | Lighthouse Laboratory Plymouth | Wellcome Sanger Institute for the COVID-19 Genomics UK (COG-UK) Consortium | Lighthouse Laboratory Plymouth and Alex Alderton, Roberto Amato, Jeffrey Barrett, Sonia Goncalves, Ewan Harrison, David K. Jackson, Ian Johnston, Dominic Kwiatkowski, Cordelia Langford, John Sillitoe on behalf of the Wellcome Sanger Institute COVID-19 Surveillance Team |
| EPI_ISL_3139479 | Lighthouse Lab in Glasgow | Wellcome Sanger Institute for the COVID-19 Genomics UK (COG-UK) Consortium | Harper VanSteenhouse, Yumi Kasai, David Gray, Carol Clugston, Anna Dominiczak and Alex Alderton, Roberto Amato, Jeffrey Barrett, Sonia Goncalves, Ewan Harrison, David K. Jackson, Ian Johnston, Dominic Kwiatkowski, Cordelia Langford, John Sillitoe on behalf of the Wellcome Sanger Institute COVID-19 Surveillance Team |
| EPI_ISL_3139480 | Lighthouse Laboratory Plymouth | Wellcome Sanger Institute for the COVID-19 Genomics UK (COG-UK) Consortium | Lighthouse Laboratory Plymouth and Alex Alderton, Roberto Amato, Jeffrey Barrett, Sonia Goncalves, Ewan Harrison, David K. Jackson, Ian Johnston, Dominic Kwiatkowski, Cordelia Langford, John Sillitoe on behalf of the Wellcome Sanger Institute COVID-19 Surveillance Team |
| EPI_ISL_3139481 | Lighthouse Lab in Glasgow | Wellcome Sanger Institute for the COVID-19 Genomics UK (COG-UK) Consortium | Harper VanSteenhouse, Yumi Kasai, David Gray, Carol Clugston, Anna Dominiczak and Alex Alderton, Roberto Amato, Jeffrey Barrett, Sonia Goncalves, Ewan Harrison, David K. Jackson, Ian Johnston, Dominic Kwiatkowski, Cordelia Langford, John Sillitoe on behalf of the Wellcome Sanger Institute COVID-19 Surveillance Team |
| EPI_ISL_3139482 | Lighthouse Laboratory Plymouth | Wellcome Sanger Institute for the COVID-19 Genomics UK (COG-UK) Consortium | Lighthouse Laboratory Plymouth and Alex Alderton, Roberto Amato, Jeffrey Barrett, Sonia Goncalves, Ewan Harrison, David K. Jackson, Ian Johnston, Dominic Kwiatkowski, Cordelia Langford, John Sillitoe on behalf of the Wellcome Sanger Institute COVID-19 Surveillance Team |
| EPI_ISL_3139483, EPI_ISL_3139484, EPI_ISL_3139485, EPI_ISL_3139486 | Lighthouse Lab in Glasgow | Wellcome Sanger Institute for the COVID-19 Genomics UK (COG-UK) Consortium | Harper VanSteenhouse, Yumi Kasai, David Gray, Carol Clugston, Anna Dominiczak and Alex Alderton, Roberto Amato, Jeffrey Barrett, Sonia Goncalves, Ewan Harrison, David K. Jackson, Ian Johnston, Dominic Kwiatkowski, Cordelia Langford, John Sillitoe on behalf of the Wellcome Sanger Institute COVID-19 Surveillance Team |
| EPI_ISL_3139487 | Lighthouse Laboratory Plymouth | Wellcome Sanger Institute for the COVID-19 Genomics UK (COG-UK) Consortium | Lighthouse Laboratory Plymouth and Alex Alderton, Roberto Amato, Jeffrey Barrett, Sonia Goncalves, Ewan Harrison, David K. Jackson, Ian Johnston, Dominic Kwiatkowski, Cordelia Langford, John Sillitoe on behalf of the Wellcome Sanger Institute COVID-19 Surveillance Team |
| EPI_ISL_3139488 | Lighthouse Lab in Glasgow | Wellcome Sanger Institute for the COVID-19 Genomics UK (COG-UK) Consortium | Harper VanSteenhouse, Yumi Kasai, David Gray, Carol Clugston, Anna Dominiczak and Alex Alderton, Roberto Amato, Jeffrey Barrett, Sonia Goncalves, Ewan Harrison, David K. Jackson, Ian Johnston, Dominic Kwiatkowski, Cordelia Langford, John Sillitoe on behalf of the Wellcome Sanger Institute COVID-19 Surveillance Team |
| EPI_ISL_3139489, EPI_ISL_3139490, EPI_ISL_3139491 | Lighthouse Laboratory Plymouth | Wellcome Sanger Institute for the COVID-19 Genomics UK (COG-UK) Consortium | Lighthouse Laboratory Plymouth and Alex Alderton, Roberto Amato, Jeffrey Barrett, Sonia Goncalves, Ewan Harrison, David K. Jackson, Ian Johnston, Dominic Kwiatkowski, Cordelia Langford, John Sillitoe on behalf of the Wellcome Sanger Institute COVID-19 Surveillance Team |
| EPI_ISL_3139492 | Lighthouse Lab in Milton Keynes | Wellcome Sanger Institute for the COVID-19 Genomics UK (COG-UK) Consortium | The Lighthouse Lab in Milton Keynes and Alex Alderton, Roberto Amato, Jeffrey Barrett, Sonia Goncalves, Ewan Harrison, David K. Jackson, Ian Johnston, Dominic Kwiatkowski, Cordelia Langford, John Sillitoe on behalf of the Wellcome Sanger Institute COVID-19 Surveillance Team |
| EPI_ISL_3139493, EPI_ISL_3139494 | Lighthouse Laboratory Plymouth | Wellcome Sanger Institute for the COVID-19 Genomics UK (COG-UK) Consortium | Lighthouse Laboratory Plymouth and Alex Alderton, Roberto Amato, Jeffrey Barrett, Sonia Goncalves, Ewan Harrison, David K. Jackson, Ian Johnston, Dominic Kwiatkowski, Cordelia Langford, John Sillitoe on behalf of the Wellcome Sanger Institute COVID-19 Surveillance Team |
| EPI_ISL_3139495 | Lighthouse Lab in Milton Keynes | Wellcome Sanger Institute for the COVID-19 Genomics UK (COG-UK) Consortium | The Lighthouse Lab in Milton Keynes and Alex Alderton, Roberto Amato, Jeffrey Barrett, Sonia Goncalves, Ewan Harrison, David K. Jackson, Ian Johnston, Dominic Kwiatkowski, Cordelia Langford, John Sillitoe on behalf of the Wellcome Sanger Institute COVID-19 Surveillance Team |
| EPI_ISL_3139496 | Lighthouse Lab in Glasgow | Wellcome Sanger Institute for the COVID-19 Genomics UK (COG-UK) Consortium | Harper VanSteenhouse, Yumi Kasai, David Gray, Carol Clugston, Anna Dominiczak and Alex Alderton, Roberto Amato, Jeffrey Barrett, Sonia Goncalves, Ewan Harrison, David K. Jackson, Ian Johnston, Dominic Kwiatkowski, Cordelia Langford, John Sillitoe on behalf of the Wellcome Sanger Institute COVID-19 Surveillance Team |
| EPI_ISL_3139497 | Lighthouse Laboratory Plymouth | Wellcome Sanger Institute for the COVID-19 Genomics UK (COG-UK) Consortium | Lighthouse Laboratory Plymouth and Alex Alderton, Roberto Amato, Jeffrey Barrett, Sonia Goncalves, Ewan Harrison, David K. Jackson, Ian Johnston, Dominic Kwiatkowski, Cordelia Langford, John Sillitoe on behalf of the Wellcome Sanger Institute COVID-19 Surveillance Team |
| EPI_ISL_3139498 | Lighthouse Lab in Glasgow | Wellcome Sanger Institute for the COVID-19 Genomics UK (COG-UK) Consortium | Harper VanSteenhouse, Yumi Kasai, David Gray, Carol Clugston, Anna Dominiczak and Alex Alderton, Roberto Amato, Jeffrey Barrett, Sonia Goncalves, Ewan Harrison, David K. Jackson, Ian Johnston, Dominic Kwiatkowski, Cordelia Langford, John Sillitoe on behalf of the Wellcome Sanger Institute COVID-19 Surveillance Team |
| EPI_ISL_3139499 | Lighthouse Laboratory Plymouth | Wellcome Sanger Institute for the COVID-19 Genomics UK (COG-UK) Consortium | Lighthouse Laboratory Plymouth and Alex Alderton, Roberto Amato, Jeffrey Barrett, Sonia Goncalves, Ewan Harrison, David K. Jackson, Ian Johnston, Dominic Kwiatkowski, Cordelia Langford, John Sillitoe on behalf of the Wellcome Sanger Institute COVID-19 Surveillance Team |
| EPI_ISL_3139500 | Lighthouse Lab in Glasgow | Wellcome Sanger Institute for the COVID-19 Genomics UK (COG-UK) Consortium | Harper VanSteenhouse, Yumi Kasai, David Gray, Carol Clugston, Anna Dominiczak and Alex Alderton, Roberto Amato, Jeffrey Barrett, Sonia Goncalves, Ewan Harrison, David K. Jackson, Ian Johnston, Dominic Kwiatkowski, Cordelia Langford, John Sillitoe on behalf of the Wellcome Sanger Institute COVID-19 Surveillance Team |
| EPI_ISL_3139501 | Lighthouse Lab in Milton Keynes | Wellcome Sanger Institute for the COVID-19 Genomics UK (COG-UK) Consortium | The Lighthouse Lab in Milton Keynes and Alex Alderton, Roberto Amato, Jeffrey Barrett, Sonia Goncalves, Ewan Harrison, David K. Jackson, Ian Johnston, Dominic Kwiatkowski, Cordelia Langford, John Sillitoe on behalf of the Wellcome Sanger Institute COVID-19 Surveillance Team |
| EPI_ISL_3139502 | Lighthouse Lab in Glasgow | Wellcome Sanger Institute for the COVID-19 Genomics UK (COG-UK) Consortium | Harper VanSteenhouse, Yumi Kasai, David Gray, Carol Clugston, Anna Dominiczak and Alex Alderton, Roberto Amato, Jeffrey Barrett, Sonia Goncalves, Ewan Harrison, David K. Jackson, Ian Johnston, Dominic Kwiatkowski, Cordelia Langford, John Sillitoe on behalf of the Wellcome Sanger Institute COVID-19 Surveillance Team |
| EPI_ISL_3139503, EPI_ISL_3139504 | Lighthouse Lab in Milton Keynes | Wellcome Sanger Institute for the COVID-19 Genomics UK (COG-UK) Consortium | The Lighthouse Lab in Milton Keynes and Alex Alderton, Roberto Amato, Jeffrey Barrett, Sonia Goncalves, Ewan Harrison, David K. Jackson, Ian Johnston, Dominic Kwiatkowski, Cordelia Langford, John Sillitoe on behalf of the Wellcome Sanger Institute COVID-19 Surveillance Team |
| EPI_ISL_3139505, EPI_ISL_3139506 | Lighthouse Lab in Glasgow | Wellcome Sanger Institute for the COVID-19 Genomics UK (COG-UK) Consortium | Harper VanSteenhouse, Yumi Kasai, David Gray, Carol Clugston, Anna Dominiczak and Alex Alderton, Roberto Amato, Jeffrey Barrett, Sonia Goncalves, Ewan Harrison, David K. Jackson, Ian Johnston, Dominic Kwiatkowski, Cordelia Langford, John Sillitoe on behalf of the Wellcome Sanger Institute COVID-19 Surveillance Team |
| EPI_ISL_3139507, EPI_ISL_3139508 | Lighthouse Laboratory Plymouth | Wellcome Sanger Institute for the COVID-19 Genomics UK (COG-UK) Consortium | Lighthouse Laboratory Plymouth and Alex Alderton, Roberto Amato, Jeffrey Barrett, Sonia Goncalves, Ewan Harrison, David K. Jackson, Ian Johnston, Dominic Kwiatkowski, Cordelia Langford, John Sillitoe on behalf of the Wellcome Sanger Institute COVID-19 Surveillance Team |
| EPI_ISL_3139509 | Lighthouse Lab in Glasgow | Wellcome Sanger Institute for the COVID-19 Genomics UK (COG-UK) Consortium | Harper VanSteenhouse, Yumi Kasai, David Gray, Carol Clugston, Anna Dominiczak and Alex Alderton, Roberto Amato, Jeffrey Barrett, Sonia Goncalves, Ewan Harrison, David K. Jackson, Ian Johnston, Dominic Kwiatkowski, Cordelia Langford, John Sillitoe on behalf of the Wellcome Sanger Institute COVID-19 Surveillance Team |
| EPI_ISL_3139510 | Lighthouse Lab in Milton Keynes | Wellcome Sanger Institute for the COVID-19 Genomics UK (COG-UK) Consortium | The Lighthouse Lab in Milton Keynes and Alex Alderton, Roberto Amato, Jeffrey Barrett, Sonia Goncalves, Ewan Harrison, David K. Jackson, Ian Johnston, Dominic Kwiatkowski, Cordelia Langford, John Sillitoe on behalf of the Wellcome Sanger Institute COVID-19 Surveillance Team |
| EPI_ISL_3139511 | Lighthouse Laboratory Plymouth | Wellcome Sanger Institute for the COVID-19 Genomics UK (COG-UK) Consortium | Lighthouse Laboratory Plymouth and Alex Alderton, Roberto Amato, Jeffrey Barrett, Sonia Goncalves, Ewan Harrison, David K. Jackson, Ian Johnston, Dominic Kwiatkowski, Cordelia Langford, John Sillitoe on behalf of the Wellcome Sanger Institute COVID-19 Surveillance Team |
| EPI_ISL_3139512 | Lighthouse Lab in Milton Keynes | Wellcome Sanger Institute for the COVID-19 Genomics UK (COG-UK) Consortium | The Lighthouse Lab in Milton Keynes and Alex Alderton, Roberto Amato, Jeffrey Barrett, Sonia Goncalves, Ewan Harrison, David K. Jackson, Ian Johnston, Dominic Kwiatkowski, Cordelia Langford, John Sillitoe on behalf of the Wellcome Sanger Institute COVID-19 Surveillance Team |
| EPI_ISL_3139513 | Lighthouse Lab in Glasgow | Wellcome Sanger Institute for the COVID-19 Genomics UK (COG-UK) Consortium | Harper VanSteenhouse, Yumi Kasai, David Gray, Carol Clugston, Anna Dominiczak and Alex Alderton, Roberto Amato, Jeffrey Barrett, Sonia Goncalves, Ewan Harrison, David K. Jackson, Ian Johnston, Dominic Kwiatkowski, Cordelia Langford, John Sillitoe on behalf of the Wellcome Sanger Institute COVID-19 Surveillance Team |
| EPI_ISL_3139514, EPI_ISL_3139515, EPI_ISL_3139516 | Lighthouse Laboratory Plymouth | Wellcome Sanger Institute for the COVID-19 Genomics UK (COG-UK) Consortium | Lighthouse Laboratory Plymouth and Alex Alderton, Roberto Amato, Jeffrey Barrett, Sonia Goncalves, Ewan Harrison, David K. Jackson, Ian Johnston, Dominic Kwiatkowski, Cordelia Langford, John Sillitoe on behalf of the Wellcome Sanger Institute COVID-19 Surveillance Team |
| EPI_ISL_3139517 | Lighthouse Lab in Milton Keynes | Wellcome Sanger Institute for the COVID-19 Genomics UK (COG-UK) Consortium | The Lighthouse Lab in Milton Keynes and Alex Alderton, Roberto Amato, Jeffrey Barrett, Sonia Goncalves, Ewan Harrison, David K. Jackson, Ian Johnston, Dominic Kwiatkowski, Cordelia Langford, John Sillitoe on behalf of the Wellcome Sanger Institute COVID-19 Surveillance Team |
| EPI_ISL_3139518 | Lighthouse Laboratory Plymouth | Wellcome Sanger Institute for the COVID-19 Genomics UK (COG-UK) Consortium | Lighthouse Laboratory Plymouth and Alex Alderton, Roberto Amato, Jeffrey Barrett, Sonia Goncalves, Ewan Harrison, David K. Jackson, Ian Johnston, Dominic Kwiatkowski, Cordelia Langford, John Sillitoe on behalf of the Wellcome Sanger Institute COVID-19 Surveillance Team |
| EPI_ISL_3139519 | Lighthouse Lab in Glasgow | Wellcome Sanger Institute for the COVID-19 Genomics UK (COG-UK) Consortium | Harper VanSteenhouse, Yumi Kasai, David Gray, Carol Clugston, Anna Dominiczak and Alex Alderton, Roberto Amato, Jeffrey Barrett, Sonia Goncalves, Ewan Harrison, David K. Jackson, Ian Johnston, Dominic Kwiatkowski, Cordelia Langford, John Sillitoe on behalf of the Wellcome Sanger Institute COVID-19 Surveillance Team |
| EPI_ISL_3139520 | Lighthouse Laboratory Plymouth | Wellcome Sanger Institute for the COVID-19 Genomics UK (COG-UK) Consortium | Lighthouse Laboratory Plymouth and Alex Alderton, Roberto Amato, Jeffrey Barrett, Sonia Goncalves, Ewan Harrison, David K. Jackson, Ian Johnston, Dominic Kwiatkowski, Cordelia Langford, John Sillitoe on behalf of the Wellcome Sanger Institute COVID-19 Surveillance Team |
| EPI_ISL_3139521 | Lighthouse Lab in Glasgow | Wellcome Sanger Institute for the COVID-19 Genomics UK (COG-UK) Consortium | Harper VanSteenhouse, Yumi Kasai, David Gray, Carol Clugston, Anna Dominiczak and Alex Alderton, Roberto Amato, Jeffrey Barrett, Sonia Goncalves, Ewan Harrison, David K. Jackson, Ian Johnston, Dominic Kwiatkowski, Cordelia Langford, John Sillitoe on behalf of the Wellcome Sanger Institute COVID-19 Surveillance Team |
| EPI_ISL_3139522, EPI_ISL_3139523 | Lighthouse Lab in Milton Keynes | Wellcome Sanger Institute for the COVID-19 Genomics UK (COG-UK) Consortium | The Lighthouse Lab in Milton Keynes and Alex Alderton, Roberto Amato, Jeffrey Barrett, Sonia Goncalves, Ewan Harrison, David K. Jackson, Ian Johnston, Dominic Kwiatkowski, Cordelia Langford, John Sillitoe on behalf of the Wellcome Sanger Institute COVID-19 Surveillance Team |
| EPI_ISL_3139524, EPI_ISL_3139525 | Lighthouse Laboratory Plymouth | Wellcome Sanger Institute for the COVID-19 Genomics UK (COG-UK) Consortium | Lighthouse Laboratory Plymouth and Alex Alderton, Roberto Amato, Jeffrey Barrett, Sonia Goncalves, Ewan Harrison, David K. Jackson, Ian Johnston, Dominic Kwiatkowski, Cordelia Langford, John Sillitoe on behalf of the Wellcome Sanger Institute COVID-19 Surveillance Team |

| | | | |
|---|---|---|---|
| EPI_ISL_3139526 | Lighthouse Lab in Milton Keynes | Wellcome Sanger Institute for the COVID-19 Genomics UK (COG-UK) Consortium | The Lighthouse Lab in Milton Keynes and Alex Alderton, Roberto Amato, Jeffrey Barrett, Sonia Goncalves, Ewan Harrison, David K. Jackson, Ian Johnston, Dominic Kwiatkowski, Cordelia Langford, John Sillitoe on behalf of the Wellcome Sanger Institute COVID-19 Surveillance Team |
| EPI_ISL_3139527, EPI_ISL_3139528, EPI_ISL_3139529 | Lighthouse Laboratory Plymouth | Wellcome Sanger Institute for the COVID-19 Genomics UK (COG-UK) Consortium | Lighthouse Laboratory Plymouth and Alex Alderton, Roberto Amato, Jeffrey Barrett, Sonia Goncalves, Ewan Harrison, David K. Jackson, Ian Johnston, Dominic Kwiatkowski, Cordelia Langford, John Sillitoe on behalf of the Wellcome Sanger Institute COVID-19 Surveillance Team |
| EPI_ISL_3139530 | Lighthouse Lab in Glasgow | Wellcome Sanger Institute for the COVID-19 Genomics UK (COG-UK) Consortium | Harper VanSteenhouse, Yumi Kasai, David Gray, Carol Clugston, Anna Dominiczak and Alex Alderton, Roberto Amato, Jeffrey Barrett, Sonia Goncalves, Ewan Harrison, David K. Jackson, Ian Johnston, Dominic Kwiatkowski, Cordelia Langford, John Sillitoe on behalf of the Wellcome Sanger Institute COVID-19 Surveillance Team |
| EPI_ISL_3139531 | Lighthouse Lab in Milton Keynes | Wellcome Sanger Institute for the COVID-19 Genomics UK (COG-UK) Consortium | The Lighthouse Lab in Milton Keynes and Alex Alderton, Roberto Amato, Jeffrey Barrett, Sonia Goncalves, Ewan Harrison, David K. Jackson, Ian Johnston, Dominic Kwiatkowski, Cordelia Langford, John Sillitoe on behalf of the Wellcome Sanger Institute COVID-19 Surveillance Team |
| EPI_ISL_3139532 | Lighthouse Lab in Glasgow | Wellcome Sanger Institute for the COVID-19 Genomics UK (COG-UK) Consortium | Harper VanSteenhouse, Yumi Kasai, David Gray, Carol Clugston, Anna Dominiczak and Alex Alderton, Roberto Amato, Jeffrey Barrett, Sonia Goncalves, Ewan Harrison, David K. Jackson, Ian Johnston, Dominic Kwiatkowski, Cordelia Langford, John Sillitoe on behalf of the Wellcome Sanger Institute COVID-19 Surveillance Team |
| EPI_ISL_3139533 | Lighthouse Lab in Milton Keynes | Wellcome Sanger Institute for the COVID-19 Genomics UK (COG-UK) Consortium | The Lighthouse Lab in Milton Keynes and Alex Alderton, Roberto Amato, Jeffrey Barrett, Sonia Goncalves, Ewan Harrison, David K. Jackson, Ian Johnston, Dominic Kwiatkowski, Cordelia Langford, John Sillitoe on behalf of the Wellcome Sanger Institute COVID-19 Surveillance Team |
| EPI_ISL_3139534 | Lighthouse Laboratory Plymouth | Wellcome Sanger Institute for the COVID-19 Genomics UK (COG-UK) Consortium | Lighthouse Laboratory Plymouth and Alex Alderton, Roberto Amato, Jeffrey Barrett, Sonia Goncalves, Ewan Harrison, David K. Jackson, Ian Johnston, Dominic Kwiatkowski, Cordelia Langford, John Sillitoe on behalf of the Wellcome Sanger Institute COVID-19 Surveillance Team |
| EPI_ISL_3139535 | Lighthouse Lab in Milton Keynes | Wellcome Sanger Institute for the COVID-19 Genomics UK (COG-UK) Consortium | The Lighthouse Lab in Milton Keynes and Alex Alderton, Roberto Amato, Jeffrey Barrett, Sonia Goncalves, Ewan Harrison, David K. Jackson, Ian Johnston, Dominic Kwiatkowski, Cordelia Langford, John Sillitoe on behalf of the Wellcome Sanger Institute COVID-19 Surveillance Team |
| EPI_ISL_3139536 | Lighthouse Lab in Glasgow | Wellcome Sanger Institute for the COVID-19 Genomics UK (COG-UK) Consortium | Harper VanSteenhouse, Yumi Kasai, David Gray, Carol Clugston, Anna Dominiczak and Alex Alderton, Roberto Amato, Jeffrey Barrett, Sonia Goncalves, Ewan Harrison, David K. Jackson, Ian Johnston, Dominic Kwiatkowski, Cordelia Langford, John Sillitoe on behalf of the Wellcome Sanger Institute COVID-19 Surveillance Team |
| EPI_ISL_3139537 | Lighthouse Lab in Milton Keynes | Wellcome Sanger Institute for the COVID-19 Genomics UK (COG-UK) Consortium | The Lighthouse Lab in Milton Keynes and Alex Alderton, Roberto Amato, Jeffrey Barrett, Sonia Goncalves, Ewan Harrison, David K. Jackson, Ian Johnston, Dominic Kwiatkowski, Cordelia Langford, John Sillitoe on behalf of the Wellcome Sanger Institute COVID-19 Surveillance Team |
| EPI_ISL_3139538, EPI_ISL_3139539 | Lighthouse Laboratory Plymouth | Wellcome Sanger Institute for the COVID-19 Genomics UK (COG-UK) Consortium | Lighthouse Laboratory Plymouth and Alex Alderton, Roberto Amato, Jeffrey Barrett, Sonia Goncalves, Ewan Harrison, David K. Jackson, Ian Johnston, Dominic Kwiatkowski, Cordelia Langford, John Sillitoe on behalf of the Wellcome Sanger Institute COVID-19 Surveillance Team |
| EPI_ISL_3139540, EPI_ISL_3139541, EPI_ISL_3139542 | Lighthouse Lab in Glasgow | Wellcome Sanger Institute for the COVID-19 Genomics UK (COG-UK) Consortium | Harper VanSteenhouse, Yumi Kasai, David Gray, Carol Clugston, Anna Dominiczak and Alex Alderton, Roberto Amato, Jeffrey Barrett, Sonia Goncalves, Ewan Harrison, David K. Jackson, Ian Johnston, Dominic Kwiatkowski, Cordelia Langford, John Sillitoe on behalf of the Wellcome Sanger Institute COVID-19 Surveillance Team |
| EPI_ISL_3139543 | Lighthouse Lab in Milton Keynes | Wellcome Sanger Institute for the COVID-19 Genomics UK (COG-UK) Consortium | The Lighthouse Lab in Milton Keynes and Alex Alderton, Roberto Amato, Jeffrey Barrett, Sonia Goncalves, Ewan Harrison, David K. Jackson, Ian Johnston, Dominic Kwiatkowski, Cordelia Langford, John Sillitoe on behalf of the Wellcome Sanger Institute COVID-19 Surveillance Team |
| EPI_ISL_3139544 | Lighthouse Lab in Glasgow | Wellcome Sanger Institute for the COVID-19 Genomics UK (COG-UK) Consortium | Harper VanSteenhouse, Yumi Kasai, David Gray, Carol Clugston, Anna Dominiczak and Alex Alderton, Roberto Amato, Jeffrey Barrett, Sonia Goncalves, Ewan Harrison, David K. Jackson, Ian Johnston, Dominic Kwiatkowski, Cordelia Langford, John Sillitoe on behalf of the Wellcome Sanger Institute COVID-19 Surveillance Team |
| EPI_ISL_3139545, EPI_ISL_3139546 | Lighthouse Laboratory Plymouth | Wellcome Sanger Institute for the COVID-19 Genomics UK (COG-UK) Consortium | Lighthouse Laboratory Plymouth and Alex Alderton, Roberto Amato, Jeffrey Barrett, Sonia Goncalves, Ewan Harrison, David K. Jackson, Ian Johnston, Dominic Kwiatkowski, Cordelia Langford, John Sillitoe on behalf of the Wellcome Sanger Institute COVID-19 Surveillance Team |
| EPI_ISL_3139547, EPI_ISL_3139548, EPI_ISL_3139549 | Lighthouse Lab in Glasgow | Wellcome Sanger Institute for the COVID-19 Genomics UK (COG-UK) Consortium | Harper VanSteenhouse, Yumi Kasai, David Gray, Carol Clugston, Anna Dominiczak and Alex Alderton, Roberto Amato, Jeffrey Barrett, Sonia Goncalves, Ewan Harrison, David K. Jackson, Ian Johnston, Dominic Kwiatkowski, Cordelia Langford, John Sillitoe on behalf of the Wellcome Sanger Institute COVID-19 Surveillance Team |
| EPI_ISL_3139550, EPI_ISL_3139551 | Lighthouse Laboratory Plymouth | Wellcome Sanger Institute for the COVID-19 Genomics UK (COG-UK) Consortium | Lighthouse Laboratory Plymouth and Alex Alderton, Roberto Amato, Jeffrey Barrett, Sonia Goncalves, Ewan Harrison, David K. Jackson, Ian Johnston, Dominic Kwiatkowski, Cordelia Langford, John Sillitoe on behalf of the Wellcome Sanger Institute COVID-19 Surveillance Team |
| EPI_ISL_3139552 | Lighthouse Lab in Milton Keynes | Wellcome Sanger Institute for the COVID-19 Genomics UK (COG-UK) Consortium | The Lighthouse Lab in Milton Keynes and Alex Alderton, Roberto Amato, Jeffrey Barrett, Sonia Goncalves, Ewan Harrison, David K. Jackson, Ian Johnston, Dominic Kwiatkowski, Cordelia Langford, John Sillitoe on behalf of the Wellcome Sanger Institute COVID-19 Surveillance Team |
| EPI_ISL_3139553, EPI_ISL_3139554 | Lighthouse Laboratory Plymouth | Wellcome Sanger Institute for the COVID-19 Genomics UK (COG-UK) Consortium | Lighthouse Laboratory Plymouth and Alex Alderton, Roberto Amato, Jeffrey Barrett, Sonia Goncalves, Ewan Harrison, David K. Jackson, Ian Johnston, Dominic Kwiatkowski, Cordelia Langford, John Sillitoe on behalf of the Wellcome Sanger Institute COVID-19 Surveillance Team |
| EPI_ISL_3139555 | Lighthouse Lab in Glasgow | Wellcome Sanger Institute for the COVID-19 Genomics UK (COG-UK) Consortium | Harper VanSteenhouse, Yumi Kasai, David Gray, Carol Clugston, Anna Dominiczak and Alex Alderton, Roberto Amato, Jeffrey Barrett, Sonia Goncalves, Ewan Harrison, David K. Jackson, Ian Johnston, Dominic Kwiatkowski, Cordelia Langford, John Sillitoe on behalf of the Wellcome Sanger Institute COVID-19 Surveillance Team |
| EPI_ISL_3139556 | Lighthouse Laboratory Plymouth | Wellcome Sanger Institute for the COVID-19 Genomics UK (COG-UK) Consortium | Lighthouse Laboratory Plymouth and Alex Alderton, Roberto Amato, Jeffrey Barrett, Sonia Goncalves, Ewan Harrison, David K. Jackson, Ian Johnston, Dominic Kwiatkowski, Cordelia Langford, John Sillitoe on behalf of the Wellcome Sanger Institute COVID-19 Surveillance Team |
| EPI_ISL_3139557 | Lighthouse Lab in Milton Keynes | Wellcome Sanger Institute for the COVID-19 Genomics UK (COG-UK) Consortium | The Lighthouse Lab in Milton Keynes and Alex Alderton, Roberto Amato, Jeffrey Barrett, Sonia Goncalves, Ewan Harrison, David K. Jackson, Ian Johnston, Dominic Kwiatkowski, Cordelia Langford, John Sillitoe on behalf of the Wellcome Sanger Institute COVID-19 Surveillance Team |
| EPI_ISL_3139558 | Lighthouse Lab in Glasgow | Wellcome Sanger Institute for the COVID-19 Genomics UK (COG-UK) Consortium | Harper VanSteenhouse, Yumi Kasai, David Gray, Carol Clugston, Anna Dominiczak and Alex Alderton, Roberto Amato, Jeffrey Barrett, Sonia Goncalves, Ewan Harrison, David K. Jackson, Ian Johnston, Dominic Kwiatkowski, Cordelia Langford, John Sillitoe on behalf of the Wellcome Sanger Institute COVID-19 Surveillance Team |
| EPI_ISL_3139559 | Lighthouse Laboratory Plymouth | Wellcome Sanger Institute for the COVID-19 Genomics UK (COG-UK) Consortium | Lighthouse Laboratory Plymouth and Alex Alderton, Roberto Amato, Jeffrey Barrett, Sonia Goncalves, Ewan Harrison, David K. Jackson, Ian Johnston, Dominic Kwiatkowski, Cordelia Langford, John Sillitoe on behalf of the Wellcome Sanger Institute COVID-19 Surveillance Team |
| EPI_ISL_3139560 | Lighthouse Lab in Milton Keynes | Wellcome Sanger Institute for the COVID-19 Genomics UK (COG-UK) Consortium | The Lighthouse Lab in Milton Keynes and Alex Alderton, Roberto Amato, Jeffrey Barrett, Sonia Goncalves, Ewan Harrison, David K. Jackson, Ian Johnston, Dominic Kwiatkowski, Cordelia Langford, John Sillitoe on behalf of the Wellcome Sanger Institute COVID-19 Surveillance Team |
| EPI_ISL_3139561 | Lighthouse Laboratory Plymouth | Wellcome Sanger Institute for the COVID-19 Genomics UK (COG-UK) Consortium | Lighthouse Laboratory Plymouth and Alex Alderton, Roberto Amato, Jeffrey Barrett, Sonia Goncalves, Ewan Harrison, David K. Jackson, Ian Johnston, Dominic Kwiatkowski, Cordelia Langford, John Sillitoe on behalf of the Wellcome Sanger Institute COVID-19 Surveillance Team |
| EPI_ISL_3139562 | Lighthouse Lab in Milton Keynes | Wellcome Sanger Institute for the COVID-19 Genomics UK (COG-UK) Consortium | The Lighthouse Lab in Milton Keynes and Alex Alderton, Roberto Amato, Jeffrey Barrett, Sonia Goncalves, Ewan Harrison, David K. Jackson, Ian Johnston, Dominic Kwiatkowski, Cordelia Langford, John Sillitoe on behalf of the Wellcome Sanger Institute COVID-19 Surveillance Team |
| EPI_ISL_3139563 | Lighthouse Laboratory Plymouth | Wellcome Sanger Institute for the COVID-19 Genomics UK (COG-UK) Consortium | Lighthouse Laboratory Plymouth and Alex Alderton, Roberto Amato, Jeffrey Barrett, Sonia Goncalves, Ewan Harrison, David K. Jackson, Ian Johnston, Dominic Kwiatkowski, Cordelia Langford, John Sillitoe on behalf of the Wellcome Sanger Institute COVID-19 Surveillance Team |
| EPI_ISL_3139564 | Lighthouse Lab in Glasgow | Wellcome Sanger Institute for the COVID-19 Genomics UK (COG-UK) Consortium | Harper VanSteenhouse, Yumi Kasai, David Gray, Carol Clugston, Anna Dominiczak and Alex Alderton, Roberto Amato, Jeffrey Barrett, Sonia Goncalves, Ewan Harrison, David K. Jackson, Ian Johnston, Dominic Kwiatkowski, Cordelia Langford, John Sillitoe on behalf of the Wellcome Sanger Institute COVID-19 Surveillance Team |
| EPI_ISL_3139565 | Lighthouse Lab in Milton Keynes | Wellcome Sanger Institute for the COVID-19 Genomics UK (COG-UK) Consortium | The Lighthouse Lab in Milton Keynes and Alex Alderton, Roberto Amato, Jeffrey Barrett, Sonia Goncalves, Ewan Harrison, David K. Jackson, Ian Johnston, Dominic Kwiatkowski, Cordelia Langford, John Sillitoe on behalf of the Wellcome Sanger Institute COVID-19 Surveillance Team |
| EPI_ISL_3139566 | Lighthouse Lab in Glasgow | Wellcome Sanger Institute for the COVID-19 Genomics UK (COG-UK) Consortium | Harper VanSteenhouse, Yumi Kasai, David Gray, Carol Clugston, Anna Dominiczak and Alex Alderton, Roberto Amato, Jeffrey Barrett, Sonia Goncalves, Ewan Harrison, David K. Jackson, Ian Johnston, Dominic Kwiatkowski, Cordelia Langford, John Sillitoe on behalf of the Wellcome Sanger Institute COVID-19 Surveillance Team |
| EPI_ISL_3139567 | Lighthouse Laboratory Plymouth | Wellcome Sanger Institute for the COVID-19 Genomics UK (COG-UK) Consortium | Lighthouse Laboratory Plymouth and Alex Alderton, Roberto Amato, Jeffrey Barrett, Sonia Goncalves, Ewan Harrison, David K. Jackson, Ian Johnston, Dominic Kwiatkowski, Cordelia Langford, John Sillitoe on behalf of the Wellcome Sanger Institute COVID-19 Surveillance Team |
| EPI_ISL_3139568 | Lighthouse Lab in Milton Keynes | Wellcome Sanger Institute for the COVID-19 Genomics UK (COG-UK) Consortium | The Lighthouse Lab in Milton Keynes and Alex Alderton, Roberto Amato, Jeffrey Barrett, Sonia Goncalves, Ewan Harrison, David K. Jackson, Ian Johnston, Dominic Kwiatkowski, Cordelia Langford, John Sillitoe on behalf of the Wellcome Sanger Institute COVID-19 Surveillance Team |
| EPI_ISL_3139569 | Lighthouse Lab in Glasgow | Wellcome Sanger Institute for the COVID-19 Genomics UK (COG-UK) Consortium | Harper VanSteenhouse, Yumi Kasai, David Gray, Carol Clugston, Anna Dominiczak and Alex Alderton, Roberto Amato, Jeffrey Barrett, Sonia Goncalves, Ewan Harrison, David K. Jackson, Ian Johnston, Dominic Kwiatkowski, Cordelia Langford, John Sillitoe on behalf of the Wellcome Sanger Institute COVID-19 Surveillance Team |
| EPI_ISL_3139570, EPI_ISL_3139571 | Lighthouse Laboratory Plymouth | Wellcome Sanger Institute for the COVID-19 Genomics UK (COG-UK) Consortium | Lighthouse Laboratory Plymouth and Alex Alderton, Roberto Amato, Jeffrey Barrett, Sonia Goncalves, Ewan Harrison, David K. Jackson, Ian Johnston, Dominic Kwiatkowski, Cordelia Langford, John Sillitoe on behalf of the Wellcome Sanger Institute COVID-19 Surveillance Team |
| EPI_ISL_3139572 | Lighthouse Lab in Glasgow | Wellcome Sanger Institute for the COVID-19 Genomics UK (COG-UK) Consortium | Harper VanSteenhouse, Yumi Kasai, David Gray, Carol Clugston, Anna Dominiczak and Alex Alderton, Roberto Amato, Jeffrey Barrett, Sonia Goncalves, Ewan Harrison, David K. Jackson, Ian Johnston, Dominic Kwiatkowski, Cordelia Langford, John Sillitoe on behalf of the Wellcome Sanger Institute COVID-19 Surveillance Team |
| EPI_ISL_3139573, EPI_ISL_3139574, EPI_ISL_3139575, EPI_ISL_3139576 | Lighthouse Laboratory Plymouth | Wellcome Sanger Institute for the COVID-19 Genomics UK (COG-UK) Consortium | Lighthouse Laboratory Plymouth and Alex Alderton, Roberto Amato, Jeffrey Barrett, Sonia Goncalves, Ewan Harrison, David K. Jackson, Ian Johnston, Dominic Kwiatkowski, Cordelia Langford, John Sillitoe on behalf of the Wellcome Sanger Institute COVID-19 Surveillance Team |
| EPI_ISL_3139577 | Lighthouse Lab in Milton Keynes | Wellcome Sanger Institute for the COVID-19 Genomics UK (COG-UK) Consortium | The Lighthouse Lab in Milton Keynes and Alex Alderton, Roberto Amato, Jeffrey Barrett, Sonia Goncalves, Ewan Harrison, David K. Jackson, Ian Johnston, Dominic Kwiatkowski, Cordelia Langford, John Sillitoe on behalf of the Wellcome Sanger Institute COVID-19 Surveillance Team |
| EPI_ISL_3139578 | Lighthouse Laboratory Plymouth | Wellcome Sanger Institute for the COVID-19 Genomics UK (COG-UK) Consortium | Lighthouse Laboratory Plymouth and Alex Alderton, Roberto Amato, Jeffrey Barrett, Sonia Goncalves, Ewan Harrison, David K. Jackson, Ian Johnston, Dominic Kwiatkowski, Cordelia Langford, John Sillitoe on behalf of the Wellcome Sanger Institute COVID-19 Surveillance Team |
| EPI_ISL_3139579, EPI_ISL_3139580, EPI_ISL_3139581 | Berkshire and Surrey Pathology Services Lighthouse Laboratory | Wellcome Sanger Institute for the COVID-19 Genomics UK (COG-UK) Consortium | Berkshire and Surrey Pathology Services Lighthouse Laboratory and Alex Alderton, Roberto Amato, Jeffrey Barrett, Sonia Goncalves, Ewan Harrison, David K. Jackson, Ian Johnston, Dominic Kwiatkowski, Cordelia Langford, John Sillitoe on behalf of the Wellcome Sanger Institute COVID-19 Surveillance Team |
| EPI_ISL_3139582 | Lighthouse Laboratory Plymouth | Wellcome Sanger Institute for the COVID-19 Genomics UK (COG-UK) Consortium | Lighthouse Laboratory Plymouth and Alex Alderton, Roberto Amato, Jeffrey Barrett, Sonia Goncalves, Ewan Harrison, David K. Jackson, Ian Johnston, Dominic Kwiatkowski, Cordelia Langford, John Sillitoe on behalf of the Wellcome Sanger Institute COVID-19 Surveillance Team |
| EPI_ISL_3139583, EPI_ISL_3139584 | Berkshire and Surrey Pathology Services Lighthouse Laboratory | Wellcome Sanger Institute for the COVID-19 Genomics UK (COG-UK) Consortium | Berkshire and Surrey Pathology Services Lighthouse Laboratory and Alex Alderton, Roberto Amato, Jeffrey Barrett, Sonia Goncalves, Ewan Harrison, David K. Jackson, Ian Johnston, Dominic Kwiatkowski, Cordelia Langford, John Sillitoe on behalf of the Wellcome Sanger Institute COVID-19 Surveillance Team |
| EPI_ISL_3139585, | Lighthouse Lab in Glasgow | Wellcome Sanger Institute for the COVID-19 Genomics UK (COG-UK) | Harper VanSteenhouse, Yumi Kasai, David Gray, Carol Clugston, Anna Dominiczak and Alex Alderton, Roberto Amato, Jeffrey Barrett, Sonia Goncalves, Ewan Harrison, David K. Jackson, Ian Johnston, Dominic Kwiatkowski, Cordelia Langford, |

| | | | |
|---|---|---|---|
| EPI_ISL_3139586, EPI_ISL_3139587, EPI_ISL_3139588 | | Consortium | John Sillitoe on behalf of the Wellcome Sanger Institute COVID-19 Surveillance Team |
| EPI_ISL_3139589 | Lighthouse Laboratory Plymouth | Wellcome Sanger Institute for the COVID-19 Genomics UK (COG-UK) Consortium | Lighthouse Laboratory Plymouth and Alex Alderton, Roberto Amato, Jeffrey Barrett, Sonia Goncalves, Ewan Harrison, David K. Jackson, Ian Johnston, Dominic Kwiatkowski, Cordelia Langford, John Sillitoe on behalf of the Wellcome Sanger Institute COVID-19 Surveillance Team |
| EPI_ISL_3139590, EPI_ISL_3139591, EPI_ISL_3139592, EPI_ISL_3139593 | Lighthouse Lab in Glasgow | Wellcome Sanger Institute for the COVID-19 Genomics UK (COG-UK) Consortium | Harper VanSteenhouse, Yumi Kasai, David Gray, Carol Clugston, Anna Dominiczak and Alex Alderton, Roberto Amato, Jeffrey Barrett, Sonia Goncalves, Ewan Harrison, David K. Jackson, Ian Johnston, Dominic Kwiatkowski, Cordelia Langford, John Sillitoe on behalf of the Wellcome Sanger Institute COVID-19 Surveillance Team |
| EPI_ISL_3139594 | Berkshire and Surrey Pathology Services Lighthouse Laboratory | Wellcome Sanger Institute for the COVID-19 Genomics UK (COG-UK) Consortium | Berkshire and Surrey Pathology Services Lighthouse Laboratory and Alex Alderton, Roberto Amato, Jeffrey Barrett, Sonia Goncalves, Ewan Harrison, David K. Jackson, Ian Johnston, Dominic Kwiatkowski, Cordelia Langford, John Sillitoe on behalf of the Wellcome Sanger Institute COVID-19 Surveillance Team |
| EPI_ISL_3139595 | Lighthouse Lab in Glasgow | Wellcome Sanger Institute for the COVID-19 Genomics UK (COG-UK) Consortium | Harper VanSteenhouse, Yumi Kasai, David Gray, Carol Clugston, Anna Dominiczak and Alex Alderton, Roberto Amato, Jeffrey Barrett, Sonia Goncalves, Ewan Harrison, David K. Jackson, Ian Johnston, Dominic Kwiatkowski, Cordelia Langford, John Sillitoe on behalf of the Wellcome Sanger Institute COVID-19 Surveillance Team |
| EPI_ISL_3139596, EPI_ISL_3139597 | Berkshire and Surrey Pathology Services Lighthouse Laboratory | Wellcome Sanger Institute for the COVID-19 Genomics UK (COG-UK) Consortium | Berkshire and Surrey Pathology Services Lighthouse Laboratory and Alex Alderton, Roberto Amato, Jeffrey Barrett, Sonia Goncalves, Ewan Harrison, David K. Jackson, Ian Johnston, Dominic Kwiatkowski, Cordelia Langford, John Sillitoe on behalf of the Wellcome Sanger Institute COVID-19 Surveillance Team |
| EPI_ISL_3139598, EPI_ISL_3139599, EPI_ISL_3139600, EPI_ISL_3139601, EPI_ISL_3139602, EPI_ISL_3139603, EPI_ISL_3139604, EPI_ISL_3139605 | Lighthouse Lab in Glasgow | Wellcome Sanger Institute for the COVID-19 Genomics UK (COG-UK) Consortium | Harper VanSteenhouse, Yumi Kasai, David Gray, Carol Clugston, Anna Dominiczak and Alex Alderton, Roberto Amato, Jeffrey Barrett, Sonia Goncalves, Ewan Harrison, David K. Jackson, Ian Johnston, Dominic Kwiatkowski, Cordelia Langford, John Sillitoe on behalf of the Wellcome Sanger Institute COVID-19 Surveillance Team |
| EPI_ISL_3139606 | Berkshire and Surrey Pathology Services Lighthouse Laboratory | Wellcome Sanger Institute for the COVID-19 Genomics UK (COG-UK) Consortium | Berkshire and Surrey Pathology Services Lighthouse Laboratory and Alex Alderton, Roberto Amato, Jeffrey Barrett, Sonia Goncalves, Ewan Harrison, David K. Jackson, Ian Johnston, Dominic Kwiatkowski, Cordelia Langford, John Sillitoe on behalf of the Wellcome Sanger Institute COVID-19 Surveillance Team |
| EPI_ISL_3139607, EPI_ISL_3139608 | Lighthouse Lab in Glasgow | Wellcome Sanger Institute for the COVID-19 Genomics UK (COG-UK) Consortium | Harper VanSteenhouse, Yumi Kasai, David Gray, Carol Clugston, Anna Dominiczak and Alex Alderton, Roberto Amato, Jeffrey Barrett, Sonia Goncalves, Ewan Harrison, David K. Jackson, Ian Johnston, Dominic Kwiatkowski, Cordelia Langford, John Sillitoe on behalf of the Wellcome Sanger Institute COVID-19 Surveillance Team |
| EPI_ISL_3139609 | Berkshire and Surrey Pathology Services Lighthouse Laboratory | Wellcome Sanger Institute for the COVID-19 Genomics UK (COG-UK) Consortium | Berkshire and Surrey Pathology Services Lighthouse Laboratory and Alex Alderton, Roberto Amato, Jeffrey Barrett, Sonia Goncalves, Ewan Harrison, David K. Jackson, Ian Johnston, Dominic Kwiatkowski, Cordelia Langford, John Sillitoe on behalf of the Wellcome Sanger Institute COVID-19 Surveillance Team |
| EPI_ISL_3139610, EPI_ISL_3139611, EPI_ISL_3139612 | Lighthouse Lab in Glasgow | Wellcome Sanger Institute for the COVID-19 Genomics UK (COG-UK) Consortium | Harper VanSteenhouse, Yumi Kasai, David Gray, Carol Clugston, Anna Dominiczak and Alex Alderton, Roberto Amato, Jeffrey Barrett, Sonia Goncalves, Ewan Harrison, David K. Jackson, Ian Johnston, Dominic Kwiatkowski, Cordelia Langford, John Sillitoe on behalf of the Wellcome Sanger Institute COVID-19 Surveillance Team |
| EPI_ISL_3139613 | Lighthouse Laboratory Plymouth | Wellcome Sanger Institute for the COVID-19 Genomics UK (COG-UK) Consortium | Lighthouse Laboratory Plymouth and Alex Alderton, Roberto Amato, Jeffrey Barrett, Sonia Goncalves, Ewan Harrison, David K. Jackson, Ian Johnston, Dominic Kwiatkowski, Cordelia Langford, John Sillitoe on behalf of the Wellcome Sanger Institute COVID-19 Surveillance Team |
| EPI_ISL_3139614 | Lighthouse Lab in Glasgow | Wellcome Sanger Institute for the COVID-19 Genomics UK (COG-UK) Consortium | Harper VanSteenhouse, Yumi Kasai, David Gray, Carol Clugston, Anna Dominiczak and Alex Alderton, Roberto Amato, Jeffrey Barrett, Sonia Goncalves, Ewan Harrison, David K. Jackson, Ian Johnston, Dominic Kwiatkowski, Cordelia Langford, John Sillitoe on behalf of the Wellcome Sanger Institute COVID-19 Surveillance Team |
| EPI_ISL_3139615, EPI_ISL_3139616, EPI_ISL_3139617 | Lighthouse Lab in Milton Keynes | Wellcome Sanger Institute for the COVID-19 Genomics UK (COG-UK) Consortium | The Lighthouse Lab in Milton Keynes and Alex Alderton, Roberto Amato, Jeffrey Barrett, Sonia Goncalves, Ewan Harrison, David K. Jackson, Ian Johnston, Dominic Kwiatkowski, Cordelia Langford, John Sillitoe on behalf of the Wellcome Sanger Institute COVID-19 Surveillance Team |
| EPI_ISL_3139618 | Lighthouse Laboratory Plymouth | Wellcome Sanger Institute for the COVID-19 Genomics UK (COG-UK) Consortium | Lighthouse Laboratory Plymouth and Alex Alderton, Roberto Amato, Jeffrey Barrett, Sonia Goncalves, Ewan Harrison, David K. Jackson, Ian Johnston, Dominic Kwiatkowski, Cordelia Langford, John Sillitoe on behalf of the Wellcome Sanger Institute COVID-19 Surveillance Team |
| EPI_ISL_3139619 | Lighthouse Lab in Alderley Park | Wellcome Sanger Institute for the COVID-19 Genomics UK (COG-UK) Consortium | Jacquelyn Wynn, Mairead Hyland, The Lighthouse Lab in Alderley Park and Alex Alderton, Roberto Amato, Jeffrey Barrett, Sonia Goncalves, Ewan Harrison, David K. Jackson, Ian Johnston, Dominic Kwiatkowski, Cordelia Langford, John Sillitoe on behalf of the Wellcome Sanger Institute COVID-19 Surveillance Team |
| EPI_ISL_3139620 | Berkshire and Surrey Pathology Services Lighthouse Laboratory | Wellcome Sanger Institute for the COVID-19 Genomics UK (COG-UK) Consortium | Berkshire and Surrey Pathology Services Lighthouse Laboratory and Alex Alderton, Roberto Amato, Jeffrey Barrett, Sonia Goncalves, Ewan Harrison, David K. Jackson, Ian Johnston, Dominic Kwiatkowski, Cordelia Langford, John Sillitoe on behalf of the Wellcome Sanger Institute COVID-19 Surveillance Team |
| EPI_ISL_3139621 | Lighthouse Lab in Alderley Park | Wellcome Sanger Institute for the COVID-19 Genomics UK (COG-UK) Consortium | Jacquelyn Wynn, Mairead Hyland, The Lighthouse Lab in Alderley Park and Alex Alderton, Roberto Amato, Jeffrey Barrett, Sonia Goncalves, Ewan Harrison, David K. Jackson, Ian Johnston, Dominic Kwiatkowski, Cordelia Langford, John Sillitoe on behalf of the Wellcome Sanger Institute COVID-19 Surveillance Team |
| EPI_ISL_3139622, EPI_ISL_3139623 | Lighthouse Laboratory Plymouth | Wellcome Sanger Institute for the COVID-19 Genomics UK (COG-UK) Consortium | Lighthouse Laboratory Plymouth and Alex Alderton, Roberto Amato, Jeffrey Barrett, Sonia Goncalves, Ewan Harrison, David K. Jackson, Ian Johnston, Dominic Kwiatkowski, Cordelia Langford, John Sillitoe on behalf of the Wellcome Sanger Institute COVID-19 Surveillance Team |
| EPI_ISL_3139624 | Lighthouse Lab in Alderley Park | Wellcome Sanger Institute for the COVID-19 Genomics UK (COG-UK) Consortium | Jacquelyn Wynn, Mairead Hyland, The Lighthouse Lab in Alderley Park and Alex Alderton, Roberto Amato, Jeffrey Barrett, Sonia Goncalves, Ewan Harrison, David K. Jackson, Ian Johnston, Dominic Kwiatkowski, Cordelia Langford, John Sillitoe on behalf of the Wellcome Sanger Institute COVID-19 Surveillance Team |
| EPI_ISL_3139625 | Lighthouse Laboratory Plymouth | Wellcome Sanger Institute for the COVID-19 Genomics UK (COG-UK) Consortium | Lighthouse Laboratory Plymouth and Alex Alderton, Roberto Amato, Jeffrey Barrett, Sonia Goncalves, Ewan Harrison, David K. Jackson, Ian Johnston, Dominic Kwiatkowski, Cordelia Langford, John Sillitoe on behalf of the Wellcome Sanger Institute COVID-19 Surveillance Team |
| EPI_ISL_3139626 | Lighthouse Lab in Alderley Park | Wellcome Sanger Institute for the COVID-19 Genomics UK (COG-UK) Consortium | Jacquelyn Wynn, Mairead Hyland, The Lighthouse Lab in Alderley Park and Alex Alderton, Roberto Amato, Jeffrey Barrett, Sonia Goncalves, Ewan Harrison, David K. Jackson, Ian Johnston, Dominic Kwiatkowski, Cordelia Langford, John Sillitoe on behalf of the Wellcome Sanger Institute COVID-19 Surveillance Team |
| EPI_ISL_3139627, EPI_ISL_3139628 | Lighthouse Lab in Milton Keynes | Wellcome Sanger Institute for the COVID-19 Genomics UK (COG-UK) Consortium | The Lighthouse Lab in Milton Keynes and Alex Alderton, Roberto Amato, Jeffrey Barrett, Sonia Goncalves, Ewan Harrison, David K. Jackson, Ian Johnston, Dominic Kwiatkowski, Cordelia Langford, John Sillitoe on behalf of the Wellcome Sanger Institute COVID-19 Surveillance Team |
| EPI_ISL_3139629 | Lighthouse Lab in Alderley Park | Wellcome Sanger Institute for the COVID-19 Genomics UK (COG-UK) Consortium | Jacquelyn Wynn, Mairead Hyland, The Lighthouse Lab in Alderley Park and Alex Alderton, Roberto Amato, Jeffrey Barrett, Sonia Goncalves, Ewan Harrison, David K. Jackson, Ian Johnston, Dominic Kwiatkowski, Cordelia Langford, John Sillitoe on behalf of the Wellcome Sanger Institute COVID-19 Surveillance Team |
| EPI_ISL_3139630, EPI_ISL_3139631 | Lighthouse Lab in Milton Keynes | Wellcome Sanger Institute for the COVID-19 Genomics UK (COG-UK) Consortium | The Lighthouse Lab in Milton Keynes and Alex Alderton, Roberto Amato, Jeffrey Barrett, Sonia Goncalves, Ewan Harrison, David K. Jackson, Ian Johnston, Dominic Kwiatkowski, Cordelia Langford, John Sillitoe on behalf of the Wellcome Sanger Institute COVID-19 Surveillance Team |
| EPI_ISL_3139632, EPI_ISL_3139633 | Lighthouse Lab in Alderley Park | Wellcome Sanger Institute for the COVID-19 Genomics UK (COG-UK) Consortium | Jacquelyn Wynn, Mairead Hyland, The Lighthouse Lab in Alderley Park and Alex Alderton, Roberto Amato, Jeffrey Barrett, Sonia Goncalves, Ewan Harrison, David K. Jackson, Ian Johnston, Dominic Kwiatkowski, Cordelia Langford, John Sillitoe on behalf of the Wellcome Sanger Institute COVID-19 Surveillance Team |
| EPI_ISL_3139634 | Lighthouse Laboratory Plymouth | Wellcome Sanger Institute for the COVID-19 Genomics UK (COG-UK) Consortium | Lighthouse Laboratory Plymouth and Alex Alderton, Roberto Amato, Jeffrey Barrett, Sonia Goncalves, Ewan Harrison, David K. Jackson, Ian Johnston, Dominic Kwiatkowski, Cordelia Langford, John Sillitoe on behalf of the Wellcome Sanger Institute COVID-19 Surveillance Team |
| EPI_ISL_3139635, EPI_ISL_3139636 | Berkshire and Surrey Pathology Services Lighthouse Laboratory | Wellcome Sanger Institute for the COVID-19 Genomics UK (COG-UK) Consortium | Berkshire and Surrey Pathology Services Lighthouse Laboratory and Alex Alderton, Roberto Amato, Jeffrey Barrett, Sonia Goncalves, Ewan Harrison, David K. Jackson, Ian Johnston, Dominic Kwiatkowski, Cordelia Langford, John Sillitoe on behalf of the Wellcome Sanger Institute COVID-19 Surveillance Team |
| EPI_ISL_3139637, EPI_ISL_3139638 | Lighthouse Laboratory Plymouth | Wellcome Sanger Institute for the COVID-19 Genomics UK (COG-UK) Consortium | Lighthouse Laboratory Plymouth and Alex Alderton, Roberto Amato, Jeffrey Barrett, Sonia Goncalves, Ewan Harrison, David K. Jackson, Ian Johnston, Dominic Kwiatkowski, Cordelia Langford, John Sillitoe on behalf of the Wellcome Sanger Institute COVID-19 Surveillance Team |
| EPI_ISL_3139639 | Lighthouse Lab in Milton Keynes | Wellcome Sanger Institute for the COVID-19 Genomics UK (COG-UK) Consortium | The Lighthouse Lab in Milton Keynes and Alex Alderton, Roberto Amato, Jeffrey Barrett, Sonia Goncalves, Ewan Harrison, David K. Jackson, Ian Johnston, Dominic Kwiatkowski, Cordelia Langford, John Sillitoe on behalf of the Wellcome Sanger Institute COVID-19 Surveillance Team |
| EPI_ISL_3139640 | Berkshire and Surrey Pathology Services Lighthouse Laboratory | Wellcome Sanger Institute for the COVID-19 Genomics UK (COG-UK) Consortium | Berkshire and Surrey Pathology Services Lighthouse Laboratory and Alex Alderton, Roberto Amato, Jeffrey Barrett, Sonia Goncalves, Ewan Harrison, David K. Jackson, Ian Johnston, Dominic Kwiatkowski, Cordelia Langford, John Sillitoe on behalf of the Wellcome Sanger Institute COVID-19 Surveillance Team |
| EPI_ISL_3139641 | Lighthouse Laboratory Plymouth | Wellcome Sanger Institute for the COVID-19 Genomics UK (COG-UK) Consortium | Lighthouse Laboratory Plymouth and Alex Alderton, Roberto Amato, Jeffrey Barrett, Sonia Goncalves, Ewan Harrison, David K. Jackson, Ian Johnston, Dominic Kwiatkowski, Cordelia Langford, John Sillitoe on behalf of the Wellcome Sanger Institute COVID-19 Surveillance Team |
| EPI_ISL_3139642 | Berkshire and Surrey Pathology Services Lighthouse Laboratory | Wellcome Sanger Institute for the COVID-19 Genomics UK (COG-UK) Consortium | Berkshire and Surrey Pathology Services Lighthouse Laboratory and Alex Alderton, Roberto Amato, Jeffrey Barrett, Sonia Goncalves, Ewan Harrison, David K. Jackson, Ian Johnston, Dominic Kwiatkowski, Cordelia Langford, John Sillitoe on behalf of the Wellcome Sanger Institute COVID-19 Surveillance Team |
| EPI_ISL_3139643 | Lighthouse Lab in Milton Keynes | Wellcome Sanger Institute for the COVID-19 Genomics UK (COG-UK) Consortium | The Lighthouse Lab in Milton Keynes and Alex Alderton, Roberto Amato, Jeffrey Barrett, Sonia Goncalves, Ewan Harrison, David K. Jackson, Ian Johnston, Dominic Kwiatkowski, Cordelia Langford, John Sillitoe on behalf of the Wellcome Sanger Institute COVID-19 Surveillance Team |
| EPI_ISL_3139644 | Berkshire and Surrey Pathology Services Lighthouse Laboratory | Wellcome Sanger Institute for the COVID-19 Genomics UK (COG-UK) Consortium | Berkshire and Surrey Pathology Services Lighthouse Laboratory and Alex Alderton, Roberto Amato, Jeffrey Barrett, Sonia Goncalves, Ewan Harrison, David K. Jackson, Ian Johnston, Dominic Kwiatkowski, Cordelia Langford, John Sillitoe on behalf of the Wellcome Sanger Institute COVID-19 Surveillance Team |
| EPI_ISL_3139645, EPI_ISL_3139646 | Lighthouse Lab in Alderley Park | Wellcome Sanger Institute for the COVID-19 Genomics UK (COG-UK) Consortium | Jacquelyn Wynn, Mairead Hyland, The Lighthouse Lab in Alderley Park and Alex Alderton, Roberto Amato, Jeffrey Barrett, Sonia Goncalves, Ewan Harrison, David K. Jackson, Ian Johnston, Dominic Kwiatkowski, Cordelia Langford, John Sillitoe on behalf of the Wellcome Sanger Institute COVID-19 Surveillance Team |
| EPI_ISL_3139647 | Lighthouse Laboratory Plymouth | Wellcome Sanger Institute for the COVID-19 Genomics UK (COG-UK) Consortium | Lighthouse Laboratory Plymouth and Alex Alderton, Roberto Amato, Jeffrey Barrett, Sonia Goncalves, Ewan Harrison, David K. Jackson, Ian Johnston, Dominic Kwiatkowski, Cordelia Langford, John Sillitoe on behalf of the Wellcome Sanger Institute COVID-19 Surveillance Team |
| EPI_ISL_3139648 | Lighthouse Lab in Alderley Park | Wellcome Sanger Institute for the COVID-19 Genomics UK (COG-UK) Consortium | Jacquelyn Wynn, Mairead Hyland, The Lighthouse Lab in Alderley Park and Alex Alderton, Roberto Amato, Jeffrey Barrett, Sonia Goncalves, Ewan Harrison, David K. Jackson, Ian Johnston, Dominic Kwiatkowski, Cordelia Langford, John Sillitoe on behalf of the Wellcome Sanger Institute COVID-19 Surveillance Team |
| EPI_ISL_3139649 | Berkshire and Surrey Pathology Services Lighthouse Laboratory | Wellcome Sanger Institute for the COVID-19 Genomics UK (COG-UK) Consortium | Berkshire and Surrey Pathology Services Lighthouse Laboratory and Alex Alderton, Roberto Amato, Jeffrey Barrett, Sonia Goncalves, Ewan Harrison, David K. Jackson, Ian Johnston, Dominic Kwiatkowski, Cordelia Langford, John Sillitoe on behalf of the Wellcome Sanger Institute COVID-19 Surveillance Team |
| EPI_ISL_3139650 | Lighthouse Lab in Alderley Park | Wellcome Sanger Institute for the COVID-19 Genomics UK (COG-UK) Consortium | Jacquelyn Wynn, Mairead Hyland, The Lighthouse Lab in Alderley Park and Alex Alderton, Roberto Amato, Jeffrey Barrett, Sonia Goncalves, Ewan Harrison, David K. Jackson, Ian Johnston, Dominic Kwiatkowski, Cordelia Langford, John Sillitoe on behalf of the Wellcome Sanger Institute COVID-19 Surveillance Team |
| EPI_ISL_3139651, | Berkshire and Surrey Pathology Services Lighthouse Laboratory | Wellcome Sanger Institute for the COVID-19 Genomics UK (COG-UK) | Berkshire and Surrey Pathology Services Lighthouse Laboratory and Alex Alderton, Roberto Amato, Jeffrey Barrett, Sonia Goncalves, Ewan Harrison, David K. Jackson, Ian Johnston, Dominic Kwiatkowski, Cordelia Langford, John Sillitoe on behalf |

| | | | |
|---|---|---|---|
| EPI_ISL_3139652 | | Consortium | of the Wellcome Sanger Institute COVID-19 Surveillance Team |
| EPI_ISL_3139653 | Lighthouse Lab in Milton Keynes | Wellcome Sanger Institute for the COVID-19 Genomics UK (COG-UK) Consortium | The Lighthouse Lab in Milton Keynes and Alex Alderton, Roberto Amato, Jeffrey Barrett, Sonia Goncalves, Ewan Harrison, David K. Jackson, Ian Johnston, Dominic Kwiatkowski, Cordelia Langford, John Sillitoe on behalf of the Wellcome Sanger Institute COVID-19 Surveillance Team |
| EPI_ISL_3139654 | Lighthouse Lab in Alderley Park | Wellcome Sanger Institute for the COVID-19 Genomics UK (COG-UK) Consortium | Jacquelyn Wynn, Mairead Hyland, The Lighthouse Lab in Alderley Park and Alex Alderton, Roberto Amato, Jeffrey Barrett, Sonia Goncalves, Ewan Harrison, David K. Jackson, Ian Johnston, Dominic Kwiatkowski, Cordelia Langford, John Sillitoe on behalf of the Wellcome Sanger Institute COVID-19 Surveillance Team |
| EPI_ISL_3139655 | Lighthouse Laboratory Plymouth | Wellcome Sanger Institute for the COVID-19 Genomics UK (COG-UK) Consortium | Lighthouse Laboratory Plymouth and Alex Alderton, Roberto Amato, Jeffrey Barrett, Sonia Goncalves, Ewan Harrison, David K. Jackson, Ian Johnston, Dominic Kwiatkowski, Cordelia Langford, John Sillitoe on behalf of the Wellcome Sanger Institute COVID-19 Surveillance Team |
| EPI_ISL_3139656, EPI_ISL_3139657 | Lighthouse Lab in Alderley Park | Wellcome Sanger Institute for the COVID-19 Genomics UK (COG-UK) Consortium | Jacquelyn Wynn, Mairead Hyland, The Lighthouse Lab in Alderley Park and Alex Alderton, Roberto Amato, Jeffrey Barrett, Sonia Goncalves, Ewan Harrison, David K. Jackson, Ian Johnston, Dominic Kwiatkowski, Cordelia Langford, John Sillitoe on behalf of the Wellcome Sanger Institute COVID-19 Surveillance Team |
| EPI_ISL_3139658 | Lighthouse Laboratory Plymouth | Wellcome Sanger Institute for the COVID-19 Genomics UK (COG-UK) Consortium | Lighthouse Laboratory Plymouth and Alex Alderton, Roberto Amato, Jeffrey Barrett, Sonia Goncalves, Ewan Harrison, David K. Jackson, Ian Johnston, Dominic Kwiatkowski, Cordelia Langford, John Sillitoe on behalf of the Wellcome Sanger Institute COVID-19 Surveillance Team |
| EPI_ISL_3139659, EPI_ISL_3139660 | Lighthouse Lab in Alderley Park | Wellcome Sanger Institute for the COVID-19 Genomics UK (COG-UK) Consortium | Jacquelyn Wynn, Mairead Hyland, The Lighthouse Lab in Alderley Park and Alex Alderton, Roberto Amato, Jeffrey Barrett, Sonia Goncalves, Ewan Harrison, David K. Jackson, Ian Johnston, Dominic Kwiatkowski, Cordelia Langford, John Sillitoe on behalf of the Wellcome Sanger Institute COVID-19 Surveillance Team |
| EPI_ISL_3139661 | Berkshire and Surrey Pathology Services Lighthouse Laboratory | Wellcome Sanger Institute for the COVID-19 Genomics UK (COG-UK) Consortium | Berkshire and Surrey Pathology Services Lighthouse Laboratory and Alex Alderton, Roberto Amato, Jeffrey Barrett, Sonia Goncalves, Ewan Harrison, David K. Jackson, Ian Johnston, Dominic Kwiatkowski, Cordelia Langford, John Sillitoe on behalf of the Wellcome Sanger Institute COVID-19 Surveillance Team |
| EPI_ISL_3139662 | Lighthouse Lab in Alderley Park | Wellcome Sanger Institute for the COVID-19 Genomics UK (COG-UK) Consortium | Jacquelyn Wynn, Mairead Hyland, The Lighthouse Lab in Alderley Park and Alex Alderton, Roberto Amato, Jeffrey Barrett, Sonia Goncalves, Ewan Harrison, David K. Jackson, Ian Johnston, Dominic Kwiatkowski, Cordelia Langford, John Sillitoe on behalf of the Wellcome Sanger Institute COVID-19 Surveillance Team |
| EPI_ISL_3139663 | Lighthouse Lab in Milton Keynes | Wellcome Sanger Institute for the COVID-19 Genomics UK (COG-UK) Consortium | The Lighthouse Lab in Milton Keynes and Alex Alderton, Roberto Amato, Jeffrey Barrett, Sonia Goncalves, Ewan Harrison, David K. Jackson, Ian Johnston, Dominic Kwiatkowski, Cordelia Langford, John Sillitoe on behalf of the Wellcome Sanger Institute COVID-19 Surveillance Team |
| EPI_ISL_3139664 | Lighthouse Lab in Alderley Park | Wellcome Sanger Institute for the COVID-19 Genomics UK (COG-UK) Consortium | Jacquelyn Wynn, Mairead Hyland, The Lighthouse Lab in Alderley Park and Alex Alderton, Roberto Amato, Jeffrey Barrett, Sonia Goncalves, Ewan Harrison, David K. Jackson, Ian Johnston, Dominic Kwiatkowski, Cordelia Langford, John Sillitoe on behalf of the Wellcome Sanger Institute COVID-19 Surveillance Team |
| EPI_ISL_3139665, EPI_ISL_3139666 | Berkshire and Surrey Pathology Services Lighthouse Laboratory | Wellcome Sanger Institute for the COVID-19 Genomics UK (COG-UK) Consortium | Berkshire and Surrey Pathology Services Lighthouse Laboratory and Alex Alderton, Roberto Amato, Jeffrey Barrett, Sonia Goncalves, Ewan Harrison, David K. Jackson, Ian Johnston, Dominic Kwiatkowski, Cordelia Langford, John Sillitoe on behalf of the Wellcome Sanger Institute COVID-19 Surveillance Team |
| EPI_ISL_3139667 | Lighthouse Lab in Milton Keynes | Wellcome Sanger Institute for the COVID-19 Genomics UK (COG-UK) Consortium | The Lighthouse Lab in Milton Keynes and Alex Alderton, Roberto Amato, Jeffrey Barrett, Sonia Goncalves, Ewan Harrison, David K. Jackson, Ian Johnston, Dominic Kwiatkowski, Cordelia Langford, John Sillitoe on behalf of the Wellcome Sanger Institute COVID-19 Surveillance Team |
| EPI_ISL_3139668 | Lighthouse Laboratory Plymouth | Wellcome Sanger Institute for the COVID-19 Genomics UK (COG-UK) Consortium | Lighthouse Laboratory Plymouth and Alex Alderton, Roberto Amato, Jeffrey Barrett, Sonia Goncalves, Ewan Harrison, David K. Jackson, Ian Johnston, Dominic Kwiatkowski, Cordelia Langford, John Sillitoe on behalf of the Wellcome Sanger Institute COVID-19 Surveillance Team |
| EPI_ISL_3139669, EPI_ISL_3139670 | Berkshire and Surrey Pathology Services Lighthouse Laboratory | Wellcome Sanger Institute for the COVID-19 Genomics UK (COG-UK) Consortium | Berkshire and Surrey Pathology Services Lighthouse Laboratory and Alex Alderton, Roberto Amato, Jeffrey Barrett, Sonia Goncalves, Ewan Harrison, David K. Jackson, Ian Johnston, Dominic Kwiatkowski, Cordelia Langford, John Sillitoe on behalf of the Wellcome Sanger Institute COVID-19 Surveillance Team |
| EPI_ISL_3139671 | Lighthouse Lab in Alderley Park | Wellcome Sanger Institute for the COVID-19 Genomics UK (COG-UK) Consortium | Jacquelyn Wynn, Mairead Hyland, The Lighthouse Lab in Alderley Park and Alex Alderton, Roberto Amato, Jeffrey Barrett, Sonia Goncalves, Ewan Harrison, David K. Jackson, Ian Johnston, Dominic Kwiatkowski, Cordelia Langford, John Sillitoe on behalf of the Wellcome Sanger Institute COVID-19 Surveillance Team |
| EPI_ISL_3139672 | Berkshire and Surrey Pathology Services Lighthouse Laboratory | Wellcome Sanger Institute for the COVID-19 Genomics UK (COG-UK) Consortium | Berkshire and Surrey Pathology Services Lighthouse Laboratory and Alex Alderton, Roberto Amato, Jeffrey Barrett, Sonia Goncalves, Ewan Harrison, David K. Jackson, Ian Johnston, Dominic Kwiatkowski, Cordelia Langford, John Sillitoe on behalf of the Wellcome Sanger Institute COVID-19 Surveillance Team |
| EPI_ISL_3139673 | Lighthouse Laboratory Plymouth | Wellcome Sanger Institute for the COVID-19 Genomics UK (COG-UK) Consortium | Lighthouse Laboratory Plymouth and Alex Alderton, Roberto Amato, Jeffrey Barrett, Sonia Goncalves, Ewan Harrison, David K. Jackson, Ian Johnston, Dominic Kwiatkowski, Cordelia Langford, John Sillitoe on behalf of the Wellcome Sanger Institute COVID-19 Surveillance Team |
| EPI_ISL_3139674 | Berkshire and Surrey Pathology Services Lighthouse Laboratory | Wellcome Sanger Institute for the COVID-19 Genomics UK (COG-UK) Consortium | Berkshire and Surrey Pathology Services Lighthouse Laboratory and Alex Alderton, Roberto Amato, Jeffrey Barrett, Sonia Goncalves, Ewan Harrison, David K. Jackson, Ian Johnston, Dominic Kwiatkowski, Cordelia Langford, John Sillitoe on behalf of the Wellcome Sanger Institute COVID-19 Surveillance Team |
| EPI_ISL_3139675 | Lighthouse Laboratory Plymouth | Wellcome Sanger Institute for the COVID-19 Genomics UK (COG-UK) Consortium | Lighthouse Laboratory Plymouth and Alex Alderton, Roberto Amato, Jeffrey Barrett, Sonia Goncalves, Ewan Harrison, David K. Jackson, Ian Johnston, Dominic Kwiatkowski, Cordelia Langford, John Sillitoe on behalf of the Wellcome Sanger Institute COVID-19 Surveillance Team |
| EPI_ISL_3139676 | Lighthouse Lab in Alderley Park | Wellcome Sanger Institute for the COVID-19 Genomics UK (COG-UK) Consortium | Jacquelyn Wynn, Mairead Hyland, The Lighthouse Lab in Alderley Park and Alex Alderton, Roberto Amato, Jeffrey Barrett, Sonia Goncalves, Ewan Harrison, David K. Jackson, Ian Johnston, Dominic Kwiatkowski, Cordelia Langford, John Sillitoe on behalf of the Wellcome Sanger Institute COVID-19 Surveillance Team |
| EPI_ISL_3139677 | Lighthouse Laboratory Plymouth | Wellcome Sanger Institute for the COVID-19 Genomics UK (COG-UK) Consortium | Lighthouse Laboratory Plymouth and Alex Alderton, Roberto Amato, Jeffrey Barrett, Sonia Goncalves, Ewan Harrison, David K. Jackson, Ian Johnston, Dominic Kwiatkowski, Cordelia Langford, John Sillitoe on behalf of the Wellcome Sanger Institute COVID-19 Surveillance Team |
| EPI_ISL_3139678 | Lighthouse Lab in Alderley Park | Wellcome Sanger Institute for the COVID-19 Genomics UK (COG-UK) Consortium | Jacquelyn Wynn, Mairead Hyland, The Lighthouse Lab in Alderley Park and Alex Alderton, Roberto Amato, Jeffrey Barrett, Sonia Goncalves, Ewan Harrison, David K. Jackson, Ian Johnston, Dominic Kwiatkowski, Cordelia Langford, John Sillitoe on behalf of the Wellcome Sanger Institute COVID-19 Surveillance Team |
| EPI_ISL_3139679 | Lighthouse Lab in Milton Keynes | Wellcome Sanger Institute for the COVID-19 Genomics UK (COG-UK) Consortium | The Lighthouse Lab in Milton Keynes and Alex Alderton, Roberto Amato, Jeffrey Barrett, Sonia Goncalves, Ewan Harrison, David K. Jackson, Ian Johnston, Dominic Kwiatkowski, Cordelia Langford, John Sillitoe on behalf of the Wellcome Sanger Institute COVID-19 Surveillance Team |
| EPI_ISL_3139680, EPI_ISL_3139681 | Lighthouse Laboratory Plymouth | Wellcome Sanger Institute for the COVID-19 Genomics UK (COG-UK) Consortium | Lighthouse Laboratory Plymouth and Alex Alderton, Roberto Amato, Jeffrey Barrett, Sonia Goncalves, Ewan Harrison, David K. Jackson, Ian Johnston, Dominic Kwiatkowski, Cordelia Langford, John Sillitoe on behalf of the Wellcome Sanger Institute COVID-19 Surveillance Team |
| EPI_ISL_3139682 | Lighthouse Lab in Milton Keynes | Wellcome Sanger Institute for the COVID-19 Genomics UK (COG-UK) Consortium | The Lighthouse Lab in Milton Keynes and Alex Alderton, Roberto Amato, Jeffrey Barrett, Sonia Goncalves, Ewan Harrison, David K. Jackson, Ian Johnston, Dominic Kwiatkowski, Cordelia Langford, John Sillitoe on behalf of the Wellcome Sanger Institute COVID-19 Surveillance Team |
| EPI_ISL_3139683 | Lighthouse Laboratory Plymouth | Wellcome Sanger Institute for the COVID-19 Genomics UK (COG-UK) Consortium | Lighthouse Laboratory Plymouth and Alex Alderton, Roberto Amato, Jeffrey Barrett, Sonia Goncalves, Ewan Harrison, David K. Jackson, Ian Johnston, Dominic Kwiatkowski, Cordelia Langford, John Sillitoe on behalf of the Wellcome Sanger Institute COVID-19 Surveillance Team |
| EPI_ISL_3139684, EPI_ISL_3139685 | Berkshire and Surrey Pathology Services Lighthouse Laboratory | Wellcome Sanger Institute for the COVID-19 Genomics UK (COG-UK) Consortium | Berkshire and Surrey Pathology Services Lighthouse Laboratory and Alex Alderton, Roberto Amato, Jeffrey Barrett, Sonia Goncalves, Ewan Harrison, David K. Jackson, Ian Johnston, Dominic Kwiatkowski, Cordelia Langford, John Sillitoe on behalf of the Wellcome Sanger Institute COVID-19 Surveillance Team |
| EPI_ISL_3139686 | Lighthouse Laboratory Plymouth | Wellcome Sanger Institute for the COVID-19 Genomics UK (COG-UK) Consortium | Lighthouse Laboratory Plymouth and Alex Alderton, Roberto Amato, Jeffrey Barrett, Sonia Goncalves, Ewan Harrison, David K. Jackson, Ian Johnston, Dominic Kwiatkowski, Cordelia Langford, John Sillitoe on behalf of the Wellcome Sanger Institute COVID-19 Surveillance Team |
| EPI_ISL_3139687 | Lighthouse Lab in Milton Keynes | Wellcome Sanger Institute for the COVID-19 Genomics UK (COG-UK) Consortium | The Lighthouse Lab in Milton Keynes and Alex Alderton, Roberto Amato, Jeffrey Barrett, Sonia Goncalves, Ewan Harrison, David K. Jackson, Ian Johnston, Dominic Kwiatkowski, Cordelia Langford, John Sillitoe on behalf of the Wellcome Sanger Institute COVID-19 Surveillance Team |
| EPI_ISL_3139688 | Berkshire and Surrey Pathology Services Lighthouse Laboratory | Wellcome Sanger Institute for the COVID-19 Genomics UK (COG-UK) Consortium | Berkshire and Surrey Pathology Services Lighthouse Laboratory and Alex Alderton, Roberto Amato, Jeffrey Barrett, Sonia Goncalves, Ewan Harrison, David K. Jackson, Ian Johnston, Dominic Kwiatkowski, Cordelia Langford, John Sillitoe on behalf of the Wellcome Sanger Institute COVID-19 Surveillance Team |
| EPI_ISL_3139689 | Lighthouse Lab in Alderley Park | Wellcome Sanger Institute for the COVID-19 Genomics UK (COG-UK) Consortium | Jacquelyn Wynn, Mairead Hyland, The Lighthouse Lab in Alderley Park and Alex Alderton, Roberto Amato, Jeffrey Barrett, Sonia Goncalves, Ewan Harrison, David K. Jackson, Ian Johnston, Dominic Kwiatkowski, Cordelia Langford, John Sillitoe on behalf of the Wellcome Sanger Institute COVID-19 Surveillance Team |
| EPI_ISL_3139690 | Lighthouse Lab in Milton Keynes | Wellcome Sanger Institute for the COVID-19 Genomics UK (COG-UK) Consortium | The Lighthouse Lab in Milton Keynes and Alex Alderton, Roberto Amato, Jeffrey Barrett, Sonia Goncalves, Ewan Harrison, David K. Jackson, Ian Johnston, Dominic Kwiatkowski, Cordelia Langford, John Sillitoe on behalf of the Wellcome Sanger Institute COVID-19 Surveillance Team |
| EPI_ISL_3139691 | Berkshire and Surrey Pathology Services Lighthouse Laboratory | Wellcome Sanger Institute for the COVID-19 Genomics UK (COG-UK) Consortium | Berkshire and Surrey Pathology Services Lighthouse Laboratory and Alex Alderton, Roberto Amato, Jeffrey Barrett, Sonia Goncalves, Ewan Harrison, David K. Jackson, Ian Johnston, Dominic Kwiatkowski, Cordelia Langford, John Sillitoe on behalf of the Wellcome Sanger Institute COVID-19 Surveillance Team |
| EPI_ISL_3139692 | Lighthouse Lab in Milton Keynes | Wellcome Sanger Institute for the COVID-19 Genomics UK (COG-UK) Consortium | The Lighthouse Lab in Milton Keynes and Alex Alderton, Roberto Amato, Jeffrey Barrett, Sonia Goncalves, Ewan Harrison, David K. Jackson, Ian Johnston, Dominic Kwiatkowski, Cordelia Langford, John Sillitoe on behalf of the Wellcome Sanger Institute COVID-19 Surveillance Team |
| EPI_ISL_3139693 | Lighthouse Laboratory Plymouth | Wellcome Sanger Institute for the COVID-19 Genomics UK (COG-UK) Consortium | Lighthouse Laboratory Plymouth and Alex Alderton, Roberto Amato, Jeffrey Barrett, Sonia Goncalves, Ewan Harrison, David K. Jackson, Ian Johnston, Dominic Kwiatkowski, Cordelia Langford, John Sillitoe on behalf of the Wellcome Sanger Institute COVID-19 Surveillance Team |
| EPI_ISL_3139694 | Berkshire and Surrey Pathology Services Lighthouse Laboratory | Wellcome Sanger Institute for the COVID-19 Genomics UK (COG-UK) Consortium | Berkshire and Surrey Pathology Services Lighthouse Laboratory and Alex Alderton, Roberto Amato, Jeffrey Barrett, Sonia Goncalves, Ewan Harrison, David K. Jackson, Ian Johnston, Dominic Kwiatkowski, Cordelia Langford, John Sillitoe on behalf of the Wellcome Sanger Institute COVID-19 Surveillance Team |
| EPI_ISL_3139695 | Lighthouse Laboratory Plymouth | Wellcome Sanger Institute for the COVID-19 Genomics UK (COG-UK) Consortium | Lighthouse Laboratory Plymouth and Alex Alderton, Roberto Amato, Jeffrey Barrett, Sonia Goncalves, Ewan Harrison, David K. Jackson, Ian Johnston, Dominic Kwiatkowski, Cordelia Langford, John Sillitoe on behalf of the Wellcome Sanger Institute COVID-19 Surveillance Team |
| EPI_ISL_3139696, EPI_ISL_3139697 | Lighthouse Lab in Alderley Park | Wellcome Sanger Institute for the COVID-19 Genomics UK (COG-UK) Consortium | Jacquelyn Wynn, Mairead Hyland, The Lighthouse Lab in Alderley Park and Alex Alderton, Roberto Amato, Jeffrey Barrett, Sonia Goncalves, Ewan Harrison, David K. Jackson, Ian Johnston, Dominic Kwiatkowski, Cordelia Langford, John Sillitoe on behalf of the Wellcome Sanger Institute COVID-19 Surveillance Team |
| EPI_ISL_3139698 | Berkshire and Surrey Pathology Services Lighthouse Laboratory | Wellcome Sanger Institute for the COVID-19 Genomics UK (COG-UK) Consortium | Berkshire and Surrey Pathology Services Lighthouse Laboratory and Alex Alderton, Roberto Amato, Jeffrey Barrett, Sonia Goncalves, Ewan Harrison, David K. Jackson, Ian Johnston, Dominic Kwiatkowski, Cordelia Langford, John Sillitoe on behalf of the Wellcome Sanger Institute COVID-19 Surveillance Team |
| EPI_ISL_3139699 | Lighthouse Laboratory Plymouth | Wellcome Sanger Institute for the COVID-19 Genomics UK (COG-UK) Consortium | Lighthouse Laboratory Plymouth and Alex Alderton, Roberto Amato, Jeffrey Barrett, Sonia Goncalves, Ewan Harrison, David K. Jackson, Ian Johnston, Dominic Kwiatkowski, Cordelia Langford, John Sillitoe on behalf of the Wellcome Sanger Institute COVID-19 Surveillance Team |
| EPI_ISL_3139700 | Lighthouse Lab in Alderley Park | Wellcome Sanger Institute for the COVID-19 Genomics UK (COG-UK) Consortium | Jacquelyn Wynn, Mairead Hyland, The Lighthouse Lab in Alderley Park and Alex Alderton, Roberto Amato, Jeffrey Barrett, Sonia Goncalves, Ewan Harrison, David K. Jackson, Ian Johnston, Dominic Kwiatkowski, Cordelia Langford, John Sillitoe on behalf of the Wellcome Sanger Institute COVID-19 Surveillance Team |
| EPI_ISL_3139701 | Lighthouse Laboratory Plymouth | Wellcome Sanger Institute for the COVID-19 Genomics UK (COG-UK) Consortium | Lighthouse Laboratory Plymouth and Alex Alderton, Roberto Amato, Jeffrey Barrett, Sonia Goncalves, Ewan Harrison, David K. Jackson, Ian Johnston, Dominic Kwiatkowski, Cordelia Langford, John Sillitoe on behalf of the Wellcome Sanger Institute COVID-19 Surveillance Team |
| EPI_ISL_3139702 | Lighthouse Lab in Milton Keynes | Wellcome Sanger Institute for the COVID-19 Genomics UK (COG-UK) Consortium | The Lighthouse Lab in Milton Keynes and Alex Alderton, Roberto Amato, Jeffrey Barrett, Sonia Goncalves, Ewan Harrison, David K. Jackson, Ian Johnston, Dominic Kwiatkowski, Cordelia Langford, John Sillitoe on behalf of the Wellcome Sanger Institute COVID-19 Surveillance Team |
| EPI_ISL_3139703 | Lighthouse Laboratory Plymouth | Wellcome Sanger Institute for the COVID-19 Genomics UK (COG-UK) Consortium | Lighthouse Laboratory Plymouth and Alex Alderton, Roberto Amato, Jeffrey Barrett, Sonia Goncalves, Ewan Harrison, David K. Jackson, Ian Johnston, Dominic Kwiatkowski, Cordelia Langford, John Sillitoe on behalf of the Wellcome Sanger Institute COVID-19 Surveillance Team |
| EPI_ISL_3139704 | Berkshire and Surrey Pathology Services Lighthouse Laboratory | Wellcome Sanger Institute for the COVID-19 Genomics UK (COG-UK) | Berkshire and Surrey Pathology Services Lighthouse Laboratory and Alex Alderton, Roberto Amato, Jeffrey Barrett, Sonia Goncalves, Ewan Harrison, David K. Jackson, Ian Johnston, Dominic Kwiatkowski, Cordelia Langford, John Sillitoe on behalf |

| | | | |
|---|---|---|---|
| | | Consortium | of the Wellcome Sanger Institute COVID-19 Surveillance Team |
| EPI_ISL_3139705, EPI_ISL_3139706 | Lighthouse Laboratory Plymouth | Wellcome Sanger Institute for the COVID-19 Genomics UK (COG-UK) Consortium | Lighthouse Laboratory Plymouth and Alex Alderton, Roberto Amato, Jeffrey Barrett, Sonia Goncalves, Ewan Harrison, David K. Jackson, Ian Johnston, Dominic Kwiatkowski, Cordelia Langford, John Sillitoe on behalf of the Wellcome Sanger Institute COVID-19 Surveillance Team |
| EPI_ISL_3139707 | Lighthouse Lab in Alderley Park | Wellcome Sanger Institute for the COVID-19 Genomics UK (COG-UK) Consortium | Jacquelyn Wynn, Mairead Hyland, The Lighthouse Lab in Alderley Park and Alex Alderton, Roberto Amato, Jeffrey Barrett, Sonia Goncalves, Ewan Harrison, David K. Jackson, Ian Johnston, Dominic Kwiatkowski, Cordelia Langford, John Sillitoe on behalf of the Wellcome Sanger Institute COVID-19 Surveillance Team |
| EPI_ISL_3139708 | Lighthouse Lab in Milton Keynes | Wellcome Sanger Institute for the COVID-19 Genomics UK (COG-UK) Consortium | The Lighthouse Lab in Milton Keynes and Alex Alderton, Roberto Amato, Jeffrey Barrett, Sonia Goncalves, Ewan Harrison, David K. Jackson, Ian Johnston, Dominic Kwiatkowski, Cordelia Langford, John Sillitoe on behalf of the Wellcome Sanger Institute COVID-19 Surveillance Team |
| EPI_ISL_3139709 | Berkshire and Surrey Pathology Services Lighthouse Laboratory | Wellcome Sanger Institute for the COVID-19 Genomics UK (COG-UK) Consortium | Berkshire and Surrey Pathology Services Lighthouse Laboratory and Alex Alderton, Roberto Amato, Jeffrey Barrett, Sonia Goncalves, Ewan Harrison, David K. Jackson, Ian Johnston, Dominic Kwiatkowski, Cordelia Langford, John Sillitoe on behalf of the Wellcome Sanger Institute COVID-19 Surveillance Team |
| EPI_ISL_3139710 | Lighthouse Lab in Milton Keynes | Wellcome Sanger Institute for the COVID-19 Genomics UK (COG-UK) Consortium | The Lighthouse Lab in Milton Keynes and Alex Alderton, Roberto Amato, Jeffrey Barrett, Sonia Goncalves, Ewan Harrison, David K. Jackson, Ian Johnston, Dominic Kwiatkowski, Cordelia Langford, John Sillitoe on behalf of the Wellcome Sanger Institute COVID-19 Surveillance Team |
| EPI_ISL_3139711 | Lighthouse Lab in Alderley Park | Wellcome Sanger Institute for the COVID-19 Genomics UK (COG-UK) Consortium | Jacquelyn Wynn, Mairead Hyland, The Lighthouse Lab in Alderley Park and Alex Alderton, Roberto Amato, Jeffrey Barrett, Sonia Goncalves, Ewan Harrison, David K. Jackson, Ian Johnston, Dominic Kwiatkowski, Cordelia Langford, John Sillitoe on behalf of the Wellcome Sanger Institute COVID-19 Surveillance Team |
| EPI_ISL_3139712, EPI_ISL_3139713 | Lighthouse Lab in Milton Keynes | Wellcome Sanger Institute for the COVID-19 Genomics UK (COG-UK) Consortium | The Lighthouse Lab in Milton Keynes and Alex Alderton, Roberto Amato, Jeffrey Barrett, Sonia Goncalves, Ewan Harrison, David K. Jackson, Ian Johnston, Dominic Kwiatkowski, Cordelia Langford, John Sillitoe on behalf of the Wellcome Sanger Institute COVID-19 Surveillance Team |
| EPI_ISL_3139714 | Berkshire and Surrey Pathology Services Lighthouse Laboratory | Wellcome Sanger Institute for the COVID-19 Genomics UK (COG-UK) Consortium | Berkshire and Surrey Pathology Services Lighthouse Laboratory and Alex Alderton, Roberto Amato, Jeffrey Barrett, Sonia Goncalves, Ewan Harrison, David K. Jackson, Ian Johnston, Dominic Kwiatkowski, Cordelia Langford, John Sillitoe on behalf of the Wellcome Sanger Institute COVID-19 Surveillance Team |
| EPI_ISL_3139715, EPI_ISL_3139716, EPI_ISL_3139717, EPI_ISL_3139718 | Lighthouse Laboratory Plymouth | Wellcome Sanger Institute for the COVID-19 Genomics UK (COG-UK) Consortium | Lighthouse Laboratory Plymouth and Alex Alderton, Roberto Amato, Jeffrey Barrett, Sonia Goncalves, Ewan Harrison, David K. Jackson, Ian Johnston, Dominic Kwiatkowski, Cordelia Langford, John Sillitoe on behalf of the Wellcome Sanger Institute COVID-19 Surveillance Team |
| EPI_ISL_3139719 | Berkshire and Surrey Pathology Services Lighthouse Laboratory | Wellcome Sanger Institute for the COVID-19 Genomics UK (COG-UK) Consortium | Berkshire and Surrey Pathology Services Lighthouse Laboratory and Alex Alderton, Roberto Amato, Jeffrey Barrett, Sonia Goncalves, Ewan Harrison, David K. Jackson, Ian Johnston, Dominic Kwiatkowski, Cordelia Langford, John Sillitoe on behalf of the Wellcome Sanger Institute COVID-19 Surveillance Team |
| EPI_ISL_3139720 | Lighthouse Lab in Alderley Park | Wellcome Sanger Institute for the COVID-19 Genomics UK (COG-UK) Consortium | Jacquelyn Wynn, Mairead Hyland, The Lighthouse Lab in Alderley Park and Alex Alderton, Roberto Amato, Jeffrey Barrett, Sonia Goncalves, Ewan Harrison, David K. Jackson, Ian Johnston, Dominic Kwiatkowski, Cordelia Langford, John Sillitoe on behalf of the Wellcome Sanger Institute COVID-19 Surveillance Team |
| EPI_ISL_3139721 | Lighthouse Lab in Milton Keynes | Wellcome Sanger Institute for the COVID-19 Genomics UK (COG-UK) Consortium | The Lighthouse Lab in Milton Keynes and Alex Alderton, Roberto Amato, Jeffrey Barrett, Sonia Goncalves, Ewan Harrison, David K. Jackson, Ian Johnston, Dominic Kwiatkowski, Cordelia Langford, John Sillitoe on behalf of the Wellcome Sanger Institute COVID-19 Surveillance Team |
| EPI_ISL_3139722, EPI_ISL_3139723 | Lighthouse Laboratory Plymouth | Wellcome Sanger Institute for the COVID-19 Genomics UK (COG-UK) Consortium | Lighthouse Laboratory Plymouth and Alex Alderton, Roberto Amato, Jeffrey Barrett, Sonia Goncalves, Ewan Harrison, David K. Jackson, Ian Johnston, Dominic Kwiatkowski, Cordelia Langford, John Sillitoe on behalf of the Wellcome Sanger Institute COVID-19 Surveillance Team |
| EPI_ISL_3139724 | Berkshire and Surrey Pathology Services Lighthouse Laboratory | Wellcome Sanger Institute for the COVID-19 Genomics UK (COG-UK) Consortium | Berkshire and Surrey Pathology Services Lighthouse Laboratory and Alex Alderton, Roberto Amato, Jeffrey Barrett, Sonia Goncalves, Ewan Harrison, David K. Jackson, Ian Johnston, Dominic Kwiatkowski, Cordelia Langford, John Sillitoe on behalf of the Wellcome Sanger Institute COVID-19 Surveillance Team |
| EPI_ISL_3139725 | Lighthouse Laboratory Plymouth | Wellcome Sanger Institute for the COVID-19 Genomics UK (COG-UK) Consortium | Lighthouse Laboratory Plymouth and Alex Alderton, Roberto Amato, Jeffrey Barrett, Sonia Goncalves, Ewan Harrison, David K. Jackson, Ian Johnston, Dominic Kwiatkowski, Cordelia Langford, John Sillitoe on behalf of the Wellcome Sanger Institute COVID-19 Surveillance Team |
| EPI_ISL_3139726 | Lighthouse Lab in Milton Keynes | Wellcome Sanger Institute for the COVID-19 Genomics UK (COG-UK) Consortium | The Lighthouse Lab in Milton Keynes and Alex Alderton, Roberto Amato, Jeffrey Barrett, Sonia Goncalves, Ewan Harrison, David K. Jackson, Ian Johnston, Dominic Kwiatkowski, Cordelia Langford, John Sillitoe on behalf of the Wellcome Sanger Institute COVID-19 Surveillance Team |
| EPI_ISL_3139727 | Berkshire and Surrey Pathology Services Lighthouse Laboratory | Wellcome Sanger Institute for the COVID-19 Genomics UK (COG-UK) Consortium | Berkshire and Surrey Pathology Services Lighthouse Laboratory and Alex Alderton, Roberto Amato, Jeffrey Barrett, Sonia Goncalves, Ewan Harrison, David K. Jackson, Ian Johnston, Dominic Kwiatkowski, Cordelia Langford, John Sillitoe on behalf of the Wellcome Sanger Institute COVID-19 Surveillance Team |
| EPI_ISL_3139728 | Lighthouse Lab in Alderley Park | Wellcome Sanger Institute for the COVID-19 Genomics UK (COG-UK) Consortium | Jacquelyn Wynn, Mairead Hyland, The Lighthouse Lab in Alderley Park and Alex Alderton, Roberto Amato, Jeffrey Barrett, Sonia Goncalves, Ewan Harrison, David K. Jackson, Ian Johnston, Dominic Kwiatkowski, Cordelia Langford, John Sillitoe on behalf of the Wellcome Sanger Institute COVID-19 Surveillance Team |
| EPI_ISL_3139729 | Berkshire and Surrey Pathology Services Lighthouse Laboratory | Wellcome Sanger Institute for the COVID-19 Genomics UK (COG-UK) Consortium | Berkshire and Surrey Pathology Services Lighthouse Laboratory and Alex Alderton, Roberto Amato, Jeffrey Barrett, Sonia Goncalves, Ewan Harrison, David K. Jackson, Ian Johnston, Dominic Kwiatkowski, Cordelia Langford, John Sillitoe on behalf of the Wellcome Sanger Institute COVID-19 Surveillance Team |
| EPI_ISL_3139730, EPI_ISL_3139731, EPI_ISL_3139732 | Lighthouse Laboratory Plymouth | Wellcome Sanger Institute for the COVID-19 Genomics UK (COG-UK) Consortium | Lighthouse Laboratory Plymouth and Alex Alderton, Roberto Amato, Jeffrey Barrett, Sonia Goncalves, Ewan Harrison, David K. Jackson, Ian Johnston, Dominic Kwiatkowski, Cordelia Langford, John Sillitoe on behalf of the Wellcome Sanger Institute COVID-19 Surveillance Team |
| EPI_ISL_3139733 | Lighthouse Lab in Milton Keynes | Wellcome Sanger Institute for the COVID-19 Genomics UK (COG-UK) Consortium | The Lighthouse Lab in Milton Keynes and Alex Alderton, Roberto Amato, Jeffrey Barrett, Sonia Goncalves, Ewan Harrison, David K. Jackson, Ian Johnston, Dominic Kwiatkowski, Cordelia Langford, John Sillitoe on behalf of the Wellcome Sanger Institute COVID-19 Surveillance Team |
| EPI_ISL_3139734 | Lighthouse Laboratory Plymouth | Wellcome Sanger Institute for the COVID-19 Genomics UK (COG-UK) Consortium | Lighthouse Laboratory Plymouth and Alex Alderton, Roberto Amato, Jeffrey Barrett, Sonia Goncalves, Ewan Harrison, David K. Jackson, Ian Johnston, Dominic Kwiatkowski, Cordelia Langford, John Sillitoe on behalf of the Wellcome Sanger Institute COVID-19 Surveillance Team |
| EPI_ISL_3139735 | Lighthouse Lab in Alderley Park | Wellcome Sanger Institute for the COVID-19 Genomics UK (COG-UK) Consortium | Jacquelyn Wynn, Mairead Hyland, The Lighthouse Lab in Alderley Park and Alex Alderton, Roberto Amato, Jeffrey Barrett, Sonia Goncalves, Ewan Harrison, David K. Jackson, Ian Johnston, Dominic Kwiatkowski, Cordelia Langford, John Sillitoe on behalf of the Wellcome Sanger Institute COVID-19 Surveillance Team |
| EPI_ISL_3139736 | Berkshire and Surrey Pathology Services Lighthouse Laboratory | Wellcome Sanger Institute for the COVID-19 Genomics UK (COG-UK) Consortium | Berkshire and Surrey Pathology Services Lighthouse Laboratory and Alex Alderton, Roberto Amato, Jeffrey Barrett, Sonia Goncalves, Ewan Harrison, David K. Jackson, Ian Johnston, Dominic Kwiatkowski, Cordelia Langford, John Sillitoe on behalf of the Wellcome Sanger Institute COVID-19 Surveillance Team |
| EPI_ISL_3139737 | Lighthouse Lab in Milton Keynes | Wellcome Sanger Institute for the COVID-19 Genomics UK (COG-UK) Consortium | The Lighthouse Lab in Milton Keynes and Alex Alderton, Roberto Amato, Jeffrey Barrett, Sonia Goncalves, Ewan Harrison, David K. Jackson, Ian Johnston, Dominic Kwiatkowski, Cordelia Langford, John Sillitoe on behalf of the Wellcome Sanger Institute COVID-19 Surveillance Team |
| EPI_ISL_3139738 | Berkshire and Surrey Pathology Services Lighthouse Laboratory | Wellcome Sanger Institute for the COVID-19 Genomics UK (COG-UK) Consortium | Berkshire and Surrey Pathology Services Lighthouse Laboratory and Alex Alderton, Roberto Amato, Jeffrey Barrett, Sonia Goncalves, Ewan Harrison, David K. Jackson, Ian Johnston, Dominic Kwiatkowski, Cordelia Langford, John Sillitoe on behalf of the Wellcome Sanger Institute COVID-19 Surveillance Team |
| EPI_ISL_3139739 | Lighthouse Laboratory Plymouth | Wellcome Sanger Institute for the COVID-19 Genomics UK (COG-UK) Consortium | Lighthouse Laboratory Plymouth and Alex Alderton, Roberto Amato, Jeffrey Barrett, Sonia Goncalves, Ewan Harrison, David K. Jackson, Ian Johnston, Dominic Kwiatkowski, Cordelia Langford, John Sillitoe on behalf of the Wellcome Sanger Institute COVID-19 Surveillance Team |
| EPI_ISL_3139740 | Berkshire and Surrey Pathology Services Lighthouse Laboratory | Wellcome Sanger Institute for the COVID-19 Genomics UK (COG-UK) Consortium | Berkshire and Surrey Pathology Services Lighthouse Laboratory and Alex Alderton, Roberto Amato, Jeffrey Barrett, Sonia Goncalves, Ewan Harrison, David K. Jackson, Ian Johnston, Dominic Kwiatkowski, Cordelia Langford, John Sillitoe on behalf of the Wellcome Sanger Institute COVID-19 Surveillance Team |
| EPI_ISL_3139741 | Lighthouse Lab in Alderley Park | Wellcome Sanger Institute for the COVID-19 Genomics UK (COG-UK) Consortium | Jacquelyn Wynn, Mairead Hyland, The Lighthouse Lab in Alderley Park and Alex Alderton, Roberto Amato, Jeffrey Barrett, Sonia Goncalves, Ewan Harrison, David K. Jackson, Ian Johnston, Dominic Kwiatkowski, Cordelia Langford, John Sillitoe on behalf of the Wellcome Sanger Institute COVID-19 Surveillance Team |
| EPI_ISL_3139742 | Lighthouse Lab in Milton Keynes | Wellcome Sanger Institute for the COVID-19 Genomics UK (COG-UK) Consortium | The Lighthouse Lab in Milton Keynes and Alex Alderton, Roberto Amato, Jeffrey Barrett, Sonia Goncalves, Ewan Harrison, David K. Jackson, Ian Johnston, Dominic Kwiatkowski, Cordelia Langford, John Sillitoe on behalf of the Wellcome Sanger Institute COVID-19 Surveillance Team |
| EPI_ISL_3139743, EPI_ISL_3139744, EPI_ISL_3139745, EPI_ISL_3139746 | Lighthouse Laboratory Plymouth | Wellcome Sanger Institute for the COVID-19 Genomics UK (COG-UK) Consortium | Lighthouse Laboratory Plymouth and Alex Alderton, Roberto Amato, Jeffrey Barrett, Sonia Goncalves, Ewan Harrison, David K. Jackson, Ian Johnston, Dominic Kwiatkowski, Cordelia Langford, John Sillitoe on behalf of the Wellcome Sanger Institute COVID-19 Surveillance Team |
| EPI_ISL_3139747, EPI_ISL_3139748 | Lighthouse Lab in Milton Keynes | Wellcome Sanger Institute for the COVID-19 Genomics UK (COG-UK) Consortium | The Lighthouse Lab in Milton Keynes and Alex Alderton, Roberto Amato, Jeffrey Barrett, Sonia Goncalves, Ewan Harrison, David K. Jackson, Ian Johnston, Dominic Kwiatkowski, Cordelia Langford, John Sillitoe on behalf of the Wellcome Sanger Institute COVID-19 Surveillance Team |
| EPI_ISL_3139749 | Lighthouse Laboratory Plymouth | Wellcome Sanger Institute for the COVID-19 Genomics UK (COG-UK) Consortium | Lighthouse Laboratory Plymouth and Alex Alderton, Roberto Amato, Jeffrey Barrett, Sonia Goncalves, Ewan Harrison, David K. Jackson, Ian Johnston, Dominic Kwiatkowski, Cordelia Langford, John Sillitoe on behalf of the Wellcome Sanger Institute COVID-19 Surveillance Team |
| EPI_ISL_3139750 | Lighthouse Lab in Milton Keynes | Wellcome Sanger Institute for the COVID-19 Genomics UK (COG-UK) Consortium | The Lighthouse Lab in Milton Keynes and Alex Alderton, Roberto Amato, Jeffrey Barrett, Sonia Goncalves, Ewan Harrison, David K. Jackson, Ian Johnston, Dominic Kwiatkowski, Cordelia Langford, John Sillitoe on behalf of the Wellcome Sanger Institute COVID-19 Surveillance Team |
| EPI_ISL_3139751 | Lighthouse Laboratory Plymouth | Wellcome Sanger Institute for the COVID-19 Genomics UK (COG-UK) Consortium | Lighthouse Laboratory Plymouth and Alex Alderton, Roberto Amato, Jeffrey Barrett, Sonia Goncalves, Ewan Harrison, David K. Jackson, Ian Johnston, Dominic Kwiatkowski, Cordelia Langford, John Sillitoe on behalf of the Wellcome Sanger Institute COVID-19 Surveillance Team |
| EPI_ISL_3139752 | Lighthouse Lab in Alderley Park | Wellcome Sanger Institute for the COVID-19 Genomics UK (COG-UK) Consortium | Jacquelyn Wynn, Mairead Hyland, The Lighthouse Lab in Alderley Park and Alex Alderton, Roberto Amato, Jeffrey Barrett, Sonia Goncalves, Ewan Harrison, David K. Jackson, Ian Johnston, Dominic Kwiatkowski, Cordelia Langford, John Sillitoe on behalf of the Wellcome Sanger Institute COVID-19 Surveillance Team |
| EPI_ISL_3139753 | Berkshire and Surrey Pathology Services Lighthouse Laboratory | Wellcome Sanger Institute for the COVID-19 Genomics UK (COG-UK) Consortium | Berkshire and Surrey Pathology Services Lighthouse Laboratory and Alex Alderton, Roberto Amato, Jeffrey Barrett, Sonia Goncalves, Ewan Harrison, David K. Jackson, Ian Johnston, Dominic Kwiatkowski, Cordelia Langford, John Sillitoe on behalf of the Wellcome Sanger Institute COVID-19 Surveillance Team |
| EPI_ISL_3139754 | Lighthouse Lab in Alderley Park | Wellcome Sanger Institute for the COVID-19 Genomics UK (COG-UK) Consortium | Jacquelyn Wynn, Mairead Hyland, The Lighthouse Lab in Alderley Park and Alex Alderton, Roberto Amato, Jeffrey Barrett, Sonia Goncalves, Ewan Harrison, David K. Jackson, Ian Johnston, Dominic Kwiatkowski, Cordelia Langford, John Sillitoe on behalf of the Wellcome Sanger Institute COVID-19 Surveillance Team |
| EPI_ISL_3139755 | Lighthouse Lab in Milton Keynes | Wellcome Sanger Institute for the COVID-19 Genomics UK (COG-UK) Consortium | The Lighthouse Lab in Milton Keynes and Alex Alderton, Roberto Amato, Jeffrey Barrett, Sonia Goncalves, Ewan Harrison, David K. Jackson, Ian Johnston, Dominic Kwiatkowski, Cordelia Langford, John Sillitoe on behalf of the Wellcome Sanger Institute COVID-19 Surveillance Team |
| EPI_ISL_3139756 | Lighthouse Laboratory Plymouth | Wellcome Sanger Institute for the COVID-19 Genomics UK (COG-UK) Consortium | Lighthouse Laboratory Plymouth and Alex Alderton, Roberto Amato, Jeffrey Barrett, Sonia Goncalves, Ewan Harrison, David K. Jackson, Ian Johnston, Dominic Kwiatkowski, Cordelia Langford, John Sillitoe on behalf of the Wellcome Sanger Institute COVID-19 Surveillance Team |
| EPI_ISL_3139757 | Lighthouse Lab in Alderley Park | Wellcome Sanger Institute for the COVID-19 Genomics UK (COG-UK) Consortium | Jacquelyn Wynn, Mairead Hyland, The Lighthouse Lab in Alderley Park and Alex Alderton, Roberto Amato, Jeffrey Barrett, Sonia Goncalves, Ewan Harrison, David K. Jackson, Ian Johnston, Dominic Kwiatkowski, Cordelia Langford, John Sillitoe on behalf of the Wellcome Sanger Institute COVID-19 Surveillance Team |
| EPI_ISL_3139758 | Lighthouse Lab in Milton Keynes | Wellcome Sanger Institute for the COVID-19 Genomics UK (COG-UK) Consortium | The Lighthouse Lab in Milton Keynes and Alex Alderton, Roberto Amato, Jeffrey Barrett, Sonia Goncalves, Ewan Harrison, David K. Jackson, Ian Johnston, Dominic Kwiatkowski, Cordelia Langford, John Sillitoe on behalf of the Wellcome Sanger Institute COVID-19 Surveillance Team |
| EPI_ISL_3139759 | Lighthouse Lab in Alderley Park | Wellcome Sanger Institute for the COVID-19 Genomics UK (COG-UK) | Jacquelyn Wynn, Mairead Hyland, The Lighthouse Lab in Alderley Park and Alex Alderton, Roberto Amato, Jeffrey Barrett, Sonia Goncalves, Ewan Harrison, David K. Jackson, Ian Johnston, Dominic Kwiatkowski, Cordelia Langford, John Sillitoe on |

| | | | |
|---|---|---|---|
| | | Consortium | behalf of the Wellcome Sanger Institute COVID-19 Surveillance Team |
| EPI_ISL_3139760 | Lighthouse Lab in Milton Keynes | Wellcome Sanger Institute for the COVID-19 Genomics UK (COG-UK) Consortium | The Lighthouse Lab in Milton Keynes and Alex Alderton, Roberto Amato, Jeffrey Barrett, Sonia Goncalves, Ewan Harrison, David K. Jackson, Ian Johnston, Dominic Kwiatkowski, Cordelia Langford, John Sillitoe on behalf of the Wellcome Sanger Institute COVID-19 Surveillance Team |
| EPI_ISL_3139761 | Berkshire and Surrey Pathology Services Lighthouse Laboratory | Wellcome Sanger Institute for the COVID-19 Genomics UK (COG-UK) Consortium | Berkshire and Surrey Pathology Services Lighthouse Laboratory and Alex Alderton, Roberto Amato, Jeffrey Barrett, Sonia Goncalves, Ewan Harrison, David K. Jackson, Ian Johnston, Dominic Kwiatkowski, Cordelia Langford, John Sillitoe on behalf of the Wellcome Sanger Institute COVID-19 Surveillance Team |
| EPI_ISL_3139762 | Lighthouse Lab in Alderley Park | Wellcome Sanger Institute for the COVID-19 Genomics UK (COG-UK) Consortium | Jacquelyn Wynn, Mairead Hyland, The Lighthouse Lab in Alderley Park and Alex Alderton, Roberto Amato, Jeffrey Barrett, Sonia Goncalves, Ewan Harrison, David K. Jackson, Ian Johnston, Dominic Kwiatkowski, Cordelia Langford, John Sillitoe on behalf of the Wellcome Sanger Institute COVID-19 Surveillance Team |
| EPI_ISL_3139763 | Lighthouse Lab in Milton Keynes | Wellcome Sanger Institute for the COVID-19 Genomics UK (COG-UK) Consortium | The Lighthouse Lab in Milton Keynes and Alex Alderton, Roberto Amato, Jeffrey Barrett, Sonia Goncalves, Ewan Harrison, David K. Jackson, Ian Johnston, Dominic Kwiatkowski, Cordelia Langford, John Sillitoe on behalf of the Wellcome Sanger Institute COVID-19 Surveillance Team |
| EPI_ISL_3139764 | Berkshire and Surrey Pathology Services Lighthouse Laboratory | Wellcome Sanger Institute for the COVID-19 Genomics UK (COG-UK) Consortium | Berkshire and Surrey Pathology Services Lighthouse Laboratory and Alex Alderton, Roberto Amato, Jeffrey Barrett, Sonia Goncalves, Ewan Harrison, David K. Jackson, Ian Johnston, Dominic Kwiatkowski, Cordelia Langford, John Sillitoe on behalf of the Wellcome Sanger Institute COVID-19 Surveillance Team |
| EPI_ISL_3139765 | Lighthouse Lab in Milton Keynes | Wellcome Sanger Institute for the COVID-19 Genomics UK (COG-UK) Consortium | The Lighthouse Lab in Milton Keynes and Alex Alderton, Roberto Amato, Jeffrey Barrett, Sonia Goncalves, Ewan Harrison, David K. Jackson, Ian Johnston, Dominic Kwiatkowski, Cordelia Langford, John Sillitoe on behalf of the Wellcome Sanger Institute COVID-19 Surveillance Team |
| EPI_ISL_3139766, EPI_ISL_3139767, EPI_ISL_3139768 | Lighthouse Lab in Alderley Park | Wellcome Sanger Institute for the COVID-19 Genomics UK (COG-UK) Consortium | Jacquelyn Wynn, Mairead Hyland, The Lighthouse Lab in Alderley Park and Alex Alderton, Roberto Amato, Jeffrey Barrett, Sonia Goncalves, Ewan Harrison, David K. Jackson, Ian Johnston, Dominic Kwiatkowski, Cordelia Langford, John Sillitoe on behalf of the Wellcome Sanger Institute COVID-19 Surveillance Team |
| EPI_ISL_3139769 | Lighthouse Laboratory Plymouth | Wellcome Sanger Institute for the COVID-19 Genomics UK (COG-UK) Consortium | Lighthouse Laboratory Plymouth and Alex Alderton, Roberto Amato, Jeffrey Barrett, Sonia Goncalves, Ewan Harrison, David K. Jackson, Ian Johnston, Dominic Kwiatkowski, Cordelia Langford, John Sillitoe on behalf of the Wellcome Sanger Institute COVID-19 Surveillance Team |
| EPI_ISL_3139770 | Lighthouse Lab in Alderley Park | Wellcome Sanger Institute for the COVID-19 Genomics UK (COG-UK) Consortium | Jacquelyn Wynn, Mairead Hyland, The Lighthouse Lab in Alderley Park and Alex Alderton, Roberto Amato, Jeffrey Barrett, Sonia Goncalves, Ewan Harrison, David K. Jackson, Ian Johnston, Dominic Kwiatkowski, Cordelia Langford, John Sillitoe on behalf of the Wellcome Sanger Institute COVID-19 Surveillance Team |
| EPI_ISL_3139771 | Lighthouse Lab in Milton Keynes | Wellcome Sanger Institute for the COVID-19 Genomics UK (COG-UK) Consortium | The Lighthouse Lab in Milton Keynes and Alex Alderton, Roberto Amato, Jeffrey Barrett, Sonia Goncalves, Ewan Harrison, David K. Jackson, Ian Johnston, Dominic Kwiatkowski, Cordelia Langford, John Sillitoe on behalf of the Wellcome Sanger Institute COVID-19 Surveillance Team |
| EPI_ISL_3139772 | Lighthouse Laboratory Plymouth | Wellcome Sanger Institute for the COVID-19 Genomics UK (COG-UK) Consortium | Lighthouse Laboratory Plymouth and Alex Alderton, Roberto Amato, Jeffrey Barrett, Sonia Goncalves, Ewan Harrison, David K. Jackson, Ian Johnston, Dominic Kwiatkowski, Cordelia Langford, John Sillitoe on behalf of the Wellcome Sanger Institute COVID-19 Surveillance Team |
| EPI_ISL_3139773 | Lighthouse Lab in Alderley Park | Wellcome Sanger Institute for the COVID-19 Genomics UK (COG-UK) Consortium | Jacquelyn Wynn, Mairead Hyland, The Lighthouse Lab in Alderley Park and Alex Alderton, Roberto Amato, Jeffrey Barrett, Sonia Goncalves, Ewan Harrison, David K. Jackson, Ian Johnston, Dominic Kwiatkowski, Cordelia Langford, John Sillitoe on behalf of the Wellcome Sanger Institute COVID-19 Surveillance Team |
| EPI_ISL_3139774, EPI_ISL_3139775 | Lighthouse Lab in Milton Keynes | Wellcome Sanger Institute for the COVID-19 Genomics UK (COG-UK) Consortium | The Lighthouse Lab in Milton Keynes and Alex Alderton, Roberto Amato, Jeffrey Barrett, Sonia Goncalves, Ewan Harrison, David K. Jackson, Ian Johnston, Dominic Kwiatkowski, Cordelia Langford, John Sillitoe on behalf of the Wellcome Sanger Institute COVID-19 Surveillance Team |
| EPI_ISL_3139776 | Lighthouse Laboratory Plymouth | Wellcome Sanger Institute for the COVID-19 Genomics UK (COG-UK) Consortium | Lighthouse Laboratory Plymouth and Alex Alderton, Roberto Amato, Jeffrey Barrett, Sonia Goncalves, Ewan Harrison, David K. Jackson, Ian Johnston, Dominic Kwiatkowski, Cordelia Langford, John Sillitoe on behalf of the Wellcome Sanger Institute COVID-19 Surveillance Team |
| EPI_ISL_3139777 | Lighthouse Lab in Alderley Park | Wellcome Sanger Institute for the COVID-19 Genomics UK (COG-UK) Consortium | Jacquelyn Wynn, Mairead Hyland, The Lighthouse Lab in Alderley Park and Alex Alderton, Roberto Amato, Jeffrey Barrett, Sonia Goncalves, Ewan Harrison, David K. Jackson, Ian Johnston, Dominic Kwiatkowski, Cordelia Langford, John Sillitoe on behalf of the Wellcome Sanger Institute COVID-19 Surveillance Team |
| EPI_ISL_3139778 | Lighthouse Lab in Milton Keynes | Wellcome Sanger Institute for the COVID-19 Genomics UK (COG-UK) Consortium | The Lighthouse Lab in Milton Keynes and Alex Alderton, Roberto Amato, Jeffrey Barrett, Sonia Goncalves, Ewan Harrison, David K. Jackson, Ian Johnston, Dominic Kwiatkowski, Cordelia Langford, John Sillitoe on behalf of the Wellcome Sanger Institute COVID-19 Surveillance Team |
| EPI_ISL_3139779 | Lighthouse Lab in Alderley Park | Wellcome Sanger Institute for the COVID-19 Genomics UK (COG-UK) Consortium | Jacquelyn Wynn, Mairead Hyland, The Lighthouse Lab in Alderley Park and Alex Alderton, Roberto Amato, Jeffrey Barrett, Sonia Goncalves, Ewan Harrison, David K. Jackson, Ian Johnston, Dominic Kwiatkowski, Cordelia Langford, John Sillitoe on behalf of the Wellcome Sanger Institute COVID-19 Surveillance Team |
| EPI_ISL_3139780 | Lighthouse Lab in Milton Keynes | Wellcome Sanger Institute for the COVID-19 Genomics UK (COG-UK) Consortium | The Lighthouse Lab in Milton Keynes and Alex Alderton, Roberto Amato, Jeffrey Barrett, Sonia Goncalves, Ewan Harrison, David K. Jackson, Ian Johnston, Dominic Kwiatkowski, Cordelia Langford, John Sillitoe on behalf of the Wellcome Sanger Institute COVID-19 Surveillance Team |
| EPI_ISL_3139781 | Lighthouse Laboratory Plymouth | Wellcome Sanger Institute for the COVID-19 Genomics UK (COG-UK) Consortium | Lighthouse Laboratory Plymouth and Alex Alderton, Roberto Amato, Jeffrey Barrett, Sonia Goncalves, Ewan Harrison, David K. Jackson, Ian Johnston, Dominic Kwiatkowski, Cordelia Langford, John Sillitoe on behalf of the Wellcome Sanger Institute COVID-19 Surveillance Team |
| EPI_ISL_3139782 | Lighthouse Lab in Alderley Park | Wellcome Sanger Institute for the COVID-19 Genomics UK (COG-UK) Consortium | Jacquelyn Wynn, Mairead Hyland, The Lighthouse Lab in Alderley Park and Alex Alderton, Roberto Amato, Jeffrey Barrett, Sonia Goncalves, Ewan Harrison, David K. Jackson, Ian Johnston, Dominic Kwiatkowski, Cordelia Langford, John Sillitoe on behalf of the Wellcome Sanger Institute COVID-19 Surveillance Team |
| EPI_ISL_3139783, EPI_ISL_3139784 | Berkshire and Surrey Pathology Services Lighthouse Laboratory | Wellcome Sanger Institute for the COVID-19 Genomics UK (COG-UK) Consortium | Berkshire and Surrey Pathology Services Lighthouse Laboratory and Alex Alderton, Roberto Amato, Jeffrey Barrett, Sonia Goncalves, Ewan Harrison, David K. Jackson, Ian Johnston, Dominic Kwiatkowski, Cordelia Langford, John Sillitoe on behalf of the Wellcome Sanger Institute COVID-19 Surveillance Team |
| EPI_ISL_3139785, EPI_ISL_3139786 | Lighthouse Lab in Alderley Park | Wellcome Sanger Institute for the COVID-19 Genomics UK (COG-UK) Consortium | Jacquelyn Wynn, Mairead Hyland, The Lighthouse Lab in Alderley Park and Alex Alderton, Roberto Amato, Jeffrey Barrett, Sonia Goncalves, Ewan Harrison, David K. Jackson, Ian Johnston, Dominic Kwiatkowski, Cordelia Langford, John Sillitoe on behalf of the Wellcome Sanger Institute COVID-19 Surveillance Team |
| EPI_ISL_3139787 | Lighthouse Lab in Milton Keynes | Wellcome Sanger Institute for the COVID-19 Genomics UK (COG-UK) Consortium | The Lighthouse Lab in Milton Keynes and Alex Alderton, Roberto Amato, Jeffrey Barrett, Sonia Goncalves, Ewan Harrison, David K. Jackson, Ian Johnston, Dominic Kwiatkowski, Cordelia Langford, John Sillitoe on behalf of the Wellcome Sanger Institute COVID-19 Surveillance Team |
| EPI_ISL_3139788, EPI_ISL_3139789 | Lighthouse Laboratory Plymouth | Wellcome Sanger Institute for the COVID-19 Genomics UK (COG-UK) Consortium | Lighthouse Laboratory Plymouth and Alex Alderton, Roberto Amato, Jeffrey Barrett, Sonia Goncalves, Ewan Harrison, David K. Jackson, Ian Johnston, Dominic Kwiatkowski, Cordelia Langford, John Sillitoe on behalf of the Wellcome Sanger Institute COVID-19 Surveillance Team |
| EPI_ISL_3139790 | Lighthouse Lab in Alderley Park | Wellcome Sanger Institute for the COVID-19 Genomics UK (COG-UK) Consortium | Jacquelyn Wynn, Mairead Hyland, The Lighthouse Lab in Alderley Park and Alex Alderton, Roberto Amato, Jeffrey Barrett, Sonia Goncalves, Ewan Harrison, David K. Jackson, Ian Johnston, Dominic Kwiatkowski, Cordelia Langford, John Sillitoe on behalf of the Wellcome Sanger Institute COVID-19 Surveillance Team |
| EPI_ISL_3139791 | Lighthouse Lab in Milton Keynes | Wellcome Sanger Institute for the COVID-19 Genomics UK (COG-UK) Consortium | The Lighthouse Lab in Milton Keynes and Alex Alderton, Roberto Amato, Jeffrey Barrett, Sonia Goncalves, Ewan Harrison, David K. Jackson, Ian Johnston, Dominic Kwiatkowski, Cordelia Langford, John Sillitoe on behalf of the Wellcome Sanger Institute COVID-19 Surveillance Team |
| EPI_ISL_3139792, EPI_ISL_3139793, EPI_ISL_3139794, EPI_ISL_3139795 | Lighthouse Lab in Alderley Park | Wellcome Sanger Institute for the COVID-19 Genomics UK (COG-UK) Consortium | Jacquelyn Wynn, Mairead Hyland, The Lighthouse Lab in Alderley Park and Alex Alderton, Roberto Amato, Jeffrey Barrett, Sonia Goncalves, Ewan Harrison, David K. Jackson, Ian Johnston, Dominic Kwiatkowski, Cordelia Langford, John Sillitoe on behalf of the Wellcome Sanger Institute COVID-19 Surveillance Team |
| EPI_ISL_3139796 | Lighthouse Lab in Milton Keynes | Wellcome Sanger Institute for the COVID-19 Genomics UK (COG-UK) Consortium | The Lighthouse Lab in Milton Keynes and Alex Alderton, Roberto Amato, Jeffrey Barrett, Sonia Goncalves, Ewan Harrison, David K. Jackson, Ian Johnston, Dominic Kwiatkowski, Cordelia Langford, John Sillitoe on behalf of the Wellcome Sanger Institute COVID-19 Surveillance Team |
| EPI_ISL_3139797 | Lighthouse Lab in Alderley Park | Wellcome Sanger Institute for the COVID-19 Genomics UK (COG-UK) Consortium | Jacquelyn Wynn, Mairead Hyland, The Lighthouse Lab in Alderley Park and Alex Alderton, Roberto Amato, Jeffrey Barrett, Sonia Goncalves, Ewan Harrison, David K. Jackson, Ian Johnston, Dominic Kwiatkowski, Cordelia Langford, John Sillitoe on behalf of the Wellcome Sanger Institute COVID-19 Surveillance Team |
| EPI_ISL_3139798, EPI_ISL_3139799 | Lighthouse Lab in Milton Keynes | Wellcome Sanger Institute for the COVID-19 Genomics UK (COG-UK) Consortium | The Lighthouse Lab in Milton Keynes and Alex Alderton, Roberto Amato, Jeffrey Barrett, Sonia Goncalves, Ewan Harrison, David K. Jackson, Ian Johnston, Dominic Kwiatkowski, Cordelia Langford, John Sillitoe on behalf of the Wellcome Sanger Institute COVID-19 Surveillance Team |
| EPI_ISL_3139800 | Berkshire and Surrey Pathology Services Lighthouse Laboratory | Wellcome Sanger Institute for the COVID-19 Genomics UK (COG-UK) Consortium | Berkshire and Surrey Pathology Services Lighthouse Laboratory and Alex Alderton, Roberto Amato, Jeffrey Barrett, Sonia Goncalves, Ewan Harrison, David K. Jackson, Ian Johnston, Dominic Kwiatkowski, Cordelia Langford, John Sillitoe on behalf of the Wellcome Sanger Institute COVID-19 Surveillance Team |
| EPI_ISL_3139801 | Lighthouse Lab in Milton Keynes | Wellcome Sanger Institute for the COVID-19 Genomics UK (COG-UK) Consortium | The Lighthouse Lab in Milton Keynes and Alex Alderton, Roberto Amato, Jeffrey Barrett, Sonia Goncalves, Ewan Harrison, David K. Jackson, Ian Johnston, Dominic Kwiatkowski, Cordelia Langford, John Sillitoe on behalf of the Wellcome Sanger Institute COVID-19 Surveillance Team |
| EPI_ISL_3139802 | Lighthouse Lab in Alderley Park | Wellcome Sanger Institute for the COVID-19 Genomics UK (COG-UK) Consortium | Jacquelyn Wynn, Mairead Hyland, The Lighthouse Lab in Alderley Park and Alex Alderton, Roberto Amato, Jeffrey Barrett, Sonia Goncalves, Ewan Harrison, David K. Jackson, Ian Johnston, Dominic Kwiatkowski, Cordelia Langford, John Sillitoe on behalf of the Wellcome Sanger Institute COVID-19 Surveillance Team |
| EPI_ISL_3139803 | Berkshire and Surrey Pathology Services Lighthouse Laboratory | Wellcome Sanger Institute for the COVID-19 Genomics UK (COG-UK) Consortium | Berkshire and Surrey Pathology Services Lighthouse Laboratory and Alex Alderton, Roberto Amato, Jeffrey Barrett, Sonia Goncalves, Ewan Harrison, David K. Jackson, Ian Johnston, Dominic Kwiatkowski, Cordelia Langford, John Sillitoe on behalf of the Wellcome Sanger Institute COVID-19 Surveillance Team |
| EPI_ISL_3139804, EPI_ISL_3139805 | Lighthouse Lab in Milton Keynes | Wellcome Sanger Institute for the COVID-19 Genomics UK (COG-UK) Consortium | The Lighthouse Lab in Milton Keynes and Alex Alderton, Roberto Amato, Jeffrey Barrett, Sonia Goncalves, Ewan Harrison, David K. Jackson, Ian Johnston, Dominic Kwiatkowski, Cordelia Langford, John Sillitoe on behalf of the Wellcome Sanger Institute COVID-19 Surveillance Team |
| EPI_ISL_3139806 | Lighthouse Laboratory Plymouth | Wellcome Sanger Institute for the COVID-19 Genomics UK (COG-UK) Consortium | Lighthouse Laboratory Plymouth and Alex Alderton, Roberto Amato, Jeffrey Barrett, Sonia Goncalves, Ewan Harrison, David K. Jackson, Ian Johnston, Dominic Kwiatkowski, Cordelia Langford, John Sillitoe on behalf of the Wellcome Sanger Institute COVID-19 Surveillance Team |
| EPI_ISL_3139807 | Lighthouse Lab in Alderley Park | Wellcome Sanger Institute for the COVID-19 Genomics UK (COG-UK) Consortium | Jacquelyn Wynn, Mairead Hyland, The Lighthouse Lab in Alderley Park and Alex Alderton, Roberto Amato, Jeffrey Barrett, Sonia Goncalves, Ewan Harrison, David K. Jackson, Ian Johnston, Dominic Kwiatkowski, Cordelia Langford, John Sillitoe on behalf of the Wellcome Sanger Institute COVID-19 Surveillance Team |
| EPI_ISL_3139808, EPI_ISL_3139809 | Lighthouse Lab in Milton Keynes | Wellcome Sanger Institute for the COVID-19 Genomics UK (COG-UK) Consortium | The Lighthouse Lab in Milton Keynes and Alex Alderton, Roberto Amato, Jeffrey Barrett, Sonia Goncalves, Ewan Harrison, David K. Jackson, Ian Johnston, Dominic Kwiatkowski, Cordelia Langford, John Sillitoe on behalf of the Wellcome Sanger Institute COVID-19 Surveillance Team |
| EPI_ISL_3139810, EPI_ISL_3139811 | Berkshire and Surrey Pathology Services Lighthouse Laboratory | Wellcome Sanger Institute for the COVID-19 Genomics UK (COG-UK) Consortium | Berkshire and Surrey Pathology Services Lighthouse Laboratory and Alex Alderton, Roberto Amato, Jeffrey Barrett, Sonia Goncalves, Ewan Harrison, David K. Jackson, Ian Johnston, Dominic Kwiatkowski, Cordelia Langford, John Sillitoe on behalf of the Wellcome Sanger Institute COVID-19 Surveillance Team |
| EPI_ISL_3139812 | Lighthouse Lab in Milton Keynes | Wellcome Sanger Institute for the COVID-19 Genomics UK (COG-UK) Consortium | The Lighthouse Lab in Milton Keynes and Alex Alderton, Roberto Amato, Jeffrey Barrett, Sonia Goncalves, Ewan Harrison, David K. Jackson, Ian Johnston, Dominic Kwiatkowski, Cordelia Langford, John Sillitoe on behalf of the Wellcome Sanger Institute COVID-19 Surveillance Team |
| EPI_ISL_3139813 | Lighthouse Laboratory Plymouth | Wellcome Sanger Institute for the COVID-19 Genomics UK (COG-UK) Consortium | Lighthouse Laboratory Plymouth and Alex Alderton, Roberto Amato, Jeffrey Barrett, Sonia Goncalves, Ewan Harrison, David K. Jackson, Ian Johnston, Dominic Kwiatkowski, Cordelia Langford, John Sillitoe on behalf of the Wellcome Sanger Institute COVID-19 Surveillance Team |
| EPI_ISL_3139814 | Lighthouse Lab in Milton Keynes | Wellcome Sanger Institute for the COVID-19 Genomics UK (COG-UK) Consortium | The Lighthouse Lab in Milton Keynes and Alex Alderton, Roberto Amato, Jeffrey Barrett, Sonia Goncalves, Ewan Harrison, David K. Jackson, Ian Johnston, Dominic Kwiatkowski, Cordelia Langford, John Sillitoe on behalf of the Wellcome Sanger Institute COVID-19 Surveillance Team |
| EPI_ISL_3139815, EPI_ISL_3139816 | Lighthouse Laboratory Plymouth | Wellcome Sanger Institute for the COVID-19 Genomics UK (COG-UK) Consortium | Lighthouse Laboratory Plymouth and Alex Alderton, Roberto Amato, Jeffrey Barrett, Sonia Goncalves, Ewan Harrison, David K. Jackson, Ian Johnston, Dominic Kwiatkowski, Cordelia Langford, John Sillitoe on behalf of the Wellcome Sanger Institute COVID-19 Surveillance Team |
| EPI_ISL_3139817 | Lighthouse Lab in Milton Keynes | Wellcome Sanger Institute for the COVID-19 Genomics UK (COG-UK) | The Lighthouse Lab in Milton Keynes and Alex Alderton, Roberto Amato, Jeffrey Barrett, Sonia Goncalves, Ewan Harrison, David K. Jackson, Ian Johnston, Dominic Kwiatkowski, Cordelia Langford, John Sillitoe on behalf of the Wellcome Sanger |

| | | | |
|---|---|---|---|
| | | Consortium | Institute COVID-19 Surveillance Team |
| EPI_ISL_3139818 | Lighthouse Laboratory Plymouth | Wellcome Sanger Institute for the COVID-19 Genomics UK (COG-UK) Consortium | Lighthouse Laboratory Plymouth and Alex Alderton, Roberto Amato, Jeffrey Barrett, Sonia Goncalves, Ewan Harrison, David K. Jackson, Ian Johnston, Dominic Kwiatkowski, Cordelia Langford, John Sillitoe on behalf of the Wellcome Sanger Institute COVID-19 Surveillance Team |
| EPI_ISL_3139819 | Berkshire and Surrey Pathology Services Lighthouse Laboratory | Wellcome Sanger Institute for the COVID-19 Genomics UK (COG-UK) Consortium | Berkshire and Surrey Pathology Services Lighthouse Laboratory and Alex Alderton, Roberto Amato, Jeffrey Barrett, Sonia Goncalves, Ewan Harrison, David K. Jackson, Ian Johnston, Dominic Kwiatkowski, Cordelia Langford, John Sillitoe on behalf of the Wellcome Sanger Institute COVID-19 Surveillance Team |
| EPI_ISL_3139820 | Lighthouse Laboratory Plymouth | Wellcome Sanger Institute for the COVID-19 Genomics UK (COG-UK) Consortium | Lighthouse Laboratory Plymouth and Alex Alderton, Roberto Amato, Jeffrey Barrett, Sonia Goncalves, Ewan Harrison, David K. Jackson, Ian Johnston, Dominic Kwiatkowski, Cordelia Langford, John Sillitoe on behalf of the Wellcome Sanger Institute COVID-19 Surveillance Team |
| EPI_ISL_3139821 | Berkshire and Surrey Pathology Services Lighthouse Laboratory | Wellcome Sanger Institute for the COVID-19 Genomics UK (COG-UK) Consortium | Berkshire and Surrey Pathology Services Lighthouse Laboratory and Alex Alderton, Roberto Amato, Jeffrey Barrett, Sonia Goncalves, Ewan Harrison, David K. Jackson, Ian Johnston, Dominic Kwiatkowski, Cordelia Langford, John Sillitoe on behalf of the Wellcome Sanger Institute COVID-19 Surveillance Team |
| EPI_ISL_3139822 | Lighthouse Lab in Alderley Park | Wellcome Sanger Institute for the COVID-19 Genomics UK (COG-UK) Consortium | Jacquelyn Wynn, Mairead Hyland, The Lighthouse Lab in Alderley Park and Alex Alderton, Roberto Amato, Jeffrey Barrett, Sonia Goncalves, Ewan Harrison, David K. Jackson, Ian Johnston, Dominic Kwiatkowski, Cordelia Langford, John Sillitoe on behalf of the Wellcome Sanger Institute COVID-19 Surveillance Team |
| EPI_ISL_3139823, EPI_ISL_3139824 | Lighthouse Laboratory Plymouth | Wellcome Sanger Institute for the COVID-19 Genomics UK (COG-UK) Consortium | Lighthouse Laboratory Plymouth and Alex Alderton, Roberto Amato, Jeffrey Barrett, Sonia Goncalves, Ewan Harrison, David K. Jackson, Ian Johnston, Dominic Kwiatkowski, Cordelia Langford, John Sillitoe on behalf of the Wellcome Sanger Institute COVID-19 Surveillance Team |
| EPI_ISL_3139825 | Lighthouse Lab in Alderley Park | Wellcome Sanger Institute for the COVID-19 Genomics UK (COG-UK) Consortium | Jacquelyn Wynn, Mairead Hyland, The Lighthouse Lab in Alderley Park and Alex Alderton, Roberto Amato, Jeffrey Barrett, Sonia Goncalves, Ewan Harrison, David K. Jackson, Ian Johnston, Dominic Kwiatkowski, Cordelia Langford, John Sillitoe on behalf of the Wellcome Sanger Institute COVID-19 Surveillance Team |
| EPI_ISL_3139826 | Lighthouse Lab in Milton Keynes | Wellcome Sanger Institute for the COVID-19 Genomics UK (COG-UK) Consortium | The Lighthouse Lab in Milton Keynes and Alex Alderton, Roberto Amato, Jeffrey Barrett, Sonia Goncalves, Ewan Harrison, David K. Jackson, Ian Johnston, Dominic Kwiatkowski, Cordelia Langford, John Sillitoe on behalf of the Wellcome Sanger Institute COVID-19 Surveillance Team |
| EPI_ISL_3139827, EPI_ISL_3139828 | Lighthouse Lab in Alderley Park | Wellcome Sanger Institute for the COVID-19 Genomics UK (COG-UK) Consortium | Jacquelyn Wynn, Mairead Hyland, The Lighthouse Lab in Alderley Park and Alex Alderton, Roberto Amato, Jeffrey Barrett, Sonia Goncalves, Ewan Harrison, David K. Jackson, Ian Johnston, Dominic Kwiatkowski, Cordelia Langford, John Sillitoe on behalf of the Wellcome Sanger Institute COVID-19 Surveillance Team |
| EPI_ISL_3139829 | Lighthouse Laboratory Plymouth | Wellcome Sanger Institute for the COVID-19 Genomics UK (COG-UK) Consortium | Lighthouse Laboratory Plymouth and Alex Alderton, Roberto Amato, Jeffrey Barrett, Sonia Goncalves, Ewan Harrison, David K. Jackson, Ian Johnston, Dominic Kwiatkowski, Cordelia Langford, John Sillitoe on behalf of the Wellcome Sanger Institute COVID-19 Surveillance Team |
| EPI_ISL_3139830 | Lighthouse Lab in Milton Keynes | Wellcome Sanger Institute for the COVID-19 Genomics UK (COG-UK) Consortium | The Lighthouse Lab in Milton Keynes and Alex Alderton, Roberto Amato, Jeffrey Barrett, Sonia Goncalves, Ewan Harrison, David K. Jackson, Ian Johnston, Dominic Kwiatkowski, Cordelia Langford, John Sillitoe on behalf of the Wellcome Sanger Institute COVID-19 Surveillance Team |
| EPI_ISL_3139831 | Berkshire and Surrey Pathology Services Lighthouse Laboratory | Wellcome Sanger Institute for the COVID-19 Genomics UK (COG-UK) Consortium | Berkshire and Surrey Pathology Services Lighthouse Laboratory and Alex Alderton, Roberto Amato, Jeffrey Barrett, Sonia Goncalves, Ewan Harrison, David K. Jackson, Ian Johnston, Dominic Kwiatkowski, Cordelia Langford, John Sillitoe on behalf of the Wellcome Sanger Institute COVID-19 Surveillance Team |
| EPI_ISL_3139832 | Lighthouse Lab in Milton Keynes | Wellcome Sanger Institute for the COVID-19 Genomics UK (COG-UK) Consortium | The Lighthouse Lab in Milton Keynes and Alex Alderton, Roberto Amato, Jeffrey Barrett, Sonia Goncalves, Ewan Harrison, David K. Jackson, Ian Johnston, Dominic Kwiatkowski, Cordelia Langford, John Sillitoe on behalf of the Wellcome Sanger Institute COVID-19 Surveillance Team |
| EPI_ISL_3139833 | Lighthouse Lab in Alderley Park | Wellcome Sanger Institute for the COVID-19 Genomics UK (COG-UK) Consortium | Jacquelyn Wynn, Mairead Hyland, The Lighthouse Lab in Alderley Park and Alex Alderton, Roberto Amato, Jeffrey Barrett, Sonia Goncalves, Ewan Harrison, David K. Jackson, Ian Johnston, Dominic Kwiatkowski, Cordelia Langford, John Sillitoe on behalf of the Wellcome Sanger Institute COVID-19 Surveillance Team |
| EPI_ISL_3139834 | Lighthouse Lab in Milton Keynes | Wellcome Sanger Institute for the COVID-19 Genomics UK (COG-UK) Consortium | The Lighthouse Lab in Milton Keynes and Alex Alderton, Roberto Amato, Jeffrey Barrett, Sonia Goncalves, Ewan Harrison, David K. Jackson, Ian Johnston, Dominic Kwiatkowski, Cordelia Langford, John Sillitoe on behalf of the Wellcome Sanger Institute COVID-19 Surveillance Team |
| EPI_ISL_3139835, EPI_ISL_3139836 | Lighthouse Laboratory Plymouth | Wellcome Sanger Institute for the COVID-19 Genomics UK (COG-UK) Consortium | Lighthouse Laboratory Plymouth and Alex Alderton, Roberto Amato, Jeffrey Barrett, Sonia Goncalves, Ewan Harrison, David K. Jackson, Ian Johnston, Dominic Kwiatkowski, Cordelia Langford, John Sillitoe on behalf of the Wellcome Sanger Institute COVID-19 Surveillance Team |
| EPI_ISL_3139837 | Lighthouse Lab in Milton Keynes | Wellcome Sanger Institute for the COVID-19 Genomics UK (COG-UK) Consortium | The Lighthouse Lab in Milton Keynes and Alex Alderton, Roberto Amato, Jeffrey Barrett, Sonia Goncalves, Ewan Harrison, David K. Jackson, Ian Johnston, Dominic Kwiatkowski, Cordelia Langford, John Sillitoe on behalf of the Wellcome Sanger Institute COVID-19 Surveillance Team |
| EPI_ISL_3139838 | Lighthouse Laboratory Plymouth | Wellcome Sanger Institute for the COVID-19 Genomics UK (COG-UK) Consortium | Lighthouse Laboratory Plymouth and Alex Alderton, Roberto Amato, Jeffrey Barrett, Sonia Goncalves, Ewan Harrison, David K. Jackson, Ian Johnston, Dominic Kwiatkowski, Cordelia Langford, John Sillitoe on behalf of the Wellcome Sanger Institute COVID-19 Surveillance Team |
| EPI_ISL_3139839 | Lighthouse Lab in Milton Keynes | Wellcome Sanger Institute for the COVID-19 Genomics UK (COG-UK) Consortium | The Lighthouse Lab in Milton Keynes and Alex Alderton, Roberto Amato, Jeffrey Barrett, Sonia Goncalves, Ewan Harrison, David K. Jackson, Ian Johnston, Dominic Kwiatkowski, Cordelia Langford, John Sillitoe on behalf of the Wellcome Sanger Institute COVID-19 Surveillance Team |
| EPI_ISL_3139840, EPI_ISL_3139841 | Lighthouse Laboratory Plymouth | Wellcome Sanger Institute for the COVID-19 Genomics UK (COG-UK) Consortium | Lighthouse Laboratory Plymouth and Alex Alderton, Roberto Amato, Jeffrey Barrett, Sonia Goncalves, Ewan Harrison, David K. Jackson, Ian Johnston, Dominic Kwiatkowski, Cordelia Langford, John Sillitoe on behalf of the Wellcome Sanger Institute COVID-19 Surveillance Team |
| EPI_ISL_3139842 | Lighthouse Lab in Milton Keynes | Wellcome Sanger Institute for the COVID-19 Genomics UK (COG-UK) Consortium | The Lighthouse Lab in Milton Keynes and Alex Alderton, Roberto Amato, Jeffrey Barrett, Sonia Goncalves, Ewan Harrison, David K. Jackson, Ian Johnston, Dominic Kwiatkowski, Cordelia Langford, John Sillitoe on behalf of the Wellcome Sanger Institute COVID-19 Surveillance Team |
| EPI_ISL_3139843 | Berkshire and Surrey Pathology Services Lighthouse Laboratory | Wellcome Sanger Institute for the COVID-19 Genomics UK (COG-UK) Consortium | Berkshire and Surrey Pathology Services Lighthouse Laboratory and Alex Alderton, Roberto Amato, Jeffrey Barrett, Sonia Goncalves, Ewan Harrison, David K. Jackson, Ian Johnston, Dominic Kwiatkowski, Cordelia Langford, John Sillitoe on behalf of the Wellcome Sanger Institute COVID-19 Surveillance Team |
| EPI_ISL_3139844 | Lighthouse Lab in Alderley Park | Wellcome Sanger Institute for the COVID-19 Genomics UK (COG-UK) Consortium | Jacquelyn Wynn, Mairead Hyland, The Lighthouse Lab in Alderley Park and Alex Alderton, Roberto Amato, Jeffrey Barrett, Sonia Goncalves, Ewan Harrison, David K. Jackson, Ian Johnston, Dominic Kwiatkowski, Cordelia Langford, John Sillitoe on behalf of the Wellcome Sanger Institute COVID-19 Surveillance Team |
| EPI_ISL_3139845 | Lighthouse Laboratory Plymouth | Wellcome Sanger Institute for the COVID-19 Genomics UK (COG-UK) Consortium | Lighthouse Laboratory Plymouth and Alex Alderton, Roberto Amato, Jeffrey Barrett, Sonia Goncalves, Ewan Harrison, David K. Jackson, Ian Johnston, Dominic Kwiatkowski, Cordelia Langford, John Sillitoe on behalf of the Wellcome Sanger Institute COVID-19 Surveillance Team |
| EPI_ISL_3139846, EPI_ISL_3139847 | Berkshire and Surrey Pathology Services Lighthouse Laboratory | Wellcome Sanger Institute for the COVID-19 Genomics UK (COG-UK) Consortium | Berkshire and Surrey Pathology Services Lighthouse Laboratory and Alex Alderton, Roberto Amato, Jeffrey Barrett, Sonia Goncalves, Ewan Harrison, David K. Jackson, Ian Johnston, Dominic Kwiatkowski, Cordelia Langford, John Sillitoe on behalf of the Wellcome Sanger Institute COVID-19 Surveillance Team |
| EPI_ISL_3139848 | Lighthouse Lab in Alderley Park | Wellcome Sanger Institute for the COVID-19 Genomics UK (COG-UK) Consortium | Jacquelyn Wynn, Mairead Hyland, The Lighthouse Lab in Alderley Park and Alex Alderton, Roberto Amato, Jeffrey Barrett, Sonia Goncalves, Ewan Harrison, David K. Jackson, Ian Johnston, Dominic Kwiatkowski, Cordelia Langford, John Sillitoe on behalf of the Wellcome Sanger Institute COVID-19 Surveillance Team |
| EPI_ISL_3139849, EPI_ISL_3139850 | Lighthouse Lab in Milton Keynes | Wellcome Sanger Institute for the COVID-19 Genomics UK (COG-UK) Consortium | The Lighthouse Lab in Milton Keynes and Alex Alderton, Roberto Amato, Jeffrey Barrett, Sonia Goncalves, Ewan Harrison, David K. Jackson, Ian Johnston, Dominic Kwiatkowski, Cordelia Langford, John Sillitoe on behalf of the Wellcome Sanger Institute COVID-19 Surveillance Team |
| EPI_ISL_3139851, EPI_ISL_3139852 | Lighthouse Lab in Alderley Park | Wellcome Sanger Institute for the COVID-19 Genomics UK (COG-UK) Consortium | Jacquelyn Wynn, Mairead Hyland, The Lighthouse Lab in Alderley Park and Alex Alderton, Roberto Amato, Jeffrey Barrett, Sonia Goncalves, Ewan Harrison, David K. Jackson, Ian Johnston, Dominic Kwiatkowski, Cordelia Langford, John Sillitoe on behalf of the Wellcome Sanger Institute COVID-19 Surveillance Team |
| EPI_ISL_3139853, EPI_ISL_3139854 | Berkshire and Surrey Pathology Services Lighthouse Laboratory | Wellcome Sanger Institute for the COVID-19 Genomics UK (COG-UK) Consortium | Berkshire and Surrey Pathology Services Lighthouse Laboratory and Alex Alderton, Roberto Amato, Jeffrey Barrett, Sonia Goncalves, Ewan Harrison, David K. Jackson, Ian Johnston, Dominic Kwiatkowski, Cordelia Langford, John Sillitoe on behalf of the Wellcome Sanger Institute COVID-19 Surveillance Team |
| EPI_ISL_3139855 | Lighthouse Lab in Alderley Park | Wellcome Sanger Institute for the COVID-19 Genomics UK (COG-UK) Consortium | Jacquelyn Wynn, Mairead Hyland, The Lighthouse Lab in Alderley Park and Alex Alderton, Roberto Amato, Jeffrey Barrett, Sonia Goncalves, Ewan Harrison, David K. Jackson, Ian Johnston, Dominic Kwiatkowski, Cordelia Langford, John Sillitoe on behalf of the Wellcome Sanger Institute COVID-19 Surveillance Team |
| EPI_ISL_3139856, EPI_ISL_3139857 | Lighthouse Lab in Milton Keynes | Wellcome Sanger Institute for the COVID-19 Genomics UK (COG-UK) Consortium | The Lighthouse Lab in Milton Keynes and Alex Alderton, Roberto Amato, Jeffrey Barrett, Sonia Goncalves, Ewan Harrison, David K. Jackson, Ian Johnston, Dominic Kwiatkowski, Cordelia Langford, John Sillitoe on behalf of the Wellcome Sanger Institute COVID-19 Surveillance Team |
| EPI_ISL_3139858, EPI_ISL_3139859 | Lighthouse Laboratory Plymouth | Wellcome Sanger Institute for the COVID-19 Genomics UK (COG-UK) Consortium | Lighthouse Laboratory Plymouth and Alex Alderton, Roberto Amato, Jeffrey Barrett, Sonia Goncalves, Ewan Harrison, David K. Jackson, Ian Johnston, Dominic Kwiatkowski, Cordelia Langford, John Sillitoe on behalf of the Wellcome Sanger Institute COVID-19 Surveillance Team |
| EPI_ISL_3139860 | Berkshire and Surrey Pathology Services Lighthouse Laboratory | Wellcome Sanger Institute for the COVID-19 Genomics UK (COG-UK) Consortium | Berkshire and Surrey Pathology Services Lighthouse Laboratory and Alex Alderton, Roberto Amato, Jeffrey Barrett, Sonia Goncalves, Ewan Harrison, David K. Jackson, Ian Johnston, Dominic Kwiatkowski, Cordelia Langford, John Sillitoe on behalf of the Wellcome Sanger Institute COVID-19 Surveillance Team |
| EPI_ISL_3139861 | Lighthouse Lab in Alderley Park | Wellcome Sanger Institute for the COVID-19 Genomics UK (COG-UK) Consortium | Jacquelyn Wynn, Mairead Hyland, The Lighthouse Lab in Alderley Park and Alex Alderton, Roberto Amato, Jeffrey Barrett, Sonia Goncalves, Ewan Harrison, David K. Jackson, Ian Johnston, Dominic Kwiatkowski, Cordelia Langford, John Sillitoe on behalf of the Wellcome Sanger Institute COVID-19 Surveillance Team |
| EPI_ISL_3139862 | Lighthouse Laboratory Plymouth | Wellcome Sanger Institute for the COVID-19 Genomics UK (COG-UK) Consortium | Lighthouse Laboratory Plymouth and Alex Alderton, Roberto Amato, Jeffrey Barrett, Sonia Goncalves, Ewan Harrison, David K. Jackson, Ian Johnston, Dominic Kwiatkowski, Cordelia Langford, John Sillitoe on behalf of the Wellcome Sanger Institute COVID-19 Surveillance Team |
| EPI_ISL_3139863, EPI_ISL_3139864 | Lighthouse Lab in Milton Keynes | Wellcome Sanger Institute for the COVID-19 Genomics UK (COG-UK) Consortium | The Lighthouse Lab in Milton Keynes and Alex Alderton, Roberto Amato, Jeffrey Barrett, Sonia Goncalves, Ewan Harrison, David K. Jackson, Ian Johnston, Dominic Kwiatkowski, Cordelia Langford, John Sillitoe on behalf of the Wellcome Sanger Institute COVID-19 Surveillance Team |
| EPI_ISL_3139865, EPI_ISL_3139866 | Lighthouse Lab in Alderley Park | Wellcome Sanger Institute for the COVID-19 Genomics UK (COG-UK) Consortium | Jacquelyn Wynn, Mairead Hyland, The Lighthouse Lab in Alderley Park and Alex Alderton, Roberto Amato, Jeffrey Barrett, Sonia Goncalves, Ewan Harrison, David K. Jackson, Ian Johnston, Dominic Kwiatkowski, Cordelia Langford, John Sillitoe on behalf of the Wellcome Sanger Institute COVID-19 Surveillance Team |
| EPI_ISL_3139867 | Lighthouse Laboratory Plymouth | Wellcome Sanger Institute for the COVID-19 Genomics UK (COG-UK) Consortium | Lighthouse Laboratory Plymouth and Alex Alderton, Roberto Amato, Jeffrey Barrett, Sonia Goncalves, Ewan Harrison, David K. Jackson, Ian Johnston, Dominic Kwiatkowski, Cordelia Langford, John Sillitoe on behalf of the Wellcome Sanger Institute COVID-19 Surveillance Team |
| EPI_ISL_3139868 | Lighthouse Lab in Alderley Park | Wellcome Sanger Institute for the COVID-19 Genomics UK (COG-UK) Consortium | Jacquelyn Wynn, Mairead Hyland, The Lighthouse Lab in Alderley Park and Alex Alderton, Roberto Amato, Jeffrey Barrett, Sonia Goncalves, Ewan Harrison, David K. Jackson, Ian Johnston, Dominic Kwiatkowski, Cordelia Langford, John Sillitoe on behalf of the Wellcome Sanger Institute COVID-19 Surveillance Team |
| EPI_ISL_3139869 | Lighthouse Lab in Milton Keynes | Wellcome Sanger Institute for the COVID-19 Genomics UK (COG-UK) Consortium | The Lighthouse Lab in Milton Keynes and Alex Alderton, Roberto Amato, Jeffrey Barrett, Sonia Goncalves, Ewan Harrison, David K. Jackson, Ian Johnston, Dominic Kwiatkowski, Cordelia Langford, John Sillitoe on behalf of the Wellcome Sanger Institute COVID-19 Surveillance Team |
| EPI_ISL_3139870, EPI_ISL_3139871 | Lighthouse Lab in Alderley Park | Wellcome Sanger Institute for the COVID-19 Genomics UK (COG-UK) Consortium | Jacquelyn Wynn, Mairead Hyland, The Lighthouse Lab in Alderley Park and Alex Alderton, Roberto Amato, Jeffrey Barrett, Sonia Goncalves, Ewan Harrison, David K. Jackson, Ian Johnston, Dominic Kwiatkowski, Cordelia Langford, John Sillitoe on behalf of the Wellcome Sanger Institute COVID-19 Surveillance Team |
| EPI_ISL_3139872 | Berkshire and Surrey Pathology Services Lighthouse Laboratory | Wellcome Sanger Institute for the COVID-19 Genomics UK (COG-UK) Consortium | Berkshire and Surrey Pathology Services Lighthouse Laboratory and Alex Alderton, Roberto Amato, Jeffrey Barrett, Sonia Goncalves, Ewan Harrison, David K. Jackson, Ian Johnston, Dominic Kwiatkowski, Cordelia Langford, John Sillitoe on behalf of the Wellcome Sanger Institute COVID-19 Surveillance Team |
| EPI_ISL_3139873 | Lighthouse Lab in Alderley Park | Wellcome Sanger Institute for the COVID-19 Genomics UK (COG-UK) Consortium | Jacquelyn Wynn, Mairead Hyland, The Lighthouse Lab in Alderley Park and Alex Alderton, Roberto Amato, Jeffrey Barrett, Sonia Goncalves, Ewan Harrison, David K. Jackson, Ian Johnston, Dominic Kwiatkowski, Cordelia Langford, John Sillitoe on behalf of the Wellcome Sanger Institute COVID-19 Surveillance Team |
| EPI_ISL_3139874 | Lighthouse Lab in Milton Keynes | Wellcome Sanger Institute for the COVID-19 Genomics UK (COG-UK) Consortium | The Lighthouse Lab in Milton Keynes and Alex Alderton, Roberto Amato, Jeffrey Barrett, Sonia Goncalves, Ewan Harrison, David K. Jackson, Ian Johnston, Dominic Kwiatkowski, Cordelia Langford, John Sillitoe on behalf of the Wellcome Sanger Institute COVID-19 Surveillance Team |
| EPI_ISL_3139875 | Berkshire and Surrey Pathology Services Lighthouse Laboratory | Wellcome Sanger Institute for the COVID-19 Genomics UK (COG-UK) | Berkshire and Surrey Pathology Services Lighthouse Laboratory and Alex Alderton, Roberto Amato, Jeffrey Barrett, Sonia Goncalves, Ewan Harrison, David K. Jackson, Ian Johnston, Dominic Kwiatkowski, Cordelia Langford, John Sillitoe on behalf |

| | | | |
|---|---|---|---|
| | | Consortium | of the Wellcome Sanger Institute COVID-19 Surveillance Team |
| EPI_ISL_3139876 | Lighthouse Laboratory Plymouth | Wellcome Sanger Institute for the COVID-19 Genomics UK (COG-UK) Consortium | Lighthouse Laboratory Plymouth and Alex Alderton, Roberto Amato, Jeffrey Barrett, Sonia Goncalves, Ewan Harrison, David K. Jackson, Ian Johnston, Dominic Kwiatkowski, Cordelia Langford, John Sillitoe on behalf of the Wellcome Sanger Institute COVID-19 Surveillance Team |
| EPI_ISL_3139877 | Lighthouse Lab in Alderley Park | Wellcome Sanger Institute for the COVID-19 Genomics UK (COG-UK) Consortium | Jacquelyn Wynn, Mairead Hyland, The Lighthouse Lab in Alderley Park and Alex Alderton, Roberto Amato, Jeffrey Barrett, Sonia Goncalves, Ewan Harrison, David K. Jackson, Ian Johnston, Dominic Kwiatkowski, Cordelia Langford, John Sillitoe on behalf of the Wellcome Sanger Institute COVID-19 Surveillance Team |
| EPI_ISL_3139878, EPI_ISL_3139879 | Lighthouse Lab in Milton Keynes | Wellcome Sanger Institute for the COVID-19 Genomics UK (COG-UK) Consortium | The Lighthouse Lab in Milton Keynes and Alex Alderton, Roberto Amato, Jeffrey Barrett, Sonia Goncalves, Ewan Harrison, David K. Jackson, Ian Johnston, Dominic Kwiatkowski, Cordelia Langford, John Sillitoe on behalf of the Wellcome Sanger Institute COVID-19 Surveillance Team |
| EPI_ISL_3139880 | Lighthouse Lab in Alderley Park | Wellcome Sanger Institute for the COVID-19 Genomics UK (COG-UK) Consortium | Jacquelyn Wynn, Mairead Hyland, The Lighthouse Lab in Alderley Park and Alex Alderton, Roberto Amato, Jeffrey Barrett, Sonia Goncalves, Ewan Harrison, David K. Jackson, Ian Johnston, Dominic Kwiatkowski, Cordelia Langford, John Sillitoe on behalf of the Wellcome Sanger Institute COVID-19 Surveillance Team |
| EPI_ISL_3139881 | Lighthouse Lab in Milton Keynes | Wellcome Sanger Institute for the COVID-19 Genomics UK (COG-UK) Consortium | The Lighthouse Lab in Milton Keynes and Alex Alderton, Roberto Amato, Jeffrey Barrett, Sonia Goncalves, Ewan Harrison, David K. Jackson, Ian Johnston, Dominic Kwiatkowski, Cordelia Langford, John Sillitoe on behalf of the Wellcome Sanger Institute COVID-19 Surveillance Team |
| EPI_ISL_3139882, EPI_ISL_3139883 | Lighthouse Lab in Alderley Park | Wellcome Sanger Institute for the COVID-19 Genomics UK (COG-UK) Consortium | Jacquelyn Wynn, Mairead Hyland, The Lighthouse Lab in Alderley Park and Alex Alderton, Roberto Amato, Jeffrey Barrett, Sonia Goncalves, Ewan Harrison, David K. Jackson, Ian Johnston, Dominic Kwiatkowski, Cordelia Langford, John Sillitoe on behalf of the Wellcome Sanger Institute COVID-19 Surveillance Team |
| EPI_ISL_3139884 | Lighthouse Laboratory Plymouth | Wellcome Sanger Institute for the COVID-19 Genomics UK (COG-UK) Consortium | Lighthouse Laboratory Plymouth and Alex Alderton, Roberto Amato, Jeffrey Barrett, Sonia Goncalves, Ewan Harrison, David K. Jackson, Ian Johnston, Dominic Kwiatkowski, Cordelia Langford, John Sillitoe on behalf of the Wellcome Sanger Institute COVID-19 Surveillance Team |
| EPI_ISL_3139885 | Berkshire and Surrey Pathology Services Lighthouse Laboratory | Wellcome Sanger Institute for the COVID-19 Genomics UK (COG-UK) Consortium | Berkshire and Surrey Pathology Services Lighthouse Laboratory and Alex Alderton, Roberto Amato, Jeffrey Barrett, Sonia Goncalves, Ewan Harrison, David K. Jackson, Ian Johnston, Dominic Kwiatkowski, Cordelia Langford, John Sillitoe on behalf of the Wellcome Sanger Institute COVID-19 Surveillance Team |
| EPI_ISL_3139886 | Lighthouse Lab in Alderley Park | Wellcome Sanger Institute for the COVID-19 Genomics UK (COG-UK) Consortium | Jacquelyn Wynn, Mairead Hyland, The Lighthouse Lab in Alderley Park and Alex Alderton, Roberto Amato, Jeffrey Barrett, Sonia Goncalves, Ewan Harrison, David K. Jackson, Ian Johnston, Dominic Kwiatkowski, Cordelia Langford, John Sillitoe on behalf of the Wellcome Sanger Institute COVID-19 Surveillance Team |
| EPI_ISL_3139887 | Berkshire and Surrey Pathology Services Lighthouse Laboratory | Wellcome Sanger Institute for the COVID-19 Genomics UK (COG-UK) Consortium | Berkshire and Surrey Pathology Services Lighthouse Laboratory and Alex Alderton, Roberto Amato, Jeffrey Barrett, Sonia Goncalves, Ewan Harrison, David K. Jackson, Ian Johnston, Dominic Kwiatkowski, Cordelia Langford, John Sillitoe on behalf of the Wellcome Sanger Institute COVID-19 Surveillance Team |
| EPI_ISL_3139888, EPI_ISL_3139889, EPI_ISL_3139890 | Lighthouse Lab in Milton Keynes | Wellcome Sanger Institute for the COVID-19 Genomics UK (COG-UK) Consortium | The Lighthouse Lab in Milton Keynes and Alex Alderton, Roberto Amato, Jeffrey Barrett, Sonia Goncalves, Ewan Harrison, David K. Jackson, Ian Johnston, Dominic Kwiatkowski, Cordelia Langford, John Sillitoe on behalf of the Wellcome Sanger Institute COVID-19 Surveillance Team |
| EPI_ISL_3139891 | Lighthouse Lab in Alderley Park | Wellcome Sanger Institute for the COVID-19 Genomics UK (COG-UK) Consortium | Jacquelyn Wynn, Mairead Hyland, The Lighthouse Lab in Alderley Park and Alex Alderton, Roberto Amato, Jeffrey Barrett, Sonia Goncalves, Ewan Harrison, David K. Jackson, Ian Johnston, Dominic Kwiatkowski, Cordelia Langford, John Sillitoe on behalf of the Wellcome Sanger Institute COVID-19 Surveillance Team |
| EPI_ISL_3139892 | Lighthouse Lab in Milton Keynes | Wellcome Sanger Institute for the COVID-19 Genomics UK (COG-UK) Consortium | The Lighthouse Lab in Milton Keynes and Alex Alderton, Roberto Amato, Jeffrey Barrett, Sonia Goncalves, Ewan Harrison, David K. Jackson, Ian Johnston, Dominic Kwiatkowski, Cordelia Langford, John Sillitoe on behalf of the Wellcome Sanger Institute COVID-19 Surveillance Team |
| EPI_ISL_3139893 | Lighthouse Lab in Alderley Park | Wellcome Sanger Institute for the COVID-19 Genomics UK (COG-UK) Consortium | Jacquelyn Wynn, Mairead Hyland, The Lighthouse Lab in Alderley Park and Alex Alderton, Roberto Amato, Jeffrey Barrett, Sonia Goncalves, Ewan Harrison, David K. Jackson, Ian Johnston, Dominic Kwiatkowski, Cordelia Langford, John Sillitoe on behalf of the Wellcome Sanger Institute COVID-19 Surveillance Team |
| EPI_ISL_3139894, EPI_ISL_3139895 | Berkshire and Surrey Pathology Services Lighthouse Laboratory | Wellcome Sanger Institute for the COVID-19 Genomics UK (COG-UK) Consortium | Berkshire and Surrey Pathology Services Lighthouse Laboratory and Alex Alderton, Roberto Amato, Jeffrey Barrett, Sonia Goncalves, Ewan Harrison, David K. Jackson, Ian Johnston, Dominic Kwiatkowski, Cordelia Langford, John Sillitoe on behalf of the Wellcome Sanger Institute COVID-19 Surveillance Team |
| EPI_ISL_3139896 | Lighthouse Lab in Alderley Park | Wellcome Sanger Institute for the COVID-19 Genomics UK (COG-UK) Consortium | Jacquelyn Wynn, Mairead Hyland, The Lighthouse Lab in Alderley Park and Alex Alderton, Roberto Amato, Jeffrey Barrett, Sonia Goncalves, Ewan Harrison, David K. Jackson, Ian Johnston, Dominic Kwiatkowski, Cordelia Langford, John Sillitoe on behalf of the Wellcome Sanger Institute COVID-19 Surveillance Team |
| EPI_ISL_3139897 | Berkshire and Surrey Pathology Services Lighthouse Laboratory | Wellcome Sanger Institute for the COVID-19 Genomics UK (COG-UK) Consortium | Berkshire and Surrey Pathology Services Lighthouse Laboratory and Alex Alderton, Roberto Amato, Jeffrey Barrett, Sonia Goncalves, Ewan Harrison, David K. Jackson, Ian Johnston, Dominic Kwiatkowski, Cordelia Langford, John Sillitoe on behalf of the Wellcome Sanger Institute COVID-19 Surveillance Team |
| EPI_ISL_3139898 | Lighthouse Lab in Alderley Park | Wellcome Sanger Institute for the COVID-19 Genomics UK (COG-UK) Consortium | Jacquelyn Wynn, Mairead Hyland, The Lighthouse Lab in Alderley Park and Alex Alderton, Roberto Amato, Jeffrey Barrett, Sonia Goncalves, Ewan Harrison, David K. Jackson, Ian Johnston, Dominic Kwiatkowski, Cordelia Langford, John Sillitoe on behalf of the Wellcome Sanger Institute COVID-19 Surveillance Team |
| EPI_ISL_3139899, EPI_ISL_3139900 | Berkshire and Surrey Pathology Services Lighthouse Laboratory | Wellcome Sanger Institute for the COVID-19 Genomics UK (COG-UK) Consortium | Berkshire and Surrey Pathology Services Lighthouse Laboratory and Alex Alderton, Roberto Amato, Jeffrey Barrett, Sonia Goncalves, Ewan Harrison, David K. Jackson, Ian Johnston, Dominic Kwiatkowski, Cordelia Langford, John Sillitoe on behalf of the Wellcome Sanger Institute COVID-19 Surveillance Team |
| EPI_ISL_3139901 | Lighthouse Lab in Alderley Park | Wellcome Sanger Institute for the COVID-19 Genomics UK (COG-UK) Consortium | Jacquelyn Wynn, Mairead Hyland, The Lighthouse Lab in Alderley Park and Alex Alderton, Roberto Amato, Jeffrey Barrett, Sonia Goncalves, Ewan Harrison, David K. Jackson, Ian Johnston, Dominic Kwiatkowski, Cordelia Langford, John Sillitoe on behalf of the Wellcome Sanger Institute COVID-19 Surveillance Team |
| EPI_ISL_3139902 | Lighthouse Lab in Milton Keynes | Wellcome Sanger Institute for the COVID-19 Genomics UK (COG-UK) Consortium | The Lighthouse Lab in Milton Keynes and Alex Alderton, Roberto Amato, Jeffrey Barrett, Sonia Goncalves, Ewan Harrison, David K. Jackson, Ian Johnston, Dominic Kwiatkowski, Cordelia Langford, John Sillitoe on behalf of the Wellcome Sanger Institute COVID-19 Surveillance Team |
| EPI_ISL_3139903 | Berkshire and Surrey Pathology Services Lighthouse Laboratory | Wellcome Sanger Institute for the COVID-19 Genomics UK (COG-UK) Consortium | Berkshire and Surrey Pathology Services Lighthouse Laboratory and Alex Alderton, Roberto Amato, Jeffrey Barrett, Sonia Goncalves, Ewan Harrison, David K. Jackson, Ian Johnston, Dominic Kwiatkowski, Cordelia Langford, John Sillitoe on behalf of the Wellcome Sanger Institute COVID-19 Surveillance Team |
| EPI_ISL_3139904 | Lighthouse Lab in Milton Keynes | Wellcome Sanger Institute for the COVID-19 Genomics UK (COG-UK) Consortium | The Lighthouse Lab in Milton Keynes and Alex Alderton, Roberto Amato, Jeffrey Barrett, Sonia Goncalves, Ewan Harrison, David K. Jackson, Ian Johnston, Dominic Kwiatkowski, Cordelia Langford, John Sillitoe on behalf of the Wellcome Sanger Institute COVID-19 Surveillance Team |
| EPI_ISL_3139905 | Berkshire and Surrey Pathology Services Lighthouse Laboratory | Wellcome Sanger Institute for the COVID-19 Genomics UK (COG-UK) Consortium | Berkshire and Surrey Pathology Services Lighthouse Laboratory and Alex Alderton, Roberto Amato, Jeffrey Barrett, Sonia Goncalves, Ewan Harrison, David K. Jackson, Ian Johnston, Dominic Kwiatkowski, Cordelia Langford, John Sillitoe on behalf of the Wellcome Sanger Institute COVID-19 Surveillance Team |
| EPI_ISL_3139906 | Lighthouse Lab in Alderley Park | Wellcome Sanger Institute for the COVID-19 Genomics UK (COG-UK) Consortium | Jacquelyn Wynn, Mairead Hyland, The Lighthouse Lab in Alderley Park and Alex Alderton, Roberto Amato, Jeffrey Barrett, Sonia Goncalves, Ewan Harrison, David K. Jackson, Ian Johnston, Dominic Kwiatkowski, Cordelia Langford, John Sillitoe on behalf of the Wellcome Sanger Institute COVID-19 Surveillance Team |
| EPI_ISL_3139907, EPI_ISL_3139908 | Lighthouse Laboratory Plymouth | Wellcome Sanger Institute for the COVID-19 Genomics UK (COG-UK) Consortium | Lighthouse Laboratory Plymouth and Alex Alderton, Roberto Amato, Jeffrey Barrett, Sonia Goncalves, Ewan Harrison, David K. Jackson, Ian Johnston, Dominic Kwiatkowski, Cordelia Langford, John Sillitoe on behalf of the Wellcome Sanger Institute COVID-19 Surveillance Team |
| EPI_ISL_3139909 | Lighthouse Lab in Alderley Park | Wellcome Sanger Institute for the COVID-19 Genomics UK (COG-UK) Consortium | Jacquelyn Wynn, Mairead Hyland, The Lighthouse Lab in Alderley Park and Alex Alderton, Roberto Amato, Jeffrey Barrett, Sonia Goncalves, Ewan Harrison, David K. Jackson, Ian Johnston, Dominic Kwiatkowski, Cordelia Langford, John Sillitoe on behalf of the Wellcome Sanger Institute COVID-19 Surveillance Team |
| EPI_ISL_3139910 | Berkshire and Surrey Pathology Services Lighthouse Laboratory | Wellcome Sanger Institute for the COVID-19 Genomics UK (COG-UK) Consortium | Berkshire and Surrey Pathology Services Lighthouse Laboratory and Alex Alderton, Roberto Amato, Jeffrey Barrett, Sonia Goncalves, Ewan Harrison, David K. Jackson, Ian Johnston, Dominic Kwiatkowski, Cordelia Langford, John Sillitoe on behalf of the Wellcome Sanger Institute COVID-19 Surveillance Team |
| EPI_ISL_3139911, EPI_ISL_3139912, EPI_ISL_3139913, EPI_ISL_3139914 | Lighthouse Lab in Milton Keynes | Wellcome Sanger Institute for the COVID-19 Genomics UK (COG-UK) Consortium | The Lighthouse Lab in Milton Keynes and Alex Alderton, Roberto Amato, Jeffrey Barrett, Sonia Goncalves, Ewan Harrison, David K. Jackson, Ian Johnston, Dominic Kwiatkowski, Cordelia Langford, John Sillitoe on behalf of the Wellcome Sanger Institute COVID-19 Surveillance Team |
| EPI_ISL_3139915, EPI_ISL_3139916, EPI_ISL_3139917, EPI_ISL_3139918 | Lighthouse Lab in Alderley Park | Wellcome Sanger Institute for the COVID-19 Genomics UK (COG-UK) Consortium | Jacquelyn Wynn, Mairead Hyland, The Lighthouse Lab in Alderley Park and Alex Alderton, Roberto Amato, Jeffrey Barrett, Sonia Goncalves, Ewan Harrison, David K. Jackson, Ian Johnston, Dominic Kwiatkowski, Cordelia Langford, John Sillitoe on behalf of the Wellcome Sanger Institute COVID-19 Surveillance Team |
| EPI_ISL_3139919 | Berkshire and Surrey Pathology Services Lighthouse Laboratory | Wellcome Sanger Institute for the COVID-19 Genomics UK (COG-UK) Consortium | Berkshire and Surrey Pathology Services Lighthouse Laboratory and Alex Alderton, Roberto Amato, Jeffrey Barrett, Sonia Goncalves, Ewan Harrison, David K. Jackson, Ian Johnston, Dominic Kwiatkowski, Cordelia Langford, John Sillitoe on behalf of the Wellcome Sanger Institute COVID-19 Surveillance Team |
| EPI_ISL_3139920 | Lighthouse Laboratory Plymouth | Wellcome Sanger Institute for the COVID-19 Genomics UK (COG-UK) Consortium | Lighthouse Laboratory Plymouth and Alex Alderton, Roberto Amato, Jeffrey Barrett, Sonia Goncalves, Ewan Harrison, David K. Jackson, Ian Johnston, Dominic Kwiatkowski, Cordelia Langford, John Sillitoe on behalf of the Wellcome Sanger Institute COVID-19 Surveillance Team |
| EPI_ISL_3139921 | Lighthouse Lab in Alderley Park | Wellcome Sanger Institute for the COVID-19 Genomics UK (COG-UK) Consortium | Jacquelyn Wynn, Mairead Hyland, The Lighthouse Lab in Alderley Park and Alex Alderton, Roberto Amato, Jeffrey Barrett, Sonia Goncalves, Ewan Harrison, David K. Jackson, Ian Johnston, Dominic Kwiatkowski, Cordelia Langford, John Sillitoe on behalf of the Wellcome Sanger Institute COVID-19 Surveillance Team |
| EPI_ISL_3139922, EPI_ISL_3139923 | Lighthouse Lab in Milton Keynes | Wellcome Sanger Institute for the COVID-19 Genomics UK (COG-UK) Consortium | The Lighthouse Lab in Milton Keynes and Alex Alderton, Roberto Amato, Jeffrey Barrett, Sonia Goncalves, Ewan Harrison, David K. Jackson, Ian Johnston, Dominic Kwiatkowski, Cordelia Langford, John Sillitoe on behalf of the Wellcome Sanger Institute COVID-19 Surveillance Team |
| EPI_ISL_3139924, EPI_ISL_3139925 | Lighthouse Lab in Glasgow | Wellcome Sanger Institute for the COVID-19 Genomics UK (COG-UK) Consortium | Harper VanSteenhouse, Yumi Kasai, David Gray, Carol Clugston, Anna Dominiczak and Alex Alderton, Roberto Amato, Jeffrey Barrett, Sonia Goncalves, Ewan Harrison, David K. Jackson, Ian Johnston, Dominic Kwiatkowski, Cordelia Langford, John Sillitoe on behalf of the Wellcome Sanger Institute COVID-19 Surveillance Team |
| EPI_ISL_3139926 | Lighthouse Laboratory Plymouth | Wellcome Sanger Institute for the COVID-19 Genomics UK (COG-UK) Consortium | Lighthouse Laboratory Plymouth and Alex Alderton, Roberto Amato, Jeffrey Barrett, Sonia Goncalves, Ewan Harrison, David K. Jackson, Ian Johnston, Dominic Kwiatkowski, Cordelia Langford, John Sillitoe on behalf of the Wellcome Sanger Institute COVID-19 Surveillance Team |
| EPI_ISL_3139927 | Lighthouse Lab in Glasgow | Wellcome Sanger Institute for the COVID-19 Genomics UK (COG-UK) Consortium | Harper VanSteenhouse, Yumi Kasai, David Gray, Carol Clugston, Anna Dominiczak and Alex Alderton, Roberto Amato, Jeffrey Barrett, Sonia Goncalves, Ewan Harrison, David K. Jackson, Ian Johnston, Dominic Kwiatkowski, Cordelia Langford, John Sillitoe on behalf of the Wellcome Sanger Institute COVID-19 Surveillance Team |
| EPI_ISL_3139928 | Berkshire and Surrey Pathology Services Lighthouse Laboratory | Wellcome Sanger Institute for the COVID-19 Genomics UK (COG-UK) Consortium | Berkshire and Surrey Pathology Services Lighthouse Laboratory and Alex Alderton, Roberto Amato, Jeffrey Barrett, Sonia Goncalves, Ewan Harrison, David K. Jackson, Ian Johnston, Dominic Kwiatkowski, Cordelia Langford, John Sillitoe on behalf of the Wellcome Sanger Institute COVID-19 Surveillance Team |
| EPI_ISL_3139929 | Lighthouse Lab in Glasgow | Wellcome Sanger Institute for the COVID-19 Genomics UK (COG-UK) Consortium | Harper VanSteenhouse, Yumi Kasai, David Gray, Carol Clugston, Anna Dominiczak and Alex Alderton, Roberto Amato, Jeffrey Barrett, Sonia Goncalves, Ewan Harrison, David K. Jackson, Ian Johnston, Dominic Kwiatkowski, Cordelia Langford, John Sillitoe on behalf of the Wellcome Sanger Institute COVID-19 Surveillance Team |
| EPI_ISL_3139930 | Berkshire and Surrey Pathology Services Lighthouse Laboratory | Wellcome Sanger Institute for the COVID-19 Genomics UK (COG-UK) Consortium | Berkshire and Surrey Pathology Services Lighthouse Laboratory and Alex Alderton, Roberto Amato, Jeffrey Barrett, Sonia Goncalves, Ewan Harrison, David K. Jackson, Ian Johnston, Dominic Kwiatkowski, Cordelia Langford, John Sillitoe on behalf of the Wellcome Sanger Institute COVID-19 Surveillance Team |
| EPI_ISL_3139931 | Lighthouse Lab in Glasgow | Wellcome Sanger Institute for the COVID-19 Genomics UK (COG-UK) Consortium | Harper VanSteenhouse, Yumi Kasai, David Gray, Carol Clugston, Anna Dominiczak and Alex Alderton, Roberto Amato, Jeffrey Barrett, Sonia Goncalves, Ewan Harrison, David K. Jackson, Ian Johnston, Dominic Kwiatkowski, Cordelia Langford, John Sillitoe on behalf of the Wellcome Sanger Institute COVID-19 Surveillance Team |
| EPI_ISL_3139932 | Berkshire and Surrey Pathology Services Lighthouse Laboratory | Wellcome Sanger Institute for the COVID-19 Genomics UK (COG-UK) Consortium | Berkshire and Surrey Pathology Services Lighthouse Laboratory and Alex Alderton, Roberto Amato, Jeffrey Barrett, Sonia Goncalves, Ewan Harrison, David K. Jackson, Ian Johnston, Dominic Kwiatkowski, Cordelia Langford, John Sillitoe on behalf of the Wellcome Sanger Institute COVID-19 Surveillance Team |
| EPI_ISL_3139933 | Lighthouse Laboratory Plymouth | Wellcome Sanger Institute for the COVID-19 Genomics UK (COG-UK) | Lighthouse Laboratory Plymouth and Alex Alderton, Roberto Amato, Jeffrey Barrett, Sonia Goncalves, Ewan Harrison, David K. Jackson, Ian Johnston, Dominic Kwiatkowski, Cordelia Langford, John Sillitoe on behalf of the Wellcome Sanger |

| | | | |
|---|---|---|---|
| | | Consortium | Institute COVID-19 Surveillance Team |
| EPI_ISL_3139934 | Berkshire and Surrey Pathology Services Lighthouse Laboratory | Wellcome Sanger Institute for the COVID-19 Genomics UK (COG-UK) Consortium | Berkshire and Surrey Pathology Services Lighthouse Laboratory and Alex Alderton, Roberto Amato, Jeffrey Barrett, Sonia Goncalves, Ewan Harrison, David K. Jackson, Ian Johnston, Dominic Kwiatkowski, Cordelia Langford, John Sillitoe on behalf of the Wellcome Sanger Institute COVID-19 Surveillance Team |
| EPI_ISL_3139935 | Lighthouse Laboratory Plymouth | Wellcome Sanger Institute for the COVID-19 Genomics UK (COG-UK) Consortium | Lighthouse Laboratory Plymouth and Alex Alderton, Roberto Amato, Jeffrey Barrett, Sonia Goncalves, Ewan Harrison, David K. Jackson, Ian Johnston, Dominic Kwiatkowski, Cordelia Langford, John Sillitoe on behalf of the Wellcome Sanger Institute COVID-19 Surveillance Team |
| EPI_ISL_3139936 | Lighthouse Lab in Glasgow | Wellcome Sanger Institute for the COVID-19 Genomics UK (COG-UK) Consortium | Harper VanSteenhouse, Yumi Kasai, David Gray, Carol Clugston, Anna Dominiczak and Alex Alderton, Roberto Amato, Jeffrey Barrett, Sonia Goncalves, Ewan Harrison, David K. Jackson, Ian Johnston, Dominic Kwiatkowski, Cordelia Langford, John Sillitoe on behalf of the Wellcome Sanger Institute COVID-19 Surveillance Team |
| EPI_ISL_3139937 | Berkshire and Surrey Pathology Services Lighthouse Laboratory | Wellcome Sanger Institute for the COVID-19 Genomics UK (COG-UK) Consortium | Berkshire and Surrey Pathology Services Lighthouse Laboratory and Alex Alderton, Roberto Amato, Jeffrey Barrett, Sonia Goncalves, Ewan Harrison, David K. Jackson, Ian Johnston, Dominic Kwiatkowski, Cordelia Langford, John Sillitoe on behalf of the Wellcome Sanger Institute COVID-19 Surveillance Team |
| EPI_ISL_3139938 | Lighthouse Lab in Glasgow | Wellcome Sanger Institute for the COVID-19 Genomics UK (COG-UK) Consortium | Harper VanSteenhouse, Yumi Kasai, David Gray, Carol Clugston, Anna Dominiczak and Alex Alderton, Roberto Amato, Jeffrey Barrett, Sonia Goncalves, Ewan Harrison, David K. Jackson, Ian Johnston, Dominic Kwiatkowski, Cordelia Langford, John Sillitoe on behalf of the Wellcome Sanger Institute COVID-19 Surveillance Team |
| EPI_ISL_3139939 | Berkshire and Surrey Pathology Services Lighthouse Laboratory | Wellcome Sanger Institute for the COVID-19 Genomics UK (COG-UK) Consortium | Berkshire and Surrey Pathology Services Lighthouse Laboratory and Alex Alderton, Roberto Amato, Jeffrey Barrett, Sonia Goncalves, Ewan Harrison, David K. Jackson, Ian Johnston, Dominic Kwiatkowski, Cordelia Langford, John Sillitoe on behalf of the Wellcome Sanger Institute COVID-19 Surveillance Team |
| EPI_ISL_3139940 | Lighthouse Laboratory Plymouth | Wellcome Sanger Institute for the COVID-19 Genomics UK (COG-UK) Consortium | Lighthouse Laboratory Plymouth and Alex Alderton, Roberto Amato, Jeffrey Barrett, Sonia Goncalves, Ewan Harrison, David K. Jackson, Ian Johnston, Dominic Kwiatkowski, Cordelia Langford, John Sillitoe on behalf of the Wellcome Sanger Institute COVID-19 Surveillance Team |
| EPI_ISL_3139941, EPI_ISL_3139942 | Berkshire and Surrey Pathology Services Lighthouse Laboratory | Wellcome Sanger Institute for the COVID-19 Genomics UK (COG-UK) Consortium | Berkshire and Surrey Pathology Services Lighthouse Laboratory and Alex Alderton, Roberto Amato, Jeffrey Barrett, Sonia Goncalves, Ewan Harrison, David K. Jackson, Ian Johnston, Dominic Kwiatkowski, Cordelia Langford, John Sillitoe on behalf of the Wellcome Sanger Institute COVID-19 Surveillance Team |
| EPI_ISL_3139943 | Lighthouse Laboratory Plymouth | Wellcome Sanger Institute for the COVID-19 Genomics UK (COG-UK) Consortium | Lighthouse Laboratory Plymouth and Alex Alderton, Roberto Amato, Jeffrey Barrett, Sonia Goncalves, Ewan Harrison, David K. Jackson, Ian Johnston, Dominic Kwiatkowski, Cordelia Langford, John Sillitoe on behalf of the Wellcome Sanger Institute COVID-19 Surveillance Team |
| EPI_ISL_3139944 | Berkshire and Surrey Pathology Services Lighthouse Laboratory | Wellcome Sanger Institute for the COVID-19 Genomics UK (COG-UK) Consortium | Berkshire and Surrey Pathology Services Lighthouse Laboratory and Alex Alderton, Roberto Amato, Jeffrey Barrett, Sonia Goncalves, Ewan Harrison, David K. Jackson, Ian Johnston, Dominic Kwiatkowski, Cordelia Langford, John Sillitoe on behalf of the Wellcome Sanger Institute COVID-19 Surveillance Team |
| EPI_ISL_3139945, EPI_ISL_3139946 | Lighthouse Lab in Glasgow | Wellcome Sanger Institute for the COVID-19 Genomics UK (COG-UK) Consortium | Harper VanSteenhouse, Yumi Kasai, David Gray, Carol Clugston, Anna Dominiczak and Alex Alderton, Roberto Amato, Jeffrey Barrett, Sonia Goncalves, Ewan Harrison, David K. Jackson, Ian Johnston, Dominic Kwiatkowski, Cordelia Langford, John Sillitoe on behalf of the Wellcome Sanger Institute COVID-19 Surveillance Team |
| EPI_ISL_3139947 | Lighthouse Laboratory Plymouth | Wellcome Sanger Institute for the COVID-19 Genomics UK (COG-UK) Consortium | Lighthouse Laboratory Plymouth and Alex Alderton, Roberto Amato, Jeffrey Barrett, Sonia Goncalves, Ewan Harrison, David K. Jackson, Ian Johnston, Dominic Kwiatkowski, Cordelia Langford, John Sillitoe on behalf of the Wellcome Sanger Institute COVID-19 Surveillance Team |
| EPI_ISL_3139948 | Berkshire and Surrey Pathology Services Lighthouse Laboratory | Wellcome Sanger Institute for the COVID-19 Genomics UK (COG-UK) Consortium | Berkshire and Surrey Pathology Services Lighthouse Laboratory and Alex Alderton, Roberto Amato, Jeffrey Barrett, Sonia Goncalves, Ewan Harrison, David K. Jackson, Ian Johnston, Dominic Kwiatkowski, Cordelia Langford, John Sillitoe on behalf of the Wellcome Sanger Institute COVID-19 Surveillance Team |
| EPI_ISL_3139949, EPI_ISL_3139950 | Lighthouse Lab in Glasgow | Wellcome Sanger Institute for the COVID-19 Genomics UK (COG-UK) Consortium | Harper VanSteenhouse, Yumi Kasai, David Gray, Carol Clugston, Anna Dominiczak and Alex Alderton, Roberto Amato, Jeffrey Barrett, Sonia Goncalves, Ewan Harrison, David K. Jackson, Ian Johnston, Dominic Kwiatkowski, Cordelia Langford, John Sillitoe on behalf of the Wellcome Sanger Institute COVID-19 Surveillance Team |
| EPI_ISL_3139951, EPI_ISL_3139952, EPI_ISL_3139953 | Berkshire and Surrey Pathology Services Lighthouse Laboratory | Wellcome Sanger Institute for the COVID-19 Genomics UK (COG-UK) Consortium | Berkshire and Surrey Pathology Services Lighthouse Laboratory and Alex Alderton, Roberto Amato, Jeffrey Barrett, Sonia Goncalves, Ewan Harrison, David K. Jackson, Ian Johnston, Dominic Kwiatkowski, Cordelia Langford, John Sillitoe on behalf of the Wellcome Sanger Institute COVID-19 Surveillance Team |
| EPI_ISL_3139954, EPI_ISL_3139955 | Lighthouse Lab in Glasgow | Wellcome Sanger Institute for the COVID-19 Genomics UK (COG-UK) Consortium | Harper VanSteenhouse, Yumi Kasai, David Gray, Carol Clugston, Anna Dominiczak and Alex Alderton, Roberto Amato, Jeffrey Barrett, Sonia Goncalves, Ewan Harrison, David K. Jackson, Ian Johnston, Dominic Kwiatkowski, Cordelia Langford, John Sillitoe on behalf of the Wellcome Sanger Institute COVID-19 Surveillance Team |
| EPI_ISL_3139956 | Berkshire and Surrey Pathology Services Lighthouse Laboratory | Wellcome Sanger Institute for the COVID-19 Genomics UK (COG-UK) Consortium | Berkshire and Surrey Pathology Services Lighthouse Laboratory and Alex Alderton, Roberto Amato, Jeffrey Barrett, Sonia Goncalves, Ewan Harrison, David K. Jackson, Ian Johnston, Dominic Kwiatkowski, Cordelia Langford, John Sillitoe on behalf of the Wellcome Sanger Institute COVID-19 Surveillance Team |
| EPI_ISL_3139957 | Lighthouse Laboratory Plymouth | Wellcome Sanger Institute for the COVID-19 Genomics UK (COG-UK) Consortium | Lighthouse Laboratory Plymouth and Alex Alderton, Roberto Amato, Jeffrey Barrett, Sonia Goncalves, Ewan Harrison, David K. Jackson, Ian Johnston, Dominic Kwiatkowski, Cordelia Langford, John Sillitoe on behalf of the Wellcome Sanger Institute COVID-19 Surveillance Team |
| EPI_ISL_3139958 | Berkshire and Surrey Pathology Services Lighthouse Laboratory | Wellcome Sanger Institute for the COVID-19 Genomics UK (COG-UK) Consortium | Berkshire and Surrey Pathology Services Lighthouse Laboratory and Alex Alderton, Roberto Amato, Jeffrey Barrett, Sonia Goncalves, Ewan Harrison, David K. Jackson, Ian Johnston, Dominic Kwiatkowski, Cordelia Langford, John Sillitoe on behalf of the Wellcome Sanger Institute COVID-19 Surveillance Team |
| EPI_ISL_3139959, EPI_ISL_3139960 | Lighthouse Lab in Glasgow | Wellcome Sanger Institute for the COVID-19 Genomics UK (COG-UK) Consortium | Harper VanSteenhouse, Yumi Kasai, David Gray, Carol Clugston, Anna Dominiczak and Alex Alderton, Roberto Amato, Jeffrey Barrett, Sonia Goncalves, Ewan Harrison, David K. Jackson, Ian Johnston, Dominic Kwiatkowski, Cordelia Langford, John Sillitoe on behalf of the Wellcome Sanger Institute COVID-19 Surveillance Team |
| EPI_ISL_3139961 | Lighthouse Laboratory Plymouth | Wellcome Sanger Institute for the COVID-19 Genomics UK (COG-UK) Consortium | Lighthouse Laboratory Plymouth and Alex Alderton, Roberto Amato, Jeffrey Barrett, Sonia Goncalves, Ewan Harrison, David K. Jackson, Ian Johnston, Dominic Kwiatkowski, Cordelia Langford, John Sillitoe on behalf of the Wellcome Sanger Institute COVID-19 Surveillance Team |
| EPI_ISL_3139962 | Berkshire and Surrey Pathology Services Lighthouse Laboratory | Wellcome Sanger Institute for the COVID-19 Genomics UK (COG-UK) Consortium | Berkshire and Surrey Pathology Services Lighthouse Laboratory and Alex Alderton, Roberto Amato, Jeffrey Barrett, Sonia Goncalves, Ewan Harrison, David K. Jackson, Ian Johnston, Dominic Kwiatkowski, Cordelia Langford, John Sillitoe on behalf of the Wellcome Sanger Institute COVID-19 Surveillance Team |
| EPI_ISL_3139963, EPI_ISL_3139964 | Lighthouse Laboratory Plymouth | Wellcome Sanger Institute for the COVID-19 Genomics UK (COG-UK) Consortium | Lighthouse Laboratory Plymouth and Alex Alderton, Roberto Amato, Jeffrey Barrett, Sonia Goncalves, Ewan Harrison, David K. Jackson, Ian Johnston, Dominic Kwiatkowski, Cordelia Langford, John Sillitoe on behalf of the Wellcome Sanger Institute COVID-19 Surveillance Team |
| EPI_ISL_3139965 | Berkshire and Surrey Pathology Services Lighthouse Laboratory | Wellcome Sanger Institute for the COVID-19 Genomics UK (COG-UK) Consortium | Berkshire and Surrey Pathology Services Lighthouse Laboratory and Alex Alderton, Roberto Amato, Jeffrey Barrett, Sonia Goncalves, Ewan Harrison, David K. Jackson, Ian Johnston, Dominic Kwiatkowski, Cordelia Langford, John Sillitoe on behalf of the Wellcome Sanger Institute COVID-19 Surveillance Team |
| EPI_ISL_3139966 | Lighthouse Laboratory Plymouth | Wellcome Sanger Institute for the COVID-19 Genomics UK (COG-UK) Consortium | Lighthouse Laboratory Plymouth and Alex Alderton, Roberto Amato, Jeffrey Barrett, Sonia Goncalves, Ewan Harrison, David K. Jackson, Ian Johnston, Dominic Kwiatkowski, Cordelia Langford, John Sillitoe on behalf of the Wellcome Sanger Institute COVID-19 Surveillance Team |
| EPI_ISL_3139967, EPI_ISL_3139968 | Berkshire and Surrey Pathology Services Lighthouse Laboratory | Wellcome Sanger Institute for the COVID-19 Genomics UK (COG-UK) Consortium | Berkshire and Surrey Pathology Services Lighthouse Laboratory and Alex Alderton, Roberto Amato, Jeffrey Barrett, Sonia Goncalves, Ewan Harrison, David K. Jackson, Ian Johnston, Dominic Kwiatkowski, Cordelia Langford, John Sillitoe on behalf of the Wellcome Sanger Institute COVID-19 Surveillance Team |
| EPI_ISL_3139969 | Lighthouse Laboratory Plymouth | Wellcome Sanger Institute for the COVID-19 Genomics UK (COG-UK) Consortium | Lighthouse Laboratory Plymouth and Alex Alderton, Roberto Amato, Jeffrey Barrett, Sonia Goncalves, Ewan Harrison, David K. Jackson, Ian Johnston, Dominic Kwiatkowski, Cordelia Langford, John Sillitoe on behalf of the Wellcome Sanger Institute COVID-19 Surveillance Team |
| EPI_ISL_3139970 | Lighthouse Lab in Glasgow | Wellcome Sanger Institute for the COVID-19 Genomics UK (COG-UK) Consortium | Harper VanSteenhouse, Yumi Kasai, David Gray, Carol Clugston, Anna Dominiczak and Alex Alderton, Roberto Amato, Jeffrey Barrett, Sonia Goncalves, Ewan Harrison, David K. Jackson, Ian Johnston, Dominic Kwiatkowski, Cordelia Langford, John Sillitoe on behalf of the Wellcome Sanger Institute COVID-19 Surveillance Team |
| EPI_ISL_3139971 | Berkshire and Surrey Pathology Services Lighthouse Laboratory | Wellcome Sanger Institute for the COVID-19 Genomics UK (COG-UK) Consortium | Berkshire and Surrey Pathology Services Lighthouse Laboratory and Alex Alderton, Roberto Amato, Jeffrey Barrett, Sonia Goncalves, Ewan Harrison, David K. Jackson, Ian Johnston, Dominic Kwiatkowski, Cordelia Langford, John Sillitoe on behalf of the Wellcome Sanger Institute COVID-19 Surveillance Team |
| EPI_ISL_3139972 | Lighthouse Lab in Glasgow | Wellcome Sanger Institute for the COVID-19 Genomics UK (COG-UK) Consortium | Harper VanSteenhouse, Yumi Kasai, David Gray, Carol Clugston, Anna Dominiczak and Alex Alderton, Roberto Amato, Jeffrey Barrett, Sonia Goncalves, Ewan Harrison, David K. Jackson, Ian Johnston, Dominic Kwiatkowski, Cordelia Langford, John Sillitoe on behalf of the Wellcome Sanger Institute COVID-19 Surveillance Team |
| EPI_ISL_3139973 | Berkshire and Surrey Pathology Services Lighthouse Laboratory | Wellcome Sanger Institute for the COVID-19 Genomics UK (COG-UK) Consortium | Berkshire and Surrey Pathology Services Lighthouse Laboratory and Alex Alderton, Roberto Amato, Jeffrey Barrett, Sonia Goncalves, Ewan Harrison, David K. Jackson, Ian Johnston, Dominic Kwiatkowski, Cordelia Langford, John Sillitoe on behalf of the Wellcome Sanger Institute COVID-19 Surveillance Team |
| EPI_ISL_3139974 | Lighthouse Lab in Glasgow | Wellcome Sanger Institute for the COVID-19 Genomics UK (COG-UK) Consortium | Harper VanSteenhouse, Yumi Kasai, David Gray, Carol Clugston, Anna Dominiczak and Alex Alderton, Roberto Amato, Jeffrey Barrett, Sonia Goncalves, Ewan Harrison, David K. Jackson, Ian Johnston, Dominic Kwiatkowski, Cordelia Langford, John Sillitoe on behalf of the Wellcome Sanger Institute COVID-19 Surveillance Team |
| EPI_ISL_3139975, EPI_ISL_3139976, EPI_ISL_3139977, EPI_ISL_3139978 | Berkshire and Surrey Pathology Services Lighthouse Laboratory | Wellcome Sanger Institute for the COVID-19 Genomics UK (COG-UK) Consortium | Berkshire and Surrey Pathology Services Lighthouse Laboratory and Alex Alderton, Roberto Amato, Jeffrey Barrett, Sonia Goncalves, Ewan Harrison, David K. Jackson, Ian Johnston, Dominic Kwiatkowski, Cordelia Langford, John Sillitoe on behalf of the Wellcome Sanger Institute COVID-19 Surveillance Team |
| EPI_ISL_3139979 | Lighthouse Laboratory Plymouth | Wellcome Sanger Institute for the COVID-19 Genomics UK (COG-UK) Consortium | Lighthouse Laboratory Plymouth and Alex Alderton, Roberto Amato, Jeffrey Barrett, Sonia Goncalves, Ewan Harrison, David K. Jackson, Ian Johnston, Dominic Kwiatkowski, Cordelia Langford, John Sillitoe on behalf of the Wellcome Sanger Institute COVID-19 Surveillance Team |
| EPI_ISL_3139980 | Lighthouse Lab in Glasgow | Wellcome Sanger Institute for the COVID-19 Genomics UK (COG-UK) Consortium | Harper VanSteenhouse, Yumi Kasai, David Gray, Carol Clugston, Anna Dominiczak and Alex Alderton, Roberto Amato, Jeffrey Barrett, Sonia Goncalves, Ewan Harrison, David K. Jackson, Ian Johnston, Dominic Kwiatkowski, Cordelia Langford, John Sillitoe on behalf of the Wellcome Sanger Institute COVID-19 Surveillance Team |
| EPI_ISL_3139981 | Lighthouse Laboratory Plymouth | Wellcome Sanger Institute for the COVID-19 Genomics UK (COG-UK) Consortium | Lighthouse Laboratory Plymouth and Alex Alderton, Roberto Amato, Jeffrey Barrett, Sonia Goncalves, Ewan Harrison, David K. Jackson, Ian Johnston, Dominic Kwiatkowski, Cordelia Langford, John Sillitoe on behalf of the Wellcome Sanger Institute COVID-19 Surveillance Team |
| EPI_ISL_3139982 | Lighthouse Lab in Glasgow | Wellcome Sanger Institute for the COVID-19 Genomics UK (COG-UK) Consortium | Harper VanSteenhouse, Yumi Kasai, David Gray, Carol Clugston, Anna Dominiczak and Alex Alderton, Roberto Amato, Jeffrey Barrett, Sonia Goncalves, Ewan Harrison, David K. Jackson, Ian Johnston, Dominic Kwiatkowski, Cordelia Langford, John Sillitoe on behalf of the Wellcome Sanger Institute COVID-19 Surveillance Team |
| EPI_ISL_3139983 | Lighthouse Laboratory Plymouth | Wellcome Sanger Institute for the COVID-19 Genomics UK (COG-UK) Consortium | Lighthouse Laboratory Plymouth and Alex Alderton, Roberto Amato, Jeffrey Barrett, Sonia Goncalves, Ewan Harrison, David K. Jackson, Ian Johnston, Dominic Kwiatkowski, Cordelia Langford, John Sillitoe on behalf of the Wellcome Sanger Institute COVID-19 Surveillance Team |
| EPI_ISL_3139984 | Berkshire and Surrey Pathology Services Lighthouse Laboratory | Wellcome Sanger Institute for the COVID-19 Genomics UK (COG-UK) Consortium | Berkshire and Surrey Pathology Services Lighthouse Laboratory and Alex Alderton, Roberto Amato, Jeffrey Barrett, Sonia Goncalves, Ewan Harrison, David K. Jackson, Ian Johnston, Dominic Kwiatkowski, Cordelia Langford, John Sillitoe on behalf of the Wellcome Sanger Institute COVID-19 Surveillance Team |
| EPI_ISL_3139985, EPI_ISL_3139986 | Lighthouse Lab in Glasgow | Wellcome Sanger Institute for the COVID-19 Genomics UK (COG-UK) Consortium | Harper VanSteenhouse, Yumi Kasai, David Gray, Carol Clugston, Anna Dominiczak and Alex Alderton, Roberto Amato, Jeffrey Barrett, Sonia Goncalves, Ewan Harrison, David K. Jackson, Ian Johnston, Dominic Kwiatkowski, Cordelia Langford, John Sillitoe on behalf of the Wellcome Sanger Institute COVID-19 Surveillance Team |
| EPI_ISL_3139987 | Lighthouse Laboratory Plymouth | Wellcome Sanger Institute for the COVID-19 Genomics UK (COG-UK) Consortium | Lighthouse Laboratory Plymouth and Alex Alderton, Roberto Amato, Jeffrey Barrett, Sonia Goncalves, Ewan Harrison, David K. Jackson, Ian Johnston, Dominic Kwiatkowski, Cordelia Langford, John Sillitoe on behalf of the Wellcome Sanger Institute COVID-19 Surveillance Team |
| EPI_ISL_3139988 | Lighthouse Lab in Glasgow | Wellcome Sanger Institute for the COVID-19 Genomics UK (COG-UK) Consortium | Harper VanSteenhouse, Yumi Kasai, David Gray, Carol Clugston, Anna Dominiczak and Alex Alderton, Roberto Amato, Jeffrey Barrett, Sonia Goncalves, Ewan Harrison, David K. Jackson, Ian Johnston, Dominic Kwiatkowski, Cordelia Langford, John Sillitoe on behalf of the Wellcome Sanger Institute COVID-19 Surveillance Team |
| EPI_ISL_3139989 | Lighthouse Laboratory Plymouth | Wellcome Sanger Institute for the COVID-19 Genomics UK (COG-UK) Consortium | Lighthouse Laboratory Plymouth and Alex Alderton, Roberto Amato, Jeffrey Barrett, Sonia Goncalves, Ewan Harrison, David K. Jackson, Ian Johnston, Dominic Kwiatkowski, Cordelia Langford, John Sillitoe on behalf of the Wellcome Sanger Institute COVID-19 Surveillance Team |
| EPI_ISL_3139990 | Berkshire and Surrey Pathology Services Lighthouse Laboratory | Wellcome Sanger Institute for the COVID-19 Genomics UK (COG-UK) | Berkshire and Surrey Pathology Services Lighthouse Laboratory and Alex Alderton, Roberto Amato, Jeffrey Barrett, Sonia Goncalves, Ewan Harrison, David K. Jackson, Ian Johnston, Dominic Kwiatkowski, Cordelia Langford, John Sillitoe on behalf |

| | | | |
|---|---|---|---|
| | | Consortium | of the Wellcome Sanger Institute COVID-19 Surveillance Team |
| EPI_ISL_3139991 | Lighthouse Lab in Glasgow | Wellcome Sanger Institute for the COVID-19 Genomics UK (COG-UK) Consortium | Harper VanSteenhouse, Yumi Kasai, David Gray, Carol Clugston, Anna Dominiczak and Alex Alderton, Roberto Amato, Jeffrey Barrett, Sonia Goncalves, Ewan Harrison, David K. Jackson, Ian Johnston, Dominic Kwiatkowski, Cordelia Langford, John Sillitoe on behalf of the Wellcome Sanger Institute COVID-19 Surveillance Team |
| EPI_ISL_3139992 | Lighthouse Laboratory Plymouth | Wellcome Sanger Institute for the COVID-19 Genomics UK (COG-UK) Consortium | Lighthouse Laboratory Plymouth and Alex Alderton, Roberto Amato, Jeffrey Barrett, Sonia Goncalves, Ewan Harrison, David K. Jackson, Ian Johnston, Dominic Kwiatkowski, Cordelia Langford, John Sillitoe on behalf of the Wellcome Sanger Institute COVID-19 Surveillance Team |
| EPI_ISL_3139993, EPI_ISL_3139994, EPI_ISL_3139995, EPI_ISL_3139996 | Berkshire and Surrey Pathology Services Lighthouse Laboratory | Wellcome Sanger Institute for the COVID-19 Genomics UK (COG-UK) Consortium | Berkshire and Surrey Pathology Services Lighthouse Laboratory and Alex Alderton, Roberto Amato, Jeffrey Barrett, Sonia Goncalves, Ewan Harrison, David K. Jackson, Ian Johnston, Dominic Kwiatkowski, Cordelia Langford, John Sillitoe on behalf of the Wellcome Sanger Institute COVID-19 Surveillance Team |
| EPI_ISL_3139997, EPI_ISL_3139998, EPI_ISL_3139999, EPI_ISL_3140000 | Lighthouse Laboratory Plymouth | Wellcome Sanger Institute for the COVID-19 Genomics UK (COG-UK) Consortium | Lighthouse Laboratory Plymouth and Alex Alderton, Roberto Amato, Jeffrey Barrett, Sonia Goncalves, Ewan Harrison, David K. Jackson, Ian Johnston, Dominic Kwiatkowski, Cordelia Langford, John Sillitoe on behalf of the Wellcome Sanger Institute COVID-19 Surveillance Team |
| EPI_ISL_3140001 | Berkshire and Surrey Pathology Services Lighthouse Laboratory | Wellcome Sanger Institute for the COVID-19 Genomics UK (COG-UK) Consortium | Berkshire and Surrey Pathology Services Lighthouse Laboratory and Alex Alderton, Roberto Amato, Jeffrey Barrett, Sonia Goncalves, Ewan Harrison, David K. Jackson, Ian Johnston, Dominic Kwiatkowski, Cordelia Langford, John Sillitoe on behalf of the Wellcome Sanger Institute COVID-19 Surveillance Team |
| EPI_ISL_3140002 | Lighthouse Laboratory Plymouth | Wellcome Sanger Institute for the COVID-19 Genomics UK (COG-UK) Consortium | Lighthouse Laboratory Plymouth and Alex Alderton, Roberto Amato, Jeffrey Barrett, Sonia Goncalves, Ewan Harrison, David K. Jackson, Ian Johnston, Dominic Kwiatkowski, Cordelia Langford, John Sillitoe on behalf of the Wellcome Sanger Institute COVID-19 Surveillance Team |
| EPI_ISL_3140003, EPI_ISL_3140004 | Berkshire and Surrey Pathology Services Lighthouse Laboratory | Wellcome Sanger Institute for the COVID-19 Genomics UK (COG-UK) Consortium | Berkshire and Surrey Pathology Services Lighthouse Laboratory and Alex Alderton, Roberto Amato, Jeffrey Barrett, Sonia Goncalves, Ewan Harrison, David K. Jackson, Ian Johnston, Dominic Kwiatkowski, Cordelia Langford, John Sillitoe on behalf of the Wellcome Sanger Institute COVID-19 Surveillance Team |
| EPI_ISL_3140005 | Lighthouse Lab in Glasgow | Wellcome Sanger Institute for the COVID-19 Genomics UK (COG-UK) Consortium | Harper VanSteenhouse, Yumi Kasai, David Gray, Carol Clugston, Anna Dominiczak and Alex Alderton, Roberto Amato, Jeffrey Barrett, Sonia Goncalves, Ewan Harrison, David K. Jackson, Ian Johnston, Dominic Kwiatkowski, Cordelia Langford, John Sillitoe on behalf of the Wellcome Sanger Institute COVID-19 Surveillance Team |
| EPI_ISL_3140006, EPI_ISL_3140007, EPI_ISL_3140008 | Berkshire and Surrey Pathology Services Lighthouse Laboratory | Wellcome Sanger Institute for the COVID-19 Genomics UK (COG-UK) Consortium | Berkshire and Surrey Pathology Services Lighthouse Laboratory and Alex Alderton, Roberto Amato, Jeffrey Barrett, Sonia Goncalves, Ewan Harrison, David K. Jackson, Ian Johnston, Dominic Kwiatkowski, Cordelia Langford, John Sillitoe on behalf of the Wellcome Sanger Institute COVID-19 Surveillance Team |
| EPI_ISL_3140009 | Lighthouse Laboratory Plymouth | Wellcome Sanger Institute for the COVID-19 Genomics UK (COG-UK) Consortium | Lighthouse Laboratory Plymouth and Alex Alderton, Roberto Amato, Jeffrey Barrett, Sonia Goncalves, Ewan Harrison, David K. Jackson, Ian Johnston, Dominic Kwiatkowski, Cordelia Langford, John Sillitoe on behalf of the Wellcome Sanger Institute COVID-19 Surveillance Team |
| EPI_ISL_3140010, EPI_ISL_3140011 | Lighthouse Lab in Glasgow | Wellcome Sanger Institute for the COVID-19 Genomics UK (COG-UK) Consortium | Harper VanSteenhouse, Yumi Kasai, David Gray, Carol Clugston, Anna Dominiczak and Alex Alderton, Roberto Amato, Jeffrey Barrett, Sonia Goncalves, Ewan Harrison, David K. Jackson, Ian Johnston, Dominic Kwiatkowski, Cordelia Langford, John Sillitoe on behalf of the Wellcome Sanger Institute COVID-19 Surveillance Team |
| EPI_ISL_3140012 | Lighthouse Laboratory Plymouth | Wellcome Sanger Institute for the COVID-19 Genomics UK (COG-UK) Consortium | Lighthouse Laboratory Plymouth and Alex Alderton, Roberto Amato, Jeffrey Barrett, Sonia Goncalves, Ewan Harrison, David K. Jackson, Ian Johnston, Dominic Kwiatkowski, Cordelia Langford, John Sillitoe on behalf of the Wellcome Sanger Institute COVID-19 Surveillance Team |
| EPI_ISL_3140013, EPI_ISL_3140014 | Lighthouse Lab in Glasgow | Wellcome Sanger Institute for the COVID-19 Genomics UK (COG-UK) Consortium | Harper VanSteenhouse, Yumi Kasai, David Gray, Carol Clugston, Anna Dominiczak and Alex Alderton, Roberto Amato, Jeffrey Barrett, Sonia Goncalves, Ewan Harrison, David K. Jackson, Ian Johnston, Dominic Kwiatkowski, Cordelia Langford, John Sillitoe on behalf of the Wellcome Sanger Institute COVID-19 Surveillance Team |
| EPI_ISL_3140015, EPI_ISL_3140016, EPI_ISL_3140017 | Berkshire and Surrey Pathology Services Lighthouse Laboratory | Wellcome Sanger Institute for the COVID-19 Genomics UK (COG-UK) Consortium | Berkshire and Surrey Pathology Services Lighthouse Laboratory and Alex Alderton, Roberto Amato, Jeffrey Barrett, Sonia Goncalves, Ewan Harrison, David K. Jackson, Ian Johnston, Dominic Kwiatkowski, Cordelia Langford, John Sillitoe on behalf of the Wellcome Sanger Institute COVID-19 Surveillance Team |
| EPI_ISL_3140018 | Lighthouse Lab in Glasgow | Wellcome Sanger Institute for the COVID-19 Genomics UK (COG-UK) Consortium | Harper VanSteenhouse, Yumi Kasai, David Gray, Carol Clugston, Anna Dominiczak and Alex Alderton, Roberto Amato, Jeffrey Barrett, Sonia Goncalves, Ewan Harrison, David K. Jackson, Ian Johnston, Dominic Kwiatkowski, Cordelia Langford, John Sillitoe on behalf of the Wellcome Sanger Institute COVID-19 Surveillance Team |
| EPI_ISL_3140019, EPI_ISL_3140020, EPI_ISL_3140021, EPI_ISL_3140022 | Berkshire and Surrey Pathology Services Lighthouse Laboratory | Wellcome Sanger Institute for the COVID-19 Genomics UK (COG-UK) Consortium | Berkshire and Surrey Pathology Services Lighthouse Laboratory and Alex Alderton, Roberto Amato, Jeffrey Barrett, Sonia Goncalves, Ewan Harrison, David K. Jackson, Ian Johnston, Dominic Kwiatkowski, Cordelia Langford, John Sillitoe on behalf of the Wellcome Sanger Institute COVID-19 Surveillance Team |
| EPI_ISL_3140023 | Lighthouse Lab in Glasgow | Wellcome Sanger Institute for the COVID-19 Genomics UK (COG-UK) Consortium | Harper VanSteenhouse, Yumi Kasai, David Gray, Carol Clugston, Anna Dominiczak and Alex Alderton, Roberto Amato, Jeffrey Barrett, Sonia Goncalves, Ewan Harrison, David K. Jackson, Ian Johnston, Dominic Kwiatkowski, Cordelia Langford, John Sillitoe on behalf of the Wellcome Sanger Institute COVID-19 Surveillance Team |
| EPI_ISL_3140024, EPI_ISL_3140025, EPI_ISL_3140026 | Lighthouse Laboratory Plymouth | Wellcome Sanger Institute for the COVID-19 Genomics UK (COG-UK) Consortium | Lighthouse Laboratory Plymouth and Alex Alderton, Roberto Amato, Jeffrey Barrett, Sonia Goncalves, Ewan Harrison, David K. Jackson, Ian Johnston, Dominic Kwiatkowski, Cordelia Langford, John Sillitoe on behalf of the Wellcome Sanger Institute COVID-19 Surveillance Team |
| EPI_ISL_3140027 | Berkshire and Surrey Pathology Services Lighthouse Laboratory | Wellcome Sanger Institute for the COVID-19 Genomics UK (COG-UK) Consortium | Berkshire and Surrey Pathology Services Lighthouse Laboratory and Alex Alderton, Roberto Amato, Jeffrey Barrett, Sonia Goncalves, Ewan Harrison, David K. Jackson, Ian Johnston, Dominic Kwiatkowski, Cordelia Langford, John Sillitoe on behalf of the Wellcome Sanger Institute COVID-19 Surveillance Team |
| EPI_ISL_3140028, EPI_ISL_3140029, EPI_ISL_3140030 | Lighthouse Lab in Glasgow | Wellcome Sanger Institute for the COVID-19 Genomics UK (COG-UK) Consortium | Harper VanSteenhouse, Yumi Kasai, David Gray, Carol Clugston, Anna Dominiczak and Alex Alderton, Roberto Amato, Jeffrey Barrett, Sonia Goncalves, Ewan Harrison, David K. Jackson, Ian Johnston, Dominic Kwiatkowski, Cordelia Langford, John Sillitoe on behalf of the Wellcome Sanger Institute COVID-19 Surveillance Team |
| EPI_ISL_3140031 | Lighthouse Laboratory Plymouth | Wellcome Sanger Institute for the COVID-19 Genomics UK (COG-UK) Consortium | Lighthouse Laboratory Plymouth and Alex Alderton, Roberto Amato, Jeffrey Barrett, Sonia Goncalves, Ewan Harrison, David K. Jackson, Ian Johnston, Dominic Kwiatkowski, Cordelia Langford, John Sillitoe on behalf of the Wellcome Sanger Institute COVID-19 Surveillance Team |
| EPI_ISL_3140032, EPI_ISL_3140033, EPI_ISL_3140034 | Lighthouse Lab in Glasgow | Wellcome Sanger Institute for the COVID-19 Genomics UK (COG-UK) Consortium | Harper VanSteenhouse, Yumi Kasai, David Gray, Carol Clugston, Anna Dominiczak and Alex Alderton, Roberto Amato, Jeffrey Barrett, Sonia Goncalves, Ewan Harrison, David K. Jackson, Ian Johnston, Dominic Kwiatkowski, Cordelia Langford, John Sillitoe on behalf of the Wellcome Sanger Institute COVID-19 Surveillance Team |
| EPI_ISL_3140035, EPI_ISL_3140036 | Berkshire and Surrey Pathology Services Lighthouse Laboratory | Wellcome Sanger Institute for the COVID-19 Genomics UK (COG-UK) Consortium | Berkshire and Surrey Pathology Services Lighthouse Laboratory and Alex Alderton, Roberto Amato, Jeffrey Barrett, Sonia Goncalves, Ewan Harrison, David K. Jackson, Ian Johnston, Dominic Kwiatkowski, Cordelia Langford, John Sillitoe on behalf of the Wellcome Sanger Institute COVID-19 Surveillance Team |
| EPI_ISL_3140037 | Lighthouse Laboratory Plymouth | Wellcome Sanger Institute for the COVID-19 Genomics UK (COG-UK) Consortium | Lighthouse Laboratory Plymouth and Alex Alderton, Roberto Amato, Jeffrey Barrett, Sonia Goncalves, Ewan Harrison, David K. Jackson, Ian Johnston, Dominic Kwiatkowski, Cordelia Langford, John Sillitoe on behalf of the Wellcome Sanger Institute COVID-19 Surveillance Team |
| EPI_ISL_3140038, EPI_ISL_3140039 | Lighthouse Lab in Glasgow | Wellcome Sanger Institute for the COVID-19 Genomics UK (COG-UK) Consortium | Harper VanSteenhouse, Yumi Kasai, David Gray, Carol Clugston, Anna Dominiczak and Alex Alderton, Roberto Amato, Jeffrey Barrett, Sonia Goncalves, Ewan Harrison, David K. Jackson, Ian Johnston, Dominic Kwiatkowski, Cordelia Langford, John Sillitoe on behalf of the Wellcome Sanger Institute COVID-19 Surveillance Team |
| EPI_ISL_3140040 | Lighthouse Laboratory Plymouth | Wellcome Sanger Institute for the COVID-19 Genomics UK (COG-UK) Consortium | Lighthouse Laboratory Plymouth and Alex Alderton, Roberto Amato, Jeffrey Barrett, Sonia Goncalves, Ewan Harrison, David K. Jackson, Ian Johnston, Dominic Kwiatkowski, Cordelia Langford, John Sillitoe on behalf of the Wellcome Sanger Institute COVID-19 Surveillance Team |
| EPI_ISL_3140041 | Berkshire and Surrey Pathology Services Lighthouse Laboratory | Wellcome Sanger Institute for the COVID-19 Genomics UK (COG-UK) Consortium | Berkshire and Surrey Pathology Services Lighthouse Laboratory and Alex Alderton, Roberto Amato, Jeffrey Barrett, Sonia Goncalves, Ewan Harrison, David K. Jackson, Ian Johnston, Dominic Kwiatkowski, Cordelia Langford, John Sillitoe on behalf of the Wellcome Sanger Institute COVID-19 Surveillance Team |
| EPI_ISL_3140042 | Lighthouse Laboratory Plymouth | Wellcome Sanger Institute for the COVID-19 Genomics UK (COG-UK) Consortium | Lighthouse Laboratory Plymouth and Alex Alderton, Roberto Amato, Jeffrey Barrett, Sonia Goncalves, Ewan Harrison, David K. Jackson, Ian Johnston, Dominic Kwiatkowski, Cordelia Langford, John Sillitoe on behalf of the Wellcome Sanger Institute COVID-19 Surveillance Team |
| EPI_ISL_3140043 | Berkshire and Surrey Pathology Services Lighthouse Laboratory | Wellcome Sanger Institute for the COVID-19 Genomics UK (COG-UK) Consortium | Berkshire and Surrey Pathology Services Lighthouse Laboratory and Alex Alderton, Roberto Amato, Jeffrey Barrett, Sonia Goncalves, Ewan Harrison, David K. Jackson, Ian Johnston, Dominic Kwiatkowski, Cordelia Langford, John Sillitoe on behalf of the Wellcome Sanger Institute COVID-19 Surveillance Team |
| EPI_ISL_3140044, EPI_ISL_3140045 | Lighthouse Laboratory Plymouth | Wellcome Sanger Institute for the COVID-19 Genomics UK (COG-UK) Consortium | Lighthouse Laboratory Plymouth and Alex Alderton, Roberto Amato, Jeffrey Barrett, Sonia Goncalves, Ewan Harrison, David K. Jackson, Ian Johnston, Dominic Kwiatkowski, Cordelia Langford, John Sillitoe on behalf of the Wellcome Sanger Institute COVID-19 Surveillance Team |
| EPI_ISL_3140046, EPI_ISL_3140047, EPI_ISL_3140048 | Berkshire and Surrey Pathology Services Lighthouse Laboratory | Wellcome Sanger Institute for the COVID-19 Genomics UK (COG-UK) Consortium | Berkshire and Surrey Pathology Services Lighthouse Laboratory and Alex Alderton, Roberto Amato, Jeffrey Barrett, Sonia Goncalves, Ewan Harrison, David K. Jackson, Ian Johnston, Dominic Kwiatkowski, Cordelia Langford, John Sillitoe on behalf of the Wellcome Sanger Institute COVID-19 Surveillance Team |
| EPI_ISL_3140049, EPI_ISL_3140050, EPI_ISL_3140051 | Lighthouse Lab in Glasgow | Wellcome Sanger Institute for the COVID-19 Genomics UK (COG-UK) Consortium | Harper VanSteenhouse, Yumi Kasai, David Gray, Carol Clugston, Anna Dominiczak and Alex Alderton, Roberto Amato, Jeffrey Barrett, Sonia Goncalves, Ewan Harrison, David K. Jackson, Ian Johnston, Dominic Kwiatkowski, Cordelia Langford, John Sillitoe on behalf of the Wellcome Sanger Institute COVID-19 Surveillance Team |
| EPI_ISL_3140052 | Lighthouse Laboratory Plymouth | Wellcome Sanger Institute for the COVID-19 Genomics UK (COG-UK) Consortium | Lighthouse Laboratory Plymouth and Alex Alderton, Roberto Amato, Jeffrey Barrett, Sonia Goncalves, Ewan Harrison, David K. Jackson, Ian Johnston, Dominic Kwiatkowski, Cordelia Langford, John Sillitoe on behalf of the Wellcome Sanger Institute COVID-19 Surveillance Team |
| EPI_ISL_3140053, EPI_ISL_3140054, EPI_ISL_3140055, EPI_ISL_3140056, EPI_ISL_3140057, EPI_ISL_3140058 | Berkshire and Surrey Pathology Services Lighthouse Laboratory | Wellcome Sanger Institute for the COVID-19 Genomics UK (COG-UK) Consortium | Berkshire and Surrey Pathology Services Lighthouse Laboratory and Alex Alderton, Roberto Amato, Jeffrey Barrett, Sonia Goncalves, Ewan Harrison, David K. Jackson, Ian Johnston, Dominic Kwiatkowski, Cordelia Langford, John Sillitoe on behalf of the Wellcome Sanger Institute COVID-19 Surveillance Team |
| EPI_ISL_3140059 | Lighthouse Laboratory Plymouth | Wellcome Sanger Institute for the COVID-19 Genomics UK (COG-UK) Consortium | Lighthouse Laboratory Plymouth and Alex Alderton, Roberto Amato, Jeffrey Barrett, Sonia Goncalves, Ewan Harrison, David K. Jackson, Ian Johnston, Dominic Kwiatkowski, Cordelia Langford, John Sillitoe on behalf of the Wellcome Sanger Institute COVID-19 Surveillance Team |
| EPI_ISL_3140060 | Lighthouse Lab in Glasgow | Wellcome Sanger Institute for the COVID-19 Genomics UK (COG-UK) Consortium | Harper VanSteenhouse, Yumi Kasai, David Gray, Carol Clugston, Anna Dominiczak and Alex Alderton, Roberto Amato, Jeffrey Barrett, Sonia Goncalves, Ewan Harrison, David K. Jackson, Ian Johnston, Dominic Kwiatkowski, Cordelia Langford, John Sillitoe on behalf of the Wellcome Sanger Institute COVID-19 Surveillance Team |
| EPI_ISL_3140061, EPI_ISL_3140062, EPI_ISL_3140063 | Berkshire and Surrey Pathology Services Lighthouse Laboratory | Wellcome Sanger Institute for the COVID-19 Genomics UK (COG-UK) Consortium | Berkshire and Surrey Pathology Services Lighthouse Laboratory and Alex Alderton, Roberto Amato, Jeffrey Barrett, Sonia Goncalves, Ewan Harrison, David K. Jackson, Ian Johnston, Dominic Kwiatkowski, Cordelia Langford, John Sillitoe on behalf of the Wellcome Sanger Institute COVID-19 Surveillance Team |

| | | | |
|---|---|---|---|
| EPI_ISL_3140064 | Lighthouse Laboratory Plymouth | Wellcome Sanger Institute for the COVID-19 Genomics UK (COG-UK) Consortium | Lighthouse Laboratory Plymouth and Alex Alderton, Roberto Amato, Jeffrey Barrett, Sonia Goncalves, Ewan Harrison, David K. Jackson, Ian Johnston, Dominic Kwiatkowski, Cordelia Langford, John Sillitoe on behalf of the Wellcome Sanger Institute COVID-19 Surveillance Team |
| EPI_ISL_3140065 | Lighthouse Lab in Glasgow | Wellcome Sanger Institute for the COVID-19 Genomics UK (COG-UK) Consortium | Harper VanSteenhouse, Yumi Kasai, David Gray, Carol Clugston, Anna Dominiczak and Alex Alderton, Roberto Amato, Jeffrey Barrett, Sonia Goncalves, Ewan Harrison, David K. Jackson, Ian Johnston, Dominic Kwiatkowski, Cordelia Langford, John Sillitoe on behalf of the Wellcome Sanger Institute COVID-19 Surveillance Team |
| EPI_ISL_3140066 | Berkshire and Surrey Pathology Services Lighthouse Laboratory | Wellcome Sanger Institute for the COVID-19 Genomics UK (COG-UK) Consortium | Berkshire and Surrey Pathology Services Lighthouse Laboratory and Alex Alderton, Roberto Amato, Jeffrey Barrett, Sonia Goncalves, Ewan Harrison, David K. Jackson, Ian Johnston, Dominic Kwiatkowski, Cordelia Langford, John Sillitoe on behalf of the Wellcome Sanger Institute COVID-19 Surveillance Team |
| EPI_ISL_3140067 | Lighthouse Laboratory Plymouth | Wellcome Sanger Institute for the COVID-19 Genomics UK (COG-UK) Consortium | Lighthouse Laboratory Plymouth and Alex Alderton, Roberto Amato, Jeffrey Barrett, Sonia Goncalves, Ewan Harrison, David K. Jackson, Ian Johnston, Dominic Kwiatkowski, Cordelia Langford, John Sillitoe on behalf of the Wellcome Sanger Institute COVID-19 Surveillance Team |
| EPI_ISL_3140068, EPI_ISL_3140069 | Berkshire and Surrey Pathology Services Lighthouse Laboratory | Wellcome Sanger Institute for the COVID-19 Genomics UK (COG-UK) Consortium | Berkshire and Surrey Pathology Services Lighthouse Laboratory and Alex Alderton, Roberto Amato, Jeffrey Barrett, Sonia Goncalves, Ewan Harrison, David K. Jackson, Ian Johnston, Dominic Kwiatkowski, Cordelia Langford, John Sillitoe on behalf of the Wellcome Sanger Institute COVID-19 Surveillance Team |
| EPI_ISL_3140070 | Lighthouse Lab in Glasgow | Wellcome Sanger Institute for the COVID-19 Genomics UK (COG-UK) Consortium | Harper VanSteenhouse, Yumi Kasai, David Gray, Carol Clugston, Anna Dominiczak and Alex Alderton, Roberto Amato, Jeffrey Barrett, Sonia Goncalves, Ewan Harrison, David K. Jackson, Ian Johnston, Dominic Kwiatkowski, Cordelia Langford, John Sillitoe on behalf of the Wellcome Sanger Institute COVID-19 Surveillance Team |
| EPI_ISL_3140071 | Lighthouse Laboratory Plymouth | Wellcome Sanger Institute for the COVID-19 Genomics UK (COG-UK) Consortium | Lighthouse Laboratory Plymouth and Alex Alderton, Roberto Amato, Jeffrey Barrett, Sonia Goncalves, Ewan Harrison, David K. Jackson, Ian Johnston, Dominic Kwiatkowski, Cordelia Langford, John Sillitoe on behalf of the Wellcome Sanger Institute COVID-19 Surveillance Team |
| EPI_ISL_3140072 | Lighthouse Lab in Glasgow | Wellcome Sanger Institute for the COVID-19 Genomics UK (COG-UK) Consortium | Harper VanSteenhouse, Yumi Kasai, David Gray, Carol Clugston, Anna Dominiczak and Alex Alderton, Roberto Amato, Jeffrey Barrett, Sonia Goncalves, Ewan Harrison, David K. Jackson, Ian Johnston, Dominic Kwiatkowski, Cordelia Langford, John Sillitoe on behalf of the Wellcome Sanger Institute COVID-19 Surveillance Team |
| EPI_ISL_3140073 | Lighthouse Laboratory Plymouth | Wellcome Sanger Institute for the COVID-19 Genomics UK (COG-UK) Consortium | Lighthouse Laboratory Plymouth and Alex Alderton, Roberto Amato, Jeffrey Barrett, Sonia Goncalves, Ewan Harrison, David K. Jackson, Ian Johnston, Dominic Kwiatkowski, Cordelia Langford, John Sillitoe on behalf of the Wellcome Sanger Institute COVID-19 Surveillance Team |
| EPI_ISL_3140074 | Lighthouse Lab in Glasgow | Wellcome Sanger Institute for the COVID-19 Genomics UK (COG-UK) Consortium | Harper VanSteenhouse, Yumi Kasai, David Gray, Carol Clugston, Anna Dominiczak and Alex Alderton, Roberto Amato, Jeffrey Barrett, Sonia Goncalves, Ewan Harrison, David K. Jackson, Ian Johnston, Dominic Kwiatkowski, Cordelia Langford, John Sillitoe on behalf of the Wellcome Sanger Institute COVID-19 Surveillance Team |
| EPI_ISL_3140075, EPI_ISL_3140076, EPI_ISL_3140077, EPI_ISL_3140078 | Berkshire and Surrey Pathology Services Lighthouse Laboratory | Wellcome Sanger Institute for the COVID-19 Genomics UK (COG-UK) Consortium | Berkshire and Surrey Pathology Services Lighthouse Laboratory and Alex Alderton, Roberto Amato, Jeffrey Barrett, Sonia Goncalves, Ewan Harrison, David K. Jackson, Ian Johnston, Dominic Kwiatkowski, Cordelia Langford, John Sillitoe on behalf of the Wellcome Sanger Institute COVID-19 Surveillance Team |
| EPI_ISL_3140079 | Lighthouse Lab in Glasgow | Wellcome Sanger Institute for the COVID-19 Genomics UK (COG-UK) Consortium | Harper VanSteenhouse, Yumi Kasai, David Gray, Carol Clugston, Anna Dominiczak and Alex Alderton, Roberto Amato, Jeffrey Barrett, Sonia Goncalves, Ewan Harrison, David K. Jackson, Ian Johnston, Dominic Kwiatkowski, Cordelia Langford, John Sillitoe on behalf of the Wellcome Sanger Institute COVID-19 Surveillance Team |
| EPI_ISL_3140080, EPI_ISL_3140081, EPI_ISL_3140082, EPI_ISL_3140083 | Berkshire and Surrey Pathology Services Lighthouse Laboratory | Wellcome Sanger Institute for the COVID-19 Genomics UK (COG-UK) Consortium | Berkshire and Surrey Pathology Services Lighthouse Laboratory and Alex Alderton, Roberto Amato, Jeffrey Barrett, Sonia Goncalves, Ewan Harrison, David K. Jackson, Ian Johnston, Dominic Kwiatkowski, Cordelia Langford, John Sillitoe on behalf of the Wellcome Sanger Institute COVID-19 Surveillance Team |
| EPI_ISL_3140084, EPI_ISL_3140085 | Lighthouse Lab in Glasgow | Wellcome Sanger Institute for the COVID-19 Genomics UK (COG-UK) Consortium | Harper VanSteenhouse, Yumi Kasai, David Gray, Carol Clugston, Anna Dominiczak and Alex Alderton, Roberto Amato, Jeffrey Barrett, Sonia Goncalves, Ewan Harrison, David K. Jackson, Ian Johnston, Dominic Kwiatkowski, Cordelia Langford, John Sillitoe on behalf of the Wellcome Sanger Institute COVID-19 Surveillance Team |
| EPI_ISL_3140086 | Berkshire and Surrey Pathology Services Lighthouse Laboratory | Wellcome Sanger Institute for the COVID-19 Genomics UK (COG-UK) Consortium | Berkshire and Surrey Pathology Services Lighthouse Laboratory and Alex Alderton, Roberto Amato, Jeffrey Barrett, Sonia Goncalves, Ewan Harrison, David K. Jackson, Ian Johnston, Dominic Kwiatkowski, Cordelia Langford, John Sillitoe on behalf of the Wellcome Sanger Institute COVID-19 Surveillance Team |
| EPI_ISL_3140087, EPI_ISL_3140088 | Lighthouse Lab in Glasgow | Wellcome Sanger Institute for the COVID-19 Genomics UK (COG-UK) Consortium | Harper VanSteenhouse, Yumi Kasai, David Gray, Carol Clugston, Anna Dominiczak and Alex Alderton, Roberto Amato, Jeffrey Barrett, Sonia Goncalves, Ewan Harrison, David K. Jackson, Ian Johnston, Dominic Kwiatkowski, Cordelia Langford, John Sillitoe on behalf of the Wellcome Sanger Institute COVID-19 Surveillance Team |
| EPI_ISL_3140089 | Lighthouse Laboratory Plymouth | Wellcome Sanger Institute for the COVID-19 Genomics UK (COG-UK) Consortium | Lighthouse Laboratory Plymouth and Alex Alderton, Roberto Amato, Jeffrey Barrett, Sonia Goncalves, Ewan Harrison, David K. Jackson, Ian Johnston, Dominic Kwiatkowski, Cordelia Langford, John Sillitoe on behalf of the Wellcome Sanger Institute COVID-19 Surveillance Team |
| EPI_ISL_3140090 | Berkshire and Surrey Pathology Services Lighthouse Laboratory | Wellcome Sanger Institute for the COVID-19 Genomics UK (COG-UK) Consortium | Berkshire and Surrey Pathology Services Lighthouse Laboratory and Alex Alderton, Roberto Amato, Jeffrey Barrett, Sonia Goncalves, Ewan Harrison, David K. Jackson, Ian Johnston, Dominic Kwiatkowski, Cordelia Langford, John Sillitoe on behalf of the Wellcome Sanger Institute COVID-19 Surveillance Team |
| EPI_ISL_3140091 | Lighthouse Lab in Glasgow | Wellcome Sanger Institute for the COVID-19 Genomics UK (COG-UK) Consortium | Harper VanSteenhouse, Yumi Kasai, David Gray, Carol Clugston, Anna Dominiczak and Alex Alderton, Roberto Amato, Jeffrey Barrett, Sonia Goncalves, Ewan Harrison, David K. Jackson, Ian Johnston, Dominic Kwiatkowski, Cordelia Langford, John Sillitoe on behalf of the Wellcome Sanger Institute COVID-19 Surveillance Team |
| EPI_ISL_3140092 | Lighthouse Laboratory Plymouth | Wellcome Sanger Institute for the COVID-19 Genomics UK (COG-UK) Consortium | Lighthouse Laboratory Plymouth and Alex Alderton, Roberto Amato, Jeffrey Barrett, Sonia Goncalves, Ewan Harrison, David K. Jackson, Ian Johnston, Dominic Kwiatkowski, Cordelia Langford, John Sillitoe on behalf of the Wellcome Sanger Institute COVID-19 Surveillance Team |
| EPI_ISL_3140093 | Lighthouse Lab in Glasgow | Wellcome Sanger Institute for the COVID-19 Genomics UK (COG-UK) Consortium | Harper VanSteenhouse, Yumi Kasai, David Gray, Carol Clugston, Anna Dominiczak and Alex Alderton, Roberto Amato, Jeffrey Barrett, Sonia Goncalves, Ewan Harrison, David K. Jackson, Ian Johnston, Dominic Kwiatkowski, Cordelia Langford, John Sillitoe on behalf of the Wellcome Sanger Institute COVID-19 Surveillance Team |
| EPI_ISL_3140094, EPI_ISL_3140095 | Berkshire and Surrey Pathology Services Lighthouse Laboratory | Wellcome Sanger Institute for the COVID-19 Genomics UK (COG-UK) Consortium | Berkshire and Surrey Pathology Services Lighthouse Laboratory and Alex Alderton, Roberto Amato, Jeffrey Barrett, Sonia Goncalves, Ewan Harrison, David K. Jackson, Ian Johnston, Dominic Kwiatkowski, Cordelia Langford, John Sillitoe on behalf of the Wellcome Sanger Institute COVID-19 Surveillance Team |
| EPI_ISL_3140096, EPI_ISL_3140097 | Lighthouse Lab in Glasgow | Wellcome Sanger Institute for the COVID-19 Genomics UK (COG-UK) Consortium | Harper VanSteenhouse, Yumi Kasai, David Gray, Carol Clugston, Anna Dominiczak and Alex Alderton, Roberto Amato, Jeffrey Barrett, Sonia Goncalves, Ewan Harrison, David K. Jackson, Ian Johnston, Dominic Kwiatkowski, Cordelia Langford, John Sillitoe on behalf of the Wellcome Sanger Institute COVID-19 Surveillance Team |
| EPI_ISL_3140098, EPI_ISL_3140099, EPI_ISL_3140100, EPI_ISL_3140101 | Lighthouse Laboratory Plymouth | Wellcome Sanger Institute for the COVID-19 Genomics UK (COG-UK) Consortium | Lighthouse Laboratory Plymouth and Alex Alderton, Roberto Amato, Jeffrey Barrett, Sonia Goncalves, Ewan Harrison, David K. Jackson, Ian Johnston, Dominic Kwiatkowski, Cordelia Langford, John Sillitoe on behalf of the Wellcome Sanger Institute COVID-19 Surveillance Team |
| EPI_ISL_3140102 | Berkshire and Surrey Pathology Services Lighthouse Laboratory | Wellcome Sanger Institute for the COVID-19 Genomics UK (COG-UK) Consortium | Berkshire and Surrey Pathology Services Lighthouse Laboratory and Alex Alderton, Roberto Amato, Jeffrey Barrett, Sonia Goncalves, Ewan Harrison, David K. Jackson, Ian Johnston, Dominic Kwiatkowski, Cordelia Langford, John Sillitoe on behalf of the Wellcome Sanger Institute COVID-19 Surveillance Team |
| EPI_ISL_3140103, EPI_ISL_3140104 | Lighthouse Laboratory Plymouth | Wellcome Sanger Institute for the COVID-19 Genomics UK (COG-UK) Consortium | Lighthouse Laboratory Plymouth and Alex Alderton, Roberto Amato, Jeffrey Barrett, Sonia Goncalves, Ewan Harrison, David K. Jackson, Ian Johnston, Dominic Kwiatkowski, Cordelia Langford, John Sillitoe on behalf of the Wellcome Sanger Institute COVID-19 Surveillance Team |
| EPI_ISL_3140105 | Lighthouse Lab in Glasgow | Wellcome Sanger Institute for the COVID-19 Genomics UK (COG-UK) Consortium | Harper VanSteenhouse, Yumi Kasai, David Gray, Carol Clugston, Anna Dominiczak and Alex Alderton, Roberto Amato, Jeffrey Barrett, Sonia Goncalves, Ewan Harrison, David K. Jackson, Ian Johnston, Dominic Kwiatkowski, Cordelia Langford, John Sillitoe on behalf of the Wellcome Sanger Institute COVID-19 Surveillance Team |
| EPI_ISL_3140106, EPI_ISL_3140107 | Berkshire and Surrey Pathology Services Lighthouse Laboratory | Wellcome Sanger Institute for the COVID-19 Genomics UK (COG-UK) Consortium | Berkshire and Surrey Pathology Services Lighthouse Laboratory and Alex Alderton, Roberto Amato, Jeffrey Barrett, Sonia Goncalves, Ewan Harrison, David K. Jackson, Ian Johnston, Dominic Kwiatkowski, Cordelia Langford, John Sillitoe on behalf of the Wellcome Sanger Institute COVID-19 Surveillance Team |
| EPI_ISL_3140108, EPI_ISL_3140109, EPI_ISL_3140110 | Lighthouse Lab in Glasgow | Wellcome Sanger Institute for the COVID-19 Genomics UK (COG-UK) Consortium | Harper VanSteenhouse, Yumi Kasai, David Gray, Carol Clugston, Anna Dominiczak and Alex Alderton, Roberto Amato, Jeffrey Barrett, Sonia Goncalves, Ewan Harrison, David K. Jackson, Ian Johnston, Dominic Kwiatkowski, Cordelia Langford, John Sillitoe on behalf of the Wellcome Sanger Institute COVID-19 Surveillance Team |
| EPI_ISL_3140111 | Berkshire and Surrey Pathology Services Lighthouse Laboratory | Wellcome Sanger Institute for the COVID-19 Genomics UK (COG-UK) Consortium | Berkshire and Surrey Pathology Services Lighthouse Laboratory and Alex Alderton, Roberto Amato, Jeffrey Barrett, Sonia Goncalves, Ewan Harrison, David K. Jackson, Ian Johnston, Dominic Kwiatkowski, Cordelia Langford, John Sillitoe on behalf of the Wellcome Sanger Institute COVID-19 Surveillance Team |
| EPI_ISL_3140112 | Lighthouse Lab in Glasgow | Wellcome Sanger Institute for the COVID-19 Genomics UK (COG-UK) Consortium | Harper VanSteenhouse, Yumi Kasai, David Gray, Carol Clugston, Anna Dominiczak and Alex Alderton, Roberto Amato, Jeffrey Barrett, Sonia Goncalves, Ewan Harrison, David K. Jackson, Ian Johnston, Dominic Kwiatkowski, Cordelia Langford, John Sillitoe on behalf of the Wellcome Sanger Institute COVID-19 Surveillance Team |
| EPI_ISL_3140113 | Berkshire and Surrey Pathology Services Lighthouse Laboratory | Wellcome Sanger Institute for the COVID-19 Genomics UK (COG-UK) Consortium | Berkshire and Surrey Pathology Services Lighthouse Laboratory and Alex Alderton, Roberto Amato, Jeffrey Barrett, Sonia Goncalves, Ewan Harrison, David K. Jackson, Ian Johnston, Dominic Kwiatkowski, Cordelia Langford, John Sillitoe on behalf of the Wellcome Sanger Institute COVID-19 Surveillance Team |
| EPI_ISL_3140114 | Lighthouse Laboratory Plymouth | Wellcome Sanger Institute for the COVID-19 Genomics UK (COG-UK) Consortium | Lighthouse Laboratory Plymouth and Alex Alderton, Roberto Amato, Jeffrey Barrett, Sonia Goncalves, Ewan Harrison, David K. Jackson, Ian Johnston, Dominic Kwiatkowski, Cordelia Langford, John Sillitoe on behalf of the Wellcome Sanger Institute COVID-19 Surveillance Team |
| EPI_ISL_3140115, EPI_ISL_3140116 | Berkshire and Surrey Pathology Services Lighthouse Laboratory | Wellcome Sanger Institute for the COVID-19 Genomics UK (COG-UK) Consortium | Berkshire and Surrey Pathology Services Lighthouse Laboratory and Alex Alderton, Roberto Amato, Jeffrey Barrett, Sonia Goncalves, Ewan Harrison, David K. Jackson, Ian Johnston, Dominic Kwiatkowski, Cordelia Langford, John Sillitoe on behalf of the Wellcome Sanger Institute COVID-19 Surveillance Team |
| EPI_ISL_3140117 | Lighthouse Lab in Glasgow | Wellcome Sanger Institute for the COVID-19 Genomics UK (COG-UK) Consortium | Harper VanSteenhouse, Yumi Kasai, David Gray, Carol Clugston, Anna Dominiczak and Alex Alderton, Roberto Amato, Jeffrey Barrett, Sonia Goncalves, Ewan Harrison, David K. Jackson, Ian Johnston, Dominic Kwiatkowski, Cordelia Langford, John Sillitoe on behalf of the Wellcome Sanger Institute COVID-19 Surveillance Team |
| EPI_ISL_3140118, EPI_ISL_3140119 | Lighthouse Laboratory Plymouth | Wellcome Sanger Institute for the COVID-19 Genomics UK (COG-UK) Consortium | Lighthouse Laboratory Plymouth and Alex Alderton, Roberto Amato, Jeffrey Barrett, Sonia Goncalves, Ewan Harrison, David K. Jackson, Ian Johnston, Dominic Kwiatkowski, Cordelia Langford, John Sillitoe on behalf of the Wellcome Sanger Institute COVID-19 Surveillance Team |
| EPI_ISL_3140120 | Berkshire and Surrey Pathology Services Lighthouse Laboratory | Wellcome Sanger Institute for the COVID-19 Genomics UK (COG-UK) Consortium | Berkshire and Surrey Pathology Services Lighthouse Laboratory and Alex Alderton, Roberto Amato, Jeffrey Barrett, Sonia Goncalves, Ewan Harrison, David K. Jackson, Ian Johnston, Dominic Kwiatkowski, Cordelia Langford, John Sillitoe on behalf of the Wellcome Sanger Institute COVID-19 Surveillance Team |
| EPI_ISL_3140121 | Lighthouse Laboratory Plymouth | Wellcome Sanger Institute for the COVID-19 Genomics UK (COG-UK) Consortium | Lighthouse Laboratory Plymouth and Alex Alderton, Roberto Amato, Jeffrey Barrett, Sonia Goncalves, Ewan Harrison, David K. Jackson, Ian Johnston, Dominic Kwiatkowski, Cordelia Langford, John Sillitoe on behalf of the Wellcome Sanger Institute COVID-19 Surveillance Team |
| EPI_ISL_3140122 | Berkshire and Surrey Pathology Services Lighthouse Laboratory | Wellcome Sanger Institute for the COVID-19 Genomics UK (COG-UK) Consortium | Berkshire and Surrey Pathology Services Lighthouse Laboratory and Alex Alderton, Roberto Amato, Jeffrey Barrett, Sonia Goncalves, Ewan Harrison, David K. Jackson, Ian Johnston, Dominic Kwiatkowski, Cordelia Langford, John Sillitoe on behalf of the Wellcome Sanger Institute COVID-19 Surveillance Team |
| EPI_ISL_3140123, EPI_ISL_3140124 | Lighthouse Laboratory Plymouth | Wellcome Sanger Institute for the COVID-19 Genomics UK (COG-UK) Consortium | Lighthouse Laboratory Plymouth and Alex Alderton, Roberto Amato, Jeffrey Barrett, Sonia Goncalves, Ewan Harrison, David K. Jackson, Ian Johnston, Dominic Kwiatkowski, Cordelia Langford, John Sillitoe on behalf of the Wellcome Sanger Institute COVID-19 Surveillance Team |
| EPI_ISL_3140125, EPI_ISL_3140126 | Berkshire and Surrey Pathology Services Lighthouse Laboratory | Wellcome Sanger Institute for the COVID-19 Genomics UK (COG-UK) Consortium | Berkshire and Surrey Pathology Services Lighthouse Laboratory and Alex Alderton, Roberto Amato, Jeffrey Barrett, Sonia Goncalves, Ewan Harrison, David K. Jackson, Ian Johnston, Dominic Kwiatkowski, Cordelia Langford, John Sillitoe on behalf of the Wellcome Sanger Institute COVID-19 Surveillance Team |
| EPI_ISL_3140127, EPI_ISL_3140128 | Lighthouse Laboratory Plymouth | Wellcome Sanger Institute for the COVID-19 Genomics UK (COG-UK) Consortium | Lighthouse Laboratory Plymouth and Alex Alderton, Roberto Amato, Jeffrey Barrett, Sonia Goncalves, Ewan Harrison, David K. Jackson, Ian Johnston, Dominic Kwiatkowski, Cordelia Langford, John Sillitoe on behalf of the Wellcome Sanger Institute COVID-19 Surveillance Team |
| EPI_ISL_3140129 | Berkshire and Surrey Pathology Services Lighthouse Laboratory | Wellcome Sanger Institute for the COVID-19 Genomics UK (COG-UK) | Berkshire and Surrey Pathology Services Lighthouse Laboratory and Alex Alderton, Roberto Amato, Jeffrey Barrett, Sonia Goncalves, Ewan Harrison, David K. Jackson, Ian Johnston, Dominic Kwiatkowski, Cordelia Langford, John Sillitoe on behalf |

| | | | |
|---|---|---|---|
| | | Consortium | of the Wellcome Sanger Institute COVID-19 Surveillance Team |
| EPI_ISL_3140130, EPI_ISL_3140131 | Lighthouse Laboratory Plymouth | Wellcome Sanger Institute for the COVID-19 Genomics UK (COG-UK) Consortium | Lighthouse Laboratory Plymouth and Alex Alderton, Roberto Amato, Jeffrey Barrett, Sonia Goncalves, Ewan Harrison, David K. Jackson, Ian Johnston, Dominic Kwiatkowski, Cordelia Langford, John Sillitoe on behalf of the Wellcome Sanger Institute COVID-19 Surveillance Team |
| EPI_ISL_3140132, EPI_ISL_3140133, EPI_ISL_3140134 | Lighthouse Lab in Glasgow | Wellcome Sanger Institute for the COVID-19 Genomics UK (COG-UK) Consortium | Harper VanSteenhouse, Yumi Kasai, David Gray, Carol Clugston, Anna Dominiczak and Alex Alderton, Roberto Amato, Jeffrey Barrett, Sonia Goncalves, Ewan Harrison, David K. Jackson, Ian Johnston, Dominic Kwiatkowski, Cordelia Langford, John Sillitoe on behalf of the Wellcome Sanger Institute COVID-19 Surveillance Team |
| EPI_ISL_3140135 | Lighthouse Laboratory Plymouth | Wellcome Sanger Institute for the COVID-19 Genomics UK (COG-UK) Consortium | Lighthouse Laboratory Plymouth and Alex Alderton, Roberto Amato, Jeffrey Barrett, Sonia Goncalves, Ewan Harrison, David K. Jackson, Ian Johnston, Dominic Kwiatkowski, Cordelia Langford, John Sillitoe on behalf of the Wellcome Sanger Institute COVID-19 Surveillance Team |
| EPI_ISL_3140136, EPI_ISL_3140137 | Lighthouse Lab in Glasgow | Wellcome Sanger Institute for the COVID-19 Genomics UK (COG-UK) Consortium | Harper VanSteenhouse, Yumi Kasai, David Gray, Carol Clugston, Anna Dominiczak and Alex Alderton, Roberto Amato, Jeffrey Barrett, Sonia Goncalves, Ewan Harrison, David K. Jackson, Ian Johnston, Dominic Kwiatkowski, Cordelia Langford, John Sillitoe on behalf of the Wellcome Sanger Institute COVID-19 Surveillance Team |
| EPI_ISL_3140138 | Lighthouse Laboratory Plymouth | Wellcome Sanger Institute for the COVID-19 Genomics UK (COG-UK) Consortium | Lighthouse Laboratory Plymouth and Alex Alderton, Roberto Amato, Jeffrey Barrett, Sonia Goncalves, Ewan Harrison, David K. Jackson, Ian Johnston, Dominic Kwiatkowski, Cordelia Langford, John Sillitoe on behalf of the Wellcome Sanger Institute COVID-19 Surveillance Team |
| EPI_ISL_3140139, EPI_ISL_3140140 | Berkshire and Surrey Pathology Services Lighthouse Laboratory | Wellcome Sanger Institute for the COVID-19 Genomics UK (COG-UK) Consortium | Berkshire and Surrey Pathology Services Lighthouse Laboratory and Alex Alderton, Roberto Amato, Jeffrey Barrett, Sonia Goncalves, Ewan Harrison, David K. Jackson, Ian Johnston, Dominic Kwiatkowski, Cordelia Langford, John Sillitoe on behalf of the Wellcome Sanger Institute COVID-19 Surveillance Team |
| EPI_ISL_3140141 | Lighthouse Lab in Glasgow | Wellcome Sanger Institute for the COVID-19 Genomics UK (COG-UK) Consortium | Harper VanSteenhouse, Yumi Kasai, David Gray, Carol Clugston, Anna Dominiczak and Alex Alderton, Roberto Amato, Jeffrey Barrett, Sonia Goncalves, Ewan Harrison, David K. Jackson, Ian Johnston, Dominic Kwiatkowski, Cordelia Langford, John Sillitoe on behalf of the Wellcome Sanger Institute COVID-19 Surveillance Team |
| EPI_ISL_3140142, EPI_ISL_3140143, EPI_ISL_3140144, EPI_ISL_3140145 | Berkshire and Surrey Pathology Services Lighthouse Laboratory | Wellcome Sanger Institute for the COVID-19 Genomics UK (COG-UK) Consortium | Berkshire and Surrey Pathology Services Lighthouse Laboratory and Alex Alderton, Roberto Amato, Jeffrey Barrett, Sonia Goncalves, Ewan Harrison, David K. Jackson, Ian Johnston, Dominic Kwiatkowski, Cordelia Langford, John Sillitoe on behalf of the Wellcome Sanger Institute COVID-19 Surveillance Team |
| EPI_ISL_3140146 | Lighthouse Lab in Glasgow | Wellcome Sanger Institute for the COVID-19 Genomics UK (COG-UK) Consortium | Harper VanSteenhouse, Yumi Kasai, David Gray, Carol Clugston, Anna Dominiczak and Alex Alderton, Roberto Amato, Jeffrey Barrett, Sonia Goncalves, Ewan Harrison, David K. Jackson, Ian Johnston, Dominic Kwiatkowski, Cordelia Langford, John Sillitoe on behalf of the Wellcome Sanger Institute COVID-19 Surveillance Team |
| EPI_ISL_3140147 | Berkshire and Surrey Pathology Services Lighthouse Laboratory | Wellcome Sanger Institute for the COVID-19 Genomics UK (COG-UK) Consortium | Berkshire and Surrey Pathology Services Lighthouse Laboratory and Alex Alderton, Roberto Amato, Jeffrey Barrett, Sonia Goncalves, Ewan Harrison, David K. Jackson, Ian Johnston, Dominic Kwiatkowski, Cordelia Langford, John Sillitoe on behalf of the Wellcome Sanger Institute COVID-19 Surveillance Team |
| EPI_ISL_3140148 | Lighthouse Laboratory Plymouth | Wellcome Sanger Institute for the COVID-19 Genomics UK (COG-UK) Consortium | Lighthouse Laboratory Plymouth and Alex Alderton, Roberto Amato, Jeffrey Barrett, Sonia Goncalves, Ewan Harrison, David K. Jackson, Ian Johnston, Dominic Kwiatkowski, Cordelia Langford, John Sillitoe on behalf of the Wellcome Sanger Institute COVID-19 Surveillance Team |
| EPI_ISL_3140149, EPI_ISL_3140150 | Berkshire and Surrey Pathology Services Lighthouse Laboratory | Wellcome Sanger Institute for the COVID-19 Genomics UK (COG-UK) Consortium | Berkshire and Surrey Pathology Services Lighthouse Laboratory and Alex Alderton, Roberto Amato, Jeffrey Barrett, Sonia Goncalves, Ewan Harrison, David K. Jackson, Ian Johnston, Dominic Kwiatkowski, Cordelia Langford, John Sillitoe on behalf of the Wellcome Sanger Institute COVID-19 Surveillance Team |
| EPI_ISL_3140151, EPI_ISL_3140152, EPI_ISL_3140153 | Lighthouse Lab in Glasgow | Wellcome Sanger Institute for the COVID-19 Genomics UK (COG-UK) Consortium | Harper VanSteenhouse, Yumi Kasai, David Gray, Carol Clugston, Anna Dominiczak and Alex Alderton, Roberto Amato, Jeffrey Barrett, Sonia Goncalves, Ewan Harrison, David K. Jackson, Ian Johnston, Dominic Kwiatkowski, Cordelia Langford, John Sillitoe on behalf of the Wellcome Sanger Institute COVID-19 Surveillance Team |
| EPI_ISL_3140154 | Lighthouse Laboratory Plymouth | Wellcome Sanger Institute for the COVID-19 Genomics UK (COG-UK) Consortium | Lighthouse Laboratory Plymouth and Alex Alderton, Roberto Amato, Jeffrey Barrett, Sonia Goncalves, Ewan Harrison, David K. Jackson, Ian Johnston, Dominic Kwiatkowski, Cordelia Langford, John Sillitoe on behalf of the Wellcome Sanger Institute COVID-19 Surveillance Team |
| EPI_ISL_3140155, EPI_ISL_3140156 | Lighthouse Lab in Glasgow | Wellcome Sanger Institute for the COVID-19 Genomics UK (COG-UK) Consortium | Harper VanSteenhouse, Yumi Kasai, David Gray, Carol Clugston, Anna Dominiczak and Alex Alderton, Roberto Amato, Jeffrey Barrett, Sonia Goncalves, Ewan Harrison, David K. Jackson, Ian Johnston, Dominic Kwiatkowski, Cordelia Langford, John Sillitoe on behalf of the Wellcome Sanger Institute COVID-19 Surveillance Team |
| EPI_ISL_3140157 | Lighthouse Laboratory Plymouth | Wellcome Sanger Institute for the COVID-19 Genomics UK (COG-UK) Consortium | Lighthouse Laboratory Plymouth and Alex Alderton, Roberto Amato, Jeffrey Barrett, Sonia Goncalves, Ewan Harrison, David K. Jackson, Ian Johnston, Dominic Kwiatkowski, Cordelia Langford, John Sillitoe on behalf of the Wellcome Sanger Institute COVID-19 Surveillance Team |
| EPI_ISL_3140158 | Lighthouse Lab in Glasgow | Wellcome Sanger Institute for the COVID-19 Genomics UK (COG-UK) Consortium | Harper VanSteenhouse, Yumi Kasai, David Gray, Carol Clugston, Anna Dominiczak and Alex Alderton, Roberto Amato, Jeffrey Barrett, Sonia Goncalves, Ewan Harrison, David K. Jackson, Ian Johnston, Dominic Kwiatkowski, Cordelia Langford, John Sillitoe on behalf of the Wellcome Sanger Institute COVID-19 Surveillance Team |
| EPI_ISL_3140159 | Lighthouse Laboratory Plymouth | Wellcome Sanger Institute for the COVID-19 Genomics UK (COG-UK) Consortium | Lighthouse Laboratory Plymouth and Alex Alderton, Roberto Amato, Jeffrey Barrett, Sonia Goncalves, Ewan Harrison, David K. Jackson, Ian Johnston, Dominic Kwiatkowski, Cordelia Langford, John Sillitoe on behalf of the Wellcome Sanger Institute COVID-19 Surveillance Team |
| EPI_ISL_3140160 | Berkshire and Surrey Pathology Services Lighthouse Laboratory | Wellcome Sanger Institute for the COVID-19 Genomics UK (COG-UK) Consortium | Berkshire and Surrey Pathology Services Lighthouse Laboratory and Alex Alderton, Roberto Amato, Jeffrey Barrett, Sonia Goncalves, Ewan Harrison, David K. Jackson, Ian Johnston, Dominic Kwiatkowski, Cordelia Langford, John Sillitoe on behalf of the Wellcome Sanger Institute COVID-19 Surveillance Team |
| EPI_ISL_3140161 | Lighthouse Laboratory Plymouth | Wellcome Sanger Institute for the COVID-19 Genomics UK (COG-UK) Consortium | Lighthouse Laboratory Plymouth and Alex Alderton, Roberto Amato, Jeffrey Barrett, Sonia Goncalves, Ewan Harrison, David K. Jackson, Ian Johnston, Dominic Kwiatkowski, Cordelia Langford, John Sillitoe on behalf of the Wellcome Sanger Institute COVID-19 Surveillance Team |
| EPI_ISL_3140162 | Lighthouse Lab in Glasgow | Wellcome Sanger Institute for the COVID-19 Genomics UK (COG-UK) Consortium | Harper VanSteenhouse, Yumi Kasai, David Gray, Carol Clugston, Anna Dominiczak and Alex Alderton, Roberto Amato, Jeffrey Barrett, Sonia Goncalves, Ewan Harrison, David K. Jackson, Ian Johnston, Dominic Kwiatkowski, Cordelia Langford, John Sillitoe on behalf of the Wellcome Sanger Institute COVID-19 Surveillance Team |
| EPI_ISL_3140163, EPI_ISL_3140164 | Berkshire and Surrey Pathology Services Lighthouse Laboratory | Wellcome Sanger Institute for the COVID-19 Genomics UK (COG-UK) Consortium | Berkshire and Surrey Pathology Services Lighthouse Laboratory and Alex Alderton, Roberto Amato, Jeffrey Barrett, Sonia Goncalves, Ewan Harrison, David K. Jackson, Ian Johnston, Dominic Kwiatkowski, Cordelia Langford, John Sillitoe on behalf of the Wellcome Sanger Institute COVID-19 Surveillance Team |
| EPI_ISL_3140165, EPI_ISL_3140166 | Lighthouse Lab in Glasgow | Wellcome Sanger Institute for the COVID-19 Genomics UK (COG-UK) Consortium | Harper VanSteenhouse, Yumi Kasai, David Gray, Carol Clugston, Anna Dominiczak and Alex Alderton, Roberto Amato, Jeffrey Barrett, Sonia Goncalves, Ewan Harrison, David K. Jackson, Ian Johnston, Dominic Kwiatkowski, Cordelia Langford, John Sillitoe on behalf of the Wellcome Sanger Institute COVID-19 Surveillance Team |
| EPI_ISL_3140167 | Lighthouse Laboratory Plymouth | Wellcome Sanger Institute for the COVID-19 Genomics UK (COG-UK) Consortium | Lighthouse Laboratory Plymouth and Alex Alderton, Roberto Amato, Jeffrey Barrett, Sonia Goncalves, Ewan Harrison, David K. Jackson, Ian Johnston, Dominic Kwiatkowski, Cordelia Langford, John Sillitoe on behalf of the Wellcome Sanger Institute COVID-19 Surveillance Team |
| EPI_ISL_3140168 | Berkshire and Surrey Pathology Services Lighthouse Laboratory | Wellcome Sanger Institute for the COVID-19 Genomics UK (COG-UK) Consortium | Berkshire and Surrey Pathology Services Lighthouse Laboratory and Alex Alderton, Roberto Amato, Jeffrey Barrett, Sonia Goncalves, Ewan Harrison, David K. Jackson, Ian Johnston, Dominic Kwiatkowski, Cordelia Langford, John Sillitoe on behalf of the Wellcome Sanger Institute COVID-19 Surveillance Team |
| EPI_ISL_3140169 | Lighthouse Lab in Glasgow | Wellcome Sanger Institute for the COVID-19 Genomics UK (COG-UK) Consortium | Harper VanSteenhouse, Yumi Kasai, David Gray, Carol Clugston, Anna Dominiczak and Alex Alderton, Roberto Amato, Jeffrey Barrett, Sonia Goncalves, Ewan Harrison, David K. Jackson, Ian Johnston, Dominic Kwiatkowski, Cordelia Langford, John Sillitoe on behalf of the Wellcome Sanger Institute COVID-19 Surveillance Team |
| EPI_ISL_3140170 | Berkshire and Surrey Pathology Services Lighthouse Laboratory | Wellcome Sanger Institute for the COVID-19 Genomics UK (COG-UK) Consortium | Berkshire and Surrey Pathology Services Lighthouse Laboratory and Alex Alderton, Roberto Amato, Jeffrey Barrett, Sonia Goncalves, Ewan Harrison, David K. Jackson, Ian Johnston, Dominic Kwiatkowski, Cordelia Langford, John Sillitoe on behalf of the Wellcome Sanger Institute COVID-19 Surveillance Team |
| EPI_ISL_3140171 | Lighthouse Lab in Glasgow | Wellcome Sanger Institute for the COVID-19 Genomics UK (COG-UK) Consortium | Harper VanSteenhouse, Yumi Kasai, David Gray, Carol Clugston, Anna Dominiczak and Alex Alderton, Roberto Amato, Jeffrey Barrett, Sonia Goncalves, Ewan Harrison, David K. Jackson, Ian Johnston, Dominic Kwiatkowski, Cordelia Langford, John Sillitoe on behalf of the Wellcome Sanger Institute COVID-19 Surveillance Team |
| EPI_ISL_3140172, EPI_ISL_3140173 | Lighthouse Laboratory Plymouth | Wellcome Sanger Institute for the COVID-19 Genomics UK (COG-UK) Consortium | Lighthouse Laboratory Plymouth and Alex Alderton, Roberto Amato, Jeffrey Barrett, Sonia Goncalves, Ewan Harrison, David K. Jackson, Ian Johnston, Dominic Kwiatkowski, Cordelia Langford, John Sillitoe on behalf of the Wellcome Sanger Institute COVID-19 Surveillance Team |
| EPI_ISL_3140174 | Berkshire and Surrey Pathology Services Lighthouse Laboratory | Wellcome Sanger Institute for the COVID-19 Genomics UK (COG-UK) Consortium | Berkshire and Surrey Pathology Services Lighthouse Laboratory and Alex Alderton, Roberto Amato, Jeffrey Barrett, Sonia Goncalves, Ewan Harrison, David K. Jackson, Ian Johnston, Dominic Kwiatkowski, Cordelia Langford, John Sillitoe on behalf of the Wellcome Sanger Institute COVID-19 Surveillance Team |
| EPI_ISL_3140175, EPI_ISL_3140176 | Lighthouse Lab in Glasgow | Wellcome Sanger Institute for the COVID-19 Genomics UK (COG-UK) Consortium | Harper VanSteenhouse, Yumi Kasai, David Gray, Carol Clugston, Anna Dominiczak and Alex Alderton, Roberto Amato, Jeffrey Barrett, Sonia Goncalves, Ewan Harrison, David K. Jackson, Ian Johnston, Dominic Kwiatkowski, Cordelia Langford, John Sillitoe on behalf of the Wellcome Sanger Institute COVID-19 Surveillance Team |
| EPI_ISL_3140177, EPI_ISL_3140178 | Lighthouse Laboratory Plymouth | Wellcome Sanger Institute for the COVID-19 Genomics UK (COG-UK) Consortium | Lighthouse Laboratory Plymouth and Alex Alderton, Roberto Amato, Jeffrey Barrett, Sonia Goncalves, Ewan Harrison, David K. Jackson, Ian Johnston, Dominic Kwiatkowski, Cordelia Langford, John Sillitoe on behalf of the Wellcome Sanger Institute COVID-19 Surveillance Team |
| EPI_ISL_3140179 | Berkshire and Surrey Pathology Services Lighthouse Laboratory | Wellcome Sanger Institute for the COVID-19 Genomics UK (COG-UK) Consortium | Berkshire and Surrey Pathology Services Lighthouse Laboratory and Alex Alderton, Roberto Amato, Jeffrey Barrett, Sonia Goncalves, Ewan Harrison, David K. Jackson, Ian Johnston, Dominic Kwiatkowski, Cordelia Langford, John Sillitoe on behalf of the Wellcome Sanger Institute COVID-19 Surveillance Team |
| EPI_ISL_3140180, EPI_ISL_3140181 | Lighthouse Lab in Glasgow | Wellcome Sanger Institute for the COVID-19 Genomics UK (COG-UK) Consortium | Harper VanSteenhouse, Yumi Kasai, David Gray, Carol Clugston, Anna Dominiczak and Alex Alderton, Roberto Amato, Jeffrey Barrett, Sonia Goncalves, Ewan Harrison, David K. Jackson, Ian Johnston, Dominic Kwiatkowski, Cordelia Langford, John Sillitoe on behalf of the Wellcome Sanger Institute COVID-19 Surveillance Team |
| EPI_ISL_3140182 | Berkshire and Surrey Pathology Services Lighthouse Laboratory | Wellcome Sanger Institute for the COVID-19 Genomics UK (COG-UK) Consortium | Berkshire and Surrey Pathology Services Lighthouse Laboratory and Alex Alderton, Roberto Amato, Jeffrey Barrett, Sonia Goncalves, Ewan Harrison, David K. Jackson, Ian Johnston, Dominic Kwiatkowski, Cordelia Langford, John Sillitoe on behalf of the Wellcome Sanger Institute COVID-19 Surveillance Team |
| EPI_ISL_3140183 | Lighthouse Laboratory Plymouth | Wellcome Sanger Institute for the COVID-19 Genomics UK (COG-UK) Consortium | Lighthouse Laboratory Plymouth and Alex Alderton, Roberto Amato, Jeffrey Barrett, Sonia Goncalves, Ewan Harrison, David K. Jackson, Ian Johnston, Dominic Kwiatkowski, Cordelia Langford, John Sillitoe on behalf of the Wellcome Sanger Institute COVID-19 Surveillance Team |
| EPI_ISL_3140184 | Lighthouse Lab in Glasgow | Wellcome Sanger Institute for the COVID-19 Genomics UK (COG-UK) Consortium | Harper VanSteenhouse, Yumi Kasai, David Gray, Carol Clugston, Anna Dominiczak and Alex Alderton, Roberto Amato, Jeffrey Barrett, Sonia Goncalves, Ewan Harrison, David K. Jackson, Ian Johnston, Dominic Kwiatkowski, Cordelia Langford, John Sillitoe on behalf of the Wellcome Sanger Institute COVID-19 Surveillance Team |
| EPI_ISL_3140185, EPI_ISL_3140186, EPI_ISL_3140187 | Berkshire and Surrey Pathology Services Lighthouse Laboratory | Wellcome Sanger Institute for the COVID-19 Genomics UK (COG-UK) Consortium | Berkshire and Surrey Pathology Services Lighthouse Laboratory and Alex Alderton, Roberto Amato, Jeffrey Barrett, Sonia Goncalves, Ewan Harrison, David K. Jackson, Ian Johnston, Dominic Kwiatkowski, Cordelia Langford, John Sillitoe on behalf of the Wellcome Sanger Institute COVID-19 Surveillance Team |
| EPI_ISL_3140188, EPI_ISL_3140189 | Lighthouse Lab in Glasgow | Wellcome Sanger Institute for the COVID-19 Genomics UK (COG-UK) Consortium | Harper VanSteenhouse, Yumi Kasai, David Gray, Carol Clugston, Anna Dominiczak and Alex Alderton, Roberto Amato, Jeffrey Barrett, Sonia Goncalves, Ewan Harrison, David K. Jackson, Ian Johnston, Dominic Kwiatkowski, Cordelia Langford, John Sillitoe on behalf of the Wellcome Sanger Institute COVID-19 Surveillance Team |
| EPI_ISL_3140190, EPI_ISL_3140191 | Berkshire and Surrey Pathology Services Lighthouse Laboratory | Wellcome Sanger Institute for the COVID-19 Genomics UK (COG-UK) Consortium | Berkshire and Surrey Pathology Services Lighthouse Laboratory and Alex Alderton, Roberto Amato, Jeffrey Barrett, Sonia Goncalves, Ewan Harrison, David K. Jackson, Ian Johnston, Dominic Kwiatkowski, Cordelia Langford, John Sillitoe on behalf of the Wellcome Sanger Institute COVID-19 Surveillance Team |
| EPI_ISL_3140192 | Lighthouse Laboratory Plymouth | Wellcome Sanger Institute for the COVID-19 Genomics UK (COG-UK) Consortium | Lighthouse Laboratory Plymouth and Alex Alderton, Roberto Amato, Jeffrey Barrett, Sonia Goncalves, Ewan Harrison, David K. Jackson, Ian Johnston, Dominic Kwiatkowski, Cordelia Langford, John Sillitoe on behalf of the Wellcome Sanger Institute COVID-19 Surveillance Team |
| EPI_ISL_3140193 | Lighthouse Lab in Alderley Park | Wellcome Sanger Institute for the COVID-19 Genomics UK (COG-UK) Consortium | Jacquelyn Wynn, Mairead Hyland, The Lighthouse Lab in Alderley Park and Alex Alderton, Roberto Amato, Jeffrey Barrett, Sonia Goncalves, Ewan Harrison, David K. Jackson, Ian Johnston, Dominic Kwiatkowski, Cordelia Langford, John Sillitoe on behalf of the Wellcome Sanger Institute COVID-19 Surveillance Team |
| EPI_ISL_3140194 | Lighthouse Lab in Milton Keynes | Wellcome Sanger Institute for the COVID-19 Genomics UK (COG-UK) | The Lighthouse Lab in Milton Keynes and Alex Alderton, Roberto Amato, Jeffrey Barrett, Sonia Goncalves, Ewan Harrison, David K. Jackson, Ian Johnston, Dominic Kwiatkowski, Cordelia Langford, John Sillitoe on behalf of the Wellcome Sanger |

| | | | |
|---|---|---|---|
| | | Consortium | Institute COVID-19 Surveillance Team |
| EPI_ISL_3140195 | Lighthouse Lab in Glasgow | Wellcome Sanger Institute for the COVID-19 Genomics UK (COG-UK) Consortium | Harper VanSteenhouse, Yumi Kasai, David Gray, Carol Clugston, Anna Dominiczak and Alex Alderton, Roberto Amato, Jeffrey Barrett, Sonia Goncalves, Ewan Harrison, David K. Jackson, Ian Johnston, Dominic Kwiatkowski, Cordelia Langford, John Sillitoe on behalf of the Wellcome Sanger Institute COVID-19 Surveillance Team |
| EPI_ISL_3140196 | Lighthouse Lab in Alderley Park | Wellcome Sanger Institute for the COVID-19 Genomics UK (COG-UK) Consortium | Jacquelyn Wynn, Mairead Hyland, The Lighthouse Lab in Alderley Park and Alex Alderton, Roberto Amato, Jeffrey Barrett, Sonia Goncalves, Ewan Harrison, David K. Jackson, Ian Johnston, Dominic Kwiatkowski, Cordelia Langford, John Sillitoe on behalf of the Wellcome Sanger Institute COVID-19 Surveillance Team |
| EPI_ISL_3140197 | Lighthouse Lab in Milton Keynes | Wellcome Sanger Institute for the COVID-19 Genomics UK (COG-UK) Consortium | The Lighthouse Lab in Milton Keynes and Alex Alderton, Roberto Amato, Jeffrey Barrett, Sonia Goncalves, Ewan Harrison, David K. Jackson, Ian Johnston, Dominic Kwiatkowski, Cordelia Langford, John Sillitoe on behalf of the Wellcome Sanger Institute COVID-19 Surveillance Team |
| EPI_ISL_3140198, EPI_ISL_3140199 | Lighthouse Lab in Glasgow | Wellcome Sanger Institute for the COVID-19 Genomics UK (COG-UK) Consortium | Harper VanSteenhouse, Yumi Kasai, David Gray, Carol Clugston, Anna Dominiczak and Alex Alderton, Roberto Amato, Jeffrey Barrett, Sonia Goncalves, Ewan Harrison, David K. Jackson, Ian Johnston, Dominic Kwiatkowski, Cordelia Langford, John Sillitoe on behalf of the Wellcome Sanger Institute COVID-19 Surveillance Team |
| EPI_ISL_3140200 | Lighthouse Lab in Milton Keynes | Wellcome Sanger Institute for the COVID-19 Genomics UK (COG-UK) Consortium | The Lighthouse Lab in Milton Keynes and Alex Alderton, Roberto Amato, Jeffrey Barrett, Sonia Goncalves, Ewan Harrison, David K. Jackson, Ian Johnston, Dominic Kwiatkowski, Cordelia Langford, John Sillitoe on behalf of the Wellcome Sanger Institute COVID-19 Surveillance Team |
| EPI_ISL_3140201 | Lighthouse Lab in Alderley Park | Wellcome Sanger Institute for the COVID-19 Genomics UK (COG-UK) Consortium | Jacquelyn Wynn, Mairead Hyland, The Lighthouse Lab in Alderley Park and Alex Alderton, Roberto Amato, Jeffrey Barrett, Sonia Goncalves, Ewan Harrison, David K. Jackson, Ian Johnston, Dominic Kwiatkowski, Cordelia Langford, John Sillitoe on behalf of the Wellcome Sanger Institute COVID-19 Surveillance Team |
| EPI_ISL_3140202, EPI_ISL_3140203, EPI_ISL_3140204 | Lighthouse Lab in Milton Keynes | Wellcome Sanger Institute for the COVID-19 Genomics UK (COG-UK) Consortium | The Lighthouse Lab in Milton Keynes and Alex Alderton, Roberto Amato, Jeffrey Barrett, Sonia Goncalves, Ewan Harrison, David K. Jackson, Ian Johnston, Dominic Kwiatkowski, Cordelia Langford, John Sillitoe on behalf of the Wellcome Sanger Institute COVID-19 Surveillance Team |
| EPI_ISL_3140205 | Lighthouse Lab in Alderley Park | Wellcome Sanger Institute for the COVID-19 Genomics UK (COG-UK) Consortium | Jacquelyn Wynn, Mairead Hyland, The Lighthouse Lab in Alderley Park and Alex Alderton, Roberto Amato, Jeffrey Barrett, Sonia Goncalves, Ewan Harrison, David K. Jackson, Ian Johnston, Dominic Kwiatkowski, Cordelia Langford, John Sillitoe on behalf of the Wellcome Sanger Institute COVID-19 Surveillance Team |
| EPI_ISL_3140206 | Lighthouse Lab in Glasgow | Wellcome Sanger Institute for the COVID-19 Genomics UK (COG-UK) Consortium | Harper VanSteenhouse, Yumi Kasai, David Gray, Carol Clugston, Anna Dominiczak and Alex Alderton, Roberto Amato, Jeffrey Barrett, Sonia Goncalves, Ewan Harrison, David K. Jackson, Ian Johnston, Dominic Kwiatkowski, Cordelia Langford, John Sillitoe on behalf of the Wellcome Sanger Institute COVID-19 Surveillance Team |
| EPI_ISL_3140207, EPI_ISL_3140208, EPI_ISL_3140209 | Lighthouse Lab in Milton Keynes | Wellcome Sanger Institute for the COVID-19 Genomics UK (COG-UK) Consortium | The Lighthouse Lab in Milton Keynes and Alex Alderton, Roberto Amato, Jeffrey Barrett, Sonia Goncalves, Ewan Harrison, David K. Jackson, Ian Johnston, Dominic Kwiatkowski, Cordelia Langford, John Sillitoe on behalf of the Wellcome Sanger Institute COVID-19 Surveillance Team |
| EPI_ISL_3140210, EPI_ISL_3140211 | Lighthouse Lab in Glasgow | Wellcome Sanger Institute for the COVID-19 Genomics UK (COG-UK) Consortium | Harper VanSteenhouse, Yumi Kasai, David Gray, Carol Clugston, Anna Dominiczak and Alex Alderton, Roberto Amato, Jeffrey Barrett, Sonia Goncalves, Ewan Harrison, David K. Jackson, Ian Johnston, Dominic Kwiatkowski, Cordelia Langford, John Sillitoe on behalf of the Wellcome Sanger Institute COVID-19 Surveillance Team |
| EPI_ISL_3140212 | Lighthouse Lab in Alderley Park | Wellcome Sanger Institute for the COVID-19 Genomics UK (COG-UK) Consortium | Jacquelyn Wynn, Mairead Hyland, The Lighthouse Lab in Alderley Park and Alex Alderton, Roberto Amato, Jeffrey Barrett, Sonia Goncalves, Ewan Harrison, David K. Jackson, Ian Johnston, Dominic Kwiatkowski, Cordelia Langford, John Sillitoe on behalf of the Wellcome Sanger Institute COVID-19 Surveillance Team |
| EPI_ISL_3140213, EPI_ISL_3140214 | Lighthouse Lab in Milton Keynes | Wellcome Sanger Institute for the COVID-19 Genomics UK (COG-UK) Consortium | The Lighthouse Lab in Milton Keynes and Alex Alderton, Roberto Amato, Jeffrey Barrett, Sonia Goncalves, Ewan Harrison, David K. Jackson, Ian Johnston, Dominic Kwiatkowski, Cordelia Langford, John Sillitoe on behalf of the Wellcome Sanger Institute COVID-19 Surveillance Team |
| EPI_ISL_3140215 | Lighthouse Lab in Glasgow | Wellcome Sanger Institute for the COVID-19 Genomics UK (COG-UK) Consortium | Harper VanSteenhouse, Yumi Kasai, David Gray, Carol Clugston, Anna Dominiczak and Alex Alderton, Roberto Amato, Jeffrey Barrett, Sonia Goncalves, Ewan Harrison, David K. Jackson, Ian Johnston, Dominic Kwiatkowski, Cordelia Langford, John Sillitoe on behalf of the Wellcome Sanger Institute COVID-19 Surveillance Team |
| EPI_ISL_3140216 | Lighthouse Lab in Milton Keynes | Wellcome Sanger Institute for the COVID-19 Genomics UK (COG-UK) Consortium | The Lighthouse Lab in Milton Keynes and Alex Alderton, Roberto Amato, Jeffrey Barrett, Sonia Goncalves, Ewan Harrison, David K. Jackson, Ian Johnston, Dominic Kwiatkowski, Cordelia Langford, John Sillitoe on behalf of the Wellcome Sanger Institute COVID-19 Surveillance Team |
| EPI_ISL_3140217 | Lighthouse Lab in Alderley Park | Wellcome Sanger Institute for the COVID-19 Genomics UK (COG-UK) Consortium | Jacquelyn Wynn, Mairead Hyland, The Lighthouse Lab in Alderley Park and Alex Alderton, Roberto Amato, Jeffrey Barrett, Sonia Goncalves, Ewan Harrison, David K. Jackson, Ian Johnston, Dominic Kwiatkowski, Cordelia Langford, John Sillitoe on behalf of the Wellcome Sanger Institute COVID-19 Surveillance Team |
| EPI_ISL_3140218 | Lighthouse Lab in Glasgow | Wellcome Sanger Institute for the COVID-19 Genomics UK (COG-UK) Consortium | Harper VanSteenhouse, Yumi Kasai, David Gray, Carol Clugston, Anna Dominiczak and Alex Alderton, Roberto Amato, Jeffrey Barrett, Sonia Goncalves, Ewan Harrison, David K. Jackson, Ian Johnston, Dominic Kwiatkowski, Cordelia Langford, John Sillitoe on behalf of the Wellcome Sanger Institute COVID-19 Surveillance Team |
| EPI_ISL_3140219, EPI_ISL_3140220 | Lighthouse Lab in Milton Keynes | Wellcome Sanger Institute for the COVID-19 Genomics UK (COG-UK) Consortium | The Lighthouse Lab in Milton Keynes and Alex Alderton, Roberto Amato, Jeffrey Barrett, Sonia Goncalves, Ewan Harrison, David K. Jackson, Ian Johnston, Dominic Kwiatkowski, Cordelia Langford, John Sillitoe on behalf of the Wellcome Sanger Institute COVID-19 Surveillance Team |
| EPI_ISL_3140221 | Lighthouse Lab in Glasgow | Wellcome Sanger Institute for the COVID-19 Genomics UK (COG-UK) Consortium | Harper VanSteenhouse, Yumi Kasai, David Gray, Carol Clugston, Anna Dominiczak and Alex Alderton, Roberto Amato, Jeffrey Barrett, Sonia Goncalves, Ewan Harrison, David K. Jackson, Ian Johnston, Dominic Kwiatkowski, Cordelia Langford, John Sillitoe on behalf of the Wellcome Sanger Institute COVID-19 Surveillance Team |
| EPI_ISL_3140222, EPI_ISL_3140223 | Lighthouse Lab in Alderley Park | Wellcome Sanger Institute for the COVID-19 Genomics UK (COG-UK) Consortium | Jacquelyn Wynn, Mairead Hyland, The Lighthouse Lab in Alderley Park and Alex Alderton, Roberto Amato, Jeffrey Barrett, Sonia Goncalves, Ewan Harrison, David K. Jackson, Ian Johnston, Dominic Kwiatkowski, Cordelia Langford, John Sillitoe on behalf of the Wellcome Sanger Institute COVID-19 Surveillance Team |
| EPI_ISL_3140224, EPI_ISL_3140225, EPI_ISL_3140226, EPI_ISL_3140227 | Lighthouse Lab in Milton Keynes | Wellcome Sanger Institute for the COVID-19 Genomics UK (COG-UK) Consortium | The Lighthouse Lab in Milton Keynes and Alex Alderton, Roberto Amato, Jeffrey Barrett, Sonia Goncalves, Ewan Harrison, David K. Jackson, Ian Johnston, Dominic Kwiatkowski, Cordelia Langford, John Sillitoe on behalf of the Wellcome Sanger Institute COVID-19 Surveillance Team |
| EPI_ISL_3140228 | Lighthouse Lab in Alderley Park | Wellcome Sanger Institute for the COVID-19 Genomics UK (COG-UK) Consortium | Jacquelyn Wynn, Mairead Hyland, The Lighthouse Lab in Alderley Park and Alex Alderton, Roberto Amato, Jeffrey Barrett, Sonia Goncalves, Ewan Harrison, David K. Jackson, Ian Johnston, Dominic Kwiatkowski, Cordelia Langford, John Sillitoe on behalf of the Wellcome Sanger Institute COVID-19 Surveillance Team |
| EPI_ISL_3140229, EPI_ISL_3140230 | Lighthouse Lab in Milton Keynes | Wellcome Sanger Institute for the COVID-19 Genomics UK (COG-UK) Consortium | The Lighthouse Lab in Milton Keynes and Alex Alderton, Roberto Amato, Jeffrey Barrett, Sonia Goncalves, Ewan Harrison, David K. Jackson, Ian Johnston, Dominic Kwiatkowski, Cordelia Langford, John Sillitoe on behalf of the Wellcome Sanger Institute COVID-19 Surveillance Team |
| EPI_ISL_3140231 | Lighthouse Lab in Glasgow | Wellcome Sanger Institute for the COVID-19 Genomics UK (COG-UK) Consortium | Harper VanSteenhouse, Yumi Kasai, David Gray, Carol Clugston, Anna Dominiczak and Alex Alderton, Roberto Amato, Jeffrey Barrett, Sonia Goncalves, Ewan Harrison, David K. Jackson, Ian Johnston, Dominic Kwiatkowski, Cordelia Langford, John Sillitoe on behalf of the Wellcome Sanger Institute COVID-19 Surveillance Team |
| EPI_ISL_3140232 | Lighthouse Lab in Alderley Park | Wellcome Sanger Institute for the COVID-19 Genomics UK (COG-UK) Consortium | Jacquelyn Wynn, Mairead Hyland, The Lighthouse Lab in Alderley Park and Alex Alderton, Roberto Amato, Jeffrey Barrett, Sonia Goncalves, Ewan Harrison, David K. Jackson, Ian Johnston, Dominic Kwiatkowski, Cordelia Langford, John Sillitoe on behalf of the Wellcome Sanger Institute COVID-19 Surveillance Team |
| EPI_ISL_3140233, EPI_ISL_3140234 | Lighthouse Lab in Milton Keynes | Wellcome Sanger Institute for the COVID-19 Genomics UK (COG-UK) Consortium | The Lighthouse Lab in Milton Keynes and Alex Alderton, Roberto Amato, Jeffrey Barrett, Sonia Goncalves, Ewan Harrison, David K. Jackson, Ian Johnston, Dominic Kwiatkowski, Cordelia Langford, John Sillitoe on behalf of the Wellcome Sanger Institute COVID-19 Surveillance Team |
| EPI_ISL_3140235 | Lighthouse Lab in Glasgow | Wellcome Sanger Institute for the COVID-19 Genomics UK (COG-UK) Consortium | Harper VanSteenhouse, Yumi Kasai, David Gray, Carol Clugston, Anna Dominiczak and Alex Alderton, Roberto Amato, Jeffrey Barrett, Sonia Goncalves, Ewan Harrison, David K. Jackson, Ian Johnston, Dominic Kwiatkowski, Cordelia Langford, John Sillitoe on behalf of the Wellcome Sanger Institute COVID-19 Surveillance Team |
| EPI_ISL_3140236 | Lighthouse Lab in Alderley Park | Wellcome Sanger Institute for the COVID-19 Genomics UK (COG-UK) Consortium | Jacquelyn Wynn, Mairead Hyland, The Lighthouse Lab in Alderley Park and Alex Alderton, Roberto Amato, Jeffrey Barrett, Sonia Goncalves, Ewan Harrison, David K. Jackson, Ian Johnston, Dominic Kwiatkowski, Cordelia Langford, John Sillitoe on behalf of the Wellcome Sanger Institute COVID-19 Surveillance Team |
| EPI_ISL_3140237 | Lighthouse Lab in Glasgow | Wellcome Sanger Institute for the COVID-19 Genomics UK (COG-UK) Consortium | Harper VanSteenhouse, Yumi Kasai, David Gray, Carol Clugston, Anna Dominiczak and Alex Alderton, Roberto Amato, Jeffrey Barrett, Sonia Goncalves, Ewan Harrison, David K. Jackson, Ian Johnston, Dominic Kwiatkowski, Cordelia Langford, John Sillitoe on behalf of the Wellcome Sanger Institute COVID-19 Surveillance Team |
| EPI_ISL_3140238 | Lighthouse Lab in Milton Keynes | Wellcome Sanger Institute for the COVID-19 Genomics UK (COG-UK) Consortium | The Lighthouse Lab in Milton Keynes and Alex Alderton, Roberto Amato, Jeffrey Barrett, Sonia Goncalves, Ewan Harrison, David K. Jackson, Ian Johnston, Dominic Kwiatkowski, Cordelia Langford, John Sillitoe on behalf of the Wellcome Sanger Institute COVID-19 Surveillance Team |
| EPI_ISL_3140239 | Lighthouse Lab in Glasgow | Wellcome Sanger Institute for the COVID-19 Genomics UK (COG-UK) Consortium | Harper VanSteenhouse, Yumi Kasai, David Gray, Carol Clugston, Anna Dominiczak and Alex Alderton, Roberto Amato, Jeffrey Barrett, Sonia Goncalves, Ewan Harrison, David K. Jackson, Ian Johnston, Dominic Kwiatkowski, Cordelia Langford, John Sillitoe on behalf of the Wellcome Sanger Institute COVID-19 Surveillance Team |
| EPI_ISL_3140240, EPI_ISL_3140241 | Lighthouse Lab in Alderley Park | Wellcome Sanger Institute for the COVID-19 Genomics UK (COG-UK) Consortium | Jacquelyn Wynn, Mairead Hyland, The Lighthouse Lab in Alderley Park and Alex Alderton, Roberto Amato, Jeffrey Barrett, Sonia Goncalves, Ewan Harrison, David K. Jackson, Ian Johnston, Dominic Kwiatkowski, Cordelia Langford, John Sillitoe on behalf of the Wellcome Sanger Institute COVID-19 Surveillance Team |
| EPI_ISL_3140242 | Lighthouse Lab in Glasgow | Wellcome Sanger Institute for the COVID-19 Genomics UK (COG-UK) Consortium | Harper VanSteenhouse, Yumi Kasai, David Gray, Carol Clugston, Anna Dominiczak and Alex Alderton, Roberto Amato, Jeffrey Barrett, Sonia Goncalves, Ewan Harrison, David K. Jackson, Ian Johnston, Dominic Kwiatkowski, Cordelia Langford, John Sillitoe on behalf of the Wellcome Sanger Institute COVID-19 Surveillance Team |
| EPI_ISL_3140243 | Lighthouse Lab in Milton Keynes | Wellcome Sanger Institute for the COVID-19 Genomics UK (COG-UK) Consortium | The Lighthouse Lab in Milton Keynes and Alex Alderton, Roberto Amato, Jeffrey Barrett, Sonia Goncalves, Ewan Harrison, David K. Jackson, Ian Johnston, Dominic Kwiatkowski, Cordelia Langford, John Sillitoe on behalf of the Wellcome Sanger Institute COVID-19 Surveillance Team |
| EPI_ISL_3140244, EPI_ISL_3140245 | Lighthouse Lab in Glasgow | Wellcome Sanger Institute for the COVID-19 Genomics UK (COG-UK) Consortium | Harper VanSteenhouse, Yumi Kasai, David Gray, Carol Clugston, Anna Dominiczak and Alex Alderton, Roberto Amato, Jeffrey Barrett, Sonia Goncalves, Ewan Harrison, David K. Jackson, Ian Johnston, Dominic Kwiatkowski, Cordelia Langford, John Sillitoe on behalf of the Wellcome Sanger Institute COVID-19 Surveillance Team |
| EPI_ISL_3140246 | Lighthouse Lab in Milton Keynes | Wellcome Sanger Institute for the COVID-19 Genomics UK (COG-UK) Consortium | The Lighthouse Lab in Milton Keynes and Alex Alderton, Roberto Amato, Jeffrey Barrett, Sonia Goncalves, Ewan Harrison, David K. Jackson, Ian Johnston, Dominic Kwiatkowski, Cordelia Langford, John Sillitoe on behalf of the Wellcome Sanger Institute COVID-19 Surveillance Team |
| EPI_ISL_3140247 | Lighthouse Lab in Glasgow | Wellcome Sanger Institute for the COVID-19 Genomics UK (COG-UK) Consortium | Harper VanSteenhouse, Yumi Kasai, David Gray, Carol Clugston, Anna Dominiczak and Alex Alderton, Roberto Amato, Jeffrey Barrett, Sonia Goncalves, Ewan Harrison, David K. Jackson, Ian Johnston, Dominic Kwiatkowski, Cordelia Langford, John Sillitoe on behalf of the Wellcome Sanger Institute COVID-19 Surveillance Team |
| EPI_ISL_3140248 | Lighthouse Lab in Milton Keynes | Wellcome Sanger Institute for the COVID-19 Genomics UK (COG-UK) Consortium | The Lighthouse Lab in Milton Keynes and Alex Alderton, Roberto Amato, Jeffrey Barrett, Sonia Goncalves, Ewan Harrison, David K. Jackson, Ian Johnston, Dominic Kwiatkowski, Cordelia Langford, John Sillitoe on behalf of the Wellcome Sanger Institute COVID-19 Surveillance Team |
| EPI_ISL_3140249 | Lighthouse Lab in Alderley Park | Wellcome Sanger Institute for the COVID-19 Genomics UK (COG-UK) Consortium | Jacquelyn Wynn, Mairead Hyland, The Lighthouse Lab in Alderley Park and Alex Alderton, Roberto Amato, Jeffrey Barrett, Sonia Goncalves, Ewan Harrison, David K. Jackson, Ian Johnston, Dominic Kwiatkowski, Cordelia Langford, John Sillitoe on behalf of the Wellcome Sanger Institute COVID-19 Surveillance Team |
| EPI_ISL_3140250 | Lighthouse Lab in Milton Keynes | Wellcome Sanger Institute for the COVID-19 Genomics UK (COG-UK) Consortium | The Lighthouse Lab in Milton Keynes and Alex Alderton, Roberto Amato, Jeffrey Barrett, Sonia Goncalves, Ewan Harrison, David K. Jackson, Ian Johnston, Dominic Kwiatkowski, Cordelia Langford, John Sillitoe on behalf of the Wellcome Sanger Institute COVID-19 Surveillance Team |
| EPI_ISL_3140251 | Lighthouse Lab in Alderley Park | Wellcome Sanger Institute for the COVID-19 Genomics UK (COG-UK) Consortium | Jacquelyn Wynn, Mairead Hyland, The Lighthouse Lab in Alderley Park and Alex Alderton, Roberto Amato, Jeffrey Barrett, Sonia Goncalves, Ewan Harrison, David K. Jackson, Ian Johnston, Dominic Kwiatkowski, Cordelia Langford, John Sillitoe on behalf of the Wellcome Sanger Institute COVID-19 Surveillance Team |
| EPI_ISL_3140252 | Lighthouse Lab in Glasgow | Wellcome Sanger Institute for the COVID-19 Genomics UK (COG-UK) Consortium | Harper VanSteenhouse, Yumi Kasai, David Gray, Carol Clugston, Anna Dominiczak and Alex Alderton, Roberto Amato, Jeffrey Barrett, Sonia Goncalves, Ewan Harrison, David K. Jackson, Ian Johnston, Dominic Kwiatkowski, Cordelia Langford, John Sillitoe on behalf of the Wellcome Sanger Institute COVID-19 Surveillance Team |
| EPI_ISL_3140253, EPI_ISL_3140254 | Lighthouse Lab in Alderley Park | Wellcome Sanger Institute for the COVID-19 Genomics UK (COG-UK) Consortium | Jacquelyn Wynn, Mairead Hyland, The Lighthouse Lab in Alderley Park and Alex Alderton, Roberto Amato, Jeffrey Barrett, Sonia Goncalves, Ewan Harrison, David K. Jackson, Ian Johnston, Dominic Kwiatkowski, Cordelia Langford, John Sillitoe on behalf of the Wellcome Sanger Institute COVID-19 Surveillance Team |

| | | | |
|---|---|---|---|
| EPI_ISL_3140255, EPI_ISL_3140256 | Lighthouse Lab in Milton Keynes | Wellcome Sanger Institute for the COVID-19 Genomics UK (COG-UK) Consortium | The Lighthouse Lab in Milton Keynes and Alex Alderton, Roberto Amato, Jeffrey Barrett, Sonia Goncalves, Ewan Harrison, David K. Jackson, Ian Johnston, Dominic Kwiatkowski, Cordelia Langford, John Sillitoe on behalf of the Wellcome Sanger Institute COVID-19 Surveillance Team |
| EPI_ISL_3140257 | Lighthouse Lab in Glasgow | Wellcome Sanger Institute for the COVID-19 Genomics UK (COG-UK) Consortium | Harper VanSteenhouse, Yumi Kasai, David Gray, Carol Clugston, Anna Dominiczak and Alex Alderton, Roberto Amato, Jeffrey Barrett, Sonia Goncalves, Ewan Harrison, David K. Jackson, Ian Johnston, Dominic Kwiatkowski, Cordelia Langford, John Sillitoe on behalf of the Wellcome Sanger Institute COVID-19 Surveillance Team |
| EPI_ISL_3140258 | Lighthouse Lab in Milton Keynes | Wellcome Sanger Institute for the COVID-19 Genomics UK (COG-UK) Consortium | The Lighthouse Lab in Milton Keynes and Alex Alderton, Roberto Amato, Jeffrey Barrett, Sonia Goncalves, Ewan Harrison, David K. Jackson, Ian Johnston, Dominic Kwiatkowski, Cordelia Langford, John Sillitoe on behalf of the Wellcome Sanger Institute COVID-19 Surveillance Team |
| EPI_ISL_3140259 | Lighthouse Lab in Alderley Park | Wellcome Sanger Institute for the COVID-19 Genomics UK (COG-UK) Consortium | Jacquelyn Wynn, Mairead Hyland, The Lighthouse Lab in Alderley Park and Alex Alderton, Roberto Amato, Jeffrey Barrett, Sonia Goncalves, Ewan Harrison, David K. Jackson, Ian Johnston, Dominic Kwiatkowski, Cordelia Langford, John Sillitoe on behalf of the Wellcome Sanger Institute COVID-19 Surveillance Team |
| EPI_ISL_3140260, EPI_ISL_3140261, EPI_ISL_3140262 | Lighthouse Lab in Milton Keynes | Wellcome Sanger Institute for the COVID-19 Genomics UK (COG-UK) Consortium | The Lighthouse Lab in Milton Keynes and Alex Alderton, Roberto Amato, Jeffrey Barrett, Sonia Goncalves, Ewan Harrison, David K. Jackson, Ian Johnston, Dominic Kwiatkowski, Cordelia Langford, John Sillitoe on behalf of the Wellcome Sanger Institute COVID-19 Surveillance Team |
| EPI_ISL_3140263 | Lighthouse Lab in Glasgow | Wellcome Sanger Institute for the COVID-19 Genomics UK (COG-UK) Consortium | Harper VanSteenhouse, Yumi Kasai, David Gray, Carol Clugston, Anna Dominiczak and Alex Alderton, Roberto Amato, Jeffrey Barrett, Sonia Goncalves, Ewan Harrison, David K. Jackson, Ian Johnston, Dominic Kwiatkowski, Cordelia Langford, John Sillitoe on behalf of the Wellcome Sanger Institute COVID-19 Surveillance Team |
| EPI_ISL_3140264 | Lighthouse Lab in Milton Keynes | Wellcome Sanger Institute for the COVID-19 Genomics UK (COG-UK) Consortium | The Lighthouse Lab in Milton Keynes and Alex Alderton, Roberto Amato, Jeffrey Barrett, Sonia Goncalves, Ewan Harrison, David K. Jackson, Ian Johnston, Dominic Kwiatkowski, Cordelia Langford, John Sillitoe on behalf of the Wellcome Sanger Institute COVID-19 Surveillance Team |
| EPI_ISL_3140265, EPI_ISL_3140266 | Lighthouse Lab in Alderley Park | Wellcome Sanger Institute for the COVID-19 Genomics UK (COG-UK) Consortium | Jacquelyn Wynn, Mairead Hyland, The Lighthouse Lab in Alderley Park and Alex Alderton, Roberto Amato, Jeffrey Barrett, Sonia Goncalves, Ewan Harrison, David K. Jackson, Ian Johnston, Dominic Kwiatkowski, Cordelia Langford, John Sillitoe on behalf of the Wellcome Sanger Institute COVID-19 Surveillance Team |
| EPI_ISL_3140267, EPI_ISL_3140268 | Lighthouse Lab in Milton Keynes | Wellcome Sanger Institute for the COVID-19 Genomics UK (COG-UK) Consortium | The Lighthouse Lab in Milton Keynes and Alex Alderton, Roberto Amato, Jeffrey Barrett, Sonia Goncalves, Ewan Harrison, David K. Jackson, Ian Johnston, Dominic Kwiatkowski, Cordelia Langford, John Sillitoe on behalf of the Wellcome Sanger Institute COVID-19 Surveillance Team |
| EPI_ISL_3140269, EPI_ISL_3140270, EPI_ISL_3140271 | Lighthouse Lab in Glasgow | Wellcome Sanger Institute for the COVID-19 Genomics UK (COG-UK) Consortium | Harper VanSteenhouse, Yumi Kasai, David Gray, Carol Clugston, Anna Dominiczak and Alex Alderton, Roberto Amato, Jeffrey Barrett, Sonia Goncalves, Ewan Harrison, David K. Jackson, Ian Johnston, Dominic Kwiatkowski, Cordelia Langford, John Sillitoe on behalf of the Wellcome Sanger Institute COVID-19 Surveillance Team |
| EPI_ISL_3140272, EPI_ISL_3140273 | Lighthouse Lab in Milton Keynes | Wellcome Sanger Institute for the COVID-19 Genomics UK (COG-UK) Consortium | The Lighthouse Lab in Milton Keynes and Alex Alderton, Roberto Amato, Jeffrey Barrett, Sonia Goncalves, Ewan Harrison, David K. Jackson, Ian Johnston, Dominic Kwiatkowski, Cordelia Langford, John Sillitoe on behalf of the Wellcome Sanger Institute COVID-19 Surveillance Team |
| EPI_ISL_3140274 | Lighthouse Lab in Glasgow | Wellcome Sanger Institute for the COVID-19 Genomics UK (COG-UK) Consortium | Harper VanSteenhouse, Yumi Kasai, David Gray, Carol Clugston, Anna Dominiczak and Alex Alderton, Roberto Amato, Jeffrey Barrett, Sonia Goncalves, Ewan Harrison, David K. Jackson, Ian Johnston, Dominic Kwiatkowski, Cordelia Langford, John Sillitoe on behalf of the Wellcome Sanger Institute COVID-19 Surveillance Team |
| EPI_ISL_3140275 | Lighthouse Lab in Milton Keynes | Wellcome Sanger Institute for the COVID-19 Genomics UK (COG-UK) Consortium | The Lighthouse Lab in Milton Keynes and Alex Alderton, Roberto Amato, Jeffrey Barrett, Sonia Goncalves, Ewan Harrison, David K. Jackson, Ian Johnston, Dominic Kwiatkowski, Cordelia Langford, John Sillitoe on behalf of the Wellcome Sanger Institute COVID-19 Surveillance Team |
| EPI_ISL_3140276 | Lighthouse Lab in Alderley Park | Wellcome Sanger Institute for the COVID-19 Genomics UK (COG-UK) Consortium | Jacquelyn Wynn, Mairead Hyland, The Lighthouse Lab in Alderley Park and Alex Alderton, Roberto Amato, Jeffrey Barrett, Sonia Goncalves, Ewan Harrison, David K. Jackson, Ian Johnston, Dominic Kwiatkowski, Cordelia Langford, John Sillitoe on behalf of the Wellcome Sanger Institute COVID-19 Surveillance Team |
| EPI_ISL_3140277 | Lighthouse Lab in Glasgow | Wellcome Sanger Institute for the COVID-19 Genomics UK (COG-UK) Consortium | Harper VanSteenhouse, Yumi Kasai, David Gray, Carol Clugston, Anna Dominiczak and Alex Alderton, Roberto Amato, Jeffrey Barrett, Sonia Goncalves, Ewan Harrison, David K. Jackson, Ian Johnston, Dominic Kwiatkowski, Cordelia Langford, John Sillitoe on behalf of the Wellcome Sanger Institute COVID-19 Surveillance Team |
| EPI_ISL_3140278, EPI_ISL_3140279, EPI_ISL_3140280, EPI_ISL_3140281, EPI_ISL_3140282 | Lighthouse Lab in Milton Keynes | Wellcome Sanger Institute for the COVID-19 Genomics UK (COG-UK) Consortium | The Lighthouse Lab in Milton Keynes and Alex Alderton, Roberto Amato, Jeffrey Barrett, Sonia Goncalves, Ewan Harrison, David K. Jackson, Ian Johnston, Dominic Kwiatkowski, Cordelia Langford, John Sillitoe on behalf of the Wellcome Sanger Institute COVID-19 Surveillance Team |
| EPI_ISL_3140283 | Lighthouse Lab in Glasgow | Wellcome Sanger Institute for the COVID-19 Genomics UK (COG-UK) Consortium | Harper VanSteenhouse, Yumi Kasai, David Gray, Carol Clugston, Anna Dominiczak and Alex Alderton, Roberto Amato, Jeffrey Barrett, Sonia Goncalves, Ewan Harrison, David K. Jackson, Ian Johnston, Dominic Kwiatkowski, Cordelia Langford, John Sillitoe on behalf of the Wellcome Sanger Institute COVID-19 Surveillance Team |
| EPI_ISL_3140284 | Lighthouse Lab in Milton Keynes | Wellcome Sanger Institute for the COVID-19 Genomics UK (COG-UK) Consortium | The Lighthouse Lab in Milton Keynes and Alex Alderton, Roberto Amato, Jeffrey Barrett, Sonia Goncalves, Ewan Harrison, David K. Jackson, Ian Johnston, Dominic Kwiatkowski, Cordelia Langford, John Sillitoe on behalf of the Wellcome Sanger Institute COVID-19 Surveillance Team |
| EPI_ISL_3140285 | Lighthouse Lab in Glasgow | Wellcome Sanger Institute for the COVID-19 Genomics UK (COG-UK) Consortium | Harper VanSteenhouse, Yumi Kasai, David Gray, Carol Clugston, Anna Dominiczak and Alex Alderton, Roberto Amato, Jeffrey Barrett, Sonia Goncalves, Ewan Harrison, David K. Jackson, Ian Johnston, Dominic Kwiatkowski, Cordelia Langford, John Sillitoe on behalf of the Wellcome Sanger Institute COVID-19 Surveillance Team |
| EPI_ISL_3140286, EPI_ISL_3140287 | Lighthouse Lab in Milton Keynes | Wellcome Sanger Institute for the COVID-19 Genomics UK (COG-UK) Consortium | The Lighthouse Lab in Milton Keynes and Alex Alderton, Roberto Amato, Jeffrey Barrett, Sonia Goncalves, Ewan Harrison, David K. Jackson, Ian Johnston, Dominic Kwiatkowski, Cordelia Langford, John Sillitoe on behalf of the Wellcome Sanger Institute COVID-19 Surveillance Team |
| EPI_ISL_3140288 | Lighthouse Lab in Alderley Park | Wellcome Sanger Institute for the COVID-19 Genomics UK (COG-UK) Consortium | Jacquelyn Wynn, Mairead Hyland, The Lighthouse Lab in Alderley Park and Alex Alderton, Roberto Amato, Jeffrey Barrett, Sonia Goncalves, Ewan Harrison, David K. Jackson, Ian Johnston, Dominic Kwiatkowski, Cordelia Langford, John Sillitoe on behalf of the Wellcome Sanger Institute COVID-19 Surveillance Team |
| EPI_ISL_3140289, EPI_ISL_3140290, EPI_ISL_3140291 | Lighthouse Lab in Milton Keynes | Wellcome Sanger Institute for the COVID-19 Genomics UK (COG-UK) Consortium | The Lighthouse Lab in Milton Keynes and Alex Alderton, Roberto Amato, Jeffrey Barrett, Sonia Goncalves, Ewan Harrison, David K. Jackson, Ian Johnston, Dominic Kwiatkowski, Cordelia Langford, John Sillitoe on behalf of the Wellcome Sanger Institute COVID-19 Surveillance Team |
| EPI_ISL_3140292 | Lighthouse Lab in Glasgow | Wellcome Sanger Institute for the COVID-19 Genomics UK (COG-UK) Consortium | Harper VanSteenhouse, Yumi Kasai, David Gray, Carol Clugston, Anna Dominiczak and Alex Alderton, Roberto Amato, Jeffrey Barrett, Sonia Goncalves, Ewan Harrison, David K. Jackson, Ian Johnston, Dominic Kwiatkowski, Cordelia Langford, John Sillitoe on behalf of the Wellcome Sanger Institute COVID-19 Surveillance Team |
| EPI_ISL_3140293 | Lighthouse Lab in Alderley Park | Wellcome Sanger Institute for the COVID-19 Genomics UK (COG-UK) Consortium | Jacquelyn Wynn, Mairead Hyland, The Lighthouse Lab in Alderley Park and Alex Alderton, Roberto Amato, Jeffrey Barrett, Sonia Goncalves, Ewan Harrison, David K. Jackson, Ian Johnston, Dominic Kwiatkowski, Cordelia Langford, John Sillitoe on behalf of the Wellcome Sanger Institute COVID-19 Surveillance Team |
| EPI_ISL_3140294 | Lighthouse Lab in Milton Keynes | Wellcome Sanger Institute for the COVID-19 Genomics UK (COG-UK) Consortium | The Lighthouse Lab in Milton Keynes and Alex Alderton, Roberto Amato, Jeffrey Barrett, Sonia Goncalves, Ewan Harrison, David K. Jackson, Ian Johnston, Dominic Kwiatkowski, Cordelia Langford, John Sillitoe on behalf of the Wellcome Sanger Institute COVID-19 Surveillance Team |
| EPI_ISL_3140295, EPI_ISL_3140296 | Lighthouse Lab in Glasgow | Wellcome Sanger Institute for the COVID-19 Genomics UK (COG-UK) Consortium | Harper VanSteenhouse, Yumi Kasai, David Gray, Carol Clugston, Anna Dominiczak and Alex Alderton, Roberto Amato, Jeffrey Barrett, Sonia Goncalves, Ewan Harrison, David K. Jackson, Ian Johnston, Dominic Kwiatkowski, Cordelia Langford, John Sillitoe on behalf of the Wellcome Sanger Institute COVID-19 Surveillance Team |
| EPI_ISL_3140297, EPI_ISL_3140298, EPI_ISL_3140299 | Lighthouse Lab in Milton Keynes | Wellcome Sanger Institute for the COVID-19 Genomics UK (COG-UK) Consortium | The Lighthouse Lab in Milton Keynes and Alex Alderton, Roberto Amato, Jeffrey Barrett, Sonia Goncalves, Ewan Harrison, David K. Jackson, Ian Johnston, Dominic Kwiatkowski, Cordelia Langford, John Sillitoe on behalf of the Wellcome Sanger Institute COVID-19 Surveillance Team |
| EPI_ISL_3140300 | Lighthouse Lab in Alderley Park | Wellcome Sanger Institute for the COVID-19 Genomics UK (COG-UK) Consortium | Jacquelyn Wynn, Mairead Hyland, The Lighthouse Lab in Alderley Park and Alex Alderton, Roberto Amato, Jeffrey Barrett, Sonia Goncalves, Ewan Harrison, David K. Jackson, Ian Johnston, Dominic Kwiatkowski, Cordelia Langford, John Sillitoe on behalf of the Wellcome Sanger Institute COVID-19 Surveillance Team |
| EPI_ISL_3140301 | Lighthouse Lab in Glasgow | Wellcome Sanger Institute for the COVID-19 Genomics UK (COG-UK) Consortium | Harper VanSteenhouse, Yumi Kasai, David Gray, Carol Clugston, Anna Dominiczak and Alex Alderton, Roberto Amato, Jeffrey Barrett, Sonia Goncalves, Ewan Harrison, David K. Jackson, Ian Johnston, Dominic Kwiatkowski, Cordelia Langford, John Sillitoe on behalf of the Wellcome Sanger Institute COVID-19 Surveillance Team |
| EPI_ISL_3140302 | Lighthouse Lab in Milton Keynes | Wellcome Sanger Institute for the COVID-19 Genomics UK (COG-UK) Consortium | The Lighthouse Lab in Milton Keynes and Alex Alderton, Roberto Amato, Jeffrey Barrett, Sonia Goncalves, Ewan Harrison, David K. Jackson, Ian Johnston, Dominic Kwiatkowski, Cordelia Langford, John Sillitoe on behalf of the Wellcome Sanger Institute COVID-19 Surveillance Team |
| EPI_ISL_3140303 | Lighthouse Lab in Alderley Park | Wellcome Sanger Institute for the COVID-19 Genomics UK (COG-UK) Consortium | Jacquelyn Wynn, Mairead Hyland, The Lighthouse Lab in Alderley Park and Alex Alderton, Roberto Amato, Jeffrey Barrett, Sonia Goncalves, Ewan Harrison, David K. Jackson, Ian Johnston, Dominic Kwiatkowski, Cordelia Langford, John Sillitoe on behalf of the Wellcome Sanger Institute COVID-19 Surveillance Team |
| EPI_ISL_3140304 | Lighthouse Lab in Milton Keynes | Wellcome Sanger Institute for the COVID-19 Genomics UK (COG-UK) Consortium | The Lighthouse Lab in Milton Keynes and Alex Alderton, Roberto Amato, Jeffrey Barrett, Sonia Goncalves, Ewan Harrison, David K. Jackson, Ian Johnston, Dominic Kwiatkowski, Cordelia Langford, John Sillitoe on behalf of the Wellcome Sanger Institute COVID-19 Surveillance Team |
| EPI_ISL_3140305 | Lighthouse Lab in Alderley Park | Wellcome Sanger Institute for the COVID-19 Genomics UK (COG-UK) Consortium | Jacquelyn Wynn, Mairead Hyland, The Lighthouse Lab in Alderley Park and Alex Alderton, Roberto Amato, Jeffrey Barrett, Sonia Goncalves, Ewan Harrison, David K. Jackson, Ian Johnston, Dominic Kwiatkowski, Cordelia Langford, John Sillitoe on behalf of the Wellcome Sanger Institute COVID-19 Surveillance Team |
| EPI_ISL_3140306 | Lighthouse Lab in Milton Keynes | Wellcome Sanger Institute for the COVID-19 Genomics UK (COG-UK) Consortium | The Lighthouse Lab in Milton Keynes and Alex Alderton, Roberto Amato, Jeffrey Barrett, Sonia Goncalves, Ewan Harrison, David K. Jackson, Ian Johnston, Dominic Kwiatkowski, Cordelia Langford, John Sillitoe on behalf of the Wellcome Sanger Institute COVID-19 Surveillance Team |
| EPI_ISL_3140307 | Lighthouse Lab in Glasgow | Wellcome Sanger Institute for the COVID-19 Genomics UK (COG-UK) Consortium | Harper VanSteenhouse, Yumi Kasai, David Gray, Carol Clugston, Anna Dominiczak and Alex Alderton, Roberto Amato, Jeffrey Barrett, Sonia Goncalves, Ewan Harrison, David K. Jackson, Ian Johnston, Dominic Kwiatkowski, Cordelia Langford, John Sillitoe on behalf of the Wellcome Sanger Institute COVID-19 Surveillance Team |
| EPI_ISL_3140308, EPI_ISL_3140309 | Lighthouse Lab in Milton Keynes | Wellcome Sanger Institute for the COVID-19 Genomics UK (COG-UK) Consortium | The Lighthouse Lab in Milton Keynes and Alex Alderton, Roberto Amato, Jeffrey Barrett, Sonia Goncalves, Ewan Harrison, David K. Jackson, Ian Johnston, Dominic Kwiatkowski, Cordelia Langford, John Sillitoe on behalf of the Wellcome Sanger Institute COVID-19 Surveillance Team |
| EPI_ISL_3140310 | Lighthouse Lab in Alderley Park | Wellcome Sanger Institute for the COVID-19 Genomics UK (COG-UK) Consortium | Jacquelyn Wynn, Mairead Hyland, The Lighthouse Lab in Alderley Park and Alex Alderton, Roberto Amato, Jeffrey Barrett, Sonia Goncalves, Ewan Harrison, David K. Jackson, Ian Johnston, Dominic Kwiatkowski, Cordelia Langford, John Sillitoe on behalf of the Wellcome Sanger Institute COVID-19 Surveillance Team |
| EPI_ISL_3140311 | Lighthouse Lab in Glasgow | Wellcome Sanger Institute for the COVID-19 Genomics UK (COG-UK) Consortium | Harper VanSteenhouse, Yumi Kasai, David Gray, Carol Clugston, Anna Dominiczak and Alex Alderton, Roberto Amato, Jeffrey Barrett, Sonia Goncalves, Ewan Harrison, David K. Jackson, Ian Johnston, Dominic Kwiatkowski, Cordelia Langford, John Sillitoe on behalf of the Wellcome Sanger Institute COVID-19 Surveillance Team |
| EPI_ISL_3140312 | Lighthouse Lab in Alderley Park | Wellcome Sanger Institute for the COVID-19 Genomics UK (COG-UK) Consortium | Jacquelyn Wynn, Mairead Hyland, The Lighthouse Lab in Alderley Park and Alex Alderton, Roberto Amato, Jeffrey Barrett, Sonia Goncalves, Ewan Harrison, David K. Jackson, Ian Johnston, Dominic Kwiatkowski, Cordelia Langford, John Sillitoe on behalf of the Wellcome Sanger Institute COVID-19 Surveillance Team |
| EPI_ISL_3140313 | Lighthouse Lab in Glasgow | Wellcome Sanger Institute for the COVID-19 Genomics UK (COG-UK) Consortium | Harper VanSteenhouse, Yumi Kasai, David Gray, Carol Clugston, Anna Dominiczak and Alex Alderton, Roberto Amato, Jeffrey Barrett, Sonia Goncalves, Ewan Harrison, David K. Jackson, Ian Johnston, Dominic Kwiatkowski, Cordelia Langford, John Sillitoe on behalf of the Wellcome Sanger Institute COVID-19 Surveillance Team |
| EPI_ISL_3140314, EPI_ISL_3140315, EPI_ISL_3140316 | Lighthouse Lab in Milton Keynes | Wellcome Sanger Institute for the COVID-19 Genomics UK (COG-UK) Consortium | The Lighthouse Lab in Milton Keynes and Alex Alderton, Roberto Amato, Jeffrey Barrett, Sonia Goncalves, Ewan Harrison, David K. Jackson, Ian Johnston, Dominic Kwiatkowski, Cordelia Langford, John Sillitoe on behalf of the Wellcome Sanger Institute COVID-19 Surveillance Team |

| | | | |
|---|---|---|---|
| EPI_ISL_3140317 | Lighthouse Lab in Alderley Park | Wellcome Sanger Institute for the COVID-19 Genomics UK (COG-UK) Consortium | Jacquelyn Wynn, Mairead Hyland, The Lighthouse Lab in Alderley Park and Alex Alderton, Roberto Amato, Jeffrey Barrett, Sonia Goncalves, Ewan Harrison, David K. Jackson, Ian Johnston, Dominic Kwiatkowski, Cordelia Langford, John Sillitoe on behalf of the Wellcome Sanger Institute COVID-19 Surveillance Team |
| EPI_ISL_3140318, EPI_ISL_3140319 | Lighthouse Lab in Milton Keynes | Wellcome Sanger Institute for the COVID-19 Genomics UK (COG-UK) Consortium | The Lighthouse Lab in Milton Keynes and Alex Alderton, Roberto Amato, Jeffrey Barrett, Sonia Goncalves, Ewan Harrison, David K. Jackson, Ian Johnston, Dominic Kwiatkowski, Cordelia Langford, John Sillitoe on behalf of the Wellcome Sanger Institute COVID-19 Surveillance Team |
| EPI_ISL_3140320 | Lighthouse Lab in Alderley Park | Wellcome Sanger Institute for the COVID-19 Genomics UK (COG-UK) Consortium | Jacquelyn Wynn, Mairead Hyland, The Lighthouse Lab in Alderley Park and Alex Alderton, Roberto Amato, Jeffrey Barrett, Sonia Goncalves, Ewan Harrison, David K. Jackson, Ian Johnston, Dominic Kwiatkowski, Cordelia Langford, John Sillitoe on behalf of the Wellcome Sanger Institute COVID-19 Surveillance Team |
| EPI_ISL_3140321, EPI_ISL_3140322, EPI_ISL_3140323 | Lighthouse Lab in Glasgow | Wellcome Sanger Institute for the COVID-19 Genomics UK (COG-UK) Consortium | Harper VanSteenhouse, Yumi Kasai, David Gray, Carol Clugston, Anna Dominiczak and Alex Alderton, Roberto Amato, Jeffrey Barrett, Sonia Goncalves, Ewan Harrison, David K. Jackson, Ian Johnston, Dominic Kwiatkowski, Cordelia Langford, John Sillitoe on behalf of the Wellcome Sanger Institute COVID-19 Surveillance Team |
| EPI_ISL_3140324 | Lighthouse Lab in Milton Keynes | Wellcome Sanger Institute for the COVID-19 Genomics UK (COG-UK) Consortium | The Lighthouse Lab in Milton Keynes and Alex Alderton, Roberto Amato, Jeffrey Barrett, Sonia Goncalves, Ewan Harrison, David K. Jackson, Ian Johnston, Dominic Kwiatkowski, Cordelia Langford, John Sillitoe on behalf of the Wellcome Sanger Institute COVID-19 Surveillance Team |
| EPI_ISL_3140325 | Lighthouse Lab in Alderley Park | Wellcome Sanger Institute for the COVID-19 Genomics UK (COG-UK) Consortium | Jacquelyn Wynn, Mairead Hyland, The Lighthouse Lab in Alderley Park and Alex Alderton, Roberto Amato, Jeffrey Barrett, Sonia Goncalves, Ewan Harrison, David K. Jackson, Ian Johnston, Dominic Kwiatkowski, Cordelia Langford, John Sillitoe on behalf of the Wellcome Sanger Institute COVID-19 Surveillance Team |
| EPI_ISL_3140326 | Lighthouse Lab in Milton Keynes | Wellcome Sanger Institute for the COVID-19 Genomics UK (COG-UK) Consortium | The Lighthouse Lab in Milton Keynes and Alex Alderton, Roberto Amato, Jeffrey Barrett, Sonia Goncalves, Ewan Harrison, David K. Jackson, Ian Johnston, Dominic Kwiatkowski, Cordelia Langford, John Sillitoe on behalf of the Wellcome Sanger Institute COVID-19 Surveillance Team |
| EPI_ISL_3140327 | Lighthouse Lab in Glasgow | Wellcome Sanger Institute for the COVID-19 Genomics UK (COG-UK) Consortium | Harper VanSteenhouse, Yumi Kasai, David Gray, Carol Clugston, Anna Dominiczak and Alex Alderton, Roberto Amato, Jeffrey Barrett, Sonia Goncalves, Ewan Harrison, David K. Jackson, Ian Johnston, Dominic Kwiatkowski, Cordelia Langford, John Sillitoe on behalf of the Wellcome Sanger Institute COVID-19 Surveillance Team |
| EPI_ISL_3140328 | Lighthouse Lab in Milton Keynes | Wellcome Sanger Institute for the COVID-19 Genomics UK (COG-UK) Consortium | The Lighthouse Lab in Milton Keynes and Alex Alderton, Roberto Amato, Jeffrey Barrett, Sonia Goncalves, Ewan Harrison, David K. Jackson, Ian Johnston, Dominic Kwiatkowski, Cordelia Langford, John Sillitoe on behalf of the Wellcome Sanger Institute COVID-19 Surveillance Team |
| EPI_ISL_3140329 | Lighthouse Lab in Glasgow | Wellcome Sanger Institute for the COVID-19 Genomics UK (COG-UK) Consortium | Harper VanSteenhouse, Yumi Kasai, David Gray, Carol Clugston, Anna Dominiczak and Alex Alderton, Roberto Amato, Jeffrey Barrett, Sonia Goncalves, Ewan Harrison, David K. Jackson, Ian Johnston, Dominic Kwiatkowski, Cordelia Langford, John Sillitoe on behalf of the Wellcome Sanger Institute COVID-19 Surveillance Team |
| EPI_ISL_3140330 | Lighthouse Lab in Milton Keynes | Wellcome Sanger Institute for the COVID-19 Genomics UK (COG-UK) Consortium | The Lighthouse Lab in Milton Keynes and Alex Alderton, Roberto Amato, Jeffrey Barrett, Sonia Goncalves, Ewan Harrison, David K. Jackson, Ian Johnston, Dominic Kwiatkowski, Cordelia Langford, John Sillitoe on behalf of the Wellcome Sanger Institute COVID-19 Surveillance Team |
| EPI_ISL_3140331 | Lighthouse Lab in Alderley Park | Wellcome Sanger Institute for the COVID-19 Genomics UK (COG-UK) Consortium | Jacquelyn Wynn, Mairead Hyland, The Lighthouse Lab in Alderley Park and Alex Alderton, Roberto Amato, Jeffrey Barrett, Sonia Goncalves, Ewan Harrison, David K. Jackson, Ian Johnston, Dominic Kwiatkowski, Cordelia Langford, John Sillitoe on behalf of the Wellcome Sanger Institute COVID-19 Surveillance Team |
| EPI_ISL_3140332 | Lighthouse Lab in Milton Keynes | Wellcome Sanger Institute for the COVID-19 Genomics UK (COG-UK) Consortium | The Lighthouse Lab in Milton Keynes and Alex Alderton, Roberto Amato, Jeffrey Barrett, Sonia Goncalves, Ewan Harrison, David K. Jackson, Ian Johnston, Dominic Kwiatkowski, Cordelia Langford, John Sillitoe on behalf of the Wellcome Sanger Institute COVID-19 Surveillance Team |
| EPI_ISL_3140333, EPI_ISL_3140334 | Lighthouse Lab in Alderley Park | Wellcome Sanger Institute for the COVID-19 Genomics UK (COG-UK) Consortium | Jacquelyn Wynn, Mairead Hyland, The Lighthouse Lab in Alderley Park and Alex Alderton, Roberto Amato, Jeffrey Barrett, Sonia Goncalves, Ewan Harrison, David K. Jackson, Ian Johnston, Dominic Kwiatkowski, Cordelia Langford, John Sillitoe on behalf of the Wellcome Sanger Institute COVID-19 Surveillance Team |
| EPI_ISL_3140335 | Lighthouse Lab in Milton Keynes | Wellcome Sanger Institute for the COVID-19 Genomics UK (COG-UK) Consortium | The Lighthouse Lab in Milton Keynes and Alex Alderton, Roberto Amato, Jeffrey Barrett, Sonia Goncalves, Ewan Harrison, David K. Jackson, Ian Johnston, Dominic Kwiatkowski, Cordelia Langford, John Sillitoe on behalf of the Wellcome Sanger Institute COVID-19 Surveillance Team |
| EPI_ISL_3140336 | Lighthouse Lab in Alderley Park | Wellcome Sanger Institute for the COVID-19 Genomics UK (COG-UK) Consortium | Jacquelyn Wynn, Mairead Hyland, The Lighthouse Lab in Alderley Park and Alex Alderton, Roberto Amato, Jeffrey Barrett, Sonia Goncalves, Ewan Harrison, David K. Jackson, Ian Johnston, Dominic Kwiatkowski, Cordelia Langford, John Sillitoe on behalf of the Wellcome Sanger Institute COVID-19 Surveillance Team |
| EPI_ISL_3140337 | Lighthouse Lab in Milton Keynes | Wellcome Sanger Institute for the COVID-19 Genomics UK (COG-UK) Consortium | The Lighthouse Lab in Milton Keynes and Alex Alderton, Roberto Amato, Jeffrey Barrett, Sonia Goncalves, Ewan Harrison, David K. Jackson, Ian Johnston, Dominic Kwiatkowski, Cordelia Langford, John Sillitoe on behalf of the Wellcome Sanger Institute COVID-19 Surveillance Team |
| EPI_ISL_3140338 | Lighthouse Lab in Alderley Park | Wellcome Sanger Institute for the COVID-19 Genomics UK (COG-UK) Consortium | Jacquelyn Wynn, Mairead Hyland, The Lighthouse Lab in Alderley Park and Alex Alderton, Roberto Amato, Jeffrey Barrett, Sonia Goncalves, Ewan Harrison, David K. Jackson, Ian Johnston, Dominic Kwiatkowski, Cordelia Langford, John Sillitoe on behalf of the Wellcome Sanger Institute COVID-19 Surveillance Team |
| EPI_ISL_3140339 | Lighthouse Lab in Milton Keynes | Wellcome Sanger Institute for the COVID-19 Genomics UK (COG-UK) Consortium | The Lighthouse Lab in Milton Keynes and Alex Alderton, Roberto Amato, Jeffrey Barrett, Sonia Goncalves, Ewan Harrison, David K. Jackson, Ian Johnston, Dominic Kwiatkowski, Cordelia Langford, John Sillitoe on behalf of the Wellcome Sanger Institute COVID-19 Surveillance Team |
| EPI_ISL_3140340, EPI_ISL_3140341 | Lighthouse Lab in Alderley Park | Wellcome Sanger Institute for the COVID-19 Genomics UK (COG-UK) Consortium | Jacquelyn Wynn, Mairead Hyland, The Lighthouse Lab in Alderley Park and Alex Alderton, Roberto Amato, Jeffrey Barrett, Sonia Goncalves, Ewan Harrison, David K. Jackson, Ian Johnston, Dominic Kwiatkowski, Cordelia Langford, John Sillitoe on behalf of the Wellcome Sanger Institute COVID-19 Surveillance Team |
| EPI_ISL_3140342, EPI_ISL_3140343 | Lighthouse Lab in Milton Keynes | Wellcome Sanger Institute for the COVID-19 Genomics UK (COG-UK) Consortium | The Lighthouse Lab in Milton Keynes and Alex Alderton, Roberto Amato, Jeffrey Barrett, Sonia Goncalves, Ewan Harrison, David K. Jackson, Ian Johnston, Dominic Kwiatkowski, Cordelia Langford, John Sillitoe on behalf of the Wellcome Sanger Institute COVID-19 Surveillance Team |
| EPI_ISL_3140344, EPI_ISL_3140345, EPI_ISL_3140346 | Lighthouse Lab in Glasgow | Wellcome Sanger Institute for the COVID-19 Genomics UK (COG-UK) Consortium | Harper VanSteenhouse, Yumi Kasai, David Gray, Carol Clugston, Anna Dominiczak and Alex Alderton, Roberto Amato, Jeffrey Barrett, Sonia Goncalves, Ewan Harrison, David K. Jackson, Ian Johnston, Dominic Kwiatkowski, Cordelia Langford, John Sillitoe on behalf of the Wellcome Sanger Institute COVID-19 Surveillance Team |
| EPI_ISL_3140347 | Lighthouse Lab in Alderley Park | Wellcome Sanger Institute for the COVID-19 Genomics UK (COG-UK) Consortium | Jacquelyn Wynn, Mairead Hyland, The Lighthouse Lab in Alderley Park and Alex Alderton, Roberto Amato, Jeffrey Barrett, Sonia Goncalves, Ewan Harrison, David K. Jackson, Ian Johnston, Dominic Kwiatkowski, Cordelia Langford, John Sillitoe on behalf of the Wellcome Sanger Institute COVID-19 Surveillance Team |
| EPI_ISL_3140348 | Lighthouse Lab in Glasgow | Wellcome Sanger Institute for the COVID-19 Genomics UK (COG-UK) Consortium | Harper VanSteenhouse, Yumi Kasai, David Gray, Carol Clugston, Anna Dominiczak and Alex Alderton, Roberto Amato, Jeffrey Barrett, Sonia Goncalves, Ewan Harrison, David K. Jackson, Ian Johnston, Dominic Kwiatkowski, Cordelia Langford, John Sillitoe on behalf of the Wellcome Sanger Institute COVID-19 Surveillance Team |
| EPI_ISL_3140349 | Lighthouse Lab in Alderley Park | Wellcome Sanger Institute for the COVID-19 Genomics UK (COG-UK) Consortium | Jacquelyn Wynn, Mairead Hyland, The Lighthouse Lab in Alderley Park and Alex Alderton, Roberto Amato, Jeffrey Barrett, Sonia Goncalves, Ewan Harrison, David K. Jackson, Ian Johnston, Dominic Kwiatkowski, Cordelia Langford, John Sillitoe on behalf of the Wellcome Sanger Institute COVID-19 Surveillance Team |
| EPI_ISL_3140350 | Lighthouse Lab in Milton Keynes | Wellcome Sanger Institute for the COVID-19 Genomics UK (COG-UK) Consortium | The Lighthouse Lab in Milton Keynes and Alex Alderton, Roberto Amato, Jeffrey Barrett, Sonia Goncalves, Ewan Harrison, David K. Jackson, Ian Johnston, Dominic Kwiatkowski, Cordelia Langford, John Sillitoe on behalf of the Wellcome Sanger Institute COVID-19 Surveillance Team |
| EPI_ISL_3140351, EPI_ISL_3140352 | Lighthouse Lab in Alderley Park | Wellcome Sanger Institute for the COVID-19 Genomics UK (COG-UK) Consortium | Jacquelyn Wynn, Mairead Hyland, The Lighthouse Lab in Alderley Park and Alex Alderton, Roberto Amato, Jeffrey Barrett, Sonia Goncalves, Ewan Harrison, David K. Jackson, Ian Johnston, Dominic Kwiatkowski, Cordelia Langford, John Sillitoe on behalf of the Wellcome Sanger Institute COVID-19 Surveillance Team |
| EPI_ISL_3140353, EPI_ISL_3140354, EPI_ISL_3140355, EPI_ISL_3140356, EPI_ISL_3140357 | Lighthouse Lab in Milton Keynes | Wellcome Sanger Institute for the COVID-19 Genomics UK (COG-UK) Consortium | The Lighthouse Lab in Milton Keynes and Alex Alderton, Roberto Amato, Jeffrey Barrett, Sonia Goncalves, Ewan Harrison, David K. Jackson, Ian Johnston, Dominic Kwiatkowski, Cordelia Langford, John Sillitoe on behalf of the Wellcome Sanger Institute COVID-19 Surveillance Team |
| EPI_ISL_3140358 | Lighthouse Lab in Alderley Park | Wellcome Sanger Institute for the COVID-19 Genomics UK (COG-UK) Consortium | Jacquelyn Wynn, Mairead Hyland, The Lighthouse Lab in Alderley Park and Alex Alderton, Roberto Amato, Jeffrey Barrett, Sonia Goncalves, Ewan Harrison, David K. Jackson, Ian Johnston, Dominic Kwiatkowski, Cordelia Langford, John Sillitoe on behalf of the Wellcome Sanger Institute COVID-19 Surveillance Team |
| EPI_ISL_3140359 | Lighthouse Lab in Milton Keynes | Wellcome Sanger Institute for the COVID-19 Genomics UK (COG-UK) Consortium | The Lighthouse Lab in Milton Keynes and Alex Alderton, Roberto Amato, Jeffrey Barrett, Sonia Goncalves, Ewan Harrison, David K. Jackson, Ian Johnston, Dominic Kwiatkowski, Cordelia Langford, John Sillitoe on behalf of the Wellcome Sanger Institute COVID-19 Surveillance Team |
| EPI_ISL_3140360 | Lighthouse Lab in Glasgow | Wellcome Sanger Institute for the COVID-19 Genomics UK (COG-UK) Consortium | Harper VanSteenhouse, Yumi Kasai, David Gray, Carol Clugston, Anna Dominiczak and Alex Alderton, Roberto Amato, Jeffrey Barrett, Sonia Goncalves, Ewan Harrison, David K. Jackson, Ian Johnston, Dominic Kwiatkowski, Cordelia Langford, John Sillitoe on behalf of the Wellcome Sanger Institute COVID-19 Surveillance Team |
| EPI_ISL_3140361 | Lighthouse Lab in Alderley Park | Wellcome Sanger Institute for the COVID-19 Genomics UK (COG-UK) Consortium | Jacquelyn Wynn, Mairead Hyland, The Lighthouse Lab in Alderley Park and Alex Alderton, Roberto Amato, Jeffrey Barrett, Sonia Goncalves, Ewan Harrison, David K. Jackson, Ian Johnston, Dominic Kwiatkowski, Cordelia Langford, John Sillitoe on behalf of the Wellcome Sanger Institute COVID-19 Surveillance Team |
| EPI_ISL_3140362, EPI_ISL_3140363 | Lighthouse Lab in Milton Keynes | Wellcome Sanger Institute for the COVID-19 Genomics UK (COG-UK) Consortium | The Lighthouse Lab in Milton Keynes and Alex Alderton, Roberto Amato, Jeffrey Barrett, Sonia Goncalves, Ewan Harrison, David K. Jackson, Ian Johnston, Dominic Kwiatkowski, Cordelia Langford, John Sillitoe on behalf of the Wellcome Sanger Institute COVID-19 Surveillance Team |
| EPI_ISL_3140364 | Lighthouse Lab in Glasgow | Wellcome Sanger Institute for the COVID-19 Genomics UK (COG-UK) Consortium | Harper VanSteenhouse, Yumi Kasai, David Gray, Carol Clugston, Anna Dominiczak and Alex Alderton, Roberto Amato, Jeffrey Barrett, Sonia Goncalves, Ewan Harrison, David K. Jackson, Ian Johnston, Dominic Kwiatkowski, Cordelia Langford, John Sillitoe on behalf of the Wellcome Sanger Institute COVID-19 Surveillance Team |
| EPI_ISL_3140365 | Lighthouse Lab in Milton Keynes | Wellcome Sanger Institute for the COVID-19 Genomics UK (COG-UK) Consortium | The Lighthouse Lab in Milton Keynes and Alex Alderton, Roberto Amato, Jeffrey Barrett, Sonia Goncalves, Ewan Harrison, David K. Jackson, Ian Johnston, Dominic Kwiatkowski, Cordelia Langford, John Sillitoe on behalf of the Wellcome Sanger Institute COVID-19 Surveillance Team |
| EPI_ISL_3140366 | Lighthouse Lab in Alderley Park | Wellcome Sanger Institute for the COVID-19 Genomics UK (COG-UK) Consortium | Jacquelyn Wynn, Mairead Hyland, The Lighthouse Lab in Alderley Park and Alex Alderton, Roberto Amato, Jeffrey Barrett, Sonia Goncalves, Ewan Harrison, David K. Jackson, Ian Johnston, Dominic Kwiatkowski, Cordelia Langford, John Sillitoe on behalf of the Wellcome Sanger Institute COVID-19 Surveillance Team |
| EPI_ISL_3140367 | Lighthouse Lab in Glasgow | Wellcome Sanger Institute for the COVID-19 Genomics UK (COG-UK) Consortium | Harper VanSteenhouse, Yumi Kasai, David Gray, Carol Clugston, Anna Dominiczak and Alex Alderton, Roberto Amato, Jeffrey Barrett, Sonia Goncalves, Ewan Harrison, David K. Jackson, Ian Johnston, Dominic Kwiatkowski, Cordelia Langford, John Sillitoe on behalf of the Wellcome Sanger Institute COVID-19 Surveillance Team |
| EPI_ISL_3140368, EPI_ISL_3140369 | Lighthouse Lab in Milton Keynes | Wellcome Sanger Institute for the COVID-19 Genomics UK (COG-UK) Consortium | The Lighthouse Lab in Milton Keynes and Alex Alderton, Roberto Amato, Jeffrey Barrett, Sonia Goncalves, Ewan Harrison, David K. Jackson, Ian Johnston, Dominic Kwiatkowski, Cordelia Langford, John Sillitoe on behalf of the Wellcome Sanger Institute COVID-19 Surveillance Team |
| EPI_ISL_3140370 | Lighthouse Lab in Glasgow | Wellcome Sanger Institute for the COVID-19 Genomics UK (COG-UK) Consortium | Harper VanSteenhouse, Yumi Kasai, David Gray, Carol Clugston, Anna Dominiczak and Alex Alderton, Roberto Amato, Jeffrey Barrett, Sonia Goncalves, Ewan Harrison, David K. Jackson, Ian Johnston, Dominic Kwiatkowski, Cordelia Langford, John Sillitoe on behalf of the Wellcome Sanger Institute COVID-19 Surveillance Team |
| EPI_ISL_3140371 | Lighthouse Lab in Milton Keynes | Wellcome Sanger Institute for the COVID-19 Genomics UK (COG-UK) Consortium | The Lighthouse Lab in Milton Keynes and Alex Alderton, Roberto Amato, Jeffrey Barrett, Sonia Goncalves, Ewan Harrison, David K. Jackson, Ian Johnston, Dominic Kwiatkowski, Cordelia Langford, John Sillitoe on behalf of the Wellcome Sanger Institute COVID-19 Surveillance Team |
| EPI_ISL_3140372 | Lighthouse Lab in Glasgow | Wellcome Sanger Institute for the COVID-19 Genomics UK (COG-UK) Consortium | Harper VanSteenhouse, Yumi Kasai, David Gray, Carol Clugston, Anna Dominiczak and Alex Alderton, Roberto Amato, Jeffrey Barrett, Sonia Goncalves, Ewan Harrison, David K. Jackson, Ian Johnston, Dominic Kwiatkowski, Cordelia Langford, John Sillitoe on behalf of the Wellcome Sanger Institute COVID-19 Surveillance Team |
| EPI_ISL_3140373 | Lighthouse Lab in Milton Keynes | Wellcome Sanger Institute for the COVID-19 Genomics UK (COG-UK) Consortium | The Lighthouse Lab in Milton Keynes and Alex Alderton, Roberto Amato, Jeffrey Barrett, Sonia Goncalves, Ewan Harrison, David K. Jackson, Ian Johnston, Dominic Kwiatkowski, Cordelia Langford, John Sillitoe on behalf of the Wellcome Sanger Institute COVID-19 Surveillance Team |
| EPI_ISL_3140374 | Lighthouse Lab in Alderley Park | Wellcome Sanger Institute for the COVID-19 Genomics UK (COG-UK) Consortium | Jacquelyn Wynn, Mairead Hyland, The Lighthouse Lab in Alderley Park and Alex Alderton, Roberto Amato, Jeffrey Barrett, Sonia Goncalves, Ewan Harrison, David K. Jackson, Ian Johnston, Dominic Kwiatkowski, Cordelia Langford, John Sillitoe on behalf of the Wellcome Sanger Institute COVID-19 Surveillance Team |

| | | | |
|---|---|---|---|
| EPI_ISL_3140375, EPI_ISL_3140376 | Lighthouse Lab in Milton Keynes | Wellcome Sanger Institute for the COVID-19 Genomics UK (COG-UK) Consortium | The Lighthouse Lab in Milton Keynes and Alex Alderton, Roberto Amato, Jeffrey Barrett, Sonia Goncalves, Ewan Harrison, David K. Jackson, Ian Johnston, Dominic Kwiatkowski, Cordelia Langford, John Sillitoe on behalf of the Wellcome Sanger Institute COVID-19 Surveillance Team |
| EPI_ISL_3140377, EPI_ISL_3140378 | Lighthouse Lab in Glasgow | Wellcome Sanger Institute for the COVID-19 Genomics UK (COG-UK) Consortium | Harper VanSteenhouse, Yumi Kasai, David Gray, Carol Clugston, Anna Dominiczak and Alex Alderton, Roberto Amato, Jeffrey Barrett, Sonia Goncalves, Ewan Harrison, David K. Jackson, Ian Johnston, Dominic Kwiatkowski, Cordelia Langford, John Sillitoe on behalf of the Wellcome Sanger Institute COVID-19 Surveillance Team |
| EPI_ISL_3140379, EPI_ISL_3140380 | Lighthouse Lab in Milton Keynes | Wellcome Sanger Institute for the COVID-19 Genomics UK (COG-UK) Consortium | The Lighthouse Lab in Milton Keynes and Alex Alderton, Roberto Amato, Jeffrey Barrett, Sonia Goncalves, Ewan Harrison, David K. Jackson, Ian Johnston, Dominic Kwiatkowski, Cordelia Langford, John Sillitoe on behalf of the Wellcome Sanger Institute COVID-19 Surveillance Team |
| EPI_ISL_3140381 | Lighthouse Lab in Glasgow | Wellcome Sanger Institute for the COVID-19 Genomics UK (COG-UK) Consortium | Harper VanSteenhouse, Yumi Kasai, David Gray, Carol Clugston, Anna Dominiczak and Alex Alderton, Roberto Amato, Jeffrey Barrett, Sonia Goncalves, Ewan Harrison, David K. Jackson, Ian Johnston, Dominic Kwiatkowski, Cordelia Langford, John Sillitoe on behalf of the Wellcome Sanger Institute COVID-19 Surveillance Team |
| EPI_ISL_3140382, EPI_ISL_3140383 | Lighthouse Lab in Alderley Park | Wellcome Sanger Institute for the COVID-19 Genomics UK (COG-UK) Consortium | Jacquelyn Wynn, Mairead Hyland, The Lighthouse Lab in Alderley Park and Alex Alderton, Roberto Amato, Jeffrey Barrett, Sonia Goncalves, Ewan Harrison, David K. Jackson, Ian Johnston, Dominic Kwiatkowski, Cordelia Langford, John Sillitoe on behalf of the Wellcome Sanger Institute COVID-19 Surveillance Team |
| EPI_ISL_3140384, EPI_ISL_3140385 | Lighthouse Lab in Glasgow | Wellcome Sanger Institute for the COVID-19 Genomics UK (COG-UK) Consortium | Harper VanSteenhouse, Yumi Kasai, David Gray, Carol Clugston, Anna Dominiczak and Alex Alderton, Roberto Amato, Jeffrey Barrett, Sonia Goncalves, Ewan Harrison, David K. Jackson, Ian Johnston, Dominic Kwiatkowski, Cordelia Langford, John Sillitoe on behalf of the Wellcome Sanger Institute COVID-19 Surveillance Team |
| EPI_ISL_3140386, EPI_ISL_3140387 | Lighthouse Lab in Alderley Park | Wellcome Sanger Institute for the COVID-19 Genomics UK (COG-UK) Consortium | Jacquelyn Wynn, Mairead Hyland, The Lighthouse Lab in Alderley Park and Alex Alderton, Roberto Amato, Jeffrey Barrett, Sonia Goncalves, Ewan Harrison, David K. Jackson, Ian Johnston, Dominic Kwiatkowski, Cordelia Langford, John Sillitoe on behalf of the Wellcome Sanger Institute COVID-19 Surveillance Team |
| EPI_ISL_3140388, EPI_ISL_3140389 | Lighthouse Lab in Milton Keynes | Wellcome Sanger Institute for the COVID-19 Genomics UK (COG-UK) Consortium | The Lighthouse Lab in Milton Keynes and Alex Alderton, Roberto Amato, Jeffrey Barrett, Sonia Goncalves, Ewan Harrison, David K. Jackson, Ian Johnston, Dominic Kwiatkowski, Cordelia Langford, John Sillitoe on behalf of the Wellcome Sanger Institute COVID-19 Surveillance Team |
| EPI_ISL_3140390, EPI_ISL_3140391 | Lighthouse Lab in Glasgow | Wellcome Sanger Institute for the COVID-19 Genomics UK (COG-UK) Consortium | Harper VanSteenhouse, Yumi Kasai, David Gray, Carol Clugston, Anna Dominiczak and Alex Alderton, Roberto Amato, Jeffrey Barrett, Sonia Goncalves, Ewan Harrison, David K. Jackson, Ian Johnston, Dominic Kwiatkowski, Cordelia Langford, John Sillitoe on behalf of the Wellcome Sanger Institute COVID-19 Surveillance Team |
| EPI_ISL_3140392 | Lighthouse Lab in Alderley Park | Wellcome Sanger Institute for the COVID-19 Genomics UK (COG-UK) Consortium | Jacquelyn Wynn, Mairead Hyland, The Lighthouse Lab in Alderley Park and Alex Alderton, Roberto Amato, Jeffrey Barrett, Sonia Goncalves, Ewan Harrison, David K. Jackson, Ian Johnston, Dominic Kwiatkowski, Cordelia Langford, John Sillitoe on behalf of the Wellcome Sanger Institute COVID-19 Surveillance Team |
| EPI_ISL_3140393 | Lighthouse Lab in Milton Keynes | Wellcome Sanger Institute for the COVID-19 Genomics UK (COG-UK) Consortium | The Lighthouse Lab in Milton Keynes and Alex Alderton, Roberto Amato, Jeffrey Barrett, Sonia Goncalves, Ewan Harrison, David K. Jackson, Ian Johnston, Dominic Kwiatkowski, Cordelia Langford, John Sillitoe on behalf of the Wellcome Sanger Institute COVID-19 Surveillance Team |
| EPI_ISL_3140394, EPI_ISL_3140395, EPI_ISL_3140396 | Lighthouse Lab in Alderley Park | Wellcome Sanger Institute for the COVID-19 Genomics UK (COG-UK) Consortium | Jacquelyn Wynn, Mairead Hyland, The Lighthouse Lab in Alderley Park and Alex Alderton, Roberto Amato, Jeffrey Barrett, Sonia Goncalves, Ewan Harrison, David K. Jackson, Ian Johnston, Dominic Kwiatkowski, Cordelia Langford, John Sillitoe on behalf of the Wellcome Sanger Institute COVID-19 Surveillance Team |
| EPI_ISL_3140397 | Lighthouse Lab in Milton Keynes | Wellcome Sanger Institute for the COVID-19 Genomics UK (COG-UK) Consortium | The Lighthouse Lab in Milton Keynes and Alex Alderton, Roberto Amato, Jeffrey Barrett, Sonia Goncalves, Ewan Harrison, David K. Jackson, Ian Johnston, Dominic Kwiatkowski, Cordelia Langford, John Sillitoe on behalf of the Wellcome Sanger Institute COVID-19 Surveillance Team |
| EPI_ISL_3140398 | Lighthouse Lab in Glasgow | Wellcome Sanger Institute for the COVID-19 Genomics UK (COG-UK) Consortium | Harper VanSteenhouse, Yumi Kasai, David Gray, Carol Clugston, Anna Dominiczak and Alex Alderton, Roberto Amato, Jeffrey Barrett, Sonia Goncalves, Ewan Harrison, David K. Jackson, Ian Johnston, Dominic Kwiatkowski, Cordelia Langford, John Sillitoe on behalf of the Wellcome Sanger Institute COVID-19 Surveillance Team |
| EPI_ISL_3140399 | Lighthouse Lab in Milton Keynes | Wellcome Sanger Institute for the COVID-19 Genomics UK (COG-UK) Consortium | The Lighthouse Lab in Milton Keynes and Alex Alderton, Roberto Amato, Jeffrey Barrett, Sonia Goncalves, Ewan Harrison, David K. Jackson, Ian Johnston, Dominic Kwiatkowski, Cordelia Langford, John Sillitoe on behalf of the Wellcome Sanger Institute COVID-19 Surveillance Team |
| EPI_ISL_3140400 | Lighthouse Lab in Alderley Park | Wellcome Sanger Institute for the COVID-19 Genomics UK (COG-UK) Consortium | Jacquelyn Wynn, Mairead Hyland, The Lighthouse Lab in Alderley Park and Alex Alderton, Roberto Amato, Jeffrey Barrett, Sonia Goncalves, Ewan Harrison, David K. Jackson, Ian Johnston, Dominic Kwiatkowski, Cordelia Langford, John Sillitoe on behalf of the Wellcome Sanger Institute COVID-19 Surveillance Team |
| EPI_ISL_3140401 | Lighthouse Lab in Glasgow | Wellcome Sanger Institute for the COVID-19 Genomics UK (COG-UK) Consortium | Harper VanSteenhouse, Yumi Kasai, David Gray, Carol Clugston, Anna Dominiczak and Alex Alderton, Roberto Amato, Jeffrey Barrett, Sonia Goncalves, Ewan Harrison, David K. Jackson, Ian Johnston, Dominic Kwiatkowski, Cordelia Langford, John Sillitoe on behalf of the Wellcome Sanger Institute COVID-19 Surveillance Team |
| EPI_ISL_3140402, EPI_ISL_3140403, EPI_ISL_3140404, EPI_ISL_3140405, EPI_ISL_3140406 | Lighthouse Lab in Milton Keynes | Wellcome Sanger Institute for the COVID-19 Genomics UK (COG-UK) Consortium | The Lighthouse Lab in Milton Keynes and Alex Alderton, Roberto Amato, Jeffrey Barrett, Sonia Goncalves, Ewan Harrison, David K. Jackson, Ian Johnston, Dominic Kwiatkowski, Cordelia Langford, John Sillitoe on behalf of the Wellcome Sanger Institute COVID-19 Surveillance Team |
| EPI_ISL_3140407 | Lighthouse Lab in Alderley Park | Wellcome Sanger Institute for the COVID-19 Genomics UK (COG-UK) Consortium | Jacquelyn Wynn, Mairead Hyland, The Lighthouse Lab in Alderley Park and Alex Alderton, Roberto Amato, Jeffrey Barrett, Sonia Goncalves, Ewan Harrison, David K. Jackson, Ian Johnston, Dominic Kwiatkowski, Cordelia Langford, John Sillitoe on behalf of the Wellcome Sanger Institute COVID-19 Surveillance Team |
| EPI_ISL_3140408, EPI_ISL_3140409, EPI_ISL_3140410, EPI_ISL_3140411 | Lighthouse Lab in Milton Keynes | Wellcome Sanger Institute for the COVID-19 Genomics UK (COG-UK) Consortium | The Lighthouse Lab in Milton Keynes and Alex Alderton, Roberto Amato, Jeffrey Barrett, Sonia Goncalves, Ewan Harrison, David K. Jackson, Ian Johnston, Dominic Kwiatkowski, Cordelia Langford, John Sillitoe on behalf of the Wellcome Sanger Institute COVID-19 Surveillance Team |
| EPI_ISL_3140412 | Lighthouse Lab in Glasgow | Wellcome Sanger Institute for the COVID-19 Genomics UK (COG-UK) Consortium | Harper VanSteenhouse, Yumi Kasai, David Gray, Carol Clugston, Anna Dominiczak and Alex Alderton, Roberto Amato, Jeffrey Barrett, Sonia Goncalves, Ewan Harrison, David K. Jackson, Ian Johnston, Dominic Kwiatkowski, Cordelia Langford, John Sillitoe on behalf of the Wellcome Sanger Institute COVID-19 Surveillance Team |
| EPI_ISL_3140413 | Lighthouse Lab in Milton Keynes | Wellcome Sanger Institute for the COVID-19 Genomics UK (COG-UK) Consortium | The Lighthouse Lab in Milton Keynes and Alex Alderton, Roberto Amato, Jeffrey Barrett, Sonia Goncalves, Ewan Harrison, David K. Jackson, Ian Johnston, Dominic Kwiatkowski, Cordelia Langford, John Sillitoe on behalf of the Wellcome Sanger Institute COVID-19 Surveillance Team |
| EPI_ISL_3140414 | Lighthouse Lab in Glasgow | Wellcome Sanger Institute for the COVID-19 Genomics UK (COG-UK) Consortium | Harper VanSteenhouse, Yumi Kasai, David Gray, Carol Clugston, Anna Dominiczak and Alex Alderton, Roberto Amato, Jeffrey Barrett, Sonia Goncalves, Ewan Harrison, David K. Jackson, Ian Johnston, Dominic Kwiatkowski, Cordelia Langford, John Sillitoe on behalf of the Wellcome Sanger Institute COVID-19 Surveillance Team |
| EPI_ISL_3140415, EPI_ISL_3140416, EPI_ISL_3140417, EPI_ISL_3140418 | Lighthouse Lab in Milton Keynes | Wellcome Sanger Institute for the COVID-19 Genomics UK (COG-UK) Consortium | The Lighthouse Lab in Milton Keynes and Alex Alderton, Roberto Amato, Jeffrey Barrett, Sonia Goncalves, Ewan Harrison, David K. Jackson, Ian Johnston, Dominic Kwiatkowski, Cordelia Langford, John Sillitoe on behalf of the Wellcome Sanger Institute COVID-19 Surveillance Team |
| EPI_ISL_3140419, EPI_ISL_3140420, EPI_ISL_3140421 | Lighthouse Lab in Alderley Park | Wellcome Sanger Institute for the COVID-19 Genomics UK (COG-UK) Consortium | Jacquelyn Wynn, Mairead Hyland, The Lighthouse Lab in Alderley Park and Alex Alderton, Roberto Amato, Jeffrey Barrett, Sonia Goncalves, Ewan Harrison, David K. Jackson, Ian Johnston, Dominic Kwiatkowski, Cordelia Langford, John Sillitoe on behalf of the Wellcome Sanger Institute COVID-19 Surveillance Team |
| EPI_ISL_3140422 | Lighthouse Lab in Glasgow | Wellcome Sanger Institute for the COVID-19 Genomics UK (COG-UK) Consortium | Harper VanSteenhouse, Yumi Kasai, David Gray, Carol Clugston, Anna Dominiczak and Alex Alderton, Roberto Amato, Jeffrey Barrett, Sonia Goncalves, Ewan Harrison, David K. Jackson, Ian Johnston, Dominic Kwiatkowski, Cordelia Langford, John Sillitoe on behalf of the Wellcome Sanger Institute COVID-19 Surveillance Team |
| EPI_ISL_3140423 | Lighthouse Lab in Alderley Park | Wellcome Sanger Institute for the COVID-19 Genomics UK (COG-UK) Consortium | Jacquelyn Wynn, Mairead Hyland, The Lighthouse Lab in Alderley Park and Alex Alderton, Roberto Amato, Jeffrey Barrett, Sonia Goncalves, Ewan Harrison, David K. Jackson, Ian Johnston, Dominic Kwiatkowski, Cordelia Langford, John Sillitoe on behalf of the Wellcome Sanger Institute COVID-19 Surveillance Team |
| EPI_ISL_3140424 | Lighthouse Lab in Milton Keynes | Wellcome Sanger Institute for the COVID-19 Genomics UK (COG-UK) Consortium | The Lighthouse Lab in Milton Keynes and Alex Alderton, Roberto Amato, Jeffrey Barrett, Sonia Goncalves, Ewan Harrison, David K. Jackson, Ian Johnston, Dominic Kwiatkowski, Cordelia Langford, John Sillitoe on behalf of the Wellcome Sanger Institute COVID-19 Surveillance Team |
| EPI_ISL_3140425 | Lighthouse Lab in Glasgow | Wellcome Sanger Institute for the COVID-19 Genomics UK (COG-UK) Consortium | Harper VanSteenhouse, Yumi Kasai, David Gray, Carol Clugston, Anna Dominiczak and Alex Alderton, Roberto Amato, Jeffrey Barrett, Sonia Goncalves, Ewan Harrison, David K. Jackson, Ian Johnston, Dominic Kwiatkowski, Cordelia Langford, John Sillitoe on behalf of the Wellcome Sanger Institute COVID-19 Surveillance Team |
| EPI_ISL_3140426, EPI_ISL_3140427 | Lighthouse Lab in Alderley Park | Wellcome Sanger Institute for the COVID-19 Genomics UK (COG-UK) Consortium | Jacquelyn Wynn, Mairead Hyland, The Lighthouse Lab in Alderley Park and Alex Alderton, Roberto Amato, Jeffrey Barrett, Sonia Goncalves, Ewan Harrison, David K. Jackson, Ian Johnston, Dominic Kwiatkowski, Cordelia Langford, John Sillitoe on behalf of the Wellcome Sanger Institute COVID-19 Surveillance Team |
| EPI_ISL_3140428 | Lighthouse Lab in Milton Keynes | Wellcome Sanger Institute for the COVID-19 Genomics UK (COG-UK) Consortium | The Lighthouse Lab in Milton Keynes and Alex Alderton, Roberto Amato, Jeffrey Barrett, Sonia Goncalves, Ewan Harrison, David K. Jackson, Ian Johnston, Dominic Kwiatkowski, Cordelia Langford, John Sillitoe on behalf of the Wellcome Sanger Institute COVID-19 Surveillance Team |
| EPI_ISL_3140429 | Lighthouse Lab in Alderley Park | Wellcome Sanger Institute for the COVID-19 Genomics UK (COG-UK) Consortium | Jacquelyn Wynn, Mairead Hyland, The Lighthouse Lab in Alderley Park and Alex Alderton, Roberto Amato, Jeffrey Barrett, Sonia Goncalves, Ewan Harrison, David K. Jackson, Ian Johnston, Dominic Kwiatkowski, Cordelia Langford, John Sillitoe on behalf of the Wellcome Sanger Institute COVID-19 Surveillance Team |
| EPI_ISL_3140430 | Lighthouse Lab in Glasgow | Wellcome Sanger Institute for the COVID-19 Genomics UK (COG-UK) Consortium | Harper VanSteenhouse, Yumi Kasai, David Gray, Carol Clugston, Anna Dominiczak and Alex Alderton, Roberto Amato, Jeffrey Barrett, Sonia Goncalves, Ewan Harrison, David K. Jackson, Ian Johnston, Dominic Kwiatkowski, Cordelia Langford, John Sillitoe on behalf of the Wellcome Sanger Institute COVID-19 Surveillance Team |
| EPI_ISL_3140431, EPI_ISL_3140432 | Lighthouse Lab in Milton Keynes | Wellcome Sanger Institute for the COVID-19 Genomics UK (COG-UK) Consortium | The Lighthouse Lab in Milton Keynes and Alex Alderton, Roberto Amato, Jeffrey Barrett, Sonia Goncalves, Ewan Harrison, David K. Jackson, Ian Johnston, Dominic Kwiatkowski, Cordelia Langford, John Sillitoe on behalf of the Wellcome Sanger Institute COVID-19 Surveillance Team |
| EPI_ISL_3140433 | Lighthouse Lab in Glasgow | Wellcome Sanger Institute for the COVID-19 Genomics UK (COG-UK) Consortium | Harper VanSteenhouse, Yumi Kasai, David Gray, Carol Clugston, Anna Dominiczak and Alex Alderton, Roberto Amato, Jeffrey Barrett, Sonia Goncalves, Ewan Harrison, David K. Jackson, Ian Johnston, Dominic Kwiatkowski, Cordelia Langford, John Sillitoe on behalf of the Wellcome Sanger Institute COVID-19 Surveillance Team |
| EPI_ISL_3140434, EPI_ISL_3140435 | Lighthouse Lab in Milton Keynes | Wellcome Sanger Institute for the COVID-19 Genomics UK (COG-UK) Consortium | The Lighthouse Lab in Milton Keynes and Alex Alderton, Roberto Amato, Jeffrey Barrett, Sonia Goncalves, Ewan Harrison, David K. Jackson, Ian Johnston, Dominic Kwiatkowski, Cordelia Langford, John Sillitoe on behalf of the Wellcome Sanger Institute COVID-19 Surveillance Team |
| EPI_ISL_3140436, EPI_ISL_3140437, EPI_ISL_3140438 | Lighthouse Lab in Glasgow | Wellcome Sanger Institute for the COVID-19 Genomics UK (COG-UK) Consortium | Harper VanSteenhouse, Yumi Kasai, David Gray, Carol Clugston, Anna Dominiczak and Alex Alderton, Roberto Amato, Jeffrey Barrett, Sonia Goncalves, Ewan Harrison, David K. Jackson, Ian Johnston, Dominic Kwiatkowski, Cordelia Langford, John Sillitoe on behalf of the Wellcome Sanger Institute COVID-19 Surveillance Team |
| EPI_ISL_3140439, EPI_ISL_3140440, EPI_ISL_3140441, EPI_ISL_3140442, EPI_ISL_3140443, EPI_ISL_3140444, EPI_ISL_3140445 | Lighthouse Lab in Milton Keynes | Wellcome Sanger Institute for the COVID-19 Genomics UK (COG-UK) Consortium | The Lighthouse Lab in Milton Keynes and Alex Alderton, Roberto Amato, Jeffrey Barrett, Sonia Goncalves, Ewan Harrison, David K. Jackson, Ian Johnston, Dominic Kwiatkowski, Cordelia Langford, John Sillitoe on behalf of the Wellcome Sanger Institute COVID-19 Surveillance Team |
| EPI_ISL_3140446, EPI_ISL_3140447 | Lighthouse Lab in Glasgow | Wellcome Sanger Institute for the COVID-19 Genomics UK (COG-UK) Consortium | Harper VanSteenhouse, Yumi Kasai, David Gray, Carol Clugston, Anna Dominiczak and Alex Alderton, Roberto Amato, Jeffrey Barrett, Sonia Goncalves, Ewan Harrison, David K. Jackson, Ian Johnston, Dominic Kwiatkowski, Cordelia Langford, John Sillitoe on behalf of the Wellcome Sanger Institute COVID-19 Surveillance Team |

| | | | |
|---|---|---|---|
| EPI_ISL_3140448 | Lighthouse Lab in Milton Keynes | Wellcome Sanger Institute for the COVID-19 Genomics UK (COG-UK) Consortium | The Lighthouse Lab in Milton Keynes and Alex Alderton, Roberto Amato, Jeffrey Barrett, Sonia Goncalves, Ewan Harrison, David K. Jackson, Ian Johnston, Dominic Kwiatkowski, Cordelia Langford, John Sillitoe on behalf of the Wellcome Sanger Institute COVID-19 Surveillance Team |
| EPI_ISL_3140449 | Lighthouse Lab in Alderley Park | Wellcome Sanger Institute for the COVID-19 Genomics UK (COG-UK) Consortium | Jacquelyn Wynn, Mairead Hyland, The Lighthouse Lab in Alderley Park and Alex Alderton, Roberto Amato, Jeffrey Barrett, Sonia Goncalves, Ewan Harrison, David K. Jackson, Ian Johnston, Dominic Kwiatkowski, Cordelia Langford, John Sillitoe on behalf of the Wellcome Sanger Institute COVID-19 Surveillance Team |
| EPI_ISL_3140450 | Lighthouse Lab in Milton Keynes | Wellcome Sanger Institute for the COVID-19 Genomics UK (COG-UK) Consortium | The Lighthouse Lab in Milton Keynes and Alex Alderton, Roberto Amato, Jeffrey Barrett, Sonia Goncalves, Ewan Harrison, David K. Jackson, Ian Johnston, Dominic Kwiatkowski, Cordelia Langford, John Sillitoe on behalf of the Wellcome Sanger Institute COVID-19 Surveillance Team |
| EPI_ISL_3140451, EPI_ISL_3140452 | Lighthouse Lab in Glasgow | Wellcome Sanger Institute for the COVID-19 Genomics UK (COG-UK) Consortium | Harper VanSteenhouse, Yumi Kasai, David Gray, Carol Clugston, Anna Dominiczak and Alex Alderton, Roberto Amato, Jeffrey Barrett, Sonia Goncalves, Ewan Harrison, David K. Jackson, Ian Johnston, Dominic Kwiatkowski, Cordelia Langford, John Sillitoe on behalf of the Wellcome Sanger Institute COVID-19 Surveillance Team |
| EPI_ISL_3140453, EPI_ISL_3140454, EPI_ISL_3140455, EPI_ISL_3140456 | Lighthouse Lab in Milton Keynes | Wellcome Sanger Institute for the COVID-19 Genomics UK (COG-UK) Consortium | The Lighthouse Lab in Milton Keynes and Alex Alderton, Roberto Amato, Jeffrey Barrett, Sonia Goncalves, Ewan Harrison, David K. Jackson, Ian Johnston, Dominic Kwiatkowski, Cordelia Langford, John Sillitoe on behalf of the Wellcome Sanger Institute COVID-19 Surveillance Team |
| EPI_ISL_3140457, EPI_ISL_3140458, EPI_ISL_3140459 | Lighthouse Lab in Glasgow | Wellcome Sanger Institute for the COVID-19 Genomics UK (COG-UK) Consortium | Harper VanSteenhouse, Yumi Kasai, David Gray, Carol Clugston, Anna Dominiczak and Alex Alderton, Roberto Amato, Jeffrey Barrett, Sonia Goncalves, Ewan Harrison, David K. Jackson, Ian Johnston, Dominic Kwiatkowski, Cordelia Langford, John Sillitoe on behalf of the Wellcome Sanger Institute COVID-19 Surveillance Team |
| EPI_ISL_3140460, EPI_ISL_3140461 | Lighthouse Lab in Milton Keynes | Wellcome Sanger Institute for the COVID-19 Genomics UK (COG-UK) Consortium | The Lighthouse Lab in Milton Keynes and Alex Alderton, Roberto Amato, Jeffrey Barrett, Sonia Goncalves, Ewan Harrison, David K. Jackson, Ian Johnston, Dominic Kwiatkowski, Cordelia Langford, John Sillitoe on behalf of the Wellcome Sanger Institute COVID-19 Surveillance Team |
| EPI_ISL_3140462, EPI_ISL_3140463, EPI_ISL_3140464 | Lighthouse Lab in Alderley Park | Wellcome Sanger Institute for the COVID-19 Genomics UK (COG-UK) Consortium | Jacquelyn Wynn, Mairead Hyland, The Lighthouse Lab in Alderley Park and Alex Alderton, Roberto Amato, Jeffrey Barrett, Sonia Goncalves, Ewan Harrison, David K. Jackson, Ian Johnston, Dominic Kwiatkowski, Cordelia Langford, John Sillitoe on behalf of the Wellcome Sanger Institute COVID-19 Surveillance Team |
| EPI_ISL_3140465 | Lighthouse Lab in Glasgow | Wellcome Sanger Institute for the COVID-19 Genomics UK (COG-UK) Consortium | Harper VanSteenhouse, Yumi Kasai, David Gray, Carol Clugston, Anna Dominiczak and Alex Alderton, Roberto Amato, Jeffrey Barrett, Sonia Goncalves, Ewan Harrison, David K. Jackson, Ian Johnston, Dominic Kwiatkowski, Cordelia Langford, John Sillitoe on behalf of the Wellcome Sanger Institute COVID-19 Surveillance Team |
| EPI_ISL_3140466 | Lighthouse Lab in Milton Keynes | Wellcome Sanger Institute for the COVID-19 Genomics UK (COG-UK) Consortium | The Lighthouse Lab in Milton Keynes and Alex Alderton, Roberto Amato, Jeffrey Barrett, Sonia Goncalves, Ewan Harrison, David K. Jackson, Ian Johnston, Dominic Kwiatkowski, Cordelia Langford, John Sillitoe on behalf of the Wellcome Sanger Institute COVID-19 Surveillance Team |
| EPI_ISL_3140467 | Lighthouse Lab in Alderley Park | Wellcome Sanger Institute for the COVID-19 Genomics UK (COG-UK) Consortium | Jacquelyn Wynn, Mairead Hyland, The Lighthouse Lab in Alderley Park and Alex Alderton, Roberto Amato, Jeffrey Barrett, Sonia Goncalves, Ewan Harrison, David K. Jackson, Ian Johnston, Dominic Kwiatkowski, Cordelia Langford, John Sillitoe on behalf of the Wellcome Sanger Institute COVID-19 Surveillance Team |
| EPI_ISL_3140468, EPI_ISL_3140469, EPI_ISL_3140470, EPI_ISL_3140471 | Lighthouse Lab in Glasgow | Wellcome Sanger Institute for the COVID-19 Genomics UK (COG-UK) Consortium | Harper VanSteenhouse, Yumi Kasai, David Gray, Carol Clugston, Anna Dominiczak and Alex Alderton, Roberto Amato, Jeffrey Barrett, Sonia Goncalves, Ewan Harrison, David K. Jackson, Ian Johnston, Dominic Kwiatkowski, Cordelia Langford, John Sillitoe on behalf of the Wellcome Sanger Institute COVID-19 Surveillance Team |
| EPI_ISL_3140472, EPI_ISL_3140473 | Lighthouse Lab in Milton Keynes | Wellcome Sanger Institute for the COVID-19 Genomics UK (COG-UK) Consortium | The Lighthouse Lab in Milton Keynes and Alex Alderton, Roberto Amato, Jeffrey Barrett, Sonia Goncalves, Ewan Harrison, David K. Jackson, Ian Johnston, Dominic Kwiatkowski, Cordelia Langford, John Sillitoe on behalf of the Wellcome Sanger Institute COVID-19 Surveillance Team |
| EPI_ISL_3140474, EPI_ISL_3140475, EPI_ISL_3140476 | Lighthouse Lab in Alderley Park | Wellcome Sanger Institute for the COVID-19 Genomics UK (COG-UK) Consortium | Jacquelyn Wynn, Mairead Hyland, The Lighthouse Lab in Alderley Park and Alex Alderton, Roberto Amato, Jeffrey Barrett, Sonia Goncalves, Ewan Harrison, David K. Jackson, Ian Johnston, Dominic Kwiatkowski, Cordelia Langford, John Sillitoe on behalf of the Wellcome Sanger Institute COVID-19 Surveillance Team |
| EPI_ISL_3140477 | Lighthouse Lab in Milton Keynes | Wellcome Sanger Institute for the COVID-19 Genomics UK (COG-UK) Consortium | The Lighthouse Lab in Milton Keynes and Alex Alderton, Roberto Amato, Jeffrey Barrett, Sonia Goncalves, Ewan Harrison, David K. Jackson, Ian Johnston, Dominic Kwiatkowski, Cordelia Langford, John Sillitoe on behalf of the Wellcome Sanger Institute COVID-19 Surveillance Team |
| EPI_ISL_3140478 | Lighthouse Lab in Alderley Park | Wellcome Sanger Institute for the COVID-19 Genomics UK (COG-UK) Consortium | Jacquelyn Wynn, Mairead Hyland, The Lighthouse Lab in Alderley Park and Alex Alderton, Roberto Amato, Jeffrey Barrett, Sonia Goncalves, Ewan Harrison, David K. Jackson, Ian Johnston, Dominic Kwiatkowski, Cordelia Langford, John Sillitoe on behalf of the Wellcome Sanger Institute COVID-19 Surveillance Team |
| EPI_ISL_3140479, EPI_ISL_3140480 | Lighthouse Lab in Milton Keynes | Wellcome Sanger Institute for the COVID-19 Genomics UK (COG-UK) Consortium | The Lighthouse Lab in Milton Keynes and Alex Alderton, Roberto Amato, Jeffrey Barrett, Sonia Goncalves, Ewan Harrison, David K. Jackson, Ian Johnston, Dominic Kwiatkowski, Cordelia Langford, John Sillitoe on behalf of the Wellcome Sanger Institute COVID-19 Surveillance Team |
| EPI_ISL_3140481 | Lighthouse Lab in Alderley Park | Wellcome Sanger Institute for the COVID-19 Genomics UK (COG-UK) Consortium | Jacquelyn Wynn, Mairead Hyland, The Lighthouse Lab in Alderley Park and Alex Alderton, Roberto Amato, Jeffrey Barrett, Sonia Goncalves, Ewan Harrison, David K. Jackson, Ian Johnston, Dominic Kwiatkowski, Cordelia Langford, John Sillitoe on behalf of the Wellcome Sanger Institute COVID-19 Surveillance Team |
| EPI_ISL_3140482, EPI_ISL_3140483, EPI_ISL_3140484, EPI_ISL_3140485, EPI_ISL_3140486, EPI_ISL_3140487 | Lighthouse Lab in Milton Keynes | Wellcome Sanger Institute for the COVID-19 Genomics UK (COG-UK) Consortium | The Lighthouse Lab in Milton Keynes and Alex Alderton, Roberto Amato, Jeffrey Barrett, Sonia Goncalves, Ewan Harrison, David K. Jackson, Ian Johnston, Dominic Kwiatkowski, Cordelia Langford, John Sillitoe on behalf of the Wellcome Sanger Institute COVID-19 Surveillance Team |
| EPI_ISL_3140488 | Lighthouse Lab in Alderley Park | Wellcome Sanger Institute for the COVID-19 Genomics UK (COG-UK) Consortium | Jacquelyn Wynn, Mairead Hyland, The Lighthouse Lab in Alderley Park and Alex Alderton, Roberto Amato, Jeffrey Barrett, Sonia Goncalves, Ewan Harrison, David K. Jackson, Ian Johnston, Dominic Kwiatkowski, Cordelia Langford, John Sillitoe on behalf of the Wellcome Sanger Institute COVID-19 Surveillance Team |
| EPI_ISL_3140489 | Lighthouse Lab in Milton Keynes | Wellcome Sanger Institute for the COVID-19 Genomics UK (COG-UK) Consortium | The Lighthouse Lab in Milton Keynes and Alex Alderton, Roberto Amato, Jeffrey Barrett, Sonia Goncalves, Ewan Harrison, David K. Jackson, Ian Johnston, Dominic Kwiatkowski, Cordelia Langford, John Sillitoe on behalf of the Wellcome Sanger Institute COVID-19 Surveillance Team |
| EPI_ISL_3140490 | Lighthouse Lab in Glasgow | Wellcome Sanger Institute for the COVID-19 Genomics UK (COG-UK) Consortium | Harper VanSteenhouse, Yumi Kasai, David Gray, Carol Clugston, Anna Dominiczak and Alex Alderton, Roberto Amato, Jeffrey Barrett, Sonia Goncalves, Ewan Harrison, David K. Jackson, Ian Johnston, Dominic Kwiatkowski, Cordelia Langford, John Sillitoe on behalf of the Wellcome Sanger Institute COVID-19 Surveillance Team |
| EPI_ISL_3140491 | Lighthouse Lab in Milton Keynes | Wellcome Sanger Institute for the COVID-19 Genomics UK (COG-UK) Consortium | The Lighthouse Lab in Milton Keynes and Alex Alderton, Roberto Amato, Jeffrey Barrett, Sonia Goncalves, Ewan Harrison, David K. Jackson, Ian Johnston, Dominic Kwiatkowski, Cordelia Langford, John Sillitoe on behalf of the Wellcome Sanger Institute COVID-19 Surveillance Team |
| EPI_ISL_3140492 | Lighthouse Lab in Glasgow | Wellcome Sanger Institute for the COVID-19 Genomics UK (COG-UK) Consortium | Harper VanSteenhouse, Yumi Kasai, David Gray, Carol Clugston, Anna Dominiczak and Alex Alderton, Roberto Amato, Jeffrey Barrett, Sonia Goncalves, Ewan Harrison, David K. Jackson, Ian Johnston, Dominic Kwiatkowski, Cordelia Langford, John Sillitoe on behalf of the Wellcome Sanger Institute COVID-19 Surveillance Team |
| EPI_ISL_3140493, EPI_ISL_3140494 | Lighthouse Lab in Milton Keynes | Wellcome Sanger Institute for the COVID-19 Genomics UK (COG-UK) Consortium | The Lighthouse Lab in Milton Keynes and Alex Alderton, Roberto Amato, Jeffrey Barrett, Sonia Goncalves, Ewan Harrison, David K. Jackson, Ian Johnston, Dominic Kwiatkowski, Cordelia Langford, John Sillitoe on behalf of the Wellcome Sanger Institute COVID-19 Surveillance Team |
| EPI_ISL_3140495, EPI_ISL_3140496 | Lighthouse Lab in Alderley Park | Wellcome Sanger Institute for the COVID-19 Genomics UK (COG-UK) Consortium | Jacquelyn Wynn, Mairead Hyland, The Lighthouse Lab in Alderley Park and Alex Alderton, Roberto Amato, Jeffrey Barrett, Sonia Goncalves, Ewan Harrison, David K. Jackson, Ian Johnston, Dominic Kwiatkowski, Cordelia Langford, John Sillitoe on behalf of the Wellcome Sanger Institute COVID-19 Surveillance Team |
| EPI_ISL_3140497, EPI_ISL_3140498, EPI_ISL_3140499 | Lighthouse Lab in Milton Keynes | Wellcome Sanger Institute for the COVID-19 Genomics UK (COG-UK) Consortium | The Lighthouse Lab in Milton Keynes and Alex Alderton, Roberto Amato, Jeffrey Barrett, Sonia Goncalves, Ewan Harrison, David K. Jackson, Ian Johnston, Dominic Kwiatkowski, Cordelia Langford, John Sillitoe on behalf of the Wellcome Sanger Institute COVID-19 Surveillance Team |
| EPI_ISL_3140500 | Lighthouse Lab in Glasgow | Wellcome Sanger Institute for the COVID-19 Genomics UK (COG-UK) Consortium | Harper VanSteenhouse, Yumi Kasai, David Gray, Carol Clugston, Anna Dominiczak and Alex Alderton, Roberto Amato, Jeffrey Barrett, Sonia Goncalves, Ewan Harrison, David K. Jackson, Ian Johnston, Dominic Kwiatkowski, Cordelia Langford, John Sillitoe on behalf of the Wellcome Sanger Institute COVID-19 Surveillance Team |
| EPI_ISL_3140501 | Lighthouse Lab in Milton Keynes | Wellcome Sanger Institute for the COVID-19 Genomics UK (COG-UK) Consortium | The Lighthouse Lab in Milton Keynes and Alex Alderton, Roberto Amato, Jeffrey Barrett, Sonia Goncalves, Ewan Harrison, David K. Jackson, Ian Johnston, Dominic Kwiatkowski, Cordelia Langford, John Sillitoe on behalf of the Wellcome Sanger Institute COVID-19 Surveillance Team |
| EPI_ISL_3140502, EPI_ISL_3140503 | Lighthouse Lab in Alderley Park | Wellcome Sanger Institute for the COVID-19 Genomics UK (COG-UK) Consortium | Jacquelyn Wynn, Mairead Hyland, The Lighthouse Lab in Alderley Park and Alex Alderton, Roberto Amato, Jeffrey Barrett, Sonia Goncalves, Ewan Harrison, David K. Jackson, Ian Johnston, Dominic Kwiatkowski, Cordelia Langford, John Sillitoe on behalf of the Wellcome Sanger Institute COVID-19 Surveillance Team |
| EPI_ISL_3140504, EPI_ISL_3140505, EPI_ISL_3140506, EPI_ISL_3140507, EPI_ISL_3140508 | Lighthouse Lab in Milton Keynes | Wellcome Sanger Institute for the COVID-19 Genomics UK (COG-UK) Consortium | The Lighthouse Lab in Milton Keynes and Alex Alderton, Roberto Amato, Jeffrey Barrett, Sonia Goncalves, Ewan Harrison, David K. Jackson, Ian Johnston, Dominic Kwiatkowski, Cordelia Langford, John Sillitoe on behalf of the Wellcome Sanger Institute COVID-19 Surveillance Team |
| EPI_ISL_3140509 | Lighthouse Lab in Glasgow | Wellcome Sanger Institute for the COVID-19 Genomics UK (COG-UK) Consortium | Harper VanSteenhouse, Yumi Kasai, David Gray, Carol Clugston, Anna Dominiczak and Alex Alderton, Roberto Amato, Jeffrey Barrett, Sonia Goncalves, Ewan Harrison, David K. Jackson, Ian Johnston, Dominic Kwiatkowski, Cordelia Langford, John Sillitoe on behalf of the Wellcome Sanger Institute COVID-19 Surveillance Team |
| EPI_ISL_3140510 | Lighthouse Lab in Milton Keynes | Wellcome Sanger Institute for the COVID-19 Genomics UK (COG-UK) Consortium | The Lighthouse Lab in Milton Keynes and Alex Alderton, Roberto Amato, Jeffrey Barrett, Sonia Goncalves, Ewan Harrison, David K. Jackson, Ian Johnston, Dominic Kwiatkowski, Cordelia Langford, John Sillitoe on behalf of the Wellcome Sanger Institute COVID-19 Surveillance Team |
| EPI_ISL_3140511 | Lighthouse Lab in Alderley Park | Wellcome Sanger Institute for the COVID-19 Genomics UK (COG-UK) Consortium | Jacquelyn Wynn, Mairead Hyland, The Lighthouse Lab in Alderley Park and Alex Alderton, Roberto Amato, Jeffrey Barrett, Sonia Goncalves, Ewan Harrison, David K. Jackson, Ian Johnston, Dominic Kwiatkowski, Cordelia Langford, John Sillitoe on behalf of the Wellcome Sanger Institute COVID-19 Surveillance Team |
| EPI_ISL_3140512 | Lighthouse Lab in Milton Keynes | Wellcome Sanger Institute for the COVID-19 Genomics UK (COG-UK) Consortium | The Lighthouse Lab in Milton Keynes and Alex Alderton, Roberto Amato, Jeffrey Barrett, Sonia Goncalves, Ewan Harrison, David K. Jackson, Ian Johnston, Dominic Kwiatkowski, Cordelia Langford, John Sillitoe on behalf of the Wellcome Sanger Institute COVID-19 Surveillance Team |
| EPI_ISL_3140513 | Lighthouse Lab in Glasgow | Wellcome Sanger Institute for the COVID-19 Genomics UK (COG-UK) Consortium | Harper VanSteenhouse, Yumi Kasai, David Gray, Carol Clugston, Anna Dominiczak and Alex Alderton, Roberto Amato, Jeffrey Barrett, Sonia Goncalves, Ewan Harrison, David K. Jackson, Ian Johnston, Dominic Kwiatkowski, Cordelia Langford, John Sillitoe on behalf of the Wellcome Sanger Institute COVID-19 Surveillance Team |
| EPI_ISL_3140514 | Lighthouse Lab in Milton Keynes | Wellcome Sanger Institute for the COVID-19 Genomics UK (COG-UK) Consortium | The Lighthouse Lab in Milton Keynes and Alex Alderton, Roberto Amato, Jeffrey Barrett, Sonia Goncalves, Ewan Harrison, David K. Jackson, Ian Johnston, Dominic Kwiatkowski, Cordelia Langford, John Sillitoe on behalf of the Wellcome Sanger Institute COVID-19 Surveillance Team |
| EPI_ISL_3140515 | Lighthouse Lab in Alderley Park | Wellcome Sanger Institute for the COVID-19 Genomics UK (COG-UK) Consortium | Jacquelyn Wynn, Mairead Hyland, The Lighthouse Lab in Alderley Park and Alex Alderton, Roberto Amato, Jeffrey Barrett, Sonia Goncalves, Ewan Harrison, David K. Jackson, Ian Johnston, Dominic Kwiatkowski, Cordelia Langford, John Sillitoe on behalf of the Wellcome Sanger Institute COVID-19 Surveillance Team |
| EPI_ISL_3140516, EPI_ISL_3140517, | Lighthouse Lab in Milton Keynes | Wellcome Sanger Institute for the COVID-19 Genomics UK (COG-UK) Consortium | The Lighthouse Lab in Milton Keynes and Alex Alderton, Roberto Amato, Jeffrey Barrett, Sonia Goncalves, Ewan Harrison, David K. Jackson, Ian Johnston, Dominic Kwiatkowski, Cordelia Langford, John Sillitoe on behalf of the Wellcome Sanger Institute COVID-19 Surveillance Team |

| | | | |
|---|---|---|---|
| EPI_ISL_3140518, EPI_ISL_3140519 | | | |
| EPI_ISL_3140520, EPI_ISL_3140521 | Lighthouse Lab in Alderley Park | Wellcome Sanger Institute for the COVID-19 Genomics UK (COG-UK) Consortium | Jacquelyn Wynn, Mairead Hyland, The Lighthouse Lab in Alderley Park and Alex Alderton, Roberto Amato, Jeffrey Barrett, Sonia Goncalves, Ewan Harrison, David K. Jackson, Ian Johnston, Dominic Kwiatkowski, Cordelia Langford, John Sillitoe on behalf of the Wellcome Sanger Institute COVID-19 Surveillance Team |
| EPI_ISL_3140522 | Lighthouse Lab in Glasgow | Wellcome Sanger Institute for the COVID-19 Genomics UK (COG-UK) Consortium | Harper VanSteenhouse, Yumi Kasai, David Gray, Carol Clugston, Anna Dominiczak and Alex Alderton, Roberto Amato, Jeffrey Barrett, Sonia Goncalves, Ewan Harrison, David K. Jackson, Ian Johnston, Dominic Kwiatkowski, Cordelia Langford, John Sillitoe on behalf of the Wellcome Sanger Institute COVID-19 Surveillance Team |
| EPI_ISL_3140523 | Lighthouse Lab in Alderley Park | Wellcome Sanger Institute for the COVID-19 Genomics UK (COG-UK) Consortium | Jacquelyn Wynn, Mairead Hyland, The Lighthouse Lab in Alderley Park and Alex Alderton, Roberto Amato, Jeffrey Barrett, Sonia Goncalves, Ewan Harrison, David K. Jackson, Ian Johnston, Dominic Kwiatkowski, Cordelia Langford, John Sillitoe on behalf of the Wellcome Sanger Institute COVID-19 Surveillance Team |
| EPI_ISL_3140524, EPI_ISL_3140525, EPI_ISL_3140526 | Lighthouse Lab in Milton Keynes | Wellcome Sanger Institute for the COVID-19 Genomics UK (COG-UK) Consortium | The Lighthouse Lab in Milton Keynes and Alex Alderton, Roberto Amato, Jeffrey Barrett, Sonia Goncalves, Ewan Harrison, David K. Jackson, Ian Johnston, Dominic Kwiatkowski, Cordelia Langford, John Sillitoe on behalf of the Wellcome Sanger Institute COVID-19 Surveillance Team |
| EPI_ISL_3140527 | Lighthouse Lab in Alderley Park | Wellcome Sanger Institute for the COVID-19 Genomics UK (COG-UK) Consortium | Jacquelyn Wynn, Mairead Hyland, The Lighthouse Lab in Alderley Park and Alex Alderton, Roberto Amato, Jeffrey Barrett, Sonia Goncalves, Ewan Harrison, David K. Jackson, Ian Johnston, Dominic Kwiatkowski, Cordelia Langford, John Sillitoe on behalf of the Wellcome Sanger Institute COVID-19 Surveillance Team |
| EPI_ISL_3140528 | Lighthouse Lab in Milton Keynes | Wellcome Sanger Institute for the COVID-19 Genomics UK (COG-UK) Consortium | The Lighthouse Lab in Milton Keynes and Alex Alderton, Roberto Amato, Jeffrey Barrett, Sonia Goncalves, Ewan Harrison, David K. Jackson, Ian Johnston, Dominic Kwiatkowski, Cordelia Langford, John Sillitoe on behalf of the Wellcome Sanger Institute COVID-19 Surveillance Team |
| EPI_ISL_3140529 | Lighthouse Lab in Alderley Park | Wellcome Sanger Institute for the COVID-19 Genomics UK (COG-UK) Consortium | Jacquelyn Wynn, Mairead Hyland, The Lighthouse Lab in Alderley Park and Alex Alderton, Roberto Amato, Jeffrey Barrett, Sonia Goncalves, Ewan Harrison, David K. Jackson, Ian Johnston, Dominic Kwiatkowski, Cordelia Langford, John Sillitoe on behalf of the Wellcome Sanger Institute COVID-19 Surveillance Team |
| EPI_ISL_3140530 | Lighthouse Lab in Glasgow | Wellcome Sanger Institute for the COVID-19 Genomics UK (COG-UK) Consortium | Harper VanSteenhouse, Yumi Kasai, David Gray, Carol Clugston, Anna Dominiczak and Alex Alderton, Roberto Amato, Jeffrey Barrett, Sonia Goncalves, Ewan Harrison, David K. Jackson, Ian Johnston, Dominic Kwiatkowski, Cordelia Langford, John Sillitoe on behalf of the Wellcome Sanger Institute COVID-19 Surveillance Team |
| EPI_ISL_3140531, EPI_ISL_3140532 | Berkshire and Surrey Pathology Services Lighthouse Laboratory | Wellcome Sanger Institute for the COVID-19 Genomics UK (COG-UK) Consortium | Berkshire and Surrey Pathology Services Lighthouse Laboratory and Alex Alderton, Roberto Amato, Jeffrey Barrett, Sonia Goncalves, Ewan Harrison, David K. Jackson, Ian Johnston, Dominic Kwiatkowski, Cordelia Langford, John Sillitoe on behalf of the Wellcome Sanger Institute COVID-19 Surveillance Team |
| EPI_ISL_3140533 | Lighthouse Lab in Milton Keynes | Wellcome Sanger Institute for the COVID-19 Genomics UK (COG-UK) Consortium | The Lighthouse Lab in Milton Keynes and Alex Alderton, Roberto Amato, Jeffrey Barrett, Sonia Goncalves, Ewan Harrison, David K. Jackson, Ian Johnston, Dominic Kwiatkowski, Cordelia Langford, John Sillitoe on behalf of the Wellcome Sanger Institute COVID-19 Surveillance Team |
| EPI_ISL_3140534 | Lighthouse Lab in Alderley Park | Wellcome Sanger Institute for the COVID-19 Genomics UK (COG-UK) Consortium | Jacquelyn Wynn, Mairead Hyland, The Lighthouse Lab in Alderley Park and Alex Alderton, Roberto Amato, Jeffrey Barrett, Sonia Goncalves, Ewan Harrison, David K. Jackson, Ian Johnston, Dominic Kwiatkowski, Cordelia Langford, John Sillitoe on behalf of the Wellcome Sanger Institute COVID-19 Surveillance Team |
| EPI_ISL_3140535 | Lighthouse Lab in Glasgow | Wellcome Sanger Institute for the COVID-19 Genomics UK (COG-UK) Consortium | Harper VanSteenhouse, Yumi Kasai, David Gray, Carol Clugston, Anna Dominiczak and Alex Alderton, Roberto Amato, Jeffrey Barrett, Sonia Goncalves, Ewan Harrison, David K. Jackson, Ian Johnston, Dominic Kwiatkowski, Cordelia Langford, John Sillitoe on behalf of the Wellcome Sanger Institute COVID-19 Surveillance Team |
| EPI_ISL_3140536 | Berkshire and Surrey Pathology Services Lighthouse Laboratory | Wellcome Sanger Institute for the COVID-19 Genomics UK (COG-UK) Consortium | Berkshire and Surrey Pathology Services Lighthouse Laboratory and Alex Alderton, Roberto Amato, Jeffrey Barrett, Sonia Goncalves, Ewan Harrison, David K. Jackson, Ian Johnston, Dominic Kwiatkowski, Cordelia Langford, John Sillitoe on behalf of the Wellcome Sanger Institute COVID-19 Surveillance Team |
| EPI_ISL_3140537, EPI_ISL_3140538 | Lighthouse Lab in Glasgow | Wellcome Sanger Institute for the COVID-19 Genomics UK (COG-UK) Consortium | Harper VanSteenhouse, Yumi Kasai, David Gray, Carol Clugston, Anna Dominiczak and Alex Alderton, Roberto Amato, Jeffrey Barrett, Sonia Goncalves, Ewan Harrison, David K. Jackson, Ian Johnston, Dominic Kwiatkowski, Cordelia Langford, John Sillitoe on behalf of the Wellcome Sanger Institute COVID-19 Surveillance Team |
| EPI_ISL_3140539, EPI_ISL_3140540 | Berkshire and Surrey Pathology Services Lighthouse Laboratory | Wellcome Sanger Institute for the COVID-19 Genomics UK (COG-UK) Consortium | Berkshire and Surrey Pathology Services Lighthouse Laboratory and Alex Alderton, Roberto Amato, Jeffrey Barrett, Sonia Goncalves, Ewan Harrison, David K. Jackson, Ian Johnston, Dominic Kwiatkowski, Cordelia Langford, John Sillitoe on behalf of the Wellcome Sanger Institute COVID-19 Surveillance Team |
| EPI_ISL_3140541 | Lighthouse Lab in Milton Keynes | Wellcome Sanger Institute for the COVID-19 Genomics UK (COG-UK) Consortium | The Lighthouse Lab in Milton Keynes and Alex Alderton, Roberto Amato, Jeffrey Barrett, Sonia Goncalves, Ewan Harrison, David K. Jackson, Ian Johnston, Dominic Kwiatkowski, Cordelia Langford, John Sillitoe on behalf of the Wellcome Sanger Institute COVID-19 Surveillance Team |
| EPI_ISL_3140542, EPI_ISL_3140543 | Lighthouse Lab in Alderley Park | Wellcome Sanger Institute for the COVID-19 Genomics UK (COG-UK) Consortium | Jacquelyn Wynn, Mairead Hyland, The Lighthouse Lab in Alderley Park and Alex Alderton, Roberto Amato, Jeffrey Barrett, Sonia Goncalves, Ewan Harrison, David K. Jackson, Ian Johnston, Dominic Kwiatkowski, Cordelia Langford, John Sillitoe on behalf of the Wellcome Sanger Institute COVID-19 Surveillance Team |
| EPI_ISL_3140544 | Lighthouse Lab in Glasgow | Wellcome Sanger Institute for the COVID-19 Genomics UK (COG-UK) Consortium | Harper VanSteenhouse, Yumi Kasai, David Gray, Carol Clugston, Anna Dominiczak and Alex Alderton, Roberto Amato, Jeffrey Barrett, Sonia Goncalves, Ewan Harrison, David K. Jackson, Ian Johnston, Dominic Kwiatkowski, Cordelia Langford, John Sillitoe on behalf of the Wellcome Sanger Institute COVID-19 Surveillance Team |
| EPI_ISL_3140545 | Lighthouse Lab in Alderley Park | Wellcome Sanger Institute for the COVID-19 Genomics UK (COG-UK) Consortium | Jacquelyn Wynn, Mairead Hyland, The Lighthouse Lab in Alderley Park and Alex Alderton, Roberto Amato, Jeffrey Barrett, Sonia Goncalves, Ewan Harrison, David K. Jackson, Ian Johnston, Dominic Kwiatkowski, Cordelia Langford, John Sillitoe on behalf of the Wellcome Sanger Institute COVID-19 Surveillance Team |
| EPI_ISL_3140546, EPI_ISL_3140547 | Lighthouse Lab in Glasgow | Wellcome Sanger Institute for the COVID-19 Genomics UK (COG-UK) Consortium | Harper VanSteenhouse, Yumi Kasai, David Gray, Carol Clugston, Anna Dominiczak and Alex Alderton, Roberto Amato, Jeffrey Barrett, Sonia Goncalves, Ewan Harrison, David K. Jackson, Ian Johnston, Dominic Kwiatkowski, Cordelia Langford, John Sillitoe on behalf of the Wellcome Sanger Institute COVID-19 Surveillance Team |
| EPI_ISL_3140548 | Lighthouse Lab in Milton Keynes | Wellcome Sanger Institute for the COVID-19 Genomics UK (COG-UK) Consortium | The Lighthouse Lab in Milton Keynes and Alex Alderton, Roberto Amato, Jeffrey Barrett, Sonia Goncalves, Ewan Harrison, David K. Jackson, Ian Johnston, Dominic Kwiatkowski, Cordelia Langford, John Sillitoe on behalf of the Wellcome Sanger Institute COVID-19 Surveillance Team |
| EPI_ISL_3140549, EPI_ISL_3140550, EPI_ISL_3140551, EPI_ISL_3140552 | Lighthouse Lab in Alderley Park | Wellcome Sanger Institute for the COVID-19 Genomics UK (COG-UK) Consortium | Jacquelyn Wynn, Mairead Hyland, The Lighthouse Lab in Alderley Park and Alex Alderton, Roberto Amato, Jeffrey Barrett, Sonia Goncalves, Ewan Harrison, David K. Jackson, Ian Johnston, Dominic Kwiatkowski, Cordelia Langford, John Sillitoe on behalf of the Wellcome Sanger Institute COVID-19 Surveillance Team |
| EPI_ISL_3140553 | Lighthouse Lab in Milton Keynes | Wellcome Sanger Institute for the COVID-19 Genomics UK (COG-UK) Consortium | The Lighthouse Lab in Milton Keynes and Alex Alderton, Roberto Amato, Jeffrey Barrett, Sonia Goncalves, Ewan Harrison, David K. Jackson, Ian Johnston, Dominic Kwiatkowski, Cordelia Langford, John Sillitoe on behalf of the Wellcome Sanger Institute COVID-19 Surveillance Team |
| EPI_ISL_3140554 | Lighthouse Lab in Glasgow | Wellcome Sanger Institute for the COVID-19 Genomics UK (COG-UK) Consortium | Harper VanSteenhouse, Yumi Kasai, David Gray, Carol Clugston, Anna Dominiczak and Alex Alderton, Roberto Amato, Jeffrey Barrett, Sonia Goncalves, Ewan Harrison, David K. Jackson, Ian Johnston, Dominic Kwiatkowski, Cordelia Langford, John Sillitoe on behalf of the Wellcome Sanger Institute COVID-19 Surveillance Team |
| EPI_ISL_3140555 | Lighthouse Lab in Alderley Park | Wellcome Sanger Institute for the COVID-19 Genomics UK (COG-UK) Consortium | Jacquelyn Wynn, Mairead Hyland, The Lighthouse Lab in Alderley Park and Alex Alderton, Roberto Amato, Jeffrey Barrett, Sonia Goncalves, Ewan Harrison, David K. Jackson, Ian Johnston, Dominic Kwiatkowski, Cordelia Langford, John Sillitoe on behalf of the Wellcome Sanger Institute COVID-19 Surveillance Team |
| EPI_ISL_3140556 | Lighthouse Lab in Milton Keynes | Wellcome Sanger Institute for the COVID-19 Genomics UK (COG-UK) Consortium | The Lighthouse Lab in Milton Keynes and Alex Alderton, Roberto Amato, Jeffrey Barrett, Sonia Goncalves, Ewan Harrison, David K. Jackson, Ian Johnston, Dominic Kwiatkowski, Cordelia Langford, John Sillitoe on behalf of the Wellcome Sanger Institute COVID-19 Surveillance Team |
| EPI_ISL_3140557, EPI_ISL_3140558 | Lighthouse Lab in Alderley Park | Wellcome Sanger Institute for the COVID-19 Genomics UK (COG-UK) Consortium | Jacquelyn Wynn, Mairead Hyland, The Lighthouse Lab in Alderley Park and Alex Alderton, Roberto Amato, Jeffrey Barrett, Sonia Goncalves, Ewan Harrison, David K. Jackson, Ian Johnston, Dominic Kwiatkowski, Cordelia Langford, John Sillitoe on behalf of the Wellcome Sanger Institute COVID-19 Surveillance Team |
| EPI_ISL_3140559 | Lighthouse Lab in Glasgow | Wellcome Sanger Institute for the COVID-19 Genomics UK (COG-UK) Consortium | Harper VanSteenhouse, Yumi Kasai, David Gray, Carol Clugston, Anna Dominiczak and Alex Alderton, Roberto Amato, Jeffrey Barrett, Sonia Goncalves, Ewan Harrison, David K. Jackson, Ian Johnston, Dominic Kwiatkowski, Cordelia Langford, John Sillitoe on behalf of the Wellcome Sanger Institute COVID-19 Surveillance Team |
| EPI_ISL_3140560 | Lighthouse Lab in Alderley Park | Wellcome Sanger Institute for the COVID-19 Genomics UK (COG-UK) Consortium | Jacquelyn Wynn, Mairead Hyland, The Lighthouse Lab in Alderley Park and Alex Alderton, Roberto Amato, Jeffrey Barrett, Sonia Goncalves, Ewan Harrison, David K. Jackson, Ian Johnston, Dominic Kwiatkowski, Cordelia Langford, John Sillitoe on behalf of the Wellcome Sanger Institute COVID-19 Surveillance Team |
| EPI_ISL_3140561 | Lighthouse Lab in Glasgow | Wellcome Sanger Institute for the COVID-19 Genomics UK (COG-UK) Consortium | Harper VanSteenhouse, Yumi Kasai, David Gray, Carol Clugston, Anna Dominiczak and Alex Alderton, Roberto Amato, Jeffrey Barrett, Sonia Goncalves, Ewan Harrison, David K. Jackson, Ian Johnston, Dominic Kwiatkowski, Cordelia Langford, John Sillitoe on behalf of the Wellcome Sanger Institute COVID-19 Surveillance Team |
| EPI_ISL_3140562 | Lighthouse Lab in Milton Keynes | Wellcome Sanger Institute for the COVID-19 Genomics UK (COG-UK) Consortium | The Lighthouse Lab in Milton Keynes and Alex Alderton, Roberto Amato, Jeffrey Barrett, Sonia Goncalves, Ewan Harrison, David K. Jackson, Ian Johnston, Dominic Kwiatkowski, Cordelia Langford, John Sillitoe on behalf of the Wellcome Sanger Institute COVID-19 Surveillance Team |
| EPI_ISL_3140563, EPI_ISL_3140564 | Lighthouse Lab in Glasgow | Wellcome Sanger Institute for the COVID-19 Genomics UK (COG-UK) Consortium | Harper VanSteenhouse, Yumi Kasai, David Gray, Carol Clugston, Anna Dominiczak and Alex Alderton, Roberto Amato, Jeffrey Barrett, Sonia Goncalves, Ewan Harrison, David K. Jackson, Ian Johnston, Dominic Kwiatkowski, Cordelia Langford, John Sillitoe on behalf of the Wellcome Sanger Institute COVID-19 Surveillance Team |
| EPI_ISL_3140565 | Lighthouse Lab in Alderley Park | Wellcome Sanger Institute for the COVID-19 Genomics UK (COG-UK) Consortium | Jacquelyn Wynn, Mairead Hyland, The Lighthouse Lab in Alderley Park and Alex Alderton, Roberto Amato, Jeffrey Barrett, Sonia Goncalves, Ewan Harrison, David K. Jackson, Ian Johnston, Dominic Kwiatkowski, Cordelia Langford, John Sillitoe on behalf of the Wellcome Sanger Institute COVID-19 Surveillance Team |
| EPI_ISL_3140566, EPI_ISL_3140567 | Berkshire and Surrey Pathology Services Lighthouse Laboratory | Wellcome Sanger Institute for the COVID-19 Genomics UK (COG-UK) Consortium | Berkshire and Surrey Pathology Services Lighthouse Laboratory and Alex Alderton, Roberto Amato, Jeffrey Barrett, Sonia Goncalves, Ewan Harrison, David K. Jackson, Ian Johnston, Dominic Kwiatkowski, Cordelia Langford, John Sillitoe on behalf of the Wellcome Sanger Institute COVID-19 Surveillance Team |
| EPI_ISL_3140568, EPI_ISL_3140569 | Lighthouse Lab in Milton Keynes | Wellcome Sanger Institute for the COVID-19 Genomics UK (COG-UK) Consortium | The Lighthouse Lab in Milton Keynes and Alex Alderton, Roberto Amato, Jeffrey Barrett, Sonia Goncalves, Ewan Harrison, David K. Jackson, Ian Johnston, Dominic Kwiatkowski, Cordelia Langford, John Sillitoe on behalf of the Wellcome Sanger Institute COVID-19 Surveillance Team |
| EPI_ISL_3140570, EPI_ISL_3140571 | Lighthouse Lab in Alderley Park | Wellcome Sanger Institute for the COVID-19 Genomics UK (COG-UK) Consortium | Jacquelyn Wynn, Mairead Hyland, The Lighthouse Lab in Alderley Park and Alex Alderton, Roberto Amato, Jeffrey Barrett, Sonia Goncalves, Ewan Harrison, David K. Jackson, Ian Johnston, Dominic Kwiatkowski, Cordelia Langford, John Sillitoe on behalf of the Wellcome Sanger Institute COVID-19 Surveillance Team |
| EPI_ISL_3140572, EPI_ISL_3140573 | Lighthouse Lab in Glasgow | Wellcome Sanger Institute for the COVID-19 Genomics UK (COG-UK) Consortium | Harper VanSteenhouse, Yumi Kasai, David Gray, Carol Clugston, Anna Dominiczak and Alex Alderton, Roberto Amato, Jeffrey Barrett, Sonia Goncalves, Ewan Harrison, David K. Jackson, Ian Johnston, Dominic Kwiatkowski, Cordelia Langford, John Sillitoe on behalf of the Wellcome Sanger Institute COVID-19 Surveillance Team |
| EPI_ISL_3140574 | Lighthouse Lab in Alderley Park | Wellcome Sanger Institute for the COVID-19 Genomics UK (COG-UK) Consortium | Jacquelyn Wynn, Mairead Hyland, The Lighthouse Lab in Alderley Park and Alex Alderton, Roberto Amato, Jeffrey Barrett, Sonia Goncalves, Ewan Harrison, David K. Jackson, Ian Johnston, Dominic Kwiatkowski, Cordelia Langford, John Sillitoe on behalf of the Wellcome Sanger Institute COVID-19 Surveillance Team |
| EPI_ISL_3140575 | Berkshire and Surrey Pathology Services Lighthouse Laboratory | Wellcome Sanger Institute for the COVID-19 Genomics UK (COG-UK) Consortium | Berkshire and Surrey Pathology Services Lighthouse Laboratory and Alex Alderton, Roberto Amato, Jeffrey Barrett, Sonia Goncalves, Ewan Harrison, David K. Jackson, Ian Johnston, Dominic Kwiatkowski, Cordelia Langford, John Sillitoe on behalf of the Wellcome Sanger Institute COVID-19 Surveillance Team |
| EPI_ISL_3140576 | Lighthouse Lab in Alderley Park | Wellcome Sanger Institute for the COVID-19 Genomics UK (COG-UK) Consortium | Jacquelyn Wynn, Mairead Hyland, The Lighthouse Lab in Alderley Park and Alex Alderton, Roberto Amato, Jeffrey Barrett, Sonia Goncalves, Ewan Harrison, David K. Jackson, Ian Johnston, Dominic Kwiatkowski, Cordelia Langford, John Sillitoe on behalf of the Wellcome Sanger Institute COVID-19 Surveillance Team |
| EPI_ISL_3140577 | Lighthouse Lab in Glasgow | Wellcome Sanger Institute for the COVID-19 Genomics UK (COG-UK) Consortium | Harper VanSteenhouse, Yumi Kasai, David Gray, Carol Clugston, Anna Dominiczak and Alex Alderton, Roberto Amato, Jeffrey Barrett, Sonia Goncalves, Ewan Harrison, David K. Jackson, Ian Johnston, Dominic Kwiatkowski, Cordelia Langford, John Sillitoe on behalf of the Wellcome Sanger Institute COVID-19 Surveillance Team |
| EPI_ISL_3140578 | Lighthouse Lab in Alderley Park | Wellcome Sanger Institute for the COVID-19 Genomics UK (COG-UK) Consortium | Jacquelyn Wynn, Mairead Hyland, The Lighthouse Lab in Alderley Park and Alex Alderton, Roberto Amato, Jeffrey Barrett, Sonia Goncalves, Ewan Harrison, David K. Jackson, Ian Johnston, Dominic Kwiatkowski, Cordelia Langford, John Sillitoe on behalf of the Wellcome Sanger Institute COVID-19 Surveillance Team |
| EPI_ISL_3140579 | Lighthouse Lab in Glasgow | Wellcome Sanger Institute for the COVID-19 Genomics UK (COG-UK) Consortium | Harper VanSteenhouse, Yumi Kasai, David Gray, Carol Clugston, Anna Dominiczak and Alex Alderton, Roberto Amato, Jeffrey Barrett, Sonia Goncalves, Ewan Harrison, David K. Jackson, Ian Johnston, Dominic Kwiatkowski, Cordelia Langford, John Sillitoe on behalf of the Wellcome Sanger Institute COVID-19 Surveillance Team |

| | | | |
|---|---|---|---|
| EPI_ISL_3140580, EPI_ISL_3140581 | Berkshire and Surrey Pathology Services Lighthouse Laboratory | Wellcome Sanger Institute for the COVID-19 Genomics UK (COG-UK) Consortium | Berkshire and Surrey Pathology Services Lighthouse Laboratory and Alex Alderton, Roberto Amato, Jeffrey Barrett, Sonia Goncalves, Ewan Harrison, David K. Jackson, Ian Johnston, Dominic Kwiatkowski, Cordelia Langford, John Sillitoe on behalf of the Wellcome Sanger Institute COVID-19 Surveillance Team |
| EPI_ISL_3140582, EPI_ISL_3140583 | Lighthouse Lab in Milton Keynes | Wellcome Sanger Institute for the COVID-19 Genomics UK (COG-UK) Consortium | The Lighthouse Lab in Milton Keynes and Alex Alderton, Roberto Amato, Jeffrey Barrett, Sonia Goncalves, Ewan Harrison, David K. Jackson, Ian Johnston, Dominic Kwiatkowski, Cordelia Langford, John Sillitoe on behalf of the Wellcome Sanger Institute COVID-19 Surveillance Team |
| EPI_ISL_3140584 | Berkshire and Surrey Pathology Services Lighthouse Laboratory | Wellcome Sanger Institute for the COVID-19 Genomics UK (COG-UK) Consortium | Berkshire and Surrey Pathology Services Lighthouse Laboratory and Alex Alderton, Roberto Amato, Jeffrey Barrett, Sonia Goncalves, Ewan Harrison, David K. Jackson, Ian Johnston, Dominic Kwiatkowski, Cordelia Langford, John Sillitoe on behalf of the Wellcome Sanger Institute COVID-19 Surveillance Team |
| EPI_ISL_3140585, EPI_ISL_3140586 | Lighthouse Lab in Glasgow | Wellcome Sanger Institute for the COVID-19 Genomics UK (COG-UK) Consortium | Harper VanSteenhouse, Yumi Kasai, David Gray, Carol Clugston, Anna Dominiczak and Alex Alderton, Roberto Amato, Jeffrey Barrett, Sonia Goncalves, Ewan Harrison, David K. Jackson, Ian Johnston, Dominic Kwiatkowski, Cordelia Langford, John Sillitoe on behalf of the Wellcome Sanger Institute COVID-19 Surveillance Team |
| EPI_ISL_3140587, EPI_ISL_3140588 | Lighthouse Lab in Milton Keynes | Wellcome Sanger Institute for the COVID-19 Genomics UK (COG-UK) Consortium | The Lighthouse Lab in Milton Keynes and Alex Alderton, Roberto Amato, Jeffrey Barrett, Sonia Goncalves, Ewan Harrison, David K. Jackson, Ian Johnston, Dominic Kwiatkowski, Cordelia Langford, John Sillitoe on behalf of the Wellcome Sanger Institute COVID-19 Surveillance Team |
| EPI_ISL_3140589, EPI_ISL_3140590 | Lighthouse Lab in Alderley Park | Wellcome Sanger Institute for the COVID-19 Genomics UK (COG-UK) Consortium | Jacquelyn Wynn, Mairead Hyland, The Lighthouse Lab in Alderley Park and Alex Alderton, Roberto Amato, Jeffrey Barrett, Sonia Goncalves, Ewan Harrison, David K. Jackson, Ian Johnston, Dominic Kwiatkowski, Cordelia Langford, John Sillitoe on behalf of the Wellcome Sanger Institute COVID-19 Surveillance Team |
| EPI_ISL_3140591 | Lighthouse Lab in Glasgow | Wellcome Sanger Institute for the COVID-19 Genomics UK (COG-UK) Consortium | Harper VanSteenhouse, Yumi Kasai, David Gray, Carol Clugston, Anna Dominiczak and Alex Alderton, Roberto Amato, Jeffrey Barrett, Sonia Goncalves, Ewan Harrison, David K. Jackson, Ian Johnston, Dominic Kwiatkowski, Cordelia Langford, John Sillitoe on behalf of the Wellcome Sanger Institute COVID-19 Surveillance Team |
| EPI_ISL_3140592 | Lighthouse Lab in Milton Keynes | Wellcome Sanger Institute for the COVID-19 Genomics UK (COG-UK) Consortium | The Lighthouse Lab in Milton Keynes and Alex Alderton, Roberto Amato, Jeffrey Barrett, Sonia Goncalves, Ewan Harrison, David K. Jackson, Ian Johnston, Dominic Kwiatkowski, Cordelia Langford, John Sillitoe on behalf of the Wellcome Sanger Institute COVID-19 Surveillance Team |
| EPI_ISL_3140593, EPI_ISL_3140594 | Lighthouse Lab in Alderley Park | Wellcome Sanger Institute for the COVID-19 Genomics UK (COG-UK) Consortium | Jacquelyn Wynn, Mairead Hyland, The Lighthouse Lab in Alderley Park and Alex Alderton, Roberto Amato, Jeffrey Barrett, Sonia Goncalves, Ewan Harrison, David K. Jackson, Ian Johnston, Dominic Kwiatkowski, Cordelia Langford, John Sillitoe on behalf of the Wellcome Sanger Institute COVID-19 Surveillance Team |
| EPI_ISL_3140595 | Lighthouse Lab in Milton Keynes | Wellcome Sanger Institute for the COVID-19 Genomics UK (COG-UK) Consortium | The Lighthouse Lab in Milton Keynes and Alex Alderton, Roberto Amato, Jeffrey Barrett, Sonia Goncalves, Ewan Harrison, David K. Jackson, Ian Johnston, Dominic Kwiatkowski, Cordelia Langford, John Sillitoe on behalf of the Wellcome Sanger Institute COVID-19 Surveillance Team |
| EPI_ISL_3140596, EPI_ISL_3140597, EPI_ISL_3140598 | Lighthouse Lab in Glasgow | Wellcome Sanger Institute for the COVID-19 Genomics UK (COG-UK) Consortium | Harper VanSteenhouse, Yumi Kasai, David Gray, Carol Clugston, Anna Dominiczak and Alex Alderton, Roberto Amato, Jeffrey Barrett, Sonia Goncalves, Ewan Harrison, David K. Jackson, Ian Johnston, Dominic Kwiatkowski, Cordelia Langford, John Sillitoe on behalf of the Wellcome Sanger Institute COVID-19 Surveillance Team |
| EPI_ISL_3140599, EPI_ISL_3140600 | Lighthouse Lab in Alderley Park | Wellcome Sanger Institute for the COVID-19 Genomics UK (COG-UK) Consortium | Jacquelyn Wynn, Mairead Hyland, The Lighthouse Lab in Alderley Park and Alex Alderton, Roberto Amato, Jeffrey Barrett, Sonia Goncalves, Ewan Harrison, David K. Jackson, Ian Johnston, Dominic Kwiatkowski, Cordelia Langford, John Sillitoe on behalf of the Wellcome Sanger Institute COVID-19 Surveillance Team |
| EPI_ISL_3140601 | Lighthouse Lab in Glasgow | Wellcome Sanger Institute for the COVID-19 Genomics UK (COG-UK) Consortium | Harper VanSteenhouse, Yumi Kasai, David Gray, Carol Clugston, Anna Dominiczak and Alex Alderton, Roberto Amato, Jeffrey Barrett, Sonia Goncalves, Ewan Harrison, David K. Jackson, Ian Johnston, Dominic Kwiatkowski, Cordelia Langford, John Sillitoe on behalf of the Wellcome Sanger Institute COVID-19 Surveillance Team |
| EPI_ISL_3140602 | Berkshire and Surrey Pathology Services Lighthouse Laboratory | Wellcome Sanger Institute for the COVID-19 Genomics UK (COG-UK) Consortium | Berkshire and Surrey Pathology Services Lighthouse Laboratory and Alex Alderton, Roberto Amato, Jeffrey Barrett, Sonia Goncalves, Ewan Harrison, David K. Jackson, Ian Johnston, Dominic Kwiatkowski, Cordelia Langford, John Sillitoe on behalf of the Wellcome Sanger Institute COVID-19 Surveillance Team |
| EPI_ISL_3140603 | Lighthouse Lab in Glasgow | Wellcome Sanger Institute for the COVID-19 Genomics UK (COG-UK) Consortium | Harper VanSteenhouse, Yumi Kasai, David Gray, Carol Clugston, Anna Dominiczak and Alex Alderton, Roberto Amato, Jeffrey Barrett, Sonia Goncalves, Ewan Harrison, David K. Jackson, Ian Johnston, Dominic Kwiatkowski, Cordelia Langford, John Sillitoe on behalf of the Wellcome Sanger Institute COVID-19 Surveillance Team |
| EPI_ISL_3140604 | Berkshire and Surrey Pathology Services Lighthouse Laboratory | Wellcome Sanger Institute for the COVID-19 Genomics UK (COG-UK) Consortium | Berkshire and Surrey Pathology Services Lighthouse Laboratory and Alex Alderton, Roberto Amato, Jeffrey Barrett, Sonia Goncalves, Ewan Harrison, David K. Jackson, Ian Johnston, Dominic Kwiatkowski, Cordelia Langford, John Sillitoe on behalf of the Wellcome Sanger Institute COVID-19 Surveillance Team |
| EPI_ISL_3140605 | Lighthouse Lab in Milton Keynes | Wellcome Sanger Institute for the COVID-19 Genomics UK (COG-UK) Consortium | The Lighthouse Lab in Milton Keynes and Alex Alderton, Roberto Amato, Jeffrey Barrett, Sonia Goncalves, Ewan Harrison, David K. Jackson, Ian Johnston, Dominic Kwiatkowski, Cordelia Langford, John Sillitoe on behalf of the Wellcome Sanger Institute COVID-19 Surveillance Team |
| EPI_ISL_3140606, EPI_ISL_3140607 | Lighthouse Lab in Glasgow | Wellcome Sanger Institute for the COVID-19 Genomics UK (COG-UK) Consortium | Harper VanSteenhouse, Yumi Kasai, David Gray, Carol Clugston, Anna Dominiczak and Alex Alderton, Roberto Amato, Jeffrey Barrett, Sonia Goncalves, Ewan Harrison, David K. Jackson, Ian Johnston, Dominic Kwiatkowski, Cordelia Langford, John Sillitoe on behalf of the Wellcome Sanger Institute COVID-19 Surveillance Team |
| EPI_ISL_3140608 | Lighthouse Lab in Milton Keynes | Wellcome Sanger Institute for the COVID-19 Genomics UK (COG-UK) Consortium | The Lighthouse Lab in Milton Keynes and Alex Alderton, Roberto Amato, Jeffrey Barrett, Sonia Goncalves, Ewan Harrison, David K. Jackson, Ian Johnston, Dominic Kwiatkowski, Cordelia Langford, John Sillitoe on behalf of the Wellcome Sanger Institute COVID-19 Surveillance Team |
| EPI_ISL_3140609 | Lighthouse Lab in Glasgow | Wellcome Sanger Institute for the COVID-19 Genomics UK (COG-UK) Consortium | Harper VanSteenhouse, Yumi Kasai, David Gray, Carol Clugston, Anna Dominiczak and Alex Alderton, Roberto Amato, Jeffrey Barrett, Sonia Goncalves, Ewan Harrison, David K. Jackson, Ian Johnston, Dominic Kwiatkowski, Cordelia Langford, John Sillitoe on behalf of the Wellcome Sanger Institute COVID-19 Surveillance Team |
| EPI_ISL_3140610 | Lighthouse Lab in Milton Keynes | Wellcome Sanger Institute for the COVID-19 Genomics UK (COG-UK) Consortium | The Lighthouse Lab in Milton Keynes and Alex Alderton, Roberto Amato, Jeffrey Barrett, Sonia Goncalves, Ewan Harrison, David K. Jackson, Ian Johnston, Dominic Kwiatkowski, Cordelia Langford, John Sillitoe on behalf of the Wellcome Sanger Institute COVID-19 Surveillance Team |
| EPI_ISL_3140611 | Lighthouse Lab in Alderley Park | Wellcome Sanger Institute for the COVID-19 Genomics UK (COG-UK) Consortium | Jacquelyn Wynn, Mairead Hyland, The Lighthouse Lab in Alderley Park and Alex Alderton, Roberto Amato, Jeffrey Barrett, Sonia Goncalves, Ewan Harrison, David K. Jackson, Ian Johnston, Dominic Kwiatkowski, Cordelia Langford, John Sillitoe on behalf of the Wellcome Sanger Institute COVID-19 Surveillance Team |
| EPI_ISL_3140612 | Lighthouse Lab in Milton Keynes | Wellcome Sanger Institute for the COVID-19 Genomics UK (COG-UK) Consortium | The Lighthouse Lab in Milton Keynes and Alex Alderton, Roberto Amato, Jeffrey Barrett, Sonia Goncalves, Ewan Harrison, David K. Jackson, Ian Johnston, Dominic Kwiatkowski, Cordelia Langford, John Sillitoe on behalf of the Wellcome Sanger Institute COVID-19 Surveillance Team |
| EPI_ISL_3140613 | Berkshire and Surrey Pathology Services Lighthouse Laboratory | Wellcome Sanger Institute for the COVID-19 Genomics UK (COG-UK) Consortium | Berkshire and Surrey Pathology Services Lighthouse Laboratory and Alex Alderton, Roberto Amato, Jeffrey Barrett, Sonia Goncalves, Ewan Harrison, David K. Jackson, Ian Johnston, Dominic Kwiatkowski, Cordelia Langford, John Sillitoe on behalf of the Wellcome Sanger Institute COVID-19 Surveillance Team |
| EPI_ISL_3140614 | Lighthouse Lab in Alderley Park | Wellcome Sanger Institute for the COVID-19 Genomics UK (COG-UK) Consortium | Jacquelyn Wynn, Mairead Hyland, The Lighthouse Lab in Alderley Park and Alex Alderton, Roberto Amato, Jeffrey Barrett, Sonia Goncalves, Ewan Harrison, David K. Jackson, Ian Johnston, Dominic Kwiatkowski, Cordelia Langford, John Sillitoe on behalf of the Wellcome Sanger Institute COVID-19 Surveillance Team |
| EPI_ISL_3140615 | Lighthouse Lab in Glasgow | Wellcome Sanger Institute for the COVID-19 Genomics UK (COG-UK) Consortium | Harper VanSteenhouse, Yumi Kasai, David Gray, Carol Clugston, Anna Dominiczak and Alex Alderton, Roberto Amato, Jeffrey Barrett, Sonia Goncalves, Ewan Harrison, David K. Jackson, Ian Johnston, Dominic Kwiatkowski, Cordelia Langford, John Sillitoe on behalf of the Wellcome Sanger Institute COVID-19 Surveillance Team |
| EPI_ISL_3140616 | Berkshire and Surrey Pathology Services Lighthouse Laboratory | Wellcome Sanger Institute for the COVID-19 Genomics UK (COG-UK) Consortium | Berkshire and Surrey Pathology Services Lighthouse Laboratory and Alex Alderton, Roberto Amato, Jeffrey Barrett, Sonia Goncalves, Ewan Harrison, David K. Jackson, Ian Johnston, Dominic Kwiatkowski, Cordelia Langford, John Sillitoe on behalf of the Wellcome Sanger Institute COVID-19 Surveillance Team |
| EPI_ISL_3140617 | Lighthouse Lab in Milton Keynes | Wellcome Sanger Institute for the COVID-19 Genomics UK (COG-UK) Consortium | The Lighthouse Lab in Milton Keynes and Alex Alderton, Roberto Amato, Jeffrey Barrett, Sonia Goncalves, Ewan Harrison, David K. Jackson, Ian Johnston, Dominic Kwiatkowski, Cordelia Langford, John Sillitoe on behalf of the Wellcome Sanger Institute COVID-19 Surveillance Team |
| EPI_ISL_3140618, EPI_ISL_3140619 | Lighthouse Lab in Alderley Park | Wellcome Sanger Institute for the COVID-19 Genomics UK (COG-UK) Consortium | Jacquelyn Wynn, Mairead Hyland, The Lighthouse Lab in Alderley Park and Alex Alderton, Roberto Amato, Jeffrey Barrett, Sonia Goncalves, Ewan Harrison, David K. Jackson, Ian Johnston, Dominic Kwiatkowski, Cordelia Langford, John Sillitoe on behalf of the Wellcome Sanger Institute COVID-19 Surveillance Team |
| EPI_ISL_3140620 | Berkshire and Surrey Pathology Services Lighthouse Laboratory | Wellcome Sanger Institute for the COVID-19 Genomics UK (COG-UK) Consortium | Berkshire and Surrey Pathology Services Lighthouse Laboratory and Alex Alderton, Roberto Amato, Jeffrey Barrett, Sonia Goncalves, Ewan Harrison, David K. Jackson, Ian Johnston, Dominic Kwiatkowski, Cordelia Langford, John Sillitoe on behalf of the Wellcome Sanger Institute COVID-19 Surveillance Team |
| EPI_ISL_3140621 | Lighthouse Lab in Alderley Park | Wellcome Sanger Institute for the COVID-19 Genomics UK (COG-UK) Consortium | Jacquelyn Wynn, Mairead Hyland, The Lighthouse Lab in Alderley Park and Alex Alderton, Roberto Amato, Jeffrey Barrett, Sonia Goncalves, Ewan Harrison, David K. Jackson, Ian Johnston, Dominic Kwiatkowski, Cordelia Langford, John Sillitoe on behalf of the Wellcome Sanger Institute COVID-19 Surveillance Team |
| EPI_ISL_3140622 | Lighthouse Lab in Milton Keynes | Wellcome Sanger Institute for the COVID-19 Genomics UK (COG-UK) Consortium | The Lighthouse Lab in Milton Keynes and Alex Alderton, Roberto Amato, Jeffrey Barrett, Sonia Goncalves, Ewan Harrison, David K. Jackson, Ian Johnston, Dominic Kwiatkowski, Cordelia Langford, John Sillitoe on behalf of the Wellcome Sanger Institute COVID-19 Surveillance Team |
| EPI_ISL_3140623 | Lighthouse Lab in Alderley Park | Wellcome Sanger Institute for the COVID-19 Genomics UK (COG-UK) Consortium | Jacquelyn Wynn, Mairead Hyland, The Lighthouse Lab in Alderley Park and Alex Alderton, Roberto Amato, Jeffrey Barrett, Sonia Goncalves, Ewan Harrison, David K. Jackson, Ian Johnston, Dominic Kwiatkowski, Cordelia Langford, John Sillitoe on behalf of the Wellcome Sanger Institute COVID-19 Surveillance Team |
| EPI_ISL_3140624 | Berkshire and Surrey Pathology Services Lighthouse Laboratory | Wellcome Sanger Institute for the COVID-19 Genomics UK (COG-UK) Consortium | Berkshire and Surrey Pathology Services Lighthouse Laboratory and Alex Alderton, Roberto Amato, Jeffrey Barrett, Sonia Goncalves, Ewan Harrison, David K. Jackson, Ian Johnston, Dominic Kwiatkowski, Cordelia Langford, John Sillitoe on behalf of the Wellcome Sanger Institute COVID-19 Surveillance Team |
| EPI_ISL_3140625 | Lighthouse Lab in Milton Keynes | Wellcome Sanger Institute for the COVID-19 Genomics UK (COG-UK) Consortium | The Lighthouse Lab in Milton Keynes and Alex Alderton, Roberto Amato, Jeffrey Barrett, Sonia Goncalves, Ewan Harrison, David K. Jackson, Ian Johnston, Dominic Kwiatkowski, Cordelia Langford, John Sillitoe on behalf of the Wellcome Sanger Institute COVID-19 Surveillance Team |
| EPI_ISL_3140626 | Berkshire and Surrey Pathology Services Lighthouse Laboratory | Wellcome Sanger Institute for the COVID-19 Genomics UK (COG-UK) Consortium | Berkshire and Surrey Pathology Services Lighthouse Laboratory and Alex Alderton, Roberto Amato, Jeffrey Barrett, Sonia Goncalves, Ewan Harrison, David K. Jackson, Ian Johnston, Dominic Kwiatkowski, Cordelia Langford, John Sillitoe on behalf of the Wellcome Sanger Institute COVID-19 Surveillance Team |
| EPI_ISL_3140627 | Lighthouse Lab in Milton Keynes | Wellcome Sanger Institute for the COVID-19 Genomics UK (COG-UK) Consortium | The Lighthouse Lab in Milton Keynes and Alex Alderton, Roberto Amato, Jeffrey Barrett, Sonia Goncalves, Ewan Harrison, David K. Jackson, Ian Johnston, Dominic Kwiatkowski, Cordelia Langford, John Sillitoe on behalf of the Wellcome Sanger Institute COVID-19 Surveillance Team |
| EPI_ISL_3140628, EPI_ISL_3140629 | Lighthouse Lab in Glasgow | Wellcome Sanger Institute for the COVID-19 Genomics UK (COG-UK) Consortium | Harper VanSteenhouse, Yumi Kasai, David Gray, Carol Clugston, Anna Dominiczak and Alex Alderton, Roberto Amato, Jeffrey Barrett, Sonia Goncalves, Ewan Harrison, David K. Jackson, Ian Johnston, Dominic Kwiatkowski, Cordelia Langford, John Sillitoe on behalf of the Wellcome Sanger Institute COVID-19 Surveillance Team |
| EPI_ISL_3140630 | Lighthouse Lab in Alderley Park | Wellcome Sanger Institute for the COVID-19 Genomics UK (COG-UK) Consortium | Jacquelyn Wynn, Mairead Hyland, The Lighthouse Lab in Alderley Park and Alex Alderton, Roberto Amato, Jeffrey Barrett, Sonia Goncalves, Ewan Harrison, David K. Jackson, Ian Johnston, Dominic Kwiatkowski, Cordelia Langford, John Sillitoe on behalf of the Wellcome Sanger Institute COVID-19 Surveillance Team |
| EPI_ISL_3140631 | Lighthouse Lab in Glasgow | Wellcome Sanger Institute for the COVID-19 Genomics UK (COG-UK) Consortium | Harper VanSteenhouse, Yumi Kasai, David Gray, Carol Clugston, Anna Dominiczak and Alex Alderton, Roberto Amato, Jeffrey Barrett, Sonia Goncalves, Ewan Harrison, David K. Jackson, Ian Johnston, Dominic Kwiatkowski, Cordelia Langford, John Sillitoe on behalf of the Wellcome Sanger Institute COVID-19 Surveillance Team |
| EPI_ISL_3140632 | Berkshire and Surrey Pathology Services Lighthouse Laboratory | Wellcome Sanger Institute for the COVID-19 Genomics UK (COG-UK) Consortium | Berkshire and Surrey Pathology Services Lighthouse Laboratory and Alex Alderton, Roberto Amato, Jeffrey Barrett, Sonia Goncalves, Ewan Harrison, David K. Jackson, Ian Johnston, Dominic Kwiatkowski, Cordelia Langford, John Sillitoe on behalf of the Wellcome Sanger Institute COVID-19 Surveillance Team |
| EPI_ISL_3140633 | Lighthouse Lab in Milton Keynes | Wellcome Sanger Institute for the COVID-19 Genomics UK (COG-UK) Consortium | The Lighthouse Lab in Milton Keynes and Alex Alderton, Roberto Amato, Jeffrey Barrett, Sonia Goncalves, Ewan Harrison, David K. Jackson, Ian Johnston, Dominic Kwiatkowski, Cordelia Langford, John Sillitoe on behalf of the Wellcome Sanger Institute COVID-19 Surveillance Team |
| EPI_ISL_3140634 | Lighthouse Lab in Alderley Park | Wellcome Sanger Institute for the COVID-19 Genomics UK (COG-UK) Consortium | Jacquelyn Wynn, Mairead Hyland, The Lighthouse Lab in Alderley Park and Alex Alderton, Roberto Amato, Jeffrey Barrett, Sonia Goncalves, Ewan Harrison, David K. Jackson, Ian Johnston, Dominic Kwiatkowski, Cordelia Langford, John Sillitoe on behalf of the Wellcome Sanger Institute COVID-19 Surveillance Team |
| EPI_ISL_3140635 | Lighthouse Lab in Milton Keynes | Wellcome Sanger Institute for the COVID-19 Genomics UK (COG-UK) Consortium | The Lighthouse Lab in Milton Keynes and Alex Alderton, Roberto Amato, Jeffrey Barrett, Sonia Goncalves, Ewan Harrison, David K. Jackson, Ian Johnston, Dominic Kwiatkowski, Cordelia Langford, John Sillitoe on behalf of the Wellcome Sanger Institute COVID-19 Surveillance Team |
| EPI_ISL_3140636 | Lighthouse Lab in Alderley Park | Wellcome Sanger Institute for the COVID-19 Genomics UK (COG-UK) | Jacquelyn Wynn, Mairead Hyland, The Lighthouse Lab in Alderley Park and Alex Alderton, Roberto Amato, Jeffrey Barrett, Sonia Goncalves, Ewan Harrison, David K. Jackson, Ian Johnston, Dominic Kwiatkowski, Cordelia Langford, John Sillitoe on |

| | | | |
|---|---|---|---|
| | | Consortium | behalf of the Wellcome Sanger Institute COVID-19 Surveillance Team |
| EPI_ISL_3140637 | Lighthouse Lab in Milton Keynes | Wellcome Sanger Institute for the COVID-19 Genomics UK (COG-UK) Consortium | The Lighthouse Lab in Milton Keynes and Alex Alderton, Roberto Amato, Jeffrey Barrett, Sonia Goncalves, Ewan Harrison, David K. Jackson, Ian Johnston, Dominic Kwiatkowski, Cordelia Langford, John Sillitoe on behalf of the Wellcome Sanger Institute COVID-19 Surveillance Team |
| EPI_ISL_3140638, EPI_ISL_3140639 | Lighthouse Lab in Alderley Park | Wellcome Sanger Institute for the COVID-19 Genomics UK (COG-UK) Consortium | Jacquelyn Wynn, Mairead Hyland, The Lighthouse Lab in Alderley Park and Alex Alderton, Roberto Amato, Jeffrey Barrett, Sonia Goncalves, Ewan Harrison, David K. Jackson, Ian Johnston, Dominic Kwiatkowski, Cordelia Langford, John Sillitoe on behalf of the Wellcome Sanger Institute COVID-19 Surveillance Team |
| EPI_ISL_3140640 | Lighthouse Lab in Milton Keynes | Wellcome Sanger Institute for the COVID-19 Genomics UK (COG-UK) Consortium | The Lighthouse Lab in Milton Keynes and Alex Alderton, Roberto Amato, Jeffrey Barrett, Sonia Goncalves, Ewan Harrison, David K. Jackson, Ian Johnston, Dominic Kwiatkowski, Cordelia Langford, John Sillitoe on behalf of the Wellcome Sanger Institute COVID-19 Surveillance Team |
| EPI_ISL_3140641, EPI_ISL_3140642 | Lighthouse Lab in Alderley Park | Wellcome Sanger Institute for the COVID-19 Genomics UK (COG-UK) Consortium | Jacquelyn Wynn, Mairead Hyland, The Lighthouse Lab in Alderley Park and Alex Alderton, Roberto Amato, Jeffrey Barrett, Sonia Goncalves, Ewan Harrison, David K. Jackson, Ian Johnston, Dominic Kwiatkowski, Cordelia Langford, John Sillitoe on behalf of the Wellcome Sanger Institute COVID-19 Surveillance Team |
| EPI_ISL_3140643, EPI_ISL_3140644, EPI_ISL_3140645 | Lighthouse Lab in Milton Keynes | Wellcome Sanger Institute for the COVID-19 Genomics UK (COG-UK) Consortium | The Lighthouse Lab in Milton Keynes and Alex Alderton, Roberto Amato, Jeffrey Barrett, Sonia Goncalves, Ewan Harrison, David K. Jackson, Ian Johnston, Dominic Kwiatkowski, Cordelia Langford, John Sillitoe on behalf of the Wellcome Sanger Institute COVID-19 Surveillance Team |
| EPI_ISL_3140646 | Lighthouse Lab in Glasgow | Wellcome Sanger Institute for the COVID-19 Genomics UK (COG-UK) Consortium | Harper VanSteenhouse, Yumi Kasai, David Gray, Carol Clugston, Anna Dominiczak and Alex Alderton, Roberto Amato, Jeffrey Barrett, Sonia Goncalves, Ewan Harrison, David K. Jackson, Ian Johnston, Dominic Kwiatkowski, Cordelia Langford, John Sillitoe on behalf of the Wellcome Sanger Institute COVID-19 Surveillance Team |
| EPI_ISL_3140647, EPI_ISL_3140648 | Lighthouse Lab in Milton Keynes | Wellcome Sanger Institute for the COVID-19 Genomics UK (COG-UK) Consortium | The Lighthouse Lab in Milton Keynes and Alex Alderton, Roberto Amato, Jeffrey Barrett, Sonia Goncalves, Ewan Harrison, David K. Jackson, Ian Johnston, Dominic Kwiatkowski, Cordelia Langford, John Sillitoe on behalf of the Wellcome Sanger Institute COVID-19 Surveillance Team |
| EPI_ISL_3140649 | Lighthouse Lab in Alderley Park | Wellcome Sanger Institute for the COVID-19 Genomics UK (COG-UK) Consortium | Jacquelyn Wynn, Mairead Hyland, The Lighthouse Lab in Alderley Park and Alex Alderton, Roberto Amato, Jeffrey Barrett, Sonia Goncalves, Ewan Harrison, David K. Jackson, Ian Johnston, Dominic Kwiatkowski, Cordelia Langford, John Sillitoe on behalf of the Wellcome Sanger Institute COVID-19 Surveillance Team |
| EPI_ISL_3140650, EPI_ISL_3140651, EPI_ISL_3140652 | Lighthouse Lab in Milton Keynes | Wellcome Sanger Institute for the COVID-19 Genomics UK (COG-UK) Consortium | The Lighthouse Lab in Milton Keynes and Alex Alderton, Roberto Amato, Jeffrey Barrett, Sonia Goncalves, Ewan Harrison, David K. Jackson, Ian Johnston, Dominic Kwiatkowski, Cordelia Langford, John Sillitoe on behalf of the Wellcome Sanger Institute COVID-19 Surveillance Team |
| EPI_ISL_3140653 | Lighthouse Lab in Alderley Park | Wellcome Sanger Institute for the COVID-19 Genomics UK (COG-UK) Consortium | Jacquelyn Wynn, Mairead Hyland, The Lighthouse Lab in Alderley Park and Alex Alderton, Roberto Amato, Jeffrey Barrett, Sonia Goncalves, Ewan Harrison, David K. Jackson, Ian Johnston, Dominic Kwiatkowski, Cordelia Langford, John Sillitoe on behalf of the Wellcome Sanger Institute COVID-19 Surveillance Team |
| EPI_ISL_3140654, EPI_ISL_3140655 | Lighthouse Lab in Milton Keynes | Wellcome Sanger Institute for the COVID-19 Genomics UK (COG-UK) Consortium | The Lighthouse Lab in Milton Keynes and Alex Alderton, Roberto Amato, Jeffrey Barrett, Sonia Goncalves, Ewan Harrison, David K. Jackson, Ian Johnston, Dominic Kwiatkowski, Cordelia Langford, John Sillitoe on behalf of the Wellcome Sanger Institute COVID-19 Surveillance Team |
| EPI_ISL_3140656 | Berkshire and Surrey Pathology Services Lighthouse Laboratory | Wellcome Sanger Institute for the COVID-19 Genomics UK (COG-UK) Consortium | Berkshire and Surrey Pathology Services Lighthouse Laboratory and Alex Alderton, Roberto Amato, Jeffrey Barrett, Sonia Goncalves, Ewan Harrison, David K. Jackson, Ian Johnston, Dominic Kwiatkowski, Cordelia Langford, John Sillitoe on behalf of the Wellcome Sanger Institute COVID-19 Surveillance Team |
| EPI_ISL_3140657, EPI_ISL_3140658 | Lighthouse Lab in Glasgow | Wellcome Sanger Institute for the COVID-19 Genomics UK (COG-UK) Consortium | Harper VanSteenhouse, Yumi Kasai, David Gray, Carol Clugston, Anna Dominiczak and Alex Alderton, Roberto Amato, Jeffrey Barrett, Sonia Goncalves, Ewan Harrison, David K. Jackson, Ian Johnston, Dominic Kwiatkowski, Cordelia Langford, John Sillitoe on behalf of the Wellcome Sanger Institute COVID-19 Surveillance Team |
| EPI_ISL_3140659 | Berkshire and Surrey Pathology Services Lighthouse Laboratory | Wellcome Sanger Institute for the COVID-19 Genomics UK (COG-UK) Consortium | Berkshire and Surrey Pathology Services Lighthouse Laboratory and Alex Alderton, Roberto Amato, Jeffrey Barrett, Sonia Goncalves, Ewan Harrison, David K. Jackson, Ian Johnston, Dominic Kwiatkowski, Cordelia Langford, John Sillitoe on behalf of the Wellcome Sanger Institute COVID-19 Surveillance Team |
| EPI_ISL_3140660 | Lighthouse Lab in Glasgow | Wellcome Sanger Institute for the COVID-19 Genomics UK (COG-UK) Consortium | Harper VanSteenhouse, Yumi Kasai, David Gray, Carol Clugston, Anna Dominiczak and Alex Alderton, Roberto Amato, Jeffrey Barrett, Sonia Goncalves, Ewan Harrison, David K. Jackson, Ian Johnston, Dominic Kwiatkowski, Cordelia Langford, John Sillitoe on behalf of the Wellcome Sanger Institute COVID-19 Surveillance Team |
| EPI_ISL_3140661 | Lighthouse Lab in Milton Keynes | Wellcome Sanger Institute for the COVID-19 Genomics UK (COG-UK) Consortium | The Lighthouse Lab in Milton Keynes and Alex Alderton, Roberto Amato, Jeffrey Barrett, Sonia Goncalves, Ewan Harrison, David K. Jackson, Ian Johnston, Dominic Kwiatkowski, Cordelia Langford, John Sillitoe on behalf of the Wellcome Sanger Institute COVID-19 Surveillance Team |
| EPI_ISL_3140662, EPI_ISL_3140663 | Berkshire and Surrey Pathology Services Lighthouse Laboratory | Wellcome Sanger Institute for the COVID-19 Genomics UK (COG-UK) Consortium | Berkshire and Surrey Pathology Services Lighthouse Laboratory and Alex Alderton, Roberto Amato, Jeffrey Barrett, Sonia Goncalves, Ewan Harrison, David K. Jackson, Ian Johnston, Dominic Kwiatkowski, Cordelia Langford, John Sillitoe on behalf of the Wellcome Sanger Institute COVID-19 Surveillance Team |
| EPI_ISL_3140664 | Lighthouse Lab in Milton Keynes | Wellcome Sanger Institute for the COVID-19 Genomics UK (COG-UK) Consortium | The Lighthouse Lab in Milton Keynes and Alex Alderton, Roberto Amato, Jeffrey Barrett, Sonia Goncalves, Ewan Harrison, David K. Jackson, Ian Johnston, Dominic Kwiatkowski, Cordelia Langford, John Sillitoe on behalf of the Wellcome Sanger Institute COVID-19 Surveillance Team |
| EPI_ISL_3140665, EPI_ISL_3140666 | Berkshire and Surrey Pathology Services Lighthouse Laboratory | Wellcome Sanger Institute for the COVID-19 Genomics UK (COG-UK) Consortium | Berkshire and Surrey Pathology Services Lighthouse Laboratory and Alex Alderton, Roberto Amato, Jeffrey Barrett, Sonia Goncalves, Ewan Harrison, David K. Jackson, Ian Johnston, Dominic Kwiatkowski, Cordelia Langford, John Sillitoe on behalf of the Wellcome Sanger Institute COVID-19 Surveillance Team |
| EPI_ISL_3140667 | Lighthouse Lab in Alderley Park | Wellcome Sanger Institute for the COVID-19 Genomics UK (COG-UK) Consortium | Jacquelyn Wynn, Mairead Hyland, The Lighthouse Lab in Alderley Park and Alex Alderton, Roberto Amato, Jeffrey Barrett, Sonia Goncalves, Ewan Harrison, David K. Jackson, Ian Johnston, Dominic Kwiatkowski, Cordelia Langford, John Sillitoe on behalf of the Wellcome Sanger Institute COVID-19 Surveillance Team |
| EPI_ISL_3140668 | Lighthouse Lab in Milton Keynes | Wellcome Sanger Institute for the COVID-19 Genomics UK (COG-UK) Consortium | The Lighthouse Lab in Milton Keynes and Alex Alderton, Roberto Amato, Jeffrey Barrett, Sonia Goncalves, Ewan Harrison, David K. Jackson, Ian Johnston, Dominic Kwiatkowski, Cordelia Langford, John Sillitoe on behalf of the Wellcome Sanger Institute COVID-19 Surveillance Team |
| EPI_ISL_3140669 | Lighthouse Lab in Glasgow | Wellcome Sanger Institute for the COVID-19 Genomics UK (COG-UK) Consortium | Harper VanSteenhouse, Yumi Kasai, David Gray, Carol Clugston, Anna Dominiczak and Alex Alderton, Roberto Amato, Jeffrey Barrett, Sonia Goncalves, Ewan Harrison, David K. Jackson, Ian Johnston, Dominic Kwiatkowski, Cordelia Langford, John Sillitoe on behalf of the Wellcome Sanger Institute COVID-19 Surveillance Team |
| EPI_ISL_3140670 | Lighthouse Lab in Milton Keynes | Wellcome Sanger Institute for the COVID-19 Genomics UK (COG-UK) Consortium | The Lighthouse Lab in Milton Keynes and Alex Alderton, Roberto Amato, Jeffrey Barrett, Sonia Goncalves, Ewan Harrison, David K. Jackson, Ian Johnston, Dominic Kwiatkowski, Cordelia Langford, John Sillitoe on behalf of the Wellcome Sanger Institute COVID-19 Surveillance Team |
| EPI_ISL_3140671 | Berkshire and Surrey Pathology Services Lighthouse Laboratory | Wellcome Sanger Institute for the COVID-19 Genomics UK (COG-UK) Consortium | Berkshire and Surrey Pathology Services Lighthouse Laboratory and Alex Alderton, Roberto Amato, Jeffrey Barrett, Sonia Goncalves, Ewan Harrison, David K. Jackson, Ian Johnston, Dominic Kwiatkowski, Cordelia Langford, John Sillitoe on behalf of the Wellcome Sanger Institute COVID-19 Surveillance Team |
| EPI_ISL_3140672, EPI_ISL_3140673 | Lighthouse Lab in Glasgow | Wellcome Sanger Institute for the COVID-19 Genomics UK (COG-UK) Consortium | Harper VanSteenhouse, Yumi Kasai, David Gray, Carol Clugston, Anna Dominiczak and Alex Alderton, Roberto Amato, Jeffrey Barrett, Sonia Goncalves, Ewan Harrison, David K. Jackson, Ian Johnston, Dominic Kwiatkowski, Cordelia Langford, John Sillitoe on behalf of the Wellcome Sanger Institute COVID-19 Surveillance Team |
| EPI_ISL_3140674 | Berkshire and Surrey Pathology Services Lighthouse Laboratory | Wellcome Sanger Institute for the COVID-19 Genomics UK (COG-UK) Consortium | Berkshire and Surrey Pathology Services Lighthouse Laboratory and Alex Alderton, Roberto Amato, Jeffrey Barrett, Sonia Goncalves, Ewan Harrison, David K. Jackson, Ian Johnston, Dominic Kwiatkowski, Cordelia Langford, John Sillitoe on behalf of the Wellcome Sanger Institute COVID-19 Surveillance Team |
| EPI_ISL_3140675 | Lighthouse Lab in Alderley Park | Wellcome Sanger Institute for the COVID-19 Genomics UK (COG-UK) Consortium | Jacquelyn Wynn, Mairead Hyland, The Lighthouse Lab in Alderley Park and Alex Alderton, Roberto Amato, Jeffrey Barrett, Sonia Goncalves, Ewan Harrison, David K. Jackson, Ian Johnston, Dominic Kwiatkowski, Cordelia Langford, John Sillitoe on behalf of the Wellcome Sanger Institute COVID-19 Surveillance Team |
| EPI_ISL_3140676, EPI_ISL_3140677, EPI_ISL_3140678 | Lighthouse Lab in Glasgow | Wellcome Sanger Institute for the COVID-19 Genomics UK (COG-UK) Consortium | Harper VanSteenhouse, Yumi Kasai, David Gray, Carol Clugston, Anna Dominiczak and Alex Alderton, Roberto Amato, Jeffrey Barrett, Sonia Goncalves, Ewan Harrison, David K. Jackson, Ian Johnston, Dominic Kwiatkowski, Cordelia Langford, John Sillitoe on behalf of the Wellcome Sanger Institute COVID-19 Surveillance Team |
| EPI_ISL_3140679, EPI_ISL_3140680 | Berkshire and Surrey Pathology Services Lighthouse Laboratory | Wellcome Sanger Institute for the COVID-19 Genomics UK (COG-UK) Consortium | Berkshire and Surrey Pathology Services Lighthouse Laboratory and Alex Alderton, Roberto Amato, Jeffrey Barrett, Sonia Goncalves, Ewan Harrison, David K. Jackson, Ian Johnston, Dominic Kwiatkowski, Cordelia Langford, John Sillitoe on behalf of the Wellcome Sanger Institute COVID-19 Surveillance Team |
| EPI_ISL_3140681, EPI_ISL_3140682 | Lighthouse Lab in Glasgow | Wellcome Sanger Institute for the COVID-19 Genomics UK (COG-UK) Consortium | Harper VanSteenhouse, Yumi Kasai, David Gray, Carol Clugston, Anna Dominiczak and Alex Alderton, Roberto Amato, Jeffrey Barrett, Sonia Goncalves, Ewan Harrison, David K. Jackson, Ian Johnston, Dominic Kwiatkowski, Cordelia Langford, John Sillitoe on behalf of the Wellcome Sanger Institute COVID-19 Surveillance Team |
| EPI_ISL_3140683 | Lighthouse Lab in Milton Keynes | Wellcome Sanger Institute for the COVID-19 Genomics UK (COG-UK) Consortium | The Lighthouse Lab in Milton Keynes and Alex Alderton, Roberto Amato, Jeffrey Barrett, Sonia Goncalves, Ewan Harrison, David K. Jackson, Ian Johnston, Dominic Kwiatkowski, Cordelia Langford, John Sillitoe on behalf of the Wellcome Sanger Institute COVID-19 Surveillance Team |
| EPI_ISL_3140684 | Lighthouse Lab in Glasgow | Wellcome Sanger Institute for the COVID-19 Genomics UK (COG-UK) Consortium | Harper VanSteenhouse, Yumi Kasai, David Gray, Carol Clugston, Anna Dominiczak and Alex Alderton, Roberto Amato, Jeffrey Barrett, Sonia Goncalves, Ewan Harrison, David K. Jackson, Ian Johnston, Dominic Kwiatkowski, Cordelia Langford, John Sillitoe on behalf of the Wellcome Sanger Institute COVID-19 Surveillance Team |
| EPI_ISL_3140685, EPI_ISL_3140686, EPI_ISL_3140687 | Berkshire and Surrey Pathology Services Lighthouse Laboratory | Wellcome Sanger Institute for the COVID-19 Genomics UK (COG-UK) Consortium | Berkshire and Surrey Pathology Services Lighthouse Laboratory and Alex Alderton, Roberto Amato, Jeffrey Barrett, Sonia Goncalves, Ewan Harrison, David K. Jackson, Ian Johnston, Dominic Kwiatkowski, Cordelia Langford, John Sillitoe on behalf of the Wellcome Sanger Institute COVID-19 Surveillance Team |
| EPI_ISL_3140688 | Lighthouse Lab in Alderley Park | Wellcome Sanger Institute for the COVID-19 Genomics UK (COG-UK) Consortium | Jacquelyn Wynn, Mairead Hyland, The Lighthouse Lab in Alderley Park and Alex Alderton, Roberto Amato, Jeffrey Barrett, Sonia Goncalves, Ewan Harrison, David K. Jackson, Ian Johnston, Dominic Kwiatkowski, Cordelia Langford, John Sillitoe on behalf of the Wellcome Sanger Institute COVID-19 Surveillance Team |
| EPI_ISL_3140689 | Berkshire and Surrey Pathology Services Lighthouse Laboratory | Wellcome Sanger Institute for the COVID-19 Genomics UK (COG-UK) Consortium | Berkshire and Surrey Pathology Services Lighthouse Laboratory and Alex Alderton, Roberto Amato, Jeffrey Barrett, Sonia Goncalves, Ewan Harrison, David K. Jackson, Ian Johnston, Dominic Kwiatkowski, Cordelia Langford, John Sillitoe on behalf of the Wellcome Sanger Institute COVID-19 Surveillance Team |
| EPI_ISL_3140690 | Lighthouse Lab in Milton Keynes | Wellcome Sanger Institute for the COVID-19 Genomics UK (COG-UK) Consortium | The Lighthouse Lab in Milton Keynes and Alex Alderton, Roberto Amato, Jeffrey Barrett, Sonia Goncalves, Ewan Harrison, David K. Jackson, Ian Johnston, Dominic Kwiatkowski, Cordelia Langford, John Sillitoe on behalf of the Wellcome Sanger Institute COVID-19 Surveillance Team |
| EPI_ISL_3140691 | Lighthouse Lab in Glasgow | Wellcome Sanger Institute for the COVID-19 Genomics UK (COG-UK) Consortium | Harper VanSteenhouse, Yumi Kasai, David Gray, Carol Clugston, Anna Dominiczak and Alex Alderton, Roberto Amato, Jeffrey Barrett, Sonia Goncalves, Ewan Harrison, David K. Jackson, Ian Johnston, Dominic Kwiatkowski, Cordelia Langford, John Sillitoe on behalf of the Wellcome Sanger Institute COVID-19 Surveillance Team |
| EPI_ISL_3140692 | Lighthouse Lab in Milton Keynes | Wellcome Sanger Institute for the COVID-19 Genomics UK (COG-UK) Consortium | The Lighthouse Lab in Milton Keynes and Alex Alderton, Roberto Amato, Jeffrey Barrett, Sonia Goncalves, Ewan Harrison, David K. Jackson, Ian Johnston, Dominic Kwiatkowski, Cordelia Langford, John Sillitoe on behalf of the Wellcome Sanger Institute COVID-19 Surveillance Team |
| EPI_ISL_3140693 | Lighthouse Lab in Glasgow | Wellcome Sanger Institute for the COVID-19 Genomics UK (COG-UK) Consortium | Harper VanSteenhouse, Yumi Kasai, David Gray, Carol Clugston, Anna Dominiczak and Alex Alderton, Roberto Amato, Jeffrey Barrett, Sonia Goncalves, Ewan Harrison, David K. Jackson, Ian Johnston, Dominic Kwiatkowski, Cordelia Langford, John Sillitoe on behalf of the Wellcome Sanger Institute COVID-19 Surveillance Team |
| EPI_ISL_3140694 | Berkshire and Surrey Pathology Services Lighthouse Laboratory | Wellcome Sanger Institute for the COVID-19 Genomics UK (COG-UK) Consortium | Berkshire and Surrey Pathology Services Lighthouse Laboratory and Alex Alderton, Roberto Amato, Jeffrey Barrett, Sonia Goncalves, Ewan Harrison, David K. Jackson, Ian Johnston, Dominic Kwiatkowski, Cordelia Langford, John Sillitoe on behalf of the Wellcome Sanger Institute COVID-19 Surveillance Team |
| EPI_ISL_3140695, EPI_ISL_3140696 | Lighthouse Lab in Alderley Park | Wellcome Sanger Institute for the COVID-19 Genomics UK (COG-UK) Consortium | Jacquelyn Wynn, Mairead Hyland, The Lighthouse Lab in Alderley Park and Alex Alderton, Roberto Amato, Jeffrey Barrett, Sonia Goncalves, Ewan Harrison, David K. Jackson, Ian Johnston, Dominic Kwiatkowski, Cordelia Langford, John Sillitoe on behalf of the Wellcome Sanger Institute COVID-19 Surveillance Team |
| EPI_ISL_3140697 | Berkshire and Surrey Pathology Services Lighthouse Laboratory | Wellcome Sanger Institute for the COVID-19 Genomics UK (COG-UK) Consortium | Berkshire and Surrey Pathology Services Lighthouse Laboratory and Alex Alderton, Roberto Amato, Jeffrey Barrett, Sonia Goncalves, Ewan Harrison, David K. Jackson, Ian Johnston, Dominic Kwiatkowski, Cordelia Langford, John Sillitoe on behalf of the Wellcome Sanger Institute COVID-19 Surveillance Team |
| EPI_ISL_3140698 | Lighthouse Lab in Glasgow | Wellcome Sanger Institute for the COVID-19 Genomics UK (COG-UK) Consortium | Harper VanSteenhouse, Yumi Kasai, David Gray, Carol Clugston, Anna Dominiczak and Alex Alderton, Roberto Amato, Jeffrey Barrett, Sonia Goncalves, Ewan Harrison, David K. Jackson, Ian Johnston, Dominic Kwiatkowski, Cordelia Langford, John Sillitoe on behalf of the Wellcome Sanger Institute COVID-19 Surveillance Team |

| | | | |
|---|---|---|---|
| EPI_ISL_3140699 | Lighthouse Lab in Alderley Park | Wellcome Sanger Institute for the COVID-19 Genomics UK (COG-UK) Consortium | Jacquelyn Wynn, Mairead Hyland, The Lighthouse Lab in Alderley Park and Alex Alderton, Roberto Amato, Jeffrey Barrett, Sonia Goncalves, Ewan Harrison, David K. Jackson, Ian Johnston, Dominic Kwiatkowski, Cordelia Langford, John Sillitoe on behalf of the Wellcome Sanger Institute COVID-19 Surveillance Team |
| EPI_ISL_3140700 | Berkshire and Surrey Pathology Services Lighthouse Laboratory | Wellcome Sanger Institute for the COVID-19 Genomics UK (COG-UK) Consortium | Berkshire and Surrey Pathology Services Lighthouse Laboratory and Alex Alderton, Roberto Amato, Jeffrey Barrett, Sonia Goncalves, Ewan Harrison, David K. Jackson, Ian Johnston, Dominic Kwiatkowski, Cordelia Langford, John Sillitoe on behalf of the Wellcome Sanger Institute COVID-19 Surveillance Team |
| EPI_ISL_3140701 | Lighthouse Lab in Alderley Park | Wellcome Sanger Institute for the COVID-19 Genomics UK (COG-UK) Consortium | Jacquelyn Wynn, Mairead Hyland, The Lighthouse Lab in Alderley Park and Alex Alderton, Roberto Amato, Jeffrey Barrett, Sonia Goncalves, Ewan Harrison, David K. Jackson, Ian Johnston, Dominic Kwiatkowski, Cordelia Langford, John Sillitoe on behalf of the Wellcome Sanger Institute COVID-19 Surveillance Team |
| EPI_ISL_3140702 | Berkshire and Surrey Pathology Services Lighthouse Laboratory | Wellcome Sanger Institute for the COVID-19 Genomics UK (COG-UK) Consortium | Berkshire and Surrey Pathology Services Lighthouse Laboratory and Alex Alderton, Roberto Amato, Jeffrey Barrett, Sonia Goncalves, Ewan Harrison, David K. Jackson, Ian Johnston, Dominic Kwiatkowski, Cordelia Langford, John Sillitoe on behalf of the Wellcome Sanger Institute COVID-19 Surveillance Team |
| EPI_ISL_3140703 | Lighthouse Lab in Milton Keynes | Wellcome Sanger Institute for the COVID-19 Genomics UK (COG-UK) Consortium | The Lighthouse Lab in Milton Keynes and Alex Alderton, Roberto Amato, Jeffrey Barrett, Sonia Goncalves, Ewan Harrison, David K. Jackson, Ian Johnston, Dominic Kwiatkowski, Cordelia Langford, John Sillitoe on behalf of the Wellcome Sanger Institute COVID-19 Surveillance Team |
| EPI_ISL_3140704 | Lighthouse Lab in Glasgow | Wellcome Sanger Institute for the COVID-19 Genomics UK (COG-UK) Consortium | Harper VanSteenhouse, Yumi Kasai, David Gray, Carol Clugston, Anna Dominiczak and Alex Alderton, Roberto Amato, Jeffrey Barrett, Sonia Goncalves, Ewan Harrison, David K. Jackson, Ian Johnston, Dominic Kwiatkowski, Cordelia Langford, John Sillitoe on behalf of the Wellcome Sanger Institute COVID-19 Surveillance Team |
| EPI_ISL_3140705, EPI_ISL_3140706 | Lighthouse Lab in Alderley Park | Wellcome Sanger Institute for the COVID-19 Genomics UK (COG-UK) Consortium | Jacquelyn Wynn, Mairead Hyland, The Lighthouse Lab in Alderley Park and Alex Alderton, Roberto Amato, Jeffrey Barrett, Sonia Goncalves, Ewan Harrison, David K. Jackson, Ian Johnston, Dominic Kwiatkowski, Cordelia Langford, John Sillitoe on behalf of the Wellcome Sanger Institute COVID-19 Surveillance Team |
| EPI_ISL_3140707, EPI_ISL_3140708, EPI_ISL_3140709 | Berkshire and Surrey Pathology Services Lighthouse Laboratory | Wellcome Sanger Institute for the COVID-19 Genomics UK (COG-UK) Consortium | Berkshire and Surrey Pathology Services Lighthouse Laboratory and Alex Alderton, Roberto Amato, Jeffrey Barrett, Sonia Goncalves, Ewan Harrison, David K. Jackson, Ian Johnston, Dominic Kwiatkowski, Cordelia Langford, John Sillitoe on behalf of the Wellcome Sanger Institute COVID-19 Surveillance Team |
| EPI_ISL_3140710 | Lighthouse Lab in Milton Keynes | Wellcome Sanger Institute for the COVID-19 Genomics UK (COG-UK) Consortium | The Lighthouse Lab in Milton Keynes and Alex Alderton, Roberto Amato, Jeffrey Barrett, Sonia Goncalves, Ewan Harrison, David K. Jackson, Ian Johnston, Dominic Kwiatkowski, Cordelia Langford, John Sillitoe on behalf of the Wellcome Sanger Institute COVID-19 Surveillance Team |
| EPI_ISL_3140711 | Lighthouse Lab in Glasgow | Wellcome Sanger Institute for the COVID-19 Genomics UK (COG-UK) Consortium | Harper VanSteenhouse, Yumi Kasai, David Gray, Carol Clugston, Anna Dominiczak and Alex Alderton, Roberto Amato, Jeffrey Barrett, Sonia Goncalves, Ewan Harrison, David K. Jackson, Ian Johnston, Dominic Kwiatkowski, Cordelia Langford, John Sillitoe on behalf of the Wellcome Sanger Institute COVID-19 Surveillance Team |
| EPI_ISL_3140712 | Berkshire and Surrey Pathology Services Lighthouse Laboratory | Wellcome Sanger Institute for the COVID-19 Genomics UK (COG-UK) Consortium | Berkshire and Surrey Pathology Services Lighthouse Laboratory and Alex Alderton, Roberto Amato, Jeffrey Barrett, Sonia Goncalves, Ewan Harrison, David K. Jackson, Ian Johnston, Dominic Kwiatkowski, Cordelia Langford, John Sillitoe on behalf of the Wellcome Sanger Institute COVID-19 Surveillance Team |
| EPI_ISL_3140713 | Lighthouse Lab in Alderley Park | Wellcome Sanger Institute for the COVID-19 Genomics UK (COG-UK) Consortium | Jacquelyn Wynn, Mairead Hyland, The Lighthouse Lab in Alderley Park and Alex Alderton, Roberto Amato, Jeffrey Barrett, Sonia Goncalves, Ewan Harrison, David K. Jackson, Ian Johnston, Dominic Kwiatkowski, Cordelia Langford, John Sillitoe on behalf of the Wellcome Sanger Institute COVID-19 Surveillance Team |
| EPI_ISL_3140714, EPI_ISL_3140715, EPI_ISL_3140716 | Lighthouse Lab in Glasgow | Wellcome Sanger Institute for the COVID-19 Genomics UK (COG-UK) Consortium | Harper VanSteenhouse, Yumi Kasai, David Gray, Carol Clugston, Anna Dominiczak and Alex Alderton, Roberto Amato, Jeffrey Barrett, Sonia Goncalves, Ewan Harrison, David K. Jackson, Ian Johnston, Dominic Kwiatkowski, Cordelia Langford, John Sillitoe on behalf of the Wellcome Sanger Institute COVID-19 Surveillance Team |
| EPI_ISL_3140717, EPI_ISL_3140718 | Lighthouse Lab in Alderley Park | Wellcome Sanger Institute for the COVID-19 Genomics UK (COG-UK) Consortium | Jacquelyn Wynn, Mairead Hyland, The Lighthouse Lab in Alderley Park and Alex Alderton, Roberto Amato, Jeffrey Barrett, Sonia Goncalves, Ewan Harrison, David K. Jackson, Ian Johnston, Dominic Kwiatkowski, Cordelia Langford, John Sillitoe on behalf of the Wellcome Sanger Institute COVID-19 Surveillance Team |
| EPI_ISL_3140719 | Berkshire and Surrey Pathology Services Lighthouse Laboratory | Wellcome Sanger Institute for the COVID-19 Genomics UK (COG-UK) Consortium | Berkshire and Surrey Pathology Services Lighthouse Laboratory and Alex Alderton, Roberto Amato, Jeffrey Barrett, Sonia Goncalves, Ewan Harrison, David K. Jackson, Ian Johnston, Dominic Kwiatkowski, Cordelia Langford, John Sillitoe on behalf of the Wellcome Sanger Institute COVID-19 Surveillance Team |
| EPI_ISL_3140720 | Lighthouse Lab in Glasgow | Wellcome Sanger Institute for the COVID-19 Genomics UK (COG-UK) Consortium | Harper VanSteenhouse, Yumi Kasai, David Gray, Carol Clugston, Anna Dominiczak and Alex Alderton, Roberto Amato, Jeffrey Barrett, Sonia Goncalves, Ewan Harrison, David K. Jackson, Ian Johnston, Dominic Kwiatkowski, Cordelia Langford, John Sillitoe on behalf of the Wellcome Sanger Institute COVID-19 Surveillance Team |
| EPI_ISL_3140721 | Berkshire and Surrey Pathology Services Lighthouse Laboratory | Wellcome Sanger Institute for the COVID-19 Genomics UK (COG-UK) Consortium | Berkshire and Surrey Pathology Services Lighthouse Laboratory and Alex Alderton, Roberto Amato, Jeffrey Barrett, Sonia Goncalves, Ewan Harrison, David K. Jackson, Ian Johnston, Dominic Kwiatkowski, Cordelia Langford, John Sillitoe on behalf of the Wellcome Sanger Institute COVID-19 Surveillance Team |
| EPI_ISL_3140722, EPI_ISL_3140723, EPI_ISL_3140724 | Lighthouse Lab in Alderley Park | Wellcome Sanger Institute for the COVID-19 Genomics UK (COG-UK) Consortium | Jacquelyn Wynn, Mairead Hyland, The Lighthouse Lab in Alderley Park and Alex Alderton, Roberto Amato, Jeffrey Barrett, Sonia Goncalves, Ewan Harrison, David K. Jackson, Ian Johnston, Dominic Kwiatkowski, Cordelia Langford, John Sillitoe on behalf of the Wellcome Sanger Institute COVID-19 Surveillance Team |
| EPI_ISL_3140725, EPI_ISL_3140726 | Lighthouse Lab in Milton Keynes | Wellcome Sanger Institute for the COVID-19 Genomics UK (COG-UK) Consortium | The Lighthouse Lab in Milton Keynes and Alex Alderton, Roberto Amato, Jeffrey Barrett, Sonia Goncalves, Ewan Harrison, David K. Jackson, Ian Johnston, Dominic Kwiatkowski, Cordelia Langford, John Sillitoe on behalf of the Wellcome Sanger Institute COVID-19 Surveillance Team |
| EPI_ISL_3140727 | Lighthouse Lab in Glasgow | Wellcome Sanger Institute for the COVID-19 Genomics UK (COG-UK) Consortium | Harper VanSteenhouse, Yumi Kasai, David Gray, Carol Clugston, Anna Dominiczak and Alex Alderton, Roberto Amato, Jeffrey Barrett, Sonia Goncalves, Ewan Harrison, David K. Jackson, Ian Johnston, Dominic Kwiatkowski, Cordelia Langford, John Sillitoe on behalf of the Wellcome Sanger Institute COVID-19 Surveillance Team |
| EPI_ISL_3140728 | Berkshire and Surrey Pathology Services Lighthouse Laboratory | Wellcome Sanger Institute for the COVID-19 Genomics UK (COG-UK) Consortium | Berkshire and Surrey Pathology Services Lighthouse Laboratory and Alex Alderton, Roberto Amato, Jeffrey Barrett, Sonia Goncalves, Ewan Harrison, David K. Jackson, Ian Johnston, Dominic Kwiatkowski, Cordelia Langford, John Sillitoe on behalf of the Wellcome Sanger Institute COVID-19 Surveillance Team |
| EPI_ISL_3140729 | Lighthouse Lab in Glasgow | Wellcome Sanger Institute for the COVID-19 Genomics UK (COG-UK) Consortium | Harper VanSteenhouse, Yumi Kasai, David Gray, Carol Clugston, Anna Dominiczak and Alex Alderton, Roberto Amato, Jeffrey Barrett, Sonia Goncalves, Ewan Harrison, David K. Jackson, Ian Johnston, Dominic Kwiatkowski, Cordelia Langford, John Sillitoe on behalf of the Wellcome Sanger Institute COVID-19 Surveillance Team |
| EPI_ISL_3140730 | Berkshire and Surrey Pathology Services Lighthouse Laboratory | Wellcome Sanger Institute for the COVID-19 Genomics UK (COG-UK) Consortium | Berkshire and Surrey Pathology Services Lighthouse Laboratory and Alex Alderton, Roberto Amato, Jeffrey Barrett, Sonia Goncalves, Ewan Harrison, David K. Jackson, Ian Johnston, Dominic Kwiatkowski, Cordelia Langford, John Sillitoe on behalf of the Wellcome Sanger Institute COVID-19 Surveillance Team |
| EPI_ISL_3140731 | Lighthouse Lab in Alderley Park | Wellcome Sanger Institute for the COVID-19 Genomics UK (COG-UK) Consortium | Jacquelyn Wynn, Mairead Hyland, The Lighthouse Lab in Alderley Park and Alex Alderton, Roberto Amato, Jeffrey Barrett, Sonia Goncalves, Ewan Harrison, David K. Jackson, Ian Johnston, Dominic Kwiatkowski, Cordelia Langford, John Sillitoe on behalf of the Wellcome Sanger Institute COVID-19 Surveillance Team |
| EPI_ISL_3140732 | Lighthouse Lab in Milton Keynes | Wellcome Sanger Institute for the COVID-19 Genomics UK (COG-UK) Consortium | The Lighthouse Lab in Milton Keynes and Alex Alderton, Roberto Amato, Jeffrey Barrett, Sonia Goncalves, Ewan Harrison, David K. Jackson, Ian Johnston, Dominic Kwiatkowski, Cordelia Langford, John Sillitoe on behalf of the Wellcome Sanger Institute COVID-19 Surveillance Team |
| EPI_ISL_3140733, EPI_ISL_3140734, EPI_ISL_3140735, EPI_ISL_3140736 | Berkshire and Surrey Pathology Services Lighthouse Laboratory | Wellcome Sanger Institute for the COVID-19 Genomics UK (COG-UK) Consortium | Berkshire and Surrey Pathology Services Lighthouse Laboratory and Alex Alderton, Roberto Amato, Jeffrey Barrett, Sonia Goncalves, Ewan Harrison, David K. Jackson, Ian Johnston, Dominic Kwiatkowski, Cordelia Langford, John Sillitoe on behalf of the Wellcome Sanger Institute COVID-19 Surveillance Team |
| EPI_ISL_3140737 | Lighthouse Lab in Alderley Park | Wellcome Sanger Institute for the COVID-19 Genomics UK (COG-UK) Consortium | Jacquelyn Wynn, Mairead Hyland, The Lighthouse Lab in Alderley Park and Alex Alderton, Roberto Amato, Jeffrey Barrett, Sonia Goncalves, Ewan Harrison, David K. Jackson, Ian Johnston, Dominic Kwiatkowski, Cordelia Langford, John Sillitoe on behalf of the Wellcome Sanger Institute COVID-19 Surveillance Team |
| EPI_ISL_3140738 | Berkshire and Surrey Pathology Services Lighthouse Laboratory | Wellcome Sanger Institute for the COVID-19 Genomics UK (COG-UK) Consortium | Berkshire and Surrey Pathology Services Lighthouse Laboratory and Alex Alderton, Roberto Amato, Jeffrey Barrett, Sonia Goncalves, Ewan Harrison, David K. Jackson, Ian Johnston, Dominic Kwiatkowski, Cordelia Langford, John Sillitoe on behalf of the Wellcome Sanger Institute COVID-19 Surveillance Team |
| EPI_ISL_3140739 | Lighthouse Lab in Alderley Park | Wellcome Sanger Institute for the COVID-19 Genomics UK (COG-UK) Consortium | Jacquelyn Wynn, Mairead Hyland, The Lighthouse Lab in Alderley Park and Alex Alderton, Roberto Amato, Jeffrey Barrett, Sonia Goncalves, Ewan Harrison, David K. Jackson, Ian Johnston, Dominic Kwiatkowski, Cordelia Langford, John Sillitoe on behalf of the Wellcome Sanger Institute COVID-19 Surveillance Team |
| EPI_ISL_3140740 | Lighthouse Lab in Glasgow | Wellcome Sanger Institute for the COVID-19 Genomics UK (COG-UK) Consortium | Harper VanSteenhouse, Yumi Kasai, David Gray, Carol Clugston, Anna Dominiczak and Alex Alderton, Roberto Amato, Jeffrey Barrett, Sonia Goncalves, Ewan Harrison, David K. Jackson, Ian Johnston, Dominic Kwiatkowski, Cordelia Langford, John Sillitoe on behalf of the Wellcome Sanger Institute COVID-19 Surveillance Team |
| EPI_ISL_3140741, EPI_ISL_3140742 | Lighthouse Lab in Alderley Park | Wellcome Sanger Institute for the COVID-19 Genomics UK (COG-UK) Consortium | Jacquelyn Wynn, Mairead Hyland, The Lighthouse Lab in Alderley Park and Alex Alderton, Roberto Amato, Jeffrey Barrett, Sonia Goncalves, Ewan Harrison, David K. Jackson, Ian Johnston, Dominic Kwiatkowski, Cordelia Langford, John Sillitoe on behalf of the Wellcome Sanger Institute COVID-19 Surveillance Team |
| EPI_ISL_3140743 | Berkshire and Surrey Pathology Services Lighthouse Laboratory | Wellcome Sanger Institute for the COVID-19 Genomics UK (COG-UK) Consortium | Berkshire and Surrey Pathology Services Lighthouse Laboratory and Alex Alderton, Roberto Amato, Jeffrey Barrett, Sonia Goncalves, Ewan Harrison, David K. Jackson, Ian Johnston, Dominic Kwiatkowski, Cordelia Langford, John Sillitoe on behalf of the Wellcome Sanger Institute COVID-19 Surveillance Team |
| EPI_ISL_3140744 | Lighthouse Lab in Alderley Park | Wellcome Sanger Institute for the COVID-19 Genomics UK (COG-UK) Consortium | Jacquelyn Wynn, Mairead Hyland, The Lighthouse Lab in Alderley Park and Alex Alderton, Roberto Amato, Jeffrey Barrett, Sonia Goncalves, Ewan Harrison, David K. Jackson, Ian Johnston, Dominic Kwiatkowski, Cordelia Langford, John Sillitoe on behalf of the Wellcome Sanger Institute COVID-19 Surveillance Team |
| EPI_ISL_3140745, EPI_ISL_3140746, EPI_ISL_3140747, EPI_ISL_3140748 | Lighthouse Lab in Milton Keynes | Wellcome Sanger Institute for the COVID-19 Genomics UK (COG-UK) Consortium | The Lighthouse Lab in Milton Keynes and Alex Alderton, Roberto Amato, Jeffrey Barrett, Sonia Goncalves, Ewan Harrison, David K. Jackson, Ian Johnston, Dominic Kwiatkowski, Cordelia Langford, John Sillitoe on behalf of the Wellcome Sanger Institute COVID-19 Surveillance Team |
| EPI_ISL_3140749 | Lighthouse Lab in Glasgow | Wellcome Sanger Institute for the COVID-19 Genomics UK (COG-UK) Consortium | Harper VanSteenhouse, Yumi Kasai, David Gray, Carol Clugston, Anna Dominiczak and Alex Alderton, Roberto Amato, Jeffrey Barrett, Sonia Goncalves, Ewan Harrison, David K. Jackson, Ian Johnston, Dominic Kwiatkowski, Cordelia Langford, John Sillitoe on behalf of the Wellcome Sanger Institute COVID-19 Surveillance Team |
| EPI_ISL_3140750 | Lighthouse Lab in Milton Keynes | Wellcome Sanger Institute for the COVID-19 Genomics UK (COG-UK) Consortium | The Lighthouse Lab in Milton Keynes and Alex Alderton, Roberto Amato, Jeffrey Barrett, Sonia Goncalves, Ewan Harrison, David K. Jackson, Ian Johnston, Dominic Kwiatkowski, Cordelia Langford, John Sillitoe on behalf of the Wellcome Sanger Institute COVID-19 Surveillance Team |
| EPI_ISL_3140751 | Lighthouse Lab in Alderley Park | Wellcome Sanger Institute for the COVID-19 Genomics UK (COG-UK) Consortium | Jacquelyn Wynn, Mairead Hyland, The Lighthouse Lab in Alderley Park and Alex Alderton, Roberto Amato, Jeffrey Barrett, Sonia Goncalves, Ewan Harrison, David K. Jackson, Ian Johnston, Dominic Kwiatkowski, Cordelia Langford, John Sillitoe on behalf of the Wellcome Sanger Institute COVID-19 Surveillance Team |
| EPI_ISL_3140752 | Lighthouse Lab in Glasgow | Wellcome Sanger Institute for the COVID-19 Genomics UK (COG-UK) Consortium | Harper VanSteenhouse, Yumi Kasai, David Gray, Carol Clugston, Anna Dominiczak and Alex Alderton, Roberto Amato, Jeffrey Barrett, Sonia Goncalves, Ewan Harrison, David K. Jackson, Ian Johnston, Dominic Kwiatkowski, Cordelia Langford, John Sillitoe on behalf of the Wellcome Sanger Institute COVID-19 Surveillance Team |
| EPI_ISL_3140753 | Lighthouse Lab in Alderley Park | Wellcome Sanger Institute for the COVID-19 Genomics UK (COG-UK) Consortium | Jacquelyn Wynn, Mairead Hyland, The Lighthouse Lab in Alderley Park and Alex Alderton, Roberto Amato, Jeffrey Barrett, Sonia Goncalves, Ewan Harrison, David K. Jackson, Ian Johnston, Dominic Kwiatkowski, Cordelia Langford, John Sillitoe on behalf of the Wellcome Sanger Institute COVID-19 Surveillance Team |
| EPI_ISL_3140754 | Berkshire and Surrey Pathology Services Lighthouse Laboratory | Wellcome Sanger Institute for the COVID-19 Genomics UK (COG-UK) Consortium | Berkshire and Surrey Pathology Services Lighthouse Laboratory and Alex Alderton, Roberto Amato, Jeffrey Barrett, Sonia Goncalves, Ewan Harrison, David K. Jackson, Ian Johnston, Dominic Kwiatkowski, Cordelia Langford, John Sillitoe on behalf of the Wellcome Sanger Institute COVID-19 Surveillance Team |
| EPI_ISL_3140755 | Lighthouse Lab in Milton Keynes | Wellcome Sanger Institute for the COVID-19 Genomics UK (COG-UK) Consortium | The Lighthouse Lab in Milton Keynes and Alex Alderton, Roberto Amato, Jeffrey Barrett, Sonia Goncalves, Ewan Harrison, David K. Jackson, Ian Johnston, Dominic Kwiatkowski, Cordelia Langford, John Sillitoe on behalf of the Wellcome Sanger Institute COVID-19 Surveillance Team |
| EPI_ISL_3140756 | Lighthouse Lab in Glasgow | Wellcome Sanger Institute for the COVID-19 Genomics UK (COG-UK) Consortium | Harper VanSteenhouse, Yumi Kasai, David Gray, Carol Clugston, Anna Dominiczak and Alex Alderton, Roberto Amato, Jeffrey Barrett, Sonia Goncalves, Ewan Harrison, David K. Jackson, Ian Johnston, Dominic Kwiatkowski, Cordelia Langford, John Sillitoe on behalf of the Wellcome Sanger Institute COVID-19 Surveillance Team |

| | | | |
|---|---|---|---|
| EPI_ISL_3140757, EPI_ISL_3140758 | Lighthouse Lab in Milton Keynes | Wellcome Sanger Institute for the COVID-19 Genomics UK (COG-UK) Consortium | The Lighthouse Lab in Milton Keynes and Alex Alderton, Roberto Amato, Jeffrey Barrett, Sonia Goncalves, Ewan Harrison, David K. Jackson, Ian Johnston, Dominic Kwiatkowski, Cordelia Langford, John Sillitoe on behalf of the Wellcome Sanger Institute COVID-19 Surveillance Team |
| EPI_ISL_3140759 | Berkshire and Surrey Pathology Services Lighthouse Laboratory | Wellcome Sanger Institute for the COVID-19 Genomics UK (COG-UK) Consortium | Berkshire and Surrey Pathology Services Lighthouse Laboratory and Alex Alderton, Roberto Amato, Jeffrey Barrett, Sonia Goncalves, Ewan Harrison, David K. Jackson, Ian Johnston, Dominic Kwiatkowski, Cordelia Langford, John Sillitoe on behalf of the Wellcome Sanger Institute COVID-19 Surveillance Team |
| EPI_ISL_3140760 | Lighthouse Lab in Alderley Park | Wellcome Sanger Institute for the COVID-19 Genomics UK (COG-UK) Consortium | Jacquelyn Wynn, Mairead Hyland, The Lighthouse Lab in Alderley Park and Alex Alderton, Roberto Amato, Jeffrey Barrett, Sonia Goncalves, Ewan Harrison, David K. Jackson, Ian Johnston, Dominic Kwiatkowski, Cordelia Langford, John Sillitoe on behalf of the Wellcome Sanger Institute COVID-19 Surveillance Team |
| EPI_ISL_3140761 | Berkshire and Surrey Pathology Services Lighthouse Laboratory | Wellcome Sanger Institute for the COVID-19 Genomics UK (COG-UK) Consortium | Berkshire and Surrey Pathology Services Lighthouse Laboratory and Alex Alderton, Roberto Amato, Jeffrey Barrett, Sonia Goncalves, Ewan Harrison, David K. Jackson, Ian Johnston, Dominic Kwiatkowski, Cordelia Langford, John Sillitoe on behalf of the Wellcome Sanger Institute COVID-19 Surveillance Team |
| EPI_ISL_3140762 | Lighthouse Lab in Glasgow | Wellcome Sanger Institute for the COVID-19 Genomics UK (COG-UK) Consortium | Harper VanSteenhouse, Yumi Kasai, David Gray, Carol Clugston, Anna Dominiczak and Alex Alderton, Roberto Amato, Jeffrey Barrett, Sonia Goncalves, Ewan Harrison, David K. Jackson, Ian Johnston, Dominic Kwiatkowski, Cordelia Langford, John Sillitoe on behalf of the Wellcome Sanger Institute COVID-19 Surveillance Team |
| EPI_ISL_3140763 | Lighthouse Lab in Alderley Park | Wellcome Sanger Institute for the COVID-19 Genomics UK (COG-UK) Consortium | Jacquelyn Wynn, Mairead Hyland, The Lighthouse Lab in Alderley Park and Alex Alderton, Roberto Amato, Jeffrey Barrett, Sonia Goncalves, Ewan Harrison, David K. Jackson, Ian Johnston, Dominic Kwiatkowski, Cordelia Langford, John Sillitoe on behalf of the Wellcome Sanger Institute COVID-19 Surveillance Team |
| EPI_ISL_3140764 | Lighthouse Lab in Glasgow | Wellcome Sanger Institute for the COVID-19 Genomics UK (COG-UK) Consortium | Harper VanSteenhouse, Yumi Kasai, David Gray, Carol Clugston, Anna Dominiczak and Alex Alderton, Roberto Amato, Jeffrey Barrett, Sonia Goncalves, Ewan Harrison, David K. Jackson, Ian Johnston, Dominic Kwiatkowski, Cordelia Langford, John Sillitoe on behalf of the Wellcome Sanger Institute COVID-19 Surveillance Team |
| EPI_ISL_3140765 | Lighthouse Lab in Milton Keynes | Wellcome Sanger Institute for the COVID-19 Genomics UK (COG-UK) Consortium | The Lighthouse Lab in Milton Keynes and Alex Alderton, Roberto Amato, Jeffrey Barrett, Sonia Goncalves, Ewan Harrison, David K. Jackson, Ian Johnston, Dominic Kwiatkowski, Cordelia Langford, John Sillitoe on behalf of the Wellcome Sanger Institute COVID-19 Surveillance Team |
| EPI_ISL_3140766 | Lighthouse Lab in Glasgow | Wellcome Sanger Institute for the COVID-19 Genomics UK (COG-UK) Consortium | Harper VanSteenhouse, Yumi Kasai, David Gray, Carol Clugston, Anna Dominiczak and Alex Alderton, Roberto Amato, Jeffrey Barrett, Sonia Goncalves, Ewan Harrison, David K. Jackson, Ian Johnston, Dominic Kwiatkowski, Cordelia Langford, John Sillitoe on behalf of the Wellcome Sanger Institute COVID-19 Surveillance Team |
| EPI_ISL_3140767 | Lighthouse Lab in Milton Keynes | Wellcome Sanger Institute for the COVID-19 Genomics UK (COG-UK) Consortium | The Lighthouse Lab in Milton Keynes and Alex Alderton, Roberto Amato, Jeffrey Barrett, Sonia Goncalves, Ewan Harrison, David K. Jackson, Ian Johnston, Dominic Kwiatkowski, Cordelia Langford, John Sillitoe on behalf of the Wellcome Sanger Institute COVID-19 Surveillance Team |
| EPI_ISL_3140768 | Lighthouse Lab in Glasgow | Wellcome Sanger Institute for the COVID-19 Genomics UK (COG-UK) Consortium | Harper VanSteenhouse, Yumi Kasai, David Gray, Carol Clugston, Anna Dominiczak and Alex Alderton, Roberto Amato, Jeffrey Barrett, Sonia Goncalves, Ewan Harrison, David K. Jackson, Ian Johnston, Dominic Kwiatkowski, Cordelia Langford, John Sillitoe on behalf of the Wellcome Sanger Institute COVID-19 Surveillance Team |
| EPI_ISL_3140769, EPI_ISL_3140770 | Lighthouse Lab in Milton Keynes | Wellcome Sanger Institute for the COVID-19 Genomics UK (COG-UK) Consortium | The Lighthouse Lab in Milton Keynes and Alex Alderton, Roberto Amato, Jeffrey Barrett, Sonia Goncalves, Ewan Harrison, David K. Jackson, Ian Johnston, Dominic Kwiatkowski, Cordelia Langford, John Sillitoe on behalf of the Wellcome Sanger Institute COVID-19 Surveillance Team |
| EPI_ISL_3140771 | Berkshire and Surrey Pathology Services Lighthouse Laboratory | Wellcome Sanger Institute for the COVID-19 Genomics UK (COG-UK) Consortium | Berkshire and Surrey Pathology Services Lighthouse Laboratory and Alex Alderton, Roberto Amato, Jeffrey Barrett, Sonia Goncalves, Ewan Harrison, David K. Jackson, Ian Johnston, Dominic Kwiatkowski, Cordelia Langford, John Sillitoe on behalf of the Wellcome Sanger Institute COVID-19 Surveillance Team |
| EPI_ISL_3140772 | Lighthouse Lab in Milton Keynes | Wellcome Sanger Institute for the COVID-19 Genomics UK (COG-UK) Consortium | The Lighthouse Lab in Milton Keynes and Alex Alderton, Roberto Amato, Jeffrey Barrett, Sonia Goncalves, Ewan Harrison, David K. Jackson, Ian Johnston, Dominic Kwiatkowski, Cordelia Langford, John Sillitoe on behalf of the Wellcome Sanger Institute COVID-19 Surveillance Team |
| EPI_ISL_3140773 | Lighthouse Lab in Alderley Park | Wellcome Sanger Institute for the COVID-19 Genomics UK (COG-UK) Consortium | Jacquelyn Wynn, Mairead Hyland, The Lighthouse Lab in Alderley Park and Alex Alderton, Roberto Amato, Jeffrey Barrett, Sonia Goncalves, Ewan Harrison, David K. Jackson, Ian Johnston, Dominic Kwiatkowski, Cordelia Langford, John Sillitoe on behalf of the Wellcome Sanger Institute COVID-19 Surveillance Team |
| EPI_ISL_3140774 | Berkshire and Surrey Pathology Services Lighthouse Laboratory | Wellcome Sanger Institute for the COVID-19 Genomics UK (COG-UK) Consortium | Berkshire and Surrey Pathology Services Lighthouse Laboratory and Alex Alderton, Roberto Amato, Jeffrey Barrett, Sonia Goncalves, Ewan Harrison, David K. Jackson, Ian Johnston, Dominic Kwiatkowski, Cordelia Langford, John Sillitoe on behalf of the Wellcome Sanger Institute COVID-19 Surveillance Team |
| EPI_ISL_3140775 | Lighthouse Lab in Milton Keynes | Wellcome Sanger Institute for the COVID-19 Genomics UK (COG-UK) Consortium | The Lighthouse Lab in Milton Keynes and Alex Alderton, Roberto Amato, Jeffrey Barrett, Sonia Goncalves, Ewan Harrison, David K. Jackson, Ian Johnston, Dominic Kwiatkowski, Cordelia Langford, John Sillitoe on behalf of the Wellcome Sanger Institute COVID-19 Surveillance Team |
| EPI_ISL_3140776, EPI_ISL_3140777 | Lighthouse Lab in Glasgow | Wellcome Sanger Institute for the COVID-19 Genomics UK (COG-UK) Consortium | Harper VanSteenhouse, Yumi Kasai, David Gray, Carol Clugston, Anna Dominiczak and Alex Alderton, Roberto Amato, Jeffrey Barrett, Sonia Goncalves, Ewan Harrison, David K. Jackson, Ian Johnston, Dominic Kwiatkowski, Cordelia Langford, John Sillitoe on behalf of the Wellcome Sanger Institute COVID-19 Surveillance Team |
| EPI_ISL_3140778 | Lighthouse Lab in Alderley Park | Wellcome Sanger Institute for the COVID-19 Genomics UK (COG-UK) Consortium | Jacquelyn Wynn, Mairead Hyland, The Lighthouse Lab in Alderley Park and Alex Alderton, Roberto Amato, Jeffrey Barrett, Sonia Goncalves, Ewan Harrison, David K. Jackson, Ian Johnston, Dominic Kwiatkowski, Cordelia Langford, John Sillitoe on behalf of the Wellcome Sanger Institute COVID-19 Surveillance Team |
| EPI_ISL_3140779, EPI_ISL_3140780, EPI_ISL_3140781, EPI_ISL_3140782 | Berkshire and Surrey Pathology Services Lighthouse Laboratory | Wellcome Sanger Institute for the COVID-19 Genomics UK (COG-UK) Consortium | Berkshire and Surrey Pathology Services Lighthouse Laboratory and Alex Alderton, Roberto Amato, Jeffrey Barrett, Sonia Goncalves, Ewan Harrison, David K. Jackson, Ian Johnston, Dominic Kwiatkowski, Cordelia Langford, John Sillitoe on behalf of the Wellcome Sanger Institute COVID-19 Surveillance Team |
| EPI_ISL_3140783 | Lighthouse Lab in Glasgow | Wellcome Sanger Institute for the COVID-19 Genomics UK (COG-UK) Consortium | Harper VanSteenhouse, Yumi Kasai, David Gray, Carol Clugston, Anna Dominiczak and Alex Alderton, Roberto Amato, Jeffrey Barrett, Sonia Goncalves, Ewan Harrison, David K. Jackson, Ian Johnston, Dominic Kwiatkowski, Cordelia Langford, John Sillitoe on behalf of the Wellcome Sanger Institute COVID-19 Surveillance Team |
| EPI_ISL_3140784, EPI_ISL_3140785 | Lighthouse Lab in Milton Keynes | Wellcome Sanger Institute for the COVID-19 Genomics UK (COG-UK) Consortium | The Lighthouse Lab in Milton Keynes and Alex Alderton, Roberto Amato, Jeffrey Barrett, Sonia Goncalves, Ewan Harrison, David K. Jackson, Ian Johnston, Dominic Kwiatkowski, Cordelia Langford, John Sillitoe on behalf of the Wellcome Sanger Institute COVID-19 Surveillance Team |
| EPI_ISL_3140786 | Berkshire and Surrey Pathology Services Lighthouse Laboratory | Wellcome Sanger Institute for the COVID-19 Genomics UK (COG-UK) Consortium | Berkshire and Surrey Pathology Services Lighthouse Laboratory and Alex Alderton, Roberto Amato, Jeffrey Barrett, Sonia Goncalves, Ewan Harrison, David K. Jackson, Ian Johnston, Dominic Kwiatkowski, Cordelia Langford, John Sillitoe on behalf of the Wellcome Sanger Institute COVID-19 Surveillance Team |
| EPI_ISL_3140787 | Lighthouse Lab in Milton Keynes | Wellcome Sanger Institute for the COVID-19 Genomics UK (COG-UK) Consortium | The Lighthouse Lab in Milton Keynes and Alex Alderton, Roberto Amato, Jeffrey Barrett, Sonia Goncalves, Ewan Harrison, David K. Jackson, Ian Johnston, Dominic Kwiatkowski, Cordelia Langford, John Sillitoe on behalf of the Wellcome Sanger Institute COVID-19 Surveillance Team |
| EPI_ISL_3140788 | Lighthouse Lab in Alderley Park | Wellcome Sanger Institute for the COVID-19 Genomics UK (COG-UK) Consortium | Jacquelyn Wynn, Mairead Hyland, The Lighthouse Lab in Alderley Park and Alex Alderton, Roberto Amato, Jeffrey Barrett, Sonia Goncalves, Ewan Harrison, David K. Jackson, Ian Johnston, Dominic Kwiatkowski, Cordelia Langford, John Sillitoe on behalf of the Wellcome Sanger Institute COVID-19 Surveillance Team |
| EPI_ISL_3140789 | Lighthouse Lab in Milton Keynes | Wellcome Sanger Institute for the COVID-19 Genomics UK (COG-UK) Consortium | The Lighthouse Lab in Milton Keynes and Alex Alderton, Roberto Amato, Jeffrey Barrett, Sonia Goncalves, Ewan Harrison, David K. Jackson, Ian Johnston, Dominic Kwiatkowski, Cordelia Langford, John Sillitoe on behalf of the Wellcome Sanger Institute COVID-19 Surveillance Team |
| EPI_ISL_3140790 | Lighthouse Lab in Alderley Park | Wellcome Sanger Institute for the COVID-19 Genomics UK (COG-UK) Consortium | Jacquelyn Wynn, Mairead Hyland, The Lighthouse Lab in Alderley Park and Alex Alderton, Roberto Amato, Jeffrey Barrett, Sonia Goncalves, Ewan Harrison, David K. Jackson, Ian Johnston, Dominic Kwiatkowski, Cordelia Langford, John Sillitoe on behalf of the Wellcome Sanger Institute COVID-19 Surveillance Team |
| EPI_ISL_3140791, EPI_ISL_3140792 | Lighthouse Lab in Glasgow | Wellcome Sanger Institute for the COVID-19 Genomics UK (COG-UK) Consortium | Harper VanSteenhouse, Yumi Kasai, David Gray, Carol Clugston, Anna Dominiczak and Alex Alderton, Roberto Amato, Jeffrey Barrett, Sonia Goncalves, Ewan Harrison, David K. Jackson, Ian Johnston, Dominic Kwiatkowski, Cordelia Langford, John Sillitoe on behalf of the Wellcome Sanger Institute COVID-19 Surveillance Team |
| EPI_ISL_3140793, EPI_ISL_3140794, EPI_ISL_3140795 | Lighthouse Lab in Alderley Park | Wellcome Sanger Institute for the COVID-19 Genomics UK (COG-UK) Consortium | Jacquelyn Wynn, Mairead Hyland, The Lighthouse Lab in Alderley Park and Alex Alderton, Roberto Amato, Jeffrey Barrett, Sonia Goncalves, Ewan Harrison, David K. Jackson, Ian Johnston, Dominic Kwiatkowski, Cordelia Langford, John Sillitoe on behalf of the Wellcome Sanger Institute COVID-19 Surveillance Team |
| EPI_ISL_3140796 | Lighthouse Lab in Glasgow | Wellcome Sanger Institute for the COVID-19 Genomics UK (COG-UK) Consortium | Harper VanSteenhouse, Yumi Kasai, David Gray, Carol Clugston, Anna Dominiczak and Alex Alderton, Roberto Amato, Jeffrey Barrett, Sonia Goncalves, Ewan Harrison, David K. Jackson, Ian Johnston, Dominic Kwiatkowski, Cordelia Langford, John Sillitoe on behalf of the Wellcome Sanger Institute COVID-19 Surveillance Team |
| EPI_ISL_3140797, EPI_ISL_3140798 | Lighthouse Lab in Alderley Park | Wellcome Sanger Institute for the COVID-19 Genomics UK (COG-UK) Consortium | Jacquelyn Wynn, Mairead Hyland, The Lighthouse Lab in Alderley Park and Alex Alderton, Roberto Amato, Jeffrey Barrett, Sonia Goncalves, Ewan Harrison, David K. Jackson, Ian Johnston, Dominic Kwiatkowski, Cordelia Langford, John Sillitoe on behalf of the Wellcome Sanger Institute COVID-19 Surveillance Team |
| EPI_ISL_3140799, EPI_ISL_3140800 | Lighthouse Lab in Milton Keynes | Wellcome Sanger Institute for the COVID-19 Genomics UK (COG-UK) Consortium | The Lighthouse Lab in Milton Keynes and Alex Alderton, Roberto Amato, Jeffrey Barrett, Sonia Goncalves, Ewan Harrison, David K. Jackson, Ian Johnston, Dominic Kwiatkowski, Cordelia Langford, John Sillitoe on behalf of the Wellcome Sanger Institute COVID-19 Surveillance Team |
| EPI_ISL_3140801 | Lighthouse Lab in Alderley Park | Wellcome Sanger Institute for the COVID-19 Genomics UK (COG-UK) Consortium | Jacquelyn Wynn, Mairead Hyland, The Lighthouse Lab in Alderley Park and Alex Alderton, Roberto Amato, Jeffrey Barrett, Sonia Goncalves, Ewan Harrison, David K. Jackson, Ian Johnston, Dominic Kwiatkowski, Cordelia Langford, John Sillitoe on behalf of the Wellcome Sanger Institute COVID-19 Surveillance Team |
| EPI_ISL_3140802, EPI_ISL_3140803 | Lighthouse Lab in Glasgow | Wellcome Sanger Institute for the COVID-19 Genomics UK (COG-UK) Consortium | Harper VanSteenhouse, Yumi Kasai, David Gray, Carol Clugston, Anna Dominiczak and Alex Alderton, Roberto Amato, Jeffrey Barrett, Sonia Goncalves, Ewan Harrison, David K. Jackson, Ian Johnston, Dominic Kwiatkowski, Cordelia Langford, John Sillitoe on behalf of the Wellcome Sanger Institute COVID-19 Surveillance Team |
| EPI_ISL_3140804 | Berkshire and Surrey Pathology Services Lighthouse Laboratory | Wellcome Sanger Institute for the COVID-19 Genomics UK (COG-UK) Consortium | Berkshire and Surrey Pathology Services Lighthouse Laboratory and Alex Alderton, Roberto Amato, Jeffrey Barrett, Sonia Goncalves, Ewan Harrison, David K. Jackson, Ian Johnston, Dominic Kwiatkowski, Cordelia Langford, John Sillitoe on behalf of the Wellcome Sanger Institute COVID-19 Surveillance Team |
| EPI_ISL_3140805, EPI_ISL_3140806 | Lighthouse Lab in Alderley Park | Wellcome Sanger Institute for the COVID-19 Genomics UK (COG-UK) Consortium | Jacquelyn Wynn, Mairead Hyland, The Lighthouse Lab in Alderley Park and Alex Alderton, Roberto Amato, Jeffrey Barrett, Sonia Goncalves, Ewan Harrison, David K. Jackson, Ian Johnston, Dominic Kwiatkowski, Cordelia Langford, John Sillitoe on behalf of the Wellcome Sanger Institute COVID-19 Surveillance Team |
| EPI_ISL_3140807, EPI_ISL_3140808 | Berkshire and Surrey Pathology Services Lighthouse Laboratory | Wellcome Sanger Institute for the COVID-19 Genomics UK (COG-UK) Consortium | Berkshire and Surrey Pathology Services Lighthouse Laboratory and Alex Alderton, Roberto Amato, Jeffrey Barrett, Sonia Goncalves, Ewan Harrison, David K. Jackson, Ian Johnston, Dominic Kwiatkowski, Cordelia Langford, John Sillitoe on behalf of the Wellcome Sanger Institute COVID-19 Surveillance Team |
| EPI_ISL_3140809 | Lighthouse Lab in Glasgow | Wellcome Sanger Institute for the COVID-19 Genomics UK (COG-UK) Consortium | Harper VanSteenhouse, Yumi Kasai, David Gray, Carol Clugston, Anna Dominiczak and Alex Alderton, Roberto Amato, Jeffrey Barrett, Sonia Goncalves, Ewan Harrison, David K. Jackson, Ian Johnston, Dominic Kwiatkowski, Cordelia Langford, John Sillitoe on behalf of the Wellcome Sanger Institute COVID-19 Surveillance Team |
| EPI_ISL_3140810 | Berkshire and Surrey Pathology Services Lighthouse Laboratory | Wellcome Sanger Institute for the COVID-19 Genomics UK (COG-UK) Consortium | Berkshire and Surrey Pathology Services Lighthouse Laboratory and Alex Alderton, Roberto Amato, Jeffrey Barrett, Sonia Goncalves, Ewan Harrison, David K. Jackson, Ian Johnston, Dominic Kwiatkowski, Cordelia Langford, John Sillitoe on behalf of the Wellcome Sanger Institute COVID-19 Surveillance Team |
| EPI_ISL_3140811 | Lighthouse Lab in Alderley Park | Wellcome Sanger Institute for the COVID-19 Genomics UK (COG-UK) Consortium | Jacquelyn Wynn, Mairead Hyland, The Lighthouse Lab in Alderley Park and Alex Alderton, Roberto Amato, Jeffrey Barrett, Sonia Goncalves, Ewan Harrison, David K. Jackson, Ian Johnston, Dominic Kwiatkowski, Cordelia Langford, John Sillitoe on behalf of the Wellcome Sanger Institute COVID-19 Surveillance Team |
| EPI_ISL_3140812 | Lighthouse Lab in Glasgow | Wellcome Sanger Institute for the COVID-19 Genomics UK (COG-UK) Consortium | Harper VanSteenhouse, Yumi Kasai, David Gray, Carol Clugston, Anna Dominiczak and Alex Alderton, Roberto Amato, Jeffrey Barrett, Sonia Goncalves, Ewan Harrison, David K. Jackson, Ian Johnston, Dominic Kwiatkowski, Cordelia Langford, John Sillitoe on behalf of the Wellcome Sanger Institute COVID-19 Surveillance Team |
| EPI_ISL_3140813, EPI_ISL_3140814 | Lighthouse Lab in Alderley Park | Wellcome Sanger Institute for the COVID-19 Genomics UK (COG-UK) Consortium | Jacquelyn Wynn, Mairead Hyland, The Lighthouse Lab in Alderley Park and Alex Alderton, Roberto Amato, Jeffrey Barrett, Sonia Goncalves, Ewan Harrison, David K. Jackson, Ian Johnston, Dominic Kwiatkowski, Cordelia Langford, John Sillitoe on behalf of the Wellcome Sanger Institute COVID-19 Surveillance Team |
| EPI_ISL_3140815 | Berkshire and Surrey Pathology Services Lighthouse Laboratory | Wellcome Sanger Institute for the COVID-19 Genomics UK (COG-UK) Consortium | Berkshire and Surrey Pathology Services Lighthouse Laboratory and Alex Alderton, Roberto Amato, Jeffrey Barrett, Sonia Goncalves, Ewan Harrison, David K. Jackson, Ian Johnston, Dominic Kwiatkowski, Cordelia Langford, John Sillitoe on behalf of the Wellcome Sanger Institute COVID-19 Surveillance Team |
| EPI_ISL_3140816, EPI_ISL_3140817 | Lighthouse Lab in Milton Keynes | Wellcome Sanger Institute for the COVID-19 Genomics UK (COG-UK) Consortium | The Lighthouse Lab in Milton Keynes and Alex Alderton, Roberto Amato, Jeffrey Barrett, Sonia Goncalves, Ewan Harrison, David K. Jackson, Ian Johnston, Dominic Kwiatkowski, Cordelia Langford, John Sillitoe on behalf of the Wellcome Sanger Institute COVID-19 Surveillance Team |

| | | | |
|---|---|---|---|
| EPI_ISL_3140818 | Lighthouse Lab in Alderley Park | Wellcome Sanger Institute for the COVID-19 Genomics UK (COG-UK) Consortium | Jacquelyn Wynn, Mairead Hyland, The Lighthouse Lab in Alderley Park and Alex Alderton, Roberto Amato, Jeffrey Barrett, Sonia Goncalves, Ewan Harrison, David K. Jackson, Ian Johnston, Dominic Kwiatkowski, Cordelia Langford, John Sillitoe on behalf of the Wellcome Sanger Institute COVID-19 Surveillance Team |
| EPI_ISL_3140819 | Berkshire and Surrey Pathology Services Lighthouse Laboratory | Wellcome Sanger Institute for the COVID-19 Genomics UK (COG-UK) Consortium | Berkshire and Surrey Pathology Services Lighthouse Laboratory and Alex Alderton, Roberto Amato, Jeffrey Barrett, Sonia Goncalves, Ewan Harrison, David K. Jackson, Ian Johnston, Dominic Kwiatkowski, Cordelia Langford, John Sillitoe on behalf of the Wellcome Sanger Institute COVID-19 Surveillance Team |
| EPI_ISL_3140820, EPI_ISL_3140821, EPI_ISL_3140822 | Lighthouse Lab in Milton Keynes | Wellcome Sanger Institute for the COVID-19 Genomics UK (COG-UK) Consortium | The Lighthouse Lab in Milton Keynes and Alex Alderton, Roberto Amato, Jeffrey Barrett, Sonia Goncalves, Ewan Harrison, David K. Jackson, Ian Johnston, Dominic Kwiatkowski, Cordelia Langford, John Sillitoe on behalf of the Wellcome Sanger Institute COVID-19 Surveillance Team |
| EPI_ISL_3140823 | Berkshire and Surrey Pathology Services Lighthouse Laboratory | Wellcome Sanger Institute for the COVID-19 Genomics UK (COG-UK) Consortium | Berkshire and Surrey Pathology Services Lighthouse Laboratory and Alex Alderton, Roberto Amato, Jeffrey Barrett, Sonia Goncalves, Ewan Harrison, David K. Jackson, Ian Johnston, Dominic Kwiatkowski, Cordelia Langford, John Sillitoe on behalf of the Wellcome Sanger Institute COVID-19 Surveillance Team |
| EPI_ISL_3140824 | Lighthouse Lab in Alderley Park | Wellcome Sanger Institute for the COVID-19 Genomics UK (COG-UK) Consortium | Jacquelyn Wynn, Mairead Hyland, The Lighthouse Lab in Alderley Park and Alex Alderton, Roberto Amato, Jeffrey Barrett, Sonia Goncalves, Ewan Harrison, David K. Jackson, Ian Johnston, Dominic Kwiatkowski, Cordelia Langford, John Sillitoe on behalf of the Wellcome Sanger Institute COVID-19 Surveillance Team |
| EPI_ISL_3140825 | Lighthouse Lab in Milton Keynes | Wellcome Sanger Institute for the COVID-19 Genomics UK (COG-UK) Consortium | The Lighthouse Lab in Milton Keynes and Alex Alderton, Roberto Amato, Jeffrey Barrett, Sonia Goncalves, Ewan Harrison, David K. Jackson, Ian Johnston, Dominic Kwiatkowski, Cordelia Langford, John Sillitoe on behalf of the Wellcome Sanger Institute COVID-19 Surveillance Team |
| EPI_ISL_3140826 | Berkshire and Surrey Pathology Services Lighthouse Laboratory | Wellcome Sanger Institute for the COVID-19 Genomics UK (COG-UK) Consortium | Berkshire and Surrey Pathology Services Lighthouse Laboratory and Alex Alderton, Roberto Amato, Jeffrey Barrett, Sonia Goncalves, Ewan Harrison, David K. Jackson, Ian Johnston, Dominic Kwiatkowski, Cordelia Langford, John Sillitoe on behalf of the Wellcome Sanger Institute COVID-19 Surveillance Team |
| EPI_ISL_3140827 | Lighthouse Lab in Glasgow | Wellcome Sanger Institute for the COVID-19 Genomics UK (COG-UK) Consortium | Harper VanSteenhouse, Yumi Kasai, David Gray, Carol Clugston, Anna Dominiczak and Alex Alderton, Roberto Amato, Jeffrey Barrett, Sonia Goncalves, Ewan Harrison, David K. Jackson, Ian Johnston, Dominic Kwiatkowski, Cordelia Langford, John Sillitoe on behalf of the Wellcome Sanger Institute COVID-19 Surveillance Team |
| EPI_ISL_3140828 | Lighthouse Lab in Milton Keynes | Wellcome Sanger Institute for the COVID-19 Genomics UK (COG-UK) Consortium | The Lighthouse Lab in Milton Keynes and Alex Alderton, Roberto Amato, Jeffrey Barrett, Sonia Goncalves, Ewan Harrison, David K. Jackson, Ian Johnston, Dominic Kwiatkowski, Cordelia Langford, John Sillitoe on behalf of the Wellcome Sanger Institute COVID-19 Surveillance Team |
| EPI_ISL_3140829, EPI_ISL_3140830 | Berkshire and Surrey Pathology Services Lighthouse Laboratory | Wellcome Sanger Institute for the COVID-19 Genomics UK (COG-UK) Consortium | Berkshire and Surrey Pathology Services Lighthouse Laboratory and Alex Alderton, Roberto Amato, Jeffrey Barrett, Sonia Goncalves, Ewan Harrison, David K. Jackson, Ian Johnston, Dominic Kwiatkowski, Cordelia Langford, John Sillitoe on behalf of the Wellcome Sanger Institute COVID-19 Surveillance Team |
| EPI_ISL_3140831 | Lighthouse Lab in Alderley Park | Wellcome Sanger Institute for the COVID-19 Genomics UK (COG-UK) Consortium | Jacquelyn Wynn, Mairead Hyland, The Lighthouse Lab in Alderley Park and Alex Alderton, Roberto Amato, Jeffrey Barrett, Sonia Goncalves, Ewan Harrison, David K. Jackson, Ian Johnston, Dominic Kwiatkowski, Cordelia Langford, John Sillitoe on behalf of the Wellcome Sanger Institute COVID-19 Surveillance Team |
| EPI_ISL_3140832 | Lighthouse Lab in Glasgow | Wellcome Sanger Institute for the COVID-19 Genomics UK (COG-UK) Consortium | Harper VanSteenhouse, Yumi Kasai, David Gray, Carol Clugston, Anna Dominiczak and Alex Alderton, Roberto Amato, Jeffrey Barrett, Sonia Goncalves, Ewan Harrison, David K. Jackson, Ian Johnston, Dominic Kwiatkowski, Cordelia Langford, John Sillitoe on behalf of the Wellcome Sanger Institute COVID-19 Surveillance Team |
| EPI_ISL_3140833 | Lighthouse Lab in Alderley Park | Wellcome Sanger Institute for the COVID-19 Genomics UK (COG-UK) Consortium | Jacquelyn Wynn, Mairead Hyland, The Lighthouse Lab in Alderley Park and Alex Alderton, Roberto Amato, Jeffrey Barrett, Sonia Goncalves, Ewan Harrison, David K. Jackson, Ian Johnston, Dominic Kwiatkowski, Cordelia Langford, John Sillitoe on behalf of the Wellcome Sanger Institute COVID-19 Surveillance Team |
| EPI_ISL_3140834 | Lighthouse Lab in Milton Keynes | Wellcome Sanger Institute for the COVID-19 Genomics UK (COG-UK) Consortium | The Lighthouse Lab in Milton Keynes and Alex Alderton, Roberto Amato, Jeffrey Barrett, Sonia Goncalves, Ewan Harrison, David K. Jackson, Ian Johnston, Dominic Kwiatkowski, Cordelia Langford, John Sillitoe on behalf of the Wellcome Sanger Institute COVID-19 Surveillance Team |
| EPI_ISL_3140835 | Lighthouse Lab in Glasgow | Wellcome Sanger Institute for the COVID-19 Genomics UK (COG-UK) Consortium | Harper VanSteenhouse, Yumi Kasai, David Gray, Carol Clugston, Anna Dominiczak and Alex Alderton, Roberto Amato, Jeffrey Barrett, Sonia Goncalves, Ewan Harrison, David K. Jackson, Ian Johnston, Dominic Kwiatkowski, Cordelia Langford, John Sillitoe on behalf of the Wellcome Sanger Institute COVID-19 Surveillance Team |
| EPI_ISL_3140836 | Lighthouse Lab in Milton Keynes | Wellcome Sanger Institute for the COVID-19 Genomics UK (COG-UK) Consortium | The Lighthouse Lab in Milton Keynes and Alex Alderton, Roberto Amato, Jeffrey Barrett, Sonia Goncalves, Ewan Harrison, David K. Jackson, Ian Johnston, Dominic Kwiatkowski, Cordelia Langford, John Sillitoe on behalf of the Wellcome Sanger Institute COVID-19 Surveillance Team |
| EPI_ISL_3140837 | Lighthouse Lab in Alderley Park | Wellcome Sanger Institute for the COVID-19 Genomics UK (COG-UK) Consortium | Jacquelyn Wynn, Mairead Hyland, The Lighthouse Lab in Alderley Park and Alex Alderton, Roberto Amato, Jeffrey Barrett, Sonia Goncalves, Ewan Harrison, David K. Jackson, Ian Johnston, Dominic Kwiatkowski, Cordelia Langford, John Sillitoe on behalf of the Wellcome Sanger Institute COVID-19 Surveillance Team |
| EPI_ISL_3140838 | Lighthouse Lab in Milton Keynes | Wellcome Sanger Institute for the COVID-19 Genomics UK (COG-UK) Consortium | The Lighthouse Lab in Milton Keynes and Alex Alderton, Roberto Amato, Jeffrey Barrett, Sonia Goncalves, Ewan Harrison, David K. Jackson, Ian Johnston, Dominic Kwiatkowski, Cordelia Langford, John Sillitoe on behalf of the Wellcome Sanger Institute COVID-19 Surveillance Team |
| EPI_ISL_3140839, EPI_ISL_3140840, EPI_ISL_3140841, EPI_ISL_3140842 | Lighthouse Lab in Glasgow | Wellcome Sanger Institute for the COVID-19 Genomics UK (COG-UK) Consortium | Harper VanSteenhouse, Yumi Kasai, David Gray, Carol Clugston, Anna Dominiczak and Alex Alderton, Roberto Amato, Jeffrey Barrett, Sonia Goncalves, Ewan Harrison, David K. Jackson, Ian Johnston, Dominic Kwiatkowski, Cordelia Langford, John Sillitoe on behalf of the Wellcome Sanger Institute COVID-19 Surveillance Team |
| EPI_ISL_3140843 | Lighthouse Lab in Alderley Park | Wellcome Sanger Institute for the COVID-19 Genomics UK (COG-UK) Consortium | Jacquelyn Wynn, Mairead Hyland, The Lighthouse Lab in Alderley Park and Alex Alderton, Roberto Amato, Jeffrey Barrett, Sonia Goncalves, Ewan Harrison, David K. Jackson, Ian Johnston, Dominic Kwiatkowski, Cordelia Langford, John Sillitoe on behalf of the Wellcome Sanger Institute COVID-19 Surveillance Team |
| EPI_ISL_3140844 | Lighthouse Lab in Glasgow | Wellcome Sanger Institute for the COVID-19 Genomics UK (COG-UK) Consortium | Harper VanSteenhouse, Yumi Kasai, David Gray, Carol Clugston, Anna Dominiczak and Alex Alderton, Roberto Amato, Jeffrey Barrett, Sonia Goncalves, Ewan Harrison, David K. Jackson, Ian Johnston, Dominic Kwiatkowski, Cordelia Langford, John Sillitoe on behalf of the Wellcome Sanger Institute COVID-19 Surveillance Team |
| EPI_ISL_3140845 | Lighthouse Lab in Alderley Park | Wellcome Sanger Institute for the COVID-19 Genomics UK (COG-UK) Consortium | Jacquelyn Wynn, Mairead Hyland, The Lighthouse Lab in Alderley Park and Alex Alderton, Roberto Amato, Jeffrey Barrett, Sonia Goncalves, Ewan Harrison, David K. Jackson, Ian Johnston, Dominic Kwiatkowski, Cordelia Langford, John Sillitoe on behalf of the Wellcome Sanger Institute COVID-19 Surveillance Team |
| EPI_ISL_3140846 | Lighthouse Lab in Milton Keynes | Wellcome Sanger Institute for the COVID-19 Genomics UK (COG-UK) Consortium | The Lighthouse Lab in Milton Keynes and Alex Alderton, Roberto Amato, Jeffrey Barrett, Sonia Goncalves, Ewan Harrison, David K. Jackson, Ian Johnston, Dominic Kwiatkowski, Cordelia Langford, John Sillitoe on behalf of the Wellcome Sanger Institute COVID-19 Surveillance Team |
| EPI_ISL_3140847 | Lighthouse Lab in Glasgow | Wellcome Sanger Institute for the COVID-19 Genomics UK (COG-UK) Consortium | Harper VanSteenhouse, Yumi Kasai, David Gray, Carol Clugston, Anna Dominiczak and Alex Alderton, Roberto Amato, Jeffrey Barrett, Sonia Goncalves, Ewan Harrison, David K. Jackson, Ian Johnston, Dominic Kwiatkowski, Cordelia Langford, John Sillitoe on behalf of the Wellcome Sanger Institute COVID-19 Surveillance Team |
| EPI_ISL_3140848 | Lighthouse Lab in Alderley Park | Wellcome Sanger Institute for the COVID-19 Genomics UK (COG-UK) Consortium | Jacquelyn Wynn, Mairead Hyland, The Lighthouse Lab in Alderley Park and Alex Alderton, Roberto Amato, Jeffrey Barrett, Sonia Goncalves, Ewan Harrison, David K. Jackson, Ian Johnston, Dominic Kwiatkowski, Cordelia Langford, John Sillitoe on behalf of the Wellcome Sanger Institute COVID-19 Surveillance Team |
| EPI_ISL_3140849 | Lighthouse Lab in Milton Keynes | Wellcome Sanger Institute for the COVID-19 Genomics UK (COG-UK) Consortium | The Lighthouse Lab in Milton Keynes and Alex Alderton, Roberto Amato, Jeffrey Barrett, Sonia Goncalves, Ewan Harrison, David K. Jackson, Ian Johnston, Dominic Kwiatkowski, Cordelia Langford, John Sillitoe on behalf of the Wellcome Sanger Institute COVID-19 Surveillance Team |
| EPI_ISL_3140850 | Lighthouse Lab in Glasgow | Wellcome Sanger Institute for the COVID-19 Genomics UK (COG-UK) Consortium | Harper VanSteenhouse, Yumi Kasai, David Gray, Carol Clugston, Anna Dominiczak and Alex Alderton, Roberto Amato, Jeffrey Barrett, Sonia Goncalves, Ewan Harrison, David K. Jackson, Ian Johnston, Dominic Kwiatkowski, Cordelia Langford, John Sillitoe on behalf of the Wellcome Sanger Institute COVID-19 Surveillance Team |
| EPI_ISL_3140851 | Lighthouse Lab in Alderley Park | Wellcome Sanger Institute for the COVID-19 Genomics UK (COG-UK) Consortium | Jacquelyn Wynn, Mairead Hyland, The Lighthouse Lab in Alderley Park and Alex Alderton, Roberto Amato, Jeffrey Barrett, Sonia Goncalves, Ewan Harrison, David K. Jackson, Ian Johnston, Dominic Kwiatkowski, Cordelia Langford, John Sillitoe on behalf of the Wellcome Sanger Institute COVID-19 Surveillance Team |
| EPI_ISL_3140852, EPI_ISL_3140853 | Berkshire and Surrey Pathology Services Lighthouse Laboratory | Wellcome Sanger Institute for the COVID-19 Genomics UK (COG-UK) Consortium | Berkshire and Surrey Pathology Services Lighthouse Laboratory and Alex Alderton, Roberto Amato, Jeffrey Barrett, Sonia Goncalves, Ewan Harrison, David K. Jackson, Ian Johnston, Dominic Kwiatkowski, Cordelia Langford, John Sillitoe on behalf of the Wellcome Sanger Institute COVID-19 Surveillance Team |
| EPI_ISL_3140854 | Lighthouse Lab in Alderley Park | Wellcome Sanger Institute for the COVID-19 Genomics UK (COG-UK) Consortium | Jacquelyn Wynn, Mairead Hyland, The Lighthouse Lab in Alderley Park and Alex Alderton, Roberto Amato, Jeffrey Barrett, Sonia Goncalves, Ewan Harrison, David K. Jackson, Ian Johnston, Dominic Kwiatkowski, Cordelia Langford, John Sillitoe on behalf of the Wellcome Sanger Institute COVID-19 Surveillance Team |
| EPI_ISL_3140855, EPI_ISL_3140856, EPI_ISL_3140857, EPI_ISL_3140858, EPI_ISL_3140859, EPI_ISL_3140860, EPI_ISL_3140861, EPI_ISL_3140862, EPI_ISL_3140863, EPI_ISL_3140864, EPI_ISL_3140865, EPI_ISL_3140866, EPI_ISL_3140867, EPI_ISL_3140868, EPI_ISL_3140869, EPI_ISL_3140870, EPI_ISL_3140871, EPI_ISL_3140872, EPI_ISL_3140873, EPI_ISL_3140874, EPI_ISL_3140875, EPI_ISL_3140876, EPI_ISL_3140877, EPI_ISL_3140878, EPI_ISL_3140879, EPI_ISL_3140880, EPI_ISL_3140881 | | | |
| see above | Lighthouse Lab in Glasgow | Wellcome Sanger Institute for the COVID-19 Genomics UK (COG-UK) Consortium | Harper VanSteenhouse, Yumi Kasai, David Gray, Carol Clugston, Anna Dominiczak and Alex Alderton, Roberto Amato, Jeffrey Barrett, Sonia Goncalves, Ewan Harrison, David K. Jackson, Ian Johnston, Dominic Kwiatkowski, Cordelia Langford, John Sillitoe on behalf of the Wellcome Sanger Institute COVID-19 Surveillance Team |
| EPI_ISL_3140882 | Berkshire and Surrey Pathology Services Lighthouse Laboratory | Wellcome Sanger Institute for the COVID-19 Genomics UK (COG-UK) Consortium | Berkshire and Surrey Pathology Services Lighthouse Laboratory and Alex Alderton, Roberto Amato, Jeffrey Barrett, Sonia Goncalves, Ewan Harrison, David K. Jackson, Ian Johnston, Dominic Kwiatkowski, Cordelia Langford, John Sillitoe on behalf of the Wellcome Sanger Institute COVID-19 Surveillance Team |
| EPI_ISL_3140883, EPI_ISL_3140884, EPI_ISL_3140885, EPI_ISL_3140886, EPI_ISL_3140887, EPI_ISL_3140888, EPI_ISL_3140889, EPI_ISL_3140890, EPI_ISL_3140891, EPI_ISL_3140892, EPI_ISL_3140893, EPI_ISL_3140894, EPI_ISL_3140895, EPI_ISL_3140896, EPI_ISL_3140897, EPI_ISL_3140898, EPI_ISL_3140899, EPI_ISL_3140900, EPI_ISL_3140901, EPI_ISL_3140902, EPI_ISL_3140903, EPI_ISL_3140904, EPI_ISL_3140905, EPI_ISL_3140906 | | | |
| see above | Lighthouse Lab in Glasgow | Wellcome Sanger Institute for the COVID-19 Genomics UK (COG-UK) Consortium | Harper VanSteenhouse, Yumi Kasai, David Gray, Carol Clugston, Anna Dominiczak and Alex Alderton, Roberto Amato, Jeffrey Barrett, Sonia Goncalves, Ewan Harrison, David K. Jackson, Ian Johnston, Dominic Kwiatkowski, Cordelia Langford, John Sillitoe on behalf of the Wellcome Sanger Institute COVID-19 Surveillance Team |
| EPI_ISL_3140907 | Berkshire and Surrey Pathology Services Lighthouse Laboratory | Wellcome Sanger Institute for the COVID-19 Genomics UK (COG-UK) Consortium | Berkshire and Surrey Pathology Services Lighthouse Laboratory and Alex Alderton, Roberto Amato, Jeffrey Barrett, Sonia Goncalves, Ewan Harrison, David K. Jackson, Ian Johnston, Dominic Kwiatkowski, Cordelia Langford, John Sillitoe on behalf of the Wellcome Sanger Institute COVID-19 Surveillance Team |
| EPI_ISL_3140908, EPI_ISL_3140909, EPI_ISL_3140910, EPI_ISL_3140911, EPI_ISL_3140912, EPI_ISL_3140913, EPI_ISL_3140914, EPI_ISL_3140915, EPI_ISL_3140916, EPI_ISL_3140917, EPI_ISL_3140918, EPI_ISL_3140919, EPI_ISL_3140920, EPI_ISL_3140921, EPI_ISL_3140922, EPI_ISL_3140923, EPI_ISL_3140924, EPI_ISL_3140925, EPI_ISL_3140926, EPI_ISL_3140927, EPI_ISL_3140928, EPI_ISL_3140929, EPI_ISL_3140930, EPI_ISL_3140931, EPI_ISL_3140932, EPI_ISL_3140933, EPI_ISL_3140934 | | | |
| see above | Lighthouse Lab in Glasgow | Wellcome Sanger Institute for the COVID-19 Genomics UK (COG-UK) Consortium | Harper VanSteenhouse, Yumi Kasai, David Gray, Carol Clugston, Anna Dominiczak and Alex Alderton, Roberto Amato, Jeffrey Barrett, Sonia Goncalves, Ewan Harrison, David K. Jackson, Ian Johnston, Dominic Kwiatkowski, Cordelia Langford, John Sillitoe on behalf of the Wellcome Sanger Institute COVID-19 Surveillance Team |
| EPI_ISL_3140935 | Berkshire and Surrey Pathology Services Lighthouse Laboratory | Wellcome Sanger Institute for the COVID-19 Genomics UK (COG-UK) Consortium | Berkshire and Surrey Pathology Services Lighthouse Laboratory and Alex Alderton, Roberto Amato, Jeffrey Barrett, Sonia Goncalves, Ewan Harrison, David K. Jackson, Ian Johnston, Dominic Kwiatkowski, Cordelia Langford, John Sillitoe on behalf of the Wellcome Sanger Institute COVID-19 Surveillance Team |
| EPI_ISL_3140936, EPI_ISL_3140937, EPI_ISL_3140938, EPI_ISL_3140939, EPI_ISL_3140940, EPI_ISL_3140941, EPI_ISL_3140942, EPI_ISL_3140943, EPI_ISL_3140944, EPI_ISL_3140945, EPI_ISL_3140946, EPI_ISL_3140947, EPI_ISL_3140948, EPI_ISL_3140949, EPI_ISL_3140950, EPI_ISL_3140951, EPI_ISL_3140952, EPI_ISL_3140953, EPI_ISL_3140954, EPI_ISL_3140955, EPI_ISL_3140956, EPI_ISL_3140957, EPI_ISL_3140958, EPI_ISL_3140959, EPI_ISL_3140960, EPI_ISL_3140961, EPI_ISL_3140962, EPI_ISL_3140963, EPI_ISL_3140964, EPI_ISL_3140965, EPI_ISL_3140966, EPI_ISL_3140967, EPI_ISL_3140968, EPI_ISL_3140969, EPI_ISL_3140970, EPI_ISL_3140971, EPI_ISL_3140972, EPI_ISL_3140973, EPI_ISL_3140974, EPI_ISL_3140976, EPI_ISL_3140977, EPI_ISL_3140978, EPI_ISL_3140979, EPI_ISL_3140980, EPI_ISL_3140981, EPI_ISL_3140982, EPI_ISL_3140983, EPI_ISL_3140984, EPI_ISL_3140985, EPI_ISL_3140986, EPI_ISL_3140987, EPI_ISL_3140988, EPI_ISL_3140989, EPI_ISL_3140990, EPI_ISL_3140991, EPI_ISL_3140992, EPI_ISL_3140993, EPI_ISL_3140994, EPI_ISL_3140995, EPI_ISL_3140996, EPI_ISL_3140997, EPI_ISL_3140998, EPI_ISL_3140999, EPI_ISL_3141000, EPI_ISL_3141001, EPI_ISL_3141002, EPI_ISL_3141003, EPI_ISL_3141004, EPI_ISL_3141005, EPI_ISL_3141006, EPI_ISL_3141007, EPI_ISL_3141008, EPI_ISL_3141009, EPI_ISL_3141010 | | | |
| see above | Lighthouse Lab in Glasgow | Wellcome Sanger Institute for the COVID-19 Genomics UK (COG-UK) Consortium | Harper VanSteenhouse, Yumi Kasai, David Gray, Carol Clugston, Anna Dominiczak and Alex Alderton, Roberto Amato, Jeffrey Barrett, Sonia Goncalves, Ewan Harrison, David K. Jackson, Ian Johnston, Dominic Kwiatkowski, Cordelia Langford, John Sillitoe on behalf of the Wellcome Sanger Institute COVID-19 Surveillance Team |
| EPI_ISL_3141011 | Berkshire and Surrey Pathology Services Lighthouse Laboratory | Wellcome Sanger Institute for the COVID-19 Genomics UK (COG-UK) Consortium | Berkshire and Surrey Pathology Services Lighthouse Laboratory and Alex Alderton, Roberto Amato, Jeffrey Barrett, Sonia Goncalves, Ewan Harrison, David K. Jackson, Ian Johnston, Dominic Kwiatkowski, Cordelia Langford, John Sillitoe on behalf of the Wellcome Sanger Institute COVID-19 Surveillance Team |
| EPI_ISL_3141012, EPI_ISL_3141013, EPI_ISL_3141014, EPI_ISL_3141015, EPI_ISL_3141016, EPI_ISL_3141017, EPI_ISL_3141018, EPI_ISL_3141019, EPI_ISL_3141020, EPI_ISL_3141021, EPI_ISL_3141022, EPI_ISL_3141023, EPI_ISL_3141024, EPI_ISL_3141025, EPI_ISL_3141026, EPI_ISL_3141027, EPI_ISL_3141028, EPI_ISL_3141029, EPI_ISL_3141030, EPI_ISL_3141031, EPI_ISL_3141032, EPI_ISL_3141033, EPI_ISL_3141034, EPI_ISL_3141035, EPI_ISL_3141036, EPI_ISL_3141037, EPI_ISL_3141038, EPI_ISL_3141039, EPI_ISL_3141040, EPI_ISL_3141041, EPI_ISL_3141042, EPI_ISL_3141043, EPI_ISL_3141044, EPI_ISL_3141045, EPI_ISL_3141046, EPI_ISL_3141047, EPI_ISL_3141048, EPI_ISL_3141049, EPI_ISL_3141050, EPI_ISL_3141051, EPI_ISL_3141052, EPI_ISL_3141053 | | | |

| | | | |
|---|---|---|---|
| see above | Lighthouse Lab in Glasgow | Wellcome Sanger Institute for the COVID-19 Genomics UK (COG-UK) Consortium | Harper VanSteenhouse, Yumi Kasai, David Gray, Carol Clugston, Anna Dominiczak and Alex Alderton, Roberto Amato, Jeffrey Barrett, Sonia Goncalves, Ewan Harrison, David K. Jackson, Ian Johnston, Dominic Kwiatkowski, Cordelia Langford, John Sillitoe on behalf of the Wellcome Sanger Institute COVID-19 Surveillance Team |
| EPI_ISL_3141055 | Lighthouse Lab in Alderley Park | Wellcome Sanger Institute for the COVID-19 Genomics UK (COG-UK) Consortium | Jacquelyn Wynn, Mairead Hyland, The Lighthouse Lab in Alderley Park and Alex Alderton, Roberto Amato, Jeffrey Barrett, Sonia Goncalves, Ewan Harrison, David K. Jackson, Ian Johnston, Dominic Kwiatkowski, Cordelia Langford, John Sillitoe on behalf of the Wellcome Sanger Institute COVID-19 Surveillance Team |
| EPI_ISL_3141056, EPI_ISL_3141057, EPI_ISL_3141058, EPI_ISL_3141059, EPI_ISL_3141060, EPI_ISL_3141061, EPI_ISL_3141062, EPI_ISL_3141063, EPI_ISL_3141064, EPI_ISL_3141065, EPI_ISL_3141066, EPI_ISL_3141067, EPI_ISL_3141068 | | | |
| see above | Lighthouse Lab in Glasgow | Wellcome Sanger Institute for the COVID-19 Genomics UK (COG-UK) Consortium | Harper VanSteenhouse, Yumi Kasai, David Gray, Carol Clugston, Anna Dominiczak and Alex Alderton, Roberto Amato, Jeffrey Barrett, Sonia Goncalves, Ewan Harrison, David K. Jackson, Ian Johnston, Dominic Kwiatkowski, Cordelia Langford, John Sillitoe on behalf of the Wellcome Sanger Institute COVID-19 Surveillance Team |
| EPI_ISL_3141069 | Lighthouse Lab in Alderley Park | Wellcome Sanger Institute for the COVID-19 Genomics UK (COG-UK) Consortium | Jacquelyn Wynn, Mairead Hyland, The Lighthouse Lab in Alderley Park and Alex Alderton, Roberto Amato, Jeffrey Barrett, Sonia Goncalves, Ewan Harrison, David K. Jackson, Ian Johnston, Dominic Kwiatkowski, Cordelia Langford, John Sillitoe on behalf of the Wellcome Sanger Institute COVID-19 Surveillance Team |
| EPI_ISL_3141070, EPI_ISL_3141071, EPI_ISL_3141072, EPI_ISL_3141073, EPI_ISL_3141074 | Lighthouse Lab in Glasgow | Wellcome Sanger Institute for the COVID-19 Genomics UK (COG-UK) Consortium | Harper VanSteenhouse, Yumi Kasai, David Gray, Carol Clugston, Anna Dominiczak and Alex Alderton, Roberto Amato, Jeffrey Barrett, Sonia Goncalves, Ewan Harrison, David K. Jackson, Ian Johnston, Dominic Kwiatkowski, Cordelia Langford, John Sillitoe on behalf of the Wellcome Sanger Institute COVID-19 Surveillance Team |
| EPI_ISL_3141075 | Berkshire and Surrey Pathology Services Lighthouse Laboratory | Wellcome Sanger Institute for the COVID-19 Genomics UK (COG-UK) Consortium | Berkshire and Surrey Pathology Services Lighthouse Laboratory and Alex Alderton, Roberto Amato, Jeffrey Barrett, Sonia Goncalves, Ewan Harrison, David K. Jackson, Ian Johnston, Dominic Kwiatkowski, Cordelia Langford, John Sillitoe on behalf of the Wellcome Sanger Institute COVID-19 Surveillance Team |
| EPI_ISL_3141076 | Lighthouse Lab in Alderley Park | Wellcome Sanger Institute for the COVID-19 Genomics UK (COG-UK) Consortium | Jacquelyn Wynn, Mairead Hyland, The Lighthouse Lab in Alderley Park and Alex Alderton, Roberto Amato, Jeffrey Barrett, Sonia Goncalves, Ewan Harrison, David K. Jackson, Ian Johnston, Dominic Kwiatkowski, Cordelia Langford, John Sillitoe on behalf of the Wellcome Sanger Institute COVID-19 Surveillance Team |
| EPI_ISL_3141077, EPI_ISL_3141078, EPI_ISL_3141079, EPI_ISL_3141080, EPI_ISL_3141081, EPI_ISL_3141082, EPI_ISL_3141083 | Lighthouse Lab in Glasgow | Wellcome Sanger Institute for the COVID-19 Genomics UK (COG-UK) Consortium | Harper VanSteenhouse, Yumi Kasai, David Gray, Carol Clugston, Anna Dominiczak and Alex Alderton, Roberto Amato, Jeffrey Barrett, Sonia Goncalves, Ewan Harrison, David K. Jackson, Ian Johnston, Dominic Kwiatkowski, Cordelia Langford, John Sillitoe on behalf of the Wellcome Sanger Institute COVID-19 Surveillance Team |
| EPI_ISL_3141084 | Lighthouse Lab in Alderley Park | Wellcome Sanger Institute for the COVID-19 Genomics UK (COG-UK) Consortium | Jacquelyn Wynn, Mairead Hyland, The Lighthouse Lab in Alderley Park and Alex Alderton, Roberto Amato, Jeffrey Barrett, Sonia Goncalves, Ewan Harrison, David K. Jackson, Ian Johnston, Dominic Kwiatkowski, Cordelia Langford, John Sillitoe on behalf of the Wellcome Sanger Institute COVID-19 Surveillance Team |
| EPI_ISL_3141085 | Lighthouse Lab in Glasgow | Wellcome Sanger Institute for the COVID-19 Genomics UK (COG-UK) Consortium | Harper VanSteenhouse, Yumi Kasai, David Gray, Carol Clugston, Anna Dominiczak and Alex Alderton, Roberto Amato, Jeffrey Barrett, Sonia Goncalves, Ewan Harrison, David K. Jackson, Ian Johnston, Dominic Kwiatkowski, Cordelia Langford, John Sillitoe on behalf of the Wellcome Sanger Institute COVID-19 Surveillance Team |
| EPI_ISL_3141086 | Berkshire and Surrey Pathology Services Lighthouse Laboratory | Wellcome Sanger Institute for the COVID-19 Genomics UK (COG-UK) Consortium | Berkshire and Surrey Pathology Services Lighthouse Laboratory and Alex Alderton, Roberto Amato, Jeffrey Barrett, Sonia Goncalves, Ewan Harrison, David K. Jackson, Ian Johnston, Dominic Kwiatkowski, Cordelia Langford, John Sillitoe on behalf of the Wellcome Sanger Institute COVID-19 Surveillance Team |
| EPI_ISL_3141087 | Lighthouse Lab in Glasgow | Wellcome Sanger Institute for the COVID-19 Genomics UK (COG-UK) Consortium | Harper VanSteenhouse, Yumi Kasai, David Gray, Carol Clugston, Anna Dominiczak and Alex Alderton, Roberto Amato, Jeffrey Barrett, Sonia Goncalves, Ewan Harrison, David K. Jackson, Ian Johnston, Dominic Kwiatkowski, Cordelia Langford, John Sillitoe on behalf of the Wellcome Sanger Institute COVID-19 Surveillance Team |
| EPI_ISL_3141088 | Lighthouse Lab in Alderley Park | Wellcome Sanger Institute for the COVID-19 Genomics UK (COG-UK) Consortium | Jacquelyn Wynn, Mairead Hyland, The Lighthouse Lab in Alderley Park and Alex Alderton, Roberto Amato, Jeffrey Barrett, Sonia Goncalves, Ewan Harrison, David K. Jackson, Ian Johnston, Dominic Kwiatkowski, Cordelia Langford, John Sillitoe on behalf of the Wellcome Sanger Institute COVID-19 Surveillance Team |
| EPI_ISL_3141089, EPI_ISL_3141090, EPI_ISL_3141091, EPI_ISL_3141092, EPI_ISL_3141093, EPI_ISL_3141094, EPI_ISL_3141095, EPI_ISL_3141096 | Lighthouse Lab in Glasgow | Wellcome Sanger Institute for the COVID-19 Genomics UK (COG-UK) Consortium | Harper VanSteenhouse, Yumi Kasai, David Gray, Carol Clugston, Anna Dominiczak and Alex Alderton, Roberto Amato, Jeffrey Barrett, Sonia Goncalves, Ewan Harrison, David K. Jackson, Ian Johnston, Dominic Kwiatkowski, Cordelia Langford, John Sillitoe on behalf of the Wellcome Sanger Institute COVID-19 Surveillance Team |
| EPI_ISL_3141097 | Lighthouse Lab in Alderley Park | Wellcome Sanger Institute for the COVID-19 Genomics UK (COG-UK) Consortium | Jacquelyn Wynn, Mairead Hyland, The Lighthouse Lab in Alderley Park and Alex Alderton, Roberto Amato, Jeffrey Barrett, Sonia Goncalves, Ewan Harrison, David K. Jackson, Ian Johnston, Dominic Kwiatkowski, Cordelia Langford, John Sillitoe on behalf of the Wellcome Sanger Institute COVID-19 Surveillance Team |
| EPI_ISL_3141098, EPI_ISL_3141099, EPI_ISL_3141100, EPI_ISL_3141101, EPI_ISL_3141102, EPI_ISL_3141103, EPI_ISL_3141104, EPI_ISL_3141105, EPI_ISL_3141106, EPI_ISL_3141107, EPI_ISL_3141108, EPI_ISL_3141109, EPI_ISL_3141110, EPI_ISL_3141111, EPI_ISL_3141112, EPI_ISL_3141113, EPI_ISL_3141114, EPI_ISL_3141115, EPI_ISL_3141116, EPI_ISL_3141117, EPI_ISL_3141118, EPI_ISL_3141119, EPI_ISL_3141120, EPI_ISL_3141121, EPI_ISL_3141122, EPI_ISL_3141123, EPI_ISL_3141124, EPI_ISL_3141125, EPI_ISL_3141126, EPI_ISL_3141127, EPI_ISL_3141128, EPI_ISL_3141129, EPI_ISL_3141130, EPI_ISL_3141131, EPI_ISL_3141132, EPI_ISL_3141133, EPI_ISL_3141134, EPI_ISL_3141135, EPI_ISL_3141136, EPI_ISL_3141137, EPI_ISL_3141138, EPI_ISL_3141139, EPI_ISL_3141140, EPI_ISL_3141141, EPI_ISL_3141142, EPI_ISL_3141143, EPI_ISL_3141144, EPI_ISL_3141145, EPI_ISL_3141146, EPI_ISL_3141147 | | | |
| see above | Lighthouse Lab in Glasgow | Wellcome Sanger Institute for the COVID-19 Genomics UK (COG-UK) Consortium | Harper VanSteenhouse, Yumi Kasai, David Gray, Carol Clugston, Anna Dominiczak and Alex Alderton, Roberto Amato, Jeffrey Barrett, Sonia Goncalves, Ewan Harrison, David K. Jackson, Ian Johnston, Dominic Kwiatkowski, Cordelia Langford, John Sillitoe on behalf of the Wellcome Sanger Institute COVID-19 Surveillance Team |
| EPI_ISL_3141148 | Lighthouse Lab in Milton Keynes | Wellcome Sanger Institute for the COVID-19 Genomics UK (COG-UK) Consortium | The Lighthouse Lab in Milton Keynes and Alex Alderton, Roberto Amato, Jeffrey Barrett, Sonia Goncalves, Ewan Harrison, David K. Jackson, Ian Johnston, Dominic Kwiatkowski, Cordelia Langford, John Sillitoe on behalf of the Wellcome Sanger Institute COVID-19 Surveillance Team |
| EPI_ISL_3141149, EPI_ISL_3141150, EPI_ISL_3141151, EPI_ISL_3141152, EPI_ISL_3141153, EPI_ISL_3141154, EPI_ISL_3141155, EPI_ISL_3141156, EPI_ISL_3141157, EPI_ISL_3141158, EPI_ISL_3141159, EPI_ISL_3141160, EPI_ISL_3141161, EPI_ISL_3141162, EPI_ISL_3141163, EPI_ISL_3141164, EPI_ISL_3141165, EPI_ISL_3141166, EPI_ISL_3141167, EPI_ISL_3141168, EPI_ISL_3141169, EPI_ISL_3141170, EPI_ISL_3141171, EPI_ISL_3141172, EPI_ISL_3141173, EPI_ISL_3141174, EPI_ISL_3141175, EPI_ISL_3141176, EPI_ISL_3141177, EPI_ISL_3141178, EPI_ISL_3141179, EPI_ISL_3141180, EPI_ISL_3141181, EPI_ISL_3141182, EPI_ISL_3141183, EPI_ISL_3141184, EPI_ISL_3141185, EPI_ISL_3141186, EPI_ISL_3141187, EPI_ISL_3141188, EPI_ISL_3141189, EPI_ISL_3141190, EPI_ISL_3141191, EPI_ISL_3141192, EPI_ISL_3141193, EPI_ISL_3141194, EPI_ISL_3141195, EPI_ISL_3141196, EPI_ISL_3141197, EPI_ISL_3141198, EPI_ISL_3141199, EPI_ISL_3141200, EPI_ISL_3141201, EPI_ISL_3141202, EPI_ISL_3141203, EPI_ISL_3141204, EPI_ISL_3141205, EPI_ISL_3141206, EPI_ISL_3141207, EPI_ISL_3141208, EPI_ISL_3141209, EPI_ISL_3141210, EPI_ISL_3141211, EPI_ISL_3141212, EPI_ISL_3141213, EPI_ISL_3141214, EPI_ISL_3141215, EPI_ISL_3141216, EPI_ISL_3141217, EPI_ISL_3141218, EPI_ISL_3141219, EPI_ISL_3141220, EPI_ISL_3141221, EPI_ISL_3141222, EPI_ISL_3141223, EPI_ISL_3141224, EPI_ISL_3141225, EPI_ISL_3141226, EPI_ISL_3141227, EPI_ISL_3141228, EPI_ISL_3141229, EPI_ISL_3141230, EPI_ISL_3141231, EPI_ISL_3141232, EPI_ISL_3141233, EPI_ISL_3141234, EPI_ISL_3141235, EPI_ISL_3141236, EPI_ISL_3141237, EPI_ISL_3141238, EPI_ISL_3141239, EPI_ISL_3141240, EPI_ISL_3141241, EPI_ISL_3141242, EPI_ISL_3141243, EPI_ISL_3141244, EPI_ISL_3141245, EPI_ISL_3141246, EPI_ISL_3141247, EPI_ISL_3141248, EPI_ISL_3141249, EPI_ISL_3141250, EPI_ISL_3141251, EPI_ISL_3141252, EPI_ISL_3141253, EPI_ISL_3141254, EPI_ISL_3141255, EPI_ISL_3141256, EPI_ISL_3141257, EPI_ISL_3141258, EPI_ISL_3141259, EPI_ISL_3141260, EPI_ISL_3141261, EPI_ISL_3141262, EPI_ISL_3141263, EPI_ISL_3141264, EPI_ISL_3141265, EPI_ISL_3141266, EPI_ISL_3141267, EPI_ISL_3141268, EPI_ISL_3141269, EPI_ISL_3141270, EPI_ISL_3141271, EPI_ISL_3141272, EPI_ISL_3141273, EPI_ISL_3141274, EPI_ISL_3141275, EPI_ISL_3141276, EPI_ISL_3141277, EPI_ISL_3141278, EPI_ISL_3141279, EPI_ISL_3141280, EPI_ISL_3141281, EPI_ISL_3141282, EPI_ISL_3141283, EPI_ISL_3141284, EPI_ISL_3141285, EPI_ISL_3141286, EPI_ISL_3141287, EPI_ISL_3141288, EPI_ISL_3141289, EPI_ISL_3141290, EPI_ISL_3141291, EPI_ISL_3141292, EPI_ISL_3141293, EPI_ISL_3141294, EPI_ISL_3141295, EPI_ISL_3141296, EPI_ISL_3141297, EPI_ISL_3141298, EPI_ISL_3141299, EPI_ISL_3141300, EPI_ISL_3141301, EPI_ISL_3141302, EPI_ISL_3141303, EPI_ISL_3141304, EPI_ISL_3141305, EPI_ISL_3141306, EPI_ISL_3141307, EPI_ISL_3141308, EPI_ISL_3141309, EPI_ISL_3141310, EPI_ISL_3141311, EPI_ISL_3141312, EPI_ISL_3141313, EPI_ISL_3141314, EPI_ISL_3141315, EPI_ISL_3141316, EPI_ISL_3141317, EPI_ISL_3141318, EPI_ISL_3141319, EPI_ISL_3141320, EPI_ISL_3141321, EPI_ISL_3141322, EPI_ISL_3141323, EPI_ISL_3141324, EPI_ISL_3141325, EPI_ISL_3141326, EPI_ISL_3141327, EPI_ISL_3141328, EPI_ISL_3141329, EPI_ISL_3141330, EPI_ISL_3141331, EPI_ISL_3141332, EPI_ISL_3141333, EPI_ISL_3141334, EPI_ISL_3141336, EPI_ISL_3141337, EPI_ISL_3141338, EPI_ISL_3141339, EPI_ISL_3141340, EPI_ISL_3141341, EPI_ISL_3141342, EPI_ISL_3141343, EPI_ISL_3141344, EPI_ISL_3141345, EPI_ISL_3141346, EPI_ISL_3141347, EPI_ISL_3141348, EPI_ISL_3141349, EPI_ISL_3141350, EPI_ISL_3141351, EPI_ISL_3141352, EPI_ISL_3141353, EPI_ISL_3141354, EPI_ISL_3141355, EPI_ISL_3141356, EPI_ISL_3141357, EPI_ISL_3141358, EPI_ISL_3141359, EPI_ISL_3141360, EPI_ISL_3141361, EPI_ISL_3141362, EPI_ISL_3141363, EPI_ISL_3141364, EPI_ISL_3141365, EPI_ISL_3141366, EPI_ISL_3141367, EPI_ISL_3141368, EPI_ISL_3141369, EPI_ISL_3141370, EPI_ISL_3141371, EPI_ISL_3141372, EPI_ISL_3141373, EPI_ISL_3141374, EPI_ISL_3141375, EPI_ISL_3141376, EPI_ISL_3141377, EPI_ISL_3141378, EPI_ISL_3141379, EPI_ISL_3141380, EPI_ISL_3141381, EPI_ISL_3141382, EPI_ISL_3141383, EPI_ISL_3141384, EPI_ISL_3141385, EPI_ISL_3141386, EPI_ISL_3141387, EPI_ISL_3141388, EPI_ISL_3141389, EPI_ISL_3141390, EPI_ISL_3141391, EPI_ISL_3141392, EPI_ISL_3141393, EPI_ISL_3141394, EPI_ISL_3141395, EPI_ISL_3141396, EPI_ISL_3141397, EPI_ISL_3141398, EPI_ISL_3141399, EPI_ISL_3141400, EPI_ISL_3141401, EPI_ISL_3141402, EPI_ISL_3141403, EPI_ISL_3141404, EPI_ISL_3141405, EPI_ISL_3141406, EPI_ISL_3141407, EPI_ISL_3141408, EPI_ISL_3141409, EPI_ISL_3141410, EPI_ISL_3141411, EPI_ISL_3141412, EPI_ISL_3141413, EPI_ISL_3141414, EPI_ISL_3141415, EPI_ISL_3141416, EPI_ISL_3141417, EPI_ISL_3141418, EPI_ISL_3141419, EPI_ISL_3141420, EPI_ISL_3141421, EPI_ISL_3141422, EPI_ISL_3141423, EPI_ISL_3141424, EPI_ISL_3141425, EPI_ISL_3141426, EPI_ISL_3141427, EPI_ISL_3141428, EPI_ISL_3141429, EPI_ISL_3141430, EPI_ISL_3141431, EPI_ISL_3141432, EPI_ISL_3141433, EPI_ISL_3141434, EPI_ISL_3141435, EPI_ISL_3141436, EPI_ISL_3141437, EPI_ISL_3141438, EPI_ISL_3141439, EPI_ISL_3141440, EPI_ISL_3141441, EPI_ISL_3141442, EPI_ISL_3141443, EPI_ISL_3141444, EPI_ISL_3141445, EPI_ISL_3141446, EPI_ISL_3141447, EPI_ISL_3141448, EPI_ISL_3141449, EPI_ISL_3141450, EPI_ISL_3141451, EPI_ISL_3141452, EPI_ISL_3141453, EPI_ISL_3141454, EPI_ISL_3141455, EPI_ISL_3141456, EPI_ISL_3141457, EPI_ISL_3141458, EPI_ISL_3141459, EPI_ISL_3141460, EPI_ISL_3141461, EPI_ISL_3141462, EPI_ISL_3141463, EPI_ISL_3141464, EPI_ISL_3141465, EPI_ISL_3141466, EPI_ISL_3141467, EPI_ISL_3141468, EPI_ISL_3141469, EPI_ISL_3141470, EPI_ISL_3141471, EPI_ISL_3141472, EPI_ISL_3141473, EPI_ISL_3141474, EPI_ISL_3141475, EPI_ISL_3141476, EPI_ISL_3141477, EPI_ISL_3141478, EPI_ISL_3141479, EPI_ISL_3141480, EPI_ISL_3141481, EPI_ISL_3141482, EPI_ISL_3141483, EPI_ISL_3141484, EPI_ISL_3141485, EPI_ISL_3141486, EPI_ISL_3141488, EPI_ISL_3141489, EPI_ISL_3141490, EPI_ISL_3141491, EPI_ISL_3141492, EPI_ISL_3141493, EPI_ISL_3141494, EPI_ISL_3141495, EPI_ISL_3141496, EPI_ISL_3141497, EPI_ISL_3141498, EPI_ISL_3141499, EPI_ISL_3141500, EPI_ISL_3141501, EPI_ISL_3141502, EPI_ISL_3141503, EPI_ISL_3141504, EPI_ISL_3141505, EPI_ISL_3141506, EPI_ISL_3141507, EPI_ISL_3141508, EPI_ISL_3141509, EPI_ISL_3141510, EPI_ISL_3141511, EPI_ISL_3141512, EPI_ISL_3141513, EPI_ISL_3141514, EPI_ISL_3141515, EPI_ISL_3141516, EPI_ISL_3141517, EPI_ISL_3141518, EPI_ISL_3141519, EPI_ISL_3141520 | | | |
| see above | Lighthouse Lab in Glasgow | Wellcome Sanger Institute for the COVID-19 Genomics UK (COG-UK) Consortium | Harper VanSteenhouse, Yumi Kasai, David Gray, Carol Clugston, Anna Dominiczak and Alex Alderton, Roberto Amato, Jeffrey Barrett, Sonia Goncalves, Ewan Harrison, David K. Jackson, Ian Johnston, Dominic Kwiatkowski, Cordelia Langford, John Sillitoe on behalf of the Wellcome Sanger Institute COVID-19 Surveillance Team |
| EPI_ISL_3141521 | Lighthouse Lab in Milton Keynes | Wellcome Sanger Institute for the COVID-19 Genomics UK (COG-UK) Consortium | The Lighthouse Lab in Milton Keynes and Alex Alderton, Roberto Amato, Jeffrey Barrett, Sonia Goncalves, Ewan Harrison, David K. Jackson, Ian Johnston, Dominic Kwiatkowski, Cordelia Langford, John Sillitoe on behalf of the Wellcome Sanger Institute COVID-19 Surveillance Team |
| EPI_ISL_3141522, EPI_ISL_3141523, EPI_ISL_3141526, EPI_ISL_3141527 | Lighthouse Lab in Glasgow | Wellcome Sanger Institute for the COVID-19 Genomics UK (COG-UK) Consortium | Harper VanSteenhouse, Yumi Kasai, David Gray, Carol Clugston, Anna Dominiczak and Alex Alderton, Roberto Amato, Jeffrey Barrett, Sonia Goncalves, Ewan Harrison, David K. Jackson, Ian Johnston, Dominic Kwiatkowski, Cordelia Langford, John Sillitoe on behalf of the Wellcome Sanger Institute COVID-19 Surveillance Team |
| EPI_ISL_3141529, EPI_ISL_3141530, EPI_ISL_3141531, EPI_ISL_3141532, EPI_ISL_3141533, EPI_ISL_3141534, EPI_ISL_3141535, EPI_ISL_3141536, EPI_ISL_3141537, EPI_ISL_3141538, EPI_ISL_3141539, EPI_ISL_3141540, EPI_ISL_3141541, EPI_ISL_3141542, EPI_ISL_3141543, EPI_ISL_3141544, EPI_ISL_3141545, EPI_ISL_3141546, EPI_ISL_3141547, EPI_ISL_3141548, EPI_ISL_3141549, EPI_ISL_3141550, EPI_ISL_3141551, EPI_ISL_3141552, EPI_ISL_3141553, EPI_ISL_3141554, EPI_ISL_3141555, EPI_ISL_3141556, EPI_ISL_3141557, EPI_ISL_3141558, EPI_ISL_3141559, EPI_ISL_3141560, EPI_ISL_3141561, EPI_ISL_3141562, EPI_ISL_3141563, EPI_ISL_3141564, EPI_ISL_3141566, EPI_ISL_3141567, EPI_ISL_3141568, EPI_ISL_3141569, EPI_ISL_3141570, EPI_ISL_3141571, EPI_ISL_3141572, EPI_ISL_3141573, EPI_ISL_3141574, EPI_ISL_3141575, EPI_ISL_3141576, EPI_ISL_3141577, EPI_ISL_3141578, EPI_ISL_3141579, EPI_ISL_3141580, EPI_ISL_3141581, EPI_ISL_3141582, EPI_ISL_3141583, EPI_ISL_3141584, EPI_ISL_3141585, EPI_ISL_3141586, EPI_ISL_3141587, EPI_ISL_3141588, EPI_ISL_3141589, EPI_ISL_3141590, EPI_ISL_3141591, EPI_ISL_3141592, EPI_ISL_3141593, EPI_ISL_3141594, EPI_ISL_3141595, EPI_ISL_3141596, EPI_ISL_3141597, EPI_ISL_3141598, EPI_ISL_3141599, EPI_ISL_3141600, EPI_ISL_3141601, EPI_ISL_3141602, EPI_ISL_3141603, EPI_ISL_3141604, EPI_ISL_3141605, EPI_ISL_3141606, EPI_ISL_3141607, EPI_ISL_3141608, EPI_ISL_3141609, EPI_ISL_3141610, EPI_ISL_3141611, EPI_ISL_3141613, EPI_ISL_3141614, EPI_ISL_3141615, EPI_ISL_3141616, EPI_ISL_3141617, EPI_ISL_3141618, EPI_ISL_3141619, EPI_ISL_3141620, EPI_ISL_3141621, EPI_ISL_3141622, EPI_ISL_3141623, EPI_ISL_3141624, EPI_ISL_3141625, EPI_ISL_3141626, EPI_ISL_3141627, EPI_ISL_3141628, EPI_ISL_3141629, EPI_ISL_3141630, EPI_ISL_3141632, EPI_ISL_3141633, EPI_ISL_3141634, EPI_ISL_3141635, EPI_ISL_3141636, EPI_ISL_3141637, EPI_ISL_3141638, EPI_ISL_3141639, EPI_ISL_3141640, EPI_ISL_3141641, EPI_ISL_3141642, EPI_ISL_3141643, EPI_ISL_3141644, EPI_ISL_3141645, EPI_ISL_3141646, EPI_ISL_3141647, EPI_ISL_3141648, EPI_ISL_3141649, EPI_ISL_3141650, EPI_ISL_3141651, EPI_ISL_3141652, EPI_ISL_3141653, EPI_ISL_3141654, EPI_ISL_3141655, EPI_ISL_3141656, EPI_ISL_3141657, EPI_ISL_3141658, EPI_ISL_3141659, EPI_ISL_3141660, EPI_ISL_3141661, EPI_ISL_3141662, EPI_ISL_3141663, EPI_ISL_3141664, EPI_ISL_3141665, EPI_ISL_3141666, EPI_ISL_3141667, EPI_ISL_3141668, EPI_ISL_3141669, EPI_ISL_3141670, EPI_ISL_3141671, EPI_ISL_3141672, EPI_ISL_3141673, EPI_ISL_3141674, EPI_ISL_3141675, EPI_ISL_3141676, EPI_ISL_3141677, EPI_ISL_3141678, EPI_ISL_3141679, EPI_ISL_3141680, EPI_ISL_3141681, EPI_ISL_3141682, EPI_ISL_3141683, EPI_ISL_3141684, EPI_ISL_3141685, EPI_ISL_3141686, EPI_ISL_3141687, EPI_ISL_3141688, EPI_ISL_3141689, EPI_ISL_3141690, EPI_ISL_3141691, EPI_ISL_3141692, EPI_ISL_3141693, EPI_ISL_3141694, EPI_ISL_3141695, EPI_ISL_3141696, EPI_ISL_3141697, EPI_ISL_3141698, EPI_ISL_3141699, EPI_ISL_3141700, EPI_ISL_3141701, EPI_ISL_3141702, EPI_ISL_3141703, EPI_ISL_3141704, EPI_ISL_3141705, EPI_ISL_3141706, EPI_ISL_3141707, EPI_ISL_3141708, EPI_ISL_3141709, EPI_ISL_3141710, EPI_ISL_3141711, EPI_ISL_3141712, EPI_ISL_3141713, EPI_ISL_3141714, EPI_ISL_3141715, EPI_ISL_3141716, EPI_ISL_3141717, EPI_ISL_3141718, EPI_ISL_3141719, EPI_ISL_3141720, EPI_ISL_3141721, EPI_ISL_3141722, EPI_ISL_3141723, EPI_ISL_3141724, EPI_ISL_3141725, EPI_ISL_3141726, EPI_ISL_3141727, EPI_ISL_3141728, EPI_ISL_3141729, EPI_ISL_3141730, EPI_ISL_3141731, EPI_ISL_3141732, EPI_ISL_3141733, EPI_ISL_3141734, EPI_ISL_3141735, EPI_ISL_3141736, EPI_ISL_3141737, EPI_ISL_3141738, EPI_ISL_3141739, EPI_ISL_3141740, EPI_ISL_3141741, EPI_ISL_3141742, EPI_ISL_3141743, EPI_ISL_3141744, EPI_ISL_3141745, EPI_ISL_3141747, EPI_ISL_3141749, EPI_ISL_3141750, EPI_ISL_3141751, EPI_ISL_3141752, EPI_ISL_3141753, EPI_ISL_3141754, EPI_ISL_3141755, EPI_ISL_3141756, EPI_ISL_3141757, EPI_ISL_3141758, EPI_ISL_3141759, EPI_ISL_3141760, EPI_ISL_3141761, EPI_ISL_3141762, EPI_ISL_3141763, EPI_ISL_3141764, EPI_ISL_3141765, EPI_ISL_3141766, EPI_ISL_3141767, EPI_ISL_3141768, EPI_ISL_3141770, EPI_ISL_3141771, EPI_ISL_3141772, EPI_ISL_3141773, EPI_ISL_3141774, EPI_ISL_3141775, EPI_ISL_3141776, EPI_ISL_3141777, EPI_ISL_3141778, EPI_ISL_3141779, EPI_ISL_3141780, EPI_ISL_3141781, EPI_ISL_3141782, EPI_ISL_3141783, EPI_ISL_3141784, EPI_ISL_3141785, EPI_ISL_3141786, EPI_ISL_3141787, EPI_ISL_3141788, EPI_ISL_3141789, EPI_ISL_3141790, EPI_ISL_3141791, EPI_ISL_3141792, EPI_ISL_3141793, EPI_ISL_3141794, EPI_ISL_3141795, EPI_ISL_3141796, EPI_ISL_3141797, EPI_ISL_3141798, EPI_ISL_3141799, EPI_ISL_3141800, EPI_ISL_3141801, EPI_ISL_3141802, EPI_ISL_3141803, EPI_ISL_3141804, EPI_ISL_3141805, EPI_ISL_3141806, EPI_ISL_3141807, EPI_ISL_3141808, EPI_ISL_3141809, EPI_ISL_3141810, EPI_ISL_3141811, EPI_ISL_3141812, EPI_ISL_3141813, EPI_ISL_3141814, EPI_ISL_3141815, EPI_ISL_3141817, EPI_ISL_3141819, EPI_ISL_3141820, EPI_ISL_3141821, EPI_ISL_3141822, EPI_ISL_3141823, EPI_ISL_3141824, EPI_ISL_3141825, EPI_ISL_3141826, EPI_ISL_3141827, EPI_ISL_3141828, EPI_ISL_3141829, EPI_ISL_3141830, EPI_ISL_3141831, EPI_ISL_3141832, EPI_ISL_3141833, EPI_ISL_3141834, EPI_ISL_3141835, EPI_ISL_3141836, EPI_ISL_3141837, EPI_ISL_3141838, EPI_ISL_3141839, EPI_ISL_3141840, EPI_ISL_3141841, EPI_ISL_3141842, EPI_ISL_3141843, EPI_ISL_3141844, EPI_ISL_3141845, EPI_ISL_3141846, EPI_ISL_3141847, EPI_ISL_3141848, EPI_ISL_3141849, EPI_ISL_3141850, EPI_ISL_3141851, EPI_ISL_3141852, EPI_ISL_3141853, EPI_ISL_3141854, EPI_ISL_3141855, EPI_ISL_3141857, EPI_ISL_3141858, EPI_ISL_3141859, EPI_ISL_3141861, EPI_ISL_3141862, EPI_ISL_3141863, EPI_ISL_3141864, EPI_ISL_3141865, EPI_ISL_3141866, EPI_ISL_3141867, EPI_ISL_3141868, EPI_ISL_3141869, EPI_ISL_3141870, EPI_ISL_3141871, EPI_ISL_3141872, EPI_ISL_3141873, EPI_ISL_3141874, EPI_ISL_3141875, EPI_ISL_3141876, EPI_ISL_3141877, EPI_ISL_3141878, EPI_ISL_3141879, EPI_ISL_3141880, EPI_ISL_3141881, EPI_ISL_3141882, EPI_ISL_3141883, EPI_ISL_3141884, EPI_ISL_3141885, EPI_ISL_3141886, EPI_ISL_3141887, EPI_ISL_3141888, EPI_ISL_3141889, EPI_ISL_3141890, EPI_ISL_3141891, EPI_ISL_3141892, EPI_ISL_3141893, EPI_ISL_3141894, EPI_ISL_3141895, EPI_ISL_3141896, EPI_ISL_3141897, EPI_ISL_3141898, EPI_ISL_3141899, EPI_ISL_3141900, EPI_ISL_3141901, EPI_ISL_3141902, EPI_ISL_3141903, EPI_ISL_3141904, EPI_ISL_3141905, EPI_ISL_3141906, EPI_ISL_3141907, EPI_ISL_3141908, EPI_ISL_3141909, EPI_ISL_3141910, EPI_ISL_3141911, EPI_ISL_3141912, EPI_ISL_3141913, EPI_ISL_3141914, EPI_ISL_3141915, EPI_ISL_3141916, EPI_ISL_3141917, EPI_ISL_3141918, EPI_ISL_3141919, EPI_ISL_3141920, EPI_ISL_3141921, EPI_ISL_3141922, EPI_ISL_3141923, EPI_ISL_3141924, EPI_ISL_3141925, EPI_ISL_3141926, EPI_ISL_3141927, EPI_ISL_3141928, EPI_ISL_3141929, EPI_ISL_3141930, EPI_ISL_3141931, EPI_ISL_3141933, EPI_ISL_3141934, EPI_ISL_3141935, EPI_ISL_3141936, EPI_ISL_3141937, EPI_ISL_3141938, EPI_ISL_3141939, EPI_ISL_3141940, EPI_ISL_3141941, EPI_ISL_3141942, EPI_ISL_3141943, EPI_ISL_3141944, EPI_ISL_3141945, EPI_ISL_3141946, EPI_ISL_3141947, EPI_ISL_3141948, EPI_ISL_3141949, EPI_ISL_3141950, EPI_ISL_3141951, EPI_ISL_3141952, EPI_ISL_3141953, EPI_ISL_3141954, EPI_ISL_3141955, EPI_ISL_3141956, EPI_ISL_3141957, EPI_ISL_3141958, EPI_ISL_3141959, EPI_ISL_3141960, EPI_ISL_3141961, EPI_ISL_3141962, EPI_ISL_3141963, EPI_ISL_3141964, EPI_ISL_3141965, EPI_ISL_3141966, EPI_ISL_3141967, EPI_ISL_3141968, EPI_ISL_3141969, EPI_ISL_3141970, EPI_ISL_3141971, EPI_ISL_3141972, EPI_ISL_3141973, EPI_ISL_3141974, EPI_ISL_3141975, EPI_ISL_3141976, EPI_ISL_3141977, EPI_ISL_3141978, EPI_ISL_3141979, EPI_ISL_3141980, EPI_ISL_3141981, EPI_ISL_3141982, EPI_ISL_3141983, EPI_ISL_3141984, EPI_ISL_3141985, EPI_ISL_3141987, EPI_ISL_3141988, EPI_ISL_3141989, EPI_ISL_3141990, EPI_ISL_3141991, EPI_ISL_3141992, EPI_ISL_3141993, EPI_ISL_3141994, EPI_ISL_3141995, EPI_ISL_3141996, EPI_ISL_3141997, EPI_ISL_3141998, EPI_ISL_3141999, EPI_ISL_3142000, EPI_ISL_3142001, EPI_ISL_3142002, EPI_ISL_3142003, EPI_ISL_3142004, EPI_ISL_3142005, EPI_ISL_3142006, EPI_ISL_3142007, EPI_ISL_3142008, EPI_ISL_3142009, EPI_ISL_3142010, EPI_ISL_3142011, EPI_ISL_3142012, EPI_ISL_3142013, EPI_ISL_3142014, EPI_ISL_3142015, EPI_ISL_3142016, EPI_ISL_3142017, EPI_ISL_3142018, EPI_ISL_3142019, EPI_ISL_3142020, EPI_ISL_3142021, EPI_ISL_3142022, EPI_ISL_3142023, EPI_ISL_3142024, | | | |

| | | | |
|---|---|---|---|
| EPI_ISL_3142025, EPI_ISL_3142026, EPI_ISL_3142027, EPI_ISL_3142028, EPI_ISL_3142029, EPI_ISL_3142030, EPI_ISL_3142031, EPI_ISL_3142032, EPI_ISL_3142033, EPI_ISL_3142034, EPI_ISL_3142035, EPI_ISL_3142036, EPI_ISL_3142037, EPI_ISL_3142038, EPI_ISL_3142039, EPI_ISL_3142040, EPI_ISL_3142041, EPI_ISL_3142042, EPI_ISL_3142043, EPI_ISL_3142044, EPI_ISL_3142045, EPI_ISL_3142046, EPI_ISL_3142047, EPI_ISL_3142048, EPI_ISL_3142049, EPI_ISL_3142050, EPI_ISL_3142051, EPI_ISL_3142052, EPI_ISL_3142053, EPI_ISL_3142054, EPI_ISL_3142055, EPI_ISL_3142056, EPI_ISL_3142057, EPI_ISL_3142058, EPI_ISL_3142059, EPI_ISL_3142060, EPI_ISL_3142061, EPI_ISL_3142062, EPI_ISL_3142063, EPI_ISL_3142064, EPI_ISL_3142065, EPI_ISL_3142066, EPI_ISL_3142067, EPI_ISL_3142068, EPI_ISL_3142069, EPI_ISL_3142070, EPI_ISL_3142071, EPI_ISL_3142072, EPI_ISL_3142073, EPI_ISL_3142074, EPI_ISL_3142075, EPI_ISL_3142076, EPI_ISL_3142077, EPI_ISL_3142078, EPI_ISL_3142080, EPI_ISL_3142081, EPI_ISL_3142082, EPI_ISL_3142083, EPI_ISL_3142085, EPI_ISL_3142086, EPI_ISL_3142087, EPI_ISL_3142088, EPI_ISL_3142089, EPI_ISL_3142090, EPI_ISL_3142091, EPI_ISL_3142092, EPI_ISL_3142093, EPI_ISL_3142094, EPI_ISL_3142095, EPI_ISL_3142096, EPI_ISL_3142097, EPI_ISL_3142098, EPI_ISL_3142099, EPI_ISL_3142100, EPI_ISL_3142101, EPI_ISL_3142102, EPI_ISL_3142103, EPI_ISL_3142104, EPI_ISL_3142105, EPI_ISL_3142106, EPI_ISL_3142107, EPI_ISL_3142108, EPI_ISL_3142109, EPI_ISL_3142110, EPI_ISL_3142111, EPI_ISL_3142112, EPI_ISL_3142113, EPI_ISL_3142114, EPI_ISL_3142115, EPI_ISL_3142116, EPI_ISL_3142117, EPI_ISL_3142118, EPI_ISL_3142119, EPI_ISL_3142121, EPI_ISL_3142122, EPI_ISL_3142123, EPI_ISL_3142124, EPI_ISL_3142125, EPI_ISL_3142126, EPI_ISL_3142127, EPI_ISL_3142128, EPI_ISL_3142129, EPI_ISL_3142130, EPI_ISL_3142131, EPI_ISL_3142132, EPI_ISL_3142133, EPI_ISL_3142134, EPI_ISL_3142135, EPI_ISL_3142136, EPI_ISL_3142137, EPI_ISL_3142138, EPI_ISL_3142139, EPI_ISL_3142140, EPI_ISL_3142142, EPI_ISL_3142143, EPI_ISL_3142144, EPI_ISL_3142145, EPI_ISL_3142146, EPI_ISL_3142147, EPI_ISL_3142148, EPI_ISL_3142149, EPI_ISL_3142150, EPI_ISL_3142151, EPI_ISL_3142152, EPI_ISL_3142153, EPI_ISL_3142154, EPI_ISL_3142155, EPI_ISL_3142156, EPI_ISL_3142157, EPI_ISL_3142158, EPI_ISL_3142159, EPI_ISL_3142160, EPI_ISL_3142161, EPI_ISL_3142162, EPI_ISL_3142163, EPI_ISL_3142164, EPI_ISL_3142165, EPI_ISL_3142166, EPI_ISL_3142167, EPI_ISL_3142168, EPI_ISL_3142169, EPI_ISL_3142170, EPI_ISL_3142171, EPI_ISL_3142172, EPI_ISL_3142173, EPI_ISL_3142174, EPI_ISL_3142175, EPI_ISL_3142176, EPI_ISL_3142177, EPI_ISL_3142178, EPI_ISL_3142179, EPI_ISL_3142180, EPI_ISL_3142181, EPI_ISL_3142182, EPI_ISL_3142183, EPI_ISL_3142184, EPI_ISL_3142185, EPI_ISL_3142186, EPI_ISL_3142187, EPI_ISL_3142188, EPI_ISL_3142189, EPI_ISL_3142190, EPI_ISL_3142191, EPI_ISL_3142192, EPI_ISL_3142193, EPI_ISL_3142194, EPI_ISL_3142195, EPI_ISL_3142196, EPI_ISL_3142197, EPI_ISL_3142198, EPI_ISL_3142199, EPI_ISL_3142200, EPI_ISL_3142201, EPI_ISL_3142202, EPI_ISL_3142203, EPI_ISL_3142204, EPI_ISL_3142205, EPI_ISL_3142206, EPI_ISL_3142207, EPI_ISL_3142208, EPI_ISL_3142209, EPI_ISL_3142210, EPI_ISL_3142211, EPI_ISL_3142212, EPI_ISL_3142213, EPI_ISL_3142214, EPI_ISL_3142215, EPI_ISL_3142216, EPI_ISL_3142217, EPI_ISL_3142218, EPI_ISL_3142219, EPI_ISL_3142220, EPI_ISL_3142221, EPI_ISL_3142222, EPI_ISL_3142223, EPI_ISL_3142224, EPI_ISL_3142225, EPI_ISL_3142226, EPI_ISL_3142227, EPI_ISL_3142228, EPI_ISL_3142229, EPI_ISL_3142230, EPI_ISL_3142231, EPI_ISL_3142232, EPI_ISL_3142233, EPI_ISL_3142234, EPI_ISL_3142235, EPI_ISL_3142236, EPI_ISL_3142237, EPI_ISL_3142238, EPI_ISL_3142239, EPI_ISL_3142240, EPI_ISL_3142241, EPI_ISL_3142242, EPI_ISL_3142243, EPI_ISL_3142244, EPI_ISL_3142245, EPI_ISL_3142246, EPI_ISL_3142247, EPI_ISL_3142248, EPI_ISL_3142249, EPI_ISL_3142250, EPI_ISL_3142251, EPI_ISL_3142253, EPI_ISL_3142254, EPI_ISL_3142255, EPI_ISL_3142256, EPI_ISL_3142257, EPI_ISL_3142258, EPI_ISL_3142259, EPI_ISL_3142260, EPI_ISL_3142261, EPI_ISL_3142262, EPI_ISL_3142263, EPI_ISL_3142264, EPI_ISL_3142265, EPI_ISL_3142266, EPI_ISL_3142267, EPI_ISL_3142268, EPI_ISL_3142269, EPI_ISL_3142270, EPI_ISL_3142271, EPI_ISL_3142272, EPI_ISL_3142273, EPI_ISL_3142274, EPI_ISL_3142275, EPI_ISL_3142276, EPI_ISL_3142277, EPI_ISL_3142278, EPI_ISL_3142279, EPI_ISL_3142280, EPI_ISL_3142281, EPI_ISL_3142282, EPI_ISL_3142283, EPI_ISL_3142284, EPI_ISL_3142285, EPI_ISL_3142286, EPI_ISL_3142287, EPI_ISL_3142288, EPI_ISL_3142289, EPI_ISL_3142290, EPI_ISL_3142291, EPI_ISL_3142292, EPI_ISL_3142293, EPI_ISL_3142294, EPI_ISL_3142295, EPI_ISL_3142296, EPI_ISL_3142297, EPI_ISL_3142298, EPI_ISL_3142299, EPI_ISL_3142300, EPI_ISL_3142302, EPI_ISL_3142303, EPI_ISL_3142304, EPI_ISL_3142305, EPI_ISL_3142306, EPI_ISL_3142307, EPI_ISL_3142308, EPI_ISL_3142309, EPI_ISL_3142310, EPI_ISL_3142311, EPI_ISL_3142312, EPI_ISL_3142313, EPI_ISL_3142314, EPI_ISL_3142316, EPI_ISL_3142317, EPI_ISL_3142318, EPI_ISL_3142319, EPI_ISL_3142320, EPI_ISL_3142321, EPI_ISL_3142322, EPI_ISL_3142323, EPI_ISL_3142324, EPI_ISL_3142325, EPI_ISL_3142326, EPI_ISL_3142327, EPI_ISL_3142328, EPI_ISL_3142329, EPI_ISL_3142330, EPI_ISL_3142331, EPI_ISL_3142332, EPI_ISL_3142333, EPI_ISL_3142334, EPI_ISL_3142335, EPI_ISL_3142336, EPI_ISL_3142337, EPI_ISL_3142338, EPI_ISL_3142339, EPI_ISL_3142340, EPI_ISL_3142341, EPI_ISL_3142342, EPI_ISL_3142343, EPI_ISL_3142344, EPI_ISL_3142345, EPI_ISL_3142346, EPI_ISL_3142348, EPI_ISL_3142349, EPI_ISL_3142350, EPI_ISL_3142351, EPI_ISL_3142352, EPI_ISL_3142353, EPI_ISL_3142354, EPI_ISL_3142355, EPI_ISL_3142356, EPI_ISL_3142357, EPI_ISL_3142358, EPI_ISL_3142359, EPI_ISL_3142360, EPI_ISL_3142361, EPI_ISL_3142362, EPI_ISL_3142363, EPI_ISL_3142364, EPI_ISL_3142365, EPI_ISL_3142366, EPI_ISL_3142367, EPI_ISL_3142368, EPI_ISL_3142369, EPI_ISL_3142370, EPI_ISL_3142371, EPI_ISL_3142372, EPI_ISL_3142373, EPI_ISL_3142374, EPI_ISL_3142376, EPI_ISL_3142377, EPI_ISL_3142378, EPI_ISL_3142379, EPI_ISL_3142380, EPI_ISL_3142381, EPI_ISL_3142382, EPI_ISL_3142383, EPI_ISL_3142384, EPI_ISL_3142385, EPI_ISL_3142386, EPI_ISL_3142387, EPI_ISL_3142388, EPI_ISL_3142389, EPI_ISL_3142390, EPI_ISL_3142392, EPI_ISL_3142393, EPI_ISL_3142394, EPI_ISL_3142395, EPI_ISL_3142396, EPI_ISL_3142397, EPI_ISL_3142398, EPI_ISL_3142399, EPI_ISL_3142400, EPI_ISL_3142401, EPI_ISL_3142402, EPI_ISL_3142404, EPI_ISL_3142406, EPI_ISL_3142408, EPI_ISL_3142410, EPI_ISL_3142411, EPI_ISL_3142412, EPI_ISL_3142413, EPI_ISL_3142414, EPI_ISL_3142415, EPI_ISL_3142416, EPI_ISL_3142417, EPI_ISL_3142418, EPI_ISL_3142419, EPI_ISL_3142420, EPI_ISL_3142421, EPI_ISL_3142422, EPI_ISL_3142423, EPI_ISL_3142424, EPI_ISL_3142425, EPI_ISL_3142426, EPI_ISL_3142427, EPI_ISL_3142428, EPI_ISL_3142429, EPI_ISL_3142430, EPI_ISL_3142431, EPI_ISL_3142432, EPI_ISL_3142433, EPI_ISL_3142434, EPI_ISL_3142435, EPI_ISL_3142436, EPI_ISL_3142437, EPI_ISL_3142438, EPI_ISL_3142439, EPI_ISL_3142440, EPI_ISL_3142441, EPI_ISL_3142442, EPI_ISL_3142443, EPI_ISL_3142444, EPI_ISL_3142445, EPI_ISL_3142446, EPI_ISL_3142447, EPI_ISL_3142448, EPI_ISL_3142449, EPI_ISL_3142450, EPI_ISL_3142451, EPI_ISL_3142452, EPI_ISL_3142453, EPI_ISL_3142454, EPI_ISL_3142455, EPI_ISL_3142457, EPI_ISL_3142458, EPI_ISL_3142459, EPI_ISL_3142460, EPI_ISL_3142461, EPI_ISL_3142462, EPI_ISL_3142463, EPI_ISL_3142464, EPI_ISL_3142465, EPI_ISL_3142466, EPI_ISL_3142467, EPI_ISL_3142468, EPI_ISL_3142469, EPI_ISL_3142470, EPI_ISL_3142471, EPI_ISL_3142473, EPI_ISL_3142475, EPI_ISL_3142476, EPI_ISL_3142477, EPI_ISL_3142478, EPI_ISL_3142479, EPI_ISL_3142480, EPI_ISL_3142481, EPI_ISL_3142482, EPI_ISL_3142483, EPI_ISL_3142484, EPI_ISL_3142485, EPI_ISL_3142486, EPI_ISL_3142487, EPI_ISL_3142489, EPI_ISL_3142490, EPI_ISL_3142491, EPI_ISL_3142492, EPI_ISL_3142493, EPI_ISL_3142494, EPI_ISL_3142495, EPI_ISL_3142496, EPI_ISL_3142497, EPI_ISL_3142498, EPI_ISL_3142499, EPI_ISL_3142501, EPI_ISL_3142503, EPI_ISL_3142504, EPI_ISL_3142505, EPI_ISL_3142506, EPI_ISL_3142507, EPI_ISL_3142508, EPI_ISL_3142510, EPI_ISL_3142511, EPI_ISL_3142512, EPI_ISL_3142513, EPI_ISL_3142514, EPI_ISL_3142515, EPI_ISL_3142516, EPI_ISL_3142517, EPI_ISL_3142518, EPI_ISL_3142519, EPI_ISL_3142520, EPI_ISL_3142521, EPI_ISL_3142522, EPI_ISL_3142523, EPI_ISL_3142524, EPI_ISL_3142525, EPI_ISL_3142526, EPI_ISL_3142527, EPI_ISL_3142528, EPI_ISL_3142529, EPI_ISL_3142530, EPI_ISL_3142531, EPI_ISL_3142532, EPI_ISL_3142533, EPI_ISL_3142534, EPI_ISL_3142535, EPI_ISL_3142536, EPI_ISL_3142537, EPI_ISL_3142538, EPI_ISL_3142539, EPI_ISL_3142540, EPI_ISL_3142542, EPI_ISL_3142543, EPI_ISL_3142544, EPI_ISL_3142545, EPI_ISL_3142546, EPI_ISL_3142547, EPI_ISL_3142548, EPI_ISL_3142549, EPI_ISL_3142550, EPI_ISL_3142551, EPI_ISL_3142552, EPI_ISL_3142553, EPI_ISL_3142554, EPI_ISL_3142555, EPI_ISL_3142556, EPI_ISL_3142557, EPI_ISL_3142558, EPI_ISL_3142559, EPI_ISL_3142560, EPI_ISL_3142561, EPI_ISL_3142562, EPI_ISL_3142563, EPI_ISL_3142564, EPI_ISL_3142565, EPI_ISL_3142566, EPI_ISL_3142567, EPI_ISL_3142568, EPI_ISL_3142569, EPI_ISL_3142571, EPI_ISL_3142572, EPI_ISL_3142573, EPI_ISL_3142574, EPI_ISL_3142575, EPI_ISL_3142576, EPI_ISL_3142577, EPI_ISL_3142578, EPI_ISL_3142579, EPI_ISL_3142580, EPI_ISL_3142581, EPI_ISL_3142582, EPI_ISL_3142583, EPI_ISL_3142584, EPI_ISL_3142585, EPI_ISL_3142586, EPI_ISL_3142587, EPI_ISL_3142588, EPI_ISL_3142589, EPI_ISL_3142590, EPI_ISL_3142591, EPI_ISL_3142592, EPI_ISL_3142593, EPI_ISL_3142594, EPI_ISL_3142595, EPI_ISL_3142596, EPI_ISL_3142598, EPI_ISL_3142599, EPI_ISL_3142600, EPI_ISL_3142601, EPI_ISL_3142602, EPI_ISL_3142603, EPI_ISL_3142604, EPI_ISL_3142605, EPI_ISL_3142607, EPI_ISL_3142608, EPI_ISL_3142609, EPI_ISL_3142610, EPI_ISL_3142611, EPI_ISL_3142612, EPI_ISL_3142613, EPI_ISL_3142614, EPI_ISL_3142615, EPI_ISL_3142616, EPI_ISL_3142617, EPI_ISL_3142618, EPI_ISL_3142619, EPI_ISL_3142620, EPI_ISL_3142621, EPI_ISL_3142622, EPI_ISL_3142623, EPI_ISL_3142624, EPI_ISL_3142625, EPI_ISL_3142626, EPI_ISL_3142627, EPI_ISL_3142628, EPI_ISL_3142629, EPI_ISL_3142630, EPI_ISL_3142631, EPI_ISL_3142632, EPI_ISL_3142634, EPI_ISL_3142635, EPI_ISL_3142636, EPI_ISL_3142637, EPI_ISL_3142638, EPI_ISL_3142639, EPI_ISL_3142640, EPI_ISL_3142641, EPI_ISL_3142642, EPI_ISL_3142643, EPI_ISL_3142644, EPI_ISL_3142645, EPI_ISL_3142646, EPI_ISL_3142647, EPI_ISL_3142648, EPI_ISL_3142649, EPI_ISL_3142650, EPI_ISL_3142651, EPI_ISL_3142652, EPI_ISL_3142653, EPI_ISL_3142654, EPI_ISL_3142655, EPI_ISL_3142656, EPI_ISL_3142657, EPI_ISL_3142658, EPI_ISL_3142659, EPI_ISL_3142660, EPI_ISL_3142661, EPI_ISL_3142662, EPI_ISL_3142663, EPI_ISL_3142664, EPI_ISL_3142665, EPI_ISL_3142666, EPI_ISL_3142667, EPI_ISL_3142668, EPI_ISL_3142670, EPI_ISL_3142671, EPI_ISL_3142672, EPI_ISL_3142673, EPI_ISL_3142674, EPI_ISL_3142675, EPI_ISL_3142676, EPI_ISL_3142677, EPI_ISL_3142678, EPI_ISL_3142679, EPI_ISL_3142680, EPI_ISL_3142681, EPI_ISL_3142682, EPI_ISL_3142683, EPI_ISL_3142684, EPI_ISL_3142685, EPI_ISL_3142686, EPI_ISL_3142687, EPI_ISL_3142688, EPI_ISL_3142689, EPI_ISL_3142690, EPI_ISL_3142691, EPI_ISL_3142692, EPI_ISL_3142693, EPI_ISL_3142694, EPI_ISL_3142695, EPI_ISL_3142696, EPI_ISL_3142697, EPI_ISL_3142698, EPI_ISL_3142699, EPI_ISL_3142700, EPI_ISL_3142701, EPI_ISL_3142702, EPI_ISL_3142703, EPI_ISL_3142704, EPI_ISL_3142705, EPI_ISL_3142706, EPI_ISL_3142707, EPI_ISL_3142708, EPI_ISL_3142709, EPI_ISL_3142710, EPI_ISL_3142711, EPI_ISL_3142712, EPI_ISL_3142713, EPI_ISL_3142714, EPI_ISL_3142715, EPI_ISL_3142716, EPI_ISL_3142717, EPI_ISL_3142718, EPI_ISL_3142719, EPI_ISL_3142720, EPI_ISL_3142721, EPI_ISL_3142722, EPI_ISL_3142723, EPI_ISL_3142724 | | | |
| see above | Department of Bacteria, Parasites and Fungi, Statens Serum Institut, Copenhagen, Denmark | Statens Serum Institut Bioinformatics and Microbial Genomics | Danish Covid-19 Genome Consortium |
| EPI_ISL_3142950, EPI_ISL_3142953, EPI_ISL_3142954 | Molecular & Genomic Pathology Laboratory, Thomas Jefferson University Hospital | Molecular & Genomic Pathology Laboratory, Thomas Jefferson University Hospital | Nitika Badjatia, Shannon Mahoney, Run Jin, Zi-xuan Wang |
| EPI_ISL_3144439, EPI_ISL_3144443, EPI_ISL_3144445, EPI_ISL_3144446, EPI_ISL_3144447, EPI_ISL_3144448, EPI_ISL_3144449, EPI_ISL_3144450, EPI_ISL_3144451, EPI_ISL_3144452, EPI_ISL_3144453, EPI_ISL_3144454, EPI_ISL_3144455, EPI_ISL_3144456, EPI_ISL_3144457, EPI_ISL_3144458, EPI_ISL_3144459, EPI_ISL_3144460, EPI_ISL_3144461, EPI_ISL_3144462 | | | |
| see above | ASL BARI | University of Bari Biomedical Sciences and Human Oncology | Maria Chironna |
| EPI_ISL_3144463, EPI_ISL_3144465, EPI_ISL_3144466, EPI_ISL_3144468 | Tivoli Fondazione Policlinico Universitario Agostino Gemelli IRCCS | Fondazione Policlinico Universitario Agostino Gemelli IRCCS | Istituto di Microbiologia e Virologia |
| EPI_ISL_3144469, EPI_ISL_3144470, EPI_ISL_3144471, EPI_ISL_3144472, EPI_ISL_3144473, EPI_ISL_3144474, EPI_ISL_3144475, EPI_ISL_3144477, EPI_ISL_3144478, EPI_ISL_3144479, EPI_ISL_3144480, EPI_ISL_3144481, EPI_ISL_3144482, EPI_ISL_3144483, EPI_ISL_3144484, EPI_ISL_3144485, EPI_ISL_3144486, EPI_ISL_3144488 | | | |
| see above | Medical Laboratories Duesseldorf | Center of Medical Microbiology, Virology, and Hospital Hygiene, University of Duesseldorf | Maximilian Damagnez;Alexander Dilthey;Angelika Helmer;Torsten Houwaart;Lisanna Hülse;Christian Lange;Malte Kohns Vasconcelos;Nadine Lübke;Jessica Nicolai;Klaus Pfeffer;Daniel Strelow;Jörg Timm;Andreas Walker;Tobias Wienemann |
| EPI_ISL_3144539, EPI_ISL_3144540, EPI_ISL_3144541, EPI_ISL_3144542, EPI_ISL_3144543, EPI_ISL_3144544, EPI_ISL_3144545, EPI_ISL_3144546, EPI_ISL_3144547, EPI_ISL_3144548, EPI_ISL_3144549, EPI_ISL_3144550, EPI_ISL_3144551, EPI_ISL_3144552, EPI_ISL_3144553, EPI_ISL_3144554, EPI_ISL_3144555, EPI_ISL_3144556, EPI_ISL_3144557, EPI_ISL_3144558, EPI_ISL_3144559, EPI_ISL_3144560, EPI_ISL_3144561, EPI_ISL_3144562, EPI_ISL_3144563, EPI_ISL_3144564, EPI_ISL_3144565, EPI_ISL_3144566, EPI_ISL_3144567, EPI_ISL_3144568, EPI_ISL_3144569, EPI_ISL_3144570, EPI_ISL_3144571, EPI_ISL_3144572, EPI_ISL_3144573, EPI_ISL_3144574, EPI_ISL_3144575, EPI_ISL_3144576, EPI_ISL_3144577, EPI_ISL_3144578, EPI_ISL_3144579, EPI_ISL_3144580, EPI_ISL_3144581, EPI_ISL_3144582, EPI_ISL_3144583, EPI_ISL_3144584, EPI_ISL_3144585, EPI_ISL_3144586, EPI_ISL_3144587, EPI_ISL_3144588, EPI_ISL_3144589, EPI_ISL_3144590, EPI_ISL_3144591, EPI_ISL_3144592, EPI_ISL_3144593, EPI_ISL_3144594, EPI_ISL_3144595, EPI_ISL_3144596, EPI_ISL_3144597, EPI_ISL_3144598, EPI_ISL_3144599, EPI_ISL_3144600, EPI_ISL_3144601, EPI_ISL_3144602, EPI_ISL_3144603, EPI_ISL_3144604, EPI_ISL_3144605, EPI_ISL_3144606, EPI_ISL_3144607, EPI_ISL_3144608, EPI_ISL_3144609, EPI_ISL_3144610, EPI_ISL_3144611, EPI_ISL_3144612, EPI_ISL_3144613, EPI_ISL_3144614, EPI_ISL_3144615, EPI_ISL_3144616, EPI_ISL_3144617, EPI_ISL_3144618, EPI_ISL_3144619, EPI_ISL_3144620, EPI_ISL_3144621, EPI_ISL_3144622, EPI_ISL_3144623, EPI_ISL_3144624, EPI_ISL_3144625, EPI_ISL_3144626, EPI_ISL_3144627, EPI_ISL_3144628, EPI_ISL_3144629, EPI_ISL_3144630, EPI_ISL_3144631, EPI_ISL_3144632, EPI_ISL_3144633, EPI_ISL_3144634, EPI_ISL_3144635, EPI_ISL_3144636, EPI_ISL_3144637, EPI_ISL_3144638, EPI_ISL_3144639, EPI_ISL_3144640, EPI_ISL_3144641, EPI_ISL_3144642, EPI_ISL_3144643, EPI_ISL_3144645, EPI_ISL_3144646, EPI_ISL_3144647, EPI_ISL_3144648, EPI_ISL_3144649, EPI_ISL_3144650, EPI_ISL_3144651, EPI_ISL_3144652, EPI_ISL_3144653, EPI_ISL_3144654, EPI_ISL_3144655, EPI_ISL_3144656, EPI_ISL_3144657, EPI_ISL_3144658, EPI_ISL_3144659, EPI_ISL_3144660, EPI_ISL_3144661, EPI_ISL_3144662, EPI_ISL_3144663, EPI_ISL_3144664, EPI_ISL_3144665, EPI_ISL_3144666, EPI_ISL_3144667, EPI_ISL_3144668, EPI_ISL_3144669, EPI_ISL_3144670, EPI_ISL_3144671, EPI_ISL_3144672, EPI_ISL_3144681, EPI_ISL_3144687, EPI_ISL_3144688, EPI_ISL_3144689, EPI_ISL_3144690, EPI_ISL_3144691, EPI_ISL_3144695, EPI_ISL_3144696, EPI_ISL_3144697, EPI_ISL_3144699, EPI_ISL_3144702, EPI_ISL_3144704, EPI_ISL_3144705, EPI_ISL_3144706, EPI_ISL_3144708, EPI_ISL_3144709, EPI_ISL_3144710, EPI_ISL_3144711, EPI_ISL_3144713, EPI_ISL_3144714, EPI_ISL_3144715, EPI_ISL_3144717, EPI_ISL_3144762, EPI_ISL_3144765, EPI_ISL_3144767, EPI_ISL_3144768, EPI_ISL_3144769, EPI_ISL_3144770, EPI_ISL_3144771, EPI_ISL_3144773, EPI_ISL_3144774, EPI_ISL_3144775, EPI_ISL_3144776, EPI_ISL_3144777, EPI_ISL_3144778, EPI_ISL_3144780, EPI_ISL_3144781, EPI_ISL_3144782, EPI_ISL_3144783, EPI_ISL_3144784, EPI_ISL_3144785, EPI_ISL_3144787, EPI_ISL_3144793, EPI_ISL_3144794, EPI_ISL_3144795, EPI_ISL_3144796, EPI_ISL_3144797, EPI_ISL_3144798, EPI_ISL_3144799, EPI_ISL_3144800, EPI_ISL_3144801, EPI_ISL_3144802, EPI_ISL_3144803, EPI_ISL_3144804, EPI_ISL_3144805, EPI_ISL_3144806, EPI_ISL_3144807, EPI_ISL_3144808, EPI_ISL_3144809, EPI_ISL_3144810, EPI_ISL_3144811, EPI_ISL_3144813, EPI_ISL_3144814, EPI_ISL_3144815, EPI_ISL_3144816, EPI_ISL_3144817, EPI_ISL_3144819, EPI_ISL_3144822, EPI_ISL_3144823, EPI_ISL_3144825, EPI_ISL_3144827, EPI_ISL_3144831, EPI_ISL_3144834, EPI_ISL_3144836, EPI_ISL_3144837, EPI_ISL_3144838, EPI_ISL_3144839, EPI_ISL_3144840, EPI_ISL_3144845, EPI_ISL_3144846, EPI_ISL_3144849, EPI_ISL_3144850 | | | |
| see above | Utah Public Health Laboratory | Utah Public Health Laboratory | Olinto Linares-Perdomo, Erin L. Young, Kelly F. Oakeson |
| EPI_ISL_3144855, EPI_ISL_3144856, EPI_ISL_3144857, EPI_ISL_3144858, EPI_ISL_3144859, EPI_ISL_3144860, EPI_ISL_3144861, EPI_ISL_3144862, EPI_ISL_3144863, EPI_ISL_3144864, EPI_ISL_3144865, EPI_ISL_3144866, EPI_ISL_3144867, EPI_ISL_3144868, EPI_ISL_3144869, EPI_ISL_3144870, EPI_ISL_3144871, EPI_ISL_3144872, EPI_ISL_3144873, EPI_ISL_3144874, EPI_ISL_3144875, EPI_ISL_3144877, EPI_ISL_3144878, EPI_ISL_3144879, EPI_ISL_3144880, EPI_ISL_3144881, EPI_ISL_3144882, EPI_ISL_3144883, EPI_ISL_3144884, EPI_ISL_3144886, EPI_ISL_3144887, EPI_ISL_3144888, EPI_ISL_3144889, EPI_ISL_3144891 | | | |
| see above | Dept. of Microbiology and Infection Control, Akershus University Hospital HF | Dept. of Microbiology and Infection Control, Akershus University Hospital HF | Hege Vangstein Aamot, Alexander Hesselberg Løvestad |
| EPI_ISL_3144946 | Center for Virology | Center for Virology | Jeremy V. Camp, Monika Redlberger-Fritz, Stephan W. Aberle |
| EPI_ISL_3145006, EPI_ISL_3145020, EPI_ISL_3145021, EPI_ISL_3145023, EPI_ISL_3145025, EPI_ISL_3145419, EPI_ISL_3145447, EPI_ISL_3145456, EPI_ISL_3145555, EPI_ISL_3145556, EPI_ISL_3145557, EPI_ISL_3145558, EPI_ISL_3145559, EPI_ISL_3145560, EPI_ISL_3145561, EPI_ISL_3145562, EPI_ISL_3145563, EPI_ISL_3145564, EPI_ISL_3145565, EPI_ISL_3145566, EPI_ISL_3145567, EPI_ISL_3145568, EPI_ISL_3145569, EPI_ISL_3145570, EPI_ISL_3145571, EPI_ISL_3145572, EPI_ISL_3145573, EPI_ISL_3145574, EPI_ISL_3145575, EPI_ISL_3145576, EPI_ISL_3145577, EPI_ISL_3145578, EPI_ISL_3145579, EPI_ISL_3145580, EPI_ISL_3145581, EPI_ISL_3145582, EPI_ISL_3145583, EPI_ISL_3145584, EPI_ISL_3145585, EPI_ISL_3145586, EPI_ISL_3145702, EPI_ISL_3145703, EPI_ISL_3145704, EPI_ISL_3145705, EPI_ISL_3145706, EPI_ISL_3145707, EPI_ISL_3145708, EPI_ISL_3145709, EPI_ISL_3145710, EPI_ISL_3145711, EPI_ISL_3145712, EPI_ISL_3145713, EPI_ISL_3145714, EPI_ISL_3145715, EPI_ISL_3145716, EPI_ISL_3145717, EPI_ISL_3145718, EPI_ISL_3145719, EPI_ISL_3145720, EPI_ISL_3145721, EPI_ISL_3145722, EPI_ISL_3145723, EPI_ISL_3145724, EPI_ISL_3145725, EPI_ISL_3145726, EPI_ISL_3145727, EPI_ISL_3145728, EPI_ISL_3145729, EPI_ISL_3145730, EPI_ISL_3145731, EPI_ISL_3145732, EPI_ISL_3145733, EPI_ISL_3145734, EPI_ISL_3145735, EPI_ISL_3145736, EPI_ISL_3145737, EPI_ISL_3145738, EPI_ISL_3145739, EPI_ISL_3145740, EPI_ISL_3145741, EPI_ISL_3145742, EPI_ISL_3145743, EPI_ISL_3145744, EPI_ISL_3145745, EPI_ISL_3145746, EPI_ISL_3145747, EPI_ISL_3145748, EPI_ISL_3145749, EPI_ISL_3145750, EPI_ISL_3145751, EPI_ISL_3145752, EPI_ISL_3145753, EPI_ISL_3145754, EPI_ISL_3145755, EPI_ISL_3145756, EPI_ISL_3145757, EPI_ISL_3145758, EPI_ISL_3145759, EPI_ISL_3145760, EPI_ISL_3145761, EPI_ISL_3145762, EPI_ISL_3145763, EPI_ISL_3145764, EPI_ISL_3145765, EPI_ISL_3145766, EPI_ISL_3145767, EPI_ISL_3145768, EPI_ISL_3145769, EPI_ISL_3145770, EPI_ISL_3145771, EPI_ISL_3145772, EPI_ISL_3145773, EPI_ISL_3145774, EPI_ISL_3145775, EPI_ISL_3145776, EPI_ISL_3145777, EPI_ISL_3145778, EPI_ISL_3145779, EPI_ISL_3145780, EPI_ISL_3145781, EPI_ISL_3145782, EPI_ISL_3145783, EPI_ISL_3145784, EPI_ISL_3145785, EPI_ISL_3145786, EPI_ISL_3145787, EPI_ISL_3145788, EPI_ISL_3145789, EPI_ISL_3145790, EPI_ISL_3145791, EPI_ISL_3145792, EPI_ISL_3145793, EPI_ISL_3145794, EPI_ISL_3145795, EPI_ISL_3145796, EPI_ISL_3145797, EPI_ISL_3145798, EPI_ISL_3145799, EPI_ISL_3145800, EPI_ISL_3145801, EPI_ISL_3145802, EPI_ISL_3145803, EPI_ISL_3145804, EPI_ISL_3145805, EPI_ISL_3145806, EPI_ISL_3145807, EPI_ISL_3145808, EPI_ISL_3145809, EPI_ISL_3145810, EPI_ISL_3145811, EPI_ISL_3145812, EPI_ISL_3145813, EPI_ISL_3145814, EPI_ISL_3145815, EPI_ISL_3145816, EPI_ISL_3145817, EPI_ISL_3145818, EPI_ISL_3145819, EPI_ISL_3145820, EPI_ISL_3145821, EPI_ISL_3145822, EPI_ISL_3145823, EPI_ISL_3145824, EPI_ISL_3145825, EPI_ISL_3145826, EPI_ISL_3145827, EPI_ISL_3145828, EPI_ISL_3145829, EPI_ISL_3145830, EPI_ISL_3145831, EPI_ISL_3145832, EPI_ISL_3145833, EPI_ISL_3145834, EPI_ISL_3145835, EPI_ISL_3145836, EPI_ISL_3145837, EPI_ISL_3145838, EPI_ISL_3145839, EPI_ISL_3145840, EPI_ISL_3145841, EPI_ISL_3145842, EPI_ISL_3145843, EPI_ISL_3145844, EPI_ISL_3145845, EPI_ISL_3145846, EPI_ISL_3145847, EPI_ISL_3145848, EPI_ISL_3145849, EPI_ISL_3145850, EPI_ISL_3145851, EPI_ISL_3145852, EPI_ISL_3145853, EPI_ISL_3145854, EPI_ISL_3145855, EPI_ISL_3145856, EPI_ISL_3145857, EPI_ISL_3145858, EPI_ISL_3145859, EPI_ISL_3145860, EPI_ISL_3145861, EPI_ISL_3145862, EPI_ISL_3145863, EPI_ISL_3145864, EPI_ISL_3145865, EPI_ISL_3145866, EPI_ISL_3145867, EPI_ISL_3145868, EPI_ISL_3145869, EPI_ISL_3145870, EPI_ISL_3145871, EPI_ISL_3145872, EPI_ISL_3145873, EPI_ISL_3145874, EPI_ISL_3145875, EPI_ISL_3145876, EPI_ISL_3145877, EPI_ISL_3145878, EPI_ISL_3145879, EPI_ISL_3145880, EPI_ISL_3145881, EPI_ISL_3145882, EPI_ISL_3145883, EPI_ISL_3145884, EPI_ISL_3145885, EPI_ISL_3145886, EPI_ISL_3145887, EPI_ISL_3145888, EPI_ISL_3145892, EPI_ISL_3145894, EPI_ISL_3145895, EPI_ISL_3145896, EPI_ISL_3145897, EPI_ISL_3145898, EPI_ISL_3145906, EPI_ISL_3146261, EPI_ISL_3146263, EPI_ISL_3146265, EPI_ISL_3146266, EPI_ISL_3146267, EPI_ISL_3146271, EPI_ISL_3146275, EPI_ISL_3146279, EPI_ISL_3146280, EPI_ISL_3146282, EPI_ISL_3146283, EPI_ISL_3146355, EPI_ISL_3146356, EPI_ISL_3146357, EPI_ISL_3146358, EPI_ISL_3146359, EPI_ISL_3146360, EPI_ISL_3146361, EPI_ISL_3146362, EPI_ISL_3146363, EPI_ISL_3146364, EPI_ISL_3146365, EPI_ISL_3146366, EPI_ISL_3146367, EPI_ISL_3146368, EPI_ISL_3146369, EPI_ISL_3146370, EPI_ISL_3146371, EPI_ISL_3146372, EPI_ISL_3146373, EPI_ISL_3146374, EPI_ISL_3146375, EPI_ISL_3146376, EPI_ISL_3146377, EPI_ISL_3146378, EPI_ISL_3146379, EPI_ISL_3146380, EPI_ISL_3146381, EPI_ISL_3146382, EPI_ISL_3146383, EPI_ISL_3146384, EPI_ISL_3146385, EPI_ISL_3146386, EPI_ISL_3146387, EPI_ISL_3146388, EPI_ISL_3146389, EPI_ISL_3146390, EPI_ISL_3146392, EPI_ISL_3146393, EPI_ISL_3146394, EPI_ISL_3146395, EPI_ISL_3146396, EPI_ISL_3146397, EPI_ISL_3146398, EPI_ISL_3146399, EPI_ISL_3146400, EPI_ISL_3146401, EPI_ISL_3146402, EPI_ISL_3146403, EPI_ISL_3146404, EPI_ISL_3146405, EPI_ISL_3146406, EPI_ISL_3146407, EPI_ISL_3146408, EPI_ISL_3146409, EPI_ISL_3146410, EPI_ISL_3146411, EPI_ISL_3146412, EPI_ISL_3146413, EPI_ISL_3146414, EPI_ISL_3146415, EPI_ISL_3146416, EPI_ISL_3146417, EPI_ISL_3146418, EPI_ISL_3146421, EPI_ISL_3146422, EPI_ISL_3146423, EPI_ISL_3146424, EPI_ISL_3146426, EPI_ISL_3146427, EPI_ISL_3146428, EPI_ISL_3146429, EPI_ISL_3146430, EPI_ISL_3146431, EPI_ISL_3146432, EPI_ISL_3146433, EPI_ISL_3146434, EPI_ISL_3146435, EPI_ISL_3146436, EPI_ISL_3146437, EPI_ISL_3146438, EPI_ISL_3146439, EPI_ISL_3146440, EPI_ISL_3146441, EPI_ISL_3146442, EPI_ISL_3146443, EPI_ISL_3146444, EPI_ISL_3146445, EPI_ISL_3146446, EPI_ISL_3146447, EPI_ISL_3146448, EPI_ISL_3146449, EPI_ISL_3146450, EPI_ISL_3146451, EPI_ISL_3146452, EPI_ISL_3146453, EPI_ISL_3146454, EPI_ISL_3146455, EPI_ISL_3146456, EPI_ISL_3146457, EPI_ISL_3146458, EPI_ISL_3146459, EPI_ISL_3146462, EPI_ISL_3146465, EPI_ISL_3146470, EPI_ISL_3146471, EPI_ISL_3146477, EPI_ISL_3146479, EPI_ISL_3146480, EPI_ISL_3146482, EPI_ISL_3146495, EPI_ISL_3146496, EPI_ISL_3146499, EPI_ISL_3146500, EPI_ISL_3146501, EPI_ISL_3146504, EPI_ISL_3146505, EPI_ISL_3146506, EPI_ISL_3146509, EPI_ISL_3146510, EPI_ISL_3146512, EPI_ISL_3146515, EPI_ISL_3146516, EPI_ISL_3146518, EPI_ISL_3146519, EPI_ISL_3146520, EPI_ISL_3146522, EPI_ISL_3146523, EPI_ISL_3146526, EPI_ISL_3146527, EPI_ISL_3146529, EPI_ISL_3146530, EPI_ISL_3146532, EPI_ISL_3146535, EPI_ISL_3146538, EPI_ISL_3146544, EPI_ISL_3146546, EPI_ISL_3146549, EPI_ISL_3146551, EPI_ISL_3146552, EPI_ISL_3146553 | | | |
| see above | BioneXt Lab | Laboratoire national de sante, Microbiology, Microbial Genomics Platform | Anke Wienecke-Baldacchino, Catherine Ragimbeau,Jessica Tapp, Fatu Djabi, Lise Pignon, Raoul Salmon,Virginie Jover, Elodie Solarino, Thibault Ferrandon, Tamir Abdelrahman |
| EPI_ISL_3146555, EPI_ISL_3146556, EPI_ISL_3146557, EPI_ISL_3146558, EPI_ISL_3146559, EPI_ISL_3146560, EPI_ISL_3146561, EPI_ISL_3146562, EPI_ISL_3146563, EPI_ISL_3146564, EPI_ISL_3146565, EPI_ISL_3146566, EPI_ISL_3146568, EPI_ISL_3146569 | | | |
| see above | Department of Microbiology, Yamanashi Institute for Public Health and Environment | Genome Analysis Center, Yamanashi Central Hospital | Yosuke Hirotsu |
| EPI_ISL_3146571, EPI_ISL_3146572, EPI_ISL_3146573, EPI_ISL_3146574, EPI_ISL_3146575, EPI_ISL_3146576, EPI_ISL_3146577, EPI_ISL_3146578, EPI_ISL_3146579, EPI_ISL_3146580, EPI_ISL_3146581, EPI_ISL_3146582, EPI_ISL_3146583, EPI_ISL_3146584, EPI_ISL_3146585, EPI_ISL_3146586, EPI_ISL_3146587, EPI_ISL_3146588, EPI_ISL_3146589, EPI_ISL_3146590, EPI_ISL_3146591, EPI_ISL_3146592, EPI_ISL_3146593, EPI_ISL_3146594, EPI_ISL_3146595, EPI_ISL_3146596, EPI_ISL_3146597, EPI_ISL_3146598, EPI_ISL_3146599, EPI_ISL_3146600, EPI_ISL_3146601, EPI_ISL_3146602, EPI_ISL_3146603, EPI_ISL_3146604, EPI_ISL_3146605, EPI_ISL_3146606, EPI_ISL_3146607, EPI_ISL_3146608, EPI_ISL_3146609, EPI_ISL_3146610, EPI_ISL_3146611, EPI_ISL_3146612, EPI_ISL_3146613, EPI_ISL_3146614, EPI_ISL_3146615, EPI_ISL_3146616, EPI_ISL_3146617, EPI_ISL_3146618, EPI_ISL_3146619, EPI_ISL_3146620, EPI_ISL_3146621, EPI_ISL_3146622, EPI_ISL_3146623, EPI_ISL_3146624, EPI_ISL_3146625, EPI_ISL_3146626, EPI_ISL_3146627, EPI_ISL_3146628, EPI_ISL_3146629, EPI_ISL_3146630, EPI_ISL_3146631, EPI_ISL_3146632, EPI_ISL_3146633, EPI_ISL_3146634, EPI_ISL_3146635, EPI_ISL_3146636 | | | |
| see above | Laboratorio di Microbiologia | Laboratorio di Microbiologia | Martinetti Lucchini Gladys, Valeria Spina |
| EPI_ISL_3146738, EPI_ISL_3146740, EPI_ISL_3146743, EPI_ISL_3146745, EPI_ISL_3146747, EPI_ISL_3146749, EPI_ISL_3146752, EPI_ISL_3146754, EPI_ISL_3146756, EPI_ISL_3146758, EPI_ISL_3146763, EPI_ISL_3146766, EPI_ISL_3146768, EPI_ISL_3146770, EPI_ISL_3146773, EPI_ISL_3146775, EPI_ISL_3146779, EPI_ISL_3146781, EPI_ISL_3146784, EPI_ISL_3146786, EPI_ISL_3146789, EPI_ISL_3146791, EPI_ISL_3146794, EPI_ISL_3146796, EPI_ISL_3146799, EPI_ISL_3146801 | | | |
| see above | IN State Department of Health Laboratory Services | IN State Department of Health Laboratory Services | Cassandra Campion, Jamie Yeadon, Brian Pope, Lixia Liu, Kyle Brownlee, Melissa Hindenlang, Mark Glazier |
| EPI_ISL_3147181, EPI_ISL_3147184 | Laboratoires Reunis | Laboratoire national de sante, Microbiology, Microbial Genomics Platform | Anke Wienecke-Baldacchino, Catherine Ragimbeau,Jessica Tapp, Fatu Djabi, Lise Pignon, Raoul Salmon, Virginie Jover, Elodie Solarino, Bernard Weber, Tamir Abdelrahman |
| EPI_ISL_3147187 | H⌠pitaux Robert Schuman | Laboratoire national de sante, Microbiology, Microbial Genomics Platform | Anke Wienecke-Baldacchino, Catherine Ragimbeau,Jessica Tapp, Fatu Djabi, Lise Pignon, Raoul Salmon, Virginie Jover, Elodie Solarino, Alain Hakim, Nawfal Faik, Tamir Abdelrahman |
| EPI_ISL_3147188, EPI_ISL_3147195, EPI_ISL_3147198, EPI_ISL_3147200, EPI_ISL_3147201 | Laboratoires Reunis | Laboratoire national de sante, Microbiology, Microbial Genomics Platform | Anke Wienecke-Baldacchino, Catherine Ragimbeau,Jessica Tapp, Fatu Djabi, Lise Pignon, Raoul Salmon, Virginie Jover, Elodie Solarino, Bernard Weber, Tamir Abdelrahman |

| | | | |
|---|---|---|---|
| EPI_ISL_3147206 | Hospital Center Luxembourg | Laboratoire national de sante, Microbiology, Microbial Genomics Platform | Anke Wienecke-Baldacchino, Catherine Ragimbeau,Jessica Tapp, Fatu Djabi, Lise Pignon, Raoul Salmon, Virginie Jover, Elodie Solarino, Michel Kohnen, Jean-Hugues Francois, Tamir Abdelrahman |
| EPI_ISL_3147208, EPI_ISL_3147217, EPI_ISL_3147218, EPI_ISL_3147219, EPI_ISL_3147220, EPI_ISL_3147222, EPI_ISL_3147225, EPI_ISL_3147228 | Laboratoires Reunis | Laboratoire national de sante, Microbiology, Microbial Genomics Platform | Anke Wienecke-Baldacchino, Catherine Ragimbeau,Jessica Tapp, Fatu Djabi, Lise Pignon, Raoul Salmon, Virginie Jover, Elodie Solarino, Bernard Weber, Tamir Abdelrahman |
| EPI_ISL_3147233 | Laboratoire national de sante, Microbiology, Virology | Laboratoire national de sante, Microbiology, Microbial Genomics Platform | Anke Wienecke-Baldacchino, Catherine Ragimbeau,Jessica Tapp, Fatu Djabi, Lise Pignon, Raoul Salmon, Virginie Jover, Elodie Solarino, Trung Nguyen Nguyen, Tamir Abdelrahman |
| EPI_ISL_3147237, EPI_ISL_3147239, EPI_ISL_3147242, EPI_ISL_3147247, EPI_ISL_3147248 | Laboratoires Reunis | Laboratoire national de sante, Microbiology, Microbial Genomics Platform | Anke Wienecke-Baldacchino, Catherine Ragimbeau,Jessica Tapp, Fatu Djabi, Lise Pignon, Raoul Salmon, Virginie Jover, Elodie Solarino, Bernard Weber, Tamir Abdelrahman |
| EPI_ISL_3147250 | Hospital Center Luxembourg | Laboratoire national de sante, Microbiology, Microbial Genomics Platform | Anke Wienecke-Baldacchino, Catherine Ragimbeau,Jessica Tapp, Fatu Djabi, Lise Pignon, Raoul Salmon, Virginie Jover, Elodie Solarino, Michel Kohnen, Jean-Hugues Francois, Tamir Abdelrahman |
| EPI_ISL_3147253, EPI_ISL_3147254, EPI_ISL_3147256 | Laboratoires Reunis | Laboratoire national de sante, Microbiology, Microbial Genomics Platform | Anke Wienecke-Baldacchino, Catherine Ragimbeau,Jessica Tapp, Fatu Djabi, Lise Pignon, Raoul Salmon, Virginie Jover, Elodie Solarino, Bernard Weber, Tamir Abdelrahman |
| EPI_ISL_3147258 | Laboratoire national de sante, Microbiology, Virology | Laboratoire national de sante, Microbiology, Microbial Genomics Platform | Anke Wienecke-Baldacchino, Catherine Ragimbeau,Jessica Tapp, Fatu Djabi, Lise Pignon, Raoul Salmon, Virginie Jover, Elodie Solarino, Trung Nguyen Nguyen, Tamir Abdelrahman |
| EPI_ISL_3147260, EPI_ISL_3147261 | Hospital Center Luxembourg | Laboratoire national de sante, Microbiology, Microbial Genomics Platform | Anke Wienecke-Baldacchino, Catherine Ragimbeau,Jessica Tapp, Fatu Djabi, Lise Pignon, Raoul Salmon, Virginie Jover, Elodie Solarino, Michel Kohnen, Jean-Hugues Francois, Tamir Abdelrahman |
| EPI_ISL_3147265 | Laboratoire national de sante, Microbiology, Virology | Laboratoire national de sante, Microbiology, Microbial Genomics Platform | Anke Wienecke-Baldacchino, Catherine Ragimbeau,Jessica Tapp, Fatu Djabi, Lise Pignon, Raoul Salmon, Virginie Jover, Elodie Solarino, Trung Nguyen Nguyen, Tamir Abdelrahman |
| EPI_ISL_3147266 | Hospital Center Luxembourg | Laboratoire national de sante, Microbiology, Microbial Genomics Platform | Anke Wienecke-Baldacchino, Catherine Ragimbeau,Jessica Tapp, Fatu Djabi, Lise Pignon, Raoul Salmon, Virginie Jover, Elodie Solarino, Michel Kohnen, Jean-Hugues Francois, Tamir Abdelrahman |
| EPI_ISL_3147273 | Laboratoires Reunis | Laboratoire national de sante, Microbiology, Microbial Genomics Platform | Anke Wienecke-Baldacchino, Catherine Ragimbeau,Jessica Tapp, Fatu Djabi, Lise Pignon, Raoul Salmon, Virginie Jover, Elodie Solarino, Bernard Weber, Tamir Abdelrahman |
| EPI_ISL_3147275 | Hospital Center Emile Mayrisch | Laboratoire national de sante, Microbiology, Microbial Genomics Platform | Anke Wienecke-Baldacchino, Catherine Ragimbeau,Jessica Tapp, Fatu Djabi, Lise Pignon, Raoul Salmon, Virginie Jover, Elodie Solarino, Cynthia Oxacelay, Tamir Abdelrahman |
| EPI_ISL_3147281, EPI_ISL_3147282 | Laboratoires Reunis | Laboratoire national de sante, Microbiology, Microbial Genomics Platform | Anke Wienecke-Baldacchino, Catherine Ragimbeau,Jessica Tapp, Fatu Djabi, Lise Pignon, Raoul Salmon, Virginie Jover, Elodie Solarino, Bernard Weber, Tamir Abdelrahman |
| EPI_ISL_3147286 | Hospital Center Luxembourg | Laboratoire national de sante, Microbiology, Microbial Genomics Platform | Anke Wienecke-Baldacchino, Catherine Ragimbeau,Jessica Tapp, Fatu Djabi, Lise Pignon, Raoul Salmon, Virginie Jover, Elodie Solarino, Michel Kohnen, Jean-Hugues Francois, Tamir Abdelrahman |
| EPI_ISL_3147287 | Laboratoires Reunis | Laboratoire national de sante, Microbiology, Microbial Genomics Platform | Anke Wienecke-Baldacchino, Catherine Ragimbeau,Jessica Tapp, Fatu Djabi, Lise Pignon, Raoul Salmon, Virginie Jover, Elodie Solarino, Bernard Weber, Tamir Abdelrahman |
| EPI_ISL_3147288 | Hospital Center Luxembourg | Laboratoire national de sante, Microbiology, Microbial Genomics Platform | Anke Wienecke-Baldacchino, Catherine Ragimbeau,Jessica Tapp, Fatu Djabi, Lise Pignon, Raoul Salmon, Virginie Jover, Elodie Solarino, Michel Kohnen, Jean-Hugues Francois, Tamir Abdelrahman |
| EPI_ISL_3147292, EPI_ISL_3147294, EPI_ISL_3147297, EPI_ISL_3147299 | Laboratoires Reunis | Laboratoire national de sante, Microbiology, Microbial Genomics Platform | Anke Wienecke-Baldacchino, Catherine Ragimbeau,Jessica Tapp, Fatu Djabi, Lise Pignon, Raoul Salmon, Virginie Jover, Elodie Solarino, Bernard Weber, Tamir Abdelrahman |
| EPI_ISL_3147304 | Laboratoire national de sante, Microbiology, Virology | Laboratoire national de sante, Microbiology, Microbial Genomics Platform | Anke Wienecke-Baldacchino, Catherine Ragimbeau,Jessica Tapp, Fatu Djabi, Lise Pignon, Raoul Salmon, Virginie Jover, Elodie Solarino, Trung Nguyen Nguyen, Tamir Abdelrahman |
| EPI_ISL_3147305, EPI_ISL_3147308, EPI_ISL_3147309 | Laboratoires Reunis | Laboratoire national de sante, Microbiology, Microbial Genomics Platform | Anke Wienecke-Baldacchino, Catherine Ragimbeau,Jessica Tapp, Fatu Djabi, Lise Pignon, Raoul Salmon, Virginie Jover, Elodie Solarino, Bernard Weber, Tamir Abdelrahman |
| EPI_ISL_3147314, EPI_ISL_3147315 | Laboratoire national de sante, Microbiology, Virology | Laboratoire national de sante, Microbiology, Microbial Genomics Platform | Anke Wienecke-Baldacchino, Catherine Ragimbeau,Jessica Tapp, Fatu Djabi, Lise Pignon, Raoul Salmon, Virginie Jover, Elodie Solarino, Trung Nguyen Nguyen, Tamir Abdelrahman |
| EPI_ISL_3147322 | Hospital Center Luxembourg | Laboratoire national de sante, Microbiology, Microbial Genomics Platform | Anke Wienecke-Baldacchino, Catherine Ragimbeau,Jessica Tapp, Fatu Djabi, Lise Pignon, Raoul Salmon, Virginie Jover, Elodie Solarino, Michel Kohnen, Jean-Hugues Francois, Tamir Abdelrahman |
| EPI_ISL_3147328 | Laboratoire national de sante, Microbiology, Virology | Laboratoire national de sante, Microbiology, Microbial Genomics Platform | Anke Wienecke-Baldacchino, Catherine Ragimbeau,Jessica Tapp, Fatu Djabi, Lise Pignon, Raoul Salmon, Virginie Jover, Elodie Solarino, Trung Nguyen Nguyen, Tamir Abdelrahman |
| EPI_ISL_3147334 | Laboratoires Reunis | Laboratoire national de sante, Microbiology, Microbial Genomics Platform | Anke Wienecke-Baldacchino, Catherine Ragimbeau,Jessica Tapp, Fatu Djabi, Lise Pignon, Raoul Salmon, Virginie Jover, Elodie Solarino, Bernard Weber, Tamir Abdelrahman |
| EPI_ISL_3147335, EPI_ISL_3147337 | Hospital Center Luxembourg | Laboratoire national de sante, Microbiology, Microbial Genomics Platform | Anke Wienecke-Baldacchino, Catherine Ragimbeau,Jessica Tapp, Fatu Djabi, Lise Pignon, Raoul Salmon, Virginie Jover, Elodie Solarino, Michel Kohnen, Jean-Hugues Francois, Tamir Abdelrahman |
| EPI_ISL_3147338 | Laboratoire national de sante, Microbiology, Virology | Laboratoire national de sante, Microbiology, Microbial Genomics Platform | Anke Wienecke-Baldacchino, Catherine Ragimbeau,Jessica Tapp, Fatu Djabi, Lise Pignon, Raoul Salmon, Virginie Jover, Elodie Solarino, Trung Nguyen Nguyen, Tamir Abdelrahman |
| EPI_ISL_3147339 | Hospital Center Luxembourg | Laboratoire national de sante, Microbiology, Microbial Genomics Platform | Anke Wienecke-Baldacchino, Catherine Ragimbeau,Jessica Tapp, Fatu Djabi, Lise Pignon, Raoul Salmon, Virginie Jover, Elodie Solarino, Michel Kohnen, Jean-Hugues Francois, Tamir Abdelrahman |
| EPI_ISL_3147340 | Hospital Center Emile Mayrisch | Laboratoire national de sante, Microbiology, Microbial Genomics Platform | Anke Wienecke-Baldacchino, Catherine Ragimbeau,Jessica Tapp, Fatu Djabi, Lise Pignon, Raoul Salmon, Virginie Jover, Elodie Solarino, Cynthia Oxacelay, Tamir Abdelrahman |
| EPI_ISL_3147341 | Laboratoire national de sante, Microbiology, Virology | Laboratoire national de sante, Microbiology, Microbial Genomics Platform | Anke Wienecke-Baldacchino, Catherine Ragimbeau,Jessica Tapp, Fatu Djabi, Lise Pignon, Raoul Salmon, Virginie Jover, Elodie Solarino, Trung Nguyen Nguyen, Tamir Abdelrahman |
| EPI_ISL_3147342 | Hospital Center Emile Mayrisch | Laboratoire national de sante, Microbiology, Microbial Genomics Platform | Anke Wienecke-Baldacchino, Catherine Ragimbeau,Jessica Tapp, Fatu Djabi, Lise Pignon, Raoul Salmon, Virginie Jover, Elodie Solarino, Cynthia Oxacelay, Tamir Abdelrahman |
| EPI_ISL_3147343 | Hospital Center Luxembourg | Laboratoire national de sante, Microbiology, Microbial Genomics Platform | Anke Wienecke-Baldacchino, Catherine Ragimbeau,Jessica Tapp, Fatu Djabi, Lise Pignon, Raoul Salmon, Virginie Jover, Elodie Solarino, Michel Kohnen, Jean-Hugues Francois, Tamir Abdelrahman |
| EPI_ISL_3147348 | H⌈pitaux Robert Schuman | Laboratoire national de sante, Microbiology, Microbial Genomics Platform | Anke Wienecke-Baldacchino, Catherine Ragimbeau,Jessica Tapp, Fatu Djabi, Lise Pignon, Raoul Salmon, Virginie Jover, Elodie Solarino, Alain Hakim, Nawfal Faik, Tamir Abdelrahman |
| EPI_ISL_3147349, EPI_ISL_3147350, EPI_ISL_3147351 | Laboratoire national de sante, Microbiology, Virology | Laboratoire national de sante, Microbiology, Microbial Genomics Platform | Anke Wienecke-Baldacchino, Catherine Ragimbeau,Jessica Tapp, Fatu Djabi, Lise Pignon, Raoul Salmon, Virginie Jover, Elodie Solarino, Trung Nguyen Nguyen, Tamir Abdelrahman |
| EPI_ISL_3147352 | Laboratoires Reunis | Laboratoire national de sante, Microbiology, Microbial Genomics Platform | Anke Wienecke-Baldacchino, Catherine Ragimbeau,Jessica Tapp, Fatu Djabi, Lise Pignon, Raoul Salmon, Virginie Jover, Elodie Solarino, Bernard Weber, Tamir Abdelrahman |
| EPI_ISL_3147353 | Hospital Center Luxembourg | Laboratoire national de sante, Microbiology, Microbial Genomics Platform | Anke Wienecke-Baldacchino, Catherine Ragimbeau,Jessica Tapp, Fatu Djabi, Lise Pignon, Raoul Salmon, Virginie Jover, Elodie Solarino, Michel Kohnen, Jean-Hugues Francois, Tamir Abdelrahman |
| EPI_ISL_3147354, EPI_ISL_3147355 | Laboratoire national de sante, Microbiology, Virology | Laboratoire national de sante, Microbiology, Microbial Genomics Platform | Anke Wienecke-Baldacchino, Catherine Ragimbeau,Jessica Tapp, Fatu Djabi, Lise Pignon, Raoul Salmon, Virginie Jover, Elodie Solarino, Trung Nguyen Nguyen, Tamir Abdelrahman |
| EPI_ISL_3147356 | Hospital Center Emile Mayrisch | Laboratoire national de sante, Microbiology, Microbial Genomics Platform | Anke Wienecke-Baldacchino, Catherine Ragimbeau,Jessica Tapp, Fatu Djabi, Lise Pignon, Raoul Salmon, Virginie Jover, Elodie Solarino, Cynthia Oxacelay, Tamir Abdelrahman |
| EPI_ISL_3147357 | Laboratoires Reunis | Laboratoire national de sante, Microbiology, Microbial Genomics Platform | Anke Wienecke-Baldacchino, Catherine Ragimbeau,Jessica Tapp, Fatu Djabi, Lise Pignon, Raoul Salmon, Virginie Jover, Elodie Solarino, Bernard Weber, Tamir Abdelrahman |
| EPI_ISL_3147391 | Laboratoire national de sante, Microbiology, Virology | Laboratoire national de sante, Microbiology, Microbial Genomics Platform | Anke Wienecke-Baldacchino, Catherine Ragimbeau,Jessica Tapp, Fatu Djabi, Lise Pignon, Raoul Salmon, Virginie Jover, Elodie Solarino, Trung Nguyen Nguyen, Tamir Abdelrahman |
| EPI_ISL_3147392 | Hospital Center Luxembourg | Laboratoire national de sante, Microbiology, Microbial Genomics Platform | Anke Wienecke-Baldacchino, Catherine Ragimbeau,Jessica Tapp, Fatu Djabi, Lise Pignon, Raoul Salmon, Virginie Jover, Elodie Solarino, Michel Kohnen, Jean-Hugues Francois, Tamir Abdelrahman |

| | | | |
|---|---|---|---|
| EPI_ISL_3147393 | Hospital Center Emile Mayrisch | Laboratoire national de sante, Microbiology, Microbial Genomics Platform | Anke Wienecke-Baldacchino, Catherine Ragimbeau,Jessica Tapp, Fatu Djabi, Lise Pignon, Raoul Salmon, Virginie Jover, Elodie Solarino, Cynthia Oxacelay, Tamir Abdelrahman |
| EPI_ISL_3147394, EPI_ISL_3147395, EPI_ISL_3147397, EPI_ISL_3147398, EPI_ISL_3147399 | Laboratoires Reunis | Laboratoire national de sante, Microbiology, Microbial Genomics Platform | Anke Wienecke-Baldacchino, Catherine Ragimbeau,Jessica Tapp, Fatu Djabi, Lise Pignon, Raoul Salmon, Virginie Jover, Elodie Solarino, Bernard Weber, Tamir Abdelrahman |
| EPI_ISL_3147400 | Hospital Center Emile Mayrisch | Laboratoire national de sante, Microbiology, Microbial Genomics Platform | Anke Wienecke-Baldacchino, Catherine Ragimbeau,Jessica Tapp, Fatu Djabi, Lise Pignon, Raoul Salmon, Virginie Jover, Elodie Solarino, Cynthia Oxacelay, Tamir Abdelrahman |
| EPI_ISL_3147401, EPI_ISL_3147403, EPI_ISL_3147404 | Laboratoires Reunis | Laboratoire national de sante, Microbiology, Microbial Genomics Platform | Anke Wienecke-Baldacchino, Catherine Ragimbeau,Jessica Tapp, Fatu Djabi, Lise Pignon, Raoul Salmon, Virginie Jover, Elodie Solarino, Bernard Weber, Tamir Abdelrahman |
| EPI_ISL_3147410 | Hospital Center Luxembourg | Laboratoire national de sante, Microbiology, Microbial Genomics Platform | Anke Wienecke-Baldacchino, Catherine Ragimbeau,Jessica Tapp, Fatu Djabi, Lise Pignon, Raoul Salmon, Virginie Jover, Elodie Solarino, Michel Kohnen, Jean-Hugues Francois, Tamir Abdelrahman |
| EPI_ISL_3147415 | Laboratoires Reunis | Laboratoire national de sante, Microbiology, Microbial Genomics Platform | Anke Wienecke-Baldacchino, Catherine Ragimbeau,Jessica Tapp, Fatu Djabi, Lise Pignon, Raoul Salmon, Virginie Jover, Elodie Solarino, Bernard Weber, Tamir Abdelrahman |
| EPI_ISL_3147417 | Laboratoire national de sante, Microbiology, Virology | Laboratoire national de sante, Microbiology, Microbial Genomics Platform | Anke Wienecke-Baldacchino, Catherine Ragimbeau,Jessica Tapp, Fatu Djabi, Lise Pignon, Raoul Salmon, Virginie Jover, Elodie Solarino, Trung Nguyen Nguyen, Tamir Abdelrahman |
| EPI_ISL_3147419, EPI_ISL_3147420, EPI_ISL_3147422, EPI_ISL_3147426, EPI_ISL_3147428, EPI_ISL_3147430, EPI_ISL_3147432 | Laboratoires Reunis | Laboratoire national de sante, Microbiology, Microbial Genomics Platform | Anke Wienecke-Baldacchino, Catherine Ragimbeau,Jessica Tapp, Fatu Djabi, Lise Pignon, Raoul Salmon, Virginie Jover, Elodie Solarino, Bernard Weber, Tamir Abdelrahman |
| EPI_ISL_3147439 | Laboratoire national de sante, Microbiology, Virology | Laboratoire national de sante, Microbiology, Microbial Genomics Platform | Anke Wienecke-Baldacchino, Catherine Ragimbeau,Jessica Tapp, Fatu Djabi, Lise Pignon, Raoul Salmon, Virginie Jover, Elodie Solarino, Trung Nguyen Nguyen, Tamir Abdelrahman |
| EPI_ISL_3147440 | H⌠pitaux Robert Schuman | Laboratoire national de sante, Microbiology, Microbial Genomics Platform | Anke Wienecke-Baldacchino, Catherine Ragimbeau,Jessica Tapp, Fatu Djabi, Lise Pignon, Raoul Salmon, Virginie Jover, Elodie Solarino, Alain Hakim, Nawfal Faik, Tamir Abdelrahman |
| EPI_ISL_3147441, EPI_ISL_3147445, EPI_ISL_3147447 | Laboratoires Reunis | Laboratoire national de sante, Microbiology, Microbial Genomics Platform | Anke Wienecke-Baldacchino, Catherine Ragimbeau,Jessica Tapp, Fatu Djabi, Lise Pignon, Raoul Salmon, Virginie Jover, Elodie Solarino, Bernard Weber, Tamir Abdelrahman |
| EPI_ISL_3147448 | Hospital Center Luxembourg | Laboratoire national de sante, Microbiology, Microbial Genomics Platform | Anke Wienecke-Baldacchino, Catherine Ragimbeau,Jessica Tapp, Fatu Djabi, Lise Pignon, Raoul Salmon, Virginie Jover, Elodie Solarino, Michel Kohnen, Jean-Hugues Francois, Tamir Abdelrahman |
| EPI_ISL_3147453 | Laboratoires Reunis | Laboratoire national de sante, Microbiology, Microbial Genomics Platform | Anke Wienecke-Baldacchino, Catherine Ragimbeau,Jessica Tapp, Fatu Djabi, Lise Pignon, Raoul Salmon, Virginie Jover, Elodie Solarino, Bernard Weber, Tamir Abdelrahman |
| EPI_ISL_3147454 | Hospital Center Luxembourg | Laboratoire national de sante, Microbiology, Microbial Genomics Platform | Anke Wienecke-Baldacchino, Catherine Ragimbeau,Jessica Tapp, Fatu Djabi, Lise Pignon, Raoul Salmon, Virginie Jover, Elodie Solarino, Michel Kohnen, Jean-Hugues Francois, Tamir Abdelrahman |
| EPI_ISL_3147455, EPI_ISL_3147463, EPI_ISL_3147464 | Laboratoires Reunis | Laboratoire national de sante, Microbiology, Microbial Genomics Platform | Anke Wienecke-Baldacchino, Catherine Ragimbeau,Jessica Tapp, Fatu Djabi, Lise Pignon, Raoul Salmon, Virginie Jover, Elodie Solarino, Bernard Weber, Tamir Abdelrahman |
| EPI_ISL_3147465 | Laboratoire national de sante, Microbiology, Virology | Laboratoire national de sante, Microbiology, Microbial Genomics Platform | Anke Wienecke-Baldacchino, Catherine Ragimbeau,Jessica Tapp, Fatu Djabi, Lise Pignon, Raoul Salmon, Virginie Jover, Elodie Solarino, Trung Nguyen Nguyen, Tamir Abdelrahman |
| EPI_ISL_3147466, EPI_ISL_3147468, EPI_ISL_3147474 | Laboratoires Reunis | Laboratoire national de sante, Microbiology, Microbial Genomics Platform | Anke Wienecke-Baldacchino, Catherine Ragimbeau,Jessica Tapp, Fatu Djabi, Lise Pignon, Raoul Salmon, Virginie Jover, Elodie Solarino, Bernard Weber, Tamir Abdelrahman |
| EPI_ISL_3147604 | Hospital Center Emile Mayrisch | Laboratoire national de sante, Microbiology, Microbial Genomics Platform | Anke Wienecke-Baldacchino, Catherine Ragimbeau,Jessica Tapp, Fatu Djabi, Lise Pignon, Raoul Salmon, Virginie Jover, Elodie Solarino, Cynthia Oxacelay, Tamir Abdelrahman |
| EPI_ISL_3147607 | Laboratoires Reunis | Laboratoire national de sante, Microbiology, Microbial Genomics Platform | Anke Wienecke-Baldacchino, Catherine Ragimbeau,Jessica Tapp, Fatu Djabi, Lise Pignon, Raoul Salmon, Virginie Jover, Elodie Solarino, Bernard Weber, Tamir Abdelrahman |
| EPI_ISL_3147608 | H⌠pitaux Robert Schuman | Laboratoire national de sante, Microbiology, Microbial Genomics Platform | Anke Wienecke-Baldacchino, Catherine Ragimbeau,Jessica Tapp, Fatu Djabi, Lise Pignon, Raoul Salmon, Virginie Jover, Elodie Solarino, Alain Hakim, Nawfal Faik, Tamir Abdelrahman |
| EPI_ISL_3147621, EPI_ISL_3147622, EPI_ISL_3147623 | Hospital Center Emile Mayrisch | Laboratoire national de sante, Microbiology, Microbial Genomics Platform | Anke Wienecke-Baldacchino, Catherine Ragimbeau,Jessica Tapp, Fatu Djabi, Lise Pignon, Raoul Salmon, Virginie Jover, Elodie Solarino, Cynthia Oxacelay, Tamir Abdelrahman |
| EPI_ISL_3147624, EPI_ISL_3147625 | Hospital Center Luxembourg | Laboratoire national de sante, Microbiology, Microbial Genomics Platform | Anke Wienecke-Baldacchino, Catherine Ragimbeau,Jessica Tapp, Fatu Djabi, Lise Pignon, Raoul Salmon, Virginie Jover, Elodie Solarino, Michel Kohnen, Jean-Hugues Francois, Tamir Abdelrahman |
| EPI_ISL_3147626 | Laboratoire national de sante, Microbiology, Virology | Laboratoire national de sante, Microbiology, Microbial Genomics Platform | Anke Wienecke-Baldacchino, Catherine Ragimbeau,Jessica Tapp, Fatu Djabi, Lise Pignon, Raoul Salmon, Virginie Jover, Elodie Solarino, Trung Nguyen Nguyen, Tamir Abdelrahman |
| EPI_ISL_3147627 | Hospital Center Emile Mayrisch | Laboratoire national de sante, Microbiology, Microbial Genomics Platform | Anke Wienecke-Baldacchino, Catherine Ragimbeau,Jessica Tapp, Fatu Djabi, Lise Pignon, Raoul Salmon, Virginie Jover, Elodie Solarino, Cynthia Oxacelay, Tamir Abdelrahman |
| EPI_ISL_3147628 | H⌠pitaux Robert Schuman | Laboratoire national de sante, Microbiology, Microbial Genomics Platform | Anke Wienecke-Baldacchino, Catherine Ragimbeau,Jessica Tapp, Fatu Djabi, Lise Pignon, Raoul Salmon, Virginie Jover, Elodie Solarino, Alain Hakim, Nawfal Faik, Tamir Abdelrahman |
| EPI_ISL_3147629 | Hospital Center Emile Mayrisch | Laboratoire national de sante, Microbiology, Microbial Genomics Platform | Anke Wienecke-Baldacchino, Catherine Ragimbeau,Jessica Tapp, Fatu Djabi, Lise Pignon, Raoul Salmon, Virginie Jover, Elodie Solarino, Cynthia Oxacelay, Tamir Abdelrahman |
| EPI_ISL_3147630 | Laboratoire national de sante, Microbiology, Virology | Laboratoire national de sante, Microbiology, Microbial Genomics Platform | Anke Wienecke-Baldacchino, Catherine Ragimbeau,Jessica Tapp, Fatu Djabi, Lise Pignon, Raoul Salmon, Virginie Jover, Elodie Solarino, Trung Nguyen Nguyen, Tamir Abdelrahman |
| EPI_ISL_3147631 | Hospital Center Emile Mayrisch | Laboratoire national de sante, Microbiology, Microbial Genomics Platform | Anke Wienecke-Baldacchino, Catherine Ragimbeau,Jessica Tapp, Fatu Djabi, Lise Pignon, Raoul Salmon, Virginie Jover, Elodie Solarino, Cynthia Oxacelay, Tamir Abdelrahman |
| EPI_ISL_3147632 | Hospital Center Luxembourg | Laboratoire national de sante, Microbiology, Microbial Genomics Platform | Anke Wienecke-Baldacchino, Catherine Ragimbeau,Jessica Tapp, Fatu Djabi, Lise Pignon, Raoul Salmon, Virginie Jover, Elodie Solarino, Michel Kohnen, Jean-Hugues Francois, Tamir Abdelrahman |
| EPI_ISL_3147633 | Laboratoire national de sante, Microbiology, Virology | Laboratoire national de sante, Microbiology, Microbial Genomics Platform | Anke Wienecke-Baldacchino, Catherine Ragimbeau,Jessica Tapp, Fatu Djabi, Lise Pignon, Raoul Salmon, Virginie Jover, Elodie Solarino, Trung Nguyen Nguyen, Tamir Abdelrahman |
| EPI_ISL_3147634 | Hospital Center Emile Mayrisch | Laboratoire national de sante, Microbiology, Microbial Genomics Platform | Anke Wienecke-Baldacchino, Catherine Ragimbeau,Jessica Tapp, Fatu Djabi, Lise Pignon, Raoul Salmon, Virginie Jover, Elodie Solarino, Cynthia Oxacelay, Tamir Abdelrahman |
| EPI_ISL_3147635 | Hospital Center Luxembourg | Laboratoire national de sante, Microbiology, Microbial Genomics Platform | Anke Wienecke-Baldacchino, Catherine Ragimbeau,Jessica Tapp, Fatu Djabi, Lise Pignon, Raoul Salmon, Virginie Jover, Elodie Solarino, Michel Kohnen, Jean-Hugues Francois, Tamir Abdelrahman |
| EPI_ISL_3147636, EPI_ISL_3147637 | Laboratoire national de sante, Microbiology, Virology | Laboratoire national de sante, Microbiology, Microbial Genomics Platform | Anke Wienecke-Baldacchino, Catherine Ragimbeau,Jessica Tapp, Fatu Djabi, Lise Pignon, Raoul Salmon, Virginie Jover, Elodie Solarino, Trung Nguyen Nguyen, Tamir Abdelrahman |
| EPI_ISL_3147638 | Hospital Center Emile Mayrisch | Laboratoire national de sante, Microbiology, Microbial Genomics Platform | Anke Wienecke-Baldacchino, Catherine Ragimbeau,Jessica Tapp, Fatu Djabi, Lise Pignon, Raoul Salmon, Virginie Jover, Elodie Solarino, Cynthia Oxacelay, Tamir Abdelrahman |
| EPI_ISL_3147639 | Laboratoire national de sante, Microbiology, Virology | Laboratoire national de sante, Microbiology, Microbial Genomics Platform | Anke Wienecke-Baldacchino, Catherine Ragimbeau,Jessica Tapp, Fatu Djabi, Lise Pignon, Raoul Salmon, Virginie Jover, Elodie Solarino, Trung Nguyen Nguyen, Tamir Abdelrahman |
| EPI_ISL_3147640, EPI_ISL_3147641 | Hospital Center Emile Mayrisch | Laboratoire national de sante, Microbiology, Microbial Genomics Platform | Anke Wienecke-Baldacchino, Catherine Ragimbeau,Jessica Tapp, Fatu Djabi, Lise Pignon, Raoul Salmon, Virginie Jover, Elodie Solarino, Cynthia Oxacelay, Tamir Abdelrahman |
| EPI_ISL_3147642 | Laboratoires Reunis | Laboratoire national de sante, Microbiology, Microbial Genomics Platform | Anke Wienecke-Baldacchino, Catherine Ragimbeau,Jessica Tapp, Fatu Djabi, Lise Pignon, Raoul Salmon, Virginie Jover, Elodie Solarino, Bernard Weber, Tamir Abdelrahman |
| EPI_ISL_3147643 | Hospital Center Emile Mayrisch | Laboratoire national de sante, Microbiology, Microbial Genomics Platform | Anke Wienecke-Baldacchino, Catherine Ragimbeau,Jessica Tapp, Fatu Djabi, Lise Pignon, Raoul Salmon, Virginie Jover, Elodie Solarino, Cynthia Oxacelay, Tamir Abdelrahman |
| EPI_ISL_3147644 | Hospital Center Luxembourg | Laboratoire national de sante, Microbiology, Microbial Genomics Platform | Anke Wienecke-Baldacchino, Catherine Ragimbeau,Jessica Tapp, Fatu Djabi, Lise Pignon, Raoul Salmon, Virginie Jover, Elodie Solarino, Michel Kohnen, Jean-Hugues Francois, Tamir Abdelrahman |
| EPI_ISL_3147645 | Laboratoires Reunis | Laboratoire national de sante, Microbiology, Microbial Genomics | Anke Wienecke-Baldacchino, Catherine Ragimbeau,Jessica Tapp, Fatu Djabi, Lise Pignon, Raoul Salmon, Virginie Jover, Elodie Solarino, Bernard Weber, Tamir Abdelrahman |

| | | | |
|---|---|---|---|
| | | Platform | |
| EPI_ISL_3147646 | Laboratoire national de sante, Microbiology, Virology | Laboratoire national de sante, Microbiology, Microbial Genomics Platform | Anke Wienecke-Baldacchino, Catherine Ragimbeau,Jessica Tapp, Fatu Djabi, Lise Pignon, Raoul Salmon, Virginie Jover, Elodie Solarino, Trung Nguyen Nguyen, Tamir Abdelrahman |
| EPI_ISL_3147647 | Hospital Center Emile Mayrisch | Laboratoire national de sante, Microbiology, Microbial Genomics Platform | Anke Wienecke-Baldacchino, Catherine Ragimbeau,Jessica Tapp, Fatu Djabi, Lise Pignon, Raoul Salmon, Virginie Jover, Elodie Solarino, Cynthia Oxacelay, Tamir Abdelrahman |
| EPI_ISL_3147648, EPI_ISL_3147649 | Laboratoires Reunis | Laboratoire national de sante, Microbiology, Microbial Genomics Platform | Anke Wienecke-Baldacchino, Catherine Ragimbeau,Jessica Tapp, Fatu Djabi, Lise Pignon, Raoul Salmon, Virginie Jover, Elodie Solarino, Bernard Weber, Tamir Abdelrahman |
| EPI_ISL_3147650 | Hospital Center Emile Mayrisch | Laboratoire national de sante, Microbiology, Microbial Genomics Platform | Anke Wienecke-Baldacchino, Catherine Ragimbeau,Jessica Tapp, Fatu Djabi, Lise Pignon, Raoul Salmon, Virginie Jover, Elodie Solarino, Cynthia Oxacelay, Tamir Abdelrahman |
| EPI_ISL_3147651 | Laboratoires Reunis | Laboratoire national de sante, Microbiology, Microbial Genomics Platform | Anke Wienecke-Baldacchino, Catherine Ragimbeau,Jessica Tapp, Fatu Djabi, Lise Pignon, Raoul Salmon, Virginie Jover, Elodie Solarino, Bernard Weber, Tamir Abdelrahman |
| EPI_ISL_3147652 | Hospital Center Emile Mayrisch | Laboratoire national de sante, Microbiology, Microbial Genomics Platform | Anke Wienecke-Baldacchino, Catherine Ragimbeau,Jessica Tapp, Fatu Djabi, Lise Pignon, Raoul Salmon, Virginie Jover, Elodie Solarino, Cynthia Oxacelay, Tamir Abdelrahman |
| EPI_ISL_3147653, EPI_ISL_3147654, EPI_ISL_3147655 | Hospital Center Luxembourg | Laboratoire national de sante, Microbiology, Microbial Genomics Platform | Anke Wienecke-Baldacchino, Catherine Ragimbeau,Jessica Tapp, Fatu Djabi, Lise Pignon, Raoul Salmon, Virginie Jover, Elodie Solarino, Michel Kohnen, Jean-Hugues Francois, Tamir Abdelrahman |
| EPI_ISL_3147656, EPI_ISL_3147657 | Hospital Center Emile Mayrisch | Laboratoire national de sante, Microbiology, Microbial Genomics Platform | Anke Wienecke-Baldacchino, Catherine Ragimbeau,Jessica Tapp, Fatu Djabi, Lise Pignon, Raoul Salmon, Virginie Jover, Elodie Solarino, Cynthia Oxacelay, Tamir Abdelrahman |
| EPI_ISL_3147658, EPI_ISL_3147659 | Laboratoire national de sante, Microbiology, Virology | Laboratoire national de sante, Microbiology, Microbial Genomics Platform | Anke Wienecke-Baldacchino, Catherine Ragimbeau,Jessica Tapp, Fatu Djabi, Lise Pignon, Raoul Salmon, Virginie Jover, Elodie Solarino, Trung Nguyen Nguyen, Tamir Abdelrahman |
| EPI_ISL_3147660 | Laboratoires Reunis | Laboratoire national de sante, Microbiology, Microbial Genomics Platform | Anke Wienecke-Baldacchino, Catherine Ragimbeau,Jessica Tapp, Fatu Djabi, Lise Pignon, Raoul Salmon, Virginie Jover, Elodie Solarino, Bernard Weber, Tamir Abdelrahman |
| EPI_ISL_3147661 | H⌠pitaux Robert Schuman | Laboratoire national de sante, Microbiology, Microbial Genomics Platform | Anke Wienecke-Baldacchino, Catherine Ragimbeau,Jessica Tapp, Fatu Djabi, Lise Pignon, Raoul Salmon, Virginie Jover, Elodie Solarino, Alain Hakim, Nawfal Faik, Tamir Abdelrahman |
| EPI_ISL_3147662, EPI_ISL_3147663 | Laboratoires Reunis | Laboratoire national de sante, Microbiology, Microbial Genomics Platform | Anke Wienecke-Baldacchino, Catherine Ragimbeau,Jessica Tapp, Fatu Djabi, Lise Pignon, Raoul Salmon, Virginie Jover, Elodie Solarino, Bernard Weber, Tamir Abdelrahman |
| EPI_ISL_3147664, EPI_ISL_3147665 | Hospital Center Emile Mayrisch | Laboratoire national de sante, Microbiology, Microbial Genomics Platform | Anke Wienecke-Baldacchino, Catherine Ragimbeau,Jessica Tapp, Fatu Djabi, Lise Pignon, Raoul Salmon, Virginie Jover, Elodie Solarino, Cynthia Oxacelay, Tamir Abdelrahman |
| EPI_ISL_3147666 | Hospital Center Luxembourg | Laboratoire national de sante, Microbiology, Microbial Genomics Platform | Anke Wienecke-Baldacchino, Catherine Ragimbeau,Jessica Tapp, Fatu Djabi, Lise Pignon, Raoul Salmon, Virginie Jover, Elodie Solarino, Michel Kohnen, Jean-Hugues Francois, Tamir Abdelrahman |
| EPI_ISL_3147667, EPI_ISL_3147668 | Hospital Center Emile Mayrisch | Laboratoire national de sante, Microbiology, Microbial Genomics Platform | Anke Wienecke-Baldacchino, Catherine Ragimbeau,Jessica Tapp, Fatu Djabi, Lise Pignon, Raoul Salmon, Virginie Jover, Elodie Solarino, Cynthia Oxacelay, Tamir Abdelrahman |
| EPI_ISL_3147669, EPI_ISL_3147670 | Laboratoire national de sante, Microbiology, Virology | Laboratoire national de sante, Microbiology, Microbial Genomics Platform | Anke Wienecke-Baldacchino, Catherine Ragimbeau,Jessica Tapp, Fatu Djabi, Lise Pignon, Raoul Salmon, Virginie Jover, Elodie Solarino, Trung Nguyen Nguyen, Tamir Abdelrahman |
| EPI_ISL_3147671 | Laboratoires Reunis | Laboratoire national de sante, Microbiology, Microbial Genomics Platform | Anke Wienecke-Baldacchino, Catherine Ragimbeau,Jessica Tapp, Fatu Djabi, Lise Pignon, Raoul Salmon, Virginie Jover, Elodie Solarino, Bernard Weber, Tamir Abdelrahman |
| EPI_ISL_3147672 | Laboratoire national de sante, Microbiology, Virology | Laboratoire national de sante, Microbiology, Microbial Genomics Platform | Anke Wienecke-Baldacchino, Catherine Ragimbeau,Jessica Tapp, Fatu Djabi, Lise Pignon, Raoul Salmon, Virginie Jover, Elodie Solarino, Trung Nguyen Nguyen, Tamir Abdelrahman |
| EPI_ISL_3147673 | Hospital Center Luxembourg | Laboratoire national de sante, Microbiology, Microbial Genomics Platform | Anke Wienecke-Baldacchino, Catherine Ragimbeau,Jessica Tapp, Fatu Djabi, Lise Pignon, Raoul Salmon, Virginie Jover, Elodie Solarino, Michel Kohnen, Jean-Hugues Francois, Tamir Abdelrahman |
| EPI_ISL_3147674 | Hospital Center Emile Mayrisch | Laboratoire national de sante, Microbiology, Microbial Genomics Platform | Anke Wienecke-Baldacchino, Catherine Ragimbeau,Jessica Tapp, Fatu Djabi, Lise Pignon, Raoul Salmon, Virginie Jover, Elodie Solarino, Cynthia Oxacelay, Tamir Abdelrahman |
| EPI_ISL_3147675 | H⌠pitaux Robert Schuman | Laboratoire national de sante, Microbiology, Microbial Genomics Platform | Anke Wienecke-Baldacchino, Catherine Ragimbeau,Jessica Tapp, Fatu Djabi, Lise Pignon, Raoul Salmon, Virginie Jover, Elodie Solarino, Alain Hakim, Nawfal Faik, Tamir Abdelrahman |
| EPI_ISL_3147676 | Laboratoire national de sante, Microbiology, Virology | Laboratoire national de sante, Microbiology, Microbial Genomics Platform | Anke Wienecke-Baldacchino, Catherine Ragimbeau,Jessica Tapp, Fatu Djabi, Lise Pignon, Raoul Salmon, Virginie Jover, Elodie Solarino, Trung Nguyen Nguyen, Tamir Abdelrahman |
| EPI_ISL_3147677, EPI_ISL_3147678 | Centre Hospitalier du Nord | Laboratoire national de sante, Microbiology, Microbial Genomics Platform | Anke Wienecke-Baldacchino, Catherine Ragimbeau,Jessica Tapp, Fatu Djabi, Lise Pignon, Raoul Salmon, Virginie Jover, Elodie Solarino, Fatiha Boulmerka, Jacqueline Parmentier, Tamir Abdelrahman |
| EPI_ISL_3147679 | Laboratoires Reunis | Laboratoire national de sante, Microbiology, Microbial Genomics Platform | Anke Wienecke-Baldacchino, Catherine Ragimbeau,Jessica Tapp, Fatu Djabi, Lise Pignon, Raoul Salmon, Virginie Jover, Elodie Solarino, Bernard Weber, Tamir Abdelrahman |
| EPI_ISL_3147680 | Centre Hospitalier du Nord | Laboratoire national de sante, Microbiology, Microbial Genomics Platform | Anke Wienecke-Baldacchino, Catherine Ragimbeau,Jessica Tapp, Fatu Djabi, Lise Pignon, Raoul Salmon, Virginie Jover, Elodie Solarino, Fatiha Boulmerka, Jacqueline Parmentier, Tamir Abdelrahman |
| EPI_ISL_3147681, EPI_ISL_3147682 | Laboratoire national de sante, Microbiology, Virology | Laboratoire national de sante, Microbiology, Microbial Genomics Platform | Anke Wienecke-Baldacchino, Catherine Ragimbeau,Jessica Tapp, Fatu Djabi, Lise Pignon, Raoul Salmon, Virginie Jover, Elodie Solarino, Trung Nguyen Nguyen, Tamir Abdelrahman |
| EPI_ISL_3147683 | Hospital Center Luxembourg | Laboratoire national de sante, Microbiology, Microbial Genomics Platform | Anke Wienecke-Baldacchino, Catherine Ragimbeau,Jessica Tapp, Fatu Djabi, Lise Pignon, Raoul Salmon, Virginie Jover, Elodie Solarino, Michel Kohnen, Jean-Hugues Francois, Tamir Abdelrahman |
| EPI_ISL_3147684, EPI_ISL_3147685, EPI_ISL_3147686 | Laboratoires Reunis | Laboratoire national de sante, Microbiology, Microbial Genomics Platform | Anke Wienecke-Baldacchino, Catherine Ragimbeau,Jessica Tapp, Fatu Djabi, Lise Pignon, Raoul Salmon, Virginie Jover, Elodie Solarino, Bernard Weber, Tamir Abdelrahman |
| EPI_ISL_3147687 | Centre Hospitalier du Nord | Laboratoire national de sante, Microbiology, Microbial Genomics Platform | Anke Wienecke-Baldacchino, Catherine Ragimbeau,Jessica Tapp, Fatu Djabi, Lise Pignon, Raoul Salmon, Virginie Jover, Elodie Solarino, Fatiha Boulmerka, Jacqueline Parmentier, Tamir Abdelrahman |
| EPI_ISL_3147688, EPI_ISL_3147689, EPI_ISL_3147690 | Laboratoires Reunis | Laboratoire national de sante, Microbiology, Microbial Genomics Platform | Anke Wienecke-Baldacchino, Catherine Ragimbeau,Jessica Tapp, Fatu Djabi, Lise Pignon, Raoul Salmon, Virginie Jover, Elodie Solarino, Bernard Weber, Tamir Abdelrahman |
| EPI_ISL_3147691, EPI_ISL_3147692 | Hospital Center Luxembourg | Laboratoire national de sante, Microbiology, Microbial Genomics Platform | Anke Wienecke-Baldacchino, Catherine Ragimbeau,Jessica Tapp, Fatu Djabi, Lise Pignon, Raoul Salmon, Virginie Jover, Elodie Solarino, Michel Kohnen, Jean-Hugues Francois, Tamir Abdelrahman |
| EPI_ISL_3147693, EPI_ISL_3147694, EPI_ISL_3147695 | Laboratoires Reunis | Laboratoire national de sante, Microbiology, Microbial Genomics Platform | Anke Wienecke-Baldacchino, Catherine Ragimbeau,Jessica Tapp, Fatu Djabi, Lise Pignon, Raoul Salmon, Virginie Jover, Elodie Solarino, Bernard Weber, Tamir Abdelrahman |
| EPI_ISL_3147696 | Hospital Center Luxembourg | Laboratoire national de sante, Microbiology, Microbial Genomics Platform | Anke Wienecke-Baldacchino, Catherine Ragimbeau,Jessica Tapp, Fatu Djabi, Lise Pignon, Raoul Salmon, Virginie Jover, Elodie Solarino, Michel Kohnen, Jean-Hugues Francois, Tamir Abdelrahman |
| EPI_ISL_3147697 | Laboratoire national de sante, Microbiology, Virology | Laboratoire national de sante, Microbiology, Microbial Genomics Platform | Anke Wienecke-Baldacchino, Catherine Ragimbeau,Jessica Tapp, Fatu Djabi, Lise Pignon, Raoul Salmon, Virginie Jover, Elodie Solarino, Trung Nguyen Nguyen, Tamir Abdelrahman |
| EPI_ISL_3147698 | Hospital Center Emile Mayrisch | Laboratoire national de sante, Microbiology, Microbial Genomics Platform | Anke Wienecke-Baldacchino, Catherine Ragimbeau,Jessica Tapp, Fatu Djabi, Lise Pignon, Raoul Salmon, Virginie Jover, Elodie Solarino, Cynthia Oxacelay, Tamir Abdelrahman |
| EPI_ISL_3147699 | Laboratoire national de sante, Microbiology, Virology | Laboratoire national de sante, Microbiology, Microbial Genomics Platform | Anke Wienecke-Baldacchino, Catherine Ragimbeau,Jessica Tapp, Fatu Djabi, Lise Pignon, Raoul Salmon, Virginie Jover, Elodie Solarino, Trung Nguyen Nguyen, Tamir Abdelrahman |
| EPI_ISL_3147700 | H⌠pitaux Robert Schuman | Laboratoire national de sante, Microbiology, Microbial Genomics Platform | Anke Wienecke-Baldacchino, Catherine Ragimbeau,Jessica Tapp, Fatu Djabi, Lise Pignon, Raoul Salmon, Virginie Jover, Elodie Solarino, Alain Hakim, Nawfal Faik, Tamir Abdelrahman |
| EPI_ISL_3147701 | Hospital Center Emile Mayrisch | Laboratoire national de sante, Microbiology, Microbial Genomics Platform | Anke Wienecke-Baldacchino, Catherine Ragimbeau,Jessica Tapp, Fatu Djabi, Lise Pignon, Raoul Salmon, Virginie Jover, Elodie Solarino, Cynthia Oxacelay, Tamir Abdelrahman |
| EPI_ISL_3147702 | Hospital Center Luxembourg | Laboratoire national de sante, Microbiology, Microbial Genomics Platform | Anke Wienecke-Baldacchino, Catherine Ragimbeau,Jessica Tapp, Fatu Djabi, Lise Pignon, Raoul Salmon, Virginie Jover, Elodie Solarino, Michel Kohnen, Jean-Hugues Francois, Tamir Abdelrahman |
| EPI_ISL_3147703 | H⌠pitaux Robert Schuman | Laboratoire national de sante, Microbiology, Microbial Genomics Platform | Anke Wienecke-Baldacchino, Catherine Ragimbeau,Jessica Tapp, Fatu Djabi, Lise Pignon, Raoul Salmon, Virginie Jover, Elodie Solarino, Alain Hakim, Nawfal Faik, Tamir Abdelrahman |
| EPI_ISL_3147704 | Hospital Center Emile Mayrisch | Laboratoire national de sante, Microbiology, Microbial Genomics Platform | Anke Wienecke-Baldacchino, Catherine Ragimbeau,Jessica Tapp, Fatu Djabi, Lise Pignon, Raoul Salmon, Virginie Jover, Elodie Solarino, Cynthia Oxacelay, Tamir Abdelrahman |
| EPI_ISL_3147705 | Centre Hospitalier du Nord | Laboratoire national de sante, Microbiology, Microbial Genomics Platform | Anke Wienecke-Baldacchino, Catherine Ragimbeau,Jessica Tapp, Fatu Djabi, Lise Pignon, Raoul Salmon, Virginie Jover, Elodie Solarino, Fatiha Boulmerka, Jacqueline Parmentier, Tamir Abdelrahman |

EPI_ISL_3147715, EPI_ISL_3147724, EPI_ISL_3147738, EPI_ISL_3147742, EPI_ISL_3147747, EPI_ISL_3147748, EPI_ISL_3147750, EPI_ISL_3147751, EPI_ISL_3147760, EPI_ISL_3147762, EPI_ISL_3147770, EPI_ISL_3147774, EPI_ISL_3147784, EPI_ISL_3147790, EPI_ISL_3147794, EPI_ISL_3147795, EPI_ISL_3147799, EPI_ISL_3147804, EPI_ISL_3147811, EPI_ISL_3147834, EPI_ISL_3147835, EPI_ISL_3147836,
EPI_ISL_3147837, EPI_ISL_3147841, EPI_ISL_3147842, EPI_ISL_3147843, EPI_ISL_3147844, EPI_ISL_3147847, EPI_ISL_3147848, EPI_ISL_3147849, EPI_ISL_3147852, EPI_ISL_3147853, EPI_ISL_3147854, EPI_ISL_3147855, EPI_ISL_3147856, EPI_ISL_3147857, EPI_ISL_3147859, EPI_ISL_3147860, EPI_ISL_3147861, EPI_ISL_3147862, EPI_ISL_3147863, EPI_ISL_3147864, EPI_ISL_3147865, EPI_ISL_3147866,

| | | | |
|---|---|---|---|
| EPI_ISL_3147867, EPI_ISL_3147868, EPI_ISL_3147869, EPI_ISL_3147870, EPI_ISL_3148064, EPI_ISL_3148067, EPI_ISL_3148075, EPI_ISL_3148083, EPI_ISL_3148092, EPI_ISL_3148099, EPI_ISL_3148103, EPI_ISL_3148106, EPI_ISL_3148114, EPI_ISL_3148119, EPI_ISL_3148130, EPI_ISL_3148299, EPI_ISL_3148300, EPI_ISL_3148301, EPI_ISL_3148302, EPI_ISL_3148310, EPI_ISL_3148311, EPI_ISL_3148358, EPI_ISL_3148359, EPI_ISL_3148360, EPI_ISL_3148361, EPI_ISL_3148362, EPI_ISL_3148363, EPI_ISL_3148364, EPI_ISL_3148365, EPI_ISL_3148366, EPI_ISL_3148367, EPI_ISL_3148368, EPI_ISL_3148369, EPI_ISL_3148370, EPI_ISL_3148371, EPI_ISL_3148372, EPI_ISL_3148373, EPI_ISL_3148374, EPI_ISL_3148375, EPI_ISL_3148376, EPI_ISL_3148377, EPI_ISL_3148378, EPI_ISL_3148379, EPI_ISL_3148380, EPI_ISL_3148381, EPI_ISL_3148382, EPI_ISL_3148383, EPI_ISL_3148384, EPI_ISL_3148385, EPI_ISL_3148386, EPI_ISL_3148387, EPI_ISL_3148388, EPI_ISL_3148389, EPI_ISL_3148390, EPI_ISL_3148391, EPI_ISL_3148392, EPI_ISL_3148393, EPI_ISL_3148394, EPI_ISL_3148395, EPI_ISL_3148396, EPI_ISL_3148397, EPI_ISL_3148398, EPI_ISL_3148399, EPI_ISL_3148400, EPI_ISL_3148401, EPI_ISL_3148402, EPI_ISL_3148403, EPI_ISL_3148404, EPI_ISL_3148405, EPI_ISL_3148406, EPI_ISL_3148407, EPI_ISL_3148408, EPI_ISL_3148409, EPI_ISL_3148410, EPI_ISL_3148411, EPI_ISL_3148412, EPI_ISL_3148413, EPI_ISL_3148414, EPI_ISL_3148415, EPI_ISL_3148416, EPI_ISL_3148417, EPI_ISL_3148418, EPI_ISL_3148419, EPI_ISL_3148420, EPI_ISL_3148421, EPI_ISL_3148422, EPI_ISL_3148423, EPI_ISL_3148424, EPI_ISL_3148425, EPI_ISL_3148426, EPI_ISL_3148427, EPI_ISL_3148428, EPI_ISL_3148429, EPI_ISL_3148430, EPI_ISL_3148431, EPI_ISL_3148432, EPI_ISL_3148433, EPI_ISL_3148434, EPI_ISL_3148435, EPI_ISL_3148436, EPI_ISL_3148437, EPI_ISL_3148438, EPI_ISL_3148439, EPI_ISL_3148440 | | | |
| see above | Laboratoires d'analyses medicales - Ketterthill | Laboratoire national de sante, Microbiology, Microbial Genomics Platform | Anke Wienecke-Baldacchino, Catherine Ragimbeau,Jessica Tapp, Fatu Djabi, Lise Pignon, Raoul Salmon, Virginie Jover, Elodie Solarino, Serge Vedy, Caroline Scheiber, Tamir Abdelrahman |
| EPI_ISL_3148580 | Dutch COVID-19 response team | Erasmus Medical Center | Bas Oude Munnink, Reina Sikkema, David Nieuwenhuijse, Irina Chestakova, Anne van der Linden, Marjan Boter, Emmanuelle Munger, Corine GeurtsvanKessel, Annemiek van der Eijk, Richard Molenkamp, Marion Koopmans, on behalf of the Dutch national COVID-19 response team. |
| EPI_ISL_3148713, EPI_ISL_3148714, EPI_ISL_3148716, EPI_ISL_3148717, EPI_ISL_3148719, EPI_ISL_3148720, EPI_ISL_3148722, EPI_ISL_3148723, EPI_ISL_3148725, EPI_ISL_3148726, EPI_ISL_3148728, EPI_ISL_3148729, EPI_ISL_3148731, EPI_ISL_3148732 | | | |
| see above | SYNLAB | GIGA Medical Genomics | Keith Durkin, Maria Artesi, Bouchra Boujemla, Nathalie Renotte, Nadine Cambisano, Sébastien Bontems, Cécile Meex, Claire Gourzonès, Olivier Ek, Laurent Gillet, Marie-Pierre Hayette, Vincent Bours |
| EPI_ISL_3148734 | Department of Clinical Microbiology | GIGA Medical Genomics | Keith Durkin, Maria Artesi, Bouchra Boujemla, Nathalie Renotte, Nadine Cambisano, Sébastien Bontems, Cécile Meex, Claire Gourzonès, Olivier Ek, Laurent Gillet, Marie-Pierre Hayette, Vincent Bours |
| EPI_ISL_3148735, EPI_ISL_3148737, EPI_ISL_3148738, EPI_ISL_3148740, EPI_ISL_3148742, EPI_ISL_3148743, EPI_ISL_3148745, EPI_ISL_3148746, EPI_ISL_3148747 | SYNLAB | GIGA Medical Genomics | Keith Durkin, Maria Artesi, Bouchra Boujemla, Nathalie Renotte, Nadine Cambisano, Sébastien Bontems, Cécile Meex, Claire Gourzonès, Olivier Ek, Laurent Gillet, Marie-Pierre Hayette, Vincent Bours |
| EPI_ISL_3148749, EPI_ISL_3148750, EPI_ISL_3148752, EPI_ISL_3148753, EPI_ISL_3148755, EPI_ISL_3148756, EPI_ISL_3148758, EPI_ISL_3148759, EPI_ISL_3148761 | Department of Clinical Microbiology | GIGA Medical Genomics | Keith Durkin, Maria Artesi, Bouchra Boujemla, Nathalie Renotte, Nadine Cambisano, Sébastien Bontems, Cécile Meex, Claire Gourzonès, Olivier Ek, Laurent Gillet, Marie-Pierre Hayette, Vincent Bours |
| EPI_ISL_3148762, EPI_ISL_3148764 | SYNLAB | GIGA Medical Genomics | Keith Durkin, Maria Artesi, Bouchra Boujemla, Nathalie Renotte, Nadine Cambisano, Sébastien Bontems, Cécile Meex, Claire Gourzonès, Olivier Ek, Laurent Gillet, Marie-Pierre Hayette, Vincent Bours |
| EPI_ISL_3148765 | Clinique André Renard Herstal | GIGA Medical Genomics | Keith Durkin, Maria Artesi, Bouchra Boujemla, Nathalie Renotte, Nadine Cambisano, Sébastien Bontems, Cécile Meex, Claire Gourzonès, Olivier Ek, Laurent Gillet, Marie-Pierre Hayette, Vincent Bours |
| EPI_ISL_3148767, EPI_ISL_3148768, EPI_ISL_3148770, EPI_ISL_3148771, EPI_ISL_3148773, EPI_ISL_3148774, EPI_ISL_3148775, EPI_ISL_3148777 | Department of Clinical Microbiology | GIGA Medical Genomics | Keith Durkin, Maria Artesi, Bouchra Boujemla, Nathalie Renotte, Nadine Cambisano, Sébastien Bontems, Cécile Meex, Claire Gourzonès, Olivier Ek, Laurent Gillet, Marie-Pierre Hayette, Vincent Bours |
| EPI_ISL_3148778, EPI_ISL_3148780, EPI_ISL_3148781, EPI_ISL_3148782, EPI_ISL_3148784, EPI_ISL_3148786 | SYNLAB | GIGA Medical Genomics | Keith Durkin, Maria Artesi, Bouchra Boujemla, Nathalie Renotte, Nadine Cambisano, Sébastien Bontems, Cécile Meex, Claire Gourzonès, Olivier Ek, Laurent Gillet, Marie-Pierre Hayette, Vincent Bours |
| EPI_ISL_3148787 | Clinique St Joseph Saint Vith | GIGA Medical Genomics | Keith Durkin, Maria Artesi, Bouchra Boujemla, Nathalie Renotte, Nadine Cambisano, Sébastien Bontems, Cécile Meex, Claire Gourzonès, Olivier Ek, Laurent Gillet, Marie-Pierre Hayette, Vincent Bours |
| EPI_ISL_3148789 | Department of Clinical Microbiology | GIGA Medical Genomics | Keith Durkin, Maria Artesi, Bouchra Boujemla, Nathalie Renotte, Nadine Cambisano, Sébastien Bontems, Cécile Meex, Claire Gourzonès, Olivier Ek, Laurent Gillet, Marie-Pierre Hayette, Vincent Bours |
| EPI_ISL_3148790, EPI_ISL_3148791, EPI_ISL_3148793, EPI_ISL_3148794, EPI_ISL_3148796, EPI_ISL_3148797, EPI_ISL_3148799 | SYNLAB | GIGA Medical Genomics | Keith Durkin, Maria Artesi, Bouchra Boujemla, Nathalie Renotte, Nadine Cambisano, Sébastien Bontems, Cécile Meex, Claire Gourzonès, Olivier Ek, Laurent Gillet, Marie-Pierre Hayette, Vincent Bours |
| EPI_ISL_3148800 | C.H.C MontLégia | GIGA Medical Genomics | Keith Durkin, Maria Artesi, Bouchra Boujemla, Nathalie Renotte, Nadine Cambisano, Sébastien Bontems, Cécile Meex, Claire Gourzonès, Olivier Ek, Laurent Gillet, Marie-Pierre Hayette, Vincent Bours |
| EPI_ISL_3148802 | SYNLAB | GIGA Medical Genomics | Keith Durkin, Maria Artesi, Bouchra Boujemla, Nathalie Renotte, Nadine Cambisano, Sébastien Bontems, Cécile Meex, Claire Gourzonès, Olivier Ek, Laurent Gillet, Marie-Pierre Hayette, Vincent Bours |
| EPI_ISL_3148803 | Department of Clinical Microbiology | GIGA Medical Genomics | Keith Durkin, Maria Artesi, Bouchra Boujemla, Nathalie Renotte, Nadine Cambisano, Sébastien Bontems, Cécile Meex, Claire Gourzonès, Olivier Ek, Laurent Gillet, Marie-Pierre Hayette, Vincent Bours |
| EPI_ISL_3148804 | SYNLAB | GIGA Medical Genomics | Keith Durkin, Maria Artesi, Bouchra Boujemla, Nathalie Renotte, Nadine Cambisano, Sébastien Bontems, Cécile Meex, Claire Gourzonès, Olivier Ek, Laurent Gillet, Marie-Pierre Hayette, Vincent Bours |
| EPI_ISL_3148809, EPI_ISL_3148815, EPI_ISL_3148816, EPI_ISL_3148817, EPI_ISL_3148819 | Department of Clinical Microbiology | GIGA Medical Genomics | Keith Durkin, Maria Artesi, Bouchra Boujemla, Nathalie Renotte, Nadine Cambisano, Sébastien Bontems, Cécile Meex, Claire Gourzonès, Olivier Ek, Laurent Gillet, Marie-Pierre Hayette, Vincent Bours |
| EPI_ISL_3148820 | C.H.C MontLégia | GIGA Medical Genomics | Keith Durkin, Maria Artesi, Bouchra Boujemla, Nathalie Renotte, Nadine Cambisano, Sébastien Bontems, Cécile Meex, Claire Gourzonès, Olivier Ek, Laurent Gillet, Marie-Pierre Hayette, Vincent Bours |
| EPI_ISL_3148822, EPI_ISL_3148823, EPI_ISL_3148824 | SYNLAB | GIGA Medical Genomics | Keith Durkin, Maria Artesi, Bouchra Boujemla, Nathalie Renotte, Nadine Cambisano, Sébastien Bontems, Cécile Meex, Claire Gourzonès, Olivier Ek, Laurent Gillet, Marie-Pierre Hayette, Vincent Bours |
| EPI_ISL_3148826, EPI_ISL_3148827 | Department of Clinical Microbiology | GIGA Medical Genomics | Keith Durkin, Maria Artesi, Bouchra Boujemla, Nathalie Renotte, Nadine Cambisano, Sébastien Bontems, Cécile Meex, Claire Gourzonès, Olivier Ek, Laurent Gillet, Marie-Pierre Hayette, Vincent Bours |
| EPI_ISL_3148829, EPI_ISL_3148830, EPI_ISL_3148832, EPI_ISL_3148834 | SYNLAB | GIGA Medical Genomics | Keith Durkin, Maria Artesi, Bouchra Boujemla, Nathalie Renotte, Nadine Cambisano, Sébastien Bontems, Cécile Meex, Claire Gourzonès, Olivier Ek, Laurent Gillet, Marie-Pierre Hayette, Vincent Bours |
| EPI_ISL_3148836 | Department of Clinical Microbiology | GIGA Medical Genomics | Keith Durkin, Maria Artesi, Bouchra Boujemla, Nathalie Renotte, Nadine Cambisano, Sébastien Bontems, Cécile Meex, Claire Gourzonès, Olivier Ek, Laurent Gillet, Marie-Pierre Hayette, Vincent Bours |
| EPI_ISL_3148876 | Tivoli Fondazione Policlinico Universitario Agostino Gemelli IRCCS | Fondazione Policlinico Universitario Agostino Gemelli | Istituto di Microbiologia e Virologia |
| EPI_ISL_3148885, EPI_ISL_3148887, EPI_ISL_3148889, EPI_ISL_3148891, EPI_ISL_3148893, EPI_ISL_3148896, EPI_ISL_3148898, EPI_ISL_3148900, EPI_ISL_3148902, EPI_ISL_3148904, EPI_ISL_3148906, EPI_ISL_3148908, EPI_ISL_3148910, EPI_ISL_3148913, EPI_ISL_3148915, EPI_ISL_3148917, EPI_ISL_3148919, EPI_ISL_3148922, EPI_ISL_3148924, EPI_ISL_3148926, EPI_ISL_3148928 | | | |
| see above | Ochsner Health | Bioinfoexperts, LLC | Amy Feehan, Judy Crabtree, Rebecca Rose, Ben Lain, Chris Huston, Samual Moot, David J. Nolan, Tessa LaFleur, Julia-Garcia-Diaz, Lucio Miele, Susanna L. Lamers |
| EPI_ISL_3148944 | URMC LABS | Wadsworth Center, New York State Department of Health | Kirsten St. George, Daryl M. Lamson, Alexis Russell, Matthew Shudt, Melissa A Leisner, Jonathan Plitnick, Catharine Prussing, Navjot Singh, John Kelly, Erasmus Schneider, Erica Lasek-Nesselquist |
| EPI_ISL_3148947 | BOSTON HEART DIAGNOSTICS CORP | Wadsworth Center, New York State Department of Health | Kirsten St. George, Daryl M. Lamson, Alexis Russell, Matthew Shudt, Melissa A Leisner, Jonathan Plitnick, Catharine Prussing, Navjot Singh, John Kelly, Erasmus Schneider, Erica Lasek-Nesselquist |
| EPI_ISL_3148948, EPI_ISL_3148949, EPI_ISL_3148950, EPI_ISL_3148951, EPI_ISL_3148952, EPI_ISL_3148954, EPI_ISL_3148955, EPI_ISL_3148963 | URMC LABS | Wadsworth Center, New York State Department of Health | Kirsten St. George, Daryl M. Lamson, Alexis Russell, Matthew Shudt, Melissa A Leisner, Jonathan Plitnick, Catharine Prussing, Navjot Singh, John Kelly, Erasmus Schneider, Erica Lasek-Nesselquist |
| EPI_ISL_3148965 | Saratoga Hospital Laboratory | Wadsworth Center, New York State Department of Health | Kirsten St. George, Daryl M. Lamson, Alexis Russell, Matthew Shudt, Melissa A Leisner, Jonathan Plitnick, Catharine Prussing, Navjot Singh, John Kelly, Erasmus Schneider, Erica Lasek-Nesselquist |
| EPI_ISL_3148966, EPI_ISL_3148967, EPI_ISL_3148968, EPI_ISL_3148969, | URMC LABS | Wadsworth Center, New York State Department of Health | Kirsten St. George, Daryl M. Lamson, Alexis Russell, Matthew Shudt, Melissa A Leisner, Jonathan Plitnick, Catharine Prussing, Navjot Singh, John Kelly, Erasmus Schneider, Erica Lasek-Nesselquist |

| | | | |
|---|---|---|---|
| EPI_ISL_3148970, EPI_ISL_3148971 | | | |
| EPI_ISL_3148972 | Wadsworth Center, New York State Department of Health | Wadsworth Center, New York State Department of Health | Kirsten St. George, Daryl M. Lamson, Alexis Russell, Matthew Shudt, Melissa A Leisner, Jonathan Plitnick, Catharine Prussing, Navjot Singh, John Kelly, Erasmus Schneider, Erica Lasek-Nesselquist |
| EPI_ISL_3148973 | ALBANY MEDICAL CENTER | Wadsworth Center, New York State Department of Health | Kirsten St. George, Daryl M. Lamson, Alexis Russell, Matthew Shudt, Melissa A Leisner, Jonathan Plitnick, Catharine Prussing, Navjot Singh, John Kelly, Erasmus Schneider, Erica Lasek-Nesselquist |
| EPI_ISL_3148974 | Wadsworth Center, New York State Department of Health | Wadsworth Center, New York State Department of Health | Kirsten St. George, Daryl M. Lamson, Alexis Russell, Matthew Shudt, Melissa A Leisner, Jonathan Plitnick, Catharine Prussing, Navjot Singh, John Kelly, Erasmus Schneider, Erica Lasek-Nesselquist |
| EPI_ISL_3148975 | Saratoga Hospital Laboratory | Wadsworth Center, New York State Department of Health | Kirsten St. George, Daryl M. Lamson, Alexis Russell, Matthew Shudt, Melissa A Leisner, Jonathan Plitnick, Catharine Prussing, Navjot Singh, John Kelly, Erasmus Schneider, Erica Lasek-Nesselquist |
| EPI_ISL_3148976 | BOSTON HEART DIAGNOSTICS CORP | Wadsworth Center, New York State Department of Health | Kirsten St. George, Daryl M. Lamson, Alexis Russell, Matthew Shudt, Melissa A Leisner, Jonathan Plitnick, Catharine Prussing, Navjot Singh, John Kelly, Erasmus Schneider, Erica Lasek-Nesselquist |
| EPI_ISL_3148977, EPI_ISL_3148978 | ALBANY MEDICAL CENTER | Wadsworth Center, New York State Department of Health | Kirsten St. George, Daryl M. Lamson, Alexis Russell, Matthew Shudt, Melissa A Leisner, Jonathan Plitnick, Catharine Prussing, Navjot Singh, John Kelly, Erasmus Schneider, Erica Lasek-Nesselquist |
| EPI_ISL_3148979, EPI_ISL_3148980, EPI_ISL_3148981, EPI_ISL_3148982, EPI_ISL_3148983, EPI_ISL_3148984, EPI_ISL_3148985, EPI_ISL_3148986, EPI_ISL_3148987, EPI_ISL_3148988, EPI_ISL_3148989, EPI_ISL_3148990 | | | |
| see above | URMC LABS | Wadsworth Center, New York State Department of Health | Kirsten St. George, Daryl M. Lamson, Alexis Russell, Matthew Shudt, Melissa A Leisner, Jonathan Plitnick, Catharine Prussing, Navjot Singh, John Kelly, Erasmus Schneider, Erica Lasek-Nesselquist |
| EPI_ISL_3148994, EPI_ISL_3148996, EPI_ISL_3148997, EPI_ISL_3148998, EPI_ISL_3148999, EPI_ISL_3149000, EPI_ISL_3149001, EPI_ISL_3149002, EPI_ISL_3149003, EPI_ISL_3149004, EPI_ISL_3149005, EPI_ISL_3149006, EPI_ISL_3149007, EPI_ISL_3149008, EPI_ISL_3149009, EPI_ISL_3149010, EPI_ISL_3149011, EPI_ISL_3149012, EPI_ISL_3149014, EPI_ISL_3149016, EPI_ISL_3149023, EPI_ISL_3149025, EPI_ISL_3149026, EPI_ISL_3149027, EPI_ISL_3149030, EPI_ISL_3149033, EPI_ISL_3149034, EPI_ISL_3149036, EPI_ISL_3149039, EPI_ISL_3149040, EPI_ISL_3149041, EPI_ISL_3149043, EPI_ISL_3149044, EPI_ISL_3149045, EPI_ISL_3149046, EPI_ISL_3149047, EPI_ISL_3149048, EPI_ISL_3149049, EPI_ISL_3149050, EPI_ISL_3149051, EPI_ISL_3149052, EPI_ISL_3149053, EPI_ISL_3149054, EPI_ISL_3149055, EPI_ISL_3149056, EPI_ISL_3149058, EPI_ISL_3149059, EPI_ISL_3149060, EPI_ISL_3149061, EPI_ISL_3149062, EPI_ISL_3149064, EPI_ISL_3149065, EPI_ISL_3149066, EPI_ISL_3149067, EPI_ISL_3149068, EPI_ISL_3149069, EPI_ISL_3149070 | | | |
| see above | Texas Children's Hospital | Texas Children's Microbiome Center | Ruth Ann Luna, Jennifer K. Spinler, James Dunn, James Versalovic, Ila Singh |
| EPI_ISL_3149241, EPI_ISL_3149242, EPI_ISL_3149243, EPI_ISL_3149244, EPI_ISL_3149245, EPI_ISL_3149246, EPI_ISL_3149247, EPI_ISL_3149248, EPI_ISL_3149249, EPI_ISL_3149250, EPI_ISL_3149251, EPI_ISL_3149252, EPI_ISL_3149253, EPI_ISL_3149254, EPI_ISL_3149255, EPI_ISL_3149256, EPI_ISL_3149257, EPI_ISL_3149258, EPI_ISL_3149259, EPI_ISL_3149260 | | | |
| see above | State Hygienic Laboratory at the University of Iowa | State Hygienic Laboratory at the University of Iowa | Valerie Reeb, Erik Twait, Anna Yakos, Mariah Knutson, Kris Eveland, Kristen Zanon, Jeff, Benfer, Davis Rieckenberg, Wes Hottel, Alankar Kampoowale |
| EPI_ISL_3149261 | University of Iowa Hospitals and Clinics | State Hygienic Laboratory at the University of Iowa | Valerie Reeb, Erik Twait, Anna Yakos, Mariah Knutson, Kris Eveland, Kristen Zanon, Jeff, Benfer, Davis Rieckenberg, Wes Hottel, Alankar Kampoowale |
| EPI_ISL_3149262, EPI_ISL_3149263, EPI_ISL_3149265 | State Hygienic Laboratory at the University of Iowa | State Hygienic Laboratory at the University of Iowa | Valerie Reeb, Erik Twait, Anna Yakos, Mariah Knutson, Kris Eveland, Kristen Zanon, Jeff, Benfer, Davis Rieckenberg, Wes Hottel, Alankar Kampoowale |
| EPI_ISL_3149266, EPI_ISL_3149267, EPI_ISL_3149268, EPI_ISL_3149269, EPI_ISL_3149270, EPI_ISL_3149271, EPI_ISL_3149272, EPI_ISL_3149273, EPI_ISL_3149274, EPI_ISL_3149275, EPI_ISL_3149276, EPI_ISL_3149277, EPI_ISL_3149278, EPI_ISL_3149279, EPI_ISL_3149280, EPI_ISL_3149281, EPI_ISL_3149282, EPI_ISL_3149283, EPI_ISL_3149284, EPI_ISL_3149285, EPI_ISL_3149286, EPI_ISL_3149287, EPI_ISL_3149288, EPI_ISL_3149289, EPI_ISL_3149290, EPI_ISL_3149291, EPI_ISL_3149292, EPI_ISL_3149293, EPI_ISL_3149294, EPI_ISL_3149295, EPI_ISL_3149296, EPI_ISL_3149297 | | | |
| see above | Kansas Health and Environmental Lab | Kansas Health and Environmental Lab | Mike Grose, Katherine Wiggins, Jonathan Barnell, Ben Olsen, and Phil Adam |
| EPI_ISL_3149298 | LEBOWAKGOMO | National Institute for Communicable Diseases of the National Health Laboratory Service | Amoako DG, Everatt J, Scheepers C, Mohale T, Ntuli N, Mahlangu B, Mnguni A, Ismail A, Bhiman JN |
| EPI_ISL_3149308, EPI_ISL_3149310, EPI_ISL_3149311 | MANKWENG LABORATORY | National Institute for Communicable Diseases of the National Health Laboratory Service | Amoako DG, Everatt J, Scheepers C, Mohale T, Ntuli N, Mahlangu B, Mnguni A, Ismail A, Bhiman JN |
| EPI_ISL_3149322 | LANIIA | LANIIA-Nayarit | Consorcio Mexicano de Vigilancia Genómica (CoViGen-Mex). Authors (in alphabetical order): Julio Elias Alvarado-Yaah, Carlos F. Arias, Santiago Ávila-Ríos, Eduardo Becerril-Vargas, Víctor Hugo Borja-Aburto, Celia Boukadida, Cristóbal Cháidez-Quiróz, Juan Bautista Chale-Dzul, Ricardo Ciria Merce, Andreu Comas-García, Célida Duque Molina, Julissa Enciso-Ibarra, José Antonio Enciso-Moreno, Gloria Elena Espinosa-Ayala, Fernando Fontove-Herrera, Daniel Fregoso-Rueda, Victor Eduardo García-Arias, Alejandra García-Gasca, Bruno Gómez-Gil, Jean Pierre González, Irvin González-López, Concepción Grajales-Muñiz, Ricardo Grande, Rosa María Gutiérrez Rios, Jesús Hernández, Alejandra Hernández-Terán, Alfredo Herrera-Estrella, Carla Ivón Herrera-Najera, Pavel Isa, Verónica Mata-Haro, Daniel Lira Morales, Susana Lopez, Antonio Loza Román, Brenda Irasema Maldonado-Meza, Bernardo Martínez-Miguel, José Arturo Martínez-Orozco, Célida Martínez-Rodríguez, Margarita Matías-Florentino, Fidencio Mejía-Nepomuceno, María Guadalupe de Jesús Mireles-Rivera, Gloria María Molina-Salinas, Hector Montoya-Fuentes, , Mario Mújica-Sánchez, José Esteban Muñoz-Medina, José de Jesús Nuñez-Contreras, Alicia Ocaña-Mondragón, Luis Alberto Ochoa-Carrera, Hector Esteban Paz-Juárez, Marissa Perez-Garcia, Francisco Pulido, Helen Haydee Fernanda Ramírez-Plascencia, Jorge Salas-Hernández, Angel Gustavo Salas-Lais, Alejandro Sánchez-Flores, Clara Esperanza Santacruz-Tinoco, María Guadalupe Santiago-Mauricio, Selene Zárate, Nelly Sélem-Mojica, Blanca Taboada, Gloria Vazquez, Joel Armando Vázquez-Pérez. LANNIA Nayarit: Manuel Ivan Girón Pérez, Carlos Eduardo Covantes-Rosales, Daniel Alberto Girón Pérez, Ulises Mercado. |
| EPI_ISL_3149323 | Hospital Sharp | Microbial Genomics Laboratory | Consorcio Mexicano de Vigilancia Genómica (CoViGen-Mex). Authors (in alphabetical order): Julio Elias Alvarado-Yaah, Carlos F. Arias, Santiago Ávila-Ríos, Eduardo Becerril-Vargas, Víctor Hugo Borja-Aburto, Celia Boukadida, Cristóbal Cháidez-Quiróz, Juan Bautista Chale-Dzul, Ricardo Ciria Merce, Andreu Comas-García, Célida Duque Molina, Julissa Enciso-Ibarra, José Antonio Enciso-Moreno, Gloria Elena Espinosa-Ayala, Fernando Fontove-Herrera, Daniel Fregoso-Rueda, Victor Eduardo García-Arias, Alejandra García-Gasca, Bruno Gómez-Gil, Jean Pierre González, Irvin González-López, Concepción Grajales-Muñiz, Ricardo Grande, Rosa María Gutiérrez Rios, Jesús Hernández, Alejandra Hernández-Terán, Alfredo Herrera-Estrella, Carla Ivón Herrera-Najera, Pavel Isa, Verónica Mata-Haro, Daniel Lira Morales, Susana Lopez, Antonio Loza Román, Brenda Irasema Maldonado-Meza, Bernardo Martínez-Miguel, José Arturo Martínez-Orozco, Célida Martínez-Rodríguez, Margarita Matías-Florentino, Fidencio Mejía-Nepomuceno, María Guadalupe de Jesús Mireles-Rivera, Gloria María Molina-Salinas, Hector Montoya-Fuentes, , Mario Mújica-Sánchez, José Esteban Muñoz-Medina, José de Jesús Nuñez-Contreras, Alicia Ocaña-Mondragón, Luis Alberto Ochoa-Carrera, Hector Esteban Paz-Juárez, Marissa Perez-Garcia, Francisco Pulido, Helen Haydee Fernanda Ramírez-Plascencia, Jorge Salas-Hernández, Angel Gustavo Salas-Lais, Alejandro Sánchez-Flores, Clara Esperanza Santacruz-Tinoco, María Guadalupe Santiago-Mauricio, Selene Zárate, Nelly Sélem-Mojica, Blanca Taboada, Gloria Vazquez, Joel Armando Vázquez-Pérez. |
| EPI_ISL_3149324 | LANIIA | LANIIA-Nayarit | Consorcio Mexicano de Vigilancia Genómica (CoViGen-Mex). Authors (in alphabetical order): Julio Elias Alvarado-Yaah, Carlos F. Arias, Santiago Ávila-Ríos, Eduardo Becerril-Vargas, Víctor Hugo Borja-Aburto, Celia Boukadida, Cristóbal Cháidez-Quiróz, Juan Bautista Chale-Dzul, Ricardo Ciria Merce, Andreu Comas-García, Célida Duque Molina, Julissa Enciso-Ibarra, José Antonio Enciso-Moreno, Gloria Elena Espinosa-Ayala, Fernando Fontove-Herrera, Daniel Fregoso-Rueda, Victor Eduardo García-Arias, Alejandra García-Gasca, Bruno Gómez-Gil, Jean Pierre González, Irvin González-López, Concepción Grajales-Muñiz, Ricardo Grande, Rosa María Gutiérrez Rios, Jesús Hernández, Alejandra Hernández-Terán, Alfredo Herrera-Estrella, Carla Ivón Herrera-Najera, Pavel Isa, Verónica Mata-Haro, Daniel Lira Morales, Susana Lopez, Antonio Loza Román, Brenda Irasema Maldonado-Meza, Bernardo Martínez-Miguel, José Arturo Martínez-Orozco, Célida Martínez-Rodríguez, Margarita Matías-Florentino, Fidencio Mejía-Nepomuceno, María Guadalupe de Jesús Mireles-Rivera, Gloria María Molina-Salinas, Hector Montoya-Fuentes, , Mario Mújica-Sánchez, José Esteban Muñoz-Medina, José de Jesús Nuñez-Contreras, Alicia Ocaña-Mondragón, Luis Alberto Ochoa-Carrera, Hector Esteban Paz-Juárez, Marissa Perez-Garcia, Francisco Pulido, Helen Haydee Fernanda Ramírez-Plascencia, Jorge Salas-Hernández, Angel Gustavo Salas-Lais, Alejandro Sánchez-Flores, Clara Esperanza Santacruz-Tinoco, María Guadalupe Santiago-Mauricio, Selene Zárate, Nelly Sélem-Mojica, Blanca Taboada, Gloria Vazquez, Joel Armando Vázquez-Pérez. LANNIA Nayarit: Manuel Ivan Girón Pérez, Carlos Eduardo Covantes-Rosales, Daniel Alberto Girón Pérez, Ulises Mercado. |
| EPI_ISL_3149325, EPI_ISL_3149326 | Hospital Margarita Maza de Juárez | Microbial Genomics Laboratory | Consorcio Mexicano de Vigilancia Genómica (CoViGen-Mex). Authors (in alphabetical order): Julio Elias Alvarado-Yaah, Carlos F. Arias, Santiago Ávila-Ríos, Eduardo Becerril-Vargas, Víctor Hugo Borja-Aburto, Celia Boukadida, Cristóbal Cháidez-Quiróz, Juan Bautista Chale-Dzul, Ricardo Ciria Merce, Andreu Comas-García, Célida Duque Molina, Julissa Enciso-Ibarra, José Antonio Enciso-Moreno, Gloria Elena Espinosa-Ayala, Fernando Fontove-Herrera, Daniel Fregoso-Rueda, Victor Eduardo García-Arias, Alejandra García-Gasca, Bruno Gómez-Gil, Jean Pierre González, Irvin González-López, Concepción Grajales-Muñiz, Ricardo Grande, Rosa María Gutiérrez Rios, Jesús Hernández, Alejandra Hernández-Terán, Alfredo Herrera-Estrella, Carla Ivón Herrera-Najera, Pavel Isa, Verónica Mata-Haro, Daniel Lira Morales, Susana Lopez, Antonio Loza Román, Brenda Irasema Maldonado-Meza, Bernardo Martínez-Miguel, José Arturo Martínez-Orozco, Célida Martínez-Rodríguez, Margarita Matías-Florentino, Fidencio Mejía-Nepomuceno, María Guadalupe de Jesús Mireles-Rivera, Gloria María Molina-Salinas, Hector Montoya-Fuentes, , Mario Mújica-Sánchez, José Esteban Muñoz-Medina, José de Jesús Nuñez-Contreras, Alicia Ocaña-Mondragón, Luis Alberto Ochoa-Carrera, Hector Esteban Paz-Juárez, Marissa Perez-Garcia, Francisco Pulido, Helen Haydee Fernanda Ramírez-Plascencia, Jorge Salas-Hernández, Angel Gustavo Salas-Lais, Alejandro Sánchez-Flores, Clara Esperanza Santacruz-Tinoco, María Guadalupe Santiago-Mauricio, Selene Zárate, Nelly Sélem-Mojica, Blanca Taboada, Gloria Vazquez, Joel Armando Vázquez-Pérez. |
| EPI_ISL_3149327 | Hospital Sharp | Microbial Genomics Laboratory | Consorcio Mexicano de Vigilancia Genómica (CoViGen-Mex). Authors (in alphabetical order): Julio Elias Alvarado-Yaah, Carlos F. Arias, Santiago Ávila-Ríos, Eduardo Becerril-Vargas, Víctor Hugo Borja-Aburto, Celia Boukadida, Cristóbal Cháidez-Quiróz, Juan Bautista Chale-Dzul, Ricardo Ciria Merce, Andreu Comas-García, Célida Duque Molina, Julissa Enciso-Ibarra, José Antonio Enciso-Moreno, Gloria Elena Espinosa-Ayala, Fernando Fontove-Herrera, Daniel Fregoso-Rueda, Victor Eduardo García-Arias, Alejandra García-Gasca, Bruno Gómez-Gil, Jean Pierre González, Irvin González-López, Concepción Grajales-Muñiz, Ricardo Grande, Rosa María Gutiérrez Rios, Jesús Hernández, Alejandra Hernández-Terán, Alfredo Herrera-Estrella, Carla Ivón Herrera-Najera, Pavel Isa, Verónica Mata-Haro, Daniel Lira Morales, Susana Lopez, Antonio Loza Román, Brenda Irasema Maldonado-Meza, Bernardo Martínez-Miguel, José Arturo Martínez-Orozco, Célida Martínez-Rodríguez, Margarita Matías-Florentino, Fidencio Mejía-Nepomuceno, María Guadalupe de Jesús Mireles-Rivera, Gloria María Molina-Salinas, Hector Montoya-Fuentes, , Mario Mújica-Sánchez, José Esteban Muñoz-Medina, José de Jesús Nuñez-Contreras, Alicia Ocaña-Mondragón, Luis Alberto Ochoa-Carrera, Hector Esteban Paz-Juárez, Marissa Perez-Garcia, Francisco Pulido, Helen Haydee Fernanda Ramírez-Plascencia, Jorge Salas-Hernández, Angel Gustavo Salas-Lais, Alejandro Sánchez-Flores, Clara Esperanza Santacruz-Tinoco, María Guadalupe Santiago-Mauricio, Selene Zárate, Nelly Sélem-Mojica, Blanca Taboada, Gloria Vazquez, Joel Armando Vázquez-Pérez. |
| EPI_ISL_3149328, EPI_ISL_3149329 | Imágenes Diagnósticas de Mazatlán | Microbial Genomics Laboratory | Consorcio Mexicano de Vigilancia Genómica (CoViGen-Mex). Authors (in alphabetical order): Julio Elias Alvarado-Yaah, Carlos F. Arias, Santiago Ávila-Ríos, Eduardo Becerril-Vargas, Víctor Hugo Borja-Aburto, Celia Boukadida, Cristóbal Cháidez-Quiróz, Juan Bautista Chale-Dzul, Ricardo Ciria Merce, Andreu Comas-García, Célida Duque Molina, Julissa Enciso-Ibarra, José Antonio Enciso-Moreno, Gloria Elena Espinosa-Ayala, Fernando Fontove-Herrera, Daniel Fregoso-Rueda, Victor Eduardo García-Arias, Alejandra García-Gasca, Bruno Gómez-Gil, Jean Pierre González, Irvin González-López, Concepción Grajales-Muñiz, Ricardo Grande, Rosa María Gutiérrez Rios, Jesús Hernández, Alejandra Hernández-Terán, Alfredo Herrera-Estrella, Carla Ivón Herrera-Najera, Pavel Isa, Verónica Mata-Haro, Daniel Lira Morales, Susana Lopez, Antonio Loza Román, Brenda Irasema Maldonado-Meza, Bernardo Martínez-Miguel, José Arturo Martínez-Orozco, Célida Martínez-Rodríguez, Margarita Matías-Florentino, Fidencio Mejía-Nepomuceno, María Guadalupe de Jesús Mireles-Rivera, Gloria María Molina-Salinas, Hector Montoya-Fuentes, , Mario Mújica-Sánchez, José Esteban Muñoz-Medina, José de Jesús Nuñez-Contreras, Alicia Ocaña-Mondragón, Luis Alberto Ochoa-Carrera, Hector Esteban Paz-Juárez, Marissa Perez-Garcia, Francisco Pulido, Helen Haydee Fernanda Ramírez-Plascencia, Jorge Salas-Hernández, Angel Gustavo Salas-Lais, Alejandro Sánchez-Flores, Clara Esperanza Santacruz-Tinoco, María Guadalupe Santiago-Mauricio, Selene Zárate, Nelly Sélem-Mojica, Blanca Taboada, Gloria Vazquez, Joel Armando Vázquez-Pérez. |
| EPI_ISL_3149330, EPI_ISL_3149331 | Hospital Margarita Maza de Juárez | Microbial Genomics Laboratory | Consorcio Mexicano de Vigilancia Genómica (CoViGen-Mex). Authors (in alphabetical order): Julio Elias Alvarado-Yaah, Carlos F. Arias, Santiago Ávila-Ríos, Eduardo Becerril-Vargas, Víctor Hugo Borja-Aburto, Celia Boukadida, Cristóbal Cháidez-Quiróz, Juan Bautista Chale-Dzul, Ricardo Ciria Merce, Andreu Comas-García, Célida Duque Molina, Julissa Enciso-Ibarra, José Antonio Enciso-Moreno, Gloria Elena Espinosa-Ayala, Fernando Fontove-Herrera, Daniel Fregoso-Rueda, Victor Eduardo García-Arias, Alejandra García-Gasca, Bruno Gómez-Gil, Jean Pierre González, Irvin González-López, Concepción Grajales-Muñiz, Ricardo Grande, Rosa María Gutiérrez Rios, Jesús Hernández, Alejandra Hernández-Terán, Alfredo Herrera-Estrella, Carla Ivón Herrera-Najera, Pavel Isa, Verónica Mata-Haro, Daniel Lira Morales, Susana Lopez, Antonio Loza Román, Brenda Irasema Maldonado-Meza, Bernardo Martínez-Miguel, José Arturo Martínez-Orozco, Célida Martínez-Rodríguez, Margarita Matías-Florentino, Fidencio Mejía-Nepomuceno, María Guadalupe de Jesús Mireles-Rivera, Gloria María Molina-Salinas, Hector Montoya-Fuentes, , Mario Mújica-Sánchez, José Esteban Muñoz-Medina, José de Jesús Nuñez-Contreras, Alicia Ocaña-Mondragón, Luis Alberto Ochoa-Carrera, Hector Esteban Paz-Juárez, Marissa Perez-Garcia, Francisco Pulido, Helen Haydee Fernanda Ramírez-Plascencia, Jorge Salas-Hernández, Angel Gustavo Salas-Lais, Alejandro Sánchez-Flores, Clara Esperanza Santacruz-Tinoco, María Guadalupe Santiago-Mauricio, Selene Zárate, Nelly Sélem-Mojica, Blanca Taboada, Gloria Vazquez, Joel Armando Vázquez-Pérez. |
| EPI_ISL_3149332 | Hospital Sharp | Microbial Genomics Laboratory | Consorcio Mexicano de Vigilancia Genómica (CoViGen-Mex). Authors (in alphabetical order): Julio Elias Alvarado-Yaah, Carlos F. Arias, Santiago Ávila-Ríos, Eduardo Becerril-Vargas, Víctor Hugo Borja-Aburto, Celia Boukadida, Cristóbal Cháidez-Quiróz, Juan Bautista Chale-Dzul, Ricardo Ciria Merce, Andreu Comas-García, Célida Duque Molina, Julissa Enciso-Ibarra, José Antonio Enciso-Moreno, Gloria Elena Espinosa-Ayala, Fernando Fontove-Herrera, Daniel Fregoso-Rueda, Victor Eduardo García-Arias, Alejandra García-Gasca, Bruno Gómez-Gil, Jean Pierre González, Irvin González-López, Concepción Grajales-Muñiz, Ricardo Grande, Rosa María Gutiérrez Rios, Jesús Hernández, Alejandra Hernández-Terán, Alfredo Herrera-Estrella, Carla Ivón Herrera-Najera, Pavel Isa, Verónica Mata-Haro, Daniel Lira Morales, Susana Lopez, Antonio Loza Román, Brenda Irasema Maldonado-Meza, Bernardo Martínez-Miguel, José Arturo Martínez-Orozco, Célida Martínez-Rodríguez, Margarita Matías-Florentino, Fidencio Mejía-Nepomuceno, María Guadalupe de Jesús Mireles-Rivera, Gloria María Molina-Salinas, Hector Montoya-Fuentes, , Mario Mújica-Sánchez, José Esteban Muñoz-Medina, José de Jesús Nuñez-Contreras, Alicia Ocaña-Mondragón, Luis Alberto Ochoa-Carrera, Hector Esteban Paz-Juárez, Marissa Perez-Garcia, Francisco Pulido, Helen Haydee Fernanda Ramírez-Plascencia, Jorge Salas-Hernández, Angel Gustavo Salas-Lais, Alejandro Sánchez-Flores, Clara Esperanza Santacruz-Tinoco, María Guadalupe Santiago-Mauricio, Selene Zárate, Nelly Sélem-Mojica, Blanca Taboada, Gloria Vazquez, Joel Armando Vázquez-Pérez. |
| EPI_ISL_3149333, EPI_ISL_3149334, EPI_ISL_3149335, EPI_ISL_3149336, EPI_ISL_3149337, EPI_ISL_3149338, EPI_ISL_3149339, EPI_ISL_3149340, EPI_ISL_3149341, EPI_ISL_3149342 | LANIIA | LANIIA-Nayarit | Consorcio Mexicano de Vigilancia Genómica (CoViGen-Mex). Authors (in alphabetical order): Julio Elias Alvarado-Yaah, Carlos F. Arias, Santiago Ávila-Ríos, Eduardo Becerril-Vargas, Víctor Hugo Borja-Aburto, Celia Boukadida, Cristóbal Cháidez-Quiróz, Juan Bautista Chale-Dzul, Ricardo Ciria Merce, Andreu Comas-García, Célida Duque Molina, Julissa Enciso-Ibarra, José Antonio Enciso-Moreno, Gloria Elena Espinosa-Ayala, Fernando Fontove-Herrera, Daniel Fregoso-Rueda, Victor Eduardo García-Arias, Alejandra García-Gasca, Bruno Gómez-Gil, Jean Pierre González, Irvin González-López, Concepción Grajales-Muñiz, Ricardo Grande, Rosa María Gutiérrez Rios, Jesús Hernández, Alejandra Hernández-Terán, Alfredo Herrera-Estrella, Carla Ivón Herrera-Najera, Pavel Isa, Verónica Mata-Haro, Daniel Lira Morales, Susana Lopez, Antonio Loza Román, Brenda Irasema Maldonado-Meza, Bernardo Martínez-Miguel, José Arturo Martínez-Orozco, Célida Martínez-Rodríguez, Margarita Matías-Florentino, Fidencio Mejía-Nepomuceno, María Guadalupe de Jesús Mireles-Rivera, Gloria María Molina-Salinas, Hector Montoya-Fuentes, , Mario Mújica-Sánchez, José Esteban Muñoz-Medina, José de Jesús Nuñez-Contreras, Alicia Ocaña-Mondragón, Luis Alberto Ochoa-Carrera, Hector Esteban Paz-Juárez, Marissa Perez-Garcia, Francisco Pulido, Helen Haydee Fernanda Ramírez-Plascencia, Jorge Salas-Hernández, Angel Gustavo Salas-Lais, Alejandro Sánchez-Flores, Clara Esperanza Santacruz-Tinoco, María Guadalupe Santiago-Mauricio, Selene Zárate, Nelly Sélem-Mojica, Blanca Taboada, Gloria Vazquez, Joel Armando Vázquez-Pérez. LANNIA Nayarit: Manuel Ivan Girón Pérez, Carlos Eduardo Covantes-Rosales, Daniel Alberto Girón Pérez, Ulises Mercado. |
| EPI_ISL_3149344, EPI_ISL_3149345, EPI_ISL_3149346, EPI_ISL_3149347, EPI_ISL_3149348, EPI_ISL_3149349, EPI_ISL_3149350, EPI_ISL_3149351, EPI_ISL_3149352, EPI_ISL_3149353, EPI_ISL_3149354, EPI_ISL_3149355, EPI_ISL_3149356, EPI_ISL_3149357, EPI_ISL_3149358, EPI_ISL_3149359, EPI_ISL_3149360, EPI_ISL_3149361, EPI_ISL_3149362, EPI_ISL_3149363, EPI_ISL_3149364, EPI_ISL_3149365, | | | |

| | | | |
|---|---|---|---|
| EPI_ISL_3149367, EPI_ISL_3149368, EPI_ISL_3149369, EPI_ISL_3149370, EPI_ISL_3149371, EPI_ISL_3149372 | | | |
| see above | MRC/UVRI & LSHTM Uganda Research Unit, Central Public Health Laboratories | MRC/UVRI & LSHTM Uganda Research Unit, Central Public Health Laboratories | Matthew Cotten, Dan Lule Bugembe, My V.T. Phan, Pontiano Kaleebu, Isaac Sseeewanyana, Patrick Semanda, Susan Nabadda, Hellen Nansumba |
| EPI_ISL_3149373, EPI_ISL_3149374, EPI_ISL_3149375, EPI_ISL_3149376, EPI_ISL_3149377, EPI_ISL_3149378, EPI_ISL_3149379, EPI_ISL_3149380, EPI_ISL_3149381, EPI_ISL_3149382, EPI_ISL_3149383, EPI_ISL_3149384, EPI_ISL_3149385, EPI_ISL_3149386, EPI_ISL_3149387, EPI_ISL_3149388, EPI_ISL_3149389, EPI_ISL_3149390, EPI_ISL_3149391, EPI_ISL_3149392, EPI_ISL_3149393, EPI_ISL_3149394, EPI_ISL_3149395, EPI_ISL_3149396, EPI_ISL_3149397, EPI_ISL_3149398, EPI_ISL_3149399, EPI_ISL_3149400, EPI_ISL_3149401, EPI_ISL_3149402, EPI_ISL_3149403 | | | |
| see above | MRC/UVRI & LSHTM Uganda Research Unit , Rakai Health Sciences Program | MRC/UVRI & LSHTM Uganda Research Unit , Rakai Health Sciences Program | Matthew Cotten Dan Lule Bugembe, My V.T. Phan, Pontiano Kaleebu, Charles Ssuuna, Ronald Moses Galiwango, Steven J Reynolds, |
| EPI_ISL_3149404, EPI_ISL_3149405, EPI_ISL_3149406, EPI_ISL_3149407, EPI_ISL_3149409, EPI_ISL_3149410, EPI_ISL_3149411, EPI_ISL_3149412, EPI_ISL_3149413, EPI_ISL_3149414, EPI_ISL_3149415, EPI_ISL_3149416, EPI_ISL_3149417, EPI_ISL_3149418, EPI_ISL_3149419, EPI_ISL_3149420, EPI_ISL_3149421, EPI_ISL_3149423, EPI_ISL_3149424, EPI_ISL_3149425, EPI_ISL_3149426, EPI_ISL_3149427, EPI_ISL_3149428, EPI_ISL_3149429, EPI_ISL_3149430, EPI_ISL_3149431, EPI_ISL_3149432, EPI_ISL_3149434, EPI_ISL_3149436, EPI_ISL_3149437, EPI_ISL_3149438, EPI_ISL_3149440, EPI_ISL_3149441, EPI_ISL_3149442, EPI_ISL_3149443, EPI_ISL_3149444, EPI_ISL_3149445 | | | |
| see above | MRC/UVRI & LSHTM Uganda Research Unit, Uganda Virus Research Institute | MRC/UVRI & LSHTM Uganda Research Unit, Uganda Virus Research Institute | Matthew Cotten, Dan Lule Bugembe, My V.T Phan, Pontiano Kaleebu, John Kayiwa, |
| EPI_ISL_3149448 | Wyoming Public Health Laboratory | Wyoming Public Health Laboratory | Jim Mildenberger, Taylor Fearing, Channing Weber, Ashley Norberg, Chayse Rowley, Marley Goetz, Brian Dominguez, Elliot Thomasson, Sam Britz, Cari Sloma, Robert Petit, and Rob Christensen |
| EPI_ISL_3149473, EPI_ISL_3149475, EPI_ISL_3149477, EPI_ISL_3149478, EPI_ISL_3149480, EPI_ISL_3149481, EPI_ISL_3149482, EPI_ISL_3149483, EPI_ISL_3149484, EPI_ISL_3149502, EPI_ISL_3149503, EPI_ISL_3149510, EPI_ISL_3149515, EPI_ISL_3149516, EPI_ISL_3149521, EPI_ISL_3149522, EPI_ISL_3149524, EPI_ISL_3149526, EPI_ISL_3149527, EPI_ISL_3149528, EPI_ISL_3149529, EPI_ISL_3149530, EPI_ISL_3149531, EPI_ISL_3149532, EPI_ISL_3149533, EPI_ISL_3149534, EPI_ISL_3149535, EPI_ISL_3149536, EPI_ISL_3149537, EPI_ISL_3149538, EPI_ISL_3149539, EPI_ISL_3149540, EPI_ISL_3149541, EPI_ISL_3149542, EPI_ISL_3149543 | | | |
| see above | Colorado Department of Public Health and Environment | Colorado Department of Public Health and Environment | Laura Bankers, Molly C. Hetherington-Rauth, Diana Ir, Alexandria Rossheim, Michael A. Martin, Mandy Waters, Shannon R. Matzinger, Sarah Elizabeth Totten, Emily A. Travanty |
| EPI_ISL_3149547, EPI_ISL_3149548, EPI_ISL_3149549, EPI_ISL_3149550, EPI_ISL_3149551, EPI_ISL_3149552, EPI_ISL_3149553, EPI_ISL_3149554, EPI_ISL_3149555, EPI_ISL_3149556, EPI_ISL_3149557, EPI_ISL_3149558, EPI_ISL_3149559, EPI_ISL_3149560, EPI_ISL_3149562, EPI_ISL_3149563, EPI_ISL_3149564, EPI_ISL_3149565, EPI_ISL_3149566, EPI_ISL_3149567, EPI_ISL_3149569, EPI_ISL_3149570, EPI_ISL_3149571, EPI_ISL_3149572, EPI_ISL_3149573, EPI_ISL_3149574, EPI_ISL_3149575, EPI_ISL_3149577, EPI_ISL_3149578, EPI_ISL_3149579, EPI_ISL_3149581, EPI_ISL_3149582, EPI_ISL_3149583, EPI_ISL_3149584, EPI_ISL_3149585, EPI_ISL_3149586, EPI_ISL_3149587, EPI_ISL_3149588, EPI_ISL_3149589, EPI_ISL_3149590, EPI_ISL_3149591, EPI_ISL_3149592, EPI_ISL_3149593, EPI_ISL_3149594, EPI_ISL_3149595, EPI_ISL_3149596, EPI_ISL_3149597, EPI_ISL_3149598, EPI_ISL_3149599, EPI_ISL_3149600, EPI_ISL_3149601, EPI_ISL_3149602, EPI_ISL_3149603, EPI_ISL_3149604, EPI_ISL_3149605, EPI_ISL_3149606, EPI_ISL_3149607, EPI_ISL_3149609, EPI_ISL_3149611, EPI_ISL_3149613, EPI_ISL_3149614, EPI_ISL_3149615, EPI_ISL_3149616, EPI_ISL_3149617, EPI_ISL_3149618, EPI_ISL_3149619, EPI_ISL_3149620, EPI_ISL_3149621, EPI_ISL_3149622, EPI_ISL_3149623, EPI_ISL_3149624, EPI_ISL_3149625, EPI_ISL_3149626, EPI_ISL_3149627, EPI_ISL_3149628, EPI_ISL_3149629, EPI_ISL_3149630, EPI_ISL_3149631, EPI_ISL_3149632, EPI_ISL_3149633, EPI_ISL_3149634, EPI_ISL_3149635, EPI_ISL_3149636, EPI_ISL_3149637, EPI_ISL_3149638, EPI_ISL_3149639, EPI_ISL_3149640, EPI_ISL_3149641, EPI_ISL_3149642, EPI_ISL_3149643, EPI_ISL_3149644, EPI_ISL_3149646, EPI_ISL_3149647, EPI_ISL_3149648, EPI_ISL_3149649, EPI_ISL_3149650, EPI_ISL_3149651, EPI_ISL_3149653, EPI_ISL_3149654, EPI_ISL_3149655, EPI_ISL_3149656, EPI_ISL_3149657, EPI_ISL_3149658, EPI_ISL_3149659, EPI_ISL_3149660, EPI_ISL_3149661, EPI_ISL_3149662, EPI_ISL_3149663, EPI_ISL_3149664, EPI_ISL_3149666, EPI_ISL_3149667, EPI_ISL_3149668, EPI_ISL_3149669, EPI_ISL_3149670, EPI_ISL_3149671, EPI_ISL_3149672, EPI_ISL_3149673, EPI_ISL_3149674, EPI_ISL_3149675, EPI_ISL_3149677, EPI_ISL_3149678, EPI_ISL_3149679, EPI_ISL_3149680, EPI_ISL_3149681, EPI_ISL_3149682, EPI_ISL_3149683, EPI_ISL_3149684, EPI_ISL_3149685, EPI_ISL_3149686, EPI_ISL_3149687, EPI_ISL_3149688, EPI_ISL_3149689, EPI_ISL_3149690, EPI_ISL_3149693, EPI_ISL_3149694, EPI_ISL_3149695, EPI_ISL_3149696, EPI_ISL_3149697, EPI_ISL_3149698, EPI_ISL_3149699, EPI_ISL_3149700, EPI_ISL_3149701, EPI_ISL_3149702, EPI_ISL_3149704, EPI_ISL_3149705, EPI_ISL_3149706, EPI_ISL_3149707, EPI_ISL_3149708, EPI_ISL_3149709, EPI_ISL_3149710, EPI_ISL_3149711, EPI_ISL_3149712 | | | |
| see above | UW Virology Lab | UW Virology Lab | Pavitra Roychoudhury, Shah Mohamed Bakhash, Ricardo Perez, Hong Xie, Tien V. Nguyen, Maria Lukes, Sean Ellis, Nathan Breit, Robert J. Livingston, Meei-Li Huang, Keith R Jerome, Patrick Mathias, Alexander Greninger |
| EPI_ISL_3149721 | IN State Department of Health Laboratory Services | IN State Department of Health Laboratory Services | Cassandra Campion, Jamie Yeadon, Brian Pope, Lixia Liu, Kyle Brownlee, Melissa Hindenlang, Mark Glazier |
| EPI_ISL_3149750, EPI_ISL_3149751, EPI_ISL_3149752 | Public Health Authority of the Slovak Republic | Laboratory of Genomics and Bioinformatics, Comenius University Science Park | Tatiana Sedláčková, Diana Rusňáková, Miroslav Böhmer, Anna Gičová, Jaroslav Budiš, Tomáš Szemes, Jakub Styk |
| EPI_ISL_3149753, EPI_ISL_3149754, EPI_ISL_3149755, EPI_ISL_3149758, EPI_ISL_3149759, EPI_ISL_3149761, EPI_ISL_3149762, EPI_ISL_3149763, EPI_ISL_3149764, EPI_ISL_3149765, EPI_ISL_3149766, EPI_ISL_3149767, EPI_ISL_3149768, EPI_ISL_3149769, EPI_ISL_3149770, EPI_ISL_3149771, EPI_ISL_3149773, EPI_ISL_3149774, EPI_ISL_3149775, EPI_ISL_3149776, EPI_ISL_3149777, EPI_ISL_3149778, EPI_ISL_3149779, EPI_ISL_3149781, EPI_ISL_3149782, EPI_ISL_3149783, EPI_ISL_3149784, EPI_ISL_3149785, EPI_ISL_3149786, EPI_ISL_3149787, EPI_ISL_3149789, EPI_ISL_3149790, EPI_ISL_3149791, EPI_ISL_3149792, EPI_ISL_3149793, EPI_ISL_3149794, EPI_ISL_3149796, EPI_ISL_3149797, EPI_ISL_3149798, EPI_ISL_3149800, EPI_ISL_3149801, EPI_ISL_3149802, EPI_ISL_3149803, EPI_ISL_3149806, EPI_ISL_3149807, EPI_ISL_3149808, EPI_ISL_3149809, EPI_ISL_3149810, EPI_ISL_3149811, EPI_ISL_3149815, EPI_ISL_3149816, EPI_ISL_3149817, EPI_ISL_3149819, EPI_ISL_3149820, EPI_ISL_3149821, EPI_ISL_3149822, EPI_ISL_3149823, EPI_ISL_3149825, EPI_ISL_3149828, EPI_ISL_3149829, EPI_ISL_3149830, EPI_ISL_3149831, EPI_ISL_3149832, EPI_ISL_3149833, EPI_ISL_3149834, EPI_ISL_3149836, EPI_ISL_3149838, EPI_ISL_3149839, EPI_ISL_3149841, EPI_ISL_3149842, EPI_ISL_3149844, EPI_ISL_3149845, EPI_ISL_3149846, EPI_ISL_3149853, EPI_ISL_3149856, EPI_ISL_3149859, EPI_ISL_3149868, EPI_ISL_3149876, EPI_ISL_3149878, EPI_ISL_3149879, EPI_ISL_3149894, EPI_ISL_3149899, EPI_ISL_3149917, EPI_ISL_3149925, EPI_ISL_3149933, EPI_ISL_3149942, EPI_ISL_3149945, EPI_ISL_3149948, EPI_ISL_3149949, EPI_ISL_3149951, EPI_ISL_3149958, EPI_ISL_3149965 | | | |
| see above | Arizona State University | Arizona State University | Ajeet Bains, LaRinda A. Holland, Matthew F. Smith, Peter T. Skidmore, Rabia Maqsood, Nicholas J. Mellor, Joy M. Blain, Valerie Harris, Joshua LaBaer, Vel Murugan, Efrem S. Lim |
| EPI_ISL_3149971 | Genomik Solidaritas Indonesia Laboratorium | Genomik Solidaritas Indonesia Laboratorium | Carissa Sintca Wijaya, Reinhart Gabriel, Vania Gavrila Wikasa, Gracia Felias Enos Korompis, Annisa Muthiah Sukirman, Anuraj Shankar, Louisa Markus, Meutia Ayuputeri Kumaheri, Ariel Pradipta |
| EPI_ISL_3149972, EPI_ISL_3149973 | UCSC Genomics Institute | UCSC Genomics Institute | Mark Akeson, Ikenna Anigbogu, Eric Beraut, Isabel Bjork, Molly Cassatt-Johnstone, Terren Chang, Russell Corbett-Detig, Namrita Dhillon, Maximilian Haeussler, David Haussler, Angie Hinrichs, Miten Jain, Joshua Kapp, A. Marm Kilpatrick, Jakob McBroome, Hugh Olsen, Jeremy Sanford, Beth Shapiro, Michael Stone, Bryan Thornlow, Yatish Turakhia, Ciara Wanket |
| EPI_ISL_3150042, EPI_ISL_3150043, EPI_ISL_3150044, EPI_ISL_3150045, EPI_ISL_3150046, EPI_ISL_3150047, EPI_ISL_3150048, EPI_ISL_3150049, EPI_ISL_3150050, EPI_ISL_3150051, EPI_ISL_3150052, EPI_ISL_3150053, EPI_ISL_3150054, EPI_ISL_3150056, EPI_ISL_3150057, EPI_ISL_3150059, EPI_ISL_3150060, EPI_ISL_3150061, EPI_ISL_3150062, EPI_ISL_3150063, EPI_ISL_3150064, EPI_ISL_3150065, EPI_ISL_3150066, EPI_ISL_3150067, EPI_ISL_3150068, EPI_ISL_3150069, EPI_ISL_3150070, EPI_ISL_3150071, EPI_ISL_3150072, EPI_ISL_3150073, EPI_ISL_3150074, EPI_ISL_3150075, EPI_ISL_3150076, EPI_ISL_3150077, EPI_ISL_3150078, EPI_ISL_3150079, EPI_ISL_3150080, EPI_ISL_3150081, EPI_ISL_3150082, EPI_ISL_3150083, EPI_ISL_3150084, EPI_ISL_3150085, EPI_ISL_3150086, EPI_ISL_3150087, EPI_ISL_3150088, EPI_ISL_3150089, EPI_ISL_3150090, EPI_ISL_3150091, EPI_ISL_3150092, EPI_ISL_3150093, EPI_ISL_3150094, EPI_ISL_3150095, EPI_ISL_3150096, EPI_ISL_3150097, EPI_ISL_3150098, EPI_ISL_3150099, EPI_ISL_3150100, EPI_ISL_3150101, EPI_ISL_3150102, EPI_ISL_3150103, EPI_ISL_3150104, EPI_ISL_3150105, EPI_ISL_3150106, EPI_ISL_3150107, EPI_ISL_3150108, EPI_ISL_3150109, EPI_ISL_3150112, EPI_ISL_3150113, EPI_ISL_3150114, EPI_ISL_3150115, EPI_ISL_3150116, EPI_ISL_3150117, EPI_ISL_3150118, EPI_ISL_3150119, EPI_ISL_3150120, EPI_ISL_3150121, EPI_ISL_3150122, EPI_ISL_3150123, EPI_ISL_3150124, EPI_ISL_3150125, EPI_ISL_3150126, EPI_ISL_3150127, EPI_ISL_3150128, EPI_ISL_3150129, EPI_ISL_3150130, EPI_ISL_3150131, EPI_ISL_3150132, EPI_ISL_3150133, EPI_ISL_3150134, EPI_ISL_3150135, EPI_ISL_3150149, EPI_ISL_3150150, EPI_ISL_3150151, EPI_ISL_3150152, EPI_ISL_3150153, EPI_ISL_3150154 | | | |
| see above | KEMRI-Wellcome Trust Research Programme,Kilifi | KEMRI-Wellcome Trust Research Programme,Kilifi | Githinji G.,Ochola I.,Agoti C.,Mohamed K.S.,de Laurent Z.,Mburu M.W.,Morobe J.,Lambisia A., Makori T., Edidah O., Ndwiga L. |
| EPI_ISL_3150156, EPI_ISL_3150157, EPI_ISL_3150158, EPI_ISL_3150159, EPI_ISL_3150160, EPI_ISL_3150166, EPI_ISL_3150167, EPI_ISL_3150168, EPI_ISL_3150169, EPI_ISL_3150170, EPI_ISL_3150171, EPI_ISL_3150172, EPI_ISL_3150173, EPI_ISL_3150174, EPI_ISL_3150175, EPI_ISL_3150178, EPI_ISL_3150179, EPI_ISL_3150180, EPI_ISL_3150181, EPI_ISL_3150182, EPI_ISL_3150183, EPI_ISL_3150184, EPI_ISL_3150185, EPI_ISL_3150186, EPI_ISL_3150187, EPI_ISL_3150188, EPI_ISL_3150189, EPI_ISL_3150190, EPI_ISL_3150191, EPI_ISL_3150192, EPI_ISL_3150193, EPI_ISL_3150194, EPI_ISL_3150195, EPI_ISL_3150196 | | | |
| see above | Indian Council of Medical Research-National Institute of Virology, Microbial Containment Complex | Indian Council of Medical Research-National Institute of Virology, Microbial Containment Complex | Pragya D Yadav |
| EPI_ISL_3150198, EPI_ISL_3150199, EPI_ISL_3150200, EPI_ISL_3150201 | PathWest Laboratory Medicine WA | PathWest Laboratory Medicine WA Microbial Surveillance Unit | PathWest Laboratory Medicine WA Microbial Surveillance Unit |
| EPI_ISL_3150204, EPI_ISL_3150205, EPI_ISL_3150206, EPI_ISL_3150207 | Tivoli Fondazione Policlinico Universitario Agostino Gemelli IRCCS | Fondazione Policlinico Universitario Agostino Gemelli | Istituto di Microbiologia e Virologia |
| EPI_ISL_3150208 | State Hygienic Laboratory at the University of Iowa | State Hygienic Laboratory at the University of Iowa | Valerie Reeb, Erik Twait, Anna Yakos, Mariah Knutson, Kris Eveland, Kristen Zanon, Jeff, Benfer, Davis Rieckenberg, Wes Hottel, Alankar Kampoowale |
| EPI_ISL_3150209 | University of Iowa Hospitals and Clinics | State Hygienic Laboratory at the University of Iowa | Valerie Reeb, Erik Twait, Anna Yakos, Mariah Knutson, Kris Eveland, Kristen Zanon, Jeff, Benfer, Davis Rieckenberg, Wes Hottel, Alankar Kampoowale |
| EPI_ISL_3150210 | Tivoli Fondazione Policlinico Universitario Agostino Gemelli IRCCS | Fondazione Policlinico Universitario Agostino Gemelli IRCCS | Istituto di Microbiologia e Virologia |
| EPI_ISL_3150211 | FYR Diagnostics | FYR Diagnostics | Blohm, Tre; Ghai, Vikas |
| EPI_ISL_3150212 | Tivoli Fondazione Policlinico Universitario Agostino Gemelli IRCCS | Fondazione Policlinico Universitario Agostino Gemelli IRCCS | Istituto di Microbiologia e Virologia |
| EPI_ISL_3150224, EPI_ISL_3150236 | FYR Diagnostics | FYR Diagnostics | Blohm, Tre; Ghai, Vikas |
| EPI_ISL_3150237, EPI_ISL_3150238 | Tivoli Fondazione Policlinico Universitario Agostino Gemelli IRCCS | Fondazione Policlinico Universitario Agostino Gemelli IRCCS | Istituto di Microbiologia e Virologia |
| EPI_ISL_3150733, EPI_ISL_3150734, EPI_ISL_3150735, EPI_ISL_3150736, EPI_ISL_3150737, EPI_ISL_3150740, EPI_ISL_3150741, EPI_ISL_3150742, EPI_ISL_3150743, EPI_ISL_3150744, EPI_ISL_3150745, EPI_ISL_3150746, EPI_ISL_3150747, EPI_ISL_3150748, EPI_ISL_3150749, EPI_ISL_3150750, EPI_ISL_3150751, EPI_ISL_3150752, EPI_ISL_3150753, EPI_ISL_3150754, EPI_ISL_3150755, EPI_ISL_3150756 | | | |
| see above | Wyoming Public Health Laboratory | Wyoming Public Health Laboratory | Jim Mildenberger, Taylor Fearing, Channing Weber, Ashley Norberg, Chayse Rowley, Marley Goetz, Brian Dominguez, Elliot Thomasson, Sam Britz, Cari Sloma, Robert Petit, and Rob Christensen |
| EPI_ISL_3150757, EPI_ISL_3150758, EPI_ISL_3150759, EPI_ISL_3150760, EPI_ISL_3150761, EPI_ISL_3150762, EPI_ISL_3150764, EPI_ISL_3150765, EPI_ISL_3150766, EPI_ISL_3150767, EPI_ISL_3150768, EPI_ISL_3150769, EPI_ISL_3150770, EPI_ISL_3150771, EPI_ISL_3150772, EPI_ISL_3150773, EPI_ISL_3150774, EPI_ISL_3150775, EPI_ISL_3150776, EPI_ISL_3150777, EPI_ISL_3150778, EPI_ISL_3150779, EPI_ISL_3150780, EPI_ISL_3150781, EPI_ISL_3150782, EPI_ISL_3150783, EPI_ISL_3150784, EPI_ISL_3150785, EPI_ISL_3150786, EPI_ISL_3150787, EPI_ISL_3150788, EPI_ISL_3150789, EPI_ISL_3150790, EPI_ISL_3150791, EPI_ISL_3150792, EPI_ISL_3150793, EPI_ISL_3150794, EPI_ISL_3150795, EPI_ISL_3150796, EPI_ISL_3150797, EPI_ISL_3150798, EPI_ISL_3150799, EPI_ISL_3150800, EPI_ISL_3150801, EPI_ISL_3150802, EPI_ISL_3150803, EPI_ISL_3150804, EPI_ISL_3150805, EPI_ISL_3150806, EPI_ISL_3150807, EPI_ISL_3150808, EPI_ISL_3150809, EPI_ISL_3150810, EPI_ISL_3150811 | | | |
| see above | Nebraska Public Health Laboratory | NPHL COVID-19 Response Team | NPHL COVID-19 Response Team |
| EPI_ISL_3150813, EPI_ISL_3150814, EPI_ISL_3150815, EPI_ISL_3150816, EPI_ISL_3150817, EPI_ISL_3150818, EPI_ISL_3150819, EPI_ISL_3150820, EPI_ISL_3150821, EPI_ISL_3150822, EPI_ISL_3150823, EPI_ISL_3150824, EPI_ISL_3150825, EPI_ISL_3150826, EPI_ISL_3150827, EPI_ISL_3150828, EPI_ISL_3150829, EPI_ISL_3150830, EPI_ISL_3150831, EPI_ISL_3150832, EPI_ISL_3150833, EPI_ISL_3150834, EPI_ISL_3150835, EPI_ISL_3150836, EPI_ISL_3150837, EPI_ISL_3150838, EPI_ISL_3150839, EPI_ISL_3150840, EPI_ISL_3150841, EPI_ISL_3150842, EPI_ISL_3150843, EPI_ISL_3150844, EPI_ISL_3150845, EPI_ISL_3150846, EPI_ISL_3150847, EPI_ISL_3150848, EPI_ISL_3150849, EPI_ISL_3150850, EPI_ISL_3150851, EPI_ISL_3150852, EPI_ISL_3150853, EPI_ISL_3150854, EPI_ISL_3150855, EPI_ISL_3150856, EPI_ISL_3150857, EPI_ISL_3150858, EPI_ISL_3150859, EPI_ISL_3150860, EPI_ISL_3150861, EPI_ISL_3150862, EPI_ISL_3150863, EPI_ISL_3150864, EPI_ISL_3150865, EPI_ISL_3150866, EPI_ISL_3150867, EPI_ISL_3150868, EPI_ISL_3150869, EPI_ISL_3150870, EPI_ISL_3150871, EPI_ISL_3150872, EPI_ISL_3150873, EPI_ISL_3150874, EPI_ISL_3150875, EPI_ISL_3150876, EPI_ISL_3150877, EPI_ISL_3150878, EPI_ISL_3150879, EPI_ISL_3150880, EPI_ISL_3150881, EPI_ISL_3150882, EPI_ISL_3150883, EPI_ISL_3150884, EPI_ISL_3150885, EPI_ISL_3150886, EPI_ISL_3150887, EPI_ISL_3150888 | | | |
| see above | Kansas Health and Environmental Lab | Kansas Health and Environmental Lab | Mike Grose, Katherine Wiggins, Jonathan Barnell, Ben Olsen, and Phil Adam |
| EPI_ISL_3150889, EPI_ISL_3150890 | Tivoli Fondazione Policlinico Universitario Agostino Gemelli IRCCS | Fondazione Policlinico Universitario Agostino Gemelli IRCCS | Istituto di Microbiologia e Virologia |
| EPI_ISL_3150891, EPI_ISL_3150893, EPI_ISL_3150894, EPI_ISL_3150895, EPI_ISL_3150896, EPI_ISL_3150897, EPI_ISL_3150898, EPI_ISL_3150899, EPI_ISL_3150900, EPI_ISL_3150901, EPI_ISL_3150902, EPI_ISL_3150904, EPI_ISL_3150905, EPI_ISL_3150906, EPI_ISL_3150907, EPI_ISL_3150908, EPI_ISL_3150909, EPI_ISL_3150910, EPI_ISL_3150911, EPI_ISL_3150912, EPI_ISL_3150913, EPI_ISL_3150914, EPI_ISL_3150915, EPI_ISL_3150916, EPI_ISL_3150917, EPI_ISL_3150918, EPI_ISL_3150919, EPI_ISL_3150920, EPI_ISL_3150921, EPI_ISL_3150922, EPI_ISL_3150923, EPI_ISL_3150924, EPI_ISL_3150925, EPI_ISL_3150926, EPI_ISL_3150927 | | | |
| see above | Kansas Health and Environmental Lab | Kansas Health and Environmental Lab | Mike Grose, Katherine Wiggins, Jonathan Barnell, Ben Olsen, and Phil Adam |
| EPI_ISL_3150929 | Tivoli Fondazione Policlinico Universitario Agostino Gemelli IRCCS | Fondazione Policlinico Universitario Agostino Gemelli IRCCS | Istituto di Microbiologia e Virologia |
| EPI_ISL_3150930, EPI_ISL_3150931, EPI_ISL_3150932, EPI_ISL_3150933, EPI_ISL_3150934, EPI_ISL_3150935, EPI_ISL_3150936, EPI_ISL_3150937, EPI_ISL_3150938, EPI_ISL_3150939, EPI_ISL_3150940, EPI_ISL_3150941, EPI_ISL_3150942, EPI_ISL_3150943, EPI_ISL_3150944, EPI_ISL_3150947, EPI_ISL_3150948, EPI_ISL_3150949, EPI_ISL_3150950, EPI_ISL_3150952, EPI_ISL_3150953, EPI_ISL_3150954, EPI_ISL_3150955, EPI_ISL_3150956, EPI_ISL_3150957, EPI_ISL_3150958, EPI_ISL_3150959, EPI_ISL_3150960, EPI_ISL_3150961 | | | |
| see above | MRCG at LSHTM Genomics lab | MRCG at LSHTM Genomics lab | Abdul Karim sesay, Abdoulie Kanteh, Jarra Manneh, Mariama Kujabi, Bakary Sanyang,Dabiri Damilari |
| EPI_ISL_3150962, EPI_ISL_3150963 | Tivoli Fondazione Policlinico Universitario Agostino Gemelli IRCCS | Fondazione Policlinico Universitario Agostino Gemelli IRCCS | Istituto di Microbiologia e Virologia |
| EPI_ISL_3150968, EPI_ISL_3150969, EPI_ISL_3150970, EPI_ISL_3150971, EPI_ISL_3150972, EPI_ISL_3150973, EPI_ISL_3150974, EPI_ISL_3150975, EPI_ISL_3150976, EPI_ISL_3150977, EPI_ISL_3150978, EPI_ISL_3150979, EPI_ISL_3150981, EPI_ISL_3150983 | | | |
| see above | NL-Dr. Leonard A. Miller Centre for Health Services | National Microbiology Laboratory (NML) | Anna Majer, Shari Tyson, Grace Seo, Philip Mabon, Elsie Grudeski, Rhiannon Huzarewich, Russell Mandes, Anneliese Landgraff, Jennifer Tanner, Natalie Knox, Morag Graham, Gary Van Domselaar, Robert Needle, Yang Yu, Laura Gilbert, George |

| | | | |
|---|---|---|---|
| | | | Zahariadis, Nathalie Bastien, Yan Li, Timothy Booth, Darian Hole, Madison Chapel, Kirsten Biggar, Kerri Smith, CanCOGeN's metadata curation team, Public Health Agency of Canada CanCOGeN team |
| EPI_ISL_3150984 | Tivoli Fondazione Policlinico Universitario Agostino Gemelli IRCCS | Fondazione Policlinico Universitario Agostino Gemelli IRCCS | Istituto di Microbiologia e Virologia |
| EPI_ISL_3150985 | Regions Hospital | Minnesota Department of Health, Public Health Laboratory | Alexandra Lorentz, Jacob Garfin, Matt Plumb, and Xiong Wang |
| EPI_ISL_3150986, EPI_ISL_3150987, EPI_ISL_3150989, EPI_ISL_3150990, EPI_ISL_3150991, EPI_ISL_3150992, EPI_ISL_3150993 | Minnesota Department of Health, Public Health Laboratory | Minnesota Department of Health, Public Health Laboratory | Alexandra Lorentz, Jacob Garfin, Matt Plumb, and Xiong Wang |
| EPI_ISL_3150994 | Regions Hospital | Minnesota Department of Health, Public Health Laboratory | Alexandra Lorentz, Jacob Garfin, Matt Plumb, and Xiong Wang |
| EPI_ISL_3150995 | Tivoli Fondazione Policlinico Universitario Agostino Gemelli IRCCS | Fondazione Policlinico Universitario Agostino Gemelli IRCCS | Istituto di Microbiologia e Virologia |
| EPI_ISL_3150996, EPI_ISL_3150997 | Regions Hospital | Minnesota Department of Health, Public Health Laboratory | Alexandra Lorentz, Jacob Garfin, Matt Plumb, and Xiong Wang |
| EPI_ISL_3150998 | Centracare Laboratory Services | Minnesota Department of Health, Public Health Laboratory | Alexandra Lorentz, Jacob Garfin, Matt Plumb, and Xiong Wang |
| EPI_ISL_3150999, EPI_ISL_3151000 | Regions Hospital | Minnesota Department of Health, Public Health Laboratory | Alexandra Lorentz, Jacob Garfin, Matt Plumb, and Xiong Wang |
| EPI_ISL_3151001, EPI_ISL_3151002, EPI_ISL_3151003, EPI_ISL_3151004 | Minnesota Department of Health, Public Health Laboratory | Minnesota Department of Health, Public Health Laboratory | Alexandra Lorentz, Jacob Garfin, Matt Plumb, and Xiong Wang |
| EPI_ISL_3151005, EPI_ISL_3151006, EPI_ISL_3151007, EPI_ISL_3151008, EPI_ISL_3151009, EPI_ISL_3151010, EPI_ISL_3151011, EPI_ISL_3151012, EPI_ISL_3151013 | Regions Hospital | Minnesota Department of Health, Public Health Laboratory | Alexandra Lorentz, Jacob Garfin, Matt Plumb, and Xiong Wang |
| EPI_ISL_3151014, EPI_ISL_3151015, EPI_ISL_3151016, EPI_ISL_3151017 | Centracare Laboratory Services | Minnesota Department of Health, Public Health Laboratory | Alexandra Lorentz, Jacob Garfin, Matt Plumb, and Xiong Wang |
| EPI_ISL_3151018 | Minnesota Department of Health, Public Health Laboratory | Minnesota Department of Health, Public Health Laboratory | Alexandra Lorentz, Jacob Garfin, Matt Plumb, and Xiong Wang |
| EPI_ISL_3151019, EPI_ISL_3151020, EPI_ISL_3151021, EPI_ISL_3151022, EPI_ISL_3151023 | Park Nicollet Methodist Hospital | Minnesota Department of Health, Public Health Laboratory | Alexandra Lorentz, Jacob Garfin, Matt Plumb, and Xiong Wang |
| EPI_ISL_3151024 | Minnesota Department of Health, Public Health Laboratory | Minnesota Department of Health, Public Health Laboratory | Alexandra Lorentz, Jacob Garfin, Matt Plumb, and Xiong Wang |
| EPI_ISL_3151025, EPI_ISL_3151026, EPI_ISL_3151027, EPI_ISL_3151028, EPI_ISL_3151029, EPI_ISL_3151030, EPI_ISL_3151032, EPI_ISL_3151033, EPI_ISL_3151034, EPI_ISL_3151035, EPI_ISL_3151036, EPI_ISL_3151037, EPI_ISL_3151038, EPI_ISL_3151039, EPI_ISL_3151040, EPI_ISL_3151041, EPI_ISL_3151042, EPI_ISL_3151043, EPI_ISL_3151044 | | | |
| see above | Centracare Laboratory Services | Minnesota Department of Health, Public Health Laboratory | Alexandra Lorentz, Jacob Garfin, Matt Plumb, and Xiong Wang |
| EPI_ISL_3151045, EPI_ISL_3151046, EPI_ISL_3151047, EPI_ISL_3151048 | Premier Medical Lab | Minnesota Department of Health, Public Health Laboratory | Alexandra Lorentz, Jacob Garfin, Matt Plumb, and Xiong Wang |
| EPI_ISL_3151049, EPI_ISL_3151050, EPI_ISL_3151051, EPI_ISL_3151052, EPI_ISL_3151053, EPI_ISL_3151055, EPI_ISL_3151056, EPI_ISL_3151057, EPI_ISL_3151058, EPI_ISL_3151059, EPI_ISL_3151060, EPI_ISL_3151061, EPI_ISL_3151063 | | | |
| see above | Mayo Clinic & Mayo Clinic Laboratories | Minnesota Department of Health, Public Health Laboratory | Alexandra Lorentz, Jacob Garfin, Matt Plumb, and Xiong Wang |
| EPI_ISL_3151064, EPI_ISL_3151065 | Tivoli Fondazione Policlinico Universitario Agostino Gemelli IRCCS | Fondazione Policlinico Universitario Agostino Gemelli IRCCS | Istituto di Microbiologia e Virologia |
| EPI_ISL_3151066, EPI_ISL_3151067, EPI_ISL_3151068, EPI_ISL_3151069, EPI_ISL_3151070, EPI_ISL_3151071, EPI_ISL_3151072, EPI_ISL_3151073, EPI_ISL_3151074, EPI_ISL_3151075, EPI_ISL_3151076, EPI_ISL_3151077, EPI_ISL_3151078, EPI_ISL_3151079, EPI_ISL_3151080, EPI_ISL_3151081, EPI_ISL_3151082, EPI_ISL_3151083, EPI_ISL_3151084, EPI_ISL_3151085, EPI_ISL_3151086, EPI_ISL_3151087 | | | |
| see above | Monterey County Public Health Laboratory | Monterey County Public Health Laboratory | Monterey County Public Health Laboratory |
| EPI_ISL_3151091, EPI_ISL_3151092, EPI_ISL_3151093, EPI_ISL_3151094, EPI_ISL_3151095, EPI_ISL_3151096, EPI_ISL_3151097, EPI_ISL_3151098, EPI_ISL_3151099, EPI_ISL_3151100, EPI_ISL_3151101, EPI_ISL_3151102, EPI_ISL_3151104, EPI_ISL_3151114, EPI_ISL_3151115, EPI_ISL_3151116, EPI_ISL_3151124, EPI_ISL_3151126, EPI_ISL_3151127, EPI_ISL_3151128, EPI_ISL_3151129 | | | |
| see above | Wyoming Public Health Laboratory | Wyoming Public Health Laboratory | Jim Mildenberger, Taylor Fearing, Channing Weber, Ashley Norberg, Chayse Rowley, Marley Goetz, Brian Dominguez, Elliot Thomasson, Sam Britz, Cari Sloma, Robert Petit, and Rob Christensen |
| EPI_ISL_3151150, EPI_ISL_3151151, EPI_ISL_3151152, EPI_ISL_3151153, EPI_ISL_3151154, EPI_ISL_3151155, EPI_ISL_3151156, EPI_ISL_3151159, EPI_ISL_3151160, EPI_ISL_3151161 | TXDSHS | TXDSHS | Rashmi Tuladhar, Bonnie Oh, Jenny Zhang, Maliha Rahman, Mayela Pedrueza, Anita Pokharel, Karen Bobier, Lorraine Rodriguez, Myong Koag, Chun Wang, Rachel Lee, Grace Kubin |
| EPI_ISL_3151162, EPI_ISL_3151165, EPI_ISL_3151167, EPI_ISL_3151168, EPI_ISL_3151169, EPI_ISL_3151170, EPI_ISL_3151171, EPI_ISL_3151172 | Nastavni zavod za javno zdravstvo Splitsko- Dalmatinske Županije | Hrvatski zavod za javno zdravstvo | Irena Tabain, Ivana Ferenčak |
| EPI_ISL_3151173, EPI_ISL_3151174, EPI_ISL_3151175, EPI_ISL_3151176, EPI_ISL_3151177, EPI_ISL_3151178, EPI_ISL_3151179, EPI_ISL_3151180, EPI_ISL_3151182, EPI_ISL_3151183, EPI_ISL_3151184, EPI_ISL_3151185, EPI_ISL_3151186, EPI_ISL_3151187, EPI_ISL_3151188, EPI_ISL_3151189, EPI_ISL_3151190, EPI_ISL_3151191, EPI_ISL_3151192, EPI_ISL_3151193 | | | |
| see above | Zavod za javno zdravstvo Šibensko- Kninske Županije | Hrvatski zavod za javno zdravstvo | Irena Tabain, Ivana Ferenčak |
| EPI_ISL_3151194, EPI_ISL_3151195, EPI_ISL_3151196 | Klinički bolnički centar Rijeka | Hrvatski zavod za javno zdravstvo | Irena Tabain, Ivana Ferenčak |
| EPI_ISL_3151197, EPI_ISL_3151198, EPI_ISL_3151199, EPI_ISL_3151200, EPI_ISL_3151201, EPI_ISL_3151202, EPI_ISL_3151203, EPI_ISL_3151204, EPI_ISL_3151205, EPI_ISL_3151206, EPI_ISL_3151207, EPI_ISL_3151208, EPI_ISL_3151209, EPI_ISL_3151210, EPI_ISL_3151211, EPI_ISL_3151212, EPI_ISL_3151213, EPI_ISL_3151214, EPI_ISL_3151215, EPI_ISL_3151216, EPI_ISL_3151217, EPI_ISL_3151218, EPI_ISL_3151219, EPI_ISL_3151220, EPI_ISL_3151221, EPI_ISL_3151222, EPI_ISL_3151223, EPI_ISL_3151224, EPI_ISL_3151225, EPI_ISL_3151226, EPI_ISL_3151227 | | | |
| see above | Nastavni zavod za javno zdravstvo Primorsko- Goranske Županije | Hrvatski zavod za javno zdravstvo | Irena Tabain, Ivana Ferenčak |
| EPI_ISL_3151231, EPI_ISL_3151233, EPI_ISL_3151238, EPI_ISL_3151242 | TriCore Reference Laboratories | Center for Global Health, University of New Mexico Health Sciences Center | Daryl Domman, Kurt Schwalm, Valerie Morley, Cecilia Thompson, Kendra Pesko, Karissa Culbreath, Darrell Dinwiddie |
| EPI_ISL_3151249, EPI_ISL_3151251, EPI_ISL_3151259, EPI_ISL_3151260, EPI_ISL_3151262, EPI_ISL_3151263, EPI_ISL_3151265, EPI_ISL_3151266, EPI_ISL_3151269, EPI_ISL_3151273, EPI_ISL_3151277, EPI_ISL_3151278, EPI_ISL_3151279, EPI_ISL_3151280, EPI_ISL_3151281, EPI_ISL_3151284, EPI_ISL_3151289, EPI_ISL_3151290, EPI_ISL_3151291, EPI_ISL_3151292, EPI_ISL_3151293, EPI_ISL_3151294, EPI_ISL_3151295, EPI_ISL_3151296, EPI_ISL_3151297 | | | |
| see above | Arkansas Children's Hospital | Center for Global Health, University of New Mexico Health Sciences Center | Daryl Dommman, Kurt Schwalm, Valerie Morley, Ashton Ingold, Bobby Boyanton, Joshua L. Kennedy, Darrell Dinwiddie |
| EPI_ISL_3151298, EPI_ISL_3151299, EPI_ISL_3151300, EPI_ISL_3151301, EPI_ISL_3151302, EPI_ISL_3151303, EPI_ISL_3151304, EPI_ISL_3151305, EPI_ISL_3151306, EPI_ISL_3151307, EPI_ISL_3151308, EPI_ISL_3151309, EPI_ISL_3151310, EPI_ISL_3151311, EPI_ISL_3151312, EPI_ISL_3151313, EPI_ISL_3151314, EPI_ISL_3151315, EPI_ISL_3151316, EPI_ISL_3151317, EPI_ISL_3151318, EPI_ISL_3151319, EPI_ISL_3151320, EPI_ISL_3151321, EPI_ISL_3151322, EPI_ISL_3151323, EPI_ISL_3151324, EPI_ISL_3151325, EPI_ISL_3151326, EPI_ISL_3151327, EPI_ISL_3151328, EPI_ISL_3151329, EPI_ISL_3151330, EPI_ISL_3151331, EPI_ISL_3151332, EPI_ISL_3151333, EPI_ISL_3151334, EPI_ISL_3151335, EPI_ISL_3151336, EPI_ISL_3151337, EPI_ISL_3151338, EPI_ISL_3151339, EPI_ISL_3151340, EPI_ISL_3151341, EPI_ISL_3151342, EPI_ISL_3151343, EPI_ISL_3151344, EPI_ISL_3151345, EPI_ISL_3151346, EPI_ISL_3151347, EPI_ISL_3151348, EPI_ISL_3151349, EPI_ISL_3151350, EPI_ISL_3151351, EPI_ISL_3151352, EPI_ISL_3151353, EPI_ISL_3151354, EPI_ISL_3151355, EPI_ISL_3151356, EPI_ISL_3151357, EPI_ISL_3151358, EPI_ISL_3151359, EPI_ISL_3151360, EPI_ISL_3151361, EPI_ISL_3151362, EPI_ISL_3151363, | | | |

| | | | |
|---|---|---|---|
| EPI_ISL_3151364, EPI_ISL_3151365, EPI_ISL_3151366, EPI_ISL_3151367, EPI_ISL_3151368, EPI_ISL_3151369, EPI_ISL_3151370, EPI_ISL_3151371, EPI_ISL_3151372, EPI_ISL_3151373, EPI_ISL_3151374, EPI_ISL_3151375, EPI_ISL_3151376, EPI_ISL_3151377, EPI_ISL_3151378, EPI_ISL_3151379, EPI_ISL_3151380, EPI_ISL_3151381, EPI_ISL_3151382, EPI_ISL_3151383, EPI_ISL_3151384, EPI_ISL_3151385, EPI_ISL_3151386, EPI_ISL_3151387, EPI_ISL_3151388, EPI_ISL_3151389, EPI_ISL_3151390, EPI_ISL_3151391, EPI_ISL_3151392, EPI_ISL_3151393, EPI_ISL_3151394, EPI_ISL_3151395, EPI_ISL_3151396, EPI_ISL_3151397, EPI_ISL_3151398, EPI_ISL_3151399, EPI_ISL_3151400, EPI_ISL_3151401, EPI_ISL_3151402, EPI_ISL_3151403, EPI_ISL_3151405, EPI_ISL_3151407, EPI_ISL_3151408, EPI_ISL_3151409, EPI_ISL_3151410, EPI_ISL_3151412, EPI_ISL_3151413, EPI_ISL_3151414, EPI_ISL_3151415, EPI_ISL_3151416, EPI_ISL_3151417, EPI_ISL_3151418, EPI_ISL_3151419, EPI_ISL_3151420, EPI_ISL_3151421, EPI_ISL_3151422, EPI_ISL_3151423, EPI_ISL_3151424, EPI_ISL_3151425, EPI_ISL_3151426, EPI_ISL_3151427, EPI_ISL_3151428, EPI_ISL_3151429, EPI_ISL_3151430, EPI_ISL_3151431, EPI_ISL_3151432, EPI_ISL_3151433, EPI_ISL_3151434, EPI_ISL_3151435, EPI_ISL_3151436, EPI_ISL_3151437, EPI_ISL_3151438, EPI_ISL_3151439, EPI_ISL_3151440, EPI_ISL_3151441, EPI_ISL_3151442, EPI_ISL_3151443, EPI_ISL_3151445, EPI_ISL_3151446, EPI_ISL_3151447, EPI_ISL_3151448, EPI_ISL_3151449, EPI_ISL_3151450, EPI_ISL_3151451, EPI_ISL_3151452, EPI_ISL_3151453, EPI_ISL_3151454, EPI_ISL_3151455, EPI_ISL_3151456, EPI_ISL_3151457, EPI_ISL_3151458, EPI_ISL_3151459, EPI_ISL_3151460, EPI_ISL_3151461, EPI_ISL_3151462, EPI_ISL_3151463, EPI_ISL_3151464, EPI_ISL_3151465, EPI_ISL_3151466, EPI_ISL_3151467, EPI_ISL_3151468, EPI_ISL_3151469, EPI_ISL_3151470, EPI_ISL_3151471, EPI_ISL_3151473, EPI_ISL_3151474, EPI_ISL_3151475, EPI_ISL_3151476, EPI_ISL_3151477, EPI_ISL_3151478, EPI_ISL_3151479, EPI_ISL_3151480, EPI_ISL_3151481, EPI_ISL_3151482, EPI_ISL_3151483, EPI_ISL_3151484, EPI_ISL_3151485, EPI_ISL_3151486, EPI_ISL_3151487, EPI_ISL_3151488, EPI_ISL_3151489, EPI_ISL_3151490, EPI_ISL_3151491, EPI_ISL_3151492, EPI_ISL_3151493, EPI_ISL_3151494, EPI_ISL_3151495, EPI_ISL_3151496, EPI_ISL_3151497, EPI_ISL_3151498, EPI_ISL_3151499, EPI_ISL_3151500, EPI_ISL_3151501, EPI_ISL_3151502, EPI_ISL_3151503, EPI_ISL_3151504, EPI_ISL_3151505, EPI_ISL_3151506, EPI_ISL_3151507, EPI_ISL_3151508, EPI_ISL_3151509, EPI_ISL_3151510, EPI_ISL_3151511, EPI_ISL_3151512, EPI_ISL_3151513, EPI_ISL_3151514, EPI_ISL_3151515, EPI_ISL_3151516, EPI_ISL_3151517, EPI_ISL_3151518, EPI_ISL_3151519, EPI_ISL_3151520, EPI_ISL_3151521, EPI_ISL_3151522, EPI_ISL_3151523, EPI_ISL_3151524, EPI_ISL_3151525, EPI_ISL_3151526, EPI_ISL_3151527, EPI_ISL_3151528, EPI_ISL_3151529, EPI_ISL_3151530, EPI_ISL_3151531, EPI_ISL_3151532, EPI_ISL_3151533, EPI_ISL_3151534, EPI_ISL_3151535, EPI_ISL_3151536, EPI_ISL_3151537, EPI_ISL_3151538, EPI_ISL_3151539, EPI_ISL_3151540, EPI_ISL_3151541, EPI_ISL_3151542, EPI_ISL_3151543, EPI_ISL_3151544, EPI_ISL_3151545, EPI_ISL_3151546, EPI_ISL_3151547, EPI_ISL_3151548, EPI_ISL_3151549, EPI_ISL_3151550, EPI_ISL_3151551, EPI_ISL_3151552, EPI_ISL_3151553, EPI_ISL_3151554, EPI_ISL_3151555, EPI_ISL_3151556, EPI_ISL_3151557, EPI_ISL_3151558, EPI_ISL_3151559, EPI_ISL_3151560 | | | |
| see above | University Hospitals of Geneva, Laboratory of Virology | HUG, Laboratory of Virology and the Health2030 Genome Center | Samuel Cordey, Aline Mamin, Francisco Perez, Ana Rita Goncalves, Laurent Kaiser, Lorenzo Cerutti, Henri Pegeot, Melyssa Elies, Deborah Penet, Keith Harshman, Ioannis Xenarios, Emmanouil Dermitzakis |
| EPI_ISL_3151563, EPI_ISL_3151564, EPI_ISL_3151565, EPI_ISL_3151566, EPI_ISL_3151567, EPI_ISL_3151568, EPI_ISL_3151569, EPI_ISL_3151570, EPI_ISL_3151571, EPI_ISL_3151572, EPI_ISL_3151574, EPI_ISL_3151576, EPI_ISL_3151579, EPI_ISL_3151580, EPI_ISL_3151581, EPI_ISL_3151582, EPI_ISL_3151583, EPI_ISL_3151584, EPI_ISL_3151585, EPI_ISL_3151586, EPI_ISL_3151587, EPI_ISL_3151588, EPI_ISL_3151589, EPI_ISL_3151590, EPI_ISL_3151591, EPI_ISL_3151592, EPI_ISL_3151594, EPI_ISL_3151595, EPI_ISL_3151596, EPI_ISL_3151597, EPI_ISL_3151598, EPI_ISL_3151599, EPI_ISL_3151600 | | | |
| see above | CT Department of Public Health | CT Department of Public Health | Claire_Pearson,Tu_N_Nguyen |
| EPI_ISL_3151604 | Dinkes Provinsi Kalimantan Timur | National Institute of Health Research and Development | Vivi Setiawaty, Hana Apsari Pawestri, Hartanti Dian Ikawati, Subangkit, Kartika Dewi Puspa, Arie Ardiansyah Nugraha, Fajar Nur Sulistiyohadi, Triyani Soekarso, Krisna Pangesti, Nelly Puspandari |
| EPI_ISL_3151605, EPI_ISL_3151606 | RSUD Dr. W.Z. Johanes kupang | National Institute of Health Research and Development | Vivi Setiawaty, Hana Apsari Pawestri, Hartanti Dian Ikawati, Subangkit, Kartika Dewi Puspa, Arie Ardiansyah Nugraha, Fajar Nur Sulistiyohadi, Triyani Soekarso, Krisna Pangesti, Nelly Puspandari |
| EPI_ISL_3151607 | RSUD Provinsi Papua Barat | National Institute of Health Research and Development | Vivi Setiawaty, Hana Apsari Pawestri, Hartanti Dian Ikawati, Subangkit, Kartika Dewi Puspa, Arie Ardiansyah Nugraha, Fajar Nur Sulistiyohadi, Triyani Soekarso, Krisna Pangesti, Nelly Puspandari |
| EPI_ISL_3151613 | RSUD Dr. W.Z. Johanes kupang | National Institute of Health Research and Development | Vivi Setiawaty, Hana Apsari Pawestri, Hartanti Dian Ikawati, Subangkit, Kartika Dewi Puspa, Arie Ardiansyah Nugraha, Fajar Nur Sulistiyohadi, Triyani Soekarso, Krisna Pangesti, Nelly Puspandari |
| EPI_ISL_3151614 | RSUD Provinsi Papua Barat | National Institute of Health Research and Development | Vivi Setiawaty, Hana Apsari Pawestri, Hartanti Dian Ikawati, Subangkit, Kartika Dewi Puspa, Arie Ardiansyah Nugraha, Fajar Nur Sulistiyohadi, Triyani Soekarso, Krisna Pangesti, Nelly Puspandari |
| EPI_ISL_3151615 | Dinas Kesehatan Provinsi Papua Barat | National Institute of Health Research and Development | Vivi Setiawaty, Hana Apsari Pawestri, Hartanti Dian Ikawati, Subangkit, Kartika Dewi Puspa, Arie Ardiansyah Nugraha, Fajar Nur Sulistiyohadi, Triyani Soekarso, Krisna Pangesti, Nelly Puspandari |
| EPI_ISL_3151617, EPI_ISL_3151618 | Dinkes Provinsi Kalimantan Timur | National Institute of Health Research and Development | Vivi Setiawaty, Hana Apsari Pawestri, Hartanti Dian Ikawati, Subangkit, Kartika Dewi Puspa, Arie Ardiansyah Nugraha, Fajar Nur Sulistiyohadi, Triyani Soekarso, Krisna Pangesti, Nelly Puspandari |
| EPI_ISL_3151624, EPI_ISL_3151625 | Dinkes Provinsi Kalimantan Timur | National Institute of Health Research and Development | Hana Apsari Pawestri, Hartanti Dian Ikawati, Subangkit, Kartika Dewi Puspa, Arie Ardiansyah Nugraha, Fajar Nur Sulistiyohadi, Triyani Soekarso, Krisna Pangesti, Nelly Puspandari, Vivi Setiawaty |
| EPI_ISL_3151626, EPI_ISL_3151627 | RSUD Dr. W.Z. Johanes kupang | National Institute of Health Research and Development | Hana Apsari Pawestri, Hartanti Dian Ikawati, Subangkit, Kartika Dewi Puspa, Arie Ardiansyah Nugraha, Fajar Nur Sulistiyohadi, Triyani Soekarso, Krisna Pangesti, Nelly Puspandari, Vivi Setiawaty |
| EPI_ISL_3151628 | RSUD Provinsi Papua Barat | National Institute of Health Research and Development | Hana Apsari Pawestri, Hartanti Dian Ikawati, Subangkit, Kartika Dewi Puspa, Arie Ardiansyah Nugraha, Fajar Nur Sulistiyohadi, Triyani Soekarso, Krisna Pangesti, Nelly Puspandari, Vivi Setiawaty |
| EPI_ISL_3151633 | RSUD Dr. W.Z. Johanes kupang | National Institute of Health Research and Development | Hana Apsari Pawestri, Hartanti Dian Ikawati, Subangkit, Kartika Dewi Puspa, Arie Ardiansyah Nugraha, Fajar Nur Sulistiyohadi, Triyani Soekarso, Krisna Pangesti, Nelly Puspandari, Vivi Setiawaty |
| EPI_ISL_3151634 | RSUD Provinsi Papua Barat | National Institute of Health Research and Development | Hana Apsari Pawestri, Hartanti Dian Ikawati, Subangkit, Kartika Dewi Puspa, Arie Ardiansyah Nugraha, Fajar Nur Sulistiyohadi, Triyani Soekarso, Krisna Pangesti, Nelly Puspandari, Vivi Setiawaty |
| EPI_ISL_3151635 | Dinas Kesehatan Provinsi Papua Barat | National Institute of Health Research and Development | Hana Apsari Pawestri, Hartanti Dian Ikawati, Subangkit, Kartika Dewi Puspa, Arie Ardiansyah Nugraha, Fajar Nur Sulistiyohadi, Triyani Soekarso, Krisna Pangesti, Nelly Puspandari, Vivi Setiawaty |
| EPI_ISL_3151636 | BPOM Makassar | National Institute of Health Research and Development | Hana Apsari Pawestri, Hartanti Dian Ikawati, Subangkit, Kartika Dewi Puspa, Arie Ardiansyah Nugraha, Fajar Nur Sulistiyohadi, Triyani Soekarso, Krisna Pangesti, Nelly Puspandari, Vivi Setiawaty |
| EPI_ISL_3151637, EPI_ISL_3151638 | Dinkes Provinsi Kalimantan Timur | National Institute of Health Research and Development | Hana Apsari Pawestri, Hartanti Dian Ikawati, Subangkit, Kartika Dewi Puspa, Arie Ardiansyah Nugraha, Fajar Nur Sulistiyohadi, Triyani Soekarso, Krisna Pangesti, Nelly Puspandari, Vivi Setiawaty |
| EPI_ISL_3151643 | BPOM Makassar | National Institute of Health Research and Development | Hana Apsari Pawestri, Hartanti Dian Ikawati, Subangkit, Kartika Dewi Puspa, Arie Ardiansyah Nugraha, Fajar Nur Sulistiyohadi, Triyani Soekarso, Krisna Pangesti, Nelly Puspandari, Vivi Setiawaty |
| EPI_ISL_3151644, EPI_ISL_3151645 | Dinkes Provinsi Kalimantan Timur | National Institute of Health Research and Development | Hana Apsari Pawestri, Hartanti Dian Ikawati, Subangkit, Kartika Dewi Puspa, Arie Ardiansyah Nugraha, Fajar Nur Sulistiyohadi, Triyani Soekarso, Krisna Pangesti, Nelly Puspandari, Vivi Setiawaty |
| EPI_ISL_3151646 | RSUD Dr. W.Z. Johanes kupang | National Institute of Health Research and Development | Hana Apsari Pawestri, Hartanti Dian Ikawati, Subangkit, Kartika Dewi Puspa, Arie Ardiansyah Nugraha, Fajar Nur Sulistiyohadi, Triyani Soekarso, Krisna Pangesti, Nelly Puspandari, Vivi Setiawaty |
| EPI_ISL_3151648 | RSUD Provinsi Papua Barat | National Institute of Health Research and Development | Hana Apsari Pawestri, Hartanti Dian Ikawati, Subangkit, Kartika Dewi Puspa, Arie Ardiansyah Nugraha, Fajar Nur Sulistiyohadi, Triyani Soekarso, Krisna Pangesti, Nelly Puspandari, Vivi Setiawaty |
| EPI_ISL_3151652 | RSUD Dr. W.Z. Johanes kupang | National Institute of Health Research and Development | Hana Apsari Pawestri, Hartanti Dian Ikawati, Subangkit, Kartika Dewi Puspa, Arie Ardiansyah Nugraha, Fajar Nur Sulistiyohadi, Triyani Soekarso, Krisna Pangesti, Nelly Puspandari, Vivi Setiawaty |
| EPI_ISL_3151654 | RSUD Provinsi Papua Barat | National Institute of Health Research and Development | Hana Apsari Pawestri, Hartanti Dian Ikawati, Subangkit, Kartika Dewi Puspa, Arie Ardiansyah Nugraha, Fajar Nur Sulistiyohadi, Triyani Soekarso, Krisna Pangesti, Nelly Puspandari, Vivi Setiawaty |
| EPI_ISL_3151657, EPI_ISL_3151658 | Dinkes Provinsi Kalimantan Timur | National Institute of Health Research and Development | Hana Apsari Pawestri, Hartanti Dian Ikawati, Subangkit, Kartika Dewi Puspa, Arie Ardiansyah Nugraha, Fajar Nur Sulistiyohadi, Triyani Soekarso, Krisna Pangesti, Nelly Puspandari, Vivi Setiawaty |
| EPI_ISL_3151662, EPI_ISL_3151663, EPI_ISL_3151664 | Dinkes Provinsi Kalimantan Timur | National Institute of Health Research and Development | Subangkit, Hana Apsari Pawestri, Hartanti Dian Ikawati, Kartika Dewi Puspa, Arie Ardiansyah Nugraha, Fajar Nur Sulistiyohadi, Triyani Soekarso, Krisna Pangesti, Nelly Puspandari, Vivi Setiawaty |
| EPI_ISL_3151665, EPI_ISL_3151666 | RSUD Dr. W.Z. Johanes kupang | National Institute of Health Research and Development | Subangkit, Hana Apsari Pawestri, Hartanti Dian Ikawati, Kartika Dewi Puspa, Arie Ardiansyah Nugraha, Fajar Nur Sulistiyohadi, Triyani Soekarso, Krisna Pangesti, Nelly Puspandari, Vivi Setiawaty |
| EPI_ISL_3151667, EPI_ISL_3151668 | RSUD Provinsi Papua Barat | National Institute of Health Research and Development | Subangkit, Hana Apsari Pawestri, Hartanti Dian Ikawati, Kartika Dewi Puspa, Arie Ardiansyah Nugraha, Fajar Nur Sulistiyohadi, Triyani Soekarso, Krisna Pangesti, Nelly Puspandari, Vivi Setiawaty |
| EPI_ISL_3151673 | RSUD Dr. W.Z. Johanes kupang | National Institute of Health Research and Development | Hartanti Dian Ikawati, Hana Apsari Pawestri, Subangkit, Kartika Dewi Puspa, Arie Ardiansyah Nugraha, Fajar Nur Sulistiyohadi, Triyani Soekarso, Krisna Pangesti, Nelly Puspandari, Vivi Setiawaty |
| EPI_ISL_3151674, EPI_ISL_3151675 | RSUD Provinsi Papua Barat | National Institute of Health Research and Development | Kartika Dewi Puspa, Hana Apsari Pawestri, Subangkit, Hartanti Dian Ikawati, Arie Ardiansyah Nugraha, Fajar Nur Sulistiyohadi, Triyani Soekarso, Krisna Pangesti, Nelly Puspandari, Vivi Setiawaty |
| EPI_ISL_3151676 | BPOM Makassar | National Institute of Health Research and Development | Kartika Dewi Puspa, Hana Apsari Pawestri, Subangkit, Hartanti Dian Ikawati, Arie Ardiansyah Nugraha, Fajar Nur Sulistiyohadi, Triyani Soekarso, Krisna Pangesti, Nelly Puspandari, Vivi Setiawaty |
| EPI_ISL_3151677, EPI_ISL_3151678, EPI_ISL_3151679 | Dinkes Provinsi Kalimantan Timur | National Institute of Health Research and Development | Kartika Dewi Puspa, Hana Apsari Pawestri, Subangkit, Hartanti Dian Ikawati, Arie Ardiansyah Nugraha, Fajar Nur Sulistiyohadi, Triyani Soekarso, Krisna Pangesti, Nelly Puspandari, Vivi Setiawaty |
| EPI_ISL_3151680, EPI_ISL_3151681, EPI_ISL_3151682, EPI_ISL_3151684, EPI_ISL_3151685, EPI_ISL_3151686, EPI_ISL_3151687, EPI_ISL_3151689, EPI_ISL_3151690, EPI_ISL_3151691, EPI_ISL_3151692, EPI_ISL_3151693, EPI_ISL_3151694, EPI_ISL_3151695, EPI_ISL_3151696, EPI_ISL_3151697, EPI_ISL_3151698, EPI_ISL_3151699, EPI_ISL_3151700, EPI_ISL_3151701, EPI_ISL_3151702, EPI_ISL_3151703, EPI_ISL_3151704, EPI_ISL_3151705, EPI_ISL_3151706, EPI_ISL_3151707, EPI_ISL_3151708, EPI_ISL_3151709, EPI_ISL_3151710, EPI_ISL_3151711, EPI_ISL_3151712, EPI_ISL_3151713, EPI_ISL_3151714, EPI_ISL_3151715, EPI_ISL_3151716, EPI_ISL_3151717, EPI_ISL_3151718, EPI_ISL_3151719, EPI_ISL_3151720, EPI_ISL_3151721, EPI_ISL_3151722, EPI_ISL_3151723, EPI_ISL_3151724, EPI_ISL_3151725, EPI_ISL_3151726, EPI_ISL_3151727, EPI_ISL_3151728, EPI_ISL_3151729, EPI_ISL_3151730, EPI_ISL_3151731, EPI_ISL_3151732, EPI_ISL_3151734, EPI_ISL_3151735, EPI_ISL_3151736, EPI_ISL_3151737, EPI_ISL_3151738, EPI_ISL_3151739, EPI_ISL_3151740, EPI_ISL_3151741, EPI_ISL_3151742, EPI_ISL_3151743, EPI_ISL_3151744, EPI_ISL_3151745, EPI_ISL_3151746, EPI_ISL_3151747, EPI_ISL_3151748, EPI_ISL_3151749, EPI_ISL_3151750, EPI_ISL_3151751, EPI_ISL_3151752 | | | |
| see above | NJDOH, Public Health and Environmental Laboratories | NJ_PHEL | Lindsey Bodnar, Shiv K. Verma, Ryan Pachucki, Chelsea San Filippo, Jacquelyn Deverell, Dana Woell, Allison Roder, Byeong Jeong |
| EPI_ISL_3151754, EPI_ISL_3151755, EPI_ISL_3151756, EPI_ISL_3151757, EPI_ISL_3151758, EPI_ISL_3151759, EPI_ISL_3151760, EPI_ISL_3151761, EPI_ISL_3151762, EPI_ISL_3151763, EPI_ISL_3151764, EPI_ISL_3151765, EPI_ISL_3151766, EPI_ISL_3151767, EPI_ISL_3151768, EPI_ISL_3151769, EPI_ISL_3151770, EPI_ISL_3151771, EPI_ISL_3151772, EPI_ISL_3151773 | | | |
| see above | Klinika za infektivne bolesti "Dr. Fran Mihaljević" | Hrvatski zavod za javno zdravstvo | Irena Tabain, Ivana Ferenčak |
| EPI_ISL_3151776, EPI_ISL_3151777, EPI_ISL_3151779, EPI_ISL_3151780, EPI_ISL_3151781, EPI_ISL_3151783 | Nastavni zavod za javno zdravstvo "Andrija Štampar" | Hrvatski zavod za javno zdravstvo | Irena Tabain, Ivana Ferenčak |
| EPI_ISL_3151785 | Zavod za javno zdravstvo Varaždinske županije | Hrvatski zavod za javno zdravstvo | Irena Tabain, Ivana Ferenčak |
| EPI_ISL_3151788, EPI_ISL_3151789, EPI_ISL_3151792, EPI_ISL_3151793, EPI_ISL_3151794, EPI_ISL_3151795, EPI_ISL_3151796, EPI_ISL_3151797 | Zavod za javno zdravstvo Dubrovačko-Neretvanske Županije | Hrvatski zavod za javno zdravstvo | Irena Tabain, Ivana Ferenčak |
| EPI_ISL_3151798, EPI_ISL_3151799 | Klinički bolnički centar "Sestre milosrdnice" | Hrvatski zavod za javno zdravstvo | Irena Tabain, Ivana Ferenčak |
| EPI_ISL_3151800, EPI_ISL_3151801, EPI_ISL_3151802, EPI_ISL_3151803, EPI_ISL_3151804, EPI_ISL_3151805, EPI_ISL_3151806, EPI_ISL_3151807, EPI_ISL_3151808, EPI_ISL_3151809, EPI_ISL_3151810, EPI_ISL_3151811 | | | |
| see above | Zavod za javno zdravstvo Zadarske Županije | Hrvatski zavod za javno zdravstvo | Irena Tabain, Ivana Ferenčak |
| EPI_ISL_3151812, EPI_ISL_3151813, EPI_ISL_3151814, EPI_ISL_3151815, EPI_ISL_3151816, EPI_ISL_3151817, EPI_ISL_3151818 | Zavod za javno zdravstvo Međimurske Županije | Hrvatski zavod za javno zdravstvo | Irena Tabain, Ivana Ferenčak |
| EPI_ISL_3151819, EPI_ISL_3151822, | Nastavni zavod za javno zdravstvo Splitsko- Dalmatinske Županije | Hrvatski zavod za javno zdravstvo | Irena Tabain, Ivana Ferenčak |

| | | | |
|---|---|---|---|
| EPI_ISL_3151823, EPI_ISL_3151824 | | | |
| EPI_ISL_3151827, EPI_ISL_3151828, EPI_ISL_3151829, EPI_ISL_3151830, EPI_ISL_3151831, EPI_ISL_3151832, EPI_ISL_3151833, EPI_ISL_3151834, EPI_ISL_3151835, EPI_ISL_3151836, EPI_ISL_3151837, EPI_ISL_3151838, EPI_ISL_3151839, EPI_ISL_3151840, EPI_ISL_3151841, EPI_ISL_3151842, EPI_ISL_3151843, EPI_ISL_3151844, EPI_ISL_3151845, EPI_ISL_3151846, EPI_ISL_3151847, EPI_ISL_3151848, EPI_ISL_3151849, EPI_ISL_3151850, EPI_ISL_3151851, EPI_ISL_3151852, EPI_ISL_3151853, EPI_ISL_3151854, EPI_ISL_3151855, EPI_ISL_3151856, EPI_ISL_3151857, EPI_ISL_3151858, EPI_ISL_3151859, EPI_ISL_3151860, EPI_ISL_3151861, EPI_ISL_3151862, EPI_ISL_3151863, EPI_ISL_3151864, EPI_ISL_3151865, EPI_ISL_3151866, EPI_ISL_3151867, EPI_ISL_3151868, EPI_ISL_3151869, EPI_ISL_3151870, EPI_ISL_3151871, EPI_ISL_3151872, EPI_ISL_3151873, EPI_ISL_3151874, EPI_ISL_3151875, EPI_ISL_3151876, EPI_ISL_3151877, EPI_ISL_3151878, EPI_ISL_3151879, EPI_ISL_3151880, EPI_ISL_3151881, EPI_ISL_3151882, EPI_ISL_3151883, EPI_ISL_3151884, EPI_ISL_3151885, EPI_ISL_3151886, EPI_ISL_3151887, EPI_ISL_3151888, EPI_ISL_3151889, EPI_ISL_3151890, EPI_ISL_3151891, EPI_ISL_3151892, EPI_ISL_3151893, EPI_ISL_3151894, EPI_ISL_3151895, EPI_ISL_3151896, EPI_ISL_3151897, EPI_ISL_3151898, EPI_ISL_3151899, EPI_ISL_3151900, EPI_ISL_3151901, EPI_ISL_3151902, EPI_ISL_3151903, EPI_ISL_3151904, EPI_ISL_3151905, EPI_ISL_3151906, EPI_ISL_3151907, EPI_ISL_3151908, EPI_ISL_3151909, EPI_ISL_3151910, EPI_ISL_3151911, EPI_ISL_3151912, EPI_ISL_3151913, EPI_ISL_3151914, EPI_ISL_3151915, EPI_ISL_3151916, EPI_ISL_3151917, EPI_ISL_3151918, EPI_ISL_3151919, EPI_ISL_3151920, EPI_ISL_3151921, EPI_ISL_3151922, EPI_ISL_3151923, EPI_ISL_3151924, EPI_ISL_3151925, EPI_ISL_3151926, EPI_ISL_3151927, EPI_ISL_3151928, EPI_ISL_3151929, EPI_ISL_3151930, EPI_ISL_3151931, EPI_ISL_3151932, EPI_ISL_3151933, EPI_ISL_3151934, EPI_ISL_3151935, EPI_ISL_3151936, EPI_ISL_3151937, EPI_ISL_3151938, EPI_ISL_3151939, EPI_ISL_3151940, EPI_ISL_3151941, EPI_ISL_3151942, EPI_ISL_3151943, EPI_ISL_3151944 | | | |
| see above | National Public Health Laboratory, National Centre for Infectious Diseases | National Public Health Laboratory, National Centre for Infectious Diseases | Tze Minn Mak, Katherine Ching, Zhenyang Zhou, Royce Ang, Lin Cui, Raymond Tzer Pin Lin |
| EPI_ISL_3151972, EPI_ISL_3152021 | Furst Medical Laboratory | Norwegian Institute of Public Health, Department of Virology | Kathrine Stene-Johansen, Kamilla Heddeland Instefjord, Hilde Elshaug, Garcia Llorente Ignacio, Jon Bråte, Engebretsen Serina Beate,Pedersen Benedikte Nevjen, Line Victoria Moen, Debech Nadia, Atiya R Ali,Marie Paulsen Madsen, Rasmus Riis Kopperud, Hilde Vollan, Karoline Bragstad, Olav Hungnes |
| EPI_ISL_3152117, EPI_ISL_3152118, EPI_ISL_3152119, EPI_ISL_3152120, EPI_ISL_3152121, EPI_ISL_3152122, EPI_ISL_3152123, EPI_ISL_3152124, EPI_ISL_3152125, EPI_ISL_3152126, EPI_ISL_3152127, EPI_ISL_3152128, EPI_ISL_3152129, EPI_ISL_3152130, EPI_ISL_3152131, EPI_ISL_3152132, EPI_ISL_3152133, EPI_ISL_3152134, EPI_ISL_3152136, EPI_ISL_3152137, EPI_ISL_3152138, EPI_ISL_3152139, EPI_ISL_3152140, EPI_ISL_3152141, EPI_ISL_3152142, EPI_ISL_3152143, EPI_ISL_3152144, EPI_ISL_3152145, EPI_ISL_3152146, EPI_ISL_3152147, EPI_ISL_3152148, EPI_ISL_3152149, EPI_ISL_3152150, EPI_ISL_3152151, EPI_ISL_3152152, EPI_ISL_3152153, EPI_ISL_3152154, EPI_ISL_3152155, EPI_ISL_3152157, EPI_ISL_3152158, EPI_ISL_3152159, EPI_ISL_3152160, EPI_ISL_3152161, EPI_ISL_3152162, EPI_ISL_3152163, EPI_ISL_3152164, EPI_ISL_3152165, EPI_ISL_3152166, EPI_ISL_3152167, EPI_ISL_3152168, EPI_ISL_3152169, EPI_ISL_3152170, EPI_ISL_3152171, EPI_ISL_3152172, EPI_ISL_3152173, EPI_ISL_3152174, EPI_ISL_3152175, EPI_ISL_3152176, EPI_ISL_3152177, EPI_ISL_3152178, EPI_ISL_3152179, EPI_ISL_3152180, EPI_ISL_3152181, EPI_ISL_3152182, EPI_ISL_3152183, EPI_ISL_3152184, EPI_ISL_3152185, EPI_ISL_3152186, EPI_ISL_3152187, EPI_ISL_3152188, EPI_ISL_3152189 | | | |
| see above | UMC Groningen, Clinical Virology, Department of Medical Microbiology and Infection Prevention | UMC Groningen, Clinical Virology, Department of Medical Microbiology and Infection Prevention | Hubert Niesters, Alexander Friedrich, Erley Lizarazo-Forero, Monika Fliss, Lilli Gard, Sigrid Rosema, Coretta Van Leer-Buter, Xuewei Zhou, Marjolein Knoester |
| EPI_ISL_3152190, EPI_ISL_3152191, EPI_ISL_3152193, EPI_ISL_3152194, EPI_ISL_3152195, EPI_ISL_3152197, EPI_ISL_3152198, EPI_ISL_3152199, EPI_ISL_3152201, EPI_ISL_3152202, EPI_ISL_3152203, EPI_ISL_3152204, EPI_ISL_3152205, EPI_ISL_3152206, EPI_ISL_3152207, EPI_ISL_3152209, EPI_ISL_3152210, EPI_ISL_3152211, EPI_ISL_3152212, EPI_ISL_3152213, EPI_ISL_3152214, EPI_ISL_3152215, EPI_ISL_3152216, EPI_ISL_3152217, EPI_ISL_3152218, EPI_ISL_3152219, EPI_ISL_3152220, EPI_ISL_3152221, EPI_ISL_3152222, EPI_ISL_3152223, EPI_ISL_3152224, EPI_ISL_3152225, EPI_ISL_3152226, EPI_ISL_3152227, EPI_ISL_3152229, EPI_ISL_3152230, EPI_ISL_3152232, EPI_ISL_3152233, EPI_ISL_3152234, EPI_ISL_3152235 | | | |
| see above | MD PHL | Maryland Department of Health Laboratories Administration | Eric N. Keller, Jillian Loomis, Kwang Low, Terence L. Moore, Ami Patel, and Robert Myers |
| EPI_ISL_3152241, EPI_ISL_3152242, EPI_ISL_3152243 | Arlon | Plateforme de testing Namuroise | ; Demars Aurore ; Maschietto Céline ; Nobis Chloé ; Otto Gaetan ; Denis Olivier ; Degosserie Jonathan ; Mullier François |
| EPI_ISL_3152244 | Plateforme de testing Namuroise | Plateforme de testing Namuroise | ; Demars Aurore ; Maschietto Céline ; Nobis Chloé ; Otto Gaetan ; Denis Olivier ; Degosserie Jonathan ; Mullier François |
| EPI_ISL_3152246 | Arlon | Plateforme de testing Namuroise | ; Demars Aurore ; Maschietto Céline ; Nobis Chloé ; Otto Gaetan ; Denis Olivier ; Degosserie Jonathan ; Mullier François |
| EPI_ISL_3152247, EPI_ISL_3152248, EPI_ISL_3152249 | Plateforme de testing Namuroise | Plateforme de testing Namuroise | ; Demars Aurore ; Maschietto Céline ; Nobis Chloé ; Otto Gaetan ; Denis Olivier ; Degosserie Jonathan ; Mullier François |
| EPI_ISL_3152250, EPI_ISL_3152251, EPI_ISL_3152252, EPI_ISL_3152253 | Arlon | Plateforme de testing Namuroise | ; Demars Aurore ; Maschietto Céline ; Nobis Chloé ; Otto Gaetan ; Denis Olivier ; Degosserie Jonathan ; Mullier François |
| EPI_ISL_3152254, EPI_ISL_3152255 | Plateforme de testing Namuroise | Plateforme de testing Namuroise | ; Demars Aurore ; Maschietto Céline ; Nobis Chloé ; Otto Gaetan ; Denis Olivier ; Degosserie Jonathan ; Mullier François |
| EPI_ISL_3152256, EPI_ISL_3152257 | Arlon | Plateforme de testing Namuroise | ; Demars Aurore ; Maschietto Céline ; Nobis Chloé ; Otto Gaetan ; Denis Olivier ; Degosserie Jonathan ; Mullier François |
| EPI_ISL_3152258, EPI_ISL_3152259 | Plateforme de testing Namuroise | Plateforme de testing Namuroise | ; Demars Aurore ; Maschietto Céline ; Nobis Chloé ; Otto Gaetan ; Denis Olivier ; Degosserie Jonathan ; Mullier François |
| EPI_ISL_3152260, EPI_ISL_3152261 | Arlon | Plateforme de testing Namuroise | ; Demars Aurore ; Maschietto Céline ; Nobis Chloé ; Otto Gaetan ; Denis Olivier ; Degosserie Jonathan ; Mullier François |
| EPI_ISL_3152262, EPI_ISL_3152263 | Plateforme de testing Namuroise | Plateforme de testing Namuroise | ; Demars Aurore ; Maschietto Céline ; Nobis Chloé ; Otto Gaetan ; Denis Olivier ; Degosserie Jonathan ; Mullier François |
| EPI_ISL_3152264, EPI_ISL_3152265 | Arlon | Plateforme de testing Namuroise | ; Demars Aurore ; Maschietto Céline ; Nobis Chloé ; Otto Gaetan ; Denis Olivier ; Degosserie Jonathan ; Mullier François |
| EPI_ISL_3152266, EPI_ISL_3152267 | Plateforme de testing Namuroise | Plateforme de testing Namuroise | ; Demars Aurore ; Maschietto Céline ; Nobis Chloé ; Otto Gaetan ; Denis Olivier ; Degosserie Jonathan ; Mullier François |
| EPI_ISL_3152269, EPI_ISL_3152270, EPI_ISL_3152271, EPI_ISL_3152272, EPI_ISL_3152274, EPI_ISL_3152275, EPI_ISL_3152276, EPI_ISL_3152277, EPI_ISL_3152278, EPI_ISL_3152279, EPI_ISL_3152280, EPI_ISL_3152281, EPI_ISL_3152282, EPI_ISL_3152283, EPI_ISL_3152284, EPI_ISL_3152285, EPI_ISL_3152287, EPI_ISL_3152289, EPI_ISL_3152290, EPI_ISL_3152293, EPI_ISL_3152294, EPI_ISL_3152295, EPI_ISL_3152296, EPI_ISL_3152298, EPI_ISL_3152299, EPI_ISL_3152300, EPI_ISL_3152301, EPI_ISL_3152302, EPI_ISL_3152303, EPI_ISL_3152304, EPI_ISL_3152305, EPI_ISL_3152306, EPI_ISL_3152307, EPI_ISL_3152308, EPI_ISL_3152310, EPI_ISL_3152312, EPI_ISL_3152313, EPI_ISL_3152314, EPI_ISL_3152315, EPI_ISL_3152316, EPI_ISL_3152317, EPI_ISL_3152319, EPI_ISL_3152320, EPI_ISL_3152321, EPI_ISL_3152322, EPI_ISL_3152323, EPI_ISL_3152325, EPI_ISL_3152327, EPI_ISL_3152328, EPI_ISL_3152329, EPI_ISL_3152330, EPI_ISL_3152331, EPI_ISL_3152332, EPI_ISL_3152333, EPI_ISL_3152335, EPI_ISL_3152336, EPI_ISL_3152337, EPI_ISL_3152338, EPI_ISL_3152339, EPI_ISL_3152340, EPI_ISL_3152341, EPI_ISL_3152342, EPI_ISL_3152343, EPI_ISL_3152344, EPI_ISL_3152345, EPI_ISL_3152347, EPI_ISL_3152348, EPI_ISL_3152349, EPI_ISL_3152350, EPI_ISL_3152351, EPI_ISL_3152352, EPI_ISL_3152354, EPI_ISL_3152355, EPI_ISL_3152356, EPI_ISL_3152357, EPI_ISL_3152358 | | | |
| see above | MD PHL | Maryland Department of Health Laboratories Administration | Eric N. Keller, Jillian Loomis, Kwang Low, Terence L. Moore, Ami Patel, and Robert Myers |
| EPI_ISL_3152363, EPI_ISL_3152364, EPI_ISL_3152365, EPI_ISL_3152366, EPI_ISL_3152368, EPI_ISL_3152369, EPI_ISL_3152371, EPI_ISL_3152372, EPI_ISL_3152374, EPI_ISL_3152375, EPI_ISL_3152376 | | | |
| see above | DC Public Health Lab/ Dept. of Forensic Sciences | DC Public Health Lab/ Dept. of Forensic Sciences | Janis Doss, Scott Nguyen, Elizabeth Zelaya, Sarah Scott, Connie Maza, Monica Mann, Brittany Hamilton, David Payne, Jocelyn Hauser |
| EPI_ISL_3152398, EPI_ISL_3152402, EPI_ISL_3152408 | TriCore Reference Laboratories | Center for Global Health, University of New Mexico Health Sciences Center | Daryl Domman, Kurt Schwalm, Valerie Morley, Cecilia Thompson, Kendra Pesko, Karissa Culbreath, Darrell Dinwiddie |
| EPI_ISL_3152444, EPI_ISL_3152445, EPI_ISL_3152446, EPI_ISL_3152447, EPI_ISL_3152448, EPI_ISL_3152449, EPI_ISL_3152450, EPI_ISL_3152451, EPI_ISL_3152452, EPI_ISL_3152453, EPI_ISL_3152454, EPI_ISL_3152455, EPI_ISL_3152456, EPI_ISL_3152457, EPI_ISL_3152458, EPI_ISL_3152459, EPI_ISL_3152460, EPI_ISL_3152461, EPI_ISL_3152462, EPI_ISL_3152463, EPI_ISL_3152464, EPI_ISL_3152465, EPI_ISL_3152466, EPI_ISL_3152467, EPI_ISL_3152468, EPI_ISL_3152469, EPI_ISL_3152470, EPI_ISL_3152471, EPI_ISL_3152472, EPI_ISL_3152473, EPI_ISL_3152474, EPI_ISL_3152475, EPI_ISL_3152476, EPI_ISL_3152477, EPI_ISL_3152478, EPI_ISL_3152479, EPI_ISL_3152480, EPI_ISL_3152481, EPI_ISL_3152482, EPI_ISL_3152483, EPI_ISL_3152484, EPI_ISL_3152485, EPI_ISL_3152486, EPI_ISL_3152487, EPI_ISL_3152488, EPI_ISL_3152489, EPI_ISL_3152490, EPI_ISL_3152491, EPI_ISL_3152492, EPI_ISL_3152493, EPI_ISL_3152494, EPI_ISL_3152495, EPI_ISL_3152496, EPI_ISL_3152497, EPI_ISL_3152498, EPI_ISL_3152499, EPI_ISL_3152500, EPI_ISL_3152501, EPI_ISL_3152502, EPI_ISL_3152503, EPI_ISL_3152504, EPI_ISL_3152505, EPI_ISL_3152506, EPI_ISL_3152507, EPI_ISL_3152508, EPI_ISL_3152509, EPI_ISL_3152510, EPI_ISL_3152511, EPI_ISL_3152512, EPI_ISL_3152513, EPI_ISL_3152514, EPI_ISL_3152515, EPI_ISL_3152516, EPI_ISL_3152517, EPI_ISL_3152518, EPI_ISL_3152519, EPI_ISL_3152520, EPI_ISL_3152521, EPI_ISL_3152522, EPI_ISL_3152523, EPI_ISL_3152524, EPI_ISL_3152525, EPI_ISL_3152526, EPI_ISL_3152527, EPI_ISL_3152528, EPI_ISL_3152529, EPI_ISL_3152530, EPI_ISL_3152531, EPI_ISL_3152532, EPI_ISL_3152533, EPI_ISL_3152534, EPI_ISL_3152535, EPI_ISL_3152536, EPI_ISL_3152537, EPI_ISL_3152538, EPI_ISL_3152539, EPI_ISL_3152540, EPI_ISL_3152541, EPI_ISL_3152542, EPI_ISL_3152543, EPI_ISL_3152544, EPI_ISL_3152545, EPI_ISL_3152546, EPI_ISL_3152547, EPI_ISL_3152548, EPI_ISL_3152549, EPI_ISL_3152550, EPI_ISL_3152551, EPI_ISL_3152552, EPI_ISL_3152553, EPI_ISL_3152554, EPI_ISL_3152555, EPI_ISL_3152556, EPI_ISL_3152557, EPI_ISL_3152558 | | | |
| see above | National Public Health Laboratory, National Centre for Infectious Diseases | National Public Health Laboratory, National Centre for Infectious Diseases | Tze Minn Mak, Katherine Ching, Zhenyang Zhou, Royce Ang, Lin Cui, Raymond Tzer Pin Lin |
| EPI_ISL_3152559, EPI_ISL_3152560 | Medical Microbiology Unit, Department for Laboratory Medicine, Drammen Hospital, Vestre Viken Health Trust, | Norwegian Institute of Public Health, Department of Virology | Kathrine Stene-Johansen, Kamilla Heddeland Instefjord, Hilde Elshaug, Garcia Llorente Ignacio, Jon Bråte, Engebretsen Serina Beate,Pedersen Benedikte Nevjen, Line Victoria Moen, Debech Nadia, Atiya R Ali,Marie Paulsen Madsen, Rasmus Riis Kopperud, Hilde Vollan, Karoline Bragstad, Olav Hungnes |
| EPI_ISL_3152561 | Furst Medical Laboratory | Norwegian Institute of Public Health, Department of Virology | Kathrine Stene-Johansen, Kamilla Heddeland Instefjord, Hilde Elshaug, Garcia Llorente Ignacio, Jon Bråte, Engebretsen Serina Beate,Pedersen Benedikte Nevjen, Line Victoria Moen, Debech Nadia, Atiya R Ali,Marie Paulsen Madsen, Rasmus Riis Kopperud, Hilde Vollan, Karoline Bragstad, Olav Hungnes |
| EPI_ISL_3152562 | Haukeland University Hospital, Dept. of Microbiology | Norwegian Institute of Public Health, Department of Virology | Kathrine Stene-Johansen, Kamilla Heddeland Instefjord, Hilde Elshaug, Garcia Llorente Ignacio, Jon Bråte, Engebretsen Serina Beate,Pedersen Benedikte Nevjen, Line Victoria Moen, Debech Nadia, Atiya R Ali,Marie Paulsen Madsen, Rasmus Riis Kopperud, Hilde Vollan, Karoline Bragstad, Olav Hungnes |
| EPI_ISL_3152563, EPI_ISL_3152564, EPI_ISL_3152566 | Furst Medical Laboratory | Norwegian Institute of Public Health, Department of Virology | Kathrine Stene-Johansen, Kamilla Heddeland Instefjord, Hilde Elshaug, Garcia Llorente Ignacio, Jon Bråte, Engebretsen Serina Beate,Pedersen Benedikte Nevjen, Line Victoria Moen, Debech Nadia, Atiya R Ali,Marie Paulsen Madsen, Rasmus Riis Kopperud, Hilde Vollan, Karoline Bragstad, Olav Hungnes |
| EPI_ISL_3152568 | Haukeland University Hospital, Dept. of Microbiology | Norwegian Institute of Public Health, Department of Virology | Kathrine Stene-Johansen, Kamilla Heddeland Instefjord, Hilde Elshaug, Garcia Llorente Ignacio, Jon Bråte, Engebretsen Serina Beate,Pedersen Benedikte Nevjen, Line Victoria Moen, Debech Nadia, Atiya R Ali,Marie Paulsen Madsen, Rasmus Riis Kopperud, Hilde Vollan, Karoline Bragstad, Olav Hungnes |
| EPI_ISL_3152569, EPI_ISL_3152571 | Furst Medical Laboratory | Norwegian Institute of Public Health, Department of Virology | Kathrine Stene-Johansen, Kamilla Heddeland Instefjord, Hilde Elshaug, Garcia Llorente Ignacio, Jon Bråte, Engebretsen Serina Beate,Pedersen Benedikte Nevjen, Line Victoria Moen, Debech Nadia, Atiya R Ali,Marie Paulsen Madsen, Rasmus Riis Kopperud, Hilde Vollan, Karoline Bragstad, Olav Hungnes |
| EPI_ISL_3152572 | Medical Microbiology Unit, Department for Laboratory Medicine, Drammen Hospital, Vestre Viken Health Trust, | Norwegian Institute of Public Health, Department of Virology | Kathrine Stene-Johansen, Kamilla Heddeland Instefjord, Hilde Elshaug, Garcia Llorente Ignacio, Jon Bråte, Engebretsen Serina Beate,Pedersen Benedikte Nevjen, Line Victoria Moen, Debech Nadia, Atiya R Ali,Marie Paulsen Madsen, Rasmus Riis Kopperud, Hilde Vollan, Karoline Bragstad, Olav Hungnes |
| EPI_ISL_3152573 | Nordland Hospital - Bodo, Laboratory Department, Molecular Biology Unit | Norwegian Institute of Public Health, Department of Virology | Kathrine Stene-Johansen, Kamilla Heddeland Instefjord, Hilde Elshaug, Garcia Llorente Ignacio, Jon Bråte, Engebretsen Serina Beate,Pedersen Benedikte Nevjen, Line Victoria Moen, Debech Nadia, Atiya R Ali,Marie Paulsen Madsen, Rasmus Riis Kopperud, Hilde Vollan, Karoline Bragstad, Olav Hungnes |
| EPI_ISL_3152577 | Medical Microbiology Unit, Department for Laboratory Medicine, Drammen Hospital, Vestre Viken Health Trust, | Norwegian Institute of Public Health, Department of Virology | Kathrine Stene-Johansen, Kamilla Heddeland Instefjord, Hilde Elshaug, Garcia Llorente Ignacio, Jon Bråte, Engebretsen Serina Beate,Pedersen Benedikte Nevjen, Line Victoria Moen, Debech Nadia, Atiya R Ali,Marie Paulsen Madsen, Rasmus Riis Kopperud, Hilde Vollan, Karoline Bragstad, Olav Hungnes |
| EPI_ISL_3152581 | Nordland Hospital - Bodo, Laboratory Department, Molecular Biology Unit | Norwegian Institute of Public Health, Department of Virology | Kathrine Stene-Johansen, Kamilla Heddeland Instefjord, Hilde Elshaug, Garcia Llorente Ignacio, Jon Bråte, Engebretsen Serina Beate,Pedersen Benedikte Nevjen, Line Victoria Moen, Debech Nadia, Atiya R Ali,Marie Paulsen Madsen, Rasmus Riis Kopperud, Hilde Vollan, Karoline Bragstad, Olav Hungnes |
| EPI_ISL_3152582, EPI_ISL_3152583 | Haukeland University Hospital, Dept. of Microbiology | Norwegian Institute of Public Health, Department of Virology | Kathrine Stene-Johansen, Kamilla Heddeland Instefjord, Hilde Elshaug, Garcia Llorente Ignacio, Jon Bråte, Engebretsen Serina Beate,Pedersen Benedikte Nevjen, Line Victoria Moen, Debech Nadia, Atiya R Ali,Marie Paulsen Madsen, Rasmus Riis Kopperud, Hilde Vollan, Karoline Bragstad, Olav Hungnes |
| EPI_ISL_3152584 | Ostfold Hospital Trust - Kalnes, Centre for Laboratory Medicine, Section for gene technology and infection serology | Norwegian Institute of Public Health, Department of Virology | Kathrine Stene-Johansen, Kamilla Heddeland Instefjord, Hilde Elshaug, Garcia Llorente Ignacio, Jon Bråte, Engebretsen Serina Beate,Pedersen Benedikte Nevjen, Line Victoria Moen, Debech Nadia, Atiya R Ali,Marie Paulsen Madsen, Rasmus Riis Kopperud, Hilde Vollan, Karoline Bragstad, Olav Hungnes |
| EPI_ISL_3152585 | Furst Medical Laboratory | Norwegian Institute of Public Health, Department of Virology | Kathrine Stene-Johansen, Kamilla Heddeland Instefjord, Hilde Elshaug, Garcia Llorente Ignacio, Jon Bråte, Engebretsen Serina Beate,Pedersen Benedikte Nevjen, Line Victoria Moen, Debech Nadia, Atiya R Ali,Marie Paulsen Madsen, Rasmus Riis Kopperud, Hilde Vollan, Karoline Bragstad, Olav Hungnes |
| EPI_ISL_3152586 | Ostfold Hospital Trust - Kalnes, Centre for Laboratory Medicine, | Norwegian Institute of Public Health, Department of Virology | Kathrine Stene-Johansen, Kamilla Heddeland Instefjord, Hilde Elshaug, Garcia Llorente Ignacio, Jon Bråte, Engebretsen Serina Beate,Pedersen Benedikte Nevjen, Line Victoria Moen, Debech Nadia, Atiya R Ali,Marie Paulsen Madsen, Rasmus Riis |

| | | | |
|---|---|---|---|
| | Section for gene technology and infection serology | | Kopperud, Hilde Vollan, Karoline Bragstad, Olav Hungnes |
| EPI_ISL_3152587 | Haukeland University Hospital, Dept. of Microbiology | Norwegian Institute of Public Health, Department of Virology | Kathrine Stene-Johansen, Kamilla Heddeland Instefjord, Hilde Elshaug, Garcia Llorente Ignacio, Jon Bråte, Engebretsen Serina Beate,Pedersen Benedikte Nevjen, Line Victoria Moen, Debech Nadia, Atiya R Ali,Marie Paulsen Madsen, Rasmus Riis Kopperud, Hilde Vollan, Karoline Bragstad, Olav Hungnes |
| EPI_ISL_3152589, EPI_ISL_3152591 | Medical Microbiology Unit, Department for Laboratory Medicine, Drammen Hospital, Vestre Viken Health Trust, | Norwegian Institute of Public Health, Department of Virology | Kathrine Stene-Johansen, Kamilla Heddeland Instefjord, Hilde Elshaug, Garcia Llorente Ignacio, Jon Bråte, Engebretsen Serina Beate,Pedersen Benedikte Nevjen, Line Victoria Moen, Debech Nadia, Atiya R Ali,Marie Paulsen Madsen, Rasmus Riis Kopperud, Hilde Vollan, Karoline Bragstad, Olav Hungnes |
| EPI_ISL_3152593, EPI_ISL_3152595 | Ostfold Hospital Trust - Kalnes, Centre for Laboratory Medicine, Section for gene technology and infection serology | Norwegian Institute of Public Health, Department of Virology | Kathrine Stene-Johansen, Kamilla Heddeland Instefjord, Hilde Elshaug, Garcia Llorente Ignacio, Jon Bråte, Engebretsen Serina Beate,Pedersen Benedikte Nevjen, Line Victoria Moen, Debech Nadia, Atiya R Ali,Marie Paulsen Madsen, Rasmus Riis Kopperud, Hilde Vollan, Karoline Bragstad, Olav Hungnes |
| EPI_ISL_3152596 | Haukeland University Hospital, Dept. of Microbiology | Norwegian Institute of Public Health, Department of Virology | Kathrine Stene-Johansen, Kamilla Heddeland Instefjord, Hilde Elshaug, Garcia Llorente Ignacio, Jon Bråte, Engebretsen Serina Beate,Pedersen Benedikte Nevjen, Line Victoria Moen, Debech Nadia, Atiya R Ali,Marie Paulsen Madsen, Rasmus Riis Kopperud, Hilde Vollan, Karoline Bragstad, Olav Hungnes |
| EPI_ISL_3152597 | Furst Medical Laboratory | Norwegian Institute of Public Health, Department of Virology | Kathrine Stene-Johansen, Kamilla Heddeland Instefjord, Hilde Elshaug, Garcia Llorente Ignacio, Jon Bråte, Engebretsen Serina Beate,Pedersen Benedikte Nevjen, Line Victoria Moen, Debech Nadia, Atiya R Ali,Marie Paulsen Madsen, Rasmus Riis Kopperud, Hilde Vollan, Karoline Bragstad, Olav Hungnes |
| EPI_ISL_3152599, EPI_ISL_3152600, EPI_ISL_3152601 | Medical Microbiology Unit, Department for Laboratory Medicine, Drammen Hospital, Vestre Viken Health Trust, | Norwegian Institute of Public Health, Department of Virology | Kathrine Stene-Johansen, Kamilla Heddeland Instefjord, Hilde Elshaug, Garcia Llorente Ignacio, Jon Bråte, Engebretsen Serina Beate,Pedersen Benedikte Nevjen, Line Victoria Moen, Debech Nadia, Atiya R Ali,Marie Paulsen Madsen, Rasmus Riis Kopperud, Hilde Vollan, Karoline Bragstad, Olav Hungnes |
| EPI_ISL_3152604 | Hospital of Southern Norway - Kristiansand, Department of Medical Microbiology | Norwegian Institute of Public Health, Department of Virology | Kathrine Stene-Johansen, Kamilla Heddeland Instefjord, Hilde Elshaug, Garcia Llorente Ignacio, Jon Bråte, Engebretsen Serina Beate,Pedersen Benedikte Nevjen, Line Victoria Moen, Debech Nadia, Atiya R Ali,Marie Paulsen Madsen, Rasmus Riis Kopperud, Hilde Vollan, Karoline Bragstad, Olav Hungnes |
| EPI_ISL_3152605 | Vestfold Hospital, Toensberg Department of Microbiology | Norwegian Institute of Public Health, Department of Virology | Kathrine Stene-Johansen, Kamilla Heddeland Instefjord, Hilde Elshaug, Garcia Llorente Ignacio, Jon Bråte, Engebretsen Serina Beate,Pedersen Benedikte Nevjen, Line Victoria Moen, Debech Nadia, Atiya R Ali,Marie Paulsen Madsen, Rasmus Riis Kopperud, Hilde Vollan, Karoline Bragstad, Olav Hungnes |
| EPI_ISL_3152607 | Medical Microbiology Unit, Department for Laboratory Medicine, Drammen Hospital, Vestre Viken Health Trust, | Norwegian Institute of Public Health, Department of Virology | Kathrine Stene-Johansen, Kamilla Heddeland Instefjord, Hilde Elshaug, Garcia Llorente Ignacio, Jon Bråte, Engebretsen Serina Beate,Pedersen Benedikte Nevjen, Line Victoria Moen, Debech Nadia, Atiya R Ali,Marie Paulsen Madsen, Rasmus Riis Kopperud, Hilde Vollan, Karoline Bragstad, Olav Hungnes |
| EPI_ISL_3152608 | Hospital of Southern Norway - Kristiansand, Department of Medical Microbiology | Norwegian Institute of Public Health, Department of Virology | Kathrine Stene-Johansen, Kamilla Heddeland Instefjord, Hilde Elshaug, Garcia Llorente Ignacio, Jon Bråte, Engebretsen Serina Beate,Pedersen Benedikte Nevjen, Line Victoria Moen, Debech Nadia, Atiya R Ali,Marie Paulsen Madsen, Rasmus Riis Kopperud, Hilde Vollan, Karoline Bragstad, Olav Hungnes |
| EPI_ISL_3152610, EPI_ISL_3152613, EPI_ISL_3152617 | Ostfold Hospital Trust - Kalnes, Centre for Laboratory Medicine, Section for gene technology and infection serology | Norwegian Institute of Public Health, Department of Virology | Kathrine Stene-Johansen, Kamilla Heddeland Instefjord, Hilde Elshaug, Garcia Llorente Ignacio, Jon Bråte, Engebretsen Serina Beate,Pedersen Benedikte Nevjen, Line Victoria Moen, Debech Nadia, Atiya R Ali,Marie Paulsen Madsen, Rasmus Riis Kopperud, Hilde Vollan, Karoline Bragstad, Olav Hungnes |
| EPI_ISL_3152618, EPI_ISL_3152621 | Medical Microbiology Unit, Department for Laboratory Medicine, Drammen Hospital, Vestre Viken Health Trust, | Norwegian Institute of Public Health, Department of Virology | Kathrine Stene-Johansen, Kamilla Heddeland Instefjord, Hilde Elshaug, Garcia Llorente Ignacio, Jon Bråte, Engebretsen Serina Beate,Pedersen Benedikte Nevjen, Line Victoria Moen, Debech Nadia, Atiya R Ali,Marie Paulsen Madsen, Rasmus Riis Kopperud, Hilde Vollan, Karoline Bragstad, Olav Hungnes |
| EPI_ISL_3152622 | Nordland Hospital - Bodo, Laboratory Department, Molecular Biology Unit | Norwegian Institute of Public Health, Department of Virology | Kathrine Stene-Johansen, Kamilla Heddeland Instefjord, Hilde Elshaug, Garcia Llorente Ignacio, Jon Bråte, Engebretsen Serina Beate,Pedersen Benedikte Nevjen, Line Victoria Moen, Debech Nadia, Atiya R Ali,Marie Paulsen Madsen, Rasmus Riis Kopperud, Hilde Vollan, Karoline Bragstad, Olav Hungnes |
| EPI_ISL_3152624 | Hospital of Southern Norway - Kristiansand, Department of Medical Microbiology | Norwegian Institute of Public Health, Department of Virology | Kathrine Stene-Johansen, Kamilla Heddeland Instefjord, Hilde Elshaug, Garcia Llorente Ignacio, Jon Bråte, Engebretsen Serina Beate,Pedersen Benedikte Nevjen, Line Victoria Moen, Debech Nadia, Atiya R Ali,Marie Paulsen Madsen, Rasmus Riis Kopperud, Hilde Vollan, Karoline Bragstad, Olav Hungnes |
| EPI_ISL_3152627 | Medical Microbiology Unit, Department for Laboratory Medicine, Drammen Hospital, Vestre Viken Health Trust, | Norwegian Institute of Public Health, Department of Virology | Kathrine Stene-Johansen, Kamilla Heddeland Instefjord, Hilde Elshaug, Garcia Llorente Ignacio, Jon Bråte, Engebretsen Serina Beate,Pedersen Benedikte Nevjen, Line Victoria Moen, Debech Nadia, Atiya R Ali,Marie Paulsen Madsen, Rasmus Riis Kopperud, Hilde Vollan, Karoline Bragstad, Olav Hungnes |
| EPI_ISL_3152628 | Nordland Hospital - Bodo, Laboratory Department, Molecular Biology Unit | Norwegian Institute of Public Health, Department of Virology | Kathrine Stene-Johansen, Kamilla Heddeland Instefjord, Hilde Elshaug, Garcia Llorente Ignacio, Jon Bråte, Engebretsen Serina Beate,Pedersen Benedikte Nevjen, Line Victoria Moen, Debech Nadia, Atiya R Ali,Marie Paulsen Madsen, Rasmus Riis Kopperud, Hilde Vollan, Karoline Bragstad, Olav Hungnes |
| EPI_ISL_3152630 | Medical Microbiology Unit, Department for Laboratory Medicine, Drammen Hospital, Vestre Viken Health Trust, | Norwegian Institute of Public Health, Department of Virology | Kathrine Stene-Johansen, Kamilla Heddeland Instefjord, Hilde Elshaug, Garcia Llorente Ignacio, Jon Bråte, Engebretsen Serina Beate,Pedersen Benedikte Nevjen, Line Victoria Moen, Debech Nadia, Atiya R Ali,Marie Paulsen Madsen, Rasmus Riis Kopperud, Hilde Vollan, Karoline Bragstad, Olav Hungnes |
| EPI_ISL_3152642 | Vestfold Hospital, Toensberg Department of Microbiology | Norwegian Institute of Public Health, Department of Virology | Kathrine Stene-Johansen, Kamilla Heddeland Instefjord, Hilde Elshaug, Garcia Llorente Ignacio, Jon Bråte, Engebretsen Serina Beate,Pedersen Benedikte Nevjen, Line Victoria Moen, Debech Nadia, Atiya R Ali,Marie Paulsen Madsen, Rasmus Riis Kopperud, Hilde Vollan, Karoline Bragstad, Olav Hungnes |
| EPI_ISL_3152643, EPI_ISL_3152645, EPI_ISL_3152648, EPI_ISL_3152650, EPI_ISL_3152651, EPI_ISL_3152654 | Furst Medical Laboratory | Norwegian Institute of Public Health, Department of Virology | Kathrine Stene-Johansen, Kamilla Heddeland Instefjord, Hilde Elshaug, Garcia Llorente Ignacio, Jon Bråte, Engebretsen Serina Beate,Pedersen Benedikte Nevjen, Line Victoria Moen, Debech Nadia, Atiya R Ali,Marie Paulsen Madsen, Rasmus Riis Kopperud, Hilde Vollan, Karoline Bragstad, Olav Hungnes |
| EPI_ISL_3152655, EPI_ISL_3152657 | Haukeland University Hospital, Dept. of Microbiology | Norwegian Institute of Public Health, Department of Virology | Kathrine Stene-Johansen, Kamilla Heddeland Instefjord, Hilde Elshaug, Garcia Llorente Ignacio, Jon Bråte, Engebretsen Serina Beate,Pedersen Benedikte Nevjen, Line Victoria Moen, Debech Nadia, Atiya R Ali,Marie Paulsen Madsen, Rasmus Riis Kopperud, Hilde Vollan, Karoline Bragstad, Olav Hungnes |
| EPI_ISL_3152659, EPI_ISL_3152660, EPI_ISL_3152662, EPI_ISL_3152663, EPI_ISL_3152664 | Medical Microbiology Unit, Department for Laboratory Medicine, Drammen Hospital, Vestre Viken Health Trust, | Norwegian Institute of Public Health, Department of Virology | Kathrine Stene-Johansen, Kamilla Heddeland Instefjord, Hilde Elshaug, Garcia Llorente Ignacio, Jon Bråte, Engebretsen Serina Beate,Pedersen Benedikte Nevjen, Line Victoria Moen, Debech Nadia, Atiya R Ali,Marie Paulsen Madsen, Rasmus Riis Kopperud, Hilde Vollan, Karoline Bragstad, Olav Hungnes |
| EPI_ISL_3152665 | Dept. of Medical Microbiology, Stavanger University Hospital, Helse Stavanger HF | Norwegian Institute of Public Health, Department of Virology | Kathrine Stene-Johansen, Kamilla Heddeland Instefjord, Hilde Elshaug, Garcia Llorente Ignacio, Jon Bråte, Engebretsen Serina Beate,Pedersen Benedikte Nevjen, Line Victoria Moen, Debech Nadia, Atiya R Ali,Marie Paulsen Madsen, Rasmus Riis Kopperud, Hilde Vollan, Karoline Bragstad, Olav Hungnes |
| EPI_ISL_3152666 | Medical Microbiology Unit, Department for Laboratory Medicine, Drammen Hospital, Vestre Viken Health Trust, | Norwegian Institute of Public Health, Department of Virology | Kathrine Stene-Johansen, Kamilla Heddeland Instefjord, Hilde Elshaug, Garcia Llorente Ignacio, Jon Bråte, Engebretsen Serina Beate,Pedersen Benedikte Nevjen, Line Victoria Moen, Debech Nadia, Atiya R Ali,Marie Paulsen Madsen, Rasmus Riis Kopperud, Hilde Vollan, Karoline Bragstad, Olav Hungnes |
| EPI_ISL_3152668 | Haukeland University Hospital, Dept. of Microbiology | Norwegian Institute of Public Health, Department of Virology | Kathrine Stene-Johansen, Kamilla Heddeland Instefjord, Hilde Elshaug, Garcia Llorente Ignacio, Jon Bråte, Engebretsen Serina Beate,Pedersen Benedikte Nevjen, Line Victoria Moen, Debech Nadia, Atiya R Ali,Marie Paulsen Madsen, Rasmus Riis Kopperud, Hilde Vollan, Karoline Bragstad, Olav Hungnes |
| EPI_ISL_3152669 | Dept. of Medical Microbiology, Stavanger University Hospital, Helse Stavanger HF | Norwegian Institute of Public Health, Department of Virology | Kathrine Stene-Johansen, Kamilla Heddeland Instefjord, Hilde Elshaug, Garcia Llorente Ignacio, Jon Bråte, Engebretsen Serina Beate,Pedersen Benedikte Nevjen, Line Victoria Moen, Debech Nadia, Atiya R Ali,Marie Paulsen Madsen, Rasmus Riis Kopperud, Hilde Vollan, Karoline Bragstad, Olav Hungnes |
| EPI_ISL_3152672, EPI_ISL_3152673 | Medical Microbiology Unit, Department for Laboratory Medicine, Drammen Hospital, Vestre Viken Health Trust, | Norwegian Institute of Public Health, Department of Virology | Kathrine Stene-Johansen, Kamilla Heddeland Instefjord, Hilde Elshaug, Garcia Llorente Ignacio, Jon Bråte, Engebretsen Serina Beate,Pedersen Benedikte Nevjen, Line Victoria Moen, Debech Nadia, Atiya R Ali,Marie Paulsen Madsen, Rasmus Riis Kopperud, Hilde Vollan, Karoline Bragstad, Olav Hungnes |
| EPI_ISL_3152674 | Hospital of Southern Norway - Kristiansand, Department of Medical Microbiology | Norwegian Institute of Public Health, Department of Virology | Kathrine Stene-Johansen, Kamilla Heddeland Instefjord, Hilde Elshaug, Garcia Llorente Ignacio, Jon Bråte, Engebretsen Serina Beate,Pedersen Benedikte Nevjen, Line Victoria Moen, Debech Nadia, Atiya R Ali,Marie Paulsen Madsen, Rasmus Riis Kopperud, Hilde Vollan, Karoline Bragstad, Olav Hungnes |
| EPI_ISL_3152676 | Medical Microbiology Unit, Department for Laboratory Medicine, Drammen Hospital, Vestre Viken Health Trust, | Norwegian Institute of Public Health, Department of Virology | Kathrine Stene-Johansen, Kamilla Heddeland Instefjord, Hilde Elshaug, Garcia Llorente Ignacio, Jon Bråte, Engebretsen Serina Beate,Pedersen Benedikte Nevjen, Line Victoria Moen, Debech Nadia, Atiya R Ali,Marie Paulsen Madsen, Rasmus Riis Kopperud, Hilde Vollan, Karoline Bragstad, Olav Hungnes |
| EPI_ISL_3152677 | Ostfold Hospital Trust - Kalnes, Centre for Laboratory Medicine, Section for gene technology and infection serology | Norwegian Institute of Public Health, Department of Virology | Kathrine Stene-Johansen, Kamilla Heddeland Instefjord, Hilde Elshaug, Garcia Llorente Ignacio, Jon Bråte, Engebretsen Serina Beate,Pedersen Benedikte Nevjen, Line Victoria Moen, Debech Nadia, Atiya R Ali,Marie Paulsen Madsen, Rasmus Riis Kopperud, Hilde Vollan, Karoline Bragstad, Olav Hungnes |
| EPI_ISL_3152678 | Hospital of Southern Norway - Kristiansand, Department of Medical Microbiology | Norwegian Institute of Public Health, Department of Virology | Kathrine Stene-Johansen, Kamilla Heddeland Instefjord, Hilde Elshaug, Garcia Llorente Ignacio, Jon Bråte, Engebretsen Serina Beate,Pedersen Benedikte Nevjen, Line Victoria Moen, Debech Nadia, Atiya R Ali,Marie Paulsen Madsen, Rasmus Riis Kopperud, Hilde Vollan, Karoline Bragstad, Olav Hungnes |
| EPI_ISL_3152679, EPI_ISL_3152680, EPI_ISL_3152681 | Medical Microbiology Unit, Department for Laboratory Medicine, Drammen Hospital, Vestre Viken Health Trust, | Norwegian Institute of Public Health, Department of Virology | Kathrine Stene-Johansen, Kamilla Heddeland Instefjord, Hilde Elshaug, Garcia Llorente Ignacio, Jon Bråte, Engebretsen Serina Beate,Pedersen Benedikte Nevjen, Line Victoria Moen, Debech Nadia, Atiya R Ali,Marie Paulsen Madsen, Rasmus Riis Kopperud, Hilde Vollan, Karoline Bragstad, Olav Hungnes |
| EPI_ISL_3152682 | Nordland Hospital - Bodo, Laboratory Department, Molecular Biology Unit | Norwegian Institute of Public Health, Department of Virology | Kathrine Stene-Johansen, Kamilla Heddeland Instefjord, Hilde Elshaug, Garcia Llorente Ignacio, Jon Bråte, Engebretsen Serina Beate,Pedersen Benedikte Nevjen, Line Victoria Moen, Debech Nadia, Atiya R Ali,Marie Paulsen Madsen, Rasmus Riis Kopperud, Hilde Vollan, Karoline Bragstad, Olav Hungnes |
| EPI_ISL_3152683 | Medical Microbiology Unit, Department for Laboratory Medicine, Drammen Hospital, Vestre Viken Health Trust, | Norwegian Institute of Public Health, Department of Virology | Kathrine Stene-Johansen, Kamilla Heddeland Instefjord, Hilde Elshaug, Garcia Llorente Ignacio, Jon Bråte, Engebretsen Serina Beate,Pedersen Benedikte Nevjen, Line Victoria Moen, Debech Nadia, Atiya R Ali,Marie Paulsen Madsen, Rasmus Riis Kopperud, Hilde Vollan, Karoline Bragstad, Olav Hungnes |
| EPI_ISL_3152684 | Hospital of Southern Norway - Kristiansand, Department of Medical Microbiology | Norwegian Institute of Public Health, Department of Virology | Kathrine Stene-Johansen, Kamilla Heddeland Instefjord, Hilde Elshaug, Garcia Llorente Ignacio, Jon Bråte, Engebretsen Serina Beate,Pedersen Benedikte Nevjen, Line Victoria Moen, Debech Nadia, Atiya R Ali,Marie Paulsen Madsen, Rasmus Riis Kopperud, Hilde Vollan, Karoline Bragstad, Olav Hungnes |
| EPI_ISL_3152685 | Haukeland University Hospital, Dept. of Microbiology | Norwegian Institute of Public Health, Department of Virology | Kathrine Stene-Johansen, Kamilla Heddeland Instefjord, Hilde Elshaug, Garcia Llorente Ignacio, Jon Bråte, Engebretsen Serina Beate,Pedersen Benedikte Nevjen, Line Victoria Moen, Debech Nadia, Atiya R Ali,Marie Paulsen Madsen, Rasmus Riis Kopperud, Hilde Vollan, Karoline Bragstad, Olav Hungnes |
| EPI_ISL_3152687, EPI_ISL_3152688 | Medical Microbiology Unit, Department for Laboratory Medicine, Drammen Hospital, Vestre Viken Health Trust, | Norwegian Institute of Public Health, Department of Virology | Kathrine Stene-Johansen, Kamilla Heddeland Instefjord, Hilde Elshaug, Garcia Llorente Ignacio, Jon Bråte, Engebretsen Serina Beate,Pedersen Benedikte Nevjen, Line Victoria Moen, Debech Nadia, Atiya R Ali,Marie Paulsen Madsen, Rasmus Riis Kopperud, Hilde Vollan, Karoline Bragstad, Olav Hungnes |
| EPI_ISL_3152690 | Nordland Hospital - Bodo, Laboratory Department, Molecular Biology Unit | Norwegian Institute of Public Health, Department of Virology | Kathrine Stene-Johansen, Kamilla Heddeland Instefjord, Hilde Elshaug, Garcia Llorente Ignacio, Jon Bråte, Engebretsen Serina Beate,Pedersen Benedikte Nevjen, Line Victoria Moen, Debech Nadia, Atiya R Ali,Marie Paulsen Madsen, Rasmus Riis Kopperud, Hilde Vollan, Karoline Bragstad, Olav Hungnes |
| EPI_ISL_3152691, EPI_ISL_3152692, EPI_ISL_3152693 | Furst Medical Laboratory | Norwegian Institute of Public Health, Department of Virology | Kathrine Stene-Johansen, Kamilla Heddeland Instefjord, Hilde Elshaug, Garcia Llorente Ignacio, Jon Bråte, Engebretsen Serina Beate,Pedersen Benedikte Nevjen, Line Victoria Moen, Debech Nadia, Atiya R Ali,Marie Paulsen Madsen, Rasmus Riis Kopperud, Hilde Vollan, Karoline Bragstad, Olav Hungnes |
| EPI_ISL_3152694 | Haukeland University Hospital, Dept. of Microbiology | Norwegian Institute of Public Health, Department of Virology | Kathrine Stene-Johansen, Kamilla Heddeland Instefjord, Hilde Elshaug, Garcia Llorente Ignacio, Jon Bråte, Engebretsen Serina Beate,Pedersen Benedikte Nevjen, Line Victoria Moen, Debech Nadia, Atiya R Ali,Marie Paulsen Madsen, Rasmus Riis Kopperud, Hilde Vollan, Karoline Bragstad, Olav Hungnes |
| EPI_ISL_3152695, EPI_ISL_3152696 | Medical Microbiology Unit, Department for Laboratory Medicine, Drammen Hospital, Vestre Viken Health Trust, | Norwegian Institute of Public Health, Department of Virology | Kathrine Stene-Johansen, Kamilla Heddeland Instefjord, Hilde Elshaug, Garcia Llorente Ignacio, Jon Bråte, Engebretsen Serina Beate,Pedersen Benedikte Nevjen, Line Victoria Moen, Debech Nadia, Atiya R Ali,Marie Paulsen Madsen, Rasmus Riis Kopperud, Hilde Vollan, Karoline Bragstad, Olav Hungnes |

| | | | |
|---|---|---|---|
| EPI_ISL_3152697 | Hospital of Southern Norway - Kristiansand, Department of Medical Microbiology | Norwegian Institute of Public Health, Department of Virology | Kathrine Stene-Johansen, Kamilla Heddeland Instefjord, Hilde Elshaug, Garcia Llorente Ignacio, Jon Bråte, Engebretsen Serina Beate,Pedersen Benedikte Nevjen, Line Victoria Moen, Debech Nadia, Atiya R Ali,Marie Paulsen Madsen, Rasmus Riis Kopperud, Hilde Vollan, Karoline Bragstad, Olav Hungnes |
| EPI_ISL_3152698, EPI_ISL_3152699 | Medical Microbiology Unit, Department for Laboratory Medicine, Drammen Hospital, Vestre Viken Health Trust, | Norwegian Institute of Public Health, Department of Virology | Kathrine Stene-Johansen, Kamilla Heddeland Instefjord, Hilde Elshaug, Garcia Llorente Ignacio, Jon Bråte, Engebretsen Serina Beate,Pedersen Benedikte Nevjen, Line Victoria Moen, Debech Nadia, Atiya R Ali,Marie Paulsen Madsen, Rasmus Riis Kopperud, Hilde Vollan, Karoline Bragstad, Olav Hungnes |
| EPI_ISL_3152700 | Hospital of Southern Norway - Kristiansand, Department of Medical Microbiology | Norwegian Institute of Public Health, Department of Virology | Kathrine Stene-Johansen, Kamilla Heddeland Instefjord, Hilde Elshaug, Garcia Llorente Ignacio, Jon Bråte, Engebretsen Serina Beate,Pedersen Benedikte Nevjen, Line Victoria Moen, Debech Nadia, Atiya R Ali,Marie Paulsen Madsen, Rasmus Riis Kopperud, Hilde Vollan, Karoline Bragstad, Olav Hungnes |
| EPI_ISL_3152702, EPI_ISL_3152704 | Medical Microbiology Unit, Department for Laboratory Medicine, Drammen Hospital, Vestre Viken Health Trust, | Norwegian Institute of Public Health, Department of Virology | Kathrine Stene-Johansen, Kamilla Heddeland Instefjord, Hilde Elshaug, Garcia Llorente Ignacio, Jon Bråte, Engebretsen Serina Beate,Pedersen Benedikte Nevjen, Line Victoria Moen, Debech Nadia, Atiya R Ali,Marie Paulsen Madsen, Rasmus Riis Kopperud, Hilde Vollan, Karoline Bragstad, Olav Hungnes |
| EPI_ISL_3152705 | Ostfold Hospital Trust - Kalnes, Centre for Laboratory Medicine, Section for gene technology and infection serology | Norwegian Institute of Public Health, Department of Virology | Kathrine Stene-Johansen, Kamilla Heddeland Instefjord, Hilde Elshaug, Garcia Llorente Ignacio, Jon Bråte, Engebretsen Serina Beate,Pedersen Benedikte Nevjen, Line Victoria Moen, Debech Nadia, Atiya R Ali,Marie Paulsen Madsen, Rasmus Riis Kopperud, Hilde Vollan, Karoline Bragstad, Olav Hungnes |
| EPI_ISL_3152706 | Haukeland University Hospital, Dept. of Microbiology | Norwegian Institute of Public Health, Department of Virology | Kathrine Stene-Johansen, Kamilla Heddeland Instefjord, Hilde Elshaug, Garcia Llorente Ignacio, Jon Bråte, Engebretsen Serina Beate,Pedersen Benedikte Nevjen, Line Victoria Moen, Debech Nadia, Atiya R Ali,Marie Paulsen Madsen, Rasmus Riis Kopperud, Hilde Vollan, Karoline Bragstad, Olav Hungnes |
| EPI_ISL_3152707, EPI_ISL_3152708 | Medical Microbiology Unit, Department for Laboratory Medicine, Drammen Hospital, Vestre Viken Health Trust, | Norwegian Institute of Public Health, Department of Virology | Kathrine Stene-Johansen, Kamilla Heddeland Instefjord, Hilde Elshaug, Garcia Llorente Ignacio, Jon Bråte, Engebretsen Serina Beate,Pedersen Benedikte Nevjen, Line Victoria Moen, Debech Nadia, Atiya R Ali,Marie Paulsen Madsen, Rasmus Riis Kopperud, Hilde Vollan, Karoline Bragstad, Olav Hungnes |
| EPI_ISL_3152709, EPI_ISL_3152710 | Nordland Hospital - Bodo, Laboratory Department, Molecular Biology Unit | Norwegian Institute of Public Health, Department of Virology | Kathrine Stene-Johansen, Kamilla Heddeland Instefjord, Hilde Elshaug, Garcia Llorente Ignacio, Jon Bråte, Engebretsen Serina Beate,Pedersen Benedikte Nevjen, Line Victoria Moen, Debech Nadia, Atiya R Ali,Marie Paulsen Madsen, Rasmus Riis Kopperud, Hilde Vollan, Karoline Bragstad, Olav Hungnes |
| EPI_ISL_3152711 | University Hospital of Northern Norway, Department for Microbiology and Infectious Disease Control | Norwegian Institute of Public Health, Department of Virology | Kathrine Stene-Johansen, Kamilla Heddeland Instefjord, Hilde Elshaug, Garcia Llorente Ignacio, Jon Bråte, Engebretsen Serina Beate,Pedersen Benedikte Nevjen, Line Victoria Moen, Debech Nadia, Atiya R Ali,Marie Paulsen Madsen, Rasmus Riis Kopperud, Hilde Vollan, Karoline Bragstad, Olav Hungnes |
| EPI_ISL_3152712 | Medical Microbiology Unit, Department for Laboratory Medicine, Drammen Hospital, Vestre Viken Health Trust, | Norwegian Institute of Public Health, Department of Virology | Kathrine Stene-Johansen, Kamilla Heddeland Instefjord, Hilde Elshaug, Garcia Llorente Ignacio, Jon Bråte, Engebretsen Serina Beate,Pedersen Benedikte Nevjen, Line Victoria Moen, Debech Nadia, Atiya R Ali,Marie Paulsen Madsen, Rasmus Riis Kopperud, Hilde Vollan, Karoline Bragstad, Olav Hungnes |
| EPI_ISL_3152713 | Nordland Hospital - Bodo, Laboratory Department, Molecular Biology Unit | Norwegian Institute of Public Health, Department of Virology | Kathrine Stene-Johansen, Kamilla Heddeland Instefjord, Hilde Elshaug, Garcia Llorente Ignacio, Jon Bråte, Engebretsen Serina Beate,Pedersen Benedikte Nevjen, Line Victoria Moen, Debech Nadia, Atiya R Ali,Marie Paulsen Madsen, Rasmus Riis Kopperud, Hilde Vollan, Karoline Bragstad, Olav Hungnes |
| EPI_ISL_3152714, EPI_ISL_3152715 | Haukeland University Hospital, Dept. of Microbiology | Norwegian Institute of Public Health, Department of Virology | Kathrine Stene-Johansen, Kamilla Heddeland Instefjord, Hilde Elshaug, Garcia Llorente Ignacio, Jon Bråte, Engebretsen Serina Beate,Pedersen Benedikte Nevjen, Line Victoria Moen, Debech Nadia, Atiya R Ali,Marie Paulsen Madsen, Rasmus Riis Kopperud, Hilde Vollan, Karoline Bragstad, Olav Hungnes |
| EPI_ISL_3152716, EPI_ISL_3152717 | Medical Microbiology Unit, Department for Laboratory Medicine, Drammen Hospital, Vestre Viken Health Trust, | Norwegian Institute of Public Health, Department of Virology | Kathrine Stene-Johansen, Kamilla Heddeland Instefjord, Hilde Elshaug, Garcia Llorente Ignacio, Jon Bråte, Engebretsen Serina Beate,Pedersen Benedikte Nevjen, Line Victoria Moen, Debech Nadia, Atiya R Ali,Marie Paulsen Madsen, Rasmus Riis Kopperud, Hilde Vollan, Karoline Bragstad, Olav Hungnes |
| EPI_ISL_3152718 | Hospital of Southern Norway - Kristiansand, Department of Medical Microbiology | Norwegian Institute of Public Health, Department of Virology | Kathrine Stene-Johansen, Kamilla Heddeland Instefjord, Hilde Elshaug, Garcia Llorente Ignacio, Jon Bråte, Engebretsen Serina Beate,Pedersen Benedikte Nevjen, Line Victoria Moen, Debech Nadia, Atiya R Ali,Marie Paulsen Madsen, Rasmus Riis Kopperud, Hilde Vollan, Karoline Bragstad, Olav Hungnes |

EPI_ISL_3152724, EPI_ISL_3152725, EPI_ISL_3152726, EPI_ISL_3152727, EPI_ISL_3152728, EPI_ISL_3152729, EPI_ISL_3152730, EPI_ISL_3152731, EPI_ISL_3152732, EPI_ISL_3152733, EPI_ISL_3152734, EPI_ISL_3152735, EPI_ISL_3152736, EPI_ISL_3152737, EPI_ISL_3152738, EPI_ISL_3152739, EPI_ISL_3152741, EPI_ISL_3152742, EPI_ISL_3152743, EPI_ISL_3152744, EPI_ISL_3152745, EPI_ISL_3152746, EPI_ISL_3152747, EPI_ISL_3152748, EPI_ISL_3152750, EPI_ISL_3152751, EPI_ISL_3152752, EPI_ISL_3152753, EPI_ISL_3152754, EPI_ISL_3152755, EPI_ISL_3152756, EPI_ISL_3152757, EPI_ISL_3152758, EPI_ISL_3152759, EPI_ISL_3152760, EPI_ISL_3152761, EPI_ISL_3152762, EPI_ISL_3152763, EPI_ISL_3152764, EPI_ISL_3152765, EPI_ISL_3152766, EPI_ISL_3152767, EPI_ISL_3152768, EPI_ISL_3152769, EPI_ISL_3152770, EPI_ISL_3152771, EPI_ISL_3152772, EPI_ISL_3152773, EPI_ISL_3152774, EPI_ISL_3152775, EPI_ISL_3152776, EPI_ISL_3152777, EPI_ISL_3152778, EPI_ISL_3152779, EPI_ISL_3152780, EPI_ISL_3152781, EPI_ISL_3152782, EPI_ISL_3152783, EPI_ISL_3152784, EPI_ISL_3152785, EPI_ISL_3152786, EPI_ISL_3152787, EPI_ISL_3152788, EPI_ISL_3152789, EPI_ISL_3152792, EPI_ISL_3152793, EPI_ISL_3152794, EPI_ISL_3152795, EPI_ISL_3152796, EPI_ISL_3152797, EPI_ISL_3152798, EPI_ISL_3152799, EPI_ISL_3152800, EPI_ISL_3152801, EPI_ISL_3152802, EPI_ISL_3152803, EPI_ISL_3152804, EPI_ISL_3152805

| | | | |
|---|---|---|---|
| see above | Servicio de Microbiología. Hospital General Universitario de Castellón | SeqCOVID-SPAIN consortium/IBV(CSIC) | Rosario Moreno Muñoz, María Dolores Tirado Balaguer and SeqCOVID-SPAIN consortium |

EPI_ISL_3152806, EPI_ISL_3152808, EPI_ISL_3152809, EPI_ISL_3152810, EPI_ISL_3152811, EPI_ISL_3152812, EPI_ISL_3152813, EPI_ISL_3152814, EPI_ISL_3152815, EPI_ISL_3152816, EPI_ISL_3152817, EPI_ISL_3152818, EPI_ISL_3152819, EPI_ISL_3152820, EPI_ISL_3152821, EPI_ISL_3152822, EPI_ISL_3152823, EPI_ISL_3152824, EPI_ISL_3152825, EPI_ISL_3152826, EPI_ISL_3152827, EPI_ISL_3152828, EPI_ISL_3152829

| | | | |
|---|---|---|---|
| see above | Plateforme de testing Namuroise | Plateforme de testing Namuroise | ; Demars Aurore ; Maschietto Céline ; Nobis Chloé ; Otto Gaetan ; Denis Olivier ; Degosserie Jonathan ; Mullier François |

EPI_ISL_3152903, EPI_ISL_3152904, EPI_ISL_3152905, EPI_ISL_3152906, EPI_ISL_3152907, EPI_ISL_3152908, EPI_ISL_3152909, EPI_ISL_3152910, EPI_ISL_3152911, EPI_ISL_3152912, EPI_ISL_3152913, EPI_ISL_3152914, EPI_ISL_3152915, EPI_ISL_3152916, EPI_ISL_3152917, EPI_ISL_3152918, EPI_ISL_3152919, EPI_ISL_3152920, EPI_ISL_3152921, EPI_ISL_3152922, EPI_ISL_3152923, EPI_ISL_3152924, EPI_ISL_3152926, EPI_ISL_3152927, EPI_ISL_3152928, EPI_ISL_3152929, EPI_ISL_3152930, EPI_ISL_3152931, EPI_ISL_3152932, EPI_ISL_3152933, EPI_ISL_3152934, EPI_ISL_3152935, EPI_ISL_3152936, EPI_ISL_3152937

| | | | |
|---|---|---|---|
| see above | Ramathibodi Hospital | COVID-19 Network Investigations (CONI) Alliance | Elizabeth Batty, Wasun Chantratita, Thanat Chookajorn, Stefan Fernandez, Angkana Huang, Anthony R. Jones, Khajohn Joonlasak, Chonticha Klungtong, Theerarat Kochakarn, Namfon Kotanan, Krittikorn Kumpornsin, Duangkamon Loesbanluechai, Wudtichai Manasatienkij, Bhakbhoom Panthan, Ekawat Pasomsub, Kingkan Rakmanee, Insee Sensorn, Janjira Thaipadungpanit, Arporn Wangwiwatsin, Treewat Watthanachockchai |

EPI_ISL_3152938, EPI_ISL_3152939, EPI_ISL_3152940, EPI_ISL_3152941, EPI_ISL_3152942, EPI_ISL_3152943, EPI_ISL_3152944, EPI_ISL_3152945, EPI_ISL_3152946, EPI_ISL_3152947, EPI_ISL_3152948, EPI_ISL_3152949, EPI_ISL_3152950, EPI_ISL_3152951, EPI_ISL_3152952, EPI_ISL_3152953, EPI_ISL_3152954, EPI_ISL_3152955, EPI_ISL_3152956, EPI_ISL_3152957, EPI_ISL_3152958, EPI_ISL_3152959, EPI_ISL_3152960, EPI_ISL_3152961, EPI_ISL_3152962, EPI_ISL_3152963, EPI_ISL_3152964, EPI_ISL_3152965, EPI_ISL_3152966, EPI_ISL_3152967, EPI_ISL_3152968, EPI_ISL_3152969, EPI_ISL_3152970, EPI_ISL_3152971, EPI_ISL_3152972, EPI_ISL_3152973, EPI_ISL_3152974, EPI_ISL_3152975, EPI_ISL_3152976, EPI_ISL_3152977, EPI_ISL_3152978, EPI_ISL_3152979, EPI_ISL_3152981, EPI_ISL_3152982, EPI_ISL_3152983, EPI_ISL_3152984, EPI_ISL_3152985, EPI_ISL_3152986, EPI_ISL_3152987, EPI_ISL_3152988, EPI_ISL_3152989, EPI_ISL_3152990, EPI_ISL_3152991, EPI_ISL_3152992, EPI_ISL_3152993, EPI_ISL_3152994, EPI_ISL_3152995, EPI_ISL_3152996, EPI_ISL_3152997, EPI_ISL_3152998, EPI_ISL_3152999, EPI_ISL_3153000, EPI_ISL_3153001, EPI_ISL_3153002, EPI_ISL_3153003, EPI_ISL_3153004, EPI_ISL_3153005, EPI_ISL_3153006, EPI_ISL_3153007, EPI_ISL_3153008, EPI_ISL_3153009, EPI_ISL_3153010, EPI_ISL_3153011, EPI_ISL_3153012, EPI_ISL_3153013, EPI_ISL_3153014, EPI_ISL_3153015, EPI_ISL_3153016, EPI_ISL_3153017, EPI_ISL_3153018, EPI_ISL_3153019, EPI_ISL_3153020, EPI_ISL_3153021, EPI_ISL_3153022, EPI_ISL_3153023, EPI_ISL_3153024, EPI_ISL_3153025, EPI_ISL_3153026, EPI_ISL_3153027, EPI_ISL_3153028, EPI_ISL_3153029, EPI_ISL_3153030, EPI_ISL_3153031, EPI_ISL_3153032, EPI_ISL_3153033, EPI_ISL_3153034, EPI_ISL_3153035, EPI_ISL_3153036, EPI_ISL_3153037, EPI_ISL_3153038, EPI_ISL_3153039, EPI_ISL_3153040, EPI_ISL_3153041, EPI_ISL_3153042, EPI_ISL_3153043, EPI_ISL_3153044, EPI_ISL_3153045, EPI_ISL_3153046, EPI_ISL_3153047, EPI_ISL_3153048, EPI_ISL_3153049, EPI_ISL_3153050, EPI_ISL_3153051, EPI_ISL_3153052, EPI_ISL_3153053, EPI_ISL_3153054, EPI_ISL_3153055, EPI_ISL_3153056, EPI_ISL_3153057, EPI_ISL_3153059, EPI_ISL_3153060, EPI_ISL_3153061, EPI_ISL_3153062, EPI_ISL_3153064, EPI_ISL_3153065, EPI_ISL_3153066, EPI_ISL_3153067, EPI_ISL_3153068, EPI_ISL_3153069, EPI_ISL_3153070, EPI_ISL_3153071, EPI_ISL_3153072, EPI_ISL_3153073, EPI_ISL_3153074, EPI_ISL_3153075, EPI_ISL_3153076, EPI_ISL_3153077, EPI_ISL_3153078, EPI_ISL_3153079, EPI_ISL_3153080, EPI_ISL_3153081, EPI_ISL_3153082, EPI_ISL_3153083, EPI_ISL_3153084, EPI_ISL_3153085, EPI_ISL_3153086, EPI_ISL_3153087, EPI_ISL_3153088, EPI_ISL_3153089, EPI_ISL_3153090, EPI_ISL_3153091, EPI_ISL_3153092, EPI_ISL_3153093, EPI_ISL_3153094, EPI_ISL_3153095, EPI_ISL_3153096, EPI_ISL_3153097, EPI_ISL_3153098, EPI_ISL_3153099, EPI_ISL_3153100, EPI_ISL_3153101, EPI_ISL_3153102, EPI_ISL_3153103, EPI_ISL_3153104, EPI_ISL_3153105, EPI_ISL_3153107, EPI_ISL_3153108, EPI_ISL_3153109, EPI_ISL_3153110, EPI_ISL_3153111, EPI_ISL_3153112, EPI_ISL_3153113, EPI_ISL_3153114, EPI_ISL_3153115, EPI_ISL_3153116, EPI_ISL_3153117, EPI_ISL_3153118, EPI_ISL_3153119, EPI_ISL_3153120, EPI_ISL_3153121, EPI_ISL_3153122, EPI_ISL_3153123, EPI_ISL_3153124, EPI_ISL_3153125, EPI_ISL_3153126, EPI_ISL_3153127, EPI_ISL_3153128, EPI_ISL_3153129, EPI_ISL_3153130, EPI_ISL_3153131, EPI_ISL_3153132, EPI_ISL_3153133, EPI_ISL_3153134, EPI_ISL_3153135, EPI_ISL_3153136, EPI_ISL_3153137, EPI_ISL_3153138, EPI_ISL_3153139, EPI_ISL_3153140, EPI_ISL_3153141, EPI_ISL_3153142, EPI_ISL_3153143, EPI_ISL_3153144, EPI_ISL_3153145, EPI_ISL_3153146, EPI_ISL_3153147, EPI_ISL_3153148, EPI_ISL_3153149, EPI_ISL_3153150, EPI_ISL_3153151, EPI_ISL_3153152, EPI_ISL_3153153, EPI_ISL_3153154, EPI_ISL_3153155, EPI_ISL_3153156, EPI_ISL_3153157, EPI_ISL_3153158, EPI_ISL_3153159, EPI_ISL_3153160, EPI_ISL_3153161, EPI_ISL_3153162, EPI_ISL_3153163, EPI_ISL_3153164, EPI_ISL_3153165, EPI_ISL_3153166, EPI_ISL_3153167, EPI_ISL_3153168, EPI_ISL_3153169, EPI_ISL_3153170, EPI_ISL_3153171, EPI_ISL_3153172, EPI_ISL_3153173, EPI_ISL_3153174, EPI_ISL_3153175, EPI_ISL_3153176, EPI_ISL_3153177, EPI_ISL_3153178, EPI_ISL_3153179, EPI_ISL_3153180, EPI_ISL_3153181, EPI_ISL_3153182, EPI_ISL_3153183, EPI_ISL_3153184, EPI_ISL_3153185, EPI_ISL_3153186, EPI_ISL_3153187, EPI_ISL_3153188, EPI_ISL_3153189, EPI_ISL_3153190, EPI_ISL_3153191, EPI_ISL_3153192, EPI_ISL_3153193, EPI_ISL_3153194, EPI_ISL_3153195, EPI_ISL_3153196, EPI_ISL_3153197, EPI_ISL_3153198, EPI_ISL_3153199, EPI_ISL_3153200, EPI_ISL_3153201

| | | | |
|---|---|---|---|
| see above | Broad Institute Clinical Research Sequencing Platform | Infectious Disease Program, Broad Institute of Harvard and MIT | Siddle,K.J., Adams,G., Pearlman,L., Gladden-Young,A., Vicente,G., Blumenstiel,B., DeFelice,M., Lee,M., McGovern,S., Lagerborg,K., Rudy,M., DeRuff,K., Carter,A., Normandin,E., Bauer,M., Reilly,S., Tomkins-Tinch,C., Loreth,C., Chaluvadi,S., Meldrim,J., Granger,B., Lemieux,J.E., Birren,B.W., Sabeti,P.C., Larkin,K., Dodge,S., Lennon,N., Madoff,L., Brown,C., Gallagher,G., Smole,S., Park,D.J., Gabriel,S., and MacInnis,B.L. |

EPI_ISL_3153202, EPI_ISL_3153203, EPI_ISL_3153204, EPI_ISL_3153205, EPI_ISL_3153206, EPI_ISL_3153208, EPI_ISL_3153209, EPI_ISL_3153210, EPI_ISL_3153211, EPI_ISL_3153212, EPI_ISL_3153213, EPI_ISL_3153214, EPI_ISL_3153215, EPI_ISL_3153216, EPI_ISL_3153217, EPI_ISL_3153218, EPI_ISL_3153219, EPI_ISL_3153220, EPI_ISL_3153221, EPI_ISL_3153222, EPI_ISL_3153223

| | | | |
|---|---|---|---|
| see above | A.S.L. CUNEO 1 | Fondazione del Piemonte per l'Oncologia IRCCS | Antonino Sottile, Silvia Brossa, Paola Marino, Giorgio Giardina |
| EPI_ISL_3153224 | A.S.L. TO3 | Fondazione del Piemonte per l'Oncologia IRCCS | Antonino Sottile, Silvia Brossa, Paola Marino, Giorgio Giardina |
| EPI_ISL_3153225 | A.S.L. CUNEO 1 | Fondazione del Piemonte per l'Oncologia IRCCS | Antonino Sottile, Silvia Brossa, Paola Marino, Giorgio Giardina |
| EPI_ISL_3153226 | A.S.L. TO4 | Fondazione del Piemonte per l'Oncologia IRCCS | Antonino Sottile, Silvia Brossa, Paola Marino, Giorgio Giardina |
| EPI_ISL_3153227 | A.S.L. CUNEO 1 | Fondazione del Piemonte per l'Oncologia IRCCS | Antonino Sottile, Silvia Brossa, Paola Marino, Giorgio Giardina |
| EPI_ISL_3153228, EPI_ISL_3153230, EPI_ISL_3153232 | A.S.L. TO4 | Fondazione del Piemonte per l'Oncologia IRCCS | Antonino Sottile, Silvia Brossa, Paola Marino, Giorgio Giardina |
| EPI_ISL_3153234 | A.S.L. CUNEO 2 | Fondazione del Piemonte per l'Oncologia IRCCS | Antonino Sottile, Silvia Brossa, Paola Marino, Giorgio Giardina |
| EPI_ISL_3153235 | A.S.L. TO4 | Fondazione del Piemonte per l'Oncologia IRCCS | Antonino Sottile, Silvia Brossa, Paola Marino, Giorgio Giardina |
| EPI_ISL_3153236, EPI_ISL_3153237 | A.S.L. CUNEO 2 | Fondazione del Piemonte per l'Oncologia IRCCS | Antonino Sottile, Silvia Brossa, Paola Marino, Giorgio Giardina |
| EPI_ISL_3153238 | A.S.L. TO4 | Fondazione del Piemonte per l'Oncologia IRCCS | Antonino Sottile, Silvia Brossa, Paola Marino, Giorgio Giardina |
| EPI_ISL_3153241 | EHNV | Laboratory of genomics and metagenomics | Trestan Pillonel, Damien Jacot, Sébastien Aeby, Gilbert Greub, Claire Bertelli |
| EPI_ISL_3153242 | VIDYMED LAUSANNE | Laboratory of genomics and metagenomics | Trestan Pillonel, Damien Jacot, Sébastien Aeby, Gilbert Greub, Claire Bertelli |
| EPI_ISL_3153243, EPI_ISL_3153244, EPI_ISL_3153245, EPI_ISL_3153246, EPI_ISL_3153247, EPI_ISL_3153248 | CHUV | Laboratory of genomics and metagenomics | Trestan Pillonel, Damien Jacot, Sébastien Aeby, Gilbert Greub, Claire Bertelli |
| EPI_ISL_3153249, EPI_ISL_3153250, EPI_ISL_3153251 | VIDYMED LAUSANNE | Laboratory of genomics and metagenomics | Trestan Pillonel, Damien Jacot, Sébastien Aeby, Gilbert Greub, Claire Bertelli |
| EPI_ISL_3153252 | CHUV | Laboratory of genomics and metagenomics | Trestan Pillonel, Damien Jacot, Sébastien Aeby, Gilbert Greub, Claire Bertelli |
| EPI_ISL_3153253, EPI_ISL_3153254, EPI_ISL_3153255, EPI_ISL_3153256, EPI_ISL_3153257, EPI_ISL_3153258, EPI_ISL_3153259, EPI_ISL_3153260 | VIDYMED LAUSANNE | Laboratory of genomics and metagenomics | Trestan Pillonel, Damien Jacot, Sébastien Aeby, Gilbert Greub, Claire Bertelli |
| EPI_ISL_3153261 | EHC MORGES | Laboratory of genomics and metagenomics | Trestan Pillonel, Damien Jacot, Sébastien Aeby, Gilbert Greub, Claire Bertelli |
| EPI_ISL_3153262, EPI_ISL_3153263 | VIDYMED LAUSANNE | Laboratory of genomics and metagenomics | Trestan Pillonel, Damien Jacot, Sébastien Aeby, Gilbert Greub, Claire Bertelli |

| | | | |
|---|---|---|---|
| EPI_ISL_3153264, EPI_ISL_3153265 | EHNV | Laboratory of genomics and metagenomics | Trestan Pillonel, Damien Jacot, Sébastien Aeby, Gilbert Greub, Claire Bertelli |
| EPI_ISL_3153266 | CHUV | Laboratory of genomics and metagenomics | Trestan Pillonel, Damien Jacot, Sébastien Aeby, Gilbert Greub, Claire Bertelli |
| EPI_ISL_3153267, EPI_ISL_3153268, EPI_ISL_3153269, EPI_ISL_3153270, EPI_ISL_3153271, EPI_ISL_3153272, EPI_ISL_3153273, EPI_ISL_3153274, EPI_ISL_3153275, EPI_ISL_3153276, EPI_ISL_3153277, EPI_ISL_3153278, EPI_ISL_3153280, EPI_ISL_3153281, EPI_ISL_3153282 | | | |
| see above | EHNV | Laboratory of genomics and metagenomics | Trestan Pillonel, Damien Jacot, Sébastien Aeby, Gilbert Greub, Claire Bertelli |
| EPI_ISL_3153283, EPI_ISL_3153284 | CHUV | Laboratory of genomics and metagenomics | Trestan Pillonel, Damien Jacot, Sébastien Aeby, Gilbert Greub, Claire Bertelli |
| EPI_ISL_3153285, EPI_ISL_3153286, EPI_ISL_3153287 | VIDYMED LAUSANNE | Laboratory of genomics and metagenomics | Trestan Pillonel, Damien Jacot, Sébastien Aeby, Gilbert Greub, Claire Bertelli |
| EPI_ISL_3153288 | UNILABS - COPPET | Laboratory of genomics and metagenomics | Trestan Pillonel, Damien Jacot, Sébastien Aeby, Gilbert Greub, Claire Bertelli |
| EPI_ISL_3153290 | VIDYMED LAUSANNE | Laboratory of genomics and metagenomics | Trestan Pillonel, Damien Jacot, Sébastien Aeby, Gilbert Greub, Claire Bertelli |
| EPI_ISL_3153291 | CHUV | Laboratory of genomics and metagenomics | Trestan Pillonel, Damien Jacot, Sébastien Aeby, Gilbert Greub, Claire Bertelli |
| EPI_ISL_3153292 | VIDYMED LAUSANNE | Laboratory of genomics and metagenomics | Trestan Pillonel, Damien Jacot, Sébastien Aeby, Gilbert Greub, Claire Bertelli |
| EPI_ISL_3153293 | EHC MORGES | Laboratory of genomics and metagenomics | Trestan Pillonel, Damien Jacot, Sébastien Aeby, Gilbert Greub, Claire Bertelli |
| EPI_ISL_3153294, EPI_ISL_3153295, EPI_ISL_3153296, EPI_ISL_3153297 | VIDYMED LAUSANNE | Laboratory of genomics and metagenomics | Trestan Pillonel, Damien Jacot, Sébastien Aeby, Gilbert Greub, Claire Bertelli |
| EPI_ISL_3153298, EPI_ISL_3153299, EPI_ISL_3153300 | CHUV | Laboratory of genomics and metagenomics | Trestan Pillonel, Damien Jacot, Sébastien Aeby, Gilbert Greub, Claire Bertelli |
| EPI_ISL_3153301 | VIDYMED LAUSANNE | Laboratory of genomics and metagenomics | Trestan Pillonel, Damien Jacot, Sébastien Aeby, Gilbert Greub, Claire Bertelli |
| EPI_ISL_3153302 | CHUV | Laboratory of genomics and metagenomics | Trestan Pillonel, Damien Jacot, Sébastien Aeby, Gilbert Greub, Claire Bertelli |
| EPI_ISL_3153303 | VIDYMED LAUSANNE | Laboratory of genomics and metagenomics | Trestan Pillonel, Damien Jacot, Sébastien Aeby, Gilbert Greub, Claire Bertelli |
| EPI_ISL_3153304 | CHUV | Laboratory of genomics and metagenomics | Trestan Pillonel, Damien Jacot, Sébastien Aeby, Gilbert Greub, Claire Bertelli |
| EPI_ISL_3153305 | VIDYMED LAUSANNE | Laboratory of genomics and metagenomics | Trestan Pillonel, Damien Jacot, Sébastien Aeby, Gilbert Greub, Claire Bertelli |
| EPI_ISL_3153306, EPI_ISL_3153307, EPI_ISL_3153308, EPI_ISL_3153309, EPI_ISL_3153310 | CHUV | Laboratory of genomics and metagenomics | Trestan Pillonel, Damien Jacot, Sébastien Aeby, Gilbert Greub, Claire Bertelli |
| EPI_ISL_3153311 | VIDYMED LAUSANNE | Laboratory of genomics and metagenomics | Trestan Pillonel, Damien Jacot, Sébastien Aeby, Gilbert Greub, Claire Bertelli |
| EPI_ISL_3153312, EPI_ISL_3153313 | CHUV | Laboratory of genomics and metagenomics | Trestan Pillonel, Damien Jacot, Sébastien Aeby, Gilbert Greub, Claire Bertelli |
| EPI_ISL_3153314 | VIDYMED LAUSANNE | Laboratory of genomics and metagenomics | Trestan Pillonel, Damien Jacot, Sébastien Aeby, Gilbert Greub, Claire Bertelli |
| EPI_ISL_3153315 | TN DOH Lab Services | TN DOH Lab Services | Karen Beasley-Maynard, Brian Selinsky |
| EPI_ISL_3153330, EPI_ISL_3153350, EPI_ISL_3153365, EPI_ISL_3153373 | Labo Analyses Med | National Reference Center for Viruses of Respiratory Infections, Institut Pasteur, Paris | Marion Barbet, Sylvie Behillil, Méline Bizard, Angela Brisebarre, Camille Capel, Vincent Enouf, Louise Lefrançois, Frédéric Lemoine, Christophe Malabat, Corinne Maufrais, Hub de Bioinformatique et Biostatistique, Etienne Simon-Lorière, Maud Vanpeene, Sylvie Van der Werf ,Marie-HéLèNe Glaudon Louveau De La Guigneraye |
| EPI_ISL_3153374 | Hospital | National Reference Center for Viruses of Respiratory Infections, Institut Pasteur, Paris | Marion Barbet, Sylvie Behillil, Méline Bizard, Angela Brisebarre, Camille Capel, Vincent Enouf, Louise Lefrançois, Frédéric Lemoine, Christophe Malabat, Corinne Maufrais, Hub de Bioinformatique et Biostatistique, Etienne Simon-Lorière, Maud Vanpeene, Sylvie Van der Werf ,Marie-HéLèNe Glaudon Louveau De La Guigneraye |
| EPI_ISL_3153375 | Labo Analyses Med | National Reference Center for Viruses of Respiratory Infections, Institut Pasteur, Paris | Marion Barbet, Sylvie Behillil, Méline Bizard, Angela Brisebarre, Camille Capel, Vincent Enouf, Louise Lefrançois, Frédéric Lemoine, Christophe Malabat, Corinne Maufrais, Hub de Bioinformatique et Biostatistique, Etienne Simon-Lorière, Maud Vanpeene, Sylvie Van der Werf ,Marie-HéLèNe Glaudon Louveau De La Guigneraye |
| EPI_ISL_3153376, EPI_ISL_3153379 | Hospital | National Reference Center for Viruses of Respiratory Infections, Institut Pasteur, Paris | Marion Barbet, Sylvie Behillil, Méline Bizard, Angela Brisebarre, Camille Capel, Vincent Enouf, Louise Lefrançois, Frédéric Lemoine, Christophe Malabat, Corinne Maufrais, Hub de Bioinformatique et Biostatistique, Etienne Simon-Lorière, Maud Vanpeene, Sylvie Van der Werf ,Marie-HéLèNe Glaudon Louveau De La Guigneraye |
| EPI_ISL_3153380 | Labo Analyses Med | National Reference Center for Viruses of Respiratory Infections, Institut Pasteur, Paris | Marion Barbet, Sylvie Behillil, Méline Bizard, Angela Brisebarre, Camille Capel, Vincent Enouf, Louise Lefrançois, Frédéric Lemoine, Christophe Malabat, Corinne Maufrais, Hub de Bioinformatique et Biostatistique, Etienne Simon-Lorière, Maud Vanpeene, Sylvie Van der Werf ,Marie-HéLèNe Glaudon Louveau De La Guigneraye |
| EPI_ISL_3153381 | Hospital | National Reference Center for Viruses of Respiratory Infections, Institut Pasteur, Paris | Marion Barbet, Sylvie Behillil, Méline Bizard, Angela Brisebarre, Camille Capel, Vincent Enouf, Louise Lefrançois, Frédéric Lemoine, Christophe Malabat, Corinne Maufrais, Hub de Bioinformatique et Biostatistique, Etienne Simon-Lorière, Maud Vanpeene, Sylvie Van der Werf ,Marie-HéLèNe Glaudon Louveau De La Guigneraye |
| EPI_ISL_3153382, EPI_ISL_3153384, EPI_ISL_3153385 | Labo Analyses Med | National Reference Center for Viruses of Respiratory Infections, Institut Pasteur, Paris | Marion Barbet, Sylvie Behillil, Méline Bizard, Angela Brisebarre, Camille Capel, Vincent Enouf, Louise Lefrançois, Frédéric Lemoine, Christophe Malabat, Corinne Maufrais, Hub de Bioinformatique et Biostatistique, Etienne Simon-Lorière, Maud Vanpeene, Sylvie Van der Werf ,Marie-HéLèNe Glaudon Louveau De La Guigneraye |
| EPI_ISL_3153386, EPI_ISL_3153387 | Hospital | National Reference Center for Viruses of Respiratory Infections, Institut Pasteur, Paris | Marion Barbet, Sylvie Behillil, Méline Bizard, Angela Brisebarre, Camille Capel, Vincent Enouf, Louise Lefrançois, Frédéric Lemoine, Christophe Malabat, Corinne Maufrais, Hub de Bioinformatique et Biostatistique, Etienne Simon-Lorière, Maud Vanpeene, Sylvie Van der Werf ,Marie-HéLèNe Glaudon Louveau De La Guigneraye |
| EPI_ISL_3153388, EPI_ISL_3153389, EPI_ISL_3153390, EPI_ISL_3153391, EPI_ISL_3153392, EPI_ISL_3153393, EPI_ISL_3153394, EPI_ISL_3153395, EPI_ISL_3153396, EPI_ISL_3153397, EPI_ISL_3153398, EPI_ISL_3153399, EPI_ISL_3153400, EPI_ISL_3153401, EPI_ISL_3153402, EPI_ISL_3153403, EPI_ISL_3153404, EPI_ISL_3153405, EPI_ISL_3153406, EPI_ISL_3153407, EPI_ISL_3153408, EPI_ISL_3153409, EPI_ISL_3153410, EPI_ISL_3153411 | | | |
| see above | Labo Analyses Med | National Reference Center for Viruses of Respiratory Infections, Institut Pasteur, Paris | Marion Barbet, Sylvie Behillil, Méline Bizard, Angela Brisebarre, Camille Capel, Vincent Enouf, Louise Lefrançois, Frédéric Lemoine, Christophe Malabat, Corinne Maufrais, Hub de Bioinformatique et Biostatistique, Etienne Simon-Lorière, Maud Vanpeene, Sylvie Van der Werf ,Marie-HéLèNe Glaudon Louveau De La Guigneraye |
| EPI_ISL_3153412, EPI_ISL_3153413, EPI_ISL_3153414, EPI_ISL_3153415, EPI_ISL_3153416 | Hospital | National Reference Center for Viruses of Respiratory Infections, Institut Pasteur, Paris | Marion Barbet, Sylvie Behillil, Méline Bizard, Angela Brisebarre, Camille Capel, Vincent Enouf, Louise Lefrançois, Frédéric Lemoine, Christophe Malabat, Corinne Maufrais, Hub de Bioinformatique et Biostatistique, Etienne Simon-Lorière, Maud Vanpeene, Sylvie Van der Werf ,Marie-HéLèNe Glaudon Louveau De La Guigneraye |
| EPI_ISL_3153417, EPI_ISL_3153418, EPI_ISL_3153419 | Labo Analyses Med | National Reference Center for Viruses of Respiratory Infections, Institut Pasteur, Paris | Marion Barbet, Sylvie Behillil, Méline Bizard, Angela Brisebarre, Camille Capel, Vincent Enouf, Louise Lefrançois, Frédéric Lemoine, Christophe Malabat, Corinne Maufrais, Hub de Bioinformatique et Biostatistique, Etienne Simon-Lorière, Maud Vanpeene, Sylvie Van der Werf ,Marie-HéLèNe Glaudon Louveau De La Guigneraye |
| EPI_ISL_3153420, EPI_ISL_3153421, EPI_ISL_3153422 | Hospital | National Reference Center for Viruses of Respiratory Infections, Institut Pasteur, Paris | Marion Barbet, Sylvie Behillil, Méline Bizard, Angela Brisebarre, Camille Capel, Vincent Enouf, Louise Lefrançois, Frédéric Lemoine, Christophe Malabat, Corinne Maufrais, Hub de Bioinformatique et Biostatistique, Etienne Simon-Lorière, Maud Vanpeene, Sylvie Van der Werf ,Marie-HéLèNe Glaudon Louveau De La Guigneraye |
| EPI_ISL_3153423, EPI_ISL_3153424, EPI_ISL_3153425, EPI_ISL_3153426, EPI_ISL_3153427, EPI_ISL_3153428, EPI_ISL_3153429, EPI_ISL_3153430, EPI_ISL_3153431 | Labo Analyses Med | National Reference Center for Viruses of Respiratory Infections, Institut Pasteur, Paris | Marion Barbet, Sylvie Behillil, Méline Bizard, Angela Brisebarre, Camille Capel, Vincent Enouf, Louise Lefrançois, Frédéric Lemoine, Christophe Malabat, Corinne Maufrais, Hub de Bioinformatique et Biostatistique, Etienne Simon-Lorière, Maud Vanpeene, Sylvie Van der Werf ,Marie-HéLèNe Glaudon Louveau De La Guigneraye |
| EPI_ISL_3153432, EPI_ISL_3153433, EPI_ISL_3153434 | Hospital | National Reference Center for Viruses of Respiratory Infections, Institut Pasteur, Paris | Marion Barbet, Sylvie Behillil, Méline Bizard, Angela Brisebarre, Camille Capel, Vincent Enouf, Louise Lefrançois, Frédéric Lemoine, Christophe Malabat, Corinne Maufrais, Hub de Bioinformatique et Biostatistique, Etienne Simon-Lorière, Maud Vanpeene, Sylvie Van der Werf ,Marie-HéLèNe Glaudon Louveau De La Guigneraye |
| EPI_ISL_3153435, EPI_ISL_3153436, EPI_ISL_3153437, EPI_ISL_3153438, EPI_ISL_3153439, EPI_ISL_3153440, EPI_ISL_3153441 | Labo Analyses Med | National Reference Center for Viruses of Respiratory Infections, Institut Pasteur, Paris | Marion Barbet, Sylvie Behillil, Méline Bizard, Angela Brisebarre, Camille Capel, Vincent Enouf, Louise Lefrançois, Frédéric Lemoine, Christophe Malabat, Corinne Maufrais, Hub de Bioinformatique et Biostatistique, Etienne Simon-Lorière, Maud Vanpeene, Sylvie Van der Werf ,Marie-HéLèNe Glaudon Louveau De La Guigneraye |
| EPI_ISL_3153442, EPI_ISL_3153443, | Hospital | National Reference Center for Viruses of Respiratory Infections, Institut Pasteur, Paris | Marion Barbet, Sylvie Behillil, Méline Bizard, Angela Brisebarre, Camille Capel, Vincent Enouf, Louise Lefrançois, Frédéric Lemoine, Christophe Malabat, Corinne Maufrais, Hub de Bioinformatique et Biostatistique, Etienne Simon-Lorière, Maud Vanpeene, Sylvie Van der Werf ,Marie-HéLèNe Glaudon Louveau De La Guigneraye |

| | | | |
|---|---|---|---|
| EPI_ISL_3153444, EPI_ISL_3153445, EPI_ISL_3153446, EPI_ISL_3153447, EPI_ISL_3153448, EPI_ISL_3153449, EPI_ISL_3153450 | | | |
| EPI_ISL_3153451, EPI_ISL_3153452, EPI_ISL_3153453, EPI_ISL_3153454, EPI_ISL_3153455, EPI_ISL_3153456, EPI_ISL_3153457, EPI_ISL_3153458, EPI_ISL_3153459, EPI_ISL_3153460, EPI_ISL_3153461, EPI_ISL_3153462, EPI_ISL_3153463, EPI_ISL_3153464, EPI_ISL_3153465, EPI_ISL_3153466, EPI_ISL_3153467, EPI_ISL_3153468, EPI_ISL_3153469, EPI_ISL_3153470, EPI_ISL_3153471, EPI_ISL_3153472, EPI_ISL_3153473, EPI_ISL_3153474, EPI_ISL_3153475, EPI_ISL_3153476, EPI_ISL_3153477, EPI_ISL_3153478, EPI_ISL_3153479, EPI_ISL_3153480, EPI_ISL_3153481, EPI_ISL_3153482, EPI_ISL_3153483, EPI_ISL_3153484, EPI_ISL_3153485, EPI_ISL_3153486, EPI_ISL_3153487, EPI_ISL_3153488, EPI_ISL_3153489, EPI_ISL_3153490, EPI_ISL_3153491, EPI_ISL_3153492, EPI_ISL_3153493, EPI_ISL_3153494, EPI_ISL_3153495, EPI_ISL_3153496, EPI_ISL_3153497, EPI_ISL_3153498, EPI_ISL_3153499, EPI_ISL_3153500, EPI_ISL_3153501, EPI_ISL_3153502, EPI_ISL_3153503, EPI_ISL_3153504, EPI_ISL_3153505, EPI_ISL_3153506, EPI_ISL_3153507, EPI_ISL_3153508, EPI_ISL_3153509, EPI_ISL_3153510, EPI_ISL_3153511, EPI_ISL_3153512, EPI_ISL_3153513, EPI_ISL_3153514, EPI_ISL_3153515, EPI_ISL_3153516, EPI_ISL_3153517, EPI_ISL_3153518, EPI_ISL_3153519, EPI_ISL_3153520, EPI_ISL_3153521, EPI_ISL_3153522, EPI_ISL_3153523, EPI_ISL_3153524, EPI_ISL_3153525, EPI_ISL_3153526, EPI_ISL_3153527, EPI_ISL_3153528, EPI_ISL_3153529, EPI_ISL_3153530, EPI_ISL_3153531, EPI_ISL_3153532, EPI_ISL_3153533, EPI_ISL_3153534, EPI_ISL_3153535, EPI_ISL_3153536, EPI_ISL_3153537 | | | |
| see above | Labo Analyses Med | National Reference Center for Viruses of Respiratory Infections, Institut Pasteur, Paris | Marion Barbet, Sylvie Behillil, Méline Bizard, Angela Brisebarre, Camille Capel, Vincent Enouf, Louise Lefrançois, Frédéric Lemoine, Christophe Malabat, Corinne Maufrais, Hub de Bioinformatique et Biostatistique, Etienne Simon-Lorière, Maud Vanpeene, Sylvie Van der Werf ,Marie-HéLèNe Glaudon Louveau De La Guigneraye |
| EPI_ISL_3153743, EPI_ISL_3154030 | CERBALLIANCE | UMR PIMIT | Dr David A Wilkinson, Dr Patrick Mavingui, Dr Camille Lebarbenchon, Magali Turpin, Marie-Alice Simbi |
| EPI_ISL_3154282, EPI_ISL_3154283, EPI_ISL_3154284, EPI_ISL_3154285, EPI_ISL_3154286 | REUNILAB | UMR PIMIT | Dr David A Wilkinson, Dr Patrick Mavingui, Dr Camille Lebarbenchon, Magali Turpin, Marie-Alice Simbi |
| EPI_ISL_3154287 | CERBALLIANCE | UMR PIMIT | Dr David A Wilkinson, Dr Patrick Mavingui, Dr Camille Lebarbenchon, Magali Turpin, Marie-Alice Simbi |
| EPI_ISL_3154288, EPI_ISL_3154289 | REUNILAB | UMR PIMIT | Dr David A Wilkinson, Dr Patrick Mavingui, Dr Camille Lebarbenchon, Magali Turpin, Marie-Alice Simbi |
| EPI_ISL_3154290, EPI_ISL_3154291, EPI_ISL_3154292, EPI_ISL_3154293, EPI_ISL_3154294, EPI_ISL_3154295, EPI_ISL_3154296, EPI_ISL_3154297, EPI_ISL_3154298, EPI_ISL_3154299, EPI_ISL_3154300, EPI_ISL_3154301, EPI_ISL_3154302, EPI_ISL_3154303, EPI_ISL_3154304, EPI_ISL_3154305, EPI_ISL_3154306, EPI_ISL_3154307, EPI_ISL_3154308, EPI_ISL_3154309 | | | |
| see above | CERBALLIANCE | UMR PIMIT | Dr David A Wilkinson, Dr Patrick Mavingui, Dr Camille Lebarbenchon, Magali Turpin, Marie-Alice Simbi |
| EPI_ISL_3154378, EPI_ISL_3154379, EPI_ISL_3154380, EPI_ISL_3154381, EPI_ISL_3154382, EPI_ISL_3154383, EPI_ISL_3154384, EPI_ISL_3154385, EPI_ISL_3154386, EPI_ISL_3154387, EPI_ISL_3154388, EPI_ISL_3154389, EPI_ISL_3154390, EPI_ISL_3154391, EPI_ISL_3154392, EPI_ISL_3154393, EPI_ISL_3154394, EPI_ISL_3154395, EPI_ISL_3154396, EPI_ISL_3154397, EPI_ISL_3154398, EPI_ISL_3154399, EPI_ISL_3154400, EPI_ISL_3154401, EPI_ISL_3154402, EPI_ISL_3154403, EPI_ISL_3154404, EPI_ISL_3154405, EPI_ISL_3154406, EPI_ISL_3154411, EPI_ISL_3154415, EPI_ISL_3154416, EPI_ISL_3154417, EPI_ISL_3154418, EPI_ISL_3154419, EPI_ISL_3154420, EPI_ISL_3154421, EPI_ISL_3154422, EPI_ISL_3154423, EPI_ISL_3154424, EPI_ISL_3154425, EPI_ISL_3154426, EPI_ISL_3154427, EPI_ISL_3154428, EPI_ISL_3154429, EPI_ISL_3154430, EPI_ISL_3154431, EPI_ISL_3154432, EPI_ISL_3154433, EPI_ISL_3154434, EPI_ISL_3154435, EPI_ISL_3154436, EPI_ISL_3154437, EPI_ISL_3154438, EPI_ISL_3154439, EPI_ISL_3154440, EPI_ISL_3154441, EPI_ISL_3154442, EPI_ISL_3154443, EPI_ISL_3154444, EPI_ISL_3154445, EPI_ISL_3154446, EPI_ISL_3154447, EPI_ISL_3154448 | | | |
| see above | Hospital Universitari Vall d'Hebron - Vall d'Hebron Institut de Recerca | Hospital Universitari Vall d'Hebron - Vall d'Hebron Institut de Recerca | Cristina Andrés, Maria Piñana, Alejandra González-Sánchez, Damir Garcia-Cehic, Ariadna Rando, Juliana Esperalba, Maria Gema Codina, Carla Castillo, Karen García, Rodrigo Vásquez, Maria Carmen Martin, Tomàs Pumarola, Josep Quer, Andrés Antón |
| EPI_ISL_3154449, EPI_ISL_3154450, EPI_ISL_3154451, EPI_ISL_3154452, EPI_ISL_3154453, EPI_ISL_3154454, EPI_ISL_3154455, EPI_ISL_3154456, EPI_ISL_3154457, EPI_ISL_3154458, EPI_ISL_3154459, EPI_ISL_3154460, EPI_ISL_3154461, EPI_ISL_3154462, EPI_ISL_3154463, EPI_ISL_3154464, EPI_ISL_3154465, EPI_ISL_3154466, EPI_ISL_3154467, EPI_ISL_3154468, EPI_ISL_3154469, EPI_ISL_3154470, EPI_ISL_3154471, EPI_ISL_3154472, EPI_ISL_3154473, EPI_ISL_3154474, EPI_ISL_3154475, EPI_ISL_3154476, EPI_ISL_3154477, EPI_ISL_3154478, EPI_ISL_3154479, EPI_ISL_3154480, EPI_ISL_3154481, EPI_ISL_3154482, EPI_ISL_3154483, EPI_ISL_3154484, EPI_ISL_3154485, EPI_ISL_3154486, EPI_ISL_3154487, EPI_ISL_3154488, EPI_ISL_3154489, EPI_ISL_3154490, EPI_ISL_3154491, EPI_ISL_3154492, EPI_ISL_3154493, EPI_ISL_3154494, EPI_ISL_3154495, EPI_ISL_3154496, EPI_ISL_3154497, EPI_ISL_3154498, EPI_ISL_3154499, EPI_ISL_3154500, EPI_ISL_3154501, EPI_ISL_3154502, EPI_ISL_3154503, EPI_ISL_3154504, EPI_ISL_3154505, EPI_ISL_3154506, EPI_ISL_3154507, EPI_ISL_3154513, EPI_ISL_3154514, EPI_ISL_3154516, EPI_ISL_3154518, EPI_ISL_3154520, EPI_ISL_3154521, EPI_ISL_3154522, EPI_ISL_3154574, EPI_ISL_3154590, EPI_ISL_3154591, EPI_ISL_3154592, EPI_ISL_3154593, EPI_ISL_3154594, EPI_ISL_3154595, EPI_ISL_3154597, EPI_ISL_3154598, EPI_ISL_3154599, EPI_ISL_3154600, EPI_ISL_3154601, EPI_ISL_3154602, EPI_ISL_3154603, EPI_ISL_3154604, EPI_ISL_3154605, EPI_ISL_3154606, EPI_ISL_3154607, EPI_ISL_3154608, EPI_ISL_3154609, EPI_ISL_3154610, EPI_ISL_3154611, EPI_ISL_3154612, EPI_ISL_3154613, EPI_ISL_3154614, EPI_ISL_3154615, EPI_ISL_3154616, EPI_ISL_3154617, EPI_ISL_3154618, EPI_ISL_3154619, EPI_ISL_3154620, EPI_ISL_3154621, EPI_ISL_3154622, EPI_ISL_3154623, EPI_ISL_3154624, EPI_ISL_3154625, EPI_ISL_3154626, EPI_ISL_3154627, EPI_ISL_3154628, EPI_ISL_3154629, EPI_ISL_3154630, EPI_ISL_3154631, EPI_ISL_3154632, EPI_ISL_3154633, EPI_ISL_3154634, EPI_ISL_3154635, EPI_ISL_3154636, EPI_ISL_3154637, EPI_ISL_3154638, EPI_ISL_3154639, EPI_ISL_3154640, EPI_ISL_3154641, EPI_ISL_3154642, EPI_ISL_3154643, EPI_ISL_3154644, EPI_ISL_3154645, EPI_ISL_3154646, EPI_ISL_3154647, EPI_ISL_3154648, EPI_ISL_3154649, EPI_ISL_3154650, EPI_ISL_3154651, EPI_ISL_3154652, EPI_ISL_3154653, EPI_ISL_3154654, EPI_ISL_3154655, EPI_ISL_3154656, EPI_ISL_3154657, EPI_ISL_3154658, EPI_ISL_3154659, EPI_ISL_3154660, EPI_ISL_3154661, EPI_ISL_3154662, EPI_ISL_3154663, EPI_ISL_3154664, EPI_ISL_3154665, EPI_ISL_3154666, EPI_ISL_3154667, EPI_ISL_3154668, EPI_ISL_3154669, EPI_ISL_3154670, EPI_ISL_3154671, EPI_ISL_3154672, EPI_ISL_3154673, EPI_ISL_3154674, EPI_ISL_3154675, EPI_ISL_3154676, EPI_ISL_3154677, EPI_ISL_3154678, EPI_ISL_3154679, EPI_ISL_3154680, EPI_ISL_3154681, EPI_ISL_3154682, EPI_ISL_3154683, EPI_ISL_3154684, EPI_ISL_3154685, EPI_ISL_3154686, EPI_ISL_3154687, EPI_ISL_3154688, EPI_ISL_3154689, EPI_ISL_3154690, EPI_ISL_3154691, EPI_ISL_3154692, EPI_ISL_3154693, EPI_ISL_3154694, EPI_ISL_3154695, EPI_ISL_3154696, EPI_ISL_3154697, EPI_ISL_3154698, EPI_ISL_3154700, EPI_ISL_3154701, EPI_ISL_3154702, EPI_ISL_3154704, EPI_ISL_3154705, EPI_ISL_3154706, EPI_ISL_3154707, EPI_ISL_3154708, EPI_ISL_3154709, EPI_ISL_3154710, EPI_ISL_3154711 | | | |
| see above | National Virus Reference Laboratory | National Virus Reference Laboratory | Zoe Yandle, Charlene Bennett, Gabriel Gonzalez, Michael Carr, Jonathan Dean, Cillian F De Gascun |
| EPI_ISL_3154716, EPI_ISL_3154717, EPI_ISL_3154718 | Labo Analyses Med | National Reference Center for Viruses of Respiratory Infections, Institut Pasteur, Paris | Marion Barbet, Sylvie Behillil, Méline Bizard, Angela Brisebarre, Camille Capel, Vincent Enouf, Louise Lefrançois, Frédéric Lemoine, Christophe Malabat, Corinne Maufrais, Hub de Bioinformatique et Statistique, Etienne Simon-Lorière, Maud Vanpeene, Sylvie Van der Werf ,Guillaume Aubin (G) |
| EPI_ISL_3154728 | Hospital | National Reference Center for Viruses of Respiratory Infections, Institut Pasteur, Paris | Marion Barbet, Sylvie Behillil, Méline Bizard, Angela Brisebarre, Camille Capel, Vincent Enouf, Louise Lefrançois, Frédéric Lemoine, Christophe Malabat, Corinne Maufrais, Hub de Bioinformatique et Statistique, Etienne Simon-Lorière, Maud Vanpeene, Sylvie Van der Werf ,Guillaume Aubin (G) |
| EPI_ISL_3154729 | Labo Analyses Med | National Reference Center for Viruses of Respiratory Infections, Institut Pasteur, Paris | Marion Barbet, Sylvie Behillil, Méline Bizard, Angela Brisebarre, Camille Capel, Vincent Enouf, Louise Lefrançois, Frédéric Lemoine, Christophe Malabat, Corinne Maufrais, Hub de Bioinformatique et Statistique, Etienne Simon-Lorière, Maud Vanpeene, Sylvie Van der Werf ,Guillaume Aubin (G) |
| EPI_ISL_3154730, EPI_ISL_3154734, EPI_ISL_3154735, EPI_ISL_3154736, EPI_ISL_3154737, EPI_ISL_3154738 | Hospital | National Reference Center for Viruses of Respiratory Infections, Institut Pasteur, Paris | Marion Barbet, Sylvie Behillil, Méline Bizard, Angela Brisebarre, Camille Capel, Vincent Enouf, Louise Lefrançois, Frédéric Lemoine, Christophe Malabat, Corinne Maufrais, Hub de Bioinformatique et Statistique, Etienne Simon-Lorière, Maud Vanpeene, Sylvie Van der Werf ,Guillaume Aubin (G) |
| EPI_ISL_3154739, EPI_ISL_3154740, EPI_ISL_3154741, EPI_ISL_3154742, EPI_ISL_3154743, EPI_ISL_3154744, EPI_ISL_3154745, EPI_ISL_3154746, EPI_ISL_3154747, EPI_ISL_3154748, EPI_ISL_3154749, EPI_ISL_3154750, EPI_ISL_3154751, EPI_ISL_3154752, EPI_ISL_3154753, EPI_ISL_3154754, EPI_ISL_3154755, EPI_ISL_3154756, EPI_ISL_3154757 | | | |
| see above | Labo Analyses Med | National Reference Center for Viruses of Respiratory Infections, Institut Pasteur, Paris | Marion Barbet, Sylvie Behillil, Méline Bizard, Angela Brisebarre, Camille Capel, Vincent Enouf, Louise Lefrançois, Frédéric Lemoine, Christophe Malabat, Corinne Maufrais, Hub de Bioinformatique et Statistique, Etienne Simon-Lorière, Maud Vanpeene, Sylvie Van der Werf ,Guillaume Aubin (G) |
| EPI_ISL_3154758, EPI_ISL_3154759, EPI_ISL_3154760, EPI_ISL_3154761, EPI_ISL_3154762, EPI_ISL_3154763, EPI_ISL_3154764 | Hospital | National Reference Center for Viruses of Respiratory Infections, Institut Pasteur, Paris | Marion Barbet, Sylvie Behillil, Méline Bizard, Angela Brisebarre, Camille Capel, Vincent Enouf, Louise Lefrançois, Frédéric Lemoine, Christophe Malabat, Corinne Maufrais, Hub de Bioinformatique et Statistique, Etienne Simon-Lorière, Maud Vanpeene, Sylvie Van der Werf ,Guillaume Aubin (G) |
| EPI_ISL_3154765, EPI_ISL_3154766, EPI_ISL_3154767, EPI_ISL_3154768, EPI_ISL_3154769 | Labo Analyses Med | National Reference Center for Viruses of Respiratory Infections, Institut Pasteur, Paris | Marion Barbet, Sylvie Behillil, Méline Bizard, Angela Brisebarre, Camille Capel, Vincent Enouf, Louise Lefrançois, Frédéric Lemoine, Christophe Malabat, Corinne Maufrais, Hub de Bioinformatique et Statistique, Etienne Simon-Lorière, Maud Vanpeene, Sylvie Van der Werf ,Guillaume Aubin (G) |
| EPI_ISL_3154770, EPI_ISL_3154771, EPI_ISL_3154772, EPI_ISL_3154773, EPI_ISL_3154774, EPI_ISL_3154775, EPI_ISL_3154776, EPI_ISL_3154777 | SYNLAB | GIGA Medical Genomics | Keith Durkin, Maria Artesi, Bouchra Boujemla, Nathalie Renotte, Nadine Cambisano, Sébastien Bontems, Cécile Meex, Claire Gourzonès, Olivier Ek, Laurent Gillet, Marie-Pierre Hayette, Vincent Bours |
| EPI_ISL_3154778, EPI_ISL_3154779 | University of Liège COVID-19 testing center | GIGA Medical Genomics | Keith Durkin, Maria Artesi, Bouchra Boujemla, Nathalie Renotte, Nadine Cambisano, Sébastien Bontems, Cécile Meex, Claire Gourzonès, Olivier Ek, Laurent Gillet, Marie-Pierre Hayette, Vincent Bours |
| EPI_ISL_3154780, EPI_ISL_3154781, EPI_ISL_3154782, EPI_ISL_3154783, EPI_ISL_3154784, EPI_ISL_3154785, EPI_ISL_3154786, EPI_ISL_3154787, EPI_ISL_3154788, EPI_ISL_3154789, EPI_ISL_3154790, EPI_ISL_3154791, EPI_ISL_3154792, EPI_ISL_3154793, EPI_ISL_3154794, EPI_ISL_3154795, EPI_ISL_3154796, EPI_ISL_3154797, EPI_ISL_3154798, EPI_ISL_3154799, EPI_ISL_3154800, EPI_ISL_3154801 | | | |
| see above | SYNLAB | GIGA Medical Genomics | Keith Durkin, Maria Artesi, Bouchra Boujemla, Nathalie Renotte, Nadine Cambisano, Sébastien Bontems, Cécile Meex, Claire Gourzonès, Olivier Ek, Laurent Gillet, Marie-Pierre Hayette, Vincent Bours |
| EPI_ISL_3154802, EPI_ISL_3154803 | University of Liège COVID-19 testing center | GIGA Medical Genomics | Keith Durkin, Maria Artesi, Bouchra Boujemla, Nathalie Renotte, Nadine Cambisano, Sébastien Bontems, Cécile Meex, Claire Gourzonès, Olivier Ek, Laurent Gillet, Marie-Pierre Hayette, Vincent Bours |
| EPI_ISL_3154804, EPI_ISL_3154805 | SYNLAB | GIGA Medical Genomics | Keith Durkin, Maria Artesi, Bouchra Boujemla, Nathalie Renotte, Nadine Cambisano, Sébastien Bontems, Cécile Meex, Claire Gourzonès, Olivier Ek, Laurent Gillet, Marie-Pierre Hayette, Vincent Bours |
| EPI_ISL_3154806 | University of Liège COVID-19 testing center | GIGA Medical Genomics | Keith Durkin, Maria Artesi, Bouchra Boujemla, Nathalie Renotte, Nadine Cambisano, Sébastien Bontems, Cécile Meex, Claire Gourzonès, Olivier Ek, Laurent Gillet, Marie-Pierre Hayette, Vincent Bours |
| EPI_ISL_3154809, EPI_ISL_3154810, EPI_ISL_3154811, EPI_ISL_3154812, EPI_ISL_3154813, EPI_ISL_3154814, EPI_ISL_3154815, EPI_ISL_3154816, EPI_ISL_3154817, EPI_ISL_3154818, EPI_ISL_3154820, EPI_ISL_3154821, EPI_ISL_3154822, EPI_ISL_3154823, EPI_ISL_3154824, EPI_ISL_3154825 | | | |
| see above | SYNLAB | GIGA Medical Genomics | Keith Durkin, Maria Artesi, Bouchra Boujemla, Nathalie Renotte, Nadine Cambisano, Sébastien Bontems, Cécile Meex, Claire Gourzonès, Olivier Ek, Laurent Gillet, Marie-Pierre Hayette, Vincent Bours |
| EPI_ISL_3154826 | University of Liège COVID-19 testing center | GIGA Medical Genomics | Keith Durkin, Maria Artesi, Bouchra Boujemla, Nathalie Renotte, Nadine Cambisano, Sébastien Bontems, Cécile Meex, Claire Gourzonès, Olivier Ek, Laurent Gillet, Marie-Pierre Hayette, Vincent Bours |
| EPI_ISL_3154827, EPI_ISL_3154828, EPI_ISL_3154829, EPI_ISL_3154830 | SYNLAB | GIGA Medical Genomics | Keith Durkin, Maria Artesi, Bouchra Boujemla, Nathalie Renotte, Nadine Cambisano, Sébastien Bontems, Cécile Meex, Claire Gourzonès, Olivier Ek, Laurent Gillet, Marie-Pierre Hayette, Vincent Bours |
| EPI_ISL_3154842, EPI_ISL_3154843, EPI_ISL_3154844, EPI_ISL_3154845, | Wyoming Public Health Laboratory | Wyoming Public Health Laboratory | Jim Mildenberger, Taylor Fearing, Channing Weber, Ashley Norberg, Chayse Rowley, Marley Goetz, Brian Dominguez, Elliot Thomasson, Sam Britz, Cari Sloma, Robert Petit, and Rob Christensen |

| | | | |
|---|---|---|---|
| EPI_ISL_3154846, EPI_ISL_3154847, EPI_ISL_3154848 | | | |
| EPI_ISL_3154851, EPI_ISL_3154852, EPI_ISL_3154853, EPI_ISL_3154854, EPI_ISL_3154855, EPI_ISL_3154856, EPI_ISL_3154857, EPI_ISL_3154858, EPI_ISL_3154859, EPI_ISL_3154861, EPI_ISL_3154862, EPI_ISL_3154863, EPI_ISL_3154864, EPI_ISL_3154865, EPI_ISL_3154866, EPI_ISL_3154867, EPI_ISL_3154868, EPI_ISL_3154869, EPI_ISL_3154870, EPI_ISL_3154871 | | | |
| see above | Hospital Universitari Joan XXIII | Hospital Universitari Joan XXIII | Gemma Recio, Cristina Gutiérrez, Clara Benavent, Carla Martín, Margarida Terrón, Ester Picó, Natalia Bastón |
| EPI_ISL_3154879, EPI_ISL_3154885, EPI_ISL_3154886, EPI_ISL_3154887, EPI_ISL_3154890, EPI_ISL_3154891, EPI_ISL_3154892, EPI_ISL_3154893, EPI_ISL_3154894, EPI_ISL_3154896, EPI_ISL_3154897, EPI_ISL_3154898, EPI_ISL_3154899, EPI_ISL_3154900 | | | |
| see above | Iressef Genomics lab | IRESSEF | Souleymane MBOUP, Abdou PADANE , Khadim GUEYE, Diabou Diagne , Samba Ndiour, Papa Alassane DIAW, Birahim Piere NDIAYE, Barada CISSE, Aminata MBOUP, Moustapha MBOW, Ndeye Coumba Toure KANE, Nafisatou LEYE, Gora LO, Ambroise AHOUIDI , Astou Gaye GAYE, Aminata DIA, Yacine DIA,Seni Ndiaye |
| EPI_ISL_3154901 | Hospital | National Reference Center for Viruses of Respiratory Infections, Institut Pasteur, Paris | Marion Barbet, Sylvie Behillil, Méline Bizard, Angela Brisebarre, Camille Capel, Vincent Enouf, Louise Lefrançois, Frédéric Lemoine, Christophe Malabat, Corinne Maufrais, Adrien Pain, Etienne Simon-Lorière, Maud Vanpeene, Sylvie Van der Werf ,Laurence Louvet |
| EPI_ISL_3154904 | Labo Analyses Med | National Reference Center for Viruses of Respiratory Infections, Institut Pasteur, Paris | Marion Barbet, Sylvie Behillil, Méline Bizard, Angela Brisebarre, Camille Capel, Vincent Enouf, Louise Lefrançois, Frédéric Lemoine, Christophe Malabat, Corinne Maufrais, Hub Bioinformatique Biostatistiques, Etienne Simon-Lorière, Maud Vanpeene, Sylvie Van der Werf ,Jean-Baptiste Rabec |
| EPI_ISL_3154905, EPI_ISL_3154906, EPI_ISL_3154910, EPI_ISL_3154912, EPI_ISL_3154913 | Labo Analyses Med | National Reference Center for Viruses of Respiratory Infections, Institut Pasteur, Paris | Marion Barbet, Sylvie Behillil, Méline Bizard, Angela Brisebarre, Camille Capel, Vincent Enouf, Louise Lefrançois, Frédéric Lemoine, Christophe Malabat, Corinne Maufrais, Hub Bioinformatique Biostatistiques, Etienne Simon-Lorière, Maud Vanpeene, Sylvie Van der Werf ,Ronan Pichard |
| EPI_ISL_3154917, EPI_ISL_3154918, EPI_ISL_3154919, EPI_ISL_3154920, EPI_ISL_3154921, EPI_ISL_3154922, EPI_ISL_3154923, EPI_ISL_3154924, EPI_ISL_3154925, EPI_ISL_3154926, EPI_ISL_3154927, EPI_ISL_3154928, EPI_ISL_3154929, EPI_ISL_3154930, EPI_ISL_3154931, EPI_ISL_3154932 | | | |
| see above | Labo Analyses Med | National Reference Center for Viruses of Respiratory Infections, Institut Pasteur, Paris | Marion Barbet, Sylvie Behillil, Méline Bizard, Angela Brisebarre, Camille Capel, Vincent Enouf, Louise Lefrançois, Frédéric Lemoine, Christophe Malabat, Corinne Maufrais, Hub Bioinformatique Biostatistiques, Etienne Simon-Lorière, Maud Vanpeene, Sylvie Van der Werf ,GréGoire Potiron |
| EPI_ISL_3154933, EPI_ISL_3154934 | Hospital | National Reference Center for Viruses of Respiratory Infections, Institut Pasteur, Paris | Marion Barbet, Sylvie Behillil, Méline Bizard, Angela Brisebarre, Camille Capel, Vincent Enouf, Louise Lefrançois, Frédéric Lemoine, Christophe Malabat, Corinne Maufrais, Hub Bioinformatique Biostatistiques, Etienne Simon-Lorière, Maud Vanpeene, Sylvie Van der Werf ,Benjamin Maneglie |
| EPI_ISL_3154935, EPI_ISL_3154937, EPI_ISL_3154938, EPI_ISL_3154940, EPI_ISL_3154941 | Labo Analyses Med | National Reference Center for Viruses of Respiratory Infections, Institut Pasteur, Paris | Marion Barbet, Sylvie Behillil, Méline Bizard, Angela Brisebarre, Camille Capel, Vincent Enouf, Louise Lefrançois, Frédéric Lemoine, Christophe Malabat, Corinne Maufrais, Hub Bioinformatique Biostatistiques, Etienne Simon-Lorière, Maud Vanpeene, Sylvie Van der Werf ,Claire Felloni |
| EPI_ISL_3154942, EPI_ISL_3154944, EPI_ISL_3154945, EPI_ISL_3154946, EPI_ISL_3154947, EPI_ISL_3154948, EPI_ISL_3154949, EPI_ISL_3154950, EPI_ISL_3154951, EPI_ISL_3154952, EPI_ISL_3154953, EPI_ISL_3154954, EPI_ISL_3154955, EPI_ISL_3154956, EPI_ISL_3154957, EPI_ISL_3154958, EPI_ISL_3154959 | | | |
| see above | Labo Analyses Med | National Reference Center for Viruses of Respiratory Infections, Institut Pasteur, Paris | Marion Barbet, Sylvie Behillil, Méline Bizard, Angela Brisebarre, Camille Capel, Vincent Enouf, Louise Lefrançois, Frédéric Lemoine, Christophe Malabat, Corinne Maufrais, Hub Bioinformatique Biostatistiques, Etienne Simon-Lorière, Maud Vanpeene, Sylvie Van der Werf ,BéAtrice Ricard |
| EPI_ISL_3154960, EPI_ISL_3154961, EPI_ISL_3154962 | Labo Analyses Med | National Reference Center for Viruses of Respiratory Infections, Institut Pasteur, Paris | Marion Barbet, Sylvie Behillil, Méline Bizard, Angela Brisebarre, Camille Capel, Vincent Enouf, Louise Lefrançois, Frédéric Lemoine, Christophe Malabat, Corinne Maufrais, Hub Bioinformatique Biostatistiques, Etienne Simon-Lorière, Maud Vanpeene, Sylvie Van der Werf ,Sylvie Robert |
| EPI_ISL_3154963, EPI_ISL_3154964, EPI_ISL_3154965, EPI_ISL_3154966, EPI_ISL_3154967, EPI_ISL_3154968, EPI_ISL_3154969, EPI_ISL_3154970 | Labo Analyses Med | National Reference Center for Viruses of Respiratory Infections, Institut Pasteur, Paris | Marion Barbet, Sylvie Behillil, Méline Bizard, Angela Brisebarre, Camille Capel, Vincent Enouf, Louise Lefrançois, Frédéric Lemoine, Christophe Malabat, Corinne Maufrais, Hub Bioinformatique Biostatistiques, Etienne Simon-Lorière, Maud Vanpeene, Sylvie Van der Werf ,Eric Morin |
| EPI_ISL_3154971, EPI_ISL_3154972, EPI_ISL_3154973, EPI_ISL_3154974 | Labo Analyses Med | National Reference Center for Viruses of Respiratory Infections, Institut Pasteur, Paris | Marion Barbet, Sylvie Behillil, Méline Bizard, Angela Brisebarre, Camille Capel, Vincent Enouf, Louise Lefrançois, Frédéric Lemoine, Christophe Malabat, Corinne Maufrais, Hub Bioinformatique Biostatistiques, Etienne Simon-Lorière, Maud Vanpeene, Sylvie Van der Werf ,Sandrine Simon |
| EPI_ISL_3154975, EPI_ISL_3154976, EPI_ISL_3154977 | Hospital | National Reference Center for Viruses of Respiratory Infections, Institut Pasteur, Paris | Marion Barbet, Sylvie Behillil, Méline Bizard, Angela Brisebarre, Camille Capel, Vincent Enouf, Louise Lefrançois, Frédéric Lemoine, Christophe Malabat, Corinne Maufrais, Hub Bioinformatique Biostatistiques, Etienne Simon-Lorière, Maud Vanpeene, Sylvie Van der Werf ,Pierre Alain Billy |
| EPI_ISL_3154978, EPI_ISL_3154979, EPI_ISL_3154980, EPI_ISL_3154981, EPI_ISL_3154982, EPI_ISL_3154983 | Labo Analyses Med | National Reference Center for Viruses of Respiratory Infections, Institut Pasteur, Paris | Marion Barbet, Sylvie Behillil, Méline Bizard, Angela Brisebarre, Camille Capel, Vincent Enouf, Louise Lefrançois, Frédéric Lemoine, Christophe Malabat, Corinne Maufrais, Hub Bioinformatique Biostatistiques, Etienne Simon-Lorière, Maud Vanpeene, Sylvie Van der Werf ,Coralie Jean |
| EPI_ISL_3154984 | Labo Analyses Med | National Reference Center for Viruses of Respiratory Infections, Institut Pasteur, Paris | Marion Barbet, Sylvie Behillil, Méline Bizard, Angela Brisebarre, Camille Capel, Vincent Enouf, Louise Lefrançois, Frédéric Lemoine, Christophe Malabat, Corinne Maufrais, Hub Bioinformatique Biostatistiques, Etienne Simon-Lorière, Maud Vanpeene, Sylvie Van der Werf ,Caroline Malderet |
| EPI_ISL_3154986 | Hospital | National Reference Center for Viruses of Respiratory Infections, Institut Pasteur, Paris | Marion Barbet, Sylvie Behillil, Méline Bizard, Angela Brisebarre, Camille Capel, Vincent Enouf, Louise Lefrançois, Frédéric Lemoine, Christophe Malabat, Corinne Maufrais, Hub Bioinformatique Biostatistiques, Etienne Simon-Lorière, Maud Vanpeene, Sylvie Van der Werf ,Sophie Mignard |
| EPI_ISL_3154987, EPI_ISL_3154988, EPI_ISL_3154989 | Labo Analyses Med | National Reference Center for Viruses of Respiratory Infections, Institut Pasteur, Paris | Marion Barbet, Sylvie Behillil, Méline Bizard, Angela Brisebarre, Camille Capel, Vincent Enouf, Louise Lefrançois, Frédéric Lemoine, Christophe Malabat, Corinne Maufrais, Hub Bioinformatique Biostatistiques, Etienne Simon-Lorière, Maud Vanpeene, Sylvie Van der Werf ,BéNéDicte Roquebert |
| EPI_ISL_3154991, EPI_ISL_3154992, EPI_ISL_3154993, EPI_ISL_3154994, EPI_ISL_3154995, EPI_ISL_3154997, EPI_ISL_3154998, EPI_ISL_3154999, EPI_ISL_3155001, EPI_ISL_3155004, EPI_ISL_3155005 | | | |
| see above | Labo Analyses Med | National Reference Center for Viruses of Respiratory Infections, Institut Pasteur, Paris | Marion Barbet, Sylvie Behillil, Méline Bizard, Angela Brisebarre, Camille Capel, Vincent Enouf, Louise Lefrançois, Frédéric Lemoine, Christophe Malabat, Corinne Maufrais, Hub Bioinformatique Biostatistiques, Etienne Simon-Lorière, Maud Vanpeene, Sylvie Van der Werf ,Nabila Belhouachi |
| EPI_ISL_3155007, EPI_ISL_3155008, EPI_ISL_3155009, EPI_ISL_3155010, EPI_ISL_3155011, EPI_ISL_3155012, EPI_ISL_3155013, EPI_ISL_3155014, EPI_ISL_3155015, EPI_ISL_3155016, EPI_ISL_3155017, EPI_ISL_3155018, EPI_ISL_3155019, EPI_ISL_3155020, EPI_ISL_3155021, EPI_ISL_3155022, EPI_ISL_3155023, EPI_ISL_3155024, EPI_ISL_3155025, EPI_ISL_3155026, EPI_ISL_3155027, EPI_ISL_3155028, EPI_ISL_3155029, EPI_ISL_3155030 | | | |
| see above | Labo Analyses Med | National Reference Center for Viruses of Respiratory Infections, Institut Pasteur, Paris | Marion Barbet, Sylvie Behillil, Méline Bizard, Angela Brisebarre, Camille Capel, Vincent Enouf, Louise Lefrançois, Frédéric Lemoine, Christophe Malabat, Corinne Maufrais, Hub Bioinformatique Biostatistiques, Etienne Simon-Lorière, Maud Vanpeene, Sylvie Van der Werf ,Jean-FrançOis Comes |
| EPI_ISL_3155031, EPI_ISL_3155032, EPI_ISL_3155033, EPI_ISL_3155034, EPI_ISL_3155035, EPI_ISL_3155036, EPI_ISL_3155037 | Labo Analyses Med | National Reference Center for Viruses of Respiratory Infections, Institut Pasteur, Paris | Marion Barbet, Sylvie Behillil, Méline Bizard, Angela Brisebarre, Camille Capel, Vincent Enouf, Louise Lefrançois, Frédéric Lemoine, Christophe Malabat, Corinne Maufrais, Hub Bioinformatique Biostatistiques, Etienne Simon-Lorière, Maud Vanpeene, Sylvie Van der Werf ,C. Lecorre |
| EPI_ISL_3155038, EPI_ISL_3155039, EPI_ISL_3155040, EPI_ISL_3155041, EPI_ISL_3155042, EPI_ISL_3155043, EPI_ISL_3155044, EPI_ISL_3155045 | Hospital | National Reference Center for Viruses of Respiratory Infections, Institut Pasteur, Paris | Marion Barbet, Sylvie Behillil, Méline Bizard, Angela Brisebarre, Camille Capel, Vincent Enouf, Louise Lefrançois, Frédéric Lemoine, Christophe Malabat, Corinne Maufrais, Hub Bioinformatique Biostatistiques, Etienne Simon-Lorière, Maud Vanpeene, Sylvie Van der Werf ,Anne Cady |
| EPI_ISL_3155046, EPI_ISL_3155047, EPI_ISL_3155048, EPI_ISL_3155049, EPI_ISL_3155050, EPI_ISL_3155051, EPI_ISL_3155052 | Hospital | National Reference Center for Viruses of Respiratory Infections, Institut Pasteur, Paris | Marion Barbet, Sylvie Behillil, Méline Bizard, Angela Brisebarre, Camille Capel, Vincent Enouf, Louise Lefrançois, Frédéric Lemoine, Christophe Malabat, Corinne Maufrais, Hub Bioinformatique Biostatistiques, Etienne Simon-Lorière, Maud Vanpeene, Sylvie Van der Werf ,CéLine Bressollette |
| EPI_ISL_3155053, EPI_ISL_3155054, EPI_ISL_3155055 | Labo Analyses Med | National Reference Center for Viruses of Respiratory Infections, Institut Pasteur, Paris | Marion Barbet, Sylvie Behillil, Méline Bizard, Angela Brisebarre, Camille Capel, Vincent Enouf, Louise Lefrançois, Frédéric Lemoine, Christophe Malabat, Corinne Maufrais, Hub Bioinformatique Biostatistiques, Etienne Simon-Lorière, Maud Vanpeene, Sylvie Van der Werf ,Emmanuel Pasquier |
| EPI_ISL_3155056 | Hospital | National Reference Center for Viruses of Respiratory Infections, Institut Pasteur, Paris | Marion Barbet, Sylvie Behillil, Méline Bizard, Angela Brisebarre, Camille Capel, Vincent Enouf, Louise Lefrançois, Frédéric Lemoine, Christophe Malabat, Corinne Maufrais, Hub Bioinformatique Biostatistiques, Etienne Simon-Lorière, Maud Vanpeene, Sylvie Van der Werf ,Delphine Bridoux |
| EPI_ISL_3155057 | Armies | National Reference Center for Viruses of Respiratory Infections, | Marion Barbet, Sylvie Behillil, Méline Bizard, Angela Brisebarre, Camille Capel, Vincent Enouf, Louise Lefrançois, Frédéric Lemoine, Christophe Malabat, Corinne Maufrais, Hub Bioinformatique Biostatistiques, Etienne Simon-Lorière, Maud |

| | | | |
|---|---|---|---|
| | | Institut Pasteur, Paris | Vanpeene, Sylvie Van der Werf ,Marine Desroches |
| EPI_ISL_3155060, EPI_ISL_3155061, EPI_ISL_3155062 | Labo Analyses Med | National Reference Center for Viruses of Respiratory Infections, Institut Pasteur, Paris | Marion Barbet, Sylvie Behillil, Méline Bizard, Angela Brisebarre, Camille Capel, Vincent Enouf, Louise Lefrançois, Frédéric Lemoine, Christophe Malabat, Corinne Maufrais, Hub Bioinformatique Biostatistiques, Etienne Simon-Lorière, Maud Vanpeene, Sylvie Van der Werf ,OphéLie Said-Delattre |
| EPI_ISL_3155063, EPI_ISL_3155070, EPI_ISL_3155071, EPI_ISL_3155072 | Hospital | National Reference Center for Viruses of Respiratory Infections, Institut Pasteur, Paris | Marion Barbet, Sylvie Behillil, Méline Bizard, Angela Brisebarre, Camille Capel, Vincent Enouf, Louise Lefrançois, Frédéric Lemoine, Christophe Malabat, Corinne Maufrais, Hub Bioinformatique Biostatistiques, Etienne Simon-Lorière, Maud Vanpeene, Sylvie Van der Werf ,Nabil Gastli |
| EPI_ISL_3155075, EPI_ISL_3155077, EPI_ISL_3155078, EPI_ISL_3155079, EPI_ISL_3155080, EPI_ISL_3155081, EPI_ISL_3155082, EPI_ISL_3155083, EPI_ISL_3155084, EPI_ISL_3155085, EPI_ISL_3155086, EPI_ISL_3155087, EPI_ISL_3155088, EPI_ISL_3155089, EPI_ISL_3155090, EPI_ISL_3155091, EPI_ISL_3155092, EPI_ISL_3155093, EPI_ISL_3155094, EPI_ISL_3155095, EPI_ISL_3155096, EPI_ISL_3155097, EPI_ISL_3155098, EPI_ISL_3155099, EPI_ISL_3155100, EPI_ISL_3155101, EPI_ISL_3155102, EPI_ISL_3155103, EPI_ISL_3155104, EPI_ISL_3155105, EPI_ISL_3155107, EPI_ISL_3155108, EPI_ISL_3155109, EPI_ISL_3155110, EPI_ISL_3155111, EPI_ISL_3155112, EPI_ISL_3155113, EPI_ISL_3155114, EPI_ISL_3155115, EPI_ISL_3155116, EPI_ISL_3155117, EPI_ISL_3155118, EPI_ISL_3155119, EPI_ISL_3155120, EPI_ISL_3155122, EPI_ISL_3155125, EPI_ISL_3155126, EPI_ISL_3155128, EPI_ISL_3155129, EPI_ISL_3155130, EPI_ISL_3155131, EPI_ISL_3155132, EPI_ISL_3155133, EPI_ISL_3155134, EPI_ISL_3155135, EPI_ISL_3155136, EPI_ISL_3155137, EPI_ISL_3155138, EPI_ISL_3155139, EPI_ISL_3155140, EPI_ISL_3155141, EPI_ISL_3155142, EPI_ISL_3155143, EPI_ISL_3155144, EPI_ISL_3155145, EPI_ISL_3155146, EPI_ISL_3155147, EPI_ISL_3155148, EPI_ISL_3155149, EPI_ISL_3155150, EPI_ISL_3155151, EPI_ISL_3155152, EPI_ISL_3155153, EPI_ISL_3155154, EPI_ISL_3155155, EPI_ISL_3155156, EPI_ISL_3155158, EPI_ISL_3155159, EPI_ISL_3155160, EPI_ISL_3155161, EPI_ISL_3155162, EPI_ISL_3155163, EPI_ISL_3155164, EPI_ISL_3155165, EPI_ISL_3155166, EPI_ISL_3155167, EPI_ISL_3155168, EPI_ISL_3155169, EPI_ISL_3155170, EPI_ISL_3155171, EPI_ISL_3155173, EPI_ISL_3155176, EPI_ISL_3155177, EPI_ISL_3155178, EPI_ISL_3155179, EPI_ISL_3155180, EPI_ISL_3155181, EPI_ISL_3155182, EPI_ISL_3155183, EPI_ISL_3155184, EPI_ISL_3155185, EPI_ISL_3155186, EPI_ISL_3155187, EPI_ISL_3155188, EPI_ISL_3155189, EPI_ISL_3155190, EPI_ISL_3155192, EPI_ISL_3155193, EPI_ISL_3155194, EPI_ISL_3155195, EPI_ISL_3155196, EPI_ISL_3155197, EPI_ISL_3155198, EPI_ISL_3155199, EPI_ISL_3155200, EPI_ISL_3155201, EPI_ISL_3155202, EPI_ISL_3155203, EPI_ISL_3155204, EPI_ISL_3155205, EPI_ISL_3155206, EPI_ISL_3155207, EPI_ISL_3155208, EPI_ISL_3155209, EPI_ISL_3155210, EPI_ISL_3155211, EPI_ISL_3155212, EPI_ISL_3155213, EPI_ISL_3155214, EPI_ISL_3155215, EPI_ISL_3155216, EPI_ISL_3155218, EPI_ISL_3155219, EPI_ISL_3155220, EPI_ISL_3155221, EPI_ISL_3155222, EPI_ISL_3155223, EPI_ISL_3155224, EPI_ISL_3155225, EPI_ISL_3155226, EPI_ISL_3155227, EPI_ISL_3155228, EPI_ISL_3155229, EPI_ISL_3155230, EPI_ISL_3155231, EPI_ISL_3155232, EPI_ISL_3155233, EPI_ISL_3155234, EPI_ISL_3155235, EPI_ISL_3155236, EPI_ISL_3155237, EPI_ISL_3155238, EPI_ISL_3155239, EPI_ISL_3155240, EPI_ISL_3155241, EPI_ISL_3155242, EPI_ISL_3155243, EPI_ISL_3155244, EPI_ISL_3155245, EPI_ISL_3155246, EPI_ISL_3155247, EPI_ISL_3155248, EPI_ISL_3155249, EPI_ISL_3155250, EPI_ISL_3155251, EPI_ISL_3155252, EPI_ISL_3155253, EPI_ISL_3155254, EPI_ISL_3155255, EPI_ISL_3155256, EPI_ISL_3155257, EPI_ISL_3155258, EPI_ISL_3155259, EPI_ISL_3155260, EPI_ISL_3155261, EPI_ISL_3155262, EPI_ISL_3155263, EPI_ISL_3155264, EPI_ISL_3155265, EPI_ISL_3155266, EPI_ISL_3155267, EPI_ISL_3155268, EPI_ISL_3155269, EPI_ISL_3155270, EPI_ISL_3155271, EPI_ISL_3155272, EPI_ISL_3155273, EPI_ISL_3155274, EPI_ISL_3155275, EPI_ISL_3155276, EPI_ISL_3155277, EPI_ISL_3155278, EPI_ISL_3155279, EPI_ISL_3155280, EPI_ISL_3155281, EPI_ISL_3155282, EPI_ISL_3155283, EPI_ISL_3155284, EPI_ISL_3155285, EPI_ISL_3155286, EPI_ISL_3155287, EPI_ISL_3155288, EPI_ISL_3155289, EPI_ISL_3155290, EPI_ISL_3155291, EPI_ISL_3155292, EPI_ISL_3155293, EPI_ISL_3155294, EPI_ISL_3155295, EPI_ISL_3155296, EPI_ISL_3155297, EPI_ISL_3155298, EPI_ISL_3155299, EPI_ISL_3155300, EPI_ISL_3155301, EPI_ISL_3155303, EPI_ISL_3155304, EPI_ISL_3155305, EPI_ISL_3155306, EPI_ISL_3155307, EPI_ISL_3155308, EPI_ISL_3155309, EPI_ISL_3155310, EPI_ISL_3155311, EPI_ISL_3155312, EPI_ISL_3155313, EPI_ISL_3155314, EPI_ISL_3155315, EPI_ISL_3155316, EPI_ISL_3155317, EPI_ISL_3155319, EPI_ISL_3155320, EPI_ISL_3155321, EPI_ISL_3155322, EPI_ISL_3155324, EPI_ISL_3155325, EPI_ISL_3155326, EPI_ISL_3155327, EPI_ISL_3155328, EPI_ISL_3155329, EPI_ISL_3155330, EPI_ISL_3155331, EPI_ISL_3155332, EPI_ISL_3155333, EPI_ISL_3155334, EPI_ISL_3155335, EPI_ISL_3155336, EPI_ISL_3155337, EPI_ISL_3155338 | | | |
| see above | Clinical Microbiology, Infection Prevention and Control | Section for Molecular Diagnostics | Björn Hallström, Jonas Björkman |
| EPI_ISL_3155339 | ANOUAL | ANOUAL | Fatima Zahra El ansari, Farah Jouali, Yassine Kasmi, Mohcine Bennani Mechita, Rachid Benhida, Jamal Fekkak. |
| EPI_ISL_3155344, EPI_ISL_3155345, EPI_ISL_3155350, EPI_ISL_3155352, EPI_ISL_3155360, EPI_ISL_3155361, EPI_ISL_3155363, EPI_ISL_3155367, EPI_ISL_3155383, EPI_ISL_3155384, EPI_ISL_3155385, EPI_ISL_3155387, EPI_ISL_3155391 | | | |
| see above | Unidad de Investigación Médica de Yucatán (UIMY) | Centro de Investigación en Enfermedades Infecciosas (CIENI), Instituto Nacional de Enfermedades Respiratorias (INER) | Consorcio Mexicano de Vigilancia Genómica (CoViGen-Mex). Authors (in alphabetical order): Julio Elias Alvarado-Yaah, Carlos F. Arias, Santiago Ávila-Ríos, Eduardo Becerril-Vargas, Víctor Hugo Borja-Aburto, Celia Boukadida, Cristóbal Cháidez-Quiróz, Juan Bautista Chale-Dzul, Ricardo Ciria Merce, Andreu Comas-García, Célida Duque Molina, Julissa Enciso-Ibarra, José Antonio Enciso-Moreno, Gloria Elena Espinosa-Ayala, Fernando Fontove-Herrera, Daniel Fregoso-Rueda, Victor Eduardo García-Arias, Alejandra García-Gasca, Bruno Gómez-Gil, Jean Pierre González, Irvin González-López, Concepción Grajales-Muñiz, Ricardo Grande, Rosa María Gutiérrez Rios, Jesús Hernández, Alejandra Hernández-Terán, Alfredo Herrera-Estrella, Carla Ivón Herrera-Najera, Pavel Isa, Verónica Mata-Haro, Daniel Lira Morales, Susana Lopez, Antonio Loza Román, Brenda Irasema Maldonado-Meza, Bernardo Martínez-Miguel, José Arturo Martínez-Orozco, Célida Martínez-Rodríguez, Margarita Matías-Florentino, Fidencio Mejía-Nepomuceno, María Guadalupe de Jesús Mireles-Rivera, Gloria María Molina-Salinas, Hector Montoya-Fuentes, , Mario Mújica-Sánchez, José Esteban Muñoz-Medina, José de Jesús Nuñez-Contreras, Alicia Ocaña-Mondragón, Luis Alberto Ochoa-Carrera, Hector Esteban Paz-Juárez, Marissa Perez-Garcia, Francisco Pulido, Helen Haydee Fernanda Ramírez-Plascencia, Jorge Salas-Hernández, Angel Gustavo Salas-Lais, Alejandro Sánchez-Flores, Clara Esperanza Santacruz-Tinoco, María Guadalupe Santiago-Mauricio, Selene Zárate, Nelly Sélem-Mojica, Blanca Taboada, Kathia Elizabeth Tapia-Diaz, Gloria Vazquez, Joel Armando Vázquez-Pérez. |
| EPI_ISL_3155395, EPI_ISL_3155396, EPI_ISL_3155403, EPI_ISL_3155404, EPI_ISL_3155405, EPI_ISL_3155408, EPI_ISL_3155411, EPI_ISL_3155412, EPI_ISL_3155415, EPI_ISL_3155416, EPI_ISL_3155417, EPI_ISL_3155419, EPI_ISL_3155420, EPI_ISL_3155421, EPI_ISL_3155422, EPI_ISL_3155424, EPI_ISL_3155425, EPI_ISL_3155426, EPI_ISL_3155427, EPI_ISL_3155428, EPI_ISL_3155429, EPI_ISL_3155431, EPI_ISL_3155432, EPI_ISL_3155434, EPI_ISL_3155439, EPI_ISL_3155440 | | | |
| see above | Centro de Investigación Biomédica del Noreste (CIBIN) | Centro de Investigación en Enfermedades Infecciosas (CIENI), Instituto Nacional de Enfermedades Respiratorias (INER) | Consorcio Mexicano de Vigilancia Genómica (CoViGen-Mex). Authors (in alphabetical order): Julio Elias Alvarado-Yaah, Carlos F. Arias, Santiago Ávila-Ríos, Eduardo Becerril-Vargas, Víctor Hugo Borja-Aburto, Celia Boukadida, Cristóbal Cháidez-Quiróz, Juan Bautista Chale-Dzul, Ricardo Ciria Merce, Andreu Comas-García, Célida Duque Molina, Julissa Enciso-Ibarra, José Antonio Enciso-Moreno, Gloria Elena Espinosa-Ayala, Fernando Fontove-Herrera, Daniel Fregoso-Rueda, Victor Eduardo García-Arias, Alejandra García-Gasca, Bruno Gómez-Gil, Jean Pierre González, Irvin González-López, Concepción Grajales-Muñiz, Ricardo Grande, Rosa María Gutiérrez Rios, Jesús Hernández, Alejandra Hernández-Terán, Alfredo Herrera-Estrella, Carla Ivón Herrera-Najera, Pavel Isa, Verónica Mata-Haro, Daniel Lira Morales, Susana Lopez, Antonio Loza Román, Brenda Irasema Maldonado-Meza, Bernardo Martínez-Miguel, José Arturo Martínez-Orozco, Célida Martínez-Rodríguez, Margarita Matías-Florentino, Fidencio Mejía-Nepomuceno, María Guadalupe de Jesús Mireles-Rivera, Gloria María Molina-Salinas, Hector Montoya-Fuentes, , Mario Mújica-Sánchez, José Esteban Muñoz-Medina, José de Jesús Nuñez-Contreras, Alicia Ocaña-Mondragón, Luis Alberto Ochoa-Carrera, Hector Esteban Paz-Juárez, Marissa Perez-Garcia, Francisco Pulido, Helen Haydee Fernanda Ramírez-Plascencia, Jorge Salas-Hernández, Angel Gustavo Salas-Lais, Alejandro Sánchez-Flores, Clara Esperanza Santacruz-Tinoco, María Guadalupe Santiago-Mauricio, Selene Zárate, Nelly Sélem-Mojica, Blanca Taboada, Kathia Elizabeth Tapia-Diaz, Gloria Vazquez, Joel Armando Vázquez-Pérez. |
| EPI_ISL_3155441, EPI_ISL_3155443, EPI_ISL_3155444, EPI_ISL_3155445, EPI_ISL_3155446, EPI_ISL_3155447, EPI_ISL_3155448, EPI_ISL_3155449, EPI_ISL_3155450, EPI_ISL_3155451 | Unidad de Investigación Biomédica de Zacatecas (UIBZ) | Centro de Investigación en Enfermedades Infecciosas (CIENI), Instituto Nacional de Enfermedades Respiratorias (INER) | Consorcio Mexicano de Vigilancia Genómica (CoViGen-Mex). Authors (in alphabetical order): Julio Elias Alvarado-Yaah, Carlos F. Arias, Santiago Ávila-Ríos, Eduardo Becerril-Vargas, Víctor Hugo Borja-Aburto, Celia Boukadida, Cristóbal Cháidez-Quiróz, Juan Bautista Chale-Dzul, Ricardo Ciria Merce, Andreu Comas-García, Célida Duque Molina, Julissa Enciso-Ibarra, José Antonio Enciso-Moreno, Gloria Elena Espinosa-Ayala, Fernando Fontove-Herrera, Daniel Fregoso-Rueda, Victor Eduardo García-Arias, Alejandra García-Gasca, Bruno Gómez-Gil, Jean Pierre González, Irvin González-López, Concepción Grajales-Muñiz, Ricardo Grande, Rosa María Gutiérrez Rios, Jesús Hernández, Alejandra Hernández-Terán, Alfredo Herrera-Estrella, Carla Ivón Herrera-Najera, Pavel Isa, Verónica Mata-Haro, Daniel Lira Morales, Susana Lopez, Antonio Loza Román, Brenda Irasema Maldonado-Meza, Bernardo Martínez-Miguel, José Arturo Martínez-Orozco, Célida Martínez-Rodríguez, Margarita Matías-Florentino, Fidencio Mejía-Nepomuceno, María Guadalupe de Jesús Mireles-Rivera, Gloria María Molina-Salinas, Hector Montoya-Fuentes, , Mario Mújica-Sánchez, José Esteban Muñoz-Medina, José de Jesús Nuñez-Contreras, Alicia Ocaña-Mondragón, Luis Alberto Ochoa-Carrera, Hector Esteban Paz-Juárez, Marissa Perez-Garcia, Francisco Pulido, Helen Haydee Fernanda Ramírez-Plascencia, Jorge Salas-Hernández, Angel Gustavo Salas-Lais, Alejandro Sánchez-Flores, Clara Esperanza Santacruz-Tinoco, María Guadalupe Santiago-Mauricio, Selene Zárate, Nelly Sélem-Mojica, Blanca Taboada, Kathia Elizabeth Tapia-Diaz, Gloria Vazquez, Joel Armando Vázquez-Pérez. |
| EPI_ISL_3155452, EPI_ISL_3155455, EPI_ISL_3155456, EPI_ISL_3155457, EPI_ISL_3155458, EPI_ISL_3155459, EPI_ISL_3155460, EPI_ISL_3155463, EPI_ISL_3155464, EPI_ISL_3155466, EPI_ISL_3155467, EPI_ISL_3155468, EPI_ISL_3155469, EPI_ISL_3155470, EPI_ISL_3155473, EPI_ISL_3155474, EPI_ISL_3155476, EPI_ISL_3155477, EPI_ISL_3155478, EPI_ISL_3155479, EPI_ISL_3155480, EPI_ISL_3155481, EPI_ISL_3155483, EPI_ISL_3155484, EPI_ISL_3155485, EPI_ISL_3155490, EPI_ISL_3155491, EPI_ISL_3155493, EPI_ISL_3155495, EPI_ISL_3155496, EPI_ISL_3155497, EPI_ISL_3155498, EPI_ISL_3155499, EPI_ISL_3155501, EPI_ISL_3155502, EPI_ISL_3155503 | | | |
| see above | Centro de Investigación Biomédica de Occidente (CIBO) | Centro de Investigación en Enfermedades Infecciosas (CIENI), Instituto Nacional de Enfermedades Respiratorias (INER) | Consorcio Mexicano de Vigilancia Genómica (CoViGen-Mex). Authors (in alphabetical order): Julio Elias Alvarado-Yaah, Carlos F. Arias, Santiago Ávila-Ríos, Eduardo Becerril-Vargas, Víctor Hugo Borja-Aburto, Celia Boukadida, Cristóbal Cháidez-Quiróz, Juan Bautista Chale-Dzul, Ricardo Ciria Merce, Andreu Comas-García, Célida Duque Molina, Julissa Enciso-Ibarra, José Antonio Enciso-Moreno, Gloria Elena Espinosa-Ayala, Fernando Fontove-Herrera, Daniel Fregoso-Rueda, Victor Eduardo García-Arias, Alejandra García-Gasca, Bruno Gómez-Gil, Jean Pierre González, Irvin González-López, Concepción Grajales-Muñiz, Ricardo Grande, Rosa María Gutiérrez Rios, Jesús Hernández, Alejandra Hernández-Terán, Alfredo Herrera-Estrella, Carla Ivón Herrera-Najera, Pavel Isa, Verónica Mata-Haro, Daniel Lira Morales, Susana Lopez, Antonio Loza Román, Brenda Irasema Maldonado-Meza, Bernardo Martínez-Miguel, José Arturo Martínez-Orozco, Célida Martínez-Rodríguez, Margarita Matías-Florentino, Fidencio Mejía-Nepomuceno, María Guadalupe de Jesús Mireles-Rivera, Gloria María Molina-Salinas, Hector Montoya-Fuentes, , Mario Mújica-Sánchez, José Esteban Muñoz-Medina, José de Jesús Nuñez-Contreras, Alicia Ocaña-Mondragón, Luis Alberto Ochoa-Carrera, Hector Esteban Paz-Juárez, Marissa Perez-Garcia, Francisco Pulido, Helen Haydee Fernanda Ramírez-Plascencia, Jorge Salas-Hernández, Angel Gustavo Salas-Lais, Alejandro Sánchez-Flores, Clara Esperanza Santacruz-Tinoco, María Guadalupe Santiago-Mauricio, Selene Zárate, Nelly Sélem-Mojica, Blanca Taboada, Kathia Elizabeth Tapia-Diaz, Gloria Vazquez, Joel Armando Vázquez-Pérez. |
| EPI_ISL_3155504, EPI_ISL_3155505, EPI_ISL_3155506, EPI_ISL_3155507, EPI_ISL_3155508, EPI_ISL_3155509, EPI_ISL_3155511, EPI_ISL_3155512, EPI_ISL_3155513, EPI_ISL_3155514, EPI_ISL_3155515, EPI_ISL_3155518, EPI_ISL_3155520, EPI_ISL_3155521, EPI_ISL_3155522, EPI_ISL_3155523, EPI_ISL_3155524, EPI_ISL_3155525, EPI_ISL_3155526, EPI_ISL_3155527, EPI_ISL_3155528, EPI_ISL_3155529, EPI_ISL_3155530, EPI_ISL_3155531, EPI_ISL_3155532, EPI_ISL_3155533, EPI_ISL_3155534, EPI_ISL_3155535, EPI_ISL_3155536, EPI_ISL_3155537, EPI_ISL_3155538, EPI_ISL_3155539, EPI_ISL_3155541, EPI_ISL_3155543, EPI_ISL_3155544, EPI_ISL_3155546, EPI_ISL_3155548, EPI_ISL_3155551, EPI_ISL_3155552, EPI_ISL_3155553, EPI_ISL_3155554, EPI_ISL_3155555, EPI_ISL_3155557, EPI_ISL_3155558, EPI_ISL_3155561, EPI_ISL_3155562, EPI_ISL_3155563, EPI_ISL_3155567, EPI_ISL_3155568, EPI_ISL_3155569, EPI_ISL_3155570, EPI_ISL_3155571, EPI_ISL_3155572, EPI_ISL_3155573, EPI_ISL_3155574, EPI_ISL_3155576, EPI_ISL_3155577, EPI_ISL_3155581, EPI_ISL_3155583, EPI_ISL_3155584, EPI_ISL_3155585, EPI_ISL_3155586, EPI_ISL_3155587, EPI_ISL_3155588, EPI_ISL_3155590, EPI_ISL_3155591, EPI_ISL_3155592, EPI_ISL_3155593, EPI_ISL_3155594, EPI_ISL_3155595, EPI_ISL_3155596, EPI_ISL_3155597, EPI_ISL_3155598, EPI_ISL_3155599, EPI_ISL_3155600, EPI_ISL_3155601, EPI_ISL_3155603, EPI_ISL_3155605, EPI_ISL_3155606, EPI_ISL_3155607, EPI_ISL_3155608, EPI_ISL_3155609, EPI_ISL_3155610, EPI_ISL_3155612, EPI_ISL_3155613, EPI_ISL_3155615, EPI_ISL_3155616, EPI_ISL_3155617, EPI_ISL_3155618, EPI_ISL_3155619, EPI_ISL_3155620, EPI_ISL_3155621, EPI_ISL_3155622, EPI_ISL_3155624, EPI_ISL_3155627, EPI_ISL_3155628, EPI_ISL_3155629, EPI_ISL_3155631, EPI_ISL_3155632, EPI_ISL_3155633, EPI_ISL_3155634, EPI_ISL_3155635, EPI_ISL_3155636, EPI_ISL_3155640, EPI_ISL_3155646, EPI_ISL_3155647, EPI_ISL_3155648, EPI_ISL_3155649, EPI_ISL_3155650, EPI_ISL_3155651, EPI_ISL_3155652, EPI_ISL_3155653, EPI_ISL_3155654 | | | |
| see above | Laboratorio Central de Epidemiología (LCE) | Centro de Investigación en Enfermedades Infecciosas (CIENI), Instituto Nacional de Enfermedades Respiratorias (INER) | Consorcio Mexicano de Vigilancia Genómica (CoViGen-Mex). Authors (in alphabetical order): Julio Elias Alvarado-Yaah, Carlos F. Arias, Santiago Ávila-Ríos, Eduardo Becerril-Vargas, Víctor Hugo Borja-Aburto, Celia Boukadida, Cristóbal Cháidez-Quiróz, Juan Bautista Chale-Dzul, Ricardo Ciria Merce, Andreu Comas-García, Célida Duque Molina, Julissa Enciso-Ibarra, José Antonio Enciso-Moreno, Gloria Elena Espinosa-Ayala, Fernando Fontove-Herrera, Daniel Fregoso-Rueda, Victor Eduardo García-Arias, Alejandra García-Gasca, Bruno Gómez-Gil, Jean Pierre González, Irvin González-López, Concepción Grajales-Muñiz, Ricardo Grande, Rosa María Gutiérrez Rios, Jesús Hernández, Alejandra Hernández-Terán, Alfredo Herrera-Estrella, Carla Ivón Herrera-Najera, Pavel Isa, Verónica Mata-Haro, Daniel Lira Morales, Susana Lopez, Antonio Loza Román, Brenda Irasema Maldonado-Meza, Bernardo Martínez-Miguel, José Arturo Martínez-Orozco, Célida Martínez-Rodríguez, Margarita Matías-Florentino, Fidencio Mejía-Nepomuceno, María Guadalupe de Jesús Mireles-Rivera, Gloria María Molina-Salinas, Hector Montoya-Fuentes, , Mario Mújica-Sánchez, José Esteban Muñoz-Medina, José de Jesús Nuñez-Contreras, Alicia Ocaña-Mondragón, Luis Alberto Ochoa-Carrera, Hector Esteban Paz-Juárez, Marissa Perez-Garcia, Francisco Pulido, Helen Haydee Fernanda Ramírez-Plascencia, Jorge Salas-Hernández, Angel Gustavo Salas-Lais, Alejandro Sánchez-Flores, Clara Esperanza Santacruz-Tinoco, María Guadalupe Santiago-Mauricio, Selene Zárate, Nelly Sélem-Mojica, Blanca Taboada, Kathia Elizabeth Tapia-Diaz, Gloria Vazquez, Joel Armando Vázquez-Pérez. |
| EPI_ISL_3155655, EPI_ISL_3155656, EPI_ISL_3155657, EPI_ISL_3155658, EPI_ISL_3155661, EPI_ISL_3155662, EPI_ISL_3155663, EPI_ISL_3155664, EPI_ISL_3155665, EPI_ISL_3155666, EPI_ISL_3155667, EPI_ISL_3155668, EPI_ISL_3155669, EPI_ISL_3155670, EPI_ISL_3155671, EPI_ISL_3155672, EPI_ISL_3155673, EPI_ISL_3155674, EPI_ISL_3155675, EPI_ISL_3155676, EPI_ISL_3155677, EPI_ISL_3155679, EPI_ISL_3155680, EPI_ISL_3155682, EPI_ISL_3155683 | | | |
| see above | Montana Public Health Laboratory | Montana Public Health Laboratory | Joy Ritter, Michelle Mozer, Carrie Biskupiak, Deborah Gibson, Michael Dills |
| EPI_ISL_3155685, EPI_ISL_3155686, EPI_ISL_3155687, EPI_ISL_3155688, EPI_ISL_3155689, EPI_ISL_3155690, EPI_ISL_3155691, EPI_ISL_3155692, EPI_ISL_3155693, EPI_ISL_3155694, EPI_ISL_3155695, EPI_ISL_3155696, EPI_ISL_3155697, EPI_ISL_3155698, EPI_ISL_3155699, EPI_ISL_3155700, EPI_ISL_3155701, EPI_ISL_3155702, EPI_ISL_3155703, EPI_ISL_3155704, EPI_ISL_3155705, EPI_ISL_3155706, EPI_ISL_3155707, EPI_ISL_3155708, EPI_ISL_3155709, EPI_ISL_3155710, EPI_ISL_3155711, EPI_ISL_3155712, EPI_ISL_3155713, EPI_ISL_3155714, EPI_ISL_3155715, EPI_ISL_3155716, EPI_ISL_3155717, EPI_ISL_3155718, EPI_ISL_3155719, EPI_ISL_3155720, EPI_ISL_3155721, EPI_ISL_3155722, EPI_ISL_3155723, EPI_ISL_3155724 | | | |
| see above | Alaska State Virology Laboratory | Alaska State Virology Laboratory | Stephanie DeRonde, Elva House, Jacob Zidek, Lisa Smith, Ph.D., Jack Chen, Ph.D. |
| EPI_ISL_3155729, EPI_ISL_3155730, EPI_ISL_3155731, EPI_ISL_3155732, EPI_ISL_3155733, EPI_ISL_3155734, EPI_ISL_3155735, EPI_ISL_3155736, EPI_ISL_3155737, EPI_ISL_3155738, EPI_ISL_3155739, EPI_ISL_3155740, EPI_ISL_3155741, EPI_ISL_3155742, EPI_ISL_3155743, EPI_ISL_3155744, EPI_ISL_3155745, EPI_ISL_3155746, EPI_ISL_3155747, EPI_ISL_3155748, EPI_ISL_3155749, EPI_ISL_3155750, EPI_ISL_3155751, EPI_ISL_3155752, EPI_ISL_3155753, EPI_ISL_3155754, EPI_ISL_3155755, EPI_ISL_3155756, EPI_ISL_3155757, EPI_ISL_3155758, EPI_ISL_3155759, EPI_ISL_3155760, EPI_ISL_3155761, EPI_ISL_3155762, EPI_ISL_3155763, EPI_ISL_3155764, EPI_ISL_3155765, EPI_ISL_3155766, EPI_ISL_3155767, EPI_ISL_3155768, EPI_ISL_3155769, EPI_ISL_3155770, EPI_ISL_3155771, EPI_ISL_3155772, EPI_ISL_3155773, EPI_ISL_3155774, EPI_ISL_3155775, EPI_ISL_3155776, EPI_ISL_3155777, EPI_ISL_3155778, EPI_ISL_3155779, EPI_ISL_3155780, EPI_ISL_3155781, EPI_ISL_3155782, EPI_ISL_3155783, EPI_ISL_3155784, EPI_ISL_3155785, EPI_ISL_3155786, EPI_ISL_3155787, EPI_ISL_3155788, EPI_ISL_3155789, EPI_ISL_3155790, EPI_ISL_3155791, EPI_ISL_3155792, EPI_ISL_3155793, EPI_ISL_3155794, EPI_ISL_3155795, EPI_ISL_3155796, EPI_ISL_3155797, EPI_ISL_3155798, EPI_ISL_3155799, EPI_ISL_3155800, EPI_ISL_3155801, EPI_ISL_3155802, EPI_ISL_3155803, EPI_ISL_3155804, EPI_ISL_3155805, EPI_ISL_3155806, EPI_ISL_3155807, EPI_ISL_3155808, EPI_ISL_3155809, EPI_ISL_3155810, EPI_ISL_3155811, EPI_ISL_3155812, EPI_ISL_3155813, EPI_ISL_3155814, EPI_ISL_3155815, EPI_ISL_3155816, EPI_ISL_3155817, EPI_ISL_3155818, EPI_ISL_3155819, EPI_ISL_3155820, EPI_ISL_3155821, EPI_ISL_3155822, EPI_ISL_3155823, EPI_ISL_3155824, EPI_ISL_3155825, EPI_ISL_3155826, EPI_ISL_3155827, EPI_ISL_3155828, EPI_ISL_3155829, EPI_ISL_3155830, EPI_ISL_3155831, EPI_ISL_3155832, EPI_ISL_3155833, EPI_ISL_3155834, EPI_ISL_3155835, EPI_ISL_3155836, EPI_ISL_3155837, EPI_ISL_3155838, EPI_ISL_3155840, EPI_ISL_3155841, EPI_ISL_3155842, EPI_ISL_3155843, EPI_ISL_3155844, EPI_ISL_3155845, EPI_ISL_3155846, EPI_ISL_3155847, EPI_ISL_3155848, EPI_ISL_3155850, EPI_ISL_3155851, EPI_ISL_3155852, EPI_ISL_3155853, EPI_ISL_3155854, EPI_ISL_3155855, EPI_ISL_3155856, EPI_ISL_3155857, EPI_ISL_3155858, EPI_ISL_3155859, EPI_ISL_3155860, EPI_ISL_3155861, EPI_ISL_3155862, EPI_ISL_3155863, EPI_ISL_3155864, EPI_ISL_3155865, EPI_ISL_3155866, EPI_ISL_3155867, EPI_ISL_3155868, EPI_ISL_3155869, EPI_ISL_3155870, EPI_ISL_3155871, EPI_ISL_3155873, EPI_ISL_3155874, EPI_ISL_3155875, EPI_ISL_3155876, EPI_ISL_3155877, EPI_ISL_3155878, EPI_ISL_3155879, EPI_ISL_3155880, EPI_ISL_3155881, EPI_ISL_3155882, EPI_ISL_3155883, EPI_ISL_3155884, EPI_ISL_3155885, EPI_ISL_3155886, EPI_ISL_3155887, EPI_ISL_3155888, EPI_ISL_3155889, EPI_ISL_3155890, EPI_ISL_3155891, EPI_ISL_3155892, EPI_ISL_3155893, EPI_ISL_3155894, EPI_ISL_3155895, EPI_ISL_3155896, EPI_ISL_3155897, EPI_ISL_3155898, EPI_ISL_3155899, EPI_ISL_3155900, EPI_ISL_3155901, EPI_ISL_3155902, EPI_ISL_3155903, EPI_ISL_3155904, EPI_ISL_3155905, EPI_ISL_3155906, EPI_ISL_3155907, EPI_ISL_3155908, EPI_ISL_3155909, EPI_ISL_3155910, EPI_ISL_3155911, EPI_ISL_3155912, EPI_ISL_3155913, EPI_ISL_3155914, EPI_ISL_3155915, EPI_ISL_3155916, EPI_ISL_3155917, EPI_ISL_3155918, EPI_ISL_3155919, EPI_ISL_3155920, EPI_ISL_3155921, EPI_ISL_3155922, EPI_ISL_3155923, EPI_ISL_3155924, EPI_ISL_3155925, EPI_ISL_3155926, EPI_ISL_3155927, EPI_ISL_3155928, EPI_ISL_3155929, EPI_ISL_3155930, EPI_ISL_3155931, EPI_ISL_3155932, EPI_ISL_3155933, EPI_ISL_3155934, EPI_ISL_3155935, EPI_ISL_3155936, EPI_ISL_3155937, EPI_ISL_3155938, EPI_ISL_3155939, EPI_ISL_3155940, EPI_ISL_3155941, EPI_ISL_3155942, EPI_ISL_3155943, EPI_ISL_3155944, EPI_ISL_3155945, EPI_ISL_3155946, EPI_ISL_3155948, EPI_ISL_3155950, EPI_ISL_3155952, EPI_ISL_3155955, EPI_ISL_3155958, EPI_ISL_3155960, EPI_ISL_3155963, EPI_ISL_3155965, EPI_ISL_3155968, EPI_ISL_3155971, EPI_ISL_3155973, EPI_ISL_3155976, EPI_ISL_3155978, EPI_ISL_3155981, EPI_ISL_3155983, EPI_ISL_3155986, EPI_ISL_3155988, EPI_ISL_3155991, EPI_ISL_3155993, EPI_ISL_3155995, EPI_ISL_3155996, EPI_ISL_3155997, EPI_ISL_3155998, EPI_ISL_3156000, EPI_ISL_3156001, EPI_ISL_3156002, EPI_ISL_3156003, EPI_ISL_3156004, EPI_ISL_3156005, EPI_ISL_3156006, EPI_ISL_3156007, EPI_ISL_3156008, EPI_ISL_3156009, EPI_ISL_3156010, EPI_ISL_3156011, EPI_ISL_3156012, EPI_ISL_3156013, EPI_ISL_3156014, EPI_ISL_3156015, EPI_ISL_3156016, EPI_ISL_3156017, EPI_ISL_3156018, EPI_ISL_3156019, EPI_ISL_3156020, EPI_ISL_3156021, EPI_ISL_3156022, EPI_ISL_3156023, EPI_ISL_3156024, EPI_ISL_3156025, EPI_ISL_3156026, EPI_ISL_3156027, EPI_ISL_3156028, EPI_ISL_3156029, EPI_ISL_3156030, EPI_ISL_3156031, EPI_ISL_3156032, EPI_ISL_3156033, EPI_ISL_3156034, EPI_ISL_3156035, EPI_ISL_3156036, EPI_ISL_3156037, EPI_ISL_3156038, EPI_ISL_3156039, EPI_ISL_3156040 | | | |
| see above | Broad Institute Clinical Research Sequencing Platform | Infectious Disease Program, Broad Institute of Harvard and MIT | Siddle,K.J., Adams,G., Pearlman,L., Gladden-Young,A., Vicente,G., Blumenstiel,B., DeFelice,M., Lee,M., McGovern,S., Lagerborg,K., Rudy,M., DeRuff,K., Carter,A., Normandin,E., Bauer,M., Reilly,S., Tomkins-Tinch,C., Loreth,C., Chaluvadi,S., Meldrim,J., Granger,B., Lemieux,J.E., Birren,B.W., Sabeti,P.C., Larkin,K., Dodge,S., Lennon,N., Madoff,L., Brown,C., Gallagher,G., Smole,S., Park,D.J., Gabriel,S., and MacInnis,B.L. |
| EPI_ISL_3156045, EPI_ISL_3156046, EPI_ISL_3156047, EPI_ISL_3156048, EPI_ISL_3156049, EPI_ISL_3156050, EPI_ISL_3156051, EPI_ISL_3156052, EPI_ISL_3156053, EPI_ISL_3156054, EPI_ISL_3156055, EPI_ISL_3156057, EPI_ISL_3156058, EPI_ISL_3156059, EPI_ISL_3156060, EPI_ISL_3156061, EPI_ISL_3156062, EPI_ISL_3156063, EPI_ISL_3156064, EPI_ISL_3156065, EPI_ISL_3156066, EPI_ISL_3156067, EPI_ISL_3156068, EPI_ISL_3156069, EPI_ISL_3156070, EPI_ISL_3156071, EPI_ISL_3156072, EPI_ISL_3156073, EPI_ISL_3156074, EPI_ISL_3156075, EPI_ISL_3156076, EPI_ISL_3156077, EPI_ISL_3156078, EPI_ISL_3156080, EPI_ISL_3156081, EPI_ISL_3156082, EPI_ISL_3156083, EPI_ISL_3156084, EPI_ISL_3156085, EPI_ISL_3156087, EPI_ISL_3156088, EPI_ISL_3156089, EPI_ISL_3156090, EPI_ISL_3156091, EPI_ISL_3156092, EPI_ISL_3156093 | | | |
| see above | Microbiology Department, Laboratori Clínic Metropolitana Nord. Hospital Universitari Germans Trias i Pujol | Can Ruti SARS-CoV-2 Sequencing Hub (HUGTiP/IrsiCaixa/IGTP) | Marc Noguera-Julian, Pilar Armengol, Ignacio Blanco, Antoni E Bordoy, Francesc Catala-Moll, Pere-Joan Cardona, Maria Casadellà, Cristina Casañ, Gemma Clara, Bonaventura Clotet, Montserrat Giménez, Mercedes Guerrero, Anna Not, David Panisello, Roger Paredes, Mariona Parera, Alexia París, Verónica Saludes, Alba Sánchez, Laia Soler, Lauro Sumoy, and Elisa Martró on behalf of the Can Ruti SARS-CoV-2 Sequencing Hub |
| EPI_ISL_3156094 | Microbiologia CATLAB | Can Ruti SARS-CoV-2 Sequencing Hub (HUGTiP/IrsiCaixa/IGTP) | Marc Noguera-Julian, Pilar Armengol, Ignacio Blanco, Antoni E Bordoy, Francesc Catala-Moll, Pere-Joan Cardona, Maria Casadellà, Cristina Casañ, Gemma Clara, Bonaventura Clotet, Montserrat Giménez, Mercedes Guerrero, Anna Not, David Panisello, Roger Paredes, Mariona Parera, Alexia París, Verónica Saludes, Alba Sánchez, Laia Soler, Lauro Sumoy, and Elisa Martró on behalf of the Can Ruti SARS-CoV-2 Sequencing Hub |
| EPI_ISL_3156095, EPI_ISL_3156096, EPI_ISL_3156097, EPI_ISL_3156098, EPI_ISL_3156099, EPI_ISL_3156100, EPI_ISL_3156101, EPI_ISL_3156102, EPI_ISL_3156103, EPI_ISL_3156104, EPI_ISL_3156105, EPI_ISL_3156106, EPI_ISL_3156107, EPI_ISL_3156108, EPI_ISL_3156109, EPI_ISL_3156110, EPI_ISL_3156112, EPI_ISL_3156113, EPI_ISL_3156114, EPI_ISL_3156115, EPI_ISL_3156116, EPI_ISL_3156117, | | | |

| | | | |
|---|---|---|---|
| EPI_ISL_3156118, EPI_ISL_3156119, EPI_ISL_3156120, EPI_ISL_3156121, EPI_ISL_3156122, EPI_ISL_3156123, EPI_ISL_3156124, EPI_ISL_3156125, EPI_ISL_3156126, EPI_ISL_3156127, EPI_ISL_3156128, EPI_ISL_3156129, EPI_ISL_3156131, EPI_ISL_3156132, EPI_ISL_3156134, EPI_ISL_3156135, EPI_ISL_3156136, EPI_ISL_3156137, EPI_ISL_3156138, EPI_ISL_3156139, EPI_ISL_3156140 | | | |
| see above | Microbiology Department, Laboratori Clínic Metropolitana Nord. Hospital Universitari Germans Trias i Pujol | Can Ruti SARS-CoV-2 Sequencing Hub (HUGTiP/IrsiCaixa/IGTP) | Marc Noguera-Julian, Pilar Armengol, Ignacio Blanco, Antoni E Bordoy, Francesc Catala-Moll, Pere-Joan Cardona, Maria Casadellà, Cristina Casañ, Gemma Clara, Bonaventura Clotet, Montserrat Giménez, Mercedes Guerrero, Anna Not, David Panisello, Roger Paredes, Mariona Parera, Alexia París, Verónica Saludes, Alba Sánchez, Laia Soler, Lauro Sumoy, and Elisa Martró on behalf of the Can Ruti SARS-CoV-2 Sequencing Hub |
| EPI_ISL_3156141 | Microbiologia CATLAB | Can Ruti SARS-CoV-2 Sequencing Hub (HUGTiP/IrsiCaixa/IGTP) | Marc Noguera-Julian, Pilar Armengol, Ignacio Blanco, Antoni E Bordoy, Francesc Catala-Moll, Pere-Joan Cardona, Maria Casadellà, Cristina Casañ, Gemma Clara, Bonaventura Clotet, Montserrat Giménez, Mercedes Guerrero, Anna Not, David Panisello, Roger Paredes, Mariona Parera, Alexia París, Verónica Saludes, Alba Sánchez, Laia Soler, Lauro Sumoy, and Elisa Martró on behalf of the Can Ruti SARS-CoV-2 Sequencing Hub |
| EPI_ISL_3156142, EPI_ISL_3156143, EPI_ISL_3156144, EPI_ISL_3156145 | Microbiology Department, Laboratori Clínic Metropolitana Nord. Hospital Universitari Germans Trias i Pujol | Can Ruti SARS-CoV-2 Sequencing Hub (HUGTiP/IrsiCaixa/IGTP) | Marc Noguera-Julian, Pilar Armengol, Ignacio Blanco, Antoni E Bordoy, Francesc Catala-Moll, Pere-Joan Cardona, Maria Casadellà, Cristina Casañ, Gemma Clara, Bonaventura Clotet, Montserrat Giménez, Mercedes Guerrero, Anna Not, David Panisello, Roger Paredes, Mariona Parera, Alexia París, Verónica Saludes, Alba Sánchez, Laia Soler, Lauro Sumoy, and Elisa Martró on behalf of the Can Ruti SARS-CoV-2 Sequencing Hub |
| EPI_ISL_3156146 | Microbiologia CATLAB | Can Ruti SARS-CoV-2 Sequencing Hub (HUGTiP/IrsiCaixa/IGTP) | Marc Noguera-Julian, Pilar Armengol, Ignacio Blanco, Antoni E Bordoy, Francesc Catala-Moll, Pere-Joan Cardona, Maria Casadellà, Cristina Casañ, Gemma Clara, Bonaventura Clotet, Montserrat Giménez, Mercedes Guerrero, Anna Not, David Panisello, Roger Paredes, Mariona Parera, Alexia París, Verónica Saludes, Alba Sánchez, Laia Soler, Lauro Sumoy, and Elisa Martró on behalf of the Can Ruti SARS-CoV-2 Sequencing Hub |
| EPI_ISL_3156147, EPI_ISL_3156148, EPI_ISL_3156149 | Microbiology Department, Laboratori Clínic Metropolitana Nord. Hospital Universitari Germans Trias i Pujol | Can Ruti SARS-CoV-2 Sequencing Hub (HUGTiP/IrsiCaixa/IGTP) | Marc Noguera-Julian, Pilar Armengol, Ignacio Blanco, Antoni E Bordoy, Francesc Catala-Moll, Pere-Joan Cardona, Maria Casadellà, Cristina Casañ, Gemma Clara, Bonaventura Clotet, Montserrat Giménez, Mercedes Guerrero, Anna Not, David Panisello, Roger Paredes, Mariona Parera, Alexia París, Verónica Saludes, Alba Sánchez, Laia Soler, Lauro Sumoy, and Elisa Martró on behalf of the Can Ruti SARS-CoV-2 Sequencing Hub |
| EPI_ISL_3156150 | Microbiologia CATLAB | Can Ruti SARS-CoV-2 Sequencing Hub (HUGTiP/IrsiCaixa/IGTP) | Marc Noguera-Julian, Pilar Armengol, Ignacio Blanco, Antoni E Bordoy, Francesc Catala-Moll, Pere-Joan Cardona, Maria Casadellà, Cristina Casañ, Gemma Clara, Bonaventura Clotet, Montserrat Giménez, Mercedes Guerrero, Anna Not, David Panisello, Roger Paredes, Mariona Parera, Alexia París, Verónica Saludes, Alba Sánchez, Laia Soler, Lauro Sumoy, and Elisa Martró on behalf of the Can Ruti SARS-CoV-2 Sequencing Hub |
| EPI_ISL_3156151, EPI_ISL_3156152, EPI_ISL_3156153, EPI_ISL_3156154 | Microbiology Department, Laboratori Clínic Metropolitana Nord. Hospital Universitari Germans Trias i Pujol | Can Ruti SARS-CoV-2 Sequencing Hub (HUGTiP/IrsiCaixa/IGTP) | Marc Noguera-Julian, Pilar Armengol, Ignacio Blanco, Antoni E Bordoy, Francesc Catala-Moll, Pere-Joan Cardona, Maria Casadellà, Cristina Casañ, Gemma Clara, Bonaventura Clotet, Montserrat Giménez, Mercedes Guerrero, Anna Not, David Panisello, Roger Paredes, Mariona Parera, Alexia París, Verónica Saludes, Alba Sánchez, Laia Soler, Lauro Sumoy, and Elisa Martró on behalf of the Can Ruti SARS-CoV-2 Sequencing Hub |
| EPI_ISL_3156155 | Microbiologia CATLAB | Can Ruti SARS-CoV-2 Sequencing Hub (HUGTiP/IrsiCaixa/IGTP) | Marc Noguera-Julian, Pilar Armengol, Ignacio Blanco, Antoni E Bordoy, Francesc Catala-Moll, Pere-Joan Cardona, Maria Casadellà, Cristina Casañ, Gemma Clara, Bonaventura Clotet, Montserrat Giménez, Mercedes Guerrero, Anna Not, David Panisello, Roger Paredes, Mariona Parera, Alexia París, Verónica Saludes, Alba Sánchez, Laia Soler, Lauro Sumoy, and Elisa Martró on behalf of the Can Ruti SARS-CoV-2 Sequencing Hub |
| EPI_ISL_3156156, EPI_ISL_3156157, EPI_ISL_3156158, EPI_ISL_3156159 | Microbiology Department, Laboratori Clínic Metropolitana Nord. Hospital Universitari Germans Trias i Pujol | Can Ruti SARS-CoV-2 Sequencing Hub (HUGTiP/IrsiCaixa/IGTP) | Marc Noguera-Julian, Pilar Armengol, Ignacio Blanco, Antoni E Bordoy, Francesc Catala-Moll, Pere-Joan Cardona, Maria Casadellà, Cristina Casañ, Gemma Clara, Bonaventura Clotet, Montserrat Giménez, Mercedes Guerrero, Anna Not, David Panisello, Roger Paredes, Mariona Parera, Alexia París, Verónica Saludes, Alba Sánchez, Laia Soler, Lauro Sumoy, and Elisa Martró on behalf of the Can Ruti SARS-CoV-2 Sequencing Hub |
| EPI_ISL_3156160 | Microbiologia CATLAB | Can Ruti SARS-CoV-2 Sequencing Hub (HUGTiP/IrsiCaixa/IGTP) | Marc Noguera-Julian, Pilar Armengol, Ignacio Blanco, Antoni E Bordoy, Francesc Catala-Moll, Pere-Joan Cardona, Maria Casadellà, Cristina Casañ, Gemma Clara, Bonaventura Clotet, Montserrat Giménez, Mercedes Guerrero, Anna Not, David Panisello, Roger Paredes, Mariona Parera, Alexia París, Verónica Saludes, Alba Sánchez, Laia Soler, Lauro Sumoy, and Elisa Martró on behalf of the Can Ruti SARS-CoV-2 Sequencing Hub |
| EPI_ISL_3156161, EPI_ISL_3156162, EPI_ISL_3156163, EPI_ISL_3156164 | Microbiology Department, Laboratori Clínic Metropolitana Nord. Hospital Universitari Germans Trias i Pujol | Can Ruti SARS-CoV-2 Sequencing Hub (HUGTiP/IrsiCaixa/IGTP) | Marc Noguera-Julian, Pilar Armengol, Ignacio Blanco, Antoni E Bordoy, Francesc Catala-Moll, Pere-Joan Cardona, Maria Casadellà, Cristina Casañ, Gemma Clara, Bonaventura Clotet, Montserrat Giménez, Mercedes Guerrero, Anna Not, David Panisello, Roger Paredes, Mariona Parera, Alexia París, Verónica Saludes, Alba Sánchez, Laia Soler, Lauro Sumoy, and Elisa Martró on behalf of the Can Ruti SARS-CoV-2 Sequencing Hub |
| EPI_ISL_3156165 | Microbiologia CATLAB | Can Ruti SARS-CoV-2 Sequencing Hub (HUGTiP/IrsiCaixa/IGTP) | Marc Noguera-Julian, Pilar Armengol, Ignacio Blanco, Antoni E Bordoy, Francesc Catala-Moll, Pere-Joan Cardona, Maria Casadellà, Cristina Casañ, Gemma Clara, Bonaventura Clotet, Montserrat Giménez, Mercedes Guerrero, Anna Not, David Panisello, Roger Paredes, Mariona Parera, Alexia París, Verónica Saludes, Alba Sánchez, Laia Soler, Lauro Sumoy, and Elisa Martró on behalf of the Can Ruti SARS-CoV-2 Sequencing Hub |
| EPI_ISL_3156166, EPI_ISL_3156167, EPI_ISL_3156168 | Microbiology Department, Laboratori Clínic Metropolitana Nord. Hospital Universitari Germans Trias i Pujol | Can Ruti SARS-CoV-2 Sequencing Hub (HUGTiP/IrsiCaixa/IGTP) | Marc Noguera-Julian, Pilar Armengol, Ignacio Blanco, Antoni E Bordoy, Francesc Catala-Moll, Pere-Joan Cardona, Maria Casadellà, Cristina Casañ, Gemma Clara, Bonaventura Clotet, Montserrat Giménez, Mercedes Guerrero, Anna Not, David Panisello, Roger Paredes, Mariona Parera, Alexia París, Verónica Saludes, Alba Sánchez, Laia Soler, Lauro Sumoy, and Elisa Martró on behalf of the Can Ruti SARS-CoV-2 Sequencing Hub |
| EPI_ISL_3156169, EPI_ISL_3156170 | Microbiologia CATLAB | Can Ruti SARS-CoV-2 Sequencing Hub (HUGTiP/IrsiCaixa/IGTP) | Marc Noguera-Julian, Pilar Armengol, Ignacio Blanco, Antoni E Bordoy, Francesc Catala-Moll, Pere-Joan Cardona, Maria Casadellà, Cristina Casañ, Gemma Clara, Bonaventura Clotet, Montserrat Giménez, Mercedes Guerrero, Anna Not, David Panisello, Roger Paredes, Mariona Parera, Alexia París, Verónica Saludes, Alba Sánchez, Laia Soler, Lauro Sumoy, and Elisa Martró on behalf of the Can Ruti SARS-CoV-2 Sequencing Hub |
| EPI_ISL_3156171, EPI_ISL_3156172, EPI_ISL_3156173 | Microbiology Department, Laboratori Clínic Metropolitana Nord. Hospital Universitari Germans Trias i Pujol | Can Ruti SARS-CoV-2 Sequencing Hub (HUGTiP/IrsiCaixa/IGTP) | Marc Noguera-Julian, Pilar Armengol, Ignacio Blanco, Antoni E Bordoy, Francesc Catala-Moll, Pere-Joan Cardona, Maria Casadellà, Cristina Casañ, Gemma Clara, Bonaventura Clotet, Montserrat Giménez, Mercedes Guerrero, Anna Not, David Panisello, Roger Paredes, Mariona Parera, Alexia París, Verónica Saludes, Alba Sánchez, Laia Soler, Lauro Sumoy, and Elisa Martró on behalf of the Can Ruti SARS-CoV-2 Sequencing Hub |
| EPI_ISL_3156174, EPI_ISL_3156175 | Microbiologia CATLAB | Can Ruti SARS-CoV-2 Sequencing Hub (HUGTiP/IrsiCaixa/IGTP) | Marc Noguera-Julian, Pilar Armengol, Ignacio Blanco, Antoni E Bordoy, Francesc Catala-Moll, Pere-Joan Cardona, Maria Casadellà, Cristina Casañ, Gemma Clara, Bonaventura Clotet, Montserrat Giménez, Mercedes Guerrero, Anna Not, David Panisello, Roger Paredes, Mariona Parera, Alexia París, Verónica Saludes, Alba Sánchez, Laia Soler, Lauro Sumoy, and Elisa Martró on behalf of the Can Ruti SARS-CoV-2 Sequencing Hub |
| EPI_ISL_3156176, EPI_ISL_3156177, EPI_ISL_3156178, EPI_ISL_3156179, EPI_ISL_3156180, EPI_ISL_3156181, EPI_ISL_3156182, EPI_ISL_3156183, EPI_ISL_3156184, EPI_ISL_3156185, EPI_ISL_3156186, EPI_ISL_3156187, EPI_ISL_3156188, EPI_ISL_3156189, EPI_ISL_3156190, EPI_ISL_3156191, EPI_ISL_3156192, EPI_ISL_3156193, EPI_ISL_3156194, EPI_ISL_3156195, EPI_ISL_3156196, EPI_ISL_3156197, EPI_ISL_3156198, EPI_ISL_3156199, EPI_ISL_3156200, EPI_ISL_3156201, EPI_ISL_3156202, EPI_ISL_3156203, EPI_ISL_3156204, EPI_ISL_3156205, EPI_ISL_3156206, EPI_ISL_3156207, EPI_ISL_3156208, EPI_ISL_3156209, EPI_ISL_3156210, EPI_ISL_3156211, EPI_ISL_3156212, EPI_ISL_3156213, EPI_ISL_3156214, EPI_ISL_3156215, EPI_ISL_3156216, EPI_ISL_3156217, EPI_ISL_3156218, EPI_ISL_3156219, EPI_ISL_3156220, EPI_ISL_3156221, EPI_ISL_3156222, EPI_ISL_3156223, EPI_ISL_3156224, EPI_ISL_3156225, EPI_ISL_3156226, EPI_ISL_3156227, EPI_ISL_3156228, EPI_ISL_3156229, EPI_ISL_3156230, EPI_ISL_3156231, EPI_ISL_3156232, EPI_ISL_3156233, EPI_ISL_3156234, EPI_ISL_3156235, EPI_ISL_3156236, EPI_ISL_3156238, EPI_ISL_3156239, EPI_ISL_3156240, EPI_ISL_3156241, EPI_ISL_3156242, EPI_ISL_3156243, EPI_ISL_3156244, EPI_ISL_3156245, EPI_ISL_3156246, EPI_ISL_3156247, EPI_ISL_3156248, EPI_ISL_3156249, EPI_ISL_3156250, EPI_ISL_3156251, EPI_ISL_3156252, EPI_ISL_3156253, EPI_ISL_3156254, EPI_ISL_3156255, EPI_ISL_3156256, EPI_ISL_3156257, EPI_ISL_3156258, EPI_ISL_3156259, EPI_ISL_3156260, EPI_ISL_3156261, EPI_ISL_3156262, EPI_ISL_3156263, EPI_ISL_3156264, EPI_ISL_3156265, EPI_ISL_3156267, EPI_ISL_3156268 | | | |
| see above | Microbiology Department, Laboratori Clínic Metropolitana Nord. Hospital Universitari Germans Trias i Pujol | Can Ruti SARS-CoV-2 Sequencing Hub (HUGTiP/IrsiCaixa/IGTP) | Marc Noguera-Julian, Pilar Armengol, Ignacio Blanco, Antoni E Bordoy, Francesc Catala-Moll, Pere-Joan Cardona, Maria Casadellà, Cristina Casañ, Gemma Clara, Bonaventura Clotet, Montserrat Giménez, Mercedes Guerrero, Anna Not, David Panisello, Roger Paredes, Mariona Parera, Alexia París, Verónica Saludes, Alba Sánchez, Laia Soler, Lauro Sumoy, and Elisa Martró on behalf of the Can Ruti SARS-CoV-2 Sequencing Hub |
| EPI_ISL_3156275, EPI_ISL_3156276, EPI_ISL_3156277, EPI_ISL_3156278, EPI_ISL_3156280, EPI_ISL_3156281, EPI_ISL_3156282, EPI_ISL_3156283, EPI_ISL_3156284, EPI_ISL_3156285, EPI_ISL_3156286, EPI_ISL_3156287, EPI_ISL_3156288, EPI_ISL_3156289, EPI_ISL_3156290, EPI_ISL_3156291, EPI_ISL_3156292, EPI_ISL_3156293, EPI_ISL_3156294, EPI_ISL_3156295, EPI_ISL_3156296, EPI_ISL_3156298, EPI_ISL_3156299, EPI_ISL_3156300, EPI_ISL_3156301, EPI_ISL_3156302, EPI_ISL_3156303, EPI_ISL_3156304, EPI_ISL_3156305, EPI_ISL_3156306, EPI_ISL_3156307, EPI_ISL_3156308, EPI_ISL_3156309, EPI_ISL_3156310, EPI_ISL_3156311, EPI_ISL_3156312, EPI_ISL_3156313, EPI_ISL_3156314, EPI_ISL_3156315, EPI_ISL_3156316, EPI_ISL_3156317, EPI_ISL_3156318, EPI_ISL_3156319, EPI_ISL_3156320, EPI_ISL_3156321, EPI_ISL_3156322, EPI_ISL_3156323, EPI_ISL_3156324, EPI_ISL_3156326, EPI_ISL_3156327, EPI_ISL_3156328, EPI_ISL_3156329, EPI_ISL_3156330, EPI_ISL_3156331, EPI_ISL_3156332, EPI_ISL_3156333, EPI_ISL_3156334, EPI_ISL_3156335, EPI_ISL_3156336, EPI_ISL_3156337, EPI_ISL_3156338, EPI_ISL_3156339, EPI_ISL_3156340, EPI_ISL_3156341, EPI_ISL_3156342, EPI_ISL_3156343, EPI_ISL_3156344, EPI_ISL_3156345, EPI_ISL_3156346, EPI_ISL_3156347, EPI_ISL_3156348, EPI_ISL_3156349, EPI_ISL_3156350, EPI_ISL_3156351, EPI_ISL_3156352, EPI_ISL_3156353, EPI_ISL_3156354, EPI_ISL_3156355, EPI_ISL_3156356, EPI_ISL_3156357, EPI_ISL_3156358, EPI_ISL_3156359, EPI_ISL_3156360, EPI_ISL_3156361, EPI_ISL_3156362, EPI_ISL_3156363, EPI_ISL_3156364, EPI_ISL_3156365, EPI_ISL_3156366, EPI_ISL_3156367, EPI_ISL_3156368, EPI_ISL_3156369, EPI_ISL_3156370, EPI_ISL_3156371, EPI_ISL_3156372, EPI_ISL_3156373, EPI_ISL_3156374, EPI_ISL_3156375, EPI_ISL_3156376, EPI_ISL_3156377, EPI_ISL_3156378, EPI_ISL_3156379, EPI_ISL_3156380, EPI_ISL_3156381, EPI_ISL_3156382, EPI_ISL_3156383, EPI_ISL_3156384, EPI_ISL_3156385, EPI_ISL_3156386, EPI_ISL_3156387, EPI_ISL_3156388, EPI_ISL_3156389, EPI_ISL_3156390, EPI_ISL_3156391, EPI_ISL_3156392, EPI_ISL_3156393, EPI_ISL_3156394, EPI_ISL_3156395, EPI_ISL_3156396, EPI_ISL_3156397, EPI_ISL_3156398, EPI_ISL_3156399, EPI_ISL_3156400, EPI_ISL_3156401, EPI_ISL_3156402, EPI_ISL_3156403, EPI_ISL_3156404, EPI_ISL_3156405, EPI_ISL_3156406, EPI_ISL_3156407, EPI_ISL_3156408, EPI_ISL_3156409, EPI_ISL_3156410, EPI_ISL_3156411, EPI_ISL_3156412, EPI_ISL_3156413, EPI_ISL_3156414, EPI_ISL_3156415, EPI_ISL_3156418, EPI_ISL_3156419, EPI_ISL_3156420, EPI_ISL_3156421, EPI_ISL_3156422, EPI_ISL_3156423, EPI_ISL_3156425, EPI_ISL_3156426, EPI_ISL_3156427, EPI_ISL_3156428, EPI_ISL_3156429, EPI_ISL_3156430, EPI_ISL_3156431, EPI_ISL_3156432, EPI_ISL_3156433, EPI_ISL_3156434, EPI_ISL_3156435, EPI_ISL_3156436, EPI_ISL_3156437, EPI_ISL_3156438, EPI_ISL_3156439, EPI_ISL_3156440, EPI_ISL_3156441, EPI_ISL_3156442, EPI_ISL_3156443, EPI_ISL_3156444, EPI_ISL_3156445 | | | |
| see above | University of Mississippi Medical Center, Department of Pathology | University of Mississippi Medical Center, Molecular and Genomics Core Facility | Ashley C. Johnson, Wenjie Wu, Ithiel J. Frame, Krishna K. Ayyalasomayajula, Michael R. Garrett, D. Ashley Robinson |
| EPI_ISL_3156455, EPI_ISL_3156456, EPI_ISL_3156457, EPI_ISL_3156458, EPI_ISL_3156459, EPI_ISL_3156460, EPI_ISL_3156462, EPI_ISL_3156463, EPI_ISL_3156465, EPI_ISL_3156466, EPI_ISL_3156467, EPI_ISL_3156468, EPI_ISL_3156470, EPI_ISL_3156471, EPI_ISL_3156472, EPI_ISL_3156473, EPI_ISL_3156474, EPI_ISL_3156475, EPI_ISL_3156476, EPI_ISL_3156479, EPI_ISL_3156480, EPI_ISL_3156481, EPI_ISL_3156482, EPI_ISL_3156483, EPI_ISL_3156484, EPI_ISL_3156485, EPI_ISL_3156486, EPI_ISL_3156487, EPI_ISL_3156488, EPI_ISL_3156489, EPI_ISL_3156490, EPI_ISL_3156492, EPI_ISL_3156493, EPI_ISL_3156494, EPI_ISL_3156495, EPI_ISL_3156496, EPI_ISL_3156497, EPI_ISL_3156498, EPI_ISL_3156499, EPI_ISL_3156500, EPI_ISL_3156501, EPI_ISL_3156502, EPI_ISL_3156503, EPI_ISL_3156504, EPI_ISL_3156505, EPI_ISL_3156506, EPI_ISL_3156508, EPI_ISL_3156509, EPI_ISL_3156510, EPI_ISL_3156511, EPI_ISL_3156512, EPI_ISL_3156513, EPI_ISL_3156514, EPI_ISL_3156515, EPI_ISL_3156516, EPI_ISL_3156517, EPI_ISL_3156518, EPI_ISL_3156519, EPI_ISL_3156520, EPI_ISL_3156521, EPI_ISL_3156522, EPI_ISL_3156523, EPI_ISL_3156524, EPI_ISL_3156525, EPI_ISL_3156526, EPI_ISL_3156527, EPI_ISL_3156528, EPI_ISL_3156529, EPI_ISL_3156530, EPI_ISL_3156531, EPI_ISL_3156532, EPI_ISL_3156533, EPI_ISL_3156534, EPI_ISL_3156536, EPI_ISL_3156537, EPI_ISL_3156538, EPI_ISL_3156539, EPI_ISL_3156540, EPI_ISL_3156541, EPI_ISL_3156542, EPI_ISL_3156543, EPI_ISL_3156544, EPI_ISL_3156545, EPI_ISL_3156546 | | | |
| see above | Idaho Bureau of Laboratories | Boise VA Medical Center, PALMS | Matthew Burns, Aimee Ceniseros, Christian Loera, Dr. Christopher Ball, Ying Pei, Cheri Lamb McFarlane, James Razor, Emily Bartlette |
| EPI_ISL_3156548, EPI_ISL_3156549, EPI_ISL_3156550, EPI_ISL_3156551, EPI_ISL_3156552, EPI_ISL_3156553, EPI_ISL_3156554, EPI_ISL_3156555, EPI_ISL_3156556, EPI_ISL_3156557, EPI_ISL_3156558, EPI_ISL_3156559, EPI_ISL_3156560, EPI_ISL_3156561, EPI_ISL_3156562, EPI_ISL_3156563, EPI_ISL_3156564, EPI_ISL_3156565, EPI_ISL_3156567, EPI_ISL_3156568, EPI_ISL_3156569, EPI_ISL_3156570, EPI_ISL_3156571, EPI_ISL_3156572, EPI_ISL_3156573, EPI_ISL_3156574, EPI_ISL_3156575, EPI_ISL_3156577, EPI_ISL_3156578, EPI_ISL_3156579, EPI_ISL_3156580, EPI_ISL_3156581, EPI_ISL_3156582, EPI_ISL_3156583, EPI_ISL_3156584, EPI_ISL_3156585, EPI_ISL_3156586, EPI_ISL_3156587, EPI_ISL_3156588, EPI_ISL_3156589, EPI_ISL_3156590, EPI_ISL_3156591, EPI_ISL_3156592, EPI_ISL_3156593, EPI_ISL_3156594, EPI_ISL_3156598, EPI_ISL_3156599, EPI_ISL_3156600, EPI_ISL_3156601, EPI_ISL_3156602, EPI_ISL_3156603, EPI_ISL_3156604, EPI_ISL_3156605, EPI_ISL_3156606, EPI_ISL_3156607, EPI_ISL_3156608, EPI_ISL_3156609, EPI_ISL_3156610, EPI_ISL_3156611, EPI_ISL_3156612, EPI_ISL_3156613, EPI_ISL_3156614, EPI_ISL_3156615, EPI_ISL_3156616, EPI_ISL_3156617, EPI_ISL_3156618, EPI_ISL_3156619, EPI_ISL_3156621, EPI_ISL_3156622, EPI_ISL_3156623, EPI_ISL_3156626, EPI_ISL_3156627, EPI_ISL_3156628, EPI_ISL_3156630, EPI_ISL_3156631, EPI_ISL_3156632, EPI_ISL_3156633, EPI_ISL_3156634, EPI_ISL_3156635, EPI_ISL_3156636, EPI_ISL_3156637, EPI_ISL_3156638, EPI_ISL_3156639, EPI_ISL_3156640, EPI_ISL_3156641, EPI_ISL_3156642, EPI_ISL_3156643, EPI_ISL_3156644, EPI_ISL_3156645, EPI_ISL_3156646, EPI_ISL_3156647, EPI_ISL_3156649, EPI_ISL_3156650, EPI_ISL_3156651, EPI_ISL_3156652, EPI_ISL_3156653, EPI_ISL_3156654, EPI_ISL_3156655, EPI_ISL_3156656, EPI_ISL_3156657, EPI_ISL_3156658, EPI_ISL_3156659, EPI_ISL_3156660, EPI_ISL_3156661, EPI_ISL_3156662, EPI_ISL_3156664, EPI_ISL_3156665, EPI_ISL_3156666, EPI_ISL_3156667, EPI_ISL_3156668, EPI_ISL_3156669, EPI_ISL_3156670, EPI_ISL_3156671, EPI_ISL_3156672, EPI_ISL_3156673, EPI_ISL_3156674, EPI_ISL_3156675, EPI_ISL_3156676, EPI_ISL_3156677, EPI_ISL_3156678, EPI_ISL_3156679, EPI_ISL_3156680, EPI_ISL_3156681, EPI_ISL_3156682, EPI_ISL_3156683, EPI_ISL_3156684, EPI_ISL_3156686, EPI_ISL_3156687, EPI_ISL_3156689, EPI_ISL_3156690, EPI_ISL_3156692, EPI_ISL_3156693, EPI_ISL_3156694, EPI_ISL_3156696, EPI_ISL_3156697, EPI_ISL_3156698, EPI_ISL_3156699, EPI_ISL_3156700, EPI_ISL_3156701, EPI_ISL_3156702, EPI_ISL_3156703, EPI_ISL_3156704, EPI_ISL_3156705, EPI_ISL_3156706, EPI_ISL_3156707, EPI_ISL_3156708, EPI_ISL_3156709, EPI_ISL_3156710, EPI_ISL_3156809, EPI_ISL_3156810, EPI_ISL_3156811, EPI_ISL_3156812, EPI_ISL_3156813, EPI_ISL_3156814, EPI_ISL_3156815, EPI_ISL_3156817, EPI_ISL_3156818, EPI_ISL_3156819, EPI_ISL_3156820, EPI_ISL_3156821, EPI_ISL_3156822, EPI_ISL_3156823, EPI_ISL_3156824, EPI_ISL_3156825, EPI_ISL_3156826, EPI_ISL_3156828, EPI_ISL_3156829, EPI_ISL_3156830, EPI_ISL_3156831, EPI_ISL_3156832, EPI_ISL_3156833, EPI_ISL_3156834, EPI_ISL_3156835, EPI_ISL_3156836, EPI_ISL_3156837, EPI_ISL_3156838, EPI_ISL_3156839, EPI_ISL_3156840, EPI_ISL_3156841, EPI_ISL_3156842, EPI_ISL_3156843, EPI_ISL_3156844, EPI_ISL_3156845, EPI_ISL_3156846, EPI_ISL_3156847, EPI_ISL_3156848, EPI_ISL_3156849, EPI_ISL_3156850, EPI_ISL_3156851, EPI_ISL_3156852, EPI_ISL_3156853, EPI_ISL_3156854, EPI_ISL_3156855, EPI_ISL_3156856, EPI_ISL_3156857, EPI_ISL_3156858, EPI_ISL_3156859, EPI_ISL_3156860, EPI_ISL_3156861, EPI_ISL_3156862, EPI_ISL_3156863, EPI_ISL_3156864, EPI_ISL_3156865, EPI_ISL_3156867, EPI_ISL_3156868, EPI_ISL_3156869, EPI_ISL_3156870, EPI_ISL_3156871, EPI_ISL_3156872, EPI_ISL_3156873, EPI_ISL_3156874, EPI_ISL_3156875, EPI_ISL_3156876, EPI_ISL_3156877, EPI_ISL_3156878, EPI_ISL_3156879, EPI_ISL_3156880, EPI_ISL_3156881, EPI_ISL_3156882, EPI_ISL_3156883, EPI_ISL_3156884, EPI_ISL_3156885, EPI_ISL_3156886, EPI_ISL_3156887, EPI_ISL_3156888, EPI_ISL_3156889, EPI_ISL_3156890, EPI_ISL_3156891, EPI_ISL_3156892, EPI_ISL_3156893, EPI_ISL_3156894, EPI_ISL_3156895, EPI_ISL_3156896, EPI_ISL_3156897, EPI_ISL_3156898, EPI_ISL_3156899, EPI_ISL_3156900, EPI_ISL_3156901, EPI_ISL_3156902, EPI_ISL_3156903, EPI_ISL_3156904, EPI_ISL_3156905, EPI_ISL_3156906, EPI_ISL_3156907, EPI_ISL_3156908, EPI_ISL_3156909, EPI_ISL_3156910, EPI_ISL_3156911, EPI_ISL_3156912, EPI_ISL_3156913, EPI_ISL_3156914, EPI_ISL_3156915, EPI_ISL_3156916, EPI_ISL_3156917, EPI_ISL_3156918, EPI_ISL_3156919, EPI_ISL_3156920, EPI_ISL_3156921, EPI_ISL_3156922, EPI_ISL_3156923, EPI_ISL_3156924, EPI_ISL_3156926, EPI_ISL_3156927, EPI_ISL_3156928, EPI_ISL_3156929, EPI_ISL_3156930, EPI_ISL_3156931, EPI_ISL_3156932, EPI_ISL_3156933, EPI_ISL_3156934, EPI_ISL_3156935, EPI_ISL_3156936, EPI_ISL_3156937, EPI_ISL_3156938, EPI_ISL_3156940, EPI_ISL_3156941, EPI_ISL_3156942, EPI_ISL_3156943, EPI_ISL_3156944, EPI_ISL_3156945, EPI_ISL_3156946, EPI_ISL_3156947, EPI_ISL_3156948, EPI_ISL_3156949, EPI_ISL_3156950, EPI_ISL_3156951, EPI_ISL_3156952, EPI_ISL_3156953, EPI_ISL_3156954, EPI_ISL_3156955, EPI_ISL_3156956, EPI_ISL_3156957, EPI_ISL_3156958, EPI_ISL_3156959, EPI_ISL_3156960, EPI_ISL_3156961, EPI_ISL_3156963, EPI_ISL_3156964, EPI_ISL_3156965, EPI_ISL_3156966, EPI_ISL_3156967, EPI_ISL_3156968, EPI_ISL_3156969, EPI_ISL_3156970, EPI_ISL_3156971, EPI_ISL_3156972, EPI_ISL_3156973, EPI_ISL_3156974, EPI_ISL_3156975, EPI_ISL_3156976, EPI_ISL_3156977, EPI_ISL_3156978, EPI_ISL_3156979, EPI_ISL_3156980, EPI_ISL_3156981, EPI_ISL_3156982, EPI_ISL_3156983, EPI_ISL_3156984, EPI_ISL_3156985, EPI_ISL_3156986, EPI_ISL_3156987, EPI_ISL_3156988, EPI_ISL_3156989, EPI_ISL_3156990, EPI_ISL_3156991, EPI_ISL_3156992, EPI_ISL_3156993, EPI_ISL_3156994, EPI_ISL_3156995, EPI_ISL_3156996, EPI_ISL_3156997, EPI_ISL_3156998, EPI_ISL_3156999, EPI_ISL_3157000, EPI_ISL_3157001, EPI_ISL_3157002, EPI_ISL_3157003, EPI_ISL_3157004, EPI_ISL_3157005, EPI_ISL_3157006, EPI_ISL_3157007, EPI_ISL_3157008, EPI_ISL_3157009, EPI_ISL_3157010, EPI_ISL_3157011, EPI_ISL_3157012, EPI_ISL_3157013, EPI_ISL_3157014, EPI_ISL_3157015, EPI_ISL_3157016, EPI_ISL_3157017, EPI_ISL_3157018, EPI_ISL_3157019, EPI_ISL_3157020, EPI_ISL_3157021, EPI_ISL_3157022, EPI_ISL_3157023, EPI_ISL_3157024, EPI_ISL_3157025, EPI_ISL_3157026, EPI_ISL_3157027, EPI_ISL_3157028, EPI_ISL_3157029, EPI_ISL_3157030, EPI_ISL_3157031, EPI_ISL_3157032, EPI_ISL_3157033, EPI_ISL_3157034, EPI_ISL_3157035, EPI_ISL_3157036, EPI_ISL_3157037, EPI_ISL_3157038, EPI_ISL_3157039, EPI_ISL_3157040, EPI_ISL_3157041, EPI_ISL_3157042, EPI_ISL_3157043, EPI_ISL_3157044, EPI_ISL_3157045, EPI_ISL_3157046, EPI_ISL_3157047, EPI_ISL_3157048, EPI_ISL_3157049, EPI_ISL_3157050, EPI_ISL_3157052, EPI_ISL_3157053, EPI_ISL_3157054, EPI_ISL_3157055, EPI_ISL_3157056, EPI_ISL_3157057, EPI_ISL_3157058, EPI_ISL_3157059, EPI_ISL_3157060, EPI_ISL_3157061, EPI_ISL_3157062, EPI_ISL_3157063, EPI_ISL_3157064, EPI_ISL_3157065, EPI_ISL_3157066, EPI_ISL_3157067, EPI_ISL_3157068, EPI_ISL_3157069, EPI_ISL_3157070, EPI_ISL_3157071, EPI_ISL_3157072, EPI_ISL_3157073, EPI_ISL_3157074, EPI_ISL_3157075, EPI_ISL_3157076, EPI_ISL_3157077, EPI_ISL_3157078, EPI_ISL_3157079, EPI_ISL_3157080, EPI_ISL_3157081, EPI_ISL_3157082, EPI_ISL_3157083, EPI_ISL_3157084, EPI_ISL_3157085, EPI_ISL_3157086, EPI_ISL_3157088, EPI_ISL_3157089, EPI_ISL_3157090, EPI_ISL_3157091, EPI_ISL_3157092, EPI_ISL_3157093, EPI_ISL_3157094, EPI_ISL_3157095, EPI_ISL_3157096, EPI_ISL_3157097, EPI_ISL_3157098, EPI_ISL_3157099, EPI_ISL_3157100, EPI_ISL_3157101, EPI_ISL_3157102, EPI_ISL_3157103, EPI_ISL_3157104, EPI_ISL_3157105, EPI_ISL_3157106, EPI_ISL_3157107, EPI_ISL_3157108, EPI_ISL_3157109, EPI_ISL_3157110, EPI_ISL_3157112, EPI_ISL_3157113, EPI_ISL_3157114, EPI_ISL_3157115, EPI_ISL_3157116, EPI_ISL_3157117, EPI_ISL_3157118, EPI_ISL_3157119, EPI_ISL_3157120, EPI_ISL_3157121, EPI_ISL_3157122, EPI_ISL_3157123, EPI_ISL_3157124, EPI_ISL_3157125, EPI_ISL_3157126, EPI_ISL_3157127, EPI_ISL_3157128, EPI_ISL_3157129, EPI_ISL_3157130, EPI_ISL_3157131, EPI_ISL_3157132, EPI_ISL_3157133, EPI_ISL_3157134, EPI_ISL_3157135, EPI_ISL_3157136, EPI_ISL_3157137, EPI_ISL_3157138, EPI_ISL_3157139, EPI_ISL_3157140, EPI_ISL_3157141, EPI_ISL_3157142, EPI_ISL_3157143, EPI_ISL_3157144, EPI_ISL_3157145, EPI_ISL_3157146, EPI_ISL_3157147, EPI_ISL_3157148, EPI_ISL_3157149, EPI_ISL_3157150, EPI_ISL_3157151, EPI_ISL_3157152, EPI_ISL_3157153, EPI_ISL_3157154, EPI_ISL_3157155, EPI_ISL_3157156, EPI_ISL_3157157, EPI_ISL_3157158, EPI_ISL_3157159, EPI_ISL_3157160, EPI_ISL_3157161, EPI_ISL_3157162, EPI_ISL_3157163, EPI_ISL_3157164, EPI_ISL_3157165, EPI_ISL_3157166, EPI_ISL_3157167, EPI_ISL_3157168, EPI_ISL_3157169, EPI_ISL_3157170, EPI_ISL_3157171, EPI_ISL_3157172, EPI_ISL_3157173, EPI_ISL_3157174, EPI_ISL_3157175, EPI_ISL_3157176, EPI_ISL_3157177, EPI_ISL_3157179, EPI_ISL_3157180, EPI_ISL_3157181, EPI_ISL_3157182, EPI_ISL_3157183, EPI_ISL_3157184, EPI_ISL_3157185, EPI_ISL_3157186, EPI_ISL_3157187, EPI_ISL_3157188, EPI_ISL_3157189, EPI_ISL_3157190, EPI_ISL_3157191, EPI_ISL_3157192, EPI_ISL_3157193, EPI_ISL_3157194, EPI_ISL_3157195, EPI_ISL_3157196, EPI_ISL_3157197, EPI_ISL_3157198, EPI_ISL_3157199, EPI_ISL_3157200, EPI_ISL_3157201, EPI_ISL_3157202, EPI_ISL_3157203, EPI_ISL_3157204, EPI_ISL_3157205, EPI_ISL_3157206, EPI_ISL_3157207, EPI_ISL_3157208, EPI_ISL_3157209, EPI_ISL_3157210, EPI_ISL_3157211, EPI_ISL_3157212, EPI_ISL_3157213, EPI_ISL_3157214, EPI_ISL_3157215, EPI_ISL_3157216, EPI_ISL_3157217, EPI_ISL_3157219, EPI_ISL_3157220, EPI_ISL_3157221, EPI_ISL_3157222, EPI_ISL_3157223, EPI_ISL_3157224, EPI_ISL_3157225, EPI_ISL_3157226, EPI_ISL_3157227, EPI_ISL_3157228, EPI_ISL_3157229, EPI_ISL_3157230, EPI_ISL_3157231, EPI_ISL_3157232, EPI_ISL_3157233, EPI_ISL_3157234, EPI_ISL_3157235, EPI_ISL_3157236, EPI_ISL_3157237, EPI_ISL_3157238, EPI_ISL_3157239, EPI_ISL_3157240, EPI_ISL_3157241, EPI_ISL_3157242, EPI_ISL_3157243, EPI_ISL_3157244, EPI_ISL_3157245, EPI_ISL_3157246, EPI_ISL_3157247, EPI_ISL_3157440, EPI_ISL_3157441, EPI_ISL_3157449, EPI_ISL_3157452, EPI_ISL_3157455, EPI_ISL_3157468, EPI_ISL_3157470, EPI_ISL_3157472, EPI_ISL_3157474, EPI_ISL_3157475, EPI_ISL_3157477, EPI_ISL_3157479, EPI_ISL_3157480, EPI_ISL_3157482, EPI_ISL_3157483, EPI_ISL_3157488, EPI_ISL_3157493 | | | |
| see above | Swedish national genomic surveillance program of SARS-CoV-2 | The Public Health Agency of Sweden | Maximilian Riess, Maria Lind Karlberg, Alma Brolund, Swedish national genomic surveillance program of SARS-CoV-2 |
| EPI_ISL_3157503 | Ramathibodi Hospital | COVID-19 Network Investigations (CONI) Alliance | Elizabeth Batty, Wasun Chantratita, Thanat Chookajorn, Stefan Fernandez, Angkana Huang, Anthony R. Jones, Khajohn Joonlasak, Chonticha Klungtong, Theerarat Kochakarn, Namfon Kotanan, Krittikorn Kumpornsin, Duangkamon Loesbanluechai, Wudtichai Manasatienkij, Bhakbhoom Panthan, Ekawat Pasomsub, Kingkan Rakmanee, Insee Sensorn, Janjira Thaipadungpanit, Arporn Wangwiwatsin, Treewat Watthanachockchai |
| EPI_ISL_3157508, EPI_ISL_3157509, EPI_ISL_3157510, EPI_ISL_3157511, EPI_ISL_3157512, EPI_ISL_3157513, EPI_ISL_3157514, EPI_ISL_3157515, EPI_ISL_3157516, EPI_ISL_3157517, EPI_ISL_3157518, EPI_ISL_3157519, EPI_ISL_3157520, EPI_ISL_3157522, EPI_ISL_3157523, EPI_ISL_3157524, EPI_ISL_3157525, EPI_ISL_3157527, EPI_ISL_3157528, EPI_ISL_3157529, EPI_ISL_3157530, EPI_ISL_3157532, EPI_ISL_3157535, EPI_ISL_3157536, EPI_ISL_3157537, EPI_ISL_3157538, EPI_ISL_3157540, EPI_ISL_3157541, EPI_ISL_3157543, EPI_ISL_3157544, EPI_ISL_3157546, EPI_ISL_3157547, EPI_ISL_3157549, EPI_ISL_3157550, EPI_ISL_3157551, EPI_ISL_3157552, EPI_ISL_3157553, EPI_ISL_3157555, EPI_ISL_3157556, EPI_ISL_3157557, EPI_ISL_3157558, EPI_ISL_3157559, EPI_ISL_3157561, EPI_ISL_3157562, | | | |

| | | | |
|---|---|---|---|
| EPI_ISL_3157563, EPI_ISL_3157566, EPI_ISL_3157568, EPI_ISL_3157569, EPI_ISL_3157570, EPI_ISL_3157571, EPI_ISL_3157572, EPI_ISL_3157574, EPI_ISL_3157576, EPI_ISL_3157577, EPI_ISL_3157579, EPI_ISL_3157581, EPI_ISL_3157582, EPI_ISL_3157583, EPI_ISL_3157584, EPI_ISL_3157586, EPI_ISL_3157587, EPI_ISL_3157588, EPI_ISL_3157590, EPI_ISL_3157593, EPI_ISL_3157594, EPI_ISL_3157595, EPI_ISL_3157596, EPI_ISL_3157597, EPI_ISL_3157598, EPI_ISL_3157599, EPI_ISL_3157600, EPI_ISL_3157601, EPI_ISL_3157602, EPI_ISL_3157604, EPI_ISL_3157606, EPI_ISL_3157607, EPI_ISL_3157608, EPI_ISL_3157610, EPI_ISL_3157612, EPI_ISL_3157613, EPI_ISL_3157614, EPI_ISL_3157615, EPI_ISL_3157616, EPI_ISL_3157617, EPI_ISL_3157619, EPI_ISL_3157620, EPI_ISL_3157621, EPI_ISL_3157622, EPI_ISL_3157623, EPI_ISL_3157624, EPI_ISL_3157625, EPI_ISL_3157626, EPI_ISL_3157627, EPI_ISL_3157628, EPI_ISL_3157629, EPI_ISL_3157630, EPI_ISL_3157631, EPI_ISL_3157632, EPI_ISL_3157633, EPI_ISL_3157635, EPI_ISL_3157636, EPI_ISL_3157637, EPI_ISL_3157638, EPI_ISL_3157639, EPI_ISL_3157640, EPI_ISL_3157641, EPI_ISL_3157643, EPI_ISL_3157644, EPI_ISL_3157645, EPI_ISL_3157646, EPI_ISL_3157647, EPI_ISL_3157648, EPI_ISL_3157649, EPI_ISL_3157651, EPI_ISL_3157652, EPI_ISL_3157653, EPI_ISL_3157654, EPI_ISL_3157655, EPI_ISL_3157656, EPI_ISL_3157657, EPI_ISL_3157658, EPI_ISL_3157659, EPI_ISL_3157660, EPI_ISL_3157661, EPI_ISL_3157662, EPI_ISL_3157663, EPI_ISL_3157664, EPI_ISL_3157665, EPI_ISL_3157666, EPI_ISL_3157667, EPI_ISL_3157668, EPI_ISL_3157669, EPI_ISL_3157670, EPI_ISL_3157671, EPI_ISL_3157672, EPI_ISL_3157673, EPI_ISL_3157674, EPI_ISL_3157675, EPI_ISL_3157676, EPI_ISL_3157677, EPI_ISL_3157678, EPI_ISL_3157679, EPI_ISL_3157681, EPI_ISL_3157682, EPI_ISL_3157683, EPI_ISL_3157684, EPI_ISL_3157685, EPI_ISL_3157686, EPI_ISL_3157687, EPI_ISL_3157688, EPI_ISL_3157689, EPI_ISL_3157691, EPI_ISL_3157692, EPI_ISL_3157693, EPI_ISL_3157694, EPI_ISL_3157695, EPI_ISL_3157696, EPI_ISL_3157697, EPI_ISL_3157698, EPI_ISL_3157699, EPI_ISL_3157700, EPI_ISL_3157701, EPI_ISL_3157702, EPI_ISL_3157703, EPI_ISL_3157704, EPI_ISL_3157705, EPI_ISL_3157706, EPI_ISL_3157707, EPI_ISL_3157708, EPI_ISL_3157709, EPI_ISL_3157710, EPI_ISL_3157711, EPI_ISL_3157712, EPI_ISL_3157713, EPI_ISL_3157714, EPI_ISL_3157715, EPI_ISL_3157716, EPI_ISL_3157717, EPI_ISL_3157718, EPI_ISL_3157719, EPI_ISL_3157720, EPI_ISL_3157721, EPI_ISL_3157722, EPI_ISL_3157723, EPI_ISL_3157724, EPI_ISL_3157725, EPI_ISL_3157726, EPI_ISL_3157728, EPI_ISL_3157729, EPI_ISL_3157730, EPI_ISL_3157731, EPI_ISL_3157732, EPI_ISL_3157733, EPI_ISL_3157734, EPI_ISL_3157735, EPI_ISL_3157736, EPI_ISL_3157737, EPI_ISL_3157738, EPI_ISL_3157739, EPI_ISL_3157740, EPI_ISL_3157741, EPI_ISL_3157742, EPI_ISL_3157743, EPI_ISL_3157744, EPI_ISL_3157745, EPI_ISL_3157746, EPI_ISL_3157747, EPI_ISL_3157748, EPI_ISL_3157749, EPI_ISL_3157750, EPI_ISL_3157751, EPI_ISL_3157752, EPI_ISL_3157753, EPI_ISL_3157754, EPI_ISL_3157755, EPI_ISL_3157756, EPI_ISL_3157757, EPI_ISL_3157758, EPI_ISL_3157759, EPI_ISL_3157760, EPI_ISL_3157761, EPI_ISL_3157762, EPI_ISL_3157763, EPI_ISL_3157764, EPI_ISL_3157765, EPI_ISL_3157766, EPI_ISL_3157767, EPI_ISL_3157768, EPI_ISL_3157769, EPI_ISL_3157770, EPI_ISL_3157771, EPI_ISL_3157772, EPI_ISL_3157773, EPI_ISL_3157774, EPI_ISL_3157775, EPI_ISL_3157777, EPI_ISL_3157779, EPI_ISL_3157780, EPI_ISL_3157781, EPI_ISL_3157782, EPI_ISL_3157783, EPI_ISL_3157784, EPI_ISL_3157785, EPI_ISL_3157786, EPI_ISL_3157787, EPI_ISL_3157788, EPI_ISL_3157790, EPI_ISL_3157791, EPI_ISL_3157792, EPI_ISL_3157793, EPI_ISL_3157795, EPI_ISL_3157796, EPI_ISL_3157797, EPI_ISL_3157798, EPI_ISL_3157799, EPI_ISL_3157800, EPI_ISL_3157801, EPI_ISL_3157802, EPI_ISL_3157803, EPI_ISL_3157804, EPI_ISL_3157805, EPI_ISL_3157806, EPI_ISL_3157807, EPI_ISL_3157808, EPI_ISL_3157809, EPI_ISL_3157810, EPI_ISL_3157811, EPI_ISL_3157812, EPI_ISL_3157813, EPI_ISL_3157814, EPI_ISL_3157815, EPI_ISL_3157816, EPI_ISL_3157817, EPI_ISL_3157818, EPI_ISL_3157819, EPI_ISL_3157820, EPI_ISL_3157821, EPI_ISL_3157822, EPI_ISL_3157823, EPI_ISL_3157824, EPI_ISL_3157827, EPI_ISL_3157828, EPI_ISL_3157829, EPI_ISL_3157830, EPI_ISL_3157831, EPI_ISL_3157832, EPI_ISL_3157833, EPI_ISL_3157835, EPI_ISL_3157836, EPI_ISL_3157837, EPI_ISL_3157838, EPI_ISL_3157839, EPI_ISL_3157840, EPI_ISL_3157841, EPI_ISL_3157842, EPI_ISL_3157843, EPI_ISL_3157844, EPI_ISL_3157845, EPI_ISL_3157846, EPI_ISL_3157847, EPI_ISL_3157848, EPI_ISL_3157849, EPI_ISL_3157850, EPI_ISL_3157851, EPI_ISL_3157852, EPI_ISL_3157853, EPI_ISL_3157854, EPI_ISL_3157855, EPI_ISL_3157856, EPI_ISL_3157857, EPI_ISL_3157866, EPI_ISL_3157914 | | | |
| see above | Swedish national genomic surveillance program of SARS-CoV-2 | The Public Health Agency of Sweden | Maximilian Riess, Maria Lind Karlberg, Alma Brolund, Swedish national genomic surveillance program of SARS-CoV-2 |
| EPI_ISL_3157941 | Ramathibodi Hospital | COVID-19 Network Investigations (CONI) Alliance | Elizabeth Batty, Wasun Chantratita, Thanat Chookajorn, Stefan Fernandez, Angkana Huang, Anthony R. Jones, Khajohn Joonlasak, Chonticha Klungtong, Theerarat Kochakarn, Namfon Kotanan, Krittikorn Kumpornsin, Duangkamon Loesbanluechai, Wudtichai Manasatienkij, Bhakbhoom Panthan, Ekawat Pasomsub, Kingkan Rakmanee, Insee Sensorn, Janjira Thaipadungpanit, Arporn Wangwiwatsin, Treewat Watthanachockchai |
| EPI_ISL_3158021, EPI_ISL_3158022, EPI_ISL_3158023, EPI_ISL_3158024, EPI_ISL_3158025, EPI_ISL_3158026, EPI_ISL_3158027, EPI_ISL_3158028, EPI_ISL_3158029, EPI_ISL_3158030, EPI_ISL_3158031, EPI_ISL_3158032, EPI_ISL_3158033, EPI_ISL_3158034, EPI_ISL_3158035, EPI_ISL_3158036, EPI_ISL_3158037, EPI_ISL_3158038, EPI_ISL_3158039, EPI_ISL_3158040, EPI_ISL_3158049, EPI_ISL_3158050, EPI_ISL_3158051, EPI_ISL_3158052, EPI_ISL_3158053, EPI_ISL_3158054, EPI_ISL_3158055, EPI_ISL_3158056, EPI_ISL_3158057, EPI_ISL_3158058, EPI_ISL_3158059, EPI_ISL_3158060, EPI_ISL_3158061, EPI_ISL_3158062, EPI_ISL_3158063, EPI_ISL_3158064, EPI_ISL_3158065, EPI_ISL_3158066, EPI_ISL_3158067, EPI_ISL_3158068, EPI_ISL_3158069, EPI_ISL_3158070, EPI_ISL_3158071, EPI_ISL_3158072, EPI_ISL_3158073, EPI_ISL_3158074, EPI_ISL_3158076, EPI_ISL_3158077, EPI_ISL_3158078, EPI_ISL_3158079, EPI_ISL_3158081, EPI_ISL_3158082, EPI_ISL_3158083, EPI_ISL_3158084, EPI_ISL_3158085, EPI_ISL_3158087, EPI_ISL_3158088, EPI_ISL_3158089, EPI_ISL_3158090, EPI_ISL_3158091, EPI_ISL_3158092, EPI_ISL_3158093, EPI_ISL_3158094, EPI_ISL_3158095, EPI_ISL_3158096, EPI_ISL_3158097, EPI_ISL_3158098, EPI_ISL_3158099, EPI_ISL_3158100, EPI_ISL_3158101, EPI_ISL_3158102, EPI_ISL_3158103, EPI_ISL_3158104, EPI_ISL_3158105, EPI_ISL_3158106, EPI_ISL_3158107, EPI_ISL_3158108, EPI_ISL_3158109, EPI_ISL_3158110, EPI_ISL_3158111, EPI_ISL_3158112, EPI_ISL_3158113, EPI_ISL_3158114, EPI_ISL_3158116, EPI_ISL_3158121, EPI_ISL_3158122, EPI_ISL_3158129, EPI_ISL_3158130, EPI_ISL_3158131, EPI_ISL_3158134, EPI_ISL_3158135, EPI_ISL_3158136, EPI_ISL_3158138, EPI_ISL_3158142, EPI_ISL_3158145, EPI_ISL_3158146, EPI_ISL_3158147, EPI_ISL_3158148, EPI_ISL_3158149, EPI_ISL_3158150, EPI_ISL_3158151, EPI_ISL_3158152, EPI_ISL_3158153, EPI_ISL_3158154, EPI_ISL_3158156, EPI_ISL_3158157, EPI_ISL_3158163, EPI_ISL_3158164, EPI_ISL_3158165, EPI_ISL_3158166, EPI_ISL_3158167, EPI_ISL_3158168, EPI_ISL_3158169, EPI_ISL_3158170, EPI_ISL_3158171, EPI_ISL_3158172, EPI_ISL_3158173, EPI_ISL_3158174, EPI_ISL_3158175, EPI_ISL_3158176, EPI_ISL_3158177, EPI_ISL_3158178, EPI_ISL_3158179, EPI_ISL_3158180, EPI_ISL_3158181, EPI_ISL_3158182, EPI_ISL_3158183, EPI_ISL_3158184, EPI_ISL_3158185, EPI_ISL_3158186, EPI_ISL_3158187, EPI_ISL_3158188, EPI_ISL_3158189, EPI_ISL_3158190, EPI_ISL_3158191, EPI_ISL_3158192, EPI_ISL_3158193, EPI_ISL_3158194, EPI_ISL_3158195, EPI_ISL_3158196, EPI_ISL_3158197, EPI_ISL_3158198, EPI_ISL_3158199, EPI_ISL_3158200, EPI_ISL_3158201, EPI_ISL_3158202, EPI_ISL_3158203, EPI_ISL_3158204, EPI_ISL_3158205, EPI_ISL_3158206, EPI_ISL_3158207, EPI_ISL_3158208, EPI_ISL_3158209, EPI_ISL_3158210, EPI_ISL_3158211, EPI_ISL_3158212, EPI_ISL_3158213, EPI_ISL_3158214, EPI_ISL_3158215, EPI_ISL_3158216, EPI_ISL_3158217, EPI_ISL_3158218, EPI_ISL_3158219, EPI_ISL_3158220, EPI_ISL_3158222, EPI_ISL_3158223, EPI_ISL_3158224, EPI_ISL_3158226, EPI_ISL_3158229, EPI_ISL_3158230, EPI_ISL_3158231, EPI_ISL_3158244, EPI_ISL_3158245, EPI_ISL_3158246, EPI_ISL_3158247, EPI_ISL_3158248, EPI_ISL_3158249, EPI_ISL_3158250, EPI_ISL_3158251, EPI_ISL_3158252, EPI_ISL_3158253, EPI_ISL_3158254, EPI_ISL_3158255, EPI_ISL_3158256, EPI_ISL_3158257, EPI_ISL_3158258 | | | |
| see above | Swedish national genomic surveillance program of SARS-CoV-2 | The Public Health Agency of Sweden | Maximilian Riess, Maria Lind Karlberg, Alma Brolund, Swedish national genomic surveillance program of SARS-CoV-2 |
| EPI_ISL_3158259 | Ramathibodi Hospital | COVID-19 Network Investigations (CONI) Alliance | Elizabeth Batty, Wasun Chantratita, Thanat Chookajorn, Stefan Fernandez, Angkana Huang, Anthony R. Jones, Khajohn Joonlasak, Chonticha Klungtong, Theerarat Kochakarn, Namfon Kotanan, Krittikorn Kumpornsin, Duangkamon Loesbanluechai, Wudtichai Manasatienkij, Bhakbhoom Panthan, Ekawat Pasomsub, Kingkan Rakmanee, Insee Sensorn, Janjira Thaipadungpanit, Arporn Wangwiwatsin, Treewat Watthanachockchai |
| EPI_ISL_3158260, EPI_ISL_3158261, EPI_ISL_3158262, EPI_ISL_3158263, EPI_ISL_3158264, EPI_ISL_3158265, EPI_ISL_3158266, EPI_ISL_3158267, EPI_ISL_3158268, EPI_ISL_3158269, EPI_ISL_3158270, EPI_ISL_3158271, EPI_ISL_3158272, EPI_ISL_3158273, EPI_ISL_3158274, EPI_ISL_3158275, EPI_ISL_3158276, EPI_ISL_3158277, EPI_ISL_3158278, EPI_ISL_3158279, EPI_ISL_3158280, EPI_ISL_3158281, EPI_ISL_3158282, EPI_ISL_3158283, EPI_ISL_3158284, EPI_ISL_3158285, EPI_ISL_3158294, EPI_ISL_3158337, EPI_ISL_3158338, EPI_ISL_3158339, EPI_ISL_3158340, EPI_ISL_3158341, EPI_ISL_3158342, EPI_ISL_3158343, EPI_ISL_3158344, EPI_ISL_3158345, EPI_ISL_3158346, EPI_ISL_3158347, EPI_ISL_3158348, EPI_ISL_3158349, EPI_ISL_3158350, EPI_ISL_3158351, EPI_ISL_3158352, EPI_ISL_3158353, EPI_ISL_3158354, EPI_ISL_3158355, EPI_ISL_3158356, EPI_ISL_3158357, EPI_ISL_3158453, EPI_ISL_3158454, EPI_ISL_3158455, EPI_ISL_3158456, EPI_ISL_3158457, EPI_ISL_3158458, EPI_ISL_3158459, EPI_ISL_3158460, EPI_ISL_3158461, EPI_ISL_3158462, EPI_ISL_3158463, EPI_ISL_3158464, EPI_ISL_3158465, EPI_ISL_3158466, EPI_ISL_3158535, EPI_ISL_3158542, EPI_ISL_3158543, EPI_ISL_3158546, EPI_ISL_3158547, EPI_ISL_3158548, EPI_ISL_3158549, EPI_ISL_3158550, EPI_ISL_3158551, EPI_ISL_3158566, EPI_ISL_3158567, EPI_ISL_3158568, EPI_ISL_3158569, EPI_ISL_3158570, EPI_ISL_3158571, EPI_ISL_3158581, EPI_ISL_3158582, EPI_ISL_3158583, EPI_ISL_3158584, EPI_ISL_3158585, EPI_ISL_3158586, EPI_ISL_3158587, EPI_ISL_3158589, EPI_ISL_3158591, EPI_ISL_3158592, EPI_ISL_3158593, EPI_ISL_3158643, EPI_ISL_3158644, EPI_ISL_3158645, EPI_ISL_3158646, EPI_ISL_3158647, EPI_ISL_3158648, EPI_ISL_3158649, EPI_ISL_3158650, EPI_ISL_3158651, EPI_ISL_3158655, EPI_ISL_3158656, EPI_ISL_3158658, EPI_ISL_3158659, EPI_ISL_3158660, EPI_ISL_3158670, EPI_ISL_3158671, EPI_ISL_3158672, EPI_ISL_3158673, EPI_ISL_3158675, EPI_ISL_3158676, EPI_ISL_3158715, EPI_ISL_3158716, EPI_ISL_3158717, EPI_ISL_3158719, EPI_ISL_3158720, EPI_ISL_3158721, EPI_ISL_3158722, EPI_ISL_3158723, EPI_ISL_3158724, EPI_ISL_3158725, EPI_ISL_3158726, EPI_ISL_3158727, EPI_ISL_3158728, EPI_ISL_3158729, EPI_ISL_3158730, EPI_ISL_3158731, EPI_ISL_3158732, EPI_ISL_3158733, EPI_ISL_3158734, EPI_ISL_3158735, EPI_ISL_3158736, EPI_ISL_3158737, EPI_ISL_3158738, EPI_ISL_3158739, EPI_ISL_3158741, EPI_ISL_3158742, EPI_ISL_3158743, EPI_ISL_3158744, EPI_ISL_3158745, EPI_ISL_3158746, EPI_ISL_3158825, EPI_ISL_3158826, EPI_ISL_3158827, EPI_ISL_3158828, EPI_ISL_3158829, EPI_ISL_3158830, EPI_ISL_3158831, EPI_ISL_3158832, EPI_ISL_3158833, EPI_ISL_3158834, EPI_ISL_3158835, EPI_ISL_3158836, EPI_ISL_3158837, EPI_ISL_3158838, EPI_ISL_3158839, EPI_ISL_3158840, EPI_ISL_3158841, EPI_ISL_3158842, EPI_ISL_3158843, EPI_ISL_3158844, EPI_ISL_3158845, EPI_ISL_3158846, EPI_ISL_3158847, EPI_ISL_3158848, EPI_ISL_3158849, EPI_ISL_3158850, EPI_ISL_3158851, EPI_ISL_3158855, EPI_ISL_3158909, EPI_ISL_3158910, EPI_ISL_3158911, EPI_ISL_3158912, EPI_ISL_3158913, EPI_ISL_3158914, EPI_ISL_3158915, EPI_ISL_3158916, EPI_ISL_3158917, EPI_ISL_3158918, EPI_ISL_3158919, EPI_ISL_3158920, EPI_ISL_3158921, EPI_ISL_3158922, EPI_ISL_3158967, EPI_ISL_3158968, EPI_ISL_3158969, EPI_ISL_3158970, EPI_ISL_3158971, EPI_ISL_3158972, EPI_ISL_3158973, EPI_ISL_3158974, EPI_ISL_3158975, EPI_ISL_3158976, EPI_ISL_3158977, EPI_ISL_3158978, EPI_ISL_3158979, EPI_ISL_3158980, EPI_ISL_3158981, EPI_ISL_3158983, EPI_ISL_3158984, EPI_ISL_3158985, EPI_ISL_3158986, EPI_ISL_3159013, EPI_ISL_3159014, EPI_ISL_3159015, EPI_ISL_3159016, EPI_ISL_3159017, EPI_ISL_3159018, EPI_ISL_3159019, EPI_ISL_3159020, EPI_ISL_3159021, EPI_ISL_3159022, EPI_ISL_3159029, EPI_ISL_3159030, EPI_ISL_3159032, EPI_ISL_3159037, EPI_ISL_3159041, EPI_ISL_3159045, EPI_ISL_3159047, EPI_ISL_3159049, EPI_ISL_3159051, EPI_ISL_3159052, EPI_ISL_3159053, EPI_ISL_3159054, EPI_ISL_3159055, EPI_ISL_3159056, EPI_ISL_3159063, EPI_ISL_3159065, EPI_ISL_3159066, EPI_ISL_3159067, EPI_ISL_3159069, EPI_ISL_3159071, EPI_ISL_3159073, EPI_ISL_3159077, EPI_ISL_3159078, EPI_ISL_3159081, EPI_ISL_3159085, EPI_ISL_3159086, EPI_ISL_3159090, EPI_ISL_3159103, EPI_ISL_3159113, EPI_ISL_3159114, EPI_ISL_3159115, EPI_ISL_3159116, EPI_ISL_3159117, EPI_ISL_3159137, EPI_ISL_3159140, EPI_ISL_3159141, EPI_ISL_3159160, EPI_ISL_3159161, EPI_ISL_3159239, EPI_ISL_3159241, EPI_ISL_3159242, EPI_ISL_3159244, EPI_ISL_3159245, EPI_ISL_3159246, EPI_ISL_3159248, EPI_ISL_3159249, EPI_ISL_3159250, EPI_ISL_3159251, EPI_ISL_3159252, EPI_ISL_3159253, EPI_ISL_3159254, EPI_ISL_3159255, EPI_ISL_3159256, EPI_ISL_3159257, EPI_ISL_3159259, EPI_ISL_3159276, EPI_ISL_3159277, EPI_ISL_3159278, EPI_ISL_3159279, EPI_ISL_3159280, EPI_ISL_3159281, EPI_ISL_3159283, EPI_ISL_3159284, EPI_ISL_3159285, EPI_ISL_3159335, EPI_ISL_3159336, EPI_ISL_3159337, EPI_ISL_3159338, EPI_ISL_3159340, EPI_ISL_3159341, EPI_ISL_3159343, EPI_ISL_3159344, EPI_ISL_3159345, EPI_ISL_3159347, EPI_ISL_3159349, EPI_ISL_3159350, EPI_ISL_3159351, EPI_ISL_3159352, EPI_ISL_3159353, EPI_ISL_3159354, EPI_ISL_3159406, EPI_ISL_3159419, EPI_ISL_3159420, EPI_ISL_3159421, EPI_ISL_3159422, EPI_ISL_3159423, EPI_ISL_3159424, EPI_ISL_3159426, EPI_ISL_3159427, EPI_ISL_3159428, EPI_ISL_3159429, EPI_ISL_3159430, EPI_ISL_3159438, EPI_ISL_3159439, EPI_ISL_3159440, EPI_ISL_3159442, EPI_ISL_3159443, EPI_ISL_3159444, EPI_ISL_3159445, EPI_ISL_3159446, EPI_ISL_3159447, EPI_ISL_3159448, EPI_ISL_3159450, EPI_ISL_3159451, EPI_ISL_3159452, EPI_ISL_3159453, EPI_ISL_3159454, EPI_ISL_3159455, EPI_ISL_3159456, EPI_ISL_3159457, EPI_ISL_3159458, EPI_ISL_3159459, EPI_ISL_3159461, EPI_ISL_3159462, EPI_ISL_3159463, EPI_ISL_3159464, EPI_ISL_3159465, EPI_ISL_3159466, EPI_ISL_3159467, EPI_ISL_3159468, EPI_ISL_3159469, EPI_ISL_3159470, EPI_ISL_3159471, EPI_ISL_3159472, EPI_ISL_3159473 | | | |
| see above | Swedish national genomic surveillance program of SARS-CoV-2 | The Public Health Agency of Sweden | Maximilian Riess, Maria Lind Karlberg, Alma Brolund, Swedish national genomic surveillance program of SARS-CoV-2 |
| EPI_ISL_3159475, EPI_ISL_3159476, EPI_ISL_3159478, EPI_ISL_3159483, EPI_ISL_3159486, EPI_ISL_3159487, EPI_ISL_3159490, EPI_ISL_3159491, EPI_ISL_3159493 | Wyoming Public Health Laboratory | Wyoming Public Health Laboratory | Jim Mildenberger, Taylor Fearing, Channing Weber, Ashley Norberg, Chayse Rowley, Marley Goetz, Brian Dominguez, Elliot Thomasson, Sam Britz, Cari Sloma, Robert Petit, and Rob Christensen |
| EPI_ISL_3159496, EPI_ISL_3159497, EPI_ISL_3159498, EPI_ISL_3159499, EPI_ISL_3159500, EPI_ISL_3159502, EPI_ISL_3159503, EPI_ISL_3159504, EPI_ISL_3159505, EPI_ISL_3159506, EPI_ISL_3159507, EPI_ISL_3159508, EPI_ISL_3159510, EPI_ISL_3159511, EPI_ISL_3159512, EPI_ISL_3159513, EPI_ISL_3159514, EPI_ISL_3159515, EPI_ISL_3159516, EPI_ISL_3159517, EPI_ISL_3159518, EPI_ISL_3159519, EPI_ISL_3159520, EPI_ISL_3159521, EPI_ISL_3159522, EPI_ISL_3159523, EPI_ISL_3159524, EPI_ISL_3159525, EPI_ISL_3159526, EPI_ISL_3159527, EPI_ISL_3159528, EPI_ISL_3159529, EPI_ISL_3159530, EPI_ISL_3159531, EPI_ISL_3159532, EPI_ISL_3159533, EPI_ISL_3159534, EPI_ISL_3159535, EPI_ISL_3159536 | | | |
| see above | Alaska State Virology Laboratory | Alaska State Virology Laboratory | Stephanie DeRonde, Elva House, Jacob Zidek, Lisa Smith, Ph.D., Jack Chen, Ph.D. |
| EPI_ISL_3159554, EPI_ISL_3159573, EPI_ISL_3159574, EPI_ISL_3159575, EPI_ISL_3159576, EPI_ISL_3159577, EPI_ISL_3159578, EPI_ISL_3159579, EPI_ISL_3159581, EPI_ISL_3159582, EPI_ISL_3159590, EPI_ISL_3159596, EPI_ISL_3159597, EPI_ISL_3159598, EPI_ISL_3159607, EPI_ISL_3159608, EPI_ISL_3159618, EPI_ISL_3159623, EPI_ISL_3159635, EPI_ISL_3159641 | | | |
| see above | Colorado Department of Public Health and Environment | Colorado Department of Public Health and Environment | Laura Bankers, Molly C. Hetherington-Rauth, Diana Ir, Alexandria Rossheim, Michael Martin, Mandy Waters, Shannon R. Matzinger, Sarah Elizabeth Totten, Emily A. Travanty |
| EPI_ISL_3159657 | Eijkman Institute for Molecular Biology, National Research and Innovation Agency | Eijkman Institute for Molecular Biology, National Research and Innovation Agency | Willy Agustine, Hidayat Trimarsanto, Lydia V. Panggalo, Iskandar Adnan, Sukma Oktavianthi, Lidwina Priliani, Edison Johar, Frilasita A Yudhaputri, Muhammad Rezki Rasyak, Safarina G Malik, Khin Saw Myint, Amin Soebandrio |
| EPI_ISL_3159660 | Eijkman Institute for Molecular Biology, National Research and Innovation Agency | Eijkman Institute for Molecular Biology, National Research and Innovation Agency | Hidayat Trimarsanto, Lydia V. Panggalo, Iskandar Adnan, Sukma Oktavianthi, Lidwina Priliani, Edison Johar, Frilasita A Yudhaputri, Muhammad Rezki Rasyak, Willy Agustine, Safarina G Malik, Khin Saw Myint, Amin Soebandrio |
| EPI_ISL_3159663 | Eijkman Institute for Molecular Biology, National Research and Innovation Agency | Eijkman Institute for Molecular Biology, National Research and Innovation Agency | Lydia V. Panggalo, Iskandar Adnan, Sukma Oktavianthi, Lidwina Priliani, Edison Johar, Frilasita A Yudhaputri, Muhammad Rezki Rasyak, Willy Agustine, Hidayat Trimarsanto, Safarina G Malik, Khin Saw Myint, Amin Soebandrio |
| EPI_ISL_3159666 | Eijkman Institute for Molecular Biology, National Research and Innovation Agency | Eijkman Institute for Molecular Biology, National Research and Innovation Agency | Iskandar Adnan, Sukma Oktavianthi, Lidwina Priliani, Edison Johar, Frilasita A Yudhaputri, Muhammad Rezki Rasyak, Willy Agustine, Hidayat Trimarsanto, Lydia V. Panggalo, Safarina G Malik, Khin Saw Myint, Amin Soebandrio |
| EPI_ISL_3159669 | Eijkman Institute for Molecular Biology, National Research and Innovation Agency | Eijkman Institute for Molecular Biology, National Research and Innovation Agency | Sukma Oktavianthi, Lidwina Priliani, Edison Johar, Frilasita A Yudhaputri, Muhammad Rezki Rasyak, Willy Agustine, Hidayat Trimarsanto, Lydia V. Panggalo, Iskandar Adnan, Safarina G Malik, Khin Saw Myint, Amin Soebandrio |
| EPI_ISL_3159670, EPI_ISL_3159671 | Colorado Department of Public Health and Environment | Colorado Department of Public Health and Environment | Laura Bankers, Molly C. Hetherington-Rauth, Diana Ir, Alexandria Rossheim, Michael Martin, Mandy Waters, Shannon R. Matzinger, Sarah Elizabeth Totten, Emily A. Travanty |
| EPI_ISL_3159673 | Eijkman Institute for Molecular Biology, National Research and Innovation Agency | Eijkman Institute for Molecular Biology, National Research and Innovation Agency | Lidwina Priliani, Edison Johar, Frilasita A Yudhaputri, Muhammad Rezki Rasyak, Willy Agustine, Hidayat Trimarsanto, Lydia V. Panggalo, Iskandar Adnan, Sukma Oktavianthi, Safarina G Malik, Khin Saw Myint, Amin Soebandrio |
| EPI_ISL_3159674, EPI_ISL_3159675 | Colorado Department of Public Health and Environment | Colorado Department of Public Health and Environment | Laura Bankers, Molly C. Hetherington-Rauth, Diana Ir, Alexandria Rossheim, Michael Martin, Mandy Waters, Shannon R. Matzinger, Sarah Elizabeth Totten, Emily A. Travanty |
| EPI_ISL_3159676 | Eijkman Institute for Molecular Biology, National Research and Innovation Agency | Eijkman Institute for Molecular Biology, National Research and Innovation Agency | Edison Johar, Frilasita A Yudhaputri, Muhammad Rezki Rasyak, Willy Agustine, Hidayat Trimarsanto, Lydia V. Panggalo, Iskandar Adnan, Sukma Oktavianthi, Lidwina Priliani, Safarina G Malik, Khin Saw Myint, Amin Soebandrio |
| EPI_ISL_3159677, EPI_ISL_3159678 | Colorado Department of Public Health and Environment | Colorado Department of Public Health and Environment | Laura Bankers, Molly C. Hetherington-Rauth, Diana Ir, Alexandria Rossheim, Michael Martin, Mandy Waters, Shannon R. Matzinger, Sarah Elizabeth Totten, Emily A. Travanty |
| EPI_ISL_3159679 | Eijkman Institute for Molecular Biology, National Research and Innovation Agency | Eijkman Institute for Molecular Biology, National Research and Innovation Agency | Frilasita A Yudhaputri, Muhammad Rezki Rasyak, Willy Agustine, Hidayat Trimarsanto, Lydia V. Panggalo, Iskandar Adnan, Sukma Oktavianthi, Lidwina Priliani, Edison Johar, Safarina G Malik, Khin Saw Myint, Amin Soebandrio |
| EPI_ISL_3159680, EPI_ISL_3159681 | Colorado Department of Public Health and Environment | Colorado Department of Public Health and Environment | Laura Bankers, Molly C. Hetherington-Rauth, Diana Ir, Alexandria Rossheim, Michael Martin, Mandy Waters, Shannon R. Matzinger, Sarah Elizabeth Totten, Emily A. Travanty |
| EPI_ISL_3159682 | Eijkman Institute for Molecular Biology, National Research and Innovation Agency | Eijkman Institute for Molecular Biology, National Research and Innovation Agency | Muhammad Rezki Rasyak, Willy Agustine, Hidayat Trimarsanto, Lydia V. Panggalo, Iskandar Adnan, Sukma Oktavianthi, Lidwina Priliani, Edison Johar, Frilasita A Yudhaputri, Safarina G Malik, Khin Saw Myint, Amin Soebandrio |
| EPI_ISL_3159683, EPI_ISL_3159684 | Colorado Department of Public Health and Environment | Colorado Department of Public Health and Environment | Laura Bankers, Molly C. Hetherington-Rauth, Diana Ir, Alexandria Rossheim, Michael Martin, Mandy Waters, Shannon R. Matzinger, Sarah Elizabeth Totten, Emily A. Travanty |
| EPI_ISL_3159685 | Eijkman Institute for Molecular Biology, National Research and Innovation Agency | Eijkman Institute for Molecular Biology, National Research and Innovation Agency | Hidayat Trimarsanto, Lydia V. Panggalo, Iskandar Adnan, Sukma Oktavianthi, Lidwina Priliani, Edison Johar, Frilasita A Yudhaputri, Muhammad Rezki Rasyak, Willy Agustine, Safarina G Malik, Khin Saw Myint, Amin Soebandrio |
| EPI_ISL_3159686, EPI_ISL_3159687 | Colorado Department of Public Health and Environment | Colorado Department of Public Health and Environment | Laura Bankers, Molly C. Hetherington-Rauth, Diana Ir, Alexandria Rossheim, Michael Martin, Mandy Waters, Shannon R. Matzinger, Sarah Elizabeth Totten, Emily A. Travanty |
| EPI_ISL_3159688 | Eijkman Institute for Molecular Biology, National Research and Innovation Agency | Eijkman Institute for Molecular Biology, National Research and Innovation Agency | Sukma Oktavianthi, Lidwina Priliani, Edison Johar, Frilasita A Yudhaputri, Muhammad Rezki Rasyak, Willy Agustine, Hidayat Trimarsanto, Lydia V. Panggalo, Iskandar Adnan, Safarina G Malik, Khin Saw Myint, Amin Soebandrio |
| EPI_ISL_3159689 | Colorado Department of Public Health and Environment | Colorado Department of Public Health and Environment | Laura Bankers, Molly C. Hetherington-Rauth, Diana Ir, Alexandria Rossheim, Michael Martin, Mandy Waters, Shannon R. Matzinger, Sarah Elizabeth Totten, Emily A. Travanty |
| EPI_ISL_3159690 | Eijkman Institute for Molecular Biology, National Research and Innovation Agency | Eijkman Institute for Molecular Biology, National Research and Innovation Agency | Iskandar Adnan, Sukma Oktavianthi, Lidwina Priliani, Edison Johar, Frilasita A Yudhaputri, Muhammad Rezki Rasyak, Willy Agustine, Hidayat Trimarsanto, Lydia V. Panggalo, Safarina G Malik, Khin Saw Myint, Amin Soebandrio |

| | | | |
|---|---|---|---|
| EPI_ISL_3159691, EPI_ISL_3159692 | Colorado Department of Public Health and Environment | Colorado Department of Public Health and Environment | Laura Bankers, Molly C. Hetherington-Rauth, Diana Ir, Alexandria Rossheim, Michael Martin, Mandy Waters, Shannon R. Matzinger, Sarah Elizabeth Totten, Emily A. Travanty |
| EPI_ISL_3159693 | RS Pondok Indah Puri Indah | Eijkman Institute for Molecular Biology, National Research and Innovation Agency | Frilasita A Yudhaputri, Muhammad Rezki Rasyak, Willy Agustine, Hidayat Trimarsanto, Lydia V. Panggalo, Iskandar Adnan, Sukma Oktavianthi, Lidwina Priliani, Edison Johar, Safarina G Malik, Khin Saw Myint, Amin Soebandrio |
| EPI_ISL_3159694, EPI_ISL_3159695, EPI_ISL_3159697, EPI_ISL_3159698, EPI_ISL_3159700, EPI_ISL_3159701, EPI_ISL_3159703, EPI_ISL_3159704, EPI_ISL_3159706, EPI_ISL_3159707, EPI_ISL_3159709, EPI_ISL_3159710, EPI_ISL_3159712, EPI_ISL_3159713, EPI_ISL_3159715, EPI_ISL_3159716, EPI_ISL_3159718, EPI_ISL_3159719, EPI_ISL_3159721, EPI_ISL_3159722, EPI_ISL_3159724, EPI_ISL_3159725, EPI_ISL_3159727, EPI_ISL_3159749 | | | |
| see above | Colorado Department of Public Health and Environment | Colorado Department of Public Health and Environment | Laura Bankers, Molly C. Hetherington-Rauth, Diana Ir, Alexandria Rossheim, Michael Martin, Mandy Waters, Shannon R. Matzinger, Sarah Elizabeth Totten, Emily A. Travanty |
| EPI_ISL_3159750 | Dinas Kesehatan Provinsi Kalimantan Timur | Eijkman Institute for Molecular Biology, National Research and Innovation Agency; National Institute of Health Research and Development | Muhammad Rezki Rasyak, Willy Agustine, Hidayat Trimarsanto, Lydia V. Panggalo, Iskandar Adnan, Sukma Oktavianthi, Lidwina Priliani, Edison Johar, Frilasita A Yudhaputri, Vivi Setiawaty, Hana Apsari Pawestri, Subangkit, Safarina G Malik, Khin Saw Myint, Amin Soebandrio |
| EPI_ISL_3159753 | Dinas Kesehatan Provinsi Kalimantan Timur | Eijkman Institute for Molecular Biology, National Research and Innovation Agency; National Institute of Health Research and Development | Willy Agustine, Hidayat Trimarsanto, Lydia V. Panggalo, Iskandar Adnan, Sukma Oktavianthi, Lidwina Priliani, Edison Johar, Frilasita A Yudhaputri, Muhammad Rezki Rasyak, Vivi Setiawaty, Hana Apsari Pawestri, Subangkit, Safarina G Malik, Khin Saw Myint, Amin Soebandrio |
| EPI_ISL_3159760 | Colorado Department of Public Health and Environment | Colorado Department of Public Health and Environment | Laura Bankers, Molly C. Hetherington-Rauth, Diana Ir, Alexandria Rossheim, Michael Martin, Mandy Waters, Shannon R. Matzinger, Sarah Elizabeth Totten, Emily A. Travanty |
| EPI_ISL_3159765 | Dinas Kesehatan Provinsi Kalimantan Timur | Eijkman Institute for Molecular Biology, National Research and Innovation Agency; National Institute of Health Research and Development | Sukma Oktavianthi, Lidwina Priliani, Edison Johar, Frilasita A Yudhaputri, Muhammad Rezki Rasyak, Willy Agustine, Hidayat Trimarsanto, Lydia V. Panggalo, Iskandar Adnan, Vivi Setiawaty, Hana Apsari Pawestri, Subangkit, Safarina G Malik, Khin Saw Myint, Amin Soebandrio |
| EPI_ISL_3159767 | Colorado Department of Public Health and Environment | Colorado Department of Public Health and Environment | Laura Bankers, Molly C. Hetherington-Rauth, Diana Ir, Alexandria Rossheim, Michael Martin, Mandy Waters, Shannon R. Matzinger, Sarah Elizabeth Totten, Emily A. Travanty |
| EPI_ISL_3159768 | Dinas Kesehatan Provinsi Kalimantan Timur | Eijkman Institute for Molecular Biology, National Research and Innovation Agency; National Institute of Health Research and Development | Lidwina Priliani, Edison Johar, Frilasita A Yudhaputri, Muhammad Rezki Rasyak, Willy Agustine, Hidayat Trimarsanto, Lydia V. Panggalo, Iskandar Adnan, Sukma Oktavianthi, Vivi Setiawaty, Hana Apsari Pawestri, Subangkit, Safarina G Malik, Khin Saw Myint, Amin Soebandrio |
| EPI_ISL_3159774 | Dinas Kesehatan Provinsi Kalimantan Timur | Eijkman Institute for Molecular Biology, National Research and Innovation Agency; National Institute of Health Research and Development | Willy Agustine, Hidayat Trimarsanto, Lydia V. Panggalo, Iskandar Adnan, Sukma Oktavianthi, Lidwina Priliani, Edison Johar, Frilasita A Yudhaputri, Muhammad Rezki Rasyak, Vivi Setiawaty, Hana Apsari Pawestri, Subangkit, Safarina G Malik, Khin Saw Myint, Amin Soebandrio |
| EPI_ISL_3159777 | Dinas Kesehatan Provinsi Kalimantan Timur | Eijkman Institute for Molecular Biology, National Research and Innovation Agency; National Institute of Health Research and Development | Hidayat Trimarsanto, Lydia V. Panggalo, Iskandar Adnan, Sukma Oktavianthi, Lidwina Priliani, Edison Johar, Frilasita A Yudhaputri, Muhammad Rezki Rasyak, Willy Agustine, Vivi Setiawaty, Hana Apsari Pawestri, Subangkit, Safarina G Malik, Khin Saw Myint, Amin Soebandrio |
| EPI_ISL_3159786 | Dinas Kesehatan Provinsi Kalimantan Timur | Eijkman Institute for Molecular Biology, National Research and Innovation Agency; National Institute of Health Research and Development | Sukma Oktavianthi, Lidwina Priliani, Edison Johar, Frilasita A Yudhaputri, Muhammad Rezki Rasyak, Willy Agustine, Hidayat Trimarsanto, Lydia V. Panggalo, Iskandar Adnan, Vivi Setiawaty, Hana Apsari Pawestri, Subangkit, Safarina G Malik, Khin Saw Myint, Amin Soebandrio |
| EPI_ISL_3159790 | Colorado Department of Public Health and Environment | Colorado Department of Public Health and Environment | Laura Bankers, Molly C. Hetherington-Rauth, Diana Ir, Alexandria Rossheim, Michael Martin, Mandy Waters, Shannon R. Matzinger, Sarah Elizabeth Totten, Emily A. Travanty |
| EPI_ISL_3159791 | Dinas Kesehatan Provinsi Kalimantan Timur | Eijkman Institute for Molecular Biology, National Research and Innovation Agency; National Institute of Health Research and Development | Edison Johar, Frilasita A Yudhaputri, Muhammad Rezki Rasyak, Willy Agustine, Hidayat Trimarsanto, Lydia V. Panggalo, Iskandar Adnan, Sukma Oktavianthi, Lidwina Priliani, Vivi Setiawaty, Hana Apsari Pawestri, Subangkit, Safarina G Malik, Khin Saw Myint, Amin Soebandrio |
| EPI_ISL_3159792 | Colorado Department of Public Health and Environment | Colorado Department of Public Health and Environment | Laura Bankers, Molly C. Hetherington-Rauth, Diana Ir, Alexandria Rossheim, Michael Martin, Mandy Waters, Shannon R. Matzinger, Sarah Elizabeth Totten, Emily A. Travanty |
| EPI_ISL_3159794 | Dinas Kesehatan Provinsi Kalimantan Timur | Eijkman Institute for Molecular Biology, National Research and Innovation Agency; National Institute of Health Research and Development | Frilasita A Yudhaputri, Muhammad Rezki Rasyak, Willy Agustine, Hidayat Trimarsanto, Lydia V. Panggalo, Iskandar Adnan, Sukma Oktavianthi, Lidwina Priliani, Edison Johar, Vivi Setiawaty, Hana Apsari Pawestri, Subangkit, Safarina G Malik, Khin Saw Myint, Amin Soebandrio |
| EPI_ISL_3159797 | Dinas Kesehatan Provinsi Kalimantan Timur | Eijkman Institute for Molecular Biology, National Research and Innovation Agency; National Institute of Health Research and Development | Muhammad Rezki Rasyak, Willy Agustine, Hidayat Trimarsanto, Lydia V. Panggalo, Iskandar Adnan, Sukma Oktavianthi, Lidwina Priliani, Edison Johar, Frilasita A Yudhaputri, Vivi Setiawaty, Hana Apsari Pawestri, Subangkit, Safarina G Malik, Khin Saw Myint, Amin Soebandrio |
| EPI_ISL_3159802 | Dinas Kesehatan Provinsi Kalimantan Timur | Eijkman Institute for Molecular Biology, National Research and Innovation Agency; National Institute of Health Research and Development | Hidayat Trimarsanto, Lydia V. Panggalo, Iskandar Adnan, Sukma Oktavianthi, Lidwina Priliani, Edison Johar, Frilasita A Yudhaputri, Muhammad Rezki Rasyak, Willy Agustine, Vivi Setiawaty, Hana Apsari Pawestri, Subangkit, Safarina G Malik, Khin Saw Myint, Amin Soebandrio |
| EPI_ISL_3159805 | Dinas Kesehatan Provinsi Kalimantan Timur | Eijkman Institute for Molecular Biology, National Research and Innovation Agency; National Institute of Health Research and Development | Lydia V. Panggalo, Iskandar Adnan, Sukma Oktavianthi, Lidwina Priliani, Edison Johar, Frilasita A Yudhaputri, Muhammad Rezki Rasyak, Willy Agustine, Hidayat Trimarsanto, Vivi Setiawaty, Hana Apsari Pawestri, Subangkit, Safarina G Malik, Khin Saw Myint, Amin Soebandrio |
| EPI_ISL_3159808 | Dinas Kesehatan Provinsi Kalimantan Timur | Eijkman Institute for Molecular Biology, National Research and Innovation Agency; National Institute of Health Research and Development | Edison Johar, Frilasita A Yudhaputri, Muhammad Rezki Rasyak, Willy Agustine, Hidayat Trimarsanto, Lydia V. Panggalo, Iskandar Adnan, Sukma Oktavianthi, Lidwina Priliani, Vivi Setiawaty, Hana Apsari Pawestri, Subangkit, Safarina G Malik, Khin Saw Myint, Amin Soebandrio |
| EPI_ISL_3159811 | Dinas Kesehatan Provinsi Kalimantan Timur | Eijkman Institute for Molecular Biology, National Research and Innovation Agency; National Institute of Health Research and Development | Frilasita A Yudhaputri, Muhammad Rezki Rasyak, Willy Agustine, Hidayat Trimarsanto, Lydia V. Panggalo, Iskandar Adnan, Sukma Oktavianthi, Lidwina Priliani, Edison Johar, Vivi Setiawaty, Hana Apsari Pawestri, Subangkit, Safarina G Malik, Khin Saw Myint, Amin Soebandrio |
| EPI_ISL_3159814 | Dinas Kesehatan Provinsi Kalimantan Timur | Eijkman Institute for Molecular Biology, National Research and Innovation Agency; National Institute of Health Research and Development | Muhammad Rezki Rasyak, Willy Agustine, Hidayat Trimarsanto, Lydia V. Panggalo, Iskandar Adnan, Sukma Oktavianthi, Lidwina Priliani, Edison Johar, Frilasita A Yudhaputri, Vivi Setiawaty, Hana Apsari Pawestri, Subangkit, Safarina G Malik, Khin Saw Myint, Amin Soebandrio |
| EPI_ISL_3159820 | Dinas Kesehatan Provinsi Kalimantan Timur | Eijkman Institute for Molecular Biology, National Research and Innovation Agency; National Institute of Health Research and Development | Hidayat Trimarsanto, Lydia V. Panggalo, Iskandar Adnan, Sukma Oktavianthi, Lidwina Priliani, Edison Johar, Frilasita A Yudhaputri, Muhammad Rezki Rasyak, Willy Agustine, Vivi Setiawaty, Hana Apsari Pawestri, Subangkit, Safarina G Malik, Khin Saw Myint, Amin Soebandrio |
| EPI_ISL_3159823 | Dinas Kesehatan Provinsi Kalimantan Timur | Eijkman Institute for Molecular Biology, National Research and Innovation Agency; National Institute of Health Research and Development | Lydia V. Panggalo, Iskandar Adnan, Sukma Oktavianthi, Lidwina Priliani, Edison Johar, Frilasita A Yudhaputri, Muhammad Rezki Rasyak, Willy Agustine, Hidayat Trimarsanto, Vivi Setiawaty, Hana Apsari Pawestri, Subangkit, Safarina G Malik, Khin Saw Myint, Amin Soebandrio |
| EPI_ISL_3159826 | Dinas Kesehatan Provinsi Kalimantan Timur | Eijkman Institute for Molecular Biology, National Research and Innovation Agency; National Institute of Health Research and Development | Iskandar Adnan, Sukma Oktavianthi, Lidwina Priliani, Edison Johar, Frilasita A Yudhaputri, Muhammad Rezki Rasyak, Willy Agustine, Hidayat Trimarsanto, Lydia V. Panggalo, Vivi Setiawaty, Hana Apsari Pawestri, Subangkit, Safarina G Malik, Khin Saw Myint, Amin Soebandrio |
| EPI_ISL_3159829 | Dinas Kesehatan Provinsi Kalimantan Timur | Eijkman Institute for Molecular Biology, National Research and Innovation Agency; National Institute of Health Research and Development | Sukma Oktavianthi, Lidwina Priliani, Edison Johar, Frilasita A Yudhaputri, Muhammad Rezki Rasyak, Willy Agustine, Hidayat Trimarsanto, Lydia V. Panggalo, Iskandar Adnan, Vivi Setiawaty, Hana Apsari Pawestri, Subangkit, Safarina G Malik, Khin Saw Myint, Amin Soebandrio |
| EPI_ISL_3159833 | Dinas Kesehatan Provinsi Kalimantan Timur | Eijkman Institute for Molecular Biology, National Research and Innovation Agency; National Institute of Health Research and Development | Lidwina Priliani, Edison Johar, Frilasita A Yudhaputri, Muhammad Rezki Rasyak, Willy Agustine, Hidayat Trimarsanto, Lydia V. Panggalo, Iskandar Adnan, Sukma Oktavianthi, Vivi Setiawaty, Hana Apsari Pawestri, Subangkit, Safarina G Malik, Khin Saw Myint, Amin Soebandrio |
| EPI_ISL_3159835 | Dinas Kesehatan Provinsi Kalimantan Timur | Eijkman Institute for Molecular Biology, National Research and Innovation Agency; National Institute of Health Research and Development | Edison Johar, Frilasita A Yudhaputri, Muhammad Rezki Rasyak, Willy Agustine, Hidayat Trimarsanto, Lydia V. Panggalo, Iskandar Adnan, Sukma Oktavianthi, Lidwina Priliani, Vivi Setiawaty, Hana Apsari Pawestri, Subangkit, Safarina G Malik, Khin Saw Myint, Amin Soebandrio |
| EPI_ISL_3159838 | Dinas Kesehatan Provinsi Kalimantan Timur | Eijkman Institute for Molecular Biology, National Research and Innovation Agency; National Institute of Health Research and Development | Frilasita A Yudhaputri, Muhammad Rezki Rasyak, Willy Agustine, Hidayat Trimarsanto, Lydia V. Panggalo, Iskandar Adnan, Sukma Oktavianthi, Lidwina Priliani, Edison Johar, Vivi Setiawaty, Hana Apsari Pawestri, Subangkit, Safarina G Malik, Khin Saw Myint, Amin Soebandrio |
| EPI_ISL_3159841 | Dinas Kesehatan Provinsi Kalimantan Timur | Eijkman Institute for Molecular Biology, National Research and Innovation Agency; National Institute of Health Research and Development | Muhammad Rezki Rasyak, Willy Agustine, Hidayat Trimarsanto, Lydia V. Panggalo, Iskandar Adnan, Sukma Oktavianthi, Lidwina Priliani, Edison Johar, Frilasita A Yudhaputri, Vivi Setiawaty, Hana Apsari Pawestri, Subangkit, Safarina G Malik, Khin Saw Myint, Amin Soebandrio |
| EPI_ISL_3159844 | Dinas Kesehatan Provinsi Kalimantan Timur | Eijkman Institute for Molecular Biology, National Research and Innovation Agency; National Institute of Health Research and Development | Hidayat Trimarsanto, Lydia V. Panggalo, Iskandar Adnan, Sukma Oktavianthi, Lidwina Priliani, Edison Johar, Frilasita A Yudhaputri, Muhammad Rezki Rasyak, Willy Agustine, Vivi Setiawaty, Hana Apsari Pawestri, Subangkit, Safarina G Malik, Khin Saw Myint, Amin Soebandrio |
| EPI_ISL_3159846 | Dinas Kesehatan Provinsi Kalimantan Timur | Eijkman Institute for Molecular Biology, National Research and Innovation Agency; National Institute of Health Research and Development | Lydia V. Panggalo, Iskandar Adnan, Sukma Oktavianthi, Lidwina Priliani, Edison Johar, Frilasita A Yudhaputri, Muhammad Rezki Rasyak, Willy Agustine, Hidayat Trimarsanto, Vivi Setiawaty, Hana Apsari Pawestri, Subangkit, Safarina G Malik, Khin Saw Myint, Amin Soebandrio |
| EPI_ISL_3159849 | Dinas Kesehatan Provinsi Kalimantan Timur | Eijkman Institute for Molecular Biology, National Research and Innovation Agency; National Institute of Health Research and Development | Iskandar Adnan, Sukma Oktavianthi, Lidwina Priliani, Edison Johar, Frilasita A Yudhaputri, Muhammad Rezki Rasyak, Willy Agustine, Hidayat Trimarsanto, Lydia V. Panggalo, Vivi Setiawaty, Hana Apsari Pawestri, Subangkit, Safarina G Malik, Khin Saw Myint, Amin Soebandrio |
| EPI_ISL_3159855 | Dinas Kesehatan Provinsi Kalimantan Timur | Eijkman Institute for Molecular Biology, National Research and Innovation Agency; National Institute of Health Research and Development | Lidwina Priliani, Edison Johar, Frilasita A Yudhaputri, Muhammad Rezki Rasyak, Willy Agustine, Hidayat Trimarsanto, Lydia V. Panggalo, Iskandar Adnan, Sukma Oktavianthi, Vivi Setiawaty, Hana Apsari Pawestri, Subangkit, Safarina G Malik, Khin Saw Myint, Amin Soebandrio |
| EPI_ISL_3159858 | Dinas Kesehatan Provinsi Kalimantan Timur | Eijkman Institute for Molecular Biology, National Research and Innovation Agency; National Institute of Health Research and | Edison Johar, Frilasita A Yudhaputri, Muhammad Rezki Rasyak, Willy Agustine, Hidayat Trimarsanto, Lydia V. Panggalo, Iskandar Adnan, Sukma Oktavianthi, Lidwina Priliani, Vivi Setiawaty, Hana Apsari Pawestri, Subangkit, Safarina G Malik, Khin Saw Myint, Amin Soebandrio |

| | | | |
|---|---|---|---|
| | | Development | |
| EPI_ISL_3159864 | Dinas Kesehatan Provinsi Kalimantan Timur | Eijkman Institute for Molecular Biology, National Research and Innovation Agency; National Institute of Health Research and Development | Muhammad Rezki Rasyak, Willy Agustine, Hidayat Trimarsanto, Lydia V. Panggalo, Iskandar Adnan, Sukma Oktavianthi, Lidwina Priliani, Edison Johar, Frilasita A Yudhaputri, Vivi Setiawaty, Hana Apsari Pawestri, Subangkit, Safarina G Malik, Khin Saw Myint, Amin Soebandrio |
| EPI_ISL_3159867 | Dinas Kesehatan Provinsi Kalimantan Timur | Eijkman Institute for Molecular Biology, National Research and Innovation Agency; National Institute of Health Research and Development | Willy Agustine, Hidayat Trimarsanto, Lydia V. Panggalo, Iskandar Adnan, Sukma Oktavianthi, Lidwina Priliani, Edison Johar, Frilasita A Yudhaputri, Muhammad Rezki Rasyak, Vivi Setiawaty, Hana Apsari Pawestri, Subangkit, Safarina G Malik, Khin Saw Myint, Amin Soebandrio |
| EPI_ISL_3159870 | Dinas Kesehatan Provinsi Kalimantan Timur | Eijkman Institute for Molecular Biology, National Research and Innovation Agency; National Institute of Health Research and Development | Hidayat Trimarsanto, Lydia V. Panggalo, Iskandar Adnan, Sukma Oktavianthi, Lidwina Priliani, Edison Johar, Frilasita A Yudhaputri, Muhammad Rezki Rasyak, Willy Agustine, Vivi Setiawaty, Hana Apsari Pawestri, Subangkit, Safarina G Malik, Khin Saw Myint, Amin Soebandrio |
| EPI_ISL_3159873 | Dinas Kesehatan Provinsi Kalimantan Timur | Eijkman Institute for Molecular Biology, National Research and Innovation Agency; National Institute of Health Research and Development | Lydia V. Panggalo, Iskandar Adnan, Sukma Oktavianthi, Lidwina Priliani, Edison Johar, Frilasita A Yudhaputri, Muhammad Rezki Rasyak, Willy Agustine, Hidayat Trimarsanto, Vivi Setiawaty, Hana Apsari Pawestri, Subangkit, Safarina G Malik, Khin Saw Myint, Amin Soebandrio |
| EPI_ISL_3159876 | Dinas Kesehatan Provinsi Kalimantan Timur | Eijkman Institute for Molecular Biology, National Research and Innovation Agency; National Institute of Health Research and Development | Iskandar Adnan, Sukma Oktavianthi, Lidwina Priliani, Edison Johar, Frilasita A Yudhaputri, Muhammad Rezki Rasyak, Willy Agustine, Hidayat Trimarsanto, Lydia V. Panggalo, Vivi Setiawaty, Hana Apsari Pawestri, Subangkit, Safarina G Malik, Khin Saw Myint, Amin Soebandrio |
| EPI_ISL_3159879 | Dinas Kesehatan Provinsi Kalimantan Timur | Eijkman Institute for Molecular Biology, National Research and Innovation Agency; National Institute of Health Research and Development | Sukma Oktavianthi, Lidwina Priliani, Edison Johar, Frilasita A Yudhaputri, Muhammad Rezki Rasyak, Willy Agustine, Hidayat Trimarsanto, Lydia V. Panggalo, Iskandar Adnan, Vivi Setiawaty, Hana Apsari Pawestri, Subangkit, Safarina G Malik, Khin Saw Myint, Amin Soebandrio |
| EPI_ISL_3159882 | Dinas Kesehatan Provinsi Kalimantan Timur | Eijkman Institute for Molecular Biology, National Research and Innovation Agency; National Institute of Health Research and Development | Lidwina Priliani, Edison Johar, Frilasita A Yudhaputri, Muhammad Rezki Rasyak, Willy Agustine, Hidayat Trimarsanto, Lydia V. Panggalo, Iskandar Adnan, Sukma Oktavianthi, Vivi Setiawaty, Hana Apsari Pawestri, Subangkit, Safarina G Malik, Khin Saw Myint, Amin Soebandrio |
| EPI_ISL_3159885 | Dinas Kesehatan Provinsi Kalimantan Timur | Eijkman Institute for Molecular Biology, National Research and Innovation Agency; National Institute of Health Research and Development | Edison Johar, Frilasita A Yudhaputri, Muhammad Rezki Rasyak, Willy Agustine, Hidayat Trimarsanto, Lydia V. Panggalo, Iskandar Adnan, Sukma Oktavianthi, Lidwina Priliani, Vivi Setiawaty, Hana Apsari Pawestri, Subangkit, Safarina G Malik, Khin Saw Myint, Amin Soebandrio |
| EPI_ISL_3159889 | Dinas Kesehatan Provinsi Kalimantan Timur | Eijkman Institute for Molecular Biology, National Research and Innovation Agency; National Institute of Health Research and Development | Frilasita A Yudhaputri, Muhammad Rezki Rasyak, Willy Agustine, Hidayat Trimarsanto, Lydia V. Panggalo, Iskandar Adnan, Sukma Oktavianthi, Lidwina Priliani, Edison Johar, Vivi Setiawaty, Hana Apsari Pawestri, Subangkit, Safarina G Malik, Khin Saw Myint, Amin Soebandrio |
| EPI_ISL_3159892 | Dinas Kesehatan Provinsi Kalimantan Timur | Eijkman Institute for Molecular Biology, National Research and Innovation Agency; National Institute of Health Research and Development | Muhammad Rezki Rasyak, Willy Agustine, Hidayat Trimarsanto, Lydia V. Panggalo, Iskandar Adnan, Sukma Oktavianthi, Lidwina Priliani, Edison Johar, Frilasita A Yudhaputri, Vivi Setiawaty, Hana Apsari Pawestri, Subangkit, Safarina G Malik, Khin Saw Myint, Amin Soebandrio |
| EPI_ISL_3159895 | Dinas Kesehatan Provinsi Kalimantan Timur | Eijkman Institute for Molecular Biology, National Research and Innovation Agency; National Institute of Health Research and Development | Willy Agustine, Hidayat Trimarsanto, Lydia V. Panggalo, Iskandar Adnan, Sukma Oktavianthi, Lidwina Priliani, Edison Johar, Frilasita A Yudhaputri, Muhammad Rezki Rasyak, Vivi Setiawaty, Hana Apsari Pawestri, Subangkit, Safarina G Malik, Khin Saw Myint, Amin Soebandrio |
| EPI_ISL_3159897 | Dinas Kesehatan Provinsi Kalimantan Timur | Eijkman Institute for Molecular Biology, National Research and Innovation Agency; National Institute of Health Research and Development | Hidayat Trimarsanto, Lydia V. Panggalo, Iskandar Adnan, Sukma Oktavianthi, Lidwina Priliani, Edison Johar, Frilasita A Yudhaputri, Muhammad Rezki Rasyak, Willy Agustine, Vivi Setiawaty, Hana Apsari Pawestri, Subangkit, Safarina G Malik, Khin Saw Myint, Amin Soebandrio |
| EPI_ISL_3159901 | Dinas Kesehatan Provinsi Kalimantan Timur | Eijkman Institute for Molecular Biology, National Research and Innovation Agency; National Institute of Health Research and Development | Lydia V. Panggalo, Iskandar Adnan, Sukma Oktavianthi, Lidwina Priliani, Edison Johar, Frilasita A Yudhaputri, Muhammad Rezki Rasyak, Willy Agustine, Hidayat Trimarsanto, Vivi Setiawaty, Hana Apsari Pawestri, Subangkit, Safarina G Malik, Khin Saw Myint, Amin Soebandrio |
| EPI_ISL_3159903 | Dinas Kesehatan Provinsi Kalimantan Timur | Eijkman Institute for Molecular Biology, National Research and Innovation Agency; National Institute of Health Research and Development | Iskandar Adnan, Sukma Oktavianthi, Lidwina Priliani, Edison Johar, Frilasita A Yudhaputri, Muhammad Rezki Rasyak, Willy Agustine, Hidayat Trimarsanto, Lydia V. Panggalo, Vivi Setiawaty, Hana Apsari Pawestri, Subangkit, Safarina G Malik, Khin Saw Myint, Amin Soebandrio |
| EPI_ISL_3159906 | Dinas Kesehatan Provinsi Kalimantan Timur | Eijkman Institute for Molecular Biology, National Research and Innovation Agency; National Institute of Health Research and Development | Sukma Oktavianthi, Lidwina Priliani, Edison Johar, Frilasita A Yudhaputri, Muhammad Rezki Rasyak, Willy Agustine, Hidayat Trimarsanto, Lydia V. Panggalo, Iskandar Adnan, Vivi Setiawaty, Hana Apsari Pawestri, Subangkit, Safarina G Malik, Khin Saw Myint, Amin Soebandrio |
| EPI_ISL_3159909 | Dinas Kesehatan Provinsi Kalimantan Timur | Eijkman Institute for Molecular Biology, National Research and Innovation Agency; National Institute of Health Research and Development | Lidwina Priliani, Edison Johar, Frilasita A Yudhaputri, Muhammad Rezki Rasyak, Willy Agustine, Hidayat Trimarsanto, Lydia V. Panggalo, Iskandar Adnan, Sukma Oktavianthi, Vivi Setiawaty, Hana Apsari Pawestri, Subangkit, Safarina G Malik, Khin Saw Myint, Amin Soebandrio |
| EPI_ISL_3159912 | Dinas Kesehatan Provinsi Kalimantan Timur | Eijkman Institute for Molecular Biology, National Research and Innovation Agency; National Institute of Health Research and Development | Edison Johar, Frilasita A Yudhaputri, Muhammad Rezki Rasyak, Willy Agustine, Hidayat Trimarsanto, Lydia V. Panggalo, Iskandar Adnan, Sukma Oktavianthi, Lidwina Priliani, Vivi Setiawaty, Hana Apsari Pawestri, Subangkit, Safarina G Malik, Khin Saw Myint, Amin Soebandrio |
| EPI_ISL_3159915 | Dinas Kesehatan Provinsi Kalimantan Timur | Eijkman Institute for Molecular Biology, National Research and Innovation Agency; National Institute of Health Research and Development | Muhammad Rezki Rasyak, Willy Agustine, Hidayat Trimarsanto, Lydia V. Panggalo, Iskandar Adnan, Sukma Oktavianthi, Lidwina Priliani, Edison Johar, Frilasita A Yudhaputri, Vivi Setiawaty, Hana Apsari Pawestri, Subangkit, Safarina G Malik, Khin Saw Myint, Amin Soebandrio |
| EPI_ISL_3159918 | Dinas Kesehatan Provinsi Kalimantan Timur | Eijkman Institute for Molecular Biology, National Research and Innovation Agency; National Institute of Health Research and Development | Hidayat Trimarsanto, Lydia V. Panggalo, Iskandar Adnan, Sukma Oktavianthi, Lidwina Priliani, Edison Johar, Frilasita A Yudhaputri, Muhammad Rezki Rasyak, Willy Agustine, Vivi Setiawaty, Hana Apsari Pawestri, Subangkit, Safarina G Malik, Khin Saw Myint, Amin Soebandrio |
| EPI_ISL_3159921 | Dinas Kesehatan Provinsi Kalimantan Timur | Eijkman Institute for Molecular Biology, National Research and Innovation Agency; National Institute of Health Research and Development | Lydia V. Panggalo, Iskandar Adnan, Sukma Oktavianthi, Lidwina Priliani, Edison Johar, Frilasita A Yudhaputri, Muhammad Rezki Rasyak, Willy Agustine, Hidayat Trimarsanto, Vivi Setiawaty, Hana Apsari Pawestri, Subangkit, Safarina G Malik, Khin Saw Myint, Amin Soebandrio |
| EPI_ISL_3159924 | Dinas Kesehatan Provinsi Kalimantan Timur | Eijkman Institute for Molecular Biology, National Research and Innovation Agency; National Institute of Health Research and Development | Iskandar Adnan, Sukma Oktavianthi, Lidwina Priliani, Edison Johar, Frilasita A Yudhaputri, Muhammad Rezki Rasyak, Willy Agustine, Hidayat Trimarsanto, Lydia V. Panggalo, Vivi Setiawaty, Hana Apsari Pawestri, Subangkit, Safarina G Malik, Khin Saw Myint, Amin Soebandrio |
| EPI_ISL_3159927 | Dinas Kesehatan Provinsi Kalimantan Timur | Eijkman Institute for Molecular Biology, National Research and Innovation Agency; National Institute of Health Research and Development | Lidwina Priliani, Edison Johar, Frilasita A Yudhaputri, Muhammad Rezki Rasyak, Willy Agustine, Hidayat Trimarsanto, Lydia V. Panggalo, Iskandar Adnan, Sukma Oktavianthi, Vivi Setiawaty, Hana Apsari Pawestri, Subangkit, Safarina G Malik, Khin Saw Myint, Amin Soebandrio |
| EPI_ISL_3159930 | Dinas Kesehatan Provinsi Kalimantan Timur | Eijkman Institute for Molecular Biology, National Research and Innovation Agency; National Institute of Health Research and Development | Edison Johar, Frilasita A Yudhaputri, Muhammad Rezki Rasyak, Willy Agustine, Hidayat Trimarsanto, Lydia V. Panggalo, Iskandar Adnan, Sukma Oktavianthi, Lidwina Priliani, Vivi Setiawaty, Hana Apsari Pawestri, Subangkit, Safarina G Malik, Khin Saw Myint, Amin Soebandrio |
| EPI_ISL_3159934 | Dinas Kesehatan Provinsi Kalimantan Timur | Eijkman Institute for Molecular Biology, National Research and Innovation Agency; National Institute of Health Research and Development | Willy Agustine, Hidayat Trimarsanto, Lydia V. Panggalo, Iskandar Adnan, Sukma Oktavianthi, Lidwina Priliani, Edison Johar, Frilasita A Yudhaputri, Muhammad Rezki Rasyak, Vivi Setiawaty, Hana Apsari Pawestri, Subangkit, Safarina G Malik, Khin Saw Myint, Amin Soebandrio |
| EPI_ISL_3159937 | Dinas Kesehatan Provinsi Kalimantan Timur | Eijkman Institute for Molecular Biology, National Research and Innovation Agency; National Institute of Health Research and Development | Hidayat Trimarsanto, Lydia V. Panggalo, Iskandar Adnan, Sukma Oktavianthi, Lidwina Priliani, Edison Johar, Frilasita A Yudhaputri, Muhammad Rezki Rasyak, Willy Agustine, Vivi Setiawaty, Hana Apsari Pawestri, Subangkit, Safarina G Malik, Khin Saw Myint, Amin Soebandrio |
| EPI_ISL_3159940 | Dinas Kesehatan Provinsi Kalimantan Timur | Eijkman Institute for Molecular Biology, National Research and Innovation Agency; National Institute of Health Research and Development | Lydia V. Panggalo, Iskandar Adnan, Sukma Oktavianthi, Lidwina Priliani, Edison Johar, Frilasita A Yudhaputri, Muhammad Rezki Rasyak, Willy Agustine, Hidayat Trimarsanto, Vivi Setiawaty, Hana Apsari Pawestri, Subangkit, Safarina G Malik, Khin Saw Myint, Amin Soebandrio |
| EPI_ISL_3159943 | Dinas Kesehatan Provinsi Kalimantan Timur | Eijkman Institute for Molecular Biology, National Research and Innovation Agency; National Institute of Health Research and Development | Iskandar Adnan, Sukma Oktavianthi, Lidwina Priliani, Edison Johar, Frilasita A Yudhaputri, Muhammad Rezki Rasyak, Willy Agustine, Hidayat Trimarsanto, Lydia V. Panggalo, Vivi Setiawaty, Hana Apsari Pawestri, Subangkit, Safarina G Malik, Khin Saw Myint, Amin Soebandrio |
| EPI_ISL_3159946 | Dinas Kesehatan Provinsi Kalimantan Timur | Eijkman Institute for Molecular Biology, National Research and Innovation Agency; National Institute of Health Research and Development | Sukma Oktavianthi, Lidwina Priliani, Edison Johar, Frilasita A Yudhaputri, Muhammad Rezki Rasyak, Willy Agustine, Hidayat Trimarsanto, Lydia V. Panggalo, Iskandar Adnan, Vivi Setiawaty, Hana Apsari Pawestri, Subangkit, Safarina G Malik, Khin Saw Myint, Amin Soebandrio |
| EPI_ISL_3159950 | Dinas Kesehatan Provinsi Kalimantan Timur | Eijkman Institute for Molecular Biology, National Research and Innovation Agency; National Institute of Health Research and Development | Lidwina Priliani, Edison Johar, Frilasita A Yudhaputri, Muhammad Rezki Rasyak, Willy Agustine, Hidayat Trimarsanto, Lydia V. Panggalo, Iskandar Adnan, Sukma Oktavianthi, Vivi Setiawaty, Hana Apsari Pawestri, Subangkit, Safarina G Malik, Khin Saw Myint, Amin Soebandrio |
| EPI_ISL_3159953 | Dinas Kesehatan Provinsi Kalimantan Timur | Eijkman Institute for Molecular Biology, National Research and Innovation Agency; National Institute of Health Research and Development | Edison Johar, Frilasita A Yudhaputri, Muhammad Rezki Rasyak, Willy Agustine, Hidayat Trimarsanto, Lydia V. Panggalo, Iskandar Adnan, Sukma Oktavianthi, Lidwina Priliani, Vivi Setiawaty, Hana Apsari Pawestri, Subangkit, Safarina G Malik, Khin Saw Myint, Amin Soebandrio |
| EPI_ISL_3159956 | Dinas Kesehatan Provinsi Kalimantan Timur | Eijkman Institute for Molecular Biology, National Research and Innovation Agency; National Institute of Health Research and Development | Frilasita A Yudhaputri, Muhammad Rezki Rasyak, Willy Agustine, Hidayat Trimarsanto, Lydia V. Panggalo, Iskandar Adnan, Sukma Oktavianthi, Lidwina Priliani, Edison Johar, Vivi Setiawaty, Hana Apsari Pawestri, Subangkit, Safarina G Malik, Khin Saw Myint, Amin Soebandrio |
| EPI_ISL_3159959 | Dinas Kesehatan Provinsi Kalimantan Timur | Eijkman Institute for Molecular Biology, National Research and | Muhammad Rezki Rasyak, Willy Agustine, Hidayat Trimarsanto, Lydia V. Panggalo, Iskandar Adnan, Sukma Oktavianthi, Lidwina Priliani, Edison Johar, Frilasita A Yudhaputri, Vivi Setiawaty, Hana Apsari Pawestri, Subangkit, Safarina G Malik, |

| | | | |
|---|---|---|---|
| | | Innovation Agency; National Institute of Health Research and Development | Khin Saw Myint, Amin Soebandrio |
| EPI_ISL_3159963 | Dinas Kesehatan Provinsi Kalimantan Timur | Eijkman Institute for Molecular Biology, National Research and Innovation Agency; National Institute of Health Research and Development | Willy Agustine, Hidayat Trimarsanto, Lydia V. Panggalo, Iskandar Adnan, Sukma Oktavianthi, Lidwina Priliani, Edison Johar, Frilasita A Yudhaputri, Muhammad Rezki Rasyak, Vivi Setiawaty, Hana Apsari Pawestri, Subangkit, Safarina G Malik, Khin Saw Myint, Amin Soebandrio |
| EPI_ISL_3159966 | Dinas Kesehatan Provinsi Kalimantan Timur | Eijkman Institute for Molecular Biology, National Research and Innovation Agency; National Institute of Health Research and Development | Hidayat Trimarsanto, Lydia V. Panggalo, Iskandar Adnan, Sukma Oktavianthi, Lidwina Priliani, Edison Johar, Frilasita A Yudhaputri, Muhammad Rezki Rasyak, Willy Agustine, Vivi Setiawaty, Hana Apsari Pawestri, Subangkit, Safarina G Malik, Khin Saw Myint, Amin Soebandrio |
| EPI_ISL_3159969 | Dinas Kesehatan Provinsi Kalimantan Timur | Eijkman Institute for Molecular Biology, National Research and Innovation Agency; National Institute of Health Research and Development | Iskandar Adnan, Sukma Oktavianthi, Lidwina Priliani, Edison Johar, Frilasita A Yudhaputri, Muhammad Rezki Rasyak, Willy Agustine, Hidayat Trimarsanto, Lydia V. Panggalo, Vivi Setiawaty, Hana Apsari Pawestri, Subangkit, Safarina G Malik, Khin Saw Myint, Amin Soebandrio |
| EPI_ISL_3159972 | Dinas Kesehatan Provinsi Kalimantan Timur | Eijkman Institute for Molecular Biology, National Research and Innovation Agency; National Institute of Health Research and Development | Sukma Oktavianthi, Lidwina Priliani, Edison Johar, Frilasita A Yudhaputri, Muhammad Rezki Rasyak, Willy Agustine, Hidayat Trimarsanto, Lydia V. Panggalo, Iskandar Adnan, Vivi Setiawaty, Hana Apsari Pawestri, Subangkit, Safarina G Malik, Khin Saw Myint, Amin Soebandrio |
| EPI_ISL_3159976 | Dinas Kesehatan Provinsi Kalimantan Timur | Eijkman Institute for Molecular Biology, National Research and Innovation Agency; National Institute of Health Research and Development | Lidwina Priliani, Edison Johar, Frilasita A Yudhaputri, Muhammad Rezki Rasyak, Willy Agustine, Hidayat Trimarsanto, Lydia V. Panggalo, Iskandar Adnan, Sukma Oktavianthi, Vivi Setiawaty, Hana Apsari Pawestri, Subangkit, Safarina G Malik, Khin Saw Myint, Amin Soebandrio |
| EPI_ISL_3159979 | Dinas Kesehatan Provinsi Kalimantan Timur | Eijkman Institute for Molecular Biology, National Research and Innovation Agency; National Institute of Health Research and Development | Edison Johar, Frilasita A Yudhaputri, Muhammad Rezki Rasyak, Willy Agustine, Hidayat Trimarsanto, Lydia V. Panggalo, Iskandar Adnan, Sukma Oktavianthi, Lidwina Priliani, Vivi Setiawaty, Hana Apsari Pawestri, Subangkit, Safarina G Malik, Khin Saw Myint, Amin Soebandrio |
| EPI_ISL_3159982 | Dinas Kesehatan Provinsi Kalimantan Timur | Eijkman Institute for Molecular Biology, National Research and Innovation Agency; National Institute of Health Research and Development | Frilasita A Yudhaputri, Muhammad Rezki Rasyak, Willy Agustine, Hidayat Trimarsanto, Lydia V. Panggalo, Iskandar Adnan, Sukma Oktavianthi, Lidwina Priliani, Edison Johar, Vivi Setiawaty, Hana Apsari Pawestri, Subangkit, Safarina G Malik, Khin Saw Myint, Amin Soebandrio |
| EPI_ISL_3159983 | Colorado Department of Public Health and Environment | Colorado Department of Public Health and Environment | Laura Bankers, Molly C. Hetherington-Rauth, Diana Ir, Alexandria Rossheim, Michael Martin, Mandy Waters, Shannon R. Matzinger, Sarah Elizabeth Totten, Emily A. Travanty |
| EPI_ISL_3159985 | Dinas Kesehatan Provinsi Kalimantan Timur | Eijkman Institute for Molecular Biology, National Research and Innovation Agency; National Institute of Health Research and Development | Muhammad Rezki Rasyak, Willy Agustine, Hidayat Trimarsanto, Lydia V. Panggalo, Iskandar Adnan, Sukma Oktavianthi, Lidwina Priliani, Edison Johar, Frilasita A Yudhaputri, Vivi Setiawaty, Hana Apsari Pawestri, Subangkit, Safarina G Malik, Khin Saw Myint, Amin Soebandrio |
| EPI_ISL_3159987 | Colorado Department of Public Health and Environment | Colorado Department of Public Health and Environment | Laura Bankers, Molly C. Hetherington-Rauth, Diana Ir, Alexandria Rossheim, Michael Martin, Mandy Waters, Shannon R. Matzinger, Sarah Elizabeth Totten, Emily A. Travanty |
| EPI_ISL_3159989 | Dinas Kesehatan Provinsi Kalimantan Timur | Eijkman Institute for Molecular Biology, National Research and Innovation Agency; National Institute of Health Research and Development | Willy Agustine, Hidayat Trimarsanto, Lydia V. Panggalo, Iskandar Adnan, Sukma Oktavianthi, Lidwina Priliani, Edison Johar, Frilasita A Yudhaputri, Muhammad Rezki Rasyak, Vivi Setiawaty, Hana Apsari Pawestri, Subangkit, Safarina G Malik, Khin Saw Myint, Amin Soebandrio |
| EPI_ISL_3159990 | Colorado Department of Public Health and Environment | Colorado Department of Public Health and Environment | Laura Bankers, Molly C. Hetherington-Rauth, Diana Ir, Alexandria Rossheim, Michael Martin, Mandy Waters, Shannon R. Matzinger, Sarah Elizabeth Totten, Emily A. Travanty |
| EPI_ISL_3159992 | Dinas Kesehatan Provinsi Kalimantan Timur | Eijkman Institute for Molecular Biology, National Research and Innovation Agency; National Institute of Health Research and Development | Lydia V. Panggalo, Iskandar Adnan, Sukma Oktavianthi, Lidwina Priliani, Edison Johar, Frilasita A Yudhaputri, Muhammad Rezki Rasyak, Willy Agustine, Hidayat Trimarsanto, Vivi Setiawaty, Hana Apsari Pawestri, Subangkit, Safarina G Malik, Khin Saw Myint, Amin Soebandrio |
| EPI_ISL_3159995 | Dinas Kesehatan Provinsi Kalimantan Timur | Eijkman Institute for Molecular Biology, National Research and Innovation Agency; National Institute of Health Research and Development | Iskandar Adnan, Sukma Oktavianthi, Lidwina Priliani, Edison Johar, Frilasita A Yudhaputri, Muhammad Rezki Rasyak, Willy Agustine, Hidayat Trimarsanto, Lydia V. Panggalo, Vivi Setiawaty, Hana Apsari Pawestri, Subangkit, Safarina G Malik, Khin Saw Myint, Amin Soebandrio |
| EPI_ISL_3159998 | Dinas Kesehatan Provinsi Kalimantan Timur | Eijkman Institute for Molecular Biology, National Research and Innovation Agency; National Institute of Health Research and Development | Sukma Oktavianthi, Lidwina Priliani, Edison Johar, Frilasita A Yudhaputri, Muhammad Rezki Rasyak, Willy Agustine, Hidayat Trimarsanto, Lydia V. Panggalo, Iskandar Adnan, Vivi Setiawaty, Hana Apsari Pawestri, Subangkit, Safarina G Malik, Khin Saw Myint, Amin Soebandrio |
| EPI_ISL_3160001 | Dinas Kesehatan Provinsi Kalimantan Timur | Eijkman Institute for Molecular Biology, National Research and Innovation Agency; National Institute of Health Research and Development | Lidwina Priliani, Edison Johar, Frilasita A Yudhaputri, Muhammad Rezki Rasyak, Willy Agustine, Hidayat Trimarsanto, Lydia V. Panggalo, Iskandar Adnan, Sukma Oktavianthi, Vivi Setiawaty, Hana Apsari Pawestri, Subangkit, Safarina G Malik, Khin Saw Myint, Amin Soebandrio |
| EPI_ISL_3160004 | Dinas Kesehatan Provinsi Kalimantan Timur | Eijkman Institute for Molecular Biology, National Research and Innovation Agency; National Institute of Health Research and Development | Edison Johar, Frilasita A Yudhaputri, Muhammad Rezki Rasyak, Willy Agustine, Hidayat Trimarsanto, Lydia V. Panggalo, Iskandar Adnan, Sukma Oktavianthi, Lidwina Priliani, Vivi Setiawaty, Hana Apsari Pawestri, Subangkit, Safarina G Malik, Khin Saw Myint, Amin Soebandrio |
| EPI_ISL_3160007 | Dinas Kesehatan Provinsi Kalimantan Timur | Eijkman Institute for Molecular Biology, National Research and Innovation Agency; National Institute of Health Research and Development | Frilasita A Yudhaputri, Muhammad Rezki Rasyak, Willy Agustine, Hidayat Trimarsanto, Lydia V. Panggalo, Iskandar Adnan, Sukma Oktavianthi, Lidwina Priliani, Edison Johar, Vivi Setiawaty, Hana Apsari Pawestri, Subangkit, Safarina G Malik, Khin Saw Myint, Amin Soebandrio |
| EPI_ISL_3160008 | Colorado Department of Public Health and Environment | Colorado Department of Public Health and Environment | Laura Bankers, Molly C. Hetherington-Rauth, Diana Ir, Alexandria Rossheim, Michael Martin, Mandy Waters, Shannon R. Matzinger, Sarah Elizabeth Totten, Emily A. Travanty |
| EPI_ISL_3160010 | Dinas Kesehatan Provinsi Kalimantan Timur | Eijkman Institute for Molecular Biology, National Research and Innovation Agency; National Institute of Health Research and Development | Muhammad Rezki Rasyak, Willy Agustine, Hidayat Trimarsanto, Lydia V. Panggalo, Iskandar Adnan, Sukma Oktavianthi, Lidwina Priliani, Edison Johar, Frilasita A Yudhaputri, Vivi Setiawaty, Hana Apsari Pawestri, Subangkit, Safarina G Malik, Khin Saw Myint, Amin Soebandrio |
| EPI_ISL_3160013 | Dinas Kesehatan Provinsi Kalimantan Timur | Eijkman Institute for Molecular Biology, National Research and Innovation Agency; National Institute of Health Research and Development | Willy Agustine, Hidayat Trimarsanto, Lydia V. Panggalo, Iskandar Adnan, Sukma Oktavianthi, Lidwina Priliani, Edison Johar, Frilasita A Yudhaputri, Muhammad Rezki Rasyak, Vivi Setiawaty, Hana Apsari Pawestri, Subangkit, Safarina G Malik, Khin Saw Myint, Amin Soebandrio |
| EPI_ISL_3160016 | Dinas Kesehatan Provinsi Kalimantan Timur | Eijkman Institute for Molecular Biology, National Research and Innovation Agency; National Institute of Health Research and Development | Hidayat Trimarsanto, Lydia V. Panggalo, Iskandar Adnan, Sukma Oktavianthi, Lidwina Priliani, Edison Johar, Frilasita A Yudhaputri, Muhammad Rezki Rasyak, Willy Agustine, Vivi Setiawaty, Hana Apsari Pawestri, Subangkit, Safarina G Malik, Khin Saw Myint, Amin Soebandrio |
| EPI_ISL_3160019 | Dinas Kesehatan Provinsi Kalimantan Timur | Eijkman Institute for Molecular Biology, National Research and Innovation Agency; National Institute of Health Research and Development | Lydia V. Panggalo, Iskandar Adnan, Sukma Oktavianthi, Lidwina Priliani, Edison Johar, Frilasita A Yudhaputri, Muhammad Rezki Rasyak, Willy Agustine, Hidayat Trimarsanto, Vivi Setiawaty, Hana Apsari Pawestri, Subangkit, Safarina G Malik, Khin Saw Myint, Amin Soebandrio |
| EPI_ISL_3160022 | Dinas Kesehatan Provinsi Kalimantan Timur | Eijkman Institute for Molecular Biology, National Research and Innovation Agency; National Institute of Health Research and Development | Iskandar Adnan, Sukma Oktavianthi, Lidwina Priliani, Edison Johar, Frilasita A Yudhaputri, Muhammad Rezki Rasyak, Willy Agustine, Hidayat Trimarsanto, Lydia V. Panggalo, Vivi Setiawaty, Hana Apsari Pawestri, Subangkit, Safarina G Malik, Khin Saw Myint, Amin Soebandrio |
| EPI_ISL_3160025 | Dinas Kesehatan Provinsi Kalimantan Timur | Eijkman Institute for Molecular Biology, National Research and Innovation Agency; National Institute of Health Research and Development | Sukma Oktavianthi, Lidwina Priliani, Edison Johar, Frilasita A Yudhaputri, Muhammad Rezki Rasyak, Willy Agustine, Hidayat Trimarsanto, Lydia V. Panggalo, Iskandar Adnan, Vivi Setiawaty, Hana Apsari Pawestri, Subangkit, Safarina G Malik, Khin Saw Myint, Amin Soebandrio |
| EPI_ISL_3160028 | Dinas Kesehatan Provinsi Kalimantan Timur | Eijkman Institute for Molecular Biology, National Research and Innovation Agency; National Institute of Health Research and Development | Lidwina Priliani, Edison Johar, Frilasita A Yudhaputri, Muhammad Rezki Rasyak, Willy Agustine, Hidayat Trimarsanto, Lydia V. Panggalo, Iskandar Adnan, Sukma Oktavianthi, Vivi Setiawaty, Hana Apsari Pawestri, Subangkit, Safarina G Malik, Khin Saw Myint, Amin Soebandrio |
| EPI_ISL_3160031 | Dinas Kesehatan Provinsi Kalimantan Timur | Eijkman Institute for Molecular Biology, National Research and Innovation Agency; National Institute of Health Research and Development | Edison Johar, Frilasita A Yudhaputri, Muhammad Rezki Rasyak, Willy Agustine, Hidayat Trimarsanto, Lydia V. Panggalo, Iskandar Adnan, Sukma Oktavianthi, Lidwina Priliani, Vivi Setiawaty, Hana Apsari Pawestri, Subangkit, Safarina G Malik, Khin Saw Myint, Amin Soebandrio |
| EPI_ISL_3160034 | Dinas Kesehatan Provinsi Kalimantan Timur | Eijkman Institute for Molecular Biology, National Research and Innovation Agency; National Institute of Health Research and Development | Frilasita A Yudhaputri, Muhammad Rezki Rasyak, Willy Agustine, Hidayat Trimarsanto, Lydia V. Panggalo, Iskandar Adnan, Sukma Oktavianthi, Lidwina Priliani, Edison Johar, Vivi Setiawaty, Hana Apsari Pawestri, Subangkit, Safarina G Malik, Khin Saw Myint, Amin Soebandrio |
| EPI_ISL_3160040 | Dinas Kesehatan Provinsi Kalimantan Timur | Eijkman Institute for Molecular Biology, National Research and Innovation Agency; National Institute of Health Research and Development | Willy Agustine, Hidayat Trimarsanto, Lydia V. Panggalo, Iskandar Adnan, Sukma Oktavianthi, Lidwina Priliani, Edison Johar, Frilasita A Yudhaputri, Muhammad Rezki Rasyak, Vivi Setiawaty, Hana Apsari Pawestri, Subangkit, Safarina G Malik, Khin Saw Myint, Amin Soebandrio |
| EPI_ISL_3160043 | Dinas Kesehatan Provinsi Kalimantan Timur | Eijkman Institute for Molecular Biology, National Research and Innovation Agency; National Institute of Health Research and Development | Lydia V. Panggalo, Iskandar Adnan, Sukma Oktavianthi, Lidwina Priliani, Edison Johar, Frilasita A Yudhaputri, Muhammad Rezki Rasyak, Willy Agustine, Hidayat Trimarsanto, Vivi Setiawaty, Hana Apsari Pawestri, Subangkit, Safarina G Malik, Khin Saw Myint, Amin Soebandrio |
| EPI_ISL_3160046 | Dinas Kesehatan Provinsi Kalimantan Timur | Eijkman Institute for Molecular Biology, National Research and Innovation Agency; National Institute of Health Research and Development | Iskandar Adnan, Sukma Oktavianthi, Lidwina Priliani, Edison Johar, Frilasita A Yudhaputri, Muhammad Rezki Rasyak, Willy Agustine, Hidayat Trimarsanto, Lydia V. Panggalo, Vivi Setiawaty, Hana Apsari Pawestri, Subangkit, Safarina G Malik, Khin Saw Myint, Amin Soebandrio |
| EPI_ISL_3160049 | Dinas Kesehatan Provinsi Kalimantan Timur | Eijkman Institute for Molecular Biology, National Research and Innovation Agency; National Institute of Health Research and Development | Sukma Oktavianthi, Lidwina Priliani, Edison Johar, Frilasita A Yudhaputri, Muhammad Rezki Rasyak, Willy Agustine, Hidayat Trimarsanto, Lydia V. Panggalo, Iskandar Adnan, Vivi Setiawaty, Hana Apsari Pawestri, Subangkit, Safarina G Malik, Khin Saw Myint, Amin Soebandrio |
| EPI_ISL_3160050 | Colorado Department of Public Health and Environment | Colorado Department of Public Health and Environment | Laura Bankers, Molly C. Hetherington-Rauth, Diana Ir, Alexandria Rossheim, Michael Martin, Mandy Waters, Shannon R. Matzinger, Sarah Elizabeth Totten, Emily A. Travanty |
| EPI_ISL_3160053 | Dinas Kesehatan Provinsi Kalimantan Timur | Eijkman Institute for Molecular Biology, National Research and Innovation Agency; National Institute of Health Research and Development | Lidwina Priliani, Edison Johar, Frilasita A Yudhaputri, Muhammad Rezki Rasyak, Willy Agustine, Hidayat Trimarsanto, Lydia V. Panggalo, Iskandar Adnan, Sukma Oktavianthi, Vivi Setiawaty, Hana Apsari Pawestri, Subangkit, Safarina G Malik, Khin Saw Myint, Amin Soebandrio |

| | | | |
|---|---|---|---|
| EPI_ISL_3160055 | Dinas Kesehatan Provinsi Kalimantan Timur | Eijkman Institute for Molecular Biology, National Research and Innovation Agency; National Institute of Health Research and Development | Edison Johar, Frilasita A Yudhaputri, Muhammad Rezki Rasyak, Willy Agustine, Hidayat Trimarsanto, Lydia V. Panggalo, Iskandar Adnan, Sukma Oktavianthi, Lidwina Priliani, Vivi Setiawaty, Hana Apsari Pawestri, Subangkit, Safarina G Malik, Khin Saw Myint, Amin Soebandrio |
| EPI_ISL_3160057, EPI_ISL_3160058 | Colorado Department of Public Health and Environment | Colorado Department of Public Health and Environment | Laura Bankers, Molly C. Hetherington-Rauth, Diana Ir, Alexandria Rossheim, Michael Martin, Mandy Waters, Shannon R. Matzinger, Sarah Elizabeth Totten, Emily A. Travanty |
| EPI_ISL_3160059 | Dinas Kesehatan Provinsi Kalimantan Timur | Eijkman Institute for Molecular Biology, National Research and Innovation Agency; National Institute of Health Research and Development | Frilasita A Yudhaputri, Muhammad Rezki Rasyak, Willy Agustine, Hidayat Trimarsanto, Lydia V. Panggalo, Iskandar Adnan, Sukma Oktavianthi, Lidwina Priliani, Edison Johar, Vivi Setiawaty, Hana Apsari Pawestri, Subangkit, Safarina G Malik, Khin Saw Myint, Amin Soebandrio |
| EPI_ISL_3160062 | Dinas Kesehatan Provinsi Kalimantan Timur | Eijkman Institute for Molecular Biology, National Research and Innovation Agency; National Institute of Health Research and Development | Muhammad Rezki Rasyak, Willy Agustine, Hidayat Trimarsanto, Lydia V. Panggalo, Iskandar Adnan, Sukma Oktavianthi, Lidwina Priliani, Edison Johar, Frilasita A Yudhaputri, Vivi Setiawaty, Hana Apsari Pawestri, Subangkit, Safarina G Malik, Khin Saw Myint, Amin Soebandrio |
| EPI_ISL_3160065 | Dinas Kesehatan Provinsi Kalimantan Timur | Eijkman Institute for Molecular Biology, National Research and Innovation Agency; National Institute of Health Research and Development | Willy Agustine, Hidayat Trimarsanto, Lydia V. Panggalo, Iskandar Adnan, Sukma Oktavianthi, Lidwina Priliani, Edison Johar, Frilasita A Yudhaputri, Muhammad Rezki Rasyak, Vivi Setiawaty, Hana Apsari Pawestri, Subangkit, Safarina G Malik, Khin Saw Myint, Amin Soebandrio |
| EPI_ISL_3160067 | Colorado Department of Public Health and Environment | Colorado Department of Public Health and Environment | Laura Bankers, Molly C. Hetherington-Rauth, Diana Ir, Alexandria Rossheim, Michael Martin, Mandy Waters, Shannon R. Matzinger, Sarah Elizabeth Totten, Emily A. Travanty |
| EPI_ISL_3160068 | Dinas Kesehatan Provinsi Kalimantan Timur | Eijkman Institute for Molecular Biology, National Research and Innovation Agency; National Institute of Health Research and Development | Hidayat Trimarsanto, Lydia V. Panggalo, Iskandar Adnan, Sukma Oktavianthi, Lidwina Priliani, Edison Johar, Frilasita A Yudhaputri, Muhammad Rezki Rasyak, Willy Agustine, Vivi Setiawaty, Hana Apsari Pawestri, Subangkit, Safarina G Malik, Khin Saw Myint, Amin Soebandrio |
| EPI_ISL_3160071 | Dinas Kesehatan Provinsi Kalimantan Timur | Eijkman Institute for Molecular Biology, National Research and Innovation Agency; National Institute of Health Research and Development | Lydia V. Panggalo, Iskandar Adnan, Sukma Oktavianthi, Lidwina Priliani, Edison Johar, Frilasita A Yudhaputri, Muhammad Rezki Rasyak, Willy Agustine, Hidayat Trimarsanto, Vivi Setiawaty, Hana Apsari Pawestri, Subangkit, Safarina G Malik, Khin Saw Myint, Amin Soebandrio |
| EPI_ISL_3160074 | Dinas Kesehatan Provinsi Kalimantan Timur | Eijkman Institute for Molecular Biology, National Research and Innovation Agency; National Institute of Health Research and Development | Iskandar Adnan, Sukma Oktavianthi, Lidwina Priliani, Edison Johar, Frilasita A Yudhaputri, Muhammad Rezki Rasyak, Willy Agustine, Hidayat Trimarsanto, Lydia V. Panggalo, Vivi Setiawaty, Hana Apsari Pawestri, Subangkit, Safarina G Malik, Khin Saw Myint, Amin Soebandrio |
| EPI_ISL_3160076 | Colorado Department of Public Health and Environment | Colorado Department of Public Health and Environment | Laura Bankers, Molly C. Hetherington-Rauth, Diana Ir, Alexandria Rossheim, Michael Martin, Mandy Waters, Shannon R. Matzinger, Sarah Elizabeth Totten, Emily A. Travanty |
| EPI_ISL_3160077 | Dinas Kesehatan Provinsi Kalimantan Timur | Eijkman Institute for Molecular Biology, National Research and Innovation Agency; National Institute of Health Research and Development | Sukma Oktavianthi, Lidwina Priliani, Edison Johar, Frilasita A Yudhaputri, Muhammad Rezki Rasyak, Willy Agustine, Hidayat Trimarsanto, Lydia V. Panggalo, Iskandar Adnan, Vivi Setiawaty, Hana Apsari Pawestri, Subangkit, Safarina G Malik, Khin Saw Myint, Amin Soebandrio |
| EPI_ISL_3160080 | Dinas Kesehatan Provinsi Kalimantan Timur | Eijkman Institute for Molecular Biology, National Research and Innovation Agency; National Institute of Health Research and Development | Lidwina Priliani, Edison Johar, Frilasita A Yudhaputri, Muhammad Rezki Rasyak, Willy Agustine, Hidayat Trimarsanto, Lydia V. Panggalo, Iskandar Adnan, Sukma Oktavianthi, Vivi Setiawaty, Hana Apsari Pawestri, Subangkit, Safarina G Malik, Khin Saw Myint, Amin Soebandrio |
| EPI_ISL_3160081 | Colorado Department of Public Health and Environment | Colorado Department of Public Health and Environment | Laura Bankers, Molly C. Hetherington-Rauth, Diana Ir, Alexandria Rossheim, Michael Martin, Mandy Waters, Shannon R. Matzinger, Sarah Elizabeth Totten, Emily A. Travanty |
| EPI_ISL_3160082 | Dinas Kesehatan Provinsi Kalimantan Timur | Eijkman Institute for Molecular Biology, National Research and Innovation Agency; National Institute of Health Research and Development | Edison Johar, Frilasita A Yudhaputri, Muhammad Rezki Rasyak, Willy Agustine, Hidayat Trimarsanto, Lydia V. Panggalo, Iskandar Adnan, Sukma Oktavianthi, Lidwina Priliani, Vivi Setiawaty, Hana Apsari Pawestri, Subangkit, Safarina G Malik, Khin Saw Myint, Amin Soebandrio |
| EPI_ISL_3160085 | Dinas Kesehatan Provinsi Kalimantan Timur | Eijkman Institute for Molecular Biology, National Research and Innovation Agency; National Institute of Health Research and Development | Frilasita A Yudhaputri, Muhammad Rezki Rasyak, Willy Agustine, Hidayat Trimarsanto, Lydia V. Panggalo, Iskandar Adnan, Sukma Oktavianthi, Lidwina Priliani, Edison Johar, Vivi Setiawaty, Hana Apsari Pawestri, Subangkit, Safarina G Malik, Khin Saw Myint, Amin Soebandrio |
| EPI_ISL_3160088 | Dinas Kesehatan Provinsi Kalimantan Timur | Eijkman Institute for Molecular Biology, National Research and Innovation Agency; National Institute of Health Research and Development | Muhammad Rezki Rasyak, Willy Agustine, Hidayat Trimarsanto, Lydia V. Panggalo, Iskandar Adnan, Sukma Oktavianthi, Lidwina Priliani, Edison Johar, Frilasita A Yudhaputri, Vivi Setiawaty, Hana Apsari Pawestri, Subangkit, Safarina G Malik, Khin Saw Myint, Amin Soebandrio |
| EPI_ISL_3160089, EPI_ISL_3160090 | Colorado Department of Public Health and Environment | Colorado Department of Public Health and Environment | Laura Bankers, Molly C. Hetherington-Rauth, Diana Ir, Alexandria Rossheim, Michael Martin, Mandy Waters, Shannon R. Matzinger, Sarah Elizabeth Totten, Emily A. Travanty |
| EPI_ISL_3160091 | Dinas Kesehatan Provinsi Kalimantan Timur | Eijkman Institute for Molecular Biology, National Research and Innovation Agency; National Institute of Health Research and Development | Willy Agustine, Hidayat Trimarsanto, Lydia V. Panggalo, Iskandar Adnan, Sukma Oktavianthi, Lidwina Priliani, Edison Johar, Frilasita A Yudhaputri, Muhammad Rezki Rasyak, Vivi Setiawaty, Hana Apsari Pawestri, Subangkit, Safarina G Malik, Khin Saw Myint, Amin Soebandrio |
| EPI_ISL_3160094 | Dinas Kesehatan Provinsi Kalimantan Timur | Eijkman Institute for Molecular Biology, National Research and Innovation Agency; National Institute of Health Research and Development | Hidayat Trimarsanto, Lydia V. Panggalo, Iskandar Adnan, Sukma Oktavianthi, Lidwina Priliani, Edison Johar, Frilasita A Yudhaputri, Muhammad Rezki Rasyak, Willy Agustine, Vivi Setiawaty, Hana Apsari Pawestri, Subangkit, Safarina G Malik, Khin Saw Myint, Amin Soebandrio |
| EPI_ISL_3160097 | Dinas Kesehatan Provinsi Kalimantan Timur | Eijkman Institute for Molecular Biology, National Research and Innovation Agency; National Institute of Health Research and Development | Lydia V. Panggalo, Iskandar Adnan, Sukma Oktavianthi, Lidwina Priliani, Edison Johar, Frilasita A Yudhaputri, Muhammad Rezki Rasyak, Willy Agustine, Hidayat Trimarsanto, Vivi Setiawaty, Hana Apsari Pawestri, Subangkit, Safarina G Malik, Khin Saw Myint, Amin Soebandrio |
| EPI_ISL_3160100 | Dinas Kesehatan Provinsi Kalimantan Timur | Eijkman Institute for Molecular Biology, National Research and Innovation Agency; National Institute of Health Research and Development | Iskandar Adnan, Sukma Oktavianthi, Lidwina Priliani, Edison Johar, Frilasita A Yudhaputri, Muhammad Rezki Rasyak, Willy Agustine, Hidayat Trimarsanto, Lydia V. Panggalo, Vivi Setiawaty, Hana Apsari Pawestri, Subangkit, Safarina G Malik, Khin Saw Myint, Amin Soebandrio |
| EPI_ISL_3160103 | Dinas Kesehatan Provinsi Kalimantan Timur | Eijkman Institute for Molecular Biology, National Research and Innovation Agency; National Institute of Health Research and Development | Sukma Oktavianthi, Lidwina Priliani, Edison Johar, Frilasita A Yudhaputri, Muhammad Rezki Rasyak, Willy Agustine, Hidayat Trimarsanto, Lydia V. Panggalo, Iskandar Adnan, Vivi Setiawaty, Hana Apsari Pawestri, Subangkit, Safarina G Malik, Khin Saw Myint, Amin Soebandrio |
| EPI_ISL_3160106 | Dinas Kesehatan Provinsi Kalimantan Timur | Eijkman Institute for Molecular Biology, National Research and Innovation Agency; National Institute of Health Research and Development | Lidwina Priliani, Edison Johar, Frilasita A Yudhaputri, Muhammad Rezki Rasyak, Willy Agustine, Hidayat Trimarsanto, Lydia V. Panggalo, Iskandar Adnan, Sukma Oktavianthi, Vivi Setiawaty, Hana Apsari Pawestri, Subangkit, Safarina G Malik, Khin Saw Myint, Amin Soebandrio |
| EPI_ISL_3160112 | Dinas Kesehatan Provinsi Kalimantan Timur | Eijkman Institute for Molecular Biology, National Research and Innovation Agency; National Institute of Health Research and Development | Frilasita A Yudhaputri, Muhammad Rezki Rasyak, Willy Agustine, Hidayat Trimarsanto, Lydia V. Panggalo, Iskandar Adnan, Sukma Oktavianthi, Lidwina Priliani, Edison Johar, Vivi Setiawaty, Hana Apsari Pawestri, Subangkit, Safarina G Malik, Khin Saw Myint, Amin Soebandrio |
| EPI_ISL_3160113 | Colorado Department of Public Health and Environment | Colorado Department of Public Health and Environment | Laura Bankers, Molly C. Hetherington-Rauth, Diana Ir, Alexandria Rossheim, Michael Martin, Mandy Waters, Shannon R. Matzinger, Sarah Elizabeth Totten, Emily A. Travanty |
| EPI_ISL_3160115 | Dinas Kesehatan Provinsi Kalimantan Timur | Eijkman Institute for Molecular Biology, National Research and Innovation Agency; National Institute of Health Research and Development | Muhammad Rezki Rasyak, Willy Agustine, Hidayat Trimarsanto, Lydia V. Panggalo, Iskandar Adnan, Sukma Oktavianthi, Lidwina Priliani, Edison Johar, Frilasita A Yudhaputri, Vivi Setiawaty, Hana Apsari Pawestri, Subangkit, Safarina G Malik, Khin Saw Myint, Amin Soebandrio |
| EPI_ISL_3160118 | Dinas Kesehatan Provinsi Kalimantan Timur | Eijkman Institute for Molecular Biology, National Research and Innovation Agency; National Institute of Health Research and Development | Willy Agustine, Hidayat Trimarsanto, Lydia V. Panggalo, Iskandar Adnan, Sukma Oktavianthi, Lidwina Priliani, Edison Johar, Frilasita A Yudhaputri, Muhammad Rezki Rasyak, Vivi Setiawaty, Hana Apsari Pawestri, Subangkit, Safarina G Malik, Khin Saw Myint, Amin Soebandrio |
| EPI_ISL_3160121 | Dinas Kesehatan Provinsi Kalimantan Timur | Eijkman Institute for Molecular Biology, National Research and Innovation Agency; National Institute of Health Research and Development | Hidayat Trimarsanto, Lydia V. Panggalo, Iskandar Adnan, Sukma Oktavianthi, Lidwina Priliani, Edison Johar, Frilasita A Yudhaputri, Muhammad Rezki Rasyak, Willy Agustine, Vivi Setiawaty, Hana Apsari Pawestri, Subangkit, Safarina G Malik, Khin Saw Myint, Amin Soebandrio |
| EPI_ISL_3160123 | Colorado Department of Public Health and Environment | Colorado Department of Public Health and Environment | Laura Bankers, Molly C. Hetherington-Rauth, Diana Ir, Alexandria Rossheim, Michael Martin, Mandy Waters, Shannon R. Matzinger, Sarah Elizabeth Totten, Emily A. Travanty |
| EPI_ISL_3160124 | Dinas Kesehatan Provinsi Kalimantan Timur | Eijkman Institute for Molecular Biology, National Research and Innovation Agency; National Institute of Health Research and Development | Lydia V. Panggalo, Iskandar Adnan, Sukma Oktavianthi, Lidwina Priliani, Edison Johar, Frilasita A Yudhaputri, Muhammad Rezki Rasyak, Willy Agustine, Hidayat Trimarsanto, Vivi Setiawaty, Hana Apsari Pawestri, Subangkit, Safarina G Malik, Khin Saw Myint, Amin Soebandrio |
| EPI_ISL_3160127 | Colorado Department of Public Health and Environment | Colorado Department of Public Health and Environment | Laura Bankers, Molly C. Hetherington-Rauth, Diana Ir, Alexandria Rossheim, Michael Martin, Mandy Waters, Shannon R. Matzinger, Sarah Elizabeth Totten, Emily A. Travanty |
| EPI_ISL_3160129 | Dinas Kesehatan Provinsi Kalimantan Timur | Eijkman Institute for Molecular Biology, National Research and Innovation Agency; National Institute of Health Research and Development | Sukma Oktavianthi, Lidwina Priliani, Edison Johar, Frilasita A Yudhaputri, Muhammad Rezki Rasyak, Willy Agustine, Hidayat Trimarsanto, Lydia V. Panggalo, Iskandar Adnan, Vivi Setiawaty, Hana Apsari Pawestri, Subangkit, Safarina G Malik, Khin Saw Myint, Amin Soebandrio |
| EPI_ISL_3160132 | Dinas Kesehatan Provinsi Kalimantan Timur | Eijkman Institute for Molecular Biology, National Research and Innovation Agency; National Institute of Health Research and Development | Lidwina Priliani, Edison Johar, Frilasita A Yudhaputri, Muhammad Rezki Rasyak, Willy Agustine, Hidayat Trimarsanto, Lydia V. Panggalo, Iskandar Adnan, Sukma Oktavianthi, Vivi Setiawaty, Hana Apsari Pawestri, Subangkit, Safarina G Malik, Khin Saw Myint, Amin Soebandrio |
| EPI_ISL_3160135 | Dinas Kesehatan Provinsi Kalimantan Timur | Eijkman Institute for Molecular Biology, National Research and Innovation Agency; National Institute of Health Research and Development | Edison Johar, Frilasita A Yudhaputri, Muhammad Rezki Rasyak, Willy Agustine, Hidayat Trimarsanto, Lydia V. Panggalo, Iskandar Adnan, Sukma Oktavianthi, Lidwina Priliani, Vivi Setiawaty, Hana Apsari Pawestri, Subangkit, Safarina G Malik, Khin Saw Myint, Amin Soebandrio |
| EPI_ISL_3160138 | Dinas Kesehatan Provinsi Kalimantan Timur | Eijkman Institute for Molecular Biology, National Research and Innovation Agency; National Institute of Health Research and Development | Frilasita A Yudhaputri, Muhammad Rezki Rasyak, Willy Agustine, Hidayat Trimarsanto, Lydia V. Panggalo, Iskandar Adnan, Sukma Oktavianthi, Lidwina Priliani, Edison Johar, Vivi Setiawaty, Hana Apsari Pawestri, Subangkit, Safarina G Malik, Khin Saw Myint, Amin Soebandrio |
| EPI_ISL_3160141 | Dinas Kesehatan Provinsi Kalimantan Timur | Eijkman Institute for Molecular Biology, National Research and Innovation Agency; National Institute of Health Research and Development | Muhammad Rezki Rasyak, Willy Agustine, Hidayat Trimarsanto, Lydia V. Panggalo, Iskandar Adnan, Sukma Oktavianthi, Lidwina Priliani, Edison Johar, Frilasita A Yudhaputri, Vivi Setiawaty, Hana Apsari Pawestri, Subangkit, Safarina G Malik, Khin Saw Myint, Amin Soebandrio |

| | | | |
|---|---|---|---|
| EPI_ISL_3160142, EPI_ISL_3160143 | Colorado Department of Public Health and Environment | Colorado Department of Public Health and Environment | Laura Bankers, Molly C. Hetherington-Rauth, Diana Ir, Alexandria Rossheim, Michael Martin, Mandy Waters, Shannon R. Matzinger, Sarah Elizabeth Totten, Emily A. Travanty |
| EPI_ISL_3160144 | Dinas Kesehatan Provinsi Kalimantan Timur | Eijkman Institute for Molecular Biology, National Research and Innovation Agency; National Institute of Health Research and Development | Willy Agustine, Hidayat Trimarsanto, Lydia V. Panggalo, Iskandar Adnan, Sukma Oktavianthi, Lidwina Priliani, Edison Johar, Frilasita A Yudhaputri, Muhammad Rezki Rasyak, Vivi Setiawaty, Hana Apsari Pawestri, Subangkit, Safarina G Malik, Khin Saw Myint, Amin Soebandrio |
| EPI_ISL_3160145 | Colorado Department of Public Health and Environment | Colorado Department of Public Health and Environment | Laura Bankers, Molly C. Hetherington-Rauth, Diana Ir, Alexandria Rossheim, Michael Martin, Mandy Waters, Shannon R. Matzinger, Sarah Elizabeth Totten, Emily A. Travanty |
| EPI_ISL_3160147 | Dinas Kesehatan Provinsi Kalimantan Timur | Eijkman Institute for Molecular Biology, National Research and Innovation Agency; National Institute of Health Research and Development | Hidayat Trimarsanto, Lydia V. Panggalo, Iskandar Adnan, Sukma Oktavianthi, Lidwina Priliani, Edison Johar, Frilasita A Yudhaputri, Muhammad Rezki Rasyak, Willy Agustine, Vivi Setiawaty, Hana Apsari Pawestri, Subangkit, Safarina G Malik, Khin Saw Myint, Amin Soebandrio |
| EPI_ISL_3160149 | Colorado Department of Public Health and Environment | Colorado Department of Public Health and Environment | Laura Bankers, Molly C. Hetherington-Rauth, Diana Ir, Alexandria Rossheim, Michael Martin, Mandy Waters, Shannon R. Matzinger, Sarah Elizabeth Totten, Emily A. Travanty |
| EPI_ISL_3160150 | Dinas Kesehatan Provinsi Kalimantan Timur | Eijkman Institute for Molecular Biology, National Research and Innovation Agency; National Institute of Health Research and Development | Lydia V. Panggalo, Iskandar Adnan, Sukma Oktavianthi, Lidwina Priliani, Edison Johar, Frilasita A Yudhaputri, Muhammad Rezki Rasyak, Willy Agustine, Hidayat Trimarsanto, Vivi Setiawaty, Hana Apsari Pawestri, Subangkit, Safarina G Malik, Khin Saw Myint, Amin Soebandrio |
| EPI_ISL_3160151 | Colorado Department of Public Health and Environment | Colorado Department of Public Health and Environment | Laura Bankers, Molly C. Hetherington-Rauth, Diana Ir, Alexandria Rossheim, Michael Martin, Mandy Waters, Shannon R. Matzinger, Sarah Elizabeth Totten, Emily A. Travanty |
| EPI_ISL_3160153 | Dinas Kesehatan Provinsi Kalimantan Timur | Eijkman Institute for Molecular Biology, National Research and Innovation Agency; National Institute of Health Research and Development | Iskandar Adnan, Sukma Oktavianthi, Lidwina Priliani, Edison Johar, Frilasita A Yudhaputri, Muhammad Rezki Rasyak, Willy Agustine, Hidayat Trimarsanto, Lydia V. Panggalo, Vivi Setiawaty, Hana Apsari Pawestri, Subangkit, Safarina G Malik, Khin Saw Myint, Amin Soebandrio |
| EPI_ISL_3160156 | Dinas Kesehatan Provinsi Kalimantan Timur | Eijkman Institute for Molecular Biology, National Research and Innovation Agency; National Institute of Health Research and Development | Sukma Oktavianthi, Lidwina Priliani, Edison Johar, Frilasita A Yudhaputri, Muhammad Rezki Rasyak, Willy Agustine, Hidayat Trimarsanto, Lydia V. Panggalo, Iskandar Adnan, Vivi Setiawaty, Hana Apsari Pawestri, Subangkit, Safarina G Malik, Khin Saw Myint, Amin Soebandrio |
| EPI_ISL_3160159 | Dinas Kesehatan Provinsi Kalimantan Timur | Eijkman Institute for Molecular Biology, National Research and Innovation Agency; National Institute of Health Research and Development | Lidwina Priliani, Edison Johar, Frilasita A Yudhaputri, Muhammad Rezki Rasyak, Willy Agustine, Hidayat Trimarsanto, Lydia V. Panggalo, Iskandar Adnan, Sukma Oktavianthi, Vivi Setiawaty, Hana Apsari Pawestri, Subangkit, Safarina G Malik, Khin Saw Myint, Amin Soebandrio |
| EPI_ISL_3160162 | Dinas Kesehatan Provinsi Kalimantan Timur | Eijkman Institute for Molecular Biology, National Research and Innovation Agency; National Institute of Health Research and Development | Edison Johar, Frilasita A Yudhaputri, Muhammad Rezki Rasyak, Willy Agustine, Hidayat Trimarsanto, Lydia V. Panggalo, Iskandar Adnan, Sukma Oktavianthi, Lidwina Priliani, Vivi Setiawaty, Hana Apsari Pawestri, Subangkit, Safarina G Malik, Khin Saw Myint, Amin Soebandrio |
| EPI_ISL_3160165 | Dinas Kesehatan Provinsi Kalimantan Timur | Eijkman Institute for Molecular Biology, National Research and Innovation Agency; National Institute of Health Research and Development | Frilasita A Yudhaputri, Muhammad Rezki Rasyak, Willy Agustine, Hidayat Trimarsanto, Lydia V. Panggalo, Iskandar Adnan, Sukma Oktavianthi, Lidwina Priliani, Edison Johar, Vivi Setiawaty, Hana Apsari Pawestri, Subangkit, Safarina G Malik, Khin Saw Myint, Amin Soebandrio |
| EPI_ISL_3160168 | Dinas Kesehatan Provinsi Kalimantan Timur | Eijkman Institute for Molecular Biology, National Research and Innovation Agency; National Institute of Health Research and Development | Muhammad Rezki Rasyak, Willy Agustine, Hidayat Trimarsanto, Lydia V. Panggalo, Iskandar Adnan, Sukma Oktavianthi, Lidwina Priliani, Edison Johar, Frilasita A Yudhaputri, Vivi Setiawaty, Hana Apsari Pawestri, Subangkit, Safarina G Malik, Khin Saw Myint, Amin Soebandrio |
| EPI_ISL_3160169 | Dinas Kesehatan Provinsi Kalimantan Timur | Eijkman Institute for Molecular Biology, National Research and Innovation Agency; National Institute of Health Research and Development | Willy Agustine, Hidayat Trimarsanto, Lydia V. Panggalo, Iskandar Adnan, Sukma Oktavianthi, Lidwina Priliani, Edison Johar, Frilasita A Yudhaputri, Muhammad Rezki Rasyak, Vivi Setiawaty, Hana Apsari Pawestri, Subangkit, Safarina G Malik, Khin Saw Myint, Amin Soebandrio |
| EPI_ISL_3160170 | Dinas Kesehatan Provinsi Kalimantan Timur | Eijkman Institute for Molecular Biology, National Research and Innovation Agency; National Institute of Health Research and Development | Hidayat Trimarsanto, Lydia V. Panggalo, Iskandar Adnan, Sukma Oktavianthi, Lidwina Priliani, Edison Johar, Frilasita A Yudhaputri, Muhammad Rezki Rasyak, Willy Agustine, Vivi Setiawaty, Hana Apsari Pawestri, Subangkit, Safarina G Malik, Khin Saw Myint, Amin Soebandrio |
| EPI_ISL_3160171 | Dinas Kesehatan Provinsi Kalimantan Timur | Eijkman Institute for Molecular Biology, National Research and Innovation Agency; National Institute of Health Research and Development | Lydia V. Panggalo, Iskandar Adnan, Sukma Oktavianthi, Lidwina Priliani, Edison Johar, Frilasita A Yudhaputri, Muhammad Rezki Rasyak, Willy Agustine, Hidayat Trimarsanto, Vivi Setiawaty, Hana Apsari Pawestri, Subangkit, Safarina G Malik, Khin Saw Myint, Amin Soebandrio |
| EPI_ISL_3160172 | Dinas Kesehatan Provinsi Kalimantan Timur | Eijkman Institute for Molecular Biology, National Research and Innovation Agency; National Institute of Health Research and Development | Iskandar Adnan, Sukma Oktavianthi, Lidwina Priliani, Edison Johar, Frilasita A Yudhaputri, Muhammad Rezki Rasyak, Willy Agustine, Hidayat Trimarsanto, Lydia V. Panggalo, Vivi Setiawaty, Hana Apsari Pawestri, Subangkit, Safarina G Malik, Khin Saw Myint, Amin Soebandrio |
| EPI_ISL_3160173 | Dinas Kesehatan Provinsi Kalimantan Timur | Eijkman Institute for Molecular Biology, National Research and Innovation Agency; National Institute of Health Research and Development | Sukma Oktavianthi, Lidwina Priliani, Edison Johar, Frilasita A Yudhaputri, Muhammad Rezki Rasyak, Willy Agustine, Hidayat Trimarsanto, Lydia V. Panggalo, Iskandar Adnan, Vivi Setiawaty, Hana Apsari Pawestri, Subangkit, Safarina G Malik, Khin Saw Myint, Amin Soebandrio |
| EPI_ISL_3160174 | Dinas Kesehatan Provinsi Kalimantan Timur | Eijkman Institute for Molecular Biology, National Research and Innovation Agency; National Institute of Health Research and Development | Lidwina Priliani, Edison Johar, Frilasita A Yudhaputri, Muhammad Rezki Rasyak, Willy Agustine, Hidayat Trimarsanto, Lydia V. Panggalo, Iskandar Adnan, Sukma Oktavianthi, Vivi Setiawaty, Hana Apsari Pawestri, Subangkit, Safarina G Malik, Khin Saw Myint, Amin Soebandrio |
| EPI_ISL_3160175 | Dinas Kesehatan Provinsi Kalimantan Timur | Eijkman Institute for Molecular Biology, National Research and Innovation Agency; National Institute of Health Research and Development | Edison Johar, Frilasita A Yudhaputri, Muhammad Rezki Rasyak, Willy Agustine, Hidayat Trimarsanto, Lydia V. Panggalo, Iskandar Adnan, Sukma Oktavianthi, Lidwina Priliani, Vivi Setiawaty, Hana Apsari Pawestri, Subangkit, Safarina G Malik, Khin Saw Myint, Amin Soebandrio |
| EPI_ISL_3160176 | Dinas Kesehatan Provinsi Kalimantan Timur | Eijkman Institute for Molecular Biology, National Research and Innovation Agency; National Institute of Health Research and Development | Frilasita A Yudhaputri, Muhammad Rezki Rasyak, Willy Agustine, Hidayat Trimarsanto, Lydia V. Panggalo, Iskandar Adnan, Sukma Oktavianthi, Lidwina Priliani, Edison Johar, Vivi Setiawaty, Hana Apsari Pawestri, Subangkit, Safarina G Malik, Khin Saw Myint, Amin Soebandrio |
| EPI_ISL_3160177 | Dinas Kesehatan Provinsi Kalimantan Timur | Eijkman Institute for Molecular Biology, National Research and Innovation Agency; National Institute of Health Research and Development | Muhammad Rezki Rasyak, Willy Agustine, Hidayat Trimarsanto, Lydia V. Panggalo, Iskandar Adnan, Sukma Oktavianthi, Lidwina Priliani, Edison Johar, Frilasita A Yudhaputri, Vivi Setiawaty, Hana Apsari Pawestri, Subangkit, Safarina G Malik, Khin Saw Myint, Amin Soebandrio |
| EPI_ISL_3160178 | Dinas Kesehatan Provinsi Kalimantan Timur | Eijkman Institute for Molecular Biology, National Research and Innovation Agency; National Institute of Health Research and Development | Hidayat Trimarsanto, Lydia V. Panggalo, Iskandar Adnan, Sukma Oktavianthi, Lidwina Priliani, Edison Johar, Frilasita A Yudhaputri, Muhammad Rezki Rasyak, Willy Agustine, Vivi Setiawaty, Hana Apsari Pawestri, Subangkit, Safarina G Malik, Khin Saw Myint, Amin Soebandrio |
| EPI_ISL_3160179 | Dinas Kesehatan Provinsi Kalimantan Timur | Eijkman Institute for Molecular Biology, National Research and Innovation Agency; National Institute of Health Research and Development | Lydia V. Panggalo, Iskandar Adnan, Sukma Oktavianthi, Lidwina Priliani, Edison Johar, Frilasita A Yudhaputri, Muhammad Rezki Rasyak, Willy Agustine, Hidayat Trimarsanto, Vivi Setiawaty, Hana Apsari Pawestri, Subangkit, Safarina G Malik, Khin Saw Myint, Amin Soebandrio |
| EPI_ISL_3160180 | Dinas Kesehatan Provinsi Kalimantan Timur | Eijkman Institute for Molecular Biology, National Research and Innovation Agency; National Institute of Health Research and Development | Iskandar Adnan, Sukma Oktavianthi, Lidwina Priliani, Edison Johar, Frilasita A Yudhaputri, Muhammad Rezki Rasyak, Willy Agustine, Hidayat Trimarsanto, Lydia V. Panggalo, Vivi Setiawaty, Hana Apsari Pawestri, Subangkit, Safarina G Malik, Khin Saw Myint, Amin Soebandrio |
| EPI_ISL_3160181 | Dinas Kesehatan Provinsi Kalimantan Timur | Eijkman Institute for Molecular Biology, National Research and Innovation Agency; National Institute of Health Research and Development | Sukma Oktavianthi, Lidwina Priliani, Edison Johar, Frilasita A Yudhaputri, Muhammad Rezki Rasyak, Willy Agustine, Hidayat Trimarsanto, Lydia V. Panggalo, Iskandar Adnan, Vivi Setiawaty, Hana Apsari Pawestri, Subangkit, Safarina G Malik, Khin Saw Myint, Amin Soebandrio |
| EPI_ISL_3160182 | Dinas Kesehatan Provinsi Kalimantan Timur | Eijkman Institute for Molecular Biology, National Research and Innovation Agency; National Institute of Health Research and Development | Lidwina Priliani, Edison Johar, Frilasita A Yudhaputri, Muhammad Rezki Rasyak, Willy Agustine, Hidayat Trimarsanto, Lydia V. Panggalo, Iskandar Adnan, Sukma Oktavianthi, Vivi Setiawaty, Hana Apsari Pawestri, Subangkit, Safarina G Malik, Khin Saw Myint, Amin Soebandrio |
| EPI_ISL_3160183 | Dinas Kesehatan Provinsi Kalimantan Timur | Eijkman Institute for Molecular Biology, National Research and Innovation Agency; National Institute of Health Research and Development | Edison Johar, Frilasita A Yudhaputri, Muhammad Rezki Rasyak, Willy Agustine, Hidayat Trimarsanto, Lydia V. Panggalo, Iskandar Adnan, Sukma Oktavianthi, Lidwina Priliani, Vivi Setiawaty, Hana Apsari Pawestri, Subangkit, Safarina G Malik, Khin Saw Myint, Amin Soebandrio |
| EPI_ISL_3160184 | Dinas Kesehatan Provinsi Kalimantan Timur | Eijkman Institute for Molecular Biology, National Research and Innovation Agency; National Institute of Health Research and Development | Frilasita A Yudhaputri, Muhammad Rezki Rasyak, Willy Agustine, Hidayat Trimarsanto, Lydia V. Panggalo, Iskandar Adnan, Sukma Oktavianthi, Lidwina Priliani, Edison Johar, Vivi Setiawaty, Hana Apsari Pawestri, Subangkit, Safarina G Malik, Khin Saw Myint, Amin Soebandrio |
| EPI_ISL_3160185, EPI_ISL_3160186 | Humboldt County Public Health Laboratory | Humboldt County Public Health Laboratory | Jeremy Corrigan |
| EPI_ISL_3160187 | Dinas Kesehatan Provinsi Kalimantan Timur | Eijkman Institute for Molecular Biology, National Research and Innovation Agency; National Institute of Health Research and Development | Muhammad Rezki Rasyak, Willy Agustine, Hidayat Trimarsanto, Lydia V. Panggalo, Iskandar Adnan, Sukma Oktavianthi, Lidwina Priliani, Edison Johar, Frilasita A Yudhaputri, Vivi Setiawaty, Hana Apsari Pawestri, Subangkit, Safarina G Malik, Khin Saw Myint, Amin Soebandrio |
| EPI_ISL_3160190 | Dinas Kesehatan Provinsi Kalimantan Timur | Eijkman Institute for Molecular Biology, National Research and Innovation Agency; National Institute of Health Research and Development | Willy Agustine, Hidayat Trimarsanto, Lydia V. Panggalo, Iskandar Adnan, Sukma Oktavianthi, Lidwina Priliani, Edison Johar, Frilasita A Yudhaputri, Muhammad Rezki Rasyak, Vivi Setiawaty, Hana Apsari Pawestri, Subangkit, Safarina G Malik, Khin Saw Myint, Amin Soebandrio |
| EPI_ISL_3160191, EPI_ISL_3160192 | Humboldt County Public Health Laboratory | Humboldt County Public Health Laboratory | Jeremy Corrigan |
| EPI_ISL_3160194 | Dinas Kesehatan Provinsi Kalimantan Timur | Eijkman Institute for Molecular Biology, National Research and Innovation Agency; National Institute of Health Research and Development | Hidayat Trimarsanto, Lydia V. Panggalo, Iskandar Adnan, Sukma Oktavianthi, Lidwina Priliani, Edison Johar, Frilasita A Yudhaputri, Muhammad Rezki Rasyak, Willy Agustine, Vivi Setiawaty, Hana Apsari Pawestri, Subangkit, Safarina G Malik, Khin Saw Myint, Amin Soebandrio |
| EPI_ISL_3160195 | Humboldt County Public Health Laboratory | Humboldt County Public Health Laboratory | Jeremy Corrigan |

| | | | |
|---|---|---|---|
| EPI_ISL_3160197 | Dinas Kesehatan Provinsi Kalimantan Timur | Eijkman Institute for Molecular Biology, National Research and Innovation Agency; National Institute of Health Research and Development | Lydia V. Panggalo, Iskandar Adnan, Sukma Oktavianthi, Lidwina Priliani, Edison Johar, Frilasita A Yudhaputri, Muhammad Rezki Rasyak, Willy Agustine, Hidayat Trimarsanto, Vivi Setiawaty, Hana Apsari Pawestri, Subangkit, Safarina G Malik, Khin Saw Myint, Amin Soebandrio |
| EPI_ISL_3160200 | Dinas Kesehatan Provinsi Kalimantan Timur | Eijkman Institute for Molecular Biology, National Research and Innovation Agency; National Institute of Health Research and Development | Iskandar Adnan, Sukma Oktavianthi, Lidwina Priliani, Edison Johar, Frilasita A Yudhaputri, Muhammad Rezki Rasyak, Willy Agustine, Hidayat Trimarsanto, Lydia V. Panggalo, Vivi Setiawaty, Hana Apsari Pawestri, Subangkit, Safarina G Malik, Khin Saw Myint, Amin Soebandrio |
| EPI_ISL_3160202 | Humboldt County Public Health Laboratory | Humboldt County Public Health Laboratory | Jeremy Corrigan |
| EPI_ISL_3160203 | Dinas Kesehatan Provinsi Kalimantan Timur | Eijkman Institute for Molecular Biology, National Research and Innovation Agency; National Institute of Health Research and Development | Sukma Oktavianthi, Lidwina Priliani, Edison Johar, Frilasita A Yudhaputri, Muhammad Rezki Rasyak, Willy Agustine, Hidayat Trimarsanto, Lydia V. Panggalo, Iskandar Adnan, Vivi Setiawaty, Hana Apsari Pawestri, Subangkit, Safarina G Malik, Khin Saw Myint, Amin Soebandrio |
| EPI_ISL_3160207, EPI_ISL_3160208, EPI_ISL_3160210, EPI_ISL_3160211 | Humboldt County Public Health Laboratory | Humboldt County Public Health Laboratory | Jeremy Corrigan |
| EPI_ISL_3160254, EPI_ISL_3160256, EPI_ISL_3160257, EPI_ISL_3160258, EPI_ISL_3160260, EPI_ISL_3160261, EPI_ISL_3160262, EPI_ISL_3160264, EPI_ISL_3160265, EPI_ISL_3160266, EPI_ISL_3160278, EPI_ISL_3160279, EPI_ISL_3160280, EPI_ISL_3160282, EPI_ISL_3160283, EPI_ISL_3160285, EPI_ISL_3160286, EPI_ISL_3160288, EPI_ISL_3160289, EPI_ISL_3160290, EPI_ISL_3160292, EPI_ISL_3160293, EPI_ISL_3160295, EPI_ISL_3160296, EPI_ISL_3160298, EPI_ISL_3160299, EPI_ISL_3160301, EPI_ISL_3160302, EPI_ISL_3160304, EPI_ISL_3160305, EPI_ISL_3160306, EPI_ISL_3160308, EPI_ISL_3160309, EPI_ISL_3160311, EPI_ISL_3160312, EPI_ISL_3160313, EPI_ISL_3160315, EPI_ISL_3160316, EPI_ISL_3160318, EPI_ISL_3160319, EPI_ISL_3160321, EPI_ISL_3160322, EPI_ISL_3160323, EPI_ISL_3160325, EPI_ISL_3160326, EPI_ISL_3160328, EPI_ISL_3160329, EPI_ISL_3160331, EPI_ISL_3160332, EPI_ISL_3160334, EPI_ISL_3160335, EPI_ISL_3160337, EPI_ISL_3160338, EPI_ISL_3160340, EPI_ISL_3160341, EPI_ISL_3160342, EPI_ISL_3160344, EPI_ISL_3160345, EPI_ISL_3160347, EPI_ISL_3160348, EPI_ISL_3160350, EPI_ISL_3160351, EPI_ISL_3160352, EPI_ISL_3160354, EPI_ISL_3160355, EPI_ISL_3160357, EPI_ISL_3160358, EPI_ISL_3160360, EPI_ISL_3160361, EPI_ISL_3160363, EPI_ISL_3160364 | | | |
| see above | Pathology and Laboratory Medicine Institute, Cleveland Clinic, Ohio, USA | Pathology and Laboratory Medicine Institute, Cleveland Clinic, Ohio, USA | Jessica Spildener, Concetta Peck, Jennifer Starbuck, Joy Nakitandwe, Kristen McDonnell, David Plunkett, Zheng Jin Tu, Jay Brock, Yu-Wei Cheng, Gary Procop, Daniel Rhoads, Daniel H. Farkas, David Bosler |
| EPI_ISL_3160399 | RSUPN Dr. Cipto Mangunkusumo | Eijkman Institute for Molecular Biology, National Research and Innovation Agency | Iskandar Adnan, Sukma Oktavianthi, Lidwina Priliani, Edison Johar, Frilasita A Yudhaputri, Muhammad Rezki Rasyak, Willy Agustine, Hidayat Trimarsanto, Lydia V. Panggalo, Safarina G Malik, Khin Saw Myint, Amin Soebandrio |
| EPI_ISL_3160400, EPI_ISL_3160401, EPI_ISL_3160402, EPI_ISL_3160403, EPI_ISL_3160404, EPI_ISL_3160405, EPI_ISL_3160406, EPI_ISL_3160407, EPI_ISL_3160408, EPI_ISL_3160409 | Dianalabs SA | Genesupport | Tanguy ARAUD, Geraldine Jost, Katia Jaton, Nadia Liassine |
| EPI_ISL_3160410 | Proxilab SA | Genesupport | Tanguy ARAUD, Geraldine Jost, Katia Jaton, Nadia Liassine |
| EPI_ISL_3160411 | Dianalabs-Valais SA | Genesupport | Tanguy ARAUD, Geraldine Jost, Katia Jaton, Nadia Liassine |
| EPI_ISL_3160412, EPI_ISL_3160413, EPI_ISL_3160414, EPI_ISL_3160415 | Dianalabs SA | Genesupport | Tanguy ARAUD, Geraldine Jost, Katia Jaton, Nadia Liassine |
| EPI_ISL_3160416 | Proxilab SA | Genesupport | Tanguy ARAUD, Geraldine Jost, Katia Jaton, Nadia Liassine |
| EPI_ISL_3160417, EPI_ISL_3160418, EPI_ISL_3160419, EPI_ISL_3160420, EPI_ISL_3160421, EPI_ISL_3160422, EPI_ISL_3160423, EPI_ISL_3160424, EPI_ISL_3160425, EPI_ISL_3160426, EPI_ISL_3160427, EPI_ISL_3160428, EPI_ISL_3160429, EPI_ISL_3160432, EPI_ISL_3160433, EPI_ISL_3160434, EPI_ISL_3160435, EPI_ISL_3160436, EPI_ISL_3160437, EPI_ISL_3160438, EPI_ISL_3160439, EPI_ISL_3160440, EPI_ISL_3160441, EPI_ISL_3160442, EPI_ISL_3160443, EPI_ISL_3160444, EPI_ISL_3160445, EPI_ISL_3160446, EPI_ISL_3160447, EPI_ISL_3160448, EPI_ISL_3160449, EPI_ISL_3160450, EPI_ISL_3160451, EPI_ISL_3160452, EPI_ISL_3160453, EPI_ISL_3160454, EPI_ISL_3160455, EPI_ISL_3160456, EPI_ISL_3160457, EPI_ISL_3160458, EPI_ISL_3160459, EPI_ISL_3160460, EPI_ISL_3160461, EPI_ISL_3160462, EPI_ISL_3160463, EPI_ISL_3160464, EPI_ISL_3160465, EPI_ISL_3160466, EPI_ISL_3160467, EPI_ISL_3160468, EPI_ISL_3160469, EPI_ISL_3160470, EPI_ISL_3160471, EPI_ISL_3160472, EPI_ISL_3160473, EPI_ISL_3160474, EPI_ISL_3160475, EPI_ISL_3160476, EPI_ISL_3160477, EPI_ISL_3160478, EPI_ISL_3160479, EPI_ISL_3160480, EPI_ISL_3160481, EPI_ISL_3160482, EPI_ISL_3160483, EPI_ISL_3160484, EPI_ISL_3160485, EPI_ISL_3160486, EPI_ISL_3160487, EPI_ISL_3160488, EPI_ISL_3160489, EPI_ISL_3160490, EPI_ISL_3160491, EPI_ISL_3160492, EPI_ISL_3160493, EPI_ISL_3160494, EPI_ISL_3160495, EPI_ISL_3160496, EPI_ISL_3160497, EPI_ISL_3160498, EPI_ISL_3160499, EPI_ISL_3160500, EPI_ISL_3160501, EPI_ISL_3160502, EPI_ISL_3160503, EPI_ISL_3160504, EPI_ISL_3160505, EPI_ISL_3160506, EPI_ISL_3160507, EPI_ISL_3160508, EPI_ISL_3160509, EPI_ISL_3160510, EPI_ISL_3160511, EPI_ISL_3160512, EPI_ISL_3160513, EPI_ISL_3160514, EPI_ISL_3160515, EPI_ISL_3160516, EPI_ISL_3160517, EPI_ISL_3160518, EPI_ISL_3160519 | | | |
| see above | Dianalabs SA | Genesupport | Tanguy ARAUD, Geraldine Jost, Katia Jaton, Nadia Liassine |
| EPI_ISL_3160520, EPI_ISL_3160521, EPI_ISL_3160522, EPI_ISL_3160523, EPI_ISL_3160524, EPI_ISL_3160525, EPI_ISL_3160526 | Polyanalytic SA | Genesupport | Tanguy ARAUD, Geraldine Jost, Katia Jaton, Nadia Liassine |
| EPI_ISL_3160527, EPI_ISL_3160528, EPI_ISL_3160529, EPI_ISL_3160530, EPI_ISL_3160531, EPI_ISL_3160532, EPI_ISL_3160533 | Dianalabs-Valais SA | Genesupport | Tanguy ARAUD, Geraldine Jost, Katia Jaton, Nadia Liassine |
| EPI_ISL_3160534, EPI_ISL_3160535, EPI_ISL_3160536, EPI_ISL_3160537, EPI_ISL_3160538, EPI_ISL_3160539, EPI_ISL_3160540, EPI_ISL_3160541, EPI_ISL_3160542, EPI_ISL_3160543, EPI_ISL_3160544, EPI_ISL_3160545, EPI_ISL_3160546, EPI_ISL_3160547, EPI_ISL_3160548, EPI_ISL_3160549, EPI_ISL_3160550, EPI_ISL_3160551, EPI_ISL_3160552, EPI_ISL_3160553, EPI_ISL_3160554, EPI_ISL_3160555, EPI_ISL_3160556, EPI_ISL_3160557, EPI_ISL_3160558, EPI_ISL_3160559, EPI_ISL_3160560, EPI_ISL_3160561, EPI_ISL_3160562, EPI_ISL_3160563, EPI_ISL_3160564, EPI_ISL_3160565, EPI_ISL_3160566, EPI_ISL_3160567, EPI_ISL_3160568, EPI_ISL_3160569 | | | |
| see above | Dianalabs SA | Genesupport | Tanguy ARAUD, Geraldine Jost, Katia Jaton, Nadia Liassine |
| EPI_ISL_3160570, EPI_ISL_3160571, EPI_ISL_3160572, EPI_ISL_3160573, EPI_ISL_3160574, EPI_ISL_3160575, EPI_ISL_3160576 | Proxilab SA | Genesupport | Tanguy ARAUD, Geraldine Jost, Katia Jaton, Nadia Liassine |
| EPI_ISL_3160577, EPI_ISL_3160578, EPI_ISL_3160579, EPI_ISL_3160580, EPI_ISL_3160581, EPI_ISL_3160582, EPI_ISL_3160583, EPI_ISL_3160584, EPI_ISL_3160585, EPI_ISL_3160586, EPI_ISL_3160587, EPI_ISL_3160588, EPI_ISL_3160589, EPI_ISL_3160590, EPI_ISL_3160591, EPI_ISL_3160592, EPI_ISL_3160598, EPI_ISL_3160599, EPI_ISL_3160601, EPI_ISL_3160602, EPI_ISL_3160603, EPI_ISL_3160604, EPI_ISL_3160605, EPI_ISL_3160606, EPI_ISL_3160608, EPI_ISL_3160609, EPI_ISL_3160610, EPI_ISL_3160611, EPI_ISL_3160612, EPI_ISL_3160614, EPI_ISL_3160615, EPI_ISL_3160616, EPI_ISL_3160617, EPI_ISL_3160618, EPI_ISL_3160619, EPI_ISL_3160620, EPI_ISL_3160621, EPI_ISL_3160622, EPI_ISL_3160623, EPI_ISL_3160625, EPI_ISL_3160626, EPI_ISL_3160627, EPI_ISL_3160628, EPI_ISL_3160629, EPI_ISL_3160631 | | | |
| see above | Wyoming Public Health Laboratory | Wyoming Public Health Laboratory | Jim Mildenberger, Taylor Fearing, Channing Weber, Ashley Norberg, Chayse Rowley, Marley Goetz, Brian Dominguez, Elliot Thomasson, Sam Britz, Cari Sloma, Robert Petit, and Rob Christensen |
| EPI_ISL_3160632, EPI_ISL_3160633 | Molecular Diagnostics Laboratory | National Reference Laboratory, Nigeria Centre for Disease Control | Dr Ndodo Nnaemeka, Olusola Anuoluwapo Akanbi, Chimaobi Chukwu, Dr Omoare Adesuyi, Celestina Obiekea, Anthony Ahumibe, Catherine Okoi, Nwando Mba, Dr Chikwe Ihekweazu |
| EPI_ISL_3160634, EPI_ISL_3160635, EPI_ISL_3160637, EPI_ISL_3160638, EPI_ISL_3160639, EPI_ISL_3160640, EPI_ISL_3160641, EPI_ISL_3160642, EPI_ISL_3160643, EPI_ISL_3160644, EPI_ISL_3160645, EPI_ISL_3160646, EPI_ISL_3160647, EPI_ISL_3160650, EPI_ISL_3160651, EPI_ISL_3160653, EPI_ISL_3160654, EPI_ISL_3160656, EPI_ISL_3160658, EPI_ISL_3160659, EPI_ISL_3160660, EPI_ISL_3160661, EPI_ISL_3160662, EPI_ISL_3160663, EPI_ISL_3160664, EPI_ISL_3160665, EPI_ISL_3160667, EPI_ISL_3160668, EPI_ISL_3160669, EPI_ISL_3160670, EPI_ISL_3160671, EPI_ISL_3160672, EPI_ISL_3160673, EPI_ISL_3160674, EPI_ISL_3160675, EPI_ISL_3160677, EPI_ISL_3160678, EPI_ISL_3160679, EPI_ISL_3160680, EPI_ISL_3160681, EPI_ISL_3160682, EPI_ISL_3160683, EPI_ISL_3160688, EPI_ISL_3160690, EPI_ISL_3160691, EPI_ISL_3160693, EPI_ISL_3160695, EPI_ISL_3160696, EPI_ISL_3160697, EPI_ISL_3160698, EPI_ISL_3160699, EPI_ISL_3160700, EPI_ISL_3160702, EPI_ISL_3160703, EPI_ISL_3160704, EPI_ISL_3160705, EPI_ISL_3160706, EPI_ISL_3160707, EPI_ISL_3160709, EPI_ISL_3160710, EPI_ISL_3160711, EPI_ISL_3160712, EPI_ISL_3160713, EPI_ISL_3160714, EPI_ISL_3160717, EPI_ISL_3160719, EPI_ISL_3160720, EPI_ISL_3160721, EPI_ISL_3160722, EPI_ISL_3160723, EPI_ISL_3160724 | | | |
| see above | Salud Digna | Instituto Nacional de Medicina Genomica | Cedro-Tanda A, Escobar-Arrazola MA, Ramirez-Vega O, Abraham Campos-Romero, Moreno-Camacho José Luis, Rodriguez-Gallegos Jorge, Luna-Ruiz Marco, Rangel-DeLeon D, Munguia-Garza P, Hidalgo-Miranda A, Mendoza-Vargas A, Reyes-Grajeda JP, Herrera-Montalvo LA. |
| EPI_ISL_3160725, EPI_ISL_3160726, EPI_ISL_3160727, EPI_ISL_3160728 | INCMNSZ | Instituto Nacional de Medicina Genomica | Cedro-Tanda A, Escobar-Arrazola MA, Ramirez-Vega O, Rangel-DeLeon D, Munguia-Garza P, Hidalgo-Miranda A, Mendoza-Vargas A, Reyes-Grajeda JP, Herrera-Montalvo LA. |
| EPI_ISL_3160729, EPI_ISL_3160730, EPI_ISL_3160731, EPI_ISL_3160732, EPI_ISL_3160733, EPI_ISL_3160734, EPI_ISL_3160735, EPI_ISL_3160736, EPI_ISL_3160737, EPI_ISL_3160738, EPI_ISL_3160739, EPI_ISL_3160740, EPI_ISL_3160741, EPI_ISL_3160742, EPI_ISL_3160743, EPI_ISL_3160744, EPI_ISL_3160745, EPI_ISL_3160746, EPI_ISL_3160747, EPI_ISL_3160748, EPI_ISL_3160749, EPI_ISL_3160750, EPI_ISL_3160751, EPI_ISL_3160752, EPI_ISL_3160753, EPI_ISL_3160754 | | | |
| see above | Instituto Nacional de Medicina Genomica | Instituto Nacional de Medicina Genomica | Cedro-Tanda A, Escobar-Arrazola MA, Ramirez-Vega O, Rangel-DeLeon D, Munguia-Garza P, Hidalgo-Miranda A, Mendoza-Vargas A, Reyes-Grajeda JP, Herrera-Montalvo LA. |
| EPI_ISL_3160755, EPI_ISL_3160756, EPI_ISL_3160757, EPI_ISL_3160758, EPI_ISL_3160759, EPI_ISL_3160760, EPI_ISL_3160761, EPI_ISL_3160762, EPI_ISL_3160763, EPI_ISL_3160764, EPI_ISL_3160765 | | | |
| see above | INCMNSZ | Instituto Nacional de Medicina Genomica | Cedro-Tanda A, Escobar-Arrazola MA, Ramirez-Vega O, Rangel-DeLeon D, Munguia-Garza P, Hidalgo-Miranda A, Mendoza-Vargas A, Reyes-Grajeda JP, Herrera-Montalvo LA. |
| EPI_ISL_3160766, EPI_ISL_3160767, EPI_ISL_3160768, EPI_ISL_3160769, EPI_ISL_3160771, EPI_ISL_3160772, EPI_ISL_3160773, EPI_ISL_3160774, EPI_ISL_3160775, EPI_ISL_3160776, EPI_ISL_3160777, EPI_ISL_3160778, EPI_ISL_3160779, EPI_ISL_3160780, EPI_ISL_3160781, EPI_ISL_3160782, EPI_ISL_3160783 | | | |
| see above | Instituto Nacional de Medicina Genomica | Instituto Nacional de Medicina Genomica | Cedro-Tanda A, Escobar-Arrazola MA, Ramirez-Vega O, Rangel-DeLeon D, Munguia-Garza P, Hidalgo-Miranda A, Mendoza-Vargas A, Reyes-Grajeda JP, Herrera-Montalvo LA. |
| EPI_ISL_3160784, EPI_ISL_3160785, EPI_ISL_3160786, | INCMNSZ | Instituto Nacional de Medicina Genomica | Cedro-Tanda A, Escobar-Arrazola MA, Ramirez-Vega O, Rangel-DeLeon D, Munguia-Garza P, Hidalgo-Miranda A, Mendoza-Vargas A, Reyes-Grajeda JP, Herrera-Montalvo LA. |

| | | | |
|---|---|---|---|
| EPI_ISL_3160787, EPI_ISL_3160788, EPI_ISL_3160789, EPI_ISL_3160790, EPI_ISL_3160791, EPI_ISL_3160792 | | | |
| EPI_ISL_3160794, EPI_ISL_3160795, EPI_ISL_3160796, EPI_ISL_3160797, EPI_ISL_3160798, EPI_ISL_3160799, EPI_ISL_3160800, EPI_ISL_3160801, EPI_ISL_3160802, EPI_ISL_3160803, EPI_ISL_3160804, EPI_ISL_3160805, EPI_ISL_3160806, EPI_ISL_3160807, EPI_ISL_3160808, EPI_ISL_3160809, EPI_ISL_3160810, EPI_ISL_3160811, EPI_ISL_3160812 | | | |
| see above | Instituto Nacional de Medicina Genomica | Instituto Nacional de Medicina Genomica | Cedro-Tanda A, Escobar-Arrazola MA, Ramirez-Vega O, Rangel-DeLeon D, Munguia-Garza P, Hidalgo-Miranda A, Mendoza-Vargas A, Reyes-Grajeda JP, Herrera-Montalvo LA. |
| EPI_ISL_3160814, EPI_ISL_3160817, EPI_ISL_3160819, EPI_ISL_3160820, EPI_ISL_3160821, EPI_ISL_3160822, EPI_ISL_3160823, EPI_ISL_3160824, EPI_ISL_3160826, EPI_ISL_3160828, EPI_ISL_3160829, EPI_ISL_3160830, EPI_ISL_3160831, EPI_ISL_3160832, EPI_ISL_3160833, EPI_ISL_3160834, EPI_ISL_3160835, EPI_ISL_3160836, EPI_ISL_3160837, EPI_ISL_3160838, EPI_ISL_3160839, EPI_ISL_3160840, EPI_ISL_3160841, EPI_ISL_3160842, EPI_ISL_3160843, EPI_ISL_3160844, EPI_ISL_3160845, EPI_ISL_3160846, EPI_ISL_3160848, EPI_ISL_3160849, EPI_ISL_3160850, EPI_ISL_3160852, EPI_ISL_3160853, EPI_ISL_3160854, EPI_ISL_3160855, EPI_ISL_3160856, EPI_ISL_3160857, EPI_ISL_3160858, EPI_ISL_3160859, EPI_ISL_3160860, EPI_ISL_3160861, EPI_ISL_3160862, EPI_ISL_3160864, EPI_ISL_3160865, EPI_ISL_3160866 | | | |
| see above | Salud Digna | Instituto Nacional de Medicina Genomica | Cedro-Tanda A, Escobar-Arrazola MA, Ramirez-Vega O, Abraham Campos-Romero, Moreno-Camacho José Luis, Rodriguez-Gallegos Jorge, Luna-Ruiz Marco, Rangel-DeLeon D, Munguia-Garza P, Hidalgo-Miranda A, Mendoza-Vargas A, Reyes-Grajeda JP, Herrera-Montalvo LA. |
| EPI_ISL_3160867, EPI_ISL_3160868, EPI_ISL_3160869, EPI_ISL_3160870, EPI_ISL_3160871, EPI_ISL_3160872, EPI_ISL_3160873, EPI_ISL_3160874, EPI_ISL_3160875, EPI_ISL_3160876, EPI_ISL_3160877, EPI_ISL_3160879, EPI_ISL_3160880 | | | |
| see above | INCMNSZ | Instituto Nacional de Medicina Genomica | Cedro-Tanda A, Escobar-Arrazola MA, Ramirez-Vega O, Rangel-DeLeon D, Munguia-Garza P, Hidalgo-Miranda A, Mendoza-Vargas A, Reyes-Grajeda JP, Herrera-Montalvo LA. |
| EPI_ISL_3160881, EPI_ISL_3160882, EPI_ISL_3160883, EPI_ISL_3160884, EPI_ISL_3160885, EPI_ISL_3160886, EPI_ISL_3160887, EPI_ISL_3160888, EPI_ISL_3160889, EPI_ISL_3160890, EPI_ISL_3160891, EPI_ISL_3160892 | | | |
| see above | Instituto Nacional de Medicina Genomica | Instituto Nacional de Medicina Genomica | Cedro-Tanda A, Escobar-Arrazola MA, Ramirez-Vega O, Rangel-DeLeon D, Munguia-Garza P, Hidalgo-Miranda A, Mendoza-Vargas A, Reyes-Grajeda JP, Herrera-Montalvo LA. |
| EPI_ISL_3160893, EPI_ISL_3160894 | INCan | Instituto Nacional de Medicina Genomica | Cedro-Tanda A, Escobar-Arrazola MA, Ramirez-Vega O, Rangel-DeLeon D, Munguia-Garza P, Hidalgo-Miranda A, Mendoza-Vargas A, Reyes-Grajeda JP, Herrera-Montalvo LA. |
| EPI_ISL_3160895, EPI_ISL_3160898, EPI_ISL_3160899, EPI_ISL_3160900, EPI_ISL_3160901, EPI_ISL_3160902, EPI_ISL_3160903, EPI_ISL_3160904, EPI_ISL_3160905, EPI_ISL_3160906, EPI_ISL_3160907, EPI_ISL_3160909, EPI_ISL_3160910 | | | |
| see above | INCMNSZ | Instituto Nacional de Medicina Genomica | Cedro-Tanda A, Escobar-Arrazola MA, Ramirez-Vega O, Rangel-DeLeon D, Munguia-Garza P, Hidalgo-Miranda A, Mendoza-Vargas A, Reyes-Grajeda JP, Herrera-Montalvo LA. |
| EPI_ISL_3160912, EPI_ISL_3160914, EPI_ISL_3160915, EPI_ISL_3160916, EPI_ISL_3160917, EPI_ISL_3160918, EPI_ISL_3160919, EPI_ISL_3160920, EPI_ISL_3160921, EPI_ISL_3160922 | Instituto Nacional de Medicina Genomica | Instituto Nacional de Medicina Genomica | Cedro-Tanda A, Escobar-Arrazola MA, Ramirez-Vega O, Rangel-DeLeon D, Munguia-Garza P, Hidalgo-Miranda A, Mendoza-Vargas A, Reyes-Grajeda JP, Herrera-Montalvo LA. |
| EPI_ISL_3160925, EPI_ISL_3160926 | Salud Digna | Instituto Nacional de Medicina Genomica | Cedro-Tanda A, Escobar-Arrazola MA, Ramirez-Vega O, Abraham Campos-Romero, Moreno-Camacho José Luis, Rodriguez-Gallegos Jorge, Luna-Ruiz Marco, Rangel-DeLeon D, Munguia-Garza P, Hidalgo-Miranda A, Mendoza-Vargas A, Reyes-Grajeda JP, Herrera-Montalvo LA. |
| EPI_ISL_3160928, EPI_ISL_3160929, EPI_ISL_3160930, EPI_ISL_3160931 | Instituto Nacional de Medicina Genomica | Instituto Nacional de Medicina Genomica | Cedro-Tanda A, Escobar-Arrazola MA, Ramirez-Vega O, Rangel-DeLeon D, Munguia-Garza P, Hidalgo-Miranda A, Mendoza-Vargas A, Reyes-Grajeda JP, Herrera-Montalvo LA. |
| EPI_ISL_3160932, EPI_ISL_3160933, EPI_ISL_3160934, EPI_ISL_3160935, EPI_ISL_3160936, EPI_ISL_3160939, EPI_ISL_3160940, EPI_ISL_3160941, EPI_ISL_3160943, EPI_ISL_3160945, EPI_ISL_3160946, EPI_ISL_3160947, EPI_ISL_3160948, EPI_ISL_3160949, EPI_ISL_3160953, EPI_ISL_3160958, EPI_ISL_3160959, EPI_ISL_3160960, EPI_ISL_3160961, EPI_ISL_3160962, EPI_ISL_3160963, EPI_ISL_3160965, EPI_ISL_3160967, EPI_ISL_3160971, EPI_ISL_3160972, EPI_ISL_3160974, EPI_ISL_3160976, EPI_ISL_3160977, EPI_ISL_3160978 | | | |
| see above | Salud Digna | Instituto Nacional de Medicina Genomica | Cedro-Tanda A, Escobar-Arrazola MA, Ramirez-Vega O, Abraham Campos-Romero, Moreno-Camacho José Luis, Rodriguez-Gallegos Jorge, Luna-Ruiz Marco, Rangel-DeLeon D, Munguia-Garza P, Hidalgo-Miranda A, Mendoza-Vargas A, Reyes-Grajeda JP, Herrera-Montalvo LA. |
| EPI_ISL_3160979, EPI_ISL_3160980, EPI_ISL_3160981, EPI_ISL_3160982, EPI_ISL_3160983, EPI_ISL_3160984, EPI_ISL_3160985, EPI_ISL_3160986, EPI_ISL_3160987, EPI_ISL_3160989, EPI_ISL_3160990, EPI_ISL_3160991, EPI_ISL_3160992, EPI_ISL_3160993, EPI_ISL_3160994, EPI_ISL_3160995, EPI_ISL_3160996, EPI_ISL_3160997, EPI_ISL_3160998, EPI_ISL_3160999, EPI_ISL_3161000, EPI_ISL_3161001, EPI_ISL_3161002, EPI_ISL_3161003, EPI_ISL_3161004, EPI_ISL_3161005, EPI_ISL_3161006, EPI_ISL_3161007, EPI_ISL_3161008, EPI_ISL_3161009, EPI_ISL_3161010, EPI_ISL_3161011, EPI_ISL_3161012, EPI_ISL_3161013, EPI_ISL_3161014, EPI_ISL_3161015, EPI_ISL_3161016, EPI_ISL_3161017, EPI_ISL_3161018, EPI_ISL_3161019, EPI_ISL_3161020, EPI_ISL_3161021, EPI_ISL_3161022, EPI_ISL_3161023, EPI_ISL_3161024, EPI_ISL_3161025, EPI_ISL_3161027, EPI_ISL_3161028, EPI_ISL_3161029, EPI_ISL_3161030, EPI_ISL_3161031, EPI_ISL_3161032, EPI_ISL_3161033, EPI_ISL_3161034, EPI_ISL_3161035, EPI_ISL_3161036, EPI_ISL_3161037, EPI_ISL_3161038, EPI_ISL_3161039, EPI_ISL_3161040, EPI_ISL_3161041, EPI_ISL_3161042, EPI_ISL_3161043, EPI_ISL_3161044, EPI_ISL_3161045, EPI_ISL_3161046, EPI_ISL_3161047, EPI_ISL_3161048, EPI_ISL_3161049, EPI_ISL_3161050, EPI_ISL_3161051, EPI_ISL_3161052, EPI_ISL_3161053, EPI_ISL_3161054, EPI_ISL_3161056, EPI_ISL_3161057, EPI_ISL_3161058, EPI_ISL_3161059, EPI_ISL_3161060, EPI_ISL_3161061, EPI_ISL_3161062, EPI_ISL_3161063, EPI_ISL_3161064, EPI_ISL_3161065, EPI_ISL_3161066, EPI_ISL_3161067, EPI_ISL_3161069, EPI_ISL_3161070, EPI_ISL_3161071, EPI_ISL_3161072, EPI_ISL_3161073, EPI_ISL_3161074, EPI_ISL_3161075, EPI_ISL_3161076, EPI_ISL_3161077, EPI_ISL_3161078, EPI_ISL_3161080, EPI_ISL_3161081, EPI_ISL_3161082, EPI_ISL_3161083, EPI_ISL_3161084, EPI_ISL_3161085, EPI_ISL_3161086, EPI_ISL_3161087, EPI_ISL_3161088, EPI_ISL_3161089, EPI_ISL_3161090, EPI_ISL_3161091, EPI_ISL_3161092, EPI_ISL_3161093, EPI_ISL_3161094, EPI_ISL_3161095, EPI_ISL_3161096, EPI_ISL_3161097, EPI_ISL_3161098, EPI_ISL_3161099, EPI_ISL_3161100, EPI_ISL_3161101, EPI_ISL_3161102, EPI_ISL_3161103, EPI_ISL_3161104, EPI_ISL_3161105, EPI_ISL_3161106, EPI_ISL_3161107, EPI_ISL_3161108, EPI_ISL_3161109, EPI_ISL_3161110, EPI_ISL_3161111, EPI_ISL_3161112, EPI_ISL_3161113, EPI_ISL_3161114, EPI_ISL_3161116, EPI_ISL_3161117, EPI_ISL_3161118, EPI_ISL_3161120, EPI_ISL_3161121, EPI_ISL_3161122, EPI_ISL_3161123, EPI_ISL_3161124, EPI_ISL_3161125, EPI_ISL_3161126, EPI_ISL_3161128, EPI_ISL_3161129, EPI_ISL_3161130, EPI_ISL_3161131, EPI_ISL_3161132, EPI_ISL_3161134, EPI_ISL_3161135, EPI_ISL_3161136, EPI_ISL_3161137, EPI_ISL_3161138, EPI_ISL_3161139, EPI_ISL_3161145, EPI_ISL_3161146, EPI_ISL_3161147, EPI_ISL_3161148, EPI_ISL_3161149, EPI_ISL_3161151, EPI_ISL_3161152, EPI_ISL_3161153, EPI_ISL_3161155, EPI_ISL_3161156, EPI_ISL_3161157, EPI_ISL_3161159, EPI_ISL_3161160, EPI_ISL_3161161, EPI_ISL_3161162, EPI_ISL_3161164, EPI_ISL_3161165, EPI_ISL_3161166, EPI_ISL_3161167, EPI_ISL_3161168, EPI_ISL_3161169, EPI_ISL_3161170, EPI_ISL_3161171, EPI_ISL_3161172, EPI_ISL_3161173, EPI_ISL_3161174, EPI_ISL_3161175, EPI_ISL_3161176, EPI_ISL_3161177, EPI_ISL_3161178, EPI_ISL_3161181, EPI_ISL_3161182, EPI_ISL_3161183, EPI_ISL_3161184, EPI_ISL_3161185, EPI_ISL_3161186, EPI_ISL_3161187, EPI_ISL_3161189, EPI_ISL_3161190, EPI_ISL_3161191, EPI_ISL_3161192, EPI_ISL_3161193, EPI_ISL_3161194, EPI_ISL_3161195, EPI_ISL_3161196, EPI_ISL_3161197, EPI_ISL_3161198, EPI_ISL_3161199, EPI_ISL_3161200, EPI_ISL_3161201, EPI_ISL_3161202, EPI_ISL_3161203, EPI_ISL_3161204, EPI_ISL_3161205, EPI_ISL_3161206, EPI_ISL_3161207 | | | |
| see above | UW Virology Lab | UW Virology Lab | Pavitra Roychoudhury, Shah Mohamed Bakhash, Ricardo Perez, Hong Xie, Tien V. Nguyen, Maria Lukes, Sean Ellis, Nathan Breit, Robert J. Livingston, Meei-Li Huang, Keith R Jerome, Patrick Mathias, Alexander Greninger |
| EPI_ISL_3161208, EPI_ISL_3161209, EPI_ISL_3161210, EPI_ISL_3161211, EPI_ISL_3161212, EPI_ISL_3161213, EPI_ISL_3161214, EPI_ISL_3161215, EPI_ISL_3161216, EPI_ISL_3161217, EPI_ISL_3161218, EPI_ISL_3161219, EPI_ISL_3161220, EPI_ISL_3161221, EPI_ISL_3161222, EPI_ISL_3161223, EPI_ISL_3161224, EPI_ISL_3161225, EPI_ISL_3161226, EPI_ISL_3161227, EPI_ISL_3161228, EPI_ISL_3161229, EPI_ISL_3161230, EPI_ISL_3161231, EPI_ISL_3161232, EPI_ISL_3161233, EPI_ISL_3161234, EPI_ISL_3161235, EPI_ISL_3161236, EPI_ISL_3161237, EPI_ISL_3161238, EPI_ISL_3161239, EPI_ISL_3161240, EPI_ISL_3161241, EPI_ISL_3161242, EPI_ISL_3161243, EPI_ISL_3161244, EPI_ISL_3161245, EPI_ISL_3161246, EPI_ISL_3161247, EPI_ISL_3161248, EPI_ISL_3161249, EPI_ISL_3161250, EPI_ISL_3161251, EPI_ISL_3161252, EPI_ISL_3161253, EPI_ISL_3161254, EPI_ISL_3161255, EPI_ISL_3161256, EPI_ISL_3161257, EPI_ISL_3161258, EPI_ISL_3161259, EPI_ISL_3161260, EPI_ISL_3161261, EPI_ISL_3161262, EPI_ISL_3161263, EPI_ISL_3161264, EPI_ISL_3161265, EPI_ISL_3161266, EPI_ISL_3161267, EPI_ISL_3161268, EPI_ISL_3161269, EPI_ISL_3161270, EPI_ISL_3161271, EPI_ISL_3161272, EPI_ISL_3161273, EPI_ISL_3161274, EPI_ISL_3161275, EPI_ISL_3161277, EPI_ISL_3161278, EPI_ISL_3161279, EPI_ISL_3161280, EPI_ISL_3161281, EPI_ISL_3161283, EPI_ISL_3161284, EPI_ISL_3161285, EPI_ISL_3161286, EPI_ISL_3161287, EPI_ISL_3161288, EPI_ISL_3161289, EPI_ISL_3161290 | | | |
| see above | Missouri State Public Health Laboratory | Missouri State Public Health Laboratory | Matthew Sinn, Joshua Barry, Ashley New |
| EPI_ISL_3161297 | Omega Diagnostics at Mounes | Omega Diagnostics st Mounes | Latira Haynes-Jacob, Cynthia Corley, Vivek Khare, MD |
| EPI_ISL_3161298 | Omega Diagnostics at Mounes | Omega Diagnostics | Latira Haynes-Jacob, Cynthia Corley, Vivek Khare, MD |
| EPI_ISL_3161299, EPI_ISL_3161301, EPI_ISL_3161302, EPI_ISL_3161303, EPI_ISL_3161304, EPI_ISL_3161305 | Omega Diagnostics at Mounes | Omega Diagnostics at Mounes | Latira Haynes-Jacob, Cynthia Corley, Vivek Khare, MD |
| EPI_ISL_3161306, EPI_ISL_3161307, EPI_ISL_3161308, EPI_ISL_3161309, EPI_ISL_3161310, EPI_ISL_3161311, EPI_ISL_3161312, EPI_ISL_3161313, EPI_ISL_3161314, EPI_ISL_3161315, EPI_ISL_3161316, EPI_ISL_3161317, EPI_ISL_3161318, EPI_ISL_3161319, EPI_ISL_3161320, EPI_ISL_3161321, EPI_ISL_3161322, EPI_ISL_3161323, EPI_ISL_3161324, EPI_ISL_3161325, EPI_ISL_3161326, EPI_ISL_3161327, EPI_ISL_3161328, EPI_ISL_3161329, EPI_ISL_3161330, EPI_ISL_3161331, EPI_ISL_3161332, EPI_ISL_3161333, EPI_ISL_3161334, EPI_ISL_3161335, EPI_ISL_3161336, EPI_ISL_3161337, EPI_ISL_3161338, EPI_ISL_3161339, EPI_ISL_3161340, EPI_ISL_3161341, EPI_ISL_3161342, EPI_ISL_3161343, EPI_ISL_3161344, EPI_ISL_3161345, EPI_ISL_3161346, EPI_ISL_3161347, EPI_ISL_3161348, EPI_ISL_3161349, EPI_ISL_3161350, EPI_ISL_3161351, EPI_ISL_3161352, EPI_ISL_3161353, EPI_ISL_3161354, EPI_ISL_3161355, EPI_ISL_3161356, EPI_ISL_3161357, EPI_ISL_3161358, EPI_ISL_3161359, EPI_ISL_3161360, EPI_ISL_3161361, EPI_ISL_3161362, EPI_ISL_3161363, EPI_ISL_3161364, EPI_ISL_3161365, EPI_ISL_3161366, EPI_ISL_3161367, EPI_ISL_3161368, EPI_ISL_3161369, EPI_ISL_3161370, EPI_ISL_3161371, EPI_ISL_3161372, EPI_ISL_3161373, EPI_ISL_3161374, EPI_ISL_3161375, EPI_ISL_3161376, EPI_ISL_3161377, EPI_ISL_3161378, EPI_ISL_3161379, EPI_ISL_3161380, EPI_ISL_3161381, EPI_ISL_3161382, EPI_ISL_3161383, EPI_ISL_3161384, EPI_ISL_3161385, EPI_ISL_3161386, EPI_ISL_3161387, EPI_ISL_3161388, EPI_ISL_3161389, EPI_ISL_3161390, EPI_ISL_3161391 | | | |
| see above | Missouri State Public Health Laboratory | Missouri State Public Health Laboratory | Matthew Sinn, Joshua Barry, Ashley New |
| EPI_ISL_3161392, EPI_ISL_3161393, EPI_ISL_3161394, EPI_ISL_3161396, EPI_ISL_3161397, EPI_ISL_3161401, EPI_ISL_3161402 | Omega Diagnostics at Mounes | Omega Diagnostics at Mounes | Latira Haynes-Jacob, Cynthia Corley, Vivek Khare, MD |
| EPI_ISL_3161403 | Omega Diagnotics at Mounes | Onega Diagnostics at Mounes | Latira Haynes-Jacob, Cynthia Corley, Vivek Khare, MD |
| EPI_ISL_3161404, EPI_ISL_3161405, EPI_ISL_3161406, EPI_ISL_3161407 | Omega Diagnostics at Mounes | Omega Diagnostics at Mounes | Latira Haynes-Jacob, Cynthia Corley, Vivek Khare, MD |
| EPI_ISL_3161414 | IRCCS San Gallicano Dermatological Institute | IRCCS Regina Elena National Cancer Institute | Sabrina Strano, Sara Donzelli, Matteo Pallocca, Grazia Prignano, Fabrizio Ensoli, Fulvia Pimpinelli, Maurizio Fanciulli, Giovanni Blandino, Aldo Morrone, Gennaro Ciliberto |
| EPI_ISL_3161415 | IRCCS San Gallicano Dermatological Institute | IRCCS Regina Elena National Cancer Institute | Francesca De Nicola, Giulia Orlandi, Eleonora Sperandio, Elisabetta Trento, Fabrizio Ensoli, Fulvia Pimpinelli, Maurizio Fanciulli, Giovanni Blandino, Aldo Morrone, Gennaro Ciliberto |
| EPI_ISL_3161416 | IRCCS San Gallicano Dermatological Institute | IRCCS Regina Elena National Cancer Institute | Ludovica Ciuffreda, Sara Donzelli, Alice Massacci, Arianna Mastrofrancesco, Fabrizio Ensoli, Fulvia Pimpinelli, Maurizio Fanciulli, Giovanni Blandino, Aldo Morrone, Gennaro Ciliberto |
| EPI_ISL_3161417 | IRCCS San Gallicano Dermatological Institute | IRCCS Regina Elena National Cancer Institute | Frauke Goeman, Giulia Orlandi, Matteo Pallocca, Giovanna D'agosto, Fabrizio Ensoli, Fulvia Pimpinelli, Maurizio Fanciulli, Giovanni Blandino, Aldo Morrone, Gennaro Ciliberto |
| EPI_ISL_3161418 | IRCCS San Gallicano Dermatological Institute | IRCCS Regina Elena National Cancer Institute | Sabrina Strano, Sara Donzelli, Eleonora Sperandio, Carla Mottini, Fabrizio Ensoli, Fulvia Pimpinelli, Maurizio Fanciulli, Giovanni Blandino, Aldo Morrone, Gennaro Ciliberto |
| EPI_ISL_3161419 | IRCCS San Gallicano Dermatological Institute | IRCCS Regina Elena National Cancer Institute | Francesca De Nicola, Giulia Orlandi, Alice Massacci, Martina Pontone, Fabrizio Ensoli, Fulvia Pimpinelli, Maurizio Fanciulli, Giovanni Blandino, Aldo Morrone, Gennaro Ciliberto |
| EPI_ISL_3161420 | IRCCS San Gallicano Dermatological Institute | IRCCS Regina Elena National Cancer Institute | Ludovica Ciuffreda, Sara Donzelli, Matteo Pallocca, di Domenico Enea Gino, Fabrizio Ensoli, Fulvia Pimpinelli, Maurizio Fanciulli, Giovanni Blandino, Aldo Morrone, Gennaro Ciliberto |
| EPI_ISL_3161421 | IRCCS San Gallicano Dermatological Institute | IRCCS Regina Elena National Cancer Institute | Frauke Goeman, Giulia Orlandi, Eleonora Sperandio, Ilaria Cavallo, Fabrizio Ensoli, Fulvia Pimpinelli, Maurizio Fanciulli, Giovanni Blandino, Aldo Morrone, Gennaro Ciliberto |
| EPI_ISL_3161422 | IRCCS San Gallicano Dermatological Institute | IRCCS Regina Elena National Cancer Institute | Sabrina Strano, Sara Donzelli, Alice Massacci, Francesca Sivori, Fabrizio Ensoli, Fulvia Pimpinelli, Maurizio Fanciulli, Giovanni Blandino, Aldo Morrone, Gennaro Ciliberto |
| EPI_ISL_3161424 | IRCCS San Gallicano Dermatological Institute | IRCCS Regina Elena National Cancer Institute | Ludovica Ciuffreda, Sara Donzelli, Eleonora Sperandio, Ilaria Celesti, Fabrizio Ensoli, Fulvia Pimpinelli, Maurizio Fanciulli, Giovanni Blandino, Aldo Morrone, Gennaro Ciliberto |
| EPI_ISL_3161425 | IRCCS San Gallicano Dermatological Institute | IRCCS Regina Elena National Cancer Institute | Frauke Goeman, Giulia Orlandi, Alice Massacci, Martina Diano, Fabrizio Ensoli, Fulvia Pimpinelli, Maurizio Fanciulli, Giovanni Blandino, Aldo Morrone, Gennaro Ciliberto |

| | | | |
|---|---|---|---|
| EPI_ISL_3161426 | IRCCS San Gallicano Dermatological Institute | IRCCS Regina Elena National Cancer Institute | Sabrina Strano, Sara Donzelli, Matteo Pallocca, Antonio Federico, Fabrizio Ensoli, Fulvia Pimpinelli, Maurizio Fanciulli, Giovanni Blandino, Aldo Morrone, Gennaro Ciliberto |
| EPI_ISL_3161427 | IRCCS San Gallicano Dermatological Institute | IRCCS Regina Elena National Cancer Institute | Francesca De Nicola, Giulia Orlandi, Eleonora Sperandio, Fulvia Fraticelli, Fabrizio Ensoli, Fulvia Pimpinelli, Maurizio Fanciulli, Giovanni Blandino, Aldo Morrone, Gennaro Ciliberto |
| EPI_ISL_3161428 | IRCCS San Gallicano Dermatological Institute | IRCCS Regina Elena National Cancer Institute | Ludovica Ciuffreda, Sara Donzelli, Alice Massacci, Lorenzo Furzi, Fabrizio Ensoli, Fulvia Pimpinelli, Maurizio Fanciulli, Giovanni Blandino, Aldo Morrone, Gennaro Ciliberto |
| EPI_ISL_3161429 | IRCCS San Gallicano Dermatological Institute | IRCCS Regina Elena National Cancer Institute | Frauke Goeman, Giulia Orlandi, Matteo Pallocca, Alessia Lauretti, Fabrizio Ensoli, Fulvia Pimpinelli, Maurizio Fanciulli, Giovanni Blandino, Aldo Morrone, Gennaro Ciliberto |
| EPI_ISL_3161430 | IRCCS San Gallicano Dermatological Institute | IRCCS Regina Elena National Cancer Institute | Sabrina Strano, Sara Donzelli, Eleonora Sperandio, Francesca Maione, Fabrizio Ensoli, Fulvia Pimpinelli, Maurizio Fanciulli, Giovanni Blandino, Aldo Morrone, Gennaro Ciliberto |
| EPI_ISL_3161431 | IRCCS San Gallicano Dermatological Institute | IRCCS Regina Elena National Cancer Institute | Francesca De Nicola, Giulia Orlandi, Alice Massacci, Francisco Obregon, Fabrizio Ensoli, Fulvia Pimpinelli, Maurizio Fanciulli, Giovanni Blandino, Aldo Morrone, Gennaro Ciliberto |
| EPI_ISL_3161432 | IRCCS San Gallicano Dermatological Institute | IRCCS Regina Elena National Cancer Institute | Ludovica Ciuffreda, Sara Donzelli, Matteo Pallocca, Silvia Paluzzi, Fabrizio Ensoli, Fulvia Pimpinelli, Maurizio Fanciulli, Giovanni Blandino, Aldo Morrone, Gennaro Ciliberto |
| EPI_ISL_3161433 | IRCCS San Gallicano Dermatological Institute | IRCCS Regina Elena National Cancer Institute | Frauke Goeman, Giulia Orlandi, Eleonora Sperandio, Sara Petrolo, Fabrizio Ensoli, Fulvia Pimpinelli, Maurizio Fanciulli, Giovanni Blandino, Aldo Morrone, Gennaro Ciliberto |
| EPI_ISL_3161434 | IRCCS San Gallicano Dermatological Institute | IRCCS Regina Elena National Cancer Institute | Sabrina Strano, Sara Donzelli, Alice Massacci, Valentina Ricca, Fabrizio Ensoli, Fulvia Pimpinelli, Maurizio Fanciulli, Giovanni Blandino, Aldo Morrone, Gennaro Ciliberto |
| EPI_ISL_3161435 | IRCCS San Gallicano Dermatological Institute | IRCCS Regina Elena National Cancer Institute | Francesca De Nicola, Giulia Orlandi, Matteo Pallocca, Serena Salvo, Fabrizio Ensoli, Fulvia Pimpinelli, Maurizio Fanciulli, Giovanni Blandino, Aldo Morrone, Gennaro Ciliberto |
| EPI_ISL_3161436 | IRCCS San Gallicano Dermatological Institute | IRCCS Regina Elena National Cancer Institute | Ludovica Ciuffreda, Sara Donzelli, Eleonora Sperandio, Grazia Prignano, Fabrizio Ensoli, Fulvia Pimpinelli, Maurizio Fanciulli, Giovanni Blandino, Aldo Morrone, Gennaro Ciliberto |
| EPI_ISL_3161438 | IRCCS San Gallicano Dermatological Institute | IRCCS Regina Elena National Cancer Institute | Sabrina Strano, Sara Donzelli, Matteo Pallocca, Arianna Mastrofrancesco, Fabrizio Ensoli, Fulvia Pimpinelli, Maurizio Fanciulli, Giovanni Blandino, Aldo Morrone, Gennaro Ciliberto |
| EPI_ISL_3161439 | IRCCS San Gallicano Dermatological Institute | IRCCS Regina Elena National Cancer Institute | Francesca De Nicola, Giulia Orlandi, Eleonora Sperandio, Giovanna D'agosto, Fabrizio Ensoli, Fulvia Pimpinelli, Maurizio Fanciulli, Giovanni Blandino, Aldo Morrone, Gennaro Ciliberto |
| EPI_ISL_3161440 | IRCCS San Gallicano Dermatological Institute | IRCCS Regina Elena National Cancer Institute | Ludovica Ciuffreda, Sara Donzelli, Alice Massacci, Carla Mottini, Fabrizio Ensoli, Fulvia Pimpinelli, Maurizio Fanciulli, Giovanni Blandino, Aldo Morrone, Gennaro Ciliberto |
| EPI_ISL_3161441 | IRCCS San Gallicano Dermatological Institute | IRCCS Regina Elena National Cancer Institute | Frauke Goeman, Giulia Orlandi, Matteo Pallocca, Martina Pontone, Fabrizio Ensoli, Fulvia Pimpinelli, Maurizio Fanciulli, Giovanni Blandino, Aldo Morrone, Gennaro Ciliberto |
| EPI_ISL_3161442 | IRCCS San Gallicano Dermatological Institute | IRCCS Regina Elena National Cancer Institute | Sabrina Strano, Sara Donzelli, Eleonora Sperandio, di Domenico Enea Gino, Fabrizio Ensoli, Fulvia Pimpinelli, Maurizio Fanciulli, Giovanni Blandino, Aldo Morrone, Gennaro Ciliberto |
| EPI_ISL_3161443 | IRCCS San Gallicano Dermatological Institute | IRCCS Regina Elena National Cancer Institute | Francesca De Nicola, Giulia Orlandi, Alice Massacci, Ilaria Cavallo, Fabrizio Ensoli, Fulvia Pimpinelli, Maurizio Fanciulli, Giovanni Blandino, Aldo Morrone, Gennaro Ciliberto |
| EPI_ISL_3161444 | IRCCS San Gallicano Dermatological Institute | IRCCS Regina Elena National Cancer Institute | Ludovica Ciuffreda, Sara Donzelli, Matteo Pallocca, Francesca Sivori, Fabrizio Ensoli, Fulvia Pimpinelli, Maurizio Fanciulli, Giovanni Blandino, Aldo Morrone, Gennaro Ciliberto |
| EPI_ISL_3161445 | IRCCS San Gallicano Dermatological Institute | IRCCS Regina Elena National Cancer Institute | Frauke Goeman, Giulia Orlandi, Eleonora Sperandio, Andrea Cazzani, Fabrizio Ensoli, Fulvia Pimpinelli, Maurizio Fanciulli, Giovanni Blandino, Aldo Morrone, Gennaro Ciliberto |
| EPI_ISL_3161446 | IRCCS San Gallicano Dermatological Institute | IRCCS Regina Elena National Cancer Institute | Sabrina Strano, Sara Donzelli, Alice Massacci, Ilaria Celesti, Fabrizio Ensoli, Fulvia Pimpinelli, Maurizio Fanciulli, Giovanni Blandino, Aldo Morrone, Gennaro Ciliberto |
| EPI_ISL_3161447 | IRCCS San Gallicano Dermatological Institute | IRCCS Regina Elena National Cancer Institute | Francesca De Nicola, Giulia Orlandi, Matteo Pallocca, Martina Diano, Fabrizio Ensoli, Fulvia Pimpinelli, Maurizio Fanciulli, Giovanni Blandino, Aldo Morrone, Gennaro Ciliberto |
| EPI_ISL_3161448 | IRCCS San Gallicano Dermatological Institute | IRCCS Regina Elena National Cancer Institute | Ludovica Ciuffreda, Sara Donzelli, Eleonora Sperandio, Antonio Federico, Fabrizio Ensoli, Fulvia Pimpinelli, Maurizio Fanciulli, Giovanni Blandino, Aldo Morrone, Gennaro Ciliberto |
| EPI_ISL_3161449 | IRCCS San Gallicano Dermatological Institute | IRCCS Regina Elena National Cancer Institute | Frauke Goeman, Giulia Orlandi, Alice Massacci, Fulvia Fraticelli, Fabrizio Ensoli, Fulvia Pimpinelli, Maurizio Fanciulli, Giovanni Blandino, Aldo Morrone, Gennaro Ciliberto |
| EPI_ISL_3161450 | IRCCS San Gallicano Dermatological Institute | IRCCS Regina Elena National Cancer Institute | Sabrina Strano, Sara Donzelli, Matteo Pallocca, Lorenzo Furzi, Fabrizio Ensoli, Fulvia Pimpinelli, Maurizio Fanciulli, Giovanni Blandino, Aldo Morrone, Gennaro Ciliberto |
| EPI_ISL_3161451 | IRCCS San Gallicano Dermatological Institute | IRCCS Regina Elena National Cancer Institute | Francesca De Nicola, Giulia Orlandi, Eleonora Sperandio, Alessia Lauretti, Fabrizio Ensoli, Fulvia Pimpinelli, Maurizio Fanciulli, Giovanni Blandino, Aldo Morrone, Gennaro Ciliberto |
| EPI_ISL_3161452 | IRCCS San Gallicano Dermatological Institute | IRCCS Regina Elena National Cancer Institute | Ludovica Ciuffreda, Sara Donzelli, Alice Massacci, Francesca Maione, Fabrizio Ensoli, Fulvia Pimpinelli, Maurizio Fanciulli, Giovanni Blandino, Aldo Morrone, Gennaro Ciliberto |
| EPI_ISL_3161453 | IRCCS San Gallicano Dermatological Institute | IRCCS Regina Elena National Cancer Institute | Frauke Goeman, Giulia Orlandi, Matteo Pallocca, Francisco Obregon, Fabrizio Ensoli, Fulvia Pimpinelli, Maurizio Fanciulli, Giovanni Blandino, Aldo Morrone, Gennaro Ciliberto |
| EPI_ISL_3161454 | IRCCS San Gallicano Dermatological Institute | IRCCS Regina Elena National Cancer Institute | Sabrina Strano, Sara Donzelli, Eleonora Sperandio, Silvia Paluzzi, Fabrizio Ensoli, Fulvia Pimpinelli, Maurizio Fanciulli, Giovanni Blandino, Aldo Morrone, Gennaro Ciliberto |
| EPI_ISL_3161455 | IRCCS San Gallicano Dermatological Institute | IRCCS Regina Elena National Cancer Institute | Francesca De Nicola, Giulia Orlandi, Alice Massacci, Sara Petrolo, Fabrizio Ensoli, Fulvia Pimpinelli, Maurizio Fanciulli, Giovanni Blandino, Aldo Morrone, Gennaro Ciliberto |
| EPI_ISL_3161456 | IRCCS San Gallicano Dermatological Institute | IRCCS Regina Elena National Cancer Institute | Ludovica Ciuffreda, Sara Donzelli, Matteo Pallocca, Valentina Ricca, Fabrizio Ensoli, Fulvia Pimpinelli, Maurizio Fanciulli, Giovanni Blandino, Aldo Morrone, Gennaro Ciliberto |
| EPI_ISL_3161457 | IRCCS San Gallicano Dermatological Institute | IRCCS Regina Elena National Cancer Institute | Frauke Goeman, Giulia Orlandi, Eleonora Sperandio, Serena Salvo, Fabrizio Ensoli, Fulvia Pimpinelli, Maurizio Fanciulli, Giovanni Blandino, Aldo Morrone, Gennaro Ciliberto |
| EPI_ISL_3161458 | IRCCS San Gallicano Dermatological Institute | IRCCS Regina Elena National Cancer Institute | Sabrina Strano, Sara Donzelli, Alice Massacci, Grazia Prignano, Fabrizio Ensoli, Fulvia Pimpinelli, Maurizio Fanciulli, Giovanni Blandino, Aldo Morrone, Gennaro Ciliberto |
| EPI_ISL_3161459 | IRCCS San Gallicano Dermatological Institute | IRCCS Regina Elena National Cancer Institute | Francesca De Nicola, Giulia Orlandi, Matteo Pallocca, Elisabetta Trento, Fabrizio Ensoli, Fulvia Pimpinelli, Maurizio Fanciulli, Giovanni Blandino, Aldo Morrone, Gennaro Ciliberto |
| EPI_ISL_3161460 | IRCCS San Gallicano Dermatological Institute | IRCCS Regina Elena National Cancer Institute | Ludovica Ciuffreda, Sara Donzelli, Eleonora Sperandio, Arianna Mastrofrancesco, Fabrizio Ensoli, Fulvia Pimpinelli, Maurizio Fanciulli, Giovanni Blandino, Aldo Morrone, Gennaro Ciliberto |
| EPI_ISL_3161461 | IRCCS San Gallicano Dermatological Institute | IRCCS Regina Elena National Cancer Institute | Frauke Goeman, Giulia Orlandi, Alice Massacci, Giovanna D'agosto, Fabrizio Ensoli, Fulvia Pimpinelli, Maurizio Fanciulli, Giovanni Blandino, Aldo Morrone, Gennaro Ciliberto |
| EPI_ISL_3161462 | IRCCS San Gallicano Dermatological Institute | IRCCS Regina Elena National Cancer Institute | Sabrina Strano, Sara Donzelli, Matteo Pallocca, Carla Mottini, Fabrizio Ensoli, Fulvia Pimpinelli, Maurizio Fanciulli, Giovanni Blandino, Aldo Morrone, Gennaro Ciliberto |
| EPI_ISL_3161463 | IRCCS San Gallicano Dermatological Institute | IRCCS Regina Elena National Cancer Institute | Francesca De Nicola, Giulia Orlandi, Eleonora Sperandio, Martina Pontone, Fabrizio Ensoli, Fulvia Pimpinelli, Maurizio Fanciulli, Giovanni Blandino, Aldo Morrone, Gennaro Ciliberto |
| EPI_ISL_3161464 | IRCCS San Gallicano Dermatological Institute | IRCCS Regina Elena National Cancer Institute | Ludovica Ciuffreda, Sara Donzelli, Alice Massacci, di Domenico Enea Gino, Fabrizio Ensoli, Fulvia Pimpinelli, Maurizio Fanciulli, Giovanni Blandino, Aldo Morrone, Gennaro Ciliberto |
| EPI_ISL_3161465 | IRCCS San Gallicano Dermatological Institute | IRCCS Regina Elena National Cancer Institute | Frauke Goeman, Giulia Orlandi, Matteo Pallocca, Ilaria Cavallo, Fabrizio Ensoli, Fulvia Pimpinelli, Maurizio Fanciulli, Giovanni Blandino, Aldo Morrone, Gennaro Ciliberto |
| EPI_ISL_3161466 | IRCCS San Gallicano Dermatological Institute | IRCCS Regina Elena National Cancer Institute | Sabrina Strano, Sara Donzelli, Eleonora Sperandio, Francesca Sivori, Fabrizio Ensoli, Fulvia Pimpinelli, Maurizio Fanciulli, Giovanni Blandino, Aldo Morrone, Gennaro Ciliberto |
| EPI_ISL_3161467 | IRCCS San Gallicano Dermatological Institute | IRCCS Regina Elena National Cancer Institute | Francesca De Nicola, Giulia Orlandi, Alice Massacci, Andrea Cazzani, Fabrizio Ensoli, Fulvia Pimpinelli, Maurizio Fanciulli, Giovanni Blandino, Aldo Morrone, Gennaro Ciliberto |
| EPI_ISL_3161468 | IRCCS San Gallicano Dermatological Institute | IRCCS Regina Elena National Cancer Institute | Ludovica Ciuffreda, Sara Donzelli, Matteo Pallocca, Ilaria Celesti, Fabrizio Ensoli, Fulvia Pimpinelli, Maurizio Fanciulli, Giovanni Blandino, Aldo Morrone, Gennaro Ciliberto |
| EPI_ISL_3161469 | IRCCS San Gallicano Dermatological Institute | IRCCS Regina Elena National Cancer Institute | Frauke Goeman, Giulia Orlandi, Eleonora Sperandio, Martina Diano, Fabrizio Ensoli, Fulvia Pimpinelli, Maurizio Fanciulli, Giovanni Blandino, Aldo Morrone, Gennaro Ciliberto |
| EPI_ISL_3161470 | IRCCS San Gallicano Dermatological Institute | IRCCS Regina Elena National Cancer Institute | Sabrina Strano, Sara Donzelli, Alice Massacci, Antonio Federico, Fabrizio Ensoli, Fulvia Pimpinelli, Maurizio Fanciulli, Giovanni Blandino, Aldo Morrone, Gennaro Ciliberto |
| EPI_ISL_3161471 | IRCCS San Gallicano Dermatological Institute | IRCCS Regina Elena National Cancer Institute | Francesca De Nicola, Giulia Orlandi, Matteo Pallocca, Fulvia Fraticelli, Fabrizio Ensoli, Fulvia Pimpinelli, Maurizio Fanciulli, Giovanni Blandino, Aldo Morrone, Gennaro Ciliberto |
| EPI_ISL_3161472 | IRCCS San Gallicano Dermatological Institute | IRCCS Regina Elena National Cancer Institute | Ludovica Ciuffreda, Sara Donzelli, Eleonora Sperandio, Lorenzo Furzi, Fabrizio Ensoli, Fulvia Pimpinelli, Maurizio Fanciulli, Giovanni Blandino, Aldo Morrone, Gennaro Ciliberto |
| EPI_ISL_3161473 | IRCCS San Gallicano Dermatological Institute | IRCCS Regina Elena National Cancer Institute | Frauke Goeman, Giulia Orlandi, Alice Massacci, Alessia Lauretti, Fabrizio Ensoli, Fulvia Pimpinelli, Maurizio Fanciulli, Giovanni Blandino, Aldo Morrone, Gennaro Ciliberto |
| EPI_ISL_3161474 | IRCCS San Gallicano Dermatological Institute | IRCCS Regina Elena National Cancer Institute | Sabrina Strano, Sara Donzelli, Matteo Pallocca, Francesca Maione, Fabrizio Ensoli, Fulvia Pimpinelli, Maurizio Fanciulli, Giovanni Blandino, Aldo Morrone, Gennaro Ciliberto |
| EPI_ISL_3161475 | IRCCS San Gallicano Dermatological Institute | IRCCS Regina Elena National Cancer Institute | Francesca De Nicola, Giulia Orlandi, Eleonora Sperandio, Francisco Obregon, Fabrizio Ensoli, Fulvia Pimpinelli, Maurizio Fanciulli, Giovanni Blandino, Aldo Morrone, Gennaro Ciliberto |
| EPI_ISL_3161476 | IRCCS San Gallicano Dermatological Institute | IRCCS Regina Elena National Cancer Institute | Ludovica Ciuffreda, Sara Donzelli, Alice Massacci, Silvia Paluzzi, Fabrizio Ensoli, Fulvia Pimpinelli, Maurizio Fanciulli, Giovanni Blandino, Aldo Morrone, Gennaro Ciliberto |
| EPI_ISL_3161477 | IRCCS San Gallicano Dermatological Institute | IRCCS Regina Elena National Cancer Institute | Frauke Goeman, Giulia Orlandi, Matteo Pallocca, Sara Petrolo, Fabrizio Ensoli, Fulvia Pimpinelli, Maurizio Fanciulli, Giovanni Blandino, Aldo Morrone, Gennaro Ciliberto |
| EPI_ISL_3161478 | IRCCS San Gallicano Dermatological Institute | IRCCS Regina Elena National Cancer Institute | Sabrina Strano, Sara Donzelli, Eleonora Sperandio, Valentina Ricca, Fabrizio Ensoli, Fulvia Pimpinelli, Maurizio Fanciulli, Giovanni Blandino, Aldo Morrone, Gennaro Ciliberto |
| EPI_ISL_3161479 | IRCCS San Gallicano Dermatological Institute | IRCCS Regina Elena National Cancer Institute | Francesca De Nicola, Giulia Orlandi, Alice Massacci, Serena Salvo, Fabrizio Ensoli, Fulvia Pimpinelli, Maurizio Fanciulli, Giovanni Blandino, Aldo Morrone, Gennaro Ciliberto |
| EPI_ISL_3161480 | IRCCS San Gallicano Dermatological Institute | IRCCS Regina Elena National Cancer Institute | Ludovica Ciuffreda, Sara Donzelli, Matteo Pallocca, Grazia Prignano, Fabrizio Ensoli, Fulvia Pimpinelli, Maurizio Fanciulli, Giovanni Blandino, Aldo Morrone, Gennaro Ciliberto |
| EPI_ISL_3161481 | IRCCS San Gallicano Dermatological Institute | IRCCS Regina Elena National Cancer Institute | Frauke Goeman, Giulia Orlandi, Eleonora Sperandio, Elisabetta Trento, Fabrizio Ensoli, Fulvia Pimpinelli, Maurizio Fanciulli, Giovanni Blandino, Aldo Morrone, Gennaro Ciliberto |
| EPI_ISL_3161482 | IRCCS San Gallicano Dermatological Institute | IRCCS Regina Elena National Cancer Institute | Sabrina Strano, Sara Donzelli, Alice Massacci, Arianna Mastrofrancesco, Fabrizio Ensoli, Fulvia Pimpinelli, Maurizio Fanciulli, Giovanni Blandino, Aldo Morrone, Gennaro Ciliberto |
| EPI_ISL_3161483 | IRCCS San Gallicano Dermatological Institute | IRCCS Regina Elena National Cancer Institute | Francesca De Nicola, Giulia Orlandi, Matteo Pallocca, Giovanna D'agosto, Fabrizio Ensoli, Fulvia Pimpinelli, Maurizio Fanciulli, Giovanni Blandino, Aldo Morrone, Gennaro Ciliberto |
| EPI_ISL_3161484 | IRCCS San Gallicano Dermatological Institute | IRCCS Regina Elena National Cancer Institute | Ludovica Ciuffreda, Sara Donzelli, Eleonora Sperandio, Carla Mottini, Fabrizio Ensoli, Fulvia Pimpinelli, Maurizio Fanciulli, Giovanni Blandino, Aldo Morrone, Gennaro Ciliberto |
| EPI_ISL_3161485 | IRCCS San Gallicano Dermatological Institute | IRCCS Regina Elena National Cancer Institute | Frauke Goeman, Giulia Orlandi, Alice Massacci, Martina Pontone, Fabrizio Ensoli, Fulvia Pimpinelli, Maurizio Fanciulli, Giovanni Blandino, Aldo Morrone, Gennaro Ciliberto |
| EPI_ISL_3161487 | IRCCS San Gallicano Dermatological Institute | IRCCS Regina Elena National Cancer Institute | Francesca De Nicola, Giulia Orlandi, Eleonora Sperandio, Ilaria Cavallo, Fabrizio Ensoli, Fulvia Pimpinelli, Maurizio Fanciulli, Giovanni Blandino, Aldo Morrone, Gennaro Ciliberto |
| EPI_ISL_3161488 | IRCCS San Gallicano Dermatological Institute | IRCCS Regina Elena National Cancer Institute | Ludovica Ciuffreda, Sara Donzelli, Alice Massacci, Francesca Sivori, Fabrizio Ensoli, Fulvia Pimpinelli, Maurizio Fanciulli, Giovanni Blandino, Aldo Morrone, Gennaro Ciliberto |
| EPI_ISL_3161489 | IRCCS San Gallicano Dermatological Institute | IRCCS Regina Elena National Cancer Institute | Frauke Goeman, Giulia Orlandi, Matteo Pallocca, Andrea Cazzani, Fabrizio Ensoli, Fulvia Pimpinelli, Maurizio Fanciulli, Giovanni Blandino, Aldo Morrone, Gennaro Ciliberto |
| EPI_ISL_3161490 | IRCCS San Gallicano Dermatological Institute | IRCCS Regina Elena National Cancer Institute | Sabrina Strano, Sara Donzelli, Eleonora Sperandio, Ilaria Celesti, Fabrizio Ensoli, Fulvia Pimpinelli, Maurizio Fanciulli, Giovanni Blandino, Aldo Morrone, Gennaro Ciliberto |
| EPI_ISL_3161491 | IRCCS San Gallicano Dermatological Institute | IRCCS Regina Elena National Cancer Institute | Francesca De Nicola, Giulia Orlandi, Alice Massacci, Martina Diano, Fabrizio Ensoli, Fulvia Pimpinelli, Maurizio Fanciulli, Giovanni Blandino, Aldo Morrone, Gennaro Ciliberto |
| EPI_ISL_3161492 | IRCCS San Gallicano Dermatological Institute | IRCCS Regina Elena National Cancer Institute | Ludovica Ciuffreda, Sara Donzelli, Matteo Pallocca, Antonio Federico, Fabrizio Ensoli, Fulvia Pimpinelli, Maurizio Fanciulli, Giovanni Blandino, Aldo Morrone, Gennaro Ciliberto |
| EPI_ISL_3161493 | IRCCS San Gallicano Dermatological Institute | IRCCS Regina Elena National Cancer Institute | Frauke Goeman, Giulia Orlandi, Eleonora Sperandio, Fulvia Fraticelli, Fabrizio Ensoli, Fulvia Pimpinelli, Maurizio Fanciulli, Giovanni Blandino, Aldo Morrone, Gennaro Ciliberto |
| EPI_ISL_3161494 | IRCCS San Gallicano Dermatological Institute | IRCCS Regina Elena National Cancer Institute | Sabrina Strano, Sara Donzelli, Alice Massacci, Lorenzo Furzi, Fabrizio Ensoli, Fulvia Pimpinelli, Maurizio Fanciulli, Giovanni Blandino, Aldo Morrone, Gennaro Ciliberto |
| EPI_ISL_3161495 | IRCCS San Gallicano Dermatological Institute | IRCCS Regina Elena National Cancer Institute | Francesca De Nicola, Giulia Orlandi, Matteo Pallocca, Alessia Lauretti, Fabrizio Ensoli, Fulvia Pimpinelli, Maurizio Fanciulli, Giovanni Blandino, Aldo Morrone, Gennaro Ciliberto |
| EPI_ISL_3161496 | IRCCS San Gallicano Dermatological Institute | IRCCS Regina Elena National Cancer Institute | Ludovica Ciuffreda, Sara Donzelli, Eleonora Sperandio, Francesca Maione, Fabrizio Ensoli, Fulvia Pimpinelli, Maurizio Fanciulli, Giovanni Blandino, Aldo Morrone, Gennaro Ciliberto |
| EPI_ISL_3161497 | IRCCS San Gallicano Dermatological Institute | IRCCS Regina Elena National Cancer Institute | Frauke Goeman, Giulia Orlandi, Alice Massacci, Francisco Obregon, Fabrizio Ensoli, Fulvia Pimpinelli, Maurizio Fanciulli, Giovanni Blandino, Aldo Morrone, Gennaro Ciliberto |
| EPI_ISL_3161498 | IRCCS San Gallicano Dermatological Institute | IRCCS Regina Elena National Cancer Institute | Sabrina Strano, Sara Donzelli, Matteo Pallocca, Silvia Paluzzi, Fabrizio Ensoli, Fulvia Pimpinelli, Maurizio Fanciulli, Giovanni Blandino, Aldo Morrone, Gennaro Ciliberto |
| EPI_ISL_3161499 | IRCCS San Gallicano Dermatological Institute | IRCCS Regina Elena National Cancer Institute | Francesca De Nicola, Giulia Orlandi, Eleonora Sperandio, Sara Petrolo, Fabrizio Ensoli, Fulvia Pimpinelli, Maurizio Fanciulli, Giovanni Blandino, Aldo Morrone, Gennaro Ciliberto |
| EPI_ISL_3161500 | IRCCS San Gallicano Dermatological Institute | IRCCS Regina Elena National Cancer Institute | Ludovica Ciuffreda, Sara Donzelli, Alice Massacci, Valentina Ricca, Fabrizio Ensoli, Fulvia Pimpinelli, Maurizio Fanciulli, Giovanni Blandino, Aldo Morrone, Gennaro Ciliberto |
| EPI_ISL_3161501 | IRCCS San Gallicano Dermatological Institute | IRCCS Regina Elena National Cancer Institute | Frauke Goeman, Giulia Orlandi, Matteo Pallocca, Serena Salvo, Fabrizio Ensoli, Fulvia Pimpinelli, Maurizio Fanciulli, Giovanni Blandino, Aldo Morrone, Gennaro Ciliberto |
| EPI_ISL_3161502 | IRCCS San Gallicano Dermatological Institute | IRCCS Regina Elena National Cancer Institute | Sabrina Strano, Sara Donzelli, Eleonora Sperandio, Grazia Prignano, Fabrizio Ensoli, Fulvia Pimpinelli, Maurizio Fanciulli, Giovanni Blandino, Aldo Morrone, Gennaro Ciliberto |
| EPI_ISL_3161503 | IRCCS San Gallicano Dermatological Institute | IRCCS Regina Elena National Cancer Institute | Francesca De Nicola, Giulia Orlandi, Alice Massacci, Elisabetta Trento, Fabrizio Ensoli, Fulvia Pimpinelli, Maurizio Fanciulli, Giovanni Blandino, Aldo Morrone, Gennaro Ciliberto |

| | | | |
|---|---|---|---|
| EPI_ISL_3161504 | IRCCS San Gallicano Dermatological Institute | IRCCS Regina Elena National Cancer Institute | Ludovica Ciuffreda, Sara Donzelli, Matteo Pallocca, Arianna Mastrofrancesco, Fabrizio Ensoli, Fulvia Pimpinelli, Maurizio Fanciulli, Giovanni Blandino, Aldo Morrone, Gennaro Ciliberto |
| EPI_ISL_3161505 | IRCCS San Gallicano Dermatological Institute | IRCCS Regina Elena National Cancer Institute | Frauke Goeman, Giulia Orlandi, Eleonora Sperandio, Giovanna D'agosto, Fabrizio Ensoli, Fulvia Pimpinelli, Maurizio Fanciulli, Giovanni Blandino, Aldo Morrone, Gennaro Ciliberto |
| EPI_ISL_3161506 | IRCCS San Gallicano Dermatological Institute | IRCCS Regina Elena National Cancer Institute | Sabrina Strano, Sara Donzelli, Alice Massacci, Carla Mottini, Fabrizio Ensoli, Fulvia Pimpinelli, Maurizio Fanciulli, Giovanni Blandino, Aldo Morrone, Gennaro Ciliberto |
| EPI_ISL_3161507 | IRCCS San Gallicano Dermatological Institute | IRCCS Regina Elena National Cancer Institute | Francesca De Nicola, Giulia Orlandi, Matteo Pallocca, Martina Pontone, Fabrizio Ensoli, Fulvia Pimpinelli, Maurizio Fanciulli, Giovanni Blandino, Aldo Morrone, Gennaro Ciliberto |
| EPI_ISL_3161508 | IRCCS San Gallicano Dermatological Institute | IRCCS Regina Elena National Cancer Institute | Ludovica Ciuffreda, Sara Donzelli, Eleonora Sperandio, di Domenico Enea Gino, Fabrizio Ensoli, Fulvia Pimpinelli, Maurizio Fanciulli, Giovanni Blandino, Aldo Morrone, Gennaro Ciliberto |
| EPI_ISL_3161509 | IRCCS San Gallicano Dermatological Institute | IRCCS Regina Elena National Cancer Institute | Frauke Goeman, Giulia Orlandi, Alice Massacci, Ilaria Cavallo, Fabrizio Ensoli, Fulvia Pimpinelli, Maurizio Fanciulli, Giovanni Blandino, Aldo Morrone, Gennaro Ciliberto |
| EPI_ISL_3161510 | IRCCS San Gallicano Dermatological Institute | IRCCS Regina Elena National Cancer Institute | Sabrina Strano, Sara Donzelli, Matteo Pallocca, Francesca Sivori, Fabrizio Ensoli, Fulvia Pimpinelli, Maurizio Fanciulli, Giovanni Blandino, Aldo Morrone, Gennaro Ciliberto |
| EPI_ISL_3161511 | IRCCS San Gallicano Dermatological Institute | IRCCS Regina Elena National Cancer Institute | Francesca De Nicola, Giulia Orlandi, Eleonora Sperandio, Andrea Cazzani, Fabrizio Ensoli, Fulvia Pimpinelli, Maurizio Fanciulli, Giovanni Blandino, Aldo Morrone, Gennaro Ciliberto |
| EPI_ISL_3161512 | IRCCS San Gallicano Dermatological Institute | IRCCS Regina Elena National Cancer Institute | Ludovica Ciuffreda, Sara Donzelli, Alice Massacci, Ilaria Celesti, Fabrizio Ensoli, Fulvia Pimpinelli, Maurizio Fanciulli, Giovanni Blandino, Aldo Morrone, Gennaro Ciliberto |
| EPI_ISL_3161513 | IRCCS San Gallicano Dermatological Institute | IRCCS Regina Elena National Cancer Institute | Frauke Goeman, Giulia Orlandi, Matteo Pallocca, Martina Diano, Fabrizio Ensoli, Fulvia Pimpinelli, Maurizio Fanciulli, Giovanni Blandino, Aldo Morrone, Gennaro Ciliberto |
| EPI_ISL_3161514 | IRCCS San Gallicano Dermatological Institute | IRCCS Regina Elena National Cancer Institute | Sabrina Strano, Sara Donzelli, Eleonora Sperandio, Antonio Federico, Fabrizio Ensoli, Fulvia Pimpinelli, Maurizio Fanciulli, Giovanni Blandino, Aldo Morrone, Gennaro Ciliberto |
| EPI_ISL_3161515 | IRCCS San Gallicano Dermatological Institute | IRCCS Regina Elena National Cancer Institute | Francesca De Nicola, Giulia Orlandi, Alice Massacci, Fulvia Fraticelli, Fabrizio Ensoli, Fulvia Pimpinelli, Maurizio Fanciulli, Giovanni Blandino, Aldo Morrone, Gennaro Ciliberto |
| EPI_ISL_3161516 | IRCCS San Gallicano Dermatological Institute | IRCCS Regina Elena National Cancer Institute | Ludovica Ciuffreda, Sara Donzelli, Matteo Pallocca, Lorenzo Furzi, Fabrizio Ensoli, Fulvia Pimpinelli, Maurizio Fanciulli, Giovanni Blandino, Aldo Morrone, Gennaro Ciliberto |
| EPI_ISL_3161517 | IRCCS San Gallicano Dermatological Institute | IRCCS Regina Elena National Cancer Institute | Frauke Goeman, Giulia Orlandi, Eleonora Sperandio, Alessia Lauretti, Fabrizio Ensoli, Fulvia Pimpinelli, Maurizio Fanciulli, Giovanni Blandino, Aldo Morrone, Gennaro Ciliberto |
| EPI_ISL_3161518 | IRCCS San Gallicano Dermatological Institute | IRCCS Regina Elena National Cancer Institute | Sabrina Strano, Sara Donzelli, Alice Massacci, Francesca Maione, Fabrizio Ensoli, Fulvia Pimpinelli, Maurizio Fanciulli, Giovanni Blandino, Aldo Morrone, Gennaro Ciliberto |
| EPI_ISL_3161519 | IRCCS San Gallicano Dermatological Institute | IRCCS Regina Elena National Cancer Institute | Francesca De Nicola, Giulia Orlandi, Matteo Pallocca, Francisco Obregon, Fabrizio Ensoli, Fulvia Pimpinelli, Maurizio Fanciulli, Giovanni Blandino, Aldo Morrone, Gennaro Ciliberto |
| EPI_ISL_3161520 | IRCCS San Gallicano Dermatological Institute | IRCCS Regina Elena National Cancer Institute | Ludovica Ciuffreda, Sara Donzelli, Eleonora Sperandio, Silvia Paluzzi, Fabrizio Ensoli, Fulvia Pimpinelli, Maurizio Fanciulli, Giovanni Blandino, Aldo Morrone, Gennaro Ciliberto |
| EPI_ISL_3161521 | IRCCS San Gallicano Dermatological Institute | IRCCS Regina Elena National Cancer Institute | Frauke Goeman, Giulia Orlandi, Alice Massacci, Sara Petrolo, Fabrizio Ensoli, Fulvia Pimpinelli, Maurizio Fanciulli, Giovanni Blandino, Aldo Morrone, Gennaro Ciliberto |
| EPI_ISL_3161522 | IRCCS San Gallicano Dermatological Institute | IRCCS Regina Elena National Cancer Institute | Sabrina Strano, Sara Donzelli, Matteo Pallocca, Valentina Ricca, Fabrizio Ensoli, Fulvia Pimpinelli, Maurizio Fanciulli, Giovanni Blandino, Aldo Morrone, Gennaro Ciliberto |
| EPI_ISL_3161523 | IRCCS San Gallicano Dermatological Institute | IRCCS Regina Elena National Cancer Institute | Francesca De Nicola, Giulia Orlandi, Eleonora Sperandio, Serena Salvo, Fabrizio Ensoli, Fulvia Pimpinelli, Maurizio Fanciulli, Giovanni Blandino, Aldo Morrone, Gennaro Ciliberto |
| EPI_ISL_3161524 | IRCCS San Gallicano Dermatological Institute | IRCCS Regina Elena National Cancer Institute | Ludovica Ciuffreda, Sara Donzelli, Alice Massacci, Grazia Prignano, Fabrizio Ensoli, Fulvia Pimpinelli, Maurizio Fanciulli, Giovanni Blandino, Aldo Morrone, Gennaro Ciliberto |
| EPI_ISL_3161525 | IRCCS San Gallicano Dermatological Institute | IRCCS Regina Elena National Cancer Institute | Frauke Goeman, Giulia Orlandi, Matteo Pallocca, Elisabetta Trento, Fabrizio Ensoli, Fulvia Pimpinelli, Maurizio Fanciulli, Giovanni Blandino, Aldo Morrone, Gennaro Ciliberto |
| EPI_ISL_3161526 | IRCCS San Gallicano Dermatological Institute | IRCCS Regina Elena National Cancer Institute | Sabrina Strano, Sara Donzelli, Eleonora Sperandio, Arianna Mastrofrancesco, Fabrizio Ensoli, Fulvia Pimpinelli, Maurizio Fanciulli, Giovanni Blandino, Aldo Morrone, Gennaro Ciliberto |
| EPI_ISL_3161527 | IRCCS San Gallicano Dermatological Institute | IRCCS Regina Elena National Cancer Institute | Francesca De Nicola, Giulia Orlandi, Alice Massacci, Giovanna D'agosto, Fabrizio Ensoli, Fulvia Pimpinelli, Maurizio Fanciulli, Giovanni Blandino, Aldo Morrone, Gennaro Ciliberto |
| EPI_ISL_3161528 | IRCCS San Gallicano Dermatological Institute | IRCCS Regina Elena National Cancer Institute | Ludovica Ciuffreda, Sara Donzelli, Matteo Pallocca, Carla Mottini, Fabrizio Ensoli, Fulvia Pimpinelli, Maurizio Fanciulli, Giovanni Blandino, Aldo Morrone, Gennaro Ciliberto |
| EPI_ISL_3161530 | IRCCS San Gallicano Dermatological Institute | IRCCS Regina Elena National Cancer Institute | Sabrina Strano, Sara Donzelli, Alice Massacci, di Domenico Enea Gino, Fabrizio Ensoli, Fulvia Pimpinelli, Maurizio Fanciulli, Giovanni Blandino, Aldo Morrone, Gennaro Ciliberto |
| EPI_ISL_3161531 | IRCCS San Gallicano Dermatological Institute | IRCCS Regina Elena National Cancer Institute | Francesca De Nicola, Giulia Orlandi, Matteo Pallocca, Ilaria Cavallo, Fabrizio Ensoli, Fulvia Pimpinelli, Maurizio Fanciulli, Giovanni Blandino, Aldo Morrone, Gennaro Ciliberto |
| EPI_ISL_3161532 | IRCCS San Gallicano Dermatological Institute | IRCCS Regina Elena National Cancer Institute | Ludovica Ciuffreda, Sara Donzelli, Eleonora Sperandio, Francesca Sivori, Fabrizio Ensoli, Fulvia Pimpinelli, Maurizio Fanciulli, Giovanni Blandino, Aldo Morrone, Gennaro Ciliberto |
| EPI_ISL_3161533 | IRCCS San Gallicano Dermatological Institute | IRCCS Regina Elena National Cancer Institute | Frauke Goeman, Giulia Orlandi, Alice Massacci, Andrea Cazzani, Fabrizio Ensoli, Fulvia Pimpinelli, Maurizio Fanciulli, Giovanni Blandino, Aldo Morrone, Gennaro Ciliberto |
| EPI_ISL_3161534 | IRCCS San Gallicano Dermatological Institute | IRCCS Regina Elena National Cancer Institute | Sabrina Strano, Sara Donzelli, Matteo Pallocca, Ilaria Celesti, Fabrizio Ensoli, Fulvia Pimpinelli, Maurizio Fanciulli, Giovanni Blandino, Aldo Morrone, Gennaro Ciliberto |
| EPI_ISL_3161535 | IRCCS San Gallicano Dermatological Institute | IRCCS Regina Elena National Cancer Institute | Francesca De Nicola, Giulia Orlandi, Eleonora Sperandio, Martina Diano, Fabrizio Ensoli, Fulvia Pimpinelli, Maurizio Fanciulli, Giovanni Blandino, Aldo Morrone, Gennaro Ciliberto |
| EPI_ISL_3161536 | IRCCS San Gallicano Dermatological Institute | IRCCS Regina Elena National Cancer Institute | Ludovica Ciuffreda, Sara Donzelli, Alice Massacci, Antonio Federico, Fabrizio Ensoli, Fulvia Pimpinelli, Maurizio Fanciulli, Giovanni Blandino, Aldo Morrone, Gennaro Ciliberto |
| EPI_ISL_3161537 | IRCCS San Gallicano Dermatological Institute | IRCCS Regina Elena National Cancer Institute | Frauke Goeman, Giulia Orlandi, Matteo Pallocca, Fulvia Fraticelli, Fabrizio Ensoli, Fulvia Pimpinelli, Maurizio Fanciulli, Giovanni Blandino, Aldo Morrone, Gennaro Ciliberto |
| EPI_ISL_3161538 | IRCCS San Gallicano Dermatological Institute | IRCCS Regina Elena National Cancer Institute | Sabrina Strano, Sara Donzelli, Eleonora Sperandio, Lorenzo Furzi, Fabrizio Ensoli, Fulvia Pimpinelli, Maurizio Fanciulli, Giovanni Blandino, Aldo Morrone, Gennaro Ciliberto |
| EPI_ISL_3161539 | IRCCS San Gallicano Dermatological Institute | IRCCS Regina Elena National Cancer Institute | Francesca De Nicola, Giulia Orlandi, Alice Massacci, Alessia Lauretti, Fabrizio Ensoli, Fulvia Pimpinelli, Maurizio Fanciulli, Giovanni Blandino, Aldo Morrone, Gennaro Ciliberto |
| EPI_ISL_3161540 | IRCCS San Gallicano Dermatological Institute | IRCCS Regina Elena National Cancer Institute | Ludovica Ciuffreda, Sara Donzelli, Matteo Pallocca, Francesca Maione, Fabrizio Ensoli, Fulvia Pimpinelli, Maurizio Fanciulli, Giovanni Blandino, Aldo Morrone, Gennaro Ciliberto |
| EPI_ISL_3161541 | IRCCS San Gallicano Dermatological Institute | IRCCS Regina Elena National Cancer Institute | Frauke Goeman, Giulia Orlandi, Eleonora Sperandio, Francisco Obregon, Fabrizio Ensoli, Fulvia Pimpinelli, Maurizio Fanciulli, Giovanni Blandino, Aldo Morrone, Gennaro Ciliberto |
| EPI_ISL_3161542 | IRCCS San Gallicano Dermatological Institute | IRCCS Regina Elena National Cancer Institute | Sabrina Strano, Sara Donzelli, Alice Massacci, Silvia Paluzzi, Fabrizio Ensoli, Fulvia Pimpinelli, Maurizio Fanciulli, Giovanni Blandino, Aldo Morrone, Gennaro Ciliberto |
| EPI_ISL_3161543 | IRCCS San Gallicano Dermatological Institute | IRCCS Regina Elena National Cancer Institute | Francesca De Nicola, Giulia Orlandi, Matteo Pallocca, Sara Petrolo, Fabrizio Ensoli, Fulvia Pimpinelli, Maurizio Fanciulli, Giovanni Blandino, Aldo Morrone, Gennaro Ciliberto |
| EPI_ISL_3161544 | IRCCS San Gallicano Dermatological Institute | IRCCS Regina Elena National Cancer Institute | Ludovica Ciuffreda, Sara Donzelli, Eleonora Sperandio, Valentina Ricca, Fabrizio Ensoli, Fulvia Pimpinelli, Maurizio Fanciulli, Giovanni Blandino, Aldo Morrone, Gennaro Ciliberto |
| EPI_ISL_3161545 | IRCCS San Gallicano Dermatological Institute | IRCCS Regina Elena National Cancer Institute | Frauke Goeman, Giulia Orlandi, Alice Massacci, Serena Salvo, Fabrizio Ensoli, Fulvia Pimpinelli, Maurizio Fanciulli, Giovanni Blandino, Aldo Morrone, Gennaro Ciliberto |
| EPI_ISL_3161546 | IRCCS San Gallicano Dermatological Institute | IRCCS Regina Elena National Cancer Institute | Sabrina Strano, Sara Donzelli, Matteo Pallocca, Grazia Prignano, Fabrizio Ensoli, Fulvia Pimpinelli, Maurizio Fanciulli, Giovanni Blandino, Aldo Morrone, Gennaro Ciliberto |
| EPI_ISL_3161547 | IRCCS San Gallicano Dermatological Institute | IRCCS Regina Elena National Cancer Institute | Francesca De Nicola, Giulia Orlandi, Eleonora Sperandio, Elisabetta Trento, Fabrizio Ensoli, Fulvia Pimpinelli, Maurizio Fanciulli, Giovanni Blandino, Aldo Morrone, Gennaro Ciliberto |
| EPI_ISL_3161548 | IRCCS San Gallicano Dermatological Institute | IRCCS Regina Elena National Cancer Institute | Ludovica Ciuffreda, Sara Donzelli, Alice Massacci, Arianna Mastrofrancesco, Fabrizio Ensoli, Fulvia Pimpinelli, Maurizio Fanciulli, Giovanni Blandino, Aldo Morrone, Gennaro Ciliberto |
| EPI_ISL_3161550 | IRCCS San Gallicano Dermatological Institute | IRCCS Regina Elena National Cancer Institute | Sabrina Strano, Sara Donzelli, Eleonora Sperandio, Carla Mottini, Fabrizio Ensoli, Fulvia Pimpinelli, Maurizio Fanciulli, Giovanni Blandino, Aldo Morrone, Gennaro Ciliberto |
| EPI_ISL_3161551 | IRCCS San Gallicano Dermatological Institute | IRCCS Regina Elena National Cancer Institute | Francesca De Nicola, Giulia Orlandi, Alice Massacci, Martina Pontone, Fabrizio Ensoli, Fulvia Pimpinelli, Maurizio Fanciulli, Giovanni Blandino, Aldo Morrone, Gennaro Ciliberto |
| EPI_ISL_3161552 | IRCCS San Gallicano Dermatological Institute | IRCCS Regina Elena National Cancer Institute | Ludovica Ciuffreda, Sara Donzelli, Matteo Pallocca, di Domenico Enea Gino, Fabrizio Ensoli, Fulvia Pimpinelli, Maurizio Fanciulli, Giovanni Blandino, Aldo Morrone, Gennaro Ciliberto |
| EPI_ISL_3161555 | IRCCS San Gallicano Dermatological Institute | IRCCS Regina Elena National Cancer Institute | Francesca De Nicola, Giulia Orlandi, Matteo Pallocca, Andrea Cazzani, Fabrizio Ensoli, Fulvia Pimpinelli, Maurizio Fanciulli, Giovanni Blandino, Aldo Morrone, Gennaro Ciliberto |
| EPI_ISL_3161556 | IRCCS San Gallicano Dermatological Institute | IRCCS Regina Elena National Cancer Institute | Ludovica Ciuffreda, Sara Donzelli, Eleonora Sperandio, Ilaria Celesti, Fabrizio Ensoli, Fulvia Pimpinelli, Maurizio Fanciulli, Giovanni Blandino, Aldo Morrone, Gennaro Ciliberto |
| EPI_ISL_3161557 | IRCCS San Gallicano Dermatological Institute | IRCCS Regina Elena National Cancer Institute | Frauke Goeman, Giulia Orlandi, Alice Massacci, Martina Diano, Fabrizio Ensoli, Fulvia Pimpinelli, Maurizio Fanciulli, Giovanni Blandino, Aldo Morrone, Gennaro Ciliberto |
| EPI_ISL_3161558 | IRCCS San Gallicano Dermatological Institute | IRCCS Regina Elena National Cancer Institute | Sabrina Strano, Sara Donzelli, Matteo Pallocca, Antonio Federico, Fabrizio Ensoli, Fulvia Pimpinelli, Maurizio Fanciulli, Giovanni Blandino, Aldo Morrone, Gennaro Ciliberto |
| EPI_ISL_3161559 | IRCCS San Gallicano Dermatological Institute | IRCCS Regina Elena National Cancer Institute | Francesca De Nicola, Giulia Orlandi, Eleonora Sperandio, Fulvia Fraticelli, Fabrizio Ensoli, Fulvia Pimpinelli, Maurizio Fanciulli, Giovanni Blandino, Aldo Morrone, Gennaro Ciliberto |
| EPI_ISL_3161560 | IRCCS San Gallicano Dermatological Institute | IRCCS Regina Elena National Cancer Institute | Ludovica Ciuffreda, Sara Donzelli, Alice Massacci, Lorenzo Furzi, Fabrizio Ensoli, Fulvia Pimpinelli, Maurizio Fanciulli, Giovanni Blandino, Aldo Morrone, Gennaro Ciliberto |
| EPI_ISL_3161561 | IRCCS San Gallicano Dermatological Institute | IRCCS Regina Elena National Cancer Institute | Frauke Goeman, Giulia Orlandi, Matteo Pallocca, Alessia Lauretti, Fabrizio Ensoli, Fulvia Pimpinelli, Maurizio Fanciulli, Giovanni Blandino, Aldo Morrone, Gennaro Ciliberto |
| EPI_ISL_3161563 | IRCCS San Gallicano Dermatological Institute | IRCCS Regina Elena National Cancer Institute | Francesca De Nicola, Giulia Orlandi, Alice Massacci, Francisco Obregon, Fabrizio Ensoli, Fulvia Pimpinelli, Maurizio Fanciulli, Giovanni Blandino, Aldo Morrone, Gennaro Ciliberto |
| EPI_ISL_3161564 | IRCCS San Gallicano Dermatological Institute | IRCCS Regina Elena National Cancer Institute | Ludovica Ciuffreda, Sara Donzelli, Matteo Pallocca, Silvia Paluzzi, Fabrizio Ensoli, Fulvia Pimpinelli, Maurizio Fanciulli, Giovanni Blandino, Aldo Morrone, Gennaro Ciliberto |
| EPI_ISL_3161565 | IRCCS San Gallicano Dermatological Institute | IRCCS Regina Elena National Cancer Institute | Frauke Goeman, Giulia Orlandi, Eleonora Sperandio, Sara Petrolo, Fabrizio Ensoli, Fulvia Pimpinelli, Maurizio Fanciulli, Giovanni Blandino, Aldo Morrone, Gennaro Ciliberto |
| EPI_ISL_3161566 | IRCCS San Gallicano Dermatological Institute | IRCCS Regina Elena National Cancer Institute | Sabrina Strano, Sara Donzelli, Alice Massacci, Valentina Ricca, Fabrizio Ensoli, Fulvia Pimpinelli, Maurizio Fanciulli, Giovanni Blandino, Aldo Morrone, Gennaro Ciliberto |
| EPI_ISL_3161567 | IRCCS San Gallicano Dermatological Institute | IRCCS Regina Elena National Cancer Institute | Francesca De Nicola, Giulia Orlandi, Matteo Pallocca, Serena Salvo, Fabrizio Ensoli, Fulvia Pimpinelli, Maurizio Fanciulli, Giovanni Blandino, Aldo Morrone, Gennaro Ciliberto |
| EPI_ISL_3161568 | IRCCS San Gallicano Dermatological Institute | IRCCS Regina Elena National Cancer Institute | Ludovica Ciuffreda, Sara Donzelli, Eleonora Sperandio, Grazia Prignano, Fabrizio Ensoli, Fulvia Pimpinelli, Maurizio Fanciulli, Giovanni Blandino, Aldo Morrone, Gennaro Ciliberto |
| EPI_ISL_3161570 | IRCCS San Gallicano Dermatological Institute | IRCCS Regina Elena National Cancer Institute | Sabrina Strano, Sara Donzelli, Matteo Pallocca, Arianna Mastrofrancesco, Fabrizio Ensoli, Fulvia Pimpinelli, Maurizio Fanciulli, Giovanni Blandino, Aldo Morrone, Gennaro Ciliberto |
| EPI_ISL_3161572 | IRCCS San Gallicano Dermatological Institute | IRCCS Regina Elena National Cancer Institute | Ludovica Ciuffreda, Sara Donzelli, Alice Massacci, Carla Mottini, Fabrizio Ensoli, Fulvia Pimpinelli, Maurizio Fanciulli, Giovanni Blandino, Aldo Morrone, Gennaro Ciliberto |
| EPI_ISL_3161573 | IRCCS San Gallicano Dermatological Institute | IRCCS Regina Elena National Cancer Institute | Frauke Goeman, Giulia Orlandi, Matteo Pallocca, Martina Pontone, Fabrizio Ensoli, Fulvia Pimpinelli, Maurizio Fanciulli, Giovanni Blandino, Aldo Morrone, Gennaro Ciliberto |
| EPI_ISL_3161574 | IRCCS San Gallicano Dermatological Institute | IRCCS Regina Elena National Cancer Institute | Sabrina Strano, Sara Donzelli, Eleonora Sperandio, di Domenico Enea Gino, Fabrizio Ensoli, Fulvia Pimpinelli, Maurizio Fanciulli, Giovanni Blandino, Aldo Morrone, Gennaro Ciliberto |
| EPI_ISL_3161575 | IRCCS San Gallicano Dermatological Institute | IRCCS Regina Elena National Cancer Institute | Francesca De Nicola, Giulia Orlandi, Alice Massacci, Ilaria Cavallo, Fabrizio Ensoli, Fulvia Pimpinelli, Maurizio Fanciulli, Giovanni Blandino, Aldo Morrone, Gennaro Ciliberto |
| EPI_ISL_3161576 | IRCCS San Gallicano Dermatological Institute | IRCCS Regina Elena National Cancer Institute | Ludovica Ciuffreda, Sara Donzelli, Matteo Pallocca, Francesca Sivori, Fabrizio Ensoli, Fulvia Pimpinelli, Maurizio Fanciulli, Giovanni Blandino, Aldo Morrone, Gennaro Ciliberto |
| EPI_ISL_3161577 | IRCCS San Gallicano Dermatological Institute | IRCCS Regina Elena National Cancer Institute | Frauke Goeman, Giulia Orlandi, Eleonora Sperandio, Andrea Cazzani, Fabrizio Ensoli, Fulvia Pimpinelli, Maurizio Fanciulli, Giovanni Blandino, Aldo Morrone, Gennaro Ciliberto |
| EPI_ISL_3161578 | IRCCS San Gallicano Dermatological Institute | IRCCS Regina Elena National Cancer Institute | Sabrina Strano, Sara Donzelli, Alice Massacci, Ilaria Celesti, Fabrizio Ensoli, Fulvia Pimpinelli, Maurizio Fanciulli, Giovanni Blandino, Aldo Morrone, Gennaro Ciliberto |
| EPI_ISL_3161579 | IRCCS San Gallicano Dermatological Institute | IRCCS Regina Elena National Cancer Institute | Francesca De Nicola, Giulia Orlandi, Matteo Pallocca, Martina Diano, Fabrizio Ensoli, Fulvia Pimpinelli, Maurizio Fanciulli, Giovanni Blandino, Aldo Morrone, Gennaro Ciliberto |
| EPI_ISL_3161580 | IRCCS San Gallicano Dermatological Institute | IRCCS Regina Elena National Cancer Institute | Ludovica Ciuffreda, Sara Donzelli, Eleonora Sperandio, Antonio Federico, Fabrizio Ensoli, Fulvia Pimpinelli, Maurizio Fanciulli, Giovanni Blandino, Aldo Morrone, Gennaro Ciliberto |
| EPI_ISL_3161581 | IRCCS San Gallicano Dermatological Institute | IRCCS Regina Elena National Cancer Institute | Frauke Goeman, Giulia Orlandi, Alice Massacci, Fulvia Fraticelli, Fabrizio Ensoli, Fulvia Pimpinelli, Maurizio Fanciulli, Giovanni Blandino, Aldo Morrone, Gennaro Ciliberto |
| EPI_ISL_3161582 | IRCCS San Gallicano Dermatological Institute | IRCCS Regina Elena National Cancer Institute | Sabrina Strano, Sara Donzelli, Matteo Pallocca, Lorenzo Furzi, Fabrizio Ensoli, Fulvia Pimpinelli, Maurizio Fanciulli, Giovanni Blandino, Aldo Morrone, Gennaro Ciliberto |
| EPI_ISL_3161583 | IRCCS San Gallicano Dermatological Institute | IRCCS Regina Elena National Cancer Institute | Francesca De Nicola, Giulia Orlandi, Eleonora Sperandio, Alessia Lauretti, Fabrizio Ensoli, Fulvia Pimpinelli, Maurizio Fanciulli, Giovanni Blandino, Aldo Morrone, Gennaro Ciliberto |
| EPI_ISL_3161584 | IRCCS San Gallicano Dermatological Institute | IRCCS Regina Elena National Cancer Institute | Ludovica Ciuffreda, Sara Donzelli, Alice Massacci, Francesca Maione, Fabrizio Ensoli, Fulvia Pimpinelli, Maurizio Fanciulli, Giovanni Blandino, Aldo Morrone, Gennaro Ciliberto |
| EPI_ISL_3161585 | IRCCS San Gallicano Dermatological Institute | IRCCS Regina Elena National Cancer Institute | Frauke Goeman, Giulia Orlandi, Matteo Pallocca, Francisco Obregon, Fabrizio Ensoli, Fulvia Pimpinelli, Maurizio Fanciulli, Giovanni Blandino, Aldo Morrone, Gennaro Ciliberto |
| EPI_ISL_3161586 | IRCCS San Gallicano Dermatological Institute | IRCCS Regina Elena National Cancer Institute | Sabrina Strano, Sara Donzelli, Eleonora Sperandio, Silvia Paluzzi, Fabrizio Ensoli, Fulvia Pimpinelli, Maurizio Fanciulli, Giovanni Blandino, Aldo Morrone, Gennaro Ciliberto |

| | | | |
|---|---|---|---|
| EPI_ISL_3161587 | IRCCS San Gallicano Dermatological Institute | IRCCS Regina Elena National Cancer Institute | Francesca De Nicola, Giulia Orlandi, Alice Massacci, Sara Petrolo, Fabrizio Ensoli, Fulvia Pimpinelli, Maurizio Fanciulli, Giovanni Blandino, Aldo Morrone, Gennaro Ciliberto |
| EPI_ISL_3161588 | IRCCS San Gallicano Dermatological Institute | IRCCS Regina Elena National Cancer Institute | Ludovica Ciuffreda, Sara Donzelli, Matteo Pallocca, Valentina Ricca, Fabrizio Ensoli, Fulvia Pimpinelli, Maurizio Fanciulli, Giovanni Blandino, Aldo Morrone, Gennaro Ciliberto |
| EPI_ISL_3161589 | IRCCS San Gallicano Dermatological Institute | IRCCS Regina Elena National Cancer Institute | Frauke Goeman, Giulia Orlandi, Eleonora Sperandio, Serena Salvo, Fabrizio Ensoli, Fulvia Pimpinelli, Maurizio Fanciulli, Giovanni Blandino, Aldo Morrone, Gennaro Ciliberto |
| EPI_ISL_3161590 | IRCCS San Gallicano Dermatological Institute | IRCCS Regina Elena National Cancer Institute | Sabrina Strano, Sara Donzelli, Alice Massacci, Grazia Prignano, Fabrizio Ensoli, Fulvia Pimpinelli, Maurizio Fanciulli, Giovanni Blandino, Aldo Morrone, Gennaro Ciliberto |
| EPI_ISL_3161591 | IRCCS San Gallicano Dermatological Institute | IRCCS Regina Elena National Cancer Institute | Francesca De Nicola, Giulia Orlandi, Matteo Pallocca, Elisabetta Trento, Fabrizio Ensoli, Fulvia Pimpinelli, Maurizio Fanciulli, Giovanni Blandino, Aldo Morrone, Gennaro Ciliberto |
| EPI_ISL_3161592 | IRCCS San Gallicano Dermatological Institute | IRCCS Regina Elena National Cancer Institute | Ludovica Ciuffreda, Sara Donzelli, Eleonora Sperandio, Arianna Mastrofrancesco, Fabrizio Ensoli, Fulvia Pimpinelli, Maurizio Fanciulli, Giovanni Blandino, Aldo Morrone, Gennaro Ciliberto |
| EPI_ISL_3161593 | IRCCS San Gallicano Dermatological Institute | IRCCS Regina Elena National Cancer Institute | Frauke Goeman, Giulia Orlandi, Alice Massacci, Giovanna D’agosto, Fabrizio Ensoli, Fulvia Pimpinelli, Maurizio Fanciulli, Giovanni Blandino, Aldo Morrone, Gennaro Ciliberto |
| EPI_ISL_3161594 | IRCCS San Gallicano Dermatological Institute | IRCCS Regina Elena National Cancer Institute | Sabrina Strano, Sara Donzelli, Matteo Pallocca, Carla Mottini, Fabrizio Ensoli, Fulvia Pimpinelli, Maurizio Fanciulli, Giovanni Blandino, Aldo Morrone, Gennaro Ciliberto |
| EPI_ISL_3161595 | IRCCS San Gallicano Dermatological Institute | IRCCS Regina Elena National Cancer Institute | Francesca De Nicola, Giulia Orlandi, Eleonora Sperandio, Martina Pontone, Fabrizio Ensoli, Fulvia Pimpinelli, Maurizio Fanciulli, Giovanni Blandino, Aldo Morrone, Gennaro Ciliberto |
| EPI_ISL_3161596 | IRCCS San Gallicano Dermatological Institute | IRCCS Regina Elena National Cancer Institute | Ludovica Ciuffreda, Sara Donzelli, Alice Massacci, di Domenico Enea Gino, Fabrizio Ensoli, Fulvia Pimpinelli, Maurizio Fanciulli, Giovanni Blandino, Aldo Morrone, Gennaro Ciliberto |
| EPI_ISL_3161597 | IRCCS San Gallicano Dermatological Institute | IRCCS Regina Elena National Cancer Institute | Frauke Goeman, Giulia Orlandi, Matteo Pallocca, Ilaria Cavallo, Fabrizio Ensoli, Fulvia Pimpinelli, Maurizio Fanciulli, Giovanni Blandino, Aldo Morrone, Gennaro Ciliberto |
| EPI_ISL_3161598 | IRCCS San Gallicano Dermatological Institute | IRCCS Regina Elena National Cancer Institute | Sabrina Strano, Sara Donzelli, Eleonora Sperandio, Francesca Sivori, Fabrizio Ensoli, Fulvia Pimpinelli, Maurizio Fanciulli, Giovanni Blandino, Aldo Morrone, Gennaro Ciliberto |
| EPI_ISL_3161599 | IRCCS San Gallicano Dermatological Institute | IRCCS Regina Elena National Cancer Institute | Francesca De Nicola, Giulia Orlandi, Alice Massacci, Andrea Cazzani, Fabrizio Ensoli, Fulvia Pimpinelli, Maurizio Fanciulli, Giovanni Blandino, Aldo Morrone, Gennaro Ciliberto |
| EPI_ISL_3161600 | IRCCS San Gallicano Dermatological Institute | IRCCS Regina Elena National Cancer Institute | Ludovica Ciuffreda, Sara Donzelli, Matteo Pallocca, Ilaria Celesti, Fabrizio Ensoli, Fulvia Pimpinelli, Maurizio Fanciulli, Giovanni Blandino, Aldo Morrone, Gennaro Ciliberto |
| EPI_ISL_3161601 | IRCCS San Gallicano Dermatological Institute | IRCCS Regina Elena National Cancer Institute | Frauke Goeman, Giulia Orlandi, Eleonora Sperandio, Martina Diano, Fabrizio Ensoli, Fulvia Pimpinelli, Maurizio Fanciulli, Giovanni Blandino, Aldo Morrone, Gennaro Ciliberto |
| EPI_ISL_3161602 | IRCCS San Gallicano Dermatological Institute | IRCCS Regina Elena National Cancer Institute | Sabrina Strano, Sara Donzelli, Alice Massacci, Antonio Federico, Fabrizio Ensoli, Fulvia Pimpinelli, Maurizio Fanciulli, Giovanni Blandino, Aldo Morrone, Gennaro Ciliberto |
| EPI_ISL_3161603 | IRCCS San Gallicano Dermatological Institute | IRCCS Regina Elena National Cancer Institute | Francesca De Nicola, Giulia Orlandi, Matteo Pallocca, Fulvia Fraticelli, Fabrizio Ensoli, Fulvia Pimpinelli, Maurizio Fanciulli, Giovanni Blandino, Aldo Morrone, Gennaro Ciliberto |
| EPI_ISL_3161604 | IRCCS San Gallicano Dermatological Institute | IRCCS Regina Elena National Cancer Institute | Ludovica Ciuffreda, Sara Donzelli, Eleonora Sperandio, Lorenzo Furzi, Fabrizio Ensoli, Fulvia Pimpinelli, Maurizio Fanciulli, Giovanni Blandino, Aldo Morrone, Gennaro Ciliberto |
| EPI_ISL_3161605 | IRCCS San Gallicano Dermatological Institute | IRCCS Regina Elena National Cancer Institute | Frauke Goeman, Giulia Orlandi, Alice Massacci, Alessia Lauretti, Fabrizio Ensoli, Fulvia Pimpinelli, Maurizio Fanciulli, Giovanni Blandino, Aldo Morrone, Gennaro Ciliberto |
| EPI_ISL_3161606 | IRCCS San Gallicano Dermatological Institute | IRCCS Regina Elena National Cancer Institute | Sabrina Strano, Sara Donzelli, Matteo Pallocca, Francesca Maione, Fabrizio Ensoli, Fulvia Pimpinelli, Maurizio Fanciulli, Giovanni Blandino, Aldo Morrone, Gennaro Ciliberto |
| EPI_ISL_3161607 | IRCCS San Gallicano Dermatological Institute | IRCCS Regina Elena National Cancer Institute | Francesca De Nicola, Giulia Orlandi, Eleonora Sperandio, Francisco Obregon, Fabrizio Ensoli, Fulvia Pimpinelli, Maurizio Fanciulli, Giovanni Blandino, Aldo Morrone, Gennaro Ciliberto |
| EPI_ISL_3161608 | IRCCS San Gallicano Dermatological Institute | IRCCS Regina Elena National Cancer Institute | Ludovica Ciuffreda, Sara Donzelli, Alice Massacci, Silvia Paluzzi, Fabrizio Ensoli, Fulvia Pimpinelli, Maurizio Fanciulli, Giovanni Blandino, Aldo Morrone, Gennaro Ciliberto |
| EPI_ISL_3161609 | IRCCS San Gallicano Dermatological Institute | IRCCS Regina Elena National Cancer Institute | Frauke Goeman, Giulia Orlandi, Matteo Pallocca, Sara Petrolo, Fabrizio Ensoli, Fulvia Pimpinelli, Maurizio Fanciulli, Giovanni Blandino, Aldo Morrone, Gennaro Ciliberto |
| EPI_ISL_3161610 | IRCCS San Gallicano Dermatological Institute | IRCCS Regina Elena National Cancer Institute | Sabrina Strano, Sara Donzelli, Eleonora Sperandio, Valentina Ricca, Fabrizio Ensoli, Fulvia Pimpinelli, Maurizio Fanciulli, Giovanni Blandino, Aldo Morrone, Gennaro Ciliberto |
| EPI_ISL_3161611 | IRCCS San Gallicano Dermatological Institute | IRCCS Regina Elena National Cancer Institute | Francesca De Nicola, Giulia Orlandi, Alice Massacci, Serena Salvo, Fabrizio Ensoli, Fulvia Pimpinelli, Maurizio Fanciulli, Giovanni Blandino, Aldo Morrone, Gennaro Ciliberto |
| EPI_ISL_3161612 | IRCCS San Gallicano Dermatological Institute | IRCCS Regina Elena National Cancer Institute | Ludovica Ciuffreda, Sara Donzelli, Matteo Pallocca, Grazia Prignano, Fabrizio Ensoli, Fulvia Pimpinelli, Maurizio Fanciulli, Giovanni Blandino, Aldo Morrone, Gennaro Ciliberto |
| EPI_ISL_3161613 | IRCCS San Gallicano Dermatological Institute | IRCCS Regina Elena National Cancer Institute | Frauke Goeman, Giulia Orlandi, Eleonora Sperandio, Elisabetta Trento, Fabrizio Ensoli, Fulvia Pimpinelli, Maurizio Fanciulli, Giovanni Blandino, Aldo Morrone, Gennaro Ciliberto |
| EPI_ISL_3161614 | IRCCS San Gallicano Dermatological Institute | IRCCS Regina Elena National Cancer Institute | Sabrina Strano, Sara Donzelli, Alice Massacci, Arianna Mastrofrancesco, Fabrizio Ensoli, Fulvia Pimpinelli, Maurizio Fanciulli, Giovanni Blandino, Aldo Morrone, Gennaro Ciliberto |
| EPI_ISL_3161615 | IRCCS San Gallicano Dermatological Institute | IRCCS Regina Elena National Cancer Institute | Francesca De Nicola, Giulia Orlandi, Matteo Pallocca, Giovanna D’agosto, Fabrizio Ensoli, Fulvia Pimpinelli, Maurizio Fanciulli, Giovanni Blandino, Aldo Morrone, Gennaro Ciliberto |
| EPI_ISL_3161616 | IRCCS San Gallicano Dermatological Institute | IRCCS Regina Elena National Cancer Institute | Ludovica Ciuffreda, Sara Donzelli, Eleonora Sperandio, Carla Mottini, Fabrizio Ensoli, Fulvia Pimpinelli, Maurizio Fanciulli, Giovanni Blandino, Aldo Morrone, Gennaro Ciliberto |
| EPI_ISL_3161617 | IRCCS San Gallicano Dermatological Institute | IRCCS Regina Elena National Cancer Institute | Frauke Goeman, Giulia Orlandi, Alice Massacci, Martina Pontone, Fabrizio Ensoli, Fulvia Pimpinelli, Maurizio Fanciulli, Giovanni Blandino, Aldo Morrone, Gennaro Ciliberto |
| EPI_ISL_3161618 | IRCCS San Gallicano Dermatological Institute | IRCCS Regina Elena National Cancer Institute | Sabrina Strano, Sara Donzelli, Matteo Pallocca, di Domenico Enea Gino, Fabrizio Ensoli, Fulvia Pimpinelli, Maurizio Fanciulli, Giovanni Blandino, Aldo Morrone, Gennaro Ciliberto |
| EPI_ISL_3161619 | IRCCS San Gallicano Dermatological Institute | IRCCS Regina Elena National Cancer Institute | Francesca De Nicola, Giulia Orlandi, Eleonora Sperandio, Ilaria Cavallo, Fabrizio Ensoli, Fulvia Pimpinelli, Maurizio Fanciulli, Giovanni Blandino, Aldo Morrone, Gennaro Ciliberto |
| EPI_ISL_3161620 | IRCCS San Gallicano Dermatological Institute | IRCCS Regina Elena National Cancer Institute | Ludovica Ciuffreda, Sara Donzelli, Alice Massacci, Francesca Sivori, Fabrizio Ensoli, Fulvia Pimpinelli, Maurizio Fanciulli, Giovanni Blandino, Aldo Morrone, Gennaro Ciliberto |
| EPI_ISL_3161621 | IRCCS San Gallicano Dermatological Institute | IRCCS Regina Elena National Cancer Institute | Frauke Goeman, Giulia Orlandi, Matteo Pallocca, Andrea Cazzani, Fabrizio Ensoli, Fulvia Pimpinelli, Maurizio Fanciulli, Giovanni Blandino, Aldo Morrone, Gennaro Ciliberto |
| EPI_ISL_3161622 | IRCCS San Gallicano Dermatological Institute | IRCCS Regina Elena National Cancer Institute | Sabrina Strano, Sara Donzelli, Eleonora Sperandio, Ilaria Celesti, Fabrizio Ensoli, Fulvia Pimpinelli, Maurizio Fanciulli, Giovanni Blandino, Aldo Morrone, Gennaro Ciliberto |
| EPI_ISL_3161623 | IRCCS San Gallicano Dermatological Institute | IRCCS Regina Elena National Cancer Institute | Francesca De Nicola, Giulia Orlandi, Alice Massacci, Martina Diano, Fabrizio Ensoli, Fulvia Pimpinelli, Maurizio Fanciulli, Giovanni Blandino, Aldo Morrone, Gennaro Ciliberto |
| EPI_ISL_3161625 | IRCCS San Gallicano Dermatological Institute | IRCCS Regina Elena National Cancer Institute | Frauke Goeman, Giulia Orlandi, Eleonora Sperandio, Fulvia Fraticelli, Fabrizio Ensoli, Fulvia Pimpinelli, Maurizio Fanciulli, Giovanni Blandino, Aldo Morrone, Gennaro Ciliberto |
| EPI_ISL_3161626 | IRCCS San Gallicano Dermatological Institute | IRCCS Regina Elena National Cancer Institute | Sabrina Strano, Sara Donzelli, Alice Massacci, Lorenzo Furzi, Fabrizio Ensoli, Fulvia Pimpinelli, Maurizio Fanciulli, Giovanni Blandino, Aldo Morrone, Gennaro Ciliberto |
| EPI_ISL_3161627 | IRCCS San Gallicano Dermatological Institute | IRCCS Regina Elena National Cancer Institute | Francesca De Nicola, Giulia Orlandi, Matteo Pallocca, Alessia Lauretti, Fabrizio Ensoli, Fulvia Pimpinelli, Maurizio Fanciulli, Giovanni Blandino, Aldo Morrone, Gennaro Ciliberto |
| EPI_ISL_3161628 | IRCCS San Gallicano Dermatological Institute | IRCCS Regina Elena National Cancer Institute | Ludovica Ciuffreda, Sara Donzelli, Eleonora Sperandio, Francesca Maione, Fabrizio Ensoli, Fulvia Pimpinelli, Maurizio Fanciulli, Giovanni Blandino, Aldo Morrone, Gennaro Ciliberto |
| EPI_ISL_3161629 | IRCCS San Gallicano Dermatological Institute | IRCCS Regina Elena National Cancer Institute | Frauke Goeman, Giulia Orlandi, Alice Massacci, Francisco Obregon, Fabrizio Ensoli, Fulvia Pimpinelli, Maurizio Fanciulli, Giovanni Blandino, Aldo Morrone, Gennaro Ciliberto |
| EPI_ISL_3161630 | IRCCS San Gallicano Dermatological Institute | IRCCS Regina Elena National Cancer Institute | Sabrina Strano, Sara Donzelli, Matteo Pallocca, Silvia Paluzzi, Fabrizio Ensoli, Fulvia Pimpinelli, Maurizio Fanciulli, Giovanni Blandino, Aldo Morrone, Gennaro Ciliberto |
| EPI_ISL_3161631 | IRCCS San Gallicano Dermatological Institute | IRCCS Regina Elena National Cancer Institute | Francesca De Nicola, Giulia Orlandi, Eleonora Sperandio, Sara Petrolo, Fabrizio Ensoli, Fulvia Pimpinelli, Maurizio Fanciulli, Giovanni Blandino, Aldo Morrone, Gennaro Ciliberto |
| EPI_ISL_3161632 | IRCCS San Gallicano Dermatological Institute | IRCCS Regina Elena National Cancer Institute | Ludovica Ciuffreda, Sara Donzelli, Alice Massacci, Valentina Ricca, Fabrizio Ensoli, Fulvia Pimpinelli, Maurizio Fanciulli, Giovanni Blandino, Aldo Morrone, Gennaro Ciliberto |
| EPI_ISL_3161633 | IRCCS San Gallicano Dermatological Institute | IRCCS Regina Elena National Cancer Institute | Frauke Goeman, Giulia Orlandi, Matteo Pallocca, Serena Salvo, Fabrizio Ensoli, Fulvia Pimpinelli, Maurizio Fanciulli, Giovanni Blandino, Aldo Morrone, Gennaro Ciliberto |
| EPI_ISL_3161634 | IRCCS San Gallicano Dermatological Institute | IRCCS Regina Elena National Cancer Institute | Sabrina Strano, Sara Donzelli, Eleonora Sperandio, Grazia Prignano, Fabrizio Ensoli, Fulvia Pimpinelli, Maurizio Fanciulli, Giovanni Blandino, Aldo Morrone, Gennaro Ciliberto |
| EPI_ISL_3161635 | IRCCS San Gallicano Dermatological Institute | IRCCS Regina Elena National Cancer Institute | Francesca De Nicola, Giulia Orlandi, Alice Massacci, Elisabetta Trento, Fabrizio Ensoli, Fulvia Pimpinelli, Maurizio Fanciulli, Giovanni Blandino, Aldo Morrone, Gennaro Ciliberto |
| EPI_ISL_3161636 | IRCCS San Gallicano Dermatological Institute | IRCCS Regina Elena National Cancer Institute | Ludovica Ciuffreda, Sara Donzelli, Matteo Pallocca, Arianna Mastrofrancesco, Fabrizio Ensoli, Fulvia Pimpinelli, Maurizio Fanciulli, Giovanni Blandino, Aldo Morrone, Gennaro Ciliberto |
| EPI_ISL_3161637 | IRCCS San Gallicano Dermatological Institute | IRCCS Regina Elena National Cancer Institute | Frauke Goeman, Giulia Orlandi, Eleonora Sperandio, Giovanna D’agosto, Fabrizio Ensoli, Fulvia Pimpinelli, Maurizio Fanciulli, Giovanni Blandino, Aldo Morrone, Gennaro Ciliberto |
| EPI_ISL_3161638 | IRCCS San Gallicano Dermatological Institute | IRCCS Regina Elena National Cancer Institute | Sabrina Strano, Sara Donzelli, Alice Massacci, Carla Mottini, Fabrizio Ensoli, Fulvia Pimpinelli, Maurizio Fanciulli, Giovanni Blandino, Aldo Morrone, Gennaro Ciliberto |
| EPI_ISL_3161639 | IRCCS San Gallicano Dermatological Institute | IRCCS Regina Elena National Cancer Institute | Francesca De Nicola, Giulia Orlandi, Matteo Pallocca, Martina Pontone, Fabrizio Ensoli, Fulvia Pimpinelli, Maurizio Fanciulli, Giovanni Blandino, Aldo Morrone, Gennaro Ciliberto |
| EPI_ISL_3161640 | IRCCS San Gallicano Dermatological Institute | IRCCS Regina Elena National Cancer Institute | Ludovica Ciuffreda, Sara Donzelli, Eleonora Sperandio, di Domenico Enea Gino, Fabrizio Ensoli, Fulvia Pimpinelli, Maurizio Fanciulli, Giovanni Blandino, Aldo Morrone, Gennaro Ciliberto |
| EPI_ISL_3161641 | IRCCS San Gallicano Dermatological Institute | IRCCS Regina Elena National Cancer Institute | Frauke Goeman, Giulia Orlandi, Alice Massacci, Ilaria Cavallo, Fabrizio Ensoli, Fulvia Pimpinelli, Maurizio Fanciulli, Giovanni Blandino, Aldo Morrone, Gennaro Ciliberto |
| EPI_ISL_3161642 | IRCCS San Gallicano Dermatological Institute | IRCCS Regina Elena National Cancer Institute | Sabrina Strano, Sara Donzelli, Matteo Pallocca, Francesca Sivori, Fabrizio Ensoli, Fulvia Pimpinelli, Maurizio Fanciulli, Giovanni Blandino, Aldo Morrone, Gennaro Ciliberto |
| EPI_ISL_3161643 | IRCCS San Gallicano Dermatological Institute | IRCCS Regina Elena National Cancer Institute | Francesca De Nicola, Giulia Orlandi, Eleonora Sperandio, Andrea Cazzani, Fabrizio Ensoli, Fulvia Pimpinelli, Maurizio Fanciulli, Giovanni Blandino, Aldo Morrone, Gennaro Ciliberto |
| EPI_ISL_3161644 | IRCCS San Gallicano Dermatological Institute | IRCCS Regina Elena National Cancer Institute | Ludovica Ciuffreda, Sara Donzelli, Alice Massacci, Ilaria Celesti, Fabrizio Ensoli, Fulvia Pimpinelli, Maurizio Fanciulli, Giovanni Blandino, Aldo Morrone, Gennaro Ciliberto |
| EPI_ISL_3161645 | IRCCS San Gallicano Dermatological Institute | IRCCS Regina Elena National Cancer Institute | Frauke Goeman, Giulia Orlandi, Matteo Pallocca, Martina Diano, Fabrizio Ensoli, Fulvia Pimpinelli, Maurizio Fanciulli, Giovanni Blandino, Aldo Morrone, Gennaro Ciliberto |
| EPI_ISL_3161646 | IRCCS San Gallicano Dermatological Institute | IRCCS Regina Elena National Cancer Institute | Sabrina Strano, Sara Donzelli, Eleonora Sperandio, Antonio Federico, Fabrizio Ensoli, Fulvia Pimpinelli, Maurizio Fanciulli, Giovanni Blandino, Aldo Morrone, Gennaro Ciliberto |
| EPI_ISL_3161647 | IRCCS San Gallicano Dermatological Institute | IRCCS Regina Elena National Cancer Institute | Francesca De Nicola, Giulia Orlandi, Alice Massacci, Fulvia Fraticelli, Fabrizio Ensoli, Fulvia Pimpinelli, Maurizio Fanciulli, Giovanni Blandino, Aldo Morrone, Gennaro Ciliberto |
| EPI_ISL_3161648 | IRCCS San Gallicano Dermatological Institute | IRCCS Regina Elena National Cancer Institute | Ludovica Ciuffreda, Sara Donzelli, Matteo Pallocca, Lorenzo Furzi, Fabrizio Ensoli, Fulvia Pimpinelli, Maurizio Fanciulli, Giovanni Blandino, Aldo Morrone, Gennaro Ciliberto |
| EPI_ISL_3161649 | IRCCS San Gallicano Dermatological Institute | IRCCS Regina Elena National Cancer Institute | Frauke Goeman, Giulia Orlandi, Eleonora Sperandio, Alessia Lauretti, Fabrizio Ensoli, Fulvia Pimpinelli, Maurizio Fanciulli, Giovanni Blandino, Aldo Morrone, Gennaro Ciliberto |
| EPI_ISL_3161650 | IRCCS San Gallicano Dermatological Institute | IRCCS Regina Elena National Cancer Institute | Sabrina Strano, Sara Donzelli, Alice Massacci, Francesca Maione, Fabrizio Ensoli, Fulvia Pimpinelli, Maurizio Fanciulli, Giovanni Blandino, Aldo Morrone, Gennaro Ciliberto |
| EPI_ISL_3161651 | IRCCS San Gallicano Dermatological Institute | IRCCS Regina Elena National Cancer Institute | Francesca De Nicola, Giulia Orlandi, Matteo Pallocca, Francisco Obregon, Fabrizio Ensoli, Fulvia Pimpinelli, Maurizio Fanciulli, Giovanni Blandino, Aldo Morrone, Gennaro Ciliberto |
| EPI_ISL_3161652 | IRCCS San Gallicano Dermatological Institute | IRCCS Regina Elena National Cancer Institute | Ludovica Ciuffreda, Sara Donzelli, Eleonora Sperandio, Silvia Paluzzi, Fabrizio Ensoli, Fulvia Pimpinelli, Maurizio Fanciulli, Giovanni Blandino, Aldo Morrone, Gennaro Ciliberto |
| EPI_ISL_3161653 | IRCCS San Gallicano Dermatological Institute | IRCCS Regina Elena National Cancer Institute | Frauke Goeman, Giulia Orlandi, Alice Massacci, Sara Petrolo, Fabrizio Ensoli, Fulvia Pimpinelli, Maurizio Fanciulli, Giovanni Blandino, Aldo Morrone, Gennaro Ciliberto |
| EPI_ISL_3161654 | IRCCS San Gallicano Dermatological Institute | IRCCS Regina Elena National Cancer Institute | Sabrina Strano, Sara Donzelli, Matteo Pallocca, Valentina Ricca, Fabrizio Ensoli, Fulvia Pimpinelli, Maurizio Fanciulli, Giovanni Blandino, Aldo Morrone, Gennaro Ciliberto |
| EPI_ISL_3161655 | IRCCS San Gallicano Dermatological Institute | IRCCS Regina Elena National Cancer Institute | Francesca De Nicola, Giulia Orlandi, Eleonora Sperandio, Serena Salvo, Fabrizio Ensoli, Fulvia Pimpinelli, Maurizio Fanciulli, Giovanni Blandino, Aldo Morrone, Gennaro Ciliberto |
| EPI_ISL_3161656 | IRCCS San Gallicano Dermatological Institute | IRCCS Regina Elena National Cancer Institute | Ludovica Ciuffreda, Sara Donzelli, Alice Massacci, Grazia Prignano, Fabrizio Ensoli, Fulvia Pimpinelli, Maurizio Fanciulli, Giovanni Blandino, Aldo Morrone, Gennaro Ciliberto |
| EPI_ISL_3161657 | IRCCS San Gallicano Dermatological Institute | IRCCS Regina Elena National Cancer Institute | Frauke Goeman, Giulia Orlandi, Matteo Pallocca, Elisabetta Trento, Fabrizio Ensoli, Fulvia Pimpinelli, Maurizio Fanciulli, Giovanni Blandino, Aldo Morrone, Gennaro Ciliberto |
| EPI_ISL_3161658 | IRCCS San Gallicano Dermatological Institute | IRCCS Regina Elena National Cancer Institute | Sabrina Strano, Sara Donzelli, Eleonora Sperandio, Arianna Mastrofrancesco, Fabrizio Ensoli, Fulvia Pimpinelli, Maurizio Fanciulli, Giovanni Blandino, Aldo Morrone, Gennaro Ciliberto |
| EPI_ISL_3161659 | IRCCS San Gallicano Dermatological Institute | IRCCS Regina Elena National Cancer Institute | Francesca De Nicola, Giulia Orlandi, Alice Massacci, Giovanna D’agosto, Fabrizio Ensoli, Fulvia Pimpinelli, Maurizio Fanciulli, Giovanni Blandino, Aldo Morrone, Gennaro Ciliberto |
| EPI_ISL_3161660 | IRCCS San Gallicano Dermatological Institute | IRCCS Regina Elena National Cancer Institute | Ludovica Ciuffreda, Sara Donzelli, Matteo Pallocca, Carla Mottini, Fabrizio Ensoli, Fulvia Pimpinelli, Maurizio Fanciulli, Giovanni Blandino, Aldo Morrone, Gennaro Ciliberto |
| EPI_ISL_3161661 | IRCCS San Gallicano Dermatological Institute | IRCCS Regina Elena National Cancer Institute | Frauke Goeman, Giulia Orlandi, Eleonora Sperandio, Martina Pontone, Fabrizio Ensoli, Fulvia Pimpinelli, Maurizio Fanciulli, Giovanni Blandino, Aldo Morrone, Gennaro Ciliberto |
| EPI_ISL_3161662 | IRCCS San Gallicano Dermatological Institute | IRCCS Regina Elena National Cancer Institute | Sabrina Strano, Sara Donzelli, Alice Massacci, di Domenico Enea Gino, Fabrizio Ensoli, Fulvia Pimpinelli, Maurizio Fanciulli, Giovanni Blandino, Aldo Morrone, Gennaro Ciliberto |
| EPI_ISL_3161663 | IRCCS San Gallicano Dermatological Institute | IRCCS Regina Elena National Cancer Institute | Francesca De Nicola, Giulia Orlandi, Matteo Pallocca, Ilaria Cavallo, Fabrizio Ensoli, Fulvia Pimpinelli, Maurizio Fanciulli, Giovanni Blandino, Aldo Morrone, Gennaro Ciliberto |
| EPI_ISL_3161664 | IRCCS San Gallicano Dermatological Institute | IRCCS Regina Elena National Cancer Institute | Ludovica Ciuffreda, Sara Donzelli, Eleonora Sperandio, Francesca Sivori, Fabrizio Ensoli, Fulvia Pimpinelli, Maurizio Fanciulli, Giovanni Blandino, Aldo Morrone, Gennaro Ciliberto |

| | | | |
|---|---|---|---|
| EPI_ISL_3161665 | IRCCS San Gallicano Dermatological Institute | IRCCS Regina Elena National Cancer Institute | Frauke Goeman, Giulia Orlandi, Alice Massacci, Andrea Cazzani, Fabrizio Ensoli, Fulvia Pimpinelli, Maurizio Fanciulli, Giovanni Blandino, Aldo Morrone, Gennaro Ciliberto |
| EPI_ISL_3161666 | IRCCS San Gallicano Dermatological Institute | IRCCS Regina Elena National Cancer Institute | Sabrina Strano, Sara Donzelli, Matteo Pallocca, Ilaria Celesti, Fabrizio Ensoli, Fulvia Pimpinelli, Maurizio Fanciulli, Giovanni Blandino, Aldo Morrone, Gennaro Ciliberto |
| EPI_ISL_3161667 | IRCCS San Gallicano Dermatological Institute | IRCCS Regina Elena National Cancer Institute | Francesca De Nicola, Giulia Orlandi, Eleonora Sperandio, Martina Diano, Fabrizio Ensoli, Fulvia Pimpinelli, Maurizio Fanciulli, Giovanni Blandino, Aldo Morrone, Gennaro Ciliberto |
| EPI_ISL_3161668 | IRCCS San Gallicano Dermatological Institute | IRCCS Regina Elena National Cancer Institute | Ludovica Ciuffreda, Sara Donzelli, Alice Massacci, Antonio Federico, Fabrizio Ensoli, Fulvia Pimpinelli, Maurizio Fanciulli, Giovanni Blandino, Aldo Morrone, Gennaro Ciliberto |
| EPI_ISL_3161669 | IRCCS San Gallicano Dermatological Institute | IRCCS Regina Elena National Cancer Institute | Frauke Goeman, Giulia Orlandi, Matteo Pallocca, Fulvia Fraticelli, Fabrizio Ensoli, Fulvia Pimpinelli, Maurizio Fanciulli, Giovanni Blandino, Aldo Morrone, Gennaro Ciliberto |
| EPI_ISL_3161670 | IRCCS San Gallicano Dermatological Institute | IRCCS Regina Elena National Cancer Institute | Sabrina Strano, Sara Donzelli, Eleonora Sperandio, Lorenzo Furzi, Fabrizio Ensoli, Fulvia Pimpinelli, Maurizio Fanciulli, Giovanni Blandino, Aldo Morrone, Gennaro Ciliberto |
| EPI_ISL_3161671 | IRCCS San Gallicano Dermatological Institute | IRCCS Regina Elena National Cancer Institute | Francesca De Nicola, Giulia Orlandi, Alice Massacci, Alessia Lauretti, Fabrizio Ensoli, Fulvia Pimpinelli, Maurizio Fanciulli, Giovanni Blandino, Aldo Morrone, Gennaro Ciliberto |
| EPI_ISL_3161672 | IRCCS San Gallicano Dermatological Institute | IRCCS Regina Elena National Cancer Institute | Ludovica Ciuffreda, Sara Donzelli, Matteo Pallocca, Francesca Maione, Fabrizio Ensoli, Fulvia Pimpinelli, Maurizio Fanciulli, Giovanni Blandino, Aldo Morrone, Gennaro Ciliberto |
| EPI_ISL_3161673 | IRCCS San Gallicano Dermatological Institute | IRCCS Regina Elena National Cancer Institute | Frauke Goeman, Giulia Orlandi, Eleonora Sperandio, Francisco Obregon, Fabrizio Ensoli, Fulvia Pimpinelli, Maurizio Fanciulli, Giovanni Blandino, Aldo Morrone, Gennaro Ciliberto |
| EPI_ISL_3161674 | IRCCS San Gallicano Dermatological Institute | IRCCS Regina Elena National Cancer Institute | Sabrina Strano, Sara Donzelli, Alice Massacci, Silvia Paluzzi, Fabrizio Ensoli, Fulvia Pimpinelli, Maurizio Fanciulli, Giovanni Blandino, Aldo Morrone, Gennaro Ciliberto |
| EPI_ISL_3161675 | IRCCS San Gallicano Dermatological Institute | IRCCS Regina Elena National Cancer Institute | Francesca De Nicola, Giulia Orlandi, Matteo Pallocca, Sara Petrolo, Fabrizio Ensoli, Fulvia Pimpinelli, Maurizio Fanciulli, Giovanni Blandino, Aldo Morrone, Gennaro Ciliberto |
| EPI_ISL_3161676 | IRCCS San Gallicano Dermatological Institute | IRCCS Regina Elena National Cancer Institute | Ludovica Ciuffreda, Sara Donzelli, Eleonora Sperandio, Valentina Ricca, Fabrizio Ensoli, Fulvia Pimpinelli, Maurizio Fanciulli, Giovanni Blandino, Aldo Morrone, Gennaro Ciliberto |
| EPI_ISL_3161677 | IRCCS San Gallicano Dermatological Institute | IRCCS Regina Elena National Cancer Institute | Frauke Goeman, Giulia Orlandi, Alice Massacci, Serena Salvo, Fabrizio Ensoli, Fulvia Pimpinelli, Maurizio Fanciulli, Giovanni Blandino, Aldo Morrone, Gennaro Ciliberto |
| EPI_ISL_3161678 | IRCCS San Gallicano Dermatological Institute | IRCCS Regina Elena National Cancer Institute | Sabrina Strano, Sara Donzelli, Matteo Pallocca, Grazia Prignano, Fabrizio Ensoli, Fulvia Pimpinelli, Maurizio Fanciulli, Giovanni Blandino, Aldo Morrone, Gennaro Ciliberto |
| EPI_ISL_3161679 | IRCCS San Gallicano Dermatological Institute | IRCCS Regina Elena National Cancer Institute | Francesca De Nicola, Giulia Orlandi, Eleonora Sperandio, Elisabetta Trento, Fabrizio Ensoli, Fulvia Pimpinelli, Maurizio Fanciulli, Giovanni Blandino, Aldo Morrone, Gennaro Ciliberto |
| EPI_ISL_3161680 | IRCCS San Gallicano Dermatological Institute | IRCCS Regina Elena National Cancer Institute | Ludovica Ciuffreda, Sara Donzelli, Alice Massacci, Arianna Mastrofrancesco, Fabrizio Ensoli, Fulvia Pimpinelli, Maurizio Fanciulli, Giovanni Blandino, Aldo Morrone, Gennaro Ciliberto |
| EPI_ISL_3161681 | IRCCS San Gallicano Dermatological Institute | IRCCS Regina Elena National Cancer Institute | Frauke Goeman, Giulia Orlandi, Matteo Pallocca, Giovanna D'agosto, Fabrizio Ensoli, Fulvia Pimpinelli, Maurizio Fanciulli, Giovanni Blandino, Aldo Morrone, Gennaro Ciliberto |
| EPI_ISL_3161682 | IRCCS San Gallicano Dermatological Institute | IRCCS Regina Elena National Cancer Institute | Sabrina Strano, Sara Donzelli, Eleonora Sperandio, Carla Mottini, Fabrizio Ensoli, Fulvia Pimpinelli, Maurizio Fanciulli, Giovanni Blandino, Aldo Morrone, Gennaro Ciliberto |
| EPI_ISL_3161683 | IRCCS San Gallicano Dermatological Institute | IRCCS Regina Elena National Cancer Institute | Francesca De Nicola, Giulia Orlandi, Alice Massacci, Martina Pontone, Fabrizio Ensoli, Fulvia Pimpinelli, Maurizio Fanciulli, Giovanni Blandino, Aldo Morrone, Gennaro Ciliberto |
| EPI_ISL_3161684 | IRCCS San Gallicano Dermatological Institute | IRCCS Regina Elena National Cancer Institute | Ludovica Ciuffreda, Sara Donzelli, Matteo Pallocca, di Domenico Enea Gino, Fabrizio Ensoli, Fulvia Pimpinelli, Maurizio Fanciulli, Giovanni Blandino, Aldo Morrone, Gennaro Ciliberto |
| EPI_ISL_3161685 | IRCCS San Gallicano Dermatological Institute | IRCCS Regina Elena National Cancer Institute | Sabrina Strano, Sara Donzelli, Alice Massacci, Francesca Sivori, Fabrizio Ensoli, Fulvia Pimpinelli, Maurizio Fanciulli, Giovanni Blandino, Aldo Morrone, Gennaro Ciliberto |
| EPI_ISL_3161686, EPI_ISL_3161687 | Instituto Adolfo Lutz - Regional de Santos | Instituto Adolfo Lutz, Interdiciplinary Procedures Center, Strategic Laboratory | Claudio Tavares Sacchi, Claudia Regina Gonçalves, Karoline Rodrigues Campos, Caio Vinicius Dias Lopes, Leonardo Jose Tadeu de Araujo |
| EPI_ISL_3161688, EPI_ISL_3161689, EPI_ISL_3161690, EPI_ISL_3161691, EPI_ISL_3161692, EPI_ISL_3161694, EPI_ISL_3161695, EPI_ISL_3161696, EPI_ISL_3161697, EPI_ISL_3161698, EPI_ISL_3161699, EPI_ISL_3161700, EPI_ISL_3161701, EPI_ISL_3161702, EPI_ISL_3161703, EPI_ISL_3161704, EPI_ISL_3161705, EPI_ISL_3161706, EPI_ISL_3161707, EPI_ISL_3161708, EPI_ISL_3161709, EPI_ISL_3161710, EPI_ISL_3161712, EPI_ISL_3161713, EPI_ISL_3161714, EPI_ISL_3161715, EPI_ISL_3161716, EPI_ISL_3161717, EPI_ISL_3161718, EPI_ISL_3161719, EPI_ISL_3161720, EPI_ISL_3161721, EPI_ISL_3161722, EPI_ISL_3161723, EPI_ISL_3161724, EPI_ISL_3161725, EPI_ISL_3161726, EPI_ISL_3161727, EPI_ISL_3161728, EPI_ISL_3161729, EPI_ISL_3161730, EPI_ISL_3161731, EPI_ISL_3161732, EPI_ISL_3161734, EPI_ISL_3161735, EPI_ISL_3161736, EPI_ISL_3161738, EPI_ISL_3161739, EPI_ISL_3161740, EPI_ISL_3161741, EPI_ISL_3161743, EPI_ISL_3161744, EPI_ISL_3161745, EPI_ISL_3161747, EPI_ISL_3161749, EPI_ISL_3161751, EPI_ISL_3161752, EPI_ISL_3161753, EPI_ISL_3161755, EPI_ISL_3161758 | | | |
| see above | UW Virology Lab | UW Virology Lab | Pavitra Roychoudhury, Shah Mohamed Bakhash, Ricardo Perez, Maria Lukes, Hong Xie, Tien V. Nguyen, Noah R. Baker, Sean Ellis, Meei-Li Huang, Keith R Jerome, Alexander Greninger |
| EPI_ISL_3161759, EPI_ISL_3161760, EPI_ISL_3161761, EPI_ISL_3161762, EPI_ISL_3161763, EPI_ISL_3161764, EPI_ISL_3161765, EPI_ISL_3161766, EPI_ISL_3161767, EPI_ISL_3161768, EPI_ISL_3161769, EPI_ISL_3161770, EPI_ISL_3161771, EPI_ISL_3161772, EPI_ISL_3161773, EPI_ISL_3161774, EPI_ISL_3161775, EPI_ISL_3161776, EPI_ISL_3161777, EPI_ISL_3161778, EPI_ISL_3161779, EPI_ISL_3161780, EPI_ISL_3161781, EPI_ISL_3161782, EPI_ISL_3161784, EPI_ISL_3161785, EPI_ISL_3161786, EPI_ISL_3161787, EPI_ISL_3161788, EPI_ISL_3161789, EPI_ISL_3161790, EPI_ISL_3161791, EPI_ISL_3161792, EPI_ISL_3161793, EPI_ISL_3161796, EPI_ISL_3161797, EPI_ISL_3161798, EPI_ISL_3161799, EPI_ISL_3161800, EPI_ISL_3161801 | | | |
| see above | NHLS_VIRO | KRISP, KZN Research Innovation and Sequencing Platform | Giandhari Jennifer, Pillay Sureshnee, Yajna Ramphal, Naidoo Yeshnee, Tshabuila Derek,Tegally Houriiyah, San James, Wilkinson Eduan, de Oliveira Tulio |
| EPI_ISL_3161806, EPI_ISL_3161807, EPI_ISL_3161808 | NIC, Viral Respiratory Unit | Virology Departement | Alia Grad, Fayez Khardine, Amel Benyahia, Fetouma Doudou, Nardjes Hihi, Mohamed Amine Beloufa, Aissam Hachid, Aicha Bensalem, Fawzi Derrar |
| EPI_ISL_3161809, EPI_ISL_3161810, EPI_ISL_3161811 | NIC, Viral Respiratory Unit | Virology Departement | Fetouma Doudou, Fayez Khardine, Mohamed Amine Beloufa, Aissam Hachid, Aicha Bensalem, Fawzi Derrar |
| EPI_ISL_3161812, EPI_ISL_3161814, EPI_ISL_3161815, EPI_ISL_3161816, EPI_ISL_3161817, EPI_ISL_3161818, EPI_ISL_3161819, EPI_ISL_3161820, EPI_ISL_3161821, EPI_ISL_3161822, EPI_ISL_3161823, EPI_ISL_3161824, EPI_ISL_3161825, EPI_ISL_3161826, EPI_ISL_3161827, EPI_ISL_3161828, EPI_ISL_3161829, EPI_ISL_3161830, EPI_ISL_3161831, EPI_ISL_3161832, EPI_ISL_3161833, EPI_ISL_3161834, EPI_ISL_3161835, EPI_ISL_3161836, EPI_ISL_3161837, EPI_ISL_3161838, EPI_ISL_3161839, EPI_ISL_3161840, EPI_ISL_3161841, EPI_ISL_3161842, EPI_ISL_3161843, EPI_ISL_3161844, EPI_ISL_3161845, EPI_ISL_3161846 | | | |
| see above | Queensland Health Forensic and Scientific Services | Queensland Health Forensic and Scientific Services | Chenwei Wang on behalf of Q-PHIRE Genomics |
| EPI_ISL_3161848 | BJ Medical College, Ahmedabad | Gujarat Biotechnology Research Centre | Twinkle Soni,Sonal Sharma ,Zuber Saiyed,Ramesh Pandit,Janvi Raval,Zarna Patel,Nitin Savaliya,Dinesh Kumar,Pranay Shah,Umang Mishra,Nitesh Shah,Chaitanya Joshi,Madhvi Joshi |
| EPI_ISL_3161849 | College of veterinary Science and A.H Dantiwada | Gujarat Biotechnology Research Centre | Sonal Sharma ,Zuber Saiyed,Ramesh Pandit,Janvi Raval,Zarna Patel,Nitin Savaliya,Dinesh Kumar,Twinkle Soni,Hemlatta Sunsara,Umang Mishra,Nitesh Shah,Chaitanya Joshi,Madhvi Joshi |
| EPI_ISL_3161850 | College of veterinary Science and A.H Dantiwada | Gujarat Biotechnology Research Centre | Zuber Saiyed,Ramesh Pandit,Janvi Raval,Zarna Patel,Nitin Savaliya,Dinesh Kumar,Twinkle Soni,Sonal Sharma ,Hemlatta Sunsara,Umang Mishra,Nitesh Shah,Chaitanya Joshi,Madhvi Joshi |
| EPI_ISL_3161851 | GMERS Medical College, Valsad | Gujarat Biotechnology Research Centre | Ramesh Pandit,Janvi Raval,Zarna Patel,Nitin Savaliya,Dinesh Kumar,Twinkle Soni,Sonal Sharma ,Zuber Saiyed,Vicky Gandhi,Umang Mishra,Nitesh Shah,Chaitanya Joshi,Madhvi Joshi |
| EPI_ISL_3161852 | GMERS Medical College, Valsad | Gujarat Biotechnology Research Centre | Janvi Raval,Zarna Patel,Nitin Savaliya,Dinesh Kumar,Twinkle Soni,Sonal Sharma ,Zuber Saiyed,Ramesh Pandit,Vicky Gandhi,Umang Mishra,Nitesh Shah,Chaitanya Joshi,Madhvi Joshi |
| EPI_ISL_3161853 | GMERS Medical College, Valsad | Gujarat Biotechnology Research Centre | Zarna Patel,Nitin Savaliya,Dinesh Kumar,Twinkle Soni,Sonal Sharma ,Zuber Saiyed,Ramesh Pandit,Janvi Raval,Vicky Gandhi,Umang Mishra,Nitesh Shah,Chaitanya Joshi,Madhvi Joshi |
| EPI_ISL_3161854 | GMERS Medical College, Valsad | Gujarat Biotechnology Research Centre | Nitin Savaliya,Dinesh Kumar,Twinkle Soni,Sonal Sharma ,Zuber Saiyed,Ramesh Pandit,Janvi Raval,Zarna Patel,Vicky Gandhi,Umang Mishra,Nitesh Shah,Chaitanya Joshi,Madhvi Joshi |
| EPI_ISL_3161855 | GMERS Medical College, Valsad | Gujarat Biotechnology Research Centre | Dinesh Kumar,Twinkle Soni,Sonal Sharma ,Zuber Saiyed,Ramesh Pandit,Janvi Raval,Zarna Patel,Nitin Savaliya,Vicky Gandhi,Umang Mishra,Nitesh Shah,Chaitanya Joshi,Madhvi Joshi |
| EPI_ISL_3161856 | GMERS Medical College, Valsad | Gujarat Biotechnology Research Centre | Twinkle Soni,Sonal Sharma ,Zuber Saiyed,Ramesh Pandit,Janvi Raval,Zarna Patel,Nitin Savaliya,Dinesh Kumar,Vicky Gandhi,Umang Mishra,Nitesh Shah,Chaitanya Joshi,Madhvi Joshi |
| EPI_ISL_3161857 | GMERS Medical College, Valsad | Gujarat Biotechnology Research Centre | Sonal Sharma ,Zuber Saiyed,Ramesh Pandit,Janvi Raval,Zarna Patel,Nitin Savaliya,Dinesh Kumar,Twinkle Soni,Vicky Gandhi,Umang Mishra,Nitesh Shah,Chaitanya Joshi,Madhvi Joshi |
| EPI_ISL_3161858 | GMERS Medical College, Valsad | Gujarat Biotechnology Research Centre | Zuber Saiyed,Ramesh Pandit,Janvi Raval,Zarna Patel,Nitin Savaliya,Dinesh Kumar,Twinkle Soni,Sonal Sharma ,Vicky Gandhi,Umang Mishra,Nitesh Shah,Chaitanya Joshi,Madhvi Joshi |
| EPI_ISL_3161859 | GMERS Medical College, Valsad | Gujarat Biotechnology Research Centre | Ramesh Pandit,Janvi Raval,Zarna Patel,Nitin Savaliya,Dinesh Kumar,Twinkle Soni,Sonal Sharma ,Zuber Saiyed,Vicky Gandhi,Umang Mishra,Nitesh Shah,Chaitanya Joshi,Madhvi Joshi |
| EPI_ISL_3161860 | GMERS Medical College, Valsad | Gujarat Biotechnology Research Centre | Janvi Raval,Zarna Patel,Nitin Savaliya,Dinesh Kumar,Twinkle Soni,Sonal Sharma ,Zuber Saiyed,Ramesh Pandit,Vicky Gandhi,Umang Mishra,Nitesh Shah,Chaitanya Joshi,Madhvi Joshi |
| EPI_ISL_3161861 | GMERS Medical College, Valsad | Gujarat Biotechnology Research Centre | Zarna Patel,Nitin Savaliya,Dinesh Kumar,Twinkle Soni,Sonal Sharma ,Zuber Saiyed,Ramesh Pandit,Janvi Raval,Vicky Gandhi,Umang Mishra,Nitesh Shah,Chaitanya Joshi,Madhvi Joshi |
| EPI_ISL_3161862 | GMERS Medical College, Valsad | Gujarat Biotechnology Research Centre | Nitin Savaliya,Dinesh Kumar,Twinkle Soni,Sonal Sharma ,Zuber Saiyed,Ramesh Pandit,Janvi Raval,Zarna Patel,Vicky Gandhi,Umang Mishra,Nitesh Shah,Chaitanya Joshi,Madhvi Joshi |
| EPI_ISL_3161863 | GMERS Medical College, Valsad | Gujarat Biotechnology Research Centre | Dinesh Kumar,Twinkle Soni,Sonal Sharma ,Zuber Saiyed,Ramesh Pandit,Janvi Raval,Zarna Patel,Nitin Savaliya,Vicky Gandhi,Umang Mishra,Nitesh Shah,Chaitanya Joshi,Madhvi Joshi |
| EPI_ISL_3161864 | GMERS Medical College, Valsad | Gujarat Biotechnology Research Centre | Twinkle Soni,Sonal Sharma ,Zuber Saiyed,Ramesh Pandit,Janvi Raval,Zarna Patel,Nitin Savaliya,Dinesh Kumar,Vicky Gandhi,Umang Mishra,Nitesh Shah,Chaitanya Joshi,Madhvi Joshi |
| EPI_ISL_3161865 | Banas Medical College & Research Institute (BMCRI), Palanpur | Gujarat Biotechnology Research Centre | Sonal Sharma ,Zuber Saiyed,Ramesh Pandit,Janvi Raval,Zarna Patel,Nitin Savaliya,Dinesh Kumar,Twinkle Soni,Anil Choudhary ,Amar Sajjan,Umang Mishra,Nitesh Shah,Chaitanya Joshi,Madhvi Joshi |
| EPI_ISL_3161866 | Banas Medical College & Research Institute (BMCRI), Palanpur | Gujarat Biotechnology Research Centre | Zuber Saiyed,Ramesh Pandit,Janvi Raval,Zarna Patel,Nitin Savaliya,Dinesh Kumar,Twinkle Soni,Sonal Sharma ,Anil Choudhary ,Amar Sajjan,Umang Mishra,Nitesh Shah,Chaitanya Joshi,Madhvi Joshi |
| EPI_ISL_3161867 | Banas Medical College & Research Institute (BMCRI), Palanpur | Gujarat Biotechnology Research Centre | Ramesh Pandit,Janvi Raval,Zarna Patel,Nitin Savaliya,Dinesh Kumar,Twinkle Soni,Sonal Sharma ,Zuber Saiyed,Anil Choudhary ,Amar Sajjan,Umang Mishra,Nitesh Shah,Chaitanya Joshi,Madhvi Joshi |
| EPI_ISL_3161868 | Banas Medical College & Research Institute (BMCRI), Palanpur | Gujarat Biotechnology Research Centre | Janvi Raval,Zarna Patel,Nitin Savaliya,Dinesh Kumar,Twinkle Soni,Sonal Sharma ,Zuber Saiyed,Ramesh Pandit,Anil Choudhary ,Amar Sajjan,Umang Mishra,Nitesh Shah,Chaitanya Joshi,Madhvi Joshi |
| EPI_ISL_3161869 | Banas Medical College & Research Institute (BMCRI), Palanpur | Gujarat Biotechnology Research Centre | Zarna Patel,Nitin Savaliya,Dinesh Kumar,Twinkle Soni,Sonal Sharma ,Zuber Saiyed,Ramesh Pandit,Janvi Raval,Anil Choudhary ,Amar Sajjan,Umang Mishra,Nitesh Shah,Chaitanya Joshi,Madhvi Joshi |
| EPI_ISL_3161870 | Banas Medical College & Research Institute (BMCRI), Palanpur | Gujarat Biotechnology Research Centre | Nitin Savaliya,Dinesh Kumar,Twinkle Soni,Sonal Sharma ,Zuber Saiyed,Ramesh Pandit,Janvi Raval,Zarna Patel,Anil Choudhary ,Amar Sajjan,Umang Mishra,Nitesh Shah,Chaitanya Joshi,Madhvi Joshi |
| EPI_ISL_3161871 | Banas Medical College & Research Institute (BMCRI), Palanpur | Gujarat Biotechnology Research Centre | Dinesh Kumar,Twinkle Soni,Sonal Sharma ,Zuber Saiyed,Ramesh Pandit,Janvi Raval,Zarna Patel,Nitin Savaliya,Anil Choudhary ,Amar Sajjan,Umang Mishra,Nitesh Shah,Chaitanya Joshi,Madhvi Joshi |
| EPI_ISL_3161872 | Banas Medical College & Research Institute (BMCRI), Palanpur | Gujarat Biotechnology Research Centre | Twinkle Soni,Sonal Sharma ,Zuber Saiyed,Ramesh Pandit,Janvi Raval,Zarna Patel,Nitin Savaliya,Dinesh Kumar,Anil Choudhary ,Amar Sajjan,Umang Mishra,Nitesh Shah,Chaitanya Joshi,Madhvi Joshi |
| EPI_ISL_3161873 | Banas Medical College & Research Institute (BMCRI), Palanpur | Gujarat Biotechnology Research Centre | Sonal Sharma ,Zuber Saiyed,Ramesh Pandit,Janvi Raval,Zarna Patel,Nitin Savaliya,Dinesh Kumar,Twinkle Soni,Anil Choudhary ,Amar Sajjan,Umang Mishra,Nitesh Shah,Chaitanya Joshi,Madhvi Joshi |
| EPI_ISL_3161874 | Banas Medical College & Research Institute (BMCRI), Palanpur | Gujarat Biotechnology Research Centre | Zuber Saiyed,Ramesh Pandit,Janvi Raval,Zarna Patel,Nitin Savaliya,Dinesh Kumar,Twinkle Soni,Sonal Sharma ,Anil Choudhary ,Amar Sajjan,Umang Mishra,Nitesh Shah,Chaitanya Joshi,Madhvi Joshi |
| EPI_ISL_3161875 | Banas Medical College & Research Institute (BMCRI), Palanpur | Gujarat Biotechnology Research Centre | Ramesh Pandit,Janvi Raval,Zarna Patel,Nitin Savaliya,Dinesh Kumar,Twinkle Soni,Sonal Sharma ,Zuber Saiyed,Anil Choudhary ,Amar Sajjan,Umang Mishra,Nitesh Shah,Chaitanya Joshi,Madhvi Joshi |
| EPI_ISL_3161876 | Banas Medical College & Research Institute (BMCRI), Palanpur | Gujarat Biotechnology Research Centre | Janvi Raval,Zarna Patel,Nitin Savaliya,Dinesh Kumar,Twinkle Soni,Sonal Sharma ,Zuber Saiyed,Ramesh Pandit,Anil Choudhary ,Amar Sajjan,Umang Mishra,Nitesh Shah,Chaitanya Joshi,Madhvi Joshi |
| EPI_ISL_3161877 | Banas Medical College & Research Institute (BMCRI), Palanpur | Gujarat Biotechnology Research Centre | Zarna Patel,Nitin Savaliya,Dinesh Kumar,Twinkle Soni,Sonal Sharma ,Zuber Saiyed,Ramesh Pandit,Janvi Raval,Anil Choudhary ,Amar Sajjan,Umang Mishra,Nitesh Shah,Chaitanya Joshi,Madhvi Joshi |
| EPI_ISL_3161878 | Banas Medical College & Research Institute (BMCRI), Palanpur | Gujarat Biotechnology Research Centre | Nitin Savaliya,Dinesh Kumar,Twinkle Soni,Sonal Sharma ,Zuber Saiyed,Ramesh Pandit,Janvi Raval,Zarna Patel,Anil Choudhary ,Amar Sajjan,Umang Mishra,Nitesh Shah,Chaitanya Joshi,Madhvi Joshi |
| EPI_ISL_3161879 | Banas Medical College & Research Institute (BMCRI), Palanpur | Gujarat Biotechnology Research Centre | Dinesh Kumar,Twinkle Soni,Sonal Sharma ,Zuber Saiyed,Ramesh Pandit,Janvi Raval,Zarna Patel,Nitin Savaliya,Anil Choudhary ,Amar Sajjan,Umang Mishra,Nitesh Shah,Chaitanya Joshi,Madhvi Joshi |
| EPI_ISL_3161880 | Banas Medical College & Research Institute (BMCRI), Palanpur | Gujarat Biotechnology Research Centre | Twinkle Soni,Sonal Sharma ,Zuber Saiyed,Ramesh Pandit,Janvi Raval,Zarna Patel,Nitin Savaliya,Dinesh Kumar,Anil Choudhary ,Amar Sajjan,Umang Mishra,Nitesh Shah,Chaitanya Joshi,Madhvi Joshi |
| EPI_ISL_3161881 | Banas Medical College & Research Institute (BMCRI), Palanpur | Gujarat Biotechnology Research Centre | Sonal Sharma ,Zuber Saiyed,Ramesh Pandit,Janvi Raval,Zarna Patel,Nitin Savaliya,Dinesh Kumar,Twinkle Soni,Anil Choudhary ,Amar Sajjan,Umang Mishra,Nitesh Shah,Chaitanya Joshi,Madhvi Joshi |
| EPI_ISL_3161882 | Banas Medical College & Research Institute (BMCRI), Palanpur | Gujarat Biotechnology Research Centre | Zuber Saiyed,Ramesh Pandit,Janvi Raval,Zarna Patel,Nitin Savaliya,Dinesh Kumar,Twinkle Soni,Sonal Sharma ,Anil Choudhary ,Amar Sajjan,Umang Mishra,Nitesh Shah,Chaitanya Joshi,Madhvi Joshi |
| EPI_ISL_3161883 | Banas Medical College & Research Institute (BMCRI), Palanpur | Gujarat Biotechnology Research Centre | Ramesh Pandit,Janvi Raval,Zarna Patel,Nitin Savaliya,Dinesh Kumar,Twinkle Soni,Sonal Sharma ,Zuber Saiyed,Anil Choudhary ,Amar Sajjan,Umang Mishra,Nitesh Shah,Chaitanya Joshi,Madhvi Joshi |
| EPI_ISL_3161884 | Banas Medical College & Research Institute (BMCRI), Palanpur | Gujarat Biotechnology Research Centre | Janvi Raval,Zarna Patel,Nitin Savaliya,Dinesh Kumar,Twinkle Soni,Sonal Sharma ,Zuber Saiyed,Ramesh Pandit,Anil Choudhary ,Amar Sajjan,Umang Mishra,Nitesh Shah,Chaitanya Joshi,Madhvi Joshi |
| EPI_ISL_3161885 | Banas Medical College & Research Institute (BMCRI), Palanpur | Gujarat Biotechnology Research Centre | Zarna Patel,Nitin Savaliya,Dinesh Kumar,Twinkle Soni,Sonal Sharma ,Zuber Saiyed,Ramesh Pandit,Janvi Raval,Anil Choudhary ,Amar Sajjan,Umang Mishra,Nitesh Shah,Chaitanya Joshi,Madhvi Joshi |
| EPI_ISL_3161886 | Banas Medical College & Research Institute (BMCRI), Palanpur | Gujarat Biotechnology Research Centre | Nitin Savaliya,Dinesh Kumar,Twinkle Soni,Sonal Sharma ,Zuber Saiyed,Ramesh Pandit,Janvi Raval,Zarna Patel,Anil Choudhary ,Amar Sajjan,Umang Mishra,Nitesh Shah,Chaitanya Joshi,Madhvi Joshi |
| EPI_ISL_3161887 | Banas Medical College & Research Institute (BMCRI), Palanpur | Gujarat Biotechnology Research Centre | Dinesh Kumar,Twinkle Soni,Sonal Sharma ,Zuber Saiyed,Ramesh Pandit,Janvi Raval,Zarna Patel,Nitin Savaliya,Anil Choudhary ,Amar Sajjan,Umang Mishra,Nitesh Shah,Chaitanya Joshi,Madhvi Joshi |

| | | | |
|---|---|---|---|
| EPI_ISL_3161888 | Banas Medical College & Research Institute (BMCRI), Palanpur | Gujarat Biotechnology Research Centre | Twinkle Soni,Sonal Sharma ,Zuber Saiyed,Ramesh Pandit,Janvi Raval,Zarna Patel,Nitin Savaliya,Dinesh Kumar,Anil Choudhary ,Amar Sajjan,Umang Mishra,Nitesh Shah,Chaitanya Joshi,Madhvi Joshi |
| EPI_ISL_3161889 | Banas Medical College & Research Institute (BMCRI), Palanpur | Gujarat Biotechnology Research Centre | Sonal Sharma ,Zuber Saiyed,Ramesh Pandit,Janvi Raval,Zarna Patel,Nitin Savaliya,Dinesh Kumar,Twinkle Soni,Anil Choudhary ,Amar Sajjan,Umang Mishra,Nitesh Shah,Chaitanya Joshi,Madhvi Joshi |
| EPI_ISL_3161890 | Banas Medical College & Research Institute (BMCRI), Palanpur | Gujarat Biotechnology Research Centre | Zuber Saiyed,Ramesh Pandit,Janvi Raval,Zarna Patel,Nitin Savaliya,Dinesh Kumar,Twinkle Soni,Sonal Sharma ,Anil Choudhary ,Amar Sajjan,Umang Mishra,Nitesh Shah,Chaitanya Joshi,Madhvi Joshi |
| EPI_ISL_3161891 | Banas Medical College & Research Institute (BMCRI), Palanpur | Gujarat Biotechnology Research Centre | Ramesh Pandit,Janvi Raval,Zarna Patel,Nitin Savaliya,Dinesh Kumar,Twinkle Soni,Sonal Sharma ,Zuber Saiyed,Anil Choudhary ,Amar Sajjan,Umang Mishra,Nitesh Shah,Chaitanya Joshi,Madhvi Joshi |
| EPI_ISL_3161892 | Banas Medical College & Research Institute (BMCRI), Palanpur | Gujarat Biotechnology Research Centre | Janvi Raval,Zarna Patel,Nitin Savaliya,Dinesh Kumar,Twinkle Soni,Sonal Sharma ,Zuber Saiyed,Ramesh Pandit,Anil Choudhary ,Amar Sajjan,Umang Mishra,Nitesh Shah,Chaitanya Joshi,Madhvi Joshi |
| EPI_ISL_3161893 | Banas Medical College & Research Institute (BMCRI), Palanpur | Gujarat Biotechnology Research Centre | Zarna Patel,Nitin Savaliya,Dinesh Kumar,Twinkle Soni,Sonal Sharma ,Zuber Saiyed,Ramesh Pandit,Janvi Raval,Anil Choudhary ,Amar Sajjan,Umang Mishra,Nitesh Shah,Chaitanya Joshi,Madhvi Joshi |
| EPI_ISL_3161894 | Banas Medical College & Research Institute (BMCRI), Palanpur | Gujarat Biotechnology Research Centre | Nitin Savaliya,Dinesh Kumar,Twinkle Soni,Sonal Sharma ,Zuber Saiyed,Ramesh Pandit,Janvi Raval,Zarna Patel,Anil Choudhary ,Amar Sajjan,Umang Mishra,Nitesh Shah,Chaitanya Joshi,Madhvi Joshi |
| EPI_ISL_3161895 | College of veterinary Science and A.H Dantiwada | Gujarat Biotechnology Research Centre | Dinesh Kumar,Twinkle Soni,Sonal Sharma ,Zuber Saiyed,Ramesh Pandit,Janvi Raval,Zarna Patel,Nitin Savaliya,Hemlatta Sunsara,Umang Mishra,Nitesh Shah,Chaitanya Joshi,Madhvi Joshi |
| EPI_ISL_3161896 | College of veterinary Science and A.H Dantiwada | Gujarat Biotechnology Research Centre | Twinkle Soni,Sonal Sharma ,Zuber Saiyed,Ramesh Pandit,Janvi Raval,Zarna Patel,Nitin Savaliya,Dinesh Kumar,Hemlatta Sunsara,Umang Mishra,Nitesh Shah,Chaitanya Joshi,Madhvi Joshi |
| EPI_ISL_3161897 | College of veterinary Science and A.H Dantiwada | Gujarat Biotechnology Research Centre | Sonal Sharma ,Zuber Saiyed,Ramesh Pandit,Janvi Raval,Zarna Patel,Nitin Savaliya,Dinesh Kumar,Twinkle Soni,Hemlatta Sunsara,Umang Mishra,Nitesh Shah,Chaitanya Joshi,Madhvi Joshi |
| EPI_ISL_3161898 | College of veterinary Science and A.H Dantiwada | Gujarat Biotechnology Research Centre | Zuber Saiyed,Ramesh Pandit,Janvi Raval,Zarna Patel,Nitin Savaliya,Dinesh Kumar,Twinkle Soni,Sonal Sharma ,Hemlatta Sunsara,Umang Mishra,Nitesh Shah,Chaitanya Joshi,Madhvi Joshi |
| EPI_ISL_3161899 | College of veterinary Science and A.H Dantiwada | Gujarat Biotechnology Research Centre | Ramesh Pandit,Janvi Raval,Zarna Patel,Nitin Savaliya,Dinesh Kumar,Twinkle Soni,Sonal Sharma ,Zuber Saiyed,Hemlatta Sunsara,Umang Mishra,Nitesh Shah,Chaitanya Joshi,Madhvi Joshi |
| EPI_ISL_3161900 | College of veterinary Science and A.H Dantiwada | Gujarat Biotechnology Research Centre | Janvi Raval,Zarna Patel,Nitin Savaliya,Dinesh Kumar,Twinkle Soni,Sonal Sharma ,Zuber Saiyed,Ramesh Pandit,Hemlatta Sunsara,Umang Mishra,Nitesh Shah,Chaitanya Joshi,Madhvi Joshi |
| EPI_ISL_3161901 | College of veterinary Science and A.H Dantiwada | Gujarat Biotechnology Research Centre | Zarna Patel,Nitin Savaliya,Dinesh Kumar,Twinkle Soni,Sonal Sharma ,Zuber Saiyed,Ramesh Pandit,Janvi Raval,Hemlatta Sunsara,Umang Mishra,Nitesh Shah,Chaitanya Joshi,Madhvi Joshi |
| EPI_ISL_3161902 | College of veterinary Science and A.H Dantiwada | Gujarat Biotechnology Research Centre | Nitin Savaliya,Dinesh Kumar,Twinkle Soni,Sonal Sharma ,Zuber Saiyed,Ramesh Pandit,Janvi Raval,Zarna Patel,Hemlatta Sunsara,Umang Mishra,Nitesh Shah,Chaitanya Joshi,Madhvi Joshi |
| EPI_ISL_3161903 | College of veterinary Science and A.H Dantiwada | Gujarat Biotechnology Research Centre | Dinesh Kumar,Twinkle Soni,Sonal Sharma ,Zuber Saiyed,Ramesh Pandit,Janvi Raval,Zarna Patel,Nitin Savaliya,Hemlatta Sunsara,Umang Mishra,Nitesh Shah,Chaitanya Joshi,Madhvi Joshi |
| EPI_ISL_3161904 | College of veterinary Science and A.H Dantiwada | Gujarat Biotechnology Research Centre | Twinkle Soni,Sonal Sharma ,Zuber Saiyed,Ramesh Pandit,Janvi Raval,Zarna Patel,Nitin Savaliya,Dinesh Kumar,Hemlatta Sunsara,Umang Mishra,Nitesh Shah,Chaitanya Joshi,Madhvi Joshi |
| EPI_ISL_3161905 | College of veterinary Science and A.H Dantiwada | Gujarat Biotechnology Research Centre | Sonal Sharma ,Zuber Saiyed,Ramesh Pandit,Janvi Raval,Zarna Patel,Nitin Savaliya,Dinesh Kumar,Twinkle Soni,Hemlatta Sunsara,Umang Mishra,Nitesh Shah,Chaitanya Joshi,Madhvi Joshi |
| EPI_ISL_3161906 | College of veterinary Science and A.H Dantiwada | Gujarat Biotechnology Research Centre | Zuber Saiyed,Ramesh Pandit,Janvi Raval,Zarna Patel,Nitin Savaliya,Dinesh Kumar,Twinkle Soni,Sonal Sharma ,Hemlatta Sunsara,Umang Mishra,Nitesh Shah,Chaitanya Joshi,Madhvi Joshi |
| EPI_ISL_3161907 | College of veterinary Science and A.H Dantiwada | Gujarat Biotechnology Research Centre | Ramesh Pandit,Janvi Raval,Zarna Patel,Nitin Savaliya,Dinesh Kumar,Twinkle Soni,Sonal Sharma ,Zuber Saiyed,Hemlatta Sunsara,Umang Mishra,Nitesh Shah,Chaitanya Joshi,Madhvi Joshi |
| EPI_ISL_3161908 | Supratech Micropath Laboratory Research Institute Pvt Ltd, Ahmedabad | Gujarat Biotechnology Research Centre | Janvi Raval,Zarna Patel,Nitin Savaliya,Dinesh Kumar,Twinkle Soni,Sonal Sharma ,Zuber Saiyed,Ramesh Pandit,Mahendra Parikh,Umang Mishra,Nitesh Shah,Chaitanya Joshi,Madhvi Joshi |
| EPI_ISL_3161909 | Neuberg Supratech Refrence Laboratory, Surat | Gujarat Biotechnology Research Centre | Zarna Patel,Nitin Savaliya,Dinesh Kumar,Twinkle Soni,Sonal Sharma ,Zuber Saiyed,Ramesh Pandit,Janvi Raval,Manthan Shah,Umang Mishra,Nitesh Shah,Chaitanya Joshi,Madhvi Joshi |
| EPI_ISL_3161910 | Neuberg Supratech Refrence Laboratory, Surat | Gujarat Biotechnology Research Centre | Nitin Savaliya,Dinesh Kumar,Twinkle Soni,Sonal Sharma ,Zuber Saiyed,Ramesh Pandit,Janvi Raval,Zarna Patel,Manthan Shah,Umang Mishra,Nitesh Shah,Chaitanya Joshi,Madhvi Joshi |
| EPI_ISL_3161911 | Neuberg Supratech Refrence Laboratory, Surat | Gujarat Biotechnology Research Centre | Dinesh Kumar,Twinkle Soni,Sonal Sharma ,Zuber Saiyed,Ramesh Pandit,Janvi Raval,Zarna Patel,Nitin Savaliya,Manthan Shah,Umang Mishra,Nitesh Shah,Chaitanya Joshi,Madhvi Joshi |
| EPI_ISL_3161912 | Neuberg Supratech Refrence Laboratory, Surat | Gujarat Biotechnology Research Centre | Twinkle Soni,Sonal Sharma ,Zuber Saiyed,Ramesh Pandit,Janvi Raval,Zarna Patel,Nitin Savaliya,Dinesh Kumar,Manthan Shah,Umang Mishra,Nitesh Shah,Chaitanya Joshi,Madhvi Joshi |

EPI_ISL_3161913, EPI_ISL_3161915, EPI_ISL_3161916, EPI_ISL_3161917, EPI_ISL_3161918, EPI_ISL_3161919, EPI_ISL_3161921, EPI_ISL_3161922, EPI_ISL_3161923, EPI_ISL_3161924, EPI_ISL_3161925, EPI_ISL_3161926, EPI_ISL_3161927, EPI_ISL_3161928, EPI_ISL_3161929, EPI_ISL_3161930, EPI_ISL_3161931, EPI_ISL_3161932, EPI_ISL_3161933, EPI_ISL_3161934, EPI_ISL_3161935, EPI_ISL_3161936, EPI_ISL_3161937, EPI_ISL_3161938, EPI_ISL_3161939, EPI_ISL_3161940, EPI_ISL_3161941, EPI_ISL_3161942, EPI_ISL_3161943, EPI_ISL_3161945, EPI_ISL_3161946, EPI_ISL_3161947, EPI_ISL_3161948, EPI_ISL_3161949, EPI_ISL_3161950, EPI_ISL_3161951, EPI_ISL_3161952, EPI_ISL_3161953, EPI_ISL_3161954

| | | | |
|---|---|---|---|
| see above | Rhode Island State Health Laboratory | Rhode Island State Health Laboratory | Ewa King, Richard C. Huard, Kristin Carpenter-Azevedo |
| EPI_ISL_3161956 | Government Medical College (GMC), Surat | Gujarat Biotechnology Research Centre | Nitin Shukla,Sonal Sharma ,Zuber Saiyed,Ramesh Pandit,Janvi Raval,Zarna Patel,Nitin Savaliya,Dinesh Kumar,Twinkle Soni,Neeta Khandelwal,Umang Mishra,Nitesh Shah,Chaitanya Joshi,Madhvi Joshi |
| EPI_ISL_3161957 | Government Medical College (GMC), Surat | Gujarat Biotechnology Research Centre | Sonal Sharma ,Zuber Saiyed,Ramesh Pandit,Janvi Raval,Zarna Patel,Nitin Savaliya,Dinesh Kumar,Twinkle Soni,Nitin Shukla,Neeta Khandelwal,Umang Mishra,Nitesh Shah,Chaitanya Joshi,Madhvi Joshi |
| EPI_ISL_3161958 | Government Medical College (GMC), Surat | Gujarat Biotechnology Research Centre | Zuber Saiyed,Ramesh Pandit,Janvi Raval,Zarna Patel,Nitin Savaliya,Dinesh Kumar,Twinkle Soni,Nitin Shukla,Sonal Sharma ,Neeta Khandelwal,Umang Mishra,Nitesh Shah,Chaitanya Joshi,Madhvi Joshi |
| EPI_ISL_3161959 | Government Medical College (GMC), Surat | Gujarat Biotechnology Research Centre | Ramesh Pandit,Janvi Raval,Zarna Patel,Nitin Savaliya,Dinesh Kumar,Twinkle Soni,Nitin Shukla,Sonal Sharma ,Zuber Saiyed,Neeta Khandelwal,Umang Mishra,Nitesh Shah,Chaitanya Joshi,Madhvi Joshi |
| EPI_ISL_3161960 | Government Medical College (GMC), Surat | Gujarat Biotechnology Research Centre | Janvi Raval,Zarna Patel,Nitin Savaliya,Dinesh Kumar,Twinkle Soni,Nitin Shukla,Sonal Sharma ,Zuber Saiyed,Ramesh Pandit,Neeta Khandelwal,Umang Mishra,Nitesh Shah,Chaitanya Joshi,Madhvi Joshi |
| EPI_ISL_3161961 | Government Medical College (GMC), Surat | Gujarat Biotechnology Research Centre | Zarna Patel,Nitin Savaliya,Dinesh Kumar,Twinkle Soni,Nitin Shukla,Sonal Sharma ,Zuber Saiyed,Ramesh Pandit,Janvi Raval,Neeta Khandelwal,Umang Mishra,Nitesh Shah,Chaitanya Joshi,Madhvi Joshi |
| EPI_ISL_3161962 | Government Medical College (GMC), Surat | Gujarat Biotechnology Research Centre | Nitin Savaliya,Dinesh Kumar,Twinkle Soni,Nitin Shukla,Sonal Sharma ,Zuber Saiyed,Ramesh Pandit,Janvi Raval,Zarna Patel,Neeta Khandelwal,Umang Mishra,Nitesh Shah,Chaitanya Joshi,Madhvi Joshi |
| EPI_ISL_3161963 | Government Medical College (GMC), Surat | Gujarat Biotechnology Research Centre | Dinesh Kumar,Twinkle Soni,Nitin Shukla,Sonal Sharma ,Zuber Saiyed,Ramesh Pandit,Janvi Raval,Zarna Patel,Nitin Savaliya,Neeta Khandelwal,Umang Mishra,Nitesh Shah,Chaitanya Joshi,Madhvi Joshi |
| EPI_ISL_3161964 | Government Medical College (GMC), Surat | Gujarat Biotechnology Research Centre | Twinkle Soni,Nitin Shukla,Sonal Sharma ,Zuber Saiyed,Ramesh Pandit,Janvi Raval,Zarna Patel,Nitin Savaliya,Dinesh Kumar,Neeta Khandelwal,Umang Mishra,Nitesh Shah,Chaitanya Joshi,Madhvi Joshi |
| EPI_ISL_3161965 | Government Medical College (GMC), Surat | Gujarat Biotechnology Research Centre | Nitin Shukla,Sonal Sharma ,Zuber Saiyed,Ramesh Pandit,Janvi Raval,Zarna Patel,Nitin Savaliya,Dinesh Kumar,Twinkle Soni,Neeta Khandelwal,Umang Mishra,Nitesh Shah,Chaitanya Joshi,Madhvi Joshi |
| EPI_ISL_3161966 | Government Medical College (GMC), Surat | Gujarat Biotechnology Research Centre | Sonal Sharma ,Zuber Saiyed,Ramesh Pandit,Janvi Raval,Zarna Patel,Nitin Savaliya,Dinesh Kumar,Twinkle Soni,Nitin Shukla,Neeta Khandelwal,Umang Mishra,Nitesh Shah,Chaitanya Joshi,Madhvi Joshi |
| EPI_ISL_3161967 | Government Medical College (GMC), Surat | Gujarat Biotechnology Research Centre | Zuber Saiyed,Ramesh Pandit,Janvi Raval,Zarna Patel,Nitin Savaliya,Dinesh Kumar,Twinkle Soni,Nitin Shukla,Sonal Sharma ,Neeta Khandelwal,Umang Mishra,Nitesh Shah,Chaitanya Joshi,Madhvi Joshi |
| EPI_ISL_3161968 | Government Medical College (GMC), Surat | Gujarat Biotechnology Research Centre | Ramesh Pandit,Janvi Raval,Zarna Patel,Nitin Savaliya,Dinesh Kumar,Twinkle Soni,Nitin Shukla,Sonal Sharma ,Zuber Saiyed,Neeta Khandelwal,Umang Mishra,Nitesh Shah,Chaitanya Joshi,Madhvi Joshi |
| EPI_ISL_3161969 | Government Medical College (GMC), Surat | Gujarat Biotechnology Research Centre | Janvi Raval,Zarna Patel,Nitin Savaliya,Dinesh Kumar,Twinkle Soni,Nitin Shukla,Sonal Sharma ,Zuber Saiyed,Ramesh Pandit,Neeta Khandelwal,Umang Mishra,Nitesh Shah,Chaitanya Joshi,Madhvi Joshi |
| EPI_ISL_3161970 | Government Medical College (GMC), Surat | Gujarat Biotechnology Research Centre | Zarna Patel,Nitin Savaliya,Dinesh Kumar,Twinkle Soni,Nitin Shukla,Sonal Sharma ,Zuber Saiyed,Ramesh Pandit,Janvi Raval,Neeta Khandelwal,Umang Mishra,Nitesh Shah,Chaitanya Joshi,Madhvi Joshi |
| EPI_ISL_3161971 | Government Medical College (GMC), Surat | Gujarat Biotechnology Research Centre | Nitin Savaliya,Dinesh Kumar,Twinkle Soni,Nitin Shukla,Sonal Sharma ,Zuber Saiyed,Ramesh Pandit,Janvi Raval,Zarna Patel,Neeta Khandelwal,Umang Mishra,Nitesh Shah,Chaitanya Joshi,Madhvi Joshi |
| EPI_ISL_3161972 | Government Medical College (GMC), Surat | Gujarat Biotechnology Research Centre | Dinesh Kumar,Twinkle Soni,Nitin Shukla,Sonal Sharma ,Zuber Saiyed,Ramesh Pandit,Janvi Raval,Zarna Patel,Nitin Savaliya,Neeta Khandelwal,Umang Mishra,Nitesh Shah,Chaitanya Joshi,Madhvi Joshi |
| EPI_ISL_3161973 | Government Medical College (GMC), Surat | Gujarat Biotechnology Research Centre | Twinkle Soni,Nitin Shukla,Sonal Sharma ,Zuber Saiyed,Ramesh Pandit,Janvi Raval,Zarna Patel,Nitin Savaliya,Dinesh Kumar,Neeta Khandelwal,Umang Mishra,Nitesh Shah,Chaitanya Joshi,Madhvi Joshi |
| EPI_ISL_3161974 | Government Medical College (GMC), Surat | Gujarat Biotechnology Research Centre | Nitin Shukla,Sonal Sharma ,Zuber Saiyed,Ramesh Pandit,Janvi Raval,Zarna Patel,Nitin Savaliya,Dinesh Kumar,Twinkle Soni,Neeta Khandelwal,Umang Mishra,Nitesh Shah,Chaitanya Joshi,Madhvi Joshi |
| EPI_ISL_3161975 | Government Medical College (GMC), Surat | Gujarat Biotechnology Research Centre | Sonal Sharma ,Zuber Saiyed,Ramesh Pandit,Janvi Raval,Zarna Patel,Nitin Savaliya,Dinesh Kumar,Twinkle Soni,Nitin Shukla,Neeta Khandelwal,Umang Mishra,Nitesh Shah,Chaitanya Joshi,Madhvi Joshi |
| EPI_ISL_3161976 | Government Medical College (GMC), Surat | Gujarat Biotechnology Research Centre | Zuber Saiyed,Ramesh Pandit,Janvi Raval,Zarna Patel,Nitin Savaliya,Dinesh Kumar,Twinkle Soni,Nitin Shukla,Sonal Sharma ,Neeta Khandelwal,Umang Mishra,Nitesh Shah,Chaitanya Joshi,Madhvi Joshi |
| EPI_ISL_3161977 | Government Medical College (GMC), Surat | Gujarat Biotechnology Research Centre | Ramesh Pandit,Janvi Raval,Zarna Patel,Nitin Savaliya,Dinesh Kumar,Twinkle Soni,Nitin Shukla,Sonal Sharma ,Zuber Saiyed,Neeta Khandelwal,Umang Mishra,Nitesh Shah,Chaitanya Joshi,Madhvi Joshi |
| EPI_ISL_3161978 | Government Medical College (GMC), Surat | Gujarat Biotechnology Research Centre | Janvi Raval,Zarna Patel,Nitin Savaliya,Dinesh Kumar,Twinkle Soni,Nitin Shukla,Sonal Sharma ,Zuber Saiyed,Ramesh Pandit,Neeta Khandelwal,Umang Mishra,Nitesh Shah,Chaitanya Joshi,Madhvi Joshi |
| EPI_ISL_3161979 | Government Medical College (GMC), Surat | Gujarat Biotechnology Research Centre | Zarna Patel,Nitin Savaliya,Dinesh Kumar,Twinkle Soni,Nitin Shukla,Sonal Sharma ,Zuber Saiyed,Ramesh Pandit,Janvi Raval,Neeta Khandelwal,Umang Mishra,Nitesh Shah,Chaitanya Joshi,Madhvi Joshi |
| EPI_ISL_3161980 | Government Medical College (GMC), Surat | Gujarat Biotechnology Research Centre | Nitin Savaliya,Dinesh Kumar,Twinkle Soni,Nitin Shukla,Sonal Sharma ,Zuber Saiyed,Ramesh Pandit,Janvi Raval,Zarna Patel,Neeta Khandelwal,Umang Mishra,Nitesh Shah,Chaitanya Joshi,Madhvi Joshi |
| EPI_ISL_3161981 | Government Medical College (GMC), Surat | Gujarat Biotechnology Research Centre | Dinesh Kumar,Twinkle Soni,Nitin Shukla,Sonal Sharma ,Zuber Saiyed,Ramesh Pandit,Janvi Raval,Zarna Patel,Nitin Savaliya,Neeta Khandelwal,Umang Mishra,Nitesh Shah,Chaitanya Joshi,Madhvi Joshi |
| EPI_ISL_3161983 | Government Medical College (GMC), Surat | Gujarat Biotechnology Research Centre | Nitin Shukla,Sonal Sharma ,Zuber Saiyed,Ramesh Pandit,Janvi Raval,Zarna Patel,Nitin Savaliya,Dinesh Kumar,Twinkle Soni,Neeta Khandelwal,Umang Mishra,Nitesh Shah,Chaitanya Joshi,Madhvi Joshi |
| EPI_ISL_3161984 | Government Medical College (GMC), Surat | Gujarat Biotechnology Research Centre | Sonal Sharma ,Zuber Saiyed,Ramesh Pandit,Janvi Raval,Zarna Patel,Nitin Savaliya,Dinesh Kumar,Twinkle Soni,Nitin Shukla,Neeta Khandelwal,Umang Mishra,Nitesh Shah,Chaitanya Joshi,Madhvi Joshi |
| EPI_ISL_3161985 | Government Medical College (GMC), Surat | Gujarat Biotechnology Research Centre | Zuber Saiyed,Ramesh Pandit,Janvi Raval,Zarna Patel,Nitin Savaliya,Dinesh Kumar,Twinkle Soni,Nitin Shukla,Sonal Sharma ,Neeta Khandelwal,Umang Mishra,Nitesh Shah,Chaitanya Joshi,Madhvi Joshi |
| EPI_ISL_3161986 | Government Medical College (GMC), Surat | Gujarat Biotechnology Research Centre | Ramesh Pandit,Janvi Raval,Zarna Patel,Nitin Savaliya,Dinesh Kumar,Twinkle Soni,Nitin Shukla,Sonal Sharma ,Zuber Saiyed,Neeta Khandelwal,Umang Mishra,Nitesh Shah,Chaitanya Joshi,Madhvi Joshi |
| EPI_ISL_3161987 | Government Medical College (GMC), Surat | Gujarat Biotechnology Research Centre | Janvi Raval,Zarna Patel,Nitin Savaliya,Dinesh Kumar,Twinkle Soni,Nitin Shukla,Sonal Sharma ,Zuber Saiyed,Ramesh Pandit,Neeta Khandelwal,Umang Mishra,Nitesh Shah,Chaitanya Joshi,Madhvi Joshi |
| EPI_ISL_3161988 | Government Medical College (GMC), Surat | Gujarat Biotechnology Research Centre | Zarna Patel,Nitin Savaliya,Dinesh Kumar,Twinkle Soni,Nitin Shukla,Sonal Sharma ,Zuber Saiyed,Ramesh Pandit,Janvi Raval,Neeta Khandelwal,Umang Mishra,Nitesh Shah,Chaitanya Joshi,Madhvi Joshi |
| EPI_ISL_3161989 | Government Medical College (GMC), Surat | Gujarat Biotechnology Research Centre | Nitin Savaliya,Dinesh Kumar,Twinkle Soni,Nitin Shukla,Sonal Sharma ,Zuber Saiyed,Ramesh Pandit,Janvi Raval,Zarna Patel,Neeta Khandelwal,Umang Mishra,Nitesh Shah,Chaitanya Joshi,Madhvi Joshi |
| EPI_ISL_3161990 | Government Medical College (GMC), Surat | Gujarat Biotechnology Research Centre | Dinesh Kumar,Twinkle Soni,Nitin Shukla,Sonal Sharma ,Zuber Saiyed,Ramesh Pandit,Janvi Raval,Zarna Patel,Nitin Savaliya,Neeta Khandelwal,Umang Mishra,Nitesh Shah,Chaitanya Joshi,Madhvi Joshi |
| EPI_ISL_3161991 | Government Medical College (GMC), Surat | Gujarat Biotechnology Research Centre | Twinkle Soni,Nitin Shukla,Sonal Sharma ,Zuber Saiyed,Ramesh Pandit,Janvi Raval,Zarna Patel,Nitin Savaliya,Dinesh Kumar,Neeta Khandelwal,Umang Mishra,Nitesh Shah,Chaitanya Joshi,Madhvi Joshi |
| EPI_ISL_3161992 | Government Medical College (GMC), Surat | Gujarat Biotechnology Research Centre | Nitin Shukla,Sonal Sharma ,Zuber Saiyed,Ramesh Pandit,Janvi Raval,Zarna Patel,Nitin Savaliya,Dinesh Kumar,Twinkle Soni,Neeta Khandelwal,Umang Mishra,Nitesh Shah,Chaitanya Joshi,Madhvi Joshi |
| EPI_ISL_3161993 | Government Medical College (GMC), Surat | Gujarat Biotechnology Research Centre | Sonal Sharma ,Zuber Saiyed,Ramesh Pandit,Janvi Raval,Zarna Patel,Nitin Savaliya,Dinesh Kumar,Twinkle Soni,Nitin Shukla,Neeta Khandelwal,Umang Mishra,Nitesh Shah,Chaitanya Joshi,Madhvi Joshi |
| EPI_ISL_3161994 | Government Medical College (GMC), Surat | Gujarat Biotechnology Research Centre | Zuber Saiyed,Ramesh Pandit,Janvi Raval,Zarna Patel,Nitin Savaliya,Dinesh Kumar,Twinkle Soni,Nitin Shukla,Sonal Sharma ,Neeta Khandelwal,Umang Mishra,Nitesh Shah,Chaitanya Joshi,Madhvi Joshi |
| EPI_ISL_3161995 | Government Medical College (GMC), Surat | Gujarat Biotechnology Research Centre | Ramesh Pandit,Janvi Raval,Zarna Patel,Nitin Savaliya,Dinesh Kumar,Twinkle Soni,Nitin Shukla,Sonal Sharma ,Zuber Saiyed,Neeta Khandelwal,Umang Mishra,Nitesh Shah,Chaitanya Joshi,Madhvi Joshi |
| EPI_ISL_3161996 | Government Medical College (GMC), Surat | Gujarat Biotechnology Research Centre | Janvi Raval,Zarna Patel,Nitin Savaliya,Dinesh Kumar,Twinkle Soni,Nitin Shukla,Sonal Sharma ,Zuber Saiyed,Ramesh Pandit,Neeta Khandelwal,Umang Mishra,Nitesh Shah,Chaitanya Joshi,Madhvi Joshi |
| EPI_ISL_3161997 | Government Medical College (GMC), Surat | Gujarat Biotechnology Research Centre | Zarna Patel,Nitin Savaliya,Dinesh Kumar,Twinkle Soni,Nitin Shukla,Sonal Sharma ,Zuber Saiyed,Ramesh Pandit,Janvi Raval,Neeta Khandelwal,Umang Mishra,Nitesh Shah,Chaitanya Joshi,Madhvi Joshi |
| EPI_ISL_3161998 | Government Medical College (GMC), Surat | Gujarat Biotechnology Research Centre | Nitin Savaliya,Dinesh Kumar,Twinkle Soni,Nitin Shukla,Sonal Sharma ,Zuber Saiyed,Ramesh Pandit,Janvi Raval,Zarna Patel,Neeta Khandelwal,Umang Mishra,Nitesh Shah,Chaitanya Joshi,Madhvi Joshi |
| EPI_ISL_3161999 | Government Medical College (GMC), Surat | Gujarat Biotechnology Research Centre | Dinesh Kumar,Twinkle Soni,Nitin Shukla,Sonal Sharma ,Zuber Saiyed,Ramesh Pandit,Janvi Raval,Zarna Patel,Nitin Savaliya,Neeta Khandelwal,Umang Mishra,Nitesh Shah,Chaitanya Joshi,Madhvi Joshi |
| EPI_ISL_3162000 | Government Medical College (GMC), Surat | Gujarat Biotechnology Research Centre | Twinkle Soni,Nitin Shukla,Sonal Sharma ,Zuber Saiyed,Ramesh Pandit,Janvi Raval,Zarna Patel,Nitin Savaliya,Dinesh Kumar,Neeta Khandelwal,Umang Mishra,Nitesh Shah,Chaitanya Joshi,Madhvi Joshi |
| EPI_ISL_3162001 | Government Medical College (GMC), Surat | Gujarat Biotechnology Research Centre | Nitin Shukla,Sonal Sharma ,Zuber Saiyed,Ramesh Pandit,Janvi Raval,Zarna Patel,Nitin Savaliya,Dinesh Kumar,Twinkle Soni,Neeta Khandelwal,Umang Mishra,Nitesh Shah,Chaitanya Joshi,Madhvi Joshi |
| EPI_ISL_3162002 | Government Medical College (GMC), Surat | Gujarat Biotechnology Research Centre | Sonal Sharma ,Zuber Saiyed,Ramesh Pandit,Janvi Raval,Zarna Patel,Nitin Savaliya,Dinesh Kumar,Twinkle Soni,Nitin Shukla,Neeta Khandelwal,Umang Mishra,Nitesh Shah,Chaitanya Joshi,Madhvi Joshi |
| EPI_ISL_3162003 | Government Medical College (GMC), Surat | Gujarat Biotechnology Research Centre | Zuber Saiyed,Ramesh Pandit,Janvi Raval,Zarna Patel,Nitin Savaliya,Dinesh Kumar,Twinkle Soni,Nitin Shukla,Sonal Sharma ,Neeta Khandelwal,Umang Mishra,Nitesh Shah,Chaitanya Joshi,Madhvi Joshi |
| EPI_ISL_3162004 | Government Medical College (GMC), Surat | Gujarat Biotechnology Research Centre | Ramesh Pandit,Janvi Raval,Zarna Patel,Nitin Savaliya,Dinesh Kumar,Twinkle Soni,Nitin Shukla,Sonal Sharma ,Zuber Saiyed,Neeta Khandelwal,Umang Mishra,Nitesh Shah,Chaitanya Joshi,Madhvi Joshi |

| | | | |
|---|---|---|---|
| EPI_ISL_3162005 | Government Medical College (GMC), Surat | Gujarat Biotechnology Research Centre | Janvi Raval,Zarna Patel,Nitin Savaliya,Dinesh Kumar,Twinkle Soni,Nitin Shukla,Sonal Sharma ,Zuber Saiyed,Ramesh Pandit,Neeta Khandelwal,Umang Mishra,Nitesh Shah,Chaitanya Joshi,Madhvi Joshi |
| EPI_ISL_3162006 | Government Medical College (GMC), Surat | Gujarat Biotechnology Research Centre | Zarna Patel,Nitin Savaliya,Dinesh Kumar,Twinkle Soni,Nitin Shukla,Sonal Sharma ,Zuber Saiyed,Ramesh Pandit,Janvi Raval,Neeta Khandelwal,Umang Mishra,Nitesh Shah,Chaitanya Joshi,Madhvi Joshi |
| EPI_ISL_3162007 | Government Medical College (GMC), Surat | Gujarat Biotechnology Research Centre | Nitin Savaliya,Dinesh Kumar,Twinkle Soni,Nitin Shukla,Sonal Sharma ,Zuber Saiyed,Ramesh Pandit,Janvi Raval,Zarna Patel,Neeta Khandelwal,Umang Mishra,Nitesh Shah,Chaitanya Joshi,Madhvi Joshi |
| EPI_ISL_3162008 | Government Medical College (GMC), Surat | Gujarat Biotechnology Research Centre | Dinesh Kumar,Twinkle Soni,Nitin Shukla,Sonal Sharma ,Zuber Saiyed,Ramesh Pandit,Janvi Raval,Zarna Patel,Nitin Savaliya,Neeta Khandelwal,Umang Mishra,Nitesh Shah,Chaitanya Joshi,Madhvi Joshi |
| EPI_ISL_3162009 | Government Medical College (GMC), Surat | Gujarat Biotechnology Research Centre | Twinkle Soni,Nitin Shukla,Sonal Sharma ,Zuber Saiyed,Ramesh Pandit,Janvi Raval,Zarna Patel,Nitin Savaliya,Dinesh Kumar,Neeta Khandelwal,Umang Mishra,Nitesh Shah,Chaitanya Joshi,Madhvi Joshi |
| EPI_ISL_3162010 | Government Medical College (GMC), Surat | Gujarat Biotechnology Research Centre | Nitin Shukla,Sonal Sharma ,Zuber Saiyed,Ramesh Pandit,Janvi Raval,Zarna Patel,Nitin Savaliya,Dinesh Kumar,Twinkle Soni,Neeta Khandelwal,Umang Mishra,Nitesh Shah,Chaitanya Joshi,Madhvi Joshi |
| EPI_ISL_3162011 | Government Medical College (GMC), Surat | Gujarat Biotechnology Research Centre | Sonal Sharma ,Zuber Saiyed,Ramesh Pandit,Janvi Raval,Zarna Patel,Nitin Savaliya,Dinesh Kumar,Twinkle Soni,Nitin Shukla,Neeta Khandelwal,Umang Mishra,Nitesh Shah,Chaitanya Joshi,Madhvi Joshi |
| EPI_ISL_3162012 | Government Medical College (GMC), Surat | Gujarat Biotechnology Research Centre | Zuber Saiyed,Ramesh Pandit,Janvi Raval,Zarna Patel,Nitin Savaliya,Dinesh Kumar,Twinkle Soni,Nitin Shukla,Sonal Sharma ,Neeta Khandelwal,Umang Mishra,Nitesh Shah,Chaitanya Joshi,Madhvi Joshi |
| EPI_ISL_3162013 | Government Medical College (GMC), Surat | Gujarat Biotechnology Research Centre | Ramesh Pandit,Janvi Raval,Zarna Patel,Nitin Savaliya,Dinesh Kumar,Twinkle Soni,Nitin Shukla,Sonal Sharma ,Zuber Saiyed,Neeta Khandelwal,Umang Mishra,Nitesh Shah,Chaitanya Joshi,Madhvi Joshi |
| EPI_ISL_3162014 | Government Medical College (GMC), Surat | Gujarat Biotechnology Research Centre | Janvi Raval,Zarna Patel,Nitin Savaliya,Dinesh Kumar,Twinkle Soni,Nitin Shukla,Sonal Sharma ,Zuber Saiyed,Ramesh Pandit,Neeta Khandelwal,Umang Mishra,Nitesh Shah,Chaitanya Joshi,Madhvi Joshi |
| EPI_ISL_3162015 | Government Medical College (GMC), Surat | Gujarat Biotechnology Research Centre | Zarna Patel,Nitin Savaliya,Dinesh Kumar,Twinkle Soni,Nitin Shukla,Sonal Sharma ,Zuber Saiyed,Ramesh Pandit,Janvi Raval,Neeta Khandelwal,Umang Mishra,Nitesh Shah,Chaitanya Joshi,Madhvi Joshi |
| EPI_ISL_3162016 | Government Medical College (GMC), Surat | Gujarat Biotechnology Research Centre | Nitin Savaliya,Dinesh Kumar,Twinkle Soni,Nitin Shukla,Sonal Sharma ,Zuber Saiyed,Ramesh Pandit,Janvi Raval,Zarna Patel,Neeta Khandelwal,Umang Mishra,Nitesh Shah,Chaitanya Joshi,Madhvi Joshi |
| EPI_ISL_3162017 | Government Medical College (GMC), Surat | Gujarat Biotechnology Research Centre | Dinesh Kumar,Twinkle Soni,Nitin Shukla,Sonal Sharma ,Zuber Saiyed,Ramesh Pandit,Janvi Raval,Zarna Patel,Nitin Savaliya,Neeta Khandelwal,Umang Mishra,Nitesh Shah,Chaitanya Joshi,Madhvi Joshi |
| EPI_ISL_3162018 | Government Medical College (GMC), Surat | Gujarat Biotechnology Research Centre | Twinkle Soni,Nitin Shukla,Sonal Sharma ,Zuber Saiyed,Ramesh Pandit,Janvi Raval,Zarna Patel,Nitin Savaliya,Dinesh Kumar,Neeta Khandelwal,Umang Mishra,Nitesh Shah,Chaitanya Joshi,Madhvi Joshi |
| EPI_ISL_3162019 | Government Medical College (GMC), Surat | Gujarat Biotechnology Research Centre | Nitin Shukla,Sonal Sharma ,Zuber Saiyed,Ramesh Pandit,Janvi Raval,Zarna Patel,Nitin Savaliya,Dinesh Kumar,Twinkle Soni,Neeta Khandelwal,Umang Mishra,Nitesh Shah,Chaitanya Joshi,Madhvi Joshi |
| EPI_ISL_3162020 | Government Medical College (GMC), Surat | Gujarat Biotechnology Research Centre | Sonal Sharma ,Zuber Saiyed,Ramesh Pandit,Janvi Raval,Zarna Patel,Nitin Savaliya,Dinesh Kumar,Twinkle Soni,Nitin Shukla,Neeta Khandelwal,Umang Mishra,Nitesh Shah,Chaitanya Joshi,Madhvi Joshi |
| EPI_ISL_3162021 | Government Medical College (GMC), Surat | Gujarat Biotechnology Research Centre | Zuber Saiyed,Ramesh Pandit,Janvi Raval,Zarna Patel,Nitin Savaliya,Dinesh Kumar,Twinkle Soni,Nitin Shukla,Sonal Sharma ,Neeta Khandelwal,Umang Mishra,Nitesh Shah,Chaitanya Joshi,Madhvi Joshi |
| EPI_ISL_3162022 | Government Medical College (GMC), Surat | Gujarat Biotechnology Research Centre | Ramesh Pandit,Janvi Raval,Zarna Patel,Nitin Savaliya,Dinesh Kumar,Twinkle Soni,Nitin Shukla,Sonal Sharma ,Zuber Saiyed,Neeta Khandelwal,Umang Mishra,Nitesh Shah,Chaitanya Joshi,Madhvi Joshi |
| EPI_ISL_3162023 | Government Medical College (GMC), Surat | Gujarat Biotechnology Research Centre | Janvi Raval,Zarna Patel,Nitin Savaliya,Dinesh Kumar,Twinkle Soni,Nitin Shukla,Sonal Sharma ,Zuber Saiyed,Ramesh Pandit,Neeta Khandelwal,Umang Mishra,Nitesh Shah,Chaitanya Joshi,Madhvi Joshi |
| EPI_ISL_3162024 | Government Medical College (GMC), Surat | Gujarat Biotechnology Research Centre | Zarna Patel,Nitin Savaliya,Dinesh Kumar,Twinkle Soni,Nitin Shukla,Sonal Sharma ,Zuber Saiyed,Ramesh Pandit,Janvi Raval,Neeta Khandelwal,Umang Mishra,Nitesh Shah,Chaitanya Joshi,Madhvi Joshi |
| EPI_ISL_3162025 | Government Medical College (GMC), Surat | Gujarat Biotechnology Research Centre | Nitin Savaliya,Dinesh Kumar,Twinkle Soni,Nitin Shukla,Sonal Sharma ,Zuber Saiyed,Ramesh Pandit,Janvi Raval,Zarna Patel,Neeta Khandelwal,Umang Mishra,Nitesh Shah,Chaitanya Joshi,Madhvi Joshi |
| EPI_ISL_3162026 | Government Medical College (GMC), Surat | Gujarat Biotechnology Research Centre | Dinesh Kumar,Twinkle Soni,Nitin Shukla,Sonal Sharma ,Zuber Saiyed,Ramesh Pandit,Janvi Raval,Zarna Patel,Nitin Savaliya,Neeta Khandelwal,Umang Mishra,Nitesh Shah,Chaitanya Joshi,Madhvi Joshi |
| EPI_ISL_3162027 | Government Medical College (GMC), Surat | Gujarat Biotechnology Research Centre | Twinkle Soni,Nitin Shukla,Sonal Sharma ,Zuber Saiyed,Ramesh Pandit,Janvi Raval,Zarna Patel,Nitin Savaliya,Dinesh Kumar,Neeta Khandelwal,Umang Mishra,Nitesh Shah,Chaitanya Joshi,Madhvi Joshi |
| EPI_ISL_3162028 | Government Medical College (GMC), Surat | Gujarat Biotechnology Research Centre | Nitin Shukla,Sonal Sharma ,Zuber Saiyed,Ramesh Pandit,Janvi Raval,Zarna Patel,Nitin Savaliya,Dinesh Kumar,Twinkle Soni,Neeta Khandelwal,Umang Mishra,Nitesh Shah,Chaitanya Joshi,Madhvi Joshi |
| EPI_ISL_3162029 | Government Medical College (GMC), Surat | Gujarat Biotechnology Research Centre | Sonal Sharma ,Zuber Saiyed,Ramesh Pandit,Janvi Raval,Zarna Patel,Nitin Savaliya,Dinesh Kumar,Twinkle Soni,Nitin Shukla,Neeta Khandelwal,Umang Mishra,Nitesh Shah,Chaitanya Joshi,Madhvi Joshi |
| EPI_ISL_3162030 | Government Medical College (GMC), Surat | Gujarat Biotechnology Research Centre | Zuber Saiyed,Ramesh Pandit,Janvi Raval,Zarna Patel,Nitin Savaliya,Dinesh Kumar,Twinkle Soni,Nitin Shukla,Sonal Sharma ,Neeta Khandelwal,Umang Mishra,Nitesh Shah,Chaitanya Joshi,Madhvi Joshi |
| EPI_ISL_3162031 | Government Medical College (GMC), Surat | Gujarat Biotechnology Research Centre | Ramesh Pandit,Janvi Raval,Zarna Patel,Nitin Savaliya,Dinesh Kumar,Twinkle Soni,Nitin Shukla,Sonal Sharma ,Zuber Saiyed,Neeta Khandelwal,Umang Mishra,Nitesh Shah,Chaitanya Joshi,Madhvi Joshi |
| EPI_ISL_3162032 | Government Medical College (GMC), Surat | Gujarat Biotechnology Research Centre | Janvi Raval,Zarna Patel,Nitin Savaliya,Dinesh Kumar,Twinkle Soni,Nitin Shukla,Sonal Sharma ,Zuber Saiyed,Ramesh Pandit,Neeta Khandelwal,Umang Mishra,Nitesh Shah,Chaitanya Joshi,Madhvi Joshi |
| EPI_ISL_3162033 | Government Medical College (GMC), Surat | Gujarat Biotechnology Research Centre | Zarna Patel,Nitin Savaliya,Dinesh Kumar,Twinkle Soni,Nitin Shukla,Sonal Sharma ,Zuber Saiyed,Ramesh Pandit,Janvi Raval,Neeta Khandelwal,Umang Mishra,Nitesh Shah,Chaitanya Joshi,Madhvi Joshi |
| EPI_ISL_3162034 | Government Medical College (GMC), Surat | Gujarat Biotechnology Research Centre | Nitin Savaliya,Dinesh Kumar,Twinkle Soni,Nitin Shukla,Sonal Sharma ,Zuber Saiyed,Ramesh Pandit,Janvi Raval,Zarna Patel,Neeta Khandelwal,Umang Mishra,Nitesh Shah,Chaitanya Joshi,Madhvi Joshi |
| EPI_ISL_3162035 | Government Medical College (GMC), Surat | Gujarat Biotechnology Research Centre | Dinesh Kumar,Twinkle Soni,Nitin Shukla,Sonal Sharma ,Zuber Saiyed,Ramesh Pandit,Janvi Raval,Zarna Patel,Nitin Savaliya,Neeta Khandelwal,Umang Mishra,Nitesh Shah,Chaitanya Joshi,Madhvi Joshi |
| EPI_ISL_3162036 | Government Medical College (GMC), Surat | Gujarat Biotechnology Research Centre | Twinkle Soni,Nitin Shukla,Sonal Sharma ,Zuber Saiyed,Ramesh Pandit,Janvi Raval,Zarna Patel,Nitin Savaliya,Dinesh Kumar,Neeta Khandelwal,Umang Mishra,Nitesh Shah,Chaitanya Joshi,Madhvi Joshi |
| EPI_ISL_3162037 | Government Medical College (GMC), Surat | Gujarat Biotechnology Research Centre | Nitin Shukla,Sonal Sharma ,Zuber Saiyed,Ramesh Pandit,Janvi Raval,Zarna Patel,Nitin Savaliya,Dinesh Kumar,Twinkle Soni,Neeta Khandelwal,Umang Mishra,Nitesh Shah,Chaitanya Joshi,Madhvi Joshi |
| EPI_ISL_3162038 | Government Medical College (GMC), Surat | Gujarat Biotechnology Research Centre | Sonal Sharma ,Zuber Saiyed,Ramesh Pandit,Janvi Raval,Zarna Patel,Nitin Savaliya,Dinesh Kumar,Twinkle Soni,Nitin Shukla,Neeta Khandelwal,Umang Mishra,Nitesh Shah,Chaitanya Joshi,Madhvi Joshi |
| EPI_ISL_3162039 | Government Medical College (GMC), Surat | Gujarat Biotechnology Research Centre | Zuber Saiyed,Ramesh Pandit,Janvi Raval,Zarna Patel,Nitin Savaliya,Dinesh Kumar,Twinkle Soni,Nitin Shukla,Sonal Sharma ,Neeta Khandelwal,Umang Mishra,Nitesh Shah,Chaitanya Joshi,Madhvi Joshi |
| EPI_ISL_3162040 | Government Medical College (GMC), Surat | Gujarat Biotechnology Research Centre | Ramesh Pandit,Janvi Raval,Zarna Patel,Nitin Savaliya,Dinesh Kumar,Twinkle Soni,Nitin Shukla,Sonal Sharma ,Zuber Saiyed,Neeta Khandelwal,Umang Mishra,Nitesh Shah,Chaitanya Joshi,Madhvi Joshi |
| EPI_ISL_3162041 | Government Medical College (GMC), Surat | Gujarat Biotechnology Research Centre | Janvi Raval,Zarna Patel,Nitin Savaliya,Dinesh Kumar,Twinkle Soni,Nitin Shukla,Sonal Sharma ,Zuber Saiyed,Ramesh Pandit,Neeta Khandelwal,Umang Mishra,Nitesh Shah,Chaitanya Joshi,Madhvi Joshi |
| EPI_ISL_3162042 | Government Medical College (GMC), Surat | Gujarat Biotechnology Research Centre | Zarna Patel,Nitin Savaliya,Dinesh Kumar,Twinkle Soni,Nitin Shukla,Sonal Sharma ,Zuber Saiyed,Ramesh Pandit,Janvi Raval,Neeta Khandelwal,Umang Mishra,Nitesh Shah,Chaitanya Joshi,Madhvi Joshi |
| EPI_ISL_3162043 | Government Medical College (GMC), Surat | Gujarat Biotechnology Research Centre | Nitin Savaliya,Dinesh Kumar,Twinkle Soni,Nitin Shukla,Sonal Sharma ,Zuber Saiyed,Ramesh Pandit,Janvi Raval,Zarna Patel,Neeta Khandelwal,Umang Mishra,Nitesh Shah,Chaitanya Joshi,Madhvi Joshi |
| EPI_ISL_3162044 | Government Medical College (GMC), Surat | Gujarat Biotechnology Research Centre | Dinesh Kumar,Twinkle Soni,Nitin Shukla,Sonal Sharma ,Zuber Saiyed,Ramesh Pandit,Janvi Raval,Zarna Patel,Nitin Savaliya,Neeta Khandelwal,Umang Mishra,Nitesh Shah,Chaitanya Joshi,Madhvi Joshi |
| EPI_ISL_3162045 | Government Medical College (GMC), Surat | Gujarat Biotechnology Research Centre | Twinkle Soni,Nitin Shukla,Sonal Sharma ,Zuber Saiyed,Ramesh Pandit,Janvi Raval,Zarna Patel,Nitin Savaliya,Dinesh Kumar,Neeta Khandelwal,Umang Mishra,Nitesh Shah,Chaitanya Joshi,Madhvi Joshi |
| EPI_ISL_3162046 | Government Medical College (GMC), Surat | Gujarat Biotechnology Research Centre | Nitin Shukla,Sonal Sharma ,Zuber Saiyed,Ramesh Pandit,Janvi Raval,Zarna Patel,Nitin Savaliya,Dinesh Kumar,Twinkle Soni,Neeta Khandelwal,Umang Mishra,Nitesh Shah,Chaitanya Joshi,Madhvi Joshi |
| EPI_ISL_3162047 | Government Medical College (GMC), Surat | Gujarat Biotechnology Research Centre | Sonal Sharma ,Zuber Saiyed,Ramesh Pandit,Janvi Raval,Zarna Patel,Nitin Savaliya,Dinesh Kumar,Twinkle Soni,Nitin Shukla,Neeta Khandelwal,Umang Mishra,Nitesh Shah,Chaitanya Joshi,Madhvi Joshi |
| EPI_ISL_3162048 | Government Medical College (GMC), Surat | Gujarat Biotechnology Research Centre | Zuber Saiyed,Ramesh Pandit,Janvi Raval,Zarna Patel,Nitin Savaliya,Dinesh Kumar,Twinkle Soni,Nitin Shukla,Sonal Sharma ,Neeta Khandelwal,Umang Mishra,Nitesh Shah,Chaitanya Joshi,Madhvi Joshi |
| EPI_ISL_3162049 | Government Medical College (GMC), Surat | Gujarat Biotechnology Research Centre | Ramesh Pandit,Janvi Raval,Zarna Patel,Nitin Savaliya,Dinesh Kumar,Twinkle Soni,Nitin Shukla,Sonal Sharma ,Zuber Saiyed,Neeta Khandelwal,Umang Mishra,Nitesh Shah,Chaitanya Joshi,Madhvi Joshi |
| EPI_ISL_3162050 | Government Medical College (GMC), Surat | Gujarat Biotechnology Research Centre | Janvi Raval,Zarna Patel,Nitin Savaliya,Dinesh Kumar,Twinkle Soni,Nitin Shukla,Sonal Sharma ,Zuber Saiyed,Ramesh Pandit,Neeta Khandelwal,Umang Mishra,Nitesh Shah,Chaitanya Joshi,Madhvi Joshi |
| EPI_ISL_3162051 | Government Medical College (GMC), Surat | Gujarat Biotechnology Research Centre | Zarna Patel,Nitin Savaliya,Dinesh Kumar,Twinkle Soni,Nitin Shukla,Sonal Sharma ,Zuber Saiyed,Ramesh Pandit,Janvi Raval,Neeta Khandelwal,Umang Mishra,Nitesh Shah,Chaitanya Joshi,Madhvi Joshi |
| EPI_ISL_3162052 | Government Medical College (GMC), Surat | Gujarat Biotechnology Research Centre | Nitin Savaliya,Dinesh Kumar,Twinkle Soni,Nitin Shukla,Sonal Sharma ,Zuber Saiyed,Ramesh Pandit,Janvi Raval,Zarna Patel,Neeta Khandelwal,Umang Mishra,Nitesh Shah,Chaitanya Joshi,Madhvi Joshi |
| EPI_ISL_3162053 | Government Medical College (GMC), Surat | Gujarat Biotechnology Research Centre | Dinesh Kumar,Twinkle Soni,Nitin Shukla,Sonal Sharma ,Zuber Saiyed,Ramesh Pandit,Janvi Raval,Zarna Patel,Nitin Savaliya,Neeta Khandelwal,Umang Mishra,Nitesh Shah,Chaitanya Joshi,Madhvi Joshi |
| EPI_ISL_3162054 | Government Medical College (GMC), Surat | Gujarat Biotechnology Research Centre | Twinkle Soni,Nitin Shukla,Sonal Sharma ,Zuber Saiyed,Ramesh Pandit,Janvi Raval,Zarna Patel,Nitin Savaliya,Dinesh Kumar,Neeta Khandelwal,Umang Mishra,Nitesh Shah,Chaitanya Joshi,Madhvi Joshi |
| EPI_ISL_3162055 | Government Medical College (GMC), Surat | Gujarat Biotechnology Research Centre | Nitin Shukla,Sonal Sharma ,Zuber Saiyed,Ramesh Pandit,Janvi Raval,Zarna Patel,Nitin Savaliya,Dinesh Kumar,Twinkle Soni,Neeta Khandelwal,Umang Mishra,Nitesh Shah,Chaitanya Joshi,Madhvi Joshi |
| EPI_ISL_3162056 | Government Medical College (GMC), Surat | Gujarat Biotechnology Research Centre | Sonal Sharma ,Zuber Saiyed,Ramesh Pandit,Janvi Raval,Zarna Patel,Nitin Savaliya,Dinesh Kumar,Twinkle Soni,Nitin Shukla,Neeta Khandelwal,Umang Mishra,Nitesh Shah,Chaitanya Joshi,Madhvi Joshi |
| EPI_ISL_3162057 | Government Medical College (GMC), Surat | Gujarat Biotechnology Research Centre | Zuber Saiyed,Ramesh Pandit,Janvi Raval,Zarna Patel,Nitin Savaliya,Dinesh Kumar,Twinkle Soni,Nitin Shukla,Sonal Sharma ,Neeta Khandelwal,Umang Mishra,Nitesh Shah,Chaitanya Joshi,Madhvi Joshi |
| EPI_ISL_3162058 | Government Medical College (GMC), Surat | Gujarat Biotechnology Research Centre | Ramesh Pandit,Janvi Raval,Zarna Patel,Nitin Savaliya,Dinesh Kumar,Twinkle Soni,Nitin Shukla,Sonal Sharma ,Zuber Saiyed,Neeta Khandelwal,Umang Mishra,Nitesh Shah,Chaitanya Joshi,Madhvi Joshi |
| EPI_ISL_3162059 | Government Medical College (GMC), Surat | Gujarat Biotechnology Research Centre | Janvi Raval,Zarna Patel,Nitin Savaliya,Dinesh Kumar,Twinkle Soni,Nitin Shukla,Sonal Sharma ,Zuber Saiyed,Ramesh Pandit,Neeta Khandelwal,Umang Mishra,Nitesh Shah,Chaitanya Joshi,Madhvi Joshi |
| EPI_ISL_3162060 | Government Medical College (GMC), Surat | Gujarat Biotechnology Research Centre | Zarna Patel,Nitin Savaliya,Dinesh Kumar,Twinkle Soni,Nitin Shukla,Sonal Sharma ,Zuber Saiyed,Ramesh Pandit,Janvi Raval,Neeta Khandelwal,Umang Mishra,Nitesh Shah,Chaitanya Joshi,Madhvi Joshi |
| EPI_ISL_3162061 | Government Medical College (GMC), Surat | Gujarat Biotechnology Research Centre | Nitin Savaliya,Dinesh Kumar,Twinkle Soni,Nitin Shukla,Sonal Sharma ,Zuber Saiyed,Ramesh Pandit,Janvi Raval,Zarna Patel,Neeta Khandelwal,Umang Mishra,Nitesh Shah,Chaitanya Joshi,Madhvi Joshi |
| EPI_ISL_3162062 | Government Medical College (GMC), Surat | Gujarat Biotechnology Research Centre | Dinesh Kumar,Twinkle Soni,Nitin Shukla,Sonal Sharma ,Zuber Saiyed,Ramesh Pandit,Janvi Raval,Zarna Patel,Nitin Savaliya,Neeta Khandelwal,Umang Mishra,Nitesh Shah,Chaitanya Joshi,Madhvi Joshi |
| EPI_ISL_3162063 | Government Medical College (GMC), Surat | Gujarat Biotechnology Research Centre | Twinkle Soni,Nitin Shukla,Sonal Sharma ,Zuber Saiyed,Ramesh Pandit,Janvi Raval,Zarna Patel,Nitin Savaliya,Dinesh Kumar,Neeta Khandelwal,Umang Mishra,Nitesh Shah,Chaitanya Joshi,Madhvi Joshi |
| EPI_ISL_3162064 | Government Medical College (GMC), Surat | Gujarat Biotechnology Research Centre | Nitin Shukla,Sonal Sharma ,Zuber Saiyed,Ramesh Pandit,Janvi Raval,Zarna Patel,Nitin Savaliya,Dinesh Kumar,Twinkle Soni,Neeta Khandelwal,Umang Mishra,Nitesh Shah,Chaitanya Joshi,Madhvi Joshi |
| EPI_ISL_3162065 | Government Medical College (GMC), Surat | Gujarat Biotechnology Research Centre | Sonal Sharma ,Zuber Saiyed,Ramesh Pandit,Janvi Raval,Zarna Patel,Nitin Savaliya,Dinesh Kumar,Twinkle Soni,Nitin Shukla,Neeta Khandelwal,Umang Mishra,Nitesh Shah,Chaitanya Joshi,Madhvi Joshi |
| EPI_ISL_3162066 | Government Medical College (GMC), Surat | Gujarat Biotechnology Research Centre | Zuber Saiyed,Ramesh Pandit,Janvi Raval,Zarna Patel,Nitin Savaliya,Dinesh Kumar,Twinkle Soni,Nitin Shukla,Sonal Sharma ,Neeta Khandelwal,Umang Mishra,Nitesh Shah,Chaitanya Joshi,Madhvi Joshi |
| EPI_ISL_3162067 | Government Medical College (GMC), Surat | Gujarat Biotechnology Research Centre | Ramesh Pandit,Janvi Raval,Zarna Patel,Nitin Savaliya,Dinesh Kumar,Twinkle Soni,Nitin Shukla,Sonal Sharma ,Zuber Saiyed,Neeta Khandelwal,Umang Mishra,Nitesh Shah,Chaitanya Joshi,Madhvi Joshi |
| EPI_ISL_3162068 | Government Medical College (GMC), Surat | Gujarat Biotechnology Research Centre | Janvi Raval,Zarna Patel,Nitin Savaliya,Dinesh Kumar,Twinkle Soni,Nitin Shukla,Sonal Sharma ,Zuber Saiyed,Ramesh Pandit,Neeta Khandelwal,Umang Mishra,Nitesh Shah,Chaitanya Joshi,Madhvi Joshi |
| EPI_ISL_3162069 | Government Medical College (GMC), Surat | Gujarat Biotechnology Research Centre | Zarna Patel,Nitin Savaliya,Dinesh Kumar,Twinkle Soni,Nitin Shukla,Sonal Sharma ,Zuber Saiyed,Ramesh Pandit,Janvi Raval,Neeta Khandelwal,Umang Mishra,Nitesh Shah,Chaitanya Joshi,Madhvi Joshi |
| EPI_ISL_3162070 | Government Medical College (GMC), Surat | Gujarat Biotechnology Research Centre | Nitin Savaliya,Dinesh Kumar,Twinkle Soni,Nitin Shukla,Sonal Sharma ,Zuber Saiyed,Ramesh Pandit,Janvi Raval,Zarna Patel,Neeta Khandelwal,Umang Mishra,Nitesh Shah,Chaitanya Joshi,Madhvi Joshi |
| EPI_ISL_3162071 | Government Medical College (GMC), Surat | Gujarat Biotechnology Research Centre | Dinesh Kumar,Twinkle Soni,Nitin Shukla,Sonal Sharma ,Zuber Saiyed,Ramesh Pandit,Janvi Raval,Zarna Patel,Nitin Savaliya,Neeta Khandelwal,Umang Mishra,Nitesh Shah,Chaitanya Joshi,Madhvi Joshi |
| EPI_ISL_3162072 | Government Medical College (GMC), Surat | Gujarat Biotechnology Research Centre | Twinkle Soni,Nitin Shukla,Sonal Sharma ,Zuber Saiyed,Ramesh Pandit,Janvi Raval,Zarna Patel,Nitin Savaliya,Dinesh Kumar,Neeta Khandelwal,Umang Mishra,Nitesh Shah,Chaitanya Joshi,Madhvi Joshi |
| EPI_ISL_3162073 | Government Medical College (GMC), Surat | Gujarat Biotechnology Research Centre | Nitin Shukla,Sonal Sharma ,Zuber Saiyed,Ramesh Pandit,Janvi Raval,Zarna Patel,Nitin Savaliya,Dinesh Kumar,Twinkle Soni,Neeta Khandelwal,Umang Mishra,Nitesh Shah,Chaitanya Joshi,Madhvi Joshi |
| EPI_ISL_3162074 | Government Medical College (GMC), Surat | Gujarat Biotechnology Research Centre | Sonal Sharma ,Zuber Saiyed,Ramesh Pandit,Janvi Raval,Zarna Patel,Nitin Savaliya,Dinesh Kumar,Twinkle Soni,Nitin Shukla,Neeta Khandelwal,Umang Mishra,Nitesh Shah,Chaitanya Joshi,Madhvi Joshi |
| EPI_ISL_3162075 | Government Medical College (GMC), Surat | Gujarat Biotechnology Research Centre | Zuber Saiyed,Ramesh Pandit,Janvi Raval,Zarna Patel,Nitin Savaliya,Dinesh Kumar,Twinkle Soni,Nitin Shukla,Sonal Sharma ,Neeta Khandelwal,Umang Mishra,Nitesh Shah,Chaitanya Joshi,Madhvi Joshi |
| EPI_ISL_3162076 | Government Medical College (GMC), Surat | Gujarat Biotechnology Research Centre | Ramesh Pandit,Janvi Raval,Zarna Patel,Nitin Savaliya,Dinesh Kumar,Twinkle Soni,Nitin Shukla,Sonal Sharma ,Zuber Saiyed,Neeta Khandelwal,Umang Mishra,Nitesh Shah,Chaitanya Joshi,Madhvi Joshi |
| EPI_ISL_3162096 | VRDL,Goverment Medical College Surat | Gujarat Biotechnology Research Centre | Zarna Patel,Nitin Savaliya,Dinesh Kumar,Twinkle Soni,Nitin Shukla,Sonal Sharma ,Zuber Saiyed,Ramesh Pandit,Janvi Raval,Neeta Khandelwal,Umang Mishra,Nitesh Shah,Chaitanya Joshi,Madhvi Joshi |
| EPI_ISL_3162110 | VRDL,Goverment Medical College, Surat | Gujarat Biotechnology Research Centre | Sonal Sharma ,Zuber Saiyed,Ramesh Pandit,Janvi Raval,Zarna Patel,Nitin Savaliya,Dinesh Kumar,Twinkle Soni,Nitin Shukla,Neeta Khandelwal,Umang Mishra,Nitesh Shah,Chaitanya Joshi,Madhvi Joshi |
| EPI_ISL_3162111 | VRDL,Goverment Medical College, Surat | Gujarat Biotechnology Research Centre | Zuber Saiyed,Ramesh Pandit,Janvi Raval,Zarna Patel,Nitin Savaliya,Dinesh Kumar,Twinkle Soni,Nitin Shukla,Sonal Sharma ,Neeta Khandelwal,Umang Mishra,Nitesh Shah,Chaitanya Joshi,Madhvi Joshi |
| EPI_ISL_3162118 | NHL Municipal Medical College, Ahmedbad | Gujarat Biotechnology Research Centre | Nitin Shukla,Sonal Sharma ,Zuber Saiyed,Ramesh Pandit,Janvi Raval,Zarna Patel,Nitin Savaliya,Dinesh Kumar,Twinkle Soni,Jayshri Pethani,Umang Mishra,Nitesh Shah,Chaitanya Joshi,Madhvi Joshi |
| EPI_ISL_3162128 | NHL Municipal Medical College, Ahmedbad | Gujarat Biotechnology Research Centre | Sonal Sharma ,Zuber Saiyed,Ramesh Pandit,Janvi Raval,Zarna Patel,Nitin Savaliya,Dinesh Kumar,Twinkle Soni,Nitin Shukla,Jayshri Pethani,Umang Mishra,Nitesh Shah,Chaitanya Joshi,Madhvi Joshi |

| | | | |
|---|---|---|---|
| EPI_ISL_3162129 | VRDL, Goverment Medical College, Surat | Gujarat Biotechnology Research Centre | Zuber Saiyed,Ramesh Pandit,Janvi Raval,Zarna Patel,Nitin Savaliya,Dinesh Kumar,Twinkle Soni,Nitin Shukla,Sonal Sharma ,Neeta Khandelwal,Umang Mishra,Nitesh Shah,Chaitanya Joshi,Madhvi Joshi |
| EPI_ISL_3162130 | VRDL, Goverment Medical College, Surat | Gujarat Biotechnology Research Centre | Ramesh Pandit,Janvi Raval,Zarna Patel,Nitin Savaliya,Dinesh Kumar,Twinkle Soni,Nitin Shukla,Sonal Sharma ,Zuber Saiyed,Neeta Khandelwal,Umang Mishra,Nitesh Shah,Chaitanya Joshi,Madhvi Joshi |
| EPI_ISL_3162132 | VRDL, Goverment Medical College, Surat | Gujarat Biotechnology Research Centre | Zarna Patel,Nitin Savaliya,Dinesh Kumar,Twinkle Soni,Nitin Shukla,Sonal Sharma ,Zuber Saiyed,Ramesh Pandit,Janvi Raval,Neeta Khandelwal,Umang Mishra,Nitesh Shah,Chaitanya Joshi,Madhvi Joshi |
| EPI_ISL_3162134 | VRDL, Goverment Medical College, Surat | Gujarat Biotechnology Research Centre | Dinesh Kumar,Twinkle Soni,Nitin Shukla,Sonal Sharma ,Zuber Saiyed,Ramesh Pandit,Janvi Raval,Zarna Patel,Nitin Savaliya,Neeta Khandelwal,Umang Mishra,Nitesh Shah,Chaitanya Joshi,Madhvi Joshi |
| EPI_ISL_3162135 | VRDL, Goverment Medical College, Surat | Gujarat Biotechnology Research Centre | Twinkle Soni,Nitin Shukla,Sonal Sharma ,Zuber Saiyed,Ramesh Pandit,Janvi Raval,Zarna Patel,Nitin Savaliya,Dinesh Kumar,Neeta Khandelwal,Umang Mishra,Nitesh Shah,Chaitanya Joshi,Madhvi Joshi |
| EPI_ISL_3162136 | VRDL, Goverment Medical College, Surat | Gujarat Biotechnology Research Centre | Nitin Shukla,Sonal Sharma ,Zuber Saiyed,Ramesh Pandit,Janvi Raval,Zarna Patel,Nitin Savaliya,Dinesh Kumar,Twinkle Soni,Neeta Khandelwal,Umang Mishra,Nitesh Shah,Chaitanya Joshi,Madhvi Joshi |
| EPI_ISL_3162137 | VRDL, Goverment Medical College, Surat | Gujarat Biotechnology Research Centre | Sonal Sharma ,Zuber Saiyed,Ramesh Pandit,Janvi Raval,Zarna Patel,Nitin Savaliya,Dinesh Kumar,Twinkle Soni,Nitin Shukla,Neeta Khandelwal,Umang Mishra,Nitesh Shah,Chaitanya Joshi,Madhvi Joshi |
| EPI_ISL_3162138 | VRDL, Goverment Medical College, Surat | Gujarat Biotechnology Research Centre | Zuber Saiyed,Ramesh Pandit,Janvi Raval,Zarna Patel,Nitin Savaliya,Dinesh Kumar,Twinkle Soni,Nitin Shukla,Sonal Sharma ,Neeta Khandelwal,Umang Mishra,Nitesh Shah,Chaitanya Joshi,Madhvi Joshi |
| EPI_ISL_3162140 | VRDL, Goverment Medical College, Surat | Gujarat Biotechnology Research Centre | Janvi Raval,Zarna Patel,Nitin Savaliya,Dinesh Kumar,Twinkle Soni,Nitin Shukla,Sonal Sharma ,Zuber Saiyed,Ramesh Pandit,Neeta Khandelwal,Umang Mishra,Nitesh Shah,Chaitanya Joshi,Madhvi Joshi |
| EPI_ISL_3162141 | VRDL, Goverment Medical College, Surat | Gujarat Biotechnology Research Centre | Zarna Patel,Nitin Savaliya,Dinesh Kumar,Twinkle Soni,Nitin Shukla,Sonal Sharma ,Zuber Saiyed,Ramesh Pandit,Janvi Raval,Neeta Khandelwal,Umang Mishra,Nitesh Shah,Chaitanya Joshi,Madhvi Joshi |
| EPI_ISL_3162143 | GMERS Medical College, Gotri | Gujarat Biotechnology Research Centre | Dinesh Kumar,Twinkle Soni,Nitin Shukla,Sonal Sharma ,Zuber Saiyed,Ramesh Pandit,Janvi Raval,Zarna Patel,Nitin Savaliya,Bithika Duttaroy,Umang Mishra,Nitesh Shah,Chaitanya Joshi,Madhvi Joshi |
| EPI_ISL_3162144 | GMERS Medical College, Gotri | Gujarat Biotechnology Research Centre | Twinkle Soni,Nitin Shukla,Sonal Sharma ,Zuber Saiyed,Ramesh Pandit,Janvi Raval,Zarna Patel,Nitin Savaliya,Dinesh Kumar,Bithika Duttaroy,Umang Mishra,Nitesh Shah,Chaitanya Joshi,Madhvi Joshi |
| EPI_ISL_3162147 | GMERS Medical College, Gotri | Gujarat Biotechnology Research Centre | Zuber Saiyed,Ramesh Pandit,Janvi Raval,Zarna Patel,Nitin Savaliya,Dinesh Kumar,Twinkle Soni,Nitin Shukla,Sonal Sharma ,Bithika Duttaroy,Umang Mishra,Nitesh Shah,Chaitanya Joshi,Madhvi Joshi |
| EPI_ISL_3162148 | GMERS Medical College, Gotri | Gujarat Biotechnology Research Centre | Ramesh Pandit,Janvi Raval,Zarna Patel,Nitin Savaliya,Dinesh Kumar,Twinkle Soni,Nitin Shukla,Sonal Sharma ,Zuber Saiyed,Bithika Duttaroy,Umang Mishra,Nitesh Shah,Chaitanya Joshi,Madhvi Joshi |
| EPI_ISL_3162149 | GMERS Medical College, Gotri | Gujarat Biotechnology Research Centre | Janvi Raval,Zarna Patel,Nitin Savaliya,Dinesh Kumar,Twinkle Soni,Nitin Shukla,Sonal Sharma ,Zuber Saiyed,Ramesh Pandit,Bithika Duttaroy,Umang Mishra,Nitesh Shah,Chaitanya Joshi,Madhvi Joshi |
| EPI_ISL_3162150 | N.H.L Municipal Medical College, Ahmedabad | Gujarat Biotechnology Research Centre | Zarna Patel,Nitin Savaliya,Dinesh Kumar,Twinkle Soni,Nitin Shukla,Sonal Sharma ,Zuber Saiyed,Ramesh Pandit,Janvi Raval,Jayshri Pethani,Umang Mishra,Nitesh Shah,Chaitanya Joshi,Madhvi Joshi |
| EPI_ISL_3162151 | N.H.L Municipal Medical College, Ahmedabad | Gujarat Biotechnology Research Centre | Nitin Savaliya,Dinesh Kumar,Twinkle Soni,Nitin Shukla,Sonal Sharma ,Zuber Saiyed,Ramesh Pandit,Janvi Raval,Zarna Patel,Jayshri Pethani,Umang Mishra,Nitesh Shah,Chaitanya Joshi,Madhvi Joshi |
| EPI_ISL_3162152 | N.H.L Municipal Medical College, Ahmedabad | Gujarat Biotechnology Research Centre | Dinesh Kumar,Twinkle Soni,Nitin Shukla,Sonal Sharma ,Zuber Saiyed,Ramesh Pandit,Janvi Raval,Zarna Patel,Nitin Savaliya,Jayshri Pethani,Umang Mishra,Nitesh Shah,Chaitanya Joshi,Madhvi Joshi |
| EPI_ISL_3162153 | N.H.L Municipal Medical College, Ahmedabad | Gujarat Biotechnology Research Centre | Twinkle Soni,Nitin Shukla,Sonal Sharma ,Zuber Saiyed,Ramesh Pandit,Janvi Raval,Zarna Patel,Nitin Savaliya,Dinesh Kumar,Jayshri Pethani,Umang Mishra,Nitesh Shah,Chaitanya Joshi,Madhvi Joshi |
| EPI_ISL_3162154 | N.H.L Municipal Medical College, Ahmedabad | Gujarat Biotechnology Research Centre | Nitin Shukla,Sonal Sharma ,Zuber Saiyed,Ramesh Pandit,Janvi Raval,Zarna Patel,Nitin Savaliya,Dinesh Kumar,Twinkle Soni,Jayshri Pethani,Umang Mishra,Nitesh Shah,Chaitanya Joshi,Madhvi Joshi |
| EPI_ISL_3162155 | N.H.L Municipal Medical College, Ahmedabad | Gujarat Biotechnology Research Centre | Sonal Sharma ,Zuber Saiyed,Ramesh Pandit,Janvi Raval,Zarna Patel,Nitin Savaliya,Dinesh Kumar,Twinkle Soni,Nitin Shukla,Jayshri Pethani,Umang Mishra,Nitesh Shah,Chaitanya Joshi,Madhvi Joshi |
| EPI_ISL_3162156 | N.H.L Municipal Medical College, Ahmedabad | Gujarat Biotechnology Research Centre | Zuber Saiyed,Ramesh Pandit,Janvi Raval,Zarna Patel,Nitin Savaliya,Dinesh Kumar,Twinkle Soni,Nitin Shukla,Sonal Sharma ,Jayshri Pethani,Umang Mishra,Nitesh Shah,Chaitanya Joshi,Madhvi Joshi |
| EPI_ISL_3162157 | N.H.L Municipal Medical College, Ahmedabad | Gujarat Biotechnology Research Centre | Ramesh Pandit,Janvi Raval,Zarna Patel,Nitin Savaliya,Dinesh Kumar,Twinkle Soni,Nitin Shukla,Sonal Sharma ,Zuber Saiyed,Jayshri Pethani,Umang Mishra,Nitesh Shah,Chaitanya Joshi,Madhvi Joshi |
| EPI_ISL_3162158 | N.H.L Municipal Medical College, Ahmedabad | Gujarat Biotechnology Research Centre | Janvi Raval,Zarna Patel,Nitin Savaliya,Dinesh Kumar,Twinkle Soni,Nitin Shukla,Sonal Sharma ,Zuber Saiyed,Ramesh Pandit,Jayshri Pethani,Umang Mishra,Nitesh Shah,Chaitanya Joshi,Madhvi Joshi |
| EPI_ISL_3162159 | N.H.L Municipal Medical College, Ahmedabad | Gujarat Biotechnology Research Centre | Zarna Patel,Nitin Savaliya,Dinesh Kumar,Twinkle Soni,Nitin Shukla,Sonal Sharma ,Zuber Saiyed,Ramesh Pandit,Janvi Raval,Jayshri Pethani,Umang Mishra,Nitesh Shah,Chaitanya Joshi,Madhvi Joshi |
| EPI_ISL_3162161 | N.H.L Municipal Medical College, Ahmedabad | Gujarat Biotechnology Research Centre | Dinesh Kumar,Twinkle Soni,Nitin Shukla,Sonal Sharma ,Zuber Saiyed,Ramesh Pandit,Janvi Raval,Zarna Patel,Nitin Savaliya,Jayshri Pethani,Umang Mishra,Nitesh Shah,Chaitanya Joshi,Madhvi Joshi |
| EPI_ISL_3162162 | N.H.L Municipal Medical College, Ahmedabad | Gujarat Biotechnology Research Centre | Twinkle Soni,Nitin Shukla,Sonal Sharma ,Zuber Saiyed,Ramesh Pandit,Janvi Raval,Zarna Patel,Nitin Savaliya,Dinesh Kumar,Jayshri Pethani,Umang Mishra,Nitesh Shah,Chaitanya Joshi,Madhvi Joshi |
| EPI_ISL_3162163 | N.H.L Municipal Medical College, Ahmedabad | Gujarat Biotechnology Research Centre | Nitin Shukla,Sonal Sharma ,Zuber Saiyed,Ramesh Pandit,Janvi Raval,Zarna Patel,Nitin Savaliya,Dinesh Kumar,Twinkle Soni,Jayshri Pethani,Umang Mishra,Nitesh Shah,Chaitanya Joshi,Madhvi Joshi |
| EPI_ISL_3162164 | N.H.L Municipal Medical College, Ahmedabad | Gujarat Biotechnology Research Centre | Sonal Sharma ,Zuber Saiyed,Ramesh Pandit,Janvi Raval,Zarna Patel,Nitin Savaliya,Dinesh Kumar,Twinkle Soni,Nitin Shukla,Jayshri Pethani,Umang Mishra,Nitesh Shah,Chaitanya Joshi,Madhvi Joshi |
| EPI_ISL_3162165 | N.H.L Municipal Medical College, Ahmedabad | Gujarat Biotechnology Research Centre | Zuber Saiyed,Ramesh Pandit,Janvi Raval,Zarna Patel,Nitin Savaliya,Dinesh Kumar,Twinkle Soni,Nitin Shukla,Sonal Sharma ,Jayshri Pethani,Umang Mishra,Nitesh Shah,Chaitanya Joshi,Madhvi Joshi |
| EPI_ISL_3162166 | N.H.L Municipal Medical College, Ahmedabad | Gujarat Biotechnology Research Centre | Ramesh Pandit,Janvi Raval,Zarna Patel,Nitin Savaliya,Dinesh Kumar,Twinkle Soni,Nitin Shukla,Sonal Sharma ,Zuber Saiyed,Jayshri Pethani,Umang Mishra,Nitesh Shah,Chaitanya Joshi,Madhvi Joshi |
| EPI_ISL_3162167 | N.H.L Municipal Medical College, Ahmedabad | Gujarat Biotechnology Research Centre | Janvi Raval,Zarna Patel,Nitin Savaliya,Dinesh Kumar,Twinkle Soni,Nitin Shukla,Sonal Sharma ,Zuber Saiyed,Ramesh Pandit,Jayshri Pethani,Umang Mishra,Nitesh Shah,Chaitanya Joshi,Madhvi Joshi |
| EPI_ISL_3162168 | N.H.L Municipal Medical College, Ahmedabad | Gujarat Biotechnology Research Centre | Zarna Patel,Nitin Savaliya,Dinesh Kumar,Twinkle Soni,Nitin Shukla,Sonal Sharma ,Zuber Saiyed,Ramesh Pandit,Janvi Raval,Jayshri Pethani,Umang Mishra,Nitesh Shah,Chaitanya Joshi,Madhvi Joshi |
| EPI_ISL_3162169 | N.H.L Municipal Medical College, Ahmedabad | Gujarat Biotechnology Research Centre | Nitin Savaliya,Dinesh Kumar,Twinkle Soni,Nitin Shukla,Sonal Sharma ,Zuber Saiyed,Ramesh Pandit,Janvi Raval,Zarna Patel,Jayshri Pethani,Umang Mishra,Nitesh Shah,Chaitanya Joshi,Madhvi Joshi |
| EPI_ISL_3162170 | N.H.L Municipal Medical College, Ahmedabad | Gujarat Biotechnology Research Centre | Dinesh Kumar,Twinkle Soni,Nitin Shukla,Sonal Sharma ,Zuber Saiyed,Ramesh Pandit,Janvi Raval,Zarna Patel,Nitin Savaliya,Jayshri Pethani,Umang Mishra,Nitesh Shah,Chaitanya Joshi,Madhvi Joshi |
| EPI_ISL_3162171 | N.H.L Municipal Medical College, Ahmedabad | Gujarat Biotechnology Research Centre | Twinkle Soni,Nitin Shukla,Sonal Sharma ,Zuber Saiyed,Ramesh Pandit,Janvi Raval,Zarna Patel,Nitin Savaliya,Dinesh Kumar,Jayshri Pethani,Umang Mishra,Nitesh Shah,Chaitanya Joshi,Madhvi Joshi |
| EPI_ISL_3162172 | N.H.L Municipal Medical College, Ahmedabad | Gujarat Biotechnology Research Centre | Nitin Shukla,Sonal Sharma ,Zuber Saiyed,Ramesh Pandit,Janvi Raval,Zarna Patel,Nitin Savaliya,Dinesh Kumar,Twinkle Soni,Jayshri Pethani,Umang Mishra,Nitesh Shah,Chaitanya Joshi,Madhvi Joshi |
| EPI_ISL_3162173 | N.H.L Municipal Medical College, Ahmedabad | Gujarat Biotechnology Research Centre | Sonal Sharma ,Zuber Saiyed,Ramesh Pandit,Janvi Raval,Zarna Patel,Nitin Savaliya,Dinesh Kumar,Twinkle Soni,Nitin Shukla,Jayshri Pethani,Umang Mishra,Nitesh Shah,Chaitanya Joshi,Madhvi Joshi |
| EPI_ISL_3162174 | N.H.L Municipal Medical College, Ahmedabad | Gujarat Biotechnology Research Centre | Zuber Saiyed,Ramesh Pandit,Janvi Raval,Zarna Patel,Nitin Savaliya,Dinesh Kumar,Twinkle Soni,Nitin Shukla,Sonal Sharma ,Jayshri Pethani,Umang Mishra,Nitesh Shah,Chaitanya Joshi,Madhvi Joshi |
| EPI_ISL_3162175 | N.H.L Municipal Medical College, Ahmedabad | Gujarat Biotechnology Research Centre | Ramesh Pandit,Janvi Raval,Zarna Patel,Nitin Savaliya,Dinesh Kumar,Twinkle Soni,Nitin Shukla,Sonal Sharma ,Zuber Saiyed,Jayshri Pethani,Umang Mishra,Nitesh Shah,Chaitanya Joshi,Madhvi Joshi |
| EPI_ISL_3162176 | N.H.L Municipal Medical College, Ahmedabad | Gujarat Biotechnology Research Centre | Janvi Raval,Zarna Patel,Nitin Savaliya,Dinesh Kumar,Twinkle Soni,Nitin Shukla,Sonal Sharma ,Zuber Saiyed,Ramesh Pandit,Jayshri Pethani,Umang Mishra,Nitesh Shah,Chaitanya Joshi,Madhvi Joshi |
| EPI_ISL_3162177 | N.H.L Municipal Medical College, Ahmedabad | Gujarat Biotechnology Research Centre | Zarna Patel,Nitin Savaliya,Dinesh Kumar,Twinkle Soni,Nitin Shukla,Sonal Sharma ,Zuber Saiyed,Ramesh Pandit,Janvi Raval,Jayshri Pethani,Umang Mishra,Nitesh Shah,Chaitanya Joshi,Madhvi Joshi |
| EPI_ISL_3162178 | N.H.L Municipal Medical College, Ahmedabad | Gujarat Biotechnology Research Centre | Nitin Savaliya,Dinesh Kumar,Twinkle Soni,Nitin Shukla,Sonal Sharma ,Zuber Saiyed,Ramesh Pandit,Janvi Raval,Zarna Patel,Jayshri Pethani,Umang Mishra,Nitesh Shah,Chaitanya Joshi,Madhvi Joshi |
| EPI_ISL_3162179 | N.H.L Municipal Medical College, Ahmedabad | Gujarat Biotechnology Research Centre | Dinesh Kumar,Twinkle Soni,Nitin Shukla,Sonal Sharma ,Zuber Saiyed,Ramesh Pandit,Janvi Raval,Zarna Patel,Nitin Savaliya,Jayshri Pethani,Umang Mishra,Nitesh Shah,Chaitanya Joshi,Madhvi Joshi |
| EPI_ISL_3162180 | N.H.L Municipal Medical College, Ahmedabad | Gujarat Biotechnology Research Centre | Twinkle Soni,Nitin Shukla,Sonal Sharma ,Zuber Saiyed,Ramesh Pandit,Janvi Raval,Zarna Patel,Nitin Savaliya,Dinesh Kumar,Jayshri Pethani,Umang Mishra,Nitesh Shah,Chaitanya Joshi,Madhvi Joshi |
| EPI_ISL_3162181 | N.H.L Municipal Medical College, Ahmedabad | Gujarat Biotechnology Research Centre | Nitin Shukla,Sonal Sharma ,Zuber Saiyed,Ramesh Pandit,Janvi Raval,Zarna Patel,Nitin Savaliya,Dinesh Kumar,Twinkle Soni,Jayshri Pethani,Umang Mishra,Nitesh Shah,Chaitanya Joshi,Madhvi Joshi |
| EPI_ISL_3162182 | N.H.L Municipal Medical College, Ahmedabad | Gujarat Biotechnology Research Centre | Sonal Sharma ,Zuber Saiyed,Ramesh Pandit,Janvi Raval,Zarna Patel,Nitin Savaliya,Dinesh Kumar,Twinkle Soni,Nitin Shukla,Jayshri Pethani,Umang Mishra,Nitesh Shah,Chaitanya Joshi,Madhvi Joshi |
| EPI_ISL_3162183 | N.H.L Municipal Medical College, Ahmedabad | Gujarat Biotechnology Research Centre | Zuber Saiyed,Ramesh Pandit,Janvi Raval,Zarna Patel,Nitin Savaliya,Dinesh Kumar,Twinkle Soni,Nitin Shukla,Sonal Sharma ,Jayshri Pethani,Umang Mishra,Nitesh Shah,Chaitanya Joshi,Madhvi Joshi |
| EPI_ISL_3162184 | N.H.L Municipal Medical College, Ahmedabad | Gujarat Biotechnology Research Centre | Ramesh Pandit,Janvi Raval,Zarna Patel,Nitin Savaliya,Dinesh Kumar,Twinkle Soni,Nitin Shukla,Sonal Sharma ,Zuber Saiyed,Jayshri Pethani,Umang Mishra,Nitesh Shah,Chaitanya Joshi,Madhvi Joshi |
| EPI_ISL_3162185 | N.H.L Municipal Medical College, Ahmedabad | Gujarat Biotechnology Research Centre | Janvi Raval,Zarna Patel,Nitin Savaliya,Dinesh Kumar,Twinkle Soni,Nitin Shukla,Sonal Sharma ,Zuber Saiyed,Ramesh Pandit,Jayshri Pethani,Umang Mishra,Nitesh Shah,Chaitanya Joshi,Madhvi Joshi |
| EPI_ISL_3162186 | Government Medical College (GMC), Surat | Gujarat Biotechnology Research Centre | Zarna Patel,Nitin Savaliya,Dinesh Kumar,Twinkle Soni,Nitin Shukla,Sonal Sharma ,Zuber Saiyed,Ramesh Pandit,Janvi Raval,Neeta Khandelwal,Umang Mishra,Nitesh Shah,Chaitanya Joshi,Madhvi Joshi |
| EPI_ISL_3162187 | Government Medical College (GMC), Surat | Gujarat Biotechnology Research Centre | Nitin Savaliya,Dinesh Kumar,Twinkle Soni,Nitin Shukla,Sonal Sharma ,Zuber Saiyed,Ramesh Pandit,Janvi Raval,Zarna Patel,Neeta Khandelwal,Umang Mishra,Nitesh Shah,Chaitanya Joshi,Madhvi Joshi |
| EPI_ISL_3162188 | Government Medical College (GMC), Surat | Gujarat Biotechnology Research Centre | Dinesh Kumar,Twinkle Soni,Nitin Shukla,Sonal Sharma ,Zuber Saiyed,Ramesh Pandit,Janvi Raval,Zarna Patel,Nitin Savaliya,Neeta Khandelwal,Umang Mishra,Nitesh Shah,Chaitanya Joshi,Madhvi Joshi |
| EPI_ISL_3162189 | Government Medical College (GMC), Surat | Gujarat Biotechnology Research Centre | Twinkle Soni,Nitin Shukla,Sonal Sharma ,Zuber Saiyed,Ramesh Pandit,Janvi Raval,Zarna Patel,Nitin Savaliya,Dinesh Kumar,Neeta Khandelwal,Umang Mishra,Nitesh Shah,Chaitanya Joshi,Madhvi Joshi |
| EPI_ISL_3162190 | Government Medical College (GMC), Surat | Gujarat Biotechnology Research Centre | Nitin Shukla,Sonal Sharma ,Zuber Saiyed,Ramesh Pandit,Janvi Raval,Zarna Patel,Nitin Savaliya,Dinesh Kumar,Twinkle Soni,Neeta Khandelwal,Umang Mishra,Nitesh Shah,Chaitanya Joshi,Madhvi Joshi |
| EPI_ISL_3162222 | Department of Medical Microbiology, Hospital Pengajar Universiti Putra Malaysia | Malaysia Genome Institute | Mohd Noor Mat Isa, Syafinaz Amin-Nordin, Irni Suhayu Sapian, Hui-Yee Chee, Yusuf Muhammad Noor, Nurhezreen Md Iqbal, Enizza Kasim, Siti Noraini Othman, Mohd Faizal Abu Bakar, Shamsidar Sopie, Azrin Ahmad, Narcisse Joseph, Muhammad MI, Avisha Richards, Nor Zahrin Hasran, Mohd Anuar Jonet, Nor Azfa Johari |
| EPI_ISL_3162224 | Pathology and Laboratory Medicine Institute, Cleveland Clinic, Ohio, USA | Pathology and Laboratory Medicine Institute, Cleveland Clinic, Ohio, USA | Jessica Spildener, Concetta Peck, Jennifer Starbuck, Joy Nakitandwe, Kristen McDonnell, David Plunkett, Zheng Jin Tu, Jay Brock, Yu-Wei Cheng, Gary Procop, Daniel Rhoads, Daniel H. Farkas, David Bosler |
| EPI_ISL_3162243 | Salud Digna | Instituto Nacional de Medicina Genomica | Cedro-Tanda A, Hidalgo-Miranda A, Cisneros-Villanueva M, Blancas S, Mendoza-Vargas A, Reyes-Grajeda JP, Abraham Campos-Romero, Moreno-Camacho José Luis, Rodriguez-Gallegos Jorge, Luna-Ruiz Marco, Gonzalez-Barrera D, Rangel-DeLeon D, Munguia-Garza P, Ramirez-Vega O, Escobar-Arrazola, M, Herrera-Montalvo LA. |
| EPI_ISL_3162253 | Instituto Nacional de Medicina Genomica | Instituto Nacional de Medicina Genomica | Cedro-Tanda A, Hidalgo-Miranda A, Cisneros-Villanueva M, Blancas S, Mendoza-Vargas A, Reyes-Grajeda JP, Abraham Campos-Romero, Moreno-Camacho José Luis, Rodriguez-Gallegos Jorge, Luna-Ruiz Marco, Gonzalez-Barrera D, Rangel-DeLeon D, Munguia-Garza P, Ramirez-Vega O, Escobar-Arrazola, M, Herrera-Montalvo LA. |
| EPI_ISL_3162262, EPI_ISL_3162263, EPI_ISL_3162265, EPI_ISL_3162266, EPI_ISL_3162268, EPI_ISL_3162270, EPI_ISL_3162271, EPI_ISL_3162272, EPI_ISL_3162273, EPI_ISL_3162277, EPI_ISL_3162278, EPI_ISL_3162283, EPI_ISL_3162284 | | | |
| see above | Botswana Harvard HIV Reference Laboratory | Botswana Harvard HIV Reference Laboratory | Sikhulile Moyo, Wonderful T. Choga, Dorcas Maruapula, Thongbotho Mphoyakgosi, Boitumelo Zuze, Botshelo Radibe, Legodile Kooepile, Ontlametse T. Bareng, Keoratile Ntshambiwa, Modisa Motswaledi, Madisa Mine, Joseph Makhema, Roger Shapiro, Shahin Lockman, Mosepele Mosepele, Simani Gaseitsiwe |
| EPI_ISL_3162315 | Dr. Yoshitaka Tamura Department of Clinical Laboratory, Osaka Habikino Medical Center | Department of Developmental Medicine, Research Institute, Osaka Women's and Children's Hospital | Itaru Yanagihara, Yukiko Nakura |
| EPI_ISL_3162316, EPI_ISL_3162317, EPI_ISL_3162318, EPI_ISL_3162319, EPI_ISL_3162320, EPI_ISL_3162321, EPI_ISL_3162322, EPI_ISL_3162323, EPI_ISL_3162324, EPI_ISL_3162325, EPI_ISL_3162326, EPI_ISL_3162327, EPI_ISL_3162328, EPI_ISL_3162329, EPI_ISL_3162330, EPI_ISL_3162332, EPI_ISL_3162333, EPI_ISL_3162334, EPI_ISL_3162335, EPI_ISL_3162336, EPI_ISL_3162337, EPI_ISL_3162338, EPI_ISL_3162339, EPI_ISL_3162340, EPI_ISL_3162341, EPI_ISL_3162342, EPI_ISL_3162343, EPI_ISL_3162344, EPI_ISL_3162345, EPI_ISL_3162346, EPI_ISL_3162347, EPI_ISL_3162348, EPI_ISL_3162351, EPI_ISL_3162352, EPI_ISL_3162353, EPI_ISL_3162354, EPI_ISL_3162355, EPI_ISL_3162356, EPI_ISL_3162357, EPI_ISL_3162358, EPI_ISL_3162359, EPI_ISL_3162360, EPI_ISL_3162361, EPI_ISL_3162362, EPI_ISL_3162363, EPI_ISL_3162364, EPI_ISL_3162365, EPI_ISL_3162366, EPI_ISL_3162367, EPI_ISL_3162368, EPI_ISL_3162369, EPI_ISL_3162370, EPI_ISL_3162371, EPI_ISL_3162372, EPI_ISL_3162373, EPI_ISL_3162374, EPI_ISL_3162375, EPI_ISL_3162376, EPI_ISL_3162377, EPI_ISL_3162378, EPI_ISL_3162379, EPI_ISL_3162380, EPI_ISL_3162381, EPI_ISL_3162382, EPI_ISL_3162383, EPI_ISL_3162384, EPI_ISL_3162387, EPI_ISL_3162388, EPI_ISL_3162389, EPI_ISL_3162390, EPI_ISL_3162391, EPI_ISL_3162393, EPI_ISL_3162394, EPI_ISL_3162395, EPI_ISL_3162396, EPI_ISL_3162397, EPI_ISL_3162399, EPI_ISL_3162400, EPI_ISL_3162401, EPI_ISL_3162402, EPI_ISL_3162403, EPI_ISL_3162404, EPI_ISL_3162405, EPI_ISL_3162406, EPI_ISL_3162407, EPI_ISL_3162408, EPI_ISL_3162409 | | | |
| see above | Illinois Department of Public Health - Springfield Lab | Illinois Department of Public Health - Springfield Lab | Bryan Sim, Marshall Smock |
| EPI_ISL_3162410, EPI_ISL_3162411, EPI_ISL_3162412, EPI_ISL_3162413, EPI_ISL_3162414, EPI_ISL_3162415, EPI_ISL_3162416, EPI_ISL_3162417, EPI_ISL_3162418, EPI_ISL_3162419, EPI_ISL_3162420, EPI_ISL_3162421, EPI_ISL_3162422, EPI_ISL_3162423, EPI_ISL_3162424 | | | |
| see above | Department of Bacteria, Parasites and Fungi, Statens Serum Institut, Copenhagen, Denmark | Statens Serum Institut Bioinformatics and Microbial Genomics | Danish Covid-19 Genome Consortium |
| EPI_ISL_3162425 | Molekylær Medicinsk Afdeling, Aarhus University Hospital, Aarhus, Denmark | Statens Serum Institut Bioinformatics and Microbial Genomics | Danish Covid-19 Genome Consortium |
| EPI_ISL_3162426, EPI_ISL_3162427, EPI_ISL_3162428, EPI_ISL_3162429, EPI_ISL_3162430, EPI_ISL_3162431, EPI_ISL_3162432, EPI_ISL_3162433, EPI_ISL_3162434, EPI_ISL_3162435, EPI_ISL_3162436, EPI_ISL_3162437, EPI_ISL_3162438, EPI_ISL_3162439, EPI_ISL_3162440, EPI_ISL_3162441, EPI_ISL_3162442, EPI_ISL_3162443, EPI_ISL_3162444, EPI_ISL_3162445, EPI_ISL_3162446, EPI_ISL_3162447, EPI_ISL_3162448, EPI_ISL_3162449, EPI_ISL_3162450, EPI_ISL_3162451, EPI_ISL_3162452, EPI_ISL_3162453, EPI_ISL_3162454, EPI_ISL_3162455, EPI_ISL_3162456, EPI_ISL_3162457, EPI_ISL_3162458, EPI_ISL_3162459, EPI_ISL_3162460, EPI_ISL_3162461, EPI_ISL_3162462, EPI_ISL_3162463, EPI_ISL_3162464, EPI_ISL_3162465, EPI_ISL_3162466, EPI_ISL_3162467, EPI_ISL_3162468, EPI_ISL_3162469, | | | |

| | | | |
|---|---|---|---|
| EPI_ISL_3162470, EPI_ISL_3162471, EPI_ISL_3162472, EPI_ISL_3162473, EPI_ISL_3162474, EPI_ISL_3162475, EPI_ISL_3162476, EPI_ISL_3162477, EPI_ISL_3162478, EPI_ISL_3162479, EPI_ISL_3162480, EPI_ISL_3162481, EPI_ISL_3162482, EPI_ISL_3162483, EPI_ISL_3162484, EPI_ISL_3162485, EPI_ISL_3162486, EPI_ISL_3162487, EPI_ISL_3162488, EPI_ISL_3162489, EPI_ISL_3162490, EPI_ISL_3162491, EPI_ISL_3162492, EPI_ISL_3162493, EPI_ISL_3162494, EPI_ISL_3162495, EPI_ISL_3162496, EPI_ISL_3162497, EPI_ISL_3162498, EPI_ISL_3162499, EPI_ISL_3162500, EPI_ISL_3162501, EPI_ISL_3162502, EPI_ISL_3162503, EPI_ISL_3162504, EPI_ISL_3162505, EPI_ISL_3162506, EPI_ISL_3162507, EPI_ISL_3162508, EPI_ISL_3162509, EPI_ISL_3162510, EPI_ISL_3162511, EPI_ISL_3162512, EPI_ISL_3162513, EPI_ISL_3162514, EPI_ISL_3162515, EPI_ISL_3162516, EPI_ISL_3162517, EPI_ISL_3162518, EPI_ISL_3162519, EPI_ISL_3162520, EPI_ISL_3162521, EPI_ISL_3162522, EPI_ISL_3162523, EPI_ISL_3162524, EPI_ISL_3162525, EPI_ISL_3162526, EPI_ISL_3162527, EPI_ISL_3162528, EPI_ISL_3162529, EPI_ISL_3162530, EPI_ISL_3162531, EPI_ISL_3162532, EPI_ISL_3162533, EPI_ISL_3162534, EPI_ISL_3162535, EPI_ISL_3162536, EPI_ISL_3162537, EPI_ISL_3162538, EPI_ISL_3162539, EPI_ISL_3162540, EPI_ISL_3162541, EPI_ISL_3162542, EPI_ISL_3162543, EPI_ISL_3162544, EPI_ISL_3162545, EPI_ISL_3162546, EPI_ISL_3162547, EPI_ISL_3162548, EPI_ISL_3162549, EPI_ISL_3162550 | | | |
| see above | Department of Bacteria, Parasites and Fungi, Statens Serum Institut, Copenhagen, Denmark | Statens Serum Institut Bioinformatics and Microbial Genomics | Danish Covid-19 Genome Consortium |
| EPI_ISL_3162551 | Molekylær Medicinsk Afdeling, Aarhus University Hospital, Aarhus, Denmark | Statens Serum Institut Bioinformatics and Microbial Genomics | Danish Covid-19 Genome Consortium |
| EPI_ISL_3162552, EPI_ISL_3162553, EPI_ISL_3162554, EPI_ISL_3162555, EPI_ISL_3162556, EPI_ISL_3162557, EPI_ISL_3162558, EPI_ISL_3162559, EPI_ISL_3162560, EPI_ISL_3162561, EPI_ISL_3162562, EPI_ISL_3162563, EPI_ISL_3162564, EPI_ISL_3162565, EPI_ISL_3162566, EPI_ISL_3162567, EPI_ISL_3162568, EPI_ISL_3162569, EPI_ISL_3162570, EPI_ISL_3162571, EPI_ISL_3162572, EPI_ISL_3162573, EPI_ISL_3162574, EPI_ISL_3162575, EPI_ISL_3162576, EPI_ISL_3162577, EPI_ISL_3162578, EPI_ISL_3162579, EPI_ISL_3162580, EPI_ISL_3162581, EPI_ISL_3162582, EPI_ISL_3162583, EPI_ISL_3162584, EPI_ISL_3162585, EPI_ISL_3162586, EPI_ISL_3162587, EPI_ISL_3162588, EPI_ISL_3162589, EPI_ISL_3162590, EPI_ISL_3162591, EPI_ISL_3162592, EPI_ISL_3162593, EPI_ISL_3162594, EPI_ISL_3162595, EPI_ISL_3162596, EPI_ISL_3162597, EPI_ISL_3162598, EPI_ISL_3162599, EPI_ISL_3162600, EPI_ISL_3162601, EPI_ISL_3162602, EPI_ISL_3162603, EPI_ISL_3162604, EPI_ISL_3162605, EPI_ISL_3162606 | | | |
| see above | Department of Bacteria, Parasites and Fungi, Statens Serum Institut, Copenhagen, Denmark | Statens Serum Institut Bioinformatics and Microbial Genomics | Danish Covid-19 Genome Consortium |
| EPI_ISL_3162607 | Department of Clinical Microbiology and Center for Genomic Medicine, Rigshospitalet, Copenhagen, Denmark | Statens Serum Institut Bioinformatics and Microbial Genomics | Danish Covid-19 Genome Consortium |
| EPI_ISL_3162608, EPI_ISL_3162609, EPI_ISL_3162610, EPI_ISL_3162611, EPI_ISL_3162612, EPI_ISL_3162613, EPI_ISL_3162614, EPI_ISL_3162615, EPI_ISL_3162616, EPI_ISL_3162617, EPI_ISL_3162618, EPI_ISL_3162619, EPI_ISL_3162620, EPI_ISL_3162621, EPI_ISL_3162622, EPI_ISL_3162623, EPI_ISL_3162624, EPI_ISL_3162625, EPI_ISL_3162626, EPI_ISL_3162627, EPI_ISL_3162628, EPI_ISL_3162629, EPI_ISL_3162630, EPI_ISL_3162631, EPI_ISL_3162632, EPI_ISL_3162633, EPI_ISL_3162634, EPI_ISL_3162635, EPI_ISL_3162636, EPI_ISL_3162637, EPI_ISL_3162638, EPI_ISL_3162639, EPI_ISL_3162640, EPI_ISL_3162641, EPI_ISL_3162642, EPI_ISL_3162643, EPI_ISL_3162644, EPI_ISL_3162645, EPI_ISL_3162646, EPI_ISL_3162647, EPI_ISL_3162648, EPI_ISL_3162649, EPI_ISL_3162650, EPI_ISL_3162651, EPI_ISL_3162652, EPI_ISL_3162653, EPI_ISL_3162654, EPI_ISL_3162655, EPI_ISL_3162656, EPI_ISL_3162657, EPI_ISL_3162658, EPI_ISL_3162659, EPI_ISL_3162660, EPI_ISL_3162661, EPI_ISL_3162662, EPI_ISL_3162663, EPI_ISL_3162664, EPI_ISL_3162665, EPI_ISL_3162666, EPI_ISL_3162667, EPI_ISL_3162668, EPI_ISL_3162669, EPI_ISL_3162670, EPI_ISL_3162671, EPI_ISL_3162672, EPI_ISL_3162673, EPI_ISL_3162674, EPI_ISL_3162675, EPI_ISL_3162676, EPI_ISL_3162677, EPI_ISL_3162678, EPI_ISL_3162679, EPI_ISL_3162680, EPI_ISL_3162681, EPI_ISL_3162682, EPI_ISL_3162683, EPI_ISL_3162684, EPI_ISL_3162685, EPI_ISL_3162686, EPI_ISL_3162687, EPI_ISL_3162688, EPI_ISL_3162689, EPI_ISL_3162690, EPI_ISL_3162691, EPI_ISL_3162692, EPI_ISL_3162694, EPI_ISL_3162695, EPI_ISL_3162696 | | | |
| see above | Department of Bacteria, Parasites and Fungi, Statens Serum Institut, Copenhagen, Denmark | Statens Serum Institut Bioinformatics and Microbial Genomics | Danish Covid-19 Genome Consortium |
| EPI_ISL_3162697 | Molekylær Medicinsk Afdeling, Aarhus University Hospital, Aarhus, Denmark | Statens Serum Institut Bioinformatics and Microbial Genomics | Danish Covid-19 Genome Consortium |
| EPI_ISL_3162698, EPI_ISL_3162699, EPI_ISL_3162700, EPI_ISL_3162701, EPI_ISL_3162702, EPI_ISL_3162703, EPI_ISL_3162704, EPI_ISL_3162705, EPI_ISL_3162706, EPI_ISL_3162707, EPI_ISL_3162708, EPI_ISL_3162709, EPI_ISL_3162710, EPI_ISL_3162711, EPI_ISL_3162712, EPI_ISL_3162713, EPI_ISL_3162714, EPI_ISL_3162715, EPI_ISL_3162716, EPI_ISL_3162717, EPI_ISL_3162718, EPI_ISL_3162719, EPI_ISL_3162720, EPI_ISL_3162721, EPI_ISL_3162722, EPI_ISL_3162723, EPI_ISL_3162724, EPI_ISL_3162725, EPI_ISL_3162727, EPI_ISL_3162728, EPI_ISL_3162729, EPI_ISL_3162730, EPI_ISL_3162731, EPI_ISL_3162732, EPI_ISL_3162733, EPI_ISL_3162734, EPI_ISL_3162735, EPI_ISL_3162736, EPI_ISL_3162737, EPI_ISL_3162738, EPI_ISL_3162739, EPI_ISL_3162740, EPI_ISL_3162741, EPI_ISL_3162742, EPI_ISL_3162743, EPI_ISL_3162744, EPI_ISL_3162745, EPI_ISL_3162746, EPI_ISL_3162747, EPI_ISL_3162748, EPI_ISL_3162749, EPI_ISL_3162750, EPI_ISL_3162751, EPI_ISL_3162752, EPI_ISL_3162753, EPI_ISL_3162754, EPI_ISL_3162755, EPI_ISL_3162756, EPI_ISL_3162757, EPI_ISL_3162758, EPI_ISL_3162759, EPI_ISL_3162760, EPI_ISL_3162761, EPI_ISL_3162762, EPI_ISL_3162763, EPI_ISL_3162764, EPI_ISL_3162765, EPI_ISL_3162766, EPI_ISL_3162767, EPI_ISL_3162768, EPI_ISL_3162769, EPI_ISL_3162770, EPI_ISL_3162771, EPI_ISL_3162772, EPI_ISL_3162773, EPI_ISL_3162774, EPI_ISL_3162775, EPI_ISL_3162776, EPI_ISL_3162777, EPI_ISL_3162778, EPI_ISL_3162779, EPI_ISL_3162780, EPI_ISL_3162781, EPI_ISL_3162782, EPI_ISL_3162783, EPI_ISL_3162784, EPI_ISL_3162785, EPI_ISL_3162786, EPI_ISL_3162787, EPI_ISL_3162788, EPI_ISL_3162789, EPI_ISL_3162790, EPI_ISL_3162791, EPI_ISL_3162792, EPI_ISL_3162793, EPI_ISL_3162794, EPI_ISL_3162795 | | | |
| see above | Department of Bacteria, Parasites and Fungi, Statens Serum Institut, Copenhagen, Denmark | Statens Serum Institut Bioinformatics and Microbial Genomics | Danish Covid-19 Genome Consortium |
| EPI_ISL_3162796 | Tokyo Metropolitan Institute of Public Health | Tokyo Metropolitan Institute of Public Health | Masaki Hayashi, Takako Yamazaki, Akane Negishi, Hirofumi Miyake, Kenshirou Kuroki, Maya Isogai, Takushi Fujiwara, Mami Nagashima, Jun Suzuki, Kenji Sadamasu |
| EPI_ISL_3162797, EPI_ISL_3162798 | Department of Bacteria, Parasites and Fungi, Statens Serum Institut, Copenhagen, Denmark | Statens Serum Institut Bioinformatics and Microbial Genomics | Danish Covid-19 Genome Consortium |
| EPI_ISL_3162799 | Tokyo Metropolitan Institute of Public Health | Tokyo Metropolitan Institute of Public Health | Masaki Hayashi, Takako Yamazaki, Akane Negishi, Hirofumi Miyake, Kenshirou Kuroki, Maya Isogai, Takushi Fujiwara, Mami Nagashima, Jun Suzuki, Kenji Sadamasu |
| EPI_ISL_3162800, EPI_ISL_3162801 | Department of Bacteria, Parasites and Fungi, Statens Serum Institut, Copenhagen, Denmark | Statens Serum Institut Bioinformatics and Microbial Genomics | Danish Covid-19 Genome Consortium |
| EPI_ISL_3162802 | Tokyo Metropolitan Institute of Public Health | Tokyo Metropolitan Institute of Public Health | Masaki Hayashi, Takako Yamazaki, Akane Negishi, Hirofumi Miyake, Kenshirou Kuroki, Maya Isogai, Takushi Fujiwara, Mami Nagashima, Jun Suzuki, Kenji Sadamasu |
| EPI_ISL_3162803 | Department of Bacteria, Parasites and Fungi, Statens Serum Institut, Copenhagen, Denmark | Statens Serum Institut Bioinformatics and Microbial Genomics | Danish Covid-19 Genome Consortium |
| EPI_ISL_3162804 | Tokyo Metropolitan Institute of Public Health | Tokyo Metropolitan Institute of Public Health | Masaki Hayashi, Takako Yamazaki, Akane Negishi, Hirofumi Miyake, Kenshirou Kuroki, Maya Isogai, Takushi Fujiwara, Mami Nagashima, Jun Suzuki, Kenji Sadamasu |
| EPI_ISL_3162805, EPI_ISL_3162806 | Department of Bacteria, Parasites and Fungi, Statens Serum Institut, Copenhagen, Denmark | Statens Serum Institut Bioinformatics and Microbial Genomics | Danish Covid-19 Genome Consortium |
| EPI_ISL_3162807 | Tokyo Metropolitan Institute of Public Health | Tokyo Metropolitan Institute of Public Health | Masaki Hayashi, Takako Yamazaki, Akane Negishi, Hirofumi Miyake, Kenshirou Kuroki, Maya Isogai, Takushi Fujiwara, Mami Nagashima, Jun Suzuki, Kenji Sadamasu |
| EPI_ISL_3162808 | Department of Bacteria, Parasites and Fungi, Statens Serum Institut, Copenhagen, Denmark | Statens Serum Institut Bioinformatics and Microbial Genomics | Danish Covid-19 Genome Consortium |
| EPI_ISL_3162809 | Tokyo Metropolitan Institute of Public Health | Tokyo Metropolitan Institute of Public Health | Masaki Hayashi, Takako Yamazaki, Akane Negishi, Hirofumi Miyake, Kenshirou Kuroki, Maya Isogai, Takushi Fujiwara, Mami Nagashima, Jun Suzuki, Kenji Sadamasu |
| EPI_ISL_3162810, EPI_ISL_3162811 | Department of Bacteria, Parasites and Fungi, Statens Serum Institut, Copenhagen, Denmark | Statens Serum Institut Bioinformatics and Microbial Genomics | Danish Covid-19 Genome Consortium |
| EPI_ISL_3162812 | Tokyo Metropolitan Institute of Public Health | Tokyo Metropolitan Institute of Public Health | Masaki Hayashi, Takako Yamazaki, Akane Negishi, Hirofumi Miyake, Kenshirou Kuroki, Maya Isogai, Takushi Fujiwara, Mami Nagashima, Jun Suzuki, Kenji Sadamasu |
| EPI_ISL_3162813 | Department of Bacteria, Parasites and Fungi, Statens Serum Institut, Copenhagen, Denmark | Statens Serum Institut Bioinformatics and Microbial Genomics | Danish Covid-19 Genome Consortium |
| EPI_ISL_3162814 | Tokyo Metropolitan Institute of Public Health | Tokyo Metropolitan Institute of Public Health | Masaki Hayashi, Takako Yamazaki, Akane Negishi, Hirofumi Miyake, Kenshirou Kuroki, Maya Isogai, Takushi Fujiwara, Mami Nagashima, Jun Suzuki, Kenji Sadamasu |
| EPI_ISL_3162815 | Department of Bacteria, Parasites and Fungi, Statens Serum Institut, Copenhagen, Denmark | Statens Serum Institut Bioinformatics and Microbial Genomics | Danish Covid-19 Genome Consortium |
| EPI_ISL_3162816 | Tokyo Metropolitan Institute of Public Health | Tokyo Metropolitan Institute of Public Health | Masaki Hayashi, Takako Yamazaki, Akane Negishi, Hirofumi Miyake, Kenshirou Kuroki, Maya Isogai, Takushi Fujiwara, Mami Nagashima, Jun Suzuki, Kenji Sadamasu |
| EPI_ISL_3162817, EPI_ISL_3162818, EPI_ISL_3162819, EPI_ISL_3162820, EPI_ISL_3162821, EPI_ISL_3162822, EPI_ISL_3162824, EPI_ISL_3162825, EPI_ISL_3162826, EPI_ISL_3162827, EPI_ISL_3162828, EPI_ISL_3162829, EPI_ISL_3162830, EPI_ISL_3162831, EPI_ISL_3162832, EPI_ISL_3162833, EPI_ISL_3162834 | | | |
| see above | Department of Bacteria, Parasites and Fungi, Statens Serum Institut, Copenhagen, Denmark | Statens Serum Institut Bioinformatics and Microbial Genomics | Danish Covid-19 Genome Consortium |
| EPI_ISL_3162835 | Department of Clinical Microbiology and Center for Genomic Medicine, Rigshospitalet, Copenhagen, Denmark | Statens Serum Institut Bioinformatics and Microbial Genomics | Danish Covid-19 Genome Consortium |
| EPI_ISL_3162836, EPI_ISL_3162837, EPI_ISL_3162838, EPI_ISL_3162839, EPI_ISL_3162840, EPI_ISL_3162841, EPI_ISL_3162842, EPI_ISL_3162843, EPI_ISL_3162844, EPI_ISL_3162845, EPI_ISL_3162846, EPI_ISL_3162847, EPI_ISL_3162848, EPI_ISL_3162849, EPI_ISL_3162850, EPI_ISL_3162851, EPI_ISL_3162852, EPI_ISL_3162853, EPI_ISL_3162854, EPI_ISL_3162855, EPI_ISL_3162856, EPI_ISL_3162857, EPI_ISL_3162858, EPI_ISL_3162859, EPI_ISL_3162860, EPI_ISL_3162861, EPI_ISL_3162862, EPI_ISL_3162863, EPI_ISL_3162864, EPI_ISL_3162865, EPI_ISL_3162866, EPI_ISL_3162867, EPI_ISL_3162868, EPI_ISL_3162869, EPI_ISL_3162870, EPI_ISL_3162871, EPI_ISL_3162872, EPI_ISL_3162873, EPI_ISL_3162874, EPI_ISL_3162875, EPI_ISL_3162876, EPI_ISL_3162877, EPI_ISL_3162878, EPI_ISL_3162879, EPI_ISL_3162880, EPI_ISL_3162881, EPI_ISL_3162882, EPI_ISL_3162883, EPI_ISL_3162884, EPI_ISL_3162885, EPI_ISL_3162886, EPI_ISL_3162887, EPI_ISL_3162888, EPI_ISL_3162889, EPI_ISL_3162890, EPI_ISL_3162891, EPI_ISL_3162892, EPI_ISL_3162893, EPI_ISL_3162894, EPI_ISL_3162895, EPI_ISL_3162896, EPI_ISL_3162897, EPI_ISL_3162898, EPI_ISL_3162899, EPI_ISL_3162900, EPI_ISL_3162901, EPI_ISL_3162902, EPI_ISL_3162903, EPI_ISL_3162904, EPI_ISL_3162905, EPI_ISL_3162906, EPI_ISL_3162907, EPI_ISL_3162908, EPI_ISL_3162909, EPI_ISL_3162910, EPI_ISL_3162912, EPI_ISL_3162913, EPI_ISL_3162914, EPI_ISL_3162915, EPI_ISL_3162916, EPI_ISL_3162917, EPI_ISL_3162918, EPI_ISL_3162919, EPI_ISL_3162920, EPI_ISL_3162921, EPI_ISL_3162922, EPI_ISL_3162923, EPI_ISL_3162924, EPI_ISL_3162925, EPI_ISL_3162926, EPI_ISL_3162927, EPI_ISL_3162928, EPI_ISL_3162929, EPI_ISL_3162930, EPI_ISL_3162931, EPI_ISL_3162932, EPI_ISL_3162933, EPI_ISL_3162934, EPI_ISL_3162935, EPI_ISL_3162936, EPI_ISL_3162937, EPI_ISL_3162938, EPI_ISL_3162939, EPI_ISL_3162940, EPI_ISL_3162941, EPI_ISL_3162942, EPI_ISL_3162943, EPI_ISL_3162944, EPI_ISL_3162945, EPI_ISL_3162946, EPI_ISL_3162947, EPI_ISL_3162948, EPI_ISL_3162949, EPI_ISL_3162950, EPI_ISL_3162951, EPI_ISL_3162952, EPI_ISL_3162953, EPI_ISL_3162954, EPI_ISL_3162955, EPI_ISL_3162956, EPI_ISL_3162957, EPI_ISL_3162958, EPI_ISL_3162959, EPI_ISL_3162960, EPI_ISL_3162961, EPI_ISL_3162962, EPI_ISL_3162963, EPI_ISL_3162964, EPI_ISL_3162965, EPI_ISL_3162966, EPI_ISL_3162967, EPI_ISL_3162968, EPI_ISL_3162969, EPI_ISL_3162970, EPI_ISL_3162971, EPI_ISL_3162972, EPI_ISL_3162973, EPI_ISL_3162974, EPI_ISL_3162975, EPI_ISL_3162976, EPI_ISL_3162977, EPI_ISL_3162978, EPI_ISL_3162979, EPI_ISL_3162980, EPI_ISL_3162981, EPI_ISL_3162982, EPI_ISL_3162983, EPI_ISL_3162984, EPI_ISL_3162985, EPI_ISL_3162986, EPI_ISL_3162987, EPI_ISL_3162988, EPI_ISL_3162989, EPI_ISL_3162990, EPI_ISL_3162991, EPI_ISL_3162992, EPI_ISL_3162993, EPI_ISL_3162994, EPI_ISL_3162995, EPI_ISL_3162996, EPI_ISL_3162997, EPI_ISL_3162998, EPI_ISL_3162999, EPI_ISL_3163000, EPI_ISL_3163001, EPI_ISL_3163002, EPI_ISL_3163003, EPI_ISL_3163004, EPI_ISL_3163005, EPI_ISL_3163006, EPI_ISL_3163007, EPI_ISL_3163008, EPI_ISL_3163009, EPI_ISL_3163010, EPI_ISL_3163011, EPI_ISL_3163012, EPI_ISL_3163013, EPI_ISL_3163014, EPI_ISL_3163015, EPI_ISL_3163016, EPI_ISL_3163017, EPI_ISL_3163018, EPI_ISL_3163019, EPI_ISL_3163020, EPI_ISL_3163021, EPI_ISL_3163022, EPI_ISL_3163023, EPI_ISL_3163024, EPI_ISL_3163025, EPI_ISL_3163026, EPI_ISL_3163027, EPI_ISL_3163028, EPI_ISL_3163029, EPI_ISL_3163030, EPI_ISL_3163031 | | | |
| see above | Department of Bacteria, Parasites and Fungi, Statens Serum Institut, Copenhagen, Denmark | Statens Serum Institut Bioinformatics and Microbial Genomics | Danish Covid-19 Genome Consortium |
| EPI_ISL_3163032 | Molekylær Medicinsk Afdeling, Aarhus University Hospital, Aarhus, Denmark | Statens Serum Institut Bioinformatics and Microbial Genomics | Danish Covid-19 Genome Consortium |
| EPI_ISL_3163033, EPI_ISL_3163034, EPI_ISL_3163035, EPI_ISL_3163036, EPI_ISL_3163037, EPI_ISL_3163038, EPI_ISL_3163039, EPI_ISL_3163040, EPI_ISL_3163041, EPI_ISL_3163042, EPI_ISL_3163043, EPI_ISL_3163044, EPI_ISL_3163045, EPI_ISL_3163046, EPI_ISL_3163047, EPI_ISL_3163048, EPI_ISL_3163049, EPI_ISL_3163050, EPI_ISL_3163051, EPI_ISL_3163052, EPI_ISL_3163053, EPI_ISL_3163054, EPI_ISL_3163055, EPI_ISL_3163056, EPI_ISL_3163057, EPI_ISL_3163058, EPI_ISL_3163059, EPI_ISL_3163060, EPI_ISL_3163061, EPI_ISL_3163062, EPI_ISL_3163063, EPI_ISL_3163064, EPI_ISL_3163065, EPI_ISL_3163066, EPI_ISL_3163067, EPI_ISL_3163068, EPI_ISL_3163069 | | | |
| see above | Department of Bacteria, Parasites and Fungi, Statens Serum Institut, Copenhagen, Denmark | Statens Serum Institut Bioinformatics and Microbial Genomics | Danish Covid-19 Genome Consortium |
| EPI_ISL_3163070 | Molekylær Medicinsk Afdeling, Aarhus University Hospital, Aarhus, Denmark | Statens Serum Institut Bioinformatics and Microbial Genomics | Danish Covid-19 Genome Consortium |
| EPI_ISL_3163071, EPI_ISL_3163072, EPI_ISL_3163073, EPI_ISL_3163074, EPI_ISL_3163075, EPI_ISL_3163076, EPI_ISL_3163077, EPI_ISL_3163078, EPI_ISL_3163079, EPI_ISL_3163080, EPI_ISL_3163081, EPI_ISL_3163082 | | | |
| see above | Department of Bacteria, Parasites and Fungi, Statens Serum Institut, Copenhagen, Denmark | Statens Serum Institut Bioinformatics and Microbial Genomics | Danish Covid-19 Genome Consortium |
| EPI_ISL_3163083 | Department of Clinical Microbiology and Center for Genomic Medicine, Rigshospitalet, Copenhagen, Denmark | Statens Serum Institut Bioinformatics and Microbial Genomics | Danish Covid-19 Genome Consortium |
| EPI_ISL_3163084, EPI_ISL_3163085, EPI_ISL_3163086, EPI_ISL_3163087, EPI_ISL_3163088, EPI_ISL_3163089, EPI_ISL_3163090, EPI_ISL_3163091, EPI_ISL_3163092, EPI_ISL_3163093, EPI_ISL_3163094, EPI_ISL_3163095, EPI_ISL_3163096, EPI_ISL_3163097, EPI_ISL_3163098, EPI_ISL_3163099, EPI_ISL_3163100, EPI_ISL_3163101, EPI_ISL_3163102, EPI_ISL_3163103, EPI_ISL_3163104, EPI_ISL_3163105, EPI_ISL_3163106, EPI_ISL_3163107, EPI_ISL_3163108, EPI_ISL_3163109, EPI_ISL_3163110, EPI_ISL_3163111, EPI_ISL_3163112, EPI_ISL_3163113, EPI_ISL_3163114, EPI_ISL_3163115, EPI_ISL_3163116, EPI_ISL_3163117, EPI_ISL_3163118, EPI_ISL_3163119, EPI_ISL_3163120, EPI_ISL_3163121, EPI_ISL_3163122, EPI_ISL_3163123, EPI_ISL_3163124, EPI_ISL_3163125, EPI_ISL_3163126, EPI_ISL_3163127, EPI_ISL_3163128, EPI_ISL_3163129, EPI_ISL_3163130, EPI_ISL_3163131, EPI_ISL_3163132, EPI_ISL_3163133, EPI_ISL_3163134, EPI_ISL_3163135, EPI_ISL_3163136, EPI_ISL_3163137, EPI_ISL_3163138, EPI_ISL_3163139, EPI_ISL_3163140, EPI_ISL_3163141, EPI_ISL_3163142, EPI_ISL_3163143, EPI_ISL_3163144, EPI_ISL_3163145, EPI_ISL_3163146, EPI_ISL_3163147, EPI_ISL_3163148, EPI_ISL_3163149, EPI_ISL_3163150, EPI_ISL_3163151, EPI_ISL_3163152, EPI_ISL_3163153, EPI_ISL_3163154, EPI_ISL_3163155, EPI_ISL_3163156, EPI_ISL_3163157, EPI_ISL_3163158, EPI_ISL_3163159, EPI_ISL_3163160, EPI_ISL_3163161, EPI_ISL_3163162, EPI_ISL_3163163, EPI_ISL_3163164, EPI_ISL_3163165, EPI_ISL_3163166, EPI_ISL_3163167, EPI_ISL_3163168, EPI_ISL_3163169, EPI_ISL_3163170, EPI_ISL_3163171, EPI_ISL_3163172, EPI_ISL_3163173, EPI_ISL_3163174, EPI_ISL_3163175, EPI_ISL_3163176, EPI_ISL_3163177, EPI_ISL_3163178, EPI_ISL_3163179, EPI_ISL_3163180, EPI_ISL_3163181, EPI_ISL_3163182, EPI_ISL_3163183, EPI_ISL_3163184, EPI_ISL_3163185, EPI_ISL_3163186, EPI_ISL_3163187, EPI_ISL_3163188, EPI_ISL_3163189, EPI_ISL_3163190, EPI_ISL_3163191, EPI_ISL_3163192, EPI_ISL_3163193, EPI_ISL_3163195, EPI_ISL_3163196, EPI_ISL_3163197, EPI_ISL_3163198, EPI_ISL_3163199, EPI_ISL_3163200, EPI_ISL_3163201, EPI_ISL_3163202, EPI_ISL_3163203, EPI_ISL_3163204, EPI_ISL_3163205, EPI_ISL_3163206, EPI_ISL_3163207, EPI_ISL_3163208, EPI_ISL_3163209, EPI_ISL_3163210, EPI_ISL_3163211, EPI_ISL_3163212, EPI_ISL_3163213, EPI_ISL_3163214, EPI_ISL_3163215, EPI_ISL_3163216, EPI_ISL_3163217 | | | |
| see above | Department of Bacteria, Parasites and Fungi, Statens Serum | Statens Serum Institut Bioinformatics and Microbial Genomics | Danish Covid-19 Genome Consortium |

| | | | |
|---|---|---|---|
| | Institut, Copenhagen, Denmark | | |
| EPI_ISL_3163218 | Molekylær Medicinsk Afdeling, Aarhus University Hospital, Aarhus, Denmark | Statens Serum Institut Bioinformatics and Microbial Genomics | Danish Covid-19 Genome Consortium |
| EPI_ISL_3163219, EPI_ISL_3163220, EPI_ISL_3163221, EPI_ISL_3163222, EPI_ISL_3163223, EPI_ISL_3163224, EPI_ISL_3163225, EPI_ISL_3163226, EPI_ISL_3163227, EPI_ISL_3163228, EPI_ISL_3163229, EPI_ISL_3163230, EPI_ISL_3163231, EPI_ISL_3163232, EPI_ISL_3163233, EPI_ISL_3163234, EPI_ISL_3163235, EPI_ISL_3163236, EPI_ISL_3163237, EPI_ISL_3163238, EPI_ISL_3163239, EPI_ISL_3163240, EPI_ISL_3163241, EPI_ISL_3163242, EPI_ISL_3163243, EPI_ISL_3163244, EPI_ISL_3163245, EPI_ISL_3163246, EPI_ISL_3163247, EPI_ISL_3163248, EPI_ISL_3163249, EPI_ISL_3163250, EPI_ISL_3163251, EPI_ISL_3163252, EPI_ISL_3163253, EPI_ISL_3163254, EPI_ISL_3163255, EPI_ISL_3163256, EPI_ISL_3163257, EPI_ISL_3163258, EPI_ISL_3163259, EPI_ISL_3163260, EPI_ISL_3163261, EPI_ISL_3163262, EPI_ISL_3163263, EPI_ISL_3163264, EPI_ISL_3163265, EPI_ISL_3163266, EPI_ISL_3163267, EPI_ISL_3163268, EPI_ISL_3163269, EPI_ISL_3163270, EPI_ISL_3163271, EPI_ISL_3163272, EPI_ISL_3163273, EPI_ISL_3163274, EPI_ISL_3163275, EPI_ISL_3163276, EPI_ISL_3163277, EPI_ISL_3163278, EPI_ISL_3163279, EPI_ISL_3163280, EPI_ISL_3163281, EPI_ISL_3163282, EPI_ISL_3163283, EPI_ISL_3163284, EPI_ISL_3163285, EPI_ISL_3163286, EPI_ISL_3163287, EPI_ISL_3163288, EPI_ISL_3163289 | | | |
| see above | Department of Bacteria, Parasites and Fungi, Statens Serum Institut, Copenhagen, Denmark | Statens Serum Institut Bioinformatics and Microbial Genomics | Danish Covid-19 Genome Consortium |
| EPI_ISL_3163290 | Molekylær Medicinsk Afdeling, Aarhus University Hospital, Aarhus, Denmark | Statens Serum Institut Bioinformatics and Microbial Genomics | Danish Covid-19 Genome Consortium |
| EPI_ISL_3163291, EPI_ISL_3163292, EPI_ISL_3163293, EPI_ISL_3163294, EPI_ISL_3163295, EPI_ISL_3163296, EPI_ISL_3163297, EPI_ISL_3163298, EPI_ISL_3163299, EPI_ISL_3163300, EPI_ISL_3163301, EPI_ISL_3163302, EPI_ISL_3163303, EPI_ISL_3163304, EPI_ISL_3163305, EPI_ISL_3163306, EPI_ISL_3163307, EPI_ISL_3163308, EPI_ISL_3163309, EPI_ISL_3163310, EPI_ISL_3163311, EPI_ISL_3163312, EPI_ISL_3163313, EPI_ISL_3163314, EPI_ISL_3163315, EPI_ISL_3163316, EPI_ISL_3163317, EPI_ISL_3163318, EPI_ISL_3163319, EPI_ISL_3163320, EPI_ISL_3163321, EPI_ISL_3163322, EPI_ISL_3163323, EPI_ISL_3163324, EPI_ISL_3163325, EPI_ISL_3163326, EPI_ISL_3163327, EPI_ISL_3163328, EPI_ISL_3163329, EPI_ISL_3163330, EPI_ISL_3163331, EPI_ISL_3163332, EPI_ISL_3163333, EPI_ISL_3163334, EPI_ISL_3163335, EPI_ISL_3163336, EPI_ISL_3163337, EPI_ISL_3163338, EPI_ISL_3163339, EPI_ISL_3163340, EPI_ISL_3163341, EPI_ISL_3163342, EPI_ISL_3163343, EPI_ISL_3163344, EPI_ISL_3163345, EPI_ISL_3163346, EPI_ISL_3163347, EPI_ISL_3163348, EPI_ISL_3163349, EPI_ISL_3163350, EPI_ISL_3163351, EPI_ISL_3163352, EPI_ISL_3163353, EPI_ISL_3163354, EPI_ISL_3163355, EPI_ISL_3163356, EPI_ISL_3163357, EPI_ISL_3163358, EPI_ISL_3163359, EPI_ISL_3163360, EPI_ISL_3163361, EPI_ISL_3163362, EPI_ISL_3163363, EPI_ISL_3163364, EPI_ISL_3163365, EPI_ISL_3163366, EPI_ISL_3163367, EPI_ISL_3163368, EPI_ISL_3163369, EPI_ISL_3163370, EPI_ISL_3163371, EPI_ISL_3163372, EPI_ISL_3163373, EPI_ISL_3163374, EPI_ISL_3163375, EPI_ISL_3163376 | | | |
| see above | Department of Bacteria, Parasites and Fungi, Statens Serum Institut, Copenhagen, Denmark | Statens Serum Institut Bioinformatics and Microbial Genomics | Danish Covid-19 Genome Consortium |
| EPI_ISL_3163377 | Molekylær Medicinsk Afdeling, Aarhus University Hospital, Aarhus, Denmark | Statens Serum Institut Bioinformatics and Microbial Genomics | Danish Covid-19 Genome Consortium |
| EPI_ISL_3163378, EPI_ISL_3163379, EPI_ISL_3163380, EPI_ISL_3163381, EPI_ISL_3163382, EPI_ISL_3163383, EPI_ISL_3163384, EPI_ISL_3163385, EPI_ISL_3163386, EPI_ISL_3163387, EPI_ISL_3163388, EPI_ISL_3163389, EPI_ISL_3163390, EPI_ISL_3163391, EPI_ISL_3163392, EPI_ISL_3163393, EPI_ISL_3163394, EPI_ISL_3163395, EPI_ISL_3163396, EPI_ISL_3163397, EPI_ISL_3163398, EPI_ISL_3163399, EPI_ISL_3163400, EPI_ISL_3163401, EPI_ISL_3163402, EPI_ISL_3163403, EPI_ISL_3163404, EPI_ISL_3163405, EPI_ISL_3163406, EPI_ISL_3163407, EPI_ISL_3163408, EPI_ISL_3163409, EPI_ISL_3163410, EPI_ISL_3163411, EPI_ISL_3163412, EPI_ISL_3163413, EPI_ISL_3163414, EPI_ISL_3163415, EPI_ISL_3163416, EPI_ISL_3163417, EPI_ISL_3163418, EPI_ISL_3163419, EPI_ISL_3163420, EPI_ISL_3163421, EPI_ISL_3163422, EPI_ISL_3163423, EPI_ISL_3163424, EPI_ISL_3163425, EPI_ISL_3163426, EPI_ISL_3163427, EPI_ISL_3163428, EPI_ISL_3163429, EPI_ISL_3163430, EPI_ISL_3163431, EPI_ISL_3163432, EPI_ISL_3163433, EPI_ISL_3163434, EPI_ISL_3163435, EPI_ISL_3163436, EPI_ISL_3163437, EPI_ISL_3163438, EPI_ISL_3163439, EPI_ISL_3163440, EPI_ISL_3163441, EPI_ISL_3163442, EPI_ISL_3163443, EPI_ISL_3163444, EPI_ISL_3163445, EPI_ISL_3163446, EPI_ISL_3163447, EPI_ISL_3163448, EPI_ISL_3163449, EPI_ISL_3163450, EPI_ISL_3163451, EPI_ISL_3163452, EPI_ISL_3163453, EPI_ISL_3163454, EPI_ISL_3163455, EPI_ISL_3163456, EPI_ISL_3163457, EPI_ISL_3163458, EPI_ISL_3163459, EPI_ISL_3163460, EPI_ISL_3163461, EPI_ISL_3163462, EPI_ISL_3163463, EPI_ISL_3163464, EPI_ISL_3163465, EPI_ISL_3163466, EPI_ISL_3163467, EPI_ISL_3163468, EPI_ISL_3163469, EPI_ISL_3163470, EPI_ISL_3163471, EPI_ISL_3163473, EPI_ISL_3163474, EPI_ISL_3163475, EPI_ISL_3163476, EPI_ISL_3163477, EPI_ISL_3163478, EPI_ISL_3163479, EPI_ISL_3163480, EPI_ISL_3163481, EPI_ISL_3163482, EPI_ISL_3163483, EPI_ISL_3163484, EPI_ISL_3163485, EPI_ISL_3163486, EPI_ISL_3163487, EPI_ISL_3163488, EPI_ISL_3163489, EPI_ISL_3163490, EPI_ISL_3163491, EPI_ISL_3163492, EPI_ISL_3163493, EPI_ISL_3163494, EPI_ISL_3163495, EPI_ISL_3163496, EPI_ISL_3163497, EPI_ISL_3163498, EPI_ISL_3163499, EPI_ISL_3163500, EPI_ISL_3163501, EPI_ISL_3163502, EPI_ISL_3163503, EPI_ISL_3163504, EPI_ISL_3163505, EPI_ISL_3163506, EPI_ISL_3163507, EPI_ISL_3163508, EPI_ISL_3163509, EPI_ISL_3163510, EPI_ISL_3163511, EPI_ISL_3163512, EPI_ISL_3163513, EPI_ISL_3163514, EPI_ISL_3163515, EPI_ISL_3163516, EPI_ISL_3163517, EPI_ISL_3163518, EPI_ISL_3163519, EPI_ISL_3163520, EPI_ISL_3163521, EPI_ISL_3163522, EPI_ISL_3163523, EPI_ISL_3163524, EPI_ISL_3163525, EPI_ISL_3163526 | | | |
| see above | Department of Bacteria, Parasites and Fungi, Statens Serum Institut, Copenhagen, Denmark | Statens Serum Institut Bioinformatics and Microbial Genomics | Danish Covid-19 Genome Consortium |
| EPI_ISL_3163527 | Molekylær Medicinsk Afdeling, Aarhus University Hospital, Aarhus, Denmark | Statens Serum Institut Bioinformatics and Microbial Genomics | Danish Covid-19 Genome Consortium |
| EPI_ISL_3163528, EPI_ISL_3163529, EPI_ISL_3163530, EPI_ISL_3163531, EPI_ISL_3163532, EPI_ISL_3163533, EPI_ISL_3163534, EPI_ISL_3163535, EPI_ISL_3163536, EPI_ISL_3163537, EPI_ISL_3163538, EPI_ISL_3163539, EPI_ISL_3163540, EPI_ISL_3163541, EPI_ISL_3163542, EPI_ISL_3163543, EPI_ISL_3163544, EPI_ISL_3163545, EPI_ISL_3163546, EPI_ISL_3163547, EPI_ISL_3163548 | | | |
| see above | Department of Bacteria, Parasites and Fungi, Statens Serum Institut, Copenhagen, Denmark | Statens Serum Institut Bioinformatics and Microbial Genomics | Danish Covid-19 Genome Consortium |
| EPI_ISL_3163549 | Molekylær Medicinsk Afdeling, Aarhus University Hospital, Aarhus, Denmark | Statens Serum Institut Bioinformatics and Microbial Genomics | Danish Covid-19 Genome Consortium |
| EPI_ISL_3163550, EPI_ISL_3163551, EPI_ISL_3163552, EPI_ISL_3163553, EPI_ISL_3163554, EPI_ISL_3163555, EPI_ISL_3163556, EPI_ISL_3163557, EPI_ISL_3163558, EPI_ISL_3163559, EPI_ISL_3163560, EPI_ISL_3163561, EPI_ISL_3163562, EPI_ISL_3163563, EPI_ISL_3163564, EPI_ISL_3163565, EPI_ISL_3163566, EPI_ISL_3163567, EPI_ISL_3163568, EPI_ISL_3163569, EPI_ISL_3163570, EPI_ISL_3163571, EPI_ISL_3163572, EPI_ISL_3163573, EPI_ISL_3163574, EPI_ISL_3163575, EPI_ISL_3163576, EPI_ISL_3163577, EPI_ISL_3163578, EPI_ISL_3163579, EPI_ISL_3163580, EPI_ISL_3163581, EPI_ISL_3163582, EPI_ISL_3163583, EPI_ISL_3163584, EPI_ISL_3163585, EPI_ISL_3163586, EPI_ISL_3163587, EPI_ISL_3163588, EPI_ISL_3163589, EPI_ISL_3163590, EPI_ISL_3163591, EPI_ISL_3163592, EPI_ISL_3163593, EPI_ISL_3163594, EPI_ISL_3163595, EPI_ISL_3163596, EPI_ISL_3163597, EPI_ISL_3163598, EPI_ISL_3163599, EPI_ISL_3163600, EPI_ISL_3163601, EPI_ISL_3163602, EPI_ISL_3163603, EPI_ISL_3163604, EPI_ISL_3163605, EPI_ISL_3163606, EPI_ISL_3163607, EPI_ISL_3163608, EPI_ISL_3163609, EPI_ISL_3163611, EPI_ISL_3163612, EPI_ISL_3163613, EPI_ISL_3163614, EPI_ISL_3163615, EPI_ISL_3163616, EPI_ISL_3163617, EPI_ISL_3163618, EPI_ISL_3163619, EPI_ISL_3163620, EPI_ISL_3163621, EPI_ISL_3163622, EPI_ISL_3163623, EPI_ISL_3163624, EPI_ISL_3163625, EPI_ISL_3163626, EPI_ISL_3163627, EPI_ISL_3163628, EPI_ISL_3163629, EPI_ISL_3163630, EPI_ISL_3163631, EPI_ISL_3163632, EPI_ISL_3163633, EPI_ISL_3163634, EPI_ISL_3163635, EPI_ISL_3163636, EPI_ISL_3163637, EPI_ISL_3163638, EPI_ISL_3163639, EPI_ISL_3163640, EPI_ISL_3163641, EPI_ISL_3163642, EPI_ISL_3163643, EPI_ISL_3163644, EPI_ISL_3163645, EPI_ISL_3163646, EPI_ISL_3163647, EPI_ISL_3163648, EPI_ISL_3163649, EPI_ISL_3163650, EPI_ISL_3163651, EPI_ISL_3163652, EPI_ISL_3163653, EPI_ISL_3163654, EPI_ISL_3163655, EPI_ISL_3163656, EPI_ISL_3163657, EPI_ISL_3163658, EPI_ISL_3163659, EPI_ISL_3163660, EPI_ISL_3163661, EPI_ISL_3163662, EPI_ISL_3163663, EPI_ISL_3163664, EPI_ISL_3163665, EPI_ISL_3163666, EPI_ISL_3163667, EPI_ISL_3163668, EPI_ISL_3163669, EPI_ISL_3163670, EPI_ISL_3163671, EPI_ISL_3163672, EPI_ISL_3163673, EPI_ISL_3163674, EPI_ISL_3163675, EPI_ISL_3163676, EPI_ISL_3163677, EPI_ISL_3163678, EPI_ISL_3163679, EPI_ISL_3163680, EPI_ISL_3163681, EPI_ISL_3163682, EPI_ISL_3163683, EPI_ISL_3163684, EPI_ISL_3163685, EPI_ISL_3163686, EPI_ISL_3163687, EPI_ISL_3163688, EPI_ISL_3163689, EPI_ISL_3163690, EPI_ISL_3163691, EPI_ISL_3163692, EPI_ISL_3163693, EPI_ISL_3163694, EPI_ISL_3163695, EPI_ISL_3163696, EPI_ISL_3163697, EPI_ISL_3163698, EPI_ISL_3163699, EPI_ISL_3163700, EPI_ISL_3163701, EPI_ISL_3163702, EPI_ISL_3163703, EPI_ISL_3163704, EPI_ISL_3163705, EPI_ISL_3163706, EPI_ISL_3163707, EPI_ISL_3163708, EPI_ISL_3163709, EPI_ISL_3163710, EPI_ISL_3163711, EPI_ISL_3163712, EPI_ISL_3163713, EPI_ISL_3163714, EPI_ISL_3163715, EPI_ISL_3163716, EPI_ISL_3163717, EPI_ISL_3163718, EPI_ISL_3163719, EPI_ISL_3163720, EPI_ISL_3163721, EPI_ISL_3163722, EPI_ISL_3163723, EPI_ISL_3163724, EPI_ISL_3163725, EPI_ISL_3163726, EPI_ISL_3163727, EPI_ISL_3163728, EPI_ISL_3163729, EPI_ISL_3163730, EPI_ISL_3163731, EPI_ISL_3163732, EPI_ISL_3163733, EPI_ISL_3163734, EPI_ISL_3163735, EPI_ISL_3163736, EPI_ISL_3163737, EPI_ISL_3163738, EPI_ISL_3163739, EPI_ISL_3163740, EPI_ISL_3163741, EPI_ISL_3163742 | | | |
| see above | Department of Bacteria, Parasites and Fungi, Statens Serum Institut, Copenhagen, Denmark | Statens Serum Institut Bioinformatics and Microbial Genomics | Danish Covid-19 Genome Consortium |
| EPI_ISL_3163743 | Molekylær Medicinsk Afdeling, Aarhus University Hospital, Aarhus, Denmark | Statens Serum Institut Bioinformatics and Microbial Genomics | Danish Covid-19 Genome Consortium |
| EPI_ISL_3163744, EPI_ISL_3163745, EPI_ISL_3163746, EPI_ISL_3163747, EPI_ISL_3163748, EPI_ISL_3163749, EPI_ISL_3163750, EPI_ISL_3163751, EPI_ISL_3163752, EPI_ISL_3163753, EPI_ISL_3163754 | | | |
| see above | Department of Bacteria, Parasites and Fungi, Statens Serum Institut, Copenhagen, Denmark | Statens Serum Institut Bioinformatics and Microbial Genomics | Danish Covid-19 Genome Consortium |
| EPI_ISL_3163755 | Department of Clinical Microbiology and Center for Genomic Medicine, Rigshospitalet, Copenhagen, Denmark | Statens Serum Institut Bioinformatics and Microbial Genomics | Danish Covid-19 Genome Consortium |
| EPI_ISL_3163756, EPI_ISL_3163757, EPI_ISL_3163758, EPI_ISL_3163759, EPI_ISL_3163760, EPI_ISL_3163761, EPI_ISL_3163762, EPI_ISL_3163763, EPI_ISL_3163764, EPI_ISL_3163765, EPI_ISL_3163767, EPI_ISL_3163768, EPI_ISL_3163769, EPI_ISL_3163770, EPI_ISL_3163771, EPI_ISL_3163772, EPI_ISL_3163773, EPI_ISL_3163774, EPI_ISL_3163775, EPI_ISL_3163776, EPI_ISL_3163777, EPI_ISL_3163778, EPI_ISL_3163779, EPI_ISL_3163780, EPI_ISL_3163781, EPI_ISL_3163782, EPI_ISL_3163783, EPI_ISL_3163784, EPI_ISL_3163785, EPI_ISL_3163786, EPI_ISL_3163787, EPI_ISL_3163788, EPI_ISL_3163789, EPI_ISL_3163790, EPI_ISL_3163791, EPI_ISL_3163792, EPI_ISL_3163793 | | | |
| see above | Department of Bacteria, Parasites and Fungi, Statens Serum Institut, Copenhagen, Denmark | Statens Serum Institut Bioinformatics and Microbial Genomics | Danish Covid-19 Genome Consortium |
| EPI_ISL_3163794, EPI_ISL_3163796 | Molekylær Medicinsk Afdeling, Aarhus University Hospital, Aarhus, Denmark | Statens Serum Institut Bioinformatics and Microbial Genomics | Danish Covid-19 Genome Consortium |
| EPI_ISL_3163797, EPI_ISL_3163798, EPI_ISL_3163799, EPI_ISL_3163800, EPI_ISL_3163801, EPI_ISL_3163802, EPI_ISL_3163803, EPI_ISL_3163804, EPI_ISL_3163805, EPI_ISL_3163806, EPI_ISL_3163807, EPI_ISL_3163808, EPI_ISL_3163809, EPI_ISL_3163810, EPI_ISL_3163811, EPI_ISL_3163812, EPI_ISL_3163813, EPI_ISL_3163814, EPI_ISL_3163815, EPI_ISL_3163816, EPI_ISL_3163817, EPI_ISL_3163818, EPI_ISL_3163819, EPI_ISL_3163820, EPI_ISL_3163821, EPI_ISL_3163822, EPI_ISL_3163823, EPI_ISL_3163824, EPI_ISL_3163825, EPI_ISL_3163826, EPI_ISL_3163827, EPI_ISL_3163828, EPI_ISL_3163829, EPI_ISL_3163830, EPI_ISL_3163831, EPI_ISL_3163832, EPI_ISL_3163833, EPI_ISL_3163834, EPI_ISL_3163835, EPI_ISL_3163836, EPI_ISL_3163837, EPI_ISL_3163838, EPI_ISL_3163839, EPI_ISL_3163840, EPI_ISL_3163841, EPI_ISL_3163842, EPI_ISL_3163843, EPI_ISL_3163844, EPI_ISL_3163845, EPI_ISL_3163846, EPI_ISL_3163847, EPI_ISL_3163848, EPI_ISL_3163849, EPI_ISL_3163850, EPI_ISL_3163851, EPI_ISL_3163852, EPI_ISL_3163853, EPI_ISL_3163854, EPI_ISL_3163855, EPI_ISL_3163856, EPI_ISL_3163857, EPI_ISL_3163858, EPI_ISL_3163859, EPI_ISL_3163860 | | | |
| see above | Department of Bacteria, Parasites and Fungi, Statens Serum Institut, Copenhagen, Denmark | Statens Serum Institut Bioinformatics and Microbial Genomics | Danish Covid-19 Genome Consortium |
| EPI_ISL_3163861 | Department of Clinical Microbiology and Center for Genomic Medicine, Rigshospitalet, Copenhagen, Denmark | Statens Serum Institut Bioinformatics and Microbial Genomics | Danish Covid-19 Genome Consortium |
| EPI_ISL_3163862, EPI_ISL_3163863, EPI_ISL_3163864, EPI_ISL_3163865, EPI_ISL_3163866, EPI_ISL_3163867, EPI_ISL_3163868, EPI_ISL_3163869, EPI_ISL_3163870, EPI_ISL_3163871, EPI_ISL_3163872, EPI_ISL_3163873, EPI_ISL_3163874, EPI_ISL_3163875, EPI_ISL_3163876, EPI_ISL_3163877, EPI_ISL_3163878, EPI_ISL_3163879, EPI_ISL_3163880, EPI_ISL_3163881, EPI_ISL_3163882, EPI_ISL_3163883, EPI_ISL_3163884, EPI_ISL_3163885, EPI_ISL_3163886, EPI_ISL_3163887, EPI_ISL_3163888, EPI_ISL_3163889, EPI_ISL_3163890, EPI_ISL_3163891, EPI_ISL_3163892, EPI_ISL_3163893, EPI_ISL_3163894, EPI_ISL_3163895, EPI_ISL_3163896, EPI_ISL_3163897, EPI_ISL_3163898, EPI_ISL_3163899, EPI_ISL_3163900, EPI_ISL_3163901 | | | |
| see above | Department of Bacteria, Parasites and Fungi, Statens Serum Institut, Copenhagen, Denmark | Statens Serum Institut Bioinformatics and Microbial Genomics | Danish Covid-19 Genome Consortium |
| EPI_ISL_3163902 | Molekylær Medicinsk Afdeling, Aarhus University Hospital, Aarhus, Denmark | Statens Serum Institut Bioinformatics and Microbial Genomics | Danish Covid-19 Genome Consortium |
| EPI_ISL_3163903, EPI_ISL_3163904, EPI_ISL_3163905, EPI_ISL_3163906, EPI_ISL_3163907, EPI_ISL_3163908, EPI_ISL_3163910, EPI_ISL_3163911, EPI_ISL_3163912, EPI_ISL_3163913, EPI_ISL_3163914, EPI_ISL_3163915, EPI_ISL_3163916, EPI_ISL_3163917, EPI_ISL_3163918, EPI_ISL_3163919, EPI_ISL_3163920, EPI_ISL_3163921, EPI_ISL_3163922, EPI_ISL_3163923, EPI_ISL_3163924, EPI_ISL_3163925, EPI_ISL_3163926, EPI_ISL_3163927, EPI_ISL_3163928, EPI_ISL_3163929, EPI_ISL_3163930, EPI_ISL_3163931, EPI_ISL_3163932, EPI_ISL_3163933 | | | |
| see above | Department of Bacteria, Parasites and Fungi, Statens Serum Institut, Copenhagen, Denmark | Statens Serum Institut Bioinformatics and Microbial Genomics | Danish Covid-19 Genome Consortium |
| EPI_ISL_3163967 | Salud Digna | Instituto Nacional de Medicina Genomica | Cedro-Tanda A, Cisneros-Villanueva M, Escobar-Arrazola MA, Ramirez-Vega O, Abraham Campos-Romero, Moreno-Camacho José Luis, Rodriguez-Gallegos Jorge, Luna-Ruiz Marco, Rangel-DeLeon D, Munguia-Garza P, Hidalgo-Miranda A, Mendoza-Vargas A, Reyes-Grajeda JP, Herrera-Montalvo LA. |
| EPI_ISL_3163968, EPI_ISL_3163969, EPI_ISL_3163970, EPI_ISL_3163972, EPI_ISL_3163973, EPI_ISL_3163974, EPI_ISL_3163975, EPI_ISL_3163976, EPI_ISL_3163977, EPI_ISL_3163978, EPI_ISL_3163979, EPI_ISL_3163980, EPI_ISL_3163981, EPI_ISL_3163982 | | | |
| see above | Microbiology Department, Laboratori Clínic Metropolitana Nord. Hospital Universitari Germans Trias i Pujol | Can Ruti SARS-CoV-2 Sequencing Hub (HUGTiP/IrsiCaixa/IGTP) | Marc Noguera-Julian, Pilar Armengol, Ignacio Blanco, Antoni E Bordoy, Francesc Catala-Moll, Pere-Joan Cardona, Maria Casadellà, Cristina Casañ, Gemma Clara, Bonaventura Clotet, Montserrat Giménez, Mercedes Guerrero, Anna Not, David Panisello, Roger Paredes, Mariona Parera, Alexia París, Verónica Saludes, Alba Sánchez, Laia Soler, Lauro Sumoy, and Elisa Martró on behalf of the Can Ruti SARS-CoV-2 Sequencing Hub |
| EPI_ISL_3163983 | Microbiologia CATLAB | Can Ruti SARS-CoV-2 Sequencing Hub (HUGTiP/IrsiCaixa/IGTP) | Marc Noguera-Julian, Pilar Armengol, Ignacio Blanco, Antoni E Bordoy, Francesc Catala-Moll, Pere-Joan Cardona, Maria Casadellà, Cristina Casañ, Gemma Clara, Bonaventura Clotet, Montserrat Giménez, Mercedes Guerrero, Anna Not, David Panisello, Roger Paredes, Mariona Parera, Alexia París, Verónica Saludes, Alba Sánchez, Laia Soler, Lauro Sumoy, and Elisa Martró on behalf of the Can Ruti SARS-CoV-2 Sequencing Hub |
| EPI_ISL_3163984, EPI_ISL_3163985, EPI_ISL_3163986, EPI_ISL_3163987, EPI_ISL_3163988, EPI_ISL_3163989, EPI_ISL_3163991, EPI_ISL_3163992, EPI_ISL_3163993 | Microbiology Department, Laboratori Clínic Metropolitana Nord. Hospital Universitari Germans Trias i Pujol | Can Ruti SARS-CoV-2 Sequencing Hub (HUGTiP/IrsiCaixa/IGTP) | Marc Noguera-Julian, Pilar Armengol, Ignacio Blanco, Antoni E Bordoy, Francesc Catala-Moll, Pere-Joan Cardona, Maria Casadellà, Cristina Casañ, Gemma Clara, Bonaventura Clotet, Montserrat Giménez, Mercedes Guerrero, Anna Not, David Panisello, Roger Paredes, Mariona Parera, Alexia París, Verónica Saludes, Alba Sánchez, Laia Soler, Lauro Sumoy, and Elisa Martró on behalf of the Can Ruti SARS-CoV-2 Sequencing Hub |
| EPI_ISL_3163994, EPI_ISL_3163995 | Microbiologia CATLAB | Can Ruti SARS-CoV-2 Sequencing Hub (HUGTiP/IrsiCaixa/IGTP) | Marc Noguera-Julian, Pilar Armengol, Ignacio Blanco, Antoni E Bordoy, Francesc Catala-Moll, Pere-Joan Cardona, Maria Casadellà, Cristina Casañ, Gemma Clara, Bonaventura Clotet, Montserrat Giménez, Mercedes Guerrero, Anna Not, David Panisello, Roger Paredes, Mariona Parera, Alexia París, Verónica Saludes, Alba Sánchez, Laia Soler, Lauro Sumoy, and Elisa Martró on behalf of the Can Ruti SARS-CoV-2 Sequencing Hub |

| | | | |
|---|---|---|---|
| EPI_ISL_3163996, EPI_ISL_3163997, EPI_ISL_3163998, EPI_ISL_3163999, EPI_ISL_3164000, EPI_ISL_3164002, EPI_ISL_3164003, EPI_ISL_3164004, EPI_ISL_3164005 | Microbiology Department, Laboratori Clínic Metropolitana Nord. Hospital Universitari Germans Trias i Pujol | Can Ruti SARS-CoV-2 Sequencing Hub (HUGTiP/IrsiCaixa/IGTP) | Marc Noguera-Julian, Pilar Armengol, Ignacio Blanco, Antoni E Bordoy, Francesc Catala-Moll, Pere-Joan Cardona, Maria Casadellà, Cristina Casañ, Gemma Clara, Bonaventura Clotet, Montserrat Giménez, Mercedes Guerrero, Anna Not, David Panisello, Roger Paredes, Mariona Parera, Alexia París, Verónica Saludes, Alba Sánchez, Laia Soler, Lauro Sumoy, and Elisa Martró on behalf of the Can Ruti SARS-CoV-2 Sequencing Hub |
| EPI_ISL_3164006, EPI_ISL_3164007 | Microbiologia CATLAB | Can Ruti SARS-CoV-2 Sequencing Hub (HUGTiP/IrsiCaixa/IGTP) | Marc Noguera-Julian, Pilar Armengol, Ignacio Blanco, Antoni E Bordoy, Francesc Catala-Moll, Pere-Joan Cardona, Maria Casadellà, Cristina Casañ, Gemma Clara, Bonaventura Clotet, Montserrat Giménez, Mercedes Guerrero, Anna Not, David Panisello, Roger Paredes, Mariona Parera, Alexia París, Verónica Saludes, Alba Sánchez, Laia Soler, Lauro Sumoy, and Elisa Martró on behalf of the Can Ruti SARS-CoV-2 Sequencing Hub |
| EPI_ISL_3164008, EPI_ISL_3164009, EPI_ISL_3164010, EPI_ISL_3164011, EPI_ISL_3164012, EPI_ISL_3164013, EPI_ISL_3164014, EPI_ISL_3164015, EPI_ISL_3164016, EPI_ISL_3164017 | Microbiology Department, Laboratori Clínic Metropolitana Nord. Hospital Universitari Germans Trias i Pujol | Can Ruti SARS-CoV-2 Sequencing Hub (HUGTiP/IrsiCaixa/IGTP) | Marc Noguera-Julian, Pilar Armengol, Ignacio Blanco, Antoni E Bordoy, Francesc Catala-Moll, Pere-Joan Cardona, Maria Casadellà, Cristina Casañ, Gemma Clara, Bonaventura Clotet, Montserrat Giménez, Mercedes Guerrero, Anna Not, David Panisello, Roger Paredes, Mariona Parera, Alexia París, Verónica Saludes, Alba Sánchez, Laia Soler, Lauro Sumoy, and Elisa Martró on behalf of the Can Ruti SARS-CoV-2 Sequencing Hub |
| EPI_ISL_3164018 | Microbiologia CATLAB | Can Ruti SARS-CoV-2 Sequencing Hub (HUGTiP/IrsiCaixa/IGTP) | Marc Noguera-Julian, Pilar Armengol, Ignacio Blanco, Antoni E Bordoy, Francesc Catala-Moll, Pere-Joan Cardona, Maria Casadellà, Cristina Casañ, Gemma Clara, Bonaventura Clotet, Montserrat Giménez, Mercedes Guerrero, Anna Not, David Panisello, Roger Paredes, Mariona Parera, Alexia París, Verónica Saludes, Alba Sánchez, Laia Soler, Lauro Sumoy, and Elisa Martró on behalf of the Can Ruti SARS-CoV-2 Sequencing Hub |
| EPI_ISL_3164019, EPI_ISL_3164020, EPI_ISL_3164021, EPI_ISL_3164022, EPI_ISL_3164023, EPI_ISL_3164024, EPI_ISL_3164025, EPI_ISL_3164026, EPI_ISL_3164027, EPI_ISL_3164028, EPI_ISL_3164029 | | | |
| see above | Microbiology Department, Laboratori Clínic Metropolitana Nord. Hospital Universitari Germans Trias i Pujol | Can Ruti SARS-CoV-2 Sequencing Hub (HUGTiP/IrsiCaixa/IGTP) | Marc Noguera-Julian, Pilar Armengol, Ignacio Blanco, Antoni E Bordoy, Francesc Catala-Moll, Pere-Joan Cardona, Maria Casadellà, Cristina Casañ, Gemma Clara, Bonaventura Clotet, Montserrat Giménez, Mercedes Guerrero, Anna Not, David Panisello, Roger Paredes, Mariona Parera, Alexia París, Verónica Saludes, Alba Sánchez, Laia Soler, Lauro Sumoy, and Elisa Martró on behalf of the Can Ruti SARS-CoV-2 Sequencing Hub |
| EPI_ISL_3164030 | Microbiologia CATLAB | Can Ruti SARS-CoV-2 Sequencing Hub (HUGTiP/IrsiCaixa/IGTP) | Marc Noguera-Julian, Pilar Armengol, Ignacio Blanco, Antoni E Bordoy, Francesc Catala-Moll, Pere-Joan Cardona, Maria Casadellà, Cristina Casañ, Gemma Clara, Bonaventura Clotet, Montserrat Giménez, Mercedes Guerrero, Anna Not, David Panisello, Roger Paredes, Mariona Parera, Alexia París, Verónica Saludes, Alba Sánchez, Laia Soler, Lauro Sumoy, and Elisa Martró on behalf of the Can Ruti SARS-CoV-2 Sequencing Hub |
| EPI_ISL_3164031, EPI_ISL_3164032, EPI_ISL_3164033, EPI_ISL_3164034, EPI_ISL_3164035, EPI_ISL_3164036, EPI_ISL_3164037, EPI_ISL_3164038, EPI_ISL_3164039, EPI_ISL_3164040 | Microbiology Department, Laboratori Clínic Metropolitana Nord. Hospital Universitari Germans Trias i Pujol | Can Ruti SARS-CoV-2 Sequencing Hub (HUGTiP/IrsiCaixa/IGTP) | Marc Noguera-Julian, Pilar Armengol, Ignacio Blanco, Antoni E Bordoy, Francesc Catala-Moll, Pere-Joan Cardona, Maria Casadellà, Cristina Casañ, Gemma Clara, Bonaventura Clotet, Montserrat Giménez, Mercedes Guerrero, Anna Not, David Panisello, Roger Paredes, Mariona Parera, Alexia París, Verónica Saludes, Alba Sánchez, Laia Soler, Lauro Sumoy, and Elisa Martró on behalf of the Can Ruti SARS-CoV-2 Sequencing Hub |
| EPI_ISL_3164041 | Microbiologia CATLAB | Can Ruti SARS-CoV-2 Sequencing Hub (HUGTiP/IrsiCaixa/IGTP) | Marc Noguera-Julian, Pilar Armengol, Ignacio Blanco, Antoni E Bordoy, Francesc Catala-Moll, Pere-Joan Cardona, Maria Casadellà, Cristina Casañ, Gemma Clara, Bonaventura Clotet, Montserrat Giménez, Mercedes Guerrero, Anna Not, David Panisello, Roger Paredes, Mariona Parera, Alexia París, Verónica Saludes, Alba Sánchez, Laia Soler, Lauro Sumoy, and Elisa Martró on behalf of the Can Ruti SARS-CoV-2 Sequencing Hub |
| EPI_ISL_3164042, EPI_ISL_3164043, EPI_ISL_3164044, EPI_ISL_3164045, EPI_ISL_3164046, EPI_ISL_3164047, EPI_ISL_3164048, EPI_ISL_3164049, EPI_ISL_3164050, EPI_ISL_3164051 | Microbiology Department, Laboratori Clínic Metropolitana Nord. Hospital Universitari Germans Trias i Pujol | Can Ruti SARS-CoV-2 Sequencing Hub (HUGTiP/IrsiCaixa/IGTP) | Marc Noguera-Julian, Pilar Armengol, Ignacio Blanco, Antoni E Bordoy, Francesc Catala-Moll, Pere-Joan Cardona, Maria Casadellà, Cristina Casañ, Gemma Clara, Bonaventura Clotet, Montserrat Giménez, Mercedes Guerrero, Anna Not, David Panisello, Roger Paredes, Mariona Parera, Alexia París, Verónica Saludes, Alba Sánchez, Laia Soler, Lauro Sumoy, and Elisa Martró on behalf of the Can Ruti SARS-CoV-2 Sequencing Hub |
| EPI_ISL_3164052 | Microbiologia CATLAB | Can Ruti SARS-CoV-2 Sequencing Hub (HUGTiP/IrsiCaixa/IGTP) | Marc Noguera-Julian, Pilar Armengol, Ignacio Blanco, Antoni E Bordoy, Francesc Catala-Moll, Pere-Joan Cardona, Maria Casadellà, Cristina Casañ, Gemma Clara, Bonaventura Clotet, Montserrat Giménez, Mercedes Guerrero, Anna Not, David Panisello, Roger Paredes, Mariona Parera, Alexia París, Verónica Saludes, Alba Sánchez, Laia Soler, Lauro Sumoy, and Elisa Martró on behalf of the Can Ruti SARS-CoV-2 Sequencing Hub |
| EPI_ISL_3164053, EPI_ISL_3164054, EPI_ISL_3164055, EPI_ISL_3164056, EPI_ISL_3164057, EPI_ISL_3164058, EPI_ISL_3164060, EPI_ISL_3164061 | Microbiology Department, Laboratori Clínic Metropolitana Nord. Hospital Universitari Germans Trias i Pujol | Can Ruti SARS-CoV-2 Sequencing Hub (HUGTiP/IrsiCaixa/IGTP) | Marc Noguera-Julian, Pilar Armengol, Ignacio Blanco, Antoni E Bordoy, Francesc Catala-Moll, Pere-Joan Cardona, Maria Casadellà, Cristina Casañ, Gemma Clara, Bonaventura Clotet, Montserrat Giménez, Mercedes Guerrero, Anna Not, David Panisello, Roger Paredes, Mariona Parera, Alexia París, Verónica Saludes, Alba Sánchez, Laia Soler, Lauro Sumoy, and Elisa Martró on behalf of the Can Ruti SARS-CoV-2 Sequencing Hub |
| EPI_ISL_3164062, EPI_ISL_3164063, EPI_ISL_3164064, EPI_ISL_3164065, EPI_ISL_3164066, EPI_ISL_3164067, EPI_ISL_3164069, EPI_ISL_3164070, EPI_ISL_3164071, EPI_ISL_3164072, EPI_ISL_3164073, EPI_ISL_3164074 | | | |
| see above | Althaia. Xarxa Assistencial Universitària de Manresa | IrsiCaixa | Marc Noguera-Julian, Mariona Parera, Maria Casadellà, Pilar Armengol, Francesc Catala-Moll, Roger Paredes, Bonaventura Clotet Gloria Trujillo, Rafel Perez Vidal, Jaume Trapé, Carolina Gonzalez-Fernandez, Miquel Micó, Antonia Flor, Roger Paredes, Eulalia Grau, Bonaventura Clotet |
| EPI_ISL_3164076, EPI_ISL_3164077 | Middlemore Hospital | Institute of Environmental Science and Research (ESR) | Rachel Boyle, SallyAnn Harbison, Olivia Stroeven, Xiaoyun Ren, Matt Storey, Nikki Freed, Muhammad Faisal, Jing Wang, Hermes Perez, Anja Werno, Antje van der Linden, Arlo Upton, Chris Mansell, David Hammer, Dragana Drinkovic, Gary McAuliffe, Hana Sofia Andersson, James Ussher, Jill Sherwood, Josh Freeman, Julia Howard, Juliet Elvy, Mary DeAlmeida, Matt Blakiston, Matthew Rogers, Max Bloomfield, Michael Addidle, Michelle Balm, Sally Roberts, Sarah Jefferies, Sharmini Muttaiyah, Susan Morpeth, Susan Taylor, Timothy Blackmore, Vani Sathyendran, Veronica Playle, Virginia Hope, Erasmus Smit, Lauren Jelly, Olin Silander, Joep de Ligt |
| EPI_ISL_3164078 | LabPLUS | Institute of Environmental Science and Research (ESR) | Rachel Boyle, SallyAnn Harbison, Olivia Stroeven, Xiaoyun Ren, Matt Storey, Nikki Freed, Muhammad Faisal, Jing Wang, Hermes Perez, Anja Werno, Antje van der Linden, Arlo Upton, Chris Mansell, David Hammer, Dragana Drinkovic, Gary McAuliffe, Hana Sofia Andersson, James Ussher, Jill Sherwood, Josh Freeman, Julia Howard, Juliet Elvy, Mary DeAlmeida, Matt Blakiston, Matthew Rogers, Max Bloomfield, Michael Addidle, Michelle Balm, Sally Roberts, Sarah Jefferies, Sharmini Muttaiyah, Susan Morpeth, Susan Taylor, Timothy Blackmore, Vani Sathyendran, Veronica Playle, Virginia Hope, Erasmus Smit, Lauren Jelly, Olin Silander, Joep de Ligt |
| EPI_ISL_3164079 | Middlemore Hospital | Institute of Environmental Science and Research (ESR) | Rachel Boyle, SallyAnn Harbison, Olivia Stroeven, Xiaoyun Ren, Matt Storey, Nikki Freed, Muhammad Faisal, Jing Wang, Hermes Perez, Anja Werno, Antje van der Linden, Arlo Upton, Chris Mansell, David Hammer, Dragana Drinkovic, Gary McAuliffe, Hana Sofia Andersson, James Ussher, Jill Sherwood, Josh Freeman, Julia Howard, Juliet Elvy, Mary DeAlmeida, Matt Blakiston, Matthew Rogers, Max Bloomfield, Michael Addidle, Michelle Balm, Sally Roberts, Sarah Jefferies, Sharmini Muttaiyah, Susan Morpeth, Susan Taylor, Timothy Blackmore, Vani Sathyendran, Veronica Playle, Virginia Hope, Erasmus Smit, Lauren Jelly, Olin Silander, Joep de Ligt |
| EPI_ISL_3164080, EPI_ISL_3164081 | LabPLUS | Institute of Environmental Science and Research (ESR) | Rachel Boyle, SallyAnn Harbison, Olivia Stroeven, Xiaoyun Ren, Matt Storey, Nikki Freed, Muhammad Faisal, Jing Wang, Hermes Perez, Anja Werno, Antje van der Linden, Arlo Upton, Chris Mansell, David Hammer, Dragana Drinkovic, Gary McAuliffe, Hana Sofia Andersson, James Ussher, Jill Sherwood, Josh Freeman, Julia Howard, Juliet Elvy, Mary DeAlmeida, Matt Blakiston, Matthew Rogers, Max Bloomfield, Michael Addidle, Michelle Balm, Sally Roberts, Sarah Jefferies, Sharmini Muttaiyah, Susan Morpeth, Susan Taylor, Timothy Blackmore, Vani Sathyendran, Veronica Playle, Virginia Hope, Erasmus Smit, Lauren Jelly, Olin Silander, Joep de Ligt |
| EPI_ISL_3164082, EPI_ISL_3164083, EPI_ISL_3164084, EPI_ISL_3164085 | Middlemore Hospital | Institute of Environmental Science and Research (ESR) | Rachel Boyle, SallyAnn Harbison, Olivia Stroeven, Xiaoyun Ren, Matt Storey, Nikki Freed, Muhammad Faisal, Jing Wang, Hermes Perez, Anja Werno, Antje van der Linden, Arlo Upton, Chris Mansell, David Hammer, Dragana Drinkovic, Gary McAuliffe, Hana Sofia Andersson, James Ussher, Jill Sherwood, Josh Freeman, Julia Howard, Juliet Elvy, Mary DeAlmeida, Matt Blakiston, Matthew Rogers, Max Bloomfield, Michael Addidle, Michelle Balm, Sally Roberts, Sarah Jefferies, Sharmini Muttaiyah, Susan Morpeth, Susan Taylor, Timothy Blackmore, Vani Sathyendran, Veronica Playle, Virginia Hope, Erasmus Smit, Lauren Jelly, Olin Silander, Joep de Ligt |
| EPI_ISL_3164086, EPI_ISL_3164087, EPI_ISL_3164088 | LabPLUS | Institute of Environmental Science and Research (ESR) | Rachel Boyle, SallyAnn Harbison, Olivia Stroeven, Xiaoyun Ren, Matt Storey, Nikki Freed, Muhammad Faisal, Jing Wang, Hermes Perez, Anja Werno, Antje van der Linden, Arlo Upton, Chris Mansell, David Hammer, Dragana Drinkovic, Gary McAuliffe, Hana Sofia Andersson, James Ussher, Jill Sherwood, Josh Freeman, Julia Howard, Juliet Elvy, Mary DeAlmeida, Matt Blakiston, Matthew Rogers, Max Bloomfield, Michael Addidle, Michelle Balm, Sally Roberts, Sarah Jefferies, Sharmini Muttaiyah, Susan Morpeth, Susan Taylor, Timothy Blackmore, Vani Sathyendran, Veronica Playle, Virginia Hope, Erasmus Smit, Lauren Jelly, Olin Silander, Joep de Ligt |
| EPI_ISL_3164089 | Middlemore Hospital | Institute of Environmental Science and Research (ESR) | Rachel Boyle, SallyAnn Harbison, Olivia Stroeven, Xiaoyun Ren, Matt Storey, Nikki Freed, Muhammad Faisal, Jing Wang, Hermes Perez, Anja Werno, Antje van der Linden, Arlo Upton, Chris Mansell, David Hammer, Dragana Drinkovic, Gary McAuliffe, Hana Sofia Andersson, James Ussher, Jill Sherwood, Josh Freeman, Julia Howard, Juliet Elvy, Mary DeAlmeida, Matt Blakiston, Matthew Rogers, Max Bloomfield, Michael Addidle, Michelle Balm, Sally Roberts, Sarah Jefferies, Sharmini Muttaiyah, Susan Morpeth, Susan Taylor, Timothy Blackmore, Vani Sathyendran, Veronica Playle, Virginia Hope, Erasmus Smit, Lauren Jelly, Olin Silander, Joep de Ligt |
| EPI_ISL_3164090, EPI_ISL_3164091 | LabPLUS | Institute of Environmental Science and Research (ESR) | Rachel Boyle, SallyAnn Harbison, Olivia Stroeven, Xiaoyun Ren, Matt Storey, Nikki Freed, Muhammad Faisal, Jing Wang, Hermes Perez, Anja Werno, Antje van der Linden, Arlo Upton, Chris Mansell, David Hammer, Dragana Drinkovic, Gary McAuliffe, Hana Sofia Andersson, James Ussher, Jill Sherwood, Josh Freeman, Julia Howard, Juliet Elvy, Mary DeAlmeida, Matt Blakiston, Matthew Rogers, Max Bloomfield, Michael Addidle, Michelle Balm, Sally Roberts, Sarah Jefferies, Sharmini Muttaiyah, Susan Morpeth, Susan Taylor, Timothy Blackmore, Vani Sathyendran, Veronica Playle, Virginia Hope, Erasmus Smit, Lauren Jelly, Olin Silander, Joep de Ligt |
| EPI_ISL_3164092 | Middlemore Hospital | Institute of Environmental Science and Research (ESR) | Rachel Boyle, SallyAnn Harbison, Olivia Stroeven, Xiaoyun Ren, Matt Storey, Nikki Freed, Muhammad Faisal, Jing Wang, Hermes Perez, Anja Werno, Antje van der Linden, Arlo Upton, Chris Mansell, David Hammer, Dragana Drinkovic, Gary McAuliffe, Hana Sofia Andersson, James Ussher, Jill Sherwood, Josh Freeman, Julia Howard, Juliet Elvy, Mary DeAlmeida, Matt Blakiston, Matthew Rogers, Max Bloomfield, Michael Addidle, Michelle Balm, Sally Roberts, Sarah Jefferies, Sharmini Muttaiyah, Susan Morpeth, Susan Taylor, Timothy Blackmore, Vani Sathyendran, Veronica Playle, Virginia Hope, Erasmus Smit, Lauren Jelly, Olin Silander, Joep de Ligt |
| EPI_ISL_3164093 | Wellington SCL (WN) | Institute of Environmental Science and Research (ESR) | Xiaoyun Ren, Matt Storey, Nikki Freed, Muhammad Faisal, Jing Wang, Hermes Perez, Anja Werno, Antje van der Linden, Arlo Upton, Chris Mansell, David Hammer, Dragana Drinkovic, Gary McAuliffe, Hana Sofia Andersson, James Ussher, Jill Sherwood, Josh Freeman, Julia Howard, Juliet Elvy, Mary DeAlmeida, Matt Blakiston, Matthew Rogers, Max Bloomfield, Michael Addidle, Michelle Balm, Sally Roberts, Sarah Jefferies, Sharmini Muttaiyah, Susan Morpeth, Susan Taylor, Timothy Blackmore, Vani Sathyendran, Veronica Playle, Virginia Hope, Erasmus Smit, Lauren Jelly, Olin Silander, Joep de Ligt |
| EPI_ISL_3164094, EPI_ISL_3164095 | Southern Community Labs Dunedin | Institute of Environmental Science and Research (ESR) | Rachel Boyle, SallyAnn Harbison, Olivia Stroeven, Xiaoyun Ren, Matt Storey, Nikki Freed, Muhammad Faisal, Jing Wang, Hermes Perez, Anja Werno, Antje van der Linden, Arlo Upton, Chris Mansell, David Hammer, Dragana Drinkovic, Gary McAuliffe, Hana Sofia Andersson, James Ussher, Jill Sherwood, Josh Freeman, Julia Howard, Juliet Elvy, Mary DeAlmeida, Matt Blakiston, Matthew Rogers, Max Bloomfield, Michael Addidle, Michelle Balm, Sally Roberts, Sarah Jefferies, Sharmini Muttaiyah, Susan Morpeth, Susan Taylor, Timothy Blackmore, Vani Sathyendran, Veronica Playle, Virginia Hope, Erasmus Smit, Lauren Jelly, Olin Silander, Joep de Ligt |
| EPI_ISL_3164096, EPI_ISL_3164097, | Wellington SCL (WN) | Institute of Environmental Science and Research (ESR) | Xiaoyun Ren, Matt Storey, Nikki Freed, Muhammad Faisal, Jing Wang, Hermes Perez, Anja Werno, Antje van der Linden, Arlo Upton, Chris Mansell, David Hammer, Dragana Drinkovic, Gary McAuliffe, Hana Sofia Andersson, James Ussher, Jill Sherwood, Josh Freeman, Julia Howard, Juliet Elvy, Mary DeAlmeida, Matt Blakiston, Matthew Rogers, Max Bloomfield, Michael Addidle, Michelle Balm, Sally Roberts, Sarah Jefferies, Sharmini Muttaiyah, Susan Morpeth, Susan Taylor, Timothy |

| | | | |
|---|---|---|---|
| EPI_ISL_3164098, EPI_ISL_3164099 | | | Blackmore, Vani Sathyendran, Veronica Playle, Virginia Hope, Erasmus Smit, Lauren Jelly, Olin Silander, Joep de Ligt |
| EPI_ISL_3164101 | LabTests | Institute of Environmental Science and Research (ESR) | Rachel Boyle, SallyAnn Harbison, Olivia Stroeven, Xiaoyun Ren, Matt Storey, Nikki Freed, Muhammad Faisal, Jing Wang, Hermes Perez, Anja Werno, Antje van der Linden, Arlo Upton, Chris Mansell, David Hammer, Dragana Drinkovic, Gary McAuliffe, Hana Sofia Andersson, James Ussher, Jill Sherwood, Josh Freeman, Julia Howard, Juliet Elvy, Mary DeAlmeida, Matt Blakiston, Matthew Rogers, Max Bloomfield, Michael Addidle, Michelle Balm, Sally Roberts, Sarah Jefferies, Sharmini Muttaiyah, Susan Morpeth, Susan Taylor, Timothy Blackmore, Vani Sathyendran, Veronica Playle, Virginia Hope, Erasmus Smit, Lauren Jelly, Olin Silander, Joep de Ligt |
| EPI_ISL_3164102, EPI_ISL_3164103, EPI_ISL_3164104, EPI_ISL_3164105, EPI_ISL_3164106, EPI_ISL_3164107, EPI_ISL_3164108, EPI_ISL_3164109, EPI_ISL_3164110 | Wellington SCL (WN) | Institute of Environmental Science and Research (ESR) | Xiaoyun Ren, Matt Storey, Nikki Freed, Muhammad Faisal, Jing Wang, Hermes Perez, Anja Werno, Antje van der Linden, Arlo Upton, Chris Mansell, David Hammer, Dragana Drinkovic, Gary McAuliffe, Hana Sofia Andersson, James Ussher, Jill Sherwood, Josh Freeman, Julia Howard, Juliet Elvy, Mary DeAlmeida, Matt Blakiston, Matthew Rogers, Max Bloomfield, Michael Addidle, Michelle Balm, Sally Roberts, Sarah Jefferies, Sharmini Muttaiyah, Susan Morpeth, Susan Taylor, Timothy Blackmore, Vani Sathyendran, Veronica Playle, Virginia Hope, Erasmus Smit, Lauren Jelly, Olin Silander, Joep de Ligt |
| EPI_ISL_3164111 | Middlemore Hospital | Institute of Environmental Science and Research (ESR) | Rachel Boyle, SallyAnn Harbison, Olivia Stroeven, Xiaoyun Ren, Matt Storey, Nikki Freed, Muhammad Faisal, Jing Wang, Hermes Perez, Anja Werno, Antje van der Linden, Arlo Upton, Chris Mansell, David Hammer, Dragana Drinkovic, Gary McAuliffe, Hana Sofia Andersson, James Ussher, Jill Sherwood, Josh Freeman, Julia Howard, Juliet Elvy, Mary DeAlmeida, Matt Blakiston, Matthew Rogers, Max Bloomfield, Michael Addidle, Michelle Balm, Sally Roberts, Sarah Jefferies, Sharmini Muttaiyah, Susan Morpeth, Susan Taylor, Timothy Blackmore, Vani Sathyendran, Veronica Playle, Virginia Hope, Erasmus Smit, Lauren Jelly, Olin Silander, Joep de Ligt |
| EPI_ISL_3164112 | Southern Community Labs Dunedin | Institute of Environmental Science and Research (ESR) | Paula scholes, Susan Lin, Xiaoyun Ren, Matt Storey, Nikki Freed, Muhammad Faisal, Jing Wang, Hermes Perez, Anja Werno, Antje van der Linden, Arlo Upton, Chris Mansell, David Hammer, Dragana Drinkovic, Gary McAuliffe, Hana Sofia Andersson, James Ussher, Jill Sherwood, Josh Freeman, Julia Howard, Juliet Elvy, Mary DeAlmeida, Matt Blakiston, Matthew Rogers, Max Bloomfield, Michael Addidle, Michelle Balm, Sally Roberts, Sarah Jefferies, Sharmini Muttaiyah, Susan Morpeth, Susan Taylor, Timothy Blackmore, Vani Sathyendran, Veronica Playle, Virginia Hope, Erasmus Smit, Lauren Jelly, Olin Silander, Joep de Ligt |
| EPI_ISL_3164113 | Southern Community Labs Dunedin | Institute of Environmental Science and Research (ESR) | Rachel Boyle, SallyAnn Harbison, Olivia Stroeven, Xiaoyun Ren, Matt Storey, Nikki Freed, Muhammad Faisal, Jing Wang, Hermes Perez, Anja Werno, Antje van der Linden, Arlo Upton, Chris Mansell, David Hammer, Dragana Drinkovic, Gary McAuliffe, Hana Sofia Andersson, James Ussher, Jill Sherwood, Josh Freeman, Julia Howard, Juliet Elvy, Mary DeAlmeida, Matt Blakiston, Matthew Rogers, Max Bloomfield, Michael Addidle, Michelle Balm, Sally Roberts, Sarah Jefferies, Sharmini Muttaiyah, Susan Morpeth, Susan Taylor, Timothy Blackmore, Vani Sathyendran, Veronica Playle, Virginia Hope, Erasmus Smit, Lauren Jelly, Olin Silander, Joep de Ligt |
| EPI_ISL_3164114, EPI_ISL_3164115, EPI_ISL_3164116, EPI_ISL_3164117, EPI_ISL_3164118, EPI_ISL_3164119 | Southern Community Labs Dunedin | Institute of Environmental Science and Research (ESR) | Paula scholes, Susan Lin, Xiaoyun Ren, Matt Storey, Nikki Freed, Muhammad Faisal, Jing Wang, Hermes Perez, Anja Werno, Antje van der Linden, Arlo Upton, Chris Mansell, David Hammer, Dragana Drinkovic, Gary McAuliffe, Hana Sofia Andersson, James Ussher, Jill Sherwood, Josh Freeman, Julia Howard, Juliet Elvy, Mary DeAlmeida, Matt Blakiston, Matthew Rogers, Max Bloomfield, Michael Addidle, Michelle Balm, Sally Roberts, Sarah Jefferies, Sharmini Muttaiyah, Susan Morpeth, Susan Taylor, Timothy Blackmore, Vani Sathyendran, Veronica Playle, Virginia Hope, Erasmus Smit, Lauren Jelly, Olin Silander, Joep de Ligt |
| EPI_ISL_3164120 | Southern Community Labs Dunedin | Institute of Environmental Science and Research (ESR) | Rachel Boyle, SallyAnn Harbison, Olivia Stroeven, Xiaoyun Ren, Matt Storey, Nikki Freed, Muhammad Faisal, Jing Wang, Hermes Perez, Anja Werno, Antje van der Linden, Arlo Upton, Chris Mansell, David Hammer, Dragana Drinkovic, Gary McAuliffe, Hana Sofia Andersson, James Ussher, Jill Sherwood, Josh Freeman, Julia Howard, Juliet Elvy, Mary DeAlmeida, Matt Blakiston, Matthew Rogers, Max Bloomfield, Michael Addidle, Michelle Balm, Sally Roberts, Sarah Jefferies, Sharmini Muttaiyah, Susan Morpeth, Susan Taylor, Timothy Blackmore, Vani Sathyendran, Veronica Playle, Virginia Hope, Erasmus Smit, Lauren Jelly, Olin Silander, Joep de Ligt |
| EPI_ISL_3164121, EPI_ISL_3164122 | Southern Community Labs Dunedin | Institute of Environmental Science and Research (ESR) | Paula scholes, Susan Lin, Xiaoyun Ren, Matt Storey, Nikki Freed, Muhammad Faisal, Jing Wang, Hermes Perez, Anja Werno, Antje van der Linden, Arlo Upton, Chris Mansell, David Hammer, Dragana Drinkovic, Gary McAuliffe, Hana Sofia Andersson, James Ussher, Jill Sherwood, Josh Freeman, Julia Howard, Juliet Elvy, Mary DeAlmeida, Matt Blakiston, Matthew Rogers, Max Bloomfield, Michael Addidle, Michelle Balm, Sally Roberts, Sarah Jefferies, Sharmini Muttaiyah, Susan Morpeth, Susan Taylor, Timothy Blackmore, Vani Sathyendran, Veronica Playle, Virginia Hope, Erasmus Smit, Lauren Jelly, Olin Silander, Joep de Ligt |
| EPI_ISL_3164123, EPI_ISL_3164124 | LabPLUS | Institute of Environmental Science and Research (ESR) | Rachel Boyle, SallyAnn Harbison, Olivia Stroeven, Xiaoyun Ren, Matt Storey, Nikki Freed, Muhammad Faisal, Jing Wang, Hermes Perez, Anja Werno, Antje van der Linden, Arlo Upton, Chris Mansell, David Hammer, Dragana Drinkovic, Gary McAuliffe, Hana Sofia Andersson, James Ussher, Jill Sherwood, Josh Freeman, Julia Howard, Juliet Elvy, Mary DeAlmeida, Matt Blakiston, Matthew Rogers, Max Bloomfield, Michael Addidle, Michelle Balm, Sally Roberts, Sarah Jefferies, Sharmini Muttaiyah, Susan Morpeth, Susan Taylor, Timothy Blackmore, Vani Sathyendran, Veronica Playle, Virginia Hope, Erasmus Smit, Lauren Jelly, Olin Silander, Joep de Ligt |
| EPI_ISL_3164136 | Osaka General Medical Center | Department of Developmental Medicine, Research Institute, Osaka Women's and Children's Hospital | Itaru Yanagihara, Yukiko Nakura |
| EPI_ISL_3164138 | National Institute of Laboratory Medicine and Referral Centre (NILMRC) | Genomic Research Laboratory, Bangabandhu Sheikh Mujib Medical University | Laila Anjuman Banu, Md.Sayedur Rahman, Sharadindu Kanti Sinha, Arifa Akram, Md Ehsanul Hoque, Md.Sharfuddin Ahmed |
| EPI_ISL_3164186, EPI_ISL_3164188, EPI_ISL_3164197, EPI_ISL_3164200, EPI_ISL_3164208, EPI_ISL_3164211, EPI_ISL_3164214 | Karolinska University Hospital Huddinge | Karolinska University Hospital | Jan Albert ,Tobias Allander ,Annelie Bjerkner ,Sandra Broddesson ,Robert Dyrdak ,Martin Ekman ,Lynda Eneh ,Lina Guerra Blomqvist ,Karolina Ininbergs ,Tanja Normark ,Isak Sylvin ,Zhibing Yun ,Martina Wahlund ,Valtteri Wirta |
| EPI_ISL_3165770 | National Institute of Laboratory Medicine and Referral Centre (NILMRC) | Genomic Research Laboratory, Bangabandhu Sheikh Mujib Medical University Shahbag, Dhaka_1000 | Laila Anjuman Banu, Md.Sayedur Rahman, Sharadindu Kanti Sinha, Arifa Akram, Md Ehsanul Hoque, Md.Sharfuddin Ahmed |
| EPI_ISL_3166181 | Osaka General Medical Center | Department of Developmental Medicine, Research Institute, Osaka Women's and Children's Hospital | Itaru Yanagihara, Yukiko Nakura |
| EPI_ISL_3166765 | Department of clinical lab., Osaka Women's and Children's Hospital | Department of Developmental Medicine, Research Institute, Osaka Women's and Children's Hospital | Itaru Yanagihara, Yukiko Nakura |
| EPI_ISL_3167131 | National Institute of Laboratory Medicine and Referral Centre (NILMRC) | Genomic Research Laboratory, Bangabandhu Sheikh Mujib Medical University Shahbag, Dhaka_1000 | Laila Anjuman Banu, Md.Sayedur Rahman, Sharadindu Kanti Sinha, Arifa Akram, Md Ehsanul Hoque, Md.Sharfuddin Ahmed |
| EPI_ISL_3167147, EPI_ISL_3167152, EPI_ISL_3167154 | NILMRC | Genomic Research Laboratory, Bangabandhu Sheikh Mujib Medical University Shahbag, Dhaka_1000 | Laila Anjuman Banu, Md.Sayedur Rahman, Sharadindu Kanti Sinha, Arifa Akram, Md Ehsanul Hoque, Md.Sharfuddin Ahmed |
| EPI_ISL_3167158, EPI_ISL_3167159, EPI_ISL_3167160, EPI_ISL_3167161, EPI_ISL_3167162, EPI_ISL_3167163, EPI_ISL_3167164, EPI_ISL_3167165 | Karolinska University Hospital Huddinge | Karolinska University Hospital | Jan Albert ,Tobias Allander ,Annelie Bjerkner ,Sandra Broddesson ,Robert Dyrdak ,Martin Ekman ,Lynda Eneh ,Lina Guerra Blomqvist ,Karolina Ininbergs ,Tanja Normark ,Isak Sylvin ,Zhibing Yun ,Martina Wahlund ,Valtteri Wirta |
| EPI_ISL_3167236, EPI_ISL_3167237, EPI_ISL_3167238 | Presidio Ospedaliero "Madonna delle Grazie" di Matera | ASM - P.O. Madonna delle Grazie - Matera | Massimo Dell'Edera |
| EPI_ISL_3167239 | Lighthouse Laboratory Plymouth | Wellcome Sanger Institute for the COVID-19 Genomics UK (COG-UK) Consortium | Lighthouse Laboratory Plymouth and Alex Alderton, Roberto Amato, Jeffrey Barrett, Sonia Goncalves, Ewan Harrison, David K. Jackson, Ian Johnston, Dominic Kwiatkowski, Cordelia Langford, John Sillitoe on behalf of the Wellcome Sanger Institute COVID-19 Surveillance Team |
| EPI_ISL_3167240, EPI_ISL_3167241 | Lighthouse Lab in Milton Keynes | Wellcome Sanger Institute for the COVID-19 Genomics UK (COG-UK) Consortium | The Lighthouse Lab in Milton Keynes and Alex Alderton, Roberto Amato, Jeffrey Barrett, Sonia Goncalves, Ewan Harrison, David K. Jackson, Ian Johnston, Dominic Kwiatkowski, Cordelia Langford, John Sillitoe on behalf of the Wellcome Sanger Institute COVID-19 Surveillance Team |
| EPI_ISL_3167242 | Lighthouse Laboratory Plymouth | Wellcome Sanger Institute for the COVID-19 Genomics UK (COG-UK) Consortium | Lighthouse Laboratory Plymouth and Alex Alderton, Roberto Amato, Jeffrey Barrett, Sonia Goncalves, Ewan Harrison, David K. Jackson, Ian Johnston, Dominic Kwiatkowski, Cordelia Langford, John Sillitoe on behalf of the Wellcome Sanger Institute COVID-19 Surveillance Team |
| EPI_ISL_3167243, EPI_ISL_3167244, EPI_ISL_3167245 | Lighthouse Lab in Glasgow | Wellcome Sanger Institute for the COVID-19 Genomics UK (COG-UK) Consortium | Harper VanSteenhouse, Yumi Kasai, David Gray, Carol Clugston, Anna Dominiczak and Alex Alderton, Roberto Amato, Jeffrey Barrett, Sonia Goncalves, Ewan Harrison, David K. Jackson, Ian Johnston, Dominic Kwiatkowski, Cordelia Langford, John Sillitoe on behalf of the Wellcome Sanger Institute COVID-19 Surveillance Team |
| EPI_ISL_3167246, EPI_ISL_3167247 | Lighthouse Laboratory Plymouth | Wellcome Sanger Institute for the COVID-19 Genomics UK (COG-UK) Consortium | Lighthouse Laboratory Plymouth and Alex Alderton, Roberto Amato, Jeffrey Barrett, Sonia Goncalves, Ewan Harrison, David K. Jackson, Ian Johnston, Dominic Kwiatkowski, Cordelia Langford, John Sillitoe on behalf of the Wellcome Sanger Institute COVID-19 Surveillance Team |
| EPI_ISL_3167248 | Lighthouse Lab in Glasgow | Wellcome Sanger Institute for the COVID-19 Genomics UK (COG-UK) Consortium | Harper VanSteenhouse, Yumi Kasai, David Gray, Carol Clugston, Anna Dominiczak and Alex Alderton, Roberto Amato, Jeffrey Barrett, Sonia Goncalves, Ewan Harrison, David K. Jackson, Ian Johnston, Dominic Kwiatkowski, Cordelia Langford, John Sillitoe on behalf of the Wellcome Sanger Institute COVID-19 Surveillance Team |
| EPI_ISL_3167249, EPI_ISL_3167250 | Lighthouse Lab in Milton Keynes | Wellcome Sanger Institute for the COVID-19 Genomics UK (COG-UK) Consortium | The Lighthouse Lab in Milton Keynes and Alex Alderton, Roberto Amato, Jeffrey Barrett, Sonia Goncalves, Ewan Harrison, David K. Jackson, Ian Johnston, Dominic Kwiatkowski, Cordelia Langford, John Sillitoe on behalf of the Wellcome Sanger Institute COVID-19 Surveillance Team |
| EPI_ISL_3167251, EPI_ISL_3167252 | Lighthouse Laboratory Plymouth | Wellcome Sanger Institute for the COVID-19 Genomics UK (COG-UK) Consortium | Lighthouse Laboratory Plymouth and Alex Alderton, Roberto Amato, Jeffrey Barrett, Sonia Goncalves, Ewan Harrison, David K. Jackson, Ian Johnston, Dominic Kwiatkowski, Cordelia Langford, John Sillitoe on behalf of the Wellcome Sanger Institute COVID-19 Surveillance Team |
| EPI_ISL_3167253 | Lighthouse Lab in Glasgow | Wellcome Sanger Institute for the COVID-19 Genomics UK (COG-UK) Consortium | Harper VanSteenhouse, Yumi Kasai, David Gray, Carol Clugston, Anna Dominiczak and Alex Alderton, Roberto Amato, Jeffrey Barrett, Sonia Goncalves, Ewan Harrison, David K. Jackson, Ian Johnston, Dominic Kwiatkowski, Cordelia Langford, John Sillitoe on behalf of the Wellcome Sanger Institute COVID-19 Surveillance Team |
| EPI_ISL_3167254 | Lighthouse Laboratory Plymouth | Wellcome Sanger Institute for the COVID-19 Genomics UK (COG-UK) Consortium | Lighthouse Laboratory Plymouth and Alex Alderton, Roberto Amato, Jeffrey Barrett, Sonia Goncalves, Ewan Harrison, David K. Jackson, Ian Johnston, Dominic Kwiatkowski, Cordelia Langford, John Sillitoe on behalf of the Wellcome Sanger Institute COVID-19 Surveillance Team |
| EPI_ISL_3167255, EPI_ISL_3167256 | Lighthouse Lab in Milton Keynes | Wellcome Sanger Institute for the COVID-19 Genomics UK (COG-UK) Consortium | The Lighthouse Lab in Milton Keynes and Alex Alderton, Roberto Amato, Jeffrey Barrett, Sonia Goncalves, Ewan Harrison, David K. Jackson, Ian Johnston, Dominic Kwiatkowski, Cordelia Langford, John Sillitoe on behalf of the Wellcome Sanger Institute COVID-19 Surveillance Team |
| EPI_ISL_3167257 | Lighthouse Lab in Glasgow | Wellcome Sanger Institute for the COVID-19 Genomics UK (COG-UK) Consortium | Harper VanSteenhouse, Yumi Kasai, David Gray, Carol Clugston, Anna Dominiczak and Alex Alderton, Roberto Amato, Jeffrey Barrett, Sonia Goncalves, Ewan Harrison, David K. Jackson, Ian Johnston, Dominic Kwiatkowski, Cordelia Langford, John Sillitoe on behalf of the Wellcome Sanger Institute COVID-19 Surveillance Team |

| | | | |
|---|---|---|---|
| EPI_ISL_3167258 | Lighthouse Laboratory Plymouth | Wellcome Sanger Institute for the COVID-19 Genomics UK (COG-UK) Consortium | Lighthouse Laboratory Plymouth and Alex Alderton, Roberto Amato, Jeffrey Barrett, Sonia Goncalves, Ewan Harrison, David K. Jackson, Ian Johnston, Dominic Kwiatkowski, Cordelia Langford, John Sillitoe on behalf of the Wellcome Sanger Institute COVID-19 Surveillance Team |
| EPI_ISL_3167259, EPI_ISL_3167260 | Lighthouse Lab in Milton Keynes | Wellcome Sanger Institute for the COVID-19 Genomics UK (COG-UK) Consortium | The Lighthouse Lab in Milton Keynes and Alex Alderton, Roberto Amato, Jeffrey Barrett, Sonia Goncalves, Ewan Harrison, David K. Jackson, Ian Johnston, Dominic Kwiatkowski, Cordelia Langford, John Sillitoe on behalf of the Wellcome Sanger Institute COVID-19 Surveillance Team |
| EPI_ISL_3167261, EPI_ISL_3167262 | Lighthouse Laboratory Plymouth | Wellcome Sanger Institute for the COVID-19 Genomics UK (COG-UK) Consortium | Lighthouse Laboratory Plymouth and Alex Alderton, Roberto Amato, Jeffrey Barrett, Sonia Goncalves, Ewan Harrison, David K. Jackson, Ian Johnston, Dominic Kwiatkowski, Cordelia Langford, John Sillitoe on behalf of the Wellcome Sanger Institute COVID-19 Surveillance Team |
| EPI_ISL_3167263 | Lighthouse Lab in Milton Keynes | Wellcome Sanger Institute for the COVID-19 Genomics UK (COG-UK) Consortium | The Lighthouse Lab in Milton Keynes and Alex Alderton, Roberto Amato, Jeffrey Barrett, Sonia Goncalves, Ewan Harrison, David K. Jackson, Ian Johnston, Dominic Kwiatkowski, Cordelia Langford, John Sillitoe on behalf of the Wellcome Sanger Institute COVID-19 Surveillance Team |
| EPI_ISL_3167264 | Lighthouse Lab in Glasgow | Wellcome Sanger Institute for the COVID-19 Genomics UK (COG-UK) Consortium | Harper VanSteenhouse, Yumi Kasai, David Gray, Carol Clugston, Anna Dominiczak and Alex Alderton, Roberto Amato, Jeffrey Barrett, Sonia Goncalves, Ewan Harrison, David K. Jackson, Ian Johnston, Dominic Kwiatkowski, Cordelia Langford, John Sillitoe on behalf of the Wellcome Sanger Institute COVID-19 Surveillance Team |
| EPI_ISL_3167265 | Lighthouse Lab in Milton Keynes | Wellcome Sanger Institute for the COVID-19 Genomics UK (COG-UK) Consortium | The Lighthouse Lab in Milton Keynes and Alex Alderton, Roberto Amato, Jeffrey Barrett, Sonia Goncalves, Ewan Harrison, David K. Jackson, Ian Johnston, Dominic Kwiatkowski, Cordelia Langford, John Sillitoe on behalf of the Wellcome Sanger Institute COVID-19 Surveillance Team |
| EPI_ISL_3167266 | Lighthouse Laboratory Plymouth | Wellcome Sanger Institute for the COVID-19 Genomics UK (COG-UK) Consortium | Lighthouse Laboratory Plymouth and Alex Alderton, Roberto Amato, Jeffrey Barrett, Sonia Goncalves, Ewan Harrison, David K. Jackson, Ian Johnston, Dominic Kwiatkowski, Cordelia Langford, John Sillitoe on behalf of the Wellcome Sanger Institute COVID-19 Surveillance Team |
| EPI_ISL_3167267, EPI_ISL_3167268 | Lighthouse Lab in Milton Keynes | Wellcome Sanger Institute for the COVID-19 Genomics UK (COG-UK) Consortium | The Lighthouse Lab in Milton Keynes and Alex Alderton, Roberto Amato, Jeffrey Barrett, Sonia Goncalves, Ewan Harrison, David K. Jackson, Ian Johnston, Dominic Kwiatkowski, Cordelia Langford, John Sillitoe on behalf of the Wellcome Sanger Institute COVID-19 Surveillance Team |
| EPI_ISL_3167269 | Lighthouse Lab in Glasgow | Wellcome Sanger Institute for the COVID-19 Genomics UK (COG-UK) Consortium | Harper VanSteenhouse, Yumi Kasai, David Gray, Carol Clugston, Anna Dominiczak and Alex Alderton, Roberto Amato, Jeffrey Barrett, Sonia Goncalves, Ewan Harrison, David K. Jackson, Ian Johnston, Dominic Kwiatkowski, Cordelia Langford, John Sillitoe on behalf of the Wellcome Sanger Institute COVID-19 Surveillance Team |
| EPI_ISL_3167270, EPI_ISL_3167271, EPI_ISL_3167272 | Lighthouse Lab in Milton Keynes | Wellcome Sanger Institute for the COVID-19 Genomics UK (COG-UK) Consortium | The Lighthouse Lab in Milton Keynes and Alex Alderton, Roberto Amato, Jeffrey Barrett, Sonia Goncalves, Ewan Harrison, David K. Jackson, Ian Johnston, Dominic Kwiatkowski, Cordelia Langford, John Sillitoe on behalf of the Wellcome Sanger Institute COVID-19 Surveillance Team |
| EPI_ISL_3167273 | Lighthouse Laboratory Plymouth | Wellcome Sanger Institute for the COVID-19 Genomics UK (COG-UK) Consortium | Lighthouse Laboratory Plymouth and Alex Alderton, Roberto Amato, Jeffrey Barrett, Sonia Goncalves, Ewan Harrison, David K. Jackson, Ian Johnston, Dominic Kwiatkowski, Cordelia Langford, John Sillitoe on behalf of the Wellcome Sanger Institute COVID-19 Surveillance Team |
| EPI_ISL_3167274 | Lighthouse Lab in Milton Keynes | Wellcome Sanger Institute for the COVID-19 Genomics UK (COG-UK) Consortium | The Lighthouse Lab in Milton Keynes and Alex Alderton, Roberto Amato, Jeffrey Barrett, Sonia Goncalves, Ewan Harrison, David K. Jackson, Ian Johnston, Dominic Kwiatkowski, Cordelia Langford, John Sillitoe on behalf of the Wellcome Sanger Institute COVID-19 Surveillance Team |
| EPI_ISL_3167275 | Lighthouse Lab in Glasgow | Wellcome Sanger Institute for the COVID-19 Genomics UK (COG-UK) Consortium | Harper VanSteenhouse, Yumi Kasai, David Gray, Carol Clugston, Anna Dominiczak and Alex Alderton, Roberto Amato, Jeffrey Barrett, Sonia Goncalves, Ewan Harrison, David K. Jackson, Ian Johnston, Dominic Kwiatkowski, Cordelia Langford, John Sillitoe on behalf of the Wellcome Sanger Institute COVID-19 Surveillance Team |
| EPI_ISL_3167276, EPI_ISL_3167277 | Lighthouse Lab in Milton Keynes | Wellcome Sanger Institute for the COVID-19 Genomics UK (COG-UK) Consortium | The Lighthouse Lab in Milton Keynes and Alex Alderton, Roberto Amato, Jeffrey Barrett, Sonia Goncalves, Ewan Harrison, David K. Jackson, Ian Johnston, Dominic Kwiatkowski, Cordelia Langford, John Sillitoe on behalf of the Wellcome Sanger Institute COVID-19 Surveillance Team |
| EPI_ISL_3167278 | Lighthouse Lab in Glasgow | Wellcome Sanger Institute for the COVID-19 Genomics UK (COG-UK) Consortium | Harper VanSteenhouse, Yumi Kasai, David Gray, Carol Clugston, Anna Dominiczak and Alex Alderton, Roberto Amato, Jeffrey Barrett, Sonia Goncalves, Ewan Harrison, David K. Jackson, Ian Johnston, Dominic Kwiatkowski, Cordelia Langford, John Sillitoe on behalf of the Wellcome Sanger Institute COVID-19 Surveillance Team |
| EPI_ISL_3167279, EPI_ISL_3167280 | Lighthouse Lab in Milton Keynes | Wellcome Sanger Institute for the COVID-19 Genomics UK (COG-UK) Consortium | The Lighthouse Lab in Milton Keynes and Alex Alderton, Roberto Amato, Jeffrey Barrett, Sonia Goncalves, Ewan Harrison, David K. Jackson, Ian Johnston, Dominic Kwiatkowski, Cordelia Langford, John Sillitoe on behalf of the Wellcome Sanger Institute COVID-19 Surveillance Team |
| EPI_ISL_3167281, EPI_ISL_3167282 | Lighthouse Laboratory Plymouth | Wellcome Sanger Institute for the COVID-19 Genomics UK (COG-UK) Consortium | Lighthouse Laboratory Plymouth and Alex Alderton, Roberto Amato, Jeffrey Barrett, Sonia Goncalves, Ewan Harrison, David K. Jackson, Ian Johnston, Dominic Kwiatkowski, Cordelia Langford, John Sillitoe on behalf of the Wellcome Sanger Institute COVID-19 Surveillance Team |
| EPI_ISL_3167283, EPI_ISL_3167284, EPI_ISL_3167285, EPI_ISL_3167286, EPI_ISL_3167287, EPI_ISL_3167288, EPI_ISL_3167289, EPI_ISL_3167290 | Lighthouse Lab in Milton Keynes | Wellcome Sanger Institute for the COVID-19 Genomics UK (COG-UK) Consortium | The Lighthouse Lab in Milton Keynes and Alex Alderton, Roberto Amato, Jeffrey Barrett, Sonia Goncalves, Ewan Harrison, David K. Jackson, Ian Johnston, Dominic Kwiatkowski, Cordelia Langford, John Sillitoe on behalf of the Wellcome Sanger Institute COVID-19 Surveillance Team |
| EPI_ISL_3167291 | Lighthouse Laboratory Plymouth | Wellcome Sanger Institute for the COVID-19 Genomics UK (COG-UK) Consortium | Lighthouse Laboratory Plymouth and Alex Alderton, Roberto Amato, Jeffrey Barrett, Sonia Goncalves, Ewan Harrison, David K. Jackson, Ian Johnston, Dominic Kwiatkowski, Cordelia Langford, John Sillitoe on behalf of the Wellcome Sanger Institute COVID-19 Surveillance Team |
| EPI_ISL_3167292, EPI_ISL_3167293 | Lighthouse Lab in Milton Keynes | Wellcome Sanger Institute for the COVID-19 Genomics UK (COG-UK) Consortium | The Lighthouse Lab in Milton Keynes and Alex Alderton, Roberto Amato, Jeffrey Barrett, Sonia Goncalves, Ewan Harrison, David K. Jackson, Ian Johnston, Dominic Kwiatkowski, Cordelia Langford, John Sillitoe on behalf of the Wellcome Sanger Institute COVID-19 Surveillance Team |
| EPI_ISL_3167294 | Lighthouse Laboratory Plymouth | Wellcome Sanger Institute for the COVID-19 Genomics UK (COG-UK) Consortium | Lighthouse Laboratory Plymouth and Alex Alderton, Roberto Amato, Jeffrey Barrett, Sonia Goncalves, Ewan Harrison, David K. Jackson, Ian Johnston, Dominic Kwiatkowski, Cordelia Langford, John Sillitoe on behalf of the Wellcome Sanger Institute COVID-19 Surveillance Team |
| EPI_ISL_3167295, EPI_ISL_3167296, EPI_ISL_3167297, EPI_ISL_3167298 | Lighthouse Lab in Milton Keynes | Wellcome Sanger Institute for the COVID-19 Genomics UK (COG-UK) Consortium | The Lighthouse Lab in Milton Keynes and Alex Alderton, Roberto Amato, Jeffrey Barrett, Sonia Goncalves, Ewan Harrison, David K. Jackson, Ian Johnston, Dominic Kwiatkowski, Cordelia Langford, John Sillitoe on behalf of the Wellcome Sanger Institute COVID-19 Surveillance Team |
| EPI_ISL_3167299 | Lighthouse Lab in Glasgow | Wellcome Sanger Institute for the COVID-19 Genomics UK (COG-UK) Consortium | Harper VanSteenhouse, Yumi Kasai, David Gray, Carol Clugston, Anna Dominiczak and Alex Alderton, Roberto Amato, Jeffrey Barrett, Sonia Goncalves, Ewan Harrison, David K. Jackson, Ian Johnston, Dominic Kwiatkowski, Cordelia Langford, John Sillitoe on behalf of the Wellcome Sanger Institute COVID-19 Surveillance Team |
| EPI_ISL_3167300 | Lighthouse Laboratory Plymouth | Wellcome Sanger Institute for the COVID-19 Genomics UK (COG-UK) Consortium | Lighthouse Laboratory Plymouth and Alex Alderton, Roberto Amato, Jeffrey Barrett, Sonia Goncalves, Ewan Harrison, David K. Jackson, Ian Johnston, Dominic Kwiatkowski, Cordelia Langford, John Sillitoe on behalf of the Wellcome Sanger Institute COVID-19 Surveillance Team |
| EPI_ISL_3167301 | Lighthouse Lab in Milton Keynes | Wellcome Sanger Institute for the COVID-19 Genomics UK (COG-UK) Consortium | The Lighthouse Lab in Milton Keynes and Alex Alderton, Roberto Amato, Jeffrey Barrett, Sonia Goncalves, Ewan Harrison, David K. Jackson, Ian Johnston, Dominic Kwiatkowski, Cordelia Langford, John Sillitoe on behalf of the Wellcome Sanger Institute COVID-19 Surveillance Team |
| EPI_ISL_3167302 | Lighthouse Laboratory Plymouth | Wellcome Sanger Institute for the COVID-19 Genomics UK (COG-UK) Consortium | Lighthouse Laboratory Plymouth and Alex Alderton, Roberto Amato, Jeffrey Barrett, Sonia Goncalves, Ewan Harrison, David K. Jackson, Ian Johnston, Dominic Kwiatkowski, Cordelia Langford, John Sillitoe on behalf of the Wellcome Sanger Institute COVID-19 Surveillance Team |
| EPI_ISL_3167303 | Lighthouse Lab in Milton Keynes | Wellcome Sanger Institute for the COVID-19 Genomics UK (COG-UK) Consortium | The Lighthouse Lab in Milton Keynes and Alex Alderton, Roberto Amato, Jeffrey Barrett, Sonia Goncalves, Ewan Harrison, David K. Jackson, Ian Johnston, Dominic Kwiatkowski, Cordelia Langford, John Sillitoe on behalf of the Wellcome Sanger Institute COVID-19 Surveillance Team |
| EPI_ISL_3167304 | Lighthouse Laboratory Plymouth | Wellcome Sanger Institute for the COVID-19 Genomics UK (COG-UK) Consortium | Lighthouse Laboratory Plymouth and Alex Alderton, Roberto Amato, Jeffrey Barrett, Sonia Goncalves, Ewan Harrison, David K. Jackson, Ian Johnston, Dominic Kwiatkowski, Cordelia Langford, John Sillitoe on behalf of the Wellcome Sanger Institute COVID-19 Surveillance Team |
| EPI_ISL_3167305, EPI_ISL_3167306 | Lighthouse Lab in Milton Keynes | Wellcome Sanger Institute for the COVID-19 Genomics UK (COG-UK) Consortium | The Lighthouse Lab in Milton Keynes and Alex Alderton, Roberto Amato, Jeffrey Barrett, Sonia Goncalves, Ewan Harrison, David K. Jackson, Ian Johnston, Dominic Kwiatkowski, Cordelia Langford, John Sillitoe on behalf of the Wellcome Sanger Institute COVID-19 Surveillance Team |
| EPI_ISL_3167307 | Lighthouse Laboratory Plymouth | Wellcome Sanger Institute for the COVID-19 Genomics UK (COG-UK) Consortium | Lighthouse Laboratory Plymouth and Alex Alderton, Roberto Amato, Jeffrey Barrett, Sonia Goncalves, Ewan Harrison, David K. Jackson, Ian Johnston, Dominic Kwiatkowski, Cordelia Langford, John Sillitoe on behalf of the Wellcome Sanger Institute COVID-19 Surveillance Team |
| EPI_ISL_3167308 | Lighthouse Lab in Glasgow | Wellcome Sanger Institute for the COVID-19 Genomics UK (COG-UK) Consortium | Harper VanSteenhouse, Yumi Kasai, David Gray, Carol Clugston, Anna Dominiczak and Alex Alderton, Roberto Amato, Jeffrey Barrett, Sonia Goncalves, Ewan Harrison, David K. Jackson, Ian Johnston, Dominic Kwiatkowski, Cordelia Langford, John Sillitoe on behalf of the Wellcome Sanger Institute COVID-19 Surveillance Team |
| EPI_ISL_3167309, EPI_ISL_3167310 | Lighthouse Laboratory Plymouth | Wellcome Sanger Institute for the COVID-19 Genomics UK (COG-UK) Consortium | Lighthouse Laboratory Plymouth and Alex Alderton, Roberto Amato, Jeffrey Barrett, Sonia Goncalves, Ewan Harrison, David K. Jackson, Ian Johnston, Dominic Kwiatkowski, Cordelia Langford, John Sillitoe on behalf of the Wellcome Sanger Institute COVID-19 Surveillance Team |
| EPI_ISL_3167311 | Lighthouse Lab in Milton Keynes | Wellcome Sanger Institute for the COVID-19 Genomics UK (COG-UK) Consortium | The Lighthouse Lab in Milton Keynes and Alex Alderton, Roberto Amato, Jeffrey Barrett, Sonia Goncalves, Ewan Harrison, David K. Jackson, Ian Johnston, Dominic Kwiatkowski, Cordelia Langford, John Sillitoe on behalf of the Wellcome Sanger Institute COVID-19 Surveillance Team |
| EPI_ISL_3167312 | Lighthouse Lab in Glasgow | Wellcome Sanger Institute for the COVID-19 Genomics UK (COG-UK) Consortium | Harper VanSteenhouse, Yumi Kasai, David Gray, Carol Clugston, Anna Dominiczak and Alex Alderton, Roberto Amato, Jeffrey Barrett, Sonia Goncalves, Ewan Harrison, David K. Jackson, Ian Johnston, Dominic Kwiatkowski, Cordelia Langford, John Sillitoe on behalf of the Wellcome Sanger Institute COVID-19 Surveillance Team |
| EPI_ISL_3167313, EPI_ISL_3167314 | Lighthouse Lab in Milton Keynes | Wellcome Sanger Institute for the COVID-19 Genomics UK (COG-UK) Consortium | The Lighthouse Lab in Milton Keynes and Alex Alderton, Roberto Amato, Jeffrey Barrett, Sonia Goncalves, Ewan Harrison, David K. Jackson, Ian Johnston, Dominic Kwiatkowski, Cordelia Langford, John Sillitoe on behalf of the Wellcome Sanger Institute COVID-19 Surveillance Team |
| EPI_ISL_3167315, EPI_ISL_3167316, EPI_ISL_3167317 | Lighthouse Lab in Glasgow | Wellcome Sanger Institute for the COVID-19 Genomics UK (COG-UK) Consortium | Harper VanSteenhouse, Yumi Kasai, David Gray, Carol Clugston, Anna Dominiczak and Alex Alderton, Roberto Amato, Jeffrey Barrett, Sonia Goncalves, Ewan Harrison, David K. Jackson, Ian Johnston, Dominic Kwiatkowski, Cordelia Langford, John Sillitoe on behalf of the Wellcome Sanger Institute COVID-19 Surveillance Team |
| EPI_ISL_3167318, EPI_ISL_3167319, EPI_ISL_3167320 | Lighthouse Lab in Milton Keynes | Wellcome Sanger Institute for the COVID-19 Genomics UK (COG-UK) Consortium | The Lighthouse Lab in Milton Keynes and Alex Alderton, Roberto Amato, Jeffrey Barrett, Sonia Goncalves, Ewan Harrison, David K. Jackson, Ian Johnston, Dominic Kwiatkowski, Cordelia Langford, John Sillitoe on behalf of the Wellcome Sanger Institute COVID-19 Surveillance Team |
| EPI_ISL_3167321 | Lighthouse Laboratory Plymouth | Wellcome Sanger Institute for the COVID-19 Genomics UK (COG-UK) Consortium | Lighthouse Laboratory Plymouth and Alex Alderton, Roberto Amato, Jeffrey Barrett, Sonia Goncalves, Ewan Harrison, David K. Jackson, Ian Johnston, Dominic Kwiatkowski, Cordelia Langford, John Sillitoe on behalf of the Wellcome Sanger Institute COVID-19 Surveillance Team |
| EPI_ISL_3167322 | Lighthouse Lab in Glasgow | Wellcome Sanger Institute for the COVID-19 Genomics UK (COG-UK) Consortium | Harper VanSteenhouse, Yumi Kasai, David Gray, Carol Clugston, Anna Dominiczak and Alex Alderton, Roberto Amato, Jeffrey Barrett, Sonia Goncalves, Ewan Harrison, David K. Jackson, Ian Johnston, Dominic Kwiatkowski, Cordelia Langford, John Sillitoe on behalf of the Wellcome Sanger Institute COVID-19 Surveillance Team |
| EPI_ISL_3167323, EPI_ISL_3167324 | Lighthouse Lab in Milton Keynes | Wellcome Sanger Institute for the COVID-19 Genomics UK (COG-UK) Consortium | The Lighthouse Lab in Milton Keynes and Alex Alderton, Roberto Amato, Jeffrey Barrett, Sonia Goncalves, Ewan Harrison, David K. Jackson, Ian Johnston, Dominic Kwiatkowski, Cordelia Langford, John Sillitoe on behalf of the Wellcome Sanger Institute COVID-19 Surveillance Team |
| EPI_ISL_3167325 | Lighthouse Laboratory Plymouth | Wellcome Sanger Institute for the COVID-19 Genomics UK (COG-UK) Consortium | Lighthouse Laboratory Plymouth and Alex Alderton, Roberto Amato, Jeffrey Barrett, Sonia Goncalves, Ewan Harrison, David K. Jackson, Ian Johnston, Dominic Kwiatkowski, Cordelia Langford, John Sillitoe on behalf of the Wellcome Sanger Institute COVID-19 Surveillance Team |

| | | | |
|---|---|---|---|
| EPI_ISL_3167326, EPI_ISL_3167327 | Lighthouse Lab in Milton Keynes | Wellcome Sanger Institute for the COVID-19 Genomics UK (COG-UK) Consortium | The Lighthouse Lab in Milton Keynes and Alex Alderton, Roberto Amato, Jeffrey Barrett, Sonia Goncalves, Ewan Harrison, David K. Jackson, Ian Johnston, Dominic Kwiatkowski, Cordelia Langford, John Sillitoe on behalf of the Wellcome Sanger Institute COVID-19 Surveillance Team |
| EPI_ISL_3167328 | Lighthouse Laboratory Plymouth | Wellcome Sanger Institute for the COVID-19 Genomics UK (COG-UK) Consortium | Lighthouse Laboratory Plymouth and Alex Alderton, Roberto Amato, Jeffrey Barrett, Sonia Goncalves, Ewan Harrison, David K. Jackson, Ian Johnston, Dominic Kwiatkowski, Cordelia Langford, John Sillitoe on behalf of the Wellcome Sanger Institute COVID-19 Surveillance Team |
| EPI_ISL_3167329 | Lighthouse Lab in Glasgow | Wellcome Sanger Institute for the COVID-19 Genomics UK (COG-UK) Consortium | Harper VanSteenhouse, Yumi Kasai, David Gray, Carol Clugston, Anna Dominiczak and Alex Alderton, Roberto Amato, Jeffrey Barrett, Sonia Goncalves, Ewan Harrison, David K. Jackson, Ian Johnston, Dominic Kwiatkowski, Cordelia Langford, John Sillitoe on behalf of the Wellcome Sanger Institute COVID-19 Surveillance Team |
| EPI_ISL_3167330 | Lighthouse Laboratory Plymouth | Wellcome Sanger Institute for the COVID-19 Genomics UK (COG-UK) Consortium | Lighthouse Laboratory Plymouth and Alex Alderton, Roberto Amato, Jeffrey Barrett, Sonia Goncalves, Ewan Harrison, David K. Jackson, Ian Johnston, Dominic Kwiatkowski, Cordelia Langford, John Sillitoe on behalf of the Wellcome Sanger Institute COVID-19 Surveillance Team |
| EPI_ISL_3167331 | Lighthouse Lab in Milton Keynes | Wellcome Sanger Institute for the COVID-19 Genomics UK (COG-UK) Consortium | The Lighthouse Lab in Milton Keynes and Alex Alderton, Roberto Amato, Jeffrey Barrett, Sonia Goncalves, Ewan Harrison, David K. Jackson, Ian Johnston, Dominic Kwiatkowski, Cordelia Langford, John Sillitoe on behalf of the Wellcome Sanger Institute COVID-19 Surveillance Team |
| EPI_ISL_3167332, EPI_ISL_3167333, EPI_ISL_3167334 | Lighthouse Laboratory Plymouth | Wellcome Sanger Institute for the COVID-19 Genomics UK (COG-UK) Consortium | Lighthouse Laboratory Plymouth and Alex Alderton, Roberto Amato, Jeffrey Barrett, Sonia Goncalves, Ewan Harrison, David K. Jackson, Ian Johnston, Dominic Kwiatkowski, Cordelia Langford, John Sillitoe on behalf of the Wellcome Sanger Institute COVID-19 Surveillance Team |
| EPI_ISL_3167335 | Lighthouse Lab in Glasgow | Wellcome Sanger Institute for the COVID-19 Genomics UK (COG-UK) Consortium | Harper VanSteenhouse, Yumi Kasai, David Gray, Carol Clugston, Anna Dominiczak and Alex Alderton, Roberto Amato, Jeffrey Barrett, Sonia Goncalves, Ewan Harrison, David K. Jackson, Ian Johnston, Dominic Kwiatkowski, Cordelia Langford, John Sillitoe on behalf of the Wellcome Sanger Institute COVID-19 Surveillance Team |
| EPI_ISL_3167336 | Lighthouse Lab in Milton Keynes | Wellcome Sanger Institute for the COVID-19 Genomics UK (COG-UK) Consortium | The Lighthouse Lab in Milton Keynes and Alex Alderton, Roberto Amato, Jeffrey Barrett, Sonia Goncalves, Ewan Harrison, David K. Jackson, Ian Johnston, Dominic Kwiatkowski, Cordelia Langford, John Sillitoe on behalf of the Wellcome Sanger Institute COVID-19 Surveillance Team |
| EPI_ISL_3167337, EPI_ISL_3167338 | Lighthouse Laboratory Plymouth | Wellcome Sanger Institute for the COVID-19 Genomics UK (COG-UK) Consortium | Lighthouse Laboratory Plymouth and Alex Alderton, Roberto Amato, Jeffrey Barrett, Sonia Goncalves, Ewan Harrison, David K. Jackson, Ian Johnston, Dominic Kwiatkowski, Cordelia Langford, John Sillitoe on behalf of the Wellcome Sanger Institute COVID-19 Surveillance Team |
| EPI_ISL_3167339, EPI_ISL_3167340, EPI_ISL_3167341, EPI_ISL_3167342 | Lighthouse Lab in Milton Keynes | Wellcome Sanger Institute for the COVID-19 Genomics UK (COG-UK) Consortium | The Lighthouse Lab in Milton Keynes and Alex Alderton, Roberto Amato, Jeffrey Barrett, Sonia Goncalves, Ewan Harrison, David K. Jackson, Ian Johnston, Dominic Kwiatkowski, Cordelia Langford, John Sillitoe on behalf of the Wellcome Sanger Institute COVID-19 Surveillance Team |
| EPI_ISL_3167343 | Lighthouse Laboratory Plymouth | Wellcome Sanger Institute for the COVID-19 Genomics UK (COG-UK) Consortium | Lighthouse Laboratory Plymouth and Alex Alderton, Roberto Amato, Jeffrey Barrett, Sonia Goncalves, Ewan Harrison, David K. Jackson, Ian Johnston, Dominic Kwiatkowski, Cordelia Langford, John Sillitoe on behalf of the Wellcome Sanger Institute COVID-19 Surveillance Team |
| EPI_ISL_3167344, EPI_ISL_3167345 | Lighthouse Lab in Milton Keynes | Wellcome Sanger Institute for the COVID-19 Genomics UK (COG-UK) Consortium | The Lighthouse Lab in Milton Keynes and Alex Alderton, Roberto Amato, Jeffrey Barrett, Sonia Goncalves, Ewan Harrison, David K. Jackson, Ian Johnston, Dominic Kwiatkowski, Cordelia Langford, John Sillitoe on behalf of the Wellcome Sanger Institute COVID-19 Surveillance Team |
| EPI_ISL_3167346 | Lighthouse Laboratory Plymouth | Wellcome Sanger Institute for the COVID-19 Genomics UK (COG-UK) Consortium | Lighthouse Laboratory Plymouth and Alex Alderton, Roberto Amato, Jeffrey Barrett, Sonia Goncalves, Ewan Harrison, David K. Jackson, Ian Johnston, Dominic Kwiatkowski, Cordelia Langford, John Sillitoe on behalf of the Wellcome Sanger Institute COVID-19 Surveillance Team |
| EPI_ISL_3167347 | Lighthouse Lab in Milton Keynes | Wellcome Sanger Institute for the COVID-19 Genomics UK (COG-UK) Consortium | The Lighthouse Lab in Milton Keynes and Alex Alderton, Roberto Amato, Jeffrey Barrett, Sonia Goncalves, Ewan Harrison, David K. Jackson, Ian Johnston, Dominic Kwiatkowski, Cordelia Langford, John Sillitoe on behalf of the Wellcome Sanger Institute COVID-19 Surveillance Team |
| EPI_ISL_3167348 | Lighthouse Laboratory Plymouth | Wellcome Sanger Institute for the COVID-19 Genomics UK (COG-UK) Consortium | Lighthouse Laboratory Plymouth and Alex Alderton, Roberto Amato, Jeffrey Barrett, Sonia Goncalves, Ewan Harrison, David K. Jackson, Ian Johnston, Dominic Kwiatkowski, Cordelia Langford, John Sillitoe on behalf of the Wellcome Sanger Institute COVID-19 Surveillance Team |
| EPI_ISL_3167349, EPI_ISL_3167350, EPI_ISL_3167351, EPI_ISL_3167352, EPI_ISL_3167353 | Lighthouse Lab in Milton Keynes | Wellcome Sanger Institute for the COVID-19 Genomics UK (COG-UK) Consortium | The Lighthouse Lab in Milton Keynes and Alex Alderton, Roberto Amato, Jeffrey Barrett, Sonia Goncalves, Ewan Harrison, David K. Jackson, Ian Johnston, Dominic Kwiatkowski, Cordelia Langford, John Sillitoe on behalf of the Wellcome Sanger Institute COVID-19 Surveillance Team |
| EPI_ISL_3167354, EPI_ISL_3167355 | Lighthouse Laboratory Plymouth | Wellcome Sanger Institute for the COVID-19 Genomics UK (COG-UK) Consortium | Lighthouse Laboratory Plymouth and Alex Alderton, Roberto Amato, Jeffrey Barrett, Sonia Goncalves, Ewan Harrison, David K. Jackson, Ian Johnston, Dominic Kwiatkowski, Cordelia Langford, John Sillitoe on behalf of the Wellcome Sanger Institute COVID-19 Surveillance Team |
| EPI_ISL_3167356 | Lighthouse Lab in Milton Keynes | Wellcome Sanger Institute for the COVID-19 Genomics UK (COG-UK) Consortium | The Lighthouse Lab in Milton Keynes and Alex Alderton, Roberto Amato, Jeffrey Barrett, Sonia Goncalves, Ewan Harrison, David K. Jackson, Ian Johnston, Dominic Kwiatkowski, Cordelia Langford, John Sillitoe on behalf of the Wellcome Sanger Institute COVID-19 Surveillance Team |
| EPI_ISL_3167357 | Lighthouse Lab in Glasgow | Wellcome Sanger Institute for the COVID-19 Genomics UK (COG-UK) Consortium | Harper VanSteenhouse, Yumi Kasai, David Gray, Carol Clugston, Anna Dominiczak and Alex Alderton, Roberto Amato, Jeffrey Barrett, Sonia Goncalves, Ewan Harrison, David K. Jackson, Ian Johnston, Dominic Kwiatkowski, Cordelia Langford, John Sillitoe on behalf of the Wellcome Sanger Institute COVID-19 Surveillance Team |
| EPI_ISL_3167358 | Lighthouse Lab in Milton Keynes | Wellcome Sanger Institute for the COVID-19 Genomics UK (COG-UK) Consortium | The Lighthouse Lab in Milton Keynes and Alex Alderton, Roberto Amato, Jeffrey Barrett, Sonia Goncalves, Ewan Harrison, David K. Jackson, Ian Johnston, Dominic Kwiatkowski, Cordelia Langford, John Sillitoe on behalf of the Wellcome Sanger Institute COVID-19 Surveillance Team |
| EPI_ISL_3167359, EPI_ISL_3167360, EPI_ISL_3167361 | Lighthouse Laboratory Plymouth | Wellcome Sanger Institute for the COVID-19 Genomics UK (COG-UK) Consortium | Lighthouse Laboratory Plymouth and Alex Alderton, Roberto Amato, Jeffrey Barrett, Sonia Goncalves, Ewan Harrison, David K. Jackson, Ian Johnston, Dominic Kwiatkowski, Cordelia Langford, John Sillitoe on behalf of the Wellcome Sanger Institute COVID-19 Surveillance Team |
| EPI_ISL_3167362 | Lighthouse Lab in Milton Keynes | Wellcome Sanger Institute for the COVID-19 Genomics UK (COG-UK) Consortium | The Lighthouse Lab in Milton Keynes and Alex Alderton, Roberto Amato, Jeffrey Barrett, Sonia Goncalves, Ewan Harrison, David K. Jackson, Ian Johnston, Dominic Kwiatkowski, Cordelia Langford, John Sillitoe on behalf of the Wellcome Sanger Institute COVID-19 Surveillance Team |
| EPI_ISL_3167363, EPI_ISL_3167364 | Lighthouse Lab in Glasgow | Wellcome Sanger Institute for the COVID-19 Genomics UK (COG-UK) Consortium | Harper VanSteenhouse, Yumi Kasai, David Gray, Carol Clugston, Anna Dominiczak and Alex Alderton, Roberto Amato, Jeffrey Barrett, Sonia Goncalves, Ewan Harrison, David K. Jackson, Ian Johnston, Dominic Kwiatkowski, Cordelia Langford, John Sillitoe on behalf of the Wellcome Sanger Institute COVID-19 Surveillance Team |
| EPI_ISL_3167365 | Lighthouse Lab in Milton Keynes | Wellcome Sanger Institute for the COVID-19 Genomics UK (COG-UK) Consortium | The Lighthouse Lab in Milton Keynes and Alex Alderton, Roberto Amato, Jeffrey Barrett, Sonia Goncalves, Ewan Harrison, David K. Jackson, Ian Johnston, Dominic Kwiatkowski, Cordelia Langford, John Sillitoe on behalf of the Wellcome Sanger Institute COVID-19 Surveillance Team |
| EPI_ISL_3167366 | Lighthouse Lab in Glasgow | Wellcome Sanger Institute for the COVID-19 Genomics UK (COG-UK) Consortium | Harper VanSteenhouse, Yumi Kasai, David Gray, Carol Clugston, Anna Dominiczak and Alex Alderton, Roberto Amato, Jeffrey Barrett, Sonia Goncalves, Ewan Harrison, David K. Jackson, Ian Johnston, Dominic Kwiatkowski, Cordelia Langford, John Sillitoe on behalf of the Wellcome Sanger Institute COVID-19 Surveillance Team |
| EPI_ISL_3167367 | Lighthouse Lab in Milton Keynes | Wellcome Sanger Institute for the COVID-19 Genomics UK (COG-UK) Consortium | The Lighthouse Lab in Milton Keynes and Alex Alderton, Roberto Amato, Jeffrey Barrett, Sonia Goncalves, Ewan Harrison, David K. Jackson, Ian Johnston, Dominic Kwiatkowski, Cordelia Langford, John Sillitoe on behalf of the Wellcome Sanger Institute COVID-19 Surveillance Team |
| EPI_ISL_3167368 | Lighthouse Laboratory Plymouth | Wellcome Sanger Institute for the COVID-19 Genomics UK (COG-UK) Consortium | Lighthouse Laboratory Plymouth and Alex Alderton, Roberto Amato, Jeffrey Barrett, Sonia Goncalves, Ewan Harrison, David K. Jackson, Ian Johnston, Dominic Kwiatkowski, Cordelia Langford, John Sillitoe on behalf of the Wellcome Sanger Institute COVID-19 Surveillance Team |
| EPI_ISL_3167369 | Lighthouse Lab in Milton Keynes | Wellcome Sanger Institute for the COVID-19 Genomics UK (COG-UK) Consortium | The Lighthouse Lab in Milton Keynes and Alex Alderton, Roberto Amato, Jeffrey Barrett, Sonia Goncalves, Ewan Harrison, David K. Jackson, Ian Johnston, Dominic Kwiatkowski, Cordelia Langford, John Sillitoe on behalf of the Wellcome Sanger Institute COVID-19 Surveillance Team |
| EPI_ISL_3167370 | Lighthouse Laboratory Plymouth | Wellcome Sanger Institute for the COVID-19 Genomics UK (COG-UK) Consortium | Lighthouse Laboratory Plymouth and Alex Alderton, Roberto Amato, Jeffrey Barrett, Sonia Goncalves, Ewan Harrison, David K. Jackson, Ian Johnston, Dominic Kwiatkowski, Cordelia Langford, John Sillitoe on behalf of the Wellcome Sanger Institute COVID-19 Surveillance Team |
| EPI_ISL_3167371, EPI_ISL_3167372, EPI_ISL_3167373 | Lighthouse Lab in Milton Keynes | Wellcome Sanger Institute for the COVID-19 Genomics UK (COG-UK) Consortium | The Lighthouse Lab in Milton Keynes and Alex Alderton, Roberto Amato, Jeffrey Barrett, Sonia Goncalves, Ewan Harrison, David K. Jackson, Ian Johnston, Dominic Kwiatkowski, Cordelia Langford, John Sillitoe on behalf of the Wellcome Sanger Institute COVID-19 Surveillance Team |
| EPI_ISL_3167374, EPI_ISL_3167375 | Lighthouse Lab in Glasgow | Wellcome Sanger Institute for the COVID-19 Genomics UK (COG-UK) Consortium | Harper VanSteenhouse, Yumi Kasai, David Gray, Carol Clugston, Anna Dominiczak and Alex Alderton, Roberto Amato, Jeffrey Barrett, Sonia Goncalves, Ewan Harrison, David K. Jackson, Ian Johnston, Dominic Kwiatkowski, Cordelia Langford, John Sillitoe on behalf of the Wellcome Sanger Institute COVID-19 Surveillance Team |
| EPI_ISL_3167376, EPI_ISL_3167377 | Lighthouse Lab in Milton Keynes | Wellcome Sanger Institute for the COVID-19 Genomics UK (COG-UK) Consortium | The Lighthouse Lab in Milton Keynes and Alex Alderton, Roberto Amato, Jeffrey Barrett, Sonia Goncalves, Ewan Harrison, David K. Jackson, Ian Johnston, Dominic Kwiatkowski, Cordelia Langford, John Sillitoe on behalf of the Wellcome Sanger Institute COVID-19 Surveillance Team |
| EPI_ISL_3167378 | Lighthouse Laboratory Plymouth | Wellcome Sanger Institute for the COVID-19 Genomics UK (COG-UK) Consortium | Lighthouse Laboratory Plymouth and Alex Alderton, Roberto Amato, Jeffrey Barrett, Sonia Goncalves, Ewan Harrison, David K. Jackson, Ian Johnston, Dominic Kwiatkowski, Cordelia Langford, John Sillitoe on behalf of the Wellcome Sanger Institute COVID-19 Surveillance Team |
| EPI_ISL_3167379, EPI_ISL_3167380, EPI_ISL_3167381 | Lighthouse Lab in Milton Keynes | Wellcome Sanger Institute for the COVID-19 Genomics UK (COG-UK) Consortium | The Lighthouse Lab in Milton Keynes and Alex Alderton, Roberto Amato, Jeffrey Barrett, Sonia Goncalves, Ewan Harrison, David K. Jackson, Ian Johnston, Dominic Kwiatkowski, Cordelia Langford, John Sillitoe on behalf of the Wellcome Sanger Institute COVID-19 Surveillance Team |
| EPI_ISL_3167382 | Lighthouse Laboratory Plymouth | Wellcome Sanger Institute for the COVID-19 Genomics UK (COG-UK) Consortium | Lighthouse Laboratory Plymouth and Alex Alderton, Roberto Amato, Jeffrey Barrett, Sonia Goncalves, Ewan Harrison, David K. Jackson, Ian Johnston, Dominic Kwiatkowski, Cordelia Langford, John Sillitoe on behalf of the Wellcome Sanger Institute COVID-19 Surveillance Team |
| EPI_ISL_3167383 | Lighthouse Lab in Milton Keynes | Wellcome Sanger Institute for the COVID-19 Genomics UK (COG-UK) Consortium | The Lighthouse Lab in Milton Keynes and Alex Alderton, Roberto Amato, Jeffrey Barrett, Sonia Goncalves, Ewan Harrison, David K. Jackson, Ian Johnston, Dominic Kwiatkowski, Cordelia Langford, John Sillitoe on behalf of the Wellcome Sanger Institute COVID-19 Surveillance Team |
| EPI_ISL_3167384 | Lighthouse Lab in Glasgow | Wellcome Sanger Institute for the COVID-19 Genomics UK (COG-UK) Consortium | Harper VanSteenhouse, Yumi Kasai, David Gray, Carol Clugston, Anna Dominiczak and Alex Alderton, Roberto Amato, Jeffrey Barrett, Sonia Goncalves, Ewan Harrison, David K. Jackson, Ian Johnston, Dominic Kwiatkowski, Cordelia Langford, John Sillitoe on behalf of the Wellcome Sanger Institute COVID-19 Surveillance Team |
| EPI_ISL_3167385, EPI_ISL_3167386, EPI_ISL_3167387 | Lighthouse Lab in Milton Keynes | Wellcome Sanger Institute for the COVID-19 Genomics UK (COG-UK) Consortium | The Lighthouse Lab in Milton Keynes and Alex Alderton, Roberto Amato, Jeffrey Barrett, Sonia Goncalves, Ewan Harrison, David K. Jackson, Ian Johnston, Dominic Kwiatkowski, Cordelia Langford, John Sillitoe on behalf of the Wellcome Sanger Institute COVID-19 Surveillance Team |
| EPI_ISL_3167388 | Lighthouse Lab in Glasgow | Wellcome Sanger Institute for the COVID-19 Genomics UK (COG-UK) Consortium | Harper VanSteenhouse, Yumi Kasai, David Gray, Carol Clugston, Anna Dominiczak and Alex Alderton, Roberto Amato, Jeffrey Barrett, Sonia Goncalves, Ewan Harrison, David K. Jackson, Ian Johnston, Dominic Kwiatkowski, Cordelia Langford, John Sillitoe on behalf of the Wellcome Sanger Institute COVID-19 Surveillance Team |
| EPI_ISL_3167389 | Lighthouse Laboratory Plymouth | Wellcome Sanger Institute for the COVID-19 Genomics UK (COG-UK) Consortium | Lighthouse Laboratory Plymouth and Alex Alderton, Roberto Amato, Jeffrey Barrett, Sonia Goncalves, Ewan Harrison, David K. Jackson, Ian Johnston, Dominic Kwiatkowski, Cordelia Langford, John Sillitoe on behalf of the Wellcome Sanger Institute COVID-19 Surveillance Team |
| EPI_ISL_3167390 | Lighthouse Lab in Milton Keynes | Wellcome Sanger Institute for the COVID-19 Genomics UK (COG-UK) | The Lighthouse Lab in Milton Keynes and Alex Alderton, Roberto Amato, Jeffrey Barrett, Sonia Goncalves, Ewan Harrison, David K. Jackson, Ian Johnston, Dominic Kwiatkowski, Cordelia Langford, John Sillitoe on behalf of the Wellcome Sanger |

| | | | |
|---|---|---|---|
| | | Consortium | Institute COVID-19 Surveillance Team |
| EPI_ISL_3167391 | Lighthouse Lab in Glasgow | Wellcome Sanger Institute for the COVID-19 Genomics UK (COG-UK) Consortium | Harper VanSteenhouse, Yumi Kasai, David Gray, Carol Clugston, Anna Dominiczak and Alex Alderton, Roberto Amato, Jeffrey Barrett, Sonia Goncalves, Ewan Harrison, David K. Jackson, Ian Johnston, Dominic Kwiatkowski, Cordelia Langford, John Sillitoe on behalf of the Wellcome Sanger Institute COVID-19 Surveillance Team |
| EPI_ISL_3167392 | Lighthouse Laboratory Plymouth | Wellcome Sanger Institute for the COVID-19 Genomics UK (COG-UK) Consortium | Lighthouse Laboratory Plymouth and Alex Alderton, Roberto Amato, Jeffrey Barrett, Sonia Goncalves, Ewan Harrison, David K. Jackson, Ian Johnston, Dominic Kwiatkowski, Cordelia Langford, John Sillitoe on behalf of the Wellcome Sanger Institute COVID-19 Surveillance Team |
| EPI_ISL_3167393 | Lighthouse Lab in Milton Keynes | Wellcome Sanger Institute for the COVID-19 Genomics UK (COG-UK) Consortium | The Lighthouse Lab in Milton Keynes and Alex Alderton, Roberto Amato, Jeffrey Barrett, Sonia Goncalves, Ewan Harrison, David K. Jackson, Ian Johnston, Dominic Kwiatkowski, Cordelia Langford, John Sillitoe on behalf of the Wellcome Sanger Institute COVID-19 Surveillance Team |
| EPI_ISL_3167394, EPI_ISL_3167395 | Lighthouse Lab in Glasgow | Wellcome Sanger Institute for the COVID-19 Genomics UK (COG-UK) Consortium | Harper VanSteenhouse, Yumi Kasai, David Gray, Carol Clugston, Anna Dominiczak and Alex Alderton, Roberto Amato, Jeffrey Barrett, Sonia Goncalves, Ewan Harrison, David K. Jackson, Ian Johnston, Dominic Kwiatkowski, Cordelia Langford, John Sillitoe on behalf of the Wellcome Sanger Institute COVID-19 Surveillance Team |
| EPI_ISL_3167396 | Lighthouse Laboratory Plymouth | Wellcome Sanger Institute for the COVID-19 Genomics UK (COG-UK) Consortium | Lighthouse Laboratory Plymouth and Alex Alderton, Roberto Amato, Jeffrey Barrett, Sonia Goncalves, Ewan Harrison, David K. Jackson, Ian Johnston, Dominic Kwiatkowski, Cordelia Langford, John Sillitoe on behalf of the Wellcome Sanger Institute COVID-19 Surveillance Team |
| EPI_ISL_3167397 | Lighthouse Lab in Milton Keynes | Wellcome Sanger Institute for the COVID-19 Genomics UK (COG-UK) Consortium | The Lighthouse Lab in Milton Keynes and Alex Alderton, Roberto Amato, Jeffrey Barrett, Sonia Goncalves, Ewan Harrison, David K. Jackson, Ian Johnston, Dominic Kwiatkowski, Cordelia Langford, John Sillitoe on behalf of the Wellcome Sanger Institute COVID-19 Surveillance Team |
| EPI_ISL_3167398 | Lighthouse Lab in Glasgow | Wellcome Sanger Institute for the COVID-19 Genomics UK (COG-UK) Consortium | Harper VanSteenhouse, Yumi Kasai, David Gray, Carol Clugston, Anna Dominiczak and Alex Alderton, Roberto Amato, Jeffrey Barrett, Sonia Goncalves, Ewan Harrison, David K. Jackson, Ian Johnston, Dominic Kwiatkowski, Cordelia Langford, John Sillitoe on behalf of the Wellcome Sanger Institute COVID-19 Surveillance Team |
| EPI_ISL_3167399 | Lighthouse Laboratory Plymouth | Wellcome Sanger Institute for the COVID-19 Genomics UK (COG-UK) Consortium | Lighthouse Laboratory Plymouth and Alex Alderton, Roberto Amato, Jeffrey Barrett, Sonia Goncalves, Ewan Harrison, David K. Jackson, Ian Johnston, Dominic Kwiatkowski, Cordelia Langford, John Sillitoe on behalf of the Wellcome Sanger Institute COVID-19 Surveillance Team |
| EPI_ISL_3167400, EPI_ISL_3167401 | Lighthouse Lab in Milton Keynes | Wellcome Sanger Institute for the COVID-19 Genomics UK (COG-UK) Consortium | The Lighthouse Lab in Milton Keynes and Alex Alderton, Roberto Amato, Jeffrey Barrett, Sonia Goncalves, Ewan Harrison, David K. Jackson, Ian Johnston, Dominic Kwiatkowski, Cordelia Langford, John Sillitoe on behalf of the Wellcome Sanger Institute COVID-19 Surveillance Team |
| EPI_ISL_3167402 | Lighthouse Lab in Glasgow | Wellcome Sanger Institute for the COVID-19 Genomics UK (COG-UK) Consortium | Harper VanSteenhouse, Yumi Kasai, David Gray, Carol Clugston, Anna Dominiczak and Alex Alderton, Roberto Amato, Jeffrey Barrett, Sonia Goncalves, Ewan Harrison, David K. Jackson, Ian Johnston, Dominic Kwiatkowski, Cordelia Langford, John Sillitoe on behalf of the Wellcome Sanger Institute COVID-19 Surveillance Team |
| EPI_ISL_3167403, EPI_ISL_3167404 | Lighthouse Lab in Milton Keynes | Wellcome Sanger Institute for the COVID-19 Genomics UK (COG-UK) Consortium | The Lighthouse Lab in Milton Keynes and Alex Alderton, Roberto Amato, Jeffrey Barrett, Sonia Goncalves, Ewan Harrison, David K. Jackson, Ian Johnston, Dominic Kwiatkowski, Cordelia Langford, John Sillitoe on behalf of the Wellcome Sanger Institute COVID-19 Surveillance Team |
| EPI_ISL_3167405 | Lighthouse Lab in Glasgow | Wellcome Sanger Institute for the COVID-19 Genomics UK (COG-UK) Consortium | Harper VanSteenhouse, Yumi Kasai, David Gray, Carol Clugston, Anna Dominiczak and Alex Alderton, Roberto Amato, Jeffrey Barrett, Sonia Goncalves, Ewan Harrison, David K. Jackson, Ian Johnston, Dominic Kwiatkowski, Cordelia Langford, John Sillitoe on behalf of the Wellcome Sanger Institute COVID-19 Surveillance Team |
| EPI_ISL_3167406 | Lighthouse Lab in Milton Keynes | Wellcome Sanger Institute for the COVID-19 Genomics UK (COG-UK) Consortium | The Lighthouse Lab in Milton Keynes and Alex Alderton, Roberto Amato, Jeffrey Barrett, Sonia Goncalves, Ewan Harrison, David K. Jackson, Ian Johnston, Dominic Kwiatkowski, Cordelia Langford, John Sillitoe on behalf of the Wellcome Sanger Institute COVID-19 Surveillance Team |
| EPI_ISL_3167407 | Lighthouse Laboratory Plymouth | Wellcome Sanger Institute for the COVID-19 Genomics UK (COG-UK) Consortium | Lighthouse Laboratory Plymouth and Alex Alderton, Roberto Amato, Jeffrey Barrett, Sonia Goncalves, Ewan Harrison, David K. Jackson, Ian Johnston, Dominic Kwiatkowski, Cordelia Langford, John Sillitoe on behalf of the Wellcome Sanger Institute COVID-19 Surveillance Team |
| EPI_ISL_3167408, EPI_ISL_3167409, EPI_ISL_3167410, EPI_ISL_3167411 | Lighthouse Lab in Milton Keynes | Wellcome Sanger Institute for the COVID-19 Genomics UK (COG-UK) Consortium | The Lighthouse Lab in Milton Keynes and Alex Alderton, Roberto Amato, Jeffrey Barrett, Sonia Goncalves, Ewan Harrison, David K. Jackson, Ian Johnston, Dominic Kwiatkowski, Cordelia Langford, John Sillitoe on behalf of the Wellcome Sanger Institute COVID-19 Surveillance Team |
| EPI_ISL_3167412 | Lighthouse Lab in Glasgow | Wellcome Sanger Institute for the COVID-19 Genomics UK (COG-UK) Consortium | Harper VanSteenhouse, Yumi Kasai, David Gray, Carol Clugston, Anna Dominiczak and Alex Alderton, Roberto Amato, Jeffrey Barrett, Sonia Goncalves, Ewan Harrison, David K. Jackson, Ian Johnston, Dominic Kwiatkowski, Cordelia Langford, John Sillitoe on behalf of the Wellcome Sanger Institute COVID-19 Surveillance Team |
| EPI_ISL_3167413 | Lighthouse Laboratory Plymouth | Wellcome Sanger Institute for the COVID-19 Genomics UK (COG-UK) Consortium | Lighthouse Laboratory Plymouth and Alex Alderton, Roberto Amato, Jeffrey Barrett, Sonia Goncalves, Ewan Harrison, David K. Jackson, Ian Johnston, Dominic Kwiatkowski, Cordelia Langford, John Sillitoe on behalf of the Wellcome Sanger Institute COVID-19 Surveillance Team |
| EPI_ISL_3167414, EPI_ISL_3167415, EPI_ISL_3167416 | Lighthouse Lab in Milton Keynes | Wellcome Sanger Institute for the COVID-19 Genomics UK (COG-UK) Consortium | The Lighthouse Lab in Milton Keynes and Alex Alderton, Roberto Amato, Jeffrey Barrett, Sonia Goncalves, Ewan Harrison, David K. Jackson, Ian Johnston, Dominic Kwiatkowski, Cordelia Langford, John Sillitoe on behalf of the Wellcome Sanger Institute COVID-19 Surveillance Team |
| EPI_ISL_3167417 | Lighthouse Lab in Glasgow | Wellcome Sanger Institute for the COVID-19 Genomics UK (COG-UK) Consortium | Harper VanSteenhouse, Yumi Kasai, David Gray, Carol Clugston, Anna Dominiczak and Alex Alderton, Roberto Amato, Jeffrey Barrett, Sonia Goncalves, Ewan Harrison, David K. Jackson, Ian Johnston, Dominic Kwiatkowski, Cordelia Langford, John Sillitoe on behalf of the Wellcome Sanger Institute COVID-19 Surveillance Team |
| EPI_ISL_3167418, EPI_ISL_3167419 | Lighthouse Lab in Milton Keynes | Wellcome Sanger Institute for the COVID-19 Genomics UK (COG-UK) Consortium | The Lighthouse Lab in Milton Keynes and Alex Alderton, Roberto Amato, Jeffrey Barrett, Sonia Goncalves, Ewan Harrison, David K. Jackson, Ian Johnston, Dominic Kwiatkowski, Cordelia Langford, John Sillitoe on behalf of the Wellcome Sanger Institute COVID-19 Surveillance Team |
| EPI_ISL_3167420 | Lighthouse Laboratory Plymouth | Wellcome Sanger Institute for the COVID-19 Genomics UK (COG-UK) Consortium | Lighthouse Laboratory Plymouth and Alex Alderton, Roberto Amato, Jeffrey Barrett, Sonia Goncalves, Ewan Harrison, David K. Jackson, Ian Johnston, Dominic Kwiatkowski, Cordelia Langford, John Sillitoe on behalf of the Wellcome Sanger Institute COVID-19 Surveillance Team |
| EPI_ISL_3167421 | Lighthouse Lab in Milton Keynes | Wellcome Sanger Institute for the COVID-19 Genomics UK (COG-UK) Consortium | The Lighthouse Lab in Milton Keynes and Alex Alderton, Roberto Amato, Jeffrey Barrett, Sonia Goncalves, Ewan Harrison, David K. Jackson, Ian Johnston, Dominic Kwiatkowski, Cordelia Langford, John Sillitoe on behalf of the Wellcome Sanger Institute COVID-19 Surveillance Team |
| EPI_ISL_3167422 | Lighthouse Lab in Glasgow | Wellcome Sanger Institute for the COVID-19 Genomics UK (COG-UK) Consortium | Harper VanSteenhouse, Yumi Kasai, David Gray, Carol Clugston, Anna Dominiczak and Alex Alderton, Roberto Amato, Jeffrey Barrett, Sonia Goncalves, Ewan Harrison, David K. Jackson, Ian Johnston, Dominic Kwiatkowski, Cordelia Langford, John Sillitoe on behalf of the Wellcome Sanger Institute COVID-19 Surveillance Team |
| EPI_ISL_3167423 | Lighthouse Lab in Milton Keynes | Wellcome Sanger Institute for the COVID-19 Genomics UK (COG-UK) Consortium | The Lighthouse Lab in Milton Keynes and Alex Alderton, Roberto Amato, Jeffrey Barrett, Sonia Goncalves, Ewan Harrison, David K. Jackson, Ian Johnston, Dominic Kwiatkowski, Cordelia Langford, John Sillitoe on behalf of the Wellcome Sanger Institute COVID-19 Surveillance Team |
| EPI_ISL_3167424, EPI_ISL_3167425 | Lighthouse Laboratory Plymouth | Wellcome Sanger Institute for the COVID-19 Genomics UK (COG-UK) Consortium | Lighthouse Laboratory Plymouth and Alex Alderton, Roberto Amato, Jeffrey Barrett, Sonia Goncalves, Ewan Harrison, David K. Jackson, Ian Johnston, Dominic Kwiatkowski, Cordelia Langford, John Sillitoe on behalf of the Wellcome Sanger Institute COVID-19 Surveillance Team |
| EPI_ISL_3167426, EPI_ISL_3167427, EPI_ISL_3167428 | Lighthouse Lab in Milton Keynes | Wellcome Sanger Institute for the COVID-19 Genomics UK (COG-UK) Consortium | The Lighthouse Lab in Milton Keynes and Alex Alderton, Roberto Amato, Jeffrey Barrett, Sonia Goncalves, Ewan Harrison, David K. Jackson, Ian Johnston, Dominic Kwiatkowski, Cordelia Langford, John Sillitoe on behalf of the Wellcome Sanger Institute COVID-19 Surveillance Team |
| EPI_ISL_3167429 | Lighthouse Laboratory Plymouth | Wellcome Sanger Institute for the COVID-19 Genomics UK (COG-UK) Consortium | Lighthouse Laboratory Plymouth and Alex Alderton, Roberto Amato, Jeffrey Barrett, Sonia Goncalves, Ewan Harrison, David K. Jackson, Ian Johnston, Dominic Kwiatkowski, Cordelia Langford, John Sillitoe on behalf of the Wellcome Sanger Institute COVID-19 Surveillance Team |
| EPI_ISL_3167430, EPI_ISL_3167431, EPI_ISL_3167432 | Lighthouse Lab in Milton Keynes | Wellcome Sanger Institute for the COVID-19 Genomics UK (COG-UK) Consortium | The Lighthouse Lab in Milton Keynes and Alex Alderton, Roberto Amato, Jeffrey Barrett, Sonia Goncalves, Ewan Harrison, David K. Jackson, Ian Johnston, Dominic Kwiatkowski, Cordelia Langford, John Sillitoe on behalf of the Wellcome Sanger Institute COVID-19 Surveillance Team |
| EPI_ISL_3167433 | Lighthouse Laboratory Plymouth | Wellcome Sanger Institute for the COVID-19 Genomics UK (COG-UK) Consortium | Lighthouse Laboratory Plymouth and Alex Alderton, Roberto Amato, Jeffrey Barrett, Sonia Goncalves, Ewan Harrison, David K. Jackson, Ian Johnston, Dominic Kwiatkowski, Cordelia Langford, John Sillitoe on behalf of the Wellcome Sanger Institute COVID-19 Surveillance Team |
| EPI_ISL_3167434 | Lighthouse Lab in Glasgow | Wellcome Sanger Institute for the COVID-19 Genomics UK (COG-UK) Consortium | Harper VanSteenhouse, Yumi Kasai, David Gray, Carol Clugston, Anna Dominiczak and Alex Alderton, Roberto Amato, Jeffrey Barrett, Sonia Goncalves, Ewan Harrison, David K. Jackson, Ian Johnston, Dominic Kwiatkowski, Cordelia Langford, John Sillitoe on behalf of the Wellcome Sanger Institute COVID-19 Surveillance Team |
| EPI_ISL_3167435, EPI_ISL_3167436 | Lighthouse Lab in Milton Keynes | Wellcome Sanger Institute for the COVID-19 Genomics UK (COG-UK) Consortium | The Lighthouse Lab in Milton Keynes and Alex Alderton, Roberto Amato, Jeffrey Barrett, Sonia Goncalves, Ewan Harrison, David K. Jackson, Ian Johnston, Dominic Kwiatkowski, Cordelia Langford, John Sillitoe on behalf of the Wellcome Sanger Institute COVID-19 Surveillance Team |
| EPI_ISL_3167437, EPI_ISL_3167438 | Lighthouse Laboratory Plymouth | Wellcome Sanger Institute for the COVID-19 Genomics UK (COG-UK) Consortium | Lighthouse Laboratory Plymouth and Alex Alderton, Roberto Amato, Jeffrey Barrett, Sonia Goncalves, Ewan Harrison, David K. Jackson, Ian Johnston, Dominic Kwiatkowski, Cordelia Langford, John Sillitoe on behalf of the Wellcome Sanger Institute COVID-19 Surveillance Team |
| EPI_ISL_3167439 | Lighthouse Lab in Milton Keynes | Wellcome Sanger Institute for the COVID-19 Genomics UK (COG-UK) Consortium | The Lighthouse Lab in Milton Keynes and Alex Alderton, Roberto Amato, Jeffrey Barrett, Sonia Goncalves, Ewan Harrison, David K. Jackson, Ian Johnston, Dominic Kwiatkowski, Cordelia Langford, John Sillitoe on behalf of the Wellcome Sanger Institute COVID-19 Surveillance Team |
| EPI_ISL_3167440, EPI_ISL_3167441, EPI_ISL_3167442 | Lighthouse Lab in Glasgow | Wellcome Sanger Institute for the COVID-19 Genomics UK (COG-UK) Consortium | Harper VanSteenhouse, Yumi Kasai, David Gray, Carol Clugston, Anna Dominiczak and Alex Alderton, Roberto Amato, Jeffrey Barrett, Sonia Goncalves, Ewan Harrison, David K. Jackson, Ian Johnston, Dominic Kwiatkowski, Cordelia Langford, John Sillitoe on behalf of the Wellcome Sanger Institute COVID-19 Surveillance Team |
| EPI_ISL_3167443 | Lighthouse Lab in Milton Keynes | Wellcome Sanger Institute for the COVID-19 Genomics UK (COG-UK) Consortium | The Lighthouse Lab in Milton Keynes and Alex Alderton, Roberto Amato, Jeffrey Barrett, Sonia Goncalves, Ewan Harrison, David K. Jackson, Ian Johnston, Dominic Kwiatkowski, Cordelia Langford, John Sillitoe on behalf of the Wellcome Sanger Institute COVID-19 Surveillance Team |
| EPI_ISL_3167444, EPI_ISL_3167445 | Lighthouse Laboratory Plymouth | Wellcome Sanger Institute for the COVID-19 Genomics UK (COG-UK) Consortium | Lighthouse Laboratory Plymouth and Alex Alderton, Roberto Amato, Jeffrey Barrett, Sonia Goncalves, Ewan Harrison, David K. Jackson, Ian Johnston, Dominic Kwiatkowski, Cordelia Langford, John Sillitoe on behalf of the Wellcome Sanger Institute COVID-19 Surveillance Team |
| EPI_ISL_3167446 | Lighthouse Lab in Milton Keynes | Wellcome Sanger Institute for the COVID-19 Genomics UK (COG-UK) Consortium | The Lighthouse Lab in Milton Keynes and Alex Alderton, Roberto Amato, Jeffrey Barrett, Sonia Goncalves, Ewan Harrison, David K. Jackson, Ian Johnston, Dominic Kwiatkowski, Cordelia Langford, John Sillitoe on behalf of the Wellcome Sanger Institute COVID-19 Surveillance Team |
| EPI_ISL_3167447 | Lighthouse Laboratory Plymouth | Wellcome Sanger Institute for the COVID-19 Genomics UK (COG-UK) Consortium | Lighthouse Laboratory Plymouth and Alex Alderton, Roberto Amato, Jeffrey Barrett, Sonia Goncalves, Ewan Harrison, David K. Jackson, Ian Johnston, Dominic Kwiatkowski, Cordelia Langford, John Sillitoe on behalf of the Wellcome Sanger Institute COVID-19 Surveillance Team |
| EPI_ISL_3167448 | Lighthouse Lab in Milton Keynes | Wellcome Sanger Institute for the COVID-19 Genomics UK (COG-UK) Consortium | The Lighthouse Lab in Milton Keynes and Alex Alderton, Roberto Amato, Jeffrey Barrett, Sonia Goncalves, Ewan Harrison, David K. Jackson, Ian Johnston, Dominic Kwiatkowski, Cordelia Langford, John Sillitoe on behalf of the Wellcome Sanger Institute COVID-19 Surveillance Team |
| EPI_ISL_3167449, EPI_ISL_3167450, EPI_ISL_3167451, EPI_ISL_3167452, EPI_ISL_3167453 | Lighthouse Lab in Glasgow | Wellcome Sanger Institute for the COVID-19 Genomics UK (COG-UK) Consortium | Harper VanSteenhouse, Yumi Kasai, David Gray, Carol Clugston, Anna Dominiczak and Alex Alderton, Roberto Amato, Jeffrey Barrett, Sonia Goncalves, Ewan Harrison, David K. Jackson, Ian Johnston, Dominic Kwiatkowski, Cordelia Langford, John Sillitoe on behalf of the Wellcome Sanger Institute COVID-19 Surveillance Team |
| EPI_ISL_3167454, EPI_ISL_3167455, | Lighthouse Lab in Milton Keynes | Wellcome Sanger Institute for the COVID-19 Genomics UK (COG-UK) Consortium | The Lighthouse Lab in Milton Keynes and Alex Alderton, Roberto Amato, Jeffrey Barrett, Sonia Goncalves, Ewan Harrison, David K. Jackson, Ian Johnston, Dominic Kwiatkowski, Cordelia Langford, John Sillitoe on behalf of the Wellcome Sanger Institute COVID-19 Surveillance Team |

| | | | |
|---|---|---|---|
| EPI_ISL_3167456, EPI_ISL_3167457 | | | |
| EPI_ISL_3167458, EPI_ISL_3167459 | Lighthouse Lab in Glasgow | Wellcome Sanger Institute for the COVID-19 Genomics UK (COG-UK) Consortium | Harper VanSteenhouse, Yumi Kasai, David Gray, Carol Clugston, Anna Dominiczak and Alex Alderton, Roberto Amato, Jeffrey Barrett, Sonia Goncalves, Ewan Harrison, David K. Jackson, Ian Johnston, Dominic Kwiatkowski, Cordelia Langford, John Sillitoe on behalf of the Wellcome Sanger Institute COVID-19 Surveillance Team |
| EPI_ISL_3167460 | Lighthouse Laboratory Plymouth | Wellcome Sanger Institute for the COVID-19 Genomics UK (COG-UK) Consortium | Lighthouse Laboratory Plymouth and Alex Alderton, Roberto Amato, Jeffrey Barrett, Sonia Goncalves, Ewan Harrison, David K. Jackson, Ian Johnston, Dominic Kwiatkowski, Cordelia Langford, John Sillitoe on behalf of the Wellcome Sanger Institute COVID-19 Surveillance Team |
| EPI_ISL_3167461, EPI_ISL_3167462 | Lighthouse Lab in Milton Keynes | Wellcome Sanger Institute for the COVID-19 Genomics UK (COG-UK) Consortium | The Lighthouse Lab in Milton Keynes and Alex Alderton, Roberto Amato, Jeffrey Barrett, Sonia Goncalves, Ewan Harrison, David K. Jackson, Ian Johnston, Dominic Kwiatkowski, Cordelia Langford, John Sillitoe on behalf of the Wellcome Sanger Institute COVID-19 Surveillance Team |
| EPI_ISL_3167463, EPI_ISL_3167464, EPI_ISL_3167465 | Lighthouse Laboratory Plymouth | Wellcome Sanger Institute for the COVID-19 Genomics UK (COG-UK) Consortium | Lighthouse Laboratory Plymouth and Alex Alderton, Roberto Amato, Jeffrey Barrett, Sonia Goncalves, Ewan Harrison, David K. Jackson, Ian Johnston, Dominic Kwiatkowski, Cordelia Langford, John Sillitoe on behalf of the Wellcome Sanger Institute COVID-19 Surveillance Team |
| EPI_ISL_3167466 | Lighthouse Lab in Glasgow | Wellcome Sanger Institute for the COVID-19 Genomics UK (COG-UK) Consortium | Harper VanSteenhouse, Yumi Kasai, David Gray, Carol Clugston, Anna Dominiczak and Alex Alderton, Roberto Amato, Jeffrey Barrett, Sonia Goncalves, Ewan Harrison, David K. Jackson, Ian Johnston, Dominic Kwiatkowski, Cordelia Langford, John Sillitoe on behalf of the Wellcome Sanger Institute COVID-19 Surveillance Team |
| EPI_ISL_3167467 | Lighthouse Lab in Milton Keynes | Wellcome Sanger Institute for the COVID-19 Genomics UK (COG-UK) Consortium | The Lighthouse Lab in Milton Keynes and Alex Alderton, Roberto Amato, Jeffrey Barrett, Sonia Goncalves, Ewan Harrison, David K. Jackson, Ian Johnston, Dominic Kwiatkowski, Cordelia Langford, John Sillitoe on behalf of the Wellcome Sanger Institute COVID-19 Surveillance Team |
| EPI_ISL_3167468 | Lighthouse Lab in Glasgow | Wellcome Sanger Institute for the COVID-19 Genomics UK (COG-UK) Consortium | Harper VanSteenhouse, Yumi Kasai, David Gray, Carol Clugston, Anna Dominiczak and Alex Alderton, Roberto Amato, Jeffrey Barrett, Sonia Goncalves, Ewan Harrison, David K. Jackson, Ian Johnston, Dominic Kwiatkowski, Cordelia Langford, John Sillitoe on behalf of the Wellcome Sanger Institute COVID-19 Surveillance Team |
| EPI_ISL_3167469, EPI_ISL_3167470, EPI_ISL_3167471, EPI_ISL_3167472 | Lighthouse Lab in Milton Keynes | Wellcome Sanger Institute for the COVID-19 Genomics UK (COG-UK) Consortium | The Lighthouse Lab in Milton Keynes and Alex Alderton, Roberto Amato, Jeffrey Barrett, Sonia Goncalves, Ewan Harrison, David K. Jackson, Ian Johnston, Dominic Kwiatkowski, Cordelia Langford, John Sillitoe on behalf of the Wellcome Sanger Institute COVID-19 Surveillance Team |
| EPI_ISL_3167473 | Lighthouse Laboratory Plymouth | Wellcome Sanger Institute for the COVID-19 Genomics UK (COG-UK) Consortium | Lighthouse Laboratory Plymouth and Alex Alderton, Roberto Amato, Jeffrey Barrett, Sonia Goncalves, Ewan Harrison, David K. Jackson, Ian Johnston, Dominic Kwiatkowski, Cordelia Langford, John Sillitoe on behalf of the Wellcome Sanger Institute COVID-19 Surveillance Team |
| EPI_ISL_3167474 | Lighthouse Lab in Glasgow | Wellcome Sanger Institute for the COVID-19 Genomics UK (COG-UK) Consortium | Harper VanSteenhouse, Yumi Kasai, David Gray, Carol Clugston, Anna Dominiczak and Alex Alderton, Roberto Amato, Jeffrey Barrett, Sonia Goncalves, Ewan Harrison, David K. Jackson, Ian Johnston, Dominic Kwiatkowski, Cordelia Langford, John Sillitoe on behalf of the Wellcome Sanger Institute COVID-19 Surveillance Team |
| EPI_ISL_3167475, EPI_ISL_3167476 | Lighthouse Lab in Milton Keynes | Wellcome Sanger Institute for the COVID-19 Genomics UK (COG-UK) Consortium | The Lighthouse Lab in Milton Keynes and Alex Alderton, Roberto Amato, Jeffrey Barrett, Sonia Goncalves, Ewan Harrison, David K. Jackson, Ian Johnston, Dominic Kwiatkowski, Cordelia Langford, John Sillitoe on behalf of the Wellcome Sanger Institute COVID-19 Surveillance Team |
| EPI_ISL_3167477, EPI_ISL_3167478 | Lighthouse Lab in Glasgow | Wellcome Sanger Institute for the COVID-19 Genomics UK (COG-UK) Consortium | Harper VanSteenhouse, Yumi Kasai, David Gray, Carol Clugston, Anna Dominiczak and Alex Alderton, Roberto Amato, Jeffrey Barrett, Sonia Goncalves, Ewan Harrison, David K. Jackson, Ian Johnston, Dominic Kwiatkowski, Cordelia Langford, John Sillitoe on behalf of the Wellcome Sanger Institute COVID-19 Surveillance Team |
| EPI_ISL_3167479, EPI_ISL_3167480 | Lighthouse Lab in Milton Keynes | Wellcome Sanger Institute for the COVID-19 Genomics UK (COG-UK) Consortium | The Lighthouse Lab in Milton Keynes and Alex Alderton, Roberto Amato, Jeffrey Barrett, Sonia Goncalves, Ewan Harrison, David K. Jackson, Ian Johnston, Dominic Kwiatkowski, Cordelia Langford, John Sillitoe on behalf of the Wellcome Sanger Institute COVID-19 Surveillance Team |
| EPI_ISL_3167481 | Lighthouse Lab in Glasgow | Wellcome Sanger Institute for the COVID-19 Genomics UK (COG-UK) Consortium | Harper VanSteenhouse, Yumi Kasai, David Gray, Carol Clugston, Anna Dominiczak and Alex Alderton, Roberto Amato, Jeffrey Barrett, Sonia Goncalves, Ewan Harrison, David K. Jackson, Ian Johnston, Dominic Kwiatkowski, Cordelia Langford, John Sillitoe on behalf of the Wellcome Sanger Institute COVID-19 Surveillance Team |
| EPI_ISL_3167482 | Lighthouse Laboratory Plymouth | Wellcome Sanger Institute for the COVID-19 Genomics UK (COG-UK) Consortium | Lighthouse Laboratory Plymouth and Alex Alderton, Roberto Amato, Jeffrey Barrett, Sonia Goncalves, Ewan Harrison, David K. Jackson, Ian Johnston, Dominic Kwiatkowski, Cordelia Langford, John Sillitoe on behalf of the Wellcome Sanger Institute COVID-19 Surveillance Team |
| EPI_ISL_3167483, EPI_ISL_3167484, EPI_ISL_3167485 | Lighthouse Lab in Milton Keynes | Wellcome Sanger Institute for the COVID-19 Genomics UK (COG-UK) Consortium | The Lighthouse Lab in Milton Keynes and Alex Alderton, Roberto Amato, Jeffrey Barrett, Sonia Goncalves, Ewan Harrison, David K. Jackson, Ian Johnston, Dominic Kwiatkowski, Cordelia Langford, John Sillitoe on behalf of the Wellcome Sanger Institute COVID-19 Surveillance Team |
| EPI_ISL_3167486 | Lighthouse Laboratory Plymouth | Wellcome Sanger Institute for the COVID-19 Genomics UK (COG-UK) Consortium | Lighthouse Laboratory Plymouth and Alex Alderton, Roberto Amato, Jeffrey Barrett, Sonia Goncalves, Ewan Harrison, David K. Jackson, Ian Johnston, Dominic Kwiatkowski, Cordelia Langford, John Sillitoe on behalf of the Wellcome Sanger Institute COVID-19 Surveillance Team |
| EPI_ISL_3167487, EPI_ISL_3167488 | Lighthouse Lab in Milton Keynes | Wellcome Sanger Institute for the COVID-19 Genomics UK (COG-UK) Consortium | The Lighthouse Lab in Milton Keynes and Alex Alderton, Roberto Amato, Jeffrey Barrett, Sonia Goncalves, Ewan Harrison, David K. Jackson, Ian Johnston, Dominic Kwiatkowski, Cordelia Langford, John Sillitoe on behalf of the Wellcome Sanger Institute COVID-19 Surveillance Team |
| EPI_ISL_3167489 | Lighthouse Laboratory Plymouth | Wellcome Sanger Institute for the COVID-19 Genomics UK (COG-UK) Consortium | Lighthouse Laboratory Plymouth and Alex Alderton, Roberto Amato, Jeffrey Barrett, Sonia Goncalves, Ewan Harrison, David K. Jackson, Ian Johnston, Dominic Kwiatkowski, Cordelia Langford, John Sillitoe on behalf of the Wellcome Sanger Institute COVID-19 Surveillance Team |
| EPI_ISL_3167490, EPI_ISL_3167491 | Lighthouse Lab in Milton Keynes | Wellcome Sanger Institute for the COVID-19 Genomics UK (COG-UK) Consortium | The Lighthouse Lab in Milton Keynes and Alex Alderton, Roberto Amato, Jeffrey Barrett, Sonia Goncalves, Ewan Harrison, David K. Jackson, Ian Johnston, Dominic Kwiatkowski, Cordelia Langford, John Sillitoe on behalf of the Wellcome Sanger Institute COVID-19 Surveillance Team |
| EPI_ISL_3167492, EPI_ISL_3167493 | Lighthouse Laboratory Plymouth | Wellcome Sanger Institute for the COVID-19 Genomics UK (COG-UK) Consortium | Lighthouse Laboratory Plymouth and Alex Alderton, Roberto Amato, Jeffrey Barrett, Sonia Goncalves, Ewan Harrison, David K. Jackson, Ian Johnston, Dominic Kwiatkowski, Cordelia Langford, John Sillitoe on behalf of the Wellcome Sanger Institute COVID-19 Surveillance Team |
| EPI_ISL_3167494 | Lighthouse Lab in Milton Keynes | Wellcome Sanger Institute for the COVID-19 Genomics UK (COG-UK) Consortium | The Lighthouse Lab in Milton Keynes and Alex Alderton, Roberto Amato, Jeffrey Barrett, Sonia Goncalves, Ewan Harrison, David K. Jackson, Ian Johnston, Dominic Kwiatkowski, Cordelia Langford, John Sillitoe on behalf of the Wellcome Sanger Institute COVID-19 Surveillance Team |
| EPI_ISL_3167495 | Lighthouse Laboratory Plymouth | Wellcome Sanger Institute for the COVID-19 Genomics UK (COG-UK) Consortium | Lighthouse Laboratory Plymouth and Alex Alderton, Roberto Amato, Jeffrey Barrett, Sonia Goncalves, Ewan Harrison, David K. Jackson, Ian Johnston, Dominic Kwiatkowski, Cordelia Langford, John Sillitoe on behalf of the Wellcome Sanger Institute COVID-19 Surveillance Team |
| EPI_ISL_3167496, EPI_ISL_3167497, EPI_ISL_3167498 | Lighthouse Lab in Milton Keynes | Wellcome Sanger Institute for the COVID-19 Genomics UK (COG-UK) Consortium | The Lighthouse Lab in Milton Keynes and Alex Alderton, Roberto Amato, Jeffrey Barrett, Sonia Goncalves, Ewan Harrison, David K. Jackson, Ian Johnston, Dominic Kwiatkowski, Cordelia Langford, John Sillitoe on behalf of the Wellcome Sanger Institute COVID-19 Surveillance Team |
| EPI_ISL_3167499 | Lighthouse Laboratory Plymouth | Wellcome Sanger Institute for the COVID-19 Genomics UK (COG-UK) Consortium | Lighthouse Laboratory Plymouth and Alex Alderton, Roberto Amato, Jeffrey Barrett, Sonia Goncalves, Ewan Harrison, David K. Jackson, Ian Johnston, Dominic Kwiatkowski, Cordelia Langford, John Sillitoe on behalf of the Wellcome Sanger Institute COVID-19 Surveillance Team |
| EPI_ISL_3167500 | Lighthouse Lab in Milton Keynes | Wellcome Sanger Institute for the COVID-19 Genomics UK (COG-UK) Consortium | The Lighthouse Lab in Milton Keynes and Alex Alderton, Roberto Amato, Jeffrey Barrett, Sonia Goncalves, Ewan Harrison, David K. Jackson, Ian Johnston, Dominic Kwiatkowski, Cordelia Langford, John Sillitoe on behalf of the Wellcome Sanger Institute COVID-19 Surveillance Team |
| EPI_ISL_3167501, EPI_ISL_3167502 | Lighthouse Laboratory Plymouth | Wellcome Sanger Institute for the COVID-19 Genomics UK (COG-UK) Consortium | Lighthouse Laboratory Plymouth and Alex Alderton, Roberto Amato, Jeffrey Barrett, Sonia Goncalves, Ewan Harrison, David K. Jackson, Ian Johnston, Dominic Kwiatkowski, Cordelia Langford, John Sillitoe on behalf of the Wellcome Sanger Institute COVID-19 Surveillance Team |
| EPI_ISL_3167503, EPI_ISL_3167504, EPI_ISL_3167505, EPI_ISL_3167506, EPI_ISL_3167507 | Lighthouse Lab in Milton Keynes | Wellcome Sanger Institute for the COVID-19 Genomics UK (COG-UK) Consortium | The Lighthouse Lab in Milton Keynes and Alex Alderton, Roberto Amato, Jeffrey Barrett, Sonia Goncalves, Ewan Harrison, David K. Jackson, Ian Johnston, Dominic Kwiatkowski, Cordelia Langford, John Sillitoe on behalf of the Wellcome Sanger Institute COVID-19 Surveillance Team |
| EPI_ISL_3167508 | Lighthouse Laboratory Plymouth | Wellcome Sanger Institute for the COVID-19 Genomics UK (COG-UK) Consortium | Lighthouse Laboratory Plymouth and Alex Alderton, Roberto Amato, Jeffrey Barrett, Sonia Goncalves, Ewan Harrison, David K. Jackson, Ian Johnston, Dominic Kwiatkowski, Cordelia Langford, John Sillitoe on behalf of the Wellcome Sanger Institute COVID-19 Surveillance Team |
| EPI_ISL_3167509, EPI_ISL_3167510 | Lighthouse Lab in Milton Keynes | Wellcome Sanger Institute for the COVID-19 Genomics UK (COG-UK) Consortium | The Lighthouse Lab in Milton Keynes and Alex Alderton, Roberto Amato, Jeffrey Barrett, Sonia Goncalves, Ewan Harrison, David K. Jackson, Ian Johnston, Dominic Kwiatkowski, Cordelia Langford, John Sillitoe on behalf of the Wellcome Sanger Institute COVID-19 Surveillance Team |
| EPI_ISL_3167511 | Lighthouse Laboratory Plymouth | Wellcome Sanger Institute for the COVID-19 Genomics UK (COG-UK) Consortium | Lighthouse Laboratory Plymouth and Alex Alderton, Roberto Amato, Jeffrey Barrett, Sonia Goncalves, Ewan Harrison, David K. Jackson, Ian Johnston, Dominic Kwiatkowski, Cordelia Langford, John Sillitoe on behalf of the Wellcome Sanger Institute COVID-19 Surveillance Team |
| EPI_ISL_3167512 | Lighthouse Lab in Milton Keynes | Wellcome Sanger Institute for the COVID-19 Genomics UK (COG-UK) Consortium | The Lighthouse Lab in Milton Keynes and Alex Alderton, Roberto Amato, Jeffrey Barrett, Sonia Goncalves, Ewan Harrison, David K. Jackson, Ian Johnston, Dominic Kwiatkowski, Cordelia Langford, John Sillitoe on behalf of the Wellcome Sanger Institute COVID-19 Surveillance Team |
| EPI_ISL_3167513, EPI_ISL_3167514 | Lighthouse Laboratory Plymouth | Wellcome Sanger Institute for the COVID-19 Genomics UK (COG-UK) Consortium | Lighthouse Laboratory Plymouth and Alex Alderton, Roberto Amato, Jeffrey Barrett, Sonia Goncalves, Ewan Harrison, David K. Jackson, Ian Johnston, Dominic Kwiatkowski, Cordelia Langford, John Sillitoe on behalf of the Wellcome Sanger Institute COVID-19 Surveillance Team |
| EPI_ISL_3167515 | Lighthouse Lab in Glasgow | Wellcome Sanger Institute for the COVID-19 Genomics UK (COG-UK) Consortium | Harper VanSteenhouse, Yumi Kasai, David Gray, Carol Clugston, Anna Dominiczak and Alex Alderton, Roberto Amato, Jeffrey Barrett, Sonia Goncalves, Ewan Harrison, David K. Jackson, Ian Johnston, Dominic Kwiatkowski, Cordelia Langford, John Sillitoe on behalf of the Wellcome Sanger Institute COVID-19 Surveillance Team |
| EPI_ISL_3167516, EPI_ISL_3167517 | Lighthouse Laboratory Plymouth | Wellcome Sanger Institute for the COVID-19 Genomics UK (COG-UK) Consortium | Lighthouse Laboratory Plymouth and Alex Alderton, Roberto Amato, Jeffrey Barrett, Sonia Goncalves, Ewan Harrison, David K. Jackson, Ian Johnston, Dominic Kwiatkowski, Cordelia Langford, John Sillitoe on behalf of the Wellcome Sanger Institute COVID-19 Surveillance Team |
| EPI_ISL_3167518, EPI_ISL_3167519 | Lighthouse Lab in Glasgow | Wellcome Sanger Institute for the COVID-19 Genomics UK (COG-UK) Consortium | Harper VanSteenhouse, Yumi Kasai, David Gray, Carol Clugston, Anna Dominiczak and Alex Alderton, Roberto Amato, Jeffrey Barrett, Sonia Goncalves, Ewan Harrison, David K. Jackson, Ian Johnston, Dominic Kwiatkowski, Cordelia Langford, John Sillitoe on behalf of the Wellcome Sanger Institute COVID-19 Surveillance Team |
| EPI_ISL_3167520 | Lighthouse Laboratory Plymouth | Wellcome Sanger Institute for the COVID-19 Genomics UK (COG-UK) Consortium | Lighthouse Laboratory Plymouth and Alex Alderton, Roberto Amato, Jeffrey Barrett, Sonia Goncalves, Ewan Harrison, David K. Jackson, Ian Johnston, Dominic Kwiatkowski, Cordelia Langford, John Sillitoe on behalf of the Wellcome Sanger Institute COVID-19 Surveillance Team |
| EPI_ISL_3167521, EPI_ISL_3167522 | Lighthouse Lab in Milton Keynes | Wellcome Sanger Institute for the COVID-19 Genomics UK (COG-UK) Consortium | The Lighthouse Lab in Milton Keynes and Alex Alderton, Roberto Amato, Jeffrey Barrett, Sonia Goncalves, Ewan Harrison, David K. Jackson, Ian Johnston, Dominic Kwiatkowski, Cordelia Langford, John Sillitoe on behalf of the Wellcome Sanger Institute COVID-19 Surveillance Team |
| EPI_ISL_3167523 | Berkshire and Surrey Pathology Services Lighthouse Laboratory | Wellcome Sanger Institute for the COVID-19 Genomics UK (COG-UK) Consortium | Berkshire and Surrey Pathology Services Lighthouse Laboratory and Alex Alderton, Roberto Amato, Jeffrey Barrett, Sonia Goncalves, Ewan Harrison, David K. Jackson, Ian Johnston, Dominic Kwiatkowski, Cordelia Langford, John Sillitoe on behalf of the Wellcome Sanger Institute COVID-19 Surveillance Team |
| EPI_ISL_3167524, EPI_ISL_3167525, EPI_ISL_3167526 | Lighthouse Laboratory Plymouth | Wellcome Sanger Institute for the COVID-19 Genomics UK (COG-UK) Consortium | Lighthouse Laboratory Plymouth and Alex Alderton, Roberto Amato, Jeffrey Barrett, Sonia Goncalves, Ewan Harrison, David K. Jackson, Ian Johnston, Dominic Kwiatkowski, Cordelia Langford, John Sillitoe on behalf of the Wellcome Sanger Institute COVID-19 Surveillance Team |

| | | | |
|---|---|---|---|
| EPI_ISL_3167527 | Lighthouse Lab in Alderley Park | Wellcome Sanger Institute for the COVID-19 Genomics UK (COG-UK) Consortium | Jacquelyn Wynn, Mairead Hyland, The Lighthouse Lab in Alderley Park and Alex Alderton, Roberto Amato, Jeffrey Barrett, Sonia Goncalves, Ewan Harrison, David K. Jackson, Ian Johnston, Dominic Kwiatkowski, Cordelia Langford, John Sillitoe on behalf of the Wellcome Sanger Institute COVID-19 Surveillance Team |
| EPI_ISL_3167528, EPI_ISL_3167529, EPI_ISL_3167530 | Berkshire and Surrey Pathology Services Lighthouse Laboratory | Wellcome Sanger Institute for the COVID-19 Genomics UK (COG-UK) Consortium | Berkshire and Surrey Pathology Services Lighthouse Laboratory and Alex Alderton, Roberto Amato, Jeffrey Barrett, Sonia Goncalves, Ewan Harrison, David K. Jackson, Ian Johnston, Dominic Kwiatkowski, Cordelia Langford, John Sillitoe on behalf of the Wellcome Sanger Institute COVID-19 Surveillance Team |
| EPI_ISL_3167531, EPI_ISL_3167532 | Lighthouse Lab in Alderley Park | Wellcome Sanger Institute for the COVID-19 Genomics UK (COG-UK) Consortium | Jacquelyn Wynn, Mairead Hyland, The Lighthouse Lab in Alderley Park and Alex Alderton, Roberto Amato, Jeffrey Barrett, Sonia Goncalves, Ewan Harrison, David K. Jackson, Ian Johnston, Dominic Kwiatkowski, Cordelia Langford, John Sillitoe on behalf of the Wellcome Sanger Institute COVID-19 Surveillance Team |
| EPI_ISL_3167533 | Berkshire and Surrey Pathology Services Lighthouse Laboratory | Wellcome Sanger Institute for the COVID-19 Genomics UK (COG-UK) Consortium | Berkshire and Surrey Pathology Services Lighthouse Laboratory and Alex Alderton, Roberto Amato, Jeffrey Barrett, Sonia Goncalves, Ewan Harrison, David K. Jackson, Ian Johnston, Dominic Kwiatkowski, Cordelia Langford, John Sillitoe on behalf of the Wellcome Sanger Institute COVID-19 Surveillance Team |
| EPI_ISL_3167534 | Lighthouse Lab in Alderley Park | Wellcome Sanger Institute for the COVID-19 Genomics UK (COG-UK) Consortium | Jacquelyn Wynn, Mairead Hyland, The Lighthouse Lab in Alderley Park and Alex Alderton, Roberto Amato, Jeffrey Barrett, Sonia Goncalves, Ewan Harrison, David K. Jackson, Ian Johnston, Dominic Kwiatkowski, Cordelia Langford, John Sillitoe on behalf of the Wellcome Sanger Institute COVID-19 Surveillance Team |
| EPI_ISL_3167535 | Berkshire and Surrey Pathology Services Lighthouse Laboratory | Wellcome Sanger Institute for the COVID-19 Genomics UK (COG-UK) Consortium | Berkshire and Surrey Pathology Services Lighthouse Laboratory and Alex Alderton, Roberto Amato, Jeffrey Barrett, Sonia Goncalves, Ewan Harrison, David K. Jackson, Ian Johnston, Dominic Kwiatkowski, Cordelia Langford, John Sillitoe on behalf of the Wellcome Sanger Institute COVID-19 Surveillance Team |
| EPI_ISL_3167536 | Lighthouse Lab in Alderley Park | Wellcome Sanger Institute for the COVID-19 Genomics UK (COG-UK) Consortium | Jacquelyn Wynn, Mairead Hyland, The Lighthouse Lab in Alderley Park and Alex Alderton, Roberto Amato, Jeffrey Barrett, Sonia Goncalves, Ewan Harrison, David K. Jackson, Ian Johnston, Dominic Kwiatkowski, Cordelia Langford, John Sillitoe on behalf of the Wellcome Sanger Institute COVID-19 Surveillance Team |
| EPI_ISL_3167537, EPI_ISL_3167538 | Lighthouse Laboratory Plymouth | Wellcome Sanger Institute for the COVID-19 Genomics UK (COG-UK) Consortium | Lighthouse Laboratory Plymouth and Alex Alderton, Roberto Amato, Jeffrey Barrett, Sonia Goncalves, Ewan Harrison, David K. Jackson, Ian Johnston, Dominic Kwiatkowski, Cordelia Langford, John Sillitoe on behalf of the Wellcome Sanger Institute COVID-19 Surveillance Team |
| EPI_ISL_3167539 | Berkshire and Surrey Pathology Services Lighthouse Laboratory | Wellcome Sanger Institute for the COVID-19 Genomics UK (COG-UK) Consortium | Berkshire and Surrey Pathology Services Lighthouse Laboratory and Alex Alderton, Roberto Amato, Jeffrey Barrett, Sonia Goncalves, Ewan Harrison, David K. Jackson, Ian Johnston, Dominic Kwiatkowski, Cordelia Langford, John Sillitoe on behalf of the Wellcome Sanger Institute COVID-19 Surveillance Team |
| EPI_ISL_3167540 | Lighthouse Laboratory Plymouth | Wellcome Sanger Institute for the COVID-19 Genomics UK (COG-UK) Consortium | Lighthouse Laboratory Plymouth and Alex Alderton, Roberto Amato, Jeffrey Barrett, Sonia Goncalves, Ewan Harrison, David K. Jackson, Ian Johnston, Dominic Kwiatkowski, Cordelia Langford, John Sillitoe on behalf of the Wellcome Sanger Institute COVID-19 Surveillance Team |
| EPI_ISL_3167541, EPI_ISL_3167542 | Berkshire and Surrey Pathology Services Lighthouse Laboratory | Wellcome Sanger Institute for the COVID-19 Genomics UK (COG-UK) Consortium | Berkshire and Surrey Pathology Services Lighthouse Laboratory and Alex Alderton, Roberto Amato, Jeffrey Barrett, Sonia Goncalves, Ewan Harrison, David K. Jackson, Ian Johnston, Dominic Kwiatkowski, Cordelia Langford, John Sillitoe on behalf of the Wellcome Sanger Institute COVID-19 Surveillance Team |
| EPI_ISL_3167543 | Lighthouse Laboratory Plymouth | Wellcome Sanger Institute for the COVID-19 Genomics UK (COG-UK) Consortium | Lighthouse Laboratory Plymouth and Alex Alderton, Roberto Amato, Jeffrey Barrett, Sonia Goncalves, Ewan Harrison, David K. Jackson, Ian Johnston, Dominic Kwiatkowski, Cordelia Langford, John Sillitoe on behalf of the Wellcome Sanger Institute COVID-19 Surveillance Team |
| EPI_ISL_3167544 | Lighthouse Lab in Alderley Park | Wellcome Sanger Institute for the COVID-19 Genomics UK (COG-UK) Consortium | Jacquelyn Wynn, Mairead Hyland, The Lighthouse Lab in Alderley Park and Alex Alderton, Roberto Amato, Jeffrey Barrett, Sonia Goncalves, Ewan Harrison, David K. Jackson, Ian Johnston, Dominic Kwiatkowski, Cordelia Langford, John Sillitoe on behalf of the Wellcome Sanger Institute COVID-19 Surveillance Team |
| EPI_ISL_3167545, EPI_ISL_3167546 | Berkshire and Surrey Pathology Services Lighthouse Laboratory | Wellcome Sanger Institute for the COVID-19 Genomics UK (COG-UK) Consortium | Berkshire and Surrey Pathology Services Lighthouse Laboratory and Alex Alderton, Roberto Amato, Jeffrey Barrett, Sonia Goncalves, Ewan Harrison, David K. Jackson, Ian Johnston, Dominic Kwiatkowski, Cordelia Langford, John Sillitoe on behalf of the Wellcome Sanger Institute COVID-19 Surveillance Team |
| EPI_ISL_3167547 | Lighthouse Lab in Alderley Park | Wellcome Sanger Institute for the COVID-19 Genomics UK (COG-UK) Consortium | Jacquelyn Wynn, Mairead Hyland, The Lighthouse Lab in Alderley Park and Alex Alderton, Roberto Amato, Jeffrey Barrett, Sonia Goncalves, Ewan Harrison, David K. Jackson, Ian Johnston, Dominic Kwiatkowski, Cordelia Langford, John Sillitoe on behalf of the Wellcome Sanger Institute COVID-19 Surveillance Team |
| EPI_ISL_3167548 | Lighthouse Laboratory Plymouth | Wellcome Sanger Institute for the COVID-19 Genomics UK (COG-UK) Consortium | Lighthouse Laboratory Plymouth and Alex Alderton, Roberto Amato, Jeffrey Barrett, Sonia Goncalves, Ewan Harrison, David K. Jackson, Ian Johnston, Dominic Kwiatkowski, Cordelia Langford, John Sillitoe on behalf of the Wellcome Sanger Institute COVID-19 Surveillance Team |
| EPI_ISL_3167549 | Lighthouse Lab in Alderley Park | Wellcome Sanger Institute for the COVID-19 Genomics UK (COG-UK) Consortium | Jacquelyn Wynn, Mairead Hyland, The Lighthouse Lab in Alderley Park and Alex Alderton, Roberto Amato, Jeffrey Barrett, Sonia Goncalves, Ewan Harrison, David K. Jackson, Ian Johnston, Dominic Kwiatkowski, Cordelia Langford, John Sillitoe on behalf of the Wellcome Sanger Institute COVID-19 Surveillance Team |
| EPI_ISL_3167550 | Berkshire and Surrey Pathology Services Lighthouse Laboratory | Wellcome Sanger Institute for the COVID-19 Genomics UK (COG-UK) Consortium | Berkshire and Surrey Pathology Services Lighthouse Laboratory and Alex Alderton, Roberto Amato, Jeffrey Barrett, Sonia Goncalves, Ewan Harrison, David K. Jackson, Ian Johnston, Dominic Kwiatkowski, Cordelia Langford, John Sillitoe on behalf of the Wellcome Sanger Institute COVID-19 Surveillance Team |
| EPI_ISL_3167551, EPI_ISL_3167552, EPI_ISL_3167553, EPI_ISL_3167554, EPI_ISL_3167555 | Lighthouse Lab in Alderley Park | Wellcome Sanger Institute for the COVID-19 Genomics UK (COG-UK) Consortium | Jacquelyn Wynn, Mairead Hyland, The Lighthouse Lab in Alderley Park and Alex Alderton, Roberto Amato, Jeffrey Barrett, Sonia Goncalves, Ewan Harrison, David K. Jackson, Ian Johnston, Dominic Kwiatkowski, Cordelia Langford, John Sillitoe on behalf of the Wellcome Sanger Institute COVID-19 Surveillance Team |
| EPI_ISL_3167556 | Lighthouse Laboratory Plymouth | Wellcome Sanger Institute for the COVID-19 Genomics UK (COG-UK) Consortium | Lighthouse Laboratory Plymouth and Alex Alderton, Roberto Amato, Jeffrey Barrett, Sonia Goncalves, Ewan Harrison, David K. Jackson, Ian Johnston, Dominic Kwiatkowski, Cordelia Langford, John Sillitoe on behalf of the Wellcome Sanger Institute COVID-19 Surveillance Team |
| EPI_ISL_3167557, EPI_ISL_3167558 | Lighthouse Lab in Alderley Park | Wellcome Sanger Institute for the COVID-19 Genomics UK (COG-UK) Consortium | Jacquelyn Wynn, Mairead Hyland, The Lighthouse Lab in Alderley Park and Alex Alderton, Roberto Amato, Jeffrey Barrett, Sonia Goncalves, Ewan Harrison, David K. Jackson, Ian Johnston, Dominic Kwiatkowski, Cordelia Langford, John Sillitoe on behalf of the Wellcome Sanger Institute COVID-19 Surveillance Team |
| EPI_ISL_3167559 | Berkshire and Surrey Pathology Services Lighthouse Laboratory | Wellcome Sanger Institute for the COVID-19 Genomics UK (COG-UK) Consortium | Berkshire and Surrey Pathology Services Lighthouse Laboratory and Alex Alderton, Roberto Amato, Jeffrey Barrett, Sonia Goncalves, Ewan Harrison, David K. Jackson, Ian Johnston, Dominic Kwiatkowski, Cordelia Langford, John Sillitoe on behalf of the Wellcome Sanger Institute COVID-19 Surveillance Team |
| EPI_ISL_3167560, EPI_ISL_3167561, EPI_ISL_3167562 | Lighthouse Lab in Alderley Park | Wellcome Sanger Institute for the COVID-19 Genomics UK (COG-UK) Consortium | Jacquelyn Wynn, Mairead Hyland, The Lighthouse Lab in Alderley Park and Alex Alderton, Roberto Amato, Jeffrey Barrett, Sonia Goncalves, Ewan Harrison, David K. Jackson, Ian Johnston, Dominic Kwiatkowski, Cordelia Langford, John Sillitoe on behalf of the Wellcome Sanger Institute COVID-19 Surveillance Team |
| EPI_ISL_3167563 | Lighthouse Laboratory Plymouth | Wellcome Sanger Institute for the COVID-19 Genomics UK (COG-UK) Consortium | Lighthouse Laboratory Plymouth and Alex Alderton, Roberto Amato, Jeffrey Barrett, Sonia Goncalves, Ewan Harrison, David K. Jackson, Ian Johnston, Dominic Kwiatkowski, Cordelia Langford, John Sillitoe on behalf of the Wellcome Sanger Institute COVID-19 Surveillance Team |
| EPI_ISL_3167564 | Berkshire and Surrey Pathology Services Lighthouse Laboratory | Wellcome Sanger Institute for the COVID-19 Genomics UK (COG-UK) Consortium | Berkshire and Surrey Pathology Services Lighthouse Laboratory and Alex Alderton, Roberto Amato, Jeffrey Barrett, Sonia Goncalves, Ewan Harrison, David K. Jackson, Ian Johnston, Dominic Kwiatkowski, Cordelia Langford, John Sillitoe on behalf of the Wellcome Sanger Institute COVID-19 Surveillance Team |
| EPI_ISL_3167565, EPI_ISL_3167566, EPI_ISL_3167567 | Lighthouse Lab in Alderley Park | Wellcome Sanger Institute for the COVID-19 Genomics UK (COG-UK) Consortium | Jacquelyn Wynn, Mairead Hyland, The Lighthouse Lab in Alderley Park and Alex Alderton, Roberto Amato, Jeffrey Barrett, Sonia Goncalves, Ewan Harrison, David K. Jackson, Ian Johnston, Dominic Kwiatkowski, Cordelia Langford, John Sillitoe on behalf of the Wellcome Sanger Institute COVID-19 Surveillance Team |
| EPI_ISL_3167568 | Berkshire and Surrey Pathology Services Lighthouse Laboratory | Wellcome Sanger Institute for the COVID-19 Genomics UK (COG-UK) Consortium | Berkshire and Surrey Pathology Services Lighthouse Laboratory and Alex Alderton, Roberto Amato, Jeffrey Barrett, Sonia Goncalves, Ewan Harrison, David K. Jackson, Ian Johnston, Dominic Kwiatkowski, Cordelia Langford, John Sillitoe on behalf of the Wellcome Sanger Institute COVID-19 Surveillance Team |
| EPI_ISL_3167569, EPI_ISL_3167570, EPI_ISL_3167571 | Lighthouse Lab in Alderley Park | Wellcome Sanger Institute for the COVID-19 Genomics UK (COG-UK) Consortium | Jacquelyn Wynn, Mairead Hyland, The Lighthouse Lab in Alderley Park and Alex Alderton, Roberto Amato, Jeffrey Barrett, Sonia Goncalves, Ewan Harrison, David K. Jackson, Ian Johnston, Dominic Kwiatkowski, Cordelia Langford, John Sillitoe on behalf of the Wellcome Sanger Institute COVID-19 Surveillance Team |
| EPI_ISL_3167572 | Lighthouse Laboratory Plymouth | Wellcome Sanger Institute for the COVID-19 Genomics UK (COG-UK) Consortium | Lighthouse Laboratory Plymouth and Alex Alderton, Roberto Amato, Jeffrey Barrett, Sonia Goncalves, Ewan Harrison, David K. Jackson, Ian Johnston, Dominic Kwiatkowski, Cordelia Langford, John Sillitoe on behalf of the Wellcome Sanger Institute COVID-19 Surveillance Team |
| EPI_ISL_3167573, EPI_ISL_3167574, EPI_ISL_3167575 | Lighthouse Lab in Alderley Park | Wellcome Sanger Institute for the COVID-19 Genomics UK (COG-UK) Consortium | Jacquelyn Wynn, Mairead Hyland, The Lighthouse Lab in Alderley Park and Alex Alderton, Roberto Amato, Jeffrey Barrett, Sonia Goncalves, Ewan Harrison, David K. Jackson, Ian Johnston, Dominic Kwiatkowski, Cordelia Langford, John Sillitoe on behalf of the Wellcome Sanger Institute COVID-19 Surveillance Team |
| EPI_ISL_3167576 | Lighthouse Laboratory Plymouth | Wellcome Sanger Institute for the COVID-19 Genomics UK (COG-UK) Consortium | Lighthouse Laboratory Plymouth and Alex Alderton, Roberto Amato, Jeffrey Barrett, Sonia Goncalves, Ewan Harrison, David K. Jackson, Ian Johnston, Dominic Kwiatkowski, Cordelia Langford, John Sillitoe on behalf of the Wellcome Sanger Institute COVID-19 Surveillance Team |
| EPI_ISL_3167577 | Lighthouse Lab in Alderley Park | Wellcome Sanger Institute for the COVID-19 Genomics UK (COG-UK) Consortium | Jacquelyn Wynn, Mairead Hyland, The Lighthouse Lab in Alderley Park and Alex Alderton, Roberto Amato, Jeffrey Barrett, Sonia Goncalves, Ewan Harrison, David K. Jackson, Ian Johnston, Dominic Kwiatkowski, Cordelia Langford, John Sillitoe on behalf of the Wellcome Sanger Institute COVID-19 Surveillance Team |
| EPI_ISL_3167578 | Berkshire and Surrey Pathology Services Lighthouse Laboratory | Wellcome Sanger Institute for the COVID-19 Genomics UK (COG-UK) Consortium | Berkshire and Surrey Pathology Services Lighthouse Laboratory and Alex Alderton, Roberto Amato, Jeffrey Barrett, Sonia Goncalves, Ewan Harrison, David K. Jackson, Ian Johnston, Dominic Kwiatkowski, Cordelia Langford, John Sillitoe on behalf of the Wellcome Sanger Institute COVID-19 Surveillance Team |
| EPI_ISL_3167579, EPI_ISL_3167580 | Lighthouse Lab in Alderley Park | Wellcome Sanger Institute for the COVID-19 Genomics UK (COG-UK) Consortium | Jacquelyn Wynn, Mairead Hyland, The Lighthouse Lab in Alderley Park and Alex Alderton, Roberto Amato, Jeffrey Barrett, Sonia Goncalves, Ewan Harrison, David K. Jackson, Ian Johnston, Dominic Kwiatkowski, Cordelia Langford, John Sillitoe on behalf of the Wellcome Sanger Institute COVID-19 Surveillance Team |
| EPI_ISL_3167581 | Berkshire and Surrey Pathology Services Lighthouse Laboratory | Wellcome Sanger Institute for the COVID-19 Genomics UK (COG-UK) Consortium | Berkshire and Surrey Pathology Services Lighthouse Laboratory and Alex Alderton, Roberto Amato, Jeffrey Barrett, Sonia Goncalves, Ewan Harrison, David K. Jackson, Ian Johnston, Dominic Kwiatkowski, Cordelia Langford, John Sillitoe on behalf of the Wellcome Sanger Institute COVID-19 Surveillance Team |
| EPI_ISL_3167582 | Lighthouse Lab in Alderley Park | Wellcome Sanger Institute for the COVID-19 Genomics UK (COG-UK) Consortium | Jacquelyn Wynn, Mairead Hyland, The Lighthouse Lab in Alderley Park and Alex Alderton, Roberto Amato, Jeffrey Barrett, Sonia Goncalves, Ewan Harrison, David K. Jackson, Ian Johnston, Dominic Kwiatkowski, Cordelia Langford, John Sillitoe on behalf of the Wellcome Sanger Institute COVID-19 Surveillance Team |
| EPI_ISL_3167583 | Berkshire and Surrey Pathology Services Lighthouse Laboratory | Wellcome Sanger Institute for the COVID-19 Genomics UK (COG-UK) Consortium | Berkshire and Surrey Pathology Services Lighthouse Laboratory and Alex Alderton, Roberto Amato, Jeffrey Barrett, Sonia Goncalves, Ewan Harrison, David K. Jackson, Ian Johnston, Dominic Kwiatkowski, Cordelia Langford, John Sillitoe on behalf of the Wellcome Sanger Institute COVID-19 Surveillance Team |
| EPI_ISL_3167584 | Lighthouse Lab in Alderley Park | Wellcome Sanger Institute for the COVID-19 Genomics UK (COG-UK) Consortium | Jacquelyn Wynn, Mairead Hyland, The Lighthouse Lab in Alderley Park and Alex Alderton, Roberto Amato, Jeffrey Barrett, Sonia Goncalves, Ewan Harrison, David K. Jackson, Ian Johnston, Dominic Kwiatkowski, Cordelia Langford, John Sillitoe on behalf of the Wellcome Sanger Institute COVID-19 Surveillance Team |
| EPI_ISL_3167585 | Lighthouse Laboratory Plymouth | Wellcome Sanger Institute for the COVID-19 Genomics UK (COG-UK) Consortium | Lighthouse Laboratory Plymouth and Alex Alderton, Roberto Amato, Jeffrey Barrett, Sonia Goncalves, Ewan Harrison, David K. Jackson, Ian Johnston, Dominic Kwiatkowski, Cordelia Langford, John Sillitoe on behalf of the Wellcome Sanger Institute COVID-19 Surveillance Team |
| EPI_ISL_3167586, EPI_ISL_3167587 | Lighthouse Lab in Alderley Park | Wellcome Sanger Institute for the COVID-19 Genomics UK (COG-UK) Consortium | Jacquelyn Wynn, Mairead Hyland, The Lighthouse Lab in Alderley Park and Alex Alderton, Roberto Amato, Jeffrey Barrett, Sonia Goncalves, Ewan Harrison, David K. Jackson, Ian Johnston, Dominic Kwiatkowski, Cordelia Langford, John Sillitoe on behalf of the Wellcome Sanger Institute COVID-19 Surveillance Team |
| EPI_ISL_3167588 | Berkshire and Surrey Pathology Services Lighthouse Laboratory | Wellcome Sanger Institute for the COVID-19 Genomics UK (COG-UK) Consortium | Berkshire and Surrey Pathology Services Lighthouse Laboratory and Alex Alderton, Roberto Amato, Jeffrey Barrett, Sonia Goncalves, Ewan Harrison, David K. Jackson, Ian Johnston, Dominic Kwiatkowski, Cordelia Langford, John Sillitoe on behalf of the Wellcome Sanger Institute COVID-19 Surveillance Team |
| EPI_ISL_3167589 | Lighthouse Lab in Alderley Park | Wellcome Sanger Institute for the COVID-19 Genomics UK (COG-UK) Consortium | Jacquelyn Wynn, Mairead Hyland, The Lighthouse Lab in Alderley Park and Alex Alderton, Roberto Amato, Jeffrey Barrett, Sonia Goncalves, Ewan Harrison, David K. Jackson, Ian Johnston, Dominic Kwiatkowski, Cordelia Langford, John Sillitoe on behalf of the Wellcome Sanger Institute COVID-19 Surveillance Team |

| | | | |
|---|---|---|---|
| EPI_ISL_3167590 | Lighthouse Laboratory Plymouth | Wellcome Sanger Institute for the COVID-19 Genomics UK (COG-UK) Consortium | Lighthouse Laboratory Plymouth and Alex Alderton, Roberto Amato, Jeffrey Barrett, Sonia Goncalves, Ewan Harrison, David K. Jackson, Ian Johnston, Dominic Kwiatkowski, Cordelia Langford, John Sillitoe on behalf of the Wellcome Sanger Institute COVID-19 Surveillance Team |
| EPI_ISL_3167591, EPI_ISL_3167592, EPI_ISL_3167593 | Lighthouse Lab in Alderley Park | Wellcome Sanger Institute for the COVID-19 Genomics UK (COG-UK) Consortium | Jacquelyn Wynn, Mairead Hyland, The Lighthouse Lab in Alderley Park and Alex Alderton, Roberto Amato, Jeffrey Barrett, Sonia Goncalves, Ewan Harrison, David K. Jackson, Ian Johnston, Dominic Kwiatkowski, Cordelia Langford, John Sillitoe on behalf of the Wellcome Sanger Institute COVID-19 Surveillance Team |
| EPI_ISL_3167594 | Berkshire and Surrey Pathology Services Lighthouse Laboratory | Wellcome Sanger Institute for the COVID-19 Genomics UK (COG-UK) Consortium | Berkshire and Surrey Pathology Services Lighthouse Laboratory and Alex Alderton, Roberto Amato, Jeffrey Barrett, Sonia Goncalves, Ewan Harrison, David K. Jackson, Ian Johnston, Dominic Kwiatkowski, Cordelia Langford, John Sillitoe on behalf of the Wellcome Sanger Institute COVID-19 Surveillance Team |
| EPI_ISL_3167595, EPI_ISL_3167596 | Lighthouse Lab in Alderley Park | Wellcome Sanger Institute for the COVID-19 Genomics UK (COG-UK) Consortium | Jacquelyn Wynn, Mairead Hyland, The Lighthouse Lab in Alderley Park and Alex Alderton, Roberto Amato, Jeffrey Barrett, Sonia Goncalves, Ewan Harrison, David K. Jackson, Ian Johnston, Dominic Kwiatkowski, Cordelia Langford, John Sillitoe on behalf of the Wellcome Sanger Institute COVID-19 Surveillance Team |
| EPI_ISL_3167597 | Lighthouse Laboratory Plymouth | Wellcome Sanger Institute for the COVID-19 Genomics UK (COG-UK) Consortium | Lighthouse Laboratory Plymouth and Alex Alderton, Roberto Amato, Jeffrey Barrett, Sonia Goncalves, Ewan Harrison, David K. Jackson, Ian Johnston, Dominic Kwiatkowski, Cordelia Langford, John Sillitoe on behalf of the Wellcome Sanger Institute COVID-19 Surveillance Team |
| EPI_ISL_3167598, EPI_ISL_3167599, EPI_ISL_3167600, EPI_ISL_3167601 | Lighthouse Lab in Alderley Park | Wellcome Sanger Institute for the COVID-19 Genomics UK (COG-UK) Consortium | Jacquelyn Wynn, Mairead Hyland, The Lighthouse Lab in Alderley Park and Alex Alderton, Roberto Amato, Jeffrey Barrett, Sonia Goncalves, Ewan Harrison, David K. Jackson, Ian Johnston, Dominic Kwiatkowski, Cordelia Langford, John Sillitoe on behalf of the Wellcome Sanger Institute COVID-19 Surveillance Team |
| EPI_ISL_3167602 | Berkshire and Surrey Pathology Services Lighthouse Laboratory | Wellcome Sanger Institute for the COVID-19 Genomics UK (COG-UK) Consortium | Berkshire and Surrey Pathology Services Lighthouse Laboratory and Alex Alderton, Roberto Amato, Jeffrey Barrett, Sonia Goncalves, Ewan Harrison, David K. Jackson, Ian Johnston, Dominic Kwiatkowski, Cordelia Langford, John Sillitoe on behalf of the Wellcome Sanger Institute COVID-19 Surveillance Team |
| EPI_ISL_3167603 | Lighthouse Lab in Alderley Park | Wellcome Sanger Institute for the COVID-19 Genomics UK (COG-UK) Consortium | Jacquelyn Wynn, Mairead Hyland, The Lighthouse Lab in Alderley Park and Alex Alderton, Roberto Amato, Jeffrey Barrett, Sonia Goncalves, Ewan Harrison, David K. Jackson, Ian Johnston, Dominic Kwiatkowski, Cordelia Langford, John Sillitoe on behalf of the Wellcome Sanger Institute COVID-19 Surveillance Team |
| EPI_ISL_3167604 | Lighthouse Laboratory Plymouth | Wellcome Sanger Institute for the COVID-19 Genomics UK (COG-UK) Consortium | Lighthouse Laboratory Plymouth and Alex Alderton, Roberto Amato, Jeffrey Barrett, Sonia Goncalves, Ewan Harrison, David K. Jackson, Ian Johnston, Dominic Kwiatkowski, Cordelia Langford, John Sillitoe on behalf of the Wellcome Sanger Institute COVID-19 Surveillance Team |
| EPI_ISL_3167605, EPI_ISL_3167606, EPI_ISL_3167607 | Lighthouse Lab in Alderley Park | Wellcome Sanger Institute for the COVID-19 Genomics UK (COG-UK) Consortium | Jacquelyn Wynn, Mairead Hyland, The Lighthouse Lab in Alderley Park and Alex Alderton, Roberto Amato, Jeffrey Barrett, Sonia Goncalves, Ewan Harrison, David K. Jackson, Ian Johnston, Dominic Kwiatkowski, Cordelia Langford, John Sillitoe on behalf of the Wellcome Sanger Institute COVID-19 Surveillance Team |
| EPI_ISL_3167608 | Berkshire and Surrey Pathology Services Lighthouse Laboratory | Wellcome Sanger Institute for the COVID-19 Genomics UK (COG-UK) Consortium | Berkshire and Surrey Pathology Services Lighthouse Laboratory and Alex Alderton, Roberto Amato, Jeffrey Barrett, Sonia Goncalves, Ewan Harrison, David K. Jackson, Ian Johnston, Dominic Kwiatkowski, Cordelia Langford, John Sillitoe on behalf of the Wellcome Sanger Institute COVID-19 Surveillance Team |
| EPI_ISL_3167609 | Lighthouse Lab in Alderley Park | Wellcome Sanger Institute for the COVID-19 Genomics UK (COG-UK) Consortium | Jacquelyn Wynn, Mairead Hyland, The Lighthouse Lab in Alderley Park and Alex Alderton, Roberto Amato, Jeffrey Barrett, Sonia Goncalves, Ewan Harrison, David K. Jackson, Ian Johnston, Dominic Kwiatkowski, Cordelia Langford, John Sillitoe on behalf of the Wellcome Sanger Institute COVID-19 Surveillance Team |
| EPI_ISL_3167610 | Berkshire and Surrey Pathology Services Lighthouse Laboratory | Wellcome Sanger Institute for the COVID-19 Genomics UK (COG-UK) Consortium | Berkshire and Surrey Pathology Services Lighthouse Laboratory and Alex Alderton, Roberto Amato, Jeffrey Barrett, Sonia Goncalves, Ewan Harrison, David K. Jackson, Ian Johnston, Dominic Kwiatkowski, Cordelia Langford, John Sillitoe on behalf of the Wellcome Sanger Institute COVID-19 Surveillance Team |
| EPI_ISL_3167611 | Lighthouse Laboratory Plymouth | Wellcome Sanger Institute for the COVID-19 Genomics UK (COG-UK) Consortium | Lighthouse Laboratory Plymouth and Alex Alderton, Roberto Amato, Jeffrey Barrett, Sonia Goncalves, Ewan Harrison, David K. Jackson, Ian Johnston, Dominic Kwiatkowski, Cordelia Langford, John Sillitoe on behalf of the Wellcome Sanger Institute COVID-19 Surveillance Team |
| EPI_ISL_3167612 | Lighthouse Lab in Alderley Park | Wellcome Sanger Institute for the COVID-19 Genomics UK (COG-UK) Consortium | Jacquelyn Wynn, Mairead Hyland, The Lighthouse Lab in Alderley Park and Alex Alderton, Roberto Amato, Jeffrey Barrett, Sonia Goncalves, Ewan Harrison, David K. Jackson, Ian Johnston, Dominic Kwiatkowski, Cordelia Langford, John Sillitoe on behalf of the Wellcome Sanger Institute COVID-19 Surveillance Team |
| EPI_ISL_3167613 | Lighthouse Laboratory Plymouth | Wellcome Sanger Institute for the COVID-19 Genomics UK (COG-UK) Consortium | Lighthouse Laboratory Plymouth and Alex Alderton, Roberto Amato, Jeffrey Barrett, Sonia Goncalves, Ewan Harrison, David K. Jackson, Ian Johnston, Dominic Kwiatkowski, Cordelia Langford, John Sillitoe on behalf of the Wellcome Sanger Institute COVID-19 Surveillance Team |
| EPI_ISL_3167614 | Lighthouse Lab in Alderley Park | Wellcome Sanger Institute for the COVID-19 Genomics UK (COG-UK) Consortium | Jacquelyn Wynn, Mairead Hyland, The Lighthouse Lab in Alderley Park and Alex Alderton, Roberto Amato, Jeffrey Barrett, Sonia Goncalves, Ewan Harrison, David K. Jackson, Ian Johnston, Dominic Kwiatkowski, Cordelia Langford, John Sillitoe on behalf of the Wellcome Sanger Institute COVID-19 Surveillance Team |
| EPI_ISL_3167615 | Berkshire and Surrey Pathology Services Lighthouse Laboratory | Wellcome Sanger Institute for the COVID-19 Genomics UK (COG-UK) Consortium | Berkshire and Surrey Pathology Services Lighthouse Laboratory and Alex Alderton, Roberto Amato, Jeffrey Barrett, Sonia Goncalves, Ewan Harrison, David K. Jackson, Ian Johnston, Dominic Kwiatkowski, Cordelia Langford, John Sillitoe on behalf of the Wellcome Sanger Institute COVID-19 Surveillance Team |
| EPI_ISL_3167616 | Lighthouse Laboratory Plymouth | Wellcome Sanger Institute for the COVID-19 Genomics UK (COG-UK) Consortium | Lighthouse Laboratory Plymouth and Alex Alderton, Roberto Amato, Jeffrey Barrett, Sonia Goncalves, Ewan Harrison, David K. Jackson, Ian Johnston, Dominic Kwiatkowski, Cordelia Langford, John Sillitoe on behalf of the Wellcome Sanger Institute COVID-19 Surveillance Team |
| EPI_ISL_3167617 | Lighthouse Lab in Alderley Park | Wellcome Sanger Institute for the COVID-19 Genomics UK (COG-UK) Consortium | Jacquelyn Wynn, Mairead Hyland, The Lighthouse Lab in Alderley Park and Alex Alderton, Roberto Amato, Jeffrey Barrett, Sonia Goncalves, Ewan Harrison, David K. Jackson, Ian Johnston, Dominic Kwiatkowski, Cordelia Langford, John Sillitoe on behalf of the Wellcome Sanger Institute COVID-19 Surveillance Team |
| EPI_ISL_3167618, EPI_ISL_3167619 | Berkshire and Surrey Pathology Services Lighthouse Laboratory | Wellcome Sanger Institute for the COVID-19 Genomics UK (COG-UK) Consortium | Berkshire and Surrey Pathology Services Lighthouse Laboratory and Alex Alderton, Roberto Amato, Jeffrey Barrett, Sonia Goncalves, Ewan Harrison, David K. Jackson, Ian Johnston, Dominic Kwiatkowski, Cordelia Langford, John Sillitoe on behalf of the Wellcome Sanger Institute COVID-19 Surveillance Team |
| EPI_ISL_3167620 | Lighthouse Laboratory Plymouth | Wellcome Sanger Institute for the COVID-19 Genomics UK (COG-UK) Consortium | Lighthouse Laboratory Plymouth and Alex Alderton, Roberto Amato, Jeffrey Barrett, Sonia Goncalves, Ewan Harrison, David K. Jackson, Ian Johnston, Dominic Kwiatkowski, Cordelia Langford, John Sillitoe on behalf of the Wellcome Sanger Institute COVID-19 Surveillance Team |
| EPI_ISL_3167621 | Lighthouse Lab in Alderley Park | Wellcome Sanger Institute for the COVID-19 Genomics UK (COG-UK) Consortium | Jacquelyn Wynn, Mairead Hyland, The Lighthouse Lab in Alderley Park and Alex Alderton, Roberto Amato, Jeffrey Barrett, Sonia Goncalves, Ewan Harrison, David K. Jackson, Ian Johnston, Dominic Kwiatkowski, Cordelia Langford, John Sillitoe on behalf of the Wellcome Sanger Institute COVID-19 Surveillance Team |
| EPI_ISL_3167622 | Berkshire and Surrey Pathology Services Lighthouse Laboratory | Wellcome Sanger Institute for the COVID-19 Genomics UK (COG-UK) Consortium | Berkshire and Surrey Pathology Services Lighthouse Laboratory and Alex Alderton, Roberto Amato, Jeffrey Barrett, Sonia Goncalves, Ewan Harrison, David K. Jackson, Ian Johnston, Dominic Kwiatkowski, Cordelia Langford, John Sillitoe on behalf of the Wellcome Sanger Institute COVID-19 Surveillance Team |
| EPI_ISL_3167623, EPI_ISL_3167624, EPI_ISL_3167625, EPI_ISL_3167626, EPI_ISL_3167627, EPI_ISL_3167628, EPI_ISL_3167629, EPI_ISL_3167630 | Lighthouse Lab in Alderley Park | Wellcome Sanger Institute for the COVID-19 Genomics UK (COG-UK) Consortium | Jacquelyn Wynn, Mairead Hyland, The Lighthouse Lab in Alderley Park and Alex Alderton, Roberto Amato, Jeffrey Barrett, Sonia Goncalves, Ewan Harrison, David K. Jackson, Ian Johnston, Dominic Kwiatkowski, Cordelia Langford, John Sillitoe on behalf of the Wellcome Sanger Institute COVID-19 Surveillance Team |
| EPI_ISL_3167631 | Berkshire and Surrey Pathology Services Lighthouse Laboratory | Wellcome Sanger Institute for the COVID-19 Genomics UK (COG-UK) Consortium | Berkshire and Surrey Pathology Services Lighthouse Laboratory and Alex Alderton, Roberto Amato, Jeffrey Barrett, Sonia Goncalves, Ewan Harrison, David K. Jackson, Ian Johnston, Dominic Kwiatkowski, Cordelia Langford, John Sillitoe on behalf of the Wellcome Sanger Institute COVID-19 Surveillance Team |
| EPI_ISL_3167632 | Lighthouse Lab in Alderley Park | Wellcome Sanger Institute for the COVID-19 Genomics UK (COG-UK) Consortium | Jacquelyn Wynn, Mairead Hyland, The Lighthouse Lab in Alderley Park and Alex Alderton, Roberto Amato, Jeffrey Barrett, Sonia Goncalves, Ewan Harrison, David K. Jackson, Ian Johnston, Dominic Kwiatkowski, Cordelia Langford, John Sillitoe on behalf of the Wellcome Sanger Institute COVID-19 Surveillance Team |
| EPI_ISL_3167633 | Lighthouse Laboratory Plymouth | Wellcome Sanger Institute for the COVID-19 Genomics UK (COG-UK) Consortium | Lighthouse Laboratory Plymouth and Alex Alderton, Roberto Amato, Jeffrey Barrett, Sonia Goncalves, Ewan Harrison, David K. Jackson, Ian Johnston, Dominic Kwiatkowski, Cordelia Langford, John Sillitoe on behalf of the Wellcome Sanger Institute COVID-19 Surveillance Team |
| EPI_ISL_3167634, EPI_ISL_3167635 | Berkshire and Surrey Pathology Services Lighthouse Laboratory | Wellcome Sanger Institute for the COVID-19 Genomics UK (COG-UK) Consortium | Berkshire and Surrey Pathology Services Lighthouse Laboratory and Alex Alderton, Roberto Amato, Jeffrey Barrett, Sonia Goncalves, Ewan Harrison, David K. Jackson, Ian Johnston, Dominic Kwiatkowski, Cordelia Langford, John Sillitoe on behalf of the Wellcome Sanger Institute COVID-19 Surveillance Team |
| EPI_ISL_3167636, EPI_ISL_3167637 | Lighthouse Lab in Alderley Park | Wellcome Sanger Institute for the COVID-19 Genomics UK (COG-UK) Consortium | Jacquelyn Wynn, Mairead Hyland, The Lighthouse Lab in Alderley Park and Alex Alderton, Roberto Amato, Jeffrey Barrett, Sonia Goncalves, Ewan Harrison, David K. Jackson, Ian Johnston, Dominic Kwiatkowski, Cordelia Langford, John Sillitoe on behalf of the Wellcome Sanger Institute COVID-19 Surveillance Team |
| EPI_ISL_3167638, EPI_ISL_3167639 | Berkshire and Surrey Pathology Services Lighthouse Laboratory | Wellcome Sanger Institute for the COVID-19 Genomics UK (COG-UK) Consortium | Berkshire and Surrey Pathology Services Lighthouse Laboratory and Alex Alderton, Roberto Amato, Jeffrey Barrett, Sonia Goncalves, Ewan Harrison, David K. Jackson, Ian Johnston, Dominic Kwiatkowski, Cordelia Langford, John Sillitoe on behalf of the Wellcome Sanger Institute COVID-19 Surveillance Team |
| EPI_ISL_3167640 | Lighthouse Laboratory Plymouth | Wellcome Sanger Institute for the COVID-19 Genomics UK (COG-UK) Consortium | Lighthouse Laboratory Plymouth and Alex Alderton, Roberto Amato, Jeffrey Barrett, Sonia Goncalves, Ewan Harrison, David K. Jackson, Ian Johnston, Dominic Kwiatkowski, Cordelia Langford, John Sillitoe on behalf of the Wellcome Sanger Institute COVID-19 Surveillance Team |
| EPI_ISL_3167641 | Lighthouse Lab in Alderley Park | Wellcome Sanger Institute for the COVID-19 Genomics UK (COG-UK) Consortium | Jacquelyn Wynn, Mairead Hyland, The Lighthouse Lab in Alderley Park and Alex Alderton, Roberto Amato, Jeffrey Barrett, Sonia Goncalves, Ewan Harrison, David K. Jackson, Ian Johnston, Dominic Kwiatkowski, Cordelia Langford, John Sillitoe on behalf of the Wellcome Sanger Institute COVID-19 Surveillance Team |
| EPI_ISL_3167642 | Lighthouse Laboratory Plymouth | Wellcome Sanger Institute for the COVID-19 Genomics UK (COG-UK) Consortium | Lighthouse Laboratory Plymouth and Alex Alderton, Roberto Amato, Jeffrey Barrett, Sonia Goncalves, Ewan Harrison, David K. Jackson, Ian Johnston, Dominic Kwiatkowski, Cordelia Langford, John Sillitoe on behalf of the Wellcome Sanger Institute COVID-19 Surveillance Team |
| EPI_ISL_3167643 | Lighthouse Lab in Alderley Park | Wellcome Sanger Institute for the COVID-19 Genomics UK (COG-UK) Consortium | Jacquelyn Wynn, Mairead Hyland, The Lighthouse Lab in Alderley Park and Alex Alderton, Roberto Amato, Jeffrey Barrett, Sonia Goncalves, Ewan Harrison, David K. Jackson, Ian Johnston, Dominic Kwiatkowski, Cordelia Langford, John Sillitoe on behalf of the Wellcome Sanger Institute COVID-19 Surveillance Team |
| EPI_ISL_3167644 | Lighthouse Laboratory Plymouth | Wellcome Sanger Institute for the COVID-19 Genomics UK (COG-UK) Consortium | Lighthouse Laboratory Plymouth and Alex Alderton, Roberto Amato, Jeffrey Barrett, Sonia Goncalves, Ewan Harrison, David K. Jackson, Ian Johnston, Dominic Kwiatkowski, Cordelia Langford, John Sillitoe on behalf of the Wellcome Sanger Institute COVID-19 Surveillance Team |
| EPI_ISL_3167645, EPI_ISL_3167646, EPI_ISL_3167647, EPI_ISL_3167648, EPI_ISL_3167649 | Lighthouse Lab in Alderley Park | Wellcome Sanger Institute for the COVID-19 Genomics UK (COG-UK) Consortium | Jacquelyn Wynn, Mairead Hyland, The Lighthouse Lab in Alderley Park and Alex Alderton, Roberto Amato, Jeffrey Barrett, Sonia Goncalves, Ewan Harrison, David K. Jackson, Ian Johnston, Dominic Kwiatkowski, Cordelia Langford, John Sillitoe on behalf of the Wellcome Sanger Institute COVID-19 Surveillance Team |
| EPI_ISL_3167650 | Lighthouse Laboratory Plymouth | Wellcome Sanger Institute for the COVID-19 Genomics UK (COG-UK) Consortium | Lighthouse Laboratory Plymouth and Alex Alderton, Roberto Amato, Jeffrey Barrett, Sonia Goncalves, Ewan Harrison, David K. Jackson, Ian Johnston, Dominic Kwiatkowski, Cordelia Langford, John Sillitoe on behalf of the Wellcome Sanger Institute COVID-19 Surveillance Team |
| EPI_ISL_3167651 | Lighthouse Lab in Alderley Park | Wellcome Sanger Institute for the COVID-19 Genomics UK (COG-UK) Consortium | Jacquelyn Wynn, Mairead Hyland, The Lighthouse Lab in Alderley Park and Alex Alderton, Roberto Amato, Jeffrey Barrett, Sonia Goncalves, Ewan Harrison, David K. Jackson, Ian Johnston, Dominic Kwiatkowski, Cordelia Langford, John Sillitoe on behalf of the Wellcome Sanger Institute COVID-19 Surveillance Team |
| EPI_ISL_3167652, EPI_ISL_3167653, | Lighthouse Laboratory Plymouth | Wellcome Sanger Institute for the COVID-19 Genomics UK (COG-UK) Consortium | Lighthouse Laboratory Plymouth and Alex Alderton, Roberto Amato, Jeffrey Barrett, Sonia Goncalves, Ewan Harrison, David K. Jackson, Ian Johnston, Dominic Kwiatkowski, Cordelia Langford, John Sillitoe on behalf of the Wellcome Sanger Institute COVID-19 Surveillance Team |

| | | | |
|---|---|---|---|
| EPI_ISL_3167654, EPI_ISL_3167655 | | | |
| EPI_ISL_3167656, EPI_ISL_3167657, EPI_ISL_3167658 | Lighthouse Lab in Alderley Park | Wellcome Sanger Institute for the COVID-19 Genomics UK (COG-UK) Consortium | Jacquelyn Wynn, Mairead Hyland, The Lighthouse Lab in Alderley Park and Alex Alderton, Roberto Amato, Jeffrey Barrett, Sonia Goncalves, Ewan Harrison, David K. Jackson, Ian Johnston, Dominic Kwiatkowski, Cordelia Langford, John Sillitoe on behalf of the Wellcome Sanger Institute COVID-19 Surveillance Team |
| EPI_ISL_3167659 | Lighthouse Laboratory Plymouth | Wellcome Sanger Institute for the COVID-19 Genomics UK (COG-UK) Consortium | Lighthouse Laboratory Plymouth and Alex Alderton, Roberto Amato, Jeffrey Barrett, Sonia Goncalves, Ewan Harrison, David K. Jackson, Ian Johnston, Dominic Kwiatkowski, Cordelia Langford, John Sillitoe on behalf of the Wellcome Sanger Institute COVID-19 Surveillance Team |
| EPI_ISL_3167660 | Lighthouse Lab in Alderley Park | Wellcome Sanger Institute for the COVID-19 Genomics UK (COG-UK) Consortium | Jacquelyn Wynn, Mairead Hyland, The Lighthouse Lab in Alderley Park and Alex Alderton, Roberto Amato, Jeffrey Barrett, Sonia Goncalves, Ewan Harrison, David K. Jackson, Ian Johnston, Dominic Kwiatkowski, Cordelia Langford, John Sillitoe on behalf of the Wellcome Sanger Institute COVID-19 Surveillance Team |
| EPI_ISL_3167661 | Berkshire and Surrey Pathology Services Lighthouse Laboratory | Wellcome Sanger Institute for the COVID-19 Genomics UK (COG-UK) Consortium | Berkshire and Surrey Pathology Services Lighthouse Laboratory and Alex Alderton, Roberto Amato, Jeffrey Barrett, Sonia Goncalves, Ewan Harrison, David K. Jackson, Ian Johnston, Dominic Kwiatkowski, Cordelia Langford, John Sillitoe on behalf of the Wellcome Sanger Institute COVID-19 Surveillance Team |
| EPI_ISL_3167662 | Lighthouse Lab in Alderley Park | Wellcome Sanger Institute for the COVID-19 Genomics UK (COG-UK) Consortium | Jacquelyn Wynn, Mairead Hyland, The Lighthouse Lab in Alderley Park and Alex Alderton, Roberto Amato, Jeffrey Barrett, Sonia Goncalves, Ewan Harrison, David K. Jackson, Ian Johnston, Dominic Kwiatkowski, Cordelia Langford, John Sillitoe on behalf of the Wellcome Sanger Institute COVID-19 Surveillance Team |
| EPI_ISL_3167663, EPI_ISL_3167664 | Lighthouse Laboratory Plymouth | Wellcome Sanger Institute for the COVID-19 Genomics UK (COG-UK) Consortium | Lighthouse Laboratory Plymouth and Alex Alderton, Roberto Amato, Jeffrey Barrett, Sonia Goncalves, Ewan Harrison, David K. Jackson, Ian Johnston, Dominic Kwiatkowski, Cordelia Langford, John Sillitoe on behalf of the Wellcome Sanger Institute COVID-19 Surveillance Team |
| EPI_ISL_3167665 | Lighthouse Lab in Alderley Park | Wellcome Sanger Institute for the COVID-19 Genomics UK (COG-UK) Consortium | Jacquelyn Wynn, Mairead Hyland, The Lighthouse Lab in Alderley Park and Alex Alderton, Roberto Amato, Jeffrey Barrett, Sonia Goncalves, Ewan Harrison, David K. Jackson, Ian Johnston, Dominic Kwiatkowski, Cordelia Langford, John Sillitoe on behalf of the Wellcome Sanger Institute COVID-19 Surveillance Team |
| EPI_ISL_3167666 | Lighthouse Laboratory Plymouth | Wellcome Sanger Institute for the COVID-19 Genomics UK (COG-UK) Consortium | Lighthouse Laboratory Plymouth and Alex Alderton, Roberto Amato, Jeffrey Barrett, Sonia Goncalves, Ewan Harrison, David K. Jackson, Ian Johnston, Dominic Kwiatkowski, Cordelia Langford, John Sillitoe on behalf of the Wellcome Sanger Institute COVID-19 Surveillance Team |
| EPI_ISL_3167667, EPI_ISL_3167668, EPI_ISL_3167669, EPI_ISL_3167670 | Lighthouse Lab in Alderley Park | Wellcome Sanger Institute for the COVID-19 Genomics UK (COG-UK) Consortium | Jacquelyn Wynn, Mairead Hyland, The Lighthouse Lab in Alderley Park and Alex Alderton, Roberto Amato, Jeffrey Barrett, Sonia Goncalves, Ewan Harrison, David K. Jackson, Ian Johnston, Dominic Kwiatkowski, Cordelia Langford, John Sillitoe on behalf of the Wellcome Sanger Institute COVID-19 Surveillance Team |
| EPI_ISL_3167671 | Berkshire and Surrey Pathology Services Lighthouse Laboratory | Wellcome Sanger Institute for the COVID-19 Genomics UK (COG-UK) Consortium | Berkshire and Surrey Pathology Services Lighthouse Laboratory and Alex Alderton, Roberto Amato, Jeffrey Barrett, Sonia Goncalves, Ewan Harrison, David K. Jackson, Ian Johnston, Dominic Kwiatkowski, Cordelia Langford, John Sillitoe on behalf of the Wellcome Sanger Institute COVID-19 Surveillance Team |
| EPI_ISL_3167672 | Lighthouse Lab in Alderley Park | Wellcome Sanger Institute for the COVID-19 Genomics UK (COG-UK) Consortium | Jacquelyn Wynn, Mairead Hyland, The Lighthouse Lab in Alderley Park and Alex Alderton, Roberto Amato, Jeffrey Barrett, Sonia Goncalves, Ewan Harrison, David K. Jackson, Ian Johnston, Dominic Kwiatkowski, Cordelia Langford, John Sillitoe on behalf of the Wellcome Sanger Institute COVID-19 Surveillance Team |
| EPI_ISL_3167673 | Berkshire and Surrey Pathology Services Lighthouse Laboratory | Wellcome Sanger Institute for the COVID-19 Genomics UK (COG-UK) Consortium | Berkshire and Surrey Pathology Services Lighthouse Laboratory and Alex Alderton, Roberto Amato, Jeffrey Barrett, Sonia Goncalves, Ewan Harrison, David K. Jackson, Ian Johnston, Dominic Kwiatkowski, Cordelia Langford, John Sillitoe on behalf of the Wellcome Sanger Institute COVID-19 Surveillance Team |
| EPI_ISL_3167674 | Lighthouse Laboratory Plymouth | Wellcome Sanger Institute for the COVID-19 Genomics UK (COG-UK) Consortium | Lighthouse Laboratory Plymouth and Alex Alderton, Roberto Amato, Jeffrey Barrett, Sonia Goncalves, Ewan Harrison, David K. Jackson, Ian Johnston, Dominic Kwiatkowski, Cordelia Langford, John Sillitoe on behalf of the Wellcome Sanger Institute COVID-19 Surveillance Team |
| EPI_ISL_3167675 | Berkshire and Surrey Pathology Services Lighthouse Laboratory | Wellcome Sanger Institute for the COVID-19 Genomics UK (COG-UK) Consortium | Berkshire and Surrey Pathology Services Lighthouse Laboratory and Alex Alderton, Roberto Amato, Jeffrey Barrett, Sonia Goncalves, Ewan Harrison, David K. Jackson, Ian Johnston, Dominic Kwiatkowski, Cordelia Langford, John Sillitoe on behalf of the Wellcome Sanger Institute COVID-19 Surveillance Team |
| EPI_ISL_3167676, EPI_ISL_3167677 | Lighthouse Laboratory Plymouth | Wellcome Sanger Institute for the COVID-19 Genomics UK (COG-UK) Consortium | Lighthouse Laboratory Plymouth and Alex Alderton, Roberto Amato, Jeffrey Barrett, Sonia Goncalves, Ewan Harrison, David K. Jackson, Ian Johnston, Dominic Kwiatkowski, Cordelia Langford, John Sillitoe on behalf of the Wellcome Sanger Institute COVID-19 Surveillance Team |
| EPI_ISL_3167678 | Berkshire and Surrey Pathology Services Lighthouse Laboratory | Wellcome Sanger Institute for the COVID-19 Genomics UK (COG-UK) Consortium | Berkshire and Surrey Pathology Services Lighthouse Laboratory and Alex Alderton, Roberto Amato, Jeffrey Barrett, Sonia Goncalves, Ewan Harrison, David K. Jackson, Ian Johnston, Dominic Kwiatkowski, Cordelia Langford, John Sillitoe on behalf of the Wellcome Sanger Institute COVID-19 Surveillance Team |
| EPI_ISL_3167679 | Lighthouse Lab in Alderley Park | Wellcome Sanger Institute for the COVID-19 Genomics UK (COG-UK) Consortium | Jacquelyn Wynn, Mairead Hyland, The Lighthouse Lab in Alderley Park and Alex Alderton, Roberto Amato, Jeffrey Barrett, Sonia Goncalves, Ewan Harrison, David K. Jackson, Ian Johnston, Dominic Kwiatkowski, Cordelia Langford, John Sillitoe on behalf of the Wellcome Sanger Institute COVID-19 Surveillance Team |
| EPI_ISL_3167680, EPI_ISL_3167681 | Lighthouse Laboratory Plymouth | Wellcome Sanger Institute for the COVID-19 Genomics UK (COG-UK) Consortium | Lighthouse Laboratory Plymouth and Alex Alderton, Roberto Amato, Jeffrey Barrett, Sonia Goncalves, Ewan Harrison, David K. Jackson, Ian Johnston, Dominic Kwiatkowski, Cordelia Langford, John Sillitoe on behalf of the Wellcome Sanger Institute COVID-19 Surveillance Team |
| EPI_ISL_3167682, EPI_ISL_3167683 | Lighthouse Lab in Alderley Park | Wellcome Sanger Institute for the COVID-19 Genomics UK (COG-UK) Consortium | Jacquelyn Wynn, Mairead Hyland, The Lighthouse Lab in Alderley Park and Alex Alderton, Roberto Amato, Jeffrey Barrett, Sonia Goncalves, Ewan Harrison, David K. Jackson, Ian Johnston, Dominic Kwiatkowski, Cordelia Langford, John Sillitoe on behalf of the Wellcome Sanger Institute COVID-19 Surveillance Team |
| EPI_ISL_3167684 | Lighthouse Laboratory Plymouth | Wellcome Sanger Institute for the COVID-19 Genomics UK (COG-UK) Consortium | Lighthouse Laboratory Plymouth and Alex Alderton, Roberto Amato, Jeffrey Barrett, Sonia Goncalves, Ewan Harrison, David K. Jackson, Ian Johnston, Dominic Kwiatkowski, Cordelia Langford, John Sillitoe on behalf of the Wellcome Sanger Institute COVID-19 Surveillance Team |
| EPI_ISL_3167685 | Berkshire and Surrey Pathology Services Lighthouse Laboratory | Wellcome Sanger Institute for the COVID-19 Genomics UK (COG-UK) Consortium | Berkshire and Surrey Pathology Services Lighthouse Laboratory and Alex Alderton, Roberto Amato, Jeffrey Barrett, Sonia Goncalves, Ewan Harrison, David K. Jackson, Ian Johnston, Dominic Kwiatkowski, Cordelia Langford, John Sillitoe on behalf of the Wellcome Sanger Institute COVID-19 Surveillance Team |
| EPI_ISL_3167686 | Lighthouse Laboratory Plymouth | Wellcome Sanger Institute for the COVID-19 Genomics UK (COG-UK) Consortium | Lighthouse Laboratory Plymouth and Alex Alderton, Roberto Amato, Jeffrey Barrett, Sonia Goncalves, Ewan Harrison, David K. Jackson, Ian Johnston, Dominic Kwiatkowski, Cordelia Langford, John Sillitoe on behalf of the Wellcome Sanger Institute COVID-19 Surveillance Team |
| EPI_ISL_3167687 | Lighthouse Lab in Alderley Park | Wellcome Sanger Institute for the COVID-19 Genomics UK (COG-UK) Consortium | Jacquelyn Wynn, Mairead Hyland, The Lighthouse Lab in Alderley Park and Alex Alderton, Roberto Amato, Jeffrey Barrett, Sonia Goncalves, Ewan Harrison, David K. Jackson, Ian Johnston, Dominic Kwiatkowski, Cordelia Langford, John Sillitoe on behalf of the Wellcome Sanger Institute COVID-19 Surveillance Team |
| EPI_ISL_3167688 | Lighthouse Laboratory Plymouth | Wellcome Sanger Institute for the COVID-19 Genomics UK (COG-UK) Consortium | Lighthouse Laboratory Plymouth and Alex Alderton, Roberto Amato, Jeffrey Barrett, Sonia Goncalves, Ewan Harrison, David K. Jackson, Ian Johnston, Dominic Kwiatkowski, Cordelia Langford, John Sillitoe on behalf of the Wellcome Sanger Institute COVID-19 Surveillance Team |
| EPI_ISL_3167689 | Lighthouse Lab in Alderley Park | Wellcome Sanger Institute for the COVID-19 Genomics UK (COG-UK) Consortium | Jacquelyn Wynn, Mairead Hyland, The Lighthouse Lab in Alderley Park and Alex Alderton, Roberto Amato, Jeffrey Barrett, Sonia Goncalves, Ewan Harrison, David K. Jackson, Ian Johnston, Dominic Kwiatkowski, Cordelia Langford, John Sillitoe on behalf of the Wellcome Sanger Institute COVID-19 Surveillance Team |
| EPI_ISL_3167690 | Berkshire and Surrey Pathology Services Lighthouse Laboratory | Wellcome Sanger Institute for the COVID-19 Genomics UK (COG-UK) Consortium | Berkshire and Surrey Pathology Services Lighthouse Laboratory and Alex Alderton, Roberto Amato, Jeffrey Barrett, Sonia Goncalves, Ewan Harrison, David K. Jackson, Ian Johnston, Dominic Kwiatkowski, Cordelia Langford, John Sillitoe on behalf of the Wellcome Sanger Institute COVID-19 Surveillance Team |
| EPI_ISL_3167691, EPI_ISL_3167692 | Lighthouse Lab in Alderley Park | Wellcome Sanger Institute for the COVID-19 Genomics UK (COG-UK) Consortium | Jacquelyn Wynn, Mairead Hyland, The Lighthouse Lab in Alderley Park and Alex Alderton, Roberto Amato, Jeffrey Barrett, Sonia Goncalves, Ewan Harrison, David K. Jackson, Ian Johnston, Dominic Kwiatkowski, Cordelia Langford, John Sillitoe on behalf of the Wellcome Sanger Institute COVID-19 Surveillance Team |
| EPI_ISL_3167693 | Lighthouse Laboratory Plymouth | Wellcome Sanger Institute for the COVID-19 Genomics UK (COG-UK) Consortium | Lighthouse Laboratory Plymouth and Alex Alderton, Roberto Amato, Jeffrey Barrett, Sonia Goncalves, Ewan Harrison, David K. Jackson, Ian Johnston, Dominic Kwiatkowski, Cordelia Langford, John Sillitoe on behalf of the Wellcome Sanger Institute COVID-19 Surveillance Team |
| EPI_ISL_3167694, EPI_ISL_3167695 | Lighthouse Lab in Alderley Park | Wellcome Sanger Institute for the COVID-19 Genomics UK (COG-UK) Consortium | Jacquelyn Wynn, Mairead Hyland, The Lighthouse Lab in Alderley Park and Alex Alderton, Roberto Amato, Jeffrey Barrett, Sonia Goncalves, Ewan Harrison, David K. Jackson, Ian Johnston, Dominic Kwiatkowski, Cordelia Langford, John Sillitoe on behalf of the Wellcome Sanger Institute COVID-19 Surveillance Team |
| EPI_ISL_3167696, EPI_ISL_3167697 | Lighthouse Laboratory Plymouth | Wellcome Sanger Institute for the COVID-19 Genomics UK (COG-UK) Consortium | Lighthouse Laboratory Plymouth and Alex Alderton, Roberto Amato, Jeffrey Barrett, Sonia Goncalves, Ewan Harrison, David K. Jackson, Ian Johnston, Dominic Kwiatkowski, Cordelia Langford, John Sillitoe on behalf of the Wellcome Sanger Institute COVID-19 Surveillance Team |
| EPI_ISL_3167698 | Lighthouse Lab in Alderley Park | Wellcome Sanger Institute for the COVID-19 Genomics UK (COG-UK) Consortium | Jacquelyn Wynn, Mairead Hyland, The Lighthouse Lab in Alderley Park and Alex Alderton, Roberto Amato, Jeffrey Barrett, Sonia Goncalves, Ewan Harrison, David K. Jackson, Ian Johnston, Dominic Kwiatkowski, Cordelia Langford, John Sillitoe on behalf of the Wellcome Sanger Institute COVID-19 Surveillance Team |
| EPI_ISL_3167699 | Berkshire and Surrey Pathology Services Lighthouse Laboratory | Wellcome Sanger Institute for the COVID-19 Genomics UK (COG-UK) Consortium | Berkshire and Surrey Pathology Services Lighthouse Laboratory and Alex Alderton, Roberto Amato, Jeffrey Barrett, Sonia Goncalves, Ewan Harrison, David K. Jackson, Ian Johnston, Dominic Kwiatkowski, Cordelia Langford, John Sillitoe on behalf of the Wellcome Sanger Institute COVID-19 Surveillance Team |
| EPI_ISL_3167700 | Lighthouse Laboratory Plymouth | Wellcome Sanger Institute for the COVID-19 Genomics UK (COG-UK) Consortium | Lighthouse Laboratory Plymouth and Alex Alderton, Roberto Amato, Jeffrey Barrett, Sonia Goncalves, Ewan Harrison, David K. Jackson, Ian Johnston, Dominic Kwiatkowski, Cordelia Langford, John Sillitoe on behalf of the Wellcome Sanger Institute COVID-19 Surveillance Team |
| EPI_ISL_3167701 | Lighthouse Lab in Alderley Park | Wellcome Sanger Institute for the COVID-19 Genomics UK (COG-UK) Consortium | Jacquelyn Wynn, Mairead Hyland, The Lighthouse Lab in Alderley Park and Alex Alderton, Roberto Amato, Jeffrey Barrett, Sonia Goncalves, Ewan Harrison, David K. Jackson, Ian Johnston, Dominic Kwiatkowski, Cordelia Langford, John Sillitoe on behalf of the Wellcome Sanger Institute COVID-19 Surveillance Team |
| EPI_ISL_3167702 | Berkshire and Surrey Pathology Services Lighthouse Laboratory | Wellcome Sanger Institute for the COVID-19 Genomics UK (COG-UK) Consortium | Berkshire and Surrey Pathology Services Lighthouse Laboratory and Alex Alderton, Roberto Amato, Jeffrey Barrett, Sonia Goncalves, Ewan Harrison, David K. Jackson, Ian Johnston, Dominic Kwiatkowski, Cordelia Langford, John Sillitoe on behalf of the Wellcome Sanger Institute COVID-19 Surveillance Team |
| EPI_ISL_3167703, EPI_ISL_3167704, EPI_ISL_3167705 | Lighthouse Lab in Alderley Park | Wellcome Sanger Institute for the COVID-19 Genomics UK (COG-UK) Consortium | Jacquelyn Wynn, Mairead Hyland, The Lighthouse Lab in Alderley Park and Alex Alderton, Roberto Amato, Jeffrey Barrett, Sonia Goncalves, Ewan Harrison, David K. Jackson, Ian Johnston, Dominic Kwiatkowski, Cordelia Langford, John Sillitoe on behalf of the Wellcome Sanger Institute COVID-19 Surveillance Team |
| EPI_ISL_3167706, EPI_ISL_3167707 | Berkshire and Surrey Pathology Services Lighthouse Laboratory | Wellcome Sanger Institute for the COVID-19 Genomics UK (COG-UK) Consortium | Berkshire and Surrey Pathology Services Lighthouse Laboratory and Alex Alderton, Roberto Amato, Jeffrey Barrett, Sonia Goncalves, Ewan Harrison, David K. Jackson, Ian Johnston, Dominic Kwiatkowski, Cordelia Langford, John Sillitoe on behalf of the Wellcome Sanger Institute COVID-19 Surveillance Team |
| EPI_ISL_3167708 | Lighthouse Lab in Alderley Park | Wellcome Sanger Institute for the COVID-19 Genomics UK (COG-UK) Consortium | Jacquelyn Wynn, Mairead Hyland, The Lighthouse Lab in Alderley Park and Alex Alderton, Roberto Amato, Jeffrey Barrett, Sonia Goncalves, Ewan Harrison, David K. Jackson, Ian Johnston, Dominic Kwiatkowski, Cordelia Langford, John Sillitoe on behalf of the Wellcome Sanger Institute COVID-19 Surveillance Team |
| EPI_ISL_3167709 | Berkshire and Surrey Pathology Services Lighthouse Laboratory | Wellcome Sanger Institute for the COVID-19 Genomics UK (COG-UK) Consortium | Berkshire and Surrey Pathology Services Lighthouse Laboratory and Alex Alderton, Roberto Amato, Jeffrey Barrett, Sonia Goncalves, Ewan Harrison, David K. Jackson, Ian Johnston, Dominic Kwiatkowski, Cordelia Langford, John Sillitoe on behalf of the Wellcome Sanger Institute COVID-19 Surveillance Team |
| EPI_ISL_3167710, EPI_ISL_3167711, EPI_ISL_3167712 | Lighthouse Lab in Alderley Park | Wellcome Sanger Institute for the COVID-19 Genomics UK (COG-UK) Consortium | Jacquelyn Wynn, Mairead Hyland, The Lighthouse Lab in Alderley Park and Alex Alderton, Roberto Amato, Jeffrey Barrett, Sonia Goncalves, Ewan Harrison, David K. Jackson, Ian Johnston, Dominic Kwiatkowski, Cordelia Langford, John Sillitoe on behalf of the Wellcome Sanger Institute COVID-19 Surveillance Team |
| EPI_ISL_3167713, EPI_ISL_3167714 | Berkshire and Surrey Pathology Services Lighthouse Laboratory | Wellcome Sanger Institute for the COVID-19 Genomics UK (COG-UK) Consortium | Berkshire and Surrey Pathology Services Lighthouse Laboratory and Alex Alderton, Roberto Amato, Jeffrey Barrett, Sonia Goncalves, Ewan Harrison, David K. Jackson, Ian Johnston, Dominic Kwiatkowski, Cordelia Langford, John Sillitoe on behalf of the Wellcome Sanger Institute COVID-19 Surveillance Team |
| EPI_ISL_3167715 | Lighthouse Lab in Alderley Park | Wellcome Sanger Institute for the COVID-19 Genomics UK (COG-UK) Consortium | Jacquelyn Wynn, Mairead Hyland, The Lighthouse Lab in Alderley Park and Alex Alderton, Roberto Amato, Jeffrey Barrett, Sonia Goncalves, Ewan Harrison, David K. Jackson, Ian Johnston, Dominic Kwiatkowski, Cordelia Langford, John Sillitoe on behalf of the Wellcome Sanger Institute COVID-19 Surveillance Team |

| | | | |
|---|---|---|---|
| EPI_ISL_3167716 | Lighthouse Laboratory Plymouth | Wellcome Sanger Institute for the COVID-19 Genomics UK (COG-UK) Consortium | Lighthouse Laboratory Plymouth and Alex Alderton, Roberto Amato, Jeffrey Barrett, Sonia Goncalves, Ewan Harrison, David K. Jackson, Ian Johnston, Dominic Kwiatkowski, Cordelia Langford, John Sillitoe on behalf of the Wellcome Sanger Institute COVID-19 Surveillance Team |
| EPI_ISL_3167717 | Lighthouse Lab in Alderley Park | Wellcome Sanger Institute for the COVID-19 Genomics UK (COG-UK) Consortium | Jacquelyn Wynn, Mairead Hyland, The Lighthouse Lab in Alderley Park and Alex Alderton, Roberto Amato, Jeffrey Barrett, Sonia Goncalves, Ewan Harrison, David K. Jackson, Ian Johnston, Dominic Kwiatkowski, Cordelia Langford, John Sillitoe on behalf of the Wellcome Sanger Institute COVID-19 Surveillance Team |
| EPI_ISL_3167718 | Berkshire and Surrey Pathology Services Lighthouse Laboratory | Wellcome Sanger Institute for the COVID-19 Genomics UK (COG-UK) Consortium | Berkshire and Surrey Pathology Services Lighthouse Laboratory and Alex Alderton, Roberto Amato, Jeffrey Barrett, Sonia Goncalves, Ewan Harrison, David K. Jackson, Ian Johnston, Dominic Kwiatkowski, Cordelia Langford, John Sillitoe on behalf of the Wellcome Sanger Institute COVID-19 Surveillance Team |
| EPI_ISL_3167719 | Lighthouse Lab in Alderley Park | Wellcome Sanger Institute for the COVID-19 Genomics UK (COG-UK) Consortium | Jacquelyn Wynn, Mairead Hyland, The Lighthouse Lab in Alderley Park and Alex Alderton, Roberto Amato, Jeffrey Barrett, Sonia Goncalves, Ewan Harrison, David K. Jackson, Ian Johnston, Dominic Kwiatkowski, Cordelia Langford, John Sillitoe on behalf of the Wellcome Sanger Institute COVID-19 Surveillance Team |
| EPI_ISL_3167720 | Berkshire and Surrey Pathology Services Lighthouse Laboratory | Wellcome Sanger Institute for the COVID-19 Genomics UK (COG-UK) Consortium | Berkshire and Surrey Pathology Services Lighthouse Laboratory and Alex Alderton, Roberto Amato, Jeffrey Barrett, Sonia Goncalves, Ewan Harrison, David K. Jackson, Ian Johnston, Dominic Kwiatkowski, Cordelia Langford, John Sillitoe on behalf of the Wellcome Sanger Institute COVID-19 Surveillance Team |
| EPI_ISL_3167721, EPI_ISL_3167722 | Lighthouse Laboratory Plymouth | Wellcome Sanger Institute for the COVID-19 Genomics UK (COG-UK) Consortium | Lighthouse Laboratory Plymouth and Alex Alderton, Roberto Amato, Jeffrey Barrett, Sonia Goncalves, Ewan Harrison, David K. Jackson, Ian Johnston, Dominic Kwiatkowski, Cordelia Langford, John Sillitoe on behalf of the Wellcome Sanger Institute COVID-19 Surveillance Team |
| EPI_ISL_3167723 | Lighthouse Lab in Alderley Park | Wellcome Sanger Institute for the COVID-19 Genomics UK (COG-UK) Consortium | Jacquelyn Wynn, Mairead Hyland, The Lighthouse Lab in Alderley Park and Alex Alderton, Roberto Amato, Jeffrey Barrett, Sonia Goncalves, Ewan Harrison, David K. Jackson, Ian Johnston, Dominic Kwiatkowski, Cordelia Langford, John Sillitoe on behalf of the Wellcome Sanger Institute COVID-19 Surveillance Team |
| EPI_ISL_3167724, EPI_ISL_3167725 | Lighthouse Laboratory Plymouth | Wellcome Sanger Institute for the COVID-19 Genomics UK (COG-UK) Consortium | Lighthouse Laboratory Plymouth and Alex Alderton, Roberto Amato, Jeffrey Barrett, Sonia Goncalves, Ewan Harrison, David K. Jackson, Ian Johnston, Dominic Kwiatkowski, Cordelia Langford, John Sillitoe on behalf of the Wellcome Sanger Institute COVID-19 Surveillance Team |
| EPI_ISL_3167726 | Lighthouse Lab in Alderley Park | Wellcome Sanger Institute for the COVID-19 Genomics UK (COG-UK) Consortium | Jacquelyn Wynn, Mairead Hyland, The Lighthouse Lab in Alderley Park and Alex Alderton, Roberto Amato, Jeffrey Barrett, Sonia Goncalves, Ewan Harrison, David K. Jackson, Ian Johnston, Dominic Kwiatkowski, Cordelia Langford, John Sillitoe on behalf of the Wellcome Sanger Institute COVID-19 Surveillance Team |
| EPI_ISL_3167727 | Lighthouse Laboratory Plymouth | Wellcome Sanger Institute for the COVID-19 Genomics UK (COG-UK) Consortium | Lighthouse Laboratory Plymouth and Alex Alderton, Roberto Amato, Jeffrey Barrett, Sonia Goncalves, Ewan Harrison, David K. Jackson, Ian Johnston, Dominic Kwiatkowski, Cordelia Langford, John Sillitoe on behalf of the Wellcome Sanger Institute COVID-19 Surveillance Team |
| EPI_ISL_3167728, EPI_ISL_3167729 | Berkshire and Surrey Pathology Services Lighthouse Laboratory | Wellcome Sanger Institute for the COVID-19 Genomics UK (COG-UK) Consortium | Berkshire and Surrey Pathology Services Lighthouse Laboratory and Alex Alderton, Roberto Amato, Jeffrey Barrett, Sonia Goncalves, Ewan Harrison, David K. Jackson, Ian Johnston, Dominic Kwiatkowski, Cordelia Langford, John Sillitoe on behalf of the Wellcome Sanger Institute COVID-19 Surveillance Team |
| EPI_ISL_3167730 | Lighthouse Laboratory Plymouth | Wellcome Sanger Institute for the COVID-19 Genomics UK (COG-UK) Consortium | Lighthouse Laboratory Plymouth and Alex Alderton, Roberto Amato, Jeffrey Barrett, Sonia Goncalves, Ewan Harrison, David K. Jackson, Ian Johnston, Dominic Kwiatkowski, Cordelia Langford, John Sillitoe on behalf of the Wellcome Sanger Institute COVID-19 Surveillance Team |
| EPI_ISL_3167731, EPI_ISL_3167732, EPI_ISL_3167733 | Lighthouse Lab in Alderley Park | Wellcome Sanger Institute for the COVID-19 Genomics UK (COG-UK) Consortium | Jacquelyn Wynn, Mairead Hyland, The Lighthouse Lab in Alderley Park and Alex Alderton, Roberto Amato, Jeffrey Barrett, Sonia Goncalves, Ewan Harrison, David K. Jackson, Ian Johnston, Dominic Kwiatkowski, Cordelia Langford, John Sillitoe on behalf of the Wellcome Sanger Institute COVID-19 Surveillance Team |
| EPI_ISL_3167734 | Lighthouse Laboratory Plymouth | Wellcome Sanger Institute for the COVID-19 Genomics UK (COG-UK) Consortium | Lighthouse Laboratory Plymouth and Alex Alderton, Roberto Amato, Jeffrey Barrett, Sonia Goncalves, Ewan Harrison, David K. Jackson, Ian Johnston, Dominic Kwiatkowski, Cordelia Langford, John Sillitoe on behalf of the Wellcome Sanger Institute COVID-19 Surveillance Team |
| EPI_ISL_3167735, EPI_ISL_3167736 | Lighthouse Lab in Alderley Park | Wellcome Sanger Institute for the COVID-19 Genomics UK (COG-UK) Consortium | Jacquelyn Wynn, Mairead Hyland, The Lighthouse Lab in Alderley Park and Alex Alderton, Roberto Amato, Jeffrey Barrett, Sonia Goncalves, Ewan Harrison, David K. Jackson, Ian Johnston, Dominic Kwiatkowski, Cordelia Langford, John Sillitoe on behalf of the Wellcome Sanger Institute COVID-19 Surveillance Team |
| EPI_ISL_3167737, EPI_ISL_3167738 | Lighthouse Laboratory Plymouth | Wellcome Sanger Institute for the COVID-19 Genomics UK (COG-UK) Consortium | Lighthouse Laboratory Plymouth and Alex Alderton, Roberto Amato, Jeffrey Barrett, Sonia Goncalves, Ewan Harrison, David K. Jackson, Ian Johnston, Dominic Kwiatkowski, Cordelia Langford, John Sillitoe on behalf of the Wellcome Sanger Institute COVID-19 Surveillance Team |
| EPI_ISL_3167739 | Berkshire and Surrey Pathology Services Lighthouse Laboratory | Wellcome Sanger Institute for the COVID-19 Genomics UK (COG-UK) Consortium | Berkshire and Surrey Pathology Services Lighthouse Laboratory and Alex Alderton, Roberto Amato, Jeffrey Barrett, Sonia Goncalves, Ewan Harrison, David K. Jackson, Ian Johnston, Dominic Kwiatkowski, Cordelia Langford, John Sillitoe on behalf of the Wellcome Sanger Institute COVID-19 Surveillance Team |
| EPI_ISL_3167740, EPI_ISL_3167741, EPI_ISL_3167742, EPI_ISL_3167743, EPI_ISL_3167744, EPI_ISL_3167745 | Lighthouse Lab in Alderley Park | Wellcome Sanger Institute for the COVID-19 Genomics UK (COG-UK) Consortium | Jacquelyn Wynn, Mairead Hyland, The Lighthouse Lab in Alderley Park and Alex Alderton, Roberto Amato, Jeffrey Barrett, Sonia Goncalves, Ewan Harrison, David K. Jackson, Ian Johnston, Dominic Kwiatkowski, Cordelia Langford, John Sillitoe on behalf of the Wellcome Sanger Institute COVID-19 Surveillance Team |
| EPI_ISL_3167746, EPI_ISL_3167747, EPI_ISL_3167748 | Berkshire and Surrey Pathology Services Lighthouse Laboratory | Wellcome Sanger Institute for the COVID-19 Genomics UK (COG-UK) Consortium | Berkshire and Surrey Pathology Services Lighthouse Laboratory and Alex Alderton, Roberto Amato, Jeffrey Barrett, Sonia Goncalves, Ewan Harrison, David K. Jackson, Ian Johnston, Dominic Kwiatkowski, Cordelia Langford, John Sillitoe on behalf of the Wellcome Sanger Institute COVID-19 Surveillance Team |
| EPI_ISL_3167749 | Lighthouse Lab in Alderley Park | Wellcome Sanger Institute for the COVID-19 Genomics UK (COG-UK) Consortium | Jacquelyn Wynn, Mairead Hyland, The Lighthouse Lab in Alderley Park and Alex Alderton, Roberto Amato, Jeffrey Barrett, Sonia Goncalves, Ewan Harrison, David K. Jackson, Ian Johnston, Dominic Kwiatkowski, Cordelia Langford, John Sillitoe on behalf of the Wellcome Sanger Institute COVID-19 Surveillance Team |
| EPI_ISL_3167750 | Berkshire and Surrey Pathology Services Lighthouse Laboratory | Wellcome Sanger Institute for the COVID-19 Genomics UK (COG-UK) Consortium | Berkshire and Surrey Pathology Services Lighthouse Laboratory and Alex Alderton, Roberto Amato, Jeffrey Barrett, Sonia Goncalves, Ewan Harrison, David K. Jackson, Ian Johnston, Dominic Kwiatkowski, Cordelia Langford, John Sillitoe on behalf of the Wellcome Sanger Institute COVID-19 Surveillance Team |
| EPI_ISL_3167751 | Lighthouse Laboratory Plymouth | Wellcome Sanger Institute for the COVID-19 Genomics UK (COG-UK) Consortium | Lighthouse Laboratory Plymouth and Alex Alderton, Roberto Amato, Jeffrey Barrett, Sonia Goncalves, Ewan Harrison, David K. Jackson, Ian Johnston, Dominic Kwiatkowski, Cordelia Langford, John Sillitoe on behalf of the Wellcome Sanger Institute COVID-19 Surveillance Team |
| EPI_ISL_3167752, EPI_ISL_3167753 | Lighthouse Lab in Alderley Park | Wellcome Sanger Institute for the COVID-19 Genomics UK (COG-UK) Consortium | Jacquelyn Wynn, Mairead Hyland, The Lighthouse Lab in Alderley Park and Alex Alderton, Roberto Amato, Jeffrey Barrett, Sonia Goncalves, Ewan Harrison, David K. Jackson, Ian Johnston, Dominic Kwiatkowski, Cordelia Langford, John Sillitoe on behalf of the Wellcome Sanger Institute COVID-19 Surveillance Team |
| EPI_ISL_3167754 | Berkshire and Surrey Pathology Services Lighthouse Laboratory | Wellcome Sanger Institute for the COVID-19 Genomics UK (COG-UK) Consortium | Berkshire and Surrey Pathology Services Lighthouse Laboratory and Alex Alderton, Roberto Amato, Jeffrey Barrett, Sonia Goncalves, Ewan Harrison, David K. Jackson, Ian Johnston, Dominic Kwiatkowski, Cordelia Langford, John Sillitoe on behalf of the Wellcome Sanger Institute COVID-19 Surveillance Team |
| EPI_ISL_3167755 | Lighthouse Laboratory Plymouth | Wellcome Sanger Institute for the COVID-19 Genomics UK (COG-UK) Consortium | Lighthouse Laboratory Plymouth and Alex Alderton, Roberto Amato, Jeffrey Barrett, Sonia Goncalves, Ewan Harrison, David K. Jackson, Ian Johnston, Dominic Kwiatkowski, Cordelia Langford, John Sillitoe on behalf of the Wellcome Sanger Institute COVID-19 Surveillance Team |
| EPI_ISL_3167756, EPI_ISL_3167757, EPI_ISL_3167758, EPI_ISL_3167759, EPI_ISL_3167760, EPI_ISL_3167761, EPI_ISL_3167762, EPI_ISL_3167763, EPI_ISL_3167764, EPI_ISL_3167765 | Lighthouse Lab in Alderley Park | Wellcome Sanger Institute for the COVID-19 Genomics UK (COG-UK) Consortium | Jacquelyn Wynn, Mairead Hyland, The Lighthouse Lab in Alderley Park and Alex Alderton, Roberto Amato, Jeffrey Barrett, Sonia Goncalves, Ewan Harrison, David K. Jackson, Ian Johnston, Dominic Kwiatkowski, Cordelia Langford, John Sillitoe on behalf of the Wellcome Sanger Institute COVID-19 Surveillance Team |
| EPI_ISL_3167766, EPI_ISL_3167767 | Lighthouse Laboratory Plymouth | Wellcome Sanger Institute for the COVID-19 Genomics UK (COG-UK) Consortium | Lighthouse Laboratory Plymouth and Alex Alderton, Roberto Amato, Jeffrey Barrett, Sonia Goncalves, Ewan Harrison, David K. Jackson, Ian Johnston, Dominic Kwiatkowski, Cordelia Langford, John Sillitoe on behalf of the Wellcome Sanger Institute COVID-19 Surveillance Team |
| EPI_ISL_3167768 | Lighthouse Lab in Alderley Park | Wellcome Sanger Institute for the COVID-19 Genomics UK (COG-UK) Consortium | Jacquelyn Wynn, Mairead Hyland, The Lighthouse Lab in Alderley Park and Alex Alderton, Roberto Amato, Jeffrey Barrett, Sonia Goncalves, Ewan Harrison, David K. Jackson, Ian Johnston, Dominic Kwiatkowski, Cordelia Langford, John Sillitoe on behalf of the Wellcome Sanger Institute COVID-19 Surveillance Team |
| EPI_ISL_3167769 | Lighthouse Laboratory Plymouth | Wellcome Sanger Institute for the COVID-19 Genomics UK (COG-UK) Consortium | Lighthouse Laboratory Plymouth and Alex Alderton, Roberto Amato, Jeffrey Barrett, Sonia Goncalves, Ewan Harrison, David K. Jackson, Ian Johnston, Dominic Kwiatkowski, Cordelia Langford, John Sillitoe on behalf of the Wellcome Sanger Institute COVID-19 Surveillance Team |
| EPI_ISL_3167770 | Lighthouse Lab in Alderley Park | Wellcome Sanger Institute for the COVID-19 Genomics UK (COG-UK) Consortium | Jacquelyn Wynn, Mairead Hyland, The Lighthouse Lab in Alderley Park and Alex Alderton, Roberto Amato, Jeffrey Barrett, Sonia Goncalves, Ewan Harrison, David K. Jackson, Ian Johnston, Dominic Kwiatkowski, Cordelia Langford, John Sillitoe on behalf of the Wellcome Sanger Institute COVID-19 Surveillance Team |
| EPI_ISL_3167771 | Berkshire and Surrey Pathology Services Lighthouse Laboratory | Wellcome Sanger Institute for the COVID-19 Genomics UK (COG-UK) Consortium | Berkshire and Surrey Pathology Services Lighthouse Laboratory and Alex Alderton, Roberto Amato, Jeffrey Barrett, Sonia Goncalves, Ewan Harrison, David K. Jackson, Ian Johnston, Dominic Kwiatkowski, Cordelia Langford, John Sillitoe on behalf of the Wellcome Sanger Institute COVID-19 Surveillance Team |
| EPI_ISL_3167772 | Lighthouse Laboratory Plymouth | Wellcome Sanger Institute for the COVID-19 Genomics UK (COG-UK) Consortium | Lighthouse Laboratory Plymouth and Alex Alderton, Roberto Amato, Jeffrey Barrett, Sonia Goncalves, Ewan Harrison, David K. Jackson, Ian Johnston, Dominic Kwiatkowski, Cordelia Langford, John Sillitoe on behalf of the Wellcome Sanger Institute COVID-19 Surveillance Team |
| EPI_ISL_3167773 | Berkshire and Surrey Pathology Services Lighthouse Laboratory | Wellcome Sanger Institute for the COVID-19 Genomics UK (COG-UK) Consortium | Berkshire and Surrey Pathology Services Lighthouse Laboratory and Alex Alderton, Roberto Amato, Jeffrey Barrett, Sonia Goncalves, Ewan Harrison, David K. Jackson, Ian Johnston, Dominic Kwiatkowski, Cordelia Langford, John Sillitoe on behalf of the Wellcome Sanger Institute COVID-19 Surveillance Team |
| EPI_ISL_3167774, EPI_ISL_3167775 | Lighthouse Laboratory Plymouth | Wellcome Sanger Institute for the COVID-19 Genomics UK (COG-UK) Consortium | Lighthouse Laboratory Plymouth and Alex Alderton, Roberto Amato, Jeffrey Barrett, Sonia Goncalves, Ewan Harrison, David K. Jackson, Ian Johnston, Dominic Kwiatkowski, Cordelia Langford, John Sillitoe on behalf of the Wellcome Sanger Institute COVID-19 Surveillance Team |
| EPI_ISL_3167776 | Lighthouse Lab in Alderley Park | Wellcome Sanger Institute for the COVID-19 Genomics UK (COG-UK) Consortium | Jacquelyn Wynn, Mairead Hyland, The Lighthouse Lab in Alderley Park and Alex Alderton, Roberto Amato, Jeffrey Barrett, Sonia Goncalves, Ewan Harrison, David K. Jackson, Ian Johnston, Dominic Kwiatkowski, Cordelia Langford, John Sillitoe on behalf of the Wellcome Sanger Institute COVID-19 Surveillance Team |
| EPI_ISL_3167777 | Lighthouse Laboratory Plymouth | Wellcome Sanger Institute for the COVID-19 Genomics UK (COG-UK) Consortium | Lighthouse Laboratory Plymouth and Alex Alderton, Roberto Amato, Jeffrey Barrett, Sonia Goncalves, Ewan Harrison, David K. Jackson, Ian Johnston, Dominic Kwiatkowski, Cordelia Langford, John Sillitoe on behalf of the Wellcome Sanger Institute COVID-19 Surveillance Team |
| EPI_ISL_3167778 | Lighthouse Lab in Alderley Park | Wellcome Sanger Institute for the COVID-19 Genomics UK (COG-UK) Consortium | Jacquelyn Wynn, Mairead Hyland, The Lighthouse Lab in Alderley Park and Alex Alderton, Roberto Amato, Jeffrey Barrett, Sonia Goncalves, Ewan Harrison, David K. Jackson, Ian Johnston, Dominic Kwiatkowski, Cordelia Langford, John Sillitoe on behalf of the Wellcome Sanger Institute COVID-19 Surveillance Team |
| EPI_ISL_3167779 | Lighthouse Laboratory Plymouth | Wellcome Sanger Institute for the COVID-19 Genomics UK (COG-UK) Consortium | Lighthouse Laboratory Plymouth and Alex Alderton, Roberto Amato, Jeffrey Barrett, Sonia Goncalves, Ewan Harrison, David K. Jackson, Ian Johnston, Dominic Kwiatkowski, Cordelia Langford, John Sillitoe on behalf of the Wellcome Sanger Institute COVID-19 Surveillance Team |
| EPI_ISL_3167780, EPI_ISL_3167781 | Lighthouse Lab in Alderley Park | Wellcome Sanger Institute for the COVID-19 Genomics UK (COG-UK) Consortium | Jacquelyn Wynn, Mairead Hyland, The Lighthouse Lab in Alderley Park and Alex Alderton, Roberto Amato, Jeffrey Barrett, Sonia Goncalves, Ewan Harrison, David K. Jackson, Ian Johnston, Dominic Kwiatkowski, Cordelia Langford, John Sillitoe on behalf of the Wellcome Sanger Institute COVID-19 Surveillance Team |

| | | | |
|---|---|---|---|
| EPI_ISL_3167782, EPI_ISL_3167783, EPI_ISL_3167784, EPI_ISL_3167785 | Lighthouse Laboratory Plymouth | Wellcome Sanger Institute for the COVID-19 Genomics UK (COG-UK) Consortium | Lighthouse Laboratory Plymouth and Alex Alderton, Roberto Amato, Jeffrey Barrett, Sonia Goncalves, Ewan Harrison, David K. Jackson, Ian Johnston, Dominic Kwiatkowski, Cordelia Langford, John Sillitoe on behalf of the Wellcome Sanger Institute COVID-19 Surveillance Team |
| EPI_ISL_3167786 | Berkshire and Surrey Pathology Services Lighthouse Laboratory | Wellcome Sanger Institute for the COVID-19 Genomics UK (COG-UK) Consortium | Berkshire and Surrey Pathology Services Lighthouse Laboratory and Alex Alderton, Roberto Amato, Jeffrey Barrett, Sonia Goncalves, Ewan Harrison, David K. Jackson, Ian Johnston, Dominic Kwiatkowski, Cordelia Langford, John Sillitoe on behalf of the Wellcome Sanger Institute COVID-19 Surveillance Team |
| EPI_ISL_3167787 | Lighthouse Laboratory Plymouth | Wellcome Sanger Institute for the COVID-19 Genomics UK (COG-UK) Consortium | Lighthouse Laboratory Plymouth and Alex Alderton, Roberto Amato, Jeffrey Barrett, Sonia Goncalves, Ewan Harrison, David K. Jackson, Ian Johnston, Dominic Kwiatkowski, Cordelia Langford, John Sillitoe on behalf of the Wellcome Sanger Institute COVID-19 Surveillance Team |
| EPI_ISL_3167788 | Berkshire and Surrey Pathology Services Lighthouse Laboratory | Wellcome Sanger Institute for the COVID-19 Genomics UK (COG-UK) Consortium | Berkshire and Surrey Pathology Services Lighthouse Laboratory and Alex Alderton, Roberto Amato, Jeffrey Barrett, Sonia Goncalves, Ewan Harrison, David K. Jackson, Ian Johnston, Dominic Kwiatkowski, Cordelia Langford, John Sillitoe on behalf of the Wellcome Sanger Institute COVID-19 Surveillance Team |
| EPI_ISL_3167789 | Lighthouse Lab in Alderley Park | Wellcome Sanger Institute for the COVID-19 Genomics UK (COG-UK) Consortium | Jacquelyn Wynn, Mairead Hyland, The Lighthouse Lab in Alderley Park and Alex Alderton, Roberto Amato, Jeffrey Barrett, Sonia Goncalves, Ewan Harrison, David K. Jackson, Ian Johnston, Dominic Kwiatkowski, Cordelia Langford, John Sillitoe on behalf of the Wellcome Sanger Institute COVID-19 Surveillance Team |
| EPI_ISL_3167790 | Lighthouse Laboratory Plymouth | Wellcome Sanger Institute for the COVID-19 Genomics UK (COG-UK) Consortium | Lighthouse Laboratory Plymouth and Alex Alderton, Roberto Amato, Jeffrey Barrett, Sonia Goncalves, Ewan Harrison, David K. Jackson, Ian Johnston, Dominic Kwiatkowski, Cordelia Langford, John Sillitoe on behalf of the Wellcome Sanger Institute COVID-19 Surveillance Team |
| EPI_ISL_3167791 | Berkshire and Surrey Pathology Services Lighthouse Laboratory | Wellcome Sanger Institute for the COVID-19 Genomics UK (COG-UK) Consortium | Berkshire and Surrey Pathology Services Lighthouse Laboratory and Alex Alderton, Roberto Amato, Jeffrey Barrett, Sonia Goncalves, Ewan Harrison, David K. Jackson, Ian Johnston, Dominic Kwiatkowski, Cordelia Langford, John Sillitoe on behalf of the Wellcome Sanger Institute COVID-19 Surveillance Team |
| EPI_ISL_3167792, EPI_ISL_3167793 | Lighthouse Laboratory Plymouth | Wellcome Sanger Institute for the COVID-19 Genomics UK (COG-UK) Consortium | Lighthouse Laboratory Plymouth and Alex Alderton, Roberto Amato, Jeffrey Barrett, Sonia Goncalves, Ewan Harrison, David K. Jackson, Ian Johnston, Dominic Kwiatkowski, Cordelia Langford, John Sillitoe on behalf of the Wellcome Sanger Institute COVID-19 Surveillance Team |
| EPI_ISL_3167794, EPI_ISL_3167795 | Lighthouse Lab in Alderley Park | Wellcome Sanger Institute for the COVID-19 Genomics UK (COG-UK) Consortium | Jacquelyn Wynn, Mairead Hyland, The Lighthouse Lab in Alderley Park and Alex Alderton, Roberto Amato, Jeffrey Barrett, Sonia Goncalves, Ewan Harrison, David K. Jackson, Ian Johnston, Dominic Kwiatkowski, Cordelia Langford, John Sillitoe on behalf of the Wellcome Sanger Institute COVID-19 Surveillance Team |
| EPI_ISL_3167796 | Lighthouse Laboratory Plymouth | Wellcome Sanger Institute for the COVID-19 Genomics UK (COG-UK) Consortium | Lighthouse Laboratory Plymouth and Alex Alderton, Roberto Amato, Jeffrey Barrett, Sonia Goncalves, Ewan Harrison, David K. Jackson, Ian Johnston, Dominic Kwiatkowski, Cordelia Langford, John Sillitoe on behalf of the Wellcome Sanger Institute COVID-19 Surveillance Team |
| EPI_ISL_3167797, EPI_ISL_3167798, EPI_ISL_3167799 | Lighthouse Lab in Alderley Park | Wellcome Sanger Institute for the COVID-19 Genomics UK (COG-UK) Consortium | Jacquelyn Wynn, Mairead Hyland, The Lighthouse Lab in Alderley Park and Alex Alderton, Roberto Amato, Jeffrey Barrett, Sonia Goncalves, Ewan Harrison, David K. Jackson, Ian Johnston, Dominic Kwiatkowski, Cordelia Langford, John Sillitoe on behalf of the Wellcome Sanger Institute COVID-19 Surveillance Team |
| EPI_ISL_3167800, EPI_ISL_3167801 | Lighthouse Laboratory Plymouth | Wellcome Sanger Institute for the COVID-19 Genomics UK (COG-UK) Consortium | Lighthouse Laboratory Plymouth and Alex Alderton, Roberto Amato, Jeffrey Barrett, Sonia Goncalves, Ewan Harrison, David K. Jackson, Ian Johnston, Dominic Kwiatkowski, Cordelia Langford, John Sillitoe on behalf of the Wellcome Sanger Institute COVID-19 Surveillance Team |
| EPI_ISL_3167802 | Lighthouse Lab in Alderley Park | Wellcome Sanger Institute for the COVID-19 Genomics UK (COG-UK) Consortium | Jacquelyn Wynn, Mairead Hyland, The Lighthouse Lab in Alderley Park and Alex Alderton, Roberto Amato, Jeffrey Barrett, Sonia Goncalves, Ewan Harrison, David K. Jackson, Ian Johnston, Dominic Kwiatkowski, Cordelia Langford, John Sillitoe on behalf of the Wellcome Sanger Institute COVID-19 Surveillance Team |
| EPI_ISL_3167803 | Lighthouse Laboratory Plymouth | Wellcome Sanger Institute for the COVID-19 Genomics UK (COG-UK) Consortium | Lighthouse Laboratory Plymouth and Alex Alderton, Roberto Amato, Jeffrey Barrett, Sonia Goncalves, Ewan Harrison, David K. Jackson, Ian Johnston, Dominic Kwiatkowski, Cordelia Langford, John Sillitoe on behalf of the Wellcome Sanger Institute COVID-19 Surveillance Team |
| EPI_ISL_3167804, EPI_ISL_3167805, EPI_ISL_3167806 | Lighthouse Lab in Alderley Park | Wellcome Sanger Institute for the COVID-19 Genomics UK (COG-UK) Consortium | Jacquelyn Wynn, Mairead Hyland, The Lighthouse Lab in Alderley Park and Alex Alderton, Roberto Amato, Jeffrey Barrett, Sonia Goncalves, Ewan Harrison, David K. Jackson, Ian Johnston, Dominic Kwiatkowski, Cordelia Langford, John Sillitoe on behalf of the Wellcome Sanger Institute COVID-19 Surveillance Team |
| EPI_ISL_3167807 | Lighthouse Laboratory Plymouth | Wellcome Sanger Institute for the COVID-19 Genomics UK (COG-UK) Consortium | Lighthouse Laboratory Plymouth and Alex Alderton, Roberto Amato, Jeffrey Barrett, Sonia Goncalves, Ewan Harrison, David K. Jackson, Ian Johnston, Dominic Kwiatkowski, Cordelia Langford, John Sillitoe on behalf of the Wellcome Sanger Institute COVID-19 Surveillance Team |
| EPI_ISL_3167808, EPI_ISL_3167809, EPI_ISL_3167810 | Lighthouse Lab in Alderley Park | Wellcome Sanger Institute for the COVID-19 Genomics UK (COG-UK) Consortium | Jacquelyn Wynn, Mairead Hyland, The Lighthouse Lab in Alderley Park and Alex Alderton, Roberto Amato, Jeffrey Barrett, Sonia Goncalves, Ewan Harrison, David K. Jackson, Ian Johnston, Dominic Kwiatkowski, Cordelia Langford, John Sillitoe on behalf of the Wellcome Sanger Institute COVID-19 Surveillance Team |
| EPI_ISL_3167811 | Berkshire and Surrey Pathology Services Lighthouse Laboratory | Wellcome Sanger Institute for the COVID-19 Genomics UK (COG-UK) Consortium | Berkshire and Surrey Pathology Services Lighthouse Laboratory and Alex Alderton, Roberto Amato, Jeffrey Barrett, Sonia Goncalves, Ewan Harrison, David K. Jackson, Ian Johnston, Dominic Kwiatkowski, Cordelia Langford, John Sillitoe on behalf of the Wellcome Sanger Institute COVID-19 Surveillance Team |
| EPI_ISL_3167812, EPI_ISL_3167813 | Lighthouse Laboratory Plymouth | Wellcome Sanger Institute for the COVID-19 Genomics UK (COG-UK) Consortium | Lighthouse Laboratory Plymouth and Alex Alderton, Roberto Amato, Jeffrey Barrett, Sonia Goncalves, Ewan Harrison, David K. Jackson, Ian Johnston, Dominic Kwiatkowski, Cordelia Langford, John Sillitoe on behalf of the Wellcome Sanger Institute COVID-19 Surveillance Team |
| EPI_ISL_3167814, EPI_ISL_3167815 | Lighthouse Lab in Alderley Park | Wellcome Sanger Institute for the COVID-19 Genomics UK (COG-UK) Consortium | Jacquelyn Wynn, Mairead Hyland, The Lighthouse Lab in Alderley Park and Alex Alderton, Roberto Amato, Jeffrey Barrett, Sonia Goncalves, Ewan Harrison, David K. Jackson, Ian Johnston, Dominic Kwiatkowski, Cordelia Langford, John Sillitoe on behalf of the Wellcome Sanger Institute COVID-19 Surveillance Team |
| EPI_ISL_3167816 | Lighthouse Laboratory Plymouth | Wellcome Sanger Institute for the COVID-19 Genomics UK (COG-UK) Consortium | Lighthouse Laboratory Plymouth and Alex Alderton, Roberto Amato, Jeffrey Barrett, Sonia Goncalves, Ewan Harrison, David K. Jackson, Ian Johnston, Dominic Kwiatkowski, Cordelia Langford, John Sillitoe on behalf of the Wellcome Sanger Institute COVID-19 Surveillance Team |
| EPI_ISL_3167817, EPI_ISL_3167818 | Lighthouse Lab in Alderley Park | Wellcome Sanger Institute for the COVID-19 Genomics UK (COG-UK) Consortium | Jacquelyn Wynn, Mairead Hyland, The Lighthouse Lab in Alderley Park and Alex Alderton, Roberto Amato, Jeffrey Barrett, Sonia Goncalves, Ewan Harrison, David K. Jackson, Ian Johnston, Dominic Kwiatkowski, Cordelia Langford, John Sillitoe on behalf of the Wellcome Sanger Institute COVID-19 Surveillance Team |
| EPI_ISL_3167819, EPI_ISL_3167820 | Berkshire and Surrey Pathology Services Lighthouse Laboratory | Wellcome Sanger Institute for the COVID-19 Genomics UK (COG-UK) Consortium | Berkshire and Surrey Pathology Services Lighthouse Laboratory and Alex Alderton, Roberto Amato, Jeffrey Barrett, Sonia Goncalves, Ewan Harrison, David K. Jackson, Ian Johnston, Dominic Kwiatkowski, Cordelia Langford, John Sillitoe on behalf of the Wellcome Sanger Institute COVID-19 Surveillance Team |
| EPI_ISL_3167821 | Lighthouse Lab in Alderley Park | Wellcome Sanger Institute for the COVID-19 Genomics UK (COG-UK) Consortium | Jacquelyn Wynn, Mairead Hyland, The Lighthouse Lab in Alderley Park and Alex Alderton, Roberto Amato, Jeffrey Barrett, Sonia Goncalves, Ewan Harrison, David K. Jackson, Ian Johnston, Dominic Kwiatkowski, Cordelia Langford, John Sillitoe on behalf of the Wellcome Sanger Institute COVID-19 Surveillance Team |
| EPI_ISL_3167822 | Berkshire and Surrey Pathology Services Lighthouse Laboratory | Wellcome Sanger Institute for the COVID-19 Genomics UK (COG-UK) Consortium | Berkshire and Surrey Pathology Services Lighthouse Laboratory and Alex Alderton, Roberto Amato, Jeffrey Barrett, Sonia Goncalves, Ewan Harrison, David K. Jackson, Ian Johnston, Dominic Kwiatkowski, Cordelia Langford, John Sillitoe on behalf of the Wellcome Sanger Institute COVID-19 Surveillance Team |
| EPI_ISL_3167823 | Lighthouse Lab in Alderley Park | Wellcome Sanger Institute for the COVID-19 Genomics UK (COG-UK) Consortium | Jacquelyn Wynn, Mairead Hyland, The Lighthouse Lab in Alderley Park and Alex Alderton, Roberto Amato, Jeffrey Barrett, Sonia Goncalves, Ewan Harrison, David K. Jackson, Ian Johnston, Dominic Kwiatkowski, Cordelia Langford, John Sillitoe on behalf of the Wellcome Sanger Institute COVID-19 Surveillance Team |
| EPI_ISL_3167824 | Lighthouse Laboratory Plymouth | Wellcome Sanger Institute for the COVID-19 Genomics UK (COG-UK) Consortium | Lighthouse Laboratory Plymouth and Alex Alderton, Roberto Amato, Jeffrey Barrett, Sonia Goncalves, Ewan Harrison, David K. Jackson, Ian Johnston, Dominic Kwiatkowski, Cordelia Langford, John Sillitoe on behalf of the Wellcome Sanger Institute COVID-19 Surveillance Team |
| EPI_ISL_3167825, EPI_ISL_3167826 | Lighthouse Lab in Alderley Park | Wellcome Sanger Institute for the COVID-19 Genomics UK (COG-UK) Consortium | Jacquelyn Wynn, Mairead Hyland, The Lighthouse Lab in Alderley Park and Alex Alderton, Roberto Amato, Jeffrey Barrett, Sonia Goncalves, Ewan Harrison, David K. Jackson, Ian Johnston, Dominic Kwiatkowski, Cordelia Langford, John Sillitoe on behalf of the Wellcome Sanger Institute COVID-19 Surveillance Team |
| EPI_ISL_3167827 | Berkshire and Surrey Pathology Services Lighthouse Laboratory | Wellcome Sanger Institute for the COVID-19 Genomics UK (COG-UK) Consortium | Berkshire and Surrey Pathology Services Lighthouse Laboratory and Alex Alderton, Roberto Amato, Jeffrey Barrett, Sonia Goncalves, Ewan Harrison, David K. Jackson, Ian Johnston, Dominic Kwiatkowski, Cordelia Langford, John Sillitoe on behalf of the Wellcome Sanger Institute COVID-19 Surveillance Team |
| EPI_ISL_3167828, EPI_ISL_3167829 | Lighthouse Laboratory Plymouth | Wellcome Sanger Institute for the COVID-19 Genomics UK (COG-UK) Consortium | Lighthouse Laboratory Plymouth and Alex Alderton, Roberto Amato, Jeffrey Barrett, Sonia Goncalves, Ewan Harrison, David K. Jackson, Ian Johnston, Dominic Kwiatkowski, Cordelia Langford, John Sillitoe on behalf of the Wellcome Sanger Institute COVID-19 Surveillance Team |
| EPI_ISL_3167830 | Lighthouse Lab in Alderley Park | Wellcome Sanger Institute for the COVID-19 Genomics UK (COG-UK) Consortium | Jacquelyn Wynn, Mairead Hyland, The Lighthouse Lab in Alderley Park and Alex Alderton, Roberto Amato, Jeffrey Barrett, Sonia Goncalves, Ewan Harrison, David K. Jackson, Ian Johnston, Dominic Kwiatkowski, Cordelia Langford, John Sillitoe on behalf of the Wellcome Sanger Institute COVID-19 Surveillance Team |
| EPI_ISL_3167831 | Lighthouse Laboratory Plymouth | Wellcome Sanger Institute for the COVID-19 Genomics UK (COG-UK) Consortium | Lighthouse Laboratory Plymouth and Alex Alderton, Roberto Amato, Jeffrey Barrett, Sonia Goncalves, Ewan Harrison, David K. Jackson, Ian Johnston, Dominic Kwiatkowski, Cordelia Langford, John Sillitoe on behalf of the Wellcome Sanger Institute COVID-19 Surveillance Team |
| EPI_ISL_3167832 | Berkshire and Surrey Pathology Services Lighthouse Laboratory | Wellcome Sanger Institute for the COVID-19 Genomics UK (COG-UK) Consortium | Berkshire and Surrey Pathology Services Lighthouse Laboratory and Alex Alderton, Roberto Amato, Jeffrey Barrett, Sonia Goncalves, Ewan Harrison, David K. Jackson, Ian Johnston, Dominic Kwiatkowski, Cordelia Langford, John Sillitoe on behalf of the Wellcome Sanger Institute COVID-19 Surveillance Team |
| EPI_ISL_3167833 | Lighthouse Lab in Alderley Park | Wellcome Sanger Institute for the COVID-19 Genomics UK (COG-UK) Consortium | Jacquelyn Wynn, Mairead Hyland, The Lighthouse Lab in Alderley Park and Alex Alderton, Roberto Amato, Jeffrey Barrett, Sonia Goncalves, Ewan Harrison, David K. Jackson, Ian Johnston, Dominic Kwiatkowski, Cordelia Langford, John Sillitoe on behalf of the Wellcome Sanger Institute COVID-19 Surveillance Team |
| EPI_ISL_3167834 | Berkshire and Surrey Pathology Services Lighthouse Laboratory | Wellcome Sanger Institute for the COVID-19 Genomics UK (COG-UK) Consortium | Berkshire and Surrey Pathology Services Lighthouse Laboratory and Alex Alderton, Roberto Amato, Jeffrey Barrett, Sonia Goncalves, Ewan Harrison, David K. Jackson, Ian Johnston, Dominic Kwiatkowski, Cordelia Langford, John Sillitoe on behalf of the Wellcome Sanger Institute COVID-19 Surveillance Team |
| EPI_ISL_3167835 | Lighthouse Laboratory Plymouth | Wellcome Sanger Institute for the COVID-19 Genomics UK (COG-UK) Consortium | Lighthouse Laboratory Plymouth and Alex Alderton, Roberto Amato, Jeffrey Barrett, Sonia Goncalves, Ewan Harrison, David K. Jackson, Ian Johnston, Dominic Kwiatkowski, Cordelia Langford, John Sillitoe on behalf of the Wellcome Sanger Institute COVID-19 Surveillance Team |
| EPI_ISL_3167836, EPI_ISL_3167837, EPI_ISL_3167838, EPI_ISL_3167839, EPI_ISL_3167840 | Lighthouse Lab in Alderley Park | Wellcome Sanger Institute for the COVID-19 Genomics UK (COG-UK) Consortium | Jacquelyn Wynn, Mairead Hyland, The Lighthouse Lab in Alderley Park and Alex Alderton, Roberto Amato, Jeffrey Barrett, Sonia Goncalves, Ewan Harrison, David K. Jackson, Ian Johnston, Dominic Kwiatkowski, Cordelia Langford, John Sillitoe on behalf of the Wellcome Sanger Institute COVID-19 Surveillance Team |
| EPI_ISL_3167841 | Berkshire and Surrey Pathology Services Lighthouse Laboratory | Wellcome Sanger Institute for the COVID-19 Genomics UK (COG-UK) Consortium | Berkshire and Surrey Pathology Services Lighthouse Laboratory and Alex Alderton, Roberto Amato, Jeffrey Barrett, Sonia Goncalves, Ewan Harrison, David K. Jackson, Ian Johnston, Dominic Kwiatkowski, Cordelia Langford, John Sillitoe on behalf of the Wellcome Sanger Institute COVID-19 Surveillance Team |
| EPI_ISL_3167842 | Lighthouse Laboratory Plymouth | Wellcome Sanger Institute for the COVID-19 Genomics UK (COG-UK) Consortium | Lighthouse Laboratory Plymouth and Alex Alderton, Roberto Amato, Jeffrey Barrett, Sonia Goncalves, Ewan Harrison, David K. Jackson, Ian Johnston, Dominic Kwiatkowski, Cordelia Langford, John Sillitoe on behalf of the Wellcome Sanger Institute COVID-19 Surveillance Team |
| EPI_ISL_3167843 | Berkshire and Surrey Pathology Services Lighthouse Laboratory | Wellcome Sanger Institute for the COVID-19 Genomics UK (COG-UK) Consortium | Berkshire and Surrey Pathology Services Lighthouse Laboratory and Alex Alderton, Roberto Amato, Jeffrey Barrett, Sonia Goncalves, Ewan Harrison, David K. Jackson, Ian Johnston, Dominic Kwiatkowski, Cordelia Langford, John Sillitoe on behalf of the Wellcome Sanger Institute COVID-19 Surveillance Team |
| EPI_ISL_3167844, EPI_ISL_3167845, EPI_ISL_3167846 | Lighthouse Lab in Alderley Park | Wellcome Sanger Institute for the COVID-19 Genomics UK (COG-UK) Consortium | Jacquelyn Wynn, Mairead Hyland, The Lighthouse Lab in Alderley Park and Alex Alderton, Roberto Amato, Jeffrey Barrett, Sonia Goncalves, Ewan Harrison, David K. Jackson, Ian Johnston, Dominic Kwiatkowski, Cordelia Langford, John Sillitoe on behalf of the Wellcome Sanger Institute COVID-19 Surveillance Team |

| | | | |
|---|---|---|---|
| EPI_ISL_3167847 | Berkshire and Surrey Pathology Services Lighthouse Laboratory | Wellcome Sanger Institute for the COVID-19 Genomics UK (COG-UK) Consortium | Berkshire and Surrey Pathology Services Lighthouse Laboratory and Alex Alderton, Roberto Amato, Jeffrey Barrett, Sonia Goncalves, Ewan Harrison, David K. Jackson, Ian Johnston, Dominic Kwiatkowski, Cordelia Langford, John Sillitoe on behalf of the Wellcome Sanger Institute COVID-19 Surveillance Team |
| EPI_ISL_3167848, EPI_ISL_3167849, EPI_ISL_3167850, EPI_ISL_3167851, EPI_ISL_3167852, EPI_ISL_3167853 | Lighthouse Lab in Glasgow | Wellcome Sanger Institute for the COVID-19 Genomics UK (COG-UK) Consortium | Harper VanSteenhouse, Yumi Kasai, David Gray, Carol Clugston, Anna Dominiczak and Alex Alderton, Roberto Amato, Jeffrey Barrett, Sonia Goncalves, Ewan Harrison, David K. Jackson, Ian Johnston, Dominic Kwiatkowski, Cordelia Langford, John Sillitoe on behalf of the Wellcome Sanger Institute COVID-19 Surveillance Team |
| EPI_ISL_3167854 | Lighthouse Laboratory Plymouth | Wellcome Sanger Institute for the COVID-19 Genomics UK (COG-UK) Consortium | Lighthouse Laboratory Plymouth and Alex Alderton, Roberto Amato, Jeffrey Barrett, Sonia Goncalves, Ewan Harrison, David K. Jackson, Ian Johnston, Dominic Kwiatkowski, Cordelia Langford, John Sillitoe on behalf of the Wellcome Sanger Institute COVID-19 Surveillance Team |
| EPI_ISL_3167855 | Lighthouse Lab in Glasgow | Wellcome Sanger Institute for the COVID-19 Genomics UK (COG-UK) Consortium | Harper VanSteenhouse, Yumi Kasai, David Gray, Carol Clugston, Anna Dominiczak and Alex Alderton, Roberto Amato, Jeffrey Barrett, Sonia Goncalves, Ewan Harrison, David K. Jackson, Ian Johnston, Dominic Kwiatkowski, Cordelia Langford, John Sillitoe on behalf of the Wellcome Sanger Institute COVID-19 Surveillance Team |
| EPI_ISL_3167856 | Lighthouse Laboratory Plymouth | Wellcome Sanger Institute for the COVID-19 Genomics UK (COG-UK) Consortium | Lighthouse Laboratory Plymouth and Alex Alderton, Roberto Amato, Jeffrey Barrett, Sonia Goncalves, Ewan Harrison, David K. Jackson, Ian Johnston, Dominic Kwiatkowski, Cordelia Langford, John Sillitoe on behalf of the Wellcome Sanger Institute COVID-19 Surveillance Team |
| EPI_ISL_3167857 | Lighthouse Lab in Glasgow | Wellcome Sanger Institute for the COVID-19 Genomics UK (COG-UK) Consortium | Harper VanSteenhouse, Yumi Kasai, David Gray, Carol Clugston, Anna Dominiczak and Alex Alderton, Roberto Amato, Jeffrey Barrett, Sonia Goncalves, Ewan Harrison, David K. Jackson, Ian Johnston, Dominic Kwiatkowski, Cordelia Langford, John Sillitoe on behalf of the Wellcome Sanger Institute COVID-19 Surveillance Team |
| EPI_ISL_3167858 | Lighthouse Laboratory Plymouth | Wellcome Sanger Institute for the COVID-19 Genomics UK (COG-UK) Consortium | Lighthouse Laboratory Plymouth and Alex Alderton, Roberto Amato, Jeffrey Barrett, Sonia Goncalves, Ewan Harrison, David K. Jackson, Ian Johnston, Dominic Kwiatkowski, Cordelia Langford, John Sillitoe on behalf of the Wellcome Sanger Institute COVID-19 Surveillance Team |
| EPI_ISL_3167859, EPI_ISL_3167860, EPI_ISL_3167861, EPI_ISL_3167862 | Lighthouse Lab in Glasgow | Wellcome Sanger Institute for the COVID-19 Genomics UK (COG-UK) Consortium | Harper VanSteenhouse, Yumi Kasai, David Gray, Carol Clugston, Anna Dominiczak and Alex Alderton, Roberto Amato, Jeffrey Barrett, Sonia Goncalves, Ewan Harrison, David K. Jackson, Ian Johnston, Dominic Kwiatkowski, Cordelia Langford, John Sillitoe on behalf of the Wellcome Sanger Institute COVID-19 Surveillance Team |
| EPI_ISL_3167863 | Lighthouse Laboratory Plymouth | Wellcome Sanger Institute for the COVID-19 Genomics UK (COG-UK) Consortium | Lighthouse Laboratory Plymouth and Alex Alderton, Roberto Amato, Jeffrey Barrett, Sonia Goncalves, Ewan Harrison, David K. Jackson, Ian Johnston, Dominic Kwiatkowski, Cordelia Langford, John Sillitoe on behalf of the Wellcome Sanger Institute COVID-19 Surveillance Team |
| EPI_ISL_3167864 | Lighthouse Lab in Glasgow | Wellcome Sanger Institute for the COVID-19 Genomics UK (COG-UK) Consortium | Harper VanSteenhouse, Yumi Kasai, David Gray, Carol Clugston, Anna Dominiczak and Alex Alderton, Roberto Amato, Jeffrey Barrett, Sonia Goncalves, Ewan Harrison, David K. Jackson, Ian Johnston, Dominic Kwiatkowski, Cordelia Langford, John Sillitoe on behalf of the Wellcome Sanger Institute COVID-19 Surveillance Team |
| EPI_ISL_3167865 | Lighthouse Laboratory Plymouth | Wellcome Sanger Institute for the COVID-19 Genomics UK (COG-UK) Consortium | Lighthouse Laboratory Plymouth and Alex Alderton, Roberto Amato, Jeffrey Barrett, Sonia Goncalves, Ewan Harrison, David K. Jackson, Ian Johnston, Dominic Kwiatkowski, Cordelia Langford, John Sillitoe on behalf of the Wellcome Sanger Institute COVID-19 Surveillance Team |
| EPI_ISL_3167866, EPI_ISL_3167867 | Lighthouse Lab in Glasgow | Wellcome Sanger Institute for the COVID-19 Genomics UK (COG-UK) Consortium | Harper VanSteenhouse, Yumi Kasai, David Gray, Carol Clugston, Anna Dominiczak and Alex Alderton, Roberto Amato, Jeffrey Barrett, Sonia Goncalves, Ewan Harrison, David K. Jackson, Ian Johnston, Dominic Kwiatkowski, Cordelia Langford, John Sillitoe on behalf of the Wellcome Sanger Institute COVID-19 Surveillance Team |
| EPI_ISL_3167868, EPI_ISL_3167869, EPI_ISL_3167870 | Lighthouse Laboratory Plymouth | Wellcome Sanger Institute for the COVID-19 Genomics UK (COG-UK) Consortium | Lighthouse Laboratory Plymouth and Alex Alderton, Roberto Amato, Jeffrey Barrett, Sonia Goncalves, Ewan Harrison, David K. Jackson, Ian Johnston, Dominic Kwiatkowski, Cordelia Langford, John Sillitoe on behalf of the Wellcome Sanger Institute COVID-19 Surveillance Team |
| EPI_ISL_3167871, EPI_ISL_3167872 | Lighthouse Lab in Glasgow | Wellcome Sanger Institute for the COVID-19 Genomics UK (COG-UK) Consortium | Harper VanSteenhouse, Yumi Kasai, David Gray, Carol Clugston, Anna Dominiczak and Alex Alderton, Roberto Amato, Jeffrey Barrett, Sonia Goncalves, Ewan Harrison, David K. Jackson, Ian Johnston, Dominic Kwiatkowski, Cordelia Langford, John Sillitoe on behalf of the Wellcome Sanger Institute COVID-19 Surveillance Team |
| EPI_ISL_3167873 | Lighthouse Laboratory Plymouth | Wellcome Sanger Institute for the COVID-19 Genomics UK (COG-UK) Consortium | Lighthouse Laboratory Plymouth and Alex Alderton, Roberto Amato, Jeffrey Barrett, Sonia Goncalves, Ewan Harrison, David K. Jackson, Ian Johnston, Dominic Kwiatkowski, Cordelia Langford, John Sillitoe on behalf of the Wellcome Sanger Institute COVID-19 Surveillance Team |
| EPI_ISL_3167874, EPI_ISL_3167875, EPI_ISL_3167876 | Lighthouse Lab in Glasgow | Wellcome Sanger Institute for the COVID-19 Genomics UK (COG-UK) Consortium | Harper VanSteenhouse, Yumi Kasai, David Gray, Carol Clugston, Anna Dominiczak and Alex Alderton, Roberto Amato, Jeffrey Barrett, Sonia Goncalves, Ewan Harrison, David K. Jackson, Ian Johnston, Dominic Kwiatkowski, Cordelia Langford, John Sillitoe on behalf of the Wellcome Sanger Institute COVID-19 Surveillance Team |
| EPI_ISL_3167877 | Lighthouse Laboratory Plymouth | Wellcome Sanger Institute for the COVID-19 Genomics UK (COG-UK) Consortium | Lighthouse Laboratory Plymouth and Alex Alderton, Roberto Amato, Jeffrey Barrett, Sonia Goncalves, Ewan Harrison, David K. Jackson, Ian Johnston, Dominic Kwiatkowski, Cordelia Langford, John Sillitoe on behalf of the Wellcome Sanger Institute COVID-19 Surveillance Team |
| EPI_ISL_3167878, EPI_ISL_3167879, EPI_ISL_3167880, EPI_ISL_3167881, EPI_ISL_3167882, EPI_ISL_3167883 | Lighthouse Lab in Glasgow | Wellcome Sanger Institute for the COVID-19 Genomics UK (COG-UK) Consortium | Harper VanSteenhouse, Yumi Kasai, David Gray, Carol Clugston, Anna Dominiczak and Alex Alderton, Roberto Amato, Jeffrey Barrett, Sonia Goncalves, Ewan Harrison, David K. Jackson, Ian Johnston, Dominic Kwiatkowski, Cordelia Langford, John Sillitoe on behalf of the Wellcome Sanger Institute COVID-19 Surveillance Team |
| EPI_ISL_3167884 | Lighthouse Laboratory Plymouth | Wellcome Sanger Institute for the COVID-19 Genomics UK (COG-UK) Consortium | Lighthouse Laboratory Plymouth and Alex Alderton, Roberto Amato, Jeffrey Barrett, Sonia Goncalves, Ewan Harrison, David K. Jackson, Ian Johnston, Dominic Kwiatkowski, Cordelia Langford, John Sillitoe on behalf of the Wellcome Sanger Institute COVID-19 Surveillance Team |
| EPI_ISL_3167885 | Lighthouse Lab in Glasgow | Wellcome Sanger Institute for the COVID-19 Genomics UK (COG-UK) Consortium | Harper VanSteenhouse, Yumi Kasai, David Gray, Carol Clugston, Anna Dominiczak and Alex Alderton, Roberto Amato, Jeffrey Barrett, Sonia Goncalves, Ewan Harrison, David K. Jackson, Ian Johnston, Dominic Kwiatkowski, Cordelia Langford, John Sillitoe on behalf of the Wellcome Sanger Institute COVID-19 Surveillance Team |
| EPI_ISL_3167886, EPI_ISL_3167887 | Lighthouse Laboratory Plymouth | Wellcome Sanger Institute for the COVID-19 Genomics UK (COG-UK) Consortium | Lighthouse Laboratory Plymouth and Alex Alderton, Roberto Amato, Jeffrey Barrett, Sonia Goncalves, Ewan Harrison, David K. Jackson, Ian Johnston, Dominic Kwiatkowski, Cordelia Langford, John Sillitoe on behalf of the Wellcome Sanger Institute COVID-19 Surveillance Team |
| EPI_ISL_3167888, EPI_ISL_3167889, EPI_ISL_3167890, EPI_ISL_3167891, EPI_ISL_3167892, EPI_ISL_3167893, EPI_ISL_3167894, EPI_ISL_3167895, EPI_ISL_3167896, EPI_ISL_3167897 | Lighthouse Lab in Glasgow | Wellcome Sanger Institute for the COVID-19 Genomics UK (COG-UK) Consortium | Harper VanSteenhouse, Yumi Kasai, David Gray, Carol Clugston, Anna Dominiczak and Alex Alderton, Roberto Amato, Jeffrey Barrett, Sonia Goncalves, Ewan Harrison, David K. Jackson, Ian Johnston, Dominic Kwiatkowski, Cordelia Langford, John Sillitoe on behalf of the Wellcome Sanger Institute COVID-19 Surveillance Team |
| EPI_ISL_3167898 | Lighthouse Laboratory Plymouth | Wellcome Sanger Institute for the COVID-19 Genomics UK (COG-UK) Consortium | Lighthouse Laboratory Plymouth and Alex Alderton, Roberto Amato, Jeffrey Barrett, Sonia Goncalves, Ewan Harrison, David K. Jackson, Ian Johnston, Dominic Kwiatkowski, Cordelia Langford, John Sillitoe on behalf of the Wellcome Sanger Institute COVID-19 Surveillance Team |
| EPI_ISL_3167899 | Lighthouse Lab in Glasgow | Wellcome Sanger Institute for the COVID-19 Genomics UK (COG-UK) Consortium | Harper VanSteenhouse, Yumi Kasai, David Gray, Carol Clugston, Anna Dominiczak and Alex Alderton, Roberto Amato, Jeffrey Barrett, Sonia Goncalves, Ewan Harrison, David K. Jackson, Ian Johnston, Dominic Kwiatkowski, Cordelia Langford, John Sillitoe on behalf of the Wellcome Sanger Institute COVID-19 Surveillance Team |
| EPI_ISL_3167900 | Lighthouse Laboratory Plymouth | Wellcome Sanger Institute for the COVID-19 Genomics UK (COG-UK) Consortium | Lighthouse Laboratory Plymouth and Alex Alderton, Roberto Amato, Jeffrey Barrett, Sonia Goncalves, Ewan Harrison, David K. Jackson, Ian Johnston, Dominic Kwiatkowski, Cordelia Langford, John Sillitoe on behalf of the Wellcome Sanger Institute COVID-19 Surveillance Team |
| EPI_ISL_3167901, EPI_ISL_3167902, EPI_ISL_3167903, EPI_ISL_3167904, EPI_ISL_3167905, EPI_ISL_3167906 | Lighthouse Lab in Glasgow | Wellcome Sanger Institute for the COVID-19 Genomics UK (COG-UK) Consortium | Harper VanSteenhouse, Yumi Kasai, David Gray, Carol Clugston, Anna Dominiczak and Alex Alderton, Roberto Amato, Jeffrey Barrett, Sonia Goncalves, Ewan Harrison, David K. Jackson, Ian Johnston, Dominic Kwiatkowski, Cordelia Langford, John Sillitoe on behalf of the Wellcome Sanger Institute COVID-19 Surveillance Team |
| EPI_ISL_3167907, EPI_ISL_3167908 | Lighthouse Laboratory Plymouth | Wellcome Sanger Institute for the COVID-19 Genomics UK (COG-UK) Consortium | Lighthouse Laboratory Plymouth and Alex Alderton, Roberto Amato, Jeffrey Barrett, Sonia Goncalves, Ewan Harrison, David K. Jackson, Ian Johnston, Dominic Kwiatkowski, Cordelia Langford, John Sillitoe on behalf of the Wellcome Sanger Institute COVID-19 Surveillance Team |
| EPI_ISL_3167909, EPI_ISL_3167910, EPI_ISL_3167911, EPI_ISL_3167912, EPI_ISL_3167913 | Lighthouse Lab in Glasgow | Wellcome Sanger Institute for the COVID-19 Genomics UK (COG-UK) Consortium | Harper VanSteenhouse, Yumi Kasai, David Gray, Carol Clugston, Anna Dominiczak and Alex Alderton, Roberto Amato, Jeffrey Barrett, Sonia Goncalves, Ewan Harrison, David K. Jackson, Ian Johnston, Dominic Kwiatkowski, Cordelia Langford, John Sillitoe on behalf of the Wellcome Sanger Institute COVID-19 Surveillance Team |
| EPI_ISL_3167914, EPI_ISL_3167915 | Lighthouse Laboratory Plymouth | Wellcome Sanger Institute for the COVID-19 Genomics UK (COG-UK) Consortium | Lighthouse Laboratory Plymouth and Alex Alderton, Roberto Amato, Jeffrey Barrett, Sonia Goncalves, Ewan Harrison, David K. Jackson, Ian Johnston, Dominic Kwiatkowski, Cordelia Langford, John Sillitoe on behalf of the Wellcome Sanger Institute COVID-19 Surveillance Team |
| EPI_ISL_3167916, EPI_ISL_3167917, EPI_ISL_3167918, EPI_ISL_3167919, EPI_ISL_3167920, EPI_ISL_3167921 | Lighthouse Lab in Glasgow | Wellcome Sanger Institute for the COVID-19 Genomics UK (COG-UK) Consortium | Harper VanSteenhouse, Yumi Kasai, David Gray, Carol Clugston, Anna Dominiczak and Alex Alderton, Roberto Amato, Jeffrey Barrett, Sonia Goncalves, Ewan Harrison, David K. Jackson, Ian Johnston, Dominic Kwiatkowski, Cordelia Langford, John Sillitoe on behalf of the Wellcome Sanger Institute COVID-19 Surveillance Team |
| EPI_ISL_3167922, EPI_ISL_3167923 | Lighthouse Laboratory Plymouth | Wellcome Sanger Institute for the COVID-19 Genomics UK (COG-UK) Consortium | Lighthouse Laboratory Plymouth and Alex Alderton, Roberto Amato, Jeffrey Barrett, Sonia Goncalves, Ewan Harrison, David K. Jackson, Ian Johnston, Dominic Kwiatkowski, Cordelia Langford, John Sillitoe on behalf of the Wellcome Sanger Institute COVID-19 Surveillance Team |
| EPI_ISL_3167924 | Lighthouse Lab in Glasgow | Wellcome Sanger Institute for the COVID-19 Genomics UK (COG-UK) Consortium | Harper VanSteenhouse, Yumi Kasai, David Gray, Carol Clugston, Anna Dominiczak and Alex Alderton, Roberto Amato, Jeffrey Barrett, Sonia Goncalves, Ewan Harrison, David K. Jackson, Ian Johnston, Dominic Kwiatkowski, Cordelia Langford, John Sillitoe on behalf of the Wellcome Sanger Institute COVID-19 Surveillance Team |
| EPI_ISL_3167925 | Lighthouse Laboratory Plymouth | Wellcome Sanger Institute for the COVID-19 Genomics UK (COG-UK) | Lighthouse Laboratory Plymouth and Alex Alderton, Roberto Amato, Jeffrey Barrett, Sonia Goncalves, Ewan Harrison, David K. Jackson, Ian Johnston, Dominic Kwiatkowski, Cordelia Langford, John Sillitoe on behalf of the Wellcome Sanger |

| | | | |
|---|---|---|---|
| | | Consortium | Institute COVID-19 Surveillance Team |
| EPI_ISL_3167926, EPI_ISL_3167927 | Lighthouse Lab in Glasgow | Wellcome Sanger Institute for the COVID-19 Genomics UK (COG-UK) Consortium | Harper VanSteenhouse, Yumi Kasai, David Gray, Carol Clugston, Anna Dominiczak and Alex Alderton, Roberto Amato, Jeffrey Barrett, Sonia Goncalves, Ewan Harrison, David K. Jackson, Ian Johnston, Dominic Kwiatkowski, Cordelia Langford, John Sillitoe on behalf of the Wellcome Sanger Institute COVID-19 Surveillance Team |
| EPI_ISL_3167928 | Lighthouse Laboratory Plymouth | Wellcome Sanger Institute for the COVID-19 Genomics UK (COG-UK) Consortium | Lighthouse Laboratory Plymouth and Alex Alderton, Roberto Amato, Jeffrey Barrett, Sonia Goncalves, Ewan Harrison, David K. Jackson, Ian Johnston, Dominic Kwiatkowski, Cordelia Langford, John Sillitoe on behalf of the Wellcome Sanger Institute COVID-19 Surveillance Team |
| EPI_ISL_3167929, EPI_ISL_3167930, EPI_ISL_3167931, EPI_ISL_3167932, EPI_ISL_3167933, EPI_ISL_3167934, EPI_ISL_3167935 | Lighthouse Lab in Glasgow | Wellcome Sanger Institute for the COVID-19 Genomics UK (COG-UK) Consortium | Harper VanSteenhouse, Yumi Kasai, David Gray, Carol Clugston, Anna Dominiczak and Alex Alderton, Roberto Amato, Jeffrey Barrett, Sonia Goncalves, Ewan Harrison, David K. Jackson, Ian Johnston, Dominic Kwiatkowski, Cordelia Langford, John Sillitoe on behalf of the Wellcome Sanger Institute COVID-19 Surveillance Team |
| EPI_ISL_3167936 | Lighthouse Laboratory Plymouth | Wellcome Sanger Institute for the COVID-19 Genomics UK (COG-UK) Consortium | Lighthouse Laboratory Plymouth and Alex Alderton, Roberto Amato, Jeffrey Barrett, Sonia Goncalves, Ewan Harrison, David K. Jackson, Ian Johnston, Dominic Kwiatkowski, Cordelia Langford, John Sillitoe on behalf of the Wellcome Sanger Institute COVID-19 Surveillance Team |
| EPI_ISL_3167937, EPI_ISL_3167938, EPI_ISL_3167939, EPI_ISL_3167940, EPI_ISL_3167941, EPI_ISL_3167942 | Lighthouse Lab in Glasgow | Wellcome Sanger Institute for the COVID-19 Genomics UK (COG-UK) Consortium | Harper VanSteenhouse, Yumi Kasai, David Gray, Carol Clugston, Anna Dominiczak and Alex Alderton, Roberto Amato, Jeffrey Barrett, Sonia Goncalves, Ewan Harrison, David K. Jackson, Ian Johnston, Dominic Kwiatkowski, Cordelia Langford, John Sillitoe on behalf of the Wellcome Sanger Institute COVID-19 Surveillance Team |
| EPI_ISL_3167943 | Lighthouse Laboratory Plymouth | Wellcome Sanger Institute for the COVID-19 Genomics UK (COG-UK) Consortium | Lighthouse Laboratory Plymouth and Alex Alderton, Roberto Amato, Jeffrey Barrett, Sonia Goncalves, Ewan Harrison, David K. Jackson, Ian Johnston, Dominic Kwiatkowski, Cordelia Langford, John Sillitoe on behalf of the Wellcome Sanger Institute COVID-19 Surveillance Team |
| EPI_ISL_3167944, EPI_ISL_3167945, EPI_ISL_3167946, EPI_ISL_3167947, EPI_ISL_3167948, EPI_ISL_3167949, EPI_ISL_3167950, EPI_ISL_3167951, EPI_ISL_3167952, EPI_ISL_3167953, EPI_ISL_3167954, EPI_ISL_3167955, EPI_ISL_3167956 | | | |
| see above | Lighthouse Lab in Glasgow | Wellcome Sanger Institute for the COVID-19 Genomics UK (COG-UK) Consortium | Harper VanSteenhouse, Yumi Kasai, David Gray, Carol Clugston, Anna Dominiczak and Alex Alderton, Roberto Amato, Jeffrey Barrett, Sonia Goncalves, Ewan Harrison, David K. Jackson, Ian Johnston, Dominic Kwiatkowski, Cordelia Langford, John Sillitoe on behalf of the Wellcome Sanger Institute COVID-19 Surveillance Team |
| EPI_ISL_3167957, EPI_ISL_3167958, EPI_ISL_3167959 | Lighthouse Laboratory Plymouth | Wellcome Sanger Institute for the COVID-19 Genomics UK (COG-UK) Consortium | Lighthouse Laboratory Plymouth and Alex Alderton, Roberto Amato, Jeffrey Barrett, Sonia Goncalves, Ewan Harrison, David K. Jackson, Ian Johnston, Dominic Kwiatkowski, Cordelia Langford, John Sillitoe on behalf of the Wellcome Sanger Institute COVID-19 Surveillance Team |
| EPI_ISL_3167960, EPI_ISL_3167961 | Lighthouse Lab in Glasgow | Wellcome Sanger Institute for the COVID-19 Genomics UK (COG-UK) Consortium | Harper VanSteenhouse, Yumi Kasai, David Gray, Carol Clugston, Anna Dominiczak and Alex Alderton, Roberto Amato, Jeffrey Barrett, Sonia Goncalves, Ewan Harrison, David K. Jackson, Ian Johnston, Dominic Kwiatkowski, Cordelia Langford, John Sillitoe on behalf of the Wellcome Sanger Institute COVID-19 Surveillance Team |
| EPI_ISL_3167962 | Lighthouse Laboratory Plymouth | Wellcome Sanger Institute for the COVID-19 Genomics UK (COG-UK) Consortium | Lighthouse Laboratory Plymouth and Alex Alderton, Roberto Amato, Jeffrey Barrett, Sonia Goncalves, Ewan Harrison, David K. Jackson, Ian Johnston, Dominic Kwiatkowski, Cordelia Langford, John Sillitoe on behalf of the Wellcome Sanger Institute COVID-19 Surveillance Team |
| EPI_ISL_3167963, EPI_ISL_3167964, EPI_ISL_3167965 | Lighthouse Lab in Glasgow | Wellcome Sanger Institute for the COVID-19 Genomics UK (COG-UK) Consortium | Harper VanSteenhouse, Yumi Kasai, David Gray, Carol Clugston, Anna Dominiczak and Alex Alderton, Roberto Amato, Jeffrey Barrett, Sonia Goncalves, Ewan Harrison, David K. Jackson, Ian Johnston, Dominic Kwiatkowski, Cordelia Langford, John Sillitoe on behalf of the Wellcome Sanger Institute COVID-19 Surveillance Team |
| EPI_ISL_3167966 | Lighthouse Laboratory Plymouth | Wellcome Sanger Institute for the COVID-19 Genomics UK (COG-UK) Consortium | Lighthouse Laboratory Plymouth and Alex Alderton, Roberto Amato, Jeffrey Barrett, Sonia Goncalves, Ewan Harrison, David K. Jackson, Ian Johnston, Dominic Kwiatkowski, Cordelia Langford, John Sillitoe on behalf of the Wellcome Sanger Institute COVID-19 Surveillance Team |
| EPI_ISL_3167967, EPI_ISL_3167968 | Lighthouse Lab in Glasgow | Wellcome Sanger Institute for the COVID-19 Genomics UK (COG-UK) Consortium | Harper VanSteenhouse, Yumi Kasai, David Gray, Carol Clugston, Anna Dominiczak and Alex Alderton, Roberto Amato, Jeffrey Barrett, Sonia Goncalves, Ewan Harrison, David K. Jackson, Ian Johnston, Dominic Kwiatkowski, Cordelia Langford, John Sillitoe on behalf of the Wellcome Sanger Institute COVID-19 Surveillance Team |
| EPI_ISL_3167969 | Lighthouse Laboratory Plymouth | Wellcome Sanger Institute for the COVID-19 Genomics UK (COG-UK) Consortium | Lighthouse Laboratory Plymouth and Alex Alderton, Roberto Amato, Jeffrey Barrett, Sonia Goncalves, Ewan Harrison, David K. Jackson, Ian Johnston, Dominic Kwiatkowski, Cordelia Langford, John Sillitoe on behalf of the Wellcome Sanger Institute COVID-19 Surveillance Team |
| EPI_ISL_3167970 | Lighthouse Lab in Glasgow | Wellcome Sanger Institute for the COVID-19 Genomics UK (COG-UK) Consortium | Harper VanSteenhouse, Yumi Kasai, David Gray, Carol Clugston, Anna Dominiczak and Alex Alderton, Roberto Amato, Jeffrey Barrett, Sonia Goncalves, Ewan Harrison, David K. Jackson, Ian Johnston, Dominic Kwiatkowski, Cordelia Langford, John Sillitoe on behalf of the Wellcome Sanger Institute COVID-19 Surveillance Team |
| EPI_ISL_3167971 | Lighthouse Laboratory Plymouth | Wellcome Sanger Institute for the COVID-19 Genomics UK (COG-UK) Consortium | Lighthouse Laboratory Plymouth and Alex Alderton, Roberto Amato, Jeffrey Barrett, Sonia Goncalves, Ewan Harrison, David K. Jackson, Ian Johnston, Dominic Kwiatkowski, Cordelia Langford, John Sillitoe on behalf of the Wellcome Sanger Institute COVID-19 Surveillance Team |
| EPI_ISL_3167972, EPI_ISL_3167973, EPI_ISL_3167974, EPI_ISL_3167975 | Lighthouse Lab in Glasgow | Wellcome Sanger Institute for the COVID-19 Genomics UK (COG-UK) Consortium | Harper VanSteenhouse, Yumi Kasai, David Gray, Carol Clugston, Anna Dominiczak and Alex Alderton, Roberto Amato, Jeffrey Barrett, Sonia Goncalves, Ewan Harrison, David K. Jackson, Ian Johnston, Dominic Kwiatkowski, Cordelia Langford, John Sillitoe on behalf of the Wellcome Sanger Institute COVID-19 Surveillance Team |
| EPI_ISL_3167976 | Lighthouse Laboratory Plymouth | Wellcome Sanger Institute for the COVID-19 Genomics UK (COG-UK) Consortium | Lighthouse Laboratory Plymouth and Alex Alderton, Roberto Amato, Jeffrey Barrett, Sonia Goncalves, Ewan Harrison, David K. Jackson, Ian Johnston, Dominic Kwiatkowski, Cordelia Langford, John Sillitoe on behalf of the Wellcome Sanger Institute COVID-19 Surveillance Team |
| EPI_ISL_3167977, EPI_ISL_3167978 | Lighthouse Lab in Glasgow | Wellcome Sanger Institute for the COVID-19 Genomics UK (COG-UK) Consortium | Harper VanSteenhouse, Yumi Kasai, David Gray, Carol Clugston, Anna Dominiczak and Alex Alderton, Roberto Amato, Jeffrey Barrett, Sonia Goncalves, Ewan Harrison, David K. Jackson, Ian Johnston, Dominic Kwiatkowski, Cordelia Langford, John Sillitoe on behalf of the Wellcome Sanger Institute COVID-19 Surveillance Team |
| EPI_ISL_3167979, EPI_ISL_3167980 | Lighthouse Laboratory Plymouth | Wellcome Sanger Institute for the COVID-19 Genomics UK (COG-UK) Consortium | Lighthouse Laboratory Plymouth and Alex Alderton, Roberto Amato, Jeffrey Barrett, Sonia Goncalves, Ewan Harrison, David K. Jackson, Ian Johnston, Dominic Kwiatkowski, Cordelia Langford, John Sillitoe on behalf of the Wellcome Sanger Institute COVID-19 Surveillance Team |
| EPI_ISL_3167981, EPI_ISL_3167982, EPI_ISL_3167983, EPI_ISL_3167984 | Lighthouse Lab in Glasgow | Wellcome Sanger Institute for the COVID-19 Genomics UK (COG-UK) Consortium | Harper VanSteenhouse, Yumi Kasai, David Gray, Carol Clugston, Anna Dominiczak and Alex Alderton, Roberto Amato, Jeffrey Barrett, Sonia Goncalves, Ewan Harrison, David K. Jackson, Ian Johnston, Dominic Kwiatkowski, Cordelia Langford, John Sillitoe on behalf of the Wellcome Sanger Institute COVID-19 Surveillance Team |
| EPI_ISL_3167985 | Lighthouse Laboratory Plymouth | Wellcome Sanger Institute for the COVID-19 Genomics UK (COG-UK) Consortium | Lighthouse Laboratory Plymouth and Alex Alderton, Roberto Amato, Jeffrey Barrett, Sonia Goncalves, Ewan Harrison, David K. Jackson, Ian Johnston, Dominic Kwiatkowski, Cordelia Langford, John Sillitoe on behalf of the Wellcome Sanger Institute COVID-19 Surveillance Team |
| EPI_ISL_3167986 | Lighthouse Lab in Glasgow | Wellcome Sanger Institute for the COVID-19 Genomics UK (COG-UK) Consortium | Harper VanSteenhouse, Yumi Kasai, David Gray, Carol Clugston, Anna Dominiczak and Alex Alderton, Roberto Amato, Jeffrey Barrett, Sonia Goncalves, Ewan Harrison, David K. Jackson, Ian Johnston, Dominic Kwiatkowski, Cordelia Langford, John Sillitoe on behalf of the Wellcome Sanger Institute COVID-19 Surveillance Team |
| EPI_ISL_3167987, EPI_ISL_3167988 | Lighthouse Laboratory Plymouth | Wellcome Sanger Institute for the COVID-19 Genomics UK (COG-UK) Consortium | Lighthouse Laboratory Plymouth and Alex Alderton, Roberto Amato, Jeffrey Barrett, Sonia Goncalves, Ewan Harrison, David K. Jackson, Ian Johnston, Dominic Kwiatkowski, Cordelia Langford, John Sillitoe on behalf of the Wellcome Sanger Institute COVID-19 Surveillance Team |
| EPI_ISL_3167989, EPI_ISL_3167990 | Lighthouse Lab in Glasgow | Wellcome Sanger Institute for the COVID-19 Genomics UK (COG-UK) Consortium | Harper VanSteenhouse, Yumi Kasai, David Gray, Carol Clugston, Anna Dominiczak and Alex Alderton, Roberto Amato, Jeffrey Barrett, Sonia Goncalves, Ewan Harrison, David K. Jackson, Ian Johnston, Dominic Kwiatkowski, Cordelia Langford, John Sillitoe on behalf of the Wellcome Sanger Institute COVID-19 Surveillance Team |
| EPI_ISL_3167991 | Lighthouse Laboratory Plymouth | Wellcome Sanger Institute for the COVID-19 Genomics UK (COG-UK) Consortium | Lighthouse Laboratory Plymouth and Alex Alderton, Roberto Amato, Jeffrey Barrett, Sonia Goncalves, Ewan Harrison, David K. Jackson, Ian Johnston, Dominic Kwiatkowski, Cordelia Langford, John Sillitoe on behalf of the Wellcome Sanger Institute COVID-19 Surveillance Team |
| EPI_ISL_3167992, EPI_ISL_3167993, EPI_ISL_3167994, EPI_ISL_3167995, EPI_ISL_3167996, EPI_ISL_3167997, EPI_ISL_3167998, EPI_ISL_3167999 | Lighthouse Lab in Glasgow | Wellcome Sanger Institute for the COVID-19 Genomics UK (COG-UK) Consortium | Harper VanSteenhouse, Yumi Kasai, David Gray, Carol Clugston, Anna Dominiczak and Alex Alderton, Roberto Amato, Jeffrey Barrett, Sonia Goncalves, Ewan Harrison, David K. Jackson, Ian Johnston, Dominic Kwiatkowski, Cordelia Langford, John Sillitoe on behalf of the Wellcome Sanger Institute COVID-19 Surveillance Team |
| EPI_ISL_3168000, EPI_ISL_3168001, EPI_ISL_3168002 | Lighthouse Laboratory Plymouth | Wellcome Sanger Institute for the COVID-19 Genomics UK (COG-UK) Consortium | Lighthouse Laboratory Plymouth and Alex Alderton, Roberto Amato, Jeffrey Barrett, Sonia Goncalves, Ewan Harrison, David K. Jackson, Ian Johnston, Dominic Kwiatkowski, Cordelia Langford, John Sillitoe on behalf of the Wellcome Sanger Institute COVID-19 Surveillance Team |
| EPI_ISL_3168003, EPI_ISL_3168004, EPI_ISL_3168005, EPI_ISL_3168006, EPI_ISL_3168007, EPI_ISL_3168008, EPI_ISL_3168009, EPI_ISL_3168010, EPI_ISL_3168011, EPI_ISL_3168012, EPI_ISL_3168013, EPI_ISL_3168014 | | | |
| see above | Lighthouse Lab in Glasgow | Wellcome Sanger Institute for the COVID-19 Genomics UK (COG-UK) Consortium | Harper VanSteenhouse, Yumi Kasai, David Gray, Carol Clugston, Anna Dominiczak and Alex Alderton, Roberto Amato, Jeffrey Barrett, Sonia Goncalves, Ewan Harrison, David K. Jackson, Ian Johnston, Dominic Kwiatkowski, Cordelia Langford, John Sillitoe on behalf of the Wellcome Sanger Institute COVID-19 Surveillance Team |
| EPI_ISL_3168015 | Lighthouse Laboratory Plymouth | Wellcome Sanger Institute for the COVID-19 Genomics UK (COG-UK) Consortium | Lighthouse Laboratory Plymouth and Alex Alderton, Roberto Amato, Jeffrey Barrett, Sonia Goncalves, Ewan Harrison, David K. Jackson, Ian Johnston, Dominic Kwiatkowski, Cordelia Langford, John Sillitoe on behalf of the Wellcome Sanger Institute COVID-19 Surveillance Team |
| EPI_ISL_3168016, EPI_ISL_3168017 | Lighthouse Lab in Glasgow | Wellcome Sanger Institute for the COVID-19 Genomics UK (COG-UK) Consortium | Harper VanSteenhouse, Yumi Kasai, David Gray, Carol Clugston, Anna Dominiczak and Alex Alderton, Roberto Amato, Jeffrey Barrett, Sonia Goncalves, Ewan Harrison, David K. Jackson, Ian Johnston, Dominic Kwiatkowski, Cordelia Langford, John Sillitoe on behalf of the Wellcome Sanger Institute COVID-19 Surveillance Team |
| EPI_ISL_3168018, EPI_ISL_3168019, EPI_ISL_3168020, EPI_ISL_3168021, EPI_ISL_3168022, EPI_ISL_3168023, EPI_ISL_3168024, EPI_ISL_3168025 | Lighthouse Laboratory Plymouth | Wellcome Sanger Institute for the COVID-19 Genomics UK (COG-UK) Consortium | Lighthouse Laboratory Plymouth and Alex Alderton, Roberto Amato, Jeffrey Barrett, Sonia Goncalves, Ewan Harrison, David K. Jackson, Ian Johnston, Dominic Kwiatkowski, Cordelia Langford, John Sillitoe on behalf of the Wellcome Sanger Institute COVID-19 Surveillance Team |

| | | | |
|---|---|---|---|
| EPI_ISL_3168026 | Lighthouse Lab in Glasgow | Wellcome Sanger Institute for the COVID-19 Genomics UK (COG-UK) Consortium | Harper VanSteenhouse, Yumi Kasai, David Gray, Carol Clugston, Anna Dominiczak and Alex Alderton, Roberto Amato, Jeffrey Barrett, Sonia Goncalves, Ewan Harrison, David K. Jackson, Ian Johnston, Dominic Kwiatkowski, Cordelia Langford, John Sillitoe on behalf of the Wellcome Sanger Institute COVID-19 Surveillance Team |
| EPI_ISL_3168027 | Lighthouse Laboratory Plymouth | Wellcome Sanger Institute for the COVID-19 Genomics UK (COG-UK) Consortium | Lighthouse Laboratory Plymouth and Alex Alderton, Roberto Amato, Jeffrey Barrett, Sonia Goncalves, Ewan Harrison, David K. Jackson, Ian Johnston, Dominic Kwiatkowski, Cordelia Langford, John Sillitoe on behalf of the Wellcome Sanger Institute COVID-19 Surveillance Team |
| EPI_ISL_3168028 | Lighthouse Lab in Glasgow | Wellcome Sanger Institute for the COVID-19 Genomics UK (COG-UK) Consortium | Harper VanSteenhouse, Yumi Kasai, David Gray, Carol Clugston, Anna Dominiczak and Alex Alderton, Roberto Amato, Jeffrey Barrett, Sonia Goncalves, Ewan Harrison, David K. Jackson, Ian Johnston, Dominic Kwiatkowski, Cordelia Langford, John Sillitoe on behalf of the Wellcome Sanger Institute COVID-19 Surveillance Team |
| EPI_ISL_3168029 | Lighthouse Laboratory Plymouth | Wellcome Sanger Institute for the COVID-19 Genomics UK (COG-UK) Consortium | Lighthouse Laboratory Plymouth and Alex Alderton, Roberto Amato, Jeffrey Barrett, Sonia Goncalves, Ewan Harrison, David K. Jackson, Ian Johnston, Dominic Kwiatkowski, Cordelia Langford, John Sillitoe on behalf of the Wellcome Sanger Institute COVID-19 Surveillance Team |
| EPI_ISL_3168030 | Lighthouse Lab in Glasgow | Wellcome Sanger Institute for the COVID-19 Genomics UK (COG-UK) Consortium | Harper VanSteenhouse, Yumi Kasai, David Gray, Carol Clugston, Anna Dominiczak and Alex Alderton, Roberto Amato, Jeffrey Barrett, Sonia Goncalves, Ewan Harrison, David K. Jackson, Ian Johnston, Dominic Kwiatkowski, Cordelia Langford, John Sillitoe on behalf of the Wellcome Sanger Institute COVID-19 Surveillance Team |
| EPI_ISL_3168031, EPI_ISL_3168032 | Lighthouse Laboratory Plymouth | Wellcome Sanger Institute for the COVID-19 Genomics UK (COG-UK) Consortium | Lighthouse Laboratory Plymouth and Alex Alderton, Roberto Amato, Jeffrey Barrett, Sonia Goncalves, Ewan Harrison, David K. Jackson, Ian Johnston, Dominic Kwiatkowski, Cordelia Langford, John Sillitoe on behalf of the Wellcome Sanger Institute COVID-19 Surveillance Team |
| EPI_ISL_3168033, EPI_ISL_3168034, EPI_ISL_3168035 | Lighthouse Lab in Glasgow | Wellcome Sanger Institute for the COVID-19 Genomics UK (COG-UK) Consortium | Harper VanSteenhouse, Yumi Kasai, David Gray, Carol Clugston, Anna Dominiczak and Alex Alderton, Roberto Amato, Jeffrey Barrett, Sonia Goncalves, Ewan Harrison, David K. Jackson, Ian Johnston, Dominic Kwiatkowski, Cordelia Langford, John Sillitoe on behalf of the Wellcome Sanger Institute COVID-19 Surveillance Team |
| EPI_ISL_3168036 | Lighthouse Laboratory Plymouth | Wellcome Sanger Institute for the COVID-19 Genomics UK (COG-UK) Consortium | Lighthouse Laboratory Plymouth and Alex Alderton, Roberto Amato, Jeffrey Barrett, Sonia Goncalves, Ewan Harrison, David K. Jackson, Ian Johnston, Dominic Kwiatkowski, Cordelia Langford, John Sillitoe on behalf of the Wellcome Sanger Institute COVID-19 Surveillance Team |
| EPI_ISL_3168037, EPI_ISL_3168038, EPI_ISL_3168039 | Lighthouse Lab in Glasgow | Wellcome Sanger Institute for the COVID-19 Genomics UK (COG-UK) Consortium | Harper VanSteenhouse, Yumi Kasai, David Gray, Carol Clugston, Anna Dominiczak and Alex Alderton, Roberto Amato, Jeffrey Barrett, Sonia Goncalves, Ewan Harrison, David K. Jackson, Ian Johnston, Dominic Kwiatkowski, Cordelia Langford, John Sillitoe on behalf of the Wellcome Sanger Institute COVID-19 Surveillance Team |
| EPI_ISL_3168040 | Lighthouse Laboratory Plymouth | Wellcome Sanger Institute for the COVID-19 Genomics UK (COG-UK) Consortium | Lighthouse Laboratory Plymouth and Alex Alderton, Roberto Amato, Jeffrey Barrett, Sonia Goncalves, Ewan Harrison, David K. Jackson, Ian Johnston, Dominic Kwiatkowski, Cordelia Langford, John Sillitoe on behalf of the Wellcome Sanger Institute COVID-19 Surveillance Team |
| EPI_ISL_3168041 | Lighthouse Lab in Glasgow | Wellcome Sanger Institute for the COVID-19 Genomics UK (COG-UK) Consortium | Harper VanSteenhouse, Yumi Kasai, David Gray, Carol Clugston, Anna Dominiczak and Alex Alderton, Roberto Amato, Jeffrey Barrett, Sonia Goncalves, Ewan Harrison, David K. Jackson, Ian Johnston, Dominic Kwiatkowski, Cordelia Langford, John Sillitoe on behalf of the Wellcome Sanger Institute COVID-19 Surveillance Team |
| EPI_ISL_3168042 | Lighthouse Laboratory Plymouth | Wellcome Sanger Institute for the COVID-19 Genomics UK (COG-UK) Consortium | Lighthouse Laboratory Plymouth and Alex Alderton, Roberto Amato, Jeffrey Barrett, Sonia Goncalves, Ewan Harrison, David K. Jackson, Ian Johnston, Dominic Kwiatkowski, Cordelia Langford, John Sillitoe on behalf of the Wellcome Sanger Institute COVID-19 Surveillance Team |
| EPI_ISL_3168043 | Lighthouse Lab in Glasgow | Wellcome Sanger Institute for the COVID-19 Genomics UK (COG-UK) Consortium | Harper VanSteenhouse, Yumi Kasai, David Gray, Carol Clugston, Anna Dominiczak and Alex Alderton, Roberto Amato, Jeffrey Barrett, Sonia Goncalves, Ewan Harrison, David K. Jackson, Ian Johnston, Dominic Kwiatkowski, Cordelia Langford, John Sillitoe on behalf of the Wellcome Sanger Institute COVID-19 Surveillance Team |
| EPI_ISL_3168044 | Lighthouse Laboratory Plymouth | Wellcome Sanger Institute for the COVID-19 Genomics UK (COG-UK) Consortium | Lighthouse Laboratory Plymouth and Alex Alderton, Roberto Amato, Jeffrey Barrett, Sonia Goncalves, Ewan Harrison, David K. Jackson, Ian Johnston, Dominic Kwiatkowski, Cordelia Langford, John Sillitoe on behalf of the Wellcome Sanger Institute COVID-19 Surveillance Team |
| EPI_ISL_3168045, EPI_ISL_3168046, EPI_ISL_3168047, EPI_ISL_3168048, EPI_ISL_3168049, EPI_ISL_3168050 | Lighthouse Lab in Glasgow | Wellcome Sanger Institute for the COVID-19 Genomics UK (COG-UK) Consortium | Harper VanSteenhouse, Yumi Kasai, David Gray, Carol Clugston, Anna Dominiczak and Alex Alderton, Roberto Amato, Jeffrey Barrett, Sonia Goncalves, Ewan Harrison, David K. Jackson, Ian Johnston, Dominic Kwiatkowski, Cordelia Langford, John Sillitoe on behalf of the Wellcome Sanger Institute COVID-19 Surveillance Team |
| EPI_ISL_3168051, EPI_ISL_3168052, EPI_ISL_3168053 | Lighthouse Laboratory Plymouth | Wellcome Sanger Institute for the COVID-19 Genomics UK (COG-UK) Consortium | Lighthouse Laboratory Plymouth and Alex Alderton, Roberto Amato, Jeffrey Barrett, Sonia Goncalves, Ewan Harrison, David K. Jackson, Ian Johnston, Dominic Kwiatkowski, Cordelia Langford, John Sillitoe on behalf of the Wellcome Sanger Institute COVID-19 Surveillance Team |
| EPI_ISL_3168054, EPI_ISL_3168055, EPI_ISL_3168056, EPI_ISL_3168057, EPI_ISL_3168058, EPI_ISL_3168059 | Lighthouse Lab in Glasgow | Wellcome Sanger Institute for the COVID-19 Genomics UK (COG-UK) Consortium | Harper VanSteenhouse, Yumi Kasai, David Gray, Carol Clugston, Anna Dominiczak and Alex Alderton, Roberto Amato, Jeffrey Barrett, Sonia Goncalves, Ewan Harrison, David K. Jackson, Ian Johnston, Dominic Kwiatkowski, Cordelia Langford, John Sillitoe on behalf of the Wellcome Sanger Institute COVID-19 Surveillance Team |
| EPI_ISL_3168060 | Lighthouse Laboratory Plymouth | Wellcome Sanger Institute for the COVID-19 Genomics UK (COG-UK) Consortium | Lighthouse Laboratory Plymouth and Alex Alderton, Roberto Amato, Jeffrey Barrett, Sonia Goncalves, Ewan Harrison, David K. Jackson, Ian Johnston, Dominic Kwiatkowski, Cordelia Langford, John Sillitoe on behalf of the Wellcome Sanger Institute COVID-19 Surveillance Team |
| EPI_ISL_3168061, EPI_ISL_3168062, EPI_ISL_3168063, EPI_ISL_3168064, EPI_ISL_3168065, EPI_ISL_3168066, EPI_ISL_3168067, EPI_ISL_3168068, EPI_ISL_3168069, EPI_ISL_3168070 | Lighthouse Lab in Glasgow | Wellcome Sanger Institute for the COVID-19 Genomics UK (COG-UK) Consortium | Harper VanSteenhouse, Yumi Kasai, David Gray, Carol Clugston, Anna Dominiczak and Alex Alderton, Roberto Amato, Jeffrey Barrett, Sonia Goncalves, Ewan Harrison, David K. Jackson, Ian Johnston, Dominic Kwiatkowski, Cordelia Langford, John Sillitoe on behalf of the Wellcome Sanger Institute COVID-19 Surveillance Team |
| EPI_ISL_3168071, EPI_ISL_3168072 | Lighthouse Laboratory Plymouth | Wellcome Sanger Institute for the COVID-19 Genomics UK (COG-UK) Consortium | Lighthouse Laboratory Plymouth and Alex Alderton, Roberto Amato, Jeffrey Barrett, Sonia Goncalves, Ewan Harrison, David K. Jackson, Ian Johnston, Dominic Kwiatkowski, Cordelia Langford, John Sillitoe on behalf of the Wellcome Sanger Institute COVID-19 Surveillance Team |
| EPI_ISL_3168073, EPI_ISL_3168074 | Lighthouse Lab in Glasgow | Wellcome Sanger Institute for the COVID-19 Genomics UK (COG-UK) Consortium | Harper VanSteenhouse, Yumi Kasai, David Gray, Carol Clugston, Anna Dominiczak and Alex Alderton, Roberto Amato, Jeffrey Barrett, Sonia Goncalves, Ewan Harrison, David K. Jackson, Ian Johnston, Dominic Kwiatkowski, Cordelia Langford, John Sillitoe on behalf of the Wellcome Sanger Institute COVID-19 Surveillance Team |
| EPI_ISL_3168075 | Lighthouse Laboratory Plymouth | Wellcome Sanger Institute for the COVID-19 Genomics UK (COG-UK) Consortium | Lighthouse Laboratory Plymouth and Alex Alderton, Roberto Amato, Jeffrey Barrett, Sonia Goncalves, Ewan Harrison, David K. Jackson, Ian Johnston, Dominic Kwiatkowski, Cordelia Langford, John Sillitoe on behalf of the Wellcome Sanger Institute COVID-19 Surveillance Team |
| EPI_ISL_3168076, EPI_ISL_3168077 | Lighthouse Lab in Glasgow | Wellcome Sanger Institute for the COVID-19 Genomics UK (COG-UK) Consortium | Harper VanSteenhouse, Yumi Kasai, David Gray, Carol Clugston, Anna Dominiczak and Alex Alderton, Roberto Amato, Jeffrey Barrett, Sonia Goncalves, Ewan Harrison, David K. Jackson, Ian Johnston, Dominic Kwiatkowski, Cordelia Langford, John Sillitoe on behalf of the Wellcome Sanger Institute COVID-19 Surveillance Team |
| EPI_ISL_3168078 | Lighthouse Laboratory Plymouth | Wellcome Sanger Institute for the COVID-19 Genomics UK (COG-UK) Consortium | Lighthouse Laboratory Plymouth and Alex Alderton, Roberto Amato, Jeffrey Barrett, Sonia Goncalves, Ewan Harrison, David K. Jackson, Ian Johnston, Dominic Kwiatkowski, Cordelia Langford, John Sillitoe on behalf of the Wellcome Sanger Institute COVID-19 Surveillance Team |
| EPI_ISL_3168079 | Lighthouse Lab in Glasgow | Wellcome Sanger Institute for the COVID-19 Genomics UK (COG-UK) Consortium | Harper VanSteenhouse, Yumi Kasai, David Gray, Carol Clugston, Anna Dominiczak and Alex Alderton, Roberto Amato, Jeffrey Barrett, Sonia Goncalves, Ewan Harrison, David K. Jackson, Ian Johnston, Dominic Kwiatkowski, Cordelia Langford, John Sillitoe on behalf of the Wellcome Sanger Institute COVID-19 Surveillance Team |
| EPI_ISL_3168080, EPI_ISL_3168081 | Lighthouse Laboratory Plymouth | Wellcome Sanger Institute for the COVID-19 Genomics UK (COG-UK) Consortium | Lighthouse Laboratory Plymouth and Alex Alderton, Roberto Amato, Jeffrey Barrett, Sonia Goncalves, Ewan Harrison, David K. Jackson, Ian Johnston, Dominic Kwiatkowski, Cordelia Langford, John Sillitoe on behalf of the Wellcome Sanger Institute COVID-19 Surveillance Team |
| EPI_ISL_3168082 | Lighthouse Lab in Glasgow | Wellcome Sanger Institute for the COVID-19 Genomics UK (COG-UK) Consortium | Harper VanSteenhouse, Yumi Kasai, David Gray, Carol Clugston, Anna Dominiczak and Alex Alderton, Roberto Amato, Jeffrey Barrett, Sonia Goncalves, Ewan Harrison, David K. Jackson, Ian Johnston, Dominic Kwiatkowski, Cordelia Langford, John Sillitoe on behalf of the Wellcome Sanger Institute COVID-19 Surveillance Team |
| EPI_ISL_3168083, EPI_ISL_3168084, EPI_ISL_3168085 | Lighthouse Laboratory Plymouth | Wellcome Sanger Institute for the COVID-19 Genomics UK (COG-UK) Consortium | Lighthouse Laboratory Plymouth and Alex Alderton, Roberto Amato, Jeffrey Barrett, Sonia Goncalves, Ewan Harrison, David K. Jackson, Ian Johnston, Dominic Kwiatkowski, Cordelia Langford, John Sillitoe on behalf of the Wellcome Sanger Institute COVID-19 Surveillance Team |
| EPI_ISL_3168086 | Lighthouse Lab in Glasgow | Wellcome Sanger Institute for the COVID-19 Genomics UK (COG-UK) Consortium | Harper VanSteenhouse, Yumi Kasai, David Gray, Carol Clugston, Anna Dominiczak and Alex Alderton, Roberto Amato, Jeffrey Barrett, Sonia Goncalves, Ewan Harrison, David K. Jackson, Ian Johnston, Dominic Kwiatkowski, Cordelia Langford, John Sillitoe on behalf of the Wellcome Sanger Institute COVID-19 Surveillance Team |
| EPI_ISL_3168087 | Lighthouse Laboratory Plymouth | Wellcome Sanger Institute for the COVID-19 Genomics UK (COG-UK) Consortium | Lighthouse Laboratory Plymouth and Alex Alderton, Roberto Amato, Jeffrey Barrett, Sonia Goncalves, Ewan Harrison, David K. Jackson, Ian Johnston, Dominic Kwiatkowski, Cordelia Langford, John Sillitoe on behalf of the Wellcome Sanger Institute COVID-19 Surveillance Team |
| EPI_ISL_3168088 | Lighthouse Lab in Glasgow | Wellcome Sanger Institute for the COVID-19 Genomics UK (COG-UK) Consortium | Harper VanSteenhouse, Yumi Kasai, David Gray, Carol Clugston, Anna Dominiczak and Alex Alderton, Roberto Amato, Jeffrey Barrett, Sonia Goncalves, Ewan Harrison, David K. Jackson, Ian Johnston, Dominic Kwiatkowski, Cordelia Langford, John Sillitoe on behalf of the Wellcome Sanger Institute COVID-19 Surveillance Team |
| EPI_ISL_3168089, EPI_ISL_3168090 | Lighthouse Laboratory Plymouth | Wellcome Sanger Institute for the COVID-19 Genomics UK (COG-UK) Consortium | Lighthouse Laboratory Plymouth and Alex Alderton, Roberto Amato, Jeffrey Barrett, Sonia Goncalves, Ewan Harrison, David K. Jackson, Ian Johnston, Dominic Kwiatkowski, Cordelia Langford, John Sillitoe on behalf of the Wellcome Sanger Institute COVID-19 Surveillance Team |
| EPI_ISL_3168091, EPI_ISL_3168092, EPI_ISL_3168093, EPI_ISL_3168094, EPI_ISL_3168095, EPI_ISL_3168096 | Lighthouse Lab in Glasgow | Wellcome Sanger Institute for the COVID-19 Genomics UK (COG-UK) Consortium | Harper VanSteenhouse, Yumi Kasai, David Gray, Carol Clugston, Anna Dominiczak and Alex Alderton, Roberto Amato, Jeffrey Barrett, Sonia Goncalves, Ewan Harrison, David K. Jackson, Ian Johnston, Dominic Kwiatkowski, Cordelia Langford, John Sillitoe on behalf of the Wellcome Sanger Institute COVID-19 Surveillance Team |
| EPI_ISL_3168097, EPI_ISL_3168098 | Lighthouse Laboratory Plymouth | Wellcome Sanger Institute for the COVID-19 Genomics UK (COG-UK) Consortium | Lighthouse Laboratory Plymouth and Alex Alderton, Roberto Amato, Jeffrey Barrett, Sonia Goncalves, Ewan Harrison, David K. Jackson, Ian Johnston, Dominic Kwiatkowski, Cordelia Langford, John Sillitoe on behalf of the Wellcome Sanger Institute COVID-19 Surveillance Team |
| EPI_ISL_3168099 | Lighthouse Lab in Glasgow | Wellcome Sanger Institute for the COVID-19 Genomics UK (COG-UK) Consortium | Harper VanSteenhouse, Yumi Kasai, David Gray, Carol Clugston, Anna Dominiczak and Alex Alderton, Roberto Amato, Jeffrey Barrett, Sonia Goncalves, Ewan Harrison, David K. Jackson, Ian Johnston, Dominic Kwiatkowski, Cordelia Langford, John Sillitoe on behalf of the Wellcome Sanger Institute COVID-19 Surveillance Team |

| | | | |
|---|---|---|---|
| EPI_ISL_3168100 | Lighthouse Laboratory Plymouth | Wellcome Sanger Institute for the COVID-19 Genomics UK (COG-UK) Consortium | Lighthouse Laboratory Plymouth and Alex Alderton, Roberto Amato, Jeffrey Barrett, Sonia Goncalves, Ewan Harrison, David K. Jackson, Ian Johnston, Dominic Kwiatkowski, Cordelia Langford, John Sillitoe on behalf of the Wellcome Sanger Institute COVID-19 Surveillance Team |
| EPI_ISL_3168101, EPI_ISL_3168102 | Lighthouse Lab in Glasgow | Wellcome Sanger Institute for the COVID-19 Genomics UK (COG-UK) Consortium | Harper VanSteenhouse, Yumi Kasai, David Gray, Carol Clugston, Anna Dominiczak and Alex Alderton, Roberto Amato, Jeffrey Barrett, Sonia Goncalves, Ewan Harrison, David K. Jackson, Ian Johnston, Dominic Kwiatkowski, Cordelia Langford, John Sillitoe on behalf of the Wellcome Sanger Institute COVID-19 Surveillance Team |
| EPI_ISL_3168103 | Lighthouse Laboratory Plymouth | Wellcome Sanger Institute for the COVID-19 Genomics UK (COG-UK) Consortium | Lighthouse Laboratory Plymouth and Alex Alderton, Roberto Amato, Jeffrey Barrett, Sonia Goncalves, Ewan Harrison, David K. Jackson, Ian Johnston, Dominic Kwiatkowski, Cordelia Langford, John Sillitoe on behalf of the Wellcome Sanger Institute COVID-19 Surveillance Team |
| EPI_ISL_3168104, EPI_ISL_3168105 | Lighthouse Lab in Glasgow | Wellcome Sanger Institute for the COVID-19 Genomics UK (COG-UK) Consortium | Harper VanSteenhouse, Yumi Kasai, David Gray, Carol Clugston, Anna Dominiczak and Alex Alderton, Roberto Amato, Jeffrey Barrett, Sonia Goncalves, Ewan Harrison, David K. Jackson, Ian Johnston, Dominic Kwiatkowski, Cordelia Langford, John Sillitoe on behalf of the Wellcome Sanger Institute COVID-19 Surveillance Team |
| EPI_ISL_3168106 | Lighthouse Laboratory Plymouth | Wellcome Sanger Institute for the COVID-19 Genomics UK (COG-UK) Consortium | Lighthouse Laboratory Plymouth and Alex Alderton, Roberto Amato, Jeffrey Barrett, Sonia Goncalves, Ewan Harrison, David K. Jackson, Ian Johnston, Dominic Kwiatkowski, Cordelia Langford, John Sillitoe on behalf of the Wellcome Sanger Institute COVID-19 Surveillance Team |
| EPI_ISL_3168107, EPI_ISL_3168108, EPI_ISL_3168109 | Lighthouse Lab in Glasgow | Wellcome Sanger Institute for the COVID-19 Genomics UK (COG-UK) Consortium | Harper VanSteenhouse, Yumi Kasai, David Gray, Carol Clugston, Anna Dominiczak and Alex Alderton, Roberto Amato, Jeffrey Barrett, Sonia Goncalves, Ewan Harrison, David K. Jackson, Ian Johnston, Dominic Kwiatkowski, Cordelia Langford, John Sillitoe on behalf of the Wellcome Sanger Institute COVID-19 Surveillance Team |
| EPI_ISL_3168110, EPI_ISL_3168111 | Lighthouse Laboratory Plymouth | Wellcome Sanger Institute for the COVID-19 Genomics UK (COG-UK) Consortium | Lighthouse Laboratory Plymouth and Alex Alderton, Roberto Amato, Jeffrey Barrett, Sonia Goncalves, Ewan Harrison, David K. Jackson, Ian Johnston, Dominic Kwiatkowski, Cordelia Langford, John Sillitoe on behalf of the Wellcome Sanger Institute COVID-19 Surveillance Team |
| EPI_ISL_3168112, EPI_ISL_3168113 | Lighthouse Lab in Glasgow | Wellcome Sanger Institute for the COVID-19 Genomics UK (COG-UK) Consortium | Harper VanSteenhouse, Yumi Kasai, David Gray, Carol Clugston, Anna Dominiczak and Alex Alderton, Roberto Amato, Jeffrey Barrett, Sonia Goncalves, Ewan Harrison, David K. Jackson, Ian Johnston, Dominic Kwiatkowski, Cordelia Langford, John Sillitoe on behalf of the Wellcome Sanger Institute COVID-19 Surveillance Team |
| EPI_ISL_3168114, EPI_ISL_3168115, EPI_ISL_3168116, EPI_ISL_3168117, EPI_ISL_3168118, EPI_ISL_3168119, EPI_ISL_3168120, EPI_ISL_3168121 | Lighthouse Lab in Milton Keynes | Wellcome Sanger Institute for the COVID-19 Genomics UK (COG-UK) Consortium | The Lighthouse Lab in Milton Keynes and Alex Alderton, Roberto Amato, Jeffrey Barrett, Sonia Goncalves, Ewan Harrison, David K. Jackson, Ian Johnston, Dominic Kwiatkowski, Cordelia Langford, John Sillitoe on behalf of the Wellcome Sanger Institute COVID-19 Surveillance Team |
| EPI_ISL_3168122 | Lighthouse Lab in Glasgow | Wellcome Sanger Institute for the COVID-19 Genomics UK (COG-UK) Consortium | Harper VanSteenhouse, Yumi Kasai, David Gray, Carol Clugston, Anna Dominiczak and Alex Alderton, Roberto Amato, Jeffrey Barrett, Sonia Goncalves, Ewan Harrison, David K. Jackson, Ian Johnston, Dominic Kwiatkowski, Cordelia Langford, John Sillitoe on behalf of the Wellcome Sanger Institute COVID-19 Surveillance Team |
| EPI_ISL_3168123, EPI_ISL_3168124, EPI_ISL_3168125, EPI_ISL_3168126, EPI_ISL_3168127, EPI_ISL_3168128, EPI_ISL_3168129 | Lighthouse Lab in Milton Keynes | Wellcome Sanger Institute for the COVID-19 Genomics UK (COG-UK) Consortium | The Lighthouse Lab in Milton Keynes and Alex Alderton, Roberto Amato, Jeffrey Barrett, Sonia Goncalves, Ewan Harrison, David K. Jackson, Ian Johnston, Dominic Kwiatkowski, Cordelia Langford, John Sillitoe on behalf of the Wellcome Sanger Institute COVID-19 Surveillance Team |
| EPI_ISL_3168130, EPI_ISL_3168131 | Lighthouse Lab in Glasgow | Wellcome Sanger Institute for the COVID-19 Genomics UK (COG-UK) Consortium | Harper VanSteenhouse, Yumi Kasai, David Gray, Carol Clugston, Anna Dominiczak and Alex Alderton, Roberto Amato, Jeffrey Barrett, Sonia Goncalves, Ewan Harrison, David K. Jackson, Ian Johnston, Dominic Kwiatkowski, Cordelia Langford, John Sillitoe on behalf of the Wellcome Sanger Institute COVID-19 Surveillance Team |
| EPI_ISL_3168132, EPI_ISL_3168133 | Lighthouse Lab in Milton Keynes | Wellcome Sanger Institute for the COVID-19 Genomics UK (COG-UK) Consortium | The Lighthouse Lab in Milton Keynes and Alex Alderton, Roberto Amato, Jeffrey Barrett, Sonia Goncalves, Ewan Harrison, David K. Jackson, Ian Johnston, Dominic Kwiatkowski, Cordelia Langford, John Sillitoe on behalf of the Wellcome Sanger Institute COVID-19 Surveillance Team |
| EPI_ISL_3168134, EPI_ISL_3168135 | Lighthouse Lab in Glasgow | Wellcome Sanger Institute for the COVID-19 Genomics UK (COG-UK) Consortium | Harper VanSteenhouse, Yumi Kasai, David Gray, Carol Clugston, Anna Dominiczak and Alex Alderton, Roberto Amato, Jeffrey Barrett, Sonia Goncalves, Ewan Harrison, David K. Jackson, Ian Johnston, Dominic Kwiatkowski, Cordelia Langford, John Sillitoe on behalf of the Wellcome Sanger Institute COVID-19 Surveillance Team |
| EPI_ISL_3168136, EPI_ISL_3168137 | Lighthouse Lab in Milton Keynes | Wellcome Sanger Institute for the COVID-19 Genomics UK (COG-UK) Consortium | The Lighthouse Lab in Milton Keynes and Alex Alderton, Roberto Amato, Jeffrey Barrett, Sonia Goncalves, Ewan Harrison, David K. Jackson, Ian Johnston, Dominic Kwiatkowski, Cordelia Langford, John Sillitoe on behalf of the Wellcome Sanger Institute COVID-19 Surveillance Team |
| EPI_ISL_3168138 | Lighthouse Lab in Glasgow | Wellcome Sanger Institute for the COVID-19 Genomics UK (COG-UK) Consortium | Harper VanSteenhouse, Yumi Kasai, David Gray, Carol Clugston, Anna Dominiczak and Alex Alderton, Roberto Amato, Jeffrey Barrett, Sonia Goncalves, Ewan Harrison, David K. Jackson, Ian Johnston, Dominic Kwiatkowski, Cordelia Langford, John Sillitoe on behalf of the Wellcome Sanger Institute COVID-19 Surveillance Team |
| EPI_ISL_3168139, EPI_ISL_3168140, EPI_ISL_3168141, EPI_ISL_3168142, EPI_ISL_3168143, EPI_ISL_3168144, EPI_ISL_3168145, EPI_ISL_3168146, EPI_ISL_3168147, EPI_ISL_3168148 | Lighthouse Lab in Milton Keynes | Wellcome Sanger Institute for the COVID-19 Genomics UK (COG-UK) Consortium | The Lighthouse Lab in Milton Keynes and Alex Alderton, Roberto Amato, Jeffrey Barrett, Sonia Goncalves, Ewan Harrison, David K. Jackson, Ian Johnston, Dominic Kwiatkowski, Cordelia Langford, John Sillitoe on behalf of the Wellcome Sanger Institute COVID-19 Surveillance Team |
| EPI_ISL_3168149 | Lighthouse Lab in Glasgow | Wellcome Sanger Institute for the COVID-19 Genomics UK (COG-UK) Consortium | Harper VanSteenhouse, Yumi Kasai, David Gray, Carol Clugston, Anna Dominiczak and Alex Alderton, Roberto Amato, Jeffrey Barrett, Sonia Goncalves, Ewan Harrison, David K. Jackson, Ian Johnston, Dominic Kwiatkowski, Cordelia Langford, John Sillitoe on behalf of the Wellcome Sanger Institute COVID-19 Surveillance Team |
| EPI_ISL_3168150, EPI_ISL_3168151, EPI_ISL_3168152, EPI_ISL_3168153, EPI_ISL_3168154, EPI_ISL_3168155, EPI_ISL_3168156, EPI_ISL_3168157, EPI_ISL_3168158, EPI_ISL_3168159, EPI_ISL_3168160, EPI_ISL_3168161, EPI_ISL_3168162, EPI_ISL_3168163, EPI_ISL_3168164, EPI_ISL_3168165 | | | |
| see above | Lighthouse Lab in Milton Keynes | Wellcome Sanger Institute for the COVID-19 Genomics UK (COG-UK) Consortium | The Lighthouse Lab in Milton Keynes and Alex Alderton, Roberto Amato, Jeffrey Barrett, Sonia Goncalves, Ewan Harrison, David K. Jackson, Ian Johnston, Dominic Kwiatkowski, Cordelia Langford, John Sillitoe on behalf of the Wellcome Sanger Institute COVID-19 Surveillance Team |
| EPI_ISL_3168166 | Lighthouse Lab in Glasgow | Wellcome Sanger Institute for the COVID-19 Genomics UK (COG-UK) Consortium | Harper VanSteenhouse, Yumi Kasai, David Gray, Carol Clugston, Anna Dominiczak and Alex Alderton, Roberto Amato, Jeffrey Barrett, Sonia Goncalves, Ewan Harrison, David K. Jackson, Ian Johnston, Dominic Kwiatkowski, Cordelia Langford, John Sillitoe on behalf of the Wellcome Sanger Institute COVID-19 Surveillance Team |
| EPI_ISL_3168167, EPI_ISL_3168168, EPI_ISL_3168169, EPI_ISL_3168170, EPI_ISL_3168171, EPI_ISL_3168172 | Lighthouse Lab in Milton Keynes | Wellcome Sanger Institute for the COVID-19 Genomics UK (COG-UK) Consortium | The Lighthouse Lab in Milton Keynes and Alex Alderton, Roberto Amato, Jeffrey Barrett, Sonia Goncalves, Ewan Harrison, David K. Jackson, Ian Johnston, Dominic Kwiatkowski, Cordelia Langford, John Sillitoe on behalf of the Wellcome Sanger Institute COVID-19 Surveillance Team |
| EPI_ISL_3168173 | Lighthouse Lab in Glasgow | Wellcome Sanger Institute for the COVID-19 Genomics UK (COG-UK) Consortium | Harper VanSteenhouse, Yumi Kasai, David Gray, Carol Clugston, Anna Dominiczak and Alex Alderton, Roberto Amato, Jeffrey Barrett, Sonia Goncalves, Ewan Harrison, David K. Jackson, Ian Johnston, Dominic Kwiatkowski, Cordelia Langford, John Sillitoe on behalf of the Wellcome Sanger Institute COVID-19 Surveillance Team |
| EPI_ISL_3168174, EPI_ISL_3168175, EPI_ISL_3168176, EPI_ISL_3168177, EPI_ISL_3168178 | Lighthouse Lab in Milton Keynes | Wellcome Sanger Institute for the COVID-19 Genomics UK (COG-UK) Consortium | The Lighthouse Lab in Milton Keynes and Alex Alderton, Roberto Amato, Jeffrey Barrett, Sonia Goncalves, Ewan Harrison, David K. Jackson, Ian Johnston, Dominic Kwiatkowski, Cordelia Langford, John Sillitoe on behalf of the Wellcome Sanger Institute COVID-19 Surveillance Team |
| EPI_ISL_3168179 | Lighthouse Lab in Glasgow | Wellcome Sanger Institute for the COVID-19 Genomics UK (COG-UK) Consortium | Harper VanSteenhouse, Yumi Kasai, David Gray, Carol Clugston, Anna Dominiczak and Alex Alderton, Roberto Amato, Jeffrey Barrett, Sonia Goncalves, Ewan Harrison, David K. Jackson, Ian Johnston, Dominic Kwiatkowski, Cordelia Langford, John Sillitoe on behalf of the Wellcome Sanger Institute COVID-19 Surveillance Team |
| EPI_ISL_3168180 | Lighthouse Lab in Milton Keynes | Wellcome Sanger Institute for the COVID-19 Genomics UK (COG-UK) Consortium | The Lighthouse Lab in Milton Keynes and Alex Alderton, Roberto Amato, Jeffrey Barrett, Sonia Goncalves, Ewan Harrison, David K. Jackson, Ian Johnston, Dominic Kwiatkowski, Cordelia Langford, John Sillitoe on behalf of the Wellcome Sanger Institute COVID-19 Surveillance Team |
| EPI_ISL_3168181 | Lighthouse Lab in Glasgow | Wellcome Sanger Institute for the COVID-19 Genomics UK (COG-UK) Consortium | Harper VanSteenhouse, Yumi Kasai, David Gray, Carol Clugston, Anna Dominiczak and Alex Alderton, Roberto Amato, Jeffrey Barrett, Sonia Goncalves, Ewan Harrison, David K. Jackson, Ian Johnston, Dominic Kwiatkowski, Cordelia Langford, John Sillitoe on behalf of the Wellcome Sanger Institute COVID-19 Surveillance Team |
| EPI_ISL_3168182, EPI_ISL_3168183, EPI_ISL_3168184, EPI_ISL_3168185, EPI_ISL_3168186, EPI_ISL_3168187, EPI_ISL_3168188, EPI_ISL_3168189, EPI_ISL_3168190 | Lighthouse Lab in Milton Keynes | Wellcome Sanger Institute for the COVID-19 Genomics UK (COG-UK) Consortium | The Lighthouse Lab in Milton Keynes and Alex Alderton, Roberto Amato, Jeffrey Barrett, Sonia Goncalves, Ewan Harrison, David K. Jackson, Ian Johnston, Dominic Kwiatkowski, Cordelia Langford, John Sillitoe on behalf of the Wellcome Sanger Institute COVID-19 Surveillance Team |
| EPI_ISL_3168191 | Lighthouse Lab in Glasgow | Wellcome Sanger Institute for the COVID-19 Genomics UK (COG-UK) Consortium | Harper VanSteenhouse, Yumi Kasai, David Gray, Carol Clugston, Anna Dominiczak and Alex Alderton, Roberto Amato, Jeffrey Barrett, Sonia Goncalves, Ewan Harrison, David K. Jackson, Ian Johnston, Dominic Kwiatkowski, Cordelia Langford, John Sillitoe on behalf of the Wellcome Sanger Institute COVID-19 Surveillance Team |
| EPI_ISL_3168192, EPI_ISL_3168193, EPI_ISL_3168194, EPI_ISL_3168195, EPI_ISL_3168196 | Lighthouse Lab in Milton Keynes | Wellcome Sanger Institute for the COVID-19 Genomics UK (COG-UK) Consortium | The Lighthouse Lab in Milton Keynes and Alex Alderton, Roberto Amato, Jeffrey Barrett, Sonia Goncalves, Ewan Harrison, David K. Jackson, Ian Johnston, Dominic Kwiatkowski, Cordelia Langford, John Sillitoe on behalf of the Wellcome Sanger Institute COVID-19 Surveillance Team |

| | | | |
|---|---|---|---|
| EPI_ISL_3168197 | Lighthouse Lab in Glasgow | Wellcome Sanger Institute for the COVID-19 Genomics UK (COG-UK) Consortium | Harper VanSteenhouse, Yumi Kasai, David Gray, Carol Clugston, Anna Dominiczak and Alex Alderton, Roberto Amato, Jeffrey Barrett, Sonia Goncalves, Ewan Harrison, David K. Jackson, Ian Johnston, Dominic Kwiatkowski, Cordelia Langford, John Sillitoe on behalf of the Wellcome Sanger Institute COVID-19 Surveillance Team |
| EPI_ISL_3168198, EPI_ISL_3168199, EPI_ISL_3168200, EPI_ISL_3168201 | Lighthouse Lab in Milton Keynes | Wellcome Sanger Institute for the COVID-19 Genomics UK (COG-UK) Consortium | The Lighthouse Lab in Milton Keynes and Alex Alderton, Roberto Amato, Jeffrey Barrett, Sonia Goncalves, Ewan Harrison, David K. Jackson, Ian Johnston, Dominic Kwiatkowski, Cordelia Langford, John Sillitoe on behalf of the Wellcome Sanger Institute COVID-19 Surveillance Team |
| EPI_ISL_3168202, EPI_ISL_3168203 | Lighthouse Lab in Glasgow | Wellcome Sanger Institute for the COVID-19 Genomics UK (COG-UK) Consortium | Harper VanSteenhouse, Yumi Kasai, David Gray, Carol Clugston, Anna Dominiczak and Alex Alderton, Roberto Amato, Jeffrey Barrett, Sonia Goncalves, Ewan Harrison, David K. Jackson, Ian Johnston, Dominic Kwiatkowski, Cordelia Langford, John Sillitoe on behalf of the Wellcome Sanger Institute COVID-19 Surveillance Team |
| EPI_ISL_3168204, EPI_ISL_3168205 | Lighthouse Lab in Milton Keynes | Wellcome Sanger Institute for the COVID-19 Genomics UK (COG-UK) Consortium | The Lighthouse Lab in Milton Keynes and Alex Alderton, Roberto Amato, Jeffrey Barrett, Sonia Goncalves, Ewan Harrison, David K. Jackson, Ian Johnston, Dominic Kwiatkowski, Cordelia Langford, John Sillitoe on behalf of the Wellcome Sanger Institute COVID-19 Surveillance Team |
| EPI_ISL_3168206 | Lighthouse Lab in Glasgow | Wellcome Sanger Institute for the COVID-19 Genomics UK (COG-UK) Consortium | Harper VanSteenhouse, Yumi Kasai, David Gray, Carol Clugston, Anna Dominiczak and Alex Alderton, Roberto Amato, Jeffrey Barrett, Sonia Goncalves, Ewan Harrison, David K. Jackson, Ian Johnston, Dominic Kwiatkowski, Cordelia Langford, John Sillitoe on behalf of the Wellcome Sanger Institute COVID-19 Surveillance Team |
| EPI_ISL_3168207, EPI_ISL_3168208, EPI_ISL_3168209, EPI_ISL_3168210, EPI_ISL_3168211, EPI_ISL_3168212, EPI_ISL_3168213 | Lighthouse Lab in Milton Keynes | Wellcome Sanger Institute for the COVID-19 Genomics UK (COG-UK) Consortium | The Lighthouse Lab in Milton Keynes and Alex Alderton, Roberto Amato, Jeffrey Barrett, Sonia Goncalves, Ewan Harrison, David K. Jackson, Ian Johnston, Dominic Kwiatkowski, Cordelia Langford, John Sillitoe on behalf of the Wellcome Sanger Institute COVID-19 Surveillance Team |
| EPI_ISL_3168214 | Lighthouse Lab in Glasgow | Wellcome Sanger Institute for the COVID-19 Genomics UK (COG-UK) Consortium | Harper VanSteenhouse, Yumi Kasai, David Gray, Carol Clugston, Anna Dominiczak and Alex Alderton, Roberto Amato, Jeffrey Barrett, Sonia Goncalves, Ewan Harrison, David K. Jackson, Ian Johnston, Dominic Kwiatkowski, Cordelia Langford, John Sillitoe on behalf of the Wellcome Sanger Institute COVID-19 Surveillance Team |
| EPI_ISL_3168215 | Lighthouse Lab in Milton Keynes | Wellcome Sanger Institute for the COVID-19 Genomics UK (COG-UK) Consortium | The Lighthouse Lab in Milton Keynes and Alex Alderton, Roberto Amato, Jeffrey Barrett, Sonia Goncalves, Ewan Harrison, David K. Jackson, Ian Johnston, Dominic Kwiatkowski, Cordelia Langford, John Sillitoe on behalf of the Wellcome Sanger Institute COVID-19 Surveillance Team |
| EPI_ISL_3168216 | Lighthouse Lab in Glasgow | Wellcome Sanger Institute for the COVID-19 Genomics UK (COG-UK) Consortium | Harper VanSteenhouse, Yumi Kasai, David Gray, Carol Clugston, Anna Dominiczak and Alex Alderton, Roberto Amato, Jeffrey Barrett, Sonia Goncalves, Ewan Harrison, David K. Jackson, Ian Johnston, Dominic Kwiatkowski, Cordelia Langford, John Sillitoe on behalf of the Wellcome Sanger Institute COVID-19 Surveillance Team |
| EPI_ISL_3168217, EPI_ISL_3168218, EPI_ISL_3168219, EPI_ISL_3168220, EPI_ISL_3168221, EPI_ISL_3168222, EPI_ISL_3168223, EPI_ISL_3168224, EPI_ISL_3168225 | Lighthouse Lab in Milton Keynes | Wellcome Sanger Institute for the COVID-19 Genomics UK (COG-UK) Consortium | The Lighthouse Lab in Milton Keynes and Alex Alderton, Roberto Amato, Jeffrey Barrett, Sonia Goncalves, Ewan Harrison, David K. Jackson, Ian Johnston, Dominic Kwiatkowski, Cordelia Langford, John Sillitoe on behalf of the Wellcome Sanger Institute COVID-19 Surveillance Team |
| EPI_ISL_3168226 | Lighthouse Lab in Glasgow | Wellcome Sanger Institute for the COVID-19 Genomics UK (COG-UK) Consortium | Harper VanSteenhouse, Yumi Kasai, David Gray, Carol Clugston, Anna Dominiczak and Alex Alderton, Roberto Amato, Jeffrey Barrett, Sonia Goncalves, Ewan Harrison, David K. Jackson, Ian Johnston, Dominic Kwiatkowski, Cordelia Langford, John Sillitoe on behalf of the Wellcome Sanger Institute COVID-19 Surveillance Team |
| EPI_ISL_3168227, EPI_ISL_3168228, EPI_ISL_3168229, EPI_ISL_3168230, EPI_ISL_3168231 | Lighthouse Lab in Milton Keynes | Wellcome Sanger Institute for the COVID-19 Genomics UK (COG-UK) Consortium | The Lighthouse Lab in Milton Keynes and Alex Alderton, Roberto Amato, Jeffrey Barrett, Sonia Goncalves, Ewan Harrison, David K. Jackson, Ian Johnston, Dominic Kwiatkowski, Cordelia Langford, John Sillitoe on behalf of the Wellcome Sanger Institute COVID-19 Surveillance Team |
| EPI_ISL_3168232 | Lighthouse Lab in Glasgow | Wellcome Sanger Institute for the COVID-19 Genomics UK (COG-UK) Consortium | Harper VanSteenhouse, Yumi Kasai, David Gray, Carol Clugston, Anna Dominiczak and Alex Alderton, Roberto Amato, Jeffrey Barrett, Sonia Goncalves, Ewan Harrison, David K. Jackson, Ian Johnston, Dominic Kwiatkowski, Cordelia Langford, John Sillitoe on behalf of the Wellcome Sanger Institute COVID-19 Surveillance Team |
| EPI_ISL_3168233, EPI_ISL_3168234, EPI_ISL_3168235, EPI_ISL_3168236, EPI_ISL_3168237 | Lighthouse Lab in Milton Keynes | Wellcome Sanger Institute for the COVID-19 Genomics UK (COG-UK) Consortium | The Lighthouse Lab in Milton Keynes and Alex Alderton, Roberto Amato, Jeffrey Barrett, Sonia Goncalves, Ewan Harrison, David K. Jackson, Ian Johnston, Dominic Kwiatkowski, Cordelia Langford, John Sillitoe on behalf of the Wellcome Sanger Institute COVID-19 Surveillance Team |
| EPI_ISL_3168238 | Lighthouse Lab in Glasgow | Wellcome Sanger Institute for the COVID-19 Genomics UK (COG-UK) Consortium | Harper VanSteenhouse, Yumi Kasai, David Gray, Carol Clugston, Anna Dominiczak and Alex Alderton, Roberto Amato, Jeffrey Barrett, Sonia Goncalves, Ewan Harrison, David K. Jackson, Ian Johnston, Dominic Kwiatkowski, Cordelia Langford, John Sillitoe on behalf of the Wellcome Sanger Institute COVID-19 Surveillance Team |
| EPI_ISL_3168239, EPI_ISL_3168240, EPI_ISL_3168241, EPI_ISL_3168242, EPI_ISL_3168243, EPI_ISL_3168244, EPI_ISL_3168245, EPI_ISL_3168246, EPI_ISL_3168247, EPI_ISL_3168248, EPI_ISL_3168249, EPI_ISL_3168250 | | | |
| see above | Lighthouse Lab in Milton Keynes | Wellcome Sanger Institute for the COVID-19 Genomics UK (COG-UK) Consortium | The Lighthouse Lab in Milton Keynes and Alex Alderton, Roberto Amato, Jeffrey Barrett, Sonia Goncalves, Ewan Harrison, David K. Jackson, Ian Johnston, Dominic Kwiatkowski, Cordelia Langford, John Sillitoe on behalf of the Wellcome Sanger Institute COVID-19 Surveillance Team |
| EPI_ISL_3168251 | Lighthouse Lab in Glasgow | Wellcome Sanger Institute for the COVID-19 Genomics UK (COG-UK) Consortium | Harper VanSteenhouse, Yumi Kasai, David Gray, Carol Clugston, Anna Dominiczak and Alex Alderton, Roberto Amato, Jeffrey Barrett, Sonia Goncalves, Ewan Harrison, David K. Jackson, Ian Johnston, Dominic Kwiatkowski, Cordelia Langford, John Sillitoe on behalf of the Wellcome Sanger Institute COVID-19 Surveillance Team |
| EPI_ISL_3168252, EPI_ISL_3168253, EPI_ISL_3168254, EPI_ISL_3168255, EPI_ISL_3168256, EPI_ISL_3168257, EPI_ISL_3168258, EPI_ISL_3168259 | Lighthouse Lab in Milton Keynes | Wellcome Sanger Institute for the COVID-19 Genomics UK (COG-UK) Consortium | The Lighthouse Lab in Milton Keynes and Alex Alderton, Roberto Amato, Jeffrey Barrett, Sonia Goncalves, Ewan Harrison, David K. Jackson, Ian Johnston, Dominic Kwiatkowski, Cordelia Langford, John Sillitoe on behalf of the Wellcome Sanger Institute COVID-19 Surveillance Team |
| EPI_ISL_3168260 | Lighthouse Lab in Glasgow | Wellcome Sanger Institute for the COVID-19 Genomics UK (COG-UK) Consortium | Harper VanSteenhouse, Yumi Kasai, David Gray, Carol Clugston, Anna Dominiczak and Alex Alderton, Roberto Amato, Jeffrey Barrett, Sonia Goncalves, Ewan Harrison, David K. Jackson, Ian Johnston, Dominic Kwiatkowski, Cordelia Langford, John Sillitoe on behalf of the Wellcome Sanger Institute COVID-19 Surveillance Team |
| EPI_ISL_3168261, EPI_ISL_3168262 | Lighthouse Lab in Milton Keynes | Wellcome Sanger Institute for the COVID-19 Genomics UK (COG-UK) Consortium | The Lighthouse Lab in Milton Keynes and Alex Alderton, Roberto Amato, Jeffrey Barrett, Sonia Goncalves, Ewan Harrison, David K. Jackson, Ian Johnston, Dominic Kwiatkowski, Cordelia Langford, John Sillitoe on behalf of the Wellcome Sanger Institute COVID-19 Surveillance Team |
| EPI_ISL_3168263 | Lighthouse Lab in Glasgow | Wellcome Sanger Institute for the COVID-19 Genomics UK (COG-UK) Consortium | Harper VanSteenhouse, Yumi Kasai, David Gray, Carol Clugston, Anna Dominiczak and Alex Alderton, Roberto Amato, Jeffrey Barrett, Sonia Goncalves, Ewan Harrison, David K. Jackson, Ian Johnston, Dominic Kwiatkowski, Cordelia Langford, John Sillitoe on behalf of the Wellcome Sanger Institute COVID-19 Surveillance Team |
| EPI_ISL_3168264, EPI_ISL_3168265, EPI_ISL_3168266, EPI_ISL_3168267, EPI_ISL_3168268, EPI_ISL_3168269, EPI_ISL_3168270, EPI_ISL_3168271, EPI_ISL_3168272, EPI_ISL_3168273, EPI_ISL_3168274, EPI_ISL_3168275, EPI_ISL_3168276, EPI_ISL_3168277, EPI_ISL_3168278, EPI_ISL_3168279, EPI_ISL_3168280, EPI_ISL_3168281, EPI_ISL_3168282, EPI_ISL_3168283, EPI_ISL_3168284, EPI_ISL_3168285, EPI_ISL_3168286, EPI_ISL_3168287, EPI_ISL_3168288, EPI_ISL_3168289, EPI_ISL_3168290, EPI_ISL_3168291, EPI_ISL_3168292, EPI_ISL_3168293, EPI_ISL_3168294, EPI_ISL_3168295, EPI_ISL_3168296, EPI_ISL_3168297, EPI_ISL_3168298, EPI_ISL_3168299, EPI_ISL_3168300, EPI_ISL_3168301, EPI_ISL_3168302, EPI_ISL_3168303, EPI_ISL_3168304, EPI_ISL_3168305, EPI_ISL_3168306, EPI_ISL_3168307, EPI_ISL_3168308, EPI_ISL_3168309, EPI_ISL_3168310, EPI_ISL_3168311 | | | |
| see above | Lighthouse Lab in Milton Keynes | Wellcome Sanger Institute for the COVID-19 Genomics UK (COG-UK) Consortium | The Lighthouse Lab in Milton Keynes and Alex Alderton, Roberto Amato, Jeffrey Barrett, Sonia Goncalves, Ewan Harrison, David K. Jackson, Ian Johnston, Dominic Kwiatkowski, Cordelia Langford, John Sillitoe on behalf of the Wellcome Sanger Institute COVID-19 Surveillance Team |
| EPI_ISL_3168312 | Lighthouse Lab in Glasgow | Wellcome Sanger Institute for the COVID-19 Genomics UK (COG-UK) Consortium | Harper VanSteenhouse, Yumi Kasai, David Gray, Carol Clugston, Anna Dominiczak and Alex Alderton, Roberto Amato, Jeffrey Barrett, Sonia Goncalves, Ewan Harrison, David K. Jackson, Ian Johnston, Dominic Kwiatkowski, Cordelia Langford, John Sillitoe on behalf of the Wellcome Sanger Institute COVID-19 Surveillance Team |
| EPI_ISL_3168313, EPI_ISL_3168314, EPI_ISL_3168315, EPI_ISL_3168316, EPI_ISL_3168317, EPI_ISL_3168318, EPI_ISL_3168319, EPI_ISL_3168320, EPI_ISL_3168321, EPI_ISL_3168322, EPI_ISL_3168323, EPI_ISL_3168324, EPI_ISL_3168325, EPI_ISL_3168326, EPI_ISL_3168327, EPI_ISL_3168328, EPI_ISL_3168329, EPI_ISL_3168330, EPI_ISL_3168331, EPI_ISL_3168332, EPI_ISL_3168333, EPI_ISL_3168334, EPI_ISL_3168335, EPI_ISL_3168336 | | | |
| see above | Lighthouse Lab in Milton Keynes | Wellcome Sanger Institute for the COVID-19 Genomics UK (COG-UK) Consortium | The Lighthouse Lab in Milton Keynes and Alex Alderton, Roberto Amato, Jeffrey Barrett, Sonia Goncalves, Ewan Harrison, David K. Jackson, Ian Johnston, Dominic Kwiatkowski, Cordelia Langford, John Sillitoe on behalf of the Wellcome Sanger Institute COVID-19 Surveillance Team |
| EPI_ISL_3168337 | Lighthouse Lab in Glasgow | Wellcome Sanger Institute for the COVID-19 Genomics UK (COG-UK) Consortium | Harper VanSteenhouse, Yumi Kasai, David Gray, Carol Clugston, Anna Dominiczak and Alex Alderton, Roberto Amato, Jeffrey Barrett, Sonia Goncalves, Ewan Harrison, David K. Jackson, Ian Johnston, Dominic Kwiatkowski, Cordelia Langford, John Sillitoe on behalf of the Wellcome Sanger Institute COVID-19 Surveillance Team |
| EPI_ISL_3168338 | Lighthouse Lab in Milton Keynes | Wellcome Sanger Institute for the COVID-19 Genomics UK (COG-UK) Consortium | The Lighthouse Lab in Milton Keynes and Alex Alderton, Roberto Amato, Jeffrey Barrett, Sonia Goncalves, Ewan Harrison, David K. Jackson, Ian Johnston, Dominic Kwiatkowski, Cordelia Langford, John Sillitoe on behalf of the Wellcome Sanger Institute COVID-19 Surveillance Team |
| EPI_ISL_3168339 | Lighthouse Lab in Glasgow | Wellcome Sanger Institute for the COVID-19 Genomics UK (COG-UK) Consortium | Harper VanSteenhouse, Yumi Kasai, David Gray, Carol Clugston, Anna Dominiczak and Alex Alderton, Roberto Amato, Jeffrey Barrett, Sonia Goncalves, Ewan Harrison, David K. Jackson, Ian Johnston, Dominic Kwiatkowski, Cordelia Langford, John Sillitoe on behalf of the Wellcome Sanger Institute COVID-19 Surveillance Team |
| EPI_ISL_3168340, EPI_ISL_3168341, EPI_ISL_3168342, EPI_ISL_3168343, EPI_ISL_3168344, EPI_ISL_3168345, EPI_ISL_3168346, EPI_ISL_3168347, EPI_ISL_3168348, EPI_ISL_3168349, EPI_ISL_3168350, EPI_ISL_3168351, EPI_ISL_3168352, EPI_ISL_3168353, EPI_ISL_3168354, EPI_ISL_3168355, EPI_ISL_3168356, EPI_ISL_3168357 | | | |
| see above | Lighthouse Lab in Milton Keynes | Wellcome Sanger Institute for the COVID-19 Genomics UK (COG-UK) Consortium | The Lighthouse Lab in Milton Keynes and Alex Alderton, Roberto Amato, Jeffrey Barrett, Sonia Goncalves, Ewan Harrison, David K. Jackson, Ian Johnston, Dominic Kwiatkowski, Cordelia Langford, John Sillitoe on behalf of the Wellcome Sanger Institute COVID-19 Surveillance Team |
| EPI_ISL_3168358 | Lighthouse Lab in Glasgow | Wellcome Sanger Institute for the COVID-19 Genomics UK (COG-UK) Consortium | Harper VanSteenhouse, Yumi Kasai, David Gray, Carol Clugston, Anna Dominiczak and Alex Alderton, Roberto Amato, Jeffrey Barrett, Sonia Goncalves, Ewan Harrison, David K. Jackson, Ian Johnston, Dominic Kwiatkowski, Cordelia Langford, John Sillitoe on behalf of the Wellcome Sanger Institute COVID-19 Surveillance Team |
| EPI_ISL_3168359, EPI_ISL_3168360, EPI_ISL_3168361, EPI_ISL_3168362 | Lighthouse Lab in Milton Keynes | Wellcome Sanger Institute for the COVID-19 Genomics UK (COG-UK) Consortium | The Lighthouse Lab in Milton Keynes and Alex Alderton, Roberto Amato, Jeffrey Barrett, Sonia Goncalves, Ewan Harrison, David K. Jackson, Ian Johnston, Dominic Kwiatkowski, Cordelia Langford, John Sillitoe on behalf of the Wellcome Sanger Institute COVID-19 Surveillance Team |

| | | | |
|---|---|---|---|
| EPI_ISL_3168363 | Lighthouse Lab in Glasgow | Wellcome Sanger Institute for the COVID-19 Genomics UK (COG-UK) Consortium | Harper VanSteenhouse, Yumi Kasai, David Gray, Carol Clugston, Anna Dominiczak and Alex Alderton, Roberto Amato, Jeffrey Barrett, Sonia Goncalves, Ewan Harrison, David K. Jackson, Ian Johnston, Dominic Kwiatkowski, Cordelia Langford, John Sillitoe on behalf of the Wellcome Sanger Institute COVID-19 Surveillance Team |
| EPI_ISL_3168364 | Lighthouse Lab in Milton Keynes | Wellcome Sanger Institute for the COVID-19 Genomics UK (COG-UK) Consortium | The Lighthouse Lab in Milton Keynes and Alex Alderton, Roberto Amato, Jeffrey Barrett, Sonia Goncalves, Ewan Harrison, David K. Jackson, Ian Johnston, Dominic Kwiatkowski, Cordelia Langford, John Sillitoe on behalf of the Wellcome Sanger Institute COVID-19 Surveillance Team |
| EPI_ISL_3168365 | Lighthouse Lab in Glasgow | Wellcome Sanger Institute for the COVID-19 Genomics UK (COG-UK) Consortium | Harper VanSteenhouse, Yumi Kasai, David Gray, Carol Clugston, Anna Dominiczak and Alex Alderton, Roberto Amato, Jeffrey Barrett, Sonia Goncalves, Ewan Harrison, David K. Jackson, Ian Johnston, Dominic Kwiatkowski, Cordelia Langford, John Sillitoe on behalf of the Wellcome Sanger Institute COVID-19 Surveillance Team |
| EPI_ISL_3168366, EPI_ISL_3168367, EPI_ISL_3168368, EPI_ISL_3168369, EPI_ISL_3168370, EPI_ISL_3168371, EPI_ISL_3168372, EPI_ISL_3168373, EPI_ISL_3168374, EPI_ISL_3168375, EPI_ISL_3168376, EPI_ISL_3168377, EPI_ISL_3168378, EPI_ISL_3168379, EPI_ISL_3168380, EPI_ISL_3168381, EPI_ISL_3168382, EPI_ISL_3168383 | | | |
| see above | Lighthouse Lab in Milton Keynes | Wellcome Sanger Institute for the COVID-19 Genomics UK (COG-UK) Consortium | The Lighthouse Lab in Milton Keynes and Alex Alderton, Roberto Amato, Jeffrey Barrett, Sonia Goncalves, Ewan Harrison, David K. Jackson, Ian Johnston, Dominic Kwiatkowski, Cordelia Langford, John Sillitoe on behalf of the Wellcome Sanger Institute COVID-19 Surveillance Team |
| EPI_ISL_3168384 | Lighthouse Lab in Glasgow | Wellcome Sanger Institute for the COVID-19 Genomics UK (COG-UK) Consortium | Harper VanSteenhouse, Yumi Kasai, David Gray, Carol Clugston, Anna Dominiczak and Alex Alderton, Roberto Amato, Jeffrey Barrett, Sonia Goncalves, Ewan Harrison, David K. Jackson, Ian Johnston, Dominic Kwiatkowski, Cordelia Langford, John Sillitoe on behalf of the Wellcome Sanger Institute COVID-19 Surveillance Team |
| EPI_ISL_3168385, EPI_ISL_3168386, EPI_ISL_3168387, EPI_ISL_3168388, EPI_ISL_3168389, EPI_ISL_3168390, EPI_ISL_3168391, EPI_ISL_3168392 | Lighthouse Lab in Milton Keynes | Wellcome Sanger Institute for the COVID-19 Genomics UK (COG-UK) Consortium | The Lighthouse Lab in Milton Keynes and Alex Alderton, Roberto Amato, Jeffrey Barrett, Sonia Goncalves, Ewan Harrison, David K. Jackson, Ian Johnston, Dominic Kwiatkowski, Cordelia Langford, John Sillitoe on behalf of the Wellcome Sanger Institute COVID-19 Surveillance Team |
| EPI_ISL_3168393 | Berkshire and Surrey Pathology Services Lighthouse Laboratory | Wellcome Sanger Institute for the COVID-19 Genomics UK (COG-UK) Consortium | Berkshire and Surrey Pathology Services Lighthouse Laboratory and Alex Alderton, Roberto Amato, Jeffrey Barrett, Sonia Goncalves, Ewan Harrison, David K. Jackson, Ian Johnston, Dominic Kwiatkowski, Cordelia Langford, John Sillitoe on behalf of the Wellcome Sanger Institute COVID-19 Surveillance Team |
| EPI_ISL_3168394, EPI_ISL_3168395, EPI_ISL_3168396, EPI_ISL_3168397 | Lighthouse Lab in Alderley Park | Wellcome Sanger Institute for the COVID-19 Genomics UK (COG-UK) Consortium | Jacquelyn Wynn, Mairead Hyland, The Lighthouse Lab in Alderley Park and Alex Alderton, Roberto Amato, Jeffrey Barrett, Sonia Goncalves, Ewan Harrison, David K. Jackson, Ian Johnston, Dominic Kwiatkowski, Cordelia Langford, John Sillitoe on behalf of the Wellcome Sanger Institute COVID-19 Surveillance Team |
| EPI_ISL_3168398 | Lighthouse Lab in Glasgow | Wellcome Sanger Institute for the COVID-19 Genomics UK (COG-UK) Consortium | Harper VanSteenhouse, Yumi Kasai, David Gray, Carol Clugston, Anna Dominiczak and Alex Alderton, Roberto Amato, Jeffrey Barrett, Sonia Goncalves, Ewan Harrison, David K. Jackson, Ian Johnston, Dominic Kwiatkowski, Cordelia Langford, John Sillitoe on behalf of the Wellcome Sanger Institute COVID-19 Surveillance Team |
| EPI_ISL_3168399 | Lighthouse Lab in Alderley Park | Wellcome Sanger Institute for the COVID-19 Genomics UK (COG-UK) Consortium | Jacquelyn Wynn, Mairead Hyland, The Lighthouse Lab in Alderley Park and Alex Alderton, Roberto Amato, Jeffrey Barrett, Sonia Goncalves, Ewan Harrison, David K. Jackson, Ian Johnston, Dominic Kwiatkowski, Cordelia Langford, John Sillitoe on behalf of the Wellcome Sanger Institute COVID-19 Surveillance Team |
| EPI_ISL_3168400 | Berkshire and Surrey Pathology Services Lighthouse Laboratory | Wellcome Sanger Institute for the COVID-19 Genomics UK (COG-UK) Consortium | Berkshire and Surrey Pathology Services Lighthouse Laboratory and Alex Alderton, Roberto Amato, Jeffrey Barrett, Sonia Goncalves, Ewan Harrison, David K. Jackson, Ian Johnston, Dominic Kwiatkowski, Cordelia Langford, John Sillitoe on behalf of the Wellcome Sanger Institute COVID-19 Surveillance Team |
| EPI_ISL_3168401 | Lighthouse Lab in Glasgow | Wellcome Sanger Institute for the COVID-19 Genomics UK (COG-UK) Consortium | Harper VanSteenhouse, Yumi Kasai, David Gray, Carol Clugston, Anna Dominiczak and Alex Alderton, Roberto Amato, Jeffrey Barrett, Sonia Goncalves, Ewan Harrison, David K. Jackson, Ian Johnston, Dominic Kwiatkowski, Cordelia Langford, John Sillitoe on behalf of the Wellcome Sanger Institute COVID-19 Surveillance Team |
| EPI_ISL_3168402 | Berkshire and Surrey Pathology Services Lighthouse Laboratory | Wellcome Sanger Institute for the COVID-19 Genomics UK (COG-UK) Consortium | Berkshire and Surrey Pathology Services Lighthouse Laboratory and Alex Alderton, Roberto Amato, Jeffrey Barrett, Sonia Goncalves, Ewan Harrison, David K. Jackson, Ian Johnston, Dominic Kwiatkowski, Cordelia Langford, John Sillitoe on behalf of the Wellcome Sanger Institute COVID-19 Surveillance Team |
| EPI_ISL_3168403, EPI_ISL_3168404 | Lighthouse Lab in Alderley Park | Wellcome Sanger Institute for the COVID-19 Genomics UK (COG-UK) Consortium | Jacquelyn Wynn, Mairead Hyland, The Lighthouse Lab in Alderley Park and Alex Alderton, Roberto Amato, Jeffrey Barrett, Sonia Goncalves, Ewan Harrison, David K. Jackson, Ian Johnston, Dominic Kwiatkowski, Cordelia Langford, John Sillitoe on behalf of the Wellcome Sanger Institute COVID-19 Surveillance Team |
| EPI_ISL_3168405, EPI_ISL_3168406 | Berkshire and Surrey Pathology Services Lighthouse Laboratory | Wellcome Sanger Institute for the COVID-19 Genomics UK (COG-UK) Consortium | Berkshire and Surrey Pathology Services Lighthouse Laboratory and Alex Alderton, Roberto Amato, Jeffrey Barrett, Sonia Goncalves, Ewan Harrison, David K. Jackson, Ian Johnston, Dominic Kwiatkowski, Cordelia Langford, John Sillitoe on behalf of the Wellcome Sanger Institute COVID-19 Surveillance Team |
| EPI_ISL_3168407 | Lighthouse Lab in Alderley Park | Wellcome Sanger Institute for the COVID-19 Genomics UK (COG-UK) Consortium | Jacquelyn Wynn, Mairead Hyland, The Lighthouse Lab in Alderley Park and Alex Alderton, Roberto Amato, Jeffrey Barrett, Sonia Goncalves, Ewan Harrison, David K. Jackson, Ian Johnston, Dominic Kwiatkowski, Cordelia Langford, John Sillitoe on behalf of the Wellcome Sanger Institute COVID-19 Surveillance Team |
| EPI_ISL_3168408, EPI_ISL_3168409 | Lighthouse Lab in Glasgow | Wellcome Sanger Institute for the COVID-19 Genomics UK (COG-UK) Consortium | Harper VanSteenhouse, Yumi Kasai, David Gray, Carol Clugston, Anna Dominiczak and Alex Alderton, Roberto Amato, Jeffrey Barrett, Sonia Goncalves, Ewan Harrison, David K. Jackson, Ian Johnston, Dominic Kwiatkowski, Cordelia Langford, John Sillitoe on behalf of the Wellcome Sanger Institute COVID-19 Surveillance Team |
| EPI_ISL_3168410, EPI_ISL_3168411, EPI_ISL_3168412, EPI_ISL_3168413 | Lighthouse Lab in Alderley Park | Wellcome Sanger Institute for the COVID-19 Genomics UK (COG-UK) Consortium | Jacquelyn Wynn, Mairead Hyland, The Lighthouse Lab in Alderley Park and Alex Alderton, Roberto Amato, Jeffrey Barrett, Sonia Goncalves, Ewan Harrison, David K. Jackson, Ian Johnston, Dominic Kwiatkowski, Cordelia Langford, John Sillitoe on behalf of the Wellcome Sanger Institute COVID-19 Surveillance Team |
| EPI_ISL_3168414 | Berkshire and Surrey Pathology Services Lighthouse Laboratory | Wellcome Sanger Institute for the COVID-19 Genomics UK (COG-UK) Consortium | Berkshire and Surrey Pathology Services Lighthouse Laboratory and Alex Alderton, Roberto Amato, Jeffrey Barrett, Sonia Goncalves, Ewan Harrison, David K. Jackson, Ian Johnston, Dominic Kwiatkowski, Cordelia Langford, John Sillitoe on behalf of the Wellcome Sanger Institute COVID-19 Surveillance Team |
| EPI_ISL_3168415 | Lighthouse Lab in Alderley Park | Wellcome Sanger Institute for the COVID-19 Genomics UK (COG-UK) Consortium | Jacquelyn Wynn, Mairead Hyland, The Lighthouse Lab in Alderley Park and Alex Alderton, Roberto Amato, Jeffrey Barrett, Sonia Goncalves, Ewan Harrison, David K. Jackson, Ian Johnston, Dominic Kwiatkowski, Cordelia Langford, John Sillitoe on behalf of the Wellcome Sanger Institute COVID-19 Surveillance Team |
| EPI_ISL_3168416 | Lighthouse Lab in Glasgow | Wellcome Sanger Institute for the COVID-19 Genomics UK (COG-UK) Consortium | Harper VanSteenhouse, Yumi Kasai, David Gray, Carol Clugston, Anna Dominiczak and Alex Alderton, Roberto Amato, Jeffrey Barrett, Sonia Goncalves, Ewan Harrison, David K. Jackson, Ian Johnston, Dominic Kwiatkowski, Cordelia Langford, John Sillitoe on behalf of the Wellcome Sanger Institute COVID-19 Surveillance Team |
| EPI_ISL_3168417, EPI_ISL_3168418 | Lighthouse Lab in Alderley Park | Wellcome Sanger Institute for the COVID-19 Genomics UK (COG-UK) Consortium | Jacquelyn Wynn, Mairead Hyland, The Lighthouse Lab in Alderley Park and Alex Alderton, Roberto Amato, Jeffrey Barrett, Sonia Goncalves, Ewan Harrison, David K. Jackson, Ian Johnston, Dominic Kwiatkowski, Cordelia Langford, John Sillitoe on behalf of the Wellcome Sanger Institute COVID-19 Surveillance Team |
| EPI_ISL_3168419 | Lighthouse Lab in Glasgow | Wellcome Sanger Institute for the COVID-19 Genomics UK (COG-UK) Consortium | Harper VanSteenhouse, Yumi Kasai, David Gray, Carol Clugston, Anna Dominiczak and Alex Alderton, Roberto Amato, Jeffrey Barrett, Sonia Goncalves, Ewan Harrison, David K. Jackson, Ian Johnston, Dominic Kwiatkowski, Cordelia Langford, John Sillitoe on behalf of the Wellcome Sanger Institute COVID-19 Surveillance Team |
| EPI_ISL_3168420, EPI_ISL_3168421, EPI_ISL_3168422, EPI_ISL_3168423, EPI_ISL_3168424 | Lighthouse Lab in Alderley Park | Wellcome Sanger Institute for the COVID-19 Genomics UK (COG-UK) Consortium | Jacquelyn Wynn, Mairead Hyland, The Lighthouse Lab in Alderley Park and Alex Alderton, Roberto Amato, Jeffrey Barrett, Sonia Goncalves, Ewan Harrison, David K. Jackson, Ian Johnston, Dominic Kwiatkowski, Cordelia Langford, John Sillitoe on behalf of the Wellcome Sanger Institute COVID-19 Surveillance Team |
| EPI_ISL_3168425, EPI_ISL_3168426 | Lighthouse Lab in Glasgow | Wellcome Sanger Institute for the COVID-19 Genomics UK (COG-UK) Consortium | Harper VanSteenhouse, Yumi Kasai, David Gray, Carol Clugston, Anna Dominiczak and Alex Alderton, Roberto Amato, Jeffrey Barrett, Sonia Goncalves, Ewan Harrison, David K. Jackson, Ian Johnston, Dominic Kwiatkowski, Cordelia Langford, John Sillitoe on behalf of the Wellcome Sanger Institute COVID-19 Surveillance Team |
| EPI_ISL_3168427, EPI_ISL_3168428 | Lighthouse Lab in Alderley Park | Wellcome Sanger Institute for the COVID-19 Genomics UK (COG-UK) Consortium | Jacquelyn Wynn, Mairead Hyland, The Lighthouse Lab in Alderley Park and Alex Alderton, Roberto Amato, Jeffrey Barrett, Sonia Goncalves, Ewan Harrison, David K. Jackson, Ian Johnston, Dominic Kwiatkowski, Cordelia Langford, John Sillitoe on behalf of the Wellcome Sanger Institute COVID-19 Surveillance Team |
| EPI_ISL_3168429 | Berkshire and Surrey Pathology Services Lighthouse Laboratory | Wellcome Sanger Institute for the COVID-19 Genomics UK (COG-UK) Consortium | Berkshire and Surrey Pathology Services Lighthouse Laboratory and Alex Alderton, Roberto Amato, Jeffrey Barrett, Sonia Goncalves, Ewan Harrison, David K. Jackson, Ian Johnston, Dominic Kwiatkowski, Cordelia Langford, John Sillitoe on behalf of the Wellcome Sanger Institute COVID-19 Surveillance Team |
| EPI_ISL_3168430, EPI_ISL_3168431 | Lighthouse Lab in Alderley Park | Wellcome Sanger Institute for the COVID-19 Genomics UK (COG-UK) Consortium | Jacquelyn Wynn, Mairead Hyland, The Lighthouse Lab in Alderley Park and Alex Alderton, Roberto Amato, Jeffrey Barrett, Sonia Goncalves, Ewan Harrison, David K. Jackson, Ian Johnston, Dominic Kwiatkowski, Cordelia Langford, John Sillitoe on behalf of the Wellcome Sanger Institute COVID-19 Surveillance Team |
| EPI_ISL_3168432 | Lighthouse Lab in Glasgow | Wellcome Sanger Institute for the COVID-19 Genomics UK (COG-UK) Consortium | Harper VanSteenhouse, Yumi Kasai, David Gray, Carol Clugston, Anna Dominiczak and Alex Alderton, Roberto Amato, Jeffrey Barrett, Sonia Goncalves, Ewan Harrison, David K. Jackson, Ian Johnston, Dominic Kwiatkowski, Cordelia Langford, John Sillitoe on behalf of the Wellcome Sanger Institute COVID-19 Surveillance Team |
| EPI_ISL_3168433, EPI_ISL_3168434 | Lighthouse Lab in Alderley Park | Wellcome Sanger Institute for the COVID-19 Genomics UK (COG-UK) Consortium | Jacquelyn Wynn, Mairead Hyland, The Lighthouse Lab in Alderley Park and Alex Alderton, Roberto Amato, Jeffrey Barrett, Sonia Goncalves, Ewan Harrison, David K. Jackson, Ian Johnston, Dominic Kwiatkowski, Cordelia Langford, John Sillitoe on behalf of the Wellcome Sanger Institute COVID-19 Surveillance Team |
| EPI_ISL_3168435 | Lighthouse Lab in Glasgow | Wellcome Sanger Institute for the COVID-19 Genomics UK (COG-UK) Consortium | Harper VanSteenhouse, Yumi Kasai, David Gray, Carol Clugston, Anna Dominiczak and Alex Alderton, Roberto Amato, Jeffrey Barrett, Sonia Goncalves, Ewan Harrison, David K. Jackson, Ian Johnston, Dominic Kwiatkowski, Cordelia Langford, John Sillitoe on behalf of the Wellcome Sanger Institute COVID-19 Surveillance Team |
| EPI_ISL_3168436 | Lighthouse Lab in Alderley Park | Wellcome Sanger Institute for the COVID-19 Genomics UK (COG-UK) Consortium | Jacquelyn Wynn, Mairead Hyland, The Lighthouse Lab in Alderley Park and Alex Alderton, Roberto Amato, Jeffrey Barrett, Sonia Goncalves, Ewan Harrison, David K. Jackson, Ian Johnston, Dominic Kwiatkowski, Cordelia Langford, John Sillitoe on behalf of the Wellcome Sanger Institute COVID-19 Surveillance Team |
| EPI_ISL_3168437 | Lighthouse Lab in Glasgow | Wellcome Sanger Institute for the COVID-19 Genomics UK (COG-UK) Consortium | Harper VanSteenhouse, Yumi Kasai, David Gray, Carol Clugston, Anna Dominiczak and Alex Alderton, Roberto Amato, Jeffrey Barrett, Sonia Goncalves, Ewan Harrison, David K. Jackson, Ian Johnston, Dominic Kwiatkowski, Cordelia Langford, John Sillitoe on behalf of the Wellcome Sanger Institute COVID-19 Surveillance Team |
| EPI_ISL_3168438 | Lighthouse Lab in Alderley Park | Wellcome Sanger Institute for the COVID-19 Genomics UK (COG-UK) Consortium | Jacquelyn Wynn, Mairead Hyland, The Lighthouse Lab in Alderley Park and Alex Alderton, Roberto Amato, Jeffrey Barrett, Sonia Goncalves, Ewan Harrison, David K. Jackson, Ian Johnston, Dominic Kwiatkowski, Cordelia Langford, John Sillitoe on behalf of the Wellcome Sanger Institute COVID-19 Surveillance Team |
| EPI_ISL_3168439 | Berkshire and Surrey Pathology Services Lighthouse Laboratory | Wellcome Sanger Institute for the COVID-19 Genomics UK (COG-UK) Consortium | Berkshire and Surrey Pathology Services Lighthouse Laboratory and Alex Alderton, Roberto Amato, Jeffrey Barrett, Sonia Goncalves, Ewan Harrison, David K. Jackson, Ian Johnston, Dominic Kwiatkowski, Cordelia Langford, John Sillitoe on behalf of the Wellcome Sanger Institute COVID-19 Surveillance Team |
| EPI_ISL_3168440, EPI_ISL_3168441, EPI_ISL_3168442, EPI_ISL_3168443, EPI_ISL_3168444 | Lighthouse Lab in Alderley Park | Wellcome Sanger Institute for the COVID-19 Genomics UK (COG-UK) Consortium | Jacquelyn Wynn, Mairead Hyland, The Lighthouse Lab in Alderley Park and Alex Alderton, Roberto Amato, Jeffrey Barrett, Sonia Goncalves, Ewan Harrison, David K. Jackson, Ian Johnston, Dominic Kwiatkowski, Cordelia Langford, John Sillitoe on behalf of the Wellcome Sanger Institute COVID-19 Surveillance Team |
| EPI_ISL_3168445, EPI_ISL_3168446, EPI_ISL_3168447 | Berkshire and Surrey Pathology Services Lighthouse Laboratory | Wellcome Sanger Institute for the COVID-19 Genomics UK (COG-UK) Consortium | Berkshire and Surrey Pathology Services Lighthouse Laboratory and Alex Alderton, Roberto Amato, Jeffrey Barrett, Sonia Goncalves, Ewan Harrison, David K. Jackson, Ian Johnston, Dominic Kwiatkowski, Cordelia Langford, John Sillitoe on behalf of the Wellcome Sanger Institute COVID-19 Surveillance Team |
| EPI_ISL_3168448 | Lighthouse Lab in Alderley Park | Wellcome Sanger Institute for the COVID-19 Genomics UK (COG-UK) Consortium | Jacquelyn Wynn, Mairead Hyland, The Lighthouse Lab in Alderley Park and Alex Alderton, Roberto Amato, Jeffrey Barrett, Sonia Goncalves, Ewan Harrison, David K. Jackson, Ian Johnston, Dominic Kwiatkowski, Cordelia Langford, John Sillitoe on |

| | | | |
|---|---|---|---|
| | | Consortium | behalf of the Wellcome Sanger Institute COVID-19 Surveillance Team |
| EPI_ISL_3168449, EPI_ISL_3168450 | Berkshire and Surrey Pathology Services Lighthouse Laboratory | Wellcome Sanger Institute for the COVID-19 Genomics UK (COG-UK) Consortium | Berkshire and Surrey Pathology Services Lighthouse Laboratory and Alex Alderton, Roberto Amato, Jeffrey Barrett, Sonia Goncalves, Ewan Harrison, David K. Jackson, Ian Johnston, Dominic Kwiatkowski, Cordelia Langford, John Sillitoe on behalf of the Wellcome Sanger Institute COVID-19 Surveillance Team |
| EPI_ISL_3168451, EPI_ISL_3168452, EPI_ISL_3168453 | Lighthouse Lab in Alderley Park | Wellcome Sanger Institute for the COVID-19 Genomics UK (COG-UK) Consortium | Jacquelyn Wynn, Mairead Hyland, The Lighthouse Lab in Alderley Park and Alex Alderton, Roberto Amato, Jeffrey Barrett, Sonia Goncalves, Ewan Harrison, David K. Jackson, Ian Johnston, Dominic Kwiatkowski, Cordelia Langford, John Sillitoe on behalf of the Wellcome Sanger Institute COVID-19 Surveillance Team |
| EPI_ISL_3168454 | Lighthouse Lab in Glasgow | Wellcome Sanger Institute for the COVID-19 Genomics UK (COG-UK) Consortium | Harper VanSteenhouse, Yumi Kasai, David Gray, Carol Clugston, Anna Dominiczak and Alex Alderton, Roberto Amato, Jeffrey Barrett, Sonia Goncalves, Ewan Harrison, David K. Jackson, Ian Johnston, Dominic Kwiatkowski, Cordelia Langford, John Sillitoe on behalf of the Wellcome Sanger Institute COVID-19 Surveillance Team |
| EPI_ISL_3168455 | Lighthouse Lab in Alderley Park | Wellcome Sanger Institute for the COVID-19 Genomics UK (COG-UK) Consortium | Jacquelyn Wynn, Mairead Hyland, The Lighthouse Lab in Alderley Park and Alex Alderton, Roberto Amato, Jeffrey Barrett, Sonia Goncalves, Ewan Harrison, David K. Jackson, Ian Johnston, Dominic Kwiatkowski, Cordelia Langford, John Sillitoe on behalf of the Wellcome Sanger Institute COVID-19 Surveillance Team |
| EPI_ISL_3168456 | Berkshire and Surrey Pathology Services Lighthouse Laboratory | Wellcome Sanger Institute for the COVID-19 Genomics UK (COG-UK) Consortium | Berkshire and Surrey Pathology Services Lighthouse Laboratory and Alex Alderton, Roberto Amato, Jeffrey Barrett, Sonia Goncalves, Ewan Harrison, David K. Jackson, Ian Johnston, Dominic Kwiatkowski, Cordelia Langford, John Sillitoe on behalf of the Wellcome Sanger Institute COVID-19 Surveillance Team |
| EPI_ISL_3168457 | Lighthouse Lab in Alderley Park | Wellcome Sanger Institute for the COVID-19 Genomics UK (COG-UK) Consortium | Jacquelyn Wynn, Mairead Hyland, The Lighthouse Lab in Alderley Park and Alex Alderton, Roberto Amato, Jeffrey Barrett, Sonia Goncalves, Ewan Harrison, David K. Jackson, Ian Johnston, Dominic Kwiatkowski, Cordelia Langford, John Sillitoe on behalf of the Wellcome Sanger Institute COVID-19 Surveillance Team |
| EPI_ISL_3168458 | Berkshire and Surrey Pathology Services Lighthouse Laboratory | Wellcome Sanger Institute for the COVID-19 Genomics UK (COG-UK) Consortium | Berkshire and Surrey Pathology Services Lighthouse Laboratory and Alex Alderton, Roberto Amato, Jeffrey Barrett, Sonia Goncalves, Ewan Harrison, David K. Jackson, Ian Johnston, Dominic Kwiatkowski, Cordelia Langford, John Sillitoe on behalf of the Wellcome Sanger Institute COVID-19 Surveillance Team |
| EPI_ISL_3168459 | Lighthouse Lab in Glasgow | Wellcome Sanger Institute for the COVID-19 Genomics UK (COG-UK) Consortium | Harper VanSteenhouse, Yumi Kasai, David Gray, Carol Clugston, Anna Dominiczak and Alex Alderton, Roberto Amato, Jeffrey Barrett, Sonia Goncalves, Ewan Harrison, David K. Jackson, Ian Johnston, Dominic Kwiatkowski, Cordelia Langford, John Sillitoe on behalf of the Wellcome Sanger Institute COVID-19 Surveillance Team |
| EPI_ISL_3168460 | Lighthouse Lab in Alderley Park | Wellcome Sanger Institute for the COVID-19 Genomics UK (COG-UK) Consortium | Jacquelyn Wynn, Mairead Hyland, The Lighthouse Lab in Alderley Park and Alex Alderton, Roberto Amato, Jeffrey Barrett, Sonia Goncalves, Ewan Harrison, David K. Jackson, Ian Johnston, Dominic Kwiatkowski, Cordelia Langford, John Sillitoe on behalf of the Wellcome Sanger Institute COVID-19 Surveillance Team |
| EPI_ISL_3168461 | Lighthouse Lab in Glasgow | Wellcome Sanger Institute for the COVID-19 Genomics UK (COG-UK) Consortium | Harper VanSteenhouse, Yumi Kasai, David Gray, Carol Clugston, Anna Dominiczak and Alex Alderton, Roberto Amato, Jeffrey Barrett, Sonia Goncalves, Ewan Harrison, David K. Jackson, Ian Johnston, Dominic Kwiatkowski, Cordelia Langford, John Sillitoe on behalf of the Wellcome Sanger Institute COVID-19 Surveillance Team |
| EPI_ISL_3168462 | Lighthouse Lab in Alderley Park | Wellcome Sanger Institute for the COVID-19 Genomics UK (COG-UK) Consortium | Jacquelyn Wynn, Mairead Hyland, The Lighthouse Lab in Alderley Park and Alex Alderton, Roberto Amato, Jeffrey Barrett, Sonia Goncalves, Ewan Harrison, David K. Jackson, Ian Johnston, Dominic Kwiatkowski, Cordelia Langford, John Sillitoe on behalf of the Wellcome Sanger Institute COVID-19 Surveillance Team |
| EPI_ISL_3168463 | Lighthouse Lab in Glasgow | Wellcome Sanger Institute for the COVID-19 Genomics UK (COG-UK) Consortium | Harper VanSteenhouse, Yumi Kasai, David Gray, Carol Clugston, Anna Dominiczak and Alex Alderton, Roberto Amato, Jeffrey Barrett, Sonia Goncalves, Ewan Harrison, David K. Jackson, Ian Johnston, Dominic Kwiatkowski, Cordelia Langford, John Sillitoe on behalf of the Wellcome Sanger Institute COVID-19 Surveillance Team |
| EPI_ISL_3168464 | Lighthouse Lab in Alderley Park | Wellcome Sanger Institute for the COVID-19 Genomics UK (COG-UK) Consortium | Jacquelyn Wynn, Mairead Hyland, The Lighthouse Lab in Alderley Park and Alex Alderton, Roberto Amato, Jeffrey Barrett, Sonia Goncalves, Ewan Harrison, David K. Jackson, Ian Johnston, Dominic Kwiatkowski, Cordelia Langford, John Sillitoe on behalf of the Wellcome Sanger Institute COVID-19 Surveillance Team |
| EPI_ISL_3168465 | Lighthouse Lab in Glasgow | Wellcome Sanger Institute for the COVID-19 Genomics UK (COG-UK) Consortium | Harper VanSteenhouse, Yumi Kasai, David Gray, Carol Clugston, Anna Dominiczak and Alex Alderton, Roberto Amato, Jeffrey Barrett, Sonia Goncalves, Ewan Harrison, David K. Jackson, Ian Johnston, Dominic Kwiatkowski, Cordelia Langford, John Sillitoe on behalf of the Wellcome Sanger Institute COVID-19 Surveillance Team |
| EPI_ISL_3168466 | Berkshire and Surrey Pathology Services Lighthouse Laboratory | Wellcome Sanger Institute for the COVID-19 Genomics UK (COG-UK) Consortium | Berkshire and Surrey Pathology Services Lighthouse Laboratory and Alex Alderton, Roberto Amato, Jeffrey Barrett, Sonia Goncalves, Ewan Harrison, David K. Jackson, Ian Johnston, Dominic Kwiatkowski, Cordelia Langford, John Sillitoe on behalf of the Wellcome Sanger Institute COVID-19 Surveillance Team |
| EPI_ISL_3168467, EPI_ISL_3168468, EPI_ISL_3168469, EPI_ISL_3168470 | Lighthouse Lab in Alderley Park | Wellcome Sanger Institute for the COVID-19 Genomics UK (COG-UK) Consortium | Jacquelyn Wynn, Mairead Hyland, The Lighthouse Lab in Alderley Park and Alex Alderton, Roberto Amato, Jeffrey Barrett, Sonia Goncalves, Ewan Harrison, David K. Jackson, Ian Johnston, Dominic Kwiatkowski, Cordelia Langford, John Sillitoe on behalf of the Wellcome Sanger Institute COVID-19 Surveillance Team |
| EPI_ISL_3168471 | Lighthouse Lab in Glasgow | Wellcome Sanger Institute for the COVID-19 Genomics UK (COG-UK) Consortium | Harper VanSteenhouse, Yumi Kasai, David Gray, Carol Clugston, Anna Dominiczak and Alex Alderton, Roberto Amato, Jeffrey Barrett, Sonia Goncalves, Ewan Harrison, David K. Jackson, Ian Johnston, Dominic Kwiatkowski, Cordelia Langford, John Sillitoe on behalf of the Wellcome Sanger Institute COVID-19 Surveillance Team |
| EPI_ISL_3168472, EPI_ISL_3168473 | Lighthouse Lab in Alderley Park | Wellcome Sanger Institute for the COVID-19 Genomics UK (COG-UK) Consortium | Jacquelyn Wynn, Mairead Hyland, The Lighthouse Lab in Alderley Park and Alex Alderton, Roberto Amato, Jeffrey Barrett, Sonia Goncalves, Ewan Harrison, David K. Jackson, Ian Johnston, Dominic Kwiatkowski, Cordelia Langford, John Sillitoe on behalf of the Wellcome Sanger Institute COVID-19 Surveillance Team |
| EPI_ISL_3168474, EPI_ISL_3168475, EPI_ISL_3168476 | Lighthouse Lab in Glasgow | Wellcome Sanger Institute for the COVID-19 Genomics UK (COG-UK) Consortium | Harper VanSteenhouse, Yumi Kasai, David Gray, Carol Clugston, Anna Dominiczak and Alex Alderton, Roberto Amato, Jeffrey Barrett, Sonia Goncalves, Ewan Harrison, David K. Jackson, Ian Johnston, Dominic Kwiatkowski, Cordelia Langford, John Sillitoe on behalf of the Wellcome Sanger Institute COVID-19 Surveillance Team |
| EPI_ISL_3168477 | Lighthouse Lab in Alderley Park | Wellcome Sanger Institute for the COVID-19 Genomics UK (COG-UK) Consortium | Jacquelyn Wynn, Mairead Hyland, The Lighthouse Lab in Alderley Park and Alex Alderton, Roberto Amato, Jeffrey Barrett, Sonia Goncalves, Ewan Harrison, David K. Jackson, Ian Johnston, Dominic Kwiatkowski, Cordelia Langford, John Sillitoe on behalf of the Wellcome Sanger Institute COVID-19 Surveillance Team |
| EPI_ISL_3168478 | Berkshire and Surrey Pathology Services Lighthouse Laboratory | Wellcome Sanger Institute for the COVID-19 Genomics UK (COG-UK) Consortium | Berkshire and Surrey Pathology Services Lighthouse Laboratory and Alex Alderton, Roberto Amato, Jeffrey Barrett, Sonia Goncalves, Ewan Harrison, David K. Jackson, Ian Johnston, Dominic Kwiatkowski, Cordelia Langford, John Sillitoe on behalf of the Wellcome Sanger Institute COVID-19 Surveillance Team |
| EPI_ISL_3168479, EPI_ISL_3168480 | Lighthouse Lab in Alderley Park | Wellcome Sanger Institute for the COVID-19 Genomics UK (COG-UK) Consortium | Jacquelyn Wynn, Mairead Hyland, The Lighthouse Lab in Alderley Park and Alex Alderton, Roberto Amato, Jeffrey Barrett, Sonia Goncalves, Ewan Harrison, David K. Jackson, Ian Johnston, Dominic Kwiatkowski, Cordelia Langford, John Sillitoe on behalf of the Wellcome Sanger Institute COVID-19 Surveillance Team |
| EPI_ISL_3168481, EPI_ISL_3168483 | Lighthouse Lab in Glasgow | Wellcome Sanger Institute for the COVID-19 Genomics UK (COG-UK) Consortium | Harper VanSteenhouse, Yumi Kasai, David Gray, Carol Clugston, Anna Dominiczak and Alex Alderton, Roberto Amato, Jeffrey Barrett, Sonia Goncalves, Ewan Harrison, David K. Jackson, Ian Johnston, Dominic Kwiatkowski, Cordelia Langford, John Sillitoe on behalf of the Wellcome Sanger Institute COVID-19 Surveillance Team |
| EPI_ISL_3168484, EPI_ISL_3168485 | Lighthouse Lab in Alderley Park | Wellcome Sanger Institute for the COVID-19 Genomics UK (COG-UK) Consortium | Jacquelyn Wynn, Mairead Hyland, The Lighthouse Lab in Alderley Park and Alex Alderton, Roberto Amato, Jeffrey Barrett, Sonia Goncalves, Ewan Harrison, David K. Jackson, Ian Johnston, Dominic Kwiatkowski, Cordelia Langford, John Sillitoe on behalf of the Wellcome Sanger Institute COVID-19 Surveillance Team |
| EPI_ISL_3168486 | Lighthouse Lab in Glasgow | Wellcome Sanger Institute for the COVID-19 Genomics UK (COG-UK) Consortium | Harper VanSteenhouse, Yumi Kasai, David Gray, Carol Clugston, Anna Dominiczak and Alex Alderton, Roberto Amato, Jeffrey Barrett, Sonia Goncalves, Ewan Harrison, David K. Jackson, Ian Johnston, Dominic Kwiatkowski, Cordelia Langford, John Sillitoe on behalf of the Wellcome Sanger Institute COVID-19 Surveillance Team |
| EPI_ISL_3168487 | Berkshire and Surrey Pathology Services Lighthouse Laboratory | Wellcome Sanger Institute for the COVID-19 Genomics UK (COG-UK) Consortium | Berkshire and Surrey Pathology Services Lighthouse Laboratory and Alex Alderton, Roberto Amato, Jeffrey Barrett, Sonia Goncalves, Ewan Harrison, David K. Jackson, Ian Johnston, Dominic Kwiatkowski, Cordelia Langford, John Sillitoe on behalf of the Wellcome Sanger Institute COVID-19 Surveillance Team |
| EPI_ISL_3168488, EPI_ISL_3168489 | Lighthouse Lab in Alderley Park | Wellcome Sanger Institute for the COVID-19 Genomics UK (COG-UK) Consortium | Jacquelyn Wynn, Mairead Hyland, The Lighthouse Lab in Alderley Park and Alex Alderton, Roberto Amato, Jeffrey Barrett, Sonia Goncalves, Ewan Harrison, David K. Jackson, Ian Johnston, Dominic Kwiatkowski, Cordelia Langford, John Sillitoe on behalf of the Wellcome Sanger Institute COVID-19 Surveillance Team |
| EPI_ISL_3168490, EPI_ISL_3168491 | Lighthouse Lab in Glasgow | Wellcome Sanger Institute for the COVID-19 Genomics UK (COG-UK) Consortium | Harper VanSteenhouse, Yumi Kasai, David Gray, Carol Clugston, Anna Dominiczak and Alex Alderton, Roberto Amato, Jeffrey Barrett, Sonia Goncalves, Ewan Harrison, David K. Jackson, Ian Johnston, Dominic Kwiatkowski, Cordelia Langford, John Sillitoe on behalf of the Wellcome Sanger Institute COVID-19 Surveillance Team |
| EPI_ISL_3168492 | Lighthouse Lab in Alderley Park | Wellcome Sanger Institute for the COVID-19 Genomics UK (COG-UK) Consortium | Jacquelyn Wynn, Mairead Hyland, The Lighthouse Lab in Alderley Park and Alex Alderton, Roberto Amato, Jeffrey Barrett, Sonia Goncalves, Ewan Harrison, David K. Jackson, Ian Johnston, Dominic Kwiatkowski, Cordelia Langford, John Sillitoe on behalf of the Wellcome Sanger Institute COVID-19 Surveillance Team |
| EPI_ISL_3168493, EPI_ISL_3168494 | Lighthouse Lab in Glasgow | Wellcome Sanger Institute for the COVID-19 Genomics UK (COG-UK) Consortium | Harper VanSteenhouse, Yumi Kasai, David Gray, Carol Clugston, Anna Dominiczak and Alex Alderton, Roberto Amato, Jeffrey Barrett, Sonia Goncalves, Ewan Harrison, David K. Jackson, Ian Johnston, Dominic Kwiatkowski, Cordelia Langford, John Sillitoe on behalf of the Wellcome Sanger Institute COVID-19 Surveillance Team |
| EPI_ISL_3168495, EPI_ISL_3168496, EPI_ISL_3168497, EPI_ISL_3168498, EPI_ISL_3168499 | Lighthouse Lab in Alderley Park | Wellcome Sanger Institute for the COVID-19 Genomics UK (COG-UK) Consortium | Jacquelyn Wynn, Mairead Hyland, The Lighthouse Lab in Alderley Park and Alex Alderton, Roberto Amato, Jeffrey Barrett, Sonia Goncalves, Ewan Harrison, David K. Jackson, Ian Johnston, Dominic Kwiatkowski, Cordelia Langford, John Sillitoe on behalf of the Wellcome Sanger Institute COVID-19 Surveillance Team |
| EPI_ISL_3168500 | Lighthouse Lab in Glasgow | Wellcome Sanger Institute for the COVID-19 Genomics UK (COG-UK) Consortium | Harper VanSteenhouse, Yumi Kasai, David Gray, Carol Clugston, Anna Dominiczak and Alex Alderton, Roberto Amato, Jeffrey Barrett, Sonia Goncalves, Ewan Harrison, David K. Jackson, Ian Johnston, Dominic Kwiatkowski, Cordelia Langford, John Sillitoe on behalf of the Wellcome Sanger Institute COVID-19 Surveillance Team |
| EPI_ISL_3168501, EPI_ISL_3168502, EPI_ISL_3168503 | Lighthouse Lab in Alderley Park | Wellcome Sanger Institute for the COVID-19 Genomics UK (COG-UK) Consortium | Jacquelyn Wynn, Mairead Hyland, The Lighthouse Lab in Alderley Park and Alex Alderton, Roberto Amato, Jeffrey Barrett, Sonia Goncalves, Ewan Harrison, David K. Jackson, Ian Johnston, Dominic Kwiatkowski, Cordelia Langford, John Sillitoe on behalf of the Wellcome Sanger Institute COVID-19 Surveillance Team |
| EPI_ISL_3168504 | Berkshire and Surrey Pathology Services Lighthouse Laboratory | Wellcome Sanger Institute for the COVID-19 Genomics UK (COG-UK) Consortium | Berkshire and Surrey Pathology Services Lighthouse Laboratory and Alex Alderton, Roberto Amato, Jeffrey Barrett, Sonia Goncalves, Ewan Harrison, David K. Jackson, Ian Johnston, Dominic Kwiatkowski, Cordelia Langford, John Sillitoe on behalf of the Wellcome Sanger Institute COVID-19 Surveillance Team |
| EPI_ISL_3168505, EPI_ISL_3168506 | Lighthouse Lab in Alderley Park | Wellcome Sanger Institute for the COVID-19 Genomics UK (COG-UK) Consortium | Jacquelyn Wynn, Mairead Hyland, The Lighthouse Lab in Alderley Park and Alex Alderton, Roberto Amato, Jeffrey Barrett, Sonia Goncalves, Ewan Harrison, David K. Jackson, Ian Johnston, Dominic Kwiatkowski, Cordelia Langford, John Sillitoe on behalf of the Wellcome Sanger Institute COVID-19 Surveillance Team |
| EPI_ISL_3168507, EPI_ISL_3168508, EPI_ISL_3168509 | Berkshire and Surrey Pathology Services Lighthouse Laboratory | Wellcome Sanger Institute for the COVID-19 Genomics UK (COG-UK) Consortium | Berkshire and Surrey Pathology Services Lighthouse Laboratory and Alex Alderton, Roberto Amato, Jeffrey Barrett, Sonia Goncalves, Ewan Harrison, David K. Jackson, Ian Johnston, Dominic Kwiatkowski, Cordelia Langford, John Sillitoe on behalf of the Wellcome Sanger Institute COVID-19 Surveillance Team |
| EPI_ISL_3168510 | Lighthouse Lab in Alderley Park | Wellcome Sanger Institute for the COVID-19 Genomics UK (COG-UK) Consortium | Jacquelyn Wynn, Mairead Hyland, The Lighthouse Lab in Alderley Park and Alex Alderton, Roberto Amato, Jeffrey Barrett, Sonia Goncalves, Ewan Harrison, David K. Jackson, Ian Johnston, Dominic Kwiatkowski, Cordelia Langford, John Sillitoe on behalf of the Wellcome Sanger Institute COVID-19 Surveillance Team |
| EPI_ISL_3168511 | Berkshire and Surrey Pathology Services Lighthouse Laboratory | Wellcome Sanger Institute for the COVID-19 Genomics UK (COG-UK) Consortium | Berkshire and Surrey Pathology Services Lighthouse Laboratory and Alex Alderton, Roberto Amato, Jeffrey Barrett, Sonia Goncalves, Ewan Harrison, David K. Jackson, Ian Johnston, Dominic Kwiatkowski, Cordelia Langford, John Sillitoe on behalf of the Wellcome Sanger Institute COVID-19 Surveillance Team |
| EPI_ISL_3168512 | Lighthouse Lab in Alderley Park | Wellcome Sanger Institute for the COVID-19 Genomics UK (COG-UK) Consortium | Jacquelyn Wynn, Mairead Hyland, The Lighthouse Lab in Alderley Park and Alex Alderton, Roberto Amato, Jeffrey Barrett, Sonia Goncalves, Ewan Harrison, David K. Jackson, Ian Johnston, Dominic Kwiatkowski, Cordelia Langford, John Sillitoe on behalf of the Wellcome Sanger Institute COVID-19 Surveillance Team |
| EPI_ISL_3168513 | Berkshire and Surrey Pathology Services Lighthouse Laboratory | Wellcome Sanger Institute for the COVID-19 Genomics UK (COG-UK) Consortium | Berkshire and Surrey Pathology Services Lighthouse Laboratory and Alex Alderton, Roberto Amato, Jeffrey Barrett, Sonia Goncalves, Ewan Harrison, David K. Jackson, Ian Johnston, Dominic Kwiatkowski, Cordelia Langford, John Sillitoe on behalf of the Wellcome Sanger Institute COVID-19 Surveillance Team |
| EPI_ISL_3168514 | Lighthouse Lab in Alderley Park | Wellcome Sanger Institute for the COVID-19 Genomics UK (COG-UK) Consortium | Jacquelyn Wynn, Mairead Hyland, The Lighthouse Lab in Alderley Park and Alex Alderton, Roberto Amato, Jeffrey Barrett, Sonia Goncalves, Ewan Harrison, David K. Jackson, Ian Johnston, Dominic Kwiatkowski, Cordelia Langford, John Sillitoe on behalf of the Wellcome Sanger Institute COVID-19 Surveillance Team |

| | | | |
|---|---|---|---|
| EPI_ISL_3168515 | Lighthouse Lab in Glasgow | Wellcome Sanger Institute for the COVID-19 Genomics UK (COG-UK) Consortium | Harper VanSteenhouse, Yumi Kasai, David Gray, Carol Clugston, Anna Dominiczak and Alex Alderton, Roberto Amato, Jeffrey Barrett, Sonia Goncalves, Ewan Harrison, David K. Jackson, Ian Johnston, Dominic Kwiatkowski, Cordelia Langford, John Sillitoe on behalf of the Wellcome Sanger Institute COVID-19 Surveillance Team |
| EPI_ISL_3168516 | Berkshire and Surrey Pathology Services Lighthouse Laboratory | Wellcome Sanger Institute for the COVID-19 Genomics UK (COG-UK) Consortium | Berkshire and Surrey Pathology Services Lighthouse Laboratory and Alex Alderton, Roberto Amato, Jeffrey Barrett, Sonia Goncalves, Ewan Harrison, David K. Jackson, Ian Johnston, Dominic Kwiatkowski, Cordelia Langford, John Sillitoe on behalf of the Wellcome Sanger Institute COVID-19 Surveillance Team |
| EPI_ISL_3168517 | Lighthouse Lab in Glasgow | Wellcome Sanger Institute for the COVID-19 Genomics UK (COG-UK) Consortium | Harper VanSteenhouse, Yumi Kasai, David Gray, Carol Clugston, Anna Dominiczak and Alex Alderton, Roberto Amato, Jeffrey Barrett, Sonia Goncalves, Ewan Harrison, David K. Jackson, Ian Johnston, Dominic Kwiatkowski, Cordelia Langford, John Sillitoe on behalf of the Wellcome Sanger Institute COVID-19 Surveillance Team |
| EPI_ISL_3168518, EPI_ISL_3168519, EPI_ISL_3168520, EPI_ISL_3168521 | Lighthouse Lab in Alderley Park | Wellcome Sanger Institute for the COVID-19 Genomics UK (COG-UK) Consortium | Jacquelyn Wynn, Mairead Hyland, The Lighthouse Lab in Alderley Park and Alex Alderton, Roberto Amato, Jeffrey Barrett, Sonia Goncalves, Ewan Harrison, David K. Jackson, Ian Johnston, Dominic Kwiatkowski, Cordelia Langford, John Sillitoe on behalf of the Wellcome Sanger Institute COVID-19 Surveillance Team |
| EPI_ISL_3168522 | Lighthouse Lab in Glasgow | Wellcome Sanger Institute for the COVID-19 Genomics UK (COG-UK) Consortium | Harper VanSteenhouse, Yumi Kasai, David Gray, Carol Clugston, Anna Dominiczak and Alex Alderton, Roberto Amato, Jeffrey Barrett, Sonia Goncalves, Ewan Harrison, David K. Jackson, Ian Johnston, Dominic Kwiatkowski, Cordelia Langford, John Sillitoe on behalf of the Wellcome Sanger Institute COVID-19 Surveillance Team |
| EPI_ISL_3168523 | Lighthouse Lab in Alderley Park | Wellcome Sanger Institute for the COVID-19 Genomics UK (COG-UK) Consortium | Jacquelyn Wynn, Mairead Hyland, The Lighthouse Lab in Alderley Park and Alex Alderton, Roberto Amato, Jeffrey Barrett, Sonia Goncalves, Ewan Harrison, David K. Jackson, Ian Johnston, Dominic Kwiatkowski, Cordelia Langford, John Sillitoe on behalf of the Wellcome Sanger Institute COVID-19 Surveillance Team |
| EPI_ISL_3168524 | Berkshire and Surrey Pathology Services Lighthouse Laboratory | Wellcome Sanger Institute for the COVID-19 Genomics UK (COG-UK) Consortium | Berkshire and Surrey Pathology Services Lighthouse Laboratory and Alex Alderton, Roberto Amato, Jeffrey Barrett, Sonia Goncalves, Ewan Harrison, David K. Jackson, Ian Johnston, Dominic Kwiatkowski, Cordelia Langford, John Sillitoe on behalf of the Wellcome Sanger Institute COVID-19 Surveillance Team |
| EPI_ISL_3168525 | Lighthouse Lab in Glasgow | Wellcome Sanger Institute for the COVID-19 Genomics UK (COG-UK) Consortium | Harper VanSteenhouse, Yumi Kasai, David Gray, Carol Clugston, Anna Dominiczak and Alex Alderton, Roberto Amato, Jeffrey Barrett, Sonia Goncalves, Ewan Harrison, David K. Jackson, Ian Johnston, Dominic Kwiatkowski, Cordelia Langford, John Sillitoe on behalf of the Wellcome Sanger Institute COVID-19 Surveillance Team |
| EPI_ISL_3168526 | Berkshire and Surrey Pathology Services Lighthouse Laboratory | Wellcome Sanger Institute for the COVID-19 Genomics UK (COG-UK) Consortium | Berkshire and Surrey Pathology Services Lighthouse Laboratory and Alex Alderton, Roberto Amato, Jeffrey Barrett, Sonia Goncalves, Ewan Harrison, David K. Jackson, Ian Johnston, Dominic Kwiatkowski, Cordelia Langford, John Sillitoe on behalf of the Wellcome Sanger Institute COVID-19 Surveillance Team |
| EPI_ISL_3168527, EPI_ISL_3168528, EPI_ISL_3168529, EPI_ISL_3168530 | Lighthouse Lab in Alderley Park | Wellcome Sanger Institute for the COVID-19 Genomics UK (COG-UK) Consortium | Jacquelyn Wynn, Mairead Hyland, The Lighthouse Lab in Alderley Park and Alex Alderton, Roberto Amato, Jeffrey Barrett, Sonia Goncalves, Ewan Harrison, David K. Jackson, Ian Johnston, Dominic Kwiatkowski, Cordelia Langford, John Sillitoe on behalf of the Wellcome Sanger Institute COVID-19 Surveillance Team |
| EPI_ISL_3168531 | Berkshire and Surrey Pathology Services Lighthouse Laboratory | Wellcome Sanger Institute for the COVID-19 Genomics UK (COG-UK) Consortium | Berkshire and Surrey Pathology Services Lighthouse Laboratory and Alex Alderton, Roberto Amato, Jeffrey Barrett, Sonia Goncalves, Ewan Harrison, David K. Jackson, Ian Johnston, Dominic Kwiatkowski, Cordelia Langford, John Sillitoe on behalf of the Wellcome Sanger Institute COVID-19 Surveillance Team |
| EPI_ISL_3168532 | Lighthouse Lab in Glasgow | Wellcome Sanger Institute for the COVID-19 Genomics UK (COG-UK) Consortium | Harper VanSteenhouse, Yumi Kasai, David Gray, Carol Clugston, Anna Dominiczak and Alex Alderton, Roberto Amato, Jeffrey Barrett, Sonia Goncalves, Ewan Harrison, David K. Jackson, Ian Johnston, Dominic Kwiatkowski, Cordelia Langford, John Sillitoe on behalf of the Wellcome Sanger Institute COVID-19 Surveillance Team |
| EPI_ISL_3168533 | Berkshire and Surrey Pathology Services Lighthouse Laboratory | Wellcome Sanger Institute for the COVID-19 Genomics UK (COG-UK) Consortium | Berkshire and Surrey Pathology Services Lighthouse Laboratory and Alex Alderton, Roberto Amato, Jeffrey Barrett, Sonia Goncalves, Ewan Harrison, David K. Jackson, Ian Johnston, Dominic Kwiatkowski, Cordelia Langford, John Sillitoe on behalf of the Wellcome Sanger Institute COVID-19 Surveillance Team |
| EPI_ISL_3168534 | Lighthouse Lab in Alderley Park | Wellcome Sanger Institute for the COVID-19 Genomics UK (COG-UK) Consortium | Jacquelyn Wynn, Mairead Hyland, The Lighthouse Lab in Alderley Park and Alex Alderton, Roberto Amato, Jeffrey Barrett, Sonia Goncalves, Ewan Harrison, David K. Jackson, Ian Johnston, Dominic Kwiatkowski, Cordelia Langford, John Sillitoe on behalf of the Wellcome Sanger Institute COVID-19 Surveillance Team |
| EPI_ISL_3168535 | Berkshire and Surrey Pathology Services Lighthouse Laboratory | Wellcome Sanger Institute for the COVID-19 Genomics UK (COG-UK) Consortium | Berkshire and Surrey Pathology Services Lighthouse Laboratory and Alex Alderton, Roberto Amato, Jeffrey Barrett, Sonia Goncalves, Ewan Harrison, David K. Jackson, Ian Johnston, Dominic Kwiatkowski, Cordelia Langford, John Sillitoe on behalf of the Wellcome Sanger Institute COVID-19 Surveillance Team |
| EPI_ISL_3168536 | Lighthouse Lab in Alderley Park | Wellcome Sanger Institute for the COVID-19 Genomics UK (COG-UK) Consortium | Jacquelyn Wynn, Mairead Hyland, The Lighthouse Lab in Alderley Park and Alex Alderton, Roberto Amato, Jeffrey Barrett, Sonia Goncalves, Ewan Harrison, David K. Jackson, Ian Johnston, Dominic Kwiatkowski, Cordelia Langford, John Sillitoe on behalf of the Wellcome Sanger Institute COVID-19 Surveillance Team |
| EPI_ISL_3168537 | Berkshire and Surrey Pathology Services Lighthouse Laboratory | Wellcome Sanger Institute for the COVID-19 Genomics UK (COG-UK) Consortium | Berkshire and Surrey Pathology Services Lighthouse Laboratory and Alex Alderton, Roberto Amato, Jeffrey Barrett, Sonia Goncalves, Ewan Harrison, David K. Jackson, Ian Johnston, Dominic Kwiatkowski, Cordelia Langford, John Sillitoe on behalf of the Wellcome Sanger Institute COVID-19 Surveillance Team |
| EPI_ISL_3168538 | Lighthouse Lab in Glasgow | Wellcome Sanger Institute for the COVID-19 Genomics UK (COG-UK) Consortium | Harper VanSteenhouse, Yumi Kasai, David Gray, Carol Clugston, Anna Dominiczak and Alex Alderton, Roberto Amato, Jeffrey Barrett, Sonia Goncalves, Ewan Harrison, David K. Jackson, Ian Johnston, Dominic Kwiatkowski, Cordelia Langford, John Sillitoe on behalf of the Wellcome Sanger Institute COVID-19 Surveillance Team |
| EPI_ISL_3168539 | Berkshire and Surrey Pathology Services Lighthouse Laboratory | Wellcome Sanger Institute for the COVID-19 Genomics UK (COG-UK) Consortium | Berkshire and Surrey Pathology Services Lighthouse Laboratory and Alex Alderton, Roberto Amato, Jeffrey Barrett, Sonia Goncalves, Ewan Harrison, David K. Jackson, Ian Johnston, Dominic Kwiatkowski, Cordelia Langford, John Sillitoe on behalf of the Wellcome Sanger Institute COVID-19 Surveillance Team |
| EPI_ISL_3168540, EPI_ISL_3168541 | Lighthouse Lab in Alderley Park | Wellcome Sanger Institute for the COVID-19 Genomics UK (COG-UK) Consortium | Jacquelyn Wynn, Mairead Hyland, The Lighthouse Lab in Alderley Park and Alex Alderton, Roberto Amato, Jeffrey Barrett, Sonia Goncalves, Ewan Harrison, David K. Jackson, Ian Johnston, Dominic Kwiatkowski, Cordelia Langford, John Sillitoe on behalf of the Wellcome Sanger Institute COVID-19 Surveillance Team |
| EPI_ISL_3168542, EPI_ISL_3168543 | Lighthouse Lab in Glasgow | Wellcome Sanger Institute for the COVID-19 Genomics UK (COG-UK) Consortium | Harper VanSteenhouse, Yumi Kasai, David Gray, Carol Clugston, Anna Dominiczak and Alex Alderton, Roberto Amato, Jeffrey Barrett, Sonia Goncalves, Ewan Harrison, David K. Jackson, Ian Johnston, Dominic Kwiatkowski, Cordelia Langford, John Sillitoe on behalf of the Wellcome Sanger Institute COVID-19 Surveillance Team |
| EPI_ISL_3168544 | Lighthouse Lab in Alderley Park | Wellcome Sanger Institute for the COVID-19 Genomics UK (COG-UK) Consortium | Jacquelyn Wynn, Mairead Hyland, The Lighthouse Lab in Alderley Park and Alex Alderton, Roberto Amato, Jeffrey Barrett, Sonia Goncalves, Ewan Harrison, David K. Jackson, Ian Johnston, Dominic Kwiatkowski, Cordelia Langford, John Sillitoe on behalf of the Wellcome Sanger Institute COVID-19 Surveillance Team |
| EPI_ISL_3168545 | Berkshire and Surrey Pathology Services Lighthouse Laboratory | Wellcome Sanger Institute for the COVID-19 Genomics UK (COG-UK) Consortium | Berkshire and Surrey Pathology Services Lighthouse Laboratory and Alex Alderton, Roberto Amato, Jeffrey Barrett, Sonia Goncalves, Ewan Harrison, David K. Jackson, Ian Johnston, Dominic Kwiatkowski, Cordelia Langford, John Sillitoe on behalf of the Wellcome Sanger Institute COVID-19 Surveillance Team |
| EPI_ISL_3168546 | Lighthouse Lab in Glasgow | Wellcome Sanger Institute for the COVID-19 Genomics UK (COG-UK) Consortium | Harper VanSteenhouse, Yumi Kasai, David Gray, Carol Clugston, Anna Dominiczak and Alex Alderton, Roberto Amato, Jeffrey Barrett, Sonia Goncalves, Ewan Harrison, David K. Jackson, Ian Johnston, Dominic Kwiatkowski, Cordelia Langford, John Sillitoe on behalf of the Wellcome Sanger Institute COVID-19 Surveillance Team |
| EPI_ISL_3168547, EPI_ISL_3168548 | Lighthouse Lab in Alderley Park | Wellcome Sanger Institute for the COVID-19 Genomics UK (COG-UK) Consortium | Jacquelyn Wynn, Mairead Hyland, The Lighthouse Lab in Alderley Park and Alex Alderton, Roberto Amato, Jeffrey Barrett, Sonia Goncalves, Ewan Harrison, David K. Jackson, Ian Johnston, Dominic Kwiatkowski, Cordelia Langford, John Sillitoe on behalf of the Wellcome Sanger Institute COVID-19 Surveillance Team |
| EPI_ISL_3168549, EPI_ISL_3168550 | Berkshire and Surrey Pathology Services Lighthouse Laboratory | Wellcome Sanger Institute for the COVID-19 Genomics UK (COG-UK) Consortium | Berkshire and Surrey Pathology Services Lighthouse Laboratory and Alex Alderton, Roberto Amato, Jeffrey Barrett, Sonia Goncalves, Ewan Harrison, David K. Jackson, Ian Johnston, Dominic Kwiatkowski, Cordelia Langford, John Sillitoe on behalf of the Wellcome Sanger Institute COVID-19 Surveillance Team |
| EPI_ISL_3168551, EPI_ISL_3168552, EPI_ISL_3168553 | Lighthouse Lab in Alderley Park | Wellcome Sanger Institute for the COVID-19 Genomics UK (COG-UK) Consortium | Jacquelyn Wynn, Mairead Hyland, The Lighthouse Lab in Alderley Park and Alex Alderton, Roberto Amato, Jeffrey Barrett, Sonia Goncalves, Ewan Harrison, David K. Jackson, Ian Johnston, Dominic Kwiatkowski, Cordelia Langford, John Sillitoe on behalf of the Wellcome Sanger Institute COVID-19 Surveillance Team |
| EPI_ISL_3168554 | Berkshire and Surrey Pathology Services Lighthouse Laboratory | Wellcome Sanger Institute for the COVID-19 Genomics UK (COG-UK) Consortium | Berkshire and Surrey Pathology Services Lighthouse Laboratory and Alex Alderton, Roberto Amato, Jeffrey Barrett, Sonia Goncalves, Ewan Harrison, David K. Jackson, Ian Johnston, Dominic Kwiatkowski, Cordelia Langford, John Sillitoe on behalf of the Wellcome Sanger Institute COVID-19 Surveillance Team |
| EPI_ISL_3168555, EPI_ISL_3168556, EPI_ISL_3168557 | Lighthouse Lab in Alderley Park | Wellcome Sanger Institute for the COVID-19 Genomics UK (COG-UK) Consortium | Jacquelyn Wynn, Mairead Hyland, The Lighthouse Lab in Alderley Park and Alex Alderton, Roberto Amato, Jeffrey Barrett, Sonia Goncalves, Ewan Harrison, David K. Jackson, Ian Johnston, Dominic Kwiatkowski, Cordelia Langford, John Sillitoe on behalf of the Wellcome Sanger Institute COVID-19 Surveillance Team |
| EPI_ISL_3168558 | Berkshire and Surrey Pathology Services Lighthouse Laboratory | Wellcome Sanger Institute for the COVID-19 Genomics UK (COG-UK) Consortium | Berkshire and Surrey Pathology Services Lighthouse Laboratory and Alex Alderton, Roberto Amato, Jeffrey Barrett, Sonia Goncalves, Ewan Harrison, David K. Jackson, Ian Johnston, Dominic Kwiatkowski, Cordelia Langford, John Sillitoe on behalf of the Wellcome Sanger Institute COVID-19 Surveillance Team |
| EPI_ISL_3168559, EPI_ISL_3168560, EPI_ISL_3168561 | Lighthouse Lab in Alderley Park | Wellcome Sanger Institute for the COVID-19 Genomics UK (COG-UK) Consortium | Jacquelyn Wynn, Mairead Hyland, The Lighthouse Lab in Alderley Park and Alex Alderton, Roberto Amato, Jeffrey Barrett, Sonia Goncalves, Ewan Harrison, David K. Jackson, Ian Johnston, Dominic Kwiatkowski, Cordelia Langford, John Sillitoe on behalf of the Wellcome Sanger Institute COVID-19 Surveillance Team |
| EPI_ISL_3168562 | Berkshire and Surrey Pathology Services Lighthouse Laboratory | Wellcome Sanger Institute for the COVID-19 Genomics UK (COG-UK) Consortium | Berkshire and Surrey Pathology Services Lighthouse Laboratory and Alex Alderton, Roberto Amato, Jeffrey Barrett, Sonia Goncalves, Ewan Harrison, David K. Jackson, Ian Johnston, Dominic Kwiatkowski, Cordelia Langford, John Sillitoe on behalf of the Wellcome Sanger Institute COVID-19 Surveillance Team |
| EPI_ISL_3168563, EPI_ISL_3168564 | Lighthouse Lab in Alderley Park | Wellcome Sanger Institute for the COVID-19 Genomics UK (COG-UK) Consortium | Jacquelyn Wynn, Mairead Hyland, The Lighthouse Lab in Alderley Park and Alex Alderton, Roberto Amato, Jeffrey Barrett, Sonia Goncalves, Ewan Harrison, David K. Jackson, Ian Johnston, Dominic Kwiatkowski, Cordelia Langford, John Sillitoe on behalf of the Wellcome Sanger Institute COVID-19 Surveillance Team |
| EPI_ISL_3168565 | Lighthouse Lab in Glasgow | Wellcome Sanger Institute for the COVID-19 Genomics UK (COG-UK) Consortium | Harper VanSteenhouse, Yumi Kasai, David Gray, Carol Clugston, Anna Dominiczak and Alex Alderton, Roberto Amato, Jeffrey Barrett, Sonia Goncalves, Ewan Harrison, David K. Jackson, Ian Johnston, Dominic Kwiatkowski, Cordelia Langford, John Sillitoe on behalf of the Wellcome Sanger Institute COVID-19 Surveillance Team |
| EPI_ISL_3168566, EPI_ISL_3168567, EPI_ISL_3168568 | Berkshire and Surrey Pathology Services Lighthouse Laboratory | Wellcome Sanger Institute for the COVID-19 Genomics UK (COG-UK) Consortium | Berkshire and Surrey Pathology Services Lighthouse Laboratory and Alex Alderton, Roberto Amato, Jeffrey Barrett, Sonia Goncalves, Ewan Harrison, David K. Jackson, Ian Johnston, Dominic Kwiatkowski, Cordelia Langford, John Sillitoe on behalf of the Wellcome Sanger Institute COVID-19 Surveillance Team |
| EPI_ISL_3168569 | Lighthouse Lab in Glasgow | Wellcome Sanger Institute for the COVID-19 Genomics UK (COG-UK) Consortium | Harper VanSteenhouse, Yumi Kasai, David Gray, Carol Clugston, Anna Dominiczak and Alex Alderton, Roberto Amato, Jeffrey Barrett, Sonia Goncalves, Ewan Harrison, David K. Jackson, Ian Johnston, Dominic Kwiatkowski, Cordelia Langford, John Sillitoe on behalf of the Wellcome Sanger Institute COVID-19 Surveillance Team |
| EPI_ISL_3168570 | Lighthouse Lab in Alderley Park | Wellcome Sanger Institute for the COVID-19 Genomics UK (COG-UK) Consortium | Jacquelyn Wynn, Mairead Hyland, The Lighthouse Lab in Alderley Park and Alex Alderton, Roberto Amato, Jeffrey Barrett, Sonia Goncalves, Ewan Harrison, David K. Jackson, Ian Johnston, Dominic Kwiatkowski, Cordelia Langford, John Sillitoe on behalf of the Wellcome Sanger Institute COVID-19 Surveillance Team |
| EPI_ISL_3168571 | Berkshire and Surrey Pathology Services Lighthouse Laboratory | Wellcome Sanger Institute for the COVID-19 Genomics UK (COG-UK) Consortium | Berkshire and Surrey Pathology Services Lighthouse Laboratory and Alex Alderton, Roberto Amato, Jeffrey Barrett, Sonia Goncalves, Ewan Harrison, David K. Jackson, Ian Johnston, Dominic Kwiatkowski, Cordelia Langford, John Sillitoe on behalf of the Wellcome Sanger Institute COVID-19 Surveillance Team |
| EPI_ISL_3168572, EPI_ISL_3168573 | Lighthouse Lab in Alderley Park | Wellcome Sanger Institute for the COVID-19 Genomics UK (COG-UK) Consortium | Jacquelyn Wynn, Mairead Hyland, The Lighthouse Lab in Alderley Park and Alex Alderton, Roberto Amato, Jeffrey Barrett, Sonia Goncalves, Ewan Harrison, David K. Jackson, Ian Johnston, Dominic Kwiatkowski, Cordelia Langford, John Sillitoe on behalf of the Wellcome Sanger Institute COVID-19 Surveillance Team |
| EPI_ISL_3168574 | Berkshire and Surrey Pathology Services Lighthouse Laboratory | Wellcome Sanger Institute for the COVID-19 Genomics UK (COG-UK) Consortium | Berkshire and Surrey Pathology Services Lighthouse Laboratory and Alex Alderton, Roberto Amato, Jeffrey Barrett, Sonia Goncalves, Ewan Harrison, David K. Jackson, Ian Johnston, Dominic Kwiatkowski, Cordelia Langford, John Sillitoe on behalf of the Wellcome Sanger Institute COVID-19 Surveillance Team |
| EPI_ISL_3168575, EPI_ISL_3168576, EPI_ISL_3168577, EPI_ISL_3168578, | Lighthouse Lab in Alderley Park | Wellcome Sanger Institute for the COVID-19 Genomics UK (COG-UK) Consortium | Jacquelyn Wynn, Mairead Hyland, The Lighthouse Lab in Alderley Park and Alex Alderton, Roberto Amato, Jeffrey Barrett, Sonia Goncalves, Ewan Harrison, David K. Jackson, Ian Johnston, Dominic Kwiatkowski, Cordelia Langford, John Sillitoe on behalf of the Wellcome Sanger Institute COVID-19 Surveillance Team |

| | | | |
|---|---|---|---|
| EPI_ISL_3168579, EPI_ISL_3168580 | | | |
| EPI_ISL_3168581 | Berkshire and Surrey Pathology Services Lighthouse Laboratory | Wellcome Sanger Institute for the COVID-19 Genomics UK (COG-UK) Consortium | Berkshire and Surrey Pathology Services Lighthouse Laboratory and Alex Alderton, Roberto Amato, Jeffrey Barrett, Sonia Goncalves, Ewan Harrison, David K. Jackson, Ian Johnston, Dominic Kwiatkowski, Cordelia Langford, John Sillitoe on behalf of the Wellcome Sanger Institute COVID-19 Surveillance Team |
| EPI_ISL_3168582 | Lighthouse Lab in Alderley Park | Wellcome Sanger Institute for the COVID-19 Genomics UK (COG-UK) Consortium | Jacquelyn Wynn, Mairead Hyland, The Lighthouse Lab in Alderley Park and Alex Alderton, Roberto Amato, Jeffrey Barrett, Sonia Goncalves, Ewan Harrison, David K. Jackson, Ian Johnston, Dominic Kwiatkowski, Cordelia Langford, John Sillitoe on behalf of the Wellcome Sanger Institute COVID-19 Surveillance Team |
| EPI_ISL_3168583, EPI_ISL_3168584 | Lighthouse Lab in Glasgow | Wellcome Sanger Institute for the COVID-19 Genomics UK (COG-UK) Consortium | Harper VanSteenhouse, Yumi Kasai, David Gray, Carol Clugston, Anna Dominiczak and Alex Alderton, Roberto Amato, Jeffrey Barrett, Sonia Goncalves, Ewan Harrison, David K. Jackson, Ian Johnston, Dominic Kwiatkowski, Cordelia Langford, John Sillitoe on behalf of the Wellcome Sanger Institute COVID-19 Surveillance Team |
| EPI_ISL_3168585 | Berkshire and Surrey Pathology Services Lighthouse Laboratory | Wellcome Sanger Institute for the COVID-19 Genomics UK (COG-UK) Consortium | Berkshire and Surrey Pathology Services Lighthouse Laboratory and Alex Alderton, Roberto Amato, Jeffrey Barrett, Sonia Goncalves, Ewan Harrison, David K. Jackson, Ian Johnston, Dominic Kwiatkowski, Cordelia Langford, John Sillitoe on behalf of the Wellcome Sanger Institute COVID-19 Surveillance Team |
| EPI_ISL_3168586 | Lighthouse Lab in Alderley Park | Wellcome Sanger Institute for the COVID-19 Genomics UK (COG-UK) Consortium | Jacquelyn Wynn, Mairead Hyland, The Lighthouse Lab in Alderley Park and Alex Alderton, Roberto Amato, Jeffrey Barrett, Sonia Goncalves, Ewan Harrison, David K. Jackson, Ian Johnston, Dominic Kwiatkowski, Cordelia Langford, John Sillitoe on behalf of the Wellcome Sanger Institute COVID-19 Surveillance Team |
| EPI_ISL_3168587, EPI_ISL_3168588, EPI_ISL_3168589 | Lighthouse Lab in Glasgow | Wellcome Sanger Institute for the COVID-19 Genomics UK (COG-UK) Consortium | Harper VanSteenhouse, Yumi Kasai, David Gray, Carol Clugston, Anna Dominiczak and Alex Alderton, Roberto Amato, Jeffrey Barrett, Sonia Goncalves, Ewan Harrison, David K. Jackson, Ian Johnston, Dominic Kwiatkowski, Cordelia Langford, John Sillitoe on behalf of the Wellcome Sanger Institute COVID-19 Surveillance Team |
| EPI_ISL_3168590 | Lighthouse Lab in Alderley Park | Wellcome Sanger Institute for the COVID-19 Genomics UK (COG-UK) Consortium | Jacquelyn Wynn, Mairead Hyland, The Lighthouse Lab in Alderley Park and Alex Alderton, Roberto Amato, Jeffrey Barrett, Sonia Goncalves, Ewan Harrison, David K. Jackson, Ian Johnston, Dominic Kwiatkowski, Cordelia Langford, John Sillitoe on behalf of the Wellcome Sanger Institute COVID-19 Surveillance Team |
| EPI_ISL_3168591 | Lighthouse Lab in Glasgow | Wellcome Sanger Institute for the COVID-19 Genomics UK (COG-UK) Consortium | Harper VanSteenhouse, Yumi Kasai, David Gray, Carol Clugston, Anna Dominiczak and Alex Alderton, Roberto Amato, Jeffrey Barrett, Sonia Goncalves, Ewan Harrison, David K. Jackson, Ian Johnston, Dominic Kwiatkowski, Cordelia Langford, John Sillitoe on behalf of the Wellcome Sanger Institute COVID-19 Surveillance Team |
| EPI_ISL_3168592 | Berkshire and Surrey Pathology Services Lighthouse Laboratory | Wellcome Sanger Institute for the COVID-19 Genomics UK (COG-UK) Consortium | Berkshire and Surrey Pathology Services Lighthouse Laboratory and Alex Alderton, Roberto Amato, Jeffrey Barrett, Sonia Goncalves, Ewan Harrison, David K. Jackson, Ian Johnston, Dominic Kwiatkowski, Cordelia Langford, John Sillitoe on behalf of the Wellcome Sanger Institute COVID-19 Surveillance Team |
| EPI_ISL_3168593 | Lighthouse Lab in Alderley Park | Wellcome Sanger Institute for the COVID-19 Genomics UK (COG-UK) Consortium | Jacquelyn Wynn, Mairead Hyland, The Lighthouse Lab in Alderley Park and Alex Alderton, Roberto Amato, Jeffrey Barrett, Sonia Goncalves, Ewan Harrison, David K. Jackson, Ian Johnston, Dominic Kwiatkowski, Cordelia Langford, John Sillitoe on behalf of the Wellcome Sanger Institute COVID-19 Surveillance Team |
| EPI_ISL_3168594 | Berkshire and Surrey Pathology Services Lighthouse Laboratory | Wellcome Sanger Institute for the COVID-19 Genomics UK (COG-UK) Consortium | Berkshire and Surrey Pathology Services Lighthouse Laboratory and Alex Alderton, Roberto Amato, Jeffrey Barrett, Sonia Goncalves, Ewan Harrison, David K. Jackson, Ian Johnston, Dominic Kwiatkowski, Cordelia Langford, John Sillitoe on behalf of the Wellcome Sanger Institute COVID-19 Surveillance Team |
| EPI_ISL_3168595, EPI_ISL_3168596, EPI_ISL_3168597, EPI_ISL_3168598 | Lighthouse Lab in Alderley Park | Wellcome Sanger Institute for the COVID-19 Genomics UK (COG-UK) Consortium | Jacquelyn Wynn, Mairead Hyland, The Lighthouse Lab in Alderley Park and Alex Alderton, Roberto Amato, Jeffrey Barrett, Sonia Goncalves, Ewan Harrison, David K. Jackson, Ian Johnston, Dominic Kwiatkowski, Cordelia Langford, John Sillitoe on behalf of the Wellcome Sanger Institute COVID-19 Surveillance Team |
| EPI_ISL_3168599 | Berkshire and Surrey Pathology Services Lighthouse Laboratory | Wellcome Sanger Institute for the COVID-19 Genomics UK (COG-UK) Consortium | Berkshire and Surrey Pathology Services Lighthouse Laboratory and Alex Alderton, Roberto Amato, Jeffrey Barrett, Sonia Goncalves, Ewan Harrison, David K. Jackson, Ian Johnston, Dominic Kwiatkowski, Cordelia Langford, John Sillitoe on behalf of the Wellcome Sanger Institute COVID-19 Surveillance Team |
| EPI_ISL_3168600 | Lighthouse Lab in Alderley Park | Wellcome Sanger Institute for the COVID-19 Genomics UK (COG-UK) Consortium | Jacquelyn Wynn, Mairead Hyland, The Lighthouse Lab in Alderley Park and Alex Alderton, Roberto Amato, Jeffrey Barrett, Sonia Goncalves, Ewan Harrison, David K. Jackson, Ian Johnston, Dominic Kwiatkowski, Cordelia Langford, John Sillitoe on behalf of the Wellcome Sanger Institute COVID-19 Surveillance Team |
| EPI_ISL_3168601 | Lighthouse Lab in Glasgow | Wellcome Sanger Institute for the COVID-19 Genomics UK (COG-UK) Consortium | Harper VanSteenhouse, Yumi Kasai, David Gray, Carol Clugston, Anna Dominiczak and Alex Alderton, Roberto Amato, Jeffrey Barrett, Sonia Goncalves, Ewan Harrison, David K. Jackson, Ian Johnston, Dominic Kwiatkowski, Cordelia Langford, John Sillitoe on behalf of the Wellcome Sanger Institute COVID-19 Surveillance Team |
| EPI_ISL_3168602, EPI_ISL_3168603 | Lighthouse Lab in Alderley Park | Wellcome Sanger Institute for the COVID-19 Genomics UK (COG-UK) Consortium | Jacquelyn Wynn, Mairead Hyland, The Lighthouse Lab in Alderley Park and Alex Alderton, Roberto Amato, Jeffrey Barrett, Sonia Goncalves, Ewan Harrison, David K. Jackson, Ian Johnston, Dominic Kwiatkowski, Cordelia Langford, John Sillitoe on behalf of the Wellcome Sanger Institute COVID-19 Surveillance Team |
| EPI_ISL_3168604, EPI_ISL_3168605 | Berkshire and Surrey Pathology Services Lighthouse Laboratory | Wellcome Sanger Institute for the COVID-19 Genomics UK (COG-UK) Consortium | Berkshire and Surrey Pathology Services Lighthouse Laboratory and Alex Alderton, Roberto Amato, Jeffrey Barrett, Sonia Goncalves, Ewan Harrison, David K. Jackson, Ian Johnston, Dominic Kwiatkowski, Cordelia Langford, John Sillitoe on behalf of the Wellcome Sanger Institute COVID-19 Surveillance Team |
| EPI_ISL_3168606 | Lighthouse Lab in Glasgow | Wellcome Sanger Institute for the COVID-19 Genomics UK (COG-UK) Consortium | Harper VanSteenhouse, Yumi Kasai, David Gray, Carol Clugston, Anna Dominiczak and Alex Alderton, Roberto Amato, Jeffrey Barrett, Sonia Goncalves, Ewan Harrison, David K. Jackson, Ian Johnston, Dominic Kwiatkowski, Cordelia Langford, John Sillitoe on behalf of the Wellcome Sanger Institute COVID-19 Surveillance Team |
| EPI_ISL_3168607, EPI_ISL_3168608 | Lighthouse Lab in Alderley Park | Wellcome Sanger Institute for the COVID-19 Genomics UK (COG-UK) Consortium | Jacquelyn Wynn, Mairead Hyland, The Lighthouse Lab in Alderley Park and Alex Alderton, Roberto Amato, Jeffrey Barrett, Sonia Goncalves, Ewan Harrison, David K. Jackson, Ian Johnston, Dominic Kwiatkowski, Cordelia Langford, John Sillitoe on behalf of the Wellcome Sanger Institute COVID-19 Surveillance Team |
| EPI_ISL_3168609 | Berkshire and Surrey Pathology Services Lighthouse Laboratory | Wellcome Sanger Institute for the COVID-19 Genomics UK (COG-UK) Consortium | Berkshire and Surrey Pathology Services Lighthouse Laboratory and Alex Alderton, Roberto Amato, Jeffrey Barrett, Sonia Goncalves, Ewan Harrison, David K. Jackson, Ian Johnston, Dominic Kwiatkowski, Cordelia Langford, John Sillitoe on behalf of the Wellcome Sanger Institute COVID-19 Surveillance Team |
| EPI_ISL_3168610 | Lighthouse Lab in Glasgow | Wellcome Sanger Institute for the COVID-19 Genomics UK (COG-UK) Consortium | Harper VanSteenhouse, Yumi Kasai, David Gray, Carol Clugston, Anna Dominiczak and Alex Alderton, Roberto Amato, Jeffrey Barrett, Sonia Goncalves, Ewan Harrison, David K. Jackson, Ian Johnston, Dominic Kwiatkowski, Cordelia Langford, John Sillitoe on behalf of the Wellcome Sanger Institute COVID-19 Surveillance Team |
| EPI_ISL_3168611 | Lighthouse Lab in Alderley Park | Wellcome Sanger Institute for the COVID-19 Genomics UK (COG-UK) Consortium | Jacquelyn Wynn, Mairead Hyland, The Lighthouse Lab in Alderley Park and Alex Alderton, Roberto Amato, Jeffrey Barrett, Sonia Goncalves, Ewan Harrison, David K. Jackson, Ian Johnston, Dominic Kwiatkowski, Cordelia Langford, John Sillitoe on behalf of the Wellcome Sanger Institute COVID-19 Surveillance Team |
| EPI_ISL_3168612 | Lighthouse Lab in Glasgow | Wellcome Sanger Institute for the COVID-19 Genomics UK (COG-UK) Consortium | Harper VanSteenhouse, Yumi Kasai, David Gray, Carol Clugston, Anna Dominiczak and Alex Alderton, Roberto Amato, Jeffrey Barrett, Sonia Goncalves, Ewan Harrison, David K. Jackson, Ian Johnston, Dominic Kwiatkowski, Cordelia Langford, John Sillitoe on behalf of the Wellcome Sanger Institute COVID-19 Surveillance Team |
| EPI_ISL_3168613, EPI_ISL_3168614 | Lighthouse Lab in Alderley Park | Wellcome Sanger Institute for the COVID-19 Genomics UK (COG-UK) Consortium | Jacquelyn Wynn, Mairead Hyland, The Lighthouse Lab in Alderley Park and Alex Alderton, Roberto Amato, Jeffrey Barrett, Sonia Goncalves, Ewan Harrison, David K. Jackson, Ian Johnston, Dominic Kwiatkowski, Cordelia Langford, John Sillitoe on behalf of the Wellcome Sanger Institute COVID-19 Surveillance Team |
| EPI_ISL_3168615 | Berkshire and Surrey Pathology Services Lighthouse Laboratory | Wellcome Sanger Institute for the COVID-19 Genomics UK (COG-UK) Consortium | Berkshire and Surrey Pathology Services Lighthouse Laboratory and Alex Alderton, Roberto Amato, Jeffrey Barrett, Sonia Goncalves, Ewan Harrison, David K. Jackson, Ian Johnston, Dominic Kwiatkowski, Cordelia Langford, John Sillitoe on behalf of the Wellcome Sanger Institute COVID-19 Surveillance Team |
| EPI_ISL_3168616 | Lighthouse Lab in Glasgow | Wellcome Sanger Institute for the COVID-19 Genomics UK (COG-UK) Consortium | Harper VanSteenhouse, Yumi Kasai, David Gray, Carol Clugston, Anna Dominiczak and Alex Alderton, Roberto Amato, Jeffrey Barrett, Sonia Goncalves, Ewan Harrison, David K. Jackson, Ian Johnston, Dominic Kwiatkowski, Cordelia Langford, John Sillitoe on behalf of the Wellcome Sanger Institute COVID-19 Surveillance Team |
| EPI_ISL_3168617 | Berkshire and Surrey Pathology Services Lighthouse Laboratory | Wellcome Sanger Institute for the COVID-19 Genomics UK (COG-UK) Consortium | Berkshire and Surrey Pathology Services Lighthouse Laboratory and Alex Alderton, Roberto Amato, Jeffrey Barrett, Sonia Goncalves, Ewan Harrison, David K. Jackson, Ian Johnston, Dominic Kwiatkowski, Cordelia Langford, John Sillitoe on behalf of the Wellcome Sanger Institute COVID-19 Surveillance Team |
| EPI_ISL_3168618, EPI_ISL_3168619 | Lighthouse Lab in Glasgow | Wellcome Sanger Institute for the COVID-19 Genomics UK (COG-UK) Consortium | Harper VanSteenhouse, Yumi Kasai, David Gray, Carol Clugston, Anna Dominiczak and Alex Alderton, Roberto Amato, Jeffrey Barrett, Sonia Goncalves, Ewan Harrison, David K. Jackson, Ian Johnston, Dominic Kwiatkowski, Cordelia Langford, John Sillitoe on behalf of the Wellcome Sanger Institute COVID-19 Surveillance Team |
| EPI_ISL_3168620 | Lighthouse Lab in Alderley Park | Wellcome Sanger Institute for the COVID-19 Genomics UK (COG-UK) Consortium | Jacquelyn Wynn, Mairead Hyland, The Lighthouse Lab in Alderley Park and Alex Alderton, Roberto Amato, Jeffrey Barrett, Sonia Goncalves, Ewan Harrison, David K. Jackson, Ian Johnston, Dominic Kwiatkowski, Cordelia Langford, John Sillitoe on behalf of the Wellcome Sanger Institute COVID-19 Surveillance Team |
| EPI_ISL_3168621 | Lighthouse Lab in Glasgow | Wellcome Sanger Institute for the COVID-19 Genomics UK (COG-UK) Consortium | Harper VanSteenhouse, Yumi Kasai, David Gray, Carol Clugston, Anna Dominiczak and Alex Alderton, Roberto Amato, Jeffrey Barrett, Sonia Goncalves, Ewan Harrison, David K. Jackson, Ian Johnston, Dominic Kwiatkowski, Cordelia Langford, John Sillitoe on behalf of the Wellcome Sanger Institute COVID-19 Surveillance Team |
| EPI_ISL_3168622 | Lighthouse Lab in Alderley Park | Wellcome Sanger Institute for the COVID-19 Genomics UK (COG-UK) Consortium | Jacquelyn Wynn, Mairead Hyland, The Lighthouse Lab in Alderley Park and Alex Alderton, Roberto Amato, Jeffrey Barrett, Sonia Goncalves, Ewan Harrison, David K. Jackson, Ian Johnston, Dominic Kwiatkowski, Cordelia Langford, John Sillitoe on behalf of the Wellcome Sanger Institute COVID-19 Surveillance Team |
| EPI_ISL_3168623 | Lighthouse Lab in Glasgow | Wellcome Sanger Institute for the COVID-19 Genomics UK (COG-UK) Consortium | Harper VanSteenhouse, Yumi Kasai, David Gray, Carol Clugston, Anna Dominiczak and Alex Alderton, Roberto Amato, Jeffrey Barrett, Sonia Goncalves, Ewan Harrison, David K. Jackson, Ian Johnston, Dominic Kwiatkowski, Cordelia Langford, John Sillitoe on behalf of the Wellcome Sanger Institute COVID-19 Surveillance Team |
| EPI_ISL_3168624 | Berkshire and Surrey Pathology Services Lighthouse Laboratory | Wellcome Sanger Institute for the COVID-19 Genomics UK (COG-UK) Consortium | Berkshire and Surrey Pathology Services Lighthouse Laboratory and Alex Alderton, Roberto Amato, Jeffrey Barrett, Sonia Goncalves, Ewan Harrison, David K. Jackson, Ian Johnston, Dominic Kwiatkowski, Cordelia Langford, John Sillitoe on behalf of the Wellcome Sanger Institute COVID-19 Surveillance Team |
| EPI_ISL_3168625, EPI_ISL_3168626, EPI_ISL_3168627, EPI_ISL_3168628 | Lighthouse Lab in Alderley Park | Wellcome Sanger Institute for the COVID-19 Genomics UK (COG-UK) Consortium | Jacquelyn Wynn, Mairead Hyland, The Lighthouse Lab in Alderley Park and Alex Alderton, Roberto Amato, Jeffrey Barrett, Sonia Goncalves, Ewan Harrison, David K. Jackson, Ian Johnston, Dominic Kwiatkowski, Cordelia Langford, John Sillitoe on behalf of the Wellcome Sanger Institute COVID-19 Surveillance Team |
| EPI_ISL_3168629, EPI_ISL_3168630 | Lighthouse Lab in Glasgow | Wellcome Sanger Institute for the COVID-19 Genomics UK (COG-UK) Consortium | Harper VanSteenhouse, Yumi Kasai, David Gray, Carol Clugston, Anna Dominiczak and Alex Alderton, Roberto Amato, Jeffrey Barrett, Sonia Goncalves, Ewan Harrison, David K. Jackson, Ian Johnston, Dominic Kwiatkowski, Cordelia Langford, John Sillitoe on behalf of the Wellcome Sanger Institute COVID-19 Surveillance Team |
| EPI_ISL_3168631, EPI_ISL_3168632, EPI_ISL_3168633, EPI_ISL_3168634 | Lighthouse Lab in Alderley Park | Wellcome Sanger Institute for the COVID-19 Genomics UK (COG-UK) Consortium | Jacquelyn Wynn, Mairead Hyland, The Lighthouse Lab in Alderley Park and Alex Alderton, Roberto Amato, Jeffrey Barrett, Sonia Goncalves, Ewan Harrison, David K. Jackson, Ian Johnston, Dominic Kwiatkowski, Cordelia Langford, John Sillitoe on behalf of the Wellcome Sanger Institute COVID-19 Surveillance Team |
| EPI_ISL_3168635 | Berkshire and Surrey Pathology Services Lighthouse Laboratory | Wellcome Sanger Institute for the COVID-19 Genomics UK (COG-UK) Consortium | Berkshire and Surrey Pathology Services Lighthouse Laboratory and Alex Alderton, Roberto Amato, Jeffrey Barrett, Sonia Goncalves, Ewan Harrison, David K. Jackson, Ian Johnston, Dominic Kwiatkowski, Cordelia Langford, John Sillitoe on behalf of the Wellcome Sanger Institute COVID-19 Surveillance Team |
| EPI_ISL_3168636, EPI_ISL_3168637 | Lighthouse Lab in Alderley Park | Wellcome Sanger Institute for the COVID-19 Genomics UK (COG-UK) Consortium | Jacquelyn Wynn, Mairead Hyland, The Lighthouse Lab in Alderley Park and Alex Alderton, Roberto Amato, Jeffrey Barrett, Sonia Goncalves, Ewan Harrison, David K. Jackson, Ian Johnston, Dominic Kwiatkowski, Cordelia Langford, John Sillitoe on behalf of the Wellcome Sanger Institute COVID-19 Surveillance Team |
| EPI_ISL_3168638, EPI_ISL_3168639 | Lighthouse Lab in Glasgow | Wellcome Sanger Institute for the COVID-19 Genomics UK (COG-UK) Consortium | Harper VanSteenhouse, Yumi Kasai, David Gray, Carol Clugston, Anna Dominiczak and Alex Alderton, Roberto Amato, Jeffrey Barrett, Sonia Goncalves, Ewan Harrison, David K. Jackson, Ian Johnston, Dominic Kwiatkowski, Cordelia Langford, John Sillitoe on behalf of the Wellcome Sanger Institute COVID-19 Surveillance Team |
| EPI_ISL_3168640 | Lighthouse Lab in Alderley Park | Wellcome Sanger Institute for the COVID-19 Genomics UK (COG-UK) Consortium | Jacquelyn Wynn, Mairead Hyland, The Lighthouse Lab in Alderley Park and Alex Alderton, Roberto Amato, Jeffrey Barrett, Sonia Goncalves, Ewan Harrison, David K. Jackson, Ian Johnston, Dominic Kwiatkowski, Cordelia Langford, John Sillitoe on behalf of the Wellcome Sanger Institute COVID-19 Surveillance Team |
| EPI_ISL_3168641 | Lighthouse Lab in Glasgow | Wellcome Sanger Institute for the COVID-19 Genomics UK (COG-UK) Consortium | Harper VanSteenhouse, Yumi Kasai, David Gray, Carol Clugston, Anna Dominiczak and Alex Alderton, Roberto Amato, Jeffrey Barrett, Sonia Goncalves, Ewan Harrison, David K. Jackson, Ian Johnston, Dominic Kwiatkowski, Cordelia Langford, John Sillitoe on behalf of the Wellcome Sanger Institute COVID-19 Surveillance Team |
| EPI_ISL_3168642 | Lighthouse Lab in Alderley Park | Wellcome Sanger Institute for the COVID-19 Genomics UK (COG-UK) | Jacquelyn Wynn, Mairead Hyland, The Lighthouse Lab in Alderley Park and Alex Alderton, Roberto Amato, Jeffrey Barrett, Sonia Goncalves, Ewan Harrison, David K. Jackson, Ian Johnston, Dominic Kwiatkowski, Cordelia Langford, John Sillitoe on |

| | | | |
|---|---|---|---|
| | | Consortium | behalf of the Wellcome Sanger Institute COVID-19 Surveillance Team |
| EPI_ISL_3168643, EPI_ISL_3168644 | Lighthouse Lab in Glasgow | Wellcome Sanger Institute for the COVID-19 Genomics UK (COG-UK) Consortium | Harper VanSteenhouse, Yumi Kasai, David Gray, Carol Clugston, Anna Dominiczak and Alex Alderton, Roberto Amato, Jeffrey Barrett, Sonia Goncalves, Ewan Harrison, David K. Jackson, Ian Johnston, Dominic Kwiatkowski, Cordelia Langford, John Sillitoe on behalf of the Wellcome Sanger Institute COVID-19 Surveillance Team |
| EPI_ISL_3168645 | Berkshire and Surrey Pathology Services Lighthouse Laboratory | Wellcome Sanger Institute for the COVID-19 Genomics UK (COG-UK) Consortium | Berkshire and Surrey Pathology Services Lighthouse Laboratory and Alex Alderton, Roberto Amato, Jeffrey Barrett, Sonia Goncalves, Ewan Harrison, David K. Jackson, Ian Johnston, Dominic Kwiatkowski, Cordelia Langford, John Sillitoe on behalf of the Wellcome Sanger Institute COVID-19 Surveillance Team |
| EPI_ISL_3168646, EPI_ISL_3168647 | Lighthouse Lab in Alderley Park | Wellcome Sanger Institute for the COVID-19 Genomics UK (COG-UK) Consortium | Jacquelyn Wynn, Mairead Hyland, The Lighthouse Lab in Alderley Park and Alex Alderton, Roberto Amato, Jeffrey Barrett, Sonia Goncalves, Ewan Harrison, David K. Jackson, Ian Johnston, Dominic Kwiatkowski, Cordelia Langford, John Sillitoe on behalf of the Wellcome Sanger Institute COVID-19 Surveillance Team |
| EPI_ISL_3168648, EPI_ISL_3168649 | Berkshire and Surrey Pathology Services Lighthouse Laboratory | Wellcome Sanger Institute for the COVID-19 Genomics UK (COG-UK) Consortium | Berkshire and Surrey Pathology Services Lighthouse Laboratory and Alex Alderton, Roberto Amato, Jeffrey Barrett, Sonia Goncalves, Ewan Harrison, David K. Jackson, Ian Johnston, Dominic Kwiatkowski, Cordelia Langford, John Sillitoe on behalf of the Wellcome Sanger Institute COVID-19 Surveillance Team |
| EPI_ISL_3168650, EPI_ISL_3168651 | Lighthouse Lab in Alderley Park | Wellcome Sanger Institute for the COVID-19 Genomics UK (COG-UK) Consortium | Jacquelyn Wynn, Mairead Hyland, The Lighthouse Lab in Alderley Park and Alex Alderton, Roberto Amato, Jeffrey Barrett, Sonia Goncalves, Ewan Harrison, David K. Jackson, Ian Johnston, Dominic Kwiatkowski, Cordelia Langford, John Sillitoe on behalf of the Wellcome Sanger Institute COVID-19 Surveillance Team |
| EPI_ISL_3168652 | Lighthouse Lab in Glasgow | Wellcome Sanger Institute for the COVID-19 Genomics UK (COG-UK) Consortium | Harper VanSteenhouse, Yumi Kasai, David Gray, Carol Clugston, Anna Dominiczak and Alex Alderton, Roberto Amato, Jeffrey Barrett, Sonia Goncalves, Ewan Harrison, David K. Jackson, Ian Johnston, Dominic Kwiatkowski, Cordelia Langford, John Sillitoe on behalf of the Wellcome Sanger Institute COVID-19 Surveillance Team |
| EPI_ISL_3168653 | Berkshire and Surrey Pathology Services Lighthouse Laboratory | Wellcome Sanger Institute for the COVID-19 Genomics UK (COG-UK) Consortium | Berkshire and Surrey Pathology Services Lighthouse Laboratory and Alex Alderton, Roberto Amato, Jeffrey Barrett, Sonia Goncalves, Ewan Harrison, David K. Jackson, Ian Johnston, Dominic Kwiatkowski, Cordelia Langford, John Sillitoe on behalf of the Wellcome Sanger Institute COVID-19 Surveillance Team |
| EPI_ISL_3168654 | Lighthouse Lab in Alderley Park | Wellcome Sanger Institute for the COVID-19 Genomics UK (COG-UK) Consortium | Jacquelyn Wynn, Mairead Hyland, The Lighthouse Lab in Alderley Park and Alex Alderton, Roberto Amato, Jeffrey Barrett, Sonia Goncalves, Ewan Harrison, David K. Jackson, Ian Johnston, Dominic Kwiatkowski, Cordelia Langford, John Sillitoe on behalf of the Wellcome Sanger Institute COVID-19 Surveillance Team |
| EPI_ISL_3168655 | Berkshire and Surrey Pathology Services Lighthouse Laboratory | Wellcome Sanger Institute for the COVID-19 Genomics UK (COG-UK) Consortium | Berkshire and Surrey Pathology Services Lighthouse Laboratory and Alex Alderton, Roberto Amato, Jeffrey Barrett, Sonia Goncalves, Ewan Harrison, David K. Jackson, Ian Johnston, Dominic Kwiatkowski, Cordelia Langford, John Sillitoe on behalf of the Wellcome Sanger Institute COVID-19 Surveillance Team |
| EPI_ISL_3168656, EPI_ISL_3168657, EPI_ISL_3168658, EPI_ISL_3168659 | Lighthouse Lab in Glasgow | Wellcome Sanger Institute for the COVID-19 Genomics UK (COG-UK) Consortium | Harper VanSteenhouse, Yumi Kasai, David Gray, Carol Clugston, Anna Dominiczak and Alex Alderton, Roberto Amato, Jeffrey Barrett, Sonia Goncalves, Ewan Harrison, David K. Jackson, Ian Johnston, Dominic Kwiatkowski, Cordelia Langford, John Sillitoe on behalf of the Wellcome Sanger Institute COVID-19 Surveillance Team |
| EPI_ISL_3168660 | Lighthouse Lab in Alderley Park | Wellcome Sanger Institute for the COVID-19 Genomics UK (COG-UK) Consortium | Jacquelyn Wynn, Mairead Hyland, The Lighthouse Lab in Alderley Park and Alex Alderton, Roberto Amato, Jeffrey Barrett, Sonia Goncalves, Ewan Harrison, David K. Jackson, Ian Johnston, Dominic Kwiatkowski, Cordelia Langford, John Sillitoe on behalf of the Wellcome Sanger Institute COVID-19 Surveillance Team |
| EPI_ISL_3168661 | Lighthouse Lab in Glasgow | Wellcome Sanger Institute for the COVID-19 Genomics UK (COG-UK) Consortium | Harper VanSteenhouse, Yumi Kasai, David Gray, Carol Clugston, Anna Dominiczak and Alex Alderton, Roberto Amato, Jeffrey Barrett, Sonia Goncalves, Ewan Harrison, David K. Jackson, Ian Johnston, Dominic Kwiatkowski, Cordelia Langford, John Sillitoe on behalf of the Wellcome Sanger Institute COVID-19 Surveillance Team |
| EPI_ISL_3168662 | Lighthouse Lab in Alderley Park | Wellcome Sanger Institute for the COVID-19 Genomics UK (COG-UK) Consortium | Jacquelyn Wynn, Mairead Hyland, The Lighthouse Lab in Alderley Park and Alex Alderton, Roberto Amato, Jeffrey Barrett, Sonia Goncalves, Ewan Harrison, David K. Jackson, Ian Johnston, Dominic Kwiatkowski, Cordelia Langford, John Sillitoe on behalf of the Wellcome Sanger Institute COVID-19 Surveillance Team |
| EPI_ISL_3168663 | Berkshire and Surrey Pathology Services Lighthouse Laboratory | Wellcome Sanger Institute for the COVID-19 Genomics UK (COG-UK) Consortium | Berkshire and Surrey Pathology Services Lighthouse Laboratory and Alex Alderton, Roberto Amato, Jeffrey Barrett, Sonia Goncalves, Ewan Harrison, David K. Jackson, Ian Johnston, Dominic Kwiatkowski, Cordelia Langford, John Sillitoe on behalf of the Wellcome Sanger Institute COVID-19 Surveillance Team |
| EPI_ISL_3168664 | Lighthouse Lab in Glasgow | Wellcome Sanger Institute for the COVID-19 Genomics UK (COG-UK) Consortium | Harper VanSteenhouse, Yumi Kasai, David Gray, Carol Clugston, Anna Dominiczak and Alex Alderton, Roberto Amato, Jeffrey Barrett, Sonia Goncalves, Ewan Harrison, David K. Jackson, Ian Johnston, Dominic Kwiatkowski, Cordelia Langford, John Sillitoe on behalf of the Wellcome Sanger Institute COVID-19 Surveillance Team |
| EPI_ISL_3168665, EPI_ISL_3168666, EPI_ISL_3168667 | Lighthouse Lab in Alderley Park | Wellcome Sanger Institute for the COVID-19 Genomics UK (COG-UK) Consortium | Jacquelyn Wynn, Mairead Hyland, The Lighthouse Lab in Alderley Park and Alex Alderton, Roberto Amato, Jeffrey Barrett, Sonia Goncalves, Ewan Harrison, David K. Jackson, Ian Johnston, Dominic Kwiatkowski, Cordelia Langford, John Sillitoe on behalf of the Wellcome Sanger Institute COVID-19 Surveillance Team |
| EPI_ISL_3168668 | Lighthouse Lab in Glasgow | Wellcome Sanger Institute for the COVID-19 Genomics UK (COG-UK) Consortium | Harper VanSteenhouse, Yumi Kasai, David Gray, Carol Clugston, Anna Dominiczak and Alex Alderton, Roberto Amato, Jeffrey Barrett, Sonia Goncalves, Ewan Harrison, David K. Jackson, Ian Johnston, Dominic Kwiatkowski, Cordelia Langford, John Sillitoe on behalf of the Wellcome Sanger Institute COVID-19 Surveillance Team |
| EPI_ISL_3168669 | Lighthouse Lab in Alderley Park | Wellcome Sanger Institute for the COVID-19 Genomics UK (COG-UK) Consortium | Jacquelyn Wynn, Mairead Hyland, The Lighthouse Lab in Alderley Park and Alex Alderton, Roberto Amato, Jeffrey Barrett, Sonia Goncalves, Ewan Harrison, David K. Jackson, Ian Johnston, Dominic Kwiatkowski, Cordelia Langford, John Sillitoe on behalf of the Wellcome Sanger Institute COVID-19 Surveillance Team |
| EPI_ISL_3168670 | Berkshire and Surrey Pathology Services Lighthouse Laboratory | Wellcome Sanger Institute for the COVID-19 Genomics UK (COG-UK) Consortium | Berkshire and Surrey Pathology Services Lighthouse Laboratory and Alex Alderton, Roberto Amato, Jeffrey Barrett, Sonia Goncalves, Ewan Harrison, David K. Jackson, Ian Johnston, Dominic Kwiatkowski, Cordelia Langford, John Sillitoe on behalf of the Wellcome Sanger Institute COVID-19 Surveillance Team |
| EPI_ISL_3168671 | Lighthouse Lab in Alderley Park | Wellcome Sanger Institute for the COVID-19 Genomics UK (COG-UK) Consortium | Jacquelyn Wynn, Mairead Hyland, The Lighthouse Lab in Alderley Park and Alex Alderton, Roberto Amato, Jeffrey Barrett, Sonia Goncalves, Ewan Harrison, David K. Jackson, Ian Johnston, Dominic Kwiatkowski, Cordelia Langford, John Sillitoe on behalf of the Wellcome Sanger Institute COVID-19 Surveillance Team |
| EPI_ISL_3168672 | Lighthouse Lab in Glasgow | Wellcome Sanger Institute for the COVID-19 Genomics UK (COG-UK) Consortium | Harper VanSteenhouse, Yumi Kasai, David Gray, Carol Clugston, Anna Dominiczak and Alex Alderton, Roberto Amato, Jeffrey Barrett, Sonia Goncalves, Ewan Harrison, David K. Jackson, Ian Johnston, Dominic Kwiatkowski, Cordelia Langford, John Sillitoe on behalf of the Wellcome Sanger Institute COVID-19 Surveillance Team |
| EPI_ISL_3168673 | Lighthouse Lab in Alderley Park | Wellcome Sanger Institute for the COVID-19 Genomics UK (COG-UK) Consortium | Jacquelyn Wynn, Mairead Hyland, The Lighthouse Lab in Alderley Park and Alex Alderton, Roberto Amato, Jeffrey Barrett, Sonia Goncalves, Ewan Harrison, David K. Jackson, Ian Johnston, Dominic Kwiatkowski, Cordelia Langford, John Sillitoe on behalf of the Wellcome Sanger Institute COVID-19 Surveillance Team |
| EPI_ISL_3168674, EPI_ISL_3168675, EPI_ISL_3168676, EPI_ISL_3168677 | Lighthouse Lab in Glasgow | Wellcome Sanger Institute for the COVID-19 Genomics UK (COG-UK) Consortium | Harper VanSteenhouse, Yumi Kasai, David Gray, Carol Clugston, Anna Dominiczak and Alex Alderton, Roberto Amato, Jeffrey Barrett, Sonia Goncalves, Ewan Harrison, David K. Jackson, Ian Johnston, Dominic Kwiatkowski, Cordelia Langford, John Sillitoe on behalf of the Wellcome Sanger Institute COVID-19 Surveillance Team |
| EPI_ISL_3168678, EPI_ISL_3168679, EPI_ISL_3168680, EPI_ISL_3168681, EPI_ISL_3168682, EPI_ISL_3168683 | Lighthouse Lab in Alderley Park | Wellcome Sanger Institute for the COVID-19 Genomics UK (COG-UK) Consortium | Jacquelyn Wynn, Mairead Hyland, The Lighthouse Lab in Alderley Park and Alex Alderton, Roberto Amato, Jeffrey Barrett, Sonia Goncalves, Ewan Harrison, David K. Jackson, Ian Johnston, Dominic Kwiatkowski, Cordelia Langford, John Sillitoe on behalf of the Wellcome Sanger Institute COVID-19 Surveillance Team |
| EPI_ISL_3168684 | Berkshire and Surrey Pathology Services Lighthouse Laboratory | Wellcome Sanger Institute for the COVID-19 Genomics UK (COG-UK) Consortium | Berkshire and Surrey Pathology Services Lighthouse Laboratory and Alex Alderton, Roberto Amato, Jeffrey Barrett, Sonia Goncalves, Ewan Harrison, David K. Jackson, Ian Johnston, Dominic Kwiatkowski, Cordelia Langford, John Sillitoe on behalf of the Wellcome Sanger Institute COVID-19 Surveillance Team |
| EPI_ISL_3168685, EPI_ISL_3168686, EPI_ISL_3168687, EPI_ISL_3168688 | Lighthouse Lab in Alderley Park | Wellcome Sanger Institute for the COVID-19 Genomics UK (COG-UK) Consortium | Jacquelyn Wynn, Mairead Hyland, The Lighthouse Lab in Alderley Park and Alex Alderton, Roberto Amato, Jeffrey Barrett, Sonia Goncalves, Ewan Harrison, David K. Jackson, Ian Johnston, Dominic Kwiatkowski, Cordelia Langford, John Sillitoe on behalf of the Wellcome Sanger Institute COVID-19 Surveillance Team |
| EPI_ISL_3168689, EPI_ISL_3168690, EPI_ISL_3168691 | Berkshire and Surrey Pathology Services Lighthouse Laboratory | Wellcome Sanger Institute for the COVID-19 Genomics UK (COG-UK) Consortium | Berkshire and Surrey Pathology Services Lighthouse Laboratory and Alex Alderton, Roberto Amato, Jeffrey Barrett, Sonia Goncalves, Ewan Harrison, David K. Jackson, Ian Johnston, Dominic Kwiatkowski, Cordelia Langford, John Sillitoe on behalf of the Wellcome Sanger Institute COVID-19 Surveillance Team |
| EPI_ISL_3168692, EPI_ISL_3168693 | Lighthouse Lab in Alderley Park | Wellcome Sanger Institute for the COVID-19 Genomics UK (COG-UK) Consortium | Jacquelyn Wynn, Mairead Hyland, The Lighthouse Lab in Alderley Park and Alex Alderton, Roberto Amato, Jeffrey Barrett, Sonia Goncalves, Ewan Harrison, David K. Jackson, Ian Johnston, Dominic Kwiatkowski, Cordelia Langford, John Sillitoe on behalf of the Wellcome Sanger Institute COVID-19 Surveillance Team |
| EPI_ISL_3168694 | Lighthouse Lab in Glasgow | Wellcome Sanger Institute for the COVID-19 Genomics UK (COG-UK) Consortium | Harper VanSteenhouse, Yumi Kasai, David Gray, Carol Clugston, Anna Dominiczak and Alex Alderton, Roberto Amato, Jeffrey Barrett, Sonia Goncalves, Ewan Harrison, David K. Jackson, Ian Johnston, Dominic Kwiatkowski, Cordelia Langford, John Sillitoe on behalf of the Wellcome Sanger Institute COVID-19 Surveillance Team |
| EPI_ISL_3168695, EPI_ISL_3168696, EPI_ISL_3168697 | Lighthouse Lab in Alderley Park | Wellcome Sanger Institute for the COVID-19 Genomics UK (COG-UK) Consortium | Jacquelyn Wynn, Mairead Hyland, The Lighthouse Lab in Alderley Park and Alex Alderton, Roberto Amato, Jeffrey Barrett, Sonia Goncalves, Ewan Harrison, David K. Jackson, Ian Johnston, Dominic Kwiatkowski, Cordelia Langford, John Sillitoe on behalf of the Wellcome Sanger Institute COVID-19 Surveillance Team |
| EPI_ISL_3168698 | Lighthouse Lab in Glasgow | Wellcome Sanger Institute for the COVID-19 Genomics UK (COG-UK) Consortium | Harper VanSteenhouse, Yumi Kasai, David Gray, Carol Clugston, Anna Dominiczak and Alex Alderton, Roberto Amato, Jeffrey Barrett, Sonia Goncalves, Ewan Harrison, David K. Jackson, Ian Johnston, Dominic Kwiatkowski, Cordelia Langford, John Sillitoe on behalf of the Wellcome Sanger Institute COVID-19 Surveillance Team |
| EPI_ISL_3168699 | Berkshire and Surrey Pathology Services Lighthouse Laboratory | Wellcome Sanger Institute for the COVID-19 Genomics UK (COG-UK) Consortium | Berkshire and Surrey Pathology Services Lighthouse Laboratory and Alex Alderton, Roberto Amato, Jeffrey Barrett, Sonia Goncalves, Ewan Harrison, David K. Jackson, Ian Johnston, Dominic Kwiatkowski, Cordelia Langford, John Sillitoe on behalf of the Wellcome Sanger Institute COVID-19 Surveillance Team |
| EPI_ISL_3168700, EPI_ISL_3168701 | Lighthouse Lab in Alderley Park | Wellcome Sanger Institute for the COVID-19 Genomics UK (COG-UK) Consortium | Jacquelyn Wynn, Mairead Hyland, The Lighthouse Lab in Alderley Park and Alex Alderton, Roberto Amato, Jeffrey Barrett, Sonia Goncalves, Ewan Harrison, David K. Jackson, Ian Johnston, Dominic Kwiatkowski, Cordelia Langford, John Sillitoe on behalf of the Wellcome Sanger Institute COVID-19 Surveillance Team |
| EPI_ISL_3168702 | Lighthouse Lab in Glasgow | Wellcome Sanger Institute for the COVID-19 Genomics UK (COG-UK) Consortium | Harper VanSteenhouse, Yumi Kasai, David Gray, Carol Clugston, Anna Dominiczak and Alex Alderton, Roberto Amato, Jeffrey Barrett, Sonia Goncalves, Ewan Harrison, David K. Jackson, Ian Johnston, Dominic Kwiatkowski, Cordelia Langford, John Sillitoe on behalf of the Wellcome Sanger Institute COVID-19 Surveillance Team |
| EPI_ISL_3168703, EPI_ISL_3168704, EPI_ISL_3168705, EPI_ISL_3168706 | Lighthouse Lab in Alderley Park | Wellcome Sanger Institute for the COVID-19 Genomics UK (COG-UK) Consortium | Jacquelyn Wynn, Mairead Hyland, The Lighthouse Lab in Alderley Park and Alex Alderton, Roberto Amato, Jeffrey Barrett, Sonia Goncalves, Ewan Harrison, David K. Jackson, Ian Johnston, Dominic Kwiatkowski, Cordelia Langford, John Sillitoe on behalf of the Wellcome Sanger Institute COVID-19 Surveillance Team |
| EPI_ISL_3168707 | Berkshire and Surrey Pathology Services Lighthouse Laboratory | Wellcome Sanger Institute for the COVID-19 Genomics UK (COG-UK) Consortium | Berkshire and Surrey Pathology Services Lighthouse Laboratory and Alex Alderton, Roberto Amato, Jeffrey Barrett, Sonia Goncalves, Ewan Harrison, David K. Jackson, Ian Johnston, Dominic Kwiatkowski, Cordelia Langford, John Sillitoe on behalf of the Wellcome Sanger Institute COVID-19 Surveillance Team |
| EPI_ISL_3168708, EPI_ISL_3168709 | Lighthouse Lab in Alderley Park | Wellcome Sanger Institute for the COVID-19 Genomics UK (COG-UK) Consortium | Jacquelyn Wynn, Mairead Hyland, The Lighthouse Lab in Alderley Park and Alex Alderton, Roberto Amato, Jeffrey Barrett, Sonia Goncalves, Ewan Harrison, David K. Jackson, Ian Johnston, Dominic Kwiatkowski, Cordelia Langford, John Sillitoe on behalf of the Wellcome Sanger Institute COVID-19 Surveillance Team |
| EPI_ISL_3168710 | Lighthouse Lab in Glasgow | Wellcome Sanger Institute for the COVID-19 Genomics UK (COG-UK) Consortium | Harper VanSteenhouse, Yumi Kasai, David Gray, Carol Clugston, Anna Dominiczak and Alex Alderton, Roberto Amato, Jeffrey Barrett, Sonia Goncalves, Ewan Harrison, David K. Jackson, Ian Johnston, Dominic Kwiatkowski, Cordelia Langford, John Sillitoe on behalf of the Wellcome Sanger Institute COVID-19 Surveillance Team |
| EPI_ISL_3168711, | Lighthouse Laboratory Plymouth | Wellcome Sanger Institute for the COVID-19 Genomics UK (COG-UK) | Lighthouse Laboratory Plymouth and Alex Alderton, Roberto Amato, Jeffrey Barrett, Sonia Goncalves, Ewan Harrison, David K. Jackson, Ian Johnston, Dominic Kwiatkowski, Cordelia Langford, John Sillitoe on behalf of the Wellcome Sanger |

| | | | |
|---|---|---|---|
| EPI_ISL_3168712 | | Consortium | Institute COVID-19 Surveillance Team |
| EPI_ISL_3168713 | Lighthouse Lab in Glasgow | Wellcome Sanger Institute for the COVID-19 Genomics UK (COG-UK) Consortium | Harper VanSteenhouse, Yumi Kasai, David Gray, Carol Clugston, Anna Dominiczak and Alex Alderton, Roberto Amato, Jeffrey Barrett, Sonia Goncalves, Ewan Harrison, David K. Jackson, Ian Johnston, Dominic Kwiatkowski, Cordelia Langford, John Sillitoe on behalf of the Wellcome Sanger Institute COVID-19 Surveillance Team |
| EPI_ISL_3168714, EPI_ISL_3168715, EPI_ISL_3168716, EPI_ISL_3168717, EPI_ISL_3168718 | Lighthouse Laboratory Plymouth | Wellcome Sanger Institute for the COVID-19 Genomics UK (COG-UK) Consortium | Lighthouse Laboratory Plymouth and Alex Alderton, Roberto Amato, Jeffrey Barrett, Sonia Goncalves, Ewan Harrison, David K. Jackson, Ian Johnston, Dominic Kwiatkowski, Cordelia Langford, John Sillitoe on behalf of the Wellcome Sanger Institute COVID-19 Surveillance Team |
| EPI_ISL_3168719, EPI_ISL_3168720 | Lighthouse Lab in Glasgow | Wellcome Sanger Institute for the COVID-19 Genomics UK (COG-UK) Consortium | Harper VanSteenhouse, Yumi Kasai, David Gray, Carol Clugston, Anna Dominiczak and Alex Alderton, Roberto Amato, Jeffrey Barrett, Sonia Goncalves, Ewan Harrison, David K. Jackson, Ian Johnston, Dominic Kwiatkowski, Cordelia Langford, John Sillitoe on behalf of the Wellcome Sanger Institute COVID-19 Surveillance Team |
| EPI_ISL_3168721, EPI_ISL_3168722 | Lighthouse Laboratory Plymouth | Wellcome Sanger Institute for the COVID-19 Genomics UK (COG-UK) Consortium | Lighthouse Laboratory Plymouth and Alex Alderton, Roberto Amato, Jeffrey Barrett, Sonia Goncalves, Ewan Harrison, David K. Jackson, Ian Johnston, Dominic Kwiatkowski, Cordelia Langford, John Sillitoe on behalf of the Wellcome Sanger Institute COVID-19 Surveillance Team |
| EPI_ISL_3168723, EPI_ISL_3168724 | Lighthouse Lab in Glasgow | Wellcome Sanger Institute for the COVID-19 Genomics UK (COG-UK) Consortium | Harper VanSteenhouse, Yumi Kasai, David Gray, Carol Clugston, Anna Dominiczak and Alex Alderton, Roberto Amato, Jeffrey Barrett, Sonia Goncalves, Ewan Harrison, David K. Jackson, Ian Johnston, Dominic Kwiatkowski, Cordelia Langford, John Sillitoe on behalf of the Wellcome Sanger Institute COVID-19 Surveillance Team |
| EPI_ISL_3168725, EPI_ISL_3168726, EPI_ISL_3168727 | Lighthouse Laboratory Plymouth | Wellcome Sanger Institute for the COVID-19 Genomics UK (COG-UK) Consortium | Lighthouse Laboratory Plymouth and Alex Alderton, Roberto Amato, Jeffrey Barrett, Sonia Goncalves, Ewan Harrison, David K. Jackson, Ian Johnston, Dominic Kwiatkowski, Cordelia Langford, John Sillitoe on behalf of the Wellcome Sanger Institute COVID-19 Surveillance Team |
| EPI_ISL_3168728 | Lighthouse Lab in Glasgow | Wellcome Sanger Institute for the COVID-19 Genomics UK (COG-UK) Consortium | Harper VanSteenhouse, Yumi Kasai, David Gray, Carol Clugston, Anna Dominiczak and Alex Alderton, Roberto Amato, Jeffrey Barrett, Sonia Goncalves, Ewan Harrison, David K. Jackson, Ian Johnston, Dominic Kwiatkowski, Cordelia Langford, John Sillitoe on behalf of the Wellcome Sanger Institute COVID-19 Surveillance Team |
| EPI_ISL_3168729 | Lighthouse Laboratory Plymouth | Wellcome Sanger Institute for the COVID-19 Genomics UK (COG-UK) Consortium | Lighthouse Laboratory Plymouth and Alex Alderton, Roberto Amato, Jeffrey Barrett, Sonia Goncalves, Ewan Harrison, David K. Jackson, Ian Johnston, Dominic Kwiatkowski, Cordelia Langford, John Sillitoe on behalf of the Wellcome Sanger Institute COVID-19 Surveillance Team |
| EPI_ISL_3168730 | Lighthouse Lab in Glasgow | Wellcome Sanger Institute for the COVID-19 Genomics UK (COG-UK) Consortium | Harper VanSteenhouse, Yumi Kasai, David Gray, Carol Clugston, Anna Dominiczak and Alex Alderton, Roberto Amato, Jeffrey Barrett, Sonia Goncalves, Ewan Harrison, David K. Jackson, Ian Johnston, Dominic Kwiatkowski, Cordelia Langford, John Sillitoe on behalf of the Wellcome Sanger Institute COVID-19 Surveillance Team |
| EPI_ISL_3168731 | Lighthouse Laboratory Plymouth | Wellcome Sanger Institute for the COVID-19 Genomics UK (COG-UK) Consortium | Lighthouse Laboratory Plymouth and Alex Alderton, Roberto Amato, Jeffrey Barrett, Sonia Goncalves, Ewan Harrison, David K. Jackson, Ian Johnston, Dominic Kwiatkowski, Cordelia Langford, John Sillitoe on behalf of the Wellcome Sanger Institute COVID-19 Surveillance Team |
| EPI_ISL_3168732 | Lighthouse Lab in Glasgow | Wellcome Sanger Institute for the COVID-19 Genomics UK (COG-UK) Consortium | Harper VanSteenhouse, Yumi Kasai, David Gray, Carol Clugston, Anna Dominiczak and Alex Alderton, Roberto Amato, Jeffrey Barrett, Sonia Goncalves, Ewan Harrison, David K. Jackson, Ian Johnston, Dominic Kwiatkowski, Cordelia Langford, John Sillitoe on behalf of the Wellcome Sanger Institute COVID-19 Surveillance Team |
| EPI_ISL_3168733, EPI_ISL_3168734 | Lighthouse Laboratory Plymouth | Wellcome Sanger Institute for the COVID-19 Genomics UK (COG-UK) Consortium | Lighthouse Laboratory Plymouth and Alex Alderton, Roberto Amato, Jeffrey Barrett, Sonia Goncalves, Ewan Harrison, David K. Jackson, Ian Johnston, Dominic Kwiatkowski, Cordelia Langford, John Sillitoe on behalf of the Wellcome Sanger Institute COVID-19 Surveillance Team |
| EPI_ISL_3168735 | Lighthouse Lab in Glasgow | Wellcome Sanger Institute for the COVID-19 Genomics UK (COG-UK) Consortium | Harper VanSteenhouse, Yumi Kasai, David Gray, Carol Clugston, Anna Dominiczak and Alex Alderton, Roberto Amato, Jeffrey Barrett, Sonia Goncalves, Ewan Harrison, David K. Jackson, Ian Johnston, Dominic Kwiatkowski, Cordelia Langford, John Sillitoe on behalf of the Wellcome Sanger Institute COVID-19 Surveillance Team |
| EPI_ISL_3168736 | Lighthouse Laboratory Plymouth | Wellcome Sanger Institute for the COVID-19 Genomics UK (COG-UK) Consortium | Lighthouse Laboratory Plymouth and Alex Alderton, Roberto Amato, Jeffrey Barrett, Sonia Goncalves, Ewan Harrison, David K. Jackson, Ian Johnston, Dominic Kwiatkowski, Cordelia Langford, John Sillitoe on behalf of the Wellcome Sanger Institute COVID-19 Surveillance Team |
| EPI_ISL_3168737, EPI_ISL_3168738, EPI_ISL_3168739 | Lighthouse Lab in Glasgow | Wellcome Sanger Institute for the COVID-19 Genomics UK (COG-UK) Consortium | Harper VanSteenhouse, Yumi Kasai, David Gray, Carol Clugston, Anna Dominiczak and Alex Alderton, Roberto Amato, Jeffrey Barrett, Sonia Goncalves, Ewan Harrison, David K. Jackson, Ian Johnston, Dominic Kwiatkowski, Cordelia Langford, John Sillitoe on behalf of the Wellcome Sanger Institute COVID-19 Surveillance Team |
| EPI_ISL_3168740 | Lighthouse Laboratory Plymouth | Wellcome Sanger Institute for the COVID-19 Genomics UK (COG-UK) Consortium | Lighthouse Laboratory Plymouth and Alex Alderton, Roberto Amato, Jeffrey Barrett, Sonia Goncalves, Ewan Harrison, David K. Jackson, Ian Johnston, Dominic Kwiatkowski, Cordelia Langford, John Sillitoe on behalf of the Wellcome Sanger Institute COVID-19 Surveillance Team |
| EPI_ISL_3168741, EPI_ISL_3168742 | Lighthouse Lab in Glasgow | Wellcome Sanger Institute for the COVID-19 Genomics UK (COG-UK) Consortium | Harper VanSteenhouse, Yumi Kasai, David Gray, Carol Clugston, Anna Dominiczak and Alex Alderton, Roberto Amato, Jeffrey Barrett, Sonia Goncalves, Ewan Harrison, David K. Jackson, Ian Johnston, Dominic Kwiatkowski, Cordelia Langford, John Sillitoe on behalf of the Wellcome Sanger Institute COVID-19 Surveillance Team |
| EPI_ISL_3168743, EPI_ISL_3168744, EPI_ISL_3168745, EPI_ISL_3168746, EPI_ISL_3168747, EPI_ISL_3168748, EPI_ISL_3168749, EPI_ISL_3168750 | Lighthouse Laboratory Plymouth | Wellcome Sanger Institute for the COVID-19 Genomics UK (COG-UK) Consortium | Lighthouse Laboratory Plymouth and Alex Alderton, Roberto Amato, Jeffrey Barrett, Sonia Goncalves, Ewan Harrison, David K. Jackson, Ian Johnston, Dominic Kwiatkowski, Cordelia Langford, John Sillitoe on behalf of the Wellcome Sanger Institute COVID-19 Surveillance Team |
| EPI_ISL_3168751 | Lighthouse Lab in Glasgow | Wellcome Sanger Institute for the COVID-19 Genomics UK (COG-UK) Consortium | Harper VanSteenhouse, Yumi Kasai, David Gray, Carol Clugston, Anna Dominiczak and Alex Alderton, Roberto Amato, Jeffrey Barrett, Sonia Goncalves, Ewan Harrison, David K. Jackson, Ian Johnston, Dominic Kwiatkowski, Cordelia Langford, John Sillitoe on behalf of the Wellcome Sanger Institute COVID-19 Surveillance Team |
| EPI_ISL_3168752, EPI_ISL_3168753, EPI_ISL_3168754 | Lighthouse Laboratory Plymouth | Wellcome Sanger Institute for the COVID-19 Genomics UK (COG-UK) Consortium | Lighthouse Laboratory Plymouth and Alex Alderton, Roberto Amato, Jeffrey Barrett, Sonia Goncalves, Ewan Harrison, David K. Jackson, Ian Johnston, Dominic Kwiatkowski, Cordelia Langford, John Sillitoe on behalf of the Wellcome Sanger Institute COVID-19 Surveillance Team |
| EPI_ISL_3168755 | Lighthouse Lab in Glasgow | Wellcome Sanger Institute for the COVID-19 Genomics UK (COG-UK) Consortium | Harper VanSteenhouse, Yumi Kasai, David Gray, Carol Clugston, Anna Dominiczak and Alex Alderton, Roberto Amato, Jeffrey Barrett, Sonia Goncalves, Ewan Harrison, David K. Jackson, Ian Johnston, Dominic Kwiatkowski, Cordelia Langford, John Sillitoe on behalf of the Wellcome Sanger Institute COVID-19 Surveillance Team |
| EPI_ISL_3168756, EPI_ISL_3168757, EPI_ISL_3168758, EPI_ISL_3168759, EPI_ISL_3168760 | Lighthouse Laboratory Plymouth | Wellcome Sanger Institute for the COVID-19 Genomics UK (COG-UK) Consortium | Lighthouse Laboratory Plymouth and Alex Alderton, Roberto Amato, Jeffrey Barrett, Sonia Goncalves, Ewan Harrison, David K. Jackson, Ian Johnston, Dominic Kwiatkowski, Cordelia Langford, John Sillitoe on behalf of the Wellcome Sanger Institute COVID-19 Surveillance Team |
| EPI_ISL_3168761 | Lighthouse Lab in Glasgow | Wellcome Sanger Institute for the COVID-19 Genomics UK (COG-UK) Consortium | Harper VanSteenhouse, Yumi Kasai, David Gray, Carol Clugston, Anna Dominiczak and Alex Alderton, Roberto Amato, Jeffrey Barrett, Sonia Goncalves, Ewan Harrison, David K. Jackson, Ian Johnston, Dominic Kwiatkowski, Cordelia Langford, John Sillitoe on behalf of the Wellcome Sanger Institute COVID-19 Surveillance Team |
| EPI_ISL_3168762 | Lighthouse Laboratory Plymouth | Wellcome Sanger Institute for the COVID-19 Genomics UK (COG-UK) Consortium | Lighthouse Laboratory Plymouth and Alex Alderton, Roberto Amato, Jeffrey Barrett, Sonia Goncalves, Ewan Harrison, David K. Jackson, Ian Johnston, Dominic Kwiatkowski, Cordelia Langford, John Sillitoe on behalf of the Wellcome Sanger Institute COVID-19 Surveillance Team |
| EPI_ISL_3168763, EPI_ISL_3168764, EPI_ISL_3168765 | Lighthouse Lab in Glasgow | Wellcome Sanger Institute for the COVID-19 Genomics UK (COG-UK) Consortium | Harper VanSteenhouse, Yumi Kasai, David Gray, Carol Clugston, Anna Dominiczak and Alex Alderton, Roberto Amato, Jeffrey Barrett, Sonia Goncalves, Ewan Harrison, David K. Jackson, Ian Johnston, Dominic Kwiatkowski, Cordelia Langford, John Sillitoe on behalf of the Wellcome Sanger Institute COVID-19 Surveillance Team |
| EPI_ISL_3168766 | Lighthouse Laboratory Plymouth | Wellcome Sanger Institute for the COVID-19 Genomics UK (COG-UK) Consortium | Lighthouse Laboratory Plymouth and Alex Alderton, Roberto Amato, Jeffrey Barrett, Sonia Goncalves, Ewan Harrison, David K. Jackson, Ian Johnston, Dominic Kwiatkowski, Cordelia Langford, John Sillitoe on behalf of the Wellcome Sanger Institute COVID-19 Surveillance Team |
| EPI_ISL_3168767 | Lighthouse Lab in Glasgow | Wellcome Sanger Institute for the COVID-19 Genomics UK (COG-UK) Consortium | Harper VanSteenhouse, Yumi Kasai, David Gray, Carol Clugston, Anna Dominiczak and Alex Alderton, Roberto Amato, Jeffrey Barrett, Sonia Goncalves, Ewan Harrison, David K. Jackson, Ian Johnston, Dominic Kwiatkowski, Cordelia Langford, John Sillitoe on behalf of the Wellcome Sanger Institute COVID-19 Surveillance Team |
| EPI_ISL_3168768, EPI_ISL_3168769 | Lighthouse Laboratory Plymouth | Wellcome Sanger Institute for the COVID-19 Genomics UK (COG-UK) Consortium | Lighthouse Laboratory Plymouth and Alex Alderton, Roberto Amato, Jeffrey Barrett, Sonia Goncalves, Ewan Harrison, David K. Jackson, Ian Johnston, Dominic Kwiatkowski, Cordelia Langford, John Sillitoe on behalf of the Wellcome Sanger Institute COVID-19 Surveillance Team |
| EPI_ISL_3168770, EPI_ISL_3168771 | Lighthouse Lab in Glasgow | Wellcome Sanger Institute for the COVID-19 Genomics UK (COG-UK) Consortium | Harper VanSteenhouse, Yumi Kasai, David Gray, Carol Clugston, Anna Dominiczak and Alex Alderton, Roberto Amato, Jeffrey Barrett, Sonia Goncalves, Ewan Harrison, David K. Jackson, Ian Johnston, Dominic Kwiatkowski, Cordelia Langford, John Sillitoe on behalf of the Wellcome Sanger Institute COVID-19 Surveillance Team |
| EPI_ISL_3168772, EPI_ISL_3168773 | Lighthouse Laboratory Plymouth | Wellcome Sanger Institute for the COVID-19 Genomics UK (COG-UK) Consortium | Lighthouse Laboratory Plymouth and Alex Alderton, Roberto Amato, Jeffrey Barrett, Sonia Goncalves, Ewan Harrison, David K. Jackson, Ian Johnston, Dominic Kwiatkowski, Cordelia Langford, John Sillitoe on behalf of the Wellcome Sanger Institute COVID-19 Surveillance Team |
| EPI_ISL_3168774 | Lighthouse Lab in Glasgow | Wellcome Sanger Institute for the COVID-19 Genomics UK (COG-UK) Consortium | Harper VanSteenhouse, Yumi Kasai, David Gray, Carol Clugston, Anna Dominiczak and Alex Alderton, Roberto Amato, Jeffrey Barrett, Sonia Goncalves, Ewan Harrison, David K. Jackson, Ian Johnston, Dominic Kwiatkowski, Cordelia Langford, John Sillitoe on behalf of the Wellcome Sanger Institute COVID-19 Surveillance Team |
| EPI_ISL_3168775 | Lighthouse Laboratory Plymouth | Wellcome Sanger Institute for the COVID-19 Genomics UK (COG-UK) Consortium | Lighthouse Laboratory Plymouth and Alex Alderton, Roberto Amato, Jeffrey Barrett, Sonia Goncalves, Ewan Harrison, David K. Jackson, Ian Johnston, Dominic Kwiatkowski, Cordelia Langford, John Sillitoe on behalf of the Wellcome Sanger Institute COVID-19 Surveillance Team |
| EPI_ISL_3168776 | Lighthouse Lab in Glasgow | Wellcome Sanger Institute for the COVID-19 Genomics UK (COG-UK) Consortium | Harper VanSteenhouse, Yumi Kasai, David Gray, Carol Clugston, Anna Dominiczak and Alex Alderton, Roberto Amato, Jeffrey Barrett, Sonia Goncalves, Ewan Harrison, David K. Jackson, Ian Johnston, Dominic Kwiatkowski, Cordelia Langford, John Sillitoe on behalf of the Wellcome Sanger Institute COVID-19 Surveillance Team |
| EPI_ISL_3168777, EPI_ISL_3168778, EPI_ISL_3168779 | Lighthouse Laboratory Plymouth | Wellcome Sanger Institute for the COVID-19 Genomics UK (COG-UK) Consortium | Lighthouse Laboratory Plymouth and Alex Alderton, Roberto Amato, Jeffrey Barrett, Sonia Goncalves, Ewan Harrison, David K. Jackson, Ian Johnston, Dominic Kwiatkowski, Cordelia Langford, John Sillitoe on behalf of the Wellcome Sanger Institute COVID-19 Surveillance Team |
| EPI_ISL_3168780, EPI_ISL_3168781 | Lighthouse Lab in Glasgow | Wellcome Sanger Institute for the COVID-19 Genomics UK (COG-UK) Consortium | Harper VanSteenhouse, Yumi Kasai, David Gray, Carol Clugston, Anna Dominiczak and Alex Alderton, Roberto Amato, Jeffrey Barrett, Sonia Goncalves, Ewan Harrison, David K. Jackson, Ian Johnston, Dominic Kwiatkowski, Cordelia Langford, John Sillitoe on behalf of the Wellcome Sanger Institute COVID-19 Surveillance Team |
| EPI_ISL_3168782 | Lighthouse Laboratory Plymouth | Wellcome Sanger Institute for the COVID-19 Genomics UK (COG-UK) Consortium | Lighthouse Laboratory Plymouth and Alex Alderton, Roberto Amato, Jeffrey Barrett, Sonia Goncalves, Ewan Harrison, David K. Jackson, Ian Johnston, Dominic Kwiatkowski, Cordelia Langford, John Sillitoe on behalf of the Wellcome Sanger Institute COVID-19 Surveillance Team |
| EPI_ISL_3168783 | Lighthouse Lab in Glasgow | Wellcome Sanger Institute for the COVID-19 Genomics UK (COG-UK) Consortium | Harper VanSteenhouse, Yumi Kasai, David Gray, Carol Clugston, Anna Dominiczak and Alex Alderton, Roberto Amato, Jeffrey Barrett, Sonia Goncalves, Ewan Harrison, David K. Jackson, Ian Johnston, Dominic Kwiatkowski, Cordelia Langford, John Sillitoe on behalf of the Wellcome Sanger Institute COVID-19 Surveillance Team |
| EPI_ISL_3168784, | Lighthouse Laboratory Plymouth | Wellcome Sanger Institute for the COVID-19 Genomics UK (COG-UK) | Lighthouse Laboratory Plymouth and Alex Alderton, Roberto Amato, Jeffrey Barrett, Sonia Goncalves, Ewan Harrison, David K. Jackson, Ian Johnston, Dominic Kwiatkowski, Cordelia Langford, John Sillitoe on behalf of the Wellcome Sanger |

| | | | |
|---|---|---|---|
| EPI_ISL_3168785, EPI_ISL_3168786, EPI_ISL_3168787 | | Consortium | Institute COVID-19 Surveillance Team |
| EPI_ISL_3168788 | Lighthouse Lab in Glasgow | Wellcome Sanger Institute for the COVID-19 Genomics UK (COG-UK) Consortium | Harper VanSteenhouse, Yumi Kasai, David Gray, Carol Clugston, Anna Dominiczak and Alex Alderton, Roberto Amato, Jeffrey Barrett, Sonia Goncalves, Ewan Harrison, David K. Jackson, Ian Johnston, Dominic Kwiatkowski, Cordelia Langford, John Sillitoe on behalf of the Wellcome Sanger Institute COVID-19 Surveillance Team |
| EPI_ISL_3168789 | Lighthouse Laboratory Plymouth | Wellcome Sanger Institute for the COVID-19 Genomics UK (COG-UK) Consortium | Lighthouse Laboratory Plymouth and Alex Alderton, Roberto Amato, Jeffrey Barrett, Sonia Goncalves, Ewan Harrison, David K. Jackson, Ian Johnston, Dominic Kwiatkowski, Cordelia Langford, John Sillitoe on behalf of the Wellcome Sanger Institute COVID-19 Surveillance Team |
| EPI_ISL_3168790 | Lighthouse Lab in Glasgow | Wellcome Sanger Institute for the COVID-19 Genomics UK (COG-UK) Consortium | Harper VanSteenhouse, Yumi Kasai, David Gray, Carol Clugston, Anna Dominiczak and Alex Alderton, Roberto Amato, Jeffrey Barrett, Sonia Goncalves, Ewan Harrison, David K. Jackson, Ian Johnston, Dominic Kwiatkowski, Cordelia Langford, John Sillitoe on behalf of the Wellcome Sanger Institute COVID-19 Surveillance Team |
| EPI_ISL_3168791 | Lighthouse Laboratory Plymouth | Wellcome Sanger Institute for the COVID-19 Genomics UK (COG-UK) Consortium | Lighthouse Laboratory Plymouth and Alex Alderton, Roberto Amato, Jeffrey Barrett, Sonia Goncalves, Ewan Harrison, David K. Jackson, Ian Johnston, Dominic Kwiatkowski, Cordelia Langford, John Sillitoe on behalf of the Wellcome Sanger Institute COVID-19 Surveillance Team |
| EPI_ISL_3168792 | Lighthouse Lab in Glasgow | Wellcome Sanger Institute for the COVID-19 Genomics UK (COG-UK) Consortium | Harper VanSteenhouse, Yumi Kasai, David Gray, Carol Clugston, Anna Dominiczak and Alex Alderton, Roberto Amato, Jeffrey Barrett, Sonia Goncalves, Ewan Harrison, David K. Jackson, Ian Johnston, Dominic Kwiatkowski, Cordelia Langford, John Sillitoe on behalf of the Wellcome Sanger Institute COVID-19 Surveillance Team |
| EPI_ISL_3168793 | Lighthouse Laboratory Plymouth | Wellcome Sanger Institute for the COVID-19 Genomics UK (COG-UK) Consortium | Lighthouse Laboratory Plymouth and Alex Alderton, Roberto Amato, Jeffrey Barrett, Sonia Goncalves, Ewan Harrison, David K. Jackson, Ian Johnston, Dominic Kwiatkowski, Cordelia Langford, John Sillitoe on behalf of the Wellcome Sanger Institute COVID-19 Surveillance Team |
| EPI_ISL_3168794, EPI_ISL_3168795, EPI_ISL_3168796 | Lighthouse Lab in Glasgow | Wellcome Sanger Institute for the COVID-19 Genomics UK (COG-UK) Consortium | Harper VanSteenhouse, Yumi Kasai, David Gray, Carol Clugston, Anna Dominiczak and Alex Alderton, Roberto Amato, Jeffrey Barrett, Sonia Goncalves, Ewan Harrison, David K. Jackson, Ian Johnston, Dominic Kwiatkowski, Cordelia Langford, John Sillitoe on behalf of the Wellcome Sanger Institute COVID-19 Surveillance Team |
| EPI_ISL_3168797 | Lighthouse Laboratory Plymouth | Wellcome Sanger Institute for the COVID-19 Genomics UK (COG-UK) Consortium | Lighthouse Laboratory Plymouth and Alex Alderton, Roberto Amato, Jeffrey Barrett, Sonia Goncalves, Ewan Harrison, David K. Jackson, Ian Johnston, Dominic Kwiatkowski, Cordelia Langford, John Sillitoe on behalf of the Wellcome Sanger Institute COVID-19 Surveillance Team |
| EPI_ISL_3168798, EPI_ISL_3168799, EPI_ISL_3168800 | Lighthouse Lab in Glasgow | Wellcome Sanger Institute for the COVID-19 Genomics UK (COG-UK) Consortium | Harper VanSteenhouse, Yumi Kasai, David Gray, Carol Clugston, Anna Dominiczak and Alex Alderton, Roberto Amato, Jeffrey Barrett, Sonia Goncalves, Ewan Harrison, David K. Jackson, Ian Johnston, Dominic Kwiatkowski, Cordelia Langford, John Sillitoe on behalf of the Wellcome Sanger Institute COVID-19 Surveillance Team |
| EPI_ISL_3168801 | Lighthouse Laboratory Plymouth | Wellcome Sanger Institute for the COVID-19 Genomics UK (COG-UK) Consortium | Lighthouse Laboratory Plymouth and Alex Alderton, Roberto Amato, Jeffrey Barrett, Sonia Goncalves, Ewan Harrison, David K. Jackson, Ian Johnston, Dominic Kwiatkowski, Cordelia Langford, John Sillitoe on behalf of the Wellcome Sanger Institute COVID-19 Surveillance Team |
| EPI_ISL_3168802, EPI_ISL_3168803 | Lighthouse Lab in Glasgow | Wellcome Sanger Institute for the COVID-19 Genomics UK (COG-UK) Consortium | Harper VanSteenhouse, Yumi Kasai, David Gray, Carol Clugston, Anna Dominiczak and Alex Alderton, Roberto Amato, Jeffrey Barrett, Sonia Goncalves, Ewan Harrison, David K. Jackson, Ian Johnston, Dominic Kwiatkowski, Cordelia Langford, John Sillitoe on behalf of the Wellcome Sanger Institute COVID-19 Surveillance Team |
| EPI_ISL_3168804 | Lighthouse Laboratory Plymouth | Wellcome Sanger Institute for the COVID-19 Genomics UK (COG-UK) Consortium | Lighthouse Laboratory Plymouth and Alex Alderton, Roberto Amato, Jeffrey Barrett, Sonia Goncalves, Ewan Harrison, David K. Jackson, Ian Johnston, Dominic Kwiatkowski, Cordelia Langford, John Sillitoe on behalf of the Wellcome Sanger Institute COVID-19 Surveillance Team |
| EPI_ISL_3168805 | Lighthouse Lab in Glasgow | Wellcome Sanger Institute for the COVID-19 Genomics UK (COG-UK) Consortium | Harper VanSteenhouse, Yumi Kasai, David Gray, Carol Clugston, Anna Dominiczak and Alex Alderton, Roberto Amato, Jeffrey Barrett, Sonia Goncalves, Ewan Harrison, David K. Jackson, Ian Johnston, Dominic Kwiatkowski, Cordelia Langford, John Sillitoe on behalf of the Wellcome Sanger Institute COVID-19 Surveillance Team |
| EPI_ISL_3168806 | Lighthouse Laboratory Plymouth | Wellcome Sanger Institute for the COVID-19 Genomics UK (COG-UK) Consortium | Lighthouse Laboratory Plymouth and Alex Alderton, Roberto Amato, Jeffrey Barrett, Sonia Goncalves, Ewan Harrison, David K. Jackson, Ian Johnston, Dominic Kwiatkowski, Cordelia Langford, John Sillitoe on behalf of the Wellcome Sanger Institute COVID-19 Surveillance Team |
| EPI_ISL_3168807 | Lighthouse Lab in Glasgow | Wellcome Sanger Institute for the COVID-19 Genomics UK (COG-UK) Consortium | Harper VanSteenhouse, Yumi Kasai, David Gray, Carol Clugston, Anna Dominiczak and Alex Alderton, Roberto Amato, Jeffrey Barrett, Sonia Goncalves, Ewan Harrison, David K. Jackson, Ian Johnston, Dominic Kwiatkowski, Cordelia Langford, John Sillitoe on behalf of the Wellcome Sanger Institute COVID-19 Surveillance Team |
| EPI_ISL_3168808 | Lighthouse Laboratory Plymouth | Wellcome Sanger Institute for the COVID-19 Genomics UK (COG-UK) Consortium | Lighthouse Laboratory Plymouth and Alex Alderton, Roberto Amato, Jeffrey Barrett, Sonia Goncalves, Ewan Harrison, David K. Jackson, Ian Johnston, Dominic Kwiatkowski, Cordelia Langford, John Sillitoe on behalf of the Wellcome Sanger Institute COVID-19 Surveillance Team |
| EPI_ISL_3168809 | Lighthouse Lab in Glasgow | Wellcome Sanger Institute for the COVID-19 Genomics UK (COG-UK) Consortium | Harper VanSteenhouse, Yumi Kasai, David Gray, Carol Clugston, Anna Dominiczak and Alex Alderton, Roberto Amato, Jeffrey Barrett, Sonia Goncalves, Ewan Harrison, David K. Jackson, Ian Johnston, Dominic Kwiatkowski, Cordelia Langford, John Sillitoe on behalf of the Wellcome Sanger Institute COVID-19 Surveillance Team |
| EPI_ISL_3168810, EPI_ISL_3168811, EPI_ISL_3168812, EPI_ISL_3168813, EPI_ISL_3168814 | Lighthouse Laboratory Plymouth | Wellcome Sanger Institute for the COVID-19 Genomics UK (COG-UK) Consortium | Lighthouse Laboratory Plymouth and Alex Alderton, Roberto Amato, Jeffrey Barrett, Sonia Goncalves, Ewan Harrison, David K. Jackson, Ian Johnston, Dominic Kwiatkowski, Cordelia Langford, John Sillitoe on behalf of the Wellcome Sanger Institute COVID-19 Surveillance Team |
| EPI_ISL_3168815, EPI_ISL_3168816 | Lighthouse Lab in Glasgow | Wellcome Sanger Institute for the COVID-19 Genomics UK (COG-UK) Consortium | Harper VanSteenhouse, Yumi Kasai, David Gray, Carol Clugston, Anna Dominiczak and Alex Alderton, Roberto Amato, Jeffrey Barrett, Sonia Goncalves, Ewan Harrison, David K. Jackson, Ian Johnston, Dominic Kwiatkowski, Cordelia Langford, John Sillitoe on behalf of the Wellcome Sanger Institute COVID-19 Surveillance Team |
| EPI_ISL_3168817, EPI_ISL_3168818 | Lighthouse Laboratory Plymouth | Wellcome Sanger Institute for the COVID-19 Genomics UK (COG-UK) Consortium | Lighthouse Laboratory Plymouth and Alex Alderton, Roberto Amato, Jeffrey Barrett, Sonia Goncalves, Ewan Harrison, David K. Jackson, Ian Johnston, Dominic Kwiatkowski, Cordelia Langford, John Sillitoe on behalf of the Wellcome Sanger Institute COVID-19 Surveillance Team |
| EPI_ISL_3168819 | Lighthouse Lab in Glasgow | Wellcome Sanger Institute for the COVID-19 Genomics UK (COG-UK) Consortium | Harper VanSteenhouse, Yumi Kasai, David Gray, Carol Clugston, Anna Dominiczak and Alex Alderton, Roberto Amato, Jeffrey Barrett, Sonia Goncalves, Ewan Harrison, David K. Jackson, Ian Johnston, Dominic Kwiatkowski, Cordelia Langford, John Sillitoe on behalf of the Wellcome Sanger Institute COVID-19 Surveillance Team |
| EPI_ISL_3168820 | Lighthouse Laboratory Plymouth | Wellcome Sanger Institute for the COVID-19 Genomics UK (COG-UK) Consortium | Lighthouse Laboratory Plymouth and Alex Alderton, Roberto Amato, Jeffrey Barrett, Sonia Goncalves, Ewan Harrison, David K. Jackson, Ian Johnston, Dominic Kwiatkowski, Cordelia Langford, John Sillitoe on behalf of the Wellcome Sanger Institute COVID-19 Surveillance Team |
| EPI_ISL_3168821, EPI_ISL_3168822, EPI_ISL_3168823 | Lighthouse Lab in Glasgow | Wellcome Sanger Institute for the COVID-19 Genomics UK (COG-UK) Consortium | Harper VanSteenhouse, Yumi Kasai, David Gray, Carol Clugston, Anna Dominiczak and Alex Alderton, Roberto Amato, Jeffrey Barrett, Sonia Goncalves, Ewan Harrison, David K. Jackson, Ian Johnston, Dominic Kwiatkowski, Cordelia Langford, John Sillitoe on behalf of the Wellcome Sanger Institute COVID-19 Surveillance Team |
| EPI_ISL_3168824 | Lighthouse Laboratory Plymouth | Wellcome Sanger Institute for the COVID-19 Genomics UK (COG-UK) Consortium | Lighthouse Laboratory Plymouth and Alex Alderton, Roberto Amato, Jeffrey Barrett, Sonia Goncalves, Ewan Harrison, David K. Jackson, Ian Johnston, Dominic Kwiatkowski, Cordelia Langford, John Sillitoe on behalf of the Wellcome Sanger Institute COVID-19 Surveillance Team |
| EPI_ISL_3168825, EPI_ISL_3168826 | Lighthouse Lab in Glasgow | Wellcome Sanger Institute for the COVID-19 Genomics UK (COG-UK) Consortium | Harper VanSteenhouse, Yumi Kasai, David Gray, Carol Clugston, Anna Dominiczak and Alex Alderton, Roberto Amato, Jeffrey Barrett, Sonia Goncalves, Ewan Harrison, David K. Jackson, Ian Johnston, Dominic Kwiatkowski, Cordelia Langford, John Sillitoe on behalf of the Wellcome Sanger Institute COVID-19 Surveillance Team |
| EPI_ISL_3168827 | Lighthouse Laboratory Plymouth | Wellcome Sanger Institute for the COVID-19 Genomics UK (COG-UK) Consortium | Lighthouse Laboratory Plymouth and Alex Alderton, Roberto Amato, Jeffrey Barrett, Sonia Goncalves, Ewan Harrison, David K. Jackson, Ian Johnston, Dominic Kwiatkowski, Cordelia Langford, John Sillitoe on behalf of the Wellcome Sanger Institute COVID-19 Surveillance Team |
| EPI_ISL_3168828 | Lighthouse Lab in Glasgow | Wellcome Sanger Institute for the COVID-19 Genomics UK (COG-UK) Consortium | Harper VanSteenhouse, Yumi Kasai, David Gray, Carol Clugston, Anna Dominiczak and Alex Alderton, Roberto Amato, Jeffrey Barrett, Sonia Goncalves, Ewan Harrison, David K. Jackson, Ian Johnston, Dominic Kwiatkowski, Cordelia Langford, John Sillitoe on behalf of the Wellcome Sanger Institute COVID-19 Surveillance Team |
| EPI_ISL_3168829, EPI_ISL_3168830, EPI_ISL_3168831, EPI_ISL_3168832, EPI_ISL_3168833, EPI_ISL_3168834 | Lighthouse Laboratory Plymouth | Wellcome Sanger Institute for the COVID-19 Genomics UK (COG-UK) Consortium | Lighthouse Laboratory Plymouth and Alex Alderton, Roberto Amato, Jeffrey Barrett, Sonia Goncalves, Ewan Harrison, David K. Jackson, Ian Johnston, Dominic Kwiatkowski, Cordelia Langford, John Sillitoe on behalf of the Wellcome Sanger Institute COVID-19 Surveillance Team |
| EPI_ISL_3168835 | Lighthouse Lab in Glasgow | Wellcome Sanger Institute for the COVID-19 Genomics UK (COG-UK) Consortium | Harper VanSteenhouse, Yumi Kasai, David Gray, Carol Clugston, Anna Dominiczak and Alex Alderton, Roberto Amato, Jeffrey Barrett, Sonia Goncalves, Ewan Harrison, David K. Jackson, Ian Johnston, Dominic Kwiatkowski, Cordelia Langford, John Sillitoe on behalf of the Wellcome Sanger Institute COVID-19 Surveillance Team |
| EPI_ISL_3168836 | Lighthouse Laboratory Plymouth | Wellcome Sanger Institute for the COVID-19 Genomics UK (COG-UK) Consortium | Lighthouse Laboratory Plymouth and Alex Alderton, Roberto Amato, Jeffrey Barrett, Sonia Goncalves, Ewan Harrison, David K. Jackson, Ian Johnston, Dominic Kwiatkowski, Cordelia Langford, John Sillitoe on behalf of the Wellcome Sanger Institute COVID-19 Surveillance Team |
| EPI_ISL_3168837 | Lighthouse Lab in Glasgow | Wellcome Sanger Institute for the COVID-19 Genomics UK (COG-UK) Consortium | Harper VanSteenhouse, Yumi Kasai, David Gray, Carol Clugston, Anna Dominiczak and Alex Alderton, Roberto Amato, Jeffrey Barrett, Sonia Goncalves, Ewan Harrison, David K. Jackson, Ian Johnston, Dominic Kwiatkowski, Cordelia Langford, John Sillitoe on behalf of the Wellcome Sanger Institute COVID-19 Surveillance Team |
| EPI_ISL_3168838 | Lighthouse Laboratory Plymouth | Wellcome Sanger Institute for the COVID-19 Genomics UK (COG-UK) Consortium | Lighthouse Laboratory Plymouth and Alex Alderton, Roberto Amato, Jeffrey Barrett, Sonia Goncalves, Ewan Harrison, David K. Jackson, Ian Johnston, Dominic Kwiatkowski, Cordelia Langford, John Sillitoe on behalf of the Wellcome Sanger Institute COVID-19 Surveillance Team |
| EPI_ISL_3168839, EPI_ISL_3168840, EPI_ISL_3168841, EPI_ISL_3168842 | Lighthouse Lab in Glasgow | Wellcome Sanger Institute for the COVID-19 Genomics UK (COG-UK) Consortium | Harper VanSteenhouse, Yumi Kasai, David Gray, Carol Clugston, Anna Dominiczak and Alex Alderton, Roberto Amato, Jeffrey Barrett, Sonia Goncalves, Ewan Harrison, David K. Jackson, Ian Johnston, Dominic Kwiatkowski, Cordelia Langford, John Sillitoe on behalf of the Wellcome Sanger Institute COVID-19 Surveillance Team |
| EPI_ISL_3168843 | Lighthouse Laboratory Plymouth | Wellcome Sanger Institute for the COVID-19 Genomics UK (COG-UK) Consortium | Lighthouse Laboratory Plymouth and Alex Alderton, Roberto Amato, Jeffrey Barrett, Sonia Goncalves, Ewan Harrison, David K. Jackson, Ian Johnston, Dominic Kwiatkowski, Cordelia Langford, John Sillitoe on behalf of the Wellcome Sanger Institute COVID-19 Surveillance Team |
| EPI_ISL_3168844, EPI_ISL_3168845, EPI_ISL_3168846, EPI_ISL_3168847 | Lighthouse Lab in Glasgow | Wellcome Sanger Institute for the COVID-19 Genomics UK (COG-UK) Consortium | Harper VanSteenhouse, Yumi Kasai, David Gray, Carol Clugston, Anna Dominiczak and Alex Alderton, Roberto Amato, Jeffrey Barrett, Sonia Goncalves, Ewan Harrison, David K. Jackson, Ian Johnston, Dominic Kwiatkowski, Cordelia Langford, John Sillitoe on behalf of the Wellcome Sanger Institute COVID-19 Surveillance Team |
| EPI_ISL_3168848, EPI_ISL_3168849 | Lighthouse Laboratory Plymouth | Wellcome Sanger Institute for the COVID-19 Genomics UK (COG-UK) Consortium | Lighthouse Laboratory Plymouth and Alex Alderton, Roberto Amato, Jeffrey Barrett, Sonia Goncalves, Ewan Harrison, David K. Jackson, Ian Johnston, Dominic Kwiatkowski, Cordelia Langford, John Sillitoe on behalf of the Wellcome Sanger Institute COVID-19 Surveillance Team |
| EPI_ISL_3168850 | Lighthouse Lab in Glasgow | Wellcome Sanger Institute for the COVID-19 Genomics UK (COG-UK) Consortium | Harper VanSteenhouse, Yumi Kasai, David Gray, Carol Clugston, Anna Dominiczak and Alex Alderton, Roberto Amato, Jeffrey Barrett, Sonia Goncalves, Ewan Harrison, David K. Jackson, Ian Johnston, Dominic Kwiatkowski, Cordelia Langford, John Sillitoe on behalf of the Wellcome Sanger Institute COVID-19 Surveillance Team |
| EPI_ISL_3168851, EPI_ISL_3168852, | Lighthouse Laboratory Plymouth | Wellcome Sanger Institute for the COVID-19 Genomics UK (COG-UK) Consortium | Lighthouse Laboratory Plymouth and Alex Alderton, Roberto Amato, Jeffrey Barrett, Sonia Goncalves, Ewan Harrison, David K. Jackson, Ian Johnston, Dominic Kwiatkowski, Cordelia Langford, John Sillitoe on behalf of the Wellcome Sanger Institute COVID-19 Surveillance Team |

| | | | |
|---|---|---|---|
| EPI_ISL_3168853, EPI_ISL_3168854, EPI_ISL_3168855 | | | |
| EPI_ISL_3168856, EPI_ISL_3168857 | Lighthouse Lab in Glasgow | Wellcome Sanger Institute for the COVID-19 Genomics UK (COG-UK) Consortium | Harper VanSteenhouse, Yumi Kasai, David Gray, Carol Clugston, Anna Dominiczak and Alex Alderton, Roberto Amato, Jeffrey Barrett, Sonia Goncalves, Ewan Harrison, David K. Jackson, Ian Johnston, Dominic Kwiatkowski, Cordelia Langford, John Sillitoe on behalf of the Wellcome Sanger Institute COVID-19 Surveillance Team |
| EPI_ISL_3168858 | Lighthouse Laboratory Plymouth | Wellcome Sanger Institute for the COVID-19 Genomics UK (COG-UK) Consortium | Lighthouse Laboratory Plymouth and Alex Alderton, Roberto Amato, Jeffrey Barrett, Sonia Goncalves, Ewan Harrison, David K. Jackson, Ian Johnston, Dominic Kwiatkowski, Cordelia Langford, John Sillitoe on behalf of the Wellcome Sanger Institute COVID-19 Surveillance Team |
| EPI_ISL_3168859, EPI_ISL_3168860, EPI_ISL_3168861 | Lighthouse Lab in Glasgow | Wellcome Sanger Institute for the COVID-19 Genomics UK (COG-UK) Consortium | Harper VanSteenhouse, Yumi Kasai, David Gray, Carol Clugston, Anna Dominiczak and Alex Alderton, Roberto Amato, Jeffrey Barrett, Sonia Goncalves, Ewan Harrison, David K. Jackson, Ian Johnston, Dominic Kwiatkowski, Cordelia Langford, John Sillitoe on behalf of the Wellcome Sanger Institute COVID-19 Surveillance Team |
| EPI_ISL_3168862, EPI_ISL_3168863, EPI_ISL_3168864 | Lighthouse Laboratory Plymouth | Wellcome Sanger Institute for the COVID-19 Genomics UK (COG-UK) Consortium | Lighthouse Laboratory Plymouth and Alex Alderton, Roberto Amato, Jeffrey Barrett, Sonia Goncalves, Ewan Harrison, David K. Jackson, Ian Johnston, Dominic Kwiatkowski, Cordelia Langford, John Sillitoe on behalf of the Wellcome Sanger Institute COVID-19 Surveillance Team |
| EPI_ISL_3168865 | Lighthouse Lab in Glasgow | Wellcome Sanger Institute for the COVID-19 Genomics UK (COG-UK) Consortium | Harper VanSteenhouse, Yumi Kasai, David Gray, Carol Clugston, Anna Dominiczak and Alex Alderton, Roberto Amato, Jeffrey Barrett, Sonia Goncalves, Ewan Harrison, David K. Jackson, Ian Johnston, Dominic Kwiatkowski, Cordelia Langford, John Sillitoe on behalf of the Wellcome Sanger Institute COVID-19 Surveillance Team |
| EPI_ISL_3168866, EPI_ISL_3168867 | Lighthouse Laboratory Plymouth | Wellcome Sanger Institute for the COVID-19 Genomics UK (COG-UK) Consortium | Lighthouse Laboratory Plymouth and Alex Alderton, Roberto Amato, Jeffrey Barrett, Sonia Goncalves, Ewan Harrison, David K. Jackson, Ian Johnston, Dominic Kwiatkowski, Cordelia Langford, John Sillitoe on behalf of the Wellcome Sanger Institute COVID-19 Surveillance Team |
| EPI_ISL_3168868, EPI_ISL_3168869 | Lighthouse Lab in Glasgow | Wellcome Sanger Institute for the COVID-19 Genomics UK (COG-UK) Consortium | Harper VanSteenhouse, Yumi Kasai, David Gray, Carol Clugston, Anna Dominiczak and Alex Alderton, Roberto Amato, Jeffrey Barrett, Sonia Goncalves, Ewan Harrison, David K. Jackson, Ian Johnston, Dominic Kwiatkowski, Cordelia Langford, John Sillitoe on behalf of the Wellcome Sanger Institute COVID-19 Surveillance Team |
| EPI_ISL_3168870 | Lighthouse Laboratory Plymouth | Wellcome Sanger Institute for the COVID-19 Genomics UK (COG-UK) Consortium | Lighthouse Laboratory Plymouth and Alex Alderton, Roberto Amato, Jeffrey Barrett, Sonia Goncalves, Ewan Harrison, David K. Jackson, Ian Johnston, Dominic Kwiatkowski, Cordelia Langford, John Sillitoe on behalf of the Wellcome Sanger Institute COVID-19 Surveillance Team |
| EPI_ISL_3168871, EPI_ISL_3168872, EPI_ISL_3168873 | Lighthouse Lab in Glasgow | Wellcome Sanger Institute for the COVID-19 Genomics UK (COG-UK) Consortium | Harper VanSteenhouse, Yumi Kasai, David Gray, Carol Clugston, Anna Dominiczak and Alex Alderton, Roberto Amato, Jeffrey Barrett, Sonia Goncalves, Ewan Harrison, David K. Jackson, Ian Johnston, Dominic Kwiatkowski, Cordelia Langford, John Sillitoe on behalf of the Wellcome Sanger Institute COVID-19 Surveillance Team |
| EPI_ISL_3168874, EPI_ISL_3168875 | Lighthouse Laboratory Plymouth | Wellcome Sanger Institute for the COVID-19 Genomics UK (COG-UK) Consortium | Lighthouse Laboratory Plymouth and Alex Alderton, Roberto Amato, Jeffrey Barrett, Sonia Goncalves, Ewan Harrison, David K. Jackson, Ian Johnston, Dominic Kwiatkowski, Cordelia Langford, John Sillitoe on behalf of the Wellcome Sanger Institute COVID-19 Surveillance Team |
| EPI_ISL_3168876, EPI_ISL_3168877 | Lighthouse Lab in Glasgow | Wellcome Sanger Institute for the COVID-19 Genomics UK (COG-UK) Consortium | Harper VanSteenhouse, Yumi Kasai, David Gray, Carol Clugston, Anna Dominiczak and Alex Alderton, Roberto Amato, Jeffrey Barrett, Sonia Goncalves, Ewan Harrison, David K. Jackson, Ian Johnston, Dominic Kwiatkowski, Cordelia Langford, John Sillitoe on behalf of the Wellcome Sanger Institute COVID-19 Surveillance Team |
| EPI_ISL_3168878, EPI_ISL_3168879, EPI_ISL_3168880, EPI_ISL_3168881, EPI_ISL_3168882, EPI_ISL_3168883 | Lighthouse Laboratory Plymouth | Wellcome Sanger Institute for the COVID-19 Genomics UK (COG-UK) Consortium | Lighthouse Laboratory Plymouth and Alex Alderton, Roberto Amato, Jeffrey Barrett, Sonia Goncalves, Ewan Harrison, David K. Jackson, Ian Johnston, Dominic Kwiatkowski, Cordelia Langford, John Sillitoe on behalf of the Wellcome Sanger Institute COVID-19 Surveillance Team |
| EPI_ISL_3168884 | Lighthouse Lab in Glasgow | Wellcome Sanger Institute for the COVID-19 Genomics UK (COG-UK) Consortium | Harper VanSteenhouse, Yumi Kasai, David Gray, Carol Clugston, Anna Dominiczak and Alex Alderton, Roberto Amato, Jeffrey Barrett, Sonia Goncalves, Ewan Harrison, David K. Jackson, Ian Johnston, Dominic Kwiatkowski, Cordelia Langford, John Sillitoe on behalf of the Wellcome Sanger Institute COVID-19 Surveillance Team |
| EPI_ISL_3168885, EPI_ISL_3168886, EPI_ISL_3168887, EPI_ISL_3168888 | Lighthouse Laboratory Plymouth | Wellcome Sanger Institute for the COVID-19 Genomics UK (COG-UK) Consortium | Lighthouse Laboratory Plymouth and Alex Alderton, Roberto Amato, Jeffrey Barrett, Sonia Goncalves, Ewan Harrison, David K. Jackson, Ian Johnston, Dominic Kwiatkowski, Cordelia Langford, John Sillitoe on behalf of the Wellcome Sanger Institute COVID-19 Surveillance Team |
| EPI_ISL_3168889, EPI_ISL_3168890 | Lighthouse Lab in Glasgow | Wellcome Sanger Institute for the COVID-19 Genomics UK (COG-UK) Consortium | Harper VanSteenhouse, Yumi Kasai, David Gray, Carol Clugston, Anna Dominiczak and Alex Alderton, Roberto Amato, Jeffrey Barrett, Sonia Goncalves, Ewan Harrison, David K. Jackson, Ian Johnston, Dominic Kwiatkowski, Cordelia Langford, John Sillitoe on behalf of the Wellcome Sanger Institute COVID-19 Surveillance Team |
| EPI_ISL_3168891, EPI_ISL_3168892 | Lighthouse Laboratory Plymouth | Wellcome Sanger Institute for the COVID-19 Genomics UK (COG-UK) Consortium | Lighthouse Laboratory Plymouth and Alex Alderton, Roberto Amato, Jeffrey Barrett, Sonia Goncalves, Ewan Harrison, David K. Jackson, Ian Johnston, Dominic Kwiatkowski, Cordelia Langford, John Sillitoe on behalf of the Wellcome Sanger Institute COVID-19 Surveillance Team |
| EPI_ISL_3168893 | Lighthouse Lab in Glasgow | Wellcome Sanger Institute for the COVID-19 Genomics UK (COG-UK) Consortium | Harper VanSteenhouse, Yumi Kasai, David Gray, Carol Clugston, Anna Dominiczak and Alex Alderton, Roberto Amato, Jeffrey Barrett, Sonia Goncalves, Ewan Harrison, David K. Jackson, Ian Johnston, Dominic Kwiatkowski, Cordelia Langford, John Sillitoe on behalf of the Wellcome Sanger Institute COVID-19 Surveillance Team |
| EPI_ISL_3168894, EPI_ISL_3168895 | Lighthouse Laboratory Plymouth | Wellcome Sanger Institute for the COVID-19 Genomics UK (COG-UK) Consortium | Lighthouse Laboratory Plymouth and Alex Alderton, Roberto Amato, Jeffrey Barrett, Sonia Goncalves, Ewan Harrison, David K. Jackson, Ian Johnston, Dominic Kwiatkowski, Cordelia Langford, John Sillitoe on behalf of the Wellcome Sanger Institute COVID-19 Surveillance Team |
| EPI_ISL_3168896 | Lighthouse Lab in Glasgow | Wellcome Sanger Institute for the COVID-19 Genomics UK (COG-UK) Consortium | Harper VanSteenhouse, Yumi Kasai, David Gray, Carol Clugston, Anna Dominiczak and Alex Alderton, Roberto Amato, Jeffrey Barrett, Sonia Goncalves, Ewan Harrison, David K. Jackson, Ian Johnston, Dominic Kwiatkowski, Cordelia Langford, John Sillitoe on behalf of the Wellcome Sanger Institute COVID-19 Surveillance Team |
| EPI_ISL_3168897 | Lighthouse Laboratory Plymouth | Wellcome Sanger Institute for the COVID-19 Genomics UK (COG-UK) Consortium | Lighthouse Laboratory Plymouth and Alex Alderton, Roberto Amato, Jeffrey Barrett, Sonia Goncalves, Ewan Harrison, David K. Jackson, Ian Johnston, Dominic Kwiatkowski, Cordelia Langford, John Sillitoe on behalf of the Wellcome Sanger Institute COVID-19 Surveillance Team |
| EPI_ISL_3168898 | Lighthouse Lab in Glasgow | Wellcome Sanger Institute for the COVID-19 Genomics UK (COG-UK) Consortium | Harper VanSteenhouse, Yumi Kasai, David Gray, Carol Clugston, Anna Dominiczak and Alex Alderton, Roberto Amato, Jeffrey Barrett, Sonia Goncalves, Ewan Harrison, David K. Jackson, Ian Johnston, Dominic Kwiatkowski, Cordelia Langford, John Sillitoe on behalf of the Wellcome Sanger Institute COVID-19 Surveillance Team |
| EPI_ISL_3168899 | Lighthouse Laboratory Plymouth | Wellcome Sanger Institute for the COVID-19 Genomics UK (COG-UK) Consortium | Lighthouse Laboratory Plymouth and Alex Alderton, Roberto Amato, Jeffrey Barrett, Sonia Goncalves, Ewan Harrison, David K. Jackson, Ian Johnston, Dominic Kwiatkowski, Cordelia Langford, John Sillitoe on behalf of the Wellcome Sanger Institute COVID-19 Surveillance Team |
| EPI_ISL_3168900 | Lighthouse Lab in Glasgow | Wellcome Sanger Institute for the COVID-19 Genomics UK (COG-UK) Consortium | Harper VanSteenhouse, Yumi Kasai, David Gray, Carol Clugston, Anna Dominiczak and Alex Alderton, Roberto Amato, Jeffrey Barrett, Sonia Goncalves, Ewan Harrison, David K. Jackson, Ian Johnston, Dominic Kwiatkowski, Cordelia Langford, John Sillitoe on behalf of the Wellcome Sanger Institute COVID-19 Surveillance Team |
| EPI_ISL_3168901, EPI_ISL_3168902 | Lighthouse Laboratory Plymouth | Wellcome Sanger Institute for the COVID-19 Genomics UK (COG-UK) Consortium | Lighthouse Laboratory Plymouth and Alex Alderton, Roberto Amato, Jeffrey Barrett, Sonia Goncalves, Ewan Harrison, David K. Jackson, Ian Johnston, Dominic Kwiatkowski, Cordelia Langford, John Sillitoe on behalf of the Wellcome Sanger Institute COVID-19 Surveillance Team |
| EPI_ISL_3168903, EPI_ISL_3168904 | Lighthouse Lab in Glasgow | Wellcome Sanger Institute for the COVID-19 Genomics UK (COG-UK) Consortium | Harper VanSteenhouse, Yumi Kasai, David Gray, Carol Clugston, Anna Dominiczak and Alex Alderton, Roberto Amato, Jeffrey Barrett, Sonia Goncalves, Ewan Harrison, David K. Jackson, Ian Johnston, Dominic Kwiatkowski, Cordelia Langford, John Sillitoe on behalf of the Wellcome Sanger Institute COVID-19 Surveillance Team |
| EPI_ISL_3168905 | Lighthouse Laboratory Plymouth | Wellcome Sanger Institute for the COVID-19 Genomics UK (COG-UK) Consortium | Lighthouse Laboratory Plymouth and Alex Alderton, Roberto Amato, Jeffrey Barrett, Sonia Goncalves, Ewan Harrison, David K. Jackson, Ian Johnston, Dominic Kwiatkowski, Cordelia Langford, John Sillitoe on behalf of the Wellcome Sanger Institute COVID-19 Surveillance Team |
| EPI_ISL_3168906, EPI_ISL_3168907, EPI_ISL_3168908, EPI_ISL_3168909, EPI_ISL_3168910, EPI_ISL_3168911 | Lighthouse Lab in Glasgow | Wellcome Sanger Institute for the COVID-19 Genomics UK (COG-UK) Consortium | Harper VanSteenhouse, Yumi Kasai, David Gray, Carol Clugston, Anna Dominiczak and Alex Alderton, Roberto Amato, Jeffrey Barrett, Sonia Goncalves, Ewan Harrison, David K. Jackson, Ian Johnston, Dominic Kwiatkowski, Cordelia Langford, John Sillitoe on behalf of the Wellcome Sanger Institute COVID-19 Surveillance Team |
| EPI_ISL_3168912 | Lighthouse Laboratory Plymouth | Wellcome Sanger Institute for the COVID-19 Genomics UK (COG-UK) Consortium | Lighthouse Laboratory Plymouth and Alex Alderton, Roberto Amato, Jeffrey Barrett, Sonia Goncalves, Ewan Harrison, David K. Jackson, Ian Johnston, Dominic Kwiatkowski, Cordelia Langford, John Sillitoe on behalf of the Wellcome Sanger Institute COVID-19 Surveillance Team |
| EPI_ISL_3168913, EPI_ISL_3168914, EPI_ISL_3168915, EPI_ISL_3168916 | Lighthouse Lab in Glasgow | Wellcome Sanger Institute for the COVID-19 Genomics UK (COG-UK) Consortium | Harper VanSteenhouse, Yumi Kasai, David Gray, Carol Clugston, Anna Dominiczak and Alex Alderton, Roberto Amato, Jeffrey Barrett, Sonia Goncalves, Ewan Harrison, David K. Jackson, Ian Johnston, Dominic Kwiatkowski, Cordelia Langford, John Sillitoe on behalf of the Wellcome Sanger Institute COVID-19 Surveillance Team |
| EPI_ISL_3168917 | Lighthouse Laboratory Plymouth | Wellcome Sanger Institute for the COVID-19 Genomics UK (COG-UK) Consortium | Lighthouse Laboratory Plymouth and Alex Alderton, Roberto Amato, Jeffrey Barrett, Sonia Goncalves, Ewan Harrison, David K. Jackson, Ian Johnston, Dominic Kwiatkowski, Cordelia Langford, John Sillitoe on behalf of the Wellcome Sanger Institute COVID-19 Surveillance Team |
| EPI_ISL_3168918 | Lighthouse Lab in Glasgow | Wellcome Sanger Institute for the COVID-19 Genomics UK (COG-UK) Consortium | Harper VanSteenhouse, Yumi Kasai, David Gray, Carol Clugston, Anna Dominiczak and Alex Alderton, Roberto Amato, Jeffrey Barrett, Sonia Goncalves, Ewan Harrison, David K. Jackson, Ian Johnston, Dominic Kwiatkowski, Cordelia Langford, John Sillitoe on behalf of the Wellcome Sanger Institute COVID-19 Surveillance Team |
| EPI_ISL_3168919, EPI_ISL_3168920, EPI_ISL_3168921 | Lighthouse Laboratory Plymouth | Wellcome Sanger Institute for the COVID-19 Genomics UK (COG-UK) Consortium | Lighthouse Laboratory Plymouth and Alex Alderton, Roberto Amato, Jeffrey Barrett, Sonia Goncalves, Ewan Harrison, David K. Jackson, Ian Johnston, Dominic Kwiatkowski, Cordelia Langford, John Sillitoe on behalf of the Wellcome Sanger Institute COVID-19 Surveillance Team |
| EPI_ISL_3168922 | Lighthouse Lab in Glasgow | Wellcome Sanger Institute for the COVID-19 Genomics UK (COG-UK) Consortium | Harper VanSteenhouse, Yumi Kasai, David Gray, Carol Clugston, Anna Dominiczak and Alex Alderton, Roberto Amato, Jeffrey Barrett, Sonia Goncalves, Ewan Harrison, David K. Jackson, Ian Johnston, Dominic Kwiatkowski, Cordelia Langford, John Sillitoe on behalf of the Wellcome Sanger Institute COVID-19 Surveillance Team |
| EPI_ISL_3168923 | Lighthouse Laboratory Plymouth | Wellcome Sanger Institute for the COVID-19 Genomics UK (COG-UK) Consortium | Lighthouse Laboratory Plymouth and Alex Alderton, Roberto Amato, Jeffrey Barrett, Sonia Goncalves, Ewan Harrison, David K. Jackson, Ian Johnston, Dominic Kwiatkowski, Cordelia Langford, John Sillitoe on behalf of the Wellcome Sanger Institute COVID-19 Surveillance Team |
| EPI_ISL_3168924 | Lighthouse Lab in Glasgow | Wellcome Sanger Institute for the COVID-19 Genomics UK (COG-UK) Consortium | Harper VanSteenhouse, Yumi Kasai, David Gray, Carol Clugston, Anna Dominiczak and Alex Alderton, Roberto Amato, Jeffrey Barrett, Sonia Goncalves, Ewan Harrison, David K. Jackson, Ian Johnston, Dominic Kwiatkowski, Cordelia Langford, John Sillitoe on behalf of the Wellcome Sanger Institute COVID-19 Surveillance Team |
| EPI_ISL_3168925, EPI_ISL_3168926, EPI_ISL_3168927, EPI_ISL_3168928, EPI_ISL_3168929, | Lighthouse Laboratory Plymouth | Wellcome Sanger Institute for the COVID-19 Genomics UK (COG-UK) Consortium | Lighthouse Laboratory Plymouth and Alex Alderton, Roberto Amato, Jeffrey Barrett, Sonia Goncalves, Ewan Harrison, David K. Jackson, Ian Johnston, Dominic Kwiatkowski, Cordelia Langford, John Sillitoe on behalf of the Wellcome Sanger Institute COVID-19 Surveillance Team |

| | | | |
|---|---|---|---|
| EPI_ISL_3168930 | | | |
| EPI_ISL_3168931, EPI_ISL_3168932 | Lighthouse Lab in Glasgow | Wellcome Sanger Institute for the COVID-19 Genomics UK (COG-UK) Consortium | Harper VanSteenhouse, Yumi Kasai, David Gray, Carol Clugston, Anna Dominiczak and Alex Alderton, Roberto Amato, Jeffrey Barrett, Sonia Goncalves, Ewan Harrison, David K. Jackson, Ian Johnston, Dominic Kwiatkowski, Cordelia Langford, John Sillitoe on behalf of the Wellcome Sanger Institute COVID-19 Surveillance Team |
| EPI_ISL_3168933 | Lighthouse Laboratory Plymouth | Wellcome Sanger Institute for the COVID-19 Genomics UK (COG-UK) Consortium | Lighthouse Laboratory Plymouth and Alex Alderton, Roberto Amato, Jeffrey Barrett, Sonia Goncalves, Ewan Harrison, David K. Jackson, Ian Johnston, Dominic Kwiatkowski, Cordelia Langford, John Sillitoe on behalf of the Wellcome Sanger Institute COVID-19 Surveillance Team |
| EPI_ISL_3168934 | Lighthouse Lab in Glasgow | Wellcome Sanger Institute for the COVID-19 Genomics UK (COG-UK) Consortium | Harper VanSteenhouse, Yumi Kasai, David Gray, Carol Clugston, Anna Dominiczak and Alex Alderton, Roberto Amato, Jeffrey Barrett, Sonia Goncalves, Ewan Harrison, David K. Jackson, Ian Johnston, Dominic Kwiatkowski, Cordelia Langford, John Sillitoe on behalf of the Wellcome Sanger Institute COVID-19 Surveillance Team |
| EPI_ISL_3168935, EPI_ISL_3168936, EPI_ISL_3168937, EPI_ISL_3168938, EPI_ISL_3168939 | Lighthouse Laboratory Plymouth | Wellcome Sanger Institute for the COVID-19 Genomics UK (COG-UK) Consortium | Lighthouse Laboratory Plymouth and Alex Alderton, Roberto Amato, Jeffrey Barrett, Sonia Goncalves, Ewan Harrison, David K. Jackson, Ian Johnston, Dominic Kwiatkowski, Cordelia Langford, John Sillitoe on behalf of the Wellcome Sanger Institute COVID-19 Surveillance Team |
| EPI_ISL_3168940 | Lighthouse Lab in Glasgow | Wellcome Sanger Institute for the COVID-19 Genomics UK (COG-UK) Consortium | Harper VanSteenhouse, Yumi Kasai, David Gray, Carol Clugston, Anna Dominiczak and Alex Alderton, Roberto Amato, Jeffrey Barrett, Sonia Goncalves, Ewan Harrison, David K. Jackson, Ian Johnston, Dominic Kwiatkowski, Cordelia Langford, John Sillitoe on behalf of the Wellcome Sanger Institute COVID-19 Surveillance Team |
| EPI_ISL_3168941 | Lighthouse Laboratory Plymouth | Wellcome Sanger Institute for the COVID-19 Genomics UK (COG-UK) Consortium | Lighthouse Laboratory Plymouth and Alex Alderton, Roberto Amato, Jeffrey Barrett, Sonia Goncalves, Ewan Harrison, David K. Jackson, Ian Johnston, Dominic Kwiatkowski, Cordelia Langford, John Sillitoe on behalf of the Wellcome Sanger Institute COVID-19 Surveillance Team |
| EPI_ISL_3168942 | Lighthouse Lab in Glasgow | Wellcome Sanger Institute for the COVID-19 Genomics UK (COG-UK) Consortium | Harper VanSteenhouse, Yumi Kasai, David Gray, Carol Clugston, Anna Dominiczak and Alex Alderton, Roberto Amato, Jeffrey Barrett, Sonia Goncalves, Ewan Harrison, David K. Jackson, Ian Johnston, Dominic Kwiatkowski, Cordelia Langford, John Sillitoe on behalf of the Wellcome Sanger Institute COVID-19 Surveillance Team |
| EPI_ISL_3168943, EPI_ISL_3168944 | Lighthouse Laboratory Plymouth | Wellcome Sanger Institute for the COVID-19 Genomics UK (COG-UK) Consortium | Lighthouse Laboratory Plymouth and Alex Alderton, Roberto Amato, Jeffrey Barrett, Sonia Goncalves, Ewan Harrison, David K. Jackson, Ian Johnston, Dominic Kwiatkowski, Cordelia Langford, John Sillitoe on behalf of the Wellcome Sanger Institute COVID-19 Surveillance Team |
| EPI_ISL_3168945 | Lighthouse Lab in Glasgow | Wellcome Sanger Institute for the COVID-19 Genomics UK (COG-UK) Consortium | Harper VanSteenhouse, Yumi Kasai, David Gray, Carol Clugston, Anna Dominiczak and Alex Alderton, Roberto Amato, Jeffrey Barrett, Sonia Goncalves, Ewan Harrison, David K. Jackson, Ian Johnston, Dominic Kwiatkowski, Cordelia Langford, John Sillitoe on behalf of the Wellcome Sanger Institute COVID-19 Surveillance Team |
| EPI_ISL_3168946, EPI_ISL_3168947, EPI_ISL_3168948, EPI_ISL_3168949, EPI_ISL_3168950 | Lighthouse Laboratory Plymouth | Wellcome Sanger Institute for the COVID-19 Genomics UK (COG-UK) Consortium | Lighthouse Laboratory Plymouth and Alex Alderton, Roberto Amato, Jeffrey Barrett, Sonia Goncalves, Ewan Harrison, David K. Jackson, Ian Johnston, Dominic Kwiatkowski, Cordelia Langford, John Sillitoe on behalf of the Wellcome Sanger Institute COVID-19 Surveillance Team |
| EPI_ISL_3168951, EPI_ISL_3168952, EPI_ISL_3168953, EPI_ISL_3168954 | Lighthouse Lab in Glasgow | Wellcome Sanger Institute for the COVID-19 Genomics UK (COG-UK) Consortium | Harper VanSteenhouse, Yumi Kasai, David Gray, Carol Clugston, Anna Dominiczak and Alex Alderton, Roberto Amato, Jeffrey Barrett, Sonia Goncalves, Ewan Harrison, David K. Jackson, Ian Johnston, Dominic Kwiatkowski, Cordelia Langford, John Sillitoe on behalf of the Wellcome Sanger Institute COVID-19 Surveillance Team |
| EPI_ISL_3168955 | Lighthouse Laboratory Plymouth | Wellcome Sanger Institute for the COVID-19 Genomics UK (COG-UK) Consortium | Lighthouse Laboratory Plymouth and Alex Alderton, Roberto Amato, Jeffrey Barrett, Sonia Goncalves, Ewan Harrison, David K. Jackson, Ian Johnston, Dominic Kwiatkowski, Cordelia Langford, John Sillitoe on behalf of the Wellcome Sanger Institute COVID-19 Surveillance Team |
| EPI_ISL_3168956, EPI_ISL_3168957 | Lighthouse Lab in Glasgow | Wellcome Sanger Institute for the COVID-19 Genomics UK (COG-UK) Consortium | Harper VanSteenhouse, Yumi Kasai, David Gray, Carol Clugston, Anna Dominiczak and Alex Alderton, Roberto Amato, Jeffrey Barrett, Sonia Goncalves, Ewan Harrison, David K. Jackson, Ian Johnston, Dominic Kwiatkowski, Cordelia Langford, John Sillitoe on behalf of the Wellcome Sanger Institute COVID-19 Surveillance Team |
| EPI_ISL_3168958 | Lighthouse Laboratory Plymouth | Wellcome Sanger Institute for the COVID-19 Genomics UK (COG-UK) Consortium | Lighthouse Laboratory Plymouth and Alex Alderton, Roberto Amato, Jeffrey Barrett, Sonia Goncalves, Ewan Harrison, David K. Jackson, Ian Johnston, Dominic Kwiatkowski, Cordelia Langford, John Sillitoe on behalf of the Wellcome Sanger Institute COVID-19 Surveillance Team |
| EPI_ISL_3168959 | Lighthouse Lab in Glasgow | Wellcome Sanger Institute for the COVID-19 Genomics UK (COG-UK) Consortium | Harper VanSteenhouse, Yumi Kasai, David Gray, Carol Clugston, Anna Dominiczak and Alex Alderton, Roberto Amato, Jeffrey Barrett, Sonia Goncalves, Ewan Harrison, David K. Jackson, Ian Johnston, Dominic Kwiatkowski, Cordelia Langford, John Sillitoe on behalf of the Wellcome Sanger Institute COVID-19 Surveillance Team |
| EPI_ISL_3168960 | Lighthouse Laboratory Plymouth | Wellcome Sanger Institute for the COVID-19 Genomics UK (COG-UK) Consortium | Lighthouse Laboratory Plymouth and Alex Alderton, Roberto Amato, Jeffrey Barrett, Sonia Goncalves, Ewan Harrison, David K. Jackson, Ian Johnston, Dominic Kwiatkowski, Cordelia Langford, John Sillitoe on behalf of the Wellcome Sanger Institute COVID-19 Surveillance Team |
| EPI_ISL_3168961 | Lighthouse Lab in Glasgow | Wellcome Sanger Institute for the COVID-19 Genomics UK (COG-UK) Consortium | Harper VanSteenhouse, Yumi Kasai, David Gray, Carol Clugston, Anna Dominiczak and Alex Alderton, Roberto Amato, Jeffrey Barrett, Sonia Goncalves, Ewan Harrison, David K. Jackson, Ian Johnston, Dominic Kwiatkowski, Cordelia Langford, John Sillitoe on behalf of the Wellcome Sanger Institute COVID-19 Surveillance Team |
| EPI_ISL_3168962, EPI_ISL_3168963, EPI_ISL_3168964, EPI_ISL_3168965, EPI_ISL_3168966, EPI_ISL_3168967 | Lighthouse Laboratory Plymouth | Wellcome Sanger Institute for the COVID-19 Genomics UK (COG-UK) Consortium | Lighthouse Laboratory Plymouth and Alex Alderton, Roberto Amato, Jeffrey Barrett, Sonia Goncalves, Ewan Harrison, David K. Jackson, Ian Johnston, Dominic Kwiatkowski, Cordelia Langford, John Sillitoe on behalf of the Wellcome Sanger Institute COVID-19 Surveillance Team |
| EPI_ISL_3168968 | Lighthouse Lab in Glasgow | Wellcome Sanger Institute for the COVID-19 Genomics UK (COG-UK) Consortium | Harper VanSteenhouse, Yumi Kasai, David Gray, Carol Clugston, Anna Dominiczak and Alex Alderton, Roberto Amato, Jeffrey Barrett, Sonia Goncalves, Ewan Harrison, David K. Jackson, Ian Johnston, Dominic Kwiatkowski, Cordelia Langford, John Sillitoe on behalf of the Wellcome Sanger Institute COVID-19 Surveillance Team |
| EPI_ISL_3168969, EPI_ISL_3168970 | Lighthouse Laboratory Plymouth | Wellcome Sanger Institute for the COVID-19 Genomics UK (COG-UK) Consortium | Lighthouse Laboratory Plymouth and Alex Alderton, Roberto Amato, Jeffrey Barrett, Sonia Goncalves, Ewan Harrison, David K. Jackson, Ian Johnston, Dominic Kwiatkowski, Cordelia Langford, John Sillitoe on behalf of the Wellcome Sanger Institute COVID-19 Surveillance Team |
| EPI_ISL_3168971 | Lighthouse Lab in Glasgow | Wellcome Sanger Institute for the COVID-19 Genomics UK (COG-UK) Consortium | Harper VanSteenhouse, Yumi Kasai, David Gray, Carol Clugston, Anna Dominiczak and Alex Alderton, Roberto Amato, Jeffrey Barrett, Sonia Goncalves, Ewan Harrison, David K. Jackson, Ian Johnston, Dominic Kwiatkowski, Cordelia Langford, John Sillitoe on behalf of the Wellcome Sanger Institute COVID-19 Surveillance Team |
| EPI_ISL_3168972 | Lighthouse Laboratory Plymouth | Wellcome Sanger Institute for the COVID-19 Genomics UK (COG-UK) Consortium | Lighthouse Laboratory Plymouth and Alex Alderton, Roberto Amato, Jeffrey Barrett, Sonia Goncalves, Ewan Harrison, David K. Jackson, Ian Johnston, Dominic Kwiatkowski, Cordelia Langford, John Sillitoe on behalf of the Wellcome Sanger Institute COVID-19 Surveillance Team |
| EPI_ISL_3168973 | Lighthouse Lab in Glasgow | Wellcome Sanger Institute for the COVID-19 Genomics UK (COG-UK) Consortium | Harper VanSteenhouse, Yumi Kasai, David Gray, Carol Clugston, Anna Dominiczak and Alex Alderton, Roberto Amato, Jeffrey Barrett, Sonia Goncalves, Ewan Harrison, David K. Jackson, Ian Johnston, Dominic Kwiatkowski, Cordelia Langford, John Sillitoe on behalf of the Wellcome Sanger Institute COVID-19 Surveillance Team |
| EPI_ISL_3168974 | Lighthouse Laboratory Plymouth | Wellcome Sanger Institute for the COVID-19 Genomics UK (COG-UK) Consortium | Lighthouse Laboratory Plymouth and Alex Alderton, Roberto Amato, Jeffrey Barrett, Sonia Goncalves, Ewan Harrison, David K. Jackson, Ian Johnston, Dominic Kwiatkowski, Cordelia Langford, John Sillitoe on behalf of the Wellcome Sanger Institute COVID-19 Surveillance Team |
| EPI_ISL_3168975, EPI_ISL_3168976, EPI_ISL_3168977 | Lighthouse Lab in Glasgow | Wellcome Sanger Institute for the COVID-19 Genomics UK (COG-UK) Consortium | Harper VanSteenhouse, Yumi Kasai, David Gray, Carol Clugston, Anna Dominiczak and Alex Alderton, Roberto Amato, Jeffrey Barrett, Sonia Goncalves, Ewan Harrison, David K. Jackson, Ian Johnston, Dominic Kwiatkowski, Cordelia Langford, John Sillitoe on behalf of the Wellcome Sanger Institute COVID-19 Surveillance Team |
| EPI_ISL_3168978, EPI_ISL_3168979 | Lighthouse Laboratory Plymouth | Wellcome Sanger Institute for the COVID-19 Genomics UK (COG-UK) Consortium | Lighthouse Laboratory Plymouth and Alex Alderton, Roberto Amato, Jeffrey Barrett, Sonia Goncalves, Ewan Harrison, David K. Jackson, Ian Johnston, Dominic Kwiatkowski, Cordelia Langford, John Sillitoe on behalf of the Wellcome Sanger Institute COVID-19 Surveillance Team |
| EPI_ISL_3168980, EPI_ISL_3168981 | Lighthouse Lab in Glasgow | Wellcome Sanger Institute for the COVID-19 Genomics UK (COG-UK) Consortium | Harper VanSteenhouse, Yumi Kasai, David Gray, Carol Clugston, Anna Dominiczak and Alex Alderton, Roberto Amato, Jeffrey Barrett, Sonia Goncalves, Ewan Harrison, David K. Jackson, Ian Johnston, Dominic Kwiatkowski, Cordelia Langford, John Sillitoe on behalf of the Wellcome Sanger Institute COVID-19 Surveillance Team |
| EPI_ISL_3168982, EPI_ISL_3168983 | Lighthouse Laboratory Plymouth | Wellcome Sanger Institute for the COVID-19 Genomics UK (COG-UK) Consortium | Lighthouse Laboratory Plymouth and Alex Alderton, Roberto Amato, Jeffrey Barrett, Sonia Goncalves, Ewan Harrison, David K. Jackson, Ian Johnston, Dominic Kwiatkowski, Cordelia Langford, John Sillitoe on behalf of the Wellcome Sanger Institute COVID-19 Surveillance Team |
| EPI_ISL_3168984 | Lighthouse Lab in Glasgow | Wellcome Sanger Institute for the COVID-19 Genomics UK (COG-UK) Consortium | Harper VanSteenhouse, Yumi Kasai, David Gray, Carol Clugston, Anna Dominiczak and Alex Alderton, Roberto Amato, Jeffrey Barrett, Sonia Goncalves, Ewan Harrison, David K. Jackson, Ian Johnston, Dominic Kwiatkowski, Cordelia Langford, John Sillitoe on behalf of the Wellcome Sanger Institute COVID-19 Surveillance Team |
| EPI_ISL_3168985, EPI_ISL_3168986 | Lighthouse Laboratory Plymouth | Wellcome Sanger Institute for the COVID-19 Genomics UK (COG-UK) Consortium | Lighthouse Laboratory Plymouth and Alex Alderton, Roberto Amato, Jeffrey Barrett, Sonia Goncalves, Ewan Harrison, David K. Jackson, Ian Johnston, Dominic Kwiatkowski, Cordelia Langford, John Sillitoe on behalf of the Wellcome Sanger Institute COVID-19 Surveillance Team |
| EPI_ISL_3168987 | Lighthouse Lab in Glasgow | Wellcome Sanger Institute for the COVID-19 Genomics UK (COG-UK) Consortium | Harper VanSteenhouse, Yumi Kasai, David Gray, Carol Clugston, Anna Dominiczak and Alex Alderton, Roberto Amato, Jeffrey Barrett, Sonia Goncalves, Ewan Harrison, David K. Jackson, Ian Johnston, Dominic Kwiatkowski, Cordelia Langford, John Sillitoe on behalf of the Wellcome Sanger Institute COVID-19 Surveillance Team |
| EPI_ISL_3168988 | Lighthouse Laboratory Plymouth | Wellcome Sanger Institute for the COVID-19 Genomics UK (COG-UK) Consortium | Lighthouse Laboratory Plymouth and Alex Alderton, Roberto Amato, Jeffrey Barrett, Sonia Goncalves, Ewan Harrison, David K. Jackson, Ian Johnston, Dominic Kwiatkowski, Cordelia Langford, John Sillitoe on behalf of the Wellcome Sanger Institute COVID-19 Surveillance Team |
| EPI_ISL_3168989, EPI_ISL_3168990, EPI_ISL_3168991, EPI_ISL_3168992, EPI_ISL_3168993, EPI_ISL_3168994 | Lighthouse Lab in Glasgow | Wellcome Sanger Institute for the COVID-19 Genomics UK (COG-UK) Consortium | Harper VanSteenhouse, Yumi Kasai, David Gray, Carol Clugston, Anna Dominiczak and Alex Alderton, Roberto Amato, Jeffrey Barrett, Sonia Goncalves, Ewan Harrison, David K. Jackson, Ian Johnston, Dominic Kwiatkowski, Cordelia Langford, John Sillitoe on behalf of the Wellcome Sanger Institute COVID-19 Surveillance Team |
| EPI_ISL_3168995, EPI_ISL_3168996, EPI_ISL_3168997 | Lighthouse Laboratory Plymouth | Wellcome Sanger Institute for the COVID-19 Genomics UK (COG-UK) Consortium | Lighthouse Laboratory Plymouth and Alex Alderton, Roberto Amato, Jeffrey Barrett, Sonia Goncalves, Ewan Harrison, David K. Jackson, Ian Johnston, Dominic Kwiatkowski, Cordelia Langford, John Sillitoe on behalf of the Wellcome Sanger Institute COVID-19 Surveillance Team |
| EPI_ISL_3168998, EPI_ISL_3168999 | Lighthouse Lab in Glasgow | Wellcome Sanger Institute for the COVID-19 Genomics UK (COG-UK) Consortium | Harper VanSteenhouse, Yumi Kasai, David Gray, Carol Clugston, Anna Dominiczak and Alex Alderton, Roberto Amato, Jeffrey Barrett, Sonia Goncalves, Ewan Harrison, David K. Jackson, Ian Johnston, Dominic Kwiatkowski, Cordelia Langford, John Sillitoe on behalf of the Wellcome Sanger Institute COVID-19 Surveillance Team |
| EPI_ISL_3169000 | Lighthouse Laboratory Plymouth | Wellcome Sanger Institute for the COVID-19 Genomics UK (COG-UK) Consortium | Lighthouse Laboratory Plymouth and Alex Alderton, Roberto Amato, Jeffrey Barrett, Sonia Goncalves, Ewan Harrison, David K. Jackson, Ian Johnston, Dominic Kwiatkowski, Cordelia Langford, John Sillitoe on behalf of the Wellcome Sanger Institute COVID-19 Surveillance Team |
| EPI_ISL_3169001 | Berkshire and Surrey Pathology Services Lighthouse Laboratory | Wellcome Sanger Institute for the COVID-19 Genomics UK (COG-UK) Consortium | Berkshire and Surrey Pathology Services Lighthouse Laboratory and Alex Alderton, Roberto Amato, Jeffrey Barrett, Sonia Goncalves, Ewan Harrison, David K. Jackson, Ian Johnston, Dominic Kwiatkowski, Cordelia Langford, John Sillitoe on behalf of the Wellcome Sanger Institute COVID-19 Surveillance Team |

| | | | |
|---|---|---|---|
| EPI_ISL_3169002, EPI_ISL_3169003 | Lighthouse Laboratory Plymouth | Wellcome Sanger Institute for the COVID-19 Genomics UK (COG-UK) Consortium | Lighthouse Laboratory Plymouth and Alex Alderton, Roberto Amato, Jeffrey Barrett, Sonia Goncalves, Ewan Harrison, David K. Jackson, Ian Johnston, Dominic Kwiatkowski, Cordelia Langford, John Sillitoe on behalf of the Wellcome Sanger Institute COVID-19 Surveillance Team |
| EPI_ISL_3169004 | Lighthouse Lab in Glasgow | Wellcome Sanger Institute for the COVID-19 Genomics UK (COG-UK) Consortium | Harper VanSteenhouse, Yumi Kasai, David Gray, Carol Clugston, Anna Dominiczak and Alex Alderton, Roberto Amato, Jeffrey Barrett, Sonia Goncalves, Ewan Harrison, David K. Jackson, Ian Johnston, Dominic Kwiatkowski, Cordelia Langford, John Sillitoe on behalf of the Wellcome Sanger Institute COVID-19 Surveillance Team |
| EPI_ISL_3169005 | Lighthouse Laboratory Plymouth | Wellcome Sanger Institute for the COVID-19 Genomics UK (COG-UK) Consortium | Lighthouse Laboratory Plymouth and Alex Alderton, Roberto Amato, Jeffrey Barrett, Sonia Goncalves, Ewan Harrison, David K. Jackson, Ian Johnston, Dominic Kwiatkowski, Cordelia Langford, John Sillitoe on behalf of the Wellcome Sanger Institute COVID-19 Surveillance Team |
| EPI_ISL_3169006 | Lighthouse Lab in Glasgow | Wellcome Sanger Institute for the COVID-19 Genomics UK (COG-UK) Consortium | Harper VanSteenhouse, Yumi Kasai, David Gray, Carol Clugston, Anna Dominiczak and Alex Alderton, Roberto Amato, Jeffrey Barrett, Sonia Goncalves, Ewan Harrison, David K. Jackson, Ian Johnston, Dominic Kwiatkowski, Cordelia Langford, John Sillitoe on behalf of the Wellcome Sanger Institute COVID-19 Surveillance Team |
| EPI_ISL_3169007 | Lighthouse Laboratory Plymouth | Wellcome Sanger Institute for the COVID-19 Genomics UK (COG-UK) Consortium | Lighthouse Laboratory Plymouth and Alex Alderton, Roberto Amato, Jeffrey Barrett, Sonia Goncalves, Ewan Harrison, David K. Jackson, Ian Johnston, Dominic Kwiatkowski, Cordelia Langford, John Sillitoe on behalf of the Wellcome Sanger Institute COVID-19 Surveillance Team |
| EPI_ISL_3169008, EPI_ISL_3169009, EPI_ISL_3169010, EPI_ISL_3169011 | Lighthouse Lab in Glasgow | Wellcome Sanger Institute for the COVID-19 Genomics UK (COG-UK) Consortium | Harper VanSteenhouse, Yumi Kasai, David Gray, Carol Clugston, Anna Dominiczak and Alex Alderton, Roberto Amato, Jeffrey Barrett, Sonia Goncalves, Ewan Harrison, David K. Jackson, Ian Johnston, Dominic Kwiatkowski, Cordelia Langford, John Sillitoe on behalf of the Wellcome Sanger Institute COVID-19 Surveillance Team |
| EPI_ISL_3169012 | Berkshire and Surrey Pathology Services Lighthouse Laboratory | Wellcome Sanger Institute for the COVID-19 Genomics UK (COG-UK) Consortium | Berkshire and Surrey Pathology Services Lighthouse Laboratory and Alex Alderton, Roberto Amato, Jeffrey Barrett, Sonia Goncalves, Ewan Harrison, David K. Jackson, Ian Johnston, Dominic Kwiatkowski, Cordelia Langford, John Sillitoe on behalf of the Wellcome Sanger Institute COVID-19 Surveillance Team |
| EPI_ISL_3169013, EPI_ISL_3169014 | Lighthouse Lab in Glasgow | Wellcome Sanger Institute for the COVID-19 Genomics UK (COG-UK) Consortium | Harper VanSteenhouse, Yumi Kasai, David Gray, Carol Clugston, Anna Dominiczak and Alex Alderton, Roberto Amato, Jeffrey Barrett, Sonia Goncalves, Ewan Harrison, David K. Jackson, Ian Johnston, Dominic Kwiatkowski, Cordelia Langford, John Sillitoe on behalf of the Wellcome Sanger Institute COVID-19 Surveillance Team |
| EPI_ISL_3169015 | Lighthouse Laboratory Plymouth | Wellcome Sanger Institute for the COVID-19 Genomics UK (COG-UK) Consortium | Lighthouse Laboratory Plymouth and Alex Alderton, Roberto Amato, Jeffrey Barrett, Sonia Goncalves, Ewan Harrison, David K. Jackson, Ian Johnston, Dominic Kwiatkowski, Cordelia Langford, John Sillitoe on behalf of the Wellcome Sanger Institute COVID-19 Surveillance Team |
| EPI_ISL_3169016 | Berkshire and Surrey Pathology Services Lighthouse Laboratory | Wellcome Sanger Institute for the COVID-19 Genomics UK (COG-UK) Consortium | Berkshire and Surrey Pathology Services Lighthouse Laboratory and Alex Alderton, Roberto Amato, Jeffrey Barrett, Sonia Goncalves, Ewan Harrison, David K. Jackson, Ian Johnston, Dominic Kwiatkowski, Cordelia Langford, John Sillitoe on behalf of the Wellcome Sanger Institute COVID-19 Surveillance Team |
| EPI_ISL_3169017, EPI_ISL_3169018 | Lighthouse Laboratory Plymouth | Wellcome Sanger Institute for the COVID-19 Genomics UK (COG-UK) Consortium | Lighthouse Laboratory Plymouth and Alex Alderton, Roberto Amato, Jeffrey Barrett, Sonia Goncalves, Ewan Harrison, David K. Jackson, Ian Johnston, Dominic Kwiatkowski, Cordelia Langford, John Sillitoe on behalf of the Wellcome Sanger Institute COVID-19 Surveillance Team |
| EPI_ISL_3169019 | Lighthouse Lab in Glasgow | Wellcome Sanger Institute for the COVID-19 Genomics UK (COG-UK) Consortium | Harper VanSteenhouse, Yumi Kasai, David Gray, Carol Clugston, Anna Dominiczak and Alex Alderton, Roberto Amato, Jeffrey Barrett, Sonia Goncalves, Ewan Harrison, David K. Jackson, Ian Johnston, Dominic Kwiatkowski, Cordelia Langford, John Sillitoe on behalf of the Wellcome Sanger Institute COVID-19 Surveillance Team |
| EPI_ISL_3169020, EPI_ISL_3169021, EPI_ISL_3169022 | Lighthouse Laboratory Plymouth | Wellcome Sanger Institute for the COVID-19 Genomics UK (COG-UK) Consortium | Lighthouse Laboratory Plymouth and Alex Alderton, Roberto Amato, Jeffrey Barrett, Sonia Goncalves, Ewan Harrison, David K. Jackson, Ian Johnston, Dominic Kwiatkowski, Cordelia Langford, John Sillitoe on behalf of the Wellcome Sanger Institute COVID-19 Surveillance Team |
| EPI_ISL_3169023 | Lighthouse Lab in Glasgow | Wellcome Sanger Institute for the COVID-19 Genomics UK (COG-UK) Consortium | Harper VanSteenhouse, Yumi Kasai, David Gray, Carol Clugston, Anna Dominiczak and Alex Alderton, Roberto Amato, Jeffrey Barrett, Sonia Goncalves, Ewan Harrison, David K. Jackson, Ian Johnston, Dominic Kwiatkowski, Cordelia Langford, John Sillitoe on behalf of the Wellcome Sanger Institute COVID-19 Surveillance Team |
| EPI_ISL_3169024 | Berkshire and Surrey Pathology Services Lighthouse Laboratory | Wellcome Sanger Institute for the COVID-19 Genomics UK (COG-UK) Consortium | Berkshire and Surrey Pathology Services Lighthouse Laboratory and Alex Alderton, Roberto Amato, Jeffrey Barrett, Sonia Goncalves, Ewan Harrison, David K. Jackson, Ian Johnston, Dominic Kwiatkowski, Cordelia Langford, John Sillitoe on behalf of the Wellcome Sanger Institute COVID-19 Surveillance Team |
| EPI_ISL_3169025, EPI_ISL_3169026, EPI_ISL_3169027 | Lighthouse Laboratory Plymouth | Wellcome Sanger Institute for the COVID-19 Genomics UK (COG-UK) Consortium | Lighthouse Laboratory Plymouth and Alex Alderton, Roberto Amato, Jeffrey Barrett, Sonia Goncalves, Ewan Harrison, David K. Jackson, Ian Johnston, Dominic Kwiatkowski, Cordelia Langford, John Sillitoe on behalf of the Wellcome Sanger Institute COVID-19 Surveillance Team |
| EPI_ISL_3169028 | Lighthouse Lab in Glasgow | Wellcome Sanger Institute for the COVID-19 Genomics UK (COG-UK) Consortium | Harper VanSteenhouse, Yumi Kasai, David Gray, Carol Clugston, Anna Dominiczak and Alex Alderton, Roberto Amato, Jeffrey Barrett, Sonia Goncalves, Ewan Harrison, David K. Jackson, Ian Johnston, Dominic Kwiatkowski, Cordelia Langford, John Sillitoe on behalf of the Wellcome Sanger Institute COVID-19 Surveillance Team |
| EPI_ISL_3169029, EPI_ISL_3169030, EPI_ISL_3169031 | Berkshire and Surrey Pathology Services Lighthouse Laboratory | Wellcome Sanger Institute for the COVID-19 Genomics UK (COG-UK) Consortium | Berkshire and Surrey Pathology Services Lighthouse Laboratory and Alex Alderton, Roberto Amato, Jeffrey Barrett, Sonia Goncalves, Ewan Harrison, David K. Jackson, Ian Johnston, Dominic Kwiatkowski, Cordelia Langford, John Sillitoe on behalf of the Wellcome Sanger Institute COVID-19 Surveillance Team |
| EPI_ISL_3169032 | Lighthouse Lab in Glasgow | Wellcome Sanger Institute for the COVID-19 Genomics UK (COG-UK) Consortium | Harper VanSteenhouse, Yumi Kasai, David Gray, Carol Clugston, Anna Dominiczak and Alex Alderton, Roberto Amato, Jeffrey Barrett, Sonia Goncalves, Ewan Harrison, David K. Jackson, Ian Johnston, Dominic Kwiatkowski, Cordelia Langford, John Sillitoe on behalf of the Wellcome Sanger Institute COVID-19 Surveillance Team |
| EPI_ISL_3169033 | Berkshire and Surrey Pathology Services Lighthouse Laboratory | Wellcome Sanger Institute for the COVID-19 Genomics UK (COG-UK) Consortium | Berkshire and Surrey Pathology Services Lighthouse Laboratory and Alex Alderton, Roberto Amato, Jeffrey Barrett, Sonia Goncalves, Ewan Harrison, David K. Jackson, Ian Johnston, Dominic Kwiatkowski, Cordelia Langford, John Sillitoe on behalf of the Wellcome Sanger Institute COVID-19 Surveillance Team |
| EPI_ISL_3169034, EPI_ISL_3169035, EPI_ISL_3169036 | Lighthouse Lab in Glasgow | Wellcome Sanger Institute for the COVID-19 Genomics UK (COG-UK) Consortium | Harper VanSteenhouse, Yumi Kasai, David Gray, Carol Clugston, Anna Dominiczak and Alex Alderton, Roberto Amato, Jeffrey Barrett, Sonia Goncalves, Ewan Harrison, David K. Jackson, Ian Johnston, Dominic Kwiatkowski, Cordelia Langford, John Sillitoe on behalf of the Wellcome Sanger Institute COVID-19 Surveillance Team |
| EPI_ISL_3169037 | Berkshire and Surrey Pathology Services Lighthouse Laboratory | Wellcome Sanger Institute for the COVID-19 Genomics UK (COG-UK) Consortium | Berkshire and Surrey Pathology Services Lighthouse Laboratory and Alex Alderton, Roberto Amato, Jeffrey Barrett, Sonia Goncalves, Ewan Harrison, David K. Jackson, Ian Johnston, Dominic Kwiatkowski, Cordelia Langford, John Sillitoe on behalf of the Wellcome Sanger Institute COVID-19 Surveillance Team |
| EPI_ISL_3169038, EPI_ISL_3169039 | Lighthouse Laboratory Plymouth | Wellcome Sanger Institute for the COVID-19 Genomics UK (COG-UK) Consortium | Lighthouse Laboratory Plymouth and Alex Alderton, Roberto Amato, Jeffrey Barrett, Sonia Goncalves, Ewan Harrison, David K. Jackson, Ian Johnston, Dominic Kwiatkowski, Cordelia Langford, John Sillitoe on behalf of the Wellcome Sanger Institute COVID-19 Surveillance Team |
| EPI_ISL_3169040 | Lighthouse Lab in Glasgow | Wellcome Sanger Institute for the COVID-19 Genomics UK (COG-UK) Consortium | Harper VanSteenhouse, Yumi Kasai, David Gray, Carol Clugston, Anna Dominiczak and Alex Alderton, Roberto Amato, Jeffrey Barrett, Sonia Goncalves, Ewan Harrison, David K. Jackson, Ian Johnston, Dominic Kwiatkowski, Cordelia Langford, John Sillitoe on behalf of the Wellcome Sanger Institute COVID-19 Surveillance Team |
| EPI_ISL_3169041, EPI_ISL_3169042 | Berkshire and Surrey Pathology Services Lighthouse Laboratory | Wellcome Sanger Institute for the COVID-19 Genomics UK (COG-UK) Consortium | Berkshire and Surrey Pathology Services Lighthouse Laboratory and Alex Alderton, Roberto Amato, Jeffrey Barrett, Sonia Goncalves, Ewan Harrison, David K. Jackson, Ian Johnston, Dominic Kwiatkowski, Cordelia Langford, John Sillitoe on behalf of the Wellcome Sanger Institute COVID-19 Surveillance Team |
| EPI_ISL_3169043, EPI_ISL_3169044, EPI_ISL_3169045, EPI_ISL_3169046 | Lighthouse Laboratory Plymouth | Wellcome Sanger Institute for the COVID-19 Genomics UK (COG-UK) Consortium | Lighthouse Laboratory Plymouth and Alex Alderton, Roberto Amato, Jeffrey Barrett, Sonia Goncalves, Ewan Harrison, David K. Jackson, Ian Johnston, Dominic Kwiatkowski, Cordelia Langford, John Sillitoe on behalf of the Wellcome Sanger Institute COVID-19 Surveillance Team |
| EPI_ISL_3169047 | Lighthouse Lab in Glasgow | Wellcome Sanger Institute for the COVID-19 Genomics UK (COG-UK) Consortium | Harper VanSteenhouse, Yumi Kasai, David Gray, Carol Clugston, Anna Dominiczak and Alex Alderton, Roberto Amato, Jeffrey Barrett, Sonia Goncalves, Ewan Harrison, David K. Jackson, Ian Johnston, Dominic Kwiatkowski, Cordelia Langford, John Sillitoe on behalf of the Wellcome Sanger Institute COVID-19 Surveillance Team |
| EPI_ISL_3169048 | Berkshire and Surrey Pathology Services Lighthouse Laboratory | Wellcome Sanger Institute for the COVID-19 Genomics UK (COG-UK) Consortium | Berkshire and Surrey Pathology Services Lighthouse Laboratory and Alex Alderton, Roberto Amato, Jeffrey Barrett, Sonia Goncalves, Ewan Harrison, David K. Jackson, Ian Johnston, Dominic Kwiatkowski, Cordelia Langford, John Sillitoe on behalf of the Wellcome Sanger Institute COVID-19 Surveillance Team |
| EPI_ISL_3169049 | Lighthouse Laboratory Plymouth | Wellcome Sanger Institute for the COVID-19 Genomics UK (COG-UK) Consortium | Lighthouse Laboratory Plymouth and Alex Alderton, Roberto Amato, Jeffrey Barrett, Sonia Goncalves, Ewan Harrison, David K. Jackson, Ian Johnston, Dominic Kwiatkowski, Cordelia Langford, John Sillitoe on behalf of the Wellcome Sanger Institute COVID-19 Surveillance Team |
| EPI_ISL_3169050, EPI_ISL_3169051 | Berkshire and Surrey Pathology Services Lighthouse Laboratory | Wellcome Sanger Institute for the COVID-19 Genomics UK (COG-UK) Consortium | Berkshire and Surrey Pathology Services Lighthouse Laboratory and Alex Alderton, Roberto Amato, Jeffrey Barrett, Sonia Goncalves, Ewan Harrison, David K. Jackson, Ian Johnston, Dominic Kwiatkowski, Cordelia Langford, John Sillitoe on behalf of the Wellcome Sanger Institute COVID-19 Surveillance Team |
| EPI_ISL_3169052, EPI_ISL_3169053 | Lighthouse Laboratory Plymouth | Wellcome Sanger Institute for the COVID-19 Genomics UK (COG-UK) Consortium | Lighthouse Laboratory Plymouth and Alex Alderton, Roberto Amato, Jeffrey Barrett, Sonia Goncalves, Ewan Harrison, David K. Jackson, Ian Johnston, Dominic Kwiatkowski, Cordelia Langford, John Sillitoe on behalf of the Wellcome Sanger Institute COVID-19 Surveillance Team |
| EPI_ISL_3169054 | Berkshire and Surrey Pathology Services Lighthouse Laboratory | Wellcome Sanger Institute for the COVID-19 Genomics UK (COG-UK) Consortium | Berkshire and Surrey Pathology Services Lighthouse Laboratory and Alex Alderton, Roberto Amato, Jeffrey Barrett, Sonia Goncalves, Ewan Harrison, David K. Jackson, Ian Johnston, Dominic Kwiatkowski, Cordelia Langford, John Sillitoe on behalf of the Wellcome Sanger Institute COVID-19 Surveillance Team |
| EPI_ISL_3169055 | Lighthouse Laboratory Plymouth | Wellcome Sanger Institute for the COVID-19 Genomics UK (COG-UK) Consortium | Lighthouse Laboratory Plymouth and Alex Alderton, Roberto Amato, Jeffrey Barrett, Sonia Goncalves, Ewan Harrison, David K. Jackson, Ian Johnston, Dominic Kwiatkowski, Cordelia Langford, John Sillitoe on behalf of the Wellcome Sanger Institute COVID-19 Surveillance Team |
| EPI_ISL_3169056, EPI_ISL_3169057, EPI_ISL_3169058 | Berkshire and Surrey Pathology Services Lighthouse Laboratory | Wellcome Sanger Institute for the COVID-19 Genomics UK (COG-UK) Consortium | Berkshire and Surrey Pathology Services Lighthouse Laboratory and Alex Alderton, Roberto Amato, Jeffrey Barrett, Sonia Goncalves, Ewan Harrison, David K. Jackson, Ian Johnston, Dominic Kwiatkowski, Cordelia Langford, John Sillitoe on behalf of the Wellcome Sanger Institute COVID-19 Surveillance Team |
| EPI_ISL_3169059 | Lighthouse Laboratory Plymouth | Wellcome Sanger Institute for the COVID-19 Genomics UK (COG-UK) Consortium | Lighthouse Laboratory Plymouth and Alex Alderton, Roberto Amato, Jeffrey Barrett, Sonia Goncalves, Ewan Harrison, David K. Jackson, Ian Johnston, Dominic Kwiatkowski, Cordelia Langford, John Sillitoe on behalf of the Wellcome Sanger Institute COVID-19 Surveillance Team |
| EPI_ISL_3169060 | Lighthouse Lab in Glasgow | Wellcome Sanger Institute for the COVID-19 Genomics UK (COG-UK) Consortium | Harper VanSteenhouse, Yumi Kasai, David Gray, Carol Clugston, Anna Dominiczak and Alex Alderton, Roberto Amato, Jeffrey Barrett, Sonia Goncalves, Ewan Harrison, David K. Jackson, Ian Johnston, Dominic Kwiatkowski, Cordelia Langford, John Sillitoe on behalf of the Wellcome Sanger Institute COVID-19 Surveillance Team |
| EPI_ISL_3169061 | Berkshire and Surrey Pathology Services Lighthouse Laboratory | Wellcome Sanger Institute for the COVID-19 Genomics UK (COG-UK) Consortium | Berkshire and Surrey Pathology Services Lighthouse Laboratory and Alex Alderton, Roberto Amato, Jeffrey Barrett, Sonia Goncalves, Ewan Harrison, David K. Jackson, Ian Johnston, Dominic Kwiatkowski, Cordelia Langford, John Sillitoe on behalf of the Wellcome Sanger Institute COVID-19 Surveillance Team |
| EPI_ISL_3169062, EPI_ISL_3169063 | Lighthouse Laboratory Plymouth | Wellcome Sanger Institute for the COVID-19 Genomics UK (COG-UK) Consortium | Lighthouse Laboratory Plymouth and Alex Alderton, Roberto Amato, Jeffrey Barrett, Sonia Goncalves, Ewan Harrison, David K. Jackson, Ian Johnston, Dominic Kwiatkowski, Cordelia Langford, John Sillitoe on behalf of the Wellcome Sanger Institute COVID-19 Surveillance Team |
| EPI_ISL_3169064, EPI_ISL_3169065 | Lighthouse Lab in Glasgow | Wellcome Sanger Institute for the COVID-19 Genomics UK (COG-UK) Consortium | Harper VanSteenhouse, Yumi Kasai, David Gray, Carol Clugston, Anna Dominiczak and Alex Alderton, Roberto Amato, Jeffrey Barrett, Sonia Goncalves, Ewan Harrison, David K. Jackson, Ian Johnston, Dominic Kwiatkowski, Cordelia Langford, John Sillitoe on behalf of the Wellcome Sanger Institute COVID-19 Surveillance Team |
| EPI_ISL_3169066 | Lighthouse Laboratory Plymouth | Wellcome Sanger Institute for the COVID-19 Genomics UK (COG-UK) Consortium | Lighthouse Laboratory Plymouth and Alex Alderton, Roberto Amato, Jeffrey Barrett, Sonia Goncalves, Ewan Harrison, David K. Jackson, Ian Johnston, Dominic Kwiatkowski, Cordelia Langford, John Sillitoe on behalf of the Wellcome Sanger Institute COVID-19 Surveillance Team |
| EPI_ISL_3169067, EPI_ISL_3169068 | Berkshire and Surrey Pathology Services Lighthouse Laboratory | Wellcome Sanger Institute for the COVID-19 Genomics UK (COG-UK) Consortium | Berkshire and Surrey Pathology Services Lighthouse Laboratory and Alex Alderton, Roberto Amato, Jeffrey Barrett, Sonia Goncalves, Ewan Harrison, David K. Jackson, Ian Johnston, Dominic Kwiatkowski, Cordelia Langford, John Sillitoe on behalf of the Wellcome Sanger Institute COVID-19 Surveillance Team |
| EPI_ISL_3169069 | Lighthouse Lab in Glasgow | Wellcome Sanger Institute for the COVID-19 Genomics UK (COG-UK) | Harper VanSteenhouse, Yumi Kasai, David Gray, Carol Clugston, Anna Dominiczak and Alex Alderton, Roberto Amato, Jeffrey Barrett, Sonia Goncalves, Ewan Harrison, David K. Jackson, Ian Johnston, Dominic Kwiatkowski, Cordelia Langford, |

| | | | |
|---|---|---|---|
| | | Consortium | John Sillitoe on behalf of the Wellcome Sanger Institute COVID-19 Surveillance Team |
| EPI_ISL_3169070 | Berkshire and Surrey Pathology Services Lighthouse Laboratory | Wellcome Sanger Institute for the COVID-19 Genomics UK (COG-UK) Consortium | Berkshire and Surrey Pathology Services Lighthouse Laboratory and Alex Alderton, Roberto Amato, Jeffrey Barrett, Sonia Goncalves, Ewan Harrison, David K. Jackson, Ian Johnston, Dominic Kwiatkowski, Cordelia Langford, John Sillitoe on behalf of the Wellcome Sanger Institute COVID-19 Surveillance Team |
| EPI_ISL_3169071 | Lighthouse Laboratory Plymouth | Wellcome Sanger Institute for the COVID-19 Genomics UK (COG-UK) Consortium | Lighthouse Laboratory Plymouth and Alex Alderton, Roberto Amato, Jeffrey Barrett, Sonia Goncalves, Ewan Harrison, David K. Jackson, Ian Johnston, Dominic Kwiatkowski, Cordelia Langford, John Sillitoe on behalf of the Wellcome Sanger Institute COVID-19 Surveillance Team |
| EPI_ISL_3169072, EPI_ISL_3169073 | Lighthouse Lab in Glasgow | Wellcome Sanger Institute for the COVID-19 Genomics UK (COG-UK) Consortium | Harper VanSteenhouse, Yumi Kasai, David Gray, Carol Clugston, Anna Dominiczak and Alex Alderton, Roberto Amato, Jeffrey Barrett, Sonia Goncalves, Ewan Harrison, David K. Jackson, Ian Johnston, Dominic Kwiatkowski, Cordelia Langford, John Sillitoe on behalf of the Wellcome Sanger Institute COVID-19 Surveillance Team |
| EPI_ISL_3169074 | Lighthouse Laboratory Plymouth | Wellcome Sanger Institute for the COVID-19 Genomics UK (COG-UK) Consortium | Lighthouse Laboratory Plymouth and Alex Alderton, Roberto Amato, Jeffrey Barrett, Sonia Goncalves, Ewan Harrison, David K. Jackson, Ian Johnston, Dominic Kwiatkowski, Cordelia Langford, John Sillitoe on behalf of the Wellcome Sanger Institute COVID-19 Surveillance Team |
| EPI_ISL_3169075, EPI_ISL_3169076 | Lighthouse Lab in Glasgow | Wellcome Sanger Institute for the COVID-19 Genomics UK (COG-UK) Consortium | Harper VanSteenhouse, Yumi Kasai, David Gray, Carol Clugston, Anna Dominiczak and Alex Alderton, Roberto Amato, Jeffrey Barrett, Sonia Goncalves, Ewan Harrison, David K. Jackson, Ian Johnston, Dominic Kwiatkowski, Cordelia Langford, John Sillitoe on behalf of the Wellcome Sanger Institute COVID-19 Surveillance Team |
| EPI_ISL_3169077 | Lighthouse Laboratory Plymouth | Wellcome Sanger Institute for the COVID-19 Genomics UK (COG-UK) Consortium | Lighthouse Laboratory Plymouth and Alex Alderton, Roberto Amato, Jeffrey Barrett, Sonia Goncalves, Ewan Harrison, David K. Jackson, Ian Johnston, Dominic Kwiatkowski, Cordelia Langford, John Sillitoe on behalf of the Wellcome Sanger Institute COVID-19 Surveillance Team |
| EPI_ISL_3169078 | Lighthouse Lab in Glasgow | Wellcome Sanger Institute for the COVID-19 Genomics UK (COG-UK) Consortium | Harper VanSteenhouse, Yumi Kasai, David Gray, Carol Clugston, Anna Dominiczak and Alex Alderton, Roberto Amato, Jeffrey Barrett, Sonia Goncalves, Ewan Harrison, David K. Jackson, Ian Johnston, Dominic Kwiatkowski, Cordelia Langford, John Sillitoe on behalf of the Wellcome Sanger Institute COVID-19 Surveillance Team |
| EPI_ISL_3169079, EPI_ISL_3169080, EPI_ISL_3169081, EPI_ISL_3169082 | Lighthouse Laboratory Plymouth | Wellcome Sanger Institute for the COVID-19 Genomics UK (COG-UK) Consortium | Lighthouse Laboratory Plymouth and Alex Alderton, Roberto Amato, Jeffrey Barrett, Sonia Goncalves, Ewan Harrison, David K. Jackson, Ian Johnston, Dominic Kwiatkowski, Cordelia Langford, John Sillitoe on behalf of the Wellcome Sanger Institute COVID-19 Surveillance Team |
| EPI_ISL_3169083, EPI_ISL_3169084 | Lighthouse Lab in Glasgow | Wellcome Sanger Institute for the COVID-19 Genomics UK (COG-UK) Consortium | Harper VanSteenhouse, Yumi Kasai, David Gray, Carol Clugston, Anna Dominiczak and Alex Alderton, Roberto Amato, Jeffrey Barrett, Sonia Goncalves, Ewan Harrison, David K. Jackson, Ian Johnston, Dominic Kwiatkowski, Cordelia Langford, John Sillitoe on behalf of the Wellcome Sanger Institute COVID-19 Surveillance Team |
| EPI_ISL_3169085 | Lighthouse Laboratory Plymouth | Wellcome Sanger Institute for the COVID-19 Genomics UK (COG-UK) Consortium | Lighthouse Laboratory Plymouth and Alex Alderton, Roberto Amato, Jeffrey Barrett, Sonia Goncalves, Ewan Harrison, David K. Jackson, Ian Johnston, Dominic Kwiatkowski, Cordelia Langford, John Sillitoe on behalf of the Wellcome Sanger Institute COVID-19 Surveillance Team |
| EPI_ISL_3169086, EPI_ISL_3169087, EPI_ISL_3169088, EPI_ISL_3169089 | Lighthouse Lab in Glasgow | Wellcome Sanger Institute for the COVID-19 Genomics UK (COG-UK) Consortium | Harper VanSteenhouse, Yumi Kasai, David Gray, Carol Clugston, Anna Dominiczak and Alex Alderton, Roberto Amato, Jeffrey Barrett, Sonia Goncalves, Ewan Harrison, David K. Jackson, Ian Johnston, Dominic Kwiatkowski, Cordelia Langford, John Sillitoe on behalf of the Wellcome Sanger Institute COVID-19 Surveillance Team |
| EPI_ISL_3169090 | Berkshire and Surrey Pathology Services Lighthouse Laboratory | Wellcome Sanger Institute for the COVID-19 Genomics UK (COG-UK) Consortium | Berkshire and Surrey Pathology Services Lighthouse Laboratory and Alex Alderton, Roberto Amato, Jeffrey Barrett, Sonia Goncalves, Ewan Harrison, David K. Jackson, Ian Johnston, Dominic Kwiatkowski, Cordelia Langford, John Sillitoe on behalf of the Wellcome Sanger Institute COVID-19 Surveillance Team |
| EPI_ISL_3169091 | Lighthouse Lab in Glasgow | Wellcome Sanger Institute for the COVID-19 Genomics UK (COG-UK) Consortium | Harper VanSteenhouse, Yumi Kasai, David Gray, Carol Clugston, Anna Dominiczak and Alex Alderton, Roberto Amato, Jeffrey Barrett, Sonia Goncalves, Ewan Harrison, David K. Jackson, Ian Johnston, Dominic Kwiatkowski, Cordelia Langford, John Sillitoe on behalf of the Wellcome Sanger Institute COVID-19 Surveillance Team |
| EPI_ISL_3169092 | Berkshire and Surrey Pathology Services Lighthouse Laboratory | Wellcome Sanger Institute for the COVID-19 Genomics UK (COG-UK) Consortium | Berkshire and Surrey Pathology Services Lighthouse Laboratory and Alex Alderton, Roberto Amato, Jeffrey Barrett, Sonia Goncalves, Ewan Harrison, David K. Jackson, Ian Johnston, Dominic Kwiatkowski, Cordelia Langford, John Sillitoe on behalf of the Wellcome Sanger Institute COVID-19 Surveillance Team |
| EPI_ISL_3169093 | Lighthouse Lab in Glasgow | Wellcome Sanger Institute for the COVID-19 Genomics UK (COG-UK) Consortium | Harper VanSteenhouse, Yumi Kasai, David Gray, Carol Clugston, Anna Dominiczak and Alex Alderton, Roberto Amato, Jeffrey Barrett, Sonia Goncalves, Ewan Harrison, David K. Jackson, Ian Johnston, Dominic Kwiatkowski, Cordelia Langford, John Sillitoe on behalf of the Wellcome Sanger Institute COVID-19 Surveillance Team |
| EPI_ISL_3169094 | Berkshire and Surrey Pathology Services Lighthouse Laboratory | Wellcome Sanger Institute for the COVID-19 Genomics UK (COG-UK) Consortium | Berkshire and Surrey Pathology Services Lighthouse Laboratory and Alex Alderton, Roberto Amato, Jeffrey Barrett, Sonia Goncalves, Ewan Harrison, David K. Jackson, Ian Johnston, Dominic Kwiatkowski, Cordelia Langford, John Sillitoe on behalf of the Wellcome Sanger Institute COVID-19 Surveillance Team |
| EPI_ISL_3169095 | Lighthouse Lab in Glasgow | Wellcome Sanger Institute for the COVID-19 Genomics UK (COG-UK) Consortium | Harper VanSteenhouse, Yumi Kasai, David Gray, Carol Clugston, Anna Dominiczak and Alex Alderton, Roberto Amato, Jeffrey Barrett, Sonia Goncalves, Ewan Harrison, David K. Jackson, Ian Johnston, Dominic Kwiatkowski, Cordelia Langford, John Sillitoe on behalf of the Wellcome Sanger Institute COVID-19 Surveillance Team |
| EPI_ISL_3169096 | Lighthouse Laboratory Plymouth | Wellcome Sanger Institute for the COVID-19 Genomics UK (COG-UK) Consortium | Lighthouse Laboratory Plymouth and Alex Alderton, Roberto Amato, Jeffrey Barrett, Sonia Goncalves, Ewan Harrison, David K. Jackson, Ian Johnston, Dominic Kwiatkowski, Cordelia Langford, John Sillitoe on behalf of the Wellcome Sanger Institute COVID-19 Surveillance Team |
| EPI_ISL_3169097 | Lighthouse Lab in Glasgow | Wellcome Sanger Institute for the COVID-19 Genomics UK (COG-UK) Consortium | Harper VanSteenhouse, Yumi Kasai, David Gray, Carol Clugston, Anna Dominiczak and Alex Alderton, Roberto Amato, Jeffrey Barrett, Sonia Goncalves, Ewan Harrison, David K. Jackson, Ian Johnston, Dominic Kwiatkowski, Cordelia Langford, John Sillitoe on behalf of the Wellcome Sanger Institute COVID-19 Surveillance Team |
| EPI_ISL_3169098 | Berkshire and Surrey Pathology Services Lighthouse Laboratory | Wellcome Sanger Institute for the COVID-19 Genomics UK (COG-UK) Consortium | Berkshire and Surrey Pathology Services Lighthouse Laboratory and Alex Alderton, Roberto Amato, Jeffrey Barrett, Sonia Goncalves, Ewan Harrison, David K. Jackson, Ian Johnston, Dominic Kwiatkowski, Cordelia Langford, John Sillitoe on behalf of the Wellcome Sanger Institute COVID-19 Surveillance Team |
| EPI_ISL_3169099, EPI_ISL_3169100 | Lighthouse Lab in Glasgow | Wellcome Sanger Institute for the COVID-19 Genomics UK (COG-UK) Consortium | Harper VanSteenhouse, Yumi Kasai, David Gray, Carol Clugston, Anna Dominiczak and Alex Alderton, Roberto Amato, Jeffrey Barrett, Sonia Goncalves, Ewan Harrison, David K. Jackson, Ian Johnston, Dominic Kwiatkowski, Cordelia Langford, John Sillitoe on behalf of the Wellcome Sanger Institute COVID-19 Surveillance Team |
| EPI_ISL_3169101 | Berkshire and Surrey Pathology Services Lighthouse Laboratory | Wellcome Sanger Institute for the COVID-19 Genomics UK (COG-UK) Consortium | Berkshire and Surrey Pathology Services Lighthouse Laboratory and Alex Alderton, Roberto Amato, Jeffrey Barrett, Sonia Goncalves, Ewan Harrison, David K. Jackson, Ian Johnston, Dominic Kwiatkowski, Cordelia Langford, John Sillitoe on behalf of the Wellcome Sanger Institute COVID-19 Surveillance Team |
| EPI_ISL_3169102 | Lighthouse Laboratory Plymouth | Wellcome Sanger Institute for the COVID-19 Genomics UK (COG-UK) Consortium | Lighthouse Laboratory Plymouth and Alex Alderton, Roberto Amato, Jeffrey Barrett, Sonia Goncalves, Ewan Harrison, David K. Jackson, Ian Johnston, Dominic Kwiatkowski, Cordelia Langford, John Sillitoe on behalf of the Wellcome Sanger Institute COVID-19 Surveillance Team |
| EPI_ISL_3169103, EPI_ISL_3169104, EPI_ISL_3169105, EPI_ISL_3169106, EPI_ISL_3169107 | Lighthouse Lab in Glasgow | Wellcome Sanger Institute for the COVID-19 Genomics UK (COG-UK) Consortium | Harper VanSteenhouse, Yumi Kasai, David Gray, Carol Clugston, Anna Dominiczak and Alex Alderton, Roberto Amato, Jeffrey Barrett, Sonia Goncalves, Ewan Harrison, David K. Jackson, Ian Johnston, Dominic Kwiatkowski, Cordelia Langford, John Sillitoe on behalf of the Wellcome Sanger Institute COVID-19 Surveillance Team |
| EPI_ISL_3169108 | Lighthouse Laboratory Plymouth | Wellcome Sanger Institute for the COVID-19 Genomics UK (COG-UK) Consortium | Lighthouse Laboratory Plymouth and Alex Alderton, Roberto Amato, Jeffrey Barrett, Sonia Goncalves, Ewan Harrison, David K. Jackson, Ian Johnston, Dominic Kwiatkowski, Cordelia Langford, John Sillitoe on behalf of the Wellcome Sanger Institute COVID-19 Surveillance Team |
| EPI_ISL_3169109, EPI_ISL_3169110, EPI_ISL_3169111, EPI_ISL_3169112, EPI_ISL_3169113 | Berkshire and Surrey Pathology Services Lighthouse Laboratory | Wellcome Sanger Institute for the COVID-19 Genomics UK (COG-UK) Consortium | Berkshire and Surrey Pathology Services Lighthouse Laboratory and Alex Alderton, Roberto Amato, Jeffrey Barrett, Sonia Goncalves, Ewan Harrison, David K. Jackson, Ian Johnston, Dominic Kwiatkowski, Cordelia Langford, John Sillitoe on behalf of the Wellcome Sanger Institute COVID-19 Surveillance Team |
| EPI_ISL_3169114 | Lighthouse Laboratory Plymouth | Wellcome Sanger Institute for the COVID-19 Genomics UK (COG-UK) Consortium | Lighthouse Laboratory Plymouth and Alex Alderton, Roberto Amato, Jeffrey Barrett, Sonia Goncalves, Ewan Harrison, David K. Jackson, Ian Johnston, Dominic Kwiatkowski, Cordelia Langford, John Sillitoe on behalf of the Wellcome Sanger Institute COVID-19 Surveillance Team |
| EPI_ISL_3169115 | Berkshire and Surrey Pathology Services Lighthouse Laboratory | Wellcome Sanger Institute for the COVID-19 Genomics UK (COG-UK) Consortium | Berkshire and Surrey Pathology Services Lighthouse Laboratory and Alex Alderton, Roberto Amato, Jeffrey Barrett, Sonia Goncalves, Ewan Harrison, David K. Jackson, Ian Johnston, Dominic Kwiatkowski, Cordelia Langford, John Sillitoe on behalf of the Wellcome Sanger Institute COVID-19 Surveillance Team |
| EPI_ISL_3169116, EPI_ISL_3169117, EPI_ISL_3169118 | Lighthouse Laboratory Plymouth | Wellcome Sanger Institute for the COVID-19 Genomics UK (COG-UK) Consortium | Lighthouse Laboratory Plymouth and Alex Alderton, Roberto Amato, Jeffrey Barrett, Sonia Goncalves, Ewan Harrison, David K. Jackson, Ian Johnston, Dominic Kwiatkowski, Cordelia Langford, John Sillitoe on behalf of the Wellcome Sanger Institute COVID-19 Surveillance Team |
| EPI_ISL_3169119 | Lighthouse Lab in Glasgow | Wellcome Sanger Institute for the COVID-19 Genomics UK (COG-UK) Consortium | Harper VanSteenhouse, Yumi Kasai, David Gray, Carol Clugston, Anna Dominiczak and Alex Alderton, Roberto Amato, Jeffrey Barrett, Sonia Goncalves, Ewan Harrison, David K. Jackson, Ian Johnston, Dominic Kwiatkowski, Cordelia Langford, John Sillitoe on behalf of the Wellcome Sanger Institute COVID-19 Surveillance Team |
| EPI_ISL_3169120, EPI_ISL_3169121 | Berkshire and Surrey Pathology Services Lighthouse Laboratory | Wellcome Sanger Institute for the COVID-19 Genomics UK (COG-UK) Consortium | Berkshire and Surrey Pathology Services Lighthouse Laboratory and Alex Alderton, Roberto Amato, Jeffrey Barrett, Sonia Goncalves, Ewan Harrison, David K. Jackson, Ian Johnston, Dominic Kwiatkowski, Cordelia Langford, John Sillitoe on behalf of the Wellcome Sanger Institute COVID-19 Surveillance Team |
| EPI_ISL_3169122 | Lighthouse Lab in Glasgow | Wellcome Sanger Institute for the COVID-19 Genomics UK (COG-UK) Consortium | Harper VanSteenhouse, Yumi Kasai, David Gray, Carol Clugston, Anna Dominiczak and Alex Alderton, Roberto Amato, Jeffrey Barrett, Sonia Goncalves, Ewan Harrison, David K. Jackson, Ian Johnston, Dominic Kwiatkowski, Cordelia Langford, John Sillitoe on behalf of the Wellcome Sanger Institute COVID-19 Surveillance Team |
| EPI_ISL_3169123, EPI_ISL_3169124 | Lighthouse Laboratory Plymouth | Wellcome Sanger Institute for the COVID-19 Genomics UK (COG-UK) Consortium | Lighthouse Laboratory Plymouth and Alex Alderton, Roberto Amato, Jeffrey Barrett, Sonia Goncalves, Ewan Harrison, David K. Jackson, Ian Johnston, Dominic Kwiatkowski, Cordelia Langford, John Sillitoe on behalf of the Wellcome Sanger Institute COVID-19 Surveillance Team |
| EPI_ISL_3169125 | Berkshire and Surrey Pathology Services Lighthouse Laboratory | Wellcome Sanger Institute for the COVID-19 Genomics UK (COG-UK) Consortium | Berkshire and Surrey Pathology Services Lighthouse Laboratory and Alex Alderton, Roberto Amato, Jeffrey Barrett, Sonia Goncalves, Ewan Harrison, David K. Jackson, Ian Johnston, Dominic Kwiatkowski, Cordelia Langford, John Sillitoe on behalf of the Wellcome Sanger Institute COVID-19 Surveillance Team |
| EPI_ISL_3169126 | Lighthouse Lab in Glasgow | Wellcome Sanger Institute for the COVID-19 Genomics UK (COG-UK) Consortium | Harper VanSteenhouse, Yumi Kasai, David Gray, Carol Clugston, Anna Dominiczak and Alex Alderton, Roberto Amato, Jeffrey Barrett, Sonia Goncalves, Ewan Harrison, David K. Jackson, Ian Johnston, Dominic Kwiatkowski, Cordelia Langford, John Sillitoe on behalf of the Wellcome Sanger Institute COVID-19 Surveillance Team |
| EPI_ISL_3169127 | Berkshire and Surrey Pathology Services Lighthouse Laboratory | Wellcome Sanger Institute for the COVID-19 Genomics UK (COG-UK) Consortium | Berkshire and Surrey Pathology Services Lighthouse Laboratory and Alex Alderton, Roberto Amato, Jeffrey Barrett, Sonia Goncalves, Ewan Harrison, David K. Jackson, Ian Johnston, Dominic Kwiatkowski, Cordelia Langford, John Sillitoe on behalf of the Wellcome Sanger Institute COVID-19 Surveillance Team |
| EPI_ISL_3169128 | Lighthouse Laboratory Plymouth | Wellcome Sanger Institute for the COVID-19 Genomics UK (COG-UK) Consortium | Lighthouse Laboratory Plymouth and Alex Alderton, Roberto Amato, Jeffrey Barrett, Sonia Goncalves, Ewan Harrison, David K. Jackson, Ian Johnston, Dominic Kwiatkowski, Cordelia Langford, John Sillitoe on behalf of the Wellcome Sanger Institute COVID-19 Surveillance Team |
| EPI_ISL_3169129 | Berkshire and Surrey Pathology Services Lighthouse Laboratory | Wellcome Sanger Institute for the COVID-19 Genomics UK (COG-UK) Consortium | Berkshire and Surrey Pathology Services Lighthouse Laboratory and Alex Alderton, Roberto Amato, Jeffrey Barrett, Sonia Goncalves, Ewan Harrison, David K. Jackson, Ian Johnston, Dominic Kwiatkowski, Cordelia Langford, John Sillitoe on behalf of the Wellcome Sanger Institute COVID-19 Surveillance Team |
| EPI_ISL_3169130 | Lighthouse Laboratory Plymouth | Wellcome Sanger Institute for the COVID-19 Genomics UK (COG-UK) Consortium | Lighthouse Laboratory Plymouth and Alex Alderton, Roberto Amato, Jeffrey Barrett, Sonia Goncalves, Ewan Harrison, David K. Jackson, Ian Johnston, Dominic Kwiatkowski, Cordelia Langford, John Sillitoe on behalf of the Wellcome Sanger Institute COVID-19 Surveillance Team |
| EPI_ISL_3169131, EPI_ISL_3169132 | Berkshire and Surrey Pathology Services Lighthouse Laboratory | Wellcome Sanger Institute for the COVID-19 Genomics UK (COG-UK) Consortium | Berkshire and Surrey Pathology Services Lighthouse Laboratory and Alex Alderton, Roberto Amato, Jeffrey Barrett, Sonia Goncalves, Ewan Harrison, David K. Jackson, Ian Johnston, Dominic Kwiatkowski, Cordelia Langford, John Sillitoe on behalf of the Wellcome Sanger Institute COVID-19 Surveillance Team |

| | | | |
|---|---|---|---|
| EPI_ISL_3169133 | Lighthouse Lab in Glasgow | Wellcome Sanger Institute for the COVID-19 Genomics UK (COG-UK) Consortium | Harper VanSteenhouse, Yumi Kasai, David Gray, Carol Clugston, Anna Dominiczak and Alex Alderton, Roberto Amato, Jeffrey Barrett, Sonia Goncalves, Ewan Harrison, David K. Jackson, Ian Johnston, Dominic Kwiatkowski, Cordelia Langford, John Sillitoe on behalf of the Wellcome Sanger Institute COVID-19 Surveillance Team |
| EPI_ISL_3169134, EPI_ISL_3169135, EPI_ISL_3169136, EPI_ISL_3169137 | Berkshire and Surrey Pathology Services Lighthouse Laboratory | Wellcome Sanger Institute for the COVID-19 Genomics UK (COG-UK) Consortium | Berkshire and Surrey Pathology Services Lighthouse Laboratory and Alex Alderton, Roberto Amato, Jeffrey Barrett, Sonia Goncalves, Ewan Harrison, David K. Jackson, Ian Johnston, Dominic Kwiatkowski, Cordelia Langford, John Sillitoe on behalf of the Wellcome Sanger Institute COVID-19 Surveillance Team |
| EPI_ISL_3169138 | Lighthouse Laboratory Plymouth | Wellcome Sanger Institute for the COVID-19 Genomics UK (COG-UK) Consortium | Lighthouse Laboratory Plymouth and Alex Alderton, Roberto Amato, Jeffrey Barrett, Sonia Goncalves, Ewan Harrison, David K. Jackson, Ian Johnston, Dominic Kwiatkowski, Cordelia Langford, John Sillitoe on behalf of the Wellcome Sanger Institute COVID-19 Surveillance Team |
| EPI_ISL_3169139 | Berkshire and Surrey Pathology Services Lighthouse Laboratory | Wellcome Sanger Institute for the COVID-19 Genomics UK (COG-UK) Consortium | Berkshire and Surrey Pathology Services Lighthouse Laboratory and Alex Alderton, Roberto Amato, Jeffrey Barrett, Sonia Goncalves, Ewan Harrison, David K. Jackson, Ian Johnston, Dominic Kwiatkowski, Cordelia Langford, John Sillitoe on behalf of the Wellcome Sanger Institute COVID-19 Surveillance Team |
| EPI_ISL_3169140 | Lighthouse Lab in Glasgow | Wellcome Sanger Institute for the COVID-19 Genomics UK (COG-UK) Consortium | Harper VanSteenhouse, Yumi Kasai, David Gray, Carol Clugston, Anna Dominiczak and Alex Alderton, Roberto Amato, Jeffrey Barrett, Sonia Goncalves, Ewan Harrison, David K. Jackson, Ian Johnston, Dominic Kwiatkowski, Cordelia Langford, John Sillitoe on behalf of the Wellcome Sanger Institute COVID-19 Surveillance Team |
| EPI_ISL_3169141 | Berkshire and Surrey Pathology Services Lighthouse Laboratory | Wellcome Sanger Institute for the COVID-19 Genomics UK (COG-UK) Consortium | Berkshire and Surrey Pathology Services Lighthouse Laboratory and Alex Alderton, Roberto Amato, Jeffrey Barrett, Sonia Goncalves, Ewan Harrison, David K. Jackson, Ian Johnston, Dominic Kwiatkowski, Cordelia Langford, John Sillitoe on behalf of the Wellcome Sanger Institute COVID-19 Surveillance Team |
| EPI_ISL_3169142 | Lighthouse Laboratory Plymouth | Wellcome Sanger Institute for the COVID-19 Genomics UK (COG-UK) Consortium | Lighthouse Laboratory Plymouth and Alex Alderton, Roberto Amato, Jeffrey Barrett, Sonia Goncalves, Ewan Harrison, David K. Jackson, Ian Johnston, Dominic Kwiatkowski, Cordelia Langford, John Sillitoe on behalf of the Wellcome Sanger Institute COVID-19 Surveillance Team |
| EPI_ISL_3169143, EPI_ISL_3169144, EPI_ISL_3169145, EPI_ISL_3169146 | Lighthouse Lab in Glasgow | Wellcome Sanger Institute for the COVID-19 Genomics UK (COG-UK) Consortium | Harper VanSteenhouse, Yumi Kasai, David Gray, Carol Clugston, Anna Dominiczak and Alex Alderton, Roberto Amato, Jeffrey Barrett, Sonia Goncalves, Ewan Harrison, David K. Jackson, Ian Johnston, Dominic Kwiatkowski, Cordelia Langford, John Sillitoe on behalf of the Wellcome Sanger Institute COVID-19 Surveillance Team |
| EPI_ISL_3169147 | Berkshire and Surrey Pathology Services Lighthouse Laboratory | Wellcome Sanger Institute for the COVID-19 Genomics UK (COG-UK) Consortium | Berkshire and Surrey Pathology Services Lighthouse Laboratory and Alex Alderton, Roberto Amato, Jeffrey Barrett, Sonia Goncalves, Ewan Harrison, David K. Jackson, Ian Johnston, Dominic Kwiatkowski, Cordelia Langford, John Sillitoe on behalf of the Wellcome Sanger Institute COVID-19 Surveillance Team |
| EPI_ISL_3169148 | Lighthouse Laboratory Plymouth | Wellcome Sanger Institute for the COVID-19 Genomics UK (COG-UK) Consortium | Lighthouse Laboratory Plymouth and Alex Alderton, Roberto Amato, Jeffrey Barrett, Sonia Goncalves, Ewan Harrison, David K. Jackson, Ian Johnston, Dominic Kwiatkowski, Cordelia Langford, John Sillitoe on behalf of the Wellcome Sanger Institute COVID-19 Surveillance Team |
| EPI_ISL_3169149, EPI_ISL_3169150, EPI_ISL_3169151, EPI_ISL_3169152, EPI_ISL_3169153 | Lighthouse Lab in Glasgow | Wellcome Sanger Institute for the COVID-19 Genomics UK (COG-UK) Consortium | Harper VanSteenhouse, Yumi Kasai, David Gray, Carol Clugston, Anna Dominiczak and Alex Alderton, Roberto Amato, Jeffrey Barrett, Sonia Goncalves, Ewan Harrison, David K. Jackson, Ian Johnston, Dominic Kwiatkowski, Cordelia Langford, John Sillitoe on behalf of the Wellcome Sanger Institute COVID-19 Surveillance Team |
| EPI_ISL_3169154 | Berkshire and Surrey Pathology Services Lighthouse Laboratory | Wellcome Sanger Institute for the COVID-19 Genomics UK (COG-UK) Consortium | Berkshire and Surrey Pathology Services Lighthouse Laboratory and Alex Alderton, Roberto Amato, Jeffrey Barrett, Sonia Goncalves, Ewan Harrison, David K. Jackson, Ian Johnston, Dominic Kwiatkowski, Cordelia Langford, John Sillitoe on behalf of the Wellcome Sanger Institute COVID-19 Surveillance Team |
| EPI_ISL_3169155, EPI_ISL_3169156, EPI_ISL_3169157 | Lighthouse Lab in Glasgow | Wellcome Sanger Institute for the COVID-19 Genomics UK (COG-UK) Consortium | Harper VanSteenhouse, Yumi Kasai, David Gray, Carol Clugston, Anna Dominiczak and Alex Alderton, Roberto Amato, Jeffrey Barrett, Sonia Goncalves, Ewan Harrison, David K. Jackson, Ian Johnston, Dominic Kwiatkowski, Cordelia Langford, John Sillitoe on behalf of the Wellcome Sanger Institute COVID-19 Surveillance Team |
| EPI_ISL_3169158, EPI_ISL_3169159 | Berkshire and Surrey Pathology Services Lighthouse Laboratory | Wellcome Sanger Institute for the COVID-19 Genomics UK (COG-UK) Consortium | Berkshire and Surrey Pathology Services Lighthouse Laboratory and Alex Alderton, Roberto Amato, Jeffrey Barrett, Sonia Goncalves, Ewan Harrison, David K. Jackson, Ian Johnston, Dominic Kwiatkowski, Cordelia Langford, John Sillitoe on behalf of the Wellcome Sanger Institute COVID-19 Surveillance Team |
| EPI_ISL_3169160, EPI_ISL_3169161, EPI_ISL_3169162 | Lighthouse Laboratory Plymouth | Wellcome Sanger Institute for the COVID-19 Genomics UK (COG-UK) Consortium | Lighthouse Laboratory Plymouth and Alex Alderton, Roberto Amato, Jeffrey Barrett, Sonia Goncalves, Ewan Harrison, David K. Jackson, Ian Johnston, Dominic Kwiatkowski, Cordelia Langford, John Sillitoe on behalf of the Wellcome Sanger Institute COVID-19 Surveillance Team |
| EPI_ISL_3169163 | Lighthouse Lab in Glasgow | Wellcome Sanger Institute for the COVID-19 Genomics UK (COG-UK) Consortium | Harper VanSteenhouse, Yumi Kasai, David Gray, Carol Clugston, Anna Dominiczak and Alex Alderton, Roberto Amato, Jeffrey Barrett, Sonia Goncalves, Ewan Harrison, David K. Jackson, Ian Johnston, Dominic Kwiatkowski, Cordelia Langford, John Sillitoe on behalf of the Wellcome Sanger Institute COVID-19 Surveillance Team |
| EPI_ISL_3169164 | Lighthouse Laboratory Plymouth | Wellcome Sanger Institute for the COVID-19 Genomics UK (COG-UK) Consortium | Lighthouse Laboratory Plymouth and Alex Alderton, Roberto Amato, Jeffrey Barrett, Sonia Goncalves, Ewan Harrison, David K. Jackson, Ian Johnston, Dominic Kwiatkowski, Cordelia Langford, John Sillitoe on behalf of the Wellcome Sanger Institute COVID-19 Surveillance Team |
| EPI_ISL_3169165 | Lighthouse Lab in Glasgow | Wellcome Sanger Institute for the COVID-19 Genomics UK (COG-UK) Consortium | Harper VanSteenhouse, Yumi Kasai, David Gray, Carol Clugston, Anna Dominiczak and Alex Alderton, Roberto Amato, Jeffrey Barrett, Sonia Goncalves, Ewan Harrison, David K. Jackson, Ian Johnston, Dominic Kwiatkowski, Cordelia Langford, John Sillitoe on behalf of the Wellcome Sanger Institute COVID-19 Surveillance Team |
| EPI_ISL_3169166 | Berkshire and Surrey Pathology Services Lighthouse Laboratory | Wellcome Sanger Institute for the COVID-19 Genomics UK (COG-UK) Consortium | Berkshire and Surrey Pathology Services Lighthouse Laboratory and Alex Alderton, Roberto Amato, Jeffrey Barrett, Sonia Goncalves, Ewan Harrison, David K. Jackson, Ian Johnston, Dominic Kwiatkowski, Cordelia Langford, John Sillitoe on behalf of the Wellcome Sanger Institute COVID-19 Surveillance Team |
| EPI_ISL_3169167 | Lighthouse Lab in Glasgow | Wellcome Sanger Institute for the COVID-19 Genomics UK (COG-UK) Consortium | Harper VanSteenhouse, Yumi Kasai, David Gray, Carol Clugston, Anna Dominiczak and Alex Alderton, Roberto Amato, Jeffrey Barrett, Sonia Goncalves, Ewan Harrison, David K. Jackson, Ian Johnston, Dominic Kwiatkowski, Cordelia Langford, John Sillitoe on behalf of the Wellcome Sanger Institute COVID-19 Surveillance Team |
| EPI_ISL_3169168, EPI_ISL_3169169 | Berkshire and Surrey Pathology Services Lighthouse Laboratory | Wellcome Sanger Institute for the COVID-19 Genomics UK (COG-UK) Consortium | Berkshire and Surrey Pathology Services Lighthouse Laboratory and Alex Alderton, Roberto Amato, Jeffrey Barrett, Sonia Goncalves, Ewan Harrison, David K. Jackson, Ian Johnston, Dominic Kwiatkowski, Cordelia Langford, John Sillitoe on behalf of the Wellcome Sanger Institute COVID-19 Surveillance Team |
| EPI_ISL_3169170 | Lighthouse Lab in Glasgow | Wellcome Sanger Institute for the COVID-19 Genomics UK (COG-UK) Consortium | Harper VanSteenhouse, Yumi Kasai, David Gray, Carol Clugston, Anna Dominiczak and Alex Alderton, Roberto Amato, Jeffrey Barrett, Sonia Goncalves, Ewan Harrison, David K. Jackson, Ian Johnston, Dominic Kwiatkowski, Cordelia Langford, John Sillitoe on behalf of the Wellcome Sanger Institute COVID-19 Surveillance Team |
| EPI_ISL_3169171 | Berkshire and Surrey Pathology Services Lighthouse Laboratory | Wellcome Sanger Institute for the COVID-19 Genomics UK (COG-UK) Consortium | Berkshire and Surrey Pathology Services Lighthouse Laboratory and Alex Alderton, Roberto Amato, Jeffrey Barrett, Sonia Goncalves, Ewan Harrison, David K. Jackson, Ian Johnston, Dominic Kwiatkowski, Cordelia Langford, John Sillitoe on behalf of the Wellcome Sanger Institute COVID-19 Surveillance Team |
| EPI_ISL_3169172 | Lighthouse Lab in Glasgow | Wellcome Sanger Institute for the COVID-19 Genomics UK (COG-UK) Consortium | Harper VanSteenhouse, Yumi Kasai, David Gray, Carol Clugston, Anna Dominiczak and Alex Alderton, Roberto Amato, Jeffrey Barrett, Sonia Goncalves, Ewan Harrison, David K. Jackson, Ian Johnston, Dominic Kwiatkowski, Cordelia Langford, John Sillitoe on behalf of the Wellcome Sanger Institute COVID-19 Surveillance Team |
| EPI_ISL_3169173, EPI_ISL_3169174 | Lighthouse Laboratory Plymouth | Wellcome Sanger Institute for the COVID-19 Genomics UK (COG-UK) Consortium | Lighthouse Laboratory Plymouth and Alex Alderton, Roberto Amato, Jeffrey Barrett, Sonia Goncalves, Ewan Harrison, David K. Jackson, Ian Johnston, Dominic Kwiatkowski, Cordelia Langford, John Sillitoe on behalf of the Wellcome Sanger Institute COVID-19 Surveillance Team |
| EPI_ISL_3169175, EPI_ISL_3169176, EPI_ISL_3169177 | Berkshire and Surrey Pathology Services Lighthouse Laboratory | Wellcome Sanger Institute for the COVID-19 Genomics UK (COG-UK) Consortium | Berkshire and Surrey Pathology Services Lighthouse Laboratory and Alex Alderton, Roberto Amato, Jeffrey Barrett, Sonia Goncalves, Ewan Harrison, David K. Jackson, Ian Johnston, Dominic Kwiatkowski, Cordelia Langford, John Sillitoe on behalf of the Wellcome Sanger Institute COVID-19 Surveillance Team |
| EPI_ISL_3169178 | Lighthouse Laboratory Plymouth | Wellcome Sanger Institute for the COVID-19 Genomics UK (COG-UK) Consortium | Lighthouse Laboratory Plymouth and Alex Alderton, Roberto Amato, Jeffrey Barrett, Sonia Goncalves, Ewan Harrison, David K. Jackson, Ian Johnston, Dominic Kwiatkowski, Cordelia Langford, John Sillitoe on behalf of the Wellcome Sanger Institute COVID-19 Surveillance Team |
| EPI_ISL_3169179 | Lighthouse Lab in Glasgow | Wellcome Sanger Institute for the COVID-19 Genomics UK (COG-UK) Consortium | Harper VanSteenhouse, Yumi Kasai, David Gray, Carol Clugston, Anna Dominiczak and Alex Alderton, Roberto Amato, Jeffrey Barrett, Sonia Goncalves, Ewan Harrison, David K. Jackson, Ian Johnston, Dominic Kwiatkowski, Cordelia Langford, John Sillitoe on behalf of the Wellcome Sanger Institute COVID-19 Surveillance Team |
| EPI_ISL_3169180 | Berkshire and Surrey Pathology Services Lighthouse Laboratory | Wellcome Sanger Institute for the COVID-19 Genomics UK (COG-UK) Consortium | Berkshire and Surrey Pathology Services Lighthouse Laboratory and Alex Alderton, Roberto Amato, Jeffrey Barrett, Sonia Goncalves, Ewan Harrison, David K. Jackson, Ian Johnston, Dominic Kwiatkowski, Cordelia Langford, John Sillitoe on behalf of the Wellcome Sanger Institute COVID-19 Surveillance Team |
| EPI_ISL_3169181, EPI_ISL_3169182 | Lighthouse Lab in Glasgow | Wellcome Sanger Institute for the COVID-19 Genomics UK (COG-UK) Consortium | Harper VanSteenhouse, Yumi Kasai, David Gray, Carol Clugston, Anna Dominiczak and Alex Alderton, Roberto Amato, Jeffrey Barrett, Sonia Goncalves, Ewan Harrison, David K. Jackson, Ian Johnston, Dominic Kwiatkowski, Cordelia Langford, John Sillitoe on behalf of the Wellcome Sanger Institute COVID-19 Surveillance Team |
| EPI_ISL_3169183 | Lighthouse Laboratory Plymouth | Wellcome Sanger Institute for the COVID-19 Genomics UK (COG-UK) Consortium | Lighthouse Laboratory Plymouth and Alex Alderton, Roberto Amato, Jeffrey Barrett, Sonia Goncalves, Ewan Harrison, David K. Jackson, Ian Johnston, Dominic Kwiatkowski, Cordelia Langford, John Sillitoe on behalf of the Wellcome Sanger Institute COVID-19 Surveillance Team |
| EPI_ISL_3169184 | Lighthouse Lab in Glasgow | Wellcome Sanger Institute for the COVID-19 Genomics UK (COG-UK) Consortium | Harper VanSteenhouse, Yumi Kasai, David Gray, Carol Clugston, Anna Dominiczak and Alex Alderton, Roberto Amato, Jeffrey Barrett, Sonia Goncalves, Ewan Harrison, David K. Jackson, Ian Johnston, Dominic Kwiatkowski, Cordelia Langford, John Sillitoe on behalf of the Wellcome Sanger Institute COVID-19 Surveillance Team |
| EPI_ISL_3169185 | Lighthouse Laboratory Plymouth | Wellcome Sanger Institute for the COVID-19 Genomics UK (COG-UK) Consortium | Lighthouse Laboratory Plymouth and Alex Alderton, Roberto Amato, Jeffrey Barrett, Sonia Goncalves, Ewan Harrison, David K. Jackson, Ian Johnston, Dominic Kwiatkowski, Cordelia Langford, John Sillitoe on behalf of the Wellcome Sanger Institute COVID-19 Surveillance Team |
| EPI_ISL_3169186 | Lighthouse Lab in Glasgow | Wellcome Sanger Institute for the COVID-19 Genomics UK (COG-UK) Consortium | Harper VanSteenhouse, Yumi Kasai, David Gray, Carol Clugston, Anna Dominiczak and Alex Alderton, Roberto Amato, Jeffrey Barrett, Sonia Goncalves, Ewan Harrison, David K. Jackson, Ian Johnston, Dominic Kwiatkowski, Cordelia Langford, John Sillitoe on behalf of the Wellcome Sanger Institute COVID-19 Surveillance Team |
| EPI_ISL_3169187 | Berkshire and Surrey Pathology Services Lighthouse Laboratory | Wellcome Sanger Institute for the COVID-19 Genomics UK (COG-UK) Consortium | Berkshire and Surrey Pathology Services Lighthouse Laboratory and Alex Alderton, Roberto Amato, Jeffrey Barrett, Sonia Goncalves, Ewan Harrison, David K. Jackson, Ian Johnston, Dominic Kwiatkowski, Cordelia Langford, John Sillitoe on behalf of the Wellcome Sanger Institute COVID-19 Surveillance Team |
| EPI_ISL_3169188, EPI_ISL_3169189 | Lighthouse Laboratory Plymouth | Wellcome Sanger Institute for the COVID-19 Genomics UK (COG-UK) Consortium | Lighthouse Laboratory Plymouth and Alex Alderton, Roberto Amato, Jeffrey Barrett, Sonia Goncalves, Ewan Harrison, David K. Jackson, Ian Johnston, Dominic Kwiatkowski, Cordelia Langford, John Sillitoe on behalf of the Wellcome Sanger Institute COVID-19 Surveillance Team |
| EPI_ISL_3169190 | Lighthouse Lab in Glasgow | Wellcome Sanger Institute for the COVID-19 Genomics UK (COG-UK) Consortium | Harper VanSteenhouse, Yumi Kasai, David Gray, Carol Clugston, Anna Dominiczak and Alex Alderton, Roberto Amato, Jeffrey Barrett, Sonia Goncalves, Ewan Harrison, David K. Jackson, Ian Johnston, Dominic Kwiatkowski, Cordelia Langford, John Sillitoe on behalf of the Wellcome Sanger Institute COVID-19 Surveillance Team |
| EPI_ISL_3169191 | Lighthouse Laboratory Plymouth | Wellcome Sanger Institute for the COVID-19 Genomics UK (COG-UK) Consortium | Lighthouse Laboratory Plymouth and Alex Alderton, Roberto Amato, Jeffrey Barrett, Sonia Goncalves, Ewan Harrison, David K. Jackson, Ian Johnston, Dominic Kwiatkowski, Cordelia Langford, John Sillitoe on behalf of the Wellcome Sanger Institute COVID-19 Surveillance Team |
| EPI_ISL_3169192 | Lighthouse Lab in Glasgow | Wellcome Sanger Institute for the COVID-19 Genomics UK (COG-UK) Consortium | Harper VanSteenhouse, Yumi Kasai, David Gray, Carol Clugston, Anna Dominiczak and Alex Alderton, Roberto Amato, Jeffrey Barrett, Sonia Goncalves, Ewan Harrison, David K. Jackson, Ian Johnston, Dominic Kwiatkowski, Cordelia Langford, John Sillitoe on behalf of the Wellcome Sanger Institute COVID-19 Surveillance Team |
| EPI_ISL_3169193 | Berkshire and Surrey Pathology Services Lighthouse Laboratory | Wellcome Sanger Institute for the COVID-19 Genomics UK (COG-UK) Consortium | Berkshire and Surrey Pathology Services Lighthouse Laboratory and Alex Alderton, Roberto Amato, Jeffrey Barrett, Sonia Goncalves, Ewan Harrison, David K. Jackson, Ian Johnston, Dominic Kwiatkowski, Cordelia Langford, John Sillitoe on behalf of the Wellcome Sanger Institute COVID-19 Surveillance Team |
| EPI_ISL_3169194, EPI_ISL_3169195, | Lighthouse Lab in Glasgow | Wellcome Sanger Institute for the COVID-19 Genomics UK (COG-UK) Consortium | Harper VanSteenhouse, Yumi Kasai, David Gray, Carol Clugston, Anna Dominiczak and Alex Alderton, Roberto Amato, Jeffrey Barrett, Sonia Goncalves, Ewan Harrison, David K. Jackson, Ian Johnston, Dominic Kwiatkowski, Cordelia Langford, John Sillitoe on behalf of the Wellcome Sanger Institute COVID-19 Surveillance Team |

| | | | |
|---|---|---|---|
| EPI_ISL_3169196, EPI_ISL_3169197 | | | |
| EPI_ISL_3169198 | Lighthouse Laboratory Plymouth | Wellcome Sanger Institute for the COVID-19 Genomics UK (COG-UK) Consortium | Lighthouse Laboratory Plymouth and Alex Alderton, Roberto Amato, Jeffrey Barrett, Sonia Goncalves, Ewan Harrison, David K. Jackson, Ian Johnston, Dominic Kwiatkowski, Cordelia Langford, John Sillitoe on behalf of the Wellcome Sanger Institute COVID-19 Surveillance Team |
| EPI_ISL_3169199, EPI_ISL_3169200, EPI_ISL_3169201 | Berkshire and Surrey Pathology Services Lighthouse Laboratory | Wellcome Sanger Institute for the COVID-19 Genomics UK (COG-UK) Consortium | Berkshire and Surrey Pathology Services Lighthouse Laboratory and Alex Alderton, Roberto Amato, Jeffrey Barrett, Sonia Goncalves, Ewan Harrison, David K. Jackson, Ian Johnston, Dominic Kwiatkowski, Cordelia Langford, John Sillitoe on behalf of the Wellcome Sanger Institute COVID-19 Surveillance Team |
| EPI_ISL_3169202, EPI_ISL_3169203 | Lighthouse Laboratory Plymouth | Wellcome Sanger Institute for the COVID-19 Genomics UK (COG-UK) Consortium | Lighthouse Laboratory Plymouth and Alex Alderton, Roberto Amato, Jeffrey Barrett, Sonia Goncalves, Ewan Harrison, David K. Jackson, Ian Johnston, Dominic Kwiatkowski, Cordelia Langford, John Sillitoe on behalf of the Wellcome Sanger Institute COVID-19 Surveillance Team |
| EPI_ISL_3169204 | Berkshire and Surrey Pathology Services Lighthouse Laboratory | Wellcome Sanger Institute for the COVID-19 Genomics UK (COG-UK) Consortium | Berkshire and Surrey Pathology Services Lighthouse Laboratory and Alex Alderton, Roberto Amato, Jeffrey Barrett, Sonia Goncalves, Ewan Harrison, David K. Jackson, Ian Johnston, Dominic Kwiatkowski, Cordelia Langford, John Sillitoe on behalf of the Wellcome Sanger Institute COVID-19 Surveillance Team |
| EPI_ISL_3169205 | Lighthouse Laboratory Plymouth | Wellcome Sanger Institute for the COVID-19 Genomics UK (COG-UK) Consortium | Lighthouse Laboratory Plymouth and Alex Alderton, Roberto Amato, Jeffrey Barrett, Sonia Goncalves, Ewan Harrison, David K. Jackson, Ian Johnston, Dominic Kwiatkowski, Cordelia Langford, John Sillitoe on behalf of the Wellcome Sanger Institute COVID-19 Surveillance Team |
| EPI_ISL_3169206, EPI_ISL_3169207, EPI_ISL_3169208, EPI_ISL_3169209 | Berkshire and Surrey Pathology Services Lighthouse Laboratory | Wellcome Sanger Institute for the COVID-19 Genomics UK (COG-UK) Consortium | Berkshire and Surrey Pathology Services Lighthouse Laboratory and Alex Alderton, Roberto Amato, Jeffrey Barrett, Sonia Goncalves, Ewan Harrison, David K. Jackson, Ian Johnston, Dominic Kwiatkowski, Cordelia Langford, John Sillitoe on behalf of the Wellcome Sanger Institute COVID-19 Surveillance Team |
| EPI_ISL_3169210, EPI_ISL_3169211 | Lighthouse Laboratory Plymouth | Wellcome Sanger Institute for the COVID-19 Genomics UK (COG-UK) Consortium | Lighthouse Laboratory Plymouth and Alex Alderton, Roberto Amato, Jeffrey Barrett, Sonia Goncalves, Ewan Harrison, David K. Jackson, Ian Johnston, Dominic Kwiatkowski, Cordelia Langford, John Sillitoe on behalf of the Wellcome Sanger Institute COVID-19 Surveillance Team |
| EPI_ISL_3169212 | Lighthouse Lab in Glasgow | Wellcome Sanger Institute for the COVID-19 Genomics UK (COG-UK) Consortium | Harper VanSteenhouse, Yumi Kasai, David Gray, Carol Clugston, Anna Dominiczak and Alex Alderton, Roberto Amato, Jeffrey Barrett, Sonia Goncalves, Ewan Harrison, David K. Jackson, Ian Johnston, Dominic Kwiatkowski, Cordelia Langford, John Sillitoe on behalf of the Wellcome Sanger Institute COVID-19 Surveillance Team |
| EPI_ISL_3169213, EPI_ISL_3169214 | Lighthouse Laboratory Plymouth | Wellcome Sanger Institute for the COVID-19 Genomics UK (COG-UK) Consortium | Lighthouse Laboratory Plymouth and Alex Alderton, Roberto Amato, Jeffrey Barrett, Sonia Goncalves, Ewan Harrison, David K. Jackson, Ian Johnston, Dominic Kwiatkowski, Cordelia Langford, John Sillitoe on behalf of the Wellcome Sanger Institute COVID-19 Surveillance Team |
| EPI_ISL_3169215 | Lighthouse Lab in Glasgow | Wellcome Sanger Institute for the COVID-19 Genomics UK (COG-UK) Consortium | Harper VanSteenhouse, Yumi Kasai, David Gray, Carol Clugston, Anna Dominiczak and Alex Alderton, Roberto Amato, Jeffrey Barrett, Sonia Goncalves, Ewan Harrison, David K. Jackson, Ian Johnston, Dominic Kwiatkowski, Cordelia Langford, John Sillitoe on behalf of the Wellcome Sanger Institute COVID-19 Surveillance Team |
| EPI_ISL_3169216 | Berkshire and Surrey Pathology Services Lighthouse Laboratory | Wellcome Sanger Institute for the COVID-19 Genomics UK (COG-UK) Consortium | Berkshire and Surrey Pathology Services Lighthouse Laboratory and Alex Alderton, Roberto Amato, Jeffrey Barrett, Sonia Goncalves, Ewan Harrison, David K. Jackson, Ian Johnston, Dominic Kwiatkowski, Cordelia Langford, John Sillitoe on behalf of the Wellcome Sanger Institute COVID-19 Surveillance Team |
| EPI_ISL_3169217, EPI_ISL_3169218 | Lighthouse Lab in Glasgow | Wellcome Sanger Institute for the COVID-19 Genomics UK (COG-UK) Consortium | Harper VanSteenhouse, Yumi Kasai, David Gray, Carol Clugston, Anna Dominiczak and Alex Alderton, Roberto Amato, Jeffrey Barrett, Sonia Goncalves, Ewan Harrison, David K. Jackson, Ian Johnston, Dominic Kwiatkowski, Cordelia Langford, John Sillitoe on behalf of the Wellcome Sanger Institute COVID-19 Surveillance Team |
| EPI_ISL_3169219 | Berkshire and Surrey Pathology Services Lighthouse Laboratory | Wellcome Sanger Institute for the COVID-19 Genomics UK (COG-UK) Consortium | Berkshire and Surrey Pathology Services Lighthouse Laboratory and Alex Alderton, Roberto Amato, Jeffrey Barrett, Sonia Goncalves, Ewan Harrison, David K. Jackson, Ian Johnston, Dominic Kwiatkowski, Cordelia Langford, John Sillitoe on behalf of the Wellcome Sanger Institute COVID-19 Surveillance Team |
| EPI_ISL_3169220, EPI_ISL_3169221 | Lighthouse Lab in Glasgow | Wellcome Sanger Institute for the COVID-19 Genomics UK (COG-UK) Consortium | Harper VanSteenhouse, Yumi Kasai, David Gray, Carol Clugston, Anna Dominiczak and Alex Alderton, Roberto Amato, Jeffrey Barrett, Sonia Goncalves, Ewan Harrison, David K. Jackson, Ian Johnston, Dominic Kwiatkowski, Cordelia Langford, John Sillitoe on behalf of the Wellcome Sanger Institute COVID-19 Surveillance Team |
| EPI_ISL_3169222, EPI_ISL_3169223 | Berkshire and Surrey Pathology Services Lighthouse Laboratory | Wellcome Sanger Institute for the COVID-19 Genomics UK (COG-UK) Consortium | Berkshire and Surrey Pathology Services Lighthouse Laboratory and Alex Alderton, Roberto Amato, Jeffrey Barrett, Sonia Goncalves, Ewan Harrison, David K. Jackson, Ian Johnston, Dominic Kwiatkowski, Cordelia Langford, John Sillitoe on behalf of the Wellcome Sanger Institute COVID-19 Surveillance Team |
| EPI_ISL_3169224 | Lighthouse Laboratory Plymouth | Wellcome Sanger Institute for the COVID-19 Genomics UK (COG-UK) Consortium | Lighthouse Laboratory Plymouth and Alex Alderton, Roberto Amato, Jeffrey Barrett, Sonia Goncalves, Ewan Harrison, David K. Jackson, Ian Johnston, Dominic Kwiatkowski, Cordelia Langford, John Sillitoe on behalf of the Wellcome Sanger Institute COVID-19 Surveillance Team |
| EPI_ISL_3169225 | Berkshire and Surrey Pathology Services Lighthouse Laboratory | Wellcome Sanger Institute for the COVID-19 Genomics UK (COG-UK) Consortium | Berkshire and Surrey Pathology Services Lighthouse Laboratory and Alex Alderton, Roberto Amato, Jeffrey Barrett, Sonia Goncalves, Ewan Harrison, David K. Jackson, Ian Johnston, Dominic Kwiatkowski, Cordelia Langford, John Sillitoe on behalf of the Wellcome Sanger Institute COVID-19 Surveillance Team |
| EPI_ISL_3169226 | Lighthouse Laboratory Plymouth | Wellcome Sanger Institute for the COVID-19 Genomics UK (COG-UK) Consortium | Lighthouse Laboratory Plymouth and Alex Alderton, Roberto Amato, Jeffrey Barrett, Sonia Goncalves, Ewan Harrison, David K. Jackson, Ian Johnston, Dominic Kwiatkowski, Cordelia Langford, John Sillitoe on behalf of the Wellcome Sanger Institute COVID-19 Surveillance Team |
| EPI_ISL_3169227 | Berkshire and Surrey Pathology Services Lighthouse Laboratory | Wellcome Sanger Institute for the COVID-19 Genomics UK (COG-UK) Consortium | Berkshire and Surrey Pathology Services Lighthouse Laboratory and Alex Alderton, Roberto Amato, Jeffrey Barrett, Sonia Goncalves, Ewan Harrison, David K. Jackson, Ian Johnston, Dominic Kwiatkowski, Cordelia Langford, John Sillitoe on behalf of the Wellcome Sanger Institute COVID-19 Surveillance Team |
| EPI_ISL_3169228 | Lighthouse Lab in Glasgow | Wellcome Sanger Institute for the COVID-19 Genomics UK (COG-UK) Consortium | Harper VanSteenhouse, Yumi Kasai, David Gray, Carol Clugston, Anna Dominiczak and Alex Alderton, Roberto Amato, Jeffrey Barrett, Sonia Goncalves, Ewan Harrison, David K. Jackson, Ian Johnston, Dominic Kwiatkowski, Cordelia Langford, John Sillitoe on behalf of the Wellcome Sanger Institute COVID-19 Surveillance Team |
| EPI_ISL_3169229 | Lighthouse Laboratory Plymouth | Wellcome Sanger Institute for the COVID-19 Genomics UK (COG-UK) Consortium | Lighthouse Laboratory Plymouth and Alex Alderton, Roberto Amato, Jeffrey Barrett, Sonia Goncalves, Ewan Harrison, David K. Jackson, Ian Johnston, Dominic Kwiatkowski, Cordelia Langford, John Sillitoe on behalf of the Wellcome Sanger Institute COVID-19 Surveillance Team |
| EPI_ISL_3169230, EPI_ISL_3169231, EPI_ISL_3169232 | Lighthouse Lab in Glasgow | Wellcome Sanger Institute for the COVID-19 Genomics UK (COG-UK) Consortium | Harper VanSteenhouse, Yumi Kasai, David Gray, Carol Clugston, Anna Dominiczak and Alex Alderton, Roberto Amato, Jeffrey Barrett, Sonia Goncalves, Ewan Harrison, David K. Jackson, Ian Johnston, Dominic Kwiatkowski, Cordelia Langford, John Sillitoe on behalf of the Wellcome Sanger Institute COVID-19 Surveillance Team |
| EPI_ISL_3169233 | Lighthouse Laboratory Plymouth | Wellcome Sanger Institute for the COVID-19 Genomics UK (COG-UK) Consortium | Lighthouse Laboratory Plymouth and Alex Alderton, Roberto Amato, Jeffrey Barrett, Sonia Goncalves, Ewan Harrison, David K. Jackson, Ian Johnston, Dominic Kwiatkowski, Cordelia Langford, John Sillitoe on behalf of the Wellcome Sanger Institute COVID-19 Surveillance Team |
| EPI_ISL_3169234 | Berkshire and Surrey Pathology Services Lighthouse Laboratory | Wellcome Sanger Institute for the COVID-19 Genomics UK (COG-UK) Consortium | Berkshire and Surrey Pathology Services Lighthouse Laboratory and Alex Alderton, Roberto Amato, Jeffrey Barrett, Sonia Goncalves, Ewan Harrison, David K. Jackson, Ian Johnston, Dominic Kwiatkowski, Cordelia Langford, John Sillitoe on behalf of the Wellcome Sanger Institute COVID-19 Surveillance Team |
| EPI_ISL_3169235 | Lighthouse Laboratory Plymouth | Wellcome Sanger Institute for the COVID-19 Genomics UK (COG-UK) Consortium | Lighthouse Laboratory Plymouth and Alex Alderton, Roberto Amato, Jeffrey Barrett, Sonia Goncalves, Ewan Harrison, David K. Jackson, Ian Johnston, Dominic Kwiatkowski, Cordelia Langford, John Sillitoe on behalf of the Wellcome Sanger Institute COVID-19 Surveillance Team |
| EPI_ISL_3169236, EPI_ISL_3169237 | Lighthouse Lab in Glasgow | Wellcome Sanger Institute for the COVID-19 Genomics UK (COG-UK) Consortium | Harper VanSteenhouse, Yumi Kasai, David Gray, Carol Clugston, Anna Dominiczak and Alex Alderton, Roberto Amato, Jeffrey Barrett, Sonia Goncalves, Ewan Harrison, David K. Jackson, Ian Johnston, Dominic Kwiatkowski, Cordelia Langford, John Sillitoe on behalf of the Wellcome Sanger Institute COVID-19 Surveillance Team |
| EPI_ISL_3169238 | Lighthouse Laboratory Plymouth | Wellcome Sanger Institute for the COVID-19 Genomics UK (COG-UK) Consortium | Lighthouse Laboratory Plymouth and Alex Alderton, Roberto Amato, Jeffrey Barrett, Sonia Goncalves, Ewan Harrison, David K. Jackson, Ian Johnston, Dominic Kwiatkowski, Cordelia Langford, John Sillitoe on behalf of the Wellcome Sanger Institute COVID-19 Surveillance Team |
| EPI_ISL_3169239 | Lighthouse Lab in Glasgow | Wellcome Sanger Institute for the COVID-19 Genomics UK (COG-UK) Consortium | Harper VanSteenhouse, Yumi Kasai, David Gray, Carol Clugston, Anna Dominiczak and Alex Alderton, Roberto Amato, Jeffrey Barrett, Sonia Goncalves, Ewan Harrison, David K. Jackson, Ian Johnston, Dominic Kwiatkowski, Cordelia Langford, John Sillitoe on behalf of the Wellcome Sanger Institute COVID-19 Surveillance Team |
| EPI_ISL_3169240, EPI_ISL_3169241 | Lighthouse Laboratory Plymouth | Wellcome Sanger Institute for the COVID-19 Genomics UK (COG-UK) Consortium | Lighthouse Laboratory Plymouth and Alex Alderton, Roberto Amato, Jeffrey Barrett, Sonia Goncalves, Ewan Harrison, David K. Jackson, Ian Johnston, Dominic Kwiatkowski, Cordelia Langford, John Sillitoe on behalf of the Wellcome Sanger Institute COVID-19 Surveillance Team |
| EPI_ISL_3169242, EPI_ISL_3169243, EPI_ISL_3169244, EPI_ISL_3169245 | Lighthouse Lab in Glasgow | Wellcome Sanger Institute for the COVID-19 Genomics UK (COG-UK) Consortium | Harper VanSteenhouse, Yumi Kasai, David Gray, Carol Clugston, Anna Dominiczak and Alex Alderton, Roberto Amato, Jeffrey Barrett, Sonia Goncalves, Ewan Harrison, David K. Jackson, Ian Johnston, Dominic Kwiatkowski, Cordelia Langford, John Sillitoe on behalf of the Wellcome Sanger Institute COVID-19 Surveillance Team |
| EPI_ISL_3169246 | Berkshire and Surrey Pathology Services Lighthouse Laboratory | Wellcome Sanger Institute for the COVID-19 Genomics UK (COG-UK) Consortium | Berkshire and Surrey Pathology Services Lighthouse Laboratory and Alex Alderton, Roberto Amato, Jeffrey Barrett, Sonia Goncalves, Ewan Harrison, David K. Jackson, Ian Johnston, Dominic Kwiatkowski, Cordelia Langford, John Sillitoe on behalf of the Wellcome Sanger Institute COVID-19 Surveillance Team |
| EPI_ISL_3169247 | Lighthouse Laboratory Plymouth | Wellcome Sanger Institute for the COVID-19 Genomics UK (COG-UK) Consortium | Lighthouse Laboratory Plymouth and Alex Alderton, Roberto Amato, Jeffrey Barrett, Sonia Goncalves, Ewan Harrison, David K. Jackson, Ian Johnston, Dominic Kwiatkowski, Cordelia Langford, John Sillitoe on behalf of the Wellcome Sanger Institute COVID-19 Surveillance Team |
| EPI_ISL_3169248 | Berkshire and Surrey Pathology Services Lighthouse Laboratory | Wellcome Sanger Institute for the COVID-19 Genomics UK (COG-UK) Consortium | Berkshire and Surrey Pathology Services Lighthouse Laboratory and Alex Alderton, Roberto Amato, Jeffrey Barrett, Sonia Goncalves, Ewan Harrison, David K. Jackson, Ian Johnston, Dominic Kwiatkowski, Cordelia Langford, John Sillitoe on behalf of the Wellcome Sanger Institute COVID-19 Surveillance Team |
| EPI_ISL_3169249, EPI_ISL_3169250 | Lighthouse Lab in Glasgow | Wellcome Sanger Institute for the COVID-19 Genomics UK (COG-UK) Consortium | Harper VanSteenhouse, Yumi Kasai, David Gray, Carol Clugston, Anna Dominiczak and Alex Alderton, Roberto Amato, Jeffrey Barrett, Sonia Goncalves, Ewan Harrison, David K. Jackson, Ian Johnston, Dominic Kwiatkowski, Cordelia Langford, John Sillitoe on behalf of the Wellcome Sanger Institute COVID-19 Surveillance Team |
| EPI_ISL_3169251, EPI_ISL_3169252, EPI_ISL_3169253 | Lighthouse Laboratory Plymouth | Wellcome Sanger Institute for the COVID-19 Genomics UK (COG-UK) Consortium | Lighthouse Laboratory Plymouth and Alex Alderton, Roberto Amato, Jeffrey Barrett, Sonia Goncalves, Ewan Harrison, David K. Jackson, Ian Johnston, Dominic Kwiatkowski, Cordelia Langford, John Sillitoe on behalf of the Wellcome Sanger Institute COVID-19 Surveillance Team |
| EPI_ISL_3169254, EPI_ISL_3169255, EPI_ISL_3169256 | Lighthouse Lab in Glasgow | Wellcome Sanger Institute for the COVID-19 Genomics UK (COG-UK) Consortium | Harper VanSteenhouse, Yumi Kasai, David Gray, Carol Clugston, Anna Dominiczak and Alex Alderton, Roberto Amato, Jeffrey Barrett, Sonia Goncalves, Ewan Harrison, David K. Jackson, Ian Johnston, Dominic Kwiatkowski, Cordelia Langford, John Sillitoe on behalf of the Wellcome Sanger Institute COVID-19 Surveillance Team |
| EPI_ISL_3169257 | Berkshire and Surrey Pathology Services Lighthouse Laboratory | Wellcome Sanger Institute for the COVID-19 Genomics UK (COG-UK) Consortium | Berkshire and Surrey Pathology Services Lighthouse Laboratory and Alex Alderton, Roberto Amato, Jeffrey Barrett, Sonia Goncalves, Ewan Harrison, David K. Jackson, Ian Johnston, Dominic Kwiatkowski, Cordelia Langford, John Sillitoe on behalf of the Wellcome Sanger Institute COVID-19 Surveillance Team |
| EPI_ISL_3169258 | Lighthouse Lab in Glasgow | Wellcome Sanger Institute for the COVID-19 Genomics UK (COG-UK) Consortium | Harper VanSteenhouse, Yumi Kasai, David Gray, Carol Clugston, Anna Dominiczak and Alex Alderton, Roberto Amato, Jeffrey Barrett, Sonia Goncalves, Ewan Harrison, David K. Jackson, Ian Johnston, Dominic Kwiatkowski, Cordelia Langford, John Sillitoe on behalf of the Wellcome Sanger Institute COVID-19 Surveillance Team |
| EPI_ISL_3169259, EPI_ISL_3169260 | Lighthouse Laboratory Plymouth | Wellcome Sanger Institute for the COVID-19 Genomics UK (COG-UK) Consortium | Lighthouse Laboratory Plymouth and Alex Alderton, Roberto Amato, Jeffrey Barrett, Sonia Goncalves, Ewan Harrison, David K. Jackson, Ian Johnston, Dominic Kwiatkowski, Cordelia Langford, John Sillitoe on behalf of the Wellcome Sanger Institute COVID-19 Surveillance Team |
| EPI_ISL_3169261, EPI_ISL_3169262 | Lighthouse Lab in Glasgow | Wellcome Sanger Institute for the COVID-19 Genomics UK (COG-UK) Consortium | Harper VanSteenhouse, Yumi Kasai, David Gray, Carol Clugston, Anna Dominiczak and Alex Alderton, Roberto Amato, Jeffrey Barrett, Sonia Goncalves, Ewan Harrison, David K. Jackson, Ian Johnston, Dominic Kwiatkowski, Cordelia Langford, John Sillitoe on behalf of the Wellcome Sanger Institute COVID-19 Surveillance Team |
| EPI_ISL_3169263 | Berkshire and Surrey Pathology Services Lighthouse Laboratory | Wellcome Sanger Institute for the COVID-19 Genomics UK (COG-UK) | Berkshire and Surrey Pathology Services Lighthouse Laboratory and Alex Alderton, Roberto Amato, Jeffrey Barrett, Sonia Goncalves, Ewan Harrison, David K. Jackson, Ian Johnston, Dominic Kwiatkowski, Cordelia Langford, John Sillitoe on behalf |

| | | | |
|---|---|---|---|
| | | Consortium | of the Wellcome Sanger Institute COVID-19 Surveillance Team |
| EPI_ISL_3169264 | Lighthouse Lab in Glasgow | Wellcome Sanger Institute for the COVID-19 Genomics UK (COG-UK) Consortium | Harper VanSteenhouse, Yumi Kasai, David Gray, Carol Clugston, Anna Dominiczak and Alex Alderton, Roberto Amato, Jeffrey Barrett, Sonia Goncalves, Ewan Harrison, David K. Jackson, Ian Johnston, Dominic Kwiatkowski, Cordelia Langford, John Sillitoe on behalf of the Wellcome Sanger Institute COVID-19 Surveillance Team |
| EPI_ISL_3169265 | Berkshire and Surrey Pathology Services Lighthouse Laboratory | Wellcome Sanger Institute for the COVID-19 Genomics UK (COG-UK) Consortium | Berkshire and Surrey Pathology Services Lighthouse Laboratory and Alex Alderton, Roberto Amato, Jeffrey Barrett, Sonia Goncalves, Ewan Harrison, David K. Jackson, Ian Johnston, Dominic Kwiatkowski, Cordelia Langford, John Sillitoe on behalf of the Wellcome Sanger Institute COVID-19 Surveillance Team |
| EPI_ISL_3169266, EPI_ISL_3169267 | Lighthouse Laboratory Plymouth | Wellcome Sanger Institute for the COVID-19 Genomics UK (COG-UK) Consortium | Lighthouse Laboratory Plymouth and Alex Alderton, Roberto Amato, Jeffrey Barrett, Sonia Goncalves, Ewan Harrison, David K. Jackson, Ian Johnston, Dominic Kwiatkowski, Cordelia Langford, John Sillitoe on behalf of the Wellcome Sanger Institute COVID-19 Surveillance Team |
| EPI_ISL_3169268, EPI_ISL_3169269 | Lighthouse Lab in Glasgow | Wellcome Sanger Institute for the COVID-19 Genomics UK (COG-UK) Consortium | Harper VanSteenhouse, Yumi Kasai, David Gray, Carol Clugston, Anna Dominiczak and Alex Alderton, Roberto Amato, Jeffrey Barrett, Sonia Goncalves, Ewan Harrison, David K. Jackson, Ian Johnston, Dominic Kwiatkowski, Cordelia Langford, John Sillitoe on behalf of the Wellcome Sanger Institute COVID-19 Surveillance Team |
| EPI_ISL_3169270 | Berkshire and Surrey Pathology Services Lighthouse Laboratory | Wellcome Sanger Institute for the COVID-19 Genomics UK (COG-UK) Consortium | Berkshire and Surrey Pathology Services Lighthouse Laboratory and Alex Alderton, Roberto Amato, Jeffrey Barrett, Sonia Goncalves, Ewan Harrison, David K. Jackson, Ian Johnston, Dominic Kwiatkowski, Cordelia Langford, John Sillitoe on behalf of the Wellcome Sanger Institute COVID-19 Surveillance Team |
| EPI_ISL_3169271 | Lighthouse Lab in Glasgow | Wellcome Sanger Institute for the COVID-19 Genomics UK (COG-UK) Consortium | Harper VanSteenhouse, Yumi Kasai, David Gray, Carol Clugston, Anna Dominiczak and Alex Alderton, Roberto Amato, Jeffrey Barrett, Sonia Goncalves, Ewan Harrison, David K. Jackson, Ian Johnston, Dominic Kwiatkowski, Cordelia Langford, John Sillitoe on behalf of the Wellcome Sanger Institute COVID-19 Surveillance Team |
| EPI_ISL_3169272 | Berkshire and Surrey Pathology Services Lighthouse Laboratory | Wellcome Sanger Institute for the COVID-19 Genomics UK (COG-UK) Consortium | Berkshire and Surrey Pathology Services Lighthouse Laboratory and Alex Alderton, Roberto Amato, Jeffrey Barrett, Sonia Goncalves, Ewan Harrison, David K. Jackson, Ian Johnston, Dominic Kwiatkowski, Cordelia Langford, John Sillitoe on behalf of the Wellcome Sanger Institute COVID-19 Surveillance Team |
| EPI_ISL_3169273 | Lighthouse Lab in Glasgow | Wellcome Sanger Institute for the COVID-19 Genomics UK (COG-UK) Consortium | Harper VanSteenhouse, Yumi Kasai, David Gray, Carol Clugston, Anna Dominiczak and Alex Alderton, Roberto Amato, Jeffrey Barrett, Sonia Goncalves, Ewan Harrison, David K. Jackson, Ian Johnston, Dominic Kwiatkowski, Cordelia Langford, John Sillitoe on behalf of the Wellcome Sanger Institute COVID-19 Surveillance Team |
| EPI_ISL_3169274 | Lighthouse Laboratory Plymouth | Wellcome Sanger Institute for the COVID-19 Genomics UK (COG-UK) Consortium | Lighthouse Laboratory Plymouth and Alex Alderton, Roberto Amato, Jeffrey Barrett, Sonia Goncalves, Ewan Harrison, David K. Jackson, Ian Johnston, Dominic Kwiatkowski, Cordelia Langford, John Sillitoe on behalf of the Wellcome Sanger Institute COVID-19 Surveillance Team |
| EPI_ISL_3169275 | Lighthouse Lab in Glasgow | Wellcome Sanger Institute for the COVID-19 Genomics UK (COG-UK) Consortium | Harper VanSteenhouse, Yumi Kasai, David Gray, Carol Clugston, Anna Dominiczak and Alex Alderton, Roberto Amato, Jeffrey Barrett, Sonia Goncalves, Ewan Harrison, David K. Jackson, Ian Johnston, Dominic Kwiatkowski, Cordelia Langford, John Sillitoe on behalf of the Wellcome Sanger Institute COVID-19 Surveillance Team |
| EPI_ISL_3169276, EPI_ISL_3169277, EPI_ISL_3169278, EPI_ISL_3169279, EPI_ISL_3169280, EPI_ISL_3169281, EPI_ISL_3169282 | Lighthouse Laboratory Plymouth | Wellcome Sanger Institute for the COVID-19 Genomics UK (COG-UK) Consortium | Lighthouse Laboratory Plymouth and Alex Alderton, Roberto Amato, Jeffrey Barrett, Sonia Goncalves, Ewan Harrison, David K. Jackson, Ian Johnston, Dominic Kwiatkowski, Cordelia Langford, John Sillitoe on behalf of the Wellcome Sanger Institute COVID-19 Surveillance Team |
| EPI_ISL_3169283 | Lighthouse Lab in Glasgow | Wellcome Sanger Institute for the COVID-19 Genomics UK (COG-UK) Consortium | Harper VanSteenhouse, Yumi Kasai, David Gray, Carol Clugston, Anna Dominiczak and Alex Alderton, Roberto Amato, Jeffrey Barrett, Sonia Goncalves, Ewan Harrison, David K. Jackson, Ian Johnston, Dominic Kwiatkowski, Cordelia Langford, John Sillitoe on behalf of the Wellcome Sanger Institute COVID-19 Surveillance Team |
| EPI_ISL_3169284, EPI_ISL_3169285, EPI_ISL_3169286, EPI_ISL_3169287, EPI_ISL_3169288, EPI_ISL_3169289, EPI_ISL_3169290, EPI_ISL_3169291 | Lighthouse Laboratory Plymouth | Wellcome Sanger Institute for the COVID-19 Genomics UK (COG-UK) Consortium | Lighthouse Laboratory Plymouth and Alex Alderton, Roberto Amato, Jeffrey Barrett, Sonia Goncalves, Ewan Harrison, David K. Jackson, Ian Johnston, Dominic Kwiatkowski, Cordelia Langford, John Sillitoe on behalf of the Wellcome Sanger Institute COVID-19 Surveillance Team |
| EPI_ISL_3169292 | Lighthouse Lab in Glasgow | Wellcome Sanger Institute for the COVID-19 Genomics UK (COG-UK) Consortium | Harper VanSteenhouse, Yumi Kasai, David Gray, Carol Clugston, Anna Dominiczak and Alex Alderton, Roberto Amato, Jeffrey Barrett, Sonia Goncalves, Ewan Harrison, David K. Jackson, Ian Johnston, Dominic Kwiatkowski, Cordelia Langford, John Sillitoe on behalf of the Wellcome Sanger Institute COVID-19 Surveillance Team |
| EPI_ISL_3169293, EPI_ISL_3169294, EPI_ISL_3169295, EPI_ISL_3169296 | Lighthouse Laboratory Plymouth | Wellcome Sanger Institute for the COVID-19 Genomics UK (COG-UK) Consortium | Lighthouse Laboratory Plymouth and Alex Alderton, Roberto Amato, Jeffrey Barrett, Sonia Goncalves, Ewan Harrison, David K. Jackson, Ian Johnston, Dominic Kwiatkowski, Cordelia Langford, John Sillitoe on behalf of the Wellcome Sanger Institute COVID-19 Surveillance Team |
| EPI_ISL_3169297 | Lighthouse Lab in Glasgow | Wellcome Sanger Institute for the COVID-19 Genomics UK (COG-UK) Consortium | Harper VanSteenhouse, Yumi Kasai, David Gray, Carol Clugston, Anna Dominiczak and Alex Alderton, Roberto Amato, Jeffrey Barrett, Sonia Goncalves, Ewan Harrison, David K. Jackson, Ian Johnston, Dominic Kwiatkowski, Cordelia Langford, John Sillitoe on behalf of the Wellcome Sanger Institute COVID-19 Surveillance Team |
| EPI_ISL_3169298, EPI_ISL_3169299, EPI_ISL_3169300, EPI_ISL_3169301, EPI_ISL_3169302 | Lighthouse Laboratory Plymouth | Wellcome Sanger Institute for the COVID-19 Genomics UK (COG-UK) Consortium | Lighthouse Laboratory Plymouth and Alex Alderton, Roberto Amato, Jeffrey Barrett, Sonia Goncalves, Ewan Harrison, David K. Jackson, Ian Johnston, Dominic Kwiatkowski, Cordelia Langford, John Sillitoe on behalf of the Wellcome Sanger Institute COVID-19 Surveillance Team |
| EPI_ISL_3169303, EPI_ISL_3169304, EPI_ISL_3169305 | Lighthouse Lab in Glasgow | Wellcome Sanger Institute for the COVID-19 Genomics UK (COG-UK) Consortium | Harper VanSteenhouse, Yumi Kasai, David Gray, Carol Clugston, Anna Dominiczak and Alex Alderton, Roberto Amato, Jeffrey Barrett, Sonia Goncalves, Ewan Harrison, David K. Jackson, Ian Johnston, Dominic Kwiatkowski, Cordelia Langford, John Sillitoe on behalf of the Wellcome Sanger Institute COVID-19 Surveillance Team |
| EPI_ISL_3169306, EPI_ISL_3169307 | Lighthouse Laboratory Plymouth | Wellcome Sanger Institute for the COVID-19 Genomics UK (COG-UK) Consortium | Lighthouse Laboratory Plymouth and Alex Alderton, Roberto Amato, Jeffrey Barrett, Sonia Goncalves, Ewan Harrison, David K. Jackson, Ian Johnston, Dominic Kwiatkowski, Cordelia Langford, John Sillitoe on behalf of the Wellcome Sanger Institute COVID-19 Surveillance Team |
| EPI_ISL_3169308, EPI_ISL_3169309 | Lighthouse Lab in Glasgow | Wellcome Sanger Institute for the COVID-19 Genomics UK (COG-UK) Consortium | Harper VanSteenhouse, Yumi Kasai, David Gray, Carol Clugston, Anna Dominiczak and Alex Alderton, Roberto Amato, Jeffrey Barrett, Sonia Goncalves, Ewan Harrison, David K. Jackson, Ian Johnston, Dominic Kwiatkowski, Cordelia Langford, John Sillitoe on behalf of the Wellcome Sanger Institute COVID-19 Surveillance Team |
| EPI_ISL_3169310, EPI_ISL_3169311, EPI_ISL_3169312, EPI_ISL_3169313 | Lighthouse Laboratory Plymouth | Wellcome Sanger Institute for the COVID-19 Genomics UK (COG-UK) Consortium | Lighthouse Laboratory Plymouth and Alex Alderton, Roberto Amato, Jeffrey Barrett, Sonia Goncalves, Ewan Harrison, David K. Jackson, Ian Johnston, Dominic Kwiatkowski, Cordelia Langford, John Sillitoe on behalf of the Wellcome Sanger Institute COVID-19 Surveillance Team |
| EPI_ISL_3169314, EPI_ISL_3169315 | Lighthouse Lab in Glasgow | Wellcome Sanger Institute for the COVID-19 Genomics UK (COG-UK) Consortium | Harper VanSteenhouse, Yumi Kasai, David Gray, Carol Clugston, Anna Dominiczak and Alex Alderton, Roberto Amato, Jeffrey Barrett, Sonia Goncalves, Ewan Harrison, David K. Jackson, Ian Johnston, Dominic Kwiatkowski, Cordelia Langford, John Sillitoe on behalf of the Wellcome Sanger Institute COVID-19 Surveillance Team |
| EPI_ISL_3169316, EPI_ISL_3169317, EPI_ISL_3169318, EPI_ISL_3169319 | Lighthouse Laboratory Plymouth | Wellcome Sanger Institute for the COVID-19 Genomics UK (COG-UK) Consortium | Lighthouse Laboratory Plymouth and Alex Alderton, Roberto Amato, Jeffrey Barrett, Sonia Goncalves, Ewan Harrison, David K. Jackson, Ian Johnston, Dominic Kwiatkowski, Cordelia Langford, John Sillitoe on behalf of the Wellcome Sanger Institute COVID-19 Surveillance Team |
| EPI_ISL_3169320 | Lighthouse Lab in Glasgow | Wellcome Sanger Institute for the COVID-19 Genomics UK (COG-UK) Consortium | Harper VanSteenhouse, Yumi Kasai, David Gray, Carol Clugston, Anna Dominiczak and Alex Alderton, Roberto Amato, Jeffrey Barrett, Sonia Goncalves, Ewan Harrison, David K. Jackson, Ian Johnston, Dominic Kwiatkowski, Cordelia Langford, John Sillitoe on behalf of the Wellcome Sanger Institute COVID-19 Surveillance Team |
| EPI_ISL_3169321, EPI_ISL_3169322, EPI_ISL_3169323, EPI_ISL_3169324 | Lighthouse Laboratory Plymouth | Wellcome Sanger Institute for the COVID-19 Genomics UK (COG-UK) Consortium | Lighthouse Laboratory Plymouth and Alex Alderton, Roberto Amato, Jeffrey Barrett, Sonia Goncalves, Ewan Harrison, David K. Jackson, Ian Johnston, Dominic Kwiatkowski, Cordelia Langford, John Sillitoe on behalf of the Wellcome Sanger Institute COVID-19 Surveillance Team |
| EPI_ISL_3169325, EPI_ISL_3169326 | Lighthouse Lab in Glasgow | Wellcome Sanger Institute for the COVID-19 Genomics UK (COG-UK) Consortium | Harper VanSteenhouse, Yumi Kasai, David Gray, Carol Clugston, Anna Dominiczak and Alex Alderton, Roberto Amato, Jeffrey Barrett, Sonia Goncalves, Ewan Harrison, David K. Jackson, Ian Johnston, Dominic Kwiatkowski, Cordelia Langford, John Sillitoe on behalf of the Wellcome Sanger Institute COVID-19 Surveillance Team |
| EPI_ISL_3169327, EPI_ISL_3169328, EPI_ISL_3169329, EPI_ISL_3169330, EPI_ISL_3169331, EPI_ISL_3169332 | Lighthouse Laboratory Plymouth | Wellcome Sanger Institute for the COVID-19 Genomics UK (COG-UK) Consortium | Lighthouse Laboratory Plymouth and Alex Alderton, Roberto Amato, Jeffrey Barrett, Sonia Goncalves, Ewan Harrison, David K. Jackson, Ian Johnston, Dominic Kwiatkowski, Cordelia Langford, John Sillitoe on behalf of the Wellcome Sanger Institute COVID-19 Surveillance Team |
| EPI_ISL_3169333, EPI_ISL_3169334 | Lighthouse Lab in Glasgow | Wellcome Sanger Institute for the COVID-19 Genomics UK (COG-UK) Consortium | Harper VanSteenhouse, Yumi Kasai, David Gray, Carol Clugston, Anna Dominiczak and Alex Alderton, Roberto Amato, Jeffrey Barrett, Sonia Goncalves, Ewan Harrison, David K. Jackson, Ian Johnston, Dominic Kwiatkowski, Cordelia Langford, John Sillitoe on behalf of the Wellcome Sanger Institute COVID-19 Surveillance Team |
| EPI_ISL_3169335 | Lighthouse Laboratory Plymouth | Wellcome Sanger Institute for the COVID-19 Genomics UK (COG-UK) Consortium | Lighthouse Laboratory Plymouth and Alex Alderton, Roberto Amato, Jeffrey Barrett, Sonia Goncalves, Ewan Harrison, David K. Jackson, Ian Johnston, Dominic Kwiatkowski, Cordelia Langford, John Sillitoe on behalf of the Wellcome Sanger Institute COVID-19 Surveillance Team |
| EPI_ISL_3169336 | Lighthouse Lab in Glasgow | Wellcome Sanger Institute for the COVID-19 Genomics UK (COG-UK) Consortium | Harper VanSteenhouse, Yumi Kasai, David Gray, Carol Clugston, Anna Dominiczak and Alex Alderton, Roberto Amato, Jeffrey Barrett, Sonia Goncalves, Ewan Harrison, David K. Jackson, Ian Johnston, Dominic Kwiatkowski, Cordelia Langford, John Sillitoe on behalf of the Wellcome Sanger Institute COVID-19 Surveillance Team |
| EPI_ISL_3169337, EPI_ISL_3169338, EPI_ISL_3169339, EPI_ISL_3169340, EPI_ISL_3169341, EPI_ISL_3169342, EPI_ISL_3169343, EPI_ISL_3169344, EPI_ISL_3169345, EPI_ISL_3169346, EPI_ISL_3169347, EPI_ISL_3169348 | | | |
| see above | Lighthouse Laboratory Plymouth | Wellcome Sanger Institute for the COVID-19 Genomics UK (COG-UK) Consortium | Lighthouse Laboratory Plymouth and Alex Alderton, Roberto Amato, Jeffrey Barrett, Sonia Goncalves, Ewan Harrison, David K. Jackson, Ian Johnston, Dominic Kwiatkowski, Cordelia Langford, John Sillitoe on behalf of the Wellcome Sanger Institute COVID-19 Surveillance Team |
| EPI_ISL_3169349 | Lighthouse Lab in Glasgow | Wellcome Sanger Institute for the COVID-19 Genomics UK (COG-UK) Consortium | Harper VanSteenhouse, Yumi Kasai, David Gray, Carol Clugston, Anna Dominiczak and Alex Alderton, Roberto Amato, Jeffrey Barrett, Sonia Goncalves, Ewan Harrison, David K. Jackson, Ian Johnston, Dominic Kwiatkowski, Cordelia Langford, John Sillitoe on behalf of the Wellcome Sanger Institute COVID-19 Surveillance Team |
| EPI_ISL_3169350, EPI_ISL_3169351, | Lighthouse Laboratory Plymouth | Wellcome Sanger Institute for the COVID-19 Genomics UK (COG-UK) Consortium | Lighthouse Laboratory Plymouth and Alex Alderton, Roberto Amato, Jeffrey Barrett, Sonia Goncalves, Ewan Harrison, David K. Jackson, Ian Johnston, Dominic Kwiatkowski, Cordelia Langford, John Sillitoe on behalf of the Wellcome Sanger Institute COVID-19 Surveillance Team |

| | | | |
|---|---|---|---|
| EPI_ISL_3169352, EPI_ISL_3169353, EPI_ISL_3169354, EPI_ISL_3169355, EPI_ISL_3169356, EPI_ISL_3169357 | | | |
| EPI_ISL_3169358 | Lighthouse Lab in Glasgow | Wellcome Sanger Institute for the COVID-19 Genomics UK (COG-UK) Consortium | Harper VanSteenhouse, Yumi Kasai, David Gray, Carol Clugston, Anna Dominiczak and Alex Alderton, Roberto Amato, Jeffrey Barrett, Sonia Goncalves, Ewan Harrison, David K. Jackson, Ian Johnston, Dominic Kwiatkowski, Cordelia Langford, John Sillitoe on behalf of the Wellcome Sanger Institute COVID-19 Surveillance Team |
| EPI_ISL_3169359 | Lighthouse Laboratory Plymouth | Wellcome Sanger Institute for the COVID-19 Genomics UK (COG-UK) Consortium | Lighthouse Laboratory Plymouth and Alex Alderton, Roberto Amato, Jeffrey Barrett, Sonia Goncalves, Ewan Harrison, David K. Jackson, Ian Johnston, Dominic Kwiatkowski, Cordelia Langford, John Sillitoe on behalf of the Wellcome Sanger Institute COVID-19 Surveillance Team |
| EPI_ISL_3169360, EPI_ISL_3169361 | Lighthouse Lab in Glasgow | Wellcome Sanger Institute for the COVID-19 Genomics UK (COG-UK) Consortium | Harper VanSteenhouse, Yumi Kasai, David Gray, Carol Clugston, Anna Dominiczak and Alex Alderton, Roberto Amato, Jeffrey Barrett, Sonia Goncalves, Ewan Harrison, David K. Jackson, Ian Johnston, Dominic Kwiatkowski, Cordelia Langford, John Sillitoe on behalf of the Wellcome Sanger Institute COVID-19 Surveillance Team |
| EPI_ISL_3169362 | Lighthouse Laboratory Plymouth | Wellcome Sanger Institute for the COVID-19 Genomics UK (COG-UK) Consortium | Lighthouse Laboratory Plymouth and Alex Alderton, Roberto Amato, Jeffrey Barrett, Sonia Goncalves, Ewan Harrison, David K. Jackson, Ian Johnston, Dominic Kwiatkowski, Cordelia Langford, John Sillitoe on behalf of the Wellcome Sanger Institute COVID-19 Surveillance Team |
| EPI_ISL_3169363, EPI_ISL_3169364 | Lighthouse Lab in Glasgow | Wellcome Sanger Institute for the COVID-19 Genomics UK (COG-UK) Consortium | Harper VanSteenhouse, Yumi Kasai, David Gray, Carol Clugston, Anna Dominiczak and Alex Alderton, Roberto Amato, Jeffrey Barrett, Sonia Goncalves, Ewan Harrison, David K. Jackson, Ian Johnston, Dominic Kwiatkowski, Cordelia Langford, John Sillitoe on behalf of the Wellcome Sanger Institute COVID-19 Surveillance Team |
| EPI_ISL_3169365 | Lighthouse Laboratory Plymouth | Wellcome Sanger Institute for the COVID-19 Genomics UK (COG-UK) Consortium | Lighthouse Laboratory Plymouth and Alex Alderton, Roberto Amato, Jeffrey Barrett, Sonia Goncalves, Ewan Harrison, David K. Jackson, Ian Johnston, Dominic Kwiatkowski, Cordelia Langford, John Sillitoe on behalf of the Wellcome Sanger Institute COVID-19 Surveillance Team |
| EPI_ISL_3169366, EPI_ISL_3169367 | Lighthouse Lab in Glasgow | Wellcome Sanger Institute for the COVID-19 Genomics UK (COG-UK) Consortium | Harper VanSteenhouse, Yumi Kasai, David Gray, Carol Clugston, Anna Dominiczak and Alex Alderton, Roberto Amato, Jeffrey Barrett, Sonia Goncalves, Ewan Harrison, David K. Jackson, Ian Johnston, Dominic Kwiatkowski, Cordelia Langford, John Sillitoe on behalf of the Wellcome Sanger Institute COVID-19 Surveillance Team |
| EPI_ISL_3169368, EPI_ISL_3169369, EPI_ISL_3169370, EPI_ISL_3169371, EPI_ISL_3169372, EPI_ISL_3169373, EPI_ISL_3169374, EPI_ISL_3169375, EPI_ISL_3169376, EPI_ISL_3169377 | Lighthouse Laboratory Plymouth | Wellcome Sanger Institute for the COVID-19 Genomics UK (COG-UK) Consortium | Lighthouse Laboratory Plymouth and Alex Alderton, Roberto Amato, Jeffrey Barrett, Sonia Goncalves, Ewan Harrison, David K. Jackson, Ian Johnston, Dominic Kwiatkowski, Cordelia Langford, John Sillitoe on behalf of the Wellcome Sanger Institute COVID-19 Surveillance Team |
| EPI_ISL_3169378, EPI_ISL_3169379 | Lighthouse Lab in Glasgow | Wellcome Sanger Institute for the COVID-19 Genomics UK (COG-UK) Consortium | Harper VanSteenhouse, Yumi Kasai, David Gray, Carol Clugston, Anna Dominiczak and Alex Alderton, Roberto Amato, Jeffrey Barrett, Sonia Goncalves, Ewan Harrison, David K. Jackson, Ian Johnston, Dominic Kwiatkowski, Cordelia Langford, John Sillitoe on behalf of the Wellcome Sanger Institute COVID-19 Surveillance Team |
| EPI_ISL_3169380, EPI_ISL_3169381, EPI_ISL_3169382, EPI_ISL_3169383 | Lighthouse Laboratory Plymouth | Wellcome Sanger Institute for the COVID-19 Genomics UK (COG-UK) Consortium | Lighthouse Laboratory Plymouth and Alex Alderton, Roberto Amato, Jeffrey Barrett, Sonia Goncalves, Ewan Harrison, David K. Jackson, Ian Johnston, Dominic Kwiatkowski, Cordelia Langford, John Sillitoe on behalf of the Wellcome Sanger Institute COVID-19 Surveillance Team |
| EPI_ISL_3169384 | Lighthouse Lab in Glasgow | Wellcome Sanger Institute for the COVID-19 Genomics UK (COG-UK) Consortium | Harper VanSteenhouse, Yumi Kasai, David Gray, Carol Clugston, Anna Dominiczak and Alex Alderton, Roberto Amato, Jeffrey Barrett, Sonia Goncalves, Ewan Harrison, David K. Jackson, Ian Johnston, Dominic Kwiatkowski, Cordelia Langford, John Sillitoe on behalf of the Wellcome Sanger Institute COVID-19 Surveillance Team |
| EPI_ISL_3169385, EPI_ISL_3169386, EPI_ISL_3169387, EPI_ISL_3169388, EPI_ISL_3169389, EPI_ISL_3169390, EPI_ISL_3169391 | Lighthouse Laboratory Plymouth | Wellcome Sanger Institute for the COVID-19 Genomics UK (COG-UK) Consortium | Lighthouse Laboratory Plymouth and Alex Alderton, Roberto Amato, Jeffrey Barrett, Sonia Goncalves, Ewan Harrison, David K. Jackson, Ian Johnston, Dominic Kwiatkowski, Cordelia Langford, John Sillitoe on behalf of the Wellcome Sanger Institute COVID-19 Surveillance Team |
| EPI_ISL_3169392 | Lighthouse Lab in Glasgow | Wellcome Sanger Institute for the COVID-19 Genomics UK (COG-UK) Consortium | Harper VanSteenhouse, Yumi Kasai, David Gray, Carol Clugston, Anna Dominiczak and Alex Alderton, Roberto Amato, Jeffrey Barrett, Sonia Goncalves, Ewan Harrison, David K. Jackson, Ian Johnston, Dominic Kwiatkowski, Cordelia Langford, John Sillitoe on behalf of the Wellcome Sanger Institute COVID-19 Surveillance Team |
| EPI_ISL_3169393, EPI_ISL_3169394, EPI_ISL_3169395, EPI_ISL_3169396, EPI_ISL_3169397 | Lighthouse Laboratory Plymouth | Wellcome Sanger Institute for the COVID-19 Genomics UK (COG-UK) Consortium | Lighthouse Laboratory Plymouth and Alex Alderton, Roberto Amato, Jeffrey Barrett, Sonia Goncalves, Ewan Harrison, David K. Jackson, Ian Johnston, Dominic Kwiatkowski, Cordelia Langford, John Sillitoe on behalf of the Wellcome Sanger Institute COVID-19 Surveillance Team |
| EPI_ISL_3169398, EPI_ISL_3169399 | Lighthouse Lab in Glasgow | Wellcome Sanger Institute for the COVID-19 Genomics UK (COG-UK) Consortium | Harper VanSteenhouse, Yumi Kasai, David Gray, Carol Clugston, Anna Dominiczak and Alex Alderton, Roberto Amato, Jeffrey Barrett, Sonia Goncalves, Ewan Harrison, David K. Jackson, Ian Johnston, Dominic Kwiatkowski, Cordelia Langford, John Sillitoe on behalf of the Wellcome Sanger Institute COVID-19 Surveillance Team |
| EPI_ISL_3169400 | Lighthouse Laboratory Plymouth | Wellcome Sanger Institute for the COVID-19 Genomics UK (COG-UK) Consortium | Lighthouse Laboratory Plymouth and Alex Alderton, Roberto Amato, Jeffrey Barrett, Sonia Goncalves, Ewan Harrison, David K. Jackson, Ian Johnston, Dominic Kwiatkowski, Cordelia Langford, John Sillitoe on behalf of the Wellcome Sanger Institute COVID-19 Surveillance Team |
| EPI_ISL_3169401 | Lighthouse Lab in Glasgow | Wellcome Sanger Institute for the COVID-19 Genomics UK (COG-UK) Consortium | Harper VanSteenhouse, Yumi Kasai, David Gray, Carol Clugston, Anna Dominiczak and Alex Alderton, Roberto Amato, Jeffrey Barrett, Sonia Goncalves, Ewan Harrison, David K. Jackson, Ian Johnston, Dominic Kwiatkowski, Cordelia Langford, John Sillitoe on behalf of the Wellcome Sanger Institute COVID-19 Surveillance Team |
| EPI_ISL_3169402, EPI_ISL_3169403, EPI_ISL_3169404, EPI_ISL_3169405 | Lighthouse Laboratory Plymouth | Wellcome Sanger Institute for the COVID-19 Genomics UK (COG-UK) Consortium | Lighthouse Laboratory Plymouth and Alex Alderton, Roberto Amato, Jeffrey Barrett, Sonia Goncalves, Ewan Harrison, David K. Jackson, Ian Johnston, Dominic Kwiatkowski, Cordelia Langford, John Sillitoe on behalf of the Wellcome Sanger Institute COVID-19 Surveillance Team |
| EPI_ISL_3169406 | Lighthouse Lab in Glasgow | Wellcome Sanger Institute for the COVID-19 Genomics UK (COG-UK) Consortium | Harper VanSteenhouse, Yumi Kasai, David Gray, Carol Clugston, Anna Dominiczak and Alex Alderton, Roberto Amato, Jeffrey Barrett, Sonia Goncalves, Ewan Harrison, David K. Jackson, Ian Johnston, Dominic Kwiatkowski, Cordelia Langford, John Sillitoe on behalf of the Wellcome Sanger Institute COVID-19 Surveillance Team |
| EPI_ISL_3169407, EPI_ISL_3169408, EPI_ISL_3169409, EPI_ISL_3169410, EPI_ISL_3169411, EPI_ISL_3169412 | Lighthouse Laboratory Plymouth | Wellcome Sanger Institute for the COVID-19 Genomics UK (COG-UK) Consortium | Lighthouse Laboratory Plymouth and Alex Alderton, Roberto Amato, Jeffrey Barrett, Sonia Goncalves, Ewan Harrison, David K. Jackson, Ian Johnston, Dominic Kwiatkowski, Cordelia Langford, John Sillitoe on behalf of the Wellcome Sanger Institute COVID-19 Surveillance Team |
| EPI_ISL_3169413 | Lighthouse Lab in Glasgow | Wellcome Sanger Institute for the COVID-19 Genomics UK (COG-UK) Consortium | Harper VanSteenhouse, Yumi Kasai, David Gray, Carol Clugston, Anna Dominiczak and Alex Alderton, Roberto Amato, Jeffrey Barrett, Sonia Goncalves, Ewan Harrison, David K. Jackson, Ian Johnston, Dominic Kwiatkowski, Cordelia Langford, John Sillitoe on behalf of the Wellcome Sanger Institute COVID-19 Surveillance Team |
| EPI_ISL_3169414, EPI_ISL_3169415, EPI_ISL_3169416, EPI_ISL_3169417, EPI_ISL_3169418, EPI_ISL_3169419 | Lighthouse Laboratory Plymouth | Wellcome Sanger Institute for the COVID-19 Genomics UK (COG-UK) Consortium | Lighthouse Laboratory Plymouth and Alex Alderton, Roberto Amato, Jeffrey Barrett, Sonia Goncalves, Ewan Harrison, David K. Jackson, Ian Johnston, Dominic Kwiatkowski, Cordelia Langford, John Sillitoe on behalf of the Wellcome Sanger Institute COVID-19 Surveillance Team |
| EPI_ISL_3169420 | Lighthouse Lab in Glasgow | Wellcome Sanger Institute for the COVID-19 Genomics UK (COG-UK) Consortium | Harper VanSteenhouse, Yumi Kasai, David Gray, Carol Clugston, Anna Dominiczak and Alex Alderton, Roberto Amato, Jeffrey Barrett, Sonia Goncalves, Ewan Harrison, David K. Jackson, Ian Johnston, Dominic Kwiatkowski, Cordelia Langford, John Sillitoe on behalf of the Wellcome Sanger Institute COVID-19 Surveillance Team |
| EPI_ISL_3169421, EPI_ISL_3169422, EPI_ISL_3169423, EPI_ISL_3169424 | Lighthouse Laboratory Plymouth | Wellcome Sanger Institute for the COVID-19 Genomics UK (COG-UK) Consortium | Lighthouse Laboratory Plymouth and Alex Alderton, Roberto Amato, Jeffrey Barrett, Sonia Goncalves, Ewan Harrison, David K. Jackson, Ian Johnston, Dominic Kwiatkowski, Cordelia Langford, John Sillitoe on behalf of the Wellcome Sanger Institute COVID-19 Surveillance Team |
| EPI_ISL_3169425 | Lighthouse Lab in Glasgow | Wellcome Sanger Institute for the COVID-19 Genomics UK (COG-UK) Consortium | Harper VanSteenhouse, Yumi Kasai, David Gray, Carol Clugston, Anna Dominiczak and Alex Alderton, Roberto Amato, Jeffrey Barrett, Sonia Goncalves, Ewan Harrison, David K. Jackson, Ian Johnston, Dominic Kwiatkowski, Cordelia Langford, John Sillitoe on behalf of the Wellcome Sanger Institute COVID-19 Surveillance Team |
| EPI_ISL_3169426, EPI_ISL_3169427, EPI_ISL_3169428, EPI_ISL_3169429, EPI_ISL_3169430 | Lighthouse Laboratory Plymouth | Wellcome Sanger Institute for the COVID-19 Genomics UK (COG-UK) Consortium | Lighthouse Laboratory Plymouth and Alex Alderton, Roberto Amato, Jeffrey Barrett, Sonia Goncalves, Ewan Harrison, David K. Jackson, Ian Johnston, Dominic Kwiatkowski, Cordelia Langford, John Sillitoe on behalf of the Wellcome Sanger Institute COVID-19 Surveillance Team |
| EPI_ISL_3169431 | Lighthouse Lab in Glasgow | Wellcome Sanger Institute for the COVID-19 Genomics UK (COG-UK) Consortium | Harper VanSteenhouse, Yumi Kasai, David Gray, Carol Clugston, Anna Dominiczak and Alex Alderton, Roberto Amato, Jeffrey Barrett, Sonia Goncalves, Ewan Harrison, David K. Jackson, Ian Johnston, Dominic Kwiatkowski, Cordelia Langford, John Sillitoe on behalf of the Wellcome Sanger Institute COVID-19 Surveillance Team |
| EPI_ISL_3169432, EPI_ISL_3169433 | Lighthouse Laboratory Plymouth | Wellcome Sanger Institute for the COVID-19 Genomics UK (COG-UK) Consortium | Lighthouse Laboratory Plymouth and Alex Alderton, Roberto Amato, Jeffrey Barrett, Sonia Goncalves, Ewan Harrison, David K. Jackson, Ian Johnston, Dominic Kwiatkowski, Cordelia Langford, John Sillitoe on behalf of the Wellcome Sanger Institute COVID-19 Surveillance Team |
| EPI_ISL_3169434 | Lighthouse Lab in Glasgow | Wellcome Sanger Institute for the COVID-19 Genomics UK (COG-UK) Consortium | Harper VanSteenhouse, Yumi Kasai, David Gray, Carol Clugston, Anna Dominiczak and Alex Alderton, Roberto Amato, Jeffrey Barrett, Sonia Goncalves, Ewan Harrison, David K. Jackson, Ian Johnston, Dominic Kwiatkowski, Cordelia Langford, John Sillitoe on behalf of the Wellcome Sanger Institute COVID-19 Surveillance Team |

| | | | |
|---|---|---|---|
| EPI_ISL_3169435, EPI_ISL_3169436, EPI_ISL_3169437, EPI_ISL_3169438, EPI_ISL_3169439, EPI_ISL_3169440, EPI_ISL_3169441, EPI_ISL_3169442 | Lighthouse Laboratory Plymouth | Wellcome Sanger Institute for the COVID-19 Genomics UK (COG-UK) Consortium | Lighthouse Laboratory Plymouth and Alex Alderton, Roberto Amato, Jeffrey Barrett, Sonia Goncalves, Ewan Harrison, David K. Jackson, Ian Johnston, Dominic Kwiatkowski, Cordelia Langford, John Sillitoe on behalf of the Wellcome Sanger Institute COVID-19 Surveillance Team |
| EPI_ISL_3169443 | Lighthouse Lab in Glasgow | Wellcome Sanger Institute for the COVID-19 Genomics UK (COG-UK) Consortium | Harper VanSteenhouse, Yumi Kasai, David Gray, Carol Clugston, Anna Dominiczak and Alex Alderton, Roberto Amato, Jeffrey Barrett, Sonia Goncalves, Ewan Harrison, David K. Jackson, Ian Johnston, Dominic Kwiatkowski, Cordelia Langford, John Sillitoe on behalf of the Wellcome Sanger Institute COVID-19 Surveillance Team |
| EPI_ISL_3169444 | Lighthouse Laboratory Plymouth | Wellcome Sanger Institute for the COVID-19 Genomics UK (COG-UK) Consortium | Lighthouse Laboratory Plymouth and Alex Alderton, Roberto Amato, Jeffrey Barrett, Sonia Goncalves, Ewan Harrison, David K. Jackson, Ian Johnston, Dominic Kwiatkowski, Cordelia Langford, John Sillitoe on behalf of the Wellcome Sanger Institute COVID-19 Surveillance Team |
| EPI_ISL_3169445 | Lighthouse Lab in Glasgow | Wellcome Sanger Institute for the COVID-19 Genomics UK (COG-UK) Consortium | Harper VanSteenhouse, Yumi Kasai, David Gray, Carol Clugston, Anna Dominiczak and Alex Alderton, Roberto Amato, Jeffrey Barrett, Sonia Goncalves, Ewan Harrison, David K. Jackson, Ian Johnston, Dominic Kwiatkowski, Cordelia Langford, John Sillitoe on behalf of the Wellcome Sanger Institute COVID-19 Surveillance Team |
| EPI_ISL_3169446, EPI_ISL_3169447, EPI_ISL_3169448, EPI_ISL_3169449, EPI_ISL_3169450, EPI_ISL_3169451, EPI_ISL_3169452, EPI_ISL_3169453, EPI_ISL_3169454, EPI_ISL_3169455 | Lighthouse Laboratory Plymouth | Wellcome Sanger Institute for the COVID-19 Genomics UK (COG-UK) Consortium | Lighthouse Laboratory Plymouth and Alex Alderton, Roberto Amato, Jeffrey Barrett, Sonia Goncalves, Ewan Harrison, David K. Jackson, Ian Johnston, Dominic Kwiatkowski, Cordelia Langford, John Sillitoe on behalf of the Wellcome Sanger Institute COVID-19 Surveillance Team |
| EPI_ISL_3169456, EPI_ISL_3169457 | Lighthouse Lab in Glasgow | Wellcome Sanger Institute for the COVID-19 Genomics UK (COG-UK) Consortium | Harper VanSteenhouse, Yumi Kasai, David Gray, Carol Clugston, Anna Dominiczak and Alex Alderton, Roberto Amato, Jeffrey Barrett, Sonia Goncalves, Ewan Harrison, David K. Jackson, Ian Johnston, Dominic Kwiatkowski, Cordelia Langford, John Sillitoe on behalf of the Wellcome Sanger Institute COVID-19 Surveillance Team |
| EPI_ISL_3169458 | Lighthouse Laboratory Plymouth | Wellcome Sanger Institute for the COVID-19 Genomics UK (COG-UK) Consortium | Lighthouse Laboratory Plymouth and Alex Alderton, Roberto Amato, Jeffrey Barrett, Sonia Goncalves, Ewan Harrison, David K. Jackson, Ian Johnston, Dominic Kwiatkowski, Cordelia Langford, John Sillitoe on behalf of the Wellcome Sanger Institute COVID-19 Surveillance Team |
| EPI_ISL_3169459 | Lighthouse Lab in Glasgow | Wellcome Sanger Institute for the COVID-19 Genomics UK (COG-UK) Consortium | Harper VanSteenhouse, Yumi Kasai, David Gray, Carol Clugston, Anna Dominiczak and Alex Alderton, Roberto Amato, Jeffrey Barrett, Sonia Goncalves, Ewan Harrison, David K. Jackson, Ian Johnston, Dominic Kwiatkowski, Cordelia Langford, John Sillitoe on behalf of the Wellcome Sanger Institute COVID-19 Surveillance Team |
| EPI_ISL_3169460 | Lighthouse Laboratory Plymouth | Wellcome Sanger Institute for the COVID-19 Genomics UK (COG-UK) Consortium | Lighthouse Laboratory Plymouth and Alex Alderton, Roberto Amato, Jeffrey Barrett, Sonia Goncalves, Ewan Harrison, David K. Jackson, Ian Johnston, Dominic Kwiatkowski, Cordelia Langford, John Sillitoe on behalf of the Wellcome Sanger Institute COVID-19 Surveillance Team |
| EPI_ISL_3169461 | Lighthouse Lab in Glasgow | Wellcome Sanger Institute for the COVID-19 Genomics UK (COG-UK) Consortium | Harper VanSteenhouse, Yumi Kasai, David Gray, Carol Clugston, Anna Dominiczak and Alex Alderton, Roberto Amato, Jeffrey Barrett, Sonia Goncalves, Ewan Harrison, David K. Jackson, Ian Johnston, Dominic Kwiatkowski, Cordelia Langford, John Sillitoe on behalf of the Wellcome Sanger Institute COVID-19 Surveillance Team |
| EPI_ISL_3169462, EPI_ISL_3169463, EPI_ISL_3169464, EPI_ISL_3169465, EPI_ISL_3169466, EPI_ISL_3169467, EPI_ISL_3169468, EPI_ISL_3169469, EPI_ISL_3169470, EPI_ISL_3169471, EPI_ISL_3169472, EPI_ISL_3169473, EPI_ISL_3169474, EPI_ISL_3169475, EPI_ISL_3169476, EPI_ISL_3169477, EPI_ISL_3169478, EPI_ISL_3169479, EPI_ISL_3169480 | | | |
| see above | Lighthouse Laboratory Plymouth | Wellcome Sanger Institute for the COVID-19 Genomics UK (COG-UK) Consortium | Lighthouse Laboratory Plymouth and Alex Alderton, Roberto Amato, Jeffrey Barrett, Sonia Goncalves, Ewan Harrison, David K. Jackson, Ian Johnston, Dominic Kwiatkowski, Cordelia Langford, John Sillitoe on behalf of the Wellcome Sanger Institute COVID-19 Surveillance Team |
| EPI_ISL_3169481 | Lighthouse Lab in Glasgow | Wellcome Sanger Institute for the COVID-19 Genomics UK (COG-UK) Consortium | Harper VanSteenhouse, Yumi Kasai, David Gray, Carol Clugston, Anna Dominiczak and Alex Alderton, Roberto Amato, Jeffrey Barrett, Sonia Goncalves, Ewan Harrison, David K. Jackson, Ian Johnston, Dominic Kwiatkowski, Cordelia Langford, John Sillitoe on behalf of the Wellcome Sanger Institute COVID-19 Surveillance Team |
| EPI_ISL_3169482, EPI_ISL_3169483, EPI_ISL_3169484, EPI_ISL_3169485, EPI_ISL_3169486, EPI_ISL_3169487, EPI_ISL_3169488, EPI_ISL_3169489, EPI_ISL_3169490, EPI_ISL_3169491, EPI_ISL_3169492, EPI_ISL_3169493, EPI_ISL_3169494 | | | |
| see above | Lighthouse Laboratory Plymouth | Wellcome Sanger Institute for the COVID-19 Genomics UK (COG-UK) Consortium | Lighthouse Laboratory Plymouth and Alex Alderton, Roberto Amato, Jeffrey Barrett, Sonia Goncalves, Ewan Harrison, David K. Jackson, Ian Johnston, Dominic Kwiatkowski, Cordelia Langford, John Sillitoe on behalf of the Wellcome Sanger Institute COVID-19 Surveillance Team |
| EPI_ISL_3169495, EPI_ISL_3169496 | Lighthouse Lab in Glasgow | Wellcome Sanger Institute for the COVID-19 Genomics UK (COG-UK) Consortium | Harper VanSteenhouse, Yumi Kasai, David Gray, Carol Clugston, Anna Dominiczak and Alex Alderton, Roberto Amato, Jeffrey Barrett, Sonia Goncalves, Ewan Harrison, David K. Jackson, Ian Johnston, Dominic Kwiatkowski, Cordelia Langford, John Sillitoe on behalf of the Wellcome Sanger Institute COVID-19 Surveillance Team |
| EPI_ISL_3169497 | Lighthouse Laboratory Plymouth | Wellcome Sanger Institute for the COVID-19 Genomics UK (COG-UK) Consortium | Lighthouse Laboratory Plymouth and Alex Alderton, Roberto Amato, Jeffrey Barrett, Sonia Goncalves, Ewan Harrison, David K. Jackson, Ian Johnston, Dominic Kwiatkowski, Cordelia Langford, John Sillitoe on behalf of the Wellcome Sanger Institute COVID-19 Surveillance Team |
| EPI_ISL_3169498 | Lighthouse Lab in Glasgow | Wellcome Sanger Institute for the COVID-19 Genomics UK (COG-UK) Consortium | Harper VanSteenhouse, Yumi Kasai, David Gray, Carol Clugston, Anna Dominiczak and Alex Alderton, Roberto Amato, Jeffrey Barrett, Sonia Goncalves, Ewan Harrison, David K. Jackson, Ian Johnston, Dominic Kwiatkowski, Cordelia Langford, John Sillitoe on behalf of the Wellcome Sanger Institute COVID-19 Surveillance Team |
| EPI_ISL_3169499, EPI_ISL_3169500, EPI_ISL_3169501, EPI_ISL_3169502, EPI_ISL_3169503, EPI_ISL_3169504, EPI_ISL_3169505 | Lighthouse Laboratory Plymouth | Wellcome Sanger Institute for the COVID-19 Genomics UK (COG-UK) Consortium | Lighthouse Laboratory Plymouth and Alex Alderton, Roberto Amato, Jeffrey Barrett, Sonia Goncalves, Ewan Harrison, David K. Jackson, Ian Johnston, Dominic Kwiatkowski, Cordelia Langford, John Sillitoe on behalf of the Wellcome Sanger Institute COVID-19 Surveillance Team |
| EPI_ISL_3169506 | Lighthouse Lab in Glasgow | Wellcome Sanger Institute for the COVID-19 Genomics UK (COG-UK) Consortium | Harper VanSteenhouse, Yumi Kasai, David Gray, Carol Clugston, Anna Dominiczak and Alex Alderton, Roberto Amato, Jeffrey Barrett, Sonia Goncalves, Ewan Harrison, David K. Jackson, Ian Johnston, Dominic Kwiatkowski, Cordelia Langford, John Sillitoe on behalf of the Wellcome Sanger Institute COVID-19 Surveillance Team |
| EPI_ISL_3169507, EPI_ISL_3169508, EPI_ISL_3169509, EPI_ISL_3169510, EPI_ISL_3169511, EPI_ISL_3169512 | Lighthouse Laboratory Plymouth | Wellcome Sanger Institute for the COVID-19 Genomics UK (COG-UK) Consortium | Lighthouse Laboratory Plymouth and Alex Alderton, Roberto Amato, Jeffrey Barrett, Sonia Goncalves, Ewan Harrison, David K. Jackson, Ian Johnston, Dominic Kwiatkowski, Cordelia Langford, John Sillitoe on behalf of the Wellcome Sanger Institute COVID-19 Surveillance Team |
| EPI_ISL_3169513 | Lighthouse Lab in Glasgow | Wellcome Sanger Institute for the COVID-19 Genomics UK (COG-UK) Consortium | Harper VanSteenhouse, Yumi Kasai, David Gray, Carol Clugston, Anna Dominiczak and Alex Alderton, Roberto Amato, Jeffrey Barrett, Sonia Goncalves, Ewan Harrison, David K. Jackson, Ian Johnston, Dominic Kwiatkowski, Cordelia Langford, John Sillitoe on behalf of the Wellcome Sanger Institute COVID-19 Surveillance Team |
| EPI_ISL_3169514, EPI_ISL_3169515, EPI_ISL_3169516, EPI_ISL_3169517, EPI_ISL_3169518, EPI_ISL_3169519, EPI_ISL_3169520, EPI_ISL_3169521 | Lighthouse Laboratory Plymouth | Wellcome Sanger Institute for the COVID-19 Genomics UK (COG-UK) Consortium | Lighthouse Laboratory Plymouth and Alex Alderton, Roberto Amato, Jeffrey Barrett, Sonia Goncalves, Ewan Harrison, David K. Jackson, Ian Johnston, Dominic Kwiatkowski, Cordelia Langford, John Sillitoe on behalf of the Wellcome Sanger Institute COVID-19 Surveillance Team |
| EPI_ISL_3169522 | Lighthouse Lab in Glasgow | Wellcome Sanger Institute for the COVID-19 Genomics UK (COG-UK) Consortium | Harper VanSteenhouse, Yumi Kasai, David Gray, Carol Clugston, Anna Dominiczak and Alex Alderton, Roberto Amato, Jeffrey Barrett, Sonia Goncalves, Ewan Harrison, David K. Jackson, Ian Johnston, Dominic Kwiatkowski, Cordelia Langford, John Sillitoe on behalf of the Wellcome Sanger Institute COVID-19 Surveillance Team |
| EPI_ISL_3169523, EPI_ISL_3169524, EPI_ISL_3169525 | Lighthouse Laboratory Plymouth | Wellcome Sanger Institute for the COVID-19 Genomics UK (COG-UK) Consortium | Lighthouse Laboratory Plymouth and Alex Alderton, Roberto Amato, Jeffrey Barrett, Sonia Goncalves, Ewan Harrison, David K. Jackson, Ian Johnston, Dominic Kwiatkowski, Cordelia Langford, John Sillitoe on behalf of the Wellcome Sanger Institute COVID-19 Surveillance Team |
| EPI_ISL_3169526 | Lighthouse Lab in Glasgow | Wellcome Sanger Institute for the COVID-19 Genomics UK (COG-UK) Consortium | Harper VanSteenhouse, Yumi Kasai, David Gray, Carol Clugston, Anna Dominiczak and Alex Alderton, Roberto Amato, Jeffrey Barrett, Sonia Goncalves, Ewan Harrison, David K. Jackson, Ian Johnston, Dominic Kwiatkowski, Cordelia Langford, John Sillitoe on behalf of the Wellcome Sanger Institute COVID-19 Surveillance Team |
| EPI_ISL_3169527, EPI_ISL_3169528, EPI_ISL_3169529, EPI_ISL_3169530, EPI_ISL_3169531 | Lighthouse Laboratory Plymouth | Wellcome Sanger Institute for the COVID-19 Genomics UK (COG-UK) Consortium | Lighthouse Laboratory Plymouth and Alex Alderton, Roberto Amato, Jeffrey Barrett, Sonia Goncalves, Ewan Harrison, David K. Jackson, Ian Johnston, Dominic Kwiatkowski, Cordelia Langford, John Sillitoe on behalf of the Wellcome Sanger Institute COVID-19 Surveillance Team |
| EPI_ISL_3169532 | Lighthouse Lab in Glasgow | Wellcome Sanger Institute for the COVID-19 Genomics UK (COG-UK) Consortium | Harper VanSteenhouse, Yumi Kasai, David Gray, Carol Clugston, Anna Dominiczak and Alex Alderton, Roberto Amato, Jeffrey Barrett, Sonia Goncalves, Ewan Harrison, David K. Jackson, Ian Johnston, Dominic Kwiatkowski, Cordelia Langford, John Sillitoe on behalf of the Wellcome Sanger Institute COVID-19 Surveillance Team |
| EPI_ISL_3169533, EPI_ISL_3169534, EPI_ISL_3169535, EPI_ISL_3169536, EPI_ISL_3169537, EPI_ISL_3169538, EPI_ISL_3169539, EPI_ISL_3169540 | Lighthouse Laboratory Plymouth | Wellcome Sanger Institute for the COVID-19 Genomics UK (COG-UK) Consortium | Lighthouse Laboratory Plymouth and Alex Alderton, Roberto Amato, Jeffrey Barrett, Sonia Goncalves, Ewan Harrison, David K. Jackson, Ian Johnston, Dominic Kwiatkowski, Cordelia Langford, John Sillitoe on behalf of the Wellcome Sanger Institute COVID-19 Surveillance Team |
| EPI_ISL_3169541 | Lighthouse Lab in Glasgow | Wellcome Sanger Institute for the COVID-19 Genomics UK (COG-UK) Consortium | Harper VanSteenhouse, Yumi Kasai, David Gray, Carol Clugston, Anna Dominiczak and Alex Alderton, Roberto Amato, Jeffrey Barrett, Sonia Goncalves, Ewan Harrison, David K. Jackson, Ian Johnston, Dominic Kwiatkowski, Cordelia Langford, John Sillitoe on behalf of the Wellcome Sanger Institute COVID-19 Surveillance Team |

| | | | |
|---|---|---|---|
| EPI_ISL_3169542, EPI_ISL_3169543, EPI_ISL_3169544, EPI_ISL_3169545, EPI_ISL_3169546 | Lighthouse Laboratory Plymouth | Wellcome Sanger Institute for the COVID-19 Genomics UK (COG-UK) Consortium | Lighthouse Laboratory Plymouth and Alex Alderton, Roberto Amato, Jeffrey Barrett, Sonia Goncalves, Ewan Harrison, David K. Jackson, Ian Johnston, Dominic Kwiatkowski, Cordelia Langford, John Sillitoe on behalf of the Wellcome Sanger Institute COVID-19 Surveillance Team |
| EPI_ISL_3169547 | Lighthouse Lab in Glasgow | Wellcome Sanger Institute for the COVID-19 Genomics UK (COG-UK) Consortium | Harper VanSteenhouse, Yumi Kasai, David Gray, Carol Clugston, Anna Dominiczak and Alex Alderton, Roberto Amato, Jeffrey Barrett, Sonia Goncalves, Ewan Harrison, David K. Jackson, Ian Johnston, Dominic Kwiatkowski, Cordelia Langford, John Sillitoe on behalf of the Wellcome Sanger Institute COVID-19 Surveillance Team |
| EPI_ISL_3169548, EPI_ISL_3169549, EPI_ISL_3169550, EPI_ISL_3169551, EPI_ISL_3169552, EPI_ISL_3169553, EPI_ISL_3169554, EPI_ISL_3169555 | Lighthouse Laboratory Plymouth | Wellcome Sanger Institute for the COVID-19 Genomics UK (COG-UK) Consortium | Lighthouse Laboratory Plymouth and Alex Alderton, Roberto Amato, Jeffrey Barrett, Sonia Goncalves, Ewan Harrison, David K. Jackson, Ian Johnston, Dominic Kwiatkowski, Cordelia Langford, John Sillitoe on behalf of the Wellcome Sanger Institute COVID-19 Surveillance Team |
| EPI_ISL_3169556 | Lighthouse Lab in Glasgow | Wellcome Sanger Institute for the COVID-19 Genomics UK (COG-UK) Consortium | Harper VanSteenhouse, Yumi Kasai, David Gray, Carol Clugston, Anna Dominiczak and Alex Alderton, Roberto Amato, Jeffrey Barrett, Sonia Goncalves, Ewan Harrison, David K. Jackson, Ian Johnston, Dominic Kwiatkowski, Cordelia Langford, John Sillitoe on behalf of the Wellcome Sanger Institute COVID-19 Surveillance Team |
| EPI_ISL_3169557, EPI_ISL_3169558, EPI_ISL_3169559, EPI_ISL_3169560, EPI_ISL_3169561, EPI_ISL_3169562, EPI_ISL_3169563, EPI_ISL_3169564, EPI_ISL_3169565, EPI_ISL_3169566, EPI_ISL_3169567, EPI_ISL_3169568, EPI_ISL_3169569, EPI_ISL_3169570, EPI_ISL_3169571, EPI_ISL_3169572 | | | |
| see above | Lighthouse Laboratory Plymouth | Wellcome Sanger Institute for the COVID-19 Genomics UK (COG-UK) Consortium | Lighthouse Laboratory Plymouth and Alex Alderton, Roberto Amato, Jeffrey Barrett, Sonia Goncalves, Ewan Harrison, David K. Jackson, Ian Johnston, Dominic Kwiatkowski, Cordelia Langford, John Sillitoe on behalf of the Wellcome Sanger Institute COVID-19 Surveillance Team |
| EPI_ISL_3169573 | Lighthouse Lab in Glasgow | Wellcome Sanger Institute for the COVID-19 Genomics UK (COG-UK) Consortium | Harper VanSteenhouse, Yumi Kasai, David Gray, Carol Clugston, Anna Dominiczak and Alex Alderton, Roberto Amato, Jeffrey Barrett, Sonia Goncalves, Ewan Harrison, David K. Jackson, Ian Johnston, Dominic Kwiatkowski, Cordelia Langford, John Sillitoe on behalf of the Wellcome Sanger Institute COVID-19 Surveillance Team |
| EPI_ISL_3169574, EPI_ISL_3169575, EPI_ISL_3169576 | Lighthouse Laboratory Plymouth | Wellcome Sanger Institute for the COVID-19 Genomics UK (COG-UK) Consortium | Lighthouse Laboratory Plymouth and Alex Alderton, Roberto Amato, Jeffrey Barrett, Sonia Goncalves, Ewan Harrison, David K. Jackson, Ian Johnston, Dominic Kwiatkowski, Cordelia Langford, John Sillitoe on behalf of the Wellcome Sanger Institute COVID-19 Surveillance Team |
| EPI_ISL_3169577 | Lighthouse Lab in Glasgow | Wellcome Sanger Institute for the COVID-19 Genomics UK (COG-UK) Consortium | Harper VanSteenhouse, Yumi Kasai, David Gray, Carol Clugston, Anna Dominiczak and Alex Alderton, Roberto Amato, Jeffrey Barrett, Sonia Goncalves, Ewan Harrison, David K. Jackson, Ian Johnston, Dominic Kwiatkowski, Cordelia Langford, John Sillitoe on behalf of the Wellcome Sanger Institute COVID-19 Surveillance Team |
| EPI_ISL_3169578 | Lighthouse Laboratory Plymouth | Wellcome Sanger Institute for the COVID-19 Genomics UK (COG-UK) Consortium | Lighthouse Laboratory Plymouth and Alex Alderton, Roberto Amato, Jeffrey Barrett, Sonia Goncalves, Ewan Harrison, David K. Jackson, Ian Johnston, Dominic Kwiatkowski, Cordelia Langford, John Sillitoe on behalf of the Wellcome Sanger Institute COVID-19 Surveillance Team |
| EPI_ISL_3169579, EPI_ISL_3169580 | Lighthouse Lab in Glasgow | Wellcome Sanger Institute for the COVID-19 Genomics UK (COG-UK) Consortium | Harper VanSteenhouse, Yumi Kasai, David Gray, Carol Clugston, Anna Dominiczak and Alex Alderton, Roberto Amato, Jeffrey Barrett, Sonia Goncalves, Ewan Harrison, David K. Jackson, Ian Johnston, Dominic Kwiatkowski, Cordelia Langford, John Sillitoe on behalf of the Wellcome Sanger Institute COVID-19 Surveillance Team |
| EPI_ISL_3169581, EPI_ISL_3169582, EPI_ISL_3169583, EPI_ISL_3169584, EPI_ISL_3169585, EPI_ISL_3169586, EPI_ISL_3169587 | Lighthouse Laboratory Plymouth | Wellcome Sanger Institute for the COVID-19 Genomics UK (COG-UK) Consortium | Lighthouse Laboratory Plymouth and Alex Alderton, Roberto Amato, Jeffrey Barrett, Sonia Goncalves, Ewan Harrison, David K. Jackson, Ian Johnston, Dominic Kwiatkowski, Cordelia Langford, John Sillitoe on behalf of the Wellcome Sanger Institute COVID-19 Surveillance Team |
| EPI_ISL_3169588, EPI_ISL_3169589 | Lighthouse Lab in Glasgow | Wellcome Sanger Institute for the COVID-19 Genomics UK (COG-UK) Consortium | Harper VanSteenhouse, Yumi Kasai, David Gray, Carol Clugston, Anna Dominiczak and Alex Alderton, Roberto Amato, Jeffrey Barrett, Sonia Goncalves, Ewan Harrison, David K. Jackson, Ian Johnston, Dominic Kwiatkowski, Cordelia Langford, John Sillitoe on behalf of the Wellcome Sanger Institute COVID-19 Surveillance Team |
| EPI_ISL_3169590 | Berkshire and Surrey Pathology Services Lighthouse Laboratory | Wellcome Sanger Institute for the COVID-19 Genomics UK (COG-UK) Consortium | Berkshire and Surrey Pathology Services Lighthouse Laboratory and Alex Alderton, Roberto Amato, Jeffrey Barrett, Sonia Goncalves, Ewan Harrison, David K. Jackson, Ian Johnston, Dominic Kwiatkowski, Cordelia Langford, John Sillitoe on behalf of the Wellcome Sanger Institute COVID-19 Surveillance Team |
| EPI_ISL_3169591 | Lighthouse Lab in Milton Keynes | Wellcome Sanger Institute for the COVID-19 Genomics UK (COG-UK) Consortium | The Lighthouse Lab in Milton Keynes and Alex Alderton, Roberto Amato, Jeffrey Barrett, Sonia Goncalves, Ewan Harrison, David K. Jackson, Ian Johnston, Dominic Kwiatkowski, Cordelia Langford, John Sillitoe on behalf of the Wellcome Sanger Institute COVID-19 Surveillance Team |
| EPI_ISL_3169592, EPI_ISL_3169593 | Berkshire and Surrey Pathology Services Lighthouse Laboratory | Wellcome Sanger Institute for the COVID-19 Genomics UK (COG-UK) Consortium | Berkshire and Surrey Pathology Services Lighthouse Laboratory and Alex Alderton, Roberto Amato, Jeffrey Barrett, Sonia Goncalves, Ewan Harrison, David K. Jackson, Ian Johnston, Dominic Kwiatkowski, Cordelia Langford, John Sillitoe on behalf of the Wellcome Sanger Institute COVID-19 Surveillance Team |
| EPI_ISL_3169594 | Lighthouse Lab in Milton Keynes | Wellcome Sanger Institute for the COVID-19 Genomics UK (COG-UK) Consortium | The Lighthouse Lab in Milton Keynes and Alex Alderton, Roberto Amato, Jeffrey Barrett, Sonia Goncalves, Ewan Harrison, David K. Jackson, Ian Johnston, Dominic Kwiatkowski, Cordelia Langford, John Sillitoe on behalf of the Wellcome Sanger Institute COVID-19 Surveillance Team |
| EPI_ISL_3169595 | Lighthouse Lab in Glasgow | Wellcome Sanger Institute for the COVID-19 Genomics UK (COG-UK) Consortium | Harper VanSteenhouse, Yumi Kasai, David Gray, Carol Clugston, Anna Dominiczak and Alex Alderton, Roberto Amato, Jeffrey Barrett, Sonia Goncalves, Ewan Harrison, David K. Jackson, Ian Johnston, Dominic Kwiatkowski, Cordelia Langford, John Sillitoe on behalf of the Wellcome Sanger Institute COVID-19 Surveillance Team |
| EPI_ISL_3169596, EPI_ISL_3169597, EPI_ISL_3169598 | Lighthouse Lab in Milton Keynes | Wellcome Sanger Institute for the COVID-19 Genomics UK (COG-UK) Consortium | The Lighthouse Lab in Milton Keynes and Alex Alderton, Roberto Amato, Jeffrey Barrett, Sonia Goncalves, Ewan Harrison, David K. Jackson, Ian Johnston, Dominic Kwiatkowski, Cordelia Langford, John Sillitoe on behalf of the Wellcome Sanger Institute COVID-19 Surveillance Team |
| EPI_ISL_3169599, EPI_ISL_3169600 | Berkshire and Surrey Pathology Services Lighthouse Laboratory | Wellcome Sanger Institute for the COVID-19 Genomics UK (COG-UK) Consortium | Berkshire and Surrey Pathology Services Lighthouse Laboratory and Alex Alderton, Roberto Amato, Jeffrey Barrett, Sonia Goncalves, Ewan Harrison, David K. Jackson, Ian Johnston, Dominic Kwiatkowski, Cordelia Langford, John Sillitoe on behalf of the Wellcome Sanger Institute COVID-19 Surveillance Team |
| EPI_ISL_3169601 | Lighthouse Lab in Glasgow | Wellcome Sanger Institute for the COVID-19 Genomics UK (COG-UK) Consortium | Harper VanSteenhouse, Yumi Kasai, David Gray, Carol Clugston, Anna Dominiczak and Alex Alderton, Roberto Amato, Jeffrey Barrett, Sonia Goncalves, Ewan Harrison, David K. Jackson, Ian Johnston, Dominic Kwiatkowski, Cordelia Langford, John Sillitoe on behalf of the Wellcome Sanger Institute COVID-19 Surveillance Team |
| EPI_ISL_3169602, EPI_ISL_3169603 | Lighthouse Lab in Milton Keynes | Wellcome Sanger Institute for the COVID-19 Genomics UK (COG-UK) Consortium | The Lighthouse Lab in Milton Keynes and Alex Alderton, Roberto Amato, Jeffrey Barrett, Sonia Goncalves, Ewan Harrison, David K. Jackson, Ian Johnston, Dominic Kwiatkowski, Cordelia Langford, John Sillitoe on behalf of the Wellcome Sanger Institute COVID-19 Surveillance Team |
| EPI_ISL_3169604 | Berkshire and Surrey Pathology Services Lighthouse Laboratory | Wellcome Sanger Institute for the COVID-19 Genomics UK (COG-UK) Consortium | Berkshire and Surrey Pathology Services Lighthouse Laboratory and Alex Alderton, Roberto Amato, Jeffrey Barrett, Sonia Goncalves, Ewan Harrison, David K. Jackson, Ian Johnston, Dominic Kwiatkowski, Cordelia Langford, John Sillitoe on behalf of the Wellcome Sanger Institute COVID-19 Surveillance Team |
| EPI_ISL_3169605 | Lighthouse Lab in Glasgow | Wellcome Sanger Institute for the COVID-19 Genomics UK (COG-UK) Consortium | Harper VanSteenhouse, Yumi Kasai, David Gray, Carol Clugston, Anna Dominiczak and Alex Alderton, Roberto Amato, Jeffrey Barrett, Sonia Goncalves, Ewan Harrison, David K. Jackson, Ian Johnston, Dominic Kwiatkowski, Cordelia Langford, John Sillitoe on behalf of the Wellcome Sanger Institute COVID-19 Surveillance Team |
| EPI_ISL_3169606, EPI_ISL_3169607, EPI_ISL_3169608 | Berkshire and Surrey Pathology Services Lighthouse Laboratory | Wellcome Sanger Institute for the COVID-19 Genomics UK (COG-UK) Consortium | Berkshire and Surrey Pathology Services Lighthouse Laboratory and Alex Alderton, Roberto Amato, Jeffrey Barrett, Sonia Goncalves, Ewan Harrison, David K. Jackson, Ian Johnston, Dominic Kwiatkowski, Cordelia Langford, John Sillitoe on behalf of the Wellcome Sanger Institute COVID-19 Surveillance Team |
| EPI_ISL_3169609, EPI_ISL_3169610 | Lighthouse Lab in Milton Keynes | Wellcome Sanger Institute for the COVID-19 Genomics UK (COG-UK) Consortium | The Lighthouse Lab in Milton Keynes and Alex Alderton, Roberto Amato, Jeffrey Barrett, Sonia Goncalves, Ewan Harrison, David K. Jackson, Ian Johnston, Dominic Kwiatkowski, Cordelia Langford, John Sillitoe on behalf of the Wellcome Sanger Institute COVID-19 Surveillance Team |
| EPI_ISL_3169611, EPI_ISL_3169612, EPI_ISL_3169613 | Berkshire and Surrey Pathology Services Lighthouse Laboratory | Wellcome Sanger Institute for the COVID-19 Genomics UK (COG-UK) Consortium | Berkshire and Surrey Pathology Services Lighthouse Laboratory and Alex Alderton, Roberto Amato, Jeffrey Barrett, Sonia Goncalves, Ewan Harrison, David K. Jackson, Ian Johnston, Dominic Kwiatkowski, Cordelia Langford, John Sillitoe on behalf of the Wellcome Sanger Institute COVID-19 Surveillance Team |
| EPI_ISL_3169614 | Lighthouse Lab in Milton Keynes | Wellcome Sanger Institute for the COVID-19 Genomics UK (COG-UK) Consortium | The Lighthouse Lab in Milton Keynes and Alex Alderton, Roberto Amato, Jeffrey Barrett, Sonia Goncalves, Ewan Harrison, David K. Jackson, Ian Johnston, Dominic Kwiatkowski, Cordelia Langford, John Sillitoe on behalf of the Wellcome Sanger Institute COVID-19 Surveillance Team |
| EPI_ISL_3169615, EPI_ISL_3169616 | Berkshire and Surrey Pathology Services Lighthouse Laboratory | Wellcome Sanger Institute for the COVID-19 Genomics UK (COG-UK) Consortium | Berkshire and Surrey Pathology Services Lighthouse Laboratory and Alex Alderton, Roberto Amato, Jeffrey Barrett, Sonia Goncalves, Ewan Harrison, David K. Jackson, Ian Johnston, Dominic Kwiatkowski, Cordelia Langford, John Sillitoe on behalf of the Wellcome Sanger Institute COVID-19 Surveillance Team |
| EPI_ISL_3169617 | Lighthouse Lab in Milton Keynes | Wellcome Sanger Institute for the COVID-19 Genomics UK (COG-UK) Consortium | The Lighthouse Lab in Milton Keynes and Alex Alderton, Roberto Amato, Jeffrey Barrett, Sonia Goncalves, Ewan Harrison, David K. Jackson, Ian Johnston, Dominic Kwiatkowski, Cordelia Langford, John Sillitoe on behalf of the Wellcome Sanger Institute COVID-19 Surveillance Team |
| EPI_ISL_3169618, EPI_ISL_3169619, EPI_ISL_3169620 | Lighthouse Lab in Glasgow | Wellcome Sanger Institute for the COVID-19 Genomics UK (COG-UK) Consortium | Harper VanSteenhouse, Yumi Kasai, David Gray, Carol Clugston, Anna Dominiczak and Alex Alderton, Roberto Amato, Jeffrey Barrett, Sonia Goncalves, Ewan Harrison, David K. Jackson, Ian Johnston, Dominic Kwiatkowski, Cordelia Langford, John Sillitoe on behalf of the Wellcome Sanger Institute COVID-19 Surveillance Team |
| EPI_ISL_3169621, EPI_ISL_3169622 | Berkshire and Surrey Pathology Services Lighthouse Laboratory | Wellcome Sanger Institute for the COVID-19 Genomics UK (COG-UK) Consortium | Berkshire and Surrey Pathology Services Lighthouse Laboratory and Alex Alderton, Roberto Amato, Jeffrey Barrett, Sonia Goncalves, Ewan Harrison, David K. Jackson, Ian Johnston, Dominic Kwiatkowski, Cordelia Langford, John Sillitoe on behalf of the Wellcome Sanger Institute COVID-19 Surveillance Team |
| EPI_ISL_3169623, EPI_ISL_3169624 | Lighthouse Lab in Glasgow | Wellcome Sanger Institute for the COVID-19 Genomics UK (COG-UK) Consortium | Harper VanSteenhouse, Yumi Kasai, David Gray, Carol Clugston, Anna Dominiczak and Alex Alderton, Roberto Amato, Jeffrey Barrett, Sonia Goncalves, Ewan Harrison, David K. Jackson, Ian Johnston, Dominic Kwiatkowski, Cordelia Langford, John Sillitoe on behalf of the Wellcome Sanger Institute COVID-19 Surveillance Team |
| EPI_ISL_3169625, EPI_ISL_3169626 | Lighthouse Lab in Milton Keynes | Wellcome Sanger Institute for the COVID-19 Genomics UK (COG-UK) Consortium | The Lighthouse Lab in Milton Keynes and Alex Alderton, Roberto Amato, Jeffrey Barrett, Sonia Goncalves, Ewan Harrison, David K. Jackson, Ian Johnston, Dominic Kwiatkowski, Cordelia Langford, John Sillitoe on behalf of the Wellcome Sanger Institute COVID-19 Surveillance Team |
| EPI_ISL_3169627, EPI_ISL_3169628 | Berkshire and Surrey Pathology Services Lighthouse Laboratory | Wellcome Sanger Institute for the COVID-19 Genomics UK (COG-UK) Consortium | Berkshire and Surrey Pathology Services Lighthouse Laboratory and Alex Alderton, Roberto Amato, Jeffrey Barrett, Sonia Goncalves, Ewan Harrison, David K. Jackson, Ian Johnston, Dominic Kwiatkowski, Cordelia Langford, John Sillitoe on behalf of the Wellcome Sanger Institute COVID-19 Surveillance Team |
| EPI_ISL_3169629 | Lighthouse Lab in Milton Keynes | Wellcome Sanger Institute for the COVID-19 Genomics UK (COG-UK) Consortium | The Lighthouse Lab in Milton Keynes and Alex Alderton, Roberto Amato, Jeffrey Barrett, Sonia Goncalves, Ewan Harrison, David K. Jackson, Ian Johnston, Dominic Kwiatkowski, Cordelia Langford, John Sillitoe on behalf of the Wellcome Sanger Institute COVID-19 Surveillance Team |
| EPI_ISL_3169630, EPI_ISL_3169631 | Lighthouse Lab in Glasgow | Wellcome Sanger Institute for the COVID-19 Genomics UK (COG-UK) Consortium | Harper VanSteenhouse, Yumi Kasai, David Gray, Carol Clugston, Anna Dominiczak and Alex Alderton, Roberto Amato, Jeffrey Barrett, Sonia Goncalves, Ewan Harrison, David K. Jackson, Ian Johnston, Dominic Kwiatkowski, Cordelia Langford, John Sillitoe on behalf of the Wellcome Sanger Institute COVID-19 Surveillance Team |
| EPI_ISL_3169632, | Lighthouse Lab in Milton Keynes | Wellcome Sanger Institute for the COVID-19 Genomics UK (COG-UK) | The Lighthouse Lab in Milton Keynes and Alex Alderton, Roberto Amato, Jeffrey Barrett, Sonia Goncalves, Ewan Harrison, David K. Jackson, Ian Johnston, Dominic Kwiatkowski, Cordelia Langford, John Sillitoe on behalf of the Wellcome Sanger |

| | | | |
|---|---|---|---|
| EPI_ISL_3169633 | | Consortium | Institute COVID-19 Surveillance Team |
| EPI_ISL_3169634, EPI_ISL_3169635 | Lighthouse Lab in Glasgow | Wellcome Sanger Institute for the COVID-19 Genomics UK (COG-UK) Consortium | Harper VanSteenhouse, Yumi Kasai, David Gray, Carol Clugston, Anna Dominiczak and Alex Alderton, Roberto Amato, Jeffrey Barrett, Sonia Goncalves, Ewan Harrison, David K. Jackson, Ian Johnston, Dominic Kwiatkowski, Cordelia Langford, John Sillitoe on behalf of the Wellcome Sanger Institute COVID-19 Surveillance Team |
| EPI_ISL_3169636, EPI_ISL_3169637, EPI_ISL_3169638 | Berkshire and Surrey Pathology Services Lighthouse Laboratory | Wellcome Sanger Institute for the COVID-19 Genomics UK (COG-UK) Consortium | Berkshire and Surrey Pathology Services Lighthouse Laboratory and Alex Alderton, Roberto Amato, Jeffrey Barrett, Sonia Goncalves, Ewan Harrison, David K. Jackson, Ian Johnston, Dominic Kwiatkowski, Cordelia Langford, John Sillitoe on behalf of the Wellcome Sanger Institute COVID-19 Surveillance Team |
| EPI_ISL_3169639 | Lighthouse Lab in Milton Keynes | Wellcome Sanger Institute for the COVID-19 Genomics UK (COG-UK) Consortium | The Lighthouse Lab in Milton Keynes and Alex Alderton, Roberto Amato, Jeffrey Barrett, Sonia Goncalves, Ewan Harrison, David K. Jackson, Ian Johnston, Dominic Kwiatkowski, Cordelia Langford, John Sillitoe on behalf of the Wellcome Sanger Institute COVID-19 Surveillance Team |
| EPI_ISL_3169640, EPI_ISL_3169641, EPI_ISL_3169642, EPI_ISL_3169643, EPI_ISL_3169644, EPI_ISL_3169645, EPI_ISL_3169646 | Berkshire and Surrey Pathology Services Lighthouse Laboratory | Wellcome Sanger Institute for the COVID-19 Genomics UK (COG-UK) Consortium | Berkshire and Surrey Pathology Services Lighthouse Laboratory and Alex Alderton, Roberto Amato, Jeffrey Barrett, Sonia Goncalves, Ewan Harrison, David K. Jackson, Ian Johnston, Dominic Kwiatkowski, Cordelia Langford, John Sillitoe on behalf of the Wellcome Sanger Institute COVID-19 Surveillance Team |
| EPI_ISL_3169647 | Lighthouse Lab in Glasgow | Wellcome Sanger Institute for the COVID-19 Genomics UK (COG-UK) Consortium | Harper VanSteenhouse, Yumi Kasai, David Gray, Carol Clugston, Anna Dominiczak and Alex Alderton, Roberto Amato, Jeffrey Barrett, Sonia Goncalves, Ewan Harrison, David K. Jackson, Ian Johnston, Dominic Kwiatkowski, Cordelia Langford, John Sillitoe on behalf of the Wellcome Sanger Institute COVID-19 Surveillance Team |
| EPI_ISL_3169648 | Lighthouse Lab in Milton Keynes | Wellcome Sanger Institute for the COVID-19 Genomics UK (COG-UK) Consortium | The Lighthouse Lab in Milton Keynes and Alex Alderton, Roberto Amato, Jeffrey Barrett, Sonia Goncalves, Ewan Harrison, David K. Jackson, Ian Johnston, Dominic Kwiatkowski, Cordelia Langford, John Sillitoe on behalf of the Wellcome Sanger Institute COVID-19 Surveillance Team |
| EPI_ISL_3169649, EPI_ISL_3169650 | Berkshire and Surrey Pathology Services Lighthouse Laboratory | Wellcome Sanger Institute for the COVID-19 Genomics UK (COG-UK) Consortium | Berkshire and Surrey Pathology Services Lighthouse Laboratory and Alex Alderton, Roberto Amato, Jeffrey Barrett, Sonia Goncalves, Ewan Harrison, David K. Jackson, Ian Johnston, Dominic Kwiatkowski, Cordelia Langford, John Sillitoe on behalf of the Wellcome Sanger Institute COVID-19 Surveillance Team |
| EPI_ISL_3169651 | Lighthouse Lab in Glasgow | Wellcome Sanger Institute for the COVID-19 Genomics UK (COG-UK) Consortium | Harper VanSteenhouse, Yumi Kasai, David Gray, Carol Clugston, Anna Dominiczak and Alex Alderton, Roberto Amato, Jeffrey Barrett, Sonia Goncalves, Ewan Harrison, David K. Jackson, Ian Johnston, Dominic Kwiatkowski, Cordelia Langford, John Sillitoe on behalf of the Wellcome Sanger Institute COVID-19 Surveillance Team |
| EPI_ISL_3169652, EPI_ISL_3169653 | Berkshire and Surrey Pathology Services Lighthouse Laboratory | Wellcome Sanger Institute for the COVID-19 Genomics UK (COG-UK) Consortium | Berkshire and Surrey Pathology Services Lighthouse Laboratory and Alex Alderton, Roberto Amato, Jeffrey Barrett, Sonia Goncalves, Ewan Harrison, David K. Jackson, Ian Johnston, Dominic Kwiatkowski, Cordelia Langford, John Sillitoe on behalf of the Wellcome Sanger Institute COVID-19 Surveillance Team |
| EPI_ISL_3169654 | Lighthouse Lab in Glasgow | Wellcome Sanger Institute for the COVID-19 Genomics UK (COG-UK) Consortium | Harper VanSteenhouse, Yumi Kasai, David Gray, Carol Clugston, Anna Dominiczak and Alex Alderton, Roberto Amato, Jeffrey Barrett, Sonia Goncalves, Ewan Harrison, David K. Jackson, Ian Johnston, Dominic Kwiatkowski, Cordelia Langford, John Sillitoe on behalf of the Wellcome Sanger Institute COVID-19 Surveillance Team |
| EPI_ISL_3169655 | Berkshire and Surrey Pathology Services Lighthouse Laboratory | Wellcome Sanger Institute for the COVID-19 Genomics UK (COG-UK) Consortium | Berkshire and Surrey Pathology Services Lighthouse Laboratory and Alex Alderton, Roberto Amato, Jeffrey Barrett, Sonia Goncalves, Ewan Harrison, David K. Jackson, Ian Johnston, Dominic Kwiatkowski, Cordelia Langford, John Sillitoe on behalf of the Wellcome Sanger Institute COVID-19 Surveillance Team |
| EPI_ISL_3169656 | Lighthouse Lab in Milton Keynes | Wellcome Sanger Institute for the COVID-19 Genomics UK (COG-UK) Consortium | The Lighthouse Lab in Milton Keynes and Alex Alderton, Roberto Amato, Jeffrey Barrett, Sonia Goncalves, Ewan Harrison, David K. Jackson, Ian Johnston, Dominic Kwiatkowski, Cordelia Langford, John Sillitoe on behalf of the Wellcome Sanger Institute COVID-19 Surveillance Team |
| EPI_ISL_3169657 | Berkshire and Surrey Pathology Services Lighthouse Laboratory | Wellcome Sanger Institute for the COVID-19 Genomics UK (COG-UK) Consortium | Berkshire and Surrey Pathology Services Lighthouse Laboratory and Alex Alderton, Roberto Amato, Jeffrey Barrett, Sonia Goncalves, Ewan Harrison, David K. Jackson, Ian Johnston, Dominic Kwiatkowski, Cordelia Langford, John Sillitoe on behalf of the Wellcome Sanger Institute COVID-19 Surveillance Team |
| EPI_ISL_3169658 | Lighthouse Lab in Milton Keynes | Wellcome Sanger Institute for the COVID-19 Genomics UK (COG-UK) Consortium | The Lighthouse Lab in Milton Keynes and Alex Alderton, Roberto Amato, Jeffrey Barrett, Sonia Goncalves, Ewan Harrison, David K. Jackson, Ian Johnston, Dominic Kwiatkowski, Cordelia Langford, John Sillitoe on behalf of the Wellcome Sanger Institute COVID-19 Surveillance Team |
| EPI_ISL_3169659 | Berkshire and Surrey Pathology Services Lighthouse Laboratory | Wellcome Sanger Institute for the COVID-19 Genomics UK (COG-UK) Consortium | Berkshire and Surrey Pathology Services Lighthouse Laboratory and Alex Alderton, Roberto Amato, Jeffrey Barrett, Sonia Goncalves, Ewan Harrison, David K. Jackson, Ian Johnston, Dominic Kwiatkowski, Cordelia Langford, John Sillitoe on behalf of the Wellcome Sanger Institute COVID-19 Surveillance Team |
| EPI_ISL_3169660 | Lighthouse Lab in Glasgow | Wellcome Sanger Institute for the COVID-19 Genomics UK (COG-UK) Consortium | Harper VanSteenhouse, Yumi Kasai, David Gray, Carol Clugston, Anna Dominiczak and Alex Alderton, Roberto Amato, Jeffrey Barrett, Sonia Goncalves, Ewan Harrison, David K. Jackson, Ian Johnston, Dominic Kwiatkowski, Cordelia Langford, John Sillitoe on behalf of the Wellcome Sanger Institute COVID-19 Surveillance Team |
| EPI_ISL_3169661 | Lighthouse Lab in Milton Keynes | Wellcome Sanger Institute for the COVID-19 Genomics UK (COG-UK) Consortium | The Lighthouse Lab in Milton Keynes and Alex Alderton, Roberto Amato, Jeffrey Barrett, Sonia Goncalves, Ewan Harrison, David K. Jackson, Ian Johnston, Dominic Kwiatkowski, Cordelia Langford, John Sillitoe on behalf of the Wellcome Sanger Institute COVID-19 Surveillance Team |
| EPI_ISL_3169662 | Berkshire and Surrey Pathology Services Lighthouse Laboratory | Wellcome Sanger Institute for the COVID-19 Genomics UK (COG-UK) Consortium | Berkshire and Surrey Pathology Services Lighthouse Laboratory and Alex Alderton, Roberto Amato, Jeffrey Barrett, Sonia Goncalves, Ewan Harrison, David K. Jackson, Ian Johnston, Dominic Kwiatkowski, Cordelia Langford, John Sillitoe on behalf of the Wellcome Sanger Institute COVID-19 Surveillance Team |
| EPI_ISL_3169663 | Lighthouse Lab in Milton Keynes | Wellcome Sanger Institute for the COVID-19 Genomics UK (COG-UK) Consortium | The Lighthouse Lab in Milton Keynes and Alex Alderton, Roberto Amato, Jeffrey Barrett, Sonia Goncalves, Ewan Harrison, David K. Jackson, Ian Johnston, Dominic Kwiatkowski, Cordelia Langford, John Sillitoe on behalf of the Wellcome Sanger Institute COVID-19 Surveillance Team |
| EPI_ISL_3169664 | Berkshire and Surrey Pathology Services Lighthouse Laboratory | Wellcome Sanger Institute for the COVID-19 Genomics UK (COG-UK) Consortium | Berkshire and Surrey Pathology Services Lighthouse Laboratory and Alex Alderton, Roberto Amato, Jeffrey Barrett, Sonia Goncalves, Ewan Harrison, David K. Jackson, Ian Johnston, Dominic Kwiatkowski, Cordelia Langford, John Sillitoe on behalf of the Wellcome Sanger Institute COVID-19 Surveillance Team |
| EPI_ISL_3169665 | Lighthouse Lab in Milton Keynes | Wellcome Sanger Institute for the COVID-19 Genomics UK (COG-UK) Consortium | The Lighthouse Lab in Milton Keynes and Alex Alderton, Roberto Amato, Jeffrey Barrett, Sonia Goncalves, Ewan Harrison, David K. Jackson, Ian Johnston, Dominic Kwiatkowski, Cordelia Langford, John Sillitoe on behalf of the Wellcome Sanger Institute COVID-19 Surveillance Team |
| EPI_ISL_3169666, EPI_ISL_3169667, EPI_ISL_3169668 | Berkshire and Surrey Pathology Services Lighthouse Laboratory | Wellcome Sanger Institute for the COVID-19 Genomics UK (COG-UK) Consortium | Berkshire and Surrey Pathology Services Lighthouse Laboratory and Alex Alderton, Roberto Amato, Jeffrey Barrett, Sonia Goncalves, Ewan Harrison, David K. Jackson, Ian Johnston, Dominic Kwiatkowski, Cordelia Langford, John Sillitoe on behalf of the Wellcome Sanger Institute COVID-19 Surveillance Team |
| EPI_ISL_3169669 | Lighthouse Lab in Glasgow | Wellcome Sanger Institute for the COVID-19 Genomics UK (COG-UK) Consortium | Harper VanSteenhouse, Yumi Kasai, David Gray, Carol Clugston, Anna Dominiczak and Alex Alderton, Roberto Amato, Jeffrey Barrett, Sonia Goncalves, Ewan Harrison, David K. Jackson, Ian Johnston, Dominic Kwiatkowski, Cordelia Langford, John Sillitoe on behalf of the Wellcome Sanger Institute COVID-19 Surveillance Team |
| EPI_ISL_3169670, EPI_ISL_3169671, EPI_ISL_3169672 | Berkshire and Surrey Pathology Services Lighthouse Laboratory | Wellcome Sanger Institute for the COVID-19 Genomics UK (COG-UK) Consortium | Berkshire and Surrey Pathology Services Lighthouse Laboratory and Alex Alderton, Roberto Amato, Jeffrey Barrett, Sonia Goncalves, Ewan Harrison, David K. Jackson, Ian Johnston, Dominic Kwiatkowski, Cordelia Langford, John Sillitoe on behalf of the Wellcome Sanger Institute COVID-19 Surveillance Team |
| EPI_ISL_3169673 | Lighthouse Lab in Glasgow | Wellcome Sanger Institute for the COVID-19 Genomics UK (COG-UK) Consortium | Harper VanSteenhouse, Yumi Kasai, David Gray, Carol Clugston, Anna Dominiczak and Alex Alderton, Roberto Amato, Jeffrey Barrett, Sonia Goncalves, Ewan Harrison, David K. Jackson, Ian Johnston, Dominic Kwiatkowski, Cordelia Langford, John Sillitoe on behalf of the Wellcome Sanger Institute COVID-19 Surveillance Team |
| EPI_ISL_3169674 | Berkshire and Surrey Pathology Services Lighthouse Laboratory | Wellcome Sanger Institute for the COVID-19 Genomics UK (COG-UK) Consortium | Berkshire and Surrey Pathology Services Lighthouse Laboratory and Alex Alderton, Roberto Amato, Jeffrey Barrett, Sonia Goncalves, Ewan Harrison, David K. Jackson, Ian Johnston, Dominic Kwiatkowski, Cordelia Langford, John Sillitoe on behalf of the Wellcome Sanger Institute COVID-19 Surveillance Team |
| EPI_ISL_3169675, EPI_ISL_3169676 | Lighthouse Lab in Milton Keynes | Wellcome Sanger Institute for the COVID-19 Genomics UK (COG-UK) Consortium | The Lighthouse Lab in Milton Keynes and Alex Alderton, Roberto Amato, Jeffrey Barrett, Sonia Goncalves, Ewan Harrison, David K. Jackson, Ian Johnston, Dominic Kwiatkowski, Cordelia Langford, John Sillitoe on behalf of the Wellcome Sanger Institute COVID-19 Surveillance Team |
| EPI_ISL_3169677 | Berkshire and Surrey Pathology Services Lighthouse Laboratory | Wellcome Sanger Institute for the COVID-19 Genomics UK (COG-UK) Consortium | Berkshire and Surrey Pathology Services Lighthouse Laboratory and Alex Alderton, Roberto Amato, Jeffrey Barrett, Sonia Goncalves, Ewan Harrison, David K. Jackson, Ian Johnston, Dominic Kwiatkowski, Cordelia Langford, John Sillitoe on behalf of the Wellcome Sanger Institute COVID-19 Surveillance Team |
| EPI_ISL_3169678 | Lighthouse Lab in Milton Keynes | Wellcome Sanger Institute for the COVID-19 Genomics UK (COG-UK) Consortium | The Lighthouse Lab in Milton Keynes and Alex Alderton, Roberto Amato, Jeffrey Barrett, Sonia Goncalves, Ewan Harrison, David K. Jackson, Ian Johnston, Dominic Kwiatkowski, Cordelia Langford, John Sillitoe on behalf of the Wellcome Sanger Institute COVID-19 Surveillance Team |
| EPI_ISL_3169679, EPI_ISL_3169680 | Berkshire and Surrey Pathology Services Lighthouse Laboratory | Wellcome Sanger Institute for the COVID-19 Genomics UK (COG-UK) Consortium | Berkshire and Surrey Pathology Services Lighthouse Laboratory and Alex Alderton, Roberto Amato, Jeffrey Barrett, Sonia Goncalves, Ewan Harrison, David K. Jackson, Ian Johnston, Dominic Kwiatkowski, Cordelia Langford, John Sillitoe on behalf of the Wellcome Sanger Institute COVID-19 Surveillance Team |
| EPI_ISL_3169681, EPI_ISL_3169682 | Lighthouse Lab in Glasgow | Wellcome Sanger Institute for the COVID-19 Genomics UK (COG-UK) Consortium | Harper VanSteenhouse, Yumi Kasai, David Gray, Carol Clugston, Anna Dominiczak and Alex Alderton, Roberto Amato, Jeffrey Barrett, Sonia Goncalves, Ewan Harrison, David K. Jackson, Ian Johnston, Dominic Kwiatkowski, Cordelia Langford, John Sillitoe on behalf of the Wellcome Sanger Institute COVID-19 Surveillance Team |
| EPI_ISL_3169683, EPI_ISL_3169684 | Berkshire and Surrey Pathology Services Lighthouse Laboratory | Wellcome Sanger Institute for the COVID-19 Genomics UK (COG-UK) Consortium | Berkshire and Surrey Pathology Services Lighthouse Laboratory and Alex Alderton, Roberto Amato, Jeffrey Barrett, Sonia Goncalves, Ewan Harrison, David K. Jackson, Ian Johnston, Dominic Kwiatkowski, Cordelia Langford, John Sillitoe on behalf of the Wellcome Sanger Institute COVID-19 Surveillance Team |
| EPI_ISL_3169685 | Lighthouse Lab in Milton Keynes | Wellcome Sanger Institute for the COVID-19 Genomics UK (COG-UK) Consortium | The Lighthouse Lab in Milton Keynes and Alex Alderton, Roberto Amato, Jeffrey Barrett, Sonia Goncalves, Ewan Harrison, David K. Jackson, Ian Johnston, Dominic Kwiatkowski, Cordelia Langford, John Sillitoe on behalf of the Wellcome Sanger Institute COVID-19 Surveillance Team |
| EPI_ISL_3169686, EPI_ISL_3169687, EPI_ISL_3169688 | Berkshire and Surrey Pathology Services Lighthouse Laboratory | Wellcome Sanger Institute for the COVID-19 Genomics UK (COG-UK) Consortium | Berkshire and Surrey Pathology Services Lighthouse Laboratory and Alex Alderton, Roberto Amato, Jeffrey Barrett, Sonia Goncalves, Ewan Harrison, David K. Jackson, Ian Johnston, Dominic Kwiatkowski, Cordelia Langford, John Sillitoe on behalf of the Wellcome Sanger Institute COVID-19 Surveillance Team |
| EPI_ISL_3169689 | Lighthouse Lab in Milton Keynes | Wellcome Sanger Institute for the COVID-19 Genomics UK (COG-UK) Consortium | The Lighthouse Lab in Milton Keynes and Alex Alderton, Roberto Amato, Jeffrey Barrett, Sonia Goncalves, Ewan Harrison, David K. Jackson, Ian Johnston, Dominic Kwiatkowski, Cordelia Langford, John Sillitoe on behalf of the Wellcome Sanger Institute COVID-19 Surveillance Team |
| EPI_ISL_3169690, EPI_ISL_3169691, EPI_ISL_3169692 | Berkshire and Surrey Pathology Services Lighthouse Laboratory | Wellcome Sanger Institute for the COVID-19 Genomics UK (COG-UK) Consortium | Berkshire and Surrey Pathology Services Lighthouse Laboratory and Alex Alderton, Roberto Amato, Jeffrey Barrett, Sonia Goncalves, Ewan Harrison, David K. Jackson, Ian Johnston, Dominic Kwiatkowski, Cordelia Langford, John Sillitoe on behalf of the Wellcome Sanger Institute COVID-19 Surveillance Team |
| EPI_ISL_3169693 | Lighthouse Lab in Milton Keynes | Wellcome Sanger Institute for the COVID-19 Genomics UK (COG-UK) Consortium | The Lighthouse Lab in Milton Keynes and Alex Alderton, Roberto Amato, Jeffrey Barrett, Sonia Goncalves, Ewan Harrison, David K. Jackson, Ian Johnston, Dominic Kwiatkowski, Cordelia Langford, John Sillitoe on behalf of the Wellcome Sanger Institute COVID-19 Surveillance Team |
| EPI_ISL_3169694 | Lighthouse Lab in Glasgow | Wellcome Sanger Institute for the COVID-19 Genomics UK (COG-UK) Consortium | Harper VanSteenhouse, Yumi Kasai, David Gray, Carol Clugston, Anna Dominiczak and Alex Alderton, Roberto Amato, Jeffrey Barrett, Sonia Goncalves, Ewan Harrison, David K. Jackson, Ian Johnston, Dominic Kwiatkowski, Cordelia Langford, John Sillitoe on behalf of the Wellcome Sanger Institute COVID-19 Surveillance Team |
| EPI_ISL_3169695, EPI_ISL_3169696 | Berkshire and Surrey Pathology Services Lighthouse Laboratory | Wellcome Sanger Institute for the COVID-19 Genomics UK (COG-UK) Consortium | Berkshire and Surrey Pathology Services Lighthouse Laboratory and Alex Alderton, Roberto Amato, Jeffrey Barrett, Sonia Goncalves, Ewan Harrison, David K. Jackson, Ian Johnston, Dominic Kwiatkowski, Cordelia Langford, John Sillitoe on behalf of the Wellcome Sanger Institute COVID-19 Surveillance Team |
| EPI_ISL_3169697 | Lighthouse Lab in Glasgow | Wellcome Sanger Institute for the COVID-19 Genomics UK (COG-UK) Consortium | Harper VanSteenhouse, Yumi Kasai, David Gray, Carol Clugston, Anna Dominiczak and Alex Alderton, Roberto Amato, Jeffrey Barrett, Sonia Goncalves, Ewan Harrison, David K. Jackson, Ian Johnston, Dominic Kwiatkowski, Cordelia Langford, John Sillitoe on behalf of the Wellcome Sanger Institute COVID-19 Surveillance Team |

| | | | |
|---|---|---|---|
| EPI_ISL_3169698, EPI_ISL_3169699 | Berkshire and Surrey Pathology Services Lighthouse Laboratory | Wellcome Sanger Institute for the COVID-19 Genomics UK (COG-UK) Consortium | Berkshire and Surrey Pathology Services Lighthouse Laboratory and Alex Alderton, Roberto Amato, Jeffrey Barrett, Sonia Goncalves, Ewan Harrison, David K. Jackson, Ian Johnston, Dominic Kwiatkowski, Cordelia Langford, John Sillitoe on behalf of the Wellcome Sanger Institute COVID-19 Surveillance Team |
| EPI_ISL_3169700 | Lighthouse Lab in Glasgow | Wellcome Sanger Institute for the COVID-19 Genomics UK (COG-UK) Consortium | Harper VanSteenhouse, Yumi Kasai, David Gray, Carol Clugston, Anna Dominiczak and Alex Alderton, Roberto Amato, Jeffrey Barrett, Sonia Goncalves, Ewan Harrison, David K. Jackson, Ian Johnston, Dominic Kwiatkowski, Cordelia Langford, John Sillitoe on behalf of the Wellcome Sanger Institute COVID-19 Surveillance Team |
| EPI_ISL_3169701, EPI_ISL_3169702, EPI_ISL_3169703, EPI_ISL_3169704 | Berkshire and Surrey Pathology Services Lighthouse Laboratory | Wellcome Sanger Institute for the COVID-19 Genomics UK (COG-UK) Consortium | Berkshire and Surrey Pathology Services Lighthouse Laboratory and Alex Alderton, Roberto Amato, Jeffrey Barrett, Sonia Goncalves, Ewan Harrison, David K. Jackson, Ian Johnston, Dominic Kwiatkowski, Cordelia Langford, John Sillitoe on behalf of the Wellcome Sanger Institute COVID-19 Surveillance Team |
| EPI_ISL_3169705 | Lighthouse Lab in Glasgow | Wellcome Sanger Institute for the COVID-19 Genomics UK (COG-UK) Consortium | Harper VanSteenhouse, Yumi Kasai, David Gray, Carol Clugston, Anna Dominiczak and Alex Alderton, Roberto Amato, Jeffrey Barrett, Sonia Goncalves, Ewan Harrison, David K. Jackson, Ian Johnston, Dominic Kwiatkowski, Cordelia Langford, John Sillitoe on behalf of the Wellcome Sanger Institute COVID-19 Surveillance Team |
| EPI_ISL_3169706, EPI_ISL_3169707, EPI_ISL_3169708 | Berkshire and Surrey Pathology Services Lighthouse Laboratory | Wellcome Sanger Institute for the COVID-19 Genomics UK (COG-UK) Consortium | Berkshire and Surrey Pathology Services Lighthouse Laboratory and Alex Alderton, Roberto Amato, Jeffrey Barrett, Sonia Goncalves, Ewan Harrison, David K. Jackson, Ian Johnston, Dominic Kwiatkowski, Cordelia Langford, John Sillitoe on behalf of the Wellcome Sanger Institute COVID-19 Surveillance Team |
| EPI_ISL_3169709, EPI_ISL_3169710 | Lighthouse Lab in Milton Keynes | Wellcome Sanger Institute for the COVID-19 Genomics UK (COG-UK) Consortium | The Lighthouse Lab in Milton Keynes and Alex Alderton, Roberto Amato, Jeffrey Barrett, Sonia Goncalves, Ewan Harrison, David K. Jackson, Ian Johnston, Dominic Kwiatkowski, Cordelia Langford, John Sillitoe on behalf of the Wellcome Sanger Institute COVID-19 Surveillance Team |
| EPI_ISL_3169711, EPI_ISL_3169712, EPI_ISL_3169713 | Lighthouse Lab in Glasgow | Wellcome Sanger Institute for the COVID-19 Genomics UK (COG-UK) Consortium | Harper VanSteenhouse, Yumi Kasai, David Gray, Carol Clugston, Anna Dominiczak and Alex Alderton, Roberto Amato, Jeffrey Barrett, Sonia Goncalves, Ewan Harrison, David K. Jackson, Ian Johnston, Dominic Kwiatkowski, Cordelia Langford, John Sillitoe on behalf of the Wellcome Sanger Institute COVID-19 Surveillance Team |
| EPI_ISL_3169714, EPI_ISL_3169715, EPI_ISL_3169716 | Berkshire and Surrey Pathology Services Lighthouse Laboratory | Wellcome Sanger Institute for the COVID-19 Genomics UK (COG-UK) Consortium | Berkshire and Surrey Pathology Services Lighthouse Laboratory and Alex Alderton, Roberto Amato, Jeffrey Barrett, Sonia Goncalves, Ewan Harrison, David K. Jackson, Ian Johnston, Dominic Kwiatkowski, Cordelia Langford, John Sillitoe on behalf of the Wellcome Sanger Institute COVID-19 Surveillance Team |
| EPI_ISL_3169717 | Lighthouse Lab in Milton Keynes | Wellcome Sanger Institute for the COVID-19 Genomics UK (COG-UK) Consortium | The Lighthouse Lab in Milton Keynes and Alex Alderton, Roberto Amato, Jeffrey Barrett, Sonia Goncalves, Ewan Harrison, David K. Jackson, Ian Johnston, Dominic Kwiatkowski, Cordelia Langford, John Sillitoe on behalf of the Wellcome Sanger Institute COVID-19 Surveillance Team |
| EPI_ISL_3169718 | Berkshire and Surrey Pathology Services Lighthouse Laboratory | Wellcome Sanger Institute for the COVID-19 Genomics UK (COG-UK) Consortium | Berkshire and Surrey Pathology Services Lighthouse Laboratory and Alex Alderton, Roberto Amato, Jeffrey Barrett, Sonia Goncalves, Ewan Harrison, David K. Jackson, Ian Johnston, Dominic Kwiatkowski, Cordelia Langford, John Sillitoe on behalf of the Wellcome Sanger Institute COVID-19 Surveillance Team |
| EPI_ISL_3169719, EPI_ISL_3169720 | Lighthouse Lab in Glasgow | Wellcome Sanger Institute for the COVID-19 Genomics UK (COG-UK) Consortium | Harper VanSteenhouse, Yumi Kasai, David Gray, Carol Clugston, Anna Dominiczak and Alex Alderton, Roberto Amato, Jeffrey Barrett, Sonia Goncalves, Ewan Harrison, David K. Jackson, Ian Johnston, Dominic Kwiatkowski, Cordelia Langford, John Sillitoe on behalf of the Wellcome Sanger Institute COVID-19 Surveillance Team |
| EPI_ISL_3169721, EPI_ISL_3169722 | Berkshire and Surrey Pathology Services Lighthouse Laboratory | Wellcome Sanger Institute for the COVID-19 Genomics UK (COG-UK) Consortium | Berkshire and Surrey Pathology Services Lighthouse Laboratory and Alex Alderton, Roberto Amato, Jeffrey Barrett, Sonia Goncalves, Ewan Harrison, David K. Jackson, Ian Johnston, Dominic Kwiatkowski, Cordelia Langford, John Sillitoe on behalf of the Wellcome Sanger Institute COVID-19 Surveillance Team |
| EPI_ISL_3169723 | Lighthouse Lab in Glasgow | Wellcome Sanger Institute for the COVID-19 Genomics UK (COG-UK) Consortium | Harper VanSteenhouse, Yumi Kasai, David Gray, Carol Clugston, Anna Dominiczak and Alex Alderton, Roberto Amato, Jeffrey Barrett, Sonia Goncalves, Ewan Harrison, David K. Jackson, Ian Johnston, Dominic Kwiatkowski, Cordelia Langford, John Sillitoe on behalf of the Wellcome Sanger Institute COVID-19 Surveillance Team |
| EPI_ISL_3169724 | Berkshire and Surrey Pathology Services Lighthouse Laboratory | Wellcome Sanger Institute for the COVID-19 Genomics UK (COG-UK) Consortium | Berkshire and Surrey Pathology Services Lighthouse Laboratory and Alex Alderton, Roberto Amato, Jeffrey Barrett, Sonia Goncalves, Ewan Harrison, David K. Jackson, Ian Johnston, Dominic Kwiatkowski, Cordelia Langford, John Sillitoe on behalf of the Wellcome Sanger Institute COVID-19 Surveillance Team |
| EPI_ISL_3169725 | Lighthouse Lab in Milton Keynes | Wellcome Sanger Institute for the COVID-19 Genomics UK (COG-UK) Consortium | The Lighthouse Lab in Milton Keynes and Alex Alderton, Roberto Amato, Jeffrey Barrett, Sonia Goncalves, Ewan Harrison, David K. Jackson, Ian Johnston, Dominic Kwiatkowski, Cordelia Langford, John Sillitoe on behalf of the Wellcome Sanger Institute COVID-19 Surveillance Team |
| EPI_ISL_3169726 | Berkshire and Surrey Pathology Services Lighthouse Laboratory | Wellcome Sanger Institute for the COVID-19 Genomics UK (COG-UK) Consortium | Berkshire and Surrey Pathology Services Lighthouse Laboratory and Alex Alderton, Roberto Amato, Jeffrey Barrett, Sonia Goncalves, Ewan Harrison, David K. Jackson, Ian Johnston, Dominic Kwiatkowski, Cordelia Langford, John Sillitoe on behalf of the Wellcome Sanger Institute COVID-19 Surveillance Team |
| EPI_ISL_3169727 | Lighthouse Lab in Milton Keynes | Wellcome Sanger Institute for the COVID-19 Genomics UK (COG-UK) Consortium | The Lighthouse Lab in Milton Keynes and Alex Alderton, Roberto Amato, Jeffrey Barrett, Sonia Goncalves, Ewan Harrison, David K. Jackson, Ian Johnston, Dominic Kwiatkowski, Cordelia Langford, John Sillitoe on behalf of the Wellcome Sanger Institute COVID-19 Surveillance Team |
| EPI_ISL_3169728 | Lighthouse Lab in Glasgow | Wellcome Sanger Institute for the COVID-19 Genomics UK (COG-UK) Consortium | Harper VanSteenhouse, Yumi Kasai, David Gray, Carol Clugston, Anna Dominiczak and Alex Alderton, Roberto Amato, Jeffrey Barrett, Sonia Goncalves, Ewan Harrison, David K. Jackson, Ian Johnston, Dominic Kwiatkowski, Cordelia Langford, John Sillitoe on behalf of the Wellcome Sanger Institute COVID-19 Surveillance Team |
| EPI_ISL_3169729 | Berkshire and Surrey Pathology Services Lighthouse Laboratory | Wellcome Sanger Institute for the COVID-19 Genomics UK (COG-UK) Consortium | Berkshire and Surrey Pathology Services Lighthouse Laboratory and Alex Alderton, Roberto Amato, Jeffrey Barrett, Sonia Goncalves, Ewan Harrison, David K. Jackson, Ian Johnston, Dominic Kwiatkowski, Cordelia Langford, John Sillitoe on behalf of the Wellcome Sanger Institute COVID-19 Surveillance Team |
| EPI_ISL_3169730, EPI_ISL_3169731 | Lighthouse Lab in Milton Keynes | Wellcome Sanger Institute for the COVID-19 Genomics UK (COG-UK) Consortium | The Lighthouse Lab in Milton Keynes and Alex Alderton, Roberto Amato, Jeffrey Barrett, Sonia Goncalves, Ewan Harrison, David K. Jackson, Ian Johnston, Dominic Kwiatkowski, Cordelia Langford, John Sillitoe on behalf of the Wellcome Sanger Institute COVID-19 Surveillance Team |
| EPI_ISL_3169732 | Lighthouse Lab in Glasgow | Wellcome Sanger Institute for the COVID-19 Genomics UK (COG-UK) Consortium | Harper VanSteenhouse, Yumi Kasai, David Gray, Carol Clugston, Anna Dominiczak and Alex Alderton, Roberto Amato, Jeffrey Barrett, Sonia Goncalves, Ewan Harrison, David K. Jackson, Ian Johnston, Dominic Kwiatkowski, Cordelia Langford, John Sillitoe on behalf of the Wellcome Sanger Institute COVID-19 Surveillance Team |
| EPI_ISL_3169733, EPI_ISL_3169734 | Berkshire and Surrey Pathology Services Lighthouse Laboratory | Wellcome Sanger Institute for the COVID-19 Genomics UK (COG-UK) Consortium | Berkshire and Surrey Pathology Services Lighthouse Laboratory and Alex Alderton, Roberto Amato, Jeffrey Barrett, Sonia Goncalves, Ewan Harrison, David K. Jackson, Ian Johnston, Dominic Kwiatkowski, Cordelia Langford, John Sillitoe on behalf of the Wellcome Sanger Institute COVID-19 Surveillance Team |
| EPI_ISL_3169735 | Lighthouse Lab in Glasgow | Wellcome Sanger Institute for the COVID-19 Genomics UK (COG-UK) Consortium | Harper VanSteenhouse, Yumi Kasai, David Gray, Carol Clugston, Anna Dominiczak and Alex Alderton, Roberto Amato, Jeffrey Barrett, Sonia Goncalves, Ewan Harrison, David K. Jackson, Ian Johnston, Dominic Kwiatkowski, Cordelia Langford, John Sillitoe on behalf of the Wellcome Sanger Institute COVID-19 Surveillance Team |
| EPI_ISL_3169736, EPI_ISL_3169737, EPI_ISL_3169738 | Berkshire and Surrey Pathology Services Lighthouse Laboratory | Wellcome Sanger Institute for the COVID-19 Genomics UK (COG-UK) Consortium | Berkshire and Surrey Pathology Services Lighthouse Laboratory and Alex Alderton, Roberto Amato, Jeffrey Barrett, Sonia Goncalves, Ewan Harrison, David K. Jackson, Ian Johnston, Dominic Kwiatkowski, Cordelia Langford, John Sillitoe on behalf of the Wellcome Sanger Institute COVID-19 Surveillance Team |
| EPI_ISL_3169739 | Lighthouse Lab in Glasgow | Wellcome Sanger Institute for the COVID-19 Genomics UK (COG-UK) Consortium | Harper VanSteenhouse, Yumi Kasai, David Gray, Carol Clugston, Anna Dominiczak and Alex Alderton, Roberto Amato, Jeffrey Barrett, Sonia Goncalves, Ewan Harrison, David K. Jackson, Ian Johnston, Dominic Kwiatkowski, Cordelia Langford, John Sillitoe on behalf of the Wellcome Sanger Institute COVID-19 Surveillance Team |
| EPI_ISL_3169740 | Lighthouse Lab in Milton Keynes | Wellcome Sanger Institute for the COVID-19 Genomics UK (COG-UK) Consortium | The Lighthouse Lab in Milton Keynes and Alex Alderton, Roberto Amato, Jeffrey Barrett, Sonia Goncalves, Ewan Harrison, David K. Jackson, Ian Johnston, Dominic Kwiatkowski, Cordelia Langford, John Sillitoe on behalf of the Wellcome Sanger Institute COVID-19 Surveillance Team |
| EPI_ISL_3169741 | Berkshire and Surrey Pathology Services Lighthouse Laboratory | Wellcome Sanger Institute for the COVID-19 Genomics UK (COG-UK) Consortium | Berkshire and Surrey Pathology Services Lighthouse Laboratory and Alex Alderton, Roberto Amato, Jeffrey Barrett, Sonia Goncalves, Ewan Harrison, David K. Jackson, Ian Johnston, Dominic Kwiatkowski, Cordelia Langford, John Sillitoe on behalf of the Wellcome Sanger Institute COVID-19 Surveillance Team |
| EPI_ISL_3169742, EPI_ISL_3169743 | Lighthouse Lab in Milton Keynes | Wellcome Sanger Institute for the COVID-19 Genomics UK (COG-UK) Consortium | The Lighthouse Lab in Milton Keynes and Alex Alderton, Roberto Amato, Jeffrey Barrett, Sonia Goncalves, Ewan Harrison, David K. Jackson, Ian Johnston, Dominic Kwiatkowski, Cordelia Langford, John Sillitoe on behalf of the Wellcome Sanger Institute COVID-19 Surveillance Team |
| EPI_ISL_3169744 | Berkshire and Surrey Pathology Services Lighthouse Laboratory | Wellcome Sanger Institute for the COVID-19 Genomics UK (COG-UK) Consortium | Berkshire and Surrey Pathology Services Lighthouse Laboratory and Alex Alderton, Roberto Amato, Jeffrey Barrett, Sonia Goncalves, Ewan Harrison, David K. Jackson, Ian Johnston, Dominic Kwiatkowski, Cordelia Langford, John Sillitoe on behalf of the Wellcome Sanger Institute COVID-19 Surveillance Team |
| EPI_ISL_3169745 | Lighthouse Lab in Milton Keynes | Wellcome Sanger Institute for the COVID-19 Genomics UK (COG-UK) Consortium | The Lighthouse Lab in Milton Keynes and Alex Alderton, Roberto Amato, Jeffrey Barrett, Sonia Goncalves, Ewan Harrison, David K. Jackson, Ian Johnston, Dominic Kwiatkowski, Cordelia Langford, John Sillitoe on behalf of the Wellcome Sanger Institute COVID-19 Surveillance Team |
| EPI_ISL_3169746, EPI_ISL_3169747, EPI_ISL_3169748, EPI_ISL_3169749 | Berkshire and Surrey Pathology Services Lighthouse Laboratory | Wellcome Sanger Institute for the COVID-19 Genomics UK (COG-UK) Consortium | Berkshire and Surrey Pathology Services Lighthouse Laboratory and Alex Alderton, Roberto Amato, Jeffrey Barrett, Sonia Goncalves, Ewan Harrison, David K. Jackson, Ian Johnston, Dominic Kwiatkowski, Cordelia Langford, John Sillitoe on behalf of the Wellcome Sanger Institute COVID-19 Surveillance Team |
| EPI_ISL_3169750, EPI_ISL_3169751, EPI_ISL_3169752 | Lighthouse Lab in Milton Keynes | Wellcome Sanger Institute for the COVID-19 Genomics UK (COG-UK) Consortium | The Lighthouse Lab in Milton Keynes and Alex Alderton, Roberto Amato, Jeffrey Barrett, Sonia Goncalves, Ewan Harrison, David K. Jackson, Ian Johnston, Dominic Kwiatkowski, Cordelia Langford, John Sillitoe on behalf of the Wellcome Sanger Institute COVID-19 Surveillance Team |
| EPI_ISL_3169753, EPI_ISL_3169754 | Berkshire and Surrey Pathology Services Lighthouse Laboratory | Wellcome Sanger Institute for the COVID-19 Genomics UK (COG-UK) Consortium | Berkshire and Surrey Pathology Services Lighthouse Laboratory and Alex Alderton, Roberto Amato, Jeffrey Barrett, Sonia Goncalves, Ewan Harrison, David K. Jackson, Ian Johnston, Dominic Kwiatkowski, Cordelia Langford, John Sillitoe on behalf of the Wellcome Sanger Institute COVID-19 Surveillance Team |
| EPI_ISL_3169755, EPI_ISL_3169756 | Lighthouse Lab in Glasgow | Wellcome Sanger Institute for the COVID-19 Genomics UK (COG-UK) Consortium | Harper VanSteenhouse, Yumi Kasai, David Gray, Carol Clugston, Anna Dominiczak and Alex Alderton, Roberto Amato, Jeffrey Barrett, Sonia Goncalves, Ewan Harrison, David K. Jackson, Ian Johnston, Dominic Kwiatkowski, Cordelia Langford, John Sillitoe on behalf of the Wellcome Sanger Institute COVID-19 Surveillance Team |
| EPI_ISL_3169757, EPI_ISL_3169758, EPI_ISL_3169759, EPI_ISL_3169760 | Berkshire and Surrey Pathology Services Lighthouse Laboratory | Wellcome Sanger Institute for the COVID-19 Genomics UK (COG-UK) Consortium | Berkshire and Surrey Pathology Services Lighthouse Laboratory and Alex Alderton, Roberto Amato, Jeffrey Barrett, Sonia Goncalves, Ewan Harrison, David K. Jackson, Ian Johnston, Dominic Kwiatkowski, Cordelia Langford, John Sillitoe on behalf of the Wellcome Sanger Institute COVID-19 Surveillance Team |
| EPI_ISL_3169761, EPI_ISL_3169762 | Lighthouse Lab in Glasgow | Wellcome Sanger Institute for the COVID-19 Genomics UK (COG-UK) Consortium | Harper VanSteenhouse, Yumi Kasai, David Gray, Carol Clugston, Anna Dominiczak and Alex Alderton, Roberto Amato, Jeffrey Barrett, Sonia Goncalves, Ewan Harrison, David K. Jackson, Ian Johnston, Dominic Kwiatkowski, Cordelia Langford, John Sillitoe on behalf of the Wellcome Sanger Institute COVID-19 Surveillance Team |
| EPI_ISL_3169763 | Lighthouse Lab in Milton Keynes | Wellcome Sanger Institute for the COVID-19 Genomics UK (COG-UK) Consortium | The Lighthouse Lab in Milton Keynes and Alex Alderton, Roberto Amato, Jeffrey Barrett, Sonia Goncalves, Ewan Harrison, David K. Jackson, Ian Johnston, Dominic Kwiatkowski, Cordelia Langford, John Sillitoe on behalf of the Wellcome Sanger Institute COVID-19 Surveillance Team |
| EPI_ISL_3169764 | Lighthouse Lab in Glasgow | Wellcome Sanger Institute for the COVID-19 Genomics UK (COG-UK) Consortium | Harper VanSteenhouse, Yumi Kasai, David Gray, Carol Clugston, Anna Dominiczak and Alex Alderton, Roberto Amato, Jeffrey Barrett, Sonia Goncalves, Ewan Harrison, David K. Jackson, Ian Johnston, Dominic Kwiatkowski, Cordelia Langford, John Sillitoe on behalf of the Wellcome Sanger Institute COVID-19 Surveillance Team |
| EPI_ISL_3169765, EPI_ISL_3169766 | Berkshire and Surrey Pathology Services Lighthouse Laboratory | Wellcome Sanger Institute for the COVID-19 Genomics UK (COG-UK) Consortium | Berkshire and Surrey Pathology Services Lighthouse Laboratory and Alex Alderton, Roberto Amato, Jeffrey Barrett, Sonia Goncalves, Ewan Harrison, David K. Jackson, Ian Johnston, Dominic Kwiatkowski, Cordelia Langford, John Sillitoe on behalf of the Wellcome Sanger Institute COVID-19 Surveillance Team |
| EPI_ISL_3169767 | Lighthouse Lab in Milton Keynes | Wellcome Sanger Institute for the COVID-19 Genomics UK (COG-UK) Consortium | The Lighthouse Lab in Milton Keynes and Alex Alderton, Roberto Amato, Jeffrey Barrett, Sonia Goncalves, Ewan Harrison, David K. Jackson, Ian Johnston, Dominic Kwiatkowski, Cordelia Langford, John Sillitoe on behalf of the Wellcome Sanger Institute COVID-19 Surveillance Team |
| EPI_ISL_3169768, | Berkshire and Surrey Pathology Services Lighthouse Laboratory | Wellcome Sanger Institute for the COVID-19 Genomics UK (COG-UK) | Berkshire and Surrey Pathology Services Lighthouse Laboratory and Alex Alderton, Roberto Amato, Jeffrey Barrett, Sonia Goncalves, Ewan Harrison, David K. Jackson, Ian Johnston, Dominic Kwiatkowski, Cordelia Langford, John Sillitoe on behalf |

| | | | |
|---|---|---|---|
| EPI_ISL_3169769, EPI_ISL_3169770, EPI_ISL_3169771, EPI_ISL_3169772 | | Consortium | of the Wellcome Sanger Institute COVID-19 Surveillance Team |
| EPI_ISL_3169773 | Lighthouse Lab in Glasgow | Wellcome Sanger Institute for the COVID-19 Genomics UK (COG-UK) Consortium | Harper VanSteenhouse, Yumi Kasai, David Gray, Carol Clugston, Anna Dominiczak and Alex Alderton, Roberto Amato, Jeffrey Barrett, Sonia Goncalves, Ewan Harrison, David K. Jackson, Ian Johnston, Dominic Kwiatkowski, Cordelia Langford, John Sillitoe on behalf of the Wellcome Sanger Institute COVID-19 Surveillance Team |
| EPI_ISL_3169774 | Berkshire and Surrey Pathology Services Lighthouse Laboratory | Wellcome Sanger Institute for the COVID-19 Genomics UK (COG-UK) Consortium | Berkshire and Surrey Pathology Services Lighthouse Laboratory and Alex Alderton, Roberto Amato, Jeffrey Barrett, Sonia Goncalves, Ewan Harrison, David K. Jackson, Ian Johnston, Dominic Kwiatkowski, Cordelia Langford, John Sillitoe on behalf of the Wellcome Sanger Institute COVID-19 Surveillance Team |
| EPI_ISL_3169775, EPI_ISL_3169776 | Lighthouse Lab in Milton Keynes | Wellcome Sanger Institute for the COVID-19 Genomics UK (COG-UK) Consortium | The Lighthouse Lab in Milton Keynes and Alex Alderton, Roberto Amato, Jeffrey Barrett, Sonia Goncalves, Ewan Harrison, David K. Jackson, Ian Johnston, Dominic Kwiatkowski, Cordelia Langford, John Sillitoe on behalf of the Wellcome Sanger Institute COVID-19 Surveillance Team |
| EPI_ISL_3169777 | Lighthouse Lab in Glasgow | Wellcome Sanger Institute for the COVID-19 Genomics UK (COG-UK) Consortium | Harper VanSteenhouse, Yumi Kasai, David Gray, Carol Clugston, Anna Dominiczak and Alex Alderton, Roberto Amato, Jeffrey Barrett, Sonia Goncalves, Ewan Harrison, David K. Jackson, Ian Johnston, Dominic Kwiatkowski, Cordelia Langford, John Sillitoe on behalf of the Wellcome Sanger Institute COVID-19 Surveillance Team |
| EPI_ISL_3169778 | Lighthouse Lab in Milton Keynes | Wellcome Sanger Institute for the COVID-19 Genomics UK (COG-UK) Consortium | The Lighthouse Lab in Milton Keynes and Alex Alderton, Roberto Amato, Jeffrey Barrett, Sonia Goncalves, Ewan Harrison, David K. Jackson, Ian Johnston, Dominic Kwiatkowski, Cordelia Langford, John Sillitoe on behalf of the Wellcome Sanger Institute COVID-19 Surveillance Team |
| EPI_ISL_3169779, EPI_ISL_3169780 | Berkshire and Surrey Pathology Services Lighthouse Laboratory | Wellcome Sanger Institute for the COVID-19 Genomics UK (COG-UK) Consortium | Berkshire and Surrey Pathology Services Lighthouse Laboratory and Alex Alderton, Roberto Amato, Jeffrey Barrett, Sonia Goncalves, Ewan Harrison, David K. Jackson, Ian Johnston, Dominic Kwiatkowski, Cordelia Langford, John Sillitoe on behalf of the Wellcome Sanger Institute COVID-19 Surveillance Team |
| EPI_ISL_3169781 | Lighthouse Lab in Milton Keynes | Wellcome Sanger Institute for the COVID-19 Genomics UK (COG-UK) Consortium | The Lighthouse Lab in Milton Keynes and Alex Alderton, Roberto Amato, Jeffrey Barrett, Sonia Goncalves, Ewan Harrison, David K. Jackson, Ian Johnston, Dominic Kwiatkowski, Cordelia Langford, John Sillitoe on behalf of the Wellcome Sanger Institute COVID-19 Surveillance Team |
| EPI_ISL_3169782 | Berkshire and Surrey Pathology Services Lighthouse Laboratory | Wellcome Sanger Institute for the COVID-19 Genomics UK (COG-UK) Consortium | Berkshire and Surrey Pathology Services Lighthouse Laboratory and Alex Alderton, Roberto Amato, Jeffrey Barrett, Sonia Goncalves, Ewan Harrison, David K. Jackson, Ian Johnston, Dominic Kwiatkowski, Cordelia Langford, John Sillitoe on behalf of the Wellcome Sanger Institute COVID-19 Surveillance Team |
| EPI_ISL_3169784 | Lighthouse Lab in Glasgow | Wellcome Sanger Institute for the COVID-19 Genomics UK (COG-UK) Consortium | Harper VanSteenhouse, Yumi Kasai, David Gray, Carol Clugston, Anna Dominiczak and Alex Alderton, Roberto Amato, Jeffrey Barrett, Sonia Goncalves, Ewan Harrison, David K. Jackson, Ian Johnston, Dominic Kwiatkowski, Cordelia Langford, John Sillitoe on behalf of the Wellcome Sanger Institute COVID-19 Surveillance Team |
| EPI_ISL_3169785 | Berkshire and Surrey Pathology Services Lighthouse Laboratory | Wellcome Sanger Institute for the COVID-19 Genomics UK (COG-UK) Consortium | Berkshire and Surrey Pathology Services Lighthouse Laboratory and Alex Alderton, Roberto Amato, Jeffrey Barrett, Sonia Goncalves, Ewan Harrison, David K. Jackson, Ian Johnston, Dominic Kwiatkowski, Cordelia Langford, John Sillitoe on behalf of the Wellcome Sanger Institute COVID-19 Surveillance Team |
| EPI_ISL_3169786 | Lighthouse Lab in Glasgow | Wellcome Sanger Institute for the COVID-19 Genomics UK (COG-UK) Consortium | Harper VanSteenhouse, Yumi Kasai, David Gray, Carol Clugston, Anna Dominiczak and Alex Alderton, Roberto Amato, Jeffrey Barrett, Sonia Goncalves, Ewan Harrison, David K. Jackson, Ian Johnston, Dominic Kwiatkowski, Cordelia Langford, John Sillitoe on behalf of the Wellcome Sanger Institute COVID-19 Surveillance Team |
| EPI_ISL_3169787, EPI_ISL_3169788 | Berkshire and Surrey Pathology Services Lighthouse Laboratory | Wellcome Sanger Institute for the COVID-19 Genomics UK (COG-UK) Consortium | Berkshire and Surrey Pathology Services Lighthouse Laboratory and Alex Alderton, Roberto Amato, Jeffrey Barrett, Sonia Goncalves, Ewan Harrison, David K. Jackson, Ian Johnston, Dominic Kwiatkowski, Cordelia Langford, John Sillitoe on behalf of the Wellcome Sanger Institute COVID-19 Surveillance Team |
| EPI_ISL_3169789 | Lighthouse Lab in Glasgow | Wellcome Sanger Institute for the COVID-19 Genomics UK (COG-UK) Consortium | Harper VanSteenhouse, Yumi Kasai, David Gray, Carol Clugston, Anna Dominiczak and Alex Alderton, Roberto Amato, Jeffrey Barrett, Sonia Goncalves, Ewan Harrison, David K. Jackson, Ian Johnston, Dominic Kwiatkowski, Cordelia Langford, John Sillitoe on behalf of the Wellcome Sanger Institute COVID-19 Surveillance Team |
| EPI_ISL_3169790 | Berkshire and Surrey Pathology Services Lighthouse Laboratory | Wellcome Sanger Institute for the COVID-19 Genomics UK (COG-UK) Consortium | Berkshire and Surrey Pathology Services Lighthouse Laboratory and Alex Alderton, Roberto Amato, Jeffrey Barrett, Sonia Goncalves, Ewan Harrison, David K. Jackson, Ian Johnston, Dominic Kwiatkowski, Cordelia Langford, John Sillitoe on behalf of the Wellcome Sanger Institute COVID-19 Surveillance Team |
| EPI_ISL_3169791 | Lighthouse Lab in Milton Keynes | Wellcome Sanger Institute for the COVID-19 Genomics UK (COG-UK) Consortium | The Lighthouse Lab in Milton Keynes and Alex Alderton, Roberto Amato, Jeffrey Barrett, Sonia Goncalves, Ewan Harrison, David K. Jackson, Ian Johnston, Dominic Kwiatkowski, Cordelia Langford, John Sillitoe on behalf of the Wellcome Sanger Institute COVID-19 Surveillance Team |
| EPI_ISL_3169792 | Berkshire and Surrey Pathology Services Lighthouse Laboratory | Wellcome Sanger Institute for the COVID-19 Genomics UK (COG-UK) Consortium | Berkshire and Surrey Pathology Services Lighthouse Laboratory and Alex Alderton, Roberto Amato, Jeffrey Barrett, Sonia Goncalves, Ewan Harrison, David K. Jackson, Ian Johnston, Dominic Kwiatkowski, Cordelia Langford, John Sillitoe on behalf of the Wellcome Sanger Institute COVID-19 Surveillance Team |
| EPI_ISL_3169793 | Lighthouse Lab in Glasgow | Wellcome Sanger Institute for the COVID-19 Genomics UK (COG-UK) Consortium | Harper VanSteenhouse, Yumi Kasai, David Gray, Carol Clugston, Anna Dominiczak and Alex Alderton, Roberto Amato, Jeffrey Barrett, Sonia Goncalves, Ewan Harrison, David K. Jackson, Ian Johnston, Dominic Kwiatkowski, Cordelia Langford, John Sillitoe on behalf of the Wellcome Sanger Institute COVID-19 Surveillance Team |
| EPI_ISL_3169794 | Lighthouse Lab in Milton Keynes | Wellcome Sanger Institute for the COVID-19 Genomics UK (COG-UK) Consortium | The Lighthouse Lab in Milton Keynes and Alex Alderton, Roberto Amato, Jeffrey Barrett, Sonia Goncalves, Ewan Harrison, David K. Jackson, Ian Johnston, Dominic Kwiatkowski, Cordelia Langford, John Sillitoe on behalf of the Wellcome Sanger Institute COVID-19 Surveillance Team |
| EPI_ISL_3169795, EPI_ISL_3169796 | Lighthouse Lab in Glasgow | Wellcome Sanger Institute for the COVID-19 Genomics UK (COG-UK) Consortium | Harper VanSteenhouse, Yumi Kasai, David Gray, Carol Clugston, Anna Dominiczak and Alex Alderton, Roberto Amato, Jeffrey Barrett, Sonia Goncalves, Ewan Harrison, David K. Jackson, Ian Johnston, Dominic Kwiatkowski, Cordelia Langford, John Sillitoe on behalf of the Wellcome Sanger Institute COVID-19 Surveillance Team |
| EPI_ISL_3169797 | Lighthouse Lab in Milton Keynes | Wellcome Sanger Institute for the COVID-19 Genomics UK (COG-UK) Consortium | The Lighthouse Lab in Milton Keynes and Alex Alderton, Roberto Amato, Jeffrey Barrett, Sonia Goncalves, Ewan Harrison, David K. Jackson, Ian Johnston, Dominic Kwiatkowski, Cordelia Langford, John Sillitoe on behalf of the Wellcome Sanger Institute COVID-19 Surveillance Team |
| EPI_ISL_3169798 | Berkshire and Surrey Pathology Services Lighthouse Laboratory | Wellcome Sanger Institute for the COVID-19 Genomics UK (COG-UK) Consortium | Berkshire and Surrey Pathology Services Lighthouse Laboratory and Alex Alderton, Roberto Amato, Jeffrey Barrett, Sonia Goncalves, Ewan Harrison, David K. Jackson, Ian Johnston, Dominic Kwiatkowski, Cordelia Langford, John Sillitoe on behalf of the Wellcome Sanger Institute COVID-19 Surveillance Team |
| EPI_ISL_3169799 | Lighthouse Lab in Glasgow | Wellcome Sanger Institute for the COVID-19 Genomics UK (COG-UK) Consortium | Harper VanSteenhouse, Yumi Kasai, David Gray, Carol Clugston, Anna Dominiczak and Alex Alderton, Roberto Amato, Jeffrey Barrett, Sonia Goncalves, Ewan Harrison, David K. Jackson, Ian Johnston, Dominic Kwiatkowski, Cordelia Langford, John Sillitoe on behalf of the Wellcome Sanger Institute COVID-19 Surveillance Team |
| EPI_ISL_3169800, EPI_ISL_3169801 | Berkshire and Surrey Pathology Services Lighthouse Laboratory | Wellcome Sanger Institute for the COVID-19 Genomics UK (COG-UK) Consortium | Berkshire and Surrey Pathology Services Lighthouse Laboratory and Alex Alderton, Roberto Amato, Jeffrey Barrett, Sonia Goncalves, Ewan Harrison, David K. Jackson, Ian Johnston, Dominic Kwiatkowski, Cordelia Langford, John Sillitoe on behalf of the Wellcome Sanger Institute COVID-19 Surveillance Team |
| EPI_ISL_3169802 | Lighthouse Lab in Glasgow | Wellcome Sanger Institute for the COVID-19 Genomics UK (COG-UK) Consortium | Harper VanSteenhouse, Yumi Kasai, David Gray, Carol Clugston, Anna Dominiczak and Alex Alderton, Roberto Amato, Jeffrey Barrett, Sonia Goncalves, Ewan Harrison, David K. Jackson, Ian Johnston, Dominic Kwiatkowski, Cordelia Langford, John Sillitoe on behalf of the Wellcome Sanger Institute COVID-19 Surveillance Team |
| EPI_ISL_3169803 | Berkshire and Surrey Pathology Services Lighthouse Laboratory | Wellcome Sanger Institute for the COVID-19 Genomics UK (COG-UK) Consortium | Berkshire and Surrey Pathology Services Lighthouse Laboratory and Alex Alderton, Roberto Amato, Jeffrey Barrett, Sonia Goncalves, Ewan Harrison, David K. Jackson, Ian Johnston, Dominic Kwiatkowski, Cordelia Langford, John Sillitoe on behalf of the Wellcome Sanger Institute COVID-19 Surveillance Team |
| EPI_ISL_3169804, EPI_ISL_3169805 | Lighthouse Lab in Milton Keynes | Wellcome Sanger Institute for the COVID-19 Genomics UK (COG-UK) Consortium | The Lighthouse Lab in Milton Keynes and Alex Alderton, Roberto Amato, Jeffrey Barrett, Sonia Goncalves, Ewan Harrison, David K. Jackson, Ian Johnston, Dominic Kwiatkowski, Cordelia Langford, John Sillitoe on behalf of the Wellcome Sanger Institute COVID-19 Surveillance Team |
| EPI_ISL_3169806, EPI_ISL_3169807, EPI_ISL_3169808, EPI_ISL_3169809 | Berkshire and Surrey Pathology Services Lighthouse Laboratory | Wellcome Sanger Institute for the COVID-19 Genomics UK (COG-UK) Consortium | Berkshire and Surrey Pathology Services Lighthouse Laboratory and Alex Alderton, Roberto Amato, Jeffrey Barrett, Sonia Goncalves, Ewan Harrison, David K. Jackson, Ian Johnston, Dominic Kwiatkowski, Cordelia Langford, John Sillitoe on behalf of the Wellcome Sanger Institute COVID-19 Surveillance Team |
| EPI_ISL_3169810 | Lighthouse Lab in Glasgow | Wellcome Sanger Institute for the COVID-19 Genomics UK (COG-UK) Consortium | Harper VanSteenhouse, Yumi Kasai, David Gray, Carol Clugston, Anna Dominiczak and Alex Alderton, Roberto Amato, Jeffrey Barrett, Sonia Goncalves, Ewan Harrison, David K. Jackson, Ian Johnston, Dominic Kwiatkowski, Cordelia Langford, John Sillitoe on behalf of the Wellcome Sanger Institute COVID-19 Surveillance Team |
| EPI_ISL_3169811, EPI_ISL_3169812 | Berkshire and Surrey Pathology Services Lighthouse Laboratory | Wellcome Sanger Institute for the COVID-19 Genomics UK (COG-UK) Consortium | Berkshire and Surrey Pathology Services Lighthouse Laboratory and Alex Alderton, Roberto Amato, Jeffrey Barrett, Sonia Goncalves, Ewan Harrison, David K. Jackson, Ian Johnston, Dominic Kwiatkowski, Cordelia Langford, John Sillitoe on behalf of the Wellcome Sanger Institute COVID-19 Surveillance Team |
| EPI_ISL_3169813 | Lighthouse Lab in Milton Keynes | Wellcome Sanger Institute for the COVID-19 Genomics UK (COG-UK) Consortium | The Lighthouse Lab in Milton Keynes and Alex Alderton, Roberto Amato, Jeffrey Barrett, Sonia Goncalves, Ewan Harrison, David K. Jackson, Ian Johnston, Dominic Kwiatkowski, Cordelia Langford, John Sillitoe on behalf of the Wellcome Sanger Institute COVID-19 Surveillance Team |
| EPI_ISL_3169814 | Lighthouse Lab in Glasgow | Wellcome Sanger Institute for the COVID-19 Genomics UK (COG-UK) Consortium | Harper VanSteenhouse, Yumi Kasai, David Gray, Carol Clugston, Anna Dominiczak and Alex Alderton, Roberto Amato, Jeffrey Barrett, Sonia Goncalves, Ewan Harrison, David K. Jackson, Ian Johnston, Dominic Kwiatkowski, Cordelia Langford, John Sillitoe on behalf of the Wellcome Sanger Institute COVID-19 Surveillance Team |
| EPI_ISL_3169815 | Berkshire and Surrey Pathology Services Lighthouse Laboratory | Wellcome Sanger Institute for the COVID-19 Genomics UK (COG-UK) Consortium | Berkshire and Surrey Pathology Services Lighthouse Laboratory and Alex Alderton, Roberto Amato, Jeffrey Barrett, Sonia Goncalves, Ewan Harrison, David K. Jackson, Ian Johnston, Dominic Kwiatkowski, Cordelia Langford, John Sillitoe on behalf of the Wellcome Sanger Institute COVID-19 Surveillance Team |
| EPI_ISL_3169816 | Lighthouse Lab in Milton Keynes | Wellcome Sanger Institute for the COVID-19 Genomics UK (COG-UK) Consortium | The Lighthouse Lab in Milton Keynes and Alex Alderton, Roberto Amato, Jeffrey Barrett, Sonia Goncalves, Ewan Harrison, David K. Jackson, Ian Johnston, Dominic Kwiatkowski, Cordelia Langford, John Sillitoe on behalf of the Wellcome Sanger Institute COVID-19 Surveillance Team |
| EPI_ISL_3169818 | Lighthouse Lab in Glasgow | Wellcome Sanger Institute for the COVID-19 Genomics UK (COG-UK) Consortium | Harper VanSteenhouse, Yumi Kasai, David Gray, Carol Clugston, Anna Dominiczak and Alex Alderton, Roberto Amato, Jeffrey Barrett, Sonia Goncalves, Ewan Harrison, David K. Jackson, Ian Johnston, Dominic Kwiatkowski, Cordelia Langford, John Sillitoe on behalf of the Wellcome Sanger Institute COVID-19 Surveillance Team |
| EPI_ISL_3169819 | Lighthouse Lab in Milton Keynes | Wellcome Sanger Institute for the COVID-19 Genomics UK (COG-UK) Consortium | The Lighthouse Lab in Milton Keynes and Alex Alderton, Roberto Amato, Jeffrey Barrett, Sonia Goncalves, Ewan Harrison, David K. Jackson, Ian Johnston, Dominic Kwiatkowski, Cordelia Langford, John Sillitoe on behalf of the Wellcome Sanger Institute COVID-19 Surveillance Team |
| EPI_ISL_3169820 | Lighthouse Lab in Glasgow | Wellcome Sanger Institute for the COVID-19 Genomics UK (COG-UK) Consortium | Harper VanSteenhouse, Yumi Kasai, David Gray, Carol Clugston, Anna Dominiczak and Alex Alderton, Roberto Amato, Jeffrey Barrett, Sonia Goncalves, Ewan Harrison, David K. Jackson, Ian Johnston, Dominic Kwiatkowski, Cordelia Langford, John Sillitoe on behalf of the Wellcome Sanger Institute COVID-19 Surveillance Team |
| EPI_ISL_3169821, EPI_ISL_3169822, EPI_ISL_3169823 | Berkshire and Surrey Pathology Services Lighthouse Laboratory | Wellcome Sanger Institute for the COVID-19 Genomics UK (COG-UK) Consortium | Berkshire and Surrey Pathology Services Lighthouse Laboratory and Alex Alderton, Roberto Amato, Jeffrey Barrett, Sonia Goncalves, Ewan Harrison, David K. Jackson, Ian Johnston, Dominic Kwiatkowski, Cordelia Langford, John Sillitoe on behalf of the Wellcome Sanger Institute COVID-19 Surveillance Team |
| EPI_ISL_3169824 | Lighthouse Lab in Glasgow | Wellcome Sanger Institute for the COVID-19 Genomics UK (COG-UK) Consortium | Harper VanSteenhouse, Yumi Kasai, David Gray, Carol Clugston, Anna Dominiczak and Alex Alderton, Roberto Amato, Jeffrey Barrett, Sonia Goncalves, Ewan Harrison, David K. Jackson, Ian Johnston, Dominic Kwiatkowski, Cordelia Langford, John Sillitoe on behalf of the Wellcome Sanger Institute COVID-19 Surveillance Team |
| EPI_ISL_3169825 | Berkshire and Surrey Pathology Services Lighthouse Laboratory | Wellcome Sanger Institute for the COVID-19 Genomics UK (COG-UK) Consortium | Berkshire and Surrey Pathology Services Lighthouse Laboratory and Alex Alderton, Roberto Amato, Jeffrey Barrett, Sonia Goncalves, Ewan Harrison, David K. Jackson, Ian Johnston, Dominic Kwiatkowski, Cordelia Langford, John Sillitoe on behalf of the Wellcome Sanger Institute COVID-19 Surveillance Team |
| EPI_ISL_3169826 | Lighthouse Lab in Glasgow | Wellcome Sanger Institute for the COVID-19 Genomics UK (COG-UK) Consortium | Harper VanSteenhouse, Yumi Kasai, David Gray, Carol Clugston, Anna Dominiczak and Alex Alderton, Roberto Amato, Jeffrey Barrett, Sonia Goncalves, Ewan Harrison, David K. Jackson, Ian Johnston, Dominic Kwiatkowski, Cordelia Langford, John Sillitoe on behalf of the Wellcome Sanger Institute COVID-19 Surveillance Team |
| EPI_ISL_3169827, EPI_ISL_3169828 | Berkshire and Surrey Pathology Services Lighthouse Laboratory | Wellcome Sanger Institute for the COVID-19 Genomics UK (COG-UK) Consortium | Berkshire and Surrey Pathology Services Lighthouse Laboratory and Alex Alderton, Roberto Amato, Jeffrey Barrett, Sonia Goncalves, Ewan Harrison, David K. Jackson, Ian Johnston, Dominic Kwiatkowski, Cordelia Langford, John Sillitoe on behalf of the Wellcome Sanger Institute COVID-19 Surveillance Team |
| EPI_ISL_3169829, EPI_ISL_3169830 | Lighthouse Lab in Milton Keynes | Wellcome Sanger Institute for the COVID-19 Genomics UK (COG-UK) Consortium | The Lighthouse Lab in Milton Keynes and Alex Alderton, Roberto Amato, Jeffrey Barrett, Sonia Goncalves, Ewan Harrison, David K. Jackson, Ian Johnston, Dominic Kwiatkowski, Cordelia Langford, John Sillitoe on behalf of the Wellcome Sanger Institute COVID-19 Surveillance Team |

| | | | |
|---|---|---|---|
| EPI_ISL_3169831 | Berkshire and Surrey Pathology Services Lighthouse Laboratory | Wellcome Sanger Institute for the COVID-19 Genomics UK (COG-UK) Consortium | Berkshire and Surrey Pathology Services Lighthouse Laboratory and Alex Alderton, Roberto Amato, Jeffrey Barrett, Sonia Goncalves, Ewan Harrison, David K. Jackson, Ian Johnston, Dominic Kwiatkowski, Cordelia Langford, John Sillitoe on behalf of the Wellcome Sanger Institute COVID-19 Surveillance Team |
| EPI_ISL_3169832 | Lighthouse Lab in Glasgow | Wellcome Sanger Institute for the COVID-19 Genomics UK (COG-UK) Consortium | Harper VanSteenhouse, Yumi Kasai, David Gray, Carol Clugston, Anna Dominiczak and Alex Alderton, Roberto Amato, Jeffrey Barrett, Sonia Goncalves, Ewan Harrison, David K. Jackson, Ian Johnston, Dominic Kwiatkowski, Cordelia Langford, John Sillitoe on behalf of the Wellcome Sanger Institute COVID-19 Surveillance Team |
| EPI_ISL_3169833, EPI_ISL_3169834, EPI_ISL_3169835 | Berkshire and Surrey Pathology Services Lighthouse Laboratory | Wellcome Sanger Institute for the COVID-19 Genomics UK (COG-UK) Consortium | Berkshire and Surrey Pathology Services Lighthouse Laboratory and Alex Alderton, Roberto Amato, Jeffrey Barrett, Sonia Goncalves, Ewan Harrison, David K. Jackson, Ian Johnston, Dominic Kwiatkowski, Cordelia Langford, John Sillitoe on behalf of the Wellcome Sanger Institute COVID-19 Surveillance Team |
| EPI_ISL_3169836 | Lighthouse Lab in Milton Keynes | Wellcome Sanger Institute for the COVID-19 Genomics UK (COG-UK) Consortium | The Lighthouse Lab in Milton Keynes and Alex Alderton, Roberto Amato, Jeffrey Barrett, Sonia Goncalves, Ewan Harrison, David K. Jackson, Ian Johnston, Dominic Kwiatkowski, Cordelia Langford, John Sillitoe on behalf of the Wellcome Sanger Institute COVID-19 Surveillance Team |
| EPI_ISL_3169837 | Lighthouse Lab in Glasgow | Wellcome Sanger Institute for the COVID-19 Genomics UK (COG-UK) Consortium | Harper VanSteenhouse, Yumi Kasai, David Gray, Carol Clugston, Anna Dominiczak and Alex Alderton, Roberto Amato, Jeffrey Barrett, Sonia Goncalves, Ewan Harrison, David K. Jackson, Ian Johnston, Dominic Kwiatkowski, Cordelia Langford, John Sillitoe on behalf of the Wellcome Sanger Institute COVID-19 Surveillance Team |
| EPI_ISL_3169838 | Lighthouse Lab in Milton Keynes | Wellcome Sanger Institute for the COVID-19 Genomics UK (COG-UK) Consortium | The Lighthouse Lab in Milton Keynes and Alex Alderton, Roberto Amato, Jeffrey Barrett, Sonia Goncalves, Ewan Harrison, David K. Jackson, Ian Johnston, Dominic Kwiatkowski, Cordelia Langford, John Sillitoe on behalf of the Wellcome Sanger Institute COVID-19 Surveillance Team |
| EPI_ISL_3169839 | Lighthouse Lab in Glasgow | Wellcome Sanger Institute for the COVID-19 Genomics UK (COG-UK) Consortium | Harper VanSteenhouse, Yumi Kasai, David Gray, Carol Clugston, Anna Dominiczak and Alex Alderton, Roberto Amato, Jeffrey Barrett, Sonia Goncalves, Ewan Harrison, David K. Jackson, Ian Johnston, Dominic Kwiatkowski, Cordelia Langford, John Sillitoe on behalf of the Wellcome Sanger Institute COVID-19 Surveillance Team |
| EPI_ISL_3169840 | Berkshire and Surrey Pathology Services Lighthouse Laboratory | Wellcome Sanger Institute for the COVID-19 Genomics UK (COG-UK) Consortium | Berkshire and Surrey Pathology Services Lighthouse Laboratory and Alex Alderton, Roberto Amato, Jeffrey Barrett, Sonia Goncalves, Ewan Harrison, David K. Jackson, Ian Johnston, Dominic Kwiatkowski, Cordelia Langford, John Sillitoe on behalf of the Wellcome Sanger Institute COVID-19 Surveillance Team |
| EPI_ISL_3169841 | Lighthouse Lab in Milton Keynes | Wellcome Sanger Institute for the COVID-19 Genomics UK (COG-UK) Consortium | The Lighthouse Lab in Milton Keynes and Alex Alderton, Roberto Amato, Jeffrey Barrett, Sonia Goncalves, Ewan Harrison, David K. Jackson, Ian Johnston, Dominic Kwiatkowski, Cordelia Langford, John Sillitoe on behalf of the Wellcome Sanger Institute COVID-19 Surveillance Team |
| EPI_ISL_3169842 | Berkshire and Surrey Pathology Services Lighthouse Laboratory | Wellcome Sanger Institute for the COVID-19 Genomics UK (COG-UK) Consortium | Berkshire and Surrey Pathology Services Lighthouse Laboratory and Alex Alderton, Roberto Amato, Jeffrey Barrett, Sonia Goncalves, Ewan Harrison, David K. Jackson, Ian Johnston, Dominic Kwiatkowski, Cordelia Langford, John Sillitoe on behalf of the Wellcome Sanger Institute COVID-19 Surveillance Team |
| EPI_ISL_3169843, EPI_ISL_3169844 | Lighthouse Lab in Milton Keynes | Wellcome Sanger Institute for the COVID-19 Genomics UK (COG-UK) Consortium | The Lighthouse Lab in Milton Keynes and Alex Alderton, Roberto Amato, Jeffrey Barrett, Sonia Goncalves, Ewan Harrison, David K. Jackson, Ian Johnston, Dominic Kwiatkowski, Cordelia Langford, John Sillitoe on behalf of the Wellcome Sanger Institute COVID-19 Surveillance Team |
| EPI_ISL_3169845 | Berkshire and Surrey Pathology Services Lighthouse Laboratory | Wellcome Sanger Institute for the COVID-19 Genomics UK (COG-UK) Consortium | Berkshire and Surrey Pathology Services Lighthouse Laboratory and Alex Alderton, Roberto Amato, Jeffrey Barrett, Sonia Goncalves, Ewan Harrison, David K. Jackson, Ian Johnston, Dominic Kwiatkowski, Cordelia Langford, John Sillitoe on behalf of the Wellcome Sanger Institute COVID-19 Surveillance Team |
| EPI_ISL_3169846, EPI_ISL_3169847, EPI_ISL_3169848, EPI_ISL_3169849, EPI_ISL_3169850 | Lighthouse Lab in Milton Keynes | Wellcome Sanger Institute for the COVID-19 Genomics UK (COG-UK) Consortium | The Lighthouse Lab in Milton Keynes and Alex Alderton, Roberto Amato, Jeffrey Barrett, Sonia Goncalves, Ewan Harrison, David K. Jackson, Ian Johnston, Dominic Kwiatkowski, Cordelia Langford, John Sillitoe on behalf of the Wellcome Sanger Institute COVID-19 Surveillance Team |
| EPI_ISL_3169851 | Berkshire and Surrey Pathology Services Lighthouse Laboratory | Wellcome Sanger Institute for the COVID-19 Genomics UK (COG-UK) Consortium | Berkshire and Surrey Pathology Services Lighthouse Laboratory and Alex Alderton, Roberto Amato, Jeffrey Barrett, Sonia Goncalves, Ewan Harrison, David K. Jackson, Ian Johnston, Dominic Kwiatkowski, Cordelia Langford, John Sillitoe on behalf of the Wellcome Sanger Institute COVID-19 Surveillance Team |
| EPI_ISL_3169852, EPI_ISL_3169853 | Lighthouse Lab in Glasgow | Wellcome Sanger Institute for the COVID-19 Genomics UK (COG-UK) Consortium | Harper VanSteenhouse, Yumi Kasai, David Gray, Carol Clugston, Anna Dominiczak and Alex Alderton, Roberto Amato, Jeffrey Barrett, Sonia Goncalves, Ewan Harrison, David K. Jackson, Ian Johnston, Dominic Kwiatkowski, Cordelia Langford, John Sillitoe on behalf of the Wellcome Sanger Institute COVID-19 Surveillance Team |
| EPI_ISL_3169854 | Berkshire and Surrey Pathology Services Lighthouse Laboratory | Wellcome Sanger Institute for the COVID-19 Genomics UK (COG-UK) Consortium | Berkshire and Surrey Pathology Services Lighthouse Laboratory and Alex Alderton, Roberto Amato, Jeffrey Barrett, Sonia Goncalves, Ewan Harrison, David K. Jackson, Ian Johnston, Dominic Kwiatkowski, Cordelia Langford, John Sillitoe on behalf of the Wellcome Sanger Institute COVID-19 Surveillance Team |
| EPI_ISL_3169855 | Lighthouse Lab in Glasgow | Wellcome Sanger Institute for the COVID-19 Genomics UK (COG-UK) Consortium | Harper VanSteenhouse, Yumi Kasai, David Gray, Carol Clugston, Anna Dominiczak and Alex Alderton, Roberto Amato, Jeffrey Barrett, Sonia Goncalves, Ewan Harrison, David K. Jackson, Ian Johnston, Dominic Kwiatkowski, Cordelia Langford, John Sillitoe on behalf of the Wellcome Sanger Institute COVID-19 Surveillance Team |
| EPI_ISL_3169856, EPI_ISL_3169857, EPI_ISL_3169858, EPI_ISL_3169859 | Berkshire and Surrey Pathology Services Lighthouse Laboratory | Wellcome Sanger Institute for the COVID-19 Genomics UK (COG-UK) Consortium | Berkshire and Surrey Pathology Services Lighthouse Laboratory and Alex Alderton, Roberto Amato, Jeffrey Barrett, Sonia Goncalves, Ewan Harrison, David K. Jackson, Ian Johnston, Dominic Kwiatkowski, Cordelia Langford, John Sillitoe on behalf of the Wellcome Sanger Institute COVID-19 Surveillance Team |
| EPI_ISL_3169860, EPI_ISL_3169861, EPI_ISL_3169862 | Lighthouse Lab in Milton Keynes | Wellcome Sanger Institute for the COVID-19 Genomics UK (COG-UK) Consortium | The Lighthouse Lab in Milton Keynes and Alex Alderton, Roberto Amato, Jeffrey Barrett, Sonia Goncalves, Ewan Harrison, David K. Jackson, Ian Johnston, Dominic Kwiatkowski, Cordelia Langford, John Sillitoe on behalf of the Wellcome Sanger Institute COVID-19 Surveillance Team |
| EPI_ISL_3169863, EPI_ISL_3169865 | Berkshire and Surrey Pathology Services Lighthouse Laboratory | Wellcome Sanger Institute for the COVID-19 Genomics UK (COG-UK) Consortium | Berkshire and Surrey Pathology Services Lighthouse Laboratory and Alex Alderton, Roberto Amato, Jeffrey Barrett, Sonia Goncalves, Ewan Harrison, David K. Jackson, Ian Johnston, Dominic Kwiatkowski, Cordelia Langford, John Sillitoe on behalf of the Wellcome Sanger Institute COVID-19 Surveillance Team |
| EPI_ISL_3169866, EPI_ISL_3169867 | Lighthouse Lab in Milton Keynes | Wellcome Sanger Institute for the COVID-19 Genomics UK (COG-UK) Consortium | The Lighthouse Lab in Milton Keynes and Alex Alderton, Roberto Amato, Jeffrey Barrett, Sonia Goncalves, Ewan Harrison, David K. Jackson, Ian Johnston, Dominic Kwiatkowski, Cordelia Langford, John Sillitoe on behalf of the Wellcome Sanger Institute COVID-19 Surveillance Team |
| EPI_ISL_3169868 | Lighthouse Lab in Glasgow | Wellcome Sanger Institute for the COVID-19 Genomics UK (COG-UK) Consortium | Harper VanSteenhouse, Yumi Kasai, David Gray, Carol Clugston, Anna Dominiczak and Alex Alderton, Roberto Amato, Jeffrey Barrett, Sonia Goncalves, Ewan Harrison, David K. Jackson, Ian Johnston, Dominic Kwiatkowski, Cordelia Langford, John Sillitoe on behalf of the Wellcome Sanger Institute COVID-19 Surveillance Team |
| EPI_ISL_3169869, EPI_ISL_3169870 | Berkshire and Surrey Pathology Services Lighthouse Laboratory | Wellcome Sanger Institute for the COVID-19 Genomics UK (COG-UK) Consortium | Berkshire and Surrey Pathology Services Lighthouse Laboratory and Alex Alderton, Roberto Amato, Jeffrey Barrett, Sonia Goncalves, Ewan Harrison, David K. Jackson, Ian Johnston, Dominic Kwiatkowski, Cordelia Langford, John Sillitoe on behalf of the Wellcome Sanger Institute COVID-19 Surveillance Team |
| EPI_ISL_3169871 | Lighthouse Lab in Milton Keynes | Wellcome Sanger Institute for the COVID-19 Genomics UK (COG-UK) Consortium | The Lighthouse Lab in Milton Keynes and Alex Alderton, Roberto Amato, Jeffrey Barrett, Sonia Goncalves, Ewan Harrison, David K. Jackson, Ian Johnston, Dominic Kwiatkowski, Cordelia Langford, John Sillitoe on behalf of the Wellcome Sanger Institute COVID-19 Surveillance Team |
| EPI_ISL_3169872 | Lighthouse Lab in Glasgow | Wellcome Sanger Institute for the COVID-19 Genomics UK (COG-UK) Consortium | Harper VanSteenhouse, Yumi Kasai, David Gray, Carol Clugston, Anna Dominiczak and Alex Alderton, Roberto Amato, Jeffrey Barrett, Sonia Goncalves, Ewan Harrison, David K. Jackson, Ian Johnston, Dominic Kwiatkowski, Cordelia Langford, John Sillitoe on behalf of the Wellcome Sanger Institute COVID-19 Surveillance Team |
| EPI_ISL_3169873 | Berkshire and Surrey Pathology Services Lighthouse Laboratory | Wellcome Sanger Institute for the COVID-19 Genomics UK (COG-UK) Consortium | Berkshire and Surrey Pathology Services Lighthouse Laboratory and Alex Alderton, Roberto Amato, Jeffrey Barrett, Sonia Goncalves, Ewan Harrison, David K. Jackson, Ian Johnston, Dominic Kwiatkowski, Cordelia Langford, John Sillitoe on behalf of the Wellcome Sanger Institute COVID-19 Surveillance Team |
| EPI_ISL_3169874 | Lighthouse Lab in Glasgow | Wellcome Sanger Institute for the COVID-19 Genomics UK (COG-UK) Consortium | Harper VanSteenhouse, Yumi Kasai, David Gray, Carol Clugston, Anna Dominiczak and Alex Alderton, Roberto Amato, Jeffrey Barrett, Sonia Goncalves, Ewan Harrison, David K. Jackson, Ian Johnston, Dominic Kwiatkowski, Cordelia Langford, John Sillitoe on behalf of the Wellcome Sanger Institute COVID-19 Surveillance Team |
| EPI_ISL_3169875 | Berkshire and Surrey Pathology Services Lighthouse Laboratory | Wellcome Sanger Institute for the COVID-19 Genomics UK (COG-UK) Consortium | Berkshire and Surrey Pathology Services Lighthouse Laboratory and Alex Alderton, Roberto Amato, Jeffrey Barrett, Sonia Goncalves, Ewan Harrison, David K. Jackson, Ian Johnston, Dominic Kwiatkowski, Cordelia Langford, John Sillitoe on behalf of the Wellcome Sanger Institute COVID-19 Surveillance Team |
| EPI_ISL_3169876 | Lighthouse Lab in Glasgow | Wellcome Sanger Institute for the COVID-19 Genomics UK (COG-UK) Consortium | Harper VanSteenhouse, Yumi Kasai, David Gray, Carol Clugston, Anna Dominiczak and Alex Alderton, Roberto Amato, Jeffrey Barrett, Sonia Goncalves, Ewan Harrison, David K. Jackson, Ian Johnston, Dominic Kwiatkowski, Cordelia Langford, John Sillitoe on behalf of the Wellcome Sanger Institute COVID-19 Surveillance Team |
| EPI_ISL_3169877 | Lighthouse Lab in Milton Keynes | Wellcome Sanger Institute for the COVID-19 Genomics UK (COG-UK) Consortium | The Lighthouse Lab in Milton Keynes and Alex Alderton, Roberto Amato, Jeffrey Barrett, Sonia Goncalves, Ewan Harrison, David K. Jackson, Ian Johnston, Dominic Kwiatkowski, Cordelia Langford, John Sillitoe on behalf of the Wellcome Sanger Institute COVID-19 Surveillance Team |
| EPI_ISL_3169878, EPI_ISL_3169879, EPI_ISL_3169880 | Berkshire and Surrey Pathology Services Lighthouse Laboratory | Wellcome Sanger Institute for the COVID-19 Genomics UK (COG-UK) Consortium | Berkshire and Surrey Pathology Services Lighthouse Laboratory and Alex Alderton, Roberto Amato, Jeffrey Barrett, Sonia Goncalves, Ewan Harrison, David K. Jackson, Ian Johnston, Dominic Kwiatkowski, Cordelia Langford, John Sillitoe on behalf of the Wellcome Sanger Institute COVID-19 Surveillance Team |
| EPI_ISL_3169881, EPI_ISL_3169882 | Lighthouse Lab in Milton Keynes | Wellcome Sanger Institute for the COVID-19 Genomics UK (COG-UK) Consortium | The Lighthouse Lab in Milton Keynes and Alex Alderton, Roberto Amato, Jeffrey Barrett, Sonia Goncalves, Ewan Harrison, David K. Jackson, Ian Johnston, Dominic Kwiatkowski, Cordelia Langford, John Sillitoe on behalf of the Wellcome Sanger Institute COVID-19 Surveillance Team |
| EPI_ISL_3169883 | Berkshire and Surrey Pathology Services Lighthouse Laboratory | Wellcome Sanger Institute for the COVID-19 Genomics UK (COG-UK) Consortium | Berkshire and Surrey Pathology Services Lighthouse Laboratory and Alex Alderton, Roberto Amato, Jeffrey Barrett, Sonia Goncalves, Ewan Harrison, David K. Jackson, Ian Johnston, Dominic Kwiatkowski, Cordelia Langford, John Sillitoe on behalf of the Wellcome Sanger Institute COVID-19 Surveillance Team |
| EPI_ISL_3169884 | Lighthouse Lab in Glasgow | Wellcome Sanger Institute for the COVID-19 Genomics UK (COG-UK) Consortium | Harper VanSteenhouse, Yumi Kasai, David Gray, Carol Clugston, Anna Dominiczak and Alex Alderton, Roberto Amato, Jeffrey Barrett, Sonia Goncalves, Ewan Harrison, David K. Jackson, Ian Johnston, Dominic Kwiatkowski, Cordelia Langford, John Sillitoe on behalf of the Wellcome Sanger Institute COVID-19 Surveillance Team |
| EPI_ISL_3169885, EPI_ISL_3169886 | Lighthouse Lab in Milton Keynes | Wellcome Sanger Institute for the COVID-19 Genomics UK (COG-UK) Consortium | The Lighthouse Lab in Milton Keynes and Alex Alderton, Roberto Amato, Jeffrey Barrett, Sonia Goncalves, Ewan Harrison, David K. Jackson, Ian Johnston, Dominic Kwiatkowski, Cordelia Langford, John Sillitoe on behalf of the Wellcome Sanger Institute COVID-19 Surveillance Team |
| EPI_ISL_3169887, EPI_ISL_3169888 | Lighthouse Lab in Glasgow | Wellcome Sanger Institute for the COVID-19 Genomics UK (COG-UK) Consortium | Harper VanSteenhouse, Yumi Kasai, David Gray, Carol Clugston, Anna Dominiczak and Alex Alderton, Roberto Amato, Jeffrey Barrett, Sonia Goncalves, Ewan Harrison, David K. Jackson, Ian Johnston, Dominic Kwiatkowski, Cordelia Langford, John Sillitoe on behalf of the Wellcome Sanger Institute COVID-19 Surveillance Team |
| EPI_ISL_3169889, EPI_ISL_3169890 | Berkshire and Surrey Pathology Services Lighthouse Laboratory | Wellcome Sanger Institute for the COVID-19 Genomics UK (COG-UK) Consortium | Berkshire and Surrey Pathology Services Lighthouse Laboratory and Alex Alderton, Roberto Amato, Jeffrey Barrett, Sonia Goncalves, Ewan Harrison, David K. Jackson, Ian Johnston, Dominic Kwiatkowski, Cordelia Langford, John Sillitoe on behalf of the Wellcome Sanger Institute COVID-19 Surveillance Team |
| EPI_ISL_3169891, EPI_ISL_3169892 | Lighthouse Lab in Milton Keynes | Wellcome Sanger Institute for the COVID-19 Genomics UK (COG-UK) Consortium | The Lighthouse Lab in Milton Keynes and Alex Alderton, Roberto Amato, Jeffrey Barrett, Sonia Goncalves, Ewan Harrison, David K. Jackson, Ian Johnston, Dominic Kwiatkowski, Cordelia Langford, John Sillitoe on behalf of the Wellcome Sanger Institute COVID-19 Surveillance Team |
| EPI_ISL_3169893, EPI_ISL_3169894, EPI_ISL_3169895 | Berkshire and Surrey Pathology Services Lighthouse Laboratory | Wellcome Sanger Institute for the COVID-19 Genomics UK (COG-UK) Consortium | Berkshire and Surrey Pathology Services Lighthouse Laboratory and Alex Alderton, Roberto Amato, Jeffrey Barrett, Sonia Goncalves, Ewan Harrison, David K. Jackson, Ian Johnston, Dominic Kwiatkowski, Cordelia Langford, John Sillitoe on behalf of the Wellcome Sanger Institute COVID-19 Surveillance Team |
| EPI_ISL_3169896 | Lighthouse Lab in Milton Keynes | Wellcome Sanger Institute for the COVID-19 Genomics UK (COG-UK) Consortium | The Lighthouse Lab in Milton Keynes and Alex Alderton, Roberto Amato, Jeffrey Barrett, Sonia Goncalves, Ewan Harrison, David K. Jackson, Ian Johnston, Dominic Kwiatkowski, Cordelia Langford, John Sillitoe on behalf of the Wellcome Sanger Institute COVID-19 Surveillance Team |
| EPI_ISL_3169897, EPI_ISL_3169898, EPI_ISL_3169899, | Berkshire and Surrey Pathology Services Lighthouse Laboratory | Wellcome Sanger Institute for the COVID-19 Genomics UK (COG-UK) Consortium | Berkshire and Surrey Pathology Services Lighthouse Laboratory and Alex Alderton, Roberto Amato, Jeffrey Barrett, Sonia Goncalves, Ewan Harrison, David K. Jackson, Ian Johnston, Dominic Kwiatkowski, Cordelia Langford, John Sillitoe on behalf of the Wellcome Sanger Institute COVID-19 Surveillance Team |

| | | | |
|---|---|---|---|
| EPI_ISL_3169900 | | | |
| EPI_ISL_3169901, EPI_ISL_3169902 | Lighthouse Lab in Milton Keynes | Wellcome Sanger Institute for the COVID-19 Genomics UK (COG-UK) Consortium | The Lighthouse Lab in Milton Keynes and Alex Alderton, Roberto Amato, Jeffrey Barrett, Sonia Goncalves, Ewan Harrison, David K. Jackson, Ian Johnston, Dominic Kwiatkowski, Cordelia Langford, John Sillitoe on behalf of the Wellcome Sanger Institute COVID-19 Surveillance Team |
| EPI_ISL_3169903, EPI_ISL_3169904 | Lighthouse Laboratory Plymouth | Wellcome Sanger Institute for the COVID-19 Genomics UK (COG-UK) Consortium | Lighthouse Laboratory Plymouth and Alex Alderton, Roberto Amato, Jeffrey Barrett, Sonia Goncalves, Ewan Harrison, David K. Jackson, Ian Johnston, Dominic Kwiatkowski, Cordelia Langford, John Sillitoe on behalf of the Wellcome Sanger Institute COVID-19 Surveillance Team |
| EPI_ISL_3169905 | Berkshire and Surrey Pathology Services Lighthouse Laboratory | Wellcome Sanger Institute for the COVID-19 Genomics UK (COG-UK) Consortium | Berkshire and Surrey Pathology Services Lighthouse Laboratory and Alex Alderton, Roberto Amato, Jeffrey Barrett, Sonia Goncalves, Ewan Harrison, David K. Jackson, Ian Johnston, Dominic Kwiatkowski, Cordelia Langford, John Sillitoe on behalf of the Wellcome Sanger Institute COVID-19 Surveillance Team |
| EPI_ISL_3169906, EPI_ISL_3169907 | Lighthouse Lab in Alderley Park | Wellcome Sanger Institute for the COVID-19 Genomics UK (COG-UK) Consortium | Jacquelyn Wynn, Mairead Hyland, The Lighthouse Lab in Alderley Park and Alex Alderton, Roberto Amato, Jeffrey Barrett, Sonia Goncalves, Ewan Harrison, David K. Jackson, Ian Johnston, Dominic Kwiatkowski, Cordelia Langford, John Sillitoe on behalf of the Wellcome Sanger Institute COVID-19 Surveillance Team |
| EPI_ISL_3169908 | Berkshire and Surrey Pathology Services Lighthouse Laboratory | Wellcome Sanger Institute for the COVID-19 Genomics UK (COG-UK) Consortium | Berkshire and Surrey Pathology Services Lighthouse Laboratory and Alex Alderton, Roberto Amato, Jeffrey Barrett, Sonia Goncalves, Ewan Harrison, David K. Jackson, Ian Johnston, Dominic Kwiatkowski, Cordelia Langford, John Sillitoe on behalf of the Wellcome Sanger Institute COVID-19 Surveillance Team |
| EPI_ISL_3169909 | Lighthouse Laboratory Plymouth | Wellcome Sanger Institute for the COVID-19 Genomics UK (COG-UK) Consortium | Lighthouse Laboratory Plymouth and Alex Alderton, Roberto Amato, Jeffrey Barrett, Sonia Goncalves, Ewan Harrison, David K. Jackson, Ian Johnston, Dominic Kwiatkowski, Cordelia Langford, John Sillitoe on behalf of the Wellcome Sanger Institute COVID-19 Surveillance Team |
| EPI_ISL_3169910 | Lighthouse Lab in Alderley Park | Wellcome Sanger Institute for the COVID-19 Genomics UK (COG-UK) Consortium | Jacquelyn Wynn, Mairead Hyland, The Lighthouse Lab in Alderley Park and Alex Alderton, Roberto Amato, Jeffrey Barrett, Sonia Goncalves, Ewan Harrison, David K. Jackson, Ian Johnston, Dominic Kwiatkowski, Cordelia Langford, John Sillitoe on behalf of the Wellcome Sanger Institute COVID-19 Surveillance Team |
| EPI_ISL_3169911 | Berkshire and Surrey Pathology Services Lighthouse Laboratory | Wellcome Sanger Institute for the COVID-19 Genomics UK (COG-UK) Consortium | Berkshire and Surrey Pathology Services Lighthouse Laboratory and Alex Alderton, Roberto Amato, Jeffrey Barrett, Sonia Goncalves, Ewan Harrison, David K. Jackson, Ian Johnston, Dominic Kwiatkowski, Cordelia Langford, John Sillitoe on behalf of the Wellcome Sanger Institute COVID-19 Surveillance Team |
| EPI_ISL_3169912 | Lighthouse Laboratory Plymouth | Wellcome Sanger Institute for the COVID-19 Genomics UK (COG-UK) Consortium | Lighthouse Laboratory Plymouth and Alex Alderton, Roberto Amato, Jeffrey Barrett, Sonia Goncalves, Ewan Harrison, David K. Jackson, Ian Johnston, Dominic Kwiatkowski, Cordelia Langford, John Sillitoe on behalf of the Wellcome Sanger Institute COVID-19 Surveillance Team |
| EPI_ISL_3169913, EPI_ISL_3169914 | Lighthouse Lab in Alderley Park | Wellcome Sanger Institute for the COVID-19 Genomics UK (COG-UK) Consortium | Jacquelyn Wynn, Mairead Hyland, The Lighthouse Lab in Alderley Park and Alex Alderton, Roberto Amato, Jeffrey Barrett, Sonia Goncalves, Ewan Harrison, David K. Jackson, Ian Johnston, Dominic Kwiatkowski, Cordelia Langford, John Sillitoe on behalf of the Wellcome Sanger Institute COVID-19 Surveillance Team |
| EPI_ISL_3169915, EPI_ISL_3169916 | Lighthouse Laboratory Plymouth | Wellcome Sanger Institute for the COVID-19 Genomics UK (COG-UK) Consortium | Lighthouse Laboratory Plymouth and Alex Alderton, Roberto Amato, Jeffrey Barrett, Sonia Goncalves, Ewan Harrison, David K. Jackson, Ian Johnston, Dominic Kwiatkowski, Cordelia Langford, John Sillitoe on behalf of the Wellcome Sanger Institute COVID-19 Surveillance Team |
| EPI_ISL_3169917, EPI_ISL_3169918, EPI_ISL_3169919, EPI_ISL_3169920, EPI_ISL_3169921 | Berkshire and Surrey Pathology Services Lighthouse Laboratory | Wellcome Sanger Institute for the COVID-19 Genomics UK (COG-UK) Consortium | Berkshire and Surrey Pathology Services Lighthouse Laboratory and Alex Alderton, Roberto Amato, Jeffrey Barrett, Sonia Goncalves, Ewan Harrison, David K. Jackson, Ian Johnston, Dominic Kwiatkowski, Cordelia Langford, John Sillitoe on behalf of the Wellcome Sanger Institute COVID-19 Surveillance Team |
| EPI_ISL_3169922 | Lighthouse Lab in Alderley Park | Wellcome Sanger Institute for the COVID-19 Genomics UK (COG-UK) Consortium | Jacquelyn Wynn, Mairead Hyland, The Lighthouse Lab in Alderley Park and Alex Alderton, Roberto Amato, Jeffrey Barrett, Sonia Goncalves, Ewan Harrison, David K. Jackson, Ian Johnston, Dominic Kwiatkowski, Cordelia Langford, John Sillitoe on behalf of the Wellcome Sanger Institute COVID-19 Surveillance Team |
| EPI_ISL_3169923 | Berkshire and Surrey Pathology Services Lighthouse Laboratory | Wellcome Sanger Institute for the COVID-19 Genomics UK (COG-UK) Consortium | Berkshire and Surrey Pathology Services Lighthouse Laboratory and Alex Alderton, Roberto Amato, Jeffrey Barrett, Sonia Goncalves, Ewan Harrison, David K. Jackson, Ian Johnston, Dominic Kwiatkowski, Cordelia Langford, John Sillitoe on behalf of the Wellcome Sanger Institute COVID-19 Surveillance Team |
| EPI_ISL_3169924 | Lighthouse Lab in Alderley Park | Wellcome Sanger Institute for the COVID-19 Genomics UK (COG-UK) Consortium | Jacquelyn Wynn, Mairead Hyland, The Lighthouse Lab in Alderley Park and Alex Alderton, Roberto Amato, Jeffrey Barrett, Sonia Goncalves, Ewan Harrison, David K. Jackson, Ian Johnston, Dominic Kwiatkowski, Cordelia Langford, John Sillitoe on behalf of the Wellcome Sanger Institute COVID-19 Surveillance Team |
| EPI_ISL_3169925 | Berkshire and Surrey Pathology Services Lighthouse Laboratory | Wellcome Sanger Institute for the COVID-19 Genomics UK (COG-UK) Consortium | Berkshire and Surrey Pathology Services Lighthouse Laboratory and Alex Alderton, Roberto Amato, Jeffrey Barrett, Sonia Goncalves, Ewan Harrison, David K. Jackson, Ian Johnston, Dominic Kwiatkowski, Cordelia Langford, John Sillitoe on behalf of the Wellcome Sanger Institute COVID-19 Surveillance Team |
| EPI_ISL_3169926 | Lighthouse Lab in Alderley Park | Wellcome Sanger Institute for the COVID-19 Genomics UK (COG-UK) Consortium | Jacquelyn Wynn, Mairead Hyland, The Lighthouse Lab in Alderley Park and Alex Alderton, Roberto Amato, Jeffrey Barrett, Sonia Goncalves, Ewan Harrison, David K. Jackson, Ian Johnston, Dominic Kwiatkowski, Cordelia Langford, John Sillitoe on behalf of the Wellcome Sanger Institute COVID-19 Surveillance Team |
| EPI_ISL_3169927, EPI_ISL_3169928 | Berkshire and Surrey Pathology Services Lighthouse Laboratory | Wellcome Sanger Institute for the COVID-19 Genomics UK (COG-UK) Consortium | Berkshire and Surrey Pathology Services Lighthouse Laboratory and Alex Alderton, Roberto Amato, Jeffrey Barrett, Sonia Goncalves, Ewan Harrison, David K. Jackson, Ian Johnston, Dominic Kwiatkowski, Cordelia Langford, John Sillitoe on behalf of the Wellcome Sanger Institute COVID-19 Surveillance Team |
| EPI_ISL_3169929 | Lighthouse Laboratory Plymouth | Wellcome Sanger Institute for the COVID-19 Genomics UK (COG-UK) Consortium | Lighthouse Laboratory Plymouth and Alex Alderton, Roberto Amato, Jeffrey Barrett, Sonia Goncalves, Ewan Harrison, David K. Jackson, Ian Johnston, Dominic Kwiatkowski, Cordelia Langford, John Sillitoe on behalf of the Wellcome Sanger Institute COVID-19 Surveillance Team |
| EPI_ISL_3169930, EPI_ISL_3169931 | Lighthouse Lab in Alderley Park | Wellcome Sanger Institute for the COVID-19 Genomics UK (COG-UK) Consortium | Jacquelyn Wynn, Mairead Hyland, The Lighthouse Lab in Alderley Park and Alex Alderton, Roberto Amato, Jeffrey Barrett, Sonia Goncalves, Ewan Harrison, David K. Jackson, Ian Johnston, Dominic Kwiatkowski, Cordelia Langford, John Sillitoe on behalf of the Wellcome Sanger Institute COVID-19 Surveillance Team |
| EPI_ISL_3169932 | Lighthouse Laboratory Plymouth | Wellcome Sanger Institute for the COVID-19 Genomics UK (COG-UK) Consortium | Lighthouse Laboratory Plymouth and Alex Alderton, Roberto Amato, Jeffrey Barrett, Sonia Goncalves, Ewan Harrison, David K. Jackson, Ian Johnston, Dominic Kwiatkowski, Cordelia Langford, John Sillitoe on behalf of the Wellcome Sanger Institute COVID-19 Surveillance Team |
| EPI_ISL_3169933 | Lighthouse Lab in Alderley Park | Wellcome Sanger Institute for the COVID-19 Genomics UK (COG-UK) Consortium | Jacquelyn Wynn, Mairead Hyland, The Lighthouse Lab in Alderley Park and Alex Alderton, Roberto Amato, Jeffrey Barrett, Sonia Goncalves, Ewan Harrison, David K. Jackson, Ian Johnston, Dominic Kwiatkowski, Cordelia Langford, John Sillitoe on behalf of the Wellcome Sanger Institute COVID-19 Surveillance Team |
| EPI_ISL_3169934 | Berkshire and Surrey Pathology Services Lighthouse Laboratory | Wellcome Sanger Institute for the COVID-19 Genomics UK (COG-UK) Consortium | Berkshire and Surrey Pathology Services Lighthouse Laboratory and Alex Alderton, Roberto Amato, Jeffrey Barrett, Sonia Goncalves, Ewan Harrison, David K. Jackson, Ian Johnston, Dominic Kwiatkowski, Cordelia Langford, John Sillitoe on behalf of the Wellcome Sanger Institute COVID-19 Surveillance Team |
| EPI_ISL_3169935 | Lighthouse Laboratory Plymouth | Wellcome Sanger Institute for the COVID-19 Genomics UK (COG-UK) Consortium | Lighthouse Laboratory Plymouth and Alex Alderton, Roberto Amato, Jeffrey Barrett, Sonia Goncalves, Ewan Harrison, David K. Jackson, Ian Johnston, Dominic Kwiatkowski, Cordelia Langford, John Sillitoe on behalf of the Wellcome Sanger Institute COVID-19 Surveillance Team |
| EPI_ISL_3169936, EPI_ISL_3169937 | Lighthouse Lab in Alderley Park | Wellcome Sanger Institute for the COVID-19 Genomics UK (COG-UK) Consortium | Jacquelyn Wynn, Mairead Hyland, The Lighthouse Lab in Alderley Park and Alex Alderton, Roberto Amato, Jeffrey Barrett, Sonia Goncalves, Ewan Harrison, David K. Jackson, Ian Johnston, Dominic Kwiatkowski, Cordelia Langford, John Sillitoe on behalf of the Wellcome Sanger Institute COVID-19 Surveillance Team |
| EPI_ISL_3169938, EPI_ISL_3169939, EPI_ISL_3169940 | Berkshire and Surrey Pathology Services Lighthouse Laboratory | Wellcome Sanger Institute for the COVID-19 Genomics UK (COG-UK) Consortium | Berkshire and Surrey Pathology Services Lighthouse Laboratory and Alex Alderton, Roberto Amato, Jeffrey Barrett, Sonia Goncalves, Ewan Harrison, David K. Jackson, Ian Johnston, Dominic Kwiatkowski, Cordelia Langford, John Sillitoe on behalf of the Wellcome Sanger Institute COVID-19 Surveillance Team |
| EPI_ISL_3169941 | Lighthouse Lab in Alderley Park | Wellcome Sanger Institute for the COVID-19 Genomics UK (COG-UK) Consortium | Jacquelyn Wynn, Mairead Hyland, The Lighthouse Lab in Alderley Park and Alex Alderton, Roberto Amato, Jeffrey Barrett, Sonia Goncalves, Ewan Harrison, David K. Jackson, Ian Johnston, Dominic Kwiatkowski, Cordelia Langford, John Sillitoe on behalf of the Wellcome Sanger Institute COVID-19 Surveillance Team |
| EPI_ISL_3169942 | Berkshire and Surrey Pathology Services Lighthouse Laboratory | Wellcome Sanger Institute for the COVID-19 Genomics UK (COG-UK) Consortium | Berkshire and Surrey Pathology Services Lighthouse Laboratory and Alex Alderton, Roberto Amato, Jeffrey Barrett, Sonia Goncalves, Ewan Harrison, David K. Jackson, Ian Johnston, Dominic Kwiatkowski, Cordelia Langford, John Sillitoe on behalf of the Wellcome Sanger Institute COVID-19 Surveillance Team |
| EPI_ISL_3169943 | Lighthouse Lab in Alderley Park | Wellcome Sanger Institute for the COVID-19 Genomics UK (COG-UK) Consortium | Jacquelyn Wynn, Mairead Hyland, The Lighthouse Lab in Alderley Park and Alex Alderton, Roberto Amato, Jeffrey Barrett, Sonia Goncalves, Ewan Harrison, David K. Jackson, Ian Johnston, Dominic Kwiatkowski, Cordelia Langford, John Sillitoe on behalf of the Wellcome Sanger Institute COVID-19 Surveillance Team |
| EPI_ISL_3169944, EPI_ISL_3169945 | Berkshire and Surrey Pathology Services Lighthouse Laboratory | Wellcome Sanger Institute for the COVID-19 Genomics UK (COG-UK) Consortium | Berkshire and Surrey Pathology Services Lighthouse Laboratory and Alex Alderton, Roberto Amato, Jeffrey Barrett, Sonia Goncalves, Ewan Harrison, David K. Jackson, Ian Johnston, Dominic Kwiatkowski, Cordelia Langford, John Sillitoe on behalf of the Wellcome Sanger Institute COVID-19 Surveillance Team |
| EPI_ISL_3169946 | Lighthouse Lab in Alderley Park | Wellcome Sanger Institute for the COVID-19 Genomics UK (COG-UK) Consortium | Jacquelyn Wynn, Mairead Hyland, The Lighthouse Lab in Alderley Park and Alex Alderton, Roberto Amato, Jeffrey Barrett, Sonia Goncalves, Ewan Harrison, David K. Jackson, Ian Johnston, Dominic Kwiatkowski, Cordelia Langford, John Sillitoe on behalf of the Wellcome Sanger Institute COVID-19 Surveillance Team |
| EPI_ISL_3169947 | Berkshire and Surrey Pathology Services Lighthouse Laboratory | Wellcome Sanger Institute for the COVID-19 Genomics UK (COG-UK) Consortium | Berkshire and Surrey Pathology Services Lighthouse Laboratory and Alex Alderton, Roberto Amato, Jeffrey Barrett, Sonia Goncalves, Ewan Harrison, David K. Jackson, Ian Johnston, Dominic Kwiatkowski, Cordelia Langford, John Sillitoe on behalf of the Wellcome Sanger Institute COVID-19 Surveillance Team |
| EPI_ISL_3169948 | Lighthouse Lab in Alderley Park | Wellcome Sanger Institute for the COVID-19 Genomics UK (COG-UK) Consortium | Jacquelyn Wynn, Mairead Hyland, The Lighthouse Lab in Alderley Park and Alex Alderton, Roberto Amato, Jeffrey Barrett, Sonia Goncalves, Ewan Harrison, David K. Jackson, Ian Johnston, Dominic Kwiatkowski, Cordelia Langford, John Sillitoe on behalf of the Wellcome Sanger Institute COVID-19 Surveillance Team |
| EPI_ISL_3169949, EPI_ISL_3169950, EPI_ISL_3169951 | Berkshire and Surrey Pathology Services Lighthouse Laboratory | Wellcome Sanger Institute for the COVID-19 Genomics UK (COG-UK) Consortium | Berkshire and Surrey Pathology Services Lighthouse Laboratory and Alex Alderton, Roberto Amato, Jeffrey Barrett, Sonia Goncalves, Ewan Harrison, David K. Jackson, Ian Johnston, Dominic Kwiatkowski, Cordelia Langford, John Sillitoe on behalf of the Wellcome Sanger Institute COVID-19 Surveillance Team |
| EPI_ISL_3169952 | Lighthouse Laboratory Plymouth | Wellcome Sanger Institute for the COVID-19 Genomics UK (COG-UK) Consortium | Lighthouse Laboratory Plymouth and Alex Alderton, Roberto Amato, Jeffrey Barrett, Sonia Goncalves, Ewan Harrison, David K. Jackson, Ian Johnston, Dominic Kwiatkowski, Cordelia Langford, John Sillitoe on behalf of the Wellcome Sanger Institute COVID-19 Surveillance Team |
| EPI_ISL_3169953, EPI_ISL_3169954 | Berkshire and Surrey Pathology Services Lighthouse Laboratory | Wellcome Sanger Institute for the COVID-19 Genomics UK (COG-UK) Consortium | Berkshire and Surrey Pathology Services Lighthouse Laboratory and Alex Alderton, Roberto Amato, Jeffrey Barrett, Sonia Goncalves, Ewan Harrison, David K. Jackson, Ian Johnston, Dominic Kwiatkowski, Cordelia Langford, John Sillitoe on behalf of the Wellcome Sanger Institute COVID-19 Surveillance Team |
| EPI_ISL_3169955 | Lighthouse Laboratory Plymouth | Wellcome Sanger Institute for the COVID-19 Genomics UK (COG-UK) Consortium | Lighthouse Laboratory Plymouth and Alex Alderton, Roberto Amato, Jeffrey Barrett, Sonia Goncalves, Ewan Harrison, David K. Jackson, Ian Johnston, Dominic Kwiatkowski, Cordelia Langford, John Sillitoe on behalf of the Wellcome Sanger Institute COVID-19 Surveillance Team |
| EPI_ISL_3169956 | Berkshire and Surrey Pathology Services Lighthouse Laboratory | Wellcome Sanger Institute for the COVID-19 Genomics UK (COG-UK) Consortium | Berkshire and Surrey Pathology Services Lighthouse Laboratory and Alex Alderton, Roberto Amato, Jeffrey Barrett, Sonia Goncalves, Ewan Harrison, David K. Jackson, Ian Johnston, Dominic Kwiatkowski, Cordelia Langford, John Sillitoe on behalf of the Wellcome Sanger Institute COVID-19 Surveillance Team |
| EPI_ISL_3169957 | Lighthouse Lab in Alderley Park | Wellcome Sanger Institute for the COVID-19 Genomics UK (COG-UK) Consortium | Jacquelyn Wynn, Mairead Hyland, The Lighthouse Lab in Alderley Park and Alex Alderton, Roberto Amato, Jeffrey Barrett, Sonia Goncalves, Ewan Harrison, David K. Jackson, Ian Johnston, Dominic Kwiatkowski, Cordelia Langford, John Sillitoe on behalf of the Wellcome Sanger Institute COVID-19 Surveillance Team |
| EPI_ISL_3169958 | Lighthouse Laboratory Plymouth | Wellcome Sanger Institute for the COVID-19 Genomics UK (COG-UK) Consortium | Lighthouse Laboratory Plymouth and Alex Alderton, Roberto Amato, Jeffrey Barrett, Sonia Goncalves, Ewan Harrison, David K. Jackson, Ian Johnston, Dominic Kwiatkowski, Cordelia Langford, John Sillitoe on behalf of the Wellcome Sanger Institute COVID-19 Surveillance Team |
| EPI_ISL_3169959, EPI_ISL_3169960 | Berkshire and Surrey Pathology Services Lighthouse Laboratory | Wellcome Sanger Institute for the COVID-19 Genomics UK (COG-UK) Consortium | Berkshire and Surrey Pathology Services Lighthouse Laboratory and Alex Alderton, Roberto Amato, Jeffrey Barrett, Sonia Goncalves, Ewan Harrison, David K. Jackson, Ian Johnston, Dominic Kwiatkowski, Cordelia Langford, John Sillitoe on behalf of the Wellcome Sanger Institute COVID-19 Surveillance Team |
| EPI_ISL_3169961, EPI_ISL_3169962 | Lighthouse Laboratory Plymouth | Wellcome Sanger Institute for the COVID-19 Genomics UK (COG-UK) Consortium | Lighthouse Laboratory Plymouth and Alex Alderton, Roberto Amato, Jeffrey Barrett, Sonia Goncalves, Ewan Harrison, David K. Jackson, Ian Johnston, Dominic Kwiatkowski, Cordelia Langford, John Sillitoe on behalf of the Wellcome Sanger Institute COVID-19 Surveillance Team |

| | | | |
|---|---|---|---|
| EPI_ISL_3169963, EPI_ISL_3169964 | Lighthouse Lab in Alderley Park | Wellcome Sanger Institute for the COVID-19 Genomics UK (COG-UK) Consortium | Jacquelyn Wynn, Mairead Hyland, The Lighthouse Lab in Alderley Park and Alex Alderton, Roberto Amato, Jeffrey Barrett, Sonia Goncalves, Ewan Harrison, David K. Jackson, Ian Johnston, Dominic Kwiatkowski, Cordelia Langford, John Sillitoe on behalf of the Wellcome Sanger Institute COVID-19 Surveillance Team |
| EPI_ISL_3169965 | Berkshire and Surrey Pathology Services Lighthouse Laboratory | Wellcome Sanger Institute for the COVID-19 Genomics UK (COG-UK) Consortium | Berkshire and Surrey Pathology Services Lighthouse Laboratory and Alex Alderton, Roberto Amato, Jeffrey Barrett, Sonia Goncalves, Ewan Harrison, David K. Jackson, Ian Johnston, Dominic Kwiatkowski, Cordelia Langford, John Sillitoe on behalf of the Wellcome Sanger Institute COVID-19 Surveillance Team |
| EPI_ISL_3169966, EPI_ISL_3169967, EPI_ISL_3169968, EPI_ISL_3169969 | Lighthouse Laboratory Plymouth | Wellcome Sanger Institute for the COVID-19 Genomics UK (COG-UK) Consortium | Lighthouse Laboratory Plymouth and Alex Alderton, Roberto Amato, Jeffrey Barrett, Sonia Goncalves, Ewan Harrison, David K. Jackson, Ian Johnston, Dominic Kwiatkowski, Cordelia Langford, John Sillitoe on behalf of the Wellcome Sanger Institute COVID-19 Surveillance Team |
| EPI_ISL_3169970 | Berkshire and Surrey Pathology Services Lighthouse Laboratory | Wellcome Sanger Institute for the COVID-19 Genomics UK (COG-UK) Consortium | Berkshire and Surrey Pathology Services Lighthouse Laboratory and Alex Alderton, Roberto Amato, Jeffrey Barrett, Sonia Goncalves, Ewan Harrison, David K. Jackson, Ian Johnston, Dominic Kwiatkowski, Cordelia Langford, John Sillitoe on behalf of the Wellcome Sanger Institute COVID-19 Surveillance Team |
| EPI_ISL_3169971 | Lighthouse Laboratory Plymouth | Wellcome Sanger Institute for the COVID-19 Genomics UK (COG-UK) Consortium | Lighthouse Laboratory Plymouth and Alex Alderton, Roberto Amato, Jeffrey Barrett, Sonia Goncalves, Ewan Harrison, David K. Jackson, Ian Johnston, Dominic Kwiatkowski, Cordelia Langford, John Sillitoe on behalf of the Wellcome Sanger Institute COVID-19 Surveillance Team |
| EPI_ISL_3169972 | Berkshire and Surrey Pathology Services Lighthouse Laboratory | Wellcome Sanger Institute for the COVID-19 Genomics UK (COG-UK) Consortium | Berkshire and Surrey Pathology Services Lighthouse Laboratory and Alex Alderton, Roberto Amato, Jeffrey Barrett, Sonia Goncalves, Ewan Harrison, David K. Jackson, Ian Johnston, Dominic Kwiatkowski, Cordelia Langford, John Sillitoe on behalf of the Wellcome Sanger Institute COVID-19 Surveillance Team |
| EPI_ISL_3169973 | Lighthouse Laboratory Plymouth | Wellcome Sanger Institute for the COVID-19 Genomics UK (COG-UK) Consortium | Lighthouse Laboratory Plymouth and Alex Alderton, Roberto Amato, Jeffrey Barrett, Sonia Goncalves, Ewan Harrison, David K. Jackson, Ian Johnston, Dominic Kwiatkowski, Cordelia Langford, John Sillitoe on behalf of the Wellcome Sanger Institute COVID-19 Surveillance Team |
| EPI_ISL_3169974 | Berkshire and Surrey Pathology Services Lighthouse Laboratory | Wellcome Sanger Institute for the COVID-19 Genomics UK (COG-UK) Consortium | Berkshire and Surrey Pathology Services Lighthouse Laboratory and Alex Alderton, Roberto Amato, Jeffrey Barrett, Sonia Goncalves, Ewan Harrison, David K. Jackson, Ian Johnston, Dominic Kwiatkowski, Cordelia Langford, John Sillitoe on behalf of the Wellcome Sanger Institute COVID-19 Surveillance Team |
| EPI_ISL_3169975 | Lighthouse Laboratory Plymouth | Wellcome Sanger Institute for the COVID-19 Genomics UK (COG-UK) Consortium | Lighthouse Laboratory Plymouth and Alex Alderton, Roberto Amato, Jeffrey Barrett, Sonia Goncalves, Ewan Harrison, David K. Jackson, Ian Johnston, Dominic Kwiatkowski, Cordelia Langford, John Sillitoe on behalf of the Wellcome Sanger Institute COVID-19 Surveillance Team |
| EPI_ISL_3169976, EPI_ISL_3169977 | Berkshire and Surrey Pathology Services Lighthouse Laboratory | Wellcome Sanger Institute for the COVID-19 Genomics UK (COG-UK) Consortium | Berkshire and Surrey Pathology Services Lighthouse Laboratory and Alex Alderton, Roberto Amato, Jeffrey Barrett, Sonia Goncalves, Ewan Harrison, David K. Jackson, Ian Johnston, Dominic Kwiatkowski, Cordelia Langford, John Sillitoe on behalf of the Wellcome Sanger Institute COVID-19 Surveillance Team |
| EPI_ISL_3169978, EPI_ISL_3169979 | Lighthouse Lab in Alderley Park | Wellcome Sanger Institute for the COVID-19 Genomics UK (COG-UK) Consortium | Jacquelyn Wynn, Mairead Hyland, The Lighthouse Lab in Alderley Park and Alex Alderton, Roberto Amato, Jeffrey Barrett, Sonia Goncalves, Ewan Harrison, David K. Jackson, Ian Johnston, Dominic Kwiatkowski, Cordelia Langford, John Sillitoe on behalf of the Wellcome Sanger Institute COVID-19 Surveillance Team |
| EPI_ISL_3169980 | Berkshire and Surrey Pathology Services Lighthouse Laboratory | Wellcome Sanger Institute for the COVID-19 Genomics UK (COG-UK) Consortium | Berkshire and Surrey Pathology Services Lighthouse Laboratory and Alex Alderton, Roberto Amato, Jeffrey Barrett, Sonia Goncalves, Ewan Harrison, David K. Jackson, Ian Johnston, Dominic Kwiatkowski, Cordelia Langford, John Sillitoe on behalf of the Wellcome Sanger Institute COVID-19 Surveillance Team |
| EPI_ISL_3169981, EPI_ISL_3169982 | Lighthouse Laboratory Plymouth | Wellcome Sanger Institute for the COVID-19 Genomics UK (COG-UK) Consortium | Lighthouse Laboratory Plymouth and Alex Alderton, Roberto Amato, Jeffrey Barrett, Sonia Goncalves, Ewan Harrison, David K. Jackson, Ian Johnston, Dominic Kwiatkowski, Cordelia Langford, John Sillitoe on behalf of the Wellcome Sanger Institute COVID-19 Surveillance Team |
| EPI_ISL_3169983 | Berkshire and Surrey Pathology Services Lighthouse Laboratory | Wellcome Sanger Institute for the COVID-19 Genomics UK (COG-UK) Consortium | Berkshire and Surrey Pathology Services Lighthouse Laboratory and Alex Alderton, Roberto Amato, Jeffrey Barrett, Sonia Goncalves, Ewan Harrison, David K. Jackson, Ian Johnston, Dominic Kwiatkowski, Cordelia Langford, John Sillitoe on behalf of the Wellcome Sanger Institute COVID-19 Surveillance Team |
| EPI_ISL_3169984 | Lighthouse Lab in Alderley Park | Wellcome Sanger Institute for the COVID-19 Genomics UK (COG-UK) Consortium | Jacquelyn Wynn, Mairead Hyland, The Lighthouse Lab in Alderley Park and Alex Alderton, Roberto Amato, Jeffrey Barrett, Sonia Goncalves, Ewan Harrison, David K. Jackson, Ian Johnston, Dominic Kwiatkowski, Cordelia Langford, John Sillitoe on behalf of the Wellcome Sanger Institute COVID-19 Surveillance Team |
| EPI_ISL_3169985, EPI_ISL_3169986, EPI_ISL_3169987, EPI_ISL_3169988 | Berkshire and Surrey Pathology Services Lighthouse Laboratory | Wellcome Sanger Institute for the COVID-19 Genomics UK (COG-UK) Consortium | Berkshire and Surrey Pathology Services Lighthouse Laboratory and Alex Alderton, Roberto Amato, Jeffrey Barrett, Sonia Goncalves, Ewan Harrison, David K. Jackson, Ian Johnston, Dominic Kwiatkowski, Cordelia Langford, John Sillitoe on behalf of the Wellcome Sanger Institute COVID-19 Surveillance Team |
| EPI_ISL_3169989 | Lighthouse Laboratory Plymouth | Wellcome Sanger Institute for the COVID-19 Genomics UK (COG-UK) Consortium | Lighthouse Laboratory Plymouth and Alex Alderton, Roberto Amato, Jeffrey Barrett, Sonia Goncalves, Ewan Harrison, David K. Jackson, Ian Johnston, Dominic Kwiatkowski, Cordelia Langford, John Sillitoe on behalf of the Wellcome Sanger Institute COVID-19 Surveillance Team |
| EPI_ISL_3169990, EPI_ISL_3169991 | Berkshire and Surrey Pathology Services Lighthouse Laboratory | Wellcome Sanger Institute for the COVID-19 Genomics UK (COG-UK) Consortium | Berkshire and Surrey Pathology Services Lighthouse Laboratory and Alex Alderton, Roberto Amato, Jeffrey Barrett, Sonia Goncalves, Ewan Harrison, David K. Jackson, Ian Johnston, Dominic Kwiatkowski, Cordelia Langford, John Sillitoe on behalf of the Wellcome Sanger Institute COVID-19 Surveillance Team |
| EPI_ISL_3169992 | Lighthouse Laboratory Plymouth | Wellcome Sanger Institute for the COVID-19 Genomics UK (COG-UK) Consortium | Lighthouse Laboratory Plymouth and Alex Alderton, Roberto Amato, Jeffrey Barrett, Sonia Goncalves, Ewan Harrison, David K. Jackson, Ian Johnston, Dominic Kwiatkowski, Cordelia Langford, John Sillitoe on behalf of the Wellcome Sanger Institute COVID-19 Surveillance Team |
| EPI_ISL_3169993, EPI_ISL_3169994 | Berkshire and Surrey Pathology Services Lighthouse Laboratory | Wellcome Sanger Institute for the COVID-19 Genomics UK (COG-UK) Consortium | Berkshire and Surrey Pathology Services Lighthouse Laboratory and Alex Alderton, Roberto Amato, Jeffrey Barrett, Sonia Goncalves, Ewan Harrison, David K. Jackson, Ian Johnston, Dominic Kwiatkowski, Cordelia Langford, John Sillitoe on behalf of the Wellcome Sanger Institute COVID-19 Surveillance Team |
| EPI_ISL_3169995, EPI_ISL_3169996, EPI_ISL_3169997 | Lighthouse Lab in Alderley Park | Wellcome Sanger Institute for the COVID-19 Genomics UK (COG-UK) Consortium | Jacquelyn Wynn, Mairead Hyland, The Lighthouse Lab in Alderley Park and Alex Alderton, Roberto Amato, Jeffrey Barrett, Sonia Goncalves, Ewan Harrison, David K. Jackson, Ian Johnston, Dominic Kwiatkowski, Cordelia Langford, John Sillitoe on behalf of the Wellcome Sanger Institute COVID-19 Surveillance Team |
| EPI_ISL_3169998 | Berkshire and Surrey Pathology Services Lighthouse Laboratory | Wellcome Sanger Institute for the COVID-19 Genomics UK (COG-UK) Consortium | Berkshire and Surrey Pathology Services Lighthouse Laboratory and Alex Alderton, Roberto Amato, Jeffrey Barrett, Sonia Goncalves, Ewan Harrison, David K. Jackson, Ian Johnston, Dominic Kwiatkowski, Cordelia Langford, John Sillitoe on behalf of the Wellcome Sanger Institute COVID-19 Surveillance Team |
| EPI_ISL_3169999 | Lighthouse Lab in Alderley Park | Wellcome Sanger Institute for the COVID-19 Genomics UK (COG-UK) Consortium | Jacquelyn Wynn, Mairead Hyland, The Lighthouse Lab in Alderley Park and Alex Alderton, Roberto Amato, Jeffrey Barrett, Sonia Goncalves, Ewan Harrison, David K. Jackson, Ian Johnston, Dominic Kwiatkowski, Cordelia Langford, John Sillitoe on behalf of the Wellcome Sanger Institute COVID-19 Surveillance Team |
| EPI_ISL_3170000 | Lighthouse Laboratory Plymouth | Wellcome Sanger Institute for the COVID-19 Genomics UK (COG-UK) Consortium | Lighthouse Laboratory Plymouth and Alex Alderton, Roberto Amato, Jeffrey Barrett, Sonia Goncalves, Ewan Harrison, David K. Jackson, Ian Johnston, Dominic Kwiatkowski, Cordelia Langford, John Sillitoe on behalf of the Wellcome Sanger Institute COVID-19 Surveillance Team |
| EPI_ISL_3170001, EPI_ISL_3170002, EPI_ISL_3170003, EPI_ISL_3170004 | Berkshire and Surrey Pathology Services Lighthouse Laboratory | Wellcome Sanger Institute for the COVID-19 Genomics UK (COG-UK) Consortium | Berkshire and Surrey Pathology Services Lighthouse Laboratory and Alex Alderton, Roberto Amato, Jeffrey Barrett, Sonia Goncalves, Ewan Harrison, David K. Jackson, Ian Johnston, Dominic Kwiatkowski, Cordelia Langford, John Sillitoe on behalf of the Wellcome Sanger Institute COVID-19 Surveillance Team |
| EPI_ISL_3170005 | Lighthouse Laboratory Plymouth | Wellcome Sanger Institute for the COVID-19 Genomics UK (COG-UK) Consortium | Lighthouse Laboratory Plymouth and Alex Alderton, Roberto Amato, Jeffrey Barrett, Sonia Goncalves, Ewan Harrison, David K. Jackson, Ian Johnston, Dominic Kwiatkowski, Cordelia Langford, John Sillitoe on behalf of the Wellcome Sanger Institute COVID-19 Surveillance Team |
| EPI_ISL_3170006, EPI_ISL_3170007, EPI_ISL_3170008 | Berkshire and Surrey Pathology Services Lighthouse Laboratory | Wellcome Sanger Institute for the COVID-19 Genomics UK (COG-UK) Consortium | Berkshire and Surrey Pathology Services Lighthouse Laboratory and Alex Alderton, Roberto Amato, Jeffrey Barrett, Sonia Goncalves, Ewan Harrison, David K. Jackson, Ian Johnston, Dominic Kwiatkowski, Cordelia Langford, John Sillitoe on behalf of the Wellcome Sanger Institute COVID-19 Surveillance Team |
| EPI_ISL_3170009 | Lighthouse Lab in Alderley Park | Wellcome Sanger Institute for the COVID-19 Genomics UK (COG-UK) Consortium | Jacquelyn Wynn, Mairead Hyland, The Lighthouse Lab in Alderley Park and Alex Alderton, Roberto Amato, Jeffrey Barrett, Sonia Goncalves, Ewan Harrison, David K. Jackson, Ian Johnston, Dominic Kwiatkowski, Cordelia Langford, John Sillitoe on behalf of the Wellcome Sanger Institute COVID-19 Surveillance Team |
| EPI_ISL_3170010 | Lighthouse Laboratory Plymouth | Wellcome Sanger Institute for the COVID-19 Genomics UK (COG-UK) Consortium | Lighthouse Laboratory Plymouth and Alex Alderton, Roberto Amato, Jeffrey Barrett, Sonia Goncalves, Ewan Harrison, David K. Jackson, Ian Johnston, Dominic Kwiatkowski, Cordelia Langford, John Sillitoe on behalf of the Wellcome Sanger Institute COVID-19 Surveillance Team |
| EPI_ISL_3170011 | Berkshire and Surrey Pathology Services Lighthouse Laboratory | Wellcome Sanger Institute for the COVID-19 Genomics UK (COG-UK) Consortium | Berkshire and Surrey Pathology Services Lighthouse Laboratory and Alex Alderton, Roberto Amato, Jeffrey Barrett, Sonia Goncalves, Ewan Harrison, David K. Jackson, Ian Johnston, Dominic Kwiatkowski, Cordelia Langford, John Sillitoe on behalf of the Wellcome Sanger Institute COVID-19 Surveillance Team |
| EPI_ISL_3170012, EPI_ISL_3170013 | Lighthouse Laboratory Plymouth | Wellcome Sanger Institute for the COVID-19 Genomics UK (COG-UK) Consortium | Lighthouse Laboratory Plymouth and Alex Alderton, Roberto Amato, Jeffrey Barrett, Sonia Goncalves, Ewan Harrison, David K. Jackson, Ian Johnston, Dominic Kwiatkowski, Cordelia Langford, John Sillitoe on behalf of the Wellcome Sanger Institute COVID-19 Surveillance Team |
| EPI_ISL_3170014, EPI_ISL_3170015, EPI_ISL_3170016 | Berkshire and Surrey Pathology Services Lighthouse Laboratory | Wellcome Sanger Institute for the COVID-19 Genomics UK (COG-UK) Consortium | Berkshire and Surrey Pathology Services Lighthouse Laboratory and Alex Alderton, Roberto Amato, Jeffrey Barrett, Sonia Goncalves, Ewan Harrison, David K. Jackson, Ian Johnston, Dominic Kwiatkowski, Cordelia Langford, John Sillitoe on behalf of the Wellcome Sanger Institute COVID-19 Surveillance Team |
| EPI_ISL_3170017, EPI_ISL_3170018 | Lighthouse Laboratory Plymouth | Wellcome Sanger Institute for the COVID-19 Genomics UK (COG-UK) Consortium | Lighthouse Laboratory Plymouth and Alex Alderton, Roberto Amato, Jeffrey Barrett, Sonia Goncalves, Ewan Harrison, David K. Jackson, Ian Johnston, Dominic Kwiatkowski, Cordelia Langford, John Sillitoe on behalf of the Wellcome Sanger Institute COVID-19 Surveillance Team |
| EPI_ISL_3170019 | Berkshire and Surrey Pathology Services Lighthouse Laboratory | Wellcome Sanger Institute for the COVID-19 Genomics UK (COG-UK) Consortium | Berkshire and Surrey Pathology Services Lighthouse Laboratory and Alex Alderton, Roberto Amato, Jeffrey Barrett, Sonia Goncalves, Ewan Harrison, David K. Jackson, Ian Johnston, Dominic Kwiatkowski, Cordelia Langford, John Sillitoe on behalf of the Wellcome Sanger Institute COVID-19 Surveillance Team |
| EPI_ISL_3170020 | Lighthouse Laboratory Plymouth | Wellcome Sanger Institute for the COVID-19 Genomics UK (COG-UK) Consortium | Lighthouse Laboratory Plymouth and Alex Alderton, Roberto Amato, Jeffrey Barrett, Sonia Goncalves, Ewan Harrison, David K. Jackson, Ian Johnston, Dominic Kwiatkowski, Cordelia Langford, John Sillitoe on behalf of the Wellcome Sanger Institute COVID-19 Surveillance Team |
| EPI_ISL_3170021, EPI_ISL_3170022, EPI_ISL_3170023 | Berkshire and Surrey Pathology Services Lighthouse Laboratory | Wellcome Sanger Institute for the COVID-19 Genomics UK (COG-UK) Consortium | Berkshire and Surrey Pathology Services Lighthouse Laboratory and Alex Alderton, Roberto Amato, Jeffrey Barrett, Sonia Goncalves, Ewan Harrison, David K. Jackson, Ian Johnston, Dominic Kwiatkowski, Cordelia Langford, John Sillitoe on behalf of the Wellcome Sanger Institute COVID-19 Surveillance Team |
| EPI_ISL_3170024 | Lighthouse Laboratory Plymouth | Wellcome Sanger Institute for the COVID-19 Genomics UK (COG-UK) Consortium | Lighthouse Laboratory Plymouth and Alex Alderton, Roberto Amato, Jeffrey Barrett, Sonia Goncalves, Ewan Harrison, David K. Jackson, Ian Johnston, Dominic Kwiatkowski, Cordelia Langford, John Sillitoe on behalf of the Wellcome Sanger Institute COVID-19 Surveillance Team |
| EPI_ISL_3170025, EPI_ISL_3170026 | Berkshire and Surrey Pathology Services Lighthouse Laboratory | Wellcome Sanger Institute for the COVID-19 Genomics UK (COG-UK) Consortium | Berkshire and Surrey Pathology Services Lighthouse Laboratory and Alex Alderton, Roberto Amato, Jeffrey Barrett, Sonia Goncalves, Ewan Harrison, David K. Jackson, Ian Johnston, Dominic Kwiatkowski, Cordelia Langford, John Sillitoe on behalf of the Wellcome Sanger Institute COVID-19 Surveillance Team |
| EPI_ISL_3170027 | Lighthouse Laboratory Plymouth | Wellcome Sanger Institute for the COVID-19 Genomics UK (COG-UK) Consortium | Lighthouse Laboratory Plymouth and Alex Alderton, Roberto Amato, Jeffrey Barrett, Sonia Goncalves, Ewan Harrison, David K. Jackson, Ian Johnston, Dominic Kwiatkowski, Cordelia Langford, John Sillitoe on behalf of the Wellcome Sanger Institute COVID-19 Surveillance Team |
| EPI_ISL_3170028, EPI_ISL_3170029 | Berkshire and Surrey Pathology Services Lighthouse Laboratory | Wellcome Sanger Institute for the COVID-19 Genomics UK (COG-UK) Consortium | Berkshire and Surrey Pathology Services Lighthouse Laboratory and Alex Alderton, Roberto Amato, Jeffrey Barrett, Sonia Goncalves, Ewan Harrison, David K. Jackson, Ian Johnston, Dominic Kwiatkowski, Cordelia Langford, John Sillitoe on behalf of the Wellcome Sanger Institute COVID-19 Surveillance Team |
| EPI_ISL_3170030 | Lighthouse Laboratory Plymouth | Wellcome Sanger Institute for the COVID-19 Genomics UK (COG-UK) Consortium | Lighthouse Laboratory Plymouth and Alex Alderton, Roberto Amato, Jeffrey Barrett, Sonia Goncalves, Ewan Harrison, David K. Jackson, Ian Johnston, Dominic Kwiatkowski, Cordelia Langford, John Sillitoe on behalf of the Wellcome Sanger Institute COVID-19 Surveillance Team |

| | | | |
|---|---|---|---|
| EPI_ISL_3170031, EPI_ISL_3170032 | Lighthouse Lab in Alderley Park | Wellcome Sanger Institute for the COVID-19 Genomics UK (COG-UK) Consortium | Jacquelyn Wynn, Mairead Hyland, The Lighthouse Lab in Alderley Park and Alex Alderton, Roberto Amato, Jeffrey Barrett, Sonia Goncalves, Ewan Harrison, David K. Jackson, Ian Johnston, Dominic Kwiatkowski, Cordelia Langford, John Sillitoe on behalf of the Wellcome Sanger Institute COVID-19 Surveillance Team |
| EPI_ISL_3170033 | Lighthouse Laboratory Plymouth | Wellcome Sanger Institute for the COVID-19 Genomics UK (COG-UK) Consortium | Lighthouse Laboratory Plymouth and Alex Alderton, Roberto Amato, Jeffrey Barrett, Sonia Goncalves, Ewan Harrison, David K. Jackson, Ian Johnston, Dominic Kwiatkowski, Cordelia Langford, John Sillitoe on behalf of the Wellcome Sanger Institute COVID-19 Surveillance Team |
| EPI_ISL_3170034 | Berkshire and Surrey Pathology Services Lighthouse Laboratory | Wellcome Sanger Institute for the COVID-19 Genomics UK (COG-UK) Consortium | Berkshire and Surrey Pathology Services Lighthouse Laboratory and Alex Alderton, Roberto Amato, Jeffrey Barrett, Sonia Goncalves, Ewan Harrison, David K. Jackson, Ian Johnston, Dominic Kwiatkowski, Cordelia Langford, John Sillitoe on behalf of the Wellcome Sanger Institute COVID-19 Surveillance Team |
| EPI_ISL_3170035, EPI_ISL_3170036, EPI_ISL_3170037 | Lighthouse Lab in Alderley Park | Wellcome Sanger Institute for the COVID-19 Genomics UK (COG-UK) Consortium | Jacquelyn Wynn, Mairead Hyland, The Lighthouse Lab in Alderley Park and Alex Alderton, Roberto Amato, Jeffrey Barrett, Sonia Goncalves, Ewan Harrison, David K. Jackson, Ian Johnston, Dominic Kwiatkowski, Cordelia Langford, John Sillitoe on behalf of the Wellcome Sanger Institute COVID-19 Surveillance Team |
| EPI_ISL_3170038 | Berkshire and Surrey Pathology Services Lighthouse Laboratory | Wellcome Sanger Institute for the COVID-19 Genomics UK (COG-UK) Consortium | Berkshire and Surrey Pathology Services Lighthouse Laboratory and Alex Alderton, Roberto Amato, Jeffrey Barrett, Sonia Goncalves, Ewan Harrison, David K. Jackson, Ian Johnston, Dominic Kwiatkowski, Cordelia Langford, John Sillitoe on behalf of the Wellcome Sanger Institute COVID-19 Surveillance Team |
| EPI_ISL_3170039 | Lighthouse Laboratory Plymouth | Wellcome Sanger Institute for the COVID-19 Genomics UK (COG-UK) Consortium | Lighthouse Laboratory Plymouth and Alex Alderton, Roberto Amato, Jeffrey Barrett, Sonia Goncalves, Ewan Harrison, David K. Jackson, Ian Johnston, Dominic Kwiatkowski, Cordelia Langford, John Sillitoe on behalf of the Wellcome Sanger Institute COVID-19 Surveillance Team |
| EPI_ISL_3170040, EPI_ISL_3170041, EPI_ISL_3170042 | Berkshire and Surrey Pathology Services Lighthouse Laboratory | Wellcome Sanger Institute for the COVID-19 Genomics UK (COG-UK) Consortium | Berkshire and Surrey Pathology Services Lighthouse Laboratory and Alex Alderton, Roberto Amato, Jeffrey Barrett, Sonia Goncalves, Ewan Harrison, David K. Jackson, Ian Johnston, Dominic Kwiatkowski, Cordelia Langford, John Sillitoe on behalf of the Wellcome Sanger Institute COVID-19 Surveillance Team |
| EPI_ISL_3170043, EPI_ISL_3170044 | Lighthouse Laboratory Plymouth | Wellcome Sanger Institute for the COVID-19 Genomics UK (COG-UK) Consortium | Lighthouse Laboratory Plymouth and Alex Alderton, Roberto Amato, Jeffrey Barrett, Sonia Goncalves, Ewan Harrison, David K. Jackson, Ian Johnston, Dominic Kwiatkowski, Cordelia Langford, John Sillitoe on behalf of the Wellcome Sanger Institute COVID-19 Surveillance Team |
| EPI_ISL_3170045 | Lighthouse Lab in Alderley Park | Wellcome Sanger Institute for the COVID-19 Genomics UK (COG-UK) Consortium | Jacquelyn Wynn, Mairead Hyland, The Lighthouse Lab in Alderley Park and Alex Alderton, Roberto Amato, Jeffrey Barrett, Sonia Goncalves, Ewan Harrison, David K. Jackson, Ian Johnston, Dominic Kwiatkowski, Cordelia Langford, John Sillitoe on behalf of the Wellcome Sanger Institute COVID-19 Surveillance Team |
| EPI_ISL_3170046, EPI_ISL_3170047, EPI_ISL_3170048, EPI_ISL_3170049 | Berkshire and Surrey Pathology Services Lighthouse Laboratory | Wellcome Sanger Institute for the COVID-19 Genomics UK (COG-UK) Consortium | Berkshire and Surrey Pathology Services Lighthouse Laboratory and Alex Alderton, Roberto Amato, Jeffrey Barrett, Sonia Goncalves, Ewan Harrison, David K. Jackson, Ian Johnston, Dominic Kwiatkowski, Cordelia Langford, John Sillitoe on behalf of the Wellcome Sanger Institute COVID-19 Surveillance Team |
| EPI_ISL_3170050 | Lighthouse Lab in Alderley Park | Wellcome Sanger Institute for the COVID-19 Genomics UK (COG-UK) Consortium | Jacquelyn Wynn, Mairead Hyland, The Lighthouse Lab in Alderley Park and Alex Alderton, Roberto Amato, Jeffrey Barrett, Sonia Goncalves, Ewan Harrison, David K. Jackson, Ian Johnston, Dominic Kwiatkowski, Cordelia Langford, John Sillitoe on behalf of the Wellcome Sanger Institute COVID-19 Surveillance Team |
| EPI_ISL_3170051 | Berkshire and Surrey Pathology Services Lighthouse Laboratory | Wellcome Sanger Institute for the COVID-19 Genomics UK (COG-UK) Consortium | Berkshire and Surrey Pathology Services Lighthouse Laboratory and Alex Alderton, Roberto Amato, Jeffrey Barrett, Sonia Goncalves, Ewan Harrison, David K. Jackson, Ian Johnston, Dominic Kwiatkowski, Cordelia Langford, John Sillitoe on behalf of the Wellcome Sanger Institute COVID-19 Surveillance Team |
| EPI_ISL_3170052 | Lighthouse Lab in Alderley Park | Wellcome Sanger Institute for the COVID-19 Genomics UK (COG-UK) Consortium | Jacquelyn Wynn, Mairead Hyland, The Lighthouse Lab in Alderley Park and Alex Alderton, Roberto Amato, Jeffrey Barrett, Sonia Goncalves, Ewan Harrison, David K. Jackson, Ian Johnston, Dominic Kwiatkowski, Cordelia Langford, John Sillitoe on behalf of the Wellcome Sanger Institute COVID-19 Surveillance Team |
| EPI_ISL_3170053, EPI_ISL_3170054 | Lighthouse Laboratory Plymouth | Wellcome Sanger Institute for the COVID-19 Genomics UK (COG-UK) Consortium | Lighthouse Laboratory Plymouth and Alex Alderton, Roberto Amato, Jeffrey Barrett, Sonia Goncalves, Ewan Harrison, David K. Jackson, Ian Johnston, Dominic Kwiatkowski, Cordelia Langford, John Sillitoe on behalf of the Wellcome Sanger Institute COVID-19 Surveillance Team |
| EPI_ISL_3170055, EPI_ISL_3170056 | Berkshire and Surrey Pathology Services Lighthouse Laboratory | Wellcome Sanger Institute for the COVID-19 Genomics UK (COG-UK) Consortium | Berkshire and Surrey Pathology Services Lighthouse Laboratory and Alex Alderton, Roberto Amato, Jeffrey Barrett, Sonia Goncalves, Ewan Harrison, David K. Jackson, Ian Johnston, Dominic Kwiatkowski, Cordelia Langford, John Sillitoe on behalf of the Wellcome Sanger Institute COVID-19 Surveillance Team |
| EPI_ISL_3170057 | Lighthouse Laboratory Plymouth | Wellcome Sanger Institute for the COVID-19 Genomics UK (COG-UK) Consortium | Lighthouse Laboratory Plymouth and Alex Alderton, Roberto Amato, Jeffrey Barrett, Sonia Goncalves, Ewan Harrison, David K. Jackson, Ian Johnston, Dominic Kwiatkowski, Cordelia Langford, John Sillitoe on behalf of the Wellcome Sanger Institute COVID-19 Surveillance Team |
| EPI_ISL_3170058 | Lighthouse Lab in Alderley Park | Wellcome Sanger Institute for the COVID-19 Genomics UK (COG-UK) Consortium | Jacquelyn Wynn, Mairead Hyland, The Lighthouse Lab in Alderley Park and Alex Alderton, Roberto Amato, Jeffrey Barrett, Sonia Goncalves, Ewan Harrison, David K. Jackson, Ian Johnston, Dominic Kwiatkowski, Cordelia Langford, John Sillitoe on behalf of the Wellcome Sanger Institute COVID-19 Surveillance Team |
| EPI_ISL_3170059 | Berkshire and Surrey Pathology Services Lighthouse Laboratory | Wellcome Sanger Institute for the COVID-19 Genomics UK (COG-UK) Consortium | Berkshire and Surrey Pathology Services Lighthouse Laboratory and Alex Alderton, Roberto Amato, Jeffrey Barrett, Sonia Goncalves, Ewan Harrison, David K. Jackson, Ian Johnston, Dominic Kwiatkowski, Cordelia Langford, John Sillitoe on behalf of the Wellcome Sanger Institute COVID-19 Surveillance Team |
| EPI_ISL_3170060 | Lighthouse Lab in Alderley Park | Wellcome Sanger Institute for the COVID-19 Genomics UK (COG-UK) Consortium | Jacquelyn Wynn, Mairead Hyland, The Lighthouse Lab in Alderley Park and Alex Alderton, Roberto Amato, Jeffrey Barrett, Sonia Goncalves, Ewan Harrison, David K. Jackson, Ian Johnston, Dominic Kwiatkowski, Cordelia Langford, John Sillitoe on behalf of the Wellcome Sanger Institute COVID-19 Surveillance Team |
| EPI_ISL_3170061 | Berkshire and Surrey Pathology Services Lighthouse Laboratory | Wellcome Sanger Institute for the COVID-19 Genomics UK (COG-UK) Consortium | Berkshire and Surrey Pathology Services Lighthouse Laboratory and Alex Alderton, Roberto Amato, Jeffrey Barrett, Sonia Goncalves, Ewan Harrison, David K. Jackson, Ian Johnston, Dominic Kwiatkowski, Cordelia Langford, John Sillitoe on behalf of the Wellcome Sanger Institute COVID-19 Surveillance Team |
| EPI_ISL_3170062 | Lighthouse Lab in Alderley Park | Wellcome Sanger Institute for the COVID-19 Genomics UK (COG-UK) Consortium | Jacquelyn Wynn, Mairead Hyland, The Lighthouse Lab in Alderley Park and Alex Alderton, Roberto Amato, Jeffrey Barrett, Sonia Goncalves, Ewan Harrison, David K. Jackson, Ian Johnston, Dominic Kwiatkowski, Cordelia Langford, John Sillitoe on behalf of the Wellcome Sanger Institute COVID-19 Surveillance Team |
| EPI_ISL_3170063 | Berkshire and Surrey Pathology Services Lighthouse Laboratory | Wellcome Sanger Institute for the COVID-19 Genomics UK (COG-UK) Consortium | Berkshire and Surrey Pathology Services Lighthouse Laboratory and Alex Alderton, Roberto Amato, Jeffrey Barrett, Sonia Goncalves, Ewan Harrison, David K. Jackson, Ian Johnston, Dominic Kwiatkowski, Cordelia Langford, John Sillitoe on behalf of the Wellcome Sanger Institute COVID-19 Surveillance Team |
| EPI_ISL_3170064 | Lighthouse Laboratory Plymouth | Wellcome Sanger Institute for the COVID-19 Genomics UK (COG-UK) Consortium | Lighthouse Laboratory Plymouth and Alex Alderton, Roberto Amato, Jeffrey Barrett, Sonia Goncalves, Ewan Harrison, David K. Jackson, Ian Johnston, Dominic Kwiatkowski, Cordelia Langford, John Sillitoe on behalf of the Wellcome Sanger Institute COVID-19 Surveillance Team |
| EPI_ISL_3170065 | Berkshire and Surrey Pathology Services Lighthouse Laboratory | Wellcome Sanger Institute for the COVID-19 Genomics UK (COG-UK) Consortium | Berkshire and Surrey Pathology Services Lighthouse Laboratory and Alex Alderton, Roberto Amato, Jeffrey Barrett, Sonia Goncalves, Ewan Harrison, David K. Jackson, Ian Johnston, Dominic Kwiatkowski, Cordelia Langford, John Sillitoe on behalf of the Wellcome Sanger Institute COVID-19 Surveillance Team |
| EPI_ISL_3170066, EPI_ISL_3170067 | Lighthouse Lab in Alderley Park | Wellcome Sanger Institute for the COVID-19 Genomics UK (COG-UK) Consortium | Jacquelyn Wynn, Mairead Hyland, The Lighthouse Lab in Alderley Park and Alex Alderton, Roberto Amato, Jeffrey Barrett, Sonia Goncalves, Ewan Harrison, David K. Jackson, Ian Johnston, Dominic Kwiatkowski, Cordelia Langford, John Sillitoe on behalf of the Wellcome Sanger Institute COVID-19 Surveillance Team |
| EPI_ISL_3170068, EPI_ISL_3170069 | Berkshire and Surrey Pathology Services Lighthouse Laboratory | Wellcome Sanger Institute for the COVID-19 Genomics UK (COG-UK) Consortium | Berkshire and Surrey Pathology Services Lighthouse Laboratory and Alex Alderton, Roberto Amato, Jeffrey Barrett, Sonia Goncalves, Ewan Harrison, David K. Jackson, Ian Johnston, Dominic Kwiatkowski, Cordelia Langford, John Sillitoe on behalf of the Wellcome Sanger Institute COVID-19 Surveillance Team |
| EPI_ISL_3170070 | Lighthouse Laboratory Plymouth | Wellcome Sanger Institute for the COVID-19 Genomics UK (COG-UK) Consortium | Lighthouse Laboratory Plymouth and Alex Alderton, Roberto Amato, Jeffrey Barrett, Sonia Goncalves, Ewan Harrison, David K. Jackson, Ian Johnston, Dominic Kwiatkowski, Cordelia Langford, John Sillitoe on behalf of the Wellcome Sanger Institute COVID-19 Surveillance Team |
| EPI_ISL_3170071, EPI_ISL_3170072, EPI_ISL_3170073, EPI_ISL_3170074, EPI_ISL_3170075 | Berkshire and Surrey Pathology Services Lighthouse Laboratory | Wellcome Sanger Institute for the COVID-19 Genomics UK (COG-UK) Consortium | Berkshire and Surrey Pathology Services Lighthouse Laboratory and Alex Alderton, Roberto Amato, Jeffrey Barrett, Sonia Goncalves, Ewan Harrison, David K. Jackson, Ian Johnston, Dominic Kwiatkowski, Cordelia Langford, John Sillitoe on behalf of the Wellcome Sanger Institute COVID-19 Surveillance Team |
| EPI_ISL_3170076, EPI_ISL_3170077 | Lighthouse Laboratory Plymouth | Wellcome Sanger Institute for the COVID-19 Genomics UK (COG-UK) Consortium | Lighthouse Laboratory Plymouth and Alex Alderton, Roberto Amato, Jeffrey Barrett, Sonia Goncalves, Ewan Harrison, David K. Jackson, Ian Johnston, Dominic Kwiatkowski, Cordelia Langford, John Sillitoe on behalf of the Wellcome Sanger Institute COVID-19 Surveillance Team |
| EPI_ISL_3170078 | Lighthouse Lab in Alderley Park | Wellcome Sanger Institute for the COVID-19 Genomics UK (COG-UK) Consortium | Jacquelyn Wynn, Mairead Hyland, The Lighthouse Lab in Alderley Park and Alex Alderton, Roberto Amato, Jeffrey Barrett, Sonia Goncalves, Ewan Harrison, David K. Jackson, Ian Johnston, Dominic Kwiatkowski, Cordelia Langford, John Sillitoe on behalf of the Wellcome Sanger Institute COVID-19 Surveillance Team |
| EPI_ISL_3170079 | Berkshire and Surrey Pathology Services Lighthouse Laboratory | Wellcome Sanger Institute for the COVID-19 Genomics UK (COG-UK) Consortium | Berkshire and Surrey Pathology Services Lighthouse Laboratory and Alex Alderton, Roberto Amato, Jeffrey Barrett, Sonia Goncalves, Ewan Harrison, David K. Jackson, Ian Johnston, Dominic Kwiatkowski, Cordelia Langford, John Sillitoe on behalf of the Wellcome Sanger Institute COVID-19 Surveillance Team |
| EPI_ISL_3170080, EPI_ISL_3170081 | Lighthouse Lab in Alderley Park | Wellcome Sanger Institute for the COVID-19 Genomics UK (COG-UK) Consortium | Jacquelyn Wynn, Mairead Hyland, The Lighthouse Lab in Alderley Park and Alex Alderton, Roberto Amato, Jeffrey Barrett, Sonia Goncalves, Ewan Harrison, David K. Jackson, Ian Johnston, Dominic Kwiatkowski, Cordelia Langford, John Sillitoe on behalf of the Wellcome Sanger Institute COVID-19 Surveillance Team |
| EPI_ISL_3170082 | Berkshire and Surrey Pathology Services Lighthouse Laboratory | Wellcome Sanger Institute for the COVID-19 Genomics UK (COG-UK) Consortium | Berkshire and Surrey Pathology Services Lighthouse Laboratory and Alex Alderton, Roberto Amato, Jeffrey Barrett, Sonia Goncalves, Ewan Harrison, David K. Jackson, Ian Johnston, Dominic Kwiatkowski, Cordelia Langford, John Sillitoe on behalf of the Wellcome Sanger Institute COVID-19 Surveillance Team |
| EPI_ISL_3170083 | Lighthouse Lab in Alderley Park | Wellcome Sanger Institute for the COVID-19 Genomics UK (COG-UK) Consortium | Jacquelyn Wynn, Mairead Hyland, The Lighthouse Lab in Alderley Park and Alex Alderton, Roberto Amato, Jeffrey Barrett, Sonia Goncalves, Ewan Harrison, David K. Jackson, Ian Johnston, Dominic Kwiatkowski, Cordelia Langford, John Sillitoe on behalf of the Wellcome Sanger Institute COVID-19 Surveillance Team |
| EPI_ISL_3170084 | Lighthouse Laboratory Plymouth | Wellcome Sanger Institute for the COVID-19 Genomics UK (COG-UK) Consortium | Lighthouse Laboratory Plymouth and Alex Alderton, Roberto Amato, Jeffrey Barrett, Sonia Goncalves, Ewan Harrison, David K. Jackson, Ian Johnston, Dominic Kwiatkowski, Cordelia Langford, John Sillitoe on behalf of the Wellcome Sanger Institute COVID-19 Surveillance Team |
| EPI_ISL_3170085 | Berkshire and Surrey Pathology Services Lighthouse Laboratory | Wellcome Sanger Institute for the COVID-19 Genomics UK (COG-UK) Consortium | Berkshire and Surrey Pathology Services Lighthouse Laboratory and Alex Alderton, Roberto Amato, Jeffrey Barrett, Sonia Goncalves, Ewan Harrison, David K. Jackson, Ian Johnston, Dominic Kwiatkowski, Cordelia Langford, John Sillitoe on behalf of the Wellcome Sanger Institute COVID-19 Surveillance Team |
| EPI_ISL_3170086 | Lighthouse Laboratory Plymouth | Wellcome Sanger Institute for the COVID-19 Genomics UK (COG-UK) Consortium | Lighthouse Laboratory Plymouth and Alex Alderton, Roberto Amato, Jeffrey Barrett, Sonia Goncalves, Ewan Harrison, David K. Jackson, Ian Johnston, Dominic Kwiatkowski, Cordelia Langford, John Sillitoe on behalf of the Wellcome Sanger Institute COVID-19 Surveillance Team |
| EPI_ISL_3170087, EPI_ISL_3170088 | Berkshire and Surrey Pathology Services Lighthouse Laboratory | Wellcome Sanger Institute for the COVID-19 Genomics UK (COG-UK) Consortium | Berkshire and Surrey Pathology Services Lighthouse Laboratory and Alex Alderton, Roberto Amato, Jeffrey Barrett, Sonia Goncalves, Ewan Harrison, David K. Jackson, Ian Johnston, Dominic Kwiatkowski, Cordelia Langford, John Sillitoe on behalf of the Wellcome Sanger Institute COVID-19 Surveillance Team |
| EPI_ISL_3170089 | Lighthouse Laboratory Plymouth | Wellcome Sanger Institute for the COVID-19 Genomics UK (COG-UK) Consortium | Lighthouse Laboratory Plymouth and Alex Alderton, Roberto Amato, Jeffrey Barrett, Sonia Goncalves, Ewan Harrison, David K. Jackson, Ian Johnston, Dominic Kwiatkowski, Cordelia Langford, John Sillitoe on behalf of the Wellcome Sanger Institute COVID-19 Surveillance Team |
| EPI_ISL_3170090 | Berkshire and Surrey Pathology Services Lighthouse Laboratory | Wellcome Sanger Institute for the COVID-19 Genomics UK (COG-UK) Consortium | Berkshire and Surrey Pathology Services Lighthouse Laboratory and Alex Alderton, Roberto Amato, Jeffrey Barrett, Sonia Goncalves, Ewan Harrison, David K. Jackson, Ian Johnston, Dominic Kwiatkowski, Cordelia Langford, John Sillitoe on behalf of the Wellcome Sanger Institute COVID-19 Surveillance Team |
| EPI_ISL_3170091, EPI_ISL_3170092 | Lighthouse Lab in Alderley Park | Wellcome Sanger Institute for the COVID-19 Genomics UK (COG-UK) Consortium | Jacquelyn Wynn, Mairead Hyland, The Lighthouse Lab in Alderley Park and Alex Alderton, Roberto Amato, Jeffrey Barrett, Sonia Goncalves, Ewan Harrison, David K. Jackson, Ian Johnston, Dominic Kwiatkowski, Cordelia Langford, John Sillitoe on behalf of the Wellcome Sanger Institute COVID-19 Surveillance Team |
| EPI_ISL_3170093 | Berkshire and Surrey Pathology Services Lighthouse Laboratory | Wellcome Sanger Institute for the COVID-19 Genomics UK (COG-UK) Consortium | Berkshire and Surrey Pathology Services Lighthouse Laboratory and Alex Alderton, Roberto Amato, Jeffrey Barrett, Sonia Goncalves, Ewan Harrison, David K. Jackson, Ian Johnston, Dominic Kwiatkowski, Cordelia Langford, John Sillitoe on behalf of the Wellcome Sanger Institute COVID-19 Surveillance Team |

| | | | |
|---|---|---|---|
| EPI_ISL_3170094 | Lighthouse Lab in Alderley Park | Wellcome Sanger Institute for the COVID-19 Genomics UK (COG-UK) Consortium | Jacquelyn Wynn, Mairead Hyland, The Lighthouse Lab in Alderley Park and Alex Alderton, Roberto Amato, Jeffrey Barrett, Sonia Goncalves, Ewan Harrison, David K. Jackson, Ian Johnston, Dominic Kwiatkowski, Cordelia Langford, John Sillitoe on behalf of the Wellcome Sanger Institute COVID-19 Surveillance Team |
| EPI_ISL_3170095 | Lighthouse Laboratory Plymouth | Wellcome Sanger Institute for the COVID-19 Genomics UK (COG-UK) Consortium | Lighthouse Laboratory Plymouth and Alex Alderton, Roberto Amato, Jeffrey Barrett, Sonia Goncalves, Ewan Harrison, David K. Jackson, Ian Johnston, Dominic Kwiatkowski, Cordelia Langford, John Sillitoe on behalf of the Wellcome Sanger Institute COVID-19 Surveillance Team |
| EPI_ISL_3170096 | Berkshire and Surrey Pathology Services Lighthouse Laboratory | Wellcome Sanger Institute for the COVID-19 Genomics UK (COG-UK) Consortium | Berkshire and Surrey Pathology Services Lighthouse Laboratory and Alex Alderton, Roberto Amato, Jeffrey Barrett, Sonia Goncalves, Ewan Harrison, David K. Jackson, Ian Johnston, Dominic Kwiatkowski, Cordelia Langford, John Sillitoe on behalf of the Wellcome Sanger Institute COVID-19 Surveillance Team |
| EPI_ISL_3170097 | Lighthouse Lab in Alderley Park | Wellcome Sanger Institute for the COVID-19 Genomics UK (COG-UK) Consortium | Jacquelyn Wynn, Mairead Hyland, The Lighthouse Lab in Alderley Park and Alex Alderton, Roberto Amato, Jeffrey Barrett, Sonia Goncalves, Ewan Harrison, David K. Jackson, Ian Johnston, Dominic Kwiatkowski, Cordelia Langford, John Sillitoe on behalf of the Wellcome Sanger Institute COVID-19 Surveillance Team |
| EPI_ISL_3170098 | Lighthouse Laboratory Plymouth | Wellcome Sanger Institute for the COVID-19 Genomics UK (COG-UK) Consortium | Lighthouse Laboratory Plymouth and Alex Alderton, Roberto Amato, Jeffrey Barrett, Sonia Goncalves, Ewan Harrison, David K. Jackson, Ian Johnston, Dominic Kwiatkowski, Cordelia Langford, John Sillitoe on behalf of the Wellcome Sanger Institute COVID-19 Surveillance Team |
| EPI_ISL_3170099 | Berkshire and Surrey Pathology Services Lighthouse Laboratory | Wellcome Sanger Institute for the COVID-19 Genomics UK (COG-UK) Consortium | Berkshire and Surrey Pathology Services Lighthouse Laboratory and Alex Alderton, Roberto Amato, Jeffrey Barrett, Sonia Goncalves, Ewan Harrison, David K. Jackson, Ian Johnston, Dominic Kwiatkowski, Cordelia Langford, John Sillitoe on behalf of the Wellcome Sanger Institute COVID-19 Surveillance Team |
| EPI_ISL_3170100 | Lighthouse Lab in Alderley Park | Wellcome Sanger Institute for the COVID-19 Genomics UK (COG-UK) Consortium | Jacquelyn Wynn, Mairead Hyland, The Lighthouse Lab in Alderley Park and Alex Alderton, Roberto Amato, Jeffrey Barrett, Sonia Goncalves, Ewan Harrison, David K. Jackson, Ian Johnston, Dominic Kwiatkowski, Cordelia Langford, John Sillitoe on behalf of the Wellcome Sanger Institute COVID-19 Surveillance Team |
| EPI_ISL_3170101, EPI_ISL_3170102, EPI_ISL_3170103, EPI_ISL_3170104 | Berkshire and Surrey Pathology Services Lighthouse Laboratory | Wellcome Sanger Institute for the COVID-19 Genomics UK (COG-UK) Consortium | Berkshire and Surrey Pathology Services Lighthouse Laboratory and Alex Alderton, Roberto Amato, Jeffrey Barrett, Sonia Goncalves, Ewan Harrison, David K. Jackson, Ian Johnston, Dominic Kwiatkowski, Cordelia Langford, John Sillitoe on behalf of the Wellcome Sanger Institute COVID-19 Surveillance Team |
| EPI_ISL_3170105 | Lighthouse Lab in Alderley Park | Wellcome Sanger Institute for the COVID-19 Genomics UK (COG-UK) Consortium | Jacquelyn Wynn, Mairead Hyland, The Lighthouse Lab in Alderley Park and Alex Alderton, Roberto Amato, Jeffrey Barrett, Sonia Goncalves, Ewan Harrison, David K. Jackson, Ian Johnston, Dominic Kwiatkowski, Cordelia Langford, John Sillitoe on behalf of the Wellcome Sanger Institute COVID-19 Surveillance Team |
| EPI_ISL_3170106, EPI_ISL_3170107, EPI_ISL_3170108 | Berkshire and Surrey Pathology Services Lighthouse Laboratory | Wellcome Sanger Institute for the COVID-19 Genomics UK (COG-UK) Consortium | Berkshire and Surrey Pathology Services Lighthouse Laboratory and Alex Alderton, Roberto Amato, Jeffrey Barrett, Sonia Goncalves, Ewan Harrison, David K. Jackson, Ian Johnston, Dominic Kwiatkowski, Cordelia Langford, John Sillitoe on behalf of the Wellcome Sanger Institute COVID-19 Surveillance Team |
| EPI_ISL_3170109 | Lighthouse Lab in Alderley Park | Wellcome Sanger Institute for the COVID-19 Genomics UK (COG-UK) Consortium | Jacquelyn Wynn, Mairead Hyland, The Lighthouse Lab in Alderley Park and Alex Alderton, Roberto Amato, Jeffrey Barrett, Sonia Goncalves, Ewan Harrison, David K. Jackson, Ian Johnston, Dominic Kwiatkowski, Cordelia Langford, John Sillitoe on behalf of the Wellcome Sanger Institute COVID-19 Surveillance Team |
| EPI_ISL_3170110 | Lighthouse Laboratory Plymouth | Wellcome Sanger Institute for the COVID-19 Genomics UK (COG-UK) Consortium | Lighthouse Laboratory Plymouth and Alex Alderton, Roberto Amato, Jeffrey Barrett, Sonia Goncalves, Ewan Harrison, David K. Jackson, Ian Johnston, Dominic Kwiatkowski, Cordelia Langford, John Sillitoe on behalf of the Wellcome Sanger Institute COVID-19 Surveillance Team |
| EPI_ISL_3170111 | Lighthouse Lab in Alderley Park | Wellcome Sanger Institute for the COVID-19 Genomics UK (COG-UK) Consortium | Jacquelyn Wynn, Mairead Hyland, The Lighthouse Lab in Alderley Park and Alex Alderton, Roberto Amato, Jeffrey Barrett, Sonia Goncalves, Ewan Harrison, David K. Jackson, Ian Johnston, Dominic Kwiatkowski, Cordelia Langford, John Sillitoe on behalf of the Wellcome Sanger Institute COVID-19 Surveillance Team |
| EPI_ISL_3170112 | Berkshire and Surrey Pathology Services Lighthouse Laboratory | Wellcome Sanger Institute for the COVID-19 Genomics UK (COG-UK) Consortium | Berkshire and Surrey Pathology Services Lighthouse Laboratory and Alex Alderton, Roberto Amato, Jeffrey Barrett, Sonia Goncalves, Ewan Harrison, David K. Jackson, Ian Johnston, Dominic Kwiatkowski, Cordelia Langford, John Sillitoe on behalf of the Wellcome Sanger Institute COVID-19 Surveillance Team |
| EPI_ISL_3170113 | Lighthouse Laboratory Plymouth | Wellcome Sanger Institute for the COVID-19 Genomics UK (COG-UK) Consortium | Lighthouse Laboratory Plymouth and Alex Alderton, Roberto Amato, Jeffrey Barrett, Sonia Goncalves, Ewan Harrison, David K. Jackson, Ian Johnston, Dominic Kwiatkowski, Cordelia Langford, John Sillitoe on behalf of the Wellcome Sanger Institute COVID-19 Surveillance Team |
| EPI_ISL_3170114, EPI_ISL_3170115 | Berkshire and Surrey Pathology Services Lighthouse Laboratory | Wellcome Sanger Institute for the COVID-19 Genomics UK (COG-UK) Consortium | Berkshire and Surrey Pathology Services Lighthouse Laboratory and Alex Alderton, Roberto Amato, Jeffrey Barrett, Sonia Goncalves, Ewan Harrison, David K. Jackson, Ian Johnston, Dominic Kwiatkowski, Cordelia Langford, John Sillitoe on behalf of the Wellcome Sanger Institute COVID-19 Surveillance Team |
| EPI_ISL_3170116 | Lighthouse Laboratory Plymouth | Wellcome Sanger Institute for the COVID-19 Genomics UK (COG-UK) Consortium | Lighthouse Laboratory Plymouth and Alex Alderton, Roberto Amato, Jeffrey Barrett, Sonia Goncalves, Ewan Harrison, David K. Jackson, Ian Johnston, Dominic Kwiatkowski, Cordelia Langford, John Sillitoe on behalf of the Wellcome Sanger Institute COVID-19 Surveillance Team |
| EPI_ISL_3170117 | Lighthouse Lab in Alderley Park | Wellcome Sanger Institute for the COVID-19 Genomics UK (COG-UK) Consortium | Jacquelyn Wynn, Mairead Hyland, The Lighthouse Lab in Alderley Park and Alex Alderton, Roberto Amato, Jeffrey Barrett, Sonia Goncalves, Ewan Harrison, David K. Jackson, Ian Johnston, Dominic Kwiatkowski, Cordelia Langford, John Sillitoe on behalf of the Wellcome Sanger Institute COVID-19 Surveillance Team |
| EPI_ISL_3170118 | Berkshire and Surrey Pathology Services Lighthouse Laboratory | Wellcome Sanger Institute for the COVID-19 Genomics UK (COG-UK) Consortium | Berkshire and Surrey Pathology Services Lighthouse Laboratory and Alex Alderton, Roberto Amato, Jeffrey Barrett, Sonia Goncalves, Ewan Harrison, David K. Jackson, Ian Johnston, Dominic Kwiatkowski, Cordelia Langford, John Sillitoe on behalf of the Wellcome Sanger Institute COVID-19 Surveillance Team |
| EPI_ISL_3170119 | Lighthouse Laboratory Plymouth | Wellcome Sanger Institute for the COVID-19 Genomics UK (COG-UK) Consortium | Lighthouse Laboratory Plymouth and Alex Alderton, Roberto Amato, Jeffrey Barrett, Sonia Goncalves, Ewan Harrison, David K. Jackson, Ian Johnston, Dominic Kwiatkowski, Cordelia Langford, John Sillitoe on behalf of the Wellcome Sanger Institute COVID-19 Surveillance Team |
| EPI_ISL_3170120 | Berkshire and Surrey Pathology Services Lighthouse Laboratory | Wellcome Sanger Institute for the COVID-19 Genomics UK (COG-UK) Consortium | Berkshire and Surrey Pathology Services Lighthouse Laboratory and Alex Alderton, Roberto Amato, Jeffrey Barrett, Sonia Goncalves, Ewan Harrison, David K. Jackson, Ian Johnston, Dominic Kwiatkowski, Cordelia Langford, John Sillitoe on behalf of the Wellcome Sanger Institute COVID-19 Surveillance Team |
| EPI_ISL_3170121 | Lighthouse Laboratory Plymouth | Wellcome Sanger Institute for the COVID-19 Genomics UK (COG-UK) Consortium | Lighthouse Laboratory Plymouth and Alex Alderton, Roberto Amato, Jeffrey Barrett, Sonia Goncalves, Ewan Harrison, David K. Jackson, Ian Johnston, Dominic Kwiatkowski, Cordelia Langford, John Sillitoe on behalf of the Wellcome Sanger Institute COVID-19 Surveillance Team |
| EPI_ISL_3170122, EPI_ISL_3170123, EPI_ISL_3170124 | Berkshire and Surrey Pathology Services Lighthouse Laboratory | Wellcome Sanger Institute for the COVID-19 Genomics UK (COG-UK) Consortium | Berkshire and Surrey Pathology Services Lighthouse Laboratory and Alex Alderton, Roberto Amato, Jeffrey Barrett, Sonia Goncalves, Ewan Harrison, David K. Jackson, Ian Johnston, Dominic Kwiatkowski, Cordelia Langford, John Sillitoe on behalf of the Wellcome Sanger Institute COVID-19 Surveillance Team |
| EPI_ISL_3170125 | Lighthouse Lab in Alderley Park | Wellcome Sanger Institute for the COVID-19 Genomics UK (COG-UK) Consortium | Jacquelyn Wynn, Mairead Hyland, The Lighthouse Lab in Alderley Park and Alex Alderton, Roberto Amato, Jeffrey Barrett, Sonia Goncalves, Ewan Harrison, David K. Jackson, Ian Johnston, Dominic Kwiatkowski, Cordelia Langford, John Sillitoe on behalf of the Wellcome Sanger Institute COVID-19 Surveillance Team |
| EPI_ISL_3170126 | Lighthouse Laboratory Plymouth | Wellcome Sanger Institute for the COVID-19 Genomics UK (COG-UK) Consortium | Lighthouse Laboratory Plymouth and Alex Alderton, Roberto Amato, Jeffrey Barrett, Sonia Goncalves, Ewan Harrison, David K. Jackson, Ian Johnston, Dominic Kwiatkowski, Cordelia Langford, John Sillitoe on behalf of the Wellcome Sanger Institute COVID-19 Surveillance Team |
| EPI_ISL_3170127 | Lighthouse Lab in Alderley Park | Wellcome Sanger Institute for the COVID-19 Genomics UK (COG-UK) Consortium | Jacquelyn Wynn, Mairead Hyland, The Lighthouse Lab in Alderley Park and Alex Alderton, Roberto Amato, Jeffrey Barrett, Sonia Goncalves, Ewan Harrison, David K. Jackson, Ian Johnston, Dominic Kwiatkowski, Cordelia Langford, John Sillitoe on behalf of the Wellcome Sanger Institute COVID-19 Surveillance Team |
| EPI_ISL_3170128 | Lighthouse Laboratory Plymouth | Wellcome Sanger Institute for the COVID-19 Genomics UK (COG-UK) Consortium | Lighthouse Laboratory Plymouth and Alex Alderton, Roberto Amato, Jeffrey Barrett, Sonia Goncalves, Ewan Harrison, David K. Jackson, Ian Johnston, Dominic Kwiatkowski, Cordelia Langford, John Sillitoe on behalf of the Wellcome Sanger Institute COVID-19 Surveillance Team |
| EPI_ISL_3170129, EPI_ISL_3170130 | Berkshire and Surrey Pathology Services Lighthouse Laboratory | Wellcome Sanger Institute for the COVID-19 Genomics UK (COG-UK) Consortium | Berkshire and Surrey Pathology Services Lighthouse Laboratory and Alex Alderton, Roberto Amato, Jeffrey Barrett, Sonia Goncalves, Ewan Harrison, David K. Jackson, Ian Johnston, Dominic Kwiatkowski, Cordelia Langford, John Sillitoe on behalf of the Wellcome Sanger Institute COVID-19 Surveillance Team |
| EPI_ISL_3170131 | Lighthouse Lab in Alderley Park | Wellcome Sanger Institute for the COVID-19 Genomics UK (COG-UK) Consortium | Jacquelyn Wynn, Mairead Hyland, The Lighthouse Lab in Alderley Park and Alex Alderton, Roberto Amato, Jeffrey Barrett, Sonia Goncalves, Ewan Harrison, David K. Jackson, Ian Johnston, Dominic Kwiatkowski, Cordelia Langford, John Sillitoe on behalf of the Wellcome Sanger Institute COVID-19 Surveillance Team |
| EPI_ISL_3170132, EPI_ISL_3170133, EPI_ISL_3170134 | Berkshire and Surrey Pathology Services Lighthouse Laboratory | Wellcome Sanger Institute for the COVID-19 Genomics UK (COG-UK) Consortium | Berkshire and Surrey Pathology Services Lighthouse Laboratory and Alex Alderton, Roberto Amato, Jeffrey Barrett, Sonia Goncalves, Ewan Harrison, David K. Jackson, Ian Johnston, Dominic Kwiatkowski, Cordelia Langford, John Sillitoe on behalf of the Wellcome Sanger Institute COVID-19 Surveillance Team |
| EPI_ISL_3170135 | Lighthouse Lab in Alderley Park | Wellcome Sanger Institute for the COVID-19 Genomics UK (COG-UK) Consortium | Jacquelyn Wynn, Mairead Hyland, The Lighthouse Lab in Alderley Park and Alex Alderton, Roberto Amato, Jeffrey Barrett, Sonia Goncalves, Ewan Harrison, David K. Jackson, Ian Johnston, Dominic Kwiatkowski, Cordelia Langford, John Sillitoe on behalf of the Wellcome Sanger Institute COVID-19 Surveillance Team |
| EPI_ISL_3170136, EPI_ISL_3170137, EPI_ISL_3170138 | Berkshire and Surrey Pathology Services Lighthouse Laboratory | Wellcome Sanger Institute for the COVID-19 Genomics UK (COG-UK) Consortium | Berkshire and Surrey Pathology Services Lighthouse Laboratory and Alex Alderton, Roberto Amato, Jeffrey Barrett, Sonia Goncalves, Ewan Harrison, David K. Jackson, Ian Johnston, Dominic Kwiatkowski, Cordelia Langford, John Sillitoe on behalf of the Wellcome Sanger Institute COVID-19 Surveillance Team |
| EPI_ISL_3170139 | Lighthouse Lab in Alderley Park | Wellcome Sanger Institute for the COVID-19 Genomics UK (COG-UK) Consortium | Jacquelyn Wynn, Mairead Hyland, The Lighthouse Lab in Alderley Park and Alex Alderton, Roberto Amato, Jeffrey Barrett, Sonia Goncalves, Ewan Harrison, David K. Jackson, Ian Johnston, Dominic Kwiatkowski, Cordelia Langford, John Sillitoe on behalf of the Wellcome Sanger Institute COVID-19 Surveillance Team |
| EPI_ISL_3170140, EPI_ISL_3170141 | Berkshire and Surrey Pathology Services Lighthouse Laboratory | Wellcome Sanger Institute for the COVID-19 Genomics UK (COG-UK) Consortium | Berkshire and Surrey Pathology Services Lighthouse Laboratory and Alex Alderton, Roberto Amato, Jeffrey Barrett, Sonia Goncalves, Ewan Harrison, David K. Jackson, Ian Johnston, Dominic Kwiatkowski, Cordelia Langford, John Sillitoe on behalf of the Wellcome Sanger Institute COVID-19 Surveillance Team |
| EPI_ISL_3170142 | Lighthouse Laboratory Plymouth | Wellcome Sanger Institute for the COVID-19 Genomics UK (COG-UK) Consortium | Lighthouse Laboratory Plymouth and Alex Alderton, Roberto Amato, Jeffrey Barrett, Sonia Goncalves, Ewan Harrison, David K. Jackson, Ian Johnston, Dominic Kwiatkowski, Cordelia Langford, John Sillitoe on behalf of the Wellcome Sanger Institute COVID-19 Surveillance Team |
| EPI_ISL_3170143, EPI_ISL_3170144, EPI_ISL_3170145 | Berkshire and Surrey Pathology Services Lighthouse Laboratory | Wellcome Sanger Institute for the COVID-19 Genomics UK (COG-UK) Consortium | Berkshire and Surrey Pathology Services Lighthouse Laboratory and Alex Alderton, Roberto Amato, Jeffrey Barrett, Sonia Goncalves, Ewan Harrison, David K. Jackson, Ian Johnston, Dominic Kwiatkowski, Cordelia Langford, John Sillitoe on behalf of the Wellcome Sanger Institute COVID-19 Surveillance Team |
| EPI_ISL_3170146 | Lighthouse Lab in Alderley Park | Wellcome Sanger Institute for the COVID-19 Genomics UK (COG-UK) Consortium | Jacquelyn Wynn, Mairead Hyland, The Lighthouse Lab in Alderley Park and Alex Alderton, Roberto Amato, Jeffrey Barrett, Sonia Goncalves, Ewan Harrison, David K. Jackson, Ian Johnston, Dominic Kwiatkowski, Cordelia Langford, John Sillitoe on behalf of the Wellcome Sanger Institute COVID-19 Surveillance Team |
| EPI_ISL_3170147 | Berkshire and Surrey Pathology Services Lighthouse Laboratory | Wellcome Sanger Institute for the COVID-19 Genomics UK (COG-UK) Consortium | Berkshire and Surrey Pathology Services Lighthouse Laboratory and Alex Alderton, Roberto Amato, Jeffrey Barrett, Sonia Goncalves, Ewan Harrison, David K. Jackson, Ian Johnston, Dominic Kwiatkowski, Cordelia Langford, John Sillitoe on behalf of the Wellcome Sanger Institute COVID-19 Surveillance Team |
| EPI_ISL_3170148, EPI_ISL_3170149 | Lighthouse Lab in Alderley Park | Wellcome Sanger Institute for the COVID-19 Genomics UK (COG-UK) Consortium | Jacquelyn Wynn, Mairead Hyland, The Lighthouse Lab in Alderley Park and Alex Alderton, Roberto Amato, Jeffrey Barrett, Sonia Goncalves, Ewan Harrison, David K. Jackson, Ian Johnston, Dominic Kwiatkowski, Cordelia Langford, John Sillitoe on behalf of the Wellcome Sanger Institute COVID-19 Surveillance Team |
| EPI_ISL_3170150, EPI_ISL_3170151, EPI_ISL_3170152 | Berkshire and Surrey Pathology Services Lighthouse Laboratory | Wellcome Sanger Institute for the COVID-19 Genomics UK (COG-UK) Consortium | Berkshire and Surrey Pathology Services Lighthouse Laboratory and Alex Alderton, Roberto Amato, Jeffrey Barrett, Sonia Goncalves, Ewan Harrison, David K. Jackson, Ian Johnston, Dominic Kwiatkowski, Cordelia Langford, John Sillitoe on behalf of the Wellcome Sanger Institute COVID-19 Surveillance Team |
| EPI_ISL_3170153 | Lighthouse Lab in Alderley Park | Wellcome Sanger Institute for the COVID-19 Genomics UK (COG-UK) Consortium | Jacquelyn Wynn, Mairead Hyland, The Lighthouse Lab in Alderley Park and Alex Alderton, Roberto Amato, Jeffrey Barrett, Sonia Goncalves, Ewan Harrison, David K. Jackson, Ian Johnston, Dominic Kwiatkowski, Cordelia Langford, John Sillitoe on behalf of the Wellcome Sanger Institute COVID-19 Surveillance Team |
| EPI_ISL_3170154 | Berkshire and Surrey Pathology Services Lighthouse Laboratory | Wellcome Sanger Institute for the COVID-19 Genomics UK (COG-UK) Consortium | Berkshire and Surrey Pathology Services Lighthouse Laboratory and Alex Alderton, Roberto Amato, Jeffrey Barrett, Sonia Goncalves, Ewan Harrison, David K. Jackson, Ian Johnston, Dominic Kwiatkowski, Cordelia Langford, John Sillitoe on behalf of the Wellcome Sanger Institute COVID-19 Surveillance Team |

| | | | |
|---|---|---|---|
| EPI_ISL_3170155, EPI_ISL_3170156 | Lighthouse Lab in Alderley Park | Wellcome Sanger Institute for the COVID-19 Genomics UK (COG-UK) Consortium | Jacquelyn Wynn, Mairead Hyland, The Lighthouse Lab in Alderley Park and Alex Alderton, Roberto Amato, Jeffrey Barrett, Sonia Goncalves, Ewan Harrison, David K. Jackson, Ian Johnston, Dominic Kwiatkowski, Cordelia Langford, John Sillitoe on behalf of the Wellcome Sanger Institute COVID-19 Surveillance Team |
| EPI_ISL_3170157 | Berkshire and Surrey Pathology Services Lighthouse Laboratory | Wellcome Sanger Institute for the COVID-19 Genomics UK (COG-UK) Consortium | Berkshire and Surrey Pathology Services Lighthouse Laboratory and Alex Alderton, Roberto Amato, Jeffrey Barrett, Sonia Goncalves, Ewan Harrison, David K. Jackson, Ian Johnston, Dominic Kwiatkowski, Cordelia Langford, John Sillitoe on behalf of the Wellcome Sanger Institute COVID-19 Surveillance Team |
| EPI_ISL_3170158, EPI_ISL_3170160 | Lighthouse Lab in Alderley Park | Wellcome Sanger Institute for the COVID-19 Genomics UK (COG-UK) Consortium | Jacquelyn Wynn, Mairead Hyland, The Lighthouse Lab in Alderley Park and Alex Alderton, Roberto Amato, Jeffrey Barrett, Sonia Goncalves, Ewan Harrison, David K. Jackson, Ian Johnston, Dominic Kwiatkowski, Cordelia Langford, John Sillitoe on behalf of the Wellcome Sanger Institute COVID-19 Surveillance Team |
| EPI_ISL_3170161 | Berkshire and Surrey Pathology Services Lighthouse Laboratory | Wellcome Sanger Institute for the COVID-19 Genomics UK (COG-UK) Consortium | Berkshire and Surrey Pathology Services Lighthouse Laboratory and Alex Alderton, Roberto Amato, Jeffrey Barrett, Sonia Goncalves, Ewan Harrison, David K. Jackson, Ian Johnston, Dominic Kwiatkowski, Cordelia Langford, John Sillitoe on behalf of the Wellcome Sanger Institute COVID-19 Surveillance Team |
| EPI_ISL_3170162 | Lighthouse Laboratory Plymouth | Wellcome Sanger Institute for the COVID-19 Genomics UK (COG-UK) Consortium | Lighthouse Laboratory Plymouth and Alex Alderton, Roberto Amato, Jeffrey Barrett, Sonia Goncalves, Ewan Harrison, David K. Jackson, Ian Johnston, Dominic Kwiatkowski, Cordelia Langford, John Sillitoe on behalf of the Wellcome Sanger Institute COVID-19 Surveillance Team |
| EPI_ISL_3170163 | Berkshire and Surrey Pathology Services Lighthouse Laboratory | Wellcome Sanger Institute for the COVID-19 Genomics UK (COG-UK) Consortium | Berkshire and Surrey Pathology Services Lighthouse Laboratory and Alex Alderton, Roberto Amato, Jeffrey Barrett, Sonia Goncalves, Ewan Harrison, David K. Jackson, Ian Johnston, Dominic Kwiatkowski, Cordelia Langford, John Sillitoe on behalf of the Wellcome Sanger Institute COVID-19 Surveillance Team |
| EPI_ISL_3170164 | Lighthouse Laboratory Plymouth | Wellcome Sanger Institute for the COVID-19 Genomics UK (COG-UK) Consortium | Lighthouse Laboratory Plymouth and Alex Alderton, Roberto Amato, Jeffrey Barrett, Sonia Goncalves, Ewan Harrison, David K. Jackson, Ian Johnston, Dominic Kwiatkowski, Cordelia Langford, John Sillitoe on behalf of the Wellcome Sanger Institute COVID-19 Surveillance Team |
| EPI_ISL_3170165 | Berkshire and Surrey Pathology Services Lighthouse Laboratory | Wellcome Sanger Institute for the COVID-19 Genomics UK (COG-UK) Consortium | Berkshire and Surrey Pathology Services Lighthouse Laboratory and Alex Alderton, Roberto Amato, Jeffrey Barrett, Sonia Goncalves, Ewan Harrison, David K. Jackson, Ian Johnston, Dominic Kwiatkowski, Cordelia Langford, John Sillitoe on behalf of the Wellcome Sanger Institute COVID-19 Surveillance Team |
| EPI_ISL_3170166 | Lighthouse Lab in Alderley Park | Wellcome Sanger Institute for the COVID-19 Genomics UK (COG-UK) Consortium | Jacquelyn Wynn, Mairead Hyland, The Lighthouse Lab in Alderley Park and Alex Alderton, Roberto Amato, Jeffrey Barrett, Sonia Goncalves, Ewan Harrison, David K. Jackson, Ian Johnston, Dominic Kwiatkowski, Cordelia Langford, John Sillitoe on behalf of the Wellcome Sanger Institute COVID-19 Surveillance Team |
| EPI_ISL_3170167 | Lighthouse Laboratory Plymouth | Wellcome Sanger Institute for the COVID-19 Genomics UK (COG-UK) Consortium | Lighthouse Laboratory Plymouth and Alex Alderton, Roberto Amato, Jeffrey Barrett, Sonia Goncalves, Ewan Harrison, David K. Jackson, Ian Johnston, Dominic Kwiatkowski, Cordelia Langford, John Sillitoe on behalf of the Wellcome Sanger Institute COVID-19 Surveillance Team |
| EPI_ISL_3170168 | Lighthouse Lab in Alderley Park | Wellcome Sanger Institute for the COVID-19 Genomics UK (COG-UK) Consortium | Jacquelyn Wynn, Mairead Hyland, The Lighthouse Lab in Alderley Park and Alex Alderton, Roberto Amato, Jeffrey Barrett, Sonia Goncalves, Ewan Harrison, David K. Jackson, Ian Johnston, Dominic Kwiatkowski, Cordelia Langford, John Sillitoe on behalf of the Wellcome Sanger Institute COVID-19 Surveillance Team |
| EPI_ISL_3170169, EPI_ISL_3170170 | Lighthouse Laboratory Plymouth | Wellcome Sanger Institute for the COVID-19 Genomics UK (COG-UK) Consortium | Lighthouse Laboratory Plymouth and Alex Alderton, Roberto Amato, Jeffrey Barrett, Sonia Goncalves, Ewan Harrison, David K. Jackson, Ian Johnston, Dominic Kwiatkowski, Cordelia Langford, John Sillitoe on behalf of the Wellcome Sanger Institute COVID-19 Surveillance Team |
| EPI_ISL_3170171 | Berkshire and Surrey Pathology Services Lighthouse Laboratory | Wellcome Sanger Institute for the COVID-19 Genomics UK (COG-UK) Consortium | Berkshire and Surrey Pathology Services Lighthouse Laboratory and Alex Alderton, Roberto Amato, Jeffrey Barrett, Sonia Goncalves, Ewan Harrison, David K. Jackson, Ian Johnston, Dominic Kwiatkowski, Cordelia Langford, John Sillitoe on behalf of the Wellcome Sanger Institute COVID-19 Surveillance Team |
| EPI_ISL_3170172 | Lighthouse Laboratory Plymouth | Wellcome Sanger Institute for the COVID-19 Genomics UK (COG-UK) Consortium | Lighthouse Laboratory Plymouth and Alex Alderton, Roberto Amato, Jeffrey Barrett, Sonia Goncalves, Ewan Harrison, David K. Jackson, Ian Johnston, Dominic Kwiatkowski, Cordelia Langford, John Sillitoe on behalf of the Wellcome Sanger Institute COVID-19 Surveillance Team |
| EPI_ISL_3170173 | Lighthouse Lab in Alderley Park | Wellcome Sanger Institute for the COVID-19 Genomics UK (COG-UK) Consortium | Jacquelyn Wynn, Mairead Hyland, The Lighthouse Lab in Alderley Park and Alex Alderton, Roberto Amato, Jeffrey Barrett, Sonia Goncalves, Ewan Harrison, David K. Jackson, Ian Johnston, Dominic Kwiatkowski, Cordelia Langford, John Sillitoe on behalf of the Wellcome Sanger Institute COVID-19 Surveillance Team |
| EPI_ISL_3170174, EPI_ISL_3170175 | Berkshire and Surrey Pathology Services Lighthouse Laboratory | Wellcome Sanger Institute for the COVID-19 Genomics UK (COG-UK) Consortium | Berkshire and Surrey Pathology Services Lighthouse Laboratory and Alex Alderton, Roberto Amato, Jeffrey Barrett, Sonia Goncalves, Ewan Harrison, David K. Jackson, Ian Johnston, Dominic Kwiatkowski, Cordelia Langford, John Sillitoe on behalf of the Wellcome Sanger Institute COVID-19 Surveillance Team |
| EPI_ISL_3170176 | Lighthouse Laboratory Plymouth | Wellcome Sanger Institute for the COVID-19 Genomics UK (COG-UK) Consortium | Lighthouse Laboratory Plymouth and Alex Alderton, Roberto Amato, Jeffrey Barrett, Sonia Goncalves, Ewan Harrison, David K. Jackson, Ian Johnston, Dominic Kwiatkowski, Cordelia Langford, John Sillitoe on behalf of the Wellcome Sanger Institute COVID-19 Surveillance Team |
| EPI_ISL_3170177, EPI_ISL_3170178 | Berkshire and Surrey Pathology Services Lighthouse Laboratory | Wellcome Sanger Institute for the COVID-19 Genomics UK (COG-UK) Consortium | Berkshire and Surrey Pathology Services Lighthouse Laboratory and Alex Alderton, Roberto Amato, Jeffrey Barrett, Sonia Goncalves, Ewan Harrison, David K. Jackson, Ian Johnston, Dominic Kwiatkowski, Cordelia Langford, John Sillitoe on behalf of the Wellcome Sanger Institute COVID-19 Surveillance Team |
| EPI_ISL_3170179 | Lighthouse Lab in Alderley Park | Wellcome Sanger Institute for the COVID-19 Genomics UK (COG-UK) Consortium | Jacquelyn Wynn, Mairead Hyland, The Lighthouse Lab in Alderley Park and Alex Alderton, Roberto Amato, Jeffrey Barrett, Sonia Goncalves, Ewan Harrison, David K. Jackson, Ian Johnston, Dominic Kwiatkowski, Cordelia Langford, John Sillitoe on behalf of the Wellcome Sanger Institute COVID-19 Surveillance Team |
| EPI_ISL_3170180, EPI_ISL_3170181 | Berkshire and Surrey Pathology Services Lighthouse Laboratory | Wellcome Sanger Institute for the COVID-19 Genomics UK (COG-UK) Consortium | Berkshire and Surrey Pathology Services Lighthouse Laboratory and Alex Alderton, Roberto Amato, Jeffrey Barrett, Sonia Goncalves, Ewan Harrison, David K. Jackson, Ian Johnston, Dominic Kwiatkowski, Cordelia Langford, John Sillitoe on behalf of the Wellcome Sanger Institute COVID-19 Surveillance Team |
| EPI_ISL_3170182 | Lighthouse Lab in Alderley Park | Wellcome Sanger Institute for the COVID-19 Genomics UK (COG-UK) Consortium | Jacquelyn Wynn, Mairead Hyland, The Lighthouse Lab in Alderley Park and Alex Alderton, Roberto Amato, Jeffrey Barrett, Sonia Goncalves, Ewan Harrison, David K. Jackson, Ian Johnston, Dominic Kwiatkowski, Cordelia Langford, John Sillitoe on behalf of the Wellcome Sanger Institute COVID-19 Surveillance Team |
| EPI_ISL_3170183 | Berkshire and Surrey Pathology Services Lighthouse Laboratory | Wellcome Sanger Institute for the COVID-19 Genomics UK (COG-UK) Consortium | Berkshire and Surrey Pathology Services Lighthouse Laboratory and Alex Alderton, Roberto Amato, Jeffrey Barrett, Sonia Goncalves, Ewan Harrison, David K. Jackson, Ian Johnston, Dominic Kwiatkowski, Cordelia Langford, John Sillitoe on behalf of the Wellcome Sanger Institute COVID-19 Surveillance Team |
| EPI_ISL_3170184 | Lighthouse Lab in Alderley Park | Wellcome Sanger Institute for the COVID-19 Genomics UK (COG-UK) Consortium | Jacquelyn Wynn, Mairead Hyland, The Lighthouse Lab in Alderley Park and Alex Alderton, Roberto Amato, Jeffrey Barrett, Sonia Goncalves, Ewan Harrison, David K. Jackson, Ian Johnston, Dominic Kwiatkowski, Cordelia Langford, John Sillitoe on behalf of the Wellcome Sanger Institute COVID-19 Surveillance Team |
| EPI_ISL_3170185 | Berkshire and Surrey Pathology Services Lighthouse Laboratory | Wellcome Sanger Institute for the COVID-19 Genomics UK (COG-UK) Consortium | Berkshire and Surrey Pathology Services Lighthouse Laboratory and Alex Alderton, Roberto Amato, Jeffrey Barrett, Sonia Goncalves, Ewan Harrison, David K. Jackson, Ian Johnston, Dominic Kwiatkowski, Cordelia Langford, John Sillitoe on behalf of the Wellcome Sanger Institute COVID-19 Surveillance Team |
| EPI_ISL_3170186 | Lighthouse Lab in Alderley Park | Wellcome Sanger Institute for the COVID-19 Genomics UK (COG-UK) Consortium | Jacquelyn Wynn, Mairead Hyland, The Lighthouse Lab in Alderley Park and Alex Alderton, Roberto Amato, Jeffrey Barrett, Sonia Goncalves, Ewan Harrison, David K. Jackson, Ian Johnston, Dominic Kwiatkowski, Cordelia Langford, John Sillitoe on behalf of the Wellcome Sanger Institute COVID-19 Surveillance Team |
| EPI_ISL_3170187, EPI_ISL_3170188, EPI_ISL_3170189, EPI_ISL_3170190 | Berkshire and Surrey Pathology Services Lighthouse Laboratory | Wellcome Sanger Institute for the COVID-19 Genomics UK (COG-UK) Consortium | Berkshire and Surrey Pathology Services Lighthouse Laboratory and Alex Alderton, Roberto Amato, Jeffrey Barrett, Sonia Goncalves, Ewan Harrison, David K. Jackson, Ian Johnston, Dominic Kwiatkowski, Cordelia Langford, John Sillitoe on behalf of the Wellcome Sanger Institute COVID-19 Surveillance Team |
| EPI_ISL_3170191 | Lighthouse Lab in Alderley Park | Wellcome Sanger Institute for the COVID-19 Genomics UK (COG-UK) Consortium | Jacquelyn Wynn, Mairead Hyland, The Lighthouse Lab in Alderley Park and Alex Alderton, Roberto Amato, Jeffrey Barrett, Sonia Goncalves, Ewan Harrison, David K. Jackson, Ian Johnston, Dominic Kwiatkowski, Cordelia Langford, John Sillitoe on behalf of the Wellcome Sanger Institute COVID-19 Surveillance Team |
| EPI_ISL_3170192, EPI_ISL_3170193, EPI_ISL_3170194, EPI_ISL_3170195, EPI_ISL_3170196, EPI_ISL_3170197 | Berkshire and Surrey Pathology Services Lighthouse Laboratory | Wellcome Sanger Institute for the COVID-19 Genomics UK (COG-UK) Consortium | Berkshire and Surrey Pathology Services Lighthouse Laboratory and Alex Alderton, Roberto Amato, Jeffrey Barrett, Sonia Goncalves, Ewan Harrison, David K. Jackson, Ian Johnston, Dominic Kwiatkowski, Cordelia Langford, John Sillitoe on behalf of the Wellcome Sanger Institute COVID-19 Surveillance Team |
| EPI_ISL_3170198 | Lighthouse Lab in Alderley Park | Wellcome Sanger Institute for the COVID-19 Genomics UK (COG-UK) Consortium | Jacquelyn Wynn, Mairead Hyland, The Lighthouse Lab in Alderley Park and Alex Alderton, Roberto Amato, Jeffrey Barrett, Sonia Goncalves, Ewan Harrison, David K. Jackson, Ian Johnston, Dominic Kwiatkowski, Cordelia Langford, John Sillitoe on behalf of the Wellcome Sanger Institute COVID-19 Surveillance Team |
| EPI_ISL_3170199 | Lighthouse Laboratory Plymouth | Wellcome Sanger Institute for the COVID-19 Genomics UK (COG-UK) Consortium | Lighthouse Laboratory Plymouth and Alex Alderton, Roberto Amato, Jeffrey Barrett, Sonia Goncalves, Ewan Harrison, David K. Jackson, Ian Johnston, Dominic Kwiatkowski, Cordelia Langford, John Sillitoe on behalf of the Wellcome Sanger Institute COVID-19 Surveillance Team |
| EPI_ISL_3170200 | Berkshire and Surrey Pathology Services Lighthouse Laboratory | Wellcome Sanger Institute for the COVID-19 Genomics UK (COG-UK) Consortium | Berkshire and Surrey Pathology Services Lighthouse Laboratory and Alex Alderton, Roberto Amato, Jeffrey Barrett, Sonia Goncalves, Ewan Harrison, David K. Jackson, Ian Johnston, Dominic Kwiatkowski, Cordelia Langford, John Sillitoe on behalf of the Wellcome Sanger Institute COVID-19 Surveillance Team |
| EPI_ISL_3170201, EPI_ISL_3170202 | Lighthouse Laboratory Plymouth | Wellcome Sanger Institute for the COVID-19 Genomics UK (COG-UK) Consortium | Lighthouse Laboratory Plymouth and Alex Alderton, Roberto Amato, Jeffrey Barrett, Sonia Goncalves, Ewan Harrison, David K. Jackson, Ian Johnston, Dominic Kwiatkowski, Cordelia Langford, John Sillitoe on behalf of the Wellcome Sanger Institute COVID-19 Surveillance Team |
| EPI_ISL_3170203 | Lighthouse Lab in Alderley Park | Wellcome Sanger Institute for the COVID-19 Genomics UK (COG-UK) Consortium | Jacquelyn Wynn, Mairead Hyland, The Lighthouse Lab in Alderley Park and Alex Alderton, Roberto Amato, Jeffrey Barrett, Sonia Goncalves, Ewan Harrison, David K. Jackson, Ian Johnston, Dominic Kwiatkowski, Cordelia Langford, John Sillitoe on behalf of the Wellcome Sanger Institute COVID-19 Surveillance Team |
| EPI_ISL_3170204 | Lighthouse Laboratory Plymouth | Wellcome Sanger Institute for the COVID-19 Genomics UK (COG-UK) Consortium | Lighthouse Laboratory Plymouth and Alex Alderton, Roberto Amato, Jeffrey Barrett, Sonia Goncalves, Ewan Harrison, David K. Jackson, Ian Johnston, Dominic Kwiatkowski, Cordelia Langford, John Sillitoe on behalf of the Wellcome Sanger Institute COVID-19 Surveillance Team |
| EPI_ISL_3170205, EPI_ISL_3170206, EPI_ISL_3170207 | Lighthouse Lab in Alderley Park | Wellcome Sanger Institute for the COVID-19 Genomics UK (COG-UK) Consortium | Jacquelyn Wynn, Mairead Hyland, The Lighthouse Lab in Alderley Park and Alex Alderton, Roberto Amato, Jeffrey Barrett, Sonia Goncalves, Ewan Harrison, David K. Jackson, Ian Johnston, Dominic Kwiatkowski, Cordelia Langford, John Sillitoe on behalf of the Wellcome Sanger Institute COVID-19 Surveillance Team |
| EPI_ISL_3170208 | Berkshire and Surrey Pathology Services Lighthouse Laboratory | Wellcome Sanger Institute for the COVID-19 Genomics UK (COG-UK) Consortium | Berkshire and Surrey Pathology Services Lighthouse Laboratory and Alex Alderton, Roberto Amato, Jeffrey Barrett, Sonia Goncalves, Ewan Harrison, David K. Jackson, Ian Johnston, Dominic Kwiatkowski, Cordelia Langford, John Sillitoe on behalf of the Wellcome Sanger Institute COVID-19 Surveillance Team |
| EPI_ISL_3170209, EPI_ISL_3170210 | Lighthouse Lab in Alderley Park | Wellcome Sanger Institute for the COVID-19 Genomics UK (COG-UK) Consortium | Jacquelyn Wynn, Mairead Hyland, The Lighthouse Lab in Alderley Park and Alex Alderton, Roberto Amato, Jeffrey Barrett, Sonia Goncalves, Ewan Harrison, David K. Jackson, Ian Johnston, Dominic Kwiatkowski, Cordelia Langford, John Sillitoe on behalf of the Wellcome Sanger Institute COVID-19 Surveillance Team |
| EPI_ISL_3170211 | Berkshire and Surrey Pathology Services Lighthouse Laboratory | Wellcome Sanger Institute for the COVID-19 Genomics UK (COG-UK) Consortium | Berkshire and Surrey Pathology Services Lighthouse Laboratory and Alex Alderton, Roberto Amato, Jeffrey Barrett, Sonia Goncalves, Ewan Harrison, David K. Jackson, Ian Johnston, Dominic Kwiatkowski, Cordelia Langford, John Sillitoe on behalf of the Wellcome Sanger Institute COVID-19 Surveillance Team |
| EPI_ISL_3170212 | Lighthouse Laboratory Plymouth | Wellcome Sanger Institute for the COVID-19 Genomics UK (COG-UK) Consortium | Lighthouse Laboratory Plymouth and Alex Alderton, Roberto Amato, Jeffrey Barrett, Sonia Goncalves, Ewan Harrison, David K. Jackson, Ian Johnston, Dominic Kwiatkowski, Cordelia Langford, John Sillitoe on behalf of the Wellcome Sanger Institute COVID-19 Surveillance Team |
| EPI_ISL_3170213, EPI_ISL_3170214 | Berkshire and Surrey Pathology Services Lighthouse Laboratory | Wellcome Sanger Institute for the COVID-19 Genomics UK (COG-UK) Consortium | Berkshire and Surrey Pathology Services Lighthouse Laboratory and Alex Alderton, Roberto Amato, Jeffrey Barrett, Sonia Goncalves, Ewan Harrison, David K. Jackson, Ian Johnston, Dominic Kwiatkowski, Cordelia Langford, John Sillitoe on behalf of the Wellcome Sanger Institute COVID-19 Surveillance Team |
| EPI_ISL_3170215 | Lighthouse Lab in Alderley Park | Wellcome Sanger Institute for the COVID-19 Genomics UK (COG-UK) Consortium | Jacquelyn Wynn, Mairead Hyland, The Lighthouse Lab in Alderley Park and Alex Alderton, Roberto Amato, Jeffrey Barrett, Sonia Goncalves, Ewan Harrison, David K. Jackson, Ian Johnston, Dominic Kwiatkowski, Cordelia Langford, John Sillitoe on behalf of the Wellcome Sanger Institute COVID-19 Surveillance Team |
| EPI_ISL_3170216, EPI_ISL_3170217 | Berkshire and Surrey Pathology Services Lighthouse Laboratory | Wellcome Sanger Institute for the COVID-19 Genomics UK (COG-UK) Consortium | Berkshire and Surrey Pathology Services Lighthouse Laboratory and Alex Alderton, Roberto Amato, Jeffrey Barrett, Sonia Goncalves, Ewan Harrison, David K. Jackson, Ian Johnston, Dominic Kwiatkowski, Cordelia Langford, John Sillitoe on behalf of the Wellcome Sanger Institute COVID-19 Surveillance Team |
| EPI_ISL_3170218, | Lighthouse Lab in Alderley Park | Wellcome Sanger Institute for the COVID-19 Genomics UK (COG-UK) | Jacquelyn Wynn, Mairead Hyland, The Lighthouse Lab in Alderley Park and Alex Alderton, Roberto Amato, Jeffrey Barrett, Sonia Goncalves, Ewan Harrison, David K. Jackson, Ian Johnston, Dominic Kwiatkowski, Cordelia Langford, John Sillitoe on |

| | | | |
|---|---|---|---|
| EPI_ISL_3170219 | | Consortium | behalf of the Wellcome Sanger Institute COVID-19 Surveillance Team |
| EPI_ISL_3170220 | Berkshire and Surrey Pathology Services Lighthouse Laboratory | Wellcome Sanger Institute for the COVID-19 Genomics UK (COG-UK) Consortium | Berkshire and Surrey Pathology Services Lighthouse Laboratory and Alex Alderton, Roberto Amato, Jeffrey Barrett, Sonia Goncalves, Ewan Harrison, David K. Jackson, Ian Johnston, Dominic Kwiatkowski, Cordelia Langford, John Sillitoe on behalf of the Wellcome Sanger Institute COVID-19 Surveillance Team |
| EPI_ISL_3170221 | Lighthouse Lab in Alderley Park | Wellcome Sanger Institute for the COVID-19 Genomics UK (COG-UK) Consortium | Jacquelyn Wynn, Mairead Hyland, The Lighthouse Lab in Alderley Park and Alex Alderton, Roberto Amato, Jeffrey Barrett, Sonia Goncalves, Ewan Harrison, David K. Jackson, Ian Johnston, Dominic Kwiatkowski, Cordelia Langford, John Sillitoe on behalf of the Wellcome Sanger Institute COVID-19 Surveillance Team |
| EPI_ISL_3170222 | Lighthouse Laboratory Plymouth | Wellcome Sanger Institute for the COVID-19 Genomics UK (COG-UK) Consortium | Lighthouse Laboratory Plymouth and Alex Alderton, Roberto Amato, Jeffrey Barrett, Sonia Goncalves, Ewan Harrison, David K. Jackson, Ian Johnston, Dominic Kwiatkowski, Cordelia Langford, John Sillitoe on behalf of the Wellcome Sanger Institute COVID-19 Surveillance Team |
| EPI_ISL_3170223 | Berkshire and Surrey Pathology Services Lighthouse Laboratory | Wellcome Sanger Institute for the COVID-19 Genomics UK (COG-UK) Consortium | Berkshire and Surrey Pathology Services Lighthouse Laboratory and Alex Alderton, Roberto Amato, Jeffrey Barrett, Sonia Goncalves, Ewan Harrison, David K. Jackson, Ian Johnston, Dominic Kwiatkowski, Cordelia Langford, John Sillitoe on behalf of the Wellcome Sanger Institute COVID-19 Surveillance Team |
| EPI_ISL_3170224 | Lighthouse Lab in Alderley Park | Wellcome Sanger Institute for the COVID-19 Genomics UK (COG-UK) Consortium | Jacquelyn Wynn, Mairead Hyland, The Lighthouse Lab in Alderley Park and Alex Alderton, Roberto Amato, Jeffrey Barrett, Sonia Goncalves, Ewan Harrison, David K. Jackson, Ian Johnston, Dominic Kwiatkowski, Cordelia Langford, John Sillitoe on behalf of the Wellcome Sanger Institute COVID-19 Surveillance Team |
| EPI_ISL_3170225, EPI_ISL_3170226, EPI_ISL_3170227 | Berkshire and Surrey Pathology Services Lighthouse Laboratory | Wellcome Sanger Institute for the COVID-19 Genomics UK (COG-UK) Consortium | Berkshire and Surrey Pathology Services Lighthouse Laboratory and Alex Alderton, Roberto Amato, Jeffrey Barrett, Sonia Goncalves, Ewan Harrison, David K. Jackson, Ian Johnston, Dominic Kwiatkowski, Cordelia Langford, John Sillitoe on behalf of the Wellcome Sanger Institute COVID-19 Surveillance Team |
| EPI_ISL_3170228, EPI_ISL_3170229 | Lighthouse Laboratory Plymouth | Wellcome Sanger Institute for the COVID-19 Genomics UK (COG-UK) Consortium | Lighthouse Laboratory Plymouth and Alex Alderton, Roberto Amato, Jeffrey Barrett, Sonia Goncalves, Ewan Harrison, David K. Jackson, Ian Johnston, Dominic Kwiatkowski, Cordelia Langford, John Sillitoe on behalf of the Wellcome Sanger Institute COVID-19 Surveillance Team |
| EPI_ISL_3170230, EPI_ISL_3170231, EPI_ISL_3170232 | Berkshire and Surrey Pathology Services Lighthouse Laboratory | Wellcome Sanger Institute for the COVID-19 Genomics UK (COG-UK) Consortium | Berkshire and Surrey Pathology Services Lighthouse Laboratory and Alex Alderton, Roberto Amato, Jeffrey Barrett, Sonia Goncalves, Ewan Harrison, David K. Jackson, Ian Johnston, Dominic Kwiatkowski, Cordelia Langford, John Sillitoe on behalf of the Wellcome Sanger Institute COVID-19 Surveillance Team |
| EPI_ISL_3170233 | Lighthouse Laboratory Plymouth | Wellcome Sanger Institute for the COVID-19 Genomics UK (COG-UK) Consortium | Lighthouse Laboratory Plymouth and Alex Alderton, Roberto Amato, Jeffrey Barrett, Sonia Goncalves, Ewan Harrison, David K. Jackson, Ian Johnston, Dominic Kwiatkowski, Cordelia Langford, John Sillitoe on behalf of the Wellcome Sanger Institute COVID-19 Surveillance Team |
| EPI_ISL_3170234 | Lighthouse Lab in Alderley Park | Wellcome Sanger Institute for the COVID-19 Genomics UK (COG-UK) Consortium | Jacquelyn Wynn, Mairead Hyland, The Lighthouse Lab in Alderley Park and Alex Alderton, Roberto Amato, Jeffrey Barrett, Sonia Goncalves, Ewan Harrison, David K. Jackson, Ian Johnston, Dominic Kwiatkowski, Cordelia Langford, John Sillitoe on behalf of the Wellcome Sanger Institute COVID-19 Surveillance Team |
| EPI_ISL_3170235, EPI_ISL_3170236 | Berkshire and Surrey Pathology Services Lighthouse Laboratory | Wellcome Sanger Institute for the COVID-19 Genomics UK (COG-UK) Consortium | Berkshire and Surrey Pathology Services Lighthouse Laboratory and Alex Alderton, Roberto Amato, Jeffrey Barrett, Sonia Goncalves, Ewan Harrison, David K. Jackson, Ian Johnston, Dominic Kwiatkowski, Cordelia Langford, John Sillitoe on behalf of the Wellcome Sanger Institute COVID-19 Surveillance Team |
| EPI_ISL_3170237 | Lighthouse Laboratory Plymouth | Wellcome Sanger Institute for the COVID-19 Genomics UK (COG-UK) Consortium | Lighthouse Laboratory Plymouth and Alex Alderton, Roberto Amato, Jeffrey Barrett, Sonia Goncalves, Ewan Harrison, David K. Jackson, Ian Johnston, Dominic Kwiatkowski, Cordelia Langford, John Sillitoe on behalf of the Wellcome Sanger Institute COVID-19 Surveillance Team |
| EPI_ISL_3170238 | Berkshire and Surrey Pathology Services Lighthouse Laboratory | Wellcome Sanger Institute for the COVID-19 Genomics UK (COG-UK) Consortium | Berkshire and Surrey Pathology Services Lighthouse Laboratory and Alex Alderton, Roberto Amato, Jeffrey Barrett, Sonia Goncalves, Ewan Harrison, David K. Jackson, Ian Johnston, Dominic Kwiatkowski, Cordelia Langford, John Sillitoe on behalf of the Wellcome Sanger Institute COVID-19 Surveillance Team |
| EPI_ISL_3170261 | Lighthouse Laboratory Plymouth | Wellcome Sanger Institute for the COVID-19 Genomics UK (COG-UK) Consortium | Lighthouse Laboratory Plymouth and Alex Alderton, Roberto Amato, Jeffrey Barrett, Sonia Goncalves, Ewan Harrison, David K. Jackson, Ian Johnston, Dominic Kwiatkowski, Cordelia Langford, John Sillitoe on behalf of the Wellcome Sanger Institute COVID-19 Surveillance Team |

EPI_ISL_3170262, EPI_ISL_3170263, EPI_ISL_3170264, EPI_ISL_3170265, EPI_ISL_3170266, EPI_ISL_3170267, EPI_ISL_3170268, EPI_ISL_3170269, EPI_ISL_3170270, EPI_ISL_3170271, EPI_ISL_3170272, EPI_ISL_3170273, EPI_ISL_3170274, EPI_ISL_3170275, EPI_ISL_3170276, EPI_ISL_3170277, EPI_ISL_3170278, EPI_ISL_3170279, EPI_ISL_3170280, EPI_ISL_3170281, EPI_ISL_3170282, EPI_ISL_3170283, EPI_ISL_3170284, EPI_ISL_3170285, EPI_ISL_3170286, EPI_ISL_3170287, EPI_ISL_3170288, EPI_ISL_3170289, EPI_ISL_3170290, EPI_ISL_3170291, EPI_ISL_3170292, EPI_ISL_3170293, EPI_ISL_3170294, EPI_ISL_3170295, EPI_ISL_3170296, EPI_ISL_3170297, EPI_ISL_3170298, EPI_ISL_3170299, EPI_ISL_3170300, EPI_ISL_3170301, EPI_ISL_3170302, EPI_ISL_3170303, EPI_ISL_3170304, EPI_ISL_3170305, EPI_ISL_3170306, EPI_ISL_3170307, EPI_ISL_3170308, EPI_ISL_3170309, EPI_ISL_3170310, EPI_ISL_3170311, EPI_ISL_3170312, EPI_ISL_3170313, EPI_ISL_3170314, EPI_ISL_3170315, EPI_ISL_3170316, EPI_ISL_3170317, EPI_ISL_3170318, EPI_ISL_3170319, EPI_ISL_3170320, EPI_ISL_3170321, EPI_ISL_3170322, EPI_ISL_3170323, EPI_ISL_3170324, EPI_ISL_3170325, EPI_ISL_3170326, EPI_ISL_3170327, EPI_ISL_3170328, EPI_ISL_3170329, EPI_ISL_3170330, EPI_ISL_3170331, EPI_ISL_3170332, EPI_ISL_3170333, EPI_ISL_3170334, EPI_ISL_3170335, EPI_ISL_3170336, EPI_ISL_3170337, EPI_ISL_3170338, EPI_ISL_3170339, EPI_ISL_3170340, EPI_ISL_3170341, EPI_ISL_3170342, EPI_ISL_3170343, EPI_ISL_3170344, EPI_ISL_3170345, EPI_ISL_3170346, EPI_ISL_3170347, EPI_ISL_3170348, EPI_ISL_3170349, EPI_ISL_3170350, EPI_ISL_3170351, EPI_ISL_3170352, EPI_ISL_3170353, EPI_ISL_3170354, EPI_ISL_3170355, EPI_ISL_3170356, EPI_ISL_3170357, EPI_ISL_3170358, EPI_ISL_3170359, EPI_ISL_3170360, EPI_ISL_3170361, EPI_ISL_3170362, EPI_ISL_3170363, EPI_ISL_3170364, EPI_ISL_3170365, EPI_ISL_3170366, EPI_ISL_3170367, EPI_ISL_3170368, EPI_ISL_3170369, EPI_ISL_3170370, EPI_ISL_3170371, EPI_ISL_3170372, EPI_ISL_3170373, EPI_ISL_3170374, EPI_ISL_3170375, EPI_ISL_3170376, EPI_ISL_3170377, EPI_ISL_3170378, EPI_ISL_3170379, EPI_ISL_3170380, EPI_ISL_3170381, EPI_ISL_3170382, EPI_ISL_3170383, EPI_ISL_3170384, EPI_ISL_3170385, EPI_ISL_3170386, EPI_ISL_3170387, EPI_ISL_3170388, EPI_ISL_3170389, EPI_ISL_3170390, EPI_ISL_3170391, EPI_ISL_3170392, EPI_ISL_3170393, EPI_ISL_3170394, EPI_ISL_3170395, EPI_ISL_3170396, EPI_ISL_3170397, EPI_ISL_3170398, EPI_ISL_3170399, EPI_ISL_3170400, EPI_ISL_3170401, EPI_ISL_3170402, EPI_ISL_3170403, EPI_ISL_3170404, EPI_ISL_3170405, EPI_ISL_3170406, EPI_ISL_3170407, EPI_ISL_3170408, EPI_ISL_3170409, EPI_ISL_3170410, EPI_ISL_3170411, EPI_ISL_3170412, EPI_ISL_3170413, EPI_ISL_3170414, EPI_ISL_3170415, EPI_ISL_3170416, EPI_ISL_3170417, EPI_ISL_3170418, EPI_ISL_3170419, EPI_ISL_3170420, EPI_ISL_3170421, EPI_ISL_3170422, EPI_ISL_3170423, EPI_ISL_3170424, EPI_ISL_3170425, EPI_ISL_3170426, EPI_ISL_3170427, EPI_ISL_3170428, EPI_ISL_3170429, EPI_ISL_3170430, EPI_ISL_3170431, EPI_ISL_3170432, EPI_ISL_3170434, EPI_ISL_3170435, EPI_ISL_3170436, EPI_ISL_3170437, EPI_ISL_3170438, EPI_ISL_3170439, EPI_ISL_3170440, EPI_ISL_3170441, EPI_ISL_3170442, EPI_ISL_3170443, EPI_ISL_3170444, EPI_ISL_3170445, EPI_ISL_3170446, EPI_ISL_3170447, EPI_ISL_3170448, EPI_ISL_3170449, EPI_ISL_3170450, EPI_ISL_3170451, EPI_ISL_3170452, EPI_ISL_3170453, EPI_ISL_3170454, EPI_ISL_3170455, EPI_ISL_3170456, EPI_ISL_3170457, EPI_ISL_3170458, EPI_ISL_3170459, EPI_ISL_3170460, EPI_ISL_3170461, EPI_ISL_3170462, EPI_ISL_3170463, EPI_ISL_3170464, EPI_ISL_3170465, EPI_ISL_3170466, EPI_ISL_3170467, EPI_ISL_3170468, EPI_ISL_3170469, EPI_ISL_3170470, EPI_ISL_3170471, EPI_ISL_3170472, EPI_ISL_3170473, EPI_ISL_3170474, EPI_ISL_3170475, EPI_ISL_3170476, EPI_ISL_3170477, EPI_ISL_3170478, EPI_ISL_3170479, EPI_ISL_3170480, EPI_ISL_3170481, EPI_ISL_3170482, EPI_ISL_3170483, EPI_ISL_3170484, EPI_ISL_3170485, EPI_ISL_3170486, EPI_ISL_3170487, EPI_ISL_3170488, EPI_ISL_3170489, EPI_ISL_3170490, EPI_ISL_3170491, EPI_ISL_3170492, EPI_ISL_3170493, EPI_ISL_3170494, EPI_ISL_3170495, EPI_ISL_3170496, EPI_ISL_3170497, EPI_ISL_3170498, EPI_ISL_3170499, EPI_ISL_3170500, EPI_ISL_3170501, EPI_ISL_3170502, EPI_ISL_3170503, EPI_ISL_3170504, EPI_ISL_3170505, EPI_ISL_3170506, EPI_ISL_3170507, EPI_ISL_3170508, EPI_ISL_3170509, EPI_ISL_3170510, EPI_ISL_3170511, EPI_ISL_3170512, EPI_ISL_3170513, EPI_ISL_3170514, EPI_ISL_3170515, EPI_ISL_3170516, EPI_ISL_3170517, EPI_ISL_3170518, EPI_ISL_3170519, EPI_ISL_3170520, EPI_ISL_3170521, EPI_ISL_3170522, EPI_ISL_3170523, EPI_ISL_3170524, EPI_ISL_3170525, EPI_ISL_3170526, EPI_ISL_3170527, EPI_ISL_3170528, EPI_ISL_3170529, EPI_ISL_3170530, EPI_ISL_3170531, EPI_ISL_3170532, EPI_ISL_3170533, EPI_ISL_3170534, EPI_ISL_3170535, EPI_ISL_3170536, EPI_ISL_3170537, EPI_ISL_3170538, EPI_ISL_3170539, EPI_ISL_3170540, EPI_ISL_3170541, EPI_ISL_3170542, EPI_ISL_3170543, EPI_ISL_3170544, EPI_ISL_3170545, EPI_ISL_3170546, EPI_ISL_3170547, EPI_ISL_3170548, EPI_ISL_3170549, EPI_ISL_3170550, EPI_ISL_3170551, EPI_ISL_3170552, EPI_ISL_3170553, EPI_ISL_3170554, EPI_ISL_3170555, EPI_ISL_3170556, EPI_ISL_3170557, EPI_ISL_3170558, EPI_ISL_3170559, EPI_ISL_3170560, EPI_ISL_3170561, EPI_ISL_3170562, EPI_ISL_3170563, EPI_ISL_3170564, EPI_ISL_3170565, EPI_ISL_3170566, EPI_ISL_3170567, EPI_ISL_3170568, EPI_ISL_3170569, EPI_ISL_3170570, EPI_ISL_3170571, EPI_ISL_3170572, EPI_ISL_3170573, EPI_ISL_3170574, EPI_ISL_3170575, EPI_ISL_3170576, EPI_ISL_3170577, EPI_ISL_3170578, EPI_ISL_3170579, EPI_ISL_3170580, EPI_ISL_3170581, EPI_ISL_3170582, EPI_ISL_3170583, EPI_ISL_3170584, EPI_ISL_3170585, EPI_ISL_3170586, EPI_ISL_3170587, EPI_ISL_3170588, EPI_ISL_3170589, EPI_ISL_3170590, EPI_ISL_3170591, EPI_ISL_3170592, EPI_ISL_3170593, EPI_ISL_3170594, EPI_ISL_3170595, EPI_ISL_3170596, EPI_ISL_3170597, EPI_ISL_3170598, EPI_ISL_3170599, EPI_ISL_3170600, EPI_ISL_3170601, EPI_ISL_3170602, EPI_ISL_3170603, EPI_ISL_3170604, EPI_ISL_3170605, EPI_ISL_3170606, EPI_ISL_3170607, EPI_ISL_3170608, EPI_ISL_3170609, EPI_ISL_3170610, EPI_ISL_3170611, EPI_ISL_3170612, EPI_ISL_3170613, EPI_ISL_3170614, EPI_ISL_3170615, EPI_ISL_3170616

| | | | |
|---|---|---|---|
| see above | Lighthouse Lab in Milton Keynes | Wellcome Sanger Institute for the COVID-19 Genomics UK (COG-UK) Consortium | The Lighthouse Lab in Milton Keynes and Alex Alderton, Roberto Amato, Jeffrey Barrett, Sonia Goncalves, Ewan Harrison, David K. Jackson, Ian Johnston, Dominic Kwiatkowski, Cordelia Langford, John Sillitoe on behalf of the Wellcome Sanger Institute COVID-19 Surveillance Team |
| EPI_ISL_3170617 | Lighthouse Laboratory Plymouth | Wellcome Sanger Institute for the COVID-19 Genomics UK (COG-UK) Consortium | Lighthouse Laboratory Plymouth and Alex Alderton, Roberto Amato, Jeffrey Barrett, Sonia Goncalves, Ewan Harrison, David K. Jackson, Ian Johnston, Dominic Kwiatkowski, Cordelia Langford, John Sillitoe on behalf of the Wellcome Sanger Institute COVID-19 Surveillance Team |
| EPI_ISL_3170618 | Lighthouse Lab in Glasgow | Wellcome Sanger Institute for the COVID-19 Genomics UK (COG-UK) Consortium | Harper VanSteenhouse, Yumi Kasai, David Gray, Carol Clugston, Anna Dominiczak and Alex Alderton, Roberto Amato, Jeffrey Barrett, Sonia Goncalves, Ewan Harrison, David K. Jackson, Ian Johnston, Dominic Kwiatkowski, Cordelia Langford, John Sillitoe on behalf of the Wellcome Sanger Institute COVID-19 Surveillance Team |
| EPI_ISL_3170619 | Lighthouse Laboratory Plymouth | Wellcome Sanger Institute for the COVID-19 Genomics UK (COG-UK) Consortium | Lighthouse Laboratory Plymouth and Alex Alderton, Roberto Amato, Jeffrey Barrett, Sonia Goncalves, Ewan Harrison, David K. Jackson, Ian Johnston, Dominic Kwiatkowski, Cordelia Langford, John Sillitoe on behalf of the Wellcome Sanger Institute COVID-19 Surveillance Team |
| EPI_ISL_3170620 | Lighthouse Lab in Glasgow | Wellcome Sanger Institute for the COVID-19 Genomics UK (COG-UK) Consortium | Harper VanSteenhouse, Yumi Kasai, David Gray, Carol Clugston, Anna Dominiczak and Alex Alderton, Roberto Amato, Jeffrey Barrett, Sonia Goncalves, Ewan Harrison, David K. Jackson, Ian Johnston, Dominic Kwiatkowski, Cordelia Langford, John Sillitoe on behalf of the Wellcome Sanger Institute COVID-19 Surveillance Team |
| EPI_ISL_3170621 | Lighthouse Lab in Milton Keynes | Wellcome Sanger Institute for the COVID-19 Genomics UK (COG-UK) Consortium | The Lighthouse Lab in Milton Keynes and Alex Alderton, Roberto Amato, Jeffrey Barrett, Sonia Goncalves, Ewan Harrison, David K. Jackson, Ian Johnston, Dominic Kwiatkowski, Cordelia Langford, John Sillitoe on behalf of the Wellcome Sanger Institute COVID-19 Surveillance Team |
| EPI_ISL_3170622 | Lighthouse Laboratory Plymouth | Wellcome Sanger Institute for the COVID-19 Genomics UK (COG-UK) Consortium | Lighthouse Laboratory Plymouth and Alex Alderton, Roberto Amato, Jeffrey Barrett, Sonia Goncalves, Ewan Harrison, David K. Jackson, Ian Johnston, Dominic Kwiatkowski, Cordelia Langford, John Sillitoe on behalf of the Wellcome Sanger Institute COVID-19 Surveillance Team |
| EPI_ISL_3170623, EPI_ISL_3170624 | Lighthouse Lab in Milton Keynes | Wellcome Sanger Institute for the COVID-19 Genomics UK (COG-UK) Consortium | The Lighthouse Lab in Milton Keynes and Alex Alderton, Roberto Amato, Jeffrey Barrett, Sonia Goncalves, Ewan Harrison, David K. Jackson, Ian Johnston, Dominic Kwiatkowski, Cordelia Langford, John Sillitoe on behalf of the Wellcome Sanger Institute COVID-19 Surveillance Team |
| EPI_ISL_3170625 | Lighthouse Lab in Glasgow | Wellcome Sanger Institute for the COVID-19 Genomics UK (COG-UK) Consortium | Harper VanSteenhouse, Yumi Kasai, David Gray, Carol Clugston, Anna Dominiczak and Alex Alderton, Roberto Amato, Jeffrey Barrett, Sonia Goncalves, Ewan Harrison, David K. Jackson, Ian Johnston, Dominic Kwiatkowski, Cordelia Langford, John Sillitoe on behalf of the Wellcome Sanger Institute COVID-19 Surveillance Team |
| EPI_ISL_3170626, EPI_ISL_3170627 | Lighthouse Lab in Milton Keynes | Wellcome Sanger Institute for the COVID-19 Genomics UK (COG-UK) Consortium | The Lighthouse Lab in Milton Keynes and Alex Alderton, Roberto Amato, Jeffrey Barrett, Sonia Goncalves, Ewan Harrison, David K. Jackson, Ian Johnston, Dominic Kwiatkowski, Cordelia Langford, John Sillitoe on behalf of the Wellcome Sanger Institute COVID-19 Surveillance Team |
| EPI_ISL_3170628 | Lighthouse Laboratory Plymouth | Wellcome Sanger Institute for the COVID-19 Genomics UK (COG-UK) Consortium | Lighthouse Laboratory Plymouth and Alex Alderton, Roberto Amato, Jeffrey Barrett, Sonia Goncalves, Ewan Harrison, David K. Jackson, Ian Johnston, Dominic Kwiatkowski, Cordelia Langford, John Sillitoe on behalf of the Wellcome Sanger Institute COVID-19 Surveillance Team |
| EPI_ISL_3170629, EPI_ISL_3170630 | Lighthouse Lab in Milton Keynes | Wellcome Sanger Institute for the COVID-19 Genomics UK (COG-UK) Consortium | The Lighthouse Lab in Milton Keynes and Alex Alderton, Roberto Amato, Jeffrey Barrett, Sonia Goncalves, Ewan Harrison, David K. Jackson, Ian Johnston, Dominic Kwiatkowski, Cordelia Langford, John Sillitoe on behalf of the Wellcome Sanger Institute COVID-19 Surveillance Team |
| EPI_ISL_3170631 | Lighthouse Laboratory Plymouth | Wellcome Sanger Institute for the COVID-19 Genomics UK (COG-UK) Consortium | Lighthouse Laboratory Plymouth and Alex Alderton, Roberto Amato, Jeffrey Barrett, Sonia Goncalves, Ewan Harrison, David K. Jackson, Ian Johnston, Dominic Kwiatkowski, Cordelia Langford, John Sillitoe on behalf of the Wellcome Sanger Institute COVID-19 Surveillance Team |
| EPI_ISL_3170632, EPI_ISL_3170633, EPI_ISL_3170634, EPI_ISL_3170635, EPI_ISL_3170636 | Lighthouse Lab in Milton Keynes | Wellcome Sanger Institute for the COVID-19 Genomics UK (COG-UK) Consortium | The Lighthouse Lab in Milton Keynes and Alex Alderton, Roberto Amato, Jeffrey Barrett, Sonia Goncalves, Ewan Harrison, David K. Jackson, Ian Johnston, Dominic Kwiatkowski, Cordelia Langford, John Sillitoe on behalf of the Wellcome Sanger Institute COVID-19 Surveillance Team |
| EPI_ISL_3170637 | Lighthouse Laboratory Plymouth | Wellcome Sanger Institute for the COVID-19 Genomics UK (COG-UK) Consortium | Lighthouse Laboratory Plymouth and Alex Alderton, Roberto Amato, Jeffrey Barrett, Sonia Goncalves, Ewan Harrison, David K. Jackson, Ian Johnston, Dominic Kwiatkowski, Cordelia Langford, John Sillitoe on behalf of the Wellcome Sanger Institute COVID-19 Surveillance Team |
| EPI_ISL_3170638 | Lighthouse Lab in Glasgow | Wellcome Sanger Institute for the COVID-19 Genomics UK (COG-UK) Consortium | Harper VanSteenhouse, Yumi Kasai, David Gray, Carol Clugston, Anna Dominiczak and Alex Alderton, Roberto Amato, Jeffrey Barrett, Sonia Goncalves, Ewan Harrison, David K. Jackson, Ian Johnston, Dominic Kwiatkowski, Cordelia Langford, John Sillitoe on behalf of the Wellcome Sanger Institute COVID-19 Surveillance Team |
| EPI_ISL_3170639 | Lighthouse Lab in Milton Keynes | Wellcome Sanger Institute for the COVID-19 Genomics UK (COG-UK) Consortium | The Lighthouse Lab in Milton Keynes and Alex Alderton, Roberto Amato, Jeffrey Barrett, Sonia Goncalves, Ewan Harrison, David K. Jackson, Ian Johnston, Dominic Kwiatkowski, Cordelia Langford, John Sillitoe on behalf of the Wellcome Sanger Institute COVID-19 Surveillance Team |
| EPI_ISL_3170640 | Lighthouse Laboratory Plymouth | Wellcome Sanger Institute for the COVID-19 Genomics UK (COG-UK) Consortium | Lighthouse Laboratory Plymouth and Alex Alderton, Roberto Amato, Jeffrey Barrett, Sonia Goncalves, Ewan Harrison, David K. Jackson, Ian Johnston, Dominic Kwiatkowski, Cordelia Langford, John Sillitoe on behalf of the Wellcome Sanger Institute COVID-19 Surveillance Team |
| EPI_ISL_3170641 | Lighthouse Lab in Milton Keynes | Wellcome Sanger Institute for the COVID-19 Genomics UK (COG-UK) Consortium | The Lighthouse Lab in Milton Keynes and Alex Alderton, Roberto Amato, Jeffrey Barrett, Sonia Goncalves, Ewan Harrison, David K. Jackson, Ian Johnston, Dominic Kwiatkowski, Cordelia Langford, John Sillitoe on behalf of the Wellcome Sanger Institute COVID-19 Surveillance Team |
| EPI_ISL_3170642 | Lighthouse Lab in Glasgow | Wellcome Sanger Institute for the COVID-19 Genomics UK (COG-UK) Consortium | Harper VanSteenhouse, Yumi Kasai, David Gray, Carol Clugston, Anna Dominiczak and Alex Alderton, Roberto Amato, Jeffrey Barrett, Sonia Goncalves, Ewan Harrison, David K. Jackson, Ian Johnston, Dominic Kwiatkowski, Cordelia Langford, John Sillitoe on behalf of the Wellcome Sanger Institute COVID-19 Surveillance Team |
| EPI_ISL_3170643, EPI_ISL_3170644, | Lighthouse Lab in Milton Keynes | Wellcome Sanger Institute for the COVID-19 Genomics UK (COG-UK) Consortium | The Lighthouse Lab in Milton Keynes and Alex Alderton, Roberto Amato, Jeffrey Barrett, Sonia Goncalves, Ewan Harrison, David K. Jackson, Ian Johnston, Dominic Kwiatkowski, Cordelia Langford, John Sillitoe on behalf of the Wellcome Sanger Institute COVID-19 Surveillance Team |

| | | | |
|---|---|---|---|
| EPI_ISL_3170645 | | | |
| EPI_ISL_3170646 | Lighthouse Lab in Glasgow | Wellcome Sanger Institute for the COVID-19 Genomics UK (COG-UK) Consortium | Harper VanSteenhouse, Yumi Kasai, David Gray, Carol Clugston, Anna Dominiczak and Alex Alderton, Roberto Amato, Jeffrey Barrett, Sonia Goncalves, Ewan Harrison, David K. Jackson, Ian Johnston, Dominic Kwiatkowski, Cordelia Langford, John Sillitoe on behalf of the Wellcome Sanger Institute COVID-19 Surveillance Team |
| EPI_ISL_3170647 | Lighthouse Laboratory Plymouth | Wellcome Sanger Institute for the COVID-19 Genomics UK (COG-UK) Consortium | Lighthouse Laboratory Plymouth and Alex Alderton, Roberto Amato, Jeffrey Barrett, Sonia Goncalves, Ewan Harrison, David K. Jackson, Ian Johnston, Dominic Kwiatkowski, Cordelia Langford, John Sillitoe on behalf of the Wellcome Sanger Institute COVID-19 Surveillance Team |
| EPI_ISL_3170648, EPI_ISL_3170649 | Lighthouse Lab in Milton Keynes | Wellcome Sanger Institute for the COVID-19 Genomics UK (COG-UK) Consortium | The Lighthouse Lab in Milton Keynes and Alex Alderton, Roberto Amato, Jeffrey Barrett, Sonia Goncalves, Ewan Harrison, David K. Jackson, Ian Johnston, Dominic Kwiatkowski, Cordelia Langford, John Sillitoe on behalf of the Wellcome Sanger Institute COVID-19 Surveillance Team |
| EPI_ISL_3170650 | Lighthouse Laboratory Plymouth | Wellcome Sanger Institute for the COVID-19 Genomics UK (COG-UK) Consortium | Lighthouse Laboratory Plymouth and Alex Alderton, Roberto Amato, Jeffrey Barrett, Sonia Goncalves, Ewan Harrison, David K. Jackson, Ian Johnston, Dominic Kwiatkowski, Cordelia Langford, John Sillitoe on behalf of the Wellcome Sanger Institute COVID-19 Surveillance Team |
| EPI_ISL_3170651 | Lighthouse Lab in Milton Keynes | Wellcome Sanger Institute for the COVID-19 Genomics UK (COG-UK) Consortium | The Lighthouse Lab in Milton Keynes and Alex Alderton, Roberto Amato, Jeffrey Barrett, Sonia Goncalves, Ewan Harrison, David K. Jackson, Ian Johnston, Dominic Kwiatkowski, Cordelia Langford, John Sillitoe on behalf of the Wellcome Sanger Institute COVID-19 Surveillance Team |
| EPI_ISL_3170652 | Lighthouse Laboratory Plymouth | Wellcome Sanger Institute for the COVID-19 Genomics UK (COG-UK) Consortium | Lighthouse Laboratory Plymouth and Alex Alderton, Roberto Amato, Jeffrey Barrett, Sonia Goncalves, Ewan Harrison, David K. Jackson, Ian Johnston, Dominic Kwiatkowski, Cordelia Langford, John Sillitoe on behalf of the Wellcome Sanger Institute COVID-19 Surveillance Team |
| EPI_ISL_3170653 | Lighthouse Lab in Milton Keynes | Wellcome Sanger Institute for the COVID-19 Genomics UK (COG-UK) Consortium | The Lighthouse Lab in Milton Keynes and Alex Alderton, Roberto Amato, Jeffrey Barrett, Sonia Goncalves, Ewan Harrison, David K. Jackson, Ian Johnston, Dominic Kwiatkowski, Cordelia Langford, John Sillitoe on behalf of the Wellcome Sanger Institute COVID-19 Surveillance Team |
| EPI_ISL_3170654 | Lighthouse Laboratory Plymouth | Wellcome Sanger Institute for the COVID-19 Genomics UK (COG-UK) Consortium | Lighthouse Laboratory Plymouth and Alex Alderton, Roberto Amato, Jeffrey Barrett, Sonia Goncalves, Ewan Harrison, David K. Jackson, Ian Johnston, Dominic Kwiatkowski, Cordelia Langford, John Sillitoe on behalf of the Wellcome Sanger Institute COVID-19 Surveillance Team |
| EPI_ISL_3170655, EPI_ISL_3170656, EPI_ISL_3170657, EPI_ISL_3170658, EPI_ISL_3170659 | Lighthouse Lab in Milton Keynes | Wellcome Sanger Institute for the COVID-19 Genomics UK (COG-UK) Consortium | The Lighthouse Lab in Milton Keynes and Alex Alderton, Roberto Amato, Jeffrey Barrett, Sonia Goncalves, Ewan Harrison, David K. Jackson, Ian Johnston, Dominic Kwiatkowski, Cordelia Langford, John Sillitoe on behalf of the Wellcome Sanger Institute COVID-19 Surveillance Team |
| EPI_ISL_3170660 | Lighthouse Laboratory Plymouth | Wellcome Sanger Institute for the COVID-19 Genomics UK (COG-UK) Consortium | Lighthouse Laboratory Plymouth and Alex Alderton, Roberto Amato, Jeffrey Barrett, Sonia Goncalves, Ewan Harrison, David K. Jackson, Ian Johnston, Dominic Kwiatkowski, Cordelia Langford, John Sillitoe on behalf of the Wellcome Sanger Institute COVID-19 Surveillance Team |
| EPI_ISL_3170661, EPI_ISL_3170662 | Lighthouse Lab in Milton Keynes | Wellcome Sanger Institute for the COVID-19 Genomics UK (COG-UK) Consortium | The Lighthouse Lab in Milton Keynes and Alex Alderton, Roberto Amato, Jeffrey Barrett, Sonia Goncalves, Ewan Harrison, David K. Jackson, Ian Johnston, Dominic Kwiatkowski, Cordelia Langford, John Sillitoe on behalf of the Wellcome Sanger Institute COVID-19 Surveillance Team |
| EPI_ISL_3170663 | Lighthouse Lab in Glasgow | Wellcome Sanger Institute for the COVID-19 Genomics UK (COG-UK) Consortium | Harper VanSteenhouse, Yumi Kasai, David Gray, Carol Clugston, Anna Dominiczak and Alex Alderton, Roberto Amato, Jeffrey Barrett, Sonia Goncalves, Ewan Harrison, David K. Jackson, Ian Johnston, Dominic Kwiatkowski, Cordelia Langford, John Sillitoe on behalf of the Wellcome Sanger Institute COVID-19 Surveillance Team |
| EPI_ISL_3170664 | Lighthouse Lab in Milton Keynes | Wellcome Sanger Institute for the COVID-19 Genomics UK (COG-UK) Consortium | The Lighthouse Lab in Milton Keynes and Alex Alderton, Roberto Amato, Jeffrey Barrett, Sonia Goncalves, Ewan Harrison, David K. Jackson, Ian Johnston, Dominic Kwiatkowski, Cordelia Langford, John Sillitoe on behalf of the Wellcome Sanger Institute COVID-19 Surveillance Team |
| EPI_ISL_3170665 | Lighthouse Laboratory Plymouth | Wellcome Sanger Institute for the COVID-19 Genomics UK (COG-UK) Consortium | Lighthouse Laboratory Plymouth and Alex Alderton, Roberto Amato, Jeffrey Barrett, Sonia Goncalves, Ewan Harrison, David K. Jackson, Ian Johnston, Dominic Kwiatkowski, Cordelia Langford, John Sillitoe on behalf of the Wellcome Sanger Institute COVID-19 Surveillance Team |
| EPI_ISL_3170666, EPI_ISL_3170667, EPI_ISL_3170668, EPI_ISL_3170669, EPI_ISL_3170670 | Lighthouse Lab in Milton Keynes | Wellcome Sanger Institute for the COVID-19 Genomics UK (COG-UK) Consortium | The Lighthouse Lab in Milton Keynes and Alex Alderton, Roberto Amato, Jeffrey Barrett, Sonia Goncalves, Ewan Harrison, David K. Jackson, Ian Johnston, Dominic Kwiatkowski, Cordelia Langford, John Sillitoe on behalf of the Wellcome Sanger Institute COVID-19 Surveillance Team |
| EPI_ISL_3170671, EPI_ISL_3170672, EPI_ISL_3170673 | Lighthouse Laboratory Plymouth | Wellcome Sanger Institute for the COVID-19 Genomics UK (COG-UK) Consortium | Lighthouse Laboratory Plymouth and Alex Alderton, Roberto Amato, Jeffrey Barrett, Sonia Goncalves, Ewan Harrison, David K. Jackson, Ian Johnston, Dominic Kwiatkowski, Cordelia Langford, John Sillitoe on behalf of the Wellcome Sanger Institute COVID-19 Surveillance Team |
| EPI_ISL_3170674 | Lighthouse Lab in Milton Keynes | Wellcome Sanger Institute for the COVID-19 Genomics UK (COG-UK) Consortium | The Lighthouse Lab in Milton Keynes and Alex Alderton, Roberto Amato, Jeffrey Barrett, Sonia Goncalves, Ewan Harrison, David K. Jackson, Ian Johnston, Dominic Kwiatkowski, Cordelia Langford, John Sillitoe on behalf of the Wellcome Sanger Institute COVID-19 Surveillance Team |
| EPI_ISL_3170675 | Lighthouse Laboratory Plymouth | Wellcome Sanger Institute for the COVID-19 Genomics UK (COG-UK) Consortium | Lighthouse Laboratory Plymouth and Alex Alderton, Roberto Amato, Jeffrey Barrett, Sonia Goncalves, Ewan Harrison, David K. Jackson, Ian Johnston, Dominic Kwiatkowski, Cordelia Langford, John Sillitoe on behalf of the Wellcome Sanger Institute COVID-19 Surveillance Team |
| EPI_ISL_3170676 | Lighthouse Lab in Glasgow | Wellcome Sanger Institute for the COVID-19 Genomics UK (COG-UK) Consortium | Harper VanSteenhouse, Yumi Kasai, David Gray, Carol Clugston, Anna Dominiczak and Alex Alderton, Roberto Amato, Jeffrey Barrett, Sonia Goncalves, Ewan Harrison, David K. Jackson, Ian Johnston, Dominic Kwiatkowski, Cordelia Langford, John Sillitoe on behalf of the Wellcome Sanger Institute COVID-19 Surveillance Team |
| EPI_ISL_3170677, EPI_ISL_3170678 | Lighthouse Laboratory Plymouth | Wellcome Sanger Institute for the COVID-19 Genomics UK (COG-UK) Consortium | Lighthouse Laboratory Plymouth and Alex Alderton, Roberto Amato, Jeffrey Barrett, Sonia Goncalves, Ewan Harrison, David K. Jackson, Ian Johnston, Dominic Kwiatkowski, Cordelia Langford, John Sillitoe on behalf of the Wellcome Sanger Institute COVID-19 Surveillance Team |
| EPI_ISL_3170679 | Lighthouse Lab in Glasgow | Wellcome Sanger Institute for the COVID-19 Genomics UK (COG-UK) Consortium | Harper VanSteenhouse, Yumi Kasai, David Gray, Carol Clugston, Anna Dominiczak and Alex Alderton, Roberto Amato, Jeffrey Barrett, Sonia Goncalves, Ewan Harrison, David K. Jackson, Ian Johnston, Dominic Kwiatkowski, Cordelia Langford, John Sillitoe on behalf of the Wellcome Sanger Institute COVID-19 Surveillance Team |
| EPI_ISL_3170680 | Lighthouse Lab in Milton Keynes | Wellcome Sanger Institute for the COVID-19 Genomics UK (COG-UK) Consortium | The Lighthouse Lab in Milton Keynes and Alex Alderton, Roberto Amato, Jeffrey Barrett, Sonia Goncalves, Ewan Harrison, David K. Jackson, Ian Johnston, Dominic Kwiatkowski, Cordelia Langford, John Sillitoe on behalf of the Wellcome Sanger Institute COVID-19 Surveillance Team |
| EPI_ISL_3170681, EPI_ISL_3170682 | Lighthouse Lab in Glasgow | Wellcome Sanger Institute for the COVID-19 Genomics UK (COG-UK) Consortium | Harper VanSteenhouse, Yumi Kasai, David Gray, Carol Clugston, Anna Dominiczak and Alex Alderton, Roberto Amato, Jeffrey Barrett, Sonia Goncalves, Ewan Harrison, David K. Jackson, Ian Johnston, Dominic Kwiatkowski, Cordelia Langford, John Sillitoe on behalf of the Wellcome Sanger Institute COVID-19 Surveillance Team |
| EPI_ISL_3170683, EPI_ISL_3170684 | Lighthouse Lab in Milton Keynes | Wellcome Sanger Institute for the COVID-19 Genomics UK (COG-UK) Consortium | The Lighthouse Lab in Milton Keynes and Alex Alderton, Roberto Amato, Jeffrey Barrett, Sonia Goncalves, Ewan Harrison, David K. Jackson, Ian Johnston, Dominic Kwiatkowski, Cordelia Langford, John Sillitoe on behalf of the Wellcome Sanger Institute COVID-19 Surveillance Team |
| EPI_ISL_3170685 | Lighthouse Lab in Glasgow | Wellcome Sanger Institute for the COVID-19 Genomics UK (COG-UK) Consortium | Harper VanSteenhouse, Yumi Kasai, David Gray, Carol Clugston, Anna Dominiczak and Alex Alderton, Roberto Amato, Jeffrey Barrett, Sonia Goncalves, Ewan Harrison, David K. Jackson, Ian Johnston, Dominic Kwiatkowski, Cordelia Langford, John Sillitoe on behalf of the Wellcome Sanger Institute COVID-19 Surveillance Team |
| EPI_ISL_3170686, EPI_ISL_3170687, EPI_ISL_3170688, EPI_ISL_3170689 | Lighthouse Lab in Milton Keynes | Wellcome Sanger Institute for the COVID-19 Genomics UK (COG-UK) Consortium | The Lighthouse Lab in Milton Keynes and Alex Alderton, Roberto Amato, Jeffrey Barrett, Sonia Goncalves, Ewan Harrison, David K. Jackson, Ian Johnston, Dominic Kwiatkowski, Cordelia Langford, John Sillitoe on behalf of the Wellcome Sanger Institute COVID-19 Surveillance Team |
| EPI_ISL_3170690 | Lighthouse Laboratory Plymouth | Wellcome Sanger Institute for the COVID-19 Genomics UK (COG-UK) Consortium | Lighthouse Laboratory Plymouth and Alex Alderton, Roberto Amato, Jeffrey Barrett, Sonia Goncalves, Ewan Harrison, David K. Jackson, Ian Johnston, Dominic Kwiatkowski, Cordelia Langford, John Sillitoe on behalf of the Wellcome Sanger Institute COVID-19 Surveillance Team |
| EPI_ISL_3170691, EPI_ISL_3170692, EPI_ISL_3170693 | Lighthouse Lab in Milton Keynes | Wellcome Sanger Institute for the COVID-19 Genomics UK (COG-UK) Consortium | The Lighthouse Lab in Milton Keynes and Alex Alderton, Roberto Amato, Jeffrey Barrett, Sonia Goncalves, Ewan Harrison, David K. Jackson, Ian Johnston, Dominic Kwiatkowski, Cordelia Langford, John Sillitoe on behalf of the Wellcome Sanger Institute COVID-19 Surveillance Team |
| EPI_ISL_3170694, EPI_ISL_3170695 | Lighthouse Lab in Glasgow | Wellcome Sanger Institute for the COVID-19 Genomics UK (COG-UK) Consortium | Harper VanSteenhouse, Yumi Kasai, David Gray, Carol Clugston, Anna Dominiczak and Alex Alderton, Roberto Amato, Jeffrey Barrett, Sonia Goncalves, Ewan Harrison, David K. Jackson, Ian Johnston, Dominic Kwiatkowski, Cordelia Langford, John Sillitoe on behalf of the Wellcome Sanger Institute COVID-19 Surveillance Team |
| EPI_ISL_3170696, EPI_ISL_3170697 | Lighthouse Lab in Milton Keynes | Wellcome Sanger Institute for the COVID-19 Genomics UK (COG-UK) Consortium | The Lighthouse Lab in Milton Keynes and Alex Alderton, Roberto Amato, Jeffrey Barrett, Sonia Goncalves, Ewan Harrison, David K. Jackson, Ian Johnston, Dominic Kwiatkowski, Cordelia Langford, John Sillitoe on behalf of the Wellcome Sanger Institute COVID-19 Surveillance Team |
| EPI_ISL_3170698 | Lighthouse Laboratory Plymouth | Wellcome Sanger Institute for the COVID-19 Genomics UK (COG-UK) Consortium | Lighthouse Laboratory Plymouth and Alex Alderton, Roberto Amato, Jeffrey Barrett, Sonia Goncalves, Ewan Harrison, David K. Jackson, Ian Johnston, Dominic Kwiatkowski, Cordelia Langford, John Sillitoe on behalf of the Wellcome Sanger Institute COVID-19 Surveillance Team |
| EPI_ISL_3170699 | Lighthouse Lab in Milton Keynes | Wellcome Sanger Institute for the COVID-19 Genomics UK (COG-UK) Consortium | The Lighthouse Lab in Milton Keynes and Alex Alderton, Roberto Amato, Jeffrey Barrett, Sonia Goncalves, Ewan Harrison, David K. Jackson, Ian Johnston, Dominic Kwiatkowski, Cordelia Langford, John Sillitoe on behalf of the Wellcome Sanger Institute COVID-19 Surveillance Team |
| EPI_ISL_3170700 | Lighthouse Laboratory Plymouth | Wellcome Sanger Institute for the COVID-19 Genomics UK (COG-UK) Consortium | Lighthouse Laboratory Plymouth and Alex Alderton, Roberto Amato, Jeffrey Barrett, Sonia Goncalves, Ewan Harrison, David K. Jackson, Ian Johnston, Dominic Kwiatkowski, Cordelia Langford, John Sillitoe on behalf of the Wellcome Sanger Institute COVID-19 Surveillance Team |
| EPI_ISL_3170701, EPI_ISL_3170702 | Lighthouse Lab in Milton Keynes | Wellcome Sanger Institute for the COVID-19 Genomics UK (COG-UK) Consortium | The Lighthouse Lab in Milton Keynes and Alex Alderton, Roberto Amato, Jeffrey Barrett, Sonia Goncalves, Ewan Harrison, David K. Jackson, Ian Johnston, Dominic Kwiatkowski, Cordelia Langford, John Sillitoe on behalf of the Wellcome Sanger Institute COVID-19 Surveillance Team |
| EPI_ISL_3170703, EPI_ISL_3170704 | Lighthouse Lab in Glasgow | Wellcome Sanger Institute for the COVID-19 Genomics UK (COG-UK) Consortium | Harper VanSteenhouse, Yumi Kasai, David Gray, Carol Clugston, Anna Dominiczak and Alex Alderton, Roberto Amato, Jeffrey Barrett, Sonia Goncalves, Ewan Harrison, David K. Jackson, Ian Johnston, Dominic Kwiatkowski, Cordelia Langford, John Sillitoe on behalf of the Wellcome Sanger Institute COVID-19 Surveillance Team |
| EPI_ISL_3170705 | Lighthouse Lab in Milton Keynes | Wellcome Sanger Institute for the COVID-19 Genomics UK (COG-UK) Consortium | The Lighthouse Lab in Milton Keynes and Alex Alderton, Roberto Amato, Jeffrey Barrett, Sonia Goncalves, Ewan Harrison, David K. Jackson, Ian Johnston, Dominic Kwiatkowski, Cordelia Langford, John Sillitoe on behalf of the Wellcome Sanger Institute COVID-19 Surveillance Team |
| EPI_ISL_3170706 | Lighthouse Lab in Glasgow | Wellcome Sanger Institute for the COVID-19 Genomics UK (COG-UK) Consortium | Harper VanSteenhouse, Yumi Kasai, David Gray, Carol Clugston, Anna Dominiczak and Alex Alderton, Roberto Amato, Jeffrey Barrett, Sonia Goncalves, Ewan Harrison, David K. Jackson, Ian Johnston, Dominic Kwiatkowski, Cordelia Langford, John Sillitoe on behalf of the Wellcome Sanger Institute COVID-19 Surveillance Team |
| EPI_ISL_3170707, EPI_ISL_3170708, EPI_ISL_3170709, EPI_ISL_3170710 | Lighthouse Lab in Milton Keynes | Wellcome Sanger Institute for the COVID-19 Genomics UK (COG-UK) Consortium | The Lighthouse Lab in Milton Keynes and Alex Alderton, Roberto Amato, Jeffrey Barrett, Sonia Goncalves, Ewan Harrison, David K. Jackson, Ian Johnston, Dominic Kwiatkowski, Cordelia Langford, John Sillitoe on behalf of the Wellcome Sanger Institute COVID-19 Surveillance Team |
| EPI_ISL_3170711 | Lighthouse Lab in Glasgow | Wellcome Sanger Institute for the COVID-19 Genomics UK (COG-UK) Consortium | Harper VanSteenhouse, Yumi Kasai, David Gray, Carol Clugston, Anna Dominiczak and Alex Alderton, Roberto Amato, Jeffrey Barrett, Sonia Goncalves, Ewan Harrison, David K. Jackson, Ian Johnston, Dominic Kwiatkowski, Cordelia Langford, John Sillitoe on behalf of the Wellcome Sanger Institute COVID-19 Surveillance Team |
| EPI_ISL_3170712, | Lighthouse Lab in Milton Keynes | Wellcome Sanger Institute for the COVID-19 Genomics UK (COG-UK) | The Lighthouse Lab in Milton Keynes and Alex Alderton, Roberto Amato, Jeffrey Barrett, Sonia Goncalves, Ewan Harrison, David K. Jackson, Ian Johnston, Dominic Kwiatkowski, Cordelia Langford, John Sillitoe on behalf of the Wellcome Sanger |

| | | | |
|---|---|---|---|
| EPI_ISL_3170713, EPI_ISL_3170714 | | Consortium | Institute COVID-19 Surveillance Team |
| EPI_ISL_3170715 | Lighthouse Lab in Glasgow | Wellcome Sanger Institute for the COVID-19 Genomics UK (COG-UK) Consortium | Harper VanSteenhouse, Yumi Kasai, David Gray, Carol Clugston, Anna Dominiczak and Alex Alderton, Roberto Amato, Jeffrey Barrett, Sonia Goncalves, Ewan Harrison, David K. Jackson, Ian Johnston, Dominic Kwiatkowski, Cordelia Langford, John Sillitoe on behalf of the Wellcome Sanger Institute COVID-19 Surveillance Team |
| EPI_ISL_3170716, EPI_ISL_3170717 | Lighthouse Lab in Milton Keynes | Wellcome Sanger Institute for the COVID-19 Genomics UK (COG-UK) Consortium | The Lighthouse Lab in Milton Keynes and Alex Alderton, Roberto Amato, Jeffrey Barrett, Sonia Goncalves, Ewan Harrison, David K. Jackson, Ian Johnston, Dominic Kwiatkowski, Cordelia Langford, John Sillitoe on behalf of the Wellcome Sanger Institute COVID-19 Surveillance Team |
| EPI_ISL_3170718 | Lighthouse Laboratory Plymouth | Wellcome Sanger Institute for the COVID-19 Genomics UK (COG-UK) Consortium | Lighthouse Laboratory Plymouth and Alex Alderton, Roberto Amato, Jeffrey Barrett, Sonia Goncalves, Ewan Harrison, David K. Jackson, Ian Johnston, Dominic Kwiatkowski, Cordelia Langford, John Sillitoe on behalf of the Wellcome Sanger Institute COVID-19 Surveillance Team |
| EPI_ISL_3170719, EPI_ISL_3170720 | Lighthouse Lab in Milton Keynes | Wellcome Sanger Institute for the COVID-19 Genomics UK (COG-UK) Consortium | The Lighthouse Lab in Milton Keynes and Alex Alderton, Roberto Amato, Jeffrey Barrett, Sonia Goncalves, Ewan Harrison, David K. Jackson, Ian Johnston, Dominic Kwiatkowski, Cordelia Langford, John Sillitoe on behalf of the Wellcome Sanger Institute COVID-19 Surveillance Team |
| EPI_ISL_3170721 | Lighthouse Laboratory Plymouth | Wellcome Sanger Institute for the COVID-19 Genomics UK (COG-UK) Consortium | Lighthouse Laboratory Plymouth and Alex Alderton, Roberto Amato, Jeffrey Barrett, Sonia Goncalves, Ewan Harrison, David K. Jackson, Ian Johnston, Dominic Kwiatkowski, Cordelia Langford, John Sillitoe on behalf of the Wellcome Sanger Institute COVID-19 Surveillance Team |
| EPI_ISL_3170722, EPI_ISL_3170723 | Lighthouse Lab in Glasgow | Wellcome Sanger Institute for the COVID-19 Genomics UK (COG-UK) Consortium | Harper VanSteenhouse, Yumi Kasai, David Gray, Carol Clugston, Anna Dominiczak and Alex Alderton, Roberto Amato, Jeffrey Barrett, Sonia Goncalves, Ewan Harrison, David K. Jackson, Ian Johnston, Dominic Kwiatkowski, Cordelia Langford, John Sillitoe on behalf of the Wellcome Sanger Institute COVID-19 Surveillance Team |
| EPI_ISL_3170724, EPI_ISL_3170725, EPI_ISL_3170726 | Lighthouse Lab in Milton Keynes | Wellcome Sanger Institute for the COVID-19 Genomics UK (COG-UK) Consortium | The Lighthouse Lab in Milton Keynes and Alex Alderton, Roberto Amato, Jeffrey Barrett, Sonia Goncalves, Ewan Harrison, David K. Jackson, Ian Johnston, Dominic Kwiatkowski, Cordelia Langford, John Sillitoe on behalf of the Wellcome Sanger Institute COVID-19 Surveillance Team |
| EPI_ISL_3170727 | Lighthouse Laboratory Plymouth | Wellcome Sanger Institute for the COVID-19 Genomics UK (COG-UK) Consortium | Lighthouse Laboratory Plymouth and Alex Alderton, Roberto Amato, Jeffrey Barrett, Sonia Goncalves, Ewan Harrison, David K. Jackson, Ian Johnston, Dominic Kwiatkowski, Cordelia Langford, John Sillitoe on behalf of the Wellcome Sanger Institute COVID-19 Surveillance Team |
| EPI_ISL_3170728, EPI_ISL_3170729 | Lighthouse Lab in Glasgow | Wellcome Sanger Institute for the COVID-19 Genomics UK (COG-UK) Consortium | Harper VanSteenhouse, Yumi Kasai, David Gray, Carol Clugston, Anna Dominiczak and Alex Alderton, Roberto Amato, Jeffrey Barrett, Sonia Goncalves, Ewan Harrison, David K. Jackson, Ian Johnston, Dominic Kwiatkowski, Cordelia Langford, John Sillitoe on behalf of the Wellcome Sanger Institute COVID-19 Surveillance Team |
| EPI_ISL_3170730, EPI_ISL_3170731 | Lighthouse Lab in Milton Keynes | Wellcome Sanger Institute for the COVID-19 Genomics UK (COG-UK) Consortium | The Lighthouse Lab in Milton Keynes and Alex Alderton, Roberto Amato, Jeffrey Barrett, Sonia Goncalves, Ewan Harrison, David K. Jackson, Ian Johnston, Dominic Kwiatkowski, Cordelia Langford, John Sillitoe on behalf of the Wellcome Sanger Institute COVID-19 Surveillance Team |
| EPI_ISL_3170732 | Lighthouse Laboratory Plymouth | Wellcome Sanger Institute for the COVID-19 Genomics UK (COG-UK) Consortium | Lighthouse Laboratory Plymouth and Alex Alderton, Roberto Amato, Jeffrey Barrett, Sonia Goncalves, Ewan Harrison, David K. Jackson, Ian Johnston, Dominic Kwiatkowski, Cordelia Langford, John Sillitoe on behalf of the Wellcome Sanger Institute COVID-19 Surveillance Team |
| EPI_ISL_3170733, EPI_ISL_3170734 | Lighthouse Lab in Milton Keynes | Wellcome Sanger Institute for the COVID-19 Genomics UK (COG-UK) Consortium | The Lighthouse Lab in Milton Keynes and Alex Alderton, Roberto Amato, Jeffrey Barrett, Sonia Goncalves, Ewan Harrison, David K. Jackson, Ian Johnston, Dominic Kwiatkowski, Cordelia Langford, John Sillitoe on behalf of the Wellcome Sanger Institute COVID-19 Surveillance Team |
| EPI_ISL_3170735 | Lighthouse Lab in Glasgow | Wellcome Sanger Institute for the COVID-19 Genomics UK (COG-UK) Consortium | Harper VanSteenhouse, Yumi Kasai, David Gray, Carol Clugston, Anna Dominiczak and Alex Alderton, Roberto Amato, Jeffrey Barrett, Sonia Goncalves, Ewan Harrison, David K. Jackson, Ian Johnston, Dominic Kwiatkowski, Cordelia Langford, John Sillitoe on behalf of the Wellcome Sanger Institute COVID-19 Surveillance Team |
| EPI_ISL_3170736, EPI_ISL_3170737, EPI_ISL_3170738 | Lighthouse Lab in Milton Keynes | Wellcome Sanger Institute for the COVID-19 Genomics UK (COG-UK) Consortium | The Lighthouse Lab in Milton Keynes and Alex Alderton, Roberto Amato, Jeffrey Barrett, Sonia Goncalves, Ewan Harrison, David K. Jackson, Ian Johnston, Dominic Kwiatkowski, Cordelia Langford, John Sillitoe on behalf of the Wellcome Sanger Institute COVID-19 Surveillance Team |
| EPI_ISL_3170739 | Lighthouse Lab in Glasgow | Wellcome Sanger Institute for the COVID-19 Genomics UK (COG-UK) Consortium | Harper VanSteenhouse, Yumi Kasai, David Gray, Carol Clugston, Anna Dominiczak and Alex Alderton, Roberto Amato, Jeffrey Barrett, Sonia Goncalves, Ewan Harrison, David K. Jackson, Ian Johnston, Dominic Kwiatkowski, Cordelia Langford, John Sillitoe on behalf of the Wellcome Sanger Institute COVID-19 Surveillance Team |
| EPI_ISL_3170740 | Lighthouse Laboratory Plymouth | Wellcome Sanger Institute for the COVID-19 Genomics UK (COG-UK) Consortium | Lighthouse Laboratory Plymouth and Alex Alderton, Roberto Amato, Jeffrey Barrett, Sonia Goncalves, Ewan Harrison, David K. Jackson, Ian Johnston, Dominic Kwiatkowski, Cordelia Langford, John Sillitoe on behalf of the Wellcome Sanger Institute COVID-19 Surveillance Team |
| EPI_ISL_3170741, EPI_ISL_3170742 | Lighthouse Lab in Milton Keynes | Wellcome Sanger Institute for the COVID-19 Genomics UK (COG-UK) Consortium | The Lighthouse Lab in Milton Keynes and Alex Alderton, Roberto Amato, Jeffrey Barrett, Sonia Goncalves, Ewan Harrison, David K. Jackson, Ian Johnston, Dominic Kwiatkowski, Cordelia Langford, John Sillitoe on behalf of the Wellcome Sanger Institute COVID-19 Surveillance Team |
| EPI_ISL_3170743 | Lighthouse Laboratory Plymouth | Wellcome Sanger Institute for the COVID-19 Genomics UK (COG-UK) Consortium | Lighthouse Laboratory Plymouth and Alex Alderton, Roberto Amato, Jeffrey Barrett, Sonia Goncalves, Ewan Harrison, David K. Jackson, Ian Johnston, Dominic Kwiatkowski, Cordelia Langford, John Sillitoe on behalf of the Wellcome Sanger Institute COVID-19 Surveillance Team |
| EPI_ISL_3170744 | Lighthouse Lab in Milton Keynes | Wellcome Sanger Institute for the COVID-19 Genomics UK (COG-UK) Consortium | The Lighthouse Lab in Milton Keynes and Alex Alderton, Roberto Amato, Jeffrey Barrett, Sonia Goncalves, Ewan Harrison, David K. Jackson, Ian Johnston, Dominic Kwiatkowski, Cordelia Langford, John Sillitoe on behalf of the Wellcome Sanger Institute COVID-19 Surveillance Team |
| EPI_ISL_3170745 | Lighthouse Laboratory Plymouth | Wellcome Sanger Institute for the COVID-19 Genomics UK (COG-UK) Consortium | Lighthouse Laboratory Plymouth and Alex Alderton, Roberto Amato, Jeffrey Barrett, Sonia Goncalves, Ewan Harrison, David K. Jackson, Ian Johnston, Dominic Kwiatkowski, Cordelia Langford, John Sillitoe on behalf of the Wellcome Sanger Institute COVID-19 Surveillance Team |
| EPI_ISL_3170746, EPI_ISL_3170747, EPI_ISL_3170748, EPI_ISL_3170749, EPI_ISL_3170750 | Lighthouse Lab in Milton Keynes | Wellcome Sanger Institute for the COVID-19 Genomics UK (COG-UK) Consortium | The Lighthouse Lab in Milton Keynes and Alex Alderton, Roberto Amato, Jeffrey Barrett, Sonia Goncalves, Ewan Harrison, David K. Jackson, Ian Johnston, Dominic Kwiatkowski, Cordelia Langford, John Sillitoe on behalf of the Wellcome Sanger Institute COVID-19 Surveillance Team |
| EPI_ISL_3170751 | Lighthouse Lab in Glasgow | Wellcome Sanger Institute for the COVID-19 Genomics UK (COG-UK) Consortium | Harper VanSteenhouse, Yumi Kasai, David Gray, Carol Clugston, Anna Dominiczak and Alex Alderton, Roberto Amato, Jeffrey Barrett, Sonia Goncalves, Ewan Harrison, David K. Jackson, Ian Johnston, Dominic Kwiatkowski, Cordelia Langford, John Sillitoe on behalf of the Wellcome Sanger Institute COVID-19 Surveillance Team |
| EPI_ISL_3170752, EPI_ISL_3170753 | Lighthouse Lab in Milton Keynes | Wellcome Sanger Institute for the COVID-19 Genomics UK (COG-UK) Consortium | The Lighthouse Lab in Milton Keynes and Alex Alderton, Roberto Amato, Jeffrey Barrett, Sonia Goncalves, Ewan Harrison, David K. Jackson, Ian Johnston, Dominic Kwiatkowski, Cordelia Langford, John Sillitoe on behalf of the Wellcome Sanger Institute COVID-19 Surveillance Team |
| EPI_ISL_3170754 | Lighthouse Laboratory Plymouth | Wellcome Sanger Institute for the COVID-19 Genomics UK (COG-UK) Consortium | Lighthouse Laboratory Plymouth and Alex Alderton, Roberto Amato, Jeffrey Barrett, Sonia Goncalves, Ewan Harrison, David K. Jackson, Ian Johnston, Dominic Kwiatkowski, Cordelia Langford, John Sillitoe on behalf of the Wellcome Sanger Institute COVID-19 Surveillance Team |
| EPI_ISL_3170755 | Lighthouse Lab in Milton Keynes | Wellcome Sanger Institute for the COVID-19 Genomics UK (COG-UK) Consortium | The Lighthouse Lab in Milton Keynes and Alex Alderton, Roberto Amato, Jeffrey Barrett, Sonia Goncalves, Ewan Harrison, David K. Jackson, Ian Johnston, Dominic Kwiatkowski, Cordelia Langford, John Sillitoe on behalf of the Wellcome Sanger Institute COVID-19 Surveillance Team |
| EPI_ISL_3170756, EPI_ISL_3170757 | Lighthouse Lab in Glasgow | Wellcome Sanger Institute for the COVID-19 Genomics UK (COG-UK) Consortium | Harper VanSteenhouse, Yumi Kasai, David Gray, Carol Clugston, Anna Dominiczak and Alex Alderton, Roberto Amato, Jeffrey Barrett, Sonia Goncalves, Ewan Harrison, David K. Jackson, Ian Johnston, Dominic Kwiatkowski, Cordelia Langford, John Sillitoe on behalf of the Wellcome Sanger Institute COVID-19 Surveillance Team |
| EPI_ISL_3170758 | Lighthouse Laboratory Plymouth | Wellcome Sanger Institute for the COVID-19 Genomics UK (COG-UK) Consortium | Lighthouse Laboratory Plymouth and Alex Alderton, Roberto Amato, Jeffrey Barrett, Sonia Goncalves, Ewan Harrison, David K. Jackson, Ian Johnston, Dominic Kwiatkowski, Cordelia Langford, John Sillitoe on behalf of the Wellcome Sanger Institute COVID-19 Surveillance Team |
| EPI_ISL_3170759, EPI_ISL_3170760 | Lighthouse Lab in Milton Keynes | Wellcome Sanger Institute for the COVID-19 Genomics UK (COG-UK) Consortium | The Lighthouse Lab in Milton Keynes and Alex Alderton, Roberto Amato, Jeffrey Barrett, Sonia Goncalves, Ewan Harrison, David K. Jackson, Ian Johnston, Dominic Kwiatkowski, Cordelia Langford, John Sillitoe on behalf of the Wellcome Sanger Institute COVID-19 Surveillance Team |
| EPI_ISL_3170761 | Lighthouse Laboratory Plymouth | Wellcome Sanger Institute for the COVID-19 Genomics UK (COG-UK) Consortium | Lighthouse Laboratory Plymouth and Alex Alderton, Roberto Amato, Jeffrey Barrett, Sonia Goncalves, Ewan Harrison, David K. Jackson, Ian Johnston, Dominic Kwiatkowski, Cordelia Langford, John Sillitoe on behalf of the Wellcome Sanger Institute COVID-19 Surveillance Team |
| EPI_ISL_3170762 | Lighthouse Lab in Glasgow | Wellcome Sanger Institute for the COVID-19 Genomics UK (COG-UK) Consortium | Harper VanSteenhouse, Yumi Kasai, David Gray, Carol Clugston, Anna Dominiczak and Alex Alderton, Roberto Amato, Jeffrey Barrett, Sonia Goncalves, Ewan Harrison, David K. Jackson, Ian Johnston, Dominic Kwiatkowski, Cordelia Langford, John Sillitoe on behalf of the Wellcome Sanger Institute COVID-19 Surveillance Team |
| EPI_ISL_3170763 | Lighthouse Laboratory Plymouth | Wellcome Sanger Institute for the COVID-19 Genomics UK (COG-UK) Consortium | Lighthouse Laboratory Plymouth and Alex Alderton, Roberto Amato, Jeffrey Barrett, Sonia Goncalves, Ewan Harrison, David K. Jackson, Ian Johnston, Dominic Kwiatkowski, Cordelia Langford, John Sillitoe on behalf of the Wellcome Sanger Institute COVID-19 Surveillance Team |
| EPI_ISL_3170764 | Lighthouse Lab in Milton Keynes | Wellcome Sanger Institute for the COVID-19 Genomics UK (COG-UK) Consortium | The Lighthouse Lab in Milton Keynes and Alex Alderton, Roberto Amato, Jeffrey Barrett, Sonia Goncalves, Ewan Harrison, David K. Jackson, Ian Johnston, Dominic Kwiatkowski, Cordelia Langford, John Sillitoe on behalf of the Wellcome Sanger Institute COVID-19 Surveillance Team |
| EPI_ISL_3170765 | Lighthouse Lab in Glasgow | Wellcome Sanger Institute for the COVID-19 Genomics UK (COG-UK) Consortium | Harper VanSteenhouse, Yumi Kasai, David Gray, Carol Clugston, Anna Dominiczak and Alex Alderton, Roberto Amato, Jeffrey Barrett, Sonia Goncalves, Ewan Harrison, David K. Jackson, Ian Johnston, Dominic Kwiatkowski, Cordelia Langford, John Sillitoe on behalf of the Wellcome Sanger Institute COVID-19 Surveillance Team |
| EPI_ISL_3170766 | Lighthouse Laboratory Plymouth | Wellcome Sanger Institute for the COVID-19 Genomics UK (COG-UK) Consortium | Lighthouse Laboratory Plymouth and Alex Alderton, Roberto Amato, Jeffrey Barrett, Sonia Goncalves, Ewan Harrison, David K. Jackson, Ian Johnston, Dominic Kwiatkowski, Cordelia Langford, John Sillitoe on behalf of the Wellcome Sanger Institute COVID-19 Surveillance Team |
| EPI_ISL_3170767 | Lighthouse Lab in Glasgow | Wellcome Sanger Institute for the COVID-19 Genomics UK (COG-UK) Consortium | Harper VanSteenhouse, Yumi Kasai, David Gray, Carol Clugston, Anna Dominiczak and Alex Alderton, Roberto Amato, Jeffrey Barrett, Sonia Goncalves, Ewan Harrison, David K. Jackson, Ian Johnston, Dominic Kwiatkowski, Cordelia Langford, John Sillitoe on behalf of the Wellcome Sanger Institute COVID-19 Surveillance Team |
| EPI_ISL_3170768, EPI_ISL_3170769 | Lighthouse Laboratory Plymouth | Wellcome Sanger Institute for the COVID-19 Genomics UK (COG-UK) Consortium | Lighthouse Laboratory Plymouth and Alex Alderton, Roberto Amato, Jeffrey Barrett, Sonia Goncalves, Ewan Harrison, David K. Jackson, Ian Johnston, Dominic Kwiatkowski, Cordelia Langford, John Sillitoe on behalf of the Wellcome Sanger Institute COVID-19 Surveillance Team |
| EPI_ISL_3170770 | Lighthouse Lab in Glasgow | Wellcome Sanger Institute for the COVID-19 Genomics UK (COG-UK) Consortium | Harper VanSteenhouse, Yumi Kasai, David Gray, Carol Clugston, Anna Dominiczak and Alex Alderton, Roberto Amato, Jeffrey Barrett, Sonia Goncalves, Ewan Harrison, David K. Jackson, Ian Johnston, Dominic Kwiatkowski, Cordelia Langford, John Sillitoe on behalf of the Wellcome Sanger Institute COVID-19 Surveillance Team |
| EPI_ISL_3170771 | Lighthouse Lab in Milton Keynes | Wellcome Sanger Institute for the COVID-19 Genomics UK (COG-UK) Consortium | The Lighthouse Lab in Milton Keynes and Alex Alderton, Roberto Amato, Jeffrey Barrett, Sonia Goncalves, Ewan Harrison, David K. Jackson, Ian Johnston, Dominic Kwiatkowski, Cordelia Langford, John Sillitoe on behalf of the Wellcome Sanger Institute COVID-19 Surveillance Team |
| EPI_ISL_3170772 | Lighthouse Lab in Glasgow | Wellcome Sanger Institute for the COVID-19 Genomics UK (COG-UK) Consortium | Harper VanSteenhouse, Yumi Kasai, David Gray, Carol Clugston, Anna Dominiczak and Alex Alderton, Roberto Amato, Jeffrey Barrett, Sonia Goncalves, Ewan Harrison, David K. Jackson, Ian Johnston, Dominic Kwiatkowski, Cordelia Langford, John Sillitoe on behalf of the Wellcome Sanger Institute COVID-19 Surveillance Team |
| EPI_ISL_3170773 | Lighthouse Laboratory Plymouth | Wellcome Sanger Institute for the COVID-19 Genomics UK (COG-UK) Consortium | Lighthouse Laboratory Plymouth and Alex Alderton, Roberto Amato, Jeffrey Barrett, Sonia Goncalves, Ewan Harrison, David K. Jackson, Ian Johnston, Dominic Kwiatkowski, Cordelia Langford, John Sillitoe on behalf of the Wellcome Sanger Institute COVID-19 Surveillance Team |
| EPI_ISL_3170774 | Lighthouse Lab in Milton Keynes | Wellcome Sanger Institute for the COVID-19 Genomics UK (COG-UK) Consortium | The Lighthouse Lab in Milton Keynes and Alex Alderton, Roberto Amato, Jeffrey Barrett, Sonia Goncalves, Ewan Harrison, David K. Jackson, Ian Johnston, Dominic Kwiatkowski, Cordelia Langford, John Sillitoe on behalf of the Wellcome Sanger Institute COVID-19 Surveillance Team |
| EPI_ISL_3170775 | Lighthouse Laboratory Plymouth | Wellcome Sanger Institute for the COVID-19 Genomics UK (COG-UK) Consortium | Lighthouse Laboratory Plymouth and Alex Alderton, Roberto Amato, Jeffrey Barrett, Sonia Goncalves, Ewan Harrison, David K. Jackson, Ian Johnston, Dominic Kwiatkowski, Cordelia Langford, John Sillitoe on behalf of the Wellcome Sanger Institute COVID-19 Surveillance Team |

| | | | |
|---|---|---|---|
| EPI_ISL_3170776 | Lighthouse Lab in Milton Keynes | Wellcome Sanger Institute for the COVID-19 Genomics UK (COG-UK) Consortium | The Lighthouse Lab in Milton Keynes and Alex Alderton, Roberto Amato, Jeffrey Barrett, Sonia Goncalves, Ewan Harrison, David K. Jackson, Ian Johnston, Dominic Kwiatkowski, Cordelia Langford, John Sillitoe on behalf of the Wellcome Sanger Institute COVID-19 Surveillance Team |
| EPI_ISL_3170777 | Lighthouse Laboratory Plymouth | Wellcome Sanger Institute for the COVID-19 Genomics UK (COG-UK) Consortium | Lighthouse Laboratory Plymouth and Alex Alderton, Roberto Amato, Jeffrey Barrett, Sonia Goncalves, Ewan Harrison, David K. Jackson, Ian Johnston, Dominic Kwiatkowski, Cordelia Langford, John Sillitoe on behalf of the Wellcome Sanger Institute COVID-19 Surveillance Team |
| EPI_ISL_3170778 | Lighthouse Lab in Glasgow | Wellcome Sanger Institute for the COVID-19 Genomics UK (COG-UK) Consortium | Harper VanSteenhouse, Yumi Kasai, David Gray, Carol Clugston, Anna Dominiczak and Alex Alderton, Roberto Amato, Jeffrey Barrett, Sonia Goncalves, Ewan Harrison, David K. Jackson, Ian Johnston, Dominic Kwiatkowski, Cordelia Langford, John Sillitoe on behalf of the Wellcome Sanger Institute COVID-19 Surveillance Team |
| EPI_ISL_3170779, EPI_ISL_3170780, EPI_ISL_3170781 | Lighthouse Lab in Milton Keynes | Wellcome Sanger Institute for the COVID-19 Genomics UK (COG-UK) Consortium | The Lighthouse Lab in Milton Keynes and Alex Alderton, Roberto Amato, Jeffrey Barrett, Sonia Goncalves, Ewan Harrison, David K. Jackson, Ian Johnston, Dominic Kwiatkowski, Cordelia Langford, John Sillitoe on behalf of the Wellcome Sanger Institute COVID-19 Surveillance Team |
| EPI_ISL_3170782 | Lighthouse Laboratory Plymouth | Wellcome Sanger Institute for the COVID-19 Genomics UK (COG-UK) Consortium | Lighthouse Laboratory Plymouth and Alex Alderton, Roberto Amato, Jeffrey Barrett, Sonia Goncalves, Ewan Harrison, David K. Jackson, Ian Johnston, Dominic Kwiatkowski, Cordelia Langford, John Sillitoe on behalf of the Wellcome Sanger Institute COVID-19 Surveillance Team |
| EPI_ISL_3170783, EPI_ISL_3170784, EPI_ISL_3170785, EPI_ISL_3170786, EPI_ISL_3170787, EPI_ISL_3170788, EPI_ISL_3170789 | Lighthouse Lab in Milton Keynes | Wellcome Sanger Institute for the COVID-19 Genomics UK (COG-UK) Consortium | The Lighthouse Lab in Milton Keynes and Alex Alderton, Roberto Amato, Jeffrey Barrett, Sonia Goncalves, Ewan Harrison, David K. Jackson, Ian Johnston, Dominic Kwiatkowski, Cordelia Langford, John Sillitoe on behalf of the Wellcome Sanger Institute COVID-19 Surveillance Team |
| EPI_ISL_3170790, EPI_ISL_3170791, EPI_ISL_3170792 | Lighthouse Laboratory Plymouth | Wellcome Sanger Institute for the COVID-19 Genomics UK (COG-UK) Consortium | Lighthouse Laboratory Plymouth and Alex Alderton, Roberto Amato, Jeffrey Barrett, Sonia Goncalves, Ewan Harrison, David K. Jackson, Ian Johnston, Dominic Kwiatkowski, Cordelia Langford, John Sillitoe on behalf of the Wellcome Sanger Institute COVID-19 Surveillance Team |
| EPI_ISL_3170793 | Lighthouse Lab in Glasgow | Wellcome Sanger Institute for the COVID-19 Genomics UK (COG-UK) Consortium | Harper VanSteenhouse, Yumi Kasai, David Gray, Carol Clugston, Anna Dominiczak and Alex Alderton, Roberto Amato, Jeffrey Barrett, Sonia Goncalves, Ewan Harrison, David K. Jackson, Ian Johnston, Dominic Kwiatkowski, Cordelia Langford, John Sillitoe on behalf of the Wellcome Sanger Institute COVID-19 Surveillance Team |
| EPI_ISL_3170794, EPI_ISL_3170795, EPI_ISL_3170796, EPI_ISL_3170797, EPI_ISL_3170798, EPI_ISL_3170799, EPI_ISL_3170800 | Lighthouse Lab in Milton Keynes | Wellcome Sanger Institute for the COVID-19 Genomics UK (COG-UK) Consortium | The Lighthouse Lab in Milton Keynes and Alex Alderton, Roberto Amato, Jeffrey Barrett, Sonia Goncalves, Ewan Harrison, David K. Jackson, Ian Johnston, Dominic Kwiatkowski, Cordelia Langford, John Sillitoe on behalf of the Wellcome Sanger Institute COVID-19 Surveillance Team |
| EPI_ISL_3170801 | Lighthouse Lab in Glasgow | Wellcome Sanger Institute for the COVID-19 Genomics UK (COG-UK) Consortium | Harper VanSteenhouse, Yumi Kasai, David Gray, Carol Clugston, Anna Dominiczak and Alex Alderton, Roberto Amato, Jeffrey Barrett, Sonia Goncalves, Ewan Harrison, David K. Jackson, Ian Johnston, Dominic Kwiatkowski, Cordelia Langford, John Sillitoe on behalf of the Wellcome Sanger Institute COVID-19 Surveillance Team |
| EPI_ISL_3170802 | Lighthouse Lab in Milton Keynes | Wellcome Sanger Institute for the COVID-19 Genomics UK (COG-UK) Consortium | The Lighthouse Lab in Milton Keynes and Alex Alderton, Roberto Amato, Jeffrey Barrett, Sonia Goncalves, Ewan Harrison, David K. Jackson, Ian Johnston, Dominic Kwiatkowski, Cordelia Langford, John Sillitoe on behalf of the Wellcome Sanger Institute COVID-19 Surveillance Team |
| EPI_ISL_3170803 | Lighthouse Laboratory Plymouth | Wellcome Sanger Institute for the COVID-19 Genomics UK (COG-UK) Consortium | Lighthouse Laboratory Plymouth and Alex Alderton, Roberto Amato, Jeffrey Barrett, Sonia Goncalves, Ewan Harrison, David K. Jackson, Ian Johnston, Dominic Kwiatkowski, Cordelia Langford, John Sillitoe on behalf of the Wellcome Sanger Institute COVID-19 Surveillance Team |
| EPI_ISL_3170804 | Lighthouse Lab in Milton Keynes | Wellcome Sanger Institute for the COVID-19 Genomics UK (COG-UK) Consortium | The Lighthouse Lab in Milton Keynes and Alex Alderton, Roberto Amato, Jeffrey Barrett, Sonia Goncalves, Ewan Harrison, David K. Jackson, Ian Johnston, Dominic Kwiatkowski, Cordelia Langford, John Sillitoe on behalf of the Wellcome Sanger Institute COVID-19 Surveillance Team |
| EPI_ISL_3170805, EPI_ISL_3170806 | Lighthouse Laboratory Plymouth | Wellcome Sanger Institute for the COVID-19 Genomics UK (COG-UK) Consortium | Lighthouse Laboratory Plymouth and Alex Alderton, Roberto Amato, Jeffrey Barrett, Sonia Goncalves, Ewan Harrison, David K. Jackson, Ian Johnston, Dominic Kwiatkowski, Cordelia Langford, John Sillitoe on behalf of the Wellcome Sanger Institute COVID-19 Surveillance Team |
| EPI_ISL_3170807, EPI_ISL_3170808, EPI_ISL_3170809, EPI_ISL_3170810, EPI_ISL_3170811, EPI_ISL_3170812, EPI_ISL_3170813 | Lighthouse Lab in Milton Keynes | Wellcome Sanger Institute for the COVID-19 Genomics UK (COG-UK) Consortium | The Lighthouse Lab in Milton Keynes and Alex Alderton, Roberto Amato, Jeffrey Barrett, Sonia Goncalves, Ewan Harrison, David K. Jackson, Ian Johnston, Dominic Kwiatkowski, Cordelia Langford, John Sillitoe on behalf of the Wellcome Sanger Institute COVID-19 Surveillance Team |
| EPI_ISL_3170814 | Lighthouse Laboratory Plymouth | Wellcome Sanger Institute for the COVID-19 Genomics UK (COG-UK) Consortium | Lighthouse Laboratory Plymouth and Alex Alderton, Roberto Amato, Jeffrey Barrett, Sonia Goncalves, Ewan Harrison, David K. Jackson, Ian Johnston, Dominic Kwiatkowski, Cordelia Langford, John Sillitoe on behalf of the Wellcome Sanger Institute COVID-19 Surveillance Team |
| EPI_ISL_3170815 | Lighthouse Lab in Milton Keynes | Wellcome Sanger Institute for the COVID-19 Genomics UK (COG-UK) Consortium | The Lighthouse Lab in Milton Keynes and Alex Alderton, Roberto Amato, Jeffrey Barrett, Sonia Goncalves, Ewan Harrison, David K. Jackson, Ian Johnston, Dominic Kwiatkowski, Cordelia Langford, John Sillitoe on behalf of the Wellcome Sanger Institute COVID-19 Surveillance Team |
| EPI_ISL_3170816 | Lighthouse Laboratory Plymouth | Wellcome Sanger Institute for the COVID-19 Genomics UK (COG-UK) Consortium | Lighthouse Laboratory Plymouth and Alex Alderton, Roberto Amato, Jeffrey Barrett, Sonia Goncalves, Ewan Harrison, David K. Jackson, Ian Johnston, Dominic Kwiatkowski, Cordelia Langford, John Sillitoe on behalf of the Wellcome Sanger Institute COVID-19 Surveillance Team |
| EPI_ISL_3170817 | Lighthouse Lab in Glasgow | Wellcome Sanger Institute for the COVID-19 Genomics UK (COG-UK) Consortium | Harper VanSteenhouse, Yumi Kasai, David Gray, Carol Clugston, Anna Dominiczak and Alex Alderton, Roberto Amato, Jeffrey Barrett, Sonia Goncalves, Ewan Harrison, David K. Jackson, Ian Johnston, Dominic Kwiatkowski, Cordelia Langford, John Sillitoe on behalf of the Wellcome Sanger Institute COVID-19 Surveillance Team |
| EPI_ISL_3170818 | Lighthouse Lab in Milton Keynes | Wellcome Sanger Institute for the COVID-19 Genomics UK (COG-UK) Consortium | The Lighthouse Lab in Milton Keynes and Alex Alderton, Roberto Amato, Jeffrey Barrett, Sonia Goncalves, Ewan Harrison, David K. Jackson, Ian Johnston, Dominic Kwiatkowski, Cordelia Langford, John Sillitoe on behalf of the Wellcome Sanger Institute COVID-19 Surveillance Team |
| EPI_ISL_3170819 | Lighthouse Laboratory Plymouth | Wellcome Sanger Institute for the COVID-19 Genomics UK (COG-UK) Consortium | Lighthouse Laboratory Plymouth and Alex Alderton, Roberto Amato, Jeffrey Barrett, Sonia Goncalves, Ewan Harrison, David K. Jackson, Ian Johnston, Dominic Kwiatkowski, Cordelia Langford, John Sillitoe on behalf of the Wellcome Sanger Institute COVID-19 Surveillance Team |
| EPI_ISL_3170820 | Lighthouse Lab in Glasgow | Wellcome Sanger Institute for the COVID-19 Genomics UK (COG-UK) Consortium | Harper VanSteenhouse, Yumi Kasai, David Gray, Carol Clugston, Anna Dominiczak and Alex Alderton, Roberto Amato, Jeffrey Barrett, Sonia Goncalves, Ewan Harrison, David K. Jackson, Ian Johnston, Dominic Kwiatkowski, Cordelia Langford, John Sillitoe on behalf of the Wellcome Sanger Institute COVID-19 Surveillance Team |
| EPI_ISL_3170821 | Lighthouse Laboratory Plymouth | Wellcome Sanger Institute for the COVID-19 Genomics UK (COG-UK) Consortium | Lighthouse Laboratory Plymouth and Alex Alderton, Roberto Amato, Jeffrey Barrett, Sonia Goncalves, Ewan Harrison, David K. Jackson, Ian Johnston, Dominic Kwiatkowski, Cordelia Langford, John Sillitoe on behalf of the Wellcome Sanger Institute COVID-19 Surveillance Team |
| EPI_ISL_3170822, EPI_ISL_3170823, EPI_ISL_3170824 | Lighthouse Lab in Milton Keynes | Wellcome Sanger Institute for the COVID-19 Genomics UK (COG-UK) Consortium | The Lighthouse Lab in Milton Keynes and Alex Alderton, Roberto Amato, Jeffrey Barrett, Sonia Goncalves, Ewan Harrison, David K. Jackson, Ian Johnston, Dominic Kwiatkowski, Cordelia Langford, John Sillitoe on behalf of the Wellcome Sanger Institute COVID-19 Surveillance Team |
| EPI_ISL_3170825 | Lighthouse Lab in Glasgow | Wellcome Sanger Institute for the COVID-19 Genomics UK (COG-UK) Consortium | Harper VanSteenhouse, Yumi Kasai, David Gray, Carol Clugston, Anna Dominiczak and Alex Alderton, Roberto Amato, Jeffrey Barrett, Sonia Goncalves, Ewan Harrison, David K. Jackson, Ian Johnston, Dominic Kwiatkowski, Cordelia Langford, John Sillitoe on behalf of the Wellcome Sanger Institute COVID-19 Surveillance Team |
| EPI_ISL_3170826 | Lighthouse Lab in Milton Keynes | Wellcome Sanger Institute for the COVID-19 Genomics UK (COG-UK) Consortium | The Lighthouse Lab in Milton Keynes and Alex Alderton, Roberto Amato, Jeffrey Barrett, Sonia Goncalves, Ewan Harrison, David K. Jackson, Ian Johnston, Dominic Kwiatkowski, Cordelia Langford, John Sillitoe on behalf of the Wellcome Sanger Institute COVID-19 Surveillance Team |
| EPI_ISL_3170827, EPI_ISL_3170828 | Lighthouse Lab in Glasgow | Wellcome Sanger Institute for the COVID-19 Genomics UK (COG-UK) Consortium | Harper VanSteenhouse, Yumi Kasai, David Gray, Carol Clugston, Anna Dominiczak and Alex Alderton, Roberto Amato, Jeffrey Barrett, Sonia Goncalves, Ewan Harrison, David K. Jackson, Ian Johnston, Dominic Kwiatkowski, Cordelia Langford, John Sillitoe on behalf of the Wellcome Sanger Institute COVID-19 Surveillance Team |
| EPI_ISL_3170829 | Lighthouse Lab in Milton Keynes | Wellcome Sanger Institute for the COVID-19 Genomics UK (COG-UK) Consortium | The Lighthouse Lab in Milton Keynes and Alex Alderton, Roberto Amato, Jeffrey Barrett, Sonia Goncalves, Ewan Harrison, David K. Jackson, Ian Johnston, Dominic Kwiatkowski, Cordelia Langford, John Sillitoe on behalf of the Wellcome Sanger Institute COVID-19 Surveillance Team |
| EPI_ISL_3170830 | Lighthouse Lab in Glasgow | Wellcome Sanger Institute for the COVID-19 Genomics UK (COG-UK) Consortium | Harper VanSteenhouse, Yumi Kasai, David Gray, Carol Clugston, Anna Dominiczak and Alex Alderton, Roberto Amato, Jeffrey Barrett, Sonia Goncalves, Ewan Harrison, David K. Jackson, Ian Johnston, Dominic Kwiatkowski, Cordelia Langford, John Sillitoe on behalf of the Wellcome Sanger Institute COVID-19 Surveillance Team |
| EPI_ISL_3170831, EPI_ISL_3170832 | Lighthouse Laboratory Plymouth | Wellcome Sanger Institute for the COVID-19 Genomics UK (COG-UK) Consortium | Lighthouse Laboratory Plymouth and Alex Alderton, Roberto Amato, Jeffrey Barrett, Sonia Goncalves, Ewan Harrison, David K. Jackson, Ian Johnston, Dominic Kwiatkowski, Cordelia Langford, John Sillitoe on behalf of the Wellcome Sanger Institute COVID-19 Surveillance Team |
| EPI_ISL_3170833 | Lighthouse Lab in Glasgow | Wellcome Sanger Institute for the COVID-19 Genomics UK (COG-UK) Consortium | Harper VanSteenhouse, Yumi Kasai, David Gray, Carol Clugston, Anna Dominiczak and Alex Alderton, Roberto Amato, Jeffrey Barrett, Sonia Goncalves, Ewan Harrison, David K. Jackson, Ian Johnston, Dominic Kwiatkowski, Cordelia Langford, John Sillitoe on behalf of the Wellcome Sanger Institute COVID-19 Surveillance Team |
| EPI_ISL_3170834, EPI_ISL_3170835 | Lighthouse Laboratory Plymouth | Wellcome Sanger Institute for the COVID-19 Genomics UK (COG-UK) Consortium | Lighthouse Laboratory Plymouth and Alex Alderton, Roberto Amato, Jeffrey Barrett, Sonia Goncalves, Ewan Harrison, David K. Jackson, Ian Johnston, Dominic Kwiatkowski, Cordelia Langford, John Sillitoe on behalf of the Wellcome Sanger Institute COVID-19 Surveillance Team |
| EPI_ISL_3170836 | Lighthouse Lab in Milton Keynes | Wellcome Sanger Institute for the COVID-19 Genomics UK (COG-UK) Consortium | The Lighthouse Lab in Milton Keynes and Alex Alderton, Roberto Amato, Jeffrey Barrett, Sonia Goncalves, Ewan Harrison, David K. Jackson, Ian Johnston, Dominic Kwiatkowski, Cordelia Langford, John Sillitoe on behalf of the Wellcome Sanger Institute COVID-19 Surveillance Team |
| EPI_ISL_3170837 | Lighthouse Lab in Glasgow | Wellcome Sanger Institute for the COVID-19 Genomics UK (COG-UK) Consortium | Harper VanSteenhouse, Yumi Kasai, David Gray, Carol Clugston, Anna Dominiczak and Alex Alderton, Roberto Amato, Jeffrey Barrett, Sonia Goncalves, Ewan Harrison, David K. Jackson, Ian Johnston, Dominic Kwiatkowski, Cordelia Langford, John Sillitoe on behalf of the Wellcome Sanger Institute COVID-19 Surveillance Team |
| EPI_ISL_3170838, EPI_ISL_3170839, EPI_ISL_3170840, EPI_ISL_3170841 | Lighthouse Lab in Milton Keynes | Wellcome Sanger Institute for the COVID-19 Genomics UK (COG-UK) Consortium | The Lighthouse Lab in Milton Keynes and Alex Alderton, Roberto Amato, Jeffrey Barrett, Sonia Goncalves, Ewan Harrison, David K. Jackson, Ian Johnston, Dominic Kwiatkowski, Cordelia Langford, John Sillitoe on behalf of the Wellcome Sanger Institute COVID-19 Surveillance Team |
| EPI_ISL_3170842 | Lighthouse Lab in Glasgow | Wellcome Sanger Institute for the COVID-19 Genomics UK (COG-UK) Consortium | Harper VanSteenhouse, Yumi Kasai, David Gray, Carol Clugston, Anna Dominiczak and Alex Alderton, Roberto Amato, Jeffrey Barrett, Sonia Goncalves, Ewan Harrison, David K. Jackson, Ian Johnston, Dominic Kwiatkowski, Cordelia Langford, John Sillitoe on behalf of the Wellcome Sanger Institute COVID-19 Surveillance Team |
| EPI_ISL_3170843 | Lighthouse Lab in Milton Keynes | Wellcome Sanger Institute for the COVID-19 Genomics UK (COG-UK) Consortium | The Lighthouse Lab in Milton Keynes and Alex Alderton, Roberto Amato, Jeffrey Barrett, Sonia Goncalves, Ewan Harrison, David K. Jackson, Ian Johnston, Dominic Kwiatkowski, Cordelia Langford, John Sillitoe on behalf of the Wellcome Sanger Institute COVID-19 Surveillance Team |

| | | | |
|---|---|---|---|
| EPI_ISL_3170844 | Lighthouse Laboratory Plymouth | Wellcome Sanger Institute for the COVID-19 Genomics UK (COG-UK) Consortium | Lighthouse Laboratory Plymouth and Alex Alderton, Roberto Amato, Jeffrey Barrett, Sonia Goncalves, Ewan Harrison, David K. Jackson, Ian Johnston, Dominic Kwiatkowski, Cordelia Langford, John Sillitoe on behalf of the Wellcome Sanger Institute COVID-19 Surveillance Team |
| EPI_ISL_3170845 | Lighthouse Lab in Glasgow | Wellcome Sanger Institute for the COVID-19 Genomics UK (COG-UK) Consortium | Harper VanSteenhouse, Yumi Kasai, David Gray, Carol Clugston, Anna Dominiczak and Alex Alderton, Roberto Amato, Jeffrey Barrett, Sonia Goncalves, Ewan Harrison, David K. Jackson, Ian Johnston, Dominic Kwiatkowski, Cordelia Langford, John Sillitoe on behalf of the Wellcome Sanger Institute COVID-19 Surveillance Team |
| EPI_ISL_3170846 | Lighthouse Laboratory Plymouth | Wellcome Sanger Institute for the COVID-19 Genomics UK (COG-UK) Consortium | Lighthouse Laboratory Plymouth and Alex Alderton, Roberto Amato, Jeffrey Barrett, Sonia Goncalves, Ewan Harrison, David K. Jackson, Ian Johnston, Dominic Kwiatkowski, Cordelia Langford, John Sillitoe on behalf of the Wellcome Sanger Institute COVID-19 Surveillance Team |
| EPI_ISL_3170847 | Lighthouse Lab in Milton Keynes | Wellcome Sanger Institute for the COVID-19 Genomics UK (COG-UK) Consortium | The Lighthouse Lab in Milton Keynes and Alex Alderton, Roberto Amato, Jeffrey Barrett, Sonia Goncalves, Ewan Harrison, David K. Jackson, Ian Johnston, Dominic Kwiatkowski, Cordelia Langford, John Sillitoe on behalf of the Wellcome Sanger Institute COVID-19 Surveillance Team |
| EPI_ISL_3170848 | Lighthouse Laboratory Plymouth | Wellcome Sanger Institute for the COVID-19 Genomics UK (COG-UK) Consortium | Lighthouse Laboratory Plymouth and Alex Alderton, Roberto Amato, Jeffrey Barrett, Sonia Goncalves, Ewan Harrison, David K. Jackson, Ian Johnston, Dominic Kwiatkowski, Cordelia Langford, John Sillitoe on behalf of the Wellcome Sanger Institute COVID-19 Surveillance Team |
| EPI_ISL_3170849, EPI_ISL_3170850 | Lighthouse Lab in Milton Keynes | Wellcome Sanger Institute for the COVID-19 Genomics UK (COG-UK) Consortium | The Lighthouse Lab in Milton Keynes and Alex Alderton, Roberto Amato, Jeffrey Barrett, Sonia Goncalves, Ewan Harrison, David K. Jackson, Ian Johnston, Dominic Kwiatkowski, Cordelia Langford, John Sillitoe on behalf of the Wellcome Sanger Institute COVID-19 Surveillance Team |
| EPI_ISL_3170851 | Lighthouse Laboratory Plymouth | Wellcome Sanger Institute for the COVID-19 Genomics UK (COG-UK) Consortium | Lighthouse Laboratory Plymouth and Alex Alderton, Roberto Amato, Jeffrey Barrett, Sonia Goncalves, Ewan Harrison, David K. Jackson, Ian Johnston, Dominic Kwiatkowski, Cordelia Langford, John Sillitoe on behalf of the Wellcome Sanger Institute COVID-19 Surveillance Team |
| EPI_ISL_3170852, EPI_ISL_3170853, EPI_ISL_3170854, EPI_ISL_3170855, EPI_ISL_3170856 | Lighthouse Lab in Milton Keynes | Wellcome Sanger Institute for the COVID-19 Genomics UK (COG-UK) Consortium | The Lighthouse Lab in Milton Keynes and Alex Alderton, Roberto Amato, Jeffrey Barrett, Sonia Goncalves, Ewan Harrison, David K. Jackson, Ian Johnston, Dominic Kwiatkowski, Cordelia Langford, John Sillitoe on behalf of the Wellcome Sanger Institute COVID-19 Surveillance Team |
| EPI_ISL_3170857 | Lighthouse Laboratory Plymouth | Wellcome Sanger Institute for the COVID-19 Genomics UK (COG-UK) Consortium | Lighthouse Laboratory Plymouth and Alex Alderton, Roberto Amato, Jeffrey Barrett, Sonia Goncalves, Ewan Harrison, David K. Jackson, Ian Johnston, Dominic Kwiatkowski, Cordelia Langford, John Sillitoe on behalf of the Wellcome Sanger Institute COVID-19 Surveillance Team |
| EPI_ISL_3170858 | Lighthouse Lab in Milton Keynes | Wellcome Sanger Institute for the COVID-19 Genomics UK (COG-UK) Consortium | The Lighthouse Lab in Milton Keynes and Alex Alderton, Roberto Amato, Jeffrey Barrett, Sonia Goncalves, Ewan Harrison, David K. Jackson, Ian Johnston, Dominic Kwiatkowski, Cordelia Langford, John Sillitoe on behalf of the Wellcome Sanger Institute COVID-19 Surveillance Team |
| EPI_ISL_3170859, EPI_ISL_3170860 | Lighthouse Laboratory Plymouth | Wellcome Sanger Institute for the COVID-19 Genomics UK (COG-UK) Consortium | Lighthouse Laboratory Plymouth and Alex Alderton, Roberto Amato, Jeffrey Barrett, Sonia Goncalves, Ewan Harrison, David K. Jackson, Ian Johnston, Dominic Kwiatkowski, Cordelia Langford, John Sillitoe on behalf of the Wellcome Sanger Institute COVID-19 Surveillance Team |
| EPI_ISL_3170861 | Lighthouse Lab in Milton Keynes | Wellcome Sanger Institute for the COVID-19 Genomics UK (COG-UK) Consortium | The Lighthouse Lab in Milton Keynes and Alex Alderton, Roberto Amato, Jeffrey Barrett, Sonia Goncalves, Ewan Harrison, David K. Jackson, Ian Johnston, Dominic Kwiatkowski, Cordelia Langford, John Sillitoe on behalf of the Wellcome Sanger Institute COVID-19 Surveillance Team |
| EPI_ISL_3170862 | Lighthouse Laboratory Plymouth | Wellcome Sanger Institute for the COVID-19 Genomics UK (COG-UK) Consortium | Lighthouse Laboratory Plymouth and Alex Alderton, Roberto Amato, Jeffrey Barrett, Sonia Goncalves, Ewan Harrison, David K. Jackson, Ian Johnston, Dominic Kwiatkowski, Cordelia Langford, John Sillitoe on behalf of the Wellcome Sanger Institute COVID-19 Surveillance Team |
| EPI_ISL_3170863 | Lighthouse Lab in Milton Keynes | Wellcome Sanger Institute for the COVID-19 Genomics UK (COG-UK) Consortium | The Lighthouse Lab in Milton Keynes and Alex Alderton, Roberto Amato, Jeffrey Barrett, Sonia Goncalves, Ewan Harrison, David K. Jackson, Ian Johnston, Dominic Kwiatkowski, Cordelia Langford, John Sillitoe on behalf of the Wellcome Sanger Institute COVID-19 Surveillance Team |
| EPI_ISL_3170864, EPI_ISL_3170865, EPI_ISL_3170866 | Lighthouse Laboratory Plymouth | Wellcome Sanger Institute for the COVID-19 Genomics UK (COG-UK) Consortium | Lighthouse Laboratory Plymouth and Alex Alderton, Roberto Amato, Jeffrey Barrett, Sonia Goncalves, Ewan Harrison, David K. Jackson, Ian Johnston, Dominic Kwiatkowski, Cordelia Langford, John Sillitoe on behalf of the Wellcome Sanger Institute COVID-19 Surveillance Team |
| EPI_ISL_3170867 | Lighthouse Lab in Milton Keynes | Wellcome Sanger Institute for the COVID-19 Genomics UK (COG-UK) Consortium | The Lighthouse Lab in Milton Keynes and Alex Alderton, Roberto Amato, Jeffrey Barrett, Sonia Goncalves, Ewan Harrison, David K. Jackson, Ian Johnston, Dominic Kwiatkowski, Cordelia Langford, John Sillitoe on behalf of the Wellcome Sanger Institute COVID-19 Surveillance Team |
| EPI_ISL_3170868 | Lighthouse Laboratory Plymouth | Wellcome Sanger Institute for the COVID-19 Genomics UK (COG-UK) Consortium | Lighthouse Laboratory Plymouth and Alex Alderton, Roberto Amato, Jeffrey Barrett, Sonia Goncalves, Ewan Harrison, David K. Jackson, Ian Johnston, Dominic Kwiatkowski, Cordelia Langford, John Sillitoe on behalf of the Wellcome Sanger Institute COVID-19 Surveillance Team |
| EPI_ISL_3170869, EPI_ISL_3170870 | Lighthouse Lab in Milton Keynes | Wellcome Sanger Institute for the COVID-19 Genomics UK (COG-UK) Consortium | The Lighthouse Lab in Milton Keynes and Alex Alderton, Roberto Amato, Jeffrey Barrett, Sonia Goncalves, Ewan Harrison, David K. Jackson, Ian Johnston, Dominic Kwiatkowski, Cordelia Langford, John Sillitoe on behalf of the Wellcome Sanger Institute COVID-19 Surveillance Team |
| EPI_ISL_3170871 | Lighthouse Laboratory Plymouth | Wellcome Sanger Institute for the COVID-19 Genomics UK (COG-UK) Consortium | Lighthouse Laboratory Plymouth and Alex Alderton, Roberto Amato, Jeffrey Barrett, Sonia Goncalves, Ewan Harrison, David K. Jackson, Ian Johnston, Dominic Kwiatkowski, Cordelia Langford, John Sillitoe on behalf of the Wellcome Sanger Institute COVID-19 Surveillance Team |
| EPI_ISL_3170872, EPI_ISL_3170873, EPI_ISL_3170874, EPI_ISL_3170875, EPI_ISL_3170876, EPI_ISL_3170877 | Lighthouse Lab in Milton Keynes | Wellcome Sanger Institute for the COVID-19 Genomics UK (COG-UK) Consortium | The Lighthouse Lab in Milton Keynes and Alex Alderton, Roberto Amato, Jeffrey Barrett, Sonia Goncalves, Ewan Harrison, David K. Jackson, Ian Johnston, Dominic Kwiatkowski, Cordelia Langford, John Sillitoe on behalf of the Wellcome Sanger Institute COVID-19 Surveillance Team |
| EPI_ISL_3170878 | Lighthouse Lab in Glasgow | Wellcome Sanger Institute for the COVID-19 Genomics UK (COG-UK) Consortium | Harper VanSteenhouse, Yumi Kasai, David Gray, Carol Clugston, Anna Dominiczak and Alex Alderton, Roberto Amato, Jeffrey Barrett, Sonia Goncalves, Ewan Harrison, David K. Jackson, Ian Johnston, Dominic Kwiatkowski, Cordelia Langford, John Sillitoe on behalf of the Wellcome Sanger Institute COVID-19 Surveillance Team |
| EPI_ISL_3170879, EPI_ISL_3170880 | Lighthouse Lab in Milton Keynes | Wellcome Sanger Institute for the COVID-19 Genomics UK (COG-UK) Consortium | The Lighthouse Lab in Milton Keynes and Alex Alderton, Roberto Amato, Jeffrey Barrett, Sonia Goncalves, Ewan Harrison, David K. Jackson, Ian Johnston, Dominic Kwiatkowski, Cordelia Langford, John Sillitoe on behalf of the Wellcome Sanger Institute COVID-19 Surveillance Team |
| EPI_ISL_3170881 | Lighthouse Lab in Glasgow | Wellcome Sanger Institute for the COVID-19 Genomics UK (COG-UK) Consortium | Harper VanSteenhouse, Yumi Kasai, David Gray, Carol Clugston, Anna Dominiczak and Alex Alderton, Roberto Amato, Jeffrey Barrett, Sonia Goncalves, Ewan Harrison, David K. Jackson, Ian Johnston, Dominic Kwiatkowski, Cordelia Langford, John Sillitoe on behalf of the Wellcome Sanger Institute COVID-19 Surveillance Team |
| EPI_ISL_3170882, EPI_ISL_3170883, EPI_ISL_3170884 | Lighthouse Lab in Milton Keynes | Wellcome Sanger Institute for the COVID-19 Genomics UK (COG-UK) Consortium | The Lighthouse Lab in Milton Keynes and Alex Alderton, Roberto Amato, Jeffrey Barrett, Sonia Goncalves, Ewan Harrison, David K. Jackson, Ian Johnston, Dominic Kwiatkowski, Cordelia Langford, John Sillitoe on behalf of the Wellcome Sanger Institute COVID-19 Surveillance Team |
| EPI_ISL_3170885 | Lighthouse Laboratory Plymouth | Wellcome Sanger Institute for the COVID-19 Genomics UK (COG-UK) Consortium | Lighthouse Laboratory Plymouth and Alex Alderton, Roberto Amato, Jeffrey Barrett, Sonia Goncalves, Ewan Harrison, David K. Jackson, Ian Johnston, Dominic Kwiatkowski, Cordelia Langford, John Sillitoe on behalf of the Wellcome Sanger Institute COVID-19 Surveillance Team |
| EPI_ISL_3170886, EPI_ISL_3170887, EPI_ISL_3170888 | Lighthouse Lab in Milton Keynes | Wellcome Sanger Institute for the COVID-19 Genomics UK (COG-UK) Consortium | The Lighthouse Lab in Milton Keynes and Alex Alderton, Roberto Amato, Jeffrey Barrett, Sonia Goncalves, Ewan Harrison, David K. Jackson, Ian Johnston, Dominic Kwiatkowski, Cordelia Langford, John Sillitoe on behalf of the Wellcome Sanger Institute COVID-19 Surveillance Team |
| EPI_ISL_3170889 | Lighthouse Laboratory Plymouth | Wellcome Sanger Institute for the COVID-19 Genomics UK (COG-UK) Consortium | Lighthouse Laboratory Plymouth and Alex Alderton, Roberto Amato, Jeffrey Barrett, Sonia Goncalves, Ewan Harrison, David K. Jackson, Ian Johnston, Dominic Kwiatkowski, Cordelia Langford, John Sillitoe on behalf of the Wellcome Sanger Institute COVID-19 Surveillance Team |
| EPI_ISL_3170890, EPI_ISL_3170891 | Lighthouse Lab in Glasgow | Wellcome Sanger Institute for the COVID-19 Genomics UK (COG-UK) Consortium | Harper VanSteenhouse, Yumi Kasai, David Gray, Carol Clugston, Anna Dominiczak and Alex Alderton, Roberto Amato, Jeffrey Barrett, Sonia Goncalves, Ewan Harrison, David K. Jackson, Ian Johnston, Dominic Kwiatkowski, Cordelia Langford, John Sillitoe on behalf of the Wellcome Sanger Institute COVID-19 Surveillance Team |
| EPI_ISL_3170892 | Lighthouse Laboratory Plymouth | Wellcome Sanger Institute for the COVID-19 Genomics UK (COG-UK) Consortium | Lighthouse Laboratory Plymouth and Alex Alderton, Roberto Amato, Jeffrey Barrett, Sonia Goncalves, Ewan Harrison, David K. Jackson, Ian Johnston, Dominic Kwiatkowski, Cordelia Langford, John Sillitoe on behalf of the Wellcome Sanger Institute COVID-19 Surveillance Team |
| EPI_ISL_3170893 | Lighthouse Lab in Milton Keynes | Wellcome Sanger Institute for the COVID-19 Genomics UK (COG-UK) Consortium | The Lighthouse Lab in Milton Keynes and Alex Alderton, Roberto Amato, Jeffrey Barrett, Sonia Goncalves, Ewan Harrison, David K. Jackson, Ian Johnston, Dominic Kwiatkowski, Cordelia Langford, John Sillitoe on behalf of the Wellcome Sanger Institute COVID-19 Surveillance Team |
| EPI_ISL_3170894 | Lighthouse Laboratory Plymouth | Wellcome Sanger Institute for the COVID-19 Genomics UK (COG-UK) Consortium | Lighthouse Laboratory Plymouth and Alex Alderton, Roberto Amato, Jeffrey Barrett, Sonia Goncalves, Ewan Harrison, David K. Jackson, Ian Johnston, Dominic Kwiatkowski, Cordelia Langford, John Sillitoe on behalf of the Wellcome Sanger Institute COVID-19 Surveillance Team |
| EPI_ISL_3170895 | Lighthouse Lab in Milton Keynes | Wellcome Sanger Institute for the COVID-19 Genomics UK (COG-UK) Consortium | The Lighthouse Lab in Milton Keynes and Alex Alderton, Roberto Amato, Jeffrey Barrett, Sonia Goncalves, Ewan Harrison, David K. Jackson, Ian Johnston, Dominic Kwiatkowski, Cordelia Langford, John Sillitoe on behalf of the Wellcome Sanger Institute COVID-19 Surveillance Team |
| EPI_ISL_3170896, EPI_ISL_3170897 | Lighthouse Laboratory Plymouth | Wellcome Sanger Institute for the COVID-19 Genomics UK (COG-UK) Consortium | Lighthouse Laboratory Plymouth and Alex Alderton, Roberto Amato, Jeffrey Barrett, Sonia Goncalves, Ewan Harrison, David K. Jackson, Ian Johnston, Dominic Kwiatkowski, Cordelia Langford, John Sillitoe on behalf of the Wellcome Sanger Institute COVID-19 Surveillance Team |
| EPI_ISL_3170898 | Lighthouse Lab in Glasgow | Wellcome Sanger Institute for the COVID-19 Genomics UK (COG-UK) Consortium | Harper VanSteenhouse, Yumi Kasai, David Gray, Carol Clugston, Anna Dominiczak and Alex Alderton, Roberto Amato, Jeffrey Barrett, Sonia Goncalves, Ewan Harrison, David K. Jackson, Ian Johnston, Dominic Kwiatkowski, Cordelia Langford, John Sillitoe on behalf of the Wellcome Sanger Institute COVID-19 Surveillance Team |
| EPI_ISL_3170899 | Lighthouse Lab in Milton Keynes | Wellcome Sanger Institute for the COVID-19 Genomics UK (COG-UK) Consortium | The Lighthouse Lab in Milton Keynes and Alex Alderton, Roberto Amato, Jeffrey Barrett, Sonia Goncalves, Ewan Harrison, David K. Jackson, Ian Johnston, Dominic Kwiatkowski, Cordelia Langford, John Sillitoe on behalf of the Wellcome Sanger Institute COVID-19 Surveillance Team |
| EPI_ISL_3170900, EPI_ISL_3170901 | Lighthouse Lab in Glasgow | Wellcome Sanger Institute for the COVID-19 Genomics UK (COG-UK) Consortium | Harper VanSteenhouse, Yumi Kasai, David Gray, Carol Clugston, Anna Dominiczak and Alex Alderton, Roberto Amato, Jeffrey Barrett, Sonia Goncalves, Ewan Harrison, David K. Jackson, Ian Johnston, Dominic Kwiatkowski, Cordelia Langford, John Sillitoe on behalf of the Wellcome Sanger Institute COVID-19 Surveillance Team |
| EPI_ISL_3170902, EPI_ISL_3170903, EPI_ISL_3170904, EPI_ISL_3170905 | Lighthouse Lab in Milton Keynes | Wellcome Sanger Institute for the COVID-19 Genomics UK (COG-UK) Consortium | The Lighthouse Lab in Milton Keynes and Alex Alderton, Roberto Amato, Jeffrey Barrett, Sonia Goncalves, Ewan Harrison, David K. Jackson, Ian Johnston, Dominic Kwiatkowski, Cordelia Langford, John Sillitoe on behalf of the Wellcome Sanger Institute COVID-19 Surveillance Team |
| EPI_ISL_3170906, EPI_ISL_3170907, EPI_ISL_3170908, EPI_ISL_3170909, EPI_ISL_3170910 | Lighthouse Laboratory Plymouth | Wellcome Sanger Institute for the COVID-19 Genomics UK (COG-UK) Consortium | Lighthouse Laboratory Plymouth and Alex Alderton, Roberto Amato, Jeffrey Barrett, Sonia Goncalves, Ewan Harrison, David K. Jackson, Ian Johnston, Dominic Kwiatkowski, Cordelia Langford, John Sillitoe on behalf of the Wellcome Sanger Institute COVID-19 Surveillance Team |
| EPI_ISL_3170911 | Lighthouse Lab in Milton Keynes | Wellcome Sanger Institute for the COVID-19 Genomics UK (COG-UK) Consortium | The Lighthouse Lab in Milton Keynes and Alex Alderton, Roberto Amato, Jeffrey Barrett, Sonia Goncalves, Ewan Harrison, David K. Jackson, Ian Johnston, Dominic Kwiatkowski, Cordelia Langford, John Sillitoe on behalf of the Wellcome Sanger Institute COVID-19 Surveillance Team |

| | | | |
|---|---|---|---|
| EPI_ISL_3170912 | Lighthouse Laboratory Plymouth | Wellcome Sanger Institute for the COVID-19 Genomics UK (COG-UK) Consortium | Lighthouse Laboratory Plymouth and Alex Alderton, Roberto Amato, Jeffrey Barrett, Sonia Goncalves, Ewan Harrison, David K. Jackson, Ian Johnston, Dominic Kwiatkowski, Cordelia Langford, John Sillitoe on behalf of the Wellcome Sanger Institute COVID-19 Surveillance Team |
| EPI_ISL_3170913, EPI_ISL_3170914 | Lighthouse Lab in Milton Keynes | Wellcome Sanger Institute for the COVID-19 Genomics UK (COG-UK) Consortium | The Lighthouse Lab in Milton Keynes and Alex Alderton, Roberto Amato, Jeffrey Barrett, Sonia Goncalves, Ewan Harrison, David K. Jackson, Ian Johnston, Dominic Kwiatkowski, Cordelia Langford, John Sillitoe on behalf of the Wellcome Sanger Institute COVID-19 Surveillance Team |
| EPI_ISL_3170915 | Lighthouse Laboratory Plymouth | Wellcome Sanger Institute for the COVID-19 Genomics UK (COG-UK) Consortium | Lighthouse Laboratory Plymouth and Alex Alderton, Roberto Amato, Jeffrey Barrett, Sonia Goncalves, Ewan Harrison, David K. Jackson, Ian Johnston, Dominic Kwiatkowski, Cordelia Langford, John Sillitoe on behalf of the Wellcome Sanger Institute COVID-19 Surveillance Team |
| EPI_ISL_3170916, EPI_ISL_3170917, EPI_ISL_3170918, EPI_ISL_3170919, EPI_ISL_3170920, EPI_ISL_3170921, EPI_ISL_3170922 | Lighthouse Lab in Milton Keynes | Wellcome Sanger Institute for the COVID-19 Genomics UK (COG-UK) Consortium | The Lighthouse Lab in Milton Keynes and Alex Alderton, Roberto Amato, Jeffrey Barrett, Sonia Goncalves, Ewan Harrison, David K. Jackson, Ian Johnston, Dominic Kwiatkowski, Cordelia Langford, John Sillitoe on behalf of the Wellcome Sanger Institute COVID-19 Surveillance Team |
| EPI_ISL_3170923, EPI_ISL_3170924 | Lighthouse Lab in Glasgow | Wellcome Sanger Institute for the COVID-19 Genomics UK (COG-UK) Consortium | Harper VanSteenhouse, Yumi Kasai, David Gray, Carol Clugston, Anna Dominiczak and Alex Alderton, Roberto Amato, Jeffrey Barrett, Sonia Goncalves, Ewan Harrison, David K. Jackson, Ian Johnston, Dominic Kwiatkowski, Cordelia Langford, John Sillitoe on behalf of the Wellcome Sanger Institute COVID-19 Surveillance Team |
| EPI_ISL_3170925 | Lighthouse Laboratory Plymouth | Wellcome Sanger Institute for the COVID-19 Genomics UK (COG-UK) Consortium | Lighthouse Laboratory Plymouth and Alex Alderton, Roberto Amato, Jeffrey Barrett, Sonia Goncalves, Ewan Harrison, David K. Jackson, Ian Johnston, Dominic Kwiatkowski, Cordelia Langford, John Sillitoe on behalf of the Wellcome Sanger Institute COVID-19 Surveillance Team |
| EPI_ISL_3170926 | Lighthouse Lab in Glasgow | Wellcome Sanger Institute for the COVID-19 Genomics UK (COG-UK) Consortium | Harper VanSteenhouse, Yumi Kasai, David Gray, Carol Clugston, Anna Dominiczak and Alex Alderton, Roberto Amato, Jeffrey Barrett, Sonia Goncalves, Ewan Harrison, David K. Jackson, Ian Johnston, Dominic Kwiatkowski, Cordelia Langford, John Sillitoe on behalf of the Wellcome Sanger Institute COVID-19 Surveillance Team |
| EPI_ISL_3170927 | Lighthouse Lab in Milton Keynes | Wellcome Sanger Institute for the COVID-19 Genomics UK (COG-UK) Consortium | The Lighthouse Lab in Milton Keynes and Alex Alderton, Roberto Amato, Jeffrey Barrett, Sonia Goncalves, Ewan Harrison, David K. Jackson, Ian Johnston, Dominic Kwiatkowski, Cordelia Langford, John Sillitoe on behalf of the Wellcome Sanger Institute COVID-19 Surveillance Team |
| EPI_ISL_3170928 | Lighthouse Lab in Glasgow | Wellcome Sanger Institute for the COVID-19 Genomics UK (COG-UK) Consortium | Harper VanSteenhouse, Yumi Kasai, David Gray, Carol Clugston, Anna Dominiczak and Alex Alderton, Roberto Amato, Jeffrey Barrett, Sonia Goncalves, Ewan Harrison, David K. Jackson, Ian Johnston, Dominic Kwiatkowski, Cordelia Langford, John Sillitoe on behalf of the Wellcome Sanger Institute COVID-19 Surveillance Team |
| EPI_ISL_3170929, EPI_ISL_3170930, EPI_ISL_3170931, EPI_ISL_3170932, EPI_ISL_3170933, EPI_ISL_3170934, EPI_ISL_3170935, EPI_ISL_3170936, EPI_ISL_3170937, EPI_ISL_3170938, EPI_ISL_3170939, EPI_ISL_3170940, EPI_ISL_3170941, EPI_ISL_3170942, EPI_ISL_3170943, EPI_ISL_3170944, EPI_ISL_3170945, EPI_ISL_3170946, EPI_ISL_3170947, EPI_ISL_3170948, EPI_ISL_3170949, EPI_ISL_3170950, EPI_ISL_3170951, EPI_ISL_3170952, EPI_ISL_3170953, EPI_ISL_3170954, EPI_ISL_3170955, EPI_ISL_3170956, EPI_ISL_3170957, EPI_ISL_3170958, EPI_ISL_3170959, EPI_ISL_3170960, EPI_ISL_3170961, EPI_ISL_3170962, EPI_ISL_3170963, EPI_ISL_3170964, EPI_ISL_3170965, EPI_ISL_3170966, EPI_ISL_3170967, EPI_ISL_3170968, EPI_ISL_3170969, EPI_ISL_3170970, EPI_ISL_3170971, EPI_ISL_3170972, EPI_ISL_3170973, EPI_ISL_3170974, EPI_ISL_3170975, EPI_ISL_3170976, EPI_ISL_3170977, EPI_ISL_3170978, EPI_ISL_3170979, EPI_ISL_3170980, EPI_ISL_3170981, EPI_ISL_3170982, EPI_ISL_3170983, EPI_ISL_3170984, EPI_ISL_3170985, EPI_ISL_3170986, EPI_ISL_3170987, EPI_ISL_3170988, EPI_ISL_3170989, EPI_ISL_3170990, EPI_ISL_3170991, EPI_ISL_3170992, EPI_ISL_3170993, EPI_ISL_3170994, EPI_ISL_3170995, EPI_ISL_3170996, EPI_ISL_3170997, EPI_ISL_3170998, EPI_ISL_3170999, EPI_ISL_3171000, EPI_ISL_3171001, EPI_ISL_3171002, EPI_ISL_3171003, EPI_ISL_3171004, EPI_ISL_3171005, EPI_ISL_3171006, EPI_ISL_3171007, EPI_ISL_3171008, EPI_ISL_3171009, EPI_ISL_3171010, EPI_ISL_3171011, EPI_ISL_3171012, EPI_ISL_3171013, EPI_ISL_3171014, EPI_ISL_3171015, EPI_ISL_3171016, EPI_ISL_3171017, EPI_ISL_3171018, EPI_ISL_3171019, EPI_ISL_3171020, EPI_ISL_3171021, EPI_ISL_3171022, EPI_ISL_3171023, EPI_ISL_3171024, EPI_ISL_3171025, EPI_ISL_3171026, EPI_ISL_3171027, EPI_ISL_3171028, EPI_ISL_3171029, EPI_ISL_3171030, EPI_ISL_3171031, EPI_ISL_3171032, EPI_ISL_3171033, EPI_ISL_3171034, EPI_ISL_3171035, EPI_ISL_3171036, EPI_ISL_3171037, EPI_ISL_3171038, EPI_ISL_3171039, EPI_ISL_3171040, EPI_ISL_3171041, EPI_ISL_3171042, EPI_ISL_3171043, EPI_ISL_3171044, EPI_ISL_3171045, EPI_ISL_3171046, EPI_ISL_3171047, EPI_ISL_3171048, EPI_ISL_3171049, EPI_ISL_3171050, EPI_ISL_3171051, EPI_ISL_3171052, EPI_ISL_3171053, EPI_ISL_3171054, EPI_ISL_3171055, EPI_ISL_3171056, EPI_ISL_3171057, EPI_ISL_3171058, EPI_ISL_3171059, EPI_ISL_3171060, EPI_ISL_3171061, EPI_ISL_3171062, EPI_ISL_3171063, EPI_ISL_3171064, EPI_ISL_3171065, EPI_ISL_3171066, EPI_ISL_3171067, EPI_ISL_3171068, EPI_ISL_3171069, EPI_ISL_3171070, EPI_ISL_3171071, EPI_ISL_3171072, EPI_ISL_3171073, EPI_ISL_3171074, EPI_ISL_3171075, EPI_ISL_3171076, EPI_ISL_3171077, EPI_ISL_3171078, EPI_ISL_3171079, EPI_ISL_3171080, EPI_ISL_3171081, EPI_ISL_3171082, EPI_ISL_3171083, EPI_ISL_3171084, EPI_ISL_3171085, EPI_ISL_3171086, EPI_ISL_3171087, EPI_ISL_3171088, EPI_ISL_3171089, EPI_ISL_3171090, EPI_ISL_3171091, EPI_ISL_3171092, EPI_ISL_3171093, EPI_ISL_3171094, EPI_ISL_3171095, EPI_ISL_3171096, EPI_ISL_3171097, EPI_ISL_3171098, EPI_ISL_3171099, EPI_ISL_3171100, EPI_ISL_3171101, EPI_ISL_3171102, EPI_ISL_3171103, EPI_ISL_3171104, EPI_ISL_3171105, EPI_ISL_3171106, EPI_ISL_3171107, EPI_ISL_3171108, EPI_ISL_3171109, EPI_ISL_3171110, EPI_ISL_3171111, EPI_ISL_3171112, EPI_ISL_3171113, EPI_ISL_3171114, EPI_ISL_3171115, EPI_ISL_3171116, EPI_ISL_3171117, EPI_ISL_3171118, EPI_ISL_3171119, EPI_ISL_3171120, EPI_ISL_3171121, EPI_ISL_3171122, EPI_ISL_3171123, EPI_ISL_3171124, EPI_ISL_3171125, EPI_ISL_3171126, EPI_ISL_3171127, EPI_ISL_3171128, EPI_ISL_3171129, EPI_ISL_3171130, EPI_ISL_3171131, EPI_ISL_3171132, EPI_ISL_3171133, EPI_ISL_3171134, EPI_ISL_3171135, EPI_ISL_3171136, EPI_ISL_3171137, EPI_ISL_3171138, EPI_ISL_3171139, EPI_ISL_3171140, EPI_ISL_3171141, EPI_ISL_3171142, EPI_ISL_3171143, EPI_ISL_3171144, EPI_ISL_3171145, EPI_ISL_3171146, EPI_ISL_3171147, EPI_ISL_3171148, EPI_ISL_3171149, EPI_ISL_3171150, EPI_ISL_3171151, EPI_ISL_3171152, EPI_ISL_3171153, EPI_ISL_3171154, EPI_ISL_3171155, EPI_ISL_3171156, EPI_ISL_3171157, EPI_ISL_3171158, EPI_ISL_3171159, EPI_ISL_3171160, EPI_ISL_3171161, EPI_ISL_3171162, EPI_ISL_3171163, EPI_ISL_3171164, EPI_ISL_3171165, EPI_ISL_3171166, EPI_ISL_3171167, EPI_ISL_3171168, EPI_ISL_3171169, EPI_ISL_3171170, EPI_ISL_3171171, EPI_ISL_3171172, EPI_ISL_3171173, EPI_ISL_3171174, EPI_ISL_3171175, EPI_ISL_3171176, EPI_ISL_3171177, EPI_ISL_3171178, EPI_ISL_3171179, EPI_ISL_3171180, EPI_ISL_3171181, EPI_ISL_3171182, EPI_ISL_3171183, EPI_ISL_3171184, EPI_ISL_3171185, EPI_ISL_3171186, EPI_ISL_3171187, EPI_ISL_3171188, EPI_ISL_3171189, EPI_ISL_3171190, EPI_ISL_3171191, EPI_ISL_3171192, EPI_ISL_3171193, EPI_ISL_3171194, EPI_ISL_3171195, EPI_ISL_3171196, EPI_ISL_3171197, EPI_ISL_3171198, EPI_ISL_3171199, EPI_ISL_3171200, EPI_ISL_3171201, EPI_ISL_3171202, EPI_ISL_3171203, EPI_ISL_3171204, EPI_ISL_3171205, EPI_ISL_3171206, EPI_ISL_3171207, EPI_ISL_3171208, EPI_ISL_3171209, EPI_ISL_3171210, EPI_ISL_3171211, EPI_ISL_3171212, EPI_ISL_3171213, EPI_ISL_3171214, EPI_ISL_3171215, EPI_ISL_3171217, EPI_ISL_3171218, EPI_ISL_3171219, EPI_ISL_3171220, EPI_ISL_3171221, EPI_ISL_3171222, EPI_ISL_3171223, EPI_ISL_3171224, EPI_ISL_3171225, EPI_ISL_3171226, EPI_ISL_3171227, EPI_ISL_3171228, EPI_ISL_3171229, EPI_ISL_3171230, EPI_ISL_3171231, EPI_ISL_3171232, EPI_ISL_3171233, EPI_ISL_3171234, EPI_ISL_3171235, EPI_ISL_3171236, EPI_ISL_3171237, EPI_ISL_3171238, EPI_ISL_3171239, EPI_ISL_3171240, EPI_ISL_3171241, EPI_ISL_3171242, EPI_ISL_3171243, EPI_ISL_3171244, EPI_ISL_3171245, EPI_ISL_3171246, EPI_ISL_3171247, EPI_ISL_3171248, EPI_ISL_3171249, EPI_ISL_3171250, EPI_ISL_3171251, EPI_ISL_3171252, EPI_ISL_3171253, EPI_ISL_3171254, EPI_ISL_3171255, EPI_ISL_3171256, EPI_ISL_3171257, EPI_ISL_3171258, EPI_ISL_3171259, EPI_ISL_3171260, EPI_ISL_3171261, EPI_ISL_3171262, EPI_ISL_3171263, EPI_ISL_3171264, EPI_ISL_3171265, EPI_ISL_3171266, EPI_ISL_3171267 | | | |
| see above | Lighthouse Lab in Milton Keynes | Wellcome Sanger Institute for the COVID-19 Genomics UK (COG-UK) Consortium | The Lighthouse Lab in Milton Keynes and Alex Alderton, Roberto Amato, Jeffrey Barrett, Sonia Goncalves, Ewan Harrison, David K. Jackson, Ian Johnston, Dominic Kwiatkowski, Cordelia Langford, John Sillitoe on behalf of the Wellcome Sanger Institute COVID-19 Surveillance Team |
| EPI_ISL_3171268, EPI_ISL_3171269, EPI_ISL_3171270, EPI_ISL_3171271, EPI_ISL_3171272 | Berkshire and Surrey Pathology Services Lighthouse Laboratory | Wellcome Sanger Institute for the COVID-19 Genomics UK (COG-UK) Consortium | Berkshire and Surrey Pathology Services Lighthouse Laboratory and Alex Alderton, Roberto Amato, Jeffrey Barrett, Sonia Goncalves, Ewan Harrison, David K. Jackson, Ian Johnston, Dominic Kwiatkowski, Cordelia Langford, John Sillitoe on behalf of the Wellcome Sanger Institute COVID-19 Surveillance Team |
| EPI_ISL_3171273, EPI_ISL_3171274 | Lighthouse Lab in Alderley Park | Wellcome Sanger Institute for the COVID-19 Genomics UK (COG-UK) Consortium | Jacquelyn Wynn, Mairead Hyland, The Lighthouse Lab in Alderley Park and Alex Alderton, Roberto Amato, Jeffrey Barrett, Sonia Goncalves, Ewan Harrison, David K. Jackson, Ian Johnston, Dominic Kwiatkowski, Cordelia Langford, John Sillitoe on behalf of the Wellcome Sanger Institute COVID-19 Surveillance Team |
| EPI_ISL_3171275 | Lighthouse Laboratory Plymouth | Wellcome Sanger Institute for the COVID-19 Genomics UK (COG-UK) Consortium | Lighthouse Laboratory Plymouth and Alex Alderton, Roberto Amato, Jeffrey Barrett, Sonia Goncalves, Ewan Harrison, David K. Jackson, Ian Johnston, Dominic Kwiatkowski, Cordelia Langford, John Sillitoe on behalf of the Wellcome Sanger Institute COVID-19 Surveillance Team |
| EPI_ISL_3171276 | Berkshire and Surrey Pathology Services Lighthouse Laboratory | Wellcome Sanger Institute for the COVID-19 Genomics UK (COG-UK) Consortium | Berkshire and Surrey Pathology Services Lighthouse Laboratory and Alex Alderton, Roberto Amato, Jeffrey Barrett, Sonia Goncalves, Ewan Harrison, David K. Jackson, Ian Johnston, Dominic Kwiatkowski, Cordelia Langford, John Sillitoe on behalf of the Wellcome Sanger Institute COVID-19 Surveillance Team |
| EPI_ISL_3171277, EPI_ISL_3171278 | Lighthouse Lab in Alderley Park | Wellcome Sanger Institute for the COVID-19 Genomics UK (COG-UK) Consortium | Jacquelyn Wynn, Mairead Hyland, The Lighthouse Lab in Alderley Park and Alex Alderton, Roberto Amato, Jeffrey Barrett, Sonia Goncalves, Ewan Harrison, David K. Jackson, Ian Johnston, Dominic Kwiatkowski, Cordelia Langford, John Sillitoe on behalf of the Wellcome Sanger Institute COVID-19 Surveillance Team |
| EPI_ISL_3171279, EPI_ISL_3171280 | Lighthouse Laboratory Plymouth | Wellcome Sanger Institute for the COVID-19 Genomics UK (COG-UK) Consortium | Lighthouse Laboratory Plymouth and Alex Alderton, Roberto Amato, Jeffrey Barrett, Sonia Goncalves, Ewan Harrison, David K. Jackson, Ian Johnston, Dominic Kwiatkowski, Cordelia Langford, John Sillitoe on behalf of the Wellcome Sanger Institute COVID-19 Surveillance Team |
| EPI_ISL_3171281, EPI_ISL_3171282 | Berkshire and Surrey Pathology Services Lighthouse Laboratory | Wellcome Sanger Institute for the COVID-19 Genomics UK (COG-UK) Consortium | Berkshire and Surrey Pathology Services Lighthouse Laboratory and Alex Alderton, Roberto Amato, Jeffrey Barrett, Sonia Goncalves, Ewan Harrison, David K. Jackson, Ian Johnston, Dominic Kwiatkowski, Cordelia Langford, John Sillitoe on behalf of the Wellcome Sanger Institute COVID-19 Surveillance Team |
| EPI_ISL_3171283 | Lighthouse Laboratory Plymouth | Wellcome Sanger Institute for the COVID-19 Genomics UK (COG-UK) Consortium | Lighthouse Laboratory Plymouth and Alex Alderton, Roberto Amato, Jeffrey Barrett, Sonia Goncalves, Ewan Harrison, David K. Jackson, Ian Johnston, Dominic Kwiatkowski, Cordelia Langford, John Sillitoe on behalf of the Wellcome Sanger Institute COVID-19 Surveillance Team |
| EPI_ISL_3171284 | Berkshire and Surrey Pathology Services Lighthouse Laboratory | Wellcome Sanger Institute for the COVID-19 Genomics UK (COG-UK) Consortium | Berkshire and Surrey Pathology Services Lighthouse Laboratory and Alex Alderton, Roberto Amato, Jeffrey Barrett, Sonia Goncalves, Ewan Harrison, David K. Jackson, Ian Johnston, Dominic Kwiatkowski, Cordelia Langford, John Sillitoe on behalf of the Wellcome Sanger Institute COVID-19 Surveillance Team |
| EPI_ISL_3171285, EPI_ISL_3171286 | Lighthouse Laboratory Plymouth | Wellcome Sanger Institute for the COVID-19 Genomics UK (COG-UK) Consortium | Lighthouse Laboratory Plymouth and Alex Alderton, Roberto Amato, Jeffrey Barrett, Sonia Goncalves, Ewan Harrison, David K. Jackson, Ian Johnston, Dominic Kwiatkowski, Cordelia Langford, John Sillitoe on behalf of the Wellcome Sanger Institute COVID-19 Surveillance Team |
| EPI_ISL_3171287, EPI_ISL_3171288 | Berkshire and Surrey Pathology Services Lighthouse Laboratory | Wellcome Sanger Institute for the COVID-19 Genomics UK (COG-UK) Consortium | Berkshire and Surrey Pathology Services Lighthouse Laboratory and Alex Alderton, Roberto Amato, Jeffrey Barrett, Sonia Goncalves, Ewan Harrison, David K. Jackson, Ian Johnston, Dominic Kwiatkowski, Cordelia Langford, John Sillitoe on behalf of the Wellcome Sanger Institute COVID-19 Surveillance Team |
| EPI_ISL_3171289 | Lighthouse Laboratory Plymouth | Wellcome Sanger Institute for the COVID-19 Genomics UK (COG-UK) Consortium | Lighthouse Laboratory Plymouth and Alex Alderton, Roberto Amato, Jeffrey Barrett, Sonia Goncalves, Ewan Harrison, David K. Jackson, Ian Johnston, Dominic Kwiatkowski, Cordelia Langford, John Sillitoe on behalf of the Wellcome Sanger Institute COVID-19 Surveillance Team |
| EPI_ISL_3171290 | Lighthouse Lab in Alderley Park | Wellcome Sanger Institute for the COVID-19 Genomics UK (COG-UK) Consortium | Jacquelyn Wynn, Mairead Hyland, The Lighthouse Lab in Alderley Park and Alex Alderton, Roberto Amato, Jeffrey Barrett, Sonia Goncalves, Ewan Harrison, David K. Jackson, Ian Johnston, Dominic Kwiatkowski, Cordelia Langford, John Sillitoe on behalf of the Wellcome Sanger Institute COVID-19 Surveillance Team |
| EPI_ISL_3171291, EPI_ISL_3171292 | Lighthouse Laboratory Plymouth | Wellcome Sanger Institute for the COVID-19 Genomics UK (COG-UK) Consortium | Lighthouse Laboratory Plymouth and Alex Alderton, Roberto Amato, Jeffrey Barrett, Sonia Goncalves, Ewan Harrison, David K. Jackson, Ian Johnston, Dominic Kwiatkowski, Cordelia Langford, John Sillitoe on behalf of the Wellcome Sanger Institute COVID-19 Surveillance Team |
| EPI_ISL_3171293 | Berkshire and Surrey Pathology Services Lighthouse Laboratory | Wellcome Sanger Institute for the COVID-19 Genomics UK (COG-UK) Consortium | Berkshire and Surrey Pathology Services Lighthouse Laboratory and Alex Alderton, Roberto Amato, Jeffrey Barrett, Sonia Goncalves, Ewan Harrison, David K. Jackson, Ian Johnston, Dominic Kwiatkowski, Cordelia Langford, John Sillitoe on behalf of the Wellcome Sanger Institute COVID-19 Surveillance Team |
| EPI_ISL_3171294, EPI_ISL_3171295 | Lighthouse Laboratory Plymouth | Wellcome Sanger Institute for the COVID-19 Genomics UK (COG-UK) Consortium | Lighthouse Laboratory Plymouth and Alex Alderton, Roberto Amato, Jeffrey Barrett, Sonia Goncalves, Ewan Harrison, David K. Jackson, Ian Johnston, Dominic Kwiatkowski, Cordelia Langford, John Sillitoe on behalf of the Wellcome Sanger Institute COVID-19 Surveillance Team |
| EPI_ISL_3171296 | Lighthouse Lab in Alderley Park | Wellcome Sanger Institute for the COVID-19 Genomics UK (COG-UK) Consortium | Jacquelyn Wynn, Mairead Hyland, The Lighthouse Lab in Alderley Park and Alex Alderton, Roberto Amato, Jeffrey Barrett, Sonia Goncalves, Ewan Harrison, David K. Jackson, Ian Johnston, Dominic Kwiatkowski, Cordelia Langford, John Sillitoe on behalf of the Wellcome Sanger Institute COVID-19 Surveillance Team |
| EPI_ISL_3171297 | Lighthouse Laboratory Plymouth | Wellcome Sanger Institute for the COVID-19 Genomics UK (COG-UK) Consortium | Lighthouse Laboratory Plymouth and Alex Alderton, Roberto Amato, Jeffrey Barrett, Sonia Goncalves, Ewan Harrison, David K. Jackson, Ian Johnston, Dominic Kwiatkowski, Cordelia Langford, John Sillitoe on behalf of the Wellcome Sanger Institute COVID-19 Surveillance Team |
| EPI_ISL_3171298, EPI_ISL_3171299 | Berkshire and Surrey Pathology Services Lighthouse Laboratory | Wellcome Sanger Institute for the COVID-19 Genomics UK (COG-UK) Consortium | Berkshire and Surrey Pathology Services Lighthouse Laboratory and Alex Alderton, Roberto Amato, Jeffrey Barrett, Sonia Goncalves, Ewan Harrison, David K. Jackson, Ian Johnston, Dominic Kwiatkowski, Cordelia Langford, John Sillitoe on behalf of the Wellcome Sanger Institute COVID-19 Surveillance Team |
| EPI_ISL_3171300 | Lighthouse Laboratory Plymouth | Wellcome Sanger Institute for the COVID-19 Genomics UK (COG-UK) Consortium | Lighthouse Laboratory Plymouth and Alex Alderton, Roberto Amato, Jeffrey Barrett, Sonia Goncalves, Ewan Harrison, David K. Jackson, Ian Johnston, Dominic Kwiatkowski, Cordelia Langford, John Sillitoe on behalf of the Wellcome Sanger Institute COVID-19 Surveillance Team |
| EPI_ISL_3171301, EPI_ISL_3171302 | Lighthouse Lab in Alderley Park | Wellcome Sanger Institute for the COVID-19 Genomics UK (COG-UK) Consortium | Jacquelyn Wynn, Mairead Hyland, The Lighthouse Lab in Alderley Park and Alex Alderton, Roberto Amato, Jeffrey Barrett, Sonia Goncalves, Ewan Harrison, David K. Jackson, Ian Johnston, Dominic Kwiatkowski, Cordelia Langford, John Sillitoe on behalf of the Wellcome Sanger Institute COVID-19 Surveillance Team |
| EPI_ISL_3171303 | Berkshire and Surrey Pathology Services Lighthouse Laboratory | Wellcome Sanger Institute for the COVID-19 Genomics UK (COG-UK) Consortium | Berkshire and Surrey Pathology Services Lighthouse Laboratory and Alex Alderton, Roberto Amato, Jeffrey Barrett, Sonia Goncalves, Ewan Harrison, David K. Jackson, Ian Johnston, Dominic Kwiatkowski, Cordelia Langford, John Sillitoe on behalf of the Wellcome Sanger Institute COVID-19 Surveillance Team |
| EPI_ISL_3171304 | Lighthouse Laboratory Plymouth | Wellcome Sanger Institute for the COVID-19 Genomics UK (COG-UK) Consortium | Lighthouse Laboratory Plymouth and Alex Alderton, Roberto Amato, Jeffrey Barrett, Sonia Goncalves, Ewan Harrison, David K. Jackson, Ian Johnston, Dominic Kwiatkowski, Cordelia Langford, John Sillitoe on behalf of the Wellcome Sanger Institute COVID-19 Surveillance Team |
| EPI_ISL_3171305 | Lighthouse Lab in Alderley Park | Wellcome Sanger Institute for the COVID-19 Genomics UK (COG-UK) Consortium | Jacquelyn Wynn, Mairead Hyland, The Lighthouse Lab in Alderley Park and Alex Alderton, Roberto Amato, Jeffrey Barrett, Sonia Goncalves, Ewan Harrison, David K. Jackson, Ian Johnston, Dominic Kwiatkowski, Cordelia Langford, John Sillitoe on behalf of the Wellcome Sanger Institute COVID-19 Surveillance Team |
| EPI_ISL_3171306, EPI_ISL_3171307 | Lighthouse Laboratory Plymouth | Wellcome Sanger Institute for the COVID-19 Genomics UK (COG-UK) Consortium | Lighthouse Laboratory Plymouth and Alex Alderton, Roberto Amato, Jeffrey Barrett, Sonia Goncalves, Ewan Harrison, David K. Jackson, Ian Johnston, Dominic Kwiatkowski, Cordelia Langford, John Sillitoe on behalf of the Wellcome Sanger Institute COVID-19 Surveillance Team |

| | | | |
|---|---|---|---|
| EPI_ISL_3171308 | Lighthouse Lab in Alderley Park | Wellcome Sanger Institute for the COVID-19 Genomics UK (COG-UK) Consortium | Jacquelyn Wynn, Mairead Hyland, The Lighthouse Lab in Alderley Park and Alex Alderton, Roberto Amato, Jeffrey Barrett, Sonia Goncalves, Ewan Harrison, David K. Jackson, Ian Johnston, Dominic Kwiatkowski, Cordelia Langford, John Sillitoe on behalf of the Wellcome Sanger Institute COVID-19 Surveillance Team |
| EPI_ISL_3171309 | Lighthouse Laboratory Plymouth | Wellcome Sanger Institute for the COVID-19 Genomics UK (COG-UK) Consortium | Lighthouse Laboratory Plymouth and Alex Alderton, Roberto Amato, Jeffrey Barrett, Sonia Goncalves, Ewan Harrison, David K. Jackson, Ian Johnston, Dominic Kwiatkowski, Cordelia Langford, John Sillitoe on behalf of the Wellcome Sanger Institute COVID-19 Surveillance Team |
| EPI_ISL_3171310 | Lighthouse Lab in Alderley Park | Wellcome Sanger Institute for the COVID-19 Genomics UK (COG-UK) Consortium | Jacquelyn Wynn, Mairead Hyland, The Lighthouse Lab in Alderley Park and Alex Alderton, Roberto Amato, Jeffrey Barrett, Sonia Goncalves, Ewan Harrison, David K. Jackson, Ian Johnston, Dominic Kwiatkowski, Cordelia Langford, John Sillitoe on behalf of the Wellcome Sanger Institute COVID-19 Surveillance Team |
| EPI_ISL_3171311, EPI_ISL_3171312 | Lighthouse Laboratory Plymouth | Wellcome Sanger Institute for the COVID-19 Genomics UK (COG-UK) Consortium | Lighthouse Laboratory Plymouth and Alex Alderton, Roberto Amato, Jeffrey Barrett, Sonia Goncalves, Ewan Harrison, David K. Jackson, Ian Johnston, Dominic Kwiatkowski, Cordelia Langford, John Sillitoe on behalf of the Wellcome Sanger Institute COVID-19 Surveillance Team |
| EPI_ISL_3171313 | Lighthouse Lab in Alderley Park | Wellcome Sanger Institute for the COVID-19 Genomics UK (COG-UK) Consortium | Jacquelyn Wynn, Mairead Hyland, The Lighthouse Lab in Alderley Park and Alex Alderton, Roberto Amato, Jeffrey Barrett, Sonia Goncalves, Ewan Harrison, David K. Jackson, Ian Johnston, Dominic Kwiatkowski, Cordelia Langford, John Sillitoe on behalf of the Wellcome Sanger Institute COVID-19 Surveillance Team |
| EPI_ISL_3171314 | Lighthouse Laboratory Plymouth | Wellcome Sanger Institute for the COVID-19 Genomics UK (COG-UK) Consortium | Lighthouse Laboratory Plymouth and Alex Alderton, Roberto Amato, Jeffrey Barrett, Sonia Goncalves, Ewan Harrison, David K. Jackson, Ian Johnston, Dominic Kwiatkowski, Cordelia Langford, John Sillitoe on behalf of the Wellcome Sanger Institute COVID-19 Surveillance Team |
| EPI_ISL_3171315, EPI_ISL_3171316 | Berkshire and Surrey Pathology Services Lighthouse Laboratory | Wellcome Sanger Institute for the COVID-19 Genomics UK (COG-UK) Consortium | Berkshire and Surrey Pathology Services Lighthouse Laboratory and Alex Alderton, Roberto Amato, Jeffrey Barrett, Sonia Goncalves, Ewan Harrison, David K. Jackson, Ian Johnston, Dominic Kwiatkowski, Cordelia Langford, John Sillitoe on behalf of the Wellcome Sanger Institute COVID-19 Surveillance Team |
| EPI_ISL_3171317, EPI_ISL_3171318, EPI_ISL_3171319, EPI_ISL_3171320 | Lighthouse Laboratory Plymouth | Wellcome Sanger Institute for the COVID-19 Genomics UK (COG-UK) Consortium | Lighthouse Laboratory Plymouth and Alex Alderton, Roberto Amato, Jeffrey Barrett, Sonia Goncalves, Ewan Harrison, David K. Jackson, Ian Johnston, Dominic Kwiatkowski, Cordelia Langford, John Sillitoe on behalf of the Wellcome Sanger Institute COVID-19 Surveillance Team |
| EPI_ISL_3171321, EPI_ISL_3171322 | Berkshire and Surrey Pathology Services Lighthouse Laboratory | Wellcome Sanger Institute for the COVID-19 Genomics UK (COG-UK) Consortium | Berkshire and Surrey Pathology Services Lighthouse Laboratory and Alex Alderton, Roberto Amato, Jeffrey Barrett, Sonia Goncalves, Ewan Harrison, David K. Jackson, Ian Johnston, Dominic Kwiatkowski, Cordelia Langford, John Sillitoe on behalf of the Wellcome Sanger Institute COVID-19 Surveillance Team |
| EPI_ISL_3171323, EPI_ISL_3171324 | Lighthouse Laboratory Plymouth | Wellcome Sanger Institute for the COVID-19 Genomics UK (COG-UK) Consortium | Lighthouse Laboratory Plymouth and Alex Alderton, Roberto Amato, Jeffrey Barrett, Sonia Goncalves, Ewan Harrison, David K. Jackson, Ian Johnston, Dominic Kwiatkowski, Cordelia Langford, John Sillitoe on behalf of the Wellcome Sanger Institute COVID-19 Surveillance Team |
| EPI_ISL_3171325 | Lighthouse Lab in Alderley Park | Wellcome Sanger Institute for the COVID-19 Genomics UK (COG-UK) Consortium | Jacquelyn Wynn, Mairead Hyland, The Lighthouse Lab in Alderley Park and Alex Alderton, Roberto Amato, Jeffrey Barrett, Sonia Goncalves, Ewan Harrison, David K. Jackson, Ian Johnston, Dominic Kwiatkowski, Cordelia Langford, John Sillitoe on behalf of the Wellcome Sanger Institute COVID-19 Surveillance Team |
| EPI_ISL_3171326 | Berkshire and Surrey Pathology Services Lighthouse Laboratory | Wellcome Sanger Institute for the COVID-19 Genomics UK (COG-UK) Consortium | Berkshire and Surrey Pathology Services Lighthouse Laboratory and Alex Alderton, Roberto Amato, Jeffrey Barrett, Sonia Goncalves, Ewan Harrison, David K. Jackson, Ian Johnston, Dominic Kwiatkowski, Cordelia Langford, John Sillitoe on behalf of the Wellcome Sanger Institute COVID-19 Surveillance Team |
| EPI_ISL_3171327, EPI_ISL_3171328 | Lighthouse Laboratory Plymouth | Wellcome Sanger Institute for the COVID-19 Genomics UK (COG-UK) Consortium | Lighthouse Laboratory Plymouth and Alex Alderton, Roberto Amato, Jeffrey Barrett, Sonia Goncalves, Ewan Harrison, David K. Jackson, Ian Johnston, Dominic Kwiatkowski, Cordelia Langford, John Sillitoe on behalf of the Wellcome Sanger Institute COVID-19 Surveillance Team |
| EPI_ISL_3171329 | Berkshire and Surrey Pathology Services Lighthouse Laboratory | Wellcome Sanger Institute for the COVID-19 Genomics UK (COG-UK) Consortium | Berkshire and Surrey Pathology Services Lighthouse Laboratory and Alex Alderton, Roberto Amato, Jeffrey Barrett, Sonia Goncalves, Ewan Harrison, David K. Jackson, Ian Johnston, Dominic Kwiatkowski, Cordelia Langford, John Sillitoe on behalf of the Wellcome Sanger Institute COVID-19 Surveillance Team |
| EPI_ISL_3171330 | Lighthouse Lab in Alderley Park | Wellcome Sanger Institute for the COVID-19 Genomics UK (COG-UK) Consortium | Jacquelyn Wynn, Mairead Hyland, The Lighthouse Lab in Alderley Park and Alex Alderton, Roberto Amato, Jeffrey Barrett, Sonia Goncalves, Ewan Harrison, David K. Jackson, Ian Johnston, Dominic Kwiatkowski, Cordelia Langford, John Sillitoe on behalf of the Wellcome Sanger Institute COVID-19 Surveillance Team |
| EPI_ISL_3171331 | Lighthouse Laboratory Plymouth | Wellcome Sanger Institute for the COVID-19 Genomics UK (COG-UK) Consortium | Lighthouse Laboratory Plymouth and Alex Alderton, Roberto Amato, Jeffrey Barrett, Sonia Goncalves, Ewan Harrison, David K. Jackson, Ian Johnston, Dominic Kwiatkowski, Cordelia Langford, John Sillitoe on behalf of the Wellcome Sanger Institute COVID-19 Surveillance Team |
| EPI_ISL_3171332 | Lighthouse Lab in Alderley Park | Wellcome Sanger Institute for the COVID-19 Genomics UK (COG-UK) Consortium | Jacquelyn Wynn, Mairead Hyland, The Lighthouse Lab in Alderley Park and Alex Alderton, Roberto Amato, Jeffrey Barrett, Sonia Goncalves, Ewan Harrison, David K. Jackson, Ian Johnston, Dominic Kwiatkowski, Cordelia Langford, John Sillitoe on behalf of the Wellcome Sanger Institute COVID-19 Surveillance Team |
| EPI_ISL_3171333, EPI_ISL_3171334, EPI_ISL_3171335 | Lighthouse Laboratory Plymouth | Wellcome Sanger Institute for the COVID-19 Genomics UK (COG-UK) Consortium | Lighthouse Laboratory Plymouth and Alex Alderton, Roberto Amato, Jeffrey Barrett, Sonia Goncalves, Ewan Harrison, David K. Jackson, Ian Johnston, Dominic Kwiatkowski, Cordelia Langford, John Sillitoe on behalf of the Wellcome Sanger Institute COVID-19 Surveillance Team |
| EPI_ISL_3171336 | Berkshire and Surrey Pathology Services Lighthouse Laboratory | Wellcome Sanger Institute for the COVID-19 Genomics UK (COG-UK) Consortium | Berkshire and Surrey Pathology Services Lighthouse Laboratory and Alex Alderton, Roberto Amato, Jeffrey Barrett, Sonia Goncalves, Ewan Harrison, David K. Jackson, Ian Johnston, Dominic Kwiatkowski, Cordelia Langford, John Sillitoe on behalf of the Wellcome Sanger Institute COVID-19 Surveillance Team |
| EPI_ISL_3171337, EPI_ISL_3171338 | Lighthouse Laboratory Plymouth | Wellcome Sanger Institute for the COVID-19 Genomics UK (COG-UK) Consortium | Lighthouse Laboratory Plymouth and Alex Alderton, Roberto Amato, Jeffrey Barrett, Sonia Goncalves, Ewan Harrison, David K. Jackson, Ian Johnston, Dominic Kwiatkowski, Cordelia Langford, John Sillitoe on behalf of the Wellcome Sanger Institute COVID-19 Surveillance Team |
| EPI_ISL_3171339 | Lighthouse Lab in Alderley Park | Wellcome Sanger Institute for the COVID-19 Genomics UK (COG-UK) Consortium | Jacquelyn Wynn, Mairead Hyland, The Lighthouse Lab in Alderley Park and Alex Alderton, Roberto Amato, Jeffrey Barrett, Sonia Goncalves, Ewan Harrison, David K. Jackson, Ian Johnston, Dominic Kwiatkowski, Cordelia Langford, John Sillitoe on behalf of the Wellcome Sanger Institute COVID-19 Surveillance Team |
| EPI_ISL_3171340 | Berkshire and Surrey Pathology Services Lighthouse Laboratory | Wellcome Sanger Institute for the COVID-19 Genomics UK (COG-UK) Consortium | Berkshire and Surrey Pathology Services Lighthouse Laboratory and Alex Alderton, Roberto Amato, Jeffrey Barrett, Sonia Goncalves, Ewan Harrison, David K. Jackson, Ian Johnston, Dominic Kwiatkowski, Cordelia Langford, John Sillitoe on behalf of the Wellcome Sanger Institute COVID-19 Surveillance Team |
| EPI_ISL_3171341, EPI_ISL_3171342 | Lighthouse Laboratory Plymouth | Wellcome Sanger Institute for the COVID-19 Genomics UK (COG-UK) Consortium | Lighthouse Laboratory Plymouth and Alex Alderton, Roberto Amato, Jeffrey Barrett, Sonia Goncalves, Ewan Harrison, David K. Jackson, Ian Johnston, Dominic Kwiatkowski, Cordelia Langford, John Sillitoe on behalf of the Wellcome Sanger Institute COVID-19 Surveillance Team |
| EPI_ISL_3171343 | Lighthouse Lab in Alderley Park | Wellcome Sanger Institute for the COVID-19 Genomics UK (COG-UK) Consortium | Jacquelyn Wynn, Mairead Hyland, The Lighthouse Lab in Alderley Park and Alex Alderton, Roberto Amato, Jeffrey Barrett, Sonia Goncalves, Ewan Harrison, David K. Jackson, Ian Johnston, Dominic Kwiatkowski, Cordelia Langford, John Sillitoe on behalf of the Wellcome Sanger Institute COVID-19 Surveillance Team |
| EPI_ISL_3171344 | Lighthouse Laboratory Plymouth | Wellcome Sanger Institute for the COVID-19 Genomics UK (COG-UK) Consortium | Lighthouse Laboratory Plymouth and Alex Alderton, Roberto Amato, Jeffrey Barrett, Sonia Goncalves, Ewan Harrison, David K. Jackson, Ian Johnston, Dominic Kwiatkowski, Cordelia Langford, John Sillitoe on behalf of the Wellcome Sanger Institute COVID-19 Surveillance Team |
| EPI_ISL_3171345, EPI_ISL_3171346 | Berkshire and Surrey Pathology Services Lighthouse Laboratory | Wellcome Sanger Institute for the COVID-19 Genomics UK (COG-UK) Consortium | Berkshire and Surrey Pathology Services Lighthouse Laboratory and Alex Alderton, Roberto Amato, Jeffrey Barrett, Sonia Goncalves, Ewan Harrison, David K. Jackson, Ian Johnston, Dominic Kwiatkowski, Cordelia Langford, John Sillitoe on behalf of the Wellcome Sanger Institute COVID-19 Surveillance Team |
| EPI_ISL_3171347, EPI_ISL_3171348 | Lighthouse Lab in Alderley Park | Wellcome Sanger Institute for the COVID-19 Genomics UK (COG-UK) Consortium | Jacquelyn Wynn, Mairead Hyland, The Lighthouse Lab in Alderley Park and Alex Alderton, Roberto Amato, Jeffrey Barrett, Sonia Goncalves, Ewan Harrison, David K. Jackson, Ian Johnston, Dominic Kwiatkowski, Cordelia Langford, John Sillitoe on behalf of the Wellcome Sanger Institute COVID-19 Surveillance Team |
| EPI_ISL_3171349 | Berkshire and Surrey Pathology Services Lighthouse Laboratory | Wellcome Sanger Institute for the COVID-19 Genomics UK (COG-UK) Consortium | Berkshire and Surrey Pathology Services Lighthouse Laboratory and Alex Alderton, Roberto Amato, Jeffrey Barrett, Sonia Goncalves, Ewan Harrison, David K. Jackson, Ian Johnston, Dominic Kwiatkowski, Cordelia Langford, John Sillitoe on behalf of the Wellcome Sanger Institute COVID-19 Surveillance Team |
| EPI_ISL_3171350, EPI_ISL_3171351 | Lighthouse Laboratory Plymouth | Wellcome Sanger Institute for the COVID-19 Genomics UK (COG-UK) Consortium | Lighthouse Laboratory Plymouth and Alex Alderton, Roberto Amato, Jeffrey Barrett, Sonia Goncalves, Ewan Harrison, David K. Jackson, Ian Johnston, Dominic Kwiatkowski, Cordelia Langford, John Sillitoe on behalf of the Wellcome Sanger Institute COVID-19 Surveillance Team |
| EPI_ISL_3171352 | Berkshire and Surrey Pathology Services Lighthouse Laboratory | Wellcome Sanger Institute for the COVID-19 Genomics UK (COG-UK) Consortium | Berkshire and Surrey Pathology Services Lighthouse Laboratory and Alex Alderton, Roberto Amato, Jeffrey Barrett, Sonia Goncalves, Ewan Harrison, David K. Jackson, Ian Johnston, Dominic Kwiatkowski, Cordelia Langford, John Sillitoe on behalf of the Wellcome Sanger Institute COVID-19 Surveillance Team |
| EPI_ISL_3171353 | Lighthouse Laboratory Plymouth | Wellcome Sanger Institute for the COVID-19 Genomics UK (COG-UK) Consortium | Lighthouse Laboratory Plymouth and Alex Alderton, Roberto Amato, Jeffrey Barrett, Sonia Goncalves, Ewan Harrison, David K. Jackson, Ian Johnston, Dominic Kwiatkowski, Cordelia Langford, John Sillitoe on behalf of the Wellcome Sanger Institute COVID-19 Surveillance Team |
| EPI_ISL_3171354 | Lighthouse Lab in Alderley Park | Wellcome Sanger Institute for the COVID-19 Genomics UK (COG-UK) Consortium | Jacquelyn Wynn, Mairead Hyland, The Lighthouse Lab in Alderley Park and Alex Alderton, Roberto Amato, Jeffrey Barrett, Sonia Goncalves, Ewan Harrison, David K. Jackson, Ian Johnston, Dominic Kwiatkowski, Cordelia Langford, John Sillitoe on behalf of the Wellcome Sanger Institute COVID-19 Surveillance Team |
| EPI_ISL_3171355, EPI_ISL_3171356 | Lighthouse Laboratory Plymouth | Wellcome Sanger Institute for the COVID-19 Genomics UK (COG-UK) Consortium | Lighthouse Laboratory Plymouth and Alex Alderton, Roberto Amato, Jeffrey Barrett, Sonia Goncalves, Ewan Harrison, David K. Jackson, Ian Johnston, Dominic Kwiatkowski, Cordelia Langford, John Sillitoe on behalf of the Wellcome Sanger Institute COVID-19 Surveillance Team |
| EPI_ISL_3171357 | Lighthouse Lab in Alderley Park | Wellcome Sanger Institute for the COVID-19 Genomics UK (COG-UK) Consortium | Jacquelyn Wynn, Mairead Hyland, The Lighthouse Lab in Alderley Park and Alex Alderton, Roberto Amato, Jeffrey Barrett, Sonia Goncalves, Ewan Harrison, David K. Jackson, Ian Johnston, Dominic Kwiatkowski, Cordelia Langford, John Sillitoe on behalf of the Wellcome Sanger Institute COVID-19 Surveillance Team |
| EPI_ISL_3171358 | Berkshire and Surrey Pathology Services Lighthouse Laboratory | Wellcome Sanger Institute for the COVID-19 Genomics UK (COG-UK) Consortium | Berkshire and Surrey Pathology Services Lighthouse Laboratory and Alex Alderton, Roberto Amato, Jeffrey Barrett, Sonia Goncalves, Ewan Harrison, David K. Jackson, Ian Johnston, Dominic Kwiatkowski, Cordelia Langford, John Sillitoe on behalf of the Wellcome Sanger Institute COVID-19 Surveillance Team |
| EPI_ISL_3171359, EPI_ISL_3171360, EPI_ISL_3171361, EPI_ISL_3171362, EPI_ISL_3171363, EPI_ISL_3171364, EPI_ISL_3171365, EPI_ISL_3171366 | Lighthouse Laboratory Plymouth | Wellcome Sanger Institute for the COVID-19 Genomics UK (COG-UK) Consortium | Lighthouse Laboratory Plymouth and Alex Alderton, Roberto Amato, Jeffrey Barrett, Sonia Goncalves, Ewan Harrison, David K. Jackson, Ian Johnston, Dominic Kwiatkowski, Cordelia Langford, John Sillitoe on behalf of the Wellcome Sanger Institute COVID-19 Surveillance Team |
| EPI_ISL_3171367 | Berkshire and Surrey Pathology Services Lighthouse Laboratory | Wellcome Sanger Institute for the COVID-19 Genomics UK (COG-UK) Consortium | Berkshire and Surrey Pathology Services Lighthouse Laboratory and Alex Alderton, Roberto Amato, Jeffrey Barrett, Sonia Goncalves, Ewan Harrison, David K. Jackson, Ian Johnston, Dominic Kwiatkowski, Cordelia Langford, John Sillitoe on behalf of the Wellcome Sanger Institute COVID-19 Surveillance Team |
| EPI_ISL_3171368 | Lighthouse Lab in Alderley Park | Wellcome Sanger Institute for the COVID-19 Genomics UK (COG-UK) Consortium | Jacquelyn Wynn, Mairead Hyland, The Lighthouse Lab in Alderley Park and Alex Alderton, Roberto Amato, Jeffrey Barrett, Sonia Goncalves, Ewan Harrison, David K. Jackson, Ian Johnston, Dominic Kwiatkowski, Cordelia Langford, John Sillitoe on behalf of the Wellcome Sanger Institute COVID-19 Surveillance Team |
| EPI_ISL_3171369 | Berkshire and Surrey Pathology Services Lighthouse Laboratory | Wellcome Sanger Institute for the COVID-19 Genomics UK (COG-UK) Consortium | Berkshire and Surrey Pathology Services Lighthouse Laboratory and Alex Alderton, Roberto Amato, Jeffrey Barrett, Sonia Goncalves, Ewan Harrison, David K. Jackson, Ian Johnston, Dominic Kwiatkowski, Cordelia Langford, John Sillitoe on behalf of the Wellcome Sanger Institute COVID-19 Surveillance Team |
| EPI_ISL_3171370, EPI_ISL_3171371, EPI_ISL_3171372, EPI_ISL_3171373 | Lighthouse Laboratory Plymouth | Wellcome Sanger Institute for the COVID-19 Genomics UK (COG-UK) Consortium | Lighthouse Laboratory Plymouth and Alex Alderton, Roberto Amato, Jeffrey Barrett, Sonia Goncalves, Ewan Harrison, David K. Jackson, Ian Johnston, Dominic Kwiatkowski, Cordelia Langford, John Sillitoe on behalf of the Wellcome Sanger Institute COVID-19 Surveillance Team |
| EPI_ISL_3171374 | Berkshire and Surrey Pathology Services Lighthouse Laboratory | Wellcome Sanger Institute for the COVID-19 Genomics UK (COG-UK) | Berkshire and Surrey Pathology Services Lighthouse Laboratory and Alex Alderton, Roberto Amato, Jeffrey Barrett, Sonia Goncalves, Ewan Harrison, David K. Jackson, Ian Johnston, Dominic Kwiatkowski, Cordelia Langford, John Sillitoe on behalf |

| | | | |
|---|---|---|---|
| | | Consortium | of the Wellcome Sanger Institute COVID-19 Surveillance Team |
| EPI_ISL_3171375 | Lighthouse Lab in Alderley Park | Wellcome Sanger Institute for the COVID-19 Genomics UK (COG-UK) Consortium | Jacquelyn Wynn, Mairead Hyland, The Lighthouse Lab in Alderley Park and Alex Alderton, Roberto Amato, Jeffrey Barrett, Sonia Goncalves, Ewan Harrison, David K. Jackson, Ian Johnston, Dominic Kwiatkowski, Cordelia Langford, John Sillitoe on behalf of the Wellcome Sanger Institute COVID-19 Surveillance Team |
| EPI_ISL_3171376 | Berkshire and Surrey Pathology Services Lighthouse Laboratory | Wellcome Sanger Institute for the COVID-19 Genomics UK (COG-UK) Consortium | Berkshire and Surrey Pathology Services Lighthouse Laboratory and Alex Alderton, Roberto Amato, Jeffrey Barrett, Sonia Goncalves, Ewan Harrison, David K. Jackson, Ian Johnston, Dominic Kwiatkowski, Cordelia Langford, John Sillitoe on behalf of the Wellcome Sanger Institute COVID-19 Surveillance Team |
| EPI_ISL_3171377 | Lighthouse Lab in Alderley Park | Wellcome Sanger Institute for the COVID-19 Genomics UK (COG-UK) Consortium | Jacquelyn Wynn, Mairead Hyland, The Lighthouse Lab in Alderley Park and Alex Alderton, Roberto Amato, Jeffrey Barrett, Sonia Goncalves, Ewan Harrison, David K. Jackson, Ian Johnston, Dominic Kwiatkowski, Cordelia Langford, John Sillitoe on behalf of the Wellcome Sanger Institute COVID-19 Surveillance Team |
| EPI_ISL_3171378 | Lighthouse Laboratory Plymouth | Wellcome Sanger Institute for the COVID-19 Genomics UK (COG-UK) Consortium | Lighthouse Laboratory Plymouth and Alex Alderton, Roberto Amato, Jeffrey Barrett, Sonia Goncalves, Ewan Harrison, David K. Jackson, Ian Johnston, Dominic Kwiatkowski, Cordelia Langford, John Sillitoe on behalf of the Wellcome Sanger Institute COVID-19 Surveillance Team |
| EPI_ISL_3171379 | Lighthouse Lab in Alderley Park | Wellcome Sanger Institute for the COVID-19 Genomics UK (COG-UK) Consortium | Jacquelyn Wynn, Mairead Hyland, The Lighthouse Lab in Alderley Park and Alex Alderton, Roberto Amato, Jeffrey Barrett, Sonia Goncalves, Ewan Harrison, David K. Jackson, Ian Johnston, Dominic Kwiatkowski, Cordelia Langford, John Sillitoe on behalf of the Wellcome Sanger Institute COVID-19 Surveillance Team |
| EPI_ISL_3171380 | Lighthouse Laboratory Plymouth | Wellcome Sanger Institute for the COVID-19 Genomics UK (COG-UK) Consortium | Lighthouse Laboratory Plymouth and Alex Alderton, Roberto Amato, Jeffrey Barrett, Sonia Goncalves, Ewan Harrison, David K. Jackson, Ian Johnston, Dominic Kwiatkowski, Cordelia Langford, John Sillitoe on behalf of the Wellcome Sanger Institute COVID-19 Surveillance Team |
| EPI_ISL_3171381 | Lighthouse Lab in Alderley Park | Wellcome Sanger Institute for the COVID-19 Genomics UK (COG-UK) Consortium | Jacquelyn Wynn, Mairead Hyland, The Lighthouse Lab in Alderley Park and Alex Alderton, Roberto Amato, Jeffrey Barrett, Sonia Goncalves, Ewan Harrison, David K. Jackson, Ian Johnston, Dominic Kwiatkowski, Cordelia Langford, John Sillitoe on behalf of the Wellcome Sanger Institute COVID-19 Surveillance Team |
| EPI_ISL_3171382, EPI_ISL_3171383, EPI_ISL_3171384 | Lighthouse Laboratory Plymouth | Wellcome Sanger Institute for the COVID-19 Genomics UK (COG-UK) Consortium | Lighthouse Laboratory Plymouth and Alex Alderton, Roberto Amato, Jeffrey Barrett, Sonia Goncalves, Ewan Harrison, David K. Jackson, Ian Johnston, Dominic Kwiatkowski, Cordelia Langford, John Sillitoe on behalf of the Wellcome Sanger Institute COVID-19 Surveillance Team |
| EPI_ISL_3171385 | Berkshire and Surrey Pathology Services Lighthouse Laboratory | Wellcome Sanger Institute for the COVID-19 Genomics UK (COG-UK) Consortium | Berkshire and Surrey Pathology Services Lighthouse Laboratory and Alex Alderton, Roberto Amato, Jeffrey Barrett, Sonia Goncalves, Ewan Harrison, David K. Jackson, Ian Johnston, Dominic Kwiatkowski, Cordelia Langford, John Sillitoe on behalf of the Wellcome Sanger Institute COVID-19 Surveillance Team |
| EPI_ISL_3171386, EPI_ISL_3171387 | Lighthouse Lab in Alderley Park | Wellcome Sanger Institute for the COVID-19 Genomics UK (COG-UK) Consortium | Jacquelyn Wynn, Mairead Hyland, The Lighthouse Lab in Alderley Park and Alex Alderton, Roberto Amato, Jeffrey Barrett, Sonia Goncalves, Ewan Harrison, David K. Jackson, Ian Johnston, Dominic Kwiatkowski, Cordelia Langford, John Sillitoe on behalf of the Wellcome Sanger Institute COVID-19 Surveillance Team |
| EPI_ISL_3171388 | Berkshire and Surrey Pathology Services Lighthouse Laboratory | Wellcome Sanger Institute for the COVID-19 Genomics UK (COG-UK) Consortium | Berkshire and Surrey Pathology Services Lighthouse Laboratory and Alex Alderton, Roberto Amato, Jeffrey Barrett, Sonia Goncalves, Ewan Harrison, David K. Jackson, Ian Johnston, Dominic Kwiatkowski, Cordelia Langford, John Sillitoe on behalf of the Wellcome Sanger Institute COVID-19 Surveillance Team |
| EPI_ISL_3171389, EPI_ISL_3171390 | Lighthouse Laboratory Plymouth | Wellcome Sanger Institute for the COVID-19 Genomics UK (COG-UK) Consortium | Lighthouse Laboratory Plymouth and Alex Alderton, Roberto Amato, Jeffrey Barrett, Sonia Goncalves, Ewan Harrison, David K. Jackson, Ian Johnston, Dominic Kwiatkowski, Cordelia Langford, John Sillitoe on behalf of the Wellcome Sanger Institute COVID-19 Surveillance Team |
| EPI_ISL_3171391 | Berkshire and Surrey Pathology Services Lighthouse Laboratory | Wellcome Sanger Institute for the COVID-19 Genomics UK (COG-UK) Consortium | Berkshire and Surrey Pathology Services Lighthouse Laboratory and Alex Alderton, Roberto Amato, Jeffrey Barrett, Sonia Goncalves, Ewan Harrison, David K. Jackson, Ian Johnston, Dominic Kwiatkowski, Cordelia Langford, John Sillitoe on behalf of the Wellcome Sanger Institute COVID-19 Surveillance Team |
| EPI_ISL_3171392, EPI_ISL_3171393 | Lighthouse Laboratory Plymouth | Wellcome Sanger Institute for the COVID-19 Genomics UK (COG-UK) Consortium | Lighthouse Laboratory Plymouth and Alex Alderton, Roberto Amato, Jeffrey Barrett, Sonia Goncalves, Ewan Harrison, David K. Jackson, Ian Johnston, Dominic Kwiatkowski, Cordelia Langford, John Sillitoe on behalf of the Wellcome Sanger Institute COVID-19 Surveillance Team |
| EPI_ISL_3171394, EPI_ISL_3171395 | Lighthouse Lab in Alderley Park | Wellcome Sanger Institute for the COVID-19 Genomics UK (COG-UK) Consortium | Jacquelyn Wynn, Mairead Hyland, The Lighthouse Lab in Alderley Park and Alex Alderton, Roberto Amato, Jeffrey Barrett, Sonia Goncalves, Ewan Harrison, David K. Jackson, Ian Johnston, Dominic Kwiatkowski, Cordelia Langford, John Sillitoe on behalf of the Wellcome Sanger Institute COVID-19 Surveillance Team |
| EPI_ISL_3171396 | Berkshire and Surrey Pathology Services Lighthouse Laboratory | Wellcome Sanger Institute for the COVID-19 Genomics UK (COG-UK) Consortium | Berkshire and Surrey Pathology Services Lighthouse Laboratory and Alex Alderton, Roberto Amato, Jeffrey Barrett, Sonia Goncalves, Ewan Harrison, David K. Jackson, Ian Johnston, Dominic Kwiatkowski, Cordelia Langford, John Sillitoe on behalf of the Wellcome Sanger Institute COVID-19 Surveillance Team |
| EPI_ISL_3171397, EPI_ISL_3171398 | Lighthouse Laboratory Plymouth | Wellcome Sanger Institute for the COVID-19 Genomics UK (COG-UK) Consortium | Lighthouse Laboratory Plymouth and Alex Alderton, Roberto Amato, Jeffrey Barrett, Sonia Goncalves, Ewan Harrison, David K. Jackson, Ian Johnston, Dominic Kwiatkowski, Cordelia Langford, John Sillitoe on behalf of the Wellcome Sanger Institute COVID-19 Surveillance Team |
| EPI_ISL_3171399 | Lighthouse Lab in Alderley Park | Wellcome Sanger Institute for the COVID-19 Genomics UK (COG-UK) Consortium | Jacquelyn Wynn, Mairead Hyland, The Lighthouse Lab in Alderley Park and Alex Alderton, Roberto Amato, Jeffrey Barrett, Sonia Goncalves, Ewan Harrison, David K. Jackson, Ian Johnston, Dominic Kwiatkowski, Cordelia Langford, John Sillitoe on behalf of the Wellcome Sanger Institute COVID-19 Surveillance Team |
| EPI_ISL_3171400, EPI_ISL_3171401 | Lighthouse Laboratory Plymouth | Wellcome Sanger Institute for the COVID-19 Genomics UK (COG-UK) Consortium | Lighthouse Laboratory Plymouth and Alex Alderton, Roberto Amato, Jeffrey Barrett, Sonia Goncalves, Ewan Harrison, David K. Jackson, Ian Johnston, Dominic Kwiatkowski, Cordelia Langford, John Sillitoe on behalf of the Wellcome Sanger Institute COVID-19 Surveillance Team |
| EPI_ISL_3171402 | Berkshire and Surrey Pathology Services Lighthouse Laboratory | Wellcome Sanger Institute for the COVID-19 Genomics UK (COG-UK) Consortium | Berkshire and Surrey Pathology Services Lighthouse Laboratory and Alex Alderton, Roberto Amato, Jeffrey Barrett, Sonia Goncalves, Ewan Harrison, David K. Jackson, Ian Johnston, Dominic Kwiatkowski, Cordelia Langford, John Sillitoe on behalf of the Wellcome Sanger Institute COVID-19 Surveillance Team |
| EPI_ISL_3171403 | Lighthouse Laboratory Plymouth | Wellcome Sanger Institute for the COVID-19 Genomics UK (COG-UK) Consortium | Lighthouse Laboratory Plymouth and Alex Alderton, Roberto Amato, Jeffrey Barrett, Sonia Goncalves, Ewan Harrison, David K. Jackson, Ian Johnston, Dominic Kwiatkowski, Cordelia Langford, John Sillitoe on behalf of the Wellcome Sanger Institute COVID-19 Surveillance Team |
| EPI_ISL_3171404 | Lighthouse Lab in Alderley Park | Wellcome Sanger Institute for the COVID-19 Genomics UK (COG-UK) Consortium | Jacquelyn Wynn, Mairead Hyland, The Lighthouse Lab in Alderley Park and Alex Alderton, Roberto Amato, Jeffrey Barrett, Sonia Goncalves, Ewan Harrison, David K. Jackson, Ian Johnston, Dominic Kwiatkowski, Cordelia Langford, John Sillitoe on behalf of the Wellcome Sanger Institute COVID-19 Surveillance Team |
| EPI_ISL_3171405, EPI_ISL_3171406 | Lighthouse Laboratory Plymouth | Wellcome Sanger Institute for the COVID-19 Genomics UK (COG-UK) Consortium | Lighthouse Laboratory Plymouth and Alex Alderton, Roberto Amato, Jeffrey Barrett, Sonia Goncalves, Ewan Harrison, David K. Jackson, Ian Johnston, Dominic Kwiatkowski, Cordelia Langford, John Sillitoe on behalf of the Wellcome Sanger Institute COVID-19 Surveillance Team |
| EPI_ISL_3171407, EPI_ISL_3171408 | Berkshire and Surrey Pathology Services Lighthouse Laboratory | Wellcome Sanger Institute for the COVID-19 Genomics UK (COG-UK) Consortium | Berkshire and Surrey Pathology Services Lighthouse Laboratory and Alex Alderton, Roberto Amato, Jeffrey Barrett, Sonia Goncalves, Ewan Harrison, David K. Jackson, Ian Johnston, Dominic Kwiatkowski, Cordelia Langford, John Sillitoe on behalf of the Wellcome Sanger Institute COVID-19 Surveillance Team |
| EPI_ISL_3171409, EPI_ISL_3171410 | Lighthouse Lab in Alderley Park | Wellcome Sanger Institute for the COVID-19 Genomics UK (COG-UK) Consortium | Jacquelyn Wynn, Mairead Hyland, The Lighthouse Lab in Alderley Park and Alex Alderton, Roberto Amato, Jeffrey Barrett, Sonia Goncalves, Ewan Harrison, David K. Jackson, Ian Johnston, Dominic Kwiatkowski, Cordelia Langford, John Sillitoe on behalf of the Wellcome Sanger Institute COVID-19 Surveillance Team |
| EPI_ISL_3171411, EPI_ISL_3171412 | Berkshire and Surrey Pathology Services Lighthouse Laboratory | Wellcome Sanger Institute for the COVID-19 Genomics UK (COG-UK) Consortium | Berkshire and Surrey Pathology Services Lighthouse Laboratory and Alex Alderton, Roberto Amato, Jeffrey Barrett, Sonia Goncalves, Ewan Harrison, David K. Jackson, Ian Johnston, Dominic Kwiatkowski, Cordelia Langford, John Sillitoe on behalf of the Wellcome Sanger Institute COVID-19 Surveillance Team |
| EPI_ISL_3171413 | Lighthouse Laboratory Plymouth | Wellcome Sanger Institute for the COVID-19 Genomics UK (COG-UK) Consortium | Lighthouse Laboratory Plymouth and Alex Alderton, Roberto Amato, Jeffrey Barrett, Sonia Goncalves, Ewan Harrison, David K. Jackson, Ian Johnston, Dominic Kwiatkowski, Cordelia Langford, John Sillitoe on behalf of the Wellcome Sanger Institute COVID-19 Surveillance Team |
| EPI_ISL_3171414 | Berkshire and Surrey Pathology Services Lighthouse Laboratory | Wellcome Sanger Institute for the COVID-19 Genomics UK (COG-UK) Consortium | Berkshire and Surrey Pathology Services Lighthouse Laboratory and Alex Alderton, Roberto Amato, Jeffrey Barrett, Sonia Goncalves, Ewan Harrison, David K. Jackson, Ian Johnston, Dominic Kwiatkowski, Cordelia Langford, John Sillitoe on behalf of the Wellcome Sanger Institute COVID-19 Surveillance Team |
| EPI_ISL_3171415 | Lighthouse Laboratory Plymouth | Wellcome Sanger Institute for the COVID-19 Genomics UK (COG-UK) Consortium | Lighthouse Laboratory Plymouth and Alex Alderton, Roberto Amato, Jeffrey Barrett, Sonia Goncalves, Ewan Harrison, David K. Jackson, Ian Johnston, Dominic Kwiatkowski, Cordelia Langford, John Sillitoe on behalf of the Wellcome Sanger Institute COVID-19 Surveillance Team |
| EPI_ISL_3171416, EPI_ISL_3171417 | Lighthouse Lab in Alderley Park | Wellcome Sanger Institute for the COVID-19 Genomics UK (COG-UK) Consortium | Jacquelyn Wynn, Mairead Hyland, The Lighthouse Lab in Alderley Park and Alex Alderton, Roberto Amato, Jeffrey Barrett, Sonia Goncalves, Ewan Harrison, David K. Jackson, Ian Johnston, Dominic Kwiatkowski, Cordelia Langford, John Sillitoe on behalf of the Wellcome Sanger Institute COVID-19 Surveillance Team |
| EPI_ISL_3171418, EPI_ISL_3171419 | Berkshire and Surrey Pathology Services Lighthouse Laboratory | Wellcome Sanger Institute for the COVID-19 Genomics UK (COG-UK) Consortium | Berkshire and Surrey Pathology Services Lighthouse Laboratory and Alex Alderton, Roberto Amato, Jeffrey Barrett, Sonia Goncalves, Ewan Harrison, David K. Jackson, Ian Johnston, Dominic Kwiatkowski, Cordelia Langford, John Sillitoe on behalf of the Wellcome Sanger Institute COVID-19 Surveillance Team |
| EPI_ISL_3171420, EPI_ISL_3171421, EPI_ISL_3171422 | Lighthouse Laboratory Plymouth | Wellcome Sanger Institute for the COVID-19 Genomics UK (COG-UK) Consortium | Lighthouse Laboratory Plymouth and Alex Alderton, Roberto Amato, Jeffrey Barrett, Sonia Goncalves, Ewan Harrison, David K. Jackson, Ian Johnston, Dominic Kwiatkowski, Cordelia Langford, John Sillitoe on behalf of the Wellcome Sanger Institute COVID-19 Surveillance Team |
| EPI_ISL_3171423 | Lighthouse Lab in Alderley Park | Wellcome Sanger Institute for the COVID-19 Genomics UK (COG-UK) Consortium | Jacquelyn Wynn, Mairead Hyland, The Lighthouse Lab in Alderley Park and Alex Alderton, Roberto Amato, Jeffrey Barrett, Sonia Goncalves, Ewan Harrison, David K. Jackson, Ian Johnston, Dominic Kwiatkowski, Cordelia Langford, John Sillitoe on behalf of the Wellcome Sanger Institute COVID-19 Surveillance Team |
| EPI_ISL_3171424, EPI_ISL_3171425 | Lighthouse Laboratory Plymouth | Wellcome Sanger Institute for the COVID-19 Genomics UK (COG-UK) Consortium | Lighthouse Laboratory Plymouth and Alex Alderton, Roberto Amato, Jeffrey Barrett, Sonia Goncalves, Ewan Harrison, David K. Jackson, Ian Johnston, Dominic Kwiatkowski, Cordelia Langford, John Sillitoe on behalf of the Wellcome Sanger Institute COVID-19 Surveillance Team |
| EPI_ISL_3171426 | Berkshire and Surrey Pathology Services Lighthouse Laboratory | Wellcome Sanger Institute for the COVID-19 Genomics UK (COG-UK) Consortium | Berkshire and Surrey Pathology Services Lighthouse Laboratory and Alex Alderton, Roberto Amato, Jeffrey Barrett, Sonia Goncalves, Ewan Harrison, David K. Jackson, Ian Johnston, Dominic Kwiatkowski, Cordelia Langford, John Sillitoe on behalf of the Wellcome Sanger Institute COVID-19 Surveillance Team |
| EPI_ISL_3171427 | Lighthouse Laboratory Plymouth | Wellcome Sanger Institute for the COVID-19 Genomics UK (COG-UK) Consortium | Lighthouse Laboratory Plymouth and Alex Alderton, Roberto Amato, Jeffrey Barrett, Sonia Goncalves, Ewan Harrison, David K. Jackson, Ian Johnston, Dominic Kwiatkowski, Cordelia Langford, John Sillitoe on behalf of the Wellcome Sanger Institute COVID-19 Surveillance Team |
| EPI_ISL_3171428 | Berkshire and Surrey Pathology Services Lighthouse Laboratory | Wellcome Sanger Institute for the COVID-19 Genomics UK (COG-UK) Consortium | Berkshire and Surrey Pathology Services Lighthouse Laboratory and Alex Alderton, Roberto Amato, Jeffrey Barrett, Sonia Goncalves, Ewan Harrison, David K. Jackson, Ian Johnston, Dominic Kwiatkowski, Cordelia Langford, John Sillitoe on behalf of the Wellcome Sanger Institute COVID-19 Surveillance Team |
| EPI_ISL_3171429 | Lighthouse Laboratory Plymouth | Wellcome Sanger Institute for the COVID-19 Genomics UK (COG-UK) Consortium | Lighthouse Laboratory Plymouth and Alex Alderton, Roberto Amato, Jeffrey Barrett, Sonia Goncalves, Ewan Harrison, David K. Jackson, Ian Johnston, Dominic Kwiatkowski, Cordelia Langford, John Sillitoe on behalf of the Wellcome Sanger Institute COVID-19 Surveillance Team |
| EPI_ISL_3171430 | Lighthouse Lab in Alderley Park | Wellcome Sanger Institute for the COVID-19 Genomics UK (COG-UK) Consortium | Jacquelyn Wynn, Mairead Hyland, The Lighthouse Lab in Alderley Park and Alex Alderton, Roberto Amato, Jeffrey Barrett, Sonia Goncalves, Ewan Harrison, David K. Jackson, Ian Johnston, Dominic Kwiatkowski, Cordelia Langford, John Sillitoe on behalf of the Wellcome Sanger Institute COVID-19 Surveillance Team |
| EPI_ISL_3171431 | Berkshire and Surrey Pathology Services Lighthouse Laboratory | Wellcome Sanger Institute for the COVID-19 Genomics UK (COG-UK) Consortium | Berkshire and Surrey Pathology Services Lighthouse Laboratory and Alex Alderton, Roberto Amato, Jeffrey Barrett, Sonia Goncalves, Ewan Harrison, David K. Jackson, Ian Johnston, Dominic Kwiatkowski, Cordelia Langford, John Sillitoe on behalf of the Wellcome Sanger Institute COVID-19 Surveillance Team |
| EPI_ISL_3171432 | Lighthouse Lab in Alderley Park | Wellcome Sanger Institute for the COVID-19 Genomics UK (COG-UK) Consortium | Jacquelyn Wynn, Mairead Hyland, The Lighthouse Lab in Alderley Park and Alex Alderton, Roberto Amato, Jeffrey Barrett, Sonia Goncalves, Ewan Harrison, David K. Jackson, Ian Johnston, Dominic Kwiatkowski, Cordelia Langford, John Sillitoe on behalf of the Wellcome Sanger Institute COVID-19 Surveillance Team |
| EPI_ISL_3171433 | Berkshire and Surrey Pathology Services Lighthouse Laboratory | Wellcome Sanger Institute for the COVID-19 Genomics UK (COG-UK) Consortium | Berkshire and Surrey Pathology Services Lighthouse Laboratory and Alex Alderton, Roberto Amato, Jeffrey Barrett, Sonia Goncalves, Ewan Harrison, David K. Jackson, Ian Johnston, Dominic Kwiatkowski, Cordelia Langford, John Sillitoe on behalf of the Wellcome Sanger Institute COVID-19 Surveillance Team |
| EPI_ISL_3171434, EPI_ISL_3171435, EPI_ISL_3171436, EPI_ISL_3171437 | Lighthouse Laboratory Plymouth | Wellcome Sanger Institute for the COVID-19 Genomics UK (COG-UK) Consortium | Lighthouse Laboratory Plymouth and Alex Alderton, Roberto Amato, Jeffrey Barrett, Sonia Goncalves, Ewan Harrison, David K. Jackson, Ian Johnston, Dominic Kwiatkowski, Cordelia Langford, John Sillitoe on behalf of the Wellcome Sanger Institute COVID-19 Surveillance Team |

| | | | |
|---|---|---|---|
| EPI_ISL_3171438 | Lighthouse Lab in Alderley Park | Wellcome Sanger Institute for the COVID-19 Genomics UK (COG-UK) Consortium | Jacquelyn Wynn, Mairead Hyland, The Lighthouse Lab in Alderley Park and Alex Alderton, Roberto Amato, Jeffrey Barrett, Sonia Goncalves, Ewan Harrison, David K. Jackson, Ian Johnston, Dominic Kwiatkowski, Cordelia Langford, John Sillitoe on behalf of the Wellcome Sanger Institute COVID-19 Surveillance Team |
| EPI_ISL_3171439, EPI_ISL_3171440 | Lighthouse Laboratory Plymouth | Wellcome Sanger Institute for the COVID-19 Genomics UK (COG-UK) Consortium | Lighthouse Laboratory Plymouth and Alex Alderton, Roberto Amato, Jeffrey Barrett, Sonia Goncalves, Ewan Harrison, David K. Jackson, Ian Johnston, Dominic Kwiatkowski, Cordelia Langford, John Sillitoe on behalf of the Wellcome Sanger Institute COVID-19 Surveillance Team |
| EPI_ISL_3171441, EPI_ISL_3171442 | Lighthouse Lab in Alderley Park | Wellcome Sanger Institute for the COVID-19 Genomics UK (COG-UK) Consortium | Jacquelyn Wynn, Mairead Hyland, The Lighthouse Lab in Alderley Park and Alex Alderton, Roberto Amato, Jeffrey Barrett, Sonia Goncalves, Ewan Harrison, David K. Jackson, Ian Johnston, Dominic Kwiatkowski, Cordelia Langford, John Sillitoe on behalf of the Wellcome Sanger Institute COVID-19 Surveillance Team |
| EPI_ISL_3171443 | Lighthouse Laboratory Plymouth | Wellcome Sanger Institute for the COVID-19 Genomics UK (COG-UK) Consortium | Lighthouse Laboratory Plymouth and Alex Alderton, Roberto Amato, Jeffrey Barrett, Sonia Goncalves, Ewan Harrison, David K. Jackson, Ian Johnston, Dominic Kwiatkowski, Cordelia Langford, John Sillitoe on behalf of the Wellcome Sanger Institute COVID-19 Surveillance Team |
| EPI_ISL_3171444 | Berkshire and Surrey Pathology Services Lighthouse Laboratory | Wellcome Sanger Institute for the COVID-19 Genomics UK (COG-UK) Consortium | Berkshire and Surrey Pathology Services Lighthouse Laboratory and Alex Alderton, Roberto Amato, Jeffrey Barrett, Sonia Goncalves, Ewan Harrison, David K. Jackson, Ian Johnston, Dominic Kwiatkowski, Cordelia Langford, John Sillitoe on behalf of the Wellcome Sanger Institute COVID-19 Surveillance Team |
| EPI_ISL_3171445 | Lighthouse Laboratory Plymouth | Wellcome Sanger Institute for the COVID-19 Genomics UK (COG-UK) Consortium | Lighthouse Laboratory Plymouth and Alex Alderton, Roberto Amato, Jeffrey Barrett, Sonia Goncalves, Ewan Harrison, David K. Jackson, Ian Johnston, Dominic Kwiatkowski, Cordelia Langford, John Sillitoe on behalf of the Wellcome Sanger Institute COVID-19 Surveillance Team |
| EPI_ISL_3171446 | Berkshire and Surrey Pathology Services Lighthouse Laboratory | Wellcome Sanger Institute for the COVID-19 Genomics UK (COG-UK) Consortium | Berkshire and Surrey Pathology Services Lighthouse Laboratory and Alex Alderton, Roberto Amato, Jeffrey Barrett, Sonia Goncalves, Ewan Harrison, David K. Jackson, Ian Johnston, Dominic Kwiatkowski, Cordelia Langford, John Sillitoe on behalf of the Wellcome Sanger Institute COVID-19 Surveillance Team |
| EPI_ISL_3171447 | Lighthouse Laboratory Plymouth | Wellcome Sanger Institute for the COVID-19 Genomics UK (COG-UK) Consortium | Lighthouse Laboratory Plymouth and Alex Alderton, Roberto Amato, Jeffrey Barrett, Sonia Goncalves, Ewan Harrison, David K. Jackson, Ian Johnston, Dominic Kwiatkowski, Cordelia Langford, John Sillitoe on behalf of the Wellcome Sanger Institute COVID-19 Surveillance Team |
| EPI_ISL_3171448 | Lighthouse Lab in Alderley Park | Wellcome Sanger Institute for the COVID-19 Genomics UK (COG-UK) Consortium | Jacquelyn Wynn, Mairead Hyland, The Lighthouse Lab in Alderley Park and Alex Alderton, Roberto Amato, Jeffrey Barrett, Sonia Goncalves, Ewan Harrison, David K. Jackson, Ian Johnston, Dominic Kwiatkowski, Cordelia Langford, John Sillitoe on behalf of the Wellcome Sanger Institute COVID-19 Surveillance Team |
| EPI_ISL_3171449, EPI_ISL_3171450, EPI_ISL_3171451, EPI_ISL_3171452, EPI_ISL_3171453, EPI_ISL_3171454 | Lighthouse Laboratory Plymouth | Wellcome Sanger Institute for the COVID-19 Genomics UK (COG-UK) Consortium | Lighthouse Laboratory Plymouth and Alex Alderton, Roberto Amato, Jeffrey Barrett, Sonia Goncalves, Ewan Harrison, David K. Jackson, Ian Johnston, Dominic Kwiatkowski, Cordelia Langford, John Sillitoe on behalf of the Wellcome Sanger Institute COVID-19 Surveillance Team |
| EPI_ISL_3171455 | Berkshire and Surrey Pathology Services Lighthouse Laboratory | Wellcome Sanger Institute for the COVID-19 Genomics UK (COG-UK) Consortium | Berkshire and Surrey Pathology Services Lighthouse Laboratory and Alex Alderton, Roberto Amato, Jeffrey Barrett, Sonia Goncalves, Ewan Harrison, David K. Jackson, Ian Johnston, Dominic Kwiatkowski, Cordelia Langford, John Sillitoe on behalf of the Wellcome Sanger Institute COVID-19 Surveillance Team |
| EPI_ISL_3171456 | Lighthouse Laboratory Plymouth | Wellcome Sanger Institute for the COVID-19 Genomics UK (COG-UK) Consortium | Lighthouse Laboratory Plymouth and Alex Alderton, Roberto Amato, Jeffrey Barrett, Sonia Goncalves, Ewan Harrison, David K. Jackson, Ian Johnston, Dominic Kwiatkowski, Cordelia Langford, John Sillitoe on behalf of the Wellcome Sanger Institute COVID-19 Surveillance Team |
| EPI_ISL_3171457 | Lighthouse Lab in Alderley Park | Wellcome Sanger Institute for the COVID-19 Genomics UK (COG-UK) Consortium | Jacquelyn Wynn, Mairead Hyland, The Lighthouse Lab in Alderley Park and Alex Alderton, Roberto Amato, Jeffrey Barrett, Sonia Goncalves, Ewan Harrison, David K. Jackson, Ian Johnston, Dominic Kwiatkowski, Cordelia Langford, John Sillitoe on behalf of the Wellcome Sanger Institute COVID-19 Surveillance Team |
| EPI_ISL_3171458, EPI_ISL_3171459 | Lighthouse Laboratory Plymouth | Wellcome Sanger Institute for the COVID-19 Genomics UK (COG-UK) Consortium | Lighthouse Laboratory Plymouth and Alex Alderton, Roberto Amato, Jeffrey Barrett, Sonia Goncalves, Ewan Harrison, David K. Jackson, Ian Johnston, Dominic Kwiatkowski, Cordelia Langford, John Sillitoe on behalf of the Wellcome Sanger Institute COVID-19 Surveillance Team |
| EPI_ISL_3171460 | Lighthouse Lab in Alderley Park | Wellcome Sanger Institute for the COVID-19 Genomics UK (COG-UK) Consortium | Jacquelyn Wynn, Mairead Hyland, The Lighthouse Lab in Alderley Park and Alex Alderton, Roberto Amato, Jeffrey Barrett, Sonia Goncalves, Ewan Harrison, David K. Jackson, Ian Johnston, Dominic Kwiatkowski, Cordelia Langford, John Sillitoe on behalf of the Wellcome Sanger Institute COVID-19 Surveillance Team |
| EPI_ISL_3171461, EPI_ISL_3171462 | Berkshire and Surrey Pathology Services Lighthouse Laboratory | Wellcome Sanger Institute for the COVID-19 Genomics UK (COG-UK) Consortium | Berkshire and Surrey Pathology Services Lighthouse Laboratory and Alex Alderton, Roberto Amato, Jeffrey Barrett, Sonia Goncalves, Ewan Harrison, David K. Jackson, Ian Johnston, Dominic Kwiatkowski, Cordelia Langford, John Sillitoe on behalf of the Wellcome Sanger Institute COVID-19 Surveillance Team |
| EPI_ISL_3171463 | Lighthouse Lab in Alderley Park | Wellcome Sanger Institute for the COVID-19 Genomics UK (COG-UK) Consortium | Jacquelyn Wynn, Mairead Hyland, The Lighthouse Lab in Alderley Park and Alex Alderton, Roberto Amato, Jeffrey Barrett, Sonia Goncalves, Ewan Harrison, David K. Jackson, Ian Johnston, Dominic Kwiatkowski, Cordelia Langford, John Sillitoe on behalf of the Wellcome Sanger Institute COVID-19 Surveillance Team |
| EPI_ISL_3171464, EPI_ISL_3171465, EPI_ISL_3171466 | Lighthouse Laboratory Plymouth | Wellcome Sanger Institute for the COVID-19 Genomics UK (COG-UK) Consortium | Lighthouse Laboratory Plymouth and Alex Alderton, Roberto Amato, Jeffrey Barrett, Sonia Goncalves, Ewan Harrison, David K. Jackson, Ian Johnston, Dominic Kwiatkowski, Cordelia Langford, John Sillitoe on behalf of the Wellcome Sanger Institute COVID-19 Surveillance Team |
| EPI_ISL_3171467, EPI_ISL_3171468 | Lighthouse Lab in Alderley Park | Wellcome Sanger Institute for the COVID-19 Genomics UK (COG-UK) Consortium | Jacquelyn Wynn, Mairead Hyland, The Lighthouse Lab in Alderley Park and Alex Alderton, Roberto Amato, Jeffrey Barrett, Sonia Goncalves, Ewan Harrison, David K. Jackson, Ian Johnston, Dominic Kwiatkowski, Cordelia Langford, John Sillitoe on behalf of the Wellcome Sanger Institute COVID-19 Surveillance Team |
| EPI_ISL_3171469, EPI_ISL_3171470, EPI_ISL_3171471 | Lighthouse Laboratory Plymouth | Wellcome Sanger Institute for the COVID-19 Genomics UK (COG-UK) Consortium | Lighthouse Laboratory Plymouth and Alex Alderton, Roberto Amato, Jeffrey Barrett, Sonia Goncalves, Ewan Harrison, David K. Jackson, Ian Johnston, Dominic Kwiatkowski, Cordelia Langford, John Sillitoe on behalf of the Wellcome Sanger Institute COVID-19 Surveillance Team |
| EPI_ISL_3171472 | Lighthouse Lab in Alderley Park | Wellcome Sanger Institute for the COVID-19 Genomics UK (COG-UK) Consortium | Jacquelyn Wynn, Mairead Hyland, The Lighthouse Lab in Alderley Park and Alex Alderton, Roberto Amato, Jeffrey Barrett, Sonia Goncalves, Ewan Harrison, David K. Jackson, Ian Johnston, Dominic Kwiatkowski, Cordelia Langford, John Sillitoe on behalf of the Wellcome Sanger Institute COVID-19 Surveillance Team |
| EPI_ISL_3171473 | Berkshire and Surrey Pathology Services Lighthouse Laboratory | Wellcome Sanger Institute for the COVID-19 Genomics UK (COG-UK) Consortium | Berkshire and Surrey Pathology Services Lighthouse Laboratory and Alex Alderton, Roberto Amato, Jeffrey Barrett, Sonia Goncalves, Ewan Harrison, David K. Jackson, Ian Johnston, Dominic Kwiatkowski, Cordelia Langford, John Sillitoe on behalf of the Wellcome Sanger Institute COVID-19 Surveillance Team |
| EPI_ISL_3171474, EPI_ISL_3171475 | Lighthouse Lab in Alderley Park | Wellcome Sanger Institute for the COVID-19 Genomics UK (COG-UK) Consortium | Jacquelyn Wynn, Mairead Hyland, The Lighthouse Lab in Alderley Park and Alex Alderton, Roberto Amato, Jeffrey Barrett, Sonia Goncalves, Ewan Harrison, David K. Jackson, Ian Johnston, Dominic Kwiatkowski, Cordelia Langford, John Sillitoe on behalf of the Wellcome Sanger Institute COVID-19 Surveillance Team |
| EPI_ISL_3171476, EPI_ISL_3171477, EPI_ISL_3171478 | Lighthouse Laboratory Plymouth | Wellcome Sanger Institute for the COVID-19 Genomics UK (COG-UK) Consortium | Lighthouse Laboratory Plymouth and Alex Alderton, Roberto Amato, Jeffrey Barrett, Sonia Goncalves, Ewan Harrison, David K. Jackson, Ian Johnston, Dominic Kwiatkowski, Cordelia Langford, John Sillitoe on behalf of the Wellcome Sanger Institute COVID-19 Surveillance Team |
| EPI_ISL_3171479 | Lighthouse Lab in Alderley Park | Wellcome Sanger Institute for the COVID-19 Genomics UK (COG-UK) Consortium | Jacquelyn Wynn, Mairead Hyland, The Lighthouse Lab in Alderley Park and Alex Alderton, Roberto Amato, Jeffrey Barrett, Sonia Goncalves, Ewan Harrison, David K. Jackson, Ian Johnston, Dominic Kwiatkowski, Cordelia Langford, John Sillitoe on behalf of the Wellcome Sanger Institute COVID-19 Surveillance Team |
| EPI_ISL_3171480 | Berkshire and Surrey Pathology Services Lighthouse Laboratory | Wellcome Sanger Institute for the COVID-19 Genomics UK (COG-UK) Consortium | Berkshire and Surrey Pathology Services Lighthouse Laboratory and Alex Alderton, Roberto Amato, Jeffrey Barrett, Sonia Goncalves, Ewan Harrison, David K. Jackson, Ian Johnston, Dominic Kwiatkowski, Cordelia Langford, John Sillitoe on behalf of the Wellcome Sanger Institute COVID-19 Surveillance Team |
| EPI_ISL_3171481, EPI_ISL_3171482 | Lighthouse Laboratory Plymouth | Wellcome Sanger Institute for the COVID-19 Genomics UK (COG-UK) Consortium | Lighthouse Laboratory Plymouth and Alex Alderton, Roberto Amato, Jeffrey Barrett, Sonia Goncalves, Ewan Harrison, David K. Jackson, Ian Johnston, Dominic Kwiatkowski, Cordelia Langford, John Sillitoe on behalf of the Wellcome Sanger Institute COVID-19 Surveillance Team |
| EPI_ISL_3171483 | Lighthouse Lab in Alderley Park | Wellcome Sanger Institute for the COVID-19 Genomics UK (COG-UK) Consortium | Jacquelyn Wynn, Mairead Hyland, The Lighthouse Lab in Alderley Park and Alex Alderton, Roberto Amato, Jeffrey Barrett, Sonia Goncalves, Ewan Harrison, David K. Jackson, Ian Johnston, Dominic Kwiatkowski, Cordelia Langford, John Sillitoe on behalf of the Wellcome Sanger Institute COVID-19 Surveillance Team |
| EPI_ISL_3171484, EPI_ISL_3171485 | Lighthouse Laboratory Plymouth | Wellcome Sanger Institute for the COVID-19 Genomics UK (COG-UK) Consortium | Lighthouse Laboratory Plymouth and Alex Alderton, Roberto Amato, Jeffrey Barrett, Sonia Goncalves, Ewan Harrison, David K. Jackson, Ian Johnston, Dominic Kwiatkowski, Cordelia Langford, John Sillitoe on behalf of the Wellcome Sanger Institute COVID-19 Surveillance Team |
| EPI_ISL_3171486 | Lighthouse Lab in Alderley Park | Wellcome Sanger Institute for the COVID-19 Genomics UK (COG-UK) Consortium | Jacquelyn Wynn, Mairead Hyland, The Lighthouse Lab in Alderley Park and Alex Alderton, Roberto Amato, Jeffrey Barrett, Sonia Goncalves, Ewan Harrison, David K. Jackson, Ian Johnston, Dominic Kwiatkowski, Cordelia Langford, John Sillitoe on behalf of the Wellcome Sanger Institute COVID-19 Surveillance Team |
| EPI_ISL_3171487 | Berkshire and Surrey Pathology Services Lighthouse Laboratory | Wellcome Sanger Institute for the COVID-19 Genomics UK (COG-UK) Consortium | Berkshire and Surrey Pathology Services Lighthouse Laboratory and Alex Alderton, Roberto Amato, Jeffrey Barrett, Sonia Goncalves, Ewan Harrison, David K. Jackson, Ian Johnston, Dominic Kwiatkowski, Cordelia Langford, John Sillitoe on behalf of the Wellcome Sanger Institute COVID-19 Surveillance Team |
| EPI_ISL_3171488, EPI_ISL_3171489 | Lighthouse Lab in Alderley Park | Wellcome Sanger Institute for the COVID-19 Genomics UK (COG-UK) Consortium | Jacquelyn Wynn, Mairead Hyland, The Lighthouse Lab in Alderley Park and Alex Alderton, Roberto Amato, Jeffrey Barrett, Sonia Goncalves, Ewan Harrison, David K. Jackson, Ian Johnston, Dominic Kwiatkowski, Cordelia Langford, John Sillitoe on behalf of the Wellcome Sanger Institute COVID-19 Surveillance Team |
| EPI_ISL_3171490 | Berkshire and Surrey Pathology Services Lighthouse Laboratory | Wellcome Sanger Institute for the COVID-19 Genomics UK (COG-UK) Consortium | Berkshire and Surrey Pathology Services Lighthouse Laboratory and Alex Alderton, Roberto Amato, Jeffrey Barrett, Sonia Goncalves, Ewan Harrison, David K. Jackson, Ian Johnston, Dominic Kwiatkowski, Cordelia Langford, John Sillitoe on behalf of the Wellcome Sanger Institute COVID-19 Surveillance Team |
| EPI_ISL_3171491 | Lighthouse Laboratory Plymouth | Wellcome Sanger Institute for the COVID-19 Genomics UK (COG-UK) Consortium | Lighthouse Laboratory Plymouth and Alex Alderton, Roberto Amato, Jeffrey Barrett, Sonia Goncalves, Ewan Harrison, David K. Jackson, Ian Johnston, Dominic Kwiatkowski, Cordelia Langford, John Sillitoe on behalf of the Wellcome Sanger Institute COVID-19 Surveillance Team |
| EPI_ISL_3171492 | Lighthouse Lab in Alderley Park | Wellcome Sanger Institute for the COVID-19 Genomics UK (COG-UK) Consortium | Jacquelyn Wynn, Mairead Hyland, The Lighthouse Lab in Alderley Park and Alex Alderton, Roberto Amato, Jeffrey Barrett, Sonia Goncalves, Ewan Harrison, David K. Jackson, Ian Johnston, Dominic Kwiatkowski, Cordelia Langford, John Sillitoe on behalf of the Wellcome Sanger Institute COVID-19 Surveillance Team |
| EPI_ISL_3171493 | Berkshire and Surrey Pathology Services Lighthouse Laboratory | Wellcome Sanger Institute for the COVID-19 Genomics UK (COG-UK) Consortium | Berkshire and Surrey Pathology Services Lighthouse Laboratory and Alex Alderton, Roberto Amato, Jeffrey Barrett, Sonia Goncalves, Ewan Harrison, David K. Jackson, Ian Johnston, Dominic Kwiatkowski, Cordelia Langford, John Sillitoe on behalf of the Wellcome Sanger Institute COVID-19 Surveillance Team |
| EPI_ISL_3171494 | Lighthouse Laboratory Plymouth | Wellcome Sanger Institute for the COVID-19 Genomics UK (COG-UK) Consortium | Lighthouse Laboratory Plymouth and Alex Alderton, Roberto Amato, Jeffrey Barrett, Sonia Goncalves, Ewan Harrison, David K. Jackson, Ian Johnston, Dominic Kwiatkowski, Cordelia Langford, John Sillitoe on behalf of the Wellcome Sanger Institute COVID-19 Surveillance Team |
| EPI_ISL_3171495 | Lighthouse Lab in Alderley Park | Wellcome Sanger Institute for the COVID-19 Genomics UK (COG-UK) Consortium | Jacquelyn Wynn, Mairead Hyland, The Lighthouse Lab in Alderley Park and Alex Alderton, Roberto Amato, Jeffrey Barrett, Sonia Goncalves, Ewan Harrison, David K. Jackson, Ian Johnston, Dominic Kwiatkowski, Cordelia Langford, John Sillitoe on behalf of the Wellcome Sanger Institute COVID-19 Surveillance Team |
| EPI_ISL_3171496, EPI_ISL_3171497 | Lighthouse Laboratory Plymouth | Wellcome Sanger Institute for the COVID-19 Genomics UK (COG-UK) Consortium | Lighthouse Laboratory Plymouth and Alex Alderton, Roberto Amato, Jeffrey Barrett, Sonia Goncalves, Ewan Harrison, David K. Jackson, Ian Johnston, Dominic Kwiatkowski, Cordelia Langford, John Sillitoe on behalf of the Wellcome Sanger Institute COVID-19 Surveillance Team |
| EPI_ISL_3171498 | Lighthouse Lab in Alderley Park | Wellcome Sanger Institute for the COVID-19 Genomics UK (COG-UK) Consortium | Jacquelyn Wynn, Mairead Hyland, The Lighthouse Lab in Alderley Park and Alex Alderton, Roberto Amato, Jeffrey Barrett, Sonia Goncalves, Ewan Harrison, David K. Jackson, Ian Johnston, Dominic Kwiatkowski, Cordelia Langford, John Sillitoe on behalf of the Wellcome Sanger Institute COVID-19 Surveillance Team |
| EPI_ISL_3171499, EPI_ISL_3171500 | Lighthouse Laboratory Plymouth | Wellcome Sanger Institute for the COVID-19 Genomics UK (COG-UK) Consortium | Lighthouse Laboratory Plymouth and Alex Alderton, Roberto Amato, Jeffrey Barrett, Sonia Goncalves, Ewan Harrison, David K. Jackson, Ian Johnston, Dominic Kwiatkowski, Cordelia Langford, John Sillitoe on behalf of the Wellcome Sanger Institute COVID-19 Surveillance Team |
| EPI_ISL_3171501 | Berkshire and Surrey Pathology Services Lighthouse Laboratory | Wellcome Sanger Institute for the COVID-19 Genomics UK (COG-UK) Consortium | Berkshire and Surrey Pathology Services Lighthouse Laboratory and Alex Alderton, Roberto Amato, Jeffrey Barrett, Sonia Goncalves, Ewan Harrison, David K. Jackson, Ian Johnston, Dominic Kwiatkowski, Cordelia Langford, John Sillitoe on behalf of the Wellcome Sanger Institute COVID-19 Surveillance Team |

| | | | |
|---|---|---|---|
| EPI_ISL_3171502 | Lighthouse Lab in Alderley Park | Wellcome Sanger Institute for the COVID-19 Genomics UK (COG-UK) Consortium | Jacquelyn Wynn, Mairead Hyland, The Lighthouse Lab in Alderley Park and Alex Alderton, Roberto Amato, Jeffrey Barrett, Sonia Goncalves, Ewan Harrison, David K. Jackson, Ian Johnston, Dominic Kwiatkowski, Cordelia Langford, John Sillitoe on behalf of the Wellcome Sanger Institute COVID-19 Surveillance Team |
| EPI_ISL_3171503 | Lighthouse Laboratory Plymouth | Wellcome Sanger Institute for the COVID-19 Genomics UK (COG-UK) Consortium | Lighthouse Laboratory Plymouth and Alex Alderton, Roberto Amato, Jeffrey Barrett, Sonia Goncalves, Ewan Harrison, David K. Jackson, Ian Johnston, Dominic Kwiatkowski, Cordelia Langford, John Sillitoe on behalf of the Wellcome Sanger Institute COVID-19 Surveillance Team |
| EPI_ISL_3171504 | Lighthouse Lab in Alderley Park | Wellcome Sanger Institute for the COVID-19 Genomics UK (COG-UK) Consortium | Jacquelyn Wynn, Mairead Hyland, The Lighthouse Lab in Alderley Park and Alex Alderton, Roberto Amato, Jeffrey Barrett, Sonia Goncalves, Ewan Harrison, David K. Jackson, Ian Johnston, Dominic Kwiatkowski, Cordelia Langford, John Sillitoe on behalf of the Wellcome Sanger Institute COVID-19 Surveillance Team |
| EPI_ISL_3171505, EPI_ISL_3171506 | Lighthouse Laboratory Plymouth | Wellcome Sanger Institute for the COVID-19 Genomics UK (COG-UK) Consortium | Lighthouse Laboratory Plymouth and Alex Alderton, Roberto Amato, Jeffrey Barrett, Sonia Goncalves, Ewan Harrison, David K. Jackson, Ian Johnston, Dominic Kwiatkowski, Cordelia Langford, John Sillitoe on behalf of the Wellcome Sanger Institute COVID-19 Surveillance Team |
| EPI_ISL_3171507 | Berkshire and Surrey Pathology Services Lighthouse Laboratory | Wellcome Sanger Institute for the COVID-19 Genomics UK (COG-UK) Consortium | Berkshire and Surrey Pathology Services Lighthouse Laboratory and Alex Alderton, Roberto Amato, Jeffrey Barrett, Sonia Goncalves, Ewan Harrison, David K. Jackson, Ian Johnston, Dominic Kwiatkowski, Cordelia Langford, John Sillitoe on behalf of the Wellcome Sanger Institute COVID-19 Surveillance Team |
| EPI_ISL_3171508 | Lighthouse Laboratory Plymouth | Wellcome Sanger Institute for the COVID-19 Genomics UK (COG-UK) Consortium | Lighthouse Laboratory Plymouth and Alex Alderton, Roberto Amato, Jeffrey Barrett, Sonia Goncalves, Ewan Harrison, David K. Jackson, Ian Johnston, Dominic Kwiatkowski, Cordelia Langford, John Sillitoe on behalf of the Wellcome Sanger Institute COVID-19 Surveillance Team |
| EPI_ISL_3171509, EPI_ISL_3171510 | Lighthouse Lab in Alderley Park | Wellcome Sanger Institute for the COVID-19 Genomics UK (COG-UK) Consortium | Jacquelyn Wynn, Mairead Hyland, The Lighthouse Lab in Alderley Park and Alex Alderton, Roberto Amato, Jeffrey Barrett, Sonia Goncalves, Ewan Harrison, David K. Jackson, Ian Johnston, Dominic Kwiatkowski, Cordelia Langford, John Sillitoe on behalf of the Wellcome Sanger Institute COVID-19 Surveillance Team |
| EPI_ISL_3171511, EPI_ISL_3171512, EPI_ISL_3171513, EPI_ISL_3171514 | Lighthouse Laboratory Plymouth | Wellcome Sanger Institute for the COVID-19 Genomics UK (COG-UK) Consortium | Lighthouse Laboratory Plymouth and Alex Alderton, Roberto Amato, Jeffrey Barrett, Sonia Goncalves, Ewan Harrison, David K. Jackson, Ian Johnston, Dominic Kwiatkowski, Cordelia Langford, John Sillitoe on behalf of the Wellcome Sanger Institute COVID-19 Surveillance Team |
| EPI_ISL_3171515, EPI_ISL_3171516 | Lighthouse Lab in Alderley Park | Wellcome Sanger Institute for the COVID-19 Genomics UK (COG-UK) Consortium | Jacquelyn Wynn, Mairead Hyland, The Lighthouse Lab in Alderley Park and Alex Alderton, Roberto Amato, Jeffrey Barrett, Sonia Goncalves, Ewan Harrison, David K. Jackson, Ian Johnston, Dominic Kwiatkowski, Cordelia Langford, John Sillitoe on behalf of the Wellcome Sanger Institute COVID-19 Surveillance Team |
| EPI_ISL_3171517, EPI_ISL_3171518, EPI_ISL_3171519 | Berkshire and Surrey Pathology Services Lighthouse Laboratory | Wellcome Sanger Institute for the COVID-19 Genomics UK (COG-UK) Consortium | Berkshire and Surrey Pathology Services Lighthouse Laboratory and Alex Alderton, Roberto Amato, Jeffrey Barrett, Sonia Goncalves, Ewan Harrison, David K. Jackson, Ian Johnston, Dominic Kwiatkowski, Cordelia Langford, John Sillitoe on behalf of the Wellcome Sanger Institute COVID-19 Surveillance Team |
| EPI_ISL_3171520 | Lighthouse Laboratory Plymouth | Wellcome Sanger Institute for the COVID-19 Genomics UK (COG-UK) Consortium | Lighthouse Laboratory Plymouth and Alex Alderton, Roberto Amato, Jeffrey Barrett, Sonia Goncalves, Ewan Harrison, David K. Jackson, Ian Johnston, Dominic Kwiatkowski, Cordelia Langford, John Sillitoe on behalf of the Wellcome Sanger Institute COVID-19 Surveillance Team |
| EPI_ISL_3171521 | Lighthouse Lab in Alderley Park | Wellcome Sanger Institute for the COVID-19 Genomics UK (COG-UK) Consortium | Jacquelyn Wynn, Mairead Hyland, The Lighthouse Lab in Alderley Park and Alex Alderton, Roberto Amato, Jeffrey Barrett, Sonia Goncalves, Ewan Harrison, David K. Jackson, Ian Johnston, Dominic Kwiatkowski, Cordelia Langford, John Sillitoe on behalf of the Wellcome Sanger Institute COVID-19 Surveillance Team |
| EPI_ISL_3171522, EPI_ISL_3171523, EPI_ISL_3171524 | Lighthouse Laboratory Plymouth | Wellcome Sanger Institute for the COVID-19 Genomics UK (COG-UK) Consortium | Lighthouse Laboratory Plymouth and Alex Alderton, Roberto Amato, Jeffrey Barrett, Sonia Goncalves, Ewan Harrison, David K. Jackson, Ian Johnston, Dominic Kwiatkowski, Cordelia Langford, John Sillitoe on behalf of the Wellcome Sanger Institute COVID-19 Surveillance Team |
| EPI_ISL_3171525 | Lighthouse Lab in Alderley Park | Wellcome Sanger Institute for the COVID-19 Genomics UK (COG-UK) Consortium | Jacquelyn Wynn, Mairead Hyland, The Lighthouse Lab in Alderley Park and Alex Alderton, Roberto Amato, Jeffrey Barrett, Sonia Goncalves, Ewan Harrison, David K. Jackson, Ian Johnston, Dominic Kwiatkowski, Cordelia Langford, John Sillitoe on behalf of the Wellcome Sanger Institute COVID-19 Surveillance Team |
| EPI_ISL_3171526, EPI_ISL_3171527, EPI_ISL_3171528 | Berkshire and Surrey Pathology Services Lighthouse Laboratory | Wellcome Sanger Institute for the COVID-19 Genomics UK (COG-UK) Consortium | Berkshire and Surrey Pathology Services Lighthouse Laboratory and Alex Alderton, Roberto Amato, Jeffrey Barrett, Sonia Goncalves, Ewan Harrison, David K. Jackson, Ian Johnston, Dominic Kwiatkowski, Cordelia Langford, John Sillitoe on behalf of the Wellcome Sanger Institute COVID-19 Surveillance Team |
| EPI_ISL_3171529, EPI_ISL_3171530, EPI_ISL_3171531, EPI_ISL_3171532, EPI_ISL_3171533 | Lighthouse Laboratory Plymouth | Wellcome Sanger Institute for the COVID-19 Genomics UK (COG-UK) Consortium | Lighthouse Laboratory Plymouth and Alex Alderton, Roberto Amato, Jeffrey Barrett, Sonia Goncalves, Ewan Harrison, David K. Jackson, Ian Johnston, Dominic Kwiatkowski, Cordelia Langford, John Sillitoe on behalf of the Wellcome Sanger Institute COVID-19 Surveillance Team |
| EPI_ISL_3171534 | Berkshire and Surrey Pathology Services Lighthouse Laboratory | Wellcome Sanger Institute for the COVID-19 Genomics UK (COG-UK) Consortium | Berkshire and Surrey Pathology Services Lighthouse Laboratory and Alex Alderton, Roberto Amato, Jeffrey Barrett, Sonia Goncalves, Ewan Harrison, David K. Jackson, Ian Johnston, Dominic Kwiatkowski, Cordelia Langford, John Sillitoe on behalf of the Wellcome Sanger Institute COVID-19 Surveillance Team |
| EPI_ISL_3171535 | Lighthouse Lab in Alderley Park | Wellcome Sanger Institute for the COVID-19 Genomics UK (COG-UK) Consortium | Jacquelyn Wynn, Mairead Hyland, The Lighthouse Lab in Alderley Park and Alex Alderton, Roberto Amato, Jeffrey Barrett, Sonia Goncalves, Ewan Harrison, David K. Jackson, Ian Johnston, Dominic Kwiatkowski, Cordelia Langford, John Sillitoe on behalf of the Wellcome Sanger Institute COVID-19 Surveillance Team |
| EPI_ISL_3171536 | Lighthouse Laboratory Plymouth | Wellcome Sanger Institute for the COVID-19 Genomics UK (COG-UK) Consortium | Lighthouse Laboratory Plymouth and Alex Alderton, Roberto Amato, Jeffrey Barrett, Sonia Goncalves, Ewan Harrison, David K. Jackson, Ian Johnston, Dominic Kwiatkowski, Cordelia Langford, John Sillitoe on behalf of the Wellcome Sanger Institute COVID-19 Surveillance Team |
| EPI_ISL_3171537 | Lighthouse Lab in Alderley Park | Wellcome Sanger Institute for the COVID-19 Genomics UK (COG-UK) Consortium | Jacquelyn Wynn, Mairead Hyland, The Lighthouse Lab in Alderley Park and Alex Alderton, Roberto Amato, Jeffrey Barrett, Sonia Goncalves, Ewan Harrison, David K. Jackson, Ian Johnston, Dominic Kwiatkowski, Cordelia Langford, John Sillitoe on behalf of the Wellcome Sanger Institute COVID-19 Surveillance Team |
| EPI_ISL_3171538 | Lighthouse Laboratory Plymouth | Wellcome Sanger Institute for the COVID-19 Genomics UK (COG-UK) Consortium | Lighthouse Laboratory Plymouth and Alex Alderton, Roberto Amato, Jeffrey Barrett, Sonia Goncalves, Ewan Harrison, David K. Jackson, Ian Johnston, Dominic Kwiatkowski, Cordelia Langford, John Sillitoe on behalf of the Wellcome Sanger Institute COVID-19 Surveillance Team |
| EPI_ISL_3171539, EPI_ISL_3171540 | Lighthouse Lab in Alderley Park | Wellcome Sanger Institute for the COVID-19 Genomics UK (COG-UK) Consortium | Jacquelyn Wynn, Mairead Hyland, The Lighthouse Lab in Alderley Park and Alex Alderton, Roberto Amato, Jeffrey Barrett, Sonia Goncalves, Ewan Harrison, David K. Jackson, Ian Johnston, Dominic Kwiatkowski, Cordelia Langford, John Sillitoe on behalf of the Wellcome Sanger Institute COVID-19 Surveillance Team |
| EPI_ISL_3171541, EPI_ISL_3171542, EPI_ISL_3171543, EPI_ISL_3171544, EPI_ISL_3171545 | Lighthouse Laboratory Plymouth | Wellcome Sanger Institute for the COVID-19 Genomics UK (COG-UK) Consortium | Lighthouse Laboratory Plymouth and Alex Alderton, Roberto Amato, Jeffrey Barrett, Sonia Goncalves, Ewan Harrison, David K. Jackson, Ian Johnston, Dominic Kwiatkowski, Cordelia Langford, John Sillitoe on behalf of the Wellcome Sanger Institute COVID-19 Surveillance Team |
| EPI_ISL_3171546 | Lighthouse Lab in Alderley Park | Wellcome Sanger Institute for the COVID-19 Genomics UK (COG-UK) Consortium | Jacquelyn Wynn, Mairead Hyland, The Lighthouse Lab in Alderley Park and Alex Alderton, Roberto Amato, Jeffrey Barrett, Sonia Goncalves, Ewan Harrison, David K. Jackson, Ian Johnston, Dominic Kwiatkowski, Cordelia Langford, John Sillitoe on behalf of the Wellcome Sanger Institute COVID-19 Surveillance Team |
| EPI_ISL_3171547, EPI_ISL_3171548, EPI_ISL_3171549 | Lighthouse Laboratory Plymouth | Wellcome Sanger Institute for the COVID-19 Genomics UK (COG-UK) Consortium | Lighthouse Laboratory Plymouth and Alex Alderton, Roberto Amato, Jeffrey Barrett, Sonia Goncalves, Ewan Harrison, David K. Jackson, Ian Johnston, Dominic Kwiatkowski, Cordelia Langford, John Sillitoe on behalf of the Wellcome Sanger Institute COVID-19 Surveillance Team |
| EPI_ISL_3171550 | Lighthouse Lab in Alderley Park | Wellcome Sanger Institute for the COVID-19 Genomics UK (COG-UK) Consortium | Jacquelyn Wynn, Mairead Hyland, The Lighthouse Lab in Alderley Park and Alex Alderton, Roberto Amato, Jeffrey Barrett, Sonia Goncalves, Ewan Harrison, David K. Jackson, Ian Johnston, Dominic Kwiatkowski, Cordelia Langford, John Sillitoe on behalf of the Wellcome Sanger Institute COVID-19 Surveillance Team |
| EPI_ISL_3171551 | Lighthouse Laboratory Plymouth | Wellcome Sanger Institute for the COVID-19 Genomics UK (COG-UK) Consortium | Lighthouse Laboratory Plymouth and Alex Alderton, Roberto Amato, Jeffrey Barrett, Sonia Goncalves, Ewan Harrison, David K. Jackson, Ian Johnston, Dominic Kwiatkowski, Cordelia Langford, John Sillitoe on behalf of the Wellcome Sanger Institute COVID-19 Surveillance Team |
| EPI_ISL_3171552, EPI_ISL_3171553, EPI_ISL_3171554 | Berkshire and Surrey Pathology Services Lighthouse Laboratory | Wellcome Sanger Institute for the COVID-19 Genomics UK (COG-UK) Consortium | Berkshire and Surrey Pathology Services Lighthouse Laboratory and Alex Alderton, Roberto Amato, Jeffrey Barrett, Sonia Goncalves, Ewan Harrison, David K. Jackson, Ian Johnston, Dominic Kwiatkowski, Cordelia Langford, John Sillitoe on behalf of the Wellcome Sanger Institute COVID-19 Surveillance Team |
| EPI_ISL_3171555 | Lighthouse Laboratory Plymouth | Wellcome Sanger Institute for the COVID-19 Genomics UK (COG-UK) Consortium | Lighthouse Laboratory Plymouth and Alex Alderton, Roberto Amato, Jeffrey Barrett, Sonia Goncalves, Ewan Harrison, David K. Jackson, Ian Johnston, Dominic Kwiatkowski, Cordelia Langford, John Sillitoe on behalf of the Wellcome Sanger Institute COVID-19 Surveillance Team |
| EPI_ISL_3171556, EPI_ISL_3171557, EPI_ISL_3171558 | Berkshire and Surrey Pathology Services Lighthouse Laboratory | Wellcome Sanger Institute for the COVID-19 Genomics UK (COG-UK) Consortium | Berkshire and Surrey Pathology Services Lighthouse Laboratory and Alex Alderton, Roberto Amato, Jeffrey Barrett, Sonia Goncalves, Ewan Harrison, David K. Jackson, Ian Johnston, Dominic Kwiatkowski, Cordelia Langford, John Sillitoe on behalf of the Wellcome Sanger Institute COVID-19 Surveillance Team |
| EPI_ISL_3171559 | Lighthouse Lab in Alderley Park | Wellcome Sanger Institute for the COVID-19 Genomics UK (COG-UK) Consortium | Jacquelyn Wynn, Mairead Hyland, The Lighthouse Lab in Alderley Park and Alex Alderton, Roberto Amato, Jeffrey Barrett, Sonia Goncalves, Ewan Harrison, David K. Jackson, Ian Johnston, Dominic Kwiatkowski, Cordelia Langford, John Sillitoe on behalf of the Wellcome Sanger Institute COVID-19 Surveillance Team |
| EPI_ISL_3171560 | Lighthouse Laboratory Plymouth | Wellcome Sanger Institute for the COVID-19 Genomics UK (COG-UK) Consortium | Lighthouse Laboratory Plymouth and Alex Alderton, Roberto Amato, Jeffrey Barrett, Sonia Goncalves, Ewan Harrison, David K. Jackson, Ian Johnston, Dominic Kwiatkowski, Cordelia Langford, John Sillitoe on behalf of the Wellcome Sanger Institute COVID-19 Surveillance Team |
| EPI_ISL_3171561 | Berkshire and Surrey Pathology Services Lighthouse Laboratory | Wellcome Sanger Institute for the COVID-19 Genomics UK (COG-UK) Consortium | Berkshire and Surrey Pathology Services Lighthouse Laboratory and Alex Alderton, Roberto Amato, Jeffrey Barrett, Sonia Goncalves, Ewan Harrison, David K. Jackson, Ian Johnston, Dominic Kwiatkowski, Cordelia Langford, John Sillitoe on behalf of the Wellcome Sanger Institute COVID-19 Surveillance Team |
| EPI_ISL_3171562, EPI_ISL_3171563 | Lighthouse Lab in Alderley Park | Wellcome Sanger Institute for the COVID-19 Genomics UK (COG-UK) Consortium | Jacquelyn Wynn, Mairead Hyland, The Lighthouse Lab in Alderley Park and Alex Alderton, Roberto Amato, Jeffrey Barrett, Sonia Goncalves, Ewan Harrison, David K. Jackson, Ian Johnston, Dominic Kwiatkowski, Cordelia Langford, John Sillitoe on behalf of the Wellcome Sanger Institute COVID-19 Surveillance Team |
| EPI_ISL_3171564 | Lighthouse Laboratory Plymouth | Wellcome Sanger Institute for the COVID-19 Genomics UK (COG-UK) Consortium | Lighthouse Laboratory Plymouth and Alex Alderton, Roberto Amato, Jeffrey Barrett, Sonia Goncalves, Ewan Harrison, David K. Jackson, Ian Johnston, Dominic Kwiatkowski, Cordelia Langford, John Sillitoe on behalf of the Wellcome Sanger Institute COVID-19 Surveillance Team |
| EPI_ISL_3171565 | Lighthouse Lab in Alderley Park | Wellcome Sanger Institute for the COVID-19 Genomics UK (COG-UK) Consortium | Jacquelyn Wynn, Mairead Hyland, The Lighthouse Lab in Alderley Park and Alex Alderton, Roberto Amato, Jeffrey Barrett, Sonia Goncalves, Ewan Harrison, David K. Jackson, Ian Johnston, Dominic Kwiatkowski, Cordelia Langford, John Sillitoe on behalf of the Wellcome Sanger Institute COVID-19 Surveillance Team |
| EPI_ISL_3171566, EPI_ISL_3171567, EPI_ISL_3171568 | Lighthouse Laboratory Plymouth | Wellcome Sanger Institute for the COVID-19 Genomics UK (COG-UK) Consortium | Lighthouse Laboratory Plymouth and Alex Alderton, Roberto Amato, Jeffrey Barrett, Sonia Goncalves, Ewan Harrison, David K. Jackson, Ian Johnston, Dominic Kwiatkowski, Cordelia Langford, John Sillitoe on behalf of the Wellcome Sanger Institute COVID-19 Surveillance Team |
| EPI_ISL_3171569 | Berkshire and Surrey Pathology Services Lighthouse Laboratory | Wellcome Sanger Institute for the COVID-19 Genomics UK (COG-UK) Consortium | Berkshire and Surrey Pathology Services Lighthouse Laboratory and Alex Alderton, Roberto Amato, Jeffrey Barrett, Sonia Goncalves, Ewan Harrison, David K. Jackson, Ian Johnston, Dominic Kwiatkowski, Cordelia Langford, John Sillitoe on behalf of the Wellcome Sanger Institute COVID-19 Surveillance Team |
| EPI_ISL_3171570, EPI_ISL_3171571, | Lighthouse Laboratory Plymouth | Wellcome Sanger Institute for the COVID-19 Genomics UK (COG-UK) Consortium | Lighthouse Laboratory Plymouth and Alex Alderton, Roberto Amato, Jeffrey Barrett, Sonia Goncalves, Ewan Harrison, David K. Jackson, Ian Johnston, Dominic Kwiatkowski, Cordelia Langford, John Sillitoe on behalf of the Wellcome Sanger Institute COVID-19 Surveillance Team |

| | | | |
|---|---|---|---|
| EPI_ISL_3171572 | | | |
| EPI_ISL_3171573, EPI_ISL_3171574, EPI_ISL_3171575 | Berkshire and Surrey Pathology Services Lighthouse Laboratory | Wellcome Sanger Institute for the COVID-19 Genomics UK (COG-UK) Consortium | Berkshire and Surrey Pathology Services Lighthouse Laboratory and Alex Alderton, Roberto Amato, Jeffrey Barrett, Sonia Goncalves, Ewan Harrison, David K. Jackson, Ian Johnston, Dominic Kwiatkowski, Cordelia Langford, John Sillitoe on behalf of the Wellcome Sanger Institute COVID-19 Surveillance Team |
| EPI_ISL_3171576, EPI_ISL_3171577, EPI_ISL_3171578, EPI_ISL_3171579, EPI_ISL_3171580, EPI_ISL_3171581, EPI_ISL_3171582, EPI_ISL_3171583, EPI_ISL_3171584, EPI_ISL_3171585, EPI_ISL_3171586, EPI_ISL_3171587, EPI_ISL_3171588, EPI_ISL_3171589, EPI_ISL_3171590, EPI_ISL_3171591, EPI_ISL_3171592, EPI_ISL_3171593, EPI_ISL_3171594, EPI_ISL_3171595, EPI_ISL_3171596, EPI_ISL_3171597, EPI_ISL_3171598, EPI_ISL_3171599, EPI_ISL_3171600, EPI_ISL_3171601, EPI_ISL_3171602, EPI_ISL_3171603, EPI_ISL_3171604, EPI_ISL_3171605, EPI_ISL_3171606, EPI_ISL_3171607, EPI_ISL_3171608, EPI_ISL_3171609, EPI_ISL_3171610, EPI_ISL_3171611, EPI_ISL_3171612, EPI_ISL_3171613, EPI_ISL_3171614, EPI_ISL_3171615, EPI_ISL_3171616, EPI_ISL_3171617, EPI_ISL_3171618, EPI_ISL_3171619, EPI_ISL_3171620, EPI_ISL_3171621, EPI_ISL_3171622, EPI_ISL_3171623, EPI_ISL_3171624, EPI_ISL_3171625, EPI_ISL_3171626, EPI_ISL_3171627, EPI_ISL_3171628, EPI_ISL_3171629, EPI_ISL_3171630, EPI_ISL_3171631, EPI_ISL_3171632, EPI_ISL_3171633, EPI_ISL_3171634, EPI_ISL_3171635, EPI_ISL_3171636, EPI_ISL_3171637, EPI_ISL_3171638, EPI_ISL_3171639, EPI_ISL_3171640, EPI_ISL_3171641, EPI_ISL_3171642, EPI_ISL_3171643, EPI_ISL_3171644, EPI_ISL_3171645, EPI_ISL_3171646, EPI_ISL_3171647, EPI_ISL_3171648, EPI_ISL_3171649, EPI_ISL_3171650, EPI_ISL_3171651, EPI_ISL_3171652, EPI_ISL_3171653, EPI_ISL_3171654, EPI_ISL_3171655, EPI_ISL_3171656, EPI_ISL_3171657, EPI_ISL_3171658, EPI_ISL_3171659, EPI_ISL_3171660, EPI_ISL_3171661, EPI_ISL_3171662, EPI_ISL_3171663, EPI_ISL_3171664, EPI_ISL_3171665, EPI_ISL_3171666, EPI_ISL_3171667, EPI_ISL_3171668, EPI_ISL_3171669, EPI_ISL_3171670, EPI_ISL_3171671, EPI_ISL_3171672, EPI_ISL_3171673, EPI_ISL_3171674, EPI_ISL_3171675, EPI_ISL_3171676, EPI_ISL_3171677, EPI_ISL_3171678, EPI_ISL_3171679, EPI_ISL_3171680, EPI_ISL_3171681, EPI_ISL_3171682, EPI_ISL_3171683, EPI_ISL_3171684, EPI_ISL_3171685, EPI_ISL_3171686, EPI_ISL_3171687, EPI_ISL_3171688, EPI_ISL_3171689, EPI_ISL_3171690, EPI_ISL_3171691, EPI_ISL_3171692, EPI_ISL_3171693, EPI_ISL_3171694, EPI_ISL_3171695, EPI_ISL_3171696, EPI_ISL_3171697, EPI_ISL_3171698, EPI_ISL_3171699, EPI_ISL_3171700, EPI_ISL_3171701, EPI_ISL_3171702, EPI_ISL_3171703, EPI_ISL_3171704, EPI_ISL_3171705, EPI_ISL_3171706, EPI_ISL_3171707, EPI_ISL_3171708, EPI_ISL_3171709, EPI_ISL_3171710, EPI_ISL_3171711, EPI_ISL_3171712, EPI_ISL_3171713, EPI_ISL_3171714, EPI_ISL_3171715, EPI_ISL_3171716, EPI_ISL_3171717, EPI_ISL_3171718, EPI_ISL_3171719, EPI_ISL_3171720, EPI_ISL_3171721, EPI_ISL_3171722, EPI_ISL_3171723, EPI_ISL_3171724, EPI_ISL_3171725, EPI_ISL_3171726, EPI_ISL_3171727, EPI_ISL_3171728, EPI_ISL_3171729, EPI_ISL_3171730, EPI_ISL_3171731, EPI_ISL_3171732, EPI_ISL_3171733, EPI_ISL_3171734, EPI_ISL_3171735, EPI_ISL_3171736, EPI_ISL_3171737, EPI_ISL_3171738, EPI_ISL_3171739, EPI_ISL_3171740, EPI_ISL_3171741, EPI_ISL_3171742, EPI_ISL_3171743, EPI_ISL_3171744, EPI_ISL_3171745, EPI_ISL_3171746, EPI_ISL_3171747, EPI_ISL_3171748, EPI_ISL_3171749, EPI_ISL_3171750, EPI_ISL_3171751, EPI_ISL_3171752, EPI_ISL_3171753, EPI_ISL_3171754, EPI_ISL_3171755, EPI_ISL_3171756, EPI_ISL_3171757, EPI_ISL_3171758, EPI_ISL_3171759, EPI_ISL_3171760, EPI_ISL_3171761, EPI_ISL_3171762, EPI_ISL_3171763, EPI_ISL_3171764, EPI_ISL_3171765, EPI_ISL_3171766, EPI_ISL_3171767, EPI_ISL_3171768, EPI_ISL_3171769, EPI_ISL_3171770, EPI_ISL_3171771, EPI_ISL_3171772, EPI_ISL_3171773, EPI_ISL_3171774, EPI_ISL_3171775, EPI_ISL_3171776, EPI_ISL_3171777, EPI_ISL_3171778, EPI_ISL_3171779, EPI_ISL_3171780, EPI_ISL_3171781, EPI_ISL_3171782, EPI_ISL_3171783, EPI_ISL_3171784, EPI_ISL_3171785, EPI_ISL_3171786, EPI_ISL_3171787, EPI_ISL_3171788, EPI_ISL_3171789, EPI_ISL_3171790, EPI_ISL_3171791, EPI_ISL_3171792, EPI_ISL_3171793, EPI_ISL_3171794, EPI_ISL_3171795, EPI_ISL_3171796, EPI_ISL_3171797, EPI_ISL_3171798, EPI_ISL_3171799, EPI_ISL_3171800, EPI_ISL_3171801, EPI_ISL_3171802, EPI_ISL_3171803, EPI_ISL_3171804, EPI_ISL_3171805, EPI_ISL_3171806, EPI_ISL_3171807, EPI_ISL_3171808, EPI_ISL_3171809, EPI_ISL_3171810, EPI_ISL_3171811, EPI_ISL_3171812, EPI_ISL_3171813, EPI_ISL_3171814, EPI_ISL_3171815, EPI_ISL_3171816, EPI_ISL_3171817, EPI_ISL_3171818, EPI_ISL_3171819, EPI_ISL_3171820, EPI_ISL_3171821, EPI_ISL_3171822, EPI_ISL_3171823, EPI_ISL_3171824, EPI_ISL_3171825, EPI_ISL_3171826, EPI_ISL_3171827, EPI_ISL_3171828, EPI_ISL_3171829, EPI_ISL_3171830, EPI_ISL_3171831, EPI_ISL_3171832, EPI_ISL_3171833, EPI_ISL_3171834, EPI_ISL_3171835, EPI_ISL_3171836, EPI_ISL_3171837, EPI_ISL_3171838, EPI_ISL_3171839, EPI_ISL_3171840, EPI_ISL_3171841, EPI_ISL_3171842, EPI_ISL_3171843, EPI_ISL_3171844, EPI_ISL_3171845, EPI_ISL_3171846, EPI_ISL_3171847, EPI_ISL_3171848, EPI_ISL_3171849, EPI_ISL_3171850, EPI_ISL_3171851, EPI_ISL_3171852, EPI_ISL_3171853, EPI_ISL_3171854, EPI_ISL_3171855, EPI_ISL_3171856, EPI_ISL_3171857, EPI_ISL_3171858, EPI_ISL_3171859, EPI_ISL_3171860, EPI_ISL_3171861, EPI_ISL_3171862, EPI_ISL_3171863, EPI_ISL_3171864, EPI_ISL_3171865, EPI_ISL_3171866, EPI_ISL_3171867, EPI_ISL_3171868, EPI_ISL_3171869, EPI_ISL_3171870, EPI_ISL_3171871, EPI_ISL_3171872, EPI_ISL_3171873, EPI_ISL_3171874, EPI_ISL_3171875, EPI_ISL_3171876, EPI_ISL_3171877, EPI_ISL_3171878, EPI_ISL_3171879, EPI_ISL_3171880, EPI_ISL_3171881, EPI_ISL_3171882, EPI_ISL_3171883, EPI_ISL_3171884, EPI_ISL_3171885, EPI_ISL_3171886, EPI_ISL_3171887, EPI_ISL_3171888, EPI_ISL_3171889, EPI_ISL_3171890 | | | |
| see above | Lighthouse Lab in Milton Keynes | Wellcome Sanger Institute for the COVID-19 Genomics UK (COG-UK) Consortium | The Lighthouse Lab in Milton Keynes and Alex Alderton, Roberto Amato, Jeffrey Barrett, Sonia Goncalves, Ewan Harrison, David K. Jackson, Ian Johnston, Dominic Kwiatkowski, Cordelia Langford, John Sillitoe on behalf of the Wellcome Sanger Institute COVID-19 Surveillance Team |
| EPI_ISL_3171891 | Lighthouse Lab in Glasgow | Wellcome Sanger Institute for the COVID-19 Genomics UK (COG-UK) Consortium | Harper VanSteenhouse, Yumi Kasai, David Gray, Carol Clugston, Anna Dominiczak and Alex Alderton, Roberto Amato, Jeffrey Barrett, Sonia Goncalves, Ewan Harrison, David K. Jackson, Ian Johnston, Dominic Kwiatkowski, Cordelia Langford, John Sillitoe on behalf of the Wellcome Sanger Institute COVID-19 Surveillance Team |
| EPI_ISL_3171892 | Lighthouse Lab in Alderley Park | Wellcome Sanger Institute for the COVID-19 Genomics UK (COG-UK) Consortium | Jacquelyn Wynn, Mairead Hyland, The Lighthouse Lab in Alderley Park and Alex Alderton, Roberto Amato, Jeffrey Barrett, Sonia Goncalves, Ewan Harrison, David K. Jackson, Ian Johnston, Dominic Kwiatkowski, Cordelia Langford, John Sillitoe on behalf of the Wellcome Sanger Institute COVID-19 Surveillance Team |
| EPI_ISL_3171893 | Lighthouse Lab in Glasgow | Wellcome Sanger Institute for the COVID-19 Genomics UK (COG-UK) Consortium | Harper VanSteenhouse, Yumi Kasai, David Gray, Carol Clugston, Anna Dominiczak and Alex Alderton, Roberto Amato, Jeffrey Barrett, Sonia Goncalves, Ewan Harrison, David K. Jackson, Ian Johnston, Dominic Kwiatkowski, Cordelia Langford, John Sillitoe on behalf of the Wellcome Sanger Institute COVID-19 Surveillance Team |
| EPI_ISL_3171894 | Lighthouse Laboratory Plymouth | Wellcome Sanger Institute for the COVID-19 Genomics UK (COG-UK) Consortium | Lighthouse Laboratory Plymouth and Alex Alderton, Roberto Amato, Jeffrey Barrett, Sonia Goncalves, Ewan Harrison, David K. Jackson, Ian Johnston, Dominic Kwiatkowski, Cordelia Langford, John Sillitoe on behalf of the Wellcome Sanger Institute COVID-19 Surveillance Team |
| EPI_ISL_3171895 | Lighthouse Lab in Glasgow | Wellcome Sanger Institute for the COVID-19 Genomics UK (COG-UK) Consortium | Harper VanSteenhouse, Yumi Kasai, David Gray, Carol Clugston, Anna Dominiczak and Alex Alderton, Roberto Amato, Jeffrey Barrett, Sonia Goncalves, Ewan Harrison, David K. Jackson, Ian Johnston, Dominic Kwiatkowski, Cordelia Langford, John Sillitoe on behalf of the Wellcome Sanger Institute COVID-19 Surveillance Team |
| EPI_ISL_3171896, EPI_ISL_3171898 | Lighthouse Lab in Milton Keynes | Wellcome Sanger Institute for the COVID-19 Genomics UK (COG-UK) Consortium | The Lighthouse Lab in Milton Keynes and Alex Alderton, Roberto Amato, Jeffrey Barrett, Sonia Goncalves, Ewan Harrison, David K. Jackson, Ian Johnston, Dominic Kwiatkowski, Cordelia Langford, John Sillitoe on behalf of the Wellcome Sanger Institute COVID-19 Surveillance Team |
| EPI_ISL_3171899, EPI_ISL_3171900 | Lighthouse Lab in Alderley Park | Wellcome Sanger Institute for the COVID-19 Genomics UK (COG-UK) Consortium | Jacquelyn Wynn, Mairead Hyland, The Lighthouse Lab in Alderley Park and Alex Alderton, Roberto Amato, Jeffrey Barrett, Sonia Goncalves, Ewan Harrison, David K. Jackson, Ian Johnston, Dominic Kwiatkowski, Cordelia Langford, John Sillitoe on behalf of the Wellcome Sanger Institute COVID-19 Surveillance Team |
| EPI_ISL_3171901 | Lighthouse Laboratory Plymouth | Wellcome Sanger Institute for the COVID-19 Genomics UK (COG-UK) Consortium | Lighthouse Laboratory Plymouth and Alex Alderton, Roberto Amato, Jeffrey Barrett, Sonia Goncalves, Ewan Harrison, David K. Jackson, Ian Johnston, Dominic Kwiatkowski, Cordelia Langford, John Sillitoe on behalf of the Wellcome Sanger Institute COVID-19 Surveillance Team |
| EPI_ISL_3171902 | Lighthouse Lab in Milton Keynes | Wellcome Sanger Institute for the COVID-19 Genomics UK (COG-UK) Consortium | The Lighthouse Lab in Milton Keynes and Alex Alderton, Roberto Amato, Jeffrey Barrett, Sonia Goncalves, Ewan Harrison, David K. Jackson, Ian Johnston, Dominic Kwiatkowski, Cordelia Langford, John Sillitoe on behalf of the Wellcome Sanger Institute COVID-19 Surveillance Team |
| EPI_ISL_3171903 | Lighthouse Lab in Alderley Park | Wellcome Sanger Institute for the COVID-19 Genomics UK (COG-UK) Consortium | Jacquelyn Wynn, Mairead Hyland, The Lighthouse Lab in Alderley Park and Alex Alderton, Roberto Amato, Jeffrey Barrett, Sonia Goncalves, Ewan Harrison, David K. Jackson, Ian Johnston, Dominic Kwiatkowski, Cordelia Langford, John Sillitoe on behalf of the Wellcome Sanger Institute COVID-19 Surveillance Team |
| EPI_ISL_3171904, EPI_ISL_3171905 | Lighthouse Lab in Milton Keynes | Wellcome Sanger Institute for the COVID-19 Genomics UK (COG-UK) Consortium | The Lighthouse Lab in Milton Keynes and Alex Alderton, Roberto Amato, Jeffrey Barrett, Sonia Goncalves, Ewan Harrison, David K. Jackson, Ian Johnston, Dominic Kwiatkowski, Cordelia Langford, John Sillitoe on behalf of the Wellcome Sanger Institute COVID-19 Surveillance Team |
| EPI_ISL_3171906 | Lighthouse Lab in Glasgow | Wellcome Sanger Institute for the COVID-19 Genomics UK (COG-UK) Consortium | Harper VanSteenhouse, Yumi Kasai, David Gray, Carol Clugston, Anna Dominiczak and Alex Alderton, Roberto Amato, Jeffrey Barrett, Sonia Goncalves, Ewan Harrison, David K. Jackson, Ian Johnston, Dominic Kwiatkowski, Cordelia Langford, John Sillitoe on behalf of the Wellcome Sanger Institute COVID-19 Surveillance Team |
| EPI_ISL_3171907 | Lighthouse Laboratory Plymouth | Wellcome Sanger Institute for the COVID-19 Genomics UK (COG-UK) Consortium | Lighthouse Laboratory Plymouth and Alex Alderton, Roberto Amato, Jeffrey Barrett, Sonia Goncalves, Ewan Harrison, David K. Jackson, Ian Johnston, Dominic Kwiatkowski, Cordelia Langford, John Sillitoe on behalf of the Wellcome Sanger Institute COVID-19 Surveillance Team |
| EPI_ISL_3171908 | Lighthouse Lab in Glasgow | Wellcome Sanger Institute for the COVID-19 Genomics UK (COG-UK) Consortium | Harper VanSteenhouse, Yumi Kasai, David Gray, Carol Clugston, Anna Dominiczak and Alex Alderton, Roberto Amato, Jeffrey Barrett, Sonia Goncalves, Ewan Harrison, David K. Jackson, Ian Johnston, Dominic Kwiatkowski, Cordelia Langford, John Sillitoe on behalf of the Wellcome Sanger Institute COVID-19 Surveillance Team |
| EPI_ISL_3171909 | Lighthouse Lab in Alderley Park | Wellcome Sanger Institute for the COVID-19 Genomics UK (COG-UK) Consortium | Jacquelyn Wynn, Mairead Hyland, The Lighthouse Lab in Alderley Park and Alex Alderton, Roberto Amato, Jeffrey Barrett, Sonia Goncalves, Ewan Harrison, David K. Jackson, Ian Johnston, Dominic Kwiatkowski, Cordelia Langford, John Sillitoe on behalf of the Wellcome Sanger Institute COVID-19 Surveillance Team |
| EPI_ISL_3171910 | Lighthouse Lab in Milton Keynes | Wellcome Sanger Institute for the COVID-19 Genomics UK (COG-UK) Consortium | The Lighthouse Lab in Milton Keynes and Alex Alderton, Roberto Amato, Jeffrey Barrett, Sonia Goncalves, Ewan Harrison, David K. Jackson, Ian Johnston, Dominic Kwiatkowski, Cordelia Langford, John Sillitoe on behalf of the Wellcome Sanger Institute COVID-19 Surveillance Team |
| EPI_ISL_3171911 | Lighthouse Lab in Alderley Park | Wellcome Sanger Institute for the COVID-19 Genomics UK (COG-UK) Consortium | Jacquelyn Wynn, Mairead Hyland, The Lighthouse Lab in Alderley Park and Alex Alderton, Roberto Amato, Jeffrey Barrett, Sonia Goncalves, Ewan Harrison, David K. Jackson, Ian Johnston, Dominic Kwiatkowski, Cordelia Langford, John Sillitoe on behalf of the Wellcome Sanger Institute COVID-19 Surveillance Team |
| EPI_ISL_3171912 | Lighthouse Laboratory Plymouth | Wellcome Sanger Institute for the COVID-19 Genomics UK (COG-UK) Consortium | Lighthouse Laboratory Plymouth and Alex Alderton, Roberto Amato, Jeffrey Barrett, Sonia Goncalves, Ewan Harrison, David K. Jackson, Ian Johnston, Dominic Kwiatkowski, Cordelia Langford, John Sillitoe on behalf of the Wellcome Sanger Institute COVID-19 Surveillance Team |
| EPI_ISL_3171913 | Lighthouse Lab in Alderley Park | Wellcome Sanger Institute for the COVID-19 Genomics UK (COG-UK) Consortium | Jacquelyn Wynn, Mairead Hyland, The Lighthouse Lab in Alderley Park and Alex Alderton, Roberto Amato, Jeffrey Barrett, Sonia Goncalves, Ewan Harrison, David K. Jackson, Ian Johnston, Dominic Kwiatkowski, Cordelia Langford, John Sillitoe on behalf of the Wellcome Sanger Institute COVID-19 Surveillance Team |
| EPI_ISL_3171914 | Lighthouse Lab in Glasgow | Wellcome Sanger Institute for the COVID-19 Genomics UK (COG-UK) Consortium | Harper VanSteenhouse, Yumi Kasai, David Gray, Carol Clugston, Anna Dominiczak and Alex Alderton, Roberto Amato, Jeffrey Barrett, Sonia Goncalves, Ewan Harrison, David K. Jackson, Ian Johnston, Dominic Kwiatkowski, Cordelia Langford, John Sillitoe on behalf of the Wellcome Sanger Institute COVID-19 Surveillance Team |
| EPI_ISL_3171915 | Lighthouse Laboratory Plymouth | Wellcome Sanger Institute for the COVID-19 Genomics UK (COG-UK) Consortium | Lighthouse Laboratory Plymouth and Alex Alderton, Roberto Amato, Jeffrey Barrett, Sonia Goncalves, Ewan Harrison, David K. Jackson, Ian Johnston, Dominic Kwiatkowski, Cordelia Langford, John Sillitoe on behalf of the Wellcome Sanger Institute COVID-19 Surveillance Team |
| EPI_ISL_3171916 | Lighthouse Lab in Glasgow | Wellcome Sanger Institute for the COVID-19 Genomics UK (COG-UK) Consortium | Harper VanSteenhouse, Yumi Kasai, David Gray, Carol Clugston, Anna Dominiczak and Alex Alderton, Roberto Amato, Jeffrey Barrett, Sonia Goncalves, Ewan Harrison, David K. Jackson, Ian Johnston, Dominic Kwiatkowski, Cordelia Langford, John Sillitoe on behalf of the Wellcome Sanger Institute COVID-19 Surveillance Team |
| EPI_ISL_3171917 | Lighthouse Laboratory Plymouth | Wellcome Sanger Institute for the COVID-19 Genomics UK (COG-UK) Consortium | Lighthouse Laboratory Plymouth and Alex Alderton, Roberto Amato, Jeffrey Barrett, Sonia Goncalves, Ewan Harrison, David K. Jackson, Ian Johnston, Dominic Kwiatkowski, Cordelia Langford, John Sillitoe on behalf of the Wellcome Sanger Institute COVID-19 Surveillance Team |
| EPI_ISL_3171918, EPI_ISL_3171919 | Lighthouse Lab in Milton Keynes | Wellcome Sanger Institute for the COVID-19 Genomics UK (COG-UK) Consortium | The Lighthouse Lab in Milton Keynes and Alex Alderton, Roberto Amato, Jeffrey Barrett, Sonia Goncalves, Ewan Harrison, David K. Jackson, Ian Johnston, Dominic Kwiatkowski, Cordelia Langford, John Sillitoe on behalf of the Wellcome Sanger Institute COVID-19 Surveillance Team |
| EPI_ISL_3171920 | Lighthouse Laboratory Plymouth | Wellcome Sanger Institute for the COVID-19 Genomics UK (COG-UK) Consortium | Lighthouse Laboratory Plymouth and Alex Alderton, Roberto Amato, Jeffrey Barrett, Sonia Goncalves, Ewan Harrison, David K. Jackson, Ian Johnston, Dominic Kwiatkowski, Cordelia Langford, John Sillitoe on behalf of the Wellcome Sanger Institute COVID-19 Surveillance Team |
| EPI_ISL_3171921 | Lighthouse Lab in Glasgow | Wellcome Sanger Institute for the COVID-19 Genomics UK (COG-UK) Consortium | Harper VanSteenhouse, Yumi Kasai, David Gray, Carol Clugston, Anna Dominiczak and Alex Alderton, Roberto Amato, Jeffrey Barrett, Sonia Goncalves, Ewan Harrison, David K. Jackson, Ian Johnston, Dominic Kwiatkowski, Cordelia Langford, John Sillitoe on behalf of the Wellcome Sanger Institute COVID-19 Surveillance Team |
| EPI_ISL_3171922 | Lighthouse Laboratory Plymouth | Wellcome Sanger Institute for the COVID-19 Genomics UK (COG-UK) Consortium | Lighthouse Laboratory Plymouth and Alex Alderton, Roberto Amato, Jeffrey Barrett, Sonia Goncalves, Ewan Harrison, David K. Jackson, Ian Johnston, Dominic Kwiatkowski, Cordelia Langford, John Sillitoe on behalf of the Wellcome Sanger Institute COVID-19 Surveillance Team |
| EPI_ISL_3171923, EPI_ISL_3171924 | Lighthouse Lab in Alderley Park | Wellcome Sanger Institute for the COVID-19 Genomics UK (COG-UK) Consortium | Jacquelyn Wynn, Mairead Hyland, The Lighthouse Lab in Alderley Park and Alex Alderton, Roberto Amato, Jeffrey Barrett, Sonia Goncalves, Ewan Harrison, David K. Jackson, Ian Johnston, Dominic Kwiatkowski, Cordelia Langford, John Sillitoe on behalf of the Wellcome Sanger Institute COVID-19 Surveillance Team |
| EPI_ISL_3171925, EPI_ISL_3171926 | Lighthouse Laboratory Plymouth | Wellcome Sanger Institute for the COVID-19 Genomics UK (COG-UK) Consortium | Lighthouse Laboratory Plymouth and Alex Alderton, Roberto Amato, Jeffrey Barrett, Sonia Goncalves, Ewan Harrison, David K. Jackson, Ian Johnston, Dominic Kwiatkowski, Cordelia Langford, John Sillitoe on behalf of the Wellcome Sanger Institute COVID-19 Surveillance Team |
| EPI_ISL_3171927, EPI_ISL_3171928 | Lighthouse Lab in Alderley Park | Wellcome Sanger Institute for the COVID-19 Genomics UK (COG-UK) Consortium | Jacquelyn Wynn, Mairead Hyland, The Lighthouse Lab in Alderley Park and Alex Alderton, Roberto Amato, Jeffrey Barrett, Sonia Goncalves, Ewan Harrison, David K. Jackson, Ian Johnston, Dominic Kwiatkowski, Cordelia Langford, John Sillitoe on behalf of the Wellcome Sanger Institute COVID-19 Surveillance Team |
| EPI_ISL_3171929 | Lighthouse Lab in Glasgow | Wellcome Sanger Institute for the COVID-19 Genomics UK (COG-UK) Consortium | Harper VanSteenhouse, Yumi Kasai, David Gray, Carol Clugston, Anna Dominiczak and Alex Alderton, Roberto Amato, Jeffrey Barrett, Sonia Goncalves, Ewan Harrison, David K. Jackson, Ian Johnston, Dominic Kwiatkowski, Cordelia Langford, John Sillitoe on behalf of the Wellcome Sanger Institute COVID-19 Surveillance Team |
| EPI_ISL_3171930 | Lighthouse Lab in Milton Keynes | Wellcome Sanger Institute for the COVID-19 Genomics UK (COG-UK) Consortium | The Lighthouse Lab in Milton Keynes and Alex Alderton, Roberto Amato, Jeffrey Barrett, Sonia Goncalves, Ewan Harrison, David K. Jackson, Ian Johnston, Dominic Kwiatkowski, Cordelia Langford, John Sillitoe on behalf of the Wellcome Sanger Institute COVID-19 Surveillance Team |
| EPI_ISL_3171931, EPI_ISL_3171932 | Lighthouse Lab in Glasgow | Wellcome Sanger Institute for the COVID-19 Genomics UK (COG-UK) Consortium | Harper VanSteenhouse, Yumi Kasai, David Gray, Carol Clugston, Anna Dominiczak and Alex Alderton, Roberto Amato, Jeffrey Barrett, Sonia Goncalves, Ewan Harrison, David K. Jackson, Ian Johnston, Dominic Kwiatkowski, Cordelia Langford, John Sillitoe on behalf of the Wellcome Sanger Institute COVID-19 Surveillance Team |
| EPI_ISL_3171933 | Lighthouse Lab in Milton Keynes | Wellcome Sanger Institute for the COVID-19 Genomics UK (COG-UK) Consortium | The Lighthouse Lab in Milton Keynes and Alex Alderton, Roberto Amato, Jeffrey Barrett, Sonia Goncalves, Ewan Harrison, David K. Jackson, Ian Johnston, Dominic Kwiatkowski, Cordelia Langford, John Sillitoe on behalf of the Wellcome Sanger Institute COVID-19 Surveillance Team |
| EPI_ISL_3171934 | Lighthouse Laboratory Plymouth | Wellcome Sanger Institute for the COVID-19 Genomics UK (COG-UK) Consortium | Lighthouse Laboratory Plymouth and Alex Alderton, Roberto Amato, Jeffrey Barrett, Sonia Goncalves, Ewan Harrison, David K. Jackson, Ian Johnston, Dominic Kwiatkowski, Cordelia Langford, John Sillitoe on behalf of the Wellcome Sanger Institute COVID-19 Surveillance Team |
| EPI_ISL_3171935 | Lighthouse Lab in Glasgow | Wellcome Sanger Institute for the COVID-19 Genomics UK (COG-UK) | Harper VanSteenhouse, Yumi Kasai, David Gray, Carol Clugston, Anna Dominiczak and Alex Alderton, Roberto Amato, Jeffrey Barrett, Sonia Goncalves, Ewan Harrison, David K. Jackson, Ian Johnston, Dominic Kwiatkowski, Cordelia Langford, |

| | | | |
|---|---|---|---|
| | | Consortium | John Sillitoe on behalf of the Wellcome Sanger Institute COVID-19 Surveillance Team |
| EPI_ISL_3171936 | Lighthouse Lab in Milton Keynes | Wellcome Sanger Institute for the COVID-19 Genomics UK (COG-UK) Consortium | The Lighthouse Lab in Milton Keynes and Alex Alderton, Roberto Amato, Jeffrey Barrett, Sonia Goncalves, Ewan Harrison, David K. Jackson, Ian Johnston, Dominic Kwiatkowski, Cordelia Langford, John Sillitoe on behalf of the Wellcome Sanger Institute COVID-19 Surveillance Team |
| EPI_ISL_3171937 | Lighthouse Lab in Glasgow | Wellcome Sanger Institute for the COVID-19 Genomics UK (COG-UK) Consortium | Harper VanSteenhouse, Yumi Kasai, David Gray, Carol Clugston, Anna Dominiczak and Alex Alderton, Roberto Amato, Jeffrey Barrett, Sonia Goncalves, Ewan Harrison, David K. Jackson, Ian Johnston, Dominic Kwiatkowski, Cordelia Langford, John Sillitoe on behalf of the Wellcome Sanger Institute COVID-19 Surveillance Team |
| EPI_ISL_3171938 | Lighthouse Lab in Milton Keynes | Wellcome Sanger Institute for the COVID-19 Genomics UK (COG-UK) Consortium | The Lighthouse Lab in Milton Keynes and Alex Alderton, Roberto Amato, Jeffrey Barrett, Sonia Goncalves, Ewan Harrison, David K. Jackson, Ian Johnston, Dominic Kwiatkowski, Cordelia Langford, John Sillitoe on behalf of the Wellcome Sanger Institute COVID-19 Surveillance Team |
| EPI_ISL_3171939 | Lighthouse Laboratory Plymouth | Wellcome Sanger Institute for the COVID-19 Genomics UK (COG-UK) Consortium | Lighthouse Laboratory Plymouth and Alex Alderton, Roberto Amato, Jeffrey Barrett, Sonia Goncalves, Ewan Harrison, David K. Jackson, Ian Johnston, Dominic Kwiatkowski, Cordelia Langford, John Sillitoe on behalf of the Wellcome Sanger Institute COVID-19 Surveillance Team |
| EPI_ISL_3171940 | Lighthouse Lab in Alderley Park | Wellcome Sanger Institute for the COVID-19 Genomics UK (COG-UK) Consortium | Jacquelyn Wynn, Mairead Hyland, The Lighthouse Lab in Alderley Park and Alex Alderton, Roberto Amato, Jeffrey Barrett, Sonia Goncalves, Ewan Harrison, David K. Jackson, Ian Johnston, Dominic Kwiatkowski, Cordelia Langford, John Sillitoe on behalf of the Wellcome Sanger Institute COVID-19 Surveillance Team |
| EPI_ISL_3171941, EPI_ISL_3171942, EPI_ISL_3171943, EPI_ISL_3171944 | Lighthouse Laboratory Plymouth | Wellcome Sanger Institute for the COVID-19 Genomics UK (COG-UK) Consortium | Lighthouse Laboratory Plymouth and Alex Alderton, Roberto Amato, Jeffrey Barrett, Sonia Goncalves, Ewan Harrison, David K. Jackson, Ian Johnston, Dominic Kwiatkowski, Cordelia Langford, John Sillitoe on behalf of the Wellcome Sanger Institute COVID-19 Surveillance Team |
| EPI_ISL_3171945 | Lighthouse Lab in Glasgow | Wellcome Sanger Institute for the COVID-19 Genomics UK (COG-UK) Consortium | Harper VanSteenhouse, Yumi Kasai, David Gray, Carol Clugston, Anna Dominiczak and Alex Alderton, Roberto Amato, Jeffrey Barrett, Sonia Goncalves, Ewan Harrison, David K. Jackson, Ian Johnston, Dominic Kwiatkowski, Cordelia Langford, John Sillitoe on behalf of the Wellcome Sanger Institute COVID-19 Surveillance Team |
| EPI_ISL_3171946 | Lighthouse Laboratory Plymouth | Wellcome Sanger Institute for the COVID-19 Genomics UK (COG-UK) Consortium | Lighthouse Laboratory Plymouth and Alex Alderton, Roberto Amato, Jeffrey Barrett, Sonia Goncalves, Ewan Harrison, David K. Jackson, Ian Johnston, Dominic Kwiatkowski, Cordelia Langford, John Sillitoe on behalf of the Wellcome Sanger Institute COVID-19 Surveillance Team |
| EPI_ISL_3171947, EPI_ISL_3171948 | Lighthouse Lab in Milton Keynes | Wellcome Sanger Institute for the COVID-19 Genomics UK (COG-UK) Consortium | The Lighthouse Lab in Milton Keynes and Alex Alderton, Roberto Amato, Jeffrey Barrett, Sonia Goncalves, Ewan Harrison, David K. Jackson, Ian Johnston, Dominic Kwiatkowski, Cordelia Langford, John Sillitoe on behalf of the Wellcome Sanger Institute COVID-19 Surveillance Team |
| EPI_ISL_3171949 | Lighthouse Lab in Glasgow | Wellcome Sanger Institute for the COVID-19 Genomics UK (COG-UK) Consortium | Harper VanSteenhouse, Yumi Kasai, David Gray, Carol Clugston, Anna Dominiczak and Alex Alderton, Roberto Amato, Jeffrey Barrett, Sonia Goncalves, Ewan Harrison, David K. Jackson, Ian Johnston, Dominic Kwiatkowski, Cordelia Langford, John Sillitoe on behalf of the Wellcome Sanger Institute COVID-19 Surveillance Team |
| EPI_ISL_3171950 | Lighthouse Lab in Alderley Park | Wellcome Sanger Institute for the COVID-19 Genomics UK (COG-UK) Consortium | Jacquelyn Wynn, Mairead Hyland, The Lighthouse Lab in Alderley Park and Alex Alderton, Roberto Amato, Jeffrey Barrett, Sonia Goncalves, Ewan Harrison, David K. Jackson, Ian Johnston, Dominic Kwiatkowski, Cordelia Langford, John Sillitoe on behalf of the Wellcome Sanger Institute COVID-19 Surveillance Team |
| EPI_ISL_3171951 | Lighthouse Laboratory Plymouth | Wellcome Sanger Institute for the COVID-19 Genomics UK (COG-UK) Consortium | Lighthouse Laboratory Plymouth and Alex Alderton, Roberto Amato, Jeffrey Barrett, Sonia Goncalves, Ewan Harrison, David K. Jackson, Ian Johnston, Dominic Kwiatkowski, Cordelia Langford, John Sillitoe on behalf of the Wellcome Sanger Institute COVID-19 Surveillance Team |
| EPI_ISL_3171952 | Lighthouse Lab in Alderley Park | Wellcome Sanger Institute for the COVID-19 Genomics UK (COG-UK) Consortium | Jacquelyn Wynn, Mairead Hyland, The Lighthouse Lab in Alderley Park and Alex Alderton, Roberto Amato, Jeffrey Barrett, Sonia Goncalves, Ewan Harrison, David K. Jackson, Ian Johnston, Dominic Kwiatkowski, Cordelia Langford, John Sillitoe on behalf of the Wellcome Sanger Institute COVID-19 Surveillance Team |
| EPI_ISL_3171953 | Lighthouse Laboratory Plymouth | Wellcome Sanger Institute for the COVID-19 Genomics UK (COG-UK) Consortium | Lighthouse Laboratory Plymouth and Alex Alderton, Roberto Amato, Jeffrey Barrett, Sonia Goncalves, Ewan Harrison, David K. Jackson, Ian Johnston, Dominic Kwiatkowski, Cordelia Langford, John Sillitoe on behalf of the Wellcome Sanger Institute COVID-19 Surveillance Team |
| EPI_ISL_3171954 | Lighthouse Lab in Glasgow | Wellcome Sanger Institute for the COVID-19 Genomics UK (COG-UK) Consortium | Harper VanSteenhouse, Yumi Kasai, David Gray, Carol Clugston, Anna Dominiczak and Alex Alderton, Roberto Amato, Jeffrey Barrett, Sonia Goncalves, Ewan Harrison, David K. Jackson, Ian Johnston, Dominic Kwiatkowski, Cordelia Langford, John Sillitoe on behalf of the Wellcome Sanger Institute COVID-19 Surveillance Team |
| EPI_ISL_3171955 | Lighthouse Lab in Milton Keynes | Wellcome Sanger Institute for the COVID-19 Genomics UK (COG-UK) Consortium | The Lighthouse Lab in Milton Keynes and Alex Alderton, Roberto Amato, Jeffrey Barrett, Sonia Goncalves, Ewan Harrison, David K. Jackson, Ian Johnston, Dominic Kwiatkowski, Cordelia Langford, John Sillitoe on behalf of the Wellcome Sanger Institute COVID-19 Surveillance Team |
| EPI_ISL_3171956 | Lighthouse Lab in Alderley Park | Wellcome Sanger Institute for the COVID-19 Genomics UK (COG-UK) Consortium | Jacquelyn Wynn, Mairead Hyland, The Lighthouse Lab in Alderley Park and Alex Alderton, Roberto Amato, Jeffrey Barrett, Sonia Goncalves, Ewan Harrison, David K. Jackson, Ian Johnston, Dominic Kwiatkowski, Cordelia Langford, John Sillitoe on behalf of the Wellcome Sanger Institute COVID-19 Surveillance Team |
| EPI_ISL_3171957, EPI_ISL_3171958 | Lighthouse Laboratory Plymouth | Wellcome Sanger Institute for the COVID-19 Genomics UK (COG-UK) Consortium | Lighthouse Laboratory Plymouth and Alex Alderton, Roberto Amato, Jeffrey Barrett, Sonia Goncalves, Ewan Harrison, David K. Jackson, Ian Johnston, Dominic Kwiatkowski, Cordelia Langford, John Sillitoe on behalf of the Wellcome Sanger Institute COVID-19 Surveillance Team |
| EPI_ISL_3171959, EPI_ISL_3171960 | Lighthouse Lab in Glasgow | Wellcome Sanger Institute for the COVID-19 Genomics UK (COG-UK) Consortium | Harper VanSteenhouse, Yumi Kasai, David Gray, Carol Clugston, Anna Dominiczak and Alex Alderton, Roberto Amato, Jeffrey Barrett, Sonia Goncalves, Ewan Harrison, David K. Jackson, Ian Johnston, Dominic Kwiatkowski, Cordelia Langford, John Sillitoe on behalf of the Wellcome Sanger Institute COVID-19 Surveillance Team |
| EPI_ISL_3171961 | Lighthouse Lab in Milton Keynes | Wellcome Sanger Institute for the COVID-19 Genomics UK (COG-UK) Consortium | The Lighthouse Lab in Milton Keynes and Alex Alderton, Roberto Amato, Jeffrey Barrett, Sonia Goncalves, Ewan Harrison, David K. Jackson, Ian Johnston, Dominic Kwiatkowski, Cordelia Langford, John Sillitoe on behalf of the Wellcome Sanger Institute COVID-19 Surveillance Team |
| EPI_ISL_3171962 | Lighthouse Lab in Alderley Park | Wellcome Sanger Institute for the COVID-19 Genomics UK (COG-UK) Consortium | Jacquelyn Wynn, Mairead Hyland, The Lighthouse Lab in Alderley Park and Alex Alderton, Roberto Amato, Jeffrey Barrett, Sonia Goncalves, Ewan Harrison, David K. Jackson, Ian Johnston, Dominic Kwiatkowski, Cordelia Langford, John Sillitoe on behalf of the Wellcome Sanger Institute COVID-19 Surveillance Team |
| EPI_ISL_3171963, EPI_ISL_3171964 | Lighthouse Lab in Milton Keynes | Wellcome Sanger Institute for the COVID-19 Genomics UK (COG-UK) Consortium | The Lighthouse Lab in Milton Keynes and Alex Alderton, Roberto Amato, Jeffrey Barrett, Sonia Goncalves, Ewan Harrison, David K. Jackson, Ian Johnston, Dominic Kwiatkowski, Cordelia Langford, John Sillitoe on behalf of the Wellcome Sanger Institute COVID-19 Surveillance Team |
| EPI_ISL_3171965 | Lighthouse Lab in Alderley Park | Wellcome Sanger Institute for the COVID-19 Genomics UK (COG-UK) Consortium | Jacquelyn Wynn, Mairead Hyland, The Lighthouse Lab in Alderley Park and Alex Alderton, Roberto Amato, Jeffrey Barrett, Sonia Goncalves, Ewan Harrison, David K. Jackson, Ian Johnston, Dominic Kwiatkowski, Cordelia Langford, John Sillitoe on behalf of the Wellcome Sanger Institute COVID-19 Surveillance Team |
| EPI_ISL_3171966 | Lighthouse Lab in Glasgow | Wellcome Sanger Institute for the COVID-19 Genomics UK (COG-UK) Consortium | Harper VanSteenhouse, Yumi Kasai, David Gray, Carol Clugston, Anna Dominiczak and Alex Alderton, Roberto Amato, Jeffrey Barrett, Sonia Goncalves, Ewan Harrison, David K. Jackson, Ian Johnston, Dominic Kwiatkowski, Cordelia Langford, John Sillitoe on behalf of the Wellcome Sanger Institute COVID-19 Surveillance Team |
| EPI_ISL_3171967 | Lighthouse Laboratory Plymouth | Wellcome Sanger Institute for the COVID-19 Genomics UK (COG-UK) Consortium | Lighthouse Laboratory Plymouth and Alex Alderton, Roberto Amato, Jeffrey Barrett, Sonia Goncalves, Ewan Harrison, David K. Jackson, Ian Johnston, Dominic Kwiatkowski, Cordelia Langford, John Sillitoe on behalf of the Wellcome Sanger Institute COVID-19 Surveillance Team |
| EPI_ISL_3171968 | Lighthouse Lab in Glasgow | Wellcome Sanger Institute for the COVID-19 Genomics UK (COG-UK) Consortium | Harper VanSteenhouse, Yumi Kasai, David Gray, Carol Clugston, Anna Dominiczak and Alex Alderton, Roberto Amato, Jeffrey Barrett, Sonia Goncalves, Ewan Harrison, David K. Jackson, Ian Johnston, Dominic Kwiatkowski, Cordelia Langford, John Sillitoe on behalf of the Wellcome Sanger Institute COVID-19 Surveillance Team |
| EPI_ISL_3171969 | Lighthouse Laboratory Plymouth | Wellcome Sanger Institute for the COVID-19 Genomics UK (COG-UK) Consortium | Lighthouse Laboratory Plymouth and Alex Alderton, Roberto Amato, Jeffrey Barrett, Sonia Goncalves, Ewan Harrison, David K. Jackson, Ian Johnston, Dominic Kwiatkowski, Cordelia Langford, John Sillitoe on behalf of the Wellcome Sanger Institute COVID-19 Surveillance Team |
| EPI_ISL_3171970 | Lighthouse Lab in Glasgow | Wellcome Sanger Institute for the COVID-19 Genomics UK (COG-UK) Consortium | Harper VanSteenhouse, Yumi Kasai, David Gray, Carol Clugston, Anna Dominiczak and Alex Alderton, Roberto Amato, Jeffrey Barrett, Sonia Goncalves, Ewan Harrison, David K. Jackson, Ian Johnston, Dominic Kwiatkowski, Cordelia Langford, John Sillitoe on behalf of the Wellcome Sanger Institute COVID-19 Surveillance Team |
| EPI_ISL_3171971 | Lighthouse Lab in Milton Keynes | Wellcome Sanger Institute for the COVID-19 Genomics UK (COG-UK) Consortium | The Lighthouse Lab in Milton Keynes and Alex Alderton, Roberto Amato, Jeffrey Barrett, Sonia Goncalves, Ewan Harrison, David K. Jackson, Ian Johnston, Dominic Kwiatkowski, Cordelia Langford, John Sillitoe on behalf of the Wellcome Sanger Institute COVID-19 Surveillance Team |
| EPI_ISL_3171972 | Lighthouse Laboratory Plymouth | Wellcome Sanger Institute for the COVID-19 Genomics UK (COG-UK) Consortium | Lighthouse Laboratory Plymouth and Alex Alderton, Roberto Amato, Jeffrey Barrett, Sonia Goncalves, Ewan Harrison, David K. Jackson, Ian Johnston, Dominic Kwiatkowski, Cordelia Langford, John Sillitoe on behalf of the Wellcome Sanger Institute COVID-19 Surveillance Team |
| EPI_ISL_3171973, EPI_ISL_3171974 | Lighthouse Lab in Alderley Park | Wellcome Sanger Institute for the COVID-19 Genomics UK (COG-UK) Consortium | Jacquelyn Wynn, Mairead Hyland, The Lighthouse Lab in Alderley Park and Alex Alderton, Roberto Amato, Jeffrey Barrett, Sonia Goncalves, Ewan Harrison, David K. Jackson, Ian Johnston, Dominic Kwiatkowski, Cordelia Langford, John Sillitoe on behalf of the Wellcome Sanger Institute COVID-19 Surveillance Team |
| EPI_ISL_3171975, EPI_ISL_3171976, EPI_ISL_3171977 | Lighthouse Laboratory Plymouth | Wellcome Sanger Institute for the COVID-19 Genomics UK (COG-UK) Consortium | Lighthouse Laboratory Plymouth and Alex Alderton, Roberto Amato, Jeffrey Barrett, Sonia Goncalves, Ewan Harrison, David K. Jackson, Ian Johnston, Dominic Kwiatkowski, Cordelia Langford, John Sillitoe on behalf of the Wellcome Sanger Institute COVID-19 Surveillance Team |
| EPI_ISL_3171978 | Lighthouse Lab in Glasgow | Wellcome Sanger Institute for the COVID-19 Genomics UK (COG-UK) Consortium | Harper VanSteenhouse, Yumi Kasai, David Gray, Carol Clugston, Anna Dominiczak and Alex Alderton, Roberto Amato, Jeffrey Barrett, Sonia Goncalves, Ewan Harrison, David K. Jackson, Ian Johnston, Dominic Kwiatkowski, Cordelia Langford, John Sillitoe on behalf of the Wellcome Sanger Institute COVID-19 Surveillance Team |
| EPI_ISL_3171979 | Lighthouse Lab in Alderley Park | Wellcome Sanger Institute for the COVID-19 Genomics UK (COG-UK) Consortium | Jacquelyn Wynn, Mairead Hyland, The Lighthouse Lab in Alderley Park and Alex Alderton, Roberto Amato, Jeffrey Barrett, Sonia Goncalves, Ewan Harrison, David K. Jackson, Ian Johnston, Dominic Kwiatkowski, Cordelia Langford, John Sillitoe on behalf of the Wellcome Sanger Institute COVID-19 Surveillance Team |
| EPI_ISL_3171980 | Lighthouse Lab in Glasgow | Wellcome Sanger Institute for the COVID-19 Genomics UK (COG-UK) Consortium | Harper VanSteenhouse, Yumi Kasai, David Gray, Carol Clugston, Anna Dominiczak and Alex Alderton, Roberto Amato, Jeffrey Barrett, Sonia Goncalves, Ewan Harrison, David K. Jackson, Ian Johnston, Dominic Kwiatkowski, Cordelia Langford, John Sillitoe on behalf of the Wellcome Sanger Institute COVID-19 Surveillance Team |
| EPI_ISL_3171981 | Lighthouse Lab in Alderley Park | Wellcome Sanger Institute for the COVID-19 Genomics UK (COG-UK) Consortium | Jacquelyn Wynn, Mairead Hyland, The Lighthouse Lab in Alderley Park and Alex Alderton, Roberto Amato, Jeffrey Barrett, Sonia Goncalves, Ewan Harrison, David K. Jackson, Ian Johnston, Dominic Kwiatkowski, Cordelia Langford, John Sillitoe on behalf of the Wellcome Sanger Institute COVID-19 Surveillance Team |
| EPI_ISL_3171982 | Lighthouse Lab in Glasgow | Wellcome Sanger Institute for the COVID-19 Genomics UK (COG-UK) Consortium | Harper VanSteenhouse, Yumi Kasai, David Gray, Carol Clugston, Anna Dominiczak and Alex Alderton, Roberto Amato, Jeffrey Barrett, Sonia Goncalves, Ewan Harrison, David K. Jackson, Ian Johnston, Dominic Kwiatkowski, Cordelia Langford, John Sillitoe on behalf of the Wellcome Sanger Institute COVID-19 Surveillance Team |
| EPI_ISL_3171983 | Lighthouse Lab in Alderley Park | Wellcome Sanger Institute for the COVID-19 Genomics UK (COG-UK) Consortium | Jacquelyn Wynn, Mairead Hyland, The Lighthouse Lab in Alderley Park and Alex Alderton, Roberto Amato, Jeffrey Barrett, Sonia Goncalves, Ewan Harrison, David K. Jackson, Ian Johnston, Dominic Kwiatkowski, Cordelia Langford, John Sillitoe on behalf of the Wellcome Sanger Institute COVID-19 Surveillance Team |
| EPI_ISL_3171984 | Lighthouse Laboratory Plymouth | Wellcome Sanger Institute for the COVID-19 Genomics UK (COG-UK) Consortium | Lighthouse Laboratory Plymouth and Alex Alderton, Roberto Amato, Jeffrey Barrett, Sonia Goncalves, Ewan Harrison, David K. Jackson, Ian Johnston, Dominic Kwiatkowski, Cordelia Langford, John Sillitoe on behalf of the Wellcome Sanger Institute COVID-19 Surveillance Team |
| EPI_ISL_3171985 | Lighthouse Lab in Milton Keynes | Wellcome Sanger Institute for the COVID-19 Genomics UK (COG-UK) Consortium | The Lighthouse Lab in Milton Keynes and Alex Alderton, Roberto Amato, Jeffrey Barrett, Sonia Goncalves, Ewan Harrison, David K. Jackson, Ian Johnston, Dominic Kwiatkowski, Cordelia Langford, John Sillitoe on behalf of the Wellcome Sanger Institute COVID-19 Surveillance Team |
| EPI_ISL_3171986 | Lighthouse Laboratory Plymouth | Wellcome Sanger Institute for the COVID-19 Genomics UK (COG-UK) Consortium | Lighthouse Laboratory Plymouth and Alex Alderton, Roberto Amato, Jeffrey Barrett, Sonia Goncalves, Ewan Harrison, David K. Jackson, Ian Johnston, Dominic Kwiatkowski, Cordelia Langford, John Sillitoe on behalf of the Wellcome Sanger Institute COVID-19 Surveillance Team |
| EPI_ISL_3171987 | Lighthouse Lab in Glasgow | Wellcome Sanger Institute for the COVID-19 Genomics UK (COG-UK) Consortium | Harper VanSteenhouse, Yumi Kasai, David Gray, Carol Clugston, Anna Dominiczak and Alex Alderton, Roberto Amato, Jeffrey Barrett, Sonia Goncalves, Ewan Harrison, David K. Jackson, Ian Johnston, Dominic Kwiatkowski, Cordelia Langford, John Sillitoe on behalf of the Wellcome Sanger Institute COVID-19 Surveillance Team |
| EPI_ISL_3171988 | Lighthouse Lab in Alderley Park | Wellcome Sanger Institute for the COVID-19 Genomics UK (COG-UK) Consortium | Jacquelyn Wynn, Mairead Hyland, The Lighthouse Lab in Alderley Park and Alex Alderton, Roberto Amato, Jeffrey Barrett, Sonia Goncalves, Ewan Harrison, David K. Jackson, Ian Johnston, Dominic Kwiatkowski, Cordelia Langford, John Sillitoe on behalf of the Wellcome Sanger Institute COVID-19 Surveillance Team |

| | | | |
|---|---|---|---|
| EPI_ISL_3171989 | Lighthouse Lab in Milton Keynes | Wellcome Sanger Institute for the COVID-19 Genomics UK (COG-UK) Consortium | The Lighthouse Lab in Milton Keynes and Alex Alderton, Roberto Amato, Jeffrey Barrett, Sonia Goncalves, Ewan Harrison, David K. Jackson, Ian Johnston, Dominic Kwiatkowski, Cordelia Langford, John Sillitoe on behalf of the Wellcome Sanger Institute COVID-19 Surveillance Team |
| EPI_ISL_3171990 | Lighthouse Laboratory Plymouth | Wellcome Sanger Institute for the COVID-19 Genomics UK (COG-UK) Consortium | Lighthouse Laboratory Plymouth and Alex Alderton, Roberto Amato, Jeffrey Barrett, Sonia Goncalves, Ewan Harrison, David K. Jackson, Ian Johnston, Dominic Kwiatkowski, Cordelia Langford, John Sillitoe on behalf of the Wellcome Sanger Institute COVID-19 Surveillance Team |
| EPI_ISL_3171991 | Lighthouse Lab in Milton Keynes | Wellcome Sanger Institute for the COVID-19 Genomics UK (COG-UK) Consortium | The Lighthouse Lab in Milton Keynes and Alex Alderton, Roberto Amato, Jeffrey Barrett, Sonia Goncalves, Ewan Harrison, David K. Jackson, Ian Johnston, Dominic Kwiatkowski, Cordelia Langford, John Sillitoe on behalf of the Wellcome Sanger Institute COVID-19 Surveillance Team |
| EPI_ISL_3171992, EPI_ISL_3171993, EPI_ISL_3171994 | Lighthouse Lab in Alderley Park | Wellcome Sanger Institute for the COVID-19 Genomics UK (COG-UK) Consortium | Jacquelyn Wynn, Mairead Hyland, The Lighthouse Lab in Alderley Park and Alex Alderton, Roberto Amato, Jeffrey Barrett, Sonia Goncalves, Ewan Harrison, David K. Jackson, Ian Johnston, Dominic Kwiatkowski, Cordelia Langford, John Sillitoe on behalf of the Wellcome Sanger Institute COVID-19 Surveillance Team |
| EPI_ISL_3171995, EPI_ISL_3171996 | Lighthouse Lab in Glasgow | Wellcome Sanger Institute for the COVID-19 Genomics UK (COG-UK) Consortium | Harper VanSteenhouse, Yumi Kasai, David Gray, Carol Clugston, Anna Dominiczak and Alex Alderton, Roberto Amato, Jeffrey Barrett, Sonia Goncalves, Ewan Harrison, David K. Jackson, Ian Johnston, Dominic Kwiatkowski, Cordelia Langford, John Sillitoe on behalf of the Wellcome Sanger Institute COVID-19 Surveillance Team |
| EPI_ISL_3171997 | Lighthouse Lab in Alderley Park | Wellcome Sanger Institute for the COVID-19 Genomics UK (COG-UK) Consortium | Jacquelyn Wynn, Mairead Hyland, The Lighthouse Lab in Alderley Park and Alex Alderton, Roberto Amato, Jeffrey Barrett, Sonia Goncalves, Ewan Harrison, David K. Jackson, Ian Johnston, Dominic Kwiatkowski, Cordelia Langford, John Sillitoe on behalf of the Wellcome Sanger Institute COVID-19 Surveillance Team |
| EPI_ISL_3171998, EPI_ISL_3171999 | Lighthouse Lab in Milton Keynes | Wellcome Sanger Institute for the COVID-19 Genomics UK (COG-UK) Consortium | The Lighthouse Lab in Milton Keynes and Alex Alderton, Roberto Amato, Jeffrey Barrett, Sonia Goncalves, Ewan Harrison, David K. Jackson, Ian Johnston, Dominic Kwiatkowski, Cordelia Langford, John Sillitoe on behalf of the Wellcome Sanger Institute COVID-19 Surveillance Team |
| EPI_ISL_3172000 | Lighthouse Lab in Alderley Park | Wellcome Sanger Institute for the COVID-19 Genomics UK (COG-UK) Consortium | Jacquelyn Wynn, Mairead Hyland, The Lighthouse Lab in Alderley Park and Alex Alderton, Roberto Amato, Jeffrey Barrett, Sonia Goncalves, Ewan Harrison, David K. Jackson, Ian Johnston, Dominic Kwiatkowski, Cordelia Langford, John Sillitoe on behalf of the Wellcome Sanger Institute COVID-19 Surveillance Team |
| EPI_ISL_3172001 | Lighthouse Lab in Glasgow | Wellcome Sanger Institute for the COVID-19 Genomics UK (COG-UK) Consortium | Harper VanSteenhouse, Yumi Kasai, David Gray, Carol Clugston, Anna Dominiczak and Alex Alderton, Roberto Amato, Jeffrey Barrett, Sonia Goncalves, Ewan Harrison, David K. Jackson, Ian Johnston, Dominic Kwiatkowski, Cordelia Langford, John Sillitoe on behalf of the Wellcome Sanger Institute COVID-19 Surveillance Team |
| EPI_ISL_3172002 | Lighthouse Laboratory Plymouth | Wellcome Sanger Institute for the COVID-19 Genomics UK (COG-UK) Consortium | Lighthouse Laboratory Plymouth and Alex Alderton, Roberto Amato, Jeffrey Barrett, Sonia Goncalves, Ewan Harrison, David K. Jackson, Ian Johnston, Dominic Kwiatkowski, Cordelia Langford, John Sillitoe on behalf of the Wellcome Sanger Institute COVID-19 Surveillance Team |
| EPI_ISL_3172003, EPI_ISL_3172004 | Lighthouse Lab in Milton Keynes | Wellcome Sanger Institute for the COVID-19 Genomics UK (COG-UK) Consortium | The Lighthouse Lab in Milton Keynes and Alex Alderton, Roberto Amato, Jeffrey Barrett, Sonia Goncalves, Ewan Harrison, David K. Jackson, Ian Johnston, Dominic Kwiatkowski, Cordelia Langford, John Sillitoe on behalf of the Wellcome Sanger Institute COVID-19 Surveillance Team |
| EPI_ISL_3172005 | Lighthouse Lab in Alderley Park | Wellcome Sanger Institute for the COVID-19 Genomics UK (COG-UK) Consortium | Jacquelyn Wynn, Mairead Hyland, The Lighthouse Lab in Alderley Park and Alex Alderton, Roberto Amato, Jeffrey Barrett, Sonia Goncalves, Ewan Harrison, David K. Jackson, Ian Johnston, Dominic Kwiatkowski, Cordelia Langford, John Sillitoe on behalf of the Wellcome Sanger Institute COVID-19 Surveillance Team |
| EPI_ISL_3172006, EPI_ISL_3172007 | Lighthouse Laboratory Plymouth | Wellcome Sanger Institute for the COVID-19 Genomics UK (COG-UK) Consortium | Lighthouse Laboratory Plymouth and Alex Alderton, Roberto Amato, Jeffrey Barrett, Sonia Goncalves, Ewan Harrison, David K. Jackson, Ian Johnston, Dominic Kwiatkowski, Cordelia Langford, John Sillitoe on behalf of the Wellcome Sanger Institute COVID-19 Surveillance Team |
| EPI_ISL_3172008 | Lighthouse Lab in Milton Keynes | Wellcome Sanger Institute for the COVID-19 Genomics UK (COG-UK) Consortium | The Lighthouse Lab in Milton Keynes and Alex Alderton, Roberto Amato, Jeffrey Barrett, Sonia Goncalves, Ewan Harrison, David K. Jackson, Ian Johnston, Dominic Kwiatkowski, Cordelia Langford, John Sillitoe on behalf of the Wellcome Sanger Institute COVID-19 Surveillance Team |
| EPI_ISL_3172009 | Lighthouse Laboratory Plymouth | Wellcome Sanger Institute for the COVID-19 Genomics UK (COG-UK) Consortium | Lighthouse Laboratory Plymouth and Alex Alderton, Roberto Amato, Jeffrey Barrett, Sonia Goncalves, Ewan Harrison, David K. Jackson, Ian Johnston, Dominic Kwiatkowski, Cordelia Langford, John Sillitoe on behalf of the Wellcome Sanger Institute COVID-19 Surveillance Team |
| EPI_ISL_3172010, EPI_ISL_3172011 | Lighthouse Lab in Glasgow | Wellcome Sanger Institute for the COVID-19 Genomics UK (COG-UK) Consortium | Harper VanSteenhouse, Yumi Kasai, David Gray, Carol Clugston, Anna Dominiczak and Alex Alderton, Roberto Amato, Jeffrey Barrett, Sonia Goncalves, Ewan Harrison, David K. Jackson, Ian Johnston, Dominic Kwiatkowski, Cordelia Langford, John Sillitoe on behalf of the Wellcome Sanger Institute COVID-19 Surveillance Team |
| EPI_ISL_3172012 | Lighthouse Laboratory Plymouth | Wellcome Sanger Institute for the COVID-19 Genomics UK (COG-UK) Consortium | Lighthouse Laboratory Plymouth and Alex Alderton, Roberto Amato, Jeffrey Barrett, Sonia Goncalves, Ewan Harrison, David K. Jackson, Ian Johnston, Dominic Kwiatkowski, Cordelia Langford, John Sillitoe on behalf of the Wellcome Sanger Institute COVID-19 Surveillance Team |
| EPI_ISL_3172013 | Lighthouse Lab in Alderley Park | Wellcome Sanger Institute for the COVID-19 Genomics UK (COG-UK) Consortium | Jacquelyn Wynn, Mairead Hyland, The Lighthouse Lab in Alderley Park and Alex Alderton, Roberto Amato, Jeffrey Barrett, Sonia Goncalves, Ewan Harrison, David K. Jackson, Ian Johnston, Dominic Kwiatkowski, Cordelia Langford, John Sillitoe on behalf of the Wellcome Sanger Institute COVID-19 Surveillance Team |
| EPI_ISL_3172014, EPI_ISL_3172015, EPI_ISL_3172016 | Lighthouse Lab in Milton Keynes | Wellcome Sanger Institute for the COVID-19 Genomics UK (COG-UK) Consortium | The Lighthouse Lab in Milton Keynes and Alex Alderton, Roberto Amato, Jeffrey Barrett, Sonia Goncalves, Ewan Harrison, David K. Jackson, Ian Johnston, Dominic Kwiatkowski, Cordelia Langford, John Sillitoe on behalf of the Wellcome Sanger Institute COVID-19 Surveillance Team |
| EPI_ISL_3172017 | Lighthouse Lab in Alderley Park | Wellcome Sanger Institute for the COVID-19 Genomics UK (COG-UK) Consortium | Jacquelyn Wynn, Mairead Hyland, The Lighthouse Lab in Alderley Park and Alex Alderton, Roberto Amato, Jeffrey Barrett, Sonia Goncalves, Ewan Harrison, David K. Jackson, Ian Johnston, Dominic Kwiatkowski, Cordelia Langford, John Sillitoe on behalf of the Wellcome Sanger Institute COVID-19 Surveillance Team |
| EPI_ISL_3172018 | Lighthouse Lab in Milton Keynes | Wellcome Sanger Institute for the COVID-19 Genomics UK (COG-UK) Consortium | The Lighthouse Lab in Milton Keynes and Alex Alderton, Roberto Amato, Jeffrey Barrett, Sonia Goncalves, Ewan Harrison, David K. Jackson, Ian Johnston, Dominic Kwiatkowski, Cordelia Langford, John Sillitoe on behalf of the Wellcome Sanger Institute COVID-19 Surveillance Team |
| EPI_ISL_3172019 | Lighthouse Lab in Glasgow | Wellcome Sanger Institute for the COVID-19 Genomics UK (COG-UK) Consortium | Harper VanSteenhouse, Yumi Kasai, David Gray, Carol Clugston, Anna Dominiczak and Alex Alderton, Roberto Amato, Jeffrey Barrett, Sonia Goncalves, Ewan Harrison, David K. Jackson, Ian Johnston, Dominic Kwiatkowski, Cordelia Langford, John Sillitoe on behalf of the Wellcome Sanger Institute COVID-19 Surveillance Team |
| EPI_ISL_3172020 | Lighthouse Laboratory Plymouth | Wellcome Sanger Institute for the COVID-19 Genomics UK (COG-UK) Consortium | Lighthouse Laboratory Plymouth and Alex Alderton, Roberto Amato, Jeffrey Barrett, Sonia Goncalves, Ewan Harrison, David K. Jackson, Ian Johnston, Dominic Kwiatkowski, Cordelia Langford, John Sillitoe on behalf of the Wellcome Sanger Institute COVID-19 Surveillance Team |
| EPI_ISL_3172021 | Lighthouse Lab in Glasgow | Wellcome Sanger Institute for the COVID-19 Genomics UK (COG-UK) Consortium | Harper VanSteenhouse, Yumi Kasai, David Gray, Carol Clugston, Anna Dominiczak and Alex Alderton, Roberto Amato, Jeffrey Barrett, Sonia Goncalves, Ewan Harrison, David K. Jackson, Ian Johnston, Dominic Kwiatkowski, Cordelia Langford, John Sillitoe on behalf of the Wellcome Sanger Institute COVID-19 Surveillance Team |
| EPI_ISL_3172022 | Lighthouse Laboratory Plymouth | Wellcome Sanger Institute for the COVID-19 Genomics UK (COG-UK) Consortium | Lighthouse Laboratory Plymouth and Alex Alderton, Roberto Amato, Jeffrey Barrett, Sonia Goncalves, Ewan Harrison, David K. Jackson, Ian Johnston, Dominic Kwiatkowski, Cordelia Langford, John Sillitoe on behalf of the Wellcome Sanger Institute COVID-19 Surveillance Team |
| EPI_ISL_3172023, EPI_ISL_3172024 | Lighthouse Lab in Glasgow | Wellcome Sanger Institute for the COVID-19 Genomics UK (COG-UK) Consortium | Harper VanSteenhouse, Yumi Kasai, David Gray, Carol Clugston, Anna Dominiczak and Alex Alderton, Roberto Amato, Jeffrey Barrett, Sonia Goncalves, Ewan Harrison, David K. Jackson, Ian Johnston, Dominic Kwiatkowski, Cordelia Langford, John Sillitoe on behalf of the Wellcome Sanger Institute COVID-19 Surveillance Team |
| EPI_ISL_3172025 | Lighthouse Lab in Milton Keynes | Wellcome Sanger Institute for the COVID-19 Genomics UK (COG-UK) Consortium | The Lighthouse Lab in Milton Keynes and Alex Alderton, Roberto Amato, Jeffrey Barrett, Sonia Goncalves, Ewan Harrison, David K. Jackson, Ian Johnston, Dominic Kwiatkowski, Cordelia Langford, John Sillitoe on behalf of the Wellcome Sanger Institute COVID-19 Surveillance Team |
| EPI_ISL_3172026 | Lighthouse Laboratory Plymouth | Wellcome Sanger Institute for the COVID-19 Genomics UK (COG-UK) Consortium | Lighthouse Laboratory Plymouth and Alex Alderton, Roberto Amato, Jeffrey Barrett, Sonia Goncalves, Ewan Harrison, David K. Jackson, Ian Johnston, Dominic Kwiatkowski, Cordelia Langford, John Sillitoe on behalf of the Wellcome Sanger Institute COVID-19 Surveillance Team |
| EPI_ISL_3172027, EPI_ISL_3172028 | Lighthouse Lab in Glasgow | Wellcome Sanger Institute for the COVID-19 Genomics UK (COG-UK) Consortium | Harper VanSteenhouse, Yumi Kasai, David Gray, Carol Clugston, Anna Dominiczak and Alex Alderton, Roberto Amato, Jeffrey Barrett, Sonia Goncalves, Ewan Harrison, David K. Jackson, Ian Johnston, Dominic Kwiatkowski, Cordelia Langford, John Sillitoe on behalf of the Wellcome Sanger Institute COVID-19 Surveillance Team |
| EPI_ISL_3172029 | Lighthouse Lab in Alderley Park | Wellcome Sanger Institute for the COVID-19 Genomics UK (COG-UK) Consortium | Jacquelyn Wynn, Mairead Hyland, The Lighthouse Lab in Alderley Park and Alex Alderton, Roberto Amato, Jeffrey Barrett, Sonia Goncalves, Ewan Harrison, David K. Jackson, Ian Johnston, Dominic Kwiatkowski, Cordelia Langford, John Sillitoe on behalf of the Wellcome Sanger Institute COVID-19 Surveillance Team |
| EPI_ISL_3172030, EPI_ISL_3172031 | Lighthouse Lab in Milton Keynes | Wellcome Sanger Institute for the COVID-19 Genomics UK (COG-UK) Consortium | The Lighthouse Lab in Milton Keynes and Alex Alderton, Roberto Amato, Jeffrey Barrett, Sonia Goncalves, Ewan Harrison, David K. Jackson, Ian Johnston, Dominic Kwiatkowski, Cordelia Langford, John Sillitoe on behalf of the Wellcome Sanger Institute COVID-19 Surveillance Team |
| EPI_ISL_3172032 | Lighthouse Lab in Glasgow | Wellcome Sanger Institute for the COVID-19 Genomics UK (COG-UK) Consortium | Harper VanSteenhouse, Yumi Kasai, David Gray, Carol Clugston, Anna Dominiczak and Alex Alderton, Roberto Amato, Jeffrey Barrett, Sonia Goncalves, Ewan Harrison, David K. Jackson, Ian Johnston, Dominic Kwiatkowski, Cordelia Langford, John Sillitoe on behalf of the Wellcome Sanger Institute COVID-19 Surveillance Team |
| EPI_ISL_3172033 | Lighthouse Laboratory Plymouth | Wellcome Sanger Institute for the COVID-19 Genomics UK (COG-UK) Consortium | Lighthouse Laboratory Plymouth and Alex Alderton, Roberto Amato, Jeffrey Barrett, Sonia Goncalves, Ewan Harrison, David K. Jackson, Ian Johnston, Dominic Kwiatkowski, Cordelia Langford, John Sillitoe on behalf of the Wellcome Sanger Institute COVID-19 Surveillance Team |
| EPI_ISL_3172034 | Lighthouse Lab in Alderley Park | Wellcome Sanger Institute for the COVID-19 Genomics UK (COG-UK) Consortium | Jacquelyn Wynn, Mairead Hyland, The Lighthouse Lab in Alderley Park and Alex Alderton, Roberto Amato, Jeffrey Barrett, Sonia Goncalves, Ewan Harrison, David K. Jackson, Ian Johnston, Dominic Kwiatkowski, Cordelia Langford, John Sillitoe on behalf of the Wellcome Sanger Institute COVID-19 Surveillance Team |
| EPI_ISL_3172035 | Lighthouse Lab in Glasgow | Wellcome Sanger Institute for the COVID-19 Genomics UK (COG-UK) Consortium | Harper VanSteenhouse, Yumi Kasai, David Gray, Carol Clugston, Anna Dominiczak and Alex Alderton, Roberto Amato, Jeffrey Barrett, Sonia Goncalves, Ewan Harrison, David K. Jackson, Ian Johnston, Dominic Kwiatkowski, Cordelia Langford, John Sillitoe on behalf of the Wellcome Sanger Institute COVID-19 Surveillance Team |
| EPI_ISL_3172036 | Lighthouse Lab in Milton Keynes | Wellcome Sanger Institute for the COVID-19 Genomics UK (COG-UK) Consortium | The Lighthouse Lab in Milton Keynes and Alex Alderton, Roberto Amato, Jeffrey Barrett, Sonia Goncalves, Ewan Harrison, David K. Jackson, Ian Johnston, Dominic Kwiatkowski, Cordelia Langford, John Sillitoe on behalf of the Wellcome Sanger Institute COVID-19 Surveillance Team |
| EPI_ISL_3172037, EPI_ISL_3172038 | Lighthouse Laboratory Plymouth | Wellcome Sanger Institute for the COVID-19 Genomics UK (COG-UK) Consortium | Lighthouse Laboratory Plymouth and Alex Alderton, Roberto Amato, Jeffrey Barrett, Sonia Goncalves, Ewan Harrison, David K. Jackson, Ian Johnston, Dominic Kwiatkowski, Cordelia Langford, John Sillitoe on behalf of the Wellcome Sanger Institute COVID-19 Surveillance Team |
| EPI_ISL_3172039, EPI_ISL_3172040, EPI_ISL_3172041 | Lighthouse Lab in Milton Keynes | Wellcome Sanger Institute for the COVID-19 Genomics UK (COG-UK) Consortium | The Lighthouse Lab in Milton Keynes and Alex Alderton, Roberto Amato, Jeffrey Barrett, Sonia Goncalves, Ewan Harrison, David K. Jackson, Ian Johnston, Dominic Kwiatkowski, Cordelia Langford, John Sillitoe on behalf of the Wellcome Sanger Institute COVID-19 Surveillance Team |
| EPI_ISL_3172042 | Lighthouse Lab in Alderley Park | Wellcome Sanger Institute for the COVID-19 Genomics UK (COG-UK) Consortium | Jacquelyn Wynn, Mairead Hyland, The Lighthouse Lab in Alderley Park and Alex Alderton, Roberto Amato, Jeffrey Barrett, Sonia Goncalves, Ewan Harrison, David K. Jackson, Ian Johnston, Dominic Kwiatkowski, Cordelia Langford, John Sillitoe on behalf of the Wellcome Sanger Institute COVID-19 Surveillance Team |
| EPI_ISL_3172043 | Lighthouse Lab in Glasgow | Wellcome Sanger Institute for the COVID-19 Genomics UK (COG-UK) Consortium | Harper VanSteenhouse, Yumi Kasai, David Gray, Carol Clugston, Anna Dominiczak and Alex Alderton, Roberto Amato, Jeffrey Barrett, Sonia Goncalves, Ewan Harrison, David K. Jackson, Ian Johnston, Dominic Kwiatkowski, Cordelia Langford, John Sillitoe on behalf of the Wellcome Sanger Institute COVID-19 Surveillance Team |
| EPI_ISL_3172044 | Lighthouse Lab in Milton Keynes | Wellcome Sanger Institute for the COVID-19 Genomics UK (COG-UK) Consortium | The Lighthouse Lab in Milton Keynes and Alex Alderton, Roberto Amato, Jeffrey Barrett, Sonia Goncalves, Ewan Harrison, David K. Jackson, Ian Johnston, Dominic Kwiatkowski, Cordelia Langford, John Sillitoe on behalf of the Wellcome Sanger Institute COVID-19 Surveillance Team |
| EPI_ISL_3172045, EPI_ISL_3172046 | Lighthouse Lab in Alderley Park | Wellcome Sanger Institute for the COVID-19 Genomics UK (COG-UK) Consortium | Jacquelyn Wynn, Mairead Hyland, The Lighthouse Lab in Alderley Park and Alex Alderton, Roberto Amato, Jeffrey Barrett, Sonia Goncalves, Ewan Harrison, David K. Jackson, Ian Johnston, Dominic Kwiatkowski, Cordelia Langford, John Sillitoe on behalf of the Wellcome Sanger Institute COVID-19 Surveillance Team |
| EPI_ISL_3172047 | Lighthouse Lab in Milton Keynes | Wellcome Sanger Institute for the COVID-19 Genomics UK (COG-UK) Consortium | The Lighthouse Lab in Milton Keynes and Alex Alderton, Roberto Amato, Jeffrey Barrett, Sonia Goncalves, Ewan Harrison, David K. Jackson, Ian Johnston, Dominic Kwiatkowski, Cordelia Langford, John Sillitoe on behalf of the Wellcome Sanger Institute COVID-19 Surveillance Team |
| EPI_ISL_3172048 | Lighthouse Lab in Glasgow | Wellcome Sanger Institute for the COVID-19 Genomics UK (COG-UK) Consortium | Harper VanSteenhouse, Yumi Kasai, David Gray, Carol Clugston, Anna Dominiczak and Alex Alderton, Roberto Amato, Jeffrey Barrett, Sonia Goncalves, Ewan Harrison, David K. Jackson, Ian Johnston, Dominic Kwiatkowski, Cordelia Langford, John Sillitoe on behalf of the Wellcome Sanger Institute COVID-19 Surveillance Team |

| | | | |
|---|---|---|---|
| EPI_ISL_3172049 | Lighthouse Lab in Milton Keynes | Wellcome Sanger Institute for the COVID-19 Genomics UK (COG-UK) Consortium | The Lighthouse Lab in Milton Keynes and Alex Alderton, Roberto Amato, Jeffrey Barrett, Sonia Goncalves, Ewan Harrison, David K. Jackson, Ian Johnston, Dominic Kwiatkowski, Cordelia Langford, John Sillitoe on behalf of the Wellcome Sanger Institute COVID-19 Surveillance Team |
| EPI_ISL_3172050, EPI_ISL_3172051 | Lighthouse Laboratory Plymouth | Wellcome Sanger Institute for the COVID-19 Genomics UK (COG-UK) Consortium | Lighthouse Laboratory Plymouth and Alex Alderton, Roberto Amato, Jeffrey Barrett, Sonia Goncalves, Ewan Harrison, David K. Jackson, Ian Johnston, Dominic Kwiatkowski, Cordelia Langford, John Sillitoe on behalf of the Wellcome Sanger Institute COVID-19 Surveillance Team |
| EPI_ISL_3172052, EPI_ISL_3172053 | Lighthouse Lab in Milton Keynes | Wellcome Sanger Institute for the COVID-19 Genomics UK (COG-UK) Consortium | The Lighthouse Lab in Milton Keynes and Alex Alderton, Roberto Amato, Jeffrey Barrett, Sonia Goncalves, Ewan Harrison, David K. Jackson, Ian Johnston, Dominic Kwiatkowski, Cordelia Langford, John Sillitoe on behalf of the Wellcome Sanger Institute COVID-19 Surveillance Team |
| EPI_ISL_3172054 | Lighthouse Lab in Alderley Park | Wellcome Sanger Institute for the COVID-19 Genomics UK (COG-UK) Consortium | Jacquelyn Wynn, Mairead Hyland, The Lighthouse Lab in Alderley Park and Alex Alderton, Roberto Amato, Jeffrey Barrett, Sonia Goncalves, Ewan Harrison, David K. Jackson, Ian Johnston, Dominic Kwiatkowski, Cordelia Langford, John Sillitoe on behalf of the Wellcome Sanger Institute COVID-19 Surveillance Team |
| EPI_ISL_3172055 | Lighthouse Lab in Milton Keynes | Wellcome Sanger Institute for the COVID-19 Genomics UK (COG-UK) Consortium | The Lighthouse Lab in Milton Keynes and Alex Alderton, Roberto Amato, Jeffrey Barrett, Sonia Goncalves, Ewan Harrison, David K. Jackson, Ian Johnston, Dominic Kwiatkowski, Cordelia Langford, John Sillitoe on behalf of the Wellcome Sanger Institute COVID-19 Surveillance Team |
| EPI_ISL_3172056 | Lighthouse Lab in Alderley Park | Wellcome Sanger Institute for the COVID-19 Genomics UK (COG-UK) Consortium | Jacquelyn Wynn, Mairead Hyland, The Lighthouse Lab in Alderley Park and Alex Alderton, Roberto Amato, Jeffrey Barrett, Sonia Goncalves, Ewan Harrison, David K. Jackson, Ian Johnston, Dominic Kwiatkowski, Cordelia Langford, John Sillitoe on behalf of the Wellcome Sanger Institute COVID-19 Surveillance Team |
| EPI_ISL_3172057 | Lighthouse Lab in Glasgow | Wellcome Sanger Institute for the COVID-19 Genomics UK (COG-UK) Consortium | Harper VanSteenhouse, Yumi Kasai, David Gray, Carol Clugston, Anna Dominiczak and Alex Alderton, Roberto Amato, Jeffrey Barrett, Sonia Goncalves, Ewan Harrison, David K. Jackson, Ian Johnston, Dominic Kwiatkowski, Cordelia Langford, John Sillitoe on behalf of the Wellcome Sanger Institute COVID-19 Surveillance Team |
| EPI_ISL_3172058 | Lighthouse Lab in Milton Keynes | Wellcome Sanger Institute for the COVID-19 Genomics UK (COG-UK) Consortium | The Lighthouse Lab in Milton Keynes and Alex Alderton, Roberto Amato, Jeffrey Barrett, Sonia Goncalves, Ewan Harrison, David K. Jackson, Ian Johnston, Dominic Kwiatkowski, Cordelia Langford, John Sillitoe on behalf of the Wellcome Sanger Institute COVID-19 Surveillance Team |
| EPI_ISL_3172059, EPI_ISL_3172060, EPI_ISL_3172061 | Lighthouse Laboratory Plymouth | Wellcome Sanger Institute for the COVID-19 Genomics UK (COG-UK) Consortium | Lighthouse Laboratory Plymouth and Alex Alderton, Roberto Amato, Jeffrey Barrett, Sonia Goncalves, Ewan Harrison, David K. Jackson, Ian Johnston, Dominic Kwiatkowski, Cordelia Langford, John Sillitoe on behalf of the Wellcome Sanger Institute COVID-19 Surveillance Team |
| EPI_ISL_3172062 | Lighthouse Lab in Glasgow | Wellcome Sanger Institute for the COVID-19 Genomics UK (COG-UK) Consortium | Harper VanSteenhouse, Yumi Kasai, David Gray, Carol Clugston, Anna Dominiczak and Alex Alderton, Roberto Amato, Jeffrey Barrett, Sonia Goncalves, Ewan Harrison, David K. Jackson, Ian Johnston, Dominic Kwiatkowski, Cordelia Langford, John Sillitoe on behalf of the Wellcome Sanger Institute COVID-19 Surveillance Team |
| EPI_ISL_3172063 | Lighthouse Laboratory Plymouth | Wellcome Sanger Institute for the COVID-19 Genomics UK (COG-UK) Consortium | Lighthouse Laboratory Plymouth and Alex Alderton, Roberto Amato, Jeffrey Barrett, Sonia Goncalves, Ewan Harrison, David K. Jackson, Ian Johnston, Dominic Kwiatkowski, Cordelia Langford, John Sillitoe on behalf of the Wellcome Sanger Institute COVID-19 Surveillance Team |
| EPI_ISL_3172064 | Lighthouse Lab in Milton Keynes | Wellcome Sanger Institute for the COVID-19 Genomics UK (COG-UK) Consortium | The Lighthouse Lab in Milton Keynes and Alex Alderton, Roberto Amato, Jeffrey Barrett, Sonia Goncalves, Ewan Harrison, David K. Jackson, Ian Johnston, Dominic Kwiatkowski, Cordelia Langford, John Sillitoe on behalf of the Wellcome Sanger Institute COVID-19 Surveillance Team |
| EPI_ISL_3172065 | Lighthouse Lab in Alderley Park | Wellcome Sanger Institute for the COVID-19 Genomics UK (COG-UK) Consortium | Jacquelyn Wynn, Mairead Hyland, The Lighthouse Lab in Alderley Park and Alex Alderton, Roberto Amato, Jeffrey Barrett, Sonia Goncalves, Ewan Harrison, David K. Jackson, Ian Johnston, Dominic Kwiatkowski, Cordelia Langford, John Sillitoe on behalf of the Wellcome Sanger Institute COVID-19 Surveillance Team |
| EPI_ISL_3172066, EPI_ISL_3172067, EPI_ISL_3172068, EPI_ISL_3172069 | Lighthouse Lab in Glasgow | Wellcome Sanger Institute for the COVID-19 Genomics UK (COG-UK) Consortium | Harper VanSteenhouse, Yumi Kasai, David Gray, Carol Clugston, Anna Dominiczak and Alex Alderton, Roberto Amato, Jeffrey Barrett, Sonia Goncalves, Ewan Harrison, David K. Jackson, Ian Johnston, Dominic Kwiatkowski, Cordelia Langford, John Sillitoe on behalf of the Wellcome Sanger Institute COVID-19 Surveillance Team |
| EPI_ISL_3172070 | Lighthouse Laboratory Plymouth | Wellcome Sanger Institute for the COVID-19 Genomics UK (COG-UK) Consortium | Lighthouse Laboratory Plymouth and Alex Alderton, Roberto Amato, Jeffrey Barrett, Sonia Goncalves, Ewan Harrison, David K. Jackson, Ian Johnston, Dominic Kwiatkowski, Cordelia Langford, John Sillitoe on behalf of the Wellcome Sanger Institute COVID-19 Surveillance Team |
| EPI_ISL_3172071 | Lighthouse Lab in Glasgow | Wellcome Sanger Institute for the COVID-19 Genomics UK (COG-UK) Consortium | Harper VanSteenhouse, Yumi Kasai, David Gray, Carol Clugston, Anna Dominiczak and Alex Alderton, Roberto Amato, Jeffrey Barrett, Sonia Goncalves, Ewan Harrison, David K. Jackson, Ian Johnston, Dominic Kwiatkowski, Cordelia Langford, John Sillitoe on behalf of the Wellcome Sanger Institute COVID-19 Surveillance Team |
| EPI_ISL_3172072 | Lighthouse Lab in Milton Keynes | Wellcome Sanger Institute for the COVID-19 Genomics UK (COG-UK) Consortium | The Lighthouse Lab in Milton Keynes and Alex Alderton, Roberto Amato, Jeffrey Barrett, Sonia Goncalves, Ewan Harrison, David K. Jackson, Ian Johnston, Dominic Kwiatkowski, Cordelia Langford, John Sillitoe on behalf of the Wellcome Sanger Institute COVID-19 Surveillance Team |
| EPI_ISL_3172073 | Lighthouse Lab in Glasgow | Wellcome Sanger Institute for the COVID-19 Genomics UK (COG-UK) Consortium | Harper VanSteenhouse, Yumi Kasai, David Gray, Carol Clugston, Anna Dominiczak and Alex Alderton, Roberto Amato, Jeffrey Barrett, Sonia Goncalves, Ewan Harrison, David K. Jackson, Ian Johnston, Dominic Kwiatkowski, Cordelia Langford, John Sillitoe on behalf of the Wellcome Sanger Institute COVID-19 Surveillance Team |
| EPI_ISL_3172074 | Lighthouse Lab in Alderley Park | Wellcome Sanger Institute for the COVID-19 Genomics UK (COG-UK) Consortium | Jacquelyn Wynn, Mairead Hyland, The Lighthouse Lab in Alderley Park and Alex Alderton, Roberto Amato, Jeffrey Barrett, Sonia Goncalves, Ewan Harrison, David K. Jackson, Ian Johnston, Dominic Kwiatkowski, Cordelia Langford, John Sillitoe on behalf of the Wellcome Sanger Institute COVID-19 Surveillance Team |
| EPI_ISL_3172075 | Lighthouse Lab in Milton Keynes | Wellcome Sanger Institute for the COVID-19 Genomics UK (COG-UK) Consortium | The Lighthouse Lab in Milton Keynes and Alex Alderton, Roberto Amato, Jeffrey Barrett, Sonia Goncalves, Ewan Harrison, David K. Jackson, Ian Johnston, Dominic Kwiatkowski, Cordelia Langford, John Sillitoe on behalf of the Wellcome Sanger Institute COVID-19 Surveillance Team |
| EPI_ISL_3172076 | Lighthouse Lab in Glasgow | Wellcome Sanger Institute for the COVID-19 Genomics UK (COG-UK) Consortium | Harper VanSteenhouse, Yumi Kasai, David Gray, Carol Clugston, Anna Dominiczak and Alex Alderton, Roberto Amato, Jeffrey Barrett, Sonia Goncalves, Ewan Harrison, David K. Jackson, Ian Johnston, Dominic Kwiatkowski, Cordelia Langford, John Sillitoe on behalf of the Wellcome Sanger Institute COVID-19 Surveillance Team |
| EPI_ISL_3172077, EPI_ISL_3172078 | Lighthouse Laboratory Plymouth | Wellcome Sanger Institute for the COVID-19 Genomics UK (COG-UK) Consortium | Lighthouse Laboratory Plymouth and Alex Alderton, Roberto Amato, Jeffrey Barrett, Sonia Goncalves, Ewan Harrison, David K. Jackson, Ian Johnston, Dominic Kwiatkowski, Cordelia Langford, John Sillitoe on behalf of the Wellcome Sanger Institute COVID-19 Surveillance Team |
| EPI_ISL_3172079 | Lighthouse Lab in Glasgow | Wellcome Sanger Institute for the COVID-19 Genomics UK (COG-UK) Consortium | Harper VanSteenhouse, Yumi Kasai, David Gray, Carol Clugston, Anna Dominiczak and Alex Alderton, Roberto Amato, Jeffrey Barrett, Sonia Goncalves, Ewan Harrison, David K. Jackson, Ian Johnston, Dominic Kwiatkowski, Cordelia Langford, John Sillitoe on behalf of the Wellcome Sanger Institute COVID-19 Surveillance Team |
| EPI_ISL_3172080 | Lighthouse Lab in Alderley Park | Wellcome Sanger Institute for the COVID-19 Genomics UK (COG-UK) Consortium | Jacquelyn Wynn, Mairead Hyland, The Lighthouse Lab in Alderley Park and Alex Alderton, Roberto Amato, Jeffrey Barrett, Sonia Goncalves, Ewan Harrison, David K. Jackson, Ian Johnston, Dominic Kwiatkowski, Cordelia Langford, John Sillitoe on behalf of the Wellcome Sanger Institute COVID-19 Surveillance Team |
| EPI_ISL_3172081 | Lighthouse Lab in Milton Keynes | Wellcome Sanger Institute for the COVID-19 Genomics UK (COG-UK) Consortium | The Lighthouse Lab in Milton Keynes and Alex Alderton, Roberto Amato, Jeffrey Barrett, Sonia Goncalves, Ewan Harrison, David K. Jackson, Ian Johnston, Dominic Kwiatkowski, Cordelia Langford, John Sillitoe on behalf of the Wellcome Sanger Institute COVID-19 Surveillance Team |
| EPI_ISL_3172082 | Lighthouse Laboratory Plymouth | Wellcome Sanger Institute for the COVID-19 Genomics UK (COG-UK) Consortium | Lighthouse Laboratory Plymouth and Alex Alderton, Roberto Amato, Jeffrey Barrett, Sonia Goncalves, Ewan Harrison, David K. Jackson, Ian Johnston, Dominic Kwiatkowski, Cordelia Langford, John Sillitoe on behalf of the Wellcome Sanger Institute COVID-19 Surveillance Team |
| EPI_ISL_3172083 | Lighthouse Lab in Glasgow | Wellcome Sanger Institute for the COVID-19 Genomics UK (COG-UK) Consortium | Harper VanSteenhouse, Yumi Kasai, David Gray, Carol Clugston, Anna Dominiczak and Alex Alderton, Roberto Amato, Jeffrey Barrett, Sonia Goncalves, Ewan Harrison, David K. Jackson, Ian Johnston, Dominic Kwiatkowski, Cordelia Langford, John Sillitoe on behalf of the Wellcome Sanger Institute COVID-19 Surveillance Team |
| EPI_ISL_3172084, EPI_ISL_3172085 | Lighthouse Lab in Alderley Park | Wellcome Sanger Institute for the COVID-19 Genomics UK (COG-UK) Consortium | Jacquelyn Wynn, Mairead Hyland, The Lighthouse Lab in Alderley Park and Alex Alderton, Roberto Amato, Jeffrey Barrett, Sonia Goncalves, Ewan Harrison, David K. Jackson, Ian Johnston, Dominic Kwiatkowski, Cordelia Langford, John Sillitoe on behalf of the Wellcome Sanger Institute COVID-19 Surveillance Team |
| EPI_ISL_3172086 | Lighthouse Laboratory Plymouth | Wellcome Sanger Institute for the COVID-19 Genomics UK (COG-UK) Consortium | Lighthouse Laboratory Plymouth and Alex Alderton, Roberto Amato, Jeffrey Barrett, Sonia Goncalves, Ewan Harrison, David K. Jackson, Ian Johnston, Dominic Kwiatkowski, Cordelia Langford, John Sillitoe on behalf of the Wellcome Sanger Institute COVID-19 Surveillance Team |
| EPI_ISL_3172087 | Lighthouse Lab in Glasgow | Wellcome Sanger Institute for the COVID-19 Genomics UK (COG-UK) Consortium | Harper VanSteenhouse, Yumi Kasai, David Gray, Carol Clugston, Anna Dominiczak and Alex Alderton, Roberto Amato, Jeffrey Barrett, Sonia Goncalves, Ewan Harrison, David K. Jackson, Ian Johnston, Dominic Kwiatkowski, Cordelia Langford, John Sillitoe on behalf of the Wellcome Sanger Institute COVID-19 Surveillance Team |
| EPI_ISL_3172088 | Lighthouse Lab in Milton Keynes | Wellcome Sanger Institute for the COVID-19 Genomics UK (COG-UK) Consortium | The Lighthouse Lab in Milton Keynes and Alex Alderton, Roberto Amato, Jeffrey Barrett, Sonia Goncalves, Ewan Harrison, David K. Jackson, Ian Johnston, Dominic Kwiatkowski, Cordelia Langford, John Sillitoe on behalf of the Wellcome Sanger Institute COVID-19 Surveillance Team |
| EPI_ISL_3172089, EPI_ISL_3172090 | Lighthouse Laboratory Plymouth | Wellcome Sanger Institute for the COVID-19 Genomics UK (COG-UK) Consortium | Lighthouse Laboratory Plymouth and Alex Alderton, Roberto Amato, Jeffrey Barrett, Sonia Goncalves, Ewan Harrison, David K. Jackson, Ian Johnston, Dominic Kwiatkowski, Cordelia Langford, John Sillitoe on behalf of the Wellcome Sanger Institute COVID-19 Surveillance Team |
| EPI_ISL_3172091, EPI_ISL_3172092, EPI_ISL_3172093 | Lighthouse Lab in Alderley Park | Wellcome Sanger Institute for the COVID-19 Genomics UK (COG-UK) Consortium | Jacquelyn Wynn, Mairead Hyland, The Lighthouse Lab in Alderley Park and Alex Alderton, Roberto Amato, Jeffrey Barrett, Sonia Goncalves, Ewan Harrison, David K. Jackson, Ian Johnston, Dominic Kwiatkowski, Cordelia Langford, John Sillitoe on behalf of the Wellcome Sanger Institute COVID-19 Surveillance Team |
| EPI_ISL_3172094 | Lighthouse Laboratory Plymouth | Wellcome Sanger Institute for the COVID-19 Genomics UK (COG-UK) Consortium | Lighthouse Laboratory Plymouth and Alex Alderton, Roberto Amato, Jeffrey Barrett, Sonia Goncalves, Ewan Harrison, David K. Jackson, Ian Johnston, Dominic Kwiatkowski, Cordelia Langford, John Sillitoe on behalf of the Wellcome Sanger Institute COVID-19 Surveillance Team |
| EPI_ISL_3172095 | Lighthouse Lab in Alderley Park | Wellcome Sanger Institute for the COVID-19 Genomics UK (COG-UK) Consortium | Jacquelyn Wynn, Mairead Hyland, The Lighthouse Lab in Alderley Park and Alex Alderton, Roberto Amato, Jeffrey Barrett, Sonia Goncalves, Ewan Harrison, David K. Jackson, Ian Johnston, Dominic Kwiatkowski, Cordelia Langford, John Sillitoe on behalf of the Wellcome Sanger Institute COVID-19 Surveillance Team |
| EPI_ISL_3172096 | Lighthouse Lab in Glasgow | Wellcome Sanger Institute for the COVID-19 Genomics UK (COG-UK) Consortium | Harper VanSteenhouse, Yumi Kasai, David Gray, Carol Clugston, Anna Dominiczak and Alex Alderton, Roberto Amato, Jeffrey Barrett, Sonia Goncalves, Ewan Harrison, David K. Jackson, Ian Johnston, Dominic Kwiatkowski, Cordelia Langford, John Sillitoe on behalf of the Wellcome Sanger Institute COVID-19 Surveillance Team |
| EPI_ISL_3172097 | Lighthouse Lab in Alderley Park | Wellcome Sanger Institute for the COVID-19 Genomics UK (COG-UK) Consortium | Jacquelyn Wynn, Mairead Hyland, The Lighthouse Lab in Alderley Park and Alex Alderton, Roberto Amato, Jeffrey Barrett, Sonia Goncalves, Ewan Harrison, David K. Jackson, Ian Johnston, Dominic Kwiatkowski, Cordelia Langford, John Sillitoe on behalf of the Wellcome Sanger Institute COVID-19 Surveillance Team |
| EPI_ISL_3172098 | Lighthouse Lab in Glasgow | Wellcome Sanger Institute for the COVID-19 Genomics UK (COG-UK) Consortium | Harper VanSteenhouse, Yumi Kasai, David Gray, Carol Clugston, Anna Dominiczak and Alex Alderton, Roberto Amato, Jeffrey Barrett, Sonia Goncalves, Ewan Harrison, David K. Jackson, Ian Johnston, Dominic Kwiatkowski, Cordelia Langford, John Sillitoe on behalf of the Wellcome Sanger Institute COVID-19 Surveillance Team |
| EPI_ISL_3172099, EPI_ISL_3172100, EPI_ISL_3172101 | Lighthouse Lab in Alderley Park | Wellcome Sanger Institute for the COVID-19 Genomics UK (COG-UK) Consortium | Jacquelyn Wynn, Mairead Hyland, The Lighthouse Lab in Alderley Park and Alex Alderton, Roberto Amato, Jeffrey Barrett, Sonia Goncalves, Ewan Harrison, David K. Jackson, Ian Johnston, Dominic Kwiatkowski, Cordelia Langford, John Sillitoe on behalf of the Wellcome Sanger Institute COVID-19 Surveillance Team |
| EPI_ISL_3172102 | Lighthouse Laboratory Plymouth | Wellcome Sanger Institute for the COVID-19 Genomics UK (COG-UK) Consortium | Lighthouse Laboratory Plymouth and Alex Alderton, Roberto Amato, Jeffrey Barrett, Sonia Goncalves, Ewan Harrison, David K. Jackson, Ian Johnston, Dominic Kwiatkowski, Cordelia Langford, John Sillitoe on behalf of the Wellcome Sanger Institute COVID-19 Surveillance Team |
| EPI_ISL_3172103 | Lighthouse Lab in Glasgow | Wellcome Sanger Institute for the COVID-19 Genomics UK (COG-UK) Consortium | Harper VanSteenhouse, Yumi Kasai, David Gray, Carol Clugston, Anna Dominiczak and Alex Alderton, Roberto Amato, Jeffrey Barrett, Sonia Goncalves, Ewan Harrison, David K. Jackson, Ian Johnston, Dominic Kwiatkowski, Cordelia Langford, John Sillitoe on behalf of the Wellcome Sanger Institute COVID-19 Surveillance Team |
| EPI_ISL_3172104 | Lighthouse Laboratory Plymouth | Wellcome Sanger Institute for the COVID-19 Genomics UK (COG-UK) Consortium | Lighthouse Laboratory Plymouth and Alex Alderton, Roberto Amato, Jeffrey Barrett, Sonia Goncalves, Ewan Harrison, David K. Jackson, Ian Johnston, Dominic Kwiatkowski, Cordelia Langford, John Sillitoe on behalf of the Wellcome Sanger Institute COVID-19 Surveillance Team |
| EPI_ISL_3172105 | Lighthouse Lab in Glasgow | Wellcome Sanger Institute for the COVID-19 Genomics UK (COG-UK) Consortium | Harper VanSteenhouse, Yumi Kasai, David Gray, Carol Clugston, Anna Dominiczak and Alex Alderton, Roberto Amato, Jeffrey Barrett, Sonia Goncalves, Ewan Harrison, David K. Jackson, Ian Johnston, Dominic Kwiatkowski, Cordelia Langford, John Sillitoe on behalf of the Wellcome Sanger Institute COVID-19 Surveillance Team |

| | | | |
|---|---|---|---|
| EPI_ISL_3172106 | Lighthouse Lab in Alderley Park | Wellcome Sanger Institute for the COVID-19 Genomics UK (COG-UK) Consortium | Jacquelyn Wynn, Mairead Hyland, The Lighthouse Lab in Alderley Park and Alex Alderton, Roberto Amato, Jeffrey Barrett, Sonia Goncalves, Ewan Harrison, David K. Jackson, Ian Johnston, Dominic Kwiatkowski, Cordelia Langford, John Sillitoe on behalf of the Wellcome Sanger Institute COVID-19 Surveillance Team |
| EPI_ISL_3172107, EPI_ISL_3172108 | Lighthouse Laboratory Plymouth | Wellcome Sanger Institute for the COVID-19 Genomics UK (COG-UK) Consortium | Lighthouse Laboratory Plymouth and Alex Alderton, Roberto Amato, Jeffrey Barrett, Sonia Goncalves, Ewan Harrison, David K. Jackson, Ian Johnston, Dominic Kwiatkowski, Cordelia Langford, John Sillitoe on behalf of the Wellcome Sanger Institute COVID-19 Surveillance Team |
| EPI_ISL_3172109 | Lighthouse Lab in Milton Keynes | Wellcome Sanger Institute for the COVID-19 Genomics UK (COG-UK) Consortium | The Lighthouse Lab in Milton Keynes and Alex Alderton, Roberto Amato, Jeffrey Barrett, Sonia Goncalves, Ewan Harrison, David K. Jackson, Ian Johnston, Dominic Kwiatkowski, Cordelia Langford, John Sillitoe on behalf of the Wellcome Sanger Institute COVID-19 Surveillance Team |
| EPI_ISL_3172110 | Lighthouse Lab in Alderley Park | Wellcome Sanger Institute for the COVID-19 Genomics UK (COG-UK) Consortium | Jacquelyn Wynn, Mairead Hyland, The Lighthouse Lab in Alderley Park and Alex Alderton, Roberto Amato, Jeffrey Barrett, Sonia Goncalves, Ewan Harrison, David K. Jackson, Ian Johnston, Dominic Kwiatkowski, Cordelia Langford, John Sillitoe on behalf of the Wellcome Sanger Institute COVID-19 Surveillance Team |
| EPI_ISL_3172111 | Lighthouse Lab in Glasgow | Wellcome Sanger Institute for the COVID-19 Genomics UK (COG-UK) Consortium | Harper VanSteenhouse, Yumi Kasai, David Gray, Carol Clugston, Anna Dominiczak and Alex Alderton, Roberto Amato, Jeffrey Barrett, Sonia Goncalves, Ewan Harrison, David K. Jackson, Ian Johnston, Dominic Kwiatkowski, Cordelia Langford, John Sillitoe on behalf of the Wellcome Sanger Institute COVID-19 Surveillance Team |
| EPI_ISL_3172112 | Lighthouse Laboratory Plymouth | Wellcome Sanger Institute for the COVID-19 Genomics UK (COG-UK) Consortium | Lighthouse Laboratory Plymouth and Alex Alderton, Roberto Amato, Jeffrey Barrett, Sonia Goncalves, Ewan Harrison, David K. Jackson, Ian Johnston, Dominic Kwiatkowski, Cordelia Langford, John Sillitoe on behalf of the Wellcome Sanger Institute COVID-19 Surveillance Team |
| EPI_ISL_3172113 | Lighthouse Lab in Milton Keynes | Wellcome Sanger Institute for the COVID-19 Genomics UK (COG-UK) Consortium | The Lighthouse Lab in Milton Keynes and Alex Alderton, Roberto Amato, Jeffrey Barrett, Sonia Goncalves, Ewan Harrison, David K. Jackson, Ian Johnston, Dominic Kwiatkowski, Cordelia Langford, John Sillitoe on behalf of the Wellcome Sanger Institute COVID-19 Surveillance Team |
| EPI_ISL_3172114 | Lighthouse Laboratory Plymouth | Wellcome Sanger Institute for the COVID-19 Genomics UK (COG-UK) Consortium | Lighthouse Laboratory Plymouth and Alex Alderton, Roberto Amato, Jeffrey Barrett, Sonia Goncalves, Ewan Harrison, David K. Jackson, Ian Johnston, Dominic Kwiatkowski, Cordelia Langford, John Sillitoe on behalf of the Wellcome Sanger Institute COVID-19 Surveillance Team |
| EPI_ISL_3172115 | Lighthouse Lab in Alderley Park | Wellcome Sanger Institute for the COVID-19 Genomics UK (COG-UK) Consortium | Jacquelyn Wynn, Mairead Hyland, The Lighthouse Lab in Alderley Park and Alex Alderton, Roberto Amato, Jeffrey Barrett, Sonia Goncalves, Ewan Harrison, David K. Jackson, Ian Johnston, Dominic Kwiatkowski, Cordelia Langford, John Sillitoe on behalf of the Wellcome Sanger Institute COVID-19 Surveillance Team |
| EPI_ISL_3172116 | Lighthouse Lab in Milton Keynes | Wellcome Sanger Institute for the COVID-19 Genomics UK (COG-UK) Consortium | The Lighthouse Lab in Milton Keynes and Alex Alderton, Roberto Amato, Jeffrey Barrett, Sonia Goncalves, Ewan Harrison, David K. Jackson, Ian Johnston, Dominic Kwiatkowski, Cordelia Langford, John Sillitoe on behalf of the Wellcome Sanger Institute COVID-19 Surveillance Team |
| EPI_ISL_3172117, EPI_ISL_3172118 | Lighthouse Lab in Alderley Park | Wellcome Sanger Institute for the COVID-19 Genomics UK (COG-UK) Consortium | Jacquelyn Wynn, Mairead Hyland, The Lighthouse Lab in Alderley Park and Alex Alderton, Roberto Amato, Jeffrey Barrett, Sonia Goncalves, Ewan Harrison, David K. Jackson, Ian Johnston, Dominic Kwiatkowski, Cordelia Langford, John Sillitoe on behalf of the Wellcome Sanger Institute COVID-19 Surveillance Team |
| EPI_ISL_3172119 | Lighthouse Lab in Milton Keynes | Wellcome Sanger Institute for the COVID-19 Genomics UK (COG-UK) Consortium | The Lighthouse Lab in Milton Keynes and Alex Alderton, Roberto Amato, Jeffrey Barrett, Sonia Goncalves, Ewan Harrison, David K. Jackson, Ian Johnston, Dominic Kwiatkowski, Cordelia Langford, John Sillitoe on behalf of the Wellcome Sanger Institute COVID-19 Surveillance Team |
| EPI_ISL_3172120 | Lighthouse Lab in Glasgow | Wellcome Sanger Institute for the COVID-19 Genomics UK (COG-UK) Consortium | Harper VanSteenhouse, Yumi Kasai, David Gray, Carol Clugston, Anna Dominiczak and Alex Alderton, Roberto Amato, Jeffrey Barrett, Sonia Goncalves, Ewan Harrison, David K. Jackson, Ian Johnston, Dominic Kwiatkowski, Cordelia Langford, John Sillitoe on behalf of the Wellcome Sanger Institute COVID-19 Surveillance Team |
| EPI_ISL_3172121 | Lighthouse Lab in Milton Keynes | Wellcome Sanger Institute for the COVID-19 Genomics UK (COG-UK) Consortium | The Lighthouse Lab in Milton Keynes and Alex Alderton, Roberto Amato, Jeffrey Barrett, Sonia Goncalves, Ewan Harrison, David K. Jackson, Ian Johnston, Dominic Kwiatkowski, Cordelia Langford, John Sillitoe on behalf of the Wellcome Sanger Institute COVID-19 Surveillance Team |
| EPI_ISL_3172122 | Lighthouse Lab in Alderley Park | Wellcome Sanger Institute for the COVID-19 Genomics UK (COG-UK) Consortium | Jacquelyn Wynn, Mairead Hyland, The Lighthouse Lab in Alderley Park and Alex Alderton, Roberto Amato, Jeffrey Barrett, Sonia Goncalves, Ewan Harrison, David K. Jackson, Ian Johnston, Dominic Kwiatkowski, Cordelia Langford, John Sillitoe on behalf of the Wellcome Sanger Institute COVID-19 Surveillance Team |
| EPI_ISL_3172123 | Lighthouse Lab in Glasgow | Wellcome Sanger Institute for the COVID-19 Genomics UK (COG-UK) Consortium | Harper VanSteenhouse, Yumi Kasai, David Gray, Carol Clugston, Anna Dominiczak and Alex Alderton, Roberto Amato, Jeffrey Barrett, Sonia Goncalves, Ewan Harrison, David K. Jackson, Ian Johnston, Dominic Kwiatkowski, Cordelia Langford, John Sillitoe on behalf of the Wellcome Sanger Institute COVID-19 Surveillance Team |
| EPI_ISL_3172124 | Lighthouse Lab in Milton Keynes | Wellcome Sanger Institute for the COVID-19 Genomics UK (COG-UK) Consortium | The Lighthouse Lab in Milton Keynes and Alex Alderton, Roberto Amato, Jeffrey Barrett, Sonia Goncalves, Ewan Harrison, David K. Jackson, Ian Johnston, Dominic Kwiatkowski, Cordelia Langford, John Sillitoe on behalf of the Wellcome Sanger Institute COVID-19 Surveillance Team |
| EPI_ISL_3172125 | Lighthouse Laboratory Plymouth | Wellcome Sanger Institute for the COVID-19 Genomics UK (COG-UK) Consortium | Lighthouse Laboratory Plymouth and Alex Alderton, Roberto Amato, Jeffrey Barrett, Sonia Goncalves, Ewan Harrison, David K. Jackson, Ian Johnston, Dominic Kwiatkowski, Cordelia Langford, John Sillitoe on behalf of the Wellcome Sanger Institute COVID-19 Surveillance Team |
| EPI_ISL_3172126, EPI_ISL_3172127, EPI_ISL_3172128, EPI_ISL_3172129 | Lighthouse Lab in Alderley Park | Wellcome Sanger Institute for the COVID-19 Genomics UK (COG-UK) Consortium | Jacquelyn Wynn, Mairead Hyland, The Lighthouse Lab in Alderley Park and Alex Alderton, Roberto Amato, Jeffrey Barrett, Sonia Goncalves, Ewan Harrison, David K. Jackson, Ian Johnston, Dominic Kwiatkowski, Cordelia Langford, John Sillitoe on behalf of the Wellcome Sanger Institute COVID-19 Surveillance Team |
| EPI_ISL_3172130 | Lighthouse Lab in Milton Keynes | Wellcome Sanger Institute for the COVID-19 Genomics UK (COG-UK) Consortium | The Lighthouse Lab in Milton Keynes and Alex Alderton, Roberto Amato, Jeffrey Barrett, Sonia Goncalves, Ewan Harrison, David K. Jackson, Ian Johnston, Dominic Kwiatkowski, Cordelia Langford, John Sillitoe on behalf of the Wellcome Sanger Institute COVID-19 Surveillance Team |
| EPI_ISL_3172131 | Lighthouse Lab in Alderley Park | Wellcome Sanger Institute for the COVID-19 Genomics UK (COG-UK) Consortium | Jacquelyn Wynn, Mairead Hyland, The Lighthouse Lab in Alderley Park and Alex Alderton, Roberto Amato, Jeffrey Barrett, Sonia Goncalves, Ewan Harrison, David K. Jackson, Ian Johnston, Dominic Kwiatkowski, Cordelia Langford, John Sillitoe on behalf of the Wellcome Sanger Institute COVID-19 Surveillance Team |
| EPI_ISL_3172132 | Lighthouse Lab in Milton Keynes | Wellcome Sanger Institute for the COVID-19 Genomics UK (COG-UK) Consortium | The Lighthouse Lab in Milton Keynes and Alex Alderton, Roberto Amato, Jeffrey Barrett, Sonia Goncalves, Ewan Harrison, David K. Jackson, Ian Johnston, Dominic Kwiatkowski, Cordelia Langford, John Sillitoe on behalf of the Wellcome Sanger Institute COVID-19 Surveillance Team |
| EPI_ISL_3172133 | Lighthouse Laboratory Plymouth | Wellcome Sanger Institute for the COVID-19 Genomics UK (COG-UK) Consortium | Lighthouse Laboratory Plymouth and Alex Alderton, Roberto Amato, Jeffrey Barrett, Sonia Goncalves, Ewan Harrison, David K. Jackson, Ian Johnston, Dominic Kwiatkowski, Cordelia Langford, John Sillitoe on behalf of the Wellcome Sanger Institute COVID-19 Surveillance Team |
| EPI_ISL_3172134 | Lighthouse Lab in Milton Keynes | Wellcome Sanger Institute for the COVID-19 Genomics UK (COG-UK) Consortium | The Lighthouse Lab in Milton Keynes and Alex Alderton, Roberto Amato, Jeffrey Barrett, Sonia Goncalves, Ewan Harrison, David K. Jackson, Ian Johnston, Dominic Kwiatkowski, Cordelia Langford, John Sillitoe on behalf of the Wellcome Sanger Institute COVID-19 Surveillance Team |
| EPI_ISL_3172135 | Lighthouse Lab in Alderley Park | Wellcome Sanger Institute for the COVID-19 Genomics UK (COG-UK) Consortium | Jacquelyn Wynn, Mairead Hyland, The Lighthouse Lab in Alderley Park and Alex Alderton, Roberto Amato, Jeffrey Barrett, Sonia Goncalves, Ewan Harrison, David K. Jackson, Ian Johnston, Dominic Kwiatkowski, Cordelia Langford, John Sillitoe on behalf of the Wellcome Sanger Institute COVID-19 Surveillance Team |
| EPI_ISL_3172136 | Lighthouse Lab in Glasgow | Wellcome Sanger Institute for the COVID-19 Genomics UK (COG-UK) Consortium | Harper VanSteenhouse, Yumi Kasai, David Gray, Carol Clugston, Anna Dominiczak and Alex Alderton, Roberto Amato, Jeffrey Barrett, Sonia Goncalves, Ewan Harrison, David K. Jackson, Ian Johnston, Dominic Kwiatkowski, Cordelia Langford, John Sillitoe on behalf of the Wellcome Sanger Institute COVID-19 Surveillance Team |
| EPI_ISL_3172137 | Lighthouse Lab in Alderley Park | Wellcome Sanger Institute for the COVID-19 Genomics UK (COG-UK) Consortium | Jacquelyn Wynn, Mairead Hyland, The Lighthouse Lab in Alderley Park and Alex Alderton, Roberto Amato, Jeffrey Barrett, Sonia Goncalves, Ewan Harrison, David K. Jackson, Ian Johnston, Dominic Kwiatkowski, Cordelia Langford, John Sillitoe on behalf of the Wellcome Sanger Institute COVID-19 Surveillance Team |
| EPI_ISL_3172138, EPI_ISL_3172139 | Lighthouse Lab in Glasgow | Wellcome Sanger Institute for the COVID-19 Genomics UK (COG-UK) Consortium | Harper VanSteenhouse, Yumi Kasai, David Gray, Carol Clugston, Anna Dominiczak and Alex Alderton, Roberto Amato, Jeffrey Barrett, Sonia Goncalves, Ewan Harrison, David K. Jackson, Ian Johnston, Dominic Kwiatkowski, Cordelia Langford, John Sillitoe on behalf of the Wellcome Sanger Institute COVID-19 Surveillance Team |
| EPI_ISL_3172140 | Lighthouse Lab in Alderley Park | Wellcome Sanger Institute for the COVID-19 Genomics UK (COG-UK) Consortium | Jacquelyn Wynn, Mairead Hyland, The Lighthouse Lab in Alderley Park and Alex Alderton, Roberto Amato, Jeffrey Barrett, Sonia Goncalves, Ewan Harrison, David K. Jackson, Ian Johnston, Dominic Kwiatkowski, Cordelia Langford, John Sillitoe on behalf of the Wellcome Sanger Institute COVID-19 Surveillance Team |
| EPI_ISL_3172141, EPI_ISL_3172142 | Lighthouse Laboratory Plymouth | Wellcome Sanger Institute for the COVID-19 Genomics UK (COG-UK) Consortium | Lighthouse Laboratory Plymouth and Alex Alderton, Roberto Amato, Jeffrey Barrett, Sonia Goncalves, Ewan Harrison, David K. Jackson, Ian Johnston, Dominic Kwiatkowski, Cordelia Langford, John Sillitoe on behalf of the Wellcome Sanger Institute COVID-19 Surveillance Team |
| EPI_ISL_3172143 | Lighthouse Lab in Milton Keynes | Wellcome Sanger Institute for the COVID-19 Genomics UK (COG-UK) Consortium | The Lighthouse Lab in Milton Keynes and Alex Alderton, Roberto Amato, Jeffrey Barrett, Sonia Goncalves, Ewan Harrison, David K. Jackson, Ian Johnston, Dominic Kwiatkowski, Cordelia Langford, John Sillitoe on behalf of the Wellcome Sanger Institute COVID-19 Surveillance Team |
| EPI_ISL_3172144 | Lighthouse Lab in Alderley Park | Wellcome Sanger Institute for the COVID-19 Genomics UK (COG-UK) Consortium | Jacquelyn Wynn, Mairead Hyland, The Lighthouse Lab in Alderley Park and Alex Alderton, Roberto Amato, Jeffrey Barrett, Sonia Goncalves, Ewan Harrison, David K. Jackson, Ian Johnston, Dominic Kwiatkowski, Cordelia Langford, John Sillitoe on behalf of the Wellcome Sanger Institute COVID-19 Surveillance Team |
| EPI_ISL_3172145 | Lighthouse Laboratory Plymouth | Wellcome Sanger Institute for the COVID-19 Genomics UK (COG-UK) Consortium | Lighthouse Laboratory Plymouth and Alex Alderton, Roberto Amato, Jeffrey Barrett, Sonia Goncalves, Ewan Harrison, David K. Jackson, Ian Johnston, Dominic Kwiatkowski, Cordelia Langford, John Sillitoe on behalf of the Wellcome Sanger Institute COVID-19 Surveillance Team |
| EPI_ISL_3172146, EPI_ISL_3172147 | Lighthouse Lab in Alderley Park | Wellcome Sanger Institute for the COVID-19 Genomics UK (COG-UK) Consortium | Jacquelyn Wynn, Mairead Hyland, The Lighthouse Lab in Alderley Park and Alex Alderton, Roberto Amato, Jeffrey Barrett, Sonia Goncalves, Ewan Harrison, David K. Jackson, Ian Johnston, Dominic Kwiatkowski, Cordelia Langford, John Sillitoe on behalf of the Wellcome Sanger Institute COVID-19 Surveillance Team |
| EPI_ISL_3172148 | Lighthouse Lab in Glasgow | Wellcome Sanger Institute for the COVID-19 Genomics UK (COG-UK) Consortium | Harper VanSteenhouse, Yumi Kasai, David Gray, Carol Clugston, Anna Dominiczak and Alex Alderton, Roberto Amato, Jeffrey Barrett, Sonia Goncalves, Ewan Harrison, David K. Jackson, Ian Johnston, Dominic Kwiatkowski, Cordelia Langford, John Sillitoe on behalf of the Wellcome Sanger Institute COVID-19 Surveillance Team |
| EPI_ISL_3172149 | Lighthouse Laboratory Plymouth | Wellcome Sanger Institute for the COVID-19 Genomics UK (COG-UK) Consortium | Lighthouse Laboratory Plymouth and Alex Alderton, Roberto Amato, Jeffrey Barrett, Sonia Goncalves, Ewan Harrison, David K. Jackson, Ian Johnston, Dominic Kwiatkowski, Cordelia Langford, John Sillitoe on behalf of the Wellcome Sanger Institute COVID-19 Surveillance Team |
| EPI_ISL_3172150 | Lighthouse Lab in Alderley Park | Wellcome Sanger Institute for the COVID-19 Genomics UK (COG-UK) Consortium | Jacquelyn Wynn, Mairead Hyland, The Lighthouse Lab in Alderley Park and Alex Alderton, Roberto Amato, Jeffrey Barrett, Sonia Goncalves, Ewan Harrison, David K. Jackson, Ian Johnston, Dominic Kwiatkowski, Cordelia Langford, John Sillitoe on behalf of the Wellcome Sanger Institute COVID-19 Surveillance Team |
| EPI_ISL_3172151, EPI_ISL_3172152, EPI_ISL_3172153 | Lighthouse Laboratory Plymouth | Wellcome Sanger Institute for the COVID-19 Genomics UK (COG-UK) Consortium | Lighthouse Laboratory Plymouth and Alex Alderton, Roberto Amato, Jeffrey Barrett, Sonia Goncalves, Ewan Harrison, David K. Jackson, Ian Johnston, Dominic Kwiatkowski, Cordelia Langford, John Sillitoe on behalf of the Wellcome Sanger Institute COVID-19 Surveillance Team |
| EPI_ISL_3172154 | Lighthouse Lab in Glasgow | Wellcome Sanger Institute for the COVID-19 Genomics UK (COG-UK) Consortium | Harper VanSteenhouse, Yumi Kasai, David Gray, Carol Clugston, Anna Dominiczak and Alex Alderton, Roberto Amato, Jeffrey Barrett, Sonia Goncalves, Ewan Harrison, David K. Jackson, Ian Johnston, Dominic Kwiatkowski, Cordelia Langford, John Sillitoe on behalf of the Wellcome Sanger Institute COVID-19 Surveillance Team |
| EPI_ISL_3172155 | Lighthouse Laboratory Plymouth | Wellcome Sanger Institute for the COVID-19 Genomics UK (COG-UK) Consortium | Lighthouse Laboratory Plymouth and Alex Alderton, Roberto Amato, Jeffrey Barrett, Sonia Goncalves, Ewan Harrison, David K. Jackson, Ian Johnston, Dominic Kwiatkowski, Cordelia Langford, John Sillitoe on behalf of the Wellcome Sanger Institute COVID-19 Surveillance Team |
| EPI_ISL_3172156, EPI_ISL_3172157 | Lighthouse Lab in Milton Keynes | Wellcome Sanger Institute for the COVID-19 Genomics UK (COG-UK) Consortium | The Lighthouse Lab in Milton Keynes and Alex Alderton, Roberto Amato, Jeffrey Barrett, Sonia Goncalves, Ewan Harrison, David K. Jackson, Ian Johnston, Dominic Kwiatkowski, Cordelia Langford, John Sillitoe on behalf of the Wellcome Sanger Institute COVID-19 Surveillance Team |
| EPI_ISL_3172158 | Lighthouse Laboratory Plymouth | Wellcome Sanger Institute for the COVID-19 Genomics UK (COG-UK) Consortium | Lighthouse Laboratory Plymouth and Alex Alderton, Roberto Amato, Jeffrey Barrett, Sonia Goncalves, Ewan Harrison, David K. Jackson, Ian Johnston, Dominic Kwiatkowski, Cordelia Langford, John Sillitoe on behalf of the Wellcome Sanger Institute COVID-19 Surveillance Team |
| EPI_ISL_3172159 | Lighthouse Lab in Alderley Park | Wellcome Sanger Institute for the COVID-19 Genomics UK (COG-UK) Consortium | Jacquelyn Wynn, Mairead Hyland, The Lighthouse Lab in Alderley Park and Alex Alderton, Roberto Amato, Jeffrey Barrett, Sonia Goncalves, Ewan Harrison, David K. Jackson, Ian Johnston, Dominic Kwiatkowski, Cordelia Langford, John Sillitoe on behalf of the Wellcome Sanger Institute COVID-19 Surveillance Team |
| EPI_ISL_3172160 | Lighthouse Lab in Milton Keynes | Wellcome Sanger Institute for the COVID-19 Genomics UK (COG-UK) Consortium | The Lighthouse Lab in Milton Keynes and Alex Alderton, Roberto Amato, Jeffrey Barrett, Sonia Goncalves, Ewan Harrison, David K. Jackson, Ian Johnston, Dominic Kwiatkowski, Cordelia Langford, John Sillitoe on behalf of the Wellcome Sanger Institute COVID-19 Surveillance Team |

| | | | |
|---|---|---|---|
| EPI_ISL_3172161, EPI_ISL_3172162, EPI_ISL_3172163 | Lighthouse Lab in Alderley Park | Wellcome Sanger Institute for the COVID-19 Genomics UK (COG-UK) Consortium | Jacquelyn Wynn, Mairead Hyland, The Lighthouse Lab in Alderley Park and Alex Alderton, Roberto Amato, Jeffrey Barrett, Sonia Goncalves, Ewan Harrison, David K. Jackson, Ian Johnston, Dominic Kwiatkowski, Cordelia Langford, John Sillitoe on behalf of the Wellcome Sanger Institute COVID-19 Surveillance Team |
| EPI_ISL_3172164 | Lighthouse Lab in Glasgow | Wellcome Sanger Institute for the COVID-19 Genomics UK (COG-UK) Consortium | Harper VanSteenhouse, Yumi Kasai, David Gray, Carol Clugston, Anna Dominiczak and Alex Alderton, Roberto Amato, Jeffrey Barrett, Sonia Goncalves, Ewan Harrison, David K. Jackson, Ian Johnston, Dominic Kwiatkowski, Cordelia Langford, John Sillitoe on behalf of the Wellcome Sanger Institute COVID-19 Surveillance Team |
| EPI_ISL_3172165, EPI_ISL_3172166 | Lighthouse Laboratory Plymouth | Wellcome Sanger Institute for the COVID-19 Genomics UK (COG-UK) Consortium | Lighthouse Laboratory Plymouth and Alex Alderton, Roberto Amato, Jeffrey Barrett, Sonia Goncalves, Ewan Harrison, David K. Jackson, Ian Johnston, Dominic Kwiatkowski, Cordelia Langford, John Sillitoe on behalf of the Wellcome Sanger Institute COVID-19 Surveillance Team |
| EPI_ISL_3172167 | Lighthouse Lab in Glasgow | Wellcome Sanger Institute for the COVID-19 Genomics UK (COG-UK) Consortium | Harper VanSteenhouse, Yumi Kasai, David Gray, Carol Clugston, Anna Dominiczak and Alex Alderton, Roberto Amato, Jeffrey Barrett, Sonia Goncalves, Ewan Harrison, David K. Jackson, Ian Johnston, Dominic Kwiatkowski, Cordelia Langford, John Sillitoe on behalf of the Wellcome Sanger Institute COVID-19 Surveillance Team |
| EPI_ISL_3172168, EPI_ISL_3172169 | Lighthouse Lab in Alderley Park | Wellcome Sanger Institute for the COVID-19 Genomics UK (COG-UK) Consortium | Jacquelyn Wynn, Mairead Hyland, The Lighthouse Lab in Alderley Park and Alex Alderton, Roberto Amato, Jeffrey Barrett, Sonia Goncalves, Ewan Harrison, David K. Jackson, Ian Johnston, Dominic Kwiatkowski, Cordelia Langford, John Sillitoe on behalf of the Wellcome Sanger Institute COVID-19 Surveillance Team |
| EPI_ISL_3172170 | Lighthouse Lab in Milton Keynes | Wellcome Sanger Institute for the COVID-19 Genomics UK (COG-UK) Consortium | The Lighthouse Lab in Milton Keynes and Alex Alderton, Roberto Amato, Jeffrey Barrett, Sonia Goncalves, Ewan Harrison, David K. Jackson, Ian Johnston, Dominic Kwiatkowski, Cordelia Langford, John Sillitoe on behalf of the Wellcome Sanger Institute COVID-19 Surveillance Team |
| EPI_ISL_3172171, EPI_ISL_3172172 | Lighthouse Lab in Alderley Park | Wellcome Sanger Institute for the COVID-19 Genomics UK (COG-UK) Consortium | Jacquelyn Wynn, Mairead Hyland, The Lighthouse Lab in Alderley Park and Alex Alderton, Roberto Amato, Jeffrey Barrett, Sonia Goncalves, Ewan Harrison, David K. Jackson, Ian Johnston, Dominic Kwiatkowski, Cordelia Langford, John Sillitoe on behalf of the Wellcome Sanger Institute COVID-19 Surveillance Team |
| EPI_ISL_3172173 | Lighthouse Lab in Glasgow | Wellcome Sanger Institute for the COVID-19 Genomics UK (COG-UK) Consortium | Harper VanSteenhouse, Yumi Kasai, David Gray, Carol Clugston, Anna Dominiczak and Alex Alderton, Roberto Amato, Jeffrey Barrett, Sonia Goncalves, Ewan Harrison, David K. Jackson, Ian Johnston, Dominic Kwiatkowski, Cordelia Langford, John Sillitoe on behalf of the Wellcome Sanger Institute COVID-19 Surveillance Team |
| EPI_ISL_3172174 | Lighthouse Lab in Alderley Park | Wellcome Sanger Institute for the COVID-19 Genomics UK (COG-UK) Consortium | Jacquelyn Wynn, Mairead Hyland, The Lighthouse Lab in Alderley Park and Alex Alderton, Roberto Amato, Jeffrey Barrett, Sonia Goncalves, Ewan Harrison, David K. Jackson, Ian Johnston, Dominic Kwiatkowski, Cordelia Langford, John Sillitoe on behalf of the Wellcome Sanger Institute COVID-19 Surveillance Team |
| EPI_ISL_3172175 | Lighthouse Lab in Glasgow | Wellcome Sanger Institute for the COVID-19 Genomics UK (COG-UK) Consortium | Harper VanSteenhouse, Yumi Kasai, David Gray, Carol Clugston, Anna Dominiczak and Alex Alderton, Roberto Amato, Jeffrey Barrett, Sonia Goncalves, Ewan Harrison, David K. Jackson, Ian Johnston, Dominic Kwiatkowski, Cordelia Langford, John Sillitoe on behalf of the Wellcome Sanger Institute COVID-19 Surveillance Team |
| EPI_ISL_3172176 | Lighthouse Lab in Milton Keynes | Wellcome Sanger Institute for the COVID-19 Genomics UK (COG-UK) Consortium | The Lighthouse Lab in Milton Keynes and Alex Alderton, Roberto Amato, Jeffrey Barrett, Sonia Goncalves, Ewan Harrison, David K. Jackson, Ian Johnston, Dominic Kwiatkowski, Cordelia Langford, John Sillitoe on behalf of the Wellcome Sanger Institute COVID-19 Surveillance Team |
| EPI_ISL_3172177 | Lighthouse Lab in Alderley Park | Wellcome Sanger Institute for the COVID-19 Genomics UK (COG-UK) Consortium | Jacquelyn Wynn, Mairead Hyland, The Lighthouse Lab in Alderley Park and Alex Alderton, Roberto Amato, Jeffrey Barrett, Sonia Goncalves, Ewan Harrison, David K. Jackson, Ian Johnston, Dominic Kwiatkowski, Cordelia Langford, John Sillitoe on behalf of the Wellcome Sanger Institute COVID-19 Surveillance Team |
| EPI_ISL_3172178 | Lighthouse Laboratory Plymouth | Wellcome Sanger Institute for the COVID-19 Genomics UK (COG-UK) Consortium | Lighthouse Laboratory Plymouth and Alex Alderton, Roberto Amato, Jeffrey Barrett, Sonia Goncalves, Ewan Harrison, David K. Jackson, Ian Johnston, Dominic Kwiatkowski, Cordelia Langford, John Sillitoe on behalf of the Wellcome Sanger Institute COVID-19 Surveillance Team |
| EPI_ISL_3172179 | Lighthouse Lab in Glasgow | Wellcome Sanger Institute for the COVID-19 Genomics UK (COG-UK) Consortium | Harper VanSteenhouse, Yumi Kasai, David Gray, Carol Clugston, Anna Dominiczak and Alex Alderton, Roberto Amato, Jeffrey Barrett, Sonia Goncalves, Ewan Harrison, David K. Jackson, Ian Johnston, Dominic Kwiatkowski, Cordelia Langford, John Sillitoe on behalf of the Wellcome Sanger Institute COVID-19 Surveillance Team |
| EPI_ISL_3172180 | Lighthouse Laboratory Plymouth | Wellcome Sanger Institute for the COVID-19 Genomics UK (COG-UK) Consortium | Lighthouse Laboratory Plymouth and Alex Alderton, Roberto Amato, Jeffrey Barrett, Sonia Goncalves, Ewan Harrison, David K. Jackson, Ian Johnston, Dominic Kwiatkowski, Cordelia Langford, John Sillitoe on behalf of the Wellcome Sanger Institute COVID-19 Surveillance Team |
| EPI_ISL_3172181 | Lighthouse Lab in Alderley Park | Wellcome Sanger Institute for the COVID-19 Genomics UK (COG-UK) Consortium | Jacquelyn Wynn, Mairead Hyland, The Lighthouse Lab in Alderley Park and Alex Alderton, Roberto Amato, Jeffrey Barrett, Sonia Goncalves, Ewan Harrison, David K. Jackson, Ian Johnston, Dominic Kwiatkowski, Cordelia Langford, John Sillitoe on behalf of the Wellcome Sanger Institute COVID-19 Surveillance Team |
| EPI_ISL_3172182, EPI_ISL_3172183, EPI_ISL_3172184 | Lighthouse Lab in Milton Keynes | Wellcome Sanger Institute for the COVID-19 Genomics UK (COG-UK) Consortium | The Lighthouse Lab in Milton Keynes and Alex Alderton, Roberto Amato, Jeffrey Barrett, Sonia Goncalves, Ewan Harrison, David K. Jackson, Ian Johnston, Dominic Kwiatkowski, Cordelia Langford, John Sillitoe on behalf of the Wellcome Sanger Institute COVID-19 Surveillance Team |
| EPI_ISL_3172185, EPI_ISL_3172186 | Lighthouse Lab in Glasgow | Wellcome Sanger Institute for the COVID-19 Genomics UK (COG-UK) Consortium | Harper VanSteenhouse, Yumi Kasai, David Gray, Carol Clugston, Anna Dominiczak and Alex Alderton, Roberto Amato, Jeffrey Barrett, Sonia Goncalves, Ewan Harrison, David K. Jackson, Ian Johnston, Dominic Kwiatkowski, Cordelia Langford, John Sillitoe on behalf of the Wellcome Sanger Institute COVID-19 Surveillance Team |
| EPI_ISL_3172187, EPI_ISL_3172188 | Lighthouse Lab in Alderley Park | Wellcome Sanger Institute for the COVID-19 Genomics UK (COG-UK) Consortium | Jacquelyn Wynn, Mairead Hyland, The Lighthouse Lab in Alderley Park and Alex Alderton, Roberto Amato, Jeffrey Barrett, Sonia Goncalves, Ewan Harrison, David K. Jackson, Ian Johnston, Dominic Kwiatkowski, Cordelia Langford, John Sillitoe on behalf of the Wellcome Sanger Institute COVID-19 Surveillance Team |
| EPI_ISL_3172189 | Lighthouse Laboratory Plymouth | Wellcome Sanger Institute for the COVID-19 Genomics UK (COG-UK) Consortium | Lighthouse Laboratory Plymouth and Alex Alderton, Roberto Amato, Jeffrey Barrett, Sonia Goncalves, Ewan Harrison, David K. Jackson, Ian Johnston, Dominic Kwiatkowski, Cordelia Langford, John Sillitoe on behalf of the Wellcome Sanger Institute COVID-19 Surveillance Team |
| EPI_ISL_3172190 | Lighthouse Lab in Glasgow | Wellcome Sanger Institute for the COVID-19 Genomics UK (COG-UK) Consortium | Harper VanSteenhouse, Yumi Kasai, David Gray, Carol Clugston, Anna Dominiczak and Alex Alderton, Roberto Amato, Jeffrey Barrett, Sonia Goncalves, Ewan Harrison, David K. Jackson, Ian Johnston, Dominic Kwiatkowski, Cordelia Langford, John Sillitoe on behalf of the Wellcome Sanger Institute COVID-19 Surveillance Team |
| EPI_ISL_3172191 | Lighthouse Lab in Milton Keynes | Wellcome Sanger Institute for the COVID-19 Genomics UK (COG-UK) Consortium | The Lighthouse Lab in Milton Keynes and Alex Alderton, Roberto Amato, Jeffrey Barrett, Sonia Goncalves, Ewan Harrison, David K. Jackson, Ian Johnston, Dominic Kwiatkowski, Cordelia Langford, John Sillitoe on behalf of the Wellcome Sanger Institute COVID-19 Surveillance Team |
| EPI_ISL_3172192, EPI_ISL_3172193 | Lighthouse Lab in Glasgow | Wellcome Sanger Institute for the COVID-19 Genomics UK (COG-UK) Consortium | Harper VanSteenhouse, Yumi Kasai, David Gray, Carol Clugston, Anna Dominiczak and Alex Alderton, Roberto Amato, Jeffrey Barrett, Sonia Goncalves, Ewan Harrison, David K. Jackson, Ian Johnston, Dominic Kwiatkowski, Cordelia Langford, John Sillitoe on behalf of the Wellcome Sanger Institute COVID-19 Surveillance Team |
| EPI_ISL_3172194, EPI_ISL_3172195 | Lighthouse Laboratory Plymouth | Wellcome Sanger Institute for the COVID-19 Genomics UK (COG-UK) Consortium | Lighthouse Laboratory Plymouth and Alex Alderton, Roberto Amato, Jeffrey Barrett, Sonia Goncalves, Ewan Harrison, David K. Jackson, Ian Johnston, Dominic Kwiatkowski, Cordelia Langford, John Sillitoe on behalf of the Wellcome Sanger Institute COVID-19 Surveillance Team |
| EPI_ISL_3172196, EPI_ISL_3172197 | Lighthouse Lab in Milton Keynes | Wellcome Sanger Institute for the COVID-19 Genomics UK (COG-UK) Consortium | The Lighthouse Lab in Milton Keynes and Alex Alderton, Roberto Amato, Jeffrey Barrett, Sonia Goncalves, Ewan Harrison, David K. Jackson, Ian Johnston, Dominic Kwiatkowski, Cordelia Langford, John Sillitoe on behalf of the Wellcome Sanger Institute COVID-19 Surveillance Team |
| EPI_ISL_3172198 | Lighthouse Laboratory Plymouth | Wellcome Sanger Institute for the COVID-19 Genomics UK (COG-UK) Consortium | Lighthouse Laboratory Plymouth and Alex Alderton, Roberto Amato, Jeffrey Barrett, Sonia Goncalves, Ewan Harrison, David K. Jackson, Ian Johnston, Dominic Kwiatkowski, Cordelia Langford, John Sillitoe on behalf of the Wellcome Sanger Institute COVID-19 Surveillance Team |
| EPI_ISL_3172199 | Lighthouse Lab in Milton Keynes | Wellcome Sanger Institute for the COVID-19 Genomics UK (COG-UK) Consortium | The Lighthouse Lab in Milton Keynes and Alex Alderton, Roberto Amato, Jeffrey Barrett, Sonia Goncalves, Ewan Harrison, David K. Jackson, Ian Johnston, Dominic Kwiatkowski, Cordelia Langford, John Sillitoe on behalf of the Wellcome Sanger Institute COVID-19 Surveillance Team |
| EPI_ISL_3172200 | Lighthouse Laboratory Plymouth | Wellcome Sanger Institute for the COVID-19 Genomics UK (COG-UK) Consortium | Lighthouse Laboratory Plymouth and Alex Alderton, Roberto Amato, Jeffrey Barrett, Sonia Goncalves, Ewan Harrison, David K. Jackson, Ian Johnston, Dominic Kwiatkowski, Cordelia Langford, John Sillitoe on behalf of the Wellcome Sanger Institute COVID-19 Surveillance Team |
| EPI_ISL_3172201 | Lighthouse Lab in Glasgow | Wellcome Sanger Institute for the COVID-19 Genomics UK (COG-UK) Consortium | Harper VanSteenhouse, Yumi Kasai, David Gray, Carol Clugston, Anna Dominiczak and Alex Alderton, Roberto Amato, Jeffrey Barrett, Sonia Goncalves, Ewan Harrison, David K. Jackson, Ian Johnston, Dominic Kwiatkowski, Cordelia Langford, John Sillitoe on behalf of the Wellcome Sanger Institute COVID-19 Surveillance Team |
| EPI_ISL_3172202 | Lighthouse Lab in Milton Keynes | Wellcome Sanger Institute for the COVID-19 Genomics UK (COG-UK) Consortium | The Lighthouse Lab in Milton Keynes and Alex Alderton, Roberto Amato, Jeffrey Barrett, Sonia Goncalves, Ewan Harrison, David K. Jackson, Ian Johnston, Dominic Kwiatkowski, Cordelia Langford, John Sillitoe on behalf of the Wellcome Sanger Institute COVID-19 Surveillance Team |
| EPI_ISL_3172203 | Lighthouse Lab in Alderley Park | Wellcome Sanger Institute for the COVID-19 Genomics UK (COG-UK) Consortium | Jacquelyn Wynn, Mairead Hyland, The Lighthouse Lab in Alderley Park and Alex Alderton, Roberto Amato, Jeffrey Barrett, Sonia Goncalves, Ewan Harrison, David K. Jackson, Ian Johnston, Dominic Kwiatkowski, Cordelia Langford, John Sillitoe on behalf of the Wellcome Sanger Institute COVID-19 Surveillance Team |
| EPI_ISL_3172204 | Lighthouse Lab in Milton Keynes | Wellcome Sanger Institute for the COVID-19 Genomics UK (COG-UK) Consortium | The Lighthouse Lab in Milton Keynes and Alex Alderton, Roberto Amato, Jeffrey Barrett, Sonia Goncalves, Ewan Harrison, David K. Jackson, Ian Johnston, Dominic Kwiatkowski, Cordelia Langford, John Sillitoe on behalf of the Wellcome Sanger Institute COVID-19 Surveillance Team |
| EPI_ISL_3172205 | Lighthouse Laboratory Plymouth | Wellcome Sanger Institute for the COVID-19 Genomics UK (COG-UK) Consortium | Lighthouse Laboratory Plymouth and Alex Alderton, Roberto Amato, Jeffrey Barrett, Sonia Goncalves, Ewan Harrison, David K. Jackson, Ian Johnston, Dominic Kwiatkowski, Cordelia Langford, John Sillitoe on behalf of the Wellcome Sanger Institute COVID-19 Surveillance Team |
| EPI_ISL_3172206, EPI_ISL_3172207 | Lighthouse Lab in Milton Keynes | Wellcome Sanger Institute for the COVID-19 Genomics UK (COG-UK) Consortium | The Lighthouse Lab in Milton Keynes and Alex Alderton, Roberto Amato, Jeffrey Barrett, Sonia Goncalves, Ewan Harrison, David K. Jackson, Ian Johnston, Dominic Kwiatkowski, Cordelia Langford, John Sillitoe on behalf of the Wellcome Sanger Institute COVID-19 Surveillance Team |
| EPI_ISL_3172208 | Lighthouse Laboratory Plymouth | Wellcome Sanger Institute for the COVID-19 Genomics UK (COG-UK) Consortium | Lighthouse Laboratory Plymouth and Alex Alderton, Roberto Amato, Jeffrey Barrett, Sonia Goncalves, Ewan Harrison, David K. Jackson, Ian Johnston, Dominic Kwiatkowski, Cordelia Langford, John Sillitoe on behalf of the Wellcome Sanger Institute COVID-19 Surveillance Team |
| EPI_ISL_3172209, EPI_ISL_3172210, EPI_ISL_3172211, EPI_ISL_3172212, EPI_ISL_3172213, EPI_ISL_3172214, EPI_ISL_3172215 | Lighthouse Lab in Milton Keynes | Wellcome Sanger Institute for the COVID-19 Genomics UK (COG-UK) Consortium | The Lighthouse Lab in Milton Keynes and Alex Alderton, Roberto Amato, Jeffrey Barrett, Sonia Goncalves, Ewan Harrison, David K. Jackson, Ian Johnston, Dominic Kwiatkowski, Cordelia Langford, John Sillitoe on behalf of the Wellcome Sanger Institute COVID-19 Surveillance Team |
| EPI_ISL_3172216 | Lighthouse Laboratory Plymouth | Wellcome Sanger Institute for the COVID-19 Genomics UK (COG-UK) Consortium | Lighthouse Laboratory Plymouth and Alex Alderton, Roberto Amato, Jeffrey Barrett, Sonia Goncalves, Ewan Harrison, David K. Jackson, Ian Johnston, Dominic Kwiatkowski, Cordelia Langford, John Sillitoe on behalf of the Wellcome Sanger Institute COVID-19 Surveillance Team |
| EPI_ISL_3172217 | Lighthouse Lab in Milton Keynes | Wellcome Sanger Institute for the COVID-19 Genomics UK (COG-UK) Consortium | The Lighthouse Lab in Milton Keynes and Alex Alderton, Roberto Amato, Jeffrey Barrett, Sonia Goncalves, Ewan Harrison, David K. Jackson, Ian Johnston, Dominic Kwiatkowski, Cordelia Langford, John Sillitoe on behalf of the Wellcome Sanger Institute COVID-19 Surveillance Team |
| EPI_ISL_3172218 | Lighthouse Laboratory Plymouth | Wellcome Sanger Institute for the COVID-19 Genomics UK (COG-UK) Consortium | Lighthouse Laboratory Plymouth and Alex Alderton, Roberto Amato, Jeffrey Barrett, Sonia Goncalves, Ewan Harrison, David K. Jackson, Ian Johnston, Dominic Kwiatkowski, Cordelia Langford, John Sillitoe on behalf of the Wellcome Sanger Institute COVID-19 Surveillance Team |
| EPI_ISL_3172219 | Lighthouse Lab in Milton Keynes | Wellcome Sanger Institute for the COVID-19 Genomics UK (COG-UK) Consortium | The Lighthouse Lab in Milton Keynes and Alex Alderton, Roberto Amato, Jeffrey Barrett, Sonia Goncalves, Ewan Harrison, David K. Jackson, Ian Johnston, Dominic Kwiatkowski, Cordelia Langford, John Sillitoe on behalf of the Wellcome Sanger Institute COVID-19 Surveillance Team |
| EPI_ISL_3172220 | Lighthouse Laboratory Plymouth | Wellcome Sanger Institute for the COVID-19 Genomics UK (COG-UK) Consortium | Lighthouse Laboratory Plymouth and Alex Alderton, Roberto Amato, Jeffrey Barrett, Sonia Goncalves, Ewan Harrison, David K. Jackson, Ian Johnston, Dominic Kwiatkowski, Cordelia Langford, John Sillitoe on behalf of the Wellcome Sanger Institute COVID-19 Surveillance Team |
| EPI_ISL_3172221, EPI_ISL_3172222, EPI_ISL_3172223, | Lighthouse Lab in Milton Keynes | Wellcome Sanger Institute for the COVID-19 Genomics UK (COG-UK) Consortium | The Lighthouse Lab in Milton Keynes and Alex Alderton, Roberto Amato, Jeffrey Barrett, Sonia Goncalves, Ewan Harrison, David K. Jackson, Ian Johnston, Dominic Kwiatkowski, Cordelia Langford, John Sillitoe on behalf of the Wellcome Sanger Institute COVID-19 Surveillance Team |

| | | | |
|---|---|---|---|
| EPI_ISL_3172224, EPI_ISL_3172225, EPI_ISL_3172226 | | | |
| EPI_ISL_3172227 | Lighthouse Laboratory Plymouth | Wellcome Sanger Institute for the COVID-19 Genomics UK (COG-UK) Consortium | Lighthouse Laboratory Plymouth and Alex Alderton, Roberto Amato, Jeffrey Barrett, Sonia Goncalves, Ewan Harrison, David K. Jackson, Ian Johnston, Dominic Kwiatkowski, Cordelia Langford, John Sillitoe on behalf of the Wellcome Sanger Institute COVID-19 Surveillance Team |
| EPI_ISL_3172228 | Berkshire and Surrey Pathology Services Lighthouse Laboratory | Wellcome Sanger Institute for the COVID-19 Genomics UK (COG-UK) Consortium | Berkshire and Surrey Pathology Services Lighthouse Laboratory and Alex Alderton, Roberto Amato, Jeffrey Barrett, Sonia Goncalves, Ewan Harrison, David K. Jackson, Ian Johnston, Dominic Kwiatkowski, Cordelia Langford, John Sillitoe on behalf of the Wellcome Sanger Institute COVID-19 Surveillance Team |
| EPI_ISL_3172229, EPI_ISL_3172230, EPI_ISL_3172231, EPI_ISL_3172232, EPI_ISL_3172233, EPI_ISL_3172234, EPI_ISL_3172235 | Lighthouse Lab in Milton Keynes | Wellcome Sanger Institute for the COVID-19 Genomics UK (COG-UK) Consortium | The Lighthouse Lab in Milton Keynes and Alex Alderton, Roberto Amato, Jeffrey Barrett, Sonia Goncalves, Ewan Harrison, David K. Jackson, Ian Johnston, Dominic Kwiatkowski, Cordelia Langford, John Sillitoe on behalf of the Wellcome Sanger Institute COVID-19 Surveillance Team |
| EPI_ISL_3172236, EPI_ISL_3172237 | Lighthouse Laboratory Plymouth | Wellcome Sanger Institute for the COVID-19 Genomics UK (COG-UK) Consortium | Lighthouse Laboratory Plymouth and Alex Alderton, Roberto Amato, Jeffrey Barrett, Sonia Goncalves, Ewan Harrison, David K. Jackson, Ian Johnston, Dominic Kwiatkowski, Cordelia Langford, John Sillitoe on behalf of the Wellcome Sanger Institute COVID-19 Surveillance Team |
| EPI_ISL_3172238 | Lighthouse Lab in Milton Keynes | Wellcome Sanger Institute for the COVID-19 Genomics UK (COG-UK) Consortium | The Lighthouse Lab in Milton Keynes and Alex Alderton, Roberto Amato, Jeffrey Barrett, Sonia Goncalves, Ewan Harrison, David K. Jackson, Ian Johnston, Dominic Kwiatkowski, Cordelia Langford, John Sillitoe on behalf of the Wellcome Sanger Institute COVID-19 Surveillance Team |
| EPI_ISL_3172239 | Berkshire and Surrey Pathology Services Lighthouse Laboratory | Wellcome Sanger Institute for the COVID-19 Genomics UK (COG-UK) Consortium | Berkshire and Surrey Pathology Services Lighthouse Laboratory and Alex Alderton, Roberto Amato, Jeffrey Barrett, Sonia Goncalves, Ewan Harrison, David K. Jackson, Ian Johnston, Dominic Kwiatkowski, Cordelia Langford, John Sillitoe on behalf of the Wellcome Sanger Institute COVID-19 Surveillance Team |
| EPI_ISL_3172240, EPI_ISL_3172241, EPI_ISL_3172242, EPI_ISL_3172243 | Lighthouse Lab in Milton Keynes | Wellcome Sanger Institute for the COVID-19 Genomics UK (COG-UK) Consortium | The Lighthouse Lab in Milton Keynes and Alex Alderton, Roberto Amato, Jeffrey Barrett, Sonia Goncalves, Ewan Harrison, David K. Jackson, Ian Johnston, Dominic Kwiatkowski, Cordelia Langford, John Sillitoe on behalf of the Wellcome Sanger Institute COVID-19 Surveillance Team |
| EPI_ISL_3172244 | Lighthouse Laboratory Plymouth | Wellcome Sanger Institute for the COVID-19 Genomics UK (COG-UK) Consortium | Lighthouse Laboratory Plymouth and Alex Alderton, Roberto Amato, Jeffrey Barrett, Sonia Goncalves, Ewan Harrison, David K. Jackson, Ian Johnston, Dominic Kwiatkowski, Cordelia Langford, John Sillitoe on behalf of the Wellcome Sanger Institute COVID-19 Surveillance Team |
| EPI_ISL_3172245, EPI_ISL_3172246, EPI_ISL_3172247 | Lighthouse Lab in Milton Keynes | Wellcome Sanger Institute for the COVID-19 Genomics UK (COG-UK) Consortium | The Lighthouse Lab in Milton Keynes and Alex Alderton, Roberto Amato, Jeffrey Barrett, Sonia Goncalves, Ewan Harrison, David K. Jackson, Ian Johnston, Dominic Kwiatkowski, Cordelia Langford, John Sillitoe on behalf of the Wellcome Sanger Institute COVID-19 Surveillance Team |
| EPI_ISL_3172248 | Lighthouse Laboratory Plymouth | Wellcome Sanger Institute for the COVID-19 Genomics UK (COG-UK) Consortium | Lighthouse Laboratory Plymouth and Alex Alderton, Roberto Amato, Jeffrey Barrett, Sonia Goncalves, Ewan Harrison, David K. Jackson, Ian Johnston, Dominic Kwiatkowski, Cordelia Langford, John Sillitoe on behalf of the Wellcome Sanger Institute COVID-19 Surveillance Team |
| EPI_ISL_3172249, EPI_ISL_3172250, EPI_ISL_3172251 | Lighthouse Lab in Milton Keynes | Wellcome Sanger Institute for the COVID-19 Genomics UK (COG-UK) Consortium | The Lighthouse Lab in Milton Keynes and Alex Alderton, Roberto Amato, Jeffrey Barrett, Sonia Goncalves, Ewan Harrison, David K. Jackson, Ian Johnston, Dominic Kwiatkowski, Cordelia Langford, John Sillitoe on behalf of the Wellcome Sanger Institute COVID-19 Surveillance Team |
| EPI_ISL_3172252 | Berkshire and Surrey Pathology Services Lighthouse Laboratory | Wellcome Sanger Institute for the COVID-19 Genomics UK (COG-UK) Consortium | Berkshire and Surrey Pathology Services Lighthouse Laboratory and Alex Alderton, Roberto Amato, Jeffrey Barrett, Sonia Goncalves, Ewan Harrison, David K. Jackson, Ian Johnston, Dominic Kwiatkowski, Cordelia Langford, John Sillitoe on behalf of the Wellcome Sanger Institute COVID-19 Surveillance Team |
| EPI_ISL_3172253 | Lighthouse Lab in Milton Keynes | Wellcome Sanger Institute for the COVID-19 Genomics UK (COG-UK) Consortium | The Lighthouse Lab in Milton Keynes and Alex Alderton, Roberto Amato, Jeffrey Barrett, Sonia Goncalves, Ewan Harrison, David K. Jackson, Ian Johnston, Dominic Kwiatkowski, Cordelia Langford, John Sillitoe on behalf of the Wellcome Sanger Institute COVID-19 Surveillance Team |
| EPI_ISL_3172254 | Berkshire and Surrey Pathology Services Lighthouse Laboratory | Wellcome Sanger Institute for the COVID-19 Genomics UK (COG-UK) Consortium | Berkshire and Surrey Pathology Services Lighthouse Laboratory and Alex Alderton, Roberto Amato, Jeffrey Barrett, Sonia Goncalves, Ewan Harrison, David K. Jackson, Ian Johnston, Dominic Kwiatkowski, Cordelia Langford, John Sillitoe on behalf of the Wellcome Sanger Institute COVID-19 Surveillance Team |
| EPI_ISL_3172255, EPI_ISL_3172256, EPI_ISL_3172257 | Lighthouse Lab in Milton Keynes | Wellcome Sanger Institute for the COVID-19 Genomics UK (COG-UK) Consortium | The Lighthouse Lab in Milton Keynes and Alex Alderton, Roberto Amato, Jeffrey Barrett, Sonia Goncalves, Ewan Harrison, David K. Jackson, Ian Johnston, Dominic Kwiatkowski, Cordelia Langford, John Sillitoe on behalf of the Wellcome Sanger Institute COVID-19 Surveillance Team |
| EPI_ISL_3172258 | Berkshire and Surrey Pathology Services Lighthouse Laboratory | Wellcome Sanger Institute for the COVID-19 Genomics UK (COG-UK) Consortium | Berkshire and Surrey Pathology Services Lighthouse Laboratory and Alex Alderton, Roberto Amato, Jeffrey Barrett, Sonia Goncalves, Ewan Harrison, David K. Jackson, Ian Johnston, Dominic Kwiatkowski, Cordelia Langford, John Sillitoe on behalf of the Wellcome Sanger Institute COVID-19 Surveillance Team |
| EPI_ISL_3172259, EPI_ISL_3172260 | Lighthouse Lab in Milton Keynes | Wellcome Sanger Institute for the COVID-19 Genomics UK (COG-UK) Consortium | The Lighthouse Lab in Milton Keynes and Alex Alderton, Roberto Amato, Jeffrey Barrett, Sonia Goncalves, Ewan Harrison, David K. Jackson, Ian Johnston, Dominic Kwiatkowski, Cordelia Langford, John Sillitoe on behalf of the Wellcome Sanger Institute COVID-19 Surveillance Team |
| EPI_ISL_3172261 | Berkshire and Surrey Pathology Services Lighthouse Laboratory | Wellcome Sanger Institute for the COVID-19 Genomics UK (COG-UK) Consortium | Berkshire and Surrey Pathology Services Lighthouse Laboratory and Alex Alderton, Roberto Amato, Jeffrey Barrett, Sonia Goncalves, Ewan Harrison, David K. Jackson, Ian Johnston, Dominic Kwiatkowski, Cordelia Langford, John Sillitoe on behalf of the Wellcome Sanger Institute COVID-19 Surveillance Team |
| EPI_ISL_3172262, EPI_ISL_3172263, EPI_ISL_3172264 | Lighthouse Lab in Milton Keynes | Wellcome Sanger Institute for the COVID-19 Genomics UK (COG-UK) Consortium | The Lighthouse Lab in Milton Keynes and Alex Alderton, Roberto Amato, Jeffrey Barrett, Sonia Goncalves, Ewan Harrison, David K. Jackson, Ian Johnston, Dominic Kwiatkowski, Cordelia Langford, John Sillitoe on behalf of the Wellcome Sanger Institute COVID-19 Surveillance Team |
| EPI_ISL_3172265, EPI_ISL_3172266, EPI_ISL_3172267 | Lighthouse Laboratory Plymouth | Wellcome Sanger Institute for the COVID-19 Genomics UK (COG-UK) Consortium | Lighthouse Laboratory Plymouth and Alex Alderton, Roberto Amato, Jeffrey Barrett, Sonia Goncalves, Ewan Harrison, David K. Jackson, Ian Johnston, Dominic Kwiatkowski, Cordelia Langford, John Sillitoe on behalf of the Wellcome Sanger Institute COVID-19 Surveillance Team |
| EPI_ISL_3172268 | Berkshire and Surrey Pathology Services Lighthouse Laboratory | Wellcome Sanger Institute for the COVID-19 Genomics UK (COG-UK) Consortium | Berkshire and Surrey Pathology Services Lighthouse Laboratory and Alex Alderton, Roberto Amato, Jeffrey Barrett, Sonia Goncalves, Ewan Harrison, David K. Jackson, Ian Johnston, Dominic Kwiatkowski, Cordelia Langford, John Sillitoe on behalf of the Wellcome Sanger Institute COVID-19 Surveillance Team |
| EPI_ISL_3172269 | Lighthouse Lab in Milton Keynes | Wellcome Sanger Institute for the COVID-19 Genomics UK (COG-UK) Consortium | The Lighthouse Lab in Milton Keynes and Alex Alderton, Roberto Amato, Jeffrey Barrett, Sonia Goncalves, Ewan Harrison, David K. Jackson, Ian Johnston, Dominic Kwiatkowski, Cordelia Langford, John Sillitoe on behalf of the Wellcome Sanger Institute COVID-19 Surveillance Team |
| EPI_ISL_3172270 | Berkshire and Surrey Pathology Services Lighthouse Laboratory | Wellcome Sanger Institute for the COVID-19 Genomics UK (COG-UK) Consortium | Berkshire and Surrey Pathology Services Lighthouse Laboratory and Alex Alderton, Roberto Amato, Jeffrey Barrett, Sonia Goncalves, Ewan Harrison, David K. Jackson, Ian Johnston, Dominic Kwiatkowski, Cordelia Langford, John Sillitoe on behalf of the Wellcome Sanger Institute COVID-19 Surveillance Team |
| EPI_ISL_3172271 | Lighthouse Laboratory Plymouth | Wellcome Sanger Institute for the COVID-19 Genomics UK (COG-UK) Consortium | Lighthouse Laboratory Plymouth and Alex Alderton, Roberto Amato, Jeffrey Barrett, Sonia Goncalves, Ewan Harrison, David K. Jackson, Ian Johnston, Dominic Kwiatkowski, Cordelia Langford, John Sillitoe on behalf of the Wellcome Sanger Institute COVID-19 Surveillance Team |
| EPI_ISL_3172272, EPI_ISL_3172273, EPI_ISL_3172274, EPI_ISL_3172275, EPI_ISL_3172276 | Lighthouse Lab in Milton Keynes | Wellcome Sanger Institute for the COVID-19 Genomics UK (COG-UK) Consortium | The Lighthouse Lab in Milton Keynes and Alex Alderton, Roberto Amato, Jeffrey Barrett, Sonia Goncalves, Ewan Harrison, David K. Jackson, Ian Johnston, Dominic Kwiatkowski, Cordelia Langford, John Sillitoe on behalf of the Wellcome Sanger Institute COVID-19 Surveillance Team |
| EPI_ISL_3172277 | Berkshire and Surrey Pathology Services Lighthouse Laboratory | Wellcome Sanger Institute for the COVID-19 Genomics UK (COG-UK) Consortium | Berkshire and Surrey Pathology Services Lighthouse Laboratory and Alex Alderton, Roberto Amato, Jeffrey Barrett, Sonia Goncalves, Ewan Harrison, David K. Jackson, Ian Johnston, Dominic Kwiatkowski, Cordelia Langford, John Sillitoe on behalf of the Wellcome Sanger Institute COVID-19 Surveillance Team |
| EPI_ISL_3172278 | Lighthouse Lab in Milton Keynes | Wellcome Sanger Institute for the COVID-19 Genomics UK (COG-UK) Consortium | The Lighthouse Lab in Milton Keynes and Alex Alderton, Roberto Amato, Jeffrey Barrett, Sonia Goncalves, Ewan Harrison, David K. Jackson, Ian Johnston, Dominic Kwiatkowski, Cordelia Langford, John Sillitoe on behalf of the Wellcome Sanger Institute COVID-19 Surveillance Team |
| EPI_ISL_3172279 | Berkshire and Surrey Pathology Services Lighthouse Laboratory | Wellcome Sanger Institute for the COVID-19 Genomics UK (COG-UK) Consortium | Berkshire and Surrey Pathology Services Lighthouse Laboratory and Alex Alderton, Roberto Amato, Jeffrey Barrett, Sonia Goncalves, Ewan Harrison, David K. Jackson, Ian Johnston, Dominic Kwiatkowski, Cordelia Langford, John Sillitoe on behalf of the Wellcome Sanger Institute COVID-19 Surveillance Team |
| EPI_ISL_3172280 | Lighthouse Lab in Milton Keynes | Wellcome Sanger Institute for the COVID-19 Genomics UK (COG-UK) Consortium | The Lighthouse Lab in Milton Keynes and Alex Alderton, Roberto Amato, Jeffrey Barrett, Sonia Goncalves, Ewan Harrison, David K. Jackson, Ian Johnston, Dominic Kwiatkowski, Cordelia Langford, John Sillitoe on behalf of the Wellcome Sanger Institute COVID-19 Surveillance Team |
| EPI_ISL_3172281 | Lighthouse Laboratory Plymouth | Wellcome Sanger Institute for the COVID-19 Genomics UK (COG-UK) Consortium | Lighthouse Laboratory Plymouth and Alex Alderton, Roberto Amato, Jeffrey Barrett, Sonia Goncalves, Ewan Harrison, David K. Jackson, Ian Johnston, Dominic Kwiatkowski, Cordelia Langford, John Sillitoe on behalf of the Wellcome Sanger Institute COVID-19 Surveillance Team |
| EPI_ISL_3172282 | Lighthouse Lab in Milton Keynes | Wellcome Sanger Institute for the COVID-19 Genomics UK (COG-UK) Consortium | The Lighthouse Lab in Milton Keynes and Alex Alderton, Roberto Amato, Jeffrey Barrett, Sonia Goncalves, Ewan Harrison, David K. Jackson, Ian Johnston, Dominic Kwiatkowski, Cordelia Langford, John Sillitoe on behalf of the Wellcome Sanger Institute COVID-19 Surveillance Team |
| EPI_ISL_3172283, EPI_ISL_3172284 | Berkshire and Surrey Pathology Services Lighthouse Laboratory | Wellcome Sanger Institute for the COVID-19 Genomics UK (COG-UK) Consortium | Berkshire and Surrey Pathology Services Lighthouse Laboratory and Alex Alderton, Roberto Amato, Jeffrey Barrett, Sonia Goncalves, Ewan Harrison, David K. Jackson, Ian Johnston, Dominic Kwiatkowski, Cordelia Langford, John Sillitoe on behalf of the Wellcome Sanger Institute COVID-19 Surveillance Team |
| EPI_ISL_3172285 | Lighthouse Lab in Milton Keynes | Wellcome Sanger Institute for the COVID-19 Genomics UK (COG-UK) Consortium | The Lighthouse Lab in Milton Keynes and Alex Alderton, Roberto Amato, Jeffrey Barrett, Sonia Goncalves, Ewan Harrison, David K. Jackson, Ian Johnston, Dominic Kwiatkowski, Cordelia Langford, John Sillitoe on behalf of the Wellcome Sanger Institute COVID-19 Surveillance Team |
| EPI_ISL_3172286 | Berkshire and Surrey Pathology Services Lighthouse Laboratory | Wellcome Sanger Institute for the COVID-19 Genomics UK (COG-UK) Consortium | Berkshire and Surrey Pathology Services Lighthouse Laboratory and Alex Alderton, Roberto Amato, Jeffrey Barrett, Sonia Goncalves, Ewan Harrison, David K. Jackson, Ian Johnston, Dominic Kwiatkowski, Cordelia Langford, John Sillitoe on behalf of the Wellcome Sanger Institute COVID-19 Surveillance Team |
| EPI_ISL_3172287 | Lighthouse Laboratory Plymouth | Wellcome Sanger Institute for the COVID-19 Genomics UK (COG-UK) Consortium | Lighthouse Laboratory Plymouth and Alex Alderton, Roberto Amato, Jeffrey Barrett, Sonia Goncalves, Ewan Harrison, David K. Jackson, Ian Johnston, Dominic Kwiatkowski, Cordelia Langford, John Sillitoe on behalf of the Wellcome Sanger Institute COVID-19 Surveillance Team |
| EPI_ISL_3172288, EPI_ISL_3172289 | Berkshire and Surrey Pathology Services Lighthouse Laboratory | Wellcome Sanger Institute for the COVID-19 Genomics UK (COG-UK) Consortium | Berkshire and Surrey Pathology Services Lighthouse Laboratory and Alex Alderton, Roberto Amato, Jeffrey Barrett, Sonia Goncalves, Ewan Harrison, David K. Jackson, Ian Johnston, Dominic Kwiatkowski, Cordelia Langford, John Sillitoe on behalf of the Wellcome Sanger Institute COVID-19 Surveillance Team |
| EPI_ISL_3172290 | Lighthouse Laboratory Plymouth | Wellcome Sanger Institute for the COVID-19 Genomics UK (COG-UK) Consortium | Lighthouse Laboratory Plymouth and Alex Alderton, Roberto Amato, Jeffrey Barrett, Sonia Goncalves, Ewan Harrison, David K. Jackson, Ian Johnston, Dominic Kwiatkowski, Cordelia Langford, John Sillitoe on behalf of the Wellcome Sanger Institute COVID-19 Surveillance Team |
| EPI_ISL_3172291, | Lighthouse Lab in Milton Keynes | Wellcome Sanger Institute for the COVID-19 Genomics UK (COG-UK) | The Lighthouse Lab in Milton Keynes and Alex Alderton, Roberto Amato, Jeffrey Barrett, Sonia Goncalves, Ewan Harrison, David K. Jackson, Ian Johnston, Dominic Kwiatkowski, Cordelia Langford, John Sillitoe on behalf of the Wellcome Sanger |

| | | | |
|---|---|---|---|
| EPI_ISL_3172292, EPI_ISL_3172293 | | Consortium | Institute COVID-19 Surveillance Team |
| EPI_ISL_3172294 | Berkshire and Surrey Pathology Services Lighthouse Laboratory | Wellcome Sanger Institute for the COVID-19 Genomics UK (COG-UK) Consortium | Berkshire and Surrey Pathology Services Lighthouse Laboratory and Alex Alderton, Roberto Amato, Jeffrey Barrett, Sonia Goncalves, Ewan Harrison, David K. Jackson, Ian Johnston, Dominic Kwiatkowski, Cordelia Langford, John Sillitoe on behalf of the Wellcome Sanger Institute COVID-19 Surveillance Team |
| EPI_ISL_3172295 | Lighthouse Lab in Milton Keynes | Wellcome Sanger Institute for the COVID-19 Genomics UK (COG-UK) Consortium | The Lighthouse Lab in Milton Keynes and Alex Alderton, Roberto Amato, Jeffrey Barrett, Sonia Goncalves, Ewan Harrison, David K. Jackson, Ian Johnston, Dominic Kwiatkowski, Cordelia Langford, John Sillitoe on behalf of the Wellcome Sanger Institute COVID-19 Surveillance Team |
| EPI_ISL_3172296 | Lighthouse Laboratory Plymouth | Wellcome Sanger Institute for the COVID-19 Genomics UK (COG-UK) Consortium | Lighthouse Laboratory Plymouth and Alex Alderton, Roberto Amato, Jeffrey Barrett, Sonia Goncalves, Ewan Harrison, David K. Jackson, Ian Johnston, Dominic Kwiatkowski, Cordelia Langford, John Sillitoe on behalf of the Wellcome Sanger Institute COVID-19 Surveillance Team |
| EPI_ISL_3172297 | Lighthouse Lab in Milton Keynes | Wellcome Sanger Institute for the COVID-19 Genomics UK (COG-UK) Consortium | The Lighthouse Lab in Milton Keynes and Alex Alderton, Roberto Amato, Jeffrey Barrett, Sonia Goncalves, Ewan Harrison, David K. Jackson, Ian Johnston, Dominic Kwiatkowski, Cordelia Langford, John Sillitoe on behalf of the Wellcome Sanger Institute COVID-19 Surveillance Team |
| EPI_ISL_3172298, EPI_ISL_3172299 | Berkshire and Surrey Pathology Services Lighthouse Laboratory | Wellcome Sanger Institute for the COVID-19 Genomics UK (COG-UK) Consortium | Berkshire and Surrey Pathology Services Lighthouse Laboratory and Alex Alderton, Roberto Amato, Jeffrey Barrett, Sonia Goncalves, Ewan Harrison, David K. Jackson, Ian Johnston, Dominic Kwiatkowski, Cordelia Langford, John Sillitoe on behalf of the Wellcome Sanger Institute COVID-19 Surveillance Team |
| EPI_ISL_3172300 | Lighthouse Laboratory Plymouth | Wellcome Sanger Institute for the COVID-19 Genomics UK (COG-UK) Consortium | Lighthouse Laboratory Plymouth and Alex Alderton, Roberto Amato, Jeffrey Barrett, Sonia Goncalves, Ewan Harrison, David K. Jackson, Ian Johnston, Dominic Kwiatkowski, Cordelia Langford, John Sillitoe on behalf of the Wellcome Sanger Institute COVID-19 Surveillance Team |
| EPI_ISL_3172301 | Lighthouse Lab in Milton Keynes | Wellcome Sanger Institute for the COVID-19 Genomics UK (COG-UK) Consortium | The Lighthouse Lab in Milton Keynes and Alex Alderton, Roberto Amato, Jeffrey Barrett, Sonia Goncalves, Ewan Harrison, David K. Jackson, Ian Johnston, Dominic Kwiatkowski, Cordelia Langford, John Sillitoe on behalf of the Wellcome Sanger Institute COVID-19 Surveillance Team |
| EPI_ISL_3172302 | Lighthouse Laboratory Plymouth | Wellcome Sanger Institute for the COVID-19 Genomics UK (COG-UK) Consortium | Lighthouse Laboratory Plymouth and Alex Alderton, Roberto Amato, Jeffrey Barrett, Sonia Goncalves, Ewan Harrison, David K. Jackson, Ian Johnston, Dominic Kwiatkowski, Cordelia Langford, John Sillitoe on behalf of the Wellcome Sanger Institute COVID-19 Surveillance Team |
| EPI_ISL_3172303 | Berkshire and Surrey Pathology Services Lighthouse Laboratory | Wellcome Sanger Institute for the COVID-19 Genomics UK (COG-UK) Consortium | Berkshire and Surrey Pathology Services Lighthouse Laboratory and Alex Alderton, Roberto Amato, Jeffrey Barrett, Sonia Goncalves, Ewan Harrison, David K. Jackson, Ian Johnston, Dominic Kwiatkowski, Cordelia Langford, John Sillitoe on behalf of the Wellcome Sanger Institute COVID-19 Surveillance Team |
| EPI_ISL_3172304 | Lighthouse Lab in Milton Keynes | Wellcome Sanger Institute for the COVID-19 Genomics UK (COG-UK) Consortium | The Lighthouse Lab in Milton Keynes and Alex Alderton, Roberto Amato, Jeffrey Barrett, Sonia Goncalves, Ewan Harrison, David K. Jackson, Ian Johnston, Dominic Kwiatkowski, Cordelia Langford, John Sillitoe on behalf of the Wellcome Sanger Institute COVID-19 Surveillance Team |
| EPI_ISL_3172305, EPI_ISL_3172306 | Berkshire and Surrey Pathology Services Lighthouse Laboratory | Wellcome Sanger Institute for the COVID-19 Genomics UK (COG-UK) Consortium | Berkshire and Surrey Pathology Services Lighthouse Laboratory and Alex Alderton, Roberto Amato, Jeffrey Barrett, Sonia Goncalves, Ewan Harrison, David K. Jackson, Ian Johnston, Dominic Kwiatkowski, Cordelia Langford, John Sillitoe on behalf of the Wellcome Sanger Institute COVID-19 Surveillance Team |
| EPI_ISL_3172307 | Lighthouse Lab in Milton Keynes | Wellcome Sanger Institute for the COVID-19 Genomics UK (COG-UK) Consortium | The Lighthouse Lab in Milton Keynes and Alex Alderton, Roberto Amato, Jeffrey Barrett, Sonia Goncalves, Ewan Harrison, David K. Jackson, Ian Johnston, Dominic Kwiatkowski, Cordelia Langford, John Sillitoe on behalf of the Wellcome Sanger Institute COVID-19 Surveillance Team |
| EPI_ISL_3172308 | Berkshire and Surrey Pathology Services Lighthouse Laboratory | Wellcome Sanger Institute for the COVID-19 Genomics UK (COG-UK) Consortium | Berkshire and Surrey Pathology Services Lighthouse Laboratory and Alex Alderton, Roberto Amato, Jeffrey Barrett, Sonia Goncalves, Ewan Harrison, David K. Jackson, Ian Johnston, Dominic Kwiatkowski, Cordelia Langford, John Sillitoe on behalf of the Wellcome Sanger Institute COVID-19 Surveillance Team |
| EPI_ISL_3172309 | Lighthouse Laboratory Plymouth | Wellcome Sanger Institute for the COVID-19 Genomics UK (COG-UK) Consortium | Lighthouse Laboratory Plymouth and Alex Alderton, Roberto Amato, Jeffrey Barrett, Sonia Goncalves, Ewan Harrison, David K. Jackson, Ian Johnston, Dominic Kwiatkowski, Cordelia Langford, John Sillitoe on behalf of the Wellcome Sanger Institute COVID-19 Surveillance Team |
| EPI_ISL_3172310, EPI_ISL_3172311 | Lighthouse Lab in Milton Keynes | Wellcome Sanger Institute for the COVID-19 Genomics UK (COG-UK) Consortium | The Lighthouse Lab in Milton Keynes and Alex Alderton, Roberto Amato, Jeffrey Barrett, Sonia Goncalves, Ewan Harrison, David K. Jackson, Ian Johnston, Dominic Kwiatkowski, Cordelia Langford, John Sillitoe on behalf of the Wellcome Sanger Institute COVID-19 Surveillance Team |
| EPI_ISL_3172312 | Berkshire and Surrey Pathology Services Lighthouse Laboratory | Wellcome Sanger Institute for the COVID-19 Genomics UK (COG-UK) Consortium | Berkshire and Surrey Pathology Services Lighthouse Laboratory and Alex Alderton, Roberto Amato, Jeffrey Barrett, Sonia Goncalves, Ewan Harrison, David K. Jackson, Ian Johnston, Dominic Kwiatkowski, Cordelia Langford, John Sillitoe on behalf of the Wellcome Sanger Institute COVID-19 Surveillance Team |
| EPI_ISL_3172313, EPI_ISL_3172314, EPI_ISL_3172315 | Lighthouse Lab in Milton Keynes | Wellcome Sanger Institute for the COVID-19 Genomics UK (COG-UK) Consortium | The Lighthouse Lab in Milton Keynes and Alex Alderton, Roberto Amato, Jeffrey Barrett, Sonia Goncalves, Ewan Harrison, David K. Jackson, Ian Johnston, Dominic Kwiatkowski, Cordelia Langford, John Sillitoe on behalf of the Wellcome Sanger Institute COVID-19 Surveillance Team |
| EPI_ISL_3172316 | Berkshire and Surrey Pathology Services Lighthouse Laboratory | Wellcome Sanger Institute for the COVID-19 Genomics UK (COG-UK) Consortium | Berkshire and Surrey Pathology Services Lighthouse Laboratory and Alex Alderton, Roberto Amato, Jeffrey Barrett, Sonia Goncalves, Ewan Harrison, David K. Jackson, Ian Johnston, Dominic Kwiatkowski, Cordelia Langford, John Sillitoe on behalf of the Wellcome Sanger Institute COVID-19 Surveillance Team |
| EPI_ISL_3172317, EPI_ISL_3172318, EPI_ISL_3172319, EPI_ISL_3172320, EPI_ISL_3172321 | Lighthouse Lab in Milton Keynes | Wellcome Sanger Institute for the COVID-19 Genomics UK (COG-UK) Consortium | The Lighthouse Lab in Milton Keynes and Alex Alderton, Roberto Amato, Jeffrey Barrett, Sonia Goncalves, Ewan Harrison, David K. Jackson, Ian Johnston, Dominic Kwiatkowski, Cordelia Langford, John Sillitoe on behalf of the Wellcome Sanger Institute COVID-19 Surveillance Team |
| EPI_ISL_3172322 | Berkshire and Surrey Pathology Services Lighthouse Laboratory | Wellcome Sanger Institute for the COVID-19 Genomics UK (COG-UK) Consortium | Berkshire and Surrey Pathology Services Lighthouse Laboratory and Alex Alderton, Roberto Amato, Jeffrey Barrett, Sonia Goncalves, Ewan Harrison, David K. Jackson, Ian Johnston, Dominic Kwiatkowski, Cordelia Langford, John Sillitoe on behalf of the Wellcome Sanger Institute COVID-19 Surveillance Team |
| EPI_ISL_3172323 | Lighthouse Lab in Milton Keynes | Wellcome Sanger Institute for the COVID-19 Genomics UK (COG-UK) Consortium | The Lighthouse Lab in Milton Keynes and Alex Alderton, Roberto Amato, Jeffrey Barrett, Sonia Goncalves, Ewan Harrison, David K. Jackson, Ian Johnston, Dominic Kwiatkowski, Cordelia Langford, John Sillitoe on behalf of the Wellcome Sanger Institute COVID-19 Surveillance Team |
| EPI_ISL_3172324 | Lighthouse Laboratory Plymouth | Wellcome Sanger Institute for the COVID-19 Genomics UK (COG-UK) Consortium | Lighthouse Laboratory Plymouth and Alex Alderton, Roberto Amato, Jeffrey Barrett, Sonia Goncalves, Ewan Harrison, David K. Jackson, Ian Johnston, Dominic Kwiatkowski, Cordelia Langford, John Sillitoe on behalf of the Wellcome Sanger Institute COVID-19 Surveillance Team |
| EPI_ISL_3172325 | Lighthouse Lab in Milton Keynes | Wellcome Sanger Institute for the COVID-19 Genomics UK (COG-UK) Consortium | The Lighthouse Lab in Milton Keynes and Alex Alderton, Roberto Amato, Jeffrey Barrett, Sonia Goncalves, Ewan Harrison, David K. Jackson, Ian Johnston, Dominic Kwiatkowski, Cordelia Langford, John Sillitoe on behalf of the Wellcome Sanger Institute COVID-19 Surveillance Team |
| EPI_ISL_3172326 | Lighthouse Laboratory Plymouth | Wellcome Sanger Institute for the COVID-19 Genomics UK (COG-UK) Consortium | Lighthouse Laboratory Plymouth and Alex Alderton, Roberto Amato, Jeffrey Barrett, Sonia Goncalves, Ewan Harrison, David K. Jackson, Ian Johnston, Dominic Kwiatkowski, Cordelia Langford, John Sillitoe on behalf of the Wellcome Sanger Institute COVID-19 Surveillance Team |
| EPI_ISL_3172327 | Berkshire and Surrey Pathology Services Lighthouse Laboratory | Wellcome Sanger Institute for the COVID-19 Genomics UK (COG-UK) Consortium | Berkshire and Surrey Pathology Services Lighthouse Laboratory and Alex Alderton, Roberto Amato, Jeffrey Barrett, Sonia Goncalves, Ewan Harrison, David K. Jackson, Ian Johnston, Dominic Kwiatkowski, Cordelia Langford, John Sillitoe on behalf of the Wellcome Sanger Institute COVID-19 Surveillance Team |
| EPI_ISL_3172328 | Lighthouse Laboratory Plymouth | Wellcome Sanger Institute for the COVID-19 Genomics UK (COG-UK) Consortium | Lighthouse Laboratory Plymouth and Alex Alderton, Roberto Amato, Jeffrey Barrett, Sonia Goncalves, Ewan Harrison, David K. Jackson, Ian Johnston, Dominic Kwiatkowski, Cordelia Langford, John Sillitoe on behalf of the Wellcome Sanger Institute COVID-19 Surveillance Team |
| EPI_ISL_3172329, EPI_ISL_3172330, EPI_ISL_3172331 | Lighthouse Lab in Milton Keynes | Wellcome Sanger Institute for the COVID-19 Genomics UK (COG-UK) Consortium | The Lighthouse Lab in Milton Keynes and Alex Alderton, Roberto Amato, Jeffrey Barrett, Sonia Goncalves, Ewan Harrison, David K. Jackson, Ian Johnston, Dominic Kwiatkowski, Cordelia Langford, John Sillitoe on behalf of the Wellcome Sanger Institute COVID-19 Surveillance Team |
| EPI_ISL_3172332 | Lighthouse Laboratory Plymouth | Wellcome Sanger Institute for the COVID-19 Genomics UK (COG-UK) Consortium | Lighthouse Laboratory Plymouth and Alex Alderton, Roberto Amato, Jeffrey Barrett, Sonia Goncalves, Ewan Harrison, David K. Jackson, Ian Johnston, Dominic Kwiatkowski, Cordelia Langford, John Sillitoe on behalf of the Wellcome Sanger Institute COVID-19 Surveillance Team |
| EPI_ISL_3172333, EPI_ISL_3172334 | Lighthouse Lab in Milton Keynes | Wellcome Sanger Institute for the COVID-19 Genomics UK (COG-UK) Consortium | The Lighthouse Lab in Milton Keynes and Alex Alderton, Roberto Amato, Jeffrey Barrett, Sonia Goncalves, Ewan Harrison, David K. Jackson, Ian Johnston, Dominic Kwiatkowski, Cordelia Langford, John Sillitoe on behalf of the Wellcome Sanger Institute COVID-19 Surveillance Team |
| EPI_ISL_3172335 | Lighthouse Laboratory Plymouth | Wellcome Sanger Institute for the COVID-19 Genomics UK (COG-UK) Consortium | Lighthouse Laboratory Plymouth and Alex Alderton, Roberto Amato, Jeffrey Barrett, Sonia Goncalves, Ewan Harrison, David K. Jackson, Ian Johnston, Dominic Kwiatkowski, Cordelia Langford, John Sillitoe on behalf of the Wellcome Sanger Institute COVID-19 Surveillance Team |
| EPI_ISL_3172336 | Berkshire and Surrey Pathology Services Lighthouse Laboratory | Wellcome Sanger Institute for the COVID-19 Genomics UK (COG-UK) Consortium | Berkshire and Surrey Pathology Services Lighthouse Laboratory and Alex Alderton, Roberto Amato, Jeffrey Barrett, Sonia Goncalves, Ewan Harrison, David K. Jackson, Ian Johnston, Dominic Kwiatkowski, Cordelia Langford, John Sillitoe on behalf of the Wellcome Sanger Institute COVID-19 Surveillance Team |
| EPI_ISL_3172337, EPI_ISL_3172338, EPI_ISL_3172339 | Lighthouse Lab in Milton Keynes | Wellcome Sanger Institute for the COVID-19 Genomics UK (COG-UK) Consortium | The Lighthouse Lab in Milton Keynes and Alex Alderton, Roberto Amato, Jeffrey Barrett, Sonia Goncalves, Ewan Harrison, David K. Jackson, Ian Johnston, Dominic Kwiatkowski, Cordelia Langford, John Sillitoe on behalf of the Wellcome Sanger Institute COVID-19 Surveillance Team |
| EPI_ISL_3172340 | Lighthouse Laboratory Plymouth | Wellcome Sanger Institute for the COVID-19 Genomics UK (COG-UK) Consortium | Lighthouse Laboratory Plymouth and Alex Alderton, Roberto Amato, Jeffrey Barrett, Sonia Goncalves, Ewan Harrison, David K. Jackson, Ian Johnston, Dominic Kwiatkowski, Cordelia Langford, John Sillitoe on behalf of the Wellcome Sanger Institute COVID-19 Surveillance Team |
| EPI_ISL_3172341, EPI_ISL_3172342, EPI_ISL_3172343 | Lighthouse Lab in Milton Keynes | Wellcome Sanger Institute for the COVID-19 Genomics UK (COG-UK) Consortium | The Lighthouse Lab in Milton Keynes and Alex Alderton, Roberto Amato, Jeffrey Barrett, Sonia Goncalves, Ewan Harrison, David K. Jackson, Ian Johnston, Dominic Kwiatkowski, Cordelia Langford, John Sillitoe on behalf of the Wellcome Sanger Institute COVID-19 Surveillance Team |
| EPI_ISL_3172344, EPI_ISL_3172345 | Berkshire and Surrey Pathology Services Lighthouse Laboratory | Wellcome Sanger Institute for the COVID-19 Genomics UK (COG-UK) Consortium | Berkshire and Surrey Pathology Services Lighthouse Laboratory and Alex Alderton, Roberto Amato, Jeffrey Barrett, Sonia Goncalves, Ewan Harrison, David K. Jackson, Ian Johnston, Dominic Kwiatkowski, Cordelia Langford, John Sillitoe on behalf of the Wellcome Sanger Institute COVID-19 Surveillance Team |
| EPI_ISL_3172346, EPI_ISL_3172347 | Lighthouse Lab in Milton Keynes | Wellcome Sanger Institute for the COVID-19 Genomics UK (COG-UK) Consortium | The Lighthouse Lab in Milton Keynes and Alex Alderton, Roberto Amato, Jeffrey Barrett, Sonia Goncalves, Ewan Harrison, David K. Jackson, Ian Johnston, Dominic Kwiatkowski, Cordelia Langford, John Sillitoe on behalf of the Wellcome Sanger Institute COVID-19 Surveillance Team |
| EPI_ISL_3172348, EPI_ISL_3172349, EPI_ISL_3172350 | Berkshire and Surrey Pathology Services Lighthouse Laboratory | Wellcome Sanger Institute for the COVID-19 Genomics UK (COG-UK) Consortium | Berkshire and Surrey Pathology Services Lighthouse Laboratory and Alex Alderton, Roberto Amato, Jeffrey Barrett, Sonia Goncalves, Ewan Harrison, David K. Jackson, Ian Johnston, Dominic Kwiatkowski, Cordelia Langford, John Sillitoe on behalf of the Wellcome Sanger Institute COVID-19 Surveillance Team |
| EPI_ISL_3172351, EPI_ISL_3172352 | Lighthouse Laboratory Plymouth | Wellcome Sanger Institute for the COVID-19 Genomics UK (COG-UK) Consortium | Lighthouse Laboratory Plymouth and Alex Alderton, Roberto Amato, Jeffrey Barrett, Sonia Goncalves, Ewan Harrison, David K. Jackson, Ian Johnston, Dominic Kwiatkowski, Cordelia Langford, John Sillitoe on behalf of the Wellcome Sanger Institute COVID-19 Surveillance Team |
| EPI_ISL_3172353 | Lighthouse Lab in Milton Keynes | Wellcome Sanger Institute for the COVID-19 Genomics UK (COG-UK) Consortium | The Lighthouse Lab in Milton Keynes and Alex Alderton, Roberto Amato, Jeffrey Barrett, Sonia Goncalves, Ewan Harrison, David K. Jackson, Ian Johnston, Dominic Kwiatkowski, Cordelia Langford, John Sillitoe on behalf of the Wellcome Sanger Institute COVID-19 Surveillance Team |
| EPI_ISL_3172354 | Berkshire and Surrey Pathology Services Lighthouse Laboratory | Wellcome Sanger Institute for the COVID-19 Genomics UK (COG-UK) Consortium | Berkshire and Surrey Pathology Services Lighthouse Laboratory and Alex Alderton, Roberto Amato, Jeffrey Barrett, Sonia Goncalves, Ewan Harrison, David K. Jackson, Ian Johnston, Dominic Kwiatkowski, Cordelia Langford, John Sillitoe on behalf of the Wellcome Sanger Institute COVID-19 Surveillance Team |
| EPI_ISL_3172355, | Lighthouse Laboratory Plymouth | Wellcome Sanger Institute for the COVID-19 Genomics UK (COG-UK) | Lighthouse Laboratory Plymouth and Alex Alderton, Roberto Amato, Jeffrey Barrett, Sonia Goncalves, Ewan Harrison, David K. Jackson, Ian Johnston, Dominic Kwiatkowski, Cordelia Langford, John Sillitoe on behalf of the Wellcome Sanger |

| | | | |
|---|---|---|---|
| EPI_ISL_3172356 | | Consortium | Institute COVID-19 Surveillance Team |
| EPI_ISL_3172357, EPI_ISL_3172358 | Lighthouse Lab in Milton Keynes | Wellcome Sanger Institute for the COVID-19 Genomics UK (COG-UK) Consortium | The Lighthouse Lab in Milton Keynes and Alex Alderton, Roberto Amato, Jeffrey Barrett, Sonia Goncalves, Ewan Harrison, David K. Jackson, Ian Johnston, Dominic Kwiatkowski, Cordelia Langford, John Sillitoe on behalf of the Wellcome Sanger Institute COVID-19 Surveillance Team |
| EPI_ISL_3172359 | Lighthouse Laboratory Plymouth | Wellcome Sanger Institute for the COVID-19 Genomics UK (COG-UK) Consortium | Lighthouse Laboratory Plymouth and Alex Alderton, Roberto Amato, Jeffrey Barrett, Sonia Goncalves, Ewan Harrison, David K. Jackson, Ian Johnston, Dominic Kwiatkowski, Cordelia Langford, John Sillitoe on behalf of the Wellcome Sanger Institute COVID-19 Surveillance Team |
| EPI_ISL_3172360 | Lighthouse Lab in Milton Keynes | Wellcome Sanger Institute for the COVID-19 Genomics UK (COG-UK) Consortium | The Lighthouse Lab in Milton Keynes and Alex Alderton, Roberto Amato, Jeffrey Barrett, Sonia Goncalves, Ewan Harrison, David K. Jackson, Ian Johnston, Dominic Kwiatkowski, Cordelia Langford, John Sillitoe on behalf of the Wellcome Sanger Institute COVID-19 Surveillance Team |
| EPI_ISL_3172361 | Lighthouse Laboratory Plymouth | Wellcome Sanger Institute for the COVID-19 Genomics UK (COG-UK) Consortium | Lighthouse Laboratory Plymouth and Alex Alderton, Roberto Amato, Jeffrey Barrett, Sonia Goncalves, Ewan Harrison, David K. Jackson, Ian Johnston, Dominic Kwiatkowski, Cordelia Langford, John Sillitoe on behalf of the Wellcome Sanger Institute COVID-19 Surveillance Team |
| EPI_ISL_3172362 | Lighthouse Lab in Milton Keynes | Wellcome Sanger Institute for the COVID-19 Genomics UK (COG-UK) Consortium | The Lighthouse Lab in Milton Keynes and Alex Alderton, Roberto Amato, Jeffrey Barrett, Sonia Goncalves, Ewan Harrison, David K. Jackson, Ian Johnston, Dominic Kwiatkowski, Cordelia Langford, John Sillitoe on behalf of the Wellcome Sanger Institute COVID-19 Surveillance Team |
| EPI_ISL_3172363 | Lighthouse Laboratory Plymouth | Wellcome Sanger Institute for the COVID-19 Genomics UK (COG-UK) Consortium | Lighthouse Laboratory Plymouth and Alex Alderton, Roberto Amato, Jeffrey Barrett, Sonia Goncalves, Ewan Harrison, David K. Jackson, Ian Johnston, Dominic Kwiatkowski, Cordelia Langford, John Sillitoe on behalf of the Wellcome Sanger Institute COVID-19 Surveillance Team |
| EPI_ISL_3172364, EPI_ISL_3172365, EPI_ISL_3172366 | Lighthouse Lab in Milton Keynes | Wellcome Sanger Institute for the COVID-19 Genomics UK (COG-UK) Consortium | The Lighthouse Lab in Milton Keynes and Alex Alderton, Roberto Amato, Jeffrey Barrett, Sonia Goncalves, Ewan Harrison, David K. Jackson, Ian Johnston, Dominic Kwiatkowski, Cordelia Langford, John Sillitoe on behalf of the Wellcome Sanger Institute COVID-19 Surveillance Team |
| EPI_ISL_3172367, EPI_ISL_3172368 | Lighthouse Laboratory Plymouth | Wellcome Sanger Institute for the COVID-19 Genomics UK (COG-UK) Consortium | Lighthouse Laboratory Plymouth and Alex Alderton, Roberto Amato, Jeffrey Barrett, Sonia Goncalves, Ewan Harrison, David K. Jackson, Ian Johnston, Dominic Kwiatkowski, Cordelia Langford, John Sillitoe on behalf of the Wellcome Sanger Institute COVID-19 Surveillance Team |
| EPI_ISL_3172369, EPI_ISL_3172370, EPI_ISL_3172371 | Lighthouse Lab in Milton Keynes | Wellcome Sanger Institute for the COVID-19 Genomics UK (COG-UK) Consortium | The Lighthouse Lab in Milton Keynes and Alex Alderton, Roberto Amato, Jeffrey Barrett, Sonia Goncalves, Ewan Harrison, David K. Jackson, Ian Johnston, Dominic Kwiatkowski, Cordelia Langford, John Sillitoe on behalf of the Wellcome Sanger Institute COVID-19 Surveillance Team |
| EPI_ISL_3172372 | Berkshire and Surrey Pathology Services Lighthouse Laboratory | Wellcome Sanger Institute for the COVID-19 Genomics UK (COG-UK) Consortium | Berkshire and Surrey Pathology Services Lighthouse Laboratory and Alex Alderton, Roberto Amato, Jeffrey Barrett, Sonia Goncalves, Ewan Harrison, David K. Jackson, Ian Johnston, Dominic Kwiatkowski, Cordelia Langford, John Sillitoe on behalf of the Wellcome Sanger Institute COVID-19 Surveillance Team |
| EPI_ISL_3172373, EPI_ISL_3172374 | Lighthouse Lab in Milton Keynes | Wellcome Sanger Institute for the COVID-19 Genomics UK (COG-UK) Consortium | The Lighthouse Lab in Milton Keynes and Alex Alderton, Roberto Amato, Jeffrey Barrett, Sonia Goncalves, Ewan Harrison, David K. Jackson, Ian Johnston, Dominic Kwiatkowski, Cordelia Langford, John Sillitoe on behalf of the Wellcome Sanger Institute COVID-19 Surveillance Team |
| EPI_ISL_3172375 | Berkshire and Surrey Pathology Services Lighthouse Laboratory | Wellcome Sanger Institute for the COVID-19 Genomics UK (COG-UK) Consortium | Berkshire and Surrey Pathology Services Lighthouse Laboratory and Alex Alderton, Roberto Amato, Jeffrey Barrett, Sonia Goncalves, Ewan Harrison, David K. Jackson, Ian Johnston, Dominic Kwiatkowski, Cordelia Langford, John Sillitoe on behalf of the Wellcome Sanger Institute COVID-19 Surveillance Team |
| EPI_ISL_3172376, EPI_ISL_3172377 | Lighthouse Laboratory Plymouth | Wellcome Sanger Institute for the COVID-19 Genomics UK (COG-UK) Consortium | Lighthouse Laboratory Plymouth and Alex Alderton, Roberto Amato, Jeffrey Barrett, Sonia Goncalves, Ewan Harrison, David K. Jackson, Ian Johnston, Dominic Kwiatkowski, Cordelia Langford, John Sillitoe on behalf of the Wellcome Sanger Institute COVID-19 Surveillance Team |
| EPI_ISL_3172378, EPI_ISL_3172379 | Lighthouse Lab in Milton Keynes | Wellcome Sanger Institute for the COVID-19 Genomics UK (COG-UK) Consortium | The Lighthouse Lab in Milton Keynes and Alex Alderton, Roberto Amato, Jeffrey Barrett, Sonia Goncalves, Ewan Harrison, David K. Jackson, Ian Johnston, Dominic Kwiatkowski, Cordelia Langford, John Sillitoe on behalf of the Wellcome Sanger Institute COVID-19 Surveillance Team |
| EPI_ISL_3172380 | Berkshire and Surrey Pathology Services Lighthouse Laboratory | Wellcome Sanger Institute for the COVID-19 Genomics UK (COG-UK) Consortium | Berkshire and Surrey Pathology Services Lighthouse Laboratory and Alex Alderton, Roberto Amato, Jeffrey Barrett, Sonia Goncalves, Ewan Harrison, David K. Jackson, Ian Johnston, Dominic Kwiatkowski, Cordelia Langford, John Sillitoe on behalf of the Wellcome Sanger Institute COVID-19 Surveillance Team |
| EPI_ISL_3172381, EPI_ISL_3172382, EPI_ISL_3172383 | Lighthouse Lab in Milton Keynes | Wellcome Sanger Institute for the COVID-19 Genomics UK (COG-UK) Consortium | The Lighthouse Lab in Milton Keynes and Alex Alderton, Roberto Amato, Jeffrey Barrett, Sonia Goncalves, Ewan Harrison, David K. Jackson, Ian Johnston, Dominic Kwiatkowski, Cordelia Langford, John Sillitoe on behalf of the Wellcome Sanger Institute COVID-19 Surveillance Team |
| EPI_ISL_3172384 | Berkshire and Surrey Pathology Services Lighthouse Laboratory | Wellcome Sanger Institute for the COVID-19 Genomics UK (COG-UK) Consortium | Berkshire and Surrey Pathology Services Lighthouse Laboratory and Alex Alderton, Roberto Amato, Jeffrey Barrett, Sonia Goncalves, Ewan Harrison, David K. Jackson, Ian Johnston, Dominic Kwiatkowski, Cordelia Langford, John Sillitoe on behalf of the Wellcome Sanger Institute COVID-19 Surveillance Team |
| EPI_ISL_3172385 | Lighthouse Lab in Milton Keynes | Wellcome Sanger Institute for the COVID-19 Genomics UK (COG-UK) Consortium | The Lighthouse Lab in Milton Keynes and Alex Alderton, Roberto Amato, Jeffrey Barrett, Sonia Goncalves, Ewan Harrison, David K. Jackson, Ian Johnston, Dominic Kwiatkowski, Cordelia Langford, John Sillitoe on behalf of the Wellcome Sanger Institute COVID-19 Surveillance Team |
| EPI_ISL_3172386, EPI_ISL_3172387 | Lighthouse Laboratory Plymouth | Wellcome Sanger Institute for the COVID-19 Genomics UK (COG-UK) Consortium | Lighthouse Laboratory Plymouth and Alex Alderton, Roberto Amato, Jeffrey Barrett, Sonia Goncalves, Ewan Harrison, David K. Jackson, Ian Johnston, Dominic Kwiatkowski, Cordelia Langford, John Sillitoe on behalf of the Wellcome Sanger Institute COVID-19 Surveillance Team |
| EPI_ISL_3172388 | Lighthouse Lab in Milton Keynes | Wellcome Sanger Institute for the COVID-19 Genomics UK (COG-UK) Consortium | The Lighthouse Lab in Milton Keynes and Alex Alderton, Roberto Amato, Jeffrey Barrett, Sonia Goncalves, Ewan Harrison, David K. Jackson, Ian Johnston, Dominic Kwiatkowski, Cordelia Langford, John Sillitoe on behalf of the Wellcome Sanger Institute COVID-19 Surveillance Team |
| EPI_ISL_3172389 | Berkshire and Surrey Pathology Services Lighthouse Laboratory | Wellcome Sanger Institute for the COVID-19 Genomics UK (COG-UK) Consortium | Berkshire and Surrey Pathology Services Lighthouse Laboratory and Alex Alderton, Roberto Amato, Jeffrey Barrett, Sonia Goncalves, Ewan Harrison, David K. Jackson, Ian Johnston, Dominic Kwiatkowski, Cordelia Langford, John Sillitoe on behalf of the Wellcome Sanger Institute COVID-19 Surveillance Team |
| EPI_ISL_3172390, EPI_ISL_3172391 | Lighthouse Laboratory Plymouth | Wellcome Sanger Institute for the COVID-19 Genomics UK (COG-UK) Consortium | Lighthouse Laboratory Plymouth and Alex Alderton, Roberto Amato, Jeffrey Barrett, Sonia Goncalves, Ewan Harrison, David K. Jackson, Ian Johnston, Dominic Kwiatkowski, Cordelia Langford, John Sillitoe on behalf of the Wellcome Sanger Institute COVID-19 Surveillance Team |
| EPI_ISL_3172392 | Berkshire and Surrey Pathology Services Lighthouse Laboratory | Wellcome Sanger Institute for the COVID-19 Genomics UK (COG-UK) Consortium | Berkshire and Surrey Pathology Services Lighthouse Laboratory and Alex Alderton, Roberto Amato, Jeffrey Barrett, Sonia Goncalves, Ewan Harrison, David K. Jackson, Ian Johnston, Dominic Kwiatkowski, Cordelia Langford, John Sillitoe on behalf of the Wellcome Sanger Institute COVID-19 Surveillance Team |
| EPI_ISL_3172393, EPI_ISL_3172394 | Lighthouse Lab in Milton Keynes | Wellcome Sanger Institute for the COVID-19 Genomics UK (COG-UK) Consortium | The Lighthouse Lab in Milton Keynes and Alex Alderton, Roberto Amato, Jeffrey Barrett, Sonia Goncalves, Ewan Harrison, David K. Jackson, Ian Johnston, Dominic Kwiatkowski, Cordelia Langford, John Sillitoe on behalf of the Wellcome Sanger Institute COVID-19 Surveillance Team |
| EPI_ISL_3172395 | Lighthouse Laboratory Plymouth | Wellcome Sanger Institute for the COVID-19 Genomics UK (COG-UK) Consortium | Lighthouse Laboratory Plymouth and Alex Alderton, Roberto Amato, Jeffrey Barrett, Sonia Goncalves, Ewan Harrison, David K. Jackson, Ian Johnston, Dominic Kwiatkowski, Cordelia Langford, John Sillitoe on behalf of the Wellcome Sanger Institute COVID-19 Surveillance Team |
| EPI_ISL_3172396 | Lighthouse Lab in Milton Keynes | Wellcome Sanger Institute for the COVID-19 Genomics UK (COG-UK) Consortium | The Lighthouse Lab in Milton Keynes and Alex Alderton, Roberto Amato, Jeffrey Barrett, Sonia Goncalves, Ewan Harrison, David K. Jackson, Ian Johnston, Dominic Kwiatkowski, Cordelia Langford, John Sillitoe on behalf of the Wellcome Sanger Institute COVID-19 Surveillance Team |
| EPI_ISL_3172397 | Berkshire and Surrey Pathology Services Lighthouse Laboratory | Wellcome Sanger Institute for the COVID-19 Genomics UK (COG-UK) Consortium | Berkshire and Surrey Pathology Services Lighthouse Laboratory and Alex Alderton, Roberto Amato, Jeffrey Barrett, Sonia Goncalves, Ewan Harrison, David K. Jackson, Ian Johnston, Dominic Kwiatkowski, Cordelia Langford, John Sillitoe on behalf of the Wellcome Sanger Institute COVID-19 Surveillance Team |
| EPI_ISL_3172398 | Lighthouse Lab in Milton Keynes | Wellcome Sanger Institute for the COVID-19 Genomics UK (COG-UK) Consortium | The Lighthouse Lab in Milton Keynes and Alex Alderton, Roberto Amato, Jeffrey Barrett, Sonia Goncalves, Ewan Harrison, David K. Jackson, Ian Johnston, Dominic Kwiatkowski, Cordelia Langford, John Sillitoe on behalf of the Wellcome Sanger Institute COVID-19 Surveillance Team |
| EPI_ISL_3172399 | Lighthouse Laboratory Plymouth | Wellcome Sanger Institute for the COVID-19 Genomics UK (COG-UK) Consortium | Lighthouse Laboratory Plymouth and Alex Alderton, Roberto Amato, Jeffrey Barrett, Sonia Goncalves, Ewan Harrison, David K. Jackson, Ian Johnston, Dominic Kwiatkowski, Cordelia Langford, John Sillitoe on behalf of the Wellcome Sanger Institute COVID-19 Surveillance Team |
| EPI_ISL_3172400 | Lighthouse Lab in Milton Keynes | Wellcome Sanger Institute for the COVID-19 Genomics UK (COG-UK) Consortium | The Lighthouse Lab in Milton Keynes and Alex Alderton, Roberto Amato, Jeffrey Barrett, Sonia Goncalves, Ewan Harrison, David K. Jackson, Ian Johnston, Dominic Kwiatkowski, Cordelia Langford, John Sillitoe on behalf of the Wellcome Sanger Institute COVID-19 Surveillance Team |
| EPI_ISL_3172401 | Berkshire and Surrey Pathology Services Lighthouse Laboratory | Wellcome Sanger Institute for the COVID-19 Genomics UK (COG-UK) Consortium | Berkshire and Surrey Pathology Services Lighthouse Laboratory and Alex Alderton, Roberto Amato, Jeffrey Barrett, Sonia Goncalves, Ewan Harrison, David K. Jackson, Ian Johnston, Dominic Kwiatkowski, Cordelia Langford, John Sillitoe on behalf of the Wellcome Sanger Institute COVID-19 Surveillance Team |
| EPI_ISL_3172402 | Lighthouse Lab in Milton Keynes | Wellcome Sanger Institute for the COVID-19 Genomics UK (COG-UK) Consortium | The Lighthouse Lab in Milton Keynes and Alex Alderton, Roberto Amato, Jeffrey Barrett, Sonia Goncalves, Ewan Harrison, David K. Jackson, Ian Johnston, Dominic Kwiatkowski, Cordelia Langford, John Sillitoe on behalf of the Wellcome Sanger Institute COVID-19 Surveillance Team |
| EPI_ISL_3172403 | Lighthouse Laboratory Plymouth | Wellcome Sanger Institute for the COVID-19 Genomics UK (COG-UK) Consortium | Lighthouse Laboratory Plymouth and Alex Alderton, Roberto Amato, Jeffrey Barrett, Sonia Goncalves, Ewan Harrison, David K. Jackson, Ian Johnston, Dominic Kwiatkowski, Cordelia Langford, John Sillitoe on behalf of the Wellcome Sanger Institute COVID-19 Surveillance Team |
| EPI_ISL_3172404 | Berkshire and Surrey Pathology Services Lighthouse Laboratory | Wellcome Sanger Institute for the COVID-19 Genomics UK (COG-UK) Consortium | Berkshire and Surrey Pathology Services Lighthouse Laboratory and Alex Alderton, Roberto Amato, Jeffrey Barrett, Sonia Goncalves, Ewan Harrison, David K. Jackson, Ian Johnston, Dominic Kwiatkowski, Cordelia Langford, John Sillitoe on behalf of the Wellcome Sanger Institute COVID-19 Surveillance Team |
| EPI_ISL_3172405 | Lighthouse Laboratory Plymouth | Wellcome Sanger Institute for the COVID-19 Genomics UK (COG-UK) Consortium | Lighthouse Laboratory Plymouth and Alex Alderton, Roberto Amato, Jeffrey Barrett, Sonia Goncalves, Ewan Harrison, David K. Jackson, Ian Johnston, Dominic Kwiatkowski, Cordelia Langford, John Sillitoe on behalf of the Wellcome Sanger Institute COVID-19 Surveillance Team |
| EPI_ISL_3172406 | Berkshire and Surrey Pathology Services Lighthouse Laboratory | Wellcome Sanger Institute for the COVID-19 Genomics UK (COG-UK) Consortium | Berkshire and Surrey Pathology Services Lighthouse Laboratory and Alex Alderton, Roberto Amato, Jeffrey Barrett, Sonia Goncalves, Ewan Harrison, David K. Jackson, Ian Johnston, Dominic Kwiatkowski, Cordelia Langford, John Sillitoe on behalf of the Wellcome Sanger Institute COVID-19 Surveillance Team |
| EPI_ISL_3172407 | Lighthouse Lab in Milton Keynes | Wellcome Sanger Institute for the COVID-19 Genomics UK (COG-UK) Consortium | The Lighthouse Lab in Milton Keynes and Alex Alderton, Roberto Amato, Jeffrey Barrett, Sonia Goncalves, Ewan Harrison, David K. Jackson, Ian Johnston, Dominic Kwiatkowski, Cordelia Langford, John Sillitoe on behalf of the Wellcome Sanger Institute COVID-19 Surveillance Team |
| EPI_ISL_3172408 | Lighthouse Laboratory Plymouth | Wellcome Sanger Institute for the COVID-19 Genomics UK (COG-UK) Consortium | Lighthouse Laboratory Plymouth and Alex Alderton, Roberto Amato, Jeffrey Barrett, Sonia Goncalves, Ewan Harrison, David K. Jackson, Ian Johnston, Dominic Kwiatkowski, Cordelia Langford, John Sillitoe on behalf of the Wellcome Sanger Institute COVID-19 Surveillance Team |
| EPI_ISL_3172409, EPI_ISL_3172410, EPI_ISL_3172411 | Lighthouse Lab in Milton Keynes | Wellcome Sanger Institute for the COVID-19 Genomics UK (COG-UK) Consortium | The Lighthouse Lab in Milton Keynes and Alex Alderton, Roberto Amato, Jeffrey Barrett, Sonia Goncalves, Ewan Harrison, David K. Jackson, Ian Johnston, Dominic Kwiatkowski, Cordelia Langford, John Sillitoe on behalf of the Wellcome Sanger Institute COVID-19 Surveillance Team |
| EPI_ISL_3172412 | Lighthouse Laboratory Plymouth | Wellcome Sanger Institute for the COVID-19 Genomics UK (COG-UK) Consortium | Lighthouse Laboratory Plymouth and Alex Alderton, Roberto Amato, Jeffrey Barrett, Sonia Goncalves, Ewan Harrison, David K. Jackson, Ian Johnston, Dominic Kwiatkowski, Cordelia Langford, John Sillitoe on behalf of the Wellcome Sanger Institute COVID-19 Surveillance Team |
| EPI_ISL_3172413 | Berkshire and Surrey Pathology Services Lighthouse Laboratory | Wellcome Sanger Institute for the COVID-19 Genomics UK (COG-UK) Consortium | Berkshire and Surrey Pathology Services Lighthouse Laboratory and Alex Alderton, Roberto Amato, Jeffrey Barrett, Sonia Goncalves, Ewan Harrison, David K. Jackson, Ian Johnston, Dominic Kwiatkowski, Cordelia Langford, John Sillitoe on behalf of the Wellcome Sanger Institute COVID-19 Surveillance Team |
| EPI_ISL_3172414, EPI_ISL_3172415 | Lighthouse Lab in Milton Keynes | Wellcome Sanger Institute for the COVID-19 Genomics UK (COG-UK) Consortium | The Lighthouse Lab in Milton Keynes and Alex Alderton, Roberto Amato, Jeffrey Barrett, Sonia Goncalves, Ewan Harrison, David K. Jackson, Ian Johnston, Dominic Kwiatkowski, Cordelia Langford, John Sillitoe on behalf of the Wellcome Sanger Institute COVID-19 Surveillance Team |
| EPI_ISL_3172416, | Berkshire and Surrey Pathology Services Lighthouse Laboratory | Wellcome Sanger Institute for the COVID-19 Genomics UK (COG-UK) | Berkshire and Surrey Pathology Services Lighthouse Laboratory and Alex Alderton, Roberto Amato, Jeffrey Barrett, Sonia Goncalves, Ewan Harrison, David K. Jackson, Ian Johnston, Dominic Kwiatkowski, Cordelia Langford, John Sillitoe on behalf |

| | | | |
|---|---|---|---|
| EPI_ISL_3172417 | | Consortium | of the Wellcome Sanger Institute COVID-19 Surveillance Team |
| EPI_ISL_3172418, EPI_ISL_3172419 | Lighthouse Lab in Milton Keynes | Wellcome Sanger Institute for the COVID-19 Genomics UK (COG-UK) Consortium | The Lighthouse Lab in Milton Keynes and Alex Alderton, Roberto Amato, Jeffrey Barrett, Sonia Goncalves, Ewan Harrison, David K. Jackson, Ian Johnston, Dominic Kwiatkowski, Cordelia Langford, John Sillitoe on behalf of the Wellcome Sanger Institute COVID-19 Surveillance Team |
| EPI_ISL_3172420, EPI_ISL_3172421 | Lighthouse Laboratory Plymouth | Wellcome Sanger Institute for the COVID-19 Genomics UK (COG-UK) Consortium | Lighthouse Laboratory Plymouth and Alex Alderton, Roberto Amato, Jeffrey Barrett, Sonia Goncalves, Ewan Harrison, David K. Jackson, Ian Johnston, Dominic Kwiatkowski, Cordelia Langford, John Sillitoe on behalf of the Wellcome Sanger Institute COVID-19 Surveillance Team |
| EPI_ISL_3172422 | Lighthouse Lab in Milton Keynes | Wellcome Sanger Institute for the COVID-19 Genomics UK (COG-UK) Consortium | The Lighthouse Lab in Milton Keynes and Alex Alderton, Roberto Amato, Jeffrey Barrett, Sonia Goncalves, Ewan Harrison, David K. Jackson, Ian Johnston, Dominic Kwiatkowski, Cordelia Langford, John Sillitoe on behalf of the Wellcome Sanger Institute COVID-19 Surveillance Team |
| EPI_ISL_3172423, EPI_ISL_3172424 | Lighthouse Laboratory Plymouth | Wellcome Sanger Institute for the COVID-19 Genomics UK (COG-UK) Consortium | Lighthouse Laboratory Plymouth and Alex Alderton, Roberto Amato, Jeffrey Barrett, Sonia Goncalves, Ewan Harrison, David K. Jackson, Ian Johnston, Dominic Kwiatkowski, Cordelia Langford, John Sillitoe on behalf of the Wellcome Sanger Institute COVID-19 Surveillance Team |
| EPI_ISL_3172425 | Lighthouse Lab in Milton Keynes | Wellcome Sanger Institute for the COVID-19 Genomics UK (COG-UK) Consortium | The Lighthouse Lab in Milton Keynes and Alex Alderton, Roberto Amato, Jeffrey Barrett, Sonia Goncalves, Ewan Harrison, David K. Jackson, Ian Johnston, Dominic Kwiatkowski, Cordelia Langford, John Sillitoe on behalf of the Wellcome Sanger Institute COVID-19 Surveillance Team |
| EPI_ISL_3172426 | Berkshire and Surrey Pathology Services Lighthouse Laboratory | Wellcome Sanger Institute for the COVID-19 Genomics UK (COG-UK) Consortium | Berkshire and Surrey Pathology Services Lighthouse Laboratory and Alex Alderton, Roberto Amato, Jeffrey Barrett, Sonia Goncalves, Ewan Harrison, David K. Jackson, Ian Johnston, Dominic Kwiatkowski, Cordelia Langford, John Sillitoe on behalf of the Wellcome Sanger Institute COVID-19 Surveillance Team |
| EPI_ISL_3172427, EPI_ISL_3172428 | Lighthouse Laboratory Plymouth | Wellcome Sanger Institute for the COVID-19 Genomics UK (COG-UK) Consortium | Lighthouse Laboratory Plymouth and Alex Alderton, Roberto Amato, Jeffrey Barrett, Sonia Goncalves, Ewan Harrison, David K. Jackson, Ian Johnston, Dominic Kwiatkowski, Cordelia Langford, John Sillitoe on behalf of the Wellcome Sanger Institute COVID-19 Surveillance Team |
| EPI_ISL_3172429 | Lighthouse Lab in Milton Keynes | Wellcome Sanger Institute for the COVID-19 Genomics UK (COG-UK) Consortium | The Lighthouse Lab in Milton Keynes and Alex Alderton, Roberto Amato, Jeffrey Barrett, Sonia Goncalves, Ewan Harrison, David K. Jackson, Ian Johnston, Dominic Kwiatkowski, Cordelia Langford, John Sillitoe on behalf of the Wellcome Sanger Institute COVID-19 Surveillance Team |
| EPI_ISL_3172430 | Lighthouse Laboratory Plymouth | Wellcome Sanger Institute for the COVID-19 Genomics UK (COG-UK) Consortium | Lighthouse Laboratory Plymouth and Alex Alderton, Roberto Amato, Jeffrey Barrett, Sonia Goncalves, Ewan Harrison, David K. Jackson, Ian Johnston, Dominic Kwiatkowski, Cordelia Langford, John Sillitoe on behalf of the Wellcome Sanger Institute COVID-19 Surveillance Team |
| EPI_ISL_3172431, EPI_ISL_3172432, EPI_ISL_3172433, EPI_ISL_3172434, EPI_ISL_3172435, EPI_ISL_3172436, EPI_ISL_3172437, EPI_ISL_3172438 | Lighthouse Lab in Milton Keynes | Wellcome Sanger Institute for the COVID-19 Genomics UK (COG-UK) Consortium | The Lighthouse Lab in Milton Keynes and Alex Alderton, Roberto Amato, Jeffrey Barrett, Sonia Goncalves, Ewan Harrison, David K. Jackson, Ian Johnston, Dominic Kwiatkowski, Cordelia Langford, John Sillitoe on behalf of the Wellcome Sanger Institute COVID-19 Surveillance Team |
| EPI_ISL_3172439 | Berkshire and Surrey Pathology Services Lighthouse Laboratory | Wellcome Sanger Institute for the COVID-19 Genomics UK (COG-UK) Consortium | Berkshire and Surrey Pathology Services Lighthouse Laboratory and Alex Alderton, Roberto Amato, Jeffrey Barrett, Sonia Goncalves, Ewan Harrison, David K. Jackson, Ian Johnston, Dominic Kwiatkowski, Cordelia Langford, John Sillitoe on behalf of the Wellcome Sanger Institute COVID-19 Surveillance Team |
| EPI_ISL_3172440 | Lighthouse Laboratory Plymouth | Wellcome Sanger Institute for the COVID-19 Genomics UK (COG-UK) Consortium | Lighthouse Laboratory Plymouth and Alex Alderton, Roberto Amato, Jeffrey Barrett, Sonia Goncalves, Ewan Harrison, David K. Jackson, Ian Johnston, Dominic Kwiatkowski, Cordelia Langford, John Sillitoe on behalf of the Wellcome Sanger Institute COVID-19 Surveillance Team |
| EPI_ISL_3172441 | Lighthouse Lab in Milton Keynes | Wellcome Sanger Institute for the COVID-19 Genomics UK (COG-UK) Consortium | The Lighthouse Lab in Milton Keynes and Alex Alderton, Roberto Amato, Jeffrey Barrett, Sonia Goncalves, Ewan Harrison, David K. Jackson, Ian Johnston, Dominic Kwiatkowski, Cordelia Langford, John Sillitoe on behalf of the Wellcome Sanger Institute COVID-19 Surveillance Team |
| EPI_ISL_3172442 | Lighthouse Laboratory Plymouth | Wellcome Sanger Institute for the COVID-19 Genomics UK (COG-UK) Consortium | Lighthouse Laboratory Plymouth and Alex Alderton, Roberto Amato, Jeffrey Barrett, Sonia Goncalves, Ewan Harrison, David K. Jackson, Ian Johnston, Dominic Kwiatkowski, Cordelia Langford, John Sillitoe on behalf of the Wellcome Sanger Institute COVID-19 Surveillance Team |
| EPI_ISL_3172443 | Lighthouse Lab in Milton Keynes | Wellcome Sanger Institute for the COVID-19 Genomics UK (COG-UK) Consortium | The Lighthouse Lab in Milton Keynes and Alex Alderton, Roberto Amato, Jeffrey Barrett, Sonia Goncalves, Ewan Harrison, David K. Jackson, Ian Johnston, Dominic Kwiatkowski, Cordelia Langford, John Sillitoe on behalf of the Wellcome Sanger Institute COVID-19 Surveillance Team |
| EPI_ISL_3172444 | Lighthouse Laboratory Plymouth | Wellcome Sanger Institute for the COVID-19 Genomics UK (COG-UK) Consortium | Lighthouse Laboratory Plymouth and Alex Alderton, Roberto Amato, Jeffrey Barrett, Sonia Goncalves, Ewan Harrison, David K. Jackson, Ian Johnston, Dominic Kwiatkowski, Cordelia Langford, John Sillitoe on behalf of the Wellcome Sanger Institute COVID-19 Surveillance Team |
| EPI_ISL_3172445 | Lighthouse Lab in Milton Keynes | Wellcome Sanger Institute for the COVID-19 Genomics UK (COG-UK) Consortium | The Lighthouse Lab in Milton Keynes and Alex Alderton, Roberto Amato, Jeffrey Barrett, Sonia Goncalves, Ewan Harrison, David K. Jackson, Ian Johnston, Dominic Kwiatkowski, Cordelia Langford, John Sillitoe on behalf of the Wellcome Sanger Institute COVID-19 Surveillance Team |
| EPI_ISL_3172446 | Lighthouse Laboratory Plymouth | Wellcome Sanger Institute for the COVID-19 Genomics UK (COG-UK) Consortium | Lighthouse Laboratory Plymouth and Alex Alderton, Roberto Amato, Jeffrey Barrett, Sonia Goncalves, Ewan Harrison, David K. Jackson, Ian Johnston, Dominic Kwiatkowski, Cordelia Langford, John Sillitoe on behalf of the Wellcome Sanger Institute COVID-19 Surveillance Team |
| EPI_ISL_3172447 | Berkshire and Surrey Pathology Services Lighthouse Laboratory | Wellcome Sanger Institute for the COVID-19 Genomics UK (COG-UK) Consortium | Berkshire and Surrey Pathology Services Lighthouse Laboratory and Alex Alderton, Roberto Amato, Jeffrey Barrett, Sonia Goncalves, Ewan Harrison, David K. Jackson, Ian Johnston, Dominic Kwiatkowski, Cordelia Langford, John Sillitoe on behalf of the Wellcome Sanger Institute COVID-19 Surveillance Team |
| EPI_ISL_3172448, EPI_ISL_3172449 | Lighthouse Laboratory Plymouth | Wellcome Sanger Institute for the COVID-19 Genomics UK (COG-UK) Consortium | Lighthouse Laboratory Plymouth and Alex Alderton, Roberto Amato, Jeffrey Barrett, Sonia Goncalves, Ewan Harrison, David K. Jackson, Ian Johnston, Dominic Kwiatkowski, Cordelia Langford, John Sillitoe on behalf of the Wellcome Sanger Institute COVID-19 Surveillance Team |
| EPI_ISL_3172450 | Lighthouse Lab in Milton Keynes | Wellcome Sanger Institute for the COVID-19 Genomics UK (COG-UK) Consortium | The Lighthouse Lab in Milton Keynes and Alex Alderton, Roberto Amato, Jeffrey Barrett, Sonia Goncalves, Ewan Harrison, David K. Jackson, Ian Johnston, Dominic Kwiatkowski, Cordelia Langford, John Sillitoe on behalf of the Wellcome Sanger Institute COVID-19 Surveillance Team |
| EPI_ISL_3172451, EPI_ISL_3172452, EPI_ISL_3172453 | Berkshire and Surrey Pathology Services Lighthouse Laboratory | Wellcome Sanger Institute for the COVID-19 Genomics UK (COG-UK) Consortium | Berkshire and Surrey Pathology Services Lighthouse Laboratory and Alex Alderton, Roberto Amato, Jeffrey Barrett, Sonia Goncalves, Ewan Harrison, David K. Jackson, Ian Johnston, Dominic Kwiatkowski, Cordelia Langford, John Sillitoe on behalf of the Wellcome Sanger Institute COVID-19 Surveillance Team |
| EPI_ISL_3172454 | Lighthouse Lab in Milton Keynes | Wellcome Sanger Institute for the COVID-19 Genomics UK (COG-UK) Consortium | The Lighthouse Lab in Milton Keynes and Alex Alderton, Roberto Amato, Jeffrey Barrett, Sonia Goncalves, Ewan Harrison, David K. Jackson, Ian Johnston, Dominic Kwiatkowski, Cordelia Langford, John Sillitoe on behalf of the Wellcome Sanger Institute COVID-19 Surveillance Team |
| EPI_ISL_3172455 | Berkshire and Surrey Pathology Services Lighthouse Laboratory | Wellcome Sanger Institute for the COVID-19 Genomics UK (COG-UK) Consortium | Berkshire and Surrey Pathology Services Lighthouse Laboratory and Alex Alderton, Roberto Amato, Jeffrey Barrett, Sonia Goncalves, Ewan Harrison, David K. Jackson, Ian Johnston, Dominic Kwiatkowski, Cordelia Langford, John Sillitoe on behalf of the Wellcome Sanger Institute COVID-19 Surveillance Team |
| EPI_ISL_3172456 | Lighthouse Laboratory Plymouth | Wellcome Sanger Institute for the COVID-19 Genomics UK (COG-UK) Consortium | Lighthouse Laboratory Plymouth and Alex Alderton, Roberto Amato, Jeffrey Barrett, Sonia Goncalves, Ewan Harrison, David K. Jackson, Ian Johnston, Dominic Kwiatkowski, Cordelia Langford, John Sillitoe on behalf of the Wellcome Sanger Institute COVID-19 Surveillance Team |
| EPI_ISL_3172457 | Berkshire and Surrey Pathology Services Lighthouse Laboratory | Wellcome Sanger Institute for the COVID-19 Genomics UK (COG-UK) Consortium | Berkshire and Surrey Pathology Services Lighthouse Laboratory and Alex Alderton, Roberto Amato, Jeffrey Barrett, Sonia Goncalves, Ewan Harrison, David K. Jackson, Ian Johnston, Dominic Kwiatkowski, Cordelia Langford, John Sillitoe on behalf of the Wellcome Sanger Institute COVID-19 Surveillance Team |
| EPI_ISL_3172458 | Lighthouse Lab in Milton Keynes | Wellcome Sanger Institute for the COVID-19 Genomics UK (COG-UK) Consortium | The Lighthouse Lab in Milton Keynes and Alex Alderton, Roberto Amato, Jeffrey Barrett, Sonia Goncalves, Ewan Harrison, David K. Jackson, Ian Johnston, Dominic Kwiatkowski, Cordelia Langford, John Sillitoe on behalf of the Wellcome Sanger Institute COVID-19 Surveillance Team |
| EPI_ISL_3172459 | Lighthouse Laboratory Plymouth | Wellcome Sanger Institute for the COVID-19 Genomics UK (COG-UK) Consortium | Lighthouse Laboratory Plymouth and Alex Alderton, Roberto Amato, Jeffrey Barrett, Sonia Goncalves, Ewan Harrison, David K. Jackson, Ian Johnston, Dominic Kwiatkowski, Cordelia Langford, John Sillitoe on behalf of the Wellcome Sanger Institute COVID-19 Surveillance Team |
| EPI_ISL_3172460 | Lighthouse Lab in Milton Keynes | Wellcome Sanger Institute for the COVID-19 Genomics UK (COG-UK) Consortium | The Lighthouse Lab in Milton Keynes and Alex Alderton, Roberto Amato, Jeffrey Barrett, Sonia Goncalves, Ewan Harrison, David K. Jackson, Ian Johnston, Dominic Kwiatkowski, Cordelia Langford, John Sillitoe on behalf of the Wellcome Sanger Institute COVID-19 Surveillance Team |
| EPI_ISL_3172461, EPI_ISL_3172462 | Lighthouse Laboratory Plymouth | Wellcome Sanger Institute for the COVID-19 Genomics UK (COG-UK) Consortium | Lighthouse Laboratory Plymouth and Alex Alderton, Roberto Amato, Jeffrey Barrett, Sonia Goncalves, Ewan Harrison, David K. Jackson, Ian Johnston, Dominic Kwiatkowski, Cordelia Langford, John Sillitoe on behalf of the Wellcome Sanger Institute COVID-19 Surveillance Team |
| EPI_ISL_3172463 | Berkshire and Surrey Pathology Services Lighthouse Laboratory | Wellcome Sanger Institute for the COVID-19 Genomics UK (COG-UK) Consortium | Berkshire and Surrey Pathology Services Lighthouse Laboratory and Alex Alderton, Roberto Amato, Jeffrey Barrett, Sonia Goncalves, Ewan Harrison, David K. Jackson, Ian Johnston, Dominic Kwiatkowski, Cordelia Langford, John Sillitoe on behalf of the Wellcome Sanger Institute COVID-19 Surveillance Team |
| EPI_ISL_3172464 | Lighthouse Lab in Milton Keynes | Wellcome Sanger Institute for the COVID-19 Genomics UK (COG-UK) Consortium | The Lighthouse Lab in Milton Keynes and Alex Alderton, Roberto Amato, Jeffrey Barrett, Sonia Goncalves, Ewan Harrison, David K. Jackson, Ian Johnston, Dominic Kwiatkowski, Cordelia Langford, John Sillitoe on behalf of the Wellcome Sanger Institute COVID-19 Surveillance Team |
| EPI_ISL_3172465 | Lighthouse Laboratory Plymouth | Wellcome Sanger Institute for the COVID-19 Genomics UK (COG-UK) Consortium | Lighthouse Laboratory Plymouth and Alex Alderton, Roberto Amato, Jeffrey Barrett, Sonia Goncalves, Ewan Harrison, David K. Jackson, Ian Johnston, Dominic Kwiatkowski, Cordelia Langford, John Sillitoe on behalf of the Wellcome Sanger Institute COVID-19 Surveillance Team |
| EPI_ISL_3172466, EPI_ISL_3172467 | Lighthouse Lab in Milton Keynes | Wellcome Sanger Institute for the COVID-19 Genomics UK (COG-UK) Consortium | The Lighthouse Lab in Milton Keynes and Alex Alderton, Roberto Amato, Jeffrey Barrett, Sonia Goncalves, Ewan Harrison, David K. Jackson, Ian Johnston, Dominic Kwiatkowski, Cordelia Langford, John Sillitoe on behalf of the Wellcome Sanger Institute COVID-19 Surveillance Team |
| EPI_ISL_3172468 | Lighthouse Laboratory Plymouth | Wellcome Sanger Institute for the COVID-19 Genomics UK (COG-UK) Consortium | Lighthouse Laboratory Plymouth and Alex Alderton, Roberto Amato, Jeffrey Barrett, Sonia Goncalves, Ewan Harrison, David K. Jackson, Ian Johnston, Dominic Kwiatkowski, Cordelia Langford, John Sillitoe on behalf of the Wellcome Sanger Institute COVID-19 Surveillance Team |
| EPI_ISL_3172469 | Berkshire and Surrey Pathology Services Lighthouse Laboratory | Wellcome Sanger Institute for the COVID-19 Genomics UK (COG-UK) Consortium | Berkshire and Surrey Pathology Services Lighthouse Laboratory and Alex Alderton, Roberto Amato, Jeffrey Barrett, Sonia Goncalves, Ewan Harrison, David K. Jackson, Ian Johnston, Dominic Kwiatkowski, Cordelia Langford, John Sillitoe on behalf of the Wellcome Sanger Institute COVID-19 Surveillance Team |
| EPI_ISL_3172470 | Lighthouse Laboratory Plymouth | Wellcome Sanger Institute for the COVID-19 Genomics UK (COG-UK) Consortium | Lighthouse Laboratory Plymouth and Alex Alderton, Roberto Amato, Jeffrey Barrett, Sonia Goncalves, Ewan Harrison, David K. Jackson, Ian Johnston, Dominic Kwiatkowski, Cordelia Langford, John Sillitoe on behalf of the Wellcome Sanger Institute COVID-19 Surveillance Team |
| EPI_ISL_3172471, EPI_ISL_3172472 | Lighthouse Lab in Milton Keynes | Wellcome Sanger Institute for the COVID-19 Genomics UK (COG-UK) Consortium | The Lighthouse Lab in Milton Keynes and Alex Alderton, Roberto Amato, Jeffrey Barrett, Sonia Goncalves, Ewan Harrison, David K. Jackson, Ian Johnston, Dominic Kwiatkowski, Cordelia Langford, John Sillitoe on behalf of the Wellcome Sanger Institute COVID-19 Surveillance Team |
| EPI_ISL_3172473 | Lighthouse Laboratory Plymouth | Wellcome Sanger Institute for the COVID-19 Genomics UK (COG-UK) Consortium | Lighthouse Laboratory Plymouth and Alex Alderton, Roberto Amato, Jeffrey Barrett, Sonia Goncalves, Ewan Harrison, David K. Jackson, Ian Johnston, Dominic Kwiatkowski, Cordelia Langford, John Sillitoe on behalf of the Wellcome Sanger Institute COVID-19 Surveillance Team |
| EPI_ISL_3172474, EPI_ISL_3172475 | Lighthouse Lab in Milton Keynes | Wellcome Sanger Institute for the COVID-19 Genomics UK (COG-UK) Consortium | The Lighthouse Lab in Milton Keynes and Alex Alderton, Roberto Amato, Jeffrey Barrett, Sonia Goncalves, Ewan Harrison, David K. Jackson, Ian Johnston, Dominic Kwiatkowski, Cordelia Langford, John Sillitoe on behalf of the Wellcome Sanger Institute COVID-19 Surveillance Team |
| EPI_ISL_3172476 | Berkshire and Surrey Pathology Services Lighthouse Laboratory | Wellcome Sanger Institute for the COVID-19 Genomics UK (COG-UK) Consortium | Berkshire and Surrey Pathology Services Lighthouse Laboratory and Alex Alderton, Roberto Amato, Jeffrey Barrett, Sonia Goncalves, Ewan Harrison, David K. Jackson, Ian Johnston, Dominic Kwiatkowski, Cordelia Langford, John Sillitoe on behalf of the Wellcome Sanger Institute COVID-19 Surveillance Team |
| EPI_ISL_3172477, | Lighthouse Lab in Milton Keynes | Wellcome Sanger Institute for the COVID-19 Genomics UK (COG-UK) | The Lighthouse Lab in Milton Keynes and Alex Alderton, Roberto Amato, Jeffrey Barrett, Sonia Goncalves, Ewan Harrison, David K. Jackson, Ian Johnston, Dominic Kwiatkowski, Cordelia Langford, John Sillitoe on behalf of the Wellcome Sanger |

| | | | |
|---|---|---|---|
| EPI_ISL_3172478 | | Consortium | Institute COVID-19 Surveillance Team |
| EPI_ISL_3172479 | Lighthouse Laboratory Plymouth | Wellcome Sanger Institute for the COVID-19 Genomics UK (COG-UK) Consortium | Lighthouse Laboratory Plymouth and Alex Alderton, Roberto Amato, Jeffrey Barrett, Sonia Goncalves, Ewan Harrison, David K. Jackson, Ian Johnston, Dominic Kwiatkowski, Cordelia Langford, John Sillitoe on behalf of the Wellcome Sanger Institute COVID-19 Surveillance Team |
| EPI_ISL_3172480, EPI_ISL_3172481 | Lighthouse Lab in Milton Keynes | Wellcome Sanger Institute for the COVID-19 Genomics UK (COG-UK) Consortium | The Lighthouse Lab in Milton Keynes and Alex Alderton, Roberto Amato, Jeffrey Barrett, Sonia Goncalves, Ewan Harrison, David K. Jackson, Ian Johnston, Dominic Kwiatkowski, Cordelia Langford, John Sillitoe on behalf of the Wellcome Sanger Institute COVID-19 Surveillance Team |
| EPI_ISL_3172482 | Lighthouse Laboratory Plymouth | Wellcome Sanger Institute for the COVID-19 Genomics UK (COG-UK) Consortium | Lighthouse Laboratory Plymouth and Alex Alderton, Roberto Amato, Jeffrey Barrett, Sonia Goncalves, Ewan Harrison, David K. Jackson, Ian Johnston, Dominic Kwiatkowski, Cordelia Langford, John Sillitoe on behalf of the Wellcome Sanger Institute COVID-19 Surveillance Team |
| EPI_ISL_3172483, EPI_ISL_3172484 | Berkshire and Surrey Pathology Services Lighthouse Laboratory | Wellcome Sanger Institute for the COVID-19 Genomics UK (COG-UK) Consortium | Berkshire and Surrey Pathology Services Lighthouse Laboratory and Alex Alderton, Roberto Amato, Jeffrey Barrett, Sonia Goncalves, Ewan Harrison, David K. Jackson, Ian Johnston, Dominic Kwiatkowski, Cordelia Langford, John Sillitoe on behalf of the Wellcome Sanger Institute COVID-19 Surveillance Team |
| EPI_ISL_3172485 | Lighthouse Lab in Milton Keynes | Wellcome Sanger Institute for the COVID-19 Genomics UK (COG-UK) Consortium | The Lighthouse Lab in Milton Keynes and Alex Alderton, Roberto Amato, Jeffrey Barrett, Sonia Goncalves, Ewan Harrison, David K. Jackson, Ian Johnston, Dominic Kwiatkowski, Cordelia Langford, John Sillitoe on behalf of the Wellcome Sanger Institute COVID-19 Surveillance Team |
| EPI_ISL_3172486 | Lighthouse Laboratory Plymouth | Wellcome Sanger Institute for the COVID-19 Genomics UK (COG-UK) Consortium | Lighthouse Laboratory Plymouth and Alex Alderton, Roberto Amato, Jeffrey Barrett, Sonia Goncalves, Ewan Harrison, David K. Jackson, Ian Johnston, Dominic Kwiatkowski, Cordelia Langford, John Sillitoe on behalf of the Wellcome Sanger Institute COVID-19 Surveillance Team |
| EPI_ISL_3172487 | Berkshire and Surrey Pathology Services Lighthouse Laboratory | Wellcome Sanger Institute for the COVID-19 Genomics UK (COG-UK) Consortium | Berkshire and Surrey Pathology Services Lighthouse Laboratory and Alex Alderton, Roberto Amato, Jeffrey Barrett, Sonia Goncalves, Ewan Harrison, David K. Jackson, Ian Johnston, Dominic Kwiatkowski, Cordelia Langford, John Sillitoe on behalf of the Wellcome Sanger Institute COVID-19 Surveillance Team |
| EPI_ISL_3172488, EPI_ISL_3172489 | Lighthouse Lab in Milton Keynes | Wellcome Sanger Institute for the COVID-19 Genomics UK (COG-UK) Consortium | The Lighthouse Lab in Milton Keynes and Alex Alderton, Roberto Amato, Jeffrey Barrett, Sonia Goncalves, Ewan Harrison, David K. Jackson, Ian Johnston, Dominic Kwiatkowski, Cordelia Langford, John Sillitoe on behalf of the Wellcome Sanger Institute COVID-19 Surveillance Team |
| EPI_ISL_3172490 | Berkshire and Surrey Pathology Services Lighthouse Laboratory | Wellcome Sanger Institute for the COVID-19 Genomics UK (COG-UK) Consortium | Berkshire and Surrey Pathology Services Lighthouse Laboratory and Alex Alderton, Roberto Amato, Jeffrey Barrett, Sonia Goncalves, Ewan Harrison, David K. Jackson, Ian Johnston, Dominic Kwiatkowski, Cordelia Langford, John Sillitoe on behalf of the Wellcome Sanger Institute COVID-19 Surveillance Team |
| EPI_ISL_3172491, EPI_ISL_3172492 | Lighthouse Lab in Milton Keynes | Wellcome Sanger Institute for the COVID-19 Genomics UK (COG-UK) Consortium | The Lighthouse Lab in Milton Keynes and Alex Alderton, Roberto Amato, Jeffrey Barrett, Sonia Goncalves, Ewan Harrison, David K. Jackson, Ian Johnston, Dominic Kwiatkowski, Cordelia Langford, John Sillitoe on behalf of the Wellcome Sanger Institute COVID-19 Surveillance Team |
| EPI_ISL_3172493 | Lighthouse Laboratory Plymouth | Wellcome Sanger Institute for the COVID-19 Genomics UK (COG-UK) Consortium | Lighthouse Laboratory Plymouth and Alex Alderton, Roberto Amato, Jeffrey Barrett, Sonia Goncalves, Ewan Harrison, David K. Jackson, Ian Johnston, Dominic Kwiatkowski, Cordelia Langford, John Sillitoe on behalf of the Wellcome Sanger Institute COVID-19 Surveillance Team |
| EPI_ISL_3172494 | Lighthouse Lab in Milton Keynes | Wellcome Sanger Institute for the COVID-19 Genomics UK (COG-UK) Consortium | The Lighthouse Lab in Milton Keynes and Alex Alderton, Roberto Amato, Jeffrey Barrett, Sonia Goncalves, Ewan Harrison, David K. Jackson, Ian Johnston, Dominic Kwiatkowski, Cordelia Langford, John Sillitoe on behalf of the Wellcome Sanger Institute COVID-19 Surveillance Team |
| EPI_ISL_3172495 | Lighthouse Laboratory Plymouth | Wellcome Sanger Institute for the COVID-19 Genomics UK (COG-UK) Consortium | Lighthouse Laboratory Plymouth and Alex Alderton, Roberto Amato, Jeffrey Barrett, Sonia Goncalves, Ewan Harrison, David K. Jackson, Ian Johnston, Dominic Kwiatkowski, Cordelia Langford, John Sillitoe on behalf of the Wellcome Sanger Institute COVID-19 Surveillance Team |
| EPI_ISL_3172496, EPI_ISL_3172497 | Lighthouse Lab in Milton Keynes | Wellcome Sanger Institute for the COVID-19 Genomics UK (COG-UK) Consortium | The Lighthouse Lab in Milton Keynes and Alex Alderton, Roberto Amato, Jeffrey Barrett, Sonia Goncalves, Ewan Harrison, David K. Jackson, Ian Johnston, Dominic Kwiatkowski, Cordelia Langford, John Sillitoe on behalf of the Wellcome Sanger Institute COVID-19 Surveillance Team |
| EPI_ISL_3172498 | Lighthouse Laboratory Plymouth | Wellcome Sanger Institute for the COVID-19 Genomics UK (COG-UK) Consortium | Lighthouse Laboratory Plymouth and Alex Alderton, Roberto Amato, Jeffrey Barrett, Sonia Goncalves, Ewan Harrison, David K. Jackson, Ian Johnston, Dominic Kwiatkowski, Cordelia Langford, John Sillitoe on behalf of the Wellcome Sanger Institute COVID-19 Surveillance Team |
| EPI_ISL_3172499 | Berkshire and Surrey Pathology Services Lighthouse Laboratory | Wellcome Sanger Institute for the COVID-19 Genomics UK (COG-UK) Consortium | Berkshire and Surrey Pathology Services Lighthouse Laboratory and Alex Alderton, Roberto Amato, Jeffrey Barrett, Sonia Goncalves, Ewan Harrison, David K. Jackson, Ian Johnston, Dominic Kwiatkowski, Cordelia Langford, John Sillitoe on behalf of the Wellcome Sanger Institute COVID-19 Surveillance Team |
| EPI_ISL_3172500, EPI_ISL_3172501, EPI_ISL_3172502 | Lighthouse Lab in Milton Keynes | Wellcome Sanger Institute for the COVID-19 Genomics UK (COG-UK) Consortium | The Lighthouse Lab in Milton Keynes and Alex Alderton, Roberto Amato, Jeffrey Barrett, Sonia Goncalves, Ewan Harrison, David K. Jackson, Ian Johnston, Dominic Kwiatkowski, Cordelia Langford, John Sillitoe on behalf of the Wellcome Sanger Institute COVID-19 Surveillance Team |
| EPI_ISL_3172503 | Berkshire and Surrey Pathology Services Lighthouse Laboratory | Wellcome Sanger Institute for the COVID-19 Genomics UK (COG-UK) Consortium | Berkshire and Surrey Pathology Services Lighthouse Laboratory and Alex Alderton, Roberto Amato, Jeffrey Barrett, Sonia Goncalves, Ewan Harrison, David K. Jackson, Ian Johnston, Dominic Kwiatkowski, Cordelia Langford, John Sillitoe on behalf of the Wellcome Sanger Institute COVID-19 Surveillance Team |
| EPI_ISL_3172504 | Lighthouse Lab in Milton Keynes | Wellcome Sanger Institute for the COVID-19 Genomics UK (COG-UK) Consortium | The Lighthouse Lab in Milton Keynes and Alex Alderton, Roberto Amato, Jeffrey Barrett, Sonia Goncalves, Ewan Harrison, David K. Jackson, Ian Johnston, Dominic Kwiatkowski, Cordelia Langford, John Sillitoe on behalf of the Wellcome Sanger Institute COVID-19 Surveillance Team |
| EPI_ISL_3172505 | Lighthouse Laboratory Plymouth | Wellcome Sanger Institute for the COVID-19 Genomics UK (COG-UK) Consortium | Lighthouse Laboratory Plymouth and Alex Alderton, Roberto Amato, Jeffrey Barrett, Sonia Goncalves, Ewan Harrison, David K. Jackson, Ian Johnston, Dominic Kwiatkowski, Cordelia Langford, John Sillitoe on behalf of the Wellcome Sanger Institute COVID-19 Surveillance Team |
| EPI_ISL_3172506, EPI_ISL_3172507, EPI_ISL_3172508 | Lighthouse Lab in Milton Keynes | Wellcome Sanger Institute for the COVID-19 Genomics UK (COG-UK) Consortium | The Lighthouse Lab in Milton Keynes and Alex Alderton, Roberto Amato, Jeffrey Barrett, Sonia Goncalves, Ewan Harrison, David K. Jackson, Ian Johnston, Dominic Kwiatkowski, Cordelia Langford, John Sillitoe on behalf of the Wellcome Sanger Institute COVID-19 Surveillance Team |
| EPI_ISL_3172509 | Berkshire and Surrey Pathology Services Lighthouse Laboratory | Wellcome Sanger Institute for the COVID-19 Genomics UK (COG-UK) Consortium | Berkshire and Surrey Pathology Services Lighthouse Laboratory and Alex Alderton, Roberto Amato, Jeffrey Barrett, Sonia Goncalves, Ewan Harrison, David K. Jackson, Ian Johnston, Dominic Kwiatkowski, Cordelia Langford, John Sillitoe on behalf of the Wellcome Sanger Institute COVID-19 Surveillance Team |
| EPI_ISL_3172510 | Lighthouse Laboratory Plymouth | Wellcome Sanger Institute for the COVID-19 Genomics UK (COG-UK) Consortium | Lighthouse Laboratory Plymouth and Alex Alderton, Roberto Amato, Jeffrey Barrett, Sonia Goncalves, Ewan Harrison, David K. Jackson, Ian Johnston, Dominic Kwiatkowski, Cordelia Langford, John Sillitoe on behalf of the Wellcome Sanger Institute COVID-19 Surveillance Team |
| EPI_ISL_3172511, EPI_ISL_3172512 | Lighthouse Lab in Milton Keynes | Wellcome Sanger Institute for the COVID-19 Genomics UK (COG-UK) Consortium | The Lighthouse Lab in Milton Keynes and Alex Alderton, Roberto Amato, Jeffrey Barrett, Sonia Goncalves, Ewan Harrison, David K. Jackson, Ian Johnston, Dominic Kwiatkowski, Cordelia Langford, John Sillitoe on behalf of the Wellcome Sanger Institute COVID-19 Surveillance Team |
| EPI_ISL_3172513 | Lighthouse Laboratory Plymouth | Wellcome Sanger Institute for the COVID-19 Genomics UK (COG-UK) Consortium | Lighthouse Laboratory Plymouth and Alex Alderton, Roberto Amato, Jeffrey Barrett, Sonia Goncalves, Ewan Harrison, David K. Jackson, Ian Johnston, Dominic Kwiatkowski, Cordelia Langford, John Sillitoe on behalf of the Wellcome Sanger Institute COVID-19 Surveillance Team |
| EPI_ISL_3172514 | Lighthouse Lab in Milton Keynes | Wellcome Sanger Institute for the COVID-19 Genomics UK (COG-UK) Consortium | The Lighthouse Lab in Milton Keynes and Alex Alderton, Roberto Amato, Jeffrey Barrett, Sonia Goncalves, Ewan Harrison, David K. Jackson, Ian Johnston, Dominic Kwiatkowski, Cordelia Langford, John Sillitoe on behalf of the Wellcome Sanger Institute COVID-19 Surveillance Team |
| EPI_ISL_3172515, EPI_ISL_3172516 | Berkshire and Surrey Pathology Services Lighthouse Laboratory | Wellcome Sanger Institute for the COVID-19 Genomics UK (COG-UK) Consortium | Berkshire and Surrey Pathology Services Lighthouse Laboratory and Alex Alderton, Roberto Amato, Jeffrey Barrett, Sonia Goncalves, Ewan Harrison, David K. Jackson, Ian Johnston, Dominic Kwiatkowski, Cordelia Langford, John Sillitoe on behalf of the Wellcome Sanger Institute COVID-19 Surveillance Team |
| EPI_ISL_3172517 | Lighthouse Lab in Alderley Park | Wellcome Sanger Institute for the COVID-19 Genomics UK (COG-UK) Consortium | Jacquelyn Wynn, Mairead Hyland, The Lighthouse Lab in Alderley Park and Alex Alderton, Roberto Amato, Jeffrey Barrett, Sonia Goncalves, Ewan Harrison, David K. Jackson, Ian Johnston, Dominic Kwiatkowski, Cordelia Langford, John Sillitoe on behalf of the Wellcome Sanger Institute COVID-19 Surveillance Team |
| EPI_ISL_3172518, EPI_ISL_3172519 | Berkshire and Surrey Pathology Services Lighthouse Laboratory | Wellcome Sanger Institute for the COVID-19 Genomics UK (COG-UK) Consortium | Berkshire and Surrey Pathology Services Lighthouse Laboratory and Alex Alderton, Roberto Amato, Jeffrey Barrett, Sonia Goncalves, Ewan Harrison, David K. Jackson, Ian Johnston, Dominic Kwiatkowski, Cordelia Langford, John Sillitoe on behalf of the Wellcome Sanger Institute COVID-19 Surveillance Team |
| EPI_ISL_3172520 | Lighthouse Lab in Glasgow | Wellcome Sanger Institute for the COVID-19 Genomics UK (COG-UK) Consortium | Harper VanSteenhouse, Yumi Kasai, David Gray, Carol Clugston, Anna Dominiczak and Alex Alderton, Roberto Amato, Jeffrey Barrett, Sonia Goncalves, Ewan Harrison, David K. Jackson, Ian Johnston, Dominic Kwiatkowski, Cordelia Langford, John Sillitoe on behalf of the Wellcome Sanger Institute COVID-19 Surveillance Team |
| EPI_ISL_3172521, EPI_ISL_3172522 | Berkshire and Surrey Pathology Services Lighthouse Laboratory | Wellcome Sanger Institute for the COVID-19 Genomics UK (COG-UK) Consortium | Berkshire and Surrey Pathology Services Lighthouse Laboratory and Alex Alderton, Roberto Amato, Jeffrey Barrett, Sonia Goncalves, Ewan Harrison, David K. Jackson, Ian Johnston, Dominic Kwiatkowski, Cordelia Langford, John Sillitoe on behalf of the Wellcome Sanger Institute COVID-19 Surveillance Team |
| EPI_ISL_3172523 | Lighthouse Lab in Glasgow | Wellcome Sanger Institute for the COVID-19 Genomics UK (COG-UK) Consortium | Harper VanSteenhouse, Yumi Kasai, David Gray, Carol Clugston, Anna Dominiczak and Alex Alderton, Roberto Amato, Jeffrey Barrett, Sonia Goncalves, Ewan Harrison, David K. Jackson, Ian Johnston, Dominic Kwiatkowski, Cordelia Langford, John Sillitoe on behalf of the Wellcome Sanger Institute COVID-19 Surveillance Team |
| EPI_ISL_3172524, EPI_ISL_3172525 | Lighthouse Lab in Alderley Park | Wellcome Sanger Institute for the COVID-19 Genomics UK (COG-UK) Consortium | Jacquelyn Wynn, Mairead Hyland, The Lighthouse Lab in Alderley Park and Alex Alderton, Roberto Amato, Jeffrey Barrett, Sonia Goncalves, Ewan Harrison, David K. Jackson, Ian Johnston, Dominic Kwiatkowski, Cordelia Langford, John Sillitoe on behalf of the Wellcome Sanger Institute COVID-19 Surveillance Team |
| EPI_ISL_3172526, EPI_ISL_3172527 | Lighthouse Lab in Glasgow | Wellcome Sanger Institute for the COVID-19 Genomics UK (COG-UK) Consortium | Harper VanSteenhouse, Yumi Kasai, David Gray, Carol Clugston, Anna Dominiczak and Alex Alderton, Roberto Amato, Jeffrey Barrett, Sonia Goncalves, Ewan Harrison, David K. Jackson, Ian Johnston, Dominic Kwiatkowski, Cordelia Langford, John Sillitoe on behalf of the Wellcome Sanger Institute COVID-19 Surveillance Team |
| EPI_ISL_3172528 | Berkshire and Surrey Pathology Services Lighthouse Laboratory | Wellcome Sanger Institute for the COVID-19 Genomics UK (COG-UK) Consortium | Berkshire and Surrey Pathology Services Lighthouse Laboratory and Alex Alderton, Roberto Amato, Jeffrey Barrett, Sonia Goncalves, Ewan Harrison, David K. Jackson, Ian Johnston, Dominic Kwiatkowski, Cordelia Langford, John Sillitoe on behalf of the Wellcome Sanger Institute COVID-19 Surveillance Team |
| EPI_ISL_3172529 | Lighthouse Lab in Glasgow | Wellcome Sanger Institute for the COVID-19 Genomics UK (COG-UK) Consortium | Harper VanSteenhouse, Yumi Kasai, David Gray, Carol Clugston, Anna Dominiczak and Alex Alderton, Roberto Amato, Jeffrey Barrett, Sonia Goncalves, Ewan Harrison, David K. Jackson, Ian Johnston, Dominic Kwiatkowski, Cordelia Langford, John Sillitoe on behalf of the Wellcome Sanger Institute COVID-19 Surveillance Team |
| EPI_ISL_3172530 | Lighthouse Lab in Alderley Park | Wellcome Sanger Institute for the COVID-19 Genomics UK (COG-UK) Consortium | Jacquelyn Wynn, Mairead Hyland, The Lighthouse Lab in Alderley Park and Alex Alderton, Roberto Amato, Jeffrey Barrett, Sonia Goncalves, Ewan Harrison, David K. Jackson, Ian Johnston, Dominic Kwiatkowski, Cordelia Langford, John Sillitoe on behalf of the Wellcome Sanger Institute COVID-19 Surveillance Team |
| EPI_ISL_3172531 | Lighthouse Lab in Glasgow | Wellcome Sanger Institute for the COVID-19 Genomics UK (COG-UK) Consortium | Harper VanSteenhouse, Yumi Kasai, David Gray, Carol Clugston, Anna Dominiczak and Alex Alderton, Roberto Amato, Jeffrey Barrett, Sonia Goncalves, Ewan Harrison, David K. Jackson, Ian Johnston, Dominic Kwiatkowski, Cordelia Langford, John Sillitoe on behalf of the Wellcome Sanger Institute COVID-19 Surveillance Team |
| EPI_ISL_3172532, EPI_ISL_3172533 | Lighthouse Lab in Alderley Park | Wellcome Sanger Institute for the COVID-19 Genomics UK (COG-UK) Consortium | Jacquelyn Wynn, Mairead Hyland, The Lighthouse Lab in Alderley Park and Alex Alderton, Roberto Amato, Jeffrey Barrett, Sonia Goncalves, Ewan Harrison, David K. Jackson, Ian Johnston, Dominic Kwiatkowski, Cordelia Langford, John Sillitoe on behalf of the Wellcome Sanger Institute COVID-19 Surveillance Team |
| EPI_ISL_3172534 | Berkshire and Surrey Pathology Services Lighthouse Laboratory | Wellcome Sanger Institute for the COVID-19 Genomics UK (COG-UK) Consortium | Berkshire and Surrey Pathology Services Lighthouse Laboratory and Alex Alderton, Roberto Amato, Jeffrey Barrett, Sonia Goncalves, Ewan Harrison, David K. Jackson, Ian Johnston, Dominic Kwiatkowski, Cordelia Langford, John Sillitoe on behalf of the Wellcome Sanger Institute COVID-19 Surveillance Team |
| EPI_ISL_3172535 | Lighthouse Lab in Alderley Park | Wellcome Sanger Institute for the COVID-19 Genomics UK (COG-UK) Consortium | Jacquelyn Wynn, Mairead Hyland, The Lighthouse Lab in Alderley Park and Alex Alderton, Roberto Amato, Jeffrey Barrett, Sonia Goncalves, Ewan Harrison, David K. Jackson, Ian Johnston, Dominic Kwiatkowski, Cordelia Langford, John Sillitoe on behalf of the Wellcome Sanger Institute COVID-19 Surveillance Team |
| EPI_ISL_3172536 | Lighthouse Lab in Glasgow | Wellcome Sanger Institute for the COVID-19 Genomics UK (COG-UK) Consortium | Harper VanSteenhouse, Yumi Kasai, David Gray, Carol Clugston, Anna Dominiczak and Alex Alderton, Roberto Amato, Jeffrey Barrett, Sonia Goncalves, Ewan Harrison, David K. Jackson, Ian Johnston, Dominic Kwiatkowski, Cordelia Langford, John Sillitoe on behalf of the Wellcome Sanger Institute COVID-19 Surveillance Team |
| EPI_ISL_3172537, EPI_ISL_3172538 | Berkshire and Surrey Pathology Services Lighthouse Laboratory | Wellcome Sanger Institute for the COVID-19 Genomics UK (COG-UK) Consortium | Berkshire and Surrey Pathology Services Lighthouse Laboratory and Alex Alderton, Roberto Amato, Jeffrey Barrett, Sonia Goncalves, Ewan Harrison, David K. Jackson, Ian Johnston, Dominic Kwiatkowski, Cordelia Langford, John Sillitoe on behalf of the Wellcome Sanger Institute COVID-19 Surveillance Team |
| EPI_ISL_3172539, EPI_ISL_3172540, | Lighthouse Lab in Glasgow | Wellcome Sanger Institute for the COVID-19 Genomics UK (COG-UK) Consortium | Harper VanSteenhouse, Yumi Kasai, David Gray, Carol Clugston, Anna Dominiczak and Alex Alderton, Roberto Amato, Jeffrey Barrett, Sonia Goncalves, Ewan Harrison, David K. Jackson, Ian Johnston, Dominic Kwiatkowski, Cordelia Langford, John Sillitoe on behalf of the Wellcome Sanger Institute COVID-19 Surveillance Team |

| | | | |
|---|---|---|---|
| EPI_ISL_3172541, EPI_ISL_3172542, EPI_ISL_3172543 | | | |
| EPI_ISL_3172544 | Berkshire and Surrey Pathology Services Lighthouse Laboratory | Wellcome Sanger Institute for the COVID-19 Genomics UK (COG-UK) Consortium | Berkshire and Surrey Pathology Services Lighthouse Laboratory and Alex Alderton, Roberto Amato, Jeffrey Barrett, Sonia Goncalves, Ewan Harrison, David K. Jackson, Ian Johnston, Dominic Kwiatkowski, Cordelia Langford, John Sillitoe on behalf of the Wellcome Sanger Institute COVID-19 Surveillance Team |
| EPI_ISL_3172545 | Lighthouse Lab in Glasgow | Wellcome Sanger Institute for the COVID-19 Genomics UK (COG-UK) Consortium | Harper VanSteenhouse, Yumi Kasai, David Gray, Carol Clugston, Anna Dominiczak and Alex Alderton, Roberto Amato, Jeffrey Barrett, Sonia Goncalves, Ewan Harrison, David K. Jackson, Ian Johnston, Dominic Kwiatkowski, Cordelia Langford, John Sillitoe on behalf of the Wellcome Sanger Institute COVID-19 Surveillance Team |
| EPI_ISL_3172546, EPI_ISL_3172547, EPI_ISL_3172548, EPI_ISL_3172549, EPI_ISL_3172550 | Lighthouse Lab in Alderley Park | Wellcome Sanger Institute for the COVID-19 Genomics UK (COG-UK) Consortium | Jacquelyn Wynn, Mairead Hyland, The Lighthouse Lab in Alderley Park and Alex Alderton, Roberto Amato, Jeffrey Barrett, Sonia Goncalves, Ewan Harrison, David K. Jackson, Ian Johnston, Dominic Kwiatkowski, Cordelia Langford, John Sillitoe on behalf of the Wellcome Sanger Institute COVID-19 Surveillance Team |
| EPI_ISL_3172551 | Lighthouse Lab in Glasgow | Wellcome Sanger Institute for the COVID-19 Genomics UK (COG-UK) Consortium | Harper VanSteenhouse, Yumi Kasai, David Gray, Carol Clugston, Anna Dominiczak and Alex Alderton, Roberto Amato, Jeffrey Barrett, Sonia Goncalves, Ewan Harrison, David K. Jackson, Ian Johnston, Dominic Kwiatkowski, Cordelia Langford, John Sillitoe on behalf of the Wellcome Sanger Institute COVID-19 Surveillance Team |
| EPI_ISL_3172552 | Lighthouse Lab in Alderley Park | Wellcome Sanger Institute for the COVID-19 Genomics UK (COG-UK) Consortium | Jacquelyn Wynn, Mairead Hyland, The Lighthouse Lab in Alderley Park and Alex Alderton, Roberto Amato, Jeffrey Barrett, Sonia Goncalves, Ewan Harrison, David K. Jackson, Ian Johnston, Dominic Kwiatkowski, Cordelia Langford, John Sillitoe on behalf of the Wellcome Sanger Institute COVID-19 Surveillance Team |
| EPI_ISL_3172553 | Lighthouse Lab in Glasgow | Wellcome Sanger Institute for the COVID-19 Genomics UK (COG-UK) Consortium | Harper VanSteenhouse, Yumi Kasai, David Gray, Carol Clugston, Anna Dominiczak and Alex Alderton, Roberto Amato, Jeffrey Barrett, Sonia Goncalves, Ewan Harrison, David K. Jackson, Ian Johnston, Dominic Kwiatkowski, Cordelia Langford, John Sillitoe on behalf of the Wellcome Sanger Institute COVID-19 Surveillance Team |
| EPI_ISL_3172554 | Berkshire and Surrey Pathology Services Lighthouse Laboratory | Wellcome Sanger Institute for the COVID-19 Genomics UK (COG-UK) Consortium | Berkshire and Surrey Pathology Services Lighthouse Laboratory and Alex Alderton, Roberto Amato, Jeffrey Barrett, Sonia Goncalves, Ewan Harrison, David K. Jackson, Ian Johnston, Dominic Kwiatkowski, Cordelia Langford, John Sillitoe on behalf of the Wellcome Sanger Institute COVID-19 Surveillance Team |
| EPI_ISL_3172555 | Lighthouse Lab in Glasgow | Wellcome Sanger Institute for the COVID-19 Genomics UK (COG-UK) Consortium | Harper VanSteenhouse, Yumi Kasai, David Gray, Carol Clugston, Anna Dominiczak and Alex Alderton, Roberto Amato, Jeffrey Barrett, Sonia Goncalves, Ewan Harrison, David K. Jackson, Ian Johnston, Dominic Kwiatkowski, Cordelia Langford, John Sillitoe on behalf of the Wellcome Sanger Institute COVID-19 Surveillance Team |
| EPI_ISL_3172556 | Berkshire and Surrey Pathology Services Lighthouse Laboratory | Wellcome Sanger Institute for the COVID-19 Genomics UK (COG-UK) Consortium | Berkshire and Surrey Pathology Services Lighthouse Laboratory and Alex Alderton, Roberto Amato, Jeffrey Barrett, Sonia Goncalves, Ewan Harrison, David K. Jackson, Ian Johnston, Dominic Kwiatkowski, Cordelia Langford, John Sillitoe on behalf of the Wellcome Sanger Institute COVID-19 Surveillance Team |
| EPI_ISL_3172557, EPI_ISL_3172558 | Lighthouse Lab in Glasgow | Wellcome Sanger Institute for the COVID-19 Genomics UK (COG-UK) Consortium | Harper VanSteenhouse, Yumi Kasai, David Gray, Carol Clugston, Anna Dominiczak and Alex Alderton, Roberto Amato, Jeffrey Barrett, Sonia Goncalves, Ewan Harrison, David K. Jackson, Ian Johnston, Dominic Kwiatkowski, Cordelia Langford, John Sillitoe on behalf of the Wellcome Sanger Institute COVID-19 Surveillance Team |
| EPI_ISL_3172559 | Berkshire and Surrey Pathology Services Lighthouse Laboratory | Wellcome Sanger Institute for the COVID-19 Genomics UK (COG-UK) Consortium | Berkshire and Surrey Pathology Services Lighthouse Laboratory and Alex Alderton, Roberto Amato, Jeffrey Barrett, Sonia Goncalves, Ewan Harrison, David K. Jackson, Ian Johnston, Dominic Kwiatkowski, Cordelia Langford, John Sillitoe on behalf of the Wellcome Sanger Institute COVID-19 Surveillance Team |
| EPI_ISL_3172560 | Lighthouse Lab in Glasgow | Wellcome Sanger Institute for the COVID-19 Genomics UK (COG-UK) Consortium | Harper VanSteenhouse, Yumi Kasai, David Gray, Carol Clugston, Anna Dominiczak and Alex Alderton, Roberto Amato, Jeffrey Barrett, Sonia Goncalves, Ewan Harrison, David K. Jackson, Ian Johnston, Dominic Kwiatkowski, Cordelia Langford, John Sillitoe on behalf of the Wellcome Sanger Institute COVID-19 Surveillance Team |
| EPI_ISL_3172561 | Berkshire and Surrey Pathology Services Lighthouse Laboratory | Wellcome Sanger Institute for the COVID-19 Genomics UK (COG-UK) Consortium | Berkshire and Surrey Pathology Services Lighthouse Laboratory and Alex Alderton, Roberto Amato, Jeffrey Barrett, Sonia Goncalves, Ewan Harrison, David K. Jackson, Ian Johnston, Dominic Kwiatkowski, Cordelia Langford, John Sillitoe on behalf of the Wellcome Sanger Institute COVID-19 Surveillance Team |
| EPI_ISL_3172562, EPI_ISL_3172563 | Lighthouse Lab in Alderley Park | Wellcome Sanger Institute for the COVID-19 Genomics UK (COG-UK) Consortium | Jacquelyn Wynn, Mairead Hyland, The Lighthouse Lab in Alderley Park and Alex Alderton, Roberto Amato, Jeffrey Barrett, Sonia Goncalves, Ewan Harrison, David K. Jackson, Ian Johnston, Dominic Kwiatkowski, Cordelia Langford, John Sillitoe on behalf of the Wellcome Sanger Institute COVID-19 Surveillance Team |
| EPI_ISL_3172564 | Lighthouse Lab in Glasgow | Wellcome Sanger Institute for the COVID-19 Genomics UK (COG-UK) Consortium | Harper VanSteenhouse, Yumi Kasai, David Gray, Carol Clugston, Anna Dominiczak and Alex Alderton, Roberto Amato, Jeffrey Barrett, Sonia Goncalves, Ewan Harrison, David K. Jackson, Ian Johnston, Dominic Kwiatkowski, Cordelia Langford, John Sillitoe on behalf of the Wellcome Sanger Institute COVID-19 Surveillance Team |
| EPI_ISL_3172565, EPI_ISL_3172566 | Lighthouse Lab in Alderley Park | Wellcome Sanger Institute for the COVID-19 Genomics UK (COG-UK) Consortium | Jacquelyn Wynn, Mairead Hyland, The Lighthouse Lab in Alderley Park and Alex Alderton, Roberto Amato, Jeffrey Barrett, Sonia Goncalves, Ewan Harrison, David K. Jackson, Ian Johnston, Dominic Kwiatkowski, Cordelia Langford, John Sillitoe on behalf of the Wellcome Sanger Institute COVID-19 Surveillance Team |
| EPI_ISL_3172567 | Lighthouse Lab in Glasgow | Wellcome Sanger Institute for the COVID-19 Genomics UK (COG-UK) Consortium | Harper VanSteenhouse, Yumi Kasai, David Gray, Carol Clugston, Anna Dominiczak and Alex Alderton, Roberto Amato, Jeffrey Barrett, Sonia Goncalves, Ewan Harrison, David K. Jackson, Ian Johnston, Dominic Kwiatkowski, Cordelia Langford, John Sillitoe on behalf of the Wellcome Sanger Institute COVID-19 Surveillance Team |
| EPI_ISL_3172568 | Lighthouse Lab in Alderley Park | Wellcome Sanger Institute for the COVID-19 Genomics UK (COG-UK) Consortium | Jacquelyn Wynn, Mairead Hyland, The Lighthouse Lab in Alderley Park and Alex Alderton, Roberto Amato, Jeffrey Barrett, Sonia Goncalves, Ewan Harrison, David K. Jackson, Ian Johnston, Dominic Kwiatkowski, Cordelia Langford, John Sillitoe on behalf of the Wellcome Sanger Institute COVID-19 Surveillance Team |
| EPI_ISL_3172569, EPI_ISL_3172570 | Lighthouse Lab in Glasgow | Wellcome Sanger Institute for the COVID-19 Genomics UK (COG-UK) Consortium | Harper VanSteenhouse, Yumi Kasai, David Gray, Carol Clugston, Anna Dominiczak and Alex Alderton, Roberto Amato, Jeffrey Barrett, Sonia Goncalves, Ewan Harrison, David K. Jackson, Ian Johnston, Dominic Kwiatkowski, Cordelia Langford, John Sillitoe on behalf of the Wellcome Sanger Institute COVID-19 Surveillance Team |
| EPI_ISL_3172571 | Lighthouse Lab in Alderley Park | Wellcome Sanger Institute for the COVID-19 Genomics UK (COG-UK) Consortium | Jacquelyn Wynn, Mairead Hyland, The Lighthouse Lab in Alderley Park and Alex Alderton, Roberto Amato, Jeffrey Barrett, Sonia Goncalves, Ewan Harrison, David K. Jackson, Ian Johnston, Dominic Kwiatkowski, Cordelia Langford, John Sillitoe on behalf of the Wellcome Sanger Institute COVID-19 Surveillance Team |
| EPI_ISL_3172572 | Lighthouse Lab in Glasgow | Wellcome Sanger Institute for the COVID-19 Genomics UK (COG-UK) Consortium | Harper VanSteenhouse, Yumi Kasai, David Gray, Carol Clugston, Anna Dominiczak and Alex Alderton, Roberto Amato, Jeffrey Barrett, Sonia Goncalves, Ewan Harrison, David K. Jackson, Ian Johnston, Dominic Kwiatkowski, Cordelia Langford, John Sillitoe on behalf of the Wellcome Sanger Institute COVID-19 Surveillance Team |
| EPI_ISL_3172573, EPI_ISL_3172574 | Berkshire and Surrey Pathology Services Lighthouse Laboratory | Wellcome Sanger Institute for the COVID-19 Genomics UK (COG-UK) Consortium | Berkshire and Surrey Pathology Services Lighthouse Laboratory and Alex Alderton, Roberto Amato, Jeffrey Barrett, Sonia Goncalves, Ewan Harrison, David K. Jackson, Ian Johnston, Dominic Kwiatkowski, Cordelia Langford, John Sillitoe on behalf of the Wellcome Sanger Institute COVID-19 Surveillance Team |
| EPI_ISL_3172575 | Lighthouse Lab in Glasgow | Wellcome Sanger Institute for the COVID-19 Genomics UK (COG-UK) Consortium | Harper VanSteenhouse, Yumi Kasai, David Gray, Carol Clugston, Anna Dominiczak and Alex Alderton, Roberto Amato, Jeffrey Barrett, Sonia Goncalves, Ewan Harrison, David K. Jackson, Ian Johnston, Dominic Kwiatkowski, Cordelia Langford, John Sillitoe on behalf of the Wellcome Sanger Institute COVID-19 Surveillance Team |
| EPI_ISL_3172576 | Lighthouse Lab in Alderley Park | Wellcome Sanger Institute for the COVID-19 Genomics UK (COG-UK) Consortium | Jacquelyn Wynn, Mairead Hyland, The Lighthouse Lab in Alderley Park and Alex Alderton, Roberto Amato, Jeffrey Barrett, Sonia Goncalves, Ewan Harrison, David K. Jackson, Ian Johnston, Dominic Kwiatkowski, Cordelia Langford, John Sillitoe on behalf of the Wellcome Sanger Institute COVID-19 Surveillance Team |
| EPI_ISL_3172577 | Lighthouse Lab in Glasgow | Wellcome Sanger Institute for the COVID-19 Genomics UK (COG-UK) Consortium | Harper VanSteenhouse, Yumi Kasai, David Gray, Carol Clugston, Anna Dominiczak and Alex Alderton, Roberto Amato, Jeffrey Barrett, Sonia Goncalves, Ewan Harrison, David K. Jackson, Ian Johnston, Dominic Kwiatkowski, Cordelia Langford, John Sillitoe on behalf of the Wellcome Sanger Institute COVID-19 Surveillance Team |
| EPI_ISL_3172578, EPI_ISL_3172579 | Lighthouse Lab in Alderley Park | Wellcome Sanger Institute for the COVID-19 Genomics UK (COG-UK) Consortium | Jacquelyn Wynn, Mairead Hyland, The Lighthouse Lab in Alderley Park and Alex Alderton, Roberto Amato, Jeffrey Barrett, Sonia Goncalves, Ewan Harrison, David K. Jackson, Ian Johnston, Dominic Kwiatkowski, Cordelia Langford, John Sillitoe on behalf of the Wellcome Sanger Institute COVID-19 Surveillance Team |
| EPI_ISL_3172580 | Berkshire and Surrey Pathology Services Lighthouse Laboratory | Wellcome Sanger Institute for the COVID-19 Genomics UK (COG-UK) Consortium | Berkshire and Surrey Pathology Services Lighthouse Laboratory and Alex Alderton, Roberto Amato, Jeffrey Barrett, Sonia Goncalves, Ewan Harrison, David K. Jackson, Ian Johnston, Dominic Kwiatkowski, Cordelia Langford, John Sillitoe on behalf of the Wellcome Sanger Institute COVID-19 Surveillance Team |
| EPI_ISL_3172581, EPI_ISL_3172582 | Lighthouse Lab in Alderley Park | Wellcome Sanger Institute for the COVID-19 Genomics UK (COG-UK) Consortium | Jacquelyn Wynn, Mairead Hyland, The Lighthouse Lab in Alderley Park and Alex Alderton, Roberto Amato, Jeffrey Barrett, Sonia Goncalves, Ewan Harrison, David K. Jackson, Ian Johnston, Dominic Kwiatkowski, Cordelia Langford, John Sillitoe on behalf of the Wellcome Sanger Institute COVID-19 Surveillance Team |
| EPI_ISL_3172583 | Lighthouse Lab in Glasgow | Wellcome Sanger Institute for the COVID-19 Genomics UK (COG-UK) Consortium | Harper VanSteenhouse, Yumi Kasai, David Gray, Carol Clugston, Anna Dominiczak and Alex Alderton, Roberto Amato, Jeffrey Barrett, Sonia Goncalves, Ewan Harrison, David K. Jackson, Ian Johnston, Dominic Kwiatkowski, Cordelia Langford, John Sillitoe on behalf of the Wellcome Sanger Institute COVID-19 Surveillance Team |
| EPI_ISL_3172584 | Lighthouse Lab in Alderley Park | Wellcome Sanger Institute for the COVID-19 Genomics UK (COG-UK) Consortium | Jacquelyn Wynn, Mairead Hyland, The Lighthouse Lab in Alderley Park and Alex Alderton, Roberto Amato, Jeffrey Barrett, Sonia Goncalves, Ewan Harrison, David K. Jackson, Ian Johnston, Dominic Kwiatkowski, Cordelia Langford, John Sillitoe on behalf of the Wellcome Sanger Institute COVID-19 Surveillance Team |
| EPI_ISL_3172585 | Lighthouse Lab in Glasgow | Wellcome Sanger Institute for the COVID-19 Genomics UK (COG-UK) Consortium | Harper VanSteenhouse, Yumi Kasai, David Gray, Carol Clugston, Anna Dominiczak and Alex Alderton, Roberto Amato, Jeffrey Barrett, Sonia Goncalves, Ewan Harrison, David K. Jackson, Ian Johnston, Dominic Kwiatkowski, Cordelia Langford, John Sillitoe on behalf of the Wellcome Sanger Institute COVID-19 Surveillance Team |
| EPI_ISL_3172586, EPI_ISL_3172587 | Berkshire and Surrey Pathology Services Lighthouse Laboratory | Wellcome Sanger Institute for the COVID-19 Genomics UK (COG-UK) Consortium | Berkshire and Surrey Pathology Services Lighthouse Laboratory and Alex Alderton, Roberto Amato, Jeffrey Barrett, Sonia Goncalves, Ewan Harrison, David K. Jackson, Ian Johnston, Dominic Kwiatkowski, Cordelia Langford, John Sillitoe on behalf of the Wellcome Sanger Institute COVID-19 Surveillance Team |
| EPI_ISL_3172588, EPI_ISL_3172589 | Lighthouse Lab in Glasgow | Wellcome Sanger Institute for the COVID-19 Genomics UK (COG-UK) Consortium | Harper VanSteenhouse, Yumi Kasai, David Gray, Carol Clugston, Anna Dominiczak and Alex Alderton, Roberto Amato, Jeffrey Barrett, Sonia Goncalves, Ewan Harrison, David K. Jackson, Ian Johnston, Dominic Kwiatkowski, Cordelia Langford, John Sillitoe on behalf of the Wellcome Sanger Institute COVID-19 Surveillance Team |
| EPI_ISL_3172590, EPI_ISL_3172591 | Lighthouse Lab in Alderley Park | Wellcome Sanger Institute for the COVID-19 Genomics UK (COG-UK) Consortium | Jacquelyn Wynn, Mairead Hyland, The Lighthouse Lab in Alderley Park and Alex Alderton, Roberto Amato, Jeffrey Barrett, Sonia Goncalves, Ewan Harrison, David K. Jackson, Ian Johnston, Dominic Kwiatkowski, Cordelia Langford, John Sillitoe on behalf of the Wellcome Sanger Institute COVID-19 Surveillance Team |
| EPI_ISL_3172592 | Lighthouse Lab in Glasgow | Wellcome Sanger Institute for the COVID-19 Genomics UK (COG-UK) Consortium | Harper VanSteenhouse, Yumi Kasai, David Gray, Carol Clugston, Anna Dominiczak and Alex Alderton, Roberto Amato, Jeffrey Barrett, Sonia Goncalves, Ewan Harrison, David K. Jackson, Ian Johnston, Dominic Kwiatkowski, Cordelia Langford, John Sillitoe on behalf of the Wellcome Sanger Institute COVID-19 Surveillance Team |
| EPI_ISL_3172593 | Lighthouse Lab in Alderley Park | Wellcome Sanger Institute for the COVID-19 Genomics UK (COG-UK) Consortium | Jacquelyn Wynn, Mairead Hyland, The Lighthouse Lab in Alderley Park and Alex Alderton, Roberto Amato, Jeffrey Barrett, Sonia Goncalves, Ewan Harrison, David K. Jackson, Ian Johnston, Dominic Kwiatkowski, Cordelia Langford, John Sillitoe on behalf of the Wellcome Sanger Institute COVID-19 Surveillance Team |
| EPI_ISL_3172594, EPI_ISL_3172595 | Lighthouse Lab in Glasgow | Wellcome Sanger Institute for the COVID-19 Genomics UK (COG-UK) Consortium | Harper VanSteenhouse, Yumi Kasai, David Gray, Carol Clugston, Anna Dominiczak and Alex Alderton, Roberto Amato, Jeffrey Barrett, Sonia Goncalves, Ewan Harrison, David K. Jackson, Ian Johnston, Dominic Kwiatkowski, Cordelia Langford, John Sillitoe on behalf of the Wellcome Sanger Institute COVID-19 Surveillance Team |
| EPI_ISL_3172596, EPI_ISL_3172597 | Lighthouse Lab in Alderley Park | Wellcome Sanger Institute for the COVID-19 Genomics UK (COG-UK) Consortium | Jacquelyn Wynn, Mairead Hyland, The Lighthouse Lab in Alderley Park and Alex Alderton, Roberto Amato, Jeffrey Barrett, Sonia Goncalves, Ewan Harrison, David K. Jackson, Ian Johnston, Dominic Kwiatkowski, Cordelia Langford, John Sillitoe on behalf of the Wellcome Sanger Institute COVID-19 Surveillance Team |
| EPI_ISL_3172598 | Berkshire and Surrey Pathology Services Lighthouse Laboratory | Wellcome Sanger Institute for the COVID-19 Genomics UK (COG-UK) Consortium | Berkshire and Surrey Pathology Services Lighthouse Laboratory and Alex Alderton, Roberto Amato, Jeffrey Barrett, Sonia Goncalves, Ewan Harrison, David K. Jackson, Ian Johnston, Dominic Kwiatkowski, Cordelia Langford, John Sillitoe on behalf of the Wellcome Sanger Institute COVID-19 Surveillance Team |
| EPI_ISL_3172599, EPI_ISL_3172600 | Lighthouse Lab in Alderley Park | Wellcome Sanger Institute for the COVID-19 Genomics UK (COG-UK) Consortium | Jacquelyn Wynn, Mairead Hyland, The Lighthouse Lab in Alderley Park and Alex Alderton, Roberto Amato, Jeffrey Barrett, Sonia Goncalves, Ewan Harrison, David K. Jackson, Ian Johnston, Dominic Kwiatkowski, Cordelia Langford, John Sillitoe on behalf of the Wellcome Sanger Institute COVID-19 Surveillance Team |
| EPI_ISL_3172601 | Berkshire and Surrey Pathology Services Lighthouse Laboratory | Wellcome Sanger Institute for the COVID-19 Genomics UK (COG-UK) Consortium | Berkshire and Surrey Pathology Services Lighthouse Laboratory and Alex Alderton, Roberto Amato, Jeffrey Barrett, Sonia Goncalves, Ewan Harrison, David K. Jackson, Ian Johnston, Dominic Kwiatkowski, Cordelia Langford, John Sillitoe on behalf of the Wellcome Sanger Institute COVID-19 Surveillance Team |
| EPI_ISL_3172602, EPI_ISL_3172603 | Lighthouse Lab in Glasgow | Wellcome Sanger Institute for the COVID-19 Genomics UK (COG-UK) Consortium | Harper VanSteenhouse, Yumi Kasai, David Gray, Carol Clugston, Anna Dominiczak and Alex Alderton, Roberto Amato, Jeffrey Barrett, Sonia Goncalves, Ewan Harrison, David K. Jackson, Ian Johnston, Dominic Kwiatkowski, Cordelia Langford, John Sillitoe on behalf of the Wellcome Sanger Institute COVID-19 Surveillance Team |
| EPI_ISL_3172604, | Lighthouse Lab in Alderley Park | Wellcome Sanger Institute for the COVID-19 Genomics UK (COG-UK) | Jacquelyn Wynn, Mairead Hyland, The Lighthouse Lab in Alderley Park and Alex Alderton, Roberto Amato, Jeffrey Barrett, Sonia Goncalves, Ewan Harrison, David K. Jackson, Ian Johnston, Dominic Kwiatkowski, Cordelia Langford, John Sillitoe on |

| | | | |
|---|---|---|---|
| EPI_ISL_3172605 | | Consortium | behalf of the Wellcome Sanger Institute COVID-19 Surveillance Team |
| EPI_ISL_3172606, EPI_ISL_3172607 | Berkshire and Surrey Pathology Services Lighthouse Laboratory | Wellcome Sanger Institute for the COVID-19 Genomics UK (COG-UK) Consortium | Berkshire and Surrey Pathology Services Lighthouse Laboratory and Alex Alderton, Roberto Amato, Jeffrey Barrett, Sonia Goncalves, Ewan Harrison, David K. Jackson, Ian Johnston, Dominic Kwiatkowski, Cordelia Langford, John Sillitoe on behalf of the Wellcome Sanger Institute COVID-19 Surveillance Team |
| EPI_ISL_3172608, EPI_ISL_3172609, EPI_ISL_3172610 | Lighthouse Lab in Glasgow | Wellcome Sanger Institute for the COVID-19 Genomics UK (COG-UK) Consortium | Harper VanSteenhouse, Yumi Kasai, David Gray, Carol Clugston, Anna Dominiczak and Alex Alderton, Roberto Amato, Jeffrey Barrett, Sonia Goncalves, Ewan Harrison, David K. Jackson, Ian Johnston, Dominic Kwiatkowski, Cordelia Langford, John Sillitoe on behalf of the Wellcome Sanger Institute COVID-19 Surveillance Team |
| EPI_ISL_3172611 | Lighthouse Lab in Alderley Park | Wellcome Sanger Institute for the COVID-19 Genomics UK (COG-UK) Consortium | Jacquelyn Wynn, Mairead Hyland, The Lighthouse Lab in Alderley Park and Alex Alderton, Roberto Amato, Jeffrey Barrett, Sonia Goncalves, Ewan Harrison, David K. Jackson, Ian Johnston, Dominic Kwiatkowski, Cordelia Langford, John Sillitoe on behalf of the Wellcome Sanger Institute COVID-19 Surveillance Team |
| EPI_ISL_3172612 | Lighthouse Lab in Glasgow | Wellcome Sanger Institute for the COVID-19 Genomics UK (COG-UK) Consortium | Harper VanSteenhouse, Yumi Kasai, David Gray, Carol Clugston, Anna Dominiczak and Alex Alderton, Roberto Amato, Jeffrey Barrett, Sonia Goncalves, Ewan Harrison, David K. Jackson, Ian Johnston, Dominic Kwiatkowski, Cordelia Langford, John Sillitoe on behalf of the Wellcome Sanger Institute COVID-19 Surveillance Team |
| EPI_ISL_3172613 | Berkshire and Surrey Pathology Services Lighthouse Laboratory | Wellcome Sanger Institute for the COVID-19 Genomics UK (COG-UK) Consortium | Berkshire and Surrey Pathology Services Lighthouse Laboratory and Alex Alderton, Roberto Amato, Jeffrey Barrett, Sonia Goncalves, Ewan Harrison, David K. Jackson, Ian Johnston, Dominic Kwiatkowski, Cordelia Langford, John Sillitoe on behalf of the Wellcome Sanger Institute COVID-19 Surveillance Team |
| EPI_ISL_3172614 | Lighthouse Lab in Alderley Park | Wellcome Sanger Institute for the COVID-19 Genomics UK (COG-UK) Consortium | Jacquelyn Wynn, Mairead Hyland, The Lighthouse Lab in Alderley Park and Alex Alderton, Roberto Amato, Jeffrey Barrett, Sonia Goncalves, Ewan Harrison, David K. Jackson, Ian Johnston, Dominic Kwiatkowski, Cordelia Langford, John Sillitoe on behalf of the Wellcome Sanger Institute COVID-19 Surveillance Team |
| EPI_ISL_3172615, EPI_ISL_3172616 | Lighthouse Lab in Glasgow | Wellcome Sanger Institute for the COVID-19 Genomics UK (COG-UK) Consortium | Harper VanSteenhouse, Yumi Kasai, David Gray, Carol Clugston, Anna Dominiczak and Alex Alderton, Roberto Amato, Jeffrey Barrett, Sonia Goncalves, Ewan Harrison, David K. Jackson, Ian Johnston, Dominic Kwiatkowski, Cordelia Langford, John Sillitoe on behalf of the Wellcome Sanger Institute COVID-19 Surveillance Team |
| EPI_ISL_3172617, EPI_ISL_3172618 | Lighthouse Lab in Alderley Park | Wellcome Sanger Institute for the COVID-19 Genomics UK (COG-UK) Consortium | Jacquelyn Wynn, Mairead Hyland, The Lighthouse Lab in Alderley Park and Alex Alderton, Roberto Amato, Jeffrey Barrett, Sonia Goncalves, Ewan Harrison, David K. Jackson, Ian Johnston, Dominic Kwiatkowski, Cordelia Langford, John Sillitoe on behalf of the Wellcome Sanger Institute COVID-19 Surveillance Team |
| EPI_ISL_3172619 | Lighthouse Lab in Glasgow | Wellcome Sanger Institute for the COVID-19 Genomics UK (COG-UK) Consortium | Harper VanSteenhouse, Yumi Kasai, David Gray, Carol Clugston, Anna Dominiczak and Alex Alderton, Roberto Amato, Jeffrey Barrett, Sonia Goncalves, Ewan Harrison, David K. Jackson, Ian Johnston, Dominic Kwiatkowski, Cordelia Langford, John Sillitoe on behalf of the Wellcome Sanger Institute COVID-19 Surveillance Team |
| EPI_ISL_3172620 | Berkshire and Surrey Pathology Services Lighthouse Laboratory | Wellcome Sanger Institute for the COVID-19 Genomics UK (COG-UK) Consortium | Berkshire and Surrey Pathology Services Lighthouse Laboratory and Alex Alderton, Roberto Amato, Jeffrey Barrett, Sonia Goncalves, Ewan Harrison, David K. Jackson, Ian Johnston, Dominic Kwiatkowski, Cordelia Langford, John Sillitoe on behalf of the Wellcome Sanger Institute COVID-19 Surveillance Team |
| EPI_ISL_3172621, EPI_ISL_3172622 | Lighthouse Lab in Glasgow | Wellcome Sanger Institute for the COVID-19 Genomics UK (COG-UK) Consortium | Harper VanSteenhouse, Yumi Kasai, David Gray, Carol Clugston, Anna Dominiczak and Alex Alderton, Roberto Amato, Jeffrey Barrett, Sonia Goncalves, Ewan Harrison, David K. Jackson, Ian Johnston, Dominic Kwiatkowski, Cordelia Langford, John Sillitoe on behalf of the Wellcome Sanger Institute COVID-19 Surveillance Team |
| EPI_ISL_3172623, EPI_ISL_3172624 | Lighthouse Lab in Alderley Park | Wellcome Sanger Institute for the COVID-19 Genomics UK (COG-UK) Consortium | Jacquelyn Wynn, Mairead Hyland, The Lighthouse Lab in Alderley Park and Alex Alderton, Roberto Amato, Jeffrey Barrett, Sonia Goncalves, Ewan Harrison, David K. Jackson, Ian Johnston, Dominic Kwiatkowski, Cordelia Langford, John Sillitoe on behalf of the Wellcome Sanger Institute COVID-19 Surveillance Team |
| EPI_ISL_3172625, EPI_ISL_3172626 | Lighthouse Lab in Glasgow | Wellcome Sanger Institute for the COVID-19 Genomics UK (COG-UK) Consortium | Harper VanSteenhouse, Yumi Kasai, David Gray, Carol Clugston, Anna Dominiczak and Alex Alderton, Roberto Amato, Jeffrey Barrett, Sonia Goncalves, Ewan Harrison, David K. Jackson, Ian Johnston, Dominic Kwiatkowski, Cordelia Langford, John Sillitoe on behalf of the Wellcome Sanger Institute COVID-19 Surveillance Team |
| EPI_ISL_3172627 | Berkshire and Surrey Pathology Services Lighthouse Laboratory | Wellcome Sanger Institute for the COVID-19 Genomics UK (COG-UK) Consortium | Berkshire and Surrey Pathology Services Lighthouse Laboratory and Alex Alderton, Roberto Amato, Jeffrey Barrett, Sonia Goncalves, Ewan Harrison, David K. Jackson, Ian Johnston, Dominic Kwiatkowski, Cordelia Langford, John Sillitoe on behalf of the Wellcome Sanger Institute COVID-19 Surveillance Team |
| EPI_ISL_3172628, EPI_ISL_3172629, EPI_ISL_3172630 | Lighthouse Lab in Glasgow | Wellcome Sanger Institute for the COVID-19 Genomics UK (COG-UK) Consortium | Harper VanSteenhouse, Yumi Kasai, David Gray, Carol Clugston, Anna Dominiczak and Alex Alderton, Roberto Amato, Jeffrey Barrett, Sonia Goncalves, Ewan Harrison, David K. Jackson, Ian Johnston, Dominic Kwiatkowski, Cordelia Langford, John Sillitoe on behalf of the Wellcome Sanger Institute COVID-19 Surveillance Team |
| EPI_ISL_3172631 | Lighthouse Lab in Alderley Park | Wellcome Sanger Institute for the COVID-19 Genomics UK (COG-UK) Consortium | Jacquelyn Wynn, Mairead Hyland, The Lighthouse Lab in Alderley Park and Alex Alderton, Roberto Amato, Jeffrey Barrett, Sonia Goncalves, Ewan Harrison, David K. Jackson, Ian Johnston, Dominic Kwiatkowski, Cordelia Langford, John Sillitoe on behalf of the Wellcome Sanger Institute COVID-19 Surveillance Team |
| EPI_ISL_3172632, EPI_ISL_3172633 | Lighthouse Lab in Glasgow | Wellcome Sanger Institute for the COVID-19 Genomics UK (COG-UK) Consortium | Harper VanSteenhouse, Yumi Kasai, David Gray, Carol Clugston, Anna Dominiczak and Alex Alderton, Roberto Amato, Jeffrey Barrett, Sonia Goncalves, Ewan Harrison, David K. Jackson, Ian Johnston, Dominic Kwiatkowski, Cordelia Langford, John Sillitoe on behalf of the Wellcome Sanger Institute COVID-19 Surveillance Team |
| EPI_ISL_3172634 | Berkshire and Surrey Pathology Services Lighthouse Laboratory | Wellcome Sanger Institute for the COVID-19 Genomics UK (COG-UK) Consortium | Berkshire and Surrey Pathology Services Lighthouse Laboratory and Alex Alderton, Roberto Amato, Jeffrey Barrett, Sonia Goncalves, Ewan Harrison, David K. Jackson, Ian Johnston, Dominic Kwiatkowski, Cordelia Langford, John Sillitoe on behalf of the Wellcome Sanger Institute COVID-19 Surveillance Team |
| EPI_ISL_3172635, EPI_ISL_3172636 | Lighthouse Lab in Alderley Park | Wellcome Sanger Institute for the COVID-19 Genomics UK (COG-UK) Consortium | Jacquelyn Wynn, Mairead Hyland, The Lighthouse Lab in Alderley Park and Alex Alderton, Roberto Amato, Jeffrey Barrett, Sonia Goncalves, Ewan Harrison, David K. Jackson, Ian Johnston, Dominic Kwiatkowski, Cordelia Langford, John Sillitoe on behalf of the Wellcome Sanger Institute COVID-19 Surveillance Team |
| EPI_ISL_3172637 | Berkshire and Surrey Pathology Services Lighthouse Laboratory | Wellcome Sanger Institute for the COVID-19 Genomics UK (COG-UK) Consortium | Berkshire and Surrey Pathology Services Lighthouse Laboratory and Alex Alderton, Roberto Amato, Jeffrey Barrett, Sonia Goncalves, Ewan Harrison, David K. Jackson, Ian Johnston, Dominic Kwiatkowski, Cordelia Langford, John Sillitoe on behalf of the Wellcome Sanger Institute COVID-19 Surveillance Team |
| EPI_ISL_3172638 | Lighthouse Lab in Alderley Park | Wellcome Sanger Institute for the COVID-19 Genomics UK (COG-UK) Consortium | Jacquelyn Wynn, Mairead Hyland, The Lighthouse Lab in Alderley Park and Alex Alderton, Roberto Amato, Jeffrey Barrett, Sonia Goncalves, Ewan Harrison, David K. Jackson, Ian Johnston, Dominic Kwiatkowski, Cordelia Langford, John Sillitoe on behalf of the Wellcome Sanger Institute COVID-19 Surveillance Team |
| EPI_ISL_3172639 | Berkshire and Surrey Pathology Services Lighthouse Laboratory | Wellcome Sanger Institute for the COVID-19 Genomics UK (COG-UK) Consortium | Berkshire and Surrey Pathology Services Lighthouse Laboratory and Alex Alderton, Roberto Amato, Jeffrey Barrett, Sonia Goncalves, Ewan Harrison, David K. Jackson, Ian Johnston, Dominic Kwiatkowski, Cordelia Langford, John Sillitoe on behalf of the Wellcome Sanger Institute COVID-19 Surveillance Team |
| EPI_ISL_3172640, EPI_ISL_3172641, EPI_ISL_3172642 | Lighthouse Lab in Alderley Park | Wellcome Sanger Institute for the COVID-19 Genomics UK (COG-UK) Consortium | Jacquelyn Wynn, Mairead Hyland, The Lighthouse Lab in Alderley Park and Alex Alderton, Roberto Amato, Jeffrey Barrett, Sonia Goncalves, Ewan Harrison, David K. Jackson, Ian Johnston, Dominic Kwiatkowski, Cordelia Langford, John Sillitoe on behalf of the Wellcome Sanger Institute COVID-19 Surveillance Team |
| EPI_ISL_3172643 | Lighthouse Lab in Glasgow | Wellcome Sanger Institute for the COVID-19 Genomics UK (COG-UK) Consortium | Harper VanSteenhouse, Yumi Kasai, David Gray, Carol Clugston, Anna Dominiczak and Alex Alderton, Roberto Amato, Jeffrey Barrett, Sonia Goncalves, Ewan Harrison, David K. Jackson, Ian Johnston, Dominic Kwiatkowski, Cordelia Langford, John Sillitoe on behalf of the Wellcome Sanger Institute COVID-19 Surveillance Team |
| EPI_ISL_3172644 | Berkshire and Surrey Pathology Services Lighthouse Laboratory | Wellcome Sanger Institute for the COVID-19 Genomics UK (COG-UK) Consortium | Berkshire and Surrey Pathology Services Lighthouse Laboratory and Alex Alderton, Roberto Amato, Jeffrey Barrett, Sonia Goncalves, Ewan Harrison, David K. Jackson, Ian Johnston, Dominic Kwiatkowski, Cordelia Langford, John Sillitoe on behalf of the Wellcome Sanger Institute COVID-19 Surveillance Team |
| EPI_ISL_3172645 | Lighthouse Lab in Glasgow | Wellcome Sanger Institute for the COVID-19 Genomics UK (COG-UK) Consortium | Harper VanSteenhouse, Yumi Kasai, David Gray, Carol Clugston, Anna Dominiczak and Alex Alderton, Roberto Amato, Jeffrey Barrett, Sonia Goncalves, Ewan Harrison, David K. Jackson, Ian Johnston, Dominic Kwiatkowski, Cordelia Langford, John Sillitoe on behalf of the Wellcome Sanger Institute COVID-19 Surveillance Team |
| EPI_ISL_3172646 | Berkshire and Surrey Pathology Services Lighthouse Laboratory | Wellcome Sanger Institute for the COVID-19 Genomics UK (COG-UK) Consortium | Berkshire and Surrey Pathology Services Lighthouse Laboratory and Alex Alderton, Roberto Amato, Jeffrey Barrett, Sonia Goncalves, Ewan Harrison, David K. Jackson, Ian Johnston, Dominic Kwiatkowski, Cordelia Langford, John Sillitoe on behalf of the Wellcome Sanger Institute COVID-19 Surveillance Team |
| EPI_ISL_3172647 | Lighthouse Lab in Glasgow | Wellcome Sanger Institute for the COVID-19 Genomics UK (COG-UK) Consortium | Harper VanSteenhouse, Yumi Kasai, David Gray, Carol Clugston, Anna Dominiczak and Alex Alderton, Roberto Amato, Jeffrey Barrett, Sonia Goncalves, Ewan Harrison, David K. Jackson, Ian Johnston, Dominic Kwiatkowski, Cordelia Langford, John Sillitoe on behalf of the Wellcome Sanger Institute COVID-19 Surveillance Team |
| EPI_ISL_3172648, EPI_ISL_3172649, EPI_ISL_3172650, EPI_ISL_3172651 | Lighthouse Lab in Alderley Park | Wellcome Sanger Institute for the COVID-19 Genomics UK (COG-UK) Consortium | Jacquelyn Wynn, Mairead Hyland, The Lighthouse Lab in Alderley Park and Alex Alderton, Roberto Amato, Jeffrey Barrett, Sonia Goncalves, Ewan Harrison, David K. Jackson, Ian Johnston, Dominic Kwiatkowski, Cordelia Langford, John Sillitoe on behalf of the Wellcome Sanger Institute COVID-19 Surveillance Team |
| EPI_ISL_3172652, EPI_ISL_3172653, EPI_ISL_3172654 | Lighthouse Lab in Glasgow | Wellcome Sanger Institute for the COVID-19 Genomics UK (COG-UK) Consortium | Harper VanSteenhouse, Yumi Kasai, David Gray, Carol Clugston, Anna Dominiczak and Alex Alderton, Roberto Amato, Jeffrey Barrett, Sonia Goncalves, Ewan Harrison, David K. Jackson, Ian Johnston, Dominic Kwiatkowski, Cordelia Langford, John Sillitoe on behalf of the Wellcome Sanger Institute COVID-19 Surveillance Team |
| EPI_ISL_3172655 | Lighthouse Lab in Alderley Park | Wellcome Sanger Institute for the COVID-19 Genomics UK (COG-UK) Consortium | Jacquelyn Wynn, Mairead Hyland, The Lighthouse Lab in Alderley Park and Alex Alderton, Roberto Amato, Jeffrey Barrett, Sonia Goncalves, Ewan Harrison, David K. Jackson, Ian Johnston, Dominic Kwiatkowski, Cordelia Langford, John Sillitoe on behalf of the Wellcome Sanger Institute COVID-19 Surveillance Team |
| EPI_ISL_3172656, EPI_ISL_3172657 | Lighthouse Lab in Glasgow | Wellcome Sanger Institute for the COVID-19 Genomics UK (COG-UK) Consortium | Harper VanSteenhouse, Yumi Kasai, David Gray, Carol Clugston, Anna Dominiczak and Alex Alderton, Roberto Amato, Jeffrey Barrett, Sonia Goncalves, Ewan Harrison, David K. Jackson, Ian Johnston, Dominic Kwiatkowski, Cordelia Langford, John Sillitoe on behalf of the Wellcome Sanger Institute COVID-19 Surveillance Team |
| EPI_ISL_3172658 | Berkshire and Surrey Pathology Services Lighthouse Laboratory | Wellcome Sanger Institute for the COVID-19 Genomics UK (COG-UK) Consortium | Berkshire and Surrey Pathology Services Lighthouse Laboratory and Alex Alderton, Roberto Amato, Jeffrey Barrett, Sonia Goncalves, Ewan Harrison, David K. Jackson, Ian Johnston, Dominic Kwiatkowski, Cordelia Langford, John Sillitoe on behalf of the Wellcome Sanger Institute COVID-19 Surveillance Team |
| EPI_ISL_3172659 | Lighthouse Lab in Glasgow | Wellcome Sanger Institute for the COVID-19 Genomics UK (COG-UK) Consortium | Harper VanSteenhouse, Yumi Kasai, David Gray, Carol Clugston, Anna Dominiczak and Alex Alderton, Roberto Amato, Jeffrey Barrett, Sonia Goncalves, Ewan Harrison, David K. Jackson, Ian Johnston, Dominic Kwiatkowski, Cordelia Langford, John Sillitoe on behalf of the Wellcome Sanger Institute COVID-19 Surveillance Team |
| EPI_ISL_3172660, EPI_ISL_3172661 | Lighthouse Lab in Alderley Park | Wellcome Sanger Institute for the COVID-19 Genomics UK (COG-UK) Consortium | Jacquelyn Wynn, Mairead Hyland, The Lighthouse Lab in Alderley Park and Alex Alderton, Roberto Amato, Jeffrey Barrett, Sonia Goncalves, Ewan Harrison, David K. Jackson, Ian Johnston, Dominic Kwiatkowski, Cordelia Langford, John Sillitoe on behalf of the Wellcome Sanger Institute COVID-19 Surveillance Team |
| EPI_ISL_3172662 | Lighthouse Lab in Glasgow | Wellcome Sanger Institute for the COVID-19 Genomics UK (COG-UK) Consortium | Harper VanSteenhouse, Yumi Kasai, David Gray, Carol Clugston, Anna Dominiczak and Alex Alderton, Roberto Amato, Jeffrey Barrett, Sonia Goncalves, Ewan Harrison, David K. Jackson, Ian Johnston, Dominic Kwiatkowski, Cordelia Langford, John Sillitoe on behalf of the Wellcome Sanger Institute COVID-19 Surveillance Team |
| EPI_ISL_3172663 | Berkshire and Surrey Pathology Services Lighthouse Laboratory | Wellcome Sanger Institute for the COVID-19 Genomics UK (COG-UK) Consortium | Berkshire and Surrey Pathology Services Lighthouse Laboratory and Alex Alderton, Roberto Amato, Jeffrey Barrett, Sonia Goncalves, Ewan Harrison, David K. Jackson, Ian Johnston, Dominic Kwiatkowski, Cordelia Langford, John Sillitoe on behalf of the Wellcome Sanger Institute COVID-19 Surveillance Team |
| EPI_ISL_3172664 | Lighthouse Lab in Alderley Park | Wellcome Sanger Institute for the COVID-19 Genomics UK (COG-UK) Consortium | Jacquelyn Wynn, Mairead Hyland, The Lighthouse Lab in Alderley Park and Alex Alderton, Roberto Amato, Jeffrey Barrett, Sonia Goncalves, Ewan Harrison, David K. Jackson, Ian Johnston, Dominic Kwiatkowski, Cordelia Langford, John Sillitoe on behalf of the Wellcome Sanger Institute COVID-19 Surveillance Team |
| EPI_ISL_3172665, EPI_ISL_3172666 | Berkshire and Surrey Pathology Services Lighthouse Laboratory | Wellcome Sanger Institute for the COVID-19 Genomics UK (COG-UK) Consortium | Berkshire and Surrey Pathology Services Lighthouse Laboratory and Alex Alderton, Roberto Amato, Jeffrey Barrett, Sonia Goncalves, Ewan Harrison, David K. Jackson, Ian Johnston, Dominic Kwiatkowski, Cordelia Langford, John Sillitoe on behalf of the Wellcome Sanger Institute COVID-19 Surveillance Team |
| EPI_ISL_3172667 | Lighthouse Lab in Glasgow | Wellcome Sanger Institute for the COVID-19 Genomics UK (COG-UK) Consortium | Harper VanSteenhouse, Yumi Kasai, David Gray, Carol Clugston, Anna Dominiczak and Alex Alderton, Roberto Amato, Jeffrey Barrett, Sonia Goncalves, Ewan Harrison, David K. Jackson, Ian Johnston, Dominic Kwiatkowski, Cordelia Langford, John Sillitoe on behalf of the Wellcome Sanger Institute COVID-19 Surveillance Team |
| EPI_ISL_3172668, EPI_ISL_3172669, EPI_ISL_3172670, EPI_ISL_3172671, EPI_ISL_3172672, | Lighthouse Lab in Alderley Park | Wellcome Sanger Institute for the COVID-19 Genomics UK (COG-UK) Consortium | Jacquelyn Wynn, Mairead Hyland, The Lighthouse Lab in Alderley Park and Alex Alderton, Roberto Amato, Jeffrey Barrett, Sonia Goncalves, Ewan Harrison, David K. Jackson, Ian Johnston, Dominic Kwiatkowski, Cordelia Langford, John Sillitoe on behalf of the Wellcome Sanger Institute COVID-19 Surveillance Team |

| | | | |
|---|---|---|---|
| EPI_ISL_3172673 | | | |
| EPI_ISL_3172674 | Lighthouse Lab in Glasgow | Wellcome Sanger Institute for the COVID-19 Genomics UK (COG-UK) Consortium | Harper VanSteenhouse, Yumi Kasai, David Gray, Carol Clugston, Anna Dominiczak and Alex Alderton, Roberto Amato, Jeffrey Barrett, Sonia Goncalves, Ewan Harrison, David K. Jackson, Ian Johnston, Dominic Kwiatkowski, Cordelia Langford, John Sillitoe on behalf of the Wellcome Sanger Institute COVID-19 Surveillance Team |
| EPI_ISL_3172675 | Berkshire and Surrey Pathology Services Lighthouse Laboratory | Wellcome Sanger Institute for the COVID-19 Genomics UK (COG-UK) Consortium | Berkshire and Surrey Pathology Services Lighthouse Laboratory and Alex Alderton, Roberto Amato, Jeffrey Barrett, Sonia Goncalves, Ewan Harrison, David K. Jackson, Ian Johnston, Dominic Kwiatkowski, Cordelia Langford, John Sillitoe on behalf of the Wellcome Sanger Institute COVID-19 Surveillance Team |
| EPI_ISL_3172676 | Lighthouse Lab in Alderley Park | Wellcome Sanger Institute for the COVID-19 Genomics UK (COG-UK) Consortium | Jacquelyn Wynn, Mairead Hyland, The Lighthouse Lab in Alderley Park and Alex Alderton, Roberto Amato, Jeffrey Barrett, Sonia Goncalves, Ewan Harrison, David K. Jackson, Ian Johnston, Dominic Kwiatkowski, Cordelia Langford, John Sillitoe on behalf of the Wellcome Sanger Institute COVID-19 Surveillance Team |
| EPI_ISL_3172677, EPI_ISL_3172678, EPI_ISL_3172679 | Berkshire and Surrey Pathology Services Lighthouse Laboratory | Wellcome Sanger Institute for the COVID-19 Genomics UK (COG-UK) Consortium | Berkshire and Surrey Pathology Services Lighthouse Laboratory and Alex Alderton, Roberto Amato, Jeffrey Barrett, Sonia Goncalves, Ewan Harrison, David K. Jackson, Ian Johnston, Dominic Kwiatkowski, Cordelia Langford, John Sillitoe on behalf of the Wellcome Sanger Institute COVID-19 Surveillance Team |
| EPI_ISL_3172680, EPI_ISL_3172681 | Lighthouse Lab in Glasgow | Wellcome Sanger Institute for the COVID-19 Genomics UK (COG-UK) Consortium | Harper VanSteenhouse, Yumi Kasai, David Gray, Carol Clugston, Anna Dominiczak and Alex Alderton, Roberto Amato, Jeffrey Barrett, Sonia Goncalves, Ewan Harrison, David K. Jackson, Ian Johnston, Dominic Kwiatkowski, Cordelia Langford, John Sillitoe on behalf of the Wellcome Sanger Institute COVID-19 Surveillance Team |
| EPI_ISL_3172682 | Lighthouse Lab in Alderley Park | Wellcome Sanger Institute for the COVID-19 Genomics UK (COG-UK) Consortium | Jacquelyn Wynn, Mairead Hyland, The Lighthouse Lab in Alderley Park and Alex Alderton, Roberto Amato, Jeffrey Barrett, Sonia Goncalves, Ewan Harrison, David K. Jackson, Ian Johnston, Dominic Kwiatkowski, Cordelia Langford, John Sillitoe on behalf of the Wellcome Sanger Institute COVID-19 Surveillance Team |
| EPI_ISL_3172683, EPI_ISL_3172684 | Lighthouse Lab in Glasgow | Wellcome Sanger Institute for the COVID-19 Genomics UK (COG-UK) Consortium | Harper VanSteenhouse, Yumi Kasai, David Gray, Carol Clugston, Anna Dominiczak and Alex Alderton, Roberto Amato, Jeffrey Barrett, Sonia Goncalves, Ewan Harrison, David K. Jackson, Ian Johnston, Dominic Kwiatkowski, Cordelia Langford, John Sillitoe on behalf of the Wellcome Sanger Institute COVID-19 Surveillance Team |
| EPI_ISL_3172685 | Lighthouse Lab in Alderley Park | Wellcome Sanger Institute for the COVID-19 Genomics UK (COG-UK) Consortium | Jacquelyn Wynn, Mairead Hyland, The Lighthouse Lab in Alderley Park and Alex Alderton, Roberto Amato, Jeffrey Barrett, Sonia Goncalves, Ewan Harrison, David K. Jackson, Ian Johnston, Dominic Kwiatkowski, Cordelia Langford, John Sillitoe on behalf of the Wellcome Sanger Institute COVID-19 Surveillance Team |
| EPI_ISL_3172686 | Lighthouse Lab in Glasgow | Wellcome Sanger Institute for the COVID-19 Genomics UK (COG-UK) Consortium | Harper VanSteenhouse, Yumi Kasai, David Gray, Carol Clugston, Anna Dominiczak and Alex Alderton, Roberto Amato, Jeffrey Barrett, Sonia Goncalves, Ewan Harrison, David K. Jackson, Ian Johnston, Dominic Kwiatkowski, Cordelia Langford, John Sillitoe on behalf of the Wellcome Sanger Institute COVID-19 Surveillance Team |
| EPI_ISL_3172687, EPI_ISL_3172688 | Berkshire and Surrey Pathology Services Lighthouse Laboratory | Wellcome Sanger Institute for the COVID-19 Genomics UK (COG-UK) Consortium | Berkshire and Surrey Pathology Services Lighthouse Laboratory and Alex Alderton, Roberto Amato, Jeffrey Barrett, Sonia Goncalves, Ewan Harrison, David K. Jackson, Ian Johnston, Dominic Kwiatkowski, Cordelia Langford, John Sillitoe on behalf of the Wellcome Sanger Institute COVID-19 Surveillance Team |
| EPI_ISL_3172689 | Lighthouse Lab in Glasgow | Wellcome Sanger Institute for the COVID-19 Genomics UK (COG-UK) Consortium | Harper VanSteenhouse, Yumi Kasai, David Gray, Carol Clugston, Anna Dominiczak and Alex Alderton, Roberto Amato, Jeffrey Barrett, Sonia Goncalves, Ewan Harrison, David K. Jackson, Ian Johnston, Dominic Kwiatkowski, Cordelia Langford, John Sillitoe on behalf of the Wellcome Sanger Institute COVID-19 Surveillance Team |
| EPI_ISL_3172690, EPI_ISL_3172691, EPI_ISL_3172692, EPI_ISL_3172693 | Lighthouse Lab in Alderley Park | Wellcome Sanger Institute for the COVID-19 Genomics UK (COG-UK) Consortium | Jacquelyn Wynn, Mairead Hyland, The Lighthouse Lab in Alderley Park and Alex Alderton, Roberto Amato, Jeffrey Barrett, Sonia Goncalves, Ewan Harrison, David K. Jackson, Ian Johnston, Dominic Kwiatkowski, Cordelia Langford, John Sillitoe on behalf of the Wellcome Sanger Institute COVID-19 Surveillance Team |
| EPI_ISL_3172694 | Lighthouse Lab in Glasgow | Wellcome Sanger Institute for the COVID-19 Genomics UK (COG-UK) Consortium | Harper VanSteenhouse, Yumi Kasai, David Gray, Carol Clugston, Anna Dominiczak and Alex Alderton, Roberto Amato, Jeffrey Barrett, Sonia Goncalves, Ewan Harrison, David K. Jackson, Ian Johnston, Dominic Kwiatkowski, Cordelia Langford, John Sillitoe on behalf of the Wellcome Sanger Institute COVID-19 Surveillance Team |
| EPI_ISL_3172695 | Berkshire and Surrey Pathology Services Lighthouse Laboratory | Wellcome Sanger Institute for the COVID-19 Genomics UK (COG-UK) Consortium | Berkshire and Surrey Pathology Services Lighthouse Laboratory and Alex Alderton, Roberto Amato, Jeffrey Barrett, Sonia Goncalves, Ewan Harrison, David K. Jackson, Ian Johnston, Dominic Kwiatkowski, Cordelia Langford, John Sillitoe on behalf of the Wellcome Sanger Institute COVID-19 Surveillance Team |
| EPI_ISL_3172696 | Lighthouse Lab in Glasgow | Wellcome Sanger Institute for the COVID-19 Genomics UK (COG-UK) Consortium | Harper VanSteenhouse, Yumi Kasai, David Gray, Carol Clugston, Anna Dominiczak and Alex Alderton, Roberto Amato, Jeffrey Barrett, Sonia Goncalves, Ewan Harrison, David K. Jackson, Ian Johnston, Dominic Kwiatkowski, Cordelia Langford, John Sillitoe on behalf of the Wellcome Sanger Institute COVID-19 Surveillance Team |
| EPI_ISL_3172697 | Lighthouse Lab in Alderley Park | Wellcome Sanger Institute for the COVID-19 Genomics UK (COG-UK) Consortium | Jacquelyn Wynn, Mairead Hyland, The Lighthouse Lab in Alderley Park and Alex Alderton, Roberto Amato, Jeffrey Barrett, Sonia Goncalves, Ewan Harrison, David K. Jackson, Ian Johnston, Dominic Kwiatkowski, Cordelia Langford, John Sillitoe on behalf of the Wellcome Sanger Institute COVID-19 Surveillance Team |
| EPI_ISL_3172698 | Berkshire and Surrey Pathology Services Lighthouse Laboratory | Wellcome Sanger Institute for the COVID-19 Genomics UK (COG-UK) Consortium | Berkshire and Surrey Pathology Services Lighthouse Laboratory and Alex Alderton, Roberto Amato, Jeffrey Barrett, Sonia Goncalves, Ewan Harrison, David K. Jackson, Ian Johnston, Dominic Kwiatkowski, Cordelia Langford, John Sillitoe on behalf of the Wellcome Sanger Institute COVID-19 Surveillance Team |
| EPI_ISL_3172699, EPI_ISL_3172700 | Lighthouse Lab in Glasgow | Wellcome Sanger Institute for the COVID-19 Genomics UK (COG-UK) Consortium | Harper VanSteenhouse, Yumi Kasai, David Gray, Carol Clugston, Anna Dominiczak and Alex Alderton, Roberto Amato, Jeffrey Barrett, Sonia Goncalves, Ewan Harrison, David K. Jackson, Ian Johnston, Dominic Kwiatkowski, Cordelia Langford, John Sillitoe on behalf of the Wellcome Sanger Institute COVID-19 Surveillance Team |
| EPI_ISL_3172701 | Berkshire and Surrey Pathology Services Lighthouse Laboratory | Wellcome Sanger Institute for the COVID-19 Genomics UK (COG-UK) Consortium | Berkshire and Surrey Pathology Services Lighthouse Laboratory and Alex Alderton, Roberto Amato, Jeffrey Barrett, Sonia Goncalves, Ewan Harrison, David K. Jackson, Ian Johnston, Dominic Kwiatkowski, Cordelia Langford, John Sillitoe on behalf of the Wellcome Sanger Institute COVID-19 Surveillance Team |
| EPI_ISL_3172702, EPI_ISL_3172703 | Lighthouse Lab in Glasgow | Wellcome Sanger Institute for the COVID-19 Genomics UK (COG-UK) Consortium | Harper VanSteenhouse, Yumi Kasai, David Gray, Carol Clugston, Anna Dominiczak and Alex Alderton, Roberto Amato, Jeffrey Barrett, Sonia Goncalves, Ewan Harrison, David K. Jackson, Ian Johnston, Dominic Kwiatkowski, Cordelia Langford, John Sillitoe on behalf of the Wellcome Sanger Institute COVID-19 Surveillance Team |
| EPI_ISL_3172704, EPI_ISL_3172705, EPI_ISL_3172706, EPI_ISL_3172707 | Berkshire and Surrey Pathology Services Lighthouse Laboratory | Wellcome Sanger Institute for the COVID-19 Genomics UK (COG-UK) Consortium | Berkshire and Surrey Pathology Services Lighthouse Laboratory and Alex Alderton, Roberto Amato, Jeffrey Barrett, Sonia Goncalves, Ewan Harrison, David K. Jackson, Ian Johnston, Dominic Kwiatkowski, Cordelia Langford, John Sillitoe on behalf of the Wellcome Sanger Institute COVID-19 Surveillance Team |
| EPI_ISL_3172708, EPI_ISL_3172709, EPI_ISL_3172710 | Lighthouse Lab in Glasgow | Wellcome Sanger Institute for the COVID-19 Genomics UK (COG-UK) Consortium | Harper VanSteenhouse, Yumi Kasai, David Gray, Carol Clugston, Anna Dominiczak and Alex Alderton, Roberto Amato, Jeffrey Barrett, Sonia Goncalves, Ewan Harrison, David K. Jackson, Ian Johnston, Dominic Kwiatkowski, Cordelia Langford, John Sillitoe on behalf of the Wellcome Sanger Institute COVID-19 Surveillance Team |
| EPI_ISL_3172711 | Berkshire and Surrey Pathology Services Lighthouse Laboratory | Wellcome Sanger Institute for the COVID-19 Genomics UK (COG-UK) Consortium | Berkshire and Surrey Pathology Services Lighthouse Laboratory and Alex Alderton, Roberto Amato, Jeffrey Barrett, Sonia Goncalves, Ewan Harrison, David K. Jackson, Ian Johnston, Dominic Kwiatkowski, Cordelia Langford, John Sillitoe on behalf of the Wellcome Sanger Institute COVID-19 Surveillance Team |
| EPI_ISL_3172712 | Lighthouse Lab in Alderley Park | Wellcome Sanger Institute for the COVID-19 Genomics UK (COG-UK) Consortium | Jacquelyn Wynn, Mairead Hyland, The Lighthouse Lab in Alderley Park and Alex Alderton, Roberto Amato, Jeffrey Barrett, Sonia Goncalves, Ewan Harrison, David K. Jackson, Ian Johnston, Dominic Kwiatkowski, Cordelia Langford, John Sillitoe on behalf of the Wellcome Sanger Institute COVID-19 Surveillance Team |
| EPI_ISL_3172713 | Lighthouse Lab in Glasgow | Wellcome Sanger Institute for the COVID-19 Genomics UK (COG-UK) Consortium | Harper VanSteenhouse, Yumi Kasai, David Gray, Carol Clugston, Anna Dominiczak and Alex Alderton, Roberto Amato, Jeffrey Barrett, Sonia Goncalves, Ewan Harrison, David K. Jackson, Ian Johnston, Dominic Kwiatkowski, Cordelia Langford, John Sillitoe on behalf of the Wellcome Sanger Institute COVID-19 Surveillance Team |
| EPI_ISL_3172714 | Berkshire and Surrey Pathology Services Lighthouse Laboratory | Wellcome Sanger Institute for the COVID-19 Genomics UK (COG-UK) Consortium | Berkshire and Surrey Pathology Services Lighthouse Laboratory and Alex Alderton, Roberto Amato, Jeffrey Barrett, Sonia Goncalves, Ewan Harrison, David K. Jackson, Ian Johnston, Dominic Kwiatkowski, Cordelia Langford, John Sillitoe on behalf of the Wellcome Sanger Institute COVID-19 Surveillance Team |
| EPI_ISL_3172715 | Lighthouse Lab in Glasgow | Wellcome Sanger Institute for the COVID-19 Genomics UK (COG-UK) Consortium | Harper VanSteenhouse, Yumi Kasai, David Gray, Carol Clugston, Anna Dominiczak and Alex Alderton, Roberto Amato, Jeffrey Barrett, Sonia Goncalves, Ewan Harrison, David K. Jackson, Ian Johnston, Dominic Kwiatkowski, Cordelia Langford, John Sillitoe on behalf of the Wellcome Sanger Institute COVID-19 Surveillance Team |
| EPI_ISL_3172716, EPI_ISL_3172717 | Lighthouse Lab in Alderley Park | Wellcome Sanger Institute for the COVID-19 Genomics UK (COG-UK) Consortium | Jacquelyn Wynn, Mairead Hyland, The Lighthouse Lab in Alderley Park and Alex Alderton, Roberto Amato, Jeffrey Barrett, Sonia Goncalves, Ewan Harrison, David K. Jackson, Ian Johnston, Dominic Kwiatkowski, Cordelia Langford, John Sillitoe on behalf of the Wellcome Sanger Institute COVID-19 Surveillance Team |
| EPI_ISL_3172718, EPI_ISL_3172719 | Berkshire and Surrey Pathology Services Lighthouse Laboratory | Wellcome Sanger Institute for the COVID-19 Genomics UK (COG-UK) Consortium | Berkshire and Surrey Pathology Services Lighthouse Laboratory and Alex Alderton, Roberto Amato, Jeffrey Barrett, Sonia Goncalves, Ewan Harrison, David K. Jackson, Ian Johnston, Dominic Kwiatkowski, Cordelia Langford, John Sillitoe on behalf of the Wellcome Sanger Institute COVID-19 Surveillance Team |
| EPI_ISL_3172720, EPI_ISL_3172721 | Lighthouse Lab in Alderley Park | Wellcome Sanger Institute for the COVID-19 Genomics UK (COG-UK) Consortium | Jacquelyn Wynn, Mairead Hyland, The Lighthouse Lab in Alderley Park and Alex Alderton, Roberto Amato, Jeffrey Barrett, Sonia Goncalves, Ewan Harrison, David K. Jackson, Ian Johnston, Dominic Kwiatkowski, Cordelia Langford, John Sillitoe on behalf of the Wellcome Sanger Institute COVID-19 Surveillance Team |
| EPI_ISL_3172722 | Lighthouse Lab in Glasgow | Wellcome Sanger Institute for the COVID-19 Genomics UK (COG-UK) Consortium | Harper VanSteenhouse, Yumi Kasai, David Gray, Carol Clugston, Anna Dominiczak and Alex Alderton, Roberto Amato, Jeffrey Barrett, Sonia Goncalves, Ewan Harrison, David K. Jackson, Ian Johnston, Dominic Kwiatkowski, Cordelia Langford, John Sillitoe on behalf of the Wellcome Sanger Institute COVID-19 Surveillance Team |
| EPI_ISL_3172723, EPI_ISL_3172724 | Lighthouse Lab in Alderley Park | Wellcome Sanger Institute for the COVID-19 Genomics UK (COG-UK) Consortium | Jacquelyn Wynn, Mairead Hyland, The Lighthouse Lab in Alderley Park and Alex Alderton, Roberto Amato, Jeffrey Barrett, Sonia Goncalves, Ewan Harrison, David K. Jackson, Ian Johnston, Dominic Kwiatkowski, Cordelia Langford, John Sillitoe on behalf of the Wellcome Sanger Institute COVID-19 Surveillance Team |
| EPI_ISL_3172725 | Berkshire and Surrey Pathology Services Lighthouse Laboratory | Wellcome Sanger Institute for the COVID-19 Genomics UK (COG-UK) Consortium | Berkshire and Surrey Pathology Services Lighthouse Laboratory and Alex Alderton, Roberto Amato, Jeffrey Barrett, Sonia Goncalves, Ewan Harrison, David K. Jackson, Ian Johnston, Dominic Kwiatkowski, Cordelia Langford, John Sillitoe on behalf of the Wellcome Sanger Institute COVID-19 Surveillance Team |
| EPI_ISL_3172726, EPI_ISL_3172727, EPI_ISL_3172728 | Lighthouse Lab in Glasgow | Wellcome Sanger Institute for the COVID-19 Genomics UK (COG-UK) Consortium | Harper VanSteenhouse, Yumi Kasai, David Gray, Carol Clugston, Anna Dominiczak and Alex Alderton, Roberto Amato, Jeffrey Barrett, Sonia Goncalves, Ewan Harrison, David K. Jackson, Ian Johnston, Dominic Kwiatkowski, Cordelia Langford, John Sillitoe on behalf of the Wellcome Sanger Institute COVID-19 Surveillance Team |
| EPI_ISL_3172729 | Berkshire and Surrey Pathology Services Lighthouse Laboratory | Wellcome Sanger Institute for the COVID-19 Genomics UK (COG-UK) Consortium | Berkshire and Surrey Pathology Services Lighthouse Laboratory and Alex Alderton, Roberto Amato, Jeffrey Barrett, Sonia Goncalves, Ewan Harrison, David K. Jackson, Ian Johnston, Dominic Kwiatkowski, Cordelia Langford, John Sillitoe on behalf of the Wellcome Sanger Institute COVID-19 Surveillance Team |
| EPI_ISL_3172730 | Lighthouse Lab in Glasgow | Wellcome Sanger Institute for the COVID-19 Genomics UK (COG-UK) Consortium | Harper VanSteenhouse, Yumi Kasai, David Gray, Carol Clugston, Anna Dominiczak and Alex Alderton, Roberto Amato, Jeffrey Barrett, Sonia Goncalves, Ewan Harrison, David K. Jackson, Ian Johnston, Dominic Kwiatkowski, Cordelia Langford, John Sillitoe on behalf of the Wellcome Sanger Institute COVID-19 Surveillance Team |
| EPI_ISL_3172731 | Berkshire and Surrey Pathology Services Lighthouse Laboratory | Wellcome Sanger Institute for the COVID-19 Genomics UK (COG-UK) Consortium | Berkshire and Surrey Pathology Services Lighthouse Laboratory and Alex Alderton, Roberto Amato, Jeffrey Barrett, Sonia Goncalves, Ewan Harrison, David K. Jackson, Ian Johnston, Dominic Kwiatkowski, Cordelia Langford, John Sillitoe on behalf of the Wellcome Sanger Institute COVID-19 Surveillance Team |
| EPI_ISL_3172732, EPI_ISL_3172733 | Lighthouse Lab in Alderley Park | Wellcome Sanger Institute for the COVID-19 Genomics UK (COG-UK) Consortium | Jacquelyn Wynn, Mairead Hyland, The Lighthouse Lab in Alderley Park and Alex Alderton, Roberto Amato, Jeffrey Barrett, Sonia Goncalves, Ewan Harrison, David K. Jackson, Ian Johnston, Dominic Kwiatkowski, Cordelia Langford, John Sillitoe on behalf of the Wellcome Sanger Institute COVID-19 Surveillance Team |
| EPI_ISL_3172734, EPI_ISL_3172735 | Lighthouse Lab in Glasgow | Wellcome Sanger Institute for the COVID-19 Genomics UK (COG-UK) Consortium | Harper VanSteenhouse, Yumi Kasai, David Gray, Carol Clugston, Anna Dominiczak and Alex Alderton, Roberto Amato, Jeffrey Barrett, Sonia Goncalves, Ewan Harrison, David K. Jackson, Ian Johnston, Dominic Kwiatkowski, Cordelia Langford, John Sillitoe on behalf of the Wellcome Sanger Institute COVID-19 Surveillance Team |
| EPI_ISL_3172736 | Lighthouse Lab in Alderley Park | Wellcome Sanger Institute for the COVID-19 Genomics UK (COG-UK) Consortium | Jacquelyn Wynn, Mairead Hyland, The Lighthouse Lab in Alderley Park and Alex Alderton, Roberto Amato, Jeffrey Barrett, Sonia Goncalves, Ewan Harrison, David K. Jackson, Ian Johnston, Dominic Kwiatkowski, Cordelia Langford, John Sillitoe on behalf of the Wellcome Sanger Institute COVID-19 Surveillance Team |
| EPI_ISL_3172737, EPI_ISL_3172738, EPI_ISL_3172739 | Lighthouse Lab in Glasgow | Wellcome Sanger Institute for the COVID-19 Genomics UK (COG-UK) Consortium | Harper VanSteenhouse, Yumi Kasai, David Gray, Carol Clugston, Anna Dominiczak and Alex Alderton, Roberto Amato, Jeffrey Barrett, Sonia Goncalves, Ewan Harrison, David K. Jackson, Ian Johnston, Dominic Kwiatkowski, Cordelia Langford, John Sillitoe on behalf of the Wellcome Sanger Institute COVID-19 Surveillance Team |

| | | | |
|---|---|---|---|
| EPI_ISL_3172740 | Berkshire and Surrey Pathology Services Lighthouse Laboratory | Wellcome Sanger Institute for the COVID-19 Genomics UK (COG-UK) Consortium | Berkshire and Surrey Pathology Services Lighthouse Laboratory and Alex Alderton, Roberto Amato, Jeffrey Barrett, Sonia Goncalves, Ewan Harrison, David K. Jackson, Ian Johnston, Dominic Kwiatkowski, Cordelia Langford, John Sillitoe on behalf of the Wellcome Sanger Institute COVID-19 Surveillance Team |
| EPI_ISL_3172741 | Lighthouse Lab in Alderley Park | Wellcome Sanger Institute for the COVID-19 Genomics UK (COG-UK) Consortium | Jacquelyn Wynn, Mairead Hyland, The Lighthouse Lab in Alderley Park and Alex Alderton, Roberto Amato, Jeffrey Barrett, Sonia Goncalves, Ewan Harrison, David K. Jackson, Ian Johnston, Dominic Kwiatkowski, Cordelia Langford, John Sillitoe on behalf of the Wellcome Sanger Institute COVID-19 Surveillance Team |
| EPI_ISL_3172742, EPI_ISL_3172743 | Lighthouse Lab in Glasgow | Wellcome Sanger Institute for the COVID-19 Genomics UK (COG-UK) Consortium | Harper VanSteenhouse, Yumi Kasai, David Gray, Carol Clugston, Anna Dominiczak and Alex Alderton, Roberto Amato, Jeffrey Barrett, Sonia Goncalves, Ewan Harrison, David K. Jackson, Ian Johnston, Dominic Kwiatkowski, Cordelia Langford, John Sillitoe on behalf of the Wellcome Sanger Institute COVID-19 Surveillance Team |
| EPI_ISL_3172744 | Berkshire and Surrey Pathology Services Lighthouse Laboratory | Wellcome Sanger Institute for the COVID-19 Genomics UK (COG-UK) Consortium | Berkshire and Surrey Pathology Services Lighthouse Laboratory and Alex Alderton, Roberto Amato, Jeffrey Barrett, Sonia Goncalves, Ewan Harrison, David K. Jackson, Ian Johnston, Dominic Kwiatkowski, Cordelia Langford, John Sillitoe on behalf of the Wellcome Sanger Institute COVID-19 Surveillance Team |
| EPI_ISL_3172745 | Lighthouse Lab in Alderley Park | Wellcome Sanger Institute for the COVID-19 Genomics UK (COG-UK) Consortium | Jacquelyn Wynn, Mairead Hyland, The Lighthouse Lab in Alderley Park and Alex Alderton, Roberto Amato, Jeffrey Barrett, Sonia Goncalves, Ewan Harrison, David K. Jackson, Ian Johnston, Dominic Kwiatkowski, Cordelia Langford, John Sillitoe on behalf of the Wellcome Sanger Institute COVID-19 Surveillance Team |
| EPI_ISL_3172746 | Berkshire and Surrey Pathology Services Lighthouse Laboratory | Wellcome Sanger Institute for the COVID-19 Genomics UK (COG-UK) Consortium | Berkshire and Surrey Pathology Services Lighthouse Laboratory and Alex Alderton, Roberto Amato, Jeffrey Barrett, Sonia Goncalves, Ewan Harrison, David K. Jackson, Ian Johnston, Dominic Kwiatkowski, Cordelia Langford, John Sillitoe on behalf of the Wellcome Sanger Institute COVID-19 Surveillance Team |
| EPI_ISL_3172747 | Lighthouse Lab in Alderley Park | Wellcome Sanger Institute for the COVID-19 Genomics UK (COG-UK) Consortium | Jacquelyn Wynn, Mairead Hyland, The Lighthouse Lab in Alderley Park and Alex Alderton, Roberto Amato, Jeffrey Barrett, Sonia Goncalves, Ewan Harrison, David K. Jackson, Ian Johnston, Dominic Kwiatkowski, Cordelia Langford, John Sillitoe on behalf of the Wellcome Sanger Institute COVID-19 Surveillance Team |
| EPI_ISL_3172748, EPI_ISL_3172749 | Lighthouse Lab in Glasgow | Wellcome Sanger Institute for the COVID-19 Genomics UK (COG-UK) Consortium | Harper VanSteenhouse, Yumi Kasai, David Gray, Carol Clugston, Anna Dominiczak and Alex Alderton, Roberto Amato, Jeffrey Barrett, Sonia Goncalves, Ewan Harrison, David K. Jackson, Ian Johnston, Dominic Kwiatkowski, Cordelia Langford, John Sillitoe on behalf of the Wellcome Sanger Institute COVID-19 Surveillance Team |
| EPI_ISL_3172750 | Berkshire and Surrey Pathology Services Lighthouse Laboratory | Wellcome Sanger Institute for the COVID-19 Genomics UK (COG-UK) Consortium | Berkshire and Surrey Pathology Services Lighthouse Laboratory and Alex Alderton, Roberto Amato, Jeffrey Barrett, Sonia Goncalves, Ewan Harrison, David K. Jackson, Ian Johnston, Dominic Kwiatkowski, Cordelia Langford, John Sillitoe on behalf of the Wellcome Sanger Institute COVID-19 Surveillance Team |
| EPI_ISL_3172751 | Lighthouse Lab in Alderley Park | Wellcome Sanger Institute for the COVID-19 Genomics UK (COG-UK) Consortium | Jacquelyn Wynn, Mairead Hyland, The Lighthouse Lab in Alderley Park and Alex Alderton, Roberto Amato, Jeffrey Barrett, Sonia Goncalves, Ewan Harrison, David K. Jackson, Ian Johnston, Dominic Kwiatkowski, Cordelia Langford, John Sillitoe on behalf of the Wellcome Sanger Institute COVID-19 Surveillance Team |
| EPI_ISL_3172752 | Berkshire and Surrey Pathology Services Lighthouse Laboratory | Wellcome Sanger Institute for the COVID-19 Genomics UK (COG-UK) Consortium | Berkshire and Surrey Pathology Services Lighthouse Laboratory and Alex Alderton, Roberto Amato, Jeffrey Barrett, Sonia Goncalves, Ewan Harrison, David K. Jackson, Ian Johnston, Dominic Kwiatkowski, Cordelia Langford, John Sillitoe on behalf of the Wellcome Sanger Institute COVID-19 Surveillance Team |
| EPI_ISL_3172753 | Lighthouse Lab in Glasgow | Wellcome Sanger Institute for the COVID-19 Genomics UK (COG-UK) Consortium | Harper VanSteenhouse, Yumi Kasai, David Gray, Carol Clugston, Anna Dominiczak and Alex Alderton, Roberto Amato, Jeffrey Barrett, Sonia Goncalves, Ewan Harrison, David K. Jackson, Ian Johnston, Dominic Kwiatkowski, Cordelia Langford, John Sillitoe on behalf of the Wellcome Sanger Institute COVID-19 Surveillance Team |
| EPI_ISL_3172754, EPI_ISL_3172755 | Berkshire and Surrey Pathology Services Lighthouse Laboratory | Wellcome Sanger Institute for the COVID-19 Genomics UK (COG-UK) Consortium | Berkshire and Surrey Pathology Services Lighthouse Laboratory and Alex Alderton, Roberto Amato, Jeffrey Barrett, Sonia Goncalves, Ewan Harrison, David K. Jackson, Ian Johnston, Dominic Kwiatkowski, Cordelia Langford, John Sillitoe on behalf of the Wellcome Sanger Institute COVID-19 Surveillance Team |
| EPI_ISL_3172756 | Lighthouse Lab in Glasgow | Wellcome Sanger Institute for the COVID-19 Genomics UK (COG-UK) Consortium | Harper VanSteenhouse, Yumi Kasai, David Gray, Carol Clugston, Anna Dominiczak and Alex Alderton, Roberto Amato, Jeffrey Barrett, Sonia Goncalves, Ewan Harrison, David K. Jackson, Ian Johnston, Dominic Kwiatkowski, Cordelia Langford, John Sillitoe on behalf of the Wellcome Sanger Institute COVID-19 Surveillance Team |
| EPI_ISL_3172757 | Lighthouse Lab in Alderley Park | Wellcome Sanger Institute for the COVID-19 Genomics UK (COG-UK) Consortium | Jacquelyn Wynn, Mairead Hyland, The Lighthouse Lab in Alderley Park and Alex Alderton, Roberto Amato, Jeffrey Barrett, Sonia Goncalves, Ewan Harrison, David K. Jackson, Ian Johnston, Dominic Kwiatkowski, Cordelia Langford, John Sillitoe on behalf of the Wellcome Sanger Institute COVID-19 Surveillance Team |
| EPI_ISL_3172758, EPI_ISL_3172759 | Berkshire and Surrey Pathology Services Lighthouse Laboratory | Wellcome Sanger Institute for the COVID-19 Genomics UK (COG-UK) Consortium | Berkshire and Surrey Pathology Services Lighthouse Laboratory and Alex Alderton, Roberto Amato, Jeffrey Barrett, Sonia Goncalves, Ewan Harrison, David K. Jackson, Ian Johnston, Dominic Kwiatkowski, Cordelia Langford, John Sillitoe on behalf of the Wellcome Sanger Institute COVID-19 Surveillance Team |
| EPI_ISL_3172760 | Lighthouse Lab in Alderley Park | Wellcome Sanger Institute for the COVID-19 Genomics UK (COG-UK) Consortium | Jacquelyn Wynn, Mairead Hyland, The Lighthouse Lab in Alderley Park and Alex Alderton, Roberto Amato, Jeffrey Barrett, Sonia Goncalves, Ewan Harrison, David K. Jackson, Ian Johnston, Dominic Kwiatkowski, Cordelia Langford, John Sillitoe on behalf of the Wellcome Sanger Institute COVID-19 Surveillance Team |
| EPI_ISL_3172761 | Berkshire and Surrey Pathology Services Lighthouse Laboratory | Wellcome Sanger Institute for the COVID-19 Genomics UK (COG-UK) Consortium | Berkshire and Surrey Pathology Services Lighthouse Laboratory and Alex Alderton, Roberto Amato, Jeffrey Barrett, Sonia Goncalves, Ewan Harrison, David K. Jackson, Ian Johnston, Dominic Kwiatkowski, Cordelia Langford, John Sillitoe on behalf of the Wellcome Sanger Institute COVID-19 Surveillance Team |
| EPI_ISL_3172762, EPI_ISL_3172763, EPI_ISL_3172764 | Lighthouse Lab in Glasgow | Wellcome Sanger Institute for the COVID-19 Genomics UK (COG-UK) Consortium | Harper VanSteenhouse, Yumi Kasai, David Gray, Carol Clugston, Anna Dominiczak and Alex Alderton, Roberto Amato, Jeffrey Barrett, Sonia Goncalves, Ewan Harrison, David K. Jackson, Ian Johnston, Dominic Kwiatkowski, Cordelia Langford, John Sillitoe on behalf of the Wellcome Sanger Institute COVID-19 Surveillance Team |
| EPI_ISL_3172765 | Lighthouse Lab in Alderley Park | Wellcome Sanger Institute for the COVID-19 Genomics UK (COG-UK) Consortium | Jacquelyn Wynn, Mairead Hyland, The Lighthouse Lab in Alderley Park and Alex Alderton, Roberto Amato, Jeffrey Barrett, Sonia Goncalves, Ewan Harrison, David K. Jackson, Ian Johnston, Dominic Kwiatkowski, Cordelia Langford, John Sillitoe on behalf of the Wellcome Sanger Institute COVID-19 Surveillance Team |
| EPI_ISL_3172766, EPI_ISL_3172767, EPI_ISL_3172768 | Lighthouse Lab in Glasgow | Wellcome Sanger Institute for the COVID-19 Genomics UK (COG-UK) Consortium | Harper VanSteenhouse, Yumi Kasai, David Gray, Carol Clugston, Anna Dominiczak and Alex Alderton, Roberto Amato, Jeffrey Barrett, Sonia Goncalves, Ewan Harrison, David K. Jackson, Ian Johnston, Dominic Kwiatkowski, Cordelia Langford, John Sillitoe on behalf of the Wellcome Sanger Institute COVID-19 Surveillance Team |
| EPI_ISL_3172769 | Lighthouse Lab in Alderley Park | Wellcome Sanger Institute for the COVID-19 Genomics UK (COG-UK) Consortium | Jacquelyn Wynn, Mairead Hyland, The Lighthouse Lab in Alderley Park and Alex Alderton, Roberto Amato, Jeffrey Barrett, Sonia Goncalves, Ewan Harrison, David K. Jackson, Ian Johnston, Dominic Kwiatkowski, Cordelia Langford, John Sillitoe on behalf of the Wellcome Sanger Institute COVID-19 Surveillance Team |
| EPI_ISL_3172770 | Lighthouse Lab in Glasgow | Wellcome Sanger Institute for the COVID-19 Genomics UK (COG-UK) Consortium | Harper VanSteenhouse, Yumi Kasai, David Gray, Carol Clugston, Anna Dominiczak and Alex Alderton, Roberto Amato, Jeffrey Barrett, Sonia Goncalves, Ewan Harrison, David K. Jackson, Ian Johnston, Dominic Kwiatkowski, Cordelia Langford, John Sillitoe on behalf of the Wellcome Sanger Institute COVID-19 Surveillance Team |
| EPI_ISL_3172771 | Lighthouse Lab in Milton Keynes | Wellcome Sanger Institute for the COVID-19 Genomics UK (COG-UK) Consortium | The Lighthouse Lab in Milton Keynes and Alex Alderton, Roberto Amato, Jeffrey Barrett, Sonia Goncalves, Ewan Harrison, David K. Jackson, Ian Johnston, Dominic Kwiatkowski, Cordelia Langford, John Sillitoe on behalf of the Wellcome Sanger Institute COVID-19 Surveillance Team |
| EPI_ISL_3172772 | Lighthouse Lab in Glasgow | Wellcome Sanger Institute for the COVID-19 Genomics UK (COG-UK) Consortium | Harper VanSteenhouse, Yumi Kasai, David Gray, Carol Clugston, Anna Dominiczak and Alex Alderton, Roberto Amato, Jeffrey Barrett, Sonia Goncalves, Ewan Harrison, David K. Jackson, Ian Johnston, Dominic Kwiatkowski, Cordelia Langford, John Sillitoe on behalf of the Wellcome Sanger Institute COVID-19 Surveillance Team |
| EPI_ISL_3172773 | Lighthouse Lab in Milton Keynes | Wellcome Sanger Institute for the COVID-19 Genomics UK (COG-UK) Consortium | The Lighthouse Lab in Milton Keynes and Alex Alderton, Roberto Amato, Jeffrey Barrett, Sonia Goncalves, Ewan Harrison, David K. Jackson, Ian Johnston, Dominic Kwiatkowski, Cordelia Langford, John Sillitoe on behalf of the Wellcome Sanger Institute COVID-19 Surveillance Team |
| EPI_ISL_3172774 | Lighthouse Lab in Glasgow | Wellcome Sanger Institute for the COVID-19 Genomics UK (COG-UK) Consortium | Harper VanSteenhouse, Yumi Kasai, David Gray, Carol Clugston, Anna Dominiczak and Alex Alderton, Roberto Amato, Jeffrey Barrett, Sonia Goncalves, Ewan Harrison, David K. Jackson, Ian Johnston, Dominic Kwiatkowski, Cordelia Langford, John Sillitoe on behalf of the Wellcome Sanger Institute COVID-19 Surveillance Team |
| EPI_ISL_3172775, EPI_ISL_3172776 | Lighthouse Lab in Milton Keynes | Wellcome Sanger Institute for the COVID-19 Genomics UK (COG-UK) Consortium | The Lighthouse Lab in Milton Keynes and Alex Alderton, Roberto Amato, Jeffrey Barrett, Sonia Goncalves, Ewan Harrison, David K. Jackson, Ian Johnston, Dominic Kwiatkowski, Cordelia Langford, John Sillitoe on behalf of the Wellcome Sanger Institute COVID-19 Surveillance Team |
| EPI_ISL_3172777, EPI_ISL_3172778 | Lighthouse Lab in Glasgow | Wellcome Sanger Institute for the COVID-19 Genomics UK (COG-UK) Consortium | Harper VanSteenhouse, Yumi Kasai, David Gray, Carol Clugston, Anna Dominiczak and Alex Alderton, Roberto Amato, Jeffrey Barrett, Sonia Goncalves, Ewan Harrison, David K. Jackson, Ian Johnston, Dominic Kwiatkowski, Cordelia Langford, John Sillitoe on behalf of the Wellcome Sanger Institute COVID-19 Surveillance Team |
| EPI_ISL_3172779 | Lighthouse Lab in Milton Keynes | Wellcome Sanger Institute for the COVID-19 Genomics UK (COG-UK) Consortium | The Lighthouse Lab in Milton Keynes and Alex Alderton, Roberto Amato, Jeffrey Barrett, Sonia Goncalves, Ewan Harrison, David K. Jackson, Ian Johnston, Dominic Kwiatkowski, Cordelia Langford, John Sillitoe on behalf of the Wellcome Sanger Institute COVID-19 Surveillance Team |
| EPI_ISL_3172780, EPI_ISL_3172781 | Lighthouse Lab in Glasgow | Wellcome Sanger Institute for the COVID-19 Genomics UK (COG-UK) Consortium | Harper VanSteenhouse, Yumi Kasai, David Gray, Carol Clugston, Anna Dominiczak and Alex Alderton, Roberto Amato, Jeffrey Barrett, Sonia Goncalves, Ewan Harrison, David K. Jackson, Ian Johnston, Dominic Kwiatkowski, Cordelia Langford, John Sillitoe on behalf of the Wellcome Sanger Institute COVID-19 Surveillance Team |
| EPI_ISL_3172782, EPI_ISL_3172783 | Lighthouse Lab in Milton Keynes | Wellcome Sanger Institute for the COVID-19 Genomics UK (COG-UK) Consortium | The Lighthouse Lab in Milton Keynes and Alex Alderton, Roberto Amato, Jeffrey Barrett, Sonia Goncalves, Ewan Harrison, David K. Jackson, Ian Johnston, Dominic Kwiatkowski, Cordelia Langford, John Sillitoe on behalf of the Wellcome Sanger Institute COVID-19 Surveillance Team |
| EPI_ISL_3172784, EPI_ISL_3172785 | Lighthouse Lab in Glasgow | Wellcome Sanger Institute for the COVID-19 Genomics UK (COG-UK) Consortium | Harper VanSteenhouse, Yumi Kasai, David Gray, Carol Clugston, Anna Dominiczak and Alex Alderton, Roberto Amato, Jeffrey Barrett, Sonia Goncalves, Ewan Harrison, David K. Jackson, Ian Johnston, Dominic Kwiatkowski, Cordelia Langford, John Sillitoe on behalf of the Wellcome Sanger Institute COVID-19 Surveillance Team |
| EPI_ISL_3172786, EPI_ISL_3172787, EPI_ISL_3172788, EPI_ISL_3172789 | Lighthouse Lab in Milton Keynes | Wellcome Sanger Institute for the COVID-19 Genomics UK (COG-UK) Consortium | The Lighthouse Lab in Milton Keynes and Alex Alderton, Roberto Amato, Jeffrey Barrett, Sonia Goncalves, Ewan Harrison, David K. Jackson, Ian Johnston, Dominic Kwiatkowski, Cordelia Langford, John Sillitoe on behalf of the Wellcome Sanger Institute COVID-19 Surveillance Team |
| EPI_ISL_3172790 | Lighthouse Lab in Glasgow | Wellcome Sanger Institute for the COVID-19 Genomics UK (COG-UK) Consortium | Harper VanSteenhouse, Yumi Kasai, David Gray, Carol Clugston, Anna Dominiczak and Alex Alderton, Roberto Amato, Jeffrey Barrett, Sonia Goncalves, Ewan Harrison, David K. Jackson, Ian Johnston, Dominic Kwiatkowski, Cordelia Langford, John Sillitoe on behalf of the Wellcome Sanger Institute COVID-19 Surveillance Team |
| EPI_ISL_3172791, EPI_ISL_3172792, EPI_ISL_3172793, EPI_ISL_3172794 | Lighthouse Lab in Milton Keynes | Wellcome Sanger Institute for the COVID-19 Genomics UK (COG-UK) Consortium | The Lighthouse Lab in Milton Keynes and Alex Alderton, Roberto Amato, Jeffrey Barrett, Sonia Goncalves, Ewan Harrison, David K. Jackson, Ian Johnston, Dominic Kwiatkowski, Cordelia Langford, John Sillitoe on behalf of the Wellcome Sanger Institute COVID-19 Surveillance Team |
| EPI_ISL_3172795 | Lighthouse Lab in Glasgow | Wellcome Sanger Institute for the COVID-19 Genomics UK (COG-UK) Consortium | Harper VanSteenhouse, Yumi Kasai, David Gray, Carol Clugston, Anna Dominiczak and Alex Alderton, Roberto Amato, Jeffrey Barrett, Sonia Goncalves, Ewan Harrison, David K. Jackson, Ian Johnston, Dominic Kwiatkowski, Cordelia Langford, John Sillitoe on behalf of the Wellcome Sanger Institute COVID-19 Surveillance Team |
| EPI_ISL_3172796 | Lighthouse Lab in Milton Keynes | Wellcome Sanger Institute for the COVID-19 Genomics UK (COG-UK) Consortium | The Lighthouse Lab in Milton Keynes and Alex Alderton, Roberto Amato, Jeffrey Barrett, Sonia Goncalves, Ewan Harrison, David K. Jackson, Ian Johnston, Dominic Kwiatkowski, Cordelia Langford, John Sillitoe on behalf of the Wellcome Sanger Institute COVID-19 Surveillance Team |
| EPI_ISL_3172797 | Lighthouse Lab in Glasgow | Wellcome Sanger Institute for the COVID-19 Genomics UK (COG-UK) Consortium | Harper VanSteenhouse, Yumi Kasai, David Gray, Carol Clugston, Anna Dominiczak and Alex Alderton, Roberto Amato, Jeffrey Barrett, Sonia Goncalves, Ewan Harrison, David K. Jackson, Ian Johnston, Dominic Kwiatkowski, Cordelia Langford, John Sillitoe on behalf of the Wellcome Sanger Institute COVID-19 Surveillance Team |
| EPI_ISL_3172798 | Lighthouse Lab in Milton Keynes | Wellcome Sanger Institute for the COVID-19 Genomics UK (COG-UK) Consortium | The Lighthouse Lab in Milton Keynes and Alex Alderton, Roberto Amato, Jeffrey Barrett, Sonia Goncalves, Ewan Harrison, David K. Jackson, Ian Johnston, Dominic Kwiatkowski, Cordelia Langford, John Sillitoe on behalf of the Wellcome Sanger Institute COVID-19 Surveillance Team |
| EPI_ISL_3172799, EPI_ISL_3172800 | Lighthouse Lab in Glasgow | Wellcome Sanger Institute for the COVID-19 Genomics UK (COG-UK) Consortium | Harper VanSteenhouse, Yumi Kasai, David Gray, Carol Clugston, Anna Dominiczak and Alex Alderton, Roberto Amato, Jeffrey Barrett, Sonia Goncalves, Ewan Harrison, David K. Jackson, Ian Johnston, Dominic Kwiatkowski, Cordelia Langford, John Sillitoe on behalf of the Wellcome Sanger Institute COVID-19 Surveillance Team |
| EPI_ISL_3172801 | Lighthouse Lab in Milton Keynes | Wellcome Sanger Institute for the COVID-19 Genomics UK (COG-UK) Consortium | The Lighthouse Lab in Milton Keynes and Alex Alderton, Roberto Amato, Jeffrey Barrett, Sonia Goncalves, Ewan Harrison, David K. Jackson, Ian Johnston, Dominic Kwiatkowski, Cordelia Langford, John Sillitoe on behalf of the Wellcome Sanger Institute COVID-19 Surveillance Team |
| EPI_ISL_3172802, EPI_ISL_3172803, | Lighthouse Lab in Glasgow | Wellcome Sanger Institute for the COVID-19 Genomics UK (COG-UK) Consortium | Harper VanSteenhouse, Yumi Kasai, David Gray, Carol Clugston, Anna Dominiczak and Alex Alderton, Roberto Amato, Jeffrey Barrett, Sonia Goncalves, Ewan Harrison, David K. Jackson, Ian Johnston, Dominic Kwiatkowski, Cordelia Langford, John Sillitoe on behalf of the Wellcome Sanger Institute COVID-19 Surveillance Team |

| | | | |
|---|---|---|---|
| EPI_ISL_3172804, EPI_ISL_3172805, EPI_ISL_3172806 | | | |
| EPI_ISL_3172807 | Lighthouse Lab in Milton Keynes | Wellcome Sanger Institute for the COVID-19 Genomics UK (COG-UK) Consortium | The Lighthouse Lab in Milton Keynes and Alex Alderton, Roberto Amato, Jeffrey Barrett, Sonia Goncalves, Ewan Harrison, David K. Jackson, Ian Johnston, Dominic Kwiatkowski, Cordelia Langford, John Sillitoe on behalf of the Wellcome Sanger Institute COVID-19 Surveillance Team |
| EPI_ISL_3172808 | Lighthouse Lab in Glasgow | Wellcome Sanger Institute for the COVID-19 Genomics UK (COG-UK) Consortium | Harper VanSteenhouse, Yumi Kasai, David Gray, Carol Clugston, Anna Dominiczak and Alex Alderton, Roberto Amato, Jeffrey Barrett, Sonia Goncalves, Ewan Harrison, David K. Jackson, Ian Johnston, Dominic Kwiatkowski, Cordelia Langford, John Sillitoe on behalf of the Wellcome Sanger Institute COVID-19 Surveillance Team |
| EPI_ISL_3172809 | Lighthouse Lab in Milton Keynes | Wellcome Sanger Institute for the COVID-19 Genomics UK (COG-UK) Consortium | The Lighthouse Lab in Milton Keynes and Alex Alderton, Roberto Amato, Jeffrey Barrett, Sonia Goncalves, Ewan Harrison, David K. Jackson, Ian Johnston, Dominic Kwiatkowski, Cordelia Langford, John Sillitoe on behalf of the Wellcome Sanger Institute COVID-19 Surveillance Team |
| EPI_ISL_3172810 | Lighthouse Lab in Glasgow | Wellcome Sanger Institute for the COVID-19 Genomics UK (COG-UK) Consortium | Harper VanSteenhouse, Yumi Kasai, David Gray, Carol Clugston, Anna Dominiczak and Alex Alderton, Roberto Amato, Jeffrey Barrett, Sonia Goncalves, Ewan Harrison, David K. Jackson, Ian Johnston, Dominic Kwiatkowski, Cordelia Langford, John Sillitoe on behalf of the Wellcome Sanger Institute COVID-19 Surveillance Team |
| EPI_ISL_3172811, EPI_ISL_3172812 | Lighthouse Lab in Milton Keynes | Wellcome Sanger Institute for the COVID-19 Genomics UK (COG-UK) Consortium | The Lighthouse Lab in Milton Keynes and Alex Alderton, Roberto Amato, Jeffrey Barrett, Sonia Goncalves, Ewan Harrison, David K. Jackson, Ian Johnston, Dominic Kwiatkowski, Cordelia Langford, John Sillitoe on behalf of the Wellcome Sanger Institute COVID-19 Surveillance Team |
| EPI_ISL_3172813 | Lighthouse Lab in Glasgow | Wellcome Sanger Institute for the COVID-19 Genomics UK (COG-UK) Consortium | Harper VanSteenhouse, Yumi Kasai, David Gray, Carol Clugston, Anna Dominiczak and Alex Alderton, Roberto Amato, Jeffrey Barrett, Sonia Goncalves, Ewan Harrison, David K. Jackson, Ian Johnston, Dominic Kwiatkowski, Cordelia Langford, John Sillitoe on behalf of the Wellcome Sanger Institute COVID-19 Surveillance Team |
| EPI_ISL_3172814, EPI_ISL_3172815 | Lighthouse Lab in Milton Keynes | Wellcome Sanger Institute for the COVID-19 Genomics UK (COG-UK) Consortium | The Lighthouse Lab in Milton Keynes and Alex Alderton, Roberto Amato, Jeffrey Barrett, Sonia Goncalves, Ewan Harrison, David K. Jackson, Ian Johnston, Dominic Kwiatkowski, Cordelia Langford, John Sillitoe on behalf of the Wellcome Sanger Institute COVID-19 Surveillance Team |
| EPI_ISL_3172816, EPI_ISL_3172817, EPI_ISL_3172818, EPI_ISL_3172819 | Lighthouse Lab in Glasgow | Wellcome Sanger Institute for the COVID-19 Genomics UK (COG-UK) Consortium | Harper VanSteenhouse, Yumi Kasai, David Gray, Carol Clugston, Anna Dominiczak and Alex Alderton, Roberto Amato, Jeffrey Barrett, Sonia Goncalves, Ewan Harrison, David K. Jackson, Ian Johnston, Dominic Kwiatkowski, Cordelia Langford, John Sillitoe on behalf of the Wellcome Sanger Institute COVID-19 Surveillance Team |
| EPI_ISL_3172820, EPI_ISL_3172821, EPI_ISL_3172822 | Lighthouse Lab in Milton Keynes | Wellcome Sanger Institute for the COVID-19 Genomics UK (COG-UK) Consortium | The Lighthouse Lab in Milton Keynes and Alex Alderton, Roberto Amato, Jeffrey Barrett, Sonia Goncalves, Ewan Harrison, David K. Jackson, Ian Johnston, Dominic Kwiatkowski, Cordelia Langford, John Sillitoe on behalf of the Wellcome Sanger Institute COVID-19 Surveillance Team |
| EPI_ISL_3172823 | Lighthouse Lab in Glasgow | Wellcome Sanger Institute for the COVID-19 Genomics UK (COG-UK) Consortium | Harper VanSteenhouse, Yumi Kasai, David Gray, Carol Clugston, Anna Dominiczak and Alex Alderton, Roberto Amato, Jeffrey Barrett, Sonia Goncalves, Ewan Harrison, David K. Jackson, Ian Johnston, Dominic Kwiatkowski, Cordelia Langford, John Sillitoe on behalf of the Wellcome Sanger Institute COVID-19 Surveillance Team |
| EPI_ISL_3172824, EPI_ISL_3172825 | Lighthouse Lab in Milton Keynes | Wellcome Sanger Institute for the COVID-19 Genomics UK (COG-UK) Consortium | The Lighthouse Lab in Milton Keynes and Alex Alderton, Roberto Amato, Jeffrey Barrett, Sonia Goncalves, Ewan Harrison, David K. Jackson, Ian Johnston, Dominic Kwiatkowski, Cordelia Langford, John Sillitoe on behalf of the Wellcome Sanger Institute COVID-19 Surveillance Team |
| EPI_ISL_3172826 | Lighthouse Lab in Glasgow | Wellcome Sanger Institute for the COVID-19 Genomics UK (COG-UK) Consortium | Harper VanSteenhouse, Yumi Kasai, David Gray, Carol Clugston, Anna Dominiczak and Alex Alderton, Roberto Amato, Jeffrey Barrett, Sonia Goncalves, Ewan Harrison, David K. Jackson, Ian Johnston, Dominic Kwiatkowski, Cordelia Langford, John Sillitoe on behalf of the Wellcome Sanger Institute COVID-19 Surveillance Team |
| EPI_ISL_3172827, EPI_ISL_3172828, EPI_ISL_3172829, EPI_ISL_3172830, EPI_ISL_3172831 | Lighthouse Lab in Milton Keynes | Wellcome Sanger Institute for the COVID-19 Genomics UK (COG-UK) Consortium | The Lighthouse Lab in Milton Keynes and Alex Alderton, Roberto Amato, Jeffrey Barrett, Sonia Goncalves, Ewan Harrison, David K. Jackson, Ian Johnston, Dominic Kwiatkowski, Cordelia Langford, John Sillitoe on behalf of the Wellcome Sanger Institute COVID-19 Surveillance Team |
| EPI_ISL_3172832, EPI_ISL_3172833, EPI_ISL_3172834 | Lighthouse Lab in Glasgow | Wellcome Sanger Institute for the COVID-19 Genomics UK (COG-UK) Consortium | Harper VanSteenhouse, Yumi Kasai, David Gray, Carol Clugston, Anna Dominiczak and Alex Alderton, Roberto Amato, Jeffrey Barrett, Sonia Goncalves, Ewan Harrison, David K. Jackson, Ian Johnston, Dominic Kwiatkowski, Cordelia Langford, John Sillitoe on behalf of the Wellcome Sanger Institute COVID-19 Surveillance Team |
| EPI_ISL_3172835, EPI_ISL_3172836 | Lighthouse Lab in Milton Keynes | Wellcome Sanger Institute for the COVID-19 Genomics UK (COG-UK) Consortium | The Lighthouse Lab in Milton Keynes and Alex Alderton, Roberto Amato, Jeffrey Barrett, Sonia Goncalves, Ewan Harrison, David K. Jackson, Ian Johnston, Dominic Kwiatkowski, Cordelia Langford, John Sillitoe on behalf of the Wellcome Sanger Institute COVID-19 Surveillance Team |
| EPI_ISL_3172837, EPI_ISL_3172838 | Lighthouse Lab in Glasgow | Wellcome Sanger Institute for the COVID-19 Genomics UK (COG-UK) Consortium | Harper VanSteenhouse, Yumi Kasai, David Gray, Carol Clugston, Anna Dominiczak and Alex Alderton, Roberto Amato, Jeffrey Barrett, Sonia Goncalves, Ewan Harrison, David K. Jackson, Ian Johnston, Dominic Kwiatkowski, Cordelia Langford, John Sillitoe on behalf of the Wellcome Sanger Institute COVID-19 Surveillance Team |
| EPI_ISL_3172839, EPI_ISL_3172840 | Lighthouse Lab in Milton Keynes | Wellcome Sanger Institute for the COVID-19 Genomics UK (COG-UK) Consortium | The Lighthouse Lab in Milton Keynes and Alex Alderton, Roberto Amato, Jeffrey Barrett, Sonia Goncalves, Ewan Harrison, David K. Jackson, Ian Johnston, Dominic Kwiatkowski, Cordelia Langford, John Sillitoe on behalf of the Wellcome Sanger Institute COVID-19 Surveillance Team |
| EPI_ISL_3172841, EPI_ISL_3172842 | Lighthouse Lab in Glasgow | Wellcome Sanger Institute for the COVID-19 Genomics UK (COG-UK) Consortium | Harper VanSteenhouse, Yumi Kasai, David Gray, Carol Clugston, Anna Dominiczak and Alex Alderton, Roberto Amato, Jeffrey Barrett, Sonia Goncalves, Ewan Harrison, David K. Jackson, Ian Johnston, Dominic Kwiatkowski, Cordelia Langford, John Sillitoe on behalf of the Wellcome Sanger Institute COVID-19 Surveillance Team |
| EPI_ISL_3172843 | Lighthouse Lab in Milton Keynes | Wellcome Sanger Institute for the COVID-19 Genomics UK (COG-UK) Consortium | The Lighthouse Lab in Milton Keynes and Alex Alderton, Roberto Amato, Jeffrey Barrett, Sonia Goncalves, Ewan Harrison, David K. Jackson, Ian Johnston, Dominic Kwiatkowski, Cordelia Langford, John Sillitoe on behalf of the Wellcome Sanger Institute COVID-19 Surveillance Team |
| EPI_ISL_3172844, EPI_ISL_3172845 | Lighthouse Lab in Glasgow | Wellcome Sanger Institute for the COVID-19 Genomics UK (COG-UK) Consortium | Harper VanSteenhouse, Yumi Kasai, David Gray, Carol Clugston, Anna Dominiczak and Alex Alderton, Roberto Amato, Jeffrey Barrett, Sonia Goncalves, Ewan Harrison, David K. Jackson, Ian Johnston, Dominic Kwiatkowski, Cordelia Langford, John Sillitoe on behalf of the Wellcome Sanger Institute COVID-19 Surveillance Team |
| EPI_ISL_3172846, EPI_ISL_3172847, EPI_ISL_3172848 | Lighthouse Lab in Milton Keynes | Wellcome Sanger Institute for the COVID-19 Genomics UK (COG-UK) Consortium | The Lighthouse Lab in Milton Keynes and Alex Alderton, Roberto Amato, Jeffrey Barrett, Sonia Goncalves, Ewan Harrison, David K. Jackson, Ian Johnston, Dominic Kwiatkowski, Cordelia Langford, John Sillitoe on behalf of the Wellcome Sanger Institute COVID-19 Surveillance Team |
| EPI_ISL_3172849 | Lighthouse Lab in Glasgow | Wellcome Sanger Institute for the COVID-19 Genomics UK (COG-UK) Consortium | Harper VanSteenhouse, Yumi Kasai, David Gray, Carol Clugston, Anna Dominiczak and Alex Alderton, Roberto Amato, Jeffrey Barrett, Sonia Goncalves, Ewan Harrison, David K. Jackson, Ian Johnston, Dominic Kwiatkowski, Cordelia Langford, John Sillitoe on behalf of the Wellcome Sanger Institute COVID-19 Surveillance Team |
| EPI_ISL_3172850, EPI_ISL_3172851 | Lighthouse Lab in Milton Keynes | Wellcome Sanger Institute for the COVID-19 Genomics UK (COG-UK) Consortium | The Lighthouse Lab in Milton Keynes and Alex Alderton, Roberto Amato, Jeffrey Barrett, Sonia Goncalves, Ewan Harrison, David K. Jackson, Ian Johnston, Dominic Kwiatkowski, Cordelia Langford, John Sillitoe on behalf of the Wellcome Sanger Institute COVID-19 Surveillance Team |
| EPI_ISL_3172852, EPI_ISL_3172853, EPI_ISL_3172854, EPI_ISL_3172855 | Lighthouse Lab in Glasgow | Wellcome Sanger Institute for the COVID-19 Genomics UK (COG-UK) Consortium | Harper VanSteenhouse, Yumi Kasai, David Gray, Carol Clugston, Anna Dominiczak and Alex Alderton, Roberto Amato, Jeffrey Barrett, Sonia Goncalves, Ewan Harrison, David K. Jackson, Ian Johnston, Dominic Kwiatkowski, Cordelia Langford, John Sillitoe on behalf of the Wellcome Sanger Institute COVID-19 Surveillance Team |
| EPI_ISL_3172856, EPI_ISL_3172857, EPI_ISL_3172858, EPI_ISL_3172859, EPI_ISL_3172860, EPI_ISL_3172861, EPI_ISL_3172862 | Lighthouse Lab in Milton Keynes | Wellcome Sanger Institute for the COVID-19 Genomics UK (COG-UK) Consortium | The Lighthouse Lab in Milton Keynes and Alex Alderton, Roberto Amato, Jeffrey Barrett, Sonia Goncalves, Ewan Harrison, David K. Jackson, Ian Johnston, Dominic Kwiatkowski, Cordelia Langford, John Sillitoe on behalf of the Wellcome Sanger Institute COVID-19 Surveillance Team |
| EPI_ISL_3172863 | Lighthouse Lab in Glasgow | Wellcome Sanger Institute for the COVID-19 Genomics UK (COG-UK) Consortium | Harper VanSteenhouse, Yumi Kasai, David Gray, Carol Clugston, Anna Dominiczak and Alex Alderton, Roberto Amato, Jeffrey Barrett, Sonia Goncalves, Ewan Harrison, David K. Jackson, Ian Johnston, Dominic Kwiatkowski, Cordelia Langford, John Sillitoe on behalf of the Wellcome Sanger Institute COVID-19 Surveillance Team |
| EPI_ISL_3172864, EPI_ISL_3172865 | Lighthouse Lab in Milton Keynes | Wellcome Sanger Institute for the COVID-19 Genomics UK (COG-UK) Consortium | The Lighthouse Lab in Milton Keynes and Alex Alderton, Roberto Amato, Jeffrey Barrett, Sonia Goncalves, Ewan Harrison, David K. Jackson, Ian Johnston, Dominic Kwiatkowski, Cordelia Langford, John Sillitoe on behalf of the Wellcome Sanger Institute COVID-19 Surveillance Team |
| EPI_ISL_3172866, EPI_ISL_3172867 | Lighthouse Lab in Glasgow | Wellcome Sanger Institute for the COVID-19 Genomics UK (COG-UK) Consortium | Harper VanSteenhouse, Yumi Kasai, David Gray, Carol Clugston, Anna Dominiczak and Alex Alderton, Roberto Amato, Jeffrey Barrett, Sonia Goncalves, Ewan Harrison, David K. Jackson, Ian Johnston, Dominic Kwiatkowski, Cordelia Langford, John Sillitoe on behalf of the Wellcome Sanger Institute COVID-19 Surveillance Team |
| EPI_ISL_3172868, EPI_ISL_3172869, EPI_ISL_3172870 | Lighthouse Lab in Milton Keynes | Wellcome Sanger Institute for the COVID-19 Genomics UK (COG-UK) Consortium | The Lighthouse Lab in Milton Keynes and Alex Alderton, Roberto Amato, Jeffrey Barrett, Sonia Goncalves, Ewan Harrison, David K. Jackson, Ian Johnston, Dominic Kwiatkowski, Cordelia Langford, John Sillitoe on behalf of the Wellcome Sanger Institute COVID-19 Surveillance Team |
| EPI_ISL_3172871 | Lighthouse Lab in Glasgow | Wellcome Sanger Institute for the COVID-19 Genomics UK (COG-UK) Consortium | Harper VanSteenhouse, Yumi Kasai, David Gray, Carol Clugston, Anna Dominiczak and Alex Alderton, Roberto Amato, Jeffrey Barrett, Sonia Goncalves, Ewan Harrison, David K. Jackson, Ian Johnston, Dominic Kwiatkowski, Cordelia Langford, John Sillitoe on behalf of the Wellcome Sanger Institute COVID-19 Surveillance Team |
| EPI_ISL_3172872 | Lighthouse Lab in Milton Keynes | Wellcome Sanger Institute for the COVID-19 Genomics UK (COG-UK) Consortium | The Lighthouse Lab in Milton Keynes and Alex Alderton, Roberto Amato, Jeffrey Barrett, Sonia Goncalves, Ewan Harrison, David K. Jackson, Ian Johnston, Dominic Kwiatkowski, Cordelia Langford, John Sillitoe on behalf of the Wellcome Sanger Institute COVID-19 Surveillance Team |
| EPI_ISL_3172873 | Lighthouse Lab in Glasgow | Wellcome Sanger Institute for the COVID-19 Genomics UK (COG-UK) Consortium | Harper VanSteenhouse, Yumi Kasai, David Gray, Carol Clugston, Anna Dominiczak and Alex Alderton, Roberto Amato, Jeffrey Barrett, Sonia Goncalves, Ewan Harrison, David K. Jackson, Ian Johnston, Dominic Kwiatkowski, Cordelia Langford, John Sillitoe on behalf of the Wellcome Sanger Institute COVID-19 Surveillance Team |
| EPI_ISL_3172874 | Lighthouse Lab in Milton Keynes | Wellcome Sanger Institute for the COVID-19 Genomics UK (COG-UK) Consortium | The Lighthouse Lab in Milton Keynes and Alex Alderton, Roberto Amato, Jeffrey Barrett, Sonia Goncalves, Ewan Harrison, David K. Jackson, Ian Johnston, Dominic Kwiatkowski, Cordelia Langford, John Sillitoe on behalf of the Wellcome Sanger Institute COVID-19 Surveillance Team |
| EPI_ISL_3172875 | Lighthouse Lab in Glasgow | Wellcome Sanger Institute for the COVID-19 Genomics UK (COG-UK) Consortium | Harper VanSteenhouse, Yumi Kasai, David Gray, Carol Clugston, Anna Dominiczak and Alex Alderton, Roberto Amato, Jeffrey Barrett, Sonia Goncalves, Ewan Harrison, David K. Jackson, Ian Johnston, Dominic Kwiatkowski, Cordelia Langford, John Sillitoe on behalf of the Wellcome Sanger Institute COVID-19 Surveillance Team |
| EPI_ISL_3172876, EPI_ISL_3172877, EPI_ISL_3172878 | Lighthouse Lab in Milton Keynes | Wellcome Sanger Institute for the COVID-19 Genomics UK (COG-UK) Consortium | The Lighthouse Lab in Milton Keynes and Alex Alderton, Roberto Amato, Jeffrey Barrett, Sonia Goncalves, Ewan Harrison, David K. Jackson, Ian Johnston, Dominic Kwiatkowski, Cordelia Langford, John Sillitoe on behalf of the Wellcome Sanger Institute COVID-19 Surveillance Team |
| EPI_ISL_3172879 | Lighthouse Lab in Glasgow | Wellcome Sanger Institute for the COVID-19 Genomics UK (COG-UK) Consortium | Harper VanSteenhouse, Yumi Kasai, David Gray, Carol Clugston, Anna Dominiczak and Alex Alderton, Roberto Amato, Jeffrey Barrett, Sonia Goncalves, Ewan Harrison, David K. Jackson, Ian Johnston, Dominic Kwiatkowski, Cordelia Langford, John Sillitoe on behalf of the Wellcome Sanger Institute COVID-19 Surveillance Team |
| EPI_ISL_3172880 | Lighthouse Lab in Milton Keynes | Wellcome Sanger Institute for the COVID-19 Genomics UK (COG-UK) | The Lighthouse Lab in Milton Keynes and Alex Alderton, Roberto Amato, Jeffrey Barrett, Sonia Goncalves, Ewan Harrison, David K. Jackson, Ian Johnston, Dominic Kwiatkowski, Cordelia Langford, John Sillitoe on behalf of the Wellcome Sanger |

| | | | |
|---|---|---|---|
| | | Consortium | Institute COVID-19 Surveillance Team |
| EPI_ISL_3172881, EPI_ISL_3172882, EPI_ISL_3172883 | Lighthouse Lab in Glasgow | Wellcome Sanger Institute for the COVID-19 Genomics UK (COG-UK) Consortium | Harper VanSteenhouse, Yumi Kasai, David Gray, Carol Clugston, Anna Dominiczak and Alex Alderton, Roberto Amato, Jeffrey Barrett, Sonia Goncalves, Ewan Harrison, David K. Jackson, Ian Johnston, Dominic Kwiatkowski, Cordelia Langford, John Sillitoe on behalf of the Wellcome Sanger Institute COVID-19 Surveillance Team |
| EPI_ISL_3172884 | Lighthouse Lab in Milton Keynes | Wellcome Sanger Institute for the COVID-19 Genomics UK (COG-UK) Consortium | The Lighthouse Lab in Milton Keynes and Alex Alderton, Roberto Amato, Jeffrey Barrett, Sonia Goncalves, Ewan Harrison, David K. Jackson, Ian Johnston, Dominic Kwiatkowski, Cordelia Langford, John Sillitoe on behalf of the Wellcome Sanger Institute COVID-19 Surveillance Team |
| EPI_ISL_3172885 | Lighthouse Lab in Glasgow | Wellcome Sanger Institute for the COVID-19 Genomics UK (COG-UK) Consortium | Harper VanSteenhouse, Yumi Kasai, David Gray, Carol Clugston, Anna Dominiczak and Alex Alderton, Roberto Amato, Jeffrey Barrett, Sonia Goncalves, Ewan Harrison, David K. Jackson, Ian Johnston, Dominic Kwiatkowski, Cordelia Langford, John Sillitoe on behalf of the Wellcome Sanger Institute COVID-19 Surveillance Team |
| EPI_ISL_3172886 | Lighthouse Lab in Milton Keynes | Wellcome Sanger Institute for the COVID-19 Genomics UK (COG-UK) Consortium | The Lighthouse Lab in Milton Keynes and Alex Alderton, Roberto Amato, Jeffrey Barrett, Sonia Goncalves, Ewan Harrison, David K. Jackson, Ian Johnston, Dominic Kwiatkowski, Cordelia Langford, John Sillitoe on behalf of the Wellcome Sanger Institute COVID-19 Surveillance Team |
| EPI_ISL_3172887, EPI_ISL_3172888, EPI_ISL_3172889, EPI_ISL_3172890 | Lighthouse Lab in Glasgow | Wellcome Sanger Institute for the COVID-19 Genomics UK (COG-UK) Consortium | Harper VanSteenhouse, Yumi Kasai, David Gray, Carol Clugston, Anna Dominiczak and Alex Alderton, Roberto Amato, Jeffrey Barrett, Sonia Goncalves, Ewan Harrison, David K. Jackson, Ian Johnston, Dominic Kwiatkowski, Cordelia Langford, John Sillitoe on behalf of the Wellcome Sanger Institute COVID-19 Surveillance Team |
| EPI_ISL_3172891, EPI_ISL_3172892, EPI_ISL_3172893 | Lighthouse Lab in Milton Keynes | Wellcome Sanger Institute for the COVID-19 Genomics UK (COG-UK) Consortium | The Lighthouse Lab in Milton Keynes and Alex Alderton, Roberto Amato, Jeffrey Barrett, Sonia Goncalves, Ewan Harrison, David K. Jackson, Ian Johnston, Dominic Kwiatkowski, Cordelia Langford, John Sillitoe on behalf of the Wellcome Sanger Institute COVID-19 Surveillance Team |
| EPI_ISL_3172894 | Lighthouse Lab in Glasgow | Wellcome Sanger Institute for the COVID-19 Genomics UK (COG-UK) Consortium | Harper VanSteenhouse, Yumi Kasai, David Gray, Carol Clugston, Anna Dominiczak and Alex Alderton, Roberto Amato, Jeffrey Barrett, Sonia Goncalves, Ewan Harrison, David K. Jackson, Ian Johnston, Dominic Kwiatkowski, Cordelia Langford, John Sillitoe on behalf of the Wellcome Sanger Institute COVID-19 Surveillance Team |
| EPI_ISL_3172895, EPI_ISL_3172896 | Lighthouse Lab in Milton Keynes | Wellcome Sanger Institute for the COVID-19 Genomics UK (COG-UK) Consortium | The Lighthouse Lab in Milton Keynes and Alex Alderton, Roberto Amato, Jeffrey Barrett, Sonia Goncalves, Ewan Harrison, David K. Jackson, Ian Johnston, Dominic Kwiatkowski, Cordelia Langford, John Sillitoe on behalf of the Wellcome Sanger Institute COVID-19 Surveillance Team |
| EPI_ISL_3172897, EPI_ISL_3172898 | Lighthouse Lab in Glasgow | Wellcome Sanger Institute for the COVID-19 Genomics UK (COG-UK) Consortium | Harper VanSteenhouse, Yumi Kasai, David Gray, Carol Clugston, Anna Dominiczak and Alex Alderton, Roberto Amato, Jeffrey Barrett, Sonia Goncalves, Ewan Harrison, David K. Jackson, Ian Johnston, Dominic Kwiatkowski, Cordelia Langford, John Sillitoe on behalf of the Wellcome Sanger Institute COVID-19 Surveillance Team |
| EPI_ISL_3172899, EPI_ISL_3172900 | Lighthouse Lab in Milton Keynes | Wellcome Sanger Institute for the COVID-19 Genomics UK (COG-UK) Consortium | The Lighthouse Lab in Milton Keynes and Alex Alderton, Roberto Amato, Jeffrey Barrett, Sonia Goncalves, Ewan Harrison, David K. Jackson, Ian Johnston, Dominic Kwiatkowski, Cordelia Langford, John Sillitoe on behalf of the Wellcome Sanger Institute COVID-19 Surveillance Team |
| EPI_ISL_3172901 | Lighthouse Lab in Glasgow | Wellcome Sanger Institute for the COVID-19 Genomics UK (COG-UK) Consortium | Harper VanSteenhouse, Yumi Kasai, David Gray, Carol Clugston, Anna Dominiczak and Alex Alderton, Roberto Amato, Jeffrey Barrett, Sonia Goncalves, Ewan Harrison, David K. Jackson, Ian Johnston, Dominic Kwiatkowski, Cordelia Langford, John Sillitoe on behalf of the Wellcome Sanger Institute COVID-19 Surveillance Team |
| EPI_ISL_3172902 | Lighthouse Lab in Milton Keynes | Wellcome Sanger Institute for the COVID-19 Genomics UK (COG-UK) Consortium | The Lighthouse Lab in Milton Keynes and Alex Alderton, Roberto Amato, Jeffrey Barrett, Sonia Goncalves, Ewan Harrison, David K. Jackson, Ian Johnston, Dominic Kwiatkowski, Cordelia Langford, John Sillitoe on behalf of the Wellcome Sanger Institute COVID-19 Surveillance Team |
| EPI_ISL_3172903, EPI_ISL_3172904, EPI_ISL_3172905, EPI_ISL_3172906 | Lighthouse Lab in Glasgow | Wellcome Sanger Institute for the COVID-19 Genomics UK (COG-UK) Consortium | Harper VanSteenhouse, Yumi Kasai, David Gray, Carol Clugston, Anna Dominiczak and Alex Alderton, Roberto Amato, Jeffrey Barrett, Sonia Goncalves, Ewan Harrison, David K. Jackson, Ian Johnston, Dominic Kwiatkowski, Cordelia Langford, John Sillitoe on behalf of the Wellcome Sanger Institute COVID-19 Surveillance Team |
| EPI_ISL_3172907 | Lighthouse Lab in Milton Keynes | Wellcome Sanger Institute for the COVID-19 Genomics UK (COG-UK) Consortium | The Lighthouse Lab in Milton Keynes and Alex Alderton, Roberto Amato, Jeffrey Barrett, Sonia Goncalves, Ewan Harrison, David K. Jackson, Ian Johnston, Dominic Kwiatkowski, Cordelia Langford, John Sillitoe on behalf of the Wellcome Sanger Institute COVID-19 Surveillance Team |
| EPI_ISL_3172908 | Lighthouse Lab in Glasgow | Wellcome Sanger Institute for the COVID-19 Genomics UK (COG-UK) Consortium | Harper VanSteenhouse, Yumi Kasai, David Gray, Carol Clugston, Anna Dominiczak and Alex Alderton, Roberto Amato, Jeffrey Barrett, Sonia Goncalves, Ewan Harrison, David K. Jackson, Ian Johnston, Dominic Kwiatkowski, Cordelia Langford, John Sillitoe on behalf of the Wellcome Sanger Institute COVID-19 Surveillance Team |
| EPI_ISL_3172909 | Lighthouse Lab in Milton Keynes | Wellcome Sanger Institute for the COVID-19 Genomics UK (COG-UK) Consortium | The Lighthouse Lab in Milton Keynes and Alex Alderton, Roberto Amato, Jeffrey Barrett, Sonia Goncalves, Ewan Harrison, David K. Jackson, Ian Johnston, Dominic Kwiatkowski, Cordelia Langford, John Sillitoe on behalf of the Wellcome Sanger Institute COVID-19 Surveillance Team |
| EPI_ISL_3172910 | Lighthouse Lab in Glasgow | Wellcome Sanger Institute for the COVID-19 Genomics UK (COG-UK) Consortium | Harper VanSteenhouse, Yumi Kasai, David Gray, Carol Clugston, Anna Dominiczak and Alex Alderton, Roberto Amato, Jeffrey Barrett, Sonia Goncalves, Ewan Harrison, David K. Jackson, Ian Johnston, Dominic Kwiatkowski, Cordelia Langford, John Sillitoe on behalf of the Wellcome Sanger Institute COVID-19 Surveillance Team |
| EPI_ISL_3172911, EPI_ISL_3172912 | Lighthouse Lab in Milton Keynes | Wellcome Sanger Institute for the COVID-19 Genomics UK (COG-UK) Consortium | The Lighthouse Lab in Milton Keynes and Alex Alderton, Roberto Amato, Jeffrey Barrett, Sonia Goncalves, Ewan Harrison, David K. Jackson, Ian Johnston, Dominic Kwiatkowski, Cordelia Langford, John Sillitoe on behalf of the Wellcome Sanger Institute COVID-19 Surveillance Team |
| EPI_ISL_3172913, EPI_ISL_3172914, EPI_ISL_3172915, EPI_ISL_3172916, EPI_ISL_3172917 | Lighthouse Lab in Glasgow | Wellcome Sanger Institute for the COVID-19 Genomics UK (COG-UK) Consortium | Harper VanSteenhouse, Yumi Kasai, David Gray, Carol Clugston, Anna Dominiczak and Alex Alderton, Roberto Amato, Jeffrey Barrett, Sonia Goncalves, Ewan Harrison, David K. Jackson, Ian Johnston, Dominic Kwiatkowski, Cordelia Langford, John Sillitoe on behalf of the Wellcome Sanger Institute COVID-19 Surveillance Team |
| EPI_ISL_3172918 | Lighthouse Lab in Milton Keynes | Wellcome Sanger Institute for the COVID-19 Genomics UK (COG-UK) Consortium | The Lighthouse Lab in Milton Keynes and Alex Alderton, Roberto Amato, Jeffrey Barrett, Sonia Goncalves, Ewan Harrison, David K. Jackson, Ian Johnston, Dominic Kwiatkowski, Cordelia Langford, John Sillitoe on behalf of the Wellcome Sanger Institute COVID-19 Surveillance Team |
| EPI_ISL_3172919 | Lighthouse Lab in Glasgow | Wellcome Sanger Institute for the COVID-19 Genomics UK (COG-UK) Consortium | Harper VanSteenhouse, Yumi Kasai, David Gray, Carol Clugston, Anna Dominiczak and Alex Alderton, Roberto Amato, Jeffrey Barrett, Sonia Goncalves, Ewan Harrison, David K. Jackson, Ian Johnston, Dominic Kwiatkowski, Cordelia Langford, John Sillitoe on behalf of the Wellcome Sanger Institute COVID-19 Surveillance Team |
| EPI_ISL_3172920, EPI_ISL_3172921 | Lighthouse Lab in Milton Keynes | Wellcome Sanger Institute for the COVID-19 Genomics UK (COG-UK) Consortium | The Lighthouse Lab in Milton Keynes and Alex Alderton, Roberto Amato, Jeffrey Barrett, Sonia Goncalves, Ewan Harrison, David K. Jackson, Ian Johnston, Dominic Kwiatkowski, Cordelia Langford, John Sillitoe on behalf of the Wellcome Sanger Institute COVID-19 Surveillance Team |
| EPI_ISL_3172922, EPI_ISL_3172923, EPI_ISL_3172924 | Lighthouse Lab in Glasgow | Wellcome Sanger Institute for the COVID-19 Genomics UK (COG-UK) Consortium | Harper VanSteenhouse, Yumi Kasai, David Gray, Carol Clugston, Anna Dominiczak and Alex Alderton, Roberto Amato, Jeffrey Barrett, Sonia Goncalves, Ewan Harrison, David K. Jackson, Ian Johnston, Dominic Kwiatkowski, Cordelia Langford, John Sillitoe on behalf of the Wellcome Sanger Institute COVID-19 Surveillance Team |
| EPI_ISL_3172925, EPI_ISL_3172926, EPI_ISL_3172927 | Lighthouse Lab in Milton Keynes | Wellcome Sanger Institute for the COVID-19 Genomics UK (COG-UK) Consortium | The Lighthouse Lab in Milton Keynes and Alex Alderton, Roberto Amato, Jeffrey Barrett, Sonia Goncalves, Ewan Harrison, David K. Jackson, Ian Johnston, Dominic Kwiatkowski, Cordelia Langford, John Sillitoe on behalf of the Wellcome Sanger Institute COVID-19 Surveillance Team |
| EPI_ISL_3172928, EPI_ISL_3172929, EPI_ISL_3172930, EPI_ISL_3172931, EPI_ISL_3172932, EPI_ISL_3172933, EPI_ISL_3172934 | Lighthouse Lab in Glasgow | Wellcome Sanger Institute for the COVID-19 Genomics UK (COG-UK) Consortium | Harper VanSteenhouse, Yumi Kasai, David Gray, Carol Clugston, Anna Dominiczak and Alex Alderton, Roberto Amato, Jeffrey Barrett, Sonia Goncalves, Ewan Harrison, David K. Jackson, Ian Johnston, Dominic Kwiatkowski, Cordelia Langford, John Sillitoe on behalf of the Wellcome Sanger Institute COVID-19 Surveillance Team |
| EPI_ISL_3172935 | Lighthouse Lab in Milton Keynes | Wellcome Sanger Institute for the COVID-19 Genomics UK (COG-UK) Consortium | The Lighthouse Lab in Milton Keynes and Alex Alderton, Roberto Amato, Jeffrey Barrett, Sonia Goncalves, Ewan Harrison, David K. Jackson, Ian Johnston, Dominic Kwiatkowski, Cordelia Langford, John Sillitoe on behalf of the Wellcome Sanger Institute COVID-19 Surveillance Team |
| EPI_ISL_3172936, EPI_ISL_3172937 | Lighthouse Lab in Glasgow | Wellcome Sanger Institute for the COVID-19 Genomics UK (COG-UK) Consortium | Harper VanSteenhouse, Yumi Kasai, David Gray, Carol Clugston, Anna Dominiczak and Alex Alderton, Roberto Amato, Jeffrey Barrett, Sonia Goncalves, Ewan Harrison, David K. Jackson, Ian Johnston, Dominic Kwiatkowski, Cordelia Langford, John Sillitoe on behalf of the Wellcome Sanger Institute COVID-19 Surveillance Team |
| EPI_ISL_3172938, EPI_ISL_3172939, EPI_ISL_3172940, EPI_ISL_3172941 | Lighthouse Lab in Milton Keynes | Wellcome Sanger Institute for the COVID-19 Genomics UK (COG-UK) Consortium | The Lighthouse Lab in Milton Keynes and Alex Alderton, Roberto Amato, Jeffrey Barrett, Sonia Goncalves, Ewan Harrison, David K. Jackson, Ian Johnston, Dominic Kwiatkowski, Cordelia Langford, John Sillitoe on behalf of the Wellcome Sanger Institute COVID-19 Surveillance Team |
| EPI_ISL_3172942 | Lighthouse Lab in Glasgow | Wellcome Sanger Institute for the COVID-19 Genomics UK (COG-UK) Consortium | Harper VanSteenhouse, Yumi Kasai, David Gray, Carol Clugston, Anna Dominiczak and Alex Alderton, Roberto Amato, Jeffrey Barrett, Sonia Goncalves, Ewan Harrison, David K. Jackson, Ian Johnston, Dominic Kwiatkowski, Cordelia Langford, John Sillitoe on behalf of the Wellcome Sanger Institute COVID-19 Surveillance Team |
| EPI_ISL_3172943 | Lighthouse Lab in Milton Keynes | Wellcome Sanger Institute for the COVID-19 Genomics UK (COG-UK) Consortium | The Lighthouse Lab in Milton Keynes and Alex Alderton, Roberto Amato, Jeffrey Barrett, Sonia Goncalves, Ewan Harrison, David K. Jackson, Ian Johnston, Dominic Kwiatkowski, Cordelia Langford, John Sillitoe on behalf of the Wellcome Sanger Institute COVID-19 Surveillance Team |
| EPI_ISL_3172944 | Lighthouse Lab in Glasgow | Wellcome Sanger Institute for the COVID-19 Genomics UK (COG-UK) Consortium | Harper VanSteenhouse, Yumi Kasai, David Gray, Carol Clugston, Anna Dominiczak and Alex Alderton, Roberto Amato, Jeffrey Barrett, Sonia Goncalves, Ewan Harrison, David K. Jackson, Ian Johnston, Dominic Kwiatkowski, Cordelia Langford, John Sillitoe on behalf of the Wellcome Sanger Institute COVID-19 Surveillance Team |
| EPI_ISL_3172945 | Lighthouse Lab in Milton Keynes | Wellcome Sanger Institute for the COVID-19 Genomics UK (COG-UK) Consortium | The Lighthouse Lab in Milton Keynes and Alex Alderton, Roberto Amato, Jeffrey Barrett, Sonia Goncalves, Ewan Harrison, David K. Jackson, Ian Johnston, Dominic Kwiatkowski, Cordelia Langford, John Sillitoe on behalf of the Wellcome Sanger Institute COVID-19 Surveillance Team |
| EPI_ISL_3172946, EPI_ISL_3172947 | Lighthouse Lab in Glasgow | Wellcome Sanger Institute for the COVID-19 Genomics UK (COG-UK) Consortium | Harper VanSteenhouse, Yumi Kasai, David Gray, Carol Clugston, Anna Dominiczak and Alex Alderton, Roberto Amato, Jeffrey Barrett, Sonia Goncalves, Ewan Harrison, David K. Jackson, Ian Johnston, Dominic Kwiatkowski, Cordelia Langford, John Sillitoe on behalf of the Wellcome Sanger Institute COVID-19 Surveillance Team |
| EPI_ISL_3172948, EPI_ISL_3172949, EPI_ISL_3172950 | Lighthouse Lab in Milton Keynes | Wellcome Sanger Institute for the COVID-19 Genomics UK (COG-UK) Consortium | The Lighthouse Lab in Milton Keynes and Alex Alderton, Roberto Amato, Jeffrey Barrett, Sonia Goncalves, Ewan Harrison, David K. Jackson, Ian Johnston, Dominic Kwiatkowski, Cordelia Langford, John Sillitoe on behalf of the Wellcome Sanger Institute COVID-19 Surveillance Team |
| EPI_ISL_3172951, EPI_ISL_3172952, EPI_ISL_3172953, EPI_ISL_3172954 | Lighthouse Lab in Glasgow | Wellcome Sanger Institute for the COVID-19 Genomics UK (COG-UK) Consortium | Harper VanSteenhouse, Yumi Kasai, David Gray, Carol Clugston, Anna Dominiczak and Alex Alderton, Roberto Amato, Jeffrey Barrett, Sonia Goncalves, Ewan Harrison, David K. Jackson, Ian Johnston, Dominic Kwiatkowski, Cordelia Langford, John Sillitoe on behalf of the Wellcome Sanger Institute COVID-19 Surveillance Team |
| EPI_ISL_3172955, EPI_ISL_3172956, | Lighthouse Lab in Milton Keynes | Wellcome Sanger Institute for the COVID-19 Genomics UK (COG-UK) Consortium | The Lighthouse Lab in Milton Keynes and Alex Alderton, Roberto Amato, Jeffrey Barrett, Sonia Goncalves, Ewan Harrison, David K. Jackson, Ian Johnston, Dominic Kwiatkowski, Cordelia Langford, John Sillitoe on behalf of the Wellcome Sanger Institute COVID-19 Surveillance Team |

| | | | |
|---|---|---|---|
| EPI_ISL_3172957, EPI_ISL_3172958, EPI_ISL_3172959 | | | |
| EPI_ISL_3172960 | Lighthouse Lab in Glasgow | Wellcome Sanger Institute for the COVID-19 Genomics UK (COG-UK) Consortium | Harper VanSteenhouse, Yumi Kasai, David Gray, Carol Clugston, Anna Dominiczak and Alex Alderton, Roberto Amato, Jeffrey Barrett, Sonia Goncalves, Ewan Harrison, David K. Jackson, Ian Johnston, Dominic Kwiatkowski, Cordelia Langford, John Sillitoe on behalf of the Wellcome Sanger Institute COVID-19 Surveillance Team |
| EPI_ISL_3172961, EPI_ISL_3172962, EPI_ISL_3172963 | Lighthouse Lab in Milton Keynes | Wellcome Sanger Institute for the COVID-19 Genomics UK (COG-UK) Consortium | The Lighthouse Lab in Milton Keynes and Alex Alderton, Roberto Amato, Jeffrey Barrett, Sonia Goncalves, Ewan Harrison, David K. Jackson, Ian Johnston, Dominic Kwiatkowski, Cordelia Langford, John Sillitoe on behalf of the Wellcome Sanger Institute COVID-19 Surveillance Team |
| EPI_ISL_3172964 | Lighthouse Lab in Glasgow | Wellcome Sanger Institute for the COVID-19 Genomics UK (COG-UK) Consortium | Harper VanSteenhouse, Yumi Kasai, David Gray, Carol Clugston, Anna Dominiczak and Alex Alderton, Roberto Amato, Jeffrey Barrett, Sonia Goncalves, Ewan Harrison, David K. Jackson, Ian Johnston, Dominic Kwiatkowski, Cordelia Langford, John Sillitoe on behalf of the Wellcome Sanger Institute COVID-19 Surveillance Team |
| EPI_ISL_3172965, EPI_ISL_3172966, EPI_ISL_3172967 | Lighthouse Lab in Milton Keynes | Wellcome Sanger Institute for the COVID-19 Genomics UK (COG-UK) Consortium | The Lighthouse Lab in Milton Keynes and Alex Alderton, Roberto Amato, Jeffrey Barrett, Sonia Goncalves, Ewan Harrison, David K. Jackson, Ian Johnston, Dominic Kwiatkowski, Cordelia Langford, John Sillitoe on behalf of the Wellcome Sanger Institute COVID-19 Surveillance Team |
| EPI_ISL_3172968, EPI_ISL_3172969, EPI_ISL_3172970, EPI_ISL_3172971 | Lighthouse Lab in Glasgow | Wellcome Sanger Institute for the COVID-19 Genomics UK (COG-UK) Consortium | Harper VanSteenhouse, Yumi Kasai, David Gray, Carol Clugston, Anna Dominiczak and Alex Alderton, Roberto Amato, Jeffrey Barrett, Sonia Goncalves, Ewan Harrison, David K. Jackson, Ian Johnston, Dominic Kwiatkowski, Cordelia Langford, John Sillitoe on behalf of the Wellcome Sanger Institute COVID-19 Surveillance Team |
| EPI_ISL_3172972 | Lighthouse Lab in Milton Keynes | Wellcome Sanger Institute for the COVID-19 Genomics UK (COG-UK) Consortium | The Lighthouse Lab in Milton Keynes and Alex Alderton, Roberto Amato, Jeffrey Barrett, Sonia Goncalves, Ewan Harrison, David K. Jackson, Ian Johnston, Dominic Kwiatkowski, Cordelia Langford, John Sillitoe on behalf of the Wellcome Sanger Institute COVID-19 Surveillance Team |
| EPI_ISL_3172973, EPI_ISL_3172974, EPI_ISL_3172975, EPI_ISL_3172976 | Lighthouse Lab in Glasgow | Wellcome Sanger Institute for the COVID-19 Genomics UK (COG-UK) Consortium | Harper VanSteenhouse, Yumi Kasai, David Gray, Carol Clugston, Anna Dominiczak and Alex Alderton, Roberto Amato, Jeffrey Barrett, Sonia Goncalves, Ewan Harrison, David K. Jackson, Ian Johnston, Dominic Kwiatkowski, Cordelia Langford, John Sillitoe on behalf of the Wellcome Sanger Institute COVID-19 Surveillance Team |
| EPI_ISL_3172977, EPI_ISL_3172978, EPI_ISL_3172979 | Lighthouse Lab in Milton Keynes | Wellcome Sanger Institute for the COVID-19 Genomics UK (COG-UK) Consortium | The Lighthouse Lab in Milton Keynes and Alex Alderton, Roberto Amato, Jeffrey Barrett, Sonia Goncalves, Ewan Harrison, David K. Jackson, Ian Johnston, Dominic Kwiatkowski, Cordelia Langford, John Sillitoe on behalf of the Wellcome Sanger Institute COVID-19 Surveillance Team |
| EPI_ISL_3172980 | Lighthouse Lab in Glasgow | Wellcome Sanger Institute for the COVID-19 Genomics UK (COG-UK) Consortium | Harper VanSteenhouse, Yumi Kasai, David Gray, Carol Clugston, Anna Dominiczak and Alex Alderton, Roberto Amato, Jeffrey Barrett, Sonia Goncalves, Ewan Harrison, David K. Jackson, Ian Johnston, Dominic Kwiatkowski, Cordelia Langford, John Sillitoe on behalf of the Wellcome Sanger Institute COVID-19 Surveillance Team |
| EPI_ISL_3172981 | Lighthouse Lab in Milton Keynes | Wellcome Sanger Institute for the COVID-19 Genomics UK (COG-UK) Consortium | The Lighthouse Lab in Milton Keynes and Alex Alderton, Roberto Amato, Jeffrey Barrett, Sonia Goncalves, Ewan Harrison, David K. Jackson, Ian Johnston, Dominic Kwiatkowski, Cordelia Langford, John Sillitoe on behalf of the Wellcome Sanger Institute COVID-19 Surveillance Team |
| EPI_ISL_3172982 | Lighthouse Lab in Glasgow | Wellcome Sanger Institute for the COVID-19 Genomics UK (COG-UK) Consortium | Harper VanSteenhouse, Yumi Kasai, David Gray, Carol Clugston, Anna Dominiczak and Alex Alderton, Roberto Amato, Jeffrey Barrett, Sonia Goncalves, Ewan Harrison, David K. Jackson, Ian Johnston, Dominic Kwiatkowski, Cordelia Langford, John Sillitoe on behalf of the Wellcome Sanger Institute COVID-19 Surveillance Team |
| EPI_ISL_3172983, EPI_ISL_3172984 | Lighthouse Lab in Milton Keynes | Wellcome Sanger Institute for the COVID-19 Genomics UK (COG-UK) Consortium | The Lighthouse Lab in Milton Keynes and Alex Alderton, Roberto Amato, Jeffrey Barrett, Sonia Goncalves, Ewan Harrison, David K. Jackson, Ian Johnston, Dominic Kwiatkowski, Cordelia Langford, John Sillitoe on behalf of the Wellcome Sanger Institute COVID-19 Surveillance Team |
| EPI_ISL_3172985 | Lighthouse Lab in Glasgow | Wellcome Sanger Institute for the COVID-19 Genomics UK (COG-UK) Consortium | Harper VanSteenhouse, Yumi Kasai, David Gray, Carol Clugston, Anna Dominiczak and Alex Alderton, Roberto Amato, Jeffrey Barrett, Sonia Goncalves, Ewan Harrison, David K. Jackson, Ian Johnston, Dominic Kwiatkowski, Cordelia Langford, John Sillitoe on behalf of the Wellcome Sanger Institute COVID-19 Surveillance Team |
| EPI_ISL_3172986, EPI_ISL_3172987, EPI_ISL_3172988 | Lighthouse Lab in Milton Keynes | Wellcome Sanger Institute for the COVID-19 Genomics UK (COG-UK) Consortium | The Lighthouse Lab in Milton Keynes and Alex Alderton, Roberto Amato, Jeffrey Barrett, Sonia Goncalves, Ewan Harrison, David K. Jackson, Ian Johnston, Dominic Kwiatkowski, Cordelia Langford, John Sillitoe on behalf of the Wellcome Sanger Institute COVID-19 Surveillance Team |
| EPI_ISL_3172989, EPI_ISL_3172990 | Lighthouse Lab in Glasgow | Wellcome Sanger Institute for the COVID-19 Genomics UK (COG-UK) Consortium | Harper VanSteenhouse, Yumi Kasai, David Gray, Carol Clugston, Anna Dominiczak and Alex Alderton, Roberto Amato, Jeffrey Barrett, Sonia Goncalves, Ewan Harrison, David K. Jackson, Ian Johnston, Dominic Kwiatkowski, Cordelia Langford, John Sillitoe on behalf of the Wellcome Sanger Institute COVID-19 Surveillance Team |
| EPI_ISL_3172991 | Lighthouse Lab in Milton Keynes | Wellcome Sanger Institute for the COVID-19 Genomics UK (COG-UK) Consortium | The Lighthouse Lab in Milton Keynes and Alex Alderton, Roberto Amato, Jeffrey Barrett, Sonia Goncalves, Ewan Harrison, David K. Jackson, Ian Johnston, Dominic Kwiatkowski, Cordelia Langford, John Sillitoe on behalf of the Wellcome Sanger Institute COVID-19 Surveillance Team |
| EPI_ISL_3172992, EPI_ISL_3172993 | Lighthouse Lab in Glasgow | Wellcome Sanger Institute for the COVID-19 Genomics UK (COG-UK) Consortium | Harper VanSteenhouse, Yumi Kasai, David Gray, Carol Clugston, Anna Dominiczak and Alex Alderton, Roberto Amato, Jeffrey Barrett, Sonia Goncalves, Ewan Harrison, David K. Jackson, Ian Johnston, Dominic Kwiatkowski, Cordelia Langford, John Sillitoe on behalf of the Wellcome Sanger Institute COVID-19 Surveillance Team |
| EPI_ISL_3172994, EPI_ISL_3172995, EPI_ISL_3172996 | Lighthouse Lab in Milton Keynes | Wellcome Sanger Institute for the COVID-19 Genomics UK (COG-UK) Consortium | The Lighthouse Lab in Milton Keynes and Alex Alderton, Roberto Amato, Jeffrey Barrett, Sonia Goncalves, Ewan Harrison, David K. Jackson, Ian Johnston, Dominic Kwiatkowski, Cordelia Langford, John Sillitoe on behalf of the Wellcome Sanger Institute COVID-19 Surveillance Team |
| EPI_ISL_3172997 | Lighthouse Lab in Glasgow | Wellcome Sanger Institute for the COVID-19 Genomics UK (COG-UK) Consortium | Harper VanSteenhouse, Yumi Kasai, David Gray, Carol Clugston, Anna Dominiczak and Alex Alderton, Roberto Amato, Jeffrey Barrett, Sonia Goncalves, Ewan Harrison, David K. Jackson, Ian Johnston, Dominic Kwiatkowski, Cordelia Langford, John Sillitoe on behalf of the Wellcome Sanger Institute COVID-19 Surveillance Team |
| EPI_ISL_3172998, EPI_ISL_3172999, EPI_ISL_3173000, EPI_ISL_3173001, EPI_ISL_3173002, EPI_ISL_3173003, EPI_ISL_3173004 | Lighthouse Lab in Milton Keynes | Wellcome Sanger Institute for the COVID-19 Genomics UK (COG-UK) Consortium | The Lighthouse Lab in Milton Keynes and Alex Alderton, Roberto Amato, Jeffrey Barrett, Sonia Goncalves, Ewan Harrison, David K. Jackson, Ian Johnston, Dominic Kwiatkowski, Cordelia Langford, John Sillitoe on behalf of the Wellcome Sanger Institute COVID-19 Surveillance Team |
| EPI_ISL_3173005 | Lighthouse Lab in Glasgow | Wellcome Sanger Institute for the COVID-19 Genomics UK (COG-UK) Consortium | Harper VanSteenhouse, Yumi Kasai, David Gray, Carol Clugston, Anna Dominiczak and Alex Alderton, Roberto Amato, Jeffrey Barrett, Sonia Goncalves, Ewan Harrison, David K. Jackson, Ian Johnston, Dominic Kwiatkowski, Cordelia Langford, John Sillitoe on behalf of the Wellcome Sanger Institute COVID-19 Surveillance Team |
| EPI_ISL_3173006, EPI_ISL_3173007 | Lighthouse Lab in Milton Keynes | Wellcome Sanger Institute for the COVID-19 Genomics UK (COG-UK) Consortium | The Lighthouse Lab in Milton Keynes and Alex Alderton, Roberto Amato, Jeffrey Barrett, Sonia Goncalves, Ewan Harrison, David K. Jackson, Ian Johnston, Dominic Kwiatkowski, Cordelia Langford, John Sillitoe on behalf of the Wellcome Sanger Institute COVID-19 Surveillance Team |
| EPI_ISL_3173008, EPI_ISL_3173009 | Lighthouse Lab in Glasgow | Wellcome Sanger Institute for the COVID-19 Genomics UK (COG-UK) Consortium | Harper VanSteenhouse, Yumi Kasai, David Gray, Carol Clugston, Anna Dominiczak and Alex Alderton, Roberto Amato, Jeffrey Barrett, Sonia Goncalves, Ewan Harrison, David K. Jackson, Ian Johnston, Dominic Kwiatkowski, Cordelia Langford, John Sillitoe on behalf of the Wellcome Sanger Institute COVID-19 Surveillance Team |
| EPI_ISL_3173010 | Lighthouse Lab in Milton Keynes | Wellcome Sanger Institute for the COVID-19 Genomics UK (COG-UK) Consortium | The Lighthouse Lab in Milton Keynes and Alex Alderton, Roberto Amato, Jeffrey Barrett, Sonia Goncalves, Ewan Harrison, David K. Jackson, Ian Johnston, Dominic Kwiatkowski, Cordelia Langford, John Sillitoe on behalf of the Wellcome Sanger Institute COVID-19 Surveillance Team |
| EPI_ISL_3173011 | Lighthouse Lab in Glasgow | Wellcome Sanger Institute for the COVID-19 Genomics UK (COG-UK) Consortium | Harper VanSteenhouse, Yumi Kasai, David Gray, Carol Clugston, Anna Dominiczak and Alex Alderton, Roberto Amato, Jeffrey Barrett, Sonia Goncalves, Ewan Harrison, David K. Jackson, Ian Johnston, Dominic Kwiatkowski, Cordelia Langford, John Sillitoe on behalf of the Wellcome Sanger Institute COVID-19 Surveillance Team |
| EPI_ISL_3173012, EPI_ISL_3173013 | Lighthouse Lab in Milton Keynes | Wellcome Sanger Institute for the COVID-19 Genomics UK (COG-UK) Consortium | The Lighthouse Lab in Milton Keynes and Alex Alderton, Roberto Amato, Jeffrey Barrett, Sonia Goncalves, Ewan Harrison, David K. Jackson, Ian Johnston, Dominic Kwiatkowski, Cordelia Langford, John Sillitoe on behalf of the Wellcome Sanger Institute COVID-19 Surveillance Team |
| EPI_ISL_3173014, EPI_ISL_3173015, EPI_ISL_3173016 | Lighthouse Lab in Glasgow | Wellcome Sanger Institute for the COVID-19 Genomics UK (COG-UK) Consortium | Harper VanSteenhouse, Yumi Kasai, David Gray, Carol Clugston, Anna Dominiczak and Alex Alderton, Roberto Amato, Jeffrey Barrett, Sonia Goncalves, Ewan Harrison, David K. Jackson, Ian Johnston, Dominic Kwiatkowski, Cordelia Langford, John Sillitoe on behalf of the Wellcome Sanger Institute COVID-19 Surveillance Team |
| EPI_ISL_3173017, EPI_ISL_3173018, EPI_ISL_3173019 | Lighthouse Lab in Milton Keynes | Wellcome Sanger Institute for the COVID-19 Genomics UK (COG-UK) Consortium | The Lighthouse Lab in Milton Keynes and Alex Alderton, Roberto Amato, Jeffrey Barrett, Sonia Goncalves, Ewan Harrison, David K. Jackson, Ian Johnston, Dominic Kwiatkowski, Cordelia Langford, John Sillitoe on behalf of the Wellcome Sanger Institute COVID-19 Surveillance Team |
| EPI_ISL_3173020, EPI_ISL_3173021 | Lighthouse Lab in Glasgow | Wellcome Sanger Institute for the COVID-19 Genomics UK (COG-UK) Consortium | Harper VanSteenhouse, Yumi Kasai, David Gray, Carol Clugston, Anna Dominiczak and Alex Alderton, Roberto Amato, Jeffrey Barrett, Sonia Goncalves, Ewan Harrison, David K. Jackson, Ian Johnston, Dominic Kwiatkowski, Cordelia Langford, John Sillitoe on behalf of the Wellcome Sanger Institute COVID-19 Surveillance Team |
| EPI_ISL_3173022 | Lighthouse Lab in Milton Keynes | Wellcome Sanger Institute for the COVID-19 Genomics UK (COG-UK) Consortium | The Lighthouse Lab in Milton Keynes and Alex Alderton, Roberto Amato, Jeffrey Barrett, Sonia Goncalves, Ewan Harrison, David K. Jackson, Ian Johnston, Dominic Kwiatkowski, Cordelia Langford, John Sillitoe on behalf of the Wellcome Sanger Institute COVID-19 Surveillance Team |
| EPI_ISL_3173023, EPI_ISL_3173024, EPI_ISL_3173025, EPI_ISL_3173026 | Lighthouse Lab in Glasgow | Wellcome Sanger Institute for the COVID-19 Genomics UK (COG-UK) Consortium | Harper VanSteenhouse, Yumi Kasai, David Gray, Carol Clugston, Anna Dominiczak and Alex Alderton, Roberto Amato, Jeffrey Barrett, Sonia Goncalves, Ewan Harrison, David K. Jackson, Ian Johnston, Dominic Kwiatkowski, Cordelia Langford, John Sillitoe on behalf of the Wellcome Sanger Institute COVID-19 Surveillance Team |
| EPI_ISL_3173027, EPI_ISL_3173028, EPI_ISL_3173029 | Lighthouse Lab in Milton Keynes | Wellcome Sanger Institute for the COVID-19 Genomics UK (COG-UK) Consortium | The Lighthouse Lab in Milton Keynes and Alex Alderton, Roberto Amato, Jeffrey Barrett, Sonia Goncalves, Ewan Harrison, David K. Jackson, Ian Johnston, Dominic Kwiatkowski, Cordelia Langford, John Sillitoe on behalf of the Wellcome Sanger Institute COVID-19 Surveillance Team |
| EPI_ISL_3173030, EPI_ISL_3173031 | Lighthouse Lab in Glasgow | Wellcome Sanger Institute for the COVID-19 Genomics UK (COG-UK) Consortium | Harper VanSteenhouse, Yumi Kasai, David Gray, Carol Clugston, Anna Dominiczak and Alex Alderton, Roberto Amato, Jeffrey Barrett, Sonia Goncalves, Ewan Harrison, David K. Jackson, Ian Johnston, Dominic Kwiatkowski, Cordelia Langford, John Sillitoe on behalf of the Wellcome Sanger Institute COVID-19 Surveillance Team |
| EPI_ISL_3173032 | Lighthouse Lab in Milton Keynes | Wellcome Sanger Institute for the COVID-19 Genomics UK (COG-UK) Consortium | The Lighthouse Lab in Milton Keynes and Alex Alderton, Roberto Amato, Jeffrey Barrett, Sonia Goncalves, Ewan Harrison, David K. Jackson, Ian Johnston, Dominic Kwiatkowski, Cordelia Langford, John Sillitoe on behalf of the Wellcome Sanger Institute COVID-19 Surveillance Team |
| EPI_ISL_3173033 | Lighthouse Lab in Glasgow | Wellcome Sanger Institute for the COVID-19 Genomics UK (COG-UK) Consortium | Harper VanSteenhouse, Yumi Kasai, David Gray, Carol Clugston, Anna Dominiczak and Alex Alderton, Roberto Amato, Jeffrey Barrett, Sonia Goncalves, Ewan Harrison, David K. Jackson, Ian Johnston, Dominic Kwiatkowski, Cordelia Langford, John Sillitoe on behalf of the Wellcome Sanger Institute COVID-19 Surveillance Team |
| EPI_ISL_3173034, EPI_ISL_3173035, | Lighthouse Lab in Milton Keynes | Wellcome Sanger Institute for the COVID-19 Genomics UK (COG-UK) Consortium | The Lighthouse Lab in Milton Keynes and Alex Alderton, Roberto Amato, Jeffrey Barrett, Sonia Goncalves, Ewan Harrison, David K. Jackson, Ian Johnston, Dominic Kwiatkowski, Cordelia Langford, John Sillitoe on behalf of the Wellcome Sanger Institute COVID-19 Surveillance Team |

| | | | |
|---|---|---|---|
| EPI_ISL_3173036, EPI_ISL_3173037, EPI_ISL_3173038, EPI_ISL_3173039, EPI_ISL_3173040 | | | |
| EPI_ISL_3173041 | Lighthouse Lab in Glasgow | Wellcome Sanger Institute for the COVID-19 Genomics UK (COG-UK) Consortium | Harper VanSteenhouse, Yumi Kasai, David Gray, Carol Clugston, Anna Dominiczak and Alex Alderton, Roberto Amato, Jeffrey Barrett, Sonia Goncalves, Ewan Harrison, David K. Jackson, Ian Johnston, Dominic Kwiatkowski, Cordelia Langford, John Sillitoe on behalf of the Wellcome Sanger Institute COVID-19 Surveillance Team |
| EPI_ISL_3173042 | Lighthouse Lab in Milton Keynes | Wellcome Sanger Institute for the COVID-19 Genomics UK (COG-UK) Consortium | The Lighthouse Lab in Milton Keynes and Alex Alderton, Roberto Amato, Jeffrey Barrett, Sonia Goncalves, Ewan Harrison, David K. Jackson, Ian Johnston, Dominic Kwiatkowski, Cordelia Langford, John Sillitoe on behalf of the Wellcome Sanger Institute COVID-19 Surveillance Team |
| EPI_ISL_3173043 | Lighthouse Lab in Glasgow | Wellcome Sanger Institute for the COVID-19 Genomics UK (COG-UK) Consortium | Harper VanSteenhouse, Yumi Kasai, David Gray, Carol Clugston, Anna Dominiczak and Alex Alderton, Roberto Amato, Jeffrey Barrett, Sonia Goncalves, Ewan Harrison, David K. Jackson, Ian Johnston, Dominic Kwiatkowski, Cordelia Langford, John Sillitoe on behalf of the Wellcome Sanger Institute COVID-19 Surveillance Team |
| EPI_ISL_3173044, EPI_ISL_3173045 | Lighthouse Lab in Milton Keynes | Wellcome Sanger Institute for the COVID-19 Genomics UK (COG-UK) Consortium | The Lighthouse Lab in Milton Keynes and Alex Alderton, Roberto Amato, Jeffrey Barrett, Sonia Goncalves, Ewan Harrison, David K. Jackson, Ian Johnston, Dominic Kwiatkowski, Cordelia Langford, John Sillitoe on behalf of the Wellcome Sanger Institute COVID-19 Surveillance Team |
| EPI_ISL_3173046, EPI_ISL_3173047 | Lighthouse Lab in Glasgow | Wellcome Sanger Institute for the COVID-19 Genomics UK (COG-UK) Consortium | Harper VanSteenhouse, Yumi Kasai, David Gray, Carol Clugston, Anna Dominiczak and Alex Alderton, Roberto Amato, Jeffrey Barrett, Sonia Goncalves, Ewan Harrison, David K. Jackson, Ian Johnston, Dominic Kwiatkowski, Cordelia Langford, John Sillitoe on behalf of the Wellcome Sanger Institute COVID-19 Surveillance Team |
| EPI_ISL_3173048 | Lighthouse Lab in Milton Keynes | Wellcome Sanger Institute for the COVID-19 Genomics UK (COG-UK) Consortium | The Lighthouse Lab in Milton Keynes and Alex Alderton, Roberto Amato, Jeffrey Barrett, Sonia Goncalves, Ewan Harrison, David K. Jackson, Ian Johnston, Dominic Kwiatkowski, Cordelia Langford, John Sillitoe on behalf of the Wellcome Sanger Institute COVID-19 Surveillance Team |
| EPI_ISL_3173049, EPI_ISL_3173050, EPI_ISL_3173051 | Lighthouse Lab in Glasgow | Wellcome Sanger Institute for the COVID-19 Genomics UK (COG-UK) Consortium | Harper VanSteenhouse, Yumi Kasai, David Gray, Carol Clugston, Anna Dominiczak and Alex Alderton, Roberto Amato, Jeffrey Barrett, Sonia Goncalves, Ewan Harrison, David K. Jackson, Ian Johnston, Dominic Kwiatkowski, Cordelia Langford, John Sillitoe on behalf of the Wellcome Sanger Institute COVID-19 Surveillance Team |
| EPI_ISL_3173052 | Lighthouse Lab in Milton Keynes | Wellcome Sanger Institute for the COVID-19 Genomics UK (COG-UK) Consortium | The Lighthouse Lab in Milton Keynes and Alex Alderton, Roberto Amato, Jeffrey Barrett, Sonia Goncalves, Ewan Harrison, David K. Jackson, Ian Johnston, Dominic Kwiatkowski, Cordelia Langford, John Sillitoe on behalf of the Wellcome Sanger Institute COVID-19 Surveillance Team |
| EPI_ISL_3173053, EPI_ISL_3173054, EPI_ISL_3173055, EPI_ISL_3173056 | Lighthouse Lab in Glasgow | Wellcome Sanger Institute for the COVID-19 Genomics UK (COG-UK) Consortium | Harper VanSteenhouse, Yumi Kasai, David Gray, Carol Clugston, Anna Dominiczak and Alex Alderton, Roberto Amato, Jeffrey Barrett, Sonia Goncalves, Ewan Harrison, David K. Jackson, Ian Johnston, Dominic Kwiatkowski, Cordelia Langford, John Sillitoe on behalf of the Wellcome Sanger Institute COVID-19 Surveillance Team |
| EPI_ISL_3173057, EPI_ISL_3173058, EPI_ISL_3173059 | Lighthouse Lab in Milton Keynes | Wellcome Sanger Institute for the COVID-19 Genomics UK (COG-UK) Consortium | The Lighthouse Lab in Milton Keynes and Alex Alderton, Roberto Amato, Jeffrey Barrett, Sonia Goncalves, Ewan Harrison, David K. Jackson, Ian Johnston, Dominic Kwiatkowski, Cordelia Langford, John Sillitoe on behalf of the Wellcome Sanger Institute COVID-19 Surveillance Team |
| EPI_ISL_3173060, EPI_ISL_3173061 | Lighthouse Lab in Glasgow | Wellcome Sanger Institute for the COVID-19 Genomics UK (COG-UK) Consortium | Harper VanSteenhouse, Yumi Kasai, David Gray, Carol Clugston, Anna Dominiczak and Alex Alderton, Roberto Amato, Jeffrey Barrett, Sonia Goncalves, Ewan Harrison, David K. Jackson, Ian Johnston, Dominic Kwiatkowski, Cordelia Langford, John Sillitoe on behalf of the Wellcome Sanger Institute COVID-19 Surveillance Team |
| EPI_ISL_3173062, EPI_ISL_3173063 | Lighthouse Lab in Milton Keynes | Wellcome Sanger Institute for the COVID-19 Genomics UK (COG-UK) Consortium | The Lighthouse Lab in Milton Keynes and Alex Alderton, Roberto Amato, Jeffrey Barrett, Sonia Goncalves, Ewan Harrison, David K. Jackson, Ian Johnston, Dominic Kwiatkowski, Cordelia Langford, John Sillitoe on behalf of the Wellcome Sanger Institute COVID-19 Surveillance Team |
| EPI_ISL_3173064 | Lighthouse Laboratory Plymouth | Wellcome Sanger Institute for the COVID-19 Genomics UK (COG-UK) Consortium | Lighthouse Laboratory Plymouth and Alex Alderton, Roberto Amato, Jeffrey Barrett, Sonia Goncalves, Ewan Harrison, David K. Jackson, Ian Johnston, Dominic Kwiatkowski, Cordelia Langford, John Sillitoe on behalf of the Wellcome Sanger Institute COVID-19 Surveillance Team |
| EPI_ISL_3173065 | Lighthouse Lab in Milton Keynes | Wellcome Sanger Institute for the COVID-19 Genomics UK (COG-UK) Consortium | The Lighthouse Lab in Milton Keynes and Alex Alderton, Roberto Amato, Jeffrey Barrett, Sonia Goncalves, Ewan Harrison, David K. Jackson, Ian Johnston, Dominic Kwiatkowski, Cordelia Langford, John Sillitoe on behalf of the Wellcome Sanger Institute COVID-19 Surveillance Team |
| EPI_ISL_3173066 | Lighthouse Lab in Glasgow | Wellcome Sanger Institute for the COVID-19 Genomics UK (COG-UK) Consortium | Harper VanSteenhouse, Yumi Kasai, David Gray, Carol Clugston, Anna Dominiczak and Alex Alderton, Roberto Amato, Jeffrey Barrett, Sonia Goncalves, Ewan Harrison, David K. Jackson, Ian Johnston, Dominic Kwiatkowski, Cordelia Langford, John Sillitoe on behalf of the Wellcome Sanger Institute COVID-19 Surveillance Team |
| EPI_ISL_3173067 | Lighthouse Laboratory Plymouth | Wellcome Sanger Institute for the COVID-19 Genomics UK (COG-UK) Consortium | Lighthouse Laboratory Plymouth and Alex Alderton, Roberto Amato, Jeffrey Barrett, Sonia Goncalves, Ewan Harrison, David K. Jackson, Ian Johnston, Dominic Kwiatkowski, Cordelia Langford, John Sillitoe on behalf of the Wellcome Sanger Institute COVID-19 Surveillance Team |
| EPI_ISL_3173068 | Lighthouse Lab in Milton Keynes | Wellcome Sanger Institute for the COVID-19 Genomics UK (COG-UK) Consortium | The Lighthouse Lab in Milton Keynes and Alex Alderton, Roberto Amato, Jeffrey Barrett, Sonia Goncalves, Ewan Harrison, David K. Jackson, Ian Johnston, Dominic Kwiatkowski, Cordelia Langford, John Sillitoe on behalf of the Wellcome Sanger Institute COVID-19 Surveillance Team |
| EPI_ISL_3173069 | Lighthouse Laboratory Plymouth | Wellcome Sanger Institute for the COVID-19 Genomics UK (COG-UK) Consortium | Lighthouse Laboratory Plymouth and Alex Alderton, Roberto Amato, Jeffrey Barrett, Sonia Goncalves, Ewan Harrison, David K. Jackson, Ian Johnston, Dominic Kwiatkowski, Cordelia Langford, John Sillitoe on behalf of the Wellcome Sanger Institute COVID-19 Surveillance Team |
| EPI_ISL_3173070 | Lighthouse Lab in Glasgow | Wellcome Sanger Institute for the COVID-19 Genomics UK (COG-UK) Consortium | Harper VanSteenhouse, Yumi Kasai, David Gray, Carol Clugston, Anna Dominiczak and Alex Alderton, Roberto Amato, Jeffrey Barrett, Sonia Goncalves, Ewan Harrison, David K. Jackson, Ian Johnston, Dominic Kwiatkowski, Cordelia Langford, John Sillitoe on behalf of the Wellcome Sanger Institute COVID-19 Surveillance Team |
| EPI_ISL_3173071, EPI_ISL_3173072 | Lighthouse Lab in Milton Keynes | Wellcome Sanger Institute for the COVID-19 Genomics UK (COG-UK) Consortium | The Lighthouse Lab in Milton Keynes and Alex Alderton, Roberto Amato, Jeffrey Barrett, Sonia Goncalves, Ewan Harrison, David K. Jackson, Ian Johnston, Dominic Kwiatkowski, Cordelia Langford, John Sillitoe on behalf of the Wellcome Sanger Institute COVID-19 Surveillance Team |
| EPI_ISL_3173073, EPI_ISL_3173074, EPI_ISL_3173075, EPI_ISL_3173076, EPI_ISL_3173077 | Lighthouse Laboratory Plymouth | Wellcome Sanger Institute for the COVID-19 Genomics UK (COG-UK) Consortium | Lighthouse Laboratory Plymouth and Alex Alderton, Roberto Amato, Jeffrey Barrett, Sonia Goncalves, Ewan Harrison, David K. Jackson, Ian Johnston, Dominic Kwiatkowski, Cordelia Langford, John Sillitoe on behalf of the Wellcome Sanger Institute COVID-19 Surveillance Team |
| EPI_ISL_3173078 | Lighthouse Lab in Glasgow | Wellcome Sanger Institute for the COVID-19 Genomics UK (COG-UK) Consortium | Harper VanSteenhouse, Yumi Kasai, David Gray, Carol Clugston, Anna Dominiczak and Alex Alderton, Roberto Amato, Jeffrey Barrett, Sonia Goncalves, Ewan Harrison, David K. Jackson, Ian Johnston, Dominic Kwiatkowski, Cordelia Langford, John Sillitoe on behalf of the Wellcome Sanger Institute COVID-19 Surveillance Team |
| EPI_ISL_3173079, EPI_ISL_3173080 | Lighthouse Lab in Milton Keynes | Wellcome Sanger Institute for the COVID-19 Genomics UK (COG-UK) Consortium | The Lighthouse Lab in Milton Keynes and Alex Alderton, Roberto Amato, Jeffrey Barrett, Sonia Goncalves, Ewan Harrison, David K. Jackson, Ian Johnston, Dominic Kwiatkowski, Cordelia Langford, John Sillitoe on behalf of the Wellcome Sanger Institute COVID-19 Surveillance Team |
| EPI_ISL_3173081, EPI_ISL_3173082 | Lighthouse Lab in Glasgow | Wellcome Sanger Institute for the COVID-19 Genomics UK (COG-UK) Consortium | Harper VanSteenhouse, Yumi Kasai, David Gray, Carol Clugston, Anna Dominiczak and Alex Alderton, Roberto Amato, Jeffrey Barrett, Sonia Goncalves, Ewan Harrison, David K. Jackson, Ian Johnston, Dominic Kwiatkowski, Cordelia Langford, John Sillitoe on behalf of the Wellcome Sanger Institute COVID-19 Surveillance Team |
| EPI_ISL_3173083, EPI_ISL_3173084 | Lighthouse Laboratory Plymouth | Wellcome Sanger Institute for the COVID-19 Genomics UK (COG-UK) Consortium | Lighthouse Laboratory Plymouth and Alex Alderton, Roberto Amato, Jeffrey Barrett, Sonia Goncalves, Ewan Harrison, David K. Jackson, Ian Johnston, Dominic Kwiatkowski, Cordelia Langford, John Sillitoe on behalf of the Wellcome Sanger Institute COVID-19 Surveillance Team |
| EPI_ISL_3173085, EPI_ISL_3173086 | Lighthouse Lab in Glasgow | Wellcome Sanger Institute for the COVID-19 Genomics UK (COG-UK) Consortium | Harper VanSteenhouse, Yumi Kasai, David Gray, Carol Clugston, Anna Dominiczak and Alex Alderton, Roberto Amato, Jeffrey Barrett, Sonia Goncalves, Ewan Harrison, David K. Jackson, Ian Johnston, Dominic Kwiatkowski, Cordelia Langford, John Sillitoe on behalf of the Wellcome Sanger Institute COVID-19 Surveillance Team |
| EPI_ISL_3173087 | Lighthouse Lab in Milton Keynes | Wellcome Sanger Institute for the COVID-19 Genomics UK (COG-UK) Consortium | The Lighthouse Lab in Milton Keynes and Alex Alderton, Roberto Amato, Jeffrey Barrett, Sonia Goncalves, Ewan Harrison, David K. Jackson, Ian Johnston, Dominic Kwiatkowski, Cordelia Langford, John Sillitoe on behalf of the Wellcome Sanger Institute COVID-19 Surveillance Team |
| EPI_ISL_3173088 | Lighthouse Lab in Glasgow | Wellcome Sanger Institute for the COVID-19 Genomics UK (COG-UK) Consortium | Harper VanSteenhouse, Yumi Kasai, David Gray, Carol Clugston, Anna Dominiczak and Alex Alderton, Roberto Amato, Jeffrey Barrett, Sonia Goncalves, Ewan Harrison, David K. Jackson, Ian Johnston, Dominic Kwiatkowski, Cordelia Langford, John Sillitoe on behalf of the Wellcome Sanger Institute COVID-19 Surveillance Team |
| EPI_ISL_3173089 | Lighthouse Laboratory Plymouth | Wellcome Sanger Institute for the COVID-19 Genomics UK (COG-UK) Consortium | Lighthouse Laboratory Plymouth and Alex Alderton, Roberto Amato, Jeffrey Barrett, Sonia Goncalves, Ewan Harrison, David K. Jackson, Ian Johnston, Dominic Kwiatkowski, Cordelia Langford, John Sillitoe on behalf of the Wellcome Sanger Institute COVID-19 Surveillance Team |
| EPI_ISL_3173090 | Lighthouse Lab in Glasgow | Wellcome Sanger Institute for the COVID-19 Genomics UK (COG-UK) Consortium | Harper VanSteenhouse, Yumi Kasai, David Gray, Carol Clugston, Anna Dominiczak and Alex Alderton, Roberto Amato, Jeffrey Barrett, Sonia Goncalves, Ewan Harrison, David K. Jackson, Ian Johnston, Dominic Kwiatkowski, Cordelia Langford, John Sillitoe on behalf of the Wellcome Sanger Institute COVID-19 Surveillance Team |
| EPI_ISL_3173091, EPI_ISL_3173092 | Lighthouse Laboratory Plymouth | Wellcome Sanger Institute for the COVID-19 Genomics UK (COG-UK) Consortium | Lighthouse Laboratory Plymouth and Alex Alderton, Roberto Amato, Jeffrey Barrett, Sonia Goncalves, Ewan Harrison, David K. Jackson, Ian Johnston, Dominic Kwiatkowski, Cordelia Langford, John Sillitoe on behalf of the Wellcome Sanger Institute COVID-19 Surveillance Team |
| EPI_ISL_3173093 | Lighthouse Lab in Glasgow | Wellcome Sanger Institute for the COVID-19 Genomics UK (COG-UK) Consortium | Harper VanSteenhouse, Yumi Kasai, David Gray, Carol Clugston, Anna Dominiczak and Alex Alderton, Roberto Amato, Jeffrey Barrett, Sonia Goncalves, Ewan Harrison, David K. Jackson, Ian Johnston, Dominic Kwiatkowski, Cordelia Langford, John Sillitoe on behalf of the Wellcome Sanger Institute COVID-19 Surveillance Team |
| EPI_ISL_3173094, EPI_ISL_3173095, EPI_ISL_3173096, EPI_ISL_3173097 | Lighthouse Lab in Milton Keynes | Wellcome Sanger Institute for the COVID-19 Genomics UK (COG-UK) Consortium | The Lighthouse Lab in Milton Keynes and Alex Alderton, Roberto Amato, Jeffrey Barrett, Sonia Goncalves, Ewan Harrison, David K. Jackson, Ian Johnston, Dominic Kwiatkowski, Cordelia Langford, John Sillitoe on behalf of the Wellcome Sanger Institute COVID-19 Surveillance Team |
| EPI_ISL_3173098 | Lighthouse Lab in Glasgow | Wellcome Sanger Institute for the COVID-19 Genomics UK (COG-UK) Consortium | Harper VanSteenhouse, Yumi Kasai, David Gray, Carol Clugston, Anna Dominiczak and Alex Alderton, Roberto Amato, Jeffrey Barrett, Sonia Goncalves, Ewan Harrison, David K. Jackson, Ian Johnston, Dominic Kwiatkowski, Cordelia Langford, John Sillitoe on behalf of the Wellcome Sanger Institute COVID-19 Surveillance Team |
| EPI_ISL_3173099 | Lighthouse Laboratory Plymouth | Wellcome Sanger Institute for the COVID-19 Genomics UK (COG-UK) Consortium | Lighthouse Laboratory Plymouth and Alex Alderton, Roberto Amato, Jeffrey Barrett, Sonia Goncalves, Ewan Harrison, David K. Jackson, Ian Johnston, Dominic Kwiatkowski, Cordelia Langford, John Sillitoe on behalf of the Wellcome Sanger Institute COVID-19 Surveillance Team |
| EPI_ISL_3173100, EPI_ISL_3173101, EPI_ISL_3173102 | Lighthouse Lab in Glasgow | Wellcome Sanger Institute for the COVID-19 Genomics UK (COG-UK) Consortium | Harper VanSteenhouse, Yumi Kasai, David Gray, Carol Clugston, Anna Dominiczak and Alex Alderton, Roberto Amato, Jeffrey Barrett, Sonia Goncalves, Ewan Harrison, David K. Jackson, Ian Johnston, Dominic Kwiatkowski, Cordelia Langford, John Sillitoe on behalf of the Wellcome Sanger Institute COVID-19 Surveillance Team |
| EPI_ISL_3173103, EPI_ISL_3173104 | Lighthouse Laboratory Plymouth | Wellcome Sanger Institute for the COVID-19 Genomics UK (COG-UK) Consortium | Lighthouse Laboratory Plymouth and Alex Alderton, Roberto Amato, Jeffrey Barrett, Sonia Goncalves, Ewan Harrison, David K. Jackson, Ian Johnston, Dominic Kwiatkowski, Cordelia Langford, John Sillitoe on behalf of the Wellcome Sanger Institute COVID-19 Surveillance Team |
| EPI_ISL_3173105, EPI_ISL_3173106 | Lighthouse Lab in Glasgow | Wellcome Sanger Institute for the COVID-19 Genomics UK (COG-UK) Consortium | Harper VanSteenhouse, Yumi Kasai, David Gray, Carol Clugston, Anna Dominiczak and Alex Alderton, Roberto Amato, Jeffrey Barrett, Sonia Goncalves, Ewan Harrison, David K. Jackson, Ian Johnston, Dominic Kwiatkowski, Cordelia Langford, John Sillitoe on behalf of the Wellcome Sanger Institute COVID-19 Surveillance Team |
| EPI_ISL_3173107 | Lighthouse Laboratory Plymouth | Wellcome Sanger Institute for the COVID-19 Genomics UK (COG-UK) | Lighthouse Laboratory Plymouth and Alex Alderton, Roberto Amato, Jeffrey Barrett, Sonia Goncalves, Ewan Harrison, David K. Jackson, Ian Johnston, Dominic Kwiatkowski, Cordelia Langford, John Sillitoe on behalf of the Wellcome Sanger |

| | | | |
|---|---|---|---|
| | | Consortium | Institute COVID-19 Surveillance Team |
| EPI_ISL_3173108 | Lighthouse Lab in Milton Keynes | Wellcome Sanger Institute for the COVID-19 Genomics UK (COG-UK) Consortium | The Lighthouse Lab in Milton Keynes and Alex Alderton, Roberto Amato, Jeffrey Barrett, Sonia Goncalves, Ewan Harrison, David K. Jackson, Ian Johnston, Dominic Kwiatkowski, Cordelia Langford, John Sillitoe on behalf of the Wellcome Sanger Institute COVID-19 Surveillance Team |
| EPI_ISL_3173109 | Lighthouse Lab in Glasgow | Wellcome Sanger Institute for the COVID-19 Genomics UK (COG-UK) Consortium | Harper VanSteenhouse, Yumi Kasai, David Gray, Carol Clugston, Anna Dominiczak and Alex Alderton, Roberto Amato, Jeffrey Barrett, Sonia Goncalves, Ewan Harrison, David K. Jackson, Ian Johnston, Dominic Kwiatkowski, Cordelia Langford, John Sillitoe on behalf of the Wellcome Sanger Institute COVID-19 Surveillance Team |
| EPI_ISL_3173110, EPI_ISL_3173111 | Lighthouse Laboratory Plymouth | Wellcome Sanger Institute for the COVID-19 Genomics UK (COG-UK) Consortium | Lighthouse Laboratory Plymouth and Alex Alderton, Roberto Amato, Jeffrey Barrett, Sonia Goncalves, Ewan Harrison, David K. Jackson, Ian Johnston, Dominic Kwiatkowski, Cordelia Langford, John Sillitoe on behalf of the Wellcome Sanger Institute COVID-19 Surveillance Team |
| EPI_ISL_3173112 | Lighthouse Lab in Milton Keynes | Wellcome Sanger Institute for the COVID-19 Genomics UK (COG-UK) Consortium | The Lighthouse Lab in Milton Keynes and Alex Alderton, Roberto Amato, Jeffrey Barrett, Sonia Goncalves, Ewan Harrison, David K. Jackson, Ian Johnston, Dominic Kwiatkowski, Cordelia Langford, John Sillitoe on behalf of the Wellcome Sanger Institute COVID-19 Surveillance Team |
| EPI_ISL_3173113 | Lighthouse Laboratory Plymouth | Wellcome Sanger Institute for the COVID-19 Genomics UK (COG-UK) Consortium | Lighthouse Laboratory Plymouth and Alex Alderton, Roberto Amato, Jeffrey Barrett, Sonia Goncalves, Ewan Harrison, David K. Jackson, Ian Johnston, Dominic Kwiatkowski, Cordelia Langford, John Sillitoe on behalf of the Wellcome Sanger Institute COVID-19 Surveillance Team |
| EPI_ISL_3173114, EPI_ISL_3173115, EPI_ISL_3173116 | Lighthouse Lab in Milton Keynes | Wellcome Sanger Institute for the COVID-19 Genomics UK (COG-UK) Consortium | The Lighthouse Lab in Milton Keynes and Alex Alderton, Roberto Amato, Jeffrey Barrett, Sonia Goncalves, Ewan Harrison, David K. Jackson, Ian Johnston, Dominic Kwiatkowski, Cordelia Langford, John Sillitoe on behalf of the Wellcome Sanger Institute COVID-19 Surveillance Team |
| EPI_ISL_3173117, EPI_ISL_3173118 | Lighthouse Laboratory Plymouth | Wellcome Sanger Institute for the COVID-19 Genomics UK (COG-UK) Consortium | Lighthouse Laboratory Plymouth and Alex Alderton, Roberto Amato, Jeffrey Barrett, Sonia Goncalves, Ewan Harrison, David K. Jackson, Ian Johnston, Dominic Kwiatkowski, Cordelia Langford, John Sillitoe on behalf of the Wellcome Sanger Institute COVID-19 Surveillance Team |
| EPI_ISL_3173119 | Lighthouse Lab in Milton Keynes | Wellcome Sanger Institute for the COVID-19 Genomics UK (COG-UK) Consortium | The Lighthouse Lab in Milton Keynes and Alex Alderton, Roberto Amato, Jeffrey Barrett, Sonia Goncalves, Ewan Harrison, David K. Jackson, Ian Johnston, Dominic Kwiatkowski, Cordelia Langford, John Sillitoe on behalf of the Wellcome Sanger Institute COVID-19 Surveillance Team |
| EPI_ISL_3173120 | Lighthouse Laboratory Plymouth | Wellcome Sanger Institute for the COVID-19 Genomics UK (COG-UK) Consortium | Lighthouse Laboratory Plymouth and Alex Alderton, Roberto Amato, Jeffrey Barrett, Sonia Goncalves, Ewan Harrison, David K. Jackson, Ian Johnston, Dominic Kwiatkowski, Cordelia Langford, John Sillitoe on behalf of the Wellcome Sanger Institute COVID-19 Surveillance Team |
| EPI_ISL_3173121, EPI_ISL_3173122 | Lighthouse Lab in Glasgow | Wellcome Sanger Institute for the COVID-19 Genomics UK (COG-UK) Consortium | Harper VanSteenhouse, Yumi Kasai, David Gray, Carol Clugston, Anna Dominiczak and Alex Alderton, Roberto Amato, Jeffrey Barrett, Sonia Goncalves, Ewan Harrison, David K. Jackson, Ian Johnston, Dominic Kwiatkowski, Cordelia Langford, John Sillitoe on behalf of the Wellcome Sanger Institute COVID-19 Surveillance Team |
| EPI_ISL_3173123 | Lighthouse Laboratory Plymouth | Wellcome Sanger Institute for the COVID-19 Genomics UK (COG-UK) Consortium | Lighthouse Laboratory Plymouth and Alex Alderton, Roberto Amato, Jeffrey Barrett, Sonia Goncalves, Ewan Harrison, David K. Jackson, Ian Johnston, Dominic Kwiatkowski, Cordelia Langford, John Sillitoe on behalf of the Wellcome Sanger Institute COVID-19 Surveillance Team |
| EPI_ISL_3173124 | Lighthouse Lab in Milton Keynes | Wellcome Sanger Institute for the COVID-19 Genomics UK (COG-UK) Consortium | The Lighthouse Lab in Milton Keynes and Alex Alderton, Roberto Amato, Jeffrey Barrett, Sonia Goncalves, Ewan Harrison, David K. Jackson, Ian Johnston, Dominic Kwiatkowski, Cordelia Langford, John Sillitoe on behalf of the Wellcome Sanger Institute COVID-19 Surveillance Team |
| EPI_ISL_3173125, EPI_ISL_3173126, EPI_ISL_3173127 | Lighthouse Lab in Glasgow | Wellcome Sanger Institute for the COVID-19 Genomics UK (COG-UK) Consortium | Harper VanSteenhouse, Yumi Kasai, David Gray, Carol Clugston, Anna Dominiczak and Alex Alderton, Roberto Amato, Jeffrey Barrett, Sonia Goncalves, Ewan Harrison, David K. Jackson, Ian Johnston, Dominic Kwiatkowski, Cordelia Langford, John Sillitoe on behalf of the Wellcome Sanger Institute COVID-19 Surveillance Team |
| EPI_ISL_3173128 | Lighthouse Laboratory Plymouth | Wellcome Sanger Institute for the COVID-19 Genomics UK (COG-UK) Consortium | Lighthouse Laboratory Plymouth and Alex Alderton, Roberto Amato, Jeffrey Barrett, Sonia Goncalves, Ewan Harrison, David K. Jackson, Ian Johnston, Dominic Kwiatkowski, Cordelia Langford, John Sillitoe on behalf of the Wellcome Sanger Institute COVID-19 Surveillance Team |
| EPI_ISL_3173129 | Lighthouse Lab in Glasgow | Wellcome Sanger Institute for the COVID-19 Genomics UK (COG-UK) Consortium | Harper VanSteenhouse, Yumi Kasai, David Gray, Carol Clugston, Anna Dominiczak and Alex Alderton, Roberto Amato, Jeffrey Barrett, Sonia Goncalves, Ewan Harrison, David K. Jackson, Ian Johnston, Dominic Kwiatkowski, Cordelia Langford, John Sillitoe on behalf of the Wellcome Sanger Institute COVID-19 Surveillance Team |
| EPI_ISL_3173130 | Lighthouse Lab in Milton Keynes | Wellcome Sanger Institute for the COVID-19 Genomics UK (COG-UK) Consortium | The Lighthouse Lab in Milton Keynes and Alex Alderton, Roberto Amato, Jeffrey Barrett, Sonia Goncalves, Ewan Harrison, David K. Jackson, Ian Johnston, Dominic Kwiatkowski, Cordelia Langford, John Sillitoe on behalf of the Wellcome Sanger Institute COVID-19 Surveillance Team |
| EPI_ISL_3173131, EPI_ISL_3173132, EPI_ISL_3173133, EPI_ISL_3173134 | Lighthouse Lab in Glasgow | Wellcome Sanger Institute for the COVID-19 Genomics UK (COG-UK) Consortium | Harper VanSteenhouse, Yumi Kasai, David Gray, Carol Clugston, Anna Dominiczak and Alex Alderton, Roberto Amato, Jeffrey Barrett, Sonia Goncalves, Ewan Harrison, David K. Jackson, Ian Johnston, Dominic Kwiatkowski, Cordelia Langford, John Sillitoe on behalf of the Wellcome Sanger Institute COVID-19 Surveillance Team |
| EPI_ISL_3173135 | Lighthouse Laboratory Plymouth | Wellcome Sanger Institute for the COVID-19 Genomics UK (COG-UK) Consortium | Lighthouse Laboratory Plymouth and Alex Alderton, Roberto Amato, Jeffrey Barrett, Sonia Goncalves, Ewan Harrison, David K. Jackson, Ian Johnston, Dominic Kwiatkowski, Cordelia Langford, John Sillitoe on behalf of the Wellcome Sanger Institute COVID-19 Surveillance Team |
| EPI_ISL_3173136, EPI_ISL_3173137 | Lighthouse Lab in Milton Keynes | Wellcome Sanger Institute for the COVID-19 Genomics UK (COG-UK) Consortium | The Lighthouse Lab in Milton Keynes and Alex Alderton, Roberto Amato, Jeffrey Barrett, Sonia Goncalves, Ewan Harrison, David K. Jackson, Ian Johnston, Dominic Kwiatkowski, Cordelia Langford, John Sillitoe on behalf of the Wellcome Sanger Institute COVID-19 Surveillance Team |
| EPI_ISL_3173138, EPI_ISL_3173139 | Lighthouse Laboratory Plymouth | Wellcome Sanger Institute for the COVID-19 Genomics UK (COG-UK) Consortium | Lighthouse Laboratory Plymouth and Alex Alderton, Roberto Amato, Jeffrey Barrett, Sonia Goncalves, Ewan Harrison, David K. Jackson, Ian Johnston, Dominic Kwiatkowski, Cordelia Langford, John Sillitoe on behalf of the Wellcome Sanger Institute COVID-19 Surveillance Team |
| EPI_ISL_3173140 | Lighthouse Lab in Glasgow | Wellcome Sanger Institute for the COVID-19 Genomics UK (COG-UK) Consortium | Harper VanSteenhouse, Yumi Kasai, David Gray, Carol Clugston, Anna Dominiczak and Alex Alderton, Roberto Amato, Jeffrey Barrett, Sonia Goncalves, Ewan Harrison, David K. Jackson, Ian Johnston, Dominic Kwiatkowski, Cordelia Langford, John Sillitoe on behalf of the Wellcome Sanger Institute COVID-19 Surveillance Team |
| EPI_ISL_3173141 | Lighthouse Lab in Milton Keynes | Wellcome Sanger Institute for the COVID-19 Genomics UK (COG-UK) Consortium | The Lighthouse Lab in Milton Keynes and Alex Alderton, Roberto Amato, Jeffrey Barrett, Sonia Goncalves, Ewan Harrison, David K. Jackson, Ian Johnston, Dominic Kwiatkowski, Cordelia Langford, John Sillitoe on behalf of the Wellcome Sanger Institute COVID-19 Surveillance Team |
| EPI_ISL_3173142 | Lighthouse Laboratory Plymouth | Wellcome Sanger Institute for the COVID-19 Genomics UK (COG-UK) Consortium | Lighthouse Laboratory Plymouth and Alex Alderton, Roberto Amato, Jeffrey Barrett, Sonia Goncalves, Ewan Harrison, David K. Jackson, Ian Johnston, Dominic Kwiatkowski, Cordelia Langford, John Sillitoe on behalf of the Wellcome Sanger Institute COVID-19 Surveillance Team |
| EPI_ISL_3173143, EPI_ISL_3173144, EPI_ISL_3173145, EPI_ISL_3173146 | Lighthouse Lab in Milton Keynes | Wellcome Sanger Institute for the COVID-19 Genomics UK (COG-UK) Consortium | The Lighthouse Lab in Milton Keynes and Alex Alderton, Roberto Amato, Jeffrey Barrett, Sonia Goncalves, Ewan Harrison, David K. Jackson, Ian Johnston, Dominic Kwiatkowski, Cordelia Langford, John Sillitoe on behalf of the Wellcome Sanger Institute COVID-19 Surveillance Team |
| EPI_ISL_3173147, EPI_ISL_3173148 | Lighthouse Laboratory Plymouth | Wellcome Sanger Institute for the COVID-19 Genomics UK (COG-UK) Consortium | Lighthouse Laboratory Plymouth and Alex Alderton, Roberto Amato, Jeffrey Barrett, Sonia Goncalves, Ewan Harrison, David K. Jackson, Ian Johnston, Dominic Kwiatkowski, Cordelia Langford, John Sillitoe on behalf of the Wellcome Sanger Institute COVID-19 Surveillance Team |
| EPI_ISL_3173149, EPI_ISL_3173150, EPI_ISL_3173151, EPI_ISL_3173152, EPI_ISL_3173153 | Lighthouse Lab in Milton Keynes | Wellcome Sanger Institute for the COVID-19 Genomics UK (COG-UK) Consortium | The Lighthouse Lab in Milton Keynes and Alex Alderton, Roberto Amato, Jeffrey Barrett, Sonia Goncalves, Ewan Harrison, David K. Jackson, Ian Johnston, Dominic Kwiatkowski, Cordelia Langford, John Sillitoe on behalf of the Wellcome Sanger Institute COVID-19 Surveillance Team |
| EPI_ISL_3173154 | Lighthouse Lab in Glasgow | Wellcome Sanger Institute for the COVID-19 Genomics UK (COG-UK) Consortium | Harper VanSteenhouse, Yumi Kasai, David Gray, Carol Clugston, Anna Dominiczak and Alex Alderton, Roberto Amato, Jeffrey Barrett, Sonia Goncalves, Ewan Harrison, David K. Jackson, Ian Johnston, Dominic Kwiatkowski, Cordelia Langford, John Sillitoe on behalf of the Wellcome Sanger Institute COVID-19 Surveillance Team |
| EPI_ISL_3173155 | Lighthouse Laboratory Plymouth | Wellcome Sanger Institute for the COVID-19 Genomics UK (COG-UK) Consortium | Lighthouse Laboratory Plymouth and Alex Alderton, Roberto Amato, Jeffrey Barrett, Sonia Goncalves, Ewan Harrison, David K. Jackson, Ian Johnston, Dominic Kwiatkowski, Cordelia Langford, John Sillitoe on behalf of the Wellcome Sanger Institute COVID-19 Surveillance Team |
| EPI_ISL_3173156 | Lighthouse Lab in Milton Keynes | Wellcome Sanger Institute for the COVID-19 Genomics UK (COG-UK) Consortium | The Lighthouse Lab in Milton Keynes and Alex Alderton, Roberto Amato, Jeffrey Barrett, Sonia Goncalves, Ewan Harrison, David K. Jackson, Ian Johnston, Dominic Kwiatkowski, Cordelia Langford, John Sillitoe on behalf of the Wellcome Sanger Institute COVID-19 Surveillance Team |
| EPI_ISL_3173157 | Lighthouse Lab in Glasgow | Wellcome Sanger Institute for the COVID-19 Genomics UK (COG-UK) Consortium | Harper VanSteenhouse, Yumi Kasai, David Gray, Carol Clugston, Anna Dominiczak and Alex Alderton, Roberto Amato, Jeffrey Barrett, Sonia Goncalves, Ewan Harrison, David K. Jackson, Ian Johnston, Dominic Kwiatkowski, Cordelia Langford, John Sillitoe on behalf of the Wellcome Sanger Institute COVID-19 Surveillance Team |
| EPI_ISL_3173158 | Lighthouse Lab in Milton Keynes | Wellcome Sanger Institute for the COVID-19 Genomics UK (COG-UK) Consortium | The Lighthouse Lab in Milton Keynes and Alex Alderton, Roberto Amato, Jeffrey Barrett, Sonia Goncalves, Ewan Harrison, David K. Jackson, Ian Johnston, Dominic Kwiatkowski, Cordelia Langford, John Sillitoe on behalf of the Wellcome Sanger Institute COVID-19 Surveillance Team |
| EPI_ISL_3173159 | Lighthouse Lab in Glasgow | Wellcome Sanger Institute for the COVID-19 Genomics UK (COG-UK) Consortium | Harper VanSteenhouse, Yumi Kasai, David Gray, Carol Clugston, Anna Dominiczak and Alex Alderton, Roberto Amato, Jeffrey Barrett, Sonia Goncalves, Ewan Harrison, David K. Jackson, Ian Johnston, Dominic Kwiatkowski, Cordelia Langford, John Sillitoe on behalf of the Wellcome Sanger Institute COVID-19 Surveillance Team |
| EPI_ISL_3173160 | Lighthouse Laboratory Plymouth | Wellcome Sanger Institute for the COVID-19 Genomics UK (COG-UK) Consortium | Lighthouse Laboratory Plymouth and Alex Alderton, Roberto Amato, Jeffrey Barrett, Sonia Goncalves, Ewan Harrison, David K. Jackson, Ian Johnston, Dominic Kwiatkowski, Cordelia Langford, John Sillitoe on behalf of the Wellcome Sanger Institute COVID-19 Surveillance Team |
| EPI_ISL_3173161 | Lighthouse Lab in Glasgow | Wellcome Sanger Institute for the COVID-19 Genomics UK (COG-UK) Consortium | Harper VanSteenhouse, Yumi Kasai, David Gray, Carol Clugston, Anna Dominiczak and Alex Alderton, Roberto Amato, Jeffrey Barrett, Sonia Goncalves, Ewan Harrison, David K. Jackson, Ian Johnston, Dominic Kwiatkowski, Cordelia Langford, John Sillitoe on behalf of the Wellcome Sanger Institute COVID-19 Surveillance Team |
| EPI_ISL_3173162, EPI_ISL_3173163 | Lighthouse Lab in Milton Keynes | Wellcome Sanger Institute for the COVID-19 Genomics UK (COG-UK) Consortium | The Lighthouse Lab in Milton Keynes and Alex Alderton, Roberto Amato, Jeffrey Barrett, Sonia Goncalves, Ewan Harrison, David K. Jackson, Ian Johnston, Dominic Kwiatkowski, Cordelia Langford, John Sillitoe on behalf of the Wellcome Sanger Institute COVID-19 Surveillance Team |
| EPI_ISL_3173164 | Lighthouse Lab in Glasgow | Wellcome Sanger Institute for the COVID-19 Genomics UK (COG-UK) Consortium | Harper VanSteenhouse, Yumi Kasai, David Gray, Carol Clugston, Anna Dominiczak and Alex Alderton, Roberto Amato, Jeffrey Barrett, Sonia Goncalves, Ewan Harrison, David K. Jackson, Ian Johnston, Dominic Kwiatkowski, Cordelia Langford, John Sillitoe on behalf of the Wellcome Sanger Institute COVID-19 Surveillance Team |
| EPI_ISL_3173165 | Lighthouse Lab in Milton Keynes | Wellcome Sanger Institute for the COVID-19 Genomics UK (COG-UK) Consortium | The Lighthouse Lab in Milton Keynes and Alex Alderton, Roberto Amato, Jeffrey Barrett, Sonia Goncalves, Ewan Harrison, David K. Jackson, Ian Johnston, Dominic Kwiatkowski, Cordelia Langford, John Sillitoe on behalf of the Wellcome Sanger Institute COVID-19 Surveillance Team |
| EPI_ISL_3173166 | Lighthouse Lab in Glasgow | Wellcome Sanger Institute for the COVID-19 Genomics UK (COG-UK) Consortium | Harper VanSteenhouse, Yumi Kasai, David Gray, Carol Clugston, Anna Dominiczak and Alex Alderton, Roberto Amato, Jeffrey Barrett, Sonia Goncalves, Ewan Harrison, David K. Jackson, Ian Johnston, Dominic Kwiatkowski, Cordelia Langford, John Sillitoe on behalf of the Wellcome Sanger Institute COVID-19 Surveillance Team |
| EPI_ISL_3173167 | Lighthouse Laboratory Plymouth | Wellcome Sanger Institute for the COVID-19 Genomics UK (COG-UK) Consortium | Lighthouse Laboratory Plymouth and Alex Alderton, Roberto Amato, Jeffrey Barrett, Sonia Goncalves, Ewan Harrison, David K. Jackson, Ian Johnston, Dominic Kwiatkowski, Cordelia Langford, John Sillitoe on behalf of the Wellcome Sanger Institute COVID-19 Surveillance Team |
| EPI_ISL_3173168 | Lighthouse Lab in Glasgow | Wellcome Sanger Institute for the COVID-19 Genomics UK (COG-UK) Consortium | Harper VanSteenhouse, Yumi Kasai, David Gray, Carol Clugston, Anna Dominiczak and Alex Alderton, Roberto Amato, Jeffrey Barrett, Sonia Goncalves, Ewan Harrison, David K. Jackson, Ian Johnston, Dominic Kwiatkowski, Cordelia Langford, John Sillitoe on behalf of the Wellcome Sanger Institute COVID-19 Surveillance Team |

| | | | |
|---|---|---|---|
| EPI_ISL_3173169 | Lighthouse Lab in Milton Keynes | Wellcome Sanger Institute for the COVID-19 Genomics UK (COG-UK) Consortium | The Lighthouse Lab in Milton Keynes and Alex Alderton, Roberto Amato, Jeffrey Barrett, Sonia Goncalves, Ewan Harrison, David K. Jackson, Ian Johnston, Dominic Kwiatkowski, Cordelia Langford, John Sillitoe on behalf of the Wellcome Sanger Institute COVID-19 Surveillance Team |
| EPI_ISL_3173170 | Lighthouse Laboratory Plymouth | Wellcome Sanger Institute for the COVID-19 Genomics UK (COG-UK) Consortium | Lighthouse Laboratory Plymouth and Alex Alderton, Roberto Amato, Jeffrey Barrett, Sonia Goncalves, Ewan Harrison, David K. Jackson, Ian Johnston, Dominic Kwiatkowski, Cordelia Langford, John Sillitoe on behalf of the Wellcome Sanger Institute COVID-19 Surveillance Team |
| EPI_ISL_3173171 | Lighthouse Lab in Milton Keynes | Wellcome Sanger Institute for the COVID-19 Genomics UK (COG-UK) Consortium | The Lighthouse Lab in Milton Keynes and Alex Alderton, Roberto Amato, Jeffrey Barrett, Sonia Goncalves, Ewan Harrison, David K. Jackson, Ian Johnston, Dominic Kwiatkowski, Cordelia Langford, John Sillitoe on behalf of the Wellcome Sanger Institute COVID-19 Surveillance Team |
| EPI_ISL_3173172 | Lighthouse Lab in Glasgow | Wellcome Sanger Institute for the COVID-19 Genomics UK (COG-UK) Consortium | Harper VanSteenhouse, Yumi Kasai, David Gray, Carol Clugston, Anna Dominiczak and Alex Alderton, Roberto Amato, Jeffrey Barrett, Sonia Goncalves, Ewan Harrison, David K. Jackson, Ian Johnston, Dominic Kwiatkowski, Cordelia Langford, John Sillitoe on behalf of the Wellcome Sanger Institute COVID-19 Surveillance Team |
| EPI_ISL_3173173 | Lighthouse Laboratory Plymouth | Wellcome Sanger Institute for the COVID-19 Genomics UK (COG-UK) Consortium | Lighthouse Laboratory Plymouth and Alex Alderton, Roberto Amato, Jeffrey Barrett, Sonia Goncalves, Ewan Harrison, David K. Jackson, Ian Johnston, Dominic Kwiatkowski, Cordelia Langford, John Sillitoe on behalf of the Wellcome Sanger Institute COVID-19 Surveillance Team |
| EPI_ISL_3173174, EPI_ISL_3173175 | Lighthouse Lab in Milton Keynes | Wellcome Sanger Institute for the COVID-19 Genomics UK (COG-UK) Consortium | The Lighthouse Lab in Milton Keynes and Alex Alderton, Roberto Amato, Jeffrey Barrett, Sonia Goncalves, Ewan Harrison, David K. Jackson, Ian Johnston, Dominic Kwiatkowski, Cordelia Langford, John Sillitoe on behalf of the Wellcome Sanger Institute COVID-19 Surveillance Team |
| EPI_ISL_3173176 | Lighthouse Lab in Glasgow | Wellcome Sanger Institute for the COVID-19 Genomics UK (COG-UK) Consortium | Harper VanSteenhouse, Yumi Kasai, David Gray, Carol Clugston, Anna Dominiczak and Alex Alderton, Roberto Amato, Jeffrey Barrett, Sonia Goncalves, Ewan Harrison, David K. Jackson, Ian Johnston, Dominic Kwiatkowski, Cordelia Langford, John Sillitoe on behalf of the Wellcome Sanger Institute COVID-19 Surveillance Team |
| EPI_ISL_3173177 | Lighthouse Lab in Milton Keynes | Wellcome Sanger Institute for the COVID-19 Genomics UK (COG-UK) Consortium | The Lighthouse Lab in Milton Keynes and Alex Alderton, Roberto Amato, Jeffrey Barrett, Sonia Goncalves, Ewan Harrison, David K. Jackson, Ian Johnston, Dominic Kwiatkowski, Cordelia Langford, John Sillitoe on behalf of the Wellcome Sanger Institute COVID-19 Surveillance Team |
| EPI_ISL_3173178, EPI_ISL_3173179 | Lighthouse Lab in Glasgow | Wellcome Sanger Institute for the COVID-19 Genomics UK (COG-UK) Consortium | Harper VanSteenhouse, Yumi Kasai, David Gray, Carol Clugston, Anna Dominiczak and Alex Alderton, Roberto Amato, Jeffrey Barrett, Sonia Goncalves, Ewan Harrison, David K. Jackson, Ian Johnston, Dominic Kwiatkowski, Cordelia Langford, John Sillitoe on behalf of the Wellcome Sanger Institute COVID-19 Surveillance Team |
| EPI_ISL_3173180 | Lighthouse Lab in Milton Keynes | Wellcome Sanger Institute for the COVID-19 Genomics UK (COG-UK) Consortium | The Lighthouse Lab in Milton Keynes and Alex Alderton, Roberto Amato, Jeffrey Barrett, Sonia Goncalves, Ewan Harrison, David K. Jackson, Ian Johnston, Dominic Kwiatkowski, Cordelia Langford, John Sillitoe on behalf of the Wellcome Sanger Institute COVID-19 Surveillance Team |
| EPI_ISL_3173181, EPI_ISL_3173182, EPI_ISL_3173183, EPI_ISL_3173184 | Lighthouse Laboratory Plymouth | Wellcome Sanger Institute for the COVID-19 Genomics UK (COG-UK) Consortium | Lighthouse Laboratory Plymouth and Alex Alderton, Roberto Amato, Jeffrey Barrett, Sonia Goncalves, Ewan Harrison, David K. Jackson, Ian Johnston, Dominic Kwiatkowski, Cordelia Langford, John Sillitoe on behalf of the Wellcome Sanger Institute COVID-19 Surveillance Team |
| EPI_ISL_3173185 | Lighthouse Lab in Glasgow | Wellcome Sanger Institute for the COVID-19 Genomics UK (COG-UK) Consortium | Harper VanSteenhouse, Yumi Kasai, David Gray, Carol Clugston, Anna Dominiczak and Alex Alderton, Roberto Amato, Jeffrey Barrett, Sonia Goncalves, Ewan Harrison, David K. Jackson, Ian Johnston, Dominic Kwiatkowski, Cordelia Langford, John Sillitoe on behalf of the Wellcome Sanger Institute COVID-19 Surveillance Team |
| EPI_ISL_3173186 | Lighthouse Laboratory Plymouth | Wellcome Sanger Institute for the COVID-19 Genomics UK (COG-UK) Consortium | Lighthouse Laboratory Plymouth and Alex Alderton, Roberto Amato, Jeffrey Barrett, Sonia Goncalves, Ewan Harrison, David K. Jackson, Ian Johnston, Dominic Kwiatkowski, Cordelia Langford, John Sillitoe on behalf of the Wellcome Sanger Institute COVID-19 Surveillance Team |
| EPI_ISL_3173187 | Lighthouse Lab in Glasgow | Wellcome Sanger Institute for the COVID-19 Genomics UK (COG-UK) Consortium | Harper VanSteenhouse, Yumi Kasai, David Gray, Carol Clugston, Anna Dominiczak and Alex Alderton, Roberto Amato, Jeffrey Barrett, Sonia Goncalves, Ewan Harrison, David K. Jackson, Ian Johnston, Dominic Kwiatkowski, Cordelia Langford, John Sillitoe on behalf of the Wellcome Sanger Institute COVID-19 Surveillance Team |
| EPI_ISL_3173188, EPI_ISL_3173189, EPI_ISL_3173190 | Lighthouse Laboratory Plymouth | Wellcome Sanger Institute for the COVID-19 Genomics UK (COG-UK) Consortium | Lighthouse Laboratory Plymouth and Alex Alderton, Roberto Amato, Jeffrey Barrett, Sonia Goncalves, Ewan Harrison, David K. Jackson, Ian Johnston, Dominic Kwiatkowski, Cordelia Langford, John Sillitoe on behalf of the Wellcome Sanger Institute COVID-19 Surveillance Team |
| EPI_ISL_3173191, EPI_ISL_3173192 | Lighthouse Lab in Glasgow | Wellcome Sanger Institute for the COVID-19 Genomics UK (COG-UK) Consortium | Harper VanSteenhouse, Yumi Kasai, David Gray, Carol Clugston, Anna Dominiczak and Alex Alderton, Roberto Amato, Jeffrey Barrett, Sonia Goncalves, Ewan Harrison, David K. Jackson, Ian Johnston, Dominic Kwiatkowski, Cordelia Langford, John Sillitoe on behalf of the Wellcome Sanger Institute COVID-19 Surveillance Team |
| EPI_ISL_3173193 | Lighthouse Lab in Milton Keynes | Wellcome Sanger Institute for the COVID-19 Genomics UK (COG-UK) Consortium | The Lighthouse Lab in Milton Keynes and Alex Alderton, Roberto Amato, Jeffrey Barrett, Sonia Goncalves, Ewan Harrison, David K. Jackson, Ian Johnston, Dominic Kwiatkowski, Cordelia Langford, John Sillitoe on behalf of the Wellcome Sanger Institute COVID-19 Surveillance Team |
| EPI_ISL_3173194, EPI_ISL_3173195, EPI_ISL_3173196, EPI_ISL_3173197 | Lighthouse Laboratory Plymouth | Wellcome Sanger Institute for the COVID-19 Genomics UK (COG-UK) Consortium | Lighthouse Laboratory Plymouth and Alex Alderton, Roberto Amato, Jeffrey Barrett, Sonia Goncalves, Ewan Harrison, David K. Jackson, Ian Johnston, Dominic Kwiatkowski, Cordelia Langford, John Sillitoe on behalf of the Wellcome Sanger Institute COVID-19 Surveillance Team |
| EPI_ISL_3173198, EPI_ISL_3173199 | Lighthouse Lab in Glasgow | Wellcome Sanger Institute for the COVID-19 Genomics UK (COG-UK) Consortium | Harper VanSteenhouse, Yumi Kasai, David Gray, Carol Clugston, Anna Dominiczak and Alex Alderton, Roberto Amato, Jeffrey Barrett, Sonia Goncalves, Ewan Harrison, David K. Jackson, Ian Johnston, Dominic Kwiatkowski, Cordelia Langford, John Sillitoe on behalf of the Wellcome Sanger Institute COVID-19 Surveillance Team |
| EPI_ISL_3173200 | Lighthouse Laboratory Plymouth | Wellcome Sanger Institute for the COVID-19 Genomics UK (COG-UK) Consortium | Lighthouse Laboratory Plymouth and Alex Alderton, Roberto Amato, Jeffrey Barrett, Sonia Goncalves, Ewan Harrison, David K. Jackson, Ian Johnston, Dominic Kwiatkowski, Cordelia Langford, John Sillitoe on behalf of the Wellcome Sanger Institute COVID-19 Surveillance Team |
| EPI_ISL_3173201 | Lighthouse Lab in Glasgow | Wellcome Sanger Institute for the COVID-19 Genomics UK (COG-UK) Consortium | Harper VanSteenhouse, Yumi Kasai, David Gray, Carol Clugston, Anna Dominiczak and Alex Alderton, Roberto Amato, Jeffrey Barrett, Sonia Goncalves, Ewan Harrison, David K. Jackson, Ian Johnston, Dominic Kwiatkowski, Cordelia Langford, John Sillitoe on behalf of the Wellcome Sanger Institute COVID-19 Surveillance Team |
| EPI_ISL_3173202 | Lighthouse Lab in Milton Keynes | Wellcome Sanger Institute for the COVID-19 Genomics UK (COG-UK) Consortium | The Lighthouse Lab in Milton Keynes and Alex Alderton, Roberto Amato, Jeffrey Barrett, Sonia Goncalves, Ewan Harrison, David K. Jackson, Ian Johnston, Dominic Kwiatkowski, Cordelia Langford, John Sillitoe on behalf of the Wellcome Sanger Institute COVID-19 Surveillance Team |
| EPI_ISL_3173203 | Lighthouse Laboratory Plymouth | Wellcome Sanger Institute for the COVID-19 Genomics UK (COG-UK) Consortium | Lighthouse Laboratory Plymouth and Alex Alderton, Roberto Amato, Jeffrey Barrett, Sonia Goncalves, Ewan Harrison, David K. Jackson, Ian Johnston, Dominic Kwiatkowski, Cordelia Langford, John Sillitoe on behalf of the Wellcome Sanger Institute COVID-19 Surveillance Team |
| EPI_ISL_3173204 | Lighthouse Lab in Glasgow | Wellcome Sanger Institute for the COVID-19 Genomics UK (COG-UK) Consortium | Harper VanSteenhouse, Yumi Kasai, David Gray, Carol Clugston, Anna Dominiczak and Alex Alderton, Roberto Amato, Jeffrey Barrett, Sonia Goncalves, Ewan Harrison, David K. Jackson, Ian Johnston, Dominic Kwiatkowski, Cordelia Langford, John Sillitoe on behalf of the Wellcome Sanger Institute COVID-19 Surveillance Team |
| EPI_ISL_3173205 | Lighthouse Laboratory Plymouth | Wellcome Sanger Institute for the COVID-19 Genomics UK (COG-UK) Consortium | Lighthouse Laboratory Plymouth and Alex Alderton, Roberto Amato, Jeffrey Barrett, Sonia Goncalves, Ewan Harrison, David K. Jackson, Ian Johnston, Dominic Kwiatkowski, Cordelia Langford, John Sillitoe on behalf of the Wellcome Sanger Institute COVID-19 Surveillance Team |
| EPI_ISL_3173206, EPI_ISL_3173207 | Lighthouse Lab in Milton Keynes | Wellcome Sanger Institute for the COVID-19 Genomics UK (COG-UK) Consortium | The Lighthouse Lab in Milton Keynes and Alex Alderton, Roberto Amato, Jeffrey Barrett, Sonia Goncalves, Ewan Harrison, David K. Jackson, Ian Johnston, Dominic Kwiatkowski, Cordelia Langford, John Sillitoe on behalf of the Wellcome Sanger Institute COVID-19 Surveillance Team |
| EPI_ISL_3173208, EPI_ISL_3173209, EPI_ISL_3173210 | Lighthouse Laboratory Plymouth | Wellcome Sanger Institute for the COVID-19 Genomics UK (COG-UK) Consortium | Lighthouse Laboratory Plymouth and Alex Alderton, Roberto Amato, Jeffrey Barrett, Sonia Goncalves, Ewan Harrison, David K. Jackson, Ian Johnston, Dominic Kwiatkowski, Cordelia Langford, John Sillitoe on behalf of the Wellcome Sanger Institute COVID-19 Surveillance Team |
| EPI_ISL_3173211, EPI_ISL_3173212 | Lighthouse Lab in Milton Keynes | Wellcome Sanger Institute for the COVID-19 Genomics UK (COG-UK) Consortium | The Lighthouse Lab in Milton Keynes and Alex Alderton, Roberto Amato, Jeffrey Barrett, Sonia Goncalves, Ewan Harrison, David K. Jackson, Ian Johnston, Dominic Kwiatkowski, Cordelia Langford, John Sillitoe on behalf of the Wellcome Sanger Institute COVID-19 Surveillance Team |
| EPI_ISL_3173213 | Lighthouse Lab in Glasgow | Wellcome Sanger Institute for the COVID-19 Genomics UK (COG-UK) Consortium | Harper VanSteenhouse, Yumi Kasai, David Gray, Carol Clugston, Anna Dominiczak and Alex Alderton, Roberto Amato, Jeffrey Barrett, Sonia Goncalves, Ewan Harrison, David K. Jackson, Ian Johnston, Dominic Kwiatkowski, Cordelia Langford, John Sillitoe on behalf of the Wellcome Sanger Institute COVID-19 Surveillance Team |
| EPI_ISL_3173214 | Lighthouse Laboratory Plymouth | Wellcome Sanger Institute for the COVID-19 Genomics UK (COG-UK) Consortium | Lighthouse Laboratory Plymouth and Alex Alderton, Roberto Amato, Jeffrey Barrett, Sonia Goncalves, Ewan Harrison, David K. Jackson, Ian Johnston, Dominic Kwiatkowski, Cordelia Langford, John Sillitoe on behalf of the Wellcome Sanger Institute COVID-19 Surveillance Team |
| EPI_ISL_3173215 | Lighthouse Lab in Glasgow | Wellcome Sanger Institute for the COVID-19 Genomics UK (COG-UK) Consortium | Harper VanSteenhouse, Yumi Kasai, David Gray, Carol Clugston, Anna Dominiczak and Alex Alderton, Roberto Amato, Jeffrey Barrett, Sonia Goncalves, Ewan Harrison, David K. Jackson, Ian Johnston, Dominic Kwiatkowski, Cordelia Langford, John Sillitoe on behalf of the Wellcome Sanger Institute COVID-19 Surveillance Team |
| EPI_ISL_3173216 | Lighthouse Lab in Milton Keynes | Wellcome Sanger Institute for the COVID-19 Genomics UK (COG-UK) Consortium | The Lighthouse Lab in Milton Keynes and Alex Alderton, Roberto Amato, Jeffrey Barrett, Sonia Goncalves, Ewan Harrison, David K. Jackson, Ian Johnston, Dominic Kwiatkowski, Cordelia Langford, John Sillitoe on behalf of the Wellcome Sanger Institute COVID-19 Surveillance Team |
| EPI_ISL_3173217 | Lighthouse Laboratory Plymouth | Wellcome Sanger Institute for the COVID-19 Genomics UK (COG-UK) Consortium | Lighthouse Laboratory Plymouth and Alex Alderton, Roberto Amato, Jeffrey Barrett, Sonia Goncalves, Ewan Harrison, David K. Jackson, Ian Johnston, Dominic Kwiatkowski, Cordelia Langford, John Sillitoe on behalf of the Wellcome Sanger Institute COVID-19 Surveillance Team |
| EPI_ISL_3173218, EPI_ISL_3173219 | Lighthouse Lab in Milton Keynes | Wellcome Sanger Institute for the COVID-19 Genomics UK (COG-UK) Consortium | The Lighthouse Lab in Milton Keynes and Alex Alderton, Roberto Amato, Jeffrey Barrett, Sonia Goncalves, Ewan Harrison, David K. Jackson, Ian Johnston, Dominic Kwiatkowski, Cordelia Langford, John Sillitoe on behalf of the Wellcome Sanger Institute COVID-19 Surveillance Team |
| EPI_ISL_3173220 | Lighthouse Lab in Glasgow | Wellcome Sanger Institute for the COVID-19 Genomics UK (COG-UK) Consortium | Harper VanSteenhouse, Yumi Kasai, David Gray, Carol Clugston, Anna Dominiczak and Alex Alderton, Roberto Amato, Jeffrey Barrett, Sonia Goncalves, Ewan Harrison, David K. Jackson, Ian Johnston, Dominic Kwiatkowski, Cordelia Langford, John Sillitoe on behalf of the Wellcome Sanger Institute COVID-19 Surveillance Team |
| EPI_ISL_3173221, EPI_ISL_3173222 | Lighthouse Lab in Milton Keynes | Wellcome Sanger Institute for the COVID-19 Genomics UK (COG-UK) Consortium | The Lighthouse Lab in Milton Keynes and Alex Alderton, Roberto Amato, Jeffrey Barrett, Sonia Goncalves, Ewan Harrison, David K. Jackson, Ian Johnston, Dominic Kwiatkowski, Cordelia Langford, John Sillitoe on behalf of the Wellcome Sanger Institute COVID-19 Surveillance Team |
| EPI_ISL_3173223, EPI_ISL_3173224, EPI_ISL_3173225, EPI_ISL_3173226 | Lighthouse Lab in Glasgow | Wellcome Sanger Institute for the COVID-19 Genomics UK (COG-UK) Consortium | Harper VanSteenhouse, Yumi Kasai, David Gray, Carol Clugston, Anna Dominiczak and Alex Alderton, Roberto Amato, Jeffrey Barrett, Sonia Goncalves, Ewan Harrison, David K. Jackson, Ian Johnston, Dominic Kwiatkowski, Cordelia Langford, John Sillitoe on behalf of the Wellcome Sanger Institute COVID-19 Surveillance Team |
| EPI_ISL_3173227 | Lighthouse Laboratory Plymouth | Wellcome Sanger Institute for the COVID-19 Genomics UK (COG-UK) Consortium | Lighthouse Laboratory Plymouth and Alex Alderton, Roberto Amato, Jeffrey Barrett, Sonia Goncalves, Ewan Harrison, David K. Jackson, Ian Johnston, Dominic Kwiatkowski, Cordelia Langford, John Sillitoe on behalf of the Wellcome Sanger Institute COVID-19 Surveillance Team |
| EPI_ISL_3173228 | Lighthouse Lab in Glasgow | Wellcome Sanger Institute for the COVID-19 Genomics UK (COG-UK) Consortium | Harper VanSteenhouse, Yumi Kasai, David Gray, Carol Clugston, Anna Dominiczak and Alex Alderton, Roberto Amato, Jeffrey Barrett, Sonia Goncalves, Ewan Harrison, David K. Jackson, Ian Johnston, Dominic Kwiatkowski, Cordelia Langford, John Sillitoe on behalf of the Wellcome Sanger Institute COVID-19 Surveillance Team |
| EPI_ISL_3173229, EPI_ISL_3173230 | Lighthouse Laboratory Plymouth | Wellcome Sanger Institute for the COVID-19 Genomics UK (COG-UK) Consortium | Lighthouse Laboratory Plymouth and Alex Alderton, Roberto Amato, Jeffrey Barrett, Sonia Goncalves, Ewan Harrison, David K. Jackson, Ian Johnston, Dominic Kwiatkowski, Cordelia Langford, John Sillitoe on behalf of the Wellcome Sanger Institute COVID-19 Surveillance Team |
| EPI_ISL_3173231, EPI_ISL_3173232 | Lighthouse Lab in Milton Keynes | Wellcome Sanger Institute for the COVID-19 Genomics UK (COG-UK) Consortium | The Lighthouse Lab in Milton Keynes and Alex Alderton, Roberto Amato, Jeffrey Barrett, Sonia Goncalves, Ewan Harrison, David K. Jackson, Ian Johnston, Dominic Kwiatkowski, Cordelia Langford, John Sillitoe on behalf of the Wellcome Sanger Institute COVID-19 Surveillance Team |
| EPI_ISL_3173233, EPI_ISL_3173234 | Lighthouse Lab in Glasgow | Wellcome Sanger Institute for the COVID-19 Genomics UK (COG-UK) Consortium | Harper VanSteenhouse, Yumi Kasai, David Gray, Carol Clugston, Anna Dominiczak and Alex Alderton, Roberto Amato, Jeffrey Barrett, Sonia Goncalves, Ewan Harrison, David K. Jackson, Ian Johnston, Dominic Kwiatkowski, Cordelia Langford, John Sillitoe on behalf of the Wellcome Sanger Institute COVID-19 Surveillance Team |

| | | | |
|---|---|---|---|
| EPI_ISL_3173235, EPI_ISL_3173236 | Lighthouse Laboratory Plymouth | Wellcome Sanger Institute for the COVID-19 Genomics UK (COG-UK) Consortium | Lighthouse Laboratory Plymouth and Alex Alderton, Roberto Amato, Jeffrey Barrett, Sonia Goncalves, Ewan Harrison, David K. Jackson, Ian Johnston, Dominic Kwiatkowski, Cordelia Langford, John Sillitoe on behalf of the Wellcome Sanger Institute COVID-19 Surveillance Team |
| EPI_ISL_3173237 | Lighthouse Lab in Milton Keynes | Wellcome Sanger Institute for the COVID-19 Genomics UK (COG-UK) Consortium | The Lighthouse Lab in Milton Keynes and Alex Alderton, Roberto Amato, Jeffrey Barrett, Sonia Goncalves, Ewan Harrison, David K. Jackson, Ian Johnston, Dominic Kwiatkowski, Cordelia Langford, John Sillitoe on behalf of the Wellcome Sanger Institute COVID-19 Surveillance Team |
| EPI_ISL_3173238, EPI_ISL_3173239 | Lighthouse Laboratory Plymouth | Wellcome Sanger Institute for the COVID-19 Genomics UK (COG-UK) Consortium | Lighthouse Laboratory Plymouth and Alex Alderton, Roberto Amato, Jeffrey Barrett, Sonia Goncalves, Ewan Harrison, David K. Jackson, Ian Johnston, Dominic Kwiatkowski, Cordelia Langford, John Sillitoe on behalf of the Wellcome Sanger Institute COVID-19 Surveillance Team |
| EPI_ISL_3173240 | Lighthouse Lab in Milton Keynes | Wellcome Sanger Institute for the COVID-19 Genomics UK (COG-UK) Consortium | The Lighthouse Lab in Milton Keynes and Alex Alderton, Roberto Amato, Jeffrey Barrett, Sonia Goncalves, Ewan Harrison, David K. Jackson, Ian Johnston, Dominic Kwiatkowski, Cordelia Langford, John Sillitoe on behalf of the Wellcome Sanger Institute COVID-19 Surveillance Team |
| EPI_ISL_3173241 | Lighthouse Lab in Glasgow | Wellcome Sanger Institute for the COVID-19 Genomics UK (COG-UK) Consortium | Harper VanSteenhouse, Yumi Kasai, David Gray, Carol Clugston, Anna Dominiczak and Alex Alderton, Roberto Amato, Jeffrey Barrett, Sonia Goncalves, Ewan Harrison, David K. Jackson, Ian Johnston, Dominic Kwiatkowski, Cordelia Langford, John Sillitoe on behalf of the Wellcome Sanger Institute COVID-19 Surveillance Team |
| EPI_ISL_3173242 | Lighthouse Lab in Milton Keynes | Wellcome Sanger Institute for the COVID-19 Genomics UK (COG-UK) Consortium | The Lighthouse Lab in Milton Keynes and Alex Alderton, Roberto Amato, Jeffrey Barrett, Sonia Goncalves, Ewan Harrison, David K. Jackson, Ian Johnston, Dominic Kwiatkowski, Cordelia Langford, John Sillitoe on behalf of the Wellcome Sanger Institute COVID-19 Surveillance Team |
| EPI_ISL_3173243 | Lighthouse Lab in Glasgow | Wellcome Sanger Institute for the COVID-19 Genomics UK (COG-UK) Consortium | Harper VanSteenhouse, Yumi Kasai, David Gray, Carol Clugston, Anna Dominiczak and Alex Alderton, Roberto Amato, Jeffrey Barrett, Sonia Goncalves, Ewan Harrison, David K. Jackson, Ian Johnston, Dominic Kwiatkowski, Cordelia Langford, John Sillitoe on behalf of the Wellcome Sanger Institute COVID-19 Surveillance Team |
| EPI_ISL_3173244 | Lighthouse Laboratory Plymouth | Wellcome Sanger Institute for the COVID-19 Genomics UK (COG-UK) Consortium | Lighthouse Laboratory Plymouth and Alex Alderton, Roberto Amato, Jeffrey Barrett, Sonia Goncalves, Ewan Harrison, David K. Jackson, Ian Johnston, Dominic Kwiatkowski, Cordelia Langford, John Sillitoe on behalf of the Wellcome Sanger Institute COVID-19 Surveillance Team |
| EPI_ISL_3173245, EPI_ISL_3173246 | Lighthouse Lab in Milton Keynes | Wellcome Sanger Institute for the COVID-19 Genomics UK (COG-UK) Consortium | The Lighthouse Lab in Milton Keynes and Alex Alderton, Roberto Amato, Jeffrey Barrett, Sonia Goncalves, Ewan Harrison, David K. Jackson, Ian Johnston, Dominic Kwiatkowski, Cordelia Langford, John Sillitoe on behalf of the Wellcome Sanger Institute COVID-19 Surveillance Team |
| EPI_ISL_3173247 | Lighthouse Laboratory Plymouth | Wellcome Sanger Institute for the COVID-19 Genomics UK (COG-UK) Consortium | Lighthouse Laboratory Plymouth and Alex Alderton, Roberto Amato, Jeffrey Barrett, Sonia Goncalves, Ewan Harrison, David K. Jackson, Ian Johnston, Dominic Kwiatkowski, Cordelia Langford, John Sillitoe on behalf of the Wellcome Sanger Institute COVID-19 Surveillance Team |
| EPI_ISL_3173248 | Lighthouse Lab in Glasgow | Wellcome Sanger Institute for the COVID-19 Genomics UK (COG-UK) Consortium | Harper VanSteenhouse, Yumi Kasai, David Gray, Carol Clugston, Anna Dominiczak and Alex Alderton, Roberto Amato, Jeffrey Barrett, Sonia Goncalves, Ewan Harrison, David K. Jackson, Ian Johnston, Dominic Kwiatkowski, Cordelia Langford, John Sillitoe on behalf of the Wellcome Sanger Institute COVID-19 Surveillance Team |
| EPI_ISL_3173249, EPI_ISL_3173250, EPI_ISL_3173251, EPI_ISL_3173252 | Lighthouse Laboratory Plymouth | Wellcome Sanger Institute for the COVID-19 Genomics UK (COG-UK) Consortium | Lighthouse Laboratory Plymouth and Alex Alderton, Roberto Amato, Jeffrey Barrett, Sonia Goncalves, Ewan Harrison, David K. Jackson, Ian Johnston, Dominic Kwiatkowski, Cordelia Langford, John Sillitoe on behalf of the Wellcome Sanger Institute COVID-19 Surveillance Team |
| EPI_ISL_3173253 | Lighthouse Lab in Glasgow | Wellcome Sanger Institute for the COVID-19 Genomics UK (COG-UK) Consortium | Harper VanSteenhouse, Yumi Kasai, David Gray, Carol Clugston, Anna Dominiczak and Alex Alderton, Roberto Amato, Jeffrey Barrett, Sonia Goncalves, Ewan Harrison, David K. Jackson, Ian Johnston, Dominic Kwiatkowski, Cordelia Langford, John Sillitoe on behalf of the Wellcome Sanger Institute COVID-19 Surveillance Team |
| EPI_ISL_3173254 | Lighthouse Laboratory Plymouth | Wellcome Sanger Institute for the COVID-19 Genomics UK (COG-UK) Consortium | Lighthouse Laboratory Plymouth and Alex Alderton, Roberto Amato, Jeffrey Barrett, Sonia Goncalves, Ewan Harrison, David K. Jackson, Ian Johnston, Dominic Kwiatkowski, Cordelia Langford, John Sillitoe on behalf of the Wellcome Sanger Institute COVID-19 Surveillance Team |
| EPI_ISL_3173255 | Lighthouse Lab in Milton Keynes | Wellcome Sanger Institute for the COVID-19 Genomics UK (COG-UK) Consortium | The Lighthouse Lab in Milton Keynes and Alex Alderton, Roberto Amato, Jeffrey Barrett, Sonia Goncalves, Ewan Harrison, David K. Jackson, Ian Johnston, Dominic Kwiatkowski, Cordelia Langford, John Sillitoe on behalf of the Wellcome Sanger Institute COVID-19 Surveillance Team |
| EPI_ISL_3173256 | Lighthouse Laboratory Plymouth | Wellcome Sanger Institute for the COVID-19 Genomics UK (COG-UK) Consortium | Lighthouse Laboratory Plymouth and Alex Alderton, Roberto Amato, Jeffrey Barrett, Sonia Goncalves, Ewan Harrison, David K. Jackson, Ian Johnston, Dominic Kwiatkowski, Cordelia Langford, John Sillitoe on behalf of the Wellcome Sanger Institute COVID-19 Surveillance Team |
| EPI_ISL_3173257 | Lighthouse Lab in Glasgow | Wellcome Sanger Institute for the COVID-19 Genomics UK (COG-UK) Consortium | Harper VanSteenhouse, Yumi Kasai, David Gray, Carol Clugston, Anna Dominiczak and Alex Alderton, Roberto Amato, Jeffrey Barrett, Sonia Goncalves, Ewan Harrison, David K. Jackson, Ian Johnston, Dominic Kwiatkowski, Cordelia Langford, John Sillitoe on behalf of the Wellcome Sanger Institute COVID-19 Surveillance Team |
| EPI_ISL_3173258 | Lighthouse Lab in Milton Keynes | Wellcome Sanger Institute for the COVID-19 Genomics UK (COG-UK) Consortium | The Lighthouse Lab in Milton Keynes and Alex Alderton, Roberto Amato, Jeffrey Barrett, Sonia Goncalves, Ewan Harrison, David K. Jackson, Ian Johnston, Dominic Kwiatkowski, Cordelia Langford, John Sillitoe on behalf of the Wellcome Sanger Institute COVID-19 Surveillance Team |
| EPI_ISL_3173259 | Lighthouse Laboratory Plymouth | Wellcome Sanger Institute for the COVID-19 Genomics UK (COG-UK) Consortium | Lighthouse Laboratory Plymouth and Alex Alderton, Roberto Amato, Jeffrey Barrett, Sonia Goncalves, Ewan Harrison, David K. Jackson, Ian Johnston, Dominic Kwiatkowski, Cordelia Langford, John Sillitoe on behalf of the Wellcome Sanger Institute COVID-19 Surveillance Team |
| EPI_ISL_3173260 | Lighthouse Lab in Milton Keynes | Wellcome Sanger Institute for the COVID-19 Genomics UK (COG-UK) Consortium | The Lighthouse Lab in Milton Keynes and Alex Alderton, Roberto Amato, Jeffrey Barrett, Sonia Goncalves, Ewan Harrison, David K. Jackson, Ian Johnston, Dominic Kwiatkowski, Cordelia Langford, John Sillitoe on behalf of the Wellcome Sanger Institute COVID-19 Surveillance Team |
| EPI_ISL_3173261, EPI_ISL_3173262, EPI_ISL_3173263, EPI_ISL_3173264, EPI_ISL_3173265 | Lighthouse Lab in Glasgow | Wellcome Sanger Institute for the COVID-19 Genomics UK (COG-UK) Consortium | Harper VanSteenhouse, Yumi Kasai, David Gray, Carol Clugston, Anna Dominiczak and Alex Alderton, Roberto Amato, Jeffrey Barrett, Sonia Goncalves, Ewan Harrison, David K. Jackson, Ian Johnston, Dominic Kwiatkowski, Cordelia Langford, John Sillitoe on behalf of the Wellcome Sanger Institute COVID-19 Surveillance Team |
| EPI_ISL_3173266, EPI_ISL_3173267, EPI_ISL_3173268, EPI_ISL_3173269 | Lighthouse Laboratory Plymouth | Wellcome Sanger Institute for the COVID-19 Genomics UK (COG-UK) Consortium | Lighthouse Laboratory Plymouth and Alex Alderton, Roberto Amato, Jeffrey Barrett, Sonia Goncalves, Ewan Harrison, David K. Jackson, Ian Johnston, Dominic Kwiatkowski, Cordelia Langford, John Sillitoe on behalf of the Wellcome Sanger Institute COVID-19 Surveillance Team |
| EPI_ISL_3173270 | Lighthouse Lab in Milton Keynes | Wellcome Sanger Institute for the COVID-19 Genomics UK (COG-UK) Consortium | The Lighthouse Lab in Milton Keynes and Alex Alderton, Roberto Amato, Jeffrey Barrett, Sonia Goncalves, Ewan Harrison, David K. Jackson, Ian Johnston, Dominic Kwiatkowski, Cordelia Langford, John Sillitoe on behalf of the Wellcome Sanger Institute COVID-19 Surveillance Team |
| EPI_ISL_3173271 | Lighthouse Laboratory Plymouth | Wellcome Sanger Institute for the COVID-19 Genomics UK (COG-UK) Consortium | Lighthouse Laboratory Plymouth and Alex Alderton, Roberto Amato, Jeffrey Barrett, Sonia Goncalves, Ewan Harrison, David K. Jackson, Ian Johnston, Dominic Kwiatkowski, Cordelia Langford, John Sillitoe on behalf of the Wellcome Sanger Institute COVID-19 Surveillance Team |
| EPI_ISL_3173272 | Lighthouse Lab in Milton Keynes | Wellcome Sanger Institute for the COVID-19 Genomics UK (COG-UK) Consortium | The Lighthouse Lab in Milton Keynes and Alex Alderton, Roberto Amato, Jeffrey Barrett, Sonia Goncalves, Ewan Harrison, David K. Jackson, Ian Johnston, Dominic Kwiatkowski, Cordelia Langford, John Sillitoe on behalf of the Wellcome Sanger Institute COVID-19 Surveillance Team |
| EPI_ISL_3173273 | Lighthouse Lab in Glasgow | Wellcome Sanger Institute for the COVID-19 Genomics UK (COG-UK) Consortium | Harper VanSteenhouse, Yumi Kasai, David Gray, Carol Clugston, Anna Dominiczak and Alex Alderton, Roberto Amato, Jeffrey Barrett, Sonia Goncalves, Ewan Harrison, David K. Jackson, Ian Johnston, Dominic Kwiatkowski, Cordelia Langford, John Sillitoe on behalf of the Wellcome Sanger Institute COVID-19 Surveillance Team |
| EPI_ISL_3173274 | Lighthouse Lab in Milton Keynes | Wellcome Sanger Institute for the COVID-19 Genomics UK (COG-UK) Consortium | The Lighthouse Lab in Milton Keynes and Alex Alderton, Roberto Amato, Jeffrey Barrett, Sonia Goncalves, Ewan Harrison, David K. Jackson, Ian Johnston, Dominic Kwiatkowski, Cordelia Langford, John Sillitoe on behalf of the Wellcome Sanger Institute COVID-19 Surveillance Team |
| EPI_ISL_3173275 | Lighthouse Laboratory Plymouth | Wellcome Sanger Institute for the COVID-19 Genomics UK (COG-UK) Consortium | Lighthouse Laboratory Plymouth and Alex Alderton, Roberto Amato, Jeffrey Barrett, Sonia Goncalves, Ewan Harrison, David K. Jackson, Ian Johnston, Dominic Kwiatkowski, Cordelia Langford, John Sillitoe on behalf of the Wellcome Sanger Institute COVID-19 Surveillance Team |
| EPI_ISL_3173276 | Lighthouse Lab in Milton Keynes | Wellcome Sanger Institute for the COVID-19 Genomics UK (COG-UK) Consortium | The Lighthouse Lab in Milton Keynes and Alex Alderton, Roberto Amato, Jeffrey Barrett, Sonia Goncalves, Ewan Harrison, David K. Jackson, Ian Johnston, Dominic Kwiatkowski, Cordelia Langford, John Sillitoe on behalf of the Wellcome Sanger Institute COVID-19 Surveillance Team |
| EPI_ISL_3173277 | Lighthouse Laboratory Plymouth | Wellcome Sanger Institute for the COVID-19 Genomics UK (COG-UK) Consortium | Lighthouse Laboratory Plymouth and Alex Alderton, Roberto Amato, Jeffrey Barrett, Sonia Goncalves, Ewan Harrison, David K. Jackson, Ian Johnston, Dominic Kwiatkowski, Cordelia Langford, John Sillitoe on behalf of the Wellcome Sanger Institute COVID-19 Surveillance Team |
| EPI_ISL_3173278, EPI_ISL_3173279 | Lighthouse Lab in Glasgow | Wellcome Sanger Institute for the COVID-19 Genomics UK (COG-UK) Consortium | Harper VanSteenhouse, Yumi Kasai, David Gray, Carol Clugston, Anna Dominiczak and Alex Alderton, Roberto Amato, Jeffrey Barrett, Sonia Goncalves, Ewan Harrison, David K. Jackson, Ian Johnston, Dominic Kwiatkowski, Cordelia Langford, John Sillitoe on behalf of the Wellcome Sanger Institute COVID-19 Surveillance Team |
| EPI_ISL_3173280, EPI_ISL_3173281 | Lighthouse Lab in Milton Keynes | Wellcome Sanger Institute for the COVID-19 Genomics UK (COG-UK) Consortium | The Lighthouse Lab in Milton Keynes and Alex Alderton, Roberto Amato, Jeffrey Barrett, Sonia Goncalves, Ewan Harrison, David K. Jackson, Ian Johnston, Dominic Kwiatkowski, Cordelia Langford, John Sillitoe on behalf of the Wellcome Sanger Institute COVID-19 Surveillance Team |
| EPI_ISL_3173282, EPI_ISL_3173283 | Lighthouse Laboratory Plymouth | Wellcome Sanger Institute for the COVID-19 Genomics UK (COG-UK) Consortium | Lighthouse Laboratory Plymouth and Alex Alderton, Roberto Amato, Jeffrey Barrett, Sonia Goncalves, Ewan Harrison, David K. Jackson, Ian Johnston, Dominic Kwiatkowski, Cordelia Langford, John Sillitoe on behalf of the Wellcome Sanger Institute COVID-19 Surveillance Team |
| EPI_ISL_3173284, EPI_ISL_3173285 | Lighthouse Lab in Milton Keynes | Wellcome Sanger Institute for the COVID-19 Genomics UK (COG-UK) Consortium | The Lighthouse Lab in Milton Keynes and Alex Alderton, Roberto Amato, Jeffrey Barrett, Sonia Goncalves, Ewan Harrison, David K. Jackson, Ian Johnston, Dominic Kwiatkowski, Cordelia Langford, John Sillitoe on behalf of the Wellcome Sanger Institute COVID-19 Surveillance Team |
| EPI_ISL_3173286 | Lighthouse Laboratory Plymouth | Wellcome Sanger Institute for the COVID-19 Genomics UK (COG-UK) Consortium | Lighthouse Laboratory Plymouth and Alex Alderton, Roberto Amato, Jeffrey Barrett, Sonia Goncalves, Ewan Harrison, David K. Jackson, Ian Johnston, Dominic Kwiatkowski, Cordelia Langford, John Sillitoe on behalf of the Wellcome Sanger Institute COVID-19 Surveillance Team |
| EPI_ISL_3173287 | Lighthouse Lab in Glasgow | Wellcome Sanger Institute for the COVID-19 Genomics UK (COG-UK) Consortium | Harper VanSteenhouse, Yumi Kasai, David Gray, Carol Clugston, Anna Dominiczak and Alex Alderton, Roberto Amato, Jeffrey Barrett, Sonia Goncalves, Ewan Harrison, David K. Jackson, Ian Johnston, Dominic Kwiatkowski, Cordelia Langford, John Sillitoe on behalf of the Wellcome Sanger Institute COVID-19 Surveillance Team |
| EPI_ISL_3173288 | Lighthouse Laboratory Plymouth | Wellcome Sanger Institute for the COVID-19 Genomics UK (COG-UK) Consortium | Lighthouse Laboratory Plymouth and Alex Alderton, Roberto Amato, Jeffrey Barrett, Sonia Goncalves, Ewan Harrison, David K. Jackson, Ian Johnston, Dominic Kwiatkowski, Cordelia Langford, John Sillitoe on behalf of the Wellcome Sanger Institute COVID-19 Surveillance Team |
| EPI_ISL_3173289, EPI_ISL_3173290, EPI_ISL_3173291, EPI_ISL_3173292 | Lighthouse Lab in Glasgow | Wellcome Sanger Institute for the COVID-19 Genomics UK (COG-UK) Consortium | Harper VanSteenhouse, Yumi Kasai, David Gray, Carol Clugston, Anna Dominiczak and Alex Alderton, Roberto Amato, Jeffrey Barrett, Sonia Goncalves, Ewan Harrison, David K. Jackson, Ian Johnston, Dominic Kwiatkowski, Cordelia Langford, John Sillitoe on behalf of the Wellcome Sanger Institute COVID-19 Surveillance Team |
| EPI_ISL_3173293 | Lighthouse Laboratory Plymouth | Wellcome Sanger Institute for the COVID-19 Genomics UK (COG-UK) Consortium | Lighthouse Laboratory Plymouth and Alex Alderton, Roberto Amato, Jeffrey Barrett, Sonia Goncalves, Ewan Harrison, David K. Jackson, Ian Johnston, Dominic Kwiatkowski, Cordelia Langford, John Sillitoe on behalf of the Wellcome Sanger Institute COVID-19 Surveillance Team |
| EPI_ISL_3173294 | Lighthouse Lab in Milton Keynes | Wellcome Sanger Institute for the COVID-19 Genomics UK (COG-UK) Consortium | The Lighthouse Lab in Milton Keynes and Alex Alderton, Roberto Amato, Jeffrey Barrett, Sonia Goncalves, Ewan Harrison, David K. Jackson, Ian Johnston, Dominic Kwiatkowski, Cordelia Langford, John Sillitoe on behalf of the Wellcome Sanger Institute COVID-19 Surveillance Team |
| EPI_ISL_3173295 | Lighthouse Laboratory Plymouth | Wellcome Sanger Institute for the COVID-19 Genomics UK (COG-UK) Consortium | Lighthouse Laboratory Plymouth and Alex Alderton, Roberto Amato, Jeffrey Barrett, Sonia Goncalves, Ewan Harrison, David K. Jackson, Ian Johnston, Dominic Kwiatkowski, Cordelia Langford, John Sillitoe on behalf of the Wellcome Sanger Institute COVID-19 Surveillance Team |
| EPI_ISL_3173296 | Lighthouse Lab in Milton Keynes | Wellcome Sanger Institute for the COVID-19 Genomics UK (COG-UK) | The Lighthouse Lab in Milton Keynes and Alex Alderton, Roberto Amato, Jeffrey Barrett, Sonia Goncalves, Ewan Harrison, David K. Jackson, Ian Johnston, Dominic Kwiatkowski, Cordelia Langford, John Sillitoe on behalf of the Wellcome Sanger |

| | | | |
|---|---|---|---|
| | | Consortium | Institute COVID-19 Surveillance Team |
| EPI_ISL_3173297, EPI_ISL_3173298 | Lighthouse Lab in Alderley Park | Wellcome Sanger Institute for the COVID-19 Genomics UK (COG-UK) Consortium | Jacquelyn Wynn, Mairead Hyland, The Lighthouse Lab in Alderley Park and Alex Alderton, Roberto Amato, Jeffrey Barrett, Sonia Goncalves, Ewan Harrison, David K. Jackson, Ian Johnston, Dominic Kwiatkowski, Cordelia Langford, John Sillitoe on behalf of the Wellcome Sanger Institute COVID-19 Surveillance Team |
| EPI_ISL_3173299 | Lighthouse Lab in Milton Keynes | Wellcome Sanger Institute for the COVID-19 Genomics UK (COG-UK) Consortium | The Lighthouse Lab in Milton Keynes and Alex Alderton, Roberto Amato, Jeffrey Barrett, Sonia Goncalves, Ewan Harrison, David K. Jackson, Ian Johnston, Dominic Kwiatkowski, Cordelia Langford, John Sillitoe on behalf of the Wellcome Sanger Institute COVID-19 Surveillance Team |
| EPI_ISL_3173300, EPI_ISL_3173301 | Lighthouse Lab in Alderley Park | Wellcome Sanger Institute for the COVID-19 Genomics UK (COG-UK) Consortium | Jacquelyn Wynn, Mairead Hyland, The Lighthouse Lab in Alderley Park and Alex Alderton, Roberto Amato, Jeffrey Barrett, Sonia Goncalves, Ewan Harrison, David K. Jackson, Ian Johnston, Dominic Kwiatkowski, Cordelia Langford, John Sillitoe on behalf of the Wellcome Sanger Institute COVID-19 Surveillance Team |
| EPI_ISL_3173302 | Lighthouse Lab in Milton Keynes | Wellcome Sanger Institute for the COVID-19 Genomics UK (COG-UK) Consortium | The Lighthouse Lab in Milton Keynes and Alex Alderton, Roberto Amato, Jeffrey Barrett, Sonia Goncalves, Ewan Harrison, David K. Jackson, Ian Johnston, Dominic Kwiatkowski, Cordelia Langford, John Sillitoe on behalf of the Wellcome Sanger Institute COVID-19 Surveillance Team |
| EPI_ISL_3173303, EPI_ISL_3173304, EPI_ISL_3173305, EPI_ISL_3173306, EPI_ISL_3173307 | Lighthouse Lab in Alderley Park | Wellcome Sanger Institute for the COVID-19 Genomics UK (COG-UK) Consortium | Jacquelyn Wynn, Mairead Hyland, The Lighthouse Lab in Alderley Park and Alex Alderton, Roberto Amato, Jeffrey Barrett, Sonia Goncalves, Ewan Harrison, David K. Jackson, Ian Johnston, Dominic Kwiatkowski, Cordelia Langford, John Sillitoe on behalf of the Wellcome Sanger Institute COVID-19 Surveillance Team |
| EPI_ISL_3173308 | Lighthouse Lab in Glasgow | Wellcome Sanger Institute for the COVID-19 Genomics UK (COG-UK) Consortium | Harper VanSteenhouse, Yumi Kasai, David Gray, Carol Clugston, Anna Dominiczak and Alex Alderton, Roberto Amato, Jeffrey Barrett, Sonia Goncalves, Ewan Harrison, David K. Jackson, Ian Johnston, Dominic Kwiatkowski, Cordelia Langford, John Sillitoe on behalf of the Wellcome Sanger Institute COVID-19 Surveillance Team |
| EPI_ISL_3173309, EPI_ISL_3173310 | Lighthouse Lab in Alderley Park | Wellcome Sanger Institute for the COVID-19 Genomics UK (COG-UK) Consortium | Jacquelyn Wynn, Mairead Hyland, The Lighthouse Lab in Alderley Park and Alex Alderton, Roberto Amato, Jeffrey Barrett, Sonia Goncalves, Ewan Harrison, David K. Jackson, Ian Johnston, Dominic Kwiatkowski, Cordelia Langford, John Sillitoe on behalf of the Wellcome Sanger Institute COVID-19 Surveillance Team |
| EPI_ISL_3173311, EPI_ISL_3173312 | Lighthouse Lab in Milton Keynes | Wellcome Sanger Institute for the COVID-19 Genomics UK (COG-UK) Consortium | The Lighthouse Lab in Milton Keynes and Alex Alderton, Roberto Amato, Jeffrey Barrett, Sonia Goncalves, Ewan Harrison, David K. Jackson, Ian Johnston, Dominic Kwiatkowski, Cordelia Langford, John Sillitoe on behalf of the Wellcome Sanger Institute COVID-19 Surveillance Team |
| EPI_ISL_3173313, EPI_ISL_3173314 | Lighthouse Lab in Alderley Park | Wellcome Sanger Institute for the COVID-19 Genomics UK (COG-UK) Consortium | Jacquelyn Wynn, Mairead Hyland, The Lighthouse Lab in Alderley Park and Alex Alderton, Roberto Amato, Jeffrey Barrett, Sonia Goncalves, Ewan Harrison, David K. Jackson, Ian Johnston, Dominic Kwiatkowski, Cordelia Langford, John Sillitoe on behalf of the Wellcome Sanger Institute COVID-19 Surveillance Team |
| EPI_ISL_3173315, EPI_ISL_3173316 | Lighthouse Lab in Milton Keynes | Wellcome Sanger Institute for the COVID-19 Genomics UK (COG-UK) Consortium | The Lighthouse Lab in Milton Keynes and Alex Alderton, Roberto Amato, Jeffrey Barrett, Sonia Goncalves, Ewan Harrison, David K. Jackson, Ian Johnston, Dominic Kwiatkowski, Cordelia Langford, John Sillitoe on behalf of the Wellcome Sanger Institute COVID-19 Surveillance Team |
| EPI_ISL_3173317, EPI_ISL_3173318 | Lighthouse Lab in Alderley Park | Wellcome Sanger Institute for the COVID-19 Genomics UK (COG-UK) Consortium | Jacquelyn Wynn, Mairead Hyland, The Lighthouse Lab in Alderley Park and Alex Alderton, Roberto Amato, Jeffrey Barrett, Sonia Goncalves, Ewan Harrison, David K. Jackson, Ian Johnston, Dominic Kwiatkowski, Cordelia Langford, John Sillitoe on behalf of the Wellcome Sanger Institute COVID-19 Surveillance Team |
| EPI_ISL_3173319 | Lighthouse Lab in Glasgow | Wellcome Sanger Institute for the COVID-19 Genomics UK (COG-UK) Consortium | Harper VanSteenhouse, Yumi Kasai, David Gray, Carol Clugston, Anna Dominiczak and Alex Alderton, Roberto Amato, Jeffrey Barrett, Sonia Goncalves, Ewan Harrison, David K. Jackson, Ian Johnston, Dominic Kwiatkowski, Cordelia Langford, John Sillitoe on behalf of the Wellcome Sanger Institute COVID-19 Surveillance Team |
| EPI_ISL_3173320 | Lighthouse Lab in Alderley Park | Wellcome Sanger Institute for the COVID-19 Genomics UK (COG-UK) Consortium | Jacquelyn Wynn, Mairead Hyland, The Lighthouse Lab in Alderley Park and Alex Alderton, Roberto Amato, Jeffrey Barrett, Sonia Goncalves, Ewan Harrison, David K. Jackson, Ian Johnston, Dominic Kwiatkowski, Cordelia Langford, John Sillitoe on behalf of the Wellcome Sanger Institute COVID-19 Surveillance Team |
| EPI_ISL_3173321 | Lighthouse Lab in Glasgow | Wellcome Sanger Institute for the COVID-19 Genomics UK (COG-UK) Consortium | Harper VanSteenhouse, Yumi Kasai, David Gray, Carol Clugston, Anna Dominiczak and Alex Alderton, Roberto Amato, Jeffrey Barrett, Sonia Goncalves, Ewan Harrison, David K. Jackson, Ian Johnston, Dominic Kwiatkowski, Cordelia Langford, John Sillitoe on behalf of the Wellcome Sanger Institute COVID-19 Surveillance Team |
| EPI_ISL_3173322 | Lighthouse Lab in Alderley Park | Wellcome Sanger Institute for the COVID-19 Genomics UK (COG-UK) Consortium | Jacquelyn Wynn, Mairead Hyland, The Lighthouse Lab in Alderley Park and Alex Alderton, Roberto Amato, Jeffrey Barrett, Sonia Goncalves, Ewan Harrison, David K. Jackson, Ian Johnston, Dominic Kwiatkowski, Cordelia Langford, John Sillitoe on behalf of the Wellcome Sanger Institute COVID-19 Surveillance Team |
| EPI_ISL_3173323 | Lighthouse Lab in Milton Keynes | Wellcome Sanger Institute for the COVID-19 Genomics UK (COG-UK) Consortium | The Lighthouse Lab in Milton Keynes and Alex Alderton, Roberto Amato, Jeffrey Barrett, Sonia Goncalves, Ewan Harrison, David K. Jackson, Ian Johnston, Dominic Kwiatkowski, Cordelia Langford, John Sillitoe on behalf of the Wellcome Sanger Institute COVID-19 Surveillance Team |
| EPI_ISL_3173324, EPI_ISL_3173325 | Lighthouse Lab in Alderley Park | Wellcome Sanger Institute for the COVID-19 Genomics UK (COG-UK) Consortium | Jacquelyn Wynn, Mairead Hyland, The Lighthouse Lab in Alderley Park and Alex Alderton, Roberto Amato, Jeffrey Barrett, Sonia Goncalves, Ewan Harrison, David K. Jackson, Ian Johnston, Dominic Kwiatkowski, Cordelia Langford, John Sillitoe on behalf of the Wellcome Sanger Institute COVID-19 Surveillance Team |
| EPI_ISL_3173326, EPI_ISL_3173327, EPI_ISL_3173328 | Lighthouse Lab in Milton Keynes | Wellcome Sanger Institute for the COVID-19 Genomics UK (COG-UK) Consortium | The Lighthouse Lab in Milton Keynes and Alex Alderton, Roberto Amato, Jeffrey Barrett, Sonia Goncalves, Ewan Harrison, David K. Jackson, Ian Johnston, Dominic Kwiatkowski, Cordelia Langford, John Sillitoe on behalf of the Wellcome Sanger Institute COVID-19 Surveillance Team |
| EPI_ISL_3173329 | Lighthouse Lab in Alderley Park | Wellcome Sanger Institute for the COVID-19 Genomics UK (COG-UK) Consortium | Jacquelyn Wynn, Mairead Hyland, The Lighthouse Lab in Alderley Park and Alex Alderton, Roberto Amato, Jeffrey Barrett, Sonia Goncalves, Ewan Harrison, David K. Jackson, Ian Johnston, Dominic Kwiatkowski, Cordelia Langford, John Sillitoe on behalf of the Wellcome Sanger Institute COVID-19 Surveillance Team |
| EPI_ISL_3173330 | Lighthouse Lab in Milton Keynes | Wellcome Sanger Institute for the COVID-19 Genomics UK (COG-UK) Consortium | The Lighthouse Lab in Milton Keynes and Alex Alderton, Roberto Amato, Jeffrey Barrett, Sonia Goncalves, Ewan Harrison, David K. Jackson, Ian Johnston, Dominic Kwiatkowski, Cordelia Langford, John Sillitoe on behalf of the Wellcome Sanger Institute COVID-19 Surveillance Team |
| EPI_ISL_3173331, EPI_ISL_3173332 | Lighthouse Lab in Alderley Park | Wellcome Sanger Institute for the COVID-19 Genomics UK (COG-UK) Consortium | Jacquelyn Wynn, Mairead Hyland, The Lighthouse Lab in Alderley Park and Alex Alderton, Roberto Amato, Jeffrey Barrett, Sonia Goncalves, Ewan Harrison, David K. Jackson, Ian Johnston, Dominic Kwiatkowski, Cordelia Langford, John Sillitoe on behalf of the Wellcome Sanger Institute COVID-19 Surveillance Team |
| EPI_ISL_3173333, EPI_ISL_3173334, EPI_ISL_3173335 | Lighthouse Lab in Milton Keynes | Wellcome Sanger Institute for the COVID-19 Genomics UK (COG-UK) Consortium | The Lighthouse Lab in Milton Keynes and Alex Alderton, Roberto Amato, Jeffrey Barrett, Sonia Goncalves, Ewan Harrison, David K. Jackson, Ian Johnston, Dominic Kwiatkowski, Cordelia Langford, John Sillitoe on behalf of the Wellcome Sanger Institute COVID-19 Surveillance Team |
| EPI_ISL_3173336, EPI_ISL_3173337, EPI_ISL_3173338 | Lighthouse Lab in Alderley Park | Wellcome Sanger Institute for the COVID-19 Genomics UK (COG-UK) Consortium | Jacquelyn Wynn, Mairead Hyland, The Lighthouse Lab in Alderley Park and Alex Alderton, Roberto Amato, Jeffrey Barrett, Sonia Goncalves, Ewan Harrison, David K. Jackson, Ian Johnston, Dominic Kwiatkowski, Cordelia Langford, John Sillitoe on behalf of the Wellcome Sanger Institute COVID-19 Surveillance Team |
| EPI_ISL_3173339 | Lighthouse Lab in Glasgow | Wellcome Sanger Institute for the COVID-19 Genomics UK (COG-UK) Consortium | Harper VanSteenhouse, Yumi Kasai, David Gray, Carol Clugston, Anna Dominiczak and Alex Alderton, Roberto Amato, Jeffrey Barrett, Sonia Goncalves, Ewan Harrison, David K. Jackson, Ian Johnston, Dominic Kwiatkowski, Cordelia Langford, John Sillitoe on behalf of the Wellcome Sanger Institute COVID-19 Surveillance Team |
| EPI_ISL_3173340 | Lighthouse Lab in Milton Keynes | Wellcome Sanger Institute for the COVID-19 Genomics UK (COG-UK) Consortium | The Lighthouse Lab in Milton Keynes and Alex Alderton, Roberto Amato, Jeffrey Barrett, Sonia Goncalves, Ewan Harrison, David K. Jackson, Ian Johnston, Dominic Kwiatkowski, Cordelia Langford, John Sillitoe on behalf of the Wellcome Sanger Institute COVID-19 Surveillance Team |
| EPI_ISL_3173341 | Lighthouse Lab in Glasgow | Wellcome Sanger Institute for the COVID-19 Genomics UK (COG-UK) Consortium | Harper VanSteenhouse, Yumi Kasai, David Gray, Carol Clugston, Anna Dominiczak and Alex Alderton, Roberto Amato, Jeffrey Barrett, Sonia Goncalves, Ewan Harrison, David K. Jackson, Ian Johnston, Dominic Kwiatkowski, Cordelia Langford, John Sillitoe on behalf of the Wellcome Sanger Institute COVID-19 Surveillance Team |
| EPI_ISL_3173342, EPI_ISL_3173343 | Lighthouse Lab in Milton Keynes | Wellcome Sanger Institute for the COVID-19 Genomics UK (COG-UK) Consortium | The Lighthouse Lab in Milton Keynes and Alex Alderton, Roberto Amato, Jeffrey Barrett, Sonia Goncalves, Ewan Harrison, David K. Jackson, Ian Johnston, Dominic Kwiatkowski, Cordelia Langford, John Sillitoe on behalf of the Wellcome Sanger Institute COVID-19 Surveillance Team |
| EPI_ISL_3173344 | Lighthouse Lab in Alderley Park | Wellcome Sanger Institute for the COVID-19 Genomics UK (COG-UK) Consortium | Jacquelyn Wynn, Mairead Hyland, The Lighthouse Lab in Alderley Park and Alex Alderton, Roberto Amato, Jeffrey Barrett, Sonia Goncalves, Ewan Harrison, David K. Jackson, Ian Johnston, Dominic Kwiatkowski, Cordelia Langford, John Sillitoe on behalf of the Wellcome Sanger Institute COVID-19 Surveillance Team |
| EPI_ISL_3173345, EPI_ISL_3173346 | Lighthouse Lab in Milton Keynes | Wellcome Sanger Institute for the COVID-19 Genomics UK (COG-UK) Consortium | The Lighthouse Lab in Milton Keynes and Alex Alderton, Roberto Amato, Jeffrey Barrett, Sonia Goncalves, Ewan Harrison, David K. Jackson, Ian Johnston, Dominic Kwiatkowski, Cordelia Langford, John Sillitoe on behalf of the Wellcome Sanger Institute COVID-19 Surveillance Team |
| EPI_ISL_3173347 | Lighthouse Lab in Alderley Park | Wellcome Sanger Institute for the COVID-19 Genomics UK (COG-UK) Consortium | Jacquelyn Wynn, Mairead Hyland, The Lighthouse Lab in Alderley Park and Alex Alderton, Roberto Amato, Jeffrey Barrett, Sonia Goncalves, Ewan Harrison, David K. Jackson, Ian Johnston, Dominic Kwiatkowski, Cordelia Langford, John Sillitoe on behalf of the Wellcome Sanger Institute COVID-19 Surveillance Team |
| EPI_ISL_3173348 | Lighthouse Lab in Glasgow | Wellcome Sanger Institute for the COVID-19 Genomics UK (COG-UK) Consortium | Harper VanSteenhouse, Yumi Kasai, David Gray, Carol Clugston, Anna Dominiczak and Alex Alderton, Roberto Amato, Jeffrey Barrett, Sonia Goncalves, Ewan Harrison, David K. Jackson, Ian Johnston, Dominic Kwiatkowski, Cordelia Langford, John Sillitoe on behalf of the Wellcome Sanger Institute COVID-19 Surveillance Team |
| EPI_ISL_3173349, EPI_ISL_3173350 | Lighthouse Lab in Milton Keynes | Wellcome Sanger Institute for the COVID-19 Genomics UK (COG-UK) Consortium | The Lighthouse Lab in Milton Keynes and Alex Alderton, Roberto Amato, Jeffrey Barrett, Sonia Goncalves, Ewan Harrison, David K. Jackson, Ian Johnston, Dominic Kwiatkowski, Cordelia Langford, John Sillitoe on behalf of the Wellcome Sanger Institute COVID-19 Surveillance Team |
| EPI_ISL_3173351 | Lighthouse Lab in Glasgow | Wellcome Sanger Institute for the COVID-19 Genomics UK (COG-UK) Consortium | Harper VanSteenhouse, Yumi Kasai, David Gray, Carol Clugston, Anna Dominiczak and Alex Alderton, Roberto Amato, Jeffrey Barrett, Sonia Goncalves, Ewan Harrison, David K. Jackson, Ian Johnston, Dominic Kwiatkowski, Cordelia Langford, John Sillitoe on behalf of the Wellcome Sanger Institute COVID-19 Surveillance Team |
| EPI_ISL_3173352 | Lighthouse Lab in Milton Keynes | Wellcome Sanger Institute for the COVID-19 Genomics UK (COG-UK) Consortium | The Lighthouse Lab in Milton Keynes and Alex Alderton, Roberto Amato, Jeffrey Barrett, Sonia Goncalves, Ewan Harrison, David K. Jackson, Ian Johnston, Dominic Kwiatkowski, Cordelia Langford, John Sillitoe on behalf of the Wellcome Sanger Institute COVID-19 Surveillance Team |
| EPI_ISL_3173353, EPI_ISL_3173354, EPI_ISL_3173355, EPI_ISL_3173356 | Lighthouse Lab in Alderley Park | Wellcome Sanger Institute for the COVID-19 Genomics UK (COG-UK) Consortium | Jacquelyn Wynn, Mairead Hyland, The Lighthouse Lab in Alderley Park and Alex Alderton, Roberto Amato, Jeffrey Barrett, Sonia Goncalves, Ewan Harrison, David K. Jackson, Ian Johnston, Dominic Kwiatkowski, Cordelia Langford, John Sillitoe on behalf of the Wellcome Sanger Institute COVID-19 Surveillance Team |
| EPI_ISL_3173357, EPI_ISL_3173358, EPI_ISL_3173359 | Lighthouse Lab in Milton Keynes | Wellcome Sanger Institute for the COVID-19 Genomics UK (COG-UK) Consortium | The Lighthouse Lab in Milton Keynes and Alex Alderton, Roberto Amato, Jeffrey Barrett, Sonia Goncalves, Ewan Harrison, David K. Jackson, Ian Johnston, Dominic Kwiatkowski, Cordelia Langford, John Sillitoe on behalf of the Wellcome Sanger Institute COVID-19 Surveillance Team |
| EPI_ISL_3173360 | Lighthouse Lab in Alderley Park | Wellcome Sanger Institute for the COVID-19 Genomics UK (COG-UK) Consortium | Jacquelyn Wynn, Mairead Hyland, The Lighthouse Lab in Alderley Park and Alex Alderton, Roberto Amato, Jeffrey Barrett, Sonia Goncalves, Ewan Harrison, David K. Jackson, Ian Johnston, Dominic Kwiatkowski, Cordelia Langford, John Sillitoe on behalf of the Wellcome Sanger Institute COVID-19 Surveillance Team |
| EPI_ISL_3173361 | Lighthouse Lab in Glasgow | Wellcome Sanger Institute for the COVID-19 Genomics UK (COG-UK) Consortium | Harper VanSteenhouse, Yumi Kasai, David Gray, Carol Clugston, Anna Dominiczak and Alex Alderton, Roberto Amato, Jeffrey Barrett, Sonia Goncalves, Ewan Harrison, David K. Jackson, Ian Johnston, Dominic Kwiatkowski, Cordelia Langford, John Sillitoe on behalf of the Wellcome Sanger Institute COVID-19 Surveillance Team |
| EPI_ISL_3173362 | Lighthouse Lab in Alderley Park | Wellcome Sanger Institute for the COVID-19 Genomics UK (COG-UK) Consortium | Jacquelyn Wynn, Mairead Hyland, The Lighthouse Lab in Alderley Park and Alex Alderton, Roberto Amato, Jeffrey Barrett, Sonia Goncalves, Ewan Harrison, David K. Jackson, Ian Johnston, Dominic Kwiatkowski, Cordelia Langford, John Sillitoe on behalf of the Wellcome Sanger Institute COVID-19 Surveillance Team |
| EPI_ISL_3173363 | Lighthouse Lab in Glasgow | Wellcome Sanger Institute for the COVID-19 Genomics UK (COG-UK) Consortium | Harper VanSteenhouse, Yumi Kasai, David Gray, Carol Clugston, Anna Dominiczak and Alex Alderton, Roberto Amato, Jeffrey Barrett, Sonia Goncalves, Ewan Harrison, David K. Jackson, Ian Johnston, Dominic Kwiatkowski, Cordelia Langford, John Sillitoe on behalf of the Wellcome Sanger Institute COVID-19 Surveillance Team |
| EPI_ISL_3173364 | Lighthouse Lab in Alderley Park | Wellcome Sanger Institute for the COVID-19 Genomics UK (COG-UK) Consortium | Jacquelyn Wynn, Mairead Hyland, The Lighthouse Lab in Alderley Park and Alex Alderton, Roberto Amato, Jeffrey Barrett, Sonia Goncalves, Ewan Harrison, David K. Jackson, Ian Johnston, Dominic Kwiatkowski, Cordelia Langford, John Sillitoe on behalf of the Wellcome Sanger Institute COVID-19 Surveillance Team |

| | | | |
|---|---|---|---|
| EPI_ISL_3173365 | Lighthouse Lab in Milton Keynes | Wellcome Sanger Institute for the COVID-19 Genomics UK (COG-UK) Consortium | The Lighthouse Lab in Milton Keynes and Alex Alderton, Roberto Amato, Jeffrey Barrett, Sonia Goncalves, Ewan Harrison, David K. Jackson, Ian Johnston, Dominic Kwiatkowski, Cordelia Langford, John Sillitoe on behalf of the Wellcome Sanger Institute COVID-19 Surveillance Team |
| EPI_ISL_3173366, EPI_ISL_3173367, EPI_ISL_3173368, EPI_ISL_3173369, EPI_ISL_3173370, EPI_ISL_3173371, EPI_ISL_3173372, EPI_ISL_3173373 | Lighthouse Lab in Alderley Park | Wellcome Sanger Institute for the COVID-19 Genomics UK (COG-UK) Consortium | Jacquelyn Wynn, Mairead Hyland, The Lighthouse Lab in Alderley Park and Alex Alderton, Roberto Amato, Jeffrey Barrett, Sonia Goncalves, Ewan Harrison, David K. Jackson, Ian Johnston, Dominic Kwiatkowski, Cordelia Langford, John Sillitoe on behalf of the Wellcome Sanger Institute COVID-19 Surveillance Team |
| EPI_ISL_3173374 | Lighthouse Lab in Milton Keynes | Wellcome Sanger Institute for the COVID-19 Genomics UK (COG-UK) Consortium | The Lighthouse Lab in Milton Keynes and Alex Alderton, Roberto Amato, Jeffrey Barrett, Sonia Goncalves, Ewan Harrison, David K. Jackson, Ian Johnston, Dominic Kwiatkowski, Cordelia Langford, John Sillitoe on behalf of the Wellcome Sanger Institute COVID-19 Surveillance Team |
| EPI_ISL_3173375, EPI_ISL_3173376 | Lighthouse Lab in Alderley Park | Wellcome Sanger Institute for the COVID-19 Genomics UK (COG-UK) Consortium | Jacquelyn Wynn, Mairead Hyland, The Lighthouse Lab in Alderley Park and Alex Alderton, Roberto Amato, Jeffrey Barrett, Sonia Goncalves, Ewan Harrison, David K. Jackson, Ian Johnston, Dominic Kwiatkowski, Cordelia Langford, John Sillitoe on behalf of the Wellcome Sanger Institute COVID-19 Surveillance Team |
| EPI_ISL_3173377, EPI_ISL_3173378 | Lighthouse Lab in Milton Keynes | Wellcome Sanger Institute for the COVID-19 Genomics UK (COG-UK) Consortium | The Lighthouse Lab in Milton Keynes and Alex Alderton, Roberto Amato, Jeffrey Barrett, Sonia Goncalves, Ewan Harrison, David K. Jackson, Ian Johnston, Dominic Kwiatkowski, Cordelia Langford, John Sillitoe on behalf of the Wellcome Sanger Institute COVID-19 Surveillance Team |
| EPI_ISL_3173379, EPI_ISL_3173380, EPI_ISL_3173381, EPI_ISL_3173382, EPI_ISL_3173383, EPI_ISL_3173384 | Lighthouse Lab in Alderley Park | Wellcome Sanger Institute for the COVID-19 Genomics UK (COG-UK) Consortium | Jacquelyn Wynn, Mairead Hyland, The Lighthouse Lab in Alderley Park and Alex Alderton, Roberto Amato, Jeffrey Barrett, Sonia Goncalves, Ewan Harrison, David K. Jackson, Ian Johnston, Dominic Kwiatkowski, Cordelia Langford, John Sillitoe on behalf of the Wellcome Sanger Institute COVID-19 Surveillance Team |
| EPI_ISL_3173385 | Lighthouse Lab in Milton Keynes | Wellcome Sanger Institute for the COVID-19 Genomics UK (COG-UK) Consortium | The Lighthouse Lab in Milton Keynes and Alex Alderton, Roberto Amato, Jeffrey Barrett, Sonia Goncalves, Ewan Harrison, David K. Jackson, Ian Johnston, Dominic Kwiatkowski, Cordelia Langford, John Sillitoe on behalf of the Wellcome Sanger Institute COVID-19 Surveillance Team |
| EPI_ISL_3173386 | Lighthouse Lab in Alderley Park | Wellcome Sanger Institute for the COVID-19 Genomics UK (COG-UK) Consortium | Jacquelyn Wynn, Mairead Hyland, The Lighthouse Lab in Alderley Park and Alex Alderton, Roberto Amato, Jeffrey Barrett, Sonia Goncalves, Ewan Harrison, David K. Jackson, Ian Johnston, Dominic Kwiatkowski, Cordelia Langford, John Sillitoe on behalf of the Wellcome Sanger Institute COVID-19 Surveillance Team |
| EPI_ISL_3173387 | Lighthouse Lab in Milton Keynes | Wellcome Sanger Institute for the COVID-19 Genomics UK (COG-UK) Consortium | The Lighthouse Lab in Milton Keynes and Alex Alderton, Roberto Amato, Jeffrey Barrett, Sonia Goncalves, Ewan Harrison, David K. Jackson, Ian Johnston, Dominic Kwiatkowski, Cordelia Langford, John Sillitoe on behalf of the Wellcome Sanger Institute COVID-19 Surveillance Team |
| EPI_ISL_3173388 | Lighthouse Lab in Alderley Park | Wellcome Sanger Institute for the COVID-19 Genomics UK (COG-UK) Consortium | Jacquelyn Wynn, Mairead Hyland, The Lighthouse Lab in Alderley Park and Alex Alderton, Roberto Amato, Jeffrey Barrett, Sonia Goncalves, Ewan Harrison, David K. Jackson, Ian Johnston, Dominic Kwiatkowski, Cordelia Langford, John Sillitoe on behalf of the Wellcome Sanger Institute COVID-19 Surveillance Team |
| EPI_ISL_3173389, EPI_ISL_3173390 | Lighthouse Lab in Milton Keynes | Wellcome Sanger Institute for the COVID-19 Genomics UK (COG-UK) Consortium | The Lighthouse Lab in Milton Keynes and Alex Alderton, Roberto Amato, Jeffrey Barrett, Sonia Goncalves, Ewan Harrison, David K. Jackson, Ian Johnston, Dominic Kwiatkowski, Cordelia Langford, John Sillitoe on behalf of the Wellcome Sanger Institute COVID-19 Surveillance Team |
| EPI_ISL_3173391, EPI_ISL_3173392, EPI_ISL_3173393, EPI_ISL_3173394, EPI_ISL_3173395, EPI_ISL_3173396 | Lighthouse Lab in Alderley Park | Wellcome Sanger Institute for the COVID-19 Genomics UK (COG-UK) Consortium | Jacquelyn Wynn, Mairead Hyland, The Lighthouse Lab in Alderley Park and Alex Alderton, Roberto Amato, Jeffrey Barrett, Sonia Goncalves, Ewan Harrison, David K. Jackson, Ian Johnston, Dominic Kwiatkowski, Cordelia Langford, John Sillitoe on behalf of the Wellcome Sanger Institute COVID-19 Surveillance Team |
| EPI_ISL_3173397 | Lighthouse Lab in Glasgow | Wellcome Sanger Institute for the COVID-19 Genomics UK (COG-UK) Consortium | Harper VanSteenhouse, Yumi Kasai, David Gray, Carol Clugston, Anna Dominiczak and Alex Alderton, Roberto Amato, Jeffrey Barrett, Sonia Goncalves, Ewan Harrison, David K. Jackson, Ian Johnston, Dominic Kwiatkowski, Cordelia Langford, John Sillitoe on behalf of the Wellcome Sanger Institute COVID-19 Surveillance Team |
| EPI_ISL_3173398 | Lighthouse Lab in Alderley Park | Wellcome Sanger Institute for the COVID-19 Genomics UK (COG-UK) Consortium | Jacquelyn Wynn, Mairead Hyland, The Lighthouse Lab in Alderley Park and Alex Alderton, Roberto Amato, Jeffrey Barrett, Sonia Goncalves, Ewan Harrison, David K. Jackson, Ian Johnston, Dominic Kwiatkowski, Cordelia Langford, John Sillitoe on behalf of the Wellcome Sanger Institute COVID-19 Surveillance Team |
| EPI_ISL_3173399 | Lighthouse Lab in Glasgow | Wellcome Sanger Institute for the COVID-19 Genomics UK (COG-UK) Consortium | Harper VanSteenhouse, Yumi Kasai, David Gray, Carol Clugston, Anna Dominiczak and Alex Alderton, Roberto Amato, Jeffrey Barrett, Sonia Goncalves, Ewan Harrison, David K. Jackson, Ian Johnston, Dominic Kwiatkowski, Cordelia Langford, John Sillitoe on behalf of the Wellcome Sanger Institute COVID-19 Surveillance Team |
| EPI_ISL_3173400 | Lighthouse Lab in Alderley Park | Wellcome Sanger Institute for the COVID-19 Genomics UK (COG-UK) Consortium | Jacquelyn Wynn, Mairead Hyland, The Lighthouse Lab in Alderley Park and Alex Alderton, Roberto Amato, Jeffrey Barrett, Sonia Goncalves, Ewan Harrison, David K. Jackson, Ian Johnston, Dominic Kwiatkowski, Cordelia Langford, John Sillitoe on behalf of the Wellcome Sanger Institute COVID-19 Surveillance Team |
| EPI_ISL_3173401 | Lighthouse Lab in Milton Keynes | Wellcome Sanger Institute for the COVID-19 Genomics UK (COG-UK) Consortium | The Lighthouse Lab in Milton Keynes and Alex Alderton, Roberto Amato, Jeffrey Barrett, Sonia Goncalves, Ewan Harrison, David K. Jackson, Ian Johnston, Dominic Kwiatkowski, Cordelia Langford, John Sillitoe on behalf of the Wellcome Sanger Institute COVID-19 Surveillance Team |
| EPI_ISL_3173402 | Lighthouse Lab in Alderley Park | Wellcome Sanger Institute for the COVID-19 Genomics UK (COG-UK) Consortium | Jacquelyn Wynn, Mairead Hyland, The Lighthouse Lab in Alderley Park and Alex Alderton, Roberto Amato, Jeffrey Barrett, Sonia Goncalves, Ewan Harrison, David K. Jackson, Ian Johnston, Dominic Kwiatkowski, Cordelia Langford, John Sillitoe on behalf of the Wellcome Sanger Institute COVID-19 Surveillance Team |
| EPI_ISL_3173403 | Lighthouse Lab in Milton Keynes | Wellcome Sanger Institute for the COVID-19 Genomics UK (COG-UK) Consortium | The Lighthouse Lab in Milton Keynes and Alex Alderton, Roberto Amato, Jeffrey Barrett, Sonia Goncalves, Ewan Harrison, David K. Jackson, Ian Johnston, Dominic Kwiatkowski, Cordelia Langford, John Sillitoe on behalf of the Wellcome Sanger Institute COVID-19 Surveillance Team |
| EPI_ISL_3173404 | Lighthouse Lab in Alderley Park | Wellcome Sanger Institute for the COVID-19 Genomics UK (COG-UK) Consortium | Jacquelyn Wynn, Mairead Hyland, The Lighthouse Lab in Alderley Park and Alex Alderton, Roberto Amato, Jeffrey Barrett, Sonia Goncalves, Ewan Harrison, David K. Jackson, Ian Johnston, Dominic Kwiatkowski, Cordelia Langford, John Sillitoe on behalf of the Wellcome Sanger Institute COVID-19 Surveillance Team |
| EPI_ISL_3173405 | Lighthouse Lab in Glasgow | Wellcome Sanger Institute for the COVID-19 Genomics UK (COG-UK) Consortium | Harper VanSteenhouse, Yumi Kasai, David Gray, Carol Clugston, Anna Dominiczak and Alex Alderton, Roberto Amato, Jeffrey Barrett, Sonia Goncalves, Ewan Harrison, David K. Jackson, Ian Johnston, Dominic Kwiatkowski, Cordelia Langford, John Sillitoe on behalf of the Wellcome Sanger Institute COVID-19 Surveillance Team |
| EPI_ISL_3173406 | Lighthouse Lab in Milton Keynes | Wellcome Sanger Institute for the COVID-19 Genomics UK (COG-UK) Consortium | The Lighthouse Lab in Milton Keynes and Alex Alderton, Roberto Amato, Jeffrey Barrett, Sonia Goncalves, Ewan Harrison, David K. Jackson, Ian Johnston, Dominic Kwiatkowski, Cordelia Langford, John Sillitoe on behalf of the Wellcome Sanger Institute COVID-19 Surveillance Team |
| EPI_ISL_3173407 | Lighthouse Lab in Glasgow | Wellcome Sanger Institute for the COVID-19 Genomics UK (COG-UK) Consortium | Harper VanSteenhouse, Yumi Kasai, David Gray, Carol Clugston, Anna Dominiczak and Alex Alderton, Roberto Amato, Jeffrey Barrett, Sonia Goncalves, Ewan Harrison, David K. Jackson, Ian Johnston, Dominic Kwiatkowski, Cordelia Langford, John Sillitoe on behalf of the Wellcome Sanger Institute COVID-19 Surveillance Team |
| EPI_ISL_3173408 | Lighthouse Lab in Alderley Park | Wellcome Sanger Institute for the COVID-19 Genomics UK (COG-UK) Consortium | Jacquelyn Wynn, Mairead Hyland, The Lighthouse Lab in Alderley Park and Alex Alderton, Roberto Amato, Jeffrey Barrett, Sonia Goncalves, Ewan Harrison, David K. Jackson, Ian Johnston, Dominic Kwiatkowski, Cordelia Langford, John Sillitoe on behalf of the Wellcome Sanger Institute COVID-19 Surveillance Team |
| EPI_ISL_3173409, EPI_ISL_3173410 | Lighthouse Lab in Milton Keynes | Wellcome Sanger Institute for the COVID-19 Genomics UK (COG-UK) Consortium | The Lighthouse Lab in Milton Keynes and Alex Alderton, Roberto Amato, Jeffrey Barrett, Sonia Goncalves, Ewan Harrison, David K. Jackson, Ian Johnston, Dominic Kwiatkowski, Cordelia Langford, John Sillitoe on behalf of the Wellcome Sanger Institute COVID-19 Surveillance Team |
| EPI_ISL_3173411, EPI_ISL_3173412 | Lighthouse Lab in Alderley Park | Wellcome Sanger Institute for the COVID-19 Genomics UK (COG-UK) Consortium | Jacquelyn Wynn, Mairead Hyland, The Lighthouse Lab in Alderley Park and Alex Alderton, Roberto Amato, Jeffrey Barrett, Sonia Goncalves, Ewan Harrison, David K. Jackson, Ian Johnston, Dominic Kwiatkowski, Cordelia Langford, John Sillitoe on behalf of the Wellcome Sanger Institute COVID-19 Surveillance Team |
| EPI_ISL_3173413 | Lighthouse Lab in Milton Keynes | Wellcome Sanger Institute for the COVID-19 Genomics UK (COG-UK) Consortium | The Lighthouse Lab in Milton Keynes and Alex Alderton, Roberto Amato, Jeffrey Barrett, Sonia Goncalves, Ewan Harrison, David K. Jackson, Ian Johnston, Dominic Kwiatkowski, Cordelia Langford, John Sillitoe on behalf of the Wellcome Sanger Institute COVID-19 Surveillance Team |
| EPI_ISL_3173414 | Lighthouse Lab in Glasgow | Wellcome Sanger Institute for the COVID-19 Genomics UK (COG-UK) Consortium | Harper VanSteenhouse, Yumi Kasai, David Gray, Carol Clugston, Anna Dominiczak and Alex Alderton, Roberto Amato, Jeffrey Barrett, Sonia Goncalves, Ewan Harrison, David K. Jackson, Ian Johnston, Dominic Kwiatkowski, Cordelia Langford, John Sillitoe on behalf of the Wellcome Sanger Institute COVID-19 Surveillance Team |
| EPI_ISL_3173415 | Lighthouse Lab in Milton Keynes | Wellcome Sanger Institute for the COVID-19 Genomics UK (COG-UK) Consortium | The Lighthouse Lab in Milton Keynes and Alex Alderton, Roberto Amato, Jeffrey Barrett, Sonia Goncalves, Ewan Harrison, David K. Jackson, Ian Johnston, Dominic Kwiatkowski, Cordelia Langford, John Sillitoe on behalf of the Wellcome Sanger Institute COVID-19 Surveillance Team |
| EPI_ISL_3173416 | Lighthouse Lab in Alderley Park | Wellcome Sanger Institute for the COVID-19 Genomics UK (COG-UK) Consortium | Jacquelyn Wynn, Mairead Hyland, The Lighthouse Lab in Alderley Park and Alex Alderton, Roberto Amato, Jeffrey Barrett, Sonia Goncalves, Ewan Harrison, David K. Jackson, Ian Johnston, Dominic Kwiatkowski, Cordelia Langford, John Sillitoe on behalf of the Wellcome Sanger Institute COVID-19 Surveillance Team |
| EPI_ISL_3173417 | Lighthouse Lab in Milton Keynes | Wellcome Sanger Institute for the COVID-19 Genomics UK (COG-UK) Consortium | The Lighthouse Lab in Milton Keynes and Alex Alderton, Roberto Amato, Jeffrey Barrett, Sonia Goncalves, Ewan Harrison, David K. Jackson, Ian Johnston, Dominic Kwiatkowski, Cordelia Langford, John Sillitoe on behalf of the Wellcome Sanger Institute COVID-19 Surveillance Team |
| EPI_ISL_3173418 | Lighthouse Lab in Alderley Park | Wellcome Sanger Institute for the COVID-19 Genomics UK (COG-UK) Consortium | Jacquelyn Wynn, Mairead Hyland, The Lighthouse Lab in Alderley Park and Alex Alderton, Roberto Amato, Jeffrey Barrett, Sonia Goncalves, Ewan Harrison, David K. Jackson, Ian Johnston, Dominic Kwiatkowski, Cordelia Langford, John Sillitoe on behalf of the Wellcome Sanger Institute COVID-19 Surveillance Team |
| EPI_ISL_3173419 | Lighthouse Lab in Milton Keynes | Wellcome Sanger Institute for the COVID-19 Genomics UK (COG-UK) Consortium | The Lighthouse Lab in Milton Keynes and Alex Alderton, Roberto Amato, Jeffrey Barrett, Sonia Goncalves, Ewan Harrison, David K. Jackson, Ian Johnston, Dominic Kwiatkowski, Cordelia Langford, John Sillitoe on behalf of the Wellcome Sanger Institute COVID-19 Surveillance Team |
| EPI_ISL_3173420 | Lighthouse Lab in Alderley Park | Wellcome Sanger Institute for the COVID-19 Genomics UK (COG-UK) Consortium | Jacquelyn Wynn, Mairead Hyland, The Lighthouse Lab in Alderley Park and Alex Alderton, Roberto Amato, Jeffrey Barrett, Sonia Goncalves, Ewan Harrison, David K. Jackson, Ian Johnston, Dominic Kwiatkowski, Cordelia Langford, John Sillitoe on behalf of the Wellcome Sanger Institute COVID-19 Surveillance Team |
| EPI_ISL_3173421 | Lighthouse Lab in Milton Keynes | Wellcome Sanger Institute for the COVID-19 Genomics UK (COG-UK) Consortium | The Lighthouse Lab in Milton Keynes and Alex Alderton, Roberto Amato, Jeffrey Barrett, Sonia Goncalves, Ewan Harrison, David K. Jackson, Ian Johnston, Dominic Kwiatkowski, Cordelia Langford, John Sillitoe on behalf of the Wellcome Sanger Institute COVID-19 Surveillance Team |
| EPI_ISL_3173422, EPI_ISL_3173423, EPI_ISL_3173424 | Lighthouse Lab in Alderley Park | Wellcome Sanger Institute for the COVID-19 Genomics UK (COG-UK) Consortium | Jacquelyn Wynn, Mairead Hyland, The Lighthouse Lab in Alderley Park and Alex Alderton, Roberto Amato, Jeffrey Barrett, Sonia Goncalves, Ewan Harrison, David K. Jackson, Ian Johnston, Dominic Kwiatkowski, Cordelia Langford, John Sillitoe on behalf of the Wellcome Sanger Institute COVID-19 Surveillance Team |
| EPI_ISL_3173425 | Lighthouse Lab in Glasgow | Wellcome Sanger Institute for the COVID-19 Genomics UK (COG-UK) Consortium | Harper VanSteenhouse, Yumi Kasai, David Gray, Carol Clugston, Anna Dominiczak and Alex Alderton, Roberto Amato, Jeffrey Barrett, Sonia Goncalves, Ewan Harrison, David K. Jackson, Ian Johnston, Dominic Kwiatkowski, Cordelia Langford, John Sillitoe on behalf of the Wellcome Sanger Institute COVID-19 Surveillance Team |
| EPI_ISL_3173426 | Lighthouse Lab in Alderley Park | Wellcome Sanger Institute for the COVID-19 Genomics UK (COG-UK) Consortium | Jacquelyn Wynn, Mairead Hyland, The Lighthouse Lab in Alderley Park and Alex Alderton, Roberto Amato, Jeffrey Barrett, Sonia Goncalves, Ewan Harrison, David K. Jackson, Ian Johnston, Dominic Kwiatkowski, Cordelia Langford, John Sillitoe on behalf of the Wellcome Sanger Institute COVID-19 Surveillance Team |
| EPI_ISL_3173427 | Lighthouse Lab in Milton Keynes | Wellcome Sanger Institute for the COVID-19 Genomics UK (COG-UK) Consortium | The Lighthouse Lab in Milton Keynes and Alex Alderton, Roberto Amato, Jeffrey Barrett, Sonia Goncalves, Ewan Harrison, David K. Jackson, Ian Johnston, Dominic Kwiatkowski, Cordelia Langford, John Sillitoe on behalf of the Wellcome Sanger Institute COVID-19 Surveillance Team |

| | | | |
|---|---|---|---|
| EPI_ISL_3173428, EPI_ISL_3173429 | Lighthouse Lab in Alderley Park | Wellcome Sanger Institute for the COVID-19 Genomics UK (COG-UK) Consortium | Jacquelyn Wynn, Mairead Hyland, The Lighthouse Lab in Alderley Park and Alex Alderton, Roberto Amato, Jeffrey Barrett, Sonia Goncalves, Ewan Harrison, David K. Jackson, Ian Johnston, Dominic Kwiatkowski, Cordelia Langford, John Sillitoe on behalf of the Wellcome Sanger Institute COVID-19 Surveillance Team |
| EPI_ISL_3173430, EPI_ISL_3173431 | Lighthouse Lab in Milton Keynes | Wellcome Sanger Institute for the COVID-19 Genomics UK (COG-UK) Consortium | The Lighthouse Lab in Milton Keynes and Alex Alderton, Roberto Amato, Jeffrey Barrett, Sonia Goncalves, Ewan Harrison, David K. Jackson, Ian Johnston, Dominic Kwiatkowski, Cordelia Langford, John Sillitoe on behalf of the Wellcome Sanger Institute COVID-19 Surveillance Team |
| EPI_ISL_3173432, EPI_ISL_3173433, EPI_ISL_3173434 | Lighthouse Lab in Alderley Park | Wellcome Sanger Institute for the COVID-19 Genomics UK (COG-UK) Consortium | Jacquelyn Wynn, Mairead Hyland, The Lighthouse Lab in Alderley Park and Alex Alderton, Roberto Amato, Jeffrey Barrett, Sonia Goncalves, Ewan Harrison, David K. Jackson, Ian Johnston, Dominic Kwiatkowski, Cordelia Langford, John Sillitoe on behalf of the Wellcome Sanger Institute COVID-19 Surveillance Team |
| EPI_ISL_3173435 | Lighthouse Lab in Glasgow | Wellcome Sanger Institute for the COVID-19 Genomics UK (COG-UK) Consortium | Harper VanSteenhouse, Yumi Kasai, David Gray, Carol Clugston, Anna Dominiczak and Alex Alderton, Roberto Amato, Jeffrey Barrett, Sonia Goncalves, Ewan Harrison, David K. Jackson, Ian Johnston, Dominic Kwiatkowski, Cordelia Langford, John Sillitoe on behalf of the Wellcome Sanger Institute COVID-19 Surveillance Team |
| EPI_ISL_3173436 | Lighthouse Lab in Milton Keynes | Wellcome Sanger Institute for the COVID-19 Genomics UK (COG-UK) Consortium | The Lighthouse Lab in Milton Keynes and Alex Alderton, Roberto Amato, Jeffrey Barrett, Sonia Goncalves, Ewan Harrison, David K. Jackson, Ian Johnston, Dominic Kwiatkowski, Cordelia Langford, John Sillitoe on behalf of the Wellcome Sanger Institute COVID-19 Surveillance Team |
| EPI_ISL_3173437, EPI_ISL_3173438 | Lighthouse Lab in Alderley Park | Wellcome Sanger Institute for the COVID-19 Genomics UK (COG-UK) Consortium | Jacquelyn Wynn, Mairead Hyland, The Lighthouse Lab in Alderley Park and Alex Alderton, Roberto Amato, Jeffrey Barrett, Sonia Goncalves, Ewan Harrison, David K. Jackson, Ian Johnston, Dominic Kwiatkowski, Cordelia Langford, John Sillitoe on behalf of the Wellcome Sanger Institute COVID-19 Surveillance Team |
| EPI_ISL_3173439 | Lighthouse Lab in Glasgow | Wellcome Sanger Institute for the COVID-19 Genomics UK (COG-UK) Consortium | Harper VanSteenhouse, Yumi Kasai, David Gray, Carol Clugston, Anna Dominiczak and Alex Alderton, Roberto Amato, Jeffrey Barrett, Sonia Goncalves, Ewan Harrison, David K. Jackson, Ian Johnston, Dominic Kwiatkowski, Cordelia Langford, John Sillitoe on behalf of the Wellcome Sanger Institute COVID-19 Surveillance Team |
| EPI_ISL_3173440 | Lighthouse Lab in Milton Keynes | Wellcome Sanger Institute for the COVID-19 Genomics UK (COG-UK) Consortium | The Lighthouse Lab in Milton Keynes and Alex Alderton, Roberto Amato, Jeffrey Barrett, Sonia Goncalves, Ewan Harrison, David K. Jackson, Ian Johnston, Dominic Kwiatkowski, Cordelia Langford, John Sillitoe on behalf of the Wellcome Sanger Institute COVID-19 Surveillance Team |
| EPI_ISL_3173441 | Lighthouse Lab in Alderley Park | Wellcome Sanger Institute for the COVID-19 Genomics UK (COG-UK) Consortium | Jacquelyn Wynn, Mairead Hyland, The Lighthouse Lab in Alderley Park and Alex Alderton, Roberto Amato, Jeffrey Barrett, Sonia Goncalves, Ewan Harrison, David K. Jackson, Ian Johnston, Dominic Kwiatkowski, Cordelia Langford, John Sillitoe on behalf of the Wellcome Sanger Institute COVID-19 Surveillance Team |
| EPI_ISL_3173442 | Lighthouse Lab in Milton Keynes | Wellcome Sanger Institute for the COVID-19 Genomics UK (COG-UK) Consortium | The Lighthouse Lab in Milton Keynes and Alex Alderton, Roberto Amato, Jeffrey Barrett, Sonia Goncalves, Ewan Harrison, David K. Jackson, Ian Johnston, Dominic Kwiatkowski, Cordelia Langford, John Sillitoe on behalf of the Wellcome Sanger Institute COVID-19 Surveillance Team |
| EPI_ISL_3173443 | Lighthouse Lab in Glasgow | Wellcome Sanger Institute for the COVID-19 Genomics UK (COG-UK) Consortium | Harper VanSteenhouse, Yumi Kasai, David Gray, Carol Clugston, Anna Dominiczak and Alex Alderton, Roberto Amato, Jeffrey Barrett, Sonia Goncalves, Ewan Harrison, David K. Jackson, Ian Johnston, Dominic Kwiatkowski, Cordelia Langford, John Sillitoe on behalf of the Wellcome Sanger Institute COVID-19 Surveillance Team |
| EPI_ISL_3173444 | Lighthouse Lab in Alderley Park | Wellcome Sanger Institute for the COVID-19 Genomics UK (COG-UK) Consortium | Jacquelyn Wynn, Mairead Hyland, The Lighthouse Lab in Alderley Park and Alex Alderton, Roberto Amato, Jeffrey Barrett, Sonia Goncalves, Ewan Harrison, David K. Jackson, Ian Johnston, Dominic Kwiatkowski, Cordelia Langford, John Sillitoe on behalf of the Wellcome Sanger Institute COVID-19 Surveillance Team |
| EPI_ISL_3173445 | Lighthouse Lab in Milton Keynes | Wellcome Sanger Institute for the COVID-19 Genomics UK (COG-UK) Consortium | The Lighthouse Lab in Milton Keynes and Alex Alderton, Roberto Amato, Jeffrey Barrett, Sonia Goncalves, Ewan Harrison, David K. Jackson, Ian Johnston, Dominic Kwiatkowski, Cordelia Langford, John Sillitoe on behalf of the Wellcome Sanger Institute COVID-19 Surveillance Team |
| EPI_ISL_3173446 | Lighthouse Lab in Glasgow | Wellcome Sanger Institute for the COVID-19 Genomics UK (COG-UK) Consortium | Harper VanSteenhouse, Yumi Kasai, David Gray, Carol Clugston, Anna Dominiczak and Alex Alderton, Roberto Amato, Jeffrey Barrett, Sonia Goncalves, Ewan Harrison, David K. Jackson, Ian Johnston, Dominic Kwiatkowski, Cordelia Langford, John Sillitoe on behalf of the Wellcome Sanger Institute COVID-19 Surveillance Team |
| EPI_ISL_3173447, EPI_ISL_3173448 | Lighthouse Lab in Alderley Park | Wellcome Sanger Institute for the COVID-19 Genomics UK (COG-UK) Consortium | Jacquelyn Wynn, Mairead Hyland, The Lighthouse Lab in Alderley Park and Alex Alderton, Roberto Amato, Jeffrey Barrett, Sonia Goncalves, Ewan Harrison, David K. Jackson, Ian Johnston, Dominic Kwiatkowski, Cordelia Langford, John Sillitoe on behalf of the Wellcome Sanger Institute COVID-19 Surveillance Team |
| EPI_ISL_3173449, EPI_ISL_3173450, EPI_ISL_3173451 | Lighthouse Lab in Milton Keynes | Wellcome Sanger Institute for the COVID-19 Genomics UK (COG-UK) Consortium | The Lighthouse Lab in Milton Keynes and Alex Alderton, Roberto Amato, Jeffrey Barrett, Sonia Goncalves, Ewan Harrison, David K. Jackson, Ian Johnston, Dominic Kwiatkowski, Cordelia Langford, John Sillitoe on behalf of the Wellcome Sanger Institute COVID-19 Surveillance Team |
| EPI_ISL_3173452, EPI_ISL_3173453 | Lighthouse Lab in Alderley Park | Wellcome Sanger Institute for the COVID-19 Genomics UK (COG-UK) Consortium | Jacquelyn Wynn, Mairead Hyland, The Lighthouse Lab in Alderley Park and Alex Alderton, Roberto Amato, Jeffrey Barrett, Sonia Goncalves, Ewan Harrison, David K. Jackson, Ian Johnston, Dominic Kwiatkowski, Cordelia Langford, John Sillitoe on behalf of the Wellcome Sanger Institute COVID-19 Surveillance Team |
| EPI_ISL_3173454 | Lighthouse Lab in Milton Keynes | Wellcome Sanger Institute for the COVID-19 Genomics UK (COG-UK) Consortium | The Lighthouse Lab in Milton Keynes and Alex Alderton, Roberto Amato, Jeffrey Barrett, Sonia Goncalves, Ewan Harrison, David K. Jackson, Ian Johnston, Dominic Kwiatkowski, Cordelia Langford, John Sillitoe on behalf of the Wellcome Sanger Institute COVID-19 Surveillance Team |
| EPI_ISL_3173455, EPI_ISL_3173456 | Lighthouse Lab in Alderley Park | Wellcome Sanger Institute for the COVID-19 Genomics UK (COG-UK) Consortium | Jacquelyn Wynn, Mairead Hyland, The Lighthouse Lab in Alderley Park and Alex Alderton, Roberto Amato, Jeffrey Barrett, Sonia Goncalves, Ewan Harrison, David K. Jackson, Ian Johnston, Dominic Kwiatkowski, Cordelia Langford, John Sillitoe on behalf of the Wellcome Sanger Institute COVID-19 Surveillance Team |
| EPI_ISL_3173457 | Lighthouse Lab in Milton Keynes | Wellcome Sanger Institute for the COVID-19 Genomics UK (COG-UK) Consortium | The Lighthouse Lab in Milton Keynes and Alex Alderton, Roberto Amato, Jeffrey Barrett, Sonia Goncalves, Ewan Harrison, David K. Jackson, Ian Johnston, Dominic Kwiatkowski, Cordelia Langford, John Sillitoe on behalf of the Wellcome Sanger Institute COVID-19 Surveillance Team |
| EPI_ISL_3173458, EPI_ISL_3173459, EPI_ISL_3173460, EPI_ISL_3173461, EPI_ISL_3173462 | Lighthouse Lab in Alderley Park | Wellcome Sanger Institute for the COVID-19 Genomics UK (COG-UK) Consortium | Jacquelyn Wynn, Mairead Hyland, The Lighthouse Lab in Alderley Park and Alex Alderton, Roberto Amato, Jeffrey Barrett, Sonia Goncalves, Ewan Harrison, David K. Jackson, Ian Johnston, Dominic Kwiatkowski, Cordelia Langford, John Sillitoe on behalf of the Wellcome Sanger Institute COVID-19 Surveillance Team |
| EPI_ISL_3173463, EPI_ISL_3173464 | Lighthouse Lab in Milton Keynes | Wellcome Sanger Institute for the COVID-19 Genomics UK (COG-UK) Consortium | The Lighthouse Lab in Milton Keynes and Alex Alderton, Roberto Amato, Jeffrey Barrett, Sonia Goncalves, Ewan Harrison, David K. Jackson, Ian Johnston, Dominic Kwiatkowski, Cordelia Langford, John Sillitoe on behalf of the Wellcome Sanger Institute COVID-19 Surveillance Team |
| EPI_ISL_3173465, EPI_ISL_3173466 | Lighthouse Lab in Glasgow | Wellcome Sanger Institute for the COVID-19 Genomics UK (COG-UK) Consortium | Harper VanSteenhouse, Yumi Kasai, David Gray, Carol Clugston, Anna Dominiczak and Alex Alderton, Roberto Amato, Jeffrey Barrett, Sonia Goncalves, Ewan Harrison, David K. Jackson, Ian Johnston, Dominic Kwiatkowski, Cordelia Langford, John Sillitoe on behalf of the Wellcome Sanger Institute COVID-19 Surveillance Team |
| EPI_ISL_3173467, EPI_ISL_3173468, EPI_ISL_3173469 | Lighthouse Lab in Milton Keynes | Wellcome Sanger Institute for the COVID-19 Genomics UK (COG-UK) Consortium | The Lighthouse Lab in Milton Keynes and Alex Alderton, Roberto Amato, Jeffrey Barrett, Sonia Goncalves, Ewan Harrison, David K. Jackson, Ian Johnston, Dominic Kwiatkowski, Cordelia Langford, John Sillitoe on behalf of the Wellcome Sanger Institute COVID-19 Surveillance Team |
| EPI_ISL_3173470 | Lighthouse Lab in Alderley Park | Wellcome Sanger Institute for the COVID-19 Genomics UK (COG-UK) Consortium | Jacquelyn Wynn, Mairead Hyland, The Lighthouse Lab in Alderley Park and Alex Alderton, Roberto Amato, Jeffrey Barrett, Sonia Goncalves, Ewan Harrison, David K. Jackson, Ian Johnston, Dominic Kwiatkowski, Cordelia Langford, John Sillitoe on behalf of the Wellcome Sanger Institute COVID-19 Surveillance Team |
| EPI_ISL_3173471 | Lighthouse Lab in Glasgow | Wellcome Sanger Institute for the COVID-19 Genomics UK (COG-UK) Consortium | Harper VanSteenhouse, Yumi Kasai, David Gray, Carol Clugston, Anna Dominiczak and Alex Alderton, Roberto Amato, Jeffrey Barrett, Sonia Goncalves, Ewan Harrison, David K. Jackson, Ian Johnston, Dominic Kwiatkowski, Cordelia Langford, John Sillitoe on behalf of the Wellcome Sanger Institute COVID-19 Surveillance Team |
| EPI_ISL_3173472, EPI_ISL_3173473, EPI_ISL_3173474 | Lighthouse Lab in Alderley Park | Wellcome Sanger Institute for the COVID-19 Genomics UK (COG-UK) Consortium | Jacquelyn Wynn, Mairead Hyland, The Lighthouse Lab in Alderley Park and Alex Alderton, Roberto Amato, Jeffrey Barrett, Sonia Goncalves, Ewan Harrison, David K. Jackson, Ian Johnston, Dominic Kwiatkowski, Cordelia Langford, John Sillitoe on behalf of the Wellcome Sanger Institute COVID-19 Surveillance Team |
| EPI_ISL_3173475, EPI_ISL_3173476 | Lighthouse Lab in Glasgow | Wellcome Sanger Institute for the COVID-19 Genomics UK (COG-UK) Consortium | Harper VanSteenhouse, Yumi Kasai, David Gray, Carol Clugston, Anna Dominiczak and Alex Alderton, Roberto Amato, Jeffrey Barrett, Sonia Goncalves, Ewan Harrison, David K. Jackson, Ian Johnston, Dominic Kwiatkowski, Cordelia Langford, John Sillitoe on behalf of the Wellcome Sanger Institute COVID-19 Surveillance Team |
| EPI_ISL_3173477 | Lighthouse Lab in Milton Keynes | Wellcome Sanger Institute for the COVID-19 Genomics UK (COG-UK) Consortium | The Lighthouse Lab in Milton Keynes and Alex Alderton, Roberto Amato, Jeffrey Barrett, Sonia Goncalves, Ewan Harrison, David K. Jackson, Ian Johnston, Dominic Kwiatkowski, Cordelia Langford, John Sillitoe on behalf of the Wellcome Sanger Institute COVID-19 Surveillance Team |
| EPI_ISL_3173478, EPI_ISL_3173479 | Lighthouse Lab in Alderley Park | Wellcome Sanger Institute for the COVID-19 Genomics UK (COG-UK) Consortium | Jacquelyn Wynn, Mairead Hyland, The Lighthouse Lab in Alderley Park and Alex Alderton, Roberto Amato, Jeffrey Barrett, Sonia Goncalves, Ewan Harrison, David K. Jackson, Ian Johnston, Dominic Kwiatkowski, Cordelia Langford, John Sillitoe on behalf of the Wellcome Sanger Institute COVID-19 Surveillance Team |
| EPI_ISL_3173480 | Lighthouse Lab in Milton Keynes | Wellcome Sanger Institute for the COVID-19 Genomics UK (COG-UK) Consortium | The Lighthouse Lab in Milton Keynes and Alex Alderton, Roberto Amato, Jeffrey Barrett, Sonia Goncalves, Ewan Harrison, David K. Jackson, Ian Johnston, Dominic Kwiatkowski, Cordelia Langford, John Sillitoe on behalf of the Wellcome Sanger Institute COVID-19 Surveillance Team |
| EPI_ISL_3173481, EPI_ISL_3173482, EPI_ISL_3173483 | Lighthouse Lab in Alderley Park | Wellcome Sanger Institute for the COVID-19 Genomics UK (COG-UK) Consortium | Jacquelyn Wynn, Mairead Hyland, The Lighthouse Lab in Alderley Park and Alex Alderton, Roberto Amato, Jeffrey Barrett, Sonia Goncalves, Ewan Harrison, David K. Jackson, Ian Johnston, Dominic Kwiatkowski, Cordelia Langford, John Sillitoe on behalf of the Wellcome Sanger Institute COVID-19 Surveillance Team |
| EPI_ISL_3173484 | Lighthouse Lab in Glasgow | Wellcome Sanger Institute for the COVID-19 Genomics UK (COG-UK) Consortium | Harper VanSteenhouse, Yumi Kasai, David Gray, Carol Clugston, Anna Dominiczak and Alex Alderton, Roberto Amato, Jeffrey Barrett, Sonia Goncalves, Ewan Harrison, David K. Jackson, Ian Johnston, Dominic Kwiatkowski, Cordelia Langford, John Sillitoe on behalf of the Wellcome Sanger Institute COVID-19 Surveillance Team |
| EPI_ISL_3173485, EPI_ISL_3173486, EPI_ISL_3173487 | Lighthouse Lab in Alderley Park | Wellcome Sanger Institute for the COVID-19 Genomics UK (COG-UK) Consortium | Jacquelyn Wynn, Mairead Hyland, The Lighthouse Lab in Alderley Park and Alex Alderton, Roberto Amato, Jeffrey Barrett, Sonia Goncalves, Ewan Harrison, David K. Jackson, Ian Johnston, Dominic Kwiatkowski, Cordelia Langford, John Sillitoe on behalf of the Wellcome Sanger Institute COVID-19 Surveillance Team |
| EPI_ISL_3173488 | Lighthouse Lab in Glasgow | Wellcome Sanger Institute for the COVID-19 Genomics UK (COG-UK) Consortium | Harper VanSteenhouse, Yumi Kasai, David Gray, Carol Clugston, Anna Dominiczak and Alex Alderton, Roberto Amato, Jeffrey Barrett, Sonia Goncalves, Ewan Harrison, David K. Jackson, Ian Johnston, Dominic Kwiatkowski, Cordelia Langford, John Sillitoe on behalf of the Wellcome Sanger Institute COVID-19 Surveillance Team |
| EPI_ISL_3173489, EPI_ISL_3173490, EPI_ISL_3173491, EPI_ISL_3173492, EPI_ISL_3173493 | Lighthouse Lab in Alderley Park | Wellcome Sanger Institute for the COVID-19 Genomics UK (COG-UK) Consortium | Jacquelyn Wynn, Mairead Hyland, The Lighthouse Lab in Alderley Park and Alex Alderton, Roberto Amato, Jeffrey Barrett, Sonia Goncalves, Ewan Harrison, David K. Jackson, Ian Johnston, Dominic Kwiatkowski, Cordelia Langford, John Sillitoe on behalf of the Wellcome Sanger Institute COVID-19 Surveillance Team |
| EPI_ISL_3173494 | Lighthouse Lab in Glasgow | Wellcome Sanger Institute for the COVID-19 Genomics UK (COG-UK) Consortium | Harper VanSteenhouse, Yumi Kasai, David Gray, Carol Clugston, Anna Dominiczak and Alex Alderton, Roberto Amato, Jeffrey Barrett, Sonia Goncalves, Ewan Harrison, David K. Jackson, Ian Johnston, Dominic Kwiatkowski, Cordelia Langford, John Sillitoe on behalf of the Wellcome Sanger Institute COVID-19 Surveillance Team |
| EPI_ISL_3173495, EPI_ISL_3173496 | Lighthouse Lab in Alderley Park | Wellcome Sanger Institute for the COVID-19 Genomics UK (COG-UK) Consortium | Jacquelyn Wynn, Mairead Hyland, The Lighthouse Lab in Alderley Park and Alex Alderton, Roberto Amato, Jeffrey Barrett, Sonia Goncalves, Ewan Harrison, David K. Jackson, Ian Johnston, Dominic Kwiatkowski, Cordelia Langford, John Sillitoe on behalf of the Wellcome Sanger Institute COVID-19 Surveillance Team |
| EPI_ISL_3173497 | Lighthouse Lab in Milton Keynes | Wellcome Sanger Institute for the COVID-19 Genomics UK (COG-UK) Consortium | The Lighthouse Lab in Milton Keynes and Alex Alderton, Roberto Amato, Jeffrey Barrett, Sonia Goncalves, Ewan Harrison, David K. Jackson, Ian Johnston, Dominic Kwiatkowski, Cordelia Langford, John Sillitoe on behalf of the Wellcome Sanger Institute COVID-19 Surveillance Team |
| EPI_ISL_3173498 | Lighthouse Lab in Alderley Park | Wellcome Sanger Institute for the COVID-19 Genomics UK (COG-UK) Consortium | Jacquelyn Wynn, Mairead Hyland, The Lighthouse Lab in Alderley Park and Alex Alderton, Roberto Amato, Jeffrey Barrett, Sonia Goncalves, Ewan Harrison, David K. Jackson, Ian Johnston, Dominic Kwiatkowski, Cordelia Langford, John Sillitoe on behalf of the Wellcome Sanger Institute COVID-19 Surveillance Team |

| | | | |
|---|---|---|---|
| EPI_ISL_3173499 | Lighthouse Lab in Glasgow | Wellcome Sanger Institute for the COVID-19 Genomics UK (COG-UK) Consortium | Harper VanSteenhouse, Yumi Kasai, David Gray, Carol Clugston, Anna Dominiczak and Alex Alderton, Roberto Amato, Jeffrey Barrett, Sonia Goncalves, Ewan Harrison, David K. Jackson, Ian Johnston, Dominic Kwiatkowski, Cordelia Langford, John Sillitoe on behalf of the Wellcome Sanger Institute COVID-19 Surveillance Team |
| EPI_ISL_3173500 | Lighthouse Lab in Alderley Park | Wellcome Sanger Institute for the COVID-19 Genomics UK (COG-UK) Consortium | Jacquelyn Wynn, Mairead Hyland, The Lighthouse Lab in Alderley Park and Alex Alderton, Roberto Amato, Jeffrey Barrett, Sonia Goncalves, Ewan Harrison, David K. Jackson, Ian Johnston, Dominic Kwiatkowski, Cordelia Langford, John Sillitoe on behalf of the Wellcome Sanger Institute COVID-19 Surveillance Team |
| EPI_ISL_3173501 | Lighthouse Lab in Milton Keynes | Wellcome Sanger Institute for the COVID-19 Genomics UK (COG-UK) Consortium | The Lighthouse Lab in Milton Keynes and Alex Alderton, Roberto Amato, Jeffrey Barrett, Sonia Goncalves, Ewan Harrison, David K. Jackson, Ian Johnston, Dominic Kwiatkowski, Cordelia Langford, John Sillitoe on behalf of the Wellcome Sanger Institute COVID-19 Surveillance Team |
| EPI_ISL_3173502, EPI_ISL_3173503 | Lighthouse Lab in Glasgow | Wellcome Sanger Institute for the COVID-19 Genomics UK (COG-UK) Consortium | Harper VanSteenhouse, Yumi Kasai, David Gray, Carol Clugston, Anna Dominiczak and Alex Alderton, Roberto Amato, Jeffrey Barrett, Sonia Goncalves, Ewan Harrison, David K. Jackson, Ian Johnston, Dominic Kwiatkowski, Cordelia Langford, John Sillitoe on behalf of the Wellcome Sanger Institute COVID-19 Surveillance Team |
| EPI_ISL_3173504 | Lighthouse Lab in Alderley Park | Wellcome Sanger Institute for the COVID-19 Genomics UK (COG-UK) Consortium | Jacquelyn Wynn, Mairead Hyland, The Lighthouse Lab in Alderley Park and Alex Alderton, Roberto Amato, Jeffrey Barrett, Sonia Goncalves, Ewan Harrison, David K. Jackson, Ian Johnston, Dominic Kwiatkowski, Cordelia Langford, John Sillitoe on behalf of the Wellcome Sanger Institute COVID-19 Surveillance Team |
| EPI_ISL_3173505 | Lighthouse Lab in Glasgow | Wellcome Sanger Institute for the COVID-19 Genomics UK (COG-UK) Consortium | Harper VanSteenhouse, Yumi Kasai, David Gray, Carol Clugston, Anna Dominiczak and Alex Alderton, Roberto Amato, Jeffrey Barrett, Sonia Goncalves, Ewan Harrison, David K. Jackson, Ian Johnston, Dominic Kwiatkowski, Cordelia Langford, John Sillitoe on behalf of the Wellcome Sanger Institute COVID-19 Surveillance Team |
| EPI_ISL_3173506 | Lighthouse Lab in Milton Keynes | Wellcome Sanger Institute for the COVID-19 Genomics UK (COG-UK) Consortium | The Lighthouse Lab in Milton Keynes and Alex Alderton, Roberto Amato, Jeffrey Barrett, Sonia Goncalves, Ewan Harrison, David K. Jackson, Ian Johnston, Dominic Kwiatkowski, Cordelia Langford, John Sillitoe on behalf of the Wellcome Sanger Institute COVID-19 Surveillance Team |
| EPI_ISL_3173507, EPI_ISL_3173508, EPI_ISL_3173509, EPI_ISL_3173510, EPI_ISL_3173511 | Lighthouse Lab in Alderley Park | Wellcome Sanger Institute for the COVID-19 Genomics UK (COG-UK) Consortium | Jacquelyn Wynn, Mairead Hyland, The Lighthouse Lab in Alderley Park and Alex Alderton, Roberto Amato, Jeffrey Barrett, Sonia Goncalves, Ewan Harrison, David K. Jackson, Ian Johnston, Dominic Kwiatkowski, Cordelia Langford, John Sillitoe on behalf of the Wellcome Sanger Institute COVID-19 Surveillance Team |
| EPI_ISL_3173512 | Lighthouse Lab in Glasgow | Wellcome Sanger Institute for the COVID-19 Genomics UK (COG-UK) Consortium | Harper VanSteenhouse, Yumi Kasai, David Gray, Carol Clugston, Anna Dominiczak and Alex Alderton, Roberto Amato, Jeffrey Barrett, Sonia Goncalves, Ewan Harrison, David K. Jackson, Ian Johnston, Dominic Kwiatkowski, Cordelia Langford, John Sillitoe on behalf of the Wellcome Sanger Institute COVID-19 Surveillance Team |
| EPI_ISL_3173513, EPI_ISL_3173514, EPI_ISL_3173515 | Lighthouse Lab in Alderley Park | Wellcome Sanger Institute for the COVID-19 Genomics UK (COG-UK) Consortium | Jacquelyn Wynn, Mairead Hyland, The Lighthouse Lab in Alderley Park and Alex Alderton, Roberto Amato, Jeffrey Barrett, Sonia Goncalves, Ewan Harrison, David K. Jackson, Ian Johnston, Dominic Kwiatkowski, Cordelia Langford, John Sillitoe on behalf of the Wellcome Sanger Institute COVID-19 Surveillance Team |
| EPI_ISL_3173516 | Lighthouse Lab in Glasgow | Wellcome Sanger Institute for the COVID-19 Genomics UK (COG-UK) Consortium | Harper VanSteenhouse, Yumi Kasai, David Gray, Carol Clugston, Anna Dominiczak and Alex Alderton, Roberto Amato, Jeffrey Barrett, Sonia Goncalves, Ewan Harrison, David K. Jackson, Ian Johnston, Dominic Kwiatkowski, Cordelia Langford, John Sillitoe on behalf of the Wellcome Sanger Institute COVID-19 Surveillance Team |
| EPI_ISL_3173517, EPI_ISL_3173518, EPI_ISL_3173519 | Lighthouse Lab in Alderley Park | Wellcome Sanger Institute for the COVID-19 Genomics UK (COG-UK) Consortium | Jacquelyn Wynn, Mairead Hyland, The Lighthouse Lab in Alderley Park and Alex Alderton, Roberto Amato, Jeffrey Barrett, Sonia Goncalves, Ewan Harrison, David K. Jackson, Ian Johnston, Dominic Kwiatkowski, Cordelia Langford, John Sillitoe on behalf of the Wellcome Sanger Institute COVID-19 Surveillance Team |
| EPI_ISL_3173520, EPI_ISL_3173521, EPI_ISL_3173522 | Lighthouse Lab in Glasgow | Wellcome Sanger Institute for the COVID-19 Genomics UK (COG-UK) Consortium | Harper VanSteenhouse, Yumi Kasai, David Gray, Carol Clugston, Anna Dominiczak and Alex Alderton, Roberto Amato, Jeffrey Barrett, Sonia Goncalves, Ewan Harrison, David K. Jackson, Ian Johnston, Dominic Kwiatkowski, Cordelia Langford, John Sillitoe on behalf of the Wellcome Sanger Institute COVID-19 Surveillance Team |
| EPI_ISL_3173523 | Lighthouse Lab in Alderley Park | Wellcome Sanger Institute for the COVID-19 Genomics UK (COG-UK) Consortium | Jacquelyn Wynn, Mairead Hyland, The Lighthouse Lab in Alderley Park and Alex Alderton, Roberto Amato, Jeffrey Barrett, Sonia Goncalves, Ewan Harrison, David K. Jackson, Ian Johnston, Dominic Kwiatkowski, Cordelia Langford, John Sillitoe on behalf of the Wellcome Sanger Institute COVID-19 Surveillance Team |
| EPI_ISL_3173524 | Lighthouse Lab in Milton Keynes | Wellcome Sanger Institute for the COVID-19 Genomics UK (COG-UK) Consortium | The Lighthouse Lab in Milton Keynes and Alex Alderton, Roberto Amato, Jeffrey Barrett, Sonia Goncalves, Ewan Harrison, David K. Jackson, Ian Johnston, Dominic Kwiatkowski, Cordelia Langford, John Sillitoe on behalf of the Wellcome Sanger Institute COVID-19 Surveillance Team |
| EPI_ISL_3173525 | Lighthouse Lab in Glasgow | Wellcome Sanger Institute for the COVID-19 Genomics UK (COG-UK) Consortium | Harper VanSteenhouse, Yumi Kasai, David Gray, Carol Clugston, Anna Dominiczak and Alex Alderton, Roberto Amato, Jeffrey Barrett, Sonia Goncalves, Ewan Harrison, David K. Jackson, Ian Johnston, Dominic Kwiatkowski, Cordelia Langford, John Sillitoe on behalf of the Wellcome Sanger Institute COVID-19 Surveillance Team |
| EPI_ISL_3173526 | Lighthouse Lab in Milton Keynes | Wellcome Sanger Institute for the COVID-19 Genomics UK (COG-UK) Consortium | The Lighthouse Lab in Milton Keynes and Alex Alderton, Roberto Amato, Jeffrey Barrett, Sonia Goncalves, Ewan Harrison, David K. Jackson, Ian Johnston, Dominic Kwiatkowski, Cordelia Langford, John Sillitoe on behalf of the Wellcome Sanger Institute COVID-19 Surveillance Team |
| EPI_ISL_3173527 | Lighthouse Lab in Glasgow | Wellcome Sanger Institute for the COVID-19 Genomics UK (COG-UK) Consortium | Harper VanSteenhouse, Yumi Kasai, David Gray, Carol Clugston, Anna Dominiczak and Alex Alderton, Roberto Amato, Jeffrey Barrett, Sonia Goncalves, Ewan Harrison, David K. Jackson, Ian Johnston, Dominic Kwiatkowski, Cordelia Langford, John Sillitoe on behalf of the Wellcome Sanger Institute COVID-19 Surveillance Team |
| EPI_ISL_3173528 | Lighthouse Lab in Milton Keynes | Wellcome Sanger Institute for the COVID-19 Genomics UK (COG-UK) Consortium | The Lighthouse Lab in Milton Keynes and Alex Alderton, Roberto Amato, Jeffrey Barrett, Sonia Goncalves, Ewan Harrison, David K. Jackson, Ian Johnston, Dominic Kwiatkowski, Cordelia Langford, John Sillitoe on behalf of the Wellcome Sanger Institute COVID-19 Surveillance Team |
| EPI_ISL_3173529, EPI_ISL_3173530, EPI_ISL_3173531, EPI_ISL_3173532 | Lighthouse Lab in Alderley Park | Wellcome Sanger Institute for the COVID-19 Genomics UK (COG-UK) Consortium | Jacquelyn Wynn, Mairead Hyland, The Lighthouse Lab in Alderley Park and Alex Alderton, Roberto Amato, Jeffrey Barrett, Sonia Goncalves, Ewan Harrison, David K. Jackson, Ian Johnston, Dominic Kwiatkowski, Cordelia Langford, John Sillitoe on behalf of the Wellcome Sanger Institute COVID-19 Surveillance Team |
| EPI_ISL_3173533, EPI_ISL_3173534 | Lighthouse Lab in Glasgow | Wellcome Sanger Institute for the COVID-19 Genomics UK (COG-UK) Consortium | Harper VanSteenhouse, Yumi Kasai, David Gray, Carol Clugston, Anna Dominiczak and Alex Alderton, Roberto Amato, Jeffrey Barrett, Sonia Goncalves, Ewan Harrison, David K. Jackson, Ian Johnston, Dominic Kwiatkowski, Cordelia Langford, John Sillitoe on behalf of the Wellcome Sanger Institute COVID-19 Surveillance Team |
| EPI_ISL_3173535 | Lighthouse Lab in Alderley Park | Wellcome Sanger Institute for the COVID-19 Genomics UK (COG-UK) Consortium | Jacquelyn Wynn, Mairead Hyland, The Lighthouse Lab in Alderley Park and Alex Alderton, Roberto Amato, Jeffrey Barrett, Sonia Goncalves, Ewan Harrison, David K. Jackson, Ian Johnston, Dominic Kwiatkowski, Cordelia Langford, John Sillitoe on behalf of the Wellcome Sanger Institute COVID-19 Surveillance Team |
| EPI_ISL_3173536 | Lighthouse Lab in Milton Keynes | Wellcome Sanger Institute for the COVID-19 Genomics UK (COG-UK) Consortium | The Lighthouse Lab in Milton Keynes and Alex Alderton, Roberto Amato, Jeffrey Barrett, Sonia Goncalves, Ewan Harrison, David K. Jackson, Ian Johnston, Dominic Kwiatkowski, Cordelia Langford, John Sillitoe on behalf of the Wellcome Sanger Institute COVID-19 Surveillance Team |
| EPI_ISL_3173537, EPI_ISL_3173538 | Lighthouse Lab in Alderley Park | Wellcome Sanger Institute for the COVID-19 Genomics UK (COG-UK) Consortium | Jacquelyn Wynn, Mairead Hyland, The Lighthouse Lab in Alderley Park and Alex Alderton, Roberto Amato, Jeffrey Barrett, Sonia Goncalves, Ewan Harrison, David K. Jackson, Ian Johnston, Dominic Kwiatkowski, Cordelia Langford, John Sillitoe on behalf of the Wellcome Sanger Institute COVID-19 Surveillance Team |
| EPI_ISL_3173539 | Lighthouse Lab in Milton Keynes | Wellcome Sanger Institute for the COVID-19 Genomics UK (COG-UK) Consortium | The Lighthouse Lab in Milton Keynes and Alex Alderton, Roberto Amato, Jeffrey Barrett, Sonia Goncalves, Ewan Harrison, David K. Jackson, Ian Johnston, Dominic Kwiatkowski, Cordelia Langford, John Sillitoe on behalf of the Wellcome Sanger Institute COVID-19 Surveillance Team |
| EPI_ISL_3173540 | Lighthouse Lab in Glasgow | Wellcome Sanger Institute for the COVID-19 Genomics UK (COG-UK) Consortium | Harper VanSteenhouse, Yumi Kasai, David Gray, Carol Clugston, Anna Dominiczak and Alex Alderton, Roberto Amato, Jeffrey Barrett, Sonia Goncalves, Ewan Harrison, David K. Jackson, Ian Johnston, Dominic Kwiatkowski, Cordelia Langford, John Sillitoe on behalf of the Wellcome Sanger Institute COVID-19 Surveillance Team |
| EPI_ISL_3173541, EPI_ISL_3173542 | Lighthouse Lab in Alderley Park | Wellcome Sanger Institute for the COVID-19 Genomics UK (COG-UK) Consortium | Jacquelyn Wynn, Mairead Hyland, The Lighthouse Lab in Alderley Park and Alex Alderton, Roberto Amato, Jeffrey Barrett, Sonia Goncalves, Ewan Harrison, David K. Jackson, Ian Johnston, Dominic Kwiatkowski, Cordelia Langford, John Sillitoe on behalf of the Wellcome Sanger Institute COVID-19 Surveillance Team |
| EPI_ISL_3173543 | Lighthouse Lab in Glasgow | Wellcome Sanger Institute for the COVID-19 Genomics UK (COG-UK) Consortium | Harper VanSteenhouse, Yumi Kasai, David Gray, Carol Clugston, Anna Dominiczak and Alex Alderton, Roberto Amato, Jeffrey Barrett, Sonia Goncalves, Ewan Harrison, David K. Jackson, Ian Johnston, Dominic Kwiatkowski, Cordelia Langford, John Sillitoe on behalf of the Wellcome Sanger Institute COVID-19 Surveillance Team |
| EPI_ISL_3173544, EPI_ISL_3173545 | Lighthouse Lab in Alderley Park | Wellcome Sanger Institute for the COVID-19 Genomics UK (COG-UK) Consortium | Jacquelyn Wynn, Mairead Hyland, The Lighthouse Lab in Alderley Park and Alex Alderton, Roberto Amato, Jeffrey Barrett, Sonia Goncalves, Ewan Harrison, David K. Jackson, Ian Johnston, Dominic Kwiatkowski, Cordelia Langford, John Sillitoe on behalf of the Wellcome Sanger Institute COVID-19 Surveillance Team |
| EPI_ISL_3173546 | Lighthouse Lab in Glasgow | Wellcome Sanger Institute for the COVID-19 Genomics UK (COG-UK) Consortium | Harper VanSteenhouse, Yumi Kasai, David Gray, Carol Clugston, Anna Dominiczak and Alex Alderton, Roberto Amato, Jeffrey Barrett, Sonia Goncalves, Ewan Harrison, David K. Jackson, Ian Johnston, Dominic Kwiatkowski, Cordelia Langford, John Sillitoe on behalf of the Wellcome Sanger Institute COVID-19 Surveillance Team |
| EPI_ISL_3173547 | Lighthouse Lab in Alderley Park | Wellcome Sanger Institute for the COVID-19 Genomics UK (COG-UK) Consortium | Jacquelyn Wynn, Mairead Hyland, The Lighthouse Lab in Alderley Park and Alex Alderton, Roberto Amato, Jeffrey Barrett, Sonia Goncalves, Ewan Harrison, David K. Jackson, Ian Johnston, Dominic Kwiatkowski, Cordelia Langford, John Sillitoe on behalf of the Wellcome Sanger Institute COVID-19 Surveillance Team |
| EPI_ISL_3173548 | Lighthouse Lab in Milton Keynes | Wellcome Sanger Institute for the COVID-19 Genomics UK (COG-UK) Consortium | The Lighthouse Lab in Milton Keynes and Alex Alderton, Roberto Amato, Jeffrey Barrett, Sonia Goncalves, Ewan Harrison, David K. Jackson, Ian Johnston, Dominic Kwiatkowski, Cordelia Langford, John Sillitoe on behalf of the Wellcome Sanger Institute COVID-19 Surveillance Team |
| EPI_ISL_3173549 | Lighthouse Lab in Glasgow | Wellcome Sanger Institute for the COVID-19 Genomics UK (COG-UK) Consortium | Harper VanSteenhouse, Yumi Kasai, David Gray, Carol Clugston, Anna Dominiczak and Alex Alderton, Roberto Amato, Jeffrey Barrett, Sonia Goncalves, Ewan Harrison, David K. Jackson, Ian Johnston, Dominic Kwiatkowski, Cordelia Langford, John Sillitoe on behalf of the Wellcome Sanger Institute COVID-19 Surveillance Team |
| EPI_ISL_3173550 | Lighthouse Lab in Alderley Park | Wellcome Sanger Institute for the COVID-19 Genomics UK (COG-UK) Consortium | Jacquelyn Wynn, Mairead Hyland, The Lighthouse Lab in Alderley Park and Alex Alderton, Roberto Amato, Jeffrey Barrett, Sonia Goncalves, Ewan Harrison, David K. Jackson, Ian Johnston, Dominic Kwiatkowski, Cordelia Langford, John Sillitoe on behalf of the Wellcome Sanger Institute COVID-19 Surveillance Team |
| EPI_ISL_3173551, EPI_ISL_3173552 | Lighthouse Lab in Milton Keynes | Wellcome Sanger Institute for the COVID-19 Genomics UK (COG-UK) Consortium | The Lighthouse Lab in Milton Keynes and Alex Alderton, Roberto Amato, Jeffrey Barrett, Sonia Goncalves, Ewan Harrison, David K. Jackson, Ian Johnston, Dominic Kwiatkowski, Cordelia Langford, John Sillitoe on behalf of the Wellcome Sanger Institute COVID-19 Surveillance Team |
| EPI_ISL_3173553, EPI_ISL_3173554 | Lighthouse Lab in Alderley Park | Wellcome Sanger Institute for the COVID-19 Genomics UK (COG-UK) Consortium | Jacquelyn Wynn, Mairead Hyland, The Lighthouse Lab in Alderley Park and Alex Alderton, Roberto Amato, Jeffrey Barrett, Sonia Goncalves, Ewan Harrison, David K. Jackson, Ian Johnston, Dominic Kwiatkowski, Cordelia Langford, John Sillitoe on behalf of the Wellcome Sanger Institute COVID-19 Surveillance Team |
| EPI_ISL_3173555 | Lighthouse Lab in Milton Keynes | Wellcome Sanger Institute for the COVID-19 Genomics UK (COG-UK) Consortium | The Lighthouse Lab in Milton Keynes and Alex Alderton, Roberto Amato, Jeffrey Barrett, Sonia Goncalves, Ewan Harrison, David K. Jackson, Ian Johnston, Dominic Kwiatkowski, Cordelia Langford, John Sillitoe on behalf of the Wellcome Sanger Institute COVID-19 Surveillance Team |
| EPI_ISL_3173556 | Lighthouse Lab in Alderley Park | Wellcome Sanger Institute for the COVID-19 Genomics UK (COG-UK) Consortium | Jacquelyn Wynn, Mairead Hyland, The Lighthouse Lab in Alderley Park and Alex Alderton, Roberto Amato, Jeffrey Barrett, Sonia Goncalves, Ewan Harrison, David K. Jackson, Ian Johnston, Dominic Kwiatkowski, Cordelia Langford, John Sillitoe on behalf of the Wellcome Sanger Institute COVID-19 Surveillance Team |
| EPI_ISL_3173557 | Lighthouse Lab in Milton Keynes | Wellcome Sanger Institute for the COVID-19 Genomics UK (COG-UK) Consortium | The Lighthouse Lab in Milton Keynes and Alex Alderton, Roberto Amato, Jeffrey Barrett, Sonia Goncalves, Ewan Harrison, David K. Jackson, Ian Johnston, Dominic Kwiatkowski, Cordelia Langford, John Sillitoe on behalf of the Wellcome Sanger Institute COVID-19 Surveillance Team |
| EPI_ISL_3173558, EPI_ISL_3173559, EPI_ISL_3173560, EPI_ISL_3173561, EPI_ISL_3173562, EPI_ISL_3173563, EPI_ISL_3173564, | Lighthouse Lab in Alderley Park | Wellcome Sanger Institute for the COVID-19 Genomics UK (COG-UK) Consortium | Jacquelyn Wynn, Mairead Hyland, The Lighthouse Lab in Alderley Park and Alex Alderton, Roberto Amato, Jeffrey Barrett, Sonia Goncalves, Ewan Harrison, David K. Jackson, Ian Johnston, Dominic Kwiatkowski, Cordelia Langford, John Sillitoe on behalf of the Wellcome Sanger Institute COVID-19 Surveillance Team |

| | | | |
|---|---|---|---|
| EPI_ISL_3173565, EPI_ISL_3173566 | | | |
| EPI_ISL_3173567 | Lighthouse Lab in Milton Keynes | Wellcome Sanger Institute for the COVID-19 Genomics UK (COG-UK) Consortium | The Lighthouse Lab in Milton Keynes and Alex Alderton, Roberto Amato, Jeffrey Barrett, Sonia Goncalves, Ewan Harrison, David K. Jackson, Ian Johnston, Dominic Kwiatkowski, Cordelia Langford, John Sillitoe on behalf of the Wellcome Sanger Institute COVID-19 Surveillance Team |
| EPI_ISL_3173568, EPI_ISL_3173569, EPI_ISL_3173570 | Lighthouse Lab in Alderley Park | Wellcome Sanger Institute for the COVID-19 Genomics UK (COG-UK) Consortium | Jacquelyn Wynn, Mairead Hyland, The Lighthouse Lab in Alderley Park and Alex Alderton, Roberto Amato, Jeffrey Barrett, Sonia Goncalves, Ewan Harrison, David K. Jackson, Ian Johnston, Dominic Kwiatkowski, Cordelia Langford, John Sillitoe on behalf of the Wellcome Sanger Institute COVID-19 Surveillance Team |
| EPI_ISL_3173571, EPI_ISL_3173572 | Lighthouse Lab in Glasgow | Wellcome Sanger Institute for the COVID-19 Genomics UK (COG-UK) Consortium | Harper VanSteenhouse, Yumi Kasai, David Gray, Carol Clugston, Anna Dominiczak and Alex Alderton, Roberto Amato, Jeffrey Barrett, Sonia Goncalves, Ewan Harrison, David K. Jackson, Ian Johnston, Dominic Kwiatkowski, Cordelia Langford, John Sillitoe on behalf of the Wellcome Sanger Institute COVID-19 Surveillance Team |
| EPI_ISL_3173573 | Lighthouse Lab in Milton Keynes | Wellcome Sanger Institute for the COVID-19 Genomics UK (COG-UK) Consortium | The Lighthouse Lab in Milton Keynes and Alex Alderton, Roberto Amato, Jeffrey Barrett, Sonia Goncalves, Ewan Harrison, David K. Jackson, Ian Johnston, Dominic Kwiatkowski, Cordelia Langford, John Sillitoe on behalf of the Wellcome Sanger Institute COVID-19 Surveillance Team |
| EPI_ISL_3173574 | Lighthouse Lab in Alderley Park | Wellcome Sanger Institute for the COVID-19 Genomics UK (COG-UK) Consortium | Jacquelyn Wynn, Mairead Hyland, The Lighthouse Lab in Alderley Park and Alex Alderton, Roberto Amato, Jeffrey Barrett, Sonia Goncalves, Ewan Harrison, David K. Jackson, Ian Johnston, Dominic Kwiatkowski, Cordelia Langford, John Sillitoe on behalf of the Wellcome Sanger Institute COVID-19 Surveillance Team |
| EPI_ISL_3173575 | Lighthouse Lab in Milton Keynes | Wellcome Sanger Institute for the COVID-19 Genomics UK (COG-UK) Consortium | The Lighthouse Lab in Milton Keynes and Alex Alderton, Roberto Amato, Jeffrey Barrett, Sonia Goncalves, Ewan Harrison, David K. Jackson, Ian Johnston, Dominic Kwiatkowski, Cordelia Langford, John Sillitoe on behalf of the Wellcome Sanger Institute COVID-19 Surveillance Team |
| EPI_ISL_3173576 | Lighthouse Lab in Glasgow | Wellcome Sanger Institute for the COVID-19 Genomics UK (COG-UK) Consortium | Harper VanSteenhouse, Yumi Kasai, David Gray, Carol Clugston, Anna Dominiczak and Alex Alderton, Roberto Amato, Jeffrey Barrett, Sonia Goncalves, Ewan Harrison, David K. Jackson, Ian Johnston, Dominic Kwiatkowski, Cordelia Langford, John Sillitoe on behalf of the Wellcome Sanger Institute COVID-19 Surveillance Team |
| EPI_ISL_3173577 | Lighthouse Lab in Milton Keynes | Wellcome Sanger Institute for the COVID-19 Genomics UK (COG-UK) Consortium | The Lighthouse Lab in Milton Keynes and Alex Alderton, Roberto Amato, Jeffrey Barrett, Sonia Goncalves, Ewan Harrison, David K. Jackson, Ian Johnston, Dominic Kwiatkowski, Cordelia Langford, John Sillitoe on behalf of the Wellcome Sanger Institute COVID-19 Surveillance Team |
| EPI_ISL_3173578 | Lighthouse Lab in Glasgow | Wellcome Sanger Institute for the COVID-19 Genomics UK (COG-UK) Consortium | Harper VanSteenhouse, Yumi Kasai, David Gray, Carol Clugston, Anna Dominiczak and Alex Alderton, Roberto Amato, Jeffrey Barrett, Sonia Goncalves, Ewan Harrison, David K. Jackson, Ian Johnston, Dominic Kwiatkowski, Cordelia Langford, John Sillitoe on behalf of the Wellcome Sanger Institute COVID-19 Surveillance Team |
| EPI_ISL_3173579 | Lighthouse Lab in Alderley Park | Wellcome Sanger Institute for the COVID-19 Genomics UK (COG-UK) Consortium | Jacquelyn Wynn, Mairead Hyland, The Lighthouse Lab in Alderley Park and Alex Alderton, Roberto Amato, Jeffrey Barrett, Sonia Goncalves, Ewan Harrison, David K. Jackson, Ian Johnston, Dominic Kwiatkowski, Cordelia Langford, John Sillitoe on behalf of the Wellcome Sanger Institute COVID-19 Surveillance Team |
| EPI_ISL_3173580 | Lighthouse Lab in Milton Keynes | Wellcome Sanger Institute for the COVID-19 Genomics UK (COG-UK) Consortium | The Lighthouse Lab in Milton Keynes and Alex Alderton, Roberto Amato, Jeffrey Barrett, Sonia Goncalves, Ewan Harrison, David K. Jackson, Ian Johnston, Dominic Kwiatkowski, Cordelia Langford, John Sillitoe on behalf of the Wellcome Sanger Institute COVID-19 Surveillance Team |
| EPI_ISL_3173581 | Lighthouse Lab in Alderley Park | Wellcome Sanger Institute for the COVID-19 Genomics UK (COG-UK) Consortium | Jacquelyn Wynn, Mairead Hyland, The Lighthouse Lab in Alderley Park and Alex Alderton, Roberto Amato, Jeffrey Barrett, Sonia Goncalves, Ewan Harrison, David K. Jackson, Ian Johnston, Dominic Kwiatkowski, Cordelia Langford, John Sillitoe on behalf of the Wellcome Sanger Institute COVID-19 Surveillance Team |
| EPI_ISL_3173582, EPI_ISL_3173583 | Lighthouse Lab in Milton Keynes | Wellcome Sanger Institute for the COVID-19 Genomics UK (COG-UK) Consortium | The Lighthouse Lab in Milton Keynes and Alex Alderton, Roberto Amato, Jeffrey Barrett, Sonia Goncalves, Ewan Harrison, David K. Jackson, Ian Johnston, Dominic Kwiatkowski, Cordelia Langford, John Sillitoe on behalf of the Wellcome Sanger Institute COVID-19 Surveillance Team |
| EPI_ISL_3173584 | Lighthouse Lab in Glasgow | Wellcome Sanger Institute for the COVID-19 Genomics UK (COG-UK) Consortium | Harper VanSteenhouse, Yumi Kasai, David Gray, Carol Clugston, Anna Dominiczak and Alex Alderton, Roberto Amato, Jeffrey Barrett, Sonia Goncalves, Ewan Harrison, David K. Jackson, Ian Johnston, Dominic Kwiatkowski, Cordelia Langford, John Sillitoe on behalf of the Wellcome Sanger Institute COVID-19 Surveillance Team |
| EPI_ISL_3173585 | Lighthouse Lab in Alderley Park | Wellcome Sanger Institute for the COVID-19 Genomics UK (COG-UK) Consortium | Jacquelyn Wynn, Mairead Hyland, The Lighthouse Lab in Alderley Park and Alex Alderton, Roberto Amato, Jeffrey Barrett, Sonia Goncalves, Ewan Harrison, David K. Jackson, Ian Johnston, Dominic Kwiatkowski, Cordelia Langford, John Sillitoe on behalf of the Wellcome Sanger Institute COVID-19 Surveillance Team |
| EPI_ISL_3173586, EPI_ISL_3173587 | Lighthouse Lab in Milton Keynes | Wellcome Sanger Institute for the COVID-19 Genomics UK (COG-UK) Consortium | The Lighthouse Lab in Milton Keynes and Alex Alderton, Roberto Amato, Jeffrey Barrett, Sonia Goncalves, Ewan Harrison, David K. Jackson, Ian Johnston, Dominic Kwiatkowski, Cordelia Langford, John Sillitoe on behalf of the Wellcome Sanger Institute COVID-19 Surveillance Team |
| EPI_ISL_3173588 | Lighthouse Lab in Glasgow | Wellcome Sanger Institute for the COVID-19 Genomics UK (COG-UK) Consortium | Harper VanSteenhouse, Yumi Kasai, David Gray, Carol Clugston, Anna Dominiczak and Alex Alderton, Roberto Amato, Jeffrey Barrett, Sonia Goncalves, Ewan Harrison, David K. Jackson, Ian Johnston, Dominic Kwiatkowski, Cordelia Langford, John Sillitoe on behalf of the Wellcome Sanger Institute COVID-19 Surveillance Team |
| EPI_ISL_3173589 | Lighthouse Lab in Milton Keynes | Wellcome Sanger Institute for the COVID-19 Genomics UK (COG-UK) Consortium | The Lighthouse Lab in Milton Keynes and Alex Alderton, Roberto Amato, Jeffrey Barrett, Sonia Goncalves, Ewan Harrison, David K. Jackson, Ian Johnston, Dominic Kwiatkowski, Cordelia Langford, John Sillitoe on behalf of the Wellcome Sanger Institute COVID-19 Surveillance Team |
| EPI_ISL_3173590 | Lighthouse Lab in Glasgow | Wellcome Sanger Institute for the COVID-19 Genomics UK (COG-UK) Consortium | Harper VanSteenhouse, Yumi Kasai, David Gray, Carol Clugston, Anna Dominiczak and Alex Alderton, Roberto Amato, Jeffrey Barrett, Sonia Goncalves, Ewan Harrison, David K. Jackson, Ian Johnston, Dominic Kwiatkowski, Cordelia Langford, John Sillitoe on behalf of the Wellcome Sanger Institute COVID-19 Surveillance Team |
| EPI_ISL_3173591, EPI_ISL_3173592, EPI_ISL_3173593 | Lighthouse Lab in Alderley Park | Wellcome Sanger Institute for the COVID-19 Genomics UK (COG-UK) Consortium | Jacquelyn Wynn, Mairead Hyland, The Lighthouse Lab in Alderley Park and Alex Alderton, Roberto Amato, Jeffrey Barrett, Sonia Goncalves, Ewan Harrison, David K. Jackson, Ian Johnston, Dominic Kwiatkowski, Cordelia Langford, John Sillitoe on behalf of the Wellcome Sanger Institute COVID-19 Surveillance Team |
| EPI_ISL_3173594 | Lighthouse Lab in Milton Keynes | Wellcome Sanger Institute for the COVID-19 Genomics UK (COG-UK) Consortium | The Lighthouse Lab in Milton Keynes and Alex Alderton, Roberto Amato, Jeffrey Barrett, Sonia Goncalves, Ewan Harrison, David K. Jackson, Ian Johnston, Dominic Kwiatkowski, Cordelia Langford, John Sillitoe on behalf of the Wellcome Sanger Institute COVID-19 Surveillance Team |
| EPI_ISL_3173595 | Lighthouse Lab in Alderley Park | Wellcome Sanger Institute for the COVID-19 Genomics UK (COG-UK) Consortium | Jacquelyn Wynn, Mairead Hyland, The Lighthouse Lab in Alderley Park and Alex Alderton, Roberto Amato, Jeffrey Barrett, Sonia Goncalves, Ewan Harrison, David K. Jackson, Ian Johnston, Dominic Kwiatkowski, Cordelia Langford, John Sillitoe on behalf of the Wellcome Sanger Institute COVID-19 Surveillance Team |
| EPI_ISL_3173596 | Lighthouse Lab in Milton Keynes | Wellcome Sanger Institute for the COVID-19 Genomics UK (COG-UK) Consortium | The Lighthouse Lab in Milton Keynes and Alex Alderton, Roberto Amato, Jeffrey Barrett, Sonia Goncalves, Ewan Harrison, David K. Jackson, Ian Johnston, Dominic Kwiatkowski, Cordelia Langford, John Sillitoe on behalf of the Wellcome Sanger Institute COVID-19 Surveillance Team |
| EPI_ISL_3173597, EPI_ISL_3173598 | Lighthouse Lab in Alderley Park | Wellcome Sanger Institute for the COVID-19 Genomics UK (COG-UK) Consortium | Jacquelyn Wynn, Mairead Hyland, The Lighthouse Lab in Alderley Park and Alex Alderton, Roberto Amato, Jeffrey Barrett, Sonia Goncalves, Ewan Harrison, David K. Jackson, Ian Johnston, Dominic Kwiatkowski, Cordelia Langford, John Sillitoe on behalf of the Wellcome Sanger Institute COVID-19 Surveillance Team |
| EPI_ISL_3173599 | Lighthouse Lab in Glasgow | Wellcome Sanger Institute for the COVID-19 Genomics UK (COG-UK) Consortium | Harper VanSteenhouse, Yumi Kasai, David Gray, Carol Clugston, Anna Dominiczak and Alex Alderton, Roberto Amato, Jeffrey Barrett, Sonia Goncalves, Ewan Harrison, David K. Jackson, Ian Johnston, Dominic Kwiatkowski, Cordelia Langford, John Sillitoe on behalf of the Wellcome Sanger Institute COVID-19 Surveillance Team |
| EPI_ISL_3173600 | Lighthouse Lab in Alderley Park | Wellcome Sanger Institute for the COVID-19 Genomics UK (COG-UK) Consortium | Jacquelyn Wynn, Mairead Hyland, The Lighthouse Lab in Alderley Park and Alex Alderton, Roberto Amato, Jeffrey Barrett, Sonia Goncalves, Ewan Harrison, David K. Jackson, Ian Johnston, Dominic Kwiatkowski, Cordelia Langford, John Sillitoe on behalf of the Wellcome Sanger Institute COVID-19 Surveillance Team |
| EPI_ISL_3173601, EPI_ISL_3173602 | Lighthouse Lab in Milton Keynes | Wellcome Sanger Institute for the COVID-19 Genomics UK (COG-UK) Consortium | The Lighthouse Lab in Milton Keynes and Alex Alderton, Roberto Amato, Jeffrey Barrett, Sonia Goncalves, Ewan Harrison, David K. Jackson, Ian Johnston, Dominic Kwiatkowski, Cordelia Langford, John Sillitoe on behalf of the Wellcome Sanger Institute COVID-19 Surveillance Team |
| EPI_ISL_3173603, EPI_ISL_3173604, EPI_ISL_3173605, EPI_ISL_3173606, EPI_ISL_3173607 | Lighthouse Lab in Alderley Park | Wellcome Sanger Institute for the COVID-19 Genomics UK (COG-UK) Consortium | Jacquelyn Wynn, Mairead Hyland, The Lighthouse Lab in Alderley Park and Alex Alderton, Roberto Amato, Jeffrey Barrett, Sonia Goncalves, Ewan Harrison, David K. Jackson, Ian Johnston, Dominic Kwiatkowski, Cordelia Langford, John Sillitoe on behalf of the Wellcome Sanger Institute COVID-19 Surveillance Team |
| EPI_ISL_3173608 | Lighthouse Lab in Milton Keynes | Wellcome Sanger Institute for the COVID-19 Genomics UK (COG-UK) Consortium | The Lighthouse Lab in Milton Keynes and Alex Alderton, Roberto Amato, Jeffrey Barrett, Sonia Goncalves, Ewan Harrison, David K. Jackson, Ian Johnston, Dominic Kwiatkowski, Cordelia Langford, John Sillitoe on behalf of the Wellcome Sanger Institute COVID-19 Surveillance Team |
| EPI_ISL_3173609, EPI_ISL_3173610, EPI_ISL_3173611, EPI_ISL_3173612, EPI_ISL_3173613 | Lighthouse Lab in Alderley Park | Wellcome Sanger Institute for the COVID-19 Genomics UK (COG-UK) Consortium | Jacquelyn Wynn, Mairead Hyland, The Lighthouse Lab in Alderley Park and Alex Alderton, Roberto Amato, Jeffrey Barrett, Sonia Goncalves, Ewan Harrison, David K. Jackson, Ian Johnston, Dominic Kwiatkowski, Cordelia Langford, John Sillitoe on behalf of the Wellcome Sanger Institute COVID-19 Surveillance Team |
| EPI_ISL_3173614 | Lighthouse Lab in Glasgow | Wellcome Sanger Institute for the COVID-19 Genomics UK (COG-UK) Consortium | Harper VanSteenhouse, Yumi Kasai, David Gray, Carol Clugston, Anna Dominiczak and Alex Alderton, Roberto Amato, Jeffrey Barrett, Sonia Goncalves, Ewan Harrison, David K. Jackson, Ian Johnston, Dominic Kwiatkowski, Cordelia Langford, John Sillitoe on behalf of the Wellcome Sanger Institute COVID-19 Surveillance Team |
| EPI_ISL_3173615, EPI_ISL_3173616 | Lighthouse Lab in Alderley Park | Wellcome Sanger Institute for the COVID-19 Genomics UK (COG-UK) Consortium | Jacquelyn Wynn, Mairead Hyland, The Lighthouse Lab in Alderley Park and Alex Alderton, Roberto Amato, Jeffrey Barrett, Sonia Goncalves, Ewan Harrison, David K. Jackson, Ian Johnston, Dominic Kwiatkowski, Cordelia Langford, John Sillitoe on behalf of the Wellcome Sanger Institute COVID-19 Surveillance Team |
| EPI_ISL_3173617 | Lighthouse Laboratory Plymouth | Wellcome Sanger Institute for the COVID-19 Genomics UK (COG-UK) Consortium | Lighthouse Laboratory Plymouth and Alex Alderton, Roberto Amato, Jeffrey Barrett, Sonia Goncalves, Ewan Harrison, David K. Jackson, Ian Johnston, Dominic Kwiatkowski, Cordelia Langford, John Sillitoe on behalf of the Wellcome Sanger Institute COVID-19 Surveillance Team |
| EPI_ISL_3173618 | Lighthouse Lab in Glasgow | Wellcome Sanger Institute for the COVID-19 Genomics UK (COG-UK) Consortium | Harper VanSteenhouse, Yumi Kasai, David Gray, Carol Clugston, Anna Dominiczak and Alex Alderton, Roberto Amato, Jeffrey Barrett, Sonia Goncalves, Ewan Harrison, David K. Jackson, Ian Johnston, Dominic Kwiatkowski, Cordelia Langford, John Sillitoe on behalf of the Wellcome Sanger Institute COVID-19 Surveillance Team |
| EPI_ISL_3173619 | Lighthouse Laboratory Plymouth | Wellcome Sanger Institute for the COVID-19 Genomics UK (COG-UK) Consortium | Lighthouse Laboratory Plymouth and Alex Alderton, Roberto Amato, Jeffrey Barrett, Sonia Goncalves, Ewan Harrison, David K. Jackson, Ian Johnston, Dominic Kwiatkowski, Cordelia Langford, John Sillitoe on behalf of the Wellcome Sanger Institute COVID-19 Surveillance Team |
| EPI_ISL_3173620 | Lighthouse Lab in Alderley Park | Wellcome Sanger Institute for the COVID-19 Genomics UK (COG-UK) Consortium | Jacquelyn Wynn, Mairead Hyland, The Lighthouse Lab in Alderley Park and Alex Alderton, Roberto Amato, Jeffrey Barrett, Sonia Goncalves, Ewan Harrison, David K. Jackson, Ian Johnston, Dominic Kwiatkowski, Cordelia Langford, John Sillitoe on behalf of the Wellcome Sanger Institute COVID-19 Surveillance Team |
| EPI_ISL_3173621, EPI_ISL_3173622 | Lighthouse Lab in Glasgow | Wellcome Sanger Institute for the COVID-19 Genomics UK (COG-UK) Consortium | Harper VanSteenhouse, Yumi Kasai, David Gray, Carol Clugston, Anna Dominiczak and Alex Alderton, Roberto Amato, Jeffrey Barrett, Sonia Goncalves, Ewan Harrison, David K. Jackson, Ian Johnston, Dominic Kwiatkowski, Cordelia Langford, John Sillitoe on behalf of the Wellcome Sanger Institute COVID-19 Surveillance Team |
| EPI_ISL_3173623 | Lighthouse Laboratory Plymouth | Wellcome Sanger Institute for the COVID-19 Genomics UK (COG-UK) Consortium | Lighthouse Laboratory Plymouth and Alex Alderton, Roberto Amato, Jeffrey Barrett, Sonia Goncalves, Ewan Harrison, David K. Jackson, Ian Johnston, Dominic Kwiatkowski, Cordelia Langford, John Sillitoe on behalf of the Wellcome Sanger Institute COVID-19 Surveillance Team |
| EPI_ISL_3173624 | Lighthouse Lab in Glasgow | Wellcome Sanger Institute for the COVID-19 Genomics UK (COG-UK) Consortium | Harper VanSteenhouse, Yumi Kasai, David Gray, Carol Clugston, Anna Dominiczak and Alex Alderton, Roberto Amato, Jeffrey Barrett, Sonia Goncalves, Ewan Harrison, David K. Jackson, Ian Johnston, Dominic Kwiatkowski, Cordelia Langford, John Sillitoe on behalf of the Wellcome Sanger Institute COVID-19 Surveillance Team |
| EPI_ISL_3173625 | Lighthouse Lab in Alderley Park | Wellcome Sanger Institute for the COVID-19 Genomics UK (COG-UK) Consortium | Jacquelyn Wynn, Mairead Hyland, The Lighthouse Lab in Alderley Park and Alex Alderton, Roberto Amato, Jeffrey Barrett, Sonia Goncalves, Ewan Harrison, David K. Jackson, Ian Johnston, Dominic Kwiatkowski, Cordelia Langford, John Sillitoe on behalf of the Wellcome Sanger Institute COVID-19 Surveillance Team |
| EPI_ISL_3173626 | Lighthouse Laboratory Plymouth | Wellcome Sanger Institute for the COVID-19 Genomics UK (COG-UK) Consortium | Lighthouse Laboratory Plymouth and Alex Alderton, Roberto Amato, Jeffrey Barrett, Sonia Goncalves, Ewan Harrison, David K. Jackson, Ian Johnston, Dominic Kwiatkowski, Cordelia Langford, John Sillitoe on behalf of the Wellcome Sanger Institute COVID-19 Surveillance Team |

| | | | |
|---|---|---|---|
| EPI_ISL_3173627 | Lighthouse Lab in Glasgow | Wellcome Sanger Institute for the COVID-19 Genomics UK (COG-UK) Consortium | Harper VanSteenhouse, Yumi Kasai, David Gray, Carol Clugston, Anna Dominiczak and Alex Alderton, Roberto Amato, Jeffrey Barrett, Sonia Goncalves, Ewan Harrison, David K. Jackson, Ian Johnston, Dominic Kwiatkowski, Cordelia Langford, John Sillitoe on behalf of the Wellcome Sanger Institute COVID-19 Surveillance Team |
| EPI_ISL_3173628 | Lighthouse Laboratory Plymouth | Wellcome Sanger Institute for the COVID-19 Genomics UK (COG-UK) Consortium | Lighthouse Laboratory Plymouth and Alex Alderton, Roberto Amato, Jeffrey Barrett, Sonia Goncalves, Ewan Harrison, David K. Jackson, Ian Johnston, Dominic Kwiatkowski, Cordelia Langford, John Sillitoe on behalf of the Wellcome Sanger Institute COVID-19 Surveillance Team |
| EPI_ISL_3173629 | Lighthouse Lab in Glasgow | Wellcome Sanger Institute for the COVID-19 Genomics UK (COG-UK) Consortium | Harper VanSteenhouse, Yumi Kasai, David Gray, Carol Clugston, Anna Dominiczak and Alex Alderton, Roberto Amato, Jeffrey Barrett, Sonia Goncalves, Ewan Harrison, David K. Jackson, Ian Johnston, Dominic Kwiatkowski, Cordelia Langford, John Sillitoe on behalf of the Wellcome Sanger Institute COVID-19 Surveillance Team |
| EPI_ISL_3173630, EPI_ISL_3173631, EPI_ISL_3173632 | Lighthouse Laboratory Plymouth | Wellcome Sanger Institute for the COVID-19 Genomics UK (COG-UK) Consortium | Lighthouse Laboratory Plymouth and Alex Alderton, Roberto Amato, Jeffrey Barrett, Sonia Goncalves, Ewan Harrison, David K. Jackson, Ian Johnston, Dominic Kwiatkowski, Cordelia Langford, John Sillitoe on behalf of the Wellcome Sanger Institute COVID-19 Surveillance Team |
| EPI_ISL_3173633 | Lighthouse Lab in Glasgow | Wellcome Sanger Institute for the COVID-19 Genomics UK (COG-UK) Consortium | Harper VanSteenhouse, Yumi Kasai, David Gray, Carol Clugston, Anna Dominiczak and Alex Alderton, Roberto Amato, Jeffrey Barrett, Sonia Goncalves, Ewan Harrison, David K. Jackson, Ian Johnston, Dominic Kwiatkowski, Cordelia Langford, John Sillitoe on behalf of the Wellcome Sanger Institute COVID-19 Surveillance Team |
| EPI_ISL_3173634, EPI_ISL_3173635 | Lighthouse Lab in Alderley Park | Wellcome Sanger Institute for the COVID-19 Genomics UK (COG-UK) Consortium | Jacquelyn Wynn, Mairead Hyland, The Lighthouse Lab in Alderley Park and Alex Alderton, Roberto Amato, Jeffrey Barrett, Sonia Goncalves, Ewan Harrison, David K. Jackson, Ian Johnston, Dominic Kwiatkowski, Cordelia Langford, John Sillitoe on behalf of the Wellcome Sanger Institute COVID-19 Surveillance Team |
| EPI_ISL_3173636 | Lighthouse Laboratory Plymouth | Wellcome Sanger Institute for the COVID-19 Genomics UK (COG-UK) Consortium | Lighthouse Laboratory Plymouth and Alex Alderton, Roberto Amato, Jeffrey Barrett, Sonia Goncalves, Ewan Harrison, David K. Jackson, Ian Johnston, Dominic Kwiatkowski, Cordelia Langford, John Sillitoe on behalf of the Wellcome Sanger Institute COVID-19 Surveillance Team |
| EPI_ISL_3173637 | Lighthouse Lab in Glasgow | Wellcome Sanger Institute for the COVID-19 Genomics UK (COG-UK) Consortium | Harper VanSteenhouse, Yumi Kasai, David Gray, Carol Clugston, Anna Dominiczak and Alex Alderton, Roberto Amato, Jeffrey Barrett, Sonia Goncalves, Ewan Harrison, David K. Jackson, Ian Johnston, Dominic Kwiatkowski, Cordelia Langford, John Sillitoe on behalf of the Wellcome Sanger Institute COVID-19 Surveillance Team |
| EPI_ISL_3173638 | Lighthouse Lab in Alderley Park | Wellcome Sanger Institute for the COVID-19 Genomics UK (COG-UK) Consortium | Jacquelyn Wynn, Mairead Hyland, The Lighthouse Lab in Alderley Park and Alex Alderton, Roberto Amato, Jeffrey Barrett, Sonia Goncalves, Ewan Harrison, David K. Jackson, Ian Johnston, Dominic Kwiatkowski, Cordelia Langford, John Sillitoe on behalf of the Wellcome Sanger Institute COVID-19 Surveillance Team |
| EPI_ISL_3173639 | Lighthouse Laboratory Plymouth | Wellcome Sanger Institute for the COVID-19 Genomics UK (COG-UK) Consortium | Lighthouse Laboratory Plymouth and Alex Alderton, Roberto Amato, Jeffrey Barrett, Sonia Goncalves, Ewan Harrison, David K. Jackson, Ian Johnston, Dominic Kwiatkowski, Cordelia Langford, John Sillitoe on behalf of the Wellcome Sanger Institute COVID-19 Surveillance Team |
| EPI_ISL_3173640 | Lighthouse Lab in Alderley Park | Wellcome Sanger Institute for the COVID-19 Genomics UK (COG-UK) Consortium | Jacquelyn Wynn, Mairead Hyland, The Lighthouse Lab in Alderley Park and Alex Alderton, Roberto Amato, Jeffrey Barrett, Sonia Goncalves, Ewan Harrison, David K. Jackson, Ian Johnston, Dominic Kwiatkowski, Cordelia Langford, John Sillitoe on behalf of the Wellcome Sanger Institute COVID-19 Surveillance Team |
| EPI_ISL_3173641, EPI_ISL_3173642 | Lighthouse Lab in Glasgow | Wellcome Sanger Institute for the COVID-19 Genomics UK (COG-UK) Consortium | Harper VanSteenhouse, Yumi Kasai, David Gray, Carol Clugston, Anna Dominiczak and Alex Alderton, Roberto Amato, Jeffrey Barrett, Sonia Goncalves, Ewan Harrison, David K. Jackson, Ian Johnston, Dominic Kwiatkowski, Cordelia Langford, John Sillitoe on behalf of the Wellcome Sanger Institute COVID-19 Surveillance Team |
| EPI_ISL_3173643, EPI_ISL_3173644 | Lighthouse Laboratory Plymouth | Wellcome Sanger Institute for the COVID-19 Genomics UK (COG-UK) Consortium | Lighthouse Laboratory Plymouth and Alex Alderton, Roberto Amato, Jeffrey Barrett, Sonia Goncalves, Ewan Harrison, David K. Jackson, Ian Johnston, Dominic Kwiatkowski, Cordelia Langford, John Sillitoe on behalf of the Wellcome Sanger Institute COVID-19 Surveillance Team |
| EPI_ISL_3173645 | Lighthouse Lab in Alderley Park | Wellcome Sanger Institute for the COVID-19 Genomics UK (COG-UK) Consortium | Jacquelyn Wynn, Mairead Hyland, The Lighthouse Lab in Alderley Park and Alex Alderton, Roberto Amato, Jeffrey Barrett, Sonia Goncalves, Ewan Harrison, David K. Jackson, Ian Johnston, Dominic Kwiatkowski, Cordelia Langford, John Sillitoe on behalf of the Wellcome Sanger Institute COVID-19 Surveillance Team |
| EPI_ISL_3173646, EPI_ISL_3173647 | Lighthouse Lab in Glasgow | Wellcome Sanger Institute for the COVID-19 Genomics UK (COG-UK) Consortium | Harper VanSteenhouse, Yumi Kasai, David Gray, Carol Clugston, Anna Dominiczak and Alex Alderton, Roberto Amato, Jeffrey Barrett, Sonia Goncalves, Ewan Harrison, David K. Jackson, Ian Johnston, Dominic Kwiatkowski, Cordelia Langford, John Sillitoe on behalf of the Wellcome Sanger Institute COVID-19 Surveillance Team |
| EPI_ISL_3173648 | Lighthouse Laboratory Plymouth | Wellcome Sanger Institute for the COVID-19 Genomics UK (COG-UK) Consortium | Lighthouse Laboratory Plymouth and Alex Alderton, Roberto Amato, Jeffrey Barrett, Sonia Goncalves, Ewan Harrison, David K. Jackson, Ian Johnston, Dominic Kwiatkowski, Cordelia Langford, John Sillitoe on behalf of the Wellcome Sanger Institute COVID-19 Surveillance Team |
| EPI_ISL_3173649 | Lighthouse Lab in Glasgow | Wellcome Sanger Institute for the COVID-19 Genomics UK (COG-UK) Consortium | Harper VanSteenhouse, Yumi Kasai, David Gray, Carol Clugston, Anna Dominiczak and Alex Alderton, Roberto Amato, Jeffrey Barrett, Sonia Goncalves, Ewan Harrison, David K. Jackson, Ian Johnston, Dominic Kwiatkowski, Cordelia Langford, John Sillitoe on behalf of the Wellcome Sanger Institute COVID-19 Surveillance Team |
| EPI_ISL_3173650 | Lighthouse Laboratory Plymouth | Wellcome Sanger Institute for the COVID-19 Genomics UK (COG-UK) Consortium | Lighthouse Laboratory Plymouth and Alex Alderton, Roberto Amato, Jeffrey Barrett, Sonia Goncalves, Ewan Harrison, David K. Jackson, Ian Johnston, Dominic Kwiatkowski, Cordelia Langford, John Sillitoe on behalf of the Wellcome Sanger Institute COVID-19 Surveillance Team |
| EPI_ISL_3173651, EPI_ISL_3173652 | Lighthouse Lab in Glasgow | Wellcome Sanger Institute for the COVID-19 Genomics UK (COG-UK) Consortium | Harper VanSteenhouse, Yumi Kasai, David Gray, Carol Clugston, Anna Dominiczak and Alex Alderton, Roberto Amato, Jeffrey Barrett, Sonia Goncalves, Ewan Harrison, David K. Jackson, Ian Johnston, Dominic Kwiatkowski, Cordelia Langford, John Sillitoe on behalf of the Wellcome Sanger Institute COVID-19 Surveillance Team |
| EPI_ISL_3173653 | Lighthouse Laboratory Plymouth | Wellcome Sanger Institute for the COVID-19 Genomics UK (COG-UK) Consortium | Lighthouse Laboratory Plymouth and Alex Alderton, Roberto Amato, Jeffrey Barrett, Sonia Goncalves, Ewan Harrison, David K. Jackson, Ian Johnston, Dominic Kwiatkowski, Cordelia Langford, John Sillitoe on behalf of the Wellcome Sanger Institute COVID-19 Surveillance Team |
| EPI_ISL_3173654 | Lighthouse Lab in Alderley Park | Wellcome Sanger Institute for the COVID-19 Genomics UK (COG-UK) Consortium | Jacquelyn Wynn, Mairead Hyland, The Lighthouse Lab in Alderley Park and Alex Alderton, Roberto Amato, Jeffrey Barrett, Sonia Goncalves, Ewan Harrison, David K. Jackson, Ian Johnston, Dominic Kwiatkowski, Cordelia Langford, John Sillitoe on behalf of the Wellcome Sanger Institute COVID-19 Surveillance Team |
| EPI_ISL_3173655, EPI_ISL_3173656, EPI_ISL_3173657, EPI_ISL_3173658 | Lighthouse Lab in Glasgow | Wellcome Sanger Institute for the COVID-19 Genomics UK (COG-UK) Consortium | Harper VanSteenhouse, Yumi Kasai, David Gray, Carol Clugston, Anna Dominiczak and Alex Alderton, Roberto Amato, Jeffrey Barrett, Sonia Goncalves, Ewan Harrison, David K. Jackson, Ian Johnston, Dominic Kwiatkowski, Cordelia Langford, John Sillitoe on behalf of the Wellcome Sanger Institute COVID-19 Surveillance Team |
| EPI_ISL_3173659, EPI_ISL_3173660, EPI_ISL_3173661 | Lighthouse Laboratory Plymouth | Wellcome Sanger Institute for the COVID-19 Genomics UK (COG-UK) Consortium | Lighthouse Laboratory Plymouth and Alex Alderton, Roberto Amato, Jeffrey Barrett, Sonia Goncalves, Ewan Harrison, David K. Jackson, Ian Johnston, Dominic Kwiatkowski, Cordelia Langford, John Sillitoe on behalf of the Wellcome Sanger Institute COVID-19 Surveillance Team |
| EPI_ISL_3173662, EPI_ISL_3173663 | Lighthouse Lab in Glasgow | Wellcome Sanger Institute for the COVID-19 Genomics UK (COG-UK) Consortium | Harper VanSteenhouse, Yumi Kasai, David Gray, Carol Clugston, Anna Dominiczak and Alex Alderton, Roberto Amato, Jeffrey Barrett, Sonia Goncalves, Ewan Harrison, David K. Jackson, Ian Johnston, Dominic Kwiatkowski, Cordelia Langford, John Sillitoe on behalf of the Wellcome Sanger Institute COVID-19 Surveillance Team |
| EPI_ISL_3173664, EPI_ISL_3173665, EPI_ISL_3173666 | Lighthouse Lab in Alderley Park | Wellcome Sanger Institute for the COVID-19 Genomics UK (COG-UK) Consortium | Jacquelyn Wynn, Mairead Hyland, The Lighthouse Lab in Alderley Park and Alex Alderton, Roberto Amato, Jeffrey Barrett, Sonia Goncalves, Ewan Harrison, David K. Jackson, Ian Johnston, Dominic Kwiatkowski, Cordelia Langford, John Sillitoe on behalf of the Wellcome Sanger Institute COVID-19 Surveillance Team |
| EPI_ISL_3173667 | Lighthouse Laboratory Plymouth | Wellcome Sanger Institute for the COVID-19 Genomics UK (COG-UK) Consortium | Lighthouse Laboratory Plymouth and Alex Alderton, Roberto Amato, Jeffrey Barrett, Sonia Goncalves, Ewan Harrison, David K. Jackson, Ian Johnston, Dominic Kwiatkowski, Cordelia Langford, John Sillitoe on behalf of the Wellcome Sanger Institute COVID-19 Surveillance Team |
| EPI_ISL_3173668 | Lighthouse Lab in Alderley Park | Wellcome Sanger Institute for the COVID-19 Genomics UK (COG-UK) Consortium | Jacquelyn Wynn, Mairead Hyland, The Lighthouse Lab in Alderley Park and Alex Alderton, Roberto Amato, Jeffrey Barrett, Sonia Goncalves, Ewan Harrison, David K. Jackson, Ian Johnston, Dominic Kwiatkowski, Cordelia Langford, John Sillitoe on behalf of the Wellcome Sanger Institute COVID-19 Surveillance Team |
| EPI_ISL_3173669 | Lighthouse Laboratory Plymouth | Wellcome Sanger Institute for the COVID-19 Genomics UK (COG-UK) Consortium | Lighthouse Laboratory Plymouth and Alex Alderton, Roberto Amato, Jeffrey Barrett, Sonia Goncalves, Ewan Harrison, David K. Jackson, Ian Johnston, Dominic Kwiatkowski, Cordelia Langford, John Sillitoe on behalf of the Wellcome Sanger Institute COVID-19 Surveillance Team |
| EPI_ISL_3173670 | Lighthouse Lab in Alderley Park | Wellcome Sanger Institute for the COVID-19 Genomics UK (COG-UK) Consortium | Jacquelyn Wynn, Mairead Hyland, The Lighthouse Lab in Alderley Park and Alex Alderton, Roberto Amato, Jeffrey Barrett, Sonia Goncalves, Ewan Harrison, David K. Jackson, Ian Johnston, Dominic Kwiatkowski, Cordelia Langford, John Sillitoe on behalf of the Wellcome Sanger Institute COVID-19 Surveillance Team |
| EPI_ISL_3173671 | Lighthouse Laboratory Plymouth | Wellcome Sanger Institute for the COVID-19 Genomics UK (COG-UK) Consortium | Lighthouse Laboratory Plymouth and Alex Alderton, Roberto Amato, Jeffrey Barrett, Sonia Goncalves, Ewan Harrison, David K. Jackson, Ian Johnston, Dominic Kwiatkowski, Cordelia Langford, John Sillitoe on behalf of the Wellcome Sanger Institute COVID-19 Surveillance Team |
| EPI_ISL_3173672 | Lighthouse Lab in Glasgow | Wellcome Sanger Institute for the COVID-19 Genomics UK (COG-UK) Consortium | Harper VanSteenhouse, Yumi Kasai, David Gray, Carol Clugston, Anna Dominiczak and Alex Alderton, Roberto Amato, Jeffrey Barrett, Sonia Goncalves, Ewan Harrison, David K. Jackson, Ian Johnston, Dominic Kwiatkowski, Cordelia Langford, John Sillitoe on behalf of the Wellcome Sanger Institute COVID-19 Surveillance Team |
| EPI_ISL_3173673, EPI_ISL_3173674, EPI_ISL_3173675, EPI_ISL_3173676 | Lighthouse Lab in Alderley Park | Wellcome Sanger Institute for the COVID-19 Genomics UK (COG-UK) Consortium | Jacquelyn Wynn, Mairead Hyland, The Lighthouse Lab in Alderley Park and Alex Alderton, Roberto Amato, Jeffrey Barrett, Sonia Goncalves, Ewan Harrison, David K. Jackson, Ian Johnston, Dominic Kwiatkowski, Cordelia Langford, John Sillitoe on behalf of the Wellcome Sanger Institute COVID-19 Surveillance Team |
| EPI_ISL_3173677 | Lighthouse Laboratory Plymouth | Wellcome Sanger Institute for the COVID-19 Genomics UK (COG-UK) Consortium | Lighthouse Laboratory Plymouth and Alex Alderton, Roberto Amato, Jeffrey Barrett, Sonia Goncalves, Ewan Harrison, David K. Jackson, Ian Johnston, Dominic Kwiatkowski, Cordelia Langford, John Sillitoe on behalf of the Wellcome Sanger Institute COVID-19 Surveillance Team |
| EPI_ISL_3173678 | Lighthouse Lab in Alderley Park | Wellcome Sanger Institute for the COVID-19 Genomics UK (COG-UK) Consortium | Jacquelyn Wynn, Mairead Hyland, The Lighthouse Lab in Alderley Park and Alex Alderton, Roberto Amato, Jeffrey Barrett, Sonia Goncalves, Ewan Harrison, David K. Jackson, Ian Johnston, Dominic Kwiatkowski, Cordelia Langford, John Sillitoe on behalf of the Wellcome Sanger Institute COVID-19 Surveillance Team |
| EPI_ISL_3173679 | Lighthouse Laboratory Plymouth | Wellcome Sanger Institute for the COVID-19 Genomics UK (COG-UK) Consortium | Lighthouse Laboratory Plymouth and Alex Alderton, Roberto Amato, Jeffrey Barrett, Sonia Goncalves, Ewan Harrison, David K. Jackson, Ian Johnston, Dominic Kwiatkowski, Cordelia Langford, John Sillitoe on behalf of the Wellcome Sanger Institute COVID-19 Surveillance Team |
| EPI_ISL_3173680 | Lighthouse Lab in Alderley Park | Wellcome Sanger Institute for the COVID-19 Genomics UK (COG-UK) Consortium | Jacquelyn Wynn, Mairead Hyland, The Lighthouse Lab in Alderley Park and Alex Alderton, Roberto Amato, Jeffrey Barrett, Sonia Goncalves, Ewan Harrison, David K. Jackson, Ian Johnston, Dominic Kwiatkowski, Cordelia Langford, John Sillitoe on behalf of the Wellcome Sanger Institute COVID-19 Surveillance Team |
| EPI_ISL_3173681 | Lighthouse Laboratory Plymouth | Wellcome Sanger Institute for the COVID-19 Genomics UK (COG-UK) Consortium | Lighthouse Laboratory Plymouth and Alex Alderton, Roberto Amato, Jeffrey Barrett, Sonia Goncalves, Ewan Harrison, David K. Jackson, Ian Johnston, Dominic Kwiatkowski, Cordelia Langford, John Sillitoe on behalf of the Wellcome Sanger Institute COVID-19 Surveillance Team |
| EPI_ISL_3173682 | Lighthouse Lab in Alderley Park | Wellcome Sanger Institute for the COVID-19 Genomics UK (COG-UK) Consortium | Jacquelyn Wynn, Mairead Hyland, The Lighthouse Lab in Alderley Park and Alex Alderton, Roberto Amato, Jeffrey Barrett, Sonia Goncalves, Ewan Harrison, David K. Jackson, Ian Johnston, Dominic Kwiatkowski, Cordelia Langford, John Sillitoe on behalf of the Wellcome Sanger Institute COVID-19 Surveillance Team |
| EPI_ISL_3173683 | Lighthouse Lab in Glasgow | Wellcome Sanger Institute for the COVID-19 Genomics UK (COG-UK) Consortium | Harper VanSteenhouse, Yumi Kasai, David Gray, Carol Clugston, Anna Dominiczak and Alex Alderton, Roberto Amato, Jeffrey Barrett, Sonia Goncalves, Ewan Harrison, David K. Jackson, Ian Johnston, Dominic Kwiatkowski, Cordelia Langford, John Sillitoe on behalf of the Wellcome Sanger Institute COVID-19 Surveillance Team |
| EPI_ISL_3173684, EPI_ISL_3173685, EPI_ISL_3173686 | Lighthouse Laboratory Plymouth | Wellcome Sanger Institute for the COVID-19 Genomics UK (COG-UK) Consortium | Lighthouse Laboratory Plymouth and Alex Alderton, Roberto Amato, Jeffrey Barrett, Sonia Goncalves, Ewan Harrison, David K. Jackson, Ian Johnston, Dominic Kwiatkowski, Cordelia Langford, John Sillitoe on behalf of the Wellcome Sanger Institute COVID-19 Surveillance Team |
| EPI_ISL_3173687, EPI_ISL_3173688, EPI_ISL_3173689, EPI_ISL_3173690 | Lighthouse Lab in Alderley Park | Wellcome Sanger Institute for the COVID-19 Genomics UK (COG-UK) Consortium | Jacquelyn Wynn, Mairead Hyland, The Lighthouse Lab in Alderley Park and Alex Alderton, Roberto Amato, Jeffrey Barrett, Sonia Goncalves, Ewan Harrison, David K. Jackson, Ian Johnston, Dominic Kwiatkowski, Cordelia Langford, John Sillitoe on behalf of the Wellcome Sanger Institute COVID-19 Surveillance Team |

| | | | |
|---|---|---|---|
| EPI_ISL_3173691, EPI_ISL_3173692 | Lighthouse Lab in Glasgow | Wellcome Sanger Institute for the COVID-19 Genomics UK (COG-UK) Consortium | Harper VanSteenhouse, Yumi Kasai, David Gray, Carol Clugston, Anna Dominiczak and Alex Alderton, Roberto Amato, Jeffrey Barrett, Sonia Goncalves, Ewan Harrison, David K. Jackson, Ian Johnston, Dominic Kwiatkowski, Cordelia Langford, John Sillitoe on behalf of the Wellcome Sanger Institute COVID-19 Surveillance Team |
| EPI_ISL_3173693 | Lighthouse Lab in Alderley Park | Wellcome Sanger Institute for the COVID-19 Genomics UK (COG-UK) Consortium | Jacquelyn Wynn, Mairead Hyland, The Lighthouse Lab in Alderley Park and Alex Alderton, Roberto Amato, Jeffrey Barrett, Sonia Goncalves, Ewan Harrison, David K. Jackson, Ian Johnston, Dominic Kwiatkowski, Cordelia Langford, John Sillitoe on behalf of the Wellcome Sanger Institute COVID-19 Surveillance Team |
| EPI_ISL_3173694 | Lighthouse Laboratory Plymouth | Wellcome Sanger Institute for the COVID-19 Genomics UK (COG-UK) Consortium | Lighthouse Laboratory Plymouth and Alex Alderton, Roberto Amato, Jeffrey Barrett, Sonia Goncalves, Ewan Harrison, David K. Jackson, Ian Johnston, Dominic Kwiatkowski, Cordelia Langford, John Sillitoe on behalf of the Wellcome Sanger Institute COVID-19 Surveillance Team |
| EPI_ISL_3173695 | Lighthouse Lab in Alderley Park | Wellcome Sanger Institute for the COVID-19 Genomics UK (COG-UK) Consortium | Jacquelyn Wynn, Mairead Hyland, The Lighthouse Lab in Alderley Park and Alex Alderton, Roberto Amato, Jeffrey Barrett, Sonia Goncalves, Ewan Harrison, David K. Jackson, Ian Johnston, Dominic Kwiatkowski, Cordelia Langford, John Sillitoe on behalf of the Wellcome Sanger Institute COVID-19 Surveillance Team |
| EPI_ISL_3173696, EPI_ISL_3173697 | Lighthouse Lab in Glasgow | Wellcome Sanger Institute for the COVID-19 Genomics UK (COG-UK) Consortium | Harper VanSteenhouse, Yumi Kasai, David Gray, Carol Clugston, Anna Dominiczak and Alex Alderton, Roberto Amato, Jeffrey Barrett, Sonia Goncalves, Ewan Harrison, David K. Jackson, Ian Johnston, Dominic Kwiatkowski, Cordelia Langford, John Sillitoe on behalf of the Wellcome Sanger Institute COVID-19 Surveillance Team |
| EPI_ISL_3173698 | Lighthouse Lab in Alderley Park | Wellcome Sanger Institute for the COVID-19 Genomics UK (COG-UK) Consortium | Jacquelyn Wynn, Mairead Hyland, The Lighthouse Lab in Alderley Park and Alex Alderton, Roberto Amato, Jeffrey Barrett, Sonia Goncalves, Ewan Harrison, David K. Jackson, Ian Johnston, Dominic Kwiatkowski, Cordelia Langford, John Sillitoe on behalf of the Wellcome Sanger Institute COVID-19 Surveillance Team |
| EPI_ISL_3173699 | Lighthouse Laboratory Plymouth | Wellcome Sanger Institute for the COVID-19 Genomics UK (COG-UK) Consortium | Lighthouse Laboratory Plymouth and Alex Alderton, Roberto Amato, Jeffrey Barrett, Sonia Goncalves, Ewan Harrison, David K. Jackson, Ian Johnston, Dominic Kwiatkowski, Cordelia Langford, John Sillitoe on behalf of the Wellcome Sanger Institute COVID-19 Surveillance Team |
| EPI_ISL_3173700 | Lighthouse Lab in Glasgow | Wellcome Sanger Institute for the COVID-19 Genomics UK (COG-UK) Consortium | Harper VanSteenhouse, Yumi Kasai, David Gray, Carol Clugston, Anna Dominiczak and Alex Alderton, Roberto Amato, Jeffrey Barrett, Sonia Goncalves, Ewan Harrison, David K. Jackson, Ian Johnston, Dominic Kwiatkowski, Cordelia Langford, John Sillitoe on behalf of the Wellcome Sanger Institute COVID-19 Surveillance Team |
| EPI_ISL_3173701 | Lighthouse Laboratory Plymouth | Wellcome Sanger Institute for the COVID-19 Genomics UK (COG-UK) Consortium | Lighthouse Laboratory Plymouth and Alex Alderton, Roberto Amato, Jeffrey Barrett, Sonia Goncalves, Ewan Harrison, David K. Jackson, Ian Johnston, Dominic Kwiatkowski, Cordelia Langford, John Sillitoe on behalf of the Wellcome Sanger Institute COVID-19 Surveillance Team |
| EPI_ISL_3173702, EPI_ISL_3173703 | Lighthouse Lab in Alderley Park | Wellcome Sanger Institute for the COVID-19 Genomics UK (COG-UK) Consortium | Jacquelyn Wynn, Mairead Hyland, The Lighthouse Lab in Alderley Park and Alex Alderton, Roberto Amato, Jeffrey Barrett, Sonia Goncalves, Ewan Harrison, David K. Jackson, Ian Johnston, Dominic Kwiatkowski, Cordelia Langford, John Sillitoe on behalf of the Wellcome Sanger Institute COVID-19 Surveillance Team |
| EPI_ISL_3173704 | Lighthouse Lab in Glasgow | Wellcome Sanger Institute for the COVID-19 Genomics UK (COG-UK) Consortium | Harper VanSteenhouse, Yumi Kasai, David Gray, Carol Clugston, Anna Dominiczak and Alex Alderton, Roberto Amato, Jeffrey Barrett, Sonia Goncalves, Ewan Harrison, David K. Jackson, Ian Johnston, Dominic Kwiatkowski, Cordelia Langford, John Sillitoe on behalf of the Wellcome Sanger Institute COVID-19 Surveillance Team |
| EPI_ISL_3173705 | Lighthouse Lab in Alderley Park | Wellcome Sanger Institute for the COVID-19 Genomics UK (COG-UK) Consortium | Jacquelyn Wynn, Mairead Hyland, The Lighthouse Lab in Alderley Park and Alex Alderton, Roberto Amato, Jeffrey Barrett, Sonia Goncalves, Ewan Harrison, David K. Jackson, Ian Johnston, Dominic Kwiatkowski, Cordelia Langford, John Sillitoe on behalf of the Wellcome Sanger Institute COVID-19 Surveillance Team |
| EPI_ISL_3173706 | Lighthouse Laboratory Plymouth | Wellcome Sanger Institute for the COVID-19 Genomics UK (COG-UK) Consortium | Lighthouse Laboratory Plymouth and Alex Alderton, Roberto Amato, Jeffrey Barrett, Sonia Goncalves, Ewan Harrison, David K. Jackson, Ian Johnston, Dominic Kwiatkowski, Cordelia Langford, John Sillitoe on behalf of the Wellcome Sanger Institute COVID-19 Surveillance Team |
| EPI_ISL_3173707, EPI_ISL_3173708, EPI_ISL_3173709, EPI_ISL_3173710 | Lighthouse Lab in Alderley Park | Wellcome Sanger Institute for the COVID-19 Genomics UK (COG-UK) Consortium | Jacquelyn Wynn, Mairead Hyland, The Lighthouse Lab in Alderley Park and Alex Alderton, Roberto Amato, Jeffrey Barrett, Sonia Goncalves, Ewan Harrison, David K. Jackson, Ian Johnston, Dominic Kwiatkowski, Cordelia Langford, John Sillitoe on behalf of the Wellcome Sanger Institute COVID-19 Surveillance Team |
| EPI_ISL_3173711 | Lighthouse Lab in Glasgow | Wellcome Sanger Institute for the COVID-19 Genomics UK (COG-UK) Consortium | Harper VanSteenhouse, Yumi Kasai, David Gray, Carol Clugston, Anna Dominiczak and Alex Alderton, Roberto Amato, Jeffrey Barrett, Sonia Goncalves, Ewan Harrison, David K. Jackson, Ian Johnston, Dominic Kwiatkowski, Cordelia Langford, John Sillitoe on behalf of the Wellcome Sanger Institute COVID-19 Surveillance Team |
| EPI_ISL_3173712 | Lighthouse Laboratory Plymouth | Wellcome Sanger Institute for the COVID-19 Genomics UK (COG-UK) Consortium | Lighthouse Laboratory Plymouth and Alex Alderton, Roberto Amato, Jeffrey Barrett, Sonia Goncalves, Ewan Harrison, David K. Jackson, Ian Johnston, Dominic Kwiatkowski, Cordelia Langford, John Sillitoe on behalf of the Wellcome Sanger Institute COVID-19 Surveillance Team |
| EPI_ISL_3173713, EPI_ISL_3173714 | Lighthouse Lab in Glasgow | Wellcome Sanger Institute for the COVID-19 Genomics UK (COG-UK) Consortium | Harper VanSteenhouse, Yumi Kasai, David Gray, Carol Clugston, Anna Dominiczak and Alex Alderton, Roberto Amato, Jeffrey Barrett, Sonia Goncalves, Ewan Harrison, David K. Jackson, Ian Johnston, Dominic Kwiatkowski, Cordelia Langford, John Sillitoe on behalf of the Wellcome Sanger Institute COVID-19 Surveillance Team |
| EPI_ISL_3173715 | Lighthouse Laboratory Plymouth | Wellcome Sanger Institute for the COVID-19 Genomics UK (COG-UK) Consortium | Lighthouse Laboratory Plymouth and Alex Alderton, Roberto Amato, Jeffrey Barrett, Sonia Goncalves, Ewan Harrison, David K. Jackson, Ian Johnston, Dominic Kwiatkowski, Cordelia Langford, John Sillitoe on behalf of the Wellcome Sanger Institute COVID-19 Surveillance Team |
| EPI_ISL_3173716 | Lighthouse Lab in Glasgow | Wellcome Sanger Institute for the COVID-19 Genomics UK (COG-UK) Consortium | Harper VanSteenhouse, Yumi Kasai, David Gray, Carol Clugston, Anna Dominiczak and Alex Alderton, Roberto Amato, Jeffrey Barrett, Sonia Goncalves, Ewan Harrison, David K. Jackson, Ian Johnston, Dominic Kwiatkowski, Cordelia Langford, John Sillitoe on behalf of the Wellcome Sanger Institute COVID-19 Surveillance Team |
| EPI_ISL_3173717 | Lighthouse Laboratory Plymouth | Wellcome Sanger Institute for the COVID-19 Genomics UK (COG-UK) Consortium | Lighthouse Laboratory Plymouth and Alex Alderton, Roberto Amato, Jeffrey Barrett, Sonia Goncalves, Ewan Harrison, David K. Jackson, Ian Johnston, Dominic Kwiatkowski, Cordelia Langford, John Sillitoe on behalf of the Wellcome Sanger Institute COVID-19 Surveillance Team |
| EPI_ISL_3173718 | Lighthouse Lab in Alderley Park | Wellcome Sanger Institute for the COVID-19 Genomics UK (COG-UK) Consortium | Jacquelyn Wynn, Mairead Hyland, The Lighthouse Lab in Alderley Park and Alex Alderton, Roberto Amato, Jeffrey Barrett, Sonia Goncalves, Ewan Harrison, David K. Jackson, Ian Johnston, Dominic Kwiatkowski, Cordelia Langford, John Sillitoe on behalf of the Wellcome Sanger Institute COVID-19 Surveillance Team |
| EPI_ISL_3173719 | Lighthouse Lab in Glasgow | Wellcome Sanger Institute for the COVID-19 Genomics UK (COG-UK) Consortium | Harper VanSteenhouse, Yumi Kasai, David Gray, Carol Clugston, Anna Dominiczak and Alex Alderton, Roberto Amato, Jeffrey Barrett, Sonia Goncalves, Ewan Harrison, David K. Jackson, Ian Johnston, Dominic Kwiatkowski, Cordelia Langford, John Sillitoe on behalf of the Wellcome Sanger Institute COVID-19 Surveillance Team |
| EPI_ISL_3173720, EPI_ISL_3173721 | Lighthouse Laboratory Plymouth | Wellcome Sanger Institute for the COVID-19 Genomics UK (COG-UK) Consortium | Lighthouse Laboratory Plymouth and Alex Alderton, Roberto Amato, Jeffrey Barrett, Sonia Goncalves, Ewan Harrison, David K. Jackson, Ian Johnston, Dominic Kwiatkowski, Cordelia Langford, John Sillitoe on behalf of the Wellcome Sanger Institute COVID-19 Surveillance Team |
| EPI_ISL_3173722 | Lighthouse Lab in Glasgow | Wellcome Sanger Institute for the COVID-19 Genomics UK (COG-UK) Consortium | Harper VanSteenhouse, Yumi Kasai, David Gray, Carol Clugston, Anna Dominiczak and Alex Alderton, Roberto Amato, Jeffrey Barrett, Sonia Goncalves, Ewan Harrison, David K. Jackson, Ian Johnston, Dominic Kwiatkowski, Cordelia Langford, John Sillitoe on behalf of the Wellcome Sanger Institute COVID-19 Surveillance Team |
| EPI_ISL_3173723 | Lighthouse Lab in Alderley Park | Wellcome Sanger Institute for the COVID-19 Genomics UK (COG-UK) Consortium | Jacquelyn Wynn, Mairead Hyland, The Lighthouse Lab in Alderley Park and Alex Alderton, Roberto Amato, Jeffrey Barrett, Sonia Goncalves, Ewan Harrison, David K. Jackson, Ian Johnston, Dominic Kwiatkowski, Cordelia Langford, John Sillitoe on behalf of the Wellcome Sanger Institute COVID-19 Surveillance Team |
| EPI_ISL_3173724 | Lighthouse Lab in Glasgow | Wellcome Sanger Institute for the COVID-19 Genomics UK (COG-UK) Consortium | Harper VanSteenhouse, Yumi Kasai, David Gray, Carol Clugston, Anna Dominiczak and Alex Alderton, Roberto Amato, Jeffrey Barrett, Sonia Goncalves, Ewan Harrison, David K. Jackson, Ian Johnston, Dominic Kwiatkowski, Cordelia Langford, John Sillitoe on behalf of the Wellcome Sanger Institute COVID-19 Surveillance Team |
| EPI_ISL_3173725 | Lighthouse Lab in Alderley Park | Wellcome Sanger Institute for the COVID-19 Genomics UK (COG-UK) Consortium | Jacquelyn Wynn, Mairead Hyland, The Lighthouse Lab in Alderley Park and Alex Alderton, Roberto Amato, Jeffrey Barrett, Sonia Goncalves, Ewan Harrison, David K. Jackson, Ian Johnston, Dominic Kwiatkowski, Cordelia Langford, John Sillitoe on behalf of the Wellcome Sanger Institute COVID-19 Surveillance Team |
| EPI_ISL_3173726, EPI_ISL_3173727, EPI_ISL_3173728 | Lighthouse Laboratory Plymouth | Wellcome Sanger Institute for the COVID-19 Genomics UK (COG-UK) Consortium | Lighthouse Laboratory Plymouth and Alex Alderton, Roberto Amato, Jeffrey Barrett, Sonia Goncalves, Ewan Harrison, David K. Jackson, Ian Johnston, Dominic Kwiatkowski, Cordelia Langford, John Sillitoe on behalf of the Wellcome Sanger Institute COVID-19 Surveillance Team |
| EPI_ISL_3173729, EPI_ISL_3173730 | Lighthouse Lab in Alderley Park | Wellcome Sanger Institute for the COVID-19 Genomics UK (COG-UK) Consortium | Jacquelyn Wynn, Mairead Hyland, The Lighthouse Lab in Alderley Park and Alex Alderton, Roberto Amato, Jeffrey Barrett, Sonia Goncalves, Ewan Harrison, David K. Jackson, Ian Johnston, Dominic Kwiatkowski, Cordelia Langford, John Sillitoe on behalf of the Wellcome Sanger Institute COVID-19 Surveillance Team |
| EPI_ISL_3173731 | Lighthouse Laboratory Plymouth | Wellcome Sanger Institute for the COVID-19 Genomics UK (COG-UK) Consortium | Lighthouse Laboratory Plymouth and Alex Alderton, Roberto Amato, Jeffrey Barrett, Sonia Goncalves, Ewan Harrison, David K. Jackson, Ian Johnston, Dominic Kwiatkowski, Cordelia Langford, John Sillitoe on behalf of the Wellcome Sanger Institute COVID-19 Surveillance Team |
| EPI_ISL_3173732, EPI_ISL_3173733 | Lighthouse Lab in Glasgow | Wellcome Sanger Institute for the COVID-19 Genomics UK (COG-UK) Consortium | Harper VanSteenhouse, Yumi Kasai, David Gray, Carol Clugston, Anna Dominiczak and Alex Alderton, Roberto Amato, Jeffrey Barrett, Sonia Goncalves, Ewan Harrison, David K. Jackson, Ian Johnston, Dominic Kwiatkowski, Cordelia Langford, John Sillitoe on behalf of the Wellcome Sanger Institute COVID-19 Surveillance Team |
| EPI_ISL_3173734 | Lighthouse Lab in Alderley Park | Wellcome Sanger Institute for the COVID-19 Genomics UK (COG-UK) Consortium | Jacquelyn Wynn, Mairead Hyland, The Lighthouse Lab in Alderley Park and Alex Alderton, Roberto Amato, Jeffrey Barrett, Sonia Goncalves, Ewan Harrison, David K. Jackson, Ian Johnston, Dominic Kwiatkowski, Cordelia Langford, John Sillitoe on behalf of the Wellcome Sanger Institute COVID-19 Surveillance Team |
| EPI_ISL_3173735 | Lighthouse Laboratory Plymouth | Wellcome Sanger Institute for the COVID-19 Genomics UK (COG-UK) Consortium | Lighthouse Laboratory Plymouth and Alex Alderton, Roberto Amato, Jeffrey Barrett, Sonia Goncalves, Ewan Harrison, David K. Jackson, Ian Johnston, Dominic Kwiatkowski, Cordelia Langford, John Sillitoe on behalf of the Wellcome Sanger Institute COVID-19 Surveillance Team |
| EPI_ISL_3173736, EPI_ISL_3173737 | Lighthouse Lab in Alderley Park | Wellcome Sanger Institute for the COVID-19 Genomics UK (COG-UK) Consortium | Jacquelyn Wynn, Mairead Hyland, The Lighthouse Lab in Alderley Park and Alex Alderton, Roberto Amato, Jeffrey Barrett, Sonia Goncalves, Ewan Harrison, David K. Jackson, Ian Johnston, Dominic Kwiatkowski, Cordelia Langford, John Sillitoe on behalf of the Wellcome Sanger Institute COVID-19 Surveillance Team |
| EPI_ISL_3173738 | Lighthouse Lab in Glasgow | Wellcome Sanger Institute for the COVID-19 Genomics UK (COG-UK) Consortium | Harper VanSteenhouse, Yumi Kasai, David Gray, Carol Clugston, Anna Dominiczak and Alex Alderton, Roberto Amato, Jeffrey Barrett, Sonia Goncalves, Ewan Harrison, David K. Jackson, Ian Johnston, Dominic Kwiatkowski, Cordelia Langford, John Sillitoe on behalf of the Wellcome Sanger Institute COVID-19 Surveillance Team |
| EPI_ISL_3173739 | Lighthouse Laboratory Plymouth | Wellcome Sanger Institute for the COVID-19 Genomics UK (COG-UK) Consortium | Lighthouse Laboratory Plymouth and Alex Alderton, Roberto Amato, Jeffrey Barrett, Sonia Goncalves, Ewan Harrison, David K. Jackson, Ian Johnston, Dominic Kwiatkowski, Cordelia Langford, John Sillitoe on behalf of the Wellcome Sanger Institute COVID-19 Surveillance Team |
| EPI_ISL_3173740 | Lighthouse Lab in Alderley Park | Wellcome Sanger Institute for the COVID-19 Genomics UK (COG-UK) Consortium | Jacquelyn Wynn, Mairead Hyland, The Lighthouse Lab in Alderley Park and Alex Alderton, Roberto Amato, Jeffrey Barrett, Sonia Goncalves, Ewan Harrison, David K. Jackson, Ian Johnston, Dominic Kwiatkowski, Cordelia Langford, John Sillitoe on behalf of the Wellcome Sanger Institute COVID-19 Surveillance Team |
| EPI_ISL_3173741 | Lighthouse Laboratory Plymouth | Wellcome Sanger Institute for the COVID-19 Genomics UK (COG-UK) Consortium | Lighthouse Laboratory Plymouth and Alex Alderton, Roberto Amato, Jeffrey Barrett, Sonia Goncalves, Ewan Harrison, David K. Jackson, Ian Johnston, Dominic Kwiatkowski, Cordelia Langford, John Sillitoe on behalf of the Wellcome Sanger Institute COVID-19 Surveillance Team |
| EPI_ISL_3173742 | Lighthouse Lab in Glasgow | Wellcome Sanger Institute for the COVID-19 Genomics UK (COG-UK) Consortium | Harper VanSteenhouse, Yumi Kasai, David Gray, Carol Clugston, Anna Dominiczak and Alex Alderton, Roberto Amato, Jeffrey Barrett, Sonia Goncalves, Ewan Harrison, David K. Jackson, Ian Johnston, Dominic Kwiatkowski, Cordelia Langford, John Sillitoe on behalf of the Wellcome Sanger Institute COVID-19 Surveillance Team |
| EPI_ISL_3173743 | Lighthouse Lab in Alderley Park | Wellcome Sanger Institute for the COVID-19 Genomics UK (COG-UK) Consortium | Jacquelyn Wynn, Mairead Hyland, The Lighthouse Lab in Alderley Park and Alex Alderton, Roberto Amato, Jeffrey Barrett, Sonia Goncalves, Ewan Harrison, David K. Jackson, Ian Johnston, Dominic Kwiatkowski, Cordelia Langford, John Sillitoe on behalf of the Wellcome Sanger Institute COVID-19 Surveillance Team |
| EPI_ISL_3173744 | Lighthouse Lab in Glasgow | Wellcome Sanger Institute for the COVID-19 Genomics UK (COG-UK) Consortium | Harper VanSteenhouse, Yumi Kasai, David Gray, Carol Clugston, Anna Dominiczak and Alex Alderton, Roberto Amato, Jeffrey Barrett, Sonia Goncalves, Ewan Harrison, David K. Jackson, Ian Johnston, Dominic Kwiatkowski, Cordelia Langford, John Sillitoe on behalf of the Wellcome Sanger Institute COVID-19 Surveillance Team |
| EPI_ISL_3173745, EPI_ISL_3173746 | Lighthouse Lab in Alderley Park | Wellcome Sanger Institute for the COVID-19 Genomics UK (COG-UK) Consortium | Jacquelyn Wynn, Mairead Hyland, The Lighthouse Lab in Alderley Park and Alex Alderton, Roberto Amato, Jeffrey Barrett, Sonia Goncalves, Ewan Harrison, David K. Jackson, Ian Johnston, Dominic Kwiatkowski, Cordelia Langford, John Sillitoe on behalf of the Wellcome Sanger Institute COVID-19 Surveillance Team |
| EPI_ISL_3173747, EPI_ISL_3173748, EPI_ISL_3173749 | Lighthouse Lab in Glasgow | Wellcome Sanger Institute for the COVID-19 Genomics UK (COG-UK) Consortium | Harper VanSteenhouse, Yumi Kasai, David Gray, Carol Clugston, Anna Dominiczak and Alex Alderton, Roberto Amato, Jeffrey Barrett, Sonia Goncalves, Ewan Harrison, David K. Jackson, Ian Johnston, Dominic Kwiatkowski, Cordelia Langford, John Sillitoe on behalf of the Wellcome Sanger Institute COVID-19 Surveillance Team |
| EPI_ISL_3173750, | Lighthouse Lab in Alderley Park | Wellcome Sanger Institute for the COVID-19 Genomics UK (COG-UK) | Jacquelyn Wynn, Mairead Hyland, The Lighthouse Lab in Alderley Park and Alex Alderton, Roberto Amato, Jeffrey Barrett, Sonia Goncalves, Ewan Harrison, David K. Jackson, Ian Johnston, Dominic Kwiatkowski, Cordelia Langford, John Sillitoe on |

| | | | |
|---|---|---|---|
| EPI_ISL_3173751 | | Consortium | behalf of the Wellcome Sanger Institute COVID-19 Surveillance Team |
| EPI_ISL_3173752 | Lighthouse Laboratory Plymouth | Wellcome Sanger Institute for the COVID-19 Genomics UK (COG-UK) Consortium | Lighthouse Laboratory Plymouth and Alex Alderton, Roberto Amato, Jeffrey Barrett, Sonia Goncalves, Ewan Harrison, David K. Jackson, Ian Johnston, Dominic Kwiatkowski, Cordelia Langford, John Sillitoe on behalf of the Wellcome Sanger Institute COVID-19 Surveillance Team |
| EPI_ISL_3173753 | Lighthouse Lab in Glasgow | Wellcome Sanger Institute for the COVID-19 Genomics UK (COG-UK) Consortium | Harper VanSteenhouse, Yumi Kasai, David Gray, Carol Clugston, Anna Dominiczak and Alex Alderton, Roberto Amato, Jeffrey Barrett, Sonia Goncalves, Ewan Harrison, David K. Jackson, Ian Johnston, Dominic Kwiatkowski, Cordelia Langford, John Sillitoe on behalf of the Wellcome Sanger Institute COVID-19 Surveillance Team |
| EPI_ISL_3173754 | Lighthouse Laboratory Plymouth | Wellcome Sanger Institute for the COVID-19 Genomics UK (COG-UK) Consortium | Lighthouse Laboratory Plymouth and Alex Alderton, Roberto Amato, Jeffrey Barrett, Sonia Goncalves, Ewan Harrison, David K. Jackson, Ian Johnston, Dominic Kwiatkowski, Cordelia Langford, John Sillitoe on behalf of the Wellcome Sanger Institute COVID-19 Surveillance Team |
| EPI_ISL_3173755 | Lighthouse Lab in Glasgow | Wellcome Sanger Institute for the COVID-19 Genomics UK (COG-UK) Consortium | Harper VanSteenhouse, Yumi Kasai, David Gray, Carol Clugston, Anna Dominiczak and Alex Alderton, Roberto Amato, Jeffrey Barrett, Sonia Goncalves, Ewan Harrison, David K. Jackson, Ian Johnston, Dominic Kwiatkowski, Cordelia Langford, John Sillitoe on behalf of the Wellcome Sanger Institute COVID-19 Surveillance Team |
| EPI_ISL_3173756, EPI_ISL_3173757, EPI_ISL_3173758 | Lighthouse Lab in Alderley Park | Wellcome Sanger Institute for the COVID-19 Genomics UK (COG-UK) Consortium | Jacquelyn Wynn, Mairead Hyland, The Lighthouse Lab in Alderley Park and Alex Alderton, Roberto Amato, Jeffrey Barrett, Sonia Goncalves, Ewan Harrison, David K. Jackson, Ian Johnston, Dominic Kwiatkowski, Cordelia Langford, John Sillitoe on behalf of the Wellcome Sanger Institute COVID-19 Surveillance Team |
| EPI_ISL_3173759 | Lighthouse Lab in Glasgow | Wellcome Sanger Institute for the COVID-19 Genomics UK (COG-UK) Consortium | Harper VanSteenhouse, Yumi Kasai, David Gray, Carol Clugston, Anna Dominiczak and Alex Alderton, Roberto Amato, Jeffrey Barrett, Sonia Goncalves, Ewan Harrison, David K. Jackson, Ian Johnston, Dominic Kwiatkowski, Cordelia Langford, John Sillitoe on behalf of the Wellcome Sanger Institute COVID-19 Surveillance Team |
| EPI_ISL_3173760 | Lighthouse Laboratory Plymouth | Wellcome Sanger Institute for the COVID-19 Genomics UK (COG-UK) Consortium | Lighthouse Laboratory Plymouth and Alex Alderton, Roberto Amato, Jeffrey Barrett, Sonia Goncalves, Ewan Harrison, David K. Jackson, Ian Johnston, Dominic Kwiatkowski, Cordelia Langford, John Sillitoe on behalf of the Wellcome Sanger Institute COVID-19 Surveillance Team |
| EPI_ISL_3173761 | Lighthouse Lab in Glasgow | Wellcome Sanger Institute for the COVID-19 Genomics UK (COG-UK) Consortium | Harper VanSteenhouse, Yumi Kasai, David Gray, Carol Clugston, Anna Dominiczak and Alex Alderton, Roberto Amato, Jeffrey Barrett, Sonia Goncalves, Ewan Harrison, David K. Jackson, Ian Johnston, Dominic Kwiatkowski, Cordelia Langford, John Sillitoe on behalf of the Wellcome Sanger Institute COVID-19 Surveillance Team |
| EPI_ISL_3173762 | Lighthouse Lab in Alderley Park | Wellcome Sanger Institute for the COVID-19 Genomics UK (COG-UK) Consortium | Jacquelyn Wynn, Mairead Hyland, The Lighthouse Lab in Alderley Park and Alex Alderton, Roberto Amato, Jeffrey Barrett, Sonia Goncalves, Ewan Harrison, David K. Jackson, Ian Johnston, Dominic Kwiatkowski, Cordelia Langford, John Sillitoe on behalf of the Wellcome Sanger Institute COVID-19 Surveillance Team |
| EPI_ISL_3173763 | Lighthouse Lab in Glasgow | Wellcome Sanger Institute for the COVID-19 Genomics UK (COG-UK) Consortium | Harper VanSteenhouse, Yumi Kasai, David Gray, Carol Clugston, Anna Dominiczak and Alex Alderton, Roberto Amato, Jeffrey Barrett, Sonia Goncalves, Ewan Harrison, David K. Jackson, Ian Johnston, Dominic Kwiatkowski, Cordelia Langford, John Sillitoe on behalf of the Wellcome Sanger Institute COVID-19 Surveillance Team |
| EPI_ISL_3173764 | Lighthouse Lab in Alderley Park | Wellcome Sanger Institute for the COVID-19 Genomics UK (COG-UK) Consortium | Jacquelyn Wynn, Mairead Hyland, The Lighthouse Lab in Alderley Park and Alex Alderton, Roberto Amato, Jeffrey Barrett, Sonia Goncalves, Ewan Harrison, David K. Jackson, Ian Johnston, Dominic Kwiatkowski, Cordelia Langford, John Sillitoe on behalf of the Wellcome Sanger Institute COVID-19 Surveillance Team |
| EPI_ISL_3173765 | Lighthouse Lab in Glasgow | Wellcome Sanger Institute for the COVID-19 Genomics UK (COG-UK) Consortium | Harper VanSteenhouse, Yumi Kasai, David Gray, Carol Clugston, Anna Dominiczak and Alex Alderton, Roberto Amato, Jeffrey Barrett, Sonia Goncalves, Ewan Harrison, David K. Jackson, Ian Johnston, Dominic Kwiatkowski, Cordelia Langford, John Sillitoe on behalf of the Wellcome Sanger Institute COVID-19 Surveillance Team |
| EPI_ISL_3173766 | Lighthouse Laboratory Plymouth | Wellcome Sanger Institute for the COVID-19 Genomics UK (COG-UK) Consortium | Lighthouse Laboratory Plymouth and Alex Alderton, Roberto Amato, Jeffrey Barrett, Sonia Goncalves, Ewan Harrison, David K. Jackson, Ian Johnston, Dominic Kwiatkowski, Cordelia Langford, John Sillitoe on behalf of the Wellcome Sanger Institute COVID-19 Surveillance Team |
| EPI_ISL_3173767 | Lighthouse Lab in Alderley Park | Wellcome Sanger Institute for the COVID-19 Genomics UK (COG-UK) Consortium | Jacquelyn Wynn, Mairead Hyland, The Lighthouse Lab in Alderley Park and Alex Alderton, Roberto Amato, Jeffrey Barrett, Sonia Goncalves, Ewan Harrison, David K. Jackson, Ian Johnston, Dominic Kwiatkowski, Cordelia Langford, John Sillitoe on behalf of the Wellcome Sanger Institute COVID-19 Surveillance Team |
| EPI_ISL_3173768 | Lighthouse Lab in Glasgow | Wellcome Sanger Institute for the COVID-19 Genomics UK (COG-UK) Consortium | Harper VanSteenhouse, Yumi Kasai, David Gray, Carol Clugston, Anna Dominiczak and Alex Alderton, Roberto Amato, Jeffrey Barrett, Sonia Goncalves, Ewan Harrison, David K. Jackson, Ian Johnston, Dominic Kwiatkowski, Cordelia Langford, John Sillitoe on behalf of the Wellcome Sanger Institute COVID-19 Surveillance Team |
| EPI_ISL_3173769, EPI_ISL_3173770 | Lighthouse Lab in Alderley Park | Wellcome Sanger Institute for the COVID-19 Genomics UK (COG-UK) Consortium | Jacquelyn Wynn, Mairead Hyland, The Lighthouse Lab in Alderley Park and Alex Alderton, Roberto Amato, Jeffrey Barrett, Sonia Goncalves, Ewan Harrison, David K. Jackson, Ian Johnston, Dominic Kwiatkowski, Cordelia Langford, John Sillitoe on behalf of the Wellcome Sanger Institute COVID-19 Surveillance Team |
| EPI_ISL_3173771, EPI_ISL_3173772, EPI_ISL_3173773 | Lighthouse Lab in Glasgow | Wellcome Sanger Institute for the COVID-19 Genomics UK (COG-UK) Consortium | Harper VanSteenhouse, Yumi Kasai, David Gray, Carol Clugston, Anna Dominiczak and Alex Alderton, Roberto Amato, Jeffrey Barrett, Sonia Goncalves, Ewan Harrison, David K. Jackson, Ian Johnston, Dominic Kwiatkowski, Cordelia Langford, John Sillitoe on behalf of the Wellcome Sanger Institute COVID-19 Surveillance Team |
| EPI_ISL_3173774 | Lighthouse Laboratory Plymouth | Wellcome Sanger Institute for the COVID-19 Genomics UK (COG-UK) Consortium | Lighthouse Laboratory Plymouth and Alex Alderton, Roberto Amato, Jeffrey Barrett, Sonia Goncalves, Ewan Harrison, David K. Jackson, Ian Johnston, Dominic Kwiatkowski, Cordelia Langford, John Sillitoe on behalf of the Wellcome Sanger Institute COVID-19 Surveillance Team |
| EPI_ISL_3173775, EPI_ISL_3173776 | Lighthouse Lab in Alderley Park | Wellcome Sanger Institute for the COVID-19 Genomics UK (COG-UK) Consortium | Jacquelyn Wynn, Mairead Hyland, The Lighthouse Lab in Alderley Park and Alex Alderton, Roberto Amato, Jeffrey Barrett, Sonia Goncalves, Ewan Harrison, David K. Jackson, Ian Johnston, Dominic Kwiatkowski, Cordelia Langford, John Sillitoe on behalf of the Wellcome Sanger Institute COVID-19 Surveillance Team |
| EPI_ISL_3173777, EPI_ISL_3173778 | Lighthouse Lab in Glasgow | Wellcome Sanger Institute for the COVID-19 Genomics UK (COG-UK) Consortium | Harper VanSteenhouse, Yumi Kasai, David Gray, Carol Clugston, Anna Dominiczak and Alex Alderton, Roberto Amato, Jeffrey Barrett, Sonia Goncalves, Ewan Harrison, David K. Jackson, Ian Johnston, Dominic Kwiatkowski, Cordelia Langford, John Sillitoe on behalf of the Wellcome Sanger Institute COVID-19 Surveillance Team |
| EPI_ISL_3173779 | Lighthouse Laboratory Plymouth | Wellcome Sanger Institute for the COVID-19 Genomics UK (COG-UK) Consortium | Lighthouse Laboratory Plymouth and Alex Alderton, Roberto Amato, Jeffrey Barrett, Sonia Goncalves, Ewan Harrison, David K. Jackson, Ian Johnston, Dominic Kwiatkowski, Cordelia Langford, John Sillitoe on behalf of the Wellcome Sanger Institute COVID-19 Surveillance Team |
| EPI_ISL_3173780 | Lighthouse Lab in Alderley Park | Wellcome Sanger Institute for the COVID-19 Genomics UK (COG-UK) Consortium | Jacquelyn Wynn, Mairead Hyland, The Lighthouse Lab in Alderley Park and Alex Alderton, Roberto Amato, Jeffrey Barrett, Sonia Goncalves, Ewan Harrison, David K. Jackson, Ian Johnston, Dominic Kwiatkowski, Cordelia Langford, John Sillitoe on behalf of the Wellcome Sanger Institute COVID-19 Surveillance Team |
| EPI_ISL_3173781 | Lighthouse Laboratory Plymouth | Wellcome Sanger Institute for the COVID-19 Genomics UK (COG-UK) Consortium | Lighthouse Laboratory Plymouth and Alex Alderton, Roberto Amato, Jeffrey Barrett, Sonia Goncalves, Ewan Harrison, David K. Jackson, Ian Johnston, Dominic Kwiatkowski, Cordelia Langford, John Sillitoe on behalf of the Wellcome Sanger Institute COVID-19 Surveillance Team |
| EPI_ISL_3173782 | Lighthouse Lab in Glasgow | Wellcome Sanger Institute for the COVID-19 Genomics UK (COG-UK) Consortium | Harper VanSteenhouse, Yumi Kasai, David Gray, Carol Clugston, Anna Dominiczak and Alex Alderton, Roberto Amato, Jeffrey Barrett, Sonia Goncalves, Ewan Harrison, David K. Jackson, Ian Johnston, Dominic Kwiatkowski, Cordelia Langford, John Sillitoe on behalf of the Wellcome Sanger Institute COVID-19 Surveillance Team |
| EPI_ISL_3173783 | Lighthouse Lab in Alderley Park | Wellcome Sanger Institute for the COVID-19 Genomics UK (COG-UK) Consortium | Jacquelyn Wynn, Mairead Hyland, The Lighthouse Lab in Alderley Park and Alex Alderton, Roberto Amato, Jeffrey Barrett, Sonia Goncalves, Ewan Harrison, David K. Jackson, Ian Johnston, Dominic Kwiatkowski, Cordelia Langford, John Sillitoe on behalf of the Wellcome Sanger Institute COVID-19 Surveillance Team |
| EPI_ISL_3173784 | Lighthouse Laboratory Plymouth | Wellcome Sanger Institute for the COVID-19 Genomics UK (COG-UK) Consortium | Lighthouse Laboratory Plymouth and Alex Alderton, Roberto Amato, Jeffrey Barrett, Sonia Goncalves, Ewan Harrison, David K. Jackson, Ian Johnston, Dominic Kwiatkowski, Cordelia Langford, John Sillitoe on behalf of the Wellcome Sanger Institute COVID-19 Surveillance Team |
| EPI_ISL_3173785 | Lighthouse Lab in Glasgow | Wellcome Sanger Institute for the COVID-19 Genomics UK (COG-UK) Consortium | Harper VanSteenhouse, Yumi Kasai, David Gray, Carol Clugston, Anna Dominiczak and Alex Alderton, Roberto Amato, Jeffrey Barrett, Sonia Goncalves, Ewan Harrison, David K. Jackson, Ian Johnston, Dominic Kwiatkowski, Cordelia Langford, John Sillitoe on behalf of the Wellcome Sanger Institute COVID-19 Surveillance Team |
| EPI_ISL_3173786 | Lighthouse Lab in Alderley Park | Wellcome Sanger Institute for the COVID-19 Genomics UK (COG-UK) Consortium | Jacquelyn Wynn, Mairead Hyland, The Lighthouse Lab in Alderley Park and Alex Alderton, Roberto Amato, Jeffrey Barrett, Sonia Goncalves, Ewan Harrison, David K. Jackson, Ian Johnston, Dominic Kwiatkowski, Cordelia Langford, John Sillitoe on behalf of the Wellcome Sanger Institute COVID-19 Surveillance Team |
| EPI_ISL_3173787 | Lighthouse Lab in Glasgow | Wellcome Sanger Institute for the COVID-19 Genomics UK (COG-UK) Consortium | Harper VanSteenhouse, Yumi Kasai, David Gray, Carol Clugston, Anna Dominiczak and Alex Alderton, Roberto Amato, Jeffrey Barrett, Sonia Goncalves, Ewan Harrison, David K. Jackson, Ian Johnston, Dominic Kwiatkowski, Cordelia Langford, John Sillitoe on behalf of the Wellcome Sanger Institute COVID-19 Surveillance Team |
| EPI_ISL_3173788 | Lighthouse Lab in Alderley Park | Wellcome Sanger Institute for the COVID-19 Genomics UK (COG-UK) Consortium | Jacquelyn Wynn, Mairead Hyland, The Lighthouse Lab in Alderley Park and Alex Alderton, Roberto Amato, Jeffrey Barrett, Sonia Goncalves, Ewan Harrison, David K. Jackson, Ian Johnston, Dominic Kwiatkowski, Cordelia Langford, John Sillitoe on behalf of the Wellcome Sanger Institute COVID-19 Surveillance Team |
| EPI_ISL_3173789, EPI_ISL_3173790 | Lighthouse Lab in Glasgow | Wellcome Sanger Institute for the COVID-19 Genomics UK (COG-UK) Consortium | Harper VanSteenhouse, Yumi Kasai, David Gray, Carol Clugston, Anna Dominiczak and Alex Alderton, Roberto Amato, Jeffrey Barrett, Sonia Goncalves, Ewan Harrison, David K. Jackson, Ian Johnston, Dominic Kwiatkowski, Cordelia Langford, John Sillitoe on behalf of the Wellcome Sanger Institute COVID-19 Surveillance Team |
| EPI_ISL_3173791, EPI_ISL_3173792 | Lighthouse Laboratory Plymouth | Wellcome Sanger Institute for the COVID-19 Genomics UK (COG-UK) Consortium | Lighthouse Laboratory Plymouth and Alex Alderton, Roberto Amato, Jeffrey Barrett, Sonia Goncalves, Ewan Harrison, David K. Jackson, Ian Johnston, Dominic Kwiatkowski, Cordelia Langford, John Sillitoe on behalf of the Wellcome Sanger Institute COVID-19 Surveillance Team |
| EPI_ISL_3173793, EPI_ISL_3173794 | Lighthouse Lab in Glasgow | Wellcome Sanger Institute for the COVID-19 Genomics UK (COG-UK) Consortium | Harper VanSteenhouse, Yumi Kasai, David Gray, Carol Clugston, Anna Dominiczak and Alex Alderton, Roberto Amato, Jeffrey Barrett, Sonia Goncalves, Ewan Harrison, David K. Jackson, Ian Johnston, Dominic Kwiatkowski, Cordelia Langford, John Sillitoe on behalf of the Wellcome Sanger Institute COVID-19 Surveillance Team |
| EPI_ISL_3173795, EPI_ISL_3173796 | Lighthouse Lab in Alderley Park | Wellcome Sanger Institute for the COVID-19 Genomics UK (COG-UK) Consortium | Jacquelyn Wynn, Mairead Hyland, The Lighthouse Lab in Alderley Park and Alex Alderton, Roberto Amato, Jeffrey Barrett, Sonia Goncalves, Ewan Harrison, David K. Jackson, Ian Johnston, Dominic Kwiatkowski, Cordelia Langford, John Sillitoe on behalf of the Wellcome Sanger Institute COVID-19 Surveillance Team |
| EPI_ISL_3173797, EPI_ISL_3173798, EPI_ISL_3173799 | Lighthouse Lab in Glasgow | Wellcome Sanger Institute for the COVID-19 Genomics UK (COG-UK) Consortium | Harper VanSteenhouse, Yumi Kasai, David Gray, Carol Clugston, Anna Dominiczak and Alex Alderton, Roberto Amato, Jeffrey Barrett, Sonia Goncalves, Ewan Harrison, David K. Jackson, Ian Johnston, Dominic Kwiatkowski, Cordelia Langford, John Sillitoe on behalf of the Wellcome Sanger Institute COVID-19 Surveillance Team |
| EPI_ISL_3173800, EPI_ISL_3173801, EPI_ISL_3173802 | Lighthouse Laboratory Plymouth | Wellcome Sanger Institute for the COVID-19 Genomics UK (COG-UK) Consortium | Lighthouse Laboratory Plymouth and Alex Alderton, Roberto Amato, Jeffrey Barrett, Sonia Goncalves, Ewan Harrison, David K. Jackson, Ian Johnston, Dominic Kwiatkowski, Cordelia Langford, John Sillitoe on behalf of the Wellcome Sanger Institute COVID-19 Surveillance Team |
| EPI_ISL_3173803 | Lighthouse Lab in Glasgow | Wellcome Sanger Institute for the COVID-19 Genomics UK (COG-UK) Consortium | Harper VanSteenhouse, Yumi Kasai, David Gray, Carol Clugston, Anna Dominiczak and Alex Alderton, Roberto Amato, Jeffrey Barrett, Sonia Goncalves, Ewan Harrison, David K. Jackson, Ian Johnston, Dominic Kwiatkowski, Cordelia Langford, John Sillitoe on behalf of the Wellcome Sanger Institute COVID-19 Surveillance Team |
| EPI_ISL_3173804 | Lighthouse Lab in Alderley Park | Wellcome Sanger Institute for the COVID-19 Genomics UK (COG-UK) Consortium | Jacquelyn Wynn, Mairead Hyland, The Lighthouse Lab in Alderley Park and Alex Alderton, Roberto Amato, Jeffrey Barrett, Sonia Goncalves, Ewan Harrison, David K. Jackson, Ian Johnston, Dominic Kwiatkowski, Cordelia Langford, John Sillitoe on behalf of the Wellcome Sanger Institute COVID-19 Surveillance Team |
| EPI_ISL_3173805 | Lighthouse Laboratory Plymouth | Wellcome Sanger Institute for the COVID-19 Genomics UK (COG-UK) Consortium | Lighthouse Laboratory Plymouth and Alex Alderton, Roberto Amato, Jeffrey Barrett, Sonia Goncalves, Ewan Harrison, David K. Jackson, Ian Johnston, Dominic Kwiatkowski, Cordelia Langford, John Sillitoe on behalf of the Wellcome Sanger Institute COVID-19 Surveillance Team |
| EPI_ISL_3173806 | Lighthouse Lab in Alderley Park | Wellcome Sanger Institute for the COVID-19 Genomics UK (COG-UK) Consortium | Jacquelyn Wynn, Mairead Hyland, The Lighthouse Lab in Alderley Park and Alex Alderton, Roberto Amato, Jeffrey Barrett, Sonia Goncalves, Ewan Harrison, David K. Jackson, Ian Johnston, Dominic Kwiatkowski, Cordelia Langford, John Sillitoe on behalf of the Wellcome Sanger Institute COVID-19 Surveillance Team |
| EPI_ISL_3173807 | Lighthouse Laboratory Plymouth | Wellcome Sanger Institute for the COVID-19 Genomics UK (COG-UK) Consortium | Lighthouse Laboratory Plymouth and Alex Alderton, Roberto Amato, Jeffrey Barrett, Sonia Goncalves, Ewan Harrison, David K. Jackson, Ian Johnston, Dominic Kwiatkowski, Cordelia Langford, John Sillitoe on behalf of the Wellcome Sanger Institute COVID-19 Surveillance Team |
| EPI_ISL_3173808, EPI_ISL_3173809 | Lighthouse Lab in Alderley Park | Wellcome Sanger Institute for the COVID-19 Genomics UK (COG-UK) Consortium | Jacquelyn Wynn, Mairead Hyland, The Lighthouse Lab in Alderley Park and Alex Alderton, Roberto Amato, Jeffrey Barrett, Sonia Goncalves, Ewan Harrison, David K. Jackson, Ian Johnston, Dominic Kwiatkowski, Cordelia Langford, John Sillitoe on behalf of the Wellcome Sanger Institute COVID-19 Surveillance Team |
| EPI_ISL_3173810 | Lighthouse Lab in Glasgow | Wellcome Sanger Institute for the COVID-19 Genomics UK (COG-UK) | Harper VanSteenhouse, Yumi Kasai, David Gray, Carol Clugston, Anna Dominiczak and Alex Alderton, Roberto Amato, Jeffrey Barrett, Sonia Goncalves, Ewan Harrison, David K. Jackson, Ian Johnston, Dominic Kwiatkowski, Cordelia Langford, |

| | | | |
|---|---|---|---|
| | | Consortium | John Sillitoe on behalf of the Wellcome Sanger Institute COVID-19 Surveillance Team |
| EPI_ISL_3173811 | Lighthouse Lab in Alderley Park | Wellcome Sanger Institute for the COVID-19 Genomics UK (COG-UK) Consortium | Jacquelyn Wynn, Mairead Hyland, The Lighthouse Lab in Alderley Park and Alex Alderton, Roberto Amato, Jeffrey Barrett, Sonia Goncalves, Ewan Harrison, David K. Jackson, Ian Johnston, Dominic Kwiatkowski, Cordelia Langford, John Sillitoe on behalf of the Wellcome Sanger Institute COVID-19 Surveillance Team |
| EPI_ISL_3173812, EPI_ISL_3173813 | Lighthouse Lab in Glasgow | Wellcome Sanger Institute for the COVID-19 Genomics UK (COG-UK) Consortium | Harper VanSteenhouse, Yumi Kasai, David Gray, Carol Clugston, Anna Dominiczak and Alex Alderton, Roberto Amato, Jeffrey Barrett, Sonia Goncalves, Ewan Harrison, David K. Jackson, Ian Johnston, Dominic Kwiatkowski, Cordelia Langford, John Sillitoe on behalf of the Wellcome Sanger Institute COVID-19 Surveillance Team |
| EPI_ISL_3173814, EPI_ISL_3173815 | Lighthouse Lab in Alderley Park | Wellcome Sanger Institute for the COVID-19 Genomics UK (COG-UK) Consortium | Jacquelyn Wynn, Mairead Hyland, The Lighthouse Lab in Alderley Park and Alex Alderton, Roberto Amato, Jeffrey Barrett, Sonia Goncalves, Ewan Harrison, David K. Jackson, Ian Johnston, Dominic Kwiatkowski, Cordelia Langford, John Sillitoe on behalf of the Wellcome Sanger Institute COVID-19 Surveillance Team |
| EPI_ISL_3173816 | Lighthouse Lab in Glasgow | Wellcome Sanger Institute for the COVID-19 Genomics UK (COG-UK) Consortium | Harper VanSteenhouse, Yumi Kasai, David Gray, Carol Clugston, Anna Dominiczak and Alex Alderton, Roberto Amato, Jeffrey Barrett, Sonia Goncalves, Ewan Harrison, David K. Jackson, Ian Johnston, Dominic Kwiatkowski, Cordelia Langford, John Sillitoe on behalf of the Wellcome Sanger Institute COVID-19 Surveillance Team |
| EPI_ISL_3173817 | Lighthouse Laboratory Plymouth | Wellcome Sanger Institute for the COVID-19 Genomics UK (COG-UK) Consortium | Lighthouse Laboratory Plymouth and Alex Alderton, Roberto Amato, Jeffrey Barrett, Sonia Goncalves, Ewan Harrison, David K. Jackson, Ian Johnston, Dominic Kwiatkowski, Cordelia Langford, John Sillitoe on behalf of the Wellcome Sanger Institute COVID-19 Surveillance Team |
| EPI_ISL_3173818, EPI_ISL_3173819 | Lighthouse Lab in Alderley Park | Wellcome Sanger Institute for the COVID-19 Genomics UK (COG-UK) Consortium | Jacquelyn Wynn, Mairead Hyland, The Lighthouse Lab in Alderley Park and Alex Alderton, Roberto Amato, Jeffrey Barrett, Sonia Goncalves, Ewan Harrison, David K. Jackson, Ian Johnston, Dominic Kwiatkowski, Cordelia Langford, John Sillitoe on behalf of the Wellcome Sanger Institute COVID-19 Surveillance Team |
| EPI_ISL_3173820, EPI_ISL_3173821, EPI_ISL_3173822, EPI_ISL_3173823 | Lighthouse Laboratory Plymouth | Wellcome Sanger Institute for the COVID-19 Genomics UK (COG-UK) Consortium | Lighthouse Laboratory Plymouth and Alex Alderton, Roberto Amato, Jeffrey Barrett, Sonia Goncalves, Ewan Harrison, David K. Jackson, Ian Johnston, Dominic Kwiatkowski, Cordelia Langford, John Sillitoe on behalf of the Wellcome Sanger Institute COVID-19 Surveillance Team |
| EPI_ISL_3173824, EPI_ISL_3173825 | Lighthouse Lab in Glasgow | Wellcome Sanger Institute for the COVID-19 Genomics UK (COG-UK) Consortium | Harper VanSteenhouse, Yumi Kasai, David Gray, Carol Clugston, Anna Dominiczak and Alex Alderton, Roberto Amato, Jeffrey Barrett, Sonia Goncalves, Ewan Harrison, David K. Jackson, Ian Johnston, Dominic Kwiatkowski, Cordelia Langford, John Sillitoe on behalf of the Wellcome Sanger Institute COVID-19 Surveillance Team |
| EPI_ISL_3173826 | Lighthouse Lab in Alderley Park | Wellcome Sanger Institute for the COVID-19 Genomics UK (COG-UK) Consortium | Jacquelyn Wynn, Mairead Hyland, The Lighthouse Lab in Alderley Park and Alex Alderton, Roberto Amato, Jeffrey Barrett, Sonia Goncalves, Ewan Harrison, David K. Jackson, Ian Johnston, Dominic Kwiatkowski, Cordelia Langford, John Sillitoe on behalf of the Wellcome Sanger Institute COVID-19 Surveillance Team |
| EPI_ISL_3173827, EPI_ISL_3173828 | Lighthouse Lab in Glasgow | Wellcome Sanger Institute for the COVID-19 Genomics UK (COG-UK) Consortium | Harper VanSteenhouse, Yumi Kasai, David Gray, Carol Clugston, Anna Dominiczak and Alex Alderton, Roberto Amato, Jeffrey Barrett, Sonia Goncalves, Ewan Harrison, David K. Jackson, Ian Johnston, Dominic Kwiatkowski, Cordelia Langford, John Sillitoe on behalf of the Wellcome Sanger Institute COVID-19 Surveillance Team |
| EPI_ISL_3173829 | Lighthouse Laboratory Plymouth | Wellcome Sanger Institute for the COVID-19 Genomics UK (COG-UK) Consortium | Lighthouse Laboratory Plymouth and Alex Alderton, Roberto Amato, Jeffrey Barrett, Sonia Goncalves, Ewan Harrison, David K. Jackson, Ian Johnston, Dominic Kwiatkowski, Cordelia Langford, John Sillitoe on behalf of the Wellcome Sanger Institute COVID-19 Surveillance Team |
| EPI_ISL_3173830 | Lighthouse Lab in Glasgow | Wellcome Sanger Institute for the COVID-19 Genomics UK (COG-UK) Consortium | Harper VanSteenhouse, Yumi Kasai, David Gray, Carol Clugston, Anna Dominiczak and Alex Alderton, Roberto Amato, Jeffrey Barrett, Sonia Goncalves, Ewan Harrison, David K. Jackson, Ian Johnston, Dominic Kwiatkowski, Cordelia Langford, John Sillitoe on behalf of the Wellcome Sanger Institute COVID-19 Surveillance Team |
| EPI_ISL_3173831, EPI_ISL_3173832 | Lighthouse Laboratory Plymouth | Wellcome Sanger Institute for the COVID-19 Genomics UK (COG-UK) Consortium | Lighthouse Laboratory Plymouth and Alex Alderton, Roberto Amato, Jeffrey Barrett, Sonia Goncalves, Ewan Harrison, David K. Jackson, Ian Johnston, Dominic Kwiatkowski, Cordelia Langford, John Sillitoe on behalf of the Wellcome Sanger Institute COVID-19 Surveillance Team |
| EPI_ISL_3173833 | Lighthouse Lab in Alderley Park | Wellcome Sanger Institute for the COVID-19 Genomics UK (COG-UK) Consortium | Jacquelyn Wynn, Mairead Hyland, The Lighthouse Lab in Alderley Park and Alex Alderton, Roberto Amato, Jeffrey Barrett, Sonia Goncalves, Ewan Harrison, David K. Jackson, Ian Johnston, Dominic Kwiatkowski, Cordelia Langford, John Sillitoe on behalf of the Wellcome Sanger Institute COVID-19 Surveillance Team |
| EPI_ISL_3173834, EPI_ISL_3173835, EPI_ISL_3173836 | Lighthouse Lab in Glasgow | Wellcome Sanger Institute for the COVID-19 Genomics UK (COG-UK) Consortium | Harper VanSteenhouse, Yumi Kasai, David Gray, Carol Clugston, Anna Dominiczak and Alex Alderton, Roberto Amato, Jeffrey Barrett, Sonia Goncalves, Ewan Harrison, David K. Jackson, Ian Johnston, Dominic Kwiatkowski, Cordelia Langford, John Sillitoe on behalf of the Wellcome Sanger Institute COVID-19 Surveillance Team |
| EPI_ISL_3173837 | Lighthouse Lab in Alderley Park | Wellcome Sanger Institute for the COVID-19 Genomics UK (COG-UK) Consortium | Jacquelyn Wynn, Mairead Hyland, The Lighthouse Lab in Alderley Park and Alex Alderton, Roberto Amato, Jeffrey Barrett, Sonia Goncalves, Ewan Harrison, David K. Jackson, Ian Johnston, Dominic Kwiatkowski, Cordelia Langford, John Sillitoe on behalf of the Wellcome Sanger Institute COVID-19 Surveillance Team |
| EPI_ISL_3173838 | Lighthouse Lab in Glasgow | Wellcome Sanger Institute for the COVID-19 Genomics UK (COG-UK) Consortium | Harper VanSteenhouse, Yumi Kasai, David Gray, Carol Clugston, Anna Dominiczak and Alex Alderton, Roberto Amato, Jeffrey Barrett, Sonia Goncalves, Ewan Harrison, David K. Jackson, Ian Johnston, Dominic Kwiatkowski, Cordelia Langford, John Sillitoe on behalf of the Wellcome Sanger Institute COVID-19 Surveillance Team |
| EPI_ISL_3173839 | Lighthouse Laboratory Plymouth | Wellcome Sanger Institute for the COVID-19 Genomics UK (COG-UK) Consortium | Lighthouse Laboratory Plymouth and Alex Alderton, Roberto Amato, Jeffrey Barrett, Sonia Goncalves, Ewan Harrison, David K. Jackson, Ian Johnston, Dominic Kwiatkowski, Cordelia Langford, John Sillitoe on behalf of the Wellcome Sanger Institute COVID-19 Surveillance Team |
| EPI_ISL_3173840, EPI_ISL_3173841 | Lighthouse Lab in Glasgow | Wellcome Sanger Institute for the COVID-19 Genomics UK (COG-UK) Consortium | Harper VanSteenhouse, Yumi Kasai, David Gray, Carol Clugston, Anna Dominiczak and Alex Alderton, Roberto Amato, Jeffrey Barrett, Sonia Goncalves, Ewan Harrison, David K. Jackson, Ian Johnston, Dominic Kwiatkowski, Cordelia Langford, John Sillitoe on behalf of the Wellcome Sanger Institute COVID-19 Surveillance Team |
| EPI_ISL_3173842, EPI_ISL_3173843 | Lighthouse Laboratory Plymouth | Wellcome Sanger Institute for the COVID-19 Genomics UK (COG-UK) Consortium | Lighthouse Laboratory Plymouth and Alex Alderton, Roberto Amato, Jeffrey Barrett, Sonia Goncalves, Ewan Harrison, David K. Jackson, Ian Johnston, Dominic Kwiatkowski, Cordelia Langford, John Sillitoe on behalf of the Wellcome Sanger Institute COVID-19 Surveillance Team |
| EPI_ISL_3173844 | Lighthouse Lab in Alderley Park | Wellcome Sanger Institute for the COVID-19 Genomics UK (COG-UK) Consortium | Jacquelyn Wynn, Mairead Hyland, The Lighthouse Lab in Alderley Park and Alex Alderton, Roberto Amato, Jeffrey Barrett, Sonia Goncalves, Ewan Harrison, David K. Jackson, Ian Johnston, Dominic Kwiatkowski, Cordelia Langford, John Sillitoe on behalf of the Wellcome Sanger Institute COVID-19 Surveillance Team |
| EPI_ISL_3173845, EPI_ISL_3173846, EPI_ISL_3173847, EPI_ISL_3173848 | Lighthouse Lab in Glasgow | Wellcome Sanger Institute for the COVID-19 Genomics UK (COG-UK) Consortium | Harper VanSteenhouse, Yumi Kasai, David Gray, Carol Clugston, Anna Dominiczak and Alex Alderton, Roberto Amato, Jeffrey Barrett, Sonia Goncalves, Ewan Harrison, David K. Jackson, Ian Johnston, Dominic Kwiatkowski, Cordelia Langford, John Sillitoe on behalf of the Wellcome Sanger Institute COVID-19 Surveillance Team |
| EPI_ISL_3173849 | Lighthouse Laboratory Plymouth | Wellcome Sanger Institute for the COVID-19 Genomics UK (COG-UK) Consortium | Lighthouse Laboratory Plymouth and Alex Alderton, Roberto Amato, Jeffrey Barrett, Sonia Goncalves, Ewan Harrison, David K. Jackson, Ian Johnston, Dominic Kwiatkowski, Cordelia Langford, John Sillitoe on behalf of the Wellcome Sanger Institute COVID-19 Surveillance Team |
| EPI_ISL_3173850 | Lighthouse Lab in Alderley Park | Wellcome Sanger Institute for the COVID-19 Genomics UK (COG-UK) Consortium | Jacquelyn Wynn, Mairead Hyland, The Lighthouse Lab in Alderley Park and Alex Alderton, Roberto Amato, Jeffrey Barrett, Sonia Goncalves, Ewan Harrison, David K. Jackson, Ian Johnston, Dominic Kwiatkowski, Cordelia Langford, John Sillitoe on behalf of the Wellcome Sanger Institute COVID-19 Surveillance Team |
| EPI_ISL_3173851 | Lighthouse Laboratory Plymouth | Wellcome Sanger Institute for the COVID-19 Genomics UK (COG-UK) Consortium | Lighthouse Laboratory Plymouth and Alex Alderton, Roberto Amato, Jeffrey Barrett, Sonia Goncalves, Ewan Harrison, David K. Jackson, Ian Johnston, Dominic Kwiatkowski, Cordelia Langford, John Sillitoe on behalf of the Wellcome Sanger Institute COVID-19 Surveillance Team |
| EPI_ISL_3173852, EPI_ISL_3173853 | Lighthouse Lab in Alderley Park | Wellcome Sanger Institute for the COVID-19 Genomics UK (COG-UK) Consortium | Jacquelyn Wynn, Mairead Hyland, The Lighthouse Lab in Alderley Park and Alex Alderton, Roberto Amato, Jeffrey Barrett, Sonia Goncalves, Ewan Harrison, David K. Jackson, Ian Johnston, Dominic Kwiatkowski, Cordelia Langford, John Sillitoe on behalf of the Wellcome Sanger Institute COVID-19 Surveillance Team |
| EPI_ISL_3173854 | Lighthouse Lab in Glasgow | Wellcome Sanger Institute for the COVID-19 Genomics UK (COG-UK) Consortium | Harper VanSteenhouse, Yumi Kasai, David Gray, Carol Clugston, Anna Dominiczak and Alex Alderton, Roberto Amato, Jeffrey Barrett, Sonia Goncalves, Ewan Harrison, David K. Jackson, Ian Johnston, Dominic Kwiatkowski, Cordelia Langford, John Sillitoe on behalf of the Wellcome Sanger Institute COVID-19 Surveillance Team |
| EPI_ISL_3173855, EPI_ISL_3173856 | Lighthouse Laboratory Plymouth | Wellcome Sanger Institute for the COVID-19 Genomics UK (COG-UK) Consortium | Lighthouse Laboratory Plymouth and Alex Alderton, Roberto Amato, Jeffrey Barrett, Sonia Goncalves, Ewan Harrison, David K. Jackson, Ian Johnston, Dominic Kwiatkowski, Cordelia Langford, John Sillitoe on behalf of the Wellcome Sanger Institute COVID-19 Surveillance Team |
| EPI_ISL_3173857, EPI_ISL_3173858 | Lighthouse Lab in Glasgow | Wellcome Sanger Institute for the COVID-19 Genomics UK (COG-UK) Consortium | Harper VanSteenhouse, Yumi Kasai, David Gray, Carol Clugston, Anna Dominiczak and Alex Alderton, Roberto Amato, Jeffrey Barrett, Sonia Goncalves, Ewan Harrison, David K. Jackson, Ian Johnston, Dominic Kwiatkowski, Cordelia Langford, John Sillitoe on behalf of the Wellcome Sanger Institute COVID-19 Surveillance Team |
| EPI_ISL_3173859, EPI_ISL_3173860 | Lighthouse Laboratory Plymouth | Wellcome Sanger Institute for the COVID-19 Genomics UK (COG-UK) Consortium | Lighthouse Laboratory Plymouth and Alex Alderton, Roberto Amato, Jeffrey Barrett, Sonia Goncalves, Ewan Harrison, David K. Jackson, Ian Johnston, Dominic Kwiatkowski, Cordelia Langford, John Sillitoe on behalf of the Wellcome Sanger Institute COVID-19 Surveillance Team |
| EPI_ISL_3173861 | Lighthouse Lab in Glasgow | Wellcome Sanger Institute for the COVID-19 Genomics UK (COG-UK) Consortium | Harper VanSteenhouse, Yumi Kasai, David Gray, Carol Clugston, Anna Dominiczak and Alex Alderton, Roberto Amato, Jeffrey Barrett, Sonia Goncalves, Ewan Harrison, David K. Jackson, Ian Johnston, Dominic Kwiatkowski, Cordelia Langford, John Sillitoe on behalf of the Wellcome Sanger Institute COVID-19 Surveillance Team |
| EPI_ISL_3173862 | Lighthouse Lab in Alderley Park | Wellcome Sanger Institute for the COVID-19 Genomics UK (COG-UK) Consortium | Jacquelyn Wynn, Mairead Hyland, The Lighthouse Lab in Alderley Park and Alex Alderton, Roberto Amato, Jeffrey Barrett, Sonia Goncalves, Ewan Harrison, David K. Jackson, Ian Johnston, Dominic Kwiatkowski, Cordelia Langford, John Sillitoe on behalf of the Wellcome Sanger Institute COVID-19 Surveillance Team |
| EPI_ISL_3173863 | Lighthouse Laboratory Plymouth | Wellcome Sanger Institute for the COVID-19 Genomics UK (COG-UK) Consortium | Lighthouse Laboratory Plymouth and Alex Alderton, Roberto Amato, Jeffrey Barrett, Sonia Goncalves, Ewan Harrison, David K. Jackson, Ian Johnston, Dominic Kwiatkowski, Cordelia Langford, John Sillitoe on behalf of the Wellcome Sanger Institute COVID-19 Surveillance Team |
| EPI_ISL_3173864, EPI_ISL_3173865, EPI_ISL_3173866 | Lighthouse Lab in Alderley Park | Wellcome Sanger Institute for the COVID-19 Genomics UK (COG-UK) Consortium | Jacquelyn Wynn, Mairead Hyland, The Lighthouse Lab in Alderley Park and Alex Alderton, Roberto Amato, Jeffrey Barrett, Sonia Goncalves, Ewan Harrison, David K. Jackson, Ian Johnston, Dominic Kwiatkowski, Cordelia Langford, John Sillitoe on behalf of the Wellcome Sanger Institute COVID-19 Surveillance Team |
| EPI_ISL_3173867 | Lighthouse Laboratory Plymouth | Wellcome Sanger Institute for the COVID-19 Genomics UK (COG-UK) Consortium | Lighthouse Laboratory Plymouth and Alex Alderton, Roberto Amato, Jeffrey Barrett, Sonia Goncalves, Ewan Harrison, David K. Jackson, Ian Johnston, Dominic Kwiatkowski, Cordelia Langford, John Sillitoe on behalf of the Wellcome Sanger Institute COVID-19 Surveillance Team |
| EPI_ISL_3173868 | Lighthouse Lab in Alderley Park | Wellcome Sanger Institute for the COVID-19 Genomics UK (COG-UK) Consortium | Jacquelyn Wynn, Mairead Hyland, The Lighthouse Lab in Alderley Park and Alex Alderton, Roberto Amato, Jeffrey Barrett, Sonia Goncalves, Ewan Harrison, David K. Jackson, Ian Johnston, Dominic Kwiatkowski, Cordelia Langford, John Sillitoe on behalf of the Wellcome Sanger Institute COVID-19 Surveillance Team |
| EPI_ISL_3173869, EPI_ISL_3173870, EPI_ISL_3173871, EPI_ISL_3173872 | Lighthouse Lab in Glasgow | Wellcome Sanger Institute for the COVID-19 Genomics UK (COG-UK) Consortium | Harper VanSteenhouse, Yumi Kasai, David Gray, Carol Clugston, Anna Dominiczak and Alex Alderton, Roberto Amato, Jeffrey Barrett, Sonia Goncalves, Ewan Harrison, David K. Jackson, Ian Johnston, Dominic Kwiatkowski, Cordelia Langford, John Sillitoe on behalf of the Wellcome Sanger Institute COVID-19 Surveillance Team |
| EPI_ISL_3173873 | Lighthouse Lab in Alderley Park | Wellcome Sanger Institute for the COVID-19 Genomics UK (COG-UK) Consortium | Jacquelyn Wynn, Mairead Hyland, The Lighthouse Lab in Alderley Park and Alex Alderton, Roberto Amato, Jeffrey Barrett, Sonia Goncalves, Ewan Harrison, David K. Jackson, Ian Johnston, Dominic Kwiatkowski, Cordelia Langford, John Sillitoe on behalf of the Wellcome Sanger Institute COVID-19 Surveillance Team |
| EPI_ISL_3173874 | Lighthouse Laboratory Plymouth | Wellcome Sanger Institute for the COVID-19 Genomics UK (COG-UK) Consortium | Lighthouse Laboratory Plymouth and Alex Alderton, Roberto Amato, Jeffrey Barrett, Sonia Goncalves, Ewan Harrison, David K. Jackson, Ian Johnston, Dominic Kwiatkowski, Cordelia Langford, John Sillitoe on behalf of the Wellcome Sanger Institute COVID-19 Surveillance Team |
| EPI_ISL_3173875, EPI_ISL_3173876 | Lighthouse Lab in Glasgow | Wellcome Sanger Institute for the COVID-19 Genomics UK (COG-UK) Consortium | Harper VanSteenhouse, Yumi Kasai, David Gray, Carol Clugston, Anna Dominiczak and Alex Alderton, Roberto Amato, Jeffrey Barrett, Sonia Goncalves, Ewan Harrison, David K. Jackson, Ian Johnston, Dominic Kwiatkowski, Cordelia Langford, John Sillitoe on behalf of the Wellcome Sanger Institute COVID-19 Surveillance Team |

EPI_ISL_3173877, EPI_ISL_3173878, EPI_ISL_3173879, EPI_ISL_3173880, EPI_ISL_3173881, EPI_ISL_3173882, EPI_ISL_3173883, EPI_ISL_3173884, EPI_ISL_3173885, EPI_ISL_3173886, EPI_ISL_3173887, EPI_ISL_3173888, EPI_ISL_3173889, EPI_ISL_3173890, EPI_ISL_3173891, EPI_ISL_3173892, EPI_ISL_3173893, EPI_ISL_3173894, EPI_ISL_3173895, EPI_ISL_3173896, EPI_ISL_3173897, EPI_ISL_3173898, EPI_ISL_3173899, EPI_ISL_3173900, EPI_ISL_3173901, EPI_ISL_3173902, EPI_ISL_3173903, EPI_ISL_3173904, EPI_ISL_3173905, EPI_ISL_3173906, EPI_ISL_3173907, EPI_ISL_3173908, EPI_ISL_3173909, EPI_ISL_3173910, EPI_ISL_3173911, EPI_ISL_3173912, EPI_ISL_3173913, EPI_ISL_3173914, EPI_ISL_3173915, EPI_ISL_3173916, EPI_ISL_3173917, EPI_ISL_3173918, EPI_ISL_3173919, EPI_ISL_3173920,

EPI_ISL_3173921, EPI_ISL_3173922, EPI_ISL_3173923, EPI_ISL_3173924, EPI_ISL_3173925, EPI_ISL_3173926, EPI_ISL_3173927, EPI_ISL_3173928, EPI_ISL_3173929, EPI_ISL_3173930, EPI_ISL_3173931, EPI_ISL_3173932, EPI_ISL_3173933, EPI_ISL_3173934, EPI_ISL_3173935, EPI_ISL_3173936, EPI_ISL_3173937, EPI_ISL_3173938, EPI_ISL_3173939, EPI_ISL_3173940, EPI_ISL_3173941, EPI_ISL_3173942, EPI_ISL_3173943, EPI_ISL_3173944, EPI_ISL_3173945, EPI_ISL_3173946, EPI_ISL_3173947, EPI_ISL_3173948, EPI_ISL_3173949, EPI_ISL_3173950, EPI_ISL_3173951, EPI_ISL_3173952, EPI_ISL_3173953, EPI_ISL_3173954, EPI_ISL_3173955, EPI_ISL_3173956, EPI_ISL_3173957, EPI_ISL_3173958, EPI_ISL_3173959, EPI_ISL_3173960, EPI_ISL_3173961, EPI_ISL_3173962, EPI_ISL_3173963, EPI_ISL_3173964, EPI_ISL_3173965, EPI_ISL_3173966, EPI_ISL_3173967, EPI_ISL_3173968, EPI_ISL_3173969, EPI_ISL_3173970, EPI_ISL_3173971, EPI_ISL_3173972, EPI_ISL_3173973, EPI_ISL_3173974, EPI_ISL_3173975, EPI_ISL_3173976, EPI_ISL_3173977, EPI_ISL_3173978, EPI_ISL_3173979, EPI_ISL_3173980, EPI_ISL_3173981, EPI_ISL_3173982, EPI_ISL_3173983, EPI_ISL_3173984, EPI_ISL_3173985, EPI_ISL_3173986, EPI_ISL_3173987, EPI_ISL_3173988, EPI_ISL_3173989, EPI_ISL_3173990, EPI_ISL_3173991, EPI_ISL_3173992, EPI_ISL_3173993, EPI_ISL_3173994, EPI_ISL_3173995, EPI_ISL_3173996, EPI_ISL_3173997, EPI_ISL_3173998, EPI_ISL_3173999, EPI_ISL_3174000, EPI_ISL_3174001, EPI_ISL_3174002, EPI_ISL_3174003, EPI_ISL_3174004, EPI_ISL_3174005, EPI_ISL_3174006, EPI_ISL_3174007, EPI_ISL_3174008, EPI_ISL_3174009, EPI_ISL_3174010, EPI_ISL_3174011, EPI_ISL_3174012, EPI_ISL_3174013, EPI_ISL_3174014, EPI_ISL_3174015, EPI_ISL_3174016, EPI_ISL_3174017, EPI_ISL_3174018, EPI_ISL_3174019, EPI_ISL_3174020, EPI_ISL_3174021, EPI_ISL_3174022, EPI_ISL_3174023, EPI_ISL_3174024, EPI_ISL_3174025, EPI_ISL_3174026, EPI_ISL_3174027, EPI_ISL_3174028, EPI_ISL_3174029, EPI_ISL_3174030, EPI_ISL_3174031, EPI_ISL_3174032, EPI_ISL_3174033, EPI_ISL_3174034, EPI_ISL_3174035, EPI_ISL_3174036, EPI_ISL_3174037, EPI_ISL_3174038, EPI_ISL_3174039, EPI_ISL_3174040, EPI_ISL_3174041, EPI_ISL_3174042, EPI_ISL_3174043, EPI_ISL_3174044, EPI_ISL_3174045, EPI_ISL_3174046, EPI_ISL_3174047, EPI_ISL_3174048, EPI_ISL_3174049, EPI_ISL_3174050, EPI_ISL_3174051, EPI_ISL_3174052, EPI_ISL_3174053, EPI_ISL_3174054, EPI_ISL_3174055, EPI_ISL_3174056, EPI_ISL_3174057, EPI_ISL_3174058, EPI_ISL_3174059, EPI_ISL_3174060, EPI_ISL_3174061, EPI_ISL_3174062, EPI_ISL_3174063, EPI_ISL_3174064, EPI_ISL_3174065, EPI_ISL_3174066, EPI_ISL_3174067, EPI_ISL_3174068, EPI_ISL_3174069, EPI_ISL_3174070, EPI_ISL_3174071, EPI_ISL_3174072, EPI_ISL_3174073, EPI_ISL_3174074, EPI_ISL_3174075, EPI_ISL_3174076, EPI_ISL_3174077, EPI_ISL_3174078, EPI_ISL_3174079, EPI_ISL_3174080, EPI_ISL_3174081, EPI_ISL_3174082, EPI_ISL_3174083, EPI_ISL_3174084, EPI_ISL_3174085, EPI_ISL_3174086, EPI_ISL_3174087, EPI_ISL_3174088, EPI_ISL_3174089, EPI_ISL_3174090, EPI_ISL_3174091, EPI_ISL_3174092, EPI_ISL_3174093, EPI_ISL_3174094, EPI_ISL_3174095, EPI_ISL_3174096, EPI_ISL_3174097, EPI_ISL_3174098, EPI_ISL_3174099, EPI_ISL_3174100, EPI_ISL_3174101, EPI_ISL_3174102, EPI_ISL_3174103, EPI_ISL_3174104, EPI_ISL_3174105, EPI_ISL_3174106, EPI_ISL_3174107, EPI_ISL_3174108, EPI_ISL_3174109, EPI_ISL_3174110, EPI_ISL_3174111, EPI_ISL_3174112, EPI_ISL_3174113, EPI_ISL_3174114, EPI_ISL_3174115, EPI_ISL_3174116, EPI_ISL_3174117, EPI_ISL_3174118, EPI_ISL_3174119, EPI_ISL_3174120, EPI_ISL_3174121, EPI_ISL_3174122, EPI_ISL_3174123, EPI_ISL_3174124, EPI_ISL_3174125, EPI_ISL_3174126, EPI_ISL_3174127, EPI_ISL_3174128, EPI_ISL_3174129, EPI_ISL_3174130, EPI_ISL_3174131, EPI_ISL_3174132, EPI_ISL_3174133, EPI_ISL_3174134, EPI_ISL_3174135, EPI_ISL_3174136, EPI_ISL_3174137, EPI_ISL_3174138, EPI_ISL_3174139, EPI_ISL_3174140, EPI_ISL_3174141, EPI_ISL_3174142, EPI_ISL_3174143, EPI_ISL_3174144, EPI_ISL_3174145, EPI_ISL_3174146, EPI_ISL_3174147, EPI_ISL_3174148, EPI_ISL_3174149, EPI_ISL_3174150, EPI_ISL_3174151, EPI_ISL_3174152, EPI_ISL_3174153, EPI_ISL_3174154, EPI_ISL_3174155, EPI_ISL_3174156, EPI_ISL_3174157, EPI_ISL_3174158, EPI_ISL_3174159, EPI_ISL_3174160, EPI_ISL_3174161, EPI_ISL_3174162, EPI_ISL_3174163, EPI_ISL_3174164, EPI_ISL_3174165, EPI_ISL_3174166, EPI_ISL_3174167, EPI_ISL_3174168, EPI_ISL_3174169, EPI_ISL_3174170, EPI_ISL_3174171, EPI_ISL_3174172, EPI_ISL_3174173, EPI_ISL_3174174, EPI_ISL_3174175, EPI_ISL_3174176, EPI_ISL_3174177, EPI_ISL_3174178, EPI_ISL_3174179, EPI_ISL_3174180, EPI_ISL_3174181, EPI_ISL_3174182, EPI_ISL_3174183, EPI_ISL_3174184, EPI_ISL_3174185, EPI_ISL_3174186, EPI_ISL_3174187, EPI_ISL_3174188, EPI_ISL_3174189, EPI_ISL_3174190, EPI_ISL_3174191, EPI_ISL_3174192, EPI_ISL_3174193, EPI_ISL_3174194, EPI_ISL_3174195, EPI_ISL_3174196, EPI_ISL_3174197, EPI_ISL_3174198

| | | | |
|---|---|---|---|
| see above | Lighthouse Lab in Milton Keynes | Wellcome Sanger Institute for the COVID-19 Genomics UK (COG-UK) Consortium | The Lighthouse Lab in Milton Keynes and Alex Alderton, Roberto Amato, Jeffrey Barrett, Sonia Goncalves, Ewan Harrison, David K. Jackson, Ian Johnston, Dominic Kwiatkowski, Cordelia Langford, John Sillitoe on behalf of the Wellcome Sanger Institute COVID-19 Surveillance Team |
| EPI_ISL_3174199, EPI_ISL_3174200, EPI_ISL_3174201, EPI_ISL_3174202 | Lighthouse Lab in Glasgow | Wellcome Sanger Institute for the COVID-19 Genomics UK (COG-UK) Consortium | Harper VanSteenhouse, Yumi Kasai, David Gray, Carol Clugston, Anna Dominiczak and Alex Alderton, Roberto Amato, Jeffrey Barrett, Sonia Goncalves, Ewan Harrison, David K. Jackson, Ian Johnston, Dominic Kwiatkowski, Cordelia Langford, John Sillitoe on behalf of the Wellcome Sanger Institute COVID-19 Surveillance Team |
| EPI_ISL_3174203 | Lighthouse Lab in Milton Keynes | Wellcome Sanger Institute for the COVID-19 Genomics UK (COG-UK) Consortium | The Lighthouse Lab in Milton Keynes and Alex Alderton, Roberto Amato, Jeffrey Barrett, Sonia Goncalves, Ewan Harrison, David K. Jackson, Ian Johnston, Dominic Kwiatkowski, Cordelia Langford, John Sillitoe on behalf of the Wellcome Sanger Institute COVID-19 Surveillance Team |
| EPI_ISL_3174204, EPI_ISL_3174205, EPI_ISL_3174206, EPI_ISL_3174207 | Lighthouse Lab in Glasgow | Wellcome Sanger Institute for the COVID-19 Genomics UK (COG-UK) Consortium | Harper VanSteenhouse, Yumi Kasai, David Gray, Carol Clugston, Anna Dominiczak and Alex Alderton, Roberto Amato, Jeffrey Barrett, Sonia Goncalves, Ewan Harrison, David K. Jackson, Ian Johnston, Dominic Kwiatkowski, Cordelia Langford, John Sillitoe on behalf of the Wellcome Sanger Institute COVID-19 Surveillance Team |
| EPI_ISL_3174208 | Lighthouse Lab in Milton Keynes | Wellcome Sanger Institute for the COVID-19 Genomics UK (COG-UK) Consortium | The Lighthouse Lab in Milton Keynes and Alex Alderton, Roberto Amato, Jeffrey Barrett, Sonia Goncalves, Ewan Harrison, David K. Jackson, Ian Johnston, Dominic Kwiatkowski, Cordelia Langford, John Sillitoe on behalf of the Wellcome Sanger Institute COVID-19 Surveillance Team |
| EPI_ISL_3174209 | Lighthouse Lab in Glasgow | Wellcome Sanger Institute for the COVID-19 Genomics UK (COG-UK) Consortium | Harper VanSteenhouse, Yumi Kasai, David Gray, Carol Clugston, Anna Dominiczak and Alex Alderton, Roberto Amato, Jeffrey Barrett, Sonia Goncalves, Ewan Harrison, David K. Jackson, Ian Johnston, Dominic Kwiatkowski, Cordelia Langford, John Sillitoe on behalf of the Wellcome Sanger Institute COVID-19 Surveillance Team |
| EPI_ISL_3174210 | Lighthouse Lab in Milton Keynes | Wellcome Sanger Institute for the COVID-19 Genomics UK (COG-UK) Consortium | The Lighthouse Lab in Milton Keynes and Alex Alderton, Roberto Amato, Jeffrey Barrett, Sonia Goncalves, Ewan Harrison, David K. Jackson, Ian Johnston, Dominic Kwiatkowski, Cordelia Langford, John Sillitoe on behalf of the Wellcome Sanger Institute COVID-19 Surveillance Team |
| EPI_ISL_3174211 | Lighthouse Lab in Glasgow | Wellcome Sanger Institute for the COVID-19 Genomics UK (COG-UK) Consortium | Harper VanSteenhouse, Yumi Kasai, David Gray, Carol Clugston, Anna Dominiczak and Alex Alderton, Roberto Amato, Jeffrey Barrett, Sonia Goncalves, Ewan Harrison, David K. Jackson, Ian Johnston, Dominic Kwiatkowski, Cordelia Langford, John Sillitoe on behalf of the Wellcome Sanger Institute COVID-19 Surveillance Team |
| EPI_ISL_3174212 | Lighthouse Lab in Milton Keynes | Wellcome Sanger Institute for the COVID-19 Genomics UK (COG-UK) Consortium | The Lighthouse Lab in Milton Keynes and Alex Alderton, Roberto Amato, Jeffrey Barrett, Sonia Goncalves, Ewan Harrison, David K. Jackson, Ian Johnston, Dominic Kwiatkowski, Cordelia Langford, John Sillitoe on behalf of the Wellcome Sanger Institute COVID-19 Surveillance Team |
| EPI_ISL_3174213 | Lighthouse Lab in Glasgow | Wellcome Sanger Institute for the COVID-19 Genomics UK (COG-UK) Consortium | Harper VanSteenhouse, Yumi Kasai, David Gray, Carol Clugston, Anna Dominiczak and Alex Alderton, Roberto Amato, Jeffrey Barrett, Sonia Goncalves, Ewan Harrison, David K. Jackson, Ian Johnston, Dominic Kwiatkowski, Cordelia Langford, John Sillitoe on behalf of the Wellcome Sanger Institute COVID-19 Surveillance Team |
| EPI_ISL_3174214, EPI_ISL_3174215, EPI_ISL_3174216, EPI_ISL_3174217, EPI_ISL_3174218, EPI_ISL_3174219 | Lighthouse Lab in Milton Keynes | Wellcome Sanger Institute for the COVID-19 Genomics UK (COG-UK) Consortium | The Lighthouse Lab in Milton Keynes and Alex Alderton, Roberto Amato, Jeffrey Barrett, Sonia Goncalves, Ewan Harrison, David K. Jackson, Ian Johnston, Dominic Kwiatkowski, Cordelia Langford, John Sillitoe on behalf of the Wellcome Sanger Institute COVID-19 Surveillance Team |
| EPI_ISL_3174220 | Lighthouse Lab in Glasgow | Wellcome Sanger Institute for the COVID-19 Genomics UK (COG-UK) Consortium | Harper VanSteenhouse, Yumi Kasai, David Gray, Carol Clugston, Anna Dominiczak and Alex Alderton, Roberto Amato, Jeffrey Barrett, Sonia Goncalves, Ewan Harrison, David K. Jackson, Ian Johnston, Dominic Kwiatkowski, Cordelia Langford, John Sillitoe on behalf of the Wellcome Sanger Institute COVID-19 Surveillance Team |
| EPI_ISL_3174221, EPI_ISL_3174222 | Lighthouse Lab in Milton Keynes | Wellcome Sanger Institute for the COVID-19 Genomics UK (COG-UK) Consortium | The Lighthouse Lab in Milton Keynes and Alex Alderton, Roberto Amato, Jeffrey Barrett, Sonia Goncalves, Ewan Harrison, David K. Jackson, Ian Johnston, Dominic Kwiatkowski, Cordelia Langford, John Sillitoe on behalf of the Wellcome Sanger Institute COVID-19 Surveillance Team |
| EPI_ISL_3174223, EPI_ISL_3174224 | Lighthouse Lab in Glasgow | Wellcome Sanger Institute for the COVID-19 Genomics UK (COG-UK) Consortium | Harper VanSteenhouse, Yumi Kasai, David Gray, Carol Clugston, Anna Dominiczak and Alex Alderton, Roberto Amato, Jeffrey Barrett, Sonia Goncalves, Ewan Harrison, David K. Jackson, Ian Johnston, Dominic Kwiatkowski, Cordelia Langford, John Sillitoe on behalf of the Wellcome Sanger Institute COVID-19 Surveillance Team |
| EPI_ISL_3174225 | Lighthouse Lab in Milton Keynes | Wellcome Sanger Institute for the COVID-19 Genomics UK (COG-UK) Consortium | The Lighthouse Lab in Milton Keynes and Alex Alderton, Roberto Amato, Jeffrey Barrett, Sonia Goncalves, Ewan Harrison, David K. Jackson, Ian Johnston, Dominic Kwiatkowski, Cordelia Langford, John Sillitoe on behalf of the Wellcome Sanger Institute COVID-19 Surveillance Team |
| EPI_ISL_3174226 | Lighthouse Lab in Glasgow | Wellcome Sanger Institute for the COVID-19 Genomics UK (COG-UK) Consortium | Harper VanSteenhouse, Yumi Kasai, David Gray, Carol Clugston, Anna Dominiczak and Alex Alderton, Roberto Amato, Jeffrey Barrett, Sonia Goncalves, Ewan Harrison, David K. Jackson, Ian Johnston, Dominic Kwiatkowski, Cordelia Langford, John Sillitoe on behalf of the Wellcome Sanger Institute COVID-19 Surveillance Team |
| EPI_ISL_3174227, EPI_ISL_3174228, EPI_ISL_3174229 | Lighthouse Lab in Milton Keynes | Wellcome Sanger Institute for the COVID-19 Genomics UK (COG-UK) Consortium | The Lighthouse Lab in Milton Keynes and Alex Alderton, Roberto Amato, Jeffrey Barrett, Sonia Goncalves, Ewan Harrison, David K. Jackson, Ian Johnston, Dominic Kwiatkowski, Cordelia Langford, John Sillitoe on behalf of the Wellcome Sanger Institute COVID-19 Surveillance Team |
| EPI_ISL_3174230 | Lighthouse Lab in Glasgow | Wellcome Sanger Institute for the COVID-19 Genomics UK (COG-UK) Consortium | Harper VanSteenhouse, Yumi Kasai, David Gray, Carol Clugston, Anna Dominiczak and Alex Alderton, Roberto Amato, Jeffrey Barrett, Sonia Goncalves, Ewan Harrison, David K. Jackson, Ian Johnston, Dominic Kwiatkowski, Cordelia Langford, John Sillitoe on behalf of the Wellcome Sanger Institute COVID-19 Surveillance Team |
| EPI_ISL_3174231, EPI_ISL_3174232, EPI_ISL_3174233, EPI_ISL_3174234 | Lighthouse Lab in Milton Keynes | Wellcome Sanger Institute for the COVID-19 Genomics UK (COG-UK) Consortium | The Lighthouse Lab in Milton Keynes and Alex Alderton, Roberto Amato, Jeffrey Barrett, Sonia Goncalves, Ewan Harrison, David K. Jackson, Ian Johnston, Dominic Kwiatkowski, Cordelia Langford, John Sillitoe on behalf of the Wellcome Sanger Institute COVID-19 Surveillance Team |
| EPI_ISL_3174235, EPI_ISL_3174236 | Lighthouse Lab in Glasgow | Wellcome Sanger Institute for the COVID-19 Genomics UK (COG-UK) Consortium | Harper VanSteenhouse, Yumi Kasai, David Gray, Carol Clugston, Anna Dominiczak and Alex Alderton, Roberto Amato, Jeffrey Barrett, Sonia Goncalves, Ewan Harrison, David K. Jackson, Ian Johnston, Dominic Kwiatkowski, Cordelia Langford, John Sillitoe on behalf of the Wellcome Sanger Institute COVID-19 Surveillance Team |
| EPI_ISL_3174237, EPI_ISL_3174238 | Lighthouse Lab in Milton Keynes | Wellcome Sanger Institute for the COVID-19 Genomics UK (COG-UK) Consortium | The Lighthouse Lab in Milton Keynes and Alex Alderton, Roberto Amato, Jeffrey Barrett, Sonia Goncalves, Ewan Harrison, David K. Jackson, Ian Johnston, Dominic Kwiatkowski, Cordelia Langford, John Sillitoe on behalf of the Wellcome Sanger Institute COVID-19 Surveillance Team |
| EPI_ISL_3174239 | Lighthouse Lab in Glasgow | Wellcome Sanger Institute for the COVID-19 Genomics UK (COG-UK) Consortium | Harper VanSteenhouse, Yumi Kasai, David Gray, Carol Clugston, Anna Dominiczak and Alex Alderton, Roberto Amato, Jeffrey Barrett, Sonia Goncalves, Ewan Harrison, David K. Jackson, Ian Johnston, Dominic Kwiatkowski, Cordelia Langford, John Sillitoe on behalf of the Wellcome Sanger Institute COVID-19 Surveillance Team |
| EPI_ISL_3174240 | Lighthouse Lab in Milton Keynes | Wellcome Sanger Institute for the COVID-19 Genomics UK (COG-UK) Consortium | The Lighthouse Lab in Milton Keynes and Alex Alderton, Roberto Amato, Jeffrey Barrett, Sonia Goncalves, Ewan Harrison, David K. Jackson, Ian Johnston, Dominic Kwiatkowski, Cordelia Langford, John Sillitoe on behalf of the Wellcome Sanger Institute COVID-19 Surveillance Team |
| EPI_ISL_3174241, EPI_ISL_3174242, EPI_ISL_3174243, EPI_ISL_3174244 | Lighthouse Lab in Glasgow | Wellcome Sanger Institute for the COVID-19 Genomics UK (COG-UK) Consortium | Harper VanSteenhouse, Yumi Kasai, David Gray, Carol Clugston, Anna Dominiczak and Alex Alderton, Roberto Amato, Jeffrey Barrett, Sonia Goncalves, Ewan Harrison, David K. Jackson, Ian Johnston, Dominic Kwiatkowski, Cordelia Langford, John Sillitoe on behalf of the Wellcome Sanger Institute COVID-19 Surveillance Team |
| EPI_ISL_3174245, EPI_ISL_3174246 | Lighthouse Lab in Milton Keynes | Wellcome Sanger Institute for the COVID-19 Genomics UK (COG-UK) Consortium | The Lighthouse Lab in Milton Keynes and Alex Alderton, Roberto Amato, Jeffrey Barrett, Sonia Goncalves, Ewan Harrison, David K. Jackson, Ian Johnston, Dominic Kwiatkowski, Cordelia Langford, John Sillitoe on behalf of the Wellcome Sanger Institute COVID-19 Surveillance Team |
| EPI_ISL_3174247 | Lighthouse Lab in Glasgow | Wellcome Sanger Institute for the COVID-19 Genomics UK (COG-UK) Consortium | Harper VanSteenhouse, Yumi Kasai, David Gray, Carol Clugston, Anna Dominiczak and Alex Alderton, Roberto Amato, Jeffrey Barrett, Sonia Goncalves, Ewan Harrison, David K. Jackson, Ian Johnston, Dominic Kwiatkowski, Cordelia Langford, John Sillitoe on behalf of the Wellcome Sanger Institute COVID-19 Surveillance Team |
| EPI_ISL_3174248 | Lighthouse Lab in Milton Keynes | Wellcome Sanger Institute for the COVID-19 Genomics UK (COG-UK) Consortium | The Lighthouse Lab in Milton Keynes and Alex Alderton, Roberto Amato, Jeffrey Barrett, Sonia Goncalves, Ewan Harrison, David K. Jackson, Ian Johnston, Dominic Kwiatkowski, Cordelia Langford, John Sillitoe on behalf of the Wellcome Sanger Institute COVID-19 Surveillance Team |
| EPI_ISL_3174249 | Lighthouse Lab in Glasgow | Wellcome Sanger Institute for the COVID-19 Genomics UK (COG-UK) Consortium | Harper VanSteenhouse, Yumi Kasai, David Gray, Carol Clugston, Anna Dominiczak and Alex Alderton, Roberto Amato, Jeffrey Barrett, Sonia Goncalves, Ewan Harrison, David K. Jackson, Ian Johnston, Dominic Kwiatkowski, Cordelia Langford, John Sillitoe on behalf of the Wellcome Sanger Institute COVID-19 Surveillance Team |
| EPI_ISL_3174250 | Lighthouse Lab in Milton Keynes | Wellcome Sanger Institute for the COVID-19 Genomics UK (COG-UK) Consortium | The Lighthouse Lab in Milton Keynes and Alex Alderton, Roberto Amato, Jeffrey Barrett, Sonia Goncalves, Ewan Harrison, David K. Jackson, Ian Johnston, Dominic Kwiatkowski, Cordelia Langford, John Sillitoe on behalf of the Wellcome Sanger Institute COVID-19 Surveillance Team |
| EPI_ISL_3174251 | Lighthouse Lab in Glasgow | Wellcome Sanger Institute for the COVID-19 Genomics UK (COG-UK) Consortium | Harper VanSteenhouse, Yumi Kasai, David Gray, Carol Clugston, Anna Dominiczak and Alex Alderton, Roberto Amato, Jeffrey Barrett, Sonia Goncalves, Ewan Harrison, David K. Jackson, Ian Johnston, Dominic Kwiatkowski, Cordelia Langford, John Sillitoe on behalf of the Wellcome Sanger Institute COVID-19 Surveillance Team |
| EPI_ISL_3174252, EPI_ISL_3174253, EPI_ISL_3174254 | Lighthouse Lab in Milton Keynes | Wellcome Sanger Institute for the COVID-19 Genomics UK (COG-UK) Consortium | The Lighthouse Lab in Milton Keynes and Alex Alderton, Roberto Amato, Jeffrey Barrett, Sonia Goncalves, Ewan Harrison, David K. Jackson, Ian Johnston, Dominic Kwiatkowski, Cordelia Langford, John Sillitoe on behalf of the Wellcome Sanger Institute COVID-19 Surveillance Team |
| EPI_ISL_3174255, EPI_ISL_3174256, EPI_ISL_3174257 | Lighthouse Lab in Glasgow | Wellcome Sanger Institute for the COVID-19 Genomics UK (COG-UK) Consortium | Harper VanSteenhouse, Yumi Kasai, David Gray, Carol Clugston, Anna Dominiczak and Alex Alderton, Roberto Amato, Jeffrey Barrett, Sonia Goncalves, Ewan Harrison, David K. Jackson, Ian Johnston, Dominic Kwiatkowski, Cordelia Langford, John Sillitoe on behalf of the Wellcome Sanger Institute COVID-19 Surveillance Team |
| EPI_ISL_3174258, EPI_ISL_3174259, EPI_ISL_3174260, | Lighthouse Lab in Milton Keynes | Wellcome Sanger Institute for the COVID-19 Genomics UK (COG-UK) Consortium | The Lighthouse Lab in Milton Keynes and Alex Alderton, Roberto Amato, Jeffrey Barrett, Sonia Goncalves, Ewan Harrison, David K. Jackson, Ian Johnston, Dominic Kwiatkowski, Cordelia Langford, John Sillitoe on behalf of the Wellcome Sanger Institute COVID-19 Surveillance Team |

| | | | |
|---|---|---|---|
| EPI_ISL_3174261 | | | |
| EPI_ISL_3174262 | Lighthouse Lab in Glasgow | Wellcome Sanger Institute for the COVID-19 Genomics UK (COG-UK) Consortium | Harper VanSteenhouse, Yumi Kasai, David Gray, Carol Clugston, Anna Dominiczak and Alex Alderton, Roberto Amato, Jeffrey Barrett, Sonia Goncalves, Ewan Harrison, David K. Jackson, Ian Johnston, Dominic Kwiatkowski, Cordelia Langford, John Sillitoe on behalf of the Wellcome Sanger Institute COVID-19 Surveillance Team |
| EPI_ISL_3174263 | Lighthouse Lab in Milton Keynes | Wellcome Sanger Institute for the COVID-19 Genomics UK (COG-UK) Consortium | The Lighthouse Lab in Milton Keynes and Alex Alderton, Roberto Amato, Jeffrey Barrett, Sonia Goncalves, Ewan Harrison, David K. Jackson, Ian Johnston, Dominic Kwiatkowski, Cordelia Langford, John Sillitoe on behalf of the Wellcome Sanger Institute COVID-19 Surveillance Team |
| EPI_ISL_3174264, EPI_ISL_3174265, EPI_ISL_3174266 | Lighthouse Lab in Glasgow | Wellcome Sanger Institute for the COVID-19 Genomics UK (COG-UK) Consortium | Harper VanSteenhouse, Yumi Kasai, David Gray, Carol Clugston, Anna Dominiczak and Alex Alderton, Roberto Amato, Jeffrey Barrett, Sonia Goncalves, Ewan Harrison, David K. Jackson, Ian Johnston, Dominic Kwiatkowski, Cordelia Langford, John Sillitoe on behalf of the Wellcome Sanger Institute COVID-19 Surveillance Team |
| EPI_ISL_3174267 | Lighthouse Lab in Milton Keynes | Wellcome Sanger Institute for the COVID-19 Genomics UK (COG-UK) Consortium | The Lighthouse Lab in Milton Keynes and Alex Alderton, Roberto Amato, Jeffrey Barrett, Sonia Goncalves, Ewan Harrison, David K. Jackson, Ian Johnston, Dominic Kwiatkowski, Cordelia Langford, John Sillitoe on behalf of the Wellcome Sanger Institute COVID-19 Surveillance Team |
| EPI_ISL_3174268, EPI_ISL_3174269 | Lighthouse Lab in Glasgow | Wellcome Sanger Institute for the COVID-19 Genomics UK (COG-UK) Consortium | Harper VanSteenhouse, Yumi Kasai, David Gray, Carol Clugston, Anna Dominiczak and Alex Alderton, Roberto Amato, Jeffrey Barrett, Sonia Goncalves, Ewan Harrison, David K. Jackson, Ian Johnston, Dominic Kwiatkowski, Cordelia Langford, John Sillitoe on behalf of the Wellcome Sanger Institute COVID-19 Surveillance Team |
| EPI_ISL_3174270 | Lighthouse Lab in Milton Keynes | Wellcome Sanger Institute for the COVID-19 Genomics UK (COG-UK) Consortium | The Lighthouse Lab in Milton Keynes and Alex Alderton, Roberto Amato, Jeffrey Barrett, Sonia Goncalves, Ewan Harrison, David K. Jackson, Ian Johnston, Dominic Kwiatkowski, Cordelia Langford, John Sillitoe on behalf of the Wellcome Sanger Institute COVID-19 Surveillance Team |
| EPI_ISL_3174271, EPI_ISL_3174272, EPI_ISL_3174273, EPI_ISL_3174274 | Lighthouse Lab in Glasgow | Wellcome Sanger Institute for the COVID-19 Genomics UK (COG-UK) Consortium | Harper VanSteenhouse, Yumi Kasai, David Gray, Carol Clugston, Anna Dominiczak and Alex Alderton, Roberto Amato, Jeffrey Barrett, Sonia Goncalves, Ewan Harrison, David K. Jackson, Ian Johnston, Dominic Kwiatkowski, Cordelia Langford, John Sillitoe on behalf of the Wellcome Sanger Institute COVID-19 Surveillance Team |
| EPI_ISL_3174275, EPI_ISL_3174276 | Lighthouse Lab in Milton Keynes | Wellcome Sanger Institute for the COVID-19 Genomics UK (COG-UK) Consortium | The Lighthouse Lab in Milton Keynes and Alex Alderton, Roberto Amato, Jeffrey Barrett, Sonia Goncalves, Ewan Harrison, David K. Jackson, Ian Johnston, Dominic Kwiatkowski, Cordelia Langford, John Sillitoe on behalf of the Wellcome Sanger Institute COVID-19 Surveillance Team |
| EPI_ISL_3174277, EPI_ISL_3174278, EPI_ISL_3174279 | Lighthouse Lab in Glasgow | Wellcome Sanger Institute for the COVID-19 Genomics UK (COG-UK) Consortium | Harper VanSteenhouse, Yumi Kasai, David Gray, Carol Clugston, Anna Dominiczak and Alex Alderton, Roberto Amato, Jeffrey Barrett, Sonia Goncalves, Ewan Harrison, David K. Jackson, Ian Johnston, Dominic Kwiatkowski, Cordelia Langford, John Sillitoe on behalf of the Wellcome Sanger Institute COVID-19 Surveillance Team |
| EPI_ISL_3174280, EPI_ISL_3174281, EPI_ISL_3174282, EPI_ISL_3174283, EPI_ISL_3174284, EPI_ISL_3174285, EPI_ISL_3174286 | Lighthouse Lab in Milton Keynes | Wellcome Sanger Institute for the COVID-19 Genomics UK (COG-UK) Consortium | The Lighthouse Lab in Milton Keynes and Alex Alderton, Roberto Amato, Jeffrey Barrett, Sonia Goncalves, Ewan Harrison, David K. Jackson, Ian Johnston, Dominic Kwiatkowski, Cordelia Langford, John Sillitoe on behalf of the Wellcome Sanger Institute COVID-19 Surveillance Team |
| EPI_ISL_3174287, EPI_ISL_3174288 | Lighthouse Lab in Glasgow | Wellcome Sanger Institute for the COVID-19 Genomics UK (COG-UK) Consortium | Harper VanSteenhouse, Yumi Kasai, David Gray, Carol Clugston, Anna Dominiczak and Alex Alderton, Roberto Amato, Jeffrey Barrett, Sonia Goncalves, Ewan Harrison, David K. Jackson, Ian Johnston, Dominic Kwiatkowski, Cordelia Langford, John Sillitoe on behalf of the Wellcome Sanger Institute COVID-19 Surveillance Team |
| EPI_ISL_3174289, EPI_ISL_3174290 | Lighthouse Lab in Milton Keynes | Wellcome Sanger Institute for the COVID-19 Genomics UK (COG-UK) Consortium | The Lighthouse Lab in Milton Keynes and Alex Alderton, Roberto Amato, Jeffrey Barrett, Sonia Goncalves, Ewan Harrison, David K. Jackson, Ian Johnston, Dominic Kwiatkowski, Cordelia Langford, John Sillitoe on behalf of the Wellcome Sanger Institute COVID-19 Surveillance Team |
| EPI_ISL_3174291, EPI_ISL_3174292, EPI_ISL_3174293 | Lighthouse Lab in Glasgow | Wellcome Sanger Institute for the COVID-19 Genomics UK (COG-UK) Consortium | Harper VanSteenhouse, Yumi Kasai, David Gray, Carol Clugston, Anna Dominiczak and Alex Alderton, Roberto Amato, Jeffrey Barrett, Sonia Goncalves, Ewan Harrison, David K. Jackson, Ian Johnston, Dominic Kwiatkowski, Cordelia Langford, John Sillitoe on behalf of the Wellcome Sanger Institute COVID-19 Surveillance Team |
| EPI_ISL_3174294, EPI_ISL_3174295 | Lighthouse Lab in Milton Keynes | Wellcome Sanger Institute for the COVID-19 Genomics UK (COG-UK) Consortium | The Lighthouse Lab in Milton Keynes and Alex Alderton, Roberto Amato, Jeffrey Barrett, Sonia Goncalves, Ewan Harrison, David K. Jackson, Ian Johnston, Dominic Kwiatkowski, Cordelia Langford, John Sillitoe on behalf of the Wellcome Sanger Institute COVID-19 Surveillance Team |
| EPI_ISL_3174296, EPI_ISL_3174297 | Lighthouse Lab in Glasgow | Wellcome Sanger Institute for the COVID-19 Genomics UK (COG-UK) Consortium | Harper VanSteenhouse, Yumi Kasai, David Gray, Carol Clugston, Anna Dominiczak and Alex Alderton, Roberto Amato, Jeffrey Barrett, Sonia Goncalves, Ewan Harrison, David K. Jackson, Ian Johnston, Dominic Kwiatkowski, Cordelia Langford, John Sillitoe on behalf of the Wellcome Sanger Institute COVID-19 Surveillance Team |
| EPI_ISL_3174298 | Lighthouse Lab in Milton Keynes | Wellcome Sanger Institute for the COVID-19 Genomics UK (COG-UK) Consortium | The Lighthouse Lab in Milton Keynes and Alex Alderton, Roberto Amato, Jeffrey Barrett, Sonia Goncalves, Ewan Harrison, David K. Jackson, Ian Johnston, Dominic Kwiatkowski, Cordelia Langford, John Sillitoe on behalf of the Wellcome Sanger Institute COVID-19 Surveillance Team |
| EPI_ISL_3174299 | Lighthouse Lab in Glasgow | Wellcome Sanger Institute for the COVID-19 Genomics UK (COG-UK) Consortium | Harper VanSteenhouse, Yumi Kasai, David Gray, Carol Clugston, Anna Dominiczak and Alex Alderton, Roberto Amato, Jeffrey Barrett, Sonia Goncalves, Ewan Harrison, David K. Jackson, Ian Johnston, Dominic Kwiatkowski, Cordelia Langford, John Sillitoe on behalf of the Wellcome Sanger Institute COVID-19 Surveillance Team |
| EPI_ISL_3174300 | Lighthouse Lab in Milton Keynes | Wellcome Sanger Institute for the COVID-19 Genomics UK (COG-UK) Consortium | The Lighthouse Lab in Milton Keynes and Alex Alderton, Roberto Amato, Jeffrey Barrett, Sonia Goncalves, Ewan Harrison, David K. Jackson, Ian Johnston, Dominic Kwiatkowski, Cordelia Langford, John Sillitoe on behalf of the Wellcome Sanger Institute COVID-19 Surveillance Team |
| EPI_ISL_3174301 | Lighthouse Lab in Glasgow | Wellcome Sanger Institute for the COVID-19 Genomics UK (COG-UK) Consortium | Harper VanSteenhouse, Yumi Kasai, David Gray, Carol Clugston, Anna Dominiczak and Alex Alderton, Roberto Amato, Jeffrey Barrett, Sonia Goncalves, Ewan Harrison, David K. Jackson, Ian Johnston, Dominic Kwiatkowski, Cordelia Langford, John Sillitoe on behalf of the Wellcome Sanger Institute COVID-19 Surveillance Team |
| EPI_ISL_3174302 | Lighthouse Lab in Milton Keynes | Wellcome Sanger Institute for the COVID-19 Genomics UK (COG-UK) Consortium | The Lighthouse Lab in Milton Keynes and Alex Alderton, Roberto Amato, Jeffrey Barrett, Sonia Goncalves, Ewan Harrison, David K. Jackson, Ian Johnston, Dominic Kwiatkowski, Cordelia Langford, John Sillitoe on behalf of the Wellcome Sanger Institute COVID-19 Surveillance Team |
| EPI_ISL_3174303 | Lighthouse Lab in Glasgow | Wellcome Sanger Institute for the COVID-19 Genomics UK (COG-UK) Consortium | Harper VanSteenhouse, Yumi Kasai, David Gray, Carol Clugston, Anna Dominiczak and Alex Alderton, Roberto Amato, Jeffrey Barrett, Sonia Goncalves, Ewan Harrison, David K. Jackson, Ian Johnston, Dominic Kwiatkowski, Cordelia Langford, John Sillitoe on behalf of the Wellcome Sanger Institute COVID-19 Surveillance Team |
| EPI_ISL_3174304, EPI_ISL_3174305, EPI_ISL_3174306, EPI_ISL_3174307, EPI_ISL_3174308 | Lighthouse Lab in Milton Keynes | Wellcome Sanger Institute for the COVID-19 Genomics UK (COG-UK) Consortium | The Lighthouse Lab in Milton Keynes and Alex Alderton, Roberto Amato, Jeffrey Barrett, Sonia Goncalves, Ewan Harrison, David K. Jackson, Ian Johnston, Dominic Kwiatkowski, Cordelia Langford, John Sillitoe on behalf of the Wellcome Sanger Institute COVID-19 Surveillance Team |
| EPI_ISL_3174309 | Lighthouse Lab in Glasgow | Wellcome Sanger Institute for the COVID-19 Genomics UK (COG-UK) Consortium | Harper VanSteenhouse, Yumi Kasai, David Gray, Carol Clugston, Anna Dominiczak and Alex Alderton, Roberto Amato, Jeffrey Barrett, Sonia Goncalves, Ewan Harrison, David K. Jackson, Ian Johnston, Dominic Kwiatkowski, Cordelia Langford, John Sillitoe on behalf of the Wellcome Sanger Institute COVID-19 Surveillance Team |
| EPI_ISL_3174310 | Lighthouse Lab in Milton Keynes | Wellcome Sanger Institute for the COVID-19 Genomics UK (COG-UK) Consortium | The Lighthouse Lab in Milton Keynes and Alex Alderton, Roberto Amato, Jeffrey Barrett, Sonia Goncalves, Ewan Harrison, David K. Jackson, Ian Johnston, Dominic Kwiatkowski, Cordelia Langford, John Sillitoe on behalf of the Wellcome Sanger Institute COVID-19 Surveillance Team |
| EPI_ISL_3174311 | Lighthouse Lab in Glasgow | Wellcome Sanger Institute for the COVID-19 Genomics UK (COG-UK) Consortium | Harper VanSteenhouse, Yumi Kasai, David Gray, Carol Clugston, Anna Dominiczak and Alex Alderton, Roberto Amato, Jeffrey Barrett, Sonia Goncalves, Ewan Harrison, David K. Jackson, Ian Johnston, Dominic Kwiatkowski, Cordelia Langford, John Sillitoe on behalf of the Wellcome Sanger Institute COVID-19 Surveillance Team |
| EPI_ISL_3174312, EPI_ISL_3174313, EPI_ISL_3174314, EPI_ISL_3174315 | Lighthouse Lab in Milton Keynes | Wellcome Sanger Institute for the COVID-19 Genomics UK (COG-UK) Consortium | The Lighthouse Lab in Milton Keynes and Alex Alderton, Roberto Amato, Jeffrey Barrett, Sonia Goncalves, Ewan Harrison, David K. Jackson, Ian Johnston, Dominic Kwiatkowski, Cordelia Langford, John Sillitoe on behalf of the Wellcome Sanger Institute COVID-19 Surveillance Team |
| EPI_ISL_3174316 | Lighthouse Lab in Glasgow | Wellcome Sanger Institute for the COVID-19 Genomics UK (COG-UK) Consortium | Harper VanSteenhouse, Yumi Kasai, David Gray, Carol Clugston, Anna Dominiczak and Alex Alderton, Roberto Amato, Jeffrey Barrett, Sonia Goncalves, Ewan Harrison, David K. Jackson, Ian Johnston, Dominic Kwiatkowski, Cordelia Langford, John Sillitoe on behalf of the Wellcome Sanger Institute COVID-19 Surveillance Team |
| EPI_ISL_3174317, EPI_ISL_3174318, EPI_ISL_3174319 | Lighthouse Lab in Milton Keynes | Wellcome Sanger Institute for the COVID-19 Genomics UK (COG-UK) Consortium | The Lighthouse Lab in Milton Keynes and Alex Alderton, Roberto Amato, Jeffrey Barrett, Sonia Goncalves, Ewan Harrison, David K. Jackson, Ian Johnston, Dominic Kwiatkowski, Cordelia Langford, John Sillitoe on behalf of the Wellcome Sanger Institute COVID-19 Surveillance Team |
| EPI_ISL_3174320, EPI_ISL_3174321 | Lighthouse Lab in Glasgow | Wellcome Sanger Institute for the COVID-19 Genomics UK (COG-UK) Consortium | Harper VanSteenhouse, Yumi Kasai, David Gray, Carol Clugston, Anna Dominiczak and Alex Alderton, Roberto Amato, Jeffrey Barrett, Sonia Goncalves, Ewan Harrison, David K. Jackson, Ian Johnston, Dominic Kwiatkowski, Cordelia Langford, John Sillitoe on behalf of the Wellcome Sanger Institute COVID-19 Surveillance Team |
| EPI_ISL_3174322, EPI_ISL_3174323, EPI_ISL_3174324, EPI_ISL_3174325 | Lighthouse Lab in Milton Keynes | Wellcome Sanger Institute for the COVID-19 Genomics UK (COG-UK) Consortium | The Lighthouse Lab in Milton Keynes and Alex Alderton, Roberto Amato, Jeffrey Barrett, Sonia Goncalves, Ewan Harrison, David K. Jackson, Ian Johnston, Dominic Kwiatkowski, Cordelia Langford, John Sillitoe on behalf of the Wellcome Sanger Institute COVID-19 Surveillance Team |
| EPI_ISL_3174326 | Lighthouse Lab in Glasgow | Wellcome Sanger Institute for the COVID-19 Genomics UK (COG-UK) Consortium | Harper VanSteenhouse, Yumi Kasai, David Gray, Carol Clugston, Anna Dominiczak and Alex Alderton, Roberto Amato, Jeffrey Barrett, Sonia Goncalves, Ewan Harrison, David K. Jackson, Ian Johnston, Dominic Kwiatkowski, Cordelia Langford, John Sillitoe on behalf of the Wellcome Sanger Institute COVID-19 Surveillance Team |
| EPI_ISL_3174327 | Lighthouse Lab in Milton Keynes | Wellcome Sanger Institute for the COVID-19 Genomics UK (COG-UK) Consortium | The Lighthouse Lab in Milton Keynes and Alex Alderton, Roberto Amato, Jeffrey Barrett, Sonia Goncalves, Ewan Harrison, David K. Jackson, Ian Johnston, Dominic Kwiatkowski, Cordelia Langford, John Sillitoe on behalf of the Wellcome Sanger Institute COVID-19 Surveillance Team |
| EPI_ISL_3174328, EPI_ISL_3174329, EPI_ISL_3174330 | Lighthouse Lab in Glasgow | Wellcome Sanger Institute for the COVID-19 Genomics UK (COG-UK) Consortium | Harper VanSteenhouse, Yumi Kasai, David Gray, Carol Clugston, Anna Dominiczak and Alex Alderton, Roberto Amato, Jeffrey Barrett, Sonia Goncalves, Ewan Harrison, David K. Jackson, Ian Johnston, Dominic Kwiatkowski, Cordelia Langford, John Sillitoe on behalf of the Wellcome Sanger Institute COVID-19 Surveillance Team |
| EPI_ISL_3174331 | Lighthouse Lab in Milton Keynes | Wellcome Sanger Institute for the COVID-19 Genomics UK (COG-UK) Consortium | The Lighthouse Lab in Milton Keynes and Alex Alderton, Roberto Amato, Jeffrey Barrett, Sonia Goncalves, Ewan Harrison, David K. Jackson, Ian Johnston, Dominic Kwiatkowski, Cordelia Langford, John Sillitoe on behalf of the Wellcome Sanger Institute COVID-19 Surveillance Team |
| EPI_ISL_3174332 | Lighthouse Lab in Glasgow | Wellcome Sanger Institute for the COVID-19 Genomics UK (COG-UK) Consortium | Harper VanSteenhouse, Yumi Kasai, David Gray, Carol Clugston, Anna Dominiczak and Alex Alderton, Roberto Amato, Jeffrey Barrett, Sonia Goncalves, Ewan Harrison, David K. Jackson, Ian Johnston, Dominic Kwiatkowski, Cordelia Langford, John Sillitoe on behalf of the Wellcome Sanger Institute COVID-19 Surveillance Team |
| EPI_ISL_3174333 | Lighthouse Lab in Milton Keynes | Wellcome Sanger Institute for the COVID-19 Genomics UK (COG-UK) Consortium | The Lighthouse Lab in Milton Keynes and Alex Alderton, Roberto Amato, Jeffrey Barrett, Sonia Goncalves, Ewan Harrison, David K. Jackson, Ian Johnston, Dominic Kwiatkowski, Cordelia Langford, John Sillitoe on behalf of the Wellcome Sanger Institute COVID-19 Surveillance Team |
| EPI_ISL_3174334 | Lighthouse Lab in Glasgow | Wellcome Sanger Institute for the COVID-19 Genomics UK (COG-UK) Consortium | Harper VanSteenhouse, Yumi Kasai, David Gray, Carol Clugston, Anna Dominiczak and Alex Alderton, Roberto Amato, Jeffrey Barrett, Sonia Goncalves, Ewan Harrison, David K. Jackson, Ian Johnston, Dominic Kwiatkowski, Cordelia Langford, John Sillitoe on behalf of the Wellcome Sanger Institute COVID-19 Surveillance Team |

| | | | |
|---|---|---|---|
| EPI_ISL_3174335, EPI_ISL_3174336, EPI_ISL_3174337, EPI_ISL_3174338 | Lighthouse Lab in Milton Keynes | Wellcome Sanger Institute for the COVID-19 Genomics UK (COG-UK) Consortium | The Lighthouse Lab in Milton Keynes and Alex Alderton, Roberto Amato, Jeffrey Barrett, Sonia Goncalves, Ewan Harrison, David K. Jackson, Ian Johnston, Dominic Kwiatkowski, Cordelia Langford, John Sillitoe on behalf of the Wellcome Sanger Institute COVID-19 Surveillance Team |
| EPI_ISL_3174339, EPI_ISL_3174340, EPI_ISL_3174341, EPI_ISL_3174342 | Lighthouse Lab in Glasgow | Wellcome Sanger Institute for the COVID-19 Genomics UK (COG-UK) Consortium | Harper VanSteenhouse, Yumi Kasai, David Gray, Carol Clugston, Anna Dominiczak and Alex Alderton, Roberto Amato, Jeffrey Barrett, Sonia Goncalves, Ewan Harrison, David K. Jackson, Ian Johnston, Dominic Kwiatkowski, Cordelia Langford, John Sillitoe on behalf of the Wellcome Sanger Institute COVID-19 Surveillance Team |
| EPI_ISL_3174343, EPI_ISL_3174344, EPI_ISL_3174345 | Lighthouse Lab in Milton Keynes | Wellcome Sanger Institute for the COVID-19 Genomics UK (COG-UK) Consortium | The Lighthouse Lab in Milton Keynes and Alex Alderton, Roberto Amato, Jeffrey Barrett, Sonia Goncalves, Ewan Harrison, David K. Jackson, Ian Johnston, Dominic Kwiatkowski, Cordelia Langford, John Sillitoe on behalf of the Wellcome Sanger Institute COVID-19 Surveillance Team |
| EPI_ISL_3174346 | Lighthouse Lab in Glasgow | Wellcome Sanger Institute for the COVID-19 Genomics UK (COG-UK) Consortium | Harper VanSteenhouse, Yumi Kasai, David Gray, Carol Clugston, Anna Dominiczak and Alex Alderton, Roberto Amato, Jeffrey Barrett, Sonia Goncalves, Ewan Harrison, David K. Jackson, Ian Johnston, Dominic Kwiatkowski, Cordelia Langford, John Sillitoe on behalf of the Wellcome Sanger Institute COVID-19 Surveillance Team |
| EPI_ISL_3174347 | Lighthouse Lab in Milton Keynes | Wellcome Sanger Institute for the COVID-19 Genomics UK (COG-UK) Consortium | The Lighthouse Lab in Milton Keynes and Alex Alderton, Roberto Amato, Jeffrey Barrett, Sonia Goncalves, Ewan Harrison, David K. Jackson, Ian Johnston, Dominic Kwiatkowski, Cordelia Langford, John Sillitoe on behalf of the Wellcome Sanger Institute COVID-19 Surveillance Team |
| EPI_ISL_3174348 | Lighthouse Lab in Glasgow | Wellcome Sanger Institute for the COVID-19 Genomics UK (COG-UK) Consortium | Harper VanSteenhouse, Yumi Kasai, David Gray, Carol Clugston, Anna Dominiczak and Alex Alderton, Roberto Amato, Jeffrey Barrett, Sonia Goncalves, Ewan Harrison, David K. Jackson, Ian Johnston, Dominic Kwiatkowski, Cordelia Langford, John Sillitoe on behalf of the Wellcome Sanger Institute COVID-19 Surveillance Team |
| EPI_ISL_3174349, EPI_ISL_3174350, EPI_ISL_3174351, EPI_ISL_3174352, EPI_ISL_3174353 | Lighthouse Lab in Milton Keynes | Wellcome Sanger Institute for the COVID-19 Genomics UK (COG-UK) Consortium | The Lighthouse Lab in Milton Keynes and Alex Alderton, Roberto Amato, Jeffrey Barrett, Sonia Goncalves, Ewan Harrison, David K. Jackson, Ian Johnston, Dominic Kwiatkowski, Cordelia Langford, John Sillitoe on behalf of the Wellcome Sanger Institute COVID-19 Surveillance Team |
| EPI_ISL_3174354, EPI_ISL_3174355, EPI_ISL_3174356 | Lighthouse Lab in Glasgow | Wellcome Sanger Institute for the COVID-19 Genomics UK (COG-UK) Consortium | Harper VanSteenhouse, Yumi Kasai, David Gray, Carol Clugston, Anna Dominiczak and Alex Alderton, Roberto Amato, Jeffrey Barrett, Sonia Goncalves, Ewan Harrison, David K. Jackson, Ian Johnston, Dominic Kwiatkowski, Cordelia Langford, John Sillitoe on behalf of the Wellcome Sanger Institute COVID-19 Surveillance Team |
| EPI_ISL_3174357 | Lighthouse Lab in Milton Keynes | Wellcome Sanger Institute for the COVID-19 Genomics UK (COG-UK) Consortium | The Lighthouse Lab in Milton Keynes and Alex Alderton, Roberto Amato, Jeffrey Barrett, Sonia Goncalves, Ewan Harrison, David K. Jackson, Ian Johnston, Dominic Kwiatkowski, Cordelia Langford, John Sillitoe on behalf of the Wellcome Sanger Institute COVID-19 Surveillance Team |
| EPI_ISL_3174358 | Lighthouse Lab in Glasgow | Wellcome Sanger Institute for the COVID-19 Genomics UK (COG-UK) Consortium | Harper VanSteenhouse, Yumi Kasai, David Gray, Carol Clugston, Anna Dominiczak and Alex Alderton, Roberto Amato, Jeffrey Barrett, Sonia Goncalves, Ewan Harrison, David K. Jackson, Ian Johnston, Dominic Kwiatkowski, Cordelia Langford, John Sillitoe on behalf of the Wellcome Sanger Institute COVID-19 Surveillance Team |
| EPI_ISL_3174359, EPI_ISL_3174360, EPI_ISL_3174361 | Lighthouse Lab in Milton Keynes | Wellcome Sanger Institute for the COVID-19 Genomics UK (COG-UK) Consortium | The Lighthouse Lab in Milton Keynes and Alex Alderton, Roberto Amato, Jeffrey Barrett, Sonia Goncalves, Ewan Harrison, David K. Jackson, Ian Johnston, Dominic Kwiatkowski, Cordelia Langford, John Sillitoe on behalf of the Wellcome Sanger Institute COVID-19 Surveillance Team |
| EPI_ISL_3174362, EPI_ISL_3174363 | Lighthouse Lab in Glasgow | Wellcome Sanger Institute for the COVID-19 Genomics UK (COG-UK) Consortium | Harper VanSteenhouse, Yumi Kasai, David Gray, Carol Clugston, Anna Dominiczak and Alex Alderton, Roberto Amato, Jeffrey Barrett, Sonia Goncalves, Ewan Harrison, David K. Jackson, Ian Johnston, Dominic Kwiatkowski, Cordelia Langford, John Sillitoe on behalf of the Wellcome Sanger Institute COVID-19 Surveillance Team |
| EPI_ISL_3174364 | Lighthouse Lab in Milton Keynes | Wellcome Sanger Institute for the COVID-19 Genomics UK (COG-UK) Consortium | The Lighthouse Lab in Milton Keynes and Alex Alderton, Roberto Amato, Jeffrey Barrett, Sonia Goncalves, Ewan Harrison, David K. Jackson, Ian Johnston, Dominic Kwiatkowski, Cordelia Langford, John Sillitoe on behalf of the Wellcome Sanger Institute COVID-19 Surveillance Team |
| EPI_ISL_3174365 | Lighthouse Lab in Glasgow | Wellcome Sanger Institute for the COVID-19 Genomics UK (COG-UK) Consortium | Harper VanSteenhouse, Yumi Kasai, David Gray, Carol Clugston, Anna Dominiczak and Alex Alderton, Roberto Amato, Jeffrey Barrett, Sonia Goncalves, Ewan Harrison, David K. Jackson, Ian Johnston, Dominic Kwiatkowski, Cordelia Langford, John Sillitoe on behalf of the Wellcome Sanger Institute COVID-19 Surveillance Team |
| EPI_ISL_3174366 | Lighthouse Lab in Milton Keynes | Wellcome Sanger Institute for the COVID-19 Genomics UK (COG-UK) Consortium | The Lighthouse Lab in Milton Keynes and Alex Alderton, Roberto Amato, Jeffrey Barrett, Sonia Goncalves, Ewan Harrison, David K. Jackson, Ian Johnston, Dominic Kwiatkowski, Cordelia Langford, John Sillitoe on behalf of the Wellcome Sanger Institute COVID-19 Surveillance Team |
| EPI_ISL_3174367 | Lighthouse Lab in Glasgow | Wellcome Sanger Institute for the COVID-19 Genomics UK (COG-UK) Consortium | Harper VanSteenhouse, Yumi Kasai, David Gray, Carol Clugston, Anna Dominiczak and Alex Alderton, Roberto Amato, Jeffrey Barrett, Sonia Goncalves, Ewan Harrison, David K. Jackson, Ian Johnston, Dominic Kwiatkowski, Cordelia Langford, John Sillitoe on behalf of the Wellcome Sanger Institute COVID-19 Surveillance Team |
| EPI_ISL_3174368, EPI_ISL_3174369 | Lighthouse Lab in Milton Keynes | Wellcome Sanger Institute for the COVID-19 Genomics UK (COG-UK) Consortium | The Lighthouse Lab in Milton Keynes and Alex Alderton, Roberto Amato, Jeffrey Barrett, Sonia Goncalves, Ewan Harrison, David K. Jackson, Ian Johnston, Dominic Kwiatkowski, Cordelia Langford, John Sillitoe on behalf of the Wellcome Sanger Institute COVID-19 Surveillance Team |
| EPI_ISL_3174370 | Lighthouse Lab in Glasgow | Wellcome Sanger Institute for the COVID-19 Genomics UK (COG-UK) Consortium | Harper VanSteenhouse, Yumi Kasai, David Gray, Carol Clugston, Anna Dominiczak and Alex Alderton, Roberto Amato, Jeffrey Barrett, Sonia Goncalves, Ewan Harrison, David K. Jackson, Ian Johnston, Dominic Kwiatkowski, Cordelia Langford, John Sillitoe on behalf of the Wellcome Sanger Institute COVID-19 Surveillance Team |
| EPI_ISL_3174371 | Lighthouse Lab in Milton Keynes | Wellcome Sanger Institute for the COVID-19 Genomics UK (COG-UK) Consortium | The Lighthouse Lab in Milton Keynes and Alex Alderton, Roberto Amato, Jeffrey Barrett, Sonia Goncalves, Ewan Harrison, David K. Jackson, Ian Johnston, Dominic Kwiatkowski, Cordelia Langford, John Sillitoe on behalf of the Wellcome Sanger Institute COVID-19 Surveillance Team |
| EPI_ISL_3174372, EPI_ISL_3174373 | Lighthouse Lab in Glasgow | Wellcome Sanger Institute for the COVID-19 Genomics UK (COG-UK) Consortium | Harper VanSteenhouse, Yumi Kasai, David Gray, Carol Clugston, Anna Dominiczak and Alex Alderton, Roberto Amato, Jeffrey Barrett, Sonia Goncalves, Ewan Harrison, David K. Jackson, Ian Johnston, Dominic Kwiatkowski, Cordelia Langford, John Sillitoe on behalf of the Wellcome Sanger Institute COVID-19 Surveillance Team |
| EPI_ISL_3174374, EPI_ISL_3174375, EPI_ISL_3174376, EPI_ISL_3174377, EPI_ISL_3174378, EPI_ISL_3174379, EPI_ISL_3174380, EPI_ISL_3174381, EPI_ISL_3174382 | Lighthouse Lab in Milton Keynes | Wellcome Sanger Institute for the COVID-19 Genomics UK (COG-UK) Consortium | The Lighthouse Lab in Milton Keynes and Alex Alderton, Roberto Amato, Jeffrey Barrett, Sonia Goncalves, Ewan Harrison, David K. Jackson, Ian Johnston, Dominic Kwiatkowski, Cordelia Langford, John Sillitoe on behalf of the Wellcome Sanger Institute COVID-19 Surveillance Team |
| EPI_ISL_3174383 | Lighthouse Lab in Glasgow | Wellcome Sanger Institute for the COVID-19 Genomics UK (COG-UK) Consortium | Harper VanSteenhouse, Yumi Kasai, David Gray, Carol Clugston, Anna Dominiczak and Alex Alderton, Roberto Amato, Jeffrey Barrett, Sonia Goncalves, Ewan Harrison, David K. Jackson, Ian Johnston, Dominic Kwiatkowski, Cordelia Langford, John Sillitoe on behalf of the Wellcome Sanger Institute COVID-19 Surveillance Team |
| EPI_ISL_3174384, EPI_ISL_3174385 | Lighthouse Lab in Milton Keynes | Wellcome Sanger Institute for the COVID-19 Genomics UK (COG-UK) Consortium | The Lighthouse Lab in Milton Keynes and Alex Alderton, Roberto Amato, Jeffrey Barrett, Sonia Goncalves, Ewan Harrison, David K. Jackson, Ian Johnston, Dominic Kwiatkowski, Cordelia Langford, John Sillitoe on behalf of the Wellcome Sanger Institute COVID-19 Surveillance Team |
| EPI_ISL_3174386 | Lighthouse Lab in Glasgow | Wellcome Sanger Institute for the COVID-19 Genomics UK (COG-UK) Consortium | Harper VanSteenhouse, Yumi Kasai, David Gray, Carol Clugston, Anna Dominiczak and Alex Alderton, Roberto Amato, Jeffrey Barrett, Sonia Goncalves, Ewan Harrison, David K. Jackson, Ian Johnston, Dominic Kwiatkowski, Cordelia Langford, John Sillitoe on behalf of the Wellcome Sanger Institute COVID-19 Surveillance Team |
| EPI_ISL_3174387, EPI_ISL_3174388, EPI_ISL_3174389, EPI_ISL_3174390, EPI_ISL_3174391, EPI_ISL_3174392, EPI_ISL_3174393, EPI_ISL_3174394, EPI_ISL_3174395 | Lighthouse Lab in Milton Keynes | Wellcome Sanger Institute for the COVID-19 Genomics UK (COG-UK) Consortium | The Lighthouse Lab in Milton Keynes and Alex Alderton, Roberto Amato, Jeffrey Barrett, Sonia Goncalves, Ewan Harrison, David K. Jackson, Ian Johnston, Dominic Kwiatkowski, Cordelia Langford, John Sillitoe on behalf of the Wellcome Sanger Institute COVID-19 Surveillance Team |
| EPI_ISL_3174396, EPI_ISL_3174397 | Lighthouse Lab in Glasgow | Wellcome Sanger Institute for the COVID-19 Genomics UK (COG-UK) Consortium | Harper VanSteenhouse, Yumi Kasai, David Gray, Carol Clugston, Anna Dominiczak and Alex Alderton, Roberto Amato, Jeffrey Barrett, Sonia Goncalves, Ewan Harrison, David K. Jackson, Ian Johnston, Dominic Kwiatkowski, Cordelia Langford, John Sillitoe on behalf of the Wellcome Sanger Institute COVID-19 Surveillance Team |
| EPI_ISL_3174398 | Lighthouse Lab in Milton Keynes | Wellcome Sanger Institute for the COVID-19 Genomics UK (COG-UK) Consortium | The Lighthouse Lab in Milton Keynes and Alex Alderton, Roberto Amato, Jeffrey Barrett, Sonia Goncalves, Ewan Harrison, David K. Jackson, Ian Johnston, Dominic Kwiatkowski, Cordelia Langford, John Sillitoe on behalf of the Wellcome Sanger Institute COVID-19 Surveillance Team |
| EPI_ISL_3174399, EPI_ISL_3174400 | Lighthouse Lab in Glasgow | Wellcome Sanger Institute for the COVID-19 Genomics UK (COG-UK) Consortium | Harper VanSteenhouse, Yumi Kasai, David Gray, Carol Clugston, Anna Dominiczak and Alex Alderton, Roberto Amato, Jeffrey Barrett, Sonia Goncalves, Ewan Harrison, David K. Jackson, Ian Johnston, Dominic Kwiatkowski, Cordelia Langford, John Sillitoe on behalf of the Wellcome Sanger Institute COVID-19 Surveillance Team |
| EPI_ISL_3174401, EPI_ISL_3174402, EPI_ISL_3174403 | Lighthouse Lab in Milton Keynes | Wellcome Sanger Institute for the COVID-19 Genomics UK (COG-UK) Consortium | The Lighthouse Lab in Milton Keynes and Alex Alderton, Roberto Amato, Jeffrey Barrett, Sonia Goncalves, Ewan Harrison, David K. Jackson, Ian Johnston, Dominic Kwiatkowski, Cordelia Langford, John Sillitoe on behalf of the Wellcome Sanger Institute COVID-19 Surveillance Team |
| EPI_ISL_3174404 | Lighthouse Lab in Glasgow | Wellcome Sanger Institute for the COVID-19 Genomics UK (COG-UK) Consortium | Harper VanSteenhouse, Yumi Kasai, David Gray, Carol Clugston, Anna Dominiczak and Alex Alderton, Roberto Amato, Jeffrey Barrett, Sonia Goncalves, Ewan Harrison, David K. Jackson, Ian Johnston, Dominic Kwiatkowski, Cordelia Langford, John Sillitoe on behalf of the Wellcome Sanger Institute COVID-19 Surveillance Team |
| EPI_ISL_3174405, EPI_ISL_3174406 | Lighthouse Lab in Milton Keynes | Wellcome Sanger Institute for the COVID-19 Genomics UK (COG-UK) Consortium | The Lighthouse Lab in Milton Keynes and Alex Alderton, Roberto Amato, Jeffrey Barrett, Sonia Goncalves, Ewan Harrison, David K. Jackson, Ian Johnston, Dominic Kwiatkowski, Cordelia Langford, John Sillitoe on behalf of the Wellcome Sanger Institute COVID-19 Surveillance Team |
| EPI_ISL_3174407 | Lighthouse Lab in Glasgow | Wellcome Sanger Institute for the COVID-19 Genomics UK (COG-UK) Consortium | Harper VanSteenhouse, Yumi Kasai, David Gray, Carol Clugston, Anna Dominiczak and Alex Alderton, Roberto Amato, Jeffrey Barrett, Sonia Goncalves, Ewan Harrison, David K. Jackson, Ian Johnston, Dominic Kwiatkowski, Cordelia Langford, John Sillitoe on behalf of the Wellcome Sanger Institute COVID-19 Surveillance Team |
| EPI_ISL_3174408, EPI_ISL_3174409 | Lighthouse Lab in Milton Keynes | Wellcome Sanger Institute for the COVID-19 Genomics UK (COG-UK) Consortium | The Lighthouse Lab in Milton Keynes and Alex Alderton, Roberto Amato, Jeffrey Barrett, Sonia Goncalves, Ewan Harrison, David K. Jackson, Ian Johnston, Dominic Kwiatkowski, Cordelia Langford, John Sillitoe on behalf of the Wellcome Sanger Institute COVID-19 Surveillance Team |
| EPI_ISL_3174410, EPI_ISL_3174411, EPI_ISL_3174412 | Lighthouse Lab in Glasgow | Wellcome Sanger Institute for the COVID-19 Genomics UK (COG-UK) Consortium | Harper VanSteenhouse, Yumi Kasai, David Gray, Carol Clugston, Anna Dominiczak and Alex Alderton, Roberto Amato, Jeffrey Barrett, Sonia Goncalves, Ewan Harrison, David K. Jackson, Ian Johnston, Dominic Kwiatkowski, Cordelia Langford, John Sillitoe on behalf of the Wellcome Sanger Institute COVID-19 Surveillance Team |

| | | | |
|---|---|---|---|
| EPI_ISL_3174413 | Lighthouse Lab in Milton Keynes | Wellcome Sanger Institute for the COVID-19 Genomics UK (COG-UK) Consortium | The Lighthouse Lab in Milton Keynes and Alex Alderton, Roberto Amato, Jeffrey Barrett, Sonia Goncalves, Ewan Harrison, David K. Jackson, Ian Johnston, Dominic Kwiatkowski, Cordelia Langford, John Sillitoe on behalf of the Wellcome Sanger Institute COVID-19 Surveillance Team |
| EPI_ISL_3174414 | Lighthouse Lab in Glasgow | Wellcome Sanger Institute for the COVID-19 Genomics UK (COG-UK) Consortium | Harper VanSteenhouse, Yumi Kasai, David Gray, Carol Clugston, Anna Dominiczak and Alex Alderton, Roberto Amato, Jeffrey Barrett, Sonia Goncalves, Ewan Harrison, David K. Jackson, Ian Johnston, Dominic Kwiatkowski, Cordelia Langford, John Sillitoe on behalf of the Wellcome Sanger Institute COVID-19 Surveillance Team |
| EPI_ISL_3174415 | Lighthouse Lab in Milton Keynes | Wellcome Sanger Institute for the COVID-19 Genomics UK (COG-UK) Consortium | The Lighthouse Lab in Milton Keynes and Alex Alderton, Roberto Amato, Jeffrey Barrett, Sonia Goncalves, Ewan Harrison, David K. Jackson, Ian Johnston, Dominic Kwiatkowski, Cordelia Langford, John Sillitoe on behalf of the Wellcome Sanger Institute COVID-19 Surveillance Team |
| EPI_ISL_3174416 | Lighthouse Lab in Glasgow | Wellcome Sanger Institute for the COVID-19 Genomics UK (COG-UK) Consortium | Harper VanSteenhouse, Yumi Kasai, David Gray, Carol Clugston, Anna Dominiczak and Alex Alderton, Roberto Amato, Jeffrey Barrett, Sonia Goncalves, Ewan Harrison, David K. Jackson, Ian Johnston, Dominic Kwiatkowski, Cordelia Langford, John Sillitoe on behalf of the Wellcome Sanger Institute COVID-19 Surveillance Team |
| EPI_ISL_3174417 | Lighthouse Lab in Milton Keynes | Wellcome Sanger Institute for the COVID-19 Genomics UK (COG-UK) Consortium | The Lighthouse Lab in Milton Keynes and Alex Alderton, Roberto Amato, Jeffrey Barrett, Sonia Goncalves, Ewan Harrison, David K. Jackson, Ian Johnston, Dominic Kwiatkowski, Cordelia Langford, John Sillitoe on behalf of the Wellcome Sanger Institute COVID-19 Surveillance Team |
| EPI_ISL_3174418, EPI_ISL_3174419, EPI_ISL_3174420, EPI_ISL_3174421, EPI_ISL_3174422 | Lighthouse Lab in Glasgow | Wellcome Sanger Institute for the COVID-19 Genomics UK (COG-UK) Consortium | Harper VanSteenhouse, Yumi Kasai, David Gray, Carol Clugston, Anna Dominiczak and Alex Alderton, Roberto Amato, Jeffrey Barrett, Sonia Goncalves, Ewan Harrison, David K. Jackson, Ian Johnston, Dominic Kwiatkowski, Cordelia Langford, John Sillitoe on behalf of the Wellcome Sanger Institute COVID-19 Surveillance Team |
| EPI_ISL_3174423, EPI_ISL_3174424 | Lighthouse Lab in Milton Keynes | Wellcome Sanger Institute for the COVID-19 Genomics UK (COG-UK) Consortium | The Lighthouse Lab in Milton Keynes and Alex Alderton, Roberto Amato, Jeffrey Barrett, Sonia Goncalves, Ewan Harrison, David K. Jackson, Ian Johnston, Dominic Kwiatkowski, Cordelia Langford, John Sillitoe on behalf of the Wellcome Sanger Institute COVID-19 Surveillance Team |
| EPI_ISL_3174425, EPI_ISL_3174426, EPI_ISL_3174427, EPI_ISL_3174428 | Lighthouse Lab in Glasgow | Wellcome Sanger Institute for the COVID-19 Genomics UK (COG-UK) Consortium | Harper VanSteenhouse, Yumi Kasai, David Gray, Carol Clugston, Anna Dominiczak and Alex Alderton, Roberto Amato, Jeffrey Barrett, Sonia Goncalves, Ewan Harrison, David K. Jackson, Ian Johnston, Dominic Kwiatkowski, Cordelia Langford, John Sillitoe on behalf of the Wellcome Sanger Institute COVID-19 Surveillance Team |
| EPI_ISL_3174429, EPI_ISL_3174430, EPI_ISL_3174431, EPI_ISL_3174432 | Lighthouse Lab in Milton Keynes | Wellcome Sanger Institute for the COVID-19 Genomics UK (COG-UK) Consortium | The Lighthouse Lab in Milton Keynes and Alex Alderton, Roberto Amato, Jeffrey Barrett, Sonia Goncalves, Ewan Harrison, David K. Jackson, Ian Johnston, Dominic Kwiatkowski, Cordelia Langford, John Sillitoe on behalf of the Wellcome Sanger Institute COVID-19 Surveillance Team |
| EPI_ISL_3174433, EPI_ISL_3174434 | Lighthouse Lab in Glasgow | Wellcome Sanger Institute for the COVID-19 Genomics UK (COG-UK) Consortium | Harper VanSteenhouse, Yumi Kasai, David Gray, Carol Clugston, Anna Dominiczak and Alex Alderton, Roberto Amato, Jeffrey Barrett, Sonia Goncalves, Ewan Harrison, David K. Jackson, Ian Johnston, Dominic Kwiatkowski, Cordelia Langford, John Sillitoe on behalf of the Wellcome Sanger Institute COVID-19 Surveillance Team |
| EPI_ISL_3174435, EPI_ISL_3174436, EPI_ISL_3174437, EPI_ISL_3174438 | Lighthouse Lab in Milton Keynes | Wellcome Sanger Institute for the COVID-19 Genomics UK (COG-UK) Consortium | The Lighthouse Lab in Milton Keynes and Alex Alderton, Roberto Amato, Jeffrey Barrett, Sonia Goncalves, Ewan Harrison, David K. Jackson, Ian Johnston, Dominic Kwiatkowski, Cordelia Langford, John Sillitoe on behalf of the Wellcome Sanger Institute COVID-19 Surveillance Team |
| EPI_ISL_3174439 | Lighthouse Lab in Glasgow | Wellcome Sanger Institute for the COVID-19 Genomics UK (COG-UK) Consortium | Harper VanSteenhouse, Yumi Kasai, David Gray, Carol Clugston, Anna Dominiczak and Alex Alderton, Roberto Amato, Jeffrey Barrett, Sonia Goncalves, Ewan Harrison, David K. Jackson, Ian Johnston, Dominic Kwiatkowski, Cordelia Langford, John Sillitoe on behalf of the Wellcome Sanger Institute COVID-19 Surveillance Team |
| EPI_ISL_3174440, EPI_ISL_3174441, EPI_ISL_3174442 | Lighthouse Lab in Milton Keynes | Wellcome Sanger Institute for the COVID-19 Genomics UK (COG-UK) Consortium | The Lighthouse Lab in Milton Keynes and Alex Alderton, Roberto Amato, Jeffrey Barrett, Sonia Goncalves, Ewan Harrison, David K. Jackson, Ian Johnston, Dominic Kwiatkowski, Cordelia Langford, John Sillitoe on behalf of the Wellcome Sanger Institute COVID-19 Surveillance Team |
| EPI_ISL_3174443, EPI_ISL_3174444, EPI_ISL_3174445 | Lighthouse Lab in Glasgow | Wellcome Sanger Institute for the COVID-19 Genomics UK (COG-UK) Consortium | Harper VanSteenhouse, Yumi Kasai, David Gray, Carol Clugston, Anna Dominiczak and Alex Alderton, Roberto Amato, Jeffrey Barrett, Sonia Goncalves, Ewan Harrison, David K. Jackson, Ian Johnston, Dominic Kwiatkowski, Cordelia Langford, John Sillitoe on behalf of the Wellcome Sanger Institute COVID-19 Surveillance Team |
| EPI_ISL_3174446, EPI_ISL_3174447 | Lighthouse Lab in Milton Keynes | Wellcome Sanger Institute for the COVID-19 Genomics UK (COG-UK) Consortium | The Lighthouse Lab in Milton Keynes and Alex Alderton, Roberto Amato, Jeffrey Barrett, Sonia Goncalves, Ewan Harrison, David K. Jackson, Ian Johnston, Dominic Kwiatkowski, Cordelia Langford, John Sillitoe on behalf of the Wellcome Sanger Institute COVID-19 Surveillance Team |
| EPI_ISL_3174448, EPI_ISL_3174449 | Lighthouse Lab in Glasgow | Wellcome Sanger Institute for the COVID-19 Genomics UK (COG-UK) Consortium | Harper VanSteenhouse, Yumi Kasai, David Gray, Carol Clugston, Anna Dominiczak and Alex Alderton, Roberto Amato, Jeffrey Barrett, Sonia Goncalves, Ewan Harrison, David K. Jackson, Ian Johnston, Dominic Kwiatkowski, Cordelia Langford, John Sillitoe on behalf of the Wellcome Sanger Institute COVID-19 Surveillance Team |
| EPI_ISL_3174450, EPI_ISL_3174451 | Lighthouse Lab in Milton Keynes | Wellcome Sanger Institute for the COVID-19 Genomics UK (COG-UK) Consortium | The Lighthouse Lab in Milton Keynes and Alex Alderton, Roberto Amato, Jeffrey Barrett, Sonia Goncalves, Ewan Harrison, David K. Jackson, Ian Johnston, Dominic Kwiatkowski, Cordelia Langford, John Sillitoe on behalf of the Wellcome Sanger Institute COVID-19 Surveillance Team |
| EPI_ISL_3174452, EPI_ISL_3174453 | Lighthouse Lab in Glasgow | Wellcome Sanger Institute for the COVID-19 Genomics UK (COG-UK) Consortium | Harper VanSteenhouse, Yumi Kasai, David Gray, Carol Clugston, Anna Dominiczak and Alex Alderton, Roberto Amato, Jeffrey Barrett, Sonia Goncalves, Ewan Harrison, David K. Jackson, Ian Johnston, Dominic Kwiatkowski, Cordelia Langford, John Sillitoe on behalf of the Wellcome Sanger Institute COVID-19 Surveillance Team |
| EPI_ISL_3174454, EPI_ISL_3174455 | Lighthouse Lab in Milton Keynes | Wellcome Sanger Institute for the COVID-19 Genomics UK (COG-UK) Consortium | The Lighthouse Lab in Milton Keynes and Alex Alderton, Roberto Amato, Jeffrey Barrett, Sonia Goncalves, Ewan Harrison, David K. Jackson, Ian Johnston, Dominic Kwiatkowski, Cordelia Langford, John Sillitoe on behalf of the Wellcome Sanger Institute COVID-19 Surveillance Team |
| EPI_ISL_3174456, EPI_ISL_3174457 | Lighthouse Lab in Glasgow | Wellcome Sanger Institute for the COVID-19 Genomics UK (COG-UK) Consortium | Harper VanSteenhouse, Yumi Kasai, David Gray, Carol Clugston, Anna Dominiczak and Alex Alderton, Roberto Amato, Jeffrey Barrett, Sonia Goncalves, Ewan Harrison, David K. Jackson, Ian Johnston, Dominic Kwiatkowski, Cordelia Langford, John Sillitoe on behalf of the Wellcome Sanger Institute COVID-19 Surveillance Team |
| EPI_ISL_3174458, EPI_ISL_3174459, EPI_ISL_3174460 | Lighthouse Lab in Milton Keynes | Wellcome Sanger Institute for the COVID-19 Genomics UK (COG-UK) Consortium | The Lighthouse Lab in Milton Keynes and Alex Alderton, Roberto Amato, Jeffrey Barrett, Sonia Goncalves, Ewan Harrison, David K. Jackson, Ian Johnston, Dominic Kwiatkowski, Cordelia Langford, John Sillitoe on behalf of the Wellcome Sanger Institute COVID-19 Surveillance Team |
| EPI_ISL_3174461, EPI_ISL_3174462 | Lighthouse Lab in Glasgow | Wellcome Sanger Institute for the COVID-19 Genomics UK (COG-UK) Consortium | Harper VanSteenhouse, Yumi Kasai, David Gray, Carol Clugston, Anna Dominiczak and Alex Alderton, Roberto Amato, Jeffrey Barrett, Sonia Goncalves, Ewan Harrison, David K. Jackson, Ian Johnston, Dominic Kwiatkowski, Cordelia Langford, John Sillitoe on behalf of the Wellcome Sanger Institute COVID-19 Surveillance Team |
| EPI_ISL_3174463, EPI_ISL_3174464, EPI_ISL_3174465, EPI_ISL_3174466 | Lighthouse Lab in Milton Keynes | Wellcome Sanger Institute for the COVID-19 Genomics UK (COG-UK) Consortium | The Lighthouse Lab in Milton Keynes and Alex Alderton, Roberto Amato, Jeffrey Barrett, Sonia Goncalves, Ewan Harrison, David K. Jackson, Ian Johnston, Dominic Kwiatkowski, Cordelia Langford, John Sillitoe on behalf of the Wellcome Sanger Institute COVID-19 Surveillance Team |
| EPI_ISL_3174467 | Lighthouse Lab in Glasgow | Wellcome Sanger Institute for the COVID-19 Genomics UK (COG-UK) Consortium | Harper VanSteenhouse, Yumi Kasai, David Gray, Carol Clugston, Anna Dominiczak and Alex Alderton, Roberto Amato, Jeffrey Barrett, Sonia Goncalves, Ewan Harrison, David K. Jackson, Ian Johnston, Dominic Kwiatkowski, Cordelia Langford, John Sillitoe on behalf of the Wellcome Sanger Institute COVID-19 Surveillance Team |
| EPI_ISL_3174468, EPI_ISL_3174469 | Lighthouse Lab in Milton Keynes | Wellcome Sanger Institute for the COVID-19 Genomics UK (COG-UK) Consortium | The Lighthouse Lab in Milton Keynes and Alex Alderton, Roberto Amato, Jeffrey Barrett, Sonia Goncalves, Ewan Harrison, David K. Jackson, Ian Johnston, Dominic Kwiatkowski, Cordelia Langford, John Sillitoe on behalf of the Wellcome Sanger Institute COVID-19 Surveillance Team |
| EPI_ISL_3174470 | Lighthouse Lab in Glasgow | Wellcome Sanger Institute for the COVID-19 Genomics UK (COG-UK) Consortium | Harper VanSteenhouse, Yumi Kasai, David Gray, Carol Clugston, Anna Dominiczak and Alex Alderton, Roberto Amato, Jeffrey Barrett, Sonia Goncalves, Ewan Harrison, David K. Jackson, Ian Johnston, Dominic Kwiatkowski, Cordelia Langford, John Sillitoe on behalf of the Wellcome Sanger Institute COVID-19 Surveillance Team |
| EPI_ISL_3174471, EPI_ISL_3174472, EPI_ISL_3174473, EPI_ISL_3174474 | Lighthouse Lab in Milton Keynes | Wellcome Sanger Institute for the COVID-19 Genomics UK (COG-UK) Consortium | The Lighthouse Lab in Milton Keynes and Alex Alderton, Roberto Amato, Jeffrey Barrett, Sonia Goncalves, Ewan Harrison, David K. Jackson, Ian Johnston, Dominic Kwiatkowski, Cordelia Langford, John Sillitoe on behalf of the Wellcome Sanger Institute COVID-19 Surveillance Team |
| EPI_ISL_3174475 | Lighthouse Lab in Glasgow | Wellcome Sanger Institute for the COVID-19 Genomics UK (COG-UK) Consortium | Harper VanSteenhouse, Yumi Kasai, David Gray, Carol Clugston, Anna Dominiczak and Alex Alderton, Roberto Amato, Jeffrey Barrett, Sonia Goncalves, Ewan Harrison, David K. Jackson, Ian Johnston, Dominic Kwiatkowski, Cordelia Langford, John Sillitoe on behalf of the Wellcome Sanger Institute COVID-19 Surveillance Team |
| EPI_ISL_3174476, EPI_ISL_3174477 | Lighthouse Lab in Milton Keynes | Wellcome Sanger Institute for the COVID-19 Genomics UK (COG-UK) Consortium | The Lighthouse Lab in Milton Keynes and Alex Alderton, Roberto Amato, Jeffrey Barrett, Sonia Goncalves, Ewan Harrison, David K. Jackson, Ian Johnston, Dominic Kwiatkowski, Cordelia Langford, John Sillitoe on behalf of the Wellcome Sanger Institute COVID-19 Surveillance Team |
| EPI_ISL_3174478, EPI_ISL_3174479, EPI_ISL_3174480 | Lighthouse Lab in Glasgow | Wellcome Sanger Institute for the COVID-19 Genomics UK (COG-UK) Consortium | Harper VanSteenhouse, Yumi Kasai, David Gray, Carol Clugston, Anna Dominiczak and Alex Alderton, Roberto Amato, Jeffrey Barrett, Sonia Goncalves, Ewan Harrison, David K. Jackson, Ian Johnston, Dominic Kwiatkowski, Cordelia Langford, John Sillitoe on behalf of the Wellcome Sanger Institute COVID-19 Surveillance Team |
| EPI_ISL_3174481 | Lighthouse Lab in Milton Keynes | Wellcome Sanger Institute for the COVID-19 Genomics UK (COG-UK) Consortium | The Lighthouse Lab in Milton Keynes and Alex Alderton, Roberto Amato, Jeffrey Barrett, Sonia Goncalves, Ewan Harrison, David K. Jackson, Ian Johnston, Dominic Kwiatkowski, Cordelia Langford, John Sillitoe on behalf of the Wellcome Sanger Institute COVID-19 Surveillance Team |
| EPI_ISL_3174482, EPI_ISL_3174483 | Lighthouse Lab in Glasgow | Wellcome Sanger Institute for the COVID-19 Genomics UK (COG-UK) Consortium | Harper VanSteenhouse, Yumi Kasai, David Gray, Carol Clugston, Anna Dominiczak and Alex Alderton, Roberto Amato, Jeffrey Barrett, Sonia Goncalves, Ewan Harrison, David K. Jackson, Ian Johnston, Dominic Kwiatkowski, Cordelia Langford, John Sillitoe on behalf of the Wellcome Sanger Institute COVID-19 Surveillance Team |
| EPI_ISL_3174484 | Lighthouse Lab in Milton Keynes | Wellcome Sanger Institute for the COVID-19 Genomics UK (COG-UK) Consortium | The Lighthouse Lab in Milton Keynes and Alex Alderton, Roberto Amato, Jeffrey Barrett, Sonia Goncalves, Ewan Harrison, David K. Jackson, Ian Johnston, Dominic Kwiatkowski, Cordelia Langford, John Sillitoe on behalf of the Wellcome Sanger Institute COVID-19 Surveillance Team |
| EPI_ISL_3174485 | Lighthouse Lab in Glasgow | Wellcome Sanger Institute for the COVID-19 Genomics UK (COG-UK) Consortium | Harper VanSteenhouse, Yumi Kasai, David Gray, Carol Clugston, Anna Dominiczak and Alex Alderton, Roberto Amato, Jeffrey Barrett, Sonia Goncalves, Ewan Harrison, David K. Jackson, Ian Johnston, Dominic Kwiatkowski, Cordelia Langford, John Sillitoe on behalf of the Wellcome Sanger Institute COVID-19 Surveillance Team |
| EPI_ISL_3174486 | Lighthouse Lab in Milton Keynes | Wellcome Sanger Institute for the COVID-19 Genomics UK (COG-UK) Consortium | The Lighthouse Lab in Milton Keynes and Alex Alderton, Roberto Amato, Jeffrey Barrett, Sonia Goncalves, Ewan Harrison, David K. Jackson, Ian Johnston, Dominic Kwiatkowski, Cordelia Langford, John Sillitoe on behalf of the Wellcome Sanger Institute COVID-19 Surveillance Team |
| EPI_ISL_3174487, EPI_ISL_3174488, EPI_ISL_3174489, EPI_ISL_3174490 | Lighthouse Lab in Glasgow | Wellcome Sanger Institute for the COVID-19 Genomics UK (COG-UK) Consortium | Harper VanSteenhouse, Yumi Kasai, David Gray, Carol Clugston, Anna Dominiczak and Alex Alderton, Roberto Amato, Jeffrey Barrett, Sonia Goncalves, Ewan Harrison, David K. Jackson, Ian Johnston, Dominic Kwiatkowski, Cordelia Langford, John Sillitoe on behalf of the Wellcome Sanger Institute COVID-19 Surveillance Team |
| EPI_ISL_3174491, EPI_ISL_3174492, EPI_ISL_3174493, | Lighthouse Lab in Milton Keynes | Wellcome Sanger Institute for the COVID-19 Genomics UK (COG-UK) Consortium | The Lighthouse Lab in Milton Keynes and Alex Alderton, Roberto Amato, Jeffrey Barrett, Sonia Goncalves, Ewan Harrison, David K. Jackson, Ian Johnston, Dominic Kwiatkowski, Cordelia Langford, John Sillitoe on behalf of the Wellcome Sanger Institute COVID-19 Surveillance Team |

| | | | |
|---|---|---|---|
| EPI_ISL_3174494 | | | |
| EPI_ISL_3174495, EPI_ISL_3174496, EPI_ISL_3174497, EPI_ISL_3174498 | Lighthouse Lab in Glasgow | Wellcome Sanger Institute for the COVID-19 Genomics UK (COG-UK) Consortium | Harper VanSteenhouse, Yumi Kasai, David Gray, Carol Clugston, Anna Dominiczak and Alex Alderton, Roberto Amato, Jeffrey Barrett, Sonia Goncalves, Ewan Harrison, David K. Jackson, Ian Johnston, Dominic Kwiatkowski, Cordelia Langford, John Sillitoe on behalf of the Wellcome Sanger Institute COVID-19 Surveillance Team |
| EPI_ISL_3174499 | Lighthouse Lab in Milton Keynes | Wellcome Sanger Institute for the COVID-19 Genomics UK (COG-UK) Consortium | The Lighthouse Lab in Milton Keynes and Alex Alderton, Roberto Amato, Jeffrey Barrett, Sonia Goncalves, Ewan Harrison, David K. Jackson, Ian Johnston, Dominic Kwiatkowski, Cordelia Langford, John Sillitoe on behalf of the Wellcome Sanger Institute COVID-19 Surveillance Team |
| EPI_ISL_3174500 | Lighthouse Lab in Glasgow | Wellcome Sanger Institute for the COVID-19 Genomics UK (COG-UK) Consortium | Harper VanSteenhouse, Yumi Kasai, David Gray, Carol Clugston, Anna Dominiczak and Alex Alderton, Roberto Amato, Jeffrey Barrett, Sonia Goncalves, Ewan Harrison, David K. Jackson, Ian Johnston, Dominic Kwiatkowski, Cordelia Langford, John Sillitoe on behalf of the Wellcome Sanger Institute COVID-19 Surveillance Team |
| EPI_ISL_3174501, EPI_ISL_3174502 | Lighthouse Lab in Milton Keynes | Wellcome Sanger Institute for the COVID-19 Genomics UK (COG-UK) Consortium | The Lighthouse Lab in Milton Keynes and Alex Alderton, Roberto Amato, Jeffrey Barrett, Sonia Goncalves, Ewan Harrison, David K. Jackson, Ian Johnston, Dominic Kwiatkowski, Cordelia Langford, John Sillitoe on behalf of the Wellcome Sanger Institute COVID-19 Surveillance Team |
| EPI_ISL_3174503 | Lighthouse Lab in Glasgow | Wellcome Sanger Institute for the COVID-19 Genomics UK (COG-UK) Consortium | Harper VanSteenhouse, Yumi Kasai, David Gray, Carol Clugston, Anna Dominiczak and Alex Alderton, Roberto Amato, Jeffrey Barrett, Sonia Goncalves, Ewan Harrison, David K. Jackson, Ian Johnston, Dominic Kwiatkowski, Cordelia Langford, John Sillitoe on behalf of the Wellcome Sanger Institute COVID-19 Surveillance Team |
| EPI_ISL_3174504, EPI_ISL_3174505, EPI_ISL_3174506, EPI_ISL_3174507, EPI_ISL_3174508, EPI_ISL_3174509, EPI_ISL_3174510, EPI_ISL_3174511, EPI_ISL_3174512 | Lighthouse Lab in Milton Keynes | Wellcome Sanger Institute for the COVID-19 Genomics UK (COG-UK) Consortium | The Lighthouse Lab in Milton Keynes and Alex Alderton, Roberto Amato, Jeffrey Barrett, Sonia Goncalves, Ewan Harrison, David K. Jackson, Ian Johnston, Dominic Kwiatkowski, Cordelia Langford, John Sillitoe on behalf of the Wellcome Sanger Institute COVID-19 Surveillance Team |
| EPI_ISL_3174513 | Lighthouse Lab in Glasgow | Wellcome Sanger Institute for the COVID-19 Genomics UK (COG-UK) Consortium | Harper VanSteenhouse, Yumi Kasai, David Gray, Carol Clugston, Anna Dominiczak and Alex Alderton, Roberto Amato, Jeffrey Barrett, Sonia Goncalves, Ewan Harrison, David K. Jackson, Ian Johnston, Dominic Kwiatkowski, Cordelia Langford, John Sillitoe on behalf of the Wellcome Sanger Institute COVID-19 Surveillance Team |
| EPI_ISL_3174514 | Lighthouse Lab in Alderley Park | Wellcome Sanger Institute for the COVID-19 Genomics UK (COG-UK) Consortium | Jacquelyn Wynn, Mairead Hyland, The Lighthouse Lab in Alderley Park and Alex Alderton, Roberto Amato, Jeffrey Barrett, Sonia Goncalves, Ewan Harrison, David K. Jackson, Ian Johnston, Dominic Kwiatkowski, Cordelia Langford, John Sillitoe on behalf of the Wellcome Sanger Institute COVID-19 Surveillance Team |
| EPI_ISL_3174515 | Lighthouse Lab in Milton Keynes | Wellcome Sanger Institute for the COVID-19 Genomics UK (COG-UK) Consortium | The Lighthouse Lab in Milton Keynes and Alex Alderton, Roberto Amato, Jeffrey Barrett, Sonia Goncalves, Ewan Harrison, David K. Jackson, Ian Johnston, Dominic Kwiatkowski, Cordelia Langford, John Sillitoe on behalf of the Wellcome Sanger Institute COVID-19 Surveillance Team |
| EPI_ISL_3174516, EPI_ISL_3174517 | Lighthouse Lab in Alderley Park | Wellcome Sanger Institute for the COVID-19 Genomics UK (COG-UK) Consortium | Jacquelyn Wynn, Mairead Hyland, The Lighthouse Lab in Alderley Park and Alex Alderton, Roberto Amato, Jeffrey Barrett, Sonia Goncalves, Ewan Harrison, David K. Jackson, Ian Johnston, Dominic Kwiatkowski, Cordelia Langford, John Sillitoe on behalf of the Wellcome Sanger Institute COVID-19 Surveillance Team |
| EPI_ISL_3174518 | Lighthouse Lab in Milton Keynes | Wellcome Sanger Institute for the COVID-19 Genomics UK (COG-UK) Consortium | The Lighthouse Lab in Milton Keynes and Alex Alderton, Roberto Amato, Jeffrey Barrett, Sonia Goncalves, Ewan Harrison, David K. Jackson, Ian Johnston, Dominic Kwiatkowski, Cordelia Langford, John Sillitoe on behalf of the Wellcome Sanger Institute COVID-19 Surveillance Team |
| EPI_ISL_3174519 | Lighthouse Laboratory Plymouth | Wellcome Sanger Institute for the COVID-19 Genomics UK (COG-UK) Consortium | Lighthouse Laboratory Plymouth and Alex Alderton, Roberto Amato, Jeffrey Barrett, Sonia Goncalves, Ewan Harrison, David K. Jackson, Ian Johnston, Dominic Kwiatkowski, Cordelia Langford, John Sillitoe on behalf of the Wellcome Sanger Institute COVID-19 Surveillance Team |
| EPI_ISL_3174520 | Lighthouse Lab in Milton Keynes | Wellcome Sanger Institute for the COVID-19 Genomics UK (COG-UK) Consortium | The Lighthouse Lab in Milton Keynes and Alex Alderton, Roberto Amato, Jeffrey Barrett, Sonia Goncalves, Ewan Harrison, David K. Jackson, Ian Johnston, Dominic Kwiatkowski, Cordelia Langford, John Sillitoe on behalf of the Wellcome Sanger Institute COVID-19 Surveillance Team |
| EPI_ISL_3174521, EPI_ISL_3174522 | Lighthouse Lab in Glasgow | Wellcome Sanger Institute for the COVID-19 Genomics UK (COG-UK) Consortium | Harper VanSteenhouse, Yumi Kasai, David Gray, Carol Clugston, Anna Dominiczak and Alex Alderton, Roberto Amato, Jeffrey Barrett, Sonia Goncalves, Ewan Harrison, David K. Jackson, Ian Johnston, Dominic Kwiatkowski, Cordelia Langford, John Sillitoe on behalf of the Wellcome Sanger Institute COVID-19 Surveillance Team |
| EPI_ISL_3174523 | Lighthouse Lab in Milton Keynes | Wellcome Sanger Institute for the COVID-19 Genomics UK (COG-UK) Consortium | The Lighthouse Lab in Milton Keynes and Alex Alderton, Roberto Amato, Jeffrey Barrett, Sonia Goncalves, Ewan Harrison, David K. Jackson, Ian Johnston, Dominic Kwiatkowski, Cordelia Langford, John Sillitoe on behalf of the Wellcome Sanger Institute COVID-19 Surveillance Team |
| EPI_ISL_3174524 | Lighthouse Lab in Alderley Park | Wellcome Sanger Institute for the COVID-19 Genomics UK (COG-UK) Consortium | Jacquelyn Wynn, Mairead Hyland, The Lighthouse Lab in Alderley Park and Alex Alderton, Roberto Amato, Jeffrey Barrett, Sonia Goncalves, Ewan Harrison, David K. Jackson, Ian Johnston, Dominic Kwiatkowski, Cordelia Langford, John Sillitoe on behalf of the Wellcome Sanger Institute COVID-19 Surveillance Team |
| EPI_ISL_3174525 | Lighthouse Lab in Milton Keynes | Wellcome Sanger Institute for the COVID-19 Genomics UK (COG-UK) Consortium | The Lighthouse Lab in Milton Keynes and Alex Alderton, Roberto Amato, Jeffrey Barrett, Sonia Goncalves, Ewan Harrison, David K. Jackson, Ian Johnston, Dominic Kwiatkowski, Cordelia Langford, John Sillitoe on behalf of the Wellcome Sanger Institute COVID-19 Surveillance Team |
| EPI_ISL_3174526, EPI_ISL_3174527, EPI_ISL_3174528 | Lighthouse Lab in Alderley Park | Wellcome Sanger Institute for the COVID-19 Genomics UK (COG-UK) Consortium | Jacquelyn Wynn, Mairead Hyland, The Lighthouse Lab in Alderley Park and Alex Alderton, Roberto Amato, Jeffrey Barrett, Sonia Goncalves, Ewan Harrison, David K. Jackson, Ian Johnston, Dominic Kwiatkowski, Cordelia Langford, John Sillitoe on behalf of the Wellcome Sanger Institute COVID-19 Surveillance Team |
| EPI_ISL_3174529 | Lighthouse Lab in Milton Keynes | Wellcome Sanger Institute for the COVID-19 Genomics UK (COG-UK) Consortium | The Lighthouse Lab in Milton Keynes and Alex Alderton, Roberto Amato, Jeffrey Barrett, Sonia Goncalves, Ewan Harrison, David K. Jackson, Ian Johnston, Dominic Kwiatkowski, Cordelia Langford, John Sillitoe on behalf of the Wellcome Sanger Institute COVID-19 Surveillance Team |
| EPI_ISL_3174530 | Lighthouse Lab in Glasgow | Wellcome Sanger Institute for the COVID-19 Genomics UK (COG-UK) Consortium | Harper VanSteenhouse, Yumi Kasai, David Gray, Carol Clugston, Anna Dominiczak and Alex Alderton, Roberto Amato, Jeffrey Barrett, Sonia Goncalves, Ewan Harrison, David K. Jackson, Ian Johnston, Dominic Kwiatkowski, Cordelia Langford, John Sillitoe on behalf of the Wellcome Sanger Institute COVID-19 Surveillance Team |
| EPI_ISL_3174531 | Lighthouse Lab in Milton Keynes | Wellcome Sanger Institute for the COVID-19 Genomics UK (COG-UK) Consortium | The Lighthouse Lab in Milton Keynes and Alex Alderton, Roberto Amato, Jeffrey Barrett, Sonia Goncalves, Ewan Harrison, David K. Jackson, Ian Johnston, Dominic Kwiatkowski, Cordelia Langford, John Sillitoe on behalf of the Wellcome Sanger Institute COVID-19 Surveillance Team |
| EPI_ISL_3174532 | Lighthouse Lab in Glasgow | Wellcome Sanger Institute for the COVID-19 Genomics UK (COG-UK) Consortium | Harper VanSteenhouse, Yumi Kasai, David Gray, Carol Clugston, Anna Dominiczak and Alex Alderton, Roberto Amato, Jeffrey Barrett, Sonia Goncalves, Ewan Harrison, David K. Jackson, Ian Johnston, Dominic Kwiatkowski, Cordelia Langford, John Sillitoe on behalf of the Wellcome Sanger Institute COVID-19 Surveillance Team |
| EPI_ISL_3174533 | Lighthouse Lab in Milton Keynes | Wellcome Sanger Institute for the COVID-19 Genomics UK (COG-UK) Consortium | The Lighthouse Lab in Milton Keynes and Alex Alderton, Roberto Amato, Jeffrey Barrett, Sonia Goncalves, Ewan Harrison, David K. Jackson, Ian Johnston, Dominic Kwiatkowski, Cordelia Langford, John Sillitoe on behalf of the Wellcome Sanger Institute COVID-19 Surveillance Team |
| EPI_ISL_3174534 | Lighthouse Laboratory Plymouth | Wellcome Sanger Institute for the COVID-19 Genomics UK (COG-UK) Consortium | Lighthouse Laboratory Plymouth and Alex Alderton, Roberto Amato, Jeffrey Barrett, Sonia Goncalves, Ewan Harrison, David K. Jackson, Ian Johnston, Dominic Kwiatkowski, Cordelia Langford, John Sillitoe on behalf of the Wellcome Sanger Institute COVID-19 Surveillance Team |
| EPI_ISL_3174535 | Lighthouse Lab in Alderley Park | Wellcome Sanger Institute for the COVID-19 Genomics UK (COG-UK) Consortium | Jacquelyn Wynn, Mairead Hyland, The Lighthouse Lab in Alderley Park and Alex Alderton, Roberto Amato, Jeffrey Barrett, Sonia Goncalves, Ewan Harrison, David K. Jackson, Ian Johnston, Dominic Kwiatkowski, Cordelia Langford, John Sillitoe on behalf of the Wellcome Sanger Institute COVID-19 Surveillance Team |
| EPI_ISL_3174536 | Lighthouse Laboratory Plymouth | Wellcome Sanger Institute for the COVID-19 Genomics UK (COG-UK) Consortium | Lighthouse Laboratory Plymouth and Alex Alderton, Roberto Amato, Jeffrey Barrett, Sonia Goncalves, Ewan Harrison, David K. Jackson, Ian Johnston, Dominic Kwiatkowski, Cordelia Langford, John Sillitoe on behalf of the Wellcome Sanger Institute COVID-19 Surveillance Team |
| EPI_ISL_3174537 | Lighthouse Lab in Alderley Park | Wellcome Sanger Institute for the COVID-19 Genomics UK (COG-UK) Consortium | Jacquelyn Wynn, Mairead Hyland, The Lighthouse Lab in Alderley Park and Alex Alderton, Roberto Amato, Jeffrey Barrett, Sonia Goncalves, Ewan Harrison, David K. Jackson, Ian Johnston, Dominic Kwiatkowski, Cordelia Langford, John Sillitoe on behalf of the Wellcome Sanger Institute COVID-19 Surveillance Team |
| EPI_ISL_3174538 | Lighthouse Lab in Glasgow | Wellcome Sanger Institute for the COVID-19 Genomics UK (COG-UK) Consortium | Harper VanSteenhouse, Yumi Kasai, David Gray, Carol Clugston, Anna Dominiczak and Alex Alderton, Roberto Amato, Jeffrey Barrett, Sonia Goncalves, Ewan Harrison, David K. Jackson, Ian Johnston, Dominic Kwiatkowski, Cordelia Langford, John Sillitoe on behalf of the Wellcome Sanger Institute COVID-19 Surveillance Team |
| EPI_ISL_3174539 | Lighthouse Lab in Alderley Park | Wellcome Sanger Institute for the COVID-19 Genomics UK (COG-UK) Consortium | Jacquelyn Wynn, Mairead Hyland, The Lighthouse Lab in Alderley Park and Alex Alderton, Roberto Amato, Jeffrey Barrett, Sonia Goncalves, Ewan Harrison, David K. Jackson, Ian Johnston, Dominic Kwiatkowski, Cordelia Langford, John Sillitoe on behalf of the Wellcome Sanger Institute COVID-19 Surveillance Team |
| EPI_ISL_3174540 | Lighthouse Laboratory Plymouth | Wellcome Sanger Institute for the COVID-19 Genomics UK (COG-UK) Consortium | Lighthouse Laboratory Plymouth and Alex Alderton, Roberto Amato, Jeffrey Barrett, Sonia Goncalves, Ewan Harrison, David K. Jackson, Ian Johnston, Dominic Kwiatkowski, Cordelia Langford, John Sillitoe on behalf of the Wellcome Sanger Institute COVID-19 Surveillance Team |
| EPI_ISL_3174541 | Lighthouse Lab in Milton Keynes | Wellcome Sanger Institute for the COVID-19 Genomics UK (COG-UK) Consortium | The Lighthouse Lab in Milton Keynes and Alex Alderton, Roberto Amato, Jeffrey Barrett, Sonia Goncalves, Ewan Harrison, David K. Jackson, Ian Johnston, Dominic Kwiatkowski, Cordelia Langford, John Sillitoe on behalf of the Wellcome Sanger Institute COVID-19 Surveillance Team |
| EPI_ISL_3174542 | Lighthouse Lab in Glasgow | Wellcome Sanger Institute for the COVID-19 Genomics UK (COG-UK) Consortium | Harper VanSteenhouse, Yumi Kasai, David Gray, Carol Clugston, Anna Dominiczak and Alex Alderton, Roberto Amato, Jeffrey Barrett, Sonia Goncalves, Ewan Harrison, David K. Jackson, Ian Johnston, Dominic Kwiatkowski, Cordelia Langford, John Sillitoe on behalf of the Wellcome Sanger Institute COVID-19 Surveillance Team |
| EPI_ISL_3174543, EPI_ISL_3174544 | Lighthouse Laboratory Plymouth | Wellcome Sanger Institute for the COVID-19 Genomics UK (COG-UK) Consortium | Lighthouse Laboratory Plymouth and Alex Alderton, Roberto Amato, Jeffrey Barrett, Sonia Goncalves, Ewan Harrison, David K. Jackson, Ian Johnston, Dominic Kwiatkowski, Cordelia Langford, John Sillitoe on behalf of the Wellcome Sanger Institute COVID-19 Surveillance Team |
| EPI_ISL_3174545 | Lighthouse Lab in Alderley Park | Wellcome Sanger Institute for the COVID-19 Genomics UK (COG-UK) Consortium | Jacquelyn Wynn, Mairead Hyland, The Lighthouse Lab in Alderley Park and Alex Alderton, Roberto Amato, Jeffrey Barrett, Sonia Goncalves, Ewan Harrison, David K. Jackson, Ian Johnston, Dominic Kwiatkowski, Cordelia Langford, John Sillitoe on behalf of the Wellcome Sanger Institute COVID-19 Surveillance Team |
| EPI_ISL_3174546 | Lighthouse Lab in Milton Keynes | Wellcome Sanger Institute for the COVID-19 Genomics UK (COG-UK) Consortium | The Lighthouse Lab in Milton Keynes and Alex Alderton, Roberto Amato, Jeffrey Barrett, Sonia Goncalves, Ewan Harrison, David K. Jackson, Ian Johnston, Dominic Kwiatkowski, Cordelia Langford, John Sillitoe on behalf of the Wellcome Sanger Institute COVID-19 Surveillance Team |
| EPI_ISL_3174547, EPI_ISL_3174548 | Lighthouse Lab in Glasgow | Wellcome Sanger Institute for the COVID-19 Genomics UK (COG-UK) Consortium | Harper VanSteenhouse, Yumi Kasai, David Gray, Carol Clugston, Anna Dominiczak and Alex Alderton, Roberto Amato, Jeffrey Barrett, Sonia Goncalves, Ewan Harrison, David K. Jackson, Ian Johnston, Dominic Kwiatkowski, Cordelia Langford, John Sillitoe on behalf of the Wellcome Sanger Institute COVID-19 Surveillance Team |
| EPI_ISL_3174549 | Lighthouse Laboratory Plymouth | Wellcome Sanger Institute for the COVID-19 Genomics UK (COG-UK) Consortium | Lighthouse Laboratory Plymouth and Alex Alderton, Roberto Amato, Jeffrey Barrett, Sonia Goncalves, Ewan Harrison, David K. Jackson, Ian Johnston, Dominic Kwiatkowski, Cordelia Langford, John Sillitoe on behalf of the Wellcome Sanger Institute COVID-19 Surveillance Team |
| EPI_ISL_3174550, EPI_ISL_3174551 | Lighthouse Lab in Milton Keynes | Wellcome Sanger Institute for the COVID-19 Genomics UK (COG-UK) Consortium | The Lighthouse Lab in Milton Keynes and Alex Alderton, Roberto Amato, Jeffrey Barrett, Sonia Goncalves, Ewan Harrison, David K. Jackson, Ian Johnston, Dominic Kwiatkowski, Cordelia Langford, John Sillitoe on behalf of the Wellcome Sanger Institute COVID-19 Surveillance Team |
| EPI_ISL_3174552 | Lighthouse Lab in Alderley Park | Wellcome Sanger Institute for the COVID-19 Genomics UK (COG-UK) Consortium | Jacquelyn Wynn, Mairead Hyland, The Lighthouse Lab in Alderley Park and Alex Alderton, Roberto Amato, Jeffrey Barrett, Sonia Goncalves, Ewan Harrison, David K. Jackson, Ian Johnston, Dominic Kwiatkowski, Cordelia Langford, John Sillitoe on behalf of the Wellcome Sanger Institute COVID-19 Surveillance Team |
| EPI_ISL_3174553 | Lighthouse Lab in Glasgow | Wellcome Sanger Institute for the COVID-19 Genomics UK (COG-UK) Consortium | Harper VanSteenhouse, Yumi Kasai, David Gray, Carol Clugston, Anna Dominiczak and Alex Alderton, Roberto Amato, Jeffrey Barrett, Sonia Goncalves, Ewan Harrison, David K. Jackson, Ian Johnston, Dominic Kwiatkowski, Cordelia Langford, John Sillitoe on behalf of the Wellcome Sanger Institute COVID-19 Surveillance Team |

| | | | |
|---|---|---|---|
| EPI_ISL_3174554 | Lighthouse Lab in Milton Keynes | Wellcome Sanger Institute for the COVID-19 Genomics UK (COG-UK) Consortium | The Lighthouse Lab in Milton Keynes and Alex Alderton, Roberto Amato, Jeffrey Barrett, Sonia Goncalves, Ewan Harrison, David K. Jackson, Ian Johnston, Dominic Kwiatkowski, Cordelia Langford, John Sillitoe on behalf of the Wellcome Sanger Institute COVID-19 Surveillance Team |
| EPI_ISL_3174555, EPI_ISL_3174556 | Lighthouse Lab in Glasgow | Wellcome Sanger Institute for the COVID-19 Genomics UK (COG-UK) Consortium | Harper VanSteenhouse, Yumi Kasai, David Gray, Carol Clugston, Anna Dominiczak and Alex Alderton, Roberto Amato, Jeffrey Barrett, Sonia Goncalves, Ewan Harrison, David K. Jackson, Ian Johnston, Dominic Kwiatkowski, Cordelia Langford, John Sillitoe on behalf of the Wellcome Sanger Institute COVID-19 Surveillance Team |
| EPI_ISL_3174557 | Lighthouse Laboratory Plymouth | Wellcome Sanger Institute for the COVID-19 Genomics UK (COG-UK) Consortium | Lighthouse Laboratory Plymouth and Alex Alderton, Roberto Amato, Jeffrey Barrett, Sonia Goncalves, Ewan Harrison, David K. Jackson, Ian Johnston, Dominic Kwiatkowski, Cordelia Langford, John Sillitoe on behalf of the Wellcome Sanger Institute COVID-19 Surveillance Team |
| EPI_ISL_3174558 | Lighthouse Lab in Milton Keynes | Wellcome Sanger Institute for the COVID-19 Genomics UK (COG-UK) Consortium | The Lighthouse Lab in Milton Keynes and Alex Alderton, Roberto Amato, Jeffrey Barrett, Sonia Goncalves, Ewan Harrison, David K. Jackson, Ian Johnston, Dominic Kwiatkowski, Cordelia Langford, John Sillitoe on behalf of the Wellcome Sanger Institute COVID-19 Surveillance Team |
| EPI_ISL_3174559 | Lighthouse Lab in Glasgow | Wellcome Sanger Institute for the COVID-19 Genomics UK (COG-UK) Consortium | Harper VanSteenhouse, Yumi Kasai, David Gray, Carol Clugston, Anna Dominiczak and Alex Alderton, Roberto Amato, Jeffrey Barrett, Sonia Goncalves, Ewan Harrison, David K. Jackson, Ian Johnston, Dominic Kwiatkowski, Cordelia Langford, John Sillitoe on behalf of the Wellcome Sanger Institute COVID-19 Surveillance Team |
| EPI_ISL_3174560, EPI_ISL_3174561 | Lighthouse Lab in Milton Keynes | Wellcome Sanger Institute for the COVID-19 Genomics UK (COG-UK) Consortium | The Lighthouse Lab in Milton Keynes and Alex Alderton, Roberto Amato, Jeffrey Barrett, Sonia Goncalves, Ewan Harrison, David K. Jackson, Ian Johnston, Dominic Kwiatkowski, Cordelia Langford, John Sillitoe on behalf of the Wellcome Sanger Institute COVID-19 Surveillance Team |
| EPI_ISL_3174562 | Lighthouse Lab in Glasgow | Wellcome Sanger Institute for the COVID-19 Genomics UK (COG-UK) Consortium | Harper VanSteenhouse, Yumi Kasai, David Gray, Carol Clugston, Anna Dominiczak and Alex Alderton, Roberto Amato, Jeffrey Barrett, Sonia Goncalves, Ewan Harrison, David K. Jackson, Ian Johnston, Dominic Kwiatkowski, Cordelia Langford, John Sillitoe on behalf of the Wellcome Sanger Institute COVID-19 Surveillance Team |
| EPI_ISL_3174563 | Lighthouse Laboratory Plymouth | Wellcome Sanger Institute for the COVID-19 Genomics UK (COG-UK) Consortium | Lighthouse Laboratory Plymouth and Alex Alderton, Roberto Amato, Jeffrey Barrett, Sonia Goncalves, Ewan Harrison, David K. Jackson, Ian Johnston, Dominic Kwiatkowski, Cordelia Langford, John Sillitoe on behalf of the Wellcome Sanger Institute COVID-19 Surveillance Team |
| EPI_ISL_3174564 | Lighthouse Lab in Alderley Park | Wellcome Sanger Institute for the COVID-19 Genomics UK (COG-UK) Consortium | Jacquelyn Wynn, Mairead Hyland, The Lighthouse Lab in Alderley Park and Alex Alderton, Roberto Amato, Jeffrey Barrett, Sonia Goncalves, Ewan Harrison, David K. Jackson, Ian Johnston, Dominic Kwiatkowski, Cordelia Langford, John Sillitoe on behalf of the Wellcome Sanger Institute COVID-19 Surveillance Team |
| EPI_ISL_3174565 | Lighthouse Lab in Glasgow | Wellcome Sanger Institute for the COVID-19 Genomics UK (COG-UK) Consortium | Harper VanSteenhouse, Yumi Kasai, David Gray, Carol Clugston, Anna Dominiczak and Alex Alderton, Roberto Amato, Jeffrey Barrett, Sonia Goncalves, Ewan Harrison, David K. Jackson, Ian Johnston, Dominic Kwiatkowski, Cordelia Langford, John Sillitoe on behalf of the Wellcome Sanger Institute COVID-19 Surveillance Team |
| EPI_ISL_3174566, EPI_ISL_3174567 | Lighthouse Lab in Alderley Park | Wellcome Sanger Institute for the COVID-19 Genomics UK (COG-UK) Consortium | Jacquelyn Wynn, Mairead Hyland, The Lighthouse Lab in Alderley Park and Alex Alderton, Roberto Amato, Jeffrey Barrett, Sonia Goncalves, Ewan Harrison, David K. Jackson, Ian Johnston, Dominic Kwiatkowski, Cordelia Langford, John Sillitoe on behalf of the Wellcome Sanger Institute COVID-19 Surveillance Team |
| EPI_ISL_3174568 | Lighthouse Lab in Milton Keynes | Wellcome Sanger Institute for the COVID-19 Genomics UK (COG-UK) Consortium | The Lighthouse Lab in Milton Keynes and Alex Alderton, Roberto Amato, Jeffrey Barrett, Sonia Goncalves, Ewan Harrison, David K. Jackson, Ian Johnston, Dominic Kwiatkowski, Cordelia Langford, John Sillitoe on behalf of the Wellcome Sanger Institute COVID-19 Surveillance Team |
| EPI_ISL_3174569 | Lighthouse Lab in Glasgow | Wellcome Sanger Institute for the COVID-19 Genomics UK (COG-UK) Consortium | Harper VanSteenhouse, Yumi Kasai, David Gray, Carol Clugston, Anna Dominiczak and Alex Alderton, Roberto Amato, Jeffrey Barrett, Sonia Goncalves, Ewan Harrison, David K. Jackson, Ian Johnston, Dominic Kwiatkowski, Cordelia Langford, John Sillitoe on behalf of the Wellcome Sanger Institute COVID-19 Surveillance Team |
| EPI_ISL_3174570 | Lighthouse Laboratory Plymouth | Wellcome Sanger Institute for the COVID-19 Genomics UK (COG-UK) Consortium | Lighthouse Laboratory Plymouth and Alex Alderton, Roberto Amato, Jeffrey Barrett, Sonia Goncalves, Ewan Harrison, David K. Jackson, Ian Johnston, Dominic Kwiatkowski, Cordelia Langford, John Sillitoe on behalf of the Wellcome Sanger Institute COVID-19 Surveillance Team |
| EPI_ISL_3174571 | Lighthouse Lab in Glasgow | Wellcome Sanger Institute for the COVID-19 Genomics UK (COG-UK) Consortium | Harper VanSteenhouse, Yumi Kasai, David Gray, Carol Clugston, Anna Dominiczak and Alex Alderton, Roberto Amato, Jeffrey Barrett, Sonia Goncalves, Ewan Harrison, David K. Jackson, Ian Johnston, Dominic Kwiatkowski, Cordelia Langford, John Sillitoe on behalf of the Wellcome Sanger Institute COVID-19 Surveillance Team |
| EPI_ISL_3174572 | Lighthouse Lab in Milton Keynes | Wellcome Sanger Institute for the COVID-19 Genomics UK (COG-UK) Consortium | The Lighthouse Lab in Milton Keynes and Alex Alderton, Roberto Amato, Jeffrey Barrett, Sonia Goncalves, Ewan Harrison, David K. Jackson, Ian Johnston, Dominic Kwiatkowski, Cordelia Langford, John Sillitoe on behalf of the Wellcome Sanger Institute COVID-19 Surveillance Team |
| EPI_ISL_3174573 | Lighthouse Lab in Glasgow | Wellcome Sanger Institute for the COVID-19 Genomics UK (COG-UK) Consortium | Harper VanSteenhouse, Yumi Kasai, David Gray, Carol Clugston, Anna Dominiczak and Alex Alderton, Roberto Amato, Jeffrey Barrett, Sonia Goncalves, Ewan Harrison, David K. Jackson, Ian Johnston, Dominic Kwiatkowski, Cordelia Langford, John Sillitoe on behalf of the Wellcome Sanger Institute COVID-19 Surveillance Team |
| EPI_ISL_3174574 | Lighthouse Lab in Milton Keynes | Wellcome Sanger Institute for the COVID-19 Genomics UK (COG-UK) Consortium | The Lighthouse Lab in Milton Keynes and Alex Alderton, Roberto Amato, Jeffrey Barrett, Sonia Goncalves, Ewan Harrison, David K. Jackson, Ian Johnston, Dominic Kwiatkowski, Cordelia Langford, John Sillitoe on behalf of the Wellcome Sanger Institute COVID-19 Surveillance Team |
| EPI_ISL_3174575, EPI_ISL_3174576 | Lighthouse Lab in Alderley Park | Wellcome Sanger Institute for the COVID-19 Genomics UK (COG-UK) Consortium | Jacquelyn Wynn, Mairead Hyland, The Lighthouse Lab in Alderley Park and Alex Alderton, Roberto Amato, Jeffrey Barrett, Sonia Goncalves, Ewan Harrison, David K. Jackson, Ian Johnston, Dominic Kwiatkowski, Cordelia Langford, John Sillitoe on behalf of the Wellcome Sanger Institute COVID-19 Surveillance Team |
| EPI_ISL_3174577 | Lighthouse Lab in Milton Keynes | Wellcome Sanger Institute for the COVID-19 Genomics UK (COG-UK) Consortium | The Lighthouse Lab in Milton Keynes and Alex Alderton, Roberto Amato, Jeffrey Barrett, Sonia Goncalves, Ewan Harrison, David K. Jackson, Ian Johnston, Dominic Kwiatkowski, Cordelia Langford, John Sillitoe on behalf of the Wellcome Sanger Institute COVID-19 Surveillance Team |
| EPI_ISL_3174578 | Lighthouse Lab in Alderley Park | Wellcome Sanger Institute for the COVID-19 Genomics UK (COG-UK) Consortium | Jacquelyn Wynn, Mairead Hyland, The Lighthouse Lab in Alderley Park and Alex Alderton, Roberto Amato, Jeffrey Barrett, Sonia Goncalves, Ewan Harrison, David K. Jackson, Ian Johnston, Dominic Kwiatkowski, Cordelia Langford, John Sillitoe on behalf of the Wellcome Sanger Institute COVID-19 Surveillance Team |
| EPI_ISL_3174579 | Lighthouse Lab in Milton Keynes | Wellcome Sanger Institute for the COVID-19 Genomics UK (COG-UK) Consortium | The Lighthouse Lab in Milton Keynes and Alex Alderton, Roberto Amato, Jeffrey Barrett, Sonia Goncalves, Ewan Harrison, David K. Jackson, Ian Johnston, Dominic Kwiatkowski, Cordelia Langford, John Sillitoe on behalf of the Wellcome Sanger Institute COVID-19 Surveillance Team |
| EPI_ISL_3174580 | Lighthouse Lab in Alderley Park | Wellcome Sanger Institute for the COVID-19 Genomics UK (COG-UK) Consortium | Jacquelyn Wynn, Mairead Hyland, The Lighthouse Lab in Alderley Park and Alex Alderton, Roberto Amato, Jeffrey Barrett, Sonia Goncalves, Ewan Harrison, David K. Jackson, Ian Johnston, Dominic Kwiatkowski, Cordelia Langford, John Sillitoe on behalf of the Wellcome Sanger Institute COVID-19 Surveillance Team |
| EPI_ISL_3174581 | Lighthouse Lab in Milton Keynes | Wellcome Sanger Institute for the COVID-19 Genomics UK (COG-UK) Consortium | The Lighthouse Lab in Milton Keynes and Alex Alderton, Roberto Amato, Jeffrey Barrett, Sonia Goncalves, Ewan Harrison, David K. Jackson, Ian Johnston, Dominic Kwiatkowski, Cordelia Langford, John Sillitoe on behalf of the Wellcome Sanger Institute COVID-19 Surveillance Team |
| EPI_ISL_3174582, EPI_ISL_3174583 | Lighthouse Lab in Alderley Park | Wellcome Sanger Institute for the COVID-19 Genomics UK (COG-UK) Consortium | Jacquelyn Wynn, Mairead Hyland, The Lighthouse Lab in Alderley Park and Alex Alderton, Roberto Amato, Jeffrey Barrett, Sonia Goncalves, Ewan Harrison, David K. Jackson, Ian Johnston, Dominic Kwiatkowski, Cordelia Langford, John Sillitoe on behalf of the Wellcome Sanger Institute COVID-19 Surveillance Team |
| EPI_ISL_3174584, EPI_ISL_3174585, EPI_ISL_3174586, EPI_ISL_3174587 | Lighthouse Lab in Milton Keynes | Wellcome Sanger Institute for the COVID-19 Genomics UK (COG-UK) Consortium | The Lighthouse Lab in Milton Keynes and Alex Alderton, Roberto Amato, Jeffrey Barrett, Sonia Goncalves, Ewan Harrison, David K. Jackson, Ian Johnston, Dominic Kwiatkowski, Cordelia Langford, John Sillitoe on behalf of the Wellcome Sanger Institute COVID-19 Surveillance Team |
| EPI_ISL_3174588 | Lighthouse Lab in Glasgow | Wellcome Sanger Institute for the COVID-19 Genomics UK (COG-UK) Consortium | Harper VanSteenhouse, Yumi Kasai, David Gray, Carol Clugston, Anna Dominiczak and Alex Alderton, Roberto Amato, Jeffrey Barrett, Sonia Goncalves, Ewan Harrison, David K. Jackson, Ian Johnston, Dominic Kwiatkowski, Cordelia Langford, John Sillitoe on behalf of the Wellcome Sanger Institute COVID-19 Surveillance Team |
| EPI_ISL_3174589 | Lighthouse Lab in Milton Keynes | Wellcome Sanger Institute for the COVID-19 Genomics UK (COG-UK) Consortium | The Lighthouse Lab in Milton Keynes and Alex Alderton, Roberto Amato, Jeffrey Barrett, Sonia Goncalves, Ewan Harrison, David K. Jackson, Ian Johnston, Dominic Kwiatkowski, Cordelia Langford, John Sillitoe on behalf of the Wellcome Sanger Institute COVID-19 Surveillance Team |
| EPI_ISL_3174590 | Lighthouse Laboratory Plymouth | Wellcome Sanger Institute for the COVID-19 Genomics UK (COG-UK) Consortium | Lighthouse Laboratory Plymouth and Alex Alderton, Roberto Amato, Jeffrey Barrett, Sonia Goncalves, Ewan Harrison, David K. Jackson, Ian Johnston, Dominic Kwiatkowski, Cordelia Langford, John Sillitoe on behalf of the Wellcome Sanger Institute COVID-19 Surveillance Team |
| EPI_ISL_3174591 | Lighthouse Lab in Alderley Park | Wellcome Sanger Institute for the COVID-19 Genomics UK (COG-UK) Consortium | Jacquelyn Wynn, Mairead Hyland, The Lighthouse Lab in Alderley Park and Alex Alderton, Roberto Amato, Jeffrey Barrett, Sonia Goncalves, Ewan Harrison, David K. Jackson, Ian Johnston, Dominic Kwiatkowski, Cordelia Langford, John Sillitoe on behalf of the Wellcome Sanger Institute COVID-19 Surveillance Team |
| EPI_ISL_3174592 | Lighthouse Laboratory Plymouth | Wellcome Sanger Institute for the COVID-19 Genomics UK (COG-UK) Consortium | Lighthouse Laboratory Plymouth and Alex Alderton, Roberto Amato, Jeffrey Barrett, Sonia Goncalves, Ewan Harrison, David K. Jackson, Ian Johnston, Dominic Kwiatkowski, Cordelia Langford, John Sillitoe on behalf of the Wellcome Sanger Institute COVID-19 Surveillance Team |
| EPI_ISL_3174593 | Lighthouse Lab in Alderley Park | Wellcome Sanger Institute for the COVID-19 Genomics UK (COG-UK) Consortium | Jacquelyn Wynn, Mairead Hyland, The Lighthouse Lab in Alderley Park and Alex Alderton, Roberto Amato, Jeffrey Barrett, Sonia Goncalves, Ewan Harrison, David K. Jackson, Ian Johnston, Dominic Kwiatkowski, Cordelia Langford, John Sillitoe on behalf of the Wellcome Sanger Institute COVID-19 Surveillance Team |
| EPI_ISL_3174594 | Lighthouse Lab in Milton Keynes | Wellcome Sanger Institute for the COVID-19 Genomics UK (COG-UK) Consortium | The Lighthouse Lab in Milton Keynes and Alex Alderton, Roberto Amato, Jeffrey Barrett, Sonia Goncalves, Ewan Harrison, David K. Jackson, Ian Johnston, Dominic Kwiatkowski, Cordelia Langford, John Sillitoe on behalf of the Wellcome Sanger Institute COVID-19 Surveillance Team |
| EPI_ISL_3174595, EPI_ISL_3174596 | Lighthouse Laboratory Plymouth | Wellcome Sanger Institute for the COVID-19 Genomics UK (COG-UK) Consortium | Lighthouse Laboratory Plymouth and Alex Alderton, Roberto Amato, Jeffrey Barrett, Sonia Goncalves, Ewan Harrison, David K. Jackson, Ian Johnston, Dominic Kwiatkowski, Cordelia Langford, John Sillitoe on behalf of the Wellcome Sanger Institute COVID-19 Surveillance Team |
| EPI_ISL_3174597 | Lighthouse Lab in Milton Keynes | Wellcome Sanger Institute for the COVID-19 Genomics UK (COG-UK) Consortium | The Lighthouse Lab in Milton Keynes and Alex Alderton, Roberto Amato, Jeffrey Barrett, Sonia Goncalves, Ewan Harrison, David K. Jackson, Ian Johnston, Dominic Kwiatkowski, Cordelia Langford, John Sillitoe on behalf of the Wellcome Sanger Institute COVID-19 Surveillance Team |
| EPI_ISL_3174598 | Lighthouse Lab in Alderley Park | Wellcome Sanger Institute for the COVID-19 Genomics UK (COG-UK) Consortium | Jacquelyn Wynn, Mairead Hyland, The Lighthouse Lab in Alderley Park and Alex Alderton, Roberto Amato, Jeffrey Barrett, Sonia Goncalves, Ewan Harrison, David K. Jackson, Ian Johnston, Dominic Kwiatkowski, Cordelia Langford, John Sillitoe on behalf of the Wellcome Sanger Institute COVID-19 Surveillance Team |
| EPI_ISL_3174599 | Lighthouse Lab in Glasgow | Wellcome Sanger Institute for the COVID-19 Genomics UK (COG-UK) Consortium | Harper VanSteenhouse, Yumi Kasai, David Gray, Carol Clugston, Anna Dominiczak and Alex Alderton, Roberto Amato, Jeffrey Barrett, Sonia Goncalves, Ewan Harrison, David K. Jackson, Ian Johnston, Dominic Kwiatkowski, Cordelia Langford, John Sillitoe on behalf of the Wellcome Sanger Institute COVID-19 Surveillance Team |
| EPI_ISL_3174600 | Lighthouse Lab in Alderley Park | Wellcome Sanger Institute for the COVID-19 Genomics UK (COG-UK) Consortium | Jacquelyn Wynn, Mairead Hyland, The Lighthouse Lab in Alderley Park and Alex Alderton, Roberto Amato, Jeffrey Barrett, Sonia Goncalves, Ewan Harrison, David K. Jackson, Ian Johnston, Dominic Kwiatkowski, Cordelia Langford, John Sillitoe on behalf of the Wellcome Sanger Institute COVID-19 Surveillance Team |
| EPI_ISL_3174601 | Lighthouse Lab in Milton Keynes | Wellcome Sanger Institute for the COVID-19 Genomics UK (COG-UK) Consortium | The Lighthouse Lab in Milton Keynes and Alex Alderton, Roberto Amato, Jeffrey Barrett, Sonia Goncalves, Ewan Harrison, David K. Jackson, Ian Johnston, Dominic Kwiatkowski, Cordelia Langford, John Sillitoe on behalf of the Wellcome Sanger Institute COVID-19 Surveillance Team |
| EPI_ISL_3174602 | Lighthouse Lab in Glasgow | Wellcome Sanger Institute for the COVID-19 Genomics UK (COG-UK) Consortium | Harper VanSteenhouse, Yumi Kasai, David Gray, Carol Clugston, Anna Dominiczak and Alex Alderton, Roberto Amato, Jeffrey Barrett, Sonia Goncalves, Ewan Harrison, David K. Jackson, Ian Johnston, Dominic Kwiatkowski, Cordelia Langford, John Sillitoe on behalf of the Wellcome Sanger Institute COVID-19 Surveillance Team |
| EPI_ISL_3174603, EPI_ISL_3174604, EPI_ISL_3174605 | Lighthouse Lab in Milton Keynes | Wellcome Sanger Institute for the COVID-19 Genomics UK (COG-UK) Consortium | The Lighthouse Lab in Milton Keynes and Alex Alderton, Roberto Amato, Jeffrey Barrett, Sonia Goncalves, Ewan Harrison, David K. Jackson, Ian Johnston, Dominic Kwiatkowski, Cordelia Langford, John Sillitoe on behalf of the Wellcome Sanger Institute COVID-19 Surveillance Team |
| EPI_ISL_3174606, EPI_ISL_3174607 | Lighthouse Lab in Alderley Park | Wellcome Sanger Institute for the COVID-19 Genomics UK (COG-UK) Consortium | Jacquelyn Wynn, Mairead Hyland, The Lighthouse Lab in Alderley Park and Alex Alderton, Roberto Amato, Jeffrey Barrett, Sonia Goncalves, Ewan Harrison, David K. Jackson, Ian Johnston, Dominic Kwiatkowski, Cordelia Langford, John Sillitoe on behalf of the Wellcome Sanger Institute COVID-19 Surveillance Team |
| EPI_ISL_3174608 | Lighthouse Laboratory Plymouth | Wellcome Sanger Institute for the COVID-19 Genomics UK (COG-UK) Consortium | Lighthouse Laboratory Plymouth and Alex Alderton, Roberto Amato, Jeffrey Barrett, Sonia Goncalves, Ewan Harrison, David K. Jackson, Ian Johnston, Dominic Kwiatkowski, Cordelia Langford, John Sillitoe on behalf of the Wellcome Sanger Institute COVID-19 Surveillance Team |
| EPI_ISL_3174609 | Lighthouse Lab in Glasgow | Wellcome Sanger Institute for the COVID-19 Genomics UK (COG-UK) | Harper VanSteenhouse, Yumi Kasai, David Gray, Carol Clugston, Anna Dominiczak and Alex Alderton, Roberto Amato, Jeffrey Barrett, Sonia Goncalves, Ewan Harrison, David K. Jackson, Ian Johnston, Dominic Kwiatkowski, Cordelia Langford, |

| | | | |
|---|---|---|---|
| | | Consortium | John Sillitoe on behalf of the Wellcome Sanger Institute COVID-19 Surveillance Team |
| EPI_ISL_3174610 | Lighthouse Lab in Milton Keynes | Wellcome Sanger Institute for the COVID-19 Genomics UK (COG-UK) Consortium | The Lighthouse Lab in Milton Keynes and Alex Alderton, Roberto Amato, Jeffrey Barrett, Sonia Goncalves, Ewan Harrison, David K. Jackson, Ian Johnston, Dominic Kwiatkowski, Cordelia Langford, John Sillitoe on behalf of the Wellcome Sanger Institute COVID-19 Surveillance Team |
| EPI_ISL_3174611 | Lighthouse Lab in Alderley Park | Wellcome Sanger Institute for the COVID-19 Genomics UK (COG-UK) Consortium | Jacquelyn Wynn, Mairead Hyland, The Lighthouse Lab in Alderley Park and Alex Alderton, Roberto Amato, Jeffrey Barrett, Sonia Goncalves, Ewan Harrison, David K. Jackson, Ian Johnston, Dominic Kwiatkowski, Cordelia Langford, John Sillitoe on behalf of the Wellcome Sanger Institute COVID-19 Surveillance Team |
| EPI_ISL_3174612 | Lighthouse Lab in Glasgow | Wellcome Sanger Institute for the COVID-19 Genomics UK (COG-UK) Consortium | Harper VanSteenhouse, Yumi Kasai, David Gray, Carol Clugston, Anna Dominiczak and Alex Alderton, Roberto Amato, Jeffrey Barrett, Sonia Goncalves, Ewan Harrison, David K. Jackson, Ian Johnston, Dominic Kwiatkowski, Cordelia Langford, John Sillitoe on behalf of the Wellcome Sanger Institute COVID-19 Surveillance Team |
| EPI_ISL_3174613 | Lighthouse Lab in Milton Keynes | Wellcome Sanger Institute for the COVID-19 Genomics UK (COG-UK) Consortium | The Lighthouse Lab in Milton Keynes and Alex Alderton, Roberto Amato, Jeffrey Barrett, Sonia Goncalves, Ewan Harrison, David K. Jackson, Ian Johnston, Dominic Kwiatkowski, Cordelia Langford, John Sillitoe on behalf of the Wellcome Sanger Institute COVID-19 Surveillance Team |
| EPI_ISL_3174614, EPI_ISL_3174615 | Lighthouse Laboratory Plymouth | Wellcome Sanger Institute for the COVID-19 Genomics UK (COG-UK) Consortium | Lighthouse Laboratory Plymouth and Alex Alderton, Roberto Amato, Jeffrey Barrett, Sonia Goncalves, Ewan Harrison, David K. Jackson, Ian Johnston, Dominic Kwiatkowski, Cordelia Langford, John Sillitoe on behalf of the Wellcome Sanger Institute COVID-19 Surveillance Team |
| EPI_ISL_3174616 | Lighthouse Lab in Milton Keynes | Wellcome Sanger Institute for the COVID-19 Genomics UK (COG-UK) Consortium | The Lighthouse Lab in Milton Keynes and Alex Alderton, Roberto Amato, Jeffrey Barrett, Sonia Goncalves, Ewan Harrison, David K. Jackson, Ian Johnston, Dominic Kwiatkowski, Cordelia Langford, John Sillitoe on behalf of the Wellcome Sanger Institute COVID-19 Surveillance Team |
| EPI_ISL_3174617 | Lighthouse Lab in Glasgow | Wellcome Sanger Institute for the COVID-19 Genomics UK (COG-UK) Consortium | Harper VanSteenhouse, Yumi Kasai, David Gray, Carol Clugston, Anna Dominiczak and Alex Alderton, Roberto Amato, Jeffrey Barrett, Sonia Goncalves, Ewan Harrison, David K. Jackson, Ian Johnston, Dominic Kwiatkowski, Cordelia Langford, John Sillitoe on behalf of the Wellcome Sanger Institute COVID-19 Surveillance Team |
| EPI_ISL_3174618, EPI_ISL_3174619 | Lighthouse Lab in Milton Keynes | Wellcome Sanger Institute for the COVID-19 Genomics UK (COG-UK) Consortium | The Lighthouse Lab in Milton Keynes and Alex Alderton, Roberto Amato, Jeffrey Barrett, Sonia Goncalves, Ewan Harrison, David K. Jackson, Ian Johnston, Dominic Kwiatkowski, Cordelia Langford, John Sillitoe on behalf of the Wellcome Sanger Institute COVID-19 Surveillance Team |
| EPI_ISL_3174620 | Lighthouse Laboratory Plymouth | Wellcome Sanger Institute for the COVID-19 Genomics UK (COG-UK) Consortium | Lighthouse Laboratory Plymouth and Alex Alderton, Roberto Amato, Jeffrey Barrett, Sonia Goncalves, Ewan Harrison, David K. Jackson, Ian Johnston, Dominic Kwiatkowski, Cordelia Langford, John Sillitoe on behalf of the Wellcome Sanger Institute COVID-19 Surveillance Team |
| EPI_ISL_3174621, EPI_ISL_3174622 | Lighthouse Lab in Alderley Park | Wellcome Sanger Institute for the COVID-19 Genomics UK (COG-UK) Consortium | Jacquelyn Wynn, Mairead Hyland, The Lighthouse Lab in Alderley Park and Alex Alderton, Roberto Amato, Jeffrey Barrett, Sonia Goncalves, Ewan Harrison, David K. Jackson, Ian Johnston, Dominic Kwiatkowski, Cordelia Langford, John Sillitoe on behalf of the Wellcome Sanger Institute COVID-19 Surveillance Team |
| EPI_ISL_3174623, EPI_ISL_3174624, EPI_ISL_3174625 | Lighthouse Laboratory Plymouth | Wellcome Sanger Institute for the COVID-19 Genomics UK (COG-UK) Consortium | Lighthouse Laboratory Plymouth and Alex Alderton, Roberto Amato, Jeffrey Barrett, Sonia Goncalves, Ewan Harrison, David K. Jackson, Ian Johnston, Dominic Kwiatkowski, Cordelia Langford, John Sillitoe on behalf of the Wellcome Sanger Institute COVID-19 Surveillance Team |
| EPI_ISL_3174626 | Lighthouse Lab in Alderley Park | Wellcome Sanger Institute for the COVID-19 Genomics UK (COG-UK) Consortium | Jacquelyn Wynn, Mairead Hyland, The Lighthouse Lab in Alderley Park and Alex Alderton, Roberto Amato, Jeffrey Barrett, Sonia Goncalves, Ewan Harrison, David K. Jackson, Ian Johnston, Dominic Kwiatkowski, Cordelia Langford, John Sillitoe on behalf of the Wellcome Sanger Institute COVID-19 Surveillance Team |
| EPI_ISL_3174627 | Lighthouse Lab in Glasgow | Wellcome Sanger Institute for the COVID-19 Genomics UK (COG-UK) Consortium | Harper VanSteenhouse, Yumi Kasai, David Gray, Carol Clugston, Anna Dominiczak and Alex Alderton, Roberto Amato, Jeffrey Barrett, Sonia Goncalves, Ewan Harrison, David K. Jackson, Ian Johnston, Dominic Kwiatkowski, Cordelia Langford, John Sillitoe on behalf of the Wellcome Sanger Institute COVID-19 Surveillance Team |
| EPI_ISL_3174628, EPI_ISL_3174629 | Lighthouse Lab in Alderley Park | Wellcome Sanger Institute for the COVID-19 Genomics UK (COG-UK) Consortium | Jacquelyn Wynn, Mairead Hyland, The Lighthouse Lab in Alderley Park and Alex Alderton, Roberto Amato, Jeffrey Barrett, Sonia Goncalves, Ewan Harrison, David K. Jackson, Ian Johnston, Dominic Kwiatkowski, Cordelia Langford, John Sillitoe on behalf of the Wellcome Sanger Institute COVID-19 Surveillance Team |
| EPI_ISL_3174630 | Lighthouse Lab in Milton Keynes | Wellcome Sanger Institute for the COVID-19 Genomics UK (COG-UK) Consortium | The Lighthouse Lab in Milton Keynes and Alex Alderton, Roberto Amato, Jeffrey Barrett, Sonia Goncalves, Ewan Harrison, David K. Jackson, Ian Johnston, Dominic Kwiatkowski, Cordelia Langford, John Sillitoe on behalf of the Wellcome Sanger Institute COVID-19 Surveillance Team |
| EPI_ISL_3174631, EPI_ISL_3174632 | Lighthouse Laboratory Plymouth | Wellcome Sanger Institute for the COVID-19 Genomics UK (COG-UK) Consortium | Lighthouse Laboratory Plymouth and Alex Alderton, Roberto Amato, Jeffrey Barrett, Sonia Goncalves, Ewan Harrison, David K. Jackson, Ian Johnston, Dominic Kwiatkowski, Cordelia Langford, John Sillitoe on behalf of the Wellcome Sanger Institute COVID-19 Surveillance Team |
| EPI_ISL_3174633 | Lighthouse Lab in Milton Keynes | Wellcome Sanger Institute for the COVID-19 Genomics UK (COG-UK) Consortium | The Lighthouse Lab in Milton Keynes and Alex Alderton, Roberto Amato, Jeffrey Barrett, Sonia Goncalves, Ewan Harrison, David K. Jackson, Ian Johnston, Dominic Kwiatkowski, Cordelia Langford, John Sillitoe on behalf of the Wellcome Sanger Institute COVID-19 Surveillance Team |
| EPI_ISL_3174634, EPI_ISL_3174635 | Lighthouse Lab in Alderley Park | Wellcome Sanger Institute for the COVID-19 Genomics UK (COG-UK) Consortium | Jacquelyn Wynn, Mairead Hyland, The Lighthouse Lab in Alderley Park and Alex Alderton, Roberto Amato, Jeffrey Barrett, Sonia Goncalves, Ewan Harrison, David K. Jackson, Ian Johnston, Dominic Kwiatkowski, Cordelia Langford, John Sillitoe on behalf of the Wellcome Sanger Institute COVID-19 Surveillance Team |
| EPI_ISL_3174636 | Lighthouse Lab in Milton Keynes | Wellcome Sanger Institute for the COVID-19 Genomics UK (COG-UK) Consortium | The Lighthouse Lab in Milton Keynes and Alex Alderton, Roberto Amato, Jeffrey Barrett, Sonia Goncalves, Ewan Harrison, David K. Jackson, Ian Johnston, Dominic Kwiatkowski, Cordelia Langford, John Sillitoe on behalf of the Wellcome Sanger Institute COVID-19 Surveillance Team |
| EPI_ISL_3174637 | Lighthouse Lab in Alderley Park | Wellcome Sanger Institute for the COVID-19 Genomics UK (COG-UK) Consortium | Jacquelyn Wynn, Mairead Hyland, The Lighthouse Lab in Alderley Park and Alex Alderton, Roberto Amato, Jeffrey Barrett, Sonia Goncalves, Ewan Harrison, David K. Jackson, Ian Johnston, Dominic Kwiatkowski, Cordelia Langford, John Sillitoe on behalf of the Wellcome Sanger Institute COVID-19 Surveillance Team |
| EPI_ISL_3174638 | Lighthouse Lab in Glasgow | Wellcome Sanger Institute for the COVID-19 Genomics UK (COG-UK) Consortium | Harper VanSteenhouse, Yumi Kasai, David Gray, Carol Clugston, Anna Dominiczak and Alex Alderton, Roberto Amato, Jeffrey Barrett, Sonia Goncalves, Ewan Harrison, David K. Jackson, Ian Johnston, Dominic Kwiatkowski, Cordelia Langford, John Sillitoe on behalf of the Wellcome Sanger Institute COVID-19 Surveillance Team |
| EPI_ISL_3174639 | Lighthouse Laboratory Plymouth | Wellcome Sanger Institute for the COVID-19 Genomics UK (COG-UK) Consortium | Lighthouse Laboratory Plymouth and Alex Alderton, Roberto Amato, Jeffrey Barrett, Sonia Goncalves, Ewan Harrison, David K. Jackson, Ian Johnston, Dominic Kwiatkowski, Cordelia Langford, John Sillitoe on behalf of the Wellcome Sanger Institute COVID-19 Surveillance Team |
| EPI_ISL_3174640 | Lighthouse Lab in Glasgow | Wellcome Sanger Institute for the COVID-19 Genomics UK (COG-UK) Consortium | Harper VanSteenhouse, Yumi Kasai, David Gray, Carol Clugston, Anna Dominiczak and Alex Alderton, Roberto Amato, Jeffrey Barrett, Sonia Goncalves, Ewan Harrison, David K. Jackson, Ian Johnston, Dominic Kwiatkowski, Cordelia Langford, John Sillitoe on behalf of the Wellcome Sanger Institute COVID-19 Surveillance Team |
| EPI_ISL_3174641 | Lighthouse Lab in Milton Keynes | Wellcome Sanger Institute for the COVID-19 Genomics UK (COG-UK) Consortium | The Lighthouse Lab in Milton Keynes and Alex Alderton, Roberto Amato, Jeffrey Barrett, Sonia Goncalves, Ewan Harrison, David K. Jackson, Ian Johnston, Dominic Kwiatkowski, Cordelia Langford, John Sillitoe on behalf of the Wellcome Sanger Institute COVID-19 Surveillance Team |
| EPI_ISL_3174642, EPI_ISL_3174643 | Lighthouse Laboratory Plymouth | Wellcome Sanger Institute for the COVID-19 Genomics UK (COG-UK) Consortium | Lighthouse Laboratory Plymouth and Alex Alderton, Roberto Amato, Jeffrey Barrett, Sonia Goncalves, Ewan Harrison, David K. Jackson, Ian Johnston, Dominic Kwiatkowski, Cordelia Langford, John Sillitoe on behalf of the Wellcome Sanger Institute COVID-19 Surveillance Team |
| EPI_ISL_3174644 | Lighthouse Lab in Alderley Park | Wellcome Sanger Institute for the COVID-19 Genomics UK (COG-UK) Consortium | Jacquelyn Wynn, Mairead Hyland, The Lighthouse Lab in Alderley Park and Alex Alderton, Roberto Amato, Jeffrey Barrett, Sonia Goncalves, Ewan Harrison, David K. Jackson, Ian Johnston, Dominic Kwiatkowski, Cordelia Langford, John Sillitoe on behalf of the Wellcome Sanger Institute COVID-19 Surveillance Team |
| EPI_ISL_3174645 | Lighthouse Lab in Glasgow | Wellcome Sanger Institute for the COVID-19 Genomics UK (COG-UK) Consortium | Harper VanSteenhouse, Yumi Kasai, David Gray, Carol Clugston, Anna Dominiczak and Alex Alderton, Roberto Amato, Jeffrey Barrett, Sonia Goncalves, Ewan Harrison, David K. Jackson, Ian Johnston, Dominic Kwiatkowski, Cordelia Langford, John Sillitoe on behalf of the Wellcome Sanger Institute COVID-19 Surveillance Team |
| EPI_ISL_3174646 | Lighthouse Laboratory Plymouth | Wellcome Sanger Institute for the COVID-19 Genomics UK (COG-UK) Consortium | Lighthouse Laboratory Plymouth and Alex Alderton, Roberto Amato, Jeffrey Barrett, Sonia Goncalves, Ewan Harrison, David K. Jackson, Ian Johnston, Dominic Kwiatkowski, Cordelia Langford, John Sillitoe on behalf of the Wellcome Sanger Institute COVID-19 Surveillance Team |
| EPI_ISL_3174647 | Lighthouse Lab in Glasgow | Wellcome Sanger Institute for the COVID-19 Genomics UK (COG-UK) Consortium | Harper VanSteenhouse, Yumi Kasai, David Gray, Carol Clugston, Anna Dominiczak and Alex Alderton, Roberto Amato, Jeffrey Barrett, Sonia Goncalves, Ewan Harrison, David K. Jackson, Ian Johnston, Dominic Kwiatkowski, Cordelia Langford, John Sillitoe on behalf of the Wellcome Sanger Institute COVID-19 Surveillance Team |
| EPI_ISL_3174648 | Lighthouse Lab in Alderley Park | Wellcome Sanger Institute for the COVID-19 Genomics UK (COG-UK) Consortium | Jacquelyn Wynn, Mairead Hyland, The Lighthouse Lab in Alderley Park and Alex Alderton, Roberto Amato, Jeffrey Barrett, Sonia Goncalves, Ewan Harrison, David K. Jackson, Ian Johnston, Dominic Kwiatkowski, Cordelia Langford, John Sillitoe on behalf of the Wellcome Sanger Institute COVID-19 Surveillance Team |
| EPI_ISL_3174649 | Lighthouse Laboratory Plymouth | Wellcome Sanger Institute for the COVID-19 Genomics UK (COG-UK) Consortium | Lighthouse Laboratory Plymouth and Alex Alderton, Roberto Amato, Jeffrey Barrett, Sonia Goncalves, Ewan Harrison, David K. Jackson, Ian Johnston, Dominic Kwiatkowski, Cordelia Langford, John Sillitoe on behalf of the Wellcome Sanger Institute COVID-19 Surveillance Team |
| EPI_ISL_3174650 | Lighthouse Lab in Milton Keynes | Wellcome Sanger Institute for the COVID-19 Genomics UK (COG-UK) Consortium | The Lighthouse Lab in Milton Keynes and Alex Alderton, Roberto Amato, Jeffrey Barrett, Sonia Goncalves, Ewan Harrison, David K. Jackson, Ian Johnston, Dominic Kwiatkowski, Cordelia Langford, John Sillitoe on behalf of the Wellcome Sanger Institute COVID-19 Surveillance Team |
| EPI_ISL_3174651 | Lighthouse Laboratory Plymouth | Wellcome Sanger Institute for the COVID-19 Genomics UK (COG-UK) Consortium | Lighthouse Laboratory Plymouth and Alex Alderton, Roberto Amato, Jeffrey Barrett, Sonia Goncalves, Ewan Harrison, David K. Jackson, Ian Johnston, Dominic Kwiatkowski, Cordelia Langford, John Sillitoe on behalf of the Wellcome Sanger Institute COVID-19 Surveillance Team |
| EPI_ISL_3174652, EPI_ISL_3174653 | Lighthouse Lab in Glasgow | Wellcome Sanger Institute for the COVID-19 Genomics UK (COG-UK) Consortium | Harper VanSteenhouse, Yumi Kasai, David Gray, Carol Clugston, Anna Dominiczak and Alex Alderton, Roberto Amato, Jeffrey Barrett, Sonia Goncalves, Ewan Harrison, David K. Jackson, Ian Johnston, Dominic Kwiatkowski, Cordelia Langford, John Sillitoe on behalf of the Wellcome Sanger Institute COVID-19 Surveillance Team |
| EPI_ISL_3174654 | Lighthouse Lab in Milton Keynes | Wellcome Sanger Institute for the COVID-19 Genomics UK (COG-UK) Consortium | The Lighthouse Lab in Milton Keynes and Alex Alderton, Roberto Amato, Jeffrey Barrett, Sonia Goncalves, Ewan Harrison, David K. Jackson, Ian Johnston, Dominic Kwiatkowski, Cordelia Langford, John Sillitoe on behalf of the Wellcome Sanger Institute COVID-19 Surveillance Team |
| EPI_ISL_3174655 | Lighthouse Laboratory Plymouth | Wellcome Sanger Institute for the COVID-19 Genomics UK (COG-UK) Consortium | Lighthouse Laboratory Plymouth and Alex Alderton, Roberto Amato, Jeffrey Barrett, Sonia Goncalves, Ewan Harrison, David K. Jackson, Ian Johnston, Dominic Kwiatkowski, Cordelia Langford, John Sillitoe on behalf of the Wellcome Sanger Institute COVID-19 Surveillance Team |
| EPI_ISL_3174656 | Lighthouse Lab in Milton Keynes | Wellcome Sanger Institute for the COVID-19 Genomics UK (COG-UK) Consortium | The Lighthouse Lab in Milton Keynes and Alex Alderton, Roberto Amato, Jeffrey Barrett, Sonia Goncalves, Ewan Harrison, David K. Jackson, Ian Johnston, Dominic Kwiatkowski, Cordelia Langford, John Sillitoe on behalf of the Wellcome Sanger Institute COVID-19 Surveillance Team |
| EPI_ISL_3174657 | Lighthouse Lab in Glasgow | Wellcome Sanger Institute for the COVID-19 Genomics UK (COG-UK) Consortium | Harper VanSteenhouse, Yumi Kasai, David Gray, Carol Clugston, Anna Dominiczak and Alex Alderton, Roberto Amato, Jeffrey Barrett, Sonia Goncalves, Ewan Harrison, David K. Jackson, Ian Johnston, Dominic Kwiatkowski, Cordelia Langford, John Sillitoe on behalf of the Wellcome Sanger Institute COVID-19 Surveillance Team |
| EPI_ISL_3174658 | Lighthouse Lab in Alderley Park | Wellcome Sanger Institute for the COVID-19 Genomics UK (COG-UK) Consortium | Jacquelyn Wynn, Mairead Hyland, The Lighthouse Lab in Alderley Park and Alex Alderton, Roberto Amato, Jeffrey Barrett, Sonia Goncalves, Ewan Harrison, David K. Jackson, Ian Johnston, Dominic Kwiatkowski, Cordelia Langford, John Sillitoe on behalf of the Wellcome Sanger Institute COVID-19 Surveillance Team |
| EPI_ISL_3174659 | Lighthouse Laboratory Plymouth | Wellcome Sanger Institute for the COVID-19 Genomics UK (COG-UK) Consortium | Lighthouse Laboratory Plymouth and Alex Alderton, Roberto Amato, Jeffrey Barrett, Sonia Goncalves, Ewan Harrison, David K. Jackson, Ian Johnston, Dominic Kwiatkowski, Cordelia Langford, John Sillitoe on behalf of the Wellcome Sanger Institute COVID-19 Surveillance Team |
| EPI_ISL_3174660, EPI_ISL_3174661 | Lighthouse Lab in Alderley Park | Wellcome Sanger Institute for the COVID-19 Genomics UK (COG-UK) Consortium | Jacquelyn Wynn, Mairead Hyland, The Lighthouse Lab in Alderley Park and Alex Alderton, Roberto Amato, Jeffrey Barrett, Sonia Goncalves, Ewan Harrison, David K. Jackson, Ian Johnston, Dominic Kwiatkowski, Cordelia Langford, John Sillitoe on behalf of the Wellcome Sanger Institute COVID-19 Surveillance Team |
| EPI_ISL_3174662 | Lighthouse Lab in Milton Keynes | Wellcome Sanger Institute for the COVID-19 Genomics UK (COG-UK) Consortium | The Lighthouse Lab in Milton Keynes and Alex Alderton, Roberto Amato, Jeffrey Barrett, Sonia Goncalves, Ewan Harrison, David K. Jackson, Ian Johnston, Dominic Kwiatkowski, Cordelia Langford, John Sillitoe on behalf of the Wellcome Sanger Institute COVID-19 Surveillance Team |
| EPI_ISL_3174663 | Lighthouse Laboratory Plymouth | Wellcome Sanger Institute for the COVID-19 Genomics UK (COG-UK) Consortium | Lighthouse Laboratory Plymouth and Alex Alderton, Roberto Amato, Jeffrey Barrett, Sonia Goncalves, Ewan Harrison, David K. Jackson, Ian Johnston, Dominic Kwiatkowski, Cordelia Langford, John Sillitoe on behalf of the Wellcome Sanger Institute COVID-19 Surveillance Team |
| EPI_ISL_3174664 | Lighthouse Lab in Milton Keynes | Wellcome Sanger Institute for the COVID-19 Genomics UK (COG-UK) Consortium | The Lighthouse Lab in Milton Keynes and Alex Alderton, Roberto Amato, Jeffrey Barrett, Sonia Goncalves, Ewan Harrison, David K. Jackson, Ian Johnston, Dominic Kwiatkowski, Cordelia Langford, John Sillitoe on behalf of the Wellcome Sanger Institute COVID-19 Surveillance Team |

| | | | |
|---|---|---|---|
| EPI_ISL_3174665, EPI_ISL_3174666 | Lighthouse Lab in Glasgow | Wellcome Sanger Institute for the COVID-19 Genomics UK (COG-UK) Consortium | Harper VanSteenhouse, Yumi Kasai, David Gray, Carol Clugston, Anna Dominiczak and Alex Alderton, Roberto Amato, Jeffrey Barrett, Sonia Goncalves, Ewan Harrison, David K. Jackson, Ian Johnston, Dominic Kwiatkowski, Cordelia Langford, John Sillitoe on behalf of the Wellcome Sanger Institute COVID-19 Surveillance Team |
| EPI_ISL_3174667 | Lighthouse Laboratory Plymouth | Wellcome Sanger Institute for the COVID-19 Genomics UK (COG-UK) Consortium | Lighthouse Laboratory Plymouth and Alex Alderton, Roberto Amato, Jeffrey Barrett, Sonia Goncalves, Ewan Harrison, David K. Jackson, Ian Johnston, Dominic Kwiatkowski, Cordelia Langford, John Sillitoe on behalf of the Wellcome Sanger Institute COVID-19 Surveillance Team |
| EPI_ISL_3174668, EPI_ISL_3174669 | Lighthouse Lab in Milton Keynes | Wellcome Sanger Institute for the COVID-19 Genomics UK (COG-UK) Consortium | The Lighthouse Lab in Milton Keynes and Alex Alderton, Roberto Amato, Jeffrey Barrett, Sonia Goncalves, Ewan Harrison, David K. Jackson, Ian Johnston, Dominic Kwiatkowski, Cordelia Langford, John Sillitoe on behalf of the Wellcome Sanger Institute COVID-19 Surveillance Team |
| EPI_ISL_3174670, EPI_ISL_3174671 | Lighthouse Lab in Alderley Park | Wellcome Sanger Institute for the COVID-19 Genomics UK (COG-UK) Consortium | Jacquelyn Wynn, Mairead Hyland, The Lighthouse Lab in Alderley Park and Alex Alderton, Roberto Amato, Jeffrey Barrett, Sonia Goncalves, Ewan Harrison, David K. Jackson, Ian Johnston, Dominic Kwiatkowski, Cordelia Langford, John Sillitoe on behalf of the Wellcome Sanger Institute COVID-19 Surveillance Team |
| EPI_ISL_3174672 | Lighthouse Lab in Milton Keynes | Wellcome Sanger Institute for the COVID-19 Genomics UK (COG-UK) Consortium | The Lighthouse Lab in Milton Keynes and Alex Alderton, Roberto Amato, Jeffrey Barrett, Sonia Goncalves, Ewan Harrison, David K. Jackson, Ian Johnston, Dominic Kwiatkowski, Cordelia Langford, John Sillitoe on behalf of the Wellcome Sanger Institute COVID-19 Surveillance Team |
| EPI_ISL_3174673, EPI_ISL_3174674, EPI_ISL_3174675, EPI_ISL_3174676 | Lighthouse Laboratory Plymouth | Wellcome Sanger Institute for the COVID-19 Genomics UK (COG-UK) Consortium | Lighthouse Laboratory Plymouth and Alex Alderton, Roberto Amato, Jeffrey Barrett, Sonia Goncalves, Ewan Harrison, David K. Jackson, Ian Johnston, Dominic Kwiatkowski, Cordelia Langford, John Sillitoe on behalf of the Wellcome Sanger Institute COVID-19 Surveillance Team |
| EPI_ISL_3174677 | Lighthouse Lab in Glasgow | Wellcome Sanger Institute for the COVID-19 Genomics UK (COG-UK) Consortium | Harper VanSteenhouse, Yumi Kasai, David Gray, Carol Clugston, Anna Dominiczak and Alex Alderton, Roberto Amato, Jeffrey Barrett, Sonia Goncalves, Ewan Harrison, David K. Jackson, Ian Johnston, Dominic Kwiatkowski, Cordelia Langford, John Sillitoe on behalf of the Wellcome Sanger Institute COVID-19 Surveillance Team |
| EPI_ISL_3174678, EPI_ISL_3174679 | Lighthouse Laboratory Plymouth | Wellcome Sanger Institute for the COVID-19 Genomics UK (COG-UK) Consortium | Lighthouse Laboratory Plymouth and Alex Alderton, Roberto Amato, Jeffrey Barrett, Sonia Goncalves, Ewan Harrison, David K. Jackson, Ian Johnston, Dominic Kwiatkowski, Cordelia Langford, John Sillitoe on behalf of the Wellcome Sanger Institute COVID-19 Surveillance Team |
| EPI_ISL_3174680 | Lighthouse Lab in Alderley Park | Wellcome Sanger Institute for the COVID-19 Genomics UK (COG-UK) Consortium | Jacquelyn Wynn, Mairead Hyland, The Lighthouse Lab in Alderley Park and Alex Alderton, Roberto Amato, Jeffrey Barrett, Sonia Goncalves, Ewan Harrison, David K. Jackson, Ian Johnston, Dominic Kwiatkowski, Cordelia Langford, John Sillitoe on behalf of the Wellcome Sanger Institute COVID-19 Surveillance Team |
| EPI_ISL_3174681 | Lighthouse Laboratory Plymouth | Wellcome Sanger Institute for the COVID-19 Genomics UK (COG-UK) Consortium | Lighthouse Laboratory Plymouth and Alex Alderton, Roberto Amato, Jeffrey Barrett, Sonia Goncalves, Ewan Harrison, David K. Jackson, Ian Johnston, Dominic Kwiatkowski, Cordelia Langford, John Sillitoe on behalf of the Wellcome Sanger Institute COVID-19 Surveillance Team |
| EPI_ISL_3174682 | Lighthouse Lab in Glasgow | Wellcome Sanger Institute for the COVID-19 Genomics UK (COG-UK) Consortium | Harper VanSteenhouse, Yumi Kasai, David Gray, Carol Clugston, Anna Dominiczak and Alex Alderton, Roberto Amato, Jeffrey Barrett, Sonia Goncalves, Ewan Harrison, David K. Jackson, Ian Johnston, Dominic Kwiatkowski, Cordelia Langford, John Sillitoe on behalf of the Wellcome Sanger Institute COVID-19 Surveillance Team |
| EPI_ISL_3174683 | Lighthouse Laboratory Plymouth | Wellcome Sanger Institute for the COVID-19 Genomics UK (COG-UK) Consortium | Lighthouse Laboratory Plymouth and Alex Alderton, Roberto Amato, Jeffrey Barrett, Sonia Goncalves, Ewan Harrison, David K. Jackson, Ian Johnston, Dominic Kwiatkowski, Cordelia Langford, John Sillitoe on behalf of the Wellcome Sanger Institute COVID-19 Surveillance Team |
| EPI_ISL_3174684, EPI_ISL_3174685 | Lighthouse Lab in Milton Keynes | Wellcome Sanger Institute for the COVID-19 Genomics UK (COG-UK) Consortium | The Lighthouse Lab in Milton Keynes and Alex Alderton, Roberto Amato, Jeffrey Barrett, Sonia Goncalves, Ewan Harrison, David K. Jackson, Ian Johnston, Dominic Kwiatkowski, Cordelia Langford, John Sillitoe on behalf of the Wellcome Sanger Institute COVID-19 Surveillance Team |
| EPI_ISL_3174686 | Lighthouse Lab in Alderley Park | Wellcome Sanger Institute for the COVID-19 Genomics UK (COG-UK) Consortium | Jacquelyn Wynn, Mairead Hyland, The Lighthouse Lab in Alderley Park and Alex Alderton, Roberto Amato, Jeffrey Barrett, Sonia Goncalves, Ewan Harrison, David K. Jackson, Ian Johnston, Dominic Kwiatkowski, Cordelia Langford, John Sillitoe on behalf of the Wellcome Sanger Institute COVID-19 Surveillance Team |
| EPI_ISL_3174687 | Lighthouse Laboratory Plymouth | Wellcome Sanger Institute for the COVID-19 Genomics UK (COG-UK) Consortium | Lighthouse Laboratory Plymouth and Alex Alderton, Roberto Amato, Jeffrey Barrett, Sonia Goncalves, Ewan Harrison, David K. Jackson, Ian Johnston, Dominic Kwiatkowski, Cordelia Langford, John Sillitoe on behalf of the Wellcome Sanger Institute COVID-19 Surveillance Team |
| EPI_ISL_3174688, EPI_ISL_3174689, EPI_ISL_3174690 | Lighthouse Lab in Alderley Park | Wellcome Sanger Institute for the COVID-19 Genomics UK (COG-UK) Consortium | Jacquelyn Wynn, Mairead Hyland, The Lighthouse Lab in Alderley Park and Alex Alderton, Roberto Amato, Jeffrey Barrett, Sonia Goncalves, Ewan Harrison, David K. Jackson, Ian Johnston, Dominic Kwiatkowski, Cordelia Langford, John Sillitoe on behalf of the Wellcome Sanger Institute COVID-19 Surveillance Team |
| EPI_ISL_3174691 | Lighthouse Laboratory Plymouth | Wellcome Sanger Institute for the COVID-19 Genomics UK (COG-UK) Consortium | Lighthouse Laboratory Plymouth and Alex Alderton, Roberto Amato, Jeffrey Barrett, Sonia Goncalves, Ewan Harrison, David K. Jackson, Ian Johnston, Dominic Kwiatkowski, Cordelia Langford, John Sillitoe on behalf of the Wellcome Sanger Institute COVID-19 Surveillance Team |
| EPI_ISL_3174692 | Lighthouse Lab in Glasgow | Wellcome Sanger Institute for the COVID-19 Genomics UK (COG-UK) Consortium | Harper VanSteenhouse, Yumi Kasai, David Gray, Carol Clugston, Anna Dominiczak and Alex Alderton, Roberto Amato, Jeffrey Barrett, Sonia Goncalves, Ewan Harrison, David K. Jackson, Ian Johnston, Dominic Kwiatkowski, Cordelia Langford, John Sillitoe on behalf of the Wellcome Sanger Institute COVID-19 Surveillance Team |
| EPI_ISL_3174693 | Lighthouse Lab in Alderley Park | Wellcome Sanger Institute for the COVID-19 Genomics UK (COG-UK) Consortium | Jacquelyn Wynn, Mairead Hyland, The Lighthouse Lab in Alderley Park and Alex Alderton, Roberto Amato, Jeffrey Barrett, Sonia Goncalves, Ewan Harrison, David K. Jackson, Ian Johnston, Dominic Kwiatkowski, Cordelia Langford, John Sillitoe on behalf of the Wellcome Sanger Institute COVID-19 Surveillance Team |
| EPI_ISL_3174694, EPI_ISL_3174695 | Lighthouse Laboratory Plymouth | Wellcome Sanger Institute for the COVID-19 Genomics UK (COG-UK) Consortium | Lighthouse Laboratory Plymouth and Alex Alderton, Roberto Amato, Jeffrey Barrett, Sonia Goncalves, Ewan Harrison, David K. Jackson, Ian Johnston, Dominic Kwiatkowski, Cordelia Langford, John Sillitoe on behalf of the Wellcome Sanger Institute COVID-19 Surveillance Team |
| EPI_ISL_3174696 | Lighthouse Lab in Milton Keynes | Wellcome Sanger Institute for the COVID-19 Genomics UK (COG-UK) Consortium | The Lighthouse Lab in Milton Keynes and Alex Alderton, Roberto Amato, Jeffrey Barrett, Sonia Goncalves, Ewan Harrison, David K. Jackson, Ian Johnston, Dominic Kwiatkowski, Cordelia Langford, John Sillitoe on behalf of the Wellcome Sanger Institute COVID-19 Surveillance Team |
| EPI_ISL_3174697 | Lighthouse Laboratory Plymouth | Wellcome Sanger Institute for the COVID-19 Genomics UK (COG-UK) Consortium | Lighthouse Laboratory Plymouth and Alex Alderton, Roberto Amato, Jeffrey Barrett, Sonia Goncalves, Ewan Harrison, David K. Jackson, Ian Johnston, Dominic Kwiatkowski, Cordelia Langford, John Sillitoe on behalf of the Wellcome Sanger Institute COVID-19 Surveillance Team |
| EPI_ISL_3174698, EPI_ISL_3174699 | Lighthouse Lab in Alderley Park | Wellcome Sanger Institute for the COVID-19 Genomics UK (COG-UK) Consortium | Jacquelyn Wynn, Mairead Hyland, The Lighthouse Lab in Alderley Park and Alex Alderton, Roberto Amato, Jeffrey Barrett, Sonia Goncalves, Ewan Harrison, David K. Jackson, Ian Johnston, Dominic Kwiatkowski, Cordelia Langford, John Sillitoe on behalf of the Wellcome Sanger Institute COVID-19 Surveillance Team |
| EPI_ISL_3174700, EPI_ISL_3174701 | Lighthouse Laboratory Plymouth | Wellcome Sanger Institute for the COVID-19 Genomics UK (COG-UK) Consortium | Lighthouse Laboratory Plymouth and Alex Alderton, Roberto Amato, Jeffrey Barrett, Sonia Goncalves, Ewan Harrison, David K. Jackson, Ian Johnston, Dominic Kwiatkowski, Cordelia Langford, John Sillitoe on behalf of the Wellcome Sanger Institute COVID-19 Surveillance Team |
| EPI_ISL_3174702 | Lighthouse Lab in Milton Keynes | Wellcome Sanger Institute for the COVID-19 Genomics UK (COG-UK) Consortium | The Lighthouse Lab in Milton Keynes and Alex Alderton, Roberto Amato, Jeffrey Barrett, Sonia Goncalves, Ewan Harrison, David K. Jackson, Ian Johnston, Dominic Kwiatkowski, Cordelia Langford, John Sillitoe on behalf of the Wellcome Sanger Institute COVID-19 Surveillance Team |
| EPI_ISL_3174703, EPI_ISL_3174704 | Lighthouse Lab in Alderley Park | Wellcome Sanger Institute for the COVID-19 Genomics UK (COG-UK) Consortium | Jacquelyn Wynn, Mairead Hyland, The Lighthouse Lab in Alderley Park and Alex Alderton, Roberto Amato, Jeffrey Barrett, Sonia Goncalves, Ewan Harrison, David K. Jackson, Ian Johnston, Dominic Kwiatkowski, Cordelia Langford, John Sillitoe on behalf of the Wellcome Sanger Institute COVID-19 Surveillance Team |
| EPI_ISL_3174705, EPI_ISL_3174706, EPI_ISL_3174707, EPI_ISL_3174708 | Lighthouse Lab in Milton Keynes | Wellcome Sanger Institute for the COVID-19 Genomics UK (COG-UK) Consortium | The Lighthouse Lab in Milton Keynes and Alex Alderton, Roberto Amato, Jeffrey Barrett, Sonia Goncalves, Ewan Harrison, David K. Jackson, Ian Johnston, Dominic Kwiatkowski, Cordelia Langford, John Sillitoe on behalf of the Wellcome Sanger Institute COVID-19 Surveillance Team |
| EPI_ISL_3174709 | Lighthouse Laboratory Plymouth | Wellcome Sanger Institute for the COVID-19 Genomics UK (COG-UK) Consortium | Lighthouse Laboratory Plymouth and Alex Alderton, Roberto Amato, Jeffrey Barrett, Sonia Goncalves, Ewan Harrison, David K. Jackson, Ian Johnston, Dominic Kwiatkowski, Cordelia Langford, John Sillitoe on behalf of the Wellcome Sanger Institute COVID-19 Surveillance Team |
| EPI_ISL_3174710, EPI_ISL_3174711 | Lighthouse Lab in Glasgow | Wellcome Sanger Institute for the COVID-19 Genomics UK (COG-UK) Consortium | Harper VanSteenhouse, Yumi Kasai, David Gray, Carol Clugston, Anna Dominiczak and Alex Alderton, Roberto Amato, Jeffrey Barrett, Sonia Goncalves, Ewan Harrison, David K. Jackson, Ian Johnston, Dominic Kwiatkowski, Cordelia Langford, John Sillitoe on behalf of the Wellcome Sanger Institute COVID-19 Surveillance Team |
| EPI_ISL_3174712 | Lighthouse Lab in Alderley Park | Wellcome Sanger Institute for the COVID-19 Genomics UK (COG-UK) Consortium | Jacquelyn Wynn, Mairead Hyland, The Lighthouse Lab in Alderley Park and Alex Alderton, Roberto Amato, Jeffrey Barrett, Sonia Goncalves, Ewan Harrison, David K. Jackson, Ian Johnston, Dominic Kwiatkowski, Cordelia Langford, John Sillitoe on behalf of the Wellcome Sanger Institute COVID-19 Surveillance Team |
| EPI_ISL_3174713 | Lighthouse Lab in Milton Keynes | Wellcome Sanger Institute for the COVID-19 Genomics UK (COG-UK) Consortium | The Lighthouse Lab in Milton Keynes and Alex Alderton, Roberto Amato, Jeffrey Barrett, Sonia Goncalves, Ewan Harrison, David K. Jackson, Ian Johnston, Dominic Kwiatkowski, Cordelia Langford, John Sillitoe on behalf of the Wellcome Sanger Institute COVID-19 Surveillance Team |
| EPI_ISL_3174714, EPI_ISL_3174715 | Lighthouse Laboratory Plymouth | Wellcome Sanger Institute for the COVID-19 Genomics UK (COG-UK) Consortium | Lighthouse Laboratory Plymouth and Alex Alderton, Roberto Amato, Jeffrey Barrett, Sonia Goncalves, Ewan Harrison, David K. Jackson, Ian Johnston, Dominic Kwiatkowski, Cordelia Langford, John Sillitoe on behalf of the Wellcome Sanger Institute COVID-19 Surveillance Team |
| EPI_ISL_3174716 | Lighthouse Lab in Alderley Park | Wellcome Sanger Institute for the COVID-19 Genomics UK (COG-UK) Consortium | Jacquelyn Wynn, Mairead Hyland, The Lighthouse Lab in Alderley Park and Alex Alderton, Roberto Amato, Jeffrey Barrett, Sonia Goncalves, Ewan Harrison, David K. Jackson, Ian Johnston, Dominic Kwiatkowski, Cordelia Langford, John Sillitoe on behalf of the Wellcome Sanger Institute COVID-19 Surveillance Team |
| EPI_ISL_3174717 | Lighthouse Lab in Milton Keynes | Wellcome Sanger Institute for the COVID-19 Genomics UK (COG-UK) Consortium | The Lighthouse Lab in Milton Keynes and Alex Alderton, Roberto Amato, Jeffrey Barrett, Sonia Goncalves, Ewan Harrison, David K. Jackson, Ian Johnston, Dominic Kwiatkowski, Cordelia Langford, John Sillitoe on behalf of the Wellcome Sanger Institute COVID-19 Surveillance Team |
| EPI_ISL_3174718, EPI_ISL_3174719, EPI_ISL_3174720, EPI_ISL_3174721 | Lighthouse Laboratory Plymouth | Wellcome Sanger Institute for the COVID-19 Genomics UK (COG-UK) Consortium | Lighthouse Laboratory Plymouth and Alex Alderton, Roberto Amato, Jeffrey Barrett, Sonia Goncalves, Ewan Harrison, David K. Jackson, Ian Johnston, Dominic Kwiatkowski, Cordelia Langford, John Sillitoe on behalf of the Wellcome Sanger Institute COVID-19 Surveillance Team |
| EPI_ISL_3174722 | Lighthouse Lab in Glasgow | Wellcome Sanger Institute for the COVID-19 Genomics UK (COG-UK) Consortium | Harper VanSteenhouse, Yumi Kasai, David Gray, Carol Clugston, Anna Dominiczak and Alex Alderton, Roberto Amato, Jeffrey Barrett, Sonia Goncalves, Ewan Harrison, David K. Jackson, Ian Johnston, Dominic Kwiatkowski, Cordelia Langford, John Sillitoe on behalf of the Wellcome Sanger Institute COVID-19 Surveillance Team |
| EPI_ISL_3174723 | Lighthouse Lab in Milton Keynes | Wellcome Sanger Institute for the COVID-19 Genomics UK (COG-UK) Consortium | The Lighthouse Lab in Milton Keynes and Alex Alderton, Roberto Amato, Jeffrey Barrett, Sonia Goncalves, Ewan Harrison, David K. Jackson, Ian Johnston, Dominic Kwiatkowski, Cordelia Langford, John Sillitoe on behalf of the Wellcome Sanger Institute COVID-19 Surveillance Team |
| EPI_ISL_3174724 | Lighthouse Laboratory Plymouth | Wellcome Sanger Institute for the COVID-19 Genomics UK (COG-UK) Consortium | Lighthouse Laboratory Plymouth and Alex Alderton, Roberto Amato, Jeffrey Barrett, Sonia Goncalves, Ewan Harrison, David K. Jackson, Ian Johnston, Dominic Kwiatkowski, Cordelia Langford, John Sillitoe on behalf of the Wellcome Sanger Institute COVID-19 Surveillance Team |
| EPI_ISL_3174725 | Lighthouse Lab in Glasgow | Wellcome Sanger Institute for the COVID-19 Genomics UK (COG-UK) Consortium | Harper VanSteenhouse, Yumi Kasai, David Gray, Carol Clugston, Anna Dominiczak and Alex Alderton, Roberto Amato, Jeffrey Barrett, Sonia Goncalves, Ewan Harrison, David K. Jackson, Ian Johnston, Dominic Kwiatkowski, Cordelia Langford, John Sillitoe on behalf of the Wellcome Sanger Institute COVID-19 Surveillance Team |
| EPI_ISL_3174726 | Lighthouse Laboratory Plymouth | Wellcome Sanger Institute for the COVID-19 Genomics UK (COG-UK) Consortium | Lighthouse Laboratory Plymouth and Alex Alderton, Roberto Amato, Jeffrey Barrett, Sonia Goncalves, Ewan Harrison, David K. Jackson, Ian Johnston, Dominic Kwiatkowski, Cordelia Langford, John Sillitoe on behalf of the Wellcome Sanger Institute COVID-19 Surveillance Team |
| EPI_ISL_3174727 | Lighthouse Lab in Alderley Park | Wellcome Sanger Institute for the COVID-19 Genomics UK (COG-UK) Consortium | Jacquelyn Wynn, Mairead Hyland, The Lighthouse Lab in Alderley Park and Alex Alderton, Roberto Amato, Jeffrey Barrett, Sonia Goncalves, Ewan Harrison, David K. Jackson, Ian Johnston, Dominic Kwiatkowski, Cordelia Langford, John Sillitoe on behalf of the Wellcome Sanger Institute COVID-19 Surveillance Team |
| EPI_ISL_3174728, EPI_ISL_3174729, EPI_ISL_3174730 | Lighthouse Laboratory Plymouth | Wellcome Sanger Institute for the COVID-19 Genomics UK (COG-UK) Consortium | Lighthouse Laboratory Plymouth and Alex Alderton, Roberto Amato, Jeffrey Barrett, Sonia Goncalves, Ewan Harrison, David K. Jackson, Ian Johnston, Dominic Kwiatkowski, Cordelia Langford, John Sillitoe on behalf of the Wellcome Sanger Institute COVID-19 Surveillance Team |

| | | | |
|---|---|---|---|
| EPI_ISL_3174731, EPI_ISL_3174732 | Lighthouse Lab in Alderley Park | Wellcome Sanger Institute for the COVID-19 Genomics UK (COG-UK) Consortium | Jacquelyn Wynn, Mairead Hyland, The Lighthouse Lab in Alderley Park and Alex Alderton, Roberto Amato, Jeffrey Barrett, Sonia Goncalves, Ewan Harrison, David K. Jackson, Ian Johnston, Dominic Kwiatkowski, Cordelia Langford, John Sillitoe on behalf of the Wellcome Sanger Institute COVID-19 Surveillance Team |
| EPI_ISL_3174733 | Lighthouse Lab in Milton Keynes | Wellcome Sanger Institute for the COVID-19 Genomics UK (COG-UK) Consortium | The Lighthouse Lab in Milton Keynes and Alex Alderton, Roberto Amato, Jeffrey Barrett, Sonia Goncalves, Ewan Harrison, David K. Jackson, Ian Johnston, Dominic Kwiatkowski, Cordelia Langford, John Sillitoe on behalf of the Wellcome Sanger Institute COVID-19 Surveillance Team |
| EPI_ISL_3174734 | Lighthouse Laboratory Plymouth | Wellcome Sanger Institute for the COVID-19 Genomics UK (COG-UK) Consortium | Lighthouse Laboratory Plymouth and Alex Alderton, Roberto Amato, Jeffrey Barrett, Sonia Goncalves, Ewan Harrison, David K. Jackson, Ian Johnston, Dominic Kwiatkowski, Cordelia Langford, John Sillitoe on behalf of the Wellcome Sanger Institute COVID-19 Surveillance Team |
| EPI_ISL_3174735 | Lighthouse Lab in Milton Keynes | Wellcome Sanger Institute for the COVID-19 Genomics UK (COG-UK) Consortium | The Lighthouse Lab in Milton Keynes and Alex Alderton, Roberto Amato, Jeffrey Barrett, Sonia Goncalves, Ewan Harrison, David K. Jackson, Ian Johnston, Dominic Kwiatkowski, Cordelia Langford, John Sillitoe on behalf of the Wellcome Sanger Institute COVID-19 Surveillance Team |
| EPI_ISL_3174736 | Lighthouse Laboratory Plymouth | Wellcome Sanger Institute for the COVID-19 Genomics UK (COG-UK) Consortium | Lighthouse Laboratory Plymouth and Alex Alderton, Roberto Amato, Jeffrey Barrett, Sonia Goncalves, Ewan Harrison, David K. Jackson, Ian Johnston, Dominic Kwiatkowski, Cordelia Langford, John Sillitoe on behalf of the Wellcome Sanger Institute COVID-19 Surveillance Team |
| EPI_ISL_3174737 | Lighthouse Lab in Glasgow | Wellcome Sanger Institute for the COVID-19 Genomics UK (COG-UK) Consortium | Harper VanSteenhouse, Yumi Kasai, David Gray, Carol Clugston, Anna Dominiczak and Alex Alderton, Roberto Amato, Jeffrey Barrett, Sonia Goncalves, Ewan Harrison, David K. Jackson, Ian Johnston, Dominic Kwiatkowski, Cordelia Langford, John Sillitoe on behalf of the Wellcome Sanger Institute COVID-19 Surveillance Team |
| EPI_ISL_3174738 | Lighthouse Lab in Milton Keynes | Wellcome Sanger Institute for the COVID-19 Genomics UK (COG-UK) Consortium | The Lighthouse Lab in Milton Keynes and Alex Alderton, Roberto Amato, Jeffrey Barrett, Sonia Goncalves, Ewan Harrison, David K. Jackson, Ian Johnston, Dominic Kwiatkowski, Cordelia Langford, John Sillitoe on behalf of the Wellcome Sanger Institute COVID-19 Surveillance Team |
| EPI_ISL_3174739 | Lighthouse Laboratory Plymouth | Wellcome Sanger Institute for the COVID-19 Genomics UK (COG-UK) Consortium | Lighthouse Laboratory Plymouth and Alex Alderton, Roberto Amato, Jeffrey Barrett, Sonia Goncalves, Ewan Harrison, David K. Jackson, Ian Johnston, Dominic Kwiatkowski, Cordelia Langford, John Sillitoe on behalf of the Wellcome Sanger Institute COVID-19 Surveillance Team |
| EPI_ISL_3174740 | Lighthouse Lab in Alderley Park | Wellcome Sanger Institute for the COVID-19 Genomics UK (COG-UK) Consortium | Jacquelyn Wynn, Mairead Hyland, The Lighthouse Lab in Alderley Park and Alex Alderton, Roberto Amato, Jeffrey Barrett, Sonia Goncalves, Ewan Harrison, David K. Jackson, Ian Johnston, Dominic Kwiatkowski, Cordelia Langford, John Sillitoe on behalf of the Wellcome Sanger Institute COVID-19 Surveillance Team |
| EPI_ISL_3174741 | Lighthouse Laboratory Plymouth | Wellcome Sanger Institute for the COVID-19 Genomics UK (COG-UK) Consortium | Lighthouse Laboratory Plymouth and Alex Alderton, Roberto Amato, Jeffrey Barrett, Sonia Goncalves, Ewan Harrison, David K. Jackson, Ian Johnston, Dominic Kwiatkowski, Cordelia Langford, John Sillitoe on behalf of the Wellcome Sanger Institute COVID-19 Surveillance Team |
| EPI_ISL_3174742, EPI_ISL_3174743 | Lighthouse Lab in Alderley Park | Wellcome Sanger Institute for the COVID-19 Genomics UK (COG-UK) Consortium | Jacquelyn Wynn, Mairead Hyland, The Lighthouse Lab in Alderley Park and Alex Alderton, Roberto Amato, Jeffrey Barrett, Sonia Goncalves, Ewan Harrison, David K. Jackson, Ian Johnston, Dominic Kwiatkowski, Cordelia Langford, John Sillitoe on behalf of the Wellcome Sanger Institute COVID-19 Surveillance Team |
| EPI_ISL_3174744, EPI_ISL_3174745 | Lighthouse Lab in Glasgow | Wellcome Sanger Institute for the COVID-19 Genomics UK (COG-UK) Consortium | Harper VanSteenhouse, Yumi Kasai, David Gray, Carol Clugston, Anna Dominiczak and Alex Alderton, Roberto Amato, Jeffrey Barrett, Sonia Goncalves, Ewan Harrison, David K. Jackson, Ian Johnston, Dominic Kwiatkowski, Cordelia Langford, John Sillitoe on behalf of the Wellcome Sanger Institute COVID-19 Surveillance Team |
| EPI_ISL_3174746 | Lighthouse Lab in Milton Keynes | Wellcome Sanger Institute for the COVID-19 Genomics UK (COG-UK) Consortium | The Lighthouse Lab in Milton Keynes and Alex Alderton, Roberto Amato, Jeffrey Barrett, Sonia Goncalves, Ewan Harrison, David K. Jackson, Ian Johnston, Dominic Kwiatkowski, Cordelia Langford, John Sillitoe on behalf of the Wellcome Sanger Institute COVID-19 Surveillance Team |
| EPI_ISL_3174747 | Lighthouse Lab in Alderley Park | Wellcome Sanger Institute for the COVID-19 Genomics UK (COG-UK) Consortium | Jacquelyn Wynn, Mairead Hyland, The Lighthouse Lab in Alderley Park and Alex Alderton, Roberto Amato, Jeffrey Barrett, Sonia Goncalves, Ewan Harrison, David K. Jackson, Ian Johnston, Dominic Kwiatkowski, Cordelia Langford, John Sillitoe on behalf of the Wellcome Sanger Institute COVID-19 Surveillance Team |
| EPI_ISL_3174748 | Lighthouse Lab in Glasgow | Wellcome Sanger Institute for the COVID-19 Genomics UK (COG-UK) Consortium | Harper VanSteenhouse, Yumi Kasai, David Gray, Carol Clugston, Anna Dominiczak and Alex Alderton, Roberto Amato, Jeffrey Barrett, Sonia Goncalves, Ewan Harrison, David K. Jackson, Ian Johnston, Dominic Kwiatkowski, Cordelia Langford, John Sillitoe on behalf of the Wellcome Sanger Institute COVID-19 Surveillance Team |
| EPI_ISL_3174749 | Lighthouse Lab in Alderley Park | Wellcome Sanger Institute for the COVID-19 Genomics UK (COG-UK) Consortium | Jacquelyn Wynn, Mairead Hyland, The Lighthouse Lab in Alderley Park and Alex Alderton, Roberto Amato, Jeffrey Barrett, Sonia Goncalves, Ewan Harrison, David K. Jackson, Ian Johnston, Dominic Kwiatkowski, Cordelia Langford, John Sillitoe on behalf of the Wellcome Sanger Institute COVID-19 Surveillance Team |
| EPI_ISL_3174750 | Lighthouse Laboratory Plymouth | Wellcome Sanger Institute for the COVID-19 Genomics UK (COG-UK) Consortium | Lighthouse Laboratory Plymouth and Alex Alderton, Roberto Amato, Jeffrey Barrett, Sonia Goncalves, Ewan Harrison, David K. Jackson, Ian Johnston, Dominic Kwiatkowski, Cordelia Langford, John Sillitoe on behalf of the Wellcome Sanger Institute COVID-19 Surveillance Team |
| EPI_ISL_3174751 | Lighthouse Lab in Milton Keynes | Wellcome Sanger Institute for the COVID-19 Genomics UK (COG-UK) Consortium | The Lighthouse Lab in Milton Keynes and Alex Alderton, Roberto Amato, Jeffrey Barrett, Sonia Goncalves, Ewan Harrison, David K. Jackson, Ian Johnston, Dominic Kwiatkowski, Cordelia Langford, John Sillitoe on behalf of the Wellcome Sanger Institute COVID-19 Surveillance Team |
| EPI_ISL_3174752 | Lighthouse Laboratory Plymouth | Wellcome Sanger Institute for the COVID-19 Genomics UK (COG-UK) Consortium | Lighthouse Laboratory Plymouth and Alex Alderton, Roberto Amato, Jeffrey Barrett, Sonia Goncalves, Ewan Harrison, David K. Jackson, Ian Johnston, Dominic Kwiatkowski, Cordelia Langford, John Sillitoe on behalf of the Wellcome Sanger Institute COVID-19 Surveillance Team |
| EPI_ISL_3174753 | Lighthouse Lab in Milton Keynes | Wellcome Sanger Institute for the COVID-19 Genomics UK (COG-UK) Consortium | The Lighthouse Lab in Milton Keynes and Alex Alderton, Roberto Amato, Jeffrey Barrett, Sonia Goncalves, Ewan Harrison, David K. Jackson, Ian Johnston, Dominic Kwiatkowski, Cordelia Langford, John Sillitoe on behalf of the Wellcome Sanger Institute COVID-19 Surveillance Team |
| EPI_ISL_3174754 | Lighthouse Lab in Glasgow | Wellcome Sanger Institute for the COVID-19 Genomics UK (COG-UK) Consortium | Harper VanSteenhouse, Yumi Kasai, David Gray, Carol Clugston, Anna Dominiczak and Alex Alderton, Roberto Amato, Jeffrey Barrett, Sonia Goncalves, Ewan Harrison, David K. Jackson, Ian Johnston, Dominic Kwiatkowski, Cordelia Langford, John Sillitoe on behalf of the Wellcome Sanger Institute COVID-19 Surveillance Team |
| EPI_ISL_3174755, EPI_ISL_3174756 | Lighthouse Laboratory Plymouth | Wellcome Sanger Institute for the COVID-19 Genomics UK (COG-UK) Consortium | Lighthouse Laboratory Plymouth and Alex Alderton, Roberto Amato, Jeffrey Barrett, Sonia Goncalves, Ewan Harrison, David K. Jackson, Ian Johnston, Dominic Kwiatkowski, Cordelia Langford, John Sillitoe on behalf of the Wellcome Sanger Institute COVID-19 Surveillance Team |
| EPI_ISL_3174757, EPI_ISL_3174758 | Lighthouse Lab in Milton Keynes | Wellcome Sanger Institute for the COVID-19 Genomics UK (COG-UK) Consortium | The Lighthouse Lab in Milton Keynes and Alex Alderton, Roberto Amato, Jeffrey Barrett, Sonia Goncalves, Ewan Harrison, David K. Jackson, Ian Johnston, Dominic Kwiatkowski, Cordelia Langford, John Sillitoe on behalf of the Wellcome Sanger Institute COVID-19 Surveillance Team |
| EPI_ISL_3174759 | Lighthouse Lab in Alderley Park | Wellcome Sanger Institute for the COVID-19 Genomics UK (COG-UK) Consortium | Jacquelyn Wynn, Mairead Hyland, The Lighthouse Lab in Alderley Park and Alex Alderton, Roberto Amato, Jeffrey Barrett, Sonia Goncalves, Ewan Harrison, David K. Jackson, Ian Johnston, Dominic Kwiatkowski, Cordelia Langford, John Sillitoe on behalf of the Wellcome Sanger Institute COVID-19 Surveillance Team |
| EPI_ISL_3174760 | Lighthouse Lab in Milton Keynes | Wellcome Sanger Institute for the COVID-19 Genomics UK (COG-UK) Consortium | The Lighthouse Lab in Milton Keynes and Alex Alderton, Roberto Amato, Jeffrey Barrett, Sonia Goncalves, Ewan Harrison, David K. Jackson, Ian Johnston, Dominic Kwiatkowski, Cordelia Langford, John Sillitoe on behalf of the Wellcome Sanger Institute COVID-19 Surveillance Team |
| EPI_ISL_3174761 | Lighthouse Lab in Alderley Park | Wellcome Sanger Institute for the COVID-19 Genomics UK (COG-UK) Consortium | Jacquelyn Wynn, Mairead Hyland, The Lighthouse Lab in Alderley Park and Alex Alderton, Roberto Amato, Jeffrey Barrett, Sonia Goncalves, Ewan Harrison, David K. Jackson, Ian Johnston, Dominic Kwiatkowski, Cordelia Langford, John Sillitoe on behalf of the Wellcome Sanger Institute COVID-19 Surveillance Team |
| EPI_ISL_3174762, EPI_ISL_3174763, EPI_ISL_3174764 | Lighthouse Laboratory Plymouth | Wellcome Sanger Institute for the COVID-19 Genomics UK (COG-UK) Consortium | Lighthouse Laboratory Plymouth and Alex Alderton, Roberto Amato, Jeffrey Barrett, Sonia Goncalves, Ewan Harrison, David K. Jackson, Ian Johnston, Dominic Kwiatkowski, Cordelia Langford, John Sillitoe on behalf of the Wellcome Sanger Institute COVID-19 Surveillance Team |
| EPI_ISL_3174765 | Lighthouse Lab in Glasgow | Wellcome Sanger Institute for the COVID-19 Genomics UK (COG-UK) Consortium | Harper VanSteenhouse, Yumi Kasai, David Gray, Carol Clugston, Anna Dominiczak and Alex Alderton, Roberto Amato, Jeffrey Barrett, Sonia Goncalves, Ewan Harrison, David K. Jackson, Ian Johnston, Dominic Kwiatkowski, Cordelia Langford, John Sillitoe on behalf of the Wellcome Sanger Institute COVID-19 Surveillance Team |
| EPI_ISL_3174766 | Lighthouse Lab in Alderley Park | Wellcome Sanger Institute for the COVID-19 Genomics UK (COG-UK) Consortium | Jacquelyn Wynn, Mairead Hyland, The Lighthouse Lab in Alderley Park and Alex Alderton, Roberto Amato, Jeffrey Barrett, Sonia Goncalves, Ewan Harrison, David K. Jackson, Ian Johnston, Dominic Kwiatkowski, Cordelia Langford, John Sillitoe on behalf of the Wellcome Sanger Institute COVID-19 Surveillance Team |
| EPI_ISL_3174767, EPI_ISL_3174768 | Lighthouse Lab in Milton Keynes | Wellcome Sanger Institute for the COVID-19 Genomics UK (COG-UK) Consortium | The Lighthouse Lab in Milton Keynes and Alex Alderton, Roberto Amato, Jeffrey Barrett, Sonia Goncalves, Ewan Harrison, David K. Jackson, Ian Johnston, Dominic Kwiatkowski, Cordelia Langford, John Sillitoe on behalf of the Wellcome Sanger Institute COVID-19 Surveillance Team |
| EPI_ISL_3174769, EPI_ISL_3174770, EPI_ISL_3174771 | Lighthouse Lab in Alderley Park | Wellcome Sanger Institute for the COVID-19 Genomics UK (COG-UK) Consortium | Jacquelyn Wynn, Mairead Hyland, The Lighthouse Lab in Alderley Park and Alex Alderton, Roberto Amato, Jeffrey Barrett, Sonia Goncalves, Ewan Harrison, David K. Jackson, Ian Johnston, Dominic Kwiatkowski, Cordelia Langford, John Sillitoe on behalf of the Wellcome Sanger Institute COVID-19 Surveillance Team |
| EPI_ISL_3174772 | Lighthouse Lab in Milton Keynes | Wellcome Sanger Institute for the COVID-19 Genomics UK (COG-UK) Consortium | The Lighthouse Lab in Milton Keynes and Alex Alderton, Roberto Amato, Jeffrey Barrett, Sonia Goncalves, Ewan Harrison, David K. Jackson, Ian Johnston, Dominic Kwiatkowski, Cordelia Langford, John Sillitoe on behalf of the Wellcome Sanger Institute COVID-19 Surveillance Team |
| EPI_ISL_3174773 | Lighthouse Lab in Glasgow | Wellcome Sanger Institute for the COVID-19 Genomics UK (COG-UK) Consortium | Harper VanSteenhouse, Yumi Kasai, David Gray, Carol Clugston, Anna Dominiczak and Alex Alderton, Roberto Amato, Jeffrey Barrett, Sonia Goncalves, Ewan Harrison, David K. Jackson, Ian Johnston, Dominic Kwiatkowski, Cordelia Langford, John Sillitoe on behalf of the Wellcome Sanger Institute COVID-19 Surveillance Team |
| EPI_ISL_3174774 | Lighthouse Lab in Alderley Park | Wellcome Sanger Institute for the COVID-19 Genomics UK (COG-UK) Consortium | Jacquelyn Wynn, Mairead Hyland, The Lighthouse Lab in Alderley Park and Alex Alderton, Roberto Amato, Jeffrey Barrett, Sonia Goncalves, Ewan Harrison, David K. Jackson, Ian Johnston, Dominic Kwiatkowski, Cordelia Langford, John Sillitoe on behalf of the Wellcome Sanger Institute COVID-19 Surveillance Team |
| EPI_ISL_3174775, EPI_ISL_3174776 | Lighthouse Laboratory Plymouth | Wellcome Sanger Institute for the COVID-19 Genomics UK (COG-UK) Consortium | Lighthouse Laboratory Plymouth and Alex Alderton, Roberto Amato, Jeffrey Barrett, Sonia Goncalves, Ewan Harrison, David K. Jackson, Ian Johnston, Dominic Kwiatkowski, Cordelia Langford, John Sillitoe on behalf of the Wellcome Sanger Institute COVID-19 Surveillance Team |
| EPI_ISL_3174777 | Lighthouse Lab in Glasgow | Wellcome Sanger Institute for the COVID-19 Genomics UK (COG-UK) Consortium | Harper VanSteenhouse, Yumi Kasai, David Gray, Carol Clugston, Anna Dominiczak and Alex Alderton, Roberto Amato, Jeffrey Barrett, Sonia Goncalves, Ewan Harrison, David K. Jackson, Ian Johnston, Dominic Kwiatkowski, Cordelia Langford, John Sillitoe on behalf of the Wellcome Sanger Institute COVID-19 Surveillance Team |
| EPI_ISL_3174778 | Lighthouse Lab in Milton Keynes | Wellcome Sanger Institute for the COVID-19 Genomics UK (COG-UK) Consortium | The Lighthouse Lab in Milton Keynes and Alex Alderton, Roberto Amato, Jeffrey Barrett, Sonia Goncalves, Ewan Harrison, David K. Jackson, Ian Johnston, Dominic Kwiatkowski, Cordelia Langford, John Sillitoe on behalf of the Wellcome Sanger Institute COVID-19 Surveillance Team |
| EPI_ISL_3174779 | Lighthouse Laboratory Plymouth | Wellcome Sanger Institute for the COVID-19 Genomics UK (COG-UK) Consortium | Lighthouse Laboratory Plymouth and Alex Alderton, Roberto Amato, Jeffrey Barrett, Sonia Goncalves, Ewan Harrison, David K. Jackson, Ian Johnston, Dominic Kwiatkowski, Cordelia Langford, John Sillitoe on behalf of the Wellcome Sanger Institute COVID-19 Surveillance Team |
| EPI_ISL_3174780 | Lighthouse Lab in Alderley Park | Wellcome Sanger Institute for the COVID-19 Genomics UK (COG-UK) Consortium | Jacquelyn Wynn, Mairead Hyland, The Lighthouse Lab in Alderley Park and Alex Alderton, Roberto Amato, Jeffrey Barrett, Sonia Goncalves, Ewan Harrison, David K. Jackson, Ian Johnston, Dominic Kwiatkowski, Cordelia Langford, John Sillitoe on behalf of the Wellcome Sanger Institute COVID-19 Surveillance Team |
| EPI_ISL_3174781, EPI_ISL_3174782 | Lighthouse Lab in Glasgow | Wellcome Sanger Institute for the COVID-19 Genomics UK (COG-UK) Consortium | Harper VanSteenhouse, Yumi Kasai, David Gray, Carol Clugston, Anna Dominiczak and Alex Alderton, Roberto Amato, Jeffrey Barrett, Sonia Goncalves, Ewan Harrison, David K. Jackson, Ian Johnston, Dominic Kwiatkowski, Cordelia Langford, John Sillitoe on behalf of the Wellcome Sanger Institute COVID-19 Surveillance Team |
| EPI_ISL_3174783 | Lighthouse Lab in Milton Keynes | Wellcome Sanger Institute for the COVID-19 Genomics UK (COG-UK) Consortium | The Lighthouse Lab in Milton Keynes and Alex Alderton, Roberto Amato, Jeffrey Barrett, Sonia Goncalves, Ewan Harrison, David K. Jackson, Ian Johnston, Dominic Kwiatkowski, Cordelia Langford, John Sillitoe on behalf of the Wellcome Sanger Institute COVID-19 Surveillance Team |
| EPI_ISL_3174784 | Lighthouse Lab in Glasgow | Wellcome Sanger Institute for the COVID-19 Genomics UK (COG-UK) Consortium | Harper VanSteenhouse, Yumi Kasai, David Gray, Carol Clugston, Anna Dominiczak and Alex Alderton, Roberto Amato, Jeffrey Barrett, Sonia Goncalves, Ewan Harrison, David K. Jackson, Ian Johnston, Dominic Kwiatkowski, Cordelia Langford, John Sillitoe on behalf of the Wellcome Sanger Institute COVID-19 Surveillance Team |
| EPI_ISL_3174785, EPI_ISL_3174786 | Lighthouse Lab in Milton Keynes | Wellcome Sanger Institute for the COVID-19 Genomics UK (COG-UK) Consortium | The Lighthouse Lab in Milton Keynes and Alex Alderton, Roberto Amato, Jeffrey Barrett, Sonia Goncalves, Ewan Harrison, David K. Jackson, Ian Johnston, Dominic Kwiatkowski, Cordelia Langford, John Sillitoe on behalf of the Wellcome Sanger Institute COVID-19 Surveillance Team |
| EPI_ISL_3174787, EPI_ISL_3174788, EPI_ISL_3174789 | Lighthouse Laboratory Plymouth | Wellcome Sanger Institute for the COVID-19 Genomics UK (COG-UK) Consortium | Lighthouse Laboratory Plymouth and Alex Alderton, Roberto Amato, Jeffrey Barrett, Sonia Goncalves, Ewan Harrison, David K. Jackson, Ian Johnston, Dominic Kwiatkowski, Cordelia Langford, John Sillitoe on behalf of the Wellcome Sanger Institute COVID-19 Surveillance Team |

| | | | |
|---|---|---|---|
| EPI_ISL_3174790, EPI_ISL_3174791 | Lighthouse Lab in Milton Keynes | Wellcome Sanger Institute for the COVID-19 Genomics UK (COG-UK) Consortium | The Lighthouse Lab in Milton Keynes and Alex Alderton, Roberto Amato, Jeffrey Barrett, Sonia Goncalves, Ewan Harrison, David K. Jackson, Ian Johnston, Dominic Kwiatkowski, Cordelia Langford, John Sillitoe on behalf of the Wellcome Sanger Institute COVID-19 Surveillance Team |
| EPI_ISL_3174792 | Lighthouse Lab in Alderley Park | Wellcome Sanger Institute for the COVID-19 Genomics UK (COG-UK) Consortium | Jacquelyn Wynn, Mairead Hyland, The Lighthouse Lab in Alderley Park and Alex Alderton, Roberto Amato, Jeffrey Barrett, Sonia Goncalves, Ewan Harrison, David K. Jackson, Ian Johnston, Dominic Kwiatkowski, Cordelia Langford, John Sillitoe on behalf of the Wellcome Sanger Institute COVID-19 Surveillance Team |
| EPI_ISL_3174793, EPI_ISL_3174794 | Lighthouse Laboratory Plymouth | Wellcome Sanger Institute for the COVID-19 Genomics UK (COG-UK) Consortium | Lighthouse Laboratory Plymouth and Alex Alderton, Roberto Amato, Jeffrey Barrett, Sonia Goncalves, Ewan Harrison, David K. Jackson, Ian Johnston, Dominic Kwiatkowski, Cordelia Langford, John Sillitoe on behalf of the Wellcome Sanger Institute COVID-19 Surveillance Team |
| EPI_ISL_3174795, EPI_ISL_3174796 | Lighthouse Lab in Alderley Park | Wellcome Sanger Institute for the COVID-19 Genomics UK (COG-UK) Consortium | Jacquelyn Wynn, Mairead Hyland, The Lighthouse Lab in Alderley Park and Alex Alderton, Roberto Amato, Jeffrey Barrett, Sonia Goncalves, Ewan Harrison, David K. Jackson, Ian Johnston, Dominic Kwiatkowski, Cordelia Langford, John Sillitoe on behalf of the Wellcome Sanger Institute COVID-19 Surveillance Team |
| EPI_ISL_3174797 | Lighthouse Laboratory Plymouth | Wellcome Sanger Institute for the COVID-19 Genomics UK (COG-UK) Consortium | Lighthouse Laboratory Plymouth and Alex Alderton, Roberto Amato, Jeffrey Barrett, Sonia Goncalves, Ewan Harrison, David K. Jackson, Ian Johnston, Dominic Kwiatkowski, Cordelia Langford, John Sillitoe on behalf of the Wellcome Sanger Institute COVID-19 Surveillance Team |
| EPI_ISL_3174798, EPI_ISL_3174799 | Lighthouse Lab in Glasgow | Wellcome Sanger Institute for the COVID-19 Genomics UK (COG-UK) Consortium | Harper VanSteenhouse, Yumi Kasai, David Gray, Carol Clugston, Anna Dominiczak and Alex Alderton, Roberto Amato, Jeffrey Barrett, Sonia Goncalves, Ewan Harrison, David K. Jackson, Ian Johnston, Dominic Kwiatkowski, Cordelia Langford, John Sillitoe on behalf of the Wellcome Sanger Institute COVID-19 Surveillance Team |
| EPI_ISL_3174800 | Lighthouse Lab in Milton Keynes | Wellcome Sanger Institute for the COVID-19 Genomics UK (COG-UK) Consortium | The Lighthouse Lab in Milton Keynes and Alex Alderton, Roberto Amato, Jeffrey Barrett, Sonia Goncalves, Ewan Harrison, David K. Jackson, Ian Johnston, Dominic Kwiatkowski, Cordelia Langford, John Sillitoe on behalf of the Wellcome Sanger Institute COVID-19 Surveillance Team |
| EPI_ISL_3174801 | Lighthouse Laboratory Plymouth | Wellcome Sanger Institute for the COVID-19 Genomics UK (COG-UK) Consortium | Lighthouse Laboratory Plymouth and Alex Alderton, Roberto Amato, Jeffrey Barrett, Sonia Goncalves, Ewan Harrison, David K. Jackson, Ian Johnston, Dominic Kwiatkowski, Cordelia Langford, John Sillitoe on behalf of the Wellcome Sanger Institute COVID-19 Surveillance Team |
| EPI_ISL_3174802 | Lighthouse Lab in Alderley Park | Wellcome Sanger Institute for the COVID-19 Genomics UK (COG-UK) Consortium | Jacquelyn Wynn, Mairead Hyland, The Lighthouse Lab in Alderley Park and Alex Alderton, Roberto Amato, Jeffrey Barrett, Sonia Goncalves, Ewan Harrison, David K. Jackson, Ian Johnston, Dominic Kwiatkowski, Cordelia Langford, John Sillitoe on behalf of the Wellcome Sanger Institute COVID-19 Surveillance Team |
| EPI_ISL_3174803, EPI_ISL_3174804 | Lighthouse Laboratory Plymouth | Wellcome Sanger Institute for the COVID-19 Genomics UK (COG-UK) Consortium | Lighthouse Laboratory Plymouth and Alex Alderton, Roberto Amato, Jeffrey Barrett, Sonia Goncalves, Ewan Harrison, David K. Jackson, Ian Johnston, Dominic Kwiatkowski, Cordelia Langford, John Sillitoe on behalf of the Wellcome Sanger Institute COVID-19 Surveillance Team |
| EPI_ISL_3174805 | Lighthouse Lab in Glasgow | Wellcome Sanger Institute for the COVID-19 Genomics UK (COG-UK) Consortium | Harper VanSteenhouse, Yumi Kasai, David Gray, Carol Clugston, Anna Dominiczak and Alex Alderton, Roberto Amato, Jeffrey Barrett, Sonia Goncalves, Ewan Harrison, David K. Jackson, Ian Johnston, Dominic Kwiatkowski, Cordelia Langford, John Sillitoe on behalf of the Wellcome Sanger Institute COVID-19 Surveillance Team |
| EPI_ISL_3174806, EPI_ISL_3174807 | Lighthouse Lab in Alderley Park | Wellcome Sanger Institute for the COVID-19 Genomics UK (COG-UK) Consortium | Jacquelyn Wynn, Mairead Hyland, The Lighthouse Lab in Alderley Park and Alex Alderton, Roberto Amato, Jeffrey Barrett, Sonia Goncalves, Ewan Harrison, David K. Jackson, Ian Johnston, Dominic Kwiatkowski, Cordelia Langford, John Sillitoe on behalf of the Wellcome Sanger Institute COVID-19 Surveillance Team |
| EPI_ISL_3174808 | Lighthouse Laboratory Plymouth | Wellcome Sanger Institute for the COVID-19 Genomics UK (COG-UK) Consortium | Lighthouse Laboratory Plymouth and Alex Alderton, Roberto Amato, Jeffrey Barrett, Sonia Goncalves, Ewan Harrison, David K. Jackson, Ian Johnston, Dominic Kwiatkowski, Cordelia Langford, John Sillitoe on behalf of the Wellcome Sanger Institute COVID-19 Surveillance Team |
| EPI_ISL_3174809 | Lighthouse Lab in Glasgow | Wellcome Sanger Institute for the COVID-19 Genomics UK (COG-UK) Consortium | Harper VanSteenhouse, Yumi Kasai, David Gray, Carol Clugston, Anna Dominiczak and Alex Alderton, Roberto Amato, Jeffrey Barrett, Sonia Goncalves, Ewan Harrison, David K. Jackson, Ian Johnston, Dominic Kwiatkowski, Cordelia Langford, John Sillitoe on behalf of the Wellcome Sanger Institute COVID-19 Surveillance Team |
| EPI_ISL_3174810 | Lighthouse Lab in Alderley Park | Wellcome Sanger Institute for the COVID-19 Genomics UK (COG-UK) Consortium | Jacquelyn Wynn, Mairead Hyland, The Lighthouse Lab in Alderley Park and Alex Alderton, Roberto Amato, Jeffrey Barrett, Sonia Goncalves, Ewan Harrison, David K. Jackson, Ian Johnston, Dominic Kwiatkowski, Cordelia Langford, John Sillitoe on behalf of the Wellcome Sanger Institute COVID-19 Surveillance Team |
| EPI_ISL_3174811 | Lighthouse Laboratory Plymouth | Wellcome Sanger Institute for the COVID-19 Genomics UK (COG-UK) Consortium | Lighthouse Laboratory Plymouth and Alex Alderton, Roberto Amato, Jeffrey Barrett, Sonia Goncalves, Ewan Harrison, David K. Jackson, Ian Johnston, Dominic Kwiatkowski, Cordelia Langford, John Sillitoe on behalf of the Wellcome Sanger Institute COVID-19 Surveillance Team |
| EPI_ISL_3174812 | Lighthouse Lab in Milton Keynes | Wellcome Sanger Institute for the COVID-19 Genomics UK (COG-UK) Consortium | The Lighthouse Lab in Milton Keynes and Alex Alderton, Roberto Amato, Jeffrey Barrett, Sonia Goncalves, Ewan Harrison, David K. Jackson, Ian Johnston, Dominic Kwiatkowski, Cordelia Langford, John Sillitoe on behalf of the Wellcome Sanger Institute COVID-19 Surveillance Team |
| EPI_ISL_3174813, EPI_ISL_3174814 | Lighthouse Lab in Glasgow | Wellcome Sanger Institute for the COVID-19 Genomics UK (COG-UK) Consortium | Harper VanSteenhouse, Yumi Kasai, David Gray, Carol Clugston, Anna Dominiczak and Alex Alderton, Roberto Amato, Jeffrey Barrett, Sonia Goncalves, Ewan Harrison, David K. Jackson, Ian Johnston, Dominic Kwiatkowski, Cordelia Langford, John Sillitoe on behalf of the Wellcome Sanger Institute COVID-19 Surveillance Team |
| EPI_ISL_3174815 | Lighthouse Lab in Milton Keynes | Wellcome Sanger Institute for the COVID-19 Genomics UK (COG-UK) Consortium | The Lighthouse Lab in Milton Keynes and Alex Alderton, Roberto Amato, Jeffrey Barrett, Sonia Goncalves, Ewan Harrison, David K. Jackson, Ian Johnston, Dominic Kwiatkowski, Cordelia Langford, John Sillitoe on behalf of the Wellcome Sanger Institute COVID-19 Surveillance Team |
| EPI_ISL_3174816 | Lighthouse Lab in Glasgow | Wellcome Sanger Institute for the COVID-19 Genomics UK (COG-UK) Consortium | Harper VanSteenhouse, Yumi Kasai, David Gray, Carol Clugston, Anna Dominiczak and Alex Alderton, Roberto Amato, Jeffrey Barrett, Sonia Goncalves, Ewan Harrison, David K. Jackson, Ian Johnston, Dominic Kwiatkowski, Cordelia Langford, John Sillitoe on behalf of the Wellcome Sanger Institute COVID-19 Surveillance Team |
| EPI_ISL_3174817 | Lighthouse Lab in Milton Keynes | Wellcome Sanger Institute for the COVID-19 Genomics UK (COG-UK) Consortium | The Lighthouse Lab in Milton Keynes and Alex Alderton, Roberto Amato, Jeffrey Barrett, Sonia Goncalves, Ewan Harrison, David K. Jackson, Ian Johnston, Dominic Kwiatkowski, Cordelia Langford, John Sillitoe on behalf of the Wellcome Sanger Institute COVID-19 Surveillance Team |
| EPI_ISL_3174818 | Lighthouse Lab in Alderley Park | Wellcome Sanger Institute for the COVID-19 Genomics UK (COG-UK) Consortium | Jacquelyn Wynn, Mairead Hyland, The Lighthouse Lab in Alderley Park and Alex Alderton, Roberto Amato, Jeffrey Barrett, Sonia Goncalves, Ewan Harrison, David K. Jackson, Ian Johnston, Dominic Kwiatkowski, Cordelia Langford, John Sillitoe on behalf of the Wellcome Sanger Institute COVID-19 Surveillance Team |
| EPI_ISL_3174819, EPI_ISL_3174820, EPI_ISL_3174821 | Lighthouse Lab in Glasgow | Wellcome Sanger Institute for the COVID-19 Genomics UK (COG-UK) Consortium | Harper VanSteenhouse, Yumi Kasai, David Gray, Carol Clugston, Anna Dominiczak and Alex Alderton, Roberto Amato, Jeffrey Barrett, Sonia Goncalves, Ewan Harrison, David K. Jackson, Ian Johnston, Dominic Kwiatkowski, Cordelia Langford, John Sillitoe on behalf of the Wellcome Sanger Institute COVID-19 Surveillance Team |
| EPI_ISL_3174822, EPI_ISL_3174823 | Lighthouse Lab in Milton Keynes | Wellcome Sanger Institute for the COVID-19 Genomics UK (COG-UK) Consortium | The Lighthouse Lab in Milton Keynes and Alex Alderton, Roberto Amato, Jeffrey Barrett, Sonia Goncalves, Ewan Harrison, David K. Jackson, Ian Johnston, Dominic Kwiatkowski, Cordelia Langford, John Sillitoe on behalf of the Wellcome Sanger Institute COVID-19 Surveillance Team |
| EPI_ISL_3174824, EPI_ISL_3174825 | Lighthouse Lab in Glasgow | Wellcome Sanger Institute for the COVID-19 Genomics UK (COG-UK) Consortium | Harper VanSteenhouse, Yumi Kasai, David Gray, Carol Clugston, Anna Dominiczak and Alex Alderton, Roberto Amato, Jeffrey Barrett, Sonia Goncalves, Ewan Harrison, David K. Jackson, Ian Johnston, Dominic Kwiatkowski, Cordelia Langford, John Sillitoe on behalf of the Wellcome Sanger Institute COVID-19 Surveillance Team |
| EPI_ISL_3174826, EPI_ISL_3174827 | Berkshire and Surrey Pathology Services Lighthouse Laboratory | Wellcome Sanger Institute for the COVID-19 Genomics UK (COG-UK) Consortium | Berkshire and Surrey Pathology Services Lighthouse Laboratory and Alex Alderton, Roberto Amato, Jeffrey Barrett, Sonia Goncalves, Ewan Harrison, David K. Jackson, Ian Johnston, Dominic Kwiatkowski, Cordelia Langford, John Sillitoe on behalf of the Wellcome Sanger Institute COVID-19 Surveillance Team |
| EPI_ISL_3174828, EPI_ISL_3174829 | Lighthouse Lab in Milton Keynes | Wellcome Sanger Institute for the COVID-19 Genomics UK (COG-UK) Consortium | The Lighthouse Lab in Milton Keynes and Alex Alderton, Roberto Amato, Jeffrey Barrett, Sonia Goncalves, Ewan Harrison, David K. Jackson, Ian Johnston, Dominic Kwiatkowski, Cordelia Langford, John Sillitoe on behalf of the Wellcome Sanger Institute COVID-19 Surveillance Team |
| EPI_ISL_3174830, EPI_ISL_3174831 | Lighthouse Lab in Glasgow | Wellcome Sanger Institute for the COVID-19 Genomics UK (COG-UK) Consortium | Harper VanSteenhouse, Yumi Kasai, David Gray, Carol Clugston, Anna Dominiczak and Alex Alderton, Roberto Amato, Jeffrey Barrett, Sonia Goncalves, Ewan Harrison, David K. Jackson, Ian Johnston, Dominic Kwiatkowski, Cordelia Langford, John Sillitoe on behalf of the Wellcome Sanger Institute COVID-19 Surveillance Team |
| EPI_ISL_3174832 | Lighthouse Lab in Milton Keynes | Wellcome Sanger Institute for the COVID-19 Genomics UK (COG-UK) Consortium | The Lighthouse Lab in Milton Keynes and Alex Alderton, Roberto Amato, Jeffrey Barrett, Sonia Goncalves, Ewan Harrison, David K. Jackson, Ian Johnston, Dominic Kwiatkowski, Cordelia Langford, John Sillitoe on behalf of the Wellcome Sanger Institute COVID-19 Surveillance Team |
| EPI_ISL_3174833 | Berkshire and Surrey Pathology Services Lighthouse Laboratory | Wellcome Sanger Institute for the COVID-19 Genomics UK (COG-UK) Consortium | Berkshire and Surrey Pathology Services Lighthouse Laboratory and Alex Alderton, Roberto Amato, Jeffrey Barrett, Sonia Goncalves, Ewan Harrison, David K. Jackson, Ian Johnston, Dominic Kwiatkowski, Cordelia Langford, John Sillitoe on behalf of the Wellcome Sanger Institute COVID-19 Surveillance Team |
| EPI_ISL_3174834 | Lighthouse Lab in Glasgow | Wellcome Sanger Institute for the COVID-19 Genomics UK (COG-UK) Consortium | Harper VanSteenhouse, Yumi Kasai, David Gray, Carol Clugston, Anna Dominiczak and Alex Alderton, Roberto Amato, Jeffrey Barrett, Sonia Goncalves, Ewan Harrison, David K. Jackson, Ian Johnston, Dominic Kwiatkowski, Cordelia Langford, John Sillitoe on behalf of the Wellcome Sanger Institute COVID-19 Surveillance Team |
| EPI_ISL_3174835 | Lighthouse Lab in Milton Keynes | Wellcome Sanger Institute for the COVID-19 Genomics UK (COG-UK) Consortium | The Lighthouse Lab in Milton Keynes and Alex Alderton, Roberto Amato, Jeffrey Barrett, Sonia Goncalves, Ewan Harrison, David K. Jackson, Ian Johnston, Dominic Kwiatkowski, Cordelia Langford, John Sillitoe on behalf of the Wellcome Sanger Institute COVID-19 Surveillance Team |
| EPI_ISL_3174836, EPI_ISL_3174837 | Berkshire and Surrey Pathology Services Lighthouse Laboratory | Wellcome Sanger Institute for the COVID-19 Genomics UK (COG-UK) Consortium | Berkshire and Surrey Pathology Services Lighthouse Laboratory and Alex Alderton, Roberto Amato, Jeffrey Barrett, Sonia Goncalves, Ewan Harrison, David K. Jackson, Ian Johnston, Dominic Kwiatkowski, Cordelia Langford, John Sillitoe on behalf of the Wellcome Sanger Institute COVID-19 Surveillance Team |
| EPI_ISL_3174838 | Lighthouse Lab in Glasgow | Wellcome Sanger Institute for the COVID-19 Genomics UK (COG-UK) Consortium | Harper VanSteenhouse, Yumi Kasai, David Gray, Carol Clugston, Anna Dominiczak and Alex Alderton, Roberto Amato, Jeffrey Barrett, Sonia Goncalves, Ewan Harrison, David K. Jackson, Ian Johnston, Dominic Kwiatkowski, Cordelia Langford, John Sillitoe on behalf of the Wellcome Sanger Institute COVID-19 Surveillance Team |
| EPI_ISL_3174839 | Lighthouse Lab in Milton Keynes | Wellcome Sanger Institute for the COVID-19 Genomics UK (COG-UK) Consortium | The Lighthouse Lab in Milton Keynes and Alex Alderton, Roberto Amato, Jeffrey Barrett, Sonia Goncalves, Ewan Harrison, David K. Jackson, Ian Johnston, Dominic Kwiatkowski, Cordelia Langford, John Sillitoe on behalf of the Wellcome Sanger Institute COVID-19 Surveillance Team |
| EPI_ISL_3174840, EPI_ISL_3174841 | Lighthouse Lab in Glasgow | Wellcome Sanger Institute for the COVID-19 Genomics UK (COG-UK) Consortium | Harper VanSteenhouse, Yumi Kasai, David Gray, Carol Clugston, Anna Dominiczak and Alex Alderton, Roberto Amato, Jeffrey Barrett, Sonia Goncalves, Ewan Harrison, David K. Jackson, Ian Johnston, Dominic Kwiatkowski, Cordelia Langford, John Sillitoe on behalf of the Wellcome Sanger Institute COVID-19 Surveillance Team |
| EPI_ISL_3174842 | Lighthouse Lab in Milton Keynes | Wellcome Sanger Institute for the COVID-19 Genomics UK (COG-UK) Consortium | The Lighthouse Lab in Milton Keynes and Alex Alderton, Roberto Amato, Jeffrey Barrett, Sonia Goncalves, Ewan Harrison, David K. Jackson, Ian Johnston, Dominic Kwiatkowski, Cordelia Langford, John Sillitoe on behalf of the Wellcome Sanger Institute COVID-19 Surveillance Team |
| EPI_ISL_3174843 | Lighthouse Lab in Glasgow | Wellcome Sanger Institute for the COVID-19 Genomics UK (COG-UK) Consortium | Harper VanSteenhouse, Yumi Kasai, David Gray, Carol Clugston, Anna Dominiczak and Alex Alderton, Roberto Amato, Jeffrey Barrett, Sonia Goncalves, Ewan Harrison, David K. Jackson, Ian Johnston, Dominic Kwiatkowski, Cordelia Langford, John Sillitoe on behalf of the Wellcome Sanger Institute COVID-19 Surveillance Team |
| EPI_ISL_3174844, EPI_ISL_3174845 | Berkshire and Surrey Pathology Services Lighthouse Laboratory | Wellcome Sanger Institute for the COVID-19 Genomics UK (COG-UK) Consortium | Berkshire and Surrey Pathology Services Lighthouse Laboratory and Alex Alderton, Roberto Amato, Jeffrey Barrett, Sonia Goncalves, Ewan Harrison, David K. Jackson, Ian Johnston, Dominic Kwiatkowski, Cordelia Langford, John Sillitoe on behalf of the Wellcome Sanger Institute COVID-19 Surveillance Team |
| EPI_ISL_3174846, EPI_ISL_3174847 | Lighthouse Lab in Glasgow | Wellcome Sanger Institute for the COVID-19 Genomics UK (COG-UK) Consortium | Harper VanSteenhouse, Yumi Kasai, David Gray, Carol Clugston, Anna Dominiczak and Alex Alderton, Roberto Amato, Jeffrey Barrett, Sonia Goncalves, Ewan Harrison, David K. Jackson, Ian Johnston, Dominic Kwiatkowski, Cordelia Langford, John Sillitoe on behalf of the Wellcome Sanger Institute COVID-19 Surveillance Team |
| EPI_ISL_3174848, EPI_ISL_3174849, EPI_ISL_3174850 | Lighthouse Lab in Milton Keynes | Wellcome Sanger Institute for the COVID-19 Genomics UK (COG-UK) Consortium | The Lighthouse Lab in Milton Keynes and Alex Alderton, Roberto Amato, Jeffrey Barrett, Sonia Goncalves, Ewan Harrison, David K. Jackson, Ian Johnston, Dominic Kwiatkowski, Cordelia Langford, John Sillitoe on behalf of the Wellcome Sanger Institute COVID-19 Surveillance Team |
| EPI_ISL_3174851 | Berkshire and Surrey Pathology Services Lighthouse Laboratory | Wellcome Sanger Institute for the COVID-19 Genomics UK (COG-UK) Consortium | Berkshire and Surrey Pathology Services Lighthouse Laboratory and Alex Alderton, Roberto Amato, Jeffrey Barrett, Sonia Goncalves, Ewan Harrison, David K. Jackson, Ian Johnston, Dominic Kwiatkowski, Cordelia Langford, John Sillitoe on behalf of the Wellcome Sanger Institute COVID-19 Surveillance Team |
| EPI_ISL_3174852 | Lighthouse Lab in Milton Keynes | Wellcome Sanger Institute for the COVID-19 Genomics UK (COG-UK) Consortium | The Lighthouse Lab in Milton Keynes and Alex Alderton, Roberto Amato, Jeffrey Barrett, Sonia Goncalves, Ewan Harrison, David K. Jackson, Ian Johnston, Dominic Kwiatkowski, Cordelia Langford, John Sillitoe on behalf of the Wellcome Sanger Institute COVID-19 Surveillance Team |
| EPI_ISL_3174853, EPI_ISL_3174854 | Lighthouse Lab in Glasgow | Wellcome Sanger Institute for the COVID-19 Genomics UK (COG-UK) Consortium | Harper VanSteenhouse, Yumi Kasai, David Gray, Carol Clugston, Anna Dominiczak and Alex Alderton, Roberto Amato, Jeffrey Barrett, Sonia Goncalves, Ewan Harrison, David K. Jackson, Ian Johnston, Dominic Kwiatkowski, Cordelia Langford, John Sillitoe on behalf of the Wellcome Sanger Institute COVID-19 Surveillance Team |

| | | | |
|---|---|---|---|
| EPI_ISL_3174855 | Lighthouse Lab in Milton Keynes | Wellcome Sanger Institute for the COVID-19 Genomics UK (COG-UK) Consortium | The Lighthouse Lab in Milton Keynes and Alex Alderton, Roberto Amato, Jeffrey Barrett, Sonia Goncalves, Ewan Harrison, David K. Jackson, Ian Johnston, Dominic Kwiatkowski, Cordelia Langford, John Sillitoe on behalf of the Wellcome Sanger Institute COVID-19 Surveillance Team |
| EPI_ISL_3174856, EPI_ISL_3174857, EPI_ISL_3174858 | Lighthouse Lab in Glasgow | Wellcome Sanger Institute for the COVID-19 Genomics UK (COG-UK) Consortium | Harper VanSteenhouse, Yumi Kasai, David Gray, Carol Clugston, Anna Dominiczak and Alex Alderton, Roberto Amato, Jeffrey Barrett, Sonia Goncalves, Ewan Harrison, David K. Jackson, Ian Johnston, Dominic Kwiatkowski, Cordelia Langford, John Sillitoe on behalf of the Wellcome Sanger Institute COVID-19 Surveillance Team |
| EPI_ISL_3174859 | Berkshire and Surrey Pathology Services Lighthouse Laboratory | Wellcome Sanger Institute for the COVID-19 Genomics UK (COG-UK) Consortium | Berkshire and Surrey Pathology Services Lighthouse Laboratory and Alex Alderton, Roberto Amato, Jeffrey Barrett, Sonia Goncalves, Ewan Harrison, David K. Jackson, Ian Johnston, Dominic Kwiatkowski, Cordelia Langford, John Sillitoe on behalf of the Wellcome Sanger Institute COVID-19 Surveillance Team |
| EPI_ISL_3174860 | Lighthouse Lab in Milton Keynes | Wellcome Sanger Institute for the COVID-19 Genomics UK (COG-UK) Consortium | The Lighthouse Lab in Milton Keynes and Alex Alderton, Roberto Amato, Jeffrey Barrett, Sonia Goncalves, Ewan Harrison, David K. Jackson, Ian Johnston, Dominic Kwiatkowski, Cordelia Langford, John Sillitoe on behalf of the Wellcome Sanger Institute COVID-19 Surveillance Team |
| EPI_ISL_3174861 | Berkshire and Surrey Pathology Services Lighthouse Laboratory | Wellcome Sanger Institute for the COVID-19 Genomics UK (COG-UK) Consortium | Berkshire and Surrey Pathology Services Lighthouse Laboratory and Alex Alderton, Roberto Amato, Jeffrey Barrett, Sonia Goncalves, Ewan Harrison, David K. Jackson, Ian Johnston, Dominic Kwiatkowski, Cordelia Langford, John Sillitoe on behalf of the Wellcome Sanger Institute COVID-19 Surveillance Team |
| EPI_ISL_3174862, EPI_ISL_3174863, EPI_ISL_3174864 | Lighthouse Lab in Glasgow | Wellcome Sanger Institute for the COVID-19 Genomics UK (COG-UK) Consortium | Harper VanSteenhouse, Yumi Kasai, David Gray, Carol Clugston, Anna Dominiczak and Alex Alderton, Roberto Amato, Jeffrey Barrett, Sonia Goncalves, Ewan Harrison, David K. Jackson, Ian Johnston, Dominic Kwiatkowski, Cordelia Langford, John Sillitoe on behalf of the Wellcome Sanger Institute COVID-19 Surveillance Team |
| EPI_ISL_3174865 | Berkshire and Surrey Pathology Services Lighthouse Laboratory | Wellcome Sanger Institute for the COVID-19 Genomics UK (COG-UK) Consortium | Berkshire and Surrey Pathology Services Lighthouse Laboratory and Alex Alderton, Roberto Amato, Jeffrey Barrett, Sonia Goncalves, Ewan Harrison, David K. Jackson, Ian Johnston, Dominic Kwiatkowski, Cordelia Langford, John Sillitoe on behalf of the Wellcome Sanger Institute COVID-19 Surveillance Team |
| EPI_ISL_3174866, EPI_ISL_3174867, EPI_ISL_3174868, EPI_ISL_3174869 | Lighthouse Lab in Glasgow | Wellcome Sanger Institute for the COVID-19 Genomics UK (COG-UK) Consortium | Harper VanSteenhouse, Yumi Kasai, David Gray, Carol Clugston, Anna Dominiczak and Alex Alderton, Roberto Amato, Jeffrey Barrett, Sonia Goncalves, Ewan Harrison, David K. Jackson, Ian Johnston, Dominic Kwiatkowski, Cordelia Langford, John Sillitoe on behalf of the Wellcome Sanger Institute COVID-19 Surveillance Team |
| EPI_ISL_3174870, EPI_ISL_3174871, EPI_ISL_3174872, EPI_ISL_3174873, EPI_ISL_3174874, EPI_ISL_3174875, EPI_ISL_3174876 | Berkshire and Surrey Pathology Services Lighthouse Laboratory | Wellcome Sanger Institute for the COVID-19 Genomics UK (COG-UK) Consortium | Berkshire and Surrey Pathology Services Lighthouse Laboratory and Alex Alderton, Roberto Amato, Jeffrey Barrett, Sonia Goncalves, Ewan Harrison, David K. Jackson, Ian Johnston, Dominic Kwiatkowski, Cordelia Langford, John Sillitoe on behalf of the Wellcome Sanger Institute COVID-19 Surveillance Team |
| EPI_ISL_3174877 | Lighthouse Lab in Glasgow | Wellcome Sanger Institute for the COVID-19 Genomics UK (COG-UK) Consortium | Harper VanSteenhouse, Yumi Kasai, David Gray, Carol Clugston, Anna Dominiczak and Alex Alderton, Roberto Amato, Jeffrey Barrett, Sonia Goncalves, Ewan Harrison, David K. Jackson, Ian Johnston, Dominic Kwiatkowski, Cordelia Langford, John Sillitoe on behalf of the Wellcome Sanger Institute COVID-19 Surveillance Team |
| EPI_ISL_3174878 | Lighthouse Lab in Milton Keynes | Wellcome Sanger Institute for the COVID-19 Genomics UK (COG-UK) Consortium | The Lighthouse Lab in Milton Keynes and Alex Alderton, Roberto Amato, Jeffrey Barrett, Sonia Goncalves, Ewan Harrison, David K. Jackson, Ian Johnston, Dominic Kwiatkowski, Cordelia Langford, John Sillitoe on behalf of the Wellcome Sanger Institute COVID-19 Surveillance Team |
| EPI_ISL_3174879 | Lighthouse Lab in Glasgow | Wellcome Sanger Institute for the COVID-19 Genomics UK (COG-UK) Consortium | Harper VanSteenhouse, Yumi Kasai, David Gray, Carol Clugston, Anna Dominiczak and Alex Alderton, Roberto Amato, Jeffrey Barrett, Sonia Goncalves, Ewan Harrison, David K. Jackson, Ian Johnston, Dominic Kwiatkowski, Cordelia Langford, John Sillitoe on behalf of the Wellcome Sanger Institute COVID-19 Surveillance Team |
| EPI_ISL_3174880 | Lighthouse Lab in Milton Keynes | Wellcome Sanger Institute for the COVID-19 Genomics UK (COG-UK) Consortium | The Lighthouse Lab in Milton Keynes and Alex Alderton, Roberto Amato, Jeffrey Barrett, Sonia Goncalves, Ewan Harrison, David K. Jackson, Ian Johnston, Dominic Kwiatkowski, Cordelia Langford, John Sillitoe on behalf of the Wellcome Sanger Institute COVID-19 Surveillance Team |
| EPI_ISL_3174881 | Berkshire and Surrey Pathology Services Lighthouse Laboratory | Wellcome Sanger Institute for the COVID-19 Genomics UK (COG-UK) Consortium | Berkshire and Surrey Pathology Services Lighthouse Laboratory and Alex Alderton, Roberto Amato, Jeffrey Barrett, Sonia Goncalves, Ewan Harrison, David K. Jackson, Ian Johnston, Dominic Kwiatkowski, Cordelia Langford, John Sillitoe on behalf of the Wellcome Sanger Institute COVID-19 Surveillance Team |
| EPI_ISL_3174882 | Lighthouse Lab in Glasgow | Wellcome Sanger Institute for the COVID-19 Genomics UK (COG-UK) Consortium | Harper VanSteenhouse, Yumi Kasai, David Gray, Carol Clugston, Anna Dominiczak and Alex Alderton, Roberto Amato, Jeffrey Barrett, Sonia Goncalves, Ewan Harrison, David K. Jackson, Ian Johnston, Dominic Kwiatkowski, Cordelia Langford, John Sillitoe on behalf of the Wellcome Sanger Institute COVID-19 Surveillance Team |
| EPI_ISL_3174883 | Lighthouse Lab in Milton Keynes | Wellcome Sanger Institute for the COVID-19 Genomics UK (COG-UK) Consortium | The Lighthouse Lab in Milton Keynes and Alex Alderton, Roberto Amato, Jeffrey Barrett, Sonia Goncalves, Ewan Harrison, David K. Jackson, Ian Johnston, Dominic Kwiatkowski, Cordelia Langford, John Sillitoe on behalf of the Wellcome Sanger Institute COVID-19 Surveillance Team |
| EPI_ISL_3174884 | Berkshire and Surrey Pathology Services Lighthouse Laboratory | Wellcome Sanger Institute for the COVID-19 Genomics UK (COG-UK) Consortium | Berkshire and Surrey Pathology Services Lighthouse Laboratory and Alex Alderton, Roberto Amato, Jeffrey Barrett, Sonia Goncalves, Ewan Harrison, David K. Jackson, Ian Johnston, Dominic Kwiatkowski, Cordelia Langford, John Sillitoe on behalf of the Wellcome Sanger Institute COVID-19 Surveillance Team |
| EPI_ISL_3174885 | Lighthouse Lab in Milton Keynes | Wellcome Sanger Institute for the COVID-19 Genomics UK (COG-UK) Consortium | The Lighthouse Lab in Milton Keynes and Alex Alderton, Roberto Amato, Jeffrey Barrett, Sonia Goncalves, Ewan Harrison, David K. Jackson, Ian Johnston, Dominic Kwiatkowski, Cordelia Langford, John Sillitoe on behalf of the Wellcome Sanger Institute COVID-19 Surveillance Team |
| EPI_ISL_3174886 | Lighthouse Lab in Glasgow | Wellcome Sanger Institute for the COVID-19 Genomics UK (COG-UK) Consortium | Harper VanSteenhouse, Yumi Kasai, David Gray, Carol Clugston, Anna Dominiczak and Alex Alderton, Roberto Amato, Jeffrey Barrett, Sonia Goncalves, Ewan Harrison, David K. Jackson, Ian Johnston, Dominic Kwiatkowski, Cordelia Langford, John Sillitoe on behalf of the Wellcome Sanger Institute COVID-19 Surveillance Team |
| EPI_ISL_3174887 | Berkshire and Surrey Pathology Services Lighthouse Laboratory | Wellcome Sanger Institute for the COVID-19 Genomics UK (COG-UK) Consortium | Berkshire and Surrey Pathology Services Lighthouse Laboratory and Alex Alderton, Roberto Amato, Jeffrey Barrett, Sonia Goncalves, Ewan Harrison, David K. Jackson, Ian Johnston, Dominic Kwiatkowski, Cordelia Langford, John Sillitoe on behalf of the Wellcome Sanger Institute COVID-19 Surveillance Team |
| EPI_ISL_3174888 | Lighthouse Lab in Milton Keynes | Wellcome Sanger Institute for the COVID-19 Genomics UK (COG-UK) Consortium | The Lighthouse Lab in Milton Keynes and Alex Alderton, Roberto Amato, Jeffrey Barrett, Sonia Goncalves, Ewan Harrison, David K. Jackson, Ian Johnston, Dominic Kwiatkowski, Cordelia Langford, John Sillitoe on behalf of the Wellcome Sanger Institute COVID-19 Surveillance Team |
| EPI_ISL_3174889 | Berkshire and Surrey Pathology Services Lighthouse Laboratory | Wellcome Sanger Institute for the COVID-19 Genomics UK (COG-UK) Consortium | Berkshire and Surrey Pathology Services Lighthouse Laboratory and Alex Alderton, Roberto Amato, Jeffrey Barrett, Sonia Goncalves, Ewan Harrison, David K. Jackson, Ian Johnston, Dominic Kwiatkowski, Cordelia Langford, John Sillitoe on behalf of the Wellcome Sanger Institute COVID-19 Surveillance Team |
| EPI_ISL_3174890, EPI_ISL_3174891, EPI_ISL_3174892, EPI_ISL_3174893 | Lighthouse Lab in Glasgow | Wellcome Sanger Institute for the COVID-19 Genomics UK (COG-UK) Consortium | Harper VanSteenhouse, Yumi Kasai, David Gray, Carol Clugston, Anna Dominiczak and Alex Alderton, Roberto Amato, Jeffrey Barrett, Sonia Goncalves, Ewan Harrison, David K. Jackson, Ian Johnston, Dominic Kwiatkowski, Cordelia Langford, John Sillitoe on behalf of the Wellcome Sanger Institute COVID-19 Surveillance Team |
| EPI_ISL_3174894 | Lighthouse Lab in Milton Keynes | Wellcome Sanger Institute for the COVID-19 Genomics UK (COG-UK) Consortium | The Lighthouse Lab in Milton Keynes and Alex Alderton, Roberto Amato, Jeffrey Barrett, Sonia Goncalves, Ewan Harrison, David K. Jackson, Ian Johnston, Dominic Kwiatkowski, Cordelia Langford, John Sillitoe on behalf of the Wellcome Sanger Institute COVID-19 Surveillance Team |
| EPI_ISL_3174895, EPI_ISL_3174896 | Lighthouse Lab in Glasgow | Wellcome Sanger Institute for the COVID-19 Genomics UK (COG-UK) Consortium | Harper VanSteenhouse, Yumi Kasai, David Gray, Carol Clugston, Anna Dominiczak and Alex Alderton, Roberto Amato, Jeffrey Barrett, Sonia Goncalves, Ewan Harrison, David K. Jackson, Ian Johnston, Dominic Kwiatkowski, Cordelia Langford, John Sillitoe on behalf of the Wellcome Sanger Institute COVID-19 Surveillance Team |
| EPI_ISL_3174897 | Berkshire and Surrey Pathology Services Lighthouse Laboratory | Wellcome Sanger Institute for the COVID-19 Genomics UK (COG-UK) Consortium | Berkshire and Surrey Pathology Services Lighthouse Laboratory and Alex Alderton, Roberto Amato, Jeffrey Barrett, Sonia Goncalves, Ewan Harrison, David K. Jackson, Ian Johnston, Dominic Kwiatkowski, Cordelia Langford, John Sillitoe on behalf of the Wellcome Sanger Institute COVID-19 Surveillance Team |
| EPI_ISL_3174898 | Lighthouse Lab in Milton Keynes | Wellcome Sanger Institute for the COVID-19 Genomics UK (COG-UK) Consortium | The Lighthouse Lab in Milton Keynes and Alex Alderton, Roberto Amato, Jeffrey Barrett, Sonia Goncalves, Ewan Harrison, David K. Jackson, Ian Johnston, Dominic Kwiatkowski, Cordelia Langford, John Sillitoe on behalf of the Wellcome Sanger Institute COVID-19 Surveillance Team |
| EPI_ISL_3174899 | Lighthouse Lab in Glasgow | Wellcome Sanger Institute for the COVID-19 Genomics UK (COG-UK) Consortium | Harper VanSteenhouse, Yumi Kasai, David Gray, Carol Clugston, Anna Dominiczak and Alex Alderton, Roberto Amato, Jeffrey Barrett, Sonia Goncalves, Ewan Harrison, David K. Jackson, Ian Johnston, Dominic Kwiatkowski, Cordelia Langford, John Sillitoe on behalf of the Wellcome Sanger Institute COVID-19 Surveillance Team |
| EPI_ISL_3174900 | Berkshire and Surrey Pathology Services Lighthouse Laboratory | Wellcome Sanger Institute for the COVID-19 Genomics UK (COG-UK) Consortium | Berkshire and Surrey Pathology Services Lighthouse Laboratory and Alex Alderton, Roberto Amato, Jeffrey Barrett, Sonia Goncalves, Ewan Harrison, David K. Jackson, Ian Johnston, Dominic Kwiatkowski, Cordelia Langford, John Sillitoe on behalf of the Wellcome Sanger Institute COVID-19 Surveillance Team |
| EPI_ISL_3174901 | Lighthouse Lab in Milton Keynes | Wellcome Sanger Institute for the COVID-19 Genomics UK (COG-UK) Consortium | The Lighthouse Lab in Milton Keynes and Alex Alderton, Roberto Amato, Jeffrey Barrett, Sonia Goncalves, Ewan Harrison, David K. Jackson, Ian Johnston, Dominic Kwiatkowski, Cordelia Langford, John Sillitoe on behalf of the Wellcome Sanger Institute COVID-19 Surveillance Team |
| EPI_ISL_3174902, EPI_ISL_3174903 | Lighthouse Lab in Glasgow | Wellcome Sanger Institute for the COVID-19 Genomics UK (COG-UK) Consortium | Harper VanSteenhouse, Yumi Kasai, David Gray, Carol Clugston, Anna Dominiczak and Alex Alderton, Roberto Amato, Jeffrey Barrett, Sonia Goncalves, Ewan Harrison, David K. Jackson, Ian Johnston, Dominic Kwiatkowski, Cordelia Langford, John Sillitoe on behalf of the Wellcome Sanger Institute COVID-19 Surveillance Team |
| EPI_ISL_3174904, EPI_ISL_3174905, EPI_ISL_3174906, EPI_ISL_3174907 | Lighthouse Lab in Milton Keynes | Wellcome Sanger Institute for the COVID-19 Genomics UK (COG-UK) Consortium | The Lighthouse Lab in Milton Keynes and Alex Alderton, Roberto Amato, Jeffrey Barrett, Sonia Goncalves, Ewan Harrison, David K. Jackson, Ian Johnston, Dominic Kwiatkowski, Cordelia Langford, John Sillitoe on behalf of the Wellcome Sanger Institute COVID-19 Surveillance Team |
| EPI_ISL_3174908 | Lighthouse Lab in Glasgow | Wellcome Sanger Institute for the COVID-19 Genomics UK (COG-UK) Consortium | Harper VanSteenhouse, Yumi Kasai, David Gray, Carol Clugston, Anna Dominiczak and Alex Alderton, Roberto Amato, Jeffrey Barrett, Sonia Goncalves, Ewan Harrison, David K. Jackson, Ian Johnston, Dominic Kwiatkowski, Cordelia Langford, John Sillitoe on behalf of the Wellcome Sanger Institute COVID-19 Surveillance Team |
| EPI_ISL_3174909, EPI_ISL_3174910 | Lighthouse Lab in Milton Keynes | Wellcome Sanger Institute for the COVID-19 Genomics UK (COG-UK) Consortium | The Lighthouse Lab in Milton Keynes and Alex Alderton, Roberto Amato, Jeffrey Barrett, Sonia Goncalves, Ewan Harrison, David K. Jackson, Ian Johnston, Dominic Kwiatkowski, Cordelia Langford, John Sillitoe on behalf of the Wellcome Sanger Institute COVID-19 Surveillance Team |
| EPI_ISL_3174911 | Lighthouse Lab in Glasgow | Wellcome Sanger Institute for the COVID-19 Genomics UK (COG-UK) Consortium | Harper VanSteenhouse, Yumi Kasai, David Gray, Carol Clugston, Anna Dominiczak and Alex Alderton, Roberto Amato, Jeffrey Barrett, Sonia Goncalves, Ewan Harrison, David K. Jackson, Ian Johnston, Dominic Kwiatkowski, Cordelia Langford, John Sillitoe on behalf of the Wellcome Sanger Institute COVID-19 Surveillance Team |
| EPI_ISL_3174912, EPI_ISL_3174913 | Berkshire and Surrey Pathology Services Lighthouse Laboratory | Wellcome Sanger Institute for the COVID-19 Genomics UK (COG-UK) Consortium | Berkshire and Surrey Pathology Services Lighthouse Laboratory and Alex Alderton, Roberto Amato, Jeffrey Barrett, Sonia Goncalves, Ewan Harrison, David K. Jackson, Ian Johnston, Dominic Kwiatkowski, Cordelia Langford, John Sillitoe on behalf of the Wellcome Sanger Institute COVID-19 Surveillance Team |
| EPI_ISL_3174914 | Lighthouse Lab in Glasgow | Wellcome Sanger Institute for the COVID-19 Genomics UK (COG-UK) Consortium | Harper VanSteenhouse, Yumi Kasai, David Gray, Carol Clugston, Anna Dominiczak and Alex Alderton, Roberto Amato, Jeffrey Barrett, Sonia Goncalves, Ewan Harrison, David K. Jackson, Ian Johnston, Dominic Kwiatkowski, Cordelia Langford, John Sillitoe on behalf of the Wellcome Sanger Institute COVID-19 Surveillance Team |
| EPI_ISL_3174915 | Berkshire and Surrey Pathology Services Lighthouse Laboratory | Wellcome Sanger Institute for the COVID-19 Genomics UK (COG-UK) Consortium | Berkshire and Surrey Pathology Services Lighthouse Laboratory and Alex Alderton, Roberto Amato, Jeffrey Barrett, Sonia Goncalves, Ewan Harrison, David K. Jackson, Ian Johnston, Dominic Kwiatkowski, Cordelia Langford, John Sillitoe on behalf of the Wellcome Sanger Institute COVID-19 Surveillance Team |
| EPI_ISL_3174916 | Lighthouse Lab in Glasgow | Wellcome Sanger Institute for the COVID-19 Genomics UK (COG-UK) Consortium | Harper VanSteenhouse, Yumi Kasai, David Gray, Carol Clugston, Anna Dominiczak and Alex Alderton, Roberto Amato, Jeffrey Barrett, Sonia Goncalves, Ewan Harrison, David K. Jackson, Ian Johnston, Dominic Kwiatkowski, Cordelia Langford, John Sillitoe on behalf of the Wellcome Sanger Institute COVID-19 Surveillance Team |
| EPI_ISL_3174917 | Lighthouse Lab in Milton Keynes | Wellcome Sanger Institute for the COVID-19 Genomics UK (COG-UK) | The Lighthouse Lab in Milton Keynes and Alex Alderton, Roberto Amato, Jeffrey Barrett, Sonia Goncalves, Ewan Harrison, David K. Jackson, Ian Johnston, Dominic Kwiatkowski, Cordelia Langford, John Sillitoe on behalf of the Wellcome Sanger |

| | | | |
|---|---|---|---|
| | | Consortium | Institute COVID-19 Surveillance Team |
| EPI_ISL_3174918 | Berkshire and Surrey Pathology Services Lighthouse Laboratory | Wellcome Sanger Institute for the COVID-19 Genomics UK (COG-UK) Consortium | Berkshire and Surrey Pathology Services Lighthouse Laboratory and Alex Alderton, Roberto Amato, Jeffrey Barrett, Sonia Goncalves, Ewan Harrison, David K. Jackson, Ian Johnston, Dominic Kwiatkowski, Cordelia Langford, John Sillitoe on behalf of the Wellcome Sanger Institute COVID-19 Surveillance Team |
| EPI_ISL_3174919, EPI_ISL_3174920, EPI_ISL_3174921, EPI_ISL_3174922, EPI_ISL_3174923 | Lighthouse Lab in Glasgow | Wellcome Sanger Institute for the COVID-19 Genomics UK (COG-UK) Consortium | Harper VanSteenhouse, Yumi Kasai, David Gray, Carol Clugston, Anna Dominiczak and Alex Alderton, Roberto Amato, Jeffrey Barrett, Sonia Goncalves, Ewan Harrison, David K. Jackson, Ian Johnston, Dominic Kwiatkowski, Cordelia Langford, John Sillitoe on behalf of the Wellcome Sanger Institute COVID-19 Surveillance Team |
| EPI_ISL_3174924 | Berkshire and Surrey Pathology Services Lighthouse Laboratory | Wellcome Sanger Institute for the COVID-19 Genomics UK (COG-UK) Consortium | Berkshire and Surrey Pathology Services Lighthouse Laboratory and Alex Alderton, Roberto Amato, Jeffrey Barrett, Sonia Goncalves, Ewan Harrison, David K. Jackson, Ian Johnston, Dominic Kwiatkowski, Cordelia Langford, John Sillitoe on behalf of the Wellcome Sanger Institute COVID-19 Surveillance Team |
| EPI_ISL_3174925 | Lighthouse Lab in Milton Keynes | Wellcome Sanger Institute for the COVID-19 Genomics UK (COG-UK) Consortium | The Lighthouse Lab in Milton Keynes and Alex Alderton, Roberto Amato, Jeffrey Barrett, Sonia Goncalves, Ewan Harrison, David K. Jackson, Ian Johnston, Dominic Kwiatkowski, Cordelia Langford, John Sillitoe on behalf of the Wellcome Sanger Institute COVID-19 Surveillance Team |
| EPI_ISL_3174926, EPI_ISL_3174927, EPI_ISL_3174928, EPI_ISL_3174929 | Lighthouse Lab in Glasgow | Wellcome Sanger Institute for the COVID-19 Genomics UK (COG-UK) Consortium | Harper VanSteenhouse, Yumi Kasai, David Gray, Carol Clugston, Anna Dominiczak and Alex Alderton, Roberto Amato, Jeffrey Barrett, Sonia Goncalves, Ewan Harrison, David K. Jackson, Ian Johnston, Dominic Kwiatkowski, Cordelia Langford, John Sillitoe on behalf of the Wellcome Sanger Institute COVID-19 Surveillance Team |
| EPI_ISL_3174930 | Lighthouse Lab in Milton Keynes | Wellcome Sanger Institute for the COVID-19 Genomics UK (COG-UK) Consortium | The Lighthouse Lab in Milton Keynes and Alex Alderton, Roberto Amato, Jeffrey Barrett, Sonia Goncalves, Ewan Harrison, David K. Jackson, Ian Johnston, Dominic Kwiatkowski, Cordelia Langford, John Sillitoe on behalf of the Wellcome Sanger Institute COVID-19 Surveillance Team |
| EPI_ISL_3174931 | Berkshire and Surrey Pathology Services Lighthouse Laboratory | Wellcome Sanger Institute for the COVID-19 Genomics UK (COG-UK) Consortium | Berkshire and Surrey Pathology Services Lighthouse Laboratory and Alex Alderton, Roberto Amato, Jeffrey Barrett, Sonia Goncalves, Ewan Harrison, David K. Jackson, Ian Johnston, Dominic Kwiatkowski, Cordelia Langford, John Sillitoe on behalf of the Wellcome Sanger Institute COVID-19 Surveillance Team |
| EPI_ISL_3174932 | Lighthouse Lab in Milton Keynes | Wellcome Sanger Institute for the COVID-19 Genomics UK (COG-UK) Consortium | The Lighthouse Lab in Milton Keynes and Alex Alderton, Roberto Amato, Jeffrey Barrett, Sonia Goncalves, Ewan Harrison, David K. Jackson, Ian Johnston, Dominic Kwiatkowski, Cordelia Langford, John Sillitoe on behalf of the Wellcome Sanger Institute COVID-19 Surveillance Team |
| EPI_ISL_3174933, EPI_ISL_3174934, EPI_ISL_3174935 | Lighthouse Lab in Glasgow | Wellcome Sanger Institute for the COVID-19 Genomics UK (COG-UK) Consortium | Harper VanSteenhouse, Yumi Kasai, David Gray, Carol Clugston, Anna Dominiczak and Alex Alderton, Roberto Amato, Jeffrey Barrett, Sonia Goncalves, Ewan Harrison, David K. Jackson, Ian Johnston, Dominic Kwiatkowski, Cordelia Langford, John Sillitoe on behalf of the Wellcome Sanger Institute COVID-19 Surveillance Team |
| EPI_ISL_3174936 | Lighthouse Lab in Milton Keynes | Wellcome Sanger Institute for the COVID-19 Genomics UK (COG-UK) Consortium | The Lighthouse Lab in Milton Keynes and Alex Alderton, Roberto Amato, Jeffrey Barrett, Sonia Goncalves, Ewan Harrison, David K. Jackson, Ian Johnston, Dominic Kwiatkowski, Cordelia Langford, John Sillitoe on behalf of the Wellcome Sanger Institute COVID-19 Surveillance Team |
| EPI_ISL_3174937 | Berkshire and Surrey Pathology Services Lighthouse Laboratory | Wellcome Sanger Institute for the COVID-19 Genomics UK (COG-UK) Consortium | Berkshire and Surrey Pathology Services Lighthouse Laboratory and Alex Alderton, Roberto Amato, Jeffrey Barrett, Sonia Goncalves, Ewan Harrison, David K. Jackson, Ian Johnston, Dominic Kwiatkowski, Cordelia Langford, John Sillitoe on behalf of the Wellcome Sanger Institute COVID-19 Surveillance Team |
| EPI_ISL_3174938 | Lighthouse Lab in Milton Keynes | Wellcome Sanger Institute for the COVID-19 Genomics UK (COG-UK) Consortium | The Lighthouse Lab in Milton Keynes and Alex Alderton, Roberto Amato, Jeffrey Barrett, Sonia Goncalves, Ewan Harrison, David K. Jackson, Ian Johnston, Dominic Kwiatkowski, Cordelia Langford, John Sillitoe on behalf of the Wellcome Sanger Institute COVID-19 Surveillance Team |
| EPI_ISL_3174939, EPI_ISL_3174940 | Lighthouse Lab in Glasgow | Wellcome Sanger Institute for the COVID-19 Genomics UK (COG-UK) Consortium | Harper VanSteenhouse, Yumi Kasai, David Gray, Carol Clugston, Anna Dominiczak and Alex Alderton, Roberto Amato, Jeffrey Barrett, Sonia Goncalves, Ewan Harrison, David K. Jackson, Ian Johnston, Dominic Kwiatkowski, Cordelia Langford, John Sillitoe on behalf of the Wellcome Sanger Institute COVID-19 Surveillance Team |
| EPI_ISL_3174941 | Lighthouse Lab in Milton Keynes | Wellcome Sanger Institute for the COVID-19 Genomics UK (COG-UK) Consortium | The Lighthouse Lab in Milton Keynes and Alex Alderton, Roberto Amato, Jeffrey Barrett, Sonia Goncalves, Ewan Harrison, David K. Jackson, Ian Johnston, Dominic Kwiatkowski, Cordelia Langford, John Sillitoe on behalf of the Wellcome Sanger Institute COVID-19 Surveillance Team |
| EPI_ISL_3174942 | Berkshire and Surrey Pathology Services Lighthouse Laboratory | Wellcome Sanger Institute for the COVID-19 Genomics UK (COG-UK) Consortium | Berkshire and Surrey Pathology Services Lighthouse Laboratory and Alex Alderton, Roberto Amato, Jeffrey Barrett, Sonia Goncalves, Ewan Harrison, David K. Jackson, Ian Johnston, Dominic Kwiatkowski, Cordelia Langford, John Sillitoe on behalf of the Wellcome Sanger Institute COVID-19 Surveillance Team |
| EPI_ISL_3174943 | Lighthouse Lab in Glasgow | Wellcome Sanger Institute for the COVID-19 Genomics UK (COG-UK) Consortium | Harper VanSteenhouse, Yumi Kasai, David Gray, Carol Clugston, Anna Dominiczak and Alex Alderton, Roberto Amato, Jeffrey Barrett, Sonia Goncalves, Ewan Harrison, David K. Jackson, Ian Johnston, Dominic Kwiatkowski, Cordelia Langford, John Sillitoe on behalf of the Wellcome Sanger Institute COVID-19 Surveillance Team |
| EPI_ISL_3174944 | Berkshire and Surrey Pathology Services Lighthouse Laboratory | Wellcome Sanger Institute for the COVID-19 Genomics UK (COG-UK) Consortium | Berkshire and Surrey Pathology Services Lighthouse Laboratory and Alex Alderton, Roberto Amato, Jeffrey Barrett, Sonia Goncalves, Ewan Harrison, David K. Jackson, Ian Johnston, Dominic Kwiatkowski, Cordelia Langford, John Sillitoe on behalf of the Wellcome Sanger Institute COVID-19 Surveillance Team |
| EPI_ISL_3174945, EPI_ISL_3174946, EPI_ISL_3174947, EPI_ISL_3174948, EPI_ISL_3174949 | Lighthouse Lab in Glasgow | Wellcome Sanger Institute for the COVID-19 Genomics UK (COG-UK) Consortium | Harper VanSteenhouse, Yumi Kasai, David Gray, Carol Clugston, Anna Dominiczak and Alex Alderton, Roberto Amato, Jeffrey Barrett, Sonia Goncalves, Ewan Harrison, David K. Jackson, Ian Johnston, Dominic Kwiatkowski, Cordelia Langford, John Sillitoe on behalf of the Wellcome Sanger Institute COVID-19 Surveillance Team |
| EPI_ISL_3174950, EPI_ISL_3174951 | Berkshire and Surrey Pathology Services Lighthouse Laboratory | Wellcome Sanger Institute for the COVID-19 Genomics UK (COG-UK) Consortium | Berkshire and Surrey Pathology Services Lighthouse Laboratory and Alex Alderton, Roberto Amato, Jeffrey Barrett, Sonia Goncalves, Ewan Harrison, David K. Jackson, Ian Johnston, Dominic Kwiatkowski, Cordelia Langford, John Sillitoe on behalf of the Wellcome Sanger Institute COVID-19 Surveillance Team |
| EPI_ISL_3174952, EPI_ISL_3174953 | Lighthouse Lab in Milton Keynes | Wellcome Sanger Institute for the COVID-19 Genomics UK (COG-UK) Consortium | The Lighthouse Lab in Milton Keynes and Alex Alderton, Roberto Amato, Jeffrey Barrett, Sonia Goncalves, Ewan Harrison, David K. Jackson, Ian Johnston, Dominic Kwiatkowski, Cordelia Langford, John Sillitoe on behalf of the Wellcome Sanger Institute COVID-19 Surveillance Team |
| EPI_ISL_3174954 | Berkshire and Surrey Pathology Services Lighthouse Laboratory | Wellcome Sanger Institute for the COVID-19 Genomics UK (COG-UK) Consortium | Berkshire and Surrey Pathology Services Lighthouse Laboratory and Alex Alderton, Roberto Amato, Jeffrey Barrett, Sonia Goncalves, Ewan Harrison, David K. Jackson, Ian Johnston, Dominic Kwiatkowski, Cordelia Langford, John Sillitoe on behalf of the Wellcome Sanger Institute COVID-19 Surveillance Team |
| EPI_ISL_3174955, EPI_ISL_3174956, EPI_ISL_3174957, EPI_ISL_3174958 | Lighthouse Lab in Glasgow | Wellcome Sanger Institute for the COVID-19 Genomics UK (COG-UK) Consortium | Harper VanSteenhouse, Yumi Kasai, David Gray, Carol Clugston, Anna Dominiczak and Alex Alderton, Roberto Amato, Jeffrey Barrett, Sonia Goncalves, Ewan Harrison, David K. Jackson, Ian Johnston, Dominic Kwiatkowski, Cordelia Langford, John Sillitoe on behalf of the Wellcome Sanger Institute COVID-19 Surveillance Team |
| EPI_ISL_3174959 | Berkshire and Surrey Pathology Services Lighthouse Laboratory | Wellcome Sanger Institute for the COVID-19 Genomics UK (COG-UK) Consortium | Berkshire and Surrey Pathology Services Lighthouse Laboratory and Alex Alderton, Roberto Amato, Jeffrey Barrett, Sonia Goncalves, Ewan Harrison, David K. Jackson, Ian Johnston, Dominic Kwiatkowski, Cordelia Langford, John Sillitoe on behalf of the Wellcome Sanger Institute COVID-19 Surveillance Team |
| EPI_ISL_3174960 | Lighthouse Lab in Milton Keynes | Wellcome Sanger Institute for the COVID-19 Genomics UK (COG-UK) Consortium | The Lighthouse Lab in Milton Keynes and Alex Alderton, Roberto Amato, Jeffrey Barrett, Sonia Goncalves, Ewan Harrison, David K. Jackson, Ian Johnston, Dominic Kwiatkowski, Cordelia Langford, John Sillitoe on behalf of the Wellcome Sanger Institute COVID-19 Surveillance Team |
| EPI_ISL_3174961, EPI_ISL_3174962, EPI_ISL_3174963 | Lighthouse Lab in Glasgow | Wellcome Sanger Institute for the COVID-19 Genomics UK (COG-UK) Consortium | Harper VanSteenhouse, Yumi Kasai, David Gray, Carol Clugston, Anna Dominiczak and Alex Alderton, Roberto Amato, Jeffrey Barrett, Sonia Goncalves, Ewan Harrison, David K. Jackson, Ian Johnston, Dominic Kwiatkowski, Cordelia Langford, John Sillitoe on behalf of the Wellcome Sanger Institute COVID-19 Surveillance Team |
| EPI_ISL_3174964 | Lighthouse Lab in Milton Keynes | Wellcome Sanger Institute for the COVID-19 Genomics UK (COG-UK) Consortium | The Lighthouse Lab in Milton Keynes and Alex Alderton, Roberto Amato, Jeffrey Barrett, Sonia Goncalves, Ewan Harrison, David K. Jackson, Ian Johnston, Dominic Kwiatkowski, Cordelia Langford, John Sillitoe on behalf of the Wellcome Sanger Institute COVID-19 Surveillance Team |
| EPI_ISL_3174965 | Berkshire and Surrey Pathology Services Lighthouse Laboratory | Wellcome Sanger Institute for the COVID-19 Genomics UK (COG-UK) Consortium | Berkshire and Surrey Pathology Services Lighthouse Laboratory and Alex Alderton, Roberto Amato, Jeffrey Barrett, Sonia Goncalves, Ewan Harrison, David K. Jackson, Ian Johnston, Dominic Kwiatkowski, Cordelia Langford, John Sillitoe on behalf of the Wellcome Sanger Institute COVID-19 Surveillance Team |
| EPI_ISL_3174966, EPI_ISL_3174967 | Lighthouse Lab in Glasgow | Wellcome Sanger Institute for the COVID-19 Genomics UK (COG-UK) Consortium | Harper VanSteenhouse, Yumi Kasai, David Gray, Carol Clugston, Anna Dominiczak and Alex Alderton, Roberto Amato, Jeffrey Barrett, Sonia Goncalves, Ewan Harrison, David K. Jackson, Ian Johnston, Dominic Kwiatkowski, Cordelia Langford, John Sillitoe on behalf of the Wellcome Sanger Institute COVID-19 Surveillance Team |
| EPI_ISL_3174968 | Lighthouse Lab in Milton Keynes | Wellcome Sanger Institute for the COVID-19 Genomics UK (COG-UK) Consortium | The Lighthouse Lab in Milton Keynes and Alex Alderton, Roberto Amato, Jeffrey Barrett, Sonia Goncalves, Ewan Harrison, David K. Jackson, Ian Johnston, Dominic Kwiatkowski, Cordelia Langford, John Sillitoe on behalf of the Wellcome Sanger Institute COVID-19 Surveillance Team |
| EPI_ISL_3174969, EPI_ISL_3174970 | Berkshire and Surrey Pathology Services Lighthouse Laboratory | Wellcome Sanger Institute for the COVID-19 Genomics UK (COG-UK) Consortium | Berkshire and Surrey Pathology Services Lighthouse Laboratory and Alex Alderton, Roberto Amato, Jeffrey Barrett, Sonia Goncalves, Ewan Harrison, David K. Jackson, Ian Johnston, Dominic Kwiatkowski, Cordelia Langford, John Sillitoe on behalf of the Wellcome Sanger Institute COVID-19 Surveillance Team |
| EPI_ISL_3174971 | Lighthouse Lab in Glasgow | Wellcome Sanger Institute for the COVID-19 Genomics UK (COG-UK) Consortium | Harper VanSteenhouse, Yumi Kasai, David Gray, Carol Clugston, Anna Dominiczak and Alex Alderton, Roberto Amato, Jeffrey Barrett, Sonia Goncalves, Ewan Harrison, David K. Jackson, Ian Johnston, Dominic Kwiatkowski, Cordelia Langford, John Sillitoe on behalf of the Wellcome Sanger Institute COVID-19 Surveillance Team |
| EPI_ISL_3174972, EPI_ISL_3174973 | Lighthouse Lab in Milton Keynes | Wellcome Sanger Institute for the COVID-19 Genomics UK (COG-UK) Consortium | The Lighthouse Lab in Milton Keynes and Alex Alderton, Roberto Amato, Jeffrey Barrett, Sonia Goncalves, Ewan Harrison, David K. Jackson, Ian Johnston, Dominic Kwiatkowski, Cordelia Langford, John Sillitoe on behalf of the Wellcome Sanger Institute COVID-19 Surveillance Team |
| EPI_ISL_3174974 | Lighthouse Lab in Glasgow | Wellcome Sanger Institute for the COVID-19 Genomics UK (COG-UK) Consortium | Harper VanSteenhouse, Yumi Kasai, David Gray, Carol Clugston, Anna Dominiczak and Alex Alderton, Roberto Amato, Jeffrey Barrett, Sonia Goncalves, Ewan Harrison, David K. Jackson, Ian Johnston, Dominic Kwiatkowski, Cordelia Langford, John Sillitoe on behalf of the Wellcome Sanger Institute COVID-19 Surveillance Team |
| EPI_ISL_3174975, EPI_ISL_3174976 | Berkshire and Surrey Pathology Services Lighthouse Laboratory | Wellcome Sanger Institute for the COVID-19 Genomics UK (COG-UK) Consortium | Berkshire and Surrey Pathology Services Lighthouse Laboratory and Alex Alderton, Roberto Amato, Jeffrey Barrett, Sonia Goncalves, Ewan Harrison, David K. Jackson, Ian Johnston, Dominic Kwiatkowski, Cordelia Langford, John Sillitoe on behalf of the Wellcome Sanger Institute COVID-19 Surveillance Team |
| EPI_ISL_3174977 | Lighthouse Lab in Milton Keynes | Wellcome Sanger Institute for the COVID-19 Genomics UK (COG-UK) Consortium | The Lighthouse Lab in Milton Keynes and Alex Alderton, Roberto Amato, Jeffrey Barrett, Sonia Goncalves, Ewan Harrison, David K. Jackson, Ian Johnston, Dominic Kwiatkowski, Cordelia Langford, John Sillitoe on behalf of the Wellcome Sanger Institute COVID-19 Surveillance Team |
| EPI_ISL_3174978 | Lighthouse Lab in Glasgow | Wellcome Sanger Institute for the COVID-19 Genomics UK (COG-UK) Consortium | Harper VanSteenhouse, Yumi Kasai, David Gray, Carol Clugston, Anna Dominiczak and Alex Alderton, Roberto Amato, Jeffrey Barrett, Sonia Goncalves, Ewan Harrison, David K. Jackson, Ian Johnston, Dominic Kwiatkowski, Cordelia Langford, John Sillitoe on behalf of the Wellcome Sanger Institute COVID-19 Surveillance Team |
| EPI_ISL_3174979 | Berkshire and Surrey Pathology Services Lighthouse Laboratory | Wellcome Sanger Institute for the COVID-19 Genomics UK (COG-UK) Consortium | Berkshire and Surrey Pathology Services Lighthouse Laboratory and Alex Alderton, Roberto Amato, Jeffrey Barrett, Sonia Goncalves, Ewan Harrison, David K. Jackson, Ian Johnston, Dominic Kwiatkowski, Cordelia Langford, John Sillitoe on behalf of the Wellcome Sanger Institute COVID-19 Surveillance Team |
| EPI_ISL_3174980, EPI_ISL_3174981 | Lighthouse Lab in Glasgow | Wellcome Sanger Institute for the COVID-19 Genomics UK (COG-UK) Consortium | Harper VanSteenhouse, Yumi Kasai, David Gray, Carol Clugston, Anna Dominiczak and Alex Alderton, Roberto Amato, Jeffrey Barrett, Sonia Goncalves, Ewan Harrison, David K. Jackson, Ian Johnston, Dominic Kwiatkowski, Cordelia Langford, John Sillitoe on behalf of the Wellcome Sanger Institute COVID-19 Surveillance Team |
| EPI_ISL_3174982 | Berkshire and Surrey Pathology Services Lighthouse Laboratory | Wellcome Sanger Institute for the COVID-19 Genomics UK (COG-UK) Consortium | Berkshire and Surrey Pathology Services Lighthouse Laboratory and Alex Alderton, Roberto Amato, Jeffrey Barrett, Sonia Goncalves, Ewan Harrison, David K. Jackson, Ian Johnston, Dominic Kwiatkowski, Cordelia Langford, John Sillitoe on behalf of the Wellcome Sanger Institute COVID-19 Surveillance Team |
| EPI_ISL_3174983 | Lighthouse Lab in Milton Keynes | Wellcome Sanger Institute for the COVID-19 Genomics UK (COG-UK) | The Lighthouse Lab in Milton Keynes and Alex Alderton, Roberto Amato, Jeffrey Barrett, Sonia Goncalves, Ewan Harrison, David K. Jackson, Ian Johnston, Dominic Kwiatkowski, Cordelia Langford, John Sillitoe on behalf of the Wellcome Sanger |

| | | | |
|---|---|---|---|
| | | Consortium | Institute COVID-19 Surveillance Team |
| EPI_ISL_3174984, EPI_ISL_3174985 | Lighthouse Lab in Glasgow | Wellcome Sanger Institute for the COVID-19 Genomics UK (COG-UK) Consortium | Harper VanSteenhouse, Yumi Kasai, David Gray, Carol Clugston, Anna Dominiczak and Alex Alderton, Roberto Amato, Jeffrey Barrett, Sonia Goncalves, Ewan Harrison, David K. Jackson, Ian Johnston, Dominic Kwiatkowski, Cordelia Langford, John Sillitoe on behalf of the Wellcome Sanger Institute COVID-19 Surveillance Team |
| EPI_ISL_3174986 | Lighthouse Lab in Milton Keynes | Wellcome Sanger Institute for the COVID-19 Genomics UK (COG-UK) Consortium | The Lighthouse Lab in Milton Keynes and Alex Alderton, Roberto Amato, Jeffrey Barrett, Sonia Goncalves, Ewan Harrison, David K. Jackson, Ian Johnston, Dominic Kwiatkowski, Cordelia Langford, John Sillitoe on behalf of the Wellcome Sanger Institute COVID-19 Surveillance Team |
| EPI_ISL_3174987 | Lighthouse Lab in Glasgow | Wellcome Sanger Institute for the COVID-19 Genomics UK (COG-UK) Consortium | Harper VanSteenhouse, Yumi Kasai, David Gray, Carol Clugston, Anna Dominiczak and Alex Alderton, Roberto Amato, Jeffrey Barrett, Sonia Goncalves, Ewan Harrison, David K. Jackson, Ian Johnston, Dominic Kwiatkowski, Cordelia Langford, John Sillitoe on behalf of the Wellcome Sanger Institute COVID-19 Surveillance Team |
| EPI_ISL_3174988 | Berkshire and Surrey Pathology Services Lighthouse Laboratory | Wellcome Sanger Institute for the COVID-19 Genomics UK (COG-UK) Consortium | Berkshire and Surrey Pathology Services Lighthouse Laboratory and Alex Alderton, Roberto Amato, Jeffrey Barrett, Sonia Goncalves, Ewan Harrison, David K. Jackson, Ian Johnston, Dominic Kwiatkowski, Cordelia Langford, John Sillitoe on behalf of the Wellcome Sanger Institute COVID-19 Surveillance Team |
| EPI_ISL_3174989 | Lighthouse Lab in Milton Keynes | Wellcome Sanger Institute for the COVID-19 Genomics UK (COG-UK) Consortium | The Lighthouse Lab in Milton Keynes and Alex Alderton, Roberto Amato, Jeffrey Barrett, Sonia Goncalves, Ewan Harrison, David K. Jackson, Ian Johnston, Dominic Kwiatkowski, Cordelia Langford, John Sillitoe on behalf of the Wellcome Sanger Institute COVID-19 Surveillance Team |
| EPI_ISL_3174990 | Lighthouse Lab in Glasgow | Wellcome Sanger Institute for the COVID-19 Genomics UK (COG-UK) Consortium | Harper VanSteenhouse, Yumi Kasai, David Gray, Carol Clugston, Anna Dominiczak and Alex Alderton, Roberto Amato, Jeffrey Barrett, Sonia Goncalves, Ewan Harrison, David K. Jackson, Ian Johnston, Dominic Kwiatkowski, Cordelia Langford, John Sillitoe on behalf of the Wellcome Sanger Institute COVID-19 Surveillance Team |
| EPI_ISL_3174991 | Berkshire and Surrey Pathology Services Lighthouse Laboratory | Wellcome Sanger Institute for the COVID-19 Genomics UK (COG-UK) Consortium | Berkshire and Surrey Pathology Services Lighthouse Laboratory and Alex Alderton, Roberto Amato, Jeffrey Barrett, Sonia Goncalves, Ewan Harrison, David K. Jackson, Ian Johnston, Dominic Kwiatkowski, Cordelia Langford, John Sillitoe on behalf of the Wellcome Sanger Institute COVID-19 Surveillance Team |
| EPI_ISL_3174992, EPI_ISL_3174993 | Lighthouse Lab in Glasgow | Wellcome Sanger Institute for the COVID-19 Genomics UK (COG-UK) Consortium | Harper VanSteenhouse, Yumi Kasai, David Gray, Carol Clugston, Anna Dominiczak and Alex Alderton, Roberto Amato, Jeffrey Barrett, Sonia Goncalves, Ewan Harrison, David K. Jackson, Ian Johnston, Dominic Kwiatkowski, Cordelia Langford, John Sillitoe on behalf of the Wellcome Sanger Institute COVID-19 Surveillance Team |
| EPI_ISL_3174994 | Berkshire and Surrey Pathology Services Lighthouse Laboratory | Wellcome Sanger Institute for the COVID-19 Genomics UK (COG-UK) Consortium | Berkshire and Surrey Pathology Services Lighthouse Laboratory and Alex Alderton, Roberto Amato, Jeffrey Barrett, Sonia Goncalves, Ewan Harrison, David K. Jackson, Ian Johnston, Dominic Kwiatkowski, Cordelia Langford, John Sillitoe on behalf of the Wellcome Sanger Institute COVID-19 Surveillance Team |
| EPI_ISL_3174995 | Lighthouse Lab in Glasgow | Wellcome Sanger Institute for the COVID-19 Genomics UK (COG-UK) Consortium | Harper VanSteenhouse, Yumi Kasai, David Gray, Carol Clugston, Anna Dominiczak and Alex Alderton, Roberto Amato, Jeffrey Barrett, Sonia Goncalves, Ewan Harrison, David K. Jackson, Ian Johnston, Dominic Kwiatkowski, Cordelia Langford, John Sillitoe on behalf of the Wellcome Sanger Institute COVID-19 Surveillance Team |
| EPI_ISL_3174996, EPI_ISL_3174997, EPI_ISL_3174998 | Lighthouse Lab in Milton Keynes | Wellcome Sanger Institute for the COVID-19 Genomics UK (COG-UK) Consortium | The Lighthouse Lab in Milton Keynes and Alex Alderton, Roberto Amato, Jeffrey Barrett, Sonia Goncalves, Ewan Harrison, David K. Jackson, Ian Johnston, Dominic Kwiatkowski, Cordelia Langford, John Sillitoe on behalf of the Wellcome Sanger Institute COVID-19 Surveillance Team |
| EPI_ISL_3174999 | Lighthouse Lab in Glasgow | Wellcome Sanger Institute for the COVID-19 Genomics UK (COG-UK) Consortium | Harper VanSteenhouse, Yumi Kasai, David Gray, Carol Clugston, Anna Dominiczak and Alex Alderton, Roberto Amato, Jeffrey Barrett, Sonia Goncalves, Ewan Harrison, David K. Jackson, Ian Johnston, Dominic Kwiatkowski, Cordelia Langford, John Sillitoe on behalf of the Wellcome Sanger Institute COVID-19 Surveillance Team |
| EPI_ISL_3175000 | Lighthouse Lab in Milton Keynes | Wellcome Sanger Institute for the COVID-19 Genomics UK (COG-UK) Consortium | The Lighthouse Lab in Milton Keynes and Alex Alderton, Roberto Amato, Jeffrey Barrett, Sonia Goncalves, Ewan Harrison, David K. Jackson, Ian Johnston, Dominic Kwiatkowski, Cordelia Langford, John Sillitoe on behalf of the Wellcome Sanger Institute COVID-19 Surveillance Team |
| EPI_ISL_3175001, EPI_ISL_3175002, EPI_ISL_3175003, EPI_ISL_3175004, EPI_ISL_3175005 | Lighthouse Lab in Glasgow | Wellcome Sanger Institute for the COVID-19 Genomics UK (COG-UK) Consortium | Harper VanSteenhouse, Yumi Kasai, David Gray, Carol Clugston, Anna Dominiczak and Alex Alderton, Roberto Amato, Jeffrey Barrett, Sonia Goncalves, Ewan Harrison, David K. Jackson, Ian Johnston, Dominic Kwiatkowski, Cordelia Langford, John Sillitoe on behalf of the Wellcome Sanger Institute COVID-19 Surveillance Team |
| EPI_ISL_3175006 | Berkshire and Surrey Pathology Services Lighthouse Laboratory | Wellcome Sanger Institute for the COVID-19 Genomics UK (COG-UK) Consortium | Berkshire and Surrey Pathology Services Lighthouse Laboratory and Alex Alderton, Roberto Amato, Jeffrey Barrett, Sonia Goncalves, Ewan Harrison, David K. Jackson, Ian Johnston, Dominic Kwiatkowski, Cordelia Langford, John Sillitoe on behalf of the Wellcome Sanger Institute COVID-19 Surveillance Team |
| EPI_ISL_3175007 | Lighthouse Lab in Glasgow | Wellcome Sanger Institute for the COVID-19 Genomics UK (COG-UK) Consortium | Harper VanSteenhouse, Yumi Kasai, David Gray, Carol Clugston, Anna Dominiczak and Alex Alderton, Roberto Amato, Jeffrey Barrett, Sonia Goncalves, Ewan Harrison, David K. Jackson, Ian Johnston, Dominic Kwiatkowski, Cordelia Langford, John Sillitoe on behalf of the Wellcome Sanger Institute COVID-19 Surveillance Team |
| EPI_ISL_3175008 | Berkshire and Surrey Pathology Services Lighthouse Laboratory | Wellcome Sanger Institute for the COVID-19 Genomics UK (COG-UK) Consortium | Berkshire and Surrey Pathology Services Lighthouse Laboratory and Alex Alderton, Roberto Amato, Jeffrey Barrett, Sonia Goncalves, Ewan Harrison, David K. Jackson, Ian Johnston, Dominic Kwiatkowski, Cordelia Langford, John Sillitoe on behalf of the Wellcome Sanger Institute COVID-19 Surveillance Team |
| EPI_ISL_3175009 | Lighthouse Lab in Glasgow | Wellcome Sanger Institute for the COVID-19 Genomics UK (COG-UK) Consortium | Harper VanSteenhouse, Yumi Kasai, David Gray, Carol Clugston, Anna Dominiczak and Alex Alderton, Roberto Amato, Jeffrey Barrett, Sonia Goncalves, Ewan Harrison, David K. Jackson, Ian Johnston, Dominic Kwiatkowski, Cordelia Langford, John Sillitoe on behalf of the Wellcome Sanger Institute COVID-19 Surveillance Team |
| EPI_ISL_3175010 | Lighthouse Lab in Milton Keynes | Wellcome Sanger Institute for the COVID-19 Genomics UK (COG-UK) Consortium | The Lighthouse Lab in Milton Keynes and Alex Alderton, Roberto Amato, Jeffrey Barrett, Sonia Goncalves, Ewan Harrison, David K. Jackson, Ian Johnston, Dominic Kwiatkowski, Cordelia Langford, John Sillitoe on behalf of the Wellcome Sanger Institute COVID-19 Surveillance Team |
| EPI_ISL_3175011 | Lighthouse Lab in Glasgow | Wellcome Sanger Institute for the COVID-19 Genomics UK (COG-UK) Consortium | Harper VanSteenhouse, Yumi Kasai, David Gray, Carol Clugston, Anna Dominiczak and Alex Alderton, Roberto Amato, Jeffrey Barrett, Sonia Goncalves, Ewan Harrison, David K. Jackson, Ian Johnston, Dominic Kwiatkowski, Cordelia Langford, John Sillitoe on behalf of the Wellcome Sanger Institute COVID-19 Surveillance Team |
| EPI_ISL_3175012 | Berkshire and Surrey Pathology Services Lighthouse Laboratory | Wellcome Sanger Institute for the COVID-19 Genomics UK (COG-UK) Consortium | Berkshire and Surrey Pathology Services Lighthouse Laboratory and Alex Alderton, Roberto Amato, Jeffrey Barrett, Sonia Goncalves, Ewan Harrison, David K. Jackson, Ian Johnston, Dominic Kwiatkowski, Cordelia Langford, John Sillitoe on behalf of the Wellcome Sanger Institute COVID-19 Surveillance Team |
| EPI_ISL_3175013, EPI_ISL_3175014 | Lighthouse Lab in Glasgow | Wellcome Sanger Institute for the COVID-19 Genomics UK (COG-UK) Consortium | Harper VanSteenhouse, Yumi Kasai, David Gray, Carol Clugston, Anna Dominiczak and Alex Alderton, Roberto Amato, Jeffrey Barrett, Sonia Goncalves, Ewan Harrison, David K. Jackson, Ian Johnston, Dominic Kwiatkowski, Cordelia Langford, John Sillitoe on behalf of the Wellcome Sanger Institute COVID-19 Surveillance Team |
| EPI_ISL_3175015, EPI_ISL_3175016, EPI_ISL_3175017 | Lighthouse Lab in Milton Keynes | Wellcome Sanger Institute for the COVID-19 Genomics UK (COG-UK) Consortium | The Lighthouse Lab in Milton Keynes and Alex Alderton, Roberto Amato, Jeffrey Barrett, Sonia Goncalves, Ewan Harrison, David K. Jackson, Ian Johnston, Dominic Kwiatkowski, Cordelia Langford, John Sillitoe on behalf of the Wellcome Sanger Institute COVID-19 Surveillance Team |
| EPI_ISL_3175018, EPI_ISL_3175019, EPI_ISL_3175020 | Berkshire and Surrey Pathology Services Lighthouse Laboratory | Wellcome Sanger Institute for the COVID-19 Genomics UK (COG-UK) Consortium | Berkshire and Surrey Pathology Services Lighthouse Laboratory and Alex Alderton, Roberto Amato, Jeffrey Barrett, Sonia Goncalves, Ewan Harrison, David K. Jackson, Ian Johnston, Dominic Kwiatkowski, Cordelia Langford, John Sillitoe on behalf of the Wellcome Sanger Institute COVID-19 Surveillance Team |
| EPI_ISL_3175021 | Lighthouse Lab in Milton Keynes | Wellcome Sanger Institute for the COVID-19 Genomics UK (COG-UK) Consortium | The Lighthouse Lab in Milton Keynes and Alex Alderton, Roberto Amato, Jeffrey Barrett, Sonia Goncalves, Ewan Harrison, David K. Jackson, Ian Johnston, Dominic Kwiatkowski, Cordelia Langford, John Sillitoe on behalf of the Wellcome Sanger Institute COVID-19 Surveillance Team |
| EPI_ISL_3175022, EPI_ISL_3175023 | Lighthouse Lab in Glasgow | Wellcome Sanger Institute for the COVID-19 Genomics UK (COG-UK) Consortium | Harper VanSteenhouse, Yumi Kasai, David Gray, Carol Clugston, Anna Dominiczak and Alex Alderton, Roberto Amato, Jeffrey Barrett, Sonia Goncalves, Ewan Harrison, David K. Jackson, Ian Johnston, Dominic Kwiatkowski, Cordelia Langford, John Sillitoe on behalf of the Wellcome Sanger Institute COVID-19 Surveillance Team |
| EPI_ISL_3175024 | Lighthouse Lab in Milton Keynes | Wellcome Sanger Institute for the COVID-19 Genomics UK (COG-UK) Consortium | The Lighthouse Lab in Milton Keynes and Alex Alderton, Roberto Amato, Jeffrey Barrett, Sonia Goncalves, Ewan Harrison, David K. Jackson, Ian Johnston, Dominic Kwiatkowski, Cordelia Langford, John Sillitoe on behalf of the Wellcome Sanger Institute COVID-19 Surveillance Team |
| EPI_ISL_3175025 | Lighthouse Lab in Glasgow | Wellcome Sanger Institute for the COVID-19 Genomics UK (COG-UK) Consortium | Harper VanSteenhouse, Yumi Kasai, David Gray, Carol Clugston, Anna Dominiczak and Alex Alderton, Roberto Amato, Jeffrey Barrett, Sonia Goncalves, Ewan Harrison, David K. Jackson, Ian Johnston, Dominic Kwiatkowski, Cordelia Langford, John Sillitoe on behalf of the Wellcome Sanger Institute COVID-19 Surveillance Team |
| EPI_ISL_3175026 | Lighthouse Lab in Milton Keynes | Wellcome Sanger Institute for the COVID-19 Genomics UK (COG-UK) Consortium | The Lighthouse Lab in Milton Keynes and Alex Alderton, Roberto Amato, Jeffrey Barrett, Sonia Goncalves, Ewan Harrison, David K. Jackson, Ian Johnston, Dominic Kwiatkowski, Cordelia Langford, John Sillitoe on behalf of the Wellcome Sanger Institute COVID-19 Surveillance Team |
| EPI_ISL_3175027, EPI_ISL_3175028 | Berkshire and Surrey Pathology Services Lighthouse Laboratory | Wellcome Sanger Institute for the COVID-19 Genomics UK (COG-UK) Consortium | Berkshire and Surrey Pathology Services Lighthouse Laboratory and Alex Alderton, Roberto Amato, Jeffrey Barrett, Sonia Goncalves, Ewan Harrison, David K. Jackson, Ian Johnston, Dominic Kwiatkowski, Cordelia Langford, John Sillitoe on behalf of the Wellcome Sanger Institute COVID-19 Surveillance Team |
| EPI_ISL_3175029 | Lighthouse Lab in Milton Keynes | Wellcome Sanger Institute for the COVID-19 Genomics UK (COG-UK) Consortium | The Lighthouse Lab in Milton Keynes and Alex Alderton, Roberto Amato, Jeffrey Barrett, Sonia Goncalves, Ewan Harrison, David K. Jackson, Ian Johnston, Dominic Kwiatkowski, Cordelia Langford, John Sillitoe on behalf of the Wellcome Sanger Institute COVID-19 Surveillance Team |
| EPI_ISL_3175030 | Berkshire and Surrey Pathology Services Lighthouse Laboratory | Wellcome Sanger Institute for the COVID-19 Genomics UK (COG-UK) Consortium | Berkshire and Surrey Pathology Services Lighthouse Laboratory and Alex Alderton, Roberto Amato, Jeffrey Barrett, Sonia Goncalves, Ewan Harrison, David K. Jackson, Ian Johnston, Dominic Kwiatkowski, Cordelia Langford, John Sillitoe on behalf of the Wellcome Sanger Institute COVID-19 Surveillance Team |
| EPI_ISL_3175031, EPI_ISL_3175032 | Lighthouse Lab in Glasgow | Wellcome Sanger Institute for the COVID-19 Genomics UK (COG-UK) Consortium | Harper VanSteenhouse, Yumi Kasai, David Gray, Carol Clugston, Anna Dominiczak and Alex Alderton, Roberto Amato, Jeffrey Barrett, Sonia Goncalves, Ewan Harrison, David K. Jackson, Ian Johnston, Dominic Kwiatkowski, Cordelia Langford, John Sillitoe on behalf of the Wellcome Sanger Institute COVID-19 Surveillance Team |
| EPI_ISL_3175033 | Lighthouse Lab in Milton Keynes | Wellcome Sanger Institute for the COVID-19 Genomics UK (COG-UK) Consortium | The Lighthouse Lab in Milton Keynes and Alex Alderton, Roberto Amato, Jeffrey Barrett, Sonia Goncalves, Ewan Harrison, David K. Jackson, Ian Johnston, Dominic Kwiatkowski, Cordelia Langford, John Sillitoe on behalf of the Wellcome Sanger Institute COVID-19 Surveillance Team |
| EPI_ISL_3175034, EPI_ISL_3175035, EPI_ISL_3175036 | Lighthouse Lab in Glasgow | Wellcome Sanger Institute for the COVID-19 Genomics UK (COG-UK) Consortium | Harper VanSteenhouse, Yumi Kasai, David Gray, Carol Clugston, Anna Dominiczak and Alex Alderton, Roberto Amato, Jeffrey Barrett, Sonia Goncalves, Ewan Harrison, David K. Jackson, Ian Johnston, Dominic Kwiatkowski, Cordelia Langford, John Sillitoe on behalf of the Wellcome Sanger Institute COVID-19 Surveillance Team |
| EPI_ISL_3175037 | Berkshire and Surrey Pathology Services Lighthouse Laboratory | Wellcome Sanger Institute for the COVID-19 Genomics UK (COG-UK) Consortium | Berkshire and Surrey Pathology Services Lighthouse Laboratory and Alex Alderton, Roberto Amato, Jeffrey Barrett, Sonia Goncalves, Ewan Harrison, David K. Jackson, Ian Johnston, Dominic Kwiatkowski, Cordelia Langford, John Sillitoe on behalf of the Wellcome Sanger Institute COVID-19 Surveillance Team |
| EPI_ISL_3175038, EPI_ISL_3175039 | Lighthouse Lab in Milton Keynes | Wellcome Sanger Institute for the COVID-19 Genomics UK (COG-UK) Consortium | The Lighthouse Lab in Milton Keynes and Alex Alderton, Roberto Amato, Jeffrey Barrett, Sonia Goncalves, Ewan Harrison, David K. Jackson, Ian Johnston, Dominic Kwiatkowski, Cordelia Langford, John Sillitoe on behalf of the Wellcome Sanger Institute COVID-19 Surveillance Team |
| EPI_ISL_3175040 | Berkshire and Surrey Pathology Services Lighthouse Laboratory | Wellcome Sanger Institute for the COVID-19 Genomics UK (COG-UK) Consortium | Berkshire and Surrey Pathology Services Lighthouse Laboratory and Alex Alderton, Roberto Amato, Jeffrey Barrett, Sonia Goncalves, Ewan Harrison, David K. Jackson, Ian Johnston, Dominic Kwiatkowski, Cordelia Langford, John Sillitoe on behalf of the Wellcome Sanger Institute COVID-19 Surveillance Team |
| EPI_ISL_3175041 | Lighthouse Lab in Milton Keynes | Wellcome Sanger Institute for the COVID-19 Genomics UK (COG-UK) Consortium | The Lighthouse Lab in Milton Keynes and Alex Alderton, Roberto Amato, Jeffrey Barrett, Sonia Goncalves, Ewan Harrison, David K. Jackson, Ian Johnston, Dominic Kwiatkowski, Cordelia Langford, John Sillitoe on behalf of the Wellcome Sanger Institute COVID-19 Surveillance Team |
| EPI_ISL_3175042 | Lighthouse Lab in Glasgow | Wellcome Sanger Institute for the COVID-19 Genomics UK (COG-UK) Consortium | Harper VanSteenhouse, Yumi Kasai, David Gray, Carol Clugston, Anna Dominiczak and Alex Alderton, Roberto Amato, Jeffrey Barrett, Sonia Goncalves, Ewan Harrison, David K. Jackson, Ian Johnston, Dominic Kwiatkowski, Cordelia Langford, John Sillitoe on behalf of the Wellcome Sanger Institute COVID-19 Surveillance Team |
| EPI_ISL_3175043 | Berkshire and Surrey Pathology Services Lighthouse Laboratory | Wellcome Sanger Institute for the COVID-19 Genomics UK (COG-UK) Consortium | Berkshire and Surrey Pathology Services Lighthouse Laboratory and Alex Alderton, Roberto Amato, Jeffrey Barrett, Sonia Goncalves, Ewan Harrison, David K. Jackson, Ian Johnston, Dominic Kwiatkowski, Cordelia Langford, John Sillitoe on behalf of the Wellcome Sanger Institute COVID-19 Surveillance Team |
| EPI_ISL_3175044, | Lighthouse Lab in Glasgow | Wellcome Sanger Institute for the COVID-19 Genomics UK (COG-UK) | Harper VanSteenhouse, Yumi Kasai, David Gray, Carol Clugston, Anna Dominiczak and Alex Alderton, Roberto Amato, Jeffrey Barrett, Sonia Goncalves, Ewan Harrison, David K. Jackson, Ian Johnston, Dominic Kwiatkowski, Cordelia Langford, |

| | | | |
|---|---|---|---|
| EPI_ISL_3175045, EPI_ISL_3175046, EPI_ISL_3175047, EPI_ISL_3175048 | | Consortium | John Sillitoe on behalf of the Wellcome Sanger Institute COVID-19 Surveillance Team |
| EPI_ISL_3175049 | Lighthouse Lab in Milton Keynes | Wellcome Sanger Institute for the COVID-19 Genomics UK (COG-UK) Consortium | The Lighthouse Lab in Milton Keynes and Alex Alderton, Roberto Amato, Jeffrey Barrett, Sonia Goncalves, Ewan Harrison, David K. Jackson, Ian Johnston, Dominic Kwiatkowski, Cordelia Langford, John Sillitoe on behalf of the Wellcome Sanger Institute COVID-19 Surveillance Team |
| EPI_ISL_3175050 | Lighthouse Lab in Glasgow | Wellcome Sanger Institute for the COVID-19 Genomics UK (COG-UK) Consortium | Harper VanSteenhouse, Yumi Kasai, David Gray, Carol Clugston, Anna Dominiczak and Alex Alderton, Roberto Amato, Jeffrey Barrett, Sonia Goncalves, Ewan Harrison, David K. Jackson, Ian Johnston, Dominic Kwiatkowski, Cordelia Langford, John Sillitoe on behalf of the Wellcome Sanger Institute COVID-19 Surveillance Team |
| EPI_ISL_3175051 | Berkshire and Surrey Pathology Services Lighthouse Laboratory | Wellcome Sanger Institute for the COVID-19 Genomics UK (COG-UK) Consortium | Berkshire and Surrey Pathology Services Lighthouse Laboratory and Alex Alderton, Roberto Amato, Jeffrey Barrett, Sonia Goncalves, Ewan Harrison, David K. Jackson, Ian Johnston, Dominic Kwiatkowski, Cordelia Langford, John Sillitoe on behalf of the Wellcome Sanger Institute COVID-19 Surveillance Team |
| EPI_ISL_3175052 | Lighthouse Lab in Milton Keynes | Wellcome Sanger Institute for the COVID-19 Genomics UK (COG-UK) Consortium | The Lighthouse Lab in Milton Keynes and Alex Alderton, Roberto Amato, Jeffrey Barrett, Sonia Goncalves, Ewan Harrison, David K. Jackson, Ian Johnston, Dominic Kwiatkowski, Cordelia Langford, John Sillitoe on behalf of the Wellcome Sanger Institute COVID-19 Surveillance Team |
| EPI_ISL_3175053, EPI_ISL_3175054, EPI_ISL_3175055 | Berkshire and Surrey Pathology Services Lighthouse Laboratory | Wellcome Sanger Institute for the COVID-19 Genomics UK (COG-UK) Consortium | Berkshire and Surrey Pathology Services Lighthouse Laboratory and Alex Alderton, Roberto Amato, Jeffrey Barrett, Sonia Goncalves, Ewan Harrison, David K. Jackson, Ian Johnston, Dominic Kwiatkowski, Cordelia Langford, John Sillitoe on behalf of the Wellcome Sanger Institute COVID-19 Surveillance Team |
| EPI_ISL_3175056 | Lighthouse Lab in Glasgow | Wellcome Sanger Institute for the COVID-19 Genomics UK (COG-UK) Consortium | Harper VanSteenhouse, Yumi Kasai, David Gray, Carol Clugston, Anna Dominiczak and Alex Alderton, Roberto Amato, Jeffrey Barrett, Sonia Goncalves, Ewan Harrison, David K. Jackson, Ian Johnston, Dominic Kwiatkowski, Cordelia Langford, John Sillitoe on behalf of the Wellcome Sanger Institute COVID-19 Surveillance Team |
| EPI_ISL_3175057 | Berkshire and Surrey Pathology Services Lighthouse Laboratory | Wellcome Sanger Institute for the COVID-19 Genomics UK (COG-UK) Consortium | Berkshire and Surrey Pathology Services Lighthouse Laboratory and Alex Alderton, Roberto Amato, Jeffrey Barrett, Sonia Goncalves, Ewan Harrison, David K. Jackson, Ian Johnston, Dominic Kwiatkowski, Cordelia Langford, John Sillitoe on behalf of the Wellcome Sanger Institute COVID-19 Surveillance Team |
| EPI_ISL_3175058 | Lighthouse Lab in Glasgow | Wellcome Sanger Institute for the COVID-19 Genomics UK (COG-UK) Consortium | Harper VanSteenhouse, Yumi Kasai, David Gray, Carol Clugston, Anna Dominiczak and Alex Alderton, Roberto Amato, Jeffrey Barrett, Sonia Goncalves, Ewan Harrison, David K. Jackson, Ian Johnston, Dominic Kwiatkowski, Cordelia Langford, John Sillitoe on behalf of the Wellcome Sanger Institute COVID-19 Surveillance Team |
| EPI_ISL_3175059 | Lighthouse Lab in Milton Keynes | Wellcome Sanger Institute for the COVID-19 Genomics UK (COG-UK) Consortium | The Lighthouse Lab in Milton Keynes and Alex Alderton, Roberto Amato, Jeffrey Barrett, Sonia Goncalves, Ewan Harrison, David K. Jackson, Ian Johnston, Dominic Kwiatkowski, Cordelia Langford, John Sillitoe on behalf of the Wellcome Sanger Institute COVID-19 Surveillance Team |
| EPI_ISL_3175060 | Lighthouse Lab in Glasgow | Wellcome Sanger Institute for the COVID-19 Genomics UK (COG-UK) Consortium | Harper VanSteenhouse, Yumi Kasai, David Gray, Carol Clugston, Anna Dominiczak and Alex Alderton, Roberto Amato, Jeffrey Barrett, Sonia Goncalves, Ewan Harrison, David K. Jackson, Ian Johnston, Dominic Kwiatkowski, Cordelia Langford, John Sillitoe on behalf of the Wellcome Sanger Institute COVID-19 Surveillance Team |
| EPI_ISL_3175061, EPI_ISL_3175062 | Lighthouse Lab in Milton Keynes | Wellcome Sanger Institute for the COVID-19 Genomics UK (COG-UK) Consortium | The Lighthouse Lab in Milton Keynes and Alex Alderton, Roberto Amato, Jeffrey Barrett, Sonia Goncalves, Ewan Harrison, David K. Jackson, Ian Johnston, Dominic Kwiatkowski, Cordelia Langford, John Sillitoe on behalf of the Wellcome Sanger Institute COVID-19 Surveillance Team |
| EPI_ISL_3175063 | Lighthouse Lab in Glasgow | Wellcome Sanger Institute for the COVID-19 Genomics UK (COG-UK) Consortium | Harper VanSteenhouse, Yumi Kasai, David Gray, Carol Clugston, Anna Dominiczak and Alex Alderton, Roberto Amato, Jeffrey Barrett, Sonia Goncalves, Ewan Harrison, David K. Jackson, Ian Johnston, Dominic Kwiatkowski, Cordelia Langford, John Sillitoe on behalf of the Wellcome Sanger Institute COVID-19 Surveillance Team |
| EPI_ISL_3175064 | Lighthouse Lab in Milton Keynes | Wellcome Sanger Institute for the COVID-19 Genomics UK (COG-UK) Consortium | The Lighthouse Lab in Milton Keynes and Alex Alderton, Roberto Amato, Jeffrey Barrett, Sonia Goncalves, Ewan Harrison, David K. Jackson, Ian Johnston, Dominic Kwiatkowski, Cordelia Langford, John Sillitoe on behalf of the Wellcome Sanger Institute COVID-19 Surveillance Team |
| EPI_ISL_3175065 | Berkshire and Surrey Pathology Services Lighthouse Laboratory | Wellcome Sanger Institute for the COVID-19 Genomics UK (COG-UK) Consortium | Berkshire and Surrey Pathology Services Lighthouse Laboratory and Alex Alderton, Roberto Amato, Jeffrey Barrett, Sonia Goncalves, Ewan Harrison, David K. Jackson, Ian Johnston, Dominic Kwiatkowski, Cordelia Langford, John Sillitoe on behalf of the Wellcome Sanger Institute COVID-19 Surveillance Team |
| EPI_ISL_3175066 | Lighthouse Lab in Milton Keynes | Wellcome Sanger Institute for the COVID-19 Genomics UK (COG-UK) Consortium | The Lighthouse Lab in Milton Keynes and Alex Alderton, Roberto Amato, Jeffrey Barrett, Sonia Goncalves, Ewan Harrison, David K. Jackson, Ian Johnston, Dominic Kwiatkowski, Cordelia Langford, John Sillitoe on behalf of the Wellcome Sanger Institute COVID-19 Surveillance Team |
| EPI_ISL_3175067 | Lighthouse Lab in Glasgow | Wellcome Sanger Institute for the COVID-19 Genomics UK (COG-UK) Consortium | Harper VanSteenhouse, Yumi Kasai, David Gray, Carol Clugston, Anna Dominiczak and Alex Alderton, Roberto Amato, Jeffrey Barrett, Sonia Goncalves, Ewan Harrison, David K. Jackson, Ian Johnston, Dominic Kwiatkowski, Cordelia Langford, John Sillitoe on behalf of the Wellcome Sanger Institute COVID-19 Surveillance Team |
| EPI_ISL_3175068 | Lighthouse Lab in Milton Keynes | Wellcome Sanger Institute for the COVID-19 Genomics UK (COG-UK) Consortium | The Lighthouse Lab in Milton Keynes and Alex Alderton, Roberto Amato, Jeffrey Barrett, Sonia Goncalves, Ewan Harrison, David K. Jackson, Ian Johnston, Dominic Kwiatkowski, Cordelia Langford, John Sillitoe on behalf of the Wellcome Sanger Institute COVID-19 Surveillance Team |
| EPI_ISL_3175069 | Berkshire and Surrey Pathology Services Lighthouse Laboratory | Wellcome Sanger Institute for the COVID-19 Genomics UK (COG-UK) Consortium | Berkshire and Surrey Pathology Services Lighthouse Laboratory and Alex Alderton, Roberto Amato, Jeffrey Barrett, Sonia Goncalves, Ewan Harrison, David K. Jackson, Ian Johnston, Dominic Kwiatkowski, Cordelia Langford, John Sillitoe on behalf of the Wellcome Sanger Institute COVID-19 Surveillance Team |
| EPI_ISL_3175070 | Lighthouse Lab in Milton Keynes | Wellcome Sanger Institute for the COVID-19 Genomics UK (COG-UK) Consortium | The Lighthouse Lab in Milton Keynes and Alex Alderton, Roberto Amato, Jeffrey Barrett, Sonia Goncalves, Ewan Harrison, David K. Jackson, Ian Johnston, Dominic Kwiatkowski, Cordelia Langford, John Sillitoe on behalf of the Wellcome Sanger Institute COVID-19 Surveillance Team |
| EPI_ISL_3175071, EPI_ISL_3175072 | Lighthouse Lab in Glasgow | Wellcome Sanger Institute for the COVID-19 Genomics UK (COG-UK) Consortium | Harper VanSteenhouse, Yumi Kasai, David Gray, Carol Clugston, Anna Dominiczak and Alex Alderton, Roberto Amato, Jeffrey Barrett, Sonia Goncalves, Ewan Harrison, David K. Jackson, Ian Johnston, Dominic Kwiatkowski, Cordelia Langford, John Sillitoe on behalf of the Wellcome Sanger Institute COVID-19 Surveillance Team |
| EPI_ISL_3175073 | Lighthouse Lab in Milton Keynes | Wellcome Sanger Institute for the COVID-19 Genomics UK (COG-UK) Consortium | The Lighthouse Lab in Milton Keynes and Alex Alderton, Roberto Amato, Jeffrey Barrett, Sonia Goncalves, Ewan Harrison, David K. Jackson, Ian Johnston, Dominic Kwiatkowski, Cordelia Langford, John Sillitoe on behalf of the Wellcome Sanger Institute COVID-19 Surveillance Team |
| EPI_ISL_3175074 | Berkshire and Surrey Pathology Services Lighthouse Laboratory | Wellcome Sanger Institute for the COVID-19 Genomics UK (COG-UK) Consortium | Berkshire and Surrey Pathology Services Lighthouse Laboratory and Alex Alderton, Roberto Amato, Jeffrey Barrett, Sonia Goncalves, Ewan Harrison, David K. Jackson, Ian Johnston, Dominic Kwiatkowski, Cordelia Langford, John Sillitoe on behalf of the Wellcome Sanger Institute COVID-19 Surveillance Team |
| EPI_ISL_3175075 | Lighthouse Lab in Glasgow | Wellcome Sanger Institute for the COVID-19 Genomics UK (COG-UK) Consortium | Harper VanSteenhouse, Yumi Kasai, David Gray, Carol Clugston, Anna Dominiczak and Alex Alderton, Roberto Amato, Jeffrey Barrett, Sonia Goncalves, Ewan Harrison, David K. Jackson, Ian Johnston, Dominic Kwiatkowski, Cordelia Langford, John Sillitoe on behalf of the Wellcome Sanger Institute COVID-19 Surveillance Team |
| EPI_ISL_3175076, EPI_ISL_3175077 | Lighthouse Lab in Milton Keynes | Wellcome Sanger Institute for the COVID-19 Genomics UK (COG-UK) Consortium | The Lighthouse Lab in Milton Keynes and Alex Alderton, Roberto Amato, Jeffrey Barrett, Sonia Goncalves, Ewan Harrison, David K. Jackson, Ian Johnston, Dominic Kwiatkowski, Cordelia Langford, John Sillitoe on behalf of the Wellcome Sanger Institute COVID-19 Surveillance Team |
| EPI_ISL_3175078, EPI_ISL_3175079 | Berkshire and Surrey Pathology Services Lighthouse Laboratory | Wellcome Sanger Institute for the COVID-19 Genomics UK (COG-UK) Consortium | Berkshire and Surrey Pathology Services Lighthouse Laboratory and Alex Alderton, Roberto Amato, Jeffrey Barrett, Sonia Goncalves, Ewan Harrison, David K. Jackson, Ian Johnston, Dominic Kwiatkowski, Cordelia Langford, John Sillitoe on behalf of the Wellcome Sanger Institute COVID-19 Surveillance Team |
| EPI_ISL_3175080 | Lighthouse Lab in Glasgow | Wellcome Sanger Institute for the COVID-19 Genomics UK (COG-UK) Consortium | Harper VanSteenhouse, Yumi Kasai, David Gray, Carol Clugston, Anna Dominiczak and Alex Alderton, Roberto Amato, Jeffrey Barrett, Sonia Goncalves, Ewan Harrison, David K. Jackson, Ian Johnston, Dominic Kwiatkowski, Cordelia Langford, John Sillitoe on behalf of the Wellcome Sanger Institute COVID-19 Surveillance Team |
| EPI_ISL_3175081 | Lighthouse Lab in Milton Keynes | Wellcome Sanger Institute for the COVID-19 Genomics UK (COG-UK) Consortium | The Lighthouse Lab in Milton Keynes and Alex Alderton, Roberto Amato, Jeffrey Barrett, Sonia Goncalves, Ewan Harrison, David K. Jackson, Ian Johnston, Dominic Kwiatkowski, Cordelia Langford, John Sillitoe on behalf of the Wellcome Sanger Institute COVID-19 Surveillance Team |
| EPI_ISL_3175082 | Lighthouse Lab in Glasgow | Wellcome Sanger Institute for the COVID-19 Genomics UK (COG-UK) Consortium | Harper VanSteenhouse, Yumi Kasai, David Gray, Carol Clugston, Anna Dominiczak and Alex Alderton, Roberto Amato, Jeffrey Barrett, Sonia Goncalves, Ewan Harrison, David K. Jackson, Ian Johnston, Dominic Kwiatkowski, Cordelia Langford, John Sillitoe on behalf of the Wellcome Sanger Institute COVID-19 Surveillance Team |
| EPI_ISL_3175083 | Lighthouse Lab in Milton Keynes | Wellcome Sanger Institute for the COVID-19 Genomics UK (COG-UK) Consortium | The Lighthouse Lab in Milton Keynes and Alex Alderton, Roberto Amato, Jeffrey Barrett, Sonia Goncalves, Ewan Harrison, David K. Jackson, Ian Johnston, Dominic Kwiatkowski, Cordelia Langford, John Sillitoe on behalf of the Wellcome Sanger Institute COVID-19 Surveillance Team |
| EPI_ISL_3175084 | Lighthouse Lab in Glasgow | Wellcome Sanger Institute for the COVID-19 Genomics UK (COG-UK) Consortium | Harper VanSteenhouse, Yumi Kasai, David Gray, Carol Clugston, Anna Dominiczak and Alex Alderton, Roberto Amato, Jeffrey Barrett, Sonia Goncalves, Ewan Harrison, David K. Jackson, Ian Johnston, Dominic Kwiatkowski, Cordelia Langford, John Sillitoe on behalf of the Wellcome Sanger Institute COVID-19 Surveillance Team |
| EPI_ISL_3175085, EPI_ISL_3175086, EPI_ISL_3175087 | Lighthouse Lab in Milton Keynes | Wellcome Sanger Institute for the COVID-19 Genomics UK (COG-UK) Consortium | The Lighthouse Lab in Milton Keynes and Alex Alderton, Roberto Amato, Jeffrey Barrett, Sonia Goncalves, Ewan Harrison, David K. Jackson, Ian Johnston, Dominic Kwiatkowski, Cordelia Langford, John Sillitoe on behalf of the Wellcome Sanger Institute COVID-19 Surveillance Team |
| EPI_ISL_3175088, EPI_ISL_3175089, EPI_ISL_3175090, EPI_ISL_3175091 | Lighthouse Lab in Glasgow | Wellcome Sanger Institute for the COVID-19 Genomics UK (COG-UK) Consortium | Harper VanSteenhouse, Yumi Kasai, David Gray, Carol Clugston, Anna Dominiczak and Alex Alderton, Roberto Amato, Jeffrey Barrett, Sonia Goncalves, Ewan Harrison, David K. Jackson, Ian Johnston, Dominic Kwiatkowski, Cordelia Langford, John Sillitoe on behalf of the Wellcome Sanger Institute COVID-19 Surveillance Team |
| EPI_ISL_3175092 | Berkshire and Surrey Pathology Services Lighthouse Laboratory | Wellcome Sanger Institute for the COVID-19 Genomics UK (COG-UK) Consortium | Berkshire and Surrey Pathology Services Lighthouse Laboratory and Alex Alderton, Roberto Amato, Jeffrey Barrett, Sonia Goncalves, Ewan Harrison, David K. Jackson, Ian Johnston, Dominic Kwiatkowski, Cordelia Langford, John Sillitoe on behalf of the Wellcome Sanger Institute COVID-19 Surveillance Team |
| EPI_ISL_3175093, EPI_ISL_3175094 | Lighthouse Lab in Glasgow | Wellcome Sanger Institute for the COVID-19 Genomics UK (COG-UK) Consortium | Harper VanSteenhouse, Yumi Kasai, David Gray, Carol Clugston, Anna Dominiczak and Alex Alderton, Roberto Amato, Jeffrey Barrett, Sonia Goncalves, Ewan Harrison, David K. Jackson, Ian Johnston, Dominic Kwiatkowski, Cordelia Langford, John Sillitoe on behalf of the Wellcome Sanger Institute COVID-19 Surveillance Team |
| EPI_ISL_3175095 | Lighthouse Lab in Milton Keynes | Wellcome Sanger Institute for the COVID-19 Genomics UK (COG-UK) Consortium | The Lighthouse Lab in Milton Keynes and Alex Alderton, Roberto Amato, Jeffrey Barrett, Sonia Goncalves, Ewan Harrison, David K. Jackson, Ian Johnston, Dominic Kwiatkowski, Cordelia Langford, John Sillitoe on behalf of the Wellcome Sanger Institute COVID-19 Surveillance Team |
| EPI_ISL_3175096 | Lighthouse Lab in Glasgow | Wellcome Sanger Institute for the COVID-19 Genomics UK (COG-UK) Consortium | Harper VanSteenhouse, Yumi Kasai, David Gray, Carol Clugston, Anna Dominiczak and Alex Alderton, Roberto Amato, Jeffrey Barrett, Sonia Goncalves, Ewan Harrison, David K. Jackson, Ian Johnston, Dominic Kwiatkowski, Cordelia Langford, John Sillitoe on behalf of the Wellcome Sanger Institute COVID-19 Surveillance Team |
| EPI_ISL_3175097 | Berkshire and Surrey Pathology Services Lighthouse Laboratory | Wellcome Sanger Institute for the COVID-19 Genomics UK (COG-UK) Consortium | Berkshire and Surrey Pathology Services Lighthouse Laboratory and Alex Alderton, Roberto Amato, Jeffrey Barrett, Sonia Goncalves, Ewan Harrison, David K. Jackson, Ian Johnston, Dominic Kwiatkowski, Cordelia Langford, John Sillitoe on behalf of the Wellcome Sanger Institute COVID-19 Surveillance Team |
| EPI_ISL_3175098 | Lighthouse Lab in Milton Keynes | Wellcome Sanger Institute for the COVID-19 Genomics UK (COG-UK) Consortium | The Lighthouse Lab in Milton Keynes and Alex Alderton, Roberto Amato, Jeffrey Barrett, Sonia Goncalves, Ewan Harrison, David K. Jackson, Ian Johnston, Dominic Kwiatkowski, Cordelia Langford, John Sillitoe on behalf of the Wellcome Sanger Institute COVID-19 Surveillance Team |
| EPI_ISL_3175099, EPI_ISL_3175100, EPI_ISL_3175101 | Lighthouse Lab in Glasgow | Wellcome Sanger Institute for the COVID-19 Genomics UK (COG-UK) Consortium | Harper VanSteenhouse, Yumi Kasai, David Gray, Carol Clugston, Anna Dominiczak and Alex Alderton, Roberto Amato, Jeffrey Barrett, Sonia Goncalves, Ewan Harrison, David K. Jackson, Ian Johnston, Dominic Kwiatkowski, Cordelia Langford, John Sillitoe on behalf of the Wellcome Sanger Institute COVID-19 Surveillance Team |
| EPI_ISL_3175102 | Lighthouse Lab in Milton Keynes | Wellcome Sanger Institute for the COVID-19 Genomics UK (COG-UK) Consortium | The Lighthouse Lab in Milton Keynes and Alex Alderton, Roberto Amato, Jeffrey Barrett, Sonia Goncalves, Ewan Harrison, David K. Jackson, Ian Johnston, Dominic Kwiatkowski, Cordelia Langford, John Sillitoe on behalf of the Wellcome Sanger Institute COVID-19 Surveillance Team |
| EPI_ISL_3175103, EPI_ISL_3175104 | Lighthouse Lab in Glasgow | Wellcome Sanger Institute for the COVID-19 Genomics UK (COG-UK) Consortium | Harper VanSteenhouse, Yumi Kasai, David Gray, Carol Clugston, Anna Dominiczak and Alex Alderton, Roberto Amato, Jeffrey Barrett, Sonia Goncalves, Ewan Harrison, David K. Jackson, Ian Johnston, Dominic Kwiatkowski, Cordelia Langford, John Sillitoe on behalf of the Wellcome Sanger Institute COVID-19 Surveillance Team |
| EPI_ISL_3175105 | Berkshire and Surrey Pathology Services Lighthouse Laboratory | Wellcome Sanger Institute for the COVID-19 Genomics UK (COG-UK) | Berkshire and Surrey Pathology Services Lighthouse Laboratory and Alex Alderton, Roberto Amato, Jeffrey Barrett, Sonia Goncalves, Ewan Harrison, David K. Jackson, Ian Johnston, Dominic Kwiatkowski, Cordelia Langford, John Sillitoe on behalf |

| | | | |
|---|---|---|---|
| | | Consortium | of the Wellcome Sanger Institute COVID-19 Surveillance Team |
| EPI_ISL_3175106 | Lighthouse Lab in Glasgow | Wellcome Sanger Institute for the COVID-19 Genomics UK (COG-UK) Consortium | Harper VanSteenhouse, Yumi Kasai, David Gray, Carol Clugston, Anna Dominiczak and Alex Alderton, Roberto Amato, Jeffrey Barrett, Sonia Goncalves, Ewan Harrison, David K. Jackson, Ian Johnston, Dominic Kwiatkowski, Cordelia Langford, John Sillitoe on behalf of the Wellcome Sanger Institute COVID-19 Surveillance Team |
| EPI_ISL_3175107 | Berkshire and Surrey Pathology Services Lighthouse Laboratory | Wellcome Sanger Institute for the COVID-19 Genomics UK (COG-UK) Consortium | Berkshire and Surrey Pathology Services Lighthouse Laboratory and Alex Alderton, Roberto Amato, Jeffrey Barrett, Sonia Goncalves, Ewan Harrison, David K. Jackson, Ian Johnston, Dominic Kwiatkowski, Cordelia Langford, John Sillitoe on behalf of the Wellcome Sanger Institute COVID-19 Surveillance Team |
| EPI_ISL_3175108 | Lighthouse Lab in Milton Keynes | Wellcome Sanger Institute for the COVID-19 Genomics UK (COG-UK) Consortium | The Lighthouse Lab in Milton Keynes and Alex Alderton, Roberto Amato, Jeffrey Barrett, Sonia Goncalves, Ewan Harrison, David K. Jackson, Ian Johnston, Dominic Kwiatkowski, Cordelia Langford, John Sillitoe on behalf of the Wellcome Sanger Institute COVID-19 Surveillance Team |
| EPI_ISL_3175109, EPI_ISL_3175110 | Berkshire and Surrey Pathology Services Lighthouse Laboratory | Wellcome Sanger Institute for the COVID-19 Genomics UK (COG-UK) Consortium | Berkshire and Surrey Pathology Services Lighthouse Laboratory and Alex Alderton, Roberto Amato, Jeffrey Barrett, Sonia Goncalves, Ewan Harrison, David K. Jackson, Ian Johnston, Dominic Kwiatkowski, Cordelia Langford, John Sillitoe on behalf of the Wellcome Sanger Institute COVID-19 Surveillance Team |
| EPI_ISL_3175111, EPI_ISL_3175112 | Lighthouse Lab in Glasgow | Wellcome Sanger Institute for the COVID-19 Genomics UK (COG-UK) Consortium | Harper VanSteenhouse, Yumi Kasai, David Gray, Carol Clugston, Anna Dominiczak and Alex Alderton, Roberto Amato, Jeffrey Barrett, Sonia Goncalves, Ewan Harrison, David K. Jackson, Ian Johnston, Dominic Kwiatkowski, Cordelia Langford, John Sillitoe on behalf of the Wellcome Sanger Institute COVID-19 Surveillance Team |
| EPI_ISL_3175113, EPI_ISL_3175114, EPI_ISL_3175115, EPI_ISL_3175116, EPI_ISL_3175117, EPI_ISL_3175118, EPI_ISL_3175119, EPI_ISL_3175120, EPI_ISL_3175121, EPI_ISL_3175122, EPI_ISL_3175123, EPI_ISL_3175124, EPI_ISL_3175125, EPI_ISL_3175126, EPI_ISL_3175127, EPI_ISL_3175128, EPI_ISL_3175129, EPI_ISL_3175130, EPI_ISL_3175131, EPI_ISL_3175132, EPI_ISL_3175133, EPI_ISL_3175134, EPI_ISL_3175135, EPI_ISL_3175136, EPI_ISL_3175137, EPI_ISL_3175138, EPI_ISL_3175139, EPI_ISL_3175140, EPI_ISL_3175141, EPI_ISL_3175142, EPI_ISL_3175143, EPI_ISL_3175144, EPI_ISL_3175145, EPI_ISL_3175146, EPI_ISL_3175147, EPI_ISL_3175148, EPI_ISL_3175149, EPI_ISL_3175150, EPI_ISL_3175151, EPI_ISL_3175152, EPI_ISL_3175153, EPI_ISL_3175154, EPI_ISL_3175155, EPI_ISL_3175156, EPI_ISL_3175157, EPI_ISL_3175158, EPI_ISL_3175159, EPI_ISL_3175160, EPI_ISL_3175161, EPI_ISL_3175162, EPI_ISL_3175163, EPI_ISL_3175164, EPI_ISL_3175165, EPI_ISL_3175166, EPI_ISL_3175167, EPI_ISL_3175168, EPI_ISL_3175169, EPI_ISL_3175170, EPI_ISL_3175171, EPI_ISL_3175172, EPI_ISL_3175173, EPI_ISL_3175174, EPI_ISL_3175175, EPI_ISL_3175176, EPI_ISL_3175177, EPI_ISL_3175178, EPI_ISL_3175179, EPI_ISL_3175180, EPI_ISL_3175181, EPI_ISL_3175182, EPI_ISL_3175183, EPI_ISL_3175184, EPI_ISL_3175185, EPI_ISL_3175186, EPI_ISL_3175187, EPI_ISL_3175188, EPI_ISL_3175189, EPI_ISL_3175190, EPI_ISL_3175191, EPI_ISL_3175192, EPI_ISL_3175193, EPI_ISL_3175194, EPI_ISL_3175195, EPI_ISL_3175196, EPI_ISL_3175197, EPI_ISL_3175198, EPI_ISL_3175199, EPI_ISL_3175200, EPI_ISL_3175201, EPI_ISL_3175202, EPI_ISL_3175203, EPI_ISL_3175204, EPI_ISL_3175205, EPI_ISL_3175206, EPI_ISL_3175207, EPI_ISL_3175208, EPI_ISL_3175209, EPI_ISL_3175210, EPI_ISL_3175211, EPI_ISL_3175212, EPI_ISL_3175213, EPI_ISL_3175214, EPI_ISL_3175215, EPI_ISL_3175216, EPI_ISL_3175217, EPI_ISL_3175218, EPI_ISL_3175219, EPI_ISL_3175220, EPI_ISL_3175221, EPI_ISL_3175222, EPI_ISL_3175223, EPI_ISL_3175224, EPI_ISL_3175225, EPI_ISL_3175226, EPI_ISL_3175227, EPI_ISL_3175228, EPI_ISL_3175229, EPI_ISL_3175230, EPI_ISL_3175231, EPI_ISL_3175232, EPI_ISL_3175233, EPI_ISL_3175234, EPI_ISL_3175235, EPI_ISL_3175236, EPI_ISL_3175237, EPI_ISL_3175238, EPI_ISL_3175239, EPI_ISL_3175240, EPI_ISL_3175241, EPI_ISL_3175242, EPI_ISL_3175243, EPI_ISL_3175244, EPI_ISL_3175245, EPI_ISL_3175246, EPI_ISL_3175247, EPI_ISL_3175248, EPI_ISL_3175249, EPI_ISL_3175250, EPI_ISL_3175251, EPI_ISL_3175252, EPI_ISL_3175253, EPI_ISL_3175254, EPI_ISL_3175255, EPI_ISL_3175256, EPI_ISL_3175257, EPI_ISL_3175258, EPI_ISL_3175259, EPI_ISL_3175260, EPI_ISL_3175261, EPI_ISL_3175262, EPI_ISL_3175263, EPI_ISL_3175264, EPI_ISL_3175265, EPI_ISL_3175266, EPI_ISL_3175267, EPI_ISL_3175268, EPI_ISL_3175269, EPI_ISL_3175270, EPI_ISL_3175271, EPI_ISL_3175272, EPI_ISL_3175273, EPI_ISL_3175274, EPI_ISL_3175275, EPI_ISL_3175276, EPI_ISL_3175277, EPI_ISL_3175278, EPI_ISL_3175279, EPI_ISL_3175280, EPI_ISL_3175281, EPI_ISL_3175282, EPI_ISL_3175283, EPI_ISL_3175284, EPI_ISL_3175285, EPI_ISL_3175286, EPI_ISL_3175287, EPI_ISL_3175288, EPI_ISL_3175289, EPI_ISL_3175290, EPI_ISL_3175291, EPI_ISL_3175292, EPI_ISL_3175293, EPI_ISL_3175294, EPI_ISL_3175295, EPI_ISL_3175296, EPI_ISL_3175297, EPI_ISL_3175298, EPI_ISL_3175299, EPI_ISL_3175300, EPI_ISL_3175301, EPI_ISL_3175302, EPI_ISL_3175303, EPI_ISL_3175304, EPI_ISL_3175305, EPI_ISL_3175306, EPI_ISL_3175307, EPI_ISL_3175308, EPI_ISL_3175309, EPI_ISL_3175310, EPI_ISL_3175311, EPI_ISL_3175312, EPI_ISL_3175313, EPI_ISL_3175314, EPI_ISL_3175315, EPI_ISL_3175316, EPI_ISL_3175317, EPI_ISL_3175318, EPI_ISL_3175319, EPI_ISL_3175320, EPI_ISL_3175321, EPI_ISL_3175322, EPI_ISL_3175323, EPI_ISL_3175324, EPI_ISL_3175325, EPI_ISL_3175326, EPI_ISL_3175327, EPI_ISL_3175328, EPI_ISL_3175329, EPI_ISL_3175330 | | | |
| see above | Lighthouse Lab in Milton Keynes | Wellcome Sanger Institute for the COVID-19 Genomics UK (COG-UK) Consortium | The Lighthouse Lab in Milton Keynes and Alex Alderton, Roberto Amato, Jeffrey Barrett, Sonia Goncalves, Ewan Harrison, David K. Jackson, Ian Johnston, Dominic Kwiatkowski, Cordelia Langford, John Sillitoe on behalf of the Wellcome Sanger Institute COVID-19 Surveillance Team |
| EPI_ISL_3175331, EPI_ISL_3175332, EPI_ISL_3175333, EPI_ISL_3175334 | Lighthouse Lab in Glasgow | Wellcome Sanger Institute for the COVID-19 Genomics UK (COG-UK) Consortium | Harper VanSteenhouse, Yumi Kasai, David Gray, Carol Clugston, Anna Dominiczak and Alex Alderton, Roberto Amato, Jeffrey Barrett, Sonia Goncalves, Ewan Harrison, David K. Jackson, Ian Johnston, Dominic Kwiatkowski, Cordelia Langford, John Sillitoe on behalf of the Wellcome Sanger Institute COVID-19 Surveillance Team |
| EPI_ISL_3175335, EPI_ISL_3175336 | Lighthouse Laboratory Plymouth | Wellcome Sanger Institute for the COVID-19 Genomics UK (COG-UK) Consortium | Lighthouse Laboratory Plymouth and Alex Alderton, Roberto Amato, Jeffrey Barrett, Sonia Goncalves, Ewan Harrison, David K. Jackson, Ian Johnston, Dominic Kwiatkowski, Cordelia Langford, John Sillitoe on behalf of the Wellcome Sanger Institute COVID-19 Surveillance Team |
| EPI_ISL_3175337, EPI_ISL_3175338, EPI_ISL_3175339, EPI_ISL_3175340, EPI_ISL_3175341 | Lighthouse Lab in Glasgow | Wellcome Sanger Institute for the COVID-19 Genomics UK (COG-UK) Consortium | Harper VanSteenhouse, Yumi Kasai, David Gray, Carol Clugston, Anna Dominiczak and Alex Alderton, Roberto Amato, Jeffrey Barrett, Sonia Goncalves, Ewan Harrison, David K. Jackson, Ian Johnston, Dominic Kwiatkowski, Cordelia Langford, John Sillitoe on behalf of the Wellcome Sanger Institute COVID-19 Surveillance Team |
| EPI_ISL_3175342 | Lighthouse Lab in Alderley Park | Wellcome Sanger Institute for the COVID-19 Genomics UK (COG-UK) Consortium | Jacquelyn Wynn, Mairead Hyland, The Lighthouse Lab in Alderley Park and Alex Alderton, Roberto Amato, Jeffrey Barrett, Sonia Goncalves, Ewan Harrison, David K. Jackson, Ian Johnston, Dominic Kwiatkowski, Cordelia Langford, John Sillitoe on behalf of the Wellcome Sanger Institute COVID-19 Surveillance Team |
| EPI_ISL_3175343 | Lighthouse Lab in Glasgow | Wellcome Sanger Institute for the COVID-19 Genomics UK (COG-UK) Consortium | Harper VanSteenhouse, Yumi Kasai, David Gray, Carol Clugston, Anna Dominiczak and Alex Alderton, Roberto Amato, Jeffrey Barrett, Sonia Goncalves, Ewan Harrison, David K. Jackson, Ian Johnston, Dominic Kwiatkowski, Cordelia Langford, John Sillitoe on behalf of the Wellcome Sanger Institute COVID-19 Surveillance Team |
| EPI_ISL_3175344 | Lighthouse Lab in Alderley Park | Wellcome Sanger Institute for the COVID-19 Genomics UK (COG-UK) Consortium | Jacquelyn Wynn, Mairead Hyland, The Lighthouse Lab in Alderley Park and Alex Alderton, Roberto Amato, Jeffrey Barrett, Sonia Goncalves, Ewan Harrison, David K. Jackson, Ian Johnston, Dominic Kwiatkowski, Cordelia Langford, John Sillitoe on behalf of the Wellcome Sanger Institute COVID-19 Surveillance Team |
| EPI_ISL_3175345, EPI_ISL_3175346, EPI_ISL_3175347, EPI_ISL_3175348, EPI_ISL_3175349, EPI_ISL_3175350, EPI_ISL_3175351, EPI_ISL_3175352, EPI_ISL_3175353 | Lighthouse Lab in Glasgow | Wellcome Sanger Institute for the COVID-19 Genomics UK (COG-UK) Consortium | Harper VanSteenhouse, Yumi Kasai, David Gray, Carol Clugston, Anna Dominiczak and Alex Alderton, Roberto Amato, Jeffrey Barrett, Sonia Goncalves, Ewan Harrison, David K. Jackson, Ian Johnston, Dominic Kwiatkowski, Cordelia Langford, John Sillitoe on behalf of the Wellcome Sanger Institute COVID-19 Surveillance Team |
| EPI_ISL_3175354 | Lighthouse Lab in Alderley Park | Wellcome Sanger Institute for the COVID-19 Genomics UK (COG-UK) Consortium | Jacquelyn Wynn, Mairead Hyland, The Lighthouse Lab in Alderley Park and Alex Alderton, Roberto Amato, Jeffrey Barrett, Sonia Goncalves, Ewan Harrison, David K. Jackson, Ian Johnston, Dominic Kwiatkowski, Cordelia Langford, John Sillitoe on behalf of the Wellcome Sanger Institute COVID-19 Surveillance Team |
| EPI_ISL_3175355, EPI_ISL_3175356, EPI_ISL_3175357, EPI_ISL_3175358, EPI_ISL_3175359, EPI_ISL_3175360, EPI_ISL_3175361 | Lighthouse Lab in Glasgow | Wellcome Sanger Institute for the COVID-19 Genomics UK (COG-UK) Consortium | Harper VanSteenhouse, Yumi Kasai, David Gray, Carol Clugston, Anna Dominiczak and Alex Alderton, Roberto Amato, Jeffrey Barrett, Sonia Goncalves, Ewan Harrison, David K. Jackson, Ian Johnston, Dominic Kwiatkowski, Cordelia Langford, John Sillitoe on behalf of the Wellcome Sanger Institute COVID-19 Surveillance Team |
| EPI_ISL_3175362 | Lighthouse Laboratory Plymouth | Wellcome Sanger Institute for the COVID-19 Genomics UK (COG-UK) Consortium | Lighthouse Laboratory Plymouth and Alex Alderton, Roberto Amato, Jeffrey Barrett, Sonia Goncalves, Ewan Harrison, David K. Jackson, Ian Johnston, Dominic Kwiatkowski, Cordelia Langford, John Sillitoe on behalf of the Wellcome Sanger Institute COVID-19 Surveillance Team |
| EPI_ISL_3175363, EPI_ISL_3175364, EPI_ISL_3175365, EPI_ISL_3175366, EPI_ISL_3175367 | Lighthouse Lab in Glasgow | Wellcome Sanger Institute for the COVID-19 Genomics UK (COG-UK) Consortium | Harper VanSteenhouse, Yumi Kasai, David Gray, Carol Clugston, Anna Dominiczak and Alex Alderton, Roberto Amato, Jeffrey Barrett, Sonia Goncalves, Ewan Harrison, David K. Jackson, Ian Johnston, Dominic Kwiatkowski, Cordelia Langford, John Sillitoe on behalf of the Wellcome Sanger Institute COVID-19 Surveillance Team |
| EPI_ISL_3175368, EPI_ISL_3175369, EPI_ISL_3175370, EPI_ISL_3175371 | ASL BARI | University of Bari Biomedical Sciences and Human Oncology | Maria Chironna |
| EPI_ISL_3175372, EPI_ISL_3175373, EPI_ISL_3175374, EPI_ISL_3175375, EPI_ISL_3175376, EPI_ISL_3175377, EPI_ISL_3175378, EPI_ISL_3175379, EPI_ISL_3175380, EPI_ISL_3175381, EPI_ISL_3175382, EPI_ISL_3175383, EPI_ISL_3175384, EPI_ISL_3175385, EPI_ISL_3175386, EPI_ISL_3175387, EPI_ISL_3175388, EPI_ISL_3175389, EPI_ISL_3175390, EPI_ISL_3175391, EPI_ISL_3175392, EPI_ISL_3175393, EPI_ISL_3175394, EPI_ISL_3175395, EPI_ISL_3175396, EPI_ISL_3175397, EPI_ISL_3175398, EPI_ISL_3175399, EPI_ISL_3175400, EPI_ISL_3175401, EPI_ISL_3175402, EPI_ISL_3175403, EPI_ISL_3175404, EPI_ISL_3175405, EPI_ISL_3175406, EPI_ISL_3175407, EPI_ISL_3175408, EPI_ISL_3175409, EPI_ISL_3175410, EPI_ISL_3175411, EPI_ISL_3175412, EPI_ISL_3175413, EPI_ISL_3175414, EPI_ISL_3175415, EPI_ISL_3175416, EPI_ISL_3175417, EPI_ISL_3175418, EPI_ISL_3175419, EPI_ISL_3175420, EPI_ISL_3175421, EPI_ISL_3175422, EPI_ISL_3175423, EPI_ISL_3175424, EPI_ISL_3175425, EPI_ISL_3175426, EPI_ISL_3175427, EPI_ISL_3175428, EPI_ISL_3175429, EPI_ISL_3175430, EPI_ISL_3175431, EPI_ISL_3175432, EPI_ISL_3175433, EPI_ISL_3175434, EPI_ISL_3175435, EPI_ISL_3175436, EPI_ISL_3175437, EPI_ISL_3175438, EPI_ISL_3175439, EPI_ISL_3175440, EPI_ISL_3175441, EPI_ISL_3175442, EPI_ISL_3175443, EPI_ISL_3175444, EPI_ISL_3175445, EPI_ISL_3175446, EPI_ISL_3175447, EPI_ISL_3175448, EPI_ISL_3175449, EPI_ISL_3175450, EPI_ISL_3175451, EPI_ISL_3175452, EPI_ISL_3175453, EPI_ISL_3175454, EPI_ISL_3175455, EPI_ISL_3175456, EPI_ISL_3175457, EPI_ISL_3175458, EPI_ISL_3175459, EPI_ISL_3175460, EPI_ISL_3175461, EPI_ISL_3175462, EPI_ISL_3175463, EPI_ISL_3175464, EPI_ISL_3175465, EPI_ISL_3175466, EPI_ISL_3175467, EPI_ISL_3175468, EPI_ISL_3175469, EPI_ISL_3175470, EPI_ISL_3175471, EPI_ISL_3175472, EPI_ISL_3175473, EPI_ISL_3175474, EPI_ISL_3175475, EPI_ISL_3175476, EPI_ISL_3175477, EPI_ISL_3175478, EPI_ISL_3175479, EPI_ISL_3175480, EPI_ISL_3175481, EPI_ISL_3175482, EPI_ISL_3175483, EPI_ISL_3175484, EPI_ISL_3175485, EPI_ISL_3175486, EPI_ISL_3175487, EPI_ISL_3175488, EPI_ISL_3175489, EPI_ISL_3175490, EPI_ISL_3175491, EPI_ISL_3175492, EPI_ISL_3175493, EPI_ISL_3175494, EPI_ISL_3175495, EPI_ISL_3175496, EPI_ISL_3175497, EPI_ISL_3175498, EPI_ISL_3175499, EPI_ISL_3175500, EPI_ISL_3175501, EPI_ISL_3175502, EPI_ISL_3175503, EPI_ISL_3175504, EPI_ISL_3175505, EPI_ISL_3175506, EPI_ISL_3175507, EPI_ISL_3175508, EPI_ISL_3175509, EPI_ISL_3175510, EPI_ISL_3175511, EPI_ISL_3175512, EPI_ISL_3175513, EPI_ISL_3175514, EPI_ISL_3175515, EPI_ISL_3175516, EPI_ISL_3175517, EPI_ISL_3175518, EPI_ISL_3175519, EPI_ISL_3175520, EPI_ISL_3175521, EPI_ISL_3175522, EPI_ISL_3175523, EPI_ISL_3175524, EPI_ISL_3175525, EPI_ISL_3175526, EPI_ISL_3175527, EPI_ISL_3175528, EPI_ISL_3175529, EPI_ISL_3175530, EPI_ISL_3175531, EPI_ISL_3175532, EPI_ISL_3175533, EPI_ISL_3175534, EPI_ISL_3175535, EPI_ISL_3175536, EPI_ISL_3175537, EPI_ISL_3175538, EPI_ISL_3175539, EPI_ISL_3175540, EPI_ISL_3175541, EPI_ISL_3175542, EPI_ISL_3175543, EPI_ISL_3175544, EPI_ISL_3175545, EPI_ISL_3175546, EPI_ISL_3175547, EPI_ISL_3175548, EPI_ISL_3175549, EPI_ISL_3175550, EPI_ISL_3175551, EPI_ISL_3175552, EPI_ISL_3175553, EPI_ISL_3175554, EPI_ISL_3175555, EPI_ISL_3175556, EPI_ISL_3175557, EPI_ISL_3175558, EPI_ISL_3175559, EPI_ISL_3175560, EPI_ISL_3175561, EPI_ISL_3175562, EPI_ISL_3175563, EPI_ISL_3175564, EPI_ISL_3175565, EPI_ISL_3175566, EPI_ISL_3175567, EPI_ISL_3175568, EPI_ISL_3175569, EPI_ISL_3175570, EPI_ISL_3175571, EPI_ISL_3175572, EPI_ISL_3175573, EPI_ISL_3175574, EPI_ISL_3175575, EPI_ISL_3175576, EPI_ISL_3175577, EPI_ISL_3175578, EPI_ISL_3175579, EPI_ISL_3175580, EPI_ISL_3175581, EPI_ISL_3175582, EPI_ISL_3175583, EPI_ISL_3175584, EPI_ISL_3175585, EPI_ISL_3175586, EPI_ISL_3175587, EPI_ISL_3175588, EPI_ISL_3175589, EPI_ISL_3175590, EPI_ISL_3175591, EPI_ISL_3175592, EPI_ISL_3175593, EPI_ISL_3175594, EPI_ISL_3175595, EPI_ISL_3175596, EPI_ISL_3175597, EPI_ISL_3175598, EPI_ISL_3175599, EPI_ISL_3175600, EPI_ISL_3175601, EPI_ISL_3175602, EPI_ISL_3175603, EPI_ISL_3175604, EPI_ISL_3175605, EPI_ISL_3175606, EPI_ISL_3175607, EPI_ISL_3175608, EPI_ISL_3175609, EPI_ISL_3175610, EPI_ISL_3175611, EPI_ISL_3175612, EPI_ISL_3175613, EPI_ISL_3175614, EPI_ISL_3175615, EPI_ISL_3175616, EPI_ISL_3175617, EPI_ISL_3175618, EPI_ISL_3175619, EPI_ISL_3175620, EPI_ISL_3175621, EPI_ISL_3175622, EPI_ISL_3175623, EPI_ISL_3175624, EPI_ISL_3175625, EPI_ISL_3175626, EPI_ISL_3175627, EPI_ISL_3175628, EPI_ISL_3175629, EPI_ISL_3175630, EPI_ISL_3175631, EPI_ISL_3175632, EPI_ISL_3175633, EPI_ISL_3175634, EPI_ISL_3175635, EPI_ISL_3175636, EPI_ISL_3175637, EPI_ISL_3175638, EPI_ISL_3175639, EPI_ISL_3175640, EPI_ISL_3175641, EPI_ISL_3175642, EPI_ISL_3175643, EPI_ISL_3175644, EPI_ISL_3175645, EPI_ISL_3175646, EPI_ISL_3175647, EPI_ISL_3175648, EPI_ISL_3175649, EPI_ISL_3175650, EPI_ISL_3175651, EPI_ISL_3175652, EPI_ISL_3175653, EPI_ISL_3175654 | | | |
| see above | University of Birmingham | COVID-19 Genomics UK (COG-UK) Consortium | Institute of Microbiology, University of Birmingham: Claire McMurray, Joanne Stockton, Samuel Nicholls, Radoslaw Poplawski, Will Rowe, Josh Quick, Nicholas Loman. University of Birmingham Testing Laboratory: Celina M Whalley, Andrew Bosworth, Charlotte Poxon, Kasun Wanigasooriya, Oliver Pickles, Mike Kidd, Alex Richter, Andrew D Beggs PHE Heartlands Lab: Husam Osman, Andrew Bosworth. Queen Elizabeth Hospital: Anna Casey |
| EPI_ISL_3175655, EPI_ISL_3175656, EPI_ISL_3175657, EPI_ISL_3175658, EPI_ISL_3175659, EPI_ISL_3175660, EPI_ISL_3175661, EPI_ISL_3175662, EPI_ISL_3175663, EPI_ISL_3175664, EPI_ISL_3175665, EPI_ISL_3175666, EPI_ISL_3175667, EPI_ISL_3175668, EPI_ISL_3175669, EPI_ISL_3175670, EPI_ISL_3175671, EPI_ISL_3175672, EPI_ISL_3175673, EPI_ISL_3175674, EPI_ISL_3175675, EPI_ISL_3175676, EPI_ISL_3175677, EPI_ISL_3175678, EPI_ISL_3175679, EPI_ISL_3175680, EPI_ISL_3175681, EPI_ISL_3175682, EPI_ISL_3175683, EPI_ISL_3175684, EPI_ISL_3175685, EPI_ISL_3175686, EPI_ISL_3175687, EPI_ISL_3175688, EPI_ISL_3175689, EPI_ISL_3175690, EPI_ISL_3175691, EPI_ISL_3175692, EPI_ISL_3175693, EPI_ISL_3175694, EPI_ISL_3175695, EPI_ISL_3175696, EPI_ISL_3175697, EPI_ISL_3175698, EPI_ISL_3175699, EPI_ISL_3175700, EPI_ISL_3175701, EPI_ISL_3175702, EPI_ISL_3175703, EPI_ISL_3175704, EPI_ISL_3175705, EPI_ISL_3175706, EPI_ISL_3175707, EPI_ISL_3175708, EPI_ISL_3175709, EPI_ISL_3175710, EPI_ISL_3175711, EPI_ISL_3175712, EPI_ISL_3175713, EPI_ISL_3175714, EPI_ISL_3175715, EPI_ISL_3175716, EPI_ISL_3175717, EPI_ISL_3175718, EPI_ISL_3175719, EPI_ISL_3175720, EPI_ISL_3175721, EPI_ISL_3175722, EPI_ISL_3175723, EPI_ISL_3175724, EPI_ISL_3175725, EPI_ISL_3175726, EPI_ISL_3175727, EPI_ISL_3175728, EPI_ISL_3175729, EPI_ISL_3175730, EPI_ISL_3175731, EPI_ISL_3175732, EPI_ISL_3175733, EPI_ISL_3175734, EPI_ISL_3175735, EPI_ISL_3175736, EPI_ISL_3175737, EPI_ISL_3175738, EPI_ISL_3175739, EPI_ISL_3175740, EPI_ISL_3175741, EPI_ISL_3175742, EPI_ISL_3175743, EPI_ISL_3175744, EPI_ISL_3175745, EPI_ISL_3175746, EPI_ISL_3175747, EPI_ISL_3175748, EPI_ISL_3175749, EPI_ISL_3175750, EPI_ISL_3175751, EPI_ISL_3175752, EPI_ISL_3175753, EPI_ISL_3175754, EPI_ISL_3175755, EPI_ISL_3175756, EPI_ISL_3175757, EPI_ISL_3175758, EPI_ISL_3175759, EPI_ISL_3175760, EPI_ISL_3175761, EPI_ISL_3175762, EPI_ISL_3175763, EPI_ISL_3175764, EPI_ISL_3175765, EPI_ISL_3175766, EPI_ISL_3175767, EPI_ISL_3175768, EPI_ISL_3175769, EPI_ISL_3175770, EPI_ISL_3175771, EPI_ISL_3175772, EPI_ISL_3175773, EPI_ISL_3175774, EPI_ISL_3175775, EPI_ISL_3175776, EPI_ISL_3175777, EPI_ISL_3175778, EPI_ISL_3175779, EPI_ISL_3175780, EPI_ISL_3175781, EPI_ISL_3175782, EPI_ISL_3175783, EPI_ISL_3175784, EPI_ISL_3175785, EPI_ISL_3175786, EPI_ISL_3175787, EPI_ISL_3175788, EPI_ISL_3175789, EPI_ISL_3175790, EPI_ISL_3175791, EPI_ISL_3175792, EPI_ISL_3175793, EPI_ISL_3175794, EPI_ISL_3175795, EPI_ISL_3175796, EPI_ISL_3175797, EPI_ISL_3175798, EPI_ISL_3175799, EPI_ISL_3175800, EPI_ISL_3175801, EPI_ISL_3175802, EPI_ISL_3175803, EPI_ISL_3175804, EPI_ISL_3175805, EPI_ISL_3175806, EPI_ISL_3175807, EPI_ISL_3175808, EPI_ISL_3175809, EPI_ISL_3175810, EPI_ISL_3175811, EPI_ISL_3175812, EPI_ISL_3175813, EPI_ISL_3175814, EPI_ISL_3175815, EPI_ISL_3175816, EPI_ISL_3175817, EPI_ISL_3175818, EPI_ISL_3175819, EPI_ISL_3175820, EPI_ISL_3175821, EPI_ISL_3175822, EPI_ISL_3175823, EPI_ISL_3175824, EPI_ISL_3175825, EPI_ISL_3175826, EPI_ISL_3175827, EPI_ISL_3175828, EPI_ISL_3175829, EPI_ISL_3175830, EPI_ISL_3175831, EPI_ISL_3175832, EPI_ISL_3175833, EPI_ISL_3175834, EPI_ISL_3175835, EPI_ISL_3175836, EPI_ISL_3175837, EPI_ISL_3175838, EPI_ISL_3175839, EPI_ISL_3175840, EPI_ISL_3175841, EPI_ISL_3175842, EPI_ISL_3175843, EPI_ISL_3175844, EPI_ISL_3175845, EPI_ISL_3175846, EPI_ISL_3175847, EPI_ISL_3175848, EPI_ISL_3175849, EPI_ISL_3175850, EPI_ISL_3175851, EPI_ISL_3175852, EPI_ISL_3175853, EPI_ISL_3175854, EPI_ISL_3175855, EPI_ISL_3175856, EPI_ISL_3175857, EPI_ISL_3175858, EPI_ISL_3175859, EPI_ISL_3175860, EPI_ISL_3175861, EPI_ISL_3175862, EPI_ISL_3175863, EPI_ISL_3175864, EPI_ISL_3175865, EPI_ISL_3175866, EPI_ISL_3175867, EPI_ISL_3175868, EPI_ISL_3175869, EPI_ISL_3175870, EPI_ISL_3175871, EPI_ISL_3175872, EPI_ISL_3175873, EPI_ISL_3175874, EPI_ISL_3175875, EPI_ISL_3175876, EPI_ISL_3175877, EPI_ISL_3175878, EPI_ISL_3175879, EPI_ISL_3175880, EPI_ISL_3175881, EPI_ISL_3175882, EPI_ISL_3175883, EPI_ISL_3175884, EPI_ISL_3175885, EPI_ISL_3175886, EPI_ISL_3175887, EPI_ISL_3175888, EPI_ISL_3175889, EPI_ISL_3175890, EPI_ISL_3175891, EPI_ISL_3175892, EPI_ISL_3175893, EPI_ISL_3175894, EPI_ISL_3175895, EPI_ISL_3175896, EPI_ISL_3175897, EPI_ISL_3175898, EPI_ISL_3175899, EPI_ISL_3175900, EPI_ISL_3175901, EPI_ISL_3175902, EPI_ISL_3175903, EPI_ISL_3175904, EPI_ISL_3175905, EPI_ISL_3175906, EPI_ISL_3175907, EPI_ISL_3175908, EPI_ISL_3175909, EPI_ISL_3175910, EPI_ISL_3175911, EPI_ISL_3175912, EPI_ISL_3175913, EPI_ISL_3175914, EPI_ISL_3175915, EPI_ISL_3175916, EPI_ISL_3175917, EPI_ISL_3175918, EPI_ISL_3175919, EPI_ISL_3175920, EPI_ISL_3175921, EPI_ISL_3175922, EPI_ISL_3175923, EPI_ISL_3175924, EPI_ISL_3175925, EPI_ISL_3175926, EPI_ISL_3175927, EPI_ISL_3175928, EPI_ISL_3175929, EPI_ISL_3175930, EPI_ISL_3175931, EPI_ISL_3175932, EPI_ISL_3175933, EPI_ISL_3175934, EPI_ISL_3175935, EPI_ISL_3175936, EPI_ISL_3175937, EPI_ISL_3175938, EPI_ISL_3175939, EPI_ISL_3175940, EPI_ISL_3175941, EPI_ISL_3175942, EPI_ISL_3175943, EPI_ISL_3175944, EPI_ISL_3175945, EPI_ISL_3175946, EPI_ISL_3175947, EPI_ISL_3175948, EPI_ISL_3175949, EPI_ISL_3175950, EPI_ISL_3175951, EPI_ISL_3175952, EPI_ISL_3175953, EPI_ISL_3175954, EPI_ISL_3175955, EPI_ISL_3175956, EPI_ISL_3175957, EPI_ISL_3175958, EPI_ISL_3175959, EPI_ISL_3175960, EPI_ISL_3175961, EPI_ISL_3175962, EPI_ISL_3175963, EPI_ISL_3175964, EPI_ISL_3175965, EPI_ISL_3175966, EPI_ISL_3175967, EPI_ISL_3175968, EPI_ISL_3175969, EPI_ISL_3175970, EPI_ISL_3175971, EPI_ISL_3175972, EPI_ISL_3175973, EPI_ISL_3175974, EPI_ISL_3175975, EPI_ISL_3175976, EPI_ISL_3175977, EPI_ISL_3175978, EPI_ISL_3175979, EPI_ISL_3175980, EPI_ISL_3175981, EPI_ISL_3175982, EPI_ISL_3175983, EPI_ISL_3175984, EPI_ISL_3175985, EPI_ISL_3175986, EPI_ISL_3175987, EPI_ISL_3175988, EPI_ISL_3175989, EPI_ISL_3175990, EPI_ISL_3175991, EPI_ISL_3175992, EPI_ISL_3175993, EPI_ISL_3175994, EPI_ISL_3175995, EPI_ISL_3175996, EPI_ISL_3175997, EPI_ISL_3175998, EPI_ISL_3175999, EPI_ISL_3176000, EPI_ISL_3176001, EPI_ISL_3176002, EPI_ISL_3176003, EPI_ISL_3176004, EPI_ISL_3176005, EPI_ISL_3176006, | | | |

| Accession ID | Originating lab | Submitting lab | Authors |
| --- | --- | --- | --- |
| EPI_ISL_3176007, EPI_ISL_3176008, EPI_ISL_3176009, EPI_ISL_3176010, EPI_ISL_3176011, EPI_ISL_3176012, EPI_ISL_3176013, EPI_ISL_3176014, EPI_ISL_3176015, EPI_ISL_3176016, EPI_ISL_3176017, EPI_ISL_3176018, EPI_ISL_3176019, EPI_ISL_3176020, EPI_ISL_3176021, EPI_ISL_3176022, EPI_ISL_3176023, EPI_ISL_3176024, EPI_ISL_3176025, EPI_ISL_3176026, EPI_ISL_3176027, EPI_ISL_3176028, EPI_ISL_3176029, EPI_ISL_3176030, EPI_ISL_3176031, EPI_ISL_3176032, EPI_ISL_3176033, EPI_ISL_3176034, EPI_ISL_3176035, EPI_ISL_3176036, EPI_ISL_3176037, EPI_ISL_3176038, EPI_ISL_3176039, EPI_ISL_3176040, EPI_ISL_3176041, EPI_ISL_3176042, EPI_ISL_3176043, EPI_ISL_3176044, EPI_ISL_3176045, EPI_ISL_3176046, EPI_ISL_3176047, EPI_ISL_3176048, EPI_ISL_3176049, EPI_ISL_3176050, EPI_ISL_3176051, EPI_ISL_3176052, EPI_ISL_3176053, EPI_ISL_3176054, EPI_ISL_3176055, EPI_ISL_3176056, EPI_ISL_3176057, EPI_ISL_3176058, EPI_ISL_3176059, EPI_ISL_3176060, EPI_ISL_3176061, EPI_ISL_3176062, EPI_ISL_3176063, EPI_ISL_3176064, EPI_ISL_3176065, EPI_ISL_3176066, EPI_ISL_3176067, EPI_ISL_3176068, EPI_ISL_3176069, EPI_ISL_3176070, EPI_ISL_3176071, EPI_ISL_3176072, EPI_ISL_3176073, EPI_ISL_3176074, EPI_ISL_3176075, EPI_ISL_3176076, EPI_ISL_3176077, EPI_ISL_3176078, EPI_ISL_3176079, EPI_ISL_3176080, EPI_ISL_3176081, EPI_ISL_3176082, EPI_ISL_3176083, EPI_ISL_3176084, EPI_ISL_3176085, EPI_ISL_3176086, EPI_ISL_3176087, EPI_ISL_3176088, EPI_ISL_3176089 | | | |
| see above | Department of Pathology, University of Cambridge | COVID-19 Genomics UK (COG-UK) Consortium | Aminu S. Jahun, Yasmin Chaudhry, Iliana Georgana, Myra Hosmillo, Rhys Izuagbe, William L. Hamilton, Martin D. Curran, Surendra Parmar, Ian Goodfellow |
| EPI_ISL_3176090, EPI_ISL_3176091, EPI_ISL_3176092, EPI_ISL_3176093, EPI_ISL_3176094, EPI_ISL_3176095, EPI_ISL_3176096, EPI_ISL_3176097, EPI_ISL_3176098, EPI_ISL_3176099, EPI_ISL_3176100, EPI_ISL_3176101, EPI_ISL_3176102, EPI_ISL_3176103, EPI_ISL_3176104, EPI_ISL_3176105, EPI_ISL_3176106, EPI_ISL_3176107, EPI_ISL_3176108, EPI_ISL_3176109, EPI_ISL_3176110, EPI_ISL_3176111, EPI_ISL_3176112, EPI_ISL_3176113, EPI_ISL_3176114, EPI_ISL_3176115, EPI_ISL_3176116, EPI_ISL_3176117, EPI_ISL_3176118, EPI_ISL_3176119, EPI_ISL_3176120, EPI_ISL_3176121, EPI_ISL_3176122, EPI_ISL_3176123, EPI_ISL_3176124, EPI_ISL_3176125, EPI_ISL_3176126, EPI_ISL_3176127, EPI_ISL_3176128, EPI_ISL_3176129, EPI_ISL_3176130, EPI_ISL_3176131, EPI_ISL_3176132, EPI_ISL_3176133, EPI_ISL_3176134, EPI_ISL_3176135, EPI_ISL_3176136, EPI_ISL_3176137, EPI_ISL_3176138, EPI_ISL_3176139, EPI_ISL_3176140, EPI_ISL_3176141, EPI_ISL_3176142, EPI_ISL_3176143, EPI_ISL_3176144, EPI_ISL_3176145, EPI_ISL_3176146, EPI_ISL_3176147, EPI_ISL_3176148, EPI_ISL_3176149, EPI_ISL_3176150, EPI_ISL_3176151, EPI_ISL_3176152, EPI_ISL_3176153, EPI_ISL_3176154, EPI_ISL_3176155, EPI_ISL_3176156, EPI_ISL_3176157, EPI_ISL_3176158, EPI_ISL_3176159, EPI_ISL_3176160, EPI_ISL_3176161, EPI_ISL_3176162, EPI_ISL_3176163, EPI_ISL_3176164, EPI_ISL_3176165, EPI_ISL_3176166, EPI_ISL_3176167, EPI_ISL_3176168, EPI_ISL_3176169, EPI_ISL_3176170, EPI_ISL_3176171, EPI_ISL_3176172, EPI_ISL_3176173, EPI_ISL_3176174, EPI_ISL_3176175, EPI_ISL_3176176, EPI_ISL_3176177, EPI_ISL_3176178, EPI_ISL_3176179, EPI_ISL_3176180, EPI_ISL_3176181, EPI_ISL_3176182, EPI_ISL_3176183, EPI_ISL_3176184, EPI_ISL_3176185, EPI_ISL_3176186, EPI_ISL_3176187, EPI_ISL_3176188, EPI_ISL_3176189, EPI_ISL_3176190, EPI_ISL_3176191, EPI_ISL_3176192, EPI_ISL_3176193, EPI_ISL_3176194, EPI_ISL_3176195, EPI_ISL_3176196, EPI_ISL_3176197, EPI_ISL_3176198, EPI_ISL_3176199, EPI_ISL_3176200, EPI_ISL_3176201, EPI_ISL_3176202, EPI_ISL_3176203, EPI_ISL_3176204, EPI_ISL_3176205, EPI_ISL_3176206, EPI_ISL_3176207, EPI_ISL_3176208, EPI_ISL_3176210, EPI_ISL_3176211, EPI_ISL_3176212, EPI_ISL_3176213, EPI_ISL_3176214, EPI_ISL_3176215, EPI_ISL_3176216, EPI_ISL_3176217, EPI_ISL_3176218, EPI_ISL_3176219, EPI_ISL_3176220, EPI_ISL_3176221, EPI_ISL_3176222, EPI_ISL_3176223, EPI_ISL_3176224, EPI_ISL_3176225, EPI_ISL_3176226, EPI_ISL_3176227, EPI_ISL_3176228, EPI_ISL_3176229, EPI_ISL_3176230, EPI_ISL_3176231, EPI_ISL_3176232, EPI_ISL_3176233, EPI_ISL_3176234, EPI_ISL_3176235, EPI_ISL_3176236, EPI_ISL_3176237, EPI_ISL_3176238, EPI_ISL_3176239, EPI_ISL_3176240, EPI_ISL_3176241, EPI_ISL_3176242, EPI_ISL_3176243, EPI_ISL_3176244, EPI_ISL_3176245, EPI_ISL_3176246, EPI_ISL_3176247, EPI_ISL_3176248, EPI_ISL_3176249, EPI_ISL_3176250, EPI_ISL_3176251, EPI_ISL_3176252, EPI_ISL_3176253, EPI_ISL_3176254, EPI_ISL_3176255, EPI_ISL_3176256, EPI_ISL_3176257, EPI_ISL_3176258, EPI_ISL_3176259, EPI_ISL_3176260, EPI_ISL_3176261, EPI_ISL_3176262, EPI_ISL_3176263, EPI_ISL_3176264, EPI_ISL_3176265, EPI_ISL_3176266, EPI_ISL_3176267, EPI_ISL_3176268, EPI_ISL_3176269, EPI_ISL_3176270, EPI_ISL_3176271, EPI_ISL_3176272, EPI_ISL_3176273, EPI_ISL_3176274, EPI_ISL_3176275, EPI_ISL_3176276, EPI_ISL_3176277, EPI_ISL_3176278, EPI_ISL_3176279, EPI_ISL_3176280, EPI_ISL_3176281, EPI_ISL_3176282, EPI_ISL_3176283, EPI_ISL_3176284, EPI_ISL_3176285, EPI_ISL_3176286, EPI_ISL_3176287, EPI_ISL_3176288, EPI_ISL_3176289, EPI_ISL_3176290, EPI_ISL_3176291, EPI_ISL_3176292, EPI_ISL_3176293, EPI_ISL_3176294, EPI_ISL_3176295, EPI_ISL_3176296, EPI_ISL_3176297, EPI_ISL_3176298, EPI_ISL_3176299, EPI_ISL_3176300, EPI_ISL_3176301, EPI_ISL_3176302, EPI_ISL_3176303, EPI_ISL_3176304, EPI_ISL_3176305, EPI_ISL_3176306, EPI_ISL_3176307, EPI_ISL_3176308, EPI_ISL_3176309, EPI_ISL_3176310, EPI_ISL_3176311, EPI_ISL_3176312, EPI_ISL_3176313, EPI_ISL_3176314, EPI_ISL_3176315, EPI_ISL_3176316, EPI_ISL_3176317, EPI_ISL_3176318, EPI_ISL_3176319, EPI_ISL_3176320, EPI_ISL_3176321, EPI_ISL_3176322, EPI_ISL_3176323, EPI_ISL_3176324, EPI_ISL_3176325, EPI_ISL_3176326, EPI_ISL_3176327, EPI_ISL_3176328, EPI_ISL_3176329, EPI_ISL_3176330, EPI_ISL_3176331, EPI_ISL_3176332, EPI_ISL_3176333, EPI_ISL_3176334, EPI_ISL_3176335, EPI_ISL_3176336, EPI_ISL_3176337, EPI_ISL_3176338, EPI_ISL_3176339, EPI_ISL_3176340, EPI_ISL_3176341, EPI_ISL_3176342, EPI_ISL_3176343, EPI_ISL_3176344, EPI_ISL_3176345, EPI_ISL_3176346, EPI_ISL_3176347, EPI_ISL_3176348, EPI_ISL_3176349, EPI_ISL_3176350, EPI_ISL_3176351, EPI_ISL_3176352, EPI_ISL_3176353, EPI_ISL_3176354, EPI_ISL_3176355, EPI_ISL_3176356, EPI_ISL_3176357, EPI_ISL_3176358, EPI_ISL_3176359, EPI_ISL_3176360, EPI_ISL_3176361, EPI_ISL_3176362, EPI_ISL_3176363, EPI_ISL_3176364, EPI_ISL_3176365, EPI_ISL_3176366, EPI_ISL_3176367, EPI_ISL_3176368, EPI_ISL_3176369, EPI_ISL_3176370, EPI_ISL_3176371, EPI_ISL_3176372, EPI_ISL_3176373, EPI_ISL_3176374, EPI_ISL_3176376, EPI_ISL_3176377, EPI_ISL_3176378, EPI_ISL_3176379, EPI_ISL_3176380, EPI_ISL_3176381, EPI_ISL_3176382, EPI_ISL_3176383, EPI_ISL_3176384, EPI_ISL_3176385, EPI_ISL_3176386, EPI_ISL_3176387, EPI_ISL_3176388, EPI_ISL_3176389, EPI_ISL_3176390, EPI_ISL_3176391, EPI_ISL_3176392, EPI_ISL_3176393, EPI_ISL_3176394, EPI_ISL_3176395, EPI_ISL_3176396, EPI_ISL_3176397, EPI_ISL_3176398, EPI_ISL_3176399, EPI_ISL_3176400, EPI_ISL_3176401, EPI_ISL_3176402, EPI_ISL_3176403, EPI_ISL_3176404, EPI_ISL_3176405, EPI_ISL_3176406, EPI_ISL_3176407, EPI_ISL_3176408, EPI_ISL_3176409, EPI_ISL_3176410, EPI_ISL_3176411, EPI_ISL_3176412, EPI_ISL_3176413, EPI_ISL_3176414, EPI_ISL_3176415, EPI_ISL_3176416, EPI_ISL_3176417, EPI_ISL_3176418, EPI_ISL_3176419, EPI_ISL_3176420, EPI_ISL_3176421, EPI_ISL_3176422, EPI_ISL_3176423, EPI_ISL_3176424, EPI_ISL_3176425, EPI_ISL_3176426, EPI_ISL_3176427, EPI_ISL_3176428, EPI_ISL_3176429, EPI_ISL_3176430, EPI_ISL_3176431, EPI_ISL_3176432, EPI_ISL_3176433, EPI_ISL_3176434, EPI_ISL_3176435, EPI_ISL_3176436, EPI_ISL_3176437, EPI_ISL_3176438, EPI_ISL_3176439, EPI_ISL_3176440, EPI_ISL_3176441 | | | |
| see above | West of Scotland Specialist Virology Centre, NHSGGC / MRC-University of Glasgow Centre for Virus Research | COVID-19 Genomics UK (COG-UK) Consortium | Ana da Silva Filipe, Natasha Johnson, Kathy Smollett, Daniel Mair, Stephen Carmichael, Alice Broos, Lily Tong, Jenna Nichols, Kyriaki Nomikou; Sarah McDonald; Ioulia Tsatsani, Richard Orton, Joseph Hughes, Sreenu Vattipally, David L Robertson; Alasdair MacLean, Rory Gunson; Shirif Shaaban, Matthew Holden; Rachel Blacow, Guy Mollett, Kathy Li, James Shepherd, Antonia Ho, Emma Thomson |
| EPI_ISL_3176442 | Virology Department, Royal Infirmary of Edinburgh, NHS Lothian / School of Biological Sciences, University of Edinburgh | COVID-19 Genomics UK (COG-UK) Consortium | McHugh M, Dewar R, Cotton S, Gallagher A, Maloney D, Fernandez G, O'Toole Á, Scher E, Hill V, McCrone JT, Colquhoun R, Yu X, Jackson B, Rambaut A, Templeton K |
| EPI_ISL_3176443, EPI_ISL_3176444, EPI_ISL_3176445, EPI_ISL_3176446, EPI_ISL_3176447, EPI_ISL_3176448, EPI_ISL_3176449, EPI_ISL_3176450, EPI_ISL_3176451, EPI_ISL_3176452, EPI_ISL_3176453, EPI_ISL_3176454, EPI_ISL_3176455, EPI_ISL_3176456, EPI_ISL_3176457, EPI_ISL_3176458, EPI_ISL_3176459, EPI_ISL_3176460, EPI_ISL_3176461, EPI_ISL_3176462, EPI_ISL_3176463, EPI_ISL_3176464, EPI_ISL_3176465, EPI_ISL_3176466, EPI_ISL_3176467, EPI_ISL_3176468, EPI_ISL_3176469, EPI_ISL_3176470, EPI_ISL_3176471, EPI_ISL_3176472, EPI_ISL_3176473, EPI_ISL_3176474, EPI_ISL_3176475, EPI_ISL_3176476, EPI_ISL_3176477, EPI_ISL_3176478, EPI_ISL_3176479, EPI_ISL_3176480, EPI_ISL_3176481, EPI_ISL_3176482, EPI_ISL_3176483, EPI_ISL_3176484, EPI_ISL_3176485, EPI_ISL_3176486, EPI_ISL_3176487, EPI_ISL_3176488, EPI_ISL_3176489, EPI_ISL_3176490, EPI_ISL_3176491 | | | |
| see above | The Roslin Institute and R(D)SVS, University of Edinburgh / Virology Department, Royal Infirmary of Edinburgh, NHS Lothian / School of Biological Sciences, University of Edinburgh | COVID-19 Genomics UK (COG-UK) Consortium | Warr A, Newman, C, Tait-Burkhard C, McHugh M, Dewar R, Cotton S, Gallagher A, Maloney D, Fernandez G, O'Toole Á, Scher E, Hill V, McCrone JT, Colquhoun R, Yu X, Jackson B, Rambaut A, Templeton K |
| EPI_ISL_3176492, EPI_ISL_3176493, EPI_ISL_3176494, EPI_ISL_3176495, EPI_ISL_3176496, EPI_ISL_3176497, EPI_ISL_3176498, EPI_ISL_3176499, EPI_ISL_3176500, EPI_ISL_3176501, EPI_ISL_3176502, EPI_ISL_3176503, EPI_ISL_3176504, EPI_ISL_3176505, EPI_ISL_3176506, EPI_ISL_3176507, EPI_ISL_3176508, EPI_ISL_3176509, EPI_ISL_3176510, EPI_ISL_3176511, EPI_ISL_3176512, EPI_ISL_3176513, EPI_ISL_3176514, EPI_ISL_3176515, EPI_ISL_3176516, EPI_ISL_3176517, EPI_ISL_3176518, EPI_ISL_3176519, EPI_ISL_3176520, EPI_ISL_3176521, EPI_ISL_3176522, EPI_ISL_3176523, EPI_ISL_3176524, EPI_ISL_3176525, EPI_ISL_3176526, EPI_ISL_3176527, EPI_ISL_3176528, EPI_ISL_3176529, EPI_ISL_3176530, EPI_ISL_3176531, EPI_ISL_3176532, EPI_ISL_3176533, EPI_ISL_3176534, EPI_ISL_3176535, EPI_ISL_3176536, EPI_ISL_3176537, EPI_ISL_3176538, EPI_ISL_3176539, EPI_ISL_3176540, EPI_ISL_3176541, EPI_ISL_3176542, EPI_ISL_3176543, EPI_ISL_3176544, EPI_ISL_3176545, EPI_ISL_3176546, EPI_ISL_3176547, EPI_ISL_3176548, EPI_ISL_3176549, EPI_ISL_3176550, EPI_ISL_3176551, EPI_ISL_3176552, EPI_ISL_3176553, EPI_ISL_3176554, EPI_ISL_3176555, EPI_ISL_3176556, EPI_ISL_3176557, EPI_ISL_3176558, EPI_ISL_3176559, EPI_ISL_3176560, EPI_ISL_3176561, EPI_ISL_3176562, EPI_ISL_3176563, EPI_ISL_3176564, EPI_ISL_3176565, EPI_ISL_3176566, EPI_ISL_3176567, EPI_ISL_3176568, EPI_ISL_3176569, EPI_ISL_3176570, EPI_ISL_3176571, EPI_ISL_3176572, EPI_ISL_3176573, EPI_ISL_3176574, EPI_ISL_3176575, EPI_ISL_3176576, EPI_ISL_3176577, EPI_ISL_3176578, EPI_ISL_3176579, EPI_ISL_3176580, EPI_ISL_3176581, EPI_ISL_3176582, EPI_ISL_3176583, EPI_ISL_3176584, EPI_ISL_3176585, EPI_ISL_3176586, EPI_ISL_3176587, EPI_ISL_3176588, EPI_ISL_3176589, EPI_ISL_3176590, EPI_ISL_3176591, EPI_ISL_3176592, EPI_ISL_3176593, EPI_ISL_3176594, EPI_ISL_3176595, EPI_ISL_3176596, EPI_ISL_3176597, EPI_ISL_3176598, EPI_ISL_3176599, EPI_ISL_3176600, EPI_ISL_3176601, EPI_ISL_3176602, EPI_ISL_3176603, EPI_ISL_3176604 | | | |
| see above | Virology Department, Royal Infirmary of Edinburgh, NHS Lothian / School of Biological Sciences, University of Edinburgh | COVID-19 Genomics UK (COG-UK) Consortium | McHugh M, Dewar R, Cotton S, Gallagher A, Maloney D, Fernandez G, O'Toole Á, Scher E, Hill V, McCrone JT, Colquhoun R, Yu X, Jackson B, Rambaut A, Templeton K |
| EPI_ISL_3176605, EPI_ISL_3176606, EPI_ISL_3176607, EPI_ISL_3176608, EPI_ISL_3176609, EPI_ISL_3176610, EPI_ISL_3176611, EPI_ISL_3176612, EPI_ISL_3176613, EPI_ISL_3176614, EPI_ISL_3176615, EPI_ISL_3176616, EPI_ISL_3176617, EPI_ISL_3176618, EPI_ISL_3176619, EPI_ISL_3176620, EPI_ISL_3176621, EPI_ISL_3176622, EPI_ISL_3176623, EPI_ISL_3176624, EPI_ISL_3176625, EPI_ISL_3176626, EPI_ISL_3176627, EPI_ISL_3176628, EPI_ISL_3176629, EPI_ISL_3176630, EPI_ISL_3176631, EPI_ISL_3176632, EPI_ISL_3176633, EPI_ISL_3176634, EPI_ISL_3176635, EPI_ISL_3176636, EPI_ISL_3176637, EPI_ISL_3176638, EPI_ISL_3176639, EPI_ISL_3176640, EPI_ISL_3176641, EPI_ISL_3176642, EPI_ISL_3176643, EPI_ISL_3176644, EPI_ISL_3176645, EPI_ISL_3176646, EPI_ISL_3176647, EPI_ISL_3176648, EPI_ISL_3176649, EPI_ISL_3176650, EPI_ISL_3176651, EPI_ISL_3176652, EPI_ISL_3176653, EPI_ISL_3176654, EPI_ISL_3176655, EPI_ISL_3176656, EPI_ISL_3176657, EPI_ISL_3176658, EPI_ISL_3176659, EPI_ISL_3176660, EPI_ISL_3176661, EPI_ISL_3176662, EPI_ISL_3176663, EPI_ISL_3176664, EPI_ISL_3176665, EPI_ISL_3176666, EPI_ISL_3176667, EPI_ISL_3176668, EPI_ISL_3176669, EPI_ISL_3176670, EPI_ISL_3176671, EPI_ISL_3176672, EPI_ISL_3176673, EPI_ISL_3176674, EPI_ISL_3176675, EPI_ISL_3176676, EPI_ISL_3176677, EPI_ISL_3176678, EPI_ISL_3176679, EPI_ISL_3176680, EPI_ISL_3176681, EPI_ISL_3176682, EPI_ISL_3176683, EPI_ISL_3176684, EPI_ISL_3176685, EPI_ISL_3176686, EPI_ISL_3176687, EPI_ISL_3176688, EPI_ISL_3176689, EPI_ISL_3176690, EPI_ISL_3176691, EPI_ISL_3176692, EPI_ISL_3176693, EPI_ISL_3176694, EPI_ISL_3176695, EPI_ISL_3176696, EPI_ISL_3176697, EPI_ISL_3176698, EPI_ISL_3176699, EPI_ISL_3176700, EPI_ISL_3176701, EPI_ISL_3176702, EPI_ISL_3176703, EPI_ISL_3176704, EPI_ISL_3176705, EPI_ISL_3176706, EPI_ISL_3176707, EPI_ISL_3176708, EPI_ISL_3176709, EPI_ISL_3176710, EPI_ISL_3176711, EPI_ISL_3176712, EPI_ISL_3176713, EPI_ISL_3176714, EPI_ISL_3176715, EPI_ISL_3176716, EPI_ISL_3176717, EPI_ISL_3176718, EPI_ISL_3176719, EPI_ISL_3176720, EPI_ISL_3176721, EPI_ISL_3176722, EPI_ISL_3176723, EPI_ISL_3176724, EPI_ISL_3176725, EPI_ISL_3176726, EPI_ISL_3176727, EPI_ISL_3176728, EPI_ISL_3176729, EPI_ISL_3176730, EPI_ISL_3176731, EPI_ISL_3176732, EPI_ISL_3176733, EPI_ISL_3176734, EPI_ISL_3176735, EPI_ISL_3176736, EPI_ISL_3176737, EPI_ISL_3176738, EPI_ISL_3176739, EPI_ISL_3176740, EPI_ISL_3176741, EPI_ISL_3176742, EPI_ISL_3176743, EPI_ISL_3176744, EPI_ISL_3176745, EPI_ISL_3176746, EPI_ISL_3176747, EPI_ISL_3176748, EPI_ISL_3176749, EPI_ISL_3176750, EPI_ISL_3176751, EPI_ISL_3176752, EPI_ISL_3176753, EPI_ISL_3176754, EPI_ISL_3176755, EPI_ISL_3176756, EPI_ISL_3176757, EPI_ISL_3176758, EPI_ISL_3176759, EPI_ISL_3176760, EPI_ISL_3176761, EPI_ISL_3176762, EPI_ISL_3176763, EPI_ISL_3176764, EPI_ISL_3176765, EPI_ISL_3176766, EPI_ISL_3176767, EPI_ISL_3176768, EPI_ISL_3176769, EPI_ISL_3176770, EPI_ISL_3176771, EPI_ISL_3176772, EPI_ISL_3176773, EPI_ISL_3176774, EPI_ISL_3176775, EPI_ISL_3176776, EPI_ISL_3176777, EPI_ISL_3176778, EPI_ISL_3176779, EPI_ISL_3176780, EPI_ISL_3176781, EPI_ISL_3176782, EPI_ISL_3176783, EPI_ISL_3176784, EPI_ISL_3176785, EPI_ISL_3176786, EPI_ISL_3176787, EPI_ISL_3176788, EPI_ISL_3176789, EPI_ISL_3176790, EPI_ISL_3176791, EPI_ISL_3176792, EPI_ISL_3176793, EPI_ISL_3176794, EPI_ISL_3176795, EPI_ISL_3176796, EPI_ISL_3176797, EPI_ISL_3176798, EPI_ISL_3176799, EPI_ISL_3176800, EPI_ISL_3176801, EPI_ISL_3176802, EPI_ISL_3176803, EPI_ISL_3176804, EPI_ISL_3176805, EPI_ISL_3176806, EPI_ISL_3176807, EPI_ISL_3176808, EPI_ISL_3176809, EPI_ISL_3176810, EPI_ISL_3176811, EPI_ISL_3176812, EPI_ISL_3176813, EPI_ISL_3176814, EPI_ISL_3176815, EPI_ISL_3176816, EPI_ISL_3176817, EPI_ISL_3176818, EPI_ISL_3176819, EPI_ISL_3176820, EPI_ISL_3176821, EPI_ISL_3176822, EPI_ISL_3176823, EPI_ISL_3176824, EPI_ISL_3176825, EPI_ISL_3176826, EPI_ISL_3176827, EPI_ISL_3176828, EPI_ISL_3176829, EPI_ISL_3176830, EPI_ISL_3176831, EPI_ISL_3176832, EPI_ISL_3176833, EPI_ISL_3176834, EPI_ISL_3176835, EPI_ISL_3176836, EPI_ISL_3176837, EPI_ISL_3176838, EPI_ISL_3176839, EPI_ISL_3176840, EPI_ISL_3176841, EPI_ISL_3176842, EPI_ISL_3176843 | | | |
| see above | University College London, Great Ormond Street Hospital for Children NHS Foundation Trust, Imperial College Healthcare NHS Trust | COVID-19 Genomics UK (COG-UK) Consortium | Sergi Castellano, Rachel Williams, Mark Kristiansen, Sunando Roy, Charlotte Williams, Nadua Bayzid, Marius Cotic, Helena Tutill, Patricia Dyal, Judith Breuer |
| EPI_ISL_3176844, EPI_ISL_3176845, EPI_ISL_3176846, EPI_ISL_3176847, EPI_ISL_3176848, EPI_ISL_3176849, EPI_ISL_3176850, EPI_ISL_3176851, EPI_ISL_3176852, EPI_ISL_3176853, EPI_ISL_3176854, EPI_ISL_3176855, EPI_ISL_3176856, EPI_ISL_3176857, EPI_ISL_3176858, EPI_ISL_3176859, EPI_ISL_3176860, EPI_ISL_3176861, EPI_ISL_3176862, EPI_ISL_3176863, EPI_ISL_3176864, EPI_ISL_3176865, EPI_ISL_3176866, EPI_ISL_3176867, EPI_ISL_3176868, EPI_ISL_3176869, EPI_ISL_3176870, EPI_ISL_3176871, EPI_ISL_3176872, EPI_ISL_3176873, EPI_ISL_3176874, EPI_ISL_3176875, EPI_ISL_3176876, EPI_ISL_3176877, EPI_ISL_3176878, EPI_ISL_3176879, EPI_ISL_3176880, EPI_ISL_3176881, EPI_ISL_3176882, EPI_ISL_3176883, EPI_ISL_3176884, EPI_ISL_3176885, EPI_ISL_3176886, EPI_ISL_3176887, EPI_ISL_3176888, EPI_ISL_3176889, EPI_ISL_3176890, EPI_ISL_3176891, EPI_ISL_3176892, EPI_ISL_3176893, EPI_ISL_3176894, EPI_ISL_3176895, EPI_ISL_3176896, EPI_ISL_3176897, EPI_ISL_3176898, EPI_ISL_3176899, EPI_ISL_3176900, EPI_ISL_3176901, EPI_ISL_3176902, EPI_ISL_3176903, EPI_ISL_3176904, EPI_ISL_3176905, EPI_ISL_3176906, EPI_ISL_3176907, EPI_ISL_3176908, EPI_ISL_3176909, EPI_ISL_3176910, EPI_ISL_3176911, EPI_ISL_3176912, EPI_ISL_3176913, EPI_ISL_3176914, EPI_ISL_3176915, EPI_ISL_3176916, EPI_ISL_3176917, EPI_ISL_3176918, EPI_ISL_3176919, EPI_ISL_3176920, EPI_ISL_3176921, EPI_ISL_3176922, EPI_ISL_3176923, EPI_ISL_3176924, EPI_ISL_3176925, EPI_ISL_3176926, EPI_ISL_3176927, EPI_ISL_3176928, EPI_ISL_3176929, EPI_ISL_3176930, EPI_ISL_3176931, EPI_ISL_3176932, EPI_ISL_3176933, EPI_ISL_3176934, EPI_ISL_3176935, EPI_ISL_3176936, EPI_ISL_3176937, EPI_ISL_3176938, EPI_ISL_3176939, EPI_ISL_3176940, EPI_ISL_3176941, EPI_ISL_3176942, EPI_ISL_3176943, EPI_ISL_3176944, EPI_ISL_3176945, EPI_ISL_3176946, EPI_ISL_3176947, EPI_ISL_3176948, EPI_ISL_3176949, EPI_ISL_3176950, EPI_ISL_3176951, EPI_ISL_3176952, EPI_ISL_3176953, EPI_ISL_3176954, EPI_ISL_3176955, EPI_ISL_3176956, EPI_ISL_3176957, EPI_ISL_3176958, EPI_ISL_3176959, EPI_ISL_3176960, EPI_ISL_3176961, EPI_ISL_3176962, EPI_ISL_3176963, EPI_ISL_3176964, EPI_ISL_3176965, EPI_ISL_3176966, EPI_ISL_3176967, EPI_ISL_3176968, EPI_ISL_3176969, EPI_ISL_3176970, EPI_ISL_3176971, EPI_ISL_3176972, EPI_ISL_3176973, EPI_ISL_3176974, EPI_ISL_3176975, EPI_ISL_3176976, EPI_ISL_3176977, EPI_ISL_3176978, EPI_ISL_3176979, EPI_ISL_3176980, EPI_ISL_3176981, EPI_ISL_3176982, EPI_ISL_3176983, EPI_ISL_3176984, EPI_ISL_3176985, EPI_ISL_3176986, EPI_ISL_3176987, EPI_ISL_3176988, EPI_ISL_3176989, EPI_ISL_3176990, EPI_ISL_3176991, EPI_ISL_3176992, EPI_ISL_3176993, EPI_ISL_3176994, EPI_ISL_3176995, EPI_ISL_3176996, EPI_ISL_3176997, EPI_ISL_3176998, EPI_ISL_3176999, EPI_ISL_3177000, EPI_ISL_3177001, EPI_ISL_3177002, EPI_ISL_3177003, EPI_ISL_3177004, EPI_ISL_3177005, EPI_ISL_3177006, EPI_ISL_3177007, EPI_ISL_3177008, EPI_ISL_3177009, EPI_ISL_3177010, EPI_ISL_3177011, EPI_ISL_3177012, EPI_ISL_3177013, EPI_ISL_3177014, EPI_ISL_3177015, EPI_ISL_3177016, EPI_ISL_3177017, EPI_ISL_3177018, EPI_ISL_3177019, EPI_ISL_3177020, EPI_ISL_3177021, EPI_ISL_3177022, EPI_ISL_3177023, EPI_ISL_3177024, EPI_ISL_3177025, EPI_ISL_3177026, EPI_ISL_3177027, EPI_ISL_3177028, EPI_ISL_3177029, EPI_ISL_3177030, EPI_ISL_3177031, EPI_ISL_3177032, EPI_ISL_3177033, EPI_ISL_3177034, EPI_ISL_3177035, EPI_ISL_3177037, EPI_ISL_3177039, EPI_ISL_3177041, EPI_ISL_3177043, EPI_ISL_3177045, EPI_ISL_3177046, EPI_ISL_3177048, EPI_ISL_3177050, EPI_ISL_3177052, EPI_ISL_3177054, EPI_ISL_3177056, EPI_ISL_3177058, EPI_ISL_3177060, EPI_ISL_3177062, EPI_ISL_3177064, EPI_ISL_3177066, EPI_ISL_3177067, EPI_ISL_3177069, EPI_ISL_3177071, EPI_ISL_3177073, EPI_ISL_3177075, EPI_ISL_3177077, EPI_ISL_3177078, EPI_ISL_3177080, EPI_ISL_3177082, EPI_ISL_3177083, EPI_ISL_3177084, EPI_ISL_3177085, EPI_ISL_3177086, EPI_ISL_3177087, EPI_ISL_3177088, EPI_ISL_3177089, EPI_ISL_3177090, EPI_ISL_3177091, EPI_ISL_3177092, EPI_ISL_3177093, EPI_ISL_3177094, EPI_ISL_3177095, EPI_ISL_3177096, EPI_ISL_3177097, EPI_ISL_3177098, EPI_ISL_3177099, EPI_ISL_3177100, EPI_ISL_3177101, EPI_ISL_3177102, EPI_ISL_3177103, EPI_ISL_3177104, EPI_ISL_3177105, EPI_ISL_3177106, EPI_ISL_3177107, EPI_ISL_3177108, EPI_ISL_3177109, EPI_ISL_3177110, EPI_ISL_3177111, EPI_ISL_3177112, EPI_ISL_3177113, EPI_ISL_3177114, EPI_ISL_3177115, EPI_ISL_3177116, EPI_ISL_3177117, EPI_ISL_3177118, EPI_ISL_3177119, EPI_ISL_3177120, EPI_ISL_3177121, EPI_ISL_3177122, EPI_ISL_3177123, EPI_ISL_3177124, EPI_ISL_3177125, EPI_ISL_3177126, EPI_ISL_3177127, EPI_ISL_3177128, EPI_ISL_3177129, EPI_ISL_3177130, EPI_ISL_3177131, EPI_ISL_3177132, EPI_ISL_3177133, EPI_ISL_3177134, EPI_ISL_3177135, EPI_ISL_3177136, EPI_ISL_3177137, EPI_ISL_3177138, EPI_ISL_3177139, EPI_ISL_3177140, EPI_ISL_3177141, EPI_ISL_3177142, EPI_ISL_3177143, EPI_ISL_3177144, EPI_ISL_3177145, EPI_ISL_3177146, EPI_ISL_3177147, EPI_ISL_3177148, EPI_ISL_3177149, EPI_ISL_3177150, EPI_ISL_3177151, EPI_ISL_3177152, EPI_ISL_3177153, EPI_ISL_3177154, EPI_ISL_3177155, EPI_ISL_3177156, EPI_ISL_3177157, EPI_ISL_3177158, EPI_ISL_3177159, EPI_ISL_3177160, EPI_ISL_3177161, EPI_ISL_3177162, EPI_ISL_3177163, EPI_ISL_3177164, EPI_ISL_3177165, EPI_ISL_3177166, EPI_ISL_3177167, EPI_ISL_3177168, EPI_ISL_3177169, EPI_ISL_3177170, EPI_ISL_3177171, EPI_ISL_3177172, EPI_ISL_3177173, EPI_ISL_3177174, EPI_ISL_3177175, EPI_ISL_3177176, EPI_ISL_3177177, EPI_ISL_3177178, EPI_ISL_3177179, EPI_ISL_3177180, EPI_ISL_3177181, EPI_ISL_3177182, EPI_ISL_3177183, EPI_ISL_3177184, EPI_ISL_3177185, EPI_ISL_3177186, EPI_ISL_3177187, EPI_ISL_3177188, EPI_ISL_3177189, EPI_ISL_3177190, EPI_ISL_3177191, EPI_ISL_3177192, EPI_ISL_3177193, EPI_ISL_3177194, EPI_ISL_3177195, EPI_ISL_3177196, EPI_ISL_3177197, EPI_ISL_3177198, EPI_ISL_3177199, EPI_ISL_3177200, EPI_ISL_3177201, EPI_ISL_3177202, EPI_ISL_3177203, EPI_ISL_3177204, EPI_ISL_3177205, EPI_ISL_3177206, EPI_ISL_3177207, EPI_ISL_3177208, EPI_ISL_3177209, EPI_ISL_3177210, EPI_ISL_3177211, EPI_ISL_3177212, EPI_ISL_3177213, EPI_ISL_3177214, EPI_ISL_3177215, EPI_ISL_3177216, EPI_ISL_3177217, EPI_ISL_3177218, EPI_ISL_3177219, EPI_ISL_3177220, EPI_ISL_3177221, EPI_ISL_3177222, EPI_ISL_3177223, EPI_ISL_3177224, EPI_ISL_3177225, EPI_ISL_3177226, EPI_ISL_3177227, EPI_ISL_3177228, EPI_ISL_3177229, EPI_ISL_3177230, EPI_ISL_3177231, EPI_ISL_3177232, EPI_ISL_3177233, EPI_ISL_3177234, EPI_ISL_3177235, EPI_ISL_3177236, EPI_ISL_3177237, EPI_ISL_3177238, EPI_ISL_3177239, EPI_ISL_3177240, EPI_ISL_3177241, EPI_ISL_3177242, EPI_ISL_3177243, EPI_ISL_3177244, EPI_ISL_3177245, EPI_ISL_3177246, EPI_ISL_3177247, EPI_ISL_3177248, EPI_ISL_3177249, EPI_ISL_3177250, EPI_ISL_3177251, EPI_ISL_3177252, EPI_ISL_3177253, EPI_ISL_3177254, EPI_ISL_3177255, EPI_ISL_3177256, EPI_ISL_3177257, EPI_ISL_3177258, EPI_ISL_3177259, EPI_ISL_3177260, EPI_ISL_3177261, EPI_ISL_3177262, EPI_ISL_3177263, EPI_ISL_3177264, EPI_ISL_3177265, EPI_ISL_3177266, EPI_ISL_3177267, EPI_ISL_3177268, EPI_ISL_3177269, EPI_ISL_3177270, EPI_ISL_3177271, EPI_ISL_3177272, EPI_ISL_3177273, EPI_ISL_3177274, EPI_ISL_3177275, EPI_ISL_3177276, EPI_ISL_3177277, EPI_ISL_3177278, EPI_ISL_3177279, EPI_ISL_3177280, EPI_ISL_3177281, EPI_ISL_3177282, EPI_ISL_3177283, EPI_ISL_3177284, EPI_ISL_3177285, EPI_ISL_3177286, EPI_ISL_3177287, EPI_ISL_3177288, EPI_ISL_3177289, EPI_ISL_3177290, EPI_ISL_3177291, EPI_ISL_3177292, EPI_ISL_3177293, EPI_ISL_3177294, EPI_ISL_3177295, EPI_ISL_3177296, EPI_ISL_3177297, EPI_ISL_3177298, EPI_ISL_3177299, EPI_ISL_3177300, EPI_ISL_3177301, EPI_ISL_3177302, EPI_ISL_3177303, EPI_ISL_3177304, EPI_ISL_3177305, EPI_ISL_3177306, EPI_ISL_3177307, EPI_ISL_3177308, EPI_ISL_3177309, EPI_ISL_3177310, EPI_ISL_3177311, EPI_ISL_3177312, EPI_ISL_3177313, EPI_ISL_3177314, EPI_ISL_3177315, EPI_ISL_3177316, EPI_ISL_3177317, EPI_ISL_3177318, EPI_ISL_3177319, EPI_ISL_3177320, EPI_ISL_3177321, EPI_ISL_3177322, EPI_ISL_3177323, EPI_ISL_3177324, EPI_ISL_3177325, EPI_ISL_3177326, EPI_ISL_3177327, EPI_ISL_3177328, EPI_ISL_3177329, EPI_ISL_3177330, EPI_ISL_3177331, EPI_ISL_3177332, EPI_ISL_3177333, EPI_ISL_3177334, EPI_ISL_3177335, EPI_ISL_3177336, EPI_ISL_3177337, EPI_ISL_3177338, EPI_ISL_3177339, EPI_ISL_3177340, EPI_ISL_3177341, EPI_ISL_3177342, EPI_ISL_3177343, EPI_ISL_3177344, EPI_ISL_3177345, EPI_ISL_3177346, EPI_ISL_3177347, EPI_ISL_3177348, EPI_ISL_3177349, EPI_ISL_3177350, EPI_ISL_3177351, EPI_ISL_3177352, EPI_ISL_3177353, EPI_ISL_3177354, EPI_ISL_3177355, EPI_ISL_3177356, EPI_ISL_3177357, EPI_ISL_3177358, EPI_ISL_3177359, EPI_ISL_3177360, EPI_ISL_3177361, EPI_ISL_3177362, EPI_ISL_3177363, EPI_ISL_3177364, EPI_ISL_3177365, EPI_ISL_3177366, EPI_ISL_3177367, EPI_ISL_3177368, EPI_ISL_3177369, EPI_ISL_3177370, EPI_ISL_3177371, EPI_ISL_3177372, EPI_ISL_3177373, EPI_ISL_3177374, EPI_ISL_3177375, EPI_ISL_3177376, EPI_ISL_3177377, EPI_ISL_3177378, EPI_ISL_3177379, EPI_ISL_3177380, EPI_ISL_3177381, EPI_ISL_3177382, EPI_ISL_3177383, EPI_ISL_3177384, EPI_ISL_3177385, EPI_ISL_3177386, EPI_ISL_3177387, EPI_ISL_3177388, EPI_ISL_3177389, EPI_ISL_3177390, EPI_ISL_3177391, EPI_ISL_3177392, EPI_ISL_3177393, EPI_ISL_3177394, EPI_ISL_3177395, EPI_ISL_3177396, EPI_ISL_3177397, EPI_ISL_3177398, EPI_ISL_3177399, EPI_ISL_3177400, EPI_ISL_3177401, EPI_ISL_3177402, EPI_ISL_3177403, EPI_ISL_3177404, EPI_ISL_3177405, EPI_ISL_3177406, EPI_ISL_3177407, EPI_ISL_3177408, EPI_ISL_3177409, EPI_ISL_3177410, EPI_ISL_3177411, EPI_ISL_3177412, EPI_ISL_3177413, EPI_ISL_3177414, EPI_ISL_3177415, EPI_ISL_3177416, EPI_ISL_3177417, EPI_ISL_3177418, EPI_ISL_3177419, EPI_ISL_3177420, EPI_ISL_3177421, EPI_ISL_3177422, EPI_ISL_3177423, EPI_ISL_3177424, EPI_ISL_3177425, EPI_ISL_3177426, EPI_ISL_3177427, EPI_ISL_3177428, EPI_ISL_3177429, EPI_ISL_3177430, EPI_ISL_3177431, EPI_ISL_3177432, EPI_ISL_3177433, EPI_ISL_3177434, EPI_ISL_3177435, EPI_ISL_3177436, EPI_ISL_3177437, EPI_ISL_3177438, EPI_ISL_3177439, EPI_ISL_3177440, EPI_ISL_3177441, EPI_ISL_3177442, EPI_ISL_3177443, EPI_ISL_3177444, EPI_ISL_3177445, EPI_ISL_3177446, EPI_ISL_3177447, EPI_ISL_3177448, EPI_ISL_3177449, EPI_ISL_3177450, EPI_ISL_3177451, EPI_ISL_3177452, EPI_ISL_3177453, EPI_ISL_3177454, EPI_ISL_3177455, EPI_ISL_3177456, EPI_ISL_3177457, EPI_ISL_3177458, EPI_ISL_3177459, EPI_ISL_3177460, EPI_ISL_3177461, EPI_ISL_3177462, EPI_ISL_3177463, EPI_ISL_3177464, EPI_ISL_3177465, EPI_ISL_3177466, EPI_ISL_3177467, EPI_ISL_3177468, EPI_ISL_3177469, EPI_ISL_3177470, EPI_ISL_3177471, EPI_ISL_3177472, EPI_ISL_3177473, EPI_ISL_3177474, EPI_ISL_3177475, EPI_ISL_3177476, EPI_ISL_3177477, EPI_ISL_3177478, EPI_ISL_3177479, EPI_ISL_3177480, EPI_ISL_3177481, EPI_ISL_3177482, EPI_ISL_3177483, EPI_ISL_3177484, EPI_ISL_3177485, EPI_ISL_3177486, EPI_ISL_3177487, EPI_ISL_3177488, EPI_ISL_3177489, EPI_ISL_3177490, EPI_ISL_3177491, EPI_ISL_3177492, EPI_ISL_3177493, EPI_ISL_3177494, EPI_ISL_3177495, EPI_ISL_3177496, EPI_ISL_3177497, EPI_ISL_3177498, EPI_ISL_3177499, EPI_ISL_3177500, EPI_ISL_3177501, EPI_ISL_3177502, EPI_ISL_3177503, EPI_ISL_3177504, EPI_ISL_3177505, EPI_ISL_3177506, EPI_ISL_3177507, EPI_ISL_3177508, EPI_ISL_3177509, EPI_ISL_3177510, EPI_ISL_3177511, EPI_ISL_3177512, EPI_ISL_3177513, EPI_ISL_3177514, EPI_ISL_3177515, EPI_ISL_3177516, EPI_ISL_3177517, EPI_ISL_3177518, EPI_ISL_3177519, EPI_ISL_3177520, EPI_ISL_3177521, EPI_ISL_3177522, EPI_ISL_3177523, EPI_ISL_3177524, EPI_ISL_3177525, EPI_ISL_3177526, EPI_ISL_3177527, EPI_ISL_3177528, EPI_ISL_3177529, EPI_ISL_3177530, EPI_ISL_3177531, EPI_ISL_3177532, EPI_ISL_3177533, EPI_ISL_3177534, EPI_ISL_3177535, EPI_ISL_3177536, EPI_ISL_3177537, EPI_ISL_3177538, EPI_ISL_3177539, EPI_ISL_3177540, EPI_ISL_3177541, EPI_ISL_3177542, EPI_ISL_3177543, EPI_ISL_3177544, EPI_ISL_3177545, EPI_ISL_3177546, EPI_ISL_3177547, EPI_ISL_3177548, EPI_ISL_3177549, EPI_ISL_3177550, EPI_ISL_3177551, EPI_ISL_3177552, EPI_ISL_3177553, EPI_ISL_3177554, EPI_ISL_3177555, EPI_ISL_3177556, EPI_ISL_3177557, EPI_ISL_3177558, EPI_ISL_3177559, EPI_ISL_3177560, EPI_ISL_3177561, EPI_ISL_3177562, EPI_ISL_3177563, EPI_ISL_3177564, EPI_ISL_3177565, EPI_ISL_3177566, EPI_ISL_3177567, EPI_ISL_3177568, EPI_ISL_3177569, EPI_ISL_3177570, EPI_ISL_3177571, EPI_ISL_3177572, EPI_ISL_3177573, EPI_ISL_3177574, EPI_ISL_3177575, EPI_ISL_3177576, EPI_ISL_3177577, EPI_ISL_3177578, EPI_ISL_3177579, EPI_ISL_3177580, EPI_ISL_3177581, EPI_ISL_3177582, EPI_ISL_3177583, EPI_ISL_3177584, EPI_ISL_3177585, EPI_ISL_3177586, EPI_ISL_3177587, EPI_ISL_3177588, EPI_ISL_3177589, EPI_ISL_3177590, EPI_ISL_3177591, EPI_ISL_3177592, EPI_ISL_3177593, EPI_ISL_3177594, EPI_ISL_3177595, EPI_ISL_3177596, EPI_ISL_3177597, EPI_ISL_3177598, EPI_ISL_3177599, EPI_ISL_3177600, EPI_ISL_3177601, EPI_ISL_3177602, EPI_ISL_3177603, EPI_ISL_3177604, EPI_ISL_3177605, EPI_ISL_3177606, EPI_ISL_3177607, EPI_ISL_3177608, EPI_ISL_3177609, EPI_ISL_3177610, EPI_ISL_3177611, EPI_ISL_3177612, EPI_ISL_3177613, EPI_ISL_3177614, EPI_ISL_3177615, EPI_ISL_3177616, EPI_ISL_3177617, EPI_ISL_3177618, EPI_ISL_3177619, EPI_ISL_3177620, EPI_ISL_3177621, EPI_ISL_3177622, EPI_ISL_3177623, EPI_ISL_3177624, EPI_ISL_3177625, EPI_ISL_3177626, EPI_ISL_3177627, EPI_ISL_3177628, EPI_ISL_3177629, EPI_ISL_3177630, EPI_ISL_3177631, EPI_ISL_3177632, EPI_ISL_3177633, EPI_ISL_3177634, EPI_ISL_3177635, EPI_ISL_3177636, EPI_ISL_3177637, EPI_ISL_3177638, EPI_ISL_3177639, EPI_ISL_3177640, EPI_ISL_3177641, EPI_ISL_3177642, EPI_ISL_3177643, EPI_ISL_3177644, EPI_ISL_3177645, EPI_ISL_3177646, EPI_ISL_3177647, EPI_ISL_3177648, EPI_ISL_3177649, EPI_ISL_3177650, EPI_ISL_3177651 | | | |
| see above | Northumbria University / South Tees Hospitals NHS Foundation Trust / North Cumbria Integrated Care NHS Foundation Trust / North Tees and Hartlepool NHS Foundation Trust / Newcastle Hospitals NHS Foundation Trust | COVID-19 Genomics UK (COG-UK) Consortium | Darren L Smith,Andrew Nelson,Matthew Bashton,Greg R Young,Joshua Loh,John Allan,Mohammad A Tariq,Giles S Holt,Gary Black,Wen C Yew,Lynn Dover,Paul Baker,Steve Liggett,Sarah Essex,Jane Greenaway,Debra Padgett,Clive Graham,Garren Scott,Edward Barton,Emma Swindells,Brendan Payne,Jennifer Collins,Yusri Taha,Gary Eltringham |
| EPI_ISL_3177652, EPI_ISL_3177653, EPI_ISL_3177654, EPI_ISL_3177655, EPI_ISL_3177656, EPI_ISL_3177657, EPI_ISL_3177658, EPI_ISL_3177659, EPI_ISL_3177660, EPI_ISL_3177661, EPI_ISL_3177662, EPI_ISL_3177663, EPI_ISL_3177664, EPI_ISL_3177665, EPI_ISL_3177666, EPI_ISL_3177667, EPI_ISL_3177668, EPI_ISL_3177669, EPI_ISL_3177670, EPI_ISL_3177671, EPI_ISL_3177672, EPI_ISL_3177673, EPI_ISL_3177674, EPI_ISL_3177675, EPI_ISL_3177676, EPI_ISL_3177677, EPI_ISL_3177678, EPI_ISL_3177679, EPI_ISL_3177680, EPI_ISL_3177681, EPI_ISL_3177682, EPI_ISL_3177683, EPI_ISL_3177684, EPI_ISL_3177685, EPI_ISL_3177686, EPI_ISL_3177687, EPI_ISL_3177688, EPI_ISL_3177689, EPI_ISL_3177690, EPI_ISL_3177691, EPI_ISL_3177692, EPI_ISL_3177693, EPI_ISL_3177694, EPI_ISL_3177695, EPI_ISL_3177696, EPI_ISL_3177697, EPI_ISL_3177698, EPI_ISL_3177699, EPI_ISL_3177700, EPI_ISL_3177701, EPI_ISL_3177702, EPI_ISL_3177703, EPI_ISL_3177704, EPI_ISL_3177705, EPI_ISL_3177706, EPI_ISL_3177707, EPI_ISL_3177708, EPI_ISL_3177709, EPI_ISL_3177710, EPI_ISL_3177711, EPI_ISL_3177712, EPI_ISL_3177713, EPI_ISL_3177714, EPI_ISL_3177715, EPI_ISL_3177716, EPI_ISL_3177717, EPI_ISL_3177718, EPI_ISL_3177719, EPI_ISL_3177720, EPI_ISL_3177721, EPI_ISL_3177722, EPI_ISL_3177723, EPI_ISL_3177724, EPI_ISL_3177725, EPI_ISL_3177726, EPI_ISL_3177727, EPI_ISL_3177728, EPI_ISL_3177729, EPI_ISL_3177730, EPI_ISL_3177731, EPI_ISL_3177732, EPI_ISL_3177733, EPI_ISL_3177734, EPI_ISL_3177735, EPI_ISL_3177736, EPI_ISL_3177737, EPI_ISL_3177738, EPI_ISL_3177739, EPI_ISL_3177740, EPI_ISL_3177741, EPI_ISL_3177742, EPI_ISL_3177743, EPI_ISL_3177744, EPI_ISL_3177745, EPI_ISL_3177746, EPI_ISL_3177747, EPI_ISL_3177748, EPI_ISL_3177749, EPI_ISL_3177750, EPI_ISL_3177751, EPI_ISL_3177752, EPI_ISL_3177753, EPI_ISL_3177754, EPI_ISL_3177755, EPI_ISL_3177756, EPI_ISL_3177757, EPI_ISL_3177758, EPI_ISL_3177759, EPI_ISL_3177760, EPI_ISL_3177761, EPI_ISL_3177762, EPI_ISL_3177763, EPI_ISL_3177764, EPI_ISL_3177765, EPI_ISL_3177766, EPI_ISL_3177767, EPI_ISL_3177768, EPI_ISL_3177769, EPI_ISL_3177770, EPI_ISL_3177771, EPI_ISL_3177772, EPI_ISL_3177773, EPI_ISL_3177774, EPI_ISL_3177775, EPI_ISL_3177776, EPI_ISL_3177777, EPI_ISL_3177778, EPI_ISL_3177779, EPI_ISL_3177780, EPI_ISL_3177781, EPI_ISL_3177782, EPI_ISL_3177783, EPI_ISL_3177784, EPI_ISL_3177785, EPI_ISL_3177786, EPI_ISL_3177787, EPI_ISL_3177788, EPI_ISL_3177789, EPI_ISL_3177790, EPI_ISL_3177791, EPI_ISL_3177792, EPI_ISL_3177793, EPI_ISL_3177794, EPI_ISL_3177795, EPI_ISL_3177796, EPI_ISL_3177797, EPI_ISL_3177798, EPI_ISL_3177799, EPI_ISL_3177800, EPI_ISL_3177801, EPI_ISL_3177802, EPI_ISL_3177803, EPI_ISL_3177804, EPI_ISL_3177805, EPI_ISL_3177806, EPI_ISL_3177807, EPI_ISL_3177808, EPI_ISL_3177809, EPI_ISL_3177810, EPI_ISL_3177811, EPI_ISL_3177812, EPI_ISL_3177813, EPI_ISL_3177814, EPI_ISL_3177815, EPI_ISL_3177816, EPI_ISL_3177817, EPI_ISL_3177818, EPI_ISL_3177819, EPI_ISL_3177820, EPI_ISL_3177821, EPI_ISL_3177822, EPI_ISL_3177823, EPI_ISL_3177824, EPI_ISL_3177825, EPI_ISL_3177826, EPI_ISL_3177827, EPI_ISL_3177828, EPI_ISL_3177829, EPI_ISL_3177830, EPI_ISL_3177831, EPI_ISL_3177832, EPI_ISL_3177833, EPI_ISL_3177834, EPI_ISL_3177835, EPI_ISL_3177836, EPI_ISL_3177837, EPI_ISL_3177838, EPI_ISL_3177839, EPI_ISL_3177840, EPI_ISL_3177841, EPI_ISL_3177842, EPI_ISL_3177843, EPI_ISL_3177844, EPI_ISL_3177845, EPI_ISL_3177846, EPI_ISL_3177847, EPI_ISL_3177848, EPI_ISL_3177849, EPI_ISL_3177850, EPI_ISL_3177851, EPI_ISL_3177852, EPI_ISL_3177853, EPI_ISL_3177854, EPI_ISL_3177855, EPI_ISL_3177856, EPI_ISL_3177857, EPI_ISL_3177858, EPI_ISL_3177859, EPI_ISL_3177860, EPI_ISL_3177861, EPI_ISL_3177862, EPI_ISL_3177863, EPI_ISL_3177864, EPI_ISL_3177865, EPI_ISL_3177866, EPI_ISL_3177867, EPI_ISL_3177868, EPI_ISL_3177869, EPI_ISL_3177870, EPI_ISL_3177871, EPI_ISL_3177872, EPI_ISL_3177873, EPI_ISL_3177874, EPI_ISL_3177875, EPI_ISL_3177876, EPI_ISL_3177877, EPI_ISL_3177878, EPI_ISL_3177879, EPI_ISL_3177880, EPI_ISL_3177881, EPI_ISL_3177882, EPI_ISL_3177883, EPI_ISL_3177884, EPI_ISL_3177885, EPI_ISL_3177886, EPI_ISL_3177887, EPI_ISL_3177888, EPI_ISL_3177889, EPI_ISL_3177890, EPI_ISL_3177891, EPI_ISL_3177892, EPI_ISL_3177893, EPI_ISL_3177894, EPI_ISL_3177895, EPI_ISL_3177896, EPI_ISL_3177897, EPI_ISL_3177898, EPI_ISL_3177899, EPI_ISL_3177900, EPI_ISL_3177901, EPI_ISL_3177902, EPI_ISL_3177903, EPI_ISL_3177904, EPI_ISL_3177905, EPI_ISL_3177906, EPI_ISL_3177907, EPI_ISL_3177908, EPI_ISL_3177909, EPI_ISL_3177910, EPI_ISL_3177911, EPI_ISL_3177912, EPI_ISL_3177913, EPI_ISL_3177914, EPI_ISL_3177915, | | | |

EPI_ISL_3177916, EPI_ISL_3177917, EPI_ISL_3177918, EPI_ISL_3177919, EPI_ISL_3177920, EPI_ISL_3177921, EPI_ISL_3177922, EPI_ISL_3177923, EPI_ISL_3177924, EPI_ISL_3177925, EPI_ISL_3177926, EPI_ISL_3177927, EPI_ISL_3177928, EPI_ISL_3177929, EPI_ISL_3177930, EPI_ISL_3177931, EPI_ISL_3177932, EPI_ISL_3177933, EPI_ISL_3177934, EPI_ISL_3177935, EPI_ISL_3177936, EPI_ISL_3177937, EPI_ISL_3177938, EPI_ISL_3177939, EPI_ISL_3177940, EPI_ISL_3177941, EPI_ISL_3177942, EPI_ISL_3177943, EPI_ISL_3177944, EPI_ISL_3177945, EPI_ISL_3177946, EPI_ISL_3177947, EPI_ISL_3177948, EPI_ISL_3177949, EPI_ISL_3177950, EPI_ISL_3177951, EPI_ISL_3177952, EPI_ISL_3177953, EPI_ISL_3177954, EPI_ISL_3177955, EPI_ISL_3177956, EPI_ISL_3177957, EPI_ISL_3177958, EPI_ISL_3177959, EPI_ISL_3177960, EPI_ISL_3177961, EPI_ISL_3177962, EPI_ISL_3177963, EPI_ISL_3177964, EPI_ISL_3177965, EPI_ISL_3177966, EPI_ISL_3177967, EPI_ISL_3177968, EPI_ISL_3177969, EPI_ISL_3177970, EPI_ISL_3177971, EPI_ISL_3177972, EPI_ISL_3177973, EPI_ISL_3177974, EPI_ISL_3177975, EPI_ISL_3177976, EPI_ISL_3177977, EPI_ISL_3177978, EPI_ISL_3177979, EPI_ISL_3177980, EPI_ISL_3177981, EPI_ISL_3177982, EPI_ISL_3177983, EPI_ISL_3177984, EPI_ISL_3177985, EPI_ISL_3177986, EPI_ISL_3177987, EPI_ISL_3177988, EPI_ISL_3177989, EPI_ISL_3177990, EPI_ISL_3177991, EPI_ISL_3177992, EPI_ISL_3177993, EPI_ISL_3177994, EPI_ISL_3177995, EPI_ISL_3177996, EPI_ISL_3177997, EPI_ISL_3177998, EPI_ISL_3177999, EPI_ISL_3178000, EPI_ISL_3178001, EPI_ISL_3178002, EPI_ISL_3178003, EPI_ISL_3178004, EPI_ISL_3178005, EPI_ISL_3178006, EPI_ISL_3178007, EPI_ISL_3178008, EPI_ISL_3178009, EPI_ISL_3178010, EPI_ISL_3178011, EPI_ISL_3178012, EPI_ISL_3178013, EPI_ISL_3178014, EPI_ISL_3178015, EPI_ISL_3178016, EPI_ISL_3178017, EPI_ISL_3178018, EPI_ISL_3178019, EPI_ISL_3178020, EPI_ISL_3178021, EPI_ISL_3178022, EPI_ISL_3178023, EPI_ISL_3178024, EPI_ISL_3178025, EPI_ISL_3178026, EPI_ISL_3178027, EPI_ISL_3178028, EPI_ISL_3178029, EPI_ISL_3178030, EPI_ISL_3178031, EPI_ISL_3178032, EPI_ISL_3178033, EPI_ISL_3178034, EPI_ISL_3178035, EPI_ISL_3178036, EPI_ISL_3178037, EPI_ISL_3178038, EPI_ISL_3178039, EPI_ISL_3178040, EPI_ISL_3178041, EPI_ISL_3178042, EPI_ISL_3178043, EPI_ISL_3178047, EPI_ISL_3178051, EPI_ISL_3178053, EPI_ISL_3178054, EPI_ISL_3178055, EPI_ISL_3178056, EPI_ISL_3178057, EPI_ISL_3178058, EPI_ISL_3178059, EPI_ISL_3178060, EPI_ISL_3178061, EPI_ISL_3178062, EPI_ISL_3178063, EPI_ISL_3178064, EPI_ISL_3178065, EPI_ISL_3178066, EPI_ISL_3178067, EPI_ISL_3178068, EPI_ISL_3178069, EPI_ISL_3178070, EPI_ISL_3178071, EPI_ISL_3178072, EPI_ISL_3178073, EPI_ISL_3178074, EPI_ISL_3178075, EPI_ISL_3178076, EPI_ISL_3178077, EPI_ISL_3178078, EPI_ISL_3178079, EPI_ISL_3178080, EPI_ISL_3178081, EPI_ISL_3178082, EPI_ISL_3178083, EPI_ISL_3178084, EPI_ISL_3178085, EPI_ISL_3178086, EPI_ISL_3178087, EPI_ISL_3178088, EPI_ISL_3178089, EPI_ISL_3178090, EPI_ISL_3178091, EPI_ISL_3178092, EPI_ISL_3178093, EPI_ISL_3178094, EPI_ISL_3178095, EPI_ISL_3178096, EPI_ISL_3178097, EPI_ISL_3178098, EPI_ISL_3178099, EPI_ISL_3178100, EPI_ISL_3178101, EPI_ISL_3178102, EPI_ISL_3178103, EPI_ISL_3178104, EPI_ISL_3178105, EPI_ISL_3178106, EPI_ISL_3178107, EPI_ISL_3178108, EPI_ISL_3178109, EPI_ISL_3178110, EPI_ISL_3178111, EPI_ISL_3178112, EPI_ISL_3178113, EPI_ISL_3178114, EPI_ISL_3178115, EPI_ISL_3178116, EPI_ISL_3178117, EPI_ISL_3178118, EPI_ISL_3178119, EPI_ISL_3178120, EPI_ISL_3178121, EPI_ISL_3178122, EPI_ISL_3178123, EPI_ISL_3178124, EPI_ISL_3178125, EPI_ISL_3178126, EPI_ISL_3178127, EPI_ISL_3178128, EPI_ISL_3178129, EPI_ISL_3178130, EPI_ISL_3178131, EPI_ISL_3178132, EPI_ISL_3178133, EPI_ISL_3178134, EPI_ISL_3178135, EPI_ISL_3178136, EPI_ISL_3178137, EPI_ISL_3178138, EPI_ISL_3178139, EPI_ISL_3178140, EPI_ISL_3178141, EPI_ISL_3178142, EPI_ISL_3178143, EPI_ISL_3178144, EPI_ISL_3178145, EPI_ISL_3178146, EPI_ISL_3178147, EPI_ISL_3178148, EPI_ISL_3178149, EPI_ISL_3178150, EPI_ISL_3178151, EPI_ISL_3178152, EPI_ISL_3178153, EPI_ISL_3178154, EPI_ISL_3178155, EPI_ISL_3178156, EPI_ISL_3178157, EPI_ISL_3178158, EPI_ISL_3178159, EPI_ISL_3178160, EPI_ISL_3178161, EPI_ISL_3178162, EPI_ISL_3178163, EPI_ISL_3178164, EPI_ISL_3178165, EPI_ISL_3178166, EPI_ISL_3178167, EPI_ISL_3178168, EPI_ISL_3178169, EPI_ISL_3178170, EPI_ISL_3178171, EPI_ISL_3178172, EPI_ISL_3178173, EPI_ISL_3178174, EPI_ISL_3178175, EPI_ISL_3178176, EPI_ISL_3178177, EPI_ISL_3178178, EPI_ISL_3178179, EPI_ISL_3178180, EPI_ISL_3178181, EPI_ISL_3178182, EPI_ISL_3178183, EPI_ISL_3178184, EPI_ISL_3178185, EPI_ISL_3178186, EPI_ISL_3178187, EPI_ISL_3178188, EPI_ISL_3178189, EPI_ISL_3178190, EPI_ISL_3178191, EPI_ISL_3178192, EPI_ISL_3178193, EPI_ISL_3178194, EPI_ISL_3178195, EPI_ISL_3178196, EPI_ISL_3178197, EPI_ISL_3178198, EPI_ISL_3178199, EPI_ISL_3178200, EPI_ISL_3178201, EPI_ISL_3178202, EPI_ISL_3178203, EPI_ISL_3178204, EPI_ISL_3178205, EPI_ISL_3178206, EPI_ISL_3178207, EPI_ISL_3178208, EPI_ISL_3178209, EPI_ISL_3178210, EPI_ISL_3178211, EPI_ISL_3178212, EPI_ISL_3178213, EPI_ISL_3178214, EPI_ISL_3178215, EPI_ISL_3178216, EPI_ISL_3178217, EPI_ISL_3178218, EPI_ISL_3178219, EPI_ISL_3178220, EPI_ISL_3178221, EPI_ISL_3178222, EPI_ISL_3178223, EPI_ISL_3178224, EPI_ISL_3178225, EPI_ISL_3178226, EPI_ISL_3178227, EPI_ISL_3178228, EPI_ISL_3178229, EPI_ISL_3178230, EPI_ISL_3178231, EPI_ISL_3178232, EPI_ISL_3178233, EPI_ISL_3178234, EPI_ISL_3178235, EPI_ISL_3178236, EPI_ISL_3178237, EPI_ISL_3178238, EPI_ISL_3178239, EPI_ISL_3178240, EPI_ISL_3178241, EPI_ISL_3178242, EPI_ISL_3178243, EPI_ISL_3178244, EPI_ISL_3178245, EPI_ISL_3178246, EPI_ISL_3178247, EPI_ISL_3178248, EPI_ISL_3178249, EPI_ISL_3178250, EPI_ISL_3178251, EPI_ISL_3178252, EPI_ISL_3178253, EPI_ISL_3178254, EPI_ISL_3178255, EPI_ISL_3178256, EPI_ISL_3178257, EPI_ISL_3178258, EPI_ISL_3178259, EPI_ISL_3178260, EPI_ISL_3178261, EPI_ISL_3178262, EPI_ISL_3178263, EPI_ISL_3178264, EPI_ISL_3178265, EPI_ISL_3178266, EPI_ISL_3178267, EPI_ISL_3178268, EPI_ISL_3178269, EPI_ISL_3178270, EPI_ISL_3178271, EPI_ISL_3178272, EPI_ISL_3178273, EPI_ISL_3178274, EPI_ISL_3178275, EPI_ISL_3178276, EPI_ISL_3178277, EPI_ISL_3178278, EPI_ISL_3178279, EPI_ISL_3178280, EPI_ISL_3178281, EPI_ISL_3178282, EPI_ISL_3178283, EPI_ISL_3178284, EPI_ISL_3178285, EPI_ISL_3178286, EPI_ISL_3178287, EPI_ISL_3178288, EPI_ISL_3178289, EPI_ISL_3178290, EPI_ISL_3178291, EPI_ISL_3178292, EPI_ISL_3178293, EPI_ISL_3178294, EPI_ISL_3178295, EPI_ISL_3178296, EPI_ISL_3178297, EPI_ISL_3178298, EPI_ISL_3178299, EPI_ISL_3178300, EPI_ISL_3178301, EPI_ISL_3178302, EPI_ISL_3178303, EPI_ISL_3178304, EPI_ISL_3178305, EPI_ISL_3178306, EPI_ISL_3178307, EPI_ISL_3178308, EPI_ISL_3178309, EPI_ISL_3178310, EPI_ISL_3178311, EPI_ISL_3178312, EPI_ISL_3178313, EPI_ISL_3178314, EPI_ISL_3178315, EPI_ISL_3178316, EPI_ISL_3178317, EPI_ISL_3178318, EPI_ISL_3178319, EPI_ISL_3178320, EPI_ISL_3178321, EPI_ISL_3178322, EPI_ISL_3178323, EPI_ISL_3178324, EPI_ISL_3178325, EPI_ISL_3178326, EPI_ISL_3178327, EPI_ISL_3178328, EPI_ISL_3178329, EPI_ISL_3178330, EPI_ISL_3178331, EPI_ISL_3178332, EPI_ISL_3178333, EPI_ISL_3178334, EPI_ISL_3178335, EPI_ISL_3178336, EPI_ISL_3178337, EPI_ISL_3178338, EPI_ISL_3178339, EPI_ISL_3178340, EPI_ISL_3178341

| | | | |
|---|---|---|---|
| see above | Quadram Institute Bioscience | COVID-19 Genomics UK (COG-UK) Consortium | Dave J. Baker, Gemma L. Kay, Alp Aydin, Thanh Le-Viet, Steven Rudder, Ana P. Tedim, Anastasia Kolyva, Maria Diaz, Leonardo de Oliveira Martins, Nabil-Fareed Alikhan, Lizzie Meadows, Rachael Stanley, Rose K Davidson, Ngozi Elumogo, Muhammed Yasir, Sophie J. Prosolek, Ebenezer Foster-Nyarko, Martin Lott, Christopher Jeanes, Lewis G. Spurgin, Nicholas M. Thomson, Alexander J Trotter, Rachel Gilroy, Luke Bedford, Samuel Bloomfield, Claire Stuart, Andrew Bell, Reenesh Prakash, Lindsay Coupland, Samir Dervisevic, Alison E. Mather, John Wain, Mark Webber, Andrew J. Page, Justin O'Grady |

EPI_ISL_3178342, EPI_ISL_3178343, EPI_ISL_3178344, EPI_ISL_3178345, EPI_ISL_3178346, EPI_ISL_3178347, EPI_ISL_3178348, EPI_ISL_3178349, EPI_ISL_3178350, EPI_ISL_3178351, EPI_ISL_3178352, EPI_ISL_3178353, EPI_ISL_3178354, EPI_ISL_3178355, EPI_ISL_3178356, EPI_ISL_3178357, EPI_ISL_3178358, EPI_ISL_3178359, EPI_ISL_3178360, EPI_ISL_3178361, EPI_ISL_3178362, EPI_ISL_3178363, EPI_ISL_3178364, EPI_ISL_3178365, EPI_ISL_3178366, EPI_ISL_3178367, EPI_ISL_3178368, EPI_ISL_3178369, EPI_ISL_3178370, EPI_ISL_3178371, EPI_ISL_3178372, EPI_ISL_3178373, EPI_ISL_3178374, EPI_ISL_3178375, EPI_ISL_3178376, EPI_ISL_3178377, EPI_ISL_3178378, EPI_ISL_3178379, EPI_ISL_3178380, EPI_ISL_3178381, EPI_ISL_3178382, EPI_ISL_3178383, EPI_ISL_3178384, EPI_ISL_3178385, EPI_ISL_3178386, EPI_ISL_3178387, EPI_ISL_3178388, EPI_ISL_3178389, EPI_ISL_3178390, EPI_ISL_3178391, EPI_ISL_3178392, EPI_ISL_3178393, EPI_ISL_3178394, EPI_ISL_3178395, EPI_ISL_3178396, EPI_ISL_3178397, EPI_ISL_3178398, EPI_ISL_3178399, EPI_ISL_3178400, EPI_ISL_3178401, EPI_ISL_3178402, EPI_ISL_3178403, EPI_ISL_3178404, EPI_ISL_3178405, EPI_ISL_3178406, EPI_ISL_3178407, EPI_ISL_3178408, EPI_ISL_3178409, EPI_ISL_3178410, EPI_ISL_3178411, EPI_ISL_3178412, EPI_ISL_3178413, EPI_ISL_3178414, EPI_ISL_3178415, EPI_ISL_3178416, EPI_ISL_3178417, EPI_ISL_3178418, EPI_ISL_3178419, EPI_ISL_3178420, EPI_ISL_3178421, EPI_ISL_3178422, EPI_ISL_3178423, EPI_ISL_3178424, EPI_ISL_3178425, EPI_ISL_3178426, EPI_ISL_3178427, EPI_ISL_3178428, EPI_ISL_3178429, EPI_ISL_3178430, EPI_ISL_3178431, EPI_ISL_3178432, EPI_ISL_3178433, EPI_ISL_3178434, EPI_ISL_3178435, EPI_ISL_3178436, EPI_ISL_3178437, EPI_ISL_3178438, EPI_ISL_3178439, EPI_ISL_3178440, EPI_ISL_3178441, EPI_ISL_3178442, EPI_ISL_3178443, EPI_ISL_3178444, EPI_ISL_3178445, EPI_ISL_3178446, EPI_ISL_3178447, EPI_ISL_3178448, EPI_ISL_3178449, EPI_ISL_3178450, EPI_ISL_3178451, EPI_ISL_3178452, EPI_ISL_3178453, EPI_ISL_3178454, EPI_ISL_3178455, EPI_ISL_3178456, EPI_ISL_3178457, EPI_ISL_3178458, EPI_ISL_3178459, EPI_ISL_3178460, EPI_ISL_3178461, EPI_ISL_3178462, EPI_ISL_3178463, EPI_ISL_3178464, EPI_ISL_3178465, EPI_ISL_3178466, EPI_ISL_3178467, EPI_ISL_3178468, EPI_ISL_3178469, EPI_ISL_3178470, EPI_ISL_3178471, EPI_ISL_3178472, EPI_ISL_3178473, EPI_ISL_3178474, EPI_ISL_3178475, EPI_ISL_3178476, EPI_ISL_3178477, EPI_ISL_3178478, EPI_ISL_3178479, EPI_ISL_3178480, EPI_ISL_3178481, EPI_ISL_3178482, EPI_ISL_3178483, EPI_ISL_3178484, EPI_ISL_3178485, EPI_ISL_3178486, EPI_ISL_3178487, EPI_ISL_3178488, EPI_ISL_3178489, EPI_ISL_3178490, EPI_ISL_3178491, EPI_ISL_3178492, EPI_ISL_3178493, EPI_ISL_3178494, EPI_ISL_3178495, EPI_ISL_3178496, EPI_ISL_3178497, EPI_ISL_3178498, EPI_ISL_3178499, EPI_ISL_3178500, EPI_ISL_3178501, EPI_ISL_3178502, EPI_ISL_3178503, EPI_ISL_3178504, EPI_ISL_3178505, EPI_ISL_3178506, EPI_ISL_3178507, EPI_ISL_3178508, EPI_ISL_3178509, EPI_ISL_3178510, EPI_ISL_3178511, EPI_ISL_3178512, EPI_ISL_3178513, EPI_ISL_3178514, EPI_ISL_3178515, EPI_ISL_3178516, EPI_ISL_3178517, EPI_ISL_3178518, EPI_ISL_3178519, EPI_ISL_3178520, EPI_ISL_3178521, EPI_ISL_3178522, EPI_ISL_3178523, EPI_ISL_3178524, EPI_ISL_3178525, EPI_ISL_3178526, EPI_ISL_3178527, EPI_ISL_3178528

| | | | |
|---|---|---|---|
| see above | Queens Medical Centre, Clinical Microbiology Department / DeepSeq Nottingham | COVID-19 Genomics UK (COG-UK) Consortium | Gemma Clark, Wendy Smith, Manjinder Khakh, Vicki M Fleming, Michelle M Lister, Hannah Howson-Wells, Jonathan Ball, Timothy Byaruhanga, Jayasree Dey, Emily Park, Jack Hill, Patrick McClure, Joseph Chappell, Theocharis Tsoleridis, Nadine Holmes, Matthew Carlisle, Christopher Moore, Fei Sang, Johnny Debebe, Victoria Wright, Matthew Loose |

EPI_ISL_3178530, EPI_ISL_3178531, EPI_ISL_3178534, EPI_ISL_3178535, EPI_ISL_3178538, EPI_ISL_3178539, EPI_ISL_3178540, EPI_ISL_3178542, EPI_ISL_3178543, EPI_ISL_3178544, EPI_ISL_3178545, EPI_ISL_3178546, EPI_ISL_3178547, EPI_ISL_3178549, EPI_ISL_3178550, EPI_ISL_3178552, EPI_ISL_3178555, EPI_ISL_3178556, EPI_ISL_3178557, EPI_ISL_3178559, EPI_ISL_3178560, EPI_ISL_3178562, EPI_ISL_3178566, EPI_ISL_3178567, EPI_ISL_3178568, EPI_ISL_3178570, EPI_ISL_3178573, EPI_ISL_3178574, EPI_ISL_3178575, EPI_ISL_3178576, EPI_ISL_3178577, EPI_ISL_3178578, EPI_ISL_3178581, EPI_ISL_3178582, EPI_ISL_3178584, EPI_ISL_3178585, EPI_ISL_3178586, EPI_ISL_3178587, EPI_ISL_3178589, EPI_ISL_3178591, EPI_ISL_3178592, EPI_ISL_3178593, EPI_ISL_3178595, EPI_ISL_3178596, EPI_ISL_3178598, EPI_ISL_3178599, EPI_ISL_3178600, EPI_ISL_3178601, EPI_ISL_3178602, EPI_ISL_3178604, EPI_ISL_3178605, EPI_ISL_3178606, EPI_ISL_3178609, EPI_ISL_3178610, EPI_ISL_3178611, EPI_ISL_3178613, EPI_ISL_3178615, EPI_ISL_3178616, EPI_ISL_3178617, EPI_ISL_3178618, EPI_ISL_3178619, EPI_ISL_3178620, EPI_ISL_3178621, EPI_ISL_3178622, EPI_ISL_3178623, EPI_ISL_3178624, EPI_ISL_3178625, EPI_ISL_3178626, EPI_ISL_3178629, EPI_ISL_3178630, EPI_ISL_3178631, EPI_ISL_3178632, EPI_ISL_3178633, EPI_ISL_3178634, EPI_ISL_3178635, EPI_ISL_3178636, EPI_ISL_3178637, EPI_ISL_3178638, EPI_ISL_3178639, EPI_ISL_3178640, EPI_ISL_3178641, EPI_ISL_3178643, EPI_ISL_3178644, EPI_ISL_3178645, EPI_ISL_3178646, EPI_ISL_3178647, EPI_ISL_3178648, EPI_ISL_3178649, EPI_ISL_3178650, EPI_ISL_3178651, EPI_ISL_3178652, EPI_ISL_3178653, EPI_ISL_3178655, EPI_ISL_3178656, EPI_ISL_3178657, EPI_ISL_3178658, EPI_ISL_3178659, EPI_ISL_3178660, EPI_ISL_3178661, EPI_ISL_3178662, EPI_ISL_3178663, EPI_ISL_3178664, EPI_ISL_3178665, EPI_ISL_3178666, EPI_ISL_3178667, EPI_ISL_3178668, EPI_ISL_3178669, EPI_ISL_3178670, EPI_ISL_3178671, EPI_ISL_3178672, EPI_ISL_3178674, EPI_ISL_3178675, EPI_ISL_3178676, EPI_ISL_3178677, EPI_ISL_3178679, EPI_ISL_3178680, EPI_ISL_3178681, EPI_ISL_3178682, EPI_ISL_3178683, EPI_ISL_3178684, EPI_ISL_3178685, EPI_ISL_3178687, EPI_ISL_3178688, EPI_ISL_3178689, EPI_ISL_3178690, EPI_ISL_3178691, EPI_ISL_3178692, EPI_ISL_3178694, EPI_ISL_3178695, EPI_ISL_3178696, EPI_ISL_3178697, EPI_ISL_3178698, EPI_ISL_3178699, EPI_ISL_3178700, EPI_ISL_3178701, EPI_ISL_3178702, EPI_ISL_3178703, EPI_ISL_3178704, EPI_ISL_3178705, EPI_ISL_3178707, EPI_ISL_3178708, EPI_ISL_3178709, EPI_ISL_3178710, EPI_ISL_3178711, EPI_ISL_3178713, EPI_ISL_3178714, EPI_ISL_3178715, EPI_ISL_3178716, EPI_ISL_3178717, EPI_ISL_3178718, EPI_ISL_3178720, EPI_ISL_3178721, EPI_ISL_3178722, EPI_ISL_3178723, EPI_ISL_3178724, EPI_ISL_3178725, EPI_ISL_3178726, EPI_ISL_3178727, EPI_ISL_3178728, EPI_ISL_3178732, EPI_ISL_3178733, EPI_ISL_3178734, EPI_ISL_3178736, EPI_ISL_3178737, EPI_ISL_3178739, EPI_ISL_3178740, EPI_ISL_3178741, EPI_ISL_3178742, EPI_ISL_3178743, EPI_ISL_3178744, EPI_ISL_3178745, EPI_ISL_3178746, EPI_ISL_3178749, EPI_ISL_3178750, EPI_ISL_3178751, EPI_ISL_3178752, EPI_ISL_3178753, EPI_ISL_3178754, EPI_ISL_3178755, EPI_ISL_3178756, EPI_ISL_3178757, EPI_ISL_3178758, EPI_ISL_3178759, EPI_ISL_3178760, EPI_ISL_3178761, EPI_ISL_3178763, EPI_ISL_3178764, EPI_ISL_3178765, EPI_ISL_3178766, EPI_ISL_3178767, EPI_ISL_3178768, EPI_ISL_3178769, EPI_ISL_3178770, EPI_ISL_3178771, EPI_ISL_3178772, EPI_ISL_3178774, EPI_ISL_3178776, EPI_ISL_3178777, EPI_ISL_3178779, EPI_ISL_3178780, EPI_ISL_3178782, EPI_ISL_3178783, EPI_ISL_3178784, EPI_ISL_3178787, EPI_ISL_3178788, EPI_ISL_3178789, EPI_ISL_3178790, EPI_ISL_3178791, EPI_ISL_3178794, EPI_ISL_3178795, EPI_ISL_3178796, EPI_ISL_3178797, EPI_ISL_3178798, EPI_ISL_3178799, EPI_ISL_3178801, EPI_ISL_3178802, EPI_ISL_3178803, EPI_ISL_3178805, EPI_ISL_3178806, EPI_ISL_3178807, EPI_ISL_3178809, EPI_ISL_3178810, EPI_ISL_3178811, EPI_ISL_3178813, EPI_ISL_3178814, EPI_ISL_3178815, EPI_ISL_3178816, EPI_ISL_3178817, EPI_ISL_3178818, EPI_ISL_3178819, EPI_ISL_3178820, EPI_ISL_3178821, EPI_ISL_3178824, EPI_ISL_3178825, EPI_ISL_3178826, EPI_ISL_3178831, EPI_ISL_3178832, EPI_ISL_3178833, EPI_ISL_3178834, EPI_ISL_3178835, EPI_ISL_3178836, EPI_ISL_3178837, EPI_ISL_3178838, EPI_ISL_3178839, EPI_ISL_3178840, EPI_ISL_3178841, EPI_ISL_3178842, EPI_ISL_3178844, EPI_ISL_3178845, EPI_ISL_3178846, EPI_ISL_3178847, EPI_ISL_3178848, EPI_ISL_3178849, EPI_ISL_3178852, EPI_ISL_3178853, EPI_ISL_3178855, EPI_ISL_3178856, EPI_ISL_3178857, EPI_ISL_3178858, EPI_ISL_3178859, EPI_ISL_3178860, EPI_ISL_3178861, EPI_ISL_3178862, EPI_ISL_3178863, EPI_ISL_3178864, EPI_ISL_3178865, EPI_ISL_3178866, EPI_ISL_3178867, EPI_ISL_3178868, EPI_ISL_3178870, EPI_ISL_3178871

| | | | |
|---|---|---|---|
| see above | Oxford Viromics, NDM, University of Oxford; Oxford University Hospitals; Basingstoke and North Hampshire Hospital | COVID-19 Genomics UK (COG-UK) Consortium | Tanya Golubchik, David Bonsall, George Macintyre, Amy Trebes, Mariateresa de Cesare, Catrin Moore, Alex Mobbs, Anita Justice, Robert Shaw, Monique Andersson, Timothy Peto, Emma Wise, Nathan Moore, Jessica Lynch, Nick Cortes, Matilde Mori, Stephen Kidd, David Buck, John Todd, Christophe Fraser |

EPI_ISL_3178972, EPI_ISL_3178973, EPI_ISL_3178974, EPI_ISL_3178975, EPI_ISL_3178976, EPI_ISL_3178977, EPI_ISL_3178978, EPI_ISL_3178979, EPI_ISL_3178980, EPI_ISL_3178981, EPI_ISL_3178982, EPI_ISL_3178983, EPI_ISL_3178984, EPI_ISL_3178985, EPI_ISL_3178986, EPI_ISL_3178987, EPI_ISL_3178988, EPI_ISL_3178989, EPI_ISL_3178990, EPI_ISL_3178991, EPI_ISL_3178992, EPI_ISL_3178993, EPI_ISL_3178994, EPI_ISL_3178995, EPI_ISL_3178996, EPI_ISL_3178997, EPI_ISL_3178998, EPI_ISL_3178999, EPI_ISL_3179000, EPI_ISL_3179001, EPI_ISL_3179002, EPI_ISL_3179003, EPI_ISL_3179004, EPI_ISL_3179005, EPI_ISL_3179006, EPI_ISL_3179007, EPI_ISL_3179008, EPI_ISL_3179009, EPI_ISL_3179010, EPI_ISL_3179011, EPI_ISL_3179018, EPI_ISL_3179019, EPI_ISL_3179020, EPI_ISL_3179021, EPI_ISL_3179022, EPI_ISL_3179023, EPI_ISL_3179024, EPI_ISL_3179025, EPI_ISL_3179026, EPI_ISL_3179027, EPI_ISL_3179028, EPI_ISL_3179029, EPI_ISL_3179030, EPI_ISL_3179031, EPI_ISL_3179032, EPI_ISL_3179033, EPI_ISL_3179034, EPI_ISL_3179035, EPI_ISL_3179036, EPI_ISL_3179037, EPI_ISL_3179038, EPI_ISL_3179039, EPI_ISL_3179040, EPI_ISL_3179041, EPI_ISL_3179042, EPI_ISL_3179043, EPI_ISL_3179044, EPI_ISL_3179045, EPI_ISL_3179046, EPI_ISL_3179047, EPI_ISL_3179048, EPI_ISL_3179049, EPI_ISL_3179050, EPI_ISL_3179051, EPI_ISL_3179052, EPI_ISL_3179053, EPI_ISL_3179054, EPI_ISL_3179055, EPI_ISL_3179056, EPI_ISL_3179057, EPI_ISL_3179058, EPI_ISL_3179059, EPI_ISL_3179060, EPI_ISL_3179061, EPI_ISL_3179062, EPI_ISL_3179063, EPI_ISL_3179064, EPI_ISL_3179065, EPI_ISL_3179066, EPI_ISL_3179067, EPI_ISL_3179068, EPI_ISL_3179069, EPI_ISL_3179070, EPI_ISL_3179071, EPI_ISL_3179072, EPI_ISL_3179073, EPI_ISL_3179074, EPI_ISL_3179075, EPI_ISL_3179076, EPI_ISL_3179077, EPI_ISL_3179078, EPI_ISL_3179079, EPI_ISL_3179080, EPI_ISL_3179081, EPI_ISL_3179082, EPI_ISL_3179083, EPI_ISL_3179084, EPI_ISL_3179085, EPI_ISL_3179086, EPI_ISL_3179087, EPI_ISL_3179088, EPI_ISL_3179089, EPI_ISL_3179090, EPI_ISL_3179091, EPI_ISL_3179092, EPI_ISL_3179093, EPI_ISL_3179094, EPI_ISL_3179095, EPI_ISL_3179096, EPI_ISL_3179097, EPI_ISL_3179098, EPI_ISL_3179099, EPI_ISL_3179100, EPI_ISL_3179101, EPI_ISL_3179102, EPI_ISL_3179103, EPI_ISL_3179104, EPI_ISL_3179105, EPI_ISL_3179106, EPI_ISL_3179107, EPI_ISL_3179108, EPI_ISL_3179109, EPI_ISL_3179110, EPI_ISL_3179111, EPI_ISL_3179112, EPI_ISL_3179113, EPI_ISL_3179114, EPI_ISL_3179115, EPI_ISL_3179116, EPI_ISL_3179117, EPI_ISL_3179118, EPI_ISL_3179119, EPI_ISL_3179120, EPI_ISL_3179121, EPI_ISL_3179122, EPI_ISL_3179123, EPI_ISL_3179124, EPI_ISL_3179125, EPI_ISL_3179126, EPI_ISL_3179127, EPI_ISL_3179128, EPI_ISL_3179129, EPI_ISL_3179130, EPI_ISL_3179131, EPI_ISL_3179132, EPI_ISL_3179133, EPI_ISL_3179134, EPI_ISL_3179135, EPI_ISL_3179136, EPI_ISL_3179137, EPI_ISL_3179138, EPI_ISL_3179139, EPI_ISL_3179140, EPI_ISL_3179141, EPI_ISL_3179142, EPI_ISL_3179143, EPI_ISL_3179144, EPI_ISL_3179145, EPI_ISL_3179146, EPI_ISL_3179147, EPI_ISL_3179148, EPI_ISL_3179149, EPI_ISL_3179150, EPI_ISL_3179151, EPI_ISL_3179152, EPI_ISL_3179153, EPI_ISL_3179154, EPI_ISL_3179155, EPI_ISL_3179156, EPI_ISL_3179157, EPI_ISL_3179158, EPI_ISL_3179159, EPI_ISL_3179160, EPI_ISL_3179161, EPI_ISL_3179162, EPI_ISL_3179163, EPI_ISL_3179164, EPI_ISL_3179165, EPI_ISL_3179166, EPI_ISL_3179167, EPI_ISL_3179168, EPI_ISL_3179169, EPI_ISL_3179170, EPI_ISL_3179171, EPI_ISL_3179172, EPI_ISL_3179173, EPI_ISL_3179174, EPI_ISL_3179175, EPI_ISL_3179176, EPI_ISL_3179177, EPI_ISL_3179178, EPI_ISL_3179179, EPI_ISL_3179180, EPI_ISL_3179181, EPI_ISL_3179182, EPI_ISL_3179183, EPI_ISL_3179184, EPI_ISL_3179185, EPI_ISL_3179186, EPI_ISL_3179187, EPI_ISL_3179188, EPI_ISL_3179189, EPI_ISL_3179191, EPI_ISL_3179192, EPI_ISL_3179193, EPI_ISL_3179194, EPI_ISL_3179197, EPI_ISL_3179198, EPI_ISL_3179199, EPI_ISL_3179200, EPI_ISL_3179201, EPI_ISL_3179202, EPI_ISL_3179203, EPI_ISL_3179204, EPI_ISL_3179205, EPI_ISL_3179206, EPI_ISL_3179207, EPI_ISL_3179208, EPI_ISL_3179209, EPI_ISL_3179210, EPI_ISL_3179211, EPI_ISL_3179212, EPI_ISL_3179213, EPI_ISL_3179214, EPI_ISL_3179215, EPI_ISL_3179216, EPI_ISL_3179217, EPI_ISL_3179218, EPI_ISL_3179219, EPI_ISL_3179220, EPI_ISL_3179221, EPI_ISL_3179222, EPI_ISL_3179223, EPI_ISL_3179224, EPI_ISL_3179225, EPI_ISL_3179226, EPI_ISL_3179227, EPI_ISL_3179228, EPI_ISL_3179229, EPI_ISL_3179230, EPI_ISL_3179231, EPI_ISL_3179232, EPI_ISL_3179233, EPI_ISL_3179234, EPI_ISL_3179235, EPI_ISL_3179236, EPI_ISL_3179237, EPI_ISL_3179238, EPI_ISL_3179239, EPI_ISL_3179240, EPI_ISL_3179241, EPI_ISL_3179242, EPI_ISL_3179243, EPI_ISL_3179244, EPI_ISL_3179245, EPI_ISL_3179246, EPI_ISL_3179247, EPI_ISL_3179248, EPI_ISL_3179249, EPI_ISL_3179250, EPI_ISL_3179252, EPI_ISL_3179253, EPI_ISL_3179254, EPI_ISL_3179255, EPI_ISL_3179256, EPI_ISL_3179257, EPI_ISL_3179258, EPI_ISL_3179259, EPI_ISL_3179260, EPI_ISL_3179261, EPI_ISL_3179262, EPI_ISL_3179263, EPI_ISL_3179264, EPI_ISL_3179265, EPI_ISL_3179266, EPI_ISL_3179267, EPI_ISL_3179268, EPI_ISL_3179269, EPI_ISL_3179270, EPI_ISL_3179271, EPI_ISL_3179272, EPI_ISL_3179273, EPI_ISL_3179274, EPI_ISL_3179276, EPI_ISL_3179278, EPI_ISL_3179279, EPI_ISL_3179280, EPI_ISL_3179281, EPI_ISL_3179282, EPI_ISL_3179284, EPI_ISL_3179285, EPI_ISL_3179286, EPI_ISL_3179287, EPI_ISL_3179288, EPI_ISL_3179290, EPI_ISL_3179291, EPI_ISL_3179293, EPI_ISL_3179294, EPI_ISL_3179295, EPI_ISL_3179296, EPI_ISL_3179297, EPI_ISL_3179298, EPI_ISL_3179299, EPI_ISL_3179300, EPI_ISL_3179302, EPI_ISL_3179303, EPI_ISL_3179304, EPI_ISL_3179305, EPI_ISL_3179306, EPI_ISL_3179307, EPI_ISL_3179308, EPI_ISL_3179309, EPI_ISL_3179310, EPI_ISL_3179311, EPI_ISL_3179312, EPI_ISL_3179313, EPI_ISL_3179314, EPI_ISL_3179315, EPI_ISL_3179316, EPI_ISL_3179317, EPI_ISL_3179318, EPI_ISL_3179319, EPI_ISL_3179320, EPI_ISL_3179321, EPI_ISL_3179322, EPI_ISL_3179323, EPI_ISL_3179324, EPI_ISL_3179325, EPI_ISL_3179326, EPI_ISL_3179327, EPI_ISL_3179328, EPI_ISL_3179329, EPI_ISL_3179330, EPI_ISL_3179331, EPI_ISL_3179332, EPI_ISL_3179333, EPI_ISL_3179334, EPI_ISL_3179335, EPI_ISL_3179336, EPI_ISL_3179337, EPI_ISL_3179338, EPI_ISL_3179339, EPI_ISL_3179340, EPI_ISL_3179341, EPI_ISL_3179342, EPI_ISL_3179343, EPI_ISL_3179344, EPI_ISL_3179345, EPI_ISL_3179346, EPI_ISL_3179347, EPI_ISL_3179348, EPI_ISL_3179349, EPI_ISL_3179350, EPI_ISL_3179351, EPI_ISL_3179352, EPI_ISL_3179353, EPI_ISL_3179354, EPI_ISL_3179355, EPI_ISL_3179356, EPI_ISL_3179357, EPI_ISL_3179358, EPI_ISL_3179359, EPI_ISL_3179360, EPI_ISL_3179361, EPI_ISL_3179362, EPI_ISL_3179363, EPI_ISL_3179364, EPI_ISL_3179365, EPI_ISL_3179366, EPI_ISL_3179367, EPI_ISL_3179368, EPI_ISL_3179369, EPI_ISL_3179370, EPI_ISL_3179371, EPI_ISL_3179372, EPI_ISL_3179373, EPI_ISL_3179374, EPI_ISL_3179375, EPI_ISL_3179376, EPI_ISL_3179377, EPI_ISL_3179378, EPI_ISL_3179379, EPI_ISL_3179380, EPI_ISL_3179381, EPI_ISL_3179382, EPI_ISL_3179383, EPI_ISL_3179384, EPI_ISL_3179385, EPI_ISL_3179386, EPI_ISL_3179387, EPI_ISL_3179388, EPI_ISL_3179389, EPI_ISL_3179390, EPI_ISL_3179391, EPI_ISL_3179392, EPI_ISL_3179393, EPI_ISL_3179394, EPI_ISL_3179395, EPI_ISL_3179396, EPI_ISL_3179397, EPI_ISL_3179398, EPI_ISL_3179400, EPI_ISL_3179401, EPI_ISL_3179402, EPI_ISL_3179403, EPI_ISL_3179404, EPI_ISL_3179405, EPI_ISL_3179406, EPI_ISL_3179407, EPI_ISL_3179408, EPI_ISL_3179409, EPI_ISL_3179410, EPI_ISL_3179411, EPI_ISL_3179412, EPI_ISL_3179413, EPI_ISL_3179414, EPI_ISL_3179415, EPI_ISL_3179416, EPI_ISL_3179417, EPI_ISL_3179418, EPI_ISL_3179419, EPI_ISL_3179420, EPI_ISL_3179421, EPI_ISL_3179422, EPI_ISL_3179423, EPI_ISL_3179424, EPI_ISL_3179425, EPI_ISL_3179426, EPI_ISL_3179427, EPI_ISL_3179428, EPI_ISL_3179429, EPI_ISL_3179430, EPI_ISL_3179431, EPI_ISL_3179432, EPI_ISL_3179433, EPI_ISL_3179434, EPI_ISL_3179435, EPI_ISL_3179436, EPI_ISL_3179437, EPI_ISL_3179438, EPI_ISL_3179439, EPI_ISL_3179440, EPI_ISL_3179441, EPI_ISL_3179442, EPI_ISL_3179443, EPI_ISL_3179444, EPI_ISL_3179445, EPI_ISL_3179446, EPI_ISL_3179447, EPI_ISL_3179448, EPI_ISL_3179449, EPI_ISL_3179450, EPI_ISL_3179451, EPI_ISL_3179452, EPI_ISL_3179453, EPI_ISL_3179454, EPI_ISL_3179455, EPI_ISL_3179456, EPI_ISL_3179457, EPI_ISL_3179458, EPI_ISL_3179459, EPI_ISL_3179460, EPI_ISL_3179461, EPI_ISL_3179462, EPI_ISL_3179463, EPI_ISL_3179464, EPI_ISL_3179465, EPI_ISL_3179466, EPI_ISL_3179467, EPI_ISL_3179468, EPI_ISL_3179469, EPI_ISL_3179470, EPI_ISL_3179471, EPI_ISL_3179472, EPI_ISL_3179473, EPI_ISL_3179474, EPI_ISL_3179475, EPI_ISL_3179476, EPI_ISL_3179477, EPI_ISL_3179478, EPI_ISL_3179479, EPI_ISL_3179480, EPI_ISL_3179481, EPI_ISL_3179482, EPI_ISL_3179483, EPI_ISL_3179484, EPI_ISL_3179485, EPI_ISL_3179486, EPI_ISL_3179487, EPI_ISL_3179488, EPI_ISL_3179489, EPI_ISL_3179490, EPI_ISL_3179491, EPI_ISL_3179492, EPI_ISL_3179493, EPI_ISL_3179494, EPI_ISL_3179495, EPI_ISL_3179496, EPI_ISL_3179497, EPI_ISL_3179498, EPI_ISL_3179499, EPI_ISL_3179500, EPI_ISL_3179501, EPI_ISL_3179502, EPI_ISL_3179503, EPI_ISL_3179504, EPI_ISL_3179505, EPI_ISL_3179506, EPI_ISL_3179507, EPI_ISL_3179508, EPI_ISL_3179509, EPI_ISL_3179510, EPI_ISL_3179511, EPI_ISL_3179512, EPI_ISL_3179513, EPI_ISL_3179514, EPI_ISL_3179515, EPI_ISL_3179516, EPI_ISL_3179517, EPI_ISL_3179518, EPI_ISL_3179519, EPI_ISL_3179521, EPI_ISL_3179522, EPI_ISL_3179523, EPI_ISL_3179524, EPI_ISL_3179525, EPI_ISL_3179526, EPI_ISL_3179527, EPI_ISL_3179528, EPI_ISL_3179529, EPI_ISL_3179530, EPI_ISL_3179531, EPI_ISL_3179532, EPI_ISL_3179533, EPI_ISL_3179534, EPI_ISL_3179535, EPI_ISL_3179536, EPI_ISL_3179537, EPI_ISL_3179538, EPI_ISL_3179539, EPI_ISL_3179540, EPI_ISL_3179541, EPI_ISL_3179542, EPI_ISL_3179543, EPI_ISL_3179544, EPI_ISL_3179545, EPI_ISL_3179546, EPI_ISL_3179547, EPI_ISL_3179548, EPI_ISL_3179549, EPI_ISL_3179550, EPI_ISL_3179551, EPI_ISL_3179552, EPI_ISL_3179553, EPI_ISL_3179554, EPI_ISL_3179555, EPI_ISL_3179556, EPI_ISL_3179557, EPI_ISL_3179558, EPI_ISL_3179559, EPI_ISL_3179561, EPI_ISL_3179562, EPI_ISL_3179563, EPI_ISL_3179564, EPI_ISL_3179565, EPI_ISL_3179566, EPI_ISL_3179567, EPI_ISL_3179568, EPI_ISL_3179569, EPI_ISL_3179570, EPI_ISL_3179571, EPI_ISL_3179572, EPI_ISL_3179573, EPI_ISL_3179574, EPI_ISL_3179575, EPI_ISL_3179576, EPI_ISL_3179577, EPI_ISL_3179579, EPI_ISL_3179580, EPI_ISL_3179581, EPI_ISL_3179582, EPI_ISL_3179583, EPI_ISL_3179584, EPI_ISL_3179585, EPI_ISL_3179586, EPI_ISL_3179587, EPI_ISL_3179588, EPI_ISL_3179589, EPI_ISL_3179590, EPI_ISL_3179591, EPI_ISL_3179592, EPI_ISL_3179593, EPI_ISL_3179594, EPI_ISL_3179595, EPI_ISL_3179596, EPI_ISL_3179597, EPI_ISL_3179598, EPI_ISL_3179599, EPI_ISL_3179600, EPI_ISL_3179601, EPI_ISL_3179602, EPI_ISL_3179603, EPI_ISL_3179604, EPI_ISL_3179605, EPI_ISL_3179606, EPI_ISL_3179607, EPI_ISL_3179608, EPI_ISL_3179609, EPI_ISL_3179610, EPI_ISL_3179611, EPI_ISL_3179612, EPI_ISL_3179613, EPI_ISL_3179614, EPI_ISL_3179615, EPI_ISL_3179616, EPI_ISL_3179617, EPI_ISL_3179618, EPI_ISL_3179620, EPI_ISL_3179621, EPI_ISL_3179622, EPI_ISL_3179623, EPI_ISL_3179624, EPI_ISL_3179625, EPI_ISL_3179626, EPI_ISL_3179627, EPI_ISL_3179628, EPI_ISL_3179629, EPI_ISL_3179630, EPI_ISL_3179631, EPI_ISL_3179632, EPI_ISL_3179633, EPI_ISL_3179634, EPI_ISL_3179635, EPI_ISL_3179636, EPI_ISL_3179637, EPI_ISL_3179638, EPI_ISL_3179639, EPI_ISL_3179640, EPI_ISL_3179641, EPI_ISL_3179642, EPI_ISL_3179643, EPI_ISL_3179644, EPI_ISL_3179645, EPI_ISL_3179646, EPI_ISL_3179647, EPI_ISL_3179648, EPI_ISL_3179649, EPI_ISL_3179650, EPI_ISL_3179651, EPI_ISL_3179652, EPI_ISL_3179653, EPI_ISL_3179654, EPI_ISL_3179655, EPI_ISL_3179656, EPI_ISL_3179657, EPI_ISL_3179658, EPI_ISL_3179659, EPI_ISL_3179660, EPI_ISL_3179661, EPI_ISL_3179662, EPI_ISL_3179663, EPI_ISL_3179664, EPI_ISL_3179665, EPI_ISL_3179666, EPI_ISL_3179667, EPI_ISL_3179668, EPI_ISL_3179669, EPI_ISL_3179670, EPI_ISL_3179671, EPI_ISL_3179672, EPI_ISL_3179673, EPI_ISL_3179674, EPI_ISL_3179675, EPI_ISL_3179676, EPI_ISL_3179677, EPI_ISL_3179678, EPI_ISL_3179679, EPI_ISL_3179680, EPI_ISL_3179681, EPI_ISL_3179682, EPI_ISL_3179683, EPI_ISL_3179684, EPI_ISL_3179685, EPI_ISL_3179686, EPI_ISL_3179687, EPI_ISL_3179688, EPI_ISL_3179689, EPI_ISL_3179691, EPI_ISL_3179692, EPI_ISL_3179693, EPI_ISL_3179694, EPI_ISL_3179695, EPI_ISL_3179696, EPI_ISL_3179697, EPI_ISL_3179698, EPI_ISL_3179699, EPI_ISL_3179700, EPI_ISL_3179701, EPI_ISL_3179702, EPI_ISL_3179703, EPI_ISL_3179704, EPI_ISL_3179705, EPI_ISL_3179706, EPI_ISL_3179707, EPI_ISL_3179708, EPI_ISL_3179709, EPI_ISL_3179710, EPI_ISL_3179711, EPI_ISL_3179712, EPI_ISL_3179713, EPI_ISL_3179714, EPI_ISL_3179715, EPI_ISL_3179716, EPI_ISL_3179717, EPI_ISL_3179718, EPI_ISL_3179719, EPI_ISL_3179720, EPI_ISL_3179721, EPI_ISL_3179722, EPI_ISL_3179723, EPI_ISL_3179724, EPI_ISL_3179725, EPI_ISL_3179726, EPI_ISL_3179727, EPI_ISL_3179728, EPI_ISL_3179729, EPI_ISL_3179730, EPI_ISL_3179731, EPI_ISL_3179732, EPI_ISL_3179733, EPI_ISL_3179734, EPI_ISL_3179735, EPI_ISL_3179736, EPI_ISL_3179737, EPI_ISL_3179738, EPI_ISL_3179739, EPI_ISL_3179740, EPI_ISL_3179741, EPI_ISL_3179742, EPI_ISL_3179743, EPI_ISL_3179744, EPI_ISL_3179745, EPI_ISL_3179746, EPI_ISL_3179747, EPI_ISL_3179748, EPI_ISL_3179749, EPI_ISL_3179750, EPI_ISL_3179751, EPI_ISL_3179752, EPI_ISL_3179753, EPI_ISL_3179754, EPI_ISL_3179755, EPI_ISL_3179756, EPI_ISL_3179757, EPI_ISL_3179758, EPI_ISL_3179759, EPI_ISL_3179760, EPI_ISL_3179761, EPI_ISL_3179762, EPI_ISL_3179763, EPI_ISL_3179764, EPI_ISL_3179765, EPI_ISL_3179766, EPI_ISL_3179767, EPI_ISL_3179768, EPI_ISL_3179769, EPI_ISL_3179770, EPI_ISL_3179771, EPI_ISL_3179772, EPI_ISL_3179773, EPI_ISL_3179774, EPI_ISL_3179775, EPI_ISL_3179776, EPI_ISL_3179777, EPI_ISL_3179778, EPI_ISL_3179779, EPI_ISL_3179780, EPI_ISL_3179781, EPI_ISL_3179782, EPI_ISL_3179783, EPI_ISL_3179784, EPI_ISL_3179785, EPI_ISL_3179786, EPI_ISL_3179787, EPI_ISL_3179788, EPI_ISL_3179789, EPI_ISL_3179790, EPI_ISL_3179791, EPI_ISL_3179792, EPI_ISL_3179793, EPI_ISL_3179794, EPI_ISL_3179795, EPI_ISL_3179796, EPI_ISL_3179797, EPI_ISL_3179798, EPI_ISL_3179799, EPI_ISL_3179800, EPI_ISL_3179801, EPI_ISL_3179802, EPI_ISL_3179803, EPI_ISL_3179804, EPI_ISL_3179805, EPI_ISL_3179806, EPI_ISL_3179807, EPI_ISL_3179808, EPI_ISL_3179809, EPI_ISL_3179810, EPI_ISL_3179811, EPI_ISL_3179812, EPI_ISL_3179813, EPI_ISL_3179814, EPI_ISL_3179815, EPI_ISL_3179816, EPI_ISL_3179817, EPI_ISL_3179818, EPI_ISL_3179819, EPI_ISL_3179820, EPI_ISL_3179821, EPI_ISL_3179822, EPI_ISL_3179823, EPI_ISL_3179824, EPI_ISL_3179825, EPI_ISL_3179826, EPI_ISL_3179827, EPI_ISL_3179828, EPI_ISL_3179829, EPI_ISL_3179830, EPI_ISL_3179831, EPI_ISL_3179832, EPI_ISL_3179833, EPI_ISL_3179834, EPI_ISL_3179835, EPI_ISL_3179836, EPI_ISL_3179837, EPI_ISL_3179838, EPI_ISL_3179839, EPI_ISL_3179840, EPI_ISL_3179841, EPI_ISL_3179842, EPI_ISL_3179843, EPI_ISL_3179844, EPI_ISL_3179845, EPI_ISL_3179846, EPI_ISL_3179847, EPI_ISL_3179848, EPI_ISL_3179849, EPI_ISL_3179850, EPI_ISL_3179851, EPI_ISL_3179852, EPI_ISL_3179853, EPI_ISL_3179854, EPI_ISL_3179855, EPI_ISL_3179856, EPI_ISL_3179857, EPI_ISL_3179858, EPI_ISL_3179859, EPI_ISL_3179860, EPI_ISL_3179861, EPI_ISL_3179862, EPI_ISL_3179863, EPI_ISL_3179864, EPI_ISL_3179865, EPI_ISL_3179866, EPI_ISL_3179867, EPI_ISL_3179868, EPI_ISL_3179869, EPI_ISL_3179870, EPI_ISL_3179871, EPI_ISL_3179872, EPI_ISL_3179873, EPI_ISL_3179874, EPI_ISL_3179875, EPI_ISL_3179876, EPI_ISL_3179877, EPI_ISL_3179878, EPI_ISL_3179879, EPI_ISL_3179880, EPI_ISL_3179881, EPI_ISL_3179882, EPI_ISL_3179884, EPI_ISL_3179885, EPI_ISL_3179886, EPI_ISL_3179887, EPI_ISL_3179888, EPI_ISL_3179889, EPI_ISL_3179890, EPI_ISL_3179891, EPI_ISL_3179892, EPI_ISL_3179893, EPI_ISL_3179894, EPI_ISL_3179895, EPI_ISL_3179896, EPI_ISL_3179897, EPI_ISL_3179898, EPI_ISL_3179899, EPI_ISL_3179900, EPI_ISL_3179901, EPI_ISL_3179902, EPI_ISL_3179903, EPI_ISL_3179904, EPI_ISL_3179905, EPI_ISL_3179906, EPI_ISL_3179907, EPI_ISL_3179908, EPI_ISL_3179909, EPI_ISL_3179910, EPI_ISL_3179911, EPI_ISL_3179912, EPI_ISL_3179913, EPI_ISL_3179914, EPI_ISL_3179915, EPI_ISL_3179916, EPI_ISL_3179917, EPI_ISL_3179918, EPI_ISL_3179919, EPI_ISL_3179920, EPI_ISL_3179921, EPI_ISL_3179922, EPI_ISL_3179923, EPI_ISL_3179924, EPI_ISL_3179925, EPI_ISL_3179926, EPI_ISL_3179927, EPI_ISL_3179928, EPI_ISL_3179929, EPI_ISL_3179930, EPI_ISL_3179931, EPI_ISL_3179932, EPI_ISL_3179933, EPI_ISL_3179934, EPI_ISL_3179935, EPI_ISL_3179936, EPI_ISL_3179937, EPI_ISL_3179938, EPI_ISL_3179939, EPI_ISL_3179940, EPI_ISL_3179941, EPI_ISL_3179942, EPI_ISL_3179943, EPI_ISL_3179944, EPI_ISL_3179945, EPI_ISL_3179946, EPI_ISL_3179947, EPI_ISL_3179948, EPI_ISL_3179949, EPI_ISL_3179950, EPI_ISL_3179951, EPI_ISL_3179952, EPI_ISL_3179953, EPI_ISL_3179954, EPI_ISL_3179955, EPI_ISL_3179956, EPI_ISL_3179957, EPI_ISL_3179958, EPI_ISL_3179959, EPI_ISL_3179960, EPI_ISL_3179961, EPI_ISL_3179962, EPI_ISL_3179963, EPI_ISL_3179964, EPI_ISL_3179965, EPI_ISL_3179966, EPI_ISL_3179967, EPI_ISL_3179968, EPI_ISL_3179969, EPI_ISL_3179970, EPI_ISL_3179971, EPI_ISL_3179972, EPI_ISL_3179973, EPI_ISL_3179974, EPI_ISL_3179975, EPI_ISL_3179976, EPI_ISL_3179977, EPI_ISL_3179978, EPI_ISL_3179979, EPI_ISL_3179980, EPI_ISL_3179981, EPI_ISL_3179982, EPI_ISL_3179983, EPI_ISL_3179984, EPI_ISL_3179985, EPI_ISL_3179986, EPI_ISL_3179987, EPI_ISL_3179988, EPI_ISL_3179989, EPI_ISL_3179990, EPI_ISL_3179991, EPI_ISL_3179992, EPI_ISL_3179993, EPI_ISL_3179994, EPI_ISL_3179995, EPI_ISL_3179996, EPI_ISL_3179997, EPI_ISL_3179998, EPI_ISL_3179999, EPI_ISL_3180000, EPI_ISL_3180001, EPI_ISL_3180002, EPI_ISL_3180003, EPI_ISL_3180004, EPI_ISL_3180005, EPI_ISL_3180006, EPI_ISL_3180007, EPI_ISL_3180009, EPI_ISL_3180010, EPI_ISL_3180011, EPI_ISL_3180012, EPI_ISL_3180013, EPI_ISL_3180014, EPI_ISL_3180015, EPI_ISL_3180016, EPI_ISL_3180017, EPI_ISL_3180018, EPI_ISL_3180019, EPI_ISL_3180020, EPI_ISL_3180021, EPI_ISL_3180022, EPI_ISL_3180023, EPI_ISL_3180024, EPI_ISL_3180025, EPI_ISL_3180026, EPI_ISL_3180027, EPI_ISL_3180028, EPI_ISL_3180029, EPI_ISL_3180030, EPI_ISL_3180031, EPI_ISL_3180032, EPI_ISL_3180033, EPI_ISL_3180034, EPI_ISL_3180035, EPI_ISL_3180036, EPI_ISL_3180037, EPI_ISL_3180038, EPI_ISL_3180039, EPI_ISL_3180040, EPI_ISL_3180041, EPI_ISL_3180042, EPI_ISL_3180043, EPI_ISL_3180044, EPI_ISL_3180045, EPI_ISL_3180046, EPI_ISL_3180047, EPI_ISL_3180048, EPI_ISL_3180049, EPI_ISL_3180050, EPI_ISL_3180051, EPI_ISL_3180052, EPI_ISL_3180053, EPI_ISL_3180054, EPI_ISL_3180055, EPI_ISL_3180056, EPI_ISL_3180057, EPI_ISL_3180058, EPI_ISL_3180059, EPI_ISL_3180060, EPI_ISL_3180061, EPI_ISL_3180062, EPI_ISL_3180063, EPI_ISL_3180064, EPI_ISL_3180065, EPI_ISL_3180066, EPI_ISL_3180067, EPI_ISL_3180068, EPI_ISL_3180069, EPI_ISL_3180070, EPI_ISL_3180071, EPI_ISL_3180072, EPI_ISL_3180073, EPI_ISL_3180074, EPI_ISL_3180075, EPI_ISL_3180076, EPI_ISL_3180077, EPI_ISL_3180078, EPI_ISL_3180079, EPI_ISL_3180080, EPI_ISL_3180081, EPI_ISL_3180082, EPI_ISL_3180083, EPI_ISL_3180084, EPI_ISL_3180085, EPI_ISL_3180086, EPI_ISL_3180087, EPI_ISL_3180088, EPI_ISL_3180089, EPI_ISL_3180090, EPI_ISL_3180091, EPI_ISL_3180092, EPI_ISL_3180093, EPI_ISL_3180094, EPI_ISL_3180095, EPI_ISL_3180096, EPI_ISL_3180097, EPI_ISL_3180098, EPI_ISL_3180099, EPI_ISL_3180100, EPI_ISL_3180101, EPI_ISL_3180102, EPI_ISL_3180103, EPI_ISL_3180104, EPI_ISL_3180105, EPI_ISL_3180106, EPI_ISL_3180107, EPI_ISL_3180108, EPI_ISL_3180109, EPI_ISL_3180110, EPI_ISL_3180111, EPI_ISL_3180112, EPI_ISL_3180113, EPI_ISL_3180114, EPI_ISL_3180115, EPI_ISL_3180116, EPI_ISL_3180117, EPI_ISL_3180118, EPI_ISL_3180119, EPI_ISL_3180120, EPI_ISL_3180121, EPI_ISL_3180122, EPI_ISL_3180123, EPI_ISL_3180124, EPI_ISL_3180125, EPI_ISL_3180126, EPI_ISL_3180127, EPI_ISL_3180128, EPI_ISL_3180129, EPI_ISL_3180130, EPI_ISL_3180131, EPI_ISL_3180132, EPI_ISL_3180133, EPI_ISL_3180134, EPI_ISL_3180135, EPI_ISL_3180136, EPI_ISL_3180137, EPI_ISL_3180138, EPI_ISL_3180139, EPI_ISL_3180140, EPI_ISL_3180141, EPI_ISL_3180142, EPI_ISL_3180143, EPI_ISL_3180144, EPI_ISL_3180145, EPI_ISL_3180146, EPI_ISL_3180147, EPI_ISL_3180148, EPI_ISL_3180149, EPI_ISL_3180150, EPI_ISL_3180151, EPI_ISL_3180152, EPI_ISL_3180153, EPI_ISL_3180154, EPI_ISL_3180155, EPI_ISL_3180156, EPI_ISL_3180157, EPI_ISL_3180158, EPI_ISL_3180159, EPI_ISL_3180160, EPI_ISL_3180161, EPI_ISL_3180162, EPI_ISL_3180163, EPI_ISL_3180164, EPI_ISL_3180165, EPI_ISL_3180166, EPI_ISL_3180167, EPI_ISL_3180168, EPI_ISL_3180169, EPI_ISL_3180170, EPI_ISL_3180171, EPI_ISL_3180172, EPI_ISL_3180173, EPI_ISL_3180174, EPI_ISL_3180175, EPI_ISL_3180176, EPI_ISL_3180177, EPI_ISL_3180178, EPI_ISL_3180179, EPI_ISL_3180180, EPI_ISL_3180181, EPI_ISL_3180182, EPI_ISL_3180183, EPI_ISL_3180184, EPI_ISL_3180185, EPI_ISL_3180186, EPI_ISL_3180187, EPI_ISL_3180188, EPI_ISL_3180189, EPI_ISL_3180190, EPI_ISL_3180191, EPI_ISL_3180192, EPI_ISL_3180193, EPI_ISL_3180194, EPI_ISL_3180195, EPI_ISL_3180196, EPI_ISL_3180197, EPI_ISL_3180198, EPI_ISL_3180199, EPI_ISL_3180200, EPI_ISL_3180201, EPI_ISL_3180202, EPI_ISL_3180203, EPI_ISL_3180204, EPI_ISL_3180205, EPI_ISL_3180206, EPI_ISL_3180207, EPI_ISL_3180208, EPI_ISL_3180209, EPI_ISL_3180210, EPI_ISL_3180211, EPI_ISL_3180212, EPI_ISL_3180213, EPI_ISL_3180214, EPI_ISL_3180215, EPI_ISL_3180216, EPI_ISL_3180217, EPI_ISL_3180218, EPI_ISL_3180219, EPI_ISL_3180220, EPI_ISL_3180221, EPI_ISL_3180222, EPI_ISL_3180223, EPI_ISL_3180224, EPI_ISL_3180225, EPI_ISL_3180226, EPI_ISL_3180227, EPI_ISL_3180228, EPI_ISL_3180229, EPI_ISL_3180230, EPI_ISL_3180231, EPI_ISL_3180232, EPI_ISL_3180233, EPI_ISL_3180234, EPI_ISL_3180235, EPI_ISL_3180236, EPI_ISL_3180237, EPI_ISL_3180238, EPI_ISL_3180239, EPI_ISL_3180240, EPI_ISL_3180241, EPI_ISL_3180242, EPI_ISL_3180243, EPI_ISL_3180244, EPI_ISL_3180245, EPI_ISL_3180246, EPI_ISL_3180247, EPI_ISL_3180248, EPI_ISL_3180249, EPI_ISL_3180250, EPI_ISL_3180251, EPI_ISL_3180252, EPI_ISL_3180253, EPI_ISL_3180254, EPI_ISL_3180255, EPI_ISL_3180256, EPI_ISL_3180257, EPI_ISL_3180258, EPI_ISL_3180259, EPI_ISL_3180260, EPI_ISL_3180261, EPI_ISL_3180262, EPI_ISL_3180263, EPI_ISL_3180264, EPI_ISL_3180265, EPI_ISL_3180266, EPI_ISL_3180267, EPI_ISL_3180268, EPI_ISL_3180269, EPI_ISL_3180270, EPI_ISL_3180271, EPI_ISL_3180272, EPI_ISL_3180273, EPI_ISL_3180274, EPI_ISL_3180275, EPI_ISL_3180276, EPI_ISL_3180277, EPI_ISL_3180278, EPI_ISL_3180279, EPI_ISL_3180280, EPI_ISL_3180281, EPI_ISL_3180282, EPI_ISL_3180283, EPI_ISL_3180284, EPI_ISL_3180285, EPI_ISL_3180286, EPI_ISL_3180287, EPI_ISL_3180288, EPI_ISL_3180289, EPI_ISL_3180290, EPI_ISL_3180291, EPI_ISL_3180292, EPI_ISL_3180293,

| | | | |
|---|---|---|---|
| EPI_ISL_3180294, EPI_ISL_3180295, EPI_ISL_3180296, EPI_ISL_3180297, EPI_ISL_3180298, EPI_ISL_3180299, EPI_ISL_3180300, EPI_ISL_3180301, EPI_ISL_3180302, EPI_ISL_3180303, EPI_ISL_3180304, EPI_ISL_3180305, EPI_ISL_3180306, EPI_ISL_3180307, EPI_ISL_3180308, EPI_ISL_3180309, EPI_ISL_3180310, EPI_ISL_3180311, EPI_ISL_3180312, EPI_ISL_3180313, EPI_ISL_3180314, EPI_ISL_3180315, EPI_ISL_3180316, EPI_ISL_3180317, EPI_ISL_3180318, EPI_ISL_3180319, EPI_ISL_3180320, EPI_ISL_3180321, EPI_ISL_3180322, EPI_ISL_3180323, EPI_ISL_3180324, EPI_ISL_3180325, EPI_ISL_3180326, EPI_ISL_3180327, EPI_ISL_3180328, EPI_ISL_3180329, EPI_ISL_3180330, EPI_ISL_3180331, EPI_ISL_3180332, EPI_ISL_3180333, EPI_ISL_3180335, EPI_ISL_3180336, EPI_ISL_3180337, EPI_ISL_3180338, EPI_ISL_3180339, EPI_ISL_3180340, EPI_ISL_3180341, EPI_ISL_3180342, EPI_ISL_3180343, EPI_ISL_3180344, EPI_ISL_3180345, EPI_ISL_3180346, EPI_ISL_3180347, EPI_ISL_3180348, EPI_ISL_3180349, EPI_ISL_3180350, EPI_ISL_3180351, EPI_ISL_3180352, EPI_ISL_3180353, EPI_ISL_3180354, EPI_ISL_3180355, EPI_ISL_3180356, EPI_ISL_3180357, EPI_ISL_3180358, EPI_ISL_3180359, EPI_ISL_3180360, EPI_ISL_3180361, EPI_ISL_3180362, EPI_ISL_3180363, EPI_ISL_3180364, EPI_ISL_3180365, EPI_ISL_3180366, EPI_ISL_3180367, EPI_ISL_3180368, EPI_ISL_3180369, EPI_ISL_3180370, EPI_ISL_3180371, EPI_ISL_3180372, EPI_ISL_3180373, EPI_ISL_3180374, EPI_ISL_3180375, EPI_ISL_3180376, EPI_ISL_3180377, EPI_ISL_3180378, EPI_ISL_3180379, EPI_ISL_3180380, EPI_ISL_3180381, EPI_ISL_3180382, EPI_ISL_3180383, EPI_ISL_3180384, EPI_ISL_3180385, EPI_ISL_3180386, EPI_ISL_3180387, EPI_ISL_3180388, EPI_ISL_3180389, EPI_ISL_3180390, EPI_ISL_3180391, EPI_ISL_3180392, EPI_ISL_3180393, EPI_ISL_3180394, EPI_ISL_3180395, EPI_ISL_3180396, EPI_ISL_3180397, EPI_ISL_3180398, EPI_ISL_3180399, EPI_ISL_3180400, EPI_ISL_3180401, EPI_ISL_3180402, EPI_ISL_3180403, EPI_ISL_3180404, EPI_ISL_3180405, EPI_ISL_3180406, EPI_ISL_3180407, EPI_ISL_3180408, EPI_ISL_3180409, EPI_ISL_3180410, EPI_ISL_3180411, EPI_ISL_3180412, EPI_ISL_3180413, EPI_ISL_3180414, EPI_ISL_3180415, EPI_ISL_3180416, EPI_ISL_3180417, EPI_ISL_3180418, EPI_ISL_3180419, EPI_ISL_3180420, EPI_ISL_3180421, EPI_ISL_3180422, EPI_ISL_3180423, EPI_ISL_3180424, EPI_ISL_3180425, EPI_ISL_3180426, EPI_ISL_3180427, EPI_ISL_3180428, EPI_ISL_3180429, EPI_ISL_3180430, EPI_ISL_3180431, EPI_ISL_3180432, EPI_ISL_3180433, EPI_ISL_3180434, EPI_ISL_3180435, EPI_ISL_3180436, EPI_ISL_3180437, EPI_ISL_3180438 | | | |
| see above | Respiratory Virus Unit, Microbiology Services Colindale, Public Health England | COVID-19 Genomics UK (COG-UK) Consortium | PHE Covid Sequencing Team |
| EPI_ISL_3180439, EPI_ISL_3180440, EPI_ISL_3180441, EPI_ISL_3180442, EPI_ISL_3180443, EPI_ISL_3180444, EPI_ISL_3180445, EPI_ISL_3180446, EPI_ISL_3180447, EPI_ISL_3180448, EPI_ISL_3180449, EPI_ISL_3180450, EPI_ISL_3180451, EPI_ISL_3180452, EPI_ISL_3180453, EPI_ISL_3180454, EPI_ISL_3180455, EPI_ISL_3180456, EPI_ISL_3180457, EPI_ISL_3180458, EPI_ISL_3180459, EPI_ISL_3180460, EPI_ISL_3180461, EPI_ISL_3180462, EPI_ISL_3180463, EPI_ISL_3180464, EPI_ISL_3180465, EPI_ISL_3180466, EPI_ISL_3180467, EPI_ISL_3180468, EPI_ISL_3180469, EPI_ISL_3180470, EPI_ISL_3180471, EPI_ISL_3180472, EPI_ISL_3180473, EPI_ISL_3180474, EPI_ISL_3180475 | | | |
| see above | Originating lab: Wales Specialist Virology Centre Sequencing lab: Pathogen Genomics Unit | Public Health Wales Microbiology Cardiff Wales Specialist Virology Centre | Catherine Moore, Johnathan Evans, Laura Gifford, Malorie Perry, Simon Cottrell, Angela Marchbank, Alec Birchley, Alexander Adams, Amy Gaskin, Bree Gatica-Wilcox, Jason Coombes, Joel Southgate, Lauren Gilbert, Lee Graham, Nicole Pacchiarini, Sara Kumziene-Summerhayes, Sarah Taylor, Sophie Jones, Sara Rey, Matthew Bull, Joanne Watkins, Sally Corden, Tom Connor |
| EPI_ISL_3180476, EPI_ISL_3180477, EPI_ISL_3180478, EPI_ISL_3180479, EPI_ISL_3180480, EPI_ISL_3180481, EPI_ISL_3180482, EPI_ISL_3180483, EPI_ISL_3180484, EPI_ISL_3180485, EPI_ISL_3180486, EPI_ISL_3180487, EPI_ISL_3180488, EPI_ISL_3180489, EPI_ISL_3180490, EPI_ISL_3180491, EPI_ISL_3180492, EPI_ISL_3180493, EPI_ISL_3180494, EPI_ISL_3180495, EPI_ISL_3180496, EPI_ISL_3180497, EPI_ISL_3180498, EPI_ISL_3180499, EPI_ISL_3180500, EPI_ISL_3180501, EPI_ISL_3180502, EPI_ISL_3180503, EPI_ISL_3180504, EPI_ISL_3180505, EPI_ISL_3180506, EPI_ISL_3180507, EPI_ISL_3180508, EPI_ISL_3180509, EPI_ISL_3180510, EPI_ISL_3180511, EPI_ISL_3180512, EPI_ISL_3180513, EPI_ISL_3180514, EPI_ISL_3180515, EPI_ISL_3180516, EPI_ISL_3180517, EPI_ISL_3180518, EPI_ISL_3180519, EPI_ISL_3180520, EPI_ISL_3180521, EPI_ISL_3180522, EPI_ISL_3180523, EPI_ISL_3180524, EPI_ISL_3180525, EPI_ISL_3180526, EPI_ISL_3180527, EPI_ISL_3180528, EPI_ISL_3180529, EPI_ISL_3180530, EPI_ISL_3180531, EPI_ISL_3180532, EPI_ISL_3180533, EPI_ISL_3180534, EPI_ISL_3180535, EPI_ISL_3180536, EPI_ISL_3180537, EPI_ISL_3180538, EPI_ISL_3180539, EPI_ISL_3180540, EPI_ISL_3180541, EPI_ISL_3180542, EPI_ISL_3180543, EPI_ISL_3180544, EPI_ISL_3180545, EPI_ISL_3180546, EPI_ISL_3180547, EPI_ISL_3180548, EPI_ISL_3180549, EPI_ISL_3180550, EPI_ISL_3180551, EPI_ISL_3180552, EPI_ISL_3180553, EPI_ISL_3180554, EPI_ISL_3180555, EPI_ISL_3180556, EPI_ISL_3180557, EPI_ISL_3180558, EPI_ISL_3180559, EPI_ISL_3180560, EPI_ISL_3180561, EPI_ISL_3180562, EPI_ISL_3180563, EPI_ISL_3180564, EPI_ISL_3180565, EPI_ISL_3180566, EPI_ISL_3180567, EPI_ISL_3180568, EPI_ISL_3180569, EPI_ISL_3180570, EPI_ISL_3180571, EPI_ISL_3180572, EPI_ISL_3180573, EPI_ISL_3180574, EPI_ISL_3180575, EPI_ISL_3180576, EPI_ISL_3180577, EPI_ISL_3180578, EPI_ISL_3180579, EPI_ISL_3180580, EPI_ISL_3180581, EPI_ISL_3180582, EPI_ISL_3180583, EPI_ISL_3180584, EPI_ISL_3180585, EPI_ISL_3180586, EPI_ISL_3180587, EPI_ISL_3180588, EPI_ISL_3180589, EPI_ISL_3180590, EPI_ISL_3180591, EPI_ISL_3180592, EPI_ISL_3180593, EPI_ISL_3180594, EPI_ISL_3180595, EPI_ISL_3180596, EPI_ISL_3180597, EPI_ISL_3180598, EPI_ISL_3180599, EPI_ISL_3180600, EPI_ISL_3180601, EPI_ISL_3180602, EPI_ISL_3180603, EPI_ISL_3180604, EPI_ISL_3180605, EPI_ISL_3180606, EPI_ISL_3180607, EPI_ISL_3180608, EPI_ISL_3180609, EPI_ISL_3180610, EPI_ISL_3180611, EPI_ISL_3180612, EPI_ISL_3180613, EPI_ISL_3180614, EPI_ISL_3180615, EPI_ISL_3180616, EPI_ISL_3180617, EPI_ISL_3180618, EPI_ISL_3180619, EPI_ISL_3180620, EPI_ISL_3180621, EPI_ISL_3180622, EPI_ISL_3180623, EPI_ISL_3180624, EPI_ISL_3180625, EPI_ISL_3180626, EPI_ISL_3180627, EPI_ISL_3180628, EPI_ISL_3180629, EPI_ISL_3180630, EPI_ISL_3180631, EPI_ISL_3180632, EPI_ISL_3180633, EPI_ISL_3180634, EPI_ISL_3180635, EPI_ISL_3180636, EPI_ISL_3180637, EPI_ISL_3180638, EPI_ISL_3180639, EPI_ISL_3180640, EPI_ISL_3180641, EPI_ISL_3180642, EPI_ISL_3180643, EPI_ISL_3180644, EPI_ISL_3180645, EPI_ISL_3180646, EPI_ISL_3180647, EPI_ISL_3180648, EPI_ISL_3180649, EPI_ISL_3180650, EPI_ISL_3180651, EPI_ISL_3180652, EPI_ISL_3180653, EPI_ISL_3180654, EPI_ISL_3180655, EPI_ISL_3180656, EPI_ISL_3180657, EPI_ISL_3180658, EPI_ISL_3180659, EPI_ISL_3180660, EPI_ISL_3180661, EPI_ISL_3180662, EPI_ISL_3180663, EPI_ISL_3180664, EPI_ISL_3180665, EPI_ISL_3180666, EPI_ISL_3180667, EPI_ISL_3180668, EPI_ISL_3180669, EPI_ISL_3180670, EPI_ISL_3180671, EPI_ISL_3180672, EPI_ISL_3180673, EPI_ISL_3180674, EPI_ISL_3180675, EPI_ISL_3180676, EPI_ISL_3180677, EPI_ISL_3180678, EPI_ISL_3180679, EPI_ISL_3180680, EPI_ISL_3180681, EPI_ISL_3180682, EPI_ISL_3180683, EPI_ISL_3180684, EPI_ISL_3180685, EPI_ISL_3180686, EPI_ISL_3180687, EPI_ISL_3180688, EPI_ISL_3180689, EPI_ISL_3180690, EPI_ISL_3180691, EPI_ISL_3180692, EPI_ISL_3180693, EPI_ISL_3180694, EPI_ISL_3180695, EPI_ISL_3180696, EPI_ISL_3180697, EPI_ISL_3180698, EPI_ISL_3180699, EPI_ISL_3180700, EPI_ISL_3180701, EPI_ISL_3180702, EPI_ISL_3180703, EPI_ISL_3180704, EPI_ISL_3180705, EPI_ISL_3180706, EPI_ISL_3180707, EPI_ISL_3180708, EPI_ISL_3180709, EPI_ISL_3180710, EPI_ISL_3180712, EPI_ISL_3180713, EPI_ISL_3180714, EPI_ISL_3180715, EPI_ISL_3180716, EPI_ISL_3180717, EPI_ISL_3180718, EPI_ISL_3180719, EPI_ISL_3180720, EPI_ISL_3180721, EPI_ISL_3180722, EPI_ISL_3180723, EPI_ISL_3180724, EPI_ISL_3180725, EPI_ISL_3180726, EPI_ISL_3180727, EPI_ISL_3180728, EPI_ISL_3180729, EPI_ISL_3180730, EPI_ISL_3180731, EPI_ISL_3180732, EPI_ISL_3180733, EPI_ISL_3180734, EPI_ISL_3180735, EPI_ISL_3180736, EPI_ISL_3180737, EPI_ISL_3180738, EPI_ISL_3180739, EPI_ISL_3180740, EPI_ISL_3180741, EPI_ISL_3180742, EPI_ISL_3180743, EPI_ISL_3180744, EPI_ISL_3180745, EPI_ISL_3180746, EPI_ISL_3180747, EPI_ISL_3180748, EPI_ISL_3180749, EPI_ISL_3180750, EPI_ISL_3180751, EPI_ISL_3180752, EPI_ISL_3180753, EPI_ISL_3180754, EPI_ISL_3180755, EPI_ISL_3180756, EPI_ISL_3180757, EPI_ISL_3180758, EPI_ISL_3180759, EPI_ISL_3180760, EPI_ISL_3180761, EPI_ISL_3180762, EPI_ISL_3180763, EPI_ISL_3180764, EPI_ISL_3180765, EPI_ISL_3180766, EPI_ISL_3180767, EPI_ISL_3180768, EPI_ISL_3180769, EPI_ISL_3180770, EPI_ISL_3180771, EPI_ISL_3180772, EPI_ISL_3180773, EPI_ISL_3180774, EPI_ISL_3180775, EPI_ISL_3180776, EPI_ISL_3180777, EPI_ISL_3180778, EPI_ISL_3180779, EPI_ISL_3180780, EPI_ISL_3180781, EPI_ISL_3180782, EPI_ISL_3180783, EPI_ISL_3180784, EPI_ISL_3180785, EPI_ISL_3180786, EPI_ISL_3180787, EPI_ISL_3180788, EPI_ISL_3180789, EPI_ISL_3180790, EPI_ISL_3180791, EPI_ISL_3180792, EPI_ISL_3180793, EPI_ISL_3180794, EPI_ISL_3180795, EPI_ISL_3180796, EPI_ISL_3180797, EPI_ISL_3180798, EPI_ISL_3180799, EPI_ISL_3180800, EPI_ISL_3180801, EPI_ISL_3180802, EPI_ISL_3180803, EPI_ISL_3180804, EPI_ISL_3180805, EPI_ISL_3180806, EPI_ISL_3180807, EPI_ISL_3180808, EPI_ISL_3180809, EPI_ISL_3180810, EPI_ISL_3180811, EPI_ISL_3180812, EPI_ISL_3180813, EPI_ISL_3180814, EPI_ISL_3180815, EPI_ISL_3180816, EPI_ISL_3180817, EPI_ISL_3180818, EPI_ISL_3180819, EPI_ISL_3180820, EPI_ISL_3180821, EPI_ISL_3180822, EPI_ISL_3180823, EPI_ISL_3180824, EPI_ISL_3180825, EPI_ISL_3180826, EPI_ISL_3180827, EPI_ISL_3180828, EPI_ISL_3180829, EPI_ISL_3180830, EPI_ISL_3180831, EPI_ISL_3180832, EPI_ISL_3180833, EPI_ISL_3180834, EPI_ISL_3180835, EPI_ISL_3180836, EPI_ISL_3180837, EPI_ISL_3180838, EPI_ISL_3180839, EPI_ISL_3180840, EPI_ISL_3180841, EPI_ISL_3180842, EPI_ISL_3180843, EPI_ISL_3180844, EPI_ISL_3180845, EPI_ISL_3180846, EPI_ISL_3180847, EPI_ISL_3180848, EPI_ISL_3180849, EPI_ISL_3180850, EPI_ISL_3180851, EPI_ISL_3180852, EPI_ISL_3180853, EPI_ISL_3180854, EPI_ISL_3180855, EPI_ISL_3180856, EPI_ISL_3180857, EPI_ISL_3180858, EPI_ISL_3180859, EPI_ISL_3180860, EPI_ISL_3180861, EPI_ISL_3180862, EPI_ISL_3180863, EPI_ISL_3180864, EPI_ISL_3180865, EPI_ISL_3180866, EPI_ISL_3180867, EPI_ISL_3180868, EPI_ISL_3180869, EPI_ISL_3180870, EPI_ISL_3180871, EPI_ISL_3180872, EPI_ISL_3180873, EPI_ISL_3180874, EPI_ISL_3180875, EPI_ISL_3180876, EPI_ISL_3180877, EPI_ISL_3180878, EPI_ISL_3180879, EPI_ISL_3180880, EPI_ISL_3180881, EPI_ISL_3180882, EPI_ISL_3180883, EPI_ISL_3180884, EPI_ISL_3180885, EPI_ISL_3180886, EPI_ISL_3180887, EPI_ISL_3180888, EPI_ISL_3180889, EPI_ISL_3180890, EPI_ISL_3180891, EPI_ISL_3180892, EPI_ISL_3180893, EPI_ISL_3180894, EPI_ISL_3180895, EPI_ISL_3180896, EPI_ISL_3180897, EPI_ISL_3180898, EPI_ISL_3180899, EPI_ISL_3180900, EPI_ISL_3180901, EPI_ISL_3180902, EPI_ISL_3180903, EPI_ISL_3180904, EPI_ISL_3180905, EPI_ISL_3180906, EPI_ISL_3180907, EPI_ISL_3180908, EPI_ISL_3180909, EPI_ISL_3180910, EPI_ISL_3180911, EPI_ISL_3180912, EPI_ISL_3180913, EPI_ISL_3180914, EPI_ISL_3180915, EPI_ISL_3180916, EPI_ISL_3180917, EPI_ISL_3180918, EPI_ISL_3180919, EPI_ISL_3180920, EPI_ISL_3180921, EPI_ISL_3180922, EPI_ISL_3180923, EPI_ISL_3180924, EPI_ISL_3180925, EPI_ISL_3180926, EPI_ISL_3180927, EPI_ISL_3180928, EPI_ISL_3180929, EPI_ISL_3180930, EPI_ISL_3180931, EPI_ISL_3180932, EPI_ISL_3180933, EPI_ISL_3180934, EPI_ISL_3180935, EPI_ISL_3180936, EPI_ISL_3180937, EPI_ISL_3180938, EPI_ISL_3180939, EPI_ISL_3180940, EPI_ISL_3180941, EPI_ISL_3180942, EPI_ISL_3180943, EPI_ISL_3180944, EPI_ISL_3180945, EPI_ISL_3180946, EPI_ISL_3180947, EPI_ISL_3180948, EPI_ISL_3180949, EPI_ISL_3180950, EPI_ISL_3180951, EPI_ISL_3180952, EPI_ISL_3180953, EPI_ISL_3180954, EPI_ISL_3180955, EPI_ISL_3180956, EPI_ISL_3180957, EPI_ISL_3180958, EPI_ISL_3180959, EPI_ISL_3180960, EPI_ISL_3180961, EPI_ISL_3180962, EPI_ISL_3180963, EPI_ISL_3180964, EPI_ISL_3180965, EPI_ISL_3180966, EPI_ISL_3180967, EPI_ISL_3180968, EPI_ISL_3180969, EPI_ISL_3180970, EPI_ISL_3180971, EPI_ISL_3180972, EPI_ISL_3180973, EPI_ISL_3180974, EPI_ISL_3180975, EPI_ISL_3180976, EPI_ISL_3180977, EPI_ISL_3180978, EPI_ISL_3180979, EPI_ISL_3180980, EPI_ISL_3180981, EPI_ISL_3180982, EPI_ISL_3180983, EPI_ISL_3180984, EPI_ISL_3180985, EPI_ISL_3180986, EPI_ISL_3180987, EPI_ISL_3180988, EPI_ISL_3180989, EPI_ISL_3180990, EPI_ISL_3180991, EPI_ISL_3180992, EPI_ISL_3180993, EPI_ISL_3180994, EPI_ISL_3180995, EPI_ISL_3180996, EPI_ISL_3180997, EPI_ISL_3180998, EPI_ISL_3180999, EPI_ISL_3181000, EPI_ISL_3181001, EPI_ISL_3181002, EPI_ISL_3181003, EPI_ISL_3181004, EPI_ISL_3181005, EPI_ISL_3181006, EPI_ISL_3181007, EPI_ISL_3181008, EPI_ISL_3181009, EPI_ISL_3181010, EPI_ISL_3181011, EPI_ISL_3181012, EPI_ISL_3181013, EPI_ISL_3181014, EPI_ISL_3181015, EPI_ISL_3181016, EPI_ISL_3181017, EPI_ISL_3181018, EPI_ISL_3181019, EPI_ISL_3181020, EPI_ISL_3181021, EPI_ISL_3181022, EPI_ISL_3181023, EPI_ISL_3181024, EPI_ISL_3181025, EPI_ISL_3181026, EPI_ISL_3181027, EPI_ISL_3181028, EPI_ISL_3181029, EPI_ISL_3181030, EPI_ISL_3181031, EPI_ISL_3181032, EPI_ISL_3181033, EPI_ISL_3181034, EPI_ISL_3181035, EPI_ISL_3181036, EPI_ISL_3181037, EPI_ISL_3181038, EPI_ISL_3181039, EPI_ISL_3181040, EPI_ISL_3181041, EPI_ISL_3181042, EPI_ISL_3181043, EPI_ISL_3181044, EPI_ISL_3181045, EPI_ISL_3181046, EPI_ISL_3181047, EPI_ISL_3181048, EPI_ISL_3181049, EPI_ISL_3181050, EPI_ISL_3181051, EPI_ISL_3181052, EPI_ISL_3181053, EPI_ISL_3181054, EPI_ISL_3181055, EPI_ISL_3181056, EPI_ISL_3181057, EPI_ISL_3181058, EPI_ISL_3181059, EPI_ISL_3181060, EPI_ISL_3181061, EPI_ISL_3181062, EPI_ISL_3181063, EPI_ISL_3181064, EPI_ISL_3181065, EPI_ISL_3181066, EPI_ISL_3181067, EPI_ISL_3181068, EPI_ISL_3181069, EPI_ISL_3181070, EPI_ISL_3181071, EPI_ISL_3181072, EPI_ISL_3181073, EPI_ISL_3181074, EPI_ISL_3181075, EPI_ISL_3181076, EPI_ISL_3181077, EPI_ISL_3181078, EPI_ISL_3181079, EPI_ISL_3181080, EPI_ISL_3181081, EPI_ISL_3181082, EPI_ISL_3181083, EPI_ISL_3181084, EPI_ISL_3181085, EPI_ISL_3181086, EPI_ISL_3181087, EPI_ISL_3181088, EPI_ISL_3181089, EPI_ISL_3181090, EPI_ISL_3181091, EPI_ISL_3181092, EPI_ISL_3181093, EPI_ISL_3181094, EPI_ISL_3181095, EPI_ISL_3181096, EPI_ISL_3181097, EPI_ISL_3181098, EPI_ISL_3181099, EPI_ISL_3181100, EPI_ISL_3181101, EPI_ISL_3181102, EPI_ISL_3181103, EPI_ISL_3181104, EPI_ISL_3181105, EPI_ISL_3181106, EPI_ISL_3181107, EPI_ISL_3181108, EPI_ISL_3181109, EPI_ISL_3181110, EPI_ISL_3181111, EPI_ISL_3181112, EPI_ISL_3181113, EPI_ISL_3181114, EPI_ISL_3181115, EPI_ISL_3181116, EPI_ISL_3181117, EPI_ISL_3181118, EPI_ISL_3181119, EPI_ISL_3181120, EPI_ISL_3181121, EPI_ISL_3181122, EPI_ISL_3181123, EPI_ISL_3181124, EPI_ISL_3181125, EPI_ISL_3181126, EPI_ISL_3181127, EPI_ISL_3181128, EPI_ISL_3181129, EPI_ISL_3181130, EPI_ISL_3181131, EPI_ISL_3181132, EPI_ISL_3181133, EPI_ISL_3181134, EPI_ISL_3181135, EPI_ISL_3181136, EPI_ISL_3181137, EPI_ISL_3181138, EPI_ISL_3181139, EPI_ISL_3181140, EPI_ISL_3181141, EPI_ISL_3181142, EPI_ISL_3181143, EPI_ISL_3181144, EPI_ISL_3181145, EPI_ISL_3181146, EPI_ISL_3181147, EPI_ISL_3181148, EPI_ISL_3181149, EPI_ISL_3181150, EPI_ISL_3181151, EPI_ISL_3181152, EPI_ISL_3181153, EPI_ISL_3181154, EPI_ISL_3181155, EPI_ISL_3181156, EPI_ISL_3181157, EPI_ISL_3181158, EPI_ISL_3181159, EPI_ISL_3181160, EPI_ISL_3181161, EPI_ISL_3181162, EPI_ISL_3181163, EPI_ISL_3181164, EPI_ISL_3181165, EPI_ISL_3181166, EPI_ISL_3181167, EPI_ISL_3181168, EPI_ISL_3181169, EPI_ISL_3181170, EPI_ISL_3181171, EPI_ISL_3181172, EPI_ISL_3181173, EPI_ISL_3181174, EPI_ISL_3181175, EPI_ISL_3181176, EPI_ISL_3181177, EPI_ISL_3181178, EPI_ISL_3181179, EPI_ISL_3181180, EPI_ISL_3181181, EPI_ISL_3181182, EPI_ISL_3181183, EPI_ISL_3181184, EPI_ISL_3181185, EPI_ISL_3181186, EPI_ISL_3181187, EPI_ISL_3181188, EPI_ISL_3181189, EPI_ISL_3181190, EPI_ISL_3181191, EPI_ISL_3181192, EPI_ISL_3181193, EPI_ISL_3181194, EPI_ISL_3181195, EPI_ISL_3181196, EPI_ISL_3181197, EPI_ISL_3181198, EPI_ISL_3181199, EPI_ISL_3181200, EPI_ISL_3181201, EPI_ISL_3181202, EPI_ISL_3181203, EPI_ISL_3181204, EPI_ISL_3181205, EPI_ISL_3181206, EPI_ISL_3181207, EPI_ISL_3181208, EPI_ISL_3181209, EPI_ISL_3181210, EPI_ISL_3181211, EPI_ISL_3181212, EPI_ISL_3181213, EPI_ISL_3181214, EPI_ISL_3181215, EPI_ISL_3181216, EPI_ISL_3181217, EPI_ISL_3181218, EPI_ISL_3181219, EPI_ISL_3181220, EPI_ISL_3181221, EPI_ISL_3181222, EPI_ISL_3181223, EPI_ISL_3181224, EPI_ISL_3181225, EPI_ISL_3181226, EPI_ISL_3181227, EPI_ISL_3181228, EPI_ISL_3181229, EPI_ISL_3181230, EPI_ISL_3181231, EPI_ISL_3181232, EPI_ISL_3181233, EPI_ISL_3181234, EPI_ISL_3181235, EPI_ISL_3181236, EPI_ISL_3181237, EPI_ISL_3181238, EPI_ISL_3181239, EPI_ISL_3181240, EPI_ISL_3181241, EPI_ISL_3181242, EPI_ISL_3181243, EPI_ISL_3181244, EPI_ISL_3181245, EPI_ISL_3181246, EPI_ISL_3181247, EPI_ISL_3181248, EPI_ISL_3181249, EPI_ISL_3181250, EPI_ISL_3181251, EPI_ISL_3181252, EPI_ISL_3181253, EPI_ISL_3181254, EPI_ISL_3181255, EPI_ISL_3181256, EPI_ISL_3181257, EPI_ISL_3181258, EPI_ISL_3181259, EPI_ISL_3181260, EPI_ISL_3181261, EPI_ISL_3181262, EPI_ISL_3181263, EPI_ISL_3181264, EPI_ISL_3181265, EPI_ISL_3181266, EPI_ISL_3181267, EPI_ISL_3181268, EPI_ISL_3181269, EPI_ISL_3181270, EPI_ISL_3181271, EPI_ISL_3181272, EPI_ISL_3181273, EPI_ISL_3181274, EPI_ISL_3181275, EPI_ISL_3181276, EPI_ISL_3181277, EPI_ISL_3181278, EPI_ISL_3181279, EPI_ISL_3181280, EPI_ISL_3181281, EPI_ISL_3181282, EPI_ISL_3181283, EPI_ISL_3181284, EPI_ISL_3181285, EPI_ISL_3181286, EPI_ISL_3181287, EPI_ISL_3181288, EPI_ISL_3181289, EPI_ISL_3181290, EPI_ISL_3181291, EPI_ISL_3181292, EPI_ISL_3181293, EPI_ISL_3181294, EPI_ISL_3181295, EPI_ISL_3181296, EPI_ISL_3181297, EPI_ISL_3181298, EPI_ISL_3181299, EPI_ISL_3181300, EPI_ISL_3181301, EPI_ISL_3181302, EPI_ISL_3181303, EPI_ISL_3181304, EPI_ISL_3181305, EPI_ISL_3181306, EPI_ISL_3181307, EPI_ISL_3181308, EPI_ISL_3181309, EPI_ISL_3181310, EPI_ISL_3181311, EPI_ISL_3181312, EPI_ISL_3181313, EPI_ISL_3181314, EPI_ISL_3181315, EPI_ISL_3181316, EPI_ISL_3181317, EPI_ISL_3181318, EPI_ISL_3181319, EPI_ISL_3181320, EPI_ISL_3181321, EPI_ISL_3181322, EPI_ISL_3181323, EPI_ISL_3181324, EPI_ISL_3181325, EPI_ISL_3181326, EPI_ISL_3181327, EPI_ISL_3181328, EPI_ISL_3181329, EPI_ISL_3181330, EPI_ISL_3181331, EPI_ISL_3181332, EPI_ISL_3181333, EPI_ISL_3181334, EPI_ISL_3181335, EPI_ISL_3181336, EPI_ISL_3181337, EPI_ISL_3181338, EPI_ISL_3181339, EPI_ISL_3181340, EPI_ISL_3181341, EPI_ISL_3181342, EPI_ISL_3181343, EPI_ISL_3181344, EPI_ISL_3181345, EPI_ISL_3181346, EPI_ISL_3181347, EPI_ISL_3181348, EPI_ISL_3181349, EPI_ISL_3181350, EPI_ISL_3181351, EPI_ISL_3181352, EPI_ISL_3181353, EPI_ISL_3181354, EPI_ISL_3181355, EPI_ISL_3181356, EPI_ISL_3181357, EPI_ISL_3181358, EPI_ISL_3181359, EPI_ISL_3181360, EPI_ISL_3181361, EPI_ISL_3181362, EPI_ISL_3181363, EPI_ISL_3181364, EPI_ISL_3181365, EPI_ISL_3181366, EPI_ISL_3181367, EPI_ISL_3181368, EPI_ISL_3181369, EPI_ISL_3181370, EPI_ISL_3181371, EPI_ISL_3181372, EPI_ISL_3181373, EPI_ISL_3181374, EPI_ISL_3181375, EPI_ISL_3181376, EPI_ISL_3181377, EPI_ISL_3181378, EPI_ISL_3181379, EPI_ISL_3181380, EPI_ISL_3181381, EPI_ISL_3181382, EPI_ISL_3181383, EPI_ISL_3181384, EPI_ISL_3181385, EPI_ISL_3181386, EPI_ISL_3181387, EPI_ISL_3181388, EPI_ISL_3181389, EPI_ISL_3181390, EPI_ISL_3181391, EPI_ISL_3181392, EPI_ISL_3181393, EPI_ISL_3181394, EPI_ISL_3181395, EPI_ISL_3181396, EPI_ISL_3181397, EPI_ISL_3181398, EPI_ISL_3181399, EPI_ISL_3181400, EPI_ISL_3181401, EPI_ISL_3181402, EPI_ISL_3181403, EPI_ISL_3181404, EPI_ISL_3181405, EPI_ISL_3181406, EPI_ISL_3181407, EPI_ISL_3181408, EPI_ISL_3181409, EPI_ISL_3181410, EPI_ISL_3181411, EPI_ISL_3181412, EPI_ISL_3181413, EPI_ISL_3181414, EPI_ISL_3181415, EPI_ISL_3181416, EPI_ISL_3181417, EPI_ISL_3181418, EPI_ISL_3181419, EPI_ISL_3181420, EPI_ISL_3181421, EPI_ISL_3181422, EPI_ISL_3181423, EPI_ISL_3181424, EPI_ISL_3181425, EPI_ISL_3181426, EPI_ISL_3181427, EPI_ISL_3181428, EPI_ISL_3181429, EPI_ISL_3181430, EPI_ISL_3181431, EPI_ISL_3181432, EPI_ISL_3181433, EPI_ISL_3181434, EPI_ISL_3181435, EPI_ISL_3181436, EPI_ISL_3181437, EPI_ISL_3181438, EPI_ISL_3181439, EPI_ISL_3181440, EPI_ISL_3181441, EPI_ISL_3181442, EPI_ISL_3181443, EPI_ISL_3181444, EPI_ISL_3181445, EPI_ISL_3181446, EPI_ISL_3181447, EPI_ISL_3181448, EPI_ISL_3181449, EPI_ISL_3181450, EPI_ISL_3181451, EPI_ISL_3181452, EPI_ISL_3181453, EPI_ISL_3181454, EPI_ISL_3181455, EPI_ISL_3181456 | | | |
| see above | Wales Specialist Virology Centre Sequencing lab: Pathogen Genomics Unit | Public Health Wales Microbiology Cardiff Wales Specialist Virology Centre | Catherine Moore, Johnathan Evans, Laura Gifford, Malorie Perry, Simon Cottrell, Angela Marchbank, Alec Birchley, Alexander Adams, Amy Gaskin, Bree Gatica-Wilcox, Jason Coombes, Joel Southgate, Lauren Gilbert, Lee Graham, Nicole Pacchiarini, Sara Kumziene-Summerhayes, Sarah Taylor, Sophie Jones, Sara Rey, Matthew Bull, Joanne Watkins, Sally Corden, Tom Connor |
| EPI_ISL_3181457, EPI_ISL_3181458, EPI_ISL_3181459, EPI_ISL_3181460, EPI_ISL_3181461, EPI_ISL_3181462, EPI_ISL_3181463, EPI_ISL_3181464, EPI_ISL_3181465, EPI_ISL_3181466, EPI_ISL_3181467, EPI_ISL_3181468, EPI_ISL_3181469, EPI_ISL_3181470, EPI_ISL_3181471, EPI_ISL_3181472, EPI_ISL_3181473, EPI_ISL_3181474, EPI_ISL_3181475, EPI_ISL_3181476, EPI_ISL_3181477, EPI_ISL_3181478, EPI_ISL_3181479, EPI_ISL_3181480, EPI_ISL_3181481, EPI_ISL_3181482, EPI_ISL_3181483, EPI_ISL_3181484, EPI_ISL_3181485, EPI_ISL_3181486, EPI_ISL_3181487, EPI_ISL_3181488, EPI_ISL_3181489, EPI_ISL_3181490, EPI_ISL_3181491, EPI_ISL_3181492, EPI_ISL_3181493, EPI_ISL_3181494, EPI_ISL_3181495, EPI_ISL_3181496, EPI_ISL_3181497, EPI_ISL_3181498, EPI_ISL_3181499, EPI_ISL_3181500, EPI_ISL_3181501, EPI_ISL_3181502, EPI_ISL_3181503, EPI_ISL_3181504, EPI_ISL_3181505, EPI_ISL_3181506, EPI_ISL_3181507, EPI_ISL_3181508, EPI_ISL_3181509, EPI_ISL_3181510, EPI_ISL_3181511, EPI_ISL_3181512, EPI_ISL_3181513, EPI_ISL_3181514, EPI_ISL_3181515, EPI_ISL_3181516, EPI_ISL_3181517, EPI_ISL_3181518, EPI_ISL_3181519, EPI_ISL_3181520, EPI_ISL_3181521, EPI_ISL_3181522, EPI_ISL_3181523, EPI_ISL_3181524, EPI_ISL_3181525, EPI_ISL_3181526, EPI_ISL_3181527, EPI_ISL_3181528, EPI_ISL_3181529, EPI_ISL_3181530, EPI_ISL_3181531, EPI_ISL_3181532, EPI_ISL_3181533, EPI_ISL_3181534, EPI_ISL_3181535, EPI_ISL_3181536, EPI_ISL_3181537, EPI_ISL_3181538, EPI_ISL_3181539, EPI_ISL_3181540, EPI_ISL_3181541, EPI_ISL_3181542, EPI_ISL_3181543, EPI_ISL_3181544, EPI_ISL_3181545, EPI_ISL_3181546, EPI_ISL_3181547, EPI_ISL_3181548, EPI_ISL_3181549, EPI_ISL_3181550, EPI_ISL_3181551, EPI_ISL_3181552, EPI_ISL_3181553, EPI_ISL_3181554, EPI_ISL_3181555, EPI_ISL_3181556, EPI_ISL_3181557, EPI_ISL_3181558, EPI_ISL_3181559, EPI_ISL_3181560, EPI_ISL_3181561, EPI_ISL_3181562, EPI_ISL_3181563, EPI_ISL_3181564, EPI_ISL_3181565, EPI_ISL_3181566, EPI_ISL_3181567, EPI_ISL_3181568, EPI_ISL_3181569, EPI_ISL_3181570, EPI_ISL_3181571, EPI_ISL_3181572, EPI_ISL_3181573, EPI_ISL_3181574, EPI_ISL_3181575, EPI_ISL_3181576, EPI_ISL_3181577, EPI_ISL_3181578, EPI_ISL_3181579, EPI_ISL_3181580, EPI_ISL_3181581, EPI_ISL_3181582, EPI_ISL_3181583, EPI_ISL_3181584, EPI_ISL_3181585, EPI_ISL_3181586, EPI_ISL_3181587, EPI_ISL_3181588, EPI_ISL_3181589, EPI_ISL_3181590, EPI_ISL_3181591, EPI_ISL_3181592, EPI_ISL_3181593, EPI_ISL_3181594, EPI_ISL_3181595, EPI_ISL_3181596, EPI_ISL_3181597, EPI_ISL_3181598, EPI_ISL_3181599, EPI_ISL_3181600, EPI_ISL_3181601, EPI_ISL_3181602, EPI_ISL_3181603, EPI_ISL_3181604, EPI_ISL_3181605, EPI_ISL_3181606, EPI_ISL_3181607, EPI_ISL_3181608, EPI_ISL_3181609, EPI_ISL_3181610, EPI_ISL_3181611, EPI_ISL_3181612, EPI_ISL_3181613, EPI_ISL_3181614, EPI_ISL_3181615, EPI_ISL_3181616, EPI_ISL_3181617, EPI_ISL_3181618, EPI_ISL_3181619, EPI_ISL_3181620, EPI_ISL_3181621, EPI_ISL_3181622, EPI_ISL_3181623, EPI_ISL_3181624, EPI_ISL_3181625, EPI_ISL_3181626, EPI_ISL_3181627, EPI_ISL_3181628, EPI_ISL_3181629, EPI_ISL_3181630, EPI_ISL_3181631, EPI_ISL_3181632, EPI_ISL_3181633, EPI_ISL_3181634, EPI_ISL_3181635, EPI_ISL_3181636, EPI_ISL_3181637, EPI_ISL_3181638, EPI_ISL_3181639, EPI_ISL_3181640, EPI_ISL_3181641, EPI_ISL_3181642, EPI_ISL_3181643, EPI_ISL_3181644, EPI_ISL_3181645, EPI_ISL_3181646, EPI_ISL_3181647, EPI_ISL_3181648, EPI_ISL_3181649, EPI_ISL_3181650, EPI_ISL_3181651, EPI_ISL_3181652, EPI_ISL_3181653, EPI_ISL_3181654, EPI_ISL_3181655, EPI_ISL_3181656, EPI_ISL_3181657, EPI_ISL_3181658, EPI_ISL_3181659, EPI_ISL_3181660, EPI_ISL_3181661, EPI_ISL_3181662, EPI_ISL_3181663, EPI_ISL_3181664, EPI_ISL_3181665, EPI_ISL_3181666, EPI_ISL_3181667, EPI_ISL_3181668, EPI_ISL_3181669, EPI_ISL_3181670, EPI_ISL_3181671, EPI_ISL_3181672, EPI_ISL_3181673, EPI_ISL_3181674, EPI_ISL_3181675, EPI_ISL_3181676, EPI_ISL_3181677, EPI_ISL_3181678, EPI_ISL_3181679, EPI_ISL_3181680, EPI_ISL_3181681, EPI_ISL_3181682, EPI_ISL_3181683, EPI_ISL_3181684, EPI_ISL_3181685, EPI_ISL_3181686, EPI_ISL_3181687, EPI_ISL_3181688, EPI_ISL_3181689, EPI_ISL_3181690, EPI_ISL_3181691, EPI_ISL_3181692, EPI_ISL_3181693, EPI_ISL_3181694, EPI_ISL_3181695, EPI_ISL_3181696, EPI_ISL_3181697, EPI_ISL_3181698, EPI_ISL_3181699, EPI_ISL_3181700, EPI_ISL_3181701, EPI_ISL_3181702, EPI_ISL_3181703, EPI_ISL_3181704, EPI_ISL_3181705, EPI_ISL_3181706, EPI_ISL_3181707, EPI_ISL_3181708, EPI_ISL_3181709, EPI_ISL_3181710, EPI_ISL_3181711, EPI_ISL_3181712, EPI_ISL_3181713, EPI_ISL_3181714, EPI_ISL_3181715, EPI_ISL_3181716, EPI_ISL_3181717, EPI_ISL_3181718, EPI_ISL_3181719, EPI_ISL_3181720, EPI_ISL_3181721, EPI_ISL_3181722, EPI_ISL_3181723, EPI_ISL_3181724, EPI_ISL_3181725, EPI_ISL_3181726, EPI_ISL_3181727, EPI_ISL_3181728, EPI_ISL_3181729, EPI_ISL_3181730, EPI_ISL_3181731, EPI_ISL_3181732, EPI_ISL_3181733, EPI_ISL_3181734, EPI_ISL_3181735, EPI_ISL_3181736, EPI_ISL_3181737, EPI_ISL_3181738, EPI_ISL_3181739, EPI_ISL_3181740, EPI_ISL_3181741, EPI_ISL_3181742, EPI_ISL_3181743, EPI_ISL_3181744, EPI_ISL_3181745, EPI_ISL_3181746, EPI_ISL_3181747, EPI_ISL_3181748, EPI_ISL_3181749, EPI_ISL_3181750, EPI_ISL_3181751, EPI_ISL_3181752, EPI_ISL_3181753, EPI_ISL_3181754, EPI_ISL_3181755, EPI_ISL_3181756, EPI_ISL_3181757, EPI_ISL_3181758, EPI_ISL_3181759, EPI_ISL_3181760, EPI_ISL_3181761, EPI_ISL_3181762, EPI_ISL_3181763, EPI_ISL_3181764, EPI_ISL_3181765, EPI_ISL_3181766, EPI_ISL_3181767, EPI_ISL_3181768, EPI_ISL_3181769, EPI_ISL_3181770, EPI_ISL_3181771, EPI_ISL_3181772, EPI_ISL_3181773, EPI_ISL_3181774, EPI_ISL_3181775, EPI_ISL_3181776, EPI_ISL_3181777, EPI_ISL_3181778, EPI_ISL_3181779, EPI_ISL_3181780, EPI_ISL_3181781, EPI_ISL_3181782, EPI_ISL_3181783, EPI_ISL_3181784, EPI_ISL_3181785, EPI_ISL_3181786, EPI_ISL_3181787, EPI_ISL_3181788, EPI_ISL_3181789, EPI_ISL_3181790, EPI_ISL_3181791, EPI_ISL_3181792, EPI_ISL_3181793, EPI_ISL_3181794, EPI_ISL_3181795, EPI_ISL_3181796, EPI_ISL_3181797, EPI_ISL_3181798, EPI_ISL_3181799, EPI_ISL_3181800, EPI_ISL_3181801, EPI_ISL_3181802, EPI_ISL_3181803, EPI_ISL_3181804, EPI_ISL_3181805, EPI_ISL_3181806, EPI_ISL_3181807, EPI_ISL_3181808, EPI_ISL_3181809, EPI_ISL_3181810, EPI_ISL_3181811, EPI_ISL_3181812, EPI_ISL_3181813, EPI_ISL_3181814, EPI_ISL_3181815, EPI_ISL_3181816, EPI_ISL_3181817, EPI_ISL_3181818, EPI_ISL_3181819, EPI_ISL_3181820, EPI_ISL_3181821, EPI_ISL_3181822, EPI_ISL_3181823, EPI_ISL_3181824, EPI_ISL_3181825, EPI_ISL_3181826, EPI_ISL_3181827, EPI_ISL_3181828, EPI_ISL_3181829, EPI_ISL_3181830, EPI_ISL_3181831, EPI_ISL_3181832, EPI_ISL_3181833, EPI_ISL_3181834, EPI_ISL_3181835, EPI_ISL_3181836, EPI_ISL_3181837, EPI_ISL_3181838, EPI_ISL_3181839, EPI_ISL_3181840, EPI_ISL_3181841, EPI_ISL_3181842, EPI_ISL_3181843, EPI_ISL_3181844, EPI_ISL_3181845, EPI_ISL_3181846, EPI_ISL_3181847, EPI_ISL_3181848, EPI_ISL_3181849, EPI_ISL_3181850, EPI_ISL_3181851, EPI_ISL_3181852, EPI_ISL_3181853, EPI_ISL_3181854, EPI_ISL_3181855, EPI_ISL_3181856, EPI_ISL_3181857, EPI_ISL_3181858, EPI_ISL_3181859, EPI_ISL_3181860, EPI_ISL_3181861, EPI_ISL_3181862, EPI_ISL_3181863, EPI_ISL_3181864, EPI_ISL_3181865, EPI_ISL_3181866, EPI_ISL_3181867, EPI_ISL_3181868, EPI_ISL_3181869, EPI_ISL_3181870, EPI_ISL_3181871, EPI_ISL_3181872, EPI_ISL_3181873, EPI_ISL_3181874, EPI_ISL_3181875, EPI_ISL_3181876, EPI_ISL_3181877, EPI_ISL_3181878, EPI_ISL_3181879, EPI_ISL_3181880, EPI_ISL_3181881, EPI_ISL_3181882, EPI_ISL_3181883, EPI_ISL_3181884, EPI_ISL_3181885, EPI_ISL_3181886, EPI_ISL_3181887, EPI_ISL_3181888, EPI_ISL_3181889, EPI_ISL_3181890, EPI_ISL_3181891, EPI_ISL_3181892, EPI_ISL_3181893, EPI_ISL_3181894, EPI_ISL_3181895, EPI_ISL_3181896, EPI_ISL_3181897, EPI_ISL_3181898, EPI_ISL_3181899, EPI_ISL_3181900, EPI_ISL_3181901, EPI_ISL_3181902, EPI_ISL_3181903, EPI_ISL_3181904, EPI_ISL_3181905, EPI_ISL_3181906, EPI_ISL_3181907, EPI_ISL_3181908, EPI_ISL_3181909, EPI_ISL_3181910, EPI_ISL_3181911, EPI_ISL_3181912, EPI_ISL_3181913, EPI_ISL_3181914, EPI_ISL_3181915, EPI_ISL_3181916, EPI_ISL_3181917, EPI_ISL_3181918, EPI_ISL_3181919, EPI_ISL_3181920, EPI_ISL_3181921, EPI_ISL_3181922, EPI_ISL_3181923, EPI_ISL_3181924, EPI_ISL_3181925, EPI_ISL_3181926, EPI_ISL_3181927, EPI_ISL_3181928, EPI_ISL_3181929, EPI_ISL_3181930, EPI_ISL_3181931, EPI_ISL_3181932, EPI_ISL_3181933, EPI_ISL_3181934, EPI_ISL_3181935, EPI_ISL_3181936, EPI_ISL_3181937, EPI_ISL_3181938, EPI_ISL_3181939, EPI_ISL_3181940, EPI_ISL_3181941, EPI_ISL_3181942, EPI_ISL_3181943, EPI_ISL_3181944, EPI_ISL_3181945, EPI_ISL_3181946, EPI_ISL_3181947, EPI_ISL_3181948, EPI_ISL_3181949, EPI_ISL_3181950, EPI_ISL_3181951, EPI_ISL_3181952 | | | |
| see above | Originating lab: Wales Specialist Virology Centre Sequencing lab: Pathogen Genomics Unit | Public Health Wales Microbiology Cardiff Wales Specialist Virology Centre | Catherine Moore, Johnathan Evans, Laura Gifford, Malorie Perry, Simon Cottrell, Angela Marchbank, Alec Birchley, Alexander Adams, Amy Gaskin, Bree Gatica-Wilcox, Jason Coombes, Joel Southgate, Lauren Gilbert, Lee Graham, Nicole Pacchiarini, Sara Kumziene-Summerhayes, Sarah Taylor, Sophie Jones, Sara Rey, Matthew Bull, Joanne Watkins, Sally Corden, Tom Connor |
| EPI_ISL_3181953, EPI_ISL_3181954, EPI_ISL_3181955, EPI_ISL_3181956, EPI_ISL_3181957, EPI_ISL_3181958, EPI_ISL_3181959, EPI_ISL_3181960, EPI_ISL_3181961, EPI_ISL_3181962, EPI_ISL_3181963, EPI_ISL_3181964, EPI_ISL_3181965, EPI_ISL_3181966, EPI_ISL_3181967, EPI_ISL_3181968, EPI_ISL_3181969, EPI_ISL_3181970, EPI_ISL_3181971, EPI_ISL_3181972, EPI_ISL_3181973, EPI_ISL_3181974, EPI_ISL_3181975, EPI_ISL_3181976, EPI_ISL_3181977, EPI_ISL_3181978, EPI_ISL_3181979, EPI_ISL_3181980, EPI_ISL_3181981, EPI_ISL_3181982, EPI_ISL_3181983, EPI_ISL_3181984, EPI_ISL_3181985, EPI_ISL_3181986, EPI_ISL_3181987, EPI_ISL_3181988, EPI_ISL_3181989, EPI_ISL_3181990, EPI_ISL_3181991, EPI_ISL_3181992, EPI_ISL_3181993, EPI_ISL_3181994, EPI_ISL_3181995, EPI_ISL_3181996, EPI_ISL_3181997, EPI_ISL_3181998, EPI_ISL_3181999, EPI_ISL_3182000, EPI_ISL_3182001, EPI_ISL_3182002, EPI_ISL_3182003, EPI_ISL_3182004, EPI_ISL_3182005, EPI_ISL_3182006, EPI_ISL_3182007, EPI_ISL_3182008, EPI_ISL_3182009, EPI_ISL_3182010, EPI_ISL_3182011, EPI_ISL_3182012, EPI_ISL_3182013, EPI_ISL_3182014, EPI_ISL_3182015, EPI_ISL_3182016, EPI_ISL_3182017, EPI_ISL_3182018, EPI_ISL_3182019, EPI_ISL_3182020, EPI_ISL_3182021, EPI_ISL_3182022, EPI_ISL_3182023, EPI_ISL_3182024, EPI_ISL_3182025, EPI_ISL_3182026, EPI_ISL_3182027, EPI_ISL_3182028, EPI_ISL_3182029, EPI_ISL_3182030, EPI_ISL_3182031, EPI_ISL_3182032, EPI_ISL_3182033, EPI_ISL_3182034, EPI_ISL_3182035, EPI_ISL_3182036, EPI_ISL_3182037, EPI_ISL_3182038, EPI_ISL_3182039, EPI_ISL_3182040, EPI_ISL_3182041, EPI_ISL_3182042, EPI_ISL_3182043, EPI_ISL_3182044, EPI_ISL_3182045, EPI_ISL_3182046, EPI_ISL_3182047, EPI_ISL_3182048, EPI_ISL_3182049, EPI_ISL_3182050, EPI_ISL_3182051, EPI_ISL_3182052, EPI_ISL_3182053, EPI_ISL_3182054, EPI_ISL_3182055, EPI_ISL_3182056, EPI_ISL_3182057, EPI_ISL_3182058, EPI_ISL_3182059, EPI_ISL_3182060, EPI_ISL_3182061, EPI_ISL_3182062, EPI_ISL_3182063, EPI_ISL_3182064, EPI_ISL_3182065, EPI_ISL_3182066, EPI_ISL_3182067, EPI_ISL_3182068, EPI_ISL_3182069, EPI_ISL_3182070, EPI_ISL_3182071, EPI_ISL_3182072, EPI_ISL_3182073, EPI_ISL_3182074, EPI_ISL_3182075, EPI_ISL_3182076, EPI_ISL_3182077, EPI_ISL_3182078, EPI_ISL_3182079, EPI_ISL_3182080, EPI_ISL_3182081, EPI_ISL_3182082, EPI_ISL_3182083, EPI_ISL_3182084, EPI_ISL_3182085, EPI_ISL_3182086, EPI_ISL_3182087, EPI_ISL_3182088, EPI_ISL_3182089, EPI_ISL_3182090, EPI_ISL_3182091, EPI_ISL_3182092, EPI_ISL_3182093, EPI_ISL_3182094, EPI_ISL_3182095, EPI_ISL_3182096, EPI_ISL_3182097, EPI_ISL_3182098, EPI_ISL_3182099, EPI_ISL_3182100, EPI_ISL_3182101, EPI_ISL_3182102, EPI_ISL_3182103, EPI_ISL_3182104, EPI_ISL_3182105, EPI_ISL_3182106, EPI_ISL_3182107, EPI_ISL_3182108, EPI_ISL_3182109, EPI_ISL_3182110, EPI_ISL_3182111, EPI_ISL_3182112, EPI_ISL_3182113, EPI_ISL_3182114, EPI_ISL_3182115, EPI_ISL_3182116, EPI_ISL_3182117, EPI_ISL_3182118, EPI_ISL_3182119, EPI_ISL_3182121, EPI_ISL_3182122 | | | |
| see above | Wales Specialist Virology Centre Sequencing lab: Pathogen Genomics Unit | Public Health Wales Microbiology Cardiff Wales Specialist Virology Centre | Catherine Moore, Johnathan Evans, Laura Gifford, Malorie Perry, Simon Cottrell, Angela Marchbank, Alec Birchley, Alexander Adams, Amy Gaskin, Bree Gatica-Wilcox, Jason Coombes, Joel Southgate, Lauren Gilbert, Lee Graham, Nicole Pacchiarini, Sara Kumziene-Summerhayes, Sarah Taylor, Sophie Jones, Sara Rey, Matthew Bull, Joanne Watkins, Sally Corden, Tom Connor |
| EPI_ISL_3182123 | Labor Berlin Charité Vivantes GmbH / Institut für Virologie | Charité Universitätsmedizin Berlin, Institut für Virologie/Labor Berlin | Peter Menzel, Christine Stephan, Rolf Schwarzer, Victor M Corman, Barbara Mühlemann, Terry Jones, Christian Drosten |
| EPI_ISL_3182124 | Wales Specialist Virology Centre Sequencing lab: Pathogen Genomics Unit | Public Health Wales Microbiology Cardiff Wales Specialist Virology Centre | Catherine Moore, Johnathan Evans, Laura Gifford, Malorie Perry, Simon Cottrell, Angela Marchbank, Alec Birchley, Alexander Adams, Amy Gaskin, Bree Gatica-Wilcox, Jason Coombes, Joel Southgate, Lauren Gilbert, Lee Graham, Nicole Pacchiarini, Sara Kumziene-Summerhayes, Sarah Taylor, Sophie Jones, Sara Rey, Matthew Bull, Joanne Watkins, Sally Corden, Tom Connor |
| EPI_ISL_3182125 | Labor Berlin Charité Vivantes GmbH / Institut für Virologie | Charité Universitätsmedizin Berlin, Institut für Virologie/Labor Berlin | Peter Menzel, Christine Stephan, Rolf Schwarzer, Victor M Corman, Barbara Mühlemann, Terry Jones, Christian Drosten |
| EPI_ISL_3182126 | Wales Specialist Virology Centre Sequencing lab: Pathogen Genomics Unit | Public Health Wales Microbiology Cardiff Wales Specialist Virology Centre | Catherine Moore, Johnathan Evans, Laura Gifford, Malorie Perry, Simon Cottrell, Angela Marchbank, Alec Birchley, Alexander Adams, Amy Gaskin, Bree Gatica-Wilcox, Jason Coombes, Joel Southgate, Lauren Gilbert, Lee Graham, Nicole Pacchiarini, Sara Kumziene-Summerhayes, Sarah Taylor, Sophie Jones, Sara Rey, Matthew Bull, Joanne Watkins, Sally Corden, Tom Connor |
| EPI_ISL_3182127 | Labor Berlin Charité Vivantes GmbH / Institut für Virologie | Charité Universitätsmedizin Berlin, Institut für Virologie/Labor Berlin | Peter Menzel, Christine Stephan, Rolf Schwarzer, Victor M Corman, Barbara Mühlemann, Terry Jones, Christian Drosten |
| EPI_ISL_3182128 | Wales Specialist Virology Centre Sequencing lab: Pathogen | Public Health Wales Microbiology Cardiff Wales Specialist Virology | Catherine Moore, Johnathan Evans, Laura Gifford, Malorie Perry, Simon Cottrell, Angela Marchbank, Alec Birchley, Alexander Adams, Amy Gaskin, Bree Gatica-Wilcox, Jason Coombes, Joel Southgate, Lauren Gilbert, Lee Graham, Nicole |

| | | | |
|---|---|---|---|
| | Genomics Unit | Centre | Pacchiarini, Sara Kumziene-Summerhayes, Sarah Taylor, Sophie Jones, Sara Rey, Matthew Bull, Joanne Watkins, Sally Corden, Tom Connor |
| EPI_ISL_3182129 | Labor Berlin Charité Vivantes GmbH / Institut für Virologie | Charité Universitätsmedizin Berlin, Institut für Virologie/Labor Berlin | Peter Menzel, Christine Stephan, Rolf Schwarzer, Victor M Corman, Barbara Mühlemann, Terry Jones, Christian Drosten |
| EPI_ISL_3182130 | Wales Specialist Virology Centre Sequencing lab: Pathogen Genomics Unit | Public Health Wales Microbiology Cardiff Wales Specialist Virology Centre | Catherine Moore, Johnathan Evans, Laura Gifford, Malorie Perry, Simon Cottrell, Angela Marchbank, Alec Birchley, Alexander Adams, Amy Gaskin, Bree Gatica-Wilcox, Jason Coombes, Joel Southgate, Lauren Gilbert, Lee Graham, Nicole Pacchiarini, Sara Kumziene-Summerhayes, Sarah Taylor, Sophie Jones, Sara Rey, Matthew Bull, Joanne Watkins, Sally Corden, Tom Connor |
| EPI_ISL_3182131 | Labor Berlin Charité Vivantes GmbH / Institut für Virologie | Charité Universitätsmedizin Berlin, Institut für Virologie/Labor Berlin | Peter Menzel, Christine Stephan, Rolf Schwarzer, Victor M Corman, Barbara Mühlemann, Terry Jones, Christian Drosten |
| EPI_ISL_3182132, EPI_ISL_3182133 | Wales Specialist Virology Centre Sequencing lab: Pathogen Genomics Unit | Public Health Wales Microbiology Cardiff Wales Specialist Virology Centre | Catherine Moore, Johnathan Evans, Laura Gifford, Malorie Perry, Simon Cottrell, Angela Marchbank, Alec Birchley, Alexander Adams, Amy Gaskin, Bree Gatica-Wilcox, Jason Coombes, Joel Southgate, Lauren Gilbert, Lee Graham, Nicole Pacchiarini, Sara Kumziene-Summerhayes, Sarah Taylor, Sophie Jones, Sara Rey, Matthew Bull, Joanne Watkins, Sally Corden, Tom Connor |
| EPI_ISL_3182134 | Labor Berlin Charité Vivantes GmbH / Institut für Virologie | Charité Universitätsmedizin Berlin, Institut für Virologie/Labor Berlin | Peter Menzel, Christine Stephan, Rolf Schwarzer, Victor M Corman, Barbara Mühlemann, Terry Jones, Christian Drosten |
| EPI_ISL_3182135 | Wales Specialist Virology Centre Sequencing lab: Pathogen Genomics Unit | Public Health Wales Microbiology Cardiff Wales Specialist Virology Centre | Catherine Moore, Johnathan Evans, Laura Gifford, Malorie Perry, Simon Cottrell, Angela Marchbank, Alec Birchley, Alexander Adams, Amy Gaskin, Bree Gatica-Wilcox, Jason Coombes, Joel Southgate, Lauren Gilbert, Lee Graham, Nicole Pacchiarini, Sara Kumziene-Summerhayes, Sarah Taylor, Sophie Jones, Sara Rey, Matthew Bull, Joanne Watkins, Sally Corden, Tom Connor |
| EPI_ISL_3182136 | Labor Berlin Charité Vivantes GmbH / Institut für Virologie | Charité Universitätsmedizin Berlin, Institut für Virologie/Labor Berlin | Peter Menzel, Christine Stephan, Rolf Schwarzer, Victor M Corman, Barbara Mühlemann, Terry Jones, Christian Drosten |
| EPI_ISL_3182137 | Wales Specialist Virology Centre Sequencing lab: Pathogen Genomics Unit | Public Health Wales Microbiology Cardiff Wales Specialist Virology Centre | Catherine Moore, Johnathan Evans, Laura Gifford, Malorie Perry, Simon Cottrell, Angela Marchbank, Alec Birchley, Alexander Adams, Amy Gaskin, Bree Gatica-Wilcox, Jason Coombes, Joel Southgate, Lauren Gilbert, Lee Graham, Nicole Pacchiarini, Sara Kumziene-Summerhayes, Sarah Taylor, Sophie Jones, Sara Rey, Matthew Bull, Joanne Watkins, Sally Corden, Tom Connor |
| EPI_ISL_3182138 | Labor Berlin Charité Vivantes GmbH / Institut für Virologie | Charité Universitätsmedizin Berlin, Institut für Virologie/Labor Berlin | Peter Menzel, Christine Stephan, Rolf Schwarzer, Victor M Corman, Barbara Mühlemann, Terry Jones, Christian Drosten |
| EPI_ISL_3182139, EPI_ISL_3182140, EPI_ISL_3182141, EPI_ISL_3182142 | Wales Specialist Virology Centre Sequencing lab: Pathogen Genomics Unit | Public Health Wales Microbiology Cardiff Wales Specialist Virology Centre | Catherine Moore, Johnathan Evans, Laura Gifford, Malorie Perry, Simon Cottrell, Angela Marchbank, Alec Birchley, Alexander Adams, Amy Gaskin, Bree Gatica-Wilcox, Jason Coombes, Joel Southgate, Lauren Gilbert, Lee Graham, Nicole Pacchiarini, Sara Kumziene-Summerhayes, Sarah Taylor, Sophie Jones, Sara Rey, Matthew Bull, Joanne Watkins, Sally Corden, Tom Connor |
| EPI_ISL_3182143 | Labor Berlin Charité Vivantes GmbH / Institut für Virologie | Charité Universitätsmedizin Berlin, Institut für Virologie/Labor Berlin | Peter Menzel, Christine Stephan, Rolf Schwarzer, Victor M Corman, Barbara Mühlemann, Terry Jones, Christian Drosten |
| EPI_ISL_3182144 | Wales Specialist Virology Centre Sequencing lab: Pathogen Genomics Unit | Public Health Wales Microbiology Cardiff Wales Specialist Virology Centre | Catherine Moore, Johnathan Evans, Laura Gifford, Malorie Perry, Simon Cottrell, Angela Marchbank, Alec Birchley, Alexander Adams, Amy Gaskin, Bree Gatica-Wilcox, Jason Coombes, Joel Southgate, Lauren Gilbert, Lee Graham, Nicole Pacchiarini, Sara Kumziene-Summerhayes, Sarah Taylor, Sophie Jones, Sara Rey, Matthew Bull, Joanne Watkins, Sally Corden, Tom Connor |
| EPI_ISL_3182145 | Labor Berlin Charité Vivantes GmbH / Institut für Virologie | Charité Universitätsmedizin Berlin, Institut für Virologie/Labor Berlin | Peter Menzel, Christine Stephan, Rolf Schwarzer, Victor M Corman, Barbara Mühlemann, Terry Jones, Christian Drosten |
| EPI_ISL_3182146 | Wales Specialist Virology Centre Sequencing lab: Pathogen Genomics Unit | Public Health Wales Microbiology Cardiff Wales Specialist Virology Centre | Catherine Moore, Johnathan Evans, Laura Gifford, Malorie Perry, Simon Cottrell, Angela Marchbank, Alec Birchley, Alexander Adams, Amy Gaskin, Bree Gatica-Wilcox, Jason Coombes, Joel Southgate, Lauren Gilbert, Lee Graham, Nicole Pacchiarini, Sara Kumziene-Summerhayes, Sarah Taylor, Sophie Jones, Sara Rey, Matthew Bull, Joanne Watkins, Sally Corden, Tom Connor |
| EPI_ISL_3182147 | Labor Berlin Charité Vivantes GmbH / Institut für Virologie | Charité Universitätsmedizin Berlin, Institut für Virologie/Labor Berlin | Peter Menzel, Christine Stephan, Rolf Schwarzer, Victor M Corman, Barbara Mühlemann, Terry Jones, Christian Drosten |
| EPI_ISL_3182148, EPI_ISL_3182149 | Wales Specialist Virology Centre Sequencing lab: Pathogen Genomics Unit | Public Health Wales Microbiology Cardiff Wales Specialist Virology Centre | Catherine Moore, Johnathan Evans, Laura Gifford, Malorie Perry, Simon Cottrell, Angela Marchbank, Alec Birchley, Alexander Adams, Amy Gaskin, Bree Gatica-Wilcox, Jason Coombes, Joel Southgate, Lauren Gilbert, Lee Graham, Nicole Pacchiarini, Sara Kumziene-Summerhayes, Sarah Taylor, Sophie Jones, Sara Rey, Matthew Bull, Joanne Watkins, Sally Corden, Tom Connor |
| EPI_ISL_3182150 | Labor Berlin Charité Vivantes GmbH / Institut für Virologie | Charité Universitätsmedizin Berlin, Institut für Virologie/Labor Berlin | Peter Menzel, Christine Stephan, Rolf Schwarzer, Victor M Corman, Barbara Mühlemann, Terry Jones, Christian Drosten |
| EPI_ISL_3182151 | Wales Specialist Virology Centre Sequencing lab: Pathogen Genomics Unit | Public Health Wales Microbiology Cardiff Wales Specialist Virology Centre | Catherine Moore, Johnathan Evans, Laura Gifford, Malorie Perry, Simon Cottrell, Angela Marchbank, Alec Birchley, Alexander Adams, Amy Gaskin, Bree Gatica-Wilcox, Jason Coombes, Joel Southgate, Lauren Gilbert, Lee Graham, Nicole Pacchiarini, Sara Kumziene-Summerhayes, Sarah Taylor, Sophie Jones, Sara Rey, Matthew Bull, Joanne Watkins, Sally Corden, Tom Connor |
| EPI_ISL_3182152 | Labor Berlin Charité Vivantes GmbH / Institut für Virologie | Charité Universitätsmedizin Berlin, Institut für Virologie/Labor Berlin | Peter Menzel, Christine Stephan, Rolf Schwarzer, Victor M Corman, Barbara Mühlemann, Terry Jones, Christian Drosten |
| EPI_ISL_3182153 | Wales Specialist Virology Centre Sequencing lab: Pathogen Genomics Unit | Public Health Wales Microbiology Cardiff Wales Specialist Virology Centre | Catherine Moore, Johnathan Evans, Laura Gifford, Malorie Perry, Simon Cottrell, Angela Marchbank, Alec Birchley, Alexander Adams, Amy Gaskin, Bree Gatica-Wilcox, Jason Coombes, Joel Southgate, Lauren Gilbert, Lee Graham, Nicole Pacchiarini, Sara Kumziene-Summerhayes, Sarah Taylor, Sophie Jones, Sara Rey, Matthew Bull, Joanne Watkins, Sally Corden, Tom Connor |
| EPI_ISL_3182154 | Labor Berlin Charité Vivantes GmbH / Institut für Virologie | Charité Universitätsmedizin Berlin, Institut für Virologie/Labor Berlin | Peter Menzel, Christine Stephan, Rolf Schwarzer, Victor M Corman, Barbara Mühlemann, Terry Jones, Christian Drosten |
| EPI_ISL_3182155 | Wales Specialist Virology Centre Sequencing lab: Pathogen Genomics Unit | Public Health Wales Microbiology Cardiff Wales Specialist Virology Centre | Catherine Moore, Johnathan Evans, Laura Gifford, Malorie Perry, Simon Cottrell, Angela Marchbank, Alec Birchley, Alexander Adams, Amy Gaskin, Bree Gatica-Wilcox, Jason Coombes, Joel Southgate, Lauren Gilbert, Lee Graham, Nicole Pacchiarini, Sara Kumziene-Summerhayes, Sarah Taylor, Sophie Jones, Sara Rey, Matthew Bull, Joanne Watkins, Sally Corden, Tom Connor |
| EPI_ISL_3182156 | Labor Berlin Charité Vivantes GmbH / Institut für Virologie | Charité Universitätsmedizin Berlin, Institut für Virologie/Labor Berlin | Peter Menzel, Christine Stephan, Rolf Schwarzer, Victor M Corman, Barbara Mühlemann, Terry Jones, Christian Drosten |
| EPI_ISL_3182157, EPI_ISL_3182158 | Wales Specialist Virology Centre Sequencing lab: Pathogen Genomics Unit | Public Health Wales Microbiology Cardiff Wales Specialist Virology Centre | Catherine Moore, Johnathan Evans, Laura Gifford, Malorie Perry, Simon Cottrell, Angela Marchbank, Alec Birchley, Alexander Adams, Amy Gaskin, Bree Gatica-Wilcox, Jason Coombes, Joel Southgate, Lauren Gilbert, Lee Graham, Nicole Pacchiarini, Sara Kumziene-Summerhayes, Sarah Taylor, Sophie Jones, Sara Rey, Matthew Bull, Joanne Watkins, Sally Corden, Tom Connor |
| EPI_ISL_3182159 | Labor Berlin Charité Vivantes GmbH / Institut für Virologie | Charité Universitätsmedizin Berlin, Institut für Virologie/Labor Berlin | Peter Menzel, Christine Stephan, Rolf Schwarzer, Victor M Corman, Barbara Mühlemann, Terry Jones, Christian Drosten |
| EPI_ISL_3182160 | Wales Specialist Virology Centre Sequencing lab: Pathogen Genomics Unit | Public Health Wales Microbiology Cardiff Wales Specialist Virology Centre | Catherine Moore, Johnathan Evans, Laura Gifford, Malorie Perry, Simon Cottrell, Angela Marchbank, Alec Birchley, Alexander Adams, Amy Gaskin, Bree Gatica-Wilcox, Jason Coombes, Joel Southgate, Lauren Gilbert, Lee Graham, Nicole Pacchiarini, Sara Kumziene-Summerhayes, Sarah Taylor, Sophie Jones, Sara Rey, Matthew Bull, Joanne Watkins, Sally Corden, Tom Connor |
| EPI_ISL_3182161 | Labor Berlin Charité Vivantes GmbH / Institut für Virologie | Charité Universitätsmedizin Berlin, Institut für Virologie/Labor Berlin | Peter Menzel, Christine Stephan, Rolf Schwarzer, Victor M Corman, Barbara Mühlemann, Terry Jones, Christian Drosten |
| EPI_ISL_3182162 | Wales Specialist Virology Centre Sequencing lab: Pathogen Genomics Unit | Public Health Wales Microbiology Cardiff Wales Specialist Virology Centre | Catherine Moore, Johnathan Evans, Laura Gifford, Malorie Perry, Simon Cottrell, Angela Marchbank, Alec Birchley, Alexander Adams, Amy Gaskin, Bree Gatica-Wilcox, Jason Coombes, Joel Southgate, Lauren Gilbert, Lee Graham, Nicole Pacchiarini, Sara Kumziene-Summerhayes, Sarah Taylor, Sophie Jones, Sara Rey, Matthew Bull, Joanne Watkins, Sally Corden, Tom Connor |
| EPI_ISL_3182163 | Labor Berlin Charité Vivantes GmbH / Institut für Virologie | Charité Universitätsmedizin Berlin, Institut für Virologie/Labor Berlin | Peter Menzel, Christine Stephan, Rolf Schwarzer, Victor M Corman, Barbara Mühlemann, Terry Jones, Christian Drosten |
| EPI_ISL_3182164 | Wales Specialist Virology Centre Sequencing lab: Pathogen Genomics Unit | Public Health Wales Microbiology Cardiff Wales Specialist Virology Centre | Catherine Moore, Johnathan Evans, Laura Gifford, Malorie Perry, Simon Cottrell, Angela Marchbank, Alec Birchley, Alexander Adams, Amy Gaskin, Bree Gatica-Wilcox, Jason Coombes, Joel Southgate, Lauren Gilbert, Lee Graham, Nicole Pacchiarini, Sara Kumziene-Summerhayes, Sarah Taylor, Sophie Jones, Sara Rey, Matthew Bull, Joanne Watkins, Sally Corden, Tom Connor |
| EPI_ISL_3182165 | Labor Berlin Charité Vivantes GmbH / Institut für Virologie | Charité Universitätsmedizin Berlin, Institut für Virologie/Labor Berlin | Peter Menzel, Christine Stephan, Rolf Schwarzer, Victor M Corman, Barbara Mühlemann, Terry Jones, Christian Drosten |
| EPI_ISL_3182166, EPI_ISL_3182168 | Wales Specialist Virology Centre Sequencing lab: Pathogen Genomics Unit | Public Health Wales Microbiology Cardiff Wales Specialist Virology Centre | Catherine Moore, Johnathan Evans, Laura Gifford, Malorie Perry, Simon Cottrell, Angela Marchbank, Alec Birchley, Alexander Adams, Amy Gaskin, Bree Gatica-Wilcox, Jason Coombes, Joel Southgate, Lauren Gilbert, Lee Graham, Nicole Pacchiarini, Sara Kumziene-Summerhayes, Sarah Taylor, Sophie Jones, Sara Rey, Matthew Bull, Joanne Watkins, Sally Corden, Tom Connor |
| EPI_ISL_3182169 | Labor Berlin Charité Vivantes GmbH / Institut für Virologie | Charité Universitätsmedizin Berlin, Institut für Virologie/Labor Berlin | Peter Menzel, Christine Stephan, Rolf Schwarzer, Victor M Corman, Barbara Mühlemann, Terry Jones, Christian Drosten |
| EPI_ISL_3182170, EPI_ISL_3182171 | Wales Specialist Virology Centre Sequencing lab: Pathogen Genomics Unit | Public Health Wales Microbiology Cardiff Wales Specialist Virology Centre | Catherine Moore, Johnathan Evans, Laura Gifford, Malorie Perry, Simon Cottrell, Angela Marchbank, Alec Birchley, Alexander Adams, Amy Gaskin, Bree Gatica-Wilcox, Jason Coombes, Joel Southgate, Lauren Gilbert, Lee Graham, Nicole Pacchiarini, Sara Kumziene-Summerhayes, Sarah Taylor, Sophie Jones, Sara Rey, Matthew Bull, Joanne Watkins, Sally Corden, Tom Connor |
| EPI_ISL_3182172 | Labor Berlin Charité Vivantes GmbH / Institut für Virologie | Charité Universitätsmedizin Berlin, Institut für Virologie/Labor Berlin | Peter Menzel, Christine Stephan, Rolf Schwarzer, Victor M Corman, Barbara Mühlemann, Terry Jones, Christian Drosten |
| EPI_ISL_3182173 | Wales Specialist Virology Centre Sequencing lab: Pathogen Genomics Unit | Public Health Wales Microbiology Cardiff Wales Specialist Virology Centre | Catherine Moore, Johnathan Evans, Laura Gifford, Malorie Perry, Simon Cottrell, Angela Marchbank, Alec Birchley, Alexander Adams, Amy Gaskin, Bree Gatica-Wilcox, Jason Coombes, Joel Southgate, Lauren Gilbert, Lee Graham, Nicole Pacchiarini, Sara Kumziene-Summerhayes, Sarah Taylor, Sophie Jones, Sara Rey, Matthew Bull, Joanne Watkins, Sally Corden, Tom Connor |
| EPI_ISL_3182174 | Labor Berlin Charité Vivantes GmbH / Institut für Virologie | Charité Universitätsmedizin Berlin, Institut für Virologie/Labor Berlin | Peter Menzel, Christine Stephan, Rolf Schwarzer, Victor M Corman, Barbara Mühlemann, Terry Jones, Christian Drosten |
| EPI_ISL_3182175 | Wales Specialist Virology Centre Sequencing lab: Pathogen Genomics Unit | Public Health Wales Microbiology Cardiff Wales Specialist Virology Centre | Catherine Moore, Johnathan Evans, Laura Gifford, Malorie Perry, Simon Cottrell, Angela Marchbank, Alec Birchley, Alexander Adams, Amy Gaskin, Bree Gatica-Wilcox, Jason Coombes, Joel Southgate, Lauren Gilbert, Lee Graham, Nicole Pacchiarini, Sara Kumziene-Summerhayes, Sarah Taylor, Sophie Jones, Sara Rey, Matthew Bull, Joanne Watkins, Sally Corden, Tom Connor |
| EPI_ISL_3182176 | Labor Berlin Charité Vivantes GmbH / Institut für Virologie | Charité Universitätsmedizin Berlin, Institut für Virologie/Labor Berlin | Peter Menzel, Christine Stephan, Rolf Schwarzer, Victor M Corman, Barbara Mühlemann, Terry Jones, Christian Drosten |
| EPI_ISL_3182177 | Wales Specialist Virology Centre Sequencing lab: Pathogen Genomics Unit | Public Health Wales Microbiology Cardiff Wales Specialist Virology Centre | Catherine Moore, Johnathan Evans, Laura Gifford, Malorie Perry, Simon Cottrell, Angela Marchbank, Alec Birchley, Alexander Adams, Amy Gaskin, Bree Gatica-Wilcox, Jason Coombes, Joel Southgate, Lauren Gilbert, Lee Graham, Nicole Pacchiarini, Sara Kumziene-Summerhayes, Sarah Taylor, Sophie Jones, Sara Rey, Matthew Bull, Joanne Watkins, Sally Corden, Tom Connor |
| EPI_ISL_3182178 | Labor Berlin Charité Vivantes GmbH / Institut für Virologie | Charité Universitätsmedizin Berlin, Institut für Virologie/Labor Berlin | Peter Menzel, Christine Stephan, Rolf Schwarzer, Victor M Corman, Barbara Mühlemann, Terry Jones, Christian Drosten |
| EPI_ISL_3182179 | Wales Specialist Virology Centre Sequencing lab: Pathogen Genomics Unit | Public Health Wales Microbiology Cardiff Wales Specialist Virology Centre | Catherine Moore, Johnathan Evans, Laura Gifford, Malorie Perry, Simon Cottrell, Angela Marchbank, Alec Birchley, Alexander Adams, Amy Gaskin, Bree Gatica-Wilcox, Jason Coombes, Joel Southgate, Lauren Gilbert, Lee Graham, Nicole Pacchiarini, Sara Kumziene-Summerhayes, Sarah Taylor, Sophie Jones, Sara Rey, Matthew Bull, Joanne Watkins, Sally Corden, Tom Connor |
| EPI_ISL_3182180 | Labor Berlin Charité Vivantes GmbH / Institut für Virologie | Charité Universitätsmedizin Berlin, Institut für Virologie/Labor Berlin | Peter Menzel, Christine Stephan, Rolf Schwarzer, Victor M Corman, Barbara Mühlemann, Terry Jones, Christian Drosten |
| EPI_ISL_3182181 | Wales Specialist Virology Centre Sequencing lab: Pathogen Genomics Unit | Public Health Wales Microbiology Cardiff Wales Specialist Virology Centre | Catherine Moore, Johnathan Evans, Laura Gifford, Malorie Perry, Simon Cottrell, Angela Marchbank, Alec Birchley, Alexander Adams, Amy Gaskin, Bree Gatica-Wilcox, Jason Coombes, Joel Southgate, Lauren Gilbert, Lee Graham, Nicole Pacchiarini, Sara Kumziene-Summerhayes, Sarah Taylor, Sophie Jones, Sara Rey, Matthew Bull, Joanne Watkins, Sally Corden, Tom Connor |
| EPI_ISL_3182182 | Labor Berlin Charité Vivantes GmbH / Institut für Virologie | Charité Universitätsmedizin Berlin, Institut für Virologie/Labor Berlin | Peter Menzel, Christine Stephan, Rolf Schwarzer, Victor M Corman, Barbara Mühlemann, Terry Jones, Christian Drosten |
| EPI_ISL_3182183 | Wales Specialist Virology Centre Sequencing lab: Pathogen Genomics Unit | Public Health Wales Microbiology Cardiff Wales Specialist Virology Centre | Catherine Moore, Johnathan Evans, Laura Gifford, Malorie Perry, Simon Cottrell, Angela Marchbank, Alec Birchley, Alexander Adams, Amy Gaskin, Bree Gatica-Wilcox, Jason Coombes, Joel Southgate, Lauren Gilbert, Lee Graham, Nicole Pacchiarini, Sara Kumziene-Summerhayes, Sarah Taylor, Sophie Jones, Sara Rey, Matthew Bull, Joanne Watkins, Sally Corden, Tom Connor |
| EPI_ISL_3182184 | Labor Berlin Charité Vivantes GmbH / Institut für Virologie | Charité Universitätsmedizin Berlin, Institut für Virologie/Labor Berlin | Peter Menzel, Christine Stephan, Rolf Schwarzer, Victor M Corman, Barbara Mühlemann, Terry Jones, Christian Drosten |
| EPI_ISL_3182185 | Wales Specialist Virology Centre Sequencing lab: Pathogen Genomics Unit | Public Health Wales Microbiology Cardiff Wales Specialist Virology Centre | Catherine Moore, Johnathan Evans, Laura Gifford, Malorie Perry, Simon Cottrell, Angela Marchbank, Alec Birchley, Alexander Adams, Amy Gaskin, Bree Gatica-Wilcox, Jason Coombes, Joel Southgate, Lauren Gilbert, Lee Graham, Nicole Pacchiarini, Sara Kumziene-Summerhayes, Sarah Taylor, Sophie Jones, Sara Rey, Matthew Bull, Joanne Watkins, Sally Corden, Tom Connor |
| EPI_ISL_3182186 | Labor Berlin Charité Vivantes GmbH / Institut für Virologie | Charité Universitätsmedizin Berlin, Institut für Virologie/Labor Berlin | Peter Menzel, Christine Stephan, Rolf Schwarzer, Victor M Corman, Barbara Mühlemann, Terry Jones, Christian Drosten |
| EPI_ISL_3182187, EPI_ISL_3182188 | Wales Specialist Virology Centre Sequencing lab: Pathogen Genomics Unit | Public Health Wales Microbiology Cardiff Wales Specialist Virology Centre | Catherine Moore, Johnathan Evans, Laura Gifford, Malorie Perry, Simon Cottrell, Angela Marchbank, Alec Birchley, Alexander Adams, Amy Gaskin, Bree Gatica-Wilcox, Jason Coombes, Joel Southgate, Lauren Gilbert, Lee Graham, Nicole Pacchiarini, Sara Kumziene-Summerhayes, Sarah Taylor, Sophie Jones, Sara Rey, Matthew Bull, Joanne Watkins, Sally Corden, Tom Connor |
| EPI_ISL_3182189 | Labor Berlin Charité Vivantes GmbH / Institut für Virologie | Charité Universitätsmedizin Berlin, Institut für Virologie/Labor Berlin | Peter Menzel, Christine Stephan, Rolf Schwarzer, Victor M Corman, Barbara Mühlemann, Terry Jones, Christian Drosten |
| EPI_ISL_3182190 | Wales Specialist Virology Centre Sequencing lab: Pathogen Genomics Unit | Public Health Wales Microbiology Cardiff Wales Specialist Virology Centre | Catherine Moore, Johnathan Evans, Laura Gifford, Malorie Perry, Simon Cottrell, Angela Marchbank, Alec Birchley, Alexander Adams, Amy Gaskin, Bree Gatica-Wilcox, Jason Coombes, Joel Southgate, Lauren Gilbert, Lee Graham, Nicole Pacchiarini, Sara Kumziene-Summerhayes, Sarah Taylor, Sophie Jones, Sara Rey, Matthew Bull, Joanne Watkins, Sally Corden, Tom Connor |
| EPI_ISL_3182191, EPI_ISL_3182192 | Labor Berlin Charité Vivantes GmbH / Institut für Virologie | Charité Universitätsmedizin Berlin, Institut für Virologie/Labor Berlin | Peter Menzel, Christine Stephan, Rolf Schwarzer, Victor M Corman, Barbara Mühlemann, Terry Jones, Christian Drosten |
| EPI_ISL_3182193 | Wales Specialist Virology Centre Sequencing lab: Pathogen Genomics Unit | Public Health Wales Microbiology Cardiff Wales Specialist Virology Centre | Catherine Moore, Johnathan Evans, Laura Gifford, Malorie Perry, Simon Cottrell, Angela Marchbank, Alec Birchley, Alexander Adams, Amy Gaskin, Bree Gatica-Wilcox, Jason Coombes, Joel Southgate, Lauren Gilbert, Lee Graham, Nicole Pacchiarini, Sara Kumziene-Summerhayes, Sarah Taylor, Sophie Jones, Sara Rey, Matthew Bull, Joanne Watkins, Sally Corden, Tom Connor |
| EPI_ISL_3182194 | Labor Berlin Charité Vivantes GmbH / Institut für Virologie | Charité Universitätsmedizin Berlin, Institut für Virologie/Labor Berlin | Peter Menzel, Christine Stephan, Rolf Schwarzer, Victor M Corman, Barbara Mühlemann, Terry Jones, Christian Drosten |

| | | | |
|---|---|---|---|
| EPI_ISL_3182195, EPI_ISL_3182196 | Wales Specialist Virology Centre Sequencing lab: Pathogen Genomics Unit | Public Health Wales Microbiology Cardiff Wales Specialist Virology Centre | Catherine Moore, Johnathan Evans, Laura Gifford, Malorie Perry, Simon Cottrell, Angela Marchbank, Alec Birchley, Alexander Adams, Amy Gaskin, Bree Gatica-Wilcox, Jason Coombes, Joel Southgate, Lauren Gilbert, Lee Graham, Nicole Pacchiarini, Sara Kumziene-Summerhayes, Sarah Taylor, Sophie Jones, Sara Rey, Matthew Bull, Joanne Watkins, Sally Corden, Tom Connor |
| EPI_ISL_3182197 | Labor Berlin Charité Vivantes GmbH / Institut für Virologie | Charité Universitätsmedizin Berlin, Institut für Virologie/Labor Berlin | Peter Menzel, Christine Stephan, Rolf Schwarzer, Victor M Corman, Barbara Mühlemann, Terry Jones, Christian Drosten |
| EPI_ISL_3182198 | Wales Specialist Virology Centre Sequencing lab: Pathogen Genomics Unit | Public Health Wales Microbiology Cardiff Wales Specialist Virology Centre | Catherine Moore, Johnathan Evans, Laura Gifford, Malorie Perry, Simon Cottrell, Angela Marchbank, Alec Birchley, Alexander Adams, Amy Gaskin, Bree Gatica-Wilcox, Jason Coombes, Joel Southgate, Lauren Gilbert, Lee Graham, Nicole Pacchiarini, Sara Kumziene-Summerhayes, Sarah Taylor, Sophie Jones, Sara Rey, Matthew Bull, Joanne Watkins, Sally Corden, Tom Connor |
| EPI_ISL_3182199 | Labor Berlin Charité Vivantes GmbH / Institut für Virologie | Charité Universitätsmedizin Berlin, Institut für Virologie/Labor Berlin | Peter Menzel, Christine Stephan, Rolf Schwarzer, Victor M Corman, Barbara Mühlemann, Terry Jones, Christian Drosten |
| EPI_ISL_3182200 | Wales Specialist Virology Centre Sequencing lab: Pathogen Genomics Unit | Public Health Wales Microbiology Cardiff Wales Specialist Virology Centre | Catherine Moore, Johnathan Evans, Laura Gifford, Malorie Perry, Simon Cottrell, Angela Marchbank, Alec Birchley, Alexander Adams, Amy Gaskin, Bree Gatica-Wilcox, Jason Coombes, Joel Southgate, Lauren Gilbert, Lee Graham, Nicole Pacchiarini, Sara Kumziene-Summerhayes, Sarah Taylor, Sophie Jones, Sara Rey, Matthew Bull, Joanne Watkins, Sally Corden, Tom Connor |
| EPI_ISL_3182201 | Labor Berlin Charité Vivantes GmbH / Institut für Virologie | Charité Universitätsmedizin Berlin, Institut für Virologie/Labor Berlin | Peter Menzel, Christine Stephan, Rolf Schwarzer, Victor M Corman, Barbara Mühlemann, Terry Jones, Christian Drosten |
| EPI_ISL_3182202, EPI_ISL_3182203 | Wales Specialist Virology Centre Sequencing lab: Pathogen Genomics Unit | Public Health Wales Microbiology Cardiff Wales Specialist Virology Centre | Catherine Moore, Johnathan Evans, Laura Gifford, Malorie Perry, Simon Cottrell, Angela Marchbank, Alec Birchley, Alexander Adams, Amy Gaskin, Bree Gatica-Wilcox, Jason Coombes, Joel Southgate, Lauren Gilbert, Lee Graham, Nicole Pacchiarini, Sara Kumziene-Summerhayes, Sarah Taylor, Sophie Jones, Sara Rey, Matthew Bull, Joanne Watkins, Sally Corden, Tom Connor |
| EPI_ISL_3182204 | Labor Berlin Charité Vivantes GmbH / Institut für Virologie | Charité Universitätsmedizin Berlin, Institut für Virologie/Labor Berlin | Peter Menzel, Christine Stephan, Rolf Schwarzer, Victor M Corman, Barbara Mühlemann, Terry Jones, Christian Drosten |
| EPI_ISL_3182205 | Wales Specialist Virology Centre Sequencing lab: Pathogen Genomics Unit | Public Health Wales Microbiology Cardiff Wales Specialist Virology Centre | Catherine Moore, Johnathan Evans, Laura Gifford, Malorie Perry, Simon Cottrell, Angela Marchbank, Alec Birchley, Alexander Adams, Amy Gaskin, Bree Gatica-Wilcox, Jason Coombes, Joel Southgate, Lauren Gilbert, Lee Graham, Nicole Pacchiarini, Sara Kumziene-Summerhayes, Sarah Taylor, Sophie Jones, Sara Rey, Matthew Bull, Joanne Watkins, Sally Corden, Tom Connor |
| EPI_ISL_3182206 | Labor Berlin Charité Vivantes GmbH / Institut für Virologie | Charité Universitätsmedizin Berlin, Institut für Virologie/Labor Berlin | Peter Menzel, Christine Stephan, Rolf Schwarzer, Victor M Corman, Barbara Mühlemann, Terry Jones, Christian Drosten |
| EPI_ISL_3182207 | Wales Specialist Virology Centre Sequencing lab: Pathogen Genomics Unit | Public Health Wales Microbiology Cardiff Wales Specialist Virology Centre | Catherine Moore, Johnathan Evans, Laura Gifford, Malorie Perry, Simon Cottrell, Angela Marchbank, Alec Birchley, Alexander Adams, Amy Gaskin, Bree Gatica-Wilcox, Jason Coombes, Joel Southgate, Lauren Gilbert, Lee Graham, Nicole Pacchiarini, Sara Kumziene-Summerhayes, Sarah Taylor, Sophie Jones, Sara Rey, Matthew Bull, Joanne Watkins, Sally Corden, Tom Connor |
| EPI_ISL_3182208 | Labor Berlin Charité Vivantes GmbH / Institut für Virologie | Charité Universitätsmedizin Berlin, Institut für Virologie/Labor Berlin | Peter Menzel, Christine Stephan, Rolf Schwarzer, Victor M Corman, Barbara Mühlemann, Terry Jones, Christian Drosten |
| EPI_ISL_3182209 | Wales Specialist Virology Centre Sequencing lab: Pathogen Genomics Unit | Public Health Wales Microbiology Cardiff Wales Specialist Virology Centre | Catherine Moore, Johnathan Evans, Laura Gifford, Malorie Perry, Simon Cottrell, Angela Marchbank, Alec Birchley, Alexander Adams, Amy Gaskin, Bree Gatica-Wilcox, Jason Coombes, Joel Southgate, Lauren Gilbert, Lee Graham, Nicole Pacchiarini, Sara Kumziene-Summerhayes, Sarah Taylor, Sophie Jones, Sara Rey, Matthew Bull, Joanne Watkins, Sally Corden, Tom Connor |
| EPI_ISL_3182210 | Labor Berlin Charité Vivantes GmbH / Institut für Virologie | Charité Universitätsmedizin Berlin, Institut für Virologie/Labor Berlin | Peter Menzel, Christine Stephan, Rolf Schwarzer, Victor M Corman, Barbara Mühlemann, Terry Jones, Christian Drosten |
| EPI_ISL_3182211, EPI_ISL_3182212 | Wales Specialist Virology Centre Sequencing lab: Pathogen Genomics Unit | Public Health Wales Microbiology Cardiff Wales Specialist Virology Centre | Catherine Moore, Johnathan Evans, Laura Gifford, Malorie Perry, Simon Cottrell, Angela Marchbank, Alec Birchley, Alexander Adams, Amy Gaskin, Bree Gatica-Wilcox, Jason Coombes, Joel Southgate, Lauren Gilbert, Lee Graham, Nicole Pacchiarini, Sara Kumziene-Summerhayes, Sarah Taylor, Sophie Jones, Sara Rey, Matthew Bull, Joanne Watkins, Sally Corden, Tom Connor |
| EPI_ISL_3182213 | Labor Berlin Charité Vivantes GmbH / Institut für Virologie | Charité Universitätsmedizin Berlin, Institut für Virologie/Labor Berlin | Peter Menzel, Christine Stephan, Rolf Schwarzer, Victor M Corman, Barbara Mühlemann, Terry Jones, Christian Drosten |
| EPI_ISL_3182214 | Wales Specialist Virology Centre Sequencing lab: Pathogen Genomics Unit | Public Health Wales Microbiology Cardiff Wales Specialist Virology Centre | Catherine Moore, Johnathan Evans, Laura Gifford, Malorie Perry, Simon Cottrell, Angela Marchbank, Alec Birchley, Alexander Adams, Amy Gaskin, Bree Gatica-Wilcox, Jason Coombes, Joel Southgate, Lauren Gilbert, Lee Graham, Nicole Pacchiarini, Sara Kumziene-Summerhayes, Sarah Taylor, Sophie Jones, Sara Rey, Matthew Bull, Joanne Watkins, Sally Corden, Tom Connor |
| EPI_ISL_3182215 | Labor Berlin Charité Vivantes GmbH / Institut für Virologie | Charité Universitätsmedizin Berlin, Institut für Virologie/Labor Berlin | Peter Menzel, Christine Stephan, Rolf Schwarzer, Victor M Corman, Barbara Mühlemann, Terry Jones, Christian Drosten |
| EPI_ISL_3182216 | Wales Specialist Virology Centre Sequencing lab: Pathogen Genomics Unit | Public Health Wales Microbiology Cardiff Wales Specialist Virology Centre | Catherine Moore, Johnathan Evans, Laura Gifford, Malorie Perry, Simon Cottrell, Angela Marchbank, Alec Birchley, Alexander Adams, Amy Gaskin, Bree Gatica-Wilcox, Jason Coombes, Joel Southgate, Lauren Gilbert, Lee Graham, Nicole Pacchiarini, Sara Kumziene-Summerhayes, Sarah Taylor, Sophie Jones, Sara Rey, Matthew Bull, Joanne Watkins, Sally Corden, Tom Connor |
| EPI_ISL_3182217 | Labor Berlin Charité Vivantes GmbH / Institut für Virologie | Charité Universitätsmedizin Berlin, Institut für Virologie/Labor Berlin | Peter Menzel, Christine Stephan, Rolf Schwarzer, Victor M Corman, Barbara Mühlemann, Terry Jones, Christian Drosten |
| EPI_ISL_3182218 | Wales Specialist Virology Centre Sequencing lab: Pathogen Genomics Unit | Public Health Wales Microbiology Cardiff Wales Specialist Virology Centre | Catherine Moore, Johnathan Evans, Laura Gifford, Malorie Perry, Simon Cottrell, Angela Marchbank, Alec Birchley, Alexander Adams, Amy Gaskin, Bree Gatica-Wilcox, Jason Coombes, Joel Southgate, Lauren Gilbert, Lee Graham, Nicole Pacchiarini, Sara Kumziene-Summerhayes, Sarah Taylor, Sophie Jones, Sara Rey, Matthew Bull, Joanne Watkins, Sally Corden, Tom Connor |
| EPI_ISL_3182219 | Labor Berlin Charité Vivantes GmbH / Institut für Virologie | Charité Universitätsmedizin Berlin, Institut für Virologie/Labor Berlin | Peter Menzel, Christine Stephan, Rolf Schwarzer, Victor M Corman, Barbara Mühlemann, Terry Jones, Christian Drosten |
| EPI_ISL_3182220 | Wales Specialist Virology Centre Sequencing lab: Pathogen Genomics Unit | Public Health Wales Microbiology Cardiff Wales Specialist Virology Centre | Catherine Moore, Johnathan Evans, Laura Gifford, Malorie Perry, Simon Cottrell, Angela Marchbank, Alec Birchley, Alexander Adams, Amy Gaskin, Bree Gatica-Wilcox, Jason Coombes, Joel Southgate, Lauren Gilbert, Lee Graham, Nicole Pacchiarini, Sara Kumziene-Summerhayes, Sarah Taylor, Sophie Jones, Sara Rey, Matthew Bull, Joanne Watkins, Sally Corden, Tom Connor |
| EPI_ISL_3182221 | Labor Berlin Charité Vivantes GmbH / Institut für Virologie | Charité Universitätsmedizin Berlin, Institut für Virologie/Labor Berlin | Peter Menzel, Christine Stephan, Rolf Schwarzer, Victor M Corman, Barbara Mühlemann, Terry Jones, Christian Drosten |
| EPI_ISL_3182222 | Wales Specialist Virology Centre Sequencing lab: Pathogen Genomics Unit | Public Health Wales Microbiology Cardiff Wales Specialist Virology Centre | Catherine Moore, Johnathan Evans, Laura Gifford, Malorie Perry, Simon Cottrell, Angela Marchbank, Alec Birchley, Alexander Adams, Amy Gaskin, Bree Gatica-Wilcox, Jason Coombes, Joel Southgate, Lauren Gilbert, Lee Graham, Nicole Pacchiarini, Sara Kumziene-Summerhayes, Sarah Taylor, Sophie Jones, Sara Rey, Matthew Bull, Joanne Watkins, Sally Corden, Tom Connor |
| EPI_ISL_3182223 | Labor Berlin Charité Vivantes GmbH / Institut für Virologie | Charité Universitätsmedizin Berlin, Institut für Virologie/Labor Berlin | Peter Menzel, Christine Stephan, Rolf Schwarzer, Victor M Corman, Barbara Mühlemann, Terry Jones, Christian Drosten |
| EPI_ISL_3182224 | Wales Specialist Virology Centre Sequencing lab: Pathogen Genomics Unit | Public Health Wales Microbiology Cardiff Wales Specialist Virology Centre | Catherine Moore, Johnathan Evans, Laura Gifford, Malorie Perry, Simon Cottrell, Angela Marchbank, Alec Birchley, Alexander Adams, Amy Gaskin, Bree Gatica-Wilcox, Jason Coombes, Joel Southgate, Lauren Gilbert, Lee Graham, Nicole Pacchiarini, Sara Kumziene-Summerhayes, Sarah Taylor, Sophie Jones, Sara Rey, Matthew Bull, Joanne Watkins, Sally Corden, Tom Connor |
| EPI_ISL_3182225 | Labor Berlin Charité Vivantes GmbH / Institut für Virologie | Charité Universitätsmedizin Berlin, Institut für Virologie/Labor Berlin | Peter Menzel, Christine Stephan, Rolf Schwarzer, Victor M Corman, Barbara Mühlemann, Terry Jones, Christian Drosten |
| EPI_ISL_3182226 | Wales Specialist Virology Centre Sequencing lab: Pathogen Genomics Unit | Public Health Wales Microbiology Cardiff Wales Specialist Virology Centre | Catherine Moore, Johnathan Evans, Laura Gifford, Malorie Perry, Simon Cottrell, Angela Marchbank, Alec Birchley, Alexander Adams, Amy Gaskin, Bree Gatica-Wilcox, Jason Coombes, Joel Southgate, Lauren Gilbert, Lee Graham, Nicole Pacchiarini, Sara Kumziene-Summerhayes, Sarah Taylor, Sophie Jones, Sara Rey, Matthew Bull, Joanne Watkins, Sally Corden, Tom Connor |
| EPI_ISL_3182227 | Labor Berlin Charité Vivantes GmbH / Institut für Virologie | Charité Universitätsmedizin Berlin, Institut für Virologie/Labor Berlin | Peter Menzel, Christine Stephan, Rolf Schwarzer, Victor M Corman, Barbara Mühlemann, Terry Jones, Christian Drosten |
| EPI_ISL_3182228, EPI_ISL_3182229 | Wales Specialist Virology Centre Sequencing lab: Pathogen Genomics Unit | Public Health Wales Microbiology Cardiff Wales Specialist Virology Centre | Catherine Moore, Johnathan Evans, Laura Gifford, Malorie Perry, Simon Cottrell, Angela Marchbank, Alec Birchley, Alexander Adams, Amy Gaskin, Bree Gatica-Wilcox, Jason Coombes, Joel Southgate, Lauren Gilbert, Lee Graham, Nicole Pacchiarini, Sara Kumziene-Summerhayes, Sarah Taylor, Sophie Jones, Sara Rey, Matthew Bull, Joanne Watkins, Sally Corden, Tom Connor |
| EPI_ISL_3182230 | Labor Berlin Charité Vivantes GmbH / Institut für Virologie | Charité Universitätsmedizin Berlin, Institut für Virologie/Labor Berlin | Peter Menzel, Christine Stephan, Rolf Schwarzer, Victor M Corman, Barbara Mühlemann, Terry Jones, Christian Drosten |
| EPI_ISL_3182231 | Wales Specialist Virology Centre Sequencing lab: Pathogen Genomics Unit | Public Health Wales Microbiology Cardiff Wales Specialist Virology Centre | Catherine Moore, Johnathan Evans, Laura Gifford, Malorie Perry, Simon Cottrell, Angela Marchbank, Alec Birchley, Alexander Adams, Amy Gaskin, Bree Gatica-Wilcox, Jason Coombes, Joel Southgate, Lauren Gilbert, Lee Graham, Nicole Pacchiarini, Sara Kumziene-Summerhayes, Sarah Taylor, Sophie Jones, Sara Rey, Matthew Bull, Joanne Watkins, Sally Corden, Tom Connor |
| EPI_ISL_3182232 | Labor Berlin Charité Vivantes GmbH / Institut für Virologie | Charité Universitätsmedizin Berlin, Institut für Virologie/Labor Berlin | Peter Menzel, Christine Stephan, Rolf Schwarzer, Victor M Corman, Barbara Mühlemann, Terry Jones, Christian Drosten |
| EPI_ISL_3182233 | Wales Specialist Virology Centre Sequencing lab: Pathogen Genomics Unit | Public Health Wales Microbiology Cardiff Wales Specialist Virology Centre | Catherine Moore, Johnathan Evans, Laura Gifford, Malorie Perry, Simon Cottrell, Angela Marchbank, Alec Birchley, Alexander Adams, Amy Gaskin, Bree Gatica-Wilcox, Jason Coombes, Joel Southgate, Lauren Gilbert, Lee Graham, Nicole Pacchiarini, Sara Kumziene-Summerhayes, Sarah Taylor, Sophie Jones, Sara Rey, Matthew Bull, Joanne Watkins, Sally Corden, Tom Connor |
| EPI_ISL_3182234 | Labor Berlin Charité Vivantes GmbH / Institut für Virologie | Charité Universitätsmedizin Berlin, Institut für Virologie/Labor Berlin | Peter Menzel, Christine Stephan, Rolf Schwarzer, Victor M Corman, Barbara Mühlemann, Terry Jones, Christian Drosten |
| EPI_ISL_3182235 | Wales Specialist Virology Centre Sequencing lab: Pathogen Genomics Unit | Public Health Wales Microbiology Cardiff Wales Specialist Virology Centre | Catherine Moore, Johnathan Evans, Laura Gifford, Malorie Perry, Simon Cottrell, Angela Marchbank, Alec Birchley, Alexander Adams, Amy Gaskin, Bree Gatica-Wilcox, Jason Coombes, Joel Southgate, Lauren Gilbert, Lee Graham, Nicole Pacchiarini, Sara Kumziene-Summerhayes, Sarah Taylor, Sophie Jones, Sara Rey, Matthew Bull, Joanne Watkins, Sally Corden, Tom Connor |
| EPI_ISL_3182236 | Labor Berlin Charité Vivantes GmbH / Institut für Virologie | Charité Universitätsmedizin Berlin, Institut für Virologie/Labor Berlin | Peter Menzel, Christine Stephan, Rolf Schwarzer, Victor M Corman, Barbara Mühlemann, Terry Jones, Christian Drosten |
| EPI_ISL_3182237, EPI_ISL_3182238 | Wales Specialist Virology Centre Sequencing lab: Pathogen Genomics Unit | Public Health Wales Microbiology Cardiff Wales Specialist Virology Centre | Catherine Moore, Johnathan Evans, Laura Gifford, Malorie Perry, Simon Cottrell, Angela Marchbank, Alec Birchley, Alexander Adams, Amy Gaskin, Bree Gatica-Wilcox, Jason Coombes, Joel Southgate, Lauren Gilbert, Lee Graham, Nicole Pacchiarini, Sara Kumziene-Summerhayes, Sarah Taylor, Sophie Jones, Sara Rey, Matthew Bull, Joanne Watkins, Sally Corden, Tom Connor |
| EPI_ISL_3182239 | Labor Berlin Charité Vivantes GmbH / Institut für Virologie | Charité Universitätsmedizin Berlin, Institut für Virologie/Labor Berlin | Peter Menzel, Christine Stephan, Rolf Schwarzer, Victor M Corman, Barbara Mühlemann, Terry Jones, Christian Drosten |
| EPI_ISL_3182240 | Wales Specialist Virology Centre Sequencing lab: Pathogen Genomics Unit | Public Health Wales Microbiology Cardiff Wales Specialist Virology Centre | Catherine Moore, Johnathan Evans, Laura Gifford, Malorie Perry, Simon Cottrell, Angela Marchbank, Alec Birchley, Alexander Adams, Amy Gaskin, Bree Gatica-Wilcox, Jason Coombes, Joel Southgate, Lauren Gilbert, Lee Graham, Nicole Pacchiarini, Sara Kumziene-Summerhayes, Sarah Taylor, Sophie Jones, Sara Rey, Matthew Bull, Joanne Watkins, Sally Corden, Tom Connor |
| EPI_ISL_3182241 | Labor Berlin Charité Vivantes GmbH / Institut für Virologie | Charité Universitätsmedizin Berlin, Institut für Virologie/Labor Berlin | Peter Menzel, Christine Stephan, Rolf Schwarzer, Victor M Corman, Barbara Mühlemann, Terry Jones, Christian Drosten |
| EPI_ISL_3182242, EPI_ISL_3182243 | Wales Specialist Virology Centre Sequencing lab: Pathogen Genomics Unit | Public Health Wales Microbiology Cardiff Wales Specialist Virology Centre | Catherine Moore, Johnathan Evans, Laura Gifford, Malorie Perry, Simon Cottrell, Angela Marchbank, Alec Birchley, Alexander Adams, Amy Gaskin, Bree Gatica-Wilcox, Jason Coombes, Joel Southgate, Lauren Gilbert, Lee Graham, Nicole Pacchiarini, Sara Kumziene-Summerhayes, Sarah Taylor, Sophie Jones, Sara Rey, Matthew Bull, Joanne Watkins, Sally Corden, Tom Connor |
| EPI_ISL_3182244 | Labor Berlin Charité Vivantes GmbH / Institut für Virologie | Charité Universitätsmedizin Berlin, Institut für Virologie/Labor Berlin | Peter Menzel, Christine Stephan, Rolf Schwarzer, Victor M Corman, Barbara Mühlemann, Terry Jones, Christian Drosten |
| EPI_ISL_3182245 | Wales Specialist Virology Centre Sequencing lab: Pathogen Genomics Unit | Public Health Wales Microbiology Cardiff Wales Specialist Virology Centre | Catherine Moore, Johnathan Evans, Laura Gifford, Malorie Perry, Simon Cottrell, Angela Marchbank, Alec Birchley, Alexander Adams, Amy Gaskin, Bree Gatica-Wilcox, Jason Coombes, Joel Southgate, Lauren Gilbert, Lee Graham, Nicole Pacchiarini, Sara Kumziene-Summerhayes, Sarah Taylor, Sophie Jones, Sara Rey, Matthew Bull, Joanne Watkins, Sally Corden, Tom Connor |
| EPI_ISL_3182246 | Labor Berlin Charité Vivantes GmbH / Institut für Virologie | Charité Universitätsmedizin Berlin, Institut für Virologie/Labor Berlin | Peter Menzel, Christine Stephan, Rolf Schwarzer, Victor M Corman, Barbara Mühlemann, Terry Jones, Christian Drosten |
| EPI_ISL_3182247 | Wales Specialist Virology Centre Sequencing lab: Pathogen Genomics Unit | Public Health Wales Microbiology Cardiff Wales Specialist Virology Centre | Catherine Moore, Johnathan Evans, Laura Gifford, Malorie Perry, Simon Cottrell, Angela Marchbank, Alec Birchley, Alexander Adams, Amy Gaskin, Bree Gatica-Wilcox, Jason Coombes, Joel Southgate, Lauren Gilbert, Lee Graham, Nicole Pacchiarini, Sara Kumziene-Summerhayes, Sarah Taylor, Sophie Jones, Sara Rey, Matthew Bull, Joanne Watkins, Sally Corden, Tom Connor |
| EPI_ISL_3182248 | Labor Berlin Charité Vivantes GmbH / Institut für Virologie | Charité Universitätsmedizin Berlin, Institut für Virologie/Labor Berlin | Peter Menzel, Christine Stephan, Rolf Schwarzer, Victor M Corman, Barbara Mühlemann, Terry Jones, Christian Drosten |
| EPI_ISL_3182249 | Wales Specialist Virology Centre Sequencing lab: Pathogen Genomics Unit | Public Health Wales Microbiology Cardiff Wales Specialist Virology Centre | Catherine Moore, Johnathan Evans, Laura Gifford, Malorie Perry, Simon Cottrell, Angela Marchbank, Alec Birchley, Alexander Adams, Amy Gaskin, Bree Gatica-Wilcox, Jason Coombes, Joel Southgate, Lauren Gilbert, Lee Graham, Nicole Pacchiarini, Sara Kumziene-Summerhayes, Sarah Taylor, Sophie Jones, Sara Rey, Matthew Bull, Joanne Watkins, Sally Corden, Tom Connor |
| EPI_ISL_3182250 | Labor Berlin Charité Vivantes GmbH / Institut für Virologie | Charité Universitätsmedizin Berlin, Institut für Virologie/Labor Berlin | Peter Menzel, Christine Stephan, Rolf Schwarzer, Victor M Corman, Barbara Mühlemann, Terry Jones, Christian Drosten |
| EPI_ISL_3182251, EPI_ISL_3182252, EPI_ISL_3182253, EPI_ISL_3182254, EPI_ISL_3182255, EPI_ISL_3182256, EPI_ISL_3182257, EPI_ISL_3182258, EPI_ISL_3182259, EPI_ISL_3182260, EPI_ISL_3182261, EPI_ISL_3182262, EPI_ISL_3182263, EPI_ISL_3182264, EPI_ISL_3182265, EPI_ISL_3182266, EPI_ISL_3182267, EPI_ISL_3182268, EPI_ISL_3182269, EPI_ISL_3182270, EPI_ISL_3182271, EPI_ISL_3182272, EPI_ISL_3182273, EPI_ISL_3182274, EPI_ISL_3182275, EPI_ISL_3182276, EPI_ISL_3182277, EPI_ISL_3182278, EPI_ISL_3182279, EPI_ISL_3182280, EPI_ISL_3182281, EPI_ISL_3182282, EPI_ISL_3182283, EPI_ISL_3182284, EPI_ISL_3182285, EPI_ISL_3182286, EPI_ISL_3182287, EPI_ISL_3182288, EPI_ISL_3182289, EPI_ISL_3182290, EPI_ISL_3182291, EPI_ISL_3182292, EPI_ISL_3182293, EPI_ISL_3182294, EPI_ISL_3182295, EPI_ISL_3182296, EPI_ISL_3182297, EPI_ISL_3182298, EPI_ISL_3182299, EPI_ISL_3182300, EPI_ISL_3182301, EPI_ISL_3182302, EPI_ISL_3182303, EPI_ISL_3182304, EPI_ISL_3182305, EPI_ISL_3182306, EPI_ISL_3182307, EPI_ISL_3182308, EPI_ISL_3182309, EPI_ISL_3182310, EPI_ISL_3182311, EPI_ISL_3182312, EPI_ISL_3182313, EPI_ISL_3182314, EPI_ISL_3182315, EPI_ISL_3182316, EPI_ISL_3182317, EPI_ISL_3182319, EPI_ISL_3182320, EPI_ISL_3182321, EPI_ISL_3182322, EPI_ISL_3182323, EPI_ISL_3182324, EPI_ISL_3182326, EPI_ISL_3182328, EPI_ISL_3182330, EPI_ISL_3182332, EPI_ISL_3182333, EPI_ISL_3182334, EPI_ISL_3182335, EPI_ISL_3182336, EPI_ISL_3182338, EPI_ISL_3182339, EPI_ISL_3182340, EPI_ISL_3182341, EPI_ISL_3182343, EPI_ISL_3182344, EPI_ISL_3182345, EPI_ISL_3182346, EPI_ISL_3182347, EPI_ISL_3182349, EPI_ISL_3182350, EPI_ISL_3182351, EPI_ISL_3182352 | | | |
| see above | Wales Specialist Virology Centre Sequencing lab: Pathogen Genomics Unit | Public Health Wales Microbiology Cardiff Wales Specialist Virology Centre | Catherine Moore, Johnathan Evans, Laura Gifford, Malorie Perry, Simon Cottrell, Angela Marchbank, Alec Birchley, Alexander Adams, Amy Gaskin, Bree Gatica-Wilcox, Jason Coombes, Joel Southgate, Lauren Gilbert, Lee Graham, Nicole Pacchiarini, Sara Kumziene-Summerhayes, Sarah Taylor, Sophie Jones, Sara Rey, Matthew Bull, Joanne Watkins, Sally Corden, Tom Connor |
| EPI_ISL_3182353 | Kagoshima Prefectural Institute for Environmental Research and Public Health | Kagoshima Prefectural Institute for Environmental Research and Public Health | Yuka Hamada, Takako Kiyotake, Kanji Ishitani, Mami Yamamoto, Kazuki Manabe, Naomi Shinkawa, Tsuyoshi Sekizuka, Kentaro Itokawa, Rina Tanaka, Masanori Hashino, Makoto Kuroda |
| EPI_ISL_3182354, EPI_ISL_3182355, EPI_ISL_3182356, | Wales Specialist Virology Centre Sequencing lab: Pathogen Genomics Unit | Public Health Wales Microbiology Cardiff Wales Specialist Virology Centre | Catherine Moore, Johnathan Evans, Laura Gifford, Malorie Perry, Simon Cottrell, Angela Marchbank, Alec Birchley, Alexander Adams, Amy Gaskin, Bree Gatica-Wilcox, Jason Coombes, Joel Southgate, Lauren Gilbert, Lee Graham, Nicole Pacchiarini, Sara Kumziene-Summerhayes, Sarah Taylor, Sophie Jones, Sara Rey, Matthew Bull, Joanne Watkins, Sally Corden, Tom Connor |

| | | | |
|---|---|---|---|
| EPI_ISL_3182357, EPI_ISL_3182358, EPI_ISL_3182360, EPI_ISL_3182362 | | | |
| EPI_ISL_3182363 | Department of Infectious Diseases, Kobe Institute of Health | Department of Infectious Diseases, Kobe Institute of Health | Ryohei Nomoto, Noriko Nakanishi, Tomotada Iwamoto, Tsuyoshi Sekizuka, Kentaro Itokawa, Rina Tanaka, Masanori Hashino, Makoto Kuroda |
| EPI_ISL_3182364, EPI_ISL_3182365, EPI_ISL_3182367, EPI_ISL_3182369, EPI_ISL_3182370, EPI_ISL_3182372, EPI_ISL_3182374, EPI_ISL_3182376, EPI_ISL_3182377 | Wales Specialist Virology Centre Sequencing lab: Pathogen Genomics Unit | Public Health Wales Microbiology Cardiff Wales Specialist Virology Centre | Catherine Moore, Johnathan Evans, Laura Gifford, Malorie Perry, Simon Cottrell, Angela Marchbank, Alec Birchley, Alexander Adams, Amy Gaskin, Bree Gatica-Wilcox, Jason Coombes, Joel Southgate, Lauren Gilbert, Lee Graham, Nicole Pacchiarini, Sara Kumziene-Summerhayes, Sarah Taylor, Sophie Jones, Sara Rey, Matthew Bull, Joanne Watkins, Sally Corden, Tom Connor |
| EPI_ISL_3182378 | Ibaraki Prefectural Institute of Public Health | Ibaraki Prefectural Institute of Public Health | Keiko Goto, Ryo Kashimura, Shuichi Osawa, Tsuyoshi Sekizuka, Kentaro Itokawa, Rina Tanaka, Masanori Hashino, Makoto Kuroda |
| EPI_ISL_3182379, EPI_ISL_3182381, EPI_ISL_3182383, EPI_ISL_3182384, EPI_ISL_3182386, EPI_ISL_3182388, EPI_ISL_3182390, EPI_ISL_3182392, EPI_ISL_3182393, EPI_ISL_3182395 | Wales Specialist Virology Centre Sequencing lab: Pathogen Genomics Unit | Public Health Wales Microbiology Cardiff Wales Specialist Virology Centre | Catherine Moore, Johnathan Evans, Laura Gifford, Malorie Perry, Simon Cottrell, Angela Marchbank, Alec Birchley, Alexander Adams, Amy Gaskin, Bree Gatica-Wilcox, Jason Coombes, Joel Southgate, Lauren Gilbert, Lee Graham, Nicole Pacchiarini, Sara Kumziene-Summerhayes, Sarah Taylor, Sophie Jones, Sara Rey, Matthew Bull, Joanne Watkins, Sally Corden, Tom Connor |
| EPI_ISL_3182396 | Department of Health Science, Iwate Prefectural Institute of Public Health and Environment | Iwate Prefectural Institute of Public Health and Environment | Hiroki Konno, Hiroko Kajita, Kaori Iwabuchi, Akiko Fujimori, Taihei Mitsui, Tomoko Takahashi, Tsuyoshi Sekizuka, Kentaro Itokawa, Rina Tanaka, Masanori Hashino, Makoto Kuroda |
| EPI_ISL_3182397, EPI_ISL_3182399, EPI_ISL_3182401, EPI_ISL_3182402, EPI_ISL_3182404, EPI_ISL_3182405, EPI_ISL_3182407, EPI_ISL_3182409, EPI_ISL_3182411, EPI_ISL_3182413, EPI_ISL_3182414, EPI_ISL_3182416, EPI_ISL_3182418, EPI_ISL_3182420, EPI_ISL_3182421, EPI_ISL_3182422, EPI_ISL_3182423, EPI_ISL_3182424, EPI_ISL_3182425, EPI_ISL_3182427, EPI_ISL_3182428, EPI_ISL_3182429, EPI_ISL_3182430, EPI_ISL_3182431, EPI_ISL_3182433, EPI_ISL_3182434, EPI_ISL_3182435, EPI_ISL_3182436, EPI_ISL_3182437, EPI_ISL_3182439, EPI_ISL_3182440, EPI_ISL_3182441, EPI_ISL_3182442, EPI_ISL_3182443, EPI_ISL_3182445, EPI_ISL_3182446, EPI_ISL_3182447, EPI_ISL_3182448, EPI_ISL_3182449, EPI_ISL_3182451, EPI_ISL_3182452, EPI_ISL_3182453, EPI_ISL_3182454, EPI_ISL_3182455, EPI_ISL_3182456, EPI_ISL_3182458, EPI_ISL_3182459, EPI_ISL_3182460, EPI_ISL_3182461, EPI_ISL_3182463, EPI_ISL_3182464, EPI_ISL_3182465, EPI_ISL_3182466, EPI_ISL_3182467, EPI_ISL_3182469, EPI_ISL_3182470, EPI_ISL_3182471, EPI_ISL_3182472, EPI_ISL_3182474, EPI_ISL_3182475, EPI_ISL_3182476, EPI_ISL_3182477, EPI_ISL_3182478, EPI_ISL_3182480, EPI_ISL_3182481, EPI_ISL_3182482, EPI_ISL_3182483, EPI_ISL_3182485, EPI_ISL_3182486, EPI_ISL_3182487, EPI_ISL_3182488, EPI_ISL_3182490, EPI_ISL_3182491, EPI_ISL_3182492, EPI_ISL_3182493, EPI_ISL_3182495, EPI_ISL_3182496, EPI_ISL_3182497, EPI_ISL_3182498, EPI_ISL_3182500, EPI_ISL_3182501, EPI_ISL_3182503, EPI_ISL_3182505, EPI_ISL_3182506, EPI_ISL_3182508, EPI_ISL_3182510, EPI_ISL_3182512, EPI_ISL_3182514, EPI_ISL_3182515, EPI_ISL_3182517, EPI_ISL_3182518, EPI_ISL_3182519, EPI_ISL_3182520, EPI_ISL_3182522, EPI_ISL_3182523, EPI_ISL_3182524, EPI_ISL_3182525, EPI_ISL_3182527, EPI_ISL_3182528, EPI_ISL_3182530, EPI_ISL_3182532, EPI_ISL_3182534, EPI_ISL_3182535, EPI_ISL_3182537, EPI_ISL_3182539, EPI_ISL_3182541, EPI_ISL_3182542, EPI_ISL_3182543, EPI_ISL_3182544, EPI_ISL_3182545, EPI_ISL_3182546, EPI_ISL_3182548, EPI_ISL_3182549, EPI_ISL_3182550, EPI_ISL_3182551, EPI_ISL_3182553, EPI_ISL_3182554, EPI_ISL_3182555, EPI_ISL_3182556, EPI_ISL_3182557, EPI_ISL_3182558, EPI_ISL_3182560, EPI_ISL_3182561, EPI_ISL_3182562, EPI_ISL_3182563, EPI_ISL_3182564, EPI_ISL_3182566, EPI_ISL_3182567, EPI_ISL_3182568, EPI_ISL_3182569, EPI_ISL_3182570, EPI_ISL_3182572, EPI_ISL_3182573, EPI_ISL_3182574, EPI_ISL_3182575, EPI_ISL_3182576, EPI_ISL_3182578, EPI_ISL_3182579, EPI_ISL_3182580, EPI_ISL_3182581, EPI_ISL_3182582 | | | |
| see above | Wales Specialist Virology Centre Sequencing lab: Pathogen Genomics Unit | Public Health Wales Microbiology Cardiff Wales Specialist Virology Centre | Catherine Moore, Johnathan Evans, Laura Gifford, Malorie Perry, Simon Cottrell, Angela Marchbank, Alec Birchley, Alexander Adams, Amy Gaskin, Bree Gatica-Wilcox, Jason Coombes, Joel Southgate, Lauren Gilbert, Lee Graham, Nicole Pacchiarini, Sara Kumziene-Summerhayes, Sarah Taylor, Sophie Jones, Sara Rey, Matthew Bull, Joanne Watkins, Sally Corden, Tom Connor |
| EPI_ISL_3182584, EPI_ISL_3182585, EPI_ISL_3182587, EPI_ISL_3182588, EPI_ISL_3182589, EPI_ISL_3182590, EPI_ISL_3182591, EPI_ISL_3182593, EPI_ISL_3182594, EPI_ISL_3182595, EPI_ISL_3182597, EPI_ISL_3182598, EPI_ISL_3182599 | | | |
| see above | Originating lab: Wales Specialist Virology Centre Sequencing lab: Pathogen Genomics Unit | Public Health Wales Microbiology Cardiff Wales Specialist Virology Centre | Catherine Moore, Johnathan Evans, Laura Gifford, Malorie Perry, Simon Cottrell, Angela Marchbank, Alec Birchley, Alexander Adams, Amy Gaskin, Bree Gatica-Wilcox, Jason Coombes, Joel Southgate, Lauren Gilbert, Lee Graham, Nicole Pacchiarini, Sara Kumziene-Summerhayes, Sarah Taylor, Sophie Jones, Sara Rey, Matthew Bull, Joanne Watkins, Sally Corden, Tom Connor |
| EPI_ISL_3182601, EPI_ISL_3182602, EPI_ISL_3182603, EPI_ISL_3182604, EPI_ISL_3182606, EPI_ISL_3182607, EPI_ISL_3182608, EPI_ISL_3182609, EPI_ISL_3182610, EPI_ISL_3182612, EPI_ISL_3182613, EPI_ISL_3182614, EPI_ISL_3182615, EPI_ISL_3182616, EPI_ISL_3182618, EPI_ISL_3182619, EPI_ISL_3182620, EPI_ISL_3182622, EPI_ISL_3182623, EPI_ISL_3182624, EPI_ISL_3182625, EPI_ISL_3182626, EPI_ISL_3182628, EPI_ISL_3182629, EPI_ISL_3182630, EPI_ISL_3182631, EPI_ISL_3182632, EPI_ISL_3182634, EPI_ISL_3182635, EPI_ISL_3182636, EPI_ISL_3182637, EPI_ISL_3182639, EPI_ISL_3182640, EPI_ISL_3182641, EPI_ISL_3182642, EPI_ISL_3182644, EPI_ISL_3182645, EPI_ISL_3182646, EPI_ISL_3182647, EPI_ISL_3182648, EPI_ISL_3182650, EPI_ISL_3182651, EPI_ISL_3182652, EPI_ISL_3182653, EPI_ISL_3182654, EPI_ISL_3182656, EPI_ISL_3182657, EPI_ISL_3182658, EPI_ISL_3182659, EPI_ISL_3182661, EPI_ISL_3182662, EPI_ISL_3182663, EPI_ISL_3182664, EPI_ISL_3182665, EPI_ISL_3182667, EPI_ISL_3182668, EPI_ISL_3182669, EPI_ISL_3182670, EPI_ISL_3182672, EPI_ISL_3182673, EPI_ISL_3182674, EPI_ISL_3182675, EPI_ISL_3182676, EPI_ISL_3182678, EPI_ISL_3182679, EPI_ISL_3182680, EPI_ISL_3182681, EPI_ISL_3182683, EPI_ISL_3182684, EPI_ISL_3182685, EPI_ISL_3182686, EPI_ISL_3182687, EPI_ISL_3182689, EPI_ISL_3182690, EPI_ISL_3182691, EPI_ISL_3182692, EPI_ISL_3182693, EPI_ISL_3182695, EPI_ISL_3182696, EPI_ISL_3182697, EPI_ISL_3182698, EPI_ISL_3182700, EPI_ISL_3182701, EPI_ISL_3182702, EPI_ISL_3182703, EPI_ISL_3182704, EPI_ISL_3182706, EPI_ISL_3182707, EPI_ISL_3182708, EPI_ISL_3182709, EPI_ISL_3182711, EPI_ISL_3182712, EPI_ISL_3182713, EPI_ISL_3182714, EPI_ISL_3182715, EPI_ISL_3182717, EPI_ISL_3182718, EPI_ISL_3182719, EPI_ISL_3182720, EPI_ISL_3182721, EPI_ISL_3182723, EPI_ISL_3182724, EPI_ISL_3182725, EPI_ISL_3182726, EPI_ISL_3182727, EPI_ISL_3182729, EPI_ISL_3182730, EPI_ISL_3182731, EPI_ISL_3182732, EPI_ISL_3182734, EPI_ISL_3182735, EPI_ISL_3182736, EPI_ISL_3182737, EPI_ISL_3182738, EPI_ISL_3182739, EPI_ISL_3182741, EPI_ISL_3182742, EPI_ISL_3182743, EPI_ISL_3182744, EPI_ISL_3182745, EPI_ISL_3182747, EPI_ISL_3182748, EPI_ISL_3182749, EPI_ISL_3182750, EPI_ISL_3182752, EPI_ISL_3182753, EPI_ISL_3182754, EPI_ISL_3182755, EPI_ISL_3182756, EPI_ISL_3182758, EPI_ISL_3182759, EPI_ISL_3182760, EPI_ISL_3182761, EPI_ISL_3182762, EPI_ISL_3182764, EPI_ISL_3182765, EPI_ISL_3182766, EPI_ISL_3182767, EPI_ISL_3182768, EPI_ISL_3182770, EPI_ISL_3182771, EPI_ISL_3182772, EPI_ISL_3182773, EPI_ISL_3182775, EPI_ISL_3182776, EPI_ISL_3182777, EPI_ISL_3182778, EPI_ISL_3182779, EPI_ISL_3182781, EPI_ISL_3182782, EPI_ISL_3182783, EPI_ISL_3182784, EPI_ISL_3182785, EPI_ISL_3182787, EPI_ISL_3182788, EPI_ISL_3182789, EPI_ISL_3182790, EPI_ISL_3182791, EPI_ISL_3182793, EPI_ISL_3182794, EPI_ISL_3182795, EPI_ISL_3182796, EPI_ISL_3182797, EPI_ISL_3182799, EPI_ISL_3182800, EPI_ISL_3182801, EPI_ISL_3182802, EPI_ISL_3182803, EPI_ISL_3182805, EPI_ISL_3182806, EPI_ISL_3182807, EPI_ISL_3182808, EPI_ISL_3182809, EPI_ISL_3182811, EPI_ISL_3182812, EPI_ISL_3182813, EPI_ISL_3182814, EPI_ISL_3182815, EPI_ISL_3182817, EPI_ISL_3182818, EPI_ISL_3182819, EPI_ISL_3182820, EPI_ISL_3182821, EPI_ISL_3182823, EPI_ISL_3182824, EPI_ISL_3182825, EPI_ISL_3182826, EPI_ISL_3182827, EPI_ISL_3182828, EPI_ISL_3182830, EPI_ISL_3182831, EPI_ISL_3182832, EPI_ISL_3182833, EPI_ISL_3182835, EPI_ISL_3182836, EPI_ISL_3182837, EPI_ISL_3182838, EPI_ISL_3182839, EPI_ISL_3182841, EPI_ISL_3182842, EPI_ISL_3182843, EPI_ISL_3182844, EPI_ISL_3182846, EPI_ISL_3182847, EPI_ISL_3182848, EPI_ISL_3182849, EPI_ISL_3182850, EPI_ISL_3182851, EPI_ISL_3182853, EPI_ISL_3182854, EPI_ISL_3182855, EPI_ISL_3182856, EPI_ISL_3182857, EPI_ISL_3182859, EPI_ISL_3182860, EPI_ISL_3182861, EPI_ISL_3182862, EPI_ISL_3182864, EPI_ISL_3182865, EPI_ISL_3182866, EPI_ISL_3182867, EPI_ISL_3182868, EPI_ISL_3182869, EPI_ISL_3182870, EPI_ISL_3182871, EPI_ISL_3182873, EPI_ISL_3182874, EPI_ISL_3182875, EPI_ISL_3182876, EPI_ISL_3182878, EPI_ISL_3182879, EPI_ISL_3182880, EPI_ISL_3182881, EPI_ISL_3182883, EPI_ISL_3182884, EPI_ISL_3182885, EPI_ISL_3182886, EPI_ISL_3182888, EPI_ISL_3182889, EPI_ISL_3182890, EPI_ISL_3182891, EPI_ISL_3182893, EPI_ISL_3182894, EPI_ISL_3182895, EPI_ISL_3182896, EPI_ISL_3182898, EPI_ISL_3182899, EPI_ISL_3182900, EPI_ISL_3182901, EPI_ISL_3182902, EPI_ISL_3182903, EPI_ISL_3182904, EPI_ISL_3182905, EPI_ISL_3182907, EPI_ISL_3182908, EPI_ISL_3182909, EPI_ISL_3182910, EPI_ISL_3182911, EPI_ISL_3182912, EPI_ISL_3182913, EPI_ISL_3182914, EPI_ISL_3182915, EPI_ISL_3182916, EPI_ISL_3182918, EPI_ISL_3182919, EPI_ISL_3182920, EPI_ISL_3182921, EPI_ISL_3182922, EPI_ISL_3182923, EPI_ISL_3182924, EPI_ISL_3182925, EPI_ISL_3182926, EPI_ISL_3182928, EPI_ISL_3182929, EPI_ISL_3182930 | | | |
| see above | Centre for Enzyme Innovation, University of Portsmouth / Translational Research Laboratory, Portsmouth Hospitals NHS Trust / Hampshire Hospitals NHS Foundation Trust / Bournemouth University / University Hospitals Dorset NHS Foundation Trust / Brighton and Sussex University Hospitals NHS Trust / Dartford and Gravesham NHS Trust / Isle of Wight NHS Trust / Maidstone and Tunbridge Wells NHS Trust / University Hospital Southampton NHS Foundation Trust | COVID-19 Genomics UK (COG-UK) Consortium | Samuel Robson, Angela Beckett, Salman Goudarzi, Christopher Fearn, Kate Cook, Katie Loveson, Sharon Glaysher, Scott Elliott, Kelly Bicknell, Sarah Wyllie, Allyson Lloyd, Robert Impey, Anoop Chauhan, Stephen Kidd, Nathan Moore, Nick Cortes, Claire Thomas, Anna Mantzaouratou, Sarah Buchan, Magdalena Barrow, Andrew Butt, Liz Sheridan, Jonnie Seymour, Dorian Crudgington, Ben Macklin, Mohammed Hassan-Ibrahim, Cassandra Malone, Benjamin Cogger, Kevin Tucker, Samirakhon Raupova, Rachael Jeremiah, Anibolina Castigador, Emily Macnaughton, Karen Withell, Kordo Saeed, Jacqui Prieto, Adhyana Mahanama, Buddhini Samaraweera, Siona Silvieri, Emanuela Pelosi, Eleri Wilson-Davies, Sarah Jeremiah, Helen Wheeler, Matthew Harvey, Thea Sass, Helen Umpleby, Stephen Aplin |
| EPI_ISL_3182950, EPI_ISL_3182951, EPI_ISL_3182952, EPI_ISL_3182953, EPI_ISL_3182955, EPI_ISL_3182956, EPI_ISL_3182957, EPI_ISL_3182959, EPI_ISL_3182960, EPI_ISL_3182961, EPI_ISL_3182962, EPI_ISL_3182963, EPI_ISL_3182964, EPI_ISL_3182965, EPI_ISL_3182966, EPI_ISL_3182968, EPI_ISL_3182969, EPI_ISL_3182970, EPI_ISL_3182971, EPI_ISL_3182972, EPI_ISL_3182973, EPI_ISL_3182974, EPI_ISL_3182976, EPI_ISL_3182977, EPI_ISL_3182978, EPI_ISL_3182979, EPI_ISL_3182981, EPI_ISL_3182982, EPI_ISL_3182983, EPI_ISL_3182984, EPI_ISL_3182985, EPI_ISL_3182986, EPI_ISL_3182987, EPI_ISL_3182988, EPI_ISL_3182990, EPI_ISL_3182991, EPI_ISL_3182992, EPI_ISL_3182993, EPI_ISL_3182995, EPI_ISL_3182996, EPI_ISL_3182997, EPI_ISL_3182998, EPI_ISL_3183000, EPI_ISL_3183001, EPI_ISL_3183002, EPI_ISL_3183003, EPI_ISL_3183004, EPI_ISL_3183005, EPI_ISL_3183006, EPI_ISL_3183008, EPI_ISL_3183010, EPI_ISL_3183011, EPI_ISL_3183013, EPI_ISL_3183014, EPI_ISL_3183015, EPI_ISL_3183016, EPI_ISL_3183017, EPI_ISL_3183018, EPI_ISL_3183019, EPI_ISL_3183021, EPI_ISL_3183022, EPI_ISL_3183023, EPI_ISL_3183024, EPI_ISL_3183026, EPI_ISL_3183027, EPI_ISL_3183028, EPI_ISL_3183030, EPI_ISL_3183031, EPI_ISL_3183032, EPI_ISL_3183033, EPI_ISL_3183035, EPI_ISL_3183036, EPI_ISL_3183037, EPI_ISL_3183038, EPI_ISL_3183039, EPI_ISL_3183040, EPI_ISL_3183041, EPI_ISL_3183042, EPI_ISL_3183045, EPI_ISL_3183046, EPI_ISL_3183048, EPI_ISL_3183049, EPI_ISL_3183050, EPI_ISL_3183051, EPI_ISL_3183052, EPI_ISL_3183054, EPI_ISL_3183055, EPI_ISL_3183056, EPI_ISL_3183057, EPI_ISL_3183058, EPI_ISL_3183059, EPI_ISL_3183060, EPI_ISL_3183061, EPI_ISL_3183063, EPI_ISL_3183064, EPI_ISL_3183065, EPI_ISL_3183066, EPI_ISL_3183067, EPI_ISL_3183068, EPI_ISL_3183069, EPI_ISL_3183071, EPI_ISL_3183072, EPI_ISL_3183073, EPI_ISL_3183074, EPI_ISL_3183076, EPI_ISL_3183077, EPI_ISL_3183078, EPI_ISL_3183079, EPI_ISL_3183081, EPI_ISL_3183082, EPI_ISL_3183083, EPI_ISL_3183084, EPI_ISL_3183085, EPI_ISL_3183086, EPI_ISL_3183087, EPI_ISL_3183090, EPI_ISL_3183091, EPI_ISL_3183092, EPI_ISL_3183094, EPI_ISL_3183095, EPI_ISL_3183096, EPI_ISL_3183097, EPI_ISL_3183098, EPI_ISL_3183099, EPI_ISL_3183100, EPI_ISL_3183101, EPI_ISL_3183103, EPI_ISL_3183104, EPI_ISL_3183105, EPI_ISL_3183106, EPI_ISL_3183107, EPI_ISL_3183109, EPI_ISL_3183110, EPI_ISL_3183111, EPI_ISL_3183112, EPI_ISL_3183113, EPI_ISL_3183114, EPI_ISL_3183115, EPI_ISL_3183117, EPI_ISL_3183118 | | | |
| see above | Virology Department, Sheffield Teaching Hospitals NHS Foundation Trust/Department of Infection, Immunity and Cardiovascular Disease, The Medical School, University of Sheffield | COVID-19 Genomics UK (COG-UK) Consortium | Thushan de Silva, Matthew Parker, Nikki Smith, Adri Angyal, Rebecca Brown, Luke Green, Rachel Tucker, Paul Parsons, Danielle Groves, Katie Johnson, Laura Carrilero, Alex Keeley, Dave Partridge, Matthew Wyles, Benjamin Lindsey, Mehmet Yavuz, Mohammad Raza, Cariad Evans |
| EPI_ISL_3183762, EPI_ISL_3183763, EPI_ISL_3183764 | Department of Infectious Diseases, Kobe Institute of Health | Department of Infectious Diseases, Kobe Institute of Health | Ryohei Nomoto, Noriko Nakanishi, Tomotada Iwamoto, Tsuyoshi Sekizuka, Kentaro Itokawa, Rina Tanaka, Masanori Hashino, Makoto Kuroda |
| EPI_ISL_3183766, EPI_ISL_3183775 | Gunma Prefectural Institute of Public Health and Environmental Sciences | Gunma Prefectural Institute of Public Health and Environmental Sciences | Hiroyuki Tsukagoshi, Daisuke Shinoda, Mariko Saito, Tsuyoshi Sekizuka, Kentaro Itokawa, Rina Tanaka, Masanori Hashino, Makoto Kuroda |
| EPI_ISL_3183776 | Kagoshima Prefectural Institute for Environmental Research and Public Health | Kagoshima Prefectural Institute for Environmental Research and Public Health | Yuka Hamada, Takako Kiyotake, Kanji Ishitani, Mami Yamamoto, Kazuki Manabe, Naomi Shinkawa, Tsuyoshi Sekizuka, Kentaro Itokawa, Rina Tanaka, Masanori Hashino, Makoto Kuroda |
| EPI_ISL_3183777, EPI_ISL_3183778, EPI_ISL_3183779, EPI_ISL_3183780, EPI_ISL_3183781, EPI_ISL_3183782, EPI_ISL_3183783, EPI_ISL_3183784, EPI_ISL_3183785, EPI_ISL_3183786, EPI_ISL_3183787, EPI_ISL_3183788, EPI_ISL_3183789, EPI_ISL_3183790, EPI_ISL_3183791, EPI_ISL_3183792, EPI_ISL_3183793, EPI_ISL_3183794, EPI_ISL_3183795, EPI_ISL_3183796, EPI_ISL_3183797, EPI_ISL_3183798, EPI_ISL_3183799, EPI_ISL_3183800, EPI_ISL_3183801 | | | |
| see above | Ibaraki Prefectural Institute of Public Health | Ibaraki Prefectural Institute of Public Health | Keiko Goto, Ryo Kashimura, Shuichi Osawa, Tsuyoshi Sekizuka, Kentaro Itokawa, Rina Tanaka, Masanori Hashino, Makoto Kuroda |
| EPI_ISL_3183802 | Utsunomiya City Institute of Public Health and Environment | Utsunomiya City Institute of Public Health and Environment | Akira Wakatsuki, Tsuyoshi Sekizuka, Kentaro Itokawa, Rina Tanaka, Masanori Hashino, Makoto Kuroda |
| EPI_ISL_3183803, EPI_ISL_3183804, EPI_ISL_3183805, EPI_ISL_3183806, EPI_ISL_3183807, EPI_ISL_3183808, EPI_ISL_3183809, EPI_ISL_3183810, EPI_ISL_3183811, EPI_ISL_3183812, EPI_ISL_3183813, EPI_ISL_3183814, EPI_ISL_3183815, EPI_ISL_3183816 | | | |
| see above | Medical Microbiology and Zoology Section, Okinawa Prefectural Institute of Health and Environment | Medical Microbiology and Zoology Section, Okinawa Prefectural Institute of Health and Environment | Tetsuya Kakita, Noriyuki Maeshiro, Tsuyoshi Kudeken, Yumani Kuba, Taketoshi Takara, Minoru Nidaira, Hisako Kyan, Tsuyoshi Sekizuka, Kentaro Itokawa, Rina Tanaka, Masanori Hashino, Makoto Kuroda |
| EPI_ISL_3183817 | Department of Health Science, Iwate Prefectural Institute of Public Health and Environment | Iwate Prefectural Institute of Public Health and Environment | Hiroki Konno, Hiroko Kajita, Kaori Iwabuchi, Akiko Fujimori, Taihei Mitsui, Tomoko Takahashi, Tsuyoshi Sekizuka, Kentaro Itokawa, Rina Tanaka, Masanori Hashino, Makoto Kuroda |
| EPI_ISL_3183818 | Gunma Prefectural Institute of Public Health and Environmental Sciences | Gunma Prefectural Institute of Public Health and Environmental Sciences | Hiroyuki Tsukagoshi, Daisuke Shinoda, Mariko Saito, Tsuyoshi Sekizuka, Kentaro Itokawa, Rina Tanaka, Masanori Hashino, Makoto Kuroda |
| EPI_ISL_3183819, EPI_ISL_3183820, EPI_ISL_3183821 | Hiroshima Prefectural Technology Research Institute, Public Health and Environment Center | Hiroshima Prefectural Technology Research Institute, Public Health and Environment Center | Naoki Shigemoto, Akie Nakashima, Yukie Shimazu, Hiroko Akita, Syuhei Ikeda, Kanako Masuda, Takahiro Hiratsuka, Mana Suei, Ayano Ito, Shinichi Takao, Tsuyoshi Sekizuka, Kentaro Itokawa, Rina Tanaka, Masanori Hashino, Makoto Kuroda |
| EPI_ISL_3183822, EPI_ISL_3183823, EPI_ISL_3183824, EPI_ISL_3183825, EPI_ISL_3183826, EPI_ISL_3183827 | Saitama Institute of Public Health | Saitama Institute of Public Health | Satoshi Daisaki, Eri Aonuma, Hayato Ehara, Tsuyoshi Sekizuka, Kentaro Itokawa, Rina Tanaka, Masanori Hashino, Makoto Kuroda |

| | | | |
|---|---|---|---|
| EPI_ISL_3183828 | Sapporo City Institute of Public Health | Sapporo City Institute of Public Health | Asami Ohnishi, Rie Shimazaki, Yusuke Oguchi, Masayuki Kikuchi, Tsuyoshi Sekizuka, Kentaro Itokawa, Rina Tanaka, Masanori Hashino, Makoto Kuroda |
| EPI_ISL_3183829, EPI_ISL_3183830, EPI_ISL_3183831, EPI_ISL_3183832, EPI_ISL_3183833, EPI_ISL_3183834, EPI_ISL_3183835, EPI_ISL_3183836, EPI_ISL_3183837, EPI_ISL_3183838, EPI_ISL_3183839, EPI_ISL_3183840, EPI_ISL_3183841, EPI_ISL_3183842, EPI_ISL_3183843, EPI_ISL_3183844, EPI_ISL_3183845 | | | |
| see above | Kagoshima Prefectural Institute for Environmental Research and Public Health | Kagoshima Prefectural Institute for Environmental Research and Public Health | Yuka Hamada, Takako Kiyotake, Kanji Ishitani, Mami Yamamoto, Kazuki Manabe, Naomi Shinkawa, Tsuyoshi Sekizuka, Kentaro Itokawa, Rina Tanaka, Masanori Hashino, Makoto Kuroda |
| EPI_ISL_3183950, EPI_ISL_3183951, EPI_ISL_3183953, EPI_ISL_3183955, EPI_ISL_3183958 | Genome Analysis Center, Yamanashi Central Hospital | Genome Analysis Center, Yamanashi Central Hospital | Yosuke Hirotsu |
| EPI_ISL_3183959, EPI_ISL_3183961, EPI_ISL_3183963, EPI_ISL_3183964 | Department of Microbiology, Yamanashi Institute for Public Health and Environment | Genome Analysis Center, Yamanashi Central Hospital | Yosuke Hirotsu |
| EPI_ISL_3183965 | Ender Diagnostics AG | Microbiology, Dr. Risch | Nadia Wohlwend |
| EPI_ISL_3183966, EPI_ISL_3183967, EPI_ISL_3183968, EPI_ISL_3183969, EPI_ISL_3183971, EPI_ISL_3183972, EPI_ISL_3183973 | Dr. Risch Ostschweiz AG | Microbiology, Dr. Risch | Nadia Wohlwend |
| EPI_ISL_3183974 | Ender Diagnostics AG | Microbiology, Dr. Risch | Nadia Wohlwend |
| EPI_ISL_3183975, EPI_ISL_3183976, EPI_ISL_3183977 | Dr. Risch Ostschweiz AG | Microbiology, Dr. Risch | Nadia Wohlwend |
| EPI_ISL_3183979 | Ender Diagnostics AG | Microbiology, Dr. Risch | Nadia Wohlwend |
| EPI_ISL_3183980, EPI_ISL_3183981, EPI_ISL_3183984, EPI_ISL_3183987, EPI_ISL_3183988, EPI_ISL_3183989, EPI_ISL_3183991, EPI_ISL_3183992, EPI_ISL_3183994 | Dr. Risch Ostschweiz AG | Microbiology, Dr. Risch | Nadia Wohlwend |
| EPI_ISL_3183998 | Ender Diagnostics AG | Microbiology, Dr. Risch | Nadia Wohlwend |
| EPI_ISL_3184001, EPI_ISL_3184002, EPI_ISL_3184003, EPI_ISL_3184004, EPI_ISL_3184005, EPI_ISL_3184006, EPI_ISL_3184007, EPI_ISL_3184008, EPI_ISL_3184009, EPI_ISL_3184010, EPI_ISL_3184011, EPI_ISL_3184012, EPI_ISL_3184013, EPI_ISL_3184014, EPI_ISL_3184015, EPI_ISL_3184016, EPI_ISL_3184017 | | | |
| see above | Karolinska University Hospital Huddinge | Karolinska University Hospital | Jan Albert ,Tobias Allander ,Annelie Bjerkner ,Sandra Broddesson ,Robert Dyrdak ,Martin Ekman ,Lynda Eneh ,Lina Guerra Blomqvist ,Karolina Ininbergs ,Tanja Normark ,Isak Sylvin ,Zhibing Yun ,Martina Wahlund ,Valtteri Wirta |
| EPI_ISL_3184019, EPI_ISL_3184020, EPI_ISL_3184021 | Karolinska University Hospital Solna | Karolinska University Hospital | Jan Albert ,Tobias Allander ,Annelie Bjerkner ,Sandra Broddesson ,Robert Dyrdak ,Martin Ekman ,Lynda Eneh ,Lina Guerra Blomqvist ,Karolina Ininbergs ,Tanja Normark ,Isak Sylvin ,Zhibing Yun ,Martina Wahlund ,Valtteri Wirta |
| EPI_ISL_3184022, EPI_ISL_3184023 | Unilabs | Karolinska University Hospital | Jan Albert ,Tobias Allander ,Annelie Bjerkner ,Sandra Broddesson ,Robert Dyrdak ,Martin Ekman ,Lynda Eneh ,Lina Guerra Blomqvist ,Karolina Ininbergs ,Tanja Normark ,Isak Sylvin ,Zhibing Yun ,Martina Wahlund ,Valtteri Wirta |
| EPI_ISL_3184024, EPI_ISL_3184025, EPI_ISL_3184026, EPI_ISL_3184027, EPI_ISL_3184028, EPI_ISL_3184029, EPI_ISL_3184030, EPI_ISL_3184031 | Karolinska University Hospital Solna | Karolinska University Hospital | Jan Albert ,Tobias Allander ,Annelie Bjerkner ,Sandra Broddesson ,Robert Dyrdak ,Martin Ekman ,Lynda Eneh ,Lina Guerra Blomqvist ,Karolina Ininbergs ,Tanja Normark ,Isak Sylvin ,Zhibing Yun ,Martina Wahlund ,Valtteri Wirta |
| EPI_ISL_3184032 | Synlab Medilab | Karolinska University Hospital | Jan Albert ,Tobias Allander ,Annelie Bjerkner ,Sandra Broddesson ,Robert Dyrdak ,Martin Ekman ,Lynda Eneh ,Lina Guerra Blomqvist ,Karolina Ininbergs ,Tanja Normark ,Isak Sylvin ,Zhibing Yun ,Martina Wahlund ,Valtteri Wirta |
| EPI_ISL_3184033, EPI_ISL_3184034, EPI_ISL_3184035, EPI_ISL_3184036, EPI_ISL_3184037 | Karolinska University Hospital Solna | Karolinska University Hospital | Jan Albert ,Tobias Allander ,Annelie Bjerkner ,Sandra Broddesson ,Robert Dyrdak ,Martin Ekman ,Lynda Eneh ,Lina Guerra Blomqvist ,Karolina Ininbergs ,Tanja Normark ,Isak Sylvin ,Zhibing Yun ,Martina Wahlund ,Valtteri Wirta |
| EPI_ISL_3184038 | Unilabs | Karolinska University Hospital | Jan Albert ,Tobias Allander ,Annelie Bjerkner ,Sandra Broddesson ,Robert Dyrdak ,Martin Ekman ,Lynda Eneh ,Lina Guerra Blomqvist ,Karolina Ininbergs ,Tanja Normark ,Isak Sylvin ,Zhibing Yun ,Martina Wahlund ,Valtteri Wirta |
| EPI_ISL_3184039, EPI_ISL_3184040, EPI_ISL_3184041 | Karolinska University Hospital Solna | Karolinska University Hospital | Jan Albert ,Tobias Allander ,Annelie Bjerkner ,Sandra Broddesson ,Robert Dyrdak ,Martin Ekman ,Lynda Eneh ,Lina Guerra Blomqvist ,Karolina Ininbergs ,Tanja Normark ,Isak Sylvin ,Zhibing Yun ,Martina Wahlund ,Valtteri Wirta |
| EPI_ISL_3184042 | Unilabs | Karolinska University Hospital | Jan Albert ,Tobias Allander ,Annelie Bjerkner ,Sandra Broddesson ,Robert Dyrdak ,Martin Ekman ,Lynda Eneh ,Lina Guerra Blomqvist ,Karolina Ininbergs ,Tanja Normark ,Isak Sylvin ,Zhibing Yun ,Martina Wahlund ,Valtteri Wirta |
| EPI_ISL_3184043 | Karolinska University Hospital Solna | Karolinska University Hospital | Jan Albert ,Tobias Allander ,Annelie Bjerkner ,Sandra Broddesson ,Robert Dyrdak ,Martin Ekman ,Lynda Eneh ,Lina Guerra Blomqvist ,Karolina Ininbergs ,Tanja Normark ,Isak Sylvin ,Zhibing Yun ,Martina Wahlund ,Valtteri Wirta |
| EPI_ISL_3184044 | Unilabs | Karolinska University Hospital | Jan Albert ,Tobias Allander ,Annelie Bjerkner ,Sandra Broddesson ,Robert Dyrdak ,Martin Ekman ,Lynda Eneh ,Lina Guerra Blomqvist ,Karolina Ininbergs ,Tanja Normark ,Isak Sylvin ,Zhibing Yun ,Martina Wahlund ,Valtteri Wirta |
| EPI_ISL_3184045, EPI_ISL_3184046 | Karolinska University Hospital Solna | Karolinska University Hospital | Jan Albert ,Tobias Allander ,Annelie Bjerkner ,Sandra Broddesson ,Robert Dyrdak ,Martin Ekman ,Lynda Eneh ,Lina Guerra Blomqvist ,Karolina Ininbergs ,Tanja Normark ,Isak Sylvin ,Zhibing Yun ,Martina Wahlund ,Valtteri Wirta |
| EPI_ISL_3184047, EPI_ISL_3184048 | Unilabs | Karolinska University Hospital | Jan Albert ,Tobias Allander ,Annelie Bjerkner ,Sandra Broddesson ,Robert Dyrdak ,Martin Ekman ,Lynda Eneh ,Lina Guerra Blomqvist ,Karolina Ininbergs ,Tanja Normark ,Isak Sylvin ,Zhibing Yun ,Martina Wahlund ,Valtteri Wirta |
| EPI_ISL_3184049 | Karolinska University Hospital Solna | Karolinska University Hospital | Jan Albert ,Tobias Allander ,Annelie Bjerkner ,Sandra Broddesson ,Robert Dyrdak ,Martin Ekman ,Lynda Eneh ,Lina Guerra Blomqvist ,Karolina Ininbergs ,Tanja Normark ,Isak Sylvin ,Zhibing Yun ,Martina Wahlund ,Valtteri Wirta |
| EPI_ISL_3184050 | Unilabs | Karolinska University Hospital | Jan Albert ,Tobias Allander ,Annelie Bjerkner ,Sandra Broddesson ,Robert Dyrdak ,Martin Ekman ,Lynda Eneh ,Lina Guerra Blomqvist ,Karolina Ininbergs ,Tanja Normark ,Isak Sylvin ,Zhibing Yun ,Martina Wahlund ,Valtteri Wirta |
| EPI_ISL_3184051, EPI_ISL_3184052 | Karolinska University Hospital Solna | Karolinska University Hospital | Jan Albert ,Tobias Allander ,Annelie Bjerkner ,Sandra Broddesson ,Robert Dyrdak ,Martin Ekman ,Lynda Eneh ,Lina Guerra Blomqvist ,Karolina Ininbergs ,Tanja Normark ,Isak Sylvin ,Zhibing Yun ,Martina Wahlund ,Valtteri Wirta |
| EPI_ISL_3184053 | Synlab Medilab | Karolinska University Hospital | Jan Albert ,Tobias Allander ,Annelie Bjerkner ,Sandra Broddesson ,Robert Dyrdak ,Martin Ekman ,Lynda Eneh ,Lina Guerra Blomqvist ,Karolina Ininbergs ,Tanja Normark ,Isak Sylvin ,Zhibing Yun ,Martina Wahlund ,Valtteri Wirta |
| EPI_ISL_3184054, EPI_ISL_3184055 | Karolinska University Hospital Solna | Karolinska University Hospital | Jan Albert ,Tobias Allander ,Annelie Bjerkner ,Sandra Broddesson ,Robert Dyrdak ,Martin Ekman ,Lynda Eneh ,Lina Guerra Blomqvist ,Karolina Ininbergs ,Tanja Normark ,Isak Sylvin ,Zhibing Yun ,Martina Wahlund ,Valtteri Wirta |
| EPI_ISL_3184057 | Unilabs | Karolinska University Hospital | Jan Albert ,Tobias Allander ,Annelie Bjerkner ,Sandra Broddesson ,Robert Dyrdak ,Martin Ekman ,Lynda Eneh ,Lina Guerra Blomqvist ,Karolina Ininbergs ,Tanja Normark ,Isak Sylvin ,Zhibing Yun ,Martina Wahlund ,Valtteri Wirta |
| EPI_ISL_3184058 | Karolinska University Hospital Solna | Karolinska University Hospital | Jan Albert ,Tobias Allander ,Annelie Bjerkner ,Sandra Broddesson ,Robert Dyrdak ,Martin Ekman ,Lynda Eneh ,Lina Guerra Blomqvist ,Karolina Ininbergs ,Tanja Normark ,Isak Sylvin ,Zhibing Yun ,Martina Wahlund ,Valtteri Wirta |
| EPI_ISL_3184059 | Unilabs | Karolinska University Hospital | Jan Albert ,Tobias Allander ,Annelie Bjerkner ,Sandra Broddesson ,Robert Dyrdak ,Martin Ekman ,Lynda Eneh ,Lina Guerra Blomqvist ,Karolina Ininbergs ,Tanja Normark ,Isak Sylvin ,Zhibing Yun ,Martina Wahlund ,Valtteri Wirta |
| EPI_ISL_3184061 | Karolinska University Hospital Solna | Karolinska University Hospital | Jan Albert ,Tobias Allander ,Annelie Bjerkner ,Sandra Broddesson ,Robert Dyrdak ,Martin Ekman ,Lynda Eneh ,Lina Guerra Blomqvist ,Karolina Ininbergs ,Tanja Normark ,Isak Sylvin ,Zhibing Yun ,Martina Wahlund ,Valtteri Wirta |
| EPI_ISL_3184062, EPI_ISL_3184063 | Unilabs | Karolinska University Hospital | Jan Albert ,Tobias Allander ,Annelie Bjerkner ,Sandra Broddesson ,Robert Dyrdak ,Martin Ekman ,Lynda Eneh ,Lina Guerra Blomqvist ,Karolina Ininbergs ,Tanja Normark ,Isak Sylvin ,Zhibing Yun ,Martina Wahlund ,Valtteri Wirta |
| EPI_ISL_3184064, EPI_ISL_3184065, EPI_ISL_3184066, EPI_ISL_3184067 | Karolinska University Hospital Solna | Karolinska University Hospital | Jan Albert ,Tobias Allander ,Annelie Bjerkner ,Sandra Broddesson ,Robert Dyrdak ,Martin Ekman ,Lynda Eneh ,Lina Guerra Blomqvist ,Karolina Ininbergs ,Tanja Normark ,Isak Sylvin ,Zhibing Yun ,Martina Wahlund ,Valtteri Wirta |
| EPI_ISL_3184068, EPI_ISL_3184069 | Unilabs | Karolinska University Hospital | Jan Albert ,Tobias Allander ,Annelie Bjerkner ,Sandra Broddesson ,Robert Dyrdak ,Martin Ekman ,Lynda Eneh ,Lina Guerra Blomqvist ,Karolina Ininbergs ,Tanja Normark ,Isak Sylvin ,Zhibing Yun ,Martina Wahlund ,Valtteri Wirta |
| EPI_ISL_3184070 | Karolinska University Hospital Solna | Karolinska University Hospital | Jan Albert ,Tobias Allander ,Annelie Bjerkner ,Sandra Broddesson ,Robert Dyrdak ,Martin Ekman ,Lynda Eneh ,Lina Guerra Blomqvist ,Karolina Ininbergs ,Tanja Normark ,Isak Sylvin ,Zhibing Yun ,Martina Wahlund ,Valtteri Wirta |
| EPI_ISL_3184071, EPI_ISL_3184072, EPI_ISL_3184073, EPI_ISL_3184074, EPI_ISL_3184075 | Unilabs | Karolinska University Hospital | Jan Albert ,Tobias Allander ,Annelie Bjerkner ,Sandra Broddesson ,Robert Dyrdak ,Martin Ekman ,Lynda Eneh ,Lina Guerra Blomqvist ,Karolina Ininbergs ,Tanja Normark ,Isak Sylvin ,Zhibing Yun ,Martina Wahlund ,Valtteri Wirta |

| | | | |
|---|---|---|---|
| EPI_ISL_3184076 | Karolinska University Hospital Solna | Karolinska University Hospital | Jan Albert ,Tobias Allander ,Annelie Bjerkner ,Sandra Broddesson ,Robert Dyrdak ,Martin Ekman ,Lynda Eneh ,Lina Guerra Blomqvist ,Karolina Ininbergs ,Tanja Normark ,Isak Sylvin ,Zhibing Yun ,Martina Wahlund ,Valtteri Wirta |
| EPI_ISL_3184077 | Unilabs | Karolinska University Hospital | Jan Albert ,Tobias Allander ,Annelie Bjerkner ,Sandra Broddesson ,Robert Dyrdak ,Martin Ekman ,Lynda Eneh ,Lina Guerra Blomqvist ,Karolina Ininbergs ,Tanja Normark ,Isak Sylvin ,Zhibing Yun ,Martina Wahlund ,Valtteri Wirta |
| EPI_ISL_3184079 | Karolinska University Hospital Solna | Karolinska University Hospital | Jan Albert ,Tobias Allander ,Annelie Bjerkner ,Sandra Broddesson ,Robert Dyrdak ,Martin Ekman ,Lynda Eneh ,Lina Guerra Blomqvist ,Karolina Ininbergs ,Tanja Normark ,Isak Sylvin ,Zhibing Yun ,Martina Wahlund ,Valtteri Wirta |
| EPI_ISL_3184080, EPI_ISL_3184081 | Unilabs | Karolinska University Hospital | Jan Albert ,Tobias Allander ,Annelie Bjerkner ,Sandra Broddesson ,Robert Dyrdak ,Martin Ekman ,Lynda Eneh ,Lina Guerra Blomqvist ,Karolina Ininbergs ,Tanja Normark ,Isak Sylvin ,Zhibing Yun ,Martina Wahlund ,Valtteri Wirta |
| EPI_ISL_3184082 | Karolinska University Hospital Solna | Karolinska University Hospital | Jan Albert ,Tobias Allander ,Annelie Bjerkner ,Sandra Broddesson ,Robert Dyrdak ,Martin Ekman ,Lynda Eneh ,Lina Guerra Blomqvist ,Karolina Ininbergs ,Tanja Normark ,Isak Sylvin ,Zhibing Yun ,Martina Wahlund ,Valtteri Wirta |
| EPI_ISL_3184083, EPI_ISL_3184084, EPI_ISL_3184085, EPI_ISL_3184086, EPI_ISL_3184087, EPI_ISL_3184088, EPI_ISL_3184089 | Unilabs | Karolinska University Hospital | Jan Albert ,Tobias Allander ,Annelie Bjerkner ,Sandra Broddesson ,Robert Dyrdak ,Martin Ekman ,Lynda Eneh ,Lina Guerra Blomqvist ,Karolina Ininbergs ,Tanja Normark ,Isak Sylvin ,Zhibing Yun ,Martina Wahlund ,Valtteri Wirta |
| EPI_ISL_3184090, EPI_ISL_3184091 | Synlab Medilab | Karolinska University Hospital | Jan Albert ,Tobias Allander ,Annelie Bjerkner ,Sandra Broddesson ,Robert Dyrdak ,Martin Ekman ,Lynda Eneh ,Lina Guerra Blomqvist ,Karolina Ininbergs ,Tanja Normark ,Isak Sylvin ,Zhibing Yun ,Martina Wahlund ,Valtteri Wirta |
| EPI_ISL_3184092, EPI_ISL_3184093 | Unilabs | Karolinska University Hospital | Jan Albert ,Tobias Allander ,Annelie Bjerkner ,Sandra Broddesson ,Robert Dyrdak ,Martin Ekman ,Lynda Eneh ,Lina Guerra Blomqvist ,Karolina Ininbergs ,Tanja Normark ,Isak Sylvin ,Zhibing Yun ,Martina Wahlund ,Valtteri Wirta |
| EPI_ISL_3184094 | Karolinska University Hospital Solna | Karolinska University Hospital | Jan Albert ,Tobias Allander ,Annelie Bjerkner ,Sandra Broddesson ,Robert Dyrdak ,Martin Ekman ,Lynda Eneh ,Lina Guerra Blomqvist ,Karolina Ininbergs ,Tanja Normark ,Isak Sylvin ,Zhibing Yun ,Martina Wahlund ,Valtteri Wirta |
| EPI_ISL_3184095 | Synlab Medilab | Karolinska University Hospital | Jan Albert ,Tobias Allander ,Annelie Bjerkner ,Sandra Broddesson ,Robert Dyrdak ,Martin Ekman ,Lynda Eneh ,Lina Guerra Blomqvist ,Karolina Ininbergs ,Tanja Normark ,Isak Sylvin ,Zhibing Yun ,Martina Wahlund ,Valtteri Wirta |
| EPI_ISL_3184096, EPI_ISL_3184097 | Unilabs | Karolinska University Hospital | Jan Albert ,Tobias Allander ,Annelie Bjerkner ,Sandra Broddesson ,Robert Dyrdak ,Martin Ekman ,Lynda Eneh ,Lina Guerra Blomqvist ,Karolina Ininbergs ,Tanja Normark ,Isak Sylvin ,Zhibing Yun ,Martina Wahlund ,Valtteri Wirta |
| EPI_ISL_3184098 | Synlab Medilab | Karolinska University Hospital | Jan Albert ,Tobias Allander ,Annelie Bjerkner ,Sandra Broddesson ,Robert Dyrdak ,Martin Ekman ,Lynda Eneh ,Lina Guerra Blomqvist ,Karolina Ininbergs ,Tanja Normark ,Isak Sylvin ,Zhibing Yun ,Martina Wahlund ,Valtteri Wirta |
| EPI_ISL_3184099, EPI_ISL_3184101, EPI_ISL_3184102, EPI_ISL_3184103 | Unilabs | Karolinska University Hospital | Jan Albert ,Tobias Allander ,Annelie Bjerkner ,Sandra Broddesson ,Robert Dyrdak ,Martin Ekman ,Lynda Eneh ,Lina Guerra Blomqvist ,Karolina Ininbergs ,Tanja Normark ,Isak Sylvin ,Zhibing Yun ,Martina Wahlund ,Valtteri Wirta |
| EPI_ISL_3184104 | Synlab Medilab | Karolinska University Hospital | Jan Albert ,Tobias Allander ,Annelie Bjerkner ,Sandra Broddesson ,Robert Dyrdak ,Martin Ekman ,Lynda Eneh ,Lina Guerra Blomqvist ,Karolina Ininbergs ,Tanja Normark ,Isak Sylvin ,Zhibing Yun ,Martina Wahlund ,Valtteri Wirta |
| EPI_ISL_3184105, EPI_ISL_3184106 | Unilabs | Karolinska University Hospital | Jan Albert ,Tobias Allander ,Annelie Bjerkner ,Sandra Broddesson ,Robert Dyrdak ,Martin Ekman ,Lynda Eneh ,Lina Guerra Blomqvist ,Karolina Ininbergs ,Tanja Normark ,Isak Sylvin ,Zhibing Yun ,Martina Wahlund ,Valtteri Wirta |
| EPI_ISL_3184107 | Synlab Medilab | Karolinska University Hospital | Jan Albert ,Tobias Allander ,Annelie Bjerkner ,Sandra Broddesson ,Robert Dyrdak ,Martin Ekman ,Lynda Eneh ,Lina Guerra Blomqvist ,Karolina Ininbergs ,Tanja Normark ,Isak Sylvin ,Zhibing Yun ,Martina Wahlund ,Valtteri Wirta |
| EPI_ISL_3184108, EPI_ISL_3184109, EPI_ISL_3184110 | Unilabs | Karolinska University Hospital | Jan Albert ,Tobias Allander ,Annelie Bjerkner ,Sandra Broddesson ,Robert Dyrdak ,Martin Ekman ,Lynda Eneh ,Lina Guerra Blomqvist ,Karolina Ininbergs ,Tanja Normark ,Isak Sylvin ,Zhibing Yun ,Martina Wahlund ,Valtteri Wirta |
| EPI_ISL_3184111, EPI_ISL_3184112, EPI_ISL_3184113 | Synlab Medilab | Karolinska University Hospital | Jan Albert ,Tobias Allander ,Annelie Bjerkner ,Sandra Broddesson ,Robert Dyrdak ,Martin Ekman ,Lynda Eneh ,Lina Guerra Blomqvist ,Karolina Ininbergs ,Tanja Normark ,Isak Sylvin ,Zhibing Yun ,Martina Wahlund ,Valtteri Wirta |
| EPI_ISL_3184114 | Karolinska University Hospital Solna | Karolinska University Hospital | Jan Albert ,Tobias Allander ,Annelie Bjerkner ,Sandra Broddesson ,Robert Dyrdak ,Martin Ekman ,Lynda Eneh ,Lina Guerra Blomqvist ,Karolina Ininbergs ,Tanja Normark ,Isak Sylvin ,Zhibing Yun ,Martina Wahlund ,Valtteri Wirta |
| EPI_ISL_3184115, EPI_ISL_3184116 | Unilabs | Karolinska University Hospital | Jan Albert ,Tobias Allander ,Annelie Bjerkner ,Sandra Broddesson ,Robert Dyrdak ,Martin Ekman ,Lynda Eneh ,Lina Guerra Blomqvist ,Karolina Ininbergs ,Tanja Normark ,Isak Sylvin ,Zhibing Yun ,Martina Wahlund ,Valtteri Wirta |
| EPI_ISL_3184117 | Synlab Medilab | Karolinska University Hospital | Jan Albert ,Tobias Allander ,Annelie Bjerkner ,Sandra Broddesson ,Robert Dyrdak ,Martin Ekman ,Lynda Eneh ,Lina Guerra Blomqvist ,Karolina Ininbergs ,Tanja Normark ,Isak Sylvin ,Zhibing Yun ,Martina Wahlund ,Valtteri Wirta |
| EPI_ISL_3184118 | Unilabs | Karolinska University Hospital | Jan Albert ,Tobias Allander ,Annelie Bjerkner ,Sandra Broddesson ,Robert Dyrdak ,Martin Ekman ,Lynda Eneh ,Lina Guerra Blomqvist ,Karolina Ininbergs ,Tanja Normark ,Isak Sylvin ,Zhibing Yun ,Martina Wahlund ,Valtteri Wirta |
| EPI_ISL_3184119 | Synlab Medilab | Karolinska University Hospital | Jan Albert ,Tobias Allander ,Annelie Bjerkner ,Sandra Broddesson ,Robert Dyrdak ,Martin Ekman ,Lynda Eneh ,Lina Guerra Blomqvist ,Karolina Ininbergs ,Tanja Normark ,Isak Sylvin ,Zhibing Yun ,Martina Wahlund ,Valtteri Wirta |
| EPI_ISL_3184120, EPI_ISL_3184121, EPI_ISL_3184122, EPI_ISL_3184124 | Unilabs | Karolinska University Hospital | Jan Albert ,Tobias Allander ,Annelie Bjerkner ,Sandra Broddesson ,Robert Dyrdak ,Martin Ekman ,Lynda Eneh ,Lina Guerra Blomqvist ,Karolina Ininbergs ,Tanja Normark ,Isak Sylvin ,Zhibing Yun ,Martina Wahlund ,Valtteri Wirta |
| EPI_ISL_3184125 | Synlab Medilab | Karolinska University Hospital | Jan Albert ,Tobias Allander ,Annelie Bjerkner ,Sandra Broddesson ,Robert Dyrdak ,Martin Ekman ,Lynda Eneh ,Lina Guerra Blomqvist ,Karolina Ininbergs ,Tanja Normark ,Isak Sylvin ,Zhibing Yun ,Martina Wahlund ,Valtteri Wirta |
| EPI_ISL_3184126 | Unilabs | Karolinska University Hospital | Jan Albert ,Tobias Allander ,Annelie Bjerkner ,Sandra Broddesson ,Robert Dyrdak ,Martin Ekman ,Lynda Eneh ,Lina Guerra Blomqvist ,Karolina Ininbergs ,Tanja Normark ,Isak Sylvin ,Zhibing Yun ,Martina Wahlund ,Valtteri Wirta |
| EPI_ISL_3184127, EPI_ISL_3184128, EPI_ISL_3184129, EPI_ISL_3184130 | Synlab Medilab | Karolinska University Hospital | Jan Albert ,Tobias Allander ,Annelie Bjerkner ,Sandra Broddesson ,Robert Dyrdak ,Martin Ekman ,Lynda Eneh ,Lina Guerra Blomqvist ,Karolina Ininbergs ,Tanja Normark ,Isak Sylvin ,Zhibing Yun ,Martina Wahlund ,Valtteri Wirta |
| EPI_ISL_3184131 | Unilabs | Karolinska University Hospital | Jan Albert ,Tobias Allander ,Annelie Bjerkner ,Sandra Broddesson ,Robert Dyrdak ,Martin Ekman ,Lynda Eneh ,Lina Guerra Blomqvist ,Karolina Ininbergs ,Tanja Normark ,Isak Sylvin ,Zhibing Yun ,Martina Wahlund ,Valtteri Wirta |
| EPI_ISL_3184132 | Synlab Medilab | Karolinska University Hospital | Jan Albert ,Tobias Allander ,Annelie Bjerkner ,Sandra Broddesson ,Robert Dyrdak ,Martin Ekman ,Lynda Eneh ,Lina Guerra Blomqvist ,Karolina Ininbergs ,Tanja Normark ,Isak Sylvin ,Zhibing Yun ,Martina Wahlund ,Valtteri Wirta |
| EPI_ISL_3184133 | Unilabs | Karolinska University Hospital | Jan Albert ,Tobias Allander ,Annelie Bjerkner ,Sandra Broddesson ,Robert Dyrdak ,Martin Ekman ,Lynda Eneh ,Lina Guerra Blomqvist ,Karolina Ininbergs ,Tanja Normark ,Isak Sylvin ,Zhibing Yun ,Martina Wahlund ,Valtteri Wirta |
| EPI_ISL_3184134 | Synlab Medilab | Karolinska University Hospital | Jan Albert ,Tobias Allander ,Annelie Bjerkner ,Sandra Broddesson ,Robert Dyrdak ,Martin Ekman ,Lynda Eneh ,Lina Guerra Blomqvist ,Karolina Ininbergs ,Tanja Normark ,Isak Sylvin ,Zhibing Yun ,Martina Wahlund ,Valtteri Wirta |
| EPI_ISL_3184135, EPI_ISL_3184136, EPI_ISL_3184137 | Unilabs | Karolinska University Hospital | Jan Albert ,Tobias Allander ,Annelie Bjerkner ,Sandra Broddesson ,Robert Dyrdak ,Martin Ekman ,Lynda Eneh ,Lina Guerra Blomqvist ,Karolina Ininbergs ,Tanja Normark ,Isak Sylvin ,Zhibing Yun ,Martina Wahlund ,Valtteri Wirta |
| EPI_ISL_3184138 | Synlab Medilab | Karolinska University Hospital | Jan Albert ,Tobias Allander ,Annelie Bjerkner ,Sandra Broddesson ,Robert Dyrdak ,Martin Ekman ,Lynda Eneh ,Lina Guerra Blomqvist ,Karolina Ininbergs ,Tanja Normark ,Isak Sylvin ,Zhibing Yun ,Martina Wahlund ,Valtteri Wirta |
| EPI_ISL_3184139 | Unilabs | Karolinska University Hospital | Jan Albert ,Tobias Allander ,Annelie Bjerkner ,Sandra Broddesson ,Robert Dyrdak ,Martin Ekman ,Lynda Eneh ,Lina Guerra Blomqvist ,Karolina Ininbergs ,Tanja Normark ,Isak Sylvin ,Zhibing Yun ,Martina Wahlund ,Valtteri Wirta |
| EPI_ISL_3184140, EPI_ISL_3184141 | Synlab Medilab | Karolinska University Hospital | Jan Albert ,Tobias Allander ,Annelie Bjerkner ,Sandra Broddesson ,Robert Dyrdak ,Martin Ekman ,Lynda Eneh ,Lina Guerra Blomqvist ,Karolina Ininbergs ,Tanja Normark ,Isak Sylvin ,Zhibing Yun ,Martina Wahlund ,Valtteri Wirta |
| EPI_ISL_3184142, EPI_ISL_3184143, EPI_ISL_3184144 | Unilabs | Karolinska University Hospital | Jan Albert ,Tobias Allander ,Annelie Bjerkner ,Sandra Broddesson ,Robert Dyrdak ,Martin Ekman ,Lynda Eneh ,Lina Guerra Blomqvist ,Karolina Ininbergs ,Tanja Normark ,Isak Sylvin ,Zhibing Yun ,Martina Wahlund ,Valtteri Wirta |
| EPI_ISL_3184145 | Synlab Medilab | Karolinska University Hospital | Jan Albert ,Tobias Allander ,Annelie Bjerkner ,Sandra Broddesson ,Robert Dyrdak ,Martin Ekman ,Lynda Eneh ,Lina Guerra Blomqvist ,Karolina Ininbergs ,Tanja Normark ,Isak Sylvin ,Zhibing Yun ,Martina Wahlund ,Valtteri Wirta |
| EPI_ISL_3184146, EPI_ISL_3184147 | Unilabs | Karolinska University Hospital | Jan Albert ,Tobias Allander ,Annelie Bjerkner ,Sandra Broddesson ,Robert Dyrdak ,Martin Ekman ,Lynda Eneh ,Lina Guerra Blomqvist ,Karolina Ininbergs ,Tanja Normark ,Isak Sylvin ,Zhibing Yun ,Martina Wahlund ,Valtteri Wirta |
| EPI_ISL_3184148, EPI_ISL_3184149 | Synlab Medilab | Karolinska University Hospital | Jan Albert ,Tobias Allander ,Annelie Bjerkner ,Sandra Broddesson ,Robert Dyrdak ,Martin Ekman ,Lynda Eneh ,Lina Guerra Blomqvist ,Karolina Ininbergs ,Tanja Normark ,Isak Sylvin ,Zhibing Yun ,Martina Wahlund ,Valtteri Wirta |
| EPI_ISL_3184150 | Unilabs | Karolinska University Hospital | Jan Albert ,Tobias Allander ,Annelie Bjerkner ,Sandra Broddesson ,Robert Dyrdak ,Martin Ekman ,Lynda Eneh ,Lina Guerra Blomqvist ,Karolina Ininbergs ,Tanja Normark ,Isak Sylvin ,Zhibing Yun ,Martina Wahlund ,Valtteri Wirta |
| EPI_ISL_3184151, EPI_ISL_3184152 | Synlab Medilab | Karolinska University Hospital | Jan Albert ,Tobias Allander ,Annelie Bjerkner ,Sandra Broddesson ,Robert Dyrdak ,Martin Ekman ,Lynda Eneh ,Lina Guerra Blomqvist ,Karolina Ininbergs ,Tanja Normark ,Isak Sylvin ,Zhibing Yun ,Martina Wahlund ,Valtteri Wirta |
| EPI_ISL_3184153, EPI_ISL_3184154, EPI_ISL_3184155, EPI_ISL_3184156 | Unilabs | Karolinska University Hospital | Jan Albert ,Tobias Allander ,Annelie Bjerkner ,Sandra Broddesson ,Robert Dyrdak ,Martin Ekman ,Lynda Eneh ,Lina Guerra Blomqvist ,Karolina Ininbergs ,Tanja Normark ,Isak Sylvin ,Zhibing Yun ,Martina Wahlund ,Valtteri Wirta |
| EPI_ISL_3184157, EPI_ISL_3184158, EPI_ISL_3184159 | Synlab Medilab | Karolinska University Hospital | Jan Albert ,Tobias Allander ,Annelie Bjerkner ,Sandra Broddesson ,Robert Dyrdak ,Martin Ekman ,Lynda Eneh ,Lina Guerra Blomqvist ,Karolina Ininbergs ,Tanja Normark ,Isak Sylvin ,Zhibing Yun ,Martina Wahlund ,Valtteri Wirta |
| EPI_ISL_3184160 | Unilabs | Karolinska University Hospital | Jan Albert ,Tobias Allander ,Annelie Bjerkner ,Sandra Broddesson ,Robert Dyrdak ,Martin Ekman ,Lynda Eneh ,Lina Guerra Blomqvist ,Karolina Ininbergs ,Tanja Normark ,Isak Sylvin ,Zhibing Yun ,Martina Wahlund ,Valtteri Wirta |
| EPI_ISL_3184161 | Karolinska University Hospital Solna | Karolinska University Hospital | Jan Albert ,Tobias Allander ,Annelie Bjerkner ,Sandra Broddesson ,Robert Dyrdak ,Martin Ekman ,Lynda Eneh ,Lina Guerra Blomqvist ,Karolina Ininbergs ,Tanja Normark ,Isak Sylvin ,Zhibing Yun ,Martina Wahlund ,Valtteri Wirta |
| EPI_ISL_3184162, EPI_ISL_3184163, EPI_ISL_3184164, EPI_ISL_3184165, EPI_ISL_3184166 | Synlab Medilab | Karolinska University Hospital | Jan Albert ,Tobias Allander ,Annelie Bjerkner ,Sandra Broddesson ,Robert Dyrdak ,Martin Ekman ,Lynda Eneh ,Lina Guerra Blomqvist ,Karolina Ininbergs ,Tanja Normark ,Isak Sylvin ,Zhibing Yun ,Martina Wahlund ,Valtteri Wirta |
| EPI_ISL_3184167 | Unilabs | Karolinska University Hospital | Jan Albert ,Tobias Allander ,Annelie Bjerkner ,Sandra Broddesson ,Robert Dyrdak ,Martin Ekman ,Lynda Eneh ,Lina Guerra Blomqvist ,Karolina Ininbergs ,Tanja Normark ,Isak Sylvin ,Zhibing Yun ,Martina Wahlund ,Valtteri Wirta |

| | | | |
|---|---|---|---|
| EPI_ISL_3184168, EPI_ISL_3184169 | Synlab Medilab | Karolinska University Hospital | Jan Albert ,Tobias Allander ,Annelie Bjerkner ,Sandra Broddesson ,Robert Dyrdak ,Martin Ekman ,Lynda Eneh ,Lina Guerra Blomqvist ,Karolina Ininbergs ,Tanja Normark ,Isak Sylvin ,Zhibing Yun ,Martina Wahlund ,Valtteri Wirta |
| EPI_ISL_3184170 | Unilabs | Karolinska University Hospital | Jan Albert ,Tobias Allander ,Annelie Bjerkner ,Sandra Broddesson ,Robert Dyrdak ,Martin Ekman ,Lynda Eneh ,Lina Guerra Blomqvist ,Karolina Ininbergs ,Tanja Normark ,Isak Sylvin ,Zhibing Yun ,Martina Wahlund ,Valtteri Wirta |
| EPI_ISL_3184171 | Synlab Medilab | Karolinska University Hospital | Jan Albert ,Tobias Allander ,Annelie Bjerkner ,Sandra Broddesson ,Robert Dyrdak ,Martin Ekman ,Lynda Eneh ,Lina Guerra Blomqvist ,Karolina Ininbergs ,Tanja Normark ,Isak Sylvin ,Zhibing Yun ,Martina Wahlund ,Valtteri Wirta |
| EPI_ISL_3184172, EPI_ISL_3184173, EPI_ISL_3184174, EPI_ISL_3184175, EPI_ISL_3184176, EPI_ISL_3184178, EPI_ISL_3184179, EPI_ISL_3184180, EPI_ISL_3184181 | Karolinska University Hospital Solna | Karolinska University Hospital | Jan Albert ,Tobias Allander ,Annelie Bjerkner ,Sandra Broddesson ,Robert Dyrdak ,Martin Ekman ,Lynda Eneh ,Lina Guerra Blomqvist ,Karolina Ininbergs ,Tanja Normark ,Isak Sylvin ,Zhibing Yun ,Martina Wahlund ,Valtteri Wirta |
| EPI_ISL_3184182 | Unilabs | Karolinska University Hospital | Jan Albert ,Tobias Allander ,Annelie Bjerkner ,Sandra Broddesson ,Robert Dyrdak ,Martin Ekman ,Lynda Eneh ,Lina Guerra Blomqvist ,Karolina Ininbergs ,Tanja Normark ,Isak Sylvin ,Zhibing Yun ,Martina Wahlund ,Valtteri Wirta |
| EPI_ISL_3184183, EPI_ISL_3184184, EPI_ISL_3184185 | Karolinska University Hospital Solna | Karolinska University Hospital | Jan Albert ,Tobias Allander ,Annelie Bjerkner ,Sandra Broddesson ,Robert Dyrdak ,Martin Ekman ,Lynda Eneh ,Lina Guerra Blomqvist ,Karolina Ininbergs ,Tanja Normark ,Isak Sylvin ,Zhibing Yun ,Martina Wahlund ,Valtteri Wirta |
| EPI_ISL_3184186, EPI_ISL_3184187 | Unilabs | Karolinska University Hospital | Jan Albert ,Tobias Allander ,Annelie Bjerkner ,Sandra Broddesson ,Robert Dyrdak ,Martin Ekman ,Lynda Eneh ,Lina Guerra Blomqvist ,Karolina Ininbergs ,Tanja Normark ,Isak Sylvin ,Zhibing Yun ,Martina Wahlund ,Valtteri Wirta |
| EPI_ISL_3184188, EPI_ISL_3184189, EPI_ISL_3184190, EPI_ISL_3184191, EPI_ISL_3184192, EPI_ISL_3184193, EPI_ISL_3184194, EPI_ISL_3184195, EPI_ISL_3184196, EPI_ISL_3184197, EPI_ISL_3184198, EPI_ISL_3184199, EPI_ISL_3184200, EPI_ISL_3184201, EPI_ISL_3184202, EPI_ISL_3184203, EPI_ISL_3184204, EPI_ISL_3184205, EPI_ISL_3184206, EPI_ISL_3184207 | | | |
| see above | Karolinska University Hospital Solna | Karolinska University Hospital | Jan Albert ,Tobias Allander ,Annelie Bjerkner ,Sandra Broddesson ,Robert Dyrdak ,Martin Ekman ,Lynda Eneh ,Lina Guerra Blomqvist ,Karolina Ininbergs ,Tanja Normark ,Isak Sylvin ,Zhibing Yun ,Martina Wahlund ,Valtteri Wirta |
| EPI_ISL_3184208, EPI_ISL_3184211 | Unilabs | Karolinska University Hospital | Jan Albert ,Tobias Allander ,Annelie Bjerkner ,Sandra Broddesson ,Robert Dyrdak ,Martin Ekman ,Lynda Eneh ,Lina Guerra Blomqvist ,Karolina Ininbergs ,Tanja Normark ,Isak Sylvin ,Zhibing Yun ,Martina Wahlund ,Valtteri Wirta |
| EPI_ISL_3184212, EPI_ISL_3184213, EPI_ISL_3184214, EPI_ISL_3184215, EPI_ISL_3184216, EPI_ISL_3184217 | Karolinska University Hospital Solna | Karolinska University Hospital | Jan Albert ,Tobias Allander ,Annelie Bjerkner ,Sandra Broddesson ,Robert Dyrdak ,Martin Ekman ,Lynda Eneh ,Lina Guerra Blomqvist ,Karolina Ininbergs ,Tanja Normark ,Isak Sylvin ,Zhibing Yun ,Martina Wahlund ,Valtteri Wirta |
| EPI_ISL_3184218 | Unilabs | Karolinska University Hospital | Jan Albert ,Tobias Allander ,Annelie Bjerkner ,Sandra Broddesson ,Robert Dyrdak ,Martin Ekman ,Lynda Eneh ,Lina Guerra Blomqvist ,Karolina Ininbergs ,Tanja Normark ,Isak Sylvin ,Zhibing Yun ,Martina Wahlund ,Valtteri Wirta |
| EPI_ISL_3184219, EPI_ISL_3184220, EPI_ISL_3184221, EPI_ISL_3184222, EPI_ISL_3184223, EPI_ISL_3184224 | Karolinska University Hospital Solna | Karolinska University Hospital | Jan Albert ,Tobias Allander ,Annelie Bjerkner ,Sandra Broddesson ,Robert Dyrdak ,Martin Ekman ,Lynda Eneh ,Lina Guerra Blomqvist ,Karolina Ininbergs ,Tanja Normark ,Isak Sylvin ,Zhibing Yun ,Martina Wahlund ,Valtteri Wirta |
| EPI_ISL_3184225 | Unilabs | Karolinska University Hospital | Jan Albert ,Tobias Allander ,Annelie Bjerkner ,Sandra Broddesson ,Robert Dyrdak ,Martin Ekman ,Lynda Eneh ,Lina Guerra Blomqvist ,Karolina Ininbergs ,Tanja Normark ,Isak Sylvin ,Zhibing Yun ,Martina Wahlund ,Valtteri Wirta |
| EPI_ISL_3184227 | Karolinska University Hospital Solna | Karolinska University Hospital | Jan Albert ,Tobias Allander ,Annelie Bjerkner ,Sandra Broddesson ,Robert Dyrdak ,Martin Ekman ,Lynda Eneh ,Lina Guerra Blomqvist ,Karolina Ininbergs ,Tanja Normark ,Isak Sylvin ,Zhibing Yun ,Martina Wahlund ,Valtteri Wirta |
| EPI_ISL_3184228, EPI_ISL_3184229, EPI_ISL_3184230 | Unilabs | Karolinska University Hospital | Jan Albert ,Tobias Allander ,Annelie Bjerkner ,Sandra Broddesson ,Robert Dyrdak ,Martin Ekman ,Lynda Eneh ,Lina Guerra Blomqvist ,Karolina Ininbergs ,Tanja Normark ,Isak Sylvin ,Zhibing Yun ,Martina Wahlund ,Valtteri Wirta |
| EPI_ISL_3184231, EPI_ISL_3184232, EPI_ISL_3184233, EPI_ISL_3184234, EPI_ISL_3184235, EPI_ISL_3184236, EPI_ISL_3184237 | Karolinska University Hospital Solna | Karolinska University Hospital | Jan Albert ,Tobias Allander ,Annelie Bjerkner ,Sandra Broddesson ,Robert Dyrdak ,Martin Ekman ,Lynda Eneh ,Lina Guerra Blomqvist ,Karolina Ininbergs ,Tanja Normark ,Isak Sylvin ,Zhibing Yun ,Martina Wahlund ,Valtteri Wirta |
| EPI_ISL_3184238, EPI_ISL_3184239, EPI_ISL_3184240 | Unilabs | Karolinska University Hospital | Jan Albert ,Tobias Allander ,Annelie Bjerkner ,Sandra Broddesson ,Robert Dyrdak ,Martin Ekman ,Lynda Eneh ,Lina Guerra Blomqvist ,Karolina Ininbergs ,Tanja Normark ,Isak Sylvin ,Zhibing Yun ,Martina Wahlund ,Valtteri Wirta |
| EPI_ISL_3184241 | Karolinska University Hospital Solna | Karolinska University Hospital | Jan Albert ,Tobias Allander ,Annelie Bjerkner ,Sandra Broddesson ,Robert Dyrdak ,Martin Ekman ,Lynda Eneh ,Lina Guerra Blomqvist ,Karolina Ininbergs ,Tanja Normark ,Isak Sylvin ,Zhibing Yun ,Martina Wahlund ,Valtteri Wirta |
| EPI_ISL_3184242 | Unilabs | Karolinska University Hospital | Jan Albert ,Tobias Allander ,Annelie Bjerkner ,Sandra Broddesson ,Robert Dyrdak ,Martin Ekman ,Lynda Eneh ,Lina Guerra Blomqvist ,Karolina Ininbergs ,Tanja Normark ,Isak Sylvin ,Zhibing Yun ,Martina Wahlund ,Valtteri Wirta |
| EPI_ISL_3184243 | Karolinska University Hospital Solna | Karolinska University Hospital | Jan Albert ,Tobias Allander ,Annelie Bjerkner ,Sandra Broddesson ,Robert Dyrdak ,Martin Ekman ,Lynda Eneh ,Lina Guerra Blomqvist ,Karolina Ininbergs ,Tanja Normark ,Isak Sylvin ,Zhibing Yun ,Martina Wahlund ,Valtteri Wirta |
| EPI_ISL_3184244 | Unilabs | Karolinska University Hospital | Jan Albert ,Tobias Allander ,Annelie Bjerkner ,Sandra Broddesson ,Robert Dyrdak ,Martin Ekman ,Lynda Eneh ,Lina Guerra Blomqvist ,Karolina Ininbergs ,Tanja Normark ,Isak Sylvin ,Zhibing Yun ,Martina Wahlund ,Valtteri Wirta |
| EPI_ISL_3184245 | Karolinska University Hospital Solna | Karolinska University Hospital | Jan Albert ,Tobias Allander ,Annelie Bjerkner ,Sandra Broddesson ,Robert Dyrdak ,Martin Ekman ,Lynda Eneh ,Lina Guerra Blomqvist ,Karolina Ininbergs ,Tanja Normark ,Isak Sylvin ,Zhibing Yun ,Martina Wahlund ,Valtteri Wirta |
| EPI_ISL_3184246 | Unilabs | Karolinska University Hospital | Jan Albert ,Tobias Allander ,Annelie Bjerkner ,Sandra Broddesson ,Robert Dyrdak ,Martin Ekman ,Lynda Eneh ,Lina Guerra Blomqvist ,Karolina Ininbergs ,Tanja Normark ,Isak Sylvin ,Zhibing Yun ,Martina Wahlund ,Valtteri Wirta |
| EPI_ISL_3184247, EPI_ISL_3184248 | Karolinska University Hospital Solna | Karolinska University Hospital | Jan Albert ,Tobias Allander ,Annelie Bjerkner ,Sandra Broddesson ,Robert Dyrdak ,Martin Ekman ,Lynda Eneh ,Lina Guerra Blomqvist ,Karolina Ininbergs ,Tanja Normark ,Isak Sylvin ,Zhibing Yun ,Martina Wahlund ,Valtteri Wirta |
| EPI_ISL_3184249, EPI_ISL_3184250 | Unilabs | Karolinska University Hospital | Jan Albert ,Tobias Allander ,Annelie Bjerkner ,Sandra Broddesson ,Robert Dyrdak ,Martin Ekman ,Lynda Eneh ,Lina Guerra Blomqvist ,Karolina Ininbergs ,Tanja Normark ,Isak Sylvin ,Zhibing Yun ,Martina Wahlund ,Valtteri Wirta |
| EPI_ISL_3184251, EPI_ISL_3184252, EPI_ISL_3184253 | Karolinska University Hospital Solna | Karolinska University Hospital | Jan Albert ,Tobias Allander ,Annelie Bjerkner ,Sandra Broddesson ,Robert Dyrdak ,Martin Ekman ,Lynda Eneh ,Lina Guerra Blomqvist ,Karolina Ininbergs ,Tanja Normark ,Isak Sylvin ,Zhibing Yun ,Martina Wahlund ,Valtteri Wirta |
| EPI_ISL_3184254 | Unilabs | Karolinska University Hospital | Jan Albert ,Tobias Allander ,Annelie Bjerkner ,Sandra Broddesson ,Robert Dyrdak ,Martin Ekman ,Lynda Eneh ,Lina Guerra Blomqvist ,Karolina Ininbergs ,Tanja Normark ,Isak Sylvin ,Zhibing Yun ,Martina Wahlund ,Valtteri Wirta |
| EPI_ISL_3184255 | Karolinska University Hospital Solna | Karolinska University Hospital | Jan Albert ,Tobias Allander ,Annelie Bjerkner ,Sandra Broddesson ,Robert Dyrdak ,Martin Ekman ,Lynda Eneh ,Lina Guerra Blomqvist ,Karolina Ininbergs ,Tanja Normark ,Isak Sylvin ,Zhibing Yun ,Martina Wahlund ,Valtteri Wirta |
| EPI_ISL_3184256 | Unilabs | Karolinska University Hospital | Jan Albert ,Tobias Allander ,Annelie Bjerkner ,Sandra Broddesson ,Robert Dyrdak ,Martin Ekman ,Lynda Eneh ,Lina Guerra Blomqvist ,Karolina Ininbergs ,Tanja Normark ,Isak Sylvin ,Zhibing Yun ,Martina Wahlund ,Valtteri Wirta |
| EPI_ISL_3184257 | Karolinska University Hospital Solna | Karolinska University Hospital | Jan Albert ,Tobias Allander ,Annelie Bjerkner ,Sandra Broddesson ,Robert Dyrdak ,Martin Ekman ,Lynda Eneh ,Lina Guerra Blomqvist ,Karolina Ininbergs ,Tanja Normark ,Isak Sylvin ,Zhibing Yun ,Martina Wahlund ,Valtteri Wirta |
| EPI_ISL_3184258 | Unilabs | Karolinska University Hospital | Jan Albert ,Tobias Allander ,Annelie Bjerkner ,Sandra Broddesson ,Robert Dyrdak ,Martin Ekman ,Lynda Eneh ,Lina Guerra Blomqvist ,Karolina Ininbergs ,Tanja Normark ,Isak Sylvin ,Zhibing Yun ,Martina Wahlund ,Valtteri Wirta |
| EPI_ISL_3184259, EPI_ISL_3184260, EPI_ISL_3184261, EPI_ISL_3184262 | Karolinska University Hospital Solna | Karolinska University Hospital | Jan Albert ,Tobias Allander ,Annelie Bjerkner ,Sandra Broddesson ,Robert Dyrdak ,Martin Ekman ,Lynda Eneh ,Lina Guerra Blomqvist ,Karolina Ininbergs ,Tanja Normark ,Isak Sylvin ,Zhibing Yun ,Martina Wahlund ,Valtteri Wirta |
| EPI_ISL_3184264, EPI_ISL_3184265, EPI_ISL_3184266 | Unilabs | Karolinska University Hospital | Jan Albert ,Tobias Allander ,Annelie Bjerkner ,Sandra Broddesson ,Robert Dyrdak ,Martin Ekman ,Lynda Eneh ,Lina Guerra Blomqvist ,Karolina Ininbergs ,Tanja Normark ,Isak Sylvin ,Zhibing Yun ,Martina Wahlund ,Valtteri Wirta |
| EPI_ISL_3184267 | Synlab Medilab | Karolinska University Hospital | Jan Albert ,Tobias Allander ,Annelie Bjerkner ,Sandra Broddesson ,Robert Dyrdak ,Martin Ekman ,Lynda Eneh ,Lina Guerra Blomqvist ,Karolina Ininbergs ,Tanja Normark ,Isak Sylvin ,Zhibing Yun ,Martina Wahlund ,Valtteri Wirta |
| EPI_ISL_3184268 | Karolinska University Hospital Solna | Karolinska University Hospital | Jan Albert ,Tobias Allander ,Annelie Bjerkner ,Sandra Broddesson ,Robert Dyrdak ,Martin Ekman ,Lynda Eneh ,Lina Guerra Blomqvist ,Karolina Ininbergs ,Tanja Normark ,Isak Sylvin ,Zhibing Yun ,Martina Wahlund ,Valtteri Wirta |
| EPI_ISL_3184269, EPI_ISL_3184270 | Unilabs | Karolinska University Hospital | Jan Albert ,Tobias Allander ,Annelie Bjerkner ,Sandra Broddesson ,Robert Dyrdak ,Martin Ekman ,Lynda Eneh ,Lina Guerra Blomqvist ,Karolina Ininbergs ,Tanja Normark ,Isak Sylvin ,Zhibing Yun ,Martina Wahlund ,Valtteri Wirta |
| EPI_ISL_3184271 | Karolinska University Hospital Solna | Karolinska University Hospital | Jan Albert ,Tobias Allander ,Annelie Bjerkner ,Sandra Broddesson ,Robert Dyrdak ,Martin Ekman ,Lynda Eneh ,Lina Guerra Blomqvist ,Karolina Ininbergs ,Tanja Normark ,Isak Sylvin ,Zhibing Yun ,Martina Wahlund ,Valtteri Wirta |
| EPI_ISL_3184272 | Unilabs | Karolinska University Hospital | Jan Albert ,Tobias Allander ,Annelie Bjerkner ,Sandra Broddesson ,Robert Dyrdak ,Martin Ekman ,Lynda Eneh ,Lina Guerra Blomqvist ,Karolina Ininbergs ,Tanja Normark ,Isak Sylvin ,Zhibing Yun ,Martina Wahlund ,Valtteri Wirta |
| EPI_ISL_3184273, EPI_ISL_3184274, EPI_ISL_3184275 | Karolinska University Hospital Solna | Karolinska University Hospital | Jan Albert ,Tobias Allander ,Annelie Bjerkner ,Sandra Broddesson ,Robert Dyrdak ,Martin Ekman ,Lynda Eneh ,Lina Guerra Blomqvist ,Karolina Ininbergs ,Tanja Normark ,Isak Sylvin ,Zhibing Yun ,Martina Wahlund ,Valtteri Wirta |
| EPI_ISL_3184276, EPI_ISL_3184277, EPI_ISL_3184278 | Unilabs | Karolinska University Hospital | Jan Albert ,Tobias Allander ,Annelie Bjerkner ,Sandra Broddesson ,Robert Dyrdak ,Martin Ekman ,Lynda Eneh ,Lina Guerra Blomqvist ,Karolina Ininbergs ,Tanja Normark ,Isak Sylvin ,Zhibing Yun ,Martina Wahlund ,Valtteri Wirta |
| EPI_ISL_3184279 | Karolinska University Hospital Solna | Karolinska University Hospital | Jan Albert ,Tobias Allander ,Annelie Bjerkner ,Sandra Broddesson ,Robert Dyrdak ,Martin Ekman ,Lynda Eneh ,Lina Guerra Blomqvist ,Karolina Ininbergs ,Tanja Normark ,Isak Sylvin ,Zhibing Yun ,Martina Wahlund ,Valtteri Wirta |
| EPI_ISL_3184281 | Unilabs | Karolinska University Hospital | Jan Albert ,Tobias Allander ,Annelie Bjerkner ,Sandra Broddesson ,Robert Dyrdak ,Martin Ekman ,Lynda Eneh ,Lina Guerra Blomqvist ,Karolina Ininbergs ,Tanja Normark ,Isak Sylvin ,Zhibing Yun ,Martina Wahlund ,Valtteri Wirta |
| EPI_ISL_3184282, | Karolinska University Hospital Solna | Karolinska University Hospital | Jan Albert ,Tobias Allander ,Annelie Bjerkner ,Sandra Broddesson ,Robert Dyrdak ,Martin Ekman ,Lynda Eneh ,Lina Guerra Blomqvist ,Karolina Ininbergs ,Tanja Normark ,Isak Sylvin ,Zhibing Yun ,Martina Wahlund ,Valtteri Wirta |

| | | | |
|---|---|---|---|
| EPI_ISL_3184283, EPI_ISL_3184284, EPI_ISL_3184285, EPI_ISL_3184286, EPI_ISL_3184287, EPI_ISL_3184288, EPI_ISL_3184289, EPI_ISL_3184290 | | | |
| EPI_ISL_3184291, EPI_ISL_3184292 | Unilabs | Karolinska University Hospital | Jan Albert ,Tobias Allander ,Annelie Bjerkner ,Sandra Broddesson ,Robert Dyrdak ,Martin Ekman ,Lynda Eneh ,Lina Guerra Blomqvist ,Karolina Ininbergs ,Tanja Normark ,Isak Sylvin ,Zhibing Yun ,Martina Wahlund ,Valtteri Wirta |
| EPI_ISL_3184293, EPI_ISL_3184294 | Karolinska University Hospital Solna | Karolinska University Hospital | Jan Albert ,Tobias Allander ,Annelie Bjerkner ,Sandra Broddesson ,Robert Dyrdak ,Martin Ekman ,Lynda Eneh ,Lina Guerra Blomqvist ,Karolina Ininbergs ,Tanja Normark ,Isak Sylvin ,Zhibing Yun ,Martina Wahlund ,Valtteri Wirta |
| EPI_ISL_3184295, EPI_ISL_3184296 | Unilabs | Karolinska University Hospital | Jan Albert ,Tobias Allander ,Annelie Bjerkner ,Sandra Broddesson ,Robert Dyrdak ,Martin Ekman ,Lynda Eneh ,Lina Guerra Blomqvist ,Karolina Ininbergs ,Tanja Normark ,Isak Sylvin ,Zhibing Yun ,Martina Wahlund ,Valtteri Wirta |
| EPI_ISL_3184297, EPI_ISL_3184298, EPI_ISL_3184299, EPI_ISL_3184300, EPI_ISL_3184301, EPI_ISL_3184302, EPI_ISL_3184303 | Karolinska University Hospital Solna | Karolinska University Hospital | Jan Albert ,Tobias Allander ,Annelie Bjerkner ,Sandra Broddesson ,Robert Dyrdak ,Martin Ekman ,Lynda Eneh ,Lina Guerra Blomqvist ,Karolina Ininbergs ,Tanja Normark ,Isak Sylvin ,Zhibing Yun ,Martina Wahlund ,Valtteri Wirta |
| EPI_ISL_3184304 | Unilabs | Karolinska University Hospital | Jan Albert ,Tobias Allander ,Annelie Bjerkner ,Sandra Broddesson ,Robert Dyrdak ,Martin Ekman ,Lynda Eneh ,Lina Guerra Blomqvist ,Karolina Ininbergs ,Tanja Normark ,Isak Sylvin ,Zhibing Yun ,Martina Wahlund ,Valtteri Wirta |
| EPI_ISL_3184307, EPI_ISL_3184308 | Virology Unit, AOUP | Virology Unit, AOUP | Mauro Pistello, Annalisa Capria, Pietro Giorgio Spezia, Susanna Cappellini, Francesca Pasqualetti |
| EPI_ISL_3184309 | Poliklinika Vitalis | Alea Genetic Centre | Rijad Konjhodzic, Lana Salihefendic, Dino Pecar |
| EPI_ISL_3184312 | General Hospital "Abdulah Nakas" | Alea Genetic Centre | Dino Pecar, Lana Salihefendic, Enis Kandic, Adis Kandic, Rijad Konjhodzic |
| EPI_ISL_3184314 | General Hospital "Abdulah Nakas" | Alea Genetic Centre | Lana Salihefendic, Dino Pecar, Ivana Ceko, Enis Kandic, Rijad Konjhodzic |
| EPI_ISL_3184315 | Virology Unit, AOUP | Virology Unit, AOUP | Mauro Pistello, Annalisa Capria, Pietro Giorgio Spezia, Susanna Cappellini, Francesca Pasqualetti |
| EPI_ISL_3184316 | PZU Alea dr Kandic | Alea Genetic Centre | Dino Pecar, Lana Salihefendic, Ivana Ceko, Enis Kandic, Adis Kandic, Rijad Konjhodzic |
| EPI_ISL_3184340, EPI_ISL_3184341, EPI_ISL_3184342, EPI_ISL_3184343, EPI_ISL_3184344, EPI_ISL_3184345, EPI_ISL_3184346, EPI_ISL_3184347, EPI_ISL_3184348, EPI_ISL_3184349, EPI_ISL_3184350, EPI_ISL_3184351, EPI_ISL_3184352, EPI_ISL_3184353, EPI_ISL_3184354, EPI_ISL_3184355, EPI_ISL_3184357, EPI_ISL_3184358, EPI_ISL_3184359 | | | |
| see above | Clinical Virology | Clinical Bacteriology | Tim Roloff, Fanny Wegner, Helena MB Seth-Smith, Alfredo Mari, Karoline Leuzinger, Julia Bielicki, Manuel Battegay, Hans Hirsch, Adrian Egli |
| EPI_ISL_3184360, EPI_ISL_3184361, EPI_ISL_3184362 | Synlab Suisse SA | Clinical Bacteriology | Tim Roloff, Fanny Wegner, Helena MB Seth-Smith, Alfredo Mari, Karoline Leuzinger, Julia Bielicki, Manuel Battegay, Hans Hirsch, Adrian Egli |
| EPI_ISL_3184363 | Clinical Virology | Clinical Bacteriology | Tim Roloff, Fanny Wegner, Helena MB Seth-Smith, Alfredo Mari, Karoline Leuzinger, Julia Bielicki, Manuel Battegay, Hans Hirsch, Adrian Egli |
| EPI_ISL_3184364, EPI_ISL_3184365 | Medics Labor AG | Clinical Bacteriology | Tim Roloff, Fanny Wegner, Helena MB Seth-Smith, Alfredo Mari, Karoline Leuzinger, Julia Bielicki, Manuel Battegay, Hans Hirsch, Adrian Egli |
| EPI_ISL_3184366, EPI_ISL_3184367 | Clinical Virology | Clinical Bacteriology | Tim Roloff, Fanny Wegner, Helena MB Seth-Smith, Alfredo Mari, Karoline Leuzinger, Julia Bielicki, Manuel Battegay, Hans Hirsch, Adrian Egli |
| EPI_ISL_3184368, EPI_ISL_3184369 | Rothen Medizinische Laboratorien AG | Clinical Bacteriology | Tim Roloff, Fanny Wegner, Helena MB Seth-Smith, Alfredo Mari, Karoline Leuzinger, Julia Bielicki, Ingrid Steffen, Manuel Battegay, Hans Hirsch, Adrian Egli |
| EPI_ISL_3184370 | Synlab Suisse SA | Clinical Bacteriology | Tim Roloff, Fanny Wegner, Helena MB Seth-Smith, Alfredo Mari, Karoline Leuzinger, Julia Bielicki, Manuel Battegay, Hans Hirsch, Adrian Egli |
| EPI_ISL_3184371, EPI_ISL_3184372, EPI_ISL_3184373, EPI_ISL_3184374, EPI_ISL_3184375, EPI_ISL_3184376, EPI_ISL_3184377, EPI_ISL_3184378, EPI_ISL_3184379, EPI_ISL_3184380, EPI_ISL_3184381, EPI_ISL_3184382, EPI_ISL_3184384, EPI_ISL_3184385, EPI_ISL_3184386, EPI_ISL_3184387, EPI_ISL_3184388, EPI_ISL_3184389, EPI_ISL_3184390, EPI_ISL_3184392, EPI_ISL_3184393, EPI_ISL_3184394, EPI_ISL_3184395, EPI_ISL_3184398, EPI_ISL_3184399, EPI_ISL_3184400 | | | |
| see above | IN State Department of Health Laboratory Services | IN State Department of Health Laboratory Services | Cassandra Campion, Jamie Yeadon, Brian Pope, Lixia Liu, Kyle Brownlee, Melissa Hindenlang, Mark Glazier |
| EPI_ISL_3184521 | National Public Health Laboratory/Bajhang, COVID-19 PCR Lab | National Public Health Laboratory/CSIR-Institute of Genomics and Integrative Biology | National Public Health Laboratory Team/Bajhang, COVID-19 PCR Lab Team |
| EPI_ISL_3184522 | National Public Health Laboratory/Kamalbazar, COVID-19 PCR Lab | National Public Health Laboratory/CSIR-Institute of Genomics and Integrative Biology | National Public Health Laboratory Team/Kamalbazar, COVID-19 PCR Lab Team |
| EPI_ISL_3184523, EPI_ISL_3184524 | National Public Health Laboratory/Mahakali, COVID-19 PCR Lab | National Public Health Laboratory/CSIR-Institute of Genomics and Integrative Biology | National Public Health Laboratory Team /Mahakali, COVID-19 PCR Lab Team |
| EPI_ISL_3184525, EPI_ISL_3184526 | National Public Health Laboratory/Baitadi, COVID-19 PCR Lab | National Public Health Laboratory/CSIR-Institute of Genomics and Integrative Biology | National Public Health Laboratory Team /Baitadi, COVID-19 PCR Lab Team |
| EPI_ISL_3184527 | National Public Health Laboratory/Dadeldhura, COVID-19 PCR Lab | National Public Health Laboratory/CSIR-Institute of Genomics and Integrative Biology | National Public Health Laboratory Team /Dadeldhura, COVID-19 PCR Lab Team |
| EPI_ISL_3184528 | National Public Health Laboratory/Seti Provincial, COVID-19 PCR Lab | National Public Health Laboratory/CSIR-Institute of Genomics and Integrative Biology | National Public Health Laboratory Team /Seti Provincial, COVID-19 PCR Lab Team |
| EPI_ISL_3184529 | National Public Health Laboratory/Bharatpur Hospital COVID-19 PCR Lab | National Public Health Laboratory/CSIR-Institute of Genomics and Integrative Biology | National Public Health Laboratory Team /Bharatpur Hospital COVID-19 PCR Lab Team |
| EPI_ISL_3184530 | National Public Health Laboratory/Doti COVID-19 PCR Lab | National Public Health Laboratory/CSIR-Institute of Genomics and Integrative Biology | National Public Health LaboratoryTeam/Doti COVID-19 PCR Lab Team |
| EPI_ISL_3184531, EPI_ISL_3184532, EPI_ISL_3184533 | National Public Health Laboratory/PPHL-Gandaki | National Public Health Laboratory/CSIR-Institute of Genomics and Integrative Biology | National Public Health Laboratory Team/PPHL-Gandaki Team |
| EPI_ISL_3184534 | National Public Health Laboratory/Narayani Hospital COVID-19 PCR Lab | National Public Health Laboratory/CSIR-Institute of Genomics and Integrative Biology | National Public Health Laboratory Team/Narayani Hospital COVID-19 PCR Lab Team |
| EPI_ISL_3184535 | National Public Health Laboratory/PPHL-5 | National Public Health Laboratory/CSIR-Institute of Genomics and Integrative Biology | National Public Health Laboratory Team/PPHL-5 Team |
| EPI_ISL_3184536, EPI_ISL_3184537 | National Public Health Laboratory/Chaurhajari COVID-19 PCR Lab | National Public Health Laboratory/CSIR-Institute of Genomics and Integrative Biology | National Public Health Laboratory Team/Chaurhajari COVID-19 PCR Lab Team |
| EPI_ISL_3184538 | National Public Health Laboratory/COVID-19 Testing Lab Dailekh | National Public Health Laboratory/CSIR-Institute of Genomics and Integrative Biology | National Public Health Laboratory Team /COVID-19 Testing Lab Dailekh Team |
| EPI_ISL_3184539 | National Public Health Laboratory/COVID-19 Testing Lab Dailekh | National Public Health Laboratory/CSIR-Institute of Genomics and Integrative Biology | National Public Health Laboratory Team /COVID-19 Testing Lab Dailekh Team |
| EPI_ISL_3184541 | National Public Health Laboratory/PPHL- Karnali | National Public Health Laboratory/CSIR-Institute of Genomics and Integrative Biology | National Public Health Laboratory Team/PPHL- Karnali Team |
| EPI_ISL_3184543, EPI_ISL_3184545 | National Public Health Laboratory/PPHL- Karnali | National Public Health Laboratory/CSIR-Institute of Genomics and Integrative Biology | National Public Health Laboratory Team /PPHL- Karnali Team |
| EPI_ISL_3184547, EPI_ISL_3184549 | National Public Health Laboratory/COVID-19 Testing Lab Dailekh | National Public Health Laboratory/CSIR-Institute of Genomics and Integrative Biology | National Public Health Laboratory Team /COVID-19 Testing Lab Dailekh Team |
| EPI_ISL_3184551 | National Public Health Laboratory/PPHL-Karnali | National Public Health Laboratory/CSIR-Institute of Genomics and Integrative Biology | National Public Health Laboratory Team /PPHL-Karnali Team |
| EPI_ISL_3184553 | National Public Health Laboratory/Crystal Diagnostic Pvt Ltd | National Public Health Laboratory/CSIR-Institute of Genomics and Integrative Biology | National Public Health LaboratoryTeam /Crystal Diagnostic Pvt Ltd Team |
| EPI_ISL_3184555 | National Public Health Laboratory/Crystal Diagnostic Pvt Ltd | National Public Health Laboratory/CSIR-Institute of Genomics and Integrative Biology | National Public Health Laboratory Team /Crystal Diagnostic Pvt Ltd Team |
| EPI_ISL_3184557 | National Public Health Laboratory/B.P Koirala Institute of Health Science, COVID-19 PCR Lab | National Public Health Laboratory/CSIR-Institute of Genomics and Integrative Biology | National Public Health Laboratory Team /B.P Koirala Institute of Health Science, COVID-19 PCR Lab Team |
| EPI_ISL_3184559 | National Public Health Laboratory/Molecular Microbiology Lab Grande International Hospital | National Public Health Laboratory/CSIR-Institute of Genomics and Integrative Biology | National Public Health Laboratory Team/Molecular Microbiology Lab Grande International Hospital Team |
| EPI_ISL_3184561 | National Public Health Laboratory/B&B Hospital, COVI-19 PCR | National Public Health Laboratory/CSIR-Institute of Genomics and | National Public Health Laboratory Team /B&B Hospital, COVI-19 PCR Lab Team |

| | | | |
|---|---|---|---|
| | Lab | Integrative Biology | |
| EPI_ISL_3184563, EPI_ISL_3184565 | National Public Health Laboratory/Nepal Korea Friendship Hospital, COVID-19 PCR Lab | National Public Health Laboratory/CSIR-Institute of Genomics and Integrative Biology | National Public Health Laboratory Team /Nepal Korea Friendship Hospital, COVID-19 PCR Lab Team |
| EPI_ISL_3184567, EPI_ISL_3184569, EPI_ISL_3184571, EPI_ISL_3184573, EPI_ISL_3184576, EPI_ISL_3184578, EPI_ISL_3184580, EPI_ISL_3184582, EPI_ISL_3184584 | National Public Health Laboratory | National Public Health Laboratory/CSIR-Institute of Genomics and Integrative Biology | National Public Health Laboratory Team |
| EPI_ISL_3184586 | National Public Health Laboratory/B.P Koirala Institute of Health Science, COVID-19 PCR Lab | National Public Health Laboratory/CSIR-Institute of Genomics and Integrative Biology | National Public Health Laboratory Team /B.P Koirala Institute of Health Science, COVID-19 PCR Lab Team |
| EPI_ISL_3184588, EPI_ISL_3184590, EPI_ISL_3184592 | National Public Health Laboratory | National Public Health Laboratory/CSIR-Institute of Genomics and Integrative Biology | National Public Health Laboratory Team |
| EPI_ISL_3184595 | National Public Health Laboratory/B.P Koirala Institute of Health Science, COVID-19 PCR Lab | National Public Health Laboratory/CSIR-Institute of Genomics and Integrative Biology | National Public Health Laboratory Team/B.P Koirala Institute of Health Science, COVID-19 PCR Lab Team |
| EPI_ISL_3184597 | National Public Health Laboratory/Bajhang, COVID-19 PCR Lab | National Public Health Laboratory/CSIR-Institute of Genomics and Integrative Biology | National Public Health Laboratory Team /Bajhang, COVID-19 PCR Lab Team |
| EPI_ISL_3184623, EPI_ISL_3184624, EPI_ISL_3184625, EPI_ISL_3184626, EPI_ISL_3184628, EPI_ISL_3184629, EPI_ISL_3184630, EPI_ISL_3184631, EPI_ISL_3184632, EPI_ISL_3184633, EPI_ISL_3184634, EPI_ISL_3184635, EPI_ISL_3184636, EPI_ISL_3184637, EPI_ISL_3184638, EPI_ISL_3184639, EPI_ISL_3184640, EPI_ISL_3184641, EPI_ISL_3184642, EPI_ISL_3184643, EPI_ISL_3184644, EPI_ISL_3184645, EPI_ISL_3184646, EPI_ISL_3184647, EPI_ISL_3184648, EPI_ISL_3184649, EPI_ISL_3184650, EPI_ISL_3184651, EPI_ISL_3184653, EPI_ISL_3184788, EPI_ISL_3184789, EPI_ISL_3184790, EPI_ISL_3184791, EPI_ISL_3184792, EPI_ISL_3184793, EPI_ISL_3184794, EPI_ISL_3184795, EPI_ISL_3184796, EPI_ISL_3184797, EPI_ISL_3184798, EPI_ISL_3184799, EPI_ISL_3184800, EPI_ISL_3184801, EPI_ISL_3184802 | | | |
| see above | Michigan Department of Health and Human Services, Bureau of Laboratories | Michigan Department of Health and Human Services, Bureau of Laboratories | Blankenship HM, Riner D, Soehnlen MK |
| EPI_ISL_3184881 | 4Cyte Pathology | NSW Health Pathology - Institute of Clinical Pathology and Medical Research; Westmead Hospital; University of Sydney | CIDM-PH et al. |
| EPI_ISL_3184882 | Sydney South West Pathology Service (SSWPS) - Liverpool Hospital - NSW Health Pathology | NSW Health Pathology - Institute of Clinical Pathology and Medical Research; Westmead Hospital; University of Sydney | CIDM-PH et al. |
| EPI_ISL_3184883 | Histopath | NSW Health Pathology - Institute of Clinical Pathology and Medical Research; Westmead Hospital; University of Sydney | CIDM-PH et al. |
| EPI_ISL_3184884 | Laverty Pathology | NSW Health Pathology - Institute of Clinical Pathology and Medical Research; Westmead Hospital; University of Sydney | CIDM-PH et al. |
| EPI_ISL_3184885 | Histopath | NSW Health Pathology - Institute of Clinical Pathology and Medical Research; Westmead Hospital; University of Sydney | CIDM-PH et al. |
| EPI_ISL_3184886 | St Vincent's Pathology (SydPath) | NSW Health Pathology - Institute of Clinical Pathology and Medical Research; Westmead Hospital; University of Sydney | CIDM-PH et al. |
| EPI_ISL_3184887 | Sydney South West Pathology Service (SSWPS) - Liverpool Hospital - NSW Health Pathology | NSW Health Pathology - Institute of Clinical Pathology and Medical Research; Westmead Hospital; University of Sydney | CIDM-PH et al. |
| EPI_ISL_3184888 | Douglass Hanly Moir Pathology | NSW Health Pathology - Institute of Clinical Pathology and Medical Research; Westmead Hospital; University of Sydney | CIDM-PH et al. |
| EPI_ISL_3184889 | Histopath | NSW Health Pathology - Institute of Clinical Pathology and Medical Research; Westmead Hospital; University of Sydney | CIDM-PH et al. |
| EPI_ISL_3184890 | Medlab Pathology | NSW Health Pathology - Institute of Clinical Pathology and Medical Research; Westmead Hospital; University of Sydney | CIDM-PH et al. |
| EPI_ISL_3184891 | Pathology West - NSW Health Pathology | NSW Health Pathology - Institute of Clinical Pathology and Medical Research; Westmead Hospital; University of Sydney | CIDM-PH et al. |
| EPI_ISL_3184892 | Histopath | NSW Health Pathology - Institute of Clinical Pathology and Medical Research; Westmead Hospital; University of Sydney | CIDM-PH et al. |
| EPI_ISL_3184893 | Laverty Pathology | NSW Health Pathology - Institute of Clinical Pathology and Medical Research; Westmead Hospital; University of Sydney | CIDM-PH et al. |
| EPI_ISL_3184894 | Histopath | NSW Health Pathology - Institute of Clinical Pathology and Medical Research; Westmead Hospital; University of Sydney | CIDM-PH et al. |
| EPI_ISL_3184895 | Sydney South West Pathology Service (SSWPS) - Liverpool Hospital - NSW Health Pathology | NSW Health Pathology - Institute of Clinical Pathology and Medical Research; Westmead Hospital; University of Sydney | CIDM-PH et al. |
| EPI_ISL_3184896 | Histopath | NSW Health Pathology - Institute of Clinical Pathology and Medical Research; Westmead Hospital; University of Sydney | CIDM-PH et al. |
| EPI_ISL_3184897 | Laverty Pathology | NSW Health Pathology - Institute of Clinical Pathology and Medical Research; Westmead Hospital; University of Sydney | CIDM-PH et al. |
| EPI_ISL_3184898 | Australian Clinical Labs (formerly Healthscope Pathology) | NSW Health Pathology - Institute of Clinical Pathology and Medical Research; Westmead Hospital; University of Sydney | CIDM-PH et al. |
| EPI_ISL_3184899 | 4Cyte Pathology | NSW Health Pathology - Institute of Clinical Pathology and Medical Research; Westmead Hospital; University of Sydney | CIDM-PH et al. |
| EPI_ISL_3184900 | Douglass Hanly Moir Pathology | NSW Health Pathology - Institute of Clinical Pathology and Medical Research; Westmead Hospital; University of Sydney | CIDM-PH et al. |
| EPI_ISL_3184901, EPI_ISL_3184902 | St Vincent's Pathology (SydPath) | NSW Health Pathology - Institute of Clinical Pathology and Medical Research; Westmead Hospital; University of Sydney | CIDM-PH et al. |
| EPI_ISL_3184903 | Australian Clinical Labs (formerly Healthscope Pathology) | NSW Health Pathology - Institute of Clinical Pathology and Medical Research; Westmead Hospital; University of Sydney | CIDM-PH et al. |
| EPI_ISL_3184904 | Sydney South West Pathology Service (SSWPS) - Liverpool Hospital - NSW Health Pathology | NSW Health Pathology - Institute of Clinical Pathology and Medical Research; Westmead Hospital; University of Sydney | CIDM-PH et al. |
| EPI_ISL_3184905 | 4Cyte Pathology | NSW Health Pathology - Institute of Clinical Pathology and Medical Research; Westmead Hospital; University of Sydney | CIDM-PH et al. |
| EPI_ISL_3184906 | Histopath | NSW Health Pathology - Institute of Clinical Pathology and Medical Research; Westmead Hospital; University of Sydney | CIDM-PH et al. |
| EPI_ISL_3184907 | Douglass Hanly Moir Pathology | NSW Health Pathology - Institute of Clinical Pathology and Medical Research; Westmead Hospital; University of Sydney | CIDM-PH et al. |
| EPI_ISL_3184908 | Histopath | NSW Health Pathology - Institute of Clinical Pathology and Medical Research; Westmead Hospital; University of Sydney | CIDM-PH et al. |
| EPI_ISL_3184909 | Medlab Pathology | NSW Health Pathology - Institute of Clinical Pathology and Medical Research; Westmead Hospital; University of Sydney | CIDM-PH et al. |
| EPI_ISL_3184910 | Histopath | NSW Health Pathology - Institute of Clinical Pathology and Medical Research; Westmead Hospital; University of Sydney | CIDM-PH et al. |
| EPI_ISL_3184911 | Sydney South West Pathology Service (SSWPS) - Liverpool Hospital - NSW Health Pathology | NSW Health Pathology - Institute of Clinical Pathology and Medical Research; Westmead Hospital; University of Sydney | CIDM-PH et al. |
| EPI_ISL_3184912, EPI_ISL_3184913 | Histopath | NSW Health Pathology - Institute of Clinical Pathology and Medical Research; Westmead Hospital; University of Sydney | CIDM-PH et al. |
| EPI_ISL_3184914 | Laverty Pathology | NSW Health Pathology - Institute of Clinical Pathology and Medical Research; Westmead Hospital; University of Sydney | CIDM-PH et al. |
| EPI_ISL_3184915, EPI_ISL_3184916 | St Vincent's Pathology (SydPath) | NSW Health Pathology - Institute of Clinical Pathology and Medical Research; Westmead Hospital; University of Sydney | CIDM-PH et al. |

| | | | |
|---|---|---|---|
| EPI_ISL_3184917 | Sydney South West Pathology Service (SSWPS) - Liverpool Hospital - NSW Health Pathology | NSW Health Pathology - Institute of Clinical Pathology and Medical Research; Westmead Hospital; University of Sydney | CIDM-PH et al. |
| EPI_ISL_3184918 | St Vincent's Pathology (SydPath) | NSW Health Pathology - Institute of Clinical Pathology and Medical Research; Westmead Hospital; University of Sydney | CIDM-PH et al. |
| EPI_ISL_3184919 | Laverty Pathology | NSW Health Pathology - Institute of Clinical Pathology and Medical Research; Westmead Hospital; University of Sydney | CIDM-PH et al. |
| EPI_ISL_3184920, EPI_ISL_3184921 | Histopath | NSW Health Pathology - Institute of Clinical Pathology and Medical Research; Westmead Hospital; University of Sydney | CIDM-PH et al. |
| EPI_ISL_3184922 | Sydney South West Pathology Service (SSWPS) - Liverpool Hospital - NSW Health Pathology | NSW Health Pathology - Institute of Clinical Pathology and Medical Research; Westmead Hospital; University of Sydney | CIDM-PH et al. |
| EPI_ISL_3184923 | Austech Medical Laboratories | NSW Health Pathology - Institute of Clinical Pathology and Medical Research; Westmead Hospital; University of Sydney | CIDM-PH et al. |
| EPI_ISL_3184924 | Laverty Pathology | NSW Health Pathology - Institute of Clinical Pathology and Medical Research; Westmead Hospital; University of Sydney | CIDM-PH et al. |
| EPI_ISL_3184925 | Sydney South West Pathology Service (SSWPS) - Liverpool Hospital - NSW Health Pathology | NSW Health Pathology - Institute of Clinical Pathology and Medical Research; Westmead Hospital; University of Sydney | CIDM-PH et al. |
| EPI_ISL_3184926 | St Vincent's Pathology (SydPath) | NSW Health Pathology - Institute of Clinical Pathology and Medical Research; Westmead Hospital; University of Sydney | CIDM-PH et al. |
| EPI_ISL_3184927 | Australian Clinical Labs (formerly Healthscope Pathology) | NSW Health Pathology - Institute of Clinical Pathology and Medical Research; Westmead Hospital; University of Sydney | CIDM-PH et al. |
| EPI_ISL_3184928 | Pathology West - NSW Health Pathology | NSW Health Pathology - Institute of Clinical Pathology and Medical Research; Westmead Hospital; University of Sydney | CIDM-PH et al. |
| EPI_ISL_3184929 | Sydney South West Pathology Service (SSWPS) - Liverpool Hospital - NSW Health Pathology | NSW Health Pathology - Institute of Clinical Pathology and Medical Research; Westmead Hospital; University of Sydney | CIDM-PH et al. |
| EPI_ISL_3184930 | Histopath | NSW Health Pathology - Institute of Clinical Pathology and Medical Research; Westmead Hospital; University of Sydney | CIDM-PH et al. |
| EPI_ISL_3184931 | St Vincent's Pathology (SydPath) | NSW Health Pathology - Institute of Clinical Pathology and Medical Research; Westmead Hospital; University of Sydney | CIDM-PH et al. |
| EPI_ISL_3184932 | Histopath | NSW Health Pathology - Institute of Clinical Pathology and Medical Research; Westmead Hospital; University of Sydney | CIDM-PH et al. |
| EPI_ISL_3184933 | Sydney South West Pathology Service (SSWPS) - Liverpool Hospital - NSW Health Pathology | NSW Health Pathology - Institute of Clinical Pathology and Medical Research; Westmead Hospital; University of Sydney | CIDM-PH et al. |
| EPI_ISL_3184934 | St Vincent's Pathology (SydPath) | NSW Health Pathology - Institute of Clinical Pathology and Medical Research; Westmead Hospital; University of Sydney | CIDM-PH et al. |
| EPI_ISL_3184936 | Laverty Pathology | NSW Health Pathology - Institute of Clinical Pathology and Medical Research; Westmead Hospital; University of Sydney | CIDM-PH et al. |
| EPI_ISL_3184937 | 4Cyte Pathology | NSW Health Pathology - Institute of Clinical Pathology and Medical Research; Westmead Hospital; University of Sydney | CIDM-PH et al. |
| EPI_ISL_3184938, EPI_ISL_3184939, EPI_ISL_3184940 | Histopath | NSW Health Pathology - Institute of Clinical Pathology and Medical Research; Westmead Hospital; University of Sydney | CIDM-PH et al. |
| EPI_ISL_3184941 | 4Cyte Pathology | NSW Health Pathology - Institute of Clinical Pathology and Medical Research; Westmead Hospital; University of Sydney | CIDM-PH et al. |
| EPI_ISL_3184942 | Histopath | NSW Health Pathology - Institute of Clinical Pathology and Medical Research; Westmead Hospital; University of Sydney | CIDM-PH et al. |
| EPI_ISL_3184943 | Laverty Pathology | NSW Health Pathology - Institute of Clinical Pathology and Medical Research; Westmead Hospital; University of Sydney | CIDM-PH et al. |
| EPI_ISL_3184944 | 4Cyte Pathology | NSW Health Pathology - Institute of Clinical Pathology and Medical Research; Westmead Hospital; University of Sydney | CIDM-PH et al. |
| EPI_ISL_3184945, EPI_ISL_3184946 | Histopath | NSW Health Pathology - Institute of Clinical Pathology and Medical Research; Westmead Hospital; University of Sydney | CIDM-PH et al. |
| EPI_ISL_3184947, EPI_ISL_3184948, EPI_ISL_3184949, EPI_ISL_3184950 | St Vincent's Pathology (SydPath) | NSW Health Pathology - Institute of Clinical Pathology and Medical Research; Westmead Hospital; University of Sydney | CIDM-PH et al. |
| EPI_ISL_3184951 | Histopath | NSW Health Pathology - Institute of Clinical Pathology and Medical Research; Westmead Hospital; University of Sydney | CIDM-PH et al. |
| EPI_ISL_3184952 | St Vincent's Pathology (SydPath) | NSW Health Pathology - Institute of Clinical Pathology and Medical Research; Westmead Hospital; University of Sydney | CIDM-PH et al. |
| EPI_ISL_3184953 | Histopath | NSW Health Pathology - Institute of Clinical Pathology and Medical Research; Westmead Hospital; University of Sydney | CIDM-PH et al. |
| EPI_ISL_3184954 | Australian Clinical Labs | NSW Health Pathology - Institute of Clinical Pathology and Medical Research; Westmead Hospital; University of Sydney | CIDM-PH et al. |
| EPI_ISL_3184955, EPI_ISL_3184956 | Laverty Pathology | NSW Health Pathology - Institute of Clinical Pathology and Medical Research; Westmead Hospital; University of Sydney | CIDM-PH et al. |
| EPI_ISL_3184957, EPI_ISL_3184958, EPI_ISL_3184959, EPI_ISL_3184960, EPI_ISL_3184961, EPI_ISL_3184962, EPI_ISL_3184963, EPI_ISL_3184964, EPI_ISL_3184965, EPI_ISL_3184966, EPI_ISL_3184967, EPI_ISL_3184968, EPI_ISL_3184969, EPI_ISL_3184970, EPI_ISL_3184971 | | | |
| see above | St Vincent's Pathology (SydPath) | NSW Health Pathology - Institute of Clinical Pathology and Medical Research; Westmead Hospital; University of Sydney | CIDM-PH et al. |
| EPI_ISL_3184972 | Childrens Hospital Westmead | NSW Health Pathology - Institute of Clinical Pathology and Medical Research; Westmead Hospital; University of Sydney | CIDM-PH et al. |
| EPI_ISL_3184973, EPI_ISL_3184974, EPI_ISL_3184975, EPI_ISL_3184976, EPI_ISL_3184977, EPI_ISL_3184978, EPI_ISL_3184979, EPI_ISL_3184980, EPI_ISL_3184981, EPI_ISL_3184982, EPI_ISL_3184983, EPI_ISL_3184984 | | | |
| see above | Sydney South West Pathology Service (SSWPS) - Royal Prince Alfred Hospital - NSW Health Pathology | NSW Health Pathology - Institute of Clinical Pathology and Medical Research; Westmead Hospital; University of Sydney | CIDM-PH et al. |
| EPI_ISL_3184985, EPI_ISL_3184986, EPI_ISL_3184987, EPI_ISL_3184988, EPI_ISL_3184989, EPI_ISL_3184990, EPI_ISL_3184991, EPI_ISL_3184992 | Laverty Pathology | NSW Health Pathology - Institute of Clinical Pathology and Medical Research; Westmead Hospital; University of Sydney | CIDM-PH et al. |
| EPI_ISL_3184993 | Douglass Hanly Moir Pathology | NSW Health Pathology - Institute of Clinical Pathology and Medical Research; Westmead Hospital; University of Sydney | CIDM-PH et al. |
| EPI_ISL_3184994, EPI_ISL_3184995, EPI_ISL_3184996 | Laverty Pathology | NSW Health Pathology - Institute of Clinical Pathology and Medical Research; Westmead Hospital; University of Sydney | CIDM-PH et al. |
| EPI_ISL_3184997, EPI_ISL_3184998, EPI_ISL_3184999, EPI_ISL_3185000, EPI_ISL_3185001, EPI_ISL_3185002 | 4Cyte Pathology | NSW Health Pathology - Institute of Clinical Pathology and Medical Research; Westmead Hospital; University of Sydney | CIDM-PH et al. |

| | | | |
|---|---|---|---|
| EPI_ISL_3185003, EPI_ISL_3185004, EPI_ISL_3185005, EPI_ISL_3185006, EPI_ISL_3185007 | Sydney South West Pathology Service (SSWPS) - Royal Prince Alfred Hospital - NSW Health Pathology | NSW Health Pathology - Institute of Clinical Pathology and Medical Research; Westmead Hospital; University of Sydney | CIDM-PH et al. |
| EPI_ISL_3185008 | Childrens Hospital Westmead | NSW Health Pathology - Institute of Clinical Pathology and Medical Research; Westmead Hospital; University of Sydney | CIDM-PH et al. |
| EPI_ISL_3185009 | Australian Clinical Labs | NSW Health Pathology - Institute of Clinical Pathology and Medical Research; Westmead Hospital; University of Sydney | CIDM-PH et al. |
| EPI_ISL_3185010, EPI_ISL_3185011, EPI_ISL_3185012, EPI_ISL_3185013, EPI_ISL_3185014, EPI_ISL_3185015, EPI_ISL_3185016, EPI_ISL_3185017, EPI_ISL_3185018, EPI_ISL_3185019, EPI_ISL_3185020, EPI_ISL_3185021, EPI_ISL_3185022, EPI_ISL_3185023, EPI_ISL_3185024, EPI_ISL_3185025, EPI_ISL_3185026, EPI_ISL_3185027, EPI_ISL_3185028, EPI_ISL_3185029, EPI_ISL_3185030, EPI_ISL_3185031, EPI_ISL_3185032, EPI_ISL_3185033, EPI_ISL_3185034, EPI_ISL_3185035, EPI_ISL_3185036, EPI_ISL_3185037, EPI_ISL_3185038, EPI_ISL_3185039 | | | |
| see above | Histopath | NSW Health Pathology - Institute of Clinical Pathology and Medical Research; Westmead Hospital; University of Sydney | CIDM-PH et al. |
| EPI_ISL_3185040, EPI_ISL_3185041, EPI_ISL_3185042, EPI_ISL_3185043 | Sydney South West Pathology Service (SSWPS) - Royal Prince Alfred Hospital - NSW Health Pathology | NSW Health Pathology - Institute of Clinical Pathology and Medical Research; Westmead Hospital; University of Sydney | CIDM-PH et al. |
| EPI_ISL_3185044 | Medlab Pathology | NSW Health Pathology - Institute of Clinical Pathology and Medical Research; Westmead Hospital; University of Sydney | CIDM-PH et al. |
| EPI_ISL_3185045, EPI_ISL_3185046 | Laverty Pathology | NSW Health Pathology - Institute of Clinical Pathology and Medical Research; Westmead Hospital; University of Sydney | CIDM-PH et al. |
| EPI_ISL_3185047, EPI_ISL_3185048 | St Vincent's Pathology (SydPath) | NSW Health Pathology - Institute of Clinical Pathology and Medical Research; Westmead Hospital; University of Sydney | CIDM-PH et al. |
| EPI_ISL_3185049 | Histopath | NSW Health Pathology - Institute of Clinical Pathology and Medical Research; Westmead Hospital; University of Sydney | CIDM-PH et al. |
| EPI_ISL_3185050 | Medlab Pathology | NSW Health Pathology - Institute of Clinical Pathology and Medical Research; Westmead Hospital; University of Sydney | CIDM-PH et al. |
| EPI_ISL_3185051 | Laverty Pathology | NSW Health Pathology - Institute of Clinical Pathology and Medical Research; Westmead Hospital; University of Sydney | CIDM-PH et al. |
| EPI_ISL_3185052 | Histopath | NSW Health Pathology - Institute of Clinical Pathology and Medical Research; Westmead Hospital; University of Sydney | CIDM-PH et al. |
| EPI_ISL_3185053 | 4Cyte Pathology | NSW Health Pathology - Institute of Clinical Pathology and Medical Research; Westmead Hospital; University of Sydney | CIDM-PH et al. |
| EPI_ISL_3185054 | Histopath | NSW Health Pathology - Institute of Clinical Pathology and Medical Research; Westmead Hospital; University of Sydney | CIDM-PH et al. |
| EPI_ISL_3185055 | Australian Clinical Labs (formerly Healthscope Pathology) | NSW Health Pathology - Institute of Clinical Pathology and Medical Research; Westmead Hospital; University of Sydney | CIDM-PH et al. |
| EPI_ISL_3185056, EPI_ISL_3185057 | Sydney South West Pathology Service (SSWPS) - Liverpool Hospital - NSW Health Pathology | NSW Health Pathology - Institute of Clinical Pathology and Medical Research; Westmead Hospital; University of Sydney | CIDM-PH et al. |
| EPI_ISL_3185058, EPI_ISL_3185059, EPI_ISL_3185060 | Histopath | NSW Health Pathology - Institute of Clinical Pathology and Medical Research; Westmead Hospital; University of Sydney | CIDM-PH et al. |
| EPI_ISL_3185061 | Medlab Pathology | NSW Health Pathology - Institute of Clinical Pathology and Medical Research; Westmead Hospital; University of Sydney | CIDM-PH et al. |
| EPI_ISL_3185062, EPI_ISL_3185063 | Histopath | NSW Health Pathology - Institute of Clinical Pathology and Medical Research; Westmead Hospital; University of Sydney | CIDM-PH et al. |
| EPI_ISL_3185064, EPI_ISL_3185065, EPI_ISL_3185066, EPI_ISL_3185067, EPI_ISL_3185068, EPI_ISL_3185069, EPI_ISL_3185070, EPI_ISL_3185071, EPI_ISL_3185072, EPI_ISL_3185073 | Sydney South West Pathology Service (SSWPS) - Liverpool Hospital - NSW Health Pathology | NSW Health Pathology - Institute of Clinical Pathology and Medical Research; Westmead Hospital; University of Sydney | CIDM-PH et al. |
| EPI_ISL_3185074 | Southern. IML Pathology | NSW Health Pathology - Institute of Clinical Pathology and Medical Research; Westmead Hospital; University of Sydney | CIDM-PH et al. |
| EPI_ISL_3185075, EPI_ISL_3185076, EPI_ISL_3185077, EPI_ISL_3185078, EPI_ISL_3185079, EPI_ISL_3185080, EPI_ISL_3185081, EPI_ISL_3185082, EPI_ISL_3185083, EPI_ISL_3185084, EPI_ISL_3185085, EPI_ISL_3185086, EPI_ISL_3185087, EPI_ISL_3185088, EPI_ISL_3185089, EPI_ISL_3185090, EPI_ISL_3185091, EPI_ISL_3185092, EPI_ISL_3185093, EPI_ISL_3185094, EPI_ISL_3185095, EPI_ISL_3185096, EPI_ISL_3185097, EPI_ISL_3185098, EPI_ISL_3185099, EPI_ISL_3185100, EPI_ISL_3185101, EPI_ISL_3185102 | | | |
| see above | St Vincent's Pathology (SydPath) | NSW Health Pathology - Institute of Clinical Pathology and Medical Research; Westmead Hospital; University of Sydney | CIDM-PH et al. |
| EPI_ISL_3185103, EPI_ISL_3185104, EPI_ISL_3185105, EPI_ISL_3185106, EPI_ISL_3185107, EPI_ISL_3185108, EPI_ISL_3185109, EPI_ISL_3185110, EPI_ISL_3185111, EPI_ISL_3185112 | 4Cyte Pathology | NSW Health Pathology - Institute of Clinical Pathology and Medical Research; Westmead Hospital; University of Sydney | CIDM-PH et al. |
| EPI_ISL_3185113, EPI_ISL_3185114 | Australian Clinical Labs (formerly Healthscope Pathology) | NSW Health Pathology - Institute of Clinical Pathology and Medical Research; Westmead Hospital; University of Sydney | CIDM-PH et al. |
| EPI_ISL_3185115, EPI_ISL_3185116 | Laverty Pathology | NSW Health Pathology - Institute of Clinical Pathology and Medical Research; Westmead Hospital; University of Sydney | CIDM-PH et al. |
| EPI_ISL_3185117, EPI_ISL_3185118, EPI_ISL_3185119, EPI_ISL_3185120, EPI_ISL_3185121, EPI_ISL_3185122, EPI_ISL_3185123, EPI_ISL_3185124, EPI_ISL_3185125, EPI_ISL_3185126, EPI_ISL_3185127, EPI_ISL_3185128, EPI_ISL_3185129, EPI_ISL_3185130, EPI_ISL_3185131, EPI_ISL_3185132, EPI_ISL_3185133, EPI_ISL_3185134, EPI_ISL_3185135 | | | |
| see above | St Vincent's Pathology (SydPath) | NSW Health Pathology - Institute of Clinical Pathology and Medical Research; Westmead Hospital; University of Sydney | CIDM-PH et al. |
| EPI_ISL_3185136, EPI_ISL_3185137, EPI_ISL_3185138 | Sydney South West Pathology Service (SSWPS) - Liverpool Hospital - NSW Health Pathology | NSW Health Pathology - Institute of Clinical Pathology and Medical Research; Westmead Hospital; University of Sydney | CIDM-PH et al. |
| EPI_ISL_3185139 | Sydney South West Pathology Service (SSWPS) - Concord Repatriation General Hospital - NSW Health Pathology | NSW Health Pathology - Institute of Clinical Pathology and Medical Research; Westmead Hospital; University of Sydney | CIDM-PH et al. |
| EPI_ISL_3185140, EPI_ISL_3185141, EPI_ISL_3185142, EPI_ISL_3185143, EPI_ISL_3185144, EPI_ISL_3185145, EPI_ISL_3185146, EPI_ISL_3185147, EPI_ISL_3185148, EPI_ISL_3185149, EPI_ISL_3185150, EPI_ISL_3185151, EPI_ISL_3185152 | | | |
| see above | Histopath | NSW Health Pathology - Institute of Clinical Pathology and Medical Research; Westmead Hospital; University of Sydney | CIDM-PH et al. |
| EPI_ISL_3185153, EPI_ISL_3185154 | Australian Clinical Labs (formerly Healthscope Pathology) | NSW Health Pathology - Institute of Clinical Pathology and Medical Research; Westmead Hospital; University of Sydney | CIDM-PH et al. |
| EPI_ISL_3185155, EPI_ISL_3185156, EPI_ISL_3185157, EPI_ISL_3185158, EPI_ISL_3185159, | Histopath | NSW Health Pathology - Institute of Clinical Pathology and Medical Research; Westmead Hospital; University of Sydney | CIDM-PH et al. |

| | | | |
|---|---|---|---|
| EPI_ISL_3185160, EPI_ISL_3185161, EPI_ISL_3185162 | | | |
| EPI_ISL_3185163, EPI_ISL_3185164, EPI_ISL_3185165, EPI_ISL_3185166, EPI_ISL_3185167, EPI_ISL_3185168, EPI_ISL_3185169, EPI_ISL_3185170 | 4Cyte Pathology | NSW Health Pathology - Institute of Clinical Pathology and Medical Research; Westmead Hospital; University of Sydney | CIDM-PH et al. |
| EPI_ISL_3185171, EPI_ISL_3185172, EPI_ISL_3185173, EPI_ISL_3185174, EPI_ISL_3185175, EPI_ISL_3185176, EPI_ISL_3185177 | Australian Clinical Labs (formerly Healthscope Pathology) | NSW Health Pathology - Institute of Clinical Pathology and Medical Research; Westmead Hospital; University of Sydney | CIDM-PH et al. |
| EPI_ISL_3185178, EPI_ISL_3185179, EPI_ISL_3185180, EPI_ISL_3185181, EPI_ISL_3185182, EPI_ISL_3185183, EPI_ISL_3185184, EPI_ISL_3185185 | Laverty Pathology | NSW Health Pathology - Institute of Clinical Pathology and Medical Research; Westmead Hospital; University of Sydney | CIDM-PH et al. |
| EPI_ISL_3185186, EPI_ISL_3185187, EPI_ISL_3185188, EPI_ISL_3185189, EPI_ISL_3185190, EPI_ISL_3185191, EPI_ISL_3185192 | Medlab Pathology | NSW Health Pathology - Institute of Clinical Pathology and Medical Research; Westmead Hospital; University of Sydney | CIDM-PH et al. |
| EPI_ISL_3185193, EPI_ISL_3185194, EPI_ISL_3185195, EPI_ISL_3185196, EPI_ISL_3185197 | St Vincent's Pathology (SydPath) | NSW Health Pathology - Institute of Clinical Pathology and Medical Research; Westmead Hospital; University of Sydney | CIDM-PH et al. |
| EPI_ISL_3185198 | The Children's Hospital at Westmead | NSW Health Pathology - Institute of Clinical Pathology and Medical Research; Westmead Hospital; University of Sydney | CIDM-PH et al. |
| EPI_ISL_3185199, EPI_ISL_3185200 | Sydney South West Pathology Service (SSWPS) - Concord Repatriation General Hospital - NSW Health Pathology | NSW Health Pathology - Institute of Clinical Pathology and Medical Research; Westmead Hospital; University of Sydney | CIDM-PH et al. |
| EPI_ISL_3185201, EPI_ISL_3185202, EPI_ISL_3185203 | Sydney South West Pathology Service (SSWPS) - Liverpool Hospital - NSW Health Pathology | NSW Health Pathology - Institute of Clinical Pathology and Medical Research; Westmead Hospital; University of Sydney | CIDM-PH et al. |
| EPI_ISL_3185204, EPI_ISL_3185205, EPI_ISL_3185206, EPI_ISL_3185207, EPI_ISL_3185208, EPI_ISL_3185209 | Histopath | NSW Health Pathology - Institute of Clinical Pathology and Medical Research; Westmead Hospital; University of Sydney | CIDM-PH et al. |
| EPI_ISL_3185210 | St Vincent's Pathology (SydPath) | NSW Health Pathology - Institute of Clinical Pathology and Medical Research; Westmead Hospital; University of Sydney | CIDM-PH et al. |
| EPI_ISL_3185211, EPI_ISL_3185212 | Childrens Hospital Westmead | NSW Health Pathology - Institute of Clinical Pathology and Medical Research; Westmead Hospital; University of Sydney | CIDM-PH et al. |
| EPI_ISL_3185213, EPI_ISL_3185214, EPI_ISL_3185215, EPI_ISL_3185216, EPI_ISL_3185217, EPI_ISL_3185218, EPI_ISL_3185219, EPI_ISL_3185220, EPI_ISL_3185221, EPI_ISL_3185222 | Sydney South West Pathology Service (SSWPS) - Royal Prince Alfred Hospital - NSW Health Pathology | NSW Health Pathology - Institute of Clinical Pathology and Medical Research; Westmead Hospital; University of Sydney | CIDM-PH et al. |
| EPI_ISL_3185223 | Laverty Pathology | NSW Health Pathology - Institute of Clinical Pathology and Medical Research; Westmead Hospital; University of Sydney | CIDM-PH et al. |
| EPI_ISL_3185224, EPI_ISL_3185225, EPI_ISL_3185226 | Sydney South West Pathology Service (SSWPS) - Royal Prince Alfred Hospital - NSW Health Pathology | NSW Health Pathology - Institute of Clinical Pathology and Medical Research; Westmead Hospital; University of Sydney | CIDM-PH et al. |
| EPI_ISL_3185227, EPI_ISL_3185228, EPI_ISL_3185229, EPI_ISL_3185230, EPI_ISL_3185231, EPI_ISL_3185232 | Histopath | NSW Health Pathology - Institute of Clinical Pathology and Medical Research; Westmead Hospital; University of Sydney | CIDM-PH et al. |
| EPI_ISL_3185233 | Sydney South West Pathology Service (SSWPS) - Royal Prince Alfred Hospital - NSW Health Pathology | NSW Health Pathology - Institute of Clinical Pathology and Medical Research; Westmead Hospital; University of Sydney | CIDM-PH et al. |
| EPI_ISL_3185234 | Sydney South West Pathology Service (SSWPS) - Liverpool Hospital - NSW Health Pathology | NSW Health Pathology - Institute of Clinical Pathology and Medical Research; Westmead Hospital; University of Sydney | CIDM-PH et al. |
| EPI_ISL_3185235, EPI_ISL_3185236, EPI_ISL_3185237, EPI_ISL_3185238, EPI_ISL_3185239, EPI_ISL_3185240 | St Vincent's Pathology (SydPath) | NSW Health Pathology - Institute of Clinical Pathology and Medical Research; Westmead Hospital; University of Sydney | CIDM-PH et al. |
| EPI_ISL_3185241, EPI_ISL_3185242 | Sydney South West Pathology Service (SSWPS) - Liverpool Hospital - NSW Health Pathology | NSW Health Pathology - Institute of Clinical Pathology and Medical Research; Westmead Hospital; University of Sydney | CIDM-PH et al. |
| EPI_ISL_3185243, EPI_ISL_3185244, EPI_ISL_3185245 | Histopath | NSW Health Pathology - Institute of Clinical Pathology and Medical Research; Westmead Hospital; University of Sydney | CIDM-PH et al. |
| EPI_ISL_3185246 | Laverty Pathology | NSW Health Pathology - Institute of Clinical Pathology and Medical Research; Westmead Hospital; University of Sydney | CIDM-PH et al. |
| EPI_ISL_3185247, EPI_ISL_3185248, EPI_ISL_3185249 | Medlab Pathology | NSW Health Pathology - Institute of Clinical Pathology and Medical Research; Westmead Hospital; University of Sydney | CIDM-PH et al. |
| EPI_ISL_3185250, EPI_ISL_3185251 | St Vincent's Pathology (SydPath) | NSW Health Pathology - Institute of Clinical Pathology and Medical Research; Westmead Hospital; University of Sydney | CIDM-PH et al. |
| EPI_ISL_3185252 | The Children's Hospital at Westmead | NSW Health Pathology - Institute of Clinical Pathology and Medical Research; Westmead Hospital; University of Sydney | CIDM-PH et al. |

| | | | |
|---|---|---|---|
| EPI_ISL_3185253, EPI_ISL_3185254, EPI_ISL_3185255, EPI_ISL_3185256 | Sydney South West Pathology Service (SSWPS) - Liverpool Hospital - NSW Health Pathology | NSW Health Pathology - Institute of Clinical Pathology and Medical Research; Westmead Hospital; University of Sydney | CIDM-PH et al. |
| EPI_ISL_3185257 | Histopath | NSW Health Pathology - Institute of Clinical Pathology and Medical Research; Westmead Hospital; University of Sydney | CIDM-PH et al. |
| EPI_ISL_3185258 | Sydney South West Pathology Service (SSWPS) - Royal Prince Alfred Hospital - NSW Health Pathology | NSW Health Pathology - Institute of Clinical Pathology and Medical Research; Westmead Hospital; University of Sydney | CIDM-PH et al. |
| EPI_ISL_3185461, EPI_ISL_3185463, EPI_ISL_3185465, EPI_ISL_3185485, EPI_ISL_3185543, EPI_ISL_3185544, EPI_ISL_3185597, EPI_ISL_3185599, EPI_ISL_3185600, EPI_ISL_3185659, EPI_ISL_3185660, EPI_ISL_3185732, EPI_ISL_3185733, EPI_ISL_3185778, EPI_ISL_3185809 | | | |
| see above | Genetica Molecular and Subdepartamento de Virologia ISP Chile | Instituto de Salud Publica de Chile | Karen Orostica, Constanza Campano, Barbara Parra, Loredana Arata, Gisselle Barra, Patricia Bustos, Rodrigo Fasce, Javier Tognarelli, Andres Castillo, Soledad Ulloa, Jorge Fernandez |
| EPI_ISL_3185810, EPI_ISL_3185811, EPI_ISL_3185813, EPI_ISL_3185814 | Public Health Authority of the Slovak Republic | Laboratory of Genomics and Bioinformatics, Comenius University Science Park | Tatiana Sedláčková, Diana Rusňáková, Miroslav Böhmer, Anna Gičová, Jaroslav Budiš, Tomáš Szemes, Jakub Styk |
| EPI_ISL_3185848, EPI_ISL_3185849, EPI_ISL_3185850, EPI_ISL_3185851 | Lurie Children's Hospital of Chicago | Northwestern University - Ozer Lab | Ramon Lorenzo-Redondo, Lacy M. Simons, Taylor J. Dean, Michael G. Ison, Xiaotian, Zheng, William J. Muller, Larry K. Kociolek, Judd F. Hultquist, Egon A. Ozer |
| EPI_ISL_3185854, EPI_ISL_3185855, EPI_ISL_3185856, EPI_ISL_3185857, EPI_ISL_3185858, EPI_ISL_3185859, EPI_ISL_3185860, EPI_ISL_3185861, EPI_ISL_3185862, EPI_ISL_3185863, EPI_ISL_3185865, EPI_ISL_3185866, EPI_ISL_3185867, EPI_ISL_3185868, EPI_ISL_3185869, EPI_ISL_3185870, EPI_ISL_3185871, EPI_ISL_3185872, EPI_ISL_3185873, EPI_ISL_3185875, EPI_ISL_3185876, EPI_ISL_3185877, EPI_ISL_3185878, EPI_ISL_3185879, EPI_ISL_3185880, EPI_ISL_3185881, EPI_ISL_3185882, EPI_ISL_3185883, EPI_ISL_3185884, EPI_ISL_3185885, EPI_ISL_3185886, EPI_ISL_3185887, EPI_ISL_3185889, EPI_ISL_3185891, EPI_ISL_3185892, EPI_ISL_3185893, EPI_ISL_3185894, EPI_ISL_3185895, EPI_ISL_3185896, EPI_ISL_3185897, EPI_ISL_3185898, EPI_ISL_3185899, EPI_ISL_3185900, EPI_ISL_3185901, EPI_ISL_3185902, EPI_ISL_3185903, EPI_ISL_3185904, EPI_ISL_3185906, EPI_ISL_3185907, EPI_ISL_3185908, EPI_ISL_3185909, EPI_ISL_3185910, EPI_ISL_3185912, EPI_ISL_3185913 | | | |
| see above | Northwestern Memorial Hospital | Northwestern University - Ozer Lab | Ramon Lorenzo-Redondo, Lacy M. Simons, Taylor J. Dean, Chad J. Achenbach, Lawrence J. Jennings, Chao Qi, Michael G. Ison, Judd F. Hultquist, Egon A. Ozer |
| EPI_ISL_3185936 | Hospital | National Reference Center for Viruses of Respiratory Infections, Institut Pasteur, Paris | Marion Barbet, Sylvie Behillil, Méline Bizard, Angela Brisebarre, Camille Capel, Vincent Enouf, Louise Lefrançois, Frédéric Lemoine, Christophe Malabat, Corinne Maufrais, Hub Bioinformatique Biostatistiques, Etienne Simon-Lorière, Maud Vanpeene, Sylvie Van der Werf ,CéLine Bressollette |
| EPI_ISL_3185969, EPI_ISL_3185970, EPI_ISL_3185971, EPI_ISL_3185972, EPI_ISL_3185973, EPI_ISL_3185974, EPI_ISL_3185975, EPI_ISL_3185976, EPI_ISL_3185977, EPI_ISL_3185978, EPI_ISL_3185979, EPI_ISL_3185980, EPI_ISL_3185981, EPI_ISL_3185982, EPI_ISL_3185983, EPI_ISL_3185984, EPI_ISL_3185985, EPI_ISL_3185986, EPI_ISL_3185987, EPI_ISL_3185988, EPI_ISL_3185989, EPI_ISL_3185990, EPI_ISL_3185991, EPI_ISL_3185992, EPI_ISL_3185993, EPI_ISL_3185994, EPI_ISL_3185995, EPI_ISL_3185996, EPI_ISL_3185997, EPI_ISL_3185998, EPI_ISL_3185999, EPI_ISL_3186000, EPI_ISL_3186001, EPI_ISL_3186002, EPI_ISL_3186003, EPI_ISL_3186004, EPI_ISL_3186005, EPI_ISL_3186006, EPI_ISL_3186007, EPI_ISL_3186008, EPI_ISL_3186009, EPI_ISL_3186010, EPI_ISL_3186011, EPI_ISL_3186012, EPI_ISL_3186013, EPI_ISL_3186014, EPI_ISL_3186015, EPI_ISL_3186016, EPI_ISL_3186017, EPI_ISL_3186018, EPI_ISL_3186019, EPI_ISL_3186020, EPI_ISL_3186021, EPI_ISL_3186022, EPI_ISL_3186023, EPI_ISL_3186024, EPI_ISL_3186025, EPI_ISL_3186026, EPI_ISL_3186027, EPI_ISL_3186028, EPI_ISL_3186029, EPI_ISL_3186030, EPI_ISL_3186031, EPI_ISL_3186032, EPI_ISL_3186033, EPI_ISL_3186034, EPI_ISL_3186035, EPI_ISL_3186036, EPI_ISL_3186037, EPI_ISL_3186038, EPI_ISL_3186039, EPI_ISL_3186040, EPI_ISL_3186041, EPI_ISL_3186042, EPI_ISL_3186043, EPI_ISL_3186044, EPI_ISL_3186045, EPI_ISL_3186046 | | | |
| see above | National Public Health Laboratory, National Centre for Infectious Diseases | National Public Health Laboratory, National Centre for Infectious Diseases | Tze Minn Mak, Katherine Ching, Zhenyang Zhou, Royce Ang, Lin Cui, Raymond Tzer Pin Lin |
| EPI_ISL_3186047, EPI_ISL_3186048, EPI_ISL_3186049 | Laboratorio Analisi Osp. Città di Castello - Azienda USL Umbria1 | Istituto Zooprofilattico Sperimentale dell'Abruzzo e Molise "G. Caporale" | Malagigi V, Tacconi P,Lorusso A, Marcacci M, Di Domenico M, Ancora M, Curini V, Mangone I, Rinaldi A, Delli Compagni E, Scialabba S, Di Pasquale A, Cammà C, Puglia I, Calistri P, Savini G |
| EPI_ISL_3186050, EPI_ISL_3186051, EPI_ISL_3186052 | Università degli Studi di Perugia | Istituto Zooprofilattico Sperimentale dell'Abruzzo e Molise "G. Caporale" | Mencacci A, Camilloni B, Biagetti M, Giammarioli M, Lorusso A, Marcacci M, Di Domenico M, Ancora M, Curini V, Mangone I, Rinaldi A, Delli Compagni E, Scialabba S, Di Pasquale A, Cammà C, Puglia I, Calistri P, Savini G |
| EPI_ISL_3186053, EPI_ISL_3186054, EPI_ISL_3186055, EPI_ISL_3186056 | Azienda USL Umbria 2 | Istituto Zooprofilattico Sperimentale dell'Abruzzo e Molise "G. Caporale" | Proietti A, Pistoni E, Lorusso A, Marcacci M, Di Domenico M, Ancora M, Curini V, Mangone I, Rinaldi A, Delli Compagni E, Scialabba S, Di Pasquale A, Cammà C, Puglia I, Calistri P, Savini G |
| EPI_ISL_3186057, EPI_ISL_3186058, EPI_ISL_3186059 | Azienda Ospedaliera Terni | Istituto Zooprofilattico Sperimentale dell'Abruzzo e Molise "G. Caporale" | Palumbo M, Scaccetti A, Lorusso A, Marcacci M, Di Domenico M, Ancora M, Curini V, Mangone I, Rinaldi A, Delli Compagni E, Scialabba S, Di Pasquale A, Cammà C, Puglia I, Calistri P, Savini G |
| EPI_ISL_3186075 | Histopath | NSW Health Pathology - Institute of Clinical Pathology and Medical Research; Westmead Hospital; University of Sydney | CIDM-PH et al. |
| EPI_ISL_3186076 | 4Cyte Pathology | NSW Health Pathology - Institute of Clinical Pathology and Medical Research; Westmead Hospital; University of Sydney | CIDM-PH et al. |
| EPI_ISL_3186077 | Histopath | NSW Health Pathology - Institute of Clinical Pathology and Medical Research; Westmead Hospital; University of Sydney | CIDM-PH et al. |
| EPI_ISL_3186078 | Sydney South West Pathology Service (SSWPS) - Liverpool Hospital - NSW Health Pathology | NSW Health Pathology - Institute of Clinical Pathology and Medical Research; Westmead Hospital; University of Sydney | CIDM-PH et al. |
| EPI_ISL_3186079 | St Vincent's Pathology (SydPath) | NSW Health Pathology - Institute of Clinical Pathology and Medical Research; Westmead Hospital; University of Sydney | CIDM-PH et al. |
| EPI_ISL_3186080 | 4Cyte Pathology | NSW Health Pathology - Institute of Clinical Pathology and Medical Research; Westmead Hospital; University of Sydney | CIDM-PH et al. |
| EPI_ISL_3186081 | Sydney South West Pathology Service (SSWPS) - Liverpool Hospital - NSW Health Pathology | NSW Health Pathology - Institute of Clinical Pathology and Medical Research; Westmead Hospital; University of Sydney | CIDM-PH et al. |
| EPI_ISL_3186082 | 4Cyte Pathology | NSW Health Pathology - Institute of Clinical Pathology and Medical Research; Westmead Hospital; University of Sydney | CIDM-PH et al. |
| EPI_ISL_3186083 | St Vincent's Pathology (SydPath) | NSW Health Pathology - Institute of Clinical Pathology and Medical Research; Westmead Hospital; University of Sydney | CIDM-PH et al. |
| EPI_ISL_3186084 | Laverty Pathology | NSW Health Pathology - Institute of Clinical Pathology and Medical Research; Westmead Hospital; University of Sydney | CIDM-PH et al. |
| EPI_ISL_3186085 | Histopath | NSW Health Pathology - Institute of Clinical Pathology and Medical Research; Westmead Hospital; University of Sydney | CIDM-PH et al. |
| EPI_ISL_3186086, EPI_ISL_3186087 | Laverty Pathology | NSW Health Pathology - Institute of Clinical Pathology and Medical Research; Westmead Hospital; University of Sydney | CIDM-PH et al. |
| EPI_ISL_3186088, EPI_ISL_3186089, EPI_ISL_3186090 | 4Cyte Pathology | NSW Health Pathology - Institute of Clinical Pathology and Medical Research; Westmead Hospital; University of Sydney | CIDM-PH et al. |
| EPI_ISL_3186091, EPI_ISL_3186092, EPI_ISL_3186093 | Douglass Hanly Moir Pathology | NSW Health Pathology - Institute of Clinical Pathology and Medical Research; Westmead Hospital; University of Sydney | CIDM-PH et al. |
| EPI_ISL_3186094, EPI_ISL_3186095, EPI_ISL_3186096, EPI_ISL_3186097, EPI_ISL_3186098, EPI_ISL_3186099, EPI_ISL_3186100, EPI_ISL_3186101, EPI_ISL_3186102, EPI_ISL_3186103, EPI_ISL_3186104, EPI_ISL_3186105, EPI_ISL_3186106 | | | |
| see above | Sydney South West Pathology Service (SSWPS) - Liverpool Hospital - NSW Health Pathology | NSW Health Pathology - Institute of Clinical Pathology and Medical Research; Westmead Hospital; University of Sydney | CIDM-PH et al. |
| EPI_ISL_3186107, EPI_ISL_3186108, EPI_ISL_3186109, EPI_ISL_3186110, EPI_ISL_3186111, EPI_ISL_3186112, EPI_ISL_3186113, EPI_ISL_3186114, EPI_ISL_3186115, EPI_ISL_3186116, EPI_ISL_3186117, EPI_ISL_3186118, EPI_ISL_3186119, EPI_ISL_3186120, EPI_ISL_3186121, EPI_ISL_3186122, EPI_ISL_3186123, EPI_ISL_3186124, EPI_ISL_3186125, EPI_ISL_3186126, EPI_ISL_3186127, EPI_ISL_3186128, EPI_ISL_3186129, EPI_ISL_3186130, EPI_ISL_3186131, EPI_ISL_3186132, EPI_ISL_3186133, EPI_ISL_3186134, EPI_ISL_3186135, EPI_ISL_3186136, EPI_ISL_3186137, EPI_ISL_3186138, EPI_ISL_3186139, EPI_ISL_3186140, EPI_ISL_3186141, EPI_ISL_3186142, EPI_ISL_3186143, EPI_ISL_3186144, EPI_ISL_3186145, EPI_ISL_3186146, EPI_ISL_3186147, EPI_ISL_3186148, EPI_ISL_3186149, EPI_ISL_3186150, EPI_ISL_3186151, EPI_ISL_3186152, EPI_ISL_3186153, EPI_ISL_3186154, EPI_ISL_3186155, EPI_ISL_3186156, EPI_ISL_3186157, EPI_ISL_3186158, EPI_ISL_3186159, EPI_ISL_3186160, EPI_ISL_3186161, EPI_ISL_3186162, EPI_ISL_3186163, EPI_ISL_3186164, EPI_ISL_3186165, EPI_ISL_3186166, EPI_ISL_3186167, EPI_ISL_3186168, EPI_ISL_3186169, EPI_ISL_3186170, EPI_ISL_3186171, EPI_ISL_3186172, EPI_ISL_3186173, EPI_ISL_3186174, EPI_ISL_3186175 | | | |
| see above | St Vincent's Pathology (SydPath) | NSW Health Pathology - Institute of Clinical Pathology and Medical Research; Westmead Hospital; University of Sydney | CIDM-PH et al. |
| EPI_ISL_3186176, EPI_ISL_3186177, EPI_ISL_3186178, EPI_ISL_3186179, EPI_ISL_3186180 | Histopath | NSW Health Pathology - Institute of Clinical Pathology and Medical Research; Westmead Hospital; University of Sydney | CIDM-PH et al. |
| EPI_ISL_3186181, EPI_ISL_3186182 | 4Cyte Pathology | NSW Health Pathology - Institute of Clinical Pathology and Medical Research; Westmead Hospital; University of Sydney | CIDM-PH et al. |
| EPI_ISL_3186183, EPI_ISL_3186184 | Laverty Pathology | NSW Health Pathology - Institute of Clinical Pathology and Medical Research; Westmead Hospital; University of Sydney | CIDM-PH et al. |
| EPI_ISL_3186185, EPI_ISL_3186186, | Sydney South West Pathology Service (SSWPS) - Liverpool Hospital - NSW Health Pathology | NSW Health Pathology - Institute of Clinical Pathology and Medical Research; Westmead Hospital; University of Sydney | CIDM-PH et al. |

| | | | |
|---|---|---|---|
| EPI_ISL_3186187, EPI_ISL_3186188 | | | |
| EPI_ISL_3186189, EPI_ISL_3186190, EPI_ISL_3186191, EPI_ISL_3186192, EPI_ISL_3186193, EPI_ISL_3186194, EPI_ISL_3186195, EPI_ISL_3186196, EPI_ISL_3186197, EPI_ISL_3186198, EPI_ISL_3186199, EPI_ISL_3186200, EPI_ISL_3186201, EPI_ISL_3186202 | | | |
| see above | St Vincent's Pathology (SydPath) | NSW Health Pathology - Institute of Clinical Pathology and Medical Research; Westmead Hospital; University of Sydney | CIDM-PH et al. |
| EPI_ISL_3186203, EPI_ISL_3186204, EPI_ISL_3186205, EPI_ISL_3186206, EPI_ISL_3186207, EPI_ISL_3186208, EPI_ISL_3186209, EPI_ISL_3186210, EPI_ISL_3186211, EPI_ISL_3186212, EPI_ISL_3186213 | | | |
| see above | Histopath | NSW Health Pathology - Institute of Clinical Pathology and Medical Research; Westmead Hospital; University of Sydney | CIDM-PH et al. |
| EPI_ISL_3186214, EPI_ISL_3186215, EPI_ISL_3186216 | Medlab Pathology | NSW Health Pathology - Institute of Clinical Pathology and Medical Research; Westmead Hospital; University of Sydney | CIDM-PH et al. |
| EPI_ISL_3186217 | Pathology West - NSW Health Pathology | NSW Health Pathology - Institute of Clinical Pathology and Medical Research; Westmead Hospital; University of Sydney | CIDM-PH et al. |
| EPI_ISL_3186218, EPI_ISL_3186219, EPI_ISL_3186220, EPI_ISL_3186221, EPI_ISL_3186222, EPI_ISL_3186223, EPI_ISL_3186224, EPI_ISL_3186225, EPI_ISL_3186226 | Laverty Pathology | NSW Health Pathology - Institute of Clinical Pathology and Medical Research; Westmead Hospital; University of Sydney | CIDM-PH et al. |
| EPI_ISL_3186227 | St Vincent's Pathology (SydPath) | NSW Health Pathology - Institute of Clinical Pathology and Medical Research; Westmead Hospital; University of Sydney | CIDM-PH et al. |
| EPI_ISL_3186228, EPI_ISL_3186229 | Douglass Hanly Moir Pathology | NSW Health Pathology - Institute of Clinical Pathology and Medical Research; Westmead Hospital; University of Sydney | CIDM-PH et al. |
| EPI_ISL_3186230, EPI_ISL_3186231, EPI_ISL_3186232 | Australian Clinical Labs | NSW Health Pathology - Institute of Clinical Pathology and Medical Research; Westmead Hospital; University of Sydney | CIDM-PH et al. |
| EPI_ISL_3186233, EPI_ISL_3186234, EPI_ISL_3186235 | Laverty Pathology | NSW Health Pathology - Institute of Clinical Pathology and Medical Research; Westmead Hospital; University of Sydney | CIDM-PH et al. |
| EPI_ISL_3186236 | Australian Clinical Labs | NSW Health Pathology - Institute of Clinical Pathology and Medical Research; Westmead Hospital; University of Sydney | CIDM-PH et al. |
| EPI_ISL_3186237 | Laverty Pathology | NSW Health Pathology - Institute of Clinical Pathology and Medical Research; Westmead Hospital; University of Sydney | CIDM-PH et al. |
| EPI_ISL_3186238, EPI_ISL_3186239 | 4Cyte Pathology | NSW Health Pathology - Institute of Clinical Pathology and Medical Research; Westmead Hospital; University of Sydney | CIDM-PH et al. |
| EPI_ISL_3186240, EPI_ISL_3186241, EPI_ISL_3186242, EPI_ISL_3186243 | Laverty Pathology | NSW Health Pathology - Institute of Clinical Pathology and Medical Research; Westmead Hospital; University of Sydney | CIDM-PH et al. |
| EPI_ISL_3186244, EPI_ISL_3186245, EPI_ISL_3186246, EPI_ISL_3186247 | 4Cyte Pathology | NSW Health Pathology - Institute of Clinical Pathology and Medical Research; Westmead Hospital; University of Sydney | CIDM-PH et al. |
| EPI_ISL_3186248, EPI_ISL_3186249, EPI_ISL_3186250, EPI_ISL_3186251, EPI_ISL_3186252, EPI_ISL_3186253 | Histopath | NSW Health Pathology - Institute of Clinical Pathology and Medical Research; Westmead Hospital; University of Sydney | CIDM-PH et al. |
| EPI_ISL_3186254, EPI_ISL_3186255, EPI_ISL_3186256 | 4Cyte Pathology | NSW Health Pathology - Institute of Clinical Pathology and Medical Research; Westmead Hospital; University of Sydney | CIDM-PH et al. |
| EPI_ISL_3186257, EPI_ISL_3186258 | Medlab Pathology | NSW Health Pathology - Institute of Clinical Pathology and Medical Research; Westmead Hospital; University of Sydney | CIDM-PH et al. |
| EPI_ISL_3186259, EPI_ISL_3186260, EPI_ISL_3186261 | Pathology West - NSW Health Pathology | NSW Health Pathology - Institute of Clinical Pathology and Medical Research; Westmead Hospital; University of Sydney | CIDM-PH et al. |
| EPI_ISL_3186262, EPI_ISL_3186263, EPI_ISL_3186264, EPI_ISL_3186265, EPI_ISL_3186266, EPI_ISL_3186267, EPI_ISL_3186268, EPI_ISL_3186269, EPI_ISL_3186270 | Histopath | NSW Health Pathology - Institute of Clinical Pathology and Medical Research; Westmead Hospital; University of Sydney | CIDM-PH et al. |
| EPI_ISL_3186271 | 4Cyte Pathology | NSW Health Pathology - Institute of Clinical Pathology and Medical Research; Westmead Hospital; University of Sydney | CIDM-PH et al. |
| EPI_ISL_3186272 | Sydney South West Pathology Service (SSWPS) - Liverpool Hospital - NSW Health Pathology | NSW Health Pathology - Institute of Clinical Pathology and Medical Research; Westmead Hospital; University of Sydney | CIDM-PH et al. |
| EPI_ISL_3186273 | 4Cyte Pathology | NSW Health Pathology - Institute of Clinical Pathology and Medical Research; Westmead Hospital; University of Sydney | CIDM-PH et al. |
| EPI_ISL_3186274 | Sydney South West Pathology Service (SSWPS) - Liverpool Hospital - NSW Health Pathology | NSW Health Pathology - Institute of Clinical Pathology and Medical Research; Westmead Hospital; University of Sydney | CIDM-PH et al. |
| EPI_ISL_3186275 | St Vincent's Pathology (SydPath) | NSW Health Pathology - Institute of Clinical Pathology and Medical Research; Westmead Hospital; University of Sydney | CIDM-PH et al. |
| EPI_ISL_3186276, EPI_ISL_3186277, EPI_ISL_3186278, EPI_ISL_3186279, EPI_ISL_3186280 | Histopath | NSW Health Pathology - Institute of Clinical Pathology and Medical Research; Westmead Hospital; University of Sydney | CIDM-PH et al. |
| EPI_ISL_3186281, EPI_ISL_3186282 | Laverty Pathology | NSW Health Pathology - Institute of Clinical Pathology and Medical Research; Westmead Hospital; University of Sydney | CIDM-PH et al. |
| EPI_ISL_3186283 | Histopath | NSW Health Pathology - Institute of Clinical Pathology and Medical Research; Westmead Hospital; University of Sydney | CIDM-PH et al. |
| EPI_ISL_3186284, EPI_ISL_3186285, EPI_ISL_3186286 | Laverty Pathology | NSW Health Pathology - Institute of Clinical Pathology and Medical Research; Westmead Hospital; University of Sydney | CIDM-PH et al. |
| EPI_ISL_3186287, EPI_ISL_3186288, EPI_ISL_3186289 | Histopath | NSW Health Pathology - Institute of Clinical Pathology and Medical Research; Westmead Hospital; University of Sydney | CIDM-PH et al. |
| EPI_ISL_3186290 | Australian Clinical Labs | NSW Health Pathology - Institute of Clinical Pathology and Medical | CIDM-PH et al. |

| | | | |
|---|---|---|---|
| | | Research; Westmead Hospital; University of Sydney | |
| EPI_ISL_3186291 | Pathology West - NSW Health Pathology | NSW Health Pathology - Institute of Clinical Pathology and Medical Research; Westmead Hospital; University of Sydney | CIDM-PH et al. |
| EPI_ISL_3186333, EPI_ISL_3186335, EPI_ISL_3186338, EPI_ISL_3186340, EPI_ISL_3186341, EPI_ISL_3186342, EPI_ISL_3186343, EPI_ISL_3186347, EPI_ISL_3186348, EPI_ISL_3186351, EPI_ISL_3186352, EPI_ISL_3186353, EPI_ISL_3186354, EPI_ISL_3186355, EPI_ISL_3186357, EPI_ISL_3186358, EPI_ISL_3186359, EPI_ISL_3186360, EPI_ISL_3186362, EPI_ISL_3186363, EPI_ISL_3186364, EPI_ISL_3186365, EPI_ISL_3186367, EPI_ISL_3186368, EPI_ISL_3186369, EPI_ISL_3186373, EPI_ISL_3186374, EPI_ISL_3186375, EPI_ISL_3186378, EPI_ISL_3186379, EPI_ISL_3186380, EPI_ISL_3186383, EPI_ISL_3186384, EPI_ISL_3186385, EPI_ISL_3186386, EPI_ISL_3186387, EPI_ISL_3186388, EPI_ISL_3186389, EPI_ISL_3186390, EPI_ISL_3186392, EPI_ISL_3186393, EPI_ISL_3186394, EPI_ISL_3186398, EPI_ISL_3186399, EPI_ISL_3186400, EPI_ISL_3186402, EPI_ISL_3186403, EPI_ISL_3186405, EPI_ISL_3186406, EPI_ISL_3186407, EPI_ISL_3186408, EPI_ISL_3186409, EPI_ISL_3186411, EPI_ISL_3186412, EPI_ISL_3186413, EPI_ISL_3186414, EPI_ISL_3186416, EPI_ISL_3186417, EPI_ISL_3186418, EPI_ISL_3186419, EPI_ISL_3186420, EPI_ISL_3186421, EPI_ISL_3186423, EPI_ISL_3186424, EPI_ISL_3186425, EPI_ISL_3186426, EPI_ISL_3186428, EPI_ISL_3186429, EPI_ISL_3186433, EPI_ISL_3186434, EPI_ISL_3186437, EPI_ISL_3186438, EPI_ISL_3186439, EPI_ISL_3186441, EPI_ISL_3186443, EPI_ISL_3186444, EPI_ISL_3186446, EPI_ISL_3186448, EPI_ISL_3186449, EPI_ISL_3186450, EPI_ISL_3186451, EPI_ISL_3186452, EPI_ISL_3186453, EPI_ISL_3186456 | | | |
| see above | Sweden/Sodermanland/Unilabs | Unilabs/Eskilstuna/Sweden | Emma Arvidsson |
| EPI_ISL_3186459, EPI_ISL_3186460, EPI_ISL_3186461, EPI_ISL_3186462, EPI_ISL_3186463, EPI_ISL_3186464 | Histopath | NSW Health Pathology - Institute of Clinical Pathology and Medical Research; Westmead Hospital; University of Sydney | CIDM-PH et al. |
| EPI_ISL_3186465 | 4Cyte Pathology | NSW Health Pathology - Institute of Clinical Pathology and Medical Research; Westmead Hospital; University of Sydney | CIDM-PH et al. |
| EPI_ISL_3186466, EPI_ISL_3186467 | The Children's Hospital at Westmead | NSW Health Pathology - Institute of Clinical Pathology and Medical Research; Westmead Hospital; University of Sydney | CIDM-PH et al. |
| EPI_ISL_3186468, EPI_ISL_3186469, EPI_ISL_3186470, EPI_ISL_3186471 | Laverty Pathology | NSW Health Pathology - Institute of Clinical Pathology and Medical Research; Westmead Hospital; University of Sydney | CIDM-PH et al. |
| EPI_ISL_3186472, EPI_ISL_3186473, EPI_ISL_3186474, EPI_ISL_3186475, EPI_ISL_3186476, EPI_ISL_3186477, EPI_ISL_3186478, EPI_ISL_3186479, EPI_ISL_3186480, EPI_ISL_3186481, EPI_ISL_3186482, EPI_ISL_3186483 | | | |
| see above | Sydney South West Pathology Service (SSWPS) - Liverpool Hospital - NSW Health Pathology | NSW Health Pathology - Institute of Clinical Pathology and Medical Research; Westmead Hospital; University of Sydney | CIDM-PH et al. |
| EPI_ISL_3186484, EPI_ISL_3186485, EPI_ISL_3186486, EPI_ISL_3186487 | Histopath | NSW Health Pathology - Institute of Clinical Pathology and Medical Research; Westmead Hospital; University of Sydney | CIDM-PH et al. |
| EPI_ISL_3186488, EPI_ISL_3186489 | Sydney South West Pathology Service (SSWPS) - Liverpool Hospital - NSW Health Pathology | NSW Health Pathology - Institute of Clinical Pathology and Medical Research; Westmead Hospital; University of Sydney | CIDM-PH et al. |
| EPI_ISL_3186493, EPI_ISL_3186499, EPI_ISL_3186500, EPI_ISL_3186501, EPI_ISL_3186502, EPI_ISL_3186503, EPI_ISL_3186504, EPI_ISL_3186507, EPI_ISL_3186508, EPI_ISL_3186509, EPI_ISL_3186510, EPI_ISL_3186514, EPI_ISL_3186516, EPI_ISL_3186517, EPI_ISL_3186518, EPI_ISL_3186522, EPI_ISL_3186523, EPI_ISL_3186524, EPI_ISL_3186525, EPI_ISL_3186526, EPI_ISL_3186527, EPI_ISL_3186528, EPI_ISL_3186529, EPI_ISL_3186530, EPI_ISL_3186531 | | | |
| see above | Maine Health and Environmental Testing Laboratory | Tewhey Lab, The Jackson Laboratory | Matluk,N., Dewey,H., Iosue,F., Barter,M., Lynch,R., Munger,H. and Tewhey,R. |
| EPI_ISL_3186574 | Histopath | NSW Health Pathology - Institute of Clinical Pathology and Medical Research; Westmead Hospital; University of Sydney | CIDM-PH et al. |
| EPI_ISL_3186575, EPI_ISL_3186576 | Laverty Pathology | NSW Health Pathology - Institute of Clinical Pathology and Medical Research; Westmead Hospital; University of Sydney | CIDM-PH et al. |
| EPI_ISL_3186577, EPI_ISL_3186578 | Histopath | NSW Health Pathology - Institute of Clinical Pathology and Medical Research; Westmead Hospital; University of Sydney | CIDM-PH et al. |
| EPI_ISL_3186579 | Sydney South West Pathology Service (SSWPS) - Liverpool Hospital - NSW Health Pathology | NSW Health Pathology - Institute of Clinical Pathology and Medical Research; Westmead Hospital; University of Sydney | CIDM-PH et al. |
| EPI_ISL_3186580, EPI_ISL_3186581, EPI_ISL_3186582, EPI_ISL_3186583, EPI_ISL_3186584 | St Vincent's Pathology (SydPath) | NSW Health Pathology - Institute of Clinical Pathology and Medical Research; Westmead Hospital; University of Sydney | CIDM-PH et al. |
| EPI_ISL_3186585 | Australian Clinical Labs (formerly Healthscope Pathology) | NSW Health Pathology - Institute of Clinical Pathology and Medical Research; Westmead Hospital; University of Sydney | CIDM-PH et al. |
| EPI_ISL_3186586 | Histopath | NSW Health Pathology - Institute of Clinical Pathology and Medical Research; Westmead Hospital; University of Sydney | CIDM-PH et al. |
| EPI_ISL_3186587, EPI_ISL_3186588, EPI_ISL_3186589, EPI_ISL_3186590 | Australian Clinical Labs (formerly Healthscope Pathology) | NSW Health Pathology - Institute of Clinical Pathology and Medical Research; Westmead Hospital; University of Sydney | CIDM-PH et al. |
| EPI_ISL_3186591 | The Children's Hospital at Westmead | NSW Health Pathology - Institute of Clinical Pathology and Medical Research; Westmead Hospital; University of Sydney | CIDM-PH et al. |
| EPI_ISL_3186592, EPI_ISL_3186593 | Douglass Hanly Moir Pathology | NSW Health Pathology - Institute of Clinical Pathology and Medical Research; Westmead Hospital; University of Sydney | CIDM-PH et al. |
| EPI_ISL_3186594, EPI_ISL_3186595, EPI_ISL_3186596, EPI_ISL_3186597, EPI_ISL_3186598, EPI_ISL_3186599, EPI_ISL_3186600, EPI_ISL_3186601, EPI_ISL_3186602, EPI_ISL_3186603, EPI_ISL_3186604, EPI_ISL_3186605 | | | |
| see above | Laverty Pathology | NSW Health Pathology - Institute of Clinical Pathology and Medical Research; Westmead Hospital; University of Sydney | CIDM-PH et al. |
| EPI_ISL_3186606, EPI_ISL_3186607, EPI_ISL_3186608, EPI_ISL_3186609, EPI_ISL_3186610, EPI_ISL_3186611, EPI_ISL_3186612 | Sydney South West Pathology Service (SSWPS) - Liverpool Hospital - NSW Health Pathology | NSW Health Pathology - Institute of Clinical Pathology and Medical Research; Westmead Hospital; University of Sydney | CIDM-PH et al. |
| EPI_ISL_3186613 | Medlab Pathology | NSW Health Pathology - Institute of Clinical Pathology and Medical Research; Westmead Hospital; University of Sydney | CIDM-PH et al. |
| EPI_ISL_3186614, EPI_ISL_3186615, EPI_ISL_3186616, EPI_ISL_3186617, EPI_ISL_3186618, EPI_ISL_3186619, EPI_ISL_3186620, EPI_ISL_3186621, EPI_ISL_3186622, EPI_ISL_3186623 | St Vincent's Pathology (SydPath) | NSW Health Pathology - Institute of Clinical Pathology and Medical Research; Westmead Hospital; University of Sydney | CIDM-PH et al. |
| EPI_ISL_3186624, EPI_ISL_3186625, EPI_ISL_3186626, EPI_ISL_3186627, EPI_ISL_3186628, EPI_ISL_3186629, EPI_ISL_3186630, EPI_ISL_3186631, EPI_ISL_3186632, EPI_ISL_3186633 | 4Cyte Pathology | NSW Health Pathology - Institute of Clinical Pathology and Medical Research; Westmead Hospital; University of Sydney | CIDM-PH et al. |
| EPI_ISL_3186634, EPI_ISL_3186635, EPI_ISL_3186636, EPI_ISL_3186637, EPI_ISL_3186638 | Medlab Pathology | NSW Health Pathology - Institute of Clinical Pathology and Medical Research; Westmead Hospital; University of Sydney | CIDM-PH et al. |

| | | | |
|---|---|---|---|
| EPI_ISL_3186639 | Pathology West - NSW Health Pathology | NSW Health Pathology - Institute of Clinical Pathology and Medical Research; Westmead Hospital; University of Sydney | CIDM-PH et al. |
| EPI_ISL_3186647, EPI_ISL_3186648, EPI_ISL_3186649, EPI_ISL_3186650, EPI_ISL_3186651, EPI_ISL_3186652 | Laverty Pathology | NSW Health Pathology - Institute of Clinical Pathology and Medical Research; Westmead Hospital; University of Sydney | Arnott A., Draper J., Gall M., Martinez E., Rockett R., Sintchenko V., on behalf of ICPMR |
| EPI_ISL_3186653, EPI_ISL_3186654, EPI_ISL_3186655, EPI_ISL_3186656, EPI_ISL_3186657, EPI_ISL_3186658 | Histopath | NSW Health Pathology - Institute of Clinical Pathology and Medical Research; Westmead Hospital; University of Sydney | Arnott A., Draper J., Gall M., Martinez E., Rockett R., Sintchenko V., on behalf of ICPMR |
| EPI_ISL_3186659 | 4Cyte Pathology | NSW Health Pathology - Institute of Clinical Pathology and Medical Research; Westmead Hospital; University of Sydney | Arnott A., Draper J., Gall M., Martinez E., Rockett R., Sintchenko V., on behalf of ICPMR |
| EPI_ISL_3186660 | Australian Clinical Labs (formerly Healthscope Pathology) | NSW Health Pathology - Institute of Clinical Pathology and Medical Research; Westmead Hospital; University of Sydney | Arnott A., Draper J., Gall M., Martinez E., Rockett R., Sintchenko V., on behalf of ICPMR |
| EPI_ISL_3186661, EPI_ISL_3186662, EPI_ISL_3186663, EPI_ISL_3186664, EPI_ISL_3186665, EPI_ISL_3186666, EPI_ISL_3186667, EPI_ISL_3186668, EPI_ISL_3186669, EPI_ISL_3186670 | Medlab Pathology | NSW Health Pathology - Institute of Clinical Pathology and Medical Research; Westmead Hospital; University of Sydney | Arnott A., Draper J., Gall M., Martinez E., Rockett R., Sintchenko V., on behalf of ICPMR |
| EPI_ISL_3186671 | Sydney South West Pathology Service (SSWPS) - Concord Repatriation General Hospital - NSW Health Pathology | NSW Health Pathology - Institute of Clinical Pathology and Medical Research; Westmead Hospital; University of Sydney | Arnott A., Draper J., Gall M., Martinez E., Rockett R., Sintchenko V., on behalf of ICPMR |
| EPI_ISL_3186672, EPI_ISL_3186673, EPI_ISL_3186674 | Douglass Hanly Moir Pathology | NSW Health Pathology - Institute of Clinical Pathology and Medical Research; Westmead Hospital; University of Sydney | Arnott A., Draper J., Gall M., Martinez E., Rockett R., Sintchenko V., on behalf of ICPMR |
| EPI_ISL_3186675 | Laverty Pathology | NSW Health Pathology - Institute of Clinical Pathology and Medical Research; Westmead Hospital; University of Sydney | Arnott A., Draper J., Gall M., Martinez E., Rockett R., Sintchenko V., on behalf of ICPMR |
| EPI_ISL_3186676, EPI_ISL_3186677 | Sydney South West Pathology Service (SSWPS) - Liverpool Hospital - NSW Health Pathology | NSW Health Pathology - Institute of Clinical Pathology and Medical Research; Westmead Hospital; University of Sydney | Arnott A., Draper J., Gall M., Martinez E., Rockett R., Sintchenko V., on behalf of ICPMR |
| EPI_ISL_3186678, EPI_ISL_3186679 | The Children's Hospital at Westmead | NSW Health Pathology - Institute of Clinical Pathology and Medical Research; Westmead Hospital; University of Sydney | Arnott A., Draper J., Gall M., Martinez E., Rockett R., Sintchenko V., on behalf of ICPMR |
| EPI_ISL_3186680, EPI_ISL_3186681, EPI_ISL_3186682, EPI_ISL_3186683, EPI_ISL_3186684, EPI_ISL_3186685, EPI_ISL_3186686 | St Vincent's Pathology (SydPath) | NSW Health Pathology - Institute of Clinical Pathology and Medical Research; Westmead Hospital; University of Sydney | Arnott A., Draper J., Gall M., Martinez E., Rockett R., Sintchenko V., on behalf of ICPMR |
| EPI_ISL_3186687, EPI_ISL_3186688, EPI_ISL_3186689, EPI_ISL_3186690, EPI_ISL_3186691, EPI_ISL_3186692, EPI_ISL_3186693 | Histopath | NSW Health Pathology - Institute of Clinical Pathology and Medical Research; Westmead Hospital; University of Sydney | Arnott A., Draper J., Gall M., Martinez E., Rockett R., Sintchenko V., on behalf of ICPMR |
| EPI_ISL_3186694, EPI_ISL_3186695 | 4Cyte Pathology | NSW Health Pathology - Institute of Clinical Pathology and Medical Research; Westmead Hospital; University of Sydney | Arnott A., Draper J., Gall M., Martinez E., Rockett R., Sintchenko V., on behalf of ICPMR |
| EPI_ISL_3186696 | Australian Clinical Labs (formerly Healthscope Pathology) | NSW Health Pathology - Institute of Clinical Pathology and Medical Research; Westmead Hospital; University of Sydney | Arnott A., Draper J., Gall M., Martinez E., Rockett R., Sintchenko V., on behalf of ICPMR |
| EPI_ISL_3186697 | Douglass Hanly Moir Pathology | NSW Health Pathology - Institute of Clinical Pathology and Medical Research; Westmead Hospital; University of Sydney | Arnott A., Draper J., Gall M., Martinez E., Rockett R., Sintchenko V., on behalf of ICPMR |
| EPI_ISL_3186698, EPI_ISL_3186699, EPI_ISL_3186700, EPI_ISL_3186701, EPI_ISL_3186702, EPI_ISL_3186703, EPI_ISL_3186704 | Sydney South West Pathology Service (SSWPS) - Liverpool Hospital - NSW Health Pathology | NSW Health Pathology - Institute of Clinical Pathology and Medical Research; Westmead Hospital; University of Sydney | Arnott A., Draper J., Gall M., Martinez E., Rockett R., Sintchenko V., on behalf of ICPMR |
| EPI_ISL_3186705, EPI_ISL_3186706, EPI_ISL_3186707, EPI_ISL_3186708, EPI_ISL_3186709, EPI_ISL_3186710, EPI_ISL_3186711, EPI_ISL_3186712, EPI_ISL_3186713, EPI_ISL_3186714 | Medlab Pathology | NSW Health Pathology - Institute of Clinical Pathology and Medical Research; Westmead Hospital; University of Sydney | Arnott A., Draper J., Gall M., Martinez E., Rockett R., Sintchenko V., on behalf of ICPMR |
| EPI_ISL_3186715, EPI_ISL_3186716, EPI_ISL_3186717, EPI_ISL_3186718 | St Vincent's Pathology (SydPath) | NSW Health Pathology - Institute of Clinical Pathology and Medical Research; Westmead Hospital; University of Sydney | Arnott A., Draper J., Gall M., Martinez E., Rockett R., Sintchenko V., on behalf of ICPMR |
| EPI_ISL_3186719 | Pathology West - NSW Health Pathology | NSW Health Pathology - Institute of Clinical Pathology and Medical Research; Westmead Hospital; University of Sydney | Arnott A., Draper J., Gall M., Martinez E., Rockett R., Sintchenko V., on behalf of ICPMR |
| EPI_ISL_3186720, EPI_ISL_3186721 | Medlab Pathology | NSW Health Pathology - Institute of Clinical Pathology and Medical Research; Westmead Hospital; University of Sydney | Arnott A., Draper J., Gall M., Martinez E., Rockett R., Sintchenko V., on behalf of ICPMR |
| EPI_ISL_3186722, EPI_ISL_3186723 | Histopath | NSW Health Pathology - Institute of Clinical Pathology and Medical Research; Westmead Hospital; University of Sydney | Arnott A., Draper J., Gall M., Martinez E., Rockett R., Sintchenko V., on behalf of ICPMR |
| EPI_ISL_3186724 | 4Cyte Pathology | NSW Health Pathology - Institute of Clinical Pathology and Medical Research; Westmead Hospital; University of Sydney | Arnott A., Draper J., Gall M., Martinez E., Rockett R., Sintchenko V., on behalf of ICPMR |
| EPI_ISL_3186725, EPI_ISL_3186726 | Australian Clinical Labs (formerly Healthscope Pathology) | NSW Health Pathology - Institute of Clinical Pathology and Medical Research; Westmead Hospital; University of Sydney | Arnott A., Draper J., Gall M., Martinez E., Rockett R., Sintchenko V., on behalf of ICPMR |
| EPI_ISL_3186727, EPI_ISL_3186728 | Laverty Pathology | NSW Health Pathology - Institute of Clinical Pathology and Medical Research; Westmead Hospital; University of Sydney | Arnott A., Draper J., Gall M., Martinez E., Rockett R., Sintchenko V., on behalf of ICPMR |
| EPI_ISL_3186729, EPI_ISL_3186730, EPI_ISL_3186731, EPI_ISL_3186732, EPI_ISL_3186733, EPI_ISL_3186734, EPI_ISL_3186735 | Medlab Pathology | NSW Health Pathology - Institute of Clinical Pathology and Medical Research; Westmead Hospital; University of Sydney | Arnott A., Draper J., Gall M., Martinez E., Rockett R., Sintchenko V., on behalf of ICPMR |

| | | | |
|---|---|---|---|
| EPI_ISL_3186736 | 4Cyte Pathology | NSW Health Pathology - Institute of Clinical Pathology and Medical Research; Westmead Hospital; University of Sydney | Arnott A., Draper J., Gall M., Martinez E., Rockett R., Sintchenko V., on behalf of ICPMR |
| EPI_ISL_3186737, EPI_ISL_3186738, EPI_ISL_3186739, EPI_ISL_3186740, EPI_ISL_3186741, EPI_ISL_3186742 | Medlab Pathology | NSW Health Pathology - Institute of Clinical Pathology and Medical Research; Westmead Hospital; University of Sydney | Arnott A., Draper J., Gall M., Martinez E., Rockett R., Sintchenko V., on behalf of ICPMR |
| EPI_ISL_3186743, EPI_ISL_3186744 | Sydney South West Pathology Service (SSWPS) - Liverpool Hospital - NSW Health Pathology | NSW Health Pathology - Institute of Clinical Pathology and Medical Research; Westmead Hospital; University of Sydney | Arnott A., Draper J., Gall M., Martinez E., Rockett R., Sintchenko V., on behalf of ICPMR |
| EPI_ISL_3186745, EPI_ISL_3186746, EPI_ISL_3186747 | Laverty Pathology | NSW Health Pathology - Institute of Clinical Pathology and Medical Research; Westmead Hospital; University of Sydney | Arnott A., Draper J., Gall M., Martinez E., Rockett R., Sintchenko V., on behalf of ICPMR |
| EPI_ISL_3186748 | The Children's Hospital at Westmead | NSW Health Pathology - Institute of Clinical Pathology and Medical Research; Westmead Hospital; University of Sydney | Arnott A., Draper J., Gall M., Martinez E., Rockett R., Sintchenko V., on behalf of ICPMR |
| EPI_ISL_3186749, EPI_ISL_3186750 | Laverty Pathology | NSW Health Pathology - Institute of Clinical Pathology and Medical Research; Westmead Hospital; University of Sydney | Arnott A., Draper J., Gall M., Martinez E., Rockett R., Sintchenko V., on behalf of ICPMR |
| EPI_ISL_3186751 | St Vincent's Pathology (SydPath) | NSW Health Pathology - Institute of Clinical Pathology and Medical Research; Westmead Hospital; University of Sydney | Arnott A., Draper J., Gall M., Martinez E., Rockett R., Sintchenko V., on behalf of ICPMR |
| EPI_ISL_3186752 | The Children's Hospital at Westmead | NSW Health Pathology - Institute of Clinical Pathology and Medical Research; Westmead Hospital; University of Sydney | Arnott A., Draper J., Gall M., Martinez E., Rockett R., Sintchenko V., on behalf of ICPMR |
| EPI_ISL_3186753 | Douglass Hanly Moir Pathology | NSW Health Pathology - Institute of Clinical Pathology and Medical Research; Westmead Hospital; University of Sydney | Arnott A., Draper J., Gall M., Martinez E., Rockett R., Sintchenko V., on behalf of ICPMR |
| EPI_ISL_3186754, EPI_ISL_3186755 | Laverty Pathology | NSW Health Pathology - Institute of Clinical Pathology and Medical Research; Westmead Hospital; University of Sydney | Arnott A., Draper J., Gall M., Martinez E., Rockett R., Sintchenko V., on behalf of ICPMR |
| EPI_ISL_3186756 | The Children's Hospital at Westmead | NSW Health Pathology - Institute of Clinical Pathology and Medical Research; Westmead Hospital; University of Sydney | Arnott A., Draper J., Gall M., Martinez E., Rockett R., Sintchenko V., on behalf of ICPMR |
| EPI_ISL_3186757, EPI_ISL_3186758, EPI_ISL_3186759, EPI_ISL_3186760, EPI_ISL_3186761, EPI_ISL_3186762, EPI_ISL_3186763, EPI_ISL_3186764 | Laverty Pathology | NSW Health Pathology - Institute of Clinical Pathology and Medical Research; Westmead Hospital; University of Sydney | Arnott A., Draper J., Gall M., Martinez E., Rockett R., Sintchenko V., on behalf of ICPMR |
| EPI_ISL_3186765 | Pathology North - Royal North Shore Hospital - NSW Health Pathology | NSW Health Pathology - Institute of Clinical Pathology and Medical Research; Westmead Hospital; University of Sydney | Arnott A., Draper J., Gall M., Martinez E., Rockett R., Sintchenko V., on behalf of ICPMR |
| EPI_ISL_3186766, EPI_ISL_3186767, EPI_ISL_3186768, EPI_ISL_3186769, EPI_ISL_3186770, EPI_ISL_3186771, EPI_ISL_3186772, EPI_ISL_3186773, EPI_ISL_3186774, EPI_ISL_3186775, EPI_ISL_3186776, EPI_ISL_3186777, EPI_ISL_3186778, EPI_ISL_3186779, EPI_ISL_3186780, EPI_ISL_3186781, EPI_ISL_3186782, EPI_ISL_3186783, EPI_ISL_3186784, EPI_ISL_3186785, EPI_ISL_3186786, EPI_ISL_3186787, EPI_ISL_3186788, EPI_ISL_3186789, EPI_ISL_3186790, EPI_ISL_3186791, EPI_ISL_3186792, EPI_ISL_3186793, EPI_ISL_3186794, EPI_ISL_3186795, EPI_ISL_3186796, EPI_ISL_3186797, EPI_ISL_3186798, EPI_ISL_3186799, EPI_ISL_3186800, EPI_ISL_3186801 | | | |
| see above | Histopath | NSW Health Pathology - Institute of Clinical Pathology and Medical Research; Westmead Hospital; University of Sydney | Arnott A., Draper J., Gall M., Martinez E., Rockett R., Sintchenko V., on behalf of ICPMR |
| EPI_ISL_3186802 | Sydney South West Pathology Service (SSWPS) - Liverpool Hospital - NSW Health Pathology | NSW Health Pathology - Institute of Clinical Pathology and Medical Research; Westmead Hospital; University of Sydney | Arnott A., Draper J., Gall M., Martinez E., Rockett R., Sintchenko V., on behalf of ICPMR |
| EPI_ISL_3186803, EPI_ISL_3186804, EPI_ISL_3186805, EPI_ISL_3186806, EPI_ISL_3186807, EPI_ISL_3186808, EPI_ISL_3186809, EPI_ISL_3186810, EPI_ISL_3186811, EPI_ISL_3186812, EPI_ISL_3186813, EPI_ISL_3186814, EPI_ISL_3186815, EPI_ISL_3186816, EPI_ISL_3186817, EPI_ISL_3186818, EPI_ISL_3186819, EPI_ISL_3186820, EPI_ISL_3186821, EPI_ISL_3186822, EPI_ISL_3186823, EPI_ISL_3186824, EPI_ISL_3186825, EPI_ISL_3186826, EPI_ISL_3186827, EPI_ISL_3186828, EPI_ISL_3186829, EPI_ISL_3186830, EPI_ISL_3186831, EPI_ISL_3186832, EPI_ISL_3186833, EPI_ISL_3186834, EPI_ISL_3186835 | | | |
| see above | Histopath | NSW Health Pathology - Institute of Clinical Pathology and Medical Research; Westmead Hospital; University of Sydney | Arnott A., Draper J., Gall M., Martinez E., Rockett R., Sintchenko V., on behalf of ICPMR |
| EPI_ISL_3186836, EPI_ISL_3186837, EPI_ISL_3186838, EPI_ISL_3186839, EPI_ISL_3186840, EPI_ISL_3186841, EPI_ISL_3186842, EPI_ISL_3186843 | 4Cyte Pathology | NSW Health Pathology - Institute of Clinical Pathology and Medical Research; Westmead Hospital; University of Sydney | Arnott A., Draper J., Gall M., Martinez E., Rockett R., Sintchenko V., on behalf of ICPMR |
| EPI_ISL_3186844, EPI_ISL_3186845, EPI_ISL_3186846, EPI_ISL_3186847, EPI_ISL_3186848, EPI_ISL_3186849, EPI_ISL_3186850, EPI_ISL_3186851, EPI_ISL_3186852, EPI_ISL_3186853, EPI_ISL_3186854, EPI_ISL_3186855, EPI_ISL_3186856, EPI_ISL_3186857, EPI_ISL_3186858, EPI_ISL_3186859, EPI_ISL_3186860, EPI_ISL_3186861, EPI_ISL_3186862, EPI_ISL_3186863, EPI_ISL_3186864, EPI_ISL_3186865 | | | |
| see above | Medlab Pathology | NSW Health Pathology - Institute of Clinical Pathology and Medical Research; Westmead Hospital; University of Sydney | Arnott A., Draper J., Gall M., Martinez E., Rockett R., Sintchenko V., on behalf of ICPMR |
| EPI_ISL_3186866, EPI_ISL_3186867, EPI_ISL_3186868, EPI_ISL_3186869, EPI_ISL_3186870, EPI_ISL_3186871, EPI_ISL_3186872, EPI_ISL_3186873, EPI_ISL_3186874 | 4Cyte Pathology | NSW Health Pathology - Institute of Clinical Pathology and Medical Research; Westmead Hospital; University of Sydney | Arnott A., Draper J., Gall M., Martinez E., Rockett R., Sintchenko V., on behalf of ICPMR |
| EPI_ISL_3186875, EPI_ISL_3186876 | Australian Clinical Labs (formerly Healthscope Pathology) | NSW Health Pathology - Institute of Clinical Pathology and Medical Research; Westmead Hospital; University of Sydney | Arnott A., Draper J., Gall M., Martinez E., Rockett R., Sintchenko V., on behalf of ICPMR |
| EPI_ISL_3186877, EPI_ISL_3186878, EPI_ISL_3186879, EPI_ISL_3186880, EPI_ISL_3186881 | Douglass Hanly Moir Pathology | NSW Health Pathology - Institute of Clinical Pathology and Medical Research; Westmead Hospital; University of Sydney | Arnott A., Draper J., Gall M., Martinez E., Rockett R., Sintchenko V., on behalf of ICPMR |
| EPI_ISL_3186882, EPI_ISL_3186883, EPI_ISL_3186884, EPI_ISL_3186885, EPI_ISL_3186886, EPI_ISL_3186887, EPI_ISL_3186888, EPI_ISL_3186889, EPI_ISL_3186890, EPI_ISL_3186891, EPI_ISL_3186892, EPI_ISL_3186893, EPI_ISL_3186894, EPI_ISL_3186895, EPI_ISL_3186896, EPI_ISL_3186897 | | | |
| see above | Laverty Pathology | NSW Health Pathology - Institute of Clinical Pathology and Medical Research; Westmead Hospital; University of Sydney | Arnott A., Draper J., Gall M., Martinez E., Rockett R., Sintchenko V., on behalf of ICPMR |
| EPI_ISL_3186898, EPI_ISL_3186899, EPI_ISL_3186900, EPI_ISL_3186901, EPI_ISL_3186902, EPI_ISL_3186903 | Sydney South West Pathology Service (SSWPS) - Liverpool Hospital - NSW Health Pathology | NSW Health Pathology - Institute of Clinical Pathology and Medical Research; Westmead Hospital; University of Sydney | Arnott A., Draper J., Gall M., Martinez E., Rockett R., Sintchenko V., on behalf of ICPMR |
| EPI_ISL_3186904, EPI_ISL_3186905, EPI_ISL_3186906, EPI_ISL_3186907, EPI_ISL_3186908, EPI_ISL_3186909 | The Children's Hospital at Westmead | NSW Health Pathology - Institute of Clinical Pathology and Medical Research; Westmead Hospital; University of Sydney | Arnott A., Draper J., Gall M., Martinez E., Rockett R., Sintchenko V., on behalf of ICPMR |
| EPI_ISL_3186910, EPI_ISL_3186911, EPI_ISL_3186912, EPI_ISL_3186913 | Pathology North - Royal North Shore Hospital - NSW Health Pathology | NSW Health Pathology - Institute of Clinical Pathology and Medical Research; Westmead Hospital; University of Sydney | Arnott A., Draper J., Gall M., Martinez E., Rockett R., Sintchenko V., on behalf of ICPMR |
| EPI_ISL_3186914, EPI_ISL_3186915, EPI_ISL_3186916, EPI_ISL_3186917, EPI_ISL_3186918, EPI_ISL_3186919, EPI_ISL_3186920, EPI_ISL_3186921, EPI_ISL_3186922, EPI_ISL_3186923, EPI_ISL_3186924, EPI_ISL_3186925, EPI_ISL_3186926, EPI_ISL_3186927, EPI_ISL_3186928, EPI_ISL_3186929, EPI_ISL_3186930, EPI_ISL_3186931, EPI_ISL_3186932 | | | |
| see above | St Vincent's Pathology (SydPath) | NSW Health Pathology - Institute of Clinical Pathology and Medical Research; Westmead Hospital; University of Sydney | Arnott A., Draper J., Gall M., Martinez E., Rockett R., Sintchenko V., on behalf of ICPMR |
| EPI_ISL_3186933, EPI_ISL_3186934, EPI_ISL_3186935, EPI_ISL_3186936, EPI_ISL_3186937, EPI_ISL_3186938, EPI_ISL_3186939, EPI_ISL_3186940, EPI_ISL_3186941, EPI_ISL_3186942, EPI_ISL_3186943, EPI_ISL_3186944, EPI_ISL_3186945, EPI_ISL_3186946, EPI_ISL_3186947, EPI_ISL_3186948, EPI_ISL_3186949, EPI_ISL_3186950 | | | |

| | | | |
|---|---|---|---|
| see above | Histopath | NSW Health Pathology - Institute of Clinical Pathology and Medical Research; Westmead Hospital; University of Sydney | Arnott A., Draper J., Gall M., Martinez E., Rockett R., Sintchenko V., on behalf of ICPMR |
| EPI_ISL_3186951 | Pathology West - NSW Health Pathology | NSW Health Pathology - Institute of Clinical Pathology and Medical Research; Westmead Hospital; University of Sydney | Arnott A., Draper J., Gall M., Martinez E., Rockett R., Sintchenko V., on behalf of ICPMR |
| EPI_ISL_3186952, EPI_ISL_3186953, EPI_ISL_3186954, EPI_ISL_3186955, EPI_ISL_3186956, EPI_ISL_3186957, EPI_ISL_3186958, EPI_ISL_3186959, EPI_ISL_3186960, EPI_ISL_3186961, EPI_ISL_3186962, EPI_ISL_3186963, EPI_ISL_3186964, EPI_ISL_3186965, EPI_ISL_3186966, EPI_ISL_3186967, EPI_ISL_3186968, EPI_ISL_3186969, EPI_ISL_3186970 | | | |
| see above | 4Cyte Pathology | NSW Health Pathology - Institute of Clinical Pathology and Medical Research; Westmead Hospital; University of Sydney | Arnott A., Draper J., Gall M., Martinez E., Rockett R., Sintchenko V., on behalf of ICPMR |
| EPI_ISL_3186971 | Australian Clinical Labs (formerly Healthscope Pathology) | NSW Health Pathology - Institute of Clinical Pathology and Medical Research; Westmead Hospital; University of Sydney | Arnott A., Draper J., Gall M., Martinez E., Rockett R., Sintchenko V., on behalf of ICPMR |
| EPI_ISL_3186972 | Douglass Hanly Moir Pathology | NSW Health Pathology - Institute of Clinical Pathology and Medical Research; Westmead Hospital; University of Sydney | Arnott A., Draper J., Gall M., Martinez E., Rockett R., Sintchenko V., on behalf of ICPMR |
| EPI_ISL_3186973, EPI_ISL_3186974, EPI_ISL_3186975, EPI_ISL_3186976, EPI_ISL_3186977, EPI_ISL_3186978, EPI_ISL_3186979, EPI_ISL_3186980, EPI_ISL_3186981, EPI_ISL_3186982, EPI_ISL_3186983, EPI_ISL_3186984, EPI_ISL_3186985, EPI_ISL_3186986, EPI_ISL_3186987, EPI_ISL_3186988 | | | |
| see above | Sydney South West Pathology Service (SSWPS) - Liverpool Hospital - NSW Health Pathology | NSW Health Pathology - Institute of Clinical Pathology and Medical Research; Westmead Hospital; University of Sydney | Arnott A., Draper J., Gall M., Martinez E., Rockett R., Sintchenko V., on behalf of ICPMR |
| EPI_ISL_3186989 | Sydney South West Pathology Service (SSWPS) - Concord Repatriation General Hospital - NSW Health Pathology | NSW Health Pathology - Institute of Clinical Pathology and Medical Research; Westmead Hospital; University of Sydney | Arnott A., Draper J., Gall M., Martinez E., Rockett R., Sintchenko V., on behalf of ICPMR |
| EPI_ISL_3186990, EPI_ISL_3186991, EPI_ISL_3186992, EPI_ISL_3186993, EPI_ISL_3186994, EPI_ISL_3186995 | Sydney South West Pathology Service (SSWPS) - Liverpool Hospital - NSW Health Pathology | NSW Health Pathology - Institute of Clinical Pathology and Medical Research; Westmead Hospital; University of Sydney | Arnott A., Draper J., Gall M., Martinez E., Rockett R., Sintchenko V., on behalf of ICPMR |
| EPI_ISL_3186996, EPI_ISL_3186997, EPI_ISL_3186998 | Medlab Pathology | NSW Health Pathology - Institute of Clinical Pathology and Medical Research; Westmead Hospital; University of Sydney | Arnott A., Draper J., Gall M., Martinez E., Rockett R., Sintchenko V., on behalf of ICPMR |
| EPI_ISL_3186999, EPI_ISL_3187000, EPI_ISL_3187001, EPI_ISL_3187002, EPI_ISL_3187003, EPI_ISL_3187004, EPI_ISL_3187005, EPI_ISL_3187006, EPI_ISL_3187007, EPI_ISL_3187008, EPI_ISL_3187009, EPI_ISL_3187010, EPI_ISL_3187011, EPI_ISL_3187012, EPI_ISL_3187013, EPI_ISL_3187014, EPI_ISL_3187015, EPI_ISL_3187016 | | | |
| see above | St Vincent's Pathology (SydPath) | NSW Health Pathology - Institute of Clinical Pathology and Medical Research; Westmead Hospital; University of Sydney | Arnott A., Draper J., Gall M., Martinez E., Rockett R., Sintchenko V., on behalf of ICPMR |
| EPI_ISL_3187017, EPI_ISL_3187018, EPI_ISL_3187019 | Pathology West - NSW Health Pathology | NSW Health Pathology - Institute of Clinical Pathology and Medical Research; Westmead Hospital; University of Sydney | Arnott A., Draper J., Gall M., Martinez E., Rockett R., Sintchenko V., on behalf of ICPMR |
| EPI_ISL_3187020 | Austech Medical Laboratories | NSW Health Pathology - Institute of Clinical Pathology and Medical Research; Westmead Hospital; University of Sydney | Arnott A., Draper J., Gall M., Martinez E., Rockett R., Sintchenko V., on behalf of ICPMR |
| EPI_ISL_3187021, EPI_ISL_3187022, EPI_ISL_3187023, EPI_ISL_3187024, EPI_ISL_3187025, EPI_ISL_3187026, EPI_ISL_3187027, EPI_ISL_3187028, EPI_ISL_3187029, EPI_ISL_3187030, EPI_ISL_3187031, EPI_ISL_3187032, EPI_ISL_3187033, EPI_ISL_3187034, EPI_ISL_3187035, EPI_ISL_3187036, EPI_ISL_3187037, EPI_ISL_3187038, EPI_ISL_3187039, EPI_ISL_3187040, EPI_ISL_3187041, EPI_ISL_3187042, EPI_ISL_3187043, EPI_ISL_3187044 | | | |
| see above | Histopath | NSW Health Pathology - Institute of Clinical Pathology and Medical Research; Westmead Hospital; University of Sydney | Arnott A., Draper J., Gall M., Martinez E., Rockett R., Sintchenko V., on behalf of ICPMR |
| EPI_ISL_3187045, EPI_ISL_3187046, EPI_ISL_3187047, EPI_ISL_3187048, EPI_ISL_3187049, EPI_ISL_3187050, EPI_ISL_3187051, EPI_ISL_3187052, EPI_ISL_3187053, EPI_ISL_3187054, EPI_ISL_3187055, EPI_ISL_3187056, EPI_ISL_3187057, EPI_ISL_3187058, EPI_ISL_3187059, EPI_ISL_3187060, EPI_ISL_3187061, EPI_ISL_3187062, EPI_ISL_3187063, EPI_ISL_3187064 | | | |
| see above | 4Cyte Pathology | NSW Health Pathology - Institute of Clinical Pathology and Medical Research; Westmead Hospital; University of Sydney | Arnott A., Draper J., Gall M., Martinez E., Rockett R., Sintchenko V., on behalf of ICPMR |
| EPI_ISL_3187065, EPI_ISL_3187066, EPI_ISL_3187067, EPI_ISL_3187068, EPI_ISL_3187069, EPI_ISL_3187070, EPI_ISL_3187071, EPI_ISL_3187072, EPI_ISL_3187073, EPI_ISL_3187074, EPI_ISL_3187075 | | | |
| see above | Australian Clinical Labs (formerly Healthscope Pathology) | NSW Health Pathology - Institute of Clinical Pathology and Medical Research; Westmead Hospital; University of Sydney | Arnott A., Draper J., Gall M., Martinez E., Rockett R., Sintchenko V., on behalf of ICPMR |
| EPI_ISL_3187076, EPI_ISL_3187077, EPI_ISL_3187078, EPI_ISL_3187079, EPI_ISL_3187080, EPI_ISL_3187081, EPI_ISL_3187082, EPI_ISL_3187083, EPI_ISL_3187084, EPI_ISL_3187085, EPI_ISL_3187086, EPI_ISL_3187087, EPI_ISL_3187088, EPI_ISL_3187089 | | | |
| see above | Laverty Pathology | NSW Health Pathology - Institute of Clinical Pathology and Medical Research; Westmead Hospital; University of Sydney | Arnott A., Draper J., Gall M., Martinez E., Rockett R., Sintchenko V., on behalf of ICPMR |
| EPI_ISL_3187090, EPI_ISL_3187091, EPI_ISL_3187092, EPI_ISL_3187093 | Sydney South West Pathology Service (SSWPS) - Liverpool Hospital - NSW Health Pathology | NSW Health Pathology - Institute of Clinical Pathology and Medical Research; Westmead Hospital; University of Sydney | Arnott A., Draper J., Gall M., Martinez E., Rockett R., Sintchenko V., on behalf of ICPMR |
| EPI_ISL_3187094, EPI_ISL_3187095, EPI_ISL_3187096, EPI_ISL_3187097, EPI_ISL_3187098, EPI_ISL_3187099, EPI_ISL_3187100, EPI_ISL_3187101, EPI_ISL_3187102 | Medlab Pathology | NSW Health Pathology - Institute of Clinical Pathology and Medical Research; Westmead Hospital; University of Sydney | Arnott A., Draper J., Gall M., Martinez E., Rockett R., Sintchenko V., on behalf of ICPMR |
| EPI_ISL_3187103 | Southern. IML Pathology | NSW Health Pathology - Institute of Clinical Pathology and Medical Research; Westmead Hospital; University of Sydney | Arnott A., Draper J., Gall M., Martinez E., Rockett R., Sintchenko V., on behalf of ICPMR |
| EPI_ISL_3187104, EPI_ISL_3187105, EPI_ISL_3187106, EPI_ISL_3187107, EPI_ISL_3187108 | St Vincent's Pathology (SydPath) | NSW Health Pathology - Institute of Clinical Pathology and Medical Research; Westmead Hospital; University of Sydney | Arnott A., Draper J., Gall M., Martinez E., Rockett R., Sintchenko V., on behalf of ICPMR |
| EPI_ISL_3187109, EPI_ISL_3187110, EPI_ISL_3187111, EPI_ISL_3187112, EPI_ISL_3187113, EPI_ISL_3187114 | Histopath | NSW Health Pathology - Institute of Clinical Pathology and Medical Research; Westmead Hospital; University of Sydney | Arnott A., Draper J., Gall M., Martinez E., Rockett R., Sintchenko V., on behalf of ICPMR |
| EPI_ISL_3187115, EPI_ISL_3187116, EPI_ISL_3187117, EPI_ISL_3187118, EPI_ISL_3187119, EPI_ISL_3187120, EPI_ISL_3187121, EPI_ISL_3187122, EPI_ISL_3187123, EPI_ISL_3187124, EPI_ISL_3187125, EPI_ISL_3187126, EPI_ISL_3187127, EPI_ISL_3187128, EPI_ISL_3187129, EPI_ISL_3187130, EPI_ISL_3187131 | | | |
| see above | 4Cyte Pathology | NSW Health Pathology - Institute of Clinical Pathology and Medical Research; Westmead Hospital; University of Sydney | Arnott A., Draper J., Gall M., Martinez E., Rockett R., Sintchenko V., on behalf of ICPMR |
| EPI_ISL_3187132, EPI_ISL_3187133, EPI_ISL_3187134, EPI_ISL_3187135, EPI_ISL_3187136, EPI_ISL_3187137, EPI_ISL_3187138, EPI_ISL_3187139, EPI_ISL_3187140, EPI_ISL_3187141, EPI_ISL_3187142, EPI_ISL_3187143, EPI_ISL_3187144, EPI_ISL_3187145, EPI_ISL_3187146, EPI_ISL_3187147, EPI_ISL_3187148, EPI_ISL_3187149, EPI_ISL_3187150, EPI_ISL_3187151 | | | |
| see above | Medlab Pathology | NSW Health Pathology - Institute of Clinical Pathology and Medical Research; Westmead Hospital; University of Sydney | Arnott A., Draper J., Gall M., Martinez E., Rockett R., Sintchenko V., on behalf of ICPMR |
| EPI_ISL_3187152, EPI_ISL_3187153, EPI_ISL_3187154, EPI_ISL_3187155, EPI_ISL_3187156, EPI_ISL_3187157, EPI_ISL_3187158 | Sydney South West Pathology Service (SSWPS) - Liverpool Hospital - NSW Health Pathology | NSW Health Pathology - Institute of Clinical Pathology and Medical Research; Westmead Hospital; University of Sydney | Arnott A., Draper J., Gall M., Martinez E., Rockett R., Sintchenko V., on behalf of ICPMR |
| EPI_ISL_3187159 | Pathology North - Royal North Shore Hospital - NSW Health Pathology | NSW Health Pathology - Institute of Clinical Pathology and Medical Research; Westmead Hospital; University of Sydney | Arnott A., Draper J., Gall M., Martinez E., Rockett R., Sintchenko V., on behalf of ICPMR |
| EPI_ISL_3187160, EPI_ISL_3187161, EPI_ISL_3187162, EPI_ISL_3187163, EPI_ISL_3187164, EPI_ISL_3187165 | Sydney South West Pathology Service (SSWPS) - Liverpool Hospital - NSW Health Pathology | NSW Health Pathology - Institute of Clinical Pathology and Medical Research; Westmead Hospital; University of Sydney | Arnott A., Draper J., Gall M., Martinez E., Rockett R., Sintchenko V., on behalf of ICPMR |
| EPI_ISL_3187166 | Pathology West - NSW Health Pathology | NSW Health Pathology - Institute of Clinical Pathology and Medical | Arnott A., Draper J., Gall M., Martinez E., Rockett R., Sintchenko V., on behalf of ICPMR |

| | | | |
|---|---|---|---|
| | | Research; Westmead Hospital; University of Sydney | |
| EPI_ISL_3187167, EPI_ISL_3187168, EPI_ISL_3187169 | Australian Clinical Labs (formerly Healthscope Pathology) | NSW Health Pathology - Institute of Clinical Pathology and Medical Research; Westmead Hospital; University of Sydney | Arnott A., Draper J., Gall M., Martinez E., Rockett R., Sintchenko V., on behalf of ICPMR |
| EPI_ISL_3187170, EPI_ISL_3187171, EPI_ISL_3187172, EPI_ISL_3187173, EPI_ISL_3187174 | Laverty Pathology | NSW Health Pathology - Institute of Clinical Pathology and Medical Research; Westmead Hospital; University of Sydney | Arnott A., Draper J., Gall M., Martinez E., Rockett R., Sintchenko V., on behalf of ICPMR |
| EPI_ISL_3187175 | The Children's Hospital at Westmead | NSW Health Pathology - Institute of Clinical Pathology and Medical Research; Westmead Hospital; University of Sydney | Arnott A., Draper J., Gall M., Martinez E., Rockett R., Sintchenko V., on behalf of ICPMR |
| EPI_ISL_3187176 | Australian Clinical Labs (formerly Healthscope Pathology) | NSW Health Pathology - Institute of Clinical Pathology and Medical Research; Westmead Hospital; University of Sydney | Arnott A., Draper J., Gall M., Martinez E., Rockett R., Sintchenko V., on behalf of ICPMR |
| EPI_ISL_3187177, EPI_ISL_3187178, EPI_ISL_3187179, EPI_ISL_3187180, EPI_ISL_3187181, EPI_ISL_3187182, EPI_ISL_3187183, EPI_ISL_3187184, EPI_ISL_3187185, EPI_ISL_3187186, EPI_ISL_3187187, EPI_ISL_3187188, EPI_ISL_3187189, EPI_ISL_3187190, EPI_ISL_3187191 | | | |
| see above | Laverty Pathology | NSW Health Pathology - Institute of Clinical Pathology and Medical Research; Westmead Hospital; University of Sydney | Arnott A., Draper J., Gall M., Martinez E., Rockett R., Sintchenko V., on behalf of ICPMR |
| EPI_ISL_3187192, EPI_ISL_3187193, EPI_ISL_3187194, EPI_ISL_3187195, EPI_ISL_3187196 | Medlab Pathology | NSW Health Pathology - Institute of Clinical Pathology and Medical Research; Westmead Hospital; University of Sydney | Arnott A., Draper J., Gall M., Martinez E., Rockett R., Sintchenko V., on behalf of ICPMR |
| EPI_ISL_3187197 | Laverty Pathology | NSW Health Pathology - Institute of Clinical Pathology and Medical Research; Westmead Hospital; University of Sydney | Arnott A., Draper J., Gall M., Martinez E., Rockett R., Sintchenko V., on behalf of ICPMR |
| EPI_ISL_3187198, EPI_ISL_3187199 | St Vincent's Pathology (SydPath) | NSW Health Pathology - Institute of Clinical Pathology and Medical Research; Westmead Hospital; University of Sydney | Arnott A., Draper J., Gall M., Martinez E., Rockett R., Sintchenko V., on behalf of ICPMR |
| EPI_ISL_3187200 | Laverty Pathology | NSW Health Pathology - Institute of Clinical Pathology and Medical Research; Westmead Hospital; University of Sydney | Arnott A., Draper J., Gall M., Martinez E., Rockett R., Sintchenko V., on behalf of ICPMR |
| EPI_ISL_3187201, EPI_ISL_3187202 | St Vincent's Pathology (SydPath) | NSW Health Pathology - Institute of Clinical Pathology and Medical Research; Westmead Hospital; University of Sydney | Arnott A., Draper J., Gall M., Martinez E., Rockett R., Sintchenko V., on behalf of ICPMR |
| EPI_ISL_3187203, EPI_ISL_3187204, EPI_ISL_3187205, EPI_ISL_3187206, EPI_ISL_3187207, EPI_ISL_3187208, EPI_ISL_3187209, EPI_ISL_3187210 | Laverty Pathology | NSW Health Pathology - Institute of Clinical Pathology and Medical Research; Westmead Hospital; University of Sydney | Arnott A., Draper J., Gall M., Martinez E., Rockett R., Sintchenko V., on behalf of ICPMR |
| EPI_ISL_3187211, EPI_ISL_3187212 | The Children's Hospital at Westmead | NSW Health Pathology - Institute of Clinical Pathology and Medical Research; Westmead Hospital; University of Sydney | Arnott A., Draper J., Gall M., Martinez E., Rockett R., Sintchenko V., on behalf of ICPMR |
| EPI_ISL_3187213, EPI_ISL_3187214 | Histopath | NSW Health Pathology - Institute of Clinical Pathology and Medical Research; Westmead Hospital; University of Sydney | Arnott A., Draper J., Gall M., Martinez E., Rockett R., Sintchenko V., on behalf of ICPMR |
| EPI_ISL_3187215 | 4Cyte Pathology | NSW Health Pathology - Institute of Clinical Pathology and Medical Research; Westmead Hospital; University of Sydney | Arnott A., Draper J., Gall M., Martinez E., Rockett R., Sintchenko V., on behalf of ICPMR |
| EPI_ISL_3187216 | Australian Clinical Labs (formerly Healthscope Pathology) | NSW Health Pathology - Institute of Clinical Pathology and Medical Research; Westmead Hospital; University of Sydney | Arnott A., Draper J., Gall M., Martinez E., Rockett R., Sintchenko V., on behalf of ICPMR |
| EPI_ISL_3187217 | Medlab Pathology | NSW Health Pathology - Institute of Clinical Pathology and Medical Research; Westmead Hospital; University of Sydney | Arnott A., Draper J., Gall M., Martinez E., Rockett R., Sintchenko V., on behalf of ICPMR |
| EPI_ISL_3187218 | Laverty Pathology | NSW Health Pathology - Institute of Clinical Pathology and Medical Research; Westmead Hospital; University of Sydney | Arnott A., Draper J., Gall M., Martinez E., Rockett R., Sintchenko V., on behalf of ICPMR |
| EPI_ISL_3187219, EPI_ISL_3187220 | Histopath | NSW Health Pathology - Institute of Clinical Pathology and Medical Research; Westmead Hospital; University of Sydney | Arnott A., Draper J., Gall M., Martinez E., Rockett R., Sintchenko V., on behalf of ICPMR |
| EPI_ISL_3187221 | St Vincent's Pathology (SydPath) | NSW Health Pathology - Institute of Clinical Pathology and Medical Research; Westmead Hospital; University of Sydney | Arnott A., Draper J., Gall M., Martinez E., Rockett R., Sintchenko V., on behalf of ICPMR |
| EPI_ISL_3187222 | Laverty Pathology | NSW Health Pathology - Institute of Clinical Pathology and Medical Research; Westmead Hospital; University of Sydney | Arnott A., Draper J., Gall M., Martinez E., Rockett R., Sintchenko V., on behalf of ICPMR |
| EPI_ISL_3187223, EPI_ISL_3187224 | St Vincent's Pathology (SydPath) | NSW Health Pathology - Institute of Clinical Pathology and Medical Research; Westmead Hospital; University of Sydney | Arnott A., Draper J., Gall M., Martinez E., Rockett R., Sintchenko V., on behalf of ICPMR |
| EPI_ISL_3187225 | The Children's Hospital at Westmead | NSW Health Pathology - Institute of Clinical Pathology and Medical Research; Westmead Hospital; University of Sydney | Arnott A., Draper J., Gall M., Martinez E., Rockett R., Sintchenko V., on behalf of ICPMR |
| EPI_ISL_3187226 | Laverty Pathology | NSW Health Pathology - Institute of Clinical Pathology and Medical Research; Westmead Hospital; University of Sydney | Arnott A., Draper J., Gall M., Martinez E., Rockett R., Sintchenko V., on behalf of ICPMR |
| EPI_ISL_3187227 | Australian Clinical Labs (formerly Healthscope Pathology) | NSW Health Pathology - Institute of Clinical Pathology and Medical Research; Westmead Hospital; University of Sydney | Arnott A., Draper J., Gall M., Martinez E., Rockett R., Sintchenko V., on behalf of ICPMR |
| EPI_ISL_3187228 | St Vincent's Pathology (SydPath) | NSW Health Pathology - Institute of Clinical Pathology and Medical Research; Westmead Hospital; University of Sydney | Arnott A., Draper J., Gall M., Martinez E., Rockett R., Sintchenko V., on behalf of ICPMR |
| EPI_ISL_3187229 | Histopath | NSW Health Pathology - Institute of Clinical Pathology and Medical Research; Westmead Hospital; University of Sydney | Arnott A., Draper J., Gall M., Martinez E., Rockett R., Sintchenko V., on behalf of ICPMR |
| EPI_ISL_3187787, EPI_ISL_3187860, EPI_ISL_3187875, EPI_ISL_3187877, EPI_ISL_3187892, EPI_ISL_3187902, EPI_ISL_3187910, EPI_ISL_3187916, EPI_ISL_3187928, EPI_ISL_3187934, EPI_ISL_3187936, EPI_ISL_3187938, EPI_ISL_3187946, EPI_ISL_3187956 | | | |
| see above | Pathogen Genomics Center, National Institute of Infectious Diseases | Pathogen Genomics Center, National Institute of Infectious Diseases | Tsuyoshi Sekizuka, Kentaro Itokawa, Rina Tanaka, Masanori Hashino, Makoto Kuroda |
| EPI_ISL_3187994, EPI_ISL_3188005, EPI_ISL_3188006, EPI_ISL_3188007, EPI_ISL_3188008, EPI_ISL_3188009, EPI_ISL_3188010, EPI_ISL_3188011, EPI_ISL_3188012, EPI_ISL_3188013, EPI_ISL_3188014, EPI_ISL_3188015, EPI_ISL_3188016, EPI_ISL_3188017, EPI_ISL_3188018, EPI_ISL_3188019, EPI_ISL_3188020, EPI_ISL_3188021, EPI_ISL_3188022, EPI_ISL_3188023, EPI_ISL_3188024, EPI_ISL_3188025, EPI_ISL_3188028, EPI_ISL_3188029, EPI_ISL_3188030, EPI_ISL_3188031 | | | |
| see above | SARS-CoV-2 testing team, National Institute of Infectious Diseases | Pathogen Genomics Center, National Institute of Infectious Diseases | Tsuyoshi Sekizuka, Kentaro Itokawa, Rina Tanaka, Masanori Hashino, Nozomu Hanaoka, Masumichi Saito, Naomi Nojiri, Hazuka Y Furihata, Hiromizu Takahashi, Tsuguto Fujimoto, Makoto Kuroda |
| EPI_ISL_3188230 | Phnom Penh Rapid Response Team | Virology Unit, Institut Pasteur du Cambodge | Leakhena Pum, Sophoannadedh Rath, Jurre Y Siegers, Cecile Troupin, Ly Sovann, Kraing Sidonn, Yi Sengdoeurn, Chin Savuth, Chau Darapheak, Veasna Duong, Erik A Karlsson |
| EPI_ISL_3188232, EPI_ISL_3188247, EPI_ISL_3188248 | Virology Unit, Institut Pasteur du Cambodge | Virology Unit, Institut Pasteur du Cambodge | Leakhena Pum, Sophoannadedh Rath, Jurre Y Siegers, Cecile Troupin, Ly Sovann, Kraing Sidonn, Yi Sengdoeurn, Chin Savuth, Chau Darapheak, Veasna Duong, Erik A Karlsson |
| EPI_ISL_3188253, EPI_ISL_3188254, EPI_ISL_3188256, EPI_ISL_3188257, EPI_ISL_3188258, EPI_ISL_3188259 | Pailin Rapid Response Team | Virology Unit, Institut Pasteur du Cambodge | Leakhena Pum, Sophoannadedh Rath, Jurre Y Siegers, Cecile Troupin, Ly Sovann, Kraing Sidonn, Yi Sengdoeurn, Chin Savuth, Chau Darapheak, Veasna Duong, Erik A Karlsson |
| EPI_ISL_3188262, EPI_ISL_3188263, EPI_ISL_3188264, EPI_ISL_3188265, EPI_ISL_3188266, EPI_ISL_3188268, EPI_ISL_3188269, EPI_ISL_3188270, EPI_ISL_3188271, EPI_ISL_3188272, EPI_ISL_3188273, EPI_ISL_3188274, EPI_ISL_3188275, EPI_ISL_3188276, EPI_ISL_3188277, EPI_ISL_3188278, EPI_ISL_3188279, EPI_ISL_3188280, EPI_ISL_3188281, EPI_ISL_3188282, EPI_ISL_3188284, EPI_ISL_3188285, EPI_ISL_3188286, EPI_ISL_3188287, EPI_ISL_3188288, EPI_ISL_3188289, EPI_ISL_3188290, EPI_ISL_3188292, EPI_ISL_3188293, EPI_ISL_3188294, EPI_ISL_3188295, EPI_ISL_3188296, EPI_ISL_3188297, EPI_ISL_3188298, EPI_ISL_3188299, EPI_ISL_3188300, EPI_ISL_3188301, EPI_ISL_3188302, EPI_ISL_3188303, EPI_ISL_3188304, EPI_ISL_3188305, EPI_ISL_3188306, EPI_ISL_3188307, EPI_ISL_3188308, EPI_ISL_3188309, EPI_ISL_3188310, EPI_ISL_3188311, EPI_ISL_3188313, EPI_ISL_3188314, EPI_ISL_3188315, EPI_ISL_3188316, EPI_ISL_3188317, EPI_ISL_3188319, EPI_ISL_3188321, EPI_ISL_3188322, EPI_ISL_3188323, EPI_ISL_3188324, EPI_ISL_3188325, EPI_ISL_3188326, EPI_ISL_3188327, EPI_ISL_3188328, EPI_ISL_3188329, EPI_ISL_3188330, EPI_ISL_3188331, EPI_ISL_3188332, EPI_ISL_3188333, EPI_ISL_3188334, EPI_ISL_3188335, EPI_ISL_3188336, EPI_ISL_3188337, EPI_ISL_3188340, EPI_ISL_3188341, EPI_ISL_3188342, EPI_ISL_3188343, EPI_ISL_3188344, EPI_ISL_3188345, EPI_ISL_3188346, EPI_ISL_3188347, EPI_ISL_3188348, EPI_ISL_3188349, EPI_ISL_3188350, EPI_ISL_3188351, EPI_ISL_3188352, EPI_ISL_3188353, EPI_ISL_3188354, EPI_ISL_3188355, EPI_ISL_3188356, EPI_ISL_3188358, EPI_ISL_3188359, EPI_ISL_3188360, EPI_ISL_3188361, EPI_ISL_3188362, EPI_ISL_3188363, EPI_ISL_3188365, EPI_ISL_3188367, EPI_ISL_3188368, EPI_ISL_3188369, EPI_ISL_3188370, EPI_ISL_3188372, EPI_ISL_3188373, EPI_ISL_3188374, EPI_ISL_3188375, EPI_ISL_3188376, EPI_ISL_3188377, EPI_ISL_3188378, EPI_ISL_3188379, EPI_ISL_3188380, EPI_ISL_3188381, EPI_ISL_3188382, EPI_ISL_3188383, EPI_ISL_3188384, EPI_ISL_3188385, EPI_ISL_3188386, EPI_ISL_3188388, EPI_ISL_3188389, EPI_ISL_3188390, EPI_ISL_3188391, EPI_ISL_3188392, EPI_ISL_3188393, EPI_ISL_3188394, EPI_ISL_3188395, EPI_ISL_3188396, EPI_ISL_3188397, EPI_ISL_3188398, EPI_ISL_3188399, EPI_ISL_3188400, EPI_ISL_3188401, EPI_ISL_3188403, EPI_ISL_3188404, EPI_ISL_3188405, EPI_ISL_3188406, EPI_ISL_3188407, EPI_ISL_3188408, EPI_ISL_3188409, EPI_ISL_3188410, EPI_ISL_3188411, EPI_ISL_3188412, EPI_ISL_3188413, EPI_ISL_3188414, EPI_ISL_3188415, EPI_ISL_3188416, EPI_ISL_3188417, EPI_ISL_3188418, EPI_ISL_3188419, EPI_ISL_3188420, EPI_ISL_3188421, EPI_ISL_3188422, EPI_ISL_3188423, EPI_ISL_3188424, EPI_ISL_3188425, EPI_ISL_3188426, EPI_ISL_3188427, EPI_ISL_3188428, EPI_ISL_3188430, EPI_ISL_3188431, EPI_ISL_3188433, EPI_ISL_3188435, EPI_ISL_3188436, EPI_ISL_3188437, EPI_ISL_3188438, EPI_ISL_3188439, EPI_ISL_3188440, EPI_ISL_3188442, EPI_ISL_3188443, EPI_ISL_3188444, EPI_ISL_3188445, EPI_ISL_3188446, EPI_ISL_3188447, EPI_ISL_3188448, EPI_ISL_3188449, EPI_ISL_3188450, EPI_ISL_3188451, EPI_ISL_3188452, EPI_ISL_3188453, EPI_ISL_3188454, EPI_ISL_3188455, EPI_ISL_3188456, EPI_ISL_3188457, EPI_ISL_3188458, EPI_ISL_3188459, EPI_ISL_3188460, EPI_ISL_3188461, EPI_ISL_3188462, EPI_ISL_3188463, EPI_ISL_3188464, EPI_ISL_3188465, EPI_ISL_3188466, EPI_ISL_3188467, EPI_ISL_3188468, EPI_ISL_3188469, EPI_ISL_3188470, EPI_ISL_3188471, EPI_ISL_3188472, EPI_ISL_3188473, EPI_ISL_3188474, EPI_ISL_3188475, EPI_ISL_3188476, EPI_ISL_3188477, EPI_ISL_3188478, EPI_ISL_3188479, EPI_ISL_3188480, EPI_ISL_3188481, EPI_ISL_3188482, EPI_ISL_3188483, EPI_ISL_3188484, EPI_ISL_3188485, EPI_ISL_3188486, EPI_ISL_3188487, EPI_ISL_3188488, EPI_ISL_3188489, EPI_ISL_3188490, EPI_ISL_3188492, EPI_ISL_3188493, EPI_ISL_3188494, EPI_ISL_3188495, EPI_ISL_3188496, EPI_ISL_3188497, EPI_ISL_3188498, EPI_ISL_3188499, EPI_ISL_3188500, EPI_ISL_3188501, EPI_ISL_3188502, EPI_ISL_3188503, EPI_ISL_3188504, EPI_ISL_3188506, EPI_ISL_3188509, EPI_ISL_3188510, EPI_ISL_3188511, EPI_ISL_3188512, EPI_ISL_3188513, EPI_ISL_3188514, EPI_ISL_3188515, EPI_ISL_3188516, EPI_ISL_3188517, EPI_ISL_3188518, EPI_ISL_3188519, EPI_ISL_3188520, EPI_ISL_3188521, EPI_ISL_3188522, EPI_ISL_3188523, EPI_ISL_3188524, EPI_ISL_3188525, | | | |

| | | | |
|---|---|---|---|
| EPI_ISL_3188526, EPI_ISL_3188527, EPI_ISL_3188528, EPI_ISL_3188529, EPI_ISL_3188530, EPI_ISL_3188531, EPI_ISL_3188532, EPI_ISL_3188533, EPI_ISL_3188534, EPI_ISL_3188535, EPI_ISL_3188536, EPI_ISL_3188538, EPI_ISL_3188539, EPI_ISL_3188540, EPI_ISL_3188541, EPI_ISL_3188542, EPI_ISL_3188543, EPI_ISL_3188544, EPI_ISL_3188545, EPI_ISL_3188546, EPI_ISL_3188547, EPI_ISL_3188548, EPI_ISL_3188549, EPI_ISL_3188550, EPI_ISL_3188551, EPI_ISL_3188552, EPI_ISL_3188553, EPI_ISL_3188554, EPI_ISL_3188555, EPI_ISL_3188556, EPI_ISL_3188557, EPI_ISL_3188558, EPI_ISL_3188559, EPI_ISL_3188560, EPI_ISL_3188561 | | | |
| see above | Pandemic Response Lab - NYC | Pandemic Response Lab, R&D | Henry Lee, Michael Hammerling, Melissa Hopkins, Cybill del Castillo, Shinyoung Clair Kang, William Ward, Pradeep Bugga, Sol Rey, Dylan Law, Alex Carpio, Isabel Fernandez Escapa, Haiping Hao, Jon Laurent |
| EPI_ISL_3188564, EPI_ISL_3188622 | UCC (Universidad Central del Caribe) | Grubaugh Lab - Yale School of Public Health | Joseph Fauver, Mallery Breban, Isabel Ott, Tara Alpert, Mary Petrone, Anderson Brito, Chantal Vogels, Annie Watkins, Chaney Kalinich, Jessica Rothman, Kendall Billig, Tobias Koch, Fabiola Cruz-Lopez, David Rivera Aponte, Fabiola Fontanet, Natalie Machargo, Andrea Firpo, Vianca Aponte, Ernesto Vazquez, Patricia Serrano, Tania Mitwalli, Karolane Gonzalez, Nathan Grubaugh |
| EPI_ISL_3188623, EPI_ISL_3188624, EPI_ISL_3188625, EPI_ISL_3188626, EPI_ISL_3188627, EPI_ISL_3188628, EPI_ISL_3188629, EPI_ISL_3188630, EPI_ISL_3188631, EPI_ISL_3188632, EPI_ISL_3188633, EPI_ISL_3188635, EPI_ISL_3188636, EPI_ISL_3188638, EPI_ISL_3188639, EPI_ISL_3188640, EPI_ISL_3188641, EPI_ISL_3188642, EPI_ISL_3188643, EPI_ISL_3188644, EPI_ISL_3188645, EPI_ISL_3188646, EPI_ISL_3188647, EPI_ISL_3188648, EPI_ISL_3188649, EPI_ISL_3188650, EPI_ISL_3188651, EPI_ISL_3188652, EPI_ISL_3188653, EPI_ISL_3188654, EPI_ISL_3188656, EPI_ISL_3188657, EPI_ISL_3188658, EPI_ISL_3188659, EPI_ISL_3188660, EPI_ISL_3188661, EPI_ISL_3188662, EPI_ISL_3188665, EPI_ISL_3188666, EPI_ISL_3188667, EPI_ISL_3188668, EPI_ISL_3188669, EPI_ISL_3188670 | | | |
| see above | Hospital General Universitario Gregorio Marañón | Hospital General Universitario Gregorio Marañón | Sergio Buenestado Serrano, Pedro Sola Campoy, Laura Pérez-Lago, Cristina Rodriguez-Grande, Marta Herranz Martin, Victor Manuel de la Cueva, Julia Suárez, Pilar Catalán, Patricia Muñoz, Darío García de Viedma |
| EPI_ISL_3188672, EPI_ISL_3188673, EPI_ISL_3188674, EPI_ISL_3188675, EPI_ISL_3188676, EPI_ISL_3188677, EPI_ISL_3188678, EPI_ISL_3188679, EPI_ISL_3188680, EPI_ISL_3188681, EPI_ISL_3188682, EPI_ISL_3188683, EPI_ISL_3188684, EPI_ISL_3188685, EPI_ISL_3188686, EPI_ISL_3188687, EPI_ISL_3188688, EPI_ISL_3188689, EPI_ISL_3188690, EPI_ISL_3188691, EPI_ISL_3188692, EPI_ISL_3188693, EPI_ISL_3188694, EPI_ISL_3188695, EPI_ISL_3188696, EPI_ISL_3188697, EPI_ISL_3188698, EPI_ISL_3188699, EPI_ISL_3188700, EPI_ISL_3188701, EPI_ISL_3188702, EPI_ISL_3188703, EPI_ISL_3188704, EPI_ISL_3188705, EPI_ISL_3188706, EPI_ISL_3188707, EPI_ISL_3188708, EPI_ISL_3188709, EPI_ISL_3188710, EPI_ISL_3188711, EPI_ISL_3188712, EPI_ISL_3188713, EPI_ISL_3188714, EPI_ISL_3188715, EPI_ISL_3188716, EPI_ISL_3188717, EPI_ISL_3188718, EPI_ISL_3188719, EPI_ISL_3188720, EPI_ISL_3188721, EPI_ISL_3188722, EPI_ISL_3188723, EPI_ISL_3188724, EPI_ISL_3188725, EPI_ISL_3188726, EPI_ISL_3188727, EPI_ISL_3188728, EPI_ISL_3188729, EPI_ISL_3188730, EPI_ISL_3188731, EPI_ISL_3188732, EPI_ISL_3188733, EPI_ISL_3188734, EPI_ISL_3188735, EPI_ISL_3188736, EPI_ISL_3188737, EPI_ISL_3188738, EPI_ISL_3188739, EPI_ISL_3188740, EPI_ISL_3188741, EPI_ISL_3188742, EPI_ISL_3188743, EPI_ISL_3188744, EPI_ISL_3188745, EPI_ISL_3188746, EPI_ISL_3188747, EPI_ISL_3188748, EPI_ISL_3188749, EPI_ISL_3188750, EPI_ISL_3188751, EPI_ISL_3188752, EPI_ISL_3188753, EPI_ISL_3188754, EPI_ISL_3188755, EPI_ISL_3188756, EPI_ISL_3188757, EPI_ISL_3188758, EPI_ISL_3188759, EPI_ISL_3188760, EPI_ISL_3188761, EPI_ISL_3188762, EPI_ISL_3188763, EPI_ISL_3188764, EPI_ISL_3188765, EPI_ISL_3188766, EPI_ISL_3188767, EPI_ISL_3188768, EPI_ISL_3188769, EPI_ISL_3188770, EPI_ISL_3188771, EPI_ISL_3188772, EPI_ISL_3188773, EPI_ISL_3188774, EPI_ISL_3188775, EPI_ISL_3188776, EPI_ISL_3188777, EPI_ISL_3188778, EPI_ISL_3188779, EPI_ISL_3188780, EPI_ISL_3188781, EPI_ISL_3188782, EPI_ISL_3188783, EPI_ISL_3188784, EPI_ISL_3188785, EPI_ISL_3188786, EPI_ISL_3188787, EPI_ISL_3188788, EPI_ISL_3188789, EPI_ISL_3188790, EPI_ISL_3188791, EPI_ISL_3188792, EPI_ISL_3188793, EPI_ISL_3188794, EPI_ISL_3188795, EPI_ISL_3188796, EPI_ISL_3188797, EPI_ISL_3188798, EPI_ISL_3188799, EPI_ISL_3188800, EPI_ISL_3188801, EPI_ISL_3188802, EPI_ISL_3188803, EPI_ISL_3188804, EPI_ISL_3188805, EPI_ISL_3188806, EPI_ISL_3188807, EPI_ISL_3188808, EPI_ISL_3188809, EPI_ISL_3188810, EPI_ISL_3188811, EPI_ISL_3188812 | | | |
| see above | National Public Health Laboratory, National Centre for Infectious Diseases | National Public Health Laboratory, National Centre for Infectious Diseases | Tze Minn Mak, Katherine Ching, Zhenyang Zhou, Royce Ang, Lin Cui, Raymond Tzer Pin Lin |
| EPI_ISL_3188814, EPI_ISL_3188815, EPI_ISL_3188816, EPI_ISL_3188817, EPI_ISL_3188818, EPI_ISL_3188819, EPI_ISL_3188820, EPI_ISL_3188821, EPI_ISL_3188822, EPI_ISL_3188823, EPI_ISL_3188824, EPI_ISL_3188825, EPI_ISL_3188826, EPI_ISL_3188827, EPI_ISL_3188828, EPI_ISL_3188829, EPI_ISL_3188830, EPI_ISL_3188831, EPI_ISL_3188832, EPI_ISL_3188833, EPI_ISL_3188834, EPI_ISL_3188835, EPI_ISL_3188836, EPI_ISL_3188837, EPI_ISL_3188838, EPI_ISL_3188839, EPI_ISL_3188840, EPI_ISL_3188841, EPI_ISL_3188842, EPI_ISL_3188843, EPI_ISL_3188844, EPI_ISL_3188845, EPI_ISL_3188846, EPI_ISL_3188847, EPI_ISL_3188848, EPI_ISL_3188849, EPI_ISL_3188850, EPI_ISL_3188851, EPI_ISL_3188852, EPI_ISL_3188853, EPI_ISL_3188854, EPI_ISL_3188855, EPI_ISL_3188856, EPI_ISL_3188857, EPI_ISL_3188858, EPI_ISL_3188859, EPI_ISL_3188860, EPI_ISL_3188861, EPI_ISL_3188863, EPI_ISL_3188864, EPI_ISL_3188865, EPI_ISL_3188866, EPI_ISL_3188869, EPI_ISL_3188870, EPI_ISL_3188871, EPI_ISL_3188873 | | | |
| see above | Hospital General Universitario Gregorio Marañón | Hospital General Universitario Gregorio Marañón | Sergio Buenestado Serrano, Pedro Sola Campoy, Laura Pérez-Lago, Cristina Rodriguez-Grande, Marta Herranz Martin, Victor Manuel de la Cueva, Julia Suárez, Pilar Catalán, Patricia Muñoz, Darío García de Viedma |
| EPI_ISL_3188874, EPI_ISL_3188875, EPI_ISL_3188876, EPI_ISL_3188877, EPI_ISL_3188878, EPI_ISL_3188879, EPI_ISL_3188880, EPI_ISL_3188881, EPI_ISL_3188882, EPI_ISL_3188883, EPI_ISL_3188884, EPI_ISL_3188885, EPI_ISL_3188887, EPI_ISL_3188888, EPI_ISL_3188889, EPI_ISL_3188890, EPI_ISL_3188891, EPI_ISL_3188892, EPI_ISL_3188893, EPI_ISL_3188894, EPI_ISL_3188895, EPI_ISL_3188896, EPI_ISL_3188897, EPI_ISL_3188898, EPI_ISL_3188899, EPI_ISL_3188900, EPI_ISL_3188901, EPI_ISL_3188902, EPI_ISL_3188903, EPI_ISL_3188904, EPI_ISL_3188905, EPI_ISL_3188906, EPI_ISL_3188907, EPI_ISL_3188908, EPI_ISL_3188909, EPI_ISL_3188910, EPI_ISL_3188911, EPI_ISL_3188912, EPI_ISL_3188913, EPI_ISL_3188914, EPI_ISL_3188915, EPI_ISL_3188917, EPI_ISL_3188918, EPI_ISL_3188919, EPI_ISL_3188920 | | | |
| see above | Microvida | Microvida | Suzan D. Pas, Jaco J. Verweij, Joep J. J. M. Stohr |
| EPI_ISL_3188922 | SYNLAB | GIGA Medical Genomics | Keith Durkin, Maria Artesi, Bouchra Boujemla, Nathalie Renotte, Nadine Cambisano, Sébastien Bontems, Cécile Meex, Claire Gourzonès, Olivier Ek, Laurent Gillet, Marie-Pierre Hayette, Vincent Bours |
| EPI_ISL_3188945 | NIMHANS | INSACOG-KA, NIMHANS | Chitra Pattabiraman, Anson Kunjumon George,Darshan Sreenivas, Ellango Ramasamy, Sony Sharma, Mahesh Kumar.C.S, Ananthapadmanabha Kotambail, Chetan G K, Gautham Arunachal Udupi, Anita S Desai, V Ravi |
| EPI_ISL_3188946 | RAILWAY HOSPITAL | INSACOG-KA, NIMHANS | Chitra Pattabiraman, Anson Kunjumon George,Darshan Sreenivas, Ellango Ramasamy, Sony Sharma, Mahesh Kumar.C.S, Ananthapadmanabha Kotambail, Chetan G K, Gautham Arunachal Udupi, Anita S Desai, V Ravi |
| EPI_ISL_3188947 | BBMP Urban | INSACOG-KA, NIMHANS | Chitra Pattabiraman, Anson Kunjumon George,Darshan Sreenivas, Ellango Ramasamy, Sony Sharma, Mahesh Kumar.C.S, Ananthapadmanabha Kotambail, Chetan G K, Gautham Arunachal Udupi, Anita S Desai, V Ravi |
| EPI_ISL_3188949, EPI_ISL_3188950, EPI_ISL_3188951 | DSU KALABURAGI | INSACOG-KA, NIMHANS | Chitra Pattabiraman, Anson Kunjumon George,Darshan Sreenivas, Ellango Ramasamy, Sony Sharma, Mahesh Kumar.C.S, Ananthapadmanabha Kotambail, Chetan G K, Gautham Arunachal Udupi, Anita S Desai, V Ravi |
| EPI_ISL_3188952, EPI_ISL_3188953 | DSU HASSAN | INSACOG-KA, NIMHANS | Chitra Pattabiraman, Anson Kunjumon George,Darshan Sreenivas, Ellango Ramasamy, Sony Sharma, Mahesh Kumar.C.S, Ananthapadmanabha Kotambail, Chetan G K, Gautham Arunachal Udupi, Anita S Desai, V Ravi |
| EPI_ISL_3188956, EPI_ISL_3188957, EPI_ISL_3188958 | Diagnostyka. Laboratoria Medyczne. | 1. ViroGenetics - BSL3 Laboratory of Virology, Małopolska Centre of Biotechnology, Jagiellonian University; 2. Diagtron Laboratoria Łukasz Rąbalski | Rabalski L., Gromowski,T., Rogalska-Kupiec M., Swadzba J., Wydmanski,W., Mazur-Panasiuk,N., Kowalski,M., Maciej Kosinski, Natalia Derewonko, Szulc,P., Sylwia Januszczak, Labaj,P.P., Pyrc,K. |
| EPI_ISL_3188964, EPI_ISL_3188966, EPI_ISL_3188967, EPI_ISL_3188968, EPI_ISL_3188969, EPI_ISL_3188970, EPI_ISL_3188971, EPI_ISL_3188972, EPI_ISL_3188973, EPI_ISL_3188974, EPI_ISL_3188975, EPI_ISL_3188976, EPI_ISL_3188977, EPI_ISL_3188978, EPI_ISL_3188979, EPI_ISL_3188980, EPI_ISL_3188981, EPI_ISL_3188982, EPI_ISL_3188984, EPI_ISL_3188986, EPI_ISL_3188987, EPI_ISL_3188988, EPI_ISL_3188989, EPI_ISL_3188991, EPI_ISL_3188993, EPI_ISL_3188996, EPI_ISL_3188997, EPI_ISL_3188998, EPI_ISL_3189000, EPI_ISL_3189001 | | | |
| see above | ZANGIATA INFECTIOUS CLINICAL HOSPITAL №2 | Biotechnology laboratory, Center for advanced technology | Alisher Abdullaev, Dilbar Dalimova, Abror Abdurakhimov, Zebinisa Mirakbarova, Ibragimova Shahnoza, Diyora Dalimova, Sharof Nuriddinov, Vladimir Tsoy, Elena Tsay, Gul Esonova, Shakhlo Turdikulova |
| EPI_ISL_3189002 | DSU KALABURAGI | INSACOG-KA, NIMHANS | Chitra Pattabiraman, Anson Kunjumon George,Darshan Sreenivas, Ellango Ramasamy, Sony Sharma, Mahesh Kumar.C.S, Ananthapadmanabha Kotambail, Chetan G K, Gautham Arunachal Udupi, Anita S Desai, V Ravi |
| EPI_ISL_3189003, EPI_ISL_3189004, EPI_ISL_3189005, EPI_ISL_3189006, EPI_ISL_3189007, EPI_ISL_3189008, EPI_ISL_3189009, EPI_ISL_3189010, EPI_ISL_3189011, EPI_ISL_3189012, EPI_ISL_3189013, EPI_ISL_3189014, EPI_ISL_3189015, EPI_ISL_3189016, EPI_ISL_3189017, EPI_ISL_3189018, EPI_ISL_3189019, EPI_ISL_3189020 | | | |
| see above | CHU Pontchaillou | CHU Pontchaillou | GROLHIER Claire, DENOUAL Florent, ETCHEVERRY Amandine, SASSI Mohamed, JAGLINE Steven, FEBREAU Christine, PRONIER Charlotte, GALIBERT Marie Dominique, DE TAYRAC Marie, THIBAULT Vincent |
| EPI_ISL_3189021, EPI_ISL_3189022, EPI_ISL_3189023, EPI_ISL_3189024, EPI_ISL_3189025, EPI_ISL_3189026, EPI_ISL_3189027, EPI_ISL_3189028, EPI_ISL_3189029, EPI_ISL_3189030, EPI_ISL_3189031, EPI_ISL_3189032, EPI_ISL_3189033, EPI_ISL_3189034, EPI_ISL_3189035, EPI_ISL_3189037, EPI_ISL_3189038, EPI_ISL_3189039, EPI_ISL_3189040, EPI_ISL_3189041, EPI_ISL_3189042, EPI_ISL_3189043, EPI_ISL_3189044, EPI_ISL_3189045, EPI_ISL_3189046, EPI_ISL_3189047, EPI_ISL_3189048, EPI_ISL_3189049, EPI_ISL_3189050, EPI_ISL_3189051, EPI_ISL_3189052, EPI_ISL_3189053, EPI_ISL_3189054, EPI_ISL_3189056, EPI_ISL_3189057, EPI_ISL_3189058, EPI_ISL_3189059, EPI_ISL_3189060, EPI_ISL_3189061, EPI_ISL_3189062, EPI_ISL_3189063, EPI_ISL_3189064, EPI_ISL_3189065, EPI_ISL_3189066, EPI_ISL_3189067, EPI_ISL_3189068, EPI_ISL_3189069, EPI_ISL_3189070, EPI_ISL_3189071, EPI_ISL_3189072, EPI_ISL_3189073, EPI_ISL_3189074, EPI_ISL_3189075, EPI_ISL_3189076, EPI_ISL_3189077, EPI_ISL_3189078, EPI_ISL_3189080, EPI_ISL_3189081, EPI_ISL_3189082, EPI_ISL_3189083, EPI_ISL_3189084, EPI_ISL_3189085, EPI_ISL_3189086, EPI_ISL_3189087, EPI_ISL_3189088, EPI_ISL_3189089, EPI_ISL_3189090, EPI_ISL_3189091, EPI_ISL_3189092, EPI_ISL_3189093, EPI_ISL_3189094, EPI_ISL_3189095, EPI_ISL_3189096, EPI_ISL_3189097, EPI_ISL_3189098, EPI_ISL_3189099, EPI_ISL_3189100, EPI_ISL_3189101, EPI_ISL_3189102, EPI_ISL_3189103, EPI_ISL_3189104, EPI_ISL_3189105, EPI_ISL_3189106, EPI_ISL_3189107, EPI_ISL_3189108, EPI_ISL_3189109, EPI_ISL_3189110, EPI_ISL_3189111, EPI_ISL_3189112, EPI_ISL_3189113, EPI_ISL_3189114, EPI_ISL_3189115, EPI_ISL_3189116, EPI_ISL_3189117, EPI_ISL_3189118, EPI_ISL_3189119, EPI_ISL_3189120, EPI_ISL_3189121, EPI_ISL_3189122, EPI_ISL_3189123, EPI_ISL_3189124, EPI_ISL_3189125, EPI_ISL_3189126, EPI_ISL_3189127, EPI_ISL_3189128, EPI_ISL_3189129, EPI_ISL_3189130, EPI_ISL_3189131, EPI_ISL_3189132, EPI_ISL_3189133, EPI_ISL_3189134, EPI_ISL_3189135, EPI_ISL_3189136, EPI_ISL_3189137, EPI_ISL_3189138, EPI_ISL_3189139, EPI_ISL_3189140, EPI_ISL_3189141, EPI_ISL_3189142, EPI_ISL_3189143, EPI_ISL_3189144, EPI_ISL_3189145, EPI_ISL_3189146, EPI_ISL_3189147, EPI_ISL_3189148, EPI_ISL_3189149, EPI_ISL_3189150, EPI_ISL_3189151, EPI_ISL_3189152, EPI_ISL_3189153, EPI_ISL_3189154 | | | |
| see above | INSACOG-Assam | National Institute of Biomedical Genomics – INSACOG | Arindam Maitra, Ajanta Sharma, Nidhan Kumar Biswas, Lahari Saikia, Sreedhar Chinnaswamy, Kailash Chamuah, Saumitra Das |
| EPI_ISL_3189155, EPI_ISL_3189156, EPI_ISL_3189157, EPI_ISL_3189158, EPI_ISL_3189159, EPI_ISL_3189160, EPI_ISL_3189161, EPI_ISL_3189162, EPI_ISL_3189163, EPI_ISL_3189164, EPI_ISL_3189165, EPI_ISL_3189166, EPI_ISL_3189167, EPI_ISL_3189168, EPI_ISL_3189169, EPI_ISL_3189170, EPI_ISL_3189171, EPI_ISL_3189172, EPI_ISL_3189173, EPI_ISL_3189174, EPI_ISL_3189176, EPI_ISL_3189177, EPI_ISL_3189178, EPI_ISL_3189179, EPI_ISL_3189180, EPI_ISL_3189181, EPI_ISL_3189182, EPI_ISL_3189183, EPI_ISL_3189184, EPI_ISL_3189185, EPI_ISL_3189186, EPI_ISL_3189187, EPI_ISL_3189188, EPI_ISL_3189189, EPI_ISL_3189190, EPI_ISL_3189191, EPI_ISL_3189192, EPI_ISL_3189193, EPI_ISL_3189194, EPI_ISL_3189195, EPI_ISL_3189196, EPI_ISL_3189197, EPI_ISL_3189198, EPI_ISL_3189199, EPI_ISL_3189200, EPI_ISL_3189201, EPI_ISL_3189202, EPI_ISL_3189203, EPI_ISL_3189204, EPI_ISL_3189205, EPI_ISL_3189206, EPI_ISL_3189207, EPI_ISL_3189209, EPI_ISL_3189210, EPI_ISL_3189211, EPI_ISL_3189212, EPI_ISL_3189213, EPI_ISL_3189214, EPI_ISL_3189215, EPI_ISL_3189216, EPI_ISL_3189217, EPI_ISL_3189218, EPI_ISL_3189219, EPI_ISL_3189220, EPI_ISL_3189221, EPI_ISL_3189222, EPI_ISL_3189223, EPI_ISL_3189224, EPI_ISL_3189225, EPI_ISL_3189226, EPI_ISL_3189227, EPI_ISL_3189228, EPI_ISL_3189229, EPI_ISL_3189230, EPI_ISL_3189231, EPI_ISL_3189232, EPI_ISL_3189233, EPI_ISL_3189234, EPI_ISL_3189235, EPI_ISL_3189236, EPI_ISL_3189237, EPI_ISL_3189238, EPI_ISL_3189239, EPI_ISL_3189240, EPI_ISL_3189241, EPI_ISL_3189242, EPI_ISL_3189243, EPI_ISL_3189244, EPI_ISL_3189245, EPI_ISL_3189246, EPI_ISL_3189247, EPI_ISL_3189248, EPI_ISL_3189249, EPI_ISL_3189250, EPI_ISL_3189251, EPI_ISL_3189252, EPI_ISL_3189253, EPI_ISL_3189254, EPI_ISL_3189255, EPI_ISL_3189256, EPI_ISL_3189257, EPI_ISL_3189258, EPI_ISL_3189259, EPI_ISL_3189260, EPI_ISL_3189261, EPI_ISL_3189262, EPI_ISL_3189263, EPI_ISL_3189264, EPI_ISL_3189265, EPI_ISL_3189266, EPI_ISL_3189267, EPI_ISL_3189268, EPI_ISL_3189269, EPI_ISL_3189270, EPI_ISL_3189271, EPI_ISL_3189272, EPI_ISL_3189273, EPI_ISL_3189274, EPI_ISL_3189275, EPI_ISL_3189276, EPI_ISL_3189277, EPI_ISL_3189278, EPI_ISL_3189279, EPI_ISL_3189280, EPI_ISL_3189281, EPI_ISL_3189282, EPI_ISL_3189283, EPI_ISL_3189284, EPI_ISL_3189285, EPI_ISL_3189286, EPI_ISL_3189287, EPI_ISL_3189288, EPI_ISL_3189289, EPI_ISL_3189290, EPI_ISL_3189291, EPI_ISL_3189292, EPI_ISL_3189293, EPI_ISL_3189294, EPI_ISL_3189295, EPI_ISL_3189296, EPI_ISL_3189297, EPI_ISL_3189298, EPI_ISL_3189299, EPI_ISL_3189300, EPI_ISL_3189301, EPI_ISL_3189302, EPI_ISL_3189303, EPI_ISL_3189304, EPI_ISL_3189305, EPI_ISL_3189306, EPI_ISL_3189307, EPI_ISL_3189308, EPI_ISL_3189309, EPI_ISL_3189310, EPI_ISL_3189311, EPI_ISL_3189312, EPI_ISL_3189313, EPI_ISL_3189314, EPI_ISL_3189315, EPI_ISL_3189316, EPI_ISL_3189317, EPI_ISL_3189318, EPI_ISL_3189319, EPI_ISL_3189320, EPI_ISL_3189321, EPI_ISL_3189322, EPI_ISL_3189323, EPI_ISL_3189324, EPI_ISL_3189325, EPI_ISL_3189326, EPI_ISL_3189327, EPI_ISL_3189328, EPI_ISL_3189329, EPI_ISL_3189330, EPI_ISL_3189331, EPI_ISL_3189332, EPI_ISL_3189333, EPI_ISL_3189334, EPI_ISL_3189335, EPI_ISL_3189336, EPI_ISL_3189337, EPI_ISL_3189338, EPI_ISL_3189339, EPI_ISL_3189340, EPI_ISL_3189341, EPI_ISL_3189342, EPI_ISL_3189343, EPI_ISL_3189344, EPI_ISL_3189345, EPI_ISL_3189346, EPI_ISL_3189347, EPI_ISL_3189348, EPI_ISL_3189349, EPI_ISL_3189350, EPI_ISL_3189351, EPI_ISL_3189353, EPI_ISL_3189354, EPI_ISL_3189355, EPI_ISL_3189356, EPI_ISL_3189357, EPI_ISL_3189358, EPI_ISL_3189359, EPI_ISL_3189360, EPI_ISL_3189361, EPI_ISL_3189362, EPI_ISL_3189363, EPI_ISL_3189364, EPI_ISL_3189365, EPI_ISL_3189366, EPI_ISL_3189367, EPI_ISL_3189368, EPI_ISL_3189369, EPI_ISL_3189370, EPI_ISL_3189371, EPI_ISL_3189372, EPI_ISL_3189373, EPI_ISL_3189374, EPI_ISL_3189375, EPI_ISL_3189376, EPI_ISL_3189377, EPI_ISL_3189378, EPI_ISL_3189379, EPI_ISL_3189380, EPI_ISL_3189381, EPI_ISL_3189382, EPI_ISL_3189383, EPI_ISL_3189384, EPI_ISL_3189385, EPI_ISL_3189386, EPI_ISL_3189387, EPI_ISL_3189388, EPI_ISL_3189389, EPI_ISL_3189390, EPI_ISL_3189391, EPI_ISL_3189392, EPI_ISL_3189393, EPI_ISL_3189394, EPI_ISL_3189395, EPI_ISL_3189396, EPI_ISL_3189397 | | | |
| see above | INSACOG-WB | National Institute of Biomedical Genomics – INSACOG | Arindam Maitra, Bhaswati Bandyopadhyay, Nidhan Kumar Biswas, Tamal Ghosh, Sreedhar Chinnaswamy, Ajay Chakraborti, Saumitra Das |
| EPI_ISL_3189398, EPI_ISL_3189399, EPI_ISL_3189400, EPI_ISL_3189401, EPI_ISL_3189402, EPI_ISL_3189403, EPI_ISL_3189404, EPI_ISL_3189405, EPI_ISL_3189406, EPI_ISL_3189407, EPI_ISL_3189408, EPI_ISL_3189409, EPI_ISL_3189410, EPI_ISL_3189411, EPI_ISL_3189413, EPI_ISL_3189414, EPI_ISL_3189415, EPI_ISL_3189416, EPI_ISL_3189417, EPI_ISL_3189418, EPI_ISL_3189419, EPI_ISL_3189420, EPI_ISL_3189421, EPI_ISL_3189422, EPI_ISL_3189423, EPI_ISL_3189424, EPI_ISL_3189425, EPI_ISL_3189426, EPI_ISL_3189427, EPI_ISL_3189428, EPI_ISL_3189429, EPI_ISL_3189430, EPI_ISL_3189431, EPI_ISL_3189433, EPI_ISL_3189434, EPI_ISL_3189435, EPI_ISL_3189436, EPI_ISL_3189437, EPI_ISL_3189438, EPI_ISL_3189439, EPI_ISL_3189440, EPI_ISL_3189441, EPI_ISL_3189442, EPI_ISL_3189443, EPI_ISL_3189444, EPI_ISL_3189445, EPI_ISL_3189447, EPI_ISL_3189448, EPI_ISL_3189449, EPI_ISL_3189450, EPI_ISL_3189451, EPI_ISL_3189452, EPI_ISL_3189453, EPI_ISL_3189454, EPI_ISL_3189455, EPI_ISL_3189456, EPI_ISL_3189457, EPI_ISL_3189458, EPI_ISL_3189459, EPI_ISL_3189460, EPI_ISL_3189461, EPI_ISL_3189462, EPI_ISL_3189463, EPI_ISL_3189464, EPI_ISL_3189465, EPI_ISL_3189466, EPI_ISL_3189467, EPI_ISL_3189468, EPI_ISL_3189469, EPI_ISL_3189470, EPI_ISL_3189471, EPI_ISL_3189472, EPI_ISL_3189473, EPI_ISL_3189474, EPI_ISL_3189476, EPI_ISL_3189478, EPI_ISL_3189479, EPI_ISL_3189480, EPI_ISL_3189482, EPI_ISL_3189484, EPI_ISL_3189485, EPI_ISL_3189486, EPI_ISL_3189487, EPI_ISL_3189489, EPI_ISL_3189490, EPI_ISL_3189491, EPI_ISL_3189492 | | | |
| see above | INSACOG-Mizoram | National Institute of Biomedical Genomics – INSACOG | Arindam Maitra, Swagnik Roy, Nidhan Kumar Biswas, Gracy Laldinmawii, Sreedhar Chinnaswamy, N Senthil Kumar, Saumitra Das |
| EPI_ISL_3189493, EPI_ISL_3189494, EPI_ISL_3189495, EPI_ISL_3189496, EPI_ISL_3189497, EPI_ISL_3189498, EPI_ISL_3189499, EPI_ISL_3189500, EPI_ISL_3189501, EPI_ISL_3189502, EPI_ISL_3189503, EPI_ISL_3189504, EPI_ISL_3189505, EPI_ISL_3189506, EPI_ISL_3189507, EPI_ISL_3189508, EPI_ISL_3189509, EPI_ISL_3189510, EPI_ISL_3189511, EPI_ISL_3189512, EPI_ISL_3189513, EPI_ISL_3189514, EPI_ISL_3189515, EPI_ISL_3189516, EPI_ISL_3189517, EPI_ISL_3189518, EPI_ISL_3189519, EPI_ISL_3189520, EPI_ISL_3189521, EPI_ISL_3189522, EPI_ISL_3189523, EPI_ISL_3189524, EPI_ISL_3189525, EPI_ISL_3189526, EPI_ISL_3189527, EPI_ISL_3189528, EPI_ISL_3189529, EPI_ISL_3189530, EPI_ISL_3189531, EPI_ISL_3189532, EPI_ISL_3189533, EPI_ISL_3189534, EPI_ISL_3189535, EPI_ISL_3189536, EPI_ISL_3189537, EPI_ISL_3189538, EPI_ISL_3189539, EPI_ISL_3189540, EPI_ISL_3189541, EPI_ISL_3189542, EPI_ISL_3189543, EPI_ISL_3189544, EPI_ISL_3189545, EPI_ISL_3189546, EPI_ISL_3189547, EPI_ISL_3189548, EPI_ISL_3189549, EPI_ISL_3189550, EPI_ISL_3189551, EPI_ISL_3189552, EPI_ISL_3189553, EPI_ISL_3189554, EPI_ISL_3189555, EPI_ISL_3189556, EPI_ISL_3189557, EPI_ISL_3189558, EPI_ISL_3189560, EPI_ISL_3189561, EPI_ISL_3189562, EPI_ISL_3189563, EPI_ISL_3189564, EPI_ISL_3189565, EPI_ISL_3189566, EPI_ISL_3189567, EPI_ISL_3189568, EPI_ISL_3189569, EPI_ISL_3189570, EPI_ISL_3189571, EPI_ISL_3189572, EPI_ISL_3189573, EPI_ISL_3189574, EPI_ISL_3189575, EPI_ISL_3189576, EPI_ISL_3189577, EPI_ISL_3189578, EPI_ISL_3189579, EPI_ISL_3189580, EPI_ISL_3189581, EPI_ISL_3189582, EPI_ISL_3189583, EPI_ISL_3189584, EPI_ISL_3189585, EPI_ISL_3189586, EPI_ISL_3189587, EPI_ISL_3189588, EPI_ISL_3189589, EPI_ISL_3189590, EPI_ISL_3189591, EPI_ISL_3189592 | | | |
| see above | INSACOG-NAGALAND | National Institute of Biomedical Genomics – INSACOG | Arindam Maitra, Vinotsole Khamo, Nidhan Kumar Biswas, Chenole Keppen, Sreedhar Chinnaswamy, Wetetsho Kapfo, Saumitra Das |
| EPI_ISL_3189594, EPI_ISL_3189595, EPI_ISL_3189596, EPI_ISL_3189597, EPI_ISL_3189598, EPI_ISL_3189599, EPI_ISL_3189601, EPI_ISL_3189602, EPI_ISL_3189603, EPI_ISL_3189604, EPI_ISL_3189605, EPI_ISL_3189606 | | | |
| see above | Orebro University Hospital, Dept Laboratory Medicine, Clinical Microbiology | Orebro University Hospital | Sundqvist M et al |
| EPI_ISL_3189608, EPI_ISL_3189613, EPI_ISL_3189614, EPI_ISL_3189616, EPI_ISL_3189617, EPI_ISL_3189619, EPI_ISL_3189622, EPI_ISL_3189623, EPI_ISL_3189626, EPI_ISL_3189627, EPI_ISL_3189628, EPI_ISL_3189629, EPI_ISL_3189632 | | | |
| see above | Microbiology Department, University Hospital Donostia | Microbiology Department, University Hospital Donostia | Marimon JM, Montes M, Piñeiro L, Martin-Peñaranda T, Gomez M, Sorarrain A, Cilla G. |
| EPI_ISL_3189635, EPI_ISL_3189636, EPI_ISL_3189641, EPI_ISL_3189642, EPI_ISL_3189643, EPI_ISL_3189644, EPI_ISL_3189645, EPI_ISL_3189647, EPI_ISL_3189648, EPI_ISL_3189649, EPI_ISL_3189650, EPI_ISL_3189651, EPI_ISL_3189652, EPI_ISL_3189653, EPI_ISL_3189658, EPI_ISL_3189660, EPI_ISL_3189667, EPI_ISL_3189670, EPI_ISL_3189671, EPI_ISL_3189673, EPI_ISL_3189674, EPI_ISL_3189676, EPI_ISL_3189677, EPI_ISL_3189678, EPI_ISL_3189679, EPI_ISL_3189680, EPI_ISL_3189681, EPI_ISL_3189683, EPI_ISL_3189684, EPI_ISL_3189685, EPI_ISL_3189686, EPI_ISL_3189687, EPI_ISL_3189688, EPI_ISL_3189689, EPI_ISL_3189690, EPI_ISL_3189693, EPI_ISL_3189694, EPI_ISL_3189695, EPI_ISL_3189696, EPI_ISL_3189698, EPI_ISL_3189699, EPI_ISL_3189700, EPI_ISL_3189701, EPI_ISL_3189702, EPI_ISL_3189703, EPI_ISL_3189704, EPI_ISL_3189706, EPI_ISL_3189707, EPI_ISL_3189709, EPI_ISL_3189710, EPI_ISL_3189711, EPI_ISL_3189712, EPI_ISL_3189713, EPI_ISL_3189714, EPI_ISL_3189715, EPI_ISL_3189716, EPI_ISL_3189717, EPI_ISL_3189718, EPI_ISL_3189719, EPI_ISL_3189720, EPI_ISL_3189721, EPI_ISL_3189722, EPI_ISL_3189724, EPI_ISL_3189725, EPI_ISL_3189726, EPI_ISL_3189727, EPI_ISL_3189728, EPI_ISL_3189729, EPI_ISL_3189730, EPI_ISL_3189731, EPI_ISL_3189732, EPI_ISL_3189733, EPI_ISL_3189734, EPI_ISL_3189735, EPI_ISL_3189737, EPI_ISL_3189738, EPI_ISL_3189739, EPI_ISL_3189740, EPI_ISL_3189741, EPI_ISL_3189742, EPI_ISL_3189743, EPI_ISL_3189744, EPI_ISL_3189745, EPI_ISL_3189746, EPI_ISL_3189747, EPI_ISL_3189748, EPI_ISL_3189749, EPI_ISL_3189750, EPI_ISL_3189751, EPI_ISL_3189752, EPI_ISL_3189753, EPI_ISL_3189754, EPI_ISL_3189755, EPI_ISL_3189756, EPI_ISL_3189757, EPI_ISL_3189758, EPI_ISL_3189759, EPI_ISL_3189760, EPI_ISL_3189761, EPI_ISL_3189762, EPI_ISL_3189763, EPI_ISL_3189764, EPI_ISL_3189765, EPI_ISL_3189766, EPI_ISL_3189770, EPI_ISL_3189771, EPI_ISL_3189773, EPI_ISL_3189774, EPI_ISL_3189775, EPI_ISL_3189776, EPI_ISL_3189778, EPI_ISL_3189779, EPI_ISL_3189780, EPI_ISL_3189781 | | | |
| see above | State Laboratories Division, Hawaii State Department of Health | State Laboratories Division, Hawaii State Department of Health | Pamela O'Brien, Drew Kuwazaki, Ayana Garnet, Daniel Strange, Razvan Sultana, Edward Desmond |
| EPI_ISL_3189782, EPI_ISL_3189783, EPI_ISL_3189785, EPI_ISL_3189786, EPI_ISL_3189787, EPI_ISL_3189788, EPI_ISL_3189789, EPI_ISL_3189790, EPI_ISL_3189791, EPI_ISL_3189792, EPI_ISL_3189793, EPI_ISL_3189794, EPI_ISL_3189795, EPI_ISL_3189796, EPI_ISL_3189797, EPI_ISL_3189798, EPI_ISL_3189799, EPI_ISL_3189800, EPI_ISL_3189801, EPI_ISL_3189802, EPI_ISL_3189803, EPI_ISL_3189804, EPI_ISL_3189805, EPI_ISL_3189806, EPI_ISL_3189807, EPI_ISL_3189808, EPI_ISL_3189809, EPI_ISL_3189810, EPI_ISL_3189811, EPI_ISL_3189812, EPI_ISL_3189813, EPI_ISL_3189814, EPI_ISL_3189815, EPI_ISL_3189816, EPI_ISL_3189817, EPI_ISL_3189818, EPI_ISL_3189819, EPI_ISL_3189820, EPI_ISL_3189821, EPI_ISL_3189822, EPI_ISL_3189823, EPI_ISL_3189824, EPI_ISL_3189825, EPI_ISL_3189826, EPI_ISL_3189827, EPI_ISL_3189828, EPI_ISL_3189829, EPI_ISL_3189830, EPI_ISL_3189831, EPI_ISL_3189832, EPI_ISL_3189833, EPI_ISL_3189834, EPI_ISL_3189835, EPI_ISL_3189836, EPI_ISL_3189837, EPI_ISL_3189838, EPI_ISL_3189839, EPI_ISL_3189840, EPI_ISL_3189841, EPI_ISL_3189842, EPI_ISL_3189843, EPI_ISL_3189844, EPI_ISL_3189845, EPI_ISL_3189846, EPI_ISL_3189847, EPI_ISL_3189848, EPI_ISL_3189849, EPI_ISL_3189850, EPI_ISL_3189851, EPI_ISL_3189852, EPI_ISL_3189853, EPI_ISL_3189854, EPI_ISL_3189855, EPI_ISL_3189856, EPI_ISL_3189857, EPI_ISL_3189858, EPI_ISL_3189859, EPI_ISL_3189860, EPI_ISL_3189861, EPI_ISL_3189862, EPI_ISL_3189863, EPI_ISL_3189864, EPI_ISL_3189865, EPI_ISL_3189866, EPI_ISL_3189867, EPI_ISL_3189868, EPI_ISL_3189869, EPI_ISL_3189870, EPI_ISL_3189871, EPI_ISL_3189872, EPI_ISL_3189873, EPI_ISL_3189874, EPI_ISL_3189875, EPI_ISL_3189876, EPI_ISL_3189877, EPI_ISL_3189878, EPI_ISL_3189879, EPI_ISL_3189880, EPI_ISL_3189881, EPI_ISL_3189882, EPI_ISL_3189883, EPI_ISL_3189884, EPI_ISL_3189885, EPI_ISL_3189886, EPI_ISL_3189887, EPI_ISL_3189888, EPI_ISL_3189889, EPI_ISL_3189890 | | | |
| see above | Hospital Universitari Vall d'Hebron - Vall d'Hebron Institut de Recerca | Hospital Universitari Vall d'Hebron - Vall d'Hebron Institut de Recerca | Cristina Andrés, Maria Piñana, Alejandra González-Sánchez, Damir Garcia-Cehic, Ariadna Rando, Juliana Esperalba, Maria Gema Codina, Carla Castillo, Karen García, Rodrigo Vásquez, Maria Carmen Martin, Tomàs Pumarola, Josep Quer, Andrés Antón |

| | | | |
|---|---|---|---|
| EPI_ISL_3189891, EPI_ISL_3189892, EPI_ISL_3189893, EPI_ISL_3189894, EPI_ISL_3189895, EPI_ISL_3189896, EPI_ISL_3189897, EPI_ISL_3189899, EPI_ISL_3189900, EPI_ISL_3189901, EPI_ISL_3189902, EPI_ISL_3189903, EPI_ISL_3189904, EPI_ISL_3189905, EPI_ISL_3189906, EPI_ISL_3189907, EPI_ISL_3189908, EPI_ISL_3189909, EPI_ISL_3189910, EPI_ISL_3189911, EPI_ISL_3189912, EPI_ISL_3189913, EPI_ISL_3189914, EPI_ISL_3189915, EPI_ISL_3189917, EPI_ISL_3189918, EPI_ISL_3189919, EPI_ISL_3189920, EPI_ISL_3189921, EPI_ISL_3189923, EPI_ISL_3189924, EPI_ISL_3189925, EPI_ISL_3189926, EPI_ISL_3189927, EPI_ISL_3189928, EPI_ISL_3189929, EPI_ISL_3189930, EPI_ISL_3189931, EPI_ISL_3189932, EPI_ISL_3189933, EPI_ISL_3189934, EPI_ISL_3189935, EPI_ISL_3189936, EPI_ISL_3189937, EPI_ISL_3189938, EPI_ISL_3189939 | | | |
| see above | ADPH | ADPH | Adrianna Walker |
| EPI_ISL_3189940, EPI_ISL_3189941, EPI_ISL_3189942 | Hospital Universitari Joan XXIII | Hospital Universitari Joan XXIII | Gemma Recio, Cristina Gutiérrez, Clara Benavent, Carla Martín, Margarida Terrón, Ester Picó, Natalia Bastón |
| EPI_ISL_3189943 | Viešoji įstaiga Vilniaus universiteto ligoninė Santaros klinikos | National Public Health Surveillance Laboratory | Lukas Zemaitis, Migle Gabrielaite, Jelena Razmuk, Svajune Muralyte, Ana Steponkiene, Lukas Vasionis, Danas Baksa |
| EPI_ISL_3189945, EPI_ISL_3189946 | UAB Diagnostikos laboratorija | National Public Health Surveillance Laboratory | Lukas Zemaitis, Migle Gabrielaite, Jelena Razmuk, Svajune Muralyte, Ana Steponkiene, Lukas Vasionis, Danas Baksa |
| EPI_ISL_3189948 | Viešoji įstaiga Vilniaus universiteto ligoninė Santaros klinikos | National Public Health Surveillance Laboratory | Lukas Zemaitis, Migle Gabrielaite, Jelena Razmuk, Svajune Muralyte, Ana Steponkiene, Lukas Vasionis, Danas Baksa |
| EPI_ISL_3189949 | UAB "Rezus.lt" | National Public Health Surveillance Laboratory | Lukas Zemaitis, Migle Gabrielaite, Jelena Razmuk, Svajune Muralyte, Ana Steponkiene, Lukas Vasionis, Danas Baksa |
| EPI_ISL_3189950 | Viešoji įstaiga Vilniaus universiteto ligoninė Santaros klinikos | National Public Health Surveillance Laboratory | Lukas Zemaitis, Migle Gabrielaite, Jelena Razmuk, Svajune Muralyte, Ana Steponkiene, Lukas Vasionis, Danas Baksa |
| EPI_ISL_3189951 | Viešoji įstaiga Klaipėdos universitetinė ligoninė | National Public Health Surveillance Laboratory | Lukas Zemaitis, Migle Gabrielaite, Jelena Razmuk, Svajune Muralyte, Ana Steponkiene, Lukas Vasionis, Danas Baksa |
| EPI_ISL_3189952 | UAB Diagnostikos laboratorija | National Public Health Surveillance Laboratory | Lukas Zemaitis, Migle Gabrielaite, Jelena Razmuk, Svajune Muralyte, Ana Steponkiene, Lukas Vasionis, Danas Baksa |
| EPI_ISL_3189956 | Nacionalinė visuomenės sveikatos priežiūros laboratorija | National Public Health Surveillance Laboratory | Lukas Zemaitis, Migle Gabrielaite, Jelena Razmuk, Svajune Muralyte, Ana Steponkiene, Lukas Vasionis, Danas Baksa |
| EPI_ISL_3189957 | UAB Diagnostikos laboratorija | National Public Health Surveillance Laboratory | Lukas Zemaitis, Migle Gabrielaite, Jelena Razmuk, Svajune Muralyte, Ana Steponkiene, Lukas Vasionis, Danas Baksa |
| EPI_ISL_3189958, EPI_ISL_3189959 | UAB "Rezus.lt" | National Public Health Surveillance Laboratory | Lukas Zemaitis, Migle Gabrielaite, Jelena Razmuk, Svajune Muralyte, Ana Steponkiene, Lukas Vasionis, Danas Baksa |
| EPI_ISL_3189960 | UAB Diagnostikos laboratorija | National Public Health Surveillance Laboratory | Lukas Zemaitis, Migle Gabrielaite, Jelena Razmuk, Svajune Muralyte, Ana Steponkiene, Lukas Vasionis, Danas Baksa |
| EPI_ISL_3189961 | Viešoji įstaiga Vilniaus universiteto ligoninė Santaros klinikos | National Public Health Surveillance Laboratory | Lukas Zemaitis, Migle Gabrielaite, Jelena Razmuk, Svajune Muralyte, Ana Steponkiene, Lukas Vasionis, Danas Baksa |
| EPI_ISL_3189963, EPI_ISL_3189964 | UAB "Rezus.lt" | National Public Health Surveillance Laboratory | Lukas Zemaitis, Migle Gabrielaite, Jelena Razmuk, Svajune Muralyte, Ana Steponkiene, Lukas Vasionis, Danas Baksa |
| EPI_ISL_3189965 | UAB InMedica | National Public Health Surveillance Laboratory | Lukas Zemaitis, Migle Gabrielaite, Jelena Razmuk, Svajune Muralyte, Ana Steponkiene, Lukas Vasionis, Danas Baksa |
| EPI_ISL_3189966, EPI_ISL_3189968 | UAB Diagnostikos laboratorija | National Public Health Surveillance Laboratory | Lukas Zemaitis, Migle Gabrielaite, Jelena Razmuk, Svajune Muralyte, Ana Steponkiene, Lukas Vasionis, Danas Baksa |
| EPI_ISL_3189969 | UAB "Rezus.lt" | National Public Health Surveillance Laboratory | Lukas Zemaitis, Migle Gabrielaite, Jelena Razmuk, Svajune Muralyte, Ana Steponkiene, Lukas Vasionis, Danas Baksa |
| EPI_ISL_3189970 | Viešoji įstaiga Klaipėdos universitetinė ligoninė | National Public Health Surveillance Laboratory | Lukas Zemaitis, Migle Gabrielaite, Jelena Razmuk, Svajune Muralyte, Ana Steponkiene, Lukas Vasionis, Danas Baksa |
| EPI_ISL_3189971 | UAB "Rezus.lt" | National Public Health Surveillance Laboratory | Lukas Zemaitis, Migle Gabrielaite, Jelena Razmuk, Svajune Muralyte, Ana Steponkiene, Lukas Vasionis, Danas Baksa |
| EPI_ISL_3189972, EPI_ISL_3189975 | Viešoji įstaiga Klaipėdos universitetinė ligoninė | National Public Health Surveillance Laboratory | Lukas Zemaitis, Migle Gabrielaite, Jelena Razmuk, Svajune Muralyte, Ana Steponkiene, Lukas Vasionis, Danas Baksa |
| EPI_ISL_3189978 | Viešoji įstaiga Respublikinė Šiaulių ligoninė | National Public Health Surveillance Laboratory | Lukas Zemaitis, Migle Gabrielaite, Jelena Razmuk, Svajune Muralyte, Ana Steponkiene, Lukas Vasionis, Danas Baksa |
| EPI_ISL_3189979 | UAB "Rezus.lt" | National Public Health Surveillance Laboratory | Lukas Zemaitis, Migle Gabrielaite, Jelena Razmuk, Svajune Muralyte, Ana Steponkiene, Lukas Vasionis, Danas Baksa |
| EPI_ISL_3189980 | Viešoji įstaiga Vilniaus universiteto ligoninė Santaros klinikos | National Public Health Surveillance Laboratory | Lukas Zemaitis, Migle Gabrielaite, Jelena Razmuk, Svajune Muralyte, Ana Steponkiene, Lukas Vasionis, Danas Baksa |
| EPI_ISL_3189983 | UAB InMedica | National Public Health Surveillance Laboratory | Lukas Zemaitis, Migle Gabrielaite, Jelena Razmuk, Svajune Muralyte, Ana Steponkiene, Lukas Vasionis, Danas Baksa |
| EPI_ISL_3189985 | UAB "Baltic Medics" | National Public Health Surveillance Laboratory | Lukas Zemaitis, Migle Gabrielaite, Jelena Razmuk, Svajune Muralyte, Ana Steponkiene, Lukas Vasionis, Danas Baksa |
| EPI_ISL_3189988, EPI_ISL_3189989 | UAB "Rezus.lt" | National Public Health Surveillance Laboratory | Lukas Zemaitis, Migle Gabrielaite, Jelena Razmuk, Svajune Muralyte, Ana Steponkiene, Lukas Vasionis, Danas Baksa |
| EPI_ISL_3189990 | Viešoji įstaiga Vilniaus universiteto ligoninė Santaros klinikos | National Public Health Surveillance Laboratory | Lukas Zemaitis, Migle Gabrielaite, Jelena Razmuk, Svajune Muralyte, Ana Steponkiene, Lukas Vasionis, Danas Baksa |
| EPI_ISL_3189991 | Viešoji įstaiga Klaipėdos universitetinė ligoninė | National Public Health Surveillance Laboratory | Lukas Zemaitis, Migle Gabrielaite, Jelena Razmuk, Svajune Muralyte, Ana Steponkiene, Lukas Vasionis, Danas Baksa |
| EPI_ISL_3189992 | UAB "Rezus.lt" | National Public Health Surveillance Laboratory | Lukas Zemaitis, Migle Gabrielaite, Jelena Razmuk, Svajune Muralyte, Ana Steponkiene, Lukas Vasionis, Danas Baksa |
| EPI_ISL_3189994 | UAB Diagnostikos laboratorija | National Public Health Surveillance Laboratory | Lukas Zemaitis, Migle Gabrielaite, Jelena Razmuk, Svajune Muralyte, Ana Steponkiene, Lukas Vasionis, Danas Baksa |
| EPI_ISL_3189995 | UAB "Rezus.lt" | National Public Health Surveillance Laboratory | Lukas Zemaitis, Migle Gabrielaite, Jelena Razmuk, Svajune Muralyte, Ana Steponkiene, Lukas Vasionis, Danas Baksa |
| EPI_ISL_3189996 | UAB InMedica | National Public Health Surveillance Laboratory | Lukas Zemaitis, Migle Gabrielaite, Jelena Razmuk, Svajune Muralyte, Ana Steponkiene, Lukas Vasionis, Danas Baksa |
| EPI_ISL_3189997 | UAB "Baltic Medics" | National Public Health Surveillance Laboratory | Lukas Zemaitis, Migle Gabrielaite, Jelena Razmuk, Svajune Muralyte, Ana Steponkiene, Lukas Vasionis, Danas Baksa |
| EPI_ISL_3189998 | Viešoji įstaiga Klaipėdos universitetinė ligoninė | National Public Health Surveillance Laboratory | Lukas Zemaitis, Migle Gabrielaite, Jelena Razmuk, Svajune Muralyte, Ana Steponkiene, Lukas Vasionis, Danas Baksa |
| EPI_ISL_3189999, EPI_ISL_3190000 | UAB "Rezus.lt" | National Public Health Surveillance Laboratory | Lukas Zemaitis, Migle Gabrielaite, Jelena Razmuk, Svajune Muralyte, Ana Steponkiene, Lukas Vasionis, Danas Baksa |
| EPI_ISL_3190001, EPI_ISL_3190002 | UAB Diagnostikos laboratorija | National Public Health Surveillance Laboratory | Lukas Zemaitis, Migle Gabrielaite, Jelena Razmuk, Svajune Muralyte, Ana Steponkiene, Lukas Vasionis, Danas Baksa |
| EPI_ISL_3190003 | Viešoji įstaiga Vilniaus universiteto ligoninė Santaros klinikos | National Public Health Surveillance Laboratory | Lukas Zemaitis, Migle Gabrielaite, Jelena Razmuk, Svajune Muralyte, Ana Steponkiene, Lukas Vasionis, Danas Baksa |
| EPI_ISL_3190004 | UAB Diagnostikos laboratorija | National Public Health Surveillance Laboratory | Lukas Zemaitis, Migle Gabrielaite, Jelena Razmuk, Svajune Muralyte, Ana Steponkiene, Lukas Vasionis, Danas Baksa |
| EPI_ISL_3190006 | UAB InMedica | National Public Health Surveillance Laboratory | Lukas Zemaitis, Migle Gabrielaite, Jelena Razmuk, Svajune Muralyte, Ana Steponkiene, Lukas Vasionis, Danas Baksa |
| EPI_ISL_3190007, EPI_ISL_3190008 | Viešoji įstaiga Vilniaus universiteto ligoninė Santaros klinikos | National Public Health Surveillance Laboratory | Lukas Zemaitis, Migle Gabrielaite, Jelena Razmuk, Svajune Muralyte, Ana Steponkiene, Lukas Vasionis, Danas Baksa |
| EPI_ISL_3190009 | Viešoji įstaiga Respublikinė Šiaulių ligoninė | National Public Health Surveillance Laboratory | Lukas Zemaitis, Migle Gabrielaite, Jelena Razmuk, Svajune Muralyte, Ana Steponkiene, Lukas Vasionis, Danas Baksa |
| EPI_ISL_3190010 | Nacionalinė visuomenės sveikatos priežiūros laboratorija | National Public Health Surveillance Laboratory | Lukas Zemaitis, Migle Gabrielaite, Jelena Razmuk, Svajune Muralyte, Ana Steponkiene, Lukas Vasionis, Danas Baksa |
| EPI_ISL_3190011 | UAB "Rezus.lt" | National Public Health Surveillance Laboratory | Lukas Zemaitis, Migle Gabrielaite, Jelena Razmuk, Svajune Muralyte, Ana Steponkiene, Lukas Vasionis, Danas Baksa |
| EPI_ISL_3190012, EPI_ISL_3190014, EPI_ISL_3190015 | Viešoji įstaiga Vilniaus universiteto ligoninė Santaros klinikos | National Public Health Surveillance Laboratory | Lukas Zemaitis, Migle Gabrielaite, Jelena Razmuk, Svajune Muralyte, Ana Steponkiene, Lukas Vasionis, Danas Baksa |
| EPI_ISL_3190016, EPI_ISL_3190017, EPI_ISL_3190018, EPI_ISL_3190019 | Viešoji įstaiga Klaipėdos universitetinė ligoninė | Institute of Biotechnology, Life Sciences Center, Vilnius University | Emilija Vasiliunaite; Milda Norkiene; Danguole Ziogiene; Albertas Timinskas; Alma Gedvilaite |
| EPI_ISL_3190021 | UAB "Baltic Medics" | Institute of Biotechnology, Life Sciences Center, Vilnius University | Emilija Vasiliunaite; Milda Norkiene; Danguole Ziogiene; Albertas Timinskas; Alma Gedvilaite |
| EPI_ISL_3190022 | Viešoji įstaiga Respublikinė Šiaulių ligoninė | Institute of Biotechnology, Life Sciences Center, Vilnius University | Emilija Vasiliunaite; Milda Norkiene; Danguole Ziogiene; Albertas Timinskas; Alma Gedvilaite |
| EPI_ISL_3190023, EPI_ISL_3190024, EPI_ISL_3190025 | Viešoji įstaiga Klaipėdos universitetinė ligoninė | Institute of Biotechnology, Life Sciences Center, Vilnius University | Emilija Vasiliunaite; Milda Norkiene; Danguole Ziogiene; Albertas Timinskas; Alma Gedvilaite |
| EPI_ISL_3190026 | SYNLAB Lietuva UAB | Institute of Biotechnology, Life Sciences Center, Vilnius University | Emilija Vasiliunaite; Milda Norkiene; Danguole Ziogiene; Albertas Timinskas; Alma Gedvilaite |
| EPI_ISL_3190027 | Viešoji įstaiga Respublikinė Šiaulių ligoninė | Institute of Biotechnology, Life Sciences Center, Vilnius University | Emilija Vasiliunaite; Milda Norkiene; Danguole Ziogiene; Albertas Timinskas; Alma Gedvilaite |
| EPI_ISL_3190028 | SYNLAB Lietuva UAB | Institute of Biotechnology, Life Sciences Center, Vilnius University | Emilija Vasiliunaite; Milda Norkiene; Danguole Ziogiene; Albertas Timinskas; Alma Gedvilaite |
| EPI_ISL_3190029 | UAB "Baltic Medics" | Institute of Biotechnology, Life Sciences Center, Vilnius University | Emilija Vasiliunaite; Milda Norkiene; Danguole Ziogiene; Albertas Timinskas; Alma Gedvilaite |
| EPI_ISL_3190030, EPI_ISL_3190031, EPI_ISL_3190032 | Viešoji įstaiga Klaipėdos universitetinė ligoninė | Institute of Biotechnology, Life Sciences Center, Vilnius University | Emilija Vasiliunaite; Milda Norkiene; Danguole Ziogiene; Albertas Timinskas; Alma Gedvilaite |
| EPI_ISL_3190033, EPI_ISL_3190035 | UAB "Baltic Medics" | Institute of Biotechnology, Life Sciences Center, Vilnius University | Emilija Vasiliunaite; Milda Norkiene; Danguole Ziogiene; Albertas Timinskas; Alma Gedvilaite |
| EPI_ISL_3190036, EPI_ISL_3190037 | Viešoji įstaiga Klaipėdos universitetinė ligoninė | Institute of Biotechnology, Life Sciences Center, Vilnius University | Emilija Vasiliunaite; Milda Norkiene; Danguole Ziogiene; Albertas Timinskas; Alma Gedvilaite |
| EPI_ISL_3190038 | Viešoji įstaiga Respublikinė Šiaulių ligoninė | Institute of Biotechnology, Life Sciences Center, Vilnius University | Emilija Vasiliunaite; Milda Norkiene; Danguole Ziogiene; Albertas Timinskas; Alma Gedvilaite |
| EPI_ISL_3190039 | UAB "Baltic Medics" | Institute of Biotechnology, Life Sciences Center, Vilnius University | Emilija Vasiliunaite; Milda Norkiene; Danguole Ziogiene; Albertas Timinskas; Alma Gedvilaite |
| EPI_ISL_3190040 | Nacionalinė visuomenės sveikatos priežiūros laboratorija | Institute of Biotechnology, Life Sciences Center, Vilnius University | Emilija Vasiliunaite; Milda Norkiene; Danguole Ziogiene; Albertas Timinskas; Alma Gedvilaite |

| | | | |
|---|---|---|---|
| EPI_ISL_3190041, EPI_ISL_3190042, EPI_ISL_3190043 | Viešoji įstaiga Klaipėdos universitetinė ligoninė | Institute of Biotechnology, Life Sciences Center, Vilnius University | Emilija Vasiliunaite; Milda Norkiene; Danguole Ziogiene; Albertas Timinskas; Alma Gedvilaite |
| EPI_ISL_3190044 | Viešoji įstaiga Respublikinė Šiaulių ligoninė | Institute of Biotechnology, Life Sciences Center, Vilnius University | Emilija Vasiliunaite; Milda Norkiene; Danguole Ziogiene; Albertas Timinskas; Alma Gedvilaite |
| EPI_ISL_3190046, EPI_ISL_3190047 | Viešoji įstaiga Klaipėdos universitetinė ligoninė | Institute of Biotechnology, Life Sciences Center, Vilnius University | Emilija Vasiliunaite; Milda Norkiene; Danguole Ziogiene; Albertas Timinskas; Alma Gedvilaite |
| EPI_ISL_3190048 | Viešoji įstaiga Respublikinė Šiaulių ligoninė | Institute of Biotechnology, Life Sciences Center, Vilnius University | Emilija Vasiliunaite; Milda Norkiene; Danguole Ziogiene; Albertas Timinskas; Alma Gedvilaite |
| EPI_ISL_3190050 | Nacionalinė visuomenės sveikatos priežiūros laboratorija | Institute of Biotechnology, Life Sciences Center, Vilnius University | Emilija Vasiliunaite; Milda Norkiene; Danguole Ziogiene; Albertas Timinskas; Alma Gedvilaite |
| EPI_ISL_3190051, EPI_ISL_3190053 | Viešoji įstaiga Klaipėdos universitetinė ligoninė | Institute of Biotechnology, Life Sciences Center, Vilnius University | Emilija Vasiliunaite; Milda Norkiene; Danguole Ziogiene; Albertas Timinskas; Alma Gedvilaite |
| EPI_ISL_3190054 | Nacionalinė visuomenės sveikatos priežiūros laboratorija | Institute of Biotechnology, Life Sciences Center, Vilnius University | Emilija Vasiliunaite; Milda Norkiene; Danguole Ziogiene; Albertas Timinskas; Alma Gedvilaite |
| EPI_ISL_3190073 | DC Public Health Lab/ Dept. of Forensic Sciences | DC Public Health Lab/ Dept. of Forensic Sciences | Janis Doss, Scott Nguyen, Elizabeth Zelaya, Sarah Scott, Connie Maza, Monica Mann, Brittany Hamilton, David Payne, Jocelyn Hauser |
| EPI_ISL_3190074 | TN DOH Lab Services | TN DOH Lab Services | Karen Beasley-Maynard, Brian Selinsky |
| EPI_ISL_3190085, EPI_ISL_3190086, EPI_ISL_3190087, EPI_ISL_3190088, EPI_ISL_3190089, EPI_ISL_3190090, EPI_ISL_3190091, EPI_ISL_3190092, EPI_ISL_3190093, EPI_ISL_3190094, EPI_ISL_3190095, EPI_ISL_3190096, EPI_ISL_3190097, EPI_ISL_3190098, EPI_ISL_3190099, EPI_ISL_3190100, EPI_ISL_3190101, EPI_ISL_3190102, EPI_ISL_3190103, EPI_ISL_3190104, EPI_ISL_3190105, EPI_ISL_3190106, EPI_ISL_3190107, EPI_ISL_3190108, EPI_ISL_3190109, EPI_ISL_3190110, EPI_ISL_3190111, EPI_ISL_3190112, EPI_ISL_3190113, EPI_ISL_3190114, EPI_ISL_3190115 | | | |
| see above | Kansas Health and Environmental Lab | Kansas Health and Environmental Lab | Mike Grose, Katherine Wiggins, Jonathan Barnell, Ben Olsen, and Phil Adam |
| EPI_ISL_3190116, EPI_ISL_3190117 | CH de l'Ouest Guyanais Frank Joly | Institut Pasteur de la Guyane | Anne Lavergne, Dominique Rousset, Antoine Enfissi, Arielle Salmier |
| EPI_ISL_3190124, EPI_ISL_3190126, EPI_ISL_3190128 | Laboratoire Carage | Institut Pasteur de la Guyane | Anne Lavergne, Dominique Rousset, Antoine Enfissi, Arielle Salmier |
| EPI_ISL_3190130, EPI_ISL_3190131 | CNR Institut Pasteur de la Guyane | Institut Pasteur de la Guyane | Anne Lavergne, Dominique Rousset, Antoine Enfissi, Arielle Salmier |
| EPI_ISL_3190134, EPI_ISL_3190135, EPI_ISL_3190136, EPI_ISL_3190137, EPI_ISL_3190138, EPI_ISL_3190139, EPI_ISL_3190140, EPI_ISL_3190141, EPI_ISL_3190142, EPI_ISL_3190143, EPI_ISL_3190144, EPI_ISL_3190145, EPI_ISL_3190146, EPI_ISL_3190147, EPI_ISL_3190148, EPI_ISL_3190149, EPI_ISL_3190151, EPI_ISL_3190152, EPI_ISL_3190153, EPI_ISL_3190154, EPI_ISL_3190155, EPI_ISL_3190156, EPI_ISL_3190157, EPI_ISL_3190158, EPI_ISL_3190159, EPI_ISL_3190160, EPI_ISL_3190161, EPI_ISL_3190162, EPI_ISL_3190163, EPI_ISL_3190164, EPI_ISL_3190165 | | | |
| see above | Kansas Health and Environmental Lab | Kansas Health and Environmental Lab | Mike Grose, Katherine Wiggins, Jonathan Barnell, Ben Olsen, and Phil Adam |
| EPI_ISL_3190412 | Division of Microbiology, Osaka Institute of Public Health | Division of Microbiology, Osaka Institute of Public Health | Ryuji Kawahara, Satoshi Hiroi, Saeko Morikawa, Haruyo Mori, Kazushi Motomura, Jun Ogasawara, Tsuyoshi Sekizuka, Kentaro Itokawa, Rina Tanaka, Masanori Hashino, Makoto Kuroda |
| EPI_ISL_3190852, EPI_ISL_3190853, EPI_ISL_3190858, EPI_ISL_3190859, EPI_ISL_3190860, EPI_ISL_3190861, EPI_ISL_3190862, EPI_ISL_3190863, EPI_ISL_3190864, EPI_ISL_3190865, EPI_ISL_3190866, EPI_ISL_3190867, EPI_ISL_3190868, EPI_ISL_3190869, EPI_ISL_3190870, EPI_ISL_3190871, EPI_ISL_3190872, EPI_ISL_3190873, EPI_ISL_3190874, EPI_ISL_3190875, EPI_ISL_3190876, EPI_ISL_3190877, EPI_ISL_3190878, EPI_ISL_3190879, EPI_ISL_3190880, EPI_ISL_3190881, EPI_ISL_3190882, EPI_ISL_3190883, EPI_ISL_3190884, EPI_ISL_3190885, EPI_ISL_3190886, EPI_ISL_3190887, EPI_ISL_3190888, EPI_ISL_3190889, EPI_ISL_3190890, EPI_ISL_3190891, EPI_ISL_3190892, EPI_ISL_3190893, EPI_ISL_3190894, EPI_ISL_3190895, EPI_ISL_3190896, EPI_ISL_3190897, EPI_ISL_3190898, EPI_ISL_3190899, EPI_ISL_3190900, EPI_ISL_3190901, EPI_ISL_3190902, EPI_ISL_3190903, EPI_ISL_3190904, EPI_ISL_3190905, EPI_ISL_3190906, EPI_ISL_3190907, EPI_ISL_3190908, EPI_ISL_3190909, EPI_ISL_3190910, EPI_ISL_3190911, EPI_ISL_3190912 | | | |
| see above | Pathogen Genomics Center, National Institute of Infectious Diseases | Pathogen Genomics Center, National Institute of Infectious Diseases | Tsuyoshi Sekizuka, Kentaro Itokawa, Rina Tanaka, Masanori Hashino, Makoto Kuroda |
| EPI_ISL_3190913 | SARS-CoV-2 testing team, National Institute of Infectious Diseases | Pathogen Genomics Center, National Institute of Infectious Diseases | Tsuyoshi Sekizuka, Kentaro Itokawa, Rina Tanaka, Masanori Hashino, Nozomu Hanaoka, Masumichi Saito, Naomi Nojiri, Hazuka Y Furihata, Hiromizu Takahashi, Tsuguto Fujimoto, Makoto Kuroda |
| EPI_ISL_3190992, EPI_ISL_3190994, EPI_ISL_3191042, EPI_ISL_3191044 | Idaho Bureau of Laboratories | Idaho Bureau of Laboratories | "R. Beukelman, Matthew Charles Burns, Aimee Ceniseros, Christian Loera, Robert L. Voermans, Christopher Ball" |
| EPI_ISL_3191058, EPI_ISL_3191682, EPI_ISL_3191683, EPI_ISL_3191684 | Department of Infectious Diseases, Kobe Institute of Health | Department of Infectious Diseases, Kobe Institute of Health | Ryohei Nomoto, Noriko Nakanishi, Tomotada Iwamoto, Tsuyoshi Sekizuka, Kentaro Itokawa, Rina Tanaka, Masanori Hashino, Makoto Kuroda |
| EPI_ISL_3191685, EPI_ISL_3191686, EPI_ISL_3191687, EPI_ISL_3191688 | Division of Microbiology, Osaka Institute of Public Health | Division of Microbiology, Osaka Institute of Public Health | Ryuji Kawahara, Satoshi Hiroi, Saeko Morikawa, Haruyo Mori, Kazushi Motomura, Jun Ogasawara, Tsuyoshi Sekizuka, Kentaro Itokawa, Rina Tanaka, Masanori Hashino, Makoto Kuroda |
| EPI_ISL_3191699, EPI_ISL_3191700, EPI_ISL_3191701, EPI_ISL_3191702, EPI_ISL_3191703, EPI_ISL_3191704, EPI_ISL_3191705, EPI_ISL_3191706 | Department of Infectious Diseases, Kobe Institute of Health | Department of Infectious Diseases, Kobe Institute of Health | Ryohei Nomoto, Noriko Nakanishi, Tomotada Iwamoto, Tsuyoshi Sekizuka, Kentaro Itokawa, Rina Tanaka, Masanori Hashino, Makoto Kuroda |
| EPI_ISL_3191707, EPI_ISL_3191708, EPI_ISL_3191709, EPI_ISL_3191710, EPI_ISL_3191711, EPI_ISL_3191712, EPI_ISL_3191713, EPI_ISL_3191714, EPI_ISL_3191715, EPI_ISL_3191716, EPI_ISL_3191717, EPI_ISL_3191718, EPI_ISL_3191719, EPI_ISL_3191720, EPI_ISL_3191721, EPI_ISL_3191722, EPI_ISL_3191723, EPI_ISL_3191724, EPI_ISL_3191725, EPI_ISL_3191726, EPI_ISL_3191727, EPI_ISL_3191728, EPI_ISL_3191729 | | | |
| see above | Kanagawa Prefectural Institute of Public Health | Kanagawa Prefectural Institute of Public Health | Takayuki Hishiki, Rieko Suzuki, Takako Sano, Makiko Kondo, Sumi Watanabe, Itsumi Toyokura, Junichi Sakuragi, Tomohiko Takasaki, Tsuyoshi Sekizuka, Kentaro Itokawa, Rina Tanaka, Masanori Hashino, Makoto Kuroda |
| EPI_ISL_3191730, EPI_ISL_3191731, EPI_ISL_3191732, EPI_ISL_3191733, EPI_ISL_3191734, EPI_ISL_3191735, EPI_ISL_3191736, EPI_ISL_3191737, EPI_ISL_3191738, EPI_ISL_3191739, EPI_ISL_3191740, EPI_ISL_3191741, EPI_ISL_3191742, EPI_ISL_3191743, EPI_ISL_3191744, EPI_ISL_3191745, EPI_ISL_3191746, EPI_ISL_3191747, EPI_ISL_3191748, EPI_ISL_3191749, EPI_ISL_3191750, EPI_ISL_3191751, EPI_ISL_3191752, EPI_ISL_3191753, EPI_ISL_3191754, EPI_ISL_3191755, EPI_ISL_3191756, EPI_ISL_3191757, EPI_ISL_3191758, EPI_ISL_3191759, EPI_ISL_3191760, EPI_ISL_3191761 | | | |
| see above | Chiba Prefectural Institute of Public Health | Division of Virology and Zoology, Chiba Prefectural Institute of Public Health | Masakatsu Taira, Yuki Fujinuma, Haruna Nishijima, Mari Ohta, Yuji Hanada, Mika Takeuchi, Akihiro Takemura, Shigenori Sato, Tsuyoshi Sekizuka, Kentaro Itokawa, Rina Tanaka, Masanori Hashino, Makoto Kuroda |
| EPI_ISL_3191762 | Division of Microbiology, Osaka Institute of Public Health | Division of Microbiology, Osaka Institute of Public Health | Ryuji Kawahara, Satoshi Hiroi, Saeko Morikawa, Haruyo Mori, Kazushi Motomura, Jun Ogasawara, Tsuyoshi Sekizuka, Kentaro Itokawa, Rina Tanaka, Masanori Hashino, Makoto Kuroda |
| EPI_ISL_3191763, EPI_ISL_3191764, EPI_ISL_3191765, EPI_ISL_3191766, EPI_ISL_3191767, EPI_ISL_3191768, EPI_ISL_3191769 | Tottori Prefectural Institute of Public Health and Environmental Science | Tottori Prefectural Institute of Public Health and Environmental Science | Yutaka Ueda, Rei Ootomo, Eiji Okuda, Tsuyoshi Sekizuka, Kentaro Itokawa, Rina Tanaka, Masanori Hashino, Makoto Kuroda |
| EPI_ISL_3193110, EPI_ISL_3193111, EPI_ISL_3193112, EPI_ISL_3193117, EPI_ISL_3193118, EPI_ISL_3193119, EPI_ISL_3193122, EPI_ISL_3193127, EPI_ISL_3193128, EPI_ISL_3193131, EPI_ISL_3193132, EPI_ISL_3193133, EPI_ISL_3193134, EPI_ISL_3193146, EPI_ISL_3193147, EPI_ISL_3193148, EPI_ISL_3193151, EPI_ISL_3193152, EPI_ISL_3193157, EPI_ISL_3193160, EPI_ISL_3193210, EPI_ISL_3193211, EPI_ISL_3194032, EPI_ISL_3194723, EPI_ISL_3194724, EPI_ISL_3194757, EPI_ISL_3194758, EPI_ISL_3194759, EPI_ISL_3194760 | | | |
| see above | SARS-CoV-2 testing team, National Institute of Infectious Diseases | Pathogen Genomics Center, National Institute of Infectious Diseases | Tsuyoshi Sekizuka, Kentaro Itokawa, Rina Tanaka, Masanori Hashino, Nozomu Hanaoka, Masumichi Saito, Naomi Nojiri, Hazuka Y Furihata, Hiromizu Takahashi, Tsuguto Fujimoto, Makoto Kuroda |
| EPI_ISL_3194785, EPI_ISL_3194801, EPI_ISL_3194821, EPI_ISL_3194837 | Pathogen Genomics Center, National Institute of Infectious Diseases | Pathogen Genomics Center, National Institute of Infectious Diseases | Tsuyoshi Sekizuka, Kentaro Itokawa, Rina Tanaka, Masanori Hashino, Makoto Kuroda |
| EPI_ISL_3195061, EPI_ISL_3195062, EPI_ISL_3195063, EPI_ISL_3195082, EPI_ISL_3195095, EPI_ISL_3195112, EPI_ISL_3195113, EPI_ISL_3195114, EPI_ISL_3195115, EPI_ISL_3195116, EPI_ISL_3195117 | | | |
| see above | SARS-CoV-2 testing team, National Institute of Infectious Diseases | Pathogen Genomics Center, National Institute of Infectious Diseases | Tsuyoshi Sekizuka, Kentaro Itokawa, Rina Tanaka, Masanori Hashino, Nozomu Hanaoka, Masumichi Saito, Naomi Nojiri, Hazuka Y Furihata, Hiromizu Takahashi, Tsuguto Fujimoto, Makoto Kuroda |
| EPI_ISL_3195268, EPI_ISL_3195269, EPI_ISL_3195270, EPI_ISL_3195271, EPI_ISL_3195276, EPI_ISL_3195277, EPI_ISL_3195279, EPI_ISL_3195280, EPI_ISL_3195282, EPI_ISL_3195283, EPI_ISL_3195284, EPI_ISL_3195285, EPI_ISL_3195286, EPI_ISL_3195287, EPI_ISL_3195288, EPI_ISL_3195293, EPI_ISL_3195294, EPI_ISL_3195296, EPI_ISL_3195297, EPI_ISL_3195300, EPI_ISL_3195301, EPI_ISL_3195302, EPI_ISL_3195303, EPI_ISL_3195304, EPI_ISL_3195305, EPI_ISL_3195306, EPI_ISL_3195308, EPI_ISL_3195309, EPI_ISL_3195310, EPI_ISL_3195311, EPI_ISL_3195312, EPI_ISL_3195313, EPI_ISL_3195314, EPI_ISL_3195315, EPI_ISL_3195316, EPI_ISL_3195317, EPI_ISL_3195318, EPI_ISL_3195319, EPI_ISL_3195345, EPI_ISL_3195354, EPI_ISL_3195375, EPI_ISL_3195398, EPI_ISL_3195423, EPI_ISL_3195439, EPI_ISL_3195441, EPI_ISL_3195449, EPI_ISL_3195452, EPI_ISL_3195455, EPI_ISL_3195467, EPI_ISL_3195470, EPI_ISL_3195487, EPI_ISL_3195492, EPI_ISL_3195497, EPI_ISL_3195502 | | | |
| see above | Pathogen Genomics Center, National Institute of Infectious Diseases | Pathogen Genomics Center, National Institute of Infectious Diseases | Tsuyoshi Sekizuka, Kentaro Itokawa, Rina Tanaka, Masanori Hashino, Makoto Kuroda |
| EPI_ISL_3195604, EPI_ISL_3195605, EPI_ISL_3195651, EPI_ISL_3195660, EPI_ISL_3195661, EPI_ISL_3195662, EPI_ISL_3195663, EPI_ISL_3195664, EPI_ISL_3195665, EPI_ISL_3195666, EPI_ISL_3195667, EPI_ISL_3195773, EPI_ISL_3195775, EPI_ISL_3195779, EPI_ISL_3195782, EPI_ISL_3195783, EPI_ISL_3195787, EPI_ISL_3195788, EPI_ISL_3195863, EPI_ISL_3195864, EPI_ISL_3195865, EPI_ISL_3195866, EPI_ISL_3195867, EPI_ISL_3195868, EPI_ISL_3195869, EPI_ISL_3195870, EPI_ISL_3195871, EPI_ISL_3195872, EPI_ISL_3195873 | | | |
| see above | SARS-CoV-2 testing team, National Institute of Infectious Diseases | Pathogen Genomics Center, National Institute of Infectious Diseases | Tsuyoshi Sekizuka, Kentaro Itokawa, Rina Tanaka, Masanori Hashino, Nozomu Hanaoka, Masumichi Saito, Naomi Nojiri, Hazuka Y Furihata, Hiromizu Takahashi, Tsuguto Fujimoto, Makoto Kuroda |
| EPI_ISL_3196952 | Pathogen Genomics Center, National Institute of Infectious Diseases | Pathogen Genomics Center, National Institute of Infectious Diseases | Tsuyoshi Sekizuka, Kentaro Itokawa, Rina Tanaka, Masanori Hashino, Makoto Kuroda |
| EPI_ISL_3196953, | SARS-CoV-2 testing team, National Institute of Infectious | Pathogen Genomics Center, National Institute of Infectious Diseases | Tsuyoshi Sekizuka, Kentaro Itokawa, Rina Tanaka, Masanori Hashino, Nozomu Hanaoka, Masumichi Saito, Naomi Nojiri, Hazuka Y Furihata, Hiromizu Takahashi, Tsuguto Fujimoto, Makoto Kuroda |

| | | | |
|---|---|---|---|
| EPI_ISL_3196954, EPI_ISL_3196955 | Diseases | | |
| EPI_ISL_3196960, EPI_ISL_3196961, EPI_ISL_3196962, EPI_ISL_3196963, EPI_ISL_3196964 | Pathogen Genomics Center, National Institute of Infectious Diseases | Pathogen Genomics Center, National Institute of Infectious Diseases | Tsuyoshi Sekizuka, Kentaro Itokawa, Rina Tanaka, Masanori Hashino, Makoto Kuroda |
| EPI_ISL_3196965, EPI_ISL_3196966, EPI_ISL_3196967, EPI_ISL_3196968, EPI_ISL_3196969, EPI_ISL_3196970, EPI_ISL_3196971, EPI_ISL_3196972, EPI_ISL_3196973, EPI_ISL_3196974, EPI_ISL_3196975, EPI_ISL_3196976, EPI_ISL_3196977, EPI_ISL_3196978, EPI_ISL_3196979, EPI_ISL_3196980, EPI_ISL_3196981, EPI_ISL_3196982, EPI_ISL_3196983, EPI_ISL_3196984, EPI_ISL_3196985, EPI_ISL_3196986, EPI_ISL_3196987, EPI_ISL_3196988, EPI_ISL_3196989, EPI_ISL_3196990, EPI_ISL_3196991, EPI_ISL_3196992, EPI_ISL_3196993, EPI_ISL_3196994, EPI_ISL_3196995, EPI_ISL_3196996, EPI_ISL_3196997, EPI_ISL_3196998, EPI_ISL_3196999, EPI_ISL_3197000, EPI_ISL_3197001, EPI_ISL_3197002, EPI_ISL_3197003, EPI_ISL_3197004, EPI_ISL_3197005, EPI_ISL_3197006, EPI_ISL_3197007, EPI_ISL_3197008, EPI_ISL_3197009, EPI_ISL_3197010, EPI_ISL_3197011, EPI_ISL_3197012, EPI_ISL_3197013, EPI_ISL_3197014, EPI_ISL_3197015, EPI_ISL_3197016, EPI_ISL_3197017, EPI_ISL_3197018, EPI_ISL_3197019, EPI_ISL_3197020, EPI_ISL_3197021, EPI_ISL_3197022, EPI_ISL_3197023, EPI_ISL_3197024, EPI_ISL_3197025, EPI_ISL_3197026, EPI_ISL_3197027, EPI_ISL_3197028, EPI_ISL_3197029, EPI_ISL_3197030, EPI_ISL_3197031, EPI_ISL_3197032, EPI_ISL_3197033, EPI_ISL_3197034, EPI_ISL_3198026, EPI_ISL_3198027, EPI_ISL_3198030, EPI_ISL_3198031, EPI_ISL_3198032, EPI_ISL_3198033, EPI_ISL_3198034, EPI_ISL_3198035, EPI_ISL_3198036, EPI_ISL_3198037, EPI_ISL_3198038, EPI_ISL_3198039, EPI_ISL_3199025 | | | |
| see above | SARS-CoV-2 testing team, National Institute of Infectious Diseases | Pathogen Genomics Center, National Institute of Infectious Diseases | Tsuyoshi Sekizuka, Kentaro Itokawa, Rina Tanaka, Masanori Hashino, Nozomu Hanaoka, Masumichi Saito, Naomi Nojiri, Hazuka Y Furihata, Hiromizu Takahashi, Tsuguto Fujimoto, Makoto Kuroda |
| EPI_ISL_3199031, EPI_ISL_3199032, EPI_ISL_3199033, EPI_ISL_3199034, EPI_ISL_3199035, EPI_ISL_3199036, EPI_ISL_3199037, EPI_ISL_3199038, EPI_ISL_3199039 | Pathogen Genomics Center, National Institute of Infectious Diseases | Pathogen Genomics Center, National Institute of Infectious Diseases | Tsuyoshi Sekizuka, Kentaro Itokawa, Rina Tanaka, Masanori Hashino, Makoto Kuroda |
| EPI_ISL_3199040, EPI_ISL_3199041 | SARS-CoV-2 testing team, National Institute of Infectious Diseases | Pathogen Genomics Center, National Institute of Infectious Diseases | Tsuyoshi Sekizuka, Kentaro Itokawa, Rina Tanaka, Masanori Hashino, Nozomu Hanaoka, Masumichi Saito, Naomi Nojiri, Hazuka Y Furihata, Hiromizu Takahashi, Tsuguto Fujimoto, Makoto Kuroda |
| EPI_ISL_3199739, EPI_ISL_3199740, EPI_ISL_3199741, EPI_ISL_3199742, EPI_ISL_3199743, EPI_ISL_3199744, EPI_ISL_3199745, EPI_ISL_3199746, EPI_ISL_3199747, EPI_ISL_3199748, EPI_ISL_3199749, EPI_ISL_3199750, EPI_ISL_3199751, EPI_ISL_3199752, EPI_ISL_3199753, EPI_ISL_3199754, EPI_ISL_3199755, EPI_ISL_3199756, EPI_ISL_3199757, EPI_ISL_3199758, EPI_ISL_3199759 | | | |
| see above | Hyogo Prefectural Institute of Public Health science | Pathogen Genomics Center, National Institute of Infectious Diseases | Kenichi Ogita, Tomohiro Oshibe, Miki Ogi, Denshi Takai, Etsuko Saito, Kyoko Suzuki, Masatsugu Chikahira, Yumi Akiyama, Tsuyoshi Sekizuka, Kentaro Itokawa, Rina Tanaka, Masanori Hashino, Makoto Kuroda |
| EPI_ISL_3199760, EPI_ISL_3199761, EPI_ISL_3199762, EPI_ISL_3199763, EPI_ISL_3199764, EPI_ISL_3199765 | SARS-CoV-2 testing team, National Institute of Infectious Diseases | Pathogen Genomics Center, National Institute of Infectious Diseases | Tsuyoshi Sekizuka, Kentaro Itokawa, Rina Tanaka, Masanori Hashino, Nozomu Hanaoka, Masumichi Saito, Naomi Nojiri, Hazuka Y Furihata, Hiromizu Takahashi, Tsuguto Fujimoto, Makoto Kuroda |
| EPI_ISL_3199781 | Fukuoka Institute of Health and Environmental Sciences | Fukuoka Institute of Health and Environmental Sciences | Yoshiki Etoh, Asako Nakamura, Hiroaki Shigemura, Akira Ohishi, Takayuki Kobayashi, Yuki Carl, Chiharu Katamune, Saori Ueda, Ikumi Sato, Mitsuhiro Hamasaki, Yuki Ashizuka, Tsuyoshi Sekizuka, Kentaro Itokawa, Rina Tanaka, Masanori Hashino, Makoto Kuroda |
| EPI_ISL_3199782, EPI_ISL_3199783, EPI_ISL_3199784, EPI_ISL_3199785, EPI_ISL_3199786, EPI_ISL_3199787 | Hyogo Prefectural Institute of Public Health science | Pathogen Genomics Center, National Institute of Infectious Diseases | Kenichi Ogita, Tomohiro Oshibe, Miki Ogi, Denshi Takai, Etsuko Saito, Kyoko Suzuki, Masatsugu Chikahira, Yumi Akiyama, Tsuyoshi Sekizuka, Kentaro Itokawa, Rina Tanaka, Masanori Hashino, Makoto Kuroda |
| EPI_ISL_3199788 | Pathogen Genomics Center, National Institute of Infectious Diseases | Pathogen Genomics Center, National Institute of Infectious Diseases | Tsuyoshi Sekizuka, Kentaro Itokawa, Rina Tanaka, Masanori Hashino, Makoto Kuroda |
| EPI_ISL_3199789, EPI_ISL_3199790, EPI_ISL_3199791, EPI_ISL_3199792, EPI_ISL_3199793, EPI_ISL_3199794, EPI_ISL_3199795, EPI_ISL_3199796, EPI_ISL_3199797, EPI_ISL_3199798, EPI_ISL_3199799 | | | |
| see above | SARS-CoV-2 testing team, National Institute of Infectious Diseases | Pathogen Genomics Center, National Institute of Infectious Diseases | Tsuyoshi Sekizuka, Kentaro Itokawa, Rina Tanaka, Masanori Hashino, Nozomu Hanaoka, Masumichi Saito, Naomi Nojiri, Hazuka Y Furihata, Hiromizu Takahashi, Tsuguto Fujimoto, Makoto Kuroda |
| EPI_ISL_3199805, EPI_ISL_3199806 | Hyogo Prefectural Institute of Public Health science | Pathogen Genomics Center, National Institute of Infectious Diseases | Kenichi Ogita, Tomohiro Oshibe, Miki Ogi, Denshi Takai, Etsuko Saito, Kyoko Suzuki, Masatsugu Chikahira, Yumi Akiyama, Tsuyoshi Sekizuka, Kentaro Itokawa, Rina Tanaka, Masanori Hashino, Makoto Kuroda |
| EPI_ISL_3200784 | Bunda Thamrin Hospital | Genomik Solidaritas Indonesia Laboratorium | Anna Christina Brazia, Beni Satria, Eka Airlangga, Ayodhia Pitaloka Pasaribu, Hendri Wijaya, Suwarti, Decy Subekti, Raph Hamers, Vania Gavrilla Wikasa, Carissa Sintca Wijaya, Annisa Muthiah Sukirman, Gracia Felias Enos Korompis, Anuraj Shankar, Louisa Markus, Meutia Ayuputeri Kumaheri, Ariel Pradipta |
| EPI_ISL_3200821 | Multidisciplinary Research Unit, Institute of Medical Sciences, Banaras Hindu University, Varanasi -221005 | Multidisciplinary Research Unit, Institute of Medical Sciences, Banaras Hindu University, Varanasi -221005 | Royana Singh, Priyoneel Basu, Chetan Sahni, Nitish Kumar Singh, Deepa Devadas, Sarita Chowdhary, Saurabh Singh, Sangeeta Rai, Saumya Singh, Ashish Ashish, Abhay Kumar Yadav, Manpreet Kaur, Ajay Kumar Yadav, Shivangi Mishra, Meghna Wadi, Shani Vishwakarma, Shivam Tiwari, Meghna Wadi, Nargis Khanam, Janhavi Yadav, Ragini Mishra, Surendra Pratap Mishra, Tribhuwan Mohan Mohapatra |
| EPI_ISL_3200822 | Multidisciplinary Research Unit, Institute of Medical Sciences, Banaras Hindu University, Varanasi - 221005 | Multidisciplinary Research Unit, Institute of Medical Sciences, Banaras Hindu University, Varanasi - 221005 | Royana Singh, Chetan Sahni, Priyoneel Basu, Nitish Kumar Singh, Saumya Singh, Manpreet Kaur, Deepa Devadas, Sarita Chowdhary, Saurabh Singh, Sangeeta Rai, Ashish Ashish, Abhay Kumar Yadav, Ajay Kumar Yadav, Shivangi Mishra, Meghna Wadi, Shani Vishwakarma, Shivam Tiwari, Meghna Wadi, Nargis Khanam, Janhavi Yadav, Ragini Mishra, Surendra Pratap Mishra, Tribhuwan Mohan Mohapatra |
| EPI_ISL_3200823, EPI_ISL_3200824, EPI_ISL_3200825, EPI_ISL_3200826, EPI_ISL_3200827, EPI_ISL_3200828, EPI_ISL_3200829, EPI_ISL_3200831, EPI_ISL_3200832 | Area of Virology, Serology and Virology Division (SAViD), New South Wales Health Pathology Randwick | Virology Research Laboratory; Area of Virology, Serology and Virology Division (SAViD), New South Wales Health Pathology Randwick | Foster, C.; Au, J.; Ruiz Silva, M.; Deveson, I.; Bull, R.; Van Hal, S.; Rawlinson, W. |
| EPI_ISL_3200862, EPI_ISL_3200863, EPI_ISL_3200864, EPI_ISL_3200865, EPI_ISL_3200866, EPI_ISL_3200867, EPI_ISL_3200868, EPI_ISL_3200869, EPI_ISL_3200870, EPI_ISL_3200871, EPI_ISL_3200872, EPI_ISL_3200873, EPI_ISL_3200874, EPI_ISL_3200875, EPI_ISL_3200876, EPI_ISL_3200877, EPI_ISL_3200878, EPI_ISL_3200879, EPI_ISL_3200880, EPI_ISL_3200881, EPI_ISL_3200882, EPI_ISL_3200883, EPI_ISL_3200884, EPI_ISL_3200885, EPI_ISL_3200886, EPI_ISL_3200887, EPI_ISL_3200888, EPI_ISL_3200889, EPI_ISL_3200890, EPI_ISL_3200891, EPI_ISL_3200892, EPI_ISL_3200893, EPI_ISL_3200894, EPI_ISL_3200895, EPI_ISL_3200896, EPI_ISL_3200897, EPI_ISL_3200898, EPI_ISL_3200899, EPI_ISL_3200900, EPI_ISL_3200901, EPI_ISL_3200902, EPI_ISL_3200903, EPI_ISL_3200904, EPI_ISL_3200905, EPI_ISL_3200906, EPI_ISL_3200907, EPI_ISL_3200908, EPI_ISL_3200909, EPI_ISL_3200910, EPI_ISL_3200911, EPI_ISL_3200912, EPI_ISL_3200913, EPI_ISL_3200914, EPI_ISL_3200915, EPI_ISL_3200916, EPI_ISL_3200917, EPI_ISL_3200918, EPI_ISL_3200919, EPI_ISL_3200920, EPI_ISL_3200921, EPI_ISL_3200922, EPI_ISL_3200923, EPI_ISL_3200924, EPI_ISL_3200925, EPI_ISL_3200926, EPI_ISL_3200927, EPI_ISL_3200928, EPI_ISL_3200929, EPI_ISL_3200930, EPI_ISL_3200931, EPI_ISL_3200932, EPI_ISL_3200933, EPI_ISL_3200934, EPI_ISL_3200935, EPI_ISL_3200936, EPI_ISL_3200937, EPI_ISL_3200938, EPI_ISL_3200939, EPI_ISL_3200940, EPI_ISL_3200941, EPI_ISL_3200942, EPI_ISL_3200943, EPI_ISL_3200944, EPI_ISL_3200945, EPI_ISL_3200946, EPI_ISL_3200947, EPI_ISL_3200948, EPI_ISL_3200949, EPI_ISL_3200950, EPI_ISL_3200951, EPI_ISL_3200952, EPI_ISL_3200953, EPI_ISL_3200954, EPI_ISL_3200955, EPI_ISL_3200956, EPI_ISL_3200957, EPI_ISL_3200958, EPI_ISL_3200959, EPI_ISL_3200960, EPI_ISL_3200961, EPI_ISL_3200962, EPI_ISL_3200963, EPI_ISL_3200964, EPI_ISL_3200965, EPI_ISL_3200966, EPI_ISL_3200967, EPI_ISL_3200968, EPI_ISL_3200969, EPI_ISL_3200970, EPI_ISL_3200971, EPI_ISL_3200972, EPI_ISL_3200973, EPI_ISL_3200974, EPI_ISL_3200975, EPI_ISL_3200976, EPI_ISL_3200977, EPI_ISL_3200978, EPI_ISL_3200979, EPI_ISL_3200980, EPI_ISL_3200981, EPI_ISL_3200982, EPI_ISL_3200983, EPI_ISL_3200984, EPI_ISL_3200985, EPI_ISL_3200986, EPI_ISL_3200987, EPI_ISL_3200988, EPI_ISL_3200989, EPI_ISL_3200990, EPI_ISL_3200991, EPI_ISL_3200992, EPI_ISL_3200993, EPI_ISL_3200994, EPI_ISL_3200995, EPI_ISL_3200996, EPI_ISL_3200997, EPI_ISL_3200998, EPI_ISL_3200999, EPI_ISL_3201000, EPI_ISL_3201001, EPI_ISL_3201002, EPI_ISL_3201003, EPI_ISL_3201004, EPI_ISL_3201005, EPI_ISL_3201006, EPI_ISL_3201007, EPI_ISL_3201008, EPI_ISL_3201009, EPI_ISL_3201010, EPI_ISL_3201011, EPI_ISL_3201012, EPI_ISL_3201013, EPI_ISL_3201014, EPI_ISL_3201015, EPI_ISL_3201016, EPI_ISL_3201017, EPI_ISL_3201018, EPI_ISL_3201019, EPI_ISL_3201020, EPI_ISL_3201021, EPI_ISL_3201022, EPI_ISL_3201023, EPI_ISL_3201024, EPI_ISL_3201025, EPI_ISL_3201026, EPI_ISL_3201027, EPI_ISL_3201028, EPI_ISL_3201029, EPI_ISL_3201030, EPI_ISL_3201031, EPI_ISL_3201032, EPI_ISL_3201033, EPI_ISL_3201034, EPI_ISL_3201035, EPI_ISL_3201036, EPI_ISL_3201037, EPI_ISL_3201038, EPI_ISL_3201039, EPI_ISL_3201040, EPI_ISL_3201041, EPI_ISL_3201042, EPI_ISL_3201043, EPI_ISL_3201044, EPI_ISL_3201045, EPI_ISL_3201046, EPI_ISL_3201047, EPI_ISL_3201048, EPI_ISL_3201049, EPI_ISL_3201050, EPI_ISL_3201051, EPI_ISL_3201052, EPI_ISL_3201053, EPI_ISL_3201054, EPI_ISL_3201055, EPI_ISL_3201056, EPI_ISL_3201057, EPI_ISL_3201058, EPI_ISL_3201059, EPI_ISL_3201060, EPI_ISL_3201061, EPI_ISL_3201062, EPI_ISL_3201063, EPI_ISL_3201064, EPI_ISL_3201065, EPI_ISL_3201066, EPI_ISL_3201067, EPI_ISL_3201068, EPI_ISL_3201069, EPI_ISL_3201070, EPI_ISL_3201071, EPI_ISL_3201072, EPI_ISL_3201073, EPI_ISL_3201074, EPI_ISL_3201075, EPI_ISL_3201076, EPI_ISL_3201077, EPI_ISL_3201078, EPI_ISL_3201079, EPI_ISL_3201080, EPI_ISL_3201081, EPI_ISL_3201082, EPI_ISL_3201083, EPI_ISL_3201084, EPI_ISL_3201085, EPI_ISL_3201086, EPI_ISL_3201087, EPI_ISL_3201088, EPI_ISL_3201089, EPI_ISL_3201090, EPI_ISL_3201091, EPI_ISL_3201092, EPI_ISL_3201093, EPI_ISL_3201094, EPI_ISL_3201095, EPI_ISL_3201096, EPI_ISL_3201097, EPI_ISL_3201098, EPI_ISL_3201099, EPI_ISL_3201100, EPI_ISL_3201101, EPI_ISL_3201102, EPI_ISL_3201103, EPI_ISL_3201104, EPI_ISL_3201105, EPI_ISL_3201106, EPI_ISL_3201107, EPI_ISL_3201108, EPI_ISL_3201109, EPI_ISL_3201110, EPI_ISL_3201111, EPI_ISL_3201112, EPI_ISL_3201113, EPI_ISL_3201114, EPI_ISL_3201115, EPI_ISL_3201116, EPI_ISL_3201117, EPI_ISL_3201118, EPI_ISL_3201119, EPI_ISL_3201120, EPI_ISL_3201121, EPI_ISL_3201122, EPI_ISL_3201123, EPI_ISL_3201124, EPI_ISL_3201125, EPI_ISL_3201126, EPI_ISL_3201127, EPI_ISL_3201128, EPI_ISL_3201129, EPI_ISL_3201130, EPI_ISL_3201131, EPI_ISL_3201132, EPI_ISL_3201133, EPI_ISL_3201134, EPI_ISL_3201135, EPI_ISL_3201136, EPI_ISL_3201137, EPI_ISL_3201138, EPI_ISL_3201139, EPI_ISL_3201140, EPI_ISL_3201141, EPI_ISL_3201142, EPI_ISL_3201143, EPI_ISL_3201144, EPI_ISL_3201145, EPI_ISL_3201146, EPI_ISL_3201147, EPI_ISL_3201148, EPI_ISL_3201149, EPI_ISL_3201150, EPI_ISL_3201151, EPI_ISL_3201152, EPI_ISL_3201153, EPI_ISL_3201154, EPI_ISL_3201155, EPI_ISL_3201156, EPI_ISL_3201157, EPI_ISL_3201158, EPI_ISL_3201159, EPI_ISL_3201160, EPI_ISL_3201161, EPI_ISL_3201162, EPI_ISL_3201163, EPI_ISL_3201164, EPI_ISL_3201165, EPI_ISL_3201166, EPI_ISL_3201167, EPI_ISL_3201168, EPI_ISL_3201169, EPI_ISL_3201170, EPI_ISL_3201171, EPI_ISL_3201172, EPI_ISL_3201173, EPI_ISL_3201174, EPI_ISL_3201175, EPI_ISL_3201176, EPI_ISL_3201177, EPI_ISL_3201178, EPI_ISL_3201179, EPI_ISL_3201180, EPI_ISL_3201181, EPI_ISL_3201182, EPI_ISL_3201183, EPI_ISL_3201184, EPI_ISL_3201185, EPI_ISL_3201186, EPI_ISL_3201187, EPI_ISL_3201188, EPI_ISL_3201189, EPI_ISL_3201190, EPI_ISL_3201191, EPI_ISL_3201192, EPI_ISL_3201193, EPI_ISL_3201194, EPI_ISL_3201195, EPI_ISL_3201196, EPI_ISL_3201197, EPI_ISL_3201198, EPI_ISL_3201199, EPI_ISL_3201200, EPI_ISL_3201201, EPI_ISL_3201202, EPI_ISL_3201203, EPI_ISL_3201204, EPI_ISL_3201205, EPI_ISL_3201206, EPI_ISL_3201207, EPI_ISL_3201208, EPI_ISL_3201209, EPI_ISL_3201210, EPI_ISL_3201211, EPI_ISL_3201212, EPI_ISL_3201213, EPI_ISL_3201214, EPI_ISL_3201215, EPI_ISL_3201216, EPI_ISL_3201217, EPI_ISL_3201218, EPI_ISL_3201219, EPI_ISL_3201220, EPI_ISL_3201221, EPI_ISL_3201222, EPI_ISL_3201223, EPI_ISL_3201224, EPI_ISL_3201225, EPI_ISL_3201226, EPI_ISL_3201227, EPI_ISL_3201228, EPI_ISL_3201229, EPI_ISL_3201230, EPI_ISL_3201231, EPI_ISL_3201232, EPI_ISL_3201233, EPI_ISL_3201234, EPI_ISL_3201235, EPI_ISL_3201236, EPI_ISL_3201237, EPI_ISL_3201238, EPI_ISL_3201239, EPI_ISL_3201240, EPI_ISL_3201241, EPI_ISL_3201242, EPI_ISL_3201243, EPI_ISL_3201244, EPI_ISL_3201245, EPI_ISL_3201246, EPI_ISL_3201247, EPI_ISL_3201248, EPI_ISL_3201249, EPI_ISL_3201250, EPI_ISL_3201251, EPI_ISL_3201252, EPI_ISL_3201253, EPI_ISL_3201254, EPI_ISL_3201255, EPI_ISL_3201256, EPI_ISL_3201257, EPI_ISL_3201258, EPI_ISL_3201259, EPI_ISL_3201260, EPI_ISL_3201261, EPI_ISL_3201262, EPI_ISL_3201263, EPI_ISL_3201264, EPI_ISL_3201265, EPI_ISL_3201266, EPI_ISL_3201267, EPI_ISL_3201268, EPI_ISL_3201269, EPI_ISL_3201270, EPI_ISL_3201271, EPI_ISL_3201272, EPI_ISL_3201273, EPI_ISL_3201274, EPI_ISL_3201275, EPI_ISL_3201276, EPI_ISL_3201277, EPI_ISL_3201278, EPI_ISL_3201279, EPI_ISL_3201280, EPI_ISL_3201281, EPI_ISL_3201282, EPI_ISL_3201283, EPI_ISL_3201284, EPI_ISL_3201285, EPI_ISL_3201286, EPI_ISL_3201287, EPI_ISL_3201288, EPI_ISL_3201289, EPI_ISL_3201290, EPI_ISL_3201291, EPI_ISL_3201292, EPI_ISL_3201293, EPI_ISL_3201294, EPI_ISL_3201295, EPI_ISL_3201296, EPI_ISL_3201297, EPI_ISL_3201298, EPI_ISL_3201299, EPI_ISL_3201300, EPI_ISL_3201301, EPI_ISL_3201302, EPI_ISL_3201303, EPI_ISL_3201304, EPI_ISL_3201305, EPI_ISL_3201306, EPI_ISL_3201307, EPI_ISL_3201308, EPI_ISL_3201309, EPI_ISL_3201310, EPI_ISL_3201311, EPI_ISL_3201312, EPI_ISL_3201313, EPI_ISL_3201314, EPI_ISL_3201315, EPI_ISL_3201316, EPI_ISL_3201317, EPI_ISL_3201318, EPI_ISL_3201319, EPI_ISL_3201320, EPI_ISL_3201321, EPI_ISL_3201322, EPI_ISL_3201323, EPI_ISL_3201324, EPI_ISL_3201325, EPI_ISL_3201326, EPI_ISL_3201327, EPI_ISL_3201328, EPI_ISL_3201329, EPI_ISL_3201330, EPI_ISL_3201331, EPI_ISL_3201332, EPI_ISL_3201333, EPI_ISL_3201334, EPI_ISL_3201335, EPI_ISL_3201336, EPI_ISL_3201337, EPI_ISL_3201338, EPI_ISL_3201339, EPI_ISL_3201340, EPI_ISL_3201341, EPI_ISL_3201342, EPI_ISL_3201343, EPI_ISL_3201344, EPI_ISL_3201345, EPI_ISL_3201346, EPI_ISL_3201347, EPI_ISL_3201348, EPI_ISL_3201349, EPI_ISL_3201350, EPI_ISL_3201351, EPI_ISL_3201352, EPI_ISL_3201353, EPI_ISL_3201354, EPI_ISL_3201355, EPI_ISL_3201356, EPI_ISL_3201357, EPI_ISL_3201358, EPI_ISL_3201359, EPI_ISL_3201360, EPI_ISL_3201361, EPI_ISL_3201362, EPI_ISL_3201363, EPI_ISL_3201364, EPI_ISL_3201365, EPI_ISL_3201366, EPI_ISL_3201367, EPI_ISL_3201368, EPI_ISL_3201369, EPI_ISL_3201370, EPI_ISL_3201371, EPI_ISL_3201372, EPI_ISL_3201373, EPI_ISL_3201374, EPI_ISL_3201375, EPI_ISL_3201376, EPI_ISL_3201377, EPI_ISL_3201378, EPI_ISL_3201379, EPI_ISL_3201380, EPI_ISL_3201381, EPI_ISL_3201382, EPI_ISL_3201383, EPI_ISL_3201384, EPI_ISL_3201385, EPI_ISL_3201386, EPI_ISL_3201387, EPI_ISL_3201388, EPI_ISL_3201389, EPI_ISL_3201390, EPI_ISL_3201391, EPI_ISL_3201392, EPI_ISL_3201393, EPI_ISL_3201394, EPI_ISL_3201395, EPI_ISL_3201396, EPI_ISL_3201397, EPI_ISL_3201398, EPI_ISL_3201399, EPI_ISL_3201400, EPI_ISL_3201401, EPI_ISL_3201402, EPI_ISL_3201403, EPI_ISL_3201404, EPI_ISL_3201405, EPI_ISL_3201406, EPI_ISL_3201407, EPI_ISL_3201408, EPI_ISL_3201409, EPI_ISL_3201410, EPI_ISL_3201411, EPI_ISL_3201412, EPI_ISL_3201413, EPI_ISL_3201414, EPI_ISL_3201415, EPI_ISL_3201416, EPI_ISL_3201417, EPI_ISL_3201418, EPI_ISL_3201419, EPI_ISL_3201420, EPI_ISL_3201421, EPI_ISL_3201422, EPI_ISL_3201423, EPI_ISL_3201424, EPI_ISL_3201425, EPI_ISL_3201426, EPI_ISL_3201427, EPI_ISL_3201428, EPI_ISL_3201429, EPI_ISL_3201430, EPI_ISL_3201431, EPI_ISL_3201432, EPI_ISL_3201433, EPI_ISL_3201434, EPI_ISL_3201435, EPI_ISL_3201436, EPI_ISL_3201437, EPI_ISL_3201438, EPI_ISL_3201439, EPI_ISL_3201440, EPI_ISL_3201441, EPI_ISL_3201442, EPI_ISL_3201443, EPI_ISL_3201444, EPI_ISL_3201445 | | | |
| see above | Lighthouse Lab in Milton Keynes | Wellcome Sanger Institute for the COVID-19 Genomics UK (COG-UK) Consortium | The Lighthouse Lab in Milton Keynes and Alex Alderton, Roberto Amato, Jeffrey Barrett, Sonia Goncalves, Ewan Harrison, David K. Jackson, Ian Johnston, Dominic Kwiatkowski, Cordelia Langford, John Sillitoe on behalf of the Wellcome Sanger Institute COVID-19 Surveillance Team |
| EPI_ISL_3201446, EPI_ISL_3201447 | Lighthouse Lab in Glasgow | Wellcome Sanger Institute for the COVID-19 Genomics UK (COG-UK) Consortium | Harper VanSteenhouse, Yumi Kasai, David Gray, Carol Clugston, Anna Dominiczak and Alex Alderton, Roberto Amato, Jeffrey Barrett, Sonia Goncalves, Ewan Harrison, David K. Jackson, Ian Johnston, Dominic Kwiatkowski, Cordelia Langford, John Sillitoe on behalf of the Wellcome Sanger Institute COVID-19 Surveillance Team |
| EPI_ISL_3201448, EPI_ISL_3201449, EPI_ISL_3201450 | Lighthouse Laboratory Plymouth | Wellcome Sanger Institute for the COVID-19 Genomics UK (COG-UK) Consortium | Lighthouse Laboratory Plymouth and Alex Alderton, Roberto Amato, Jeffrey Barrett, Sonia Goncalves, Ewan Harrison, David K. Jackson, Ian Johnston, Dominic Kwiatkowski, Cordelia Langford, John Sillitoe on behalf of the Wellcome Sanger Institute COVID-19 Surveillance Team |
| EPI_ISL_3201451, EPI_ISL_3201452 | Lighthouse Lab in Milton Keynes | Wellcome Sanger Institute for the COVID-19 Genomics UK (COG-UK) Consortium | The Lighthouse Lab in Milton Keynes and Alex Alderton, Roberto Amato, Jeffrey Barrett, Sonia Goncalves, Ewan Harrison, David K. Jackson, Ian Johnston, Dominic Kwiatkowski, Cordelia Langford, John Sillitoe on behalf of the Wellcome Sanger Institute COVID-19 Surveillance Team |
| EPI_ISL_3201453 | Lighthouse Lab in Glasgow | Wellcome Sanger Institute for the COVID-19 Genomics UK (COG-UK) | Harper VanSteenhouse, Yumi Kasai, David Gray, Carol Clugston, Anna Dominiczak and Alex Alderton, Roberto Amato, Jeffrey Barrett, Sonia Goncalves, Ewan Harrison, David K. Jackson, Ian Johnston, Dominic Kwiatkowski, Cordelia Langford, |

| | | | |
|---|---|---|---|
| | | Consortium | John Sillitoe on behalf of the Wellcome Sanger Institute COVID-19 Surveillance Team |
| EPI_ISL_3201454 | Lighthouse Laboratory Plymouth | Wellcome Sanger Institute for the COVID-19 Genomics UK (COG-UK) Consortium | Lighthouse Laboratory Plymouth and Alex Alderton, Roberto Amato, Jeffrey Barrett, Sonia Goncalves, Ewan Harrison, David K. Jackson, Ian Johnston, Dominic Kwiatkowski, Cordelia Langford, John Sillitoe on behalf of the Wellcome Sanger Institute COVID-19 Surveillance Team |
| EPI_ISL_3201455 | Lighthouse Lab in Milton Keynes | Wellcome Sanger Institute for the COVID-19 Genomics UK (COG-UK) Consortium | The Lighthouse Lab in Milton Keynes and Alex Alderton, Roberto Amato, Jeffrey Barrett, Sonia Goncalves, Ewan Harrison, David K. Jackson, Ian Johnston, Dominic Kwiatkowski, Cordelia Langford, John Sillitoe on behalf of the Wellcome Sanger Institute COVID-19 Surveillance Team |
| EPI_ISL_3201456 | Lighthouse Lab in Glasgow | Wellcome Sanger Institute for the COVID-19 Genomics UK (COG-UK) Consortium | Harper VanSteenhouse, Yumi Kasai, David Gray, Carol Clugston, Anna Dominiczak and Alex Alderton, Roberto Amato, Jeffrey Barrett, Sonia Goncalves, Ewan Harrison, David K. Jackson, Ian Johnston, Dominic Kwiatkowski, Cordelia Langford, John Sillitoe on behalf of the Wellcome Sanger Institute COVID-19 Surveillance Team |
| EPI_ISL_3201457 | Lighthouse Lab in Milton Keynes | Wellcome Sanger Institute for the COVID-19 Genomics UK (COG-UK) Consortium | The Lighthouse Lab in Milton Keynes and Alex Alderton, Roberto Amato, Jeffrey Barrett, Sonia Goncalves, Ewan Harrison, David K. Jackson, Ian Johnston, Dominic Kwiatkowski, Cordelia Langford, John Sillitoe on behalf of the Wellcome Sanger Institute COVID-19 Surveillance Team |
| EPI_ISL_3201458 | Lighthouse Laboratory Plymouth | Wellcome Sanger Institute for the COVID-19 Genomics UK (COG-UK) Consortium | Lighthouse Laboratory Plymouth and Alex Alderton, Roberto Amato, Jeffrey Barrett, Sonia Goncalves, Ewan Harrison, David K. Jackson, Ian Johnston, Dominic Kwiatkowski, Cordelia Langford, John Sillitoe on behalf of the Wellcome Sanger Institute COVID-19 Surveillance Team |
| EPI_ISL_3201459, EPI_ISL_3201460 | Lighthouse Lab in Milton Keynes | Wellcome Sanger Institute for the COVID-19 Genomics UK (COG-UK) Consortium | The Lighthouse Lab in Milton Keynes and Alex Alderton, Roberto Amato, Jeffrey Barrett, Sonia Goncalves, Ewan Harrison, David K. Jackson, Ian Johnston, Dominic Kwiatkowski, Cordelia Langford, John Sillitoe on behalf of the Wellcome Sanger Institute COVID-19 Surveillance Team |
| EPI_ISL_3201461, EPI_ISL_3201462 | Lighthouse Lab in Glasgow | Wellcome Sanger Institute for the COVID-19 Genomics UK (COG-UK) Consortium | Harper VanSteenhouse, Yumi Kasai, David Gray, Carol Clugston, Anna Dominiczak and Alex Alderton, Roberto Amato, Jeffrey Barrett, Sonia Goncalves, Ewan Harrison, David K. Jackson, Ian Johnston, Dominic Kwiatkowski, Cordelia Langford, John Sillitoe on behalf of the Wellcome Sanger Institute COVID-19 Surveillance Team |
| EPI_ISL_3201463, EPI_ISL_3201464, EPI_ISL_3201465, EPI_ISL_3201466 | Lighthouse Lab in Milton Keynes | Wellcome Sanger Institute for the COVID-19 Genomics UK (COG-UK) Consortium | The Lighthouse Lab in Milton Keynes and Alex Alderton, Roberto Amato, Jeffrey Barrett, Sonia Goncalves, Ewan Harrison, David K. Jackson, Ian Johnston, Dominic Kwiatkowski, Cordelia Langford, John Sillitoe on behalf of the Wellcome Sanger Institute COVID-19 Surveillance Team |
| EPI_ISL_3201467 | Lighthouse Lab in Glasgow | Wellcome Sanger Institute for the COVID-19 Genomics UK (COG-UK) Consortium | Harper VanSteenhouse, Yumi Kasai, David Gray, Carol Clugston, Anna Dominiczak and Alex Alderton, Roberto Amato, Jeffrey Barrett, Sonia Goncalves, Ewan Harrison, David K. Jackson, Ian Johnston, Dominic Kwiatkowski, Cordelia Langford, John Sillitoe on behalf of the Wellcome Sanger Institute COVID-19 Surveillance Team |
| EPI_ISL_3201468, EPI_ISL_3201469, EPI_ISL_3201470 | Lighthouse Laboratory Plymouth | Wellcome Sanger Institute for the COVID-19 Genomics UK (COG-UK) Consortium | Lighthouse Laboratory Plymouth and Alex Alderton, Roberto Amato, Jeffrey Barrett, Sonia Goncalves, Ewan Harrison, David K. Jackson, Ian Johnston, Dominic Kwiatkowski, Cordelia Langford, John Sillitoe on behalf of the Wellcome Sanger Institute COVID-19 Surveillance Team |
| EPI_ISL_3201471, EPI_ISL_3201472 | Lighthouse Lab in Glasgow | Wellcome Sanger Institute for the COVID-19 Genomics UK (COG-UK) Consortium | Harper VanSteenhouse, Yumi Kasai, David Gray, Carol Clugston, Anna Dominiczak and Alex Alderton, Roberto Amato, Jeffrey Barrett, Sonia Goncalves, Ewan Harrison, David K. Jackson, Ian Johnston, Dominic Kwiatkowski, Cordelia Langford, John Sillitoe on behalf of the Wellcome Sanger Institute COVID-19 Surveillance Team |
| EPI_ISL_3201473 | Lighthouse Lab in Milton Keynes | Wellcome Sanger Institute for the COVID-19 Genomics UK (COG-UK) Consortium | The Lighthouse Lab in Milton Keynes and Alex Alderton, Roberto Amato, Jeffrey Barrett, Sonia Goncalves, Ewan Harrison, David K. Jackson, Ian Johnston, Dominic Kwiatkowski, Cordelia Langford, John Sillitoe on behalf of the Wellcome Sanger Institute COVID-19 Surveillance Team |
| EPI_ISL_3201474 | Lighthouse Laboratory Plymouth | Wellcome Sanger Institute for the COVID-19 Genomics UK (COG-UK) Consortium | Lighthouse Laboratory Plymouth and Alex Alderton, Roberto Amato, Jeffrey Barrett, Sonia Goncalves, Ewan Harrison, David K. Jackson, Ian Johnston, Dominic Kwiatkowski, Cordelia Langford, John Sillitoe on behalf of the Wellcome Sanger Institute COVID-19 Surveillance Team |
| EPI_ISL_3201475, EPI_ISL_3201476, EPI_ISL_3201477, EPI_ISL_3201478, EPI_ISL_3201479 | Lighthouse Lab in Glasgow | Wellcome Sanger Institute for the COVID-19 Genomics UK (COG-UK) Consortium | Harper VanSteenhouse, Yumi Kasai, David Gray, Carol Clugston, Anna Dominiczak and Alex Alderton, Roberto Amato, Jeffrey Barrett, Sonia Goncalves, Ewan Harrison, David K. Jackson, Ian Johnston, Dominic Kwiatkowski, Cordelia Langford, John Sillitoe on behalf of the Wellcome Sanger Institute COVID-19 Surveillance Team |
| EPI_ISL_3201480 | Lighthouse Laboratory Plymouth | Wellcome Sanger Institute for the COVID-19 Genomics UK (COG-UK) Consortium | Lighthouse Laboratory Plymouth and Alex Alderton, Roberto Amato, Jeffrey Barrett, Sonia Goncalves, Ewan Harrison, David K. Jackson, Ian Johnston, Dominic Kwiatkowski, Cordelia Langford, John Sillitoe on behalf of the Wellcome Sanger Institute COVID-19 Surveillance Team |
| EPI_ISL_3201481 | Lighthouse Lab in Glasgow | Wellcome Sanger Institute for the COVID-19 Genomics UK (COG-UK) Consortium | Harper VanSteenhouse, Yumi Kasai, David Gray, Carol Clugston, Anna Dominiczak and Alex Alderton, Roberto Amato, Jeffrey Barrett, Sonia Goncalves, Ewan Harrison, David K. Jackson, Ian Johnston, Dominic Kwiatkowski, Cordelia Langford, John Sillitoe on behalf of the Wellcome Sanger Institute COVID-19 Surveillance Team |
| EPI_ISL_3201482 | Lighthouse Lab in Milton Keynes | Wellcome Sanger Institute for the COVID-19 Genomics UK (COG-UK) Consortium | The Lighthouse Lab in Milton Keynes and Alex Alderton, Roberto Amato, Jeffrey Barrett, Sonia Goncalves, Ewan Harrison, David K. Jackson, Ian Johnston, Dominic Kwiatkowski, Cordelia Langford, John Sillitoe on behalf of the Wellcome Sanger Institute COVID-19 Surveillance Team |
| EPI_ISL_3201483 | Lighthouse Laboratory Plymouth | Wellcome Sanger Institute for the COVID-19 Genomics UK (COG-UK) Consortium | Lighthouse Laboratory Plymouth and Alex Alderton, Roberto Amato, Jeffrey Barrett, Sonia Goncalves, Ewan Harrison, David K. Jackson, Ian Johnston, Dominic Kwiatkowski, Cordelia Langford, John Sillitoe on behalf of the Wellcome Sanger Institute COVID-19 Surveillance Team |
| EPI_ISL_3201484 | Lighthouse Lab in Glasgow | Wellcome Sanger Institute for the COVID-19 Genomics UK (COG-UK) Consortium | Harper VanSteenhouse, Yumi Kasai, David Gray, Carol Clugston, Anna Dominiczak and Alex Alderton, Roberto Amato, Jeffrey Barrett, Sonia Goncalves, Ewan Harrison, David K. Jackson, Ian Johnston, Dominic Kwiatkowski, Cordelia Langford, John Sillitoe on behalf of the Wellcome Sanger Institute COVID-19 Surveillance Team |
| EPI_ISL_3201485, EPI_ISL_3201486, EPI_ISL_3201487 | Lighthouse Laboratory Plymouth | Wellcome Sanger Institute for the COVID-19 Genomics UK (COG-UK) Consortium | Lighthouse Laboratory Plymouth and Alex Alderton, Roberto Amato, Jeffrey Barrett, Sonia Goncalves, Ewan Harrison, David K. Jackson, Ian Johnston, Dominic Kwiatkowski, Cordelia Langford, John Sillitoe on behalf of the Wellcome Sanger Institute COVID-19 Surveillance Team |
| EPI_ISL_3201488, EPI_ISL_3201489, EPI_ISL_3201490 | Lighthouse Lab in Milton Keynes | Wellcome Sanger Institute for the COVID-19 Genomics UK (COG-UK) Consortium | The Lighthouse Lab in Milton Keynes and Alex Alderton, Roberto Amato, Jeffrey Barrett, Sonia Goncalves, Ewan Harrison, David K. Jackson, Ian Johnston, Dominic Kwiatkowski, Cordelia Langford, John Sillitoe on behalf of the Wellcome Sanger Institute COVID-19 Surveillance Team |
| EPI_ISL_3201491, EPI_ISL_3201492, EPI_ISL_3201493, EPI_ISL_3201494 | Lighthouse Lab in Glasgow | Wellcome Sanger Institute for the COVID-19 Genomics UK (COG-UK) Consortium | Harper VanSteenhouse, Yumi Kasai, David Gray, Carol Clugston, Anna Dominiczak and Alex Alderton, Roberto Amato, Jeffrey Barrett, Sonia Goncalves, Ewan Harrison, David K. Jackson, Ian Johnston, Dominic Kwiatkowski, Cordelia Langford, John Sillitoe on behalf of the Wellcome Sanger Institute COVID-19 Surveillance Team |
| EPI_ISL_3201495 | Lighthouse Laboratory Plymouth | Wellcome Sanger Institute for the COVID-19 Genomics UK (COG-UK) Consortium | Lighthouse Laboratory Plymouth and Alex Alderton, Roberto Amato, Jeffrey Barrett, Sonia Goncalves, Ewan Harrison, David K. Jackson, Ian Johnston, Dominic Kwiatkowski, Cordelia Langford, John Sillitoe on behalf of the Wellcome Sanger Institute COVID-19 Surveillance Team |
| EPI_ISL_3201496 | Lighthouse Lab in Glasgow | Wellcome Sanger Institute for the COVID-19 Genomics UK (COG-UK) Consortium | Harper VanSteenhouse, Yumi Kasai, David Gray, Carol Clugston, Anna Dominiczak and Alex Alderton, Roberto Amato, Jeffrey Barrett, Sonia Goncalves, Ewan Harrison, David K. Jackson, Ian Johnston, Dominic Kwiatkowski, Cordelia Langford, John Sillitoe on behalf of the Wellcome Sanger Institute COVID-19 Surveillance Team |
| EPI_ISL_3201497 | Lighthouse Lab in Milton Keynes | Wellcome Sanger Institute for the COVID-19 Genomics UK (COG-UK) Consortium | The Lighthouse Lab in Milton Keynes and Alex Alderton, Roberto Amato, Jeffrey Barrett, Sonia Goncalves, Ewan Harrison, David K. Jackson, Ian Johnston, Dominic Kwiatkowski, Cordelia Langford, John Sillitoe on behalf of the Wellcome Sanger Institute COVID-19 Surveillance Team |
| EPI_ISL_3201498, EPI_ISL_3201499 | Lighthouse Lab in Glasgow | Wellcome Sanger Institute for the COVID-19 Genomics UK (COG-UK) Consortium | Harper VanSteenhouse, Yumi Kasai, David Gray, Carol Clugston, Anna Dominiczak and Alex Alderton, Roberto Amato, Jeffrey Barrett, Sonia Goncalves, Ewan Harrison, David K. Jackson, Ian Johnston, Dominic Kwiatkowski, Cordelia Langford, John Sillitoe on behalf of the Wellcome Sanger Institute COVID-19 Surveillance Team |
| EPI_ISL_3201500, EPI_ISL_3201501 | Lighthouse Lab in Milton Keynes | Wellcome Sanger Institute for the COVID-19 Genomics UK (COG-UK) Consortium | The Lighthouse Lab in Milton Keynes and Alex Alderton, Roberto Amato, Jeffrey Barrett, Sonia Goncalves, Ewan Harrison, David K. Jackson, Ian Johnston, Dominic Kwiatkowski, Cordelia Langford, John Sillitoe on behalf of the Wellcome Sanger Institute COVID-19 Surveillance Team |
| EPI_ISL_3201502 | Lighthouse Laboratory Plymouth | Wellcome Sanger Institute for the COVID-19 Genomics UK (COG-UK) Consortium | Lighthouse Laboratory Plymouth and Alex Alderton, Roberto Amato, Jeffrey Barrett, Sonia Goncalves, Ewan Harrison, David K. Jackson, Ian Johnston, Dominic Kwiatkowski, Cordelia Langford, John Sillitoe on behalf of the Wellcome Sanger Institute COVID-19 Surveillance Team |
| EPI_ISL_3201503 | Lighthouse Lab in Glasgow | Wellcome Sanger Institute for the COVID-19 Genomics UK (COG-UK) Consortium | Harper VanSteenhouse, Yumi Kasai, David Gray, Carol Clugston, Anna Dominiczak and Alex Alderton, Roberto Amato, Jeffrey Barrett, Sonia Goncalves, Ewan Harrison, David K. Jackson, Ian Johnston, Dominic Kwiatkowski, Cordelia Langford, John Sillitoe on behalf of the Wellcome Sanger Institute COVID-19 Surveillance Team |
| EPI_ISL_3201504, EPI_ISL_3201505, EPI_ISL_3201506 | Lighthouse Lab in Milton Keynes | Wellcome Sanger Institute for the COVID-19 Genomics UK (COG-UK) Consortium | The Lighthouse Lab in Milton Keynes and Alex Alderton, Roberto Amato, Jeffrey Barrett, Sonia Goncalves, Ewan Harrison, David K. Jackson, Ian Johnston, Dominic Kwiatkowski, Cordelia Langford, John Sillitoe on behalf of the Wellcome Sanger Institute COVID-19 Surveillance Team |
| EPI_ISL_3201507 | Lighthouse Lab in Glasgow | Wellcome Sanger Institute for the COVID-19 Genomics UK (COG-UK) Consortium | Harper VanSteenhouse, Yumi Kasai, David Gray, Carol Clugston, Anna Dominiczak and Alex Alderton, Roberto Amato, Jeffrey Barrett, Sonia Goncalves, Ewan Harrison, David K. Jackson, Ian Johnston, Dominic Kwiatkowski, Cordelia Langford, John Sillitoe on behalf of the Wellcome Sanger Institute COVID-19 Surveillance Team |
| EPI_ISL_3201508 | Lighthouse Laboratory Plymouth | Wellcome Sanger Institute for the COVID-19 Genomics UK (COG-UK) Consortium | Lighthouse Laboratory Plymouth and Alex Alderton, Roberto Amato, Jeffrey Barrett, Sonia Goncalves, Ewan Harrison, David K. Jackson, Ian Johnston, Dominic Kwiatkowski, Cordelia Langford, John Sillitoe on behalf of the Wellcome Sanger Institute COVID-19 Surveillance Team |
| EPI_ISL_3201509 | Lighthouse Lab in Milton Keynes | Wellcome Sanger Institute for the COVID-19 Genomics UK (COG-UK) Consortium | The Lighthouse Lab in Milton Keynes and Alex Alderton, Roberto Amato, Jeffrey Barrett, Sonia Goncalves, Ewan Harrison, David K. Jackson, Ian Johnston, Dominic Kwiatkowski, Cordelia Langford, John Sillitoe on behalf of the Wellcome Sanger Institute COVID-19 Surveillance Team |
| EPI_ISL_3201510 | Lighthouse Laboratory Plymouth | Wellcome Sanger Institute for the COVID-19 Genomics UK (COG-UK) Consortium | Lighthouse Laboratory Plymouth and Alex Alderton, Roberto Amato, Jeffrey Barrett, Sonia Goncalves, Ewan Harrison, David K. Jackson, Ian Johnston, Dominic Kwiatkowski, Cordelia Langford, John Sillitoe on behalf of the Wellcome Sanger Institute COVID-19 Surveillance Team |
| EPI_ISL_3201511, EPI_ISL_3201512, EPI_ISL_3201513 | Lighthouse Lab in Glasgow | Wellcome Sanger Institute for the COVID-19 Genomics UK (COG-UK) Consortium | Harper VanSteenhouse, Yumi Kasai, David Gray, Carol Clugston, Anna Dominiczak and Alex Alderton, Roberto Amato, Jeffrey Barrett, Sonia Goncalves, Ewan Harrison, David K. Jackson, Ian Johnston, Dominic Kwiatkowski, Cordelia Langford, John Sillitoe on behalf of the Wellcome Sanger Institute COVID-19 Surveillance Team |
| EPI_ISL_3201514 | Lighthouse Laboratory Plymouth | Wellcome Sanger Institute for the COVID-19 Genomics UK (COG-UK) Consortium | Lighthouse Laboratory Plymouth and Alex Alderton, Roberto Amato, Jeffrey Barrett, Sonia Goncalves, Ewan Harrison, David K. Jackson, Ian Johnston, Dominic Kwiatkowski, Cordelia Langford, John Sillitoe on behalf of the Wellcome Sanger Institute COVID-19 Surveillance Team |
| EPI_ISL_3201515, EPI_ISL_3201516 | Lighthouse Lab in Glasgow | Wellcome Sanger Institute for the COVID-19 Genomics UK (COG-UK) Consortium | Harper VanSteenhouse, Yumi Kasai, David Gray, Carol Clugston, Anna Dominiczak and Alex Alderton, Roberto Amato, Jeffrey Barrett, Sonia Goncalves, Ewan Harrison, David K. Jackson, Ian Johnston, Dominic Kwiatkowski, Cordelia Langford, John Sillitoe on behalf of the Wellcome Sanger Institute COVID-19 Surveillance Team |
| EPI_ISL_3201517 | Lighthouse Lab in Milton Keynes | Wellcome Sanger Institute for the COVID-19 Genomics UK (COG-UK) Consortium | The Lighthouse Lab in Milton Keynes and Alex Alderton, Roberto Amato, Jeffrey Barrett, Sonia Goncalves, Ewan Harrison, David K. Jackson, Ian Johnston, Dominic Kwiatkowski, Cordelia Langford, John Sillitoe on behalf of the Wellcome Sanger Institute COVID-19 Surveillance Team |
| EPI_ISL_3201518, EPI_ISL_3201519 | Lighthouse Lab in Glasgow | Wellcome Sanger Institute for the COVID-19 Genomics UK (COG-UK) Consortium | Harper VanSteenhouse, Yumi Kasai, David Gray, Carol Clugston, Anna Dominiczak and Alex Alderton, Roberto Amato, Jeffrey Barrett, Sonia Goncalves, Ewan Harrison, David K. Jackson, Ian Johnston, Dominic Kwiatkowski, Cordelia Langford, John Sillitoe on behalf of the Wellcome Sanger Institute COVID-19 Surveillance Team |
| EPI_ISL_3201520 | Lighthouse Laboratory Plymouth | Wellcome Sanger Institute for the COVID-19 Genomics UK (COG-UK) | Lighthouse Laboratory Plymouth and Alex Alderton, Roberto Amato, Jeffrey Barrett, Sonia Goncalves, Ewan Harrison, David K. Jackson, Ian Johnston, Dominic Kwiatkowski, Cordelia Langford, John Sillitoe on behalf of the Wellcome Sanger |

| | | | |
|---|---|---|---|
| | | Consortium | Institute COVID-19 Surveillance Team |
| EPI_ISL_3201521 | Lighthouse Lab in Milton Keynes | Wellcome Sanger Institute for the COVID-19 Genomics UK (COG-UK) Consortium | The Lighthouse Lab in Milton Keynes and Alex Alderton, Roberto Amato, Jeffrey Barrett, Sonia Goncalves, Ewan Harrison, David K. Jackson, Ian Johnston, Dominic Kwiatkowski, Cordelia Langford, John Sillitoe on behalf of the Wellcome Sanger Institute COVID-19 Surveillance Team |
| EPI_ISL_3201522, EPI_ISL_3201523, EPI_ISL_3201524, EPI_ISL_3201525, EPI_ISL_3201526, EPI_ISL_3201527, EPI_ISL_3201528, EPI_ISL_3201529 | Lighthouse Lab in Glasgow | Wellcome Sanger Institute for the COVID-19 Genomics UK (COG-UK) Consortium | Harper VanSteenhouse, Yumi Kasai, David Gray, Carol Clugston, Anna Dominiczak and Alex Alderton, Roberto Amato, Jeffrey Barrett, Sonia Goncalves, Ewan Harrison, David K. Jackson, Ian Johnston, Dominic Kwiatkowski, Cordelia Langford, John Sillitoe on behalf of the Wellcome Sanger Institute COVID-19 Surveillance Team |
| EPI_ISL_3201530 | Lighthouse Lab in Milton Keynes | Wellcome Sanger Institute for the COVID-19 Genomics UK (COG-UK) Consortium | The Lighthouse Lab in Milton Keynes and Alex Alderton, Roberto Amato, Jeffrey Barrett, Sonia Goncalves, Ewan Harrison, David K. Jackson, Ian Johnston, Dominic Kwiatkowski, Cordelia Langford, John Sillitoe on behalf of the Wellcome Sanger Institute COVID-19 Surveillance Team |
| EPI_ISL_3201531 | Lighthouse Laboratory Plymouth | Wellcome Sanger Institute for the COVID-19 Genomics UK (COG-UK) Consortium | Lighthouse Laboratory Plymouth and Alex Alderton, Roberto Amato, Jeffrey Barrett, Sonia Goncalves, Ewan Harrison, David K. Jackson, Ian Johnston, Dominic Kwiatkowski, Cordelia Langford, John Sillitoe on behalf of the Wellcome Sanger Institute COVID-19 Surveillance Team |
| EPI_ISL_3201532 | Lighthouse Lab in Glasgow | Wellcome Sanger Institute for the COVID-19 Genomics UK (COG-UK) Consortium | Harper VanSteenhouse, Yumi Kasai, David Gray, Carol Clugston, Anna Dominiczak and Alex Alderton, Roberto Amato, Jeffrey Barrett, Sonia Goncalves, Ewan Harrison, David K. Jackson, Ian Johnston, Dominic Kwiatkowski, Cordelia Langford, John Sillitoe on behalf of the Wellcome Sanger Institute COVID-19 Surveillance Team |
| EPI_ISL_3201533 | Lighthouse Lab in Milton Keynes | Wellcome Sanger Institute for the COVID-19 Genomics UK (COG-UK) Consortium | The Lighthouse Lab in Milton Keynes and Alex Alderton, Roberto Amato, Jeffrey Barrett, Sonia Goncalves, Ewan Harrison, David K. Jackson, Ian Johnston, Dominic Kwiatkowski, Cordelia Langford, John Sillitoe on behalf of the Wellcome Sanger Institute COVID-19 Surveillance Team |
| EPI_ISL_3201534 | Lighthouse Laboratory Plymouth | Wellcome Sanger Institute for the COVID-19 Genomics UK (COG-UK) Consortium | Lighthouse Laboratory Plymouth and Alex Alderton, Roberto Amato, Jeffrey Barrett, Sonia Goncalves, Ewan Harrison, David K. Jackson, Ian Johnston, Dominic Kwiatkowski, Cordelia Langford, John Sillitoe on behalf of the Wellcome Sanger Institute COVID-19 Surveillance Team |
| EPI_ISL_3201535 | Lighthouse Lab in Milton Keynes | Wellcome Sanger Institute for the COVID-19 Genomics UK (COG-UK) Consortium | The Lighthouse Lab in Milton Keynes and Alex Alderton, Roberto Amato, Jeffrey Barrett, Sonia Goncalves, Ewan Harrison, David K. Jackson, Ian Johnston, Dominic Kwiatkowski, Cordelia Langford, John Sillitoe on behalf of the Wellcome Sanger Institute COVID-19 Surveillance Team |
| EPI_ISL_3201536 | Lighthouse Lab in Glasgow | Wellcome Sanger Institute for the COVID-19 Genomics UK (COG-UK) Consortium | Harper VanSteenhouse, Yumi Kasai, David Gray, Carol Clugston, Anna Dominiczak and Alex Alderton, Roberto Amato, Jeffrey Barrett, Sonia Goncalves, Ewan Harrison, David K. Jackson, Ian Johnston, Dominic Kwiatkowski, Cordelia Langford, John Sillitoe on behalf of the Wellcome Sanger Institute COVID-19 Surveillance Team |
| EPI_ISL_3201537 | Lighthouse Laboratory Plymouth | Wellcome Sanger Institute for the COVID-19 Genomics UK (COG-UK) Consortium | Lighthouse Laboratory Plymouth and Alex Alderton, Roberto Amato, Jeffrey Barrett, Sonia Goncalves, Ewan Harrison, David K. Jackson, Ian Johnston, Dominic Kwiatkowski, Cordelia Langford, John Sillitoe on behalf of the Wellcome Sanger Institute COVID-19 Surveillance Team |
| EPI_ISL_3201538, EPI_ISL_3201539 | Lighthouse Lab in Milton Keynes | Wellcome Sanger Institute for the COVID-19 Genomics UK (COG-UK) Consortium | The Lighthouse Lab in Milton Keynes and Alex Alderton, Roberto Amato, Jeffrey Barrett, Sonia Goncalves, Ewan Harrison, David K. Jackson, Ian Johnston, Dominic Kwiatkowski, Cordelia Langford, John Sillitoe on behalf of the Wellcome Sanger Institute COVID-19 Surveillance Team |
| EPI_ISL_3201540 | Lighthouse Laboratory Plymouth | Wellcome Sanger Institute for the COVID-19 Genomics UK (COG-UK) Consortium | Lighthouse Laboratory Plymouth and Alex Alderton, Roberto Amato, Jeffrey Barrett, Sonia Goncalves, Ewan Harrison, David K. Jackson, Ian Johnston, Dominic Kwiatkowski, Cordelia Langford, John Sillitoe on behalf of the Wellcome Sanger Institute COVID-19 Surveillance Team |
| EPI_ISL_3201541 | Lighthouse Lab in Milton Keynes | Wellcome Sanger Institute for the COVID-19 Genomics UK (COG-UK) Consortium | The Lighthouse Lab in Milton Keynes and Alex Alderton, Roberto Amato, Jeffrey Barrett, Sonia Goncalves, Ewan Harrison, David K. Jackson, Ian Johnston, Dominic Kwiatkowski, Cordelia Langford, John Sillitoe on behalf of the Wellcome Sanger Institute COVID-19 Surveillance Team |
| EPI_ISL_3201542 | Lighthouse Laboratory Plymouth | Wellcome Sanger Institute for the COVID-19 Genomics UK (COG-UK) Consortium | Lighthouse Laboratory Plymouth and Alex Alderton, Roberto Amato, Jeffrey Barrett, Sonia Goncalves, Ewan Harrison, David K. Jackson, Ian Johnston, Dominic Kwiatkowski, Cordelia Langford, John Sillitoe on behalf of the Wellcome Sanger Institute COVID-19 Surveillance Team |
| EPI_ISL_3201543 | Lighthouse Lab in Glasgow | Wellcome Sanger Institute for the COVID-19 Genomics UK (COG-UK) Consortium | Harper VanSteenhouse, Yumi Kasai, David Gray, Carol Clugston, Anna Dominiczak and Alex Alderton, Roberto Amato, Jeffrey Barrett, Sonia Goncalves, Ewan Harrison, David K. Jackson, Ian Johnston, Dominic Kwiatkowski, Cordelia Langford, John Sillitoe on behalf of the Wellcome Sanger Institute COVID-19 Surveillance Team |
| EPI_ISL_3201544, EPI_ISL_3201545 | Lighthouse Laboratory Plymouth | Wellcome Sanger Institute for the COVID-19 Genomics UK (COG-UK) Consortium | Lighthouse Laboratory Plymouth and Alex Alderton, Roberto Amato, Jeffrey Barrett, Sonia Goncalves, Ewan Harrison, David K. Jackson, Ian Johnston, Dominic Kwiatkowski, Cordelia Langford, John Sillitoe on behalf of the Wellcome Sanger Institute COVID-19 Surveillance Team |
| EPI_ISL_3201546, EPI_ISL_3201547 | Lighthouse Lab in Glasgow | Wellcome Sanger Institute for the COVID-19 Genomics UK (COG-UK) Consortium | Harper VanSteenhouse, Yumi Kasai, David Gray, Carol Clugston, Anna Dominiczak and Alex Alderton, Roberto Amato, Jeffrey Barrett, Sonia Goncalves, Ewan Harrison, David K. Jackson, Ian Johnston, Dominic Kwiatkowski, Cordelia Langford, John Sillitoe on behalf of the Wellcome Sanger Institute COVID-19 Surveillance Team |
| EPI_ISL_3201548, EPI_ISL_3201549, EPI_ISL_3201550 | Lighthouse Lab in Milton Keynes | Wellcome Sanger Institute for the COVID-19 Genomics UK (COG-UK) Consortium | The Lighthouse Lab in Milton Keynes and Alex Alderton, Roberto Amato, Jeffrey Barrett, Sonia Goncalves, Ewan Harrison, David K. Jackson, Ian Johnston, Dominic Kwiatkowski, Cordelia Langford, John Sillitoe on behalf of the Wellcome Sanger Institute COVID-19 Surveillance Team |
| EPI_ISL_3201551, EPI_ISL_3201552 | Lighthouse Laboratory Plymouth | Wellcome Sanger Institute for the COVID-19 Genomics UK (COG-UK) Consortium | Lighthouse Laboratory Plymouth and Alex Alderton, Roberto Amato, Jeffrey Barrett, Sonia Goncalves, Ewan Harrison, David K. Jackson, Ian Johnston, Dominic Kwiatkowski, Cordelia Langford, John Sillitoe on behalf of the Wellcome Sanger Institute COVID-19 Surveillance Team |
| EPI_ISL_3201553 | Lighthouse Lab in Milton Keynes | Wellcome Sanger Institute for the COVID-19 Genomics UK (COG-UK) Consortium | The Lighthouse Lab in Milton Keynes and Alex Alderton, Roberto Amato, Jeffrey Barrett, Sonia Goncalves, Ewan Harrison, David K. Jackson, Ian Johnston, Dominic Kwiatkowski, Cordelia Langford, John Sillitoe on behalf of the Wellcome Sanger Institute COVID-19 Surveillance Team |
| EPI_ISL_3201554, EPI_ISL_3201555, EPI_ISL_3201556, EPI_ISL_3201557 | Lighthouse Lab in Glasgow | Wellcome Sanger Institute for the COVID-19 Genomics UK (COG-UK) Consortium | Harper VanSteenhouse, Yumi Kasai, David Gray, Carol Clugston, Anna Dominiczak and Alex Alderton, Roberto Amato, Jeffrey Barrett, Sonia Goncalves, Ewan Harrison, David K. Jackson, Ian Johnston, Dominic Kwiatkowski, Cordelia Langford, John Sillitoe on behalf of the Wellcome Sanger Institute COVID-19 Surveillance Team |
| EPI_ISL_3201558 | Lighthouse Lab in Milton Keynes | Wellcome Sanger Institute for the COVID-19 Genomics UK (COG-UK) Consortium | The Lighthouse Lab in Milton Keynes and Alex Alderton, Roberto Amato, Jeffrey Barrett, Sonia Goncalves, Ewan Harrison, David K. Jackson, Ian Johnston, Dominic Kwiatkowski, Cordelia Langford, John Sillitoe on behalf of the Wellcome Sanger Institute COVID-19 Surveillance Team |
| EPI_ISL_3201559, EPI_ISL_3201560 | Lighthouse Lab in Glasgow | Wellcome Sanger Institute for the COVID-19 Genomics UK (COG-UK) Consortium | Harper VanSteenhouse, Yumi Kasai, David Gray, Carol Clugston, Anna Dominiczak and Alex Alderton, Roberto Amato, Jeffrey Barrett, Sonia Goncalves, Ewan Harrison, David K. Jackson, Ian Johnston, Dominic Kwiatkowski, Cordelia Langford, John Sillitoe on behalf of the Wellcome Sanger Institute COVID-19 Surveillance Team |
| EPI_ISL_3201561 | Lighthouse Laboratory Plymouth | Wellcome Sanger Institute for the COVID-19 Genomics UK (COG-UK) Consortium | Lighthouse Laboratory Plymouth and Alex Alderton, Roberto Amato, Jeffrey Barrett, Sonia Goncalves, Ewan Harrison, David K. Jackson, Ian Johnston, Dominic Kwiatkowski, Cordelia Langford, John Sillitoe on behalf of the Wellcome Sanger Institute COVID-19 Surveillance Team |
| EPI_ISL_3201562 | Lighthouse Lab in Milton Keynes | Wellcome Sanger Institute for the COVID-19 Genomics UK (COG-UK) Consortium | The Lighthouse Lab in Milton Keynes and Alex Alderton, Roberto Amato, Jeffrey Barrett, Sonia Goncalves, Ewan Harrison, David K. Jackson, Ian Johnston, Dominic Kwiatkowski, Cordelia Langford, John Sillitoe on behalf of the Wellcome Sanger Institute COVID-19 Surveillance Team |
| EPI_ISL_3201563 | Lighthouse Lab in Glasgow | Wellcome Sanger Institute for the COVID-19 Genomics UK (COG-UK) Consortium | Harper VanSteenhouse, Yumi Kasai, David Gray, Carol Clugston, Anna Dominiczak and Alex Alderton, Roberto Amato, Jeffrey Barrett, Sonia Goncalves, Ewan Harrison, David K. Jackson, Ian Johnston, Dominic Kwiatkowski, Cordelia Langford, John Sillitoe on behalf of the Wellcome Sanger Institute COVID-19 Surveillance Team |
| EPI_ISL_3201564 | Lighthouse Lab in Milton Keynes | Wellcome Sanger Institute for the COVID-19 Genomics UK (COG-UK) Consortium | The Lighthouse Lab in Milton Keynes and Alex Alderton, Roberto Amato, Jeffrey Barrett, Sonia Goncalves, Ewan Harrison, David K. Jackson, Ian Johnston, Dominic Kwiatkowski, Cordelia Langford, John Sillitoe on behalf of the Wellcome Sanger Institute COVID-19 Surveillance Team |
| EPI_ISL_3201565 | Lighthouse Laboratory Plymouth | Wellcome Sanger Institute for the COVID-19 Genomics UK (COG-UK) Consortium | Lighthouse Laboratory Plymouth and Alex Alderton, Roberto Amato, Jeffrey Barrett, Sonia Goncalves, Ewan Harrison, David K. Jackson, Ian Johnston, Dominic Kwiatkowski, Cordelia Langford, John Sillitoe on behalf of the Wellcome Sanger Institute COVID-19 Surveillance Team |
| EPI_ISL_3201566, EPI_ISL_3201567, EPI_ISL_3201568 | Lighthouse Lab in Milton Keynes | Wellcome Sanger Institute for the COVID-19 Genomics UK (COG-UK) Consortium | The Lighthouse Lab in Milton Keynes and Alex Alderton, Roberto Amato, Jeffrey Barrett, Sonia Goncalves, Ewan Harrison, David K. Jackson, Ian Johnston, Dominic Kwiatkowski, Cordelia Langford, John Sillitoe on behalf of the Wellcome Sanger Institute COVID-19 Surveillance Team |
| EPI_ISL_3201569 | Lighthouse Laboratory Plymouth | Wellcome Sanger Institute for the COVID-19 Genomics UK (COG-UK) Consortium | Lighthouse Laboratory Plymouth and Alex Alderton, Roberto Amato, Jeffrey Barrett, Sonia Goncalves, Ewan Harrison, David K. Jackson, Ian Johnston, Dominic Kwiatkowski, Cordelia Langford, John Sillitoe on behalf of the Wellcome Sanger Institute COVID-19 Surveillance Team |
| EPI_ISL_3201570 | Lighthouse Lab in Glasgow | Wellcome Sanger Institute for the COVID-19 Genomics UK (COG-UK) Consortium | Harper VanSteenhouse, Yumi Kasai, David Gray, Carol Clugston, Anna Dominiczak and Alex Alderton, Roberto Amato, Jeffrey Barrett, Sonia Goncalves, Ewan Harrison, David K. Jackson, Ian Johnston, Dominic Kwiatkowski, Cordelia Langford, John Sillitoe on behalf of the Wellcome Sanger Institute COVID-19 Surveillance Team |
| EPI_ISL_3201571 | Lighthouse Lab in Milton Keynes | Wellcome Sanger Institute for the COVID-19 Genomics UK (COG-UK) Consortium | The Lighthouse Lab in Milton Keynes and Alex Alderton, Roberto Amato, Jeffrey Barrett, Sonia Goncalves, Ewan Harrison, David K. Jackson, Ian Johnston, Dominic Kwiatkowski, Cordelia Langford, John Sillitoe on behalf of the Wellcome Sanger Institute COVID-19 Surveillance Team |
| EPI_ISL_3201572 | Lighthouse Laboratory Plymouth | Wellcome Sanger Institute for the COVID-19 Genomics UK (COG-UK) Consortium | Lighthouse Laboratory Plymouth and Alex Alderton, Roberto Amato, Jeffrey Barrett, Sonia Goncalves, Ewan Harrison, David K. Jackson, Ian Johnston, Dominic Kwiatkowski, Cordelia Langford, John Sillitoe on behalf of the Wellcome Sanger Institute COVID-19 Surveillance Team |
| EPI_ISL_3201573, EPI_ISL_3201574, EPI_ISL_3201575 | Lighthouse Lab in Glasgow | Wellcome Sanger Institute for the COVID-19 Genomics UK (COG-UK) Consortium | Harper VanSteenhouse, Yumi Kasai, David Gray, Carol Clugston, Anna Dominiczak and Alex Alderton, Roberto Amato, Jeffrey Barrett, Sonia Goncalves, Ewan Harrison, David K. Jackson, Ian Johnston, Dominic Kwiatkowski, Cordelia Langford, John Sillitoe on behalf of the Wellcome Sanger Institute COVID-19 Surveillance Team |
| EPI_ISL_3201576, EPI_ISL_3201577 | Lighthouse Laboratory Plymouth | Wellcome Sanger Institute for the COVID-19 Genomics UK (COG-UK) Consortium | Lighthouse Laboratory Plymouth and Alex Alderton, Roberto Amato, Jeffrey Barrett, Sonia Goncalves, Ewan Harrison, David K. Jackson, Ian Johnston, Dominic Kwiatkowski, Cordelia Langford, John Sillitoe on behalf of the Wellcome Sanger Institute COVID-19 Surveillance Team |
| EPI_ISL_3201578 | Lighthouse Lab in Milton Keynes | Wellcome Sanger Institute for the COVID-19 Genomics UK (COG-UK) Consortium | The Lighthouse Lab in Milton Keynes and Alex Alderton, Roberto Amato, Jeffrey Barrett, Sonia Goncalves, Ewan Harrison, David K. Jackson, Ian Johnston, Dominic Kwiatkowski, Cordelia Langford, John Sillitoe on behalf of the Wellcome Sanger Institute COVID-19 Surveillance Team |
| EPI_ISL_3201579 | Lighthouse Lab in Glasgow | Wellcome Sanger Institute for the COVID-19 Genomics UK (COG-UK) Consortium | Harper VanSteenhouse, Yumi Kasai, David Gray, Carol Clugston, Anna Dominiczak and Alex Alderton, Roberto Amato, Jeffrey Barrett, Sonia Goncalves, Ewan Harrison, David K. Jackson, Ian Johnston, Dominic Kwiatkowski, Cordelia Langford, John Sillitoe on behalf of the Wellcome Sanger Institute COVID-19 Surveillance Team |
| EPI_ISL_3201580 | Lighthouse Lab in Milton Keynes | Wellcome Sanger Institute for the COVID-19 Genomics UK (COG-UK) Consortium | The Lighthouse Lab in Milton Keynes and Alex Alderton, Roberto Amato, Jeffrey Barrett, Sonia Goncalves, Ewan Harrison, David K. Jackson, Ian Johnston, Dominic Kwiatkowski, Cordelia Langford, John Sillitoe on behalf of the Wellcome Sanger Institute COVID-19 Surveillance Team |
| EPI_ISL_3201581 | Lighthouse Laboratory Plymouth | Wellcome Sanger Institute for the COVID-19 Genomics UK (COG-UK) Consortium | Lighthouse Laboratory Plymouth and Alex Alderton, Roberto Amato, Jeffrey Barrett, Sonia Goncalves, Ewan Harrison, David K. Jackson, Ian Johnston, Dominic Kwiatkowski, Cordelia Langford, John Sillitoe on behalf of the Wellcome Sanger Institute COVID-19 Surveillance Team |
| EPI_ISL_3201582, EPI_ISL_3201583, | Lighthouse Lab in Glasgow | Wellcome Sanger Institute for the COVID-19 Genomics UK (COG-UK) Consortium | Harper VanSteenhouse, Yumi Kasai, David Gray, Carol Clugston, Anna Dominiczak and Alex Alderton, Roberto Amato, Jeffrey Barrett, Sonia Goncalves, Ewan Harrison, David K. Jackson, Ian Johnston, Dominic Kwiatkowski, Cordelia Langford, John Sillitoe on behalf of the Wellcome Sanger Institute COVID-19 Surveillance Team |

| | | | |
|---|---|---|---|
| EPI_ISL_3201584 | | | |
| EPI_ISL_3201585 | Lighthouse Laboratory Plymouth | Wellcome Sanger Institute for the COVID-19 Genomics UK (COG-UK) Consortium | Lighthouse Laboratory Plymouth and Alex Alderton, Roberto Amato, Jeffrey Barrett, Sonia Goncalves, Ewan Harrison, David K. Jackson, Ian Johnston, Dominic Kwiatkowski, Cordelia Langford, John Sillitoe on behalf of the Wellcome Sanger Institute COVID-19 Surveillance Team |
| EPI_ISL_3201586, EPI_ISL_3201587, EPI_ISL_3201588 | Lighthouse Lab in Glasgow | Wellcome Sanger Institute for the COVID-19 Genomics UK (COG-UK) Consortium | Harper VanSteenhouse, Yumi Kasai, David Gray, Carol Clugston, Anna Dominiczak and Alex Alderton, Roberto Amato, Jeffrey Barrett, Sonia Goncalves, Ewan Harrison, David K. Jackson, Ian Johnston, Dominic Kwiatkowski, Cordelia Langford, John Sillitoe on behalf of the Wellcome Sanger Institute COVID-19 Surveillance Team |
| EPI_ISL_3201589, EPI_ISL_3201590, EPI_ISL_3201591, EPI_ISL_3201592 | Lighthouse Laboratory Plymouth | Wellcome Sanger Institute for the COVID-19 Genomics UK (COG-UK) Consortium | Lighthouse Laboratory Plymouth and Alex Alderton, Roberto Amato, Jeffrey Barrett, Sonia Goncalves, Ewan Harrison, David K. Jackson, Ian Johnston, Dominic Kwiatkowski, Cordelia Langford, John Sillitoe on behalf of the Wellcome Sanger Institute COVID-19 Surveillance Team |
| EPI_ISL_3201593, EPI_ISL_3201594 | Lighthouse Lab in Milton Keynes | Wellcome Sanger Institute for the COVID-19 Genomics UK (COG-UK) Consortium | The Lighthouse Lab in Milton Keynes and Alex Alderton, Roberto Amato, Jeffrey Barrett, Sonia Goncalves, Ewan Harrison, David K. Jackson, Ian Johnston, Dominic Kwiatkowski, Cordelia Langford, John Sillitoe on behalf of the Wellcome Sanger Institute COVID-19 Surveillance Team |
| EPI_ISL_3201595, EPI_ISL_3201596 | Lighthouse Lab in Glasgow | Wellcome Sanger Institute for the COVID-19 Genomics UK (COG-UK) Consortium | Harper VanSteenhouse, Yumi Kasai, David Gray, Carol Clugston, Anna Dominiczak and Alex Alderton, Roberto Amato, Jeffrey Barrett, Sonia Goncalves, Ewan Harrison, David K. Jackson, Ian Johnston, Dominic Kwiatkowski, Cordelia Langford, John Sillitoe on behalf of the Wellcome Sanger Institute COVID-19 Surveillance Team |
| EPI_ISL_3201597, EPI_ISL_3201598 | Lighthouse Lab in Milton Keynes | Wellcome Sanger Institute for the COVID-19 Genomics UK (COG-UK) Consortium | The Lighthouse Lab in Milton Keynes and Alex Alderton, Roberto Amato, Jeffrey Barrett, Sonia Goncalves, Ewan Harrison, David K. Jackson, Ian Johnston, Dominic Kwiatkowski, Cordelia Langford, John Sillitoe on behalf of the Wellcome Sanger Institute COVID-19 Surveillance Team |
| EPI_ISL_3201599 | Lighthouse Laboratory Plymouth | Wellcome Sanger Institute for the COVID-19 Genomics UK (COG-UK) Consortium | Lighthouse Laboratory Plymouth and Alex Alderton, Roberto Amato, Jeffrey Barrett, Sonia Goncalves, Ewan Harrison, David K. Jackson, Ian Johnston, Dominic Kwiatkowski, Cordelia Langford, John Sillitoe on behalf of the Wellcome Sanger Institute COVID-19 Surveillance Team |
| EPI_ISL_3201600 | Lighthouse Lab in Milton Keynes | Wellcome Sanger Institute for the COVID-19 Genomics UK (COG-UK) Consortium | The Lighthouse Lab in Milton Keynes and Alex Alderton, Roberto Amato, Jeffrey Barrett, Sonia Goncalves, Ewan Harrison, David K. Jackson, Ian Johnston, Dominic Kwiatkowski, Cordelia Langford, John Sillitoe on behalf of the Wellcome Sanger Institute COVID-19 Surveillance Team |
| EPI_ISL_3201601 | Lighthouse Lab in Glasgow | Wellcome Sanger Institute for the COVID-19 Genomics UK (COG-UK) Consortium | Harper VanSteenhouse, Yumi Kasai, David Gray, Carol Clugston, Anna Dominiczak and Alex Alderton, Roberto Amato, Jeffrey Barrett, Sonia Goncalves, Ewan Harrison, David K. Jackson, Ian Johnston, Dominic Kwiatkowski, Cordelia Langford, John Sillitoe on behalf of the Wellcome Sanger Institute COVID-19 Surveillance Team |
| EPI_ISL_3201602, EPI_ISL_3201603, EPI_ISL_3201604 | Lighthouse Lab in Milton Keynes | Wellcome Sanger Institute for the COVID-19 Genomics UK (COG-UK) Consortium | The Lighthouse Lab in Milton Keynes and Alex Alderton, Roberto Amato, Jeffrey Barrett, Sonia Goncalves, Ewan Harrison, David K. Jackson, Ian Johnston, Dominic Kwiatkowski, Cordelia Langford, John Sillitoe on behalf of the Wellcome Sanger Institute COVID-19 Surveillance Team |
| EPI_ISL_3201605 | Lighthouse Lab in Glasgow | Wellcome Sanger Institute for the COVID-19 Genomics UK (COG-UK) Consortium | Harper VanSteenhouse, Yumi Kasai, David Gray, Carol Clugston, Anna Dominiczak and Alex Alderton, Roberto Amato, Jeffrey Barrett, Sonia Goncalves, Ewan Harrison, David K. Jackson, Ian Johnston, Dominic Kwiatkowski, Cordelia Langford, John Sillitoe on behalf of the Wellcome Sanger Institute COVID-19 Surveillance Team |
| EPI_ISL_3201606 | Lighthouse Laboratory Plymouth | Wellcome Sanger Institute for the COVID-19 Genomics UK (COG-UK) Consortium | Lighthouse Laboratory Plymouth and Alex Alderton, Roberto Amato, Jeffrey Barrett, Sonia Goncalves, Ewan Harrison, David K. Jackson, Ian Johnston, Dominic Kwiatkowski, Cordelia Langford, John Sillitoe on behalf of the Wellcome Sanger Institute COVID-19 Surveillance Team |
| EPI_ISL_3201607 | Lighthouse Lab in Glasgow | Wellcome Sanger Institute for the COVID-19 Genomics UK (COG-UK) Consortium | Harper VanSteenhouse, Yumi Kasai, David Gray, Carol Clugston, Anna Dominiczak and Alex Alderton, Roberto Amato, Jeffrey Barrett, Sonia Goncalves, Ewan Harrison, David K. Jackson, Ian Johnston, Dominic Kwiatkowski, Cordelia Langford, John Sillitoe on behalf of the Wellcome Sanger Institute COVID-19 Surveillance Team |
| EPI_ISL_3201608 | Lighthouse Lab in Milton Keynes | Wellcome Sanger Institute for the COVID-19 Genomics UK (COG-UK) Consortium | The Lighthouse Lab in Milton Keynes and Alex Alderton, Roberto Amato, Jeffrey Barrett, Sonia Goncalves, Ewan Harrison, David K. Jackson, Ian Johnston, Dominic Kwiatkowski, Cordelia Langford, John Sillitoe on behalf of the Wellcome Sanger Institute COVID-19 Surveillance Team |
| EPI_ISL_3201609, EPI_ISL_3201610 | Lighthouse Lab in Glasgow | Wellcome Sanger Institute for the COVID-19 Genomics UK (COG-UK) Consortium | Harper VanSteenhouse, Yumi Kasai, David Gray, Carol Clugston, Anna Dominiczak and Alex Alderton, Roberto Amato, Jeffrey Barrett, Sonia Goncalves, Ewan Harrison, David K. Jackson, Ian Johnston, Dominic Kwiatkowski, Cordelia Langford, John Sillitoe on behalf of the Wellcome Sanger Institute COVID-19 Surveillance Team |
| EPI_ISL_3201611 | Lighthouse Lab in Milton Keynes | Wellcome Sanger Institute for the COVID-19 Genomics UK (COG-UK) Consortium | The Lighthouse Lab in Milton Keynes and Alex Alderton, Roberto Amato, Jeffrey Barrett, Sonia Goncalves, Ewan Harrison, David K. Jackson, Ian Johnston, Dominic Kwiatkowski, Cordelia Langford, John Sillitoe on behalf of the Wellcome Sanger Institute COVID-19 Surveillance Team |
| EPI_ISL_3201612 | Lighthouse Lab in Glasgow | Wellcome Sanger Institute for the COVID-19 Genomics UK (COG-UK) Consortium | Harper VanSteenhouse, Yumi Kasai, David Gray, Carol Clugston, Anna Dominiczak and Alex Alderton, Roberto Amato, Jeffrey Barrett, Sonia Goncalves, Ewan Harrison, David K. Jackson, Ian Johnston, Dominic Kwiatkowski, Cordelia Langford, John Sillitoe on behalf of the Wellcome Sanger Institute COVID-19 Surveillance Team |
| EPI_ISL_3201613 | Lighthouse Laboratory Plymouth | Wellcome Sanger Institute for the COVID-19 Genomics UK (COG-UK) Consortium | Lighthouse Laboratory Plymouth and Alex Alderton, Roberto Amato, Jeffrey Barrett, Sonia Goncalves, Ewan Harrison, David K. Jackson, Ian Johnston, Dominic Kwiatkowski, Cordelia Langford, John Sillitoe on behalf of the Wellcome Sanger Institute COVID-19 Surveillance Team |
| EPI_ISL_3201614 | Lighthouse Lab in Glasgow | Wellcome Sanger Institute for the COVID-19 Genomics UK (COG-UK) Consortium | Harper VanSteenhouse, Yumi Kasai, David Gray, Carol Clugston, Anna Dominiczak and Alex Alderton, Roberto Amato, Jeffrey Barrett, Sonia Goncalves, Ewan Harrison, David K. Jackson, Ian Johnston, Dominic Kwiatkowski, Cordelia Langford, John Sillitoe on behalf of the Wellcome Sanger Institute COVID-19 Surveillance Team |
| EPI_ISL_3201615 | Lighthouse Laboratory Plymouth | Wellcome Sanger Institute for the COVID-19 Genomics UK (COG-UK) Consortium | Lighthouse Laboratory Plymouth and Alex Alderton, Roberto Amato, Jeffrey Barrett, Sonia Goncalves, Ewan Harrison, David K. Jackson, Ian Johnston, Dominic Kwiatkowski, Cordelia Langford, John Sillitoe on behalf of the Wellcome Sanger Institute COVID-19 Surveillance Team |
| EPI_ISL_3201616 | Lighthouse Lab in Glasgow | Wellcome Sanger Institute for the COVID-19 Genomics UK (COG-UK) Consortium | Harper VanSteenhouse, Yumi Kasai, David Gray, Carol Clugston, Anna Dominiczak and Alex Alderton, Roberto Amato, Jeffrey Barrett, Sonia Goncalves, Ewan Harrison, David K. Jackson, Ian Johnston, Dominic Kwiatkowski, Cordelia Langford, John Sillitoe on behalf of the Wellcome Sanger Institute COVID-19 Surveillance Team |
| EPI_ISL_3201617, EPI_ISL_3201618, EPI_ISL_3201619 | Lighthouse Lab in Milton Keynes | Wellcome Sanger Institute for the COVID-19 Genomics UK (COG-UK) Consortium | The Lighthouse Lab in Milton Keynes and Alex Alderton, Roberto Amato, Jeffrey Barrett, Sonia Goncalves, Ewan Harrison, David K. Jackson, Ian Johnston, Dominic Kwiatkowski, Cordelia Langford, John Sillitoe on behalf of the Wellcome Sanger Institute COVID-19 Surveillance Team |
| EPI_ISL_3201620, EPI_ISL_3201621, EPI_ISL_3201622, EPI_ISL_3201623, EPI_ISL_3201624, EPI_ISL_3201625 | Lighthouse Lab in Glasgow | Wellcome Sanger Institute for the COVID-19 Genomics UK (COG-UK) Consortium | Harper VanSteenhouse, Yumi Kasai, David Gray, Carol Clugston, Anna Dominiczak and Alex Alderton, Roberto Amato, Jeffrey Barrett, Sonia Goncalves, Ewan Harrison, David K. Jackson, Ian Johnston, Dominic Kwiatkowski, Cordelia Langford, John Sillitoe on behalf of the Wellcome Sanger Institute COVID-19 Surveillance Team |
| EPI_ISL_3201626 | Lighthouse Lab in Milton Keynes | Wellcome Sanger Institute for the COVID-19 Genomics UK (COG-UK) Consortium | The Lighthouse Lab in Milton Keynes and Alex Alderton, Roberto Amato, Jeffrey Barrett, Sonia Goncalves, Ewan Harrison, David K. Jackson, Ian Johnston, Dominic Kwiatkowski, Cordelia Langford, John Sillitoe on behalf of the Wellcome Sanger Institute COVID-19 Surveillance Team |
| EPI_ISL_3201627, EPI_ISL_3201628 | Lighthouse Lab in Glasgow | Wellcome Sanger Institute for the COVID-19 Genomics UK (COG-UK) Consortium | Harper VanSteenhouse, Yumi Kasai, David Gray, Carol Clugston, Anna Dominiczak and Alex Alderton, Roberto Amato, Jeffrey Barrett, Sonia Goncalves, Ewan Harrison, David K. Jackson, Ian Johnston, Dominic Kwiatkowski, Cordelia Langford, John Sillitoe on behalf of the Wellcome Sanger Institute COVID-19 Surveillance Team |
| EPI_ISL_3201629 | Lighthouse Lab in Milton Keynes | Wellcome Sanger Institute for the COVID-19 Genomics UK (COG-UK) Consortium | The Lighthouse Lab in Milton Keynes and Alex Alderton, Roberto Amato, Jeffrey Barrett, Sonia Goncalves, Ewan Harrison, David K. Jackson, Ian Johnston, Dominic Kwiatkowski, Cordelia Langford, John Sillitoe on behalf of the Wellcome Sanger Institute COVID-19 Surveillance Team |
| EPI_ISL_3201630 | Lighthouse Laboratory Plymouth | Wellcome Sanger Institute for the COVID-19 Genomics UK (COG-UK) Consortium | Lighthouse Laboratory Plymouth and Alex Alderton, Roberto Amato, Jeffrey Barrett, Sonia Goncalves, Ewan Harrison, David K. Jackson, Ian Johnston, Dominic Kwiatkowski, Cordelia Langford, John Sillitoe on behalf of the Wellcome Sanger Institute COVID-19 Surveillance Team |
| EPI_ISL_3201631 | Lighthouse Lab in Milton Keynes | Wellcome Sanger Institute for the COVID-19 Genomics UK (COG-UK) Consortium | The Lighthouse Lab in Milton Keynes and Alex Alderton, Roberto Amato, Jeffrey Barrett, Sonia Goncalves, Ewan Harrison, David K. Jackson, Ian Johnston, Dominic Kwiatkowski, Cordelia Langford, John Sillitoe on behalf of the Wellcome Sanger Institute COVID-19 Surveillance Team |
| EPI_ISL_3201632 | Lighthouse Lab in Glasgow | Wellcome Sanger Institute for the COVID-19 Genomics UK (COG-UK) Consortium | Harper VanSteenhouse, Yumi Kasai, David Gray, Carol Clugston, Anna Dominiczak and Alex Alderton, Roberto Amato, Jeffrey Barrett, Sonia Goncalves, Ewan Harrison, David K. Jackson, Ian Johnston, Dominic Kwiatkowski, Cordelia Langford, John Sillitoe on behalf of the Wellcome Sanger Institute COVID-19 Surveillance Team |
| EPI_ISL_3201633, EPI_ISL_3201634 | Lighthouse Lab in Milton Keynes | Wellcome Sanger Institute for the COVID-19 Genomics UK (COG-UK) Consortium | The Lighthouse Lab in Milton Keynes and Alex Alderton, Roberto Amato, Jeffrey Barrett, Sonia Goncalves, Ewan Harrison, David K. Jackson, Ian Johnston, Dominic Kwiatkowski, Cordelia Langford, John Sillitoe on behalf of the Wellcome Sanger Institute COVID-19 Surveillance Team |
| EPI_ISL_3201635, EPI_ISL_3201636 | Lighthouse Lab in Glasgow | Wellcome Sanger Institute for the COVID-19 Genomics UK (COG-UK) Consortium | Harper VanSteenhouse, Yumi Kasai, David Gray, Carol Clugston, Anna Dominiczak and Alex Alderton, Roberto Amato, Jeffrey Barrett, Sonia Goncalves, Ewan Harrison, David K. Jackson, Ian Johnston, Dominic Kwiatkowski, Cordelia Langford, John Sillitoe on behalf of the Wellcome Sanger Institute COVID-19 Surveillance Team |
| EPI_ISL_3201637, EPI_ISL_3201638 | Lighthouse Lab in Milton Keynes | Wellcome Sanger Institute for the COVID-19 Genomics UK (COG-UK) Consortium | The Lighthouse Lab in Milton Keynes and Alex Alderton, Roberto Amato, Jeffrey Barrett, Sonia Goncalves, Ewan Harrison, David K. Jackson, Ian Johnston, Dominic Kwiatkowski, Cordelia Langford, John Sillitoe on behalf of the Wellcome Sanger Institute COVID-19 Surveillance Team |
| EPI_ISL_3201639, EPI_ISL_3201640 | Lighthouse Lab in Glasgow | Wellcome Sanger Institute for the COVID-19 Genomics UK (COG-UK) Consortium | Harper VanSteenhouse, Yumi Kasai, David Gray, Carol Clugston, Anna Dominiczak and Alex Alderton, Roberto Amato, Jeffrey Barrett, Sonia Goncalves, Ewan Harrison, David K. Jackson, Ian Johnston, Dominic Kwiatkowski, Cordelia Langford, John Sillitoe on behalf of the Wellcome Sanger Institute COVID-19 Surveillance Team |
| EPI_ISL_3201641 | Lighthouse Lab in Milton Keynes | Wellcome Sanger Institute for the COVID-19 Genomics UK (COG-UK) Consortium | The Lighthouse Lab in Milton Keynes and Alex Alderton, Roberto Amato, Jeffrey Barrett, Sonia Goncalves, Ewan Harrison, David K. Jackson, Ian Johnston, Dominic Kwiatkowski, Cordelia Langford, John Sillitoe on behalf of the Wellcome Sanger Institute COVID-19 Surveillance Team |
| EPI_ISL_3201642 | Lighthouse Laboratory Plymouth | Wellcome Sanger Institute for the COVID-19 Genomics UK (COG-UK) Consortium | Lighthouse Laboratory Plymouth and Alex Alderton, Roberto Amato, Jeffrey Barrett, Sonia Goncalves, Ewan Harrison, David K. Jackson, Ian Johnston, Dominic Kwiatkowski, Cordelia Langford, John Sillitoe on behalf of the Wellcome Sanger Institute COVID-19 Surveillance Team |
| EPI_ISL_3201643 | Lighthouse Lab in Glasgow | Wellcome Sanger Institute for the COVID-19 Genomics UK (COG-UK) Consortium | Harper VanSteenhouse, Yumi Kasai, David Gray, Carol Clugston, Anna Dominiczak and Alex Alderton, Roberto Amato, Jeffrey Barrett, Sonia Goncalves, Ewan Harrison, David K. Jackson, Ian Johnston, Dominic Kwiatkowski, Cordelia Langford, John Sillitoe on behalf of the Wellcome Sanger Institute COVID-19 Surveillance Team |
| EPI_ISL_3201644, EPI_ISL_3201645 | Lighthouse Laboratory Plymouth | Wellcome Sanger Institute for the COVID-19 Genomics UK (COG-UK) Consortium | Lighthouse Laboratory Plymouth and Alex Alderton, Roberto Amato, Jeffrey Barrett, Sonia Goncalves, Ewan Harrison, David K. Jackson, Ian Johnston, Dominic Kwiatkowski, Cordelia Langford, John Sillitoe on behalf of the Wellcome Sanger Institute COVID-19 Surveillance Team |
| EPI_ISL_3201646 | Lighthouse Lab in Glasgow | Wellcome Sanger Institute for the COVID-19 Genomics UK (COG-UK) Consortium | Harper VanSteenhouse, Yumi Kasai, David Gray, Carol Clugston, Anna Dominiczak and Alex Alderton, Roberto Amato, Jeffrey Barrett, Sonia Goncalves, Ewan Harrison, David K. Jackson, Ian Johnston, Dominic Kwiatkowski, Cordelia Langford, John Sillitoe on behalf of the Wellcome Sanger Institute COVID-19 Surveillance Team |
| EPI_ISL_3201647 | Lighthouse Laboratory Plymouth | Wellcome Sanger Institute for the COVID-19 Genomics UK (COG-UK) Consortium | Lighthouse Laboratory Plymouth and Alex Alderton, Roberto Amato, Jeffrey Barrett, Sonia Goncalves, Ewan Harrison, David K. Jackson, Ian Johnston, Dominic Kwiatkowski, Cordelia Langford, John Sillitoe on behalf of the Wellcome Sanger Institute COVID-19 Surveillance Team |
| EPI_ISL_3201648, EPI_ISL_3201649 | Lighthouse Lab in Glasgow | Wellcome Sanger Institute for the COVID-19 Genomics UK (COG-UK) Consortium | Harper VanSteenhouse, Yumi Kasai, David Gray, Carol Clugston, Anna Dominiczak and Alex Alderton, Roberto Amato, Jeffrey Barrett, Sonia Goncalves, Ewan Harrison, David K. Jackson, Ian Johnston, Dominic Kwiatkowski, Cordelia Langford, John Sillitoe on behalf of the Wellcome Sanger Institute COVID-19 Surveillance Team |
| EPI_ISL_3201650, EPI_ISL_3201651 | Lighthouse Lab in Milton Keynes | Wellcome Sanger Institute for the COVID-19 Genomics UK (COG-UK) Consortium | The Lighthouse Lab in Milton Keynes and Alex Alderton, Roberto Amato, Jeffrey Barrett, Sonia Goncalves, Ewan Harrison, David K. Jackson, Ian Johnston, Dominic Kwiatkowski, Cordelia Langford, John Sillitoe on behalf of the Wellcome Sanger Institute COVID-19 Surveillance Team |

| | | | |
|---|---|---|---|
| EPI_ISL_3201652, EPI_ISL_3201653, EPI_ISL_3201654 | Lighthouse Lab in Glasgow | Wellcome Sanger Institute for the COVID-19 Genomics UK (COG-UK) Consortium | Harper VanSteenhouse, Yumi Kasai, David Gray, Carol Clugston, Anna Dominiczak and Alex Alderton, Roberto Amato, Jeffrey Barrett, Sonia Goncalves, Ewan Harrison, David K. Jackson, Ian Johnston, Dominic Kwiatkowski, Cordelia Langford, John Sillitoe on behalf of the Wellcome Sanger Institute COVID-19 Surveillance Team |
| EPI_ISL_3201655, EPI_ISL_3201656 | Lighthouse Laboratory Plymouth | Wellcome Sanger Institute for the COVID-19 Genomics UK (COG-UK) Consortium | Lighthouse Laboratory Plymouth and Alex Alderton, Roberto Amato, Jeffrey Barrett, Sonia Goncalves, Ewan Harrison, David K. Jackson, Ian Johnston, Dominic Kwiatkowski, Cordelia Langford, John Sillitoe on behalf of the Wellcome Sanger Institute COVID-19 Surveillance Team |
| EPI_ISL_3201657, EPI_ISL_3201658, EPI_ISL_3201659, EPI_ISL_3201660 | Lighthouse Lab in Glasgow | Wellcome Sanger Institute for the COVID-19 Genomics UK (COG-UK) Consortium | Harper VanSteenhouse, Yumi Kasai, David Gray, Carol Clugston, Anna Dominiczak and Alex Alderton, Roberto Amato, Jeffrey Barrett, Sonia Goncalves, Ewan Harrison, David K. Jackson, Ian Johnston, Dominic Kwiatkowski, Cordelia Langford, John Sillitoe on behalf of the Wellcome Sanger Institute COVID-19 Surveillance Team |
| EPI_ISL_3201661 | Lighthouse Lab in Milton Keynes | Wellcome Sanger Institute for the COVID-19 Genomics UK (COG-UK) Consortium | The Lighthouse Lab in Milton Keynes and Alex Alderton, Roberto Amato, Jeffrey Barrett, Sonia Goncalves, Ewan Harrison, David K. Jackson, Ian Johnston, Dominic Kwiatkowski, Cordelia Langford, John Sillitoe on behalf of the Wellcome Sanger Institute COVID-19 Surveillance Team |
| EPI_ISL_3201662 | Lighthouse Lab in Glasgow | Wellcome Sanger Institute for the COVID-19 Genomics UK (COG-UK) Consortium | Harper VanSteenhouse, Yumi Kasai, David Gray, Carol Clugston, Anna Dominiczak and Alex Alderton, Roberto Amato, Jeffrey Barrett, Sonia Goncalves, Ewan Harrison, David K. Jackson, Ian Johnston, Dominic Kwiatkowski, Cordelia Langford, John Sillitoe on behalf of the Wellcome Sanger Institute COVID-19 Surveillance Team |
| EPI_ISL_3201663, EPI_ISL_3201664, EPI_ISL_3201665 | Lighthouse Laboratory Plymouth | Wellcome Sanger Institute for the COVID-19 Genomics UK (COG-UK) Consortium | Lighthouse Laboratory Plymouth and Alex Alderton, Roberto Amato, Jeffrey Barrett, Sonia Goncalves, Ewan Harrison, David K. Jackson, Ian Johnston, Dominic Kwiatkowski, Cordelia Langford, John Sillitoe on behalf of the Wellcome Sanger Institute COVID-19 Surveillance Team |
| EPI_ISL_3201666 | Lighthouse Lab in Glasgow | Wellcome Sanger Institute for the COVID-19 Genomics UK (COG-UK) Consortium | Harper VanSteenhouse, Yumi Kasai, David Gray, Carol Clugston, Anna Dominiczak and Alex Alderton, Roberto Amato, Jeffrey Barrett, Sonia Goncalves, Ewan Harrison, David K. Jackson, Ian Johnston, Dominic Kwiatkowski, Cordelia Langford, John Sillitoe on behalf of the Wellcome Sanger Institute COVID-19 Surveillance Team |
| EPI_ISL_3201667, EPI_ISL_3201668 | Lighthouse Lab in Milton Keynes | Wellcome Sanger Institute for the COVID-19 Genomics UK (COG-UK) Consortium | The Lighthouse Lab in Milton Keynes and Alex Alderton, Roberto Amato, Jeffrey Barrett, Sonia Goncalves, Ewan Harrison, David K. Jackson, Ian Johnston, Dominic Kwiatkowski, Cordelia Langford, John Sillitoe on behalf of the Wellcome Sanger Institute COVID-19 Surveillance Team |
| EPI_ISL_3201669, EPI_ISL_3201670 | Lighthouse Lab in Glasgow | Wellcome Sanger Institute for the COVID-19 Genomics UK (COG-UK) Consortium | Harper VanSteenhouse, Yumi Kasai, David Gray, Carol Clugston, Anna Dominiczak and Alex Alderton, Roberto Amato, Jeffrey Barrett, Sonia Goncalves, Ewan Harrison, David K. Jackson, Ian Johnston, Dominic Kwiatkowski, Cordelia Langford, John Sillitoe on behalf of the Wellcome Sanger Institute COVID-19 Surveillance Team |
| EPI_ISL_3201671 | Lighthouse Lab in Milton Keynes | Wellcome Sanger Institute for the COVID-19 Genomics UK (COG-UK) Consortium | The Lighthouse Lab in Milton Keynes and Alex Alderton, Roberto Amato, Jeffrey Barrett, Sonia Goncalves, Ewan Harrison, David K. Jackson, Ian Johnston, Dominic Kwiatkowski, Cordelia Langford, John Sillitoe on behalf of the Wellcome Sanger Institute COVID-19 Surveillance Team |
| EPI_ISL_3201672, EPI_ISL_3201673 | Lighthouse Lab in Glasgow | Wellcome Sanger Institute for the COVID-19 Genomics UK (COG-UK) Consortium | Harper VanSteenhouse, Yumi Kasai, David Gray, Carol Clugston, Anna Dominiczak and Alex Alderton, Roberto Amato, Jeffrey Barrett, Sonia Goncalves, Ewan Harrison, David K. Jackson, Ian Johnston, Dominic Kwiatkowski, Cordelia Langford, John Sillitoe on behalf of the Wellcome Sanger Institute COVID-19 Surveillance Team |
| EPI_ISL_3201674 | Lighthouse Laboratory Plymouth | Wellcome Sanger Institute for the COVID-19 Genomics UK (COG-UK) Consortium | Lighthouse Laboratory Plymouth and Alex Alderton, Roberto Amato, Jeffrey Barrett, Sonia Goncalves, Ewan Harrison, David K. Jackson, Ian Johnston, Dominic Kwiatkowski, Cordelia Langford, John Sillitoe on behalf of the Wellcome Sanger Institute COVID-19 Surveillance Team |
| EPI_ISL_3201675 | Lighthouse Lab in Glasgow | Wellcome Sanger Institute for the COVID-19 Genomics UK (COG-UK) Consortium | Harper VanSteenhouse, Yumi Kasai, David Gray, Carol Clugston, Anna Dominiczak and Alex Alderton, Roberto Amato, Jeffrey Barrett, Sonia Goncalves, Ewan Harrison, David K. Jackson, Ian Johnston, Dominic Kwiatkowski, Cordelia Langford, John Sillitoe on behalf of the Wellcome Sanger Institute COVID-19 Surveillance Team |
| EPI_ISL_3201676, EPI_ISL_3201677, EPI_ISL_3201678, EPI_ISL_3201679 | Lighthouse Laboratory Plymouth | Wellcome Sanger Institute for the COVID-19 Genomics UK (COG-UK) Consortium | Lighthouse Laboratory Plymouth and Alex Alderton, Roberto Amato, Jeffrey Barrett, Sonia Goncalves, Ewan Harrison, David K. Jackson, Ian Johnston, Dominic Kwiatkowski, Cordelia Langford, John Sillitoe on behalf of the Wellcome Sanger Institute COVID-19 Surveillance Team |
| EPI_ISL_3201680 | Lighthouse Lab in Milton Keynes | Wellcome Sanger Institute for the COVID-19 Genomics UK (COG-UK) Consortium | The Lighthouse Lab in Milton Keynes and Alex Alderton, Roberto Amato, Jeffrey Barrett, Sonia Goncalves, Ewan Harrison, David K. Jackson, Ian Johnston, Dominic Kwiatkowski, Cordelia Langford, John Sillitoe on behalf of the Wellcome Sanger Institute COVID-19 Surveillance Team |
| EPI_ISL_3201681, EPI_ISL_3201682 | Lighthouse Lab in Glasgow | Wellcome Sanger Institute for the COVID-19 Genomics UK (COG-UK) Consortium | Harper VanSteenhouse, Yumi Kasai, David Gray, Carol Clugston, Anna Dominiczak and Alex Alderton, Roberto Amato, Jeffrey Barrett, Sonia Goncalves, Ewan Harrison, David K. Jackson, Ian Johnston, Dominic Kwiatkowski, Cordelia Langford, John Sillitoe on behalf of the Wellcome Sanger Institute COVID-19 Surveillance Team |
| EPI_ISL_3201683 | Lighthouse Lab in Milton Keynes | Wellcome Sanger Institute for the COVID-19 Genomics UK (COG-UK) Consortium | The Lighthouse Lab in Milton Keynes and Alex Alderton, Roberto Amato, Jeffrey Barrett, Sonia Goncalves, Ewan Harrison, David K. Jackson, Ian Johnston, Dominic Kwiatkowski, Cordelia Langford, John Sillitoe on behalf of the Wellcome Sanger Institute COVID-19 Surveillance Team |
| EPI_ISL_3201684, EPI_ISL_3201685 | Lighthouse Lab in Glasgow | Wellcome Sanger Institute for the COVID-19 Genomics UK (COG-UK) Consortium | Harper VanSteenhouse, Yumi Kasai, David Gray, Carol Clugston, Anna Dominiczak and Alex Alderton, Roberto Amato, Jeffrey Barrett, Sonia Goncalves, Ewan Harrison, David K. Jackson, Ian Johnston, Dominic Kwiatkowski, Cordelia Langford, John Sillitoe on behalf of the Wellcome Sanger Institute COVID-19 Surveillance Team |
| EPI_ISL_3201686, EPI_ISL_3201687, EPI_ISL_3201688 | Lighthouse Laboratory Plymouth | Wellcome Sanger Institute for the COVID-19 Genomics UK (COG-UK) Consortium | Lighthouse Laboratory Plymouth and Alex Alderton, Roberto Amato, Jeffrey Barrett, Sonia Goncalves, Ewan Harrison, David K. Jackson, Ian Johnston, Dominic Kwiatkowski, Cordelia Langford, John Sillitoe on behalf of the Wellcome Sanger Institute COVID-19 Surveillance Team |
| EPI_ISL_3201689, EPI_ISL_3201690, EPI_ISL_3201691, EPI_ISL_3201692 | Lighthouse Lab in Glasgow | Wellcome Sanger Institute for the COVID-19 Genomics UK (COG-UK) Consortium | Harper VanSteenhouse, Yumi Kasai, David Gray, Carol Clugston, Anna Dominiczak and Alex Alderton, Roberto Amato, Jeffrey Barrett, Sonia Goncalves, Ewan Harrison, David K. Jackson, Ian Johnston, Dominic Kwiatkowski, Cordelia Langford, John Sillitoe on behalf of the Wellcome Sanger Institute COVID-19 Surveillance Team |
| EPI_ISL_3201693, EPI_ISL_3201694 | Lighthouse Laboratory Plymouth | Wellcome Sanger Institute for the COVID-19 Genomics UK (COG-UK) Consortium | Lighthouse Laboratory Plymouth and Alex Alderton, Roberto Amato, Jeffrey Barrett, Sonia Goncalves, Ewan Harrison, David K. Jackson, Ian Johnston, Dominic Kwiatkowski, Cordelia Langford, John Sillitoe on behalf of the Wellcome Sanger Institute COVID-19 Surveillance Team |
| EPI_ISL_3201695 | Lighthouse Lab in Glasgow | Wellcome Sanger Institute for the COVID-19 Genomics UK (COG-UK) Consortium | Harper VanSteenhouse, Yumi Kasai, David Gray, Carol Clugston, Anna Dominiczak and Alex Alderton, Roberto Amato, Jeffrey Barrett, Sonia Goncalves, Ewan Harrison, David K. Jackson, Ian Johnston, Dominic Kwiatkowski, Cordelia Langford, John Sillitoe on behalf of the Wellcome Sanger Institute COVID-19 Surveillance Team |
| EPI_ISL_3201696 | Lighthouse Laboratory Plymouth | Wellcome Sanger Institute for the COVID-19 Genomics UK (COG-UK) Consortium | Lighthouse Laboratory Plymouth and Alex Alderton, Roberto Amato, Jeffrey Barrett, Sonia Goncalves, Ewan Harrison, David K. Jackson, Ian Johnston, Dominic Kwiatkowski, Cordelia Langford, John Sillitoe on behalf of the Wellcome Sanger Institute COVID-19 Surveillance Team |
| EPI_ISL_3201697 | Lighthouse Lab in Milton Keynes | Wellcome Sanger Institute for the COVID-19 Genomics UK (COG-UK) Consortium | The Lighthouse Lab in Milton Keynes and Alex Alderton, Roberto Amato, Jeffrey Barrett, Sonia Goncalves, Ewan Harrison, David K. Jackson, Ian Johnston, Dominic Kwiatkowski, Cordelia Langford, John Sillitoe on behalf of the Wellcome Sanger Institute COVID-19 Surveillance Team |
| EPI_ISL_3201698, EPI_ISL_3201699, EPI_ISL_3201700 | Lighthouse Lab in Glasgow | Wellcome Sanger Institute for the COVID-19 Genomics UK (COG-UK) Consortium | Harper VanSteenhouse, Yumi Kasai, David Gray, Carol Clugston, Anna Dominiczak and Alex Alderton, Roberto Amato, Jeffrey Barrett, Sonia Goncalves, Ewan Harrison, David K. Jackson, Ian Johnston, Dominic Kwiatkowski, Cordelia Langford, John Sillitoe on behalf of the Wellcome Sanger Institute COVID-19 Surveillance Team |
| EPI_ISL_3201701 | Lighthouse Lab in Milton Keynes | Wellcome Sanger Institute for the COVID-19 Genomics UK (COG-UK) Consortium | The Lighthouse Lab in Milton Keynes and Alex Alderton, Roberto Amato, Jeffrey Barrett, Sonia Goncalves, Ewan Harrison, David K. Jackson, Ian Johnston, Dominic Kwiatkowski, Cordelia Langford, John Sillitoe on behalf of the Wellcome Sanger Institute COVID-19 Surveillance Team |
| EPI_ISL_3201702 | Lighthouse Lab in Glasgow | Wellcome Sanger Institute for the COVID-19 Genomics UK (COG-UK) Consortium | Harper VanSteenhouse, Yumi Kasai, David Gray, Carol Clugston, Anna Dominiczak and Alex Alderton, Roberto Amato, Jeffrey Barrett, Sonia Goncalves, Ewan Harrison, David K. Jackson, Ian Johnston, Dominic Kwiatkowski, Cordelia Langford, John Sillitoe on behalf of the Wellcome Sanger Institute COVID-19 Surveillance Team |
| EPI_ISL_3201703 | Lighthouse Lab in Milton Keynes | Wellcome Sanger Institute for the COVID-19 Genomics UK (COG-UK) Consortium | The Lighthouse Lab in Milton Keynes and Alex Alderton, Roberto Amato, Jeffrey Barrett, Sonia Goncalves, Ewan Harrison, David K. Jackson, Ian Johnston, Dominic Kwiatkowski, Cordelia Langford, John Sillitoe on behalf of the Wellcome Sanger Institute COVID-19 Surveillance Team |
| EPI_ISL_3201704, EPI_ISL_3201705 | Lighthouse Lab in Glasgow | Wellcome Sanger Institute for the COVID-19 Genomics UK (COG-UK) Consortium | Harper VanSteenhouse, Yumi Kasai, David Gray, Carol Clugston, Anna Dominiczak and Alex Alderton, Roberto Amato, Jeffrey Barrett, Sonia Goncalves, Ewan Harrison, David K. Jackson, Ian Johnston, Dominic Kwiatkowski, Cordelia Langford, John Sillitoe on behalf of the Wellcome Sanger Institute COVID-19 Surveillance Team |
| EPI_ISL_3201706 | Lighthouse Lab in Milton Keynes | Wellcome Sanger Institute for the COVID-19 Genomics UK (COG-UK) Consortium | The Lighthouse Lab in Milton Keynes and Alex Alderton, Roberto Amato, Jeffrey Barrett, Sonia Goncalves, Ewan Harrison, David K. Jackson, Ian Johnston, Dominic Kwiatkowski, Cordelia Langford, John Sillitoe on behalf of the Wellcome Sanger Institute COVID-19 Surveillance Team |
| EPI_ISL_3201707 | Lighthouse Laboratory Plymouth | Wellcome Sanger Institute for the COVID-19 Genomics UK (COG-UK) Consortium | Lighthouse Laboratory Plymouth and Alex Alderton, Roberto Amato, Jeffrey Barrett, Sonia Goncalves, Ewan Harrison, David K. Jackson, Ian Johnston, Dominic Kwiatkowski, Cordelia Langford, John Sillitoe on behalf of the Wellcome Sanger Institute COVID-19 Surveillance Team |
| EPI_ISL_3201708 | Lighthouse Lab in Glasgow | Wellcome Sanger Institute for the COVID-19 Genomics UK (COG-UK) Consortium | Harper VanSteenhouse, Yumi Kasai, David Gray, Carol Clugston, Anna Dominiczak and Alex Alderton, Roberto Amato, Jeffrey Barrett, Sonia Goncalves, Ewan Harrison, David K. Jackson, Ian Johnston, Dominic Kwiatkowski, Cordelia Langford, John Sillitoe on behalf of the Wellcome Sanger Institute COVID-19 Surveillance Team |
| EPI_ISL_3201709, EPI_ISL_3201710 | Lighthouse Laboratory Plymouth | Wellcome Sanger Institute for the COVID-19 Genomics UK (COG-UK) Consortium | Lighthouse Laboratory Plymouth and Alex Alderton, Roberto Amato, Jeffrey Barrett, Sonia Goncalves, Ewan Harrison, David K. Jackson, Ian Johnston, Dominic Kwiatkowski, Cordelia Langford, John Sillitoe on behalf of the Wellcome Sanger Institute COVID-19 Surveillance Team |
| EPI_ISL_3201711 | Lighthouse Lab in Milton Keynes | Wellcome Sanger Institute for the COVID-19 Genomics UK (COG-UK) Consortium | The Lighthouse Lab in Milton Keynes and Alex Alderton, Roberto Amato, Jeffrey Barrett, Sonia Goncalves, Ewan Harrison, David K. Jackson, Ian Johnston, Dominic Kwiatkowski, Cordelia Langford, John Sillitoe on behalf of the Wellcome Sanger Institute COVID-19 Surveillance Team |
| EPI_ISL_3201712 | Lighthouse Laboratory Plymouth | Wellcome Sanger Institute for the COVID-19 Genomics UK (COG-UK) Consortium | Lighthouse Laboratory Plymouth and Alex Alderton, Roberto Amato, Jeffrey Barrett, Sonia Goncalves, Ewan Harrison, David K. Jackson, Ian Johnston, Dominic Kwiatkowski, Cordelia Langford, John Sillitoe on behalf of the Wellcome Sanger Institute COVID-19 Surveillance Team |
| EPI_ISL_3201713, EPI_ISL_3201714, EPI_ISL_3201715 | Lighthouse Lab in Glasgow | Wellcome Sanger Institute for the COVID-19 Genomics UK (COG-UK) Consortium | Harper VanSteenhouse, Yumi Kasai, David Gray, Carol Clugston, Anna Dominiczak and Alex Alderton, Roberto Amato, Jeffrey Barrett, Sonia Goncalves, Ewan Harrison, David K. Jackson, Ian Johnston, Dominic Kwiatkowski, Cordelia Langford, John Sillitoe on behalf of the Wellcome Sanger Institute COVID-19 Surveillance Team |
| EPI_ISL_3201716 | Lighthouse Laboratory Plymouth | Wellcome Sanger Institute for the COVID-19 Genomics UK (COG-UK) Consortium | Lighthouse Laboratory Plymouth and Alex Alderton, Roberto Amato, Jeffrey Barrett, Sonia Goncalves, Ewan Harrison, David K. Jackson, Ian Johnston, Dominic Kwiatkowski, Cordelia Langford, John Sillitoe on behalf of the Wellcome Sanger Institute COVID-19 Surveillance Team |
| EPI_ISL_3201717 | Lighthouse Lab in Glasgow | Wellcome Sanger Institute for the COVID-19 Genomics UK (COG-UK) Consortium | Harper VanSteenhouse, Yumi Kasai, David Gray, Carol Clugston, Anna Dominiczak and Alex Alderton, Roberto Amato, Jeffrey Barrett, Sonia Goncalves, Ewan Harrison, David K. Jackson, Ian Johnston, Dominic Kwiatkowski, Cordelia Langford, John Sillitoe on behalf of the Wellcome Sanger Institute COVID-19 Surveillance Team |
| EPI_ISL_3201718 | Lighthouse Laboratory Plymouth | Wellcome Sanger Institute for the COVID-19 Genomics UK (COG-UK) Consortium | Lighthouse Laboratory Plymouth and Alex Alderton, Roberto Amato, Jeffrey Barrett, Sonia Goncalves, Ewan Harrison, David K. Jackson, Ian Johnston, Dominic Kwiatkowski, Cordelia Langford, John Sillitoe on behalf of the Wellcome Sanger Institute COVID-19 Surveillance Team |
| EPI_ISL_3201719 | Lighthouse Lab in Glasgow | Wellcome Sanger Institute for the COVID-19 Genomics UK (COG-UK) Consortium | Harper VanSteenhouse, Yumi Kasai, David Gray, Carol Clugston, Anna Dominiczak and Alex Alderton, Roberto Amato, Jeffrey Barrett, Sonia Goncalves, Ewan Harrison, David K. Jackson, Ian Johnston, Dominic Kwiatkowski, Cordelia Langford, John Sillitoe on behalf of the Wellcome Sanger Institute COVID-19 Surveillance Team |
| EPI_ISL_3201720 | Lighthouse Lab in Milton Keynes | Wellcome Sanger Institute for the COVID-19 Genomics UK (COG-UK) | The Lighthouse Lab in Milton Keynes and Alex Alderton, Roberto Amato, Jeffrey Barrett, Sonia Goncalves, Ewan Harrison, David K. Jackson, Ian Johnston, Dominic Kwiatkowski, Cordelia Langford, John Sillitoe on behalf of the Wellcome Sanger |

| | | | |
|---|---|---|---|
| | | Consortium | Institute COVID-19 Surveillance Team |
| EPI_ISL_3201721, EPI_ISL_3201722 | Lighthouse Lab in Glasgow | Wellcome Sanger Institute for the COVID-19 Genomics UK (COG-UK) Consortium | Harper VanSteenhouse, Yumi Kasai, David Gray, Carol Clugston, Anna Dominiczak and Alex Alderton, Roberto Amato, Jeffrey Barrett, Sonia Goncalves, Ewan Harrison, David K. Jackson, Ian Johnston, Dominic Kwiatkowski, Cordelia Langford, John Sillitoe on behalf of the Wellcome Sanger Institute COVID-19 Surveillance Team |
| EPI_ISL_3201723 | Lighthouse Laboratory Plymouth | Wellcome Sanger Institute for the COVID-19 Genomics UK (COG-UK) Consortium | Lighthouse Laboratory Plymouth and Alex Alderton, Roberto Amato, Jeffrey Barrett, Sonia Goncalves, Ewan Harrison, David K. Jackson, Ian Johnston, Dominic Kwiatkowski, Cordelia Langford, John Sillitoe on behalf of the Wellcome Sanger Institute COVID-19 Surveillance Team |
| EPI_ISL_3201724 | Lighthouse Lab in Milton Keynes | Wellcome Sanger Institute for the COVID-19 Genomics UK (COG-UK) Consortium | The Lighthouse Lab in Milton Keynes and Alex Alderton, Roberto Amato, Jeffrey Barrett, Sonia Goncalves, Ewan Harrison, David K. Jackson, Ian Johnston, Dominic Kwiatkowski, Cordelia Langford, John Sillitoe on behalf of the Wellcome Sanger Institute COVID-19 Surveillance Team |
| EPI_ISL_3201725 | Lighthouse Lab in Glasgow | Wellcome Sanger Institute for the COVID-19 Genomics UK (COG-UK) Consortium | Harper VanSteenhouse, Yumi Kasai, David Gray, Carol Clugston, Anna Dominiczak and Alex Alderton, Roberto Amato, Jeffrey Barrett, Sonia Goncalves, Ewan Harrison, David K. Jackson, Ian Johnston, Dominic Kwiatkowski, Cordelia Langford, John Sillitoe on behalf of the Wellcome Sanger Institute COVID-19 Surveillance Team |
| EPI_ISL_3201726 | Lighthouse Laboratory Plymouth | Wellcome Sanger Institute for the COVID-19 Genomics UK (COG-UK) Consortium | Lighthouse Laboratory Plymouth and Alex Alderton, Roberto Amato, Jeffrey Barrett, Sonia Goncalves, Ewan Harrison, David K. Jackson, Ian Johnston, Dominic Kwiatkowski, Cordelia Langford, John Sillitoe on behalf of the Wellcome Sanger Institute COVID-19 Surveillance Team |
| EPI_ISL_3201727, EPI_ISL_3201728 | Lighthouse Lab in Glasgow | Wellcome Sanger Institute for the COVID-19 Genomics UK (COG-UK) Consortium | Harper VanSteenhouse, Yumi Kasai, David Gray, Carol Clugston, Anna Dominiczak and Alex Alderton, Roberto Amato, Jeffrey Barrett, Sonia Goncalves, Ewan Harrison, David K. Jackson, Ian Johnston, Dominic Kwiatkowski, Cordelia Langford, John Sillitoe on behalf of the Wellcome Sanger Institute COVID-19 Surveillance Team |
| EPI_ISL_3201729 | Lighthouse Laboratory Plymouth | Wellcome Sanger Institute for the COVID-19 Genomics UK (COG-UK) Consortium | Lighthouse Laboratory Plymouth and Alex Alderton, Roberto Amato, Jeffrey Barrett, Sonia Goncalves, Ewan Harrison, David K. Jackson, Ian Johnston, Dominic Kwiatkowski, Cordelia Langford, John Sillitoe on behalf of the Wellcome Sanger Institute COVID-19 Surveillance Team |
| EPI_ISL_3201730, EPI_ISL_3201731 | Lighthouse Lab in Glasgow | Wellcome Sanger Institute for the COVID-19 Genomics UK (COG-UK) Consortium | Harper VanSteenhouse, Yumi Kasai, David Gray, Carol Clugston, Anna Dominiczak and Alex Alderton, Roberto Amato, Jeffrey Barrett, Sonia Goncalves, Ewan Harrison, David K. Jackson, Ian Johnston, Dominic Kwiatkowski, Cordelia Langford, John Sillitoe on behalf of the Wellcome Sanger Institute COVID-19 Surveillance Team |
| EPI_ISL_3201732, EPI_ISL_3201733 | Lighthouse Laboratory Plymouth | Wellcome Sanger Institute for the COVID-19 Genomics UK (COG-UK) Consortium | Lighthouse Laboratory Plymouth and Alex Alderton, Roberto Amato, Jeffrey Barrett, Sonia Goncalves, Ewan Harrison, David K. Jackson, Ian Johnston, Dominic Kwiatkowski, Cordelia Langford, John Sillitoe on behalf of the Wellcome Sanger Institute COVID-19 Surveillance Team |
| EPI_ISL_3201734 | Lighthouse Lab in Glasgow | Wellcome Sanger Institute for the COVID-19 Genomics UK (COG-UK) Consortium | Harper VanSteenhouse, Yumi Kasai, David Gray, Carol Clugston, Anna Dominiczak and Alex Alderton, Roberto Amato, Jeffrey Barrett, Sonia Goncalves, Ewan Harrison, David K. Jackson, Ian Johnston, Dominic Kwiatkowski, Cordelia Langford, John Sillitoe on behalf of the Wellcome Sanger Institute COVID-19 Surveillance Team |
| EPI_ISL_3201735 | Lighthouse Laboratory Plymouth | Wellcome Sanger Institute for the COVID-19 Genomics UK (COG-UK) Consortium | Lighthouse Laboratory Plymouth and Alex Alderton, Roberto Amato, Jeffrey Barrett, Sonia Goncalves, Ewan Harrison, David K. Jackson, Ian Johnston, Dominic Kwiatkowski, Cordelia Langford, John Sillitoe on behalf of the Wellcome Sanger Institute COVID-19 Surveillance Team |
| EPI_ISL_3201736 | Lighthouse Lab in Glasgow | Wellcome Sanger Institute for the COVID-19 Genomics UK (COG-UK) Consortium | Harper VanSteenhouse, Yumi Kasai, David Gray, Carol Clugston, Anna Dominiczak and Alex Alderton, Roberto Amato, Jeffrey Barrett, Sonia Goncalves, Ewan Harrison, David K. Jackson, Ian Johnston, Dominic Kwiatkowski, Cordelia Langford, John Sillitoe on behalf of the Wellcome Sanger Institute COVID-19 Surveillance Team |
| EPI_ISL_3201737 | Lighthouse Lab in Milton Keynes | Wellcome Sanger Institute for the COVID-19 Genomics UK (COG-UK) Consortium | The Lighthouse Lab in Milton Keynes and Alex Alderton, Roberto Amato, Jeffrey Barrett, Sonia Goncalves, Ewan Harrison, David K. Jackson, Ian Johnston, Dominic Kwiatkowski, Cordelia Langford, John Sillitoe on behalf of the Wellcome Sanger Institute COVID-19 Surveillance Team |
| EPI_ISL_3201738 | Lighthouse Laboratory Plymouth | Wellcome Sanger Institute for the COVID-19 Genomics UK (COG-UK) Consortium | Lighthouse Laboratory Plymouth and Alex Alderton, Roberto Amato, Jeffrey Barrett, Sonia Goncalves, Ewan Harrison, David K. Jackson, Ian Johnston, Dominic Kwiatkowski, Cordelia Langford, John Sillitoe on behalf of the Wellcome Sanger Institute COVID-19 Surveillance Team |
| EPI_ISL_3201739 | Lighthouse Lab in Glasgow | Wellcome Sanger Institute for the COVID-19 Genomics UK (COG-UK) Consortium | Harper VanSteenhouse, Yumi Kasai, David Gray, Carol Clugston, Anna Dominiczak and Alex Alderton, Roberto Amato, Jeffrey Barrett, Sonia Goncalves, Ewan Harrison, David K. Jackson, Ian Johnston, Dominic Kwiatkowski, Cordelia Langford, John Sillitoe on behalf of the Wellcome Sanger Institute COVID-19 Surveillance Team |
| EPI_ISL_3201740 | Lighthouse Lab in Milton Keynes | Wellcome Sanger Institute for the COVID-19 Genomics UK (COG-UK) Consortium | The Lighthouse Lab in Milton Keynes and Alex Alderton, Roberto Amato, Jeffrey Barrett, Sonia Goncalves, Ewan Harrison, David K. Jackson, Ian Johnston, Dominic Kwiatkowski, Cordelia Langford, John Sillitoe on behalf of the Wellcome Sanger Institute COVID-19 Surveillance Team |
| EPI_ISL_3201741, EPI_ISL_3201742 | Lighthouse Lab in Glasgow | Wellcome Sanger Institute for the COVID-19 Genomics UK (COG-UK) Consortium | Harper VanSteenhouse, Yumi Kasai, David Gray, Carol Clugston, Anna Dominiczak and Alex Alderton, Roberto Amato, Jeffrey Barrett, Sonia Goncalves, Ewan Harrison, David K. Jackson, Ian Johnston, Dominic Kwiatkowski, Cordelia Langford, John Sillitoe on behalf of the Wellcome Sanger Institute COVID-19 Surveillance Team |
| EPI_ISL_3201743 | Lighthouse Laboratory Plymouth | Wellcome Sanger Institute for the COVID-19 Genomics UK (COG-UK) Consortium | Lighthouse Laboratory Plymouth and Alex Alderton, Roberto Amato, Jeffrey Barrett, Sonia Goncalves, Ewan Harrison, David K. Jackson, Ian Johnston, Dominic Kwiatkowski, Cordelia Langford, John Sillitoe on behalf of the Wellcome Sanger Institute COVID-19 Surveillance Team |
| EPI_ISL_3201744, EPI_ISL_3201745 | Lighthouse Lab in Glasgow | Wellcome Sanger Institute for the COVID-19 Genomics UK (COG-UK) Consortium | Harper VanSteenhouse, Yumi Kasai, David Gray, Carol Clugston, Anna Dominiczak and Alex Alderton, Roberto Amato, Jeffrey Barrett, Sonia Goncalves, Ewan Harrison, David K. Jackson, Ian Johnston, Dominic Kwiatkowski, Cordelia Langford, John Sillitoe on behalf of the Wellcome Sanger Institute COVID-19 Surveillance Team |
| EPI_ISL_3201746 | Lighthouse Laboratory Plymouth | Wellcome Sanger Institute for the COVID-19 Genomics UK (COG-UK) Consortium | Lighthouse Laboratory Plymouth and Alex Alderton, Roberto Amato, Jeffrey Barrett, Sonia Goncalves, Ewan Harrison, David K. Jackson, Ian Johnston, Dominic Kwiatkowski, Cordelia Langford, John Sillitoe on behalf of the Wellcome Sanger Institute COVID-19 Surveillance Team |
| EPI_ISL_3201747 | Lighthouse Lab in Milton Keynes | Wellcome Sanger Institute for the COVID-19 Genomics UK (COG-UK) Consortium | The Lighthouse Lab in Milton Keynes and Alex Alderton, Roberto Amato, Jeffrey Barrett, Sonia Goncalves, Ewan Harrison, David K. Jackson, Ian Johnston, Dominic Kwiatkowski, Cordelia Langford, John Sillitoe on behalf of the Wellcome Sanger Institute COVID-19 Surveillance Team |
| EPI_ISL_3201748, EPI_ISL_3201749, EPI_ISL_3201750 | Lighthouse Lab in Glasgow | Wellcome Sanger Institute for the COVID-19 Genomics UK (COG-UK) Consortium | Harper VanSteenhouse, Yumi Kasai, David Gray, Carol Clugston, Anna Dominiczak and Alex Alderton, Roberto Amato, Jeffrey Barrett, Sonia Goncalves, Ewan Harrison, David K. Jackson, Ian Johnston, Dominic Kwiatkowski, Cordelia Langford, John Sillitoe on behalf of the Wellcome Sanger Institute COVID-19 Surveillance Team |
| EPI_ISL_3201751 | Lighthouse Laboratory Plymouth | Wellcome Sanger Institute for the COVID-19 Genomics UK (COG-UK) Consortium | Lighthouse Laboratory Plymouth and Alex Alderton, Roberto Amato, Jeffrey Barrett, Sonia Goncalves, Ewan Harrison, David K. Jackson, Ian Johnston, Dominic Kwiatkowski, Cordelia Langford, John Sillitoe on behalf of the Wellcome Sanger Institute COVID-19 Surveillance Team |
| EPI_ISL_3201752, EPI_ISL_3201753, EPI_ISL_3201754 | Lighthouse Lab in Glasgow | Wellcome Sanger Institute for the COVID-19 Genomics UK (COG-UK) Consortium | Harper VanSteenhouse, Yumi Kasai, David Gray, Carol Clugston, Anna Dominiczak and Alex Alderton, Roberto Amato, Jeffrey Barrett, Sonia Goncalves, Ewan Harrison, David K. Jackson, Ian Johnston, Dominic Kwiatkowski, Cordelia Langford, John Sillitoe on behalf of the Wellcome Sanger Institute COVID-19 Surveillance Team |
| EPI_ISL_3201755 | Lighthouse Lab in Milton Keynes | Wellcome Sanger Institute for the COVID-19 Genomics UK (COG-UK) Consortium | The Lighthouse Lab in Milton Keynes and Alex Alderton, Roberto Amato, Jeffrey Barrett, Sonia Goncalves, Ewan Harrison, David K. Jackson, Ian Johnston, Dominic Kwiatkowski, Cordelia Langford, John Sillitoe on behalf of the Wellcome Sanger Institute COVID-19 Surveillance Team |
| EPI_ISL_3201756 | Lighthouse Lab in Glasgow | Wellcome Sanger Institute for the COVID-19 Genomics UK (COG-UK) Consortium | Harper VanSteenhouse, Yumi Kasai, David Gray, Carol Clugston, Anna Dominiczak and Alex Alderton, Roberto Amato, Jeffrey Barrett, Sonia Goncalves, Ewan Harrison, David K. Jackson, Ian Johnston, Dominic Kwiatkowski, Cordelia Langford, John Sillitoe on behalf of the Wellcome Sanger Institute COVID-19 Surveillance Team |
| EPI_ISL_3201757, EPI_ISL_3201758 | Lighthouse Laboratory Plymouth | Wellcome Sanger Institute for the COVID-19 Genomics UK (COG-UK) Consortium | Lighthouse Laboratory Plymouth and Alex Alderton, Roberto Amato, Jeffrey Barrett, Sonia Goncalves, Ewan Harrison, David K. Jackson, Ian Johnston, Dominic Kwiatkowski, Cordelia Langford, John Sillitoe on behalf of the Wellcome Sanger Institute COVID-19 Surveillance Team |
| EPI_ISL_3201759, EPI_ISL_3201760, EPI_ISL_3201761, EPI_ISL_3201762, EPI_ISL_3201763 | Lighthouse Lab in Glasgow | Wellcome Sanger Institute for the COVID-19 Genomics UK (COG-UK) Consortium | Harper VanSteenhouse, Yumi Kasai, David Gray, Carol Clugston, Anna Dominiczak and Alex Alderton, Roberto Amato, Jeffrey Barrett, Sonia Goncalves, Ewan Harrison, David K. Jackson, Ian Johnston, Dominic Kwiatkowski, Cordelia Langford, John Sillitoe on behalf of the Wellcome Sanger Institute COVID-19 Surveillance Team |
| EPI_ISL_3201764 | Lighthouse Laboratory Plymouth | Wellcome Sanger Institute for the COVID-19 Genomics UK (COG-UK) Consortium | Lighthouse Laboratory Plymouth and Alex Alderton, Roberto Amato, Jeffrey Barrett, Sonia Goncalves, Ewan Harrison, David K. Jackson, Ian Johnston, Dominic Kwiatkowski, Cordelia Langford, John Sillitoe on behalf of the Wellcome Sanger Institute COVID-19 Surveillance Team |
| EPI_ISL_3201765, EPI_ISL_3201766, EPI_ISL_3201767, EPI_ISL_3201768, EPI_ISL_3201769 | Lighthouse Lab in Glasgow | Wellcome Sanger Institute for the COVID-19 Genomics UK (COG-UK) Consortium | Harper VanSteenhouse, Yumi Kasai, David Gray, Carol Clugston, Anna Dominiczak and Alex Alderton, Roberto Amato, Jeffrey Barrett, Sonia Goncalves, Ewan Harrison, David K. Jackson, Ian Johnston, Dominic Kwiatkowski, Cordelia Langford, John Sillitoe on behalf of the Wellcome Sanger Institute COVID-19 Surveillance Team |
| EPI_ISL_3201770 | Lighthouse Lab in Milton Keynes | Wellcome Sanger Institute for the COVID-19 Genomics UK (COG-UK) Consortium | The Lighthouse Lab in Milton Keynes and Alex Alderton, Roberto Amato, Jeffrey Barrett, Sonia Goncalves, Ewan Harrison, David K. Jackson, Ian Johnston, Dominic Kwiatkowski, Cordelia Langford, John Sillitoe on behalf of the Wellcome Sanger Institute COVID-19 Surveillance Team |
| EPI_ISL_3201771 | Lighthouse Laboratory Plymouth | Wellcome Sanger Institute for the COVID-19 Genomics UK (COG-UK) Consortium | Lighthouse Laboratory Plymouth and Alex Alderton, Roberto Amato, Jeffrey Barrett, Sonia Goncalves, Ewan Harrison, David K. Jackson, Ian Johnston, Dominic Kwiatkowski, Cordelia Langford, John Sillitoe on behalf of the Wellcome Sanger Institute COVID-19 Surveillance Team |
| EPI_ISL_3201772 | Lighthouse Lab in Alderley Park | Wellcome Sanger Institute for the COVID-19 Genomics UK (COG-UK) Consortium | Jacquelyn Wynn, Mairead Hyland, The Lighthouse Lab in Alderley Park and Alex Alderton, Roberto Amato, Jeffrey Barrett, Sonia Goncalves, Ewan Harrison, David K. Jackson, Ian Johnston, Dominic Kwiatkowski, Cordelia Langford, John Sillitoe on behalf of the Wellcome Sanger Institute COVID-19 Surveillance Team |
| EPI_ISL_3201773 | Lighthouse Laboratory Plymouth | Wellcome Sanger Institute for the COVID-19 Genomics UK (COG-UK) Consortium | Lighthouse Laboratory Plymouth and Alex Alderton, Roberto Amato, Jeffrey Barrett, Sonia Goncalves, Ewan Harrison, David K. Jackson, Ian Johnston, Dominic Kwiatkowski, Cordelia Langford, John Sillitoe on behalf of the Wellcome Sanger Institute COVID-19 Surveillance Team |
| EPI_ISL_3201774, EPI_ISL_3201775 | Lighthouse Lab in Milton Keynes | Wellcome Sanger Institute for the COVID-19 Genomics UK (COG-UK) Consortium | The Lighthouse Lab in Milton Keynes and Alex Alderton, Roberto Amato, Jeffrey Barrett, Sonia Goncalves, Ewan Harrison, David K. Jackson, Ian Johnston, Dominic Kwiatkowski, Cordelia Langford, John Sillitoe on behalf of the Wellcome Sanger Institute COVID-19 Surveillance Team |
| EPI_ISL_3201776 | Lighthouse Laboratory Plymouth | Wellcome Sanger Institute for the COVID-19 Genomics UK (COG-UK) Consortium | Lighthouse Laboratory Plymouth and Alex Alderton, Roberto Amato, Jeffrey Barrett, Sonia Goncalves, Ewan Harrison, David K. Jackson, Ian Johnston, Dominic Kwiatkowski, Cordelia Langford, John Sillitoe on behalf of the Wellcome Sanger Institute COVID-19 Surveillance Team |
| EPI_ISL_3201777 | Lighthouse Lab in Alderley Park | Wellcome Sanger Institute for the COVID-19 Genomics UK (COG-UK) Consortium | Jacquelyn Wynn, Mairead Hyland, The Lighthouse Lab in Alderley Park and Alex Alderton, Roberto Amato, Jeffrey Barrett, Sonia Goncalves, Ewan Harrison, David K. Jackson, Ian Johnston, Dominic Kwiatkowski, Cordelia Langford, John Sillitoe on behalf of the Wellcome Sanger Institute COVID-19 Surveillance Team |
| EPI_ISL_3201778, EPI_ISL_3201779, EPI_ISL_3201780 | Lighthouse Lab in Milton Keynes | Wellcome Sanger Institute for the COVID-19 Genomics UK (COG-UK) Consortium | The Lighthouse Lab in Milton Keynes and Alex Alderton, Roberto Amato, Jeffrey Barrett, Sonia Goncalves, Ewan Harrison, David K. Jackson, Ian Johnston, Dominic Kwiatkowski, Cordelia Langford, John Sillitoe on behalf of the Wellcome Sanger Institute COVID-19 Surveillance Team |
| EPI_ISL_3201781 | Lighthouse Laboratory Plymouth | Wellcome Sanger Institute for the COVID-19 Genomics UK (COG-UK) Consortium | Lighthouse Laboratory Plymouth and Alex Alderton, Roberto Amato, Jeffrey Barrett, Sonia Goncalves, Ewan Harrison, David K. Jackson, Ian Johnston, Dominic Kwiatkowski, Cordelia Langford, John Sillitoe on behalf of the Wellcome Sanger Institute COVID-19 Surveillance Team |
| EPI_ISL_3201782 | Lighthouse Lab in Alderley Park | Wellcome Sanger Institute for the COVID-19 Genomics UK (COG-UK) Consortium | Jacquelyn Wynn, Mairead Hyland, The Lighthouse Lab in Alderley Park and Alex Alderton, Roberto Amato, Jeffrey Barrett, Sonia Goncalves, Ewan Harrison, David K. Jackson, Ian Johnston, Dominic Kwiatkowski, Cordelia Langford, John Sillitoe on behalf of the Wellcome Sanger Institute COVID-19 Surveillance Team |
| EPI_ISL_3201783, EPI_ISL_3201784, | Lighthouse Lab in Milton Keynes | Wellcome Sanger Institute for the COVID-19 Genomics UK (COG-UK) Consortium | The Lighthouse Lab in Milton Keynes and Alex Alderton, Roberto Amato, Jeffrey Barrett, Sonia Goncalves, Ewan Harrison, David K. Jackson, Ian Johnston, Dominic Kwiatkowski, Cordelia Langford, John Sillitoe on behalf of the Wellcome Sanger Institute COVID-19 Surveillance Team |

| | | | |
|---|---|---|---|
| EPI_ISL_3201785, EPI_ISL_3201786 | | | |
| EPI_ISL_3201787 | Lighthouse Lab in Alderley Park | Wellcome Sanger Institute for the COVID-19 Genomics UK (COG-UK) Consortium | Jacquelyn Wynn, Mairead Hyland, The Lighthouse Lab in Alderley Park and Alex Alderton, Roberto Amato, Jeffrey Barrett, Sonia Goncalves, Ewan Harrison, David K. Jackson, Ian Johnston, Dominic Kwiatkowski, Cordelia Langford, John Sillitoe on behalf of the Wellcome Sanger Institute COVID-19 Surveillance Team |
| EPI_ISL_3201788 | Lighthouse Laboratory Plymouth | Wellcome Sanger Institute for the COVID-19 Genomics UK (COG-UK) Consortium | Lighthouse Laboratory Plymouth and Alex Alderton, Roberto Amato, Jeffrey Barrett, Sonia Goncalves, Ewan Harrison, David K. Jackson, Ian Johnston, Dominic Kwiatkowski, Cordelia Langford, John Sillitoe on behalf of the Wellcome Sanger Institute COVID-19 Surveillance Team |
| EPI_ISL_3201789 | Lighthouse Lab in Milton Keynes | Wellcome Sanger Institute for the COVID-19 Genomics UK (COG-UK) Consortium | The Lighthouse Lab in Milton Keynes and Alex Alderton, Roberto Amato, Jeffrey Barrett, Sonia Goncalves, Ewan Harrison, David K. Jackson, Ian Johnston, Dominic Kwiatkowski, Cordelia Langford, John Sillitoe on behalf of the Wellcome Sanger Institute COVID-19 Surveillance Team |
| EPI_ISL_3201790 | Lighthouse Laboratory Plymouth | Wellcome Sanger Institute for the COVID-19 Genomics UK (COG-UK) Consortium | Lighthouse Laboratory Plymouth and Alex Alderton, Roberto Amato, Jeffrey Barrett, Sonia Goncalves, Ewan Harrison, David K. Jackson, Ian Johnston, Dominic Kwiatkowski, Cordelia Langford, John Sillitoe on behalf of the Wellcome Sanger Institute COVID-19 Surveillance Team |
| EPI_ISL_3201791 | Lighthouse Lab in Alderley Park | Wellcome Sanger Institute for the COVID-19 Genomics UK (COG-UK) Consortium | Jacquelyn Wynn, Mairead Hyland, The Lighthouse Lab in Alderley Park and Alex Alderton, Roberto Amato, Jeffrey Barrett, Sonia Goncalves, Ewan Harrison, David K. Jackson, Ian Johnston, Dominic Kwiatkowski, Cordelia Langford, John Sillitoe on behalf of the Wellcome Sanger Institute COVID-19 Surveillance Team |
| EPI_ISL_3201792 | Lighthouse Lab in Milton Keynes | Wellcome Sanger Institute for the COVID-19 Genomics UK (COG-UK) Consortium | The Lighthouse Lab in Milton Keynes and Alex Alderton, Roberto Amato, Jeffrey Barrett, Sonia Goncalves, Ewan Harrison, David K. Jackson, Ian Johnston, Dominic Kwiatkowski, Cordelia Langford, John Sillitoe on behalf of the Wellcome Sanger Institute COVID-19 Surveillance Team |
| EPI_ISL_3201793 | Lighthouse Laboratory Plymouth | Wellcome Sanger Institute for the COVID-19 Genomics UK (COG-UK) Consortium | Lighthouse Laboratory Plymouth and Alex Alderton, Roberto Amato, Jeffrey Barrett, Sonia Goncalves, Ewan Harrison, David K. Jackson, Ian Johnston, Dominic Kwiatkowski, Cordelia Langford, John Sillitoe on behalf of the Wellcome Sanger Institute COVID-19 Surveillance Team |
| EPI_ISL_3201794 | Lighthouse Lab in Alderley Park | Wellcome Sanger Institute for the COVID-19 Genomics UK (COG-UK) Consortium | Jacquelyn Wynn, Mairead Hyland, The Lighthouse Lab in Alderley Park and Alex Alderton, Roberto Amato, Jeffrey Barrett, Sonia Goncalves, Ewan Harrison, David K. Jackson, Ian Johnston, Dominic Kwiatkowski, Cordelia Langford, John Sillitoe on behalf of the Wellcome Sanger Institute COVID-19 Surveillance Team |
| EPI_ISL_3201795 | Lighthouse Laboratory Plymouth | Wellcome Sanger Institute for the COVID-19 Genomics UK (COG-UK) Consortium | Lighthouse Laboratory Plymouth and Alex Alderton, Roberto Amato, Jeffrey Barrett, Sonia Goncalves, Ewan Harrison, David K. Jackson, Ian Johnston, Dominic Kwiatkowski, Cordelia Langford, John Sillitoe on behalf of the Wellcome Sanger Institute COVID-19 Surveillance Team |
| EPI_ISL_3201796 | Lighthouse Lab in Alderley Park | Wellcome Sanger Institute for the COVID-19 Genomics UK (COG-UK) Consortium | Jacquelyn Wynn, Mairead Hyland, The Lighthouse Lab in Alderley Park and Alex Alderton, Roberto Amato, Jeffrey Barrett, Sonia Goncalves, Ewan Harrison, David K. Jackson, Ian Johnston, Dominic Kwiatkowski, Cordelia Langford, John Sillitoe on behalf of the Wellcome Sanger Institute COVID-19 Surveillance Team |
| EPI_ISL_3201797 | Lighthouse Laboratory Plymouth | Wellcome Sanger Institute for the COVID-19 Genomics UK (COG-UK) Consortium | Lighthouse Laboratory Plymouth and Alex Alderton, Roberto Amato, Jeffrey Barrett, Sonia Goncalves, Ewan Harrison, David K. Jackson, Ian Johnston, Dominic Kwiatkowski, Cordelia Langford, John Sillitoe on behalf of the Wellcome Sanger Institute COVID-19 Surveillance Team |
| EPI_ISL_3201798 | Lighthouse Lab in Alderley Park | Wellcome Sanger Institute for the COVID-19 Genomics UK (COG-UK) Consortium | Jacquelyn Wynn, Mairead Hyland, The Lighthouse Lab in Alderley Park and Alex Alderton, Roberto Amato, Jeffrey Barrett, Sonia Goncalves, Ewan Harrison, David K. Jackson, Ian Johnston, Dominic Kwiatkowski, Cordelia Langford, John Sillitoe on behalf of the Wellcome Sanger Institute COVID-19 Surveillance Team |
| EPI_ISL_3201799 | Lighthouse Lab in Milton Keynes | Wellcome Sanger Institute for the COVID-19 Genomics UK (COG-UK) Consortium | The Lighthouse Lab in Milton Keynes and Alex Alderton, Roberto Amato, Jeffrey Barrett, Sonia Goncalves, Ewan Harrison, David K. Jackson, Ian Johnston, Dominic Kwiatkowski, Cordelia Langford, John Sillitoe on behalf of the Wellcome Sanger Institute COVID-19 Surveillance Team |
| EPI_ISL_3201800 | Lighthouse Laboratory Plymouth | Wellcome Sanger Institute for the COVID-19 Genomics UK (COG-UK) Consortium | Lighthouse Laboratory Plymouth and Alex Alderton, Roberto Amato, Jeffrey Barrett, Sonia Goncalves, Ewan Harrison, David K. Jackson, Ian Johnston, Dominic Kwiatkowski, Cordelia Langford, John Sillitoe on behalf of the Wellcome Sanger Institute COVID-19 Surveillance Team |
| EPI_ISL_3201801 | Lighthouse Lab in Alderley Park | Wellcome Sanger Institute for the COVID-19 Genomics UK (COG-UK) Consortium | Jacquelyn Wynn, Mairead Hyland, The Lighthouse Lab in Alderley Park and Alex Alderton, Roberto Amato, Jeffrey Barrett, Sonia Goncalves, Ewan Harrison, David K. Jackson, Ian Johnston, Dominic Kwiatkowski, Cordelia Langford, John Sillitoe on behalf of the Wellcome Sanger Institute COVID-19 Surveillance Team |
| EPI_ISL_3201802 | Lighthouse Laboratory Plymouth | Wellcome Sanger Institute for the COVID-19 Genomics UK (COG-UK) Consortium | Lighthouse Laboratory Plymouth and Alex Alderton, Roberto Amato, Jeffrey Barrett, Sonia Goncalves, Ewan Harrison, David K. Jackson, Ian Johnston, Dominic Kwiatkowski, Cordelia Langford, John Sillitoe on behalf of the Wellcome Sanger Institute COVID-19 Surveillance Team |
| EPI_ISL_3201803, EPI_ISL_3201804 | Lighthouse Lab in Alderley Park | Wellcome Sanger Institute for the COVID-19 Genomics UK (COG-UK) Consortium | Jacquelyn Wynn, Mairead Hyland, The Lighthouse Lab in Alderley Park and Alex Alderton, Roberto Amato, Jeffrey Barrett, Sonia Goncalves, Ewan Harrison, David K. Jackson, Ian Johnston, Dominic Kwiatkowski, Cordelia Langford, John Sillitoe on behalf of the Wellcome Sanger Institute COVID-19 Surveillance Team |
| EPI_ISL_3201805 | Lighthouse Lab in Milton Keynes | Wellcome Sanger Institute for the COVID-19 Genomics UK (COG-UK) Consortium | The Lighthouse Lab in Milton Keynes and Alex Alderton, Roberto Amato, Jeffrey Barrett, Sonia Goncalves, Ewan Harrison, David K. Jackson, Ian Johnston, Dominic Kwiatkowski, Cordelia Langford, John Sillitoe on behalf of the Wellcome Sanger Institute COVID-19 Surveillance Team |
| EPI_ISL_3201806, EPI_ISL_3201807 | Lighthouse Laboratory Plymouth | Wellcome Sanger Institute for the COVID-19 Genomics UK (COG-UK) Consortium | Lighthouse Laboratory Plymouth and Alex Alderton, Roberto Amato, Jeffrey Barrett, Sonia Goncalves, Ewan Harrison, David K. Jackson, Ian Johnston, Dominic Kwiatkowski, Cordelia Langford, John Sillitoe on behalf of the Wellcome Sanger Institute COVID-19 Surveillance Team |
| EPI_ISL_3201808 | Lighthouse Lab in Milton Keynes | Wellcome Sanger Institute for the COVID-19 Genomics UK (COG-UK) Consortium | The Lighthouse Lab in Milton Keynes and Alex Alderton, Roberto Amato, Jeffrey Barrett, Sonia Goncalves, Ewan Harrison, David K. Jackson, Ian Johnston, Dominic Kwiatkowski, Cordelia Langford, John Sillitoe on behalf of the Wellcome Sanger Institute COVID-19 Surveillance Team |
| EPI_ISL_3201809 | Lighthouse Lab in Alderley Park | Wellcome Sanger Institute for the COVID-19 Genomics UK (COG-UK) Consortium | Jacquelyn Wynn, Mairead Hyland, The Lighthouse Lab in Alderley Park and Alex Alderton, Roberto Amato, Jeffrey Barrett, Sonia Goncalves, Ewan Harrison, David K. Jackson, Ian Johnston, Dominic Kwiatkowski, Cordelia Langford, John Sillitoe on behalf of the Wellcome Sanger Institute COVID-19 Surveillance Team |
| EPI_ISL_3201810 | Lighthouse Lab in Milton Keynes | Wellcome Sanger Institute for the COVID-19 Genomics UK (COG-UK) Consortium | The Lighthouse Lab in Milton Keynes and Alex Alderton, Roberto Amato, Jeffrey Barrett, Sonia Goncalves, Ewan Harrison, David K. Jackson, Ian Johnston, Dominic Kwiatkowski, Cordelia Langford, John Sillitoe on behalf of the Wellcome Sanger Institute COVID-19 Surveillance Team |
| EPI_ISL_3201811, EPI_ISL_3201812 | Lighthouse Lab in Alderley Park | Wellcome Sanger Institute for the COVID-19 Genomics UK (COG-UK) Consortium | Jacquelyn Wynn, Mairead Hyland, The Lighthouse Lab in Alderley Park and Alex Alderton, Roberto Amato, Jeffrey Barrett, Sonia Goncalves, Ewan Harrison, David K. Jackson, Ian Johnston, Dominic Kwiatkowski, Cordelia Langford, John Sillitoe on behalf of the Wellcome Sanger Institute COVID-19 Surveillance Team |
| EPI_ISL_3201813, EPI_ISL_3201814 | Lighthouse Lab in Milton Keynes | Wellcome Sanger Institute for the COVID-19 Genomics UK (COG-UK) Consortium | The Lighthouse Lab in Milton Keynes and Alex Alderton, Roberto Amato, Jeffrey Barrett, Sonia Goncalves, Ewan Harrison, David K. Jackson, Ian Johnston, Dominic Kwiatkowski, Cordelia Langford, John Sillitoe on behalf of the Wellcome Sanger Institute COVID-19 Surveillance Team |
| EPI_ISL_3201815, EPI_ISL_3201816, EPI_ISL_3201817, EPI_ISL_3201818 | Lighthouse Laboratory Plymouth | Wellcome Sanger Institute for the COVID-19 Genomics UK (COG-UK) Consortium | Lighthouse Laboratory Plymouth and Alex Alderton, Roberto Amato, Jeffrey Barrett, Sonia Goncalves, Ewan Harrison, David K. Jackson, Ian Johnston, Dominic Kwiatkowski, Cordelia Langford, John Sillitoe on behalf of the Wellcome Sanger Institute COVID-19 Surveillance Team |
| EPI_ISL_3201819 | Lighthouse Lab in Alderley Park | Wellcome Sanger Institute for the COVID-19 Genomics UK (COG-UK) Consortium | Jacquelyn Wynn, Mairead Hyland, The Lighthouse Lab in Alderley Park and Alex Alderton, Roberto Amato, Jeffrey Barrett, Sonia Goncalves, Ewan Harrison, David K. Jackson, Ian Johnston, Dominic Kwiatkowski, Cordelia Langford, John Sillitoe on behalf of the Wellcome Sanger Institute COVID-19 Surveillance Team |
| EPI_ISL_3201820 | Lighthouse Laboratory Plymouth | Wellcome Sanger Institute for the COVID-19 Genomics UK (COG-UK) Consortium | Lighthouse Laboratory Plymouth and Alex Alderton, Roberto Amato, Jeffrey Barrett, Sonia Goncalves, Ewan Harrison, David K. Jackson, Ian Johnston, Dominic Kwiatkowski, Cordelia Langford, John Sillitoe on behalf of the Wellcome Sanger Institute COVID-19 Surveillance Team |
| EPI_ISL_3201821 | Lighthouse Lab in Alderley Park | Wellcome Sanger Institute for the COVID-19 Genomics UK (COG-UK) Consortium | Jacquelyn Wynn, Mairead Hyland, The Lighthouse Lab in Alderley Park and Alex Alderton, Roberto Amato, Jeffrey Barrett, Sonia Goncalves, Ewan Harrison, David K. Jackson, Ian Johnston, Dominic Kwiatkowski, Cordelia Langford, John Sillitoe on behalf of the Wellcome Sanger Institute COVID-19 Surveillance Team |
| EPI_ISL_3201822 | Lighthouse Laboratory Plymouth | Wellcome Sanger Institute for the COVID-19 Genomics UK (COG-UK) Consortium | Lighthouse Laboratory Plymouth and Alex Alderton, Roberto Amato, Jeffrey Barrett, Sonia Goncalves, Ewan Harrison, David K. Jackson, Ian Johnston, Dominic Kwiatkowski, Cordelia Langford, John Sillitoe on behalf of the Wellcome Sanger Institute COVID-19 Surveillance Team |
| EPI_ISL_3201823 | Lighthouse Lab in Milton Keynes | Wellcome Sanger Institute for the COVID-19 Genomics UK (COG-UK) Consortium | The Lighthouse Lab in Milton Keynes and Alex Alderton, Roberto Amato, Jeffrey Barrett, Sonia Goncalves, Ewan Harrison, David K. Jackson, Ian Johnston, Dominic Kwiatkowski, Cordelia Langford, John Sillitoe on behalf of the Wellcome Sanger Institute COVID-19 Surveillance Team |
| EPI_ISL_3201824 | Lighthouse Laboratory Plymouth | Wellcome Sanger Institute for the COVID-19 Genomics UK (COG-UK) Consortium | Lighthouse Laboratory Plymouth and Alex Alderton, Roberto Amato, Jeffrey Barrett, Sonia Goncalves, Ewan Harrison, David K. Jackson, Ian Johnston, Dominic Kwiatkowski, Cordelia Langford, John Sillitoe on behalf of the Wellcome Sanger Institute COVID-19 Surveillance Team |
| EPI_ISL_3201825 | Lighthouse Lab in Milton Keynes | Wellcome Sanger Institute for the COVID-19 Genomics UK (COG-UK) Consortium | The Lighthouse Lab in Milton Keynes and Alex Alderton, Roberto Amato, Jeffrey Barrett, Sonia Goncalves, Ewan Harrison, David K. Jackson, Ian Johnston, Dominic Kwiatkowski, Cordelia Langford, John Sillitoe on behalf of the Wellcome Sanger Institute COVID-19 Surveillance Team |
| EPI_ISL_3201826 | Lighthouse Lab in Alderley Park | Wellcome Sanger Institute for the COVID-19 Genomics UK (COG-UK) Consortium | Jacquelyn Wynn, Mairead Hyland, The Lighthouse Lab in Alderley Park and Alex Alderton, Roberto Amato, Jeffrey Barrett, Sonia Goncalves, Ewan Harrison, David K. Jackson, Ian Johnston, Dominic Kwiatkowski, Cordelia Langford, John Sillitoe on behalf of the Wellcome Sanger Institute COVID-19 Surveillance Team |
| EPI_ISL_3201827 | Lighthouse Laboratory Plymouth | Wellcome Sanger Institute for the COVID-19 Genomics UK (COG-UK) Consortium | Lighthouse Laboratory Plymouth and Alex Alderton, Roberto Amato, Jeffrey Barrett, Sonia Goncalves, Ewan Harrison, David K. Jackson, Ian Johnston, Dominic Kwiatkowski, Cordelia Langford, John Sillitoe on behalf of the Wellcome Sanger Institute COVID-19 Surveillance Team |
| EPI_ISL_3201828, EPI_ISL_3201829 | Lighthouse Lab in Milton Keynes | Wellcome Sanger Institute for the COVID-19 Genomics UK (COG-UK) Consortium | The Lighthouse Lab in Milton Keynes and Alex Alderton, Roberto Amato, Jeffrey Barrett, Sonia Goncalves, Ewan Harrison, David K. Jackson, Ian Johnston, Dominic Kwiatkowski, Cordelia Langford, John Sillitoe on behalf of the Wellcome Sanger Institute COVID-19 Surveillance Team |
| EPI_ISL_3201830 | Lighthouse Lab in Alderley Park | Wellcome Sanger Institute for the COVID-19 Genomics UK (COG-UK) Consortium | Jacquelyn Wynn, Mairead Hyland, The Lighthouse Lab in Alderley Park and Alex Alderton, Roberto Amato, Jeffrey Barrett, Sonia Goncalves, Ewan Harrison, David K. Jackson, Ian Johnston, Dominic Kwiatkowski, Cordelia Langford, John Sillitoe on behalf of the Wellcome Sanger Institute COVID-19 Surveillance Team |
| EPI_ISL_3201831 | Lighthouse Lab in Milton Keynes | Wellcome Sanger Institute for the COVID-19 Genomics UK (COG-UK) Consortium | The Lighthouse Lab in Milton Keynes and Alex Alderton, Roberto Amato, Jeffrey Barrett, Sonia Goncalves, Ewan Harrison, David K. Jackson, Ian Johnston, Dominic Kwiatkowski, Cordelia Langford, John Sillitoe on behalf of the Wellcome Sanger Institute COVID-19 Surveillance Team |
| EPI_ISL_3201832 | Lighthouse Laboratory Plymouth | Wellcome Sanger Institute for the COVID-19 Genomics UK (COG-UK) Consortium | Lighthouse Laboratory Plymouth and Alex Alderton, Roberto Amato, Jeffrey Barrett, Sonia Goncalves, Ewan Harrison, David K. Jackson, Ian Johnston, Dominic Kwiatkowski, Cordelia Langford, John Sillitoe on behalf of the Wellcome Sanger Institute COVID-19 Surveillance Team |
| EPI_ISL_3201833, EPI_ISL_3201834, EPI_ISL_3201835 | Lighthouse Lab in Alderley Park | Wellcome Sanger Institute for the COVID-19 Genomics UK (COG-UK) Consortium | Jacquelyn Wynn, Mairead Hyland, The Lighthouse Lab in Alderley Park and Alex Alderton, Roberto Amato, Jeffrey Barrett, Sonia Goncalves, Ewan Harrison, David K. Jackson, Ian Johnston, Dominic Kwiatkowski, Cordelia Langford, John Sillitoe on behalf of the Wellcome Sanger Institute COVID-19 Surveillance Team |
| EPI_ISL_3201836 | Lighthouse Lab in Milton Keynes | Wellcome Sanger Institute for the COVID-19 Genomics UK (COG-UK) Consortium | The Lighthouse Lab in Milton Keynes and Alex Alderton, Roberto Amato, Jeffrey Barrett, Sonia Goncalves, Ewan Harrison, David K. Jackson, Ian Johnston, Dominic Kwiatkowski, Cordelia Langford, John Sillitoe on behalf of the Wellcome Sanger Institute COVID-19 Surveillance Team |
| EPI_ISL_3201837 | Lighthouse Laboratory Plymouth | Wellcome Sanger Institute for the COVID-19 Genomics UK (COG-UK) Consortium | Lighthouse Laboratory Plymouth and Alex Alderton, Roberto Amato, Jeffrey Barrett, Sonia Goncalves, Ewan Harrison, David K. Jackson, Ian Johnston, Dominic Kwiatkowski, Cordelia Langford, John Sillitoe on behalf of the Wellcome Sanger Institute COVID-19 Surveillance Team |
| EPI_ISL_3201838 | Lighthouse Lab in Milton Keynes | Wellcome Sanger Institute for the COVID-19 Genomics UK (COG-UK) Consortium | The Lighthouse Lab in Milton Keynes and Alex Alderton, Roberto Amato, Jeffrey Barrett, Sonia Goncalves, Ewan Harrison, David K. Jackson, Ian Johnston, Dominic Kwiatkowski, Cordelia Langford, John Sillitoe on behalf of the Wellcome Sanger Institute COVID-19 Surveillance Team |
| EPI_ISL_3201839 | Lighthouse Lab in Alderley Park | Wellcome Sanger Institute for the COVID-19 Genomics UK (COG-UK) Consortium | Jacquelyn Wynn, Mairead Hyland, The Lighthouse Lab in Alderley Park and Alex Alderton, Roberto Amato, Jeffrey Barrett, Sonia Goncalves, Ewan Harrison, David K. Jackson, Ian Johnston, Dominic Kwiatkowski, Cordelia Langford, John Sillitoe on behalf of the Wellcome Sanger Institute COVID-19 Surveillance Team |

| | | | |
|---|---|---|---|
| EPI_ISL_3201840, EPI_ISL_3201841 | Lighthouse Lab in Milton Keynes | Wellcome Sanger Institute for the COVID-19 Genomics UK (COG-UK) Consortium | The Lighthouse Lab in Milton Keynes and Alex Alderton, Roberto Amato, Jeffrey Barrett, Sonia Goncalves, Ewan Harrison, David K. Jackson, Ian Johnston, Dominic Kwiatkowski, Cordelia Langford, John Sillitoe on behalf of the Wellcome Sanger Institute COVID-19 Surveillance Team |
| EPI_ISL_3201842, EPI_ISL_3201843 | Lighthouse Lab in Alderley Park | Wellcome Sanger Institute for the COVID-19 Genomics UK (COG-UK) Consortium | Jacquelyn Wynn, Mairead Hyland, The Lighthouse Lab in Alderley Park and Alex Alderton, Roberto Amato, Jeffrey Barrett, Sonia Goncalves, Ewan Harrison, David K. Jackson, Ian Johnston, Dominic Kwiatkowski, Cordelia Langford, John Sillitoe on behalf of the Wellcome Sanger Institute COVID-19 Surveillance Team |
| EPI_ISL_3201844, EPI_ISL_3201845 | Lighthouse Lab in Milton Keynes | Wellcome Sanger Institute for the COVID-19 Genomics UK (COG-UK) Consortium | The Lighthouse Lab in Milton Keynes and Alex Alderton, Roberto Amato, Jeffrey Barrett, Sonia Goncalves, Ewan Harrison, David K. Jackson, Ian Johnston, Dominic Kwiatkowski, Cordelia Langford, John Sillitoe on behalf of the Wellcome Sanger Institute COVID-19 Surveillance Team |
| EPI_ISL_3201846 | Lighthouse Laboratory Plymouth | Wellcome Sanger Institute for the COVID-19 Genomics UK (COG-UK) Consortium | Lighthouse Laboratory Plymouth and Alex Alderton, Roberto Amato, Jeffrey Barrett, Sonia Goncalves, Ewan Harrison, David K. Jackson, Ian Johnston, Dominic Kwiatkowski, Cordelia Langford, John Sillitoe on behalf of the Wellcome Sanger Institute COVID-19 Surveillance Team |
| EPI_ISL_3201847 | Lighthouse Lab in Alderley Park | Wellcome Sanger Institute for the COVID-19 Genomics UK (COG-UK) Consortium | Jacquelyn Wynn, Mairead Hyland, The Lighthouse Lab in Alderley Park and Alex Alderton, Roberto Amato, Jeffrey Barrett, Sonia Goncalves, Ewan Harrison, David K. Jackson, Ian Johnston, Dominic Kwiatkowski, Cordelia Langford, John Sillitoe on behalf of the Wellcome Sanger Institute COVID-19 Surveillance Team |
| EPI_ISL_3201848, EPI_ISL_3201849, EPI_ISL_3201850 | Lighthouse Lab in Milton Keynes | Wellcome Sanger Institute for the COVID-19 Genomics UK (COG-UK) Consortium | The Lighthouse Lab in Milton Keynes and Alex Alderton, Roberto Amato, Jeffrey Barrett, Sonia Goncalves, Ewan Harrison, David K. Jackson, Ian Johnston, Dominic Kwiatkowski, Cordelia Langford, John Sillitoe on behalf of the Wellcome Sanger Institute COVID-19 Surveillance Team |
| EPI_ISL_3201851 | Lighthouse Laboratory Plymouth | Wellcome Sanger Institute for the COVID-19 Genomics UK (COG-UK) Consortium | Lighthouse Laboratory Plymouth and Alex Alderton, Roberto Amato, Jeffrey Barrett, Sonia Goncalves, Ewan Harrison, David K. Jackson, Ian Johnston, Dominic Kwiatkowski, Cordelia Langford, John Sillitoe on behalf of the Wellcome Sanger Institute COVID-19 Surveillance Team |
| EPI_ISL_3201852, EPI_ISL_3201853, EPI_ISL_3201854 | Lighthouse Lab in Milton Keynes | Wellcome Sanger Institute for the COVID-19 Genomics UK (COG-UK) Consortium | The Lighthouse Lab in Milton Keynes and Alex Alderton, Roberto Amato, Jeffrey Barrett, Sonia Goncalves, Ewan Harrison, David K. Jackson, Ian Johnston, Dominic Kwiatkowski, Cordelia Langford, John Sillitoe on behalf of the Wellcome Sanger Institute COVID-19 Surveillance Team |
| EPI_ISL_3201855 | Lighthouse Laboratory Plymouth | Wellcome Sanger Institute for the COVID-19 Genomics UK (COG-UK) Consortium | Lighthouse Laboratory Plymouth and Alex Alderton, Roberto Amato, Jeffrey Barrett, Sonia Goncalves, Ewan Harrison, David K. Jackson, Ian Johnston, Dominic Kwiatkowski, Cordelia Langford, John Sillitoe on behalf of the Wellcome Sanger Institute COVID-19 Surveillance Team |
| EPI_ISL_3201856 | Lighthouse Lab in Milton Keynes | Wellcome Sanger Institute for the COVID-19 Genomics UK (COG-UK) Consortium | The Lighthouse Lab in Milton Keynes and Alex Alderton, Roberto Amato, Jeffrey Barrett, Sonia Goncalves, Ewan Harrison, David K. Jackson, Ian Johnston, Dominic Kwiatkowski, Cordelia Langford, John Sillitoe on behalf of the Wellcome Sanger Institute COVID-19 Surveillance Team |
| EPI_ISL_3201857 | Lighthouse Laboratory Plymouth | Wellcome Sanger Institute for the COVID-19 Genomics UK (COG-UK) Consortium | Lighthouse Laboratory Plymouth and Alex Alderton, Roberto Amato, Jeffrey Barrett, Sonia Goncalves, Ewan Harrison, David K. Jackson, Ian Johnston, Dominic Kwiatkowski, Cordelia Langford, John Sillitoe on behalf of the Wellcome Sanger Institute COVID-19 Surveillance Team |
| EPI_ISL_3201858 | Lighthouse Lab in Milton Keynes | Wellcome Sanger Institute for the COVID-19 Genomics UK (COG-UK) Consortium | The Lighthouse Lab in Milton Keynes and Alex Alderton, Roberto Amato, Jeffrey Barrett, Sonia Goncalves, Ewan Harrison, David K. Jackson, Ian Johnston, Dominic Kwiatkowski, Cordelia Langford, John Sillitoe on behalf of the Wellcome Sanger Institute COVID-19 Surveillance Team |
| EPI_ISL_3201859 | Lighthouse Laboratory Plymouth | Wellcome Sanger Institute for the COVID-19 Genomics UK (COG-UK) Consortium | Lighthouse Laboratory Plymouth and Alex Alderton, Roberto Amato, Jeffrey Barrett, Sonia Goncalves, Ewan Harrison, David K. Jackson, Ian Johnston, Dominic Kwiatkowski, Cordelia Langford, John Sillitoe on behalf of the Wellcome Sanger Institute COVID-19 Surveillance Team |
| EPI_ISL_3201860 | Lighthouse Lab in Milton Keynes | Wellcome Sanger Institute for the COVID-19 Genomics UK (COG-UK) Consortium | The Lighthouse Lab in Milton Keynes and Alex Alderton, Roberto Amato, Jeffrey Barrett, Sonia Goncalves, Ewan Harrison, David K. Jackson, Ian Johnston, Dominic Kwiatkowski, Cordelia Langford, John Sillitoe on behalf of the Wellcome Sanger Institute COVID-19 Surveillance Team |
| EPI_ISL_3201861, EPI_ISL_3201862 | Lighthouse Laboratory Plymouth | Wellcome Sanger Institute for the COVID-19 Genomics UK (COG-UK) Consortium | Lighthouse Laboratory Plymouth and Alex Alderton, Roberto Amato, Jeffrey Barrett, Sonia Goncalves, Ewan Harrison, David K. Jackson, Ian Johnston, Dominic Kwiatkowski, Cordelia Langford, John Sillitoe on behalf of the Wellcome Sanger Institute COVID-19 Surveillance Team |
| EPI_ISL_3201863, EPI_ISL_3201864, EPI_ISL_3201865, EPI_ISL_3201866, EPI_ISL_3201867 | Lighthouse Lab in Milton Keynes | Wellcome Sanger Institute for the COVID-19 Genomics UK (COG-UK) Consortium | The Lighthouse Lab in Milton Keynes and Alex Alderton, Roberto Amato, Jeffrey Barrett, Sonia Goncalves, Ewan Harrison, David K. Jackson, Ian Johnston, Dominic Kwiatkowski, Cordelia Langford, John Sillitoe on behalf of the Wellcome Sanger Institute COVID-19 Surveillance Team |
| EPI_ISL_3201868 | Lighthouse Lab in Alderley Park | Wellcome Sanger Institute for the COVID-19 Genomics UK (COG-UK) Consortium | Jacquelyn Wynn, Mairead Hyland, The Lighthouse Lab in Alderley Park and Alex Alderton, Roberto Amato, Jeffrey Barrett, Sonia Goncalves, Ewan Harrison, David K. Jackson, Ian Johnston, Dominic Kwiatkowski, Cordelia Langford, John Sillitoe on behalf of the Wellcome Sanger Institute COVID-19 Surveillance Team |
| EPI_ISL_3201869, EPI_ISL_3201870, EPI_ISL_3201871 | Lighthouse Lab in Milton Keynes | Wellcome Sanger Institute for the COVID-19 Genomics UK (COG-UK) Consortium | The Lighthouse Lab in Milton Keynes and Alex Alderton, Roberto Amato, Jeffrey Barrett, Sonia Goncalves, Ewan Harrison, David K. Jackson, Ian Johnston, Dominic Kwiatkowski, Cordelia Langford, John Sillitoe on behalf of the Wellcome Sanger Institute COVID-19 Surveillance Team |
| EPI_ISL_3201872 | Lighthouse Lab in Alderley Park | Wellcome Sanger Institute for the COVID-19 Genomics UK (COG-UK) Consortium | Jacquelyn Wynn, Mairead Hyland, The Lighthouse Lab in Alderley Park and Alex Alderton, Roberto Amato, Jeffrey Barrett, Sonia Goncalves, Ewan Harrison, David K. Jackson, Ian Johnston, Dominic Kwiatkowski, Cordelia Langford, John Sillitoe on behalf of the Wellcome Sanger Institute COVID-19 Surveillance Team |
| EPI_ISL_3201873 | Lighthouse Lab in Milton Keynes | Wellcome Sanger Institute for the COVID-19 Genomics UK (COG-UK) Consortium | The Lighthouse Lab in Milton Keynes and Alex Alderton, Roberto Amato, Jeffrey Barrett, Sonia Goncalves, Ewan Harrison, David K. Jackson, Ian Johnston, Dominic Kwiatkowski, Cordelia Langford, John Sillitoe on behalf of the Wellcome Sanger Institute COVID-19 Surveillance Team |
| EPI_ISL_3201874 | Lighthouse Laboratory Plymouth | Wellcome Sanger Institute for the COVID-19 Genomics UK (COG-UK) Consortium | Lighthouse Laboratory Plymouth and Alex Alderton, Roberto Amato, Jeffrey Barrett, Sonia Goncalves, Ewan Harrison, David K. Jackson, Ian Johnston, Dominic Kwiatkowski, Cordelia Langford, John Sillitoe on behalf of the Wellcome Sanger Institute COVID-19 Surveillance Team |
| EPI_ISL_3201875 | Lighthouse Lab in Alderley Park | Wellcome Sanger Institute for the COVID-19 Genomics UK (COG-UK) Consortium | Jacquelyn Wynn, Mairead Hyland, The Lighthouse Lab in Alderley Park and Alex Alderton, Roberto Amato, Jeffrey Barrett, Sonia Goncalves, Ewan Harrison, David K. Jackson, Ian Johnston, Dominic Kwiatkowski, Cordelia Langford, John Sillitoe on behalf of the Wellcome Sanger Institute COVID-19 Surveillance Team |
| EPI_ISL_3201876, EPI_ISL_3201877, EPI_ISL_3201878 | Lighthouse Lab in Milton Keynes | Wellcome Sanger Institute for the COVID-19 Genomics UK (COG-UK) Consortium | The Lighthouse Lab in Milton Keynes and Alex Alderton, Roberto Amato, Jeffrey Barrett, Sonia Goncalves, Ewan Harrison, David K. Jackson, Ian Johnston, Dominic Kwiatkowski, Cordelia Langford, John Sillitoe on behalf of the Wellcome Sanger Institute COVID-19 Surveillance Team |
| EPI_ISL_3201879 | Lighthouse Lab in Alderley Park | Wellcome Sanger Institute for the COVID-19 Genomics UK (COG-UK) Consortium | Jacquelyn Wynn, Mairead Hyland, The Lighthouse Lab in Alderley Park and Alex Alderton, Roberto Amato, Jeffrey Barrett, Sonia Goncalves, Ewan Harrison, David K. Jackson, Ian Johnston, Dominic Kwiatkowski, Cordelia Langford, John Sillitoe on behalf of the Wellcome Sanger Institute COVID-19 Surveillance Team |
| EPI_ISL_3201880 | Lighthouse Lab in Milton Keynes | Wellcome Sanger Institute for the COVID-19 Genomics UK (COG-UK) Consortium | The Lighthouse Lab in Milton Keynes and Alex Alderton, Roberto Amato, Jeffrey Barrett, Sonia Goncalves, Ewan Harrison, David K. Jackson, Ian Johnston, Dominic Kwiatkowski, Cordelia Langford, John Sillitoe on behalf of the Wellcome Sanger Institute COVID-19 Surveillance Team |
| EPI_ISL_3201881 | Lighthouse Lab in Alderley Park | Wellcome Sanger Institute for the COVID-19 Genomics UK (COG-UK) Consortium | Jacquelyn Wynn, Mairead Hyland, The Lighthouse Lab in Alderley Park and Alex Alderton, Roberto Amato, Jeffrey Barrett, Sonia Goncalves, Ewan Harrison, David K. Jackson, Ian Johnston, Dominic Kwiatkowski, Cordelia Langford, John Sillitoe on behalf of the Wellcome Sanger Institute COVID-19 Surveillance Team |
| EPI_ISL_3201882 | Lighthouse Lab in Milton Keynes | Wellcome Sanger Institute for the COVID-19 Genomics UK (COG-UK) Consortium | The Lighthouse Lab in Milton Keynes and Alex Alderton, Roberto Amato, Jeffrey Barrett, Sonia Goncalves, Ewan Harrison, David K. Jackson, Ian Johnston, Dominic Kwiatkowski, Cordelia Langford, John Sillitoe on behalf of the Wellcome Sanger Institute COVID-19 Surveillance Team |
| EPI_ISL_3201883, EPI_ISL_3201884 | Lighthouse Lab in Alderley Park | Wellcome Sanger Institute for the COVID-19 Genomics UK (COG-UK) Consortium | Jacquelyn Wynn, Mairead Hyland, The Lighthouse Lab in Alderley Park and Alex Alderton, Roberto Amato, Jeffrey Barrett, Sonia Goncalves, Ewan Harrison, David K. Jackson, Ian Johnston, Dominic Kwiatkowski, Cordelia Langford, John Sillitoe on behalf of the Wellcome Sanger Institute COVID-19 Surveillance Team |
| EPI_ISL_3201885, EPI_ISL_3201886 | Lighthouse Lab in Milton Keynes | Wellcome Sanger Institute for the COVID-19 Genomics UK (COG-UK) Consortium | The Lighthouse Lab in Milton Keynes and Alex Alderton, Roberto Amato, Jeffrey Barrett, Sonia Goncalves, Ewan Harrison, David K. Jackson, Ian Johnston, Dominic Kwiatkowski, Cordelia Langford, John Sillitoe on behalf of the Wellcome Sanger Institute COVID-19 Surveillance Team |
| EPI_ISL_3201887 | Lighthouse Laboratory Plymouth | Wellcome Sanger Institute for the COVID-19 Genomics UK (COG-UK) Consortium | Lighthouse Laboratory Plymouth and Alex Alderton, Roberto Amato, Jeffrey Barrett, Sonia Goncalves, Ewan Harrison, David K. Jackson, Ian Johnston, Dominic Kwiatkowski, Cordelia Langford, John Sillitoe on behalf of the Wellcome Sanger Institute COVID-19 Surveillance Team |
| EPI_ISL_3201888 | Lighthouse Lab in Alderley Park | Wellcome Sanger Institute for the COVID-19 Genomics UK (COG-UK) Consortium | Jacquelyn Wynn, Mairead Hyland, The Lighthouse Lab in Alderley Park and Alex Alderton, Roberto Amato, Jeffrey Barrett, Sonia Goncalves, Ewan Harrison, David K. Jackson, Ian Johnston, Dominic Kwiatkowski, Cordelia Langford, John Sillitoe on behalf of the Wellcome Sanger Institute COVID-19 Surveillance Team |
| EPI_ISL_3201889 | Lighthouse Laboratory Plymouth | Wellcome Sanger Institute for the COVID-19 Genomics UK (COG-UK) Consortium | Lighthouse Laboratory Plymouth and Alex Alderton, Roberto Amato, Jeffrey Barrett, Sonia Goncalves, Ewan Harrison, David K. Jackson, Ian Johnston, Dominic Kwiatkowski, Cordelia Langford, John Sillitoe on behalf of the Wellcome Sanger Institute COVID-19 Surveillance Team |
| EPI_ISL_3201890 | Lighthouse Lab in Milton Keynes | Wellcome Sanger Institute for the COVID-19 Genomics UK (COG-UK) Consortium | The Lighthouse Lab in Milton Keynes and Alex Alderton, Roberto Amato, Jeffrey Barrett, Sonia Goncalves, Ewan Harrison, David K. Jackson, Ian Johnston, Dominic Kwiatkowski, Cordelia Langford, John Sillitoe on behalf of the Wellcome Sanger Institute COVID-19 Surveillance Team |
| EPI_ISL_3201891 | Lighthouse Laboratory Plymouth | Wellcome Sanger Institute for the COVID-19 Genomics UK (COG-UK) Consortium | Lighthouse Laboratory Plymouth and Alex Alderton, Roberto Amato, Jeffrey Barrett, Sonia Goncalves, Ewan Harrison, David K. Jackson, Ian Johnston, Dominic Kwiatkowski, Cordelia Langford, John Sillitoe on behalf of the Wellcome Sanger Institute COVID-19 Surveillance Team |
| EPI_ISL_3201892, EPI_ISL_3201893, EPI_ISL_3201894 | Lighthouse Lab in Milton Keynes | Wellcome Sanger Institute for the COVID-19 Genomics UK (COG-UK) Consortium | The Lighthouse Lab in Milton Keynes and Alex Alderton, Roberto Amato, Jeffrey Barrett, Sonia Goncalves, Ewan Harrison, David K. Jackson, Ian Johnston, Dominic Kwiatkowski, Cordelia Langford, John Sillitoe on behalf of the Wellcome Sanger Institute COVID-19 Surveillance Team |
| EPI_ISL_3201895 | Lighthouse Lab in Alderley Park | Wellcome Sanger Institute for the COVID-19 Genomics UK (COG-UK) Consortium | Jacquelyn Wynn, Mairead Hyland, The Lighthouse Lab in Alderley Park and Alex Alderton, Roberto Amato, Jeffrey Barrett, Sonia Goncalves, Ewan Harrison, David K. Jackson, Ian Johnston, Dominic Kwiatkowski, Cordelia Langford, John Sillitoe on behalf of the Wellcome Sanger Institute COVID-19 Surveillance Team |
| EPI_ISL_3201896, EPI_ISL_3201897, EPI_ISL_3201898 | Lighthouse Lab in Milton Keynes | Wellcome Sanger Institute for the COVID-19 Genomics UK (COG-UK) Consortium | The Lighthouse Lab in Milton Keynes and Alex Alderton, Roberto Amato, Jeffrey Barrett, Sonia Goncalves, Ewan Harrison, David K. Jackson, Ian Johnston, Dominic Kwiatkowski, Cordelia Langford, John Sillitoe on behalf of the Wellcome Sanger Institute COVID-19 Surveillance Team |
| EPI_ISL_3201899 | Lighthouse Lab in Alderley Park | Wellcome Sanger Institute for the COVID-19 Genomics UK (COG-UK) Consortium | Jacquelyn Wynn, Mairead Hyland, The Lighthouse Lab in Alderley Park and Alex Alderton, Roberto Amato, Jeffrey Barrett, Sonia Goncalves, Ewan Harrison, David K. Jackson, Ian Johnston, Dominic Kwiatkowski, Cordelia Langford, John Sillitoe on behalf of the Wellcome Sanger Institute COVID-19 Surveillance Team |
| EPI_ISL_3201900 | Lighthouse Lab in Milton Keynes | Wellcome Sanger Institute for the COVID-19 Genomics UK (COG-UK) Consortium | The Lighthouse Lab in Milton Keynes and Alex Alderton, Roberto Amato, Jeffrey Barrett, Sonia Goncalves, Ewan Harrison, David K. Jackson, Ian Johnston, Dominic Kwiatkowski, Cordelia Langford, John Sillitoe on behalf of the Wellcome Sanger Institute COVID-19 Surveillance Team |
| EPI_ISL_3201901 | Lighthouse Laboratory Plymouth | Wellcome Sanger Institute for the COVID-19 Genomics UK (COG-UK) Consortium | Lighthouse Laboratory Plymouth and Alex Alderton, Roberto Amato, Jeffrey Barrett, Sonia Goncalves, Ewan Harrison, David K. Jackson, Ian Johnston, Dominic Kwiatkowski, Cordelia Langford, John Sillitoe on behalf of the Wellcome Sanger Institute COVID-19 Surveillance Team |
| EPI_ISL_3201902, EPI_ISL_3201903, EPI_ISL_3201904, EPI_ISL_3201905 | Lighthouse Lab in Milton Keynes | Wellcome Sanger Institute for the COVID-19 Genomics UK (COG-UK) Consortium | The Lighthouse Lab in Milton Keynes and Alex Alderton, Roberto Amato, Jeffrey Barrett, Sonia Goncalves, Ewan Harrison, David K. Jackson, Ian Johnston, Dominic Kwiatkowski, Cordelia Langford, John Sillitoe on behalf of the Wellcome Sanger Institute COVID-19 Surveillance Team |

| | | | |
|---|---|---|---|
| EPI_ISL_3201906 | Lighthouse Lab in Alderley Park | Wellcome Sanger Institute for the COVID-19 Genomics UK (COG-UK) Consortium | Jacquelyn Wynn, Mairead Hyland, The Lighthouse Lab in Alderley Park and Alex Alderton, Roberto Amato, Jeffrey Barrett, Sonia Goncalves, Ewan Harrison, David K. Jackson, Ian Johnston, Dominic Kwiatkowski, Cordelia Langford, John Sillitoe on behalf of the Wellcome Sanger Institute COVID-19 Surveillance Team |
| EPI_ISL_3201907 | Lighthouse Lab in Milton Keynes | Wellcome Sanger Institute for the COVID-19 Genomics UK (COG-UK) Consortium | The Lighthouse Lab in Milton Keynes and Alex Alderton, Roberto Amato, Jeffrey Barrett, Sonia Goncalves, Ewan Harrison, David K. Jackson, Ian Johnston, Dominic Kwiatkowski, Cordelia Langford, John Sillitoe on behalf of the Wellcome Sanger Institute COVID-19 Surveillance Team |
| EPI_ISL_3201908 | Lighthouse Lab in Alderley Park | Wellcome Sanger Institute for the COVID-19 Genomics UK (COG-UK) Consortium | Jacquelyn Wynn, Mairead Hyland, The Lighthouse Lab in Alderley Park and Alex Alderton, Roberto Amato, Jeffrey Barrett, Sonia Goncalves, Ewan Harrison, David K. Jackson, Ian Johnston, Dominic Kwiatkowski, Cordelia Langford, John Sillitoe on behalf of the Wellcome Sanger Institute COVID-19 Surveillance Team |
| EPI_ISL_3201909 | Lighthouse Lab in Milton Keynes | Wellcome Sanger Institute for the COVID-19 Genomics UK (COG-UK) Consortium | The Lighthouse Lab in Milton Keynes and Alex Alderton, Roberto Amato, Jeffrey Barrett, Sonia Goncalves, Ewan Harrison, David K. Jackson, Ian Johnston, Dominic Kwiatkowski, Cordelia Langford, John Sillitoe on behalf of the Wellcome Sanger Institute COVID-19 Surveillance Team |
| EPI_ISL_3201910 | Lighthouse Lab in Alderley Park | Wellcome Sanger Institute for the COVID-19 Genomics UK (COG-UK) Consortium | Jacquelyn Wynn, Mairead Hyland, The Lighthouse Lab in Alderley Park and Alex Alderton, Roberto Amato, Jeffrey Barrett, Sonia Goncalves, Ewan Harrison, David K. Jackson, Ian Johnston, Dominic Kwiatkowski, Cordelia Langford, John Sillitoe on behalf of the Wellcome Sanger Institute COVID-19 Surveillance Team |
| EPI_ISL_3201911 | Lighthouse Laboratory Plymouth | Wellcome Sanger Institute for the COVID-19 Genomics UK (COG-UK) Consortium | Lighthouse Laboratory Plymouth and Alex Alderton, Roberto Amato, Jeffrey Barrett, Sonia Goncalves, Ewan Harrison, David K. Jackson, Ian Johnston, Dominic Kwiatkowski, Cordelia Langford, John Sillitoe on behalf of the Wellcome Sanger Institute COVID-19 Surveillance Team |
| EPI_ISL_3201912, EPI_ISL_3201913, EPI_ISL_3201914, EPI_ISL_3201915 | Lighthouse Lab in Milton Keynes | Wellcome Sanger Institute for the COVID-19 Genomics UK (COG-UK) Consortium | The Lighthouse Lab in Milton Keynes and Alex Alderton, Roberto Amato, Jeffrey Barrett, Sonia Goncalves, Ewan Harrison, David K. Jackson, Ian Johnston, Dominic Kwiatkowski, Cordelia Langford, John Sillitoe on behalf of the Wellcome Sanger Institute COVID-19 Surveillance Team |
| EPI_ISL_3201916 | Lighthouse Laboratory Plymouth | Wellcome Sanger Institute for the COVID-19 Genomics UK (COG-UK) Consortium | Lighthouse Laboratory Plymouth and Alex Alderton, Roberto Amato, Jeffrey Barrett, Sonia Goncalves, Ewan Harrison, David K. Jackson, Ian Johnston, Dominic Kwiatkowski, Cordelia Langford, John Sillitoe on behalf of the Wellcome Sanger Institute COVID-19 Surveillance Team |
| EPI_ISL_3201917 | Lighthouse Lab in Alderley Park | Wellcome Sanger Institute for the COVID-19 Genomics UK (COG-UK) Consortium | Jacquelyn Wynn, Mairead Hyland, The Lighthouse Lab in Alderley Park and Alex Alderton, Roberto Amato, Jeffrey Barrett, Sonia Goncalves, Ewan Harrison, David K. Jackson, Ian Johnston, Dominic Kwiatkowski, Cordelia Langford, John Sillitoe on behalf of the Wellcome Sanger Institute COVID-19 Surveillance Team |
| EPI_ISL_3201918 | Lighthouse Laboratory Plymouth | Wellcome Sanger Institute for the COVID-19 Genomics UK (COG-UK) Consortium | Lighthouse Laboratory Plymouth and Alex Alderton, Roberto Amato, Jeffrey Barrett, Sonia Goncalves, Ewan Harrison, David K. Jackson, Ian Johnston, Dominic Kwiatkowski, Cordelia Langford, John Sillitoe on behalf of the Wellcome Sanger Institute COVID-19 Surveillance Team |
| EPI_ISL_3201919 | Lighthouse Lab in Milton Keynes | Wellcome Sanger Institute for the COVID-19 Genomics UK (COG-UK) Consortium | The Lighthouse Lab in Milton Keynes and Alex Alderton, Roberto Amato, Jeffrey Barrett, Sonia Goncalves, Ewan Harrison, David K. Jackson, Ian Johnston, Dominic Kwiatkowski, Cordelia Langford, John Sillitoe on behalf of the Wellcome Sanger Institute COVID-19 Surveillance Team |
| EPI_ISL_3201920 | Lighthouse Laboratory Plymouth | Wellcome Sanger Institute for the COVID-19 Genomics UK (COG-UK) Consortium | Lighthouse Laboratory Plymouth and Alex Alderton, Roberto Amato, Jeffrey Barrett, Sonia Goncalves, Ewan Harrison, David K. Jackson, Ian Johnston, Dominic Kwiatkowski, Cordelia Langford, John Sillitoe on behalf of the Wellcome Sanger Institute COVID-19 Surveillance Team |
| EPI_ISL_3201921 | Lighthouse Lab in Alderley Park | Wellcome Sanger Institute for the COVID-19 Genomics UK (COG-UK) Consortium | Jacquelyn Wynn, Mairead Hyland, The Lighthouse Lab in Alderley Park and Alex Alderton, Roberto Amato, Jeffrey Barrett, Sonia Goncalves, Ewan Harrison, David K. Jackson, Ian Johnston, Dominic Kwiatkowski, Cordelia Langford, John Sillitoe on behalf of the Wellcome Sanger Institute COVID-19 Surveillance Team |
| EPI_ISL_3201922, EPI_ISL_3201923, EPI_ISL_3201924, EPI_ISL_3201925 | Lighthouse Lab in Milton Keynes | Wellcome Sanger Institute for the COVID-19 Genomics UK (COG-UK) Consortium | The Lighthouse Lab in Milton Keynes and Alex Alderton, Roberto Amato, Jeffrey Barrett, Sonia Goncalves, Ewan Harrison, David K. Jackson, Ian Johnston, Dominic Kwiatkowski, Cordelia Langford, John Sillitoe on behalf of the Wellcome Sanger Institute COVID-19 Surveillance Team |
| EPI_ISL_3201926, EPI_ISL_3201927 | Lighthouse Laboratory Plymouth | Wellcome Sanger Institute for the COVID-19 Genomics UK (COG-UK) Consortium | Lighthouse Laboratory Plymouth and Alex Alderton, Roberto Amato, Jeffrey Barrett, Sonia Goncalves, Ewan Harrison, David K. Jackson, Ian Johnston, Dominic Kwiatkowski, Cordelia Langford, John Sillitoe on behalf of the Wellcome Sanger Institute COVID-19 Surveillance Team |
| EPI_ISL_3201928 | Lighthouse Lab in Milton Keynes | Wellcome Sanger Institute for the COVID-19 Genomics UK (COG-UK) Consortium | The Lighthouse Lab in Milton Keynes and Alex Alderton, Roberto Amato, Jeffrey Barrett, Sonia Goncalves, Ewan Harrison, David K. Jackson, Ian Johnston, Dominic Kwiatkowski, Cordelia Langford, John Sillitoe on behalf of the Wellcome Sanger Institute COVID-19 Surveillance Team |
| EPI_ISL_3201929 | Lighthouse Laboratory Plymouth | Wellcome Sanger Institute for the COVID-19 Genomics UK (COG-UK) Consortium | Lighthouse Laboratory Plymouth and Alex Alderton, Roberto Amato, Jeffrey Barrett, Sonia Goncalves, Ewan Harrison, David K. Jackson, Ian Johnston, Dominic Kwiatkowski, Cordelia Langford, John Sillitoe on behalf of the Wellcome Sanger Institute COVID-19 Surveillance Team |
| EPI_ISL_3201930 | Lighthouse Lab in Alderley Park | Wellcome Sanger Institute for the COVID-19 Genomics UK (COG-UK) Consortium | Jacquelyn Wynn, Mairead Hyland, The Lighthouse Lab in Alderley Park and Alex Alderton, Roberto Amato, Jeffrey Barrett, Sonia Goncalves, Ewan Harrison, David K. Jackson, Ian Johnston, Dominic Kwiatkowski, Cordelia Langford, John Sillitoe on behalf of the Wellcome Sanger Institute COVID-19 Surveillance Team |
| EPI_ISL_3201931, EPI_ISL_3201932, EPI_ISL_3201933 | Lighthouse Lab in Milton Keynes | Wellcome Sanger Institute for the COVID-19 Genomics UK (COG-UK) Consortium | The Lighthouse Lab in Milton Keynes and Alex Alderton, Roberto Amato, Jeffrey Barrett, Sonia Goncalves, Ewan Harrison, David K. Jackson, Ian Johnston, Dominic Kwiatkowski, Cordelia Langford, John Sillitoe on behalf of the Wellcome Sanger Institute COVID-19 Surveillance Team |
| EPI_ISL_3201934 | Lighthouse Laboratory Plymouth | Wellcome Sanger Institute for the COVID-19 Genomics UK (COG-UK) Consortium | Lighthouse Laboratory Plymouth and Alex Alderton, Roberto Amato, Jeffrey Barrett, Sonia Goncalves, Ewan Harrison, David K. Jackson, Ian Johnston, Dominic Kwiatkowski, Cordelia Langford, John Sillitoe on behalf of the Wellcome Sanger Institute COVID-19 Surveillance Team |
| EPI_ISL_3201935, EPI_ISL_3201936, EPI_ISL_3201937, EPI_ISL_3201938, EPI_ISL_3201939 | Lighthouse Lab in Milton Keynes | Wellcome Sanger Institute for the COVID-19 Genomics UK (COG-UK) Consortium | The Lighthouse Lab in Milton Keynes and Alex Alderton, Roberto Amato, Jeffrey Barrett, Sonia Goncalves, Ewan Harrison, David K. Jackson, Ian Johnston, Dominic Kwiatkowski, Cordelia Langford, John Sillitoe on behalf of the Wellcome Sanger Institute COVID-19 Surveillance Team |
| EPI_ISL_3201940 | Lighthouse Laboratory Plymouth | Wellcome Sanger Institute for the COVID-19 Genomics UK (COG-UK) Consortium | Lighthouse Laboratory Plymouth and Alex Alderton, Roberto Amato, Jeffrey Barrett, Sonia Goncalves, Ewan Harrison, David K. Jackson, Ian Johnston, Dominic Kwiatkowski, Cordelia Langford, John Sillitoe on behalf of the Wellcome Sanger Institute COVID-19 Surveillance Team |
| EPI_ISL_3201941, EPI_ISL_3201942 | Lighthouse Lab in Milton Keynes | Wellcome Sanger Institute for the COVID-19 Genomics UK (COG-UK) Consortium | The Lighthouse Lab in Milton Keynes and Alex Alderton, Roberto Amato, Jeffrey Barrett, Sonia Goncalves, Ewan Harrison, David K. Jackson, Ian Johnston, Dominic Kwiatkowski, Cordelia Langford, John Sillitoe on behalf of the Wellcome Sanger Institute COVID-19 Surveillance Team |
| EPI_ISL_3201943 | Lighthouse Laboratory Plymouth | Wellcome Sanger Institute for the COVID-19 Genomics UK (COG-UK) Consortium | Lighthouse Laboratory Plymouth and Alex Alderton, Roberto Amato, Jeffrey Barrett, Sonia Goncalves, Ewan Harrison, David K. Jackson, Ian Johnston, Dominic Kwiatkowski, Cordelia Langford, John Sillitoe on behalf of the Wellcome Sanger Institute COVID-19 Surveillance Team |
| EPI_ISL_3201944, EPI_ISL_3201945, EPI_ISL_3201946, EPI_ISL_3201947, EPI_ISL_3201948 | Lighthouse Lab in Alderley Park | Wellcome Sanger Institute for the COVID-19 Genomics UK (COG-UK) Consortium | Jacquelyn Wynn, Mairead Hyland, The Lighthouse Lab in Alderley Park and Alex Alderton, Roberto Amato, Jeffrey Barrett, Sonia Goncalves, Ewan Harrison, David K. Jackson, Ian Johnston, Dominic Kwiatkowski, Cordelia Langford, John Sillitoe on behalf of the Wellcome Sanger Institute COVID-19 Surveillance Team |
| EPI_ISL_3201949, EPI_ISL_3201950 | Lighthouse Laboratory Plymouth | Wellcome Sanger Institute for the COVID-19 Genomics UK (COG-UK) Consortium | Lighthouse Laboratory Plymouth and Alex Alderton, Roberto Amato, Jeffrey Barrett, Sonia Goncalves, Ewan Harrison, David K. Jackson, Ian Johnston, Dominic Kwiatkowski, Cordelia Langford, John Sillitoe on behalf of the Wellcome Sanger Institute COVID-19 Surveillance Team |
| EPI_ISL_3201951 | Lighthouse Lab in Milton Keynes | Wellcome Sanger Institute for the COVID-19 Genomics UK (COG-UK) Consortium | The Lighthouse Lab in Milton Keynes and Alex Alderton, Roberto Amato, Jeffrey Barrett, Sonia Goncalves, Ewan Harrison, David K. Jackson, Ian Johnston, Dominic Kwiatkowski, Cordelia Langford, John Sillitoe on behalf of the Wellcome Sanger Institute COVID-19 Surveillance Team |
| EPI_ISL_3201952 | Lighthouse Lab in Alderley Park | Wellcome Sanger Institute for the COVID-19 Genomics UK (COG-UK) Consortium | Jacquelyn Wynn, Mairead Hyland, The Lighthouse Lab in Alderley Park and Alex Alderton, Roberto Amato, Jeffrey Barrett, Sonia Goncalves, Ewan Harrison, David K. Jackson, Ian Johnston, Dominic Kwiatkowski, Cordelia Langford, John Sillitoe on behalf of the Wellcome Sanger Institute COVID-19 Surveillance Team |
| EPI_ISL_3201953, EPI_ISL_3201954 | Lighthouse Lab in Milton Keynes | Wellcome Sanger Institute for the COVID-19 Genomics UK (COG-UK) Consortium | The Lighthouse Lab in Milton Keynes and Alex Alderton, Roberto Amato, Jeffrey Barrett, Sonia Goncalves, Ewan Harrison, David K. Jackson, Ian Johnston, Dominic Kwiatkowski, Cordelia Langford, John Sillitoe on behalf of the Wellcome Sanger Institute COVID-19 Surveillance Team |
| EPI_ISL_3201955 | Lighthouse Laboratory Plymouth | Wellcome Sanger Institute for the COVID-19 Genomics UK (COG-UK) Consortium | Lighthouse Laboratory Plymouth and Alex Alderton, Roberto Amato, Jeffrey Barrett, Sonia Goncalves, Ewan Harrison, David K. Jackson, Ian Johnston, Dominic Kwiatkowski, Cordelia Langford, John Sillitoe on behalf of the Wellcome Sanger Institute COVID-19 Surveillance Team |
| EPI_ISL_3201956 | Lighthouse Lab in Alderley Park | Wellcome Sanger Institute for the COVID-19 Genomics UK (COG-UK) Consortium | Jacquelyn Wynn, Mairead Hyland, The Lighthouse Lab in Alderley Park and Alex Alderton, Roberto Amato, Jeffrey Barrett, Sonia Goncalves, Ewan Harrison, David K. Jackson, Ian Johnston, Dominic Kwiatkowski, Cordelia Langford, John Sillitoe on behalf of the Wellcome Sanger Institute COVID-19 Surveillance Team |
| EPI_ISL_3201957, EPI_ISL_3201958 | Lighthouse Lab in Milton Keynes | Wellcome Sanger Institute for the COVID-19 Genomics UK (COG-UK) Consortium | The Lighthouse Lab in Milton Keynes and Alex Alderton, Roberto Amato, Jeffrey Barrett, Sonia Goncalves, Ewan Harrison, David K. Jackson, Ian Johnston, Dominic Kwiatkowski, Cordelia Langford, John Sillitoe on behalf of the Wellcome Sanger Institute COVID-19 Surveillance Team |
| EPI_ISL_3201959 | Lighthouse Laboratory Plymouth | Wellcome Sanger Institute for the COVID-19 Genomics UK (COG-UK) Consortium | Lighthouse Laboratory Plymouth and Alex Alderton, Roberto Amato, Jeffrey Barrett, Sonia Goncalves, Ewan Harrison, David K. Jackson, Ian Johnston, Dominic Kwiatkowski, Cordelia Langford, John Sillitoe on behalf of the Wellcome Sanger Institute COVID-19 Surveillance Team |
| EPI_ISL_3201960 | Lighthouse Lab in Milton Keynes | Wellcome Sanger Institute for the COVID-19 Genomics UK (COG-UK) Consortium | The Lighthouse Lab in Milton Keynes and Alex Alderton, Roberto Amato, Jeffrey Barrett, Sonia Goncalves, Ewan Harrison, David K. Jackson, Ian Johnston, Dominic Kwiatkowski, Cordelia Langford, John Sillitoe on behalf of the Wellcome Sanger Institute COVID-19 Surveillance Team |
| EPI_ISL_3201961, EPI_ISL_3201962 | Lighthouse Laboratory Plymouth | Wellcome Sanger Institute for the COVID-19 Genomics UK (COG-UK) Consortium | Lighthouse Laboratory Plymouth and Alex Alderton, Roberto Amato, Jeffrey Barrett, Sonia Goncalves, Ewan Harrison, David K. Jackson, Ian Johnston, Dominic Kwiatkowski, Cordelia Langford, John Sillitoe on behalf of the Wellcome Sanger Institute COVID-19 Surveillance Team |
| EPI_ISL_3201963, EPI_ISL_3201964 | Lighthouse Lab in Milton Keynes | Wellcome Sanger Institute for the COVID-19 Genomics UK (COG-UK) Consortium | The Lighthouse Lab in Milton Keynes and Alex Alderton, Roberto Amato, Jeffrey Barrett, Sonia Goncalves, Ewan Harrison, David K. Jackson, Ian Johnston, Dominic Kwiatkowski, Cordelia Langford, John Sillitoe on behalf of the Wellcome Sanger Institute COVID-19 Surveillance Team |
| EPI_ISL_3201965 | Lighthouse Laboratory Plymouth | Wellcome Sanger Institute for the COVID-19 Genomics UK (COG-UK) Consortium | Lighthouse Laboratory Plymouth and Alex Alderton, Roberto Amato, Jeffrey Barrett, Sonia Goncalves, Ewan Harrison, David K. Jackson, Ian Johnston, Dominic Kwiatkowski, Cordelia Langford, John Sillitoe on behalf of the Wellcome Sanger Institute COVID-19 Surveillance Team |
| EPI_ISL_3201966, EPI_ISL_3201967, EPI_ISL_3201968 | Lighthouse Lab in Milton Keynes | Wellcome Sanger Institute for the COVID-19 Genomics UK (COG-UK) Consortium | The Lighthouse Lab in Milton Keynes and Alex Alderton, Roberto Amato, Jeffrey Barrett, Sonia Goncalves, Ewan Harrison, David K. Jackson, Ian Johnston, Dominic Kwiatkowski, Cordelia Langford, John Sillitoe on behalf of the Wellcome Sanger Institute COVID-19 Surveillance Team |
| EPI_ISL_3201969 | Lighthouse Lab in Alderley Park | Wellcome Sanger Institute for the COVID-19 Genomics UK (COG-UK) Consortium | Jacquelyn Wynn, Mairead Hyland, The Lighthouse Lab in Alderley Park and Alex Alderton, Roberto Amato, Jeffrey Barrett, Sonia Goncalves, Ewan Harrison, David K. Jackson, Ian Johnston, Dominic Kwiatkowski, Cordelia Langford, John Sillitoe on behalf of the Wellcome Sanger Institute COVID-19 Surveillance Team |
| EPI_ISL_3201970 | Lighthouse Lab in Milton Keynes | Wellcome Sanger Institute for the COVID-19 Genomics UK (COG-UK) Consortium | The Lighthouse Lab in Milton Keynes and Alex Alderton, Roberto Amato, Jeffrey Barrett, Sonia Goncalves, Ewan Harrison, David K. Jackson, Ian Johnston, Dominic Kwiatkowski, Cordelia Langford, John Sillitoe on behalf of the Wellcome Sanger Institute COVID-19 Surveillance Team |

| | | | |
|---|---|---|---|
| EPI_ISL_3201971 | Lighthouse Laboratory Plymouth | Wellcome Sanger Institute for the COVID-19 Genomics UK (COG-UK) Consortium | Lighthouse Laboratory Plymouth and Alex Alderton, Roberto Amato, Jeffrey Barrett, Sonia Goncalves, Ewan Harrison, David K. Jackson, Ian Johnston, Dominic Kwiatkowski, Cordelia Langford, John Sillitoe on behalf of the Wellcome Sanger Institute COVID-19 Surveillance Team |
| EPI_ISL_3201972 | Lighthouse Lab in Milton Keynes | Wellcome Sanger Institute for the COVID-19 Genomics UK (COG-UK) Consortium | The Lighthouse Lab in Milton Keynes and Alex Alderton, Roberto Amato, Jeffrey Barrett, Sonia Goncalves, Ewan Harrison, David K. Jackson, Ian Johnston, Dominic Kwiatkowski, Cordelia Langford, John Sillitoe on behalf of the Wellcome Sanger Institute COVID-19 Surveillance Team |
| EPI_ISL_3201973 | Lighthouse Lab in Alderley Park | Wellcome Sanger Institute for the COVID-19 Genomics UK (COG-UK) Consortium | Jacquelyn Wynn, Mairead Hyland, The Lighthouse Lab in Alderley Park and Alex Alderton, Roberto Amato, Jeffrey Barrett, Sonia Goncalves, Ewan Harrison, David K. Jackson, Ian Johnston, Dominic Kwiatkowski, Cordelia Langford, John Sillitoe on behalf of the Wellcome Sanger Institute COVID-19 Surveillance Team |
| EPI_ISL_3201974, EPI_ISL_3201975, EPI_ISL_3201976, EPI_ISL_3201977, EPI_ISL_3201978, EPI_ISL_3201979, EPI_ISL_3201980 | Lighthouse Lab in Milton Keynes | Wellcome Sanger Institute for the COVID-19 Genomics UK (COG-UK) Consortium | The Lighthouse Lab in Milton Keynes and Alex Alderton, Roberto Amato, Jeffrey Barrett, Sonia Goncalves, Ewan Harrison, David K. Jackson, Ian Johnston, Dominic Kwiatkowski, Cordelia Langford, John Sillitoe on behalf of the Wellcome Sanger Institute COVID-19 Surveillance Team |
| EPI_ISL_3201981, EPI_ISL_3201982 | Lighthouse Lab in Alderley Park | Wellcome Sanger Institute for the COVID-19 Genomics UK (COG-UK) Consortium | Jacquelyn Wynn, Mairead Hyland, The Lighthouse Lab in Alderley Park and Alex Alderton, Roberto Amato, Jeffrey Barrett, Sonia Goncalves, Ewan Harrison, David K. Jackson, Ian Johnston, Dominic Kwiatkowski, Cordelia Langford, John Sillitoe on behalf of the Wellcome Sanger Institute COVID-19 Surveillance Team |
| EPI_ISL_3201983 | Lighthouse Laboratory Plymouth | Wellcome Sanger Institute for the COVID-19 Genomics UK (COG-UK) Consortium | Lighthouse Laboratory Plymouth and Alex Alderton, Roberto Amato, Jeffrey Barrett, Sonia Goncalves, Ewan Harrison, David K. Jackson, Ian Johnston, Dominic Kwiatkowski, Cordelia Langford, John Sillitoe on behalf of the Wellcome Sanger Institute COVID-19 Surveillance Team |
| EPI_ISL_3201984 | Lighthouse Lab in Alderley Park | Wellcome Sanger Institute for the COVID-19 Genomics UK (COG-UK) Consortium | Jacquelyn Wynn, Mairead Hyland, The Lighthouse Lab in Alderley Park and Alex Alderton, Roberto Amato, Jeffrey Barrett, Sonia Goncalves, Ewan Harrison, David K. Jackson, Ian Johnston, Dominic Kwiatkowski, Cordelia Langford, John Sillitoe on behalf of the Wellcome Sanger Institute COVID-19 Surveillance Team |
| EPI_ISL_3201985 | Lighthouse Lab in Milton Keynes | Wellcome Sanger Institute for the COVID-19 Genomics UK (COG-UK) Consortium | The Lighthouse Lab in Milton Keynes and Alex Alderton, Roberto Amato, Jeffrey Barrett, Sonia Goncalves, Ewan Harrison, David K. Jackson, Ian Johnston, Dominic Kwiatkowski, Cordelia Langford, John Sillitoe on behalf of the Wellcome Sanger Institute COVID-19 Surveillance Team |
| EPI_ISL_3201986 | Lighthouse Laboratory Plymouth | Wellcome Sanger Institute for the COVID-19 Genomics UK (COG-UK) Consortium | Lighthouse Laboratory Plymouth and Alex Alderton, Roberto Amato, Jeffrey Barrett, Sonia Goncalves, Ewan Harrison, David K. Jackson, Ian Johnston, Dominic Kwiatkowski, Cordelia Langford, John Sillitoe on behalf of the Wellcome Sanger Institute COVID-19 Surveillance Team |
| EPI_ISL_3201987, EPI_ISL_3201988 | Lighthouse Lab in Alderley Park | Wellcome Sanger Institute for the COVID-19 Genomics UK (COG-UK) Consortium | Jacquelyn Wynn, Mairead Hyland, The Lighthouse Lab in Alderley Park and Alex Alderton, Roberto Amato, Jeffrey Barrett, Sonia Goncalves, Ewan Harrison, David K. Jackson, Ian Johnston, Dominic Kwiatkowski, Cordelia Langford, John Sillitoe on behalf of the Wellcome Sanger Institute COVID-19 Surveillance Team |
| EPI_ISL_3201989, EPI_ISL_3201990 | Lighthouse Laboratory Plymouth | Wellcome Sanger Institute for the COVID-19 Genomics UK (COG-UK) Consortium | Lighthouse Laboratory Plymouth and Alex Alderton, Roberto Amato, Jeffrey Barrett, Sonia Goncalves, Ewan Harrison, David K. Jackson, Ian Johnston, Dominic Kwiatkowski, Cordelia Langford, John Sillitoe on behalf of the Wellcome Sanger Institute COVID-19 Surveillance Team |
| EPI_ISL_3201991, EPI_ISL_3201992, EPI_ISL_3201993, EPI_ISL_3201994, EPI_ISL_3201995, EPI_ISL_3201996, EPI_ISL_3201997, EPI_ISL_3201998 | Lighthouse Lab in Milton Keynes | Wellcome Sanger Institute for the COVID-19 Genomics UK (COG-UK) Consortium | The Lighthouse Lab in Milton Keynes and Alex Alderton, Roberto Amato, Jeffrey Barrett, Sonia Goncalves, Ewan Harrison, David K. Jackson, Ian Johnston, Dominic Kwiatkowski, Cordelia Langford, John Sillitoe on behalf of the Wellcome Sanger Institute COVID-19 Surveillance Team |
| EPI_ISL_3201999 | Lighthouse Laboratory Plymouth | Wellcome Sanger Institute for the COVID-19 Genomics UK (COG-UK) Consortium | Lighthouse Laboratory Plymouth and Alex Alderton, Roberto Amato, Jeffrey Barrett, Sonia Goncalves, Ewan Harrison, David K. Jackson, Ian Johnston, Dominic Kwiatkowski, Cordelia Langford, John Sillitoe on behalf of the Wellcome Sanger Institute COVID-19 Surveillance Team |
| EPI_ISL_3202000 | Lighthouse Lab in Alderley Park | Wellcome Sanger Institute for the COVID-19 Genomics UK (COG-UK) Consortium | Jacquelyn Wynn, Mairead Hyland, The Lighthouse Lab in Alderley Park and Alex Alderton, Roberto Amato, Jeffrey Barrett, Sonia Goncalves, Ewan Harrison, David K. Jackson, Ian Johnston, Dominic Kwiatkowski, Cordelia Langford, John Sillitoe on behalf of the Wellcome Sanger Institute COVID-19 Surveillance Team |
| EPI_ISL_3202001, EPI_ISL_3202002, EPI_ISL_3202003 | Lighthouse Lab in Milton Keynes | Wellcome Sanger Institute for the COVID-19 Genomics UK (COG-UK) Consortium | The Lighthouse Lab in Milton Keynes and Alex Alderton, Roberto Amato, Jeffrey Barrett, Sonia Goncalves, Ewan Harrison, David K. Jackson, Ian Johnston, Dominic Kwiatkowski, Cordelia Langford, John Sillitoe on behalf of the Wellcome Sanger Institute COVID-19 Surveillance Team |
| EPI_ISL_3202004, EPI_ISL_3202005 | Lighthouse Lab in Alderley Park | Wellcome Sanger Institute for the COVID-19 Genomics UK (COG-UK) Consortium | Jacquelyn Wynn, Mairead Hyland, The Lighthouse Lab in Alderley Park and Alex Alderton, Roberto Amato, Jeffrey Barrett, Sonia Goncalves, Ewan Harrison, David K. Jackson, Ian Johnston, Dominic Kwiatkowski, Cordelia Langford, John Sillitoe on behalf of the Wellcome Sanger Institute COVID-19 Surveillance Team |
| EPI_ISL_3202006 | Lighthouse Lab in Milton Keynes | Wellcome Sanger Institute for the COVID-19 Genomics UK (COG-UK) Consortium | The Lighthouse Lab in Milton Keynes and Alex Alderton, Roberto Amato, Jeffrey Barrett, Sonia Goncalves, Ewan Harrison, David K. Jackson, Ian Johnston, Dominic Kwiatkowski, Cordelia Langford, John Sillitoe on behalf of the Wellcome Sanger Institute COVID-19 Surveillance Team |
| EPI_ISL_3202007 | Lighthouse Laboratory Plymouth | Wellcome Sanger Institute for the COVID-19 Genomics UK (COG-UK) Consortium | Lighthouse Laboratory Plymouth and Alex Alderton, Roberto Amato, Jeffrey Barrett, Sonia Goncalves, Ewan Harrison, David K. Jackson, Ian Johnston, Dominic Kwiatkowski, Cordelia Langford, John Sillitoe on behalf of the Wellcome Sanger Institute COVID-19 Surveillance Team |
| EPI_ISL_3202008, EPI_ISL_3202009, EPI_ISL_3202010 | Lighthouse Lab in Alderley Park | Wellcome Sanger Institute for the COVID-19 Genomics UK (COG-UK) Consortium | Jacquelyn Wynn, Mairead Hyland, The Lighthouse Lab in Alderley Park and Alex Alderton, Roberto Amato, Jeffrey Barrett, Sonia Goncalves, Ewan Harrison, David K. Jackson, Ian Johnston, Dominic Kwiatkowski, Cordelia Langford, John Sillitoe on behalf of the Wellcome Sanger Institute COVID-19 Surveillance Team |
| EPI_ISL_3202011, EPI_ISL_3202012 | Lighthouse Lab in Milton Keynes | Wellcome Sanger Institute for the COVID-19 Genomics UK (COG-UK) Consortium | The Lighthouse Lab in Milton Keynes and Alex Alderton, Roberto Amato, Jeffrey Barrett, Sonia Goncalves, Ewan Harrison, David K. Jackson, Ian Johnston, Dominic Kwiatkowski, Cordelia Langford, John Sillitoe on behalf of the Wellcome Sanger Institute COVID-19 Surveillance Team |
| EPI_ISL_3202013 | Lighthouse Laboratory Plymouth | Wellcome Sanger Institute for the COVID-19 Genomics UK (COG-UK) Consortium | Lighthouse Laboratory Plymouth and Alex Alderton, Roberto Amato, Jeffrey Barrett, Sonia Goncalves, Ewan Harrison, David K. Jackson, Ian Johnston, Dominic Kwiatkowski, Cordelia Langford, John Sillitoe on behalf of the Wellcome Sanger Institute COVID-19 Surveillance Team |
| EPI_ISL_3202014 | Lighthouse Lab in Milton Keynes | Wellcome Sanger Institute for the COVID-19 Genomics UK (COG-UK) Consortium | The Lighthouse Lab in Milton Keynes and Alex Alderton, Roberto Amato, Jeffrey Barrett, Sonia Goncalves, Ewan Harrison, David K. Jackson, Ian Johnston, Dominic Kwiatkowski, Cordelia Langford, John Sillitoe on behalf of the Wellcome Sanger Institute COVID-19 Surveillance Team |
| EPI_ISL_3202015 | Lighthouse Lab in Alderley Park | Wellcome Sanger Institute for the COVID-19 Genomics UK (COG-UK) Consortium | Jacquelyn Wynn, Mairead Hyland, The Lighthouse Lab in Alderley Park and Alex Alderton, Roberto Amato, Jeffrey Barrett, Sonia Goncalves, Ewan Harrison, David K. Jackson, Ian Johnston, Dominic Kwiatkowski, Cordelia Langford, John Sillitoe on behalf of the Wellcome Sanger Institute COVID-19 Surveillance Team |
| EPI_ISL_3202016, EPI_ISL_3202017 | Lighthouse Lab in Milton Keynes | Wellcome Sanger Institute for the COVID-19 Genomics UK (COG-UK) Consortium | The Lighthouse Lab in Milton Keynes and Alex Alderton, Roberto Amato, Jeffrey Barrett, Sonia Goncalves, Ewan Harrison, David K. Jackson, Ian Johnston, Dominic Kwiatkowski, Cordelia Langford, John Sillitoe on behalf of the Wellcome Sanger Institute COVID-19 Surveillance Team |
| EPI_ISL_3202018 | Lighthouse Lab in Alderley Park | Wellcome Sanger Institute for the COVID-19 Genomics UK (COG-UK) Consortium | Jacquelyn Wynn, Mairead Hyland, The Lighthouse Lab in Alderley Park and Alex Alderton, Roberto Amato, Jeffrey Barrett, Sonia Goncalves, Ewan Harrison, David K. Jackson, Ian Johnston, Dominic Kwiatkowski, Cordelia Langford, John Sillitoe on behalf of the Wellcome Sanger Institute COVID-19 Surveillance Team |
| EPI_ISL_3202019 | Lighthouse Laboratory Plymouth | Wellcome Sanger Institute for the COVID-19 Genomics UK (COG-UK) Consortium | Lighthouse Laboratory Plymouth and Alex Alderton, Roberto Amato, Jeffrey Barrett, Sonia Goncalves, Ewan Harrison, David K. Jackson, Ian Johnston, Dominic Kwiatkowski, Cordelia Langford, John Sillitoe on behalf of the Wellcome Sanger Institute COVID-19 Surveillance Team |
| EPI_ISL_3202020, EPI_ISL_3202021 | Lighthouse Lab in Alderley Park | Wellcome Sanger Institute for the COVID-19 Genomics UK (COG-UK) Consortium | Jacquelyn Wynn, Mairead Hyland, The Lighthouse Lab in Alderley Park and Alex Alderton, Roberto Amato, Jeffrey Barrett, Sonia Goncalves, Ewan Harrison, David K. Jackson, Ian Johnston, Dominic Kwiatkowski, Cordelia Langford, John Sillitoe on behalf of the Wellcome Sanger Institute COVID-19 Surveillance Team |
| EPI_ISL_3202022 | Lighthouse Laboratory Plymouth | Wellcome Sanger Institute for the COVID-19 Genomics UK (COG-UK) Consortium | Lighthouse Laboratory Plymouth and Alex Alderton, Roberto Amato, Jeffrey Barrett, Sonia Goncalves, Ewan Harrison, David K. Jackson, Ian Johnston, Dominic Kwiatkowski, Cordelia Langford, John Sillitoe on behalf of the Wellcome Sanger Institute COVID-19 Surveillance Team |
| EPI_ISL_3202023 | Lighthouse Lab in Milton Keynes | Wellcome Sanger Institute for the COVID-19 Genomics UK (COG-UK) Consortium | The Lighthouse Lab in Milton Keynes and Alex Alderton, Roberto Amato, Jeffrey Barrett, Sonia Goncalves, Ewan Harrison, David K. Jackson, Ian Johnston, Dominic Kwiatkowski, Cordelia Langford, John Sillitoe on behalf of the Wellcome Sanger Institute COVID-19 Surveillance Team |
| EPI_ISL_3202024 | Lighthouse Lab in Alderley Park | Wellcome Sanger Institute for the COVID-19 Genomics UK (COG-UK) Consortium | Jacquelyn Wynn, Mairead Hyland, The Lighthouse Lab in Alderley Park and Alex Alderton, Roberto Amato, Jeffrey Barrett, Sonia Goncalves, Ewan Harrison, David K. Jackson, Ian Johnston, Dominic Kwiatkowski, Cordelia Langford, John Sillitoe on behalf of the Wellcome Sanger Institute COVID-19 Surveillance Team |
| EPI_ISL_3202025, EPI_ISL_3202026, EPI_ISL_3202027 | Lighthouse Lab in Milton Keynes | Wellcome Sanger Institute for the COVID-19 Genomics UK (COG-UK) Consortium | The Lighthouse Lab in Milton Keynes and Alex Alderton, Roberto Amato, Jeffrey Barrett, Sonia Goncalves, Ewan Harrison, David K. Jackson, Ian Johnston, Dominic Kwiatkowski, Cordelia Langford, John Sillitoe on behalf of the Wellcome Sanger Institute COVID-19 Surveillance Team |
| EPI_ISL_3202028 | Lighthouse Lab in Alderley Park | Wellcome Sanger Institute for the COVID-19 Genomics UK (COG-UK) Consortium | Jacquelyn Wynn, Mairead Hyland, The Lighthouse Lab in Alderley Park and Alex Alderton, Roberto Amato, Jeffrey Barrett, Sonia Goncalves, Ewan Harrison, David K. Jackson, Ian Johnston, Dominic Kwiatkowski, Cordelia Langford, John Sillitoe on behalf of the Wellcome Sanger Institute COVID-19 Surveillance Team |
| EPI_ISL_3202029, EPI_ISL_3202030, EPI_ISL_3202031 | Lighthouse Lab in Milton Keynes | Wellcome Sanger Institute for the COVID-19 Genomics UK (COG-UK) Consortium | The Lighthouse Lab in Milton Keynes and Alex Alderton, Roberto Amato, Jeffrey Barrett, Sonia Goncalves, Ewan Harrison, David K. Jackson, Ian Johnston, Dominic Kwiatkowski, Cordelia Langford, John Sillitoe on behalf of the Wellcome Sanger Institute COVID-19 Surveillance Team |
| EPI_ISL_3202032 | Lighthouse Laboratory Plymouth | Wellcome Sanger Institute for the COVID-19 Genomics UK (COG-UK) Consortium | Lighthouse Laboratory Plymouth and Alex Alderton, Roberto Amato, Jeffrey Barrett, Sonia Goncalves, Ewan Harrison, David K. Jackson, Ian Johnston, Dominic Kwiatkowski, Cordelia Langford, John Sillitoe on behalf of the Wellcome Sanger Institute COVID-19 Surveillance Team |
| EPI_ISL_3202033 | Lighthouse Lab in Milton Keynes | Wellcome Sanger Institute for the COVID-19 Genomics UK (COG-UK) Consortium | The Lighthouse Lab in Milton Keynes and Alex Alderton, Roberto Amato, Jeffrey Barrett, Sonia Goncalves, Ewan Harrison, David K. Jackson, Ian Johnston, Dominic Kwiatkowski, Cordelia Langford, John Sillitoe on behalf of the Wellcome Sanger Institute COVID-19 Surveillance Team |
| EPI_ISL_3202034 | Lighthouse Laboratory Plymouth | Wellcome Sanger Institute for the COVID-19 Genomics UK (COG-UK) Consortium | Lighthouse Laboratory Plymouth and Alex Alderton, Roberto Amato, Jeffrey Barrett, Sonia Goncalves, Ewan Harrison, David K. Jackson, Ian Johnston, Dominic Kwiatkowski, Cordelia Langford, John Sillitoe on behalf of the Wellcome Sanger Institute COVID-19 Surveillance Team |
| EPI_ISL_3202035, EPI_ISL_3202036, EPI_ISL_3202037 | Lighthouse Lab in Milton Keynes | Wellcome Sanger Institute for the COVID-19 Genomics UK (COG-UK) Consortium | The Lighthouse Lab in Milton Keynes and Alex Alderton, Roberto Amato, Jeffrey Barrett, Sonia Goncalves, Ewan Harrison, David K. Jackson, Ian Johnston, Dominic Kwiatkowski, Cordelia Langford, John Sillitoe on behalf of the Wellcome Sanger Institute COVID-19 Surveillance Team |
| EPI_ISL_3202038, EPI_ISL_3202039 | Lighthouse Lab in Alderley Park | Wellcome Sanger Institute for the COVID-19 Genomics UK (COG-UK) Consortium | Jacquelyn Wynn, Mairead Hyland, The Lighthouse Lab in Alderley Park and Alex Alderton, Roberto Amato, Jeffrey Barrett, Sonia Goncalves, Ewan Harrison, David K. Jackson, Ian Johnston, Dominic Kwiatkowski, Cordelia Langford, John Sillitoe on behalf of the Wellcome Sanger Institute COVID-19 Surveillance Team |
| EPI_ISL_3202040 | Lighthouse Lab in Milton Keynes | Wellcome Sanger Institute for the COVID-19 Genomics UK (COG-UK) | The Lighthouse Lab in Milton Keynes and Alex Alderton, Roberto Amato, Jeffrey Barrett, Sonia Goncalves, Ewan Harrison, David K. Jackson, Ian Johnston, Dominic Kwiatkowski, Cordelia Langford, John Sillitoe on behalf of the Wellcome Sanger |

| | | | |
|---|---|---|---|
| | | Consortium | Institute COVID-19 Surveillance Team |
| EPI_ISL_3202041 | Lighthouse Laboratory Plymouth | Wellcome Sanger Institute for the COVID-19 Genomics UK (COG-UK) Consortium | Lighthouse Laboratory Plymouth and Alex Alderton, Roberto Amato, Jeffrey Barrett, Sonia Goncalves, Ewan Harrison, David K. Jackson, Ian Johnston, Dominic Kwiatkowski, Cordelia Langford, John Sillitoe on behalf of the Wellcome Sanger Institute COVID-19 Surveillance Team |
| EPI_ISL_3202042, EPI_ISL_3202043, EPI_ISL_3202044, EPI_ISL_3202045, EPI_ISL_3202046, EPI_ISL_3202047 | Lighthouse Lab in Milton Keynes | Wellcome Sanger Institute for the COVID-19 Genomics UK (COG-UK) Consortium | The Lighthouse Lab in Milton Keynes and Alex Alderton, Roberto Amato, Jeffrey Barrett, Sonia Goncalves, Ewan Harrison, David K. Jackson, Ian Johnston, Dominic Kwiatkowski, Cordelia Langford, John Sillitoe on behalf of the Wellcome Sanger Institute COVID-19 Surveillance Team |
| EPI_ISL_3202048, EPI_ISL_3202049 | Lighthouse Lab in Alderley Park | Wellcome Sanger Institute for the COVID-19 Genomics UK (COG-UK) Consortium | Jacquelyn Wynn, Mairead Hyland, The Lighthouse Lab in Alderley Park and Alex Alderton, Roberto Amato, Jeffrey Barrett, Sonia Goncalves, Ewan Harrison, David K. Jackson, Ian Johnston, Dominic Kwiatkowski, Cordelia Langford, John Sillitoe on behalf of the Wellcome Sanger Institute COVID-19 Surveillance Team |
| EPI_ISL_3202050 | Lighthouse Lab in Milton Keynes | Wellcome Sanger Institute for the COVID-19 Genomics UK (COG-UK) Consortium | The Lighthouse Lab in Milton Keynes and Alex Alderton, Roberto Amato, Jeffrey Barrett, Sonia Goncalves, Ewan Harrison, David K. Jackson, Ian Johnston, Dominic Kwiatkowski, Cordelia Langford, John Sillitoe on behalf of the Wellcome Sanger Institute COVID-19 Surveillance Team |
| EPI_ISL_3202051, EPI_ISL_3202052 | Lighthouse Laboratory Plymouth | Wellcome Sanger Institute for the COVID-19 Genomics UK (COG-UK) Consortium | Lighthouse Laboratory Plymouth and Alex Alderton, Roberto Amato, Jeffrey Barrett, Sonia Goncalves, Ewan Harrison, David K. Jackson, Ian Johnston, Dominic Kwiatkowski, Cordelia Langford, John Sillitoe on behalf of the Wellcome Sanger Institute COVID-19 Surveillance Team |
| EPI_ISL_3202053 | Lighthouse Lab in Milton Keynes | Wellcome Sanger Institute for the COVID-19 Genomics UK (COG-UK) Consortium | The Lighthouse Lab in Milton Keynes and Alex Alderton, Roberto Amato, Jeffrey Barrett, Sonia Goncalves, Ewan Harrison, David K. Jackson, Ian Johnston, Dominic Kwiatkowski, Cordelia Langford, John Sillitoe on behalf of the Wellcome Sanger Institute COVID-19 Surveillance Team |
| EPI_ISL_3202054 | Lighthouse Lab in Alderley Park | Wellcome Sanger Institute for the COVID-19 Genomics UK (COG-UK) Consortium | Jacquelyn Wynn, Mairead Hyland, The Lighthouse Lab in Alderley Park and Alex Alderton, Roberto Amato, Jeffrey Barrett, Sonia Goncalves, Ewan Harrison, David K. Jackson, Ian Johnston, Dominic Kwiatkowski, Cordelia Langford, John Sillitoe on behalf of the Wellcome Sanger Institute COVID-19 Surveillance Team |
| EPI_ISL_3202055 | Lighthouse Laboratory Plymouth | Wellcome Sanger Institute for the COVID-19 Genomics UK (COG-UK) Consortium | Lighthouse Laboratory Plymouth and Alex Alderton, Roberto Amato, Jeffrey Barrett, Sonia Goncalves, Ewan Harrison, David K. Jackson, Ian Johnston, Dominic Kwiatkowski, Cordelia Langford, John Sillitoe on behalf of the Wellcome Sanger Institute COVID-19 Surveillance Team |
| EPI_ISL_3202056 | Lighthouse Lab in Milton Keynes | Wellcome Sanger Institute for the COVID-19 Genomics UK (COG-UK) Consortium | The Lighthouse Lab in Milton Keynes and Alex Alderton, Roberto Amato, Jeffrey Barrett, Sonia Goncalves, Ewan Harrison, David K. Jackson, Ian Johnston, Dominic Kwiatkowski, Cordelia Langford, John Sillitoe on behalf of the Wellcome Sanger Institute COVID-19 Surveillance Team |
| EPI_ISL_3202057 | Lighthouse Lab in Alderley Park | Wellcome Sanger Institute for the COVID-19 Genomics UK (COG-UK) Consortium | Jacquelyn Wynn, Mairead Hyland, The Lighthouse Lab in Alderley Park and Alex Alderton, Roberto Amato, Jeffrey Barrett, Sonia Goncalves, Ewan Harrison, David K. Jackson, Ian Johnston, Dominic Kwiatkowski, Cordelia Langford, John Sillitoe on behalf of the Wellcome Sanger Institute COVID-19 Surveillance Team |
| EPI_ISL_3202058 | Lighthouse Laboratory Plymouth | Wellcome Sanger Institute for the COVID-19 Genomics UK (COG-UK) Consortium | Lighthouse Laboratory Plymouth and Alex Alderton, Roberto Amato, Jeffrey Barrett, Sonia Goncalves, Ewan Harrison, David K. Jackson, Ian Johnston, Dominic Kwiatkowski, Cordelia Langford, John Sillitoe on behalf of the Wellcome Sanger Institute COVID-19 Surveillance Team |
| EPI_ISL_3202059 | Lighthouse Lab in Milton Keynes | Wellcome Sanger Institute for the COVID-19 Genomics UK (COG-UK) Consortium | The Lighthouse Lab in Milton Keynes and Alex Alderton, Roberto Amato, Jeffrey Barrett, Sonia Goncalves, Ewan Harrison, David K. Jackson, Ian Johnston, Dominic Kwiatkowski, Cordelia Langford, John Sillitoe on behalf of the Wellcome Sanger Institute COVID-19 Surveillance Team |
| EPI_ISL_3202060 | Lighthouse Lab in Alderley Park | Wellcome Sanger Institute for the COVID-19 Genomics UK (COG-UK) Consortium | Jacquelyn Wynn, Mairead Hyland, The Lighthouse Lab in Alderley Park and Alex Alderton, Roberto Amato, Jeffrey Barrett, Sonia Goncalves, Ewan Harrison, David K. Jackson, Ian Johnston, Dominic Kwiatkowski, Cordelia Langford, John Sillitoe on behalf of the Wellcome Sanger Institute COVID-19 Surveillance Team |
| EPI_ISL_3202061 | Lighthouse Lab in Milton Keynes | Wellcome Sanger Institute for the COVID-19 Genomics UK (COG-UK) Consortium | The Lighthouse Lab in Milton Keynes and Alex Alderton, Roberto Amato, Jeffrey Barrett, Sonia Goncalves, Ewan Harrison, David K. Jackson, Ian Johnston, Dominic Kwiatkowski, Cordelia Langford, John Sillitoe on behalf of the Wellcome Sanger Institute COVID-19 Surveillance Team |
| EPI_ISL_3202062, EPI_ISL_3202063 | Lighthouse Laboratory Plymouth | Wellcome Sanger Institute for the COVID-19 Genomics UK (COG-UK) Consortium | Lighthouse Laboratory Plymouth and Alex Alderton, Roberto Amato, Jeffrey Barrett, Sonia Goncalves, Ewan Harrison, David K. Jackson, Ian Johnston, Dominic Kwiatkowski, Cordelia Langford, John Sillitoe on behalf of the Wellcome Sanger Institute COVID-19 Surveillance Team |
| EPI_ISL_3202064 | Lighthouse Lab in Milton Keynes | Wellcome Sanger Institute for the COVID-19 Genomics UK (COG-UK) Consortium | The Lighthouse Lab in Milton Keynes and Alex Alderton, Roberto Amato, Jeffrey Barrett, Sonia Goncalves, Ewan Harrison, David K. Jackson, Ian Johnston, Dominic Kwiatkowski, Cordelia Langford, John Sillitoe on behalf of the Wellcome Sanger Institute COVID-19 Surveillance Team |
| EPI_ISL_3202065 | Lighthouse Laboratory Plymouth | Wellcome Sanger Institute for the COVID-19 Genomics UK (COG-UK) Consortium | Lighthouse Laboratory Plymouth and Alex Alderton, Roberto Amato, Jeffrey Barrett, Sonia Goncalves, Ewan Harrison, David K. Jackson, Ian Johnston, Dominic Kwiatkowski, Cordelia Langford, John Sillitoe on behalf of the Wellcome Sanger Institute COVID-19 Surveillance Team |
| EPI_ISL_3202066 | Lighthouse Lab in Milton Keynes | Wellcome Sanger Institute for the COVID-19 Genomics UK (COG-UK) Consortium | The Lighthouse Lab in Milton Keynes and Alex Alderton, Roberto Amato, Jeffrey Barrett, Sonia Goncalves, Ewan Harrison, David K. Jackson, Ian Johnston, Dominic Kwiatkowski, Cordelia Langford, John Sillitoe on behalf of the Wellcome Sanger Institute COVID-19 Surveillance Team |
| EPI_ISL_3202067 | Lighthouse Laboratory Plymouth | Wellcome Sanger Institute for the COVID-19 Genomics UK (COG-UK) Consortium | Lighthouse Laboratory Plymouth and Alex Alderton, Roberto Amato, Jeffrey Barrett, Sonia Goncalves, Ewan Harrison, David K. Jackson, Ian Johnston, Dominic Kwiatkowski, Cordelia Langford, John Sillitoe on behalf of the Wellcome Sanger Institute COVID-19 Surveillance Team |
| EPI_ISL_3202068, EPI_ISL_3202069, EPI_ISL_3202070 | Lighthouse Lab in Milton Keynes | Wellcome Sanger Institute for the COVID-19 Genomics UK (COG-UK) Consortium | The Lighthouse Lab in Milton Keynes and Alex Alderton, Roberto Amato, Jeffrey Barrett, Sonia Goncalves, Ewan Harrison, David K. Jackson, Ian Johnston, Dominic Kwiatkowski, Cordelia Langford, John Sillitoe on behalf of the Wellcome Sanger Institute COVID-19 Surveillance Team |
| EPI_ISL_3202071, EPI_ISL_3202072, EPI_ISL_3202073 | Lighthouse Lab in Alderley Park | Wellcome Sanger Institute for the COVID-19 Genomics UK (COG-UK) Consortium | Jacquelyn Wynn, Mairead Hyland, The Lighthouse Lab in Alderley Park and Alex Alderton, Roberto Amato, Jeffrey Barrett, Sonia Goncalves, Ewan Harrison, David K. Jackson, Ian Johnston, Dominic Kwiatkowski, Cordelia Langford, John Sillitoe on behalf of the Wellcome Sanger Institute COVID-19 Surveillance Team |
| EPI_ISL_3202074 | Lighthouse Lab in Milton Keynes | Wellcome Sanger Institute for the COVID-19 Genomics UK (COG-UK) Consortium | The Lighthouse Lab in Milton Keynes and Alex Alderton, Roberto Amato, Jeffrey Barrett, Sonia Goncalves, Ewan Harrison, David K. Jackson, Ian Johnston, Dominic Kwiatkowski, Cordelia Langford, John Sillitoe on behalf of the Wellcome Sanger Institute COVID-19 Surveillance Team |
| EPI_ISL_3202075 | Lighthouse Laboratory Plymouth | Wellcome Sanger Institute for the COVID-19 Genomics UK (COG-UK) Consortium | Lighthouse Laboratory Plymouth and Alex Alderton, Roberto Amato, Jeffrey Barrett, Sonia Goncalves, Ewan Harrison, David K. Jackson, Ian Johnston, Dominic Kwiatkowski, Cordelia Langford, John Sillitoe on behalf of the Wellcome Sanger Institute COVID-19 Surveillance Team |
| EPI_ISL_3202076 | Lighthouse Lab in Milton Keynes | Wellcome Sanger Institute for the COVID-19 Genomics UK (COG-UK) Consortium | The Lighthouse Lab in Milton Keynes and Alex Alderton, Roberto Amato, Jeffrey Barrett, Sonia Goncalves, Ewan Harrison, David K. Jackson, Ian Johnston, Dominic Kwiatkowski, Cordelia Langford, John Sillitoe on behalf of the Wellcome Sanger Institute COVID-19 Surveillance Team |
| EPI_ISL_3202077, EPI_ISL_3202078 | Lighthouse Laboratory Plymouth | Wellcome Sanger Institute for the COVID-19 Genomics UK (COG-UK) Consortium | Lighthouse Laboratory Plymouth and Alex Alderton, Roberto Amato, Jeffrey Barrett, Sonia Goncalves, Ewan Harrison, David K. Jackson, Ian Johnston, Dominic Kwiatkowski, Cordelia Langford, John Sillitoe on behalf of the Wellcome Sanger Institute COVID-19 Surveillance Team |
| EPI_ISL_3202079, EPI_ISL_3202080 | Lighthouse Lab in Milton Keynes | Wellcome Sanger Institute for the COVID-19 Genomics UK (COG-UK) Consortium | The Lighthouse Lab in Milton Keynes and Alex Alderton, Roberto Amato, Jeffrey Barrett, Sonia Goncalves, Ewan Harrison, David K. Jackson, Ian Johnston, Dominic Kwiatkowski, Cordelia Langford, John Sillitoe on behalf of the Wellcome Sanger Institute COVID-19 Surveillance Team |
| EPI_ISL_3202081, EPI_ISL_3202082 | Lighthouse Lab in Alderley Park | Wellcome Sanger Institute for the COVID-19 Genomics UK (COG-UK) Consortium | Jacquelyn Wynn, Mairead Hyland, The Lighthouse Lab in Alderley Park and Alex Alderton, Roberto Amato, Jeffrey Barrett, Sonia Goncalves, Ewan Harrison, David K. Jackson, Ian Johnston, Dominic Kwiatkowski, Cordelia Langford, John Sillitoe on behalf of the Wellcome Sanger Institute COVID-19 Surveillance Team |
| EPI_ISL_3202083, EPI_ISL_3202084 | Lighthouse Lab in Milton Keynes | Wellcome Sanger Institute for the COVID-19 Genomics UK (COG-UK) Consortium | The Lighthouse Lab in Milton Keynes and Alex Alderton, Roberto Amato, Jeffrey Barrett, Sonia Goncalves, Ewan Harrison, David K. Jackson, Ian Johnston, Dominic Kwiatkowski, Cordelia Langford, John Sillitoe on behalf of the Wellcome Sanger Institute COVID-19 Surveillance Team |
| EPI_ISL_3202085 | Lighthouse Laboratory Plymouth | Wellcome Sanger Institute for the COVID-19 Genomics UK (COG-UK) Consortium | Lighthouse Laboratory Plymouth and Alex Alderton, Roberto Amato, Jeffrey Barrett, Sonia Goncalves, Ewan Harrison, David K. Jackson, Ian Johnston, Dominic Kwiatkowski, Cordelia Langford, John Sillitoe on behalf of the Wellcome Sanger Institute COVID-19 Surveillance Team |
| EPI_ISL_3202086 | Lighthouse Lab in Alderley Park | Wellcome Sanger Institute for the COVID-19 Genomics UK (COG-UK) Consortium | Jacquelyn Wynn, Mairead Hyland, The Lighthouse Lab in Alderley Park and Alex Alderton, Roberto Amato, Jeffrey Barrett, Sonia Goncalves, Ewan Harrison, David K. Jackson, Ian Johnston, Dominic Kwiatkowski, Cordelia Langford, John Sillitoe on behalf of the Wellcome Sanger Institute COVID-19 Surveillance Team |
| EPI_ISL_3202087, EPI_ISL_3202088 | Lighthouse Lab in Milton Keynes | Wellcome Sanger Institute for the COVID-19 Genomics UK (COG-UK) Consortium | The Lighthouse Lab in Milton Keynes and Alex Alderton, Roberto Amato, Jeffrey Barrett, Sonia Goncalves, Ewan Harrison, David K. Jackson, Ian Johnston, Dominic Kwiatkowski, Cordelia Langford, John Sillitoe on behalf of the Wellcome Sanger Institute COVID-19 Surveillance Team |
| EPI_ISL_3202089 | Lighthouse Lab in Alderley Park | Wellcome Sanger Institute for the COVID-19 Genomics UK (COG-UK) Consortium | Jacquelyn Wynn, Mairead Hyland, The Lighthouse Lab in Alderley Park and Alex Alderton, Roberto Amato, Jeffrey Barrett, Sonia Goncalves, Ewan Harrison, David K. Jackson, Ian Johnston, Dominic Kwiatkowski, Cordelia Langford, John Sillitoe on behalf of the Wellcome Sanger Institute COVID-19 Surveillance Team |
| EPI_ISL_3202090 | Lighthouse Laboratory Plymouth | Wellcome Sanger Institute for the COVID-19 Genomics UK (COG-UK) Consortium | Lighthouse Laboratory Plymouth and Alex Alderton, Roberto Amato, Jeffrey Barrett, Sonia Goncalves, Ewan Harrison, David K. Jackson, Ian Johnston, Dominic Kwiatkowski, Cordelia Langford, John Sillitoe on behalf of the Wellcome Sanger Institute COVID-19 Surveillance Team |
| EPI_ISL_3202091 | Lighthouse Lab in Alderley Park | Wellcome Sanger Institute for the COVID-19 Genomics UK (COG-UK) Consortium | Jacquelyn Wynn, Mairead Hyland, The Lighthouse Lab in Alderley Park and Alex Alderton, Roberto Amato, Jeffrey Barrett, Sonia Goncalves, Ewan Harrison, David K. Jackson, Ian Johnston, Dominic Kwiatkowski, Cordelia Langford, John Sillitoe on behalf of the Wellcome Sanger Institute COVID-19 Surveillance Team |
| EPI_ISL_3202092, EPI_ISL_3202093, EPI_ISL_3202094, EPI_ISL_3202095, EPI_ISL_3202096, EPI_ISL_3202097, EPI_ISL_3202098 | Lighthouse Lab in Milton Keynes | Wellcome Sanger Institute for the COVID-19 Genomics UK (COG-UK) Consortium | The Lighthouse Lab in Milton Keynes and Alex Alderton, Roberto Amato, Jeffrey Barrett, Sonia Goncalves, Ewan Harrison, David K. Jackson, Ian Johnston, Dominic Kwiatkowski, Cordelia Langford, John Sillitoe on behalf of the Wellcome Sanger Institute COVID-19 Surveillance Team |
| EPI_ISL_3202099, EPI_ISL_3202100, EPI_ISL_3202101, EPI_ISL_3202102, EPI_ISL_3202103, EPI_ISL_3202104 | Lighthouse Lab in Alderley Park | Wellcome Sanger Institute for the COVID-19 Genomics UK (COG-UK) Consortium | Jacquelyn Wynn, Mairead Hyland, The Lighthouse Lab in Alderley Park and Alex Alderton, Roberto Amato, Jeffrey Barrett, Sonia Goncalves, Ewan Harrison, David K. Jackson, Ian Johnston, Dominic Kwiatkowski, Cordelia Langford, John Sillitoe on behalf of the Wellcome Sanger Institute COVID-19 Surveillance Team |
| EPI_ISL_3202105 | Berkshire and Surrey Pathology Services Lighthouse Laboratory | Wellcome Sanger Institute for the COVID-19 Genomics UK (COG-UK) Consortium | Berkshire and Surrey Pathology Services Lighthouse Laboratory and Alex Alderton, Roberto Amato, Jeffrey Barrett, Sonia Goncalves, Ewan Harrison, David K. Jackson, Ian Johnston, Dominic Kwiatkowski, Cordelia Langford, John Sillitoe on behalf of the Wellcome Sanger Institute COVID-19 Surveillance Team |
| EPI_ISL_3202106 | Lighthouse Lab in Alderley Park | Wellcome Sanger Institute for the COVID-19 Genomics UK (COG-UK) Consortium | Jacquelyn Wynn, Mairead Hyland, The Lighthouse Lab in Alderley Park and Alex Alderton, Roberto Amato, Jeffrey Barrett, Sonia Goncalves, Ewan Harrison, David K. Jackson, Ian Johnston, Dominic Kwiatkowski, Cordelia Langford, John Sillitoe on behalf of the Wellcome Sanger Institute COVID-19 Surveillance Team |
| EPI_ISL_3202107 | Berkshire and Surrey Pathology Services Lighthouse Laboratory | Wellcome Sanger Institute for the COVID-19 Genomics UK (COG-UK) | Berkshire and Surrey Pathology Services Lighthouse Laboratory and Alex Alderton, Roberto Amato, Jeffrey Barrett, Sonia Goncalves, Ewan Harrison, David K. Jackson, Ian Johnston, Dominic Kwiatkowski, Cordelia Langford, John Sillitoe on behalf |

| | | | |
|---|---|---|---|
| | | Consortium | of the Wellcome Sanger Institute COVID-19 Surveillance Team |
| EPI_ISL_3202108, EPI_ISL_3202109 | Lighthouse Lab in Alderley Park | Wellcome Sanger Institute for the COVID-19 Genomics UK (COG-UK) Consortium | Jacquelyn Wynn, Mairead Hyland, The Lighthouse Lab in Alderley Park and Alex Alderton, Roberto Amato, Jeffrey Barrett, Sonia Goncalves, Ewan Harrison, David K. Jackson, Ian Johnston, Dominic Kwiatkowski, Cordelia Langford, John Sillitoe on behalf of the Wellcome Sanger Institute COVID-19 Surveillance Team |
| EPI_ISL_3202110 | Berkshire and Surrey Pathology Services Lighthouse Laboratory | Wellcome Sanger Institute for the COVID-19 Genomics UK (COG-UK) Consortium | Berkshire and Surrey Pathology Services Lighthouse Laboratory and Alex Alderton, Roberto Amato, Jeffrey Barrett, Sonia Goncalves, Ewan Harrison, David K. Jackson, Ian Johnston, Dominic Kwiatkowski, Cordelia Langford, John Sillitoe on behalf of the Wellcome Sanger Institute COVID-19 Surveillance Team |
| EPI_ISL_3202111, EPI_ISL_3202112, EPI_ISL_3202113, EPI_ISL_3202114 | Lighthouse Lab in Alderley Park | Wellcome Sanger Institute for the COVID-19 Genomics UK (COG-UK) Consortium | Jacquelyn Wynn, Mairead Hyland, The Lighthouse Lab in Alderley Park and Alex Alderton, Roberto Amato, Jeffrey Barrett, Sonia Goncalves, Ewan Harrison, David K. Jackson, Ian Johnston, Dominic Kwiatkowski, Cordelia Langford, John Sillitoe on behalf of the Wellcome Sanger Institute COVID-19 Surveillance Team |
| EPI_ISL_3202115 | Berkshire and Surrey Pathology Services Lighthouse Laboratory | Wellcome Sanger Institute for the COVID-19 Genomics UK (COG-UK) Consortium | Berkshire and Surrey Pathology Services Lighthouse Laboratory and Alex Alderton, Roberto Amato, Jeffrey Barrett, Sonia Goncalves, Ewan Harrison, David K. Jackson, Ian Johnston, Dominic Kwiatkowski, Cordelia Langford, John Sillitoe on behalf of the Wellcome Sanger Institute COVID-19 Surveillance Team |
| EPI_ISL_3202116 | Lighthouse Lab in Alderley Park | Wellcome Sanger Institute for the COVID-19 Genomics UK (COG-UK) Consortium | Jacquelyn Wynn, Mairead Hyland, The Lighthouse Lab in Alderley Park and Alex Alderton, Roberto Amato, Jeffrey Barrett, Sonia Goncalves, Ewan Harrison, David K. Jackson, Ian Johnston, Dominic Kwiatkowski, Cordelia Langford, John Sillitoe on behalf of the Wellcome Sanger Institute COVID-19 Surveillance Team |
| EPI_ISL_3202117, EPI_ISL_3202118 | Berkshire and Surrey Pathology Services Lighthouse Laboratory | Wellcome Sanger Institute for the COVID-19 Genomics UK (COG-UK) Consortium | Berkshire and Surrey Pathology Services Lighthouse Laboratory and Alex Alderton, Roberto Amato, Jeffrey Barrett, Sonia Goncalves, Ewan Harrison, David K. Jackson, Ian Johnston, Dominic Kwiatkowski, Cordelia Langford, John Sillitoe on behalf of the Wellcome Sanger Institute COVID-19 Surveillance Team |
| EPI_ISL_3202119, EPI_ISL_3202120, EPI_ISL_3202121 | Lighthouse Lab in Alderley Park | Wellcome Sanger Institute for the COVID-19 Genomics UK (COG-UK) Consortium | Jacquelyn Wynn, Mairead Hyland, The Lighthouse Lab in Alderley Park and Alex Alderton, Roberto Amato, Jeffrey Barrett, Sonia Goncalves, Ewan Harrison, David K. Jackson, Ian Johnston, Dominic Kwiatkowski, Cordelia Langford, John Sillitoe on behalf of the Wellcome Sanger Institute COVID-19 Surveillance Team |
| EPI_ISL_3202122, EPI_ISL_3202123, EPI_ISL_3202124, EPI_ISL_3202125, EPI_ISL_3202126 | Berkshire and Surrey Pathology Services Lighthouse Laboratory | Wellcome Sanger Institute for the COVID-19 Genomics UK (COG-UK) Consortium | Berkshire and Surrey Pathology Services Lighthouse Laboratory and Alex Alderton, Roberto Amato, Jeffrey Barrett, Sonia Goncalves, Ewan Harrison, David K. Jackson, Ian Johnston, Dominic Kwiatkowski, Cordelia Langford, John Sillitoe on behalf of the Wellcome Sanger Institute COVID-19 Surveillance Team |
| EPI_ISL_3202127 | Lighthouse Lab in Alderley Park | Wellcome Sanger Institute for the COVID-19 Genomics UK (COG-UK) Consortium | Jacquelyn Wynn, Mairead Hyland, The Lighthouse Lab in Alderley Park and Alex Alderton, Roberto Amato, Jeffrey Barrett, Sonia Goncalves, Ewan Harrison, David K. Jackson, Ian Johnston, Dominic Kwiatkowski, Cordelia Langford, John Sillitoe on behalf of the Wellcome Sanger Institute COVID-19 Surveillance Team |
| EPI_ISL_3202128 | Berkshire and Surrey Pathology Services Lighthouse Laboratory | Wellcome Sanger Institute for the COVID-19 Genomics UK (COG-UK) Consortium | Berkshire and Surrey Pathology Services Lighthouse Laboratory and Alex Alderton, Roberto Amato, Jeffrey Barrett, Sonia Goncalves, Ewan Harrison, David K. Jackson, Ian Johnston, Dominic Kwiatkowski, Cordelia Langford, John Sillitoe on behalf of the Wellcome Sanger Institute COVID-19 Surveillance Team |
| EPI_ISL_3202129 | Lighthouse Lab in Alderley Park | Wellcome Sanger Institute for the COVID-19 Genomics UK (COG-UK) Consortium | Jacquelyn Wynn, Mairead Hyland, The Lighthouse Lab in Alderley Park and Alex Alderton, Roberto Amato, Jeffrey Barrett, Sonia Goncalves, Ewan Harrison, David K. Jackson, Ian Johnston, Dominic Kwiatkowski, Cordelia Langford, John Sillitoe on behalf of the Wellcome Sanger Institute COVID-19 Surveillance Team |
| EPI_ISL_3202130 | Berkshire and Surrey Pathology Services Lighthouse Laboratory | Wellcome Sanger Institute for the COVID-19 Genomics UK (COG-UK) Consortium | Berkshire and Surrey Pathology Services Lighthouse Laboratory and Alex Alderton, Roberto Amato, Jeffrey Barrett, Sonia Goncalves, Ewan Harrison, David K. Jackson, Ian Johnston, Dominic Kwiatkowski, Cordelia Langford, John Sillitoe on behalf of the Wellcome Sanger Institute COVID-19 Surveillance Team |
| EPI_ISL_3202131, EPI_ISL_3202132 | Lighthouse Lab in Alderley Park | Wellcome Sanger Institute for the COVID-19 Genomics UK (COG-UK) Consortium | Jacquelyn Wynn, Mairead Hyland, The Lighthouse Lab in Alderley Park and Alex Alderton, Roberto Amato, Jeffrey Barrett, Sonia Goncalves, Ewan Harrison, David K. Jackson, Ian Johnston, Dominic Kwiatkowski, Cordelia Langford, John Sillitoe on behalf of the Wellcome Sanger Institute COVID-19 Surveillance Team |
| EPI_ISL_3202133 | Berkshire and Surrey Pathology Services Lighthouse Laboratory | Wellcome Sanger Institute for the COVID-19 Genomics UK (COG-UK) Consortium | Berkshire and Surrey Pathology Services Lighthouse Laboratory and Alex Alderton, Roberto Amato, Jeffrey Barrett, Sonia Goncalves, Ewan Harrison, David K. Jackson, Ian Johnston, Dominic Kwiatkowski, Cordelia Langford, John Sillitoe on behalf of the Wellcome Sanger Institute COVID-19 Surveillance Team |
| EPI_ISL_3202134 | Lighthouse Lab in Alderley Park | Wellcome Sanger Institute for the COVID-19 Genomics UK (COG-UK) Consortium | Jacquelyn Wynn, Mairead Hyland, The Lighthouse Lab in Alderley Park and Alex Alderton, Roberto Amato, Jeffrey Barrett, Sonia Goncalves, Ewan Harrison, David K. Jackson, Ian Johnston, Dominic Kwiatkowski, Cordelia Langford, John Sillitoe on behalf of the Wellcome Sanger Institute COVID-19 Surveillance Team |
| EPI_ISL_3202135, EPI_ISL_3202136 | Berkshire and Surrey Pathology Services Lighthouse Laboratory | Wellcome Sanger Institute for the COVID-19 Genomics UK (COG-UK) Consortium | Berkshire and Surrey Pathology Services Lighthouse Laboratory and Alex Alderton, Roberto Amato, Jeffrey Barrett, Sonia Goncalves, Ewan Harrison, David K. Jackson, Ian Johnston, Dominic Kwiatkowski, Cordelia Langford, John Sillitoe on behalf of the Wellcome Sanger Institute COVID-19 Surveillance Team |
| EPI_ISL_3202137, EPI_ISL_3202138 | Lighthouse Lab in Alderley Park | Wellcome Sanger Institute for the COVID-19 Genomics UK (COG-UK) Consortium | Jacquelyn Wynn, Mairead Hyland, The Lighthouse Lab in Alderley Park and Alex Alderton, Roberto Amato, Jeffrey Barrett, Sonia Goncalves, Ewan Harrison, David K. Jackson, Ian Johnston, Dominic Kwiatkowski, Cordelia Langford, John Sillitoe on behalf of the Wellcome Sanger Institute COVID-19 Surveillance Team |
| EPI_ISL_3202139 | Berkshire and Surrey Pathology Services Lighthouse Laboratory | Wellcome Sanger Institute for the COVID-19 Genomics UK (COG-UK) Consortium | Berkshire and Surrey Pathology Services Lighthouse Laboratory and Alex Alderton, Roberto Amato, Jeffrey Barrett, Sonia Goncalves, Ewan Harrison, David K. Jackson, Ian Johnston, Dominic Kwiatkowski, Cordelia Langford, John Sillitoe on behalf of the Wellcome Sanger Institute COVID-19 Surveillance Team |
| EPI_ISL_3202140, EPI_ISL_3202141 | Lighthouse Lab in Alderley Park | Wellcome Sanger Institute for the COVID-19 Genomics UK (COG-UK) Consortium | Jacquelyn Wynn, Mairead Hyland, The Lighthouse Lab in Alderley Park and Alex Alderton, Roberto Amato, Jeffrey Barrett, Sonia Goncalves, Ewan Harrison, David K. Jackson, Ian Johnston, Dominic Kwiatkowski, Cordelia Langford, John Sillitoe on behalf of the Wellcome Sanger Institute COVID-19 Surveillance Team |
| EPI_ISL_3202142 | Berkshire and Surrey Pathology Services Lighthouse Laboratory | Wellcome Sanger Institute for the COVID-19 Genomics UK (COG-UK) Consortium | Berkshire and Surrey Pathology Services Lighthouse Laboratory and Alex Alderton, Roberto Amato, Jeffrey Barrett, Sonia Goncalves, Ewan Harrison, David K. Jackson, Ian Johnston, Dominic Kwiatkowski, Cordelia Langford, John Sillitoe on behalf of the Wellcome Sanger Institute COVID-19 Surveillance Team |
| EPI_ISL_3202143, EPI_ISL_3202144, EPI_ISL_3202145 | Lighthouse Lab in Alderley Park | Wellcome Sanger Institute for the COVID-19 Genomics UK (COG-UK) Consortium | Jacquelyn Wynn, Mairead Hyland, The Lighthouse Lab in Alderley Park and Alex Alderton, Roberto Amato, Jeffrey Barrett, Sonia Goncalves, Ewan Harrison, David K. Jackson, Ian Johnston, Dominic Kwiatkowski, Cordelia Langford, John Sillitoe on behalf of the Wellcome Sanger Institute COVID-19 Surveillance Team |
| EPI_ISL_3202146, EPI_ISL_3202147 | Berkshire and Surrey Pathology Services Lighthouse Laboratory | Wellcome Sanger Institute for the COVID-19 Genomics UK (COG-UK) Consortium | Berkshire and Surrey Pathology Services Lighthouse Laboratory and Alex Alderton, Roberto Amato, Jeffrey Barrett, Sonia Goncalves, Ewan Harrison, David K. Jackson, Ian Johnston, Dominic Kwiatkowski, Cordelia Langford, John Sillitoe on behalf of the Wellcome Sanger Institute COVID-19 Surveillance Team |
| EPI_ISL_3202148, EPI_ISL_3202149, EPI_ISL_3202150 | Lighthouse Lab in Alderley Park | Wellcome Sanger Institute for the COVID-19 Genomics UK (COG-UK) Consortium | Jacquelyn Wynn, Mairead Hyland, The Lighthouse Lab in Alderley Park and Alex Alderton, Roberto Amato, Jeffrey Barrett, Sonia Goncalves, Ewan Harrison, David K. Jackson, Ian Johnston, Dominic Kwiatkowski, Cordelia Langford, John Sillitoe on behalf of the Wellcome Sanger Institute COVID-19 Surveillance Team |
| EPI_ISL_3202151 | Berkshire and Surrey Pathology Services Lighthouse Laboratory | Wellcome Sanger Institute for the COVID-19 Genomics UK (COG-UK) Consortium | Berkshire and Surrey Pathology Services Lighthouse Laboratory and Alex Alderton, Roberto Amato, Jeffrey Barrett, Sonia Goncalves, Ewan Harrison, David K. Jackson, Ian Johnston, Dominic Kwiatkowski, Cordelia Langford, John Sillitoe on behalf of the Wellcome Sanger Institute COVID-19 Surveillance Team |
| EPI_ISL_3202152 | Lighthouse Lab in Alderley Park | Wellcome Sanger Institute for the COVID-19 Genomics UK (COG-UK) Consortium | Jacquelyn Wynn, Mairead Hyland, The Lighthouse Lab in Alderley Park and Alex Alderton, Roberto Amato, Jeffrey Barrett, Sonia Goncalves, Ewan Harrison, David K. Jackson, Ian Johnston, Dominic Kwiatkowski, Cordelia Langford, John Sillitoe on behalf of the Wellcome Sanger Institute COVID-19 Surveillance Team |
| EPI_ISL_3202153, EPI_ISL_3202154 | Berkshire and Surrey Pathology Services Lighthouse Laboratory | Wellcome Sanger Institute for the COVID-19 Genomics UK (COG-UK) Consortium | Berkshire and Surrey Pathology Services Lighthouse Laboratory and Alex Alderton, Roberto Amato, Jeffrey Barrett, Sonia Goncalves, Ewan Harrison, David K. Jackson, Ian Johnston, Dominic Kwiatkowski, Cordelia Langford, John Sillitoe on behalf of the Wellcome Sanger Institute COVID-19 Surveillance Team |
| EPI_ISL_3202155 | Lighthouse Lab in Alderley Park | Wellcome Sanger Institute for the COVID-19 Genomics UK (COG-UK) Consortium | Jacquelyn Wynn, Mairead Hyland, The Lighthouse Lab in Alderley Park and Alex Alderton, Roberto Amato, Jeffrey Barrett, Sonia Goncalves, Ewan Harrison, David K. Jackson, Ian Johnston, Dominic Kwiatkowski, Cordelia Langford, John Sillitoe on behalf of the Wellcome Sanger Institute COVID-19 Surveillance Team |
| EPI_ISL_3202156, EPI_ISL_3202157, EPI_ISL_3202158, EPI_ISL_3202159, EPI_ISL_3202160 | Berkshire and Surrey Pathology Services Lighthouse Laboratory | Wellcome Sanger Institute for the COVID-19 Genomics UK (COG-UK) Consortium | Berkshire and Surrey Pathology Services Lighthouse Laboratory and Alex Alderton, Roberto Amato, Jeffrey Barrett, Sonia Goncalves, Ewan Harrison, David K. Jackson, Ian Johnston, Dominic Kwiatkowski, Cordelia Langford, John Sillitoe on behalf of the Wellcome Sanger Institute COVID-19 Surveillance Team |
| EPI_ISL_3202161 | Lighthouse Lab in Alderley Park | Wellcome Sanger Institute for the COVID-19 Genomics UK (COG-UK) Consortium | Jacquelyn Wynn, Mairead Hyland, The Lighthouse Lab in Alderley Park and Alex Alderton, Roberto Amato, Jeffrey Barrett, Sonia Goncalves, Ewan Harrison, David K. Jackson, Ian Johnston, Dominic Kwiatkowski, Cordelia Langford, John Sillitoe on behalf of the Wellcome Sanger Institute COVID-19 Surveillance Team |
| EPI_ISL_3202162 | Berkshire and Surrey Pathology Services Lighthouse Laboratory | Wellcome Sanger Institute for the COVID-19 Genomics UK (COG-UK) Consortium | Berkshire and Surrey Pathology Services Lighthouse Laboratory and Alex Alderton, Roberto Amato, Jeffrey Barrett, Sonia Goncalves, Ewan Harrison, David K. Jackson, Ian Johnston, Dominic Kwiatkowski, Cordelia Langford, John Sillitoe on behalf of the Wellcome Sanger Institute COVID-19 Surveillance Team |
| EPI_ISL_3202163 | Lighthouse Lab in Alderley Park | Wellcome Sanger Institute for the COVID-19 Genomics UK (COG-UK) Consortium | Jacquelyn Wynn, Mairead Hyland, The Lighthouse Lab in Alderley Park and Alex Alderton, Roberto Amato, Jeffrey Barrett, Sonia Goncalves, Ewan Harrison, David K. Jackson, Ian Johnston, Dominic Kwiatkowski, Cordelia Langford, John Sillitoe on behalf of the Wellcome Sanger Institute COVID-19 Surveillance Team |
| EPI_ISL_3202164, EPI_ISL_3202165, EPI_ISL_3202166, EPI_ISL_3202167 | Berkshire and Surrey Pathology Services Lighthouse Laboratory | Wellcome Sanger Institute for the COVID-19 Genomics UK (COG-UK) Consortium | Berkshire and Surrey Pathology Services Lighthouse Laboratory and Alex Alderton, Roberto Amato, Jeffrey Barrett, Sonia Goncalves, Ewan Harrison, David K. Jackson, Ian Johnston, Dominic Kwiatkowski, Cordelia Langford, John Sillitoe on behalf of the Wellcome Sanger Institute COVID-19 Surveillance Team |
| EPI_ISL_3202168 | Lighthouse Lab in Alderley Park | Wellcome Sanger Institute for the COVID-19 Genomics UK (COG-UK) Consortium | Jacquelyn Wynn, Mairead Hyland, The Lighthouse Lab in Alderley Park and Alex Alderton, Roberto Amato, Jeffrey Barrett, Sonia Goncalves, Ewan Harrison, David K. Jackson, Ian Johnston, Dominic Kwiatkowski, Cordelia Langford, John Sillitoe on behalf of the Wellcome Sanger Institute COVID-19 Surveillance Team |
| EPI_ISL_3202169, EPI_ISL_3202170 | Berkshire and Surrey Pathology Services Lighthouse Laboratory | Wellcome Sanger Institute for the COVID-19 Genomics UK (COG-UK) Consortium | Berkshire and Surrey Pathology Services Lighthouse Laboratory and Alex Alderton, Roberto Amato, Jeffrey Barrett, Sonia Goncalves, Ewan Harrison, David K. Jackson, Ian Johnston, Dominic Kwiatkowski, Cordelia Langford, John Sillitoe on behalf of the Wellcome Sanger Institute COVID-19 Surveillance Team |
| EPI_ISL_3202171, EPI_ISL_3202172, EPI_ISL_3202173, EPI_ISL_3202174 | Lighthouse Lab in Alderley Park | Wellcome Sanger Institute for the COVID-19 Genomics UK (COG-UK) Consortium | Jacquelyn Wynn, Mairead Hyland, The Lighthouse Lab in Alderley Park and Alex Alderton, Roberto Amato, Jeffrey Barrett, Sonia Goncalves, Ewan Harrison, David K. Jackson, Ian Johnston, Dominic Kwiatkowski, Cordelia Langford, John Sillitoe on behalf of the Wellcome Sanger Institute COVID-19 Surveillance Team |
| EPI_ISL_3202175 | Berkshire and Surrey Pathology Services Lighthouse Laboratory | Wellcome Sanger Institute for the COVID-19 Genomics UK (COG-UK) Consortium | Berkshire and Surrey Pathology Services Lighthouse Laboratory and Alex Alderton, Roberto Amato, Jeffrey Barrett, Sonia Goncalves, Ewan Harrison, David K. Jackson, Ian Johnston, Dominic Kwiatkowski, Cordelia Langford, John Sillitoe on behalf of the Wellcome Sanger Institute COVID-19 Surveillance Team |
| EPI_ISL_3202176 | Lighthouse Lab in Alderley Park | Wellcome Sanger Institute for the COVID-19 Genomics UK (COG-UK) Consortium | Jacquelyn Wynn, Mairead Hyland, The Lighthouse Lab in Alderley Park and Alex Alderton, Roberto Amato, Jeffrey Barrett, Sonia Goncalves, Ewan Harrison, David K. Jackson, Ian Johnston, Dominic Kwiatkowski, Cordelia Langford, John Sillitoe on behalf of the Wellcome Sanger Institute COVID-19 Surveillance Team |
| EPI_ISL_3202177, EPI_ISL_3202178 | Berkshire and Surrey Pathology Services Lighthouse Laboratory | Wellcome Sanger Institute for the COVID-19 Genomics UK (COG-UK) Consortium | Berkshire and Surrey Pathology Services Lighthouse Laboratory and Alex Alderton, Roberto Amato, Jeffrey Barrett, Sonia Goncalves, Ewan Harrison, David K. Jackson, Ian Johnston, Dominic Kwiatkowski, Cordelia Langford, John Sillitoe on behalf of the Wellcome Sanger Institute COVID-19 Surveillance Team |
| EPI_ISL_3202179, | Lighthouse Lab in Alderley Park | Wellcome Sanger Institute for the COVID-19 Genomics UK (COG-UK) | Jacquelyn Wynn, Mairead Hyland, The Lighthouse Lab in Alderley Park and Alex Alderton, Roberto Amato, Jeffrey Barrett, Sonia Goncalves, Ewan Harrison, David K. Jackson, Ian Johnston, Dominic Kwiatkowski, Cordelia Langford, John Sillitoe on |

| | | | |
|---|---|---|---|
| EPI_ISL_3202180, EPI_ISL_3202181 | | Consortium | behalf of the Wellcome Sanger Institute COVID-19 Surveillance Team |
| EPI_ISL_3202182 | Berkshire and Surrey Pathology Services Lighthouse Laboratory | Wellcome Sanger Institute for the COVID-19 Genomics UK (COG-UK) Consortium | Berkshire and Surrey Pathology Services Lighthouse Laboratory and Alex Alderton, Roberto Amato, Jeffrey Barrett, Sonia Goncalves, Ewan Harrison, David K. Jackson, Ian Johnston, Dominic Kwiatkowski, Cordelia Langford, John Sillitoe on behalf of the Wellcome Sanger Institute COVID-19 Surveillance Team |
| EPI_ISL_3202183 | Lighthouse Lab in Alderley Park | Wellcome Sanger Institute for the COVID-19 Genomics UK (COG-UK) Consortium | Jacquelyn Wynn, Mairead Hyland, The Lighthouse Lab in Alderley Park and Alex Alderton, Roberto Amato, Jeffrey Barrett, Sonia Goncalves, Ewan Harrison, David K. Jackson, Ian Johnston, Dominic Kwiatkowski, Cordelia Langford, John Sillitoe on behalf of the Wellcome Sanger Institute COVID-19 Surveillance Team |
| EPI_ISL_3202184, EPI_ISL_3202185 | Berkshire and Surrey Pathology Services Lighthouse Laboratory | Wellcome Sanger Institute for the COVID-19 Genomics UK (COG-UK) Consortium | Berkshire and Surrey Pathology Services Lighthouse Laboratory and Alex Alderton, Roberto Amato, Jeffrey Barrett, Sonia Goncalves, Ewan Harrison, David K. Jackson, Ian Johnston, Dominic Kwiatkowski, Cordelia Langford, John Sillitoe on behalf of the Wellcome Sanger Institute COVID-19 Surveillance Team |
| EPI_ISL_3202186, EPI_ISL_3202187, EPI_ISL_3202188, EPI_ISL_3202189 | Lighthouse Lab in Alderley Park | Wellcome Sanger Institute for the COVID-19 Genomics UK (COG-UK) Consortium | Jacquelyn Wynn, Mairead Hyland, The Lighthouse Lab in Alderley Park and Alex Alderton, Roberto Amato, Jeffrey Barrett, Sonia Goncalves, Ewan Harrison, David K. Jackson, Ian Johnston, Dominic Kwiatkowski, Cordelia Langford, John Sillitoe on behalf of the Wellcome Sanger Institute COVID-19 Surveillance Team |
| EPI_ISL_3202190, EPI_ISL_3202191 | Berkshire and Surrey Pathology Services Lighthouse Laboratory | Wellcome Sanger Institute for the COVID-19 Genomics UK (COG-UK) Consortium | Berkshire and Surrey Pathology Services Lighthouse Laboratory and Alex Alderton, Roberto Amato, Jeffrey Barrett, Sonia Goncalves, Ewan Harrison, David K. Jackson, Ian Johnston, Dominic Kwiatkowski, Cordelia Langford, John Sillitoe on behalf of the Wellcome Sanger Institute COVID-19 Surveillance Team |
| EPI_ISL_3202192 | Lighthouse Lab in Alderley Park | Wellcome Sanger Institute for the COVID-19 Genomics UK (COG-UK) Consortium | Jacquelyn Wynn, Mairead Hyland, The Lighthouse Lab in Alderley Park and Alex Alderton, Roberto Amato, Jeffrey Barrett, Sonia Goncalves, Ewan Harrison, David K. Jackson, Ian Johnston, Dominic Kwiatkowski, Cordelia Langford, John Sillitoe on behalf of the Wellcome Sanger Institute COVID-19 Surveillance Team |
| EPI_ISL_3202193 | Berkshire and Surrey Pathology Services Lighthouse Laboratory | Wellcome Sanger Institute for the COVID-19 Genomics UK (COG-UK) Consortium | Berkshire and Surrey Pathology Services Lighthouse Laboratory and Alex Alderton, Roberto Amato, Jeffrey Barrett, Sonia Goncalves, Ewan Harrison, David K. Jackson, Ian Johnston, Dominic Kwiatkowski, Cordelia Langford, John Sillitoe on behalf of the Wellcome Sanger Institute COVID-19 Surveillance Team |
| EPI_ISL_3202194, EPI_ISL_3202195, EPI_ISL_3202196 | Lighthouse Lab in Alderley Park | Wellcome Sanger Institute for the COVID-19 Genomics UK (COG-UK) Consortium | Jacquelyn Wynn, Mairead Hyland, The Lighthouse Lab in Alderley Park and Alex Alderton, Roberto Amato, Jeffrey Barrett, Sonia Goncalves, Ewan Harrison, David K. Jackson, Ian Johnston, Dominic Kwiatkowski, Cordelia Langford, John Sillitoe on behalf of the Wellcome Sanger Institute COVID-19 Surveillance Team |
| EPI_ISL_3202197, EPI_ISL_3202198, EPI_ISL_3202199, EPI_ISL_3202200 | Berkshire and Surrey Pathology Services Lighthouse Laboratory | Wellcome Sanger Institute for the COVID-19 Genomics UK (COG-UK) Consortium | Berkshire and Surrey Pathology Services Lighthouse Laboratory and Alex Alderton, Roberto Amato, Jeffrey Barrett, Sonia Goncalves, Ewan Harrison, David K. Jackson, Ian Johnston, Dominic Kwiatkowski, Cordelia Langford, John Sillitoe on behalf of the Wellcome Sanger Institute COVID-19 Surveillance Team |
| EPI_ISL_3202201 | Lighthouse Lab in Alderley Park | Wellcome Sanger Institute for the COVID-19 Genomics UK (COG-UK) Consortium | Jacquelyn Wynn, Mairead Hyland, The Lighthouse Lab in Alderley Park and Alex Alderton, Roberto Amato, Jeffrey Barrett, Sonia Goncalves, Ewan Harrison, David K. Jackson, Ian Johnston, Dominic Kwiatkowski, Cordelia Langford, John Sillitoe on behalf of the Wellcome Sanger Institute COVID-19 Surveillance Team |
| EPI_ISL_3202202 | Berkshire and Surrey Pathology Services Lighthouse Laboratory | Wellcome Sanger Institute for the COVID-19 Genomics UK (COG-UK) Consortium | Berkshire and Surrey Pathology Services Lighthouse Laboratory and Alex Alderton, Roberto Amato, Jeffrey Barrett, Sonia Goncalves, Ewan Harrison, David K. Jackson, Ian Johnston, Dominic Kwiatkowski, Cordelia Langford, John Sillitoe on behalf of the Wellcome Sanger Institute COVID-19 Surveillance Team |
| EPI_ISL_3202203, EPI_ISL_3202204, EPI_ISL_3202205 | Lighthouse Lab in Alderley Park | Wellcome Sanger Institute for the COVID-19 Genomics UK (COG-UK) Consortium | Jacquelyn Wynn, Mairead Hyland, The Lighthouse Lab in Alderley Park and Alex Alderton, Roberto Amato, Jeffrey Barrett, Sonia Goncalves, Ewan Harrison, David K. Jackson, Ian Johnston, Dominic Kwiatkowski, Cordelia Langford, John Sillitoe on behalf of the Wellcome Sanger Institute COVID-19 Surveillance Team |
| EPI_ISL_3202206 | Berkshire and Surrey Pathology Services Lighthouse Laboratory | Wellcome Sanger Institute for the COVID-19 Genomics UK (COG-UK) Consortium | Berkshire and Surrey Pathology Services Lighthouse Laboratory and Alex Alderton, Roberto Amato, Jeffrey Barrett, Sonia Goncalves, Ewan Harrison, David K. Jackson, Ian Johnston, Dominic Kwiatkowski, Cordelia Langford, John Sillitoe on behalf of the Wellcome Sanger Institute COVID-19 Surveillance Team |
| EPI_ISL_3202207, EPI_ISL_3202208 | Lighthouse Lab in Alderley Park | Wellcome Sanger Institute for the COVID-19 Genomics UK (COG-UK) Consortium | Jacquelyn Wynn, Mairead Hyland, The Lighthouse Lab in Alderley Park and Alex Alderton, Roberto Amato, Jeffrey Barrett, Sonia Goncalves, Ewan Harrison, David K. Jackson, Ian Johnston, Dominic Kwiatkowski, Cordelia Langford, John Sillitoe on behalf of the Wellcome Sanger Institute COVID-19 Surveillance Team |
| EPI_ISL_3202209 | Berkshire and Surrey Pathology Services Lighthouse Laboratory | Wellcome Sanger Institute for the COVID-19 Genomics UK (COG-UK) Consortium | Berkshire and Surrey Pathology Services Lighthouse Laboratory and Alex Alderton, Roberto Amato, Jeffrey Barrett, Sonia Goncalves, Ewan Harrison, David K. Jackson, Ian Johnston, Dominic Kwiatkowski, Cordelia Langford, John Sillitoe on behalf of the Wellcome Sanger Institute COVID-19 Surveillance Team |
| EPI_ISL_3202210 | Lighthouse Lab in Alderley Park | Wellcome Sanger Institute for the COVID-19 Genomics UK (COG-UK) Consortium | Jacquelyn Wynn, Mairead Hyland, The Lighthouse Lab in Alderley Park and Alex Alderton, Roberto Amato, Jeffrey Barrett, Sonia Goncalves, Ewan Harrison, David K. Jackson, Ian Johnston, Dominic Kwiatkowski, Cordelia Langford, John Sillitoe on behalf of the Wellcome Sanger Institute COVID-19 Surveillance Team |
| EPI_ISL_3202211 | Berkshire and Surrey Pathology Services Lighthouse Laboratory | Wellcome Sanger Institute for the COVID-19 Genomics UK (COG-UK) Consortium | Berkshire and Surrey Pathology Services Lighthouse Laboratory and Alex Alderton, Roberto Amato, Jeffrey Barrett, Sonia Goncalves, Ewan Harrison, David K. Jackson, Ian Johnston, Dominic Kwiatkowski, Cordelia Langford, John Sillitoe on behalf of the Wellcome Sanger Institute COVID-19 Surveillance Team |
| EPI_ISL_3202212, EPI_ISL_3202213, EPI_ISL_3202214 | Lighthouse Lab in Alderley Park | Wellcome Sanger Institute for the COVID-19 Genomics UK (COG-UK) Consortium | Jacquelyn Wynn, Mairead Hyland, The Lighthouse Lab in Alderley Park and Alex Alderton, Roberto Amato, Jeffrey Barrett, Sonia Goncalves, Ewan Harrison, David K. Jackson, Ian Johnston, Dominic Kwiatkowski, Cordelia Langford, John Sillitoe on behalf of the Wellcome Sanger Institute COVID-19 Surveillance Team |
| EPI_ISL_3202215 | Berkshire and Surrey Pathology Services Lighthouse Laboratory | Wellcome Sanger Institute for the COVID-19 Genomics UK (COG-UK) Consortium | Berkshire and Surrey Pathology Services Lighthouse Laboratory and Alex Alderton, Roberto Amato, Jeffrey Barrett, Sonia Goncalves, Ewan Harrison, David K. Jackson, Ian Johnston, Dominic Kwiatkowski, Cordelia Langford, John Sillitoe on behalf of the Wellcome Sanger Institute COVID-19 Surveillance Team |
| EPI_ISL_3202216, EPI_ISL_3202217 | Lighthouse Lab in Alderley Park | Wellcome Sanger Institute for the COVID-19 Genomics UK (COG-UK) Consortium | Jacquelyn Wynn, Mairead Hyland, The Lighthouse Lab in Alderley Park and Alex Alderton, Roberto Amato, Jeffrey Barrett, Sonia Goncalves, Ewan Harrison, David K. Jackson, Ian Johnston, Dominic Kwiatkowski, Cordelia Langford, John Sillitoe on behalf of the Wellcome Sanger Institute COVID-19 Surveillance Team |
| EPI_ISL_3202218 | Berkshire and Surrey Pathology Services Lighthouse Laboratory | Wellcome Sanger Institute for the COVID-19 Genomics UK (COG-UK) Consortium | Berkshire and Surrey Pathology Services Lighthouse Laboratory and Alex Alderton, Roberto Amato, Jeffrey Barrett, Sonia Goncalves, Ewan Harrison, David K. Jackson, Ian Johnston, Dominic Kwiatkowski, Cordelia Langford, John Sillitoe on behalf of the Wellcome Sanger Institute COVID-19 Surveillance Team |
| EPI_ISL_3202219 | Lighthouse Lab in Alderley Park | Wellcome Sanger Institute for the COVID-19 Genomics UK (COG-UK) Consortium | Jacquelyn Wynn, Mairead Hyland, The Lighthouse Lab in Alderley Park and Alex Alderton, Roberto Amato, Jeffrey Barrett, Sonia Goncalves, Ewan Harrison, David K. Jackson, Ian Johnston, Dominic Kwiatkowski, Cordelia Langford, John Sillitoe on behalf of the Wellcome Sanger Institute COVID-19 Surveillance Team |
| EPI_ISL_3202220, EPI_ISL_3202221 | Berkshire and Surrey Pathology Services Lighthouse Laboratory | Wellcome Sanger Institute for the COVID-19 Genomics UK (COG-UK) Consortium | Berkshire and Surrey Pathology Services Lighthouse Laboratory and Alex Alderton, Roberto Amato, Jeffrey Barrett, Sonia Goncalves, Ewan Harrison, David K. Jackson, Ian Johnston, Dominic Kwiatkowski, Cordelia Langford, John Sillitoe on behalf of the Wellcome Sanger Institute COVID-19 Surveillance Team |
| EPI_ISL_3202222, EPI_ISL_3202223 | Lighthouse Lab in Alderley Park | Wellcome Sanger Institute for the COVID-19 Genomics UK (COG-UK) Consortium | Jacquelyn Wynn, Mairead Hyland, The Lighthouse Lab in Alderley Park and Alex Alderton, Roberto Amato, Jeffrey Barrett, Sonia Goncalves, Ewan Harrison, David K. Jackson, Ian Johnston, Dominic Kwiatkowski, Cordelia Langford, John Sillitoe on behalf of the Wellcome Sanger Institute COVID-19 Surveillance Team |
| EPI_ISL_3202224, EPI_ISL_3202225 | Berkshire and Surrey Pathology Services Lighthouse Laboratory | Wellcome Sanger Institute for the COVID-19 Genomics UK (COG-UK) Consortium | Berkshire and Surrey Pathology Services Lighthouse Laboratory and Alex Alderton, Roberto Amato, Jeffrey Barrett, Sonia Goncalves, Ewan Harrison, David K. Jackson, Ian Johnston, Dominic Kwiatkowski, Cordelia Langford, John Sillitoe on behalf of the Wellcome Sanger Institute COVID-19 Surveillance Team |
| EPI_ISL_3202226, EPI_ISL_3202227 | Lighthouse Lab in Alderley Park | Wellcome Sanger Institute for the COVID-19 Genomics UK (COG-UK) Consortium | Jacquelyn Wynn, Mairead Hyland, The Lighthouse Lab in Alderley Park and Alex Alderton, Roberto Amato, Jeffrey Barrett, Sonia Goncalves, Ewan Harrison, David K. Jackson, Ian Johnston, Dominic Kwiatkowski, Cordelia Langford, John Sillitoe on behalf of the Wellcome Sanger Institute COVID-19 Surveillance Team |
| EPI_ISL_3202228 | Berkshire and Surrey Pathology Services Lighthouse Laboratory | Wellcome Sanger Institute for the COVID-19 Genomics UK (COG-UK) Consortium | Berkshire and Surrey Pathology Services Lighthouse Laboratory and Alex Alderton, Roberto Amato, Jeffrey Barrett, Sonia Goncalves, Ewan Harrison, David K. Jackson, Ian Johnston, Dominic Kwiatkowski, Cordelia Langford, John Sillitoe on behalf of the Wellcome Sanger Institute COVID-19 Surveillance Team |
| EPI_ISL_3202229, EPI_ISL_3202230, EPI_ISL_3202231, EPI_ISL_3202232, EPI_ISL_3202233, EPI_ISL_3202234 | Lighthouse Lab in Alderley Park | Wellcome Sanger Institute for the COVID-19 Genomics UK (COG-UK) Consortium | Jacquelyn Wynn, Mairead Hyland, The Lighthouse Lab in Alderley Park and Alex Alderton, Roberto Amato, Jeffrey Barrett, Sonia Goncalves, Ewan Harrison, David K. Jackson, Ian Johnston, Dominic Kwiatkowski, Cordelia Langford, John Sillitoe on behalf of the Wellcome Sanger Institute COVID-19 Surveillance Team |
| EPI_ISL_3202235 | Berkshire and Surrey Pathology Services Lighthouse Laboratory | Wellcome Sanger Institute for the COVID-19 Genomics UK (COG-UK) Consortium | Berkshire and Surrey Pathology Services Lighthouse Laboratory and Alex Alderton, Roberto Amato, Jeffrey Barrett, Sonia Goncalves, Ewan Harrison, David K. Jackson, Ian Johnston, Dominic Kwiatkowski, Cordelia Langford, John Sillitoe on behalf of the Wellcome Sanger Institute COVID-19 Surveillance Team |
| EPI_ISL_3202236, EPI_ISL_3202237, EPI_ISL_3202238 | Lighthouse Lab in Alderley Park | Wellcome Sanger Institute for the COVID-19 Genomics UK (COG-UK) Consortium | Jacquelyn Wynn, Mairead Hyland, The Lighthouse Lab in Alderley Park and Alex Alderton, Roberto Amato, Jeffrey Barrett, Sonia Goncalves, Ewan Harrison, David K. Jackson, Ian Johnston, Dominic Kwiatkowski, Cordelia Langford, John Sillitoe on behalf of the Wellcome Sanger Institute COVID-19 Surveillance Team |
| EPI_ISL_3202239 | Berkshire and Surrey Pathology Services Lighthouse Laboratory | Wellcome Sanger Institute for the COVID-19 Genomics UK (COG-UK) Consortium | Berkshire and Surrey Pathology Services Lighthouse Laboratory and Alex Alderton, Roberto Amato, Jeffrey Barrett, Sonia Goncalves, Ewan Harrison, David K. Jackson, Ian Johnston, Dominic Kwiatkowski, Cordelia Langford, John Sillitoe on behalf of the Wellcome Sanger Institute COVID-19 Surveillance Team |
| EPI_ISL_3202240 | Lighthouse Lab in Alderley Park | Wellcome Sanger Institute for the COVID-19 Genomics UK (COG-UK) Consortium | Jacquelyn Wynn, Mairead Hyland, The Lighthouse Lab in Alderley Park and Alex Alderton, Roberto Amato, Jeffrey Barrett, Sonia Goncalves, Ewan Harrison, David K. Jackson, Ian Johnston, Dominic Kwiatkowski, Cordelia Langford, John Sillitoe on behalf of the Wellcome Sanger Institute COVID-19 Surveillance Team |
| EPI_ISL_3202241 | Berkshire and Surrey Pathology Services Lighthouse Laboratory | Wellcome Sanger Institute for the COVID-19 Genomics UK (COG-UK) Consortium | Berkshire and Surrey Pathology Services Lighthouse Laboratory and Alex Alderton, Roberto Amato, Jeffrey Barrett, Sonia Goncalves, Ewan Harrison, David K. Jackson, Ian Johnston, Dominic Kwiatkowski, Cordelia Langford, John Sillitoe on behalf of the Wellcome Sanger Institute COVID-19 Surveillance Team |
| EPI_ISL_3202242 | Lighthouse Lab in Alderley Park | Wellcome Sanger Institute for the COVID-19 Genomics UK (COG-UK) Consortium | Jacquelyn Wynn, Mairead Hyland, The Lighthouse Lab in Alderley Park and Alex Alderton, Roberto Amato, Jeffrey Barrett, Sonia Goncalves, Ewan Harrison, David K. Jackson, Ian Johnston, Dominic Kwiatkowski, Cordelia Langford, John Sillitoe on behalf of the Wellcome Sanger Institute COVID-19 Surveillance Team |
| EPI_ISL_3202243 | Berkshire and Surrey Pathology Services Lighthouse Laboratory | Wellcome Sanger Institute for the COVID-19 Genomics UK (COG-UK) Consortium | Berkshire and Surrey Pathology Services Lighthouse Laboratory and Alex Alderton, Roberto Amato, Jeffrey Barrett, Sonia Goncalves, Ewan Harrison, David K. Jackson, Ian Johnston, Dominic Kwiatkowski, Cordelia Langford, John Sillitoe on behalf of the Wellcome Sanger Institute COVID-19 Surveillance Team |
| EPI_ISL_3202244 | Lighthouse Lab in Alderley Park | Wellcome Sanger Institute for the COVID-19 Genomics UK (COG-UK) Consortium | Jacquelyn Wynn, Mairead Hyland, The Lighthouse Lab in Alderley Park and Alex Alderton, Roberto Amato, Jeffrey Barrett, Sonia Goncalves, Ewan Harrison, David K. Jackson, Ian Johnston, Dominic Kwiatkowski, Cordelia Langford, John Sillitoe on behalf of the Wellcome Sanger Institute COVID-19 Surveillance Team |
| EPI_ISL_3202245 | Berkshire and Surrey Pathology Services Lighthouse Laboratory | Wellcome Sanger Institute for the COVID-19 Genomics UK (COG-UK) Consortium | Berkshire and Surrey Pathology Services Lighthouse Laboratory and Alex Alderton, Roberto Amato, Jeffrey Barrett, Sonia Goncalves, Ewan Harrison, David K. Jackson, Ian Johnston, Dominic Kwiatkowski, Cordelia Langford, John Sillitoe on behalf of the Wellcome Sanger Institute COVID-19 Surveillance Team |
| EPI_ISL_3202246, EPI_ISL_3202247 | Lighthouse Lab in Alderley Park | Wellcome Sanger Institute for the COVID-19 Genomics UK (COG-UK) Consortium | Jacquelyn Wynn, Mairead Hyland, The Lighthouse Lab in Alderley Park and Alex Alderton, Roberto Amato, Jeffrey Barrett, Sonia Goncalves, Ewan Harrison, David K. Jackson, Ian Johnston, Dominic Kwiatkowski, Cordelia Langford, John Sillitoe on behalf of the Wellcome Sanger Institute COVID-19 Surveillance Team |
| EPI_ISL_3202248 | Berkshire and Surrey Pathology Services Lighthouse Laboratory | Wellcome Sanger Institute for the COVID-19 Genomics UK (COG-UK) Consortium | Berkshire and Surrey Pathology Services Lighthouse Laboratory and Alex Alderton, Roberto Amato, Jeffrey Barrett, Sonia Goncalves, Ewan Harrison, David K. Jackson, Ian Johnston, Dominic Kwiatkowski, Cordelia Langford, John Sillitoe on behalf of the Wellcome Sanger Institute COVID-19 Surveillance Team |
| EPI_ISL_3202249 | Lighthouse Lab in Alderley Park | Wellcome Sanger Institute for the COVID-19 Genomics UK (COG-UK) | Jacquelyn Wynn, Mairead Hyland, The Lighthouse Lab in Alderley Park and Alex Alderton, Roberto Amato, Jeffrey Barrett, Sonia Goncalves, Ewan Harrison, David K. Jackson, Ian Johnston, Dominic Kwiatkowski, Cordelia Langford, John Sillitoe on |

| | | | |
|---|---|---|---|
| | | Consortium | behalf of the Wellcome Sanger Institute COVID-19 Surveillance Team |
| EPI_ISL_3202250 | Berkshire and Surrey Pathology Services Lighthouse Laboratory | Wellcome Sanger Institute for the COVID-19 Genomics UK (COG-UK) Consortium | Berkshire and Surrey Pathology Services Lighthouse Laboratory and Alex Alderton, Roberto Amato, Jeffrey Barrett, Sonia Goncalves, Ewan Harrison, David K. Jackson, Ian Johnston, Dominic Kwiatkowski, Cordelia Langford, John Sillitoe on behalf of the Wellcome Sanger Institute COVID-19 Surveillance Team |
| EPI_ISL_3202251, EPI_ISL_3202252, EPI_ISL_3202253 | Lighthouse Lab in Alderley Park | Wellcome Sanger Institute for the COVID-19 Genomics UK (COG-UK) Consortium | Jacquelyn Wynn, Mairead Hyland, The Lighthouse Lab in Alderley Park and Alex Alderton, Roberto Amato, Jeffrey Barrett, Sonia Goncalves, Ewan Harrison, David K. Jackson, Ian Johnston, Dominic Kwiatkowski, Cordelia Langford, John Sillitoe on behalf of the Wellcome Sanger Institute COVID-19 Surveillance Team |
| EPI_ISL_3202254 | Berkshire and Surrey Pathology Services Lighthouse Laboratory | Wellcome Sanger Institute for the COVID-19 Genomics UK (COG-UK) Consortium | Berkshire and Surrey Pathology Services Lighthouse Laboratory and Alex Alderton, Roberto Amato, Jeffrey Barrett, Sonia Goncalves, Ewan Harrison, David K. Jackson, Ian Johnston, Dominic Kwiatkowski, Cordelia Langford, John Sillitoe on behalf of the Wellcome Sanger Institute COVID-19 Surveillance Team |
| EPI_ISL_3202255, EPI_ISL_3202256 | Lighthouse Lab in Alderley Park | Wellcome Sanger Institute for the COVID-19 Genomics UK (COG-UK) Consortium | Jacquelyn Wynn, Mairead Hyland, The Lighthouse Lab in Alderley Park and Alex Alderton, Roberto Amato, Jeffrey Barrett, Sonia Goncalves, Ewan Harrison, David K. Jackson, Ian Johnston, Dominic Kwiatkowski, Cordelia Langford, John Sillitoe on behalf of the Wellcome Sanger Institute COVID-19 Surveillance Team |
| EPI_ISL_3202257, EPI_ISL_3202258 | Berkshire and Surrey Pathology Services Lighthouse Laboratory | Wellcome Sanger Institute for the COVID-19 Genomics UK (COG-UK) Consortium | Berkshire and Surrey Pathology Services Lighthouse Laboratory and Alex Alderton, Roberto Amato, Jeffrey Barrett, Sonia Goncalves, Ewan Harrison, David K. Jackson, Ian Johnston, Dominic Kwiatkowski, Cordelia Langford, John Sillitoe on behalf of the Wellcome Sanger Institute COVID-19 Surveillance Team |
| EPI_ISL_3202259, EPI_ISL_3202260, EPI_ISL_3202261, EPI_ISL_3202262 | Lighthouse Lab in Alderley Park | Wellcome Sanger Institute for the COVID-19 Genomics UK (COG-UK) Consortium | Jacquelyn Wynn, Mairead Hyland, The Lighthouse Lab in Alderley Park and Alex Alderton, Roberto Amato, Jeffrey Barrett, Sonia Goncalves, Ewan Harrison, David K. Jackson, Ian Johnston, Dominic Kwiatkowski, Cordelia Langford, John Sillitoe on behalf of the Wellcome Sanger Institute COVID-19 Surveillance Team |
| EPI_ISL_3202263 | Berkshire and Surrey Pathology Services Lighthouse Laboratory | Wellcome Sanger Institute for the COVID-19 Genomics UK (COG-UK) Consortium | Berkshire and Surrey Pathology Services Lighthouse Laboratory and Alex Alderton, Roberto Amato, Jeffrey Barrett, Sonia Goncalves, Ewan Harrison, David K. Jackson, Ian Johnston, Dominic Kwiatkowski, Cordelia Langford, John Sillitoe on behalf of the Wellcome Sanger Institute COVID-19 Surveillance Team |
| EPI_ISL_3202264, EPI_ISL_3202265, EPI_ISL_3202266, EPI_ISL_3202267, EPI_ISL_3202268, EPI_ISL_3202269, EPI_ISL_3202270 | Lighthouse Lab in Alderley Park | Wellcome Sanger Institute for the COVID-19 Genomics UK (COG-UK) Consortium | Jacquelyn Wynn, Mairead Hyland, The Lighthouse Lab in Alderley Park and Alex Alderton, Roberto Amato, Jeffrey Barrett, Sonia Goncalves, Ewan Harrison, David K. Jackson, Ian Johnston, Dominic Kwiatkowski, Cordelia Langford, John Sillitoe on behalf of the Wellcome Sanger Institute COVID-19 Surveillance Team |
| EPI_ISL_3202271, EPI_ISL_3202272, EPI_ISL_3202273 | Berkshire and Surrey Pathology Services Lighthouse Laboratory | Wellcome Sanger Institute for the COVID-19 Genomics UK (COG-UK) Consortium | Berkshire and Surrey Pathology Services Lighthouse Laboratory and Alex Alderton, Roberto Amato, Jeffrey Barrett, Sonia Goncalves, Ewan Harrison, David K. Jackson, Ian Johnston, Dominic Kwiatkowski, Cordelia Langford, John Sillitoe on behalf of the Wellcome Sanger Institute COVID-19 Surveillance Team |
| EPI_ISL_3202274 | Lighthouse Lab in Alderley Park | Wellcome Sanger Institute for the COVID-19 Genomics UK (COG-UK) Consortium | Jacquelyn Wynn, Mairead Hyland, The Lighthouse Lab in Alderley Park and Alex Alderton, Roberto Amato, Jeffrey Barrett, Sonia Goncalves, Ewan Harrison, David K. Jackson, Ian Johnston, Dominic Kwiatkowski, Cordelia Langford, John Sillitoe on behalf of the Wellcome Sanger Institute COVID-19 Surveillance Team |
| EPI_ISL_3202275, EPI_ISL_3202276, EPI_ISL_3202277, EPI_ISL_3202278 | Berkshire and Surrey Pathology Services Lighthouse Laboratory | Wellcome Sanger Institute for the COVID-19 Genomics UK (COG-UK) Consortium | Berkshire and Surrey Pathology Services Lighthouse Laboratory and Alex Alderton, Roberto Amato, Jeffrey Barrett, Sonia Goncalves, Ewan Harrison, David K. Jackson, Ian Johnston, Dominic Kwiatkowski, Cordelia Langford, John Sillitoe on behalf of the Wellcome Sanger Institute COVID-19 Surveillance Team |
| EPI_ISL_3202279, EPI_ISL_3202280, EPI_ISL_3202281 | Lighthouse Lab in Alderley Park | Wellcome Sanger Institute for the COVID-19 Genomics UK (COG-UK) Consortium | Jacquelyn Wynn, Mairead Hyland, The Lighthouse Lab in Alderley Park and Alex Alderton, Roberto Amato, Jeffrey Barrett, Sonia Goncalves, Ewan Harrison, David K. Jackson, Ian Johnston, Dominic Kwiatkowski, Cordelia Langford, John Sillitoe on behalf of the Wellcome Sanger Institute COVID-19 Surveillance Team |
| EPI_ISL_3202282 | Berkshire and Surrey Pathology Services Lighthouse Laboratory | Wellcome Sanger Institute for the COVID-19 Genomics UK (COG-UK) Consortium | Berkshire and Surrey Pathology Services Lighthouse Laboratory and Alex Alderton, Roberto Amato, Jeffrey Barrett, Sonia Goncalves, Ewan Harrison, David K. Jackson, Ian Johnston, Dominic Kwiatkowski, Cordelia Langford, John Sillitoe on behalf of the Wellcome Sanger Institute COVID-19 Surveillance Team |
| EPI_ISL_3202283, EPI_ISL_3202284, EPI_ISL_3202285, EPI_ISL_3202286 | Lighthouse Lab in Alderley Park | Wellcome Sanger Institute for the COVID-19 Genomics UK (COG-UK) Consortium | Jacquelyn Wynn, Mairead Hyland, The Lighthouse Lab in Alderley Park and Alex Alderton, Roberto Amato, Jeffrey Barrett, Sonia Goncalves, Ewan Harrison, David K. Jackson, Ian Johnston, Dominic Kwiatkowski, Cordelia Langford, John Sillitoe on behalf of the Wellcome Sanger Institute COVID-19 Surveillance Team |
| EPI_ISL_3202287, EPI_ISL_3202288 | Berkshire and Surrey Pathology Services Lighthouse Laboratory | Wellcome Sanger Institute for the COVID-19 Genomics UK (COG-UK) Consortium | Berkshire and Surrey Pathology Services Lighthouse Laboratory and Alex Alderton, Roberto Amato, Jeffrey Barrett, Sonia Goncalves, Ewan Harrison, David K. Jackson, Ian Johnston, Dominic Kwiatkowski, Cordelia Langford, John Sillitoe on behalf of the Wellcome Sanger Institute COVID-19 Surveillance Team |
| EPI_ISL_3202289, EPI_ISL_3202290, EPI_ISL_3202291 | Lighthouse Lab in Alderley Park | Wellcome Sanger Institute for the COVID-19 Genomics UK (COG-UK) Consortium | Jacquelyn Wynn, Mairead Hyland, The Lighthouse Lab in Alderley Park and Alex Alderton, Roberto Amato, Jeffrey Barrett, Sonia Goncalves, Ewan Harrison, David K. Jackson, Ian Johnston, Dominic Kwiatkowski, Cordelia Langford, John Sillitoe on behalf of the Wellcome Sanger Institute COVID-19 Surveillance Team |
| EPI_ISL_3202292 | Berkshire and Surrey Pathology Services Lighthouse Laboratory | Wellcome Sanger Institute for the COVID-19 Genomics UK (COG-UK) Consortium | Berkshire and Surrey Pathology Services Lighthouse Laboratory and Alex Alderton, Roberto Amato, Jeffrey Barrett, Sonia Goncalves, Ewan Harrison, David K. Jackson, Ian Johnston, Dominic Kwiatkowski, Cordelia Langford, John Sillitoe on behalf of the Wellcome Sanger Institute COVID-19 Surveillance Team |
| EPI_ISL_3202293 | Lighthouse Lab in Alderley Park | Wellcome Sanger Institute for the COVID-19 Genomics UK (COG-UK) Consortium | Jacquelyn Wynn, Mairead Hyland, The Lighthouse Lab in Alderley Park and Alex Alderton, Roberto Amato, Jeffrey Barrett, Sonia Goncalves, Ewan Harrison, David K. Jackson, Ian Johnston, Dominic Kwiatkowski, Cordelia Langford, John Sillitoe on behalf of the Wellcome Sanger Institute COVID-19 Surveillance Team |
| EPI_ISL_3202294, EPI_ISL_3202295, EPI_ISL_3202296 | Berkshire and Surrey Pathology Services Lighthouse Laboratory | Wellcome Sanger Institute for the COVID-19 Genomics UK (COG-UK) Consortium | Berkshire and Surrey Pathology Services Lighthouse Laboratory and Alex Alderton, Roberto Amato, Jeffrey Barrett, Sonia Goncalves, Ewan Harrison, David K. Jackson, Ian Johnston, Dominic Kwiatkowski, Cordelia Langford, John Sillitoe on behalf of the Wellcome Sanger Institute COVID-19 Surveillance Team |
| EPI_ISL_3202297, EPI_ISL_3202298, EPI_ISL_3202299 | Lighthouse Lab in Alderley Park | Wellcome Sanger Institute for the COVID-19 Genomics UK (COG-UK) Consortium | Jacquelyn Wynn, Mairead Hyland, The Lighthouse Lab in Alderley Park and Alex Alderton, Roberto Amato, Jeffrey Barrett, Sonia Goncalves, Ewan Harrison, David K. Jackson, Ian Johnston, Dominic Kwiatkowski, Cordelia Langford, John Sillitoe on behalf of the Wellcome Sanger Institute COVID-19 Surveillance Team |
| EPI_ISL_3202300, EPI_ISL_3202301 | Berkshire and Surrey Pathology Services Lighthouse Laboratory | Wellcome Sanger Institute for the COVID-19 Genomics UK (COG-UK) Consortium | Berkshire and Surrey Pathology Services Lighthouse Laboratory and Alex Alderton, Roberto Amato, Jeffrey Barrett, Sonia Goncalves, Ewan Harrison, David K. Jackson, Ian Johnston, Dominic Kwiatkowski, Cordelia Langford, John Sillitoe on behalf of the Wellcome Sanger Institute COVID-19 Surveillance Team |
| EPI_ISL_3202302, EPI_ISL_3202303, EPI_ISL_3202304, EPI_ISL_3202305 | Lighthouse Lab in Alderley Park | Wellcome Sanger Institute for the COVID-19 Genomics UK (COG-UK) Consortium | Jacquelyn Wynn, Mairead Hyland, The Lighthouse Lab in Alderley Park and Alex Alderton, Roberto Amato, Jeffrey Barrett, Sonia Goncalves, Ewan Harrison, David K. Jackson, Ian Johnston, Dominic Kwiatkowski, Cordelia Langford, John Sillitoe on behalf of the Wellcome Sanger Institute COVID-19 Surveillance Team |
| EPI_ISL_3202306, EPI_ISL_3202307 | Berkshire and Surrey Pathology Services Lighthouse Laboratory | Wellcome Sanger Institute for the COVID-19 Genomics UK (COG-UK) Consortium | Berkshire and Surrey Pathology Services Lighthouse Laboratory and Alex Alderton, Roberto Amato, Jeffrey Barrett, Sonia Goncalves, Ewan Harrison, David K. Jackson, Ian Johnston, Dominic Kwiatkowski, Cordelia Langford, John Sillitoe on behalf of the Wellcome Sanger Institute COVID-19 Surveillance Team |
| EPI_ISL_3202308, EPI_ISL_3202309, EPI_ISL_3202310, EPI_ISL_3202311 | Lighthouse Lab in Alderley Park | Wellcome Sanger Institute for the COVID-19 Genomics UK (COG-UK) Consortium | Jacquelyn Wynn, Mairead Hyland, The Lighthouse Lab in Alderley Park and Alex Alderton, Roberto Amato, Jeffrey Barrett, Sonia Goncalves, Ewan Harrison, David K. Jackson, Ian Johnston, Dominic Kwiatkowski, Cordelia Langford, John Sillitoe on behalf of the Wellcome Sanger Institute COVID-19 Surveillance Team |
| EPI_ISL_3202312 | Berkshire and Surrey Pathology Services Lighthouse Laboratory | Wellcome Sanger Institute for the COVID-19 Genomics UK (COG-UK) Consortium | Berkshire and Surrey Pathology Services Lighthouse Laboratory and Alex Alderton, Roberto Amato, Jeffrey Barrett, Sonia Goncalves, Ewan Harrison, David K. Jackson, Ian Johnston, Dominic Kwiatkowski, Cordelia Langford, John Sillitoe on behalf of the Wellcome Sanger Institute COVID-19 Surveillance Team |
| EPI_ISL_3202313, EPI_ISL_3202314 | Lighthouse Lab in Alderley Park | Wellcome Sanger Institute for the COVID-19 Genomics UK (COG-UK) Consortium | Jacquelyn Wynn, Mairead Hyland, The Lighthouse Lab in Alderley Park and Alex Alderton, Roberto Amato, Jeffrey Barrett, Sonia Goncalves, Ewan Harrison, David K. Jackson, Ian Johnston, Dominic Kwiatkowski, Cordelia Langford, John Sillitoe on behalf of the Wellcome Sanger Institute COVID-19 Surveillance Team |
| EPI_ISL_3202315, EPI_ISL_3202316 | Berkshire and Surrey Pathology Services Lighthouse Laboratory | Wellcome Sanger Institute for the COVID-19 Genomics UK (COG-UK) Consortium | Berkshire and Surrey Pathology Services Lighthouse Laboratory and Alex Alderton, Roberto Amato, Jeffrey Barrett, Sonia Goncalves, Ewan Harrison, David K. Jackson, Ian Johnston, Dominic Kwiatkowski, Cordelia Langford, John Sillitoe on behalf of the Wellcome Sanger Institute COVID-19 Surveillance Team |
| EPI_ISL_3202317 | Lighthouse Lab in Alderley Park | Wellcome Sanger Institute for the COVID-19 Genomics UK (COG-UK) Consortium | Jacquelyn Wynn, Mairead Hyland, The Lighthouse Lab in Alderley Park and Alex Alderton, Roberto Amato, Jeffrey Barrett, Sonia Goncalves, Ewan Harrison, David K. Jackson, Ian Johnston, Dominic Kwiatkowski, Cordelia Langford, John Sillitoe on behalf of the Wellcome Sanger Institute COVID-19 Surveillance Team |
| EPI_ISL_3202318 | Berkshire and Surrey Pathology Services Lighthouse Laboratory | Wellcome Sanger Institute for the COVID-19 Genomics UK (COG-UK) Consortium | Berkshire and Surrey Pathology Services Lighthouse Laboratory and Alex Alderton, Roberto Amato, Jeffrey Barrett, Sonia Goncalves, Ewan Harrison, David K. Jackson, Ian Johnston, Dominic Kwiatkowski, Cordelia Langford, John Sillitoe on behalf of the Wellcome Sanger Institute COVID-19 Surveillance Team |
| EPI_ISL_3202319, EPI_ISL_3202320 | Lighthouse Lab in Alderley Park | Wellcome Sanger Institute for the COVID-19 Genomics UK (COG-UK) Consortium | Jacquelyn Wynn, Mairead Hyland, The Lighthouse Lab in Alderley Park and Alex Alderton, Roberto Amato, Jeffrey Barrett, Sonia Goncalves, Ewan Harrison, David K. Jackson, Ian Johnston, Dominic Kwiatkowski, Cordelia Langford, John Sillitoe on behalf of the Wellcome Sanger Institute COVID-19 Surveillance Team |
| EPI_ISL_3202321, EPI_ISL_3202322 | Berkshire and Surrey Pathology Services Lighthouse Laboratory | Wellcome Sanger Institute for the COVID-19 Genomics UK (COG-UK) Consortium | Berkshire and Surrey Pathology Services Lighthouse Laboratory and Alex Alderton, Roberto Amato, Jeffrey Barrett, Sonia Goncalves, Ewan Harrison, David K. Jackson, Ian Johnston, Dominic Kwiatkowski, Cordelia Langford, John Sillitoe on behalf of the Wellcome Sanger Institute COVID-19 Surveillance Team |
| EPI_ISL_3202323 | Lighthouse Lab in Alderley Park | Wellcome Sanger Institute for the COVID-19 Genomics UK (COG-UK) Consortium | Jacquelyn Wynn, Mairead Hyland, The Lighthouse Lab in Alderley Park and Alex Alderton, Roberto Amato, Jeffrey Barrett, Sonia Goncalves, Ewan Harrison, David K. Jackson, Ian Johnston, Dominic Kwiatkowski, Cordelia Langford, John Sillitoe on behalf of the Wellcome Sanger Institute COVID-19 Surveillance Team |
| EPI_ISL_3202324 | Berkshire and Surrey Pathology Services Lighthouse Laboratory | Wellcome Sanger Institute for the COVID-19 Genomics UK (COG-UK) Consortium | Berkshire and Surrey Pathology Services Lighthouse Laboratory and Alex Alderton, Roberto Amato, Jeffrey Barrett, Sonia Goncalves, Ewan Harrison, David K. Jackson, Ian Johnston, Dominic Kwiatkowski, Cordelia Langford, John Sillitoe on behalf of the Wellcome Sanger Institute COVID-19 Surveillance Team |
| EPI_ISL_3202325 | Lighthouse Lab in Alderley Park | Wellcome Sanger Institute for the COVID-19 Genomics UK (COG-UK) Consortium | Jacquelyn Wynn, Mairead Hyland, The Lighthouse Lab in Alderley Park and Alex Alderton, Roberto Amato, Jeffrey Barrett, Sonia Goncalves, Ewan Harrison, David K. Jackson, Ian Johnston, Dominic Kwiatkowski, Cordelia Langford, John Sillitoe on behalf of the Wellcome Sanger Institute COVID-19 Surveillance Team |
| EPI_ISL_3202326 | Berkshire and Surrey Pathology Services Lighthouse Laboratory | Wellcome Sanger Institute for the COVID-19 Genomics UK (COG-UK) Consortium | Berkshire and Surrey Pathology Services Lighthouse Laboratory and Alex Alderton, Roberto Amato, Jeffrey Barrett, Sonia Goncalves, Ewan Harrison, David K. Jackson, Ian Johnston, Dominic Kwiatkowski, Cordelia Langford, John Sillitoe on behalf of the Wellcome Sanger Institute COVID-19 Surveillance Team |
| EPI_ISL_3202327, EPI_ISL_3202328 | Lighthouse Lab in Alderley Park | Wellcome Sanger Institute for the COVID-19 Genomics UK (COG-UK) Consortium | Jacquelyn Wynn, Mairead Hyland, The Lighthouse Lab in Alderley Park and Alex Alderton, Roberto Amato, Jeffrey Barrett, Sonia Goncalves, Ewan Harrison, David K. Jackson, Ian Johnston, Dominic Kwiatkowski, Cordelia Langford, John Sillitoe on behalf of the Wellcome Sanger Institute COVID-19 Surveillance Team |

| | | | |
|---|---|---|---|
| EPI_ISL_3202329, EPI_ISL_3202330 | Berkshire and Surrey Pathology Services Lighthouse Laboratory | Wellcome Sanger Institute for the COVID-19 Genomics UK (COG-UK) Consortium | Berkshire and Surrey Pathology Services Lighthouse Laboratory and Alex Alderton, Roberto Amato, Jeffrey Barrett, Sonia Goncalves, Ewan Harrison, David K. Jackson, Ian Johnston, Dominic Kwiatkowski, Cordelia Langford, John Sillitoe on behalf of the Wellcome Sanger Institute COVID-19 Surveillance Team |
| EPI_ISL_3202331 | Lighthouse Lab in Alderley Park | Wellcome Sanger Institute for the COVID-19 Genomics UK (COG-UK) Consortium | Jacquelyn Wynn, Mairead Hyland, The Lighthouse Lab in Alderley Park and Alex Alderton, Roberto Amato, Jeffrey Barrett, Sonia Goncalves, Ewan Harrison, David K. Jackson, Ian Johnston, Dominic Kwiatkowski, Cordelia Langford, John Sillitoe on behalf of the Wellcome Sanger Institute COVID-19 Surveillance Team |
| EPI_ISL_3202332, EPI_ISL_3202333, EPI_ISL_3202334 | Berkshire and Surrey Pathology Services Lighthouse Laboratory | Wellcome Sanger Institute for the COVID-19 Genomics UK (COG-UK) Consortium | Berkshire and Surrey Pathology Services Lighthouse Laboratory and Alex Alderton, Roberto Amato, Jeffrey Barrett, Sonia Goncalves, Ewan Harrison, David K. Jackson, Ian Johnston, Dominic Kwiatkowski, Cordelia Langford, John Sillitoe on behalf of the Wellcome Sanger Institute COVID-19 Surveillance Team |
| EPI_ISL_3202335 | Lighthouse Lab in Alderley Park | Wellcome Sanger Institute for the COVID-19 Genomics UK (COG-UK) Consortium | Jacquelyn Wynn, Mairead Hyland, The Lighthouse Lab in Alderley Park and Alex Alderton, Roberto Amato, Jeffrey Barrett, Sonia Goncalves, Ewan Harrison, David K. Jackson, Ian Johnston, Dominic Kwiatkowski, Cordelia Langford, John Sillitoe on behalf of the Wellcome Sanger Institute COVID-19 Surveillance Team |
| EPI_ISL_3202336, EPI_ISL_3202337 | Berkshire and Surrey Pathology Services Lighthouse Laboratory | Wellcome Sanger Institute for the COVID-19 Genomics UK (COG-UK) Consortium | Berkshire and Surrey Pathology Services Lighthouse Laboratory and Alex Alderton, Roberto Amato, Jeffrey Barrett, Sonia Goncalves, Ewan Harrison, David K. Jackson, Ian Johnston, Dominic Kwiatkowski, Cordelia Langford, John Sillitoe on behalf of the Wellcome Sanger Institute COVID-19 Surveillance Team |
| EPI_ISL_3202338 | Lighthouse Lab in Alderley Park | Wellcome Sanger Institute for the COVID-19 Genomics UK (COG-UK) Consortium | Jacquelyn Wynn, Mairead Hyland, The Lighthouse Lab in Alderley Park and Alex Alderton, Roberto Amato, Jeffrey Barrett, Sonia Goncalves, Ewan Harrison, David K. Jackson, Ian Johnston, Dominic Kwiatkowski, Cordelia Langford, John Sillitoe on behalf of the Wellcome Sanger Institute COVID-19 Surveillance Team |
| EPI_ISL_3202339 | Berkshire and Surrey Pathology Services Lighthouse Laboratory | Wellcome Sanger Institute for the COVID-19 Genomics UK (COG-UK) Consortium | Berkshire and Surrey Pathology Services Lighthouse Laboratory and Alex Alderton, Roberto Amato, Jeffrey Barrett, Sonia Goncalves, Ewan Harrison, David K. Jackson, Ian Johnston, Dominic Kwiatkowski, Cordelia Langford, John Sillitoe on behalf of the Wellcome Sanger Institute COVID-19 Surveillance Team |
| EPI_ISL_3202340 | Lighthouse Lab in Alderley Park | Wellcome Sanger Institute for the COVID-19 Genomics UK (COG-UK) Consortium | Jacquelyn Wynn, Mairead Hyland, The Lighthouse Lab in Alderley Park and Alex Alderton, Roberto Amato, Jeffrey Barrett, Sonia Goncalves, Ewan Harrison, David K. Jackson, Ian Johnston, Dominic Kwiatkowski, Cordelia Langford, John Sillitoe on behalf of the Wellcome Sanger Institute COVID-19 Surveillance Team |
| EPI_ISL_3202341, EPI_ISL_3202342, EPI_ISL_3202343 | Berkshire and Surrey Pathology Services Lighthouse Laboratory | Wellcome Sanger Institute for the COVID-19 Genomics UK (COG-UK) Consortium | Berkshire and Surrey Pathology Services Lighthouse Laboratory and Alex Alderton, Roberto Amato, Jeffrey Barrett, Sonia Goncalves, Ewan Harrison, David K. Jackson, Ian Johnston, Dominic Kwiatkowski, Cordelia Langford, John Sillitoe on behalf of the Wellcome Sanger Institute COVID-19 Surveillance Team |
| EPI_ISL_3202344, EPI_ISL_3202345 | Lighthouse Lab in Alderley Park | Wellcome Sanger Institute for the COVID-19 Genomics UK (COG-UK) Consortium | Jacquelyn Wynn, Mairead Hyland, The Lighthouse Lab in Alderley Park and Alex Alderton, Roberto Amato, Jeffrey Barrett, Sonia Goncalves, Ewan Harrison, David K. Jackson, Ian Johnston, Dominic Kwiatkowski, Cordelia Langford, John Sillitoe on behalf of the Wellcome Sanger Institute COVID-19 Surveillance Team |
| EPI_ISL_3202346, EPI_ISL_3202347 | Berkshire and Surrey Pathology Services Lighthouse Laboratory | Wellcome Sanger Institute for the COVID-19 Genomics UK (COG-UK) Consortium | Berkshire and Surrey Pathology Services Lighthouse Laboratory and Alex Alderton, Roberto Amato, Jeffrey Barrett, Sonia Goncalves, Ewan Harrison, David K. Jackson, Ian Johnston, Dominic Kwiatkowski, Cordelia Langford, John Sillitoe on behalf of the Wellcome Sanger Institute COVID-19 Surveillance Team |
| EPI_ISL_3202348 | Lighthouse Lab in Alderley Park | Wellcome Sanger Institute for the COVID-19 Genomics UK (COG-UK) Consortium | Jacquelyn Wynn, Mairead Hyland, The Lighthouse Lab in Alderley Park and Alex Alderton, Roberto Amato, Jeffrey Barrett, Sonia Goncalves, Ewan Harrison, David K. Jackson, Ian Johnston, Dominic Kwiatkowski, Cordelia Langford, John Sillitoe on behalf of the Wellcome Sanger Institute COVID-19 Surveillance Team |
| EPI_ISL_3202349 | Berkshire and Surrey Pathology Services Lighthouse Laboratory | Wellcome Sanger Institute for the COVID-19 Genomics UK (COG-UK) Consortium | Berkshire and Surrey Pathology Services Lighthouse Laboratory and Alex Alderton, Roberto Amato, Jeffrey Barrett, Sonia Goncalves, Ewan Harrison, David K. Jackson, Ian Johnston, Dominic Kwiatkowski, Cordelia Langford, John Sillitoe on behalf of the Wellcome Sanger Institute COVID-19 Surveillance Team |
| EPI_ISL_3202350, EPI_ISL_3202351 | Lighthouse Lab in Alderley Park | Wellcome Sanger Institute for the COVID-19 Genomics UK (COG-UK) Consortium | Jacquelyn Wynn, Mairead Hyland, The Lighthouse Lab in Alderley Park and Alex Alderton, Roberto Amato, Jeffrey Barrett, Sonia Goncalves, Ewan Harrison, David K. Jackson, Ian Johnston, Dominic Kwiatkowski, Cordelia Langford, John Sillitoe on behalf of the Wellcome Sanger Institute COVID-19 Surveillance Team |
| EPI_ISL_3202352, EPI_ISL_3202353 | Berkshire and Surrey Pathology Services Lighthouse Laboratory | Wellcome Sanger Institute for the COVID-19 Genomics UK (COG-UK) Consortium | Berkshire and Surrey Pathology Services Lighthouse Laboratory and Alex Alderton, Roberto Amato, Jeffrey Barrett, Sonia Goncalves, Ewan Harrison, David K. Jackson, Ian Johnston, Dominic Kwiatkowski, Cordelia Langford, John Sillitoe on behalf of the Wellcome Sanger Institute COVID-19 Surveillance Team |
| EPI_ISL_3202354 | Lighthouse Lab in Alderley Park | Wellcome Sanger Institute for the COVID-19 Genomics UK (COG-UK) Consortium | Jacquelyn Wynn, Mairead Hyland, The Lighthouse Lab in Alderley Park and Alex Alderton, Roberto Amato, Jeffrey Barrett, Sonia Goncalves, Ewan Harrison, David K. Jackson, Ian Johnston, Dominic Kwiatkowski, Cordelia Langford, John Sillitoe on behalf of the Wellcome Sanger Institute COVID-19 Surveillance Team |
| EPI_ISL_3202355, EPI_ISL_3202356, EPI_ISL_3202357 | Berkshire and Surrey Pathology Services Lighthouse Laboratory | Wellcome Sanger Institute for the COVID-19 Genomics UK (COG-UK) Consortium | Berkshire and Surrey Pathology Services Lighthouse Laboratory and Alex Alderton, Roberto Amato, Jeffrey Barrett, Sonia Goncalves, Ewan Harrison, David K. Jackson, Ian Johnston, Dominic Kwiatkowski, Cordelia Langford, John Sillitoe on behalf of the Wellcome Sanger Institute COVID-19 Surveillance Team |
| EPI_ISL_3202358, EPI_ISL_3202359, EPI_ISL_3202360 | Lighthouse Lab in Alderley Park | Wellcome Sanger Institute for the COVID-19 Genomics UK (COG-UK) Consortium | Jacquelyn Wynn, Mairead Hyland, The Lighthouse Lab in Alderley Park and Alex Alderton, Roberto Amato, Jeffrey Barrett, Sonia Goncalves, Ewan Harrison, David K. Jackson, Ian Johnston, Dominic Kwiatkowski, Cordelia Langford, John Sillitoe on behalf of the Wellcome Sanger Institute COVID-19 Surveillance Team |
| EPI_ISL_3202361, EPI_ISL_3202362, EPI_ISL_3202363, EPI_ISL_3202364 | Berkshire and Surrey Pathology Services Lighthouse Laboratory | Wellcome Sanger Institute for the COVID-19 Genomics UK (COG-UK) Consortium | Berkshire and Surrey Pathology Services Lighthouse Laboratory and Alex Alderton, Roberto Amato, Jeffrey Barrett, Sonia Goncalves, Ewan Harrison, David K. Jackson, Ian Johnston, Dominic Kwiatkowski, Cordelia Langford, John Sillitoe on behalf of the Wellcome Sanger Institute COVID-19 Surveillance Team |
| EPI_ISL_3202365 | Lighthouse Lab in Alderley Park | Wellcome Sanger Institute for the COVID-19 Genomics UK (COG-UK) Consortium | Jacquelyn Wynn, Mairead Hyland, The Lighthouse Lab in Alderley Park and Alex Alderton, Roberto Amato, Jeffrey Barrett, Sonia Goncalves, Ewan Harrison, David K. Jackson, Ian Johnston, Dominic Kwiatkowski, Cordelia Langford, John Sillitoe on behalf of the Wellcome Sanger Institute COVID-19 Surveillance Team |
| EPI_ISL_3202366, EPI_ISL_3202368 | Berkshire and Surrey Pathology Services Lighthouse Laboratory | Wellcome Sanger Institute for the COVID-19 Genomics UK (COG-UK) Consortium | Berkshire and Surrey Pathology Services Lighthouse Laboratory and Alex Alderton, Roberto Amato, Jeffrey Barrett, Sonia Goncalves, Ewan Harrison, David K. Jackson, Ian Johnston, Dominic Kwiatkowski, Cordelia Langford, John Sillitoe on behalf of the Wellcome Sanger Institute COVID-19 Surveillance Team |
| EPI_ISL_3202369 | Lighthouse Lab in Alderley Park | Wellcome Sanger Institute for the COVID-19 Genomics UK (COG-UK) Consortium | Jacquelyn Wynn, Mairead Hyland, The Lighthouse Lab in Alderley Park and Alex Alderton, Roberto Amato, Jeffrey Barrett, Sonia Goncalves, Ewan Harrison, David K. Jackson, Ian Johnston, Dominic Kwiatkowski, Cordelia Langford, John Sillitoe on behalf of the Wellcome Sanger Institute COVID-19 Surveillance Team |
| EPI_ISL_3202370 | Berkshire and Surrey Pathology Services Lighthouse Laboratory | Wellcome Sanger Institute for the COVID-19 Genomics UK (COG-UK) Consortium | Berkshire and Surrey Pathology Services Lighthouse Laboratory and Alex Alderton, Roberto Amato, Jeffrey Barrett, Sonia Goncalves, Ewan Harrison, David K. Jackson, Ian Johnston, Dominic Kwiatkowski, Cordelia Langford, John Sillitoe on behalf of the Wellcome Sanger Institute COVID-19 Surveillance Team |
| EPI_ISL_3202371 | Lighthouse Lab in Alderley Park | Wellcome Sanger Institute for the COVID-19 Genomics UK (COG-UK) Consortium | Jacquelyn Wynn, Mairead Hyland, The Lighthouse Lab in Alderley Park and Alex Alderton, Roberto Amato, Jeffrey Barrett, Sonia Goncalves, Ewan Harrison, David K. Jackson, Ian Johnston, Dominic Kwiatkowski, Cordelia Langford, John Sillitoe on behalf of the Wellcome Sanger Institute COVID-19 Surveillance Team |
| EPI_ISL_3202372, EPI_ISL_3202373, EPI_ISL_3202374, EPI_ISL_3202375, EPI_ISL_3202376 | Berkshire and Surrey Pathology Services Lighthouse Laboratory | Wellcome Sanger Institute for the COVID-19 Genomics UK (COG-UK) Consortium | Berkshire and Surrey Pathology Services Lighthouse Laboratory and Alex Alderton, Roberto Amato, Jeffrey Barrett, Sonia Goncalves, Ewan Harrison, David K. Jackson, Ian Johnston, Dominic Kwiatkowski, Cordelia Langford, John Sillitoe on behalf of the Wellcome Sanger Institute COVID-19 Surveillance Team |
| EPI_ISL_3202377, EPI_ISL_3202378 | Lighthouse Lab in Alderley Park | Wellcome Sanger Institute for the COVID-19 Genomics UK (COG-UK) Consortium | Jacquelyn Wynn, Mairead Hyland, The Lighthouse Lab in Alderley Park and Alex Alderton, Roberto Amato, Jeffrey Barrett, Sonia Goncalves, Ewan Harrison, David K. Jackson, Ian Johnston, Dominic Kwiatkowski, Cordelia Langford, John Sillitoe on behalf of the Wellcome Sanger Institute COVID-19 Surveillance Team |
| EPI_ISL_3202379, EPI_ISL_3202380 | Berkshire and Surrey Pathology Services Lighthouse Laboratory | Wellcome Sanger Institute for the COVID-19 Genomics UK (COG-UK) Consortium | Berkshire and Surrey Pathology Services Lighthouse Laboratory and Alex Alderton, Roberto Amato, Jeffrey Barrett, Sonia Goncalves, Ewan Harrison, David K. Jackson, Ian Johnston, Dominic Kwiatkowski, Cordelia Langford, John Sillitoe on behalf of the Wellcome Sanger Institute COVID-19 Surveillance Team |
| EPI_ISL_3202381 | Lighthouse Lab in Alderley Park | Wellcome Sanger Institute for the COVID-19 Genomics UK (COG-UK) Consortium | Jacquelyn Wynn, Mairead Hyland, The Lighthouse Lab in Alderley Park and Alex Alderton, Roberto Amato, Jeffrey Barrett, Sonia Goncalves, Ewan Harrison, David K. Jackson, Ian Johnston, Dominic Kwiatkowski, Cordelia Langford, John Sillitoe on behalf of the Wellcome Sanger Institute COVID-19 Surveillance Team |
| EPI_ISL_3202382 | Berkshire and Surrey Pathology Services Lighthouse Laboratory | Wellcome Sanger Institute for the COVID-19 Genomics UK (COG-UK) Consortium | Berkshire and Surrey Pathology Services Lighthouse Laboratory and Alex Alderton, Roberto Amato, Jeffrey Barrett, Sonia Goncalves, Ewan Harrison, David K. Jackson, Ian Johnston, Dominic Kwiatkowski, Cordelia Langford, John Sillitoe on behalf of the Wellcome Sanger Institute COVID-19 Surveillance Team |
| EPI_ISL_3202383, EPI_ISL_3202384, EPI_ISL_3202385 | Lighthouse Lab in Alderley Park | Wellcome Sanger Institute for the COVID-19 Genomics UK (COG-UK) Consortium | Jacquelyn Wynn, Mairead Hyland, The Lighthouse Lab in Alderley Park and Alex Alderton, Roberto Amato, Jeffrey Barrett, Sonia Goncalves, Ewan Harrison, David K. Jackson, Ian Johnston, Dominic Kwiatkowski, Cordelia Langford, John Sillitoe on behalf of the Wellcome Sanger Institute COVID-19 Surveillance Team |
| EPI_ISL_3202386 | Berkshire and Surrey Pathology Services Lighthouse Laboratory | Wellcome Sanger Institute for the COVID-19 Genomics UK (COG-UK) Consortium | Berkshire and Surrey Pathology Services Lighthouse Laboratory and Alex Alderton, Roberto Amato, Jeffrey Barrett, Sonia Goncalves, Ewan Harrison, David K. Jackson, Ian Johnston, Dominic Kwiatkowski, Cordelia Langford, John Sillitoe on behalf of the Wellcome Sanger Institute COVID-19 Surveillance Team |
| EPI_ISL_3202387, EPI_ISL_3202388, EPI_ISL_3202389 | Lighthouse Lab in Alderley Park | Wellcome Sanger Institute for the COVID-19 Genomics UK (COG-UK) Consortium | Jacquelyn Wynn, Mairead Hyland, The Lighthouse Lab in Alderley Park and Alex Alderton, Roberto Amato, Jeffrey Barrett, Sonia Goncalves, Ewan Harrison, David K. Jackson, Ian Johnston, Dominic Kwiatkowski, Cordelia Langford, John Sillitoe on behalf of the Wellcome Sanger Institute COVID-19 Surveillance Team |
| EPI_ISL_3202390 | Berkshire and Surrey Pathology Services Lighthouse Laboratory | Wellcome Sanger Institute for the COVID-19 Genomics UK (COG-UK) Consortium | Berkshire and Surrey Pathology Services Lighthouse Laboratory and Alex Alderton, Roberto Amato, Jeffrey Barrett, Sonia Goncalves, Ewan Harrison, David K. Jackson, Ian Johnston, Dominic Kwiatkowski, Cordelia Langford, John Sillitoe on behalf of the Wellcome Sanger Institute COVID-19 Surveillance Team |
| EPI_ISL_3202391, EPI_ISL_3202392, EPI_ISL_3202393, EPI_ISL_3202394 | Lighthouse Lab in Alderley Park | Wellcome Sanger Institute for the COVID-19 Genomics UK (COG-UK) Consortium | Jacquelyn Wynn, Mairead Hyland, The Lighthouse Lab in Alderley Park and Alex Alderton, Roberto Amato, Jeffrey Barrett, Sonia Goncalves, Ewan Harrison, David K. Jackson, Ian Johnston, Dominic Kwiatkowski, Cordelia Langford, John Sillitoe on behalf of the Wellcome Sanger Institute COVID-19 Surveillance Team |
| EPI_ISL_3202395, EPI_ISL_3202396, EPI_ISL_3202397 | Berkshire and Surrey Pathology Services Lighthouse Laboratory | Wellcome Sanger Institute for the COVID-19 Genomics UK (COG-UK) Consortium | Berkshire and Surrey Pathology Services Lighthouse Laboratory and Alex Alderton, Roberto Amato, Jeffrey Barrett, Sonia Goncalves, Ewan Harrison, David K. Jackson, Ian Johnston, Dominic Kwiatkowski, Cordelia Langford, John Sillitoe on behalf of the Wellcome Sanger Institute COVID-19 Surveillance Team |
| EPI_ISL_3202398, EPI_ISL_3202399, EPI_ISL_3202400 | Lighthouse Lab in Alderley Park | Wellcome Sanger Institute for the COVID-19 Genomics UK (COG-UK) Consortium | Jacquelyn Wynn, Mairead Hyland, The Lighthouse Lab in Alderley Park and Alex Alderton, Roberto Amato, Jeffrey Barrett, Sonia Goncalves, Ewan Harrison, David K. Jackson, Ian Johnston, Dominic Kwiatkowski, Cordelia Langford, John Sillitoe on behalf of the Wellcome Sanger Institute COVID-19 Surveillance Team |
| EPI_ISL_3202401 | Berkshire and Surrey Pathology Services Lighthouse Laboratory | Wellcome Sanger Institute for the COVID-19 Genomics UK (COG-UK) Consortium | Berkshire and Surrey Pathology Services Lighthouse Laboratory and Alex Alderton, Roberto Amato, Jeffrey Barrett, Sonia Goncalves, Ewan Harrison, David K. Jackson, Ian Johnston, Dominic Kwiatkowski, Cordelia Langford, John Sillitoe on behalf of the Wellcome Sanger Institute COVID-19 Surveillance Team |
| EPI_ISL_3202402, EPI_ISL_3202403 | Lighthouse Lab in Alderley Park | Wellcome Sanger Institute for the COVID-19 Genomics UK (COG-UK) Consortium | Jacquelyn Wynn, Mairead Hyland, The Lighthouse Lab in Alderley Park and Alex Alderton, Roberto Amato, Jeffrey Barrett, Sonia Goncalves, Ewan Harrison, David K. Jackson, Ian Johnston, Dominic Kwiatkowski, Cordelia Langford, John Sillitoe on behalf of the Wellcome Sanger Institute COVID-19 Surveillance Team |
| EPI_ISL_3202404, EPI_ISL_3202405, | Berkshire and Surrey Pathology Services Lighthouse Laboratory | Wellcome Sanger Institute for the COVID-19 Genomics UK (COG-UK) Consortium | Berkshire and Surrey Pathology Services Lighthouse Laboratory and Alex Alderton, Roberto Amato, Jeffrey Barrett, Sonia Goncalves, Ewan Harrison, David K. Jackson, Ian Johnston, Dominic Kwiatkowski, Cordelia Langford, John Sillitoe on behalf of the Wellcome Sanger Institute COVID-19 Surveillance Team |

| | | | |
|---|---|---|---|
| EPI_ISL_3202406, EPI_ISL_3202407, EPI_ISL_3202408 | | | |
| EPI_ISL_3202409, EPI_ISL_3202410 | Lighthouse Lab in Alderley Park | Wellcome Sanger Institute for the COVID-19 Genomics UK (COG-UK) Consortium | Jacquelyn Wynn, Mairead Hyland, The Lighthouse Lab in Alderley Park and Alex Alderton, Roberto Amato, Jeffrey Barrett, Sonia Goncalves, Ewan Harrison, David K. Jackson, Ian Johnston, Dominic Kwiatkowski, Cordelia Langford, John Sillitoe on behalf of the Wellcome Sanger Institute COVID-19 Surveillance Team |
| EPI_ISL_3202411, EPI_ISL_3202412, EPI_ISL_3202413 | Berkshire and Surrey Pathology Services Lighthouse Laboratory | Wellcome Sanger Institute for the COVID-19 Genomics UK (COG-UK) Consortium | Berkshire and Surrey Pathology Services Lighthouse Laboratory and Alex Alderton, Roberto Amato, Jeffrey Barrett, Sonia Goncalves, Ewan Harrison, David K. Jackson, Ian Johnston, Dominic Kwiatkowski, Cordelia Langford, John Sillitoe on behalf of the Wellcome Sanger Institute COVID-19 Surveillance Team |
| EPI_ISL_3202414, EPI_ISL_3202415 | Lighthouse Lab in Alderley Park | Wellcome Sanger Institute for the COVID-19 Genomics UK (COG-UK) Consortium | Jacquelyn Wynn, Mairead Hyland, The Lighthouse Lab in Alderley Park and Alex Alderton, Roberto Amato, Jeffrey Barrett, Sonia Goncalves, Ewan Harrison, David K. Jackson, Ian Johnston, Dominic Kwiatkowski, Cordelia Langford, John Sillitoe on behalf of the Wellcome Sanger Institute COVID-19 Surveillance Team |
| EPI_ISL_3202416, EPI_ISL_3202417 | Berkshire and Surrey Pathology Services Lighthouse Laboratory | Wellcome Sanger Institute for the COVID-19 Genomics UK (COG-UK) Consortium | Berkshire and Surrey Pathology Services Lighthouse Laboratory and Alex Alderton, Roberto Amato, Jeffrey Barrett, Sonia Goncalves, Ewan Harrison, David K. Jackson, Ian Johnston, Dominic Kwiatkowski, Cordelia Langford, John Sillitoe on behalf of the Wellcome Sanger Institute COVID-19 Surveillance Team |
| EPI_ISL_3202418, EPI_ISL_3202419, EPI_ISL_3202420, EPI_ISL_3202421 | Lighthouse Lab in Alderley Park | Wellcome Sanger Institute for the COVID-19 Genomics UK (COG-UK) Consortium | Jacquelyn Wynn, Mairead Hyland, The Lighthouse Lab in Alderley Park and Alex Alderton, Roberto Amato, Jeffrey Barrett, Sonia Goncalves, Ewan Harrison, David K. Jackson, Ian Johnston, Dominic Kwiatkowski, Cordelia Langford, John Sillitoe on behalf of the Wellcome Sanger Institute COVID-19 Surveillance Team |
| EPI_ISL_3202422, EPI_ISL_3202423, EPI_ISL_3202424 | Berkshire and Surrey Pathology Services Lighthouse Laboratory | Wellcome Sanger Institute for the COVID-19 Genomics UK (COG-UK) Consortium | Berkshire and Surrey Pathology Services Lighthouse Laboratory and Alex Alderton, Roberto Amato, Jeffrey Barrett, Sonia Goncalves, Ewan Harrison, David K. Jackson, Ian Johnston, Dominic Kwiatkowski, Cordelia Langford, John Sillitoe on behalf of the Wellcome Sanger Institute COVID-19 Surveillance Team |
| EPI_ISL_3202425, EPI_ISL_3202426, EPI_ISL_3202427, EPI_ISL_3202428, EPI_ISL_3202429, EPI_ISL_3202430 | Lighthouse Lab in Alderley Park | Wellcome Sanger Institute for the COVID-19 Genomics UK (COG-UK) Consortium | Jacquelyn Wynn, Mairead Hyland, The Lighthouse Lab in Alderley Park and Alex Alderton, Roberto Amato, Jeffrey Barrett, Sonia Goncalves, Ewan Harrison, David K. Jackson, Ian Johnston, Dominic Kwiatkowski, Cordelia Langford, John Sillitoe on behalf of the Wellcome Sanger Institute COVID-19 Surveillance Team |
| EPI_ISL_3202431 | Health Services Laboratories | Wellcome Sanger Institute for the COVID-19 Genomics UK (COG-UK) Consortium | Health Services Laboratories and Alex Alderton, Roberto Amato, Jeffrey Barrett, Sonia Goncalves, Ewan Harrison, David K. Jackson, Ian Johnston, Dominic Kwiatkowski, Cordelia Langford, John Sillitoe on behalf of the Wellcome Sanger Institute COVID-19 Surveillance Team |
| EPI_ISL_3202432 | Lighthouse Lab in Alderley Park | Wellcome Sanger Institute for the COVID-19 Genomics UK (COG-UK) Consortium | Jacquelyn Wynn, Mairead Hyland, The Lighthouse Lab in Alderley Park and Alex Alderton, Roberto Amato, Jeffrey Barrett, Sonia Goncalves, Ewan Harrison, David K. Jackson, Ian Johnston, Dominic Kwiatkowski, Cordelia Langford, John Sillitoe on behalf of the Wellcome Sanger Institute COVID-19 Surveillance Team |
| EPI_ISL_3202433 | Health Services Laboratories | Wellcome Sanger Institute for the COVID-19 Genomics UK (COG-UK) Consortium | Health Services Laboratories and Alex Alderton, Roberto Amato, Jeffrey Barrett, Sonia Goncalves, Ewan Harrison, David K. Jackson, Ian Johnston, Dominic Kwiatkowski, Cordelia Langford, John Sillitoe on behalf of the Wellcome Sanger Institute COVID-19 Surveillance Team |
| EPI_ISL_3202434 | Lighthouse Lab in Alderley Park | Wellcome Sanger Institute for the COVID-19 Genomics UK (COG-UK) Consortium | Jacquelyn Wynn, Mairead Hyland, The Lighthouse Lab in Alderley Park and Alex Alderton, Roberto Amato, Jeffrey Barrett, Sonia Goncalves, Ewan Harrison, David K. Jackson, Ian Johnston, Dominic Kwiatkowski, Cordelia Langford, John Sillitoe on behalf of the Wellcome Sanger Institute COVID-19 Surveillance Team |
| EPI_ISL_3202435 | Health Services Laboratories | Wellcome Sanger Institute for the COVID-19 Genomics UK (COG-UK) Consortium | Health Services Laboratories and Alex Alderton, Roberto Amato, Jeffrey Barrett, Sonia Goncalves, Ewan Harrison, David K. Jackson, Ian Johnston, Dominic Kwiatkowski, Cordelia Langford, John Sillitoe on behalf of the Wellcome Sanger Institute COVID-19 Surveillance Team |
| EPI_ISL_3202436 | Berkshire and Surrey Pathology Services Lighthouse Laboratory | Wellcome Sanger Institute for the COVID-19 Genomics UK (COG-UK) Consortium | Berkshire and Surrey Pathology Services Lighthouse Laboratory and Alex Alderton, Roberto Amato, Jeffrey Barrett, Sonia Goncalves, Ewan Harrison, David K. Jackson, Ian Johnston, Dominic Kwiatkowski, Cordelia Langford, John Sillitoe on behalf of the Wellcome Sanger Institute COVID-19 Surveillance Team |
| EPI_ISL_3202437 | Health Services Laboratories | Wellcome Sanger Institute for the COVID-19 Genomics UK (COG-UK) Consortium | Health Services Laboratories and Alex Alderton, Roberto Amato, Jeffrey Barrett, Sonia Goncalves, Ewan Harrison, David K. Jackson, Ian Johnston, Dominic Kwiatkowski, Cordelia Langford, John Sillitoe on behalf of the Wellcome Sanger Institute COVID-19 Surveillance Team |
| EPI_ISL_3202438 | Lighthouse Lab in Alderley Park | Wellcome Sanger Institute for the COVID-19 Genomics UK (COG-UK) Consortium | Jacquelyn Wynn, Mairead Hyland, The Lighthouse Lab in Alderley Park and Alex Alderton, Roberto Amato, Jeffrey Barrett, Sonia Goncalves, Ewan Harrison, David K. Jackson, Ian Johnston, Dominic Kwiatkowski, Cordelia Langford, John Sillitoe on behalf of the Wellcome Sanger Institute COVID-19 Surveillance Team |
| EPI_ISL_3202439 | Health Services Laboratories | Wellcome Sanger Institute for the COVID-19 Genomics UK (COG-UK) Consortium | Health Services Laboratories and Alex Alderton, Roberto Amato, Jeffrey Barrett, Sonia Goncalves, Ewan Harrison, David K. Jackson, Ian Johnston, Dominic Kwiatkowski, Cordelia Langford, John Sillitoe on behalf of the Wellcome Sanger Institute COVID-19 Surveillance Team |
| EPI_ISL_3202440 | Lighthouse Lab in Alderley Park | Wellcome Sanger Institute for the COVID-19 Genomics UK (COG-UK) Consortium | Jacquelyn Wynn, Mairead Hyland, The Lighthouse Lab in Alderley Park and Alex Alderton, Roberto Amato, Jeffrey Barrett, Sonia Goncalves, Ewan Harrison, David K. Jackson, Ian Johnston, Dominic Kwiatkowski, Cordelia Langford, John Sillitoe on behalf of the Wellcome Sanger Institute COVID-19 Surveillance Team |
| EPI_ISL_3202441 | Health Services Laboratories | Wellcome Sanger Institute for the COVID-19 Genomics UK (COG-UK) Consortium | Health Services Laboratories and Alex Alderton, Roberto Amato, Jeffrey Barrett, Sonia Goncalves, Ewan Harrison, David K. Jackson, Ian Johnston, Dominic Kwiatkowski, Cordelia Langford, John Sillitoe on behalf of the Wellcome Sanger Institute COVID-19 Surveillance Team |
| EPI_ISL_3202442, EPI_ISL_3202443, EPI_ISL_3202444, EPI_ISL_3202445 | Lighthouse Lab in Alderley Park | Wellcome Sanger Institute for the COVID-19 Genomics UK (COG-UK) Consortium | Jacquelyn Wynn, Mairead Hyland, The Lighthouse Lab in Alderley Park and Alex Alderton, Roberto Amato, Jeffrey Barrett, Sonia Goncalves, Ewan Harrison, David K. Jackson, Ian Johnston, Dominic Kwiatkowski, Cordelia Langford, John Sillitoe on behalf of the Wellcome Sanger Institute COVID-19 Surveillance Team |
| EPI_ISL_3202446 | Berkshire and Surrey Pathology Services Lighthouse Laboratory | Wellcome Sanger Institute for the COVID-19 Genomics UK (COG-UK) Consortium | Berkshire and Surrey Pathology Services Lighthouse Laboratory and Alex Alderton, Roberto Amato, Jeffrey Barrett, Sonia Goncalves, Ewan Harrison, David K. Jackson, Ian Johnston, Dominic Kwiatkowski, Cordelia Langford, John Sillitoe on behalf of the Wellcome Sanger Institute COVID-19 Surveillance Team |
| EPI_ISL_3202447, EPI_ISL_3202448 | Lighthouse Lab in Alderley Park | Wellcome Sanger Institute for the COVID-19 Genomics UK (COG-UK) Consortium | Jacquelyn Wynn, Mairead Hyland, The Lighthouse Lab in Alderley Park and Alex Alderton, Roberto Amato, Jeffrey Barrett, Sonia Goncalves, Ewan Harrison, David K. Jackson, Ian Johnston, Dominic Kwiatkowski, Cordelia Langford, John Sillitoe on behalf of the Wellcome Sanger Institute COVID-19 Surveillance Team |
| EPI_ISL_3202449 | Health Services Laboratories | Wellcome Sanger Institute for the COVID-19 Genomics UK (COG-UK) Consortium | Health Services Laboratories and Alex Alderton, Roberto Amato, Jeffrey Barrett, Sonia Goncalves, Ewan Harrison, David K. Jackson, Ian Johnston, Dominic Kwiatkowski, Cordelia Langford, John Sillitoe on behalf of the Wellcome Sanger Institute COVID-19 Surveillance Team |
| EPI_ISL_3202450, EPI_ISL_3202451 | Lighthouse Lab in Alderley Park | Wellcome Sanger Institute for the COVID-19 Genomics UK (COG-UK) Consortium | Jacquelyn Wynn, Mairead Hyland, The Lighthouse Lab in Alderley Park and Alex Alderton, Roberto Amato, Jeffrey Barrett, Sonia Goncalves, Ewan Harrison, David K. Jackson, Ian Johnston, Dominic Kwiatkowski, Cordelia Langford, John Sillitoe on behalf of the Wellcome Sanger Institute COVID-19 Surveillance Team |
| EPI_ISL_3202452 | Berkshire and Surrey Pathology Services Lighthouse Laboratory | Wellcome Sanger Institute for the COVID-19 Genomics UK (COG-UK) Consortium | Berkshire and Surrey Pathology Services Lighthouse Laboratory and Alex Alderton, Roberto Amato, Jeffrey Barrett, Sonia Goncalves, Ewan Harrison, David K. Jackson, Ian Johnston, Dominic Kwiatkowski, Cordelia Langford, John Sillitoe on behalf of the Wellcome Sanger Institute COVID-19 Surveillance Team |
| EPI_ISL_3202453 | Health Services Laboratories | Wellcome Sanger Institute for the COVID-19 Genomics UK (COG-UK) Consortium | Health Services Laboratories and Alex Alderton, Roberto Amato, Jeffrey Barrett, Sonia Goncalves, Ewan Harrison, David K. Jackson, Ian Johnston, Dominic Kwiatkowski, Cordelia Langford, John Sillitoe on behalf of the Wellcome Sanger Institute COVID-19 Surveillance Team |
| EPI_ISL_3202454, EPI_ISL_3202455 | Lighthouse Lab in Alderley Park | Wellcome Sanger Institute for the COVID-19 Genomics UK (COG-UK) Consortium | Jacquelyn Wynn, Mairead Hyland, The Lighthouse Lab in Alderley Park and Alex Alderton, Roberto Amato, Jeffrey Barrett, Sonia Goncalves, Ewan Harrison, David K. Jackson, Ian Johnston, Dominic Kwiatkowski, Cordelia Langford, John Sillitoe on behalf of the Wellcome Sanger Institute COVID-19 Surveillance Team |
| EPI_ISL_3202456 | Health Services Laboratories | Wellcome Sanger Institute for the COVID-19 Genomics UK (COG-UK) Consortium | Health Services Laboratories and Alex Alderton, Roberto Amato, Jeffrey Barrett, Sonia Goncalves, Ewan Harrison, David K. Jackson, Ian Johnston, Dominic Kwiatkowski, Cordelia Langford, John Sillitoe on behalf of the Wellcome Sanger Institute COVID-19 Surveillance Team |
| EPI_ISL_3202457, EPI_ISL_3202458, EPI_ISL_3202459, EPI_ISL_3202460 | Lighthouse Lab in Alderley Park | Wellcome Sanger Institute for the COVID-19 Genomics UK (COG-UK) Consortium | Jacquelyn Wynn, Mairead Hyland, The Lighthouse Lab in Alderley Park and Alex Alderton, Roberto Amato, Jeffrey Barrett, Sonia Goncalves, Ewan Harrison, David K. Jackson, Ian Johnston, Dominic Kwiatkowski, Cordelia Langford, John Sillitoe on behalf of the Wellcome Sanger Institute COVID-19 Surveillance Team |
| EPI_ISL_3202461 | Health Services Laboratories | Wellcome Sanger Institute for the COVID-19 Genomics UK (COG-UK) Consortium | Health Services Laboratories and Alex Alderton, Roberto Amato, Jeffrey Barrett, Sonia Goncalves, Ewan Harrison, David K. Jackson, Ian Johnston, Dominic Kwiatkowski, Cordelia Langford, John Sillitoe on behalf of the Wellcome Sanger Institute COVID-19 Surveillance Team |
| EPI_ISL_3202462 | Berkshire and Surrey Pathology Services Lighthouse Laboratory | Wellcome Sanger Institute for the COVID-19 Genomics UK (COG-UK) Consortium | Berkshire and Surrey Pathology Services Lighthouse Laboratory and Alex Alderton, Roberto Amato, Jeffrey Barrett, Sonia Goncalves, Ewan Harrison, David K. Jackson, Ian Johnston, Dominic Kwiatkowski, Cordelia Langford, John Sillitoe on behalf of the Wellcome Sanger Institute COVID-19 Surveillance Team |
| EPI_ISL_3202463 | Lighthouse Lab in Alderley Park | Wellcome Sanger Institute for the COVID-19 Genomics UK (COG-UK) Consortium | Jacquelyn Wynn, Mairead Hyland, The Lighthouse Lab in Alderley Park and Alex Alderton, Roberto Amato, Jeffrey Barrett, Sonia Goncalves, Ewan Harrison, David K. Jackson, Ian Johnston, Dominic Kwiatkowski, Cordelia Langford, John Sillitoe on behalf of the Wellcome Sanger Institute COVID-19 Surveillance Team |
| EPI_ISL_3202464, EPI_ISL_3202465 | Health Services Laboratories | Wellcome Sanger Institute for the COVID-19 Genomics UK (COG-UK) Consortium | Health Services Laboratories and Alex Alderton, Roberto Amato, Jeffrey Barrett, Sonia Goncalves, Ewan Harrison, David K. Jackson, Ian Johnston, Dominic Kwiatkowski, Cordelia Langford, John Sillitoe on behalf of the Wellcome Sanger Institute COVID-19 Surveillance Team |
| EPI_ISL_3202466 | Lighthouse Lab in Alderley Park | Wellcome Sanger Institute for the COVID-19 Genomics UK (COG-UK) Consortium | Jacquelyn Wynn, Mairead Hyland, The Lighthouse Lab in Alderley Park and Alex Alderton, Roberto Amato, Jeffrey Barrett, Sonia Goncalves, Ewan Harrison, David K. Jackson, Ian Johnston, Dominic Kwiatkowski, Cordelia Langford, John Sillitoe on behalf of the Wellcome Sanger Institute COVID-19 Surveillance Team |
| EPI_ISL_3202467 | Berkshire and Surrey Pathology Services Lighthouse Laboratory | Wellcome Sanger Institute for the COVID-19 Genomics UK (COG-UK) Consortium | Berkshire and Surrey Pathology Services Lighthouse Laboratory and Alex Alderton, Roberto Amato, Jeffrey Barrett, Sonia Goncalves, Ewan Harrison, David K. Jackson, Ian Johnston, Dominic Kwiatkowski, Cordelia Langford, John Sillitoe on behalf of the Wellcome Sanger Institute COVID-19 Surveillance Team |
| EPI_ISL_3202468 | Lighthouse Lab in Alderley Park | Wellcome Sanger Institute for the COVID-19 Genomics UK (COG-UK) Consortium | Jacquelyn Wynn, Mairead Hyland, The Lighthouse Lab in Alderley Park and Alex Alderton, Roberto Amato, Jeffrey Barrett, Sonia Goncalves, Ewan Harrison, David K. Jackson, Ian Johnston, Dominic Kwiatkowski, Cordelia Langford, John Sillitoe on behalf of the Wellcome Sanger Institute COVID-19 Surveillance Team |
| EPI_ISL_3202469, EPI_ISL_3202470 | Berkshire and Surrey Pathology Services Lighthouse Laboratory | Wellcome Sanger Institute for the COVID-19 Genomics UK (COG-UK) Consortium | Berkshire and Surrey Pathology Services Lighthouse Laboratory and Alex Alderton, Roberto Amato, Jeffrey Barrett, Sonia Goncalves, Ewan Harrison, David K. Jackson, Ian Johnston, Dominic Kwiatkowski, Cordelia Langford, John Sillitoe on behalf of the Wellcome Sanger Institute COVID-19 Surveillance Team |
| EPI_ISL_3202471 | Health Services Laboratories | Wellcome Sanger Institute for the COVID-19 Genomics UK (COG-UK) Consortium | Health Services Laboratories and Alex Alderton, Roberto Amato, Jeffrey Barrett, Sonia Goncalves, Ewan Harrison, David K. Jackson, Ian Johnston, Dominic Kwiatkowski, Cordelia Langford, John Sillitoe on behalf of the Wellcome Sanger Institute COVID-19 Surveillance Team |
| EPI_ISL_3202472 | Lighthouse Lab in Alderley Park | Wellcome Sanger Institute for the COVID-19 Genomics UK (COG-UK) Consortium | Jacquelyn Wynn, Mairead Hyland, The Lighthouse Lab in Alderley Park and Alex Alderton, Roberto Amato, Jeffrey Barrett, Sonia Goncalves, Ewan Harrison, David K. Jackson, Ian Johnston, Dominic Kwiatkowski, Cordelia Langford, John Sillitoe on behalf of the Wellcome Sanger Institute COVID-19 Surveillance Team |
| EPI_ISL_3202473, EPI_ISL_3202474, | Health Services Laboratories | Wellcome Sanger Institute for the COVID-19 Genomics UK (COG-UK) Consortium | Health Services Laboratories and Alex Alderton, Roberto Amato, Jeffrey Barrett, Sonia Goncalves, Ewan Harrison, David K. Jackson, Ian Johnston, Dominic Kwiatkowski, Cordelia Langford, John Sillitoe on behalf of the Wellcome Sanger Institute COVID-19 Surveillance Team |

| | | | |
|---|---|---|---|
| EPI_ISL_3202475 | | | |
| EPI_ISL_3202476, EPI_ISL_3202477, EPI_ISL_3202478, EPI_ISL_3202479, EPI_ISL_3202480, EPI_ISL_3202481 | Lighthouse Lab in Alderley Park | Wellcome Sanger Institute for the COVID-19 Genomics UK (COG-UK) Consortium | Jacquelyn Wynn, Mairead Hyland, The Lighthouse Lab in Alderley Park and Alex Alderton, Roberto Amato, Jeffrey Barrett, Sonia Goncalves, Ewan Harrison, David K. Jackson, Ian Johnston, Dominic Kwiatkowski, Cordelia Langford, John Sillitoe on behalf of the Wellcome Sanger Institute COVID-19 Surveillance Team |
| EPI_ISL_3202482 | Berkshire and Surrey Pathology Services Lighthouse Laboratory | Wellcome Sanger Institute for the COVID-19 Genomics UK (COG-UK) Consortium | Berkshire and Surrey Pathology Services Lighthouse Laboratory and Alex Alderton, Roberto Amato, Jeffrey Barrett, Sonia Goncalves, Ewan Harrison, David K. Jackson, Ian Johnston, Dominic Kwiatkowski, Cordelia Langford, John Sillitoe on behalf of the Wellcome Sanger Institute COVID-19 Surveillance Team |
| EPI_ISL_3202483 | Health Services Laboratories | Wellcome Sanger Institute for the COVID-19 Genomics UK (COG-UK) Consortium | Health Services Laboratories and Alex Alderton, Roberto Amato, Jeffrey Barrett, Sonia Goncalves, Ewan Harrison, David K. Jackson, Ian Johnston, Dominic Kwiatkowski, Cordelia Langford, John Sillitoe on behalf of the Wellcome Sanger Institute COVID-19 Surveillance Team |
| EPI_ISL_3202484 | Berkshire and Surrey Pathology Services Lighthouse Laboratory | Wellcome Sanger Institute for the COVID-19 Genomics UK (COG-UK) Consortium | Berkshire and Surrey Pathology Services Lighthouse Laboratory and Alex Alderton, Roberto Amato, Jeffrey Barrett, Sonia Goncalves, Ewan Harrison, David K. Jackson, Ian Johnston, Dominic Kwiatkowski, Cordelia Langford, John Sillitoe on behalf of the Wellcome Sanger Institute COVID-19 Surveillance Team |
| EPI_ISL_3202485, EPI_ISL_3202486, EPI_ISL_3202487 | Lighthouse Lab in Alderley Park | Wellcome Sanger Institute for the COVID-19 Genomics UK (COG-UK) Consortium | Jacquelyn Wynn, Mairead Hyland, The Lighthouse Lab in Alderley Park and Alex Alderton, Roberto Amato, Jeffrey Barrett, Sonia Goncalves, Ewan Harrison, David K. Jackson, Ian Johnston, Dominic Kwiatkowski, Cordelia Langford, John Sillitoe on behalf of the Wellcome Sanger Institute COVID-19 Surveillance Team |
| EPI_ISL_3202488, EPI_ISL_3202489, EPI_ISL_3202490 | Health Services Laboratories | Wellcome Sanger Institute for the COVID-19 Genomics UK (COG-UK) Consortium | Health Services Laboratories and Alex Alderton, Roberto Amato, Jeffrey Barrett, Sonia Goncalves, Ewan Harrison, David K. Jackson, Ian Johnston, Dominic Kwiatkowski, Cordelia Langford, John Sillitoe on behalf of the Wellcome Sanger Institute COVID-19 Surveillance Team |
| EPI_ISL_3202491 | Berkshire and Surrey Pathology Services Lighthouse Laboratory | Wellcome Sanger Institute for the COVID-19 Genomics UK (COG-UK) Consortium | Berkshire and Surrey Pathology Services Lighthouse Laboratory and Alex Alderton, Roberto Amato, Jeffrey Barrett, Sonia Goncalves, Ewan Harrison, David K. Jackson, Ian Johnston, Dominic Kwiatkowski, Cordelia Langford, John Sillitoe on behalf of the Wellcome Sanger Institute COVID-19 Surveillance Team |
| EPI_ISL_3202492 | Health Services Laboratories | Wellcome Sanger Institute for the COVID-19 Genomics UK (COG-UK) Consortium | Health Services Laboratories and Alex Alderton, Roberto Amato, Jeffrey Barrett, Sonia Goncalves, Ewan Harrison, David K. Jackson, Ian Johnston, Dominic Kwiatkowski, Cordelia Langford, John Sillitoe on behalf of the Wellcome Sanger Institute COVID-19 Surveillance Team |
| EPI_ISL_3202493 | Berkshire and Surrey Pathology Services Lighthouse Laboratory | Wellcome Sanger Institute for the COVID-19 Genomics UK (COG-UK) Consortium | Berkshire and Surrey Pathology Services Lighthouse Laboratory and Alex Alderton, Roberto Amato, Jeffrey Barrett, Sonia Goncalves, Ewan Harrison, David K. Jackson, Ian Johnston, Dominic Kwiatkowski, Cordelia Langford, John Sillitoe on behalf of the Wellcome Sanger Institute COVID-19 Surveillance Team |
| EPI_ISL_3202494 | Lighthouse Lab in Glasgow | Wellcome Sanger Institute for the COVID-19 Genomics UK (COG-UK) Consortium | Harper VanSteenhouse, Yumi Kasai, David Gray, Carol Clugston, Anna Dominiczak and Alex Alderton, Roberto Amato, Jeffrey Barrett, Sonia Goncalves, Ewan Harrison, David K. Jackson, Ian Johnston, Dominic Kwiatkowski, Cordelia Langford, John Sillitoe on behalf of the Wellcome Sanger Institute COVID-19 Surveillance Team |
| EPI_ISL_3202495, EPI_ISL_3202496, EPI_ISL_3202497 | Lighthouse Lab in Alderley Park | Wellcome Sanger Institute for the COVID-19 Genomics UK (COG-UK) Consortium | Jacquelyn Wynn, Mairead Hyland, The Lighthouse Lab in Alderley Park and Alex Alderton, Roberto Amato, Jeffrey Barrett, Sonia Goncalves, Ewan Harrison, David K. Jackson, Ian Johnston, Dominic Kwiatkowski, Cordelia Langford, John Sillitoe on behalf of the Wellcome Sanger Institute COVID-19 Surveillance Team |
| EPI_ISL_3202498, EPI_ISL_3202499, EPI_ISL_3202500 | Berkshire and Surrey Pathology Services Lighthouse Laboratory | Wellcome Sanger Institute for the COVID-19 Genomics UK (COG-UK) Consortium | Berkshire and Surrey Pathology Services Lighthouse Laboratory and Alex Alderton, Roberto Amato, Jeffrey Barrett, Sonia Goncalves, Ewan Harrison, David K. Jackson, Ian Johnston, Dominic Kwiatkowski, Cordelia Langford, John Sillitoe on behalf of the Wellcome Sanger Institute COVID-19 Surveillance Team |
| EPI_ISL_3202501 | Lighthouse Lab in Alderley Park | Wellcome Sanger Institute for the COVID-19 Genomics UK (COG-UK) Consortium | Jacquelyn Wynn, Mairead Hyland, The Lighthouse Lab in Alderley Park and Alex Alderton, Roberto Amato, Jeffrey Barrett, Sonia Goncalves, Ewan Harrison, David K. Jackson, Ian Johnston, Dominic Kwiatkowski, Cordelia Langford, John Sillitoe on behalf of the Wellcome Sanger Institute COVID-19 Surveillance Team |
| EPI_ISL_3202502 | Health Services Laboratories | Wellcome Sanger Institute for the COVID-19 Genomics UK (COG-UK) Consortium | Health Services Laboratories and Alex Alderton, Roberto Amato, Jeffrey Barrett, Sonia Goncalves, Ewan Harrison, David K. Jackson, Ian Johnston, Dominic Kwiatkowski, Cordelia Langford, John Sillitoe on behalf of the Wellcome Sanger Institute COVID-19 Surveillance Team |
| EPI_ISL_3202503 | Berkshire and Surrey Pathology Services Lighthouse Laboratory | Wellcome Sanger Institute for the COVID-19 Genomics UK (COG-UK) Consortium | Berkshire and Surrey Pathology Services Lighthouse Laboratory and Alex Alderton, Roberto Amato, Jeffrey Barrett, Sonia Goncalves, Ewan Harrison, David K. Jackson, Ian Johnston, Dominic Kwiatkowski, Cordelia Langford, John Sillitoe on behalf of the Wellcome Sanger Institute COVID-19 Surveillance Team |
| EPI_ISL_3202504 | Lighthouse Lab in Alderley Park | Wellcome Sanger Institute for the COVID-19 Genomics UK (COG-UK) Consortium | Jacquelyn Wynn, Mairead Hyland, The Lighthouse Lab in Alderley Park and Alex Alderton, Roberto Amato, Jeffrey Barrett, Sonia Goncalves, Ewan Harrison, David K. Jackson, Ian Johnston, Dominic Kwiatkowski, Cordelia Langford, John Sillitoe on behalf of the Wellcome Sanger Institute COVID-19 Surveillance Team |
| EPI_ISL_3202505 | Berkshire and Surrey Pathology Services Lighthouse Laboratory | Wellcome Sanger Institute for the COVID-19 Genomics UK (COG-UK) Consortium | Berkshire and Surrey Pathology Services Lighthouse Laboratory and Alex Alderton, Roberto Amato, Jeffrey Barrett, Sonia Goncalves, Ewan Harrison, David K. Jackson, Ian Johnston, Dominic Kwiatkowski, Cordelia Langford, John Sillitoe on behalf of the Wellcome Sanger Institute COVID-19 Surveillance Team |
| EPI_ISL_3202506, EPI_ISL_3202507, EPI_ISL_3202508 | Lighthouse Lab in Alderley Park | Wellcome Sanger Institute for the COVID-19 Genomics UK (COG-UK) Consortium | Jacquelyn Wynn, Mairead Hyland, The Lighthouse Lab in Alderley Park and Alex Alderton, Roberto Amato, Jeffrey Barrett, Sonia Goncalves, Ewan Harrison, David K. Jackson, Ian Johnston, Dominic Kwiatkowski, Cordelia Langford, John Sillitoe on behalf of the Wellcome Sanger Institute COVID-19 Surveillance Team |
| EPI_ISL_3202509 | Berkshire and Surrey Pathology Services Lighthouse Laboratory | Wellcome Sanger Institute for the COVID-19 Genomics UK (COG-UK) Consortium | Berkshire and Surrey Pathology Services Lighthouse Laboratory and Alex Alderton, Roberto Amato, Jeffrey Barrett, Sonia Goncalves, Ewan Harrison, David K. Jackson, Ian Johnston, Dominic Kwiatkowski, Cordelia Langford, John Sillitoe on behalf of the Wellcome Sanger Institute COVID-19 Surveillance Team |
| EPI_ISL_3202510 | Lighthouse Lab in Glasgow | Wellcome Sanger Institute for the COVID-19 Genomics UK (COG-UK) Consortium | Harper VanSteenhouse, Yumi Kasai, David Gray, Carol Clugston, Anna Dominiczak and Alex Alderton, Roberto Amato, Jeffrey Barrett, Sonia Goncalves, Ewan Harrison, David K. Jackson, Ian Johnston, Dominic Kwiatkowski, Cordelia Langford, John Sillitoe on behalf of the Wellcome Sanger Institute COVID-19 Surveillance Team |
| EPI_ISL_3202511 | Berkshire and Surrey Pathology Services Lighthouse Laboratory | Wellcome Sanger Institute for the COVID-19 Genomics UK (COG-UK) Consortium | Berkshire and Surrey Pathology Services Lighthouse Laboratory and Alex Alderton, Roberto Amato, Jeffrey Barrett, Sonia Goncalves, Ewan Harrison, David K. Jackson, Ian Johnston, Dominic Kwiatkowski, Cordelia Langford, John Sillitoe on behalf of the Wellcome Sanger Institute COVID-19 Surveillance Team |
| EPI_ISL_3202512, EPI_ISL_3202513, EPI_ISL_3202514 | Health Services Laboratories | Wellcome Sanger Institute for the COVID-19 Genomics UK (COG-UK) Consortium | Health Services Laboratories and Alex Alderton, Roberto Amato, Jeffrey Barrett, Sonia Goncalves, Ewan Harrison, David K. Jackson, Ian Johnston, Dominic Kwiatkowski, Cordelia Langford, John Sillitoe on behalf of the Wellcome Sanger Institute COVID-19 Surveillance Team |
| EPI_ISL_3202515, EPI_ISL_3202516 | Lighthouse Lab in Alderley Park | Wellcome Sanger Institute for the COVID-19 Genomics UK (COG-UK) Consortium | Jacquelyn Wynn, Mairead Hyland, The Lighthouse Lab in Alderley Park and Alex Alderton, Roberto Amato, Jeffrey Barrett, Sonia Goncalves, Ewan Harrison, David K. Jackson, Ian Johnston, Dominic Kwiatkowski, Cordelia Langford, John Sillitoe on behalf of the Wellcome Sanger Institute COVID-19 Surveillance Team |
| EPI_ISL_3202517 | Berkshire and Surrey Pathology Services Lighthouse Laboratory | Wellcome Sanger Institute for the COVID-19 Genomics UK (COG-UK) Consortium | Berkshire and Surrey Pathology Services Lighthouse Laboratory and Alex Alderton, Roberto Amato, Jeffrey Barrett, Sonia Goncalves, Ewan Harrison, David K. Jackson, Ian Johnston, Dominic Kwiatkowski, Cordelia Langford, John Sillitoe on behalf of the Wellcome Sanger Institute COVID-19 Surveillance Team |
| EPI_ISL_3202518, EPI_ISL_3202519 | Lighthouse Lab in Alderley Park | Wellcome Sanger Institute for the COVID-19 Genomics UK (COG-UK) Consortium | Jacquelyn Wynn, Mairead Hyland, The Lighthouse Lab in Alderley Park and Alex Alderton, Roberto Amato, Jeffrey Barrett, Sonia Goncalves, Ewan Harrison, David K. Jackson, Ian Johnston, Dominic Kwiatkowski, Cordelia Langford, John Sillitoe on behalf of the Wellcome Sanger Institute COVID-19 Surveillance Team |
| EPI_ISL_3202520, EPI_ISL_3202521 | Health Services Laboratories | Wellcome Sanger Institute for the COVID-19 Genomics UK (COG-UK) Consortium | Health Services Laboratories and Alex Alderton, Roberto Amato, Jeffrey Barrett, Sonia Goncalves, Ewan Harrison, David K. Jackson, Ian Johnston, Dominic Kwiatkowski, Cordelia Langford, John Sillitoe on behalf of the Wellcome Sanger Institute COVID-19 Surveillance Team |
| EPI_ISL_3202522 | Lighthouse Lab in Alderley Park | Wellcome Sanger Institute for the COVID-19 Genomics UK (COG-UK) Consortium | Jacquelyn Wynn, Mairead Hyland, The Lighthouse Lab in Alderley Park and Alex Alderton, Roberto Amato, Jeffrey Barrett, Sonia Goncalves, Ewan Harrison, David K. Jackson, Ian Johnston, Dominic Kwiatkowski, Cordelia Langford, John Sillitoe on behalf of the Wellcome Sanger Institute COVID-19 Surveillance Team |
| EPI_ISL_3202523 | Health Services Laboratories | Wellcome Sanger Institute for the COVID-19 Genomics UK (COG-UK) Consortium | Health Services Laboratories and Alex Alderton, Roberto Amato, Jeffrey Barrett, Sonia Goncalves, Ewan Harrison, David K. Jackson, Ian Johnston, Dominic Kwiatkowski, Cordelia Langford, John Sillitoe on behalf of the Wellcome Sanger Institute COVID-19 Surveillance Team |
| EPI_ISL_3202524 | Lighthouse Lab in Glasgow | Wellcome Sanger Institute for the COVID-19 Genomics UK (COG-UK) Consortium | Harper VanSteenhouse, Yumi Kasai, David Gray, Carol Clugston, Anna Dominiczak and Alex Alderton, Roberto Amato, Jeffrey Barrett, Sonia Goncalves, Ewan Harrison, David K. Jackson, Ian Johnston, Dominic Kwiatkowski, Cordelia Langford, John Sillitoe on behalf of the Wellcome Sanger Institute COVID-19 Surveillance Team |
| EPI_ISL_3202525 | Lighthouse Lab in Alderley Park | Wellcome Sanger Institute for the COVID-19 Genomics UK (COG-UK) Consortium | Jacquelyn Wynn, Mairead Hyland, The Lighthouse Lab in Alderley Park and Alex Alderton, Roberto Amato, Jeffrey Barrett, Sonia Goncalves, Ewan Harrison, David K. Jackson, Ian Johnston, Dominic Kwiatkowski, Cordelia Langford, John Sillitoe on behalf of the Wellcome Sanger Institute COVID-19 Surveillance Team |
| EPI_ISL_3202526 | Health Services Laboratories | Wellcome Sanger Institute for the COVID-19 Genomics UK (COG-UK) Consortium | Health Services Laboratories and Alex Alderton, Roberto Amato, Jeffrey Barrett, Sonia Goncalves, Ewan Harrison, David K. Jackson, Ian Johnston, Dominic Kwiatkowski, Cordelia Langford, John Sillitoe on behalf of the Wellcome Sanger Institute COVID-19 Surveillance Team |
| EPI_ISL_3202527 | Berkshire and Surrey Pathology Services Lighthouse Laboratory | Wellcome Sanger Institute for the COVID-19 Genomics UK (COG-UK) Consortium | Berkshire and Surrey Pathology Services Lighthouse Laboratory and Alex Alderton, Roberto Amato, Jeffrey Barrett, Sonia Goncalves, Ewan Harrison, David K. Jackson, Ian Johnston, Dominic Kwiatkowski, Cordelia Langford, John Sillitoe on behalf of the Wellcome Sanger Institute COVID-19 Surveillance Team |
| EPI_ISL_3202528 | Lighthouse Lab in Alderley Park | Wellcome Sanger Institute for the COVID-19 Genomics UK (COG-UK) Consortium | Jacquelyn Wynn, Mairead Hyland, The Lighthouse Lab in Alderley Park and Alex Alderton, Roberto Amato, Jeffrey Barrett, Sonia Goncalves, Ewan Harrison, David K. Jackson, Ian Johnston, Dominic Kwiatkowski, Cordelia Langford, John Sillitoe on behalf of the Wellcome Sanger Institute COVID-19 Surveillance Team |
| EPI_ISL_3202529 | Lighthouse Lab in Glasgow | Wellcome Sanger Institute for the COVID-19 Genomics UK (COG-UK) Consortium | Harper VanSteenhouse, Yumi Kasai, David Gray, Carol Clugston, Anna Dominiczak and Alex Alderton, Roberto Amato, Jeffrey Barrett, Sonia Goncalves, Ewan Harrison, David K. Jackson, Ian Johnston, Dominic Kwiatkowski, Cordelia Langford, John Sillitoe on behalf of the Wellcome Sanger Institute COVID-19 Surveillance Team |
| EPI_ISL_3202530, EPI_ISL_3202531, EPI_ISL_3202532, EPI_ISL_3202533 | Lighthouse Lab in Alderley Park | Wellcome Sanger Institute for the COVID-19 Genomics UK (COG-UK) Consortium | Jacquelyn Wynn, Mairead Hyland, The Lighthouse Lab in Alderley Park and Alex Alderton, Roberto Amato, Jeffrey Barrett, Sonia Goncalves, Ewan Harrison, David K. Jackson, Ian Johnston, Dominic Kwiatkowski, Cordelia Langford, John Sillitoe on behalf of the Wellcome Sanger Institute COVID-19 Surveillance Team |
| EPI_ISL_3202534 | Health Services Laboratories | Wellcome Sanger Institute for the COVID-19 Genomics UK (COG-UK) Consortium | Health Services Laboratories and Alex Alderton, Roberto Amato, Jeffrey Barrett, Sonia Goncalves, Ewan Harrison, David K. Jackson, Ian Johnston, Dominic Kwiatkowski, Cordelia Langford, John Sillitoe on behalf of the Wellcome Sanger Institute COVID-19 Surveillance Team |
| EPI_ISL_3202535 | Berkshire and Surrey Pathology Services Lighthouse Laboratory | Wellcome Sanger Institute for the COVID-19 Genomics UK (COG-UK) Consortium | Berkshire and Surrey Pathology Services Lighthouse Laboratory and Alex Alderton, Roberto Amato, Jeffrey Barrett, Sonia Goncalves, Ewan Harrison, David K. Jackson, Ian Johnston, Dominic Kwiatkowski, Cordelia Langford, John Sillitoe on behalf of the Wellcome Sanger Institute COVID-19 Surveillance Team |
| EPI_ISL_3202536 | Lighthouse Lab in Alderley Park | Wellcome Sanger Institute for the COVID-19 Genomics UK (COG-UK) Consortium | Jacquelyn Wynn, Mairead Hyland, The Lighthouse Lab in Alderley Park and Alex Alderton, Roberto Amato, Jeffrey Barrett, Sonia Goncalves, Ewan Harrison, David K. Jackson, Ian Johnston, Dominic Kwiatkowski, Cordelia Langford, John Sillitoe on behalf of the Wellcome Sanger Institute COVID-19 Surveillance Team |
| EPI_ISL_3202537 | Health Services Laboratories | Wellcome Sanger Institute for the COVID-19 Genomics UK (COG-UK) Consortium | Health Services Laboratories and Alex Alderton, Roberto Amato, Jeffrey Barrett, Sonia Goncalves, Ewan Harrison, David K. Jackson, Ian Johnston, Dominic Kwiatkowski, Cordelia Langford, John Sillitoe on behalf of the Wellcome Sanger Institute COVID-19 Surveillance Team |
| EPI_ISL_3202538, EPI_ISL_3202539, | Lighthouse Lab in Alderley Park | Wellcome Sanger Institute for the COVID-19 Genomics UK (COG-UK) Consortium | Jacquelyn Wynn, Mairead Hyland, The Lighthouse Lab in Alderley Park and Alex Alderton, Roberto Amato, Jeffrey Barrett, Sonia Goncalves, Ewan Harrison, David K. Jackson, Ian Johnston, Dominic Kwiatkowski, Cordelia Langford, John Sillitoe on behalf of the Wellcome Sanger Institute COVID-19 Surveillance Team |

| | | | |
|---|---|---|---|
| EPI_ISL_3202540 | | | |
| EPI_ISL_3202541 | Health Services Laboratories | Wellcome Sanger Institute for the COVID-19 Genomics UK (COG-UK) Consortium | Health Services Laboratories and Alex Alderton, Roberto Amato, Jeffrey Barrett, Sonia Goncalves, Ewan Harrison, David K. Jackson, Ian Johnston, Dominic Kwiatkowski, Cordelia Langford, John Sillitoe on behalf of the Wellcome Sanger Institute COVID-19 Surveillance Team |
| EPI_ISL_3202542 | Lighthouse Lab in Alderley Park | Wellcome Sanger Institute for the COVID-19 Genomics UK (COG-UK) Consortium | Jacquelyn Wynn, Mairead Hyland, The Lighthouse Lab in Alderley Park and Alex Alderton, Roberto Amato, Jeffrey Barrett, Sonia Goncalves, Ewan Harrison, David K. Jackson, Ian Johnston, Dominic Kwiatkowski, Cordelia Langford, John Sillitoe on behalf of the Wellcome Sanger Institute COVID-19 Surveillance Team |
| EPI_ISL_3202543 | Berkshire and Surrey Pathology Services Lighthouse Laboratory | Wellcome Sanger Institute for the COVID-19 Genomics UK (COG-UK) Consortium | Berkshire and Surrey Pathology Services Lighthouse Laboratory and Alex Alderton, Roberto Amato, Jeffrey Barrett, Sonia Goncalves, Ewan Harrison, David K. Jackson, Ian Johnston, Dominic Kwiatkowski, Cordelia Langford, John Sillitoe on behalf of the Wellcome Sanger Institute COVID-19 Surveillance Team |
| EPI_ISL_3202544 | Lighthouse Lab in Alderley Park | Wellcome Sanger Institute for the COVID-19 Genomics UK (COG-UK) Consortium | Jacquelyn Wynn, Mairead Hyland, The Lighthouse Lab in Alderley Park and Alex Alderton, Roberto Amato, Jeffrey Barrett, Sonia Goncalves, Ewan Harrison, David K. Jackson, Ian Johnston, Dominic Kwiatkowski, Cordelia Langford, John Sillitoe on behalf of the Wellcome Sanger Institute COVID-19 Surveillance Team |
| EPI_ISL_3202545 | Health Services Laboratories | Wellcome Sanger Institute for the COVID-19 Genomics UK (COG-UK) Consortium | Health Services Laboratories and Alex Alderton, Roberto Amato, Jeffrey Barrett, Sonia Goncalves, Ewan Harrison, David K. Jackson, Ian Johnston, Dominic Kwiatkowski, Cordelia Langford, John Sillitoe on behalf of the Wellcome Sanger Institute COVID-19 Surveillance Team |
| EPI_ISL_3202546 | Lighthouse Lab in Alderley Park | Wellcome Sanger Institute for the COVID-19 Genomics UK (COG-UK) Consortium | Jacquelyn Wynn, Mairead Hyland, The Lighthouse Lab in Alderley Park and Alex Alderton, Roberto Amato, Jeffrey Barrett, Sonia Goncalves, Ewan Harrison, David K. Jackson, Ian Johnston, Dominic Kwiatkowski, Cordelia Langford, John Sillitoe on behalf of the Wellcome Sanger Institute COVID-19 Surveillance Team |
| EPI_ISL_3202547 | Health Services Laboratories | Wellcome Sanger Institute for the COVID-19 Genomics UK (COG-UK) Consortium | Health Services Laboratories and Alex Alderton, Roberto Amato, Jeffrey Barrett, Sonia Goncalves, Ewan Harrison, David K. Jackson, Ian Johnston, Dominic Kwiatkowski, Cordelia Langford, John Sillitoe on behalf of the Wellcome Sanger Institute COVID-19 Surveillance Team |
| EPI_ISL_3202548 | Lighthouse Lab in Alderley Park | Wellcome Sanger Institute for the COVID-19 Genomics UK (COG-UK) Consortium | Jacquelyn Wynn, Mairead Hyland, The Lighthouse Lab in Alderley Park and Alex Alderton, Roberto Amato, Jeffrey Barrett, Sonia Goncalves, Ewan Harrison, David K. Jackson, Ian Johnston, Dominic Kwiatkowski, Cordelia Langford, John Sillitoe on behalf of the Wellcome Sanger Institute COVID-19 Surveillance Team |
| EPI_ISL_3202549, EPI_ISL_3202550 | Berkshire and Surrey Pathology Services Lighthouse Laboratory | Wellcome Sanger Institute for the COVID-19 Genomics UK (COG-UK) Consortium | Berkshire and Surrey Pathology Services Lighthouse Laboratory and Alex Alderton, Roberto Amato, Jeffrey Barrett, Sonia Goncalves, Ewan Harrison, David K. Jackson, Ian Johnston, Dominic Kwiatkowski, Cordelia Langford, John Sillitoe on behalf of the Wellcome Sanger Institute COVID-19 Surveillance Team |
| EPI_ISL_3202551 | Health Services Laboratories | Wellcome Sanger Institute for the COVID-19 Genomics UK (COG-UK) Consortium | Health Services Laboratories and Alex Alderton, Roberto Amato, Jeffrey Barrett, Sonia Goncalves, Ewan Harrison, David K. Jackson, Ian Johnston, Dominic Kwiatkowski, Cordelia Langford, John Sillitoe on behalf of the Wellcome Sanger Institute COVID-19 Surveillance Team |
| EPI_ISL_3202552, EPI_ISL_3202553, EPI_ISL_3202554 | Lighthouse Lab in Alderley Park | Wellcome Sanger Institute for the COVID-19 Genomics UK (COG-UK) Consortium | Jacquelyn Wynn, Mairead Hyland, The Lighthouse Lab in Alderley Park and Alex Alderton, Roberto Amato, Jeffrey Barrett, Sonia Goncalves, Ewan Harrison, David K. Jackson, Ian Johnston, Dominic Kwiatkowski, Cordelia Langford, John Sillitoe on behalf of the Wellcome Sanger Institute COVID-19 Surveillance Team |
| EPI_ISL_3202555, EPI_ISL_3202556 | Berkshire and Surrey Pathology Services Lighthouse Laboratory | Wellcome Sanger Institute for the COVID-19 Genomics UK (COG-UK) Consortium | Berkshire and Surrey Pathology Services Lighthouse Laboratory and Alex Alderton, Roberto Amato, Jeffrey Barrett, Sonia Goncalves, Ewan Harrison, David K. Jackson, Ian Johnston, Dominic Kwiatkowski, Cordelia Langford, John Sillitoe on behalf of the Wellcome Sanger Institute COVID-19 Surveillance Team |
| EPI_ISL_3202557 | Lighthouse Lab in Alderley Park | Wellcome Sanger Institute for the COVID-19 Genomics UK (COG-UK) Consortium | Jacquelyn Wynn, Mairead Hyland, The Lighthouse Lab in Alderley Park and Alex Alderton, Roberto Amato, Jeffrey Barrett, Sonia Goncalves, Ewan Harrison, David K. Jackson, Ian Johnston, Dominic Kwiatkowski, Cordelia Langford, John Sillitoe on behalf of the Wellcome Sanger Institute COVID-19 Surveillance Team |
| EPI_ISL_3202558 | Health Services Laboratories | Wellcome Sanger Institute for the COVID-19 Genomics UK (COG-UK) Consortium | Health Services Laboratories and Alex Alderton, Roberto Amato, Jeffrey Barrett, Sonia Goncalves, Ewan Harrison, David K. Jackson, Ian Johnston, Dominic Kwiatkowski, Cordelia Langford, John Sillitoe on behalf of the Wellcome Sanger Institute COVID-19 Surveillance Team |
| EPI_ISL_3202559, EPI_ISL_3202560, EPI_ISL_3202561 | Lighthouse Lab in Alderley Park | Wellcome Sanger Institute for the COVID-19 Genomics UK (COG-UK) Consortium | Jacquelyn Wynn, Mairead Hyland, The Lighthouse Lab in Alderley Park and Alex Alderton, Roberto Amato, Jeffrey Barrett, Sonia Goncalves, Ewan Harrison, David K. Jackson, Ian Johnston, Dominic Kwiatkowski, Cordelia Langford, John Sillitoe on behalf of the Wellcome Sanger Institute COVID-19 Surveillance Team |
| EPI_ISL_3202562 | Berkshire and Surrey Pathology Services Lighthouse Laboratory | Wellcome Sanger Institute for the COVID-19 Genomics UK (COG-UK) Consortium | Berkshire and Surrey Pathology Services Lighthouse Laboratory and Alex Alderton, Roberto Amato, Jeffrey Barrett, Sonia Goncalves, Ewan Harrison, David K. Jackson, Ian Johnston, Dominic Kwiatkowski, Cordelia Langford, John Sillitoe on behalf of the Wellcome Sanger Institute COVID-19 Surveillance Team |
| EPI_ISL_3202563 | Lighthouse Lab in Alderley Park | Wellcome Sanger Institute for the COVID-19 Genomics UK (COG-UK) Consortium | Jacquelyn Wynn, Mairead Hyland, The Lighthouse Lab in Alderley Park and Alex Alderton, Roberto Amato, Jeffrey Barrett, Sonia Goncalves, Ewan Harrison, David K. Jackson, Ian Johnston, Dominic Kwiatkowski, Cordelia Langford, John Sillitoe on behalf of the Wellcome Sanger Institute COVID-19 Surveillance Team |
| EPI_ISL_3202564 | Health Services Laboratories | Wellcome Sanger Institute for the COVID-19 Genomics UK (COG-UK) Consortium | Health Services Laboratories and Alex Alderton, Roberto Amato, Jeffrey Barrett, Sonia Goncalves, Ewan Harrison, David K. Jackson, Ian Johnston, Dominic Kwiatkowski, Cordelia Langford, John Sillitoe on behalf of the Wellcome Sanger Institute COVID-19 Surveillance Team |
| EPI_ISL_3202565, EPI_ISL_3202566 | Berkshire and Surrey Pathology Services Lighthouse Laboratory | Wellcome Sanger Institute for the COVID-19 Genomics UK (COG-UK) Consortium | Berkshire and Surrey Pathology Services Lighthouse Laboratory and Alex Alderton, Roberto Amato, Jeffrey Barrett, Sonia Goncalves, Ewan Harrison, David K. Jackson, Ian Johnston, Dominic Kwiatkowski, Cordelia Langford, John Sillitoe on behalf of the Wellcome Sanger Institute COVID-19 Surveillance Team |
| EPI_ISL_3202567, EPI_ISL_3202568, EPI_ISL_3202569, EPI_ISL_3202570 | Health Services Laboratories | Wellcome Sanger Institute for the COVID-19 Genomics UK (COG-UK) Consortium | Health Services Laboratories and Alex Alderton, Roberto Amato, Jeffrey Barrett, Sonia Goncalves, Ewan Harrison, David K. Jackson, Ian Johnston, Dominic Kwiatkowski, Cordelia Langford, John Sillitoe on behalf of the Wellcome Sanger Institute COVID-19 Surveillance Team |
| EPI_ISL_3202571 | Berkshire and Surrey Pathology Services Lighthouse Laboratory | Wellcome Sanger Institute for the COVID-19 Genomics UK (COG-UK) Consortium | Berkshire and Surrey Pathology Services Lighthouse Laboratory and Alex Alderton, Roberto Amato, Jeffrey Barrett, Sonia Goncalves, Ewan Harrison, David K. Jackson, Ian Johnston, Dominic Kwiatkowski, Cordelia Langford, John Sillitoe on behalf of the Wellcome Sanger Institute COVID-19 Surveillance Team |
| EPI_ISL_3202572, EPI_ISL_3202573 | Health Services Laboratories | Wellcome Sanger Institute for the COVID-19 Genomics UK (COG-UK) Consortium | Health Services Laboratories and Alex Alderton, Roberto Amato, Jeffrey Barrett, Sonia Goncalves, Ewan Harrison, David K. Jackson, Ian Johnston, Dominic Kwiatkowski, Cordelia Langford, John Sillitoe on behalf of the Wellcome Sanger Institute COVID-19 Surveillance Team |
| EPI_ISL_3202574 | Berkshire and Surrey Pathology Services Lighthouse Laboratory | Wellcome Sanger Institute for the COVID-19 Genomics UK (COG-UK) Consortium | Berkshire and Surrey Pathology Services Lighthouse Laboratory and Alex Alderton, Roberto Amato, Jeffrey Barrett, Sonia Goncalves, Ewan Harrison, David K. Jackson, Ian Johnston, Dominic Kwiatkowski, Cordelia Langford, John Sillitoe on behalf of the Wellcome Sanger Institute COVID-19 Surveillance Team |
| EPI_ISL_3202575, EPI_ISL_3202576, EPI_ISL_3202577 | Health Services Laboratories | Wellcome Sanger Institute for the COVID-19 Genomics UK (COG-UK) Consortium | Health Services Laboratories and Alex Alderton, Roberto Amato, Jeffrey Barrett, Sonia Goncalves, Ewan Harrison, David K. Jackson, Ian Johnston, Dominic Kwiatkowski, Cordelia Langford, John Sillitoe on behalf of the Wellcome Sanger Institute COVID-19 Surveillance Team |
| EPI_ISL_3202578 | Lighthouse Lab in Alderley Park | Wellcome Sanger Institute for the COVID-19 Genomics UK (COG-UK) Consortium | Jacquelyn Wynn, Mairead Hyland, The Lighthouse Lab in Alderley Park and Alex Alderton, Roberto Amato, Jeffrey Barrett, Sonia Goncalves, Ewan Harrison, David K. Jackson, Ian Johnston, Dominic Kwiatkowski, Cordelia Langford, John Sillitoe on behalf of the Wellcome Sanger Institute COVID-19 Surveillance Team |
| EPI_ISL_3202579, EPI_ISL_3202580 | Berkshire and Surrey Pathology Services Lighthouse Laboratory | Wellcome Sanger Institute for the COVID-19 Genomics UK (COG-UK) Consortium | Berkshire and Surrey Pathology Services Lighthouse Laboratory and Alex Alderton, Roberto Amato, Jeffrey Barrett, Sonia Goncalves, Ewan Harrison, David K. Jackson, Ian Johnston, Dominic Kwiatkowski, Cordelia Langford, John Sillitoe on behalf of the Wellcome Sanger Institute COVID-19 Surveillance Team |
| EPI_ISL_3202581 | Health Services Laboratories | Wellcome Sanger Institute for the COVID-19 Genomics UK (COG-UK) Consortium | Health Services Laboratories and Alex Alderton, Roberto Amato, Jeffrey Barrett, Sonia Goncalves, Ewan Harrison, David K. Jackson, Ian Johnston, Dominic Kwiatkowski, Cordelia Langford, John Sillitoe on behalf of the Wellcome Sanger Institute COVID-19 Surveillance Team |
| EPI_ISL_3202582, EPI_ISL_3202583, EPI_ISL_3202584, EPI_ISL_3202585, EPI_ISL_3202586, EPI_ISL_3202587, EPI_ISL_3202588, EPI_ISL_3202589 | Lighthouse Lab in Alderley Park | Wellcome Sanger Institute for the COVID-19 Genomics UK (COG-UK) Consortium | Jacquelyn Wynn, Mairead Hyland, The Lighthouse Lab in Alderley Park and Alex Alderton, Roberto Amato, Jeffrey Barrett, Sonia Goncalves, Ewan Harrison, David K. Jackson, Ian Johnston, Dominic Kwiatkowski, Cordelia Langford, John Sillitoe on behalf of the Wellcome Sanger Institute COVID-19 Surveillance Team |
| EPI_ISL_3202590 | Berkshire and Surrey Pathology Services Lighthouse Laboratory | Wellcome Sanger Institute for the COVID-19 Genomics UK (COG-UK) Consortium | Berkshire and Surrey Pathology Services Lighthouse Laboratory and Alex Alderton, Roberto Amato, Jeffrey Barrett, Sonia Goncalves, Ewan Harrison, David K. Jackson, Ian Johnston, Dominic Kwiatkowski, Cordelia Langford, John Sillitoe on behalf of the Wellcome Sanger Institute COVID-19 Surveillance Team |
| EPI_ISL_3202591, EPI_ISL_3202592, EPI_ISL_3202593, EPI_ISL_3202594, EPI_ISL_3202595, EPI_ISL_3202596, EPI_ISL_3202597 | Lighthouse Lab in Alderley Park | Wellcome Sanger Institute for the COVID-19 Genomics UK (COG-UK) Consortium | Jacquelyn Wynn, Mairead Hyland, The Lighthouse Lab in Alderley Park and Alex Alderton, Roberto Amato, Jeffrey Barrett, Sonia Goncalves, Ewan Harrison, David K. Jackson, Ian Johnston, Dominic Kwiatkowski, Cordelia Langford, John Sillitoe on behalf of the Wellcome Sanger Institute COVID-19 Surveillance Team |
| EPI_ISL_3202598 | Health Services Laboratories | Wellcome Sanger Institute for the COVID-19 Genomics UK (COG-UK) Consortium | Health Services Laboratories and Alex Alderton, Roberto Amato, Jeffrey Barrett, Sonia Goncalves, Ewan Harrison, David K. Jackson, Ian Johnston, Dominic Kwiatkowski, Cordelia Langford, John Sillitoe on behalf of the Wellcome Sanger Institute COVID-19 Surveillance Team |
| EPI_ISL_3202599, EPI_ISL_3202600, EPI_ISL_3202601, EPI_ISL_3202602 | Lighthouse Lab in Alderley Park | Wellcome Sanger Institute for the COVID-19 Genomics UK (COG-UK) Consortium | Jacquelyn Wynn, Mairead Hyland, The Lighthouse Lab in Alderley Park and Alex Alderton, Roberto Amato, Jeffrey Barrett, Sonia Goncalves, Ewan Harrison, David K. Jackson, Ian Johnston, Dominic Kwiatkowski, Cordelia Langford, John Sillitoe on behalf of the Wellcome Sanger Institute COVID-19 Surveillance Team |
| EPI_ISL_3202603, EPI_ISL_3202604 | Berkshire and Surrey Pathology Services Lighthouse Laboratory | Wellcome Sanger Institute for the COVID-19 Genomics UK (COG-UK) Consortium | Berkshire and Surrey Pathology Services Lighthouse Laboratory and Alex Alderton, Roberto Amato, Jeffrey Barrett, Sonia Goncalves, Ewan Harrison, David K. Jackson, Ian Johnston, Dominic Kwiatkowski, Cordelia Langford, John Sillitoe on behalf of the Wellcome Sanger Institute COVID-19 Surveillance Team |
| EPI_ISL_3202605, EPI_ISL_3202606 | Lighthouse Lab in Alderley Park | Wellcome Sanger Institute for the COVID-19 Genomics UK (COG-UK) Consortium | Jacquelyn Wynn, Mairead Hyland, The Lighthouse Lab in Alderley Park and Alex Alderton, Roberto Amato, Jeffrey Barrett, Sonia Goncalves, Ewan Harrison, David K. Jackson, Ian Johnston, Dominic Kwiatkowski, Cordelia Langford, John Sillitoe on behalf of the Wellcome Sanger Institute COVID-19 Surveillance Team |
| EPI_ISL_3202607 | Health Services Laboratories | Wellcome Sanger Institute for the COVID-19 Genomics UK (COG-UK) Consortium | Health Services Laboratories and Alex Alderton, Roberto Amato, Jeffrey Barrett, Sonia Goncalves, Ewan Harrison, David K. Jackson, Ian Johnston, Dominic Kwiatkowski, Cordelia Langford, John Sillitoe on behalf of the Wellcome Sanger Institute COVID-19 Surveillance Team |
| EPI_ISL_3202608, EPI_ISL_3202609, EPI_ISL_3202610, EPI_ISL_3202611, | Lighthouse Lab in Alderley Park | Wellcome Sanger Institute for the COVID-19 Genomics UK (COG-UK) Consortium | Jacquelyn Wynn, Mairead Hyland, The Lighthouse Lab in Alderley Park and Alex Alderton, Roberto Amato, Jeffrey Barrett, Sonia Goncalves, Ewan Harrison, David K. Jackson, Ian Johnston, Dominic Kwiatkowski, Cordelia Langford, John Sillitoe on behalf of the Wellcome Sanger Institute COVID-19 Surveillance Team |

| | | | |
|---|---|---|---|
| EPI_ISL_3202612, EPI_ISL_3202613 | | | |
| EPI_ISL_3202614, EPI_ISL_3202615 | Health Services Laboratories | Wellcome Sanger Institute for the COVID-19 Genomics UK (COG-UK) Consortium | Health Services Laboratories and Alex Alderton, Roberto Amato, Jeffrey Barrett, Sonia Goncalves, Ewan Harrison, David K. Jackson, Ian Johnston, Dominic Kwiatkowski, Cordelia Langford, John Sillitoe on behalf of the Wellcome Sanger Institute COVID-19 Surveillance Team |
| EPI_ISL_3202616, EPI_ISL_3202617, EPI_ISL_3202618, EPI_ISL_3202619 | Lighthouse Lab in Alderley Park | Wellcome Sanger Institute for the COVID-19 Genomics UK (COG-UK) Consortium | Jacquelyn Wynn, Mairead Hyland, The Lighthouse Lab in Alderley Park and Alex Alderton, Roberto Amato, Jeffrey Barrett, Sonia Goncalves, Ewan Harrison, David K. Jackson, Ian Johnston, Dominic Kwiatkowski, Cordelia Langford, John Sillitoe on behalf of the Wellcome Sanger Institute COVID-19 Surveillance Team |
| EPI_ISL_3202620 | Berkshire and Surrey Pathology Services Lighthouse Laboratory | Wellcome Sanger Institute for the COVID-19 Genomics UK (COG-UK) Consortium | Berkshire and Surrey Pathology Services Lighthouse Laboratory and Alex Alderton, Roberto Amato, Jeffrey Barrett, Sonia Goncalves, Ewan Harrison, David K. Jackson, Ian Johnston, Dominic Kwiatkowski, Cordelia Langford, John Sillitoe on behalf of the Wellcome Sanger Institute COVID-19 Surveillance Team |
| EPI_ISL_3202621 | Lighthouse Lab in Alderley Park | Wellcome Sanger Institute for the COVID-19 Genomics UK (COG-UK) Consortium | Jacquelyn Wynn, Mairead Hyland, The Lighthouse Lab in Alderley Park and Alex Alderton, Roberto Amato, Jeffrey Barrett, Sonia Goncalves, Ewan Harrison, David K. Jackson, Ian Johnston, Dominic Kwiatkowski, Cordelia Langford, John Sillitoe on behalf of the Wellcome Sanger Institute COVID-19 Surveillance Team |
| EPI_ISL_3202622 | Berkshire and Surrey Pathology Services Lighthouse Laboratory | Wellcome Sanger Institute for the COVID-19 Genomics UK (COG-UK) Consortium | Berkshire and Surrey Pathology Services Lighthouse Laboratory and Alex Alderton, Roberto Amato, Jeffrey Barrett, Sonia Goncalves, Ewan Harrison, David K. Jackson, Ian Johnston, Dominic Kwiatkowski, Cordelia Langford, John Sillitoe on behalf of the Wellcome Sanger Institute COVID-19 Surveillance Team |
| EPI_ISL_3202623 | Lighthouse Lab in Alderley Park | Wellcome Sanger Institute for the COVID-19 Genomics UK (COG-UK) Consortium | Jacquelyn Wynn, Mairead Hyland, The Lighthouse Lab in Alderley Park and Alex Alderton, Roberto Amato, Jeffrey Barrett, Sonia Goncalves, Ewan Harrison, David K. Jackson, Ian Johnston, Dominic Kwiatkowski, Cordelia Langford, John Sillitoe on behalf of the Wellcome Sanger Institute COVID-19 Surveillance Team |
| EPI_ISL_3202624 | Berkshire and Surrey Pathology Services Lighthouse Laboratory | Wellcome Sanger Institute for the COVID-19 Genomics UK (COG-UK) Consortium | Berkshire and Surrey Pathology Services Lighthouse Laboratory and Alex Alderton, Roberto Amato, Jeffrey Barrett, Sonia Goncalves, Ewan Harrison, David K. Jackson, Ian Johnston, Dominic Kwiatkowski, Cordelia Langford, John Sillitoe on behalf of the Wellcome Sanger Institute COVID-19 Surveillance Team |
| EPI_ISL_3202625 | Lighthouse Lab in Alderley Park | Wellcome Sanger Institute for the COVID-19 Genomics UK (COG-UK) Consortium | Jacquelyn Wynn, Mairead Hyland, The Lighthouse Lab in Alderley Park and Alex Alderton, Roberto Amato, Jeffrey Barrett, Sonia Goncalves, Ewan Harrison, David K. Jackson, Ian Johnston, Dominic Kwiatkowski, Cordelia Langford, John Sillitoe on behalf of the Wellcome Sanger Institute COVID-19 Surveillance Team |
| EPI_ISL_3202626 | Health Services Laboratories | Wellcome Sanger Institute for the COVID-19 Genomics UK (COG-UK) Consortium | Health Services Laboratories and Alex Alderton, Roberto Amato, Jeffrey Barrett, Sonia Goncalves, Ewan Harrison, David K. Jackson, Ian Johnston, Dominic Kwiatkowski, Cordelia Langford, John Sillitoe on behalf of the Wellcome Sanger Institute COVID-19 Surveillance Team |
| EPI_ISL_3202627, EPI_ISL_3202628, EPI_ISL_3202629, EPI_ISL_3202630 | Lighthouse Lab in Alderley Park | Wellcome Sanger Institute for the COVID-19 Genomics UK (COG-UK) Consortium | Jacquelyn Wynn, Mairead Hyland, The Lighthouse Lab in Alderley Park and Alex Alderton, Roberto Amato, Jeffrey Barrett, Sonia Goncalves, Ewan Harrison, David K. Jackson, Ian Johnston, Dominic Kwiatkowski, Cordelia Langford, John Sillitoe on behalf of the Wellcome Sanger Institute COVID-19 Surveillance Team |
| EPI_ISL_3202631, EPI_ISL_3202632 | Health Services Laboratories | Wellcome Sanger Institute for the COVID-19 Genomics UK (COG-UK) Consortium | Health Services Laboratories and Alex Alderton, Roberto Amato, Jeffrey Barrett, Sonia Goncalves, Ewan Harrison, David K. Jackson, Ian Johnston, Dominic Kwiatkowski, Cordelia Langford, John Sillitoe on behalf of the Wellcome Sanger Institute COVID-19 Surveillance Team |
| EPI_ISL_3202633, EPI_ISL_3202634, EPI_ISL_3202635 | Lighthouse Lab in Alderley Park | Wellcome Sanger Institute for the COVID-19 Genomics UK (COG-UK) Consortium | Jacquelyn Wynn, Mairead Hyland, The Lighthouse Lab in Alderley Park and Alex Alderton, Roberto Amato, Jeffrey Barrett, Sonia Goncalves, Ewan Harrison, David K. Jackson, Ian Johnston, Dominic Kwiatkowski, Cordelia Langford, John Sillitoe on behalf of the Wellcome Sanger Institute COVID-19 Surveillance Team |
| EPI_ISL_3202636 | Health Services Laboratories | Wellcome Sanger Institute for the COVID-19 Genomics UK (COG-UK) Consortium | Health Services Laboratories and Alex Alderton, Roberto Amato, Jeffrey Barrett, Sonia Goncalves, Ewan Harrison, David K. Jackson, Ian Johnston, Dominic Kwiatkowski, Cordelia Langford, John Sillitoe on behalf of the Wellcome Sanger Institute COVID-19 Surveillance Team |
| EPI_ISL_3202637, EPI_ISL_3202638, EPI_ISL_3202639 | Lighthouse Lab in Alderley Park | Wellcome Sanger Institute for the COVID-19 Genomics UK (COG-UK) Consortium | Jacquelyn Wynn, Mairead Hyland, The Lighthouse Lab in Alderley Park and Alex Alderton, Roberto Amato, Jeffrey Barrett, Sonia Goncalves, Ewan Harrison, David K. Jackson, Ian Johnston, Dominic Kwiatkowski, Cordelia Langford, John Sillitoe on behalf of the Wellcome Sanger Institute COVID-19 Surveillance Team |
| EPI_ISL_3202640 | Berkshire and Surrey Pathology Services Lighthouse Laboratory | Wellcome Sanger Institute for the COVID-19 Genomics UK (COG-UK) Consortium | Berkshire and Surrey Pathology Services Lighthouse Laboratory and Alex Alderton, Roberto Amato, Jeffrey Barrett, Sonia Goncalves, Ewan Harrison, David K. Jackson, Ian Johnston, Dominic Kwiatkowski, Cordelia Langford, John Sillitoe on behalf of the Wellcome Sanger Institute COVID-19 Surveillance Team |
| EPI_ISL_3202641 | Lighthouse Lab in Alderley Park | Wellcome Sanger Institute for the COVID-19 Genomics UK (COG-UK) Consortium | Jacquelyn Wynn, Mairead Hyland, The Lighthouse Lab in Alderley Park and Alex Alderton, Roberto Amato, Jeffrey Barrett, Sonia Goncalves, Ewan Harrison, David K. Jackson, Ian Johnston, Dominic Kwiatkowski, Cordelia Langford, John Sillitoe on behalf of the Wellcome Sanger Institute COVID-19 Surveillance Team |
| EPI_ISL_3202642 | Health Services Laboratories | Wellcome Sanger Institute for the COVID-19 Genomics UK (COG-UK) Consortium | Health Services Laboratories and Alex Alderton, Roberto Amato, Jeffrey Barrett, Sonia Goncalves, Ewan Harrison, David K. Jackson, Ian Johnston, Dominic Kwiatkowski, Cordelia Langford, John Sillitoe on behalf of the Wellcome Sanger Institute COVID-19 Surveillance Team |
| EPI_ISL_3202643 | Berkshire and Surrey Pathology Services Lighthouse Laboratory | Wellcome Sanger Institute for the COVID-19 Genomics UK (COG-UK) Consortium | Berkshire and Surrey Pathology Services Lighthouse Laboratory and Alex Alderton, Roberto Amato, Jeffrey Barrett, Sonia Goncalves, Ewan Harrison, David K. Jackson, Ian Johnston, Dominic Kwiatkowski, Cordelia Langford, John Sillitoe on behalf of the Wellcome Sanger Institute COVID-19 Surveillance Team |
| EPI_ISL_3202644, EPI_ISL_3202645 | Health Services Laboratories | Wellcome Sanger Institute for the COVID-19 Genomics UK (COG-UK) Consortium | Health Services Laboratories and Alex Alderton, Roberto Amato, Jeffrey Barrett, Sonia Goncalves, Ewan Harrison, David K. Jackson, Ian Johnston, Dominic Kwiatkowski, Cordelia Langford, John Sillitoe on behalf of the Wellcome Sanger Institute COVID-19 Surveillance Team |
| EPI_ISL_3202646 | Lighthouse Lab in Alderley Park | Wellcome Sanger Institute for the COVID-19 Genomics UK (COG-UK) Consortium | Jacquelyn Wynn, Mairead Hyland, The Lighthouse Lab in Alderley Park and Alex Alderton, Roberto Amato, Jeffrey Barrett, Sonia Goncalves, Ewan Harrison, David K. Jackson, Ian Johnston, Dominic Kwiatkowski, Cordelia Langford, John Sillitoe on behalf of the Wellcome Sanger Institute COVID-19 Surveillance Team |
| EPI_ISL_3202647 | Health Services Laboratories | Wellcome Sanger Institute for the COVID-19 Genomics UK (COG-UK) Consortium | Health Services Laboratories and Alex Alderton, Roberto Amato, Jeffrey Barrett, Sonia Goncalves, Ewan Harrison, David K. Jackson, Ian Johnston, Dominic Kwiatkowski, Cordelia Langford, John Sillitoe on behalf of the Wellcome Sanger Institute COVID-19 Surveillance Team |
| EPI_ISL_3202648 | Berkshire and Surrey Pathology Services Lighthouse Laboratory | Wellcome Sanger Institute for the COVID-19 Genomics UK (COG-UK) Consortium | Berkshire and Surrey Pathology Services Lighthouse Laboratory and Alex Alderton, Roberto Amato, Jeffrey Barrett, Sonia Goncalves, Ewan Harrison, David K. Jackson, Ian Johnston, Dominic Kwiatkowski, Cordelia Langford, John Sillitoe on behalf of the Wellcome Sanger Institute COVID-19 Surveillance Team |
| EPI_ISL_3202649, EPI_ISL_3202650, EPI_ISL_3202651, EPI_ISL_3202652 | Lighthouse Lab in Alderley Park | Wellcome Sanger Institute for the COVID-19 Genomics UK (COG-UK) Consortium | Jacquelyn Wynn, Mairead Hyland, The Lighthouse Lab in Alderley Park and Alex Alderton, Roberto Amato, Jeffrey Barrett, Sonia Goncalves, Ewan Harrison, David K. Jackson, Ian Johnston, Dominic Kwiatkowski, Cordelia Langford, John Sillitoe on behalf of the Wellcome Sanger Institute COVID-19 Surveillance Team |
| EPI_ISL_3202653 | Berkshire and Surrey Pathology Services Lighthouse Laboratory | Wellcome Sanger Institute for the COVID-19 Genomics UK (COG-UK) Consortium | Berkshire and Surrey Pathology Services Lighthouse Laboratory and Alex Alderton, Roberto Amato, Jeffrey Barrett, Sonia Goncalves, Ewan Harrison, David K. Jackson, Ian Johnston, Dominic Kwiatkowski, Cordelia Langford, John Sillitoe on behalf of the Wellcome Sanger Institute COVID-19 Surveillance Team |
| EPI_ISL_3202654, EPI_ISL_3202655 | Lighthouse Lab in Alderley Park | Wellcome Sanger Institute for the COVID-19 Genomics UK (COG-UK) Consortium | Jacquelyn Wynn, Mairead Hyland, The Lighthouse Lab in Alderley Park and Alex Alderton, Roberto Amato, Jeffrey Barrett, Sonia Goncalves, Ewan Harrison, David K. Jackson, Ian Johnston, Dominic Kwiatkowski, Cordelia Langford, John Sillitoe on behalf of the Wellcome Sanger Institute COVID-19 Surveillance Team |
| EPI_ISL_3202656 | Health Services Laboratories | Wellcome Sanger Institute for the COVID-19 Genomics UK (COG-UK) Consortium | Health Services Laboratories and Alex Alderton, Roberto Amato, Jeffrey Barrett, Sonia Goncalves, Ewan Harrison, David K. Jackson, Ian Johnston, Dominic Kwiatkowski, Cordelia Langford, John Sillitoe on behalf of the Wellcome Sanger Institute COVID-19 Surveillance Team |
| EPI_ISL_3202657 | Lighthouse Lab in Alderley Park | Wellcome Sanger Institute for the COVID-19 Genomics UK (COG-UK) Consortium | Jacquelyn Wynn, Mairead Hyland, The Lighthouse Lab in Alderley Park and Alex Alderton, Roberto Amato, Jeffrey Barrett, Sonia Goncalves, Ewan Harrison, David K. Jackson, Ian Johnston, Dominic Kwiatkowski, Cordelia Langford, John Sillitoe on behalf of the Wellcome Sanger Institute COVID-19 Surveillance Team |
| EPI_ISL_3202658 | Berkshire and Surrey Pathology Services Lighthouse Laboratory | Wellcome Sanger Institute for the COVID-19 Genomics UK (COG-UK) Consortium | Berkshire and Surrey Pathology Services Lighthouse Laboratory and Alex Alderton, Roberto Amato, Jeffrey Barrett, Sonia Goncalves, Ewan Harrison, David K. Jackson, Ian Johnston, Dominic Kwiatkowski, Cordelia Langford, John Sillitoe on behalf of the Wellcome Sanger Institute COVID-19 Surveillance Team |
| EPI_ISL_3202659, EPI_ISL_3202660, EPI_ISL_3202661 | Health Services Laboratories | Wellcome Sanger Institute for the COVID-19 Genomics UK (COG-UK) Consortium | Health Services Laboratories and Alex Alderton, Roberto Amato, Jeffrey Barrett, Sonia Goncalves, Ewan Harrison, David K. Jackson, Ian Johnston, Dominic Kwiatkowski, Cordelia Langford, John Sillitoe on behalf of the Wellcome Sanger Institute COVID-19 Surveillance Team |
| EPI_ISL_3202662, EPI_ISL_3202663 | Lighthouse Lab in Alderley Park | Wellcome Sanger Institute for the COVID-19 Genomics UK (COG-UK) Consortium | Jacquelyn Wynn, Mairead Hyland, The Lighthouse Lab in Alderley Park and Alex Alderton, Roberto Amato, Jeffrey Barrett, Sonia Goncalves, Ewan Harrison, David K. Jackson, Ian Johnston, Dominic Kwiatkowski, Cordelia Langford, John Sillitoe on behalf of the Wellcome Sanger Institute COVID-19 Surveillance Team |
| EPI_ISL_3202664 | Berkshire and Surrey Pathology Services Lighthouse Laboratory | Wellcome Sanger Institute for the COVID-19 Genomics UK (COG-UK) Consortium | Berkshire and Surrey Pathology Services Lighthouse Laboratory and Alex Alderton, Roberto Amato, Jeffrey Barrett, Sonia Goncalves, Ewan Harrison, David K. Jackson, Ian Johnston, Dominic Kwiatkowski, Cordelia Langford, John Sillitoe on behalf of the Wellcome Sanger Institute COVID-19 Surveillance Team |
| EPI_ISL_3202665, EPI_ISL_3202666, EPI_ISL_3202667 | Lighthouse Lab in Alderley Park | Wellcome Sanger Institute for the COVID-19 Genomics UK (COG-UK) Consortium | Jacquelyn Wynn, Mairead Hyland, The Lighthouse Lab in Alderley Park and Alex Alderton, Roberto Amato, Jeffrey Barrett, Sonia Goncalves, Ewan Harrison, David K. Jackson, Ian Johnston, Dominic Kwiatkowski, Cordelia Langford, John Sillitoe on behalf of the Wellcome Sanger Institute COVID-19 Surveillance Team |
| EPI_ISL_3202668, EPI_ISL_3202669 | Health Services Laboratories | Wellcome Sanger Institute for the COVID-19 Genomics UK (COG-UK) Consortium | Health Services Laboratories and Alex Alderton, Roberto Amato, Jeffrey Barrett, Sonia Goncalves, Ewan Harrison, David K. Jackson, Ian Johnston, Dominic Kwiatkowski, Cordelia Langford, John Sillitoe on behalf of the Wellcome Sanger Institute COVID-19 Surveillance Team |
| EPI_ISL_3202670, EPI_ISL_3202671 | Berkshire and Surrey Pathology Services Lighthouse Laboratory | Wellcome Sanger Institute for the COVID-19 Genomics UK (COG-UK) Consortium | Berkshire and Surrey Pathology Services Lighthouse Laboratory and Alex Alderton, Roberto Amato, Jeffrey Barrett, Sonia Goncalves, Ewan Harrison, David K. Jackson, Ian Johnston, Dominic Kwiatkowski, Cordelia Langford, John Sillitoe on behalf of the Wellcome Sanger Institute COVID-19 Surveillance Team |
| EPI_ISL_3202672, EPI_ISL_3202673, EPI_ISL_3202674 | Lighthouse Lab in Alderley Park | Wellcome Sanger Institute for the COVID-19 Genomics UK (COG-UK) Consortium | Jacquelyn Wynn, Mairead Hyland, The Lighthouse Lab in Alderley Park and Alex Alderton, Roberto Amato, Jeffrey Barrett, Sonia Goncalves, Ewan Harrison, David K. Jackson, Ian Johnston, Dominic Kwiatkowski, Cordelia Langford, John Sillitoe on behalf of the Wellcome Sanger Institute COVID-19 Surveillance Team |
| EPI_ISL_3202675 | Health Services Laboratories | Wellcome Sanger Institute for the COVID-19 Genomics UK (COG-UK) Consortium | Health Services Laboratories and Alex Alderton, Roberto Amato, Jeffrey Barrett, Sonia Goncalves, Ewan Harrison, David K. Jackson, Ian Johnston, Dominic Kwiatkowski, Cordelia Langford, John Sillitoe on behalf of the Wellcome Sanger Institute COVID-19 Surveillance Team |
| EPI_ISL_3202676, EPI_ISL_3202677 | Lighthouse Lab in Alderley Park | Wellcome Sanger Institute for the COVID-19 Genomics UK (COG-UK) Consortium | Jacquelyn Wynn, Mairead Hyland, The Lighthouse Lab in Alderley Park and Alex Alderton, Roberto Amato, Jeffrey Barrett, Sonia Goncalves, Ewan Harrison, David K. Jackson, Ian Johnston, Dominic Kwiatkowski, Cordelia Langford, John Sillitoe on behalf of the Wellcome Sanger Institute COVID-19 Surveillance Team |
| EPI_ISL_3202678 | Health Services Laboratories | Wellcome Sanger Institute for the COVID-19 Genomics UK (COG-UK) Consortium | Health Services Laboratories and Alex Alderton, Roberto Amato, Jeffrey Barrett, Sonia Goncalves, Ewan Harrison, David K. Jackson, Ian Johnston, Dominic Kwiatkowski, Cordelia Langford, John Sillitoe on behalf of the Wellcome Sanger Institute COVID-19 Surveillance Team |
| EPI_ISL_3202679 | Lighthouse Lab in Alderley Park | Wellcome Sanger Institute for the COVID-19 Genomics UK (COG-UK) Consortium | Jacquelyn Wynn, Mairead Hyland, The Lighthouse Lab in Alderley Park and Alex Alderton, Roberto Amato, Jeffrey Barrett, Sonia Goncalves, Ewan Harrison, David K. Jackson, Ian Johnston, Dominic Kwiatkowski, Cordelia Langford, John Sillitoe on behalf of the Wellcome Sanger Institute COVID-19 Surveillance Team |
| EPI_ISL_3202680 | Lighthouse Lab in Glasgow | Wellcome Sanger Institute for the COVID-19 Genomics UK (COG-UK) Consortium | Harper VanSteenhouse, Yumi Kasai, David Gray, Carol Clugston, Anna Dominiczak and Alex Alderton, Roberto Amato, Jeffrey Barrett, Sonia Goncalves, Ewan Harrison, David K. Jackson, Ian Johnston, Dominic Kwiatkowski, Cordelia Langford, John Sillitoe on behalf of the Wellcome Sanger Institute COVID-19 Surveillance Team |

| | | | |
|---|---|---|---|
| EPI_ISL_3202681, EPI_ISL_3202682, EPI_ISL_3202683 | Lighthouse Lab in Alderley Park | Wellcome Sanger Institute for the COVID-19 Genomics UK (COG-UK) Consortium | Jacquelyn Wynn, Mairead Hyland, The Lighthouse Lab in Alderley Park and Alex Alderton, Roberto Amato, Jeffrey Barrett, Sonia Goncalves, Ewan Harrison, David K. Jackson, Ian Johnston, Dominic Kwiatkowski, Cordelia Langford, John Sillitoe on behalf of the Wellcome Sanger Institute COVID-19 Surveillance Team |
| EPI_ISL_3202684 | Health Services Laboratories | Wellcome Sanger Institute for the COVID-19 Genomics UK (COG-UK) Consortium | Health Services Laboratories and Alex Alderton, Roberto Amato, Jeffrey Barrett, Sonia Goncalves, Ewan Harrison, David K. Jackson, Ian Johnston, Dominic Kwiatkowski, Cordelia Langford, John Sillitoe on behalf of the Wellcome Sanger Institute COVID-19 Surveillance Team |
| EPI_ISL_3202685 | Lighthouse Lab in Alderley Park | Wellcome Sanger Institute for the COVID-19 Genomics UK (COG-UK) Consortium | Jacquelyn Wynn, Mairead Hyland, The Lighthouse Lab in Alderley Park and Alex Alderton, Roberto Amato, Jeffrey Barrett, Sonia Goncalves, Ewan Harrison, David K. Jackson, Ian Johnston, Dominic Kwiatkowski, Cordelia Langford, John Sillitoe on behalf of the Wellcome Sanger Institute COVID-19 Surveillance Team |
| EPI_ISL_3202686 | Health Services Laboratories | Wellcome Sanger Institute for the COVID-19 Genomics UK (COG-UK) Consortium | Health Services Laboratories and Alex Alderton, Roberto Amato, Jeffrey Barrett, Sonia Goncalves, Ewan Harrison, David K. Jackson, Ian Johnston, Dominic Kwiatkowski, Cordelia Langford, John Sillitoe on behalf of the Wellcome Sanger Institute COVID-19 Surveillance Team |
| EPI_ISL_3202687 | Berkshire and Surrey Pathology Services Lighthouse Laboratory | Wellcome Sanger Institute for the COVID-19 Genomics UK (COG-UK) Consortium | Berkshire and Surrey Pathology Services Lighthouse Laboratory and Alex Alderton, Roberto Amato, Jeffrey Barrett, Sonia Goncalves, Ewan Harrison, David K. Jackson, Ian Johnston, Dominic Kwiatkowski, Cordelia Langford, John Sillitoe on behalf of the Wellcome Sanger Institute COVID-19 Surveillance Team |
| EPI_ISL_3202688 | Lighthouse Lab in Alderley Park | Wellcome Sanger Institute for the COVID-19 Genomics UK (COG-UK) Consortium | Jacquelyn Wynn, Mairead Hyland, The Lighthouse Lab in Alderley Park and Alex Alderton, Roberto Amato, Jeffrey Barrett, Sonia Goncalves, Ewan Harrison, David K. Jackson, Ian Johnston, Dominic Kwiatkowski, Cordelia Langford, John Sillitoe on behalf of the Wellcome Sanger Institute COVID-19 Surveillance Team |
| EPI_ISL_3202689 | Berkshire and Surrey Pathology Services Lighthouse Laboratory | Wellcome Sanger Institute for the COVID-19 Genomics UK (COG-UK) Consortium | Berkshire and Surrey Pathology Services Lighthouse Laboratory and Alex Alderton, Roberto Amato, Jeffrey Barrett, Sonia Goncalves, Ewan Harrison, David K. Jackson, Ian Johnston, Dominic Kwiatkowski, Cordelia Langford, John Sillitoe on behalf of the Wellcome Sanger Institute COVID-19 Surveillance Team |
| EPI_ISL_3202690 | Lighthouse Lab in Alderley Park | Wellcome Sanger Institute for the COVID-19 Genomics UK (COG-UK) Consortium | Jacquelyn Wynn, Mairead Hyland, The Lighthouse Lab in Alderley Park and Alex Alderton, Roberto Amato, Jeffrey Barrett, Sonia Goncalves, Ewan Harrison, David K. Jackson, Ian Johnston, Dominic Kwiatkowski, Cordelia Langford, John Sillitoe on behalf of the Wellcome Sanger Institute COVID-19 Surveillance Team |
| EPI_ISL_3202691 | Lighthouse Lab in Glasgow | Wellcome Sanger Institute for the COVID-19 Genomics UK (COG-UK) Consortium | Harper VanSteenhouse, Yumi Kasai, David Gray, Carol Clugston, Anna Dominiczak and Alex Alderton, Roberto Amato, Jeffrey Barrett, Sonia Goncalves, Ewan Harrison, David K. Jackson, Ian Johnston, Dominic Kwiatkowski, Cordelia Langford, John Sillitoe on behalf of the Wellcome Sanger Institute COVID-19 Surveillance Team |
| EPI_ISL_3202692, EPI_ISL_3202693 | Lighthouse Lab in Alderley Park | Wellcome Sanger Institute for the COVID-19 Genomics UK (COG-UK) Consortium | Jacquelyn Wynn, Mairead Hyland, The Lighthouse Lab in Alderley Park and Alex Alderton, Roberto Amato, Jeffrey Barrett, Sonia Goncalves, Ewan Harrison, David K. Jackson, Ian Johnston, Dominic Kwiatkowski, Cordelia Langford, John Sillitoe on behalf of the Wellcome Sanger Institute COVID-19 Surveillance Team |
| EPI_ISL_3202694 | Health Services Laboratories | Wellcome Sanger Institute for the COVID-19 Genomics UK (COG-UK) Consortium | Health Services Laboratories and Alex Alderton, Roberto Amato, Jeffrey Barrett, Sonia Goncalves, Ewan Harrison, David K. Jackson, Ian Johnston, Dominic Kwiatkowski, Cordelia Langford, John Sillitoe on behalf of the Wellcome Sanger Institute COVID-19 Surveillance Team |
| EPI_ISL_3202695, EPI_ISL_3202696, EPI_ISL_3202697 | Berkshire and Surrey Pathology Services Lighthouse Laboratory | Wellcome Sanger Institute for the COVID-19 Genomics UK (COG-UK) Consortium | Berkshire and Surrey Pathology Services Lighthouse Laboratory and Alex Alderton, Roberto Amato, Jeffrey Barrett, Sonia Goncalves, Ewan Harrison, David K. Jackson, Ian Johnston, Dominic Kwiatkowski, Cordelia Langford, John Sillitoe on behalf of the Wellcome Sanger Institute COVID-19 Surveillance Team |
| EPI_ISL_3202698, EPI_ISL_3202699 | Lighthouse Lab in Milton Keynes | Wellcome Sanger Institute for the COVID-19 Genomics UK (COG-UK) Consortium | The Lighthouse Lab in Milton Keynes and Alex Alderton, Roberto Amato, Jeffrey Barrett, Sonia Goncalves, Ewan Harrison, David K. Jackson, Ian Johnston, Dominic Kwiatkowski, Cordelia Langford, John Sillitoe on behalf of the Wellcome Sanger Institute COVID-19 Surveillance Team |
| EPI_ISL_3202700, EPI_ISL_3202701, EPI_ISL_3202702, EPI_ISL_3202703, EPI_ISL_3202704 | Lighthouse Laboratory Plymouth | Wellcome Sanger Institute for the COVID-19 Genomics UK (COG-UK) Consortium | Lighthouse Laboratory Plymouth and Alex Alderton, Roberto Amato, Jeffrey Barrett, Sonia Goncalves, Ewan Harrison, David K. Jackson, Ian Johnston, Dominic Kwiatkowski, Cordelia Langford, John Sillitoe on behalf of the Wellcome Sanger Institute COVID-19 Surveillance Team |
| EPI_ISL_3202705, EPI_ISL_3202706 | Lighthouse Lab in Alderley Park | Wellcome Sanger Institute for the COVID-19 Genomics UK (COG-UK) Consortium | Jacquelyn Wynn, Mairead Hyland, The Lighthouse Lab in Alderley Park and Alex Alderton, Roberto Amato, Jeffrey Barrett, Sonia Goncalves, Ewan Harrison, David K. Jackson, Ian Johnston, Dominic Kwiatkowski, Cordelia Langford, John Sillitoe on behalf of the Wellcome Sanger Institute COVID-19 Surveillance Team |
| EPI_ISL_3202707 | Lighthouse Lab in Milton Keynes | Wellcome Sanger Institute for the COVID-19 Genomics UK (COG-UK) Consortium | The Lighthouse Lab in Milton Keynes and Alex Alderton, Roberto Amato, Jeffrey Barrett, Sonia Goncalves, Ewan Harrison, David K. Jackson, Ian Johnston, Dominic Kwiatkowski, Cordelia Langford, John Sillitoe on behalf of the Wellcome Sanger Institute COVID-19 Surveillance Team |
| EPI_ISL_3202708, EPI_ISL_3202709 | Lighthouse Laboratory Plymouth | Wellcome Sanger Institute for the COVID-19 Genomics UK (COG-UK) Consortium | Lighthouse Laboratory Plymouth and Alex Alderton, Roberto Amato, Jeffrey Barrett, Sonia Goncalves, Ewan Harrison, David K. Jackson, Ian Johnston, Dominic Kwiatkowski, Cordelia Langford, John Sillitoe on behalf of the Wellcome Sanger Institute COVID-19 Surveillance Team |
| EPI_ISL_3202710, EPI_ISL_3202711, EPI_ISL_3202712 | Lighthouse Lab in Alderley Park | Wellcome Sanger Institute for the COVID-19 Genomics UK (COG-UK) Consortium | Jacquelyn Wynn, Mairead Hyland, The Lighthouse Lab in Alderley Park and Alex Alderton, Roberto Amato, Jeffrey Barrett, Sonia Goncalves, Ewan Harrison, David K. Jackson, Ian Johnston, Dominic Kwiatkowski, Cordelia Langford, John Sillitoe on behalf of the Wellcome Sanger Institute COVID-19 Surveillance Team |
| EPI_ISL_3202713 | Berkshire and Surrey Pathology Services Lighthouse Laboratory | Wellcome Sanger Institute for the COVID-19 Genomics UK (COG-UK) Consortium | Berkshire and Surrey Pathology Services Lighthouse Laboratory and Alex Alderton, Roberto Amato, Jeffrey Barrett, Sonia Goncalves, Ewan Harrison, David K. Jackson, Ian Johnston, Dominic Kwiatkowski, Cordelia Langford, John Sillitoe on behalf of the Wellcome Sanger Institute COVID-19 Surveillance Team |
| EPI_ISL_3202714 | Lighthouse Lab in Milton Keynes | Wellcome Sanger Institute for the COVID-19 Genomics UK (COG-UK) Consortium | The Lighthouse Lab in Milton Keynes and Alex Alderton, Roberto Amato, Jeffrey Barrett, Sonia Goncalves, Ewan Harrison, David K. Jackson, Ian Johnston, Dominic Kwiatkowski, Cordelia Langford, John Sillitoe on behalf of the Wellcome Sanger Institute COVID-19 Surveillance Team |
| EPI_ISL_3202715 | Lighthouse Laboratory Plymouth | Wellcome Sanger Institute for the COVID-19 Genomics UK (COG-UK) Consortium | Lighthouse Laboratory Plymouth and Alex Alderton, Roberto Amato, Jeffrey Barrett, Sonia Goncalves, Ewan Harrison, David K. Jackson, Ian Johnston, Dominic Kwiatkowski, Cordelia Langford, John Sillitoe on behalf of the Wellcome Sanger Institute COVID-19 Surveillance Team |
| EPI_ISL_3202716 | Lighthouse Lab in Alderley Park | Wellcome Sanger Institute for the COVID-19 Genomics UK (COG-UK) Consortium | Jacquelyn Wynn, Mairead Hyland, The Lighthouse Lab in Alderley Park and Alex Alderton, Roberto Amato, Jeffrey Barrett, Sonia Goncalves, Ewan Harrison, David K. Jackson, Ian Johnston, Dominic Kwiatkowski, Cordelia Langford, John Sillitoe on behalf of the Wellcome Sanger Institute COVID-19 Surveillance Team |
| EPI_ISL_3202717 | Lighthouse Lab in Milton Keynes | Wellcome Sanger Institute for the COVID-19 Genomics UK (COG-UK) Consortium | The Lighthouse Lab in Milton Keynes and Alex Alderton, Roberto Amato, Jeffrey Barrett, Sonia Goncalves, Ewan Harrison, David K. Jackson, Ian Johnston, Dominic Kwiatkowski, Cordelia Langford, John Sillitoe on behalf of the Wellcome Sanger Institute COVID-19 Surveillance Team |
| EPI_ISL_3202718 | Lighthouse Laboratory Plymouth | Wellcome Sanger Institute for the COVID-19 Genomics UK (COG-UK) Consortium | Lighthouse Laboratory Plymouth and Alex Alderton, Roberto Amato, Jeffrey Barrett, Sonia Goncalves, Ewan Harrison, David K. Jackson, Ian Johnston, Dominic Kwiatkowski, Cordelia Langford, John Sillitoe on behalf of the Wellcome Sanger Institute COVID-19 Surveillance Team |
| EPI_ISL_3202719 | Lighthouse Lab in Alderley Park | Wellcome Sanger Institute for the COVID-19 Genomics UK (COG-UK) Consortium | Jacquelyn Wynn, Mairead Hyland, The Lighthouse Lab in Alderley Park and Alex Alderton, Roberto Amato, Jeffrey Barrett, Sonia Goncalves, Ewan Harrison, David K. Jackson, Ian Johnston, Dominic Kwiatkowski, Cordelia Langford, John Sillitoe on behalf of the Wellcome Sanger Institute COVID-19 Surveillance Team |
| EPI_ISL_3202720 | Berkshire and Surrey Pathology Services Lighthouse Laboratory | Wellcome Sanger Institute for the COVID-19 Genomics UK (COG-UK) Consortium | Berkshire and Surrey Pathology Services Lighthouse Laboratory and Alex Alderton, Roberto Amato, Jeffrey Barrett, Sonia Goncalves, Ewan Harrison, David K. Jackson, Ian Johnston, Dominic Kwiatkowski, Cordelia Langford, John Sillitoe on behalf of the Wellcome Sanger Institute COVID-19 Surveillance Team |
| EPI_ISL_3202721, EPI_ISL_3202722, EPI_ISL_3202723 | Lighthouse Lab in Milton Keynes | Wellcome Sanger Institute for the COVID-19 Genomics UK (COG-UK) Consortium | The Lighthouse Lab in Milton Keynes and Alex Alderton, Roberto Amato, Jeffrey Barrett, Sonia Goncalves, Ewan Harrison, David K. Jackson, Ian Johnston, Dominic Kwiatkowski, Cordelia Langford, John Sillitoe on behalf of the Wellcome Sanger Institute COVID-19 Surveillance Team |
| EPI_ISL_3202724 | Lighthouse Laboratory Plymouth | Wellcome Sanger Institute for the COVID-19 Genomics UK (COG-UK) Consortium | Lighthouse Laboratory Plymouth and Alex Alderton, Roberto Amato, Jeffrey Barrett, Sonia Goncalves, Ewan Harrison, David K. Jackson, Ian Johnston, Dominic Kwiatkowski, Cordelia Langford, John Sillitoe on behalf of the Wellcome Sanger Institute COVID-19 Surveillance Team |
| EPI_ISL_3202725, EPI_ISL_3202726 | Lighthouse Lab in Alderley Park | Wellcome Sanger Institute for the COVID-19 Genomics UK (COG-UK) Consortium | Jacquelyn Wynn, Mairead Hyland, The Lighthouse Lab in Alderley Park and Alex Alderton, Roberto Amato, Jeffrey Barrett, Sonia Goncalves, Ewan Harrison, David K. Jackson, Ian Johnston, Dominic Kwiatkowski, Cordelia Langford, John Sillitoe on behalf of the Wellcome Sanger Institute COVID-19 Surveillance Team |
| EPI_ISL_3202727 | Berkshire and Surrey Pathology Services Lighthouse Laboratory | Wellcome Sanger Institute for the COVID-19 Genomics UK (COG-UK) Consortium | Berkshire and Surrey Pathology Services Lighthouse Laboratory and Alex Alderton, Roberto Amato, Jeffrey Barrett, Sonia Goncalves, Ewan Harrison, David K. Jackson, Ian Johnston, Dominic Kwiatkowski, Cordelia Langford, John Sillitoe on behalf of the Wellcome Sanger Institute COVID-19 Surveillance Team |
| EPI_ISL_3202728 | Lighthouse Laboratory Plymouth | Wellcome Sanger Institute for the COVID-19 Genomics UK (COG-UK) Consortium | Lighthouse Laboratory Plymouth and Alex Alderton, Roberto Amato, Jeffrey Barrett, Sonia Goncalves, Ewan Harrison, David K. Jackson, Ian Johnston, Dominic Kwiatkowski, Cordelia Langford, John Sillitoe on behalf of the Wellcome Sanger Institute COVID-19 Surveillance Team |
| EPI_ISL_3202729 | Lighthouse Lab in Alderley Park | Wellcome Sanger Institute for the COVID-19 Genomics UK (COG-UK) Consortium | Jacquelyn Wynn, Mairead Hyland, The Lighthouse Lab in Alderley Park and Alex Alderton, Roberto Amato, Jeffrey Barrett, Sonia Goncalves, Ewan Harrison, David K. Jackson, Ian Johnston, Dominic Kwiatkowski, Cordelia Langford, John Sillitoe on behalf of the Wellcome Sanger Institute COVID-19 Surveillance Team |
| EPI_ISL_3202730 | Lighthouse Lab in Milton Keynes | Wellcome Sanger Institute for the COVID-19 Genomics UK (COG-UK) Consortium | The Lighthouse Lab in Milton Keynes and Alex Alderton, Roberto Amato, Jeffrey Barrett, Sonia Goncalves, Ewan Harrison, David K. Jackson, Ian Johnston, Dominic Kwiatkowski, Cordelia Langford, John Sillitoe on behalf of the Wellcome Sanger Institute COVID-19 Surveillance Team |
| EPI_ISL_3202731, EPI_ISL_3202732, EPI_ISL_3202733 | Lighthouse Lab in Alderley Park | Wellcome Sanger Institute for the COVID-19 Genomics UK (COG-UK) Consortium | Jacquelyn Wynn, Mairead Hyland, The Lighthouse Lab in Alderley Park and Alex Alderton, Roberto Amato, Jeffrey Barrett, Sonia Goncalves, Ewan Harrison, David K. Jackson, Ian Johnston, Dominic Kwiatkowski, Cordelia Langford, John Sillitoe on behalf of the Wellcome Sanger Institute COVID-19 Surveillance Team |
| EPI_ISL_3202734 | Berkshire and Surrey Pathology Services Lighthouse Laboratory | Wellcome Sanger Institute for the COVID-19 Genomics UK (COG-UK) Consortium | Berkshire and Surrey Pathology Services Lighthouse Laboratory and Alex Alderton, Roberto Amato, Jeffrey Barrett, Sonia Goncalves, Ewan Harrison, David K. Jackson, Ian Johnston, Dominic Kwiatkowski, Cordelia Langford, John Sillitoe on behalf of the Wellcome Sanger Institute COVID-19 Surveillance Team |
| EPI_ISL_3202735 | Lighthouse Lab in Milton Keynes | Wellcome Sanger Institute for the COVID-19 Genomics UK (COG-UK) Consortium | The Lighthouse Lab in Milton Keynes and Alex Alderton, Roberto Amato, Jeffrey Barrett, Sonia Goncalves, Ewan Harrison, David K. Jackson, Ian Johnston, Dominic Kwiatkowski, Cordelia Langford, John Sillitoe on behalf of the Wellcome Sanger Institute COVID-19 Surveillance Team |
| EPI_ISL_3202736 | Berkshire and Surrey Pathology Services Lighthouse Laboratory | Wellcome Sanger Institute for the COVID-19 Genomics UK (COG-UK) Consortium | Berkshire and Surrey Pathology Services Lighthouse Laboratory and Alex Alderton, Roberto Amato, Jeffrey Barrett, Sonia Goncalves, Ewan Harrison, David K. Jackson, Ian Johnston, Dominic Kwiatkowski, Cordelia Langford, John Sillitoe on behalf of the Wellcome Sanger Institute COVID-19 Surveillance Team |
| EPI_ISL_3202737 | Lighthouse Lab in Milton Keynes | Wellcome Sanger Institute for the COVID-19 Genomics UK (COG-UK) Consortium | The Lighthouse Lab in Milton Keynes and Alex Alderton, Roberto Amato, Jeffrey Barrett, Sonia Goncalves, Ewan Harrison, David K. Jackson, Ian Johnston, Dominic Kwiatkowski, Cordelia Langford, John Sillitoe on behalf of the Wellcome Sanger Institute COVID-19 Surveillance Team |
| EPI_ISL_3202738 | Lighthouse Laboratory Plymouth | Wellcome Sanger Institute for the COVID-19 Genomics UK (COG-UK) Consortium | Lighthouse Laboratory Plymouth and Alex Alderton, Roberto Amato, Jeffrey Barrett, Sonia Goncalves, Ewan Harrison, David K. Jackson, Ian Johnston, Dominic Kwiatkowski, Cordelia Langford, John Sillitoe on behalf of the Wellcome Sanger Institute COVID-19 Surveillance Team |
| EPI_ISL_3202739 | Lighthouse Lab in Milton Keynes | Wellcome Sanger Institute for the COVID-19 Genomics UK (COG-UK) Consortium | The Lighthouse Lab in Milton Keynes and Alex Alderton, Roberto Amato, Jeffrey Barrett, Sonia Goncalves, Ewan Harrison, David K. Jackson, Ian Johnston, Dominic Kwiatkowski, Cordelia Langford, John Sillitoe on behalf of the Wellcome Sanger Institute COVID-19 Surveillance Team |
| EPI_ISL_3202740, EPI_ISL_3202741, EPI_ISL_3202742 | Berkshire and Surrey Pathology Services Lighthouse Laboratory | Wellcome Sanger Institute for the COVID-19 Genomics UK (COG-UK) Consortium | Berkshire and Surrey Pathology Services Lighthouse Laboratory and Alex Alderton, Roberto Amato, Jeffrey Barrett, Sonia Goncalves, Ewan Harrison, David K. Jackson, Ian Johnston, Dominic Kwiatkowski, Cordelia Langford, John Sillitoe on behalf of the Wellcome Sanger Institute COVID-19 Surveillance Team |

| | | | |
|---|---|---|---|
| EPI_ISL_3202743 | Lighthouse Laboratory Plymouth | Wellcome Sanger Institute for the COVID-19 Genomics UK (COG-UK) Consortium | Lighthouse Laboratory Plymouth and Alex Alderton, Roberto Amato, Jeffrey Barrett, Sonia Goncalves, Ewan Harrison, David K. Jackson, Ian Johnston, Dominic Kwiatkowski, Cordelia Langford, John Sillitoe on behalf of the Wellcome Sanger Institute COVID-19 Surveillance Team |
| EPI_ISL_3202744, EPI_ISL_3202745 | Lighthouse Lab in Alderley Park | Wellcome Sanger Institute for the COVID-19 Genomics UK (COG-UK) Consortium | Jacquelyn Wynn, Mairead Hyland, The Lighthouse Lab in Alderley Park and Alex Alderton, Roberto Amato, Jeffrey Barrett, Sonia Goncalves, Ewan Harrison, David K. Jackson, Ian Johnston, Dominic Kwiatkowski, Cordelia Langford, John Sillitoe on behalf of the Wellcome Sanger Institute COVID-19 Surveillance Team |
| EPI_ISL_3202746 | Berkshire and Surrey Pathology Services Lighthouse Laboratory | Wellcome Sanger Institute for the COVID-19 Genomics UK (COG-UK) Consortium | Berkshire and Surrey Pathology Services Lighthouse Laboratory and Alex Alderton, Roberto Amato, Jeffrey Barrett, Sonia Goncalves, Ewan Harrison, David K. Jackson, Ian Johnston, Dominic Kwiatkowski, Cordelia Langford, John Sillitoe on behalf of the Wellcome Sanger Institute COVID-19 Surveillance Team |
| EPI_ISL_3202747 | Lighthouse Lab in Alderley Park | Wellcome Sanger Institute for the COVID-19 Genomics UK (COG-UK) Consortium | Jacquelyn Wynn, Mairead Hyland, The Lighthouse Lab in Alderley Park and Alex Alderton, Roberto Amato, Jeffrey Barrett, Sonia Goncalves, Ewan Harrison, David K. Jackson, Ian Johnston, Dominic Kwiatkowski, Cordelia Langford, John Sillitoe on behalf of the Wellcome Sanger Institute COVID-19 Surveillance Team |
| EPI_ISL_3202748 | Lighthouse Laboratory Plymouth | Wellcome Sanger Institute for the COVID-19 Genomics UK (COG-UK) Consortium | Lighthouse Laboratory Plymouth and Alex Alderton, Roberto Amato, Jeffrey Barrett, Sonia Goncalves, Ewan Harrison, David K. Jackson, Ian Johnston, Dominic Kwiatkowski, Cordelia Langford, John Sillitoe on behalf of the Wellcome Sanger Institute COVID-19 Surveillance Team |
| EPI_ISL_3202749 | Berkshire and Surrey Pathology Services Lighthouse Laboratory | Wellcome Sanger Institute for the COVID-19 Genomics UK (COG-UK) Consortium | Berkshire and Surrey Pathology Services Lighthouse Laboratory and Alex Alderton, Roberto Amato, Jeffrey Barrett, Sonia Goncalves, Ewan Harrison, David K. Jackson, Ian Johnston, Dominic Kwiatkowski, Cordelia Langford, John Sillitoe on behalf of the Wellcome Sanger Institute COVID-19 Surveillance Team |
| EPI_ISL_3202750 | Lighthouse Laboratory Plymouth | Wellcome Sanger Institute for the COVID-19 Genomics UK (COG-UK) Consortium | Lighthouse Laboratory Plymouth and Alex Alderton, Roberto Amato, Jeffrey Barrett, Sonia Goncalves, Ewan Harrison, David K. Jackson, Ian Johnston, Dominic Kwiatkowski, Cordelia Langford, John Sillitoe on behalf of the Wellcome Sanger Institute COVID-19 Surveillance Team |
| EPI_ISL_3202751 | Lighthouse Lab in Milton Keynes | Wellcome Sanger Institute for the COVID-19 Genomics UK (COG-UK) Consortium | The Lighthouse Lab in Milton Keynes and Alex Alderton, Roberto Amato, Jeffrey Barrett, Sonia Goncalves, Ewan Harrison, David K. Jackson, Ian Johnston, Dominic Kwiatkowski, Cordelia Langford, John Sillitoe on behalf of the Wellcome Sanger Institute COVID-19 Surveillance Team |
| EPI_ISL_3202752, EPI_ISL_3202753 | Berkshire and Surrey Pathology Services Lighthouse Laboratory | Wellcome Sanger Institute for the COVID-19 Genomics UK (COG-UK) Consortium | Berkshire and Surrey Pathology Services Lighthouse Laboratory and Alex Alderton, Roberto Amato, Jeffrey Barrett, Sonia Goncalves, Ewan Harrison, David K. Jackson, Ian Johnston, Dominic Kwiatkowski, Cordelia Langford, John Sillitoe on behalf of the Wellcome Sanger Institute COVID-19 Surveillance Team |
| EPI_ISL_3202754, EPI_ISL_3202755 | Lighthouse Lab in Alderley Park | Wellcome Sanger Institute for the COVID-19 Genomics UK (COG-UK) Consortium | Jacquelyn Wynn, Mairead Hyland, The Lighthouse Lab in Alderley Park and Alex Alderton, Roberto Amato, Jeffrey Barrett, Sonia Goncalves, Ewan Harrison, David K. Jackson, Ian Johnston, Dominic Kwiatkowski, Cordelia Langford, John Sillitoe on behalf of the Wellcome Sanger Institute COVID-19 Surveillance Team |
| EPI_ISL_3202756 | Lighthouse Laboratory Plymouth | Wellcome Sanger Institute for the COVID-19 Genomics UK (COG-UK) Consortium | Lighthouse Laboratory Plymouth and Alex Alderton, Roberto Amato, Jeffrey Barrett, Sonia Goncalves, Ewan Harrison, David K. Jackson, Ian Johnston, Dominic Kwiatkowski, Cordelia Langford, John Sillitoe on behalf of the Wellcome Sanger Institute COVID-19 Surveillance Team |
| EPI_ISL_3202757 | Lighthouse Lab in Alderley Park | Wellcome Sanger Institute for the COVID-19 Genomics UK (COG-UK) Consortium | Jacquelyn Wynn, Mairead Hyland, The Lighthouse Lab in Alderley Park and Alex Alderton, Roberto Amato, Jeffrey Barrett, Sonia Goncalves, Ewan Harrison, David K. Jackson, Ian Johnston, Dominic Kwiatkowski, Cordelia Langford, John Sillitoe on behalf of the Wellcome Sanger Institute COVID-19 Surveillance Team |
| EPI_ISL_3202758 | Berkshire and Surrey Pathology Services Lighthouse Laboratory | Wellcome Sanger Institute for the COVID-19 Genomics UK (COG-UK) Consortium | Berkshire and Surrey Pathology Services Lighthouse Laboratory and Alex Alderton, Roberto Amato, Jeffrey Barrett, Sonia Goncalves, Ewan Harrison, David K. Jackson, Ian Johnston, Dominic Kwiatkowski, Cordelia Langford, John Sillitoe on behalf of the Wellcome Sanger Institute COVID-19 Surveillance Team |
| EPI_ISL_3202759 | Lighthouse Laboratory Plymouth | Wellcome Sanger Institute for the COVID-19 Genomics UK (COG-UK) Consortium | Lighthouse Laboratory Plymouth and Alex Alderton, Roberto Amato, Jeffrey Barrett, Sonia Goncalves, Ewan Harrison, David K. Jackson, Ian Johnston, Dominic Kwiatkowski, Cordelia Langford, John Sillitoe on behalf of the Wellcome Sanger Institute COVID-19 Surveillance Team |
| EPI_ISL_3202760 | Lighthouse Lab in Alderley Park | Wellcome Sanger Institute for the COVID-19 Genomics UK (COG-UK) Consortium | Jacquelyn Wynn, Mairead Hyland, The Lighthouse Lab in Alderley Park and Alex Alderton, Roberto Amato, Jeffrey Barrett, Sonia Goncalves, Ewan Harrison, David K. Jackson, Ian Johnston, Dominic Kwiatkowski, Cordelia Langford, John Sillitoe on behalf of the Wellcome Sanger Institute COVID-19 Surveillance Team |
| EPI_ISL_3202761 | Berkshire and Surrey Pathology Services Lighthouse Laboratory | Wellcome Sanger Institute for the COVID-19 Genomics UK (COG-UK) Consortium | Berkshire and Surrey Pathology Services Lighthouse Laboratory and Alex Alderton, Roberto Amato, Jeffrey Barrett, Sonia Goncalves, Ewan Harrison, David K. Jackson, Ian Johnston, Dominic Kwiatkowski, Cordelia Langford, John Sillitoe on behalf of the Wellcome Sanger Institute COVID-19 Surveillance Team |
| EPI_ISL_3202762 | Lighthouse Laboratory Plymouth | Wellcome Sanger Institute for the COVID-19 Genomics UK (COG-UK) Consortium | Lighthouse Laboratory Plymouth and Alex Alderton, Roberto Amato, Jeffrey Barrett, Sonia Goncalves, Ewan Harrison, David K. Jackson, Ian Johnston, Dominic Kwiatkowski, Cordelia Langford, John Sillitoe on behalf of the Wellcome Sanger Institute COVID-19 Surveillance Team |
| EPI_ISL_3202763 | Lighthouse Lab in Alderley Park | Wellcome Sanger Institute for the COVID-19 Genomics UK (COG-UK) Consortium | Jacquelyn Wynn, Mairead Hyland, The Lighthouse Lab in Alderley Park and Alex Alderton, Roberto Amato, Jeffrey Barrett, Sonia Goncalves, Ewan Harrison, David K. Jackson, Ian Johnston, Dominic Kwiatkowski, Cordelia Langford, John Sillitoe on behalf of the Wellcome Sanger Institute COVID-19 Surveillance Team |
| EPI_ISL_3202764, EPI_ISL_3202765 | Lighthouse Lab in Milton Keynes | Wellcome Sanger Institute for the COVID-19 Genomics UK (COG-UK) Consortium | The Lighthouse Lab in Milton Keynes and Alex Alderton, Roberto Amato, Jeffrey Barrett, Sonia Goncalves, Ewan Harrison, David K. Jackson, Ian Johnston, Dominic Kwiatkowski, Cordelia Langford, John Sillitoe on behalf of the Wellcome Sanger Institute COVID-19 Surveillance Team |
| EPI_ISL_3202766 | Lighthouse Lab in Alderley Park | Wellcome Sanger Institute for the COVID-19 Genomics UK (COG-UK) Consortium | Jacquelyn Wynn, Mairead Hyland, The Lighthouse Lab in Alderley Park and Alex Alderton, Roberto Amato, Jeffrey Barrett, Sonia Goncalves, Ewan Harrison, David K. Jackson, Ian Johnston, Dominic Kwiatkowski, Cordelia Langford, John Sillitoe on behalf of the Wellcome Sanger Institute COVID-19 Surveillance Team |
| EPI_ISL_3202767 | Lighthouse Lab in Milton Keynes | Wellcome Sanger Institute for the COVID-19 Genomics UK (COG-UK) Consortium | The Lighthouse Lab in Milton Keynes and Alex Alderton, Roberto Amato, Jeffrey Barrett, Sonia Goncalves, Ewan Harrison, David K. Jackson, Ian Johnston, Dominic Kwiatkowski, Cordelia Langford, John Sillitoe on behalf of the Wellcome Sanger Institute COVID-19 Surveillance Team |
| EPI_ISL_3202768 | Lighthouse Lab in Alderley Park | Wellcome Sanger Institute for the COVID-19 Genomics UK (COG-UK) Consortium | Jacquelyn Wynn, Mairead Hyland, The Lighthouse Lab in Alderley Park and Alex Alderton, Roberto Amato, Jeffrey Barrett, Sonia Goncalves, Ewan Harrison, David K. Jackson, Ian Johnston, Dominic Kwiatkowski, Cordelia Langford, John Sillitoe on behalf of the Wellcome Sanger Institute COVID-19 Surveillance Team |
| EPI_ISL_3202769 | Lighthouse Laboratory Plymouth | Wellcome Sanger Institute for the COVID-19 Genomics UK (COG-UK) Consortium | Lighthouse Laboratory Plymouth and Alex Alderton, Roberto Amato, Jeffrey Barrett, Sonia Goncalves, Ewan Harrison, David K. Jackson, Ian Johnston, Dominic Kwiatkowski, Cordelia Langford, John Sillitoe on behalf of the Wellcome Sanger Institute COVID-19 Surveillance Team |
| EPI_ISL_3202770 | Lighthouse Lab in Milton Keynes | Wellcome Sanger Institute for the COVID-19 Genomics UK (COG-UK) Consortium | The Lighthouse Lab in Milton Keynes and Alex Alderton, Roberto Amato, Jeffrey Barrett, Sonia Goncalves, Ewan Harrison, David K. Jackson, Ian Johnston, Dominic Kwiatkowski, Cordelia Langford, John Sillitoe on behalf of the Wellcome Sanger Institute COVID-19 Surveillance Team |
| EPI_ISL_3202771 | Lighthouse Lab in Alderley Park | Wellcome Sanger Institute for the COVID-19 Genomics UK (COG-UK) Consortium | Jacquelyn Wynn, Mairead Hyland, The Lighthouse Lab in Alderley Park and Alex Alderton, Roberto Amato, Jeffrey Barrett, Sonia Goncalves, Ewan Harrison, David K. Jackson, Ian Johnston, Dominic Kwiatkowski, Cordelia Langford, John Sillitoe on behalf of the Wellcome Sanger Institute COVID-19 Surveillance Team |
| EPI_ISL_3202772 | Berkshire and Surrey Pathology Services Lighthouse Laboratory | Wellcome Sanger Institute for the COVID-19 Genomics UK (COG-UK) Consortium | Berkshire and Surrey Pathology Services Lighthouse Laboratory and Alex Alderton, Roberto Amato, Jeffrey Barrett, Sonia Goncalves, Ewan Harrison, David K. Jackson, Ian Johnston, Dominic Kwiatkowski, Cordelia Langford, John Sillitoe on behalf of the Wellcome Sanger Institute COVID-19 Surveillance Team |
| EPI_ISL_3202773 | Lighthouse Laboratory Plymouth | Wellcome Sanger Institute for the COVID-19 Genomics UK (COG-UK) Consortium | Lighthouse Laboratory Plymouth and Alex Alderton, Roberto Amato, Jeffrey Barrett, Sonia Goncalves, Ewan Harrison, David K. Jackson, Ian Johnston, Dominic Kwiatkowski, Cordelia Langford, John Sillitoe on behalf of the Wellcome Sanger Institute COVID-19 Surveillance Team |
| EPI_ISL_3202774 | Lighthouse Lab in Milton Keynes | Wellcome Sanger Institute for the COVID-19 Genomics UK (COG-UK) Consortium | The Lighthouse Lab in Milton Keynes and Alex Alderton, Roberto Amato, Jeffrey Barrett, Sonia Goncalves, Ewan Harrison, David K. Jackson, Ian Johnston, Dominic Kwiatkowski, Cordelia Langford, John Sillitoe on behalf of the Wellcome Sanger Institute COVID-19 Surveillance Team |
| EPI_ISL_3202775 | Lighthouse Laboratory Plymouth | Wellcome Sanger Institute for the COVID-19 Genomics UK (COG-UK) Consortium | Lighthouse Laboratory Plymouth and Alex Alderton, Roberto Amato, Jeffrey Barrett, Sonia Goncalves, Ewan Harrison, David K. Jackson, Ian Johnston, Dominic Kwiatkowski, Cordelia Langford, John Sillitoe on behalf of the Wellcome Sanger Institute COVID-19 Surveillance Team |
| EPI_ISL_3202776 | Lighthouse Lab in Milton Keynes | Wellcome Sanger Institute for the COVID-19 Genomics UK (COG-UK) Consortium | The Lighthouse Lab in Milton Keynes and Alex Alderton, Roberto Amato, Jeffrey Barrett, Sonia Goncalves, Ewan Harrison, David K. Jackson, Ian Johnston, Dominic Kwiatkowski, Cordelia Langford, John Sillitoe on behalf of the Wellcome Sanger Institute COVID-19 Surveillance Team |
| EPI_ISL_3202777 | Lighthouse Laboratory Plymouth | Wellcome Sanger Institute for the COVID-19 Genomics UK (COG-UK) Consortium | Lighthouse Laboratory Plymouth and Alex Alderton, Roberto Amato, Jeffrey Barrett, Sonia Goncalves, Ewan Harrison, David K. Jackson, Ian Johnston, Dominic Kwiatkowski, Cordelia Langford, John Sillitoe on behalf of the Wellcome Sanger Institute COVID-19 Surveillance Team |
| EPI_ISL_3202778 | Lighthouse Lab in Alderley Park | Wellcome Sanger Institute for the COVID-19 Genomics UK (COG-UK) Consortium | Jacquelyn Wynn, Mairead Hyland, The Lighthouse Lab in Alderley Park and Alex Alderton, Roberto Amato, Jeffrey Barrett, Sonia Goncalves, Ewan Harrison, David K. Jackson, Ian Johnston, Dominic Kwiatkowski, Cordelia Langford, John Sillitoe on behalf of the Wellcome Sanger Institute COVID-19 Surveillance Team |
| EPI_ISL_3202779, EPI_ISL_3202780, EPI_ISL_3202781, EPI_ISL_3202782, EPI_ISL_3202783 | Berkshire and Surrey Pathology Services Lighthouse Laboratory | Wellcome Sanger Institute for the COVID-19 Genomics UK (COG-UK) Consortium | Berkshire and Surrey Pathology Services Lighthouse Laboratory and Alex Alderton, Roberto Amato, Jeffrey Barrett, Sonia Goncalves, Ewan Harrison, David K. Jackson, Ian Johnston, Dominic Kwiatkowski, Cordelia Langford, John Sillitoe on behalf of the Wellcome Sanger Institute COVID-19 Surveillance Team |
| EPI_ISL_3202784, EPI_ISL_3202785 | Lighthouse Lab in Alderley Park | Wellcome Sanger Institute for the COVID-19 Genomics UK (COG-UK) Consortium | Jacquelyn Wynn, Mairead Hyland, The Lighthouse Lab in Alderley Park and Alex Alderton, Roberto Amato, Jeffrey Barrett, Sonia Goncalves, Ewan Harrison, David K. Jackson, Ian Johnston, Dominic Kwiatkowski, Cordelia Langford, John Sillitoe on behalf of the Wellcome Sanger Institute COVID-19 Surveillance Team |
| EPI_ISL_3202786 | Lighthouse Laboratory Plymouth | Wellcome Sanger Institute for the COVID-19 Genomics UK (COG-UK) Consortium | Lighthouse Laboratory Plymouth and Alex Alderton, Roberto Amato, Jeffrey Barrett, Sonia Goncalves, Ewan Harrison, David K. Jackson, Ian Johnston, Dominic Kwiatkowski, Cordelia Langford, John Sillitoe on behalf of the Wellcome Sanger Institute COVID-19 Surveillance Team |
| EPI_ISL_3202787 | Lighthouse Lab in Milton Keynes | Wellcome Sanger Institute for the COVID-19 Genomics UK (COG-UK) Consortium | The Lighthouse Lab in Milton Keynes and Alex Alderton, Roberto Amato, Jeffrey Barrett, Sonia Goncalves, Ewan Harrison, David K. Jackson, Ian Johnston, Dominic Kwiatkowski, Cordelia Langford, John Sillitoe on behalf of the Wellcome Sanger Institute COVID-19 Surveillance Team |
| EPI_ISL_3202788 | Lighthouse Laboratory Plymouth | Wellcome Sanger Institute for the COVID-19 Genomics UK (COG-UK) Consortium | Lighthouse Laboratory Plymouth and Alex Alderton, Roberto Amato, Jeffrey Barrett, Sonia Goncalves, Ewan Harrison, David K. Jackson, Ian Johnston, Dominic Kwiatkowski, Cordelia Langford, John Sillitoe on behalf of the Wellcome Sanger Institute COVID-19 Surveillance Team |
| EPI_ISL_3202789 | Berkshire and Surrey Pathology Services Lighthouse Laboratory | Wellcome Sanger Institute for the COVID-19 Genomics UK (COG-UK) Consortium | Berkshire and Surrey Pathology Services Lighthouse Laboratory and Alex Alderton, Roberto Amato, Jeffrey Barrett, Sonia Goncalves, Ewan Harrison, David K. Jackson, Ian Johnston, Dominic Kwiatkowski, Cordelia Langford, John Sillitoe on behalf of the Wellcome Sanger Institute COVID-19 Surveillance Team |
| EPI_ISL_3202790 | Lighthouse Lab in Alderley Park | Wellcome Sanger Institute for the COVID-19 Genomics UK (COG-UK) Consortium | Jacquelyn Wynn, Mairead Hyland, The Lighthouse Lab in Alderley Park and Alex Alderton, Roberto Amato, Jeffrey Barrett, Sonia Goncalves, Ewan Harrison, David K. Jackson, Ian Johnston, Dominic Kwiatkowski, Cordelia Langford, John Sillitoe on behalf of the Wellcome Sanger Institute COVID-19 Surveillance Team |
| EPI_ISL_3202791, EPI_ISL_3202792 | Lighthouse Laboratory Plymouth | Wellcome Sanger Institute for the COVID-19 Genomics UK (COG-UK) Consortium | Lighthouse Laboratory Plymouth and Alex Alderton, Roberto Amato, Jeffrey Barrett, Sonia Goncalves, Ewan Harrison, David K. Jackson, Ian Johnston, Dominic Kwiatkowski, Cordelia Langford, John Sillitoe on behalf of the Wellcome Sanger Institute COVID-19 Surveillance Team |
| EPI_ISL_3202793 | Lighthouse Lab in Alderley Park | Wellcome Sanger Institute for the COVID-19 Genomics UK (COG-UK) Consortium | Jacquelyn Wynn, Mairead Hyland, The Lighthouse Lab in Alderley Park and Alex Alderton, Roberto Amato, Jeffrey Barrett, Sonia Goncalves, Ewan Harrison, David K. Jackson, Ian Johnston, Dominic Kwiatkowski, Cordelia Langford, John Sillitoe on behalf of the Wellcome Sanger Institute COVID-19 Surveillance Team |
| EPI_ISL_3202794 | Lighthouse Laboratory Plymouth | Wellcome Sanger Institute for the COVID-19 Genomics UK (COG-UK) Consortium | Lighthouse Laboratory Plymouth and Alex Alderton, Roberto Amato, Jeffrey Barrett, Sonia Goncalves, Ewan Harrison, David K. Jackson, Ian Johnston, Dominic Kwiatkowski, Cordelia Langford, John Sillitoe on behalf of the Wellcome Sanger Institute COVID-19 Surveillance Team |
| EPI_ISL_3202795 | Berkshire and Surrey Pathology Services Lighthouse Laboratory | Wellcome Sanger Institute for the COVID-19 Genomics UK (COG-UK) Consortium | Berkshire and Surrey Pathology Services Lighthouse Laboratory and Alex Alderton, Roberto Amato, Jeffrey Barrett, Sonia Goncalves, Ewan Harrison, David K. Jackson, Ian Johnston, Dominic Kwiatkowski, Cordelia Langford, John Sillitoe on behalf of the Wellcome Sanger Institute COVID-19 Surveillance Team |
| EPI_ISL_3202796, EPI_ISL_3202797, | Lighthouse Lab in Alderley Park | Wellcome Sanger Institute for the COVID-19 Genomics UK (COG-UK) Consortium | Jacquelyn Wynn, Mairead Hyland, The Lighthouse Lab in Alderley Park and Alex Alderton, Roberto Amato, Jeffrey Barrett, Sonia Goncalves, Ewan Harrison, David K. Jackson, Ian Johnston, Dominic Kwiatkowski, Cordelia Langford, John Sillitoe on behalf of the Wellcome Sanger Institute COVID-19 Surveillance Team |

| | | | |
|---|---|---|---|
| EPI_ISL_3202798, EPI_ISL_3202799 | | | |
| EPI_ISL_3202800, EPI_ISL_3202801, EPI_ISL_3202802 | Lighthouse Lab in Milton Keynes | Wellcome Sanger Institute for the COVID-19 Genomics UK (COG-UK) Consortium | The Lighthouse Lab in Milton Keynes and Alex Alderton, Roberto Amato, Jeffrey Barrett, Sonia Goncalves, Ewan Harrison, David K. Jackson, Ian Johnston, Dominic Kwiatkowski, Cordelia Langford, John Sillitoe on behalf of the Wellcome Sanger Institute COVID-19 Surveillance Team |
| EPI_ISL_3202803 | Berkshire and Surrey Pathology Services Lighthouse Laboratory | Wellcome Sanger Institute for the COVID-19 Genomics UK (COG-UK) Consortium | Berkshire and Surrey Pathology Services Lighthouse Laboratory and Alex Alderton, Roberto Amato, Jeffrey Barrett, Sonia Goncalves, Ewan Harrison, David K. Jackson, Ian Johnston, Dominic Kwiatkowski, Cordelia Langford, John Sillitoe on behalf of the Wellcome Sanger Institute COVID-19 Surveillance Team |
| EPI_ISL_3202804 | Lighthouse Lab in Milton Keynes | Wellcome Sanger Institute for the COVID-19 Genomics UK (COG-UK) Consortium | The Lighthouse Lab in Milton Keynes and Alex Alderton, Roberto Amato, Jeffrey Barrett, Sonia Goncalves, Ewan Harrison, David K. Jackson, Ian Johnston, Dominic Kwiatkowski, Cordelia Langford, John Sillitoe on behalf of the Wellcome Sanger Institute COVID-19 Surveillance Team |
| EPI_ISL_3202805 | Berkshire and Surrey Pathology Services Lighthouse Laboratory | Wellcome Sanger Institute for the COVID-19 Genomics UK (COG-UK) Consortium | Berkshire and Surrey Pathology Services Lighthouse Laboratory and Alex Alderton, Roberto Amato, Jeffrey Barrett, Sonia Goncalves, Ewan Harrison, David K. Jackson, Ian Johnston, Dominic Kwiatkowski, Cordelia Langford, John Sillitoe on behalf of the Wellcome Sanger Institute COVID-19 Surveillance Team |
| EPI_ISL_3202806, EPI_ISL_3202807 | Lighthouse Lab in Alderley Park | Wellcome Sanger Institute for the COVID-19 Genomics UK (COG-UK) Consortium | Jacquelyn Wynn, Mairead Hyland, The Lighthouse Lab in Alderley Park and Alex Alderton, Roberto Amato, Jeffrey Barrett, Sonia Goncalves, Ewan Harrison, David K. Jackson, Ian Johnston, Dominic Kwiatkowski, Cordelia Langford, John Sillitoe on behalf of the Wellcome Sanger Institute COVID-19 Surveillance Team |
| EPI_ISL_3202808 | Lighthouse Lab in Milton Keynes | Wellcome Sanger Institute for the COVID-19 Genomics UK (COG-UK) Consortium | The Lighthouse Lab in Milton Keynes and Alex Alderton, Roberto Amato, Jeffrey Barrett, Sonia Goncalves, Ewan Harrison, David K. Jackson, Ian Johnston, Dominic Kwiatkowski, Cordelia Langford, John Sillitoe on behalf of the Wellcome Sanger Institute COVID-19 Surveillance Team |
| EPI_ISL_3202809 | Berkshire and Surrey Pathology Services Lighthouse Laboratory | Wellcome Sanger Institute for the COVID-19 Genomics UK (COG-UK) Consortium | Berkshire and Surrey Pathology Services Lighthouse Laboratory and Alex Alderton, Roberto Amato, Jeffrey Barrett, Sonia Goncalves, Ewan Harrison, David K. Jackson, Ian Johnston, Dominic Kwiatkowski, Cordelia Langford, John Sillitoe on behalf of the Wellcome Sanger Institute COVID-19 Surveillance Team |
| EPI_ISL_3202810 | Lighthouse Laboratory Plymouth | Wellcome Sanger Institute for the COVID-19 Genomics UK (COG-UK) Consortium | Lighthouse Laboratory Plymouth and Alex Alderton, Roberto Amato, Jeffrey Barrett, Sonia Goncalves, Ewan Harrison, David K. Jackson, Ian Johnston, Dominic Kwiatkowski, Cordelia Langford, John Sillitoe on behalf of the Wellcome Sanger Institute COVID-19 Surveillance Team |
| EPI_ISL_3202811 | Lighthouse Lab in Milton Keynes | Wellcome Sanger Institute for the COVID-19 Genomics UK (COG-UK) Consortium | The Lighthouse Lab in Milton Keynes and Alex Alderton, Roberto Amato, Jeffrey Barrett, Sonia Goncalves, Ewan Harrison, David K. Jackson, Ian Johnston, Dominic Kwiatkowski, Cordelia Langford, John Sillitoe on behalf of the Wellcome Sanger Institute COVID-19 Surveillance Team |
| EPI_ISL_3202812 | Berkshire and Surrey Pathology Services Lighthouse Laboratory | Wellcome Sanger Institute for the COVID-19 Genomics UK (COG-UK) Consortium | Berkshire and Surrey Pathology Services Lighthouse Laboratory and Alex Alderton, Roberto Amato, Jeffrey Barrett, Sonia Goncalves, Ewan Harrison, David K. Jackson, Ian Johnston, Dominic Kwiatkowski, Cordelia Langford, John Sillitoe on behalf of the Wellcome Sanger Institute COVID-19 Surveillance Team |
| EPI_ISL_3202813 | Lighthouse Laboratory Plymouth | Wellcome Sanger Institute for the COVID-19 Genomics UK (COG-UK) Consortium | Lighthouse Laboratory Plymouth and Alex Alderton, Roberto Amato, Jeffrey Barrett, Sonia Goncalves, Ewan Harrison, David K. Jackson, Ian Johnston, Dominic Kwiatkowski, Cordelia Langford, John Sillitoe on behalf of the Wellcome Sanger Institute COVID-19 Surveillance Team |
| EPI_ISL_3202814, EPI_ISL_3202815, EPI_ISL_3202816 | Berkshire and Surrey Pathology Services Lighthouse Laboratory | Wellcome Sanger Institute for the COVID-19 Genomics UK (COG-UK) Consortium | Berkshire and Surrey Pathology Services Lighthouse Laboratory and Alex Alderton, Roberto Amato, Jeffrey Barrett, Sonia Goncalves, Ewan Harrison, David K. Jackson, Ian Johnston, Dominic Kwiatkowski, Cordelia Langford, John Sillitoe on behalf of the Wellcome Sanger Institute COVID-19 Surveillance Team |
| EPI_ISL_3202817 | Lighthouse Laboratory Plymouth | Wellcome Sanger Institute for the COVID-19 Genomics UK (COG-UK) Consortium | Lighthouse Laboratory Plymouth and Alex Alderton, Roberto Amato, Jeffrey Barrett, Sonia Goncalves, Ewan Harrison, David K. Jackson, Ian Johnston, Dominic Kwiatkowski, Cordelia Langford, John Sillitoe on behalf of the Wellcome Sanger Institute COVID-19 Surveillance Team |
| EPI_ISL_3202818 | Lighthouse Lab in Milton Keynes | Wellcome Sanger Institute for the COVID-19 Genomics UK (COG-UK) Consortium | The Lighthouse Lab in Milton Keynes and Alex Alderton, Roberto Amato, Jeffrey Barrett, Sonia Goncalves, Ewan Harrison, David K. Jackson, Ian Johnston, Dominic Kwiatkowski, Cordelia Langford, John Sillitoe on behalf of the Wellcome Sanger Institute COVID-19 Surveillance Team |
| EPI_ISL_3202819 | Berkshire and Surrey Pathology Services Lighthouse Laboratory | Wellcome Sanger Institute for the COVID-19 Genomics UK (COG-UK) Consortium | Berkshire and Surrey Pathology Services Lighthouse Laboratory and Alex Alderton, Roberto Amato, Jeffrey Barrett, Sonia Goncalves, Ewan Harrison, David K. Jackson, Ian Johnston, Dominic Kwiatkowski, Cordelia Langford, John Sillitoe on behalf of the Wellcome Sanger Institute COVID-19 Surveillance Team |
| EPI_ISL_3202820 | Lighthouse Lab in Alderley Park | Wellcome Sanger Institute for the COVID-19 Genomics UK (COG-UK) Consortium | Jacquelyn Wynn, Mairead Hyland, The Lighthouse Lab in Alderley Park and Alex Alderton, Roberto Amato, Jeffrey Barrett, Sonia Goncalves, Ewan Harrison, David K. Jackson, Ian Johnston, Dominic Kwiatkowski, Cordelia Langford, John Sillitoe on behalf of the Wellcome Sanger Institute COVID-19 Surveillance Team |
| EPI_ISL_3202821 | Lighthouse Lab in Milton Keynes | Wellcome Sanger Institute for the COVID-19 Genomics UK (COG-UK) Consortium | The Lighthouse Lab in Milton Keynes and Alex Alderton, Roberto Amato, Jeffrey Barrett, Sonia Goncalves, Ewan Harrison, David K. Jackson, Ian Johnston, Dominic Kwiatkowski, Cordelia Langford, John Sillitoe on behalf of the Wellcome Sanger Institute COVID-19 Surveillance Team |
| EPI_ISL_3202822, EPI_ISL_3202823 | Lighthouse Laboratory Plymouth | Wellcome Sanger Institute for the COVID-19 Genomics UK (COG-UK) Consortium | Lighthouse Laboratory Plymouth and Alex Alderton, Roberto Amato, Jeffrey Barrett, Sonia Goncalves, Ewan Harrison, David K. Jackson, Ian Johnston, Dominic Kwiatkowski, Cordelia Langford, John Sillitoe on behalf of the Wellcome Sanger Institute COVID-19 Surveillance Team |
| EPI_ISL_3202824 | Berkshire and Surrey Pathology Services Lighthouse Laboratory | Wellcome Sanger Institute for the COVID-19 Genomics UK (COG-UK) Consortium | Berkshire and Surrey Pathology Services Lighthouse Laboratory and Alex Alderton, Roberto Amato, Jeffrey Barrett, Sonia Goncalves, Ewan Harrison, David K. Jackson, Ian Johnston, Dominic Kwiatkowski, Cordelia Langford, John Sillitoe on behalf of the Wellcome Sanger Institute COVID-19 Surveillance Team |
| EPI_ISL_3202825, EPI_ISL_3202826 | Lighthouse Lab in Milton Keynes | Wellcome Sanger Institute for the COVID-19 Genomics UK (COG-UK) Consortium | The Lighthouse Lab in Milton Keynes and Alex Alderton, Roberto Amato, Jeffrey Barrett, Sonia Goncalves, Ewan Harrison, David K. Jackson, Ian Johnston, Dominic Kwiatkowski, Cordelia Langford, John Sillitoe on behalf of the Wellcome Sanger Institute COVID-19 Surveillance Team |
| EPI_ISL_3202827 | Berkshire and Surrey Pathology Services Lighthouse Laboratory | Wellcome Sanger Institute for the COVID-19 Genomics UK (COG-UK) Consortium | Berkshire and Surrey Pathology Services Lighthouse Laboratory and Alex Alderton, Roberto Amato, Jeffrey Barrett, Sonia Goncalves, Ewan Harrison, David K. Jackson, Ian Johnston, Dominic Kwiatkowski, Cordelia Langford, John Sillitoe on behalf of the Wellcome Sanger Institute COVID-19 Surveillance Team |
| EPI_ISL_3202828, EPI_ISL_3202829 | Lighthouse Lab in Alderley Park | Wellcome Sanger Institute for the COVID-19 Genomics UK (COG-UK) Consortium | Jacquelyn Wynn, Mairead Hyland, The Lighthouse Lab in Alderley Park and Alex Alderton, Roberto Amato, Jeffrey Barrett, Sonia Goncalves, Ewan Harrison, David K. Jackson, Ian Johnston, Dominic Kwiatkowski, Cordelia Langford, John Sillitoe on behalf of the Wellcome Sanger Institute COVID-19 Surveillance Team |
| EPI_ISL_3202830 | Berkshire and Surrey Pathology Services Lighthouse Laboratory | Wellcome Sanger Institute for the COVID-19 Genomics UK (COG-UK) Consortium | Berkshire and Surrey Pathology Services Lighthouse Laboratory and Alex Alderton, Roberto Amato, Jeffrey Barrett, Sonia Goncalves, Ewan Harrison, David K. Jackson, Ian Johnston, Dominic Kwiatkowski, Cordelia Langford, John Sillitoe on behalf of the Wellcome Sanger Institute COVID-19 Surveillance Team |
| EPI_ISL_3202831 | Lighthouse Lab in Milton Keynes | Wellcome Sanger Institute for the COVID-19 Genomics UK (COG-UK) Consortium | The Lighthouse Lab in Milton Keynes and Alex Alderton, Roberto Amato, Jeffrey Barrett, Sonia Goncalves, Ewan Harrison, David K. Jackson, Ian Johnston, Dominic Kwiatkowski, Cordelia Langford, John Sillitoe on behalf of the Wellcome Sanger Institute COVID-19 Surveillance Team |
| EPI_ISL_3202832, EPI_ISL_3202833, EPI_ISL_3202834, EPI_ISL_3202835 | Lighthouse Laboratory Plymouth | Wellcome Sanger Institute for the COVID-19 Genomics UK (COG-UK) Consortium | Lighthouse Laboratory Plymouth and Alex Alderton, Roberto Amato, Jeffrey Barrett, Sonia Goncalves, Ewan Harrison, David K. Jackson, Ian Johnston, Dominic Kwiatkowski, Cordelia Langford, John Sillitoe on behalf of the Wellcome Sanger Institute COVID-19 Surveillance Team |
| EPI_ISL_3202836 | Lighthouse Lab in Milton Keynes | Wellcome Sanger Institute for the COVID-19 Genomics UK (COG-UK) Consortium | The Lighthouse Lab in Milton Keynes and Alex Alderton, Roberto Amato, Jeffrey Barrett, Sonia Goncalves, Ewan Harrison, David K. Jackson, Ian Johnston, Dominic Kwiatkowski, Cordelia Langford, John Sillitoe on behalf of the Wellcome Sanger Institute COVID-19 Surveillance Team |
| EPI_ISL_3202837, EPI_ISL_3202838 | Berkshire and Surrey Pathology Services Lighthouse Laboratory | Wellcome Sanger Institute for the COVID-19 Genomics UK (COG-UK) Consortium | Berkshire and Surrey Pathology Services Lighthouse Laboratory and Alex Alderton, Roberto Amato, Jeffrey Barrett, Sonia Goncalves, Ewan Harrison, David K. Jackson, Ian Johnston, Dominic Kwiatkowski, Cordelia Langford, John Sillitoe on behalf of the Wellcome Sanger Institute COVID-19 Surveillance Team |
| EPI_ISL_3202839 | Lighthouse Lab in Milton Keynes | Wellcome Sanger Institute for the COVID-19 Genomics UK (COG-UK) Consortium | The Lighthouse Lab in Milton Keynes and Alex Alderton, Roberto Amato, Jeffrey Barrett, Sonia Goncalves, Ewan Harrison, David K. Jackson, Ian Johnston, Dominic Kwiatkowski, Cordelia Langford, John Sillitoe on behalf of the Wellcome Sanger Institute COVID-19 Surveillance Team |
| EPI_ISL_3202840 | Lighthouse Lab in Alderley Park | Wellcome Sanger Institute for the COVID-19 Genomics UK (COG-UK) Consortium | Jacquelyn Wynn, Mairead Hyland, The Lighthouse Lab in Alderley Park and Alex Alderton, Roberto Amato, Jeffrey Barrett, Sonia Goncalves, Ewan Harrison, David K. Jackson, Ian Johnston, Dominic Kwiatkowski, Cordelia Langford, John Sillitoe on behalf of the Wellcome Sanger Institute COVID-19 Surveillance Team |
| EPI_ISL_3202841, EPI_ISL_3202842 | Lighthouse Laboratory Plymouth | Wellcome Sanger Institute for the COVID-19 Genomics UK (COG-UK) Consortium | Lighthouse Laboratory Plymouth and Alex Alderton, Roberto Amato, Jeffrey Barrett, Sonia Goncalves, Ewan Harrison, David K. Jackson, Ian Johnston, Dominic Kwiatkowski, Cordelia Langford, John Sillitoe on behalf of the Wellcome Sanger Institute COVID-19 Surveillance Team |
| EPI_ISL_3202843 | Lighthouse Lab in Milton Keynes | Wellcome Sanger Institute for the COVID-19 Genomics UK (COG-UK) Consortium | The Lighthouse Lab in Milton Keynes and Alex Alderton, Roberto Amato, Jeffrey Barrett, Sonia Goncalves, Ewan Harrison, David K. Jackson, Ian Johnston, Dominic Kwiatkowski, Cordelia Langford, John Sillitoe on behalf of the Wellcome Sanger Institute COVID-19 Surveillance Team |
| EPI_ISL_3202844, EPI_ISL_3202845 | Lighthouse Laboratory Plymouth | Wellcome Sanger Institute for the COVID-19 Genomics UK (COG-UK) Consortium | Lighthouse Laboratory Plymouth and Alex Alderton, Roberto Amato, Jeffrey Barrett, Sonia Goncalves, Ewan Harrison, David K. Jackson, Ian Johnston, Dominic Kwiatkowski, Cordelia Langford, John Sillitoe on behalf of the Wellcome Sanger Institute COVID-19 Surveillance Team |
| EPI_ISL_3202846 | Lighthouse Lab in Milton Keynes | Wellcome Sanger Institute for the COVID-19 Genomics UK (COG-UK) Consortium | The Lighthouse Lab in Milton Keynes and Alex Alderton, Roberto Amato, Jeffrey Barrett, Sonia Goncalves, Ewan Harrison, David K. Jackson, Ian Johnston, Dominic Kwiatkowski, Cordelia Langford, John Sillitoe on behalf of the Wellcome Sanger Institute COVID-19 Surveillance Team |
| EPI_ISL_3202847, EPI_ISL_3202848 | Berkshire and Surrey Pathology Services Lighthouse Laboratory | Wellcome Sanger Institute for the COVID-19 Genomics UK (COG-UK) Consortium | Berkshire and Surrey Pathology Services Lighthouse Laboratory and Alex Alderton, Roberto Amato, Jeffrey Barrett, Sonia Goncalves, Ewan Harrison, David K. Jackson, Ian Johnston, Dominic Kwiatkowski, Cordelia Langford, John Sillitoe on behalf of the Wellcome Sanger Institute COVID-19 Surveillance Team |
| EPI_ISL_3202849 | Lighthouse Lab in Alderley Park | Wellcome Sanger Institute for the COVID-19 Genomics UK (COG-UK) Consortium | Jacquelyn Wynn, Mairead Hyland, The Lighthouse Lab in Alderley Park and Alex Alderton, Roberto Amato, Jeffrey Barrett, Sonia Goncalves, Ewan Harrison, David K. Jackson, Ian Johnston, Dominic Kwiatkowski, Cordelia Langford, John Sillitoe on behalf of the Wellcome Sanger Institute COVID-19 Surveillance Team |
| EPI_ISL_3202850 | Lighthouse Laboratory Plymouth | Wellcome Sanger Institute for the COVID-19 Genomics UK (COG-UK) Consortium | Lighthouse Laboratory Plymouth and Alex Alderton, Roberto Amato, Jeffrey Barrett, Sonia Goncalves, Ewan Harrison, David K. Jackson, Ian Johnston, Dominic Kwiatkowski, Cordelia Langford, John Sillitoe on behalf of the Wellcome Sanger Institute COVID-19 Surveillance Team |
| EPI_ISL_3202851 | Lighthouse Lab in Milton Keynes | Wellcome Sanger Institute for the COVID-19 Genomics UK (COG-UK) Consortium | The Lighthouse Lab in Milton Keynes and Alex Alderton, Roberto Amato, Jeffrey Barrett, Sonia Goncalves, Ewan Harrison, David K. Jackson, Ian Johnston, Dominic Kwiatkowski, Cordelia Langford, John Sillitoe on behalf of the Wellcome Sanger Institute COVID-19 Surveillance Team |
| EPI_ISL_3202852, EPI_ISL_3202853 | Lighthouse Lab in Alderley Park | Wellcome Sanger Institute for the COVID-19 Genomics UK (COG-UK) Consortium | Jacquelyn Wynn, Mairead Hyland, The Lighthouse Lab in Alderley Park and Alex Alderton, Roberto Amato, Jeffrey Barrett, Sonia Goncalves, Ewan Harrison, David K. Jackson, Ian Johnston, Dominic Kwiatkowski, Cordelia Langford, John Sillitoe on behalf of the Wellcome Sanger Institute COVID-19 Surveillance Team |
| EPI_ISL_3202854 | Lighthouse Lab in Milton Keynes | Wellcome Sanger Institute for the COVID-19 Genomics UK (COG-UK) Consortium | The Lighthouse Lab in Milton Keynes and Alex Alderton, Roberto Amato, Jeffrey Barrett, Sonia Goncalves, Ewan Harrison, David K. Jackson, Ian Johnston, Dominic Kwiatkowski, Cordelia Langford, John Sillitoe on behalf of the Wellcome Sanger Institute COVID-19 Surveillance Team |
| EPI_ISL_3202855 | Lighthouse Lab in Alderley Park | Wellcome Sanger Institute for the COVID-19 Genomics UK (COG-UK) Consortium | Jacquelyn Wynn, Mairead Hyland, The Lighthouse Lab in Alderley Park and Alex Alderton, Roberto Amato, Jeffrey Barrett, Sonia Goncalves, Ewan Harrison, David K. Jackson, Ian Johnston, Dominic Kwiatkowski, Cordelia Langford, John Sillitoe on behalf of the Wellcome Sanger Institute COVID-19 Surveillance Team |
| EPI_ISL_3202856 | Lighthouse Laboratory Plymouth | Wellcome Sanger Institute for the COVID-19 Genomics UK (COG-UK) Consortium | Lighthouse Laboratory Plymouth and Alex Alderton, Roberto Amato, Jeffrey Barrett, Sonia Goncalves, Ewan Harrison, David K. Jackson, Ian Johnston, Dominic Kwiatkowski, Cordelia Langford, John Sillitoe on behalf of the Wellcome Sanger Institute COVID-19 Surveillance Team |
| EPI_ISL_3202857, EPI_ISL_3202858 | Lighthouse Lab in Milton Keynes | Wellcome Sanger Institute for the COVID-19 Genomics UK (COG-UK) Consortium | The Lighthouse Lab in Milton Keynes and Alex Alderton, Roberto Amato, Jeffrey Barrett, Sonia Goncalves, Ewan Harrison, David K. Jackson, Ian Johnston, Dominic Kwiatkowski, Cordelia Langford, John Sillitoe on behalf of the Wellcome Sanger Institute COVID-19 Surveillance Team |
| EPI_ISL_3202859 | Lighthouse Lab in Alderley Park | Wellcome Sanger Institute for the COVID-19 Genomics UK (COG-UK) | Jacquelyn Wynn, Mairead Hyland, The Lighthouse Lab in Alderley Park and Alex Alderton, Roberto Amato, Jeffrey Barrett, Sonia Goncalves, Ewan Harrison, David K. Jackson, Ian Johnston, Dominic Kwiatkowski, Cordelia Langford, John Sillitoe on |

| | | | |
|---|---|---|---|
| | | Consortium | behalf of the Wellcome Sanger Institute COVID-19 Surveillance Team |
| EPI_ISL_3202860 | Berkshire and Surrey Pathology Services Lighthouse Laboratory | Wellcome Sanger Institute for the COVID-19 Genomics UK (COG-UK) Consortium | Berkshire and Surrey Pathology Services Lighthouse Laboratory and Alex Alderton, Roberto Amato, Jeffrey Barrett, Sonia Goncalves, Ewan Harrison, David K. Jackson, Ian Johnston, Dominic Kwiatkowski, Cordelia Langford, John Sillitoe on behalf of the Wellcome Sanger Institute COVID-19 Surveillance Team |
| EPI_ISL_3202861, EPI_ISL_3202862 | Lighthouse Laboratory Plymouth | Wellcome Sanger Institute for the COVID-19 Genomics UK (COG-UK) Consortium | Lighthouse Laboratory Plymouth and Alex Alderton, Roberto Amato, Jeffrey Barrett, Sonia Goncalves, Ewan Harrison, David K. Jackson, Ian Johnston, Dominic Kwiatkowski, Cordelia Langford, John Sillitoe on behalf of the Wellcome Sanger Institute COVID-19 Surveillance Team |
| EPI_ISL_3202863 | Lighthouse Lab in Alderley Park | Wellcome Sanger Institute for the COVID-19 Genomics UK (COG-UK) Consortium | Jacquelyn Wynn, Mairead Hyland, The Lighthouse Lab in Alderley Park and Alex Alderton, Roberto Amato, Jeffrey Barrett, Sonia Goncalves, Ewan Harrison, David K. Jackson, Ian Johnston, Dominic Kwiatkowski, Cordelia Langford, John Sillitoe on behalf of the Wellcome Sanger Institute COVID-19 Surveillance Team |
| EPI_ISL_3202864, EPI_ISL_3202865 | Lighthouse Lab in Milton Keynes | Wellcome Sanger Institute for the COVID-19 Genomics UK (COG-UK) Consortium | The Lighthouse Lab in Milton Keynes and Alex Alderton, Roberto Amato, Jeffrey Barrett, Sonia Goncalves, Ewan Harrison, David K. Jackson, Ian Johnston, Dominic Kwiatkowski, Cordelia Langford, John Sillitoe on behalf of the Wellcome Sanger Institute COVID-19 Surveillance Team |
| EPI_ISL_3202866 | Lighthouse Laboratory Plymouth | Wellcome Sanger Institute for the COVID-19 Genomics UK (COG-UK) Consortium | Lighthouse Laboratory Plymouth and Alex Alderton, Roberto Amato, Jeffrey Barrett, Sonia Goncalves, Ewan Harrison, David K. Jackson, Ian Johnston, Dominic Kwiatkowski, Cordelia Langford, John Sillitoe on behalf of the Wellcome Sanger Institute COVID-19 Surveillance Team |
| EPI_ISL_3202867 | Lighthouse Lab in Alderley Park | Wellcome Sanger Institute for the COVID-19 Genomics UK (COG-UK) Consortium | Jacquelyn Wynn, Mairead Hyland, The Lighthouse Lab in Alderley Park and Alex Alderton, Roberto Amato, Jeffrey Barrett, Sonia Goncalves, Ewan Harrison, David K. Jackson, Ian Johnston, Dominic Kwiatkowski, Cordelia Langford, John Sillitoe on behalf of the Wellcome Sanger Institute COVID-19 Surveillance Team |
| EPI_ISL_3202868 | Berkshire and Surrey Pathology Services Lighthouse Laboratory | Wellcome Sanger Institute for the COVID-19 Genomics UK (COG-UK) Consortium | Berkshire and Surrey Pathology Services Lighthouse Laboratory and Alex Alderton, Roberto Amato, Jeffrey Barrett, Sonia Goncalves, Ewan Harrison, David K. Jackson, Ian Johnston, Dominic Kwiatkowski, Cordelia Langford, John Sillitoe on behalf of the Wellcome Sanger Institute COVID-19 Surveillance Team |
| EPI_ISL_3202869, EPI_ISL_3202870 | Lighthouse Lab in Alderley Park | Wellcome Sanger Institute for the COVID-19 Genomics UK (COG-UK) Consortium | Jacquelyn Wynn, Mairead Hyland, The Lighthouse Lab in Alderley Park and Alex Alderton, Roberto Amato, Jeffrey Barrett, Sonia Goncalves, Ewan Harrison, David K. Jackson, Ian Johnston, Dominic Kwiatkowski, Cordelia Langford, John Sillitoe on behalf of the Wellcome Sanger Institute COVID-19 Surveillance Team |
| EPI_ISL_3202871 | Lighthouse Laboratory Plymouth | Wellcome Sanger Institute for the COVID-19 Genomics UK (COG-UK) Consortium | Lighthouse Laboratory Plymouth and Alex Alderton, Roberto Amato, Jeffrey Barrett, Sonia Goncalves, Ewan Harrison, David K. Jackson, Ian Johnston, Dominic Kwiatkowski, Cordelia Langford, John Sillitoe on behalf of the Wellcome Sanger Institute COVID-19 Surveillance Team |
| EPI_ISL_3202872 | Berkshire and Surrey Pathology Services Lighthouse Laboratory | Wellcome Sanger Institute for the COVID-19 Genomics UK (COG-UK) Consortium | Berkshire and Surrey Pathology Services Lighthouse Laboratory and Alex Alderton, Roberto Amato, Jeffrey Barrett, Sonia Goncalves, Ewan Harrison, David K. Jackson, Ian Johnston, Dominic Kwiatkowski, Cordelia Langford, John Sillitoe on behalf of the Wellcome Sanger Institute COVID-19 Surveillance Team |
| EPI_ISL_3202873, EPI_ISL_3202874 | Lighthouse Laboratory Plymouth | Wellcome Sanger Institute for the COVID-19 Genomics UK (COG-UK) Consortium | Lighthouse Laboratory Plymouth and Alex Alderton, Roberto Amato, Jeffrey Barrett, Sonia Goncalves, Ewan Harrison, David K. Jackson, Ian Johnston, Dominic Kwiatkowski, Cordelia Langford, John Sillitoe on behalf of the Wellcome Sanger Institute COVID-19 Surveillance Team |
| EPI_ISL_3202875, EPI_ISL_3202876 | Lighthouse Lab in Alderley Park | Wellcome Sanger Institute for the COVID-19 Genomics UK (COG-UK) Consortium | Jacquelyn Wynn, Mairead Hyland, The Lighthouse Lab in Alderley Park and Alex Alderton, Roberto Amato, Jeffrey Barrett, Sonia Goncalves, Ewan Harrison, David K. Jackson, Ian Johnston, Dominic Kwiatkowski, Cordelia Langford, John Sillitoe on behalf of the Wellcome Sanger Institute COVID-19 Surveillance Team |
| EPI_ISL_3202877, EPI_ISL_3202878 | Lighthouse Laboratory Plymouth | Wellcome Sanger Institute for the COVID-19 Genomics UK (COG-UK) Consortium | Lighthouse Laboratory Plymouth and Alex Alderton, Roberto Amato, Jeffrey Barrett, Sonia Goncalves, Ewan Harrison, David K. Jackson, Ian Johnston, Dominic Kwiatkowski, Cordelia Langford, John Sillitoe on behalf of the Wellcome Sanger Institute COVID-19 Surveillance Team |
| EPI_ISL_3202879, EPI_ISL_3202880 | Lighthouse Lab in Alderley Park | Wellcome Sanger Institute for the COVID-19 Genomics UK (COG-UK) Consortium | Jacquelyn Wynn, Mairead Hyland, The Lighthouse Lab in Alderley Park and Alex Alderton, Roberto Amato, Jeffrey Barrett, Sonia Goncalves, Ewan Harrison, David K. Jackson, Ian Johnston, Dominic Kwiatkowski, Cordelia Langford, John Sillitoe on behalf of the Wellcome Sanger Institute COVID-19 Surveillance Team |
| EPI_ISL_3202881 | Lighthouse Laboratory Plymouth | Wellcome Sanger Institute for the COVID-19 Genomics UK (COG-UK) Consortium | Lighthouse Laboratory Plymouth and Alex Alderton, Roberto Amato, Jeffrey Barrett, Sonia Goncalves, Ewan Harrison, David K. Jackson, Ian Johnston, Dominic Kwiatkowski, Cordelia Langford, John Sillitoe on behalf of the Wellcome Sanger Institute COVID-19 Surveillance Team |
| EPI_ISL_3202882 | Berkshire and Surrey Pathology Services Lighthouse Laboratory | Wellcome Sanger Institute for the COVID-19 Genomics UK (COG-UK) Consortium | Berkshire and Surrey Pathology Services Lighthouse Laboratory and Alex Alderton, Roberto Amato, Jeffrey Barrett, Sonia Goncalves, Ewan Harrison, David K. Jackson, Ian Johnston, Dominic Kwiatkowski, Cordelia Langford, John Sillitoe on behalf of the Wellcome Sanger Institute COVID-19 Surveillance Team |
| EPI_ISL_3202883 | Lighthouse Lab in Alderley Park | Wellcome Sanger Institute for the COVID-19 Genomics UK (COG-UK) Consortium | Jacquelyn Wynn, Mairead Hyland, The Lighthouse Lab in Alderley Park and Alex Alderton, Roberto Amato, Jeffrey Barrett, Sonia Goncalves, Ewan Harrison, David K. Jackson, Ian Johnston, Dominic Kwiatkowski, Cordelia Langford, John Sillitoe on behalf of the Wellcome Sanger Institute COVID-19 Surveillance Team |
| EPI_ISL_3202884 | Lighthouse Laboratory Plymouth | Wellcome Sanger Institute for the COVID-19 Genomics UK (COG-UK) Consortium | Lighthouse Laboratory Plymouth and Alex Alderton, Roberto Amato, Jeffrey Barrett, Sonia Goncalves, Ewan Harrison, David K. Jackson, Ian Johnston, Dominic Kwiatkowski, Cordelia Langford, John Sillitoe on behalf of the Wellcome Sanger Institute COVID-19 Surveillance Team |
| EPI_ISL_3202885 | Lighthouse Lab in Milton Keynes | Wellcome Sanger Institute for the COVID-19 Genomics UK (COG-UK) Consortium | The Lighthouse Lab in Milton Keynes and Alex Alderton, Roberto Amato, Jeffrey Barrett, Sonia Goncalves, Ewan Harrison, David K. Jackson, Ian Johnston, Dominic Kwiatkowski, Cordelia Langford, John Sillitoe on behalf of the Wellcome Sanger Institute COVID-19 Surveillance Team |
| EPI_ISL_3202886 | Lighthouse Laboratory Plymouth | Wellcome Sanger Institute for the COVID-19 Genomics UK (COG-UK) Consortium | Lighthouse Laboratory Plymouth and Alex Alderton, Roberto Amato, Jeffrey Barrett, Sonia Goncalves, Ewan Harrison, David K. Jackson, Ian Johnston, Dominic Kwiatkowski, Cordelia Langford, John Sillitoe on behalf of the Wellcome Sanger Institute COVID-19 Surveillance Team |
| EPI_ISL_3202887, EPI_ISL_3202888 | Lighthouse Lab in Alderley Park | Wellcome Sanger Institute for the COVID-19 Genomics UK (COG-UK) Consortium | Jacquelyn Wynn, Mairead Hyland, The Lighthouse Lab in Alderley Park and Alex Alderton, Roberto Amato, Jeffrey Barrett, Sonia Goncalves, Ewan Harrison, David K. Jackson, Ian Johnston, Dominic Kwiatkowski, Cordelia Langford, John Sillitoe on behalf of the Wellcome Sanger Institute COVID-19 Surveillance Team |
| EPI_ISL_3202889 | Lighthouse Laboratory Plymouth | Wellcome Sanger Institute for the COVID-19 Genomics UK (COG-UK) Consortium | Lighthouse Laboratory Plymouth and Alex Alderton, Roberto Amato, Jeffrey Barrett, Sonia Goncalves, Ewan Harrison, David K. Jackson, Ian Johnston, Dominic Kwiatkowski, Cordelia Langford, John Sillitoe on behalf of the Wellcome Sanger Institute COVID-19 Surveillance Team |
| EPI_ISL_3202890 | Lighthouse Lab in Milton Keynes | Wellcome Sanger Institute for the COVID-19 Genomics UK (COG-UK) Consortium | The Lighthouse Lab in Milton Keynes and Alex Alderton, Roberto Amato, Jeffrey Barrett, Sonia Goncalves, Ewan Harrison, David K. Jackson, Ian Johnston, Dominic Kwiatkowski, Cordelia Langford, John Sillitoe on behalf of the Wellcome Sanger Institute COVID-19 Surveillance Team |
| EPI_ISL_3202891 | Berkshire and Surrey Pathology Services Lighthouse Laboratory | Wellcome Sanger Institute for the COVID-19 Genomics UK (COG-UK) Consortium | Berkshire and Surrey Pathology Services Lighthouse Laboratory and Alex Alderton, Roberto Amato, Jeffrey Barrett, Sonia Goncalves, Ewan Harrison, David K. Jackson, Ian Johnston, Dominic Kwiatkowski, Cordelia Langford, John Sillitoe on behalf of the Wellcome Sanger Institute COVID-19 Surveillance Team |
| EPI_ISL_3202892, EPI_ISL_3202893 | Lighthouse Lab in Alderley Park | Wellcome Sanger Institute for the COVID-19 Genomics UK (COG-UK) Consortium | Jacquelyn Wynn, Mairead Hyland, The Lighthouse Lab in Alderley Park and Alex Alderton, Roberto Amato, Jeffrey Barrett, Sonia Goncalves, Ewan Harrison, David K. Jackson, Ian Johnston, Dominic Kwiatkowski, Cordelia Langford, John Sillitoe on behalf of the Wellcome Sanger Institute COVID-19 Surveillance Team |
| EPI_ISL_3202894 | Berkshire and Surrey Pathology Services Lighthouse Laboratory | Wellcome Sanger Institute for the COVID-19 Genomics UK (COG-UK) Consortium | Berkshire and Surrey Pathology Services Lighthouse Laboratory and Alex Alderton, Roberto Amato, Jeffrey Barrett, Sonia Goncalves, Ewan Harrison, David K. Jackson, Ian Johnston, Dominic Kwiatkowski, Cordelia Langford, John Sillitoe on behalf of the Wellcome Sanger Institute COVID-19 Surveillance Team |
| EPI_ISL_3202895 | Lighthouse Laboratory Plymouth | Wellcome Sanger Institute for the COVID-19 Genomics UK (COG-UK) Consortium | Lighthouse Laboratory Plymouth and Alex Alderton, Roberto Amato, Jeffrey Barrett, Sonia Goncalves, Ewan Harrison, David K. Jackson, Ian Johnston, Dominic Kwiatkowski, Cordelia Langford, John Sillitoe on behalf of the Wellcome Sanger Institute COVID-19 Surveillance Team |
| EPI_ISL_3202896 | Lighthouse Lab in Alderley Park | Wellcome Sanger Institute for the COVID-19 Genomics UK (COG-UK) Consortium | Jacquelyn Wynn, Mairead Hyland, The Lighthouse Lab in Alderley Park and Alex Alderton, Roberto Amato, Jeffrey Barrett, Sonia Goncalves, Ewan Harrison, David K. Jackson, Ian Johnston, Dominic Kwiatkowski, Cordelia Langford, John Sillitoe on behalf of the Wellcome Sanger Institute COVID-19 Surveillance Team |
| EPI_ISL_3202897, EPI_ISL_3202898 | Berkshire and Surrey Pathology Services Lighthouse Laboratory | Wellcome Sanger Institute for the COVID-19 Genomics UK (COG-UK) Consortium | Berkshire and Surrey Pathology Services Lighthouse Laboratory and Alex Alderton, Roberto Amato, Jeffrey Barrett, Sonia Goncalves, Ewan Harrison, David K. Jackson, Ian Johnston, Dominic Kwiatkowski, Cordelia Langford, John Sillitoe on behalf of the Wellcome Sanger Institute COVID-19 Surveillance Team |
| EPI_ISL_3202899 | Lighthouse Lab in Milton Keynes | Wellcome Sanger Institute for the COVID-19 Genomics UK (COG-UK) Consortium | The Lighthouse Lab in Milton Keynes and Alex Alderton, Roberto Amato, Jeffrey Barrett, Sonia Goncalves, Ewan Harrison, David K. Jackson, Ian Johnston, Dominic Kwiatkowski, Cordelia Langford, John Sillitoe on behalf of the Wellcome Sanger Institute COVID-19 Surveillance Team |
| EPI_ISL_3202900 | Lighthouse Laboratory Plymouth | Wellcome Sanger Institute for the COVID-19 Genomics UK (COG-UK) Consortium | Lighthouse Laboratory Plymouth and Alex Alderton, Roberto Amato, Jeffrey Barrett, Sonia Goncalves, Ewan Harrison, David K. Jackson, Ian Johnston, Dominic Kwiatkowski, Cordelia Langford, John Sillitoe on behalf of the Wellcome Sanger Institute COVID-19 Surveillance Team |
| EPI_ISL_3202901 | Lighthouse Lab in Milton Keynes | Wellcome Sanger Institute for the COVID-19 Genomics UK (COG-UK) Consortium | The Lighthouse Lab in Milton Keynes and Alex Alderton, Roberto Amato, Jeffrey Barrett, Sonia Goncalves, Ewan Harrison, David K. Jackson, Ian Johnston, Dominic Kwiatkowski, Cordelia Langford, John Sillitoe on behalf of the Wellcome Sanger Institute COVID-19 Surveillance Team |
| EPI_ISL_3202902 | Berkshire and Surrey Pathology Services Lighthouse Laboratory | Wellcome Sanger Institute for the COVID-19 Genomics UK (COG-UK) Consortium | Berkshire and Surrey Pathology Services Lighthouse Laboratory and Alex Alderton, Roberto Amato, Jeffrey Barrett, Sonia Goncalves, Ewan Harrison, David K. Jackson, Ian Johnston, Dominic Kwiatkowski, Cordelia Langford, John Sillitoe on behalf of the Wellcome Sanger Institute COVID-19 Surveillance Team |
| EPI_ISL_3202903 | Lighthouse Laboratory Plymouth | Wellcome Sanger Institute for the COVID-19 Genomics UK (COG-UK) Consortium | Lighthouse Laboratory Plymouth and Alex Alderton, Roberto Amato, Jeffrey Barrett, Sonia Goncalves, Ewan Harrison, David K. Jackson, Ian Johnston, Dominic Kwiatkowski, Cordelia Langford, John Sillitoe on behalf of the Wellcome Sanger Institute COVID-19 Surveillance Team |
| EPI_ISL_3202904 | Lighthouse Lab in Milton Keynes | Wellcome Sanger Institute for the COVID-19 Genomics UK (COG-UK) Consortium | The Lighthouse Lab in Milton Keynes and Alex Alderton, Roberto Amato, Jeffrey Barrett, Sonia Goncalves, Ewan Harrison, David K. Jackson, Ian Johnston, Dominic Kwiatkowski, Cordelia Langford, John Sillitoe on behalf of the Wellcome Sanger Institute COVID-19 Surveillance Team |
| EPI_ISL_3202905 | Berkshire and Surrey Pathology Services Lighthouse Laboratory | Wellcome Sanger Institute for the COVID-19 Genomics UK (COG-UK) Consortium | Berkshire and Surrey Pathology Services Lighthouse Laboratory and Alex Alderton, Roberto Amato, Jeffrey Barrett, Sonia Goncalves, Ewan Harrison, David K. Jackson, Ian Johnston, Dominic Kwiatkowski, Cordelia Langford, John Sillitoe on behalf of the Wellcome Sanger Institute COVID-19 Surveillance Team |
| EPI_ISL_3202906 | Lighthouse Lab in Alderley Park | Wellcome Sanger Institute for the COVID-19 Genomics UK (COG-UK) Consortium | Jacquelyn Wynn, Mairead Hyland, The Lighthouse Lab in Alderley Park and Alex Alderton, Roberto Amato, Jeffrey Barrett, Sonia Goncalves, Ewan Harrison, David K. Jackson, Ian Johnston, Dominic Kwiatkowski, Cordelia Langford, John Sillitoe on behalf of the Wellcome Sanger Institute COVID-19 Surveillance Team |
| EPI_ISL_3202907 | Lighthouse Lab in Milton Keynes | Wellcome Sanger Institute for the COVID-19 Genomics UK (COG-UK) Consortium | The Lighthouse Lab in Milton Keynes and Alex Alderton, Roberto Amato, Jeffrey Barrett, Sonia Goncalves, Ewan Harrison, David K. Jackson, Ian Johnston, Dominic Kwiatkowski, Cordelia Langford, John Sillitoe on behalf of the Wellcome Sanger Institute COVID-19 Surveillance Team |
| EPI_ISL_3202908, EPI_ISL_3202909, EPI_ISL_3202910 | Lighthouse Lab in Alderley Park | Wellcome Sanger Institute for the COVID-19 Genomics UK (COG-UK) Consortium | Jacquelyn Wynn, Mairead Hyland, The Lighthouse Lab in Alderley Park and Alex Alderton, Roberto Amato, Jeffrey Barrett, Sonia Goncalves, Ewan Harrison, David K. Jackson, Ian Johnston, Dominic Kwiatkowski, Cordelia Langford, John Sillitoe on behalf of the Wellcome Sanger Institute COVID-19 Surveillance Team |
| EPI_ISL_3202911 | Berkshire and Surrey Pathology Services Lighthouse Laboratory | Wellcome Sanger Institute for the COVID-19 Genomics UK (COG-UK) Consortium | Berkshire and Surrey Pathology Services Lighthouse Laboratory and Alex Alderton, Roberto Amato, Jeffrey Barrett, Sonia Goncalves, Ewan Harrison, David K. Jackson, Ian Johnston, Dominic Kwiatkowski, Cordelia Langford, John Sillitoe on behalf of the Wellcome Sanger Institute COVID-19 Surveillance Team |
| EPI_ISL_3202912, EPI_ISL_3202913 | Lighthouse Lab in Alderley Park | Wellcome Sanger Institute for the COVID-19 Genomics UK (COG-UK) Consortium | Jacquelyn Wynn, Mairead Hyland, The Lighthouse Lab in Alderley Park and Alex Alderton, Roberto Amato, Jeffrey Barrett, Sonia Goncalves, Ewan Harrison, David K. Jackson, Ian Johnston, Dominic Kwiatkowski, Cordelia Langford, John Sillitoe on behalf of the Wellcome Sanger Institute COVID-19 Surveillance Team |
| EPI_ISL_3202914 | Lighthouse Lab in Milton Keynes | Wellcome Sanger Institute for the COVID-19 Genomics UK (COG-UK) Consortium | The Lighthouse Lab in Milton Keynes and Alex Alderton, Roberto Amato, Jeffrey Barrett, Sonia Goncalves, Ewan Harrison, David K. Jackson, Ian Johnston, Dominic Kwiatkowski, Cordelia Langford, John Sillitoe on behalf of the Wellcome Sanger Institute COVID-19 Surveillance Team |
| EPI_ISL_3202915 | Lighthouse Laboratory Plymouth | Wellcome Sanger Institute for the COVID-19 Genomics UK (COG-UK) Consortium | Lighthouse Laboratory Plymouth and Alex Alderton, Roberto Amato, Jeffrey Barrett, Sonia Goncalves, Ewan Harrison, David K. Jackson, Ian Johnston, Dominic Kwiatkowski, Cordelia Langford, John Sillitoe on behalf of the Wellcome Sanger Institute COVID-19 Surveillance Team |
| EPI_ISL_3202916, EPI_ISL_3202917 | Lighthouse Lab in Alderley Park | Wellcome Sanger Institute for the COVID-19 Genomics UK (COG-UK) Consortium | Jacquelyn Wynn, Mairead Hyland, The Lighthouse Lab in Alderley Park and Alex Alderton, Roberto Amato, Jeffrey Barrett, Sonia Goncalves, Ewan Harrison, David K. Jackson, Ian Johnston, Dominic Kwiatkowski, Cordelia Langford, John Sillitoe on behalf of the Wellcome Sanger Institute COVID-19 Surveillance Team |

| | | | |
|---|---|---|---|
| EPI_ISL_3202918 | Lighthouse Lab in Milton Keynes | Wellcome Sanger Institute for the COVID-19 Genomics UK (COG-UK) Consortium | The Lighthouse Lab in Milton Keynes and Alex Alderton, Roberto Amato, Jeffrey Barrett, Sonia Goncalves, Ewan Harrison, David K. Jackson, Ian Johnston, Dominic Kwiatkowski, Cordelia Langford, John Sillitoe on behalf of the Wellcome Sanger Institute COVID-19 Surveillance Team |
| EPI_ISL_3202919 | Berkshire and Surrey Pathology Services Lighthouse Laboratory | Wellcome Sanger Institute for the COVID-19 Genomics UK (COG-UK) Consortium | Berkshire and Surrey Pathology Services Lighthouse Laboratory and Alex Alderton, Roberto Amato, Jeffrey Barrett, Sonia Goncalves, Ewan Harrison, David K. Jackson, Ian Johnston, Dominic Kwiatkowski, Cordelia Langford, John Sillitoe on behalf of the Wellcome Sanger Institute COVID-19 Surveillance Team |
| EPI_ISL_3202920, EPI_ISL_3202921 | Lighthouse Laboratory Plymouth | Wellcome Sanger Institute for the COVID-19 Genomics UK (COG-UK) Consortium | Lighthouse Laboratory Plymouth and Alex Alderton, Roberto Amato, Jeffrey Barrett, Sonia Goncalves, Ewan Harrison, David K. Jackson, Ian Johnston, Dominic Kwiatkowski, Cordelia Langford, John Sillitoe on behalf of the Wellcome Sanger Institute COVID-19 Surveillance Team |
| EPI_ISL_3202922 | Lighthouse Lab in Alderley Park | Wellcome Sanger Institute for the COVID-19 Genomics UK (COG-UK) Consortium | Jacquelyn Wynn, Mairead Hyland, The Lighthouse Lab in Alderley Park and Alex Alderton, Roberto Amato, Jeffrey Barrett, Sonia Goncalves, Ewan Harrison, David K. Jackson, Ian Johnston, Dominic Kwiatkowski, Cordelia Langford, John Sillitoe on behalf of the Wellcome Sanger Institute COVID-19 Surveillance Team |
| EPI_ISL_3202923 | Berkshire and Surrey Pathology Services Lighthouse Laboratory | Wellcome Sanger Institute for the COVID-19 Genomics UK (COG-UK) Consortium | Berkshire and Surrey Pathology Services Lighthouse Laboratory and Alex Alderton, Roberto Amato, Jeffrey Barrett, Sonia Goncalves, Ewan Harrison, David K. Jackson, Ian Johnston, Dominic Kwiatkowski, Cordelia Langford, John Sillitoe on behalf of the Wellcome Sanger Institute COVID-19 Surveillance Team |
| EPI_ISL_3202924 | Lighthouse Lab in Alderley Park | Wellcome Sanger Institute for the COVID-19 Genomics UK (COG-UK) Consortium | Jacquelyn Wynn, Mairead Hyland, The Lighthouse Lab in Alderley Park and Alex Alderton, Roberto Amato, Jeffrey Barrett, Sonia Goncalves, Ewan Harrison, David K. Jackson, Ian Johnston, Dominic Kwiatkowski, Cordelia Langford, John Sillitoe on behalf of the Wellcome Sanger Institute COVID-19 Surveillance Team |
| EPI_ISL_3202925 | Berkshire and Surrey Pathology Services Lighthouse Laboratory | Wellcome Sanger Institute for the COVID-19 Genomics UK (COG-UK) Consortium | Berkshire and Surrey Pathology Services Lighthouse Laboratory and Alex Alderton, Roberto Amato, Jeffrey Barrett, Sonia Goncalves, Ewan Harrison, David K. Jackson, Ian Johnston, Dominic Kwiatkowski, Cordelia Langford, John Sillitoe on behalf of the Wellcome Sanger Institute COVID-19 Surveillance Team |
| EPI_ISL_3202926, EPI_ISL_3202927 | Lighthouse Laboratory Plymouth | Wellcome Sanger Institute for the COVID-19 Genomics UK (COG-UK) Consortium | Lighthouse Laboratory Plymouth and Alex Alderton, Roberto Amato, Jeffrey Barrett, Sonia Goncalves, Ewan Harrison, David K. Jackson, Ian Johnston, Dominic Kwiatkowski, Cordelia Langford, John Sillitoe on behalf of the Wellcome Sanger Institute COVID-19 Surveillance Team |
| EPI_ISL_3202928 | Lighthouse Lab in Alderley Park | Wellcome Sanger Institute for the COVID-19 Genomics UK (COG-UK) Consortium | Jacquelyn Wynn, Mairead Hyland, The Lighthouse Lab in Alderley Park and Alex Alderton, Roberto Amato, Jeffrey Barrett, Sonia Goncalves, Ewan Harrison, David K. Jackson, Ian Johnston, Dominic Kwiatkowski, Cordelia Langford, John Sillitoe on behalf of the Wellcome Sanger Institute COVID-19 Surveillance Team |
| EPI_ISL_3202929, EPI_ISL_3202930 | Lighthouse Laboratory Plymouth | Wellcome Sanger Institute for the COVID-19 Genomics UK (COG-UK) Consortium | Lighthouse Laboratory Plymouth and Alex Alderton, Roberto Amato, Jeffrey Barrett, Sonia Goncalves, Ewan Harrison, David K. Jackson, Ian Johnston, Dominic Kwiatkowski, Cordelia Langford, John Sillitoe on behalf of the Wellcome Sanger Institute COVID-19 Surveillance Team |
| EPI_ISL_3202931 | Lighthouse Lab in Alderley Park | Wellcome Sanger Institute for the COVID-19 Genomics UK (COG-UK) Consortium | Jacquelyn Wynn, Mairead Hyland, The Lighthouse Lab in Alderley Park and Alex Alderton, Roberto Amato, Jeffrey Barrett, Sonia Goncalves, Ewan Harrison, David K. Jackson, Ian Johnston, Dominic Kwiatkowski, Cordelia Langford, John Sillitoe on behalf of the Wellcome Sanger Institute COVID-19 Surveillance Team |
| EPI_ISL_3202932, EPI_ISL_3202933 | Lighthouse Laboratory Plymouth | Wellcome Sanger Institute for the COVID-19 Genomics UK (COG-UK) Consortium | Lighthouse Laboratory Plymouth and Alex Alderton, Roberto Amato, Jeffrey Barrett, Sonia Goncalves, Ewan Harrison, David K. Jackson, Ian Johnston, Dominic Kwiatkowski, Cordelia Langford, John Sillitoe on behalf of the Wellcome Sanger Institute COVID-19 Surveillance Team |
| EPI_ISL_3202934, EPI_ISL_3202935, EPI_ISL_3202936 | Lighthouse Lab in Milton Keynes | Wellcome Sanger Institute for the COVID-19 Genomics UK (COG-UK) Consortium | The Lighthouse Lab in Milton Keynes and Alex Alderton, Roberto Amato, Jeffrey Barrett, Sonia Goncalves, Ewan Harrison, David K. Jackson, Ian Johnston, Dominic Kwiatkowski, Cordelia Langford, John Sillitoe on behalf of the Wellcome Sanger Institute COVID-19 Surveillance Team |
| EPI_ISL_3202937, EPI_ISL_3202938, EPI_ISL_3202939, EPI_ISL_3202940 | Lighthouse Laboratory Plymouth | Wellcome Sanger Institute for the COVID-19 Genomics UK (COG-UK) Consortium | Lighthouse Laboratory Plymouth and Alex Alderton, Roberto Amato, Jeffrey Barrett, Sonia Goncalves, Ewan Harrison, David K. Jackson, Ian Johnston, Dominic Kwiatkowski, Cordelia Langford, John Sillitoe on behalf of the Wellcome Sanger Institute COVID-19 Surveillance Team |
| EPI_ISL_3202941, EPI_ISL_3202942 | Lighthouse Lab in Alderley Park | Wellcome Sanger Institute for the COVID-19 Genomics UK (COG-UK) Consortium | Jacquelyn Wynn, Mairead Hyland, The Lighthouse Lab in Alderley Park and Alex Alderton, Roberto Amato, Jeffrey Barrett, Sonia Goncalves, Ewan Harrison, David K. Jackson, Ian Johnston, Dominic Kwiatkowski, Cordelia Langford, John Sillitoe on behalf of the Wellcome Sanger Institute COVID-19 Surveillance Team |
| EPI_ISL_3202943 | Lighthouse Laboratory Plymouth | Wellcome Sanger Institute for the COVID-19 Genomics UK (COG-UK) Consortium | Lighthouse Laboratory Plymouth and Alex Alderton, Roberto Amato, Jeffrey Barrett, Sonia Goncalves, Ewan Harrison, David K. Jackson, Ian Johnston, Dominic Kwiatkowski, Cordelia Langford, John Sillitoe on behalf of the Wellcome Sanger Institute COVID-19 Surveillance Team |
| EPI_ISL_3202944 | Lighthouse Lab in Milton Keynes | Wellcome Sanger Institute for the COVID-19 Genomics UK (COG-UK) Consortium | The Lighthouse Lab in Milton Keynes and Alex Alderton, Roberto Amato, Jeffrey Barrett, Sonia Goncalves, Ewan Harrison, David K. Jackson, Ian Johnston, Dominic Kwiatkowski, Cordelia Langford, John Sillitoe on behalf of the Wellcome Sanger Institute COVID-19 Surveillance Team |
| EPI_ISL_3202945 | Berkshire and Surrey Pathology Services Lighthouse Laboratory | Wellcome Sanger Institute for the COVID-19 Genomics UK (COG-UK) Consortium | Berkshire and Surrey Pathology Services Lighthouse Laboratory and Alex Alderton, Roberto Amato, Jeffrey Barrett, Sonia Goncalves, Ewan Harrison, David K. Jackson, Ian Johnston, Dominic Kwiatkowski, Cordelia Langford, John Sillitoe on behalf of the Wellcome Sanger Institute COVID-19 Surveillance Team |
| EPI_ISL_3202946 | Lighthouse Laboratory Plymouth | Wellcome Sanger Institute for the COVID-19 Genomics UK (COG-UK) Consortium | Lighthouse Laboratory Plymouth and Alex Alderton, Roberto Amato, Jeffrey Barrett, Sonia Goncalves, Ewan Harrison, David K. Jackson, Ian Johnston, Dominic Kwiatkowski, Cordelia Langford, John Sillitoe on behalf of the Wellcome Sanger Institute COVID-19 Surveillance Team |
| EPI_ISL_3202947 | Lighthouse Lab in Alderley Park | Wellcome Sanger Institute for the COVID-19 Genomics UK (COG-UK) Consortium | Jacquelyn Wynn, Mairead Hyland, The Lighthouse Lab in Alderley Park and Alex Alderton, Roberto Amato, Jeffrey Barrett, Sonia Goncalves, Ewan Harrison, David K. Jackson, Ian Johnston, Dominic Kwiatkowski, Cordelia Langford, John Sillitoe on behalf of the Wellcome Sanger Institute COVID-19 Surveillance Team |
| EPI_ISL_3202948 | Berkshire and Surrey Pathology Services Lighthouse Laboratory | Wellcome Sanger Institute for the COVID-19 Genomics UK (COG-UK) Consortium | Berkshire and Surrey Pathology Services Lighthouse Laboratory and Alex Alderton, Roberto Amato, Jeffrey Barrett, Sonia Goncalves, Ewan Harrison, David K. Jackson, Ian Johnston, Dominic Kwiatkowski, Cordelia Langford, John Sillitoe on behalf of the Wellcome Sanger Institute COVID-19 Surveillance Team |
| EPI_ISL_3202949, EPI_ISL_3202950 | Lighthouse Lab in Milton Keynes | Wellcome Sanger Institute for the COVID-19 Genomics UK (COG-UK) Consortium | The Lighthouse Lab in Milton Keynes and Alex Alderton, Roberto Amato, Jeffrey Barrett, Sonia Goncalves, Ewan Harrison, David K. Jackson, Ian Johnston, Dominic Kwiatkowski, Cordelia Langford, John Sillitoe on behalf of the Wellcome Sanger Institute COVID-19 Surveillance Team |
| EPI_ISL_3202951, EPI_ISL_3202952, EPI_ISL_3202953 | Berkshire and Surrey Pathology Services Lighthouse Laboratory | Wellcome Sanger Institute for the COVID-19 Genomics UK (COG-UK) Consortium | Berkshire and Surrey Pathology Services Lighthouse Laboratory and Alex Alderton, Roberto Amato, Jeffrey Barrett, Sonia Goncalves, Ewan Harrison, David K. Jackson, Ian Johnston, Dominic Kwiatkowski, Cordelia Langford, John Sillitoe on behalf of the Wellcome Sanger Institute COVID-19 Surveillance Team |
| EPI_ISL_3202954, EPI_ISL_3202955 | Lighthouse Lab in Milton Keynes | Wellcome Sanger Institute for the COVID-19 Genomics UK (COG-UK) Consortium | The Lighthouse Lab in Milton Keynes and Alex Alderton, Roberto Amato, Jeffrey Barrett, Sonia Goncalves, Ewan Harrison, David K. Jackson, Ian Johnston, Dominic Kwiatkowski, Cordelia Langford, John Sillitoe on behalf of the Wellcome Sanger Institute COVID-19 Surveillance Team |
| EPI_ISL_3202956 | Lighthouse Lab in Alderley Park | Wellcome Sanger Institute for the COVID-19 Genomics UK (COG-UK) Consortium | Jacquelyn Wynn, Mairead Hyland, The Lighthouse Lab in Alderley Park and Alex Alderton, Roberto Amato, Jeffrey Barrett, Sonia Goncalves, Ewan Harrison, David K. Jackson, Ian Johnston, Dominic Kwiatkowski, Cordelia Langford, John Sillitoe on behalf of the Wellcome Sanger Institute COVID-19 Surveillance Team |
| EPI_ISL_3202957 | Lighthouse Laboratory Plymouth | Wellcome Sanger Institute for the COVID-19 Genomics UK (COG-UK) Consortium | Lighthouse Laboratory Plymouth and Alex Alderton, Roberto Amato, Jeffrey Barrett, Sonia Goncalves, Ewan Harrison, David K. Jackson, Ian Johnston, Dominic Kwiatkowski, Cordelia Langford, John Sillitoe on behalf of the Wellcome Sanger Institute COVID-19 Surveillance Team |
| EPI_ISL_3202958, EPI_ISL_3202959, EPI_ISL_3202960 | Berkshire and Surrey Pathology Services Lighthouse Laboratory | Wellcome Sanger Institute for the COVID-19 Genomics UK (COG-UK) Consortium | Berkshire and Surrey Pathology Services Lighthouse Laboratory and Alex Alderton, Roberto Amato, Jeffrey Barrett, Sonia Goncalves, Ewan Harrison, David K. Jackson, Ian Johnston, Dominic Kwiatkowski, Cordelia Langford, John Sillitoe on behalf of the Wellcome Sanger Institute COVID-19 Surveillance Team |
| EPI_ISL_3202961 | Lighthouse Lab in Milton Keynes | Wellcome Sanger Institute for the COVID-19 Genomics UK (COG-UK) Consortium | The Lighthouse Lab in Milton Keynes and Alex Alderton, Roberto Amato, Jeffrey Barrett, Sonia Goncalves, Ewan Harrison, David K. Jackson, Ian Johnston, Dominic Kwiatkowski, Cordelia Langford, John Sillitoe on behalf of the Wellcome Sanger Institute COVID-19 Surveillance Team |
| EPI_ISL_3202962, EPI_ISL_3202963 | Lighthouse Laboratory Plymouth | Wellcome Sanger Institute for the COVID-19 Genomics UK (COG-UK) Consortium | Lighthouse Laboratory Plymouth and Alex Alderton, Roberto Amato, Jeffrey Barrett, Sonia Goncalves, Ewan Harrison, David K. Jackson, Ian Johnston, Dominic Kwiatkowski, Cordelia Langford, John Sillitoe on behalf of the Wellcome Sanger Institute COVID-19 Surveillance Team |
| EPI_ISL_3202964, EPI_ISL_3202965, EPI_ISL_3202966 | Lighthouse Lab in Milton Keynes | Wellcome Sanger Institute for the COVID-19 Genomics UK (COG-UK) Consortium | The Lighthouse Lab in Milton Keynes and Alex Alderton, Roberto Amato, Jeffrey Barrett, Sonia Goncalves, Ewan Harrison, David K. Jackson, Ian Johnston, Dominic Kwiatkowski, Cordelia Langford, John Sillitoe on behalf of the Wellcome Sanger Institute COVID-19 Surveillance Team |
| EPI_ISL_3202967 | Lighthouse Lab in Alderley Park | Wellcome Sanger Institute for the COVID-19 Genomics UK (COG-UK) Consortium | Jacquelyn Wynn, Mairead Hyland, The Lighthouse Lab in Alderley Park and Alex Alderton, Roberto Amato, Jeffrey Barrett, Sonia Goncalves, Ewan Harrison, David K. Jackson, Ian Johnston, Dominic Kwiatkowski, Cordelia Langford, John Sillitoe on behalf of the Wellcome Sanger Institute COVID-19 Surveillance Team |
| EPI_ISL_3202968, EPI_ISL_3202969 | Lighthouse Lab in Milton Keynes | Wellcome Sanger Institute for the COVID-19 Genomics UK (COG-UK) Consortium | The Lighthouse Lab in Milton Keynes and Alex Alderton, Roberto Amato, Jeffrey Barrett, Sonia Goncalves, Ewan Harrison, David K. Jackson, Ian Johnston, Dominic Kwiatkowski, Cordelia Langford, John Sillitoe on behalf of the Wellcome Sanger Institute COVID-19 Surveillance Team |
| EPI_ISL_3202970 | Berkshire and Surrey Pathology Services Lighthouse Laboratory | Wellcome Sanger Institute for the COVID-19 Genomics UK (COG-UK) Consortium | Berkshire and Surrey Pathology Services Lighthouse Laboratory and Alex Alderton, Roberto Amato, Jeffrey Barrett, Sonia Goncalves, Ewan Harrison, David K. Jackson, Ian Johnston, Dominic Kwiatkowski, Cordelia Langford, John Sillitoe on behalf of the Wellcome Sanger Institute COVID-19 Surveillance Team |
| EPI_ISL_3202971 | Lighthouse Lab in Alderley Park | Wellcome Sanger Institute for the COVID-19 Genomics UK (COG-UK) Consortium | Jacquelyn Wynn, Mairead Hyland, The Lighthouse Lab in Alderley Park and Alex Alderton, Roberto Amato, Jeffrey Barrett, Sonia Goncalves, Ewan Harrison, David K. Jackson, Ian Johnston, Dominic Kwiatkowski, Cordelia Langford, John Sillitoe on behalf of the Wellcome Sanger Institute COVID-19 Surveillance Team |
| EPI_ISL_3202972, EPI_ISL_3202973, EPI_ISL_3202974 | Lighthouse Laboratory Plymouth | Wellcome Sanger Institute for the COVID-19 Genomics UK (COG-UK) Consortium | Lighthouse Laboratory Plymouth and Alex Alderton, Roberto Amato, Jeffrey Barrett, Sonia Goncalves, Ewan Harrison, David K. Jackson, Ian Johnston, Dominic Kwiatkowski, Cordelia Langford, John Sillitoe on behalf of the Wellcome Sanger Institute COVID-19 Surveillance Team |
| EPI_ISL_3202975 | Lighthouse Lab in Milton Keynes | Wellcome Sanger Institute for the COVID-19 Genomics UK (COG-UK) Consortium | The Lighthouse Lab in Milton Keynes and Alex Alderton, Roberto Amato, Jeffrey Barrett, Sonia Goncalves, Ewan Harrison, David K. Jackson, Ian Johnston, Dominic Kwiatkowski, Cordelia Langford, John Sillitoe on behalf of the Wellcome Sanger Institute COVID-19 Surveillance Team |
| EPI_ISL_3202976 | Lighthouse Lab in Alderley Park | Wellcome Sanger Institute for the COVID-19 Genomics UK (COG-UK) Consortium | Jacquelyn Wynn, Mairead Hyland, The Lighthouse Lab in Alderley Park and Alex Alderton, Roberto Amato, Jeffrey Barrett, Sonia Goncalves, Ewan Harrison, David K. Jackson, Ian Johnston, Dominic Kwiatkowski, Cordelia Langford, John Sillitoe on behalf of the Wellcome Sanger Institute COVID-19 Surveillance Team |
| EPI_ISL_3202977 | Berkshire and Surrey Pathology Services Lighthouse Laboratory | Wellcome Sanger Institute for the COVID-19 Genomics UK (COG-UK) Consortium | Berkshire and Surrey Pathology Services Lighthouse Laboratory and Alex Alderton, Roberto Amato, Jeffrey Barrett, Sonia Goncalves, Ewan Harrison, David K. Jackson, Ian Johnston, Dominic Kwiatkowski, Cordelia Langford, John Sillitoe on behalf of the Wellcome Sanger Institute COVID-19 Surveillance Team |
| EPI_ISL_3202978 | Lighthouse Lab in Milton Keynes | Wellcome Sanger Institute for the COVID-19 Genomics UK (COG-UK) Consortium | The Lighthouse Lab in Milton Keynes and Alex Alderton, Roberto Amato, Jeffrey Barrett, Sonia Goncalves, Ewan Harrison, David K. Jackson, Ian Johnston, Dominic Kwiatkowski, Cordelia Langford, John Sillitoe on behalf of the Wellcome Sanger Institute COVID-19 Surveillance Team |
| EPI_ISL_3202979 | Berkshire and Surrey Pathology Services Lighthouse Laboratory | Wellcome Sanger Institute for the COVID-19 Genomics UK (COG-UK) Consortium | Berkshire and Surrey Pathology Services Lighthouse Laboratory and Alex Alderton, Roberto Amato, Jeffrey Barrett, Sonia Goncalves, Ewan Harrison, David K. Jackson, Ian Johnston, Dominic Kwiatkowski, Cordelia Langford, John Sillitoe on behalf of the Wellcome Sanger Institute COVID-19 Surveillance Team |
| EPI_ISL_3202980 | Lighthouse Lab in Milton Keynes | Wellcome Sanger Institute for the COVID-19 Genomics UK (COG-UK) Consortium | The Lighthouse Lab in Milton Keynes and Alex Alderton, Roberto Amato, Jeffrey Barrett, Sonia Goncalves, Ewan Harrison, David K. Jackson, Ian Johnston, Dominic Kwiatkowski, Cordelia Langford, John Sillitoe on behalf of the Wellcome Sanger Institute COVID-19 Surveillance Team |
| EPI_ISL_3202981 | Berkshire and Surrey Pathology Services Lighthouse Laboratory | Wellcome Sanger Institute for the COVID-19 Genomics UK (COG-UK) Consortium | Berkshire and Surrey Pathology Services Lighthouse Laboratory and Alex Alderton, Roberto Amato, Jeffrey Barrett, Sonia Goncalves, Ewan Harrison, David K. Jackson, Ian Johnston, Dominic Kwiatkowski, Cordelia Langford, John Sillitoe on behalf of the Wellcome Sanger Institute COVID-19 Surveillance Team |

| | | | |
|---|---|---|---|
| EPI_ISL_3202982, EPI_ISL_3202983 | Lighthouse Lab in Alderley Park | Wellcome Sanger Institute for the COVID-19 Genomics UK (COG-UK) Consortium | Jacquelyn Wynn, Mairead Hyland, The Lighthouse Lab in Alderley Park and Alex Alderton, Roberto Amato, Jeffrey Barrett, Sonia Goncalves, Ewan Harrison, David K. Jackson, Ian Johnston, Dominic Kwiatkowski, Cordelia Langford, John Sillitoe on behalf of the Wellcome Sanger Institute COVID-19 Surveillance Team |
| EPI_ISL_3202984, EPI_ISL_3202985 | Lighthouse Lab in Milton Keynes | Wellcome Sanger Institute for the COVID-19 Genomics UK (COG-UK) Consortium | The Lighthouse Lab in Milton Keynes and Alex Alderton, Roberto Amato, Jeffrey Barrett, Sonia Goncalves, Ewan Harrison, David K. Jackson, Ian Johnston, Dominic Kwiatkowski, Cordelia Langford, John Sillitoe on behalf of the Wellcome Sanger Institute COVID-19 Surveillance Team |
| EPI_ISL_3202986 | Berkshire and Surrey Pathology Services Lighthouse Laboratory | Wellcome Sanger Institute for the COVID-19 Genomics UK (COG-UK) Consortium | Berkshire and Surrey Pathology Services Lighthouse Laboratory and Alex Alderton, Roberto Amato, Jeffrey Barrett, Sonia Goncalves, Ewan Harrison, David K. Jackson, Ian Johnston, Dominic Kwiatkowski, Cordelia Langford, John Sillitoe on behalf of the Wellcome Sanger Institute COVID-19 Surveillance Team |
| EPI_ISL_3202987, EPI_ISL_3202988 | Lighthouse Lab in Milton Keynes | Wellcome Sanger Institute for the COVID-19 Genomics UK (COG-UK) Consortium | The Lighthouse Lab in Milton Keynes and Alex Alderton, Roberto Amato, Jeffrey Barrett, Sonia Goncalves, Ewan Harrison, David K. Jackson, Ian Johnston, Dominic Kwiatkowski, Cordelia Langford, John Sillitoe on behalf of the Wellcome Sanger Institute COVID-19 Surveillance Team |
| EPI_ISL_3202989 | Berkshire and Surrey Pathology Services Lighthouse Laboratory | Wellcome Sanger Institute for the COVID-19 Genomics UK (COG-UK) Consortium | Berkshire and Surrey Pathology Services Lighthouse Laboratory and Alex Alderton, Roberto Amato, Jeffrey Barrett, Sonia Goncalves, Ewan Harrison, David K. Jackson, Ian Johnston, Dominic Kwiatkowski, Cordelia Langford, John Sillitoe on behalf of the Wellcome Sanger Institute COVID-19 Surveillance Team |
| EPI_ISL_3202990 | Lighthouse Lab in Alderley Park | Wellcome Sanger Institute for the COVID-19 Genomics UK (COG-UK) Consortium | Jacquelyn Wynn, Mairead Hyland, The Lighthouse Lab in Alderley Park and Alex Alderton, Roberto Amato, Jeffrey Barrett, Sonia Goncalves, Ewan Harrison, David K. Jackson, Ian Johnston, Dominic Kwiatkowski, Cordelia Langford, John Sillitoe on behalf of the Wellcome Sanger Institute COVID-19 Surveillance Team |
| EPI_ISL_3202991 | Berkshire and Surrey Pathology Services Lighthouse Laboratory | Wellcome Sanger Institute for the COVID-19 Genomics UK (COG-UK) Consortium | Berkshire and Surrey Pathology Services Lighthouse Laboratory and Alex Alderton, Roberto Amato, Jeffrey Barrett, Sonia Goncalves, Ewan Harrison, David K. Jackson, Ian Johnston, Dominic Kwiatkowski, Cordelia Langford, John Sillitoe on behalf of the Wellcome Sanger Institute COVID-19 Surveillance Team |
| EPI_ISL_3202992 | Lighthouse Lab in Milton Keynes | Wellcome Sanger Institute for the COVID-19 Genomics UK (COG-UK) Consortium | The Lighthouse Lab in Milton Keynes and Alex Alderton, Roberto Amato, Jeffrey Barrett, Sonia Goncalves, Ewan Harrison, David K. Jackson, Ian Johnston, Dominic Kwiatkowski, Cordelia Langford, John Sillitoe on behalf of the Wellcome Sanger Institute COVID-19 Surveillance Team |
| EPI_ISL_3202993, EPI_ISL_3202994 | Lighthouse Lab in Alderley Park | Wellcome Sanger Institute for the COVID-19 Genomics UK (COG-UK) Consortium | Jacquelyn Wynn, Mairead Hyland, The Lighthouse Lab in Alderley Park and Alex Alderton, Roberto Amato, Jeffrey Barrett, Sonia Goncalves, Ewan Harrison, David K. Jackson, Ian Johnston, Dominic Kwiatkowski, Cordelia Langford, John Sillitoe on behalf of the Wellcome Sanger Institute COVID-19 Surveillance Team |
| EPI_ISL_3202995 | Lighthouse Lab in Milton Keynes | Wellcome Sanger Institute for the COVID-19 Genomics UK (COG-UK) Consortium | The Lighthouse Lab in Milton Keynes and Alex Alderton, Roberto Amato, Jeffrey Barrett, Sonia Goncalves, Ewan Harrison, David K. Jackson, Ian Johnston, Dominic Kwiatkowski, Cordelia Langford, John Sillitoe on behalf of the Wellcome Sanger Institute COVID-19 Surveillance Team |
| EPI_ISL_3202996, EPI_ISL_3202997 | Lighthouse Lab in Alderley Park | Wellcome Sanger Institute for the COVID-19 Genomics UK (COG-UK) Consortium | Jacquelyn Wynn, Mairead Hyland, The Lighthouse Lab in Alderley Park and Alex Alderton, Roberto Amato, Jeffrey Barrett, Sonia Goncalves, Ewan Harrison, David K. Jackson, Ian Johnston, Dominic Kwiatkowski, Cordelia Langford, John Sillitoe on behalf of the Wellcome Sanger Institute COVID-19 Surveillance Team |
| EPI_ISL_3202998 | Berkshire and Surrey Pathology Services Lighthouse Laboratory | Wellcome Sanger Institute for the COVID-19 Genomics UK (COG-UK) Consortium | Berkshire and Surrey Pathology Services Lighthouse Laboratory and Alex Alderton, Roberto Amato, Jeffrey Barrett, Sonia Goncalves, Ewan Harrison, David K. Jackson, Ian Johnston, Dominic Kwiatkowski, Cordelia Langford, John Sillitoe on behalf of the Wellcome Sanger Institute COVID-19 Surveillance Team |
| EPI_ISL_3202999 | Lighthouse Lab in Alderley Park | Wellcome Sanger Institute for the COVID-19 Genomics UK (COG-UK) Consortium | Jacquelyn Wynn, Mairead Hyland, The Lighthouse Lab in Alderley Park and Alex Alderton, Roberto Amato, Jeffrey Barrett, Sonia Goncalves, Ewan Harrison, David K. Jackson, Ian Johnston, Dominic Kwiatkowski, Cordelia Langford, John Sillitoe on behalf of the Wellcome Sanger Institute COVID-19 Surveillance Team |
| EPI_ISL_3203000, EPI_ISL_3203001 | Berkshire and Surrey Pathology Services Lighthouse Laboratory | Wellcome Sanger Institute for the COVID-19 Genomics UK (COG-UK) Consortium | Berkshire and Surrey Pathology Services Lighthouse Laboratory and Alex Alderton, Roberto Amato, Jeffrey Barrett, Sonia Goncalves, Ewan Harrison, David K. Jackson, Ian Johnston, Dominic Kwiatkowski, Cordelia Langford, John Sillitoe on behalf of the Wellcome Sanger Institute COVID-19 Surveillance Team |
| EPI_ISL_3203002 | Lighthouse Lab in Milton Keynes | Wellcome Sanger Institute for the COVID-19 Genomics UK (COG-UK) Consortium | The Lighthouse Lab in Milton Keynes and Alex Alderton, Roberto Amato, Jeffrey Barrett, Sonia Goncalves, Ewan Harrison, David K. Jackson, Ian Johnston, Dominic Kwiatkowski, Cordelia Langford, John Sillitoe on behalf of the Wellcome Sanger Institute COVID-19 Surveillance Team |
| EPI_ISL_3203003, EPI_ISL_3203004, EPI_ISL_3203005 | Lighthouse Lab in Alderley Park | Wellcome Sanger Institute for the COVID-19 Genomics UK (COG-UK) Consortium | Jacquelyn Wynn, Mairead Hyland, The Lighthouse Lab in Alderley Park and Alex Alderton, Roberto Amato, Jeffrey Barrett, Sonia Goncalves, Ewan Harrison, David K. Jackson, Ian Johnston, Dominic Kwiatkowski, Cordelia Langford, John Sillitoe on behalf of the Wellcome Sanger Institute COVID-19 Surveillance Team |
| EPI_ISL_3203006, EPI_ISL_3203007 | Lighthouse Laboratory Plymouth | Wellcome Sanger Institute for the COVID-19 Genomics UK (COG-UK) Consortium | Lighthouse Laboratory Plymouth and Alex Alderton, Roberto Amato, Jeffrey Barrett, Sonia Goncalves, Ewan Harrison, David K. Jackson, Ian Johnston, Dominic Kwiatkowski, Cordelia Langford, John Sillitoe on behalf of the Wellcome Sanger Institute COVID-19 Surveillance Team |
| EPI_ISL_3203008 | Lighthouse Lab in Milton Keynes | Wellcome Sanger Institute for the COVID-19 Genomics UK (COG-UK) Consortium | The Lighthouse Lab in Milton Keynes and Alex Alderton, Roberto Amato, Jeffrey Barrett, Sonia Goncalves, Ewan Harrison, David K. Jackson, Ian Johnston, Dominic Kwiatkowski, Cordelia Langford, John Sillitoe on behalf of the Wellcome Sanger Institute COVID-19 Surveillance Team |
| EPI_ISL_3203009 | Lighthouse Lab in Alderley Park | Wellcome Sanger Institute for the COVID-19 Genomics UK (COG-UK) Consortium | Jacquelyn Wynn, Mairead Hyland, The Lighthouse Lab in Alderley Park and Alex Alderton, Roberto Amato, Jeffrey Barrett, Sonia Goncalves, Ewan Harrison, David K. Jackson, Ian Johnston, Dominic Kwiatkowski, Cordelia Langford, John Sillitoe on behalf of the Wellcome Sanger Institute COVID-19 Surveillance Team |
| EPI_ISL_3203010 | Lighthouse Laboratory Plymouth | Wellcome Sanger Institute for the COVID-19 Genomics UK (COG-UK) Consortium | Lighthouse Laboratory Plymouth and Alex Alderton, Roberto Amato, Jeffrey Barrett, Sonia Goncalves, Ewan Harrison, David K. Jackson, Ian Johnston, Dominic Kwiatkowski, Cordelia Langford, John Sillitoe on behalf of the Wellcome Sanger Institute COVID-19 Surveillance Team |
| EPI_ISL_3203011 | Lighthouse Lab in Alderley Park | Wellcome Sanger Institute for the COVID-19 Genomics UK (COG-UK) Consortium | Jacquelyn Wynn, Mairead Hyland, The Lighthouse Lab in Alderley Park and Alex Alderton, Roberto Amato, Jeffrey Barrett, Sonia Goncalves, Ewan Harrison, David K. Jackson, Ian Johnston, Dominic Kwiatkowski, Cordelia Langford, John Sillitoe on behalf of the Wellcome Sanger Institute COVID-19 Surveillance Team |
| EPI_ISL_3203012 | Lighthouse Laboratory Plymouth | Wellcome Sanger Institute for the COVID-19 Genomics UK (COG-UK) Consortium | Lighthouse Laboratory Plymouth and Alex Alderton, Roberto Amato, Jeffrey Barrett, Sonia Goncalves, Ewan Harrison, David K. Jackson, Ian Johnston, Dominic Kwiatkowski, Cordelia Langford, John Sillitoe on behalf of the Wellcome Sanger Institute COVID-19 Surveillance Team |
| EPI_ISL_3203013 | Berkshire and Surrey Pathology Services Lighthouse Laboratory | Wellcome Sanger Institute for the COVID-19 Genomics UK (COG-UK) Consortium | Berkshire and Surrey Pathology Services Lighthouse Laboratory and Alex Alderton, Roberto Amato, Jeffrey Barrett, Sonia Goncalves, Ewan Harrison, David K. Jackson, Ian Johnston, Dominic Kwiatkowski, Cordelia Langford, John Sillitoe on behalf of the Wellcome Sanger Institute COVID-19 Surveillance Team |
| EPI_ISL_3203014 | Lighthouse Lab in Milton Keynes | Wellcome Sanger Institute for the COVID-19 Genomics UK (COG-UK) Consortium | The Lighthouse Lab in Milton Keynes and Alex Alderton, Roberto Amato, Jeffrey Barrett, Sonia Goncalves, Ewan Harrison, David K. Jackson, Ian Johnston, Dominic Kwiatkowski, Cordelia Langford, John Sillitoe on behalf of the Wellcome Sanger Institute COVID-19 Surveillance Team |
| EPI_ISL_3203015, EPI_ISL_3203016, EPI_ISL_3203017 | Berkshire and Surrey Pathology Services Lighthouse Laboratory | Wellcome Sanger Institute for the COVID-19 Genomics UK (COG-UK) Consortium | Berkshire and Surrey Pathology Services Lighthouse Laboratory and Alex Alderton, Roberto Amato, Jeffrey Barrett, Sonia Goncalves, Ewan Harrison, David K. Jackson, Ian Johnston, Dominic Kwiatkowski, Cordelia Langford, John Sillitoe on behalf of the Wellcome Sanger Institute COVID-19 Surveillance Team |
| EPI_ISL_3203018, EPI_ISL_3203019 | Lighthouse Lab in Alderley Park | Wellcome Sanger Institute for the COVID-19 Genomics UK (COG-UK) Consortium | Jacquelyn Wynn, Mairead Hyland, The Lighthouse Lab in Alderley Park and Alex Alderton, Roberto Amato, Jeffrey Barrett, Sonia Goncalves, Ewan Harrison, David K. Jackson, Ian Johnston, Dominic Kwiatkowski, Cordelia Langford, John Sillitoe on behalf of the Wellcome Sanger Institute COVID-19 Surveillance Team |
| EPI_ISL_3203020 | Lighthouse Laboratory Plymouth | Wellcome Sanger Institute for the COVID-19 Genomics UK (COG-UK) Consortium | Lighthouse Laboratory Plymouth and Alex Alderton, Roberto Amato, Jeffrey Barrett, Sonia Goncalves, Ewan Harrison, David K. Jackson, Ian Johnston, Dominic Kwiatkowski, Cordelia Langford, John Sillitoe on behalf of the Wellcome Sanger Institute COVID-19 Surveillance Team |
| EPI_ISL_3203021 | Lighthouse Lab in Milton Keynes | Wellcome Sanger Institute for the COVID-19 Genomics UK (COG-UK) Consortium | The Lighthouse Lab in Milton Keynes and Alex Alderton, Roberto Amato, Jeffrey Barrett, Sonia Goncalves, Ewan Harrison, David K. Jackson, Ian Johnston, Dominic Kwiatkowski, Cordelia Langford, John Sillitoe on behalf of the Wellcome Sanger Institute COVID-19 Surveillance Team |
| EPI_ISL_3203022, EPI_ISL_3203023 | Lighthouse Laboratory Plymouth | Wellcome Sanger Institute for the COVID-19 Genomics UK (COG-UK) Consortium | Lighthouse Laboratory Plymouth and Alex Alderton, Roberto Amato, Jeffrey Barrett, Sonia Goncalves, Ewan Harrison, David K. Jackson, Ian Johnston, Dominic Kwiatkowski, Cordelia Langford, John Sillitoe on behalf of the Wellcome Sanger Institute COVID-19 Surveillance Team |
| EPI_ISL_3203024 | Lighthouse Lab in Glasgow | Wellcome Sanger Institute for the COVID-19 Genomics UK (COG-UK) Consortium | Harper VanSteenhouse, Yumi Kasai, David Gray, Carol Clugston, Anna Dominiczak and Alex Alderton, Roberto Amato, Jeffrey Barrett, Sonia Goncalves, Ewan Harrison, David K. Jackson, Ian Johnston, Dominic Kwiatkowski, Cordelia Langford, John Sillitoe on behalf of the Wellcome Sanger Institute COVID-19 Surveillance Team |
| EPI_ISL_3203025, EPI_ISL_3203026 | Lighthouse Laboratory Plymouth | Wellcome Sanger Institute for the COVID-19 Genomics UK (COG-UK) Consortium | Lighthouse Laboratory Plymouth and Alex Alderton, Roberto Amato, Jeffrey Barrett, Sonia Goncalves, Ewan Harrison, David K. Jackson, Ian Johnston, Dominic Kwiatkowski, Cordelia Langford, John Sillitoe on behalf of the Wellcome Sanger Institute COVID-19 Surveillance Team |
| EPI_ISL_3203027 | Lighthouse Lab in Milton Keynes | Wellcome Sanger Institute for the COVID-19 Genomics UK (COG-UK) Consortium | The Lighthouse Lab in Milton Keynes and Alex Alderton, Roberto Amato, Jeffrey Barrett, Sonia Goncalves, Ewan Harrison, David K. Jackson, Ian Johnston, Dominic Kwiatkowski, Cordelia Langford, John Sillitoe on behalf of the Wellcome Sanger Institute COVID-19 Surveillance Team |
| EPI_ISL_3203028, EPI_ISL_3203029, EPI_ISL_3203030, EPI_ISL_3203031 | Lighthouse Laboratory Plymouth | Wellcome Sanger Institute for the COVID-19 Genomics UK (COG-UK) Consortium | Lighthouse Laboratory Plymouth and Alex Alderton, Roberto Amato, Jeffrey Barrett, Sonia Goncalves, Ewan Harrison, David K. Jackson, Ian Johnston, Dominic Kwiatkowski, Cordelia Langford, John Sillitoe on behalf of the Wellcome Sanger Institute COVID-19 Surveillance Team |
| EPI_ISL_3203032 | Lighthouse Lab in Glasgow | Wellcome Sanger Institute for the COVID-19 Genomics UK (COG-UK) Consortium | Harper VanSteenhouse, Yumi Kasai, David Gray, Carol Clugston, Anna Dominiczak and Alex Alderton, Roberto Amato, Jeffrey Barrett, Sonia Goncalves, Ewan Harrison, David K. Jackson, Ian Johnston, Dominic Kwiatkowski, Cordelia Langford, John Sillitoe on behalf of the Wellcome Sanger Institute COVID-19 Surveillance Team |
| EPI_ISL_3203033 | Lighthouse Lab in Milton Keynes | Wellcome Sanger Institute for the COVID-19 Genomics UK (COG-UK) Consortium | The Lighthouse Lab in Milton Keynes and Alex Alderton, Roberto Amato, Jeffrey Barrett, Sonia Goncalves, Ewan Harrison, David K. Jackson, Ian Johnston, Dominic Kwiatkowski, Cordelia Langford, John Sillitoe on behalf of the Wellcome Sanger Institute COVID-19 Surveillance Team |
| EPI_ISL_3203034 | Lighthouse Laboratory Plymouth | Wellcome Sanger Institute for the COVID-19 Genomics UK (COG-UK) Consortium | Lighthouse Laboratory Plymouth and Alex Alderton, Roberto Amato, Jeffrey Barrett, Sonia Goncalves, Ewan Harrison, David K. Jackson, Ian Johnston, Dominic Kwiatkowski, Cordelia Langford, John Sillitoe on behalf of the Wellcome Sanger Institute COVID-19 Surveillance Team |
| EPI_ISL_3203035 | Lighthouse Lab in Milton Keynes | Wellcome Sanger Institute for the COVID-19 Genomics UK (COG-UK) Consortium | The Lighthouse Lab in Milton Keynes and Alex Alderton, Roberto Amato, Jeffrey Barrett, Sonia Goncalves, Ewan Harrison, David K. Jackson, Ian Johnston, Dominic Kwiatkowski, Cordelia Langford, John Sillitoe on behalf of the Wellcome Sanger Institute COVID-19 Surveillance Team |
| EPI_ISL_3203036 | Lighthouse Lab in Glasgow | Wellcome Sanger Institute for the COVID-19 Genomics UK (COG-UK) Consortium | Harper VanSteenhouse, Yumi Kasai, David Gray, Carol Clugston, Anna Dominiczak and Alex Alderton, Roberto Amato, Jeffrey Barrett, Sonia Goncalves, Ewan Harrison, David K. Jackson, Ian Johnston, Dominic Kwiatkowski, Cordelia Langford, John Sillitoe on behalf of the Wellcome Sanger Institute COVID-19 Surveillance Team |
| EPI_ISL_3203037 | Lighthouse Laboratory Plymouth | Wellcome Sanger Institute for the COVID-19 Genomics UK (COG-UK) Consortium | Lighthouse Laboratory Plymouth and Alex Alderton, Roberto Amato, Jeffrey Barrett, Sonia Goncalves, Ewan Harrison, David K. Jackson, Ian Johnston, Dominic Kwiatkowski, Cordelia Langford, John Sillitoe on behalf of the Wellcome Sanger Institute COVID-19 Surveillance Team |
| EPI_ISL_3203038, EPI_ISL_3203039 | Lighthouse Lab in Milton Keynes | Wellcome Sanger Institute for the COVID-19 Genomics UK (COG-UK) Consortium | The Lighthouse Lab in Milton Keynes and Alex Alderton, Roberto Amato, Jeffrey Barrett, Sonia Goncalves, Ewan Harrison, David K. Jackson, Ian Johnston, Dominic Kwiatkowski, Cordelia Langford, John Sillitoe on behalf of the Wellcome Sanger Institute COVID-19 Surveillance Team |
| EPI_ISL_3203040 | Lighthouse Laboratory Plymouth | Wellcome Sanger Institute for the COVID-19 Genomics UK (COG-UK) Consortium | Lighthouse Laboratory Plymouth and Alex Alderton, Roberto Amato, Jeffrey Barrett, Sonia Goncalves, Ewan Harrison, David K. Jackson, Ian Johnston, Dominic Kwiatkowski, Cordelia Langford, John Sillitoe on behalf of the Wellcome Sanger Institute COVID-19 Surveillance Team |
| EPI_ISL_3203041 | Lighthouse Lab in Glasgow | Wellcome Sanger Institute for the COVID-19 Genomics UK (COG-UK) Consortium | Harper VanSteenhouse, Yumi Kasai, David Gray, Carol Clugston, Anna Dominiczak and Alex Alderton, Roberto Amato, Jeffrey Barrett, Sonia Goncalves, Ewan Harrison, David K. Jackson, Ian Johnston, Dominic Kwiatkowski, Cordelia Langford, John Sillitoe on behalf of the Wellcome Sanger Institute COVID-19 Surveillance Team |
| EPI_ISL_3203042 | Lighthouse Laboratory Plymouth | Wellcome Sanger Institute for the COVID-19 Genomics UK (COG-UK) Consortium | Lighthouse Laboratory Plymouth and Alex Alderton, Roberto Amato, Jeffrey Barrett, Sonia Goncalves, Ewan Harrison, David K. Jackson, Ian Johnston, Dominic Kwiatkowski, Cordelia Langford, John Sillitoe on behalf of the Wellcome Sanger Institute COVID-19 Surveillance Team |
| EPI_ISL_3203043 | Lighthouse Lab in Glasgow | Wellcome Sanger Institute for the COVID-19 Genomics UK (COG-UK) | Harper VanSteenhouse, Yumi Kasai, David Gray, Carol Clugston, Anna Dominiczak and Alex Alderton, Roberto Amato, Jeffrey Barrett, Sonia Goncalves, Ewan Harrison, David K. Jackson, Ian Johnston, Dominic Kwiatkowski, Cordelia Langford, |

| | | | |
|---|---|---|---|
| | | Consortium | John Sillitoe on behalf of the Wellcome Sanger Institute COVID-19 Surveillance Team |
| EPI_ISL_3203044 | Lighthouse Lab in Milton Keynes | Wellcome Sanger Institute for the COVID-19 Genomics UK (COG-UK) Consortium | The Lighthouse Lab in Milton Keynes and Alex Alderton, Roberto Amato, Jeffrey Barrett, Sonia Goncalves, Ewan Harrison, David K. Jackson, Ian Johnston, Dominic Kwiatkowski, Cordelia Langford, John Sillitoe on behalf of the Wellcome Sanger Institute COVID-19 Surveillance Team |
| EPI_ISL_3203045, EPI_ISL_3203046, EPI_ISL_3203047 | Lighthouse Laboratory Plymouth | Wellcome Sanger Institute for the COVID-19 Genomics UK (COG-UK) Consortium | Lighthouse Laboratory Plymouth and Alex Alderton, Roberto Amato, Jeffrey Barrett, Sonia Goncalves, Ewan Harrison, David K. Jackson, Ian Johnston, Dominic Kwiatkowski, Cordelia Langford, John Sillitoe on behalf of the Wellcome Sanger Institute COVID-19 Surveillance Team |
| EPI_ISL_3203048 | Lighthouse Lab in Milton Keynes | Wellcome Sanger Institute for the COVID-19 Genomics UK (COG-UK) Consortium | The Lighthouse Lab in Milton Keynes and Alex Alderton, Roberto Amato, Jeffrey Barrett, Sonia Goncalves, Ewan Harrison, David K. Jackson, Ian Johnston, Dominic Kwiatkowski, Cordelia Langford, John Sillitoe on behalf of the Wellcome Sanger Institute COVID-19 Surveillance Team |
| EPI_ISL_3203049 | Lighthouse Lab in Glasgow | Wellcome Sanger Institute for the COVID-19 Genomics UK (COG-UK) Consortium | Harper VanSteenhouse, Yumi Kasai, David Gray, Carol Clugston, Anna Dominiczak and Alex Alderton, Roberto Amato, Jeffrey Barrett, Sonia Goncalves, Ewan Harrison, David K. Jackson, Ian Johnston, Dominic Kwiatkowski, Cordelia Langford, John Sillitoe on behalf of the Wellcome Sanger Institute COVID-19 Surveillance Team |
| EPI_ISL_3203050, EPI_ISL_3203051, EPI_ISL_3203052 | Lighthouse Laboratory Plymouth | Wellcome Sanger Institute for the COVID-19 Genomics UK (COG-UK) Consortium | Lighthouse Laboratory Plymouth and Alex Alderton, Roberto Amato, Jeffrey Barrett, Sonia Goncalves, Ewan Harrison, David K. Jackson, Ian Johnston, Dominic Kwiatkowski, Cordelia Langford, John Sillitoe on behalf of the Wellcome Sanger Institute COVID-19 Surveillance Team |
| EPI_ISL_3203053 | Lighthouse Lab in Milton Keynes | Wellcome Sanger Institute for the COVID-19 Genomics UK (COG-UK) Consortium | The Lighthouse Lab in Milton Keynes and Alex Alderton, Roberto Amato, Jeffrey Barrett, Sonia Goncalves, Ewan Harrison, David K. Jackson, Ian Johnston, Dominic Kwiatkowski, Cordelia Langford, John Sillitoe on behalf of the Wellcome Sanger Institute COVID-19 Surveillance Team |
| EPI_ISL_3203054 | Lighthouse Laboratory Plymouth | Wellcome Sanger Institute for the COVID-19 Genomics UK (COG-UK) Consortium | Lighthouse Laboratory Plymouth and Alex Alderton, Roberto Amato, Jeffrey Barrett, Sonia Goncalves, Ewan Harrison, David K. Jackson, Ian Johnston, Dominic Kwiatkowski, Cordelia Langford, John Sillitoe on behalf of the Wellcome Sanger Institute COVID-19 Surveillance Team |
| EPI_ISL_3203055 | Lighthouse Lab in Milton Keynes | Wellcome Sanger Institute for the COVID-19 Genomics UK (COG-UK) Consortium | The Lighthouse Lab in Milton Keynes and Alex Alderton, Roberto Amato, Jeffrey Barrett, Sonia Goncalves, Ewan Harrison, David K. Jackson, Ian Johnston, Dominic Kwiatkowski, Cordelia Langford, John Sillitoe on behalf of the Wellcome Sanger Institute COVID-19 Surveillance Team |
| EPI_ISL_3203056 | Lighthouse Laboratory Plymouth | Wellcome Sanger Institute for the COVID-19 Genomics UK (COG-UK) Consortium | Lighthouse Laboratory Plymouth and Alex Alderton, Roberto Amato, Jeffrey Barrett, Sonia Goncalves, Ewan Harrison, David K. Jackson, Ian Johnston, Dominic Kwiatkowski, Cordelia Langford, John Sillitoe on behalf of the Wellcome Sanger Institute COVID-19 Surveillance Team |
| EPI_ISL_3203057 | Lighthouse Lab in Glasgow | Wellcome Sanger Institute for the COVID-19 Genomics UK (COG-UK) Consortium | Harper VanSteenhouse, Yumi Kasai, David Gray, Carol Clugston, Anna Dominiczak and Alex Alderton, Roberto Amato, Jeffrey Barrett, Sonia Goncalves, Ewan Harrison, David K. Jackson, Ian Johnston, Dominic Kwiatkowski, Cordelia Langford, John Sillitoe on behalf of the Wellcome Sanger Institute COVID-19 Surveillance Team |
| EPI_ISL_3203058, EPI_ISL_3203059 | Lighthouse Lab in Milton Keynes | Wellcome Sanger Institute for the COVID-19 Genomics UK (COG-UK) Consortium | The Lighthouse Lab in Milton Keynes and Alex Alderton, Roberto Amato, Jeffrey Barrett, Sonia Goncalves, Ewan Harrison, David K. Jackson, Ian Johnston, Dominic Kwiatkowski, Cordelia Langford, John Sillitoe on behalf of the Wellcome Sanger Institute COVID-19 Surveillance Team |
| EPI_ISL_3203060, EPI_ISL_3203061 | Lighthouse Laboratory Plymouth | Wellcome Sanger Institute for the COVID-19 Genomics UK (COG-UK) Consortium | Lighthouse Laboratory Plymouth and Alex Alderton, Roberto Amato, Jeffrey Barrett, Sonia Goncalves, Ewan Harrison, David K. Jackson, Ian Johnston, Dominic Kwiatkowski, Cordelia Langford, John Sillitoe on behalf of the Wellcome Sanger Institute COVID-19 Surveillance Team |
| EPI_ISL_3203062 | Lighthouse Lab in Milton Keynes | Wellcome Sanger Institute for the COVID-19 Genomics UK (COG-UK) Consortium | The Lighthouse Lab in Milton Keynes and Alex Alderton, Roberto Amato, Jeffrey Barrett, Sonia Goncalves, Ewan Harrison, David K. Jackson, Ian Johnston, Dominic Kwiatkowski, Cordelia Langford, John Sillitoe on behalf of the Wellcome Sanger Institute COVID-19 Surveillance Team |
| EPI_ISL_3203063, EPI_ISL_3203064 | Lighthouse Laboratory Plymouth | Wellcome Sanger Institute for the COVID-19 Genomics UK (COG-UK) Consortium | Lighthouse Laboratory Plymouth and Alex Alderton, Roberto Amato, Jeffrey Barrett, Sonia Goncalves, Ewan Harrison, David K. Jackson, Ian Johnston, Dominic Kwiatkowski, Cordelia Langford, John Sillitoe on behalf of the Wellcome Sanger Institute COVID-19 Surveillance Team |
| EPI_ISL_3203065 | Lighthouse Lab in Glasgow | Wellcome Sanger Institute for the COVID-19 Genomics UK (COG-UK) Consortium | Harper VanSteenhouse, Yumi Kasai, David Gray, Carol Clugston, Anna Dominiczak and Alex Alderton, Roberto Amato, Jeffrey Barrett, Sonia Goncalves, Ewan Harrison, David K. Jackson, Ian Johnston, Dominic Kwiatkowski, Cordelia Langford, John Sillitoe on behalf of the Wellcome Sanger Institute COVID-19 Surveillance Team |
| EPI_ISL_3203066, EPI_ISL_3203067 | Lighthouse Laboratory Plymouth | Wellcome Sanger Institute for the COVID-19 Genomics UK (COG-UK) Consortium | Lighthouse Laboratory Plymouth and Alex Alderton, Roberto Amato, Jeffrey Barrett, Sonia Goncalves, Ewan Harrison, David K. Jackson, Ian Johnston, Dominic Kwiatkowski, Cordelia Langford, John Sillitoe on behalf of the Wellcome Sanger Institute COVID-19 Surveillance Team |
| EPI_ISL_3203068 | Lighthouse Lab in Glasgow | Wellcome Sanger Institute for the COVID-19 Genomics UK (COG-UK) Consortium | Harper VanSteenhouse, Yumi Kasai, David Gray, Carol Clugston, Anna Dominiczak and Alex Alderton, Roberto Amato, Jeffrey Barrett, Sonia Goncalves, Ewan Harrison, David K. Jackson, Ian Johnston, Dominic Kwiatkowski, Cordelia Langford, John Sillitoe on behalf of the Wellcome Sanger Institute COVID-19 Surveillance Team |
| EPI_ISL_3203069 | Lighthouse Laboratory Plymouth | Wellcome Sanger Institute for the COVID-19 Genomics UK (COG-UK) Consortium | Lighthouse Laboratory Plymouth and Alex Alderton, Roberto Amato, Jeffrey Barrett, Sonia Goncalves, Ewan Harrison, David K. Jackson, Ian Johnston, Dominic Kwiatkowski, Cordelia Langford, John Sillitoe on behalf of the Wellcome Sanger Institute COVID-19 Surveillance Team |
| EPI_ISL_3203070 | Lighthouse Lab in Milton Keynes | Wellcome Sanger Institute for the COVID-19 Genomics UK (COG-UK) Consortium | The Lighthouse Lab in Milton Keynes and Alex Alderton, Roberto Amato, Jeffrey Barrett, Sonia Goncalves, Ewan Harrison, David K. Jackson, Ian Johnston, Dominic Kwiatkowski, Cordelia Langford, John Sillitoe on behalf of the Wellcome Sanger Institute COVID-19 Surveillance Team |
| EPI_ISL_3203071 | Lighthouse Laboratory Plymouth | Wellcome Sanger Institute for the COVID-19 Genomics UK (COG-UK) Consortium | Lighthouse Laboratory Plymouth and Alex Alderton, Roberto Amato, Jeffrey Barrett, Sonia Goncalves, Ewan Harrison, David K. Jackson, Ian Johnston, Dominic Kwiatkowski, Cordelia Langford, John Sillitoe on behalf of the Wellcome Sanger Institute COVID-19 Surveillance Team |
| EPI_ISL_3203072 | Lighthouse Lab in Milton Keynes | Wellcome Sanger Institute for the COVID-19 Genomics UK (COG-UK) Consortium | The Lighthouse Lab in Milton Keynes and Alex Alderton, Roberto Amato, Jeffrey Barrett, Sonia Goncalves, Ewan Harrison, David K. Jackson, Ian Johnston, Dominic Kwiatkowski, Cordelia Langford, John Sillitoe on behalf of the Wellcome Sanger Institute COVID-19 Surveillance Team |
| EPI_ISL_3203073 | Lighthouse Laboratory Plymouth | Wellcome Sanger Institute for the COVID-19 Genomics UK (COG-UK) Consortium | Lighthouse Laboratory Plymouth and Alex Alderton, Roberto Amato, Jeffrey Barrett, Sonia Goncalves, Ewan Harrison, David K. Jackson, Ian Johnston, Dominic Kwiatkowski, Cordelia Langford, John Sillitoe on behalf of the Wellcome Sanger Institute COVID-19 Surveillance Team |
| EPI_ISL_3203074 | Lighthouse Lab in Glasgow | Wellcome Sanger Institute for the COVID-19 Genomics UK (COG-UK) Consortium | Harper VanSteenhouse, Yumi Kasai, David Gray, Carol Clugston, Anna Dominiczak and Alex Alderton, Roberto Amato, Jeffrey Barrett, Sonia Goncalves, Ewan Harrison, David K. Jackson, Ian Johnston, Dominic Kwiatkowski, Cordelia Langford, John Sillitoe on behalf of the Wellcome Sanger Institute COVID-19 Surveillance Team |
| EPI_ISL_3203075, EPI_ISL_3203076, EPI_ISL_3203077 | Lighthouse Laboratory Plymouth | Wellcome Sanger Institute for the COVID-19 Genomics UK (COG-UK) Consortium | Lighthouse Laboratory Plymouth and Alex Alderton, Roberto Amato, Jeffrey Barrett, Sonia Goncalves, Ewan Harrison, David K. Jackson, Ian Johnston, Dominic Kwiatkowski, Cordelia Langford, John Sillitoe on behalf of the Wellcome Sanger Institute COVID-19 Surveillance Team |
| EPI_ISL_3203078 | Lighthouse Lab in Glasgow | Wellcome Sanger Institute for the COVID-19 Genomics UK (COG-UK) Consortium | Harper VanSteenhouse, Yumi Kasai, David Gray, Carol Clugston, Anna Dominiczak and Alex Alderton, Roberto Amato, Jeffrey Barrett, Sonia Goncalves, Ewan Harrison, David K. Jackson, Ian Johnston, Dominic Kwiatkowski, Cordelia Langford, John Sillitoe on behalf of the Wellcome Sanger Institute COVID-19 Surveillance Team |
| EPI_ISL_3203079 | Lighthouse Lab in Milton Keynes | Wellcome Sanger Institute for the COVID-19 Genomics UK (COG-UK) Consortium | The Lighthouse Lab in Milton Keynes and Alex Alderton, Roberto Amato, Jeffrey Barrett, Sonia Goncalves, Ewan Harrison, David K. Jackson, Ian Johnston, Dominic Kwiatkowski, Cordelia Langford, John Sillitoe on behalf of the Wellcome Sanger Institute COVID-19 Surveillance Team |
| EPI_ISL_3203080 | Lighthouse Laboratory Plymouth | Wellcome Sanger Institute for the COVID-19 Genomics UK (COG-UK) Consortium | Lighthouse Laboratory Plymouth and Alex Alderton, Roberto Amato, Jeffrey Barrett, Sonia Goncalves, Ewan Harrison, David K. Jackson, Ian Johnston, Dominic Kwiatkowski, Cordelia Langford, John Sillitoe on behalf of the Wellcome Sanger Institute COVID-19 Surveillance Team |
| EPI_ISL_3203081 | Lighthouse Lab in Glasgow | Wellcome Sanger Institute for the COVID-19 Genomics UK (COG-UK) Consortium | Harper VanSteenhouse, Yumi Kasai, David Gray, Carol Clugston, Anna Dominiczak and Alex Alderton, Roberto Amato, Jeffrey Barrett, Sonia Goncalves, Ewan Harrison, David K. Jackson, Ian Johnston, Dominic Kwiatkowski, Cordelia Langford, John Sillitoe on behalf of the Wellcome Sanger Institute COVID-19 Surveillance Team |
| EPI_ISL_3203082 | Lighthouse Laboratory Plymouth | Wellcome Sanger Institute for the COVID-19 Genomics UK (COG-UK) Consortium | Lighthouse Laboratory Plymouth and Alex Alderton, Roberto Amato, Jeffrey Barrett, Sonia Goncalves, Ewan Harrison, David K. Jackson, Ian Johnston, Dominic Kwiatkowski, Cordelia Langford, John Sillitoe on behalf of the Wellcome Sanger Institute COVID-19 Surveillance Team |
| EPI_ISL_3203083, EPI_ISL_3203084 | Lighthouse Lab in Glasgow | Wellcome Sanger Institute for the COVID-19 Genomics UK (COG-UK) Consortium | Harper VanSteenhouse, Yumi Kasai, David Gray, Carol Clugston, Anna Dominiczak and Alex Alderton, Roberto Amato, Jeffrey Barrett, Sonia Goncalves, Ewan Harrison, David K. Jackson, Ian Johnston, Dominic Kwiatkowski, Cordelia Langford, John Sillitoe on behalf of the Wellcome Sanger Institute COVID-19 Surveillance Team |
| EPI_ISL_3203085, EPI_ISL_3203086 | Lighthouse Lab in Milton Keynes | Wellcome Sanger Institute for the COVID-19 Genomics UK (COG-UK) Consortium | The Lighthouse Lab in Milton Keynes and Alex Alderton, Roberto Amato, Jeffrey Barrett, Sonia Goncalves, Ewan Harrison, David K. Jackson, Ian Johnston, Dominic Kwiatkowski, Cordelia Langford, John Sillitoe on behalf of the Wellcome Sanger Institute COVID-19 Surveillance Team |
| EPI_ISL_3203087 | Lighthouse Lab in Glasgow | Wellcome Sanger Institute for the COVID-19 Genomics UK (COG-UK) Consortium | Harper VanSteenhouse, Yumi Kasai, David Gray, Carol Clugston, Anna Dominiczak and Alex Alderton, Roberto Amato, Jeffrey Barrett, Sonia Goncalves, Ewan Harrison, David K. Jackson, Ian Johnston, Dominic Kwiatkowski, Cordelia Langford, John Sillitoe on behalf of the Wellcome Sanger Institute COVID-19 Surveillance Team |
| EPI_ISL_3203088 | Lighthouse Lab in Milton Keynes | Wellcome Sanger Institute for the COVID-19 Genomics UK (COG-UK) Consortium | The Lighthouse Lab in Milton Keynes and Alex Alderton, Roberto Amato, Jeffrey Barrett, Sonia Goncalves, Ewan Harrison, David K. Jackson, Ian Johnston, Dominic Kwiatkowski, Cordelia Langford, John Sillitoe on behalf of the Wellcome Sanger Institute COVID-19 Surveillance Team |
| EPI_ISL_3203089 | Lighthouse Lab in Glasgow | Wellcome Sanger Institute for the COVID-19 Genomics UK (COG-UK) Consortium | Harper VanSteenhouse, Yumi Kasai, David Gray, Carol Clugston, Anna Dominiczak and Alex Alderton, Roberto Amato, Jeffrey Barrett, Sonia Goncalves, Ewan Harrison, David K. Jackson, Ian Johnston, Dominic Kwiatkowski, Cordelia Langford, John Sillitoe on behalf of the Wellcome Sanger Institute COVID-19 Surveillance Team |
| EPI_ISL_3203090 | Lighthouse Laboratory Plymouth | Wellcome Sanger Institute for the COVID-19 Genomics UK (COG-UK) Consortium | Lighthouse Laboratory Plymouth and Alex Alderton, Roberto Amato, Jeffrey Barrett, Sonia Goncalves, Ewan Harrison, David K. Jackson, Ian Johnston, Dominic Kwiatkowski, Cordelia Langford, John Sillitoe on behalf of the Wellcome Sanger Institute COVID-19 Surveillance Team |
| EPI_ISL_3203091, EPI_ISL_3203092 | Lighthouse Lab in Milton Keynes | Wellcome Sanger Institute for the COVID-19 Genomics UK (COG-UK) Consortium | The Lighthouse Lab in Milton Keynes and Alex Alderton, Roberto Amato, Jeffrey Barrett, Sonia Goncalves, Ewan Harrison, David K. Jackson, Ian Johnston, Dominic Kwiatkowski, Cordelia Langford, John Sillitoe on behalf of the Wellcome Sanger Institute COVID-19 Surveillance Team |
| EPI_ISL_3203093, EPI_ISL_3203094, EPI_ISL_3203095, EPI_ISL_3203096 | Lighthouse Laboratory Plymouth | Wellcome Sanger Institute for the COVID-19 Genomics UK (COG-UK) Consortium | Lighthouse Laboratory Plymouth and Alex Alderton, Roberto Amato, Jeffrey Barrett, Sonia Goncalves, Ewan Harrison, David K. Jackson, Ian Johnston, Dominic Kwiatkowski, Cordelia Langford, John Sillitoe on behalf of the Wellcome Sanger Institute COVID-19 Surveillance Team |
| EPI_ISL_3203097 | Lighthouse Lab in Glasgow | Wellcome Sanger Institute for the COVID-19 Genomics UK (COG-UK) Consortium | Harper VanSteenhouse, Yumi Kasai, David Gray, Carol Clugston, Anna Dominiczak and Alex Alderton, Roberto Amato, Jeffrey Barrett, Sonia Goncalves, Ewan Harrison, David K. Jackson, Ian Johnston, Dominic Kwiatkowski, Cordelia Langford, John Sillitoe on behalf of the Wellcome Sanger Institute COVID-19 Surveillance Team |
| EPI_ISL_3203098, EPI_ISL_3203099 | Lighthouse Lab in Milton Keynes | Wellcome Sanger Institute for the COVID-19 Genomics UK (COG-UK) Consortium | The Lighthouse Lab in Milton Keynes and Alex Alderton, Roberto Amato, Jeffrey Barrett, Sonia Goncalves, Ewan Harrison, David K. Jackson, Ian Johnston, Dominic Kwiatkowski, Cordelia Langford, John Sillitoe on behalf of the Wellcome Sanger Institute COVID-19 Surveillance Team |
| EPI_ISL_3203100, EPI_ISL_3203101, EPI_ISL_3203102, EPI_ISL_3203103, EPI_ISL_3203104, EPI_ISL_3203105 | Lighthouse Laboratory Plymouth | Wellcome Sanger Institute for the COVID-19 Genomics UK (COG-UK) Consortium | Lighthouse Laboratory Plymouth and Alex Alderton, Roberto Amato, Jeffrey Barrett, Sonia Goncalves, Ewan Harrison, David K. Jackson, Ian Johnston, Dominic Kwiatkowski, Cordelia Langford, John Sillitoe on behalf of the Wellcome Sanger Institute COVID-19 Surveillance Team |
| EPI_ISL_3203106 | Lighthouse Lab in Milton Keynes | Wellcome Sanger Institute for the COVID-19 Genomics UK (COG-UK) Consortium | The Lighthouse Lab in Milton Keynes and Alex Alderton, Roberto Amato, Jeffrey Barrett, Sonia Goncalves, Ewan Harrison, David K. Jackson, Ian Johnston, Dominic Kwiatkowski, Cordelia Langford, John Sillitoe on behalf of the Wellcome Sanger Institute COVID-19 Surveillance Team |

| | | | |
|---|---|---|---|
| EPI_ISL_3203107 | Lighthouse Lab in Glasgow | Wellcome Sanger Institute for the COVID-19 Genomics UK (COG-UK) Consortium | Harper VanSteenhouse, Yumi Kasai, David Gray, Carol Clugston, Anna Dominiczak and Alex Alderton, Roberto Amato, Jeffrey Barrett, Sonia Goncalves, Ewan Harrison, David K. Jackson, Ian Johnston, Dominic Kwiatkowski, Cordelia Langford, John Sillitoe on behalf of the Wellcome Sanger Institute COVID-19 Surveillance Team |
| EPI_ISL_3203108 | Lighthouse Laboratory Plymouth | Wellcome Sanger Institute for the COVID-19 Genomics UK (COG-UK) Consortium | Lighthouse Laboratory Plymouth and Alex Alderton, Roberto Amato, Jeffrey Barrett, Sonia Goncalves, Ewan Harrison, David K. Jackson, Ian Johnston, Dominic Kwiatkowski, Cordelia Langford, John Sillitoe on behalf of the Wellcome Sanger Institute COVID-19 Surveillance Team |
| EPI_ISL_3203109 | Lighthouse Lab in Glasgow | Wellcome Sanger Institute for the COVID-19 Genomics UK (COG-UK) Consortium | Harper VanSteenhouse, Yumi Kasai, David Gray, Carol Clugston, Anna Dominiczak and Alex Alderton, Roberto Amato, Jeffrey Barrett, Sonia Goncalves, Ewan Harrison, David K. Jackson, Ian Johnston, Dominic Kwiatkowski, Cordelia Langford, John Sillitoe on behalf of the Wellcome Sanger Institute COVID-19 Surveillance Team |
| EPI_ISL_3203110 | Lighthouse Laboratory Plymouth | Wellcome Sanger Institute for the COVID-19 Genomics UK (COG-UK) Consortium | Lighthouse Laboratory Plymouth and Alex Alderton, Roberto Amato, Jeffrey Barrett, Sonia Goncalves, Ewan Harrison, David K. Jackson, Ian Johnston, Dominic Kwiatkowski, Cordelia Langford, John Sillitoe on behalf of the Wellcome Sanger Institute COVID-19 Surveillance Team |
| EPI_ISL_3203111 | Lighthouse Lab in Milton Keynes | Wellcome Sanger Institute for the COVID-19 Genomics UK (COG-UK) Consortium | The Lighthouse Lab in Milton Keynes and Alex Alderton, Roberto Amato, Jeffrey Barrett, Sonia Goncalves, Ewan Harrison, David K. Jackson, Ian Johnston, Dominic Kwiatkowski, Cordelia Langford, John Sillitoe on behalf of the Wellcome Sanger Institute COVID-19 Surveillance Team |
| EPI_ISL_3203112 | Lighthouse Lab in Glasgow | Wellcome Sanger Institute for the COVID-19 Genomics UK (COG-UK) Consortium | Harper VanSteenhouse, Yumi Kasai, David Gray, Carol Clugston, Anna Dominiczak and Alex Alderton, Roberto Amato, Jeffrey Barrett, Sonia Goncalves, Ewan Harrison, David K. Jackson, Ian Johnston, Dominic Kwiatkowski, Cordelia Langford, John Sillitoe on behalf of the Wellcome Sanger Institute COVID-19 Surveillance Team |
| EPI_ISL_3203113 | Lighthouse Lab in Milton Keynes | Wellcome Sanger Institute for the COVID-19 Genomics UK (COG-UK) Consortium | The Lighthouse Lab in Milton Keynes and Alex Alderton, Roberto Amato, Jeffrey Barrett, Sonia Goncalves, Ewan Harrison, David K. Jackson, Ian Johnston, Dominic Kwiatkowski, Cordelia Langford, John Sillitoe on behalf of the Wellcome Sanger Institute COVID-19 Surveillance Team |
| EPI_ISL_3203114, EPI_ISL_3203115 | Lighthouse Laboratory Plymouth | Wellcome Sanger Institute for the COVID-19 Genomics UK (COG-UK) Consortium | Lighthouse Laboratory Plymouth and Alex Alderton, Roberto Amato, Jeffrey Barrett, Sonia Goncalves, Ewan Harrison, David K. Jackson, Ian Johnston, Dominic Kwiatkowski, Cordelia Langford, John Sillitoe on behalf of the Wellcome Sanger Institute COVID-19 Surveillance Team |
| EPI_ISL_3203116 | Lighthouse Lab in Glasgow | Wellcome Sanger Institute for the COVID-19 Genomics UK (COG-UK) Consortium | Harper VanSteenhouse, Yumi Kasai, David Gray, Carol Clugston, Anna Dominiczak and Alex Alderton, Roberto Amato, Jeffrey Barrett, Sonia Goncalves, Ewan Harrison, David K. Jackson, Ian Johnston, Dominic Kwiatkowski, Cordelia Langford, John Sillitoe on behalf of the Wellcome Sanger Institute COVID-19 Surveillance Team |
| EPI_ISL_3203117 | Lighthouse Lab in Milton Keynes | Wellcome Sanger Institute for the COVID-19 Genomics UK (COG-UK) Consortium | The Lighthouse Lab in Milton Keynes and Alex Alderton, Roberto Amato, Jeffrey Barrett, Sonia Goncalves, Ewan Harrison, David K. Jackson, Ian Johnston, Dominic Kwiatkowski, Cordelia Langford, John Sillitoe on behalf of the Wellcome Sanger Institute COVID-19 Surveillance Team |
| EPI_ISL_3203118 | Lighthouse Lab in Glasgow | Wellcome Sanger Institute for the COVID-19 Genomics UK (COG-UK) Consortium | Harper VanSteenhouse, Yumi Kasai, David Gray, Carol Clugston, Anna Dominiczak and Alex Alderton, Roberto Amato, Jeffrey Barrett, Sonia Goncalves, Ewan Harrison, David K. Jackson, Ian Johnston, Dominic Kwiatkowski, Cordelia Langford, John Sillitoe on behalf of the Wellcome Sanger Institute COVID-19 Surveillance Team |
| EPI_ISL_3203119, EPI_ISL_3203120 | Lighthouse Laboratory Plymouth | Wellcome Sanger Institute for the COVID-19 Genomics UK (COG-UK) Consortium | Lighthouse Laboratory Plymouth and Alex Alderton, Roberto Amato, Jeffrey Barrett, Sonia Goncalves, Ewan Harrison, David K. Jackson, Ian Johnston, Dominic Kwiatkowski, Cordelia Langford, John Sillitoe on behalf of the Wellcome Sanger Institute COVID-19 Surveillance Team |
| EPI_ISL_3203121 | Lighthouse Lab in Milton Keynes | Wellcome Sanger Institute for the COVID-19 Genomics UK (COG-UK) Consortium | The Lighthouse Lab in Milton Keynes and Alex Alderton, Roberto Amato, Jeffrey Barrett, Sonia Goncalves, Ewan Harrison, David K. Jackson, Ian Johnston, Dominic Kwiatkowski, Cordelia Langford, John Sillitoe on behalf of the Wellcome Sanger Institute COVID-19 Surveillance Team |
| EPI_ISL_3203122 | Lighthouse Laboratory Plymouth | Wellcome Sanger Institute for the COVID-19 Genomics UK (COG-UK) Consortium | Lighthouse Laboratory Plymouth and Alex Alderton, Roberto Amato, Jeffrey Barrett, Sonia Goncalves, Ewan Harrison, David K. Jackson, Ian Johnston, Dominic Kwiatkowski, Cordelia Langford, John Sillitoe on behalf of the Wellcome Sanger Institute COVID-19 Surveillance Team |
| EPI_ISL_3203123 | Lighthouse Lab in Glasgow | Wellcome Sanger Institute for the COVID-19 Genomics UK (COG-UK) Consortium | Harper VanSteenhouse, Yumi Kasai, David Gray, Carol Clugston, Anna Dominiczak and Alex Alderton, Roberto Amato, Jeffrey Barrett, Sonia Goncalves, Ewan Harrison, David K. Jackson, Ian Johnston, Dominic Kwiatkowski, Cordelia Langford, John Sillitoe on behalf of the Wellcome Sanger Institute COVID-19 Surveillance Team |
| EPI_ISL_3203124, EPI_ISL_3203125 | Lighthouse Laboratory Plymouth | Wellcome Sanger Institute for the COVID-19 Genomics UK (COG-UK) Consortium | Lighthouse Laboratory Plymouth and Alex Alderton, Roberto Amato, Jeffrey Barrett, Sonia Goncalves, Ewan Harrison, David K. Jackson, Ian Johnston, Dominic Kwiatkowski, Cordelia Langford, John Sillitoe on behalf of the Wellcome Sanger Institute COVID-19 Surveillance Team |
| EPI_ISL_3203126 | Lighthouse Lab in Milton Keynes | Wellcome Sanger Institute for the COVID-19 Genomics UK (COG-UK) Consortium | The Lighthouse Lab in Milton Keynes and Alex Alderton, Roberto Amato, Jeffrey Barrett, Sonia Goncalves, Ewan Harrison, David K. Jackson, Ian Johnston, Dominic Kwiatkowski, Cordelia Langford, John Sillitoe on behalf of the Wellcome Sanger Institute COVID-19 Surveillance Team |
| EPI_ISL_3203127 | Lighthouse Laboratory Plymouth | Wellcome Sanger Institute for the COVID-19 Genomics UK (COG-UK) Consortium | Lighthouse Laboratory Plymouth and Alex Alderton, Roberto Amato, Jeffrey Barrett, Sonia Goncalves, Ewan Harrison, David K. Jackson, Ian Johnston, Dominic Kwiatkowski, Cordelia Langford, John Sillitoe on behalf of the Wellcome Sanger Institute COVID-19 Surveillance Team |
| EPI_ISL_3203128 | Lighthouse Lab in Milton Keynes | Wellcome Sanger Institute for the COVID-19 Genomics UK (COG-UK) Consortium | The Lighthouse Lab in Milton Keynes and Alex Alderton, Roberto Amato, Jeffrey Barrett, Sonia Goncalves, Ewan Harrison, David K. Jackson, Ian Johnston, Dominic Kwiatkowski, Cordelia Langford, John Sillitoe on behalf of the Wellcome Sanger Institute COVID-19 Surveillance Team |
| EPI_ISL_3203129 | Lighthouse Laboratory Plymouth | Wellcome Sanger Institute for the COVID-19 Genomics UK (COG-UK) Consortium | Lighthouse Laboratory Plymouth and Alex Alderton, Roberto Amato, Jeffrey Barrett, Sonia Goncalves, Ewan Harrison, David K. Jackson, Ian Johnston, Dominic Kwiatkowski, Cordelia Langford, John Sillitoe on behalf of the Wellcome Sanger Institute COVID-19 Surveillance Team |
| EPI_ISL_3203130, EPI_ISL_3203131, EPI_ISL_3203132, EPI_ISL_3203133 | Lighthouse Lab in Milton Keynes | Wellcome Sanger Institute for the COVID-19 Genomics UK (COG-UK) Consortium | The Lighthouse Lab in Milton Keynes and Alex Alderton, Roberto Amato, Jeffrey Barrett, Sonia Goncalves, Ewan Harrison, David K. Jackson, Ian Johnston, Dominic Kwiatkowski, Cordelia Langford, John Sillitoe on behalf of the Wellcome Sanger Institute COVID-19 Surveillance Team |
| EPI_ISL_3203134, EPI_ISL_3203135, EPI_ISL_3203136 | Lighthouse Laboratory Plymouth | Wellcome Sanger Institute for the COVID-19 Genomics UK (COG-UK) Consortium | Lighthouse Laboratory Plymouth and Alex Alderton, Roberto Amato, Jeffrey Barrett, Sonia Goncalves, Ewan Harrison, David K. Jackson, Ian Johnston, Dominic Kwiatkowski, Cordelia Langford, John Sillitoe on behalf of the Wellcome Sanger Institute COVID-19 Surveillance Team |
| EPI_ISL_3203137 | Lighthouse Lab in Milton Keynes | Wellcome Sanger Institute for the COVID-19 Genomics UK (COG-UK) Consortium | The Lighthouse Lab in Milton Keynes and Alex Alderton, Roberto Amato, Jeffrey Barrett, Sonia Goncalves, Ewan Harrison, David K. Jackson, Ian Johnston, Dominic Kwiatkowski, Cordelia Langford, John Sillitoe on behalf of the Wellcome Sanger Institute COVID-19 Surveillance Team |
| EPI_ISL_3203138, EPI_ISL_3203139, EPI_ISL_3203140 | Lighthouse Lab in Glasgow | Wellcome Sanger Institute for the COVID-19 Genomics UK (COG-UK) Consortium | Harper VanSteenhouse, Yumi Kasai, David Gray, Carol Clugston, Anna Dominiczak and Alex Alderton, Roberto Amato, Jeffrey Barrett, Sonia Goncalves, Ewan Harrison, David K. Jackson, Ian Johnston, Dominic Kwiatkowski, Cordelia Langford, John Sillitoe on behalf of the Wellcome Sanger Institute COVID-19 Surveillance Team |
| EPI_ISL_3203141 | Lighthouse Laboratory Plymouth | Wellcome Sanger Institute for the COVID-19 Genomics UK (COG-UK) Consortium | Lighthouse Laboratory Plymouth and Alex Alderton, Roberto Amato, Jeffrey Barrett, Sonia Goncalves, Ewan Harrison, David K. Jackson, Ian Johnston, Dominic Kwiatkowski, Cordelia Langford, John Sillitoe on behalf of the Wellcome Sanger Institute COVID-19 Surveillance Team |
| EPI_ISL_3203142 | Lighthouse Lab in Milton Keynes | Wellcome Sanger Institute for the COVID-19 Genomics UK (COG-UK) Consortium | The Lighthouse Lab in Milton Keynes and Alex Alderton, Roberto Amato, Jeffrey Barrett, Sonia Goncalves, Ewan Harrison, David K. Jackson, Ian Johnston, Dominic Kwiatkowski, Cordelia Langford, John Sillitoe on behalf of the Wellcome Sanger Institute COVID-19 Surveillance Team |
| EPI_ISL_3203143, EPI_ISL_3203144 | Lighthouse Laboratory Plymouth | Wellcome Sanger Institute for the COVID-19 Genomics UK (COG-UK) Consortium | Lighthouse Laboratory Plymouth and Alex Alderton, Roberto Amato, Jeffrey Barrett, Sonia Goncalves, Ewan Harrison, David K. Jackson, Ian Johnston, Dominic Kwiatkowski, Cordelia Langford, John Sillitoe on behalf of the Wellcome Sanger Institute COVID-19 Surveillance Team |
| EPI_ISL_3203145, EPI_ISL_3203146 | Lighthouse Lab in Milton Keynes | Wellcome Sanger Institute for the COVID-19 Genomics UK (COG-UK) Consortium | The Lighthouse Lab in Milton Keynes and Alex Alderton, Roberto Amato, Jeffrey Barrett, Sonia Goncalves, Ewan Harrison, David K. Jackson, Ian Johnston, Dominic Kwiatkowski, Cordelia Langford, John Sillitoe on behalf of the Wellcome Sanger Institute COVID-19 Surveillance Team |
| EPI_ISL_3203147, EPI_ISL_3203148 | Lighthouse Laboratory Plymouth | Wellcome Sanger Institute for the COVID-19 Genomics UK (COG-UK) Consortium | Lighthouse Laboratory Plymouth and Alex Alderton, Roberto Amato, Jeffrey Barrett, Sonia Goncalves, Ewan Harrison, David K. Jackson, Ian Johnston, Dominic Kwiatkowski, Cordelia Langford, John Sillitoe on behalf of the Wellcome Sanger Institute COVID-19 Surveillance Team |
| EPI_ISL_3203149 | Lighthouse Lab in Glasgow | Wellcome Sanger Institute for the COVID-19 Genomics UK (COG-UK) Consortium | Harper VanSteenhouse, Yumi Kasai, David Gray, Carol Clugston, Anna Dominiczak and Alex Alderton, Roberto Amato, Jeffrey Barrett, Sonia Goncalves, Ewan Harrison, David K. Jackson, Ian Johnston, Dominic Kwiatkowski, Cordelia Langford, John Sillitoe on behalf of the Wellcome Sanger Institute COVID-19 Surveillance Team |
| EPI_ISL_3203150, EPI_ISL_3203151, EPI_ISL_3203152, EPI_ISL_3203153, EPI_ISL_3203154, EPI_ISL_3203155 | Lighthouse Laboratory Plymouth | Wellcome Sanger Institute for the COVID-19 Genomics UK (COG-UK) Consortium | Lighthouse Laboratory Plymouth and Alex Alderton, Roberto Amato, Jeffrey Barrett, Sonia Goncalves, Ewan Harrison, David K. Jackson, Ian Johnston, Dominic Kwiatkowski, Cordelia Langford, John Sillitoe on behalf of the Wellcome Sanger Institute COVID-19 Surveillance Team |
| EPI_ISL_3203156 | Lighthouse Lab in Milton Keynes | Wellcome Sanger Institute for the COVID-19 Genomics UK (COG-UK) Consortium | The Lighthouse Lab in Milton Keynes and Alex Alderton, Roberto Amato, Jeffrey Barrett, Sonia Goncalves, Ewan Harrison, David K. Jackson, Ian Johnston, Dominic Kwiatkowski, Cordelia Langford, John Sillitoe on behalf of the Wellcome Sanger Institute COVID-19 Surveillance Team |
| EPI_ISL_3203157 | Lighthouse Lab in Glasgow | Wellcome Sanger Institute for the COVID-19 Genomics UK (COG-UK) Consortium | Harper VanSteenhouse, Yumi Kasai, David Gray, Carol Clugston, Anna Dominiczak and Alex Alderton, Roberto Amato, Jeffrey Barrett, Sonia Goncalves, Ewan Harrison, David K. Jackson, Ian Johnston, Dominic Kwiatkowski, Cordelia Langford, John Sillitoe on behalf of the Wellcome Sanger Institute COVID-19 Surveillance Team |
| EPI_ISL_3203158 | Lighthouse Laboratory Plymouth | Wellcome Sanger Institute for the COVID-19 Genomics UK (COG-UK) Consortium | Lighthouse Laboratory Plymouth and Alex Alderton, Roberto Amato, Jeffrey Barrett, Sonia Goncalves, Ewan Harrison, David K. Jackson, Ian Johnston, Dominic Kwiatkowski, Cordelia Langford, John Sillitoe on behalf of the Wellcome Sanger Institute COVID-19 Surveillance Team |
| EPI_ISL_3203159, EPI_ISL_3203160 | Lighthouse Lab in Glasgow | Wellcome Sanger Institute for the COVID-19 Genomics UK (COG-UK) Consortium | Harper VanSteenhouse, Yumi Kasai, David Gray, Carol Clugston, Anna Dominiczak and Alex Alderton, Roberto Amato, Jeffrey Barrett, Sonia Goncalves, Ewan Harrison, David K. Jackson, Ian Johnston, Dominic Kwiatkowski, Cordelia Langford, John Sillitoe on behalf of the Wellcome Sanger Institute COVID-19 Surveillance Team |
| EPI_ISL_3203161, EPI_ISL_3203162, EPI_ISL_3203163, EPI_ISL_3203164, EPI_ISL_3203165 | Lighthouse Laboratory Plymouth | Wellcome Sanger Institute for the COVID-19 Genomics UK (COG-UK) Consortium | Lighthouse Laboratory Plymouth and Alex Alderton, Roberto Amato, Jeffrey Barrett, Sonia Goncalves, Ewan Harrison, David K. Jackson, Ian Johnston, Dominic Kwiatkowski, Cordelia Langford, John Sillitoe on behalf of the Wellcome Sanger Institute COVID-19 Surveillance Team |
| EPI_ISL_3203166 | Lighthouse Lab in Milton Keynes | Wellcome Sanger Institute for the COVID-19 Genomics UK (COG-UK) Consortium | The Lighthouse Lab in Milton Keynes and Alex Alderton, Roberto Amato, Jeffrey Barrett, Sonia Goncalves, Ewan Harrison, David K. Jackson, Ian Johnston, Dominic Kwiatkowski, Cordelia Langford, John Sillitoe on behalf of the Wellcome Sanger Institute COVID-19 Surveillance Team |
| EPI_ISL_3203167 | Lighthouse Laboratory Plymouth | Wellcome Sanger Institute for the COVID-19 Genomics UK (COG-UK) Consortium | Lighthouse Laboratory Plymouth and Alex Alderton, Roberto Amato, Jeffrey Barrett, Sonia Goncalves, Ewan Harrison, David K. Jackson, Ian Johnston, Dominic Kwiatkowski, Cordelia Langford, John Sillitoe on behalf of the Wellcome Sanger Institute COVID-19 Surveillance Team |
| EPI_ISL_3203168 | Lighthouse Lab in Glasgow | Wellcome Sanger Institute for the COVID-19 Genomics UK (COG-UK) Consortium | Harper VanSteenhouse, Yumi Kasai, David Gray, Carol Clugston, Anna Dominiczak and Alex Alderton, Roberto Amato, Jeffrey Barrett, Sonia Goncalves, Ewan Harrison, David K. Jackson, Ian Johnston, Dominic Kwiatkowski, Cordelia Langford, John Sillitoe on behalf of the Wellcome Sanger Institute COVID-19 Surveillance Team |
| EPI_ISL_3203169 | Lighthouse Lab in Milton Keynes | Wellcome Sanger Institute for the COVID-19 Genomics UK (COG-UK) Consortium | The Lighthouse Lab in Milton Keynes and Alex Alderton, Roberto Amato, Jeffrey Barrett, Sonia Goncalves, Ewan Harrison, David K. Jackson, Ian Johnston, Dominic Kwiatkowski, Cordelia Langford, John Sillitoe on behalf of the Wellcome Sanger Institute COVID-19 Surveillance Team |
| EPI_ISL_3203170, | Lighthouse Lab in Glasgow | Wellcome Sanger Institute for the COVID-19 Genomics UK (COG-UK) | Harper VanSteenhouse, Yumi Kasai, David Gray, Carol Clugston, Anna Dominiczak and Alex Alderton, Roberto Amato, Jeffrey Barrett, Sonia Goncalves, Ewan Harrison, David K. Jackson, Ian Johnston, Dominic Kwiatkowski, Cordelia Langford, |

| | | | |
|---|---|---|---|
| EPI_ISL_3203171 | | Consortium | John Sillitoe on behalf of the Wellcome Sanger Institute COVID-19 Surveillance Team |
| EPI_ISL_3203172 | Lighthouse Laboratory Plymouth | Wellcome Sanger Institute for the COVID-19 Genomics UK (COG-UK) Consortium | Lighthouse Laboratory Plymouth and Alex Alderton, Roberto Amato, Jeffrey Barrett, Sonia Goncalves, Ewan Harrison, David K. Jackson, Ian Johnston, Dominic Kwiatkowski, Cordelia Langford, John Sillitoe on behalf of the Wellcome Sanger Institute COVID-19 Surveillance Team |
| EPI_ISL_3203173 | Lighthouse Lab in Milton Keynes | Wellcome Sanger Institute for the COVID-19 Genomics UK (COG-UK) Consortium | The Lighthouse Lab in Milton Keynes and Alex Alderton, Roberto Amato, Jeffrey Barrett, Sonia Goncalves, Ewan Harrison, David K. Jackson, Ian Johnston, Dominic Kwiatkowski, Cordelia Langford, John Sillitoe on behalf of the Wellcome Sanger Institute COVID-19 Surveillance Team |
| EPI_ISL_3203174, EPI_ISL_3203175 | Lighthouse Laboratory Plymouth | Wellcome Sanger Institute for the COVID-19 Genomics UK (COG-UK) Consortium | Lighthouse Laboratory Plymouth and Alex Alderton, Roberto Amato, Jeffrey Barrett, Sonia Goncalves, Ewan Harrison, David K. Jackson, Ian Johnston, Dominic Kwiatkowski, Cordelia Langford, John Sillitoe on behalf of the Wellcome Sanger Institute COVID-19 Surveillance Team |
| EPI_ISL_3203176, EPI_ISL_3203177 | Lighthouse Lab in Glasgow | Wellcome Sanger Institute for the COVID-19 Genomics UK (COG-UK) Consortium | Harper VanSteenhouse, Yumi Kasai, David Gray, Carol Clugston, Anna Dominiczak and Alex Alderton, Roberto Amato, Jeffrey Barrett, Sonia Goncalves, Ewan Harrison, David K. Jackson, Ian Johnston, Dominic Kwiatkowski, Cordelia Langford, John Sillitoe on behalf of the Wellcome Sanger Institute COVID-19 Surveillance Team |
| EPI_ISL_3203178 | Lighthouse Lab in Milton Keynes | Wellcome Sanger Institute for the COVID-19 Genomics UK (COG-UK) Consortium | The Lighthouse Lab in Milton Keynes and Alex Alderton, Roberto Amato, Jeffrey Barrett, Sonia Goncalves, Ewan Harrison, David K. Jackson, Ian Johnston, Dominic Kwiatkowski, Cordelia Langford, John Sillitoe on behalf of the Wellcome Sanger Institute COVID-19 Surveillance Team |
| EPI_ISL_3203179 | Lighthouse Laboratory Plymouth | Wellcome Sanger Institute for the COVID-19 Genomics UK (COG-UK) Consortium | Lighthouse Laboratory Plymouth and Alex Alderton, Roberto Amato, Jeffrey Barrett, Sonia Goncalves, Ewan Harrison, David K. Jackson, Ian Johnston, Dominic Kwiatkowski, Cordelia Langford, John Sillitoe on behalf of the Wellcome Sanger Institute COVID-19 Surveillance Team |
| EPI_ISL_3203180 | Lighthouse Lab in Glasgow | Wellcome Sanger Institute for the COVID-19 Genomics UK (COG-UK) Consortium | Harper VanSteenhouse, Yumi Kasai, David Gray, Carol Clugston, Anna Dominiczak and Alex Alderton, Roberto Amato, Jeffrey Barrett, Sonia Goncalves, Ewan Harrison, David K. Jackson, Ian Johnston, Dominic Kwiatkowski, Cordelia Langford, John Sillitoe on behalf of the Wellcome Sanger Institute COVID-19 Surveillance Team |
| EPI_ISL_3203181 | Lighthouse Laboratory Plymouth | Wellcome Sanger Institute for the COVID-19 Genomics UK (COG-UK) Consortium | Lighthouse Laboratory Plymouth and Alex Alderton, Roberto Amato, Jeffrey Barrett, Sonia Goncalves, Ewan Harrison, David K. Jackson, Ian Johnston, Dominic Kwiatkowski, Cordelia Langford, John Sillitoe on behalf of the Wellcome Sanger Institute COVID-19 Surveillance Team |
| EPI_ISL_3203182 | Lighthouse Lab in Glasgow | Wellcome Sanger Institute for the COVID-19 Genomics UK (COG-UK) Consortium | Harper VanSteenhouse, Yumi Kasai, David Gray, Carol Clugston, Anna Dominiczak and Alex Alderton, Roberto Amato, Jeffrey Barrett, Sonia Goncalves, Ewan Harrison, David K. Jackson, Ian Johnston, Dominic Kwiatkowski, Cordelia Langford, John Sillitoe on behalf of the Wellcome Sanger Institute COVID-19 Surveillance Team |
| EPI_ISL_3203183 | Lighthouse Laboratory Plymouth | Wellcome Sanger Institute for the COVID-19 Genomics UK (COG-UK) Consortium | Lighthouse Laboratory Plymouth and Alex Alderton, Roberto Amato, Jeffrey Barrett, Sonia Goncalves, Ewan Harrison, David K. Jackson, Ian Johnston, Dominic Kwiatkowski, Cordelia Langford, John Sillitoe on behalf of the Wellcome Sanger Institute COVID-19 Surveillance Team |
| EPI_ISL_3203184 | Lighthouse Lab in Milton Keynes | Wellcome Sanger Institute for the COVID-19 Genomics UK (COG-UK) Consortium | The Lighthouse Lab in Milton Keynes and Alex Alderton, Roberto Amato, Jeffrey Barrett, Sonia Goncalves, Ewan Harrison, David K. Jackson, Ian Johnston, Dominic Kwiatkowski, Cordelia Langford, John Sillitoe on behalf of the Wellcome Sanger Institute COVID-19 Surveillance Team |
| EPI_ISL_3203185, EPI_ISL_3203186 | Lighthouse Laboratory Plymouth | Wellcome Sanger Institute for the COVID-19 Genomics UK (COG-UK) Consortium | Lighthouse Laboratory Plymouth and Alex Alderton, Roberto Amato, Jeffrey Barrett, Sonia Goncalves, Ewan Harrison, David K. Jackson, Ian Johnston, Dominic Kwiatkowski, Cordelia Langford, John Sillitoe on behalf of the Wellcome Sanger Institute COVID-19 Surveillance Team |
| EPI_ISL_3203187 | Lighthouse Lab in Glasgow | Wellcome Sanger Institute for the COVID-19 Genomics UK (COG-UK) Consortium | Harper VanSteenhouse, Yumi Kasai, David Gray, Carol Clugston, Anna Dominiczak and Alex Alderton, Roberto Amato, Jeffrey Barrett, Sonia Goncalves, Ewan Harrison, David K. Jackson, Ian Johnston, Dominic Kwiatkowski, Cordelia Langford, John Sillitoe on behalf of the Wellcome Sanger Institute COVID-19 Surveillance Team |
| EPI_ISL_3203188 | Lighthouse Lab in Milton Keynes | Wellcome Sanger Institute for the COVID-19 Genomics UK (COG-UK) Consortium | The Lighthouse Lab in Milton Keynes and Alex Alderton, Roberto Amato, Jeffrey Barrett, Sonia Goncalves, Ewan Harrison, David K. Jackson, Ian Johnston, Dominic Kwiatkowski, Cordelia Langford, John Sillitoe on behalf of the Wellcome Sanger Institute COVID-19 Surveillance Team |
| EPI_ISL_3203189, EPI_ISL_3203190, EPI_ISL_3203191 | Lighthouse Laboratory Plymouth | Wellcome Sanger Institute for the COVID-19 Genomics UK (COG-UK) Consortium | Lighthouse Laboratory Plymouth and Alex Alderton, Roberto Amato, Jeffrey Barrett, Sonia Goncalves, Ewan Harrison, David K. Jackson, Ian Johnston, Dominic Kwiatkowski, Cordelia Langford, John Sillitoe on behalf of the Wellcome Sanger Institute COVID-19 Surveillance Team |
| EPI_ISL_3203192 | Lighthouse Lab in Milton Keynes | Wellcome Sanger Institute for the COVID-19 Genomics UK (COG-UK) Consortium | The Lighthouse Lab in Milton Keynes and Alex Alderton, Roberto Amato, Jeffrey Barrett, Sonia Goncalves, Ewan Harrison, David K. Jackson, Ian Johnston, Dominic Kwiatkowski, Cordelia Langford, John Sillitoe on behalf of the Wellcome Sanger Institute COVID-19 Surveillance Team |
| EPI_ISL_3203193, EPI_ISL_3203194 | Lighthouse Lab in Glasgow | Wellcome Sanger Institute for the COVID-19 Genomics UK (COG-UK) Consortium | Harper VanSteenhouse, Yumi Kasai, David Gray, Carol Clugston, Anna Dominiczak and Alex Alderton, Roberto Amato, Jeffrey Barrett, Sonia Goncalves, Ewan Harrison, David K. Jackson, Ian Johnston, Dominic Kwiatkowski, Cordelia Langford, John Sillitoe on behalf of the Wellcome Sanger Institute COVID-19 Surveillance Team |
| EPI_ISL_3203195, EPI_ISL_3203196 | Lighthouse Laboratory Plymouth | Wellcome Sanger Institute for the COVID-19 Genomics UK (COG-UK) Consortium | Lighthouse Laboratory Plymouth and Alex Alderton, Roberto Amato, Jeffrey Barrett, Sonia Goncalves, Ewan Harrison, David K. Jackson, Ian Johnston, Dominic Kwiatkowski, Cordelia Langford, John Sillitoe on behalf of the Wellcome Sanger Institute COVID-19 Surveillance Team |
| EPI_ISL_3203197 | Lighthouse Lab in Glasgow | Wellcome Sanger Institute for the COVID-19 Genomics UK (COG-UK) Consortium | Harper VanSteenhouse, Yumi Kasai, David Gray, Carol Clugston, Anna Dominiczak and Alex Alderton, Roberto Amato, Jeffrey Barrett, Sonia Goncalves, Ewan Harrison, David K. Jackson, Ian Johnston, Dominic Kwiatkowski, Cordelia Langford, John Sillitoe on behalf of the Wellcome Sanger Institute COVID-19 Surveillance Team |
| EPI_ISL_3203198 | Lighthouse Laboratory Plymouth | Wellcome Sanger Institute for the COVID-19 Genomics UK (COG-UK) Consortium | Lighthouse Laboratory Plymouth and Alex Alderton, Roberto Amato, Jeffrey Barrett, Sonia Goncalves, Ewan Harrison, David K. Jackson, Ian Johnston, Dominic Kwiatkowski, Cordelia Langford, John Sillitoe on behalf of the Wellcome Sanger Institute COVID-19 Surveillance Team |
| EPI_ISL_3203199, EPI_ISL_3203200 | Lighthouse Lab in Glasgow | Wellcome Sanger Institute for the COVID-19 Genomics UK (COG-UK) Consortium | Harper VanSteenhouse, Yumi Kasai, David Gray, Carol Clugston, Anna Dominiczak and Alex Alderton, Roberto Amato, Jeffrey Barrett, Sonia Goncalves, Ewan Harrison, David K. Jackson, Ian Johnston, Dominic Kwiatkowski, Cordelia Langford, John Sillitoe on behalf of the Wellcome Sanger Institute COVID-19 Surveillance Team |
| EPI_ISL_3203201 | Lighthouse Lab in Milton Keynes | Wellcome Sanger Institute for the COVID-19 Genomics UK (COG-UK) Consortium | The Lighthouse Lab in Milton Keynes and Alex Alderton, Roberto Amato, Jeffrey Barrett, Sonia Goncalves, Ewan Harrison, David K. Jackson, Ian Johnston, Dominic Kwiatkowski, Cordelia Langford, John Sillitoe on behalf of the Wellcome Sanger Institute COVID-19 Surveillance Team |
| EPI_ISL_3203202 | Lighthouse Laboratory Plymouth | Wellcome Sanger Institute for the COVID-19 Genomics UK (COG-UK) Consortium | Lighthouse Laboratory Plymouth and Alex Alderton, Roberto Amato, Jeffrey Barrett, Sonia Goncalves, Ewan Harrison, David K. Jackson, Ian Johnston, Dominic Kwiatkowski, Cordelia Langford, John Sillitoe on behalf of the Wellcome Sanger Institute COVID-19 Surveillance Team |
| EPI_ISL_3203203 | Lighthouse Lab in Milton Keynes | Wellcome Sanger Institute for the COVID-19 Genomics UK (COG-UK) Consortium | The Lighthouse Lab in Milton Keynes and Alex Alderton, Roberto Amato, Jeffrey Barrett, Sonia Goncalves, Ewan Harrison, David K. Jackson, Ian Johnston, Dominic Kwiatkowski, Cordelia Langford, John Sillitoe on behalf of the Wellcome Sanger Institute COVID-19 Surveillance Team |
| EPI_ISL_3203204, EPI_ISL_3203205, EPI_ISL_3203206, EPI_ISL_3203207 | Lighthouse Laboratory Plymouth | Wellcome Sanger Institute for the COVID-19 Genomics UK (COG-UK) Consortium | Lighthouse Laboratory Plymouth and Alex Alderton, Roberto Amato, Jeffrey Barrett, Sonia Goncalves, Ewan Harrison, David K. Jackson, Ian Johnston, Dominic Kwiatkowski, Cordelia Langford, John Sillitoe on behalf of the Wellcome Sanger Institute COVID-19 Surveillance Team |
| EPI_ISL_3203208 | Lighthouse Lab in Glasgow | Wellcome Sanger Institute for the COVID-19 Genomics UK (COG-UK) Consortium | Harper VanSteenhouse, Yumi Kasai, David Gray, Carol Clugston, Anna Dominiczak and Alex Alderton, Roberto Amato, Jeffrey Barrett, Sonia Goncalves, Ewan Harrison, David K. Jackson, Ian Johnston, Dominic Kwiatkowski, Cordelia Langford, John Sillitoe on behalf of the Wellcome Sanger Institute COVID-19 Surveillance Team |
| EPI_ISL_3203209 | Lighthouse Laboratory Plymouth | Wellcome Sanger Institute for the COVID-19 Genomics UK (COG-UK) Consortium | Lighthouse Laboratory Plymouth and Alex Alderton, Roberto Amato, Jeffrey Barrett, Sonia Goncalves, Ewan Harrison, David K. Jackson, Ian Johnston, Dominic Kwiatkowski, Cordelia Langford, John Sillitoe on behalf of the Wellcome Sanger Institute COVID-19 Surveillance Team |
| EPI_ISL_3203210 | Lighthouse Lab in Milton Keynes | Wellcome Sanger Institute for the COVID-19 Genomics UK (COG-UK) Consortium | The Lighthouse Lab in Milton Keynes and Alex Alderton, Roberto Amato, Jeffrey Barrett, Sonia Goncalves, Ewan Harrison, David K. Jackson, Ian Johnston, Dominic Kwiatkowski, Cordelia Langford, John Sillitoe on behalf of the Wellcome Sanger Institute COVID-19 Surveillance Team |
| EPI_ISL_3203211, EPI_ISL_3203212 | Lighthouse Lab in Glasgow | Wellcome Sanger Institute for the COVID-19 Genomics UK (COG-UK) Consortium | Harper VanSteenhouse, Yumi Kasai, David Gray, Carol Clugston, Anna Dominiczak and Alex Alderton, Roberto Amato, Jeffrey Barrett, Sonia Goncalves, Ewan Harrison, David K. Jackson, Ian Johnston, Dominic Kwiatkowski, Cordelia Langford, John Sillitoe on behalf of the Wellcome Sanger Institute COVID-19 Surveillance Team |
| EPI_ISL_3203213, EPI_ISL_3203214, EPI_ISL_3203215 | Lighthouse Lab in Milton Keynes | Wellcome Sanger Institute for the COVID-19 Genomics UK (COG-UK) Consortium | The Lighthouse Lab in Milton Keynes and Alex Alderton, Roberto Amato, Jeffrey Barrett, Sonia Goncalves, Ewan Harrison, David K. Jackson, Ian Johnston, Dominic Kwiatkowski, Cordelia Langford, John Sillitoe on behalf of the Wellcome Sanger Institute COVID-19 Surveillance Team |
| EPI_ISL_3203216 | Lighthouse Lab in Glasgow | Wellcome Sanger Institute for the COVID-19 Genomics UK (COG-UK) Consortium | Harper VanSteenhouse, Yumi Kasai, David Gray, Carol Clugston, Anna Dominiczak and Alex Alderton, Roberto Amato, Jeffrey Barrett, Sonia Goncalves, Ewan Harrison, David K. Jackson, Ian Johnston, Dominic Kwiatkowski, Cordelia Langford, John Sillitoe on behalf of the Wellcome Sanger Institute COVID-19 Surveillance Team |
| EPI_ISL_3203217, EPI_ISL_3203218, EPI_ISL_3203219 | Lighthouse Laboratory Plymouth | Wellcome Sanger Institute for the COVID-19 Genomics UK (COG-UK) Consortium | Lighthouse Laboratory Plymouth and Alex Alderton, Roberto Amato, Jeffrey Barrett, Sonia Goncalves, Ewan Harrison, David K. Jackson, Ian Johnston, Dominic Kwiatkowski, Cordelia Langford, John Sillitoe on behalf of the Wellcome Sanger Institute COVID-19 Surveillance Team |
| EPI_ISL_3203220 | Lighthouse Lab in Glasgow | Wellcome Sanger Institute for the COVID-19 Genomics UK (COG-UK) Consortium | Harper VanSteenhouse, Yumi Kasai, David Gray, Carol Clugston, Anna Dominiczak and Alex Alderton, Roberto Amato, Jeffrey Barrett, Sonia Goncalves, Ewan Harrison, David K. Jackson, Ian Johnston, Dominic Kwiatkowski, Cordelia Langford, John Sillitoe on behalf of the Wellcome Sanger Institute COVID-19 Surveillance Team |
| EPI_ISL_3203221 | Lighthouse Laboratory Plymouth | Wellcome Sanger Institute for the COVID-19 Genomics UK (COG-UK) Consortium | Lighthouse Laboratory Plymouth and Alex Alderton, Roberto Amato, Jeffrey Barrett, Sonia Goncalves, Ewan Harrison, David K. Jackson, Ian Johnston, Dominic Kwiatkowski, Cordelia Langford, John Sillitoe on behalf of the Wellcome Sanger Institute COVID-19 Surveillance Team |
| EPI_ISL_3203222 | Lighthouse Lab in Glasgow | Wellcome Sanger Institute for the COVID-19 Genomics UK (COG-UK) Consortium | Harper VanSteenhouse, Yumi Kasai, David Gray, Carol Clugston, Anna Dominiczak and Alex Alderton, Roberto Amato, Jeffrey Barrett, Sonia Goncalves, Ewan Harrison, David K. Jackson, Ian Johnston, Dominic Kwiatkowski, Cordelia Langford, John Sillitoe on behalf of the Wellcome Sanger Institute COVID-19 Surveillance Team |
| EPI_ISL_3203223, EPI_ISL_3203224, EPI_ISL_3203225, EPI_ISL_3203226 | Lighthouse Laboratory Plymouth | Wellcome Sanger Institute for the COVID-19 Genomics UK (COG-UK) Consortium | Lighthouse Laboratory Plymouth and Alex Alderton, Roberto Amato, Jeffrey Barrett, Sonia Goncalves, Ewan Harrison, David K. Jackson, Ian Johnston, Dominic Kwiatkowski, Cordelia Langford, John Sillitoe on behalf of the Wellcome Sanger Institute COVID-19 Surveillance Team |
| EPI_ISL_3203227, EPI_ISL_3203228, EPI_ISL_3203229 | Lighthouse Lab in Glasgow | Wellcome Sanger Institute for the COVID-19 Genomics UK (COG-UK) Consortium | Harper VanSteenhouse, Yumi Kasai, David Gray, Carol Clugston, Anna Dominiczak and Alex Alderton, Roberto Amato, Jeffrey Barrett, Sonia Goncalves, Ewan Harrison, David K. Jackson, Ian Johnston, Dominic Kwiatkowski, Cordelia Langford, John Sillitoe on behalf of the Wellcome Sanger Institute COVID-19 Surveillance Team |
| EPI_ISL_3203230, EPI_ISL_3203231 | Lighthouse Laboratory Plymouth | Wellcome Sanger Institute for the COVID-19 Genomics UK (COG-UK) Consortium | Lighthouse Laboratory Plymouth and Alex Alderton, Roberto Amato, Jeffrey Barrett, Sonia Goncalves, Ewan Harrison, David K. Jackson, Ian Johnston, Dominic Kwiatkowski, Cordelia Langford, John Sillitoe on behalf of the Wellcome Sanger Institute COVID-19 Surveillance Team |
| EPI_ISL_3203232 | Lighthouse Lab in Milton Keynes | Wellcome Sanger Institute for the COVID-19 Genomics UK (COG-UK) Consortium | The Lighthouse Lab in Milton Keynes and Alex Alderton, Roberto Amato, Jeffrey Barrett, Sonia Goncalves, Ewan Harrison, David K. Jackson, Ian Johnston, Dominic Kwiatkowski, Cordelia Langford, John Sillitoe on behalf of the Wellcome Sanger Institute COVID-19 Surveillance Team |
| EPI_ISL_3203233 | Lighthouse Lab in Glasgow | Wellcome Sanger Institute for the COVID-19 Genomics UK (COG-UK) Consortium | Harper VanSteenhouse, Yumi Kasai, David Gray, Carol Clugston, Anna Dominiczak and Alex Alderton, Roberto Amato, Jeffrey Barrett, Sonia Goncalves, Ewan Harrison, David K. Jackson, Ian Johnston, Dominic Kwiatkowski, Cordelia Langford, John Sillitoe on behalf of the Wellcome Sanger Institute COVID-19 Surveillance Team |
| EPI_ISL_3203234, EPI_ISL_3203235, | Lighthouse Laboratory Plymouth | Wellcome Sanger Institute for the COVID-19 Genomics UK (COG-UK) Consortium | Lighthouse Laboratory Plymouth and Alex Alderton, Roberto Amato, Jeffrey Barrett, Sonia Goncalves, Ewan Harrison, David K. Jackson, Ian Johnston, Dominic Kwiatkowski, Cordelia Langford, John Sillitoe on behalf of the Wellcome Sanger Institute COVID-19 Surveillance Team |

| | | | |
|---|---|---|---|
| EPI_ISL_3203236 | | | |
| EPI_ISL_3203237, EPI_ISL_3203238 | Lighthouse Lab in Glasgow | Wellcome Sanger Institute for the COVID-19 Genomics UK (COG-UK) Consortium | Harper VanSteenhouse, Yumi Kasai, David Gray, Carol Clugston, Anna Dominiczak and Alex Alderton, Roberto Amato, Jeffrey Barrett, Sonia Goncalves, Ewan Harrison, David K. Jackson, Ian Johnston, Dominic Kwiatkowski, Cordelia Langford, John Sillitoe on behalf of the Wellcome Sanger Institute COVID-19 Surveillance Team |
| EPI_ISL_3203239 | Lighthouse Lab in Milton Keynes | Wellcome Sanger Institute for the COVID-19 Genomics UK (COG-UK) Consortium | The Lighthouse Lab in Milton Keynes and Alex Alderton, Roberto Amato, Jeffrey Barrett, Sonia Goncalves, Ewan Harrison, David K. Jackson, Ian Johnston, Dominic Kwiatkowski, Cordelia Langford, John Sillitoe on behalf of the Wellcome Sanger Institute COVID-19 Surveillance Team |
| EPI_ISL_3203240 | Lighthouse Laboratory Plymouth | Wellcome Sanger Institute for the COVID-19 Genomics UK (COG-UK) Consortium | Lighthouse Laboratory Plymouth and Alex Alderton, Roberto Amato, Jeffrey Barrett, Sonia Goncalves, Ewan Harrison, David K. Jackson, Ian Johnston, Dominic Kwiatkowski, Cordelia Langford, John Sillitoe on behalf of the Wellcome Sanger Institute COVID-19 Surveillance Team |
| EPI_ISL_3203241 | Lighthouse Lab in Milton Keynes | Wellcome Sanger Institute for the COVID-19 Genomics UK (COG-UK) Consortium | The Lighthouse Lab in Milton Keynes and Alex Alderton, Roberto Amato, Jeffrey Barrett, Sonia Goncalves, Ewan Harrison, David K. Jackson, Ian Johnston, Dominic Kwiatkowski, Cordelia Langford, John Sillitoe on behalf of the Wellcome Sanger Institute COVID-19 Surveillance Team |
| EPI_ISL_3203242 | Lighthouse Lab in Glasgow | Wellcome Sanger Institute for the COVID-19 Genomics UK (COG-UK) Consortium | Harper VanSteenhouse, Yumi Kasai, David Gray, Carol Clugston, Anna Dominiczak and Alex Alderton, Roberto Amato, Jeffrey Barrett, Sonia Goncalves, Ewan Harrison, David K. Jackson, Ian Johnston, Dominic Kwiatkowski, Cordelia Langford, John Sillitoe on behalf of the Wellcome Sanger Institute COVID-19 Surveillance Team |
| EPI_ISL_3203243 | Lighthouse Lab in Milton Keynes | Wellcome Sanger Institute for the COVID-19 Genomics UK (COG-UK) Consortium | The Lighthouse Lab in Milton Keynes and Alex Alderton, Roberto Amato, Jeffrey Barrett, Sonia Goncalves, Ewan Harrison, David K. Jackson, Ian Johnston, Dominic Kwiatkowski, Cordelia Langford, John Sillitoe on behalf of the Wellcome Sanger Institute COVID-19 Surveillance Team |
| EPI_ISL_3203244, EPI_ISL_3203245, EPI_ISL_3203246 | Lighthouse Laboratory Plymouth | Wellcome Sanger Institute for the COVID-19 Genomics UK (COG-UK) Consortium | Lighthouse Laboratory Plymouth and Alex Alderton, Roberto Amato, Jeffrey Barrett, Sonia Goncalves, Ewan Harrison, David K. Jackson, Ian Johnston, Dominic Kwiatkowski, Cordelia Langford, John Sillitoe on behalf of the Wellcome Sanger Institute COVID-19 Surveillance Team |
| EPI_ISL_3203247 | Lighthouse Lab in Milton Keynes | Wellcome Sanger Institute for the COVID-19 Genomics UK (COG-UK) Consortium | The Lighthouse Lab in Milton Keynes and Alex Alderton, Roberto Amato, Jeffrey Barrett, Sonia Goncalves, Ewan Harrison, David K. Jackson, Ian Johnston, Dominic Kwiatkowski, Cordelia Langford, John Sillitoe on behalf of the Wellcome Sanger Institute COVID-19 Surveillance Team |
| EPI_ISL_3203248 | Lighthouse Laboratory Plymouth | Wellcome Sanger Institute for the COVID-19 Genomics UK (COG-UK) Consortium | Lighthouse Laboratory Plymouth and Alex Alderton, Roberto Amato, Jeffrey Barrett, Sonia Goncalves, Ewan Harrison, David K. Jackson, Ian Johnston, Dominic Kwiatkowski, Cordelia Langford, John Sillitoe on behalf of the Wellcome Sanger Institute COVID-19 Surveillance Team |
| EPI_ISL_3203249, EPI_ISL_3203250, EPI_ISL_3203251, EPI_ISL_3203252 | Lighthouse Lab in Milton Keynes | Wellcome Sanger Institute for the COVID-19 Genomics UK (COG-UK) Consortium | The Lighthouse Lab in Milton Keynes and Alex Alderton, Roberto Amato, Jeffrey Barrett, Sonia Goncalves, Ewan Harrison, David K. Jackson, Ian Johnston, Dominic Kwiatkowski, Cordelia Langford, John Sillitoe on behalf of the Wellcome Sanger Institute COVID-19 Surveillance Team |
| EPI_ISL_3203253, EPI_ISL_3203254 | Lighthouse Laboratory Plymouth | Wellcome Sanger Institute for the COVID-19 Genomics UK (COG-UK) Consortium | Lighthouse Laboratory Plymouth and Alex Alderton, Roberto Amato, Jeffrey Barrett, Sonia Goncalves, Ewan Harrison, David K. Jackson, Ian Johnston, Dominic Kwiatkowski, Cordelia Langford, John Sillitoe on behalf of the Wellcome Sanger Institute COVID-19 Surveillance Team |
| EPI_ISL_3203255, EPI_ISL_3203256 | Lighthouse Lab in Milton Keynes | Wellcome Sanger Institute for the COVID-19 Genomics UK (COG-UK) Consortium | The Lighthouse Lab in Milton Keynes and Alex Alderton, Roberto Amato, Jeffrey Barrett, Sonia Goncalves, Ewan Harrison, David K. Jackson, Ian Johnston, Dominic Kwiatkowski, Cordelia Langford, John Sillitoe on behalf of the Wellcome Sanger Institute COVID-19 Surveillance Team |
| EPI_ISL_3203257, EPI_ISL_3203258 | Lighthouse Laboratory Plymouth | Wellcome Sanger Institute for the COVID-19 Genomics UK (COG-UK) Consortium | Lighthouse Laboratory Plymouth and Alex Alderton, Roberto Amato, Jeffrey Barrett, Sonia Goncalves, Ewan Harrison, David K. Jackson, Ian Johnston, Dominic Kwiatkowski, Cordelia Langford, John Sillitoe on behalf of the Wellcome Sanger Institute COVID-19 Surveillance Team |
| EPI_ISL_3203259, EPI_ISL_3203260, EPI_ISL_3203261, EPI_ISL_3203262 | Lighthouse Lab in Glasgow | Wellcome Sanger Institute for the COVID-19 Genomics UK (COG-UK) Consortium | Harper VanSteenhouse, Yumi Kasai, David Gray, Carol Clugston, Anna Dominiczak and Alex Alderton, Roberto Amato, Jeffrey Barrett, Sonia Goncalves, Ewan Harrison, David K. Jackson, Ian Johnston, Dominic Kwiatkowski, Cordelia Langford, John Sillitoe on behalf of the Wellcome Sanger Institute COVID-19 Surveillance Team |
| EPI_ISL_3203263 | Lighthouse Laboratory Plymouth | Wellcome Sanger Institute for the COVID-19 Genomics UK (COG-UK) Consortium | Lighthouse Laboratory Plymouth and Alex Alderton, Roberto Amato, Jeffrey Barrett, Sonia Goncalves, Ewan Harrison, David K. Jackson, Ian Johnston, Dominic Kwiatkowski, Cordelia Langford, John Sillitoe on behalf of the Wellcome Sanger Institute COVID-19 Surveillance Team |
| EPI_ISL_3203264 | Lighthouse Lab in Milton Keynes | Wellcome Sanger Institute for the COVID-19 Genomics UK (COG-UK) Consortium | The Lighthouse Lab in Milton Keynes and Alex Alderton, Roberto Amato, Jeffrey Barrett, Sonia Goncalves, Ewan Harrison, David K. Jackson, Ian Johnston, Dominic Kwiatkowski, Cordelia Langford, John Sillitoe on behalf of the Wellcome Sanger Institute COVID-19 Surveillance Team |
| EPI_ISL_3203265 | Lighthouse Lab in Glasgow | Wellcome Sanger Institute for the COVID-19 Genomics UK (COG-UK) Consortium | Harper VanSteenhouse, Yumi Kasai, David Gray, Carol Clugston, Anna Dominiczak and Alex Alderton, Roberto Amato, Jeffrey Barrett, Sonia Goncalves, Ewan Harrison, David K. Jackson, Ian Johnston, Dominic Kwiatkowski, Cordelia Langford, John Sillitoe on behalf of the Wellcome Sanger Institute COVID-19 Surveillance Team |
| EPI_ISL_3203266 | Lighthouse Laboratory Plymouth | Wellcome Sanger Institute for the COVID-19 Genomics UK (COG-UK) Consortium | Lighthouse Laboratory Plymouth and Alex Alderton, Roberto Amato, Jeffrey Barrett, Sonia Goncalves, Ewan Harrison, David K. Jackson, Ian Johnston, Dominic Kwiatkowski, Cordelia Langford, John Sillitoe on behalf of the Wellcome Sanger Institute COVID-19 Surveillance Team |
| EPI_ISL_3203267, EPI_ISL_3203268 | Lighthouse Lab in Glasgow | Wellcome Sanger Institute for the COVID-19 Genomics UK (COG-UK) Consortium | Harper VanSteenhouse, Yumi Kasai, David Gray, Carol Clugston, Anna Dominiczak and Alex Alderton, Roberto Amato, Jeffrey Barrett, Sonia Goncalves, Ewan Harrison, David K. Jackson, Ian Johnston, Dominic Kwiatkowski, Cordelia Langford, John Sillitoe on behalf of the Wellcome Sanger Institute COVID-19 Surveillance Team |
| EPI_ISL_3203269, EPI_ISL_3203270 | Lighthouse Lab in Milton Keynes | Wellcome Sanger Institute for the COVID-19 Genomics UK (COG-UK) Consortium | The Lighthouse Lab in Milton Keynes and Alex Alderton, Roberto Amato, Jeffrey Barrett, Sonia Goncalves, Ewan Harrison, David K. Jackson, Ian Johnston, Dominic Kwiatkowski, Cordelia Langford, John Sillitoe on behalf of the Wellcome Sanger Institute COVID-19 Surveillance Team |
| EPI_ISL_3203271, EPI_ISL_3203272 | Lighthouse Lab in Glasgow | Wellcome Sanger Institute for the COVID-19 Genomics UK (COG-UK) Consortium | Harper VanSteenhouse, Yumi Kasai, David Gray, Carol Clugston, Anna Dominiczak and Alex Alderton, Roberto Amato, Jeffrey Barrett, Sonia Goncalves, Ewan Harrison, David K. Jackson, Ian Johnston, Dominic Kwiatkowski, Cordelia Langford, John Sillitoe on behalf of the Wellcome Sanger Institute COVID-19 Surveillance Team |
| EPI_ISL_3203273, EPI_ISL_3203274, EPI_ISL_3203275, EPI_ISL_3203276 | Lighthouse Laboratory Plymouth | Wellcome Sanger Institute for the COVID-19 Genomics UK (COG-UK) Consortium | Lighthouse Laboratory Plymouth and Alex Alderton, Roberto Amato, Jeffrey Barrett, Sonia Goncalves, Ewan Harrison, David K. Jackson, Ian Johnston, Dominic Kwiatkowski, Cordelia Langford, John Sillitoe on behalf of the Wellcome Sanger Institute COVID-19 Surveillance Team |
| EPI_ISL_3203277, EPI_ISL_3203278 | Lighthouse Lab in Milton Keynes | Wellcome Sanger Institute for the COVID-19 Genomics UK (COG-UK) Consortium | The Lighthouse Lab in Milton Keynes and Alex Alderton, Roberto Amato, Jeffrey Barrett, Sonia Goncalves, Ewan Harrison, David K. Jackson, Ian Johnston, Dominic Kwiatkowski, Cordelia Langford, John Sillitoe on behalf of the Wellcome Sanger Institute COVID-19 Surveillance Team |
| EPI_ISL_3203279 | Lighthouse Lab in Glasgow | Wellcome Sanger Institute for the COVID-19 Genomics UK (COG-UK) Consortium | Harper VanSteenhouse, Yumi Kasai, David Gray, Carol Clugston, Anna Dominiczak and Alex Alderton, Roberto Amato, Jeffrey Barrett, Sonia Goncalves, Ewan Harrison, David K. Jackson, Ian Johnston, Dominic Kwiatkowski, Cordelia Langford, John Sillitoe on behalf of the Wellcome Sanger Institute COVID-19 Surveillance Team |
| EPI_ISL_3203280, EPI_ISL_3203281 | Lighthouse Laboratory Plymouth | Wellcome Sanger Institute for the COVID-19 Genomics UK (COG-UK) Consortium | Lighthouse Laboratory Plymouth and Alex Alderton, Roberto Amato, Jeffrey Barrett, Sonia Goncalves, Ewan Harrison, David K. Jackson, Ian Johnston, Dominic Kwiatkowski, Cordelia Langford, John Sillitoe on behalf of the Wellcome Sanger Institute COVID-19 Surveillance Team |
| EPI_ISL_3203282, EPI_ISL_3203283 | Lighthouse Lab in Milton Keynes | Wellcome Sanger Institute for the COVID-19 Genomics UK (COG-UK) Consortium | The Lighthouse Lab in Milton Keynes and Alex Alderton, Roberto Amato, Jeffrey Barrett, Sonia Goncalves, Ewan Harrison, David K. Jackson, Ian Johnston, Dominic Kwiatkowski, Cordelia Langford, John Sillitoe on behalf of the Wellcome Sanger Institute COVID-19 Surveillance Team |
| EPI_ISL_3203284 | Lighthouse Laboratory Plymouth | Wellcome Sanger Institute for the COVID-19 Genomics UK (COG-UK) Consortium | Lighthouse Laboratory Plymouth and Alex Alderton, Roberto Amato, Jeffrey Barrett, Sonia Goncalves, Ewan Harrison, David K. Jackson, Ian Johnston, Dominic Kwiatkowski, Cordelia Langford, John Sillitoe on behalf of the Wellcome Sanger Institute COVID-19 Surveillance Team |
| EPI_ISL_3203285 | Lighthouse Lab in Milton Keynes | Wellcome Sanger Institute for the COVID-19 Genomics UK (COG-UK) Consortium | The Lighthouse Lab in Milton Keynes and Alex Alderton, Roberto Amato, Jeffrey Barrett, Sonia Goncalves, Ewan Harrison, David K. Jackson, Ian Johnston, Dominic Kwiatkowski, Cordelia Langford, John Sillitoe on behalf of the Wellcome Sanger Institute COVID-19 Surveillance Team |
| EPI_ISL_3203286 | Lighthouse Laboratory Plymouth | Wellcome Sanger Institute for the COVID-19 Genomics UK (COG-UK) Consortium | Lighthouse Laboratory Plymouth and Alex Alderton, Roberto Amato, Jeffrey Barrett, Sonia Goncalves, Ewan Harrison, David K. Jackson, Ian Johnston, Dominic Kwiatkowski, Cordelia Langford, John Sillitoe on behalf of the Wellcome Sanger Institute COVID-19 Surveillance Team |
| EPI_ISL_3203287 | Lighthouse Lab in Glasgow | Wellcome Sanger Institute for the COVID-19 Genomics UK (COG-UK) Consortium | Harper VanSteenhouse, Yumi Kasai, David Gray, Carol Clugston, Anna Dominiczak and Alex Alderton, Roberto Amato, Jeffrey Barrett, Sonia Goncalves, Ewan Harrison, David K. Jackson, Ian Johnston, Dominic Kwiatkowski, Cordelia Langford, John Sillitoe on behalf of the Wellcome Sanger Institute COVID-19 Surveillance Team |
| EPI_ISL_3203288 | Lighthouse Lab in Milton Keynes | Wellcome Sanger Institute for the COVID-19 Genomics UK (COG-UK) Consortium | The Lighthouse Lab in Milton Keynes and Alex Alderton, Roberto Amato, Jeffrey Barrett, Sonia Goncalves, Ewan Harrison, David K. Jackson, Ian Johnston, Dominic Kwiatkowski, Cordelia Langford, John Sillitoe on behalf of the Wellcome Sanger Institute COVID-19 Surveillance Team |
| EPI_ISL_3203289, EPI_ISL_3203290, EPI_ISL_3203291 | Lighthouse Laboratory Plymouth | Wellcome Sanger Institute for the COVID-19 Genomics UK (COG-UK) Consortium | Lighthouse Laboratory Plymouth and Alex Alderton, Roberto Amato, Jeffrey Barrett, Sonia Goncalves, Ewan Harrison, David K. Jackson, Ian Johnston, Dominic Kwiatkowski, Cordelia Langford, John Sillitoe on behalf of the Wellcome Sanger Institute COVID-19 Surveillance Team |
| EPI_ISL_3203292, EPI_ISL_3203293, EPI_ISL_3203294 | Lighthouse Lab in Glasgow | Wellcome Sanger Institute for the COVID-19 Genomics UK (COG-UK) Consortium | Harper VanSteenhouse, Yumi Kasai, David Gray, Carol Clugston, Anna Dominiczak and Alex Alderton, Roberto Amato, Jeffrey Barrett, Sonia Goncalves, Ewan Harrison, David K. Jackson, Ian Johnston, Dominic Kwiatkowski, Cordelia Langford, John Sillitoe on behalf of the Wellcome Sanger Institute COVID-19 Surveillance Team |
| EPI_ISL_3203295, EPI_ISL_3203296 | Lighthouse Laboratory Plymouth | Wellcome Sanger Institute for the COVID-19 Genomics UK (COG-UK) Consortium | Lighthouse Laboratory Plymouth and Alex Alderton, Roberto Amato, Jeffrey Barrett, Sonia Goncalves, Ewan Harrison, David K. Jackson, Ian Johnston, Dominic Kwiatkowski, Cordelia Langford, John Sillitoe on behalf of the Wellcome Sanger Institute COVID-19 Surveillance Team |
| EPI_ISL_3203297 | Lighthouse Lab in Glasgow | Wellcome Sanger Institute for the COVID-19 Genomics UK (COG-UK) Consortium | Harper VanSteenhouse, Yumi Kasai, David Gray, Carol Clugston, Anna Dominiczak and Alex Alderton, Roberto Amato, Jeffrey Barrett, Sonia Goncalves, Ewan Harrison, David K. Jackson, Ian Johnston, Dominic Kwiatkowski, Cordelia Langford, John Sillitoe on behalf of the Wellcome Sanger Institute COVID-19 Surveillance Team |
| EPI_ISL_3203298, EPI_ISL_3203299, EPI_ISL_3203300, EPI_ISL_3203301, EPI_ISL_3203302, EPI_ISL_3203303, EPI_ISL_3203304 | Lighthouse Laboratory Plymouth | Wellcome Sanger Institute for the COVID-19 Genomics UK (COG-UK) Consortium | Lighthouse Laboratory Plymouth and Alex Alderton, Roberto Amato, Jeffrey Barrett, Sonia Goncalves, Ewan Harrison, David K. Jackson, Ian Johnston, Dominic Kwiatkowski, Cordelia Langford, John Sillitoe on behalf of the Wellcome Sanger Institute COVID-19 Surveillance Team |
| EPI_ISL_3203305, EPI_ISL_3203306, EPI_ISL_3203307 | Lighthouse Lab in Glasgow | Wellcome Sanger Institute for the COVID-19 Genomics UK (COG-UK) Consortium | Harper VanSteenhouse, Yumi Kasai, David Gray, Carol Clugston, Anna Dominiczak and Alex Alderton, Roberto Amato, Jeffrey Barrett, Sonia Goncalves, Ewan Harrison, David K. Jackson, Ian Johnston, Dominic Kwiatkowski, Cordelia Langford, John Sillitoe on behalf of the Wellcome Sanger Institute COVID-19 Surveillance Team |
| EPI_ISL_3203308 | Lighthouse Laboratory Plymouth | Wellcome Sanger Institute for the COVID-19 Genomics UK (COG-UK) Consortium | Lighthouse Laboratory Plymouth and Alex Alderton, Roberto Amato, Jeffrey Barrett, Sonia Goncalves, Ewan Harrison, David K. Jackson, Ian Johnston, Dominic Kwiatkowski, Cordelia Langford, John Sillitoe on behalf of the Wellcome Sanger Institute COVID-19 Surveillance Team |

| | | | |
|---|---|---|---|
| EPI_ISL_3203309 | Lighthouse Lab in Milton Keynes | Wellcome Sanger Institute for the COVID-19 Genomics UK (COG-UK) Consortium | The Lighthouse Lab in Milton Keynes and Alex Alderton, Roberto Amato, Jeffrey Barrett, Sonia Goncalves, Ewan Harrison, David K. Jackson, Ian Johnston, Dominic Kwiatkowski, Cordelia Langford, John Sillitoe on behalf of the Wellcome Sanger Institute COVID-19 Surveillance Team |
| EPI_ISL_3203310 | Lighthouse Laboratory Plymouth | Wellcome Sanger Institute for the COVID-19 Genomics UK (COG-UK) Consortium | Lighthouse Laboratory Plymouth and Alex Alderton, Roberto Amato, Jeffrey Barrett, Sonia Goncalves, Ewan Harrison, David K. Jackson, Ian Johnston, Dominic Kwiatkowski, Cordelia Langford, John Sillitoe on behalf of the Wellcome Sanger Institute COVID-19 Surveillance Team |
| EPI_ISL_3203311 | Lighthouse Lab in Glasgow | Wellcome Sanger Institute for the COVID-19 Genomics UK (COG-UK) Consortium | Harper VanSteenhouse, Yumi Kasai, David Gray, Carol Clugston, Anna Dominiczak and Alex Alderton, Roberto Amato, Jeffrey Barrett, Sonia Goncalves, Ewan Harrison, David K. Jackson, Ian Johnston, Dominic Kwiatkowski, Cordelia Langford, John Sillitoe on behalf of the Wellcome Sanger Institute COVID-19 Surveillance Team |
| EPI_ISL_3203312 | Lighthouse Lab in Milton Keynes | Wellcome Sanger Institute for the COVID-19 Genomics UK (COG-UK) Consortium | The Lighthouse Lab in Milton Keynes and Alex Alderton, Roberto Amato, Jeffrey Barrett, Sonia Goncalves, Ewan Harrison, David K. Jackson, Ian Johnston, Dominic Kwiatkowski, Cordelia Langford, John Sillitoe on behalf of the Wellcome Sanger Institute COVID-19 Surveillance Team |
| EPI_ISL_3203313, EPI_ISL_3203314, EPI_ISL_3203315, EPI_ISL_3203316 | Lighthouse Laboratory Plymouth | Wellcome Sanger Institute for the COVID-19 Genomics UK (COG-UK) Consortium | Lighthouse Laboratory Plymouth and Alex Alderton, Roberto Amato, Jeffrey Barrett, Sonia Goncalves, Ewan Harrison, David K. Jackson, Ian Johnston, Dominic Kwiatkowski, Cordelia Langford, John Sillitoe on behalf of the Wellcome Sanger Institute COVID-19 Surveillance Team |
| EPI_ISL_3203317 | Lighthouse Lab in Milton Keynes | Wellcome Sanger Institute for the COVID-19 Genomics UK (COG-UK) Consortium | The Lighthouse Lab in Milton Keynes and Alex Alderton, Roberto Amato, Jeffrey Barrett, Sonia Goncalves, Ewan Harrison, David K. Jackson, Ian Johnston, Dominic Kwiatkowski, Cordelia Langford, John Sillitoe on behalf of the Wellcome Sanger Institute COVID-19 Surveillance Team |
| EPI_ISL_3203318 | Lighthouse Lab in Glasgow | Wellcome Sanger Institute for the COVID-19 Genomics UK (COG-UK) Consortium | Harper VanSteenhouse, Yumi Kasai, David Gray, Carol Clugston, Anna Dominiczak and Alex Alderton, Roberto Amato, Jeffrey Barrett, Sonia Goncalves, Ewan Harrison, David K. Jackson, Ian Johnston, Dominic Kwiatkowski, Cordelia Langford, John Sillitoe on behalf of the Wellcome Sanger Institute COVID-19 Surveillance Team |
| EPI_ISL_3203319 | Lighthouse Laboratory Plymouth | Wellcome Sanger Institute for the COVID-19 Genomics UK (COG-UK) Consortium | Lighthouse Laboratory Plymouth and Alex Alderton, Roberto Amato, Jeffrey Barrett, Sonia Goncalves, Ewan Harrison, David K. Jackson, Ian Johnston, Dominic Kwiatkowski, Cordelia Langford, John Sillitoe on behalf of the Wellcome Sanger Institute COVID-19 Surveillance Team |
| EPI_ISL_3203320 | Lighthouse Lab in Glasgow | Wellcome Sanger Institute for the COVID-19 Genomics UK (COG-UK) Consortium | Harper VanSteenhouse, Yumi Kasai, David Gray, Carol Clugston, Anna Dominiczak and Alex Alderton, Roberto Amato, Jeffrey Barrett, Sonia Goncalves, Ewan Harrison, David K. Jackson, Ian Johnston, Dominic Kwiatkowski, Cordelia Langford, John Sillitoe on behalf of the Wellcome Sanger Institute COVID-19 Surveillance Team |
| EPI_ISL_3203321 | Lighthouse Laboratory Plymouth | Wellcome Sanger Institute for the COVID-19 Genomics UK (COG-UK) Consortium | Lighthouse Laboratory Plymouth and Alex Alderton, Roberto Amato, Jeffrey Barrett, Sonia Goncalves, Ewan Harrison, David K. Jackson, Ian Johnston, Dominic Kwiatkowski, Cordelia Langford, John Sillitoe on behalf of the Wellcome Sanger Institute COVID-19 Surveillance Team |
| EPI_ISL_3203322 | Lighthouse Lab in Milton Keynes | Wellcome Sanger Institute for the COVID-19 Genomics UK (COG-UK) Consortium | The Lighthouse Lab in Milton Keynes and Alex Alderton, Roberto Amato, Jeffrey Barrett, Sonia Goncalves, Ewan Harrison, David K. Jackson, Ian Johnston, Dominic Kwiatkowski, Cordelia Langford, John Sillitoe on behalf of the Wellcome Sanger Institute COVID-19 Surveillance Team |
| EPI_ISL_3203323, EPI_ISL_3203324, EPI_ISL_3203325 | Lighthouse Lab in Glasgow | Wellcome Sanger Institute for the COVID-19 Genomics UK (COG-UK) Consortium | Harper VanSteenhouse, Yumi Kasai, David Gray, Carol Clugston, Anna Dominiczak and Alex Alderton, Roberto Amato, Jeffrey Barrett, Sonia Goncalves, Ewan Harrison, David K. Jackson, Ian Johnston, Dominic Kwiatkowski, Cordelia Langford, John Sillitoe on behalf of the Wellcome Sanger Institute COVID-19 Surveillance Team |
| EPI_ISL_3203326 | Lighthouse Laboratory Plymouth | Wellcome Sanger Institute for the COVID-19 Genomics UK (COG-UK) Consortium | Lighthouse Laboratory Plymouth and Alex Alderton, Roberto Amato, Jeffrey Barrett, Sonia Goncalves, Ewan Harrison, David K. Jackson, Ian Johnston, Dominic Kwiatkowski, Cordelia Langford, John Sillitoe on behalf of the Wellcome Sanger Institute COVID-19 Surveillance Team |
| EPI_ISL_3203327 | Lighthouse Lab in Glasgow | Wellcome Sanger Institute for the COVID-19 Genomics UK (COG-UK) Consortium | Harper VanSteenhouse, Yumi Kasai, David Gray, Carol Clugston, Anna Dominiczak and Alex Alderton, Roberto Amato, Jeffrey Barrett, Sonia Goncalves, Ewan Harrison, David K. Jackson, Ian Johnston, Dominic Kwiatkowski, Cordelia Langford, John Sillitoe on behalf of the Wellcome Sanger Institute COVID-19 Surveillance Team |
| EPI_ISL_3203328 | Lighthouse Lab in Milton Keynes | Wellcome Sanger Institute for the COVID-19 Genomics UK (COG-UK) Consortium | The Lighthouse Lab in Milton Keynes and Alex Alderton, Roberto Amato, Jeffrey Barrett, Sonia Goncalves, Ewan Harrison, David K. Jackson, Ian Johnston, Dominic Kwiatkowski, Cordelia Langford, John Sillitoe on behalf of the Wellcome Sanger Institute COVID-19 Surveillance Team |
| EPI_ISL_3203329 | Lighthouse Laboratory Plymouth | Wellcome Sanger Institute for the COVID-19 Genomics UK (COG-UK) Consortium | Lighthouse Laboratory Plymouth and Alex Alderton, Roberto Amato, Jeffrey Barrett, Sonia Goncalves, Ewan Harrison, David K. Jackson, Ian Johnston, Dominic Kwiatkowski, Cordelia Langford, John Sillitoe on behalf of the Wellcome Sanger Institute COVID-19 Surveillance Team |
| EPI_ISL_3203330 | Lighthouse Lab in Milton Keynes | Wellcome Sanger Institute for the COVID-19 Genomics UK (COG-UK) Consortium | The Lighthouse Lab in Milton Keynes and Alex Alderton, Roberto Amato, Jeffrey Barrett, Sonia Goncalves, Ewan Harrison, David K. Jackson, Ian Johnston, Dominic Kwiatkowski, Cordelia Langford, John Sillitoe on behalf of the Wellcome Sanger Institute COVID-19 Surveillance Team |
| EPI_ISL_3203331 | Lighthouse Lab in Glasgow | Wellcome Sanger Institute for the COVID-19 Genomics UK (COG-UK) Consortium | Harper VanSteenhouse, Yumi Kasai, David Gray, Carol Clugston, Anna Dominiczak and Alex Alderton, Roberto Amato, Jeffrey Barrett, Sonia Goncalves, Ewan Harrison, David K. Jackson, Ian Johnston, Dominic Kwiatkowski, Cordelia Langford, John Sillitoe on behalf of the Wellcome Sanger Institute COVID-19 Surveillance Team |
| EPI_ISL_3203332 | Lighthouse Lab in Milton Keynes | Wellcome Sanger Institute for the COVID-19 Genomics UK (COG-UK) Consortium | The Lighthouse Lab in Milton Keynes and Alex Alderton, Roberto Amato, Jeffrey Barrett, Sonia Goncalves, Ewan Harrison, David K. Jackson, Ian Johnston, Dominic Kwiatkowski, Cordelia Langford, John Sillitoe on behalf of the Wellcome Sanger Institute COVID-19 Surveillance Team |
| EPI_ISL_3203333 | Lighthouse Laboratory Plymouth | Wellcome Sanger Institute for the COVID-19 Genomics UK (COG-UK) Consortium | Lighthouse Laboratory Plymouth and Alex Alderton, Roberto Amato, Jeffrey Barrett, Sonia Goncalves, Ewan Harrison, David K. Jackson, Ian Johnston, Dominic Kwiatkowski, Cordelia Langford, John Sillitoe on behalf of the Wellcome Sanger Institute COVID-19 Surveillance Team |
| EPI_ISL_3203334 | Lighthouse Lab in Milton Keynes | Wellcome Sanger Institute for the COVID-19 Genomics UK (COG-UK) Consortium | The Lighthouse Lab in Milton Keynes and Alex Alderton, Roberto Amato, Jeffrey Barrett, Sonia Goncalves, Ewan Harrison, David K. Jackson, Ian Johnston, Dominic Kwiatkowski, Cordelia Langford, John Sillitoe on behalf of the Wellcome Sanger Institute COVID-19 Surveillance Team |
| EPI_ISL_3203335, EPI_ISL_3203336, EPI_ISL_3203337, EPI_ISL_3203338, EPI_ISL_3203339, EPI_ISL_3203340 | Lighthouse Laboratory Plymouth | Wellcome Sanger Institute for the COVID-19 Genomics UK (COG-UK) Consortium | Lighthouse Laboratory Plymouth and Alex Alderton, Roberto Amato, Jeffrey Barrett, Sonia Goncalves, Ewan Harrison, David K. Jackson, Ian Johnston, Dominic Kwiatkowski, Cordelia Langford, John Sillitoe on behalf of the Wellcome Sanger Institute COVID-19 Surveillance Team |
| EPI_ISL_3203341 | Lighthouse Lab in Milton Keynes | Wellcome Sanger Institute for the COVID-19 Genomics UK (COG-UK) Consortium | The Lighthouse Lab in Milton Keynes and Alex Alderton, Roberto Amato, Jeffrey Barrett, Sonia Goncalves, Ewan Harrison, David K. Jackson, Ian Johnston, Dominic Kwiatkowski, Cordelia Langford, John Sillitoe on behalf of the Wellcome Sanger Institute COVID-19 Surveillance Team |
| EPI_ISL_3203342 | Lighthouse Lab in Glasgow | Wellcome Sanger Institute for the COVID-19 Genomics UK (COG-UK) Consortium | Harper VanSteenhouse, Yumi Kasai, David Gray, Carol Clugston, Anna Dominiczak and Alex Alderton, Roberto Amato, Jeffrey Barrett, Sonia Goncalves, Ewan Harrison, David K. Jackson, Ian Johnston, Dominic Kwiatkowski, Cordelia Langford, John Sillitoe on behalf of the Wellcome Sanger Institute COVID-19 Surveillance Team |
| EPI_ISL_3203343 | Lighthouse Lab in Milton Keynes | Wellcome Sanger Institute for the COVID-19 Genomics UK (COG-UK) Consortium | The Lighthouse Lab in Milton Keynes and Alex Alderton, Roberto Amato, Jeffrey Barrett, Sonia Goncalves, Ewan Harrison, David K. Jackson, Ian Johnston, Dominic Kwiatkowski, Cordelia Langford, John Sillitoe on behalf of the Wellcome Sanger Institute COVID-19 Surveillance Team |
| EPI_ISL_3203344 | Lighthouse Laboratory Plymouth | Wellcome Sanger Institute for the COVID-19 Genomics UK (COG-UK) Consortium | Lighthouse Laboratory Plymouth and Alex Alderton, Roberto Amato, Jeffrey Barrett, Sonia Goncalves, Ewan Harrison, David K. Jackson, Ian Johnston, Dominic Kwiatkowski, Cordelia Langford, John Sillitoe on behalf of the Wellcome Sanger Institute COVID-19 Surveillance Team |
| EPI_ISL_3203345 | Lighthouse Lab in Milton Keynes | Wellcome Sanger Institute for the COVID-19 Genomics UK (COG-UK) Consortium | The Lighthouse Lab in Milton Keynes and Alex Alderton, Roberto Amato, Jeffrey Barrett, Sonia Goncalves, Ewan Harrison, David K. Jackson, Ian Johnston, Dominic Kwiatkowski, Cordelia Langford, John Sillitoe on behalf of the Wellcome Sanger Institute COVID-19 Surveillance Team |
| EPI_ISL_3203346, EPI_ISL_3203347, EPI_ISL_3203348 | Lighthouse Lab in Glasgow | Wellcome Sanger Institute for the COVID-19 Genomics UK (COG-UK) Consortium | Harper VanSteenhouse, Yumi Kasai, David Gray, Carol Clugston, Anna Dominiczak and Alex Alderton, Roberto Amato, Jeffrey Barrett, Sonia Goncalves, Ewan Harrison, David K. Jackson, Ian Johnston, Dominic Kwiatkowski, Cordelia Langford, John Sillitoe on behalf of the Wellcome Sanger Institute COVID-19 Surveillance Team |
| EPI_ISL_3203349, EPI_ISL_3203350, EPI_ISL_3203351 | Lighthouse Lab in Milton Keynes | Wellcome Sanger Institute for the COVID-19 Genomics UK (COG-UK) Consortium | The Lighthouse Lab in Milton Keynes and Alex Alderton, Roberto Amato, Jeffrey Barrett, Sonia Goncalves, Ewan Harrison, David K. Jackson, Ian Johnston, Dominic Kwiatkowski, Cordelia Langford, John Sillitoe on behalf of the Wellcome Sanger Institute COVID-19 Surveillance Team |
| EPI_ISL_3203352, EPI_ISL_3203353, EPI_ISL_3203354 | Lighthouse Lab in Glasgow | Wellcome Sanger Institute for the COVID-19 Genomics UK (COG-UK) Consortium | Harper VanSteenhouse, Yumi Kasai, David Gray, Carol Clugston, Anna Dominiczak and Alex Alderton, Roberto Amato, Jeffrey Barrett, Sonia Goncalves, Ewan Harrison, David K. Jackson, Ian Johnston, Dominic Kwiatkowski, Cordelia Langford, John Sillitoe on behalf of the Wellcome Sanger Institute COVID-19 Surveillance Team |
| EPI_ISL_3203355 | Lighthouse Lab in Milton Keynes | Wellcome Sanger Institute for the COVID-19 Genomics UK (COG-UK) Consortium | The Lighthouse Lab in Milton Keynes and Alex Alderton, Roberto Amato, Jeffrey Barrett, Sonia Goncalves, Ewan Harrison, David K. Jackson, Ian Johnston, Dominic Kwiatkowski, Cordelia Langford, John Sillitoe on behalf of the Wellcome Sanger Institute COVID-19 Surveillance Team |
| EPI_ISL_3203356, EPI_ISL_3203357, EPI_ISL_3203358, EPI_ISL_3203359 | Lighthouse Lab in Glasgow | Wellcome Sanger Institute for the COVID-19 Genomics UK (COG-UK) Consortium | Harper VanSteenhouse, Yumi Kasai, David Gray, Carol Clugston, Anna Dominiczak and Alex Alderton, Roberto Amato, Jeffrey Barrett, Sonia Goncalves, Ewan Harrison, David K. Jackson, Ian Johnston, Dominic Kwiatkowski, Cordelia Langford, John Sillitoe on behalf of the Wellcome Sanger Institute COVID-19 Surveillance Team |
| EPI_ISL_3203360, EPI_ISL_3203361 | Lighthouse Lab in Milton Keynes | Wellcome Sanger Institute for the COVID-19 Genomics UK (COG-UK) Consortium | The Lighthouse Lab in Milton Keynes and Alex Alderton, Roberto Amato, Jeffrey Barrett, Sonia Goncalves, Ewan Harrison, David K. Jackson, Ian Johnston, Dominic Kwiatkowski, Cordelia Langford, John Sillitoe on behalf of the Wellcome Sanger Institute COVID-19 Surveillance Team |
| EPI_ISL_3203362 | Lighthouse Lab in Glasgow | Wellcome Sanger Institute for the COVID-19 Genomics UK (COG-UK) Consortium | Harper VanSteenhouse, Yumi Kasai, David Gray, Carol Clugston, Anna Dominiczak and Alex Alderton, Roberto Amato, Jeffrey Barrett, Sonia Goncalves, Ewan Harrison, David K. Jackson, Ian Johnston, Dominic Kwiatkowski, Cordelia Langford, John Sillitoe on behalf of the Wellcome Sanger Institute COVID-19 Surveillance Team |
| EPI_ISL_3203363, EPI_ISL_3203364, EPI_ISL_3203365 | Lighthouse Lab in Milton Keynes | Wellcome Sanger Institute for the COVID-19 Genomics UK (COG-UK) Consortium | The Lighthouse Lab in Milton Keynes and Alex Alderton, Roberto Amato, Jeffrey Barrett, Sonia Goncalves, Ewan Harrison, David K. Jackson, Ian Johnston, Dominic Kwiatkowski, Cordelia Langford, John Sillitoe on behalf of the Wellcome Sanger Institute COVID-19 Surveillance Team |
| EPI_ISL_3203366 | Lighthouse Lab in Glasgow | Wellcome Sanger Institute for the COVID-19 Genomics UK (COG-UK) Consortium | Harper VanSteenhouse, Yumi Kasai, David Gray, Carol Clugston, Anna Dominiczak and Alex Alderton, Roberto Amato, Jeffrey Barrett, Sonia Goncalves, Ewan Harrison, David K. Jackson, Ian Johnston, Dominic Kwiatkowski, Cordelia Langford, John Sillitoe on behalf of the Wellcome Sanger Institute COVID-19 Surveillance Team |
| EPI_ISL_3203367, EPI_ISL_3203368 | Lighthouse Lab in Milton Keynes | Wellcome Sanger Institute for the COVID-19 Genomics UK (COG-UK) Consortium | The Lighthouse Lab in Milton Keynes and Alex Alderton, Roberto Amato, Jeffrey Barrett, Sonia Goncalves, Ewan Harrison, David K. Jackson, Ian Johnston, Dominic Kwiatkowski, Cordelia Langford, John Sillitoe on behalf of the Wellcome Sanger Institute COVID-19 Surveillance Team |
| EPI_ISL_3203369 | Lighthouse Lab in Glasgow | Wellcome Sanger Institute for the COVID-19 Genomics UK (COG-UK) Consortium | Harper VanSteenhouse, Yumi Kasai, David Gray, Carol Clugston, Anna Dominiczak and Alex Alderton, Roberto Amato, Jeffrey Barrett, Sonia Goncalves, Ewan Harrison, David K. Jackson, Ian Johnston, Dominic Kwiatkowski, Cordelia Langford, John Sillitoe on behalf of the Wellcome Sanger Institute COVID-19 Surveillance Team |
| EPI_ISL_3203370, EPI_ISL_3203371 | Lighthouse Lab in Milton Keynes | Wellcome Sanger Institute for the COVID-19 Genomics UK (COG-UK) Consortium | The Lighthouse Lab in Milton Keynes and Alex Alderton, Roberto Amato, Jeffrey Barrett, Sonia Goncalves, Ewan Harrison, David K. Jackson, Ian Johnston, Dominic Kwiatkowski, Cordelia Langford, John Sillitoe on behalf of the Wellcome Sanger Institute COVID-19 Surveillance Team |
| EPI_ISL_3203372, EPI_ISL_3203373, | Lighthouse Lab in Glasgow | Wellcome Sanger Institute for the COVID-19 Genomics UK (COG-UK) Consortium | Harper VanSteenhouse, Yumi Kasai, David Gray, Carol Clugston, Anna Dominiczak and Alex Alderton, Roberto Amato, Jeffrey Barrett, Sonia Goncalves, Ewan Harrison, David K. Jackson, Ian Johnston, Dominic Kwiatkowski, Cordelia Langford, John Sillitoe on behalf of the Wellcome Sanger Institute COVID-19 Surveillance Team |

| | | | |
|---|---|---|---|
| EPI_ISL_3203374 | | | |
| EPI_ISL_3203375 | Lighthouse Lab in Milton Keynes | Wellcome Sanger Institute for the COVID-19 Genomics UK (COG-UK) Consortium | The Lighthouse Lab in Milton Keynes and Alex Alderton, Roberto Amato, Jeffrey Barrett, Sonia Goncalves, Ewan Harrison, David K. Jackson, Ian Johnston, Dominic Kwiatkowski, Cordelia Langford, John Sillitoe on behalf of the Wellcome Sanger Institute COVID-19 Surveillance Team |
| EPI_ISL_3203376 | Lighthouse Lab in Glasgow | Wellcome Sanger Institute for the COVID-19 Genomics UK (COG-UK) Consortium | Harper VanSteenhouse, Yumi Kasai, David Gray, Carol Clugston, Anna Dominiczak and Alex Alderton, Roberto Amato, Jeffrey Barrett, Sonia Goncalves, Ewan Harrison, David K. Jackson, Ian Johnston, Dominic Kwiatkowski, Cordelia Langford, John Sillitoe on behalf of the Wellcome Sanger Institute COVID-19 Surveillance Team |
| EPI_ISL_3203377, EPI_ISL_3203378, EPI_ISL_3203379 | Lighthouse Lab in Milton Keynes | Wellcome Sanger Institute for the COVID-19 Genomics UK (COG-UK) Consortium | The Lighthouse Lab in Milton Keynes and Alex Alderton, Roberto Amato, Jeffrey Barrett, Sonia Goncalves, Ewan Harrison, David K. Jackson, Ian Johnston, Dominic Kwiatkowski, Cordelia Langford, John Sillitoe on behalf of the Wellcome Sanger Institute COVID-19 Surveillance Team |
| EPI_ISL_3203380, EPI_ISL_3203381, EPI_ISL_3203382, EPI_ISL_3203383, EPI_ISL_3203384, EPI_ISL_3203385, EPI_ISL_3203386, EPI_ISL_3203387 | Lighthouse Lab in Glasgow | Wellcome Sanger Institute for the COVID-19 Genomics UK (COG-UK) Consortium | Harper VanSteenhouse, Yumi Kasai, David Gray, Carol Clugston, Anna Dominiczak and Alex Alderton, Roberto Amato, Jeffrey Barrett, Sonia Goncalves, Ewan Harrison, David K. Jackson, Ian Johnston, Dominic Kwiatkowski, Cordelia Langford, John Sillitoe on behalf of the Wellcome Sanger Institute COVID-19 Surveillance Team |
| EPI_ISL_3203388, EPI_ISL_3203389 | Lighthouse Lab in Milton Keynes | Wellcome Sanger Institute for the COVID-19 Genomics UK (COG-UK) Consortium | The Lighthouse Lab in Milton Keynes and Alex Alderton, Roberto Amato, Jeffrey Barrett, Sonia Goncalves, Ewan Harrison, David K. Jackson, Ian Johnston, Dominic Kwiatkowski, Cordelia Langford, John Sillitoe on behalf of the Wellcome Sanger Institute COVID-19 Surveillance Team |
| EPI_ISL_3203390 | Lighthouse Lab in Glasgow | Wellcome Sanger Institute for the COVID-19 Genomics UK (COG-UK) Consortium | Harper VanSteenhouse, Yumi Kasai, David Gray, Carol Clugston, Anna Dominiczak and Alex Alderton, Roberto Amato, Jeffrey Barrett, Sonia Goncalves, Ewan Harrison, David K. Jackson, Ian Johnston, Dominic Kwiatkowski, Cordelia Langford, John Sillitoe on behalf of the Wellcome Sanger Institute COVID-19 Surveillance Team |
| EPI_ISL_3203391, EPI_ISL_3203392, EPI_ISL_3203393 | Lighthouse Lab in Milton Keynes | Wellcome Sanger Institute for the COVID-19 Genomics UK (COG-UK) Consortium | The Lighthouse Lab in Milton Keynes and Alex Alderton, Roberto Amato, Jeffrey Barrett, Sonia Goncalves, Ewan Harrison, David K. Jackson, Ian Johnston, Dominic Kwiatkowski, Cordelia Langford, John Sillitoe on behalf of the Wellcome Sanger Institute COVID-19 Surveillance Team |
| EPI_ISL_3203394, EPI_ISL_3203395 | Lighthouse Lab in Glasgow | Wellcome Sanger Institute for the COVID-19 Genomics UK (COG-UK) Consortium | Harper VanSteenhouse, Yumi Kasai, David Gray, Carol Clugston, Anna Dominiczak and Alex Alderton, Roberto Amato, Jeffrey Barrett, Sonia Goncalves, Ewan Harrison, David K. Jackson, Ian Johnston, Dominic Kwiatkowski, Cordelia Langford, John Sillitoe on behalf of the Wellcome Sanger Institute COVID-19 Surveillance Team |
| EPI_ISL_3203396 | Lighthouse Lab in Milton Keynes | Wellcome Sanger Institute for the COVID-19 Genomics UK (COG-UK) Consortium | The Lighthouse Lab in Milton Keynes and Alex Alderton, Roberto Amato, Jeffrey Barrett, Sonia Goncalves, Ewan Harrison, David K. Jackson, Ian Johnston, Dominic Kwiatkowski, Cordelia Langford, John Sillitoe on behalf of the Wellcome Sanger Institute COVID-19 Surveillance Team |
| EPI_ISL_3203397 | Lighthouse Lab in Glasgow | Wellcome Sanger Institute for the COVID-19 Genomics UK (COG-UK) Consortium | Harper VanSteenhouse, Yumi Kasai, David Gray, Carol Clugston, Anna Dominiczak and Alex Alderton, Roberto Amato, Jeffrey Barrett, Sonia Goncalves, Ewan Harrison, David K. Jackson, Ian Johnston, Dominic Kwiatkowski, Cordelia Langford, John Sillitoe on behalf of the Wellcome Sanger Institute COVID-19 Surveillance Team |
| EPI_ISL_3203398, EPI_ISL_3203399 | Lighthouse Lab in Milton Keynes | Wellcome Sanger Institute for the COVID-19 Genomics UK (COG-UK) Consortium | The Lighthouse Lab in Milton Keynes and Alex Alderton, Roberto Amato, Jeffrey Barrett, Sonia Goncalves, Ewan Harrison, David K. Jackson, Ian Johnston, Dominic Kwiatkowski, Cordelia Langford, John Sillitoe on behalf of the Wellcome Sanger Institute COVID-19 Surveillance Team |
| EPI_ISL_3203400 | Lighthouse Lab in Glasgow | Wellcome Sanger Institute for the COVID-19 Genomics UK (COG-UK) Consortium | Harper VanSteenhouse, Yumi Kasai, David Gray, Carol Clugston, Anna Dominiczak and Alex Alderton, Roberto Amato, Jeffrey Barrett, Sonia Goncalves, Ewan Harrison, David K. Jackson, Ian Johnston, Dominic Kwiatkowski, Cordelia Langford, John Sillitoe on behalf of the Wellcome Sanger Institute COVID-19 Surveillance Team |
| EPI_ISL_3203401 | Lighthouse Lab in Milton Keynes | Wellcome Sanger Institute for the COVID-19 Genomics UK (COG-UK) Consortium | The Lighthouse Lab in Milton Keynes and Alex Alderton, Roberto Amato, Jeffrey Barrett, Sonia Goncalves, Ewan Harrison, David K. Jackson, Ian Johnston, Dominic Kwiatkowski, Cordelia Langford, John Sillitoe on behalf of the Wellcome Sanger Institute COVID-19 Surveillance Team |
| EPI_ISL_3203402 | Lighthouse Lab in Glasgow | Wellcome Sanger Institute for the COVID-19 Genomics UK (COG-UK) Consortium | Harper VanSteenhouse, Yumi Kasai, David Gray, Carol Clugston, Anna Dominiczak and Alex Alderton, Roberto Amato, Jeffrey Barrett, Sonia Goncalves, Ewan Harrison, David K. Jackson, Ian Johnston, Dominic Kwiatkowski, Cordelia Langford, John Sillitoe on behalf of the Wellcome Sanger Institute COVID-19 Surveillance Team |
| EPI_ISL_3203403, EPI_ISL_3203404 | Lighthouse Lab in Milton Keynes | Wellcome Sanger Institute for the COVID-19 Genomics UK (COG-UK) Consortium | The Lighthouse Lab in Milton Keynes and Alex Alderton, Roberto Amato, Jeffrey Barrett, Sonia Goncalves, Ewan Harrison, David K. Jackson, Ian Johnston, Dominic Kwiatkowski, Cordelia Langford, John Sillitoe on behalf of the Wellcome Sanger Institute COVID-19 Surveillance Team |
| EPI_ISL_3203405, EPI_ISL_3203406, EPI_ISL_3203407, EPI_ISL_3203408, EPI_ISL_3203409 | Lighthouse Lab in Glasgow | Wellcome Sanger Institute for the COVID-19 Genomics UK (COG-UK) Consortium | Harper VanSteenhouse, Yumi Kasai, David Gray, Carol Clugston, Anna Dominiczak and Alex Alderton, Roberto Amato, Jeffrey Barrett, Sonia Goncalves, Ewan Harrison, David K. Jackson, Ian Johnston, Dominic Kwiatkowski, Cordelia Langford, John Sillitoe on behalf of the Wellcome Sanger Institute COVID-19 Surveillance Team |
| EPI_ISL_3203410, EPI_ISL_3203411 | Lighthouse Lab in Milton Keynes | Wellcome Sanger Institute for the COVID-19 Genomics UK (COG-UK) Consortium | The Lighthouse Lab in Milton Keynes and Alex Alderton, Roberto Amato, Jeffrey Barrett, Sonia Goncalves, Ewan Harrison, David K. Jackson, Ian Johnston, Dominic Kwiatkowski, Cordelia Langford, John Sillitoe on behalf of the Wellcome Sanger Institute COVID-19 Surveillance Team |
| EPI_ISL_3203412 | Lighthouse Lab in Glasgow | Wellcome Sanger Institute for the COVID-19 Genomics UK (COG-UK) Consortium | Harper VanSteenhouse, Yumi Kasai, David Gray, Carol Clugston, Anna Dominiczak and Alex Alderton, Roberto Amato, Jeffrey Barrett, Sonia Goncalves, Ewan Harrison, David K. Jackson, Ian Johnston, Dominic Kwiatkowski, Cordelia Langford, John Sillitoe on behalf of the Wellcome Sanger Institute COVID-19 Surveillance Team |
| EPI_ISL_3203413, EPI_ISL_3203414, EPI_ISL_3203415, EPI_ISL_3203416, EPI_ISL_3203417, EPI_ISL_3203418, EPI_ISL_3203419, EPI_ISL_3203420, EPI_ISL_3203421 | Lighthouse Lab in Milton Keynes | Wellcome Sanger Institute for the COVID-19 Genomics UK (COG-UK) Consortium | The Lighthouse Lab in Milton Keynes and Alex Alderton, Roberto Amato, Jeffrey Barrett, Sonia Goncalves, Ewan Harrison, David K. Jackson, Ian Johnston, Dominic Kwiatkowski, Cordelia Langford, John Sillitoe on behalf of the Wellcome Sanger Institute COVID-19 Surveillance Team |
| EPI_ISL_3203422 | Lighthouse Lab in Glasgow | Wellcome Sanger Institute for the COVID-19 Genomics UK (COG-UK) Consortium | Harper VanSteenhouse, Yumi Kasai, David Gray, Carol Clugston, Anna Dominiczak and Alex Alderton, Roberto Amato, Jeffrey Barrett, Sonia Goncalves, Ewan Harrison, David K. Jackson, Ian Johnston, Dominic Kwiatkowski, Cordelia Langford, John Sillitoe on behalf of the Wellcome Sanger Institute COVID-19 Surveillance Team |
| EPI_ISL_3203423 | Lighthouse Lab in Milton Keynes | Wellcome Sanger Institute for the COVID-19 Genomics UK (COG-UK) Consortium | The Lighthouse Lab in Milton Keynes and Alex Alderton, Roberto Amato, Jeffrey Barrett, Sonia Goncalves, Ewan Harrison, David K. Jackson, Ian Johnston, Dominic Kwiatkowski, Cordelia Langford, John Sillitoe on behalf of the Wellcome Sanger Institute COVID-19 Surveillance Team |
| EPI_ISL_3203424 | Lighthouse Lab in Glasgow | Wellcome Sanger Institute for the COVID-19 Genomics UK (COG-UK) Consortium | Harper VanSteenhouse, Yumi Kasai, David Gray, Carol Clugston, Anna Dominiczak and Alex Alderton, Roberto Amato, Jeffrey Barrett, Sonia Goncalves, Ewan Harrison, David K. Jackson, Ian Johnston, Dominic Kwiatkowski, Cordelia Langford, John Sillitoe on behalf of the Wellcome Sanger Institute COVID-19 Surveillance Team |
| EPI_ISL_3203425 | Lighthouse Lab in Milton Keynes | Wellcome Sanger Institute for the COVID-19 Genomics UK (COG-UK) Consortium | The Lighthouse Lab in Milton Keynes and Alex Alderton, Roberto Amato, Jeffrey Barrett, Sonia Goncalves, Ewan Harrison, David K. Jackson, Ian Johnston, Dominic Kwiatkowski, Cordelia Langford, John Sillitoe on behalf of the Wellcome Sanger Institute COVID-19 Surveillance Team |
| EPI_ISL_3203426 | Lighthouse Lab in Glasgow | Wellcome Sanger Institute for the COVID-19 Genomics UK (COG-UK) Consortium | Harper VanSteenhouse, Yumi Kasai, David Gray, Carol Clugston, Anna Dominiczak and Alex Alderton, Roberto Amato, Jeffrey Barrett, Sonia Goncalves, Ewan Harrison, David K. Jackson, Ian Johnston, Dominic Kwiatkowski, Cordelia Langford, John Sillitoe on behalf of the Wellcome Sanger Institute COVID-19 Surveillance Team |
| EPI_ISL_3203427 | Lighthouse Lab in Milton Keynes | Wellcome Sanger Institute for the COVID-19 Genomics UK (COG-UK) Consortium | The Lighthouse Lab in Milton Keynes and Alex Alderton, Roberto Amato, Jeffrey Barrett, Sonia Goncalves, Ewan Harrison, David K. Jackson, Ian Johnston, Dominic Kwiatkowski, Cordelia Langford, John Sillitoe on behalf of the Wellcome Sanger Institute COVID-19 Surveillance Team |
| EPI_ISL_3203428 | Lighthouse Lab in Glasgow | Wellcome Sanger Institute for the COVID-19 Genomics UK (COG-UK) Consortium | Harper VanSteenhouse, Yumi Kasai, David Gray, Carol Clugston, Anna Dominiczak and Alex Alderton, Roberto Amato, Jeffrey Barrett, Sonia Goncalves, Ewan Harrison, David K. Jackson, Ian Johnston, Dominic Kwiatkowski, Cordelia Langford, John Sillitoe on behalf of the Wellcome Sanger Institute COVID-19 Surveillance Team |
| EPI_ISL_3203429, EPI_ISL_3203430 | Lighthouse Lab in Milton Keynes | Wellcome Sanger Institute for the COVID-19 Genomics UK (COG-UK) Consortium | The Lighthouse Lab in Milton Keynes and Alex Alderton, Roberto Amato, Jeffrey Barrett, Sonia Goncalves, Ewan Harrison, David K. Jackson, Ian Johnston, Dominic Kwiatkowski, Cordelia Langford, John Sillitoe on behalf of the Wellcome Sanger Institute COVID-19 Surveillance Team |
| EPI_ISL_3203431, EPI_ISL_3203432 | Lighthouse Lab in Glasgow | Wellcome Sanger Institute for the COVID-19 Genomics UK (COG-UK) Consortium | Harper VanSteenhouse, Yumi Kasai, David Gray, Carol Clugston, Anna Dominiczak and Alex Alderton, Roberto Amato, Jeffrey Barrett, Sonia Goncalves, Ewan Harrison, David K. Jackson, Ian Johnston, Dominic Kwiatkowski, Cordelia Langford, John Sillitoe on behalf of the Wellcome Sanger Institute COVID-19 Surveillance Team |
| EPI_ISL_3203433, EPI_ISL_3203434 | Lighthouse Lab in Milton Keynes | Wellcome Sanger Institute for the COVID-19 Genomics UK (COG-UK) Consortium | The Lighthouse Lab in Milton Keynes and Alex Alderton, Roberto Amato, Jeffrey Barrett, Sonia Goncalves, Ewan Harrison, David K. Jackson, Ian Johnston, Dominic Kwiatkowski, Cordelia Langford, John Sillitoe on behalf of the Wellcome Sanger Institute COVID-19 Surveillance Team |
| EPI_ISL_3203435, EPI_ISL_3203436 | Lighthouse Lab in Glasgow | Wellcome Sanger Institute for the COVID-19 Genomics UK (COG-UK) Consortium | Harper VanSteenhouse, Yumi Kasai, David Gray, Carol Clugston, Anna Dominiczak and Alex Alderton, Roberto Amato, Jeffrey Barrett, Sonia Goncalves, Ewan Harrison, David K. Jackson, Ian Johnston, Dominic Kwiatkowski, Cordelia Langford, John Sillitoe on behalf of the Wellcome Sanger Institute COVID-19 Surveillance Team |
| EPI_ISL_3203437, EPI_ISL_3203438, EPI_ISL_3203439, EPI_ISL_3203440 | Lighthouse Lab in Milton Keynes | Wellcome Sanger Institute for the COVID-19 Genomics UK (COG-UK) Consortium | The Lighthouse Lab in Milton Keynes and Alex Alderton, Roberto Amato, Jeffrey Barrett, Sonia Goncalves, Ewan Harrison, David K. Jackson, Ian Johnston, Dominic Kwiatkowski, Cordelia Langford, John Sillitoe on behalf of the Wellcome Sanger Institute COVID-19 Surveillance Team |
| EPI_ISL_3203441, EPI_ISL_3203442 | Lighthouse Lab in Glasgow | Wellcome Sanger Institute for the COVID-19 Genomics UK (COG-UK) Consortium | Harper VanSteenhouse, Yumi Kasai, David Gray, Carol Clugston, Anna Dominiczak and Alex Alderton, Roberto Amato, Jeffrey Barrett, Sonia Goncalves, Ewan Harrison, David K. Jackson, Ian Johnston, Dominic Kwiatkowski, Cordelia Langford, John Sillitoe on behalf of the Wellcome Sanger Institute COVID-19 Surveillance Team |
| EPI_ISL_3203443 | Lighthouse Lab in Milton Keynes | Wellcome Sanger Institute for the COVID-19 Genomics UK (COG-UK) Consortium | The Lighthouse Lab in Milton Keynes and Alex Alderton, Roberto Amato, Jeffrey Barrett, Sonia Goncalves, Ewan Harrison, David K. Jackson, Ian Johnston, Dominic Kwiatkowski, Cordelia Langford, John Sillitoe on behalf of the Wellcome Sanger Institute COVID-19 Surveillance Team |
| EPI_ISL_3203444 | Lighthouse Lab in Glasgow | Wellcome Sanger Institute for the COVID-19 Genomics UK (COG-UK) Consortium | Harper VanSteenhouse, Yumi Kasai, David Gray, Carol Clugston, Anna Dominiczak and Alex Alderton, Roberto Amato, Jeffrey Barrett, Sonia Goncalves, Ewan Harrison, David K. Jackson, Ian Johnston, Dominic Kwiatkowski, Cordelia Langford, John Sillitoe on behalf of the Wellcome Sanger Institute COVID-19 Surveillance Team |
| EPI_ISL_3203445 | Lighthouse Lab in Milton Keynes | Wellcome Sanger Institute for the COVID-19 Genomics UK (COG-UK) Consortium | The Lighthouse Lab in Milton Keynes and Alex Alderton, Roberto Amato, Jeffrey Barrett, Sonia Goncalves, Ewan Harrison, David K. Jackson, Ian Johnston, Dominic Kwiatkowski, Cordelia Langford, John Sillitoe on behalf of the Wellcome Sanger Institute COVID-19 Surveillance Team |
| EPI_ISL_3203446, EPI_ISL_3203447, EPI_ISL_3203448 | Lighthouse Lab in Glasgow | Wellcome Sanger Institute for the COVID-19 Genomics UK (COG-UK) Consortium | Harper VanSteenhouse, Yumi Kasai, David Gray, Carol Clugston, Anna Dominiczak and Alex Alderton, Roberto Amato, Jeffrey Barrett, Sonia Goncalves, Ewan Harrison, David K. Jackson, Ian Johnston, Dominic Kwiatkowski, Cordelia Langford, John Sillitoe on behalf of the Wellcome Sanger Institute COVID-19 Surveillance Team |

| | | | |
|---|---|---|---|
| EPI_ISL_3203449, EPI_ISL_3203450, EPI_ISL_3203451, EPI_ISL_3203452, EPI_ISL_3203453 | Lighthouse Lab in Milton Keynes | Wellcome Sanger Institute for the COVID-19 Genomics UK (COG-UK) Consortium | The Lighthouse Lab in Milton Keynes and Alex Alderton, Roberto Amato, Jeffrey Barrett, Sonia Goncalves, Ewan Harrison, David K. Jackson, Ian Johnston, Dominic Kwiatkowski, Cordelia Langford, John Sillitoe on behalf of the Wellcome Sanger Institute COVID-19 Surveillance Team |
| EPI_ISL_3203454, EPI_ISL_3203455 | Lighthouse Lab in Glasgow | Wellcome Sanger Institute for the COVID-19 Genomics UK (COG-UK) Consortium | Harper VanSteenhouse, Yumi Kasai, David Gray, Carol Clugston, Anna Dominiczak and Alex Alderton, Roberto Amato, Jeffrey Barrett, Sonia Goncalves, Ewan Harrison, David K. Jackson, Ian Johnston, Dominic Kwiatkowski, Cordelia Langford, John Sillitoe on behalf of the Wellcome Sanger Institute COVID-19 Surveillance Team |
| EPI_ISL_3203456, EPI_ISL_3203457 | Lighthouse Lab in Milton Keynes | Wellcome Sanger Institute for the COVID-19 Genomics UK (COG-UK) Consortium | The Lighthouse Lab in Milton Keynes and Alex Alderton, Roberto Amato, Jeffrey Barrett, Sonia Goncalves, Ewan Harrison, David K. Jackson, Ian Johnston, Dominic Kwiatkowski, Cordelia Langford, John Sillitoe on behalf of the Wellcome Sanger Institute COVID-19 Surveillance Team |
| EPI_ISL_3203458 | Lighthouse Lab in Glasgow | Wellcome Sanger Institute for the COVID-19 Genomics UK (COG-UK) Consortium | Harper VanSteenhouse, Yumi Kasai, David Gray, Carol Clugston, Anna Dominiczak and Alex Alderton, Roberto Amato, Jeffrey Barrett, Sonia Goncalves, Ewan Harrison, David K. Jackson, Ian Johnston, Dominic Kwiatkowski, Cordelia Langford, John Sillitoe on behalf of the Wellcome Sanger Institute COVID-19 Surveillance Team |
| EPI_ISL_3203459, EPI_ISL_3203460, EPI_ISL_3203461 | Lighthouse Lab in Milton Keynes | Wellcome Sanger Institute for the COVID-19 Genomics UK (COG-UK) Consortium | The Lighthouse Lab in Milton Keynes and Alex Alderton, Roberto Amato, Jeffrey Barrett, Sonia Goncalves, Ewan Harrison, David K. Jackson, Ian Johnston, Dominic Kwiatkowski, Cordelia Langford, John Sillitoe on behalf of the Wellcome Sanger Institute COVID-19 Surveillance Team |
| EPI_ISL_3203462 | Lighthouse Lab in Glasgow | Wellcome Sanger Institute for the COVID-19 Genomics UK (COG-UK) Consortium | Harper VanSteenhouse, Yumi Kasai, David Gray, Carol Clugston, Anna Dominiczak and Alex Alderton, Roberto Amato, Jeffrey Barrett, Sonia Goncalves, Ewan Harrison, David K. Jackson, Ian Johnston, Dominic Kwiatkowski, Cordelia Langford, John Sillitoe on behalf of the Wellcome Sanger Institute COVID-19 Surveillance Team |
| EPI_ISL_3203463, EPI_ISL_3203464 | Lighthouse Lab in Milton Keynes | Wellcome Sanger Institute for the COVID-19 Genomics UK (COG-UK) Consortium | The Lighthouse Lab in Milton Keynes and Alex Alderton, Roberto Amato, Jeffrey Barrett, Sonia Goncalves, Ewan Harrison, David K. Jackson, Ian Johnston, Dominic Kwiatkowski, Cordelia Langford, John Sillitoe on behalf of the Wellcome Sanger Institute COVID-19 Surveillance Team |
| EPI_ISL_3203465, EPI_ISL_3203466 | Lighthouse Lab in Glasgow | Wellcome Sanger Institute for the COVID-19 Genomics UK (COG-UK) Consortium | Harper VanSteenhouse, Yumi Kasai, David Gray, Carol Clugston, Anna Dominiczak and Alex Alderton, Roberto Amato, Jeffrey Barrett, Sonia Goncalves, Ewan Harrison, David K. Jackson, Ian Johnston, Dominic Kwiatkowski, Cordelia Langford, John Sillitoe on behalf of the Wellcome Sanger Institute COVID-19 Surveillance Team |
| EPI_ISL_3203467 | Lighthouse Lab in Milton Keynes | Wellcome Sanger Institute for the COVID-19 Genomics UK (COG-UK) Consortium | The Lighthouse Lab in Milton Keynes and Alex Alderton, Roberto Amato, Jeffrey Barrett, Sonia Goncalves, Ewan Harrison, David K. Jackson, Ian Johnston, Dominic Kwiatkowski, Cordelia Langford, John Sillitoe on behalf of the Wellcome Sanger Institute COVID-19 Surveillance Team |
| EPI_ISL_3203468 | Lighthouse Lab in Glasgow | Wellcome Sanger Institute for the COVID-19 Genomics UK (COG-UK) Consortium | Harper VanSteenhouse, Yumi Kasai, David Gray, Carol Clugston, Anna Dominiczak and Alex Alderton, Roberto Amato, Jeffrey Barrett, Sonia Goncalves, Ewan Harrison, David K. Jackson, Ian Johnston, Dominic Kwiatkowski, Cordelia Langford, John Sillitoe on behalf of the Wellcome Sanger Institute COVID-19 Surveillance Team |
| EPI_ISL_3203469, EPI_ISL_3203470, EPI_ISL_3203471, EPI_ISL_3203472, EPI_ISL_3203473 | Lighthouse Lab in Milton Keynes | Wellcome Sanger Institute for the COVID-19 Genomics UK (COG-UK) Consortium | The Lighthouse Lab in Milton Keynes and Alex Alderton, Roberto Amato, Jeffrey Barrett, Sonia Goncalves, Ewan Harrison, David K. Jackson, Ian Johnston, Dominic Kwiatkowski, Cordelia Langford, John Sillitoe on behalf of the Wellcome Sanger Institute COVID-19 Surveillance Team |
| EPI_ISL_3203474 | Lighthouse Lab in Glasgow | Wellcome Sanger Institute for the COVID-19 Genomics UK (COG-UK) Consortium | Harper VanSteenhouse, Yumi Kasai, David Gray, Carol Clugston, Anna Dominiczak and Alex Alderton, Roberto Amato, Jeffrey Barrett, Sonia Goncalves, Ewan Harrison, David K. Jackson, Ian Johnston, Dominic Kwiatkowski, Cordelia Langford, John Sillitoe on behalf of the Wellcome Sanger Institute COVID-19 Surveillance Team |
| EPI_ISL_3203475 | Lighthouse Lab in Milton Keynes | Wellcome Sanger Institute for the COVID-19 Genomics UK (COG-UK) Consortium | The Lighthouse Lab in Milton Keynes and Alex Alderton, Roberto Amato, Jeffrey Barrett, Sonia Goncalves, Ewan Harrison, David K. Jackson, Ian Johnston, Dominic Kwiatkowski, Cordelia Langford, John Sillitoe on behalf of the Wellcome Sanger Institute COVID-19 Surveillance Team |
| EPI_ISL_3203476 | Lighthouse Lab in Glasgow | Wellcome Sanger Institute for the COVID-19 Genomics UK (COG-UK) Consortium | Harper VanSteenhouse, Yumi Kasai, David Gray, Carol Clugston, Anna Dominiczak and Alex Alderton, Roberto Amato, Jeffrey Barrett, Sonia Goncalves, Ewan Harrison, David K. Jackson, Ian Johnston, Dominic Kwiatkowski, Cordelia Langford, John Sillitoe on behalf of the Wellcome Sanger Institute COVID-19 Surveillance Team |
| EPI_ISL_3203477, EPI_ISL_3203478 | Lighthouse Lab in Milton Keynes | Wellcome Sanger Institute for the COVID-19 Genomics UK (COG-UK) Consortium | The Lighthouse Lab in Milton Keynes and Alex Alderton, Roberto Amato, Jeffrey Barrett, Sonia Goncalves, Ewan Harrison, David K. Jackson, Ian Johnston, Dominic Kwiatkowski, Cordelia Langford, John Sillitoe on behalf of the Wellcome Sanger Institute COVID-19 Surveillance Team |
| EPI_ISL_3203479 | Lighthouse Lab in Glasgow | Wellcome Sanger Institute for the COVID-19 Genomics UK (COG-UK) Consortium | Harper VanSteenhouse, Yumi Kasai, David Gray, Carol Clugston, Anna Dominiczak and Alex Alderton, Roberto Amato, Jeffrey Barrett, Sonia Goncalves, Ewan Harrison, David K. Jackson, Ian Johnston, Dominic Kwiatkowski, Cordelia Langford, John Sillitoe on behalf of the Wellcome Sanger Institute COVID-19 Surveillance Team |
| EPI_ISL_3203480, EPI_ISL_3203481, EPI_ISL_3203482, EPI_ISL_3203483 | Lighthouse Lab in Milton Keynes | Wellcome Sanger Institute for the COVID-19 Genomics UK (COG-UK) Consortium | The Lighthouse Lab in Milton Keynes and Alex Alderton, Roberto Amato, Jeffrey Barrett, Sonia Goncalves, Ewan Harrison, David K. Jackson, Ian Johnston, Dominic Kwiatkowski, Cordelia Langford, John Sillitoe on behalf of the Wellcome Sanger Institute COVID-19 Surveillance Team |
| EPI_ISL_3203484, EPI_ISL_3203485 | Lighthouse Lab in Glasgow | Wellcome Sanger Institute for the COVID-19 Genomics UK (COG-UK) Consortium | Harper VanSteenhouse, Yumi Kasai, David Gray, Carol Clugston, Anna Dominiczak and Alex Alderton, Roberto Amato, Jeffrey Barrett, Sonia Goncalves, Ewan Harrison, David K. Jackson, Ian Johnston, Dominic Kwiatkowski, Cordelia Langford, John Sillitoe on behalf of the Wellcome Sanger Institute COVID-19 Surveillance Team |
| EPI_ISL_3203486, EPI_ISL_3203487 | Lighthouse Lab in Milton Keynes | Wellcome Sanger Institute for the COVID-19 Genomics UK (COG-UK) Consortium | The Lighthouse Lab in Milton Keynes and Alex Alderton, Roberto Amato, Jeffrey Barrett, Sonia Goncalves, Ewan Harrison, David K. Jackson, Ian Johnston, Dominic Kwiatkowski, Cordelia Langford, John Sillitoe on behalf of the Wellcome Sanger Institute COVID-19 Surveillance Team |
| EPI_ISL_3203488 | Lighthouse Lab in Glasgow | Wellcome Sanger Institute for the COVID-19 Genomics UK (COG-UK) Consortium | Harper VanSteenhouse, Yumi Kasai, David Gray, Carol Clugston, Anna Dominiczak and Alex Alderton, Roberto Amato, Jeffrey Barrett, Sonia Goncalves, Ewan Harrison, David K. Jackson, Ian Johnston, Dominic Kwiatkowski, Cordelia Langford, John Sillitoe on behalf of the Wellcome Sanger Institute COVID-19 Surveillance Team |
| EPI_ISL_3203489 | Lighthouse Lab in Milton Keynes | Wellcome Sanger Institute for the COVID-19 Genomics UK (COG-UK) Consortium | The Lighthouse Lab in Milton Keynes and Alex Alderton, Roberto Amato, Jeffrey Barrett, Sonia Goncalves, Ewan Harrison, David K. Jackson, Ian Johnston, Dominic Kwiatkowski, Cordelia Langford, John Sillitoe on behalf of the Wellcome Sanger Institute COVID-19 Surveillance Team |
| EPI_ISL_3203490 | Lighthouse Lab in Glasgow | Wellcome Sanger Institute for the COVID-19 Genomics UK (COG-UK) Consortium | Harper VanSteenhouse, Yumi Kasai, David Gray, Carol Clugston, Anna Dominiczak and Alex Alderton, Roberto Amato, Jeffrey Barrett, Sonia Goncalves, Ewan Harrison, David K. Jackson, Ian Johnston, Dominic Kwiatkowski, Cordelia Langford, John Sillitoe on behalf of the Wellcome Sanger Institute COVID-19 Surveillance Team |
| EPI_ISL_3203491, EPI_ISL_3203492 | Lighthouse Lab in Milton Keynes | Wellcome Sanger Institute for the COVID-19 Genomics UK (COG-UK) Consortium | The Lighthouse Lab in Milton Keynes and Alex Alderton, Roberto Amato, Jeffrey Barrett, Sonia Goncalves, Ewan Harrison, David K. Jackson, Ian Johnston, Dominic Kwiatkowski, Cordelia Langford, John Sillitoe on behalf of the Wellcome Sanger Institute COVID-19 Surveillance Team |
| EPI_ISL_3203493 | Lighthouse Lab in Glasgow | Wellcome Sanger Institute for the COVID-19 Genomics UK (COG-UK) Consortium | Harper VanSteenhouse, Yumi Kasai, David Gray, Carol Clugston, Anna Dominiczak and Alex Alderton, Roberto Amato, Jeffrey Barrett, Sonia Goncalves, Ewan Harrison, David K. Jackson, Ian Johnston, Dominic Kwiatkowski, Cordelia Langford, John Sillitoe on behalf of the Wellcome Sanger Institute COVID-19 Surveillance Team |
| EPI_ISL_3203494, EPI_ISL_3203495, EPI_ISL_3203496, EPI_ISL_3203497, EPI_ISL_3203498 | Lighthouse Lab in Milton Keynes | Wellcome Sanger Institute for the COVID-19 Genomics UK (COG-UK) Consortium | The Lighthouse Lab in Milton Keynes and Alex Alderton, Roberto Amato, Jeffrey Barrett, Sonia Goncalves, Ewan Harrison, David K. Jackson, Ian Johnston, Dominic Kwiatkowski, Cordelia Langford, John Sillitoe on behalf of the Wellcome Sanger Institute COVID-19 Surveillance Team |
| EPI_ISL_3203499 | Lighthouse Lab in Glasgow | Wellcome Sanger Institute for the COVID-19 Genomics UK (COG-UK) Consortium | Harper VanSteenhouse, Yumi Kasai, David Gray, Carol Clugston, Anna Dominiczak and Alex Alderton, Roberto Amato, Jeffrey Barrett, Sonia Goncalves, Ewan Harrison, David K. Jackson, Ian Johnston, Dominic Kwiatkowski, Cordelia Langford, John Sillitoe on behalf of the Wellcome Sanger Institute COVID-19 Surveillance Team |
| EPI_ISL_3203500, EPI_ISL_3203501, EPI_ISL_3203502 | Lighthouse Lab in Milton Keynes | Wellcome Sanger Institute for the COVID-19 Genomics UK (COG-UK) Consortium | The Lighthouse Lab in Milton Keynes and Alex Alderton, Roberto Amato, Jeffrey Barrett, Sonia Goncalves, Ewan Harrison, David K. Jackson, Ian Johnston, Dominic Kwiatkowski, Cordelia Langford, John Sillitoe on behalf of the Wellcome Sanger Institute COVID-19 Surveillance Team |
| EPI_ISL_3203503, EPI_ISL_3203504, EPI_ISL_3203505 | Lighthouse Lab in Glasgow | Wellcome Sanger Institute for the COVID-19 Genomics UK (COG-UK) Consortium | Harper VanSteenhouse, Yumi Kasai, David Gray, Carol Clugston, Anna Dominiczak and Alex Alderton, Roberto Amato, Jeffrey Barrett, Sonia Goncalves, Ewan Harrison, David K. Jackson, Ian Johnston, Dominic Kwiatkowski, Cordelia Langford, John Sillitoe on behalf of the Wellcome Sanger Institute COVID-19 Surveillance Team |
| EPI_ISL_3203506 | Lighthouse Lab in Milton Keynes | Wellcome Sanger Institute for the COVID-19 Genomics UK (COG-UK) Consortium | The Lighthouse Lab in Milton Keynes and Alex Alderton, Roberto Amato, Jeffrey Barrett, Sonia Goncalves, Ewan Harrison, David K. Jackson, Ian Johnston, Dominic Kwiatkowski, Cordelia Langford, John Sillitoe on behalf of the Wellcome Sanger Institute COVID-19 Surveillance Team |
| EPI_ISL_3203507 | Lighthouse Lab in Glasgow | Wellcome Sanger Institute for the COVID-19 Genomics UK (COG-UK) Consortium | Harper VanSteenhouse, Yumi Kasai, David Gray, Carol Clugston, Anna Dominiczak and Alex Alderton, Roberto Amato, Jeffrey Barrett, Sonia Goncalves, Ewan Harrison, David K. Jackson, Ian Johnston, Dominic Kwiatkowski, Cordelia Langford, John Sillitoe on behalf of the Wellcome Sanger Institute COVID-19 Surveillance Team |
| EPI_ISL_3203508, EPI_ISL_3203509, EPI_ISL_3203510 | Lighthouse Lab in Milton Keynes | Wellcome Sanger Institute for the COVID-19 Genomics UK (COG-UK) Consortium | The Lighthouse Lab in Milton Keynes and Alex Alderton, Roberto Amato, Jeffrey Barrett, Sonia Goncalves, Ewan Harrison, David K. Jackson, Ian Johnston, Dominic Kwiatkowski, Cordelia Langford, John Sillitoe on behalf of the Wellcome Sanger Institute COVID-19 Surveillance Team |
| EPI_ISL_3203511 | Lighthouse Lab in Glasgow | Wellcome Sanger Institute for the COVID-19 Genomics UK (COG-UK) Consortium | Harper VanSteenhouse, Yumi Kasai, David Gray, Carol Clugston, Anna Dominiczak and Alex Alderton, Roberto Amato, Jeffrey Barrett, Sonia Goncalves, Ewan Harrison, David K. Jackson, Ian Johnston, Dominic Kwiatkowski, Cordelia Langford, John Sillitoe on behalf of the Wellcome Sanger Institute COVID-19 Surveillance Team |
| EPI_ISL_3203512, EPI_ISL_3203513 | Lighthouse Lab in Milton Keynes | Wellcome Sanger Institute for the COVID-19 Genomics UK (COG-UK) Consortium | The Lighthouse Lab in Milton Keynes and Alex Alderton, Roberto Amato, Jeffrey Barrett, Sonia Goncalves, Ewan Harrison, David K. Jackson, Ian Johnston, Dominic Kwiatkowski, Cordelia Langford, John Sillitoe on behalf of the Wellcome Sanger Institute COVID-19 Surveillance Team |
| EPI_ISL_3203514 | Lighthouse Lab in Glasgow | Wellcome Sanger Institute for the COVID-19 Genomics UK (COG-UK) Consortium | Harper VanSteenhouse, Yumi Kasai, David Gray, Carol Clugston, Anna Dominiczak and Alex Alderton, Roberto Amato, Jeffrey Barrett, Sonia Goncalves, Ewan Harrison, David K. Jackson, Ian Johnston, Dominic Kwiatkowski, Cordelia Langford, John Sillitoe on behalf of the Wellcome Sanger Institute COVID-19 Surveillance Team |
| EPI_ISL_3203515, EPI_ISL_3203516, EPI_ISL_3203517, EPI_ISL_3203518, EPI_ISL_3203519, EPI_ISL_3203520 | Lighthouse Lab in Milton Keynes | Wellcome Sanger Institute for the COVID-19 Genomics UK (COG-UK) Consortium | The Lighthouse Lab in Milton Keynes and Alex Alderton, Roberto Amato, Jeffrey Barrett, Sonia Goncalves, Ewan Harrison, David K. Jackson, Ian Johnston, Dominic Kwiatkowski, Cordelia Langford, John Sillitoe on behalf of the Wellcome Sanger Institute COVID-19 Surveillance Team |
| EPI_ISL_3203521 | Lighthouse Lab in Glasgow | Wellcome Sanger Institute for the COVID-19 Genomics UK (COG-UK) Consortium | Harper VanSteenhouse, Yumi Kasai, David Gray, Carol Clugston, Anna Dominiczak and Alex Alderton, Roberto Amato, Jeffrey Barrett, Sonia Goncalves, Ewan Harrison, David K. Jackson, Ian Johnston, Dominic Kwiatkowski, Cordelia Langford, John Sillitoe on behalf of the Wellcome Sanger Institute COVID-19 Surveillance Team |
| EPI_ISL_3203522, EPI_ISL_3203523 | Lighthouse Lab in Milton Keynes | Wellcome Sanger Institute for the COVID-19 Genomics UK (COG-UK) Consortium | The Lighthouse Lab in Milton Keynes and Alex Alderton, Roberto Amato, Jeffrey Barrett, Sonia Goncalves, Ewan Harrison, David K. Jackson, Ian Johnston, Dominic Kwiatkowski, Cordelia Langford, John Sillitoe on behalf of the Wellcome Sanger Institute COVID-19 Surveillance Team |
| EPI_ISL_3203524 | Lighthouse Lab in Glasgow | Wellcome Sanger Institute for the COVID-19 Genomics UK (COG-UK) | Harper VanSteenhouse, Yumi Kasai, David Gray, Carol Clugston, Anna Dominiczak and Alex Alderton, Roberto Amato, Jeffrey Barrett, Sonia Goncalves, Ewan Harrison, David K. Jackson, Ian Johnston, Dominic Kwiatkowski, Cordelia Langford, |

| | | | |
|---|---|---|---|
| | | Consortium | John Sillitoe on behalf of the Wellcome Sanger Institute COVID-19 Surveillance Team |
| EPI_ISL_3203525, EPI_ISL_3203526, EPI_ISL_3203527, EPI_ISL_3203528, EPI_ISL_3203529, EPI_ISL_3203530, EPI_ISL_3203531, EPI_ISL_3203532, EPI_ISL_3203533 | Lighthouse Lab in Milton Keynes | Wellcome Sanger Institute for the COVID-19 Genomics UK (COG-UK) Consortium | The Lighthouse Lab in Milton Keynes and Alex Alderton, Roberto Amato, Jeffrey Barrett, Sonia Goncalves, Ewan Harrison, David K. Jackson, Ian Johnston, Dominic Kwiatkowski, Cordelia Langford, John Sillitoe on behalf of the Wellcome Sanger Institute COVID-19 Surveillance Team |
| EPI_ISL_3203534 | Lighthouse Lab in Glasgow | Wellcome Sanger Institute for the COVID-19 Genomics UK (COG-UK) Consortium | Harper VanSteenhouse, Yumi Kasai, David Gray, Carol Clugston, Anna Dominiczak and Alex Alderton, Roberto Amato, Jeffrey Barrett, Sonia Goncalves, Ewan Harrison, David K. Jackson, Ian Johnston, Dominic Kwiatkowski, Cordelia Langford, John Sillitoe on behalf of the Wellcome Sanger Institute COVID-19 Surveillance Team |
| EPI_ISL_3203535, EPI_ISL_3203536, EPI_ISL_3203537, EPI_ISL_3203538, EPI_ISL_3203539, EPI_ISL_3203540, EPI_ISL_3203541, EPI_ISL_3203542, EPI_ISL_3203543 | Lighthouse Lab in Milton Keynes | Wellcome Sanger Institute for the COVID-19 Genomics UK (COG-UK) Consortium | The Lighthouse Lab in Milton Keynes and Alex Alderton, Roberto Amato, Jeffrey Barrett, Sonia Goncalves, Ewan Harrison, David K. Jackson, Ian Johnston, Dominic Kwiatkowski, Cordelia Langford, John Sillitoe on behalf of the Wellcome Sanger Institute COVID-19 Surveillance Team |
| EPI_ISL_3203544, EPI_ISL_3203545 | Lighthouse Lab in Glasgow | Wellcome Sanger Institute for the COVID-19 Genomics UK (COG-UK) Consortium | Harper VanSteenhouse, Yumi Kasai, David Gray, Carol Clugston, Anna Dominiczak and Alex Alderton, Roberto Amato, Jeffrey Barrett, Sonia Goncalves, Ewan Harrison, David K. Jackson, Ian Johnston, Dominic Kwiatkowski, Cordelia Langford, John Sillitoe on behalf of the Wellcome Sanger Institute COVID-19 Surveillance Team |
| EPI_ISL_3203546, EPI_ISL_3203547 | Lighthouse Lab in Milton Keynes | Wellcome Sanger Institute for the COVID-19 Genomics UK (COG-UK) Consortium | The Lighthouse Lab in Milton Keynes and Alex Alderton, Roberto Amato, Jeffrey Barrett, Sonia Goncalves, Ewan Harrison, David K. Jackson, Ian Johnston, Dominic Kwiatkowski, Cordelia Langford, John Sillitoe on behalf of the Wellcome Sanger Institute COVID-19 Surveillance Team |
| EPI_ISL_3203548 | Lighthouse Lab in Glasgow | Wellcome Sanger Institute for the COVID-19 Genomics UK (COG-UK) Consortium | Harper VanSteenhouse, Yumi Kasai, David Gray, Carol Clugston, Anna Dominiczak and Alex Alderton, Roberto Amato, Jeffrey Barrett, Sonia Goncalves, Ewan Harrison, David K. Jackson, Ian Johnston, Dominic Kwiatkowski, Cordelia Langford, John Sillitoe on behalf of the Wellcome Sanger Institute COVID-19 Surveillance Team |
| EPI_ISL_3203549, EPI_ISL_3203550 | Lighthouse Lab in Milton Keynes | Wellcome Sanger Institute for the COVID-19 Genomics UK (COG-UK) Consortium | The Lighthouse Lab in Milton Keynes and Alex Alderton, Roberto Amato, Jeffrey Barrett, Sonia Goncalves, Ewan Harrison, David K. Jackson, Ian Johnston, Dominic Kwiatkowski, Cordelia Langford, John Sillitoe on behalf of the Wellcome Sanger Institute COVID-19 Surveillance Team |
| EPI_ISL_3203551 | Lighthouse Lab in Glasgow | Wellcome Sanger Institute for the COVID-19 Genomics UK (COG-UK) Consortium | Harper VanSteenhouse, Yumi Kasai, David Gray, Carol Clugston, Anna Dominiczak and Alex Alderton, Roberto Amato, Jeffrey Barrett, Sonia Goncalves, Ewan Harrison, David K. Jackson, Ian Johnston, Dominic Kwiatkowski, Cordelia Langford, John Sillitoe on behalf of the Wellcome Sanger Institute COVID-19 Surveillance Team |
| EPI_ISL_3203552, EPI_ISL_3203553 | Lighthouse Lab in Milton Keynes | Wellcome Sanger Institute for the COVID-19 Genomics UK (COG-UK) Consortium | The Lighthouse Lab in Milton Keynes and Alex Alderton, Roberto Amato, Jeffrey Barrett, Sonia Goncalves, Ewan Harrison, David K. Jackson, Ian Johnston, Dominic Kwiatkowski, Cordelia Langford, John Sillitoe on behalf of the Wellcome Sanger Institute COVID-19 Surveillance Team |
| EPI_ISL_3203554, EPI_ISL_3203555, EPI_ISL_3203556 | Lighthouse Lab in Glasgow | Wellcome Sanger Institute for the COVID-19 Genomics UK (COG-UK) Consortium | Harper VanSteenhouse, Yumi Kasai, David Gray, Carol Clugston, Anna Dominiczak and Alex Alderton, Roberto Amato, Jeffrey Barrett, Sonia Goncalves, Ewan Harrison, David K. Jackson, Ian Johnston, Dominic Kwiatkowski, Cordelia Langford, John Sillitoe on behalf of the Wellcome Sanger Institute COVID-19 Surveillance Team |
| EPI_ISL_3203557, EPI_ISL_3203558 | Lighthouse Lab in Milton Keynes | Wellcome Sanger Institute for the COVID-19 Genomics UK (COG-UK) Consortium | The Lighthouse Lab in Milton Keynes and Alex Alderton, Roberto Amato, Jeffrey Barrett, Sonia Goncalves, Ewan Harrison, David K. Jackson, Ian Johnston, Dominic Kwiatkowski, Cordelia Langford, John Sillitoe on behalf of the Wellcome Sanger Institute COVID-19 Surveillance Team |
| EPI_ISL_3203559, EPI_ISL_3203560, EPI_ISL_3203561 | Lighthouse Lab in Glasgow | Wellcome Sanger Institute for the COVID-19 Genomics UK (COG-UK) Consortium | Harper VanSteenhouse, Yumi Kasai, David Gray, Carol Clugston, Anna Dominiczak and Alex Alderton, Roberto Amato, Jeffrey Barrett, Sonia Goncalves, Ewan Harrison, David K. Jackson, Ian Johnston, Dominic Kwiatkowski, Cordelia Langford, John Sillitoe on behalf of the Wellcome Sanger Institute COVID-19 Surveillance Team |
| EPI_ISL_3203562, EPI_ISL_3203563, EPI_ISL_3203564, EPI_ISL_3203565, EPI_ISL_3203566, EPI_ISL_3203567, EPI_ISL_3203568, EPI_ISL_3203569, EPI_ISL_3203570, EPI_ISL_3203571, EPI_ISL_3203572 | | | |
| see above | Lighthouse Lab in Milton Keynes | Wellcome Sanger Institute for the COVID-19 Genomics UK (COG-UK) Consortium | The Lighthouse Lab in Milton Keynes and Alex Alderton, Roberto Amato, Jeffrey Barrett, Sonia Goncalves, Ewan Harrison, David K. Jackson, Ian Johnston, Dominic Kwiatkowski, Cordelia Langford, John Sillitoe on behalf of the Wellcome Sanger Institute COVID-19 Surveillance Team |
| EPI_ISL_3203573 | Lighthouse Lab in Glasgow | Wellcome Sanger Institute for the COVID-19 Genomics UK (COG-UK) Consortium | Harper VanSteenhouse, Yumi Kasai, David Gray, Carol Clugston, Anna Dominiczak and Alex Alderton, Roberto Amato, Jeffrey Barrett, Sonia Goncalves, Ewan Harrison, David K. Jackson, Ian Johnston, Dominic Kwiatkowski, Cordelia Langford, John Sillitoe on behalf of the Wellcome Sanger Institute COVID-19 Surveillance Team |
| EPI_ISL_3203574, EPI_ISL_3203575 | Lighthouse Lab in Milton Keynes | Wellcome Sanger Institute for the COVID-19 Genomics UK (COG-UK) Consortium | The Lighthouse Lab in Milton Keynes and Alex Alderton, Roberto Amato, Jeffrey Barrett, Sonia Goncalves, Ewan Harrison, David K. Jackson, Ian Johnston, Dominic Kwiatkowski, Cordelia Langford, John Sillitoe on behalf of the Wellcome Sanger Institute COVID-19 Surveillance Team |
| EPI_ISL_3203576, EPI_ISL_3203577 | Lighthouse Lab in Glasgow | Wellcome Sanger Institute for the COVID-19 Genomics UK (COG-UK) Consortium | Harper VanSteenhouse, Yumi Kasai, David Gray, Carol Clugston, Anna Dominiczak and Alex Alderton, Roberto Amato, Jeffrey Barrett, Sonia Goncalves, Ewan Harrison, David K. Jackson, Ian Johnston, Dominic Kwiatkowski, Cordelia Langford, John Sillitoe on behalf of the Wellcome Sanger Institute COVID-19 Surveillance Team |
| EPI_ISL_3203578 | Lighthouse Lab in Milton Keynes | Wellcome Sanger Institute for the COVID-19 Genomics UK (COG-UK) Consortium | The Lighthouse Lab in Milton Keynes and Alex Alderton, Roberto Amato, Jeffrey Barrett, Sonia Goncalves, Ewan Harrison, David K. Jackson, Ian Johnston, Dominic Kwiatkowski, Cordelia Langford, John Sillitoe on behalf of the Wellcome Sanger Institute COVID-19 Surveillance Team |
| EPI_ISL_3203579 | Lighthouse Lab in Glasgow | Wellcome Sanger Institute for the COVID-19 Genomics UK (COG-UK) Consortium | Harper VanSteenhouse, Yumi Kasai, David Gray, Carol Clugston, Anna Dominiczak and Alex Alderton, Roberto Amato, Jeffrey Barrett, Sonia Goncalves, Ewan Harrison, David K. Jackson, Ian Johnston, Dominic Kwiatkowski, Cordelia Langford, John Sillitoe on behalf of the Wellcome Sanger Institute COVID-19 Surveillance Team |
| EPI_ISL_3203580, EPI_ISL_3203581, EPI_ISL_3203582, EPI_ISL_3203583, EPI_ISL_3203584 | Lighthouse Lab in Milton Keynes | Wellcome Sanger Institute for the COVID-19 Genomics UK (COG-UK) Consortium | The Lighthouse Lab in Milton Keynes and Alex Alderton, Roberto Amato, Jeffrey Barrett, Sonia Goncalves, Ewan Harrison, David K. Jackson, Ian Johnston, Dominic Kwiatkowski, Cordelia Langford, John Sillitoe on behalf of the Wellcome Sanger Institute COVID-19 Surveillance Team |
| EPI_ISL_3203585 | Lighthouse Lab in Glasgow | Wellcome Sanger Institute for the COVID-19 Genomics UK (COG-UK) Consortium | Harper VanSteenhouse, Yumi Kasai, David Gray, Carol Clugston, Anna Dominiczak and Alex Alderton, Roberto Amato, Jeffrey Barrett, Sonia Goncalves, Ewan Harrison, David K. Jackson, Ian Johnston, Dominic Kwiatkowski, Cordelia Langford, John Sillitoe on behalf of the Wellcome Sanger Institute COVID-19 Surveillance Team |
| EPI_ISL_3203586 | Lighthouse Lab in Milton Keynes | Wellcome Sanger Institute for the COVID-19 Genomics UK (COG-UK) Consortium | The Lighthouse Lab in Milton Keynes and Alex Alderton, Roberto Amato, Jeffrey Barrett, Sonia Goncalves, Ewan Harrison, David K. Jackson, Ian Johnston, Dominic Kwiatkowski, Cordelia Langford, John Sillitoe on behalf of the Wellcome Sanger Institute COVID-19 Surveillance Team |
| EPI_ISL_3203587, EPI_ISL_3203588 | Lighthouse Lab in Glasgow | Wellcome Sanger Institute for the COVID-19 Genomics UK (COG-UK) Consortium | Harper VanSteenhouse, Yumi Kasai, David Gray, Carol Clugston, Anna Dominiczak and Alex Alderton, Roberto Amato, Jeffrey Barrett, Sonia Goncalves, Ewan Harrison, David K. Jackson, Ian Johnston, Dominic Kwiatkowski, Cordelia Langford, John Sillitoe on behalf of the Wellcome Sanger Institute COVID-19 Surveillance Team |
| EPI_ISL_3203589 | Lighthouse Lab in Milton Keynes | Wellcome Sanger Institute for the COVID-19 Genomics UK (COG-UK) Consortium | The Lighthouse Lab in Milton Keynes and Alex Alderton, Roberto Amato, Jeffrey Barrett, Sonia Goncalves, Ewan Harrison, David K. Jackson, Ian Johnston, Dominic Kwiatkowski, Cordelia Langford, John Sillitoe on behalf of the Wellcome Sanger Institute COVID-19 Surveillance Team |
| EPI_ISL_3203590 | Lighthouse Lab in Glasgow | Wellcome Sanger Institute for the COVID-19 Genomics UK (COG-UK) Consortium | Harper VanSteenhouse, Yumi Kasai, David Gray, Carol Clugston, Anna Dominiczak and Alex Alderton, Roberto Amato, Jeffrey Barrett, Sonia Goncalves, Ewan Harrison, David K. Jackson, Ian Johnston, Dominic Kwiatkowski, Cordelia Langford, John Sillitoe on behalf of the Wellcome Sanger Institute COVID-19 Surveillance Team |
| EPI_ISL_3203591, EPI_ISL_3203592, EPI_ISL_3203593, EPI_ISL_3203594, EPI_ISL_3203595, EPI_ISL_3203596, EPI_ISL_3203597, EPI_ISL_3203598, EPI_ISL_3203599, EPI_ISL_3203600, EPI_ISL_3203601, EPI_ISL_3203602, EPI_ISL_3203603, EPI_ISL_3203604, EPI_ISL_3203605, EPI_ISL_3203606 | | | |
| see above | Lighthouse Lab in Milton Keynes | Wellcome Sanger Institute for the COVID-19 Genomics UK (COG-UK) Consortium | The Lighthouse Lab in Milton Keynes and Alex Alderton, Roberto Amato, Jeffrey Barrett, Sonia Goncalves, Ewan Harrison, David K. Jackson, Ian Johnston, Dominic Kwiatkowski, Cordelia Langford, John Sillitoe on behalf of the Wellcome Sanger Institute COVID-19 Surveillance Team |
| EPI_ISL_3203607 | Lighthouse Lab in Glasgow | Wellcome Sanger Institute for the COVID-19 Genomics UK (COG-UK) Consortium | Harper VanSteenhouse, Yumi Kasai, David Gray, Carol Clugston, Anna Dominiczak and Alex Alderton, Roberto Amato, Jeffrey Barrett, Sonia Goncalves, Ewan Harrison, David K. Jackson, Ian Johnston, Dominic Kwiatkowski, Cordelia Langford, John Sillitoe on behalf of the Wellcome Sanger Institute COVID-19 Surveillance Team |
| EPI_ISL_3203608, EPI_ISL_3203609 | Lighthouse Lab in Milton Keynes | Wellcome Sanger Institute for the COVID-19 Genomics UK (COG-UK) Consortium | The Lighthouse Lab in Milton Keynes and Alex Alderton, Roberto Amato, Jeffrey Barrett, Sonia Goncalves, Ewan Harrison, David K. Jackson, Ian Johnston, Dominic Kwiatkowski, Cordelia Langford, John Sillitoe on behalf of the Wellcome Sanger Institute COVID-19 Surveillance Team |
| EPI_ISL_3203610 | Lighthouse Lab in Glasgow | Wellcome Sanger Institute for the COVID-19 Genomics UK (COG-UK) Consortium | Harper VanSteenhouse, Yumi Kasai, David Gray, Carol Clugston, Anna Dominiczak and Alex Alderton, Roberto Amato, Jeffrey Barrett, Sonia Goncalves, Ewan Harrison, David K. Jackson, Ian Johnston, Dominic Kwiatkowski, Cordelia Langford, John Sillitoe on behalf of the Wellcome Sanger Institute COVID-19 Surveillance Team |
| EPI_ISL_3203611, EPI_ISL_3203612, EPI_ISL_3203613 | Lighthouse Lab in Milton Keynes | Wellcome Sanger Institute for the COVID-19 Genomics UK (COG-UK) Consortium | The Lighthouse Lab in Milton Keynes and Alex Alderton, Roberto Amato, Jeffrey Barrett, Sonia Goncalves, Ewan Harrison, David K. Jackson, Ian Johnston, Dominic Kwiatkowski, Cordelia Langford, John Sillitoe on behalf of the Wellcome Sanger Institute COVID-19 Surveillance Team |
| EPI_ISL_3203614 | Lighthouse Lab in Glasgow | Wellcome Sanger Institute for the COVID-19 Genomics UK (COG-UK) Consortium | Harper VanSteenhouse, Yumi Kasai, David Gray, Carol Clugston, Anna Dominiczak and Alex Alderton, Roberto Amato, Jeffrey Barrett, Sonia Goncalves, Ewan Harrison, David K. Jackson, Ian Johnston, Dominic Kwiatkowski, Cordelia Langford, John Sillitoe on behalf of the Wellcome Sanger Institute COVID-19 Surveillance Team |
| EPI_ISL_3203615 | Lighthouse Lab in Milton Keynes | Wellcome Sanger Institute for the COVID-19 Genomics UK (COG-UK) Consortium | The Lighthouse Lab in Milton Keynes and Alex Alderton, Roberto Amato, Jeffrey Barrett, Sonia Goncalves, Ewan Harrison, David K. Jackson, Ian Johnston, Dominic Kwiatkowski, Cordelia Langford, John Sillitoe on behalf of the Wellcome Sanger Institute COVID-19 Surveillance Team |
| EPI_ISL_3203616, EPI_ISL_3203617 | Lighthouse Lab in Glasgow | Wellcome Sanger Institute for the COVID-19 Genomics UK (COG-UK) Consortium | Harper VanSteenhouse, Yumi Kasai, David Gray, Carol Clugston, Anna Dominiczak and Alex Alderton, Roberto Amato, Jeffrey Barrett, Sonia Goncalves, Ewan Harrison, David K. Jackson, Ian Johnston, Dominic Kwiatkowski, Cordelia Langford, John Sillitoe on behalf of the Wellcome Sanger Institute COVID-19 Surveillance Team |
| EPI_ISL_3203618, EPI_ISL_3203619, EPI_ISL_3203620, EPI_ISL_3203621 | Lighthouse Lab in Milton Keynes | Wellcome Sanger Institute for the COVID-19 Genomics UK (COG-UK) Consortium | The Lighthouse Lab in Milton Keynes and Alex Alderton, Roberto Amato, Jeffrey Barrett, Sonia Goncalves, Ewan Harrison, David K. Jackson, Ian Johnston, Dominic Kwiatkowski, Cordelia Langford, John Sillitoe on behalf of the Wellcome Sanger Institute COVID-19 Surveillance Team |
| EPI_ISL_3203622, EPI_ISL_3203623, EPI_ISL_3203624, EPI_ISL_3203625, EPI_ISL_3203626, EPI_ISL_3203627, EPI_ISL_3203628, EPI_ISL_3203629, EPI_ISL_3203630, EPI_ISL_3203631, EPI_ISL_3203632, EPI_ISL_3203633, EPI_ISL_3203634, EPI_ISL_3203635, EPI_ISL_3203636, EPI_ISL_3203637, EPI_ISL_3203639, EPI_ISL_3203640, EPI_ISL_3203641, EPI_ISL_3203642, EPI_ISL_3203643, EPI_ISL_3203644, EPI_ISL_3203645, EPI_ISL_3203646, EPI_ISL_3203647, EPI_ISL_3203648, EPI_ISL_3203649, EPI_ISL_3203650, EPI_ISL_3203651, EPI_ISL_3203652, EPI_ISL_3203653, EPI_ISL_3203654, EPI_ISL_3203655, EPI_ISL_3203656, EPI_ISL_3203657, EPI_ISL_3203658, EPI_ISL_3203659, EPI_ISL_3203660, EPI_ISL_3203661, EPI_ISL_3203662, EPI_ISL_3203663, EPI_ISL_3203664, EPI_ISL_3203665, EPI_ISL_3203666, EPI_ISL_3203667, EPI_ISL_3203668, EPI_ISL_3203669, EPI_ISL_3203670, EPI_ISL_3203671, EPI_ISL_3203672, EPI_ISL_3203673, EPI_ISL_3203674, EPI_ISL_3203675, EPI_ISL_3203676, EPI_ISL_3203677, EPI_ISL_3203678, EPI_ISL_3203679, EPI_ISL_3203680, EPI_ISL_3203681, EPI_ISL_3203682, EPI_ISL_3203683, EPI_ISL_3203684, EPI_ISL_3203685, EPI_ISL_3203686, EPI_ISL_3203687, EPI_ISL_3203688, | | | |

EPI_ISL_3203689, EPI_ISL_3203691, EPI_ISL_3203692, EPI_ISL_3203693, EPI_ISL_3203694, EPI_ISL_3203695, EPI_ISL_3203696, EPI_ISL_3203697, EPI_ISL_3203698, EPI_ISL_3203699, EPI_ISL_3203700, EPI_ISL_3203701, EPI_ISL_3203702, EPI_ISL_3203703, EPI_ISL_3203704, EPI_ISL_3203705, EPI_ISL_3203706, EPI_ISL_3203707, EPI_ISL_3203708, EPI_ISL_3203709, EPI_ISL_3203710, EPI_ISL_3203711, EPI_ISL_3203712, EPI_ISL_3203713, EPI_ISL_3203714, EPI_ISL_3203715, EPI_ISL_3203716, EPI_ISL_3203717, EPI_ISL_3203718, EPI_ISL_3203719, EPI_ISL_3203720, EPI_ISL_3203721, EPI_ISL_3203722, EPI_ISL_3203723, EPI_ISL_3203724, EPI_ISL_3203725, EPI_ISL_3203726, EPI_ISL_3203727, EPI_ISL_3203728, EPI_ISL_3203729, EPI_ISL_3203730, EPI_ISL_3203731, EPI_ISL_3203732, EPI_ISL_3203733, EPI_ISL_3203734, EPI_ISL_3203735, EPI_ISL_3203736, EPI_ISL_3203737, EPI_ISL_3203738, EPI_ISL_3203739, EPI_ISL_3203740, EPI_ISL_3203741, EPI_ISL_3203742, EPI_ISL_3203743, EPI_ISL_3203744, EPI_ISL_3203745, EPI_ISL_3203746, EPI_ISL_3203747, EPI_ISL_3203748, EPI_ISL_3203749, EPI_ISL_3203750, EPI_ISL_3203751, EPI_ISL_3203752, EPI_ISL_3203753, EPI_ISL_3203754, EPI_ISL_3203755, EPI_ISL_3203756, EPI_ISL_3203757, EPI_ISL_3203758, EPI_ISL_3203759, EPI_ISL_3203760, EPI_ISL_3203761, EPI_ISL_3203762, EPI_ISL_3203763, EPI_ISL_3203764, EPI_ISL_3203765, EPI_ISL_3203766, EPI_ISL_3203767, EPI_ISL_3203768, EPI_ISL_3203769, EPI_ISL_3203771, EPI_ISL_3203772, EPI_ISL_3203773, EPI_ISL_3203774, EPI_ISL_3203775, EPI_ISL_3203776, EPI_ISL_3203777, EPI_ISL_3203778, EPI_ISL_3203779, EPI_ISL_3203780, EPI_ISL_3203781, EPI_ISL_3203782, EPI_ISL_3203783, EPI_ISL_3203784, EPI_ISL_3203785, EPI_ISL_3203786, EPI_ISL_3203787, EPI_ISL_3203788, EPI_ISL_3203789, EPI_ISL_3203790, EPI_ISL_3203791, EPI_ISL_3203792, EPI_ISL_3203793, EPI_ISL_3203794, EPI_ISL_3203795, EPI_ISL_3203796, EPI_ISL_3203797, EPI_ISL_3203798, EPI_ISL_3203799, EPI_ISL_3203800, EPI_ISL_3203801, EPI_ISL_3203802, EPI_ISL_3203803, EPI_ISL_3203804, EPI_ISL_3203805, EPI_ISL_3203806, EPI_ISL_3203807, EPI_ISL_3203808, EPI_ISL_3203809, EPI_ISL_3203810, EPI_ISL_3203811, EPI_ISL_3203812, EPI_ISL_3203813, EPI_ISL_3203814, EPI_ISL_3203815, EPI_ISL_3203816, EPI_ISL_3203817, EPI_ISL_3203818, EPI_ISL_3203819, EPI_ISL_3203820, EPI_ISL_3203821, EPI_ISL_3203822, EPI_ISL_3203823, EPI_ISL_3203824, EPI_ISL_3203825, EPI_ISL_3203826, EPI_ISL_3203827, EPI_ISL_3203828, EPI_ISL_3203829, EPI_ISL_3203830, EPI_ISL_3203831, EPI_ISL_3203832

| | | | |
|---|---|---|---|
| see above | Lighthouse Lab in Glasgow | Wellcome Sanger Institute for the COVID-19 Genomics UK (COG-UK) Consortium | Harper VanSteenhouse, Yumi Kasai, David Gray, Carol Clugston, Anna Dominiczak and Alex Alderton, Roberto Amato, Jeffrey Barrett, Sonia Goncalves, Ewan Harrison, David K. Jackson, Ian Johnston, Dominic Kwiatkowski, Cordelia Langford, John Sillitoe on behalf of the Wellcome Sanger Institute COVID-19 Surveillance Team |
| EPI_ISL_3203833, EPI_ISL_3203834 | Lighthouse Lab in Milton Keynes | Wellcome Sanger Institute for the COVID-19 Genomics UK (COG-UK) Consortium | The Lighthouse Lab in Milton Keynes and Alex Alderton, Roberto Amato, Jeffrey Barrett, Sonia Goncalves, Ewan Harrison, David K. Jackson, Ian Johnston, Dominic Kwiatkowski, Cordelia Langford, John Sillitoe on behalf of the Wellcome Sanger Institute COVID-19 Surveillance Team |
| EPI_ISL_3203835, EPI_ISL_3203836, EPI_ISL_3203837, EPI_ISL_3203838, EPI_ISL_3203839, EPI_ISL_3203840 | Lighthouse Lab in Glasgow | Wellcome Sanger Institute for the COVID-19 Genomics UK (COG-UK) Consortium | Harper VanSteenhouse, Yumi Kasai, David Gray, Carol Clugston, Anna Dominiczak and Alex Alderton, Roberto Amato, Jeffrey Barrett, Sonia Goncalves, Ewan Harrison, David K. Jackson, Ian Johnston, Dominic Kwiatkowski, Cordelia Langford, John Sillitoe on behalf of the Wellcome Sanger Institute COVID-19 Surveillance Team |
| EPI_ISL_3203841, EPI_ISL_3203842 | Lighthouse Lab in Milton Keynes | Wellcome Sanger Institute for the COVID-19 Genomics UK (COG-UK) Consortium | The Lighthouse Lab in Milton Keynes and Alex Alderton, Roberto Amato, Jeffrey Barrett, Sonia Goncalves, Ewan Harrison, David K. Jackson, Ian Johnston, Dominic Kwiatkowski, Cordelia Langford, John Sillitoe on behalf of the Wellcome Sanger Institute COVID-19 Surveillance Team |
| EPI_ISL_3203843, EPI_ISL_3203844 | Lighthouse Lab in Glasgow | Wellcome Sanger Institute for the COVID-19 Genomics UK (COG-UK) Consortium | Harper VanSteenhouse, Yumi Kasai, David Gray, Carol Clugston, Anna Dominiczak and Alex Alderton, Roberto Amato, Jeffrey Barrett, Sonia Goncalves, Ewan Harrison, David K. Jackson, Ian Johnston, Dominic Kwiatkowski, Cordelia Langford, John Sillitoe on behalf of the Wellcome Sanger Institute COVID-19 Surveillance Team |
| EPI_ISL_3203845, EPI_ISL_3203846 | Lighthouse Lab in Milton Keynes | Wellcome Sanger Institute for the COVID-19 Genomics UK (COG-UK) Consortium | The Lighthouse Lab in Milton Keynes and Alex Alderton, Roberto Amato, Jeffrey Barrett, Sonia Goncalves, Ewan Harrison, David K. Jackson, Ian Johnston, Dominic Kwiatkowski, Cordelia Langford, John Sillitoe on behalf of the Wellcome Sanger Institute COVID-19 Surveillance Team |
| EPI_ISL_3203847, EPI_ISL_3203848 | Lighthouse Lab in Glasgow | Wellcome Sanger Institute for the COVID-19 Genomics UK (COG-UK) Consortium | Harper VanSteenhouse, Yumi Kasai, David Gray, Carol Clugston, Anna Dominiczak and Alex Alderton, Roberto Amato, Jeffrey Barrett, Sonia Goncalves, Ewan Harrison, David K. Jackson, Ian Johnston, Dominic Kwiatkowski, Cordelia Langford, John Sillitoe on behalf of the Wellcome Sanger Institute COVID-19 Surveillance Team |
| EPI_ISL_3203849 | Lighthouse Lab in Milton Keynes | Wellcome Sanger Institute for the COVID-19 Genomics UK (COG-UK) Consortium | The Lighthouse Lab in Milton Keynes and Alex Alderton, Roberto Amato, Jeffrey Barrett, Sonia Goncalves, Ewan Harrison, David K. Jackson, Ian Johnston, Dominic Kwiatkowski, Cordelia Langford, John Sillitoe on behalf of the Wellcome Sanger Institute COVID-19 Surveillance Team |
| EPI_ISL_3203850 | Lighthouse Lab in Glasgow | Wellcome Sanger Institute for the COVID-19 Genomics UK (COG-UK) Consortium | Harper VanSteenhouse, Yumi Kasai, David Gray, Carol Clugston, Anna Dominiczak and Alex Alderton, Roberto Amato, Jeffrey Barrett, Sonia Goncalves, Ewan Harrison, David K. Jackson, Ian Johnston, Dominic Kwiatkowski, Cordelia Langford, John Sillitoe on behalf of the Wellcome Sanger Institute COVID-19 Surveillance Team |
| EPI_ISL_3203851, EPI_ISL_3203852 | Lighthouse Lab in Milton Keynes | Wellcome Sanger Institute for the COVID-19 Genomics UK (COG-UK) Consortium | The Lighthouse Lab in Milton Keynes and Alex Alderton, Roberto Amato, Jeffrey Barrett, Sonia Goncalves, Ewan Harrison, David K. Jackson, Ian Johnston, Dominic Kwiatkowski, Cordelia Langford, John Sillitoe on behalf of the Wellcome Sanger Institute COVID-19 Surveillance Team |
| EPI_ISL_3203853 | Lighthouse Lab in Glasgow | Wellcome Sanger Institute for the COVID-19 Genomics UK (COG-UK) Consortium | Harper VanSteenhouse, Yumi Kasai, David Gray, Carol Clugston, Anna Dominiczak and Alex Alderton, Roberto Amato, Jeffrey Barrett, Sonia Goncalves, Ewan Harrison, David K. Jackson, Ian Johnston, Dominic Kwiatkowski, Cordelia Langford, John Sillitoe on behalf of the Wellcome Sanger Institute COVID-19 Surveillance Team |
| EPI_ISL_3203854 | Lighthouse Lab in Milton Keynes | Wellcome Sanger Institute for the COVID-19 Genomics UK (COG-UK) Consortium | The Lighthouse Lab in Milton Keynes and Alex Alderton, Roberto Amato, Jeffrey Barrett, Sonia Goncalves, Ewan Harrison, David K. Jackson, Ian Johnston, Dominic Kwiatkowski, Cordelia Langford, John Sillitoe on behalf of the Wellcome Sanger Institute COVID-19 Surveillance Team |
| EPI_ISL_3203855 | Lighthouse Lab in Glasgow | Wellcome Sanger Institute for the COVID-19 Genomics UK (COG-UK) Consortium | Harper VanSteenhouse, Yumi Kasai, David Gray, Carol Clugston, Anna Dominiczak and Alex Alderton, Roberto Amato, Jeffrey Barrett, Sonia Goncalves, Ewan Harrison, David K. Jackson, Ian Johnston, Dominic Kwiatkowski, Cordelia Langford, John Sillitoe on behalf of the Wellcome Sanger Institute COVID-19 Surveillance Team |
| EPI_ISL_3203856 | Lighthouse Lab in Milton Keynes | Wellcome Sanger Institute for the COVID-19 Genomics UK (COG-UK) Consortium | The Lighthouse Lab in Milton Keynes and Alex Alderton, Roberto Amato, Jeffrey Barrett, Sonia Goncalves, Ewan Harrison, David K. Jackson, Ian Johnston, Dominic Kwiatkowski, Cordelia Langford, John Sillitoe on behalf of the Wellcome Sanger Institute COVID-19 Surveillance Team |
| EPI_ISL_3203857, EPI_ISL_3203858 | Lighthouse Lab in Glasgow | Wellcome Sanger Institute for the COVID-19 Genomics UK (COG-UK) Consortium | Harper VanSteenhouse, Yumi Kasai, David Gray, Carol Clugston, Anna Dominiczak and Alex Alderton, Roberto Amato, Jeffrey Barrett, Sonia Goncalves, Ewan Harrison, David K. Jackson, Ian Johnston, Dominic Kwiatkowski, Cordelia Langford, John Sillitoe on behalf of the Wellcome Sanger Institute COVID-19 Surveillance Team |
| EPI_ISL_3203859, EPI_ISL_3203860 | Lighthouse Lab in Milton Keynes | Wellcome Sanger Institute for the COVID-19 Genomics UK (COG-UK) Consortium | The Lighthouse Lab in Milton Keynes and Alex Alderton, Roberto Amato, Jeffrey Barrett, Sonia Goncalves, Ewan Harrison, David K. Jackson, Ian Johnston, Dominic Kwiatkowski, Cordelia Langford, John Sillitoe on behalf of the Wellcome Sanger Institute COVID-19 Surveillance Team |
| EPI_ISL_3203861, EPI_ISL_3203862, EPI_ISL_3203863 | Lighthouse Lab in Glasgow | Wellcome Sanger Institute for the COVID-19 Genomics UK (COG-UK) Consortium | Harper VanSteenhouse, Yumi Kasai, David Gray, Carol Clugston, Anna Dominiczak and Alex Alderton, Roberto Amato, Jeffrey Barrett, Sonia Goncalves, Ewan Harrison, David K. Jackson, Ian Johnston, Dominic Kwiatkowski, Cordelia Langford, John Sillitoe on behalf of the Wellcome Sanger Institute COVID-19 Surveillance Team |
| EPI_ISL_3203864 | Lighthouse Lab in Milton Keynes | Wellcome Sanger Institute for the COVID-19 Genomics UK (COG-UK) Consortium | The Lighthouse Lab in Milton Keynes and Alex Alderton, Roberto Amato, Jeffrey Barrett, Sonia Goncalves, Ewan Harrison, David K. Jackson, Ian Johnston, Dominic Kwiatkowski, Cordelia Langford, John Sillitoe on behalf of the Wellcome Sanger Institute COVID-19 Surveillance Team |
| EPI_ISL_3203865 | Lighthouse Lab in Glasgow | Wellcome Sanger Institute for the COVID-19 Genomics UK (COG-UK) Consortium | Harper VanSteenhouse, Yumi Kasai, David Gray, Carol Clugston, Anna Dominiczak and Alex Alderton, Roberto Amato, Jeffrey Barrett, Sonia Goncalves, Ewan Harrison, David K. Jackson, Ian Johnston, Dominic Kwiatkowski, Cordelia Langford, John Sillitoe on behalf of the Wellcome Sanger Institute COVID-19 Surveillance Team |
| EPI_ISL_3203866 | Lighthouse Lab in Milton Keynes | Wellcome Sanger Institute for the COVID-19 Genomics UK (COG-UK) Consortium | The Lighthouse Lab in Milton Keynes and Alex Alderton, Roberto Amato, Jeffrey Barrett, Sonia Goncalves, Ewan Harrison, David K. Jackson, Ian Johnston, Dominic Kwiatkowski, Cordelia Langford, John Sillitoe on behalf of the Wellcome Sanger Institute COVID-19 Surveillance Team |
| EPI_ISL_3203867 | Lighthouse Lab in Glasgow | Wellcome Sanger Institute for the COVID-19 Genomics UK (COG-UK) Consortium | Harper VanSteenhouse, Yumi Kasai, David Gray, Carol Clugston, Anna Dominiczak and Alex Alderton, Roberto Amato, Jeffrey Barrett, Sonia Goncalves, Ewan Harrison, David K. Jackson, Ian Johnston, Dominic Kwiatkowski, Cordelia Langford, John Sillitoe on behalf of the Wellcome Sanger Institute COVID-19 Surveillance Team |
| EPI_ISL_3203868, EPI_ISL_3203869 | Lighthouse Lab in Milton Keynes | Wellcome Sanger Institute for the COVID-19 Genomics UK (COG-UK) Consortium | The Lighthouse Lab in Milton Keynes and Alex Alderton, Roberto Amato, Jeffrey Barrett, Sonia Goncalves, Ewan Harrison, David K. Jackson, Ian Johnston, Dominic Kwiatkowski, Cordelia Langford, John Sillitoe on behalf of the Wellcome Sanger Institute COVID-19 Surveillance Team |
| EPI_ISL_3203870 | Lighthouse Lab in Glasgow | Wellcome Sanger Institute for the COVID-19 Genomics UK (COG-UK) Consortium | Harper VanSteenhouse, Yumi Kasai, David Gray, Carol Clugston, Anna Dominiczak and Alex Alderton, Roberto Amato, Jeffrey Barrett, Sonia Goncalves, Ewan Harrison, David K. Jackson, Ian Johnston, Dominic Kwiatkowski, Cordelia Langford, John Sillitoe on behalf of the Wellcome Sanger Institute COVID-19 Surveillance Team |
| EPI_ISL_3203871 | Lighthouse Lab in Milton Keynes | Wellcome Sanger Institute for the COVID-19 Genomics UK (COG-UK) Consortium | The Lighthouse Lab in Milton Keynes and Alex Alderton, Roberto Amato, Jeffrey Barrett, Sonia Goncalves, Ewan Harrison, David K. Jackson, Ian Johnston, Dominic Kwiatkowski, Cordelia Langford, John Sillitoe on behalf of the Wellcome Sanger Institute COVID-19 Surveillance Team |
| EPI_ISL_3203872, EPI_ISL_3203873 | Lighthouse Lab in Glasgow | Wellcome Sanger Institute for the COVID-19 Genomics UK (COG-UK) Consortium | Harper VanSteenhouse, Yumi Kasai, David Gray, Carol Clugston, Anna Dominiczak and Alex Alderton, Roberto Amato, Jeffrey Barrett, Sonia Goncalves, Ewan Harrison, David K. Jackson, Ian Johnston, Dominic Kwiatkowski, Cordelia Langford, John Sillitoe on behalf of the Wellcome Sanger Institute COVID-19 Surveillance Team |
| EPI_ISL_3203874 | Lighthouse Lab in Milton Keynes | Wellcome Sanger Institute for the COVID-19 Genomics UK (COG-UK) Consortium | The Lighthouse Lab in Milton Keynes and Alex Alderton, Roberto Amato, Jeffrey Barrett, Sonia Goncalves, Ewan Harrison, David K. Jackson, Ian Johnston, Dominic Kwiatkowski, Cordelia Langford, John Sillitoe on behalf of the Wellcome Sanger Institute COVID-19 Surveillance Team |
| EPI_ISL_3203875, EPI_ISL_3203876, EPI_ISL_3203877 | Lighthouse Lab in Glasgow | Wellcome Sanger Institute for the COVID-19 Genomics UK (COG-UK) Consortium | Harper VanSteenhouse, Yumi Kasai, David Gray, Carol Clugston, Anna Dominiczak and Alex Alderton, Roberto Amato, Jeffrey Barrett, Sonia Goncalves, Ewan Harrison, David K. Jackson, Ian Johnston, Dominic Kwiatkowski, Cordelia Langford, John Sillitoe on behalf of the Wellcome Sanger Institute COVID-19 Surveillance Team |
| EPI_ISL_3203878, EPI_ISL_3203879, EPI_ISL_3203880 | Lighthouse Lab in Milton Keynes | Wellcome Sanger Institute for the COVID-19 Genomics UK (COG-UK) Consortium | The Lighthouse Lab in Milton Keynes and Alex Alderton, Roberto Amato, Jeffrey Barrett, Sonia Goncalves, Ewan Harrison, David K. Jackson, Ian Johnston, Dominic Kwiatkowski, Cordelia Langford, John Sillitoe on behalf of the Wellcome Sanger Institute COVID-19 Surveillance Team |
| EPI_ISL_3203881, EPI_ISL_3203882 | Lighthouse Lab in Glasgow | Wellcome Sanger Institute for the COVID-19 Genomics UK (COG-UK) Consortium | Harper VanSteenhouse, Yumi Kasai, David Gray, Carol Clugston, Anna Dominiczak and Alex Alderton, Roberto Amato, Jeffrey Barrett, Sonia Goncalves, Ewan Harrison, David K. Jackson, Ian Johnston, Dominic Kwiatkowski, Cordelia Langford, John Sillitoe on behalf of the Wellcome Sanger Institute COVID-19 Surveillance Team |
| EPI_ISL_3203883, EPI_ISL_3203884 | Lighthouse Lab in Milton Keynes | Wellcome Sanger Institute for the COVID-19 Genomics UK (COG-UK) Consortium | The Lighthouse Lab in Milton Keynes and Alex Alderton, Roberto Amato, Jeffrey Barrett, Sonia Goncalves, Ewan Harrison, David K. Jackson, Ian Johnston, Dominic Kwiatkowski, Cordelia Langford, John Sillitoe on behalf of the Wellcome Sanger Institute COVID-19 Surveillance Team |
| EPI_ISL_3203885, EPI_ISL_3203886, EPI_ISL_3203887 | Lighthouse Lab in Glasgow | Wellcome Sanger Institute for the COVID-19 Genomics UK (COG-UK) Consortium | Harper VanSteenhouse, Yumi Kasai, David Gray, Carol Clugston, Anna Dominiczak and Alex Alderton, Roberto Amato, Jeffrey Barrett, Sonia Goncalves, Ewan Harrison, David K. Jackson, Ian Johnston, Dominic Kwiatkowski, Cordelia Langford, John Sillitoe on behalf of the Wellcome Sanger Institute COVID-19 Surveillance Team |
| EPI_ISL_3203888 | Lighthouse Lab in Milton Keynes | Wellcome Sanger Institute for the COVID-19 Genomics UK (COG-UK) Consortium | The Lighthouse Lab in Milton Keynes and Alex Alderton, Roberto Amato, Jeffrey Barrett, Sonia Goncalves, Ewan Harrison, David K. Jackson, Ian Johnston, Dominic Kwiatkowski, Cordelia Langford, John Sillitoe on behalf of the Wellcome Sanger Institute COVID-19 Surveillance Team |
| EPI_ISL_3203889 | Lighthouse Laboratory Plymouth | Wellcome Sanger Institute for the COVID-19 Genomics UK (COG-UK) Consortium | Lighthouse Laboratory Plymouth and Alex Alderton, Roberto Amato, Jeffrey Barrett, Sonia Goncalves, Ewan Harrison, David K. Jackson, Ian Johnston, Dominic Kwiatkowski, Cordelia Langford, John Sillitoe on behalf of the Wellcome Sanger Institute COVID-19 Surveillance Team |
| EPI_ISL_3203890 | Lighthouse Lab in Glasgow | Wellcome Sanger Institute for the COVID-19 Genomics UK (COG-UK) Consortium | Harper VanSteenhouse, Yumi Kasai, David Gray, Carol Clugston, Anna Dominiczak and Alex Alderton, Roberto Amato, Jeffrey Barrett, Sonia Goncalves, Ewan Harrison, David K. Jackson, Ian Johnston, Dominic Kwiatkowski, Cordelia Langford, John Sillitoe on behalf of the Wellcome Sanger Institute COVID-19 Surveillance Team |
| EPI_ISL_3203891, EPI_ISL_3203892, EPI_ISL_3203893 | Lighthouse Lab in Milton Keynes | Wellcome Sanger Institute for the COVID-19 Genomics UK (COG-UK) Consortium | The Lighthouse Lab in Milton Keynes and Alex Alderton, Roberto Amato, Jeffrey Barrett, Sonia Goncalves, Ewan Harrison, David K. Jackson, Ian Johnston, Dominic Kwiatkowski, Cordelia Langford, John Sillitoe on behalf of the Wellcome Sanger Institute COVID-19 Surveillance Team |
| EPI_ISL_3203894, EPI_ISL_3203895, EPI_ISL_3203896 | Lighthouse Lab in Glasgow | Wellcome Sanger Institute for the COVID-19 Genomics UK (COG-UK) Consortium | Harper VanSteenhouse, Yumi Kasai, David Gray, Carol Clugston, Anna Dominiczak and Alex Alderton, Roberto Amato, Jeffrey Barrett, Sonia Goncalves, Ewan Harrison, David K. Jackson, Ian Johnston, Dominic Kwiatkowski, Cordelia Langford, John Sillitoe on behalf of the Wellcome Sanger Institute COVID-19 Surveillance Team |
| EPI_ISL_3203897, EPI_ISL_3203898 | Lighthouse Lab in Milton Keynes | Wellcome Sanger Institute for the COVID-19 Genomics UK (COG-UK) Consortium | The Lighthouse Lab in Milton Keynes and Alex Alderton, Roberto Amato, Jeffrey Barrett, Sonia Goncalves, Ewan Harrison, David K. Jackson, Ian Johnston, Dominic Kwiatkowski, Cordelia Langford, John Sillitoe on behalf of the Wellcome Sanger Institute COVID-19 Surveillance Team |
| EPI_ISL_3203899 | Lighthouse Lab in Glasgow | Wellcome Sanger Institute for the COVID-19 Genomics UK (COG-UK) Consortium | Harper VanSteenhouse, Yumi Kasai, David Gray, Carol Clugston, Anna Dominiczak and Alex Alderton, Roberto Amato, Jeffrey Barrett, Sonia Goncalves, Ewan Harrison, David K. Jackson, Ian Johnston, Dominic Kwiatkowski, Cordelia Langford, John Sillitoe on behalf of the Wellcome Sanger Institute COVID-19 Surveillance Team |

| | | | |
|---|---|---|---|
| EPI_ISL_3203900 | Lighthouse Lab in Milton Keynes | Wellcome Sanger Institute for the COVID-19 Genomics UK (COG-UK) Consortium | The Lighthouse Lab in Milton Keynes and Alex Alderton, Roberto Amato, Jeffrey Barrett, Sonia Goncalves, Ewan Harrison, David K. Jackson, Ian Johnston, Dominic Kwiatkowski, Cordelia Langford, John Sillitoe on behalf of the Wellcome Sanger Institute COVID-19 Surveillance Team |
| EPI_ISL_3203901 | Lighthouse Lab in Glasgow | Wellcome Sanger Institute for the COVID-19 Genomics UK (COG-UK) Consortium | Harper VanSteenhouse, Yumi Kasai, David Gray, Carol Clugston, Anna Dominiczak and Alex Alderton, Roberto Amato, Jeffrey Barrett, Sonia Goncalves, Ewan Harrison, David K. Jackson, Ian Johnston, Dominic Kwiatkowski, Cordelia Langford, John Sillitoe on behalf of the Wellcome Sanger Institute COVID-19 Surveillance Team |
| EPI_ISL_3203902 | Lighthouse Lab in Milton Keynes | Wellcome Sanger Institute for the COVID-19 Genomics UK (COG-UK) Consortium | The Lighthouse Lab in Milton Keynes and Alex Alderton, Roberto Amato, Jeffrey Barrett, Sonia Goncalves, Ewan Harrison, David K. Jackson, Ian Johnston, Dominic Kwiatkowski, Cordelia Langford, John Sillitoe on behalf of the Wellcome Sanger Institute COVID-19 Surveillance Team |
| EPI_ISL_3203903 | Lighthouse Lab in Glasgow | Wellcome Sanger Institute for the COVID-19 Genomics UK (COG-UK) Consortium | Harper VanSteenhouse, Yumi Kasai, David Gray, Carol Clugston, Anna Dominiczak and Alex Alderton, Roberto Amato, Jeffrey Barrett, Sonia Goncalves, Ewan Harrison, David K. Jackson, Ian Johnston, Dominic Kwiatkowski, Cordelia Langford, John Sillitoe on behalf of the Wellcome Sanger Institute COVID-19 Surveillance Team |
| EPI_ISL_3203904 | Lighthouse Lab in Milton Keynes | Wellcome Sanger Institute for the COVID-19 Genomics UK (COG-UK) Consortium | The Lighthouse Lab in Milton Keynes and Alex Alderton, Roberto Amato, Jeffrey Barrett, Sonia Goncalves, Ewan Harrison, David K. Jackson, Ian Johnston, Dominic Kwiatkowski, Cordelia Langford, John Sillitoe on behalf of the Wellcome Sanger Institute COVID-19 Surveillance Team |
| EPI_ISL_3203905 | Lighthouse Lab in Glasgow | Wellcome Sanger Institute for the COVID-19 Genomics UK (COG-UK) Consortium | Harper VanSteenhouse, Yumi Kasai, David Gray, Carol Clugston, Anna Dominiczak and Alex Alderton, Roberto Amato, Jeffrey Barrett, Sonia Goncalves, Ewan Harrison, David K. Jackson, Ian Johnston, Dominic Kwiatkowski, Cordelia Langford, John Sillitoe on behalf of the Wellcome Sanger Institute COVID-19 Surveillance Team |
| EPI_ISL_3203906, EPI_ISL_3203907 | Lighthouse Lab in Milton Keynes | Wellcome Sanger Institute for the COVID-19 Genomics UK (COG-UK) Consortium | The Lighthouse Lab in Milton Keynes and Alex Alderton, Roberto Amato, Jeffrey Barrett, Sonia Goncalves, Ewan Harrison, David K. Jackson, Ian Johnston, Dominic Kwiatkowski, Cordelia Langford, John Sillitoe on behalf of the Wellcome Sanger Institute COVID-19 Surveillance Team |
| EPI_ISL_3203908, EPI_ISL_3203909 | Lighthouse Lab in Glasgow | Wellcome Sanger Institute for the COVID-19 Genomics UK (COG-UK) Consortium | Harper VanSteenhouse, Yumi Kasai, David Gray, Carol Clugston, Anna Dominiczak and Alex Alderton, Roberto Amato, Jeffrey Barrett, Sonia Goncalves, Ewan Harrison, David K. Jackson, Ian Johnston, Dominic Kwiatkowski, Cordelia Langford, John Sillitoe on behalf of the Wellcome Sanger Institute COVID-19 Surveillance Team |
| EPI_ISL_3203910 | Lighthouse Lab in Milton Keynes | Wellcome Sanger Institute for the COVID-19 Genomics UK (COG-UK) Consortium | The Lighthouse Lab in Milton Keynes and Alex Alderton, Roberto Amato, Jeffrey Barrett, Sonia Goncalves, Ewan Harrison, David K. Jackson, Ian Johnston, Dominic Kwiatkowski, Cordelia Langford, John Sillitoe on behalf of the Wellcome Sanger Institute COVID-19 Surveillance Team |
| EPI_ISL_3203911, EPI_ISL_3203912, EPI_ISL_3203913, EPI_ISL_3203914, EPI_ISL_3203915, EPI_ISL_3203916, EPI_ISL_3203917, EPI_ISL_3203918, EPI_ISL_3203919, EPI_ISL_3203920, EPI_ISL_3203921, EPI_ISL_3203922, EPI_ISL_3203923, EPI_ISL_3203924, EPI_ISL_3203925, EPI_ISL_3203926, EPI_ISL_3203927, EPI_ISL_3203928, EPI_ISL_3203929, EPI_ISL_3203930, EPI_ISL_3203931, EPI_ISL_3203932, EPI_ISL_3203933, EPI_ISL_3203934, EPI_ISL_3203935, EPI_ISL_3203936, EPI_ISL_3203937, EPI_ISL_3203938, EPI_ISL_3203939, EPI_ISL_3203940, EPI_ISL_3203941, EPI_ISL_3203942, EPI_ISL_3203943, EPI_ISL_3203944, EPI_ISL_3203945, EPI_ISL_3203946, EPI_ISL_3203947, EPI_ISL_3203948, EPI_ISL_3203949, EPI_ISL_3203950, EPI_ISL_3203951, EPI_ISL_3203952, EPI_ISL_3203953, EPI_ISL_3203954, EPI_ISL_3203955, EPI_ISL_3203956, EPI_ISL_3203957, EPI_ISL_3203958, EPI_ISL_3203959, EPI_ISL_3203960, EPI_ISL_3203961, EPI_ISL_3203962, EPI_ISL_3203963, EPI_ISL_3203964, EPI_ISL_3203965, EPI_ISL_3203966, EPI_ISL_3203967, EPI_ISL_3203968, EPI_ISL_3203969, EPI_ISL_3203970, EPI_ISL_3203971, EPI_ISL_3203972, EPI_ISL_3203973, EPI_ISL_3203974, EPI_ISL_3203975, EPI_ISL_3203976, EPI_ISL_3203977, EPI_ISL_3203978, EPI_ISL_3203979, EPI_ISL_3203980, EPI_ISL_3203981, EPI_ISL_3203982, EPI_ISL_3203983, EPI_ISL_3203984, EPI_ISL_3203985, EPI_ISL_3203986, EPI_ISL_3203987, EPI_ISL_3203988, EPI_ISL_3203989, EPI_ISL_3203990, EPI_ISL_3203991, EPI_ISL_3203992, EPI_ISL_3203993, EPI_ISL_3203994, EPI_ISL_3203995, EPI_ISL_3203996, EPI_ISL_3203997, EPI_ISL_3203998, EPI_ISL_3203999, EPI_ISL_3204000, EPI_ISL_3204001, EPI_ISL_3204002, EPI_ISL_3204003, EPI_ISL_3204004, EPI_ISL_3204005, EPI_ISL_3204006, EPI_ISL_3204007, EPI_ISL_3204008, EPI_ISL_3204009, EPI_ISL_3204010, EPI_ISL_3204011, EPI_ISL_3204012, EPI_ISL_3204013, EPI_ISL_3204014, EPI_ISL_3204015, EPI_ISL_3204016, EPI_ISL_3204017, EPI_ISL_3204018, EPI_ISL_3204019, EPI_ISL_3204020, EPI_ISL_3204021, EPI_ISL_3204022, EPI_ISL_3204023, EPI_ISL_3204024, EPI_ISL_3204025, EPI_ISL_3204026, EPI_ISL_3204027, EPI_ISL_3204028, EPI_ISL_3204029, EPI_ISL_3204030, EPI_ISL_3204031, EPI_ISL_3204032, EPI_ISL_3204033, EPI_ISL_3204034, EPI_ISL_3204035, EPI_ISL_3204036, EPI_ISL_3204037, EPI_ISL_3204038, EPI_ISL_3204039, EPI_ISL_3204040, EPI_ISL_3204041, EPI_ISL_3204042, EPI_ISL_3204043, EPI_ISL_3204044, EPI_ISL_3204045, EPI_ISL_3204046, EPI_ISL_3204047, EPI_ISL_3204048, EPI_ISL_3204049, EPI_ISL_3204050, EPI_ISL_3204051, EPI_ISL_3204052, EPI_ISL_3204053, EPI_ISL_3204054, EPI_ISL_3204055, EPI_ISL_3204056, EPI_ISL_3204057, EPI_ISL_3204058, EPI_ISL_3204059, EPI_ISL_3204060, EPI_ISL_3204061, EPI_ISL_3204062, EPI_ISL_3204063, EPI_ISL_3204064, EPI_ISL_3204065, EPI_ISL_3204066, EPI_ISL_3204067, EPI_ISL_3204068, EPI_ISL_3204069, EPI_ISL_3204070, EPI_ISL_3204071, EPI_ISL_3204072, EPI_ISL_3204073, EPI_ISL_3204074, EPI_ISL_3204075, EPI_ISL_3204076, EPI_ISL_3204077, EPI_ISL_3204078, EPI_ISL_3204079, EPI_ISL_3204080, EPI_ISL_3204081, EPI_ISL_3204082, EPI_ISL_3204083, EPI_ISL_3204084, EPI_ISL_3204085, EPI_ISL_3204086, EPI_ISL_3204087, EPI_ISL_3204088, EPI_ISL_3204089, EPI_ISL_3204090, EPI_ISL_3204091, EPI_ISL_3204092, EPI_ISL_3204093, EPI_ISL_3204094, EPI_ISL_3204095, EPI_ISL_3204096, EPI_ISL_3204097, EPI_ISL_3204098, EPI_ISL_3204099, EPI_ISL_3204100, EPI_ISL_3204101, EPI_ISL_3204102, EPI_ISL_3204103, EPI_ISL_3204104, EPI_ISL_3204105, EPI_ISL_3204106, EPI_ISL_3204107, EPI_ISL_3204108, EPI_ISL_3204109, EPI_ISL_3204110, EPI_ISL_3204111, EPI_ISL_3204112, EPI_ISL_3204113, EPI_ISL_3204114, EPI_ISL_3204115, EPI_ISL_3204116, EPI_ISL_3204117, EPI_ISL_3204118, EPI_ISL_3204119, EPI_ISL_3204120, EPI_ISL_3204121, EPI_ISL_3204122, EPI_ISL_3204123, EPI_ISL_3204124, EPI_ISL_3204125, EPI_ISL_3204126, EPI_ISL_3204127, EPI_ISL_3204128, EPI_ISL_3204129, EPI_ISL_3204130, EPI_ISL_3204131, EPI_ISL_3204132, EPI_ISL_3204133, EPI_ISL_3204134, EPI_ISL_3204135, EPI_ISL_3204136, EPI_ISL_3204137, EPI_ISL_3204138, EPI_ISL_3204139, EPI_ISL_3204140, EPI_ISL_3204141, EPI_ISL_3204142, EPI_ISL_3204143, EPI_ISL_3204144, EPI_ISL_3204145, EPI_ISL_3204146, EPI_ISL_3204147, EPI_ISL_3204148, EPI_ISL_3204149, EPI_ISL_3204150, EPI_ISL_3204151, EPI_ISL_3204152, EPI_ISL_3204153, EPI_ISL_3204154, EPI_ISL_3204155, EPI_ISL_3204156, EPI_ISL_3204157, EPI_ISL_3204158, EPI_ISL_3204159, EPI_ISL_3204160, EPI_ISL_3204161, EPI_ISL_3204162, EPI_ISL_3204163, EPI_ISL_3204164, EPI_ISL_3204165, EPI_ISL_3204166, EPI_ISL_3204167, EPI_ISL_3204168, EPI_ISL_3204169, EPI_ISL_3204170, EPI_ISL_3204171, EPI_ISL_3204172, EPI_ISL_3204173, EPI_ISL_3204174, EPI_ISL_3204175, EPI_ISL_3204176, EPI_ISL_3204177, EPI_ISL_3204178, EPI_ISL_3204179, EPI_ISL_3204180, EPI_ISL_3204181, EPI_ISL_3204182, EPI_ISL_3204183, EPI_ISL_3204184, EPI_ISL_3204185, EPI_ISL_3204186, EPI_ISL_3204187, EPI_ISL_3204188, EPI_ISL_3204189, EPI_ISL_3204190, EPI_ISL_3204191, EPI_ISL_3204192, EPI_ISL_3204193, EPI_ISL_3204194, EPI_ISL_3204195, EPI_ISL_3204196, EPI_ISL_3204197, EPI_ISL_3204198, EPI_ISL_3204199, EPI_ISL_3204200, EPI_ISL_3204201, EPI_ISL_3204202, EPI_ISL_3204203, EPI_ISL_3204204, EPI_ISL_3204205, EPI_ISL_3204206, EPI_ISL_3204207, EPI_ISL_3204208, EPI_ISL_3204209, EPI_ISL_3204210, EPI_ISL_3204211, EPI_ISL_3204212, EPI_ISL_3204213, EPI_ISL_3204214, EPI_ISL_3204215, EPI_ISL_3204216, EPI_ISL_3204217, EPI_ISL_3204218, EPI_ISL_3204219, EPI_ISL_3204220, EPI_ISL_3204221, EPI_ISL_3204222, EPI_ISL_3204223, EPI_ISL_3204224, EPI_ISL_3204225, EPI_ISL_3204226 | | | |
| see above | Lighthouse Lab in Glasgow | Wellcome Sanger Institute for the COVID-19 Genomics UK (COG-UK) Consortium | Harper VanSteenhouse, Yumi Kasai, David Gray, Carol Clugston, Anna Dominiczak and Alex Alderton, Roberto Amato, Jeffrey Barrett, Sonia Goncalves, Ewan Harrison, David K. Jackson, Ian Johnston, Dominic Kwiatkowski, Cordelia Langford, John Sillitoe on behalf of the Wellcome Sanger Institute COVID-19 Surveillance Team |
| EPI_ISL_3204227, EPI_ISL_3204228 | Lighthouse Lab in Milton Keynes | Wellcome Sanger Institute for the COVID-19 Genomics UK (COG-UK) Consortium | The Lighthouse Lab in Milton Keynes and Alex Alderton, Roberto Amato, Jeffrey Barrett, Sonia Goncalves, Ewan Harrison, David K. Jackson, Ian Johnston, Dominic Kwiatkowski, Cordelia Langford, John Sillitoe on behalf of the Wellcome Sanger Institute COVID-19 Surveillance Team |
| EPI_ISL_3204263, EPI_ISL_3204264, EPI_ISL_3204265, EPI_ISL_3204266, EPI_ISL_3204267, EPI_ISL_3204268, EPI_ISL_3204269, EPI_ISL_3204271, EPI_ISL_3204272, EPI_ISL_3204273, EPI_ISL_3204274, EPI_ISL_3204275, EPI_ISL_3204276, EPI_ISL_3204277 | | | |
| see above | LAC UCoimbra | Instituto Nacional de Saude (INSA) | Borges et al |
| EPI_ISL_3204278, EPI_ISL_3204279, EPI_ISL_3204280, EPI_ISL_3204281, EPI_ISL_3204282, EPI_ISL_3204283, EPI_ISL_3204284 | H Santarem | Instituto Nacional de Saude (INSA) | Borges et al |
| EPI_ISL_3204285, EPI_ISL_3204286, EPI_ISL_3204287, EPI_ISL_3204288, EPI_ISL_3204289, EPI_ISL_3204290, EPI_ISL_3204291, EPI_ISL_3204292, EPI_ISL_3204293, EPI_ISL_3204294, EPI_ISL_3204295, EPI_ISL_3204296, EPI_ISL_3204297, EPI_ISL_3204298, EPI_ISL_3204299, EPI_ISL_3204300, EPI_ISL_3204301, EPI_ISL_3204302, EPI_ISL_3204303 | | | |
| see above | ULS Castelo Branco | Instituto Nacional de Saude (INSA) | Borges et al |
| EPI_ISL_3204304, EPI_ISL_3204305, EPI_ISL_3204306, EPI_ISL_3204307, EPI_ISL_3204308, EPI_ISL_3204309, EPI_ISL_3204310, EPI_ISL_3204311, EPI_ISL_3204312, EPI_ISL_3204313, EPI_ISL_3204314, EPI_ISL_3204315, EPI_ISL_3204316, EPI_ISL_3204317 | | | |
| see above | ULS Litoral Alentejano | Instituto Nacional de Saude (INSA) | Borges et al |
| EPI_ISL_3204318, EPI_ISL_3204319, EPI_ISL_3204320, EPI_ISL_3204321, EPI_ISL_3204322, EPI_ISL_3204323, EPI_ISL_3204324, EPI_ISL_3204325, EPI_ISL_3204326, EPI_ISL_3204327, EPI_ISL_3204328, EPI_ISL_3204329, EPI_ISL_3204330, EPI_ISL_3204331 | | | |
| see above | H Fernando Fonseca | Instituto Nacional de Saude (INSA) | Borges et al |
| EPI_ISL_3204332, EPI_ISL_3204333, EPI_ISL_3204334, EPI_ISL_3204335, EPI_ISL_3204336, EPI_ISL_3204337, EPI_ISL_3204338, EPI_ISL_3204339, EPI_ISL_3204340, EPI_ISL_3204341, EPI_ISL_3204342, EPI_ISL_3204343, EPI_ISL_3204344, EPI_ISL_3204345, EPI_ISL_3204346 | | | |
| see above | ULS Norte Alentejano | Instituto Nacional de Saude (INSA) | Borges et al |
| EPI_ISL_3204347, EPI_ISL_3204348, EPI_ISL_3204349, EPI_ISL_3204350, EPI_ISL_3204351, EPI_ISL_3204352, EPI_ISL_3204353, EPI_ISL_3204355, EPI_ISL_3204356, EPI_ISL_3204357, EPI_ISL_3204358, EPI_ISL_3204359 | | | |
| see above | H Garcia de Orta | Instituto Nacional de Saude (INSA) | Borges et al |
| EPI_ISL_3204360, EPI_ISL_3204361, EPI_ISL_3204362, EPI_ISL_3204363, EPI_ISL_3204364, EPI_ISL_3204365, EPI_ISL_3204366, EPI_ISL_3204367, EPI_ISL_3204368, EPI_ISL_3204369, EPI_ISL_3204370, EPI_ISL_3204371, EPI_ISL_3204372, EPI_ISL_3204373, EPI_ISL_3204374, EPI_ISL_3204375, EPI_ISL_3204376, EPI_ISL_3204377, EPI_ISL_3204378, EPI_ISL_3204379, EPI_ISL_3204380, EPI_ISL_3204381, EPI_ISL_3204382, EPI_ISL_3204383, EPI_ISL_3204384, EPI_ISL_3204385, EPI_ISL_3204386, EPI_ISL_3204387, EPI_ISL_3204388, EPI_ISL_3204389, EPI_ISL_3204390, EPI_ISL_3204391, EPI_ISL_3204392, EPI_ISL_3204393, EPI_ISL_3204394, EPI_ISL_3204395, EPI_ISL_3204396, EPI_ISL_3204397, EPI_ISL_3204398, EPI_ISL_3204399, EPI_ISL_3204400, EPI_ISL_3204402, EPI_ISL_3204403, EPI_ISL_3204404, EPI_ISL_3204405, EPI_ISL_3204406, EPI_ISL_3204407, EPI_ISL_3204408, EPI_ISL_3204409 | | | |
| see above | ABC Algarve | Instituto Nacional de Saude (INSA) | Borges et al |
| EPI_ISL_3204410, EPI_ISL_3204411, EPI_ISL_3204412, EPI_ISL_3204413, EPI_ISL_3204414, EPI_ISL_3204415, EPI_ISL_3204416, EPI_ISL_3204417, EPI_ISL_3204418, EPI_ISL_3204419, EPI_ISL_3204420, EPI_ISL_3204421 | | | |
| see above | H Evora | Instituto Nacional de Saude (INSA) | Borges et al |
| EPI_ISL_3204422, EPI_ISL_3204423, EPI_ISL_3204424, EPI_ISL_3204425, EPI_ISL_3204426, EPI_ISL_3204427, EPI_ISL_3204428, EPI_ISL_3204429, EPI_ISL_3204430, EPI_ISL_3204431, EPI_ISL_3204432, EPI_ISL_3204433, EPI_ISL_3204434, EPI_ISL_3204435, EPI_ISL_3204436, EPI_ISL_3204437 | | | |
| see above | CHLC | Instituto Nacional de Saude (INSA) | Borges et al |
| EPI_ISL_3204438, EPI_ISL_3204439, EPI_ISL_3204440, EPI_ISL_3204441, EPI_ISL_3204442, EPI_ISL_3204443, EPI_ISL_3204444, EPI_ISL_3204445, EPI_ISL_3204446, EPI_ISL_3204447, EPI_ISL_3204448, EPI_ISL_3204449, EPI_ISL_3204450, EPI_ISL_3204451, EPI_ISL_3204452 | | | |
| see above | H Vila Franca Xira | Instituto Nacional de Saude (INSA) | Borges et al |
| EPI_ISL_3204453, EPI_ISL_3204454, EPI_ISL_3204455, EPI_ISL_3204456, EPI_ISL_3204457, EPI_ISL_3204458, EPI_ISL_3204459, EPI_ISL_3204460, EPI_ISL_3204461, EPI_ISL_3204462, EPI_ISL_3204463, EPI_ISL_3204464, EPI_ISL_3204465, EPI_ISL_3204466, EPI_ISL_3204467, EPI_ISL_3204468, EPI_ISL_3204469, EPI_ISL_3204470, EPI_ISL_3204471, EPI_ISL_3204472, EPI_ISL_3204473, EPI_ISL_3204474, EPI_ISL_3204475, EPI_ISL_3204476, EPI_ISL_3204477, EPI_ISL_3204478, EPI_ISL_3204479, EPI_ISL_3204480, EPI_ISL_3204481, EPI_ISL_3204482, EPI_ISL_3204483, EPI_ISL_3204484, EPI_ISL_3204485, EPI_ISL_3204486, EPI_ISL_3204487, EPI_ISL_3204488, EPI_ISL_3204489, EPI_ISL_3204490, EPI_ISL_3204491, EPI_ISL_3204492, EPI_ISL_3204493, EPI_ISL_3204494, EPI_ISL_3204495, EPI_ISL_3204496, EPI_ISL_3204497, EPI_ISL_3204498, EPI_ISL_3204499, EPI_ISL_3204500, EPI_ISL_3204501, EPI_ISL_3204502, EPI_ISL_3204503, EPI_ISL_3204504 | | | |
| see above | ARS Algarve - Laboratorio Laura Ayres | Instituto Nacional de Saude (INSA) | Borges et al |
| EPI_ISL_3204505, EPI_ISL_3204506, EPI_ISL_3204507, EPI_ISL_3204508, EPI_ISL_3204509, EPI_ISL_3204510, EPI_ISL_3204511, EPI_ISL_3204512, EPI_ISL_3204513, EPI_ISL_3204514, EPI_ISL_3204515, EPI_ISL_3204516, EPI_ISL_3204517, EPI_ISL_3204518, EPI_ISL_3204519 | | | |
| see above | Labeto - CAB - Leiria | Instituto Nacional de Saude (INSA) | Borges et al |
| EPI_ISL_3204520, EPI_ISL_3204521, EPI_ISL_3204522, EPI_ISL_3204523, EPI_ISL_3204524, EPI_ISL_3204525, EPI_ISL_3204526, EPI_ISL_3204527, EPI_ISL_3204528, EPI_ISL_3204529, EPI_ISL_3204530, EPI_ISL_3204531, EPI_ISL_3204532, EPI_ISL_3204533, EPI_ISL_3204534 | | | |
| see above | CH Porto - H Sto Antonio | Instituto Nacional de Saude (INSA) | Borges et al |
| EPI_ISL_3204536, EPI_ISL_3204537, EPI_ISL_3204539, EPI_ISL_3204540, EPI_ISL_3204541, EPI_ISL_3204542, EPI_ISL_3204543, EPI_ISL_3204544, EPI_ISL_3204545, EPI_ISL_3204546, EPI_ISL_3204547, EPI_ISL_3204548 | | | |
| see above | H Braga | Instituto Nacional de Saude (INSA) | Borges et al |
| EPI_ISL_3204549, EPI_ISL_3204550, EPI_ISL_3204551, EPI_ISL_3204552, EPI_ISL_3204553, EPI_ISL_3204554, EPI_ISL_3204555, EPI_ISL_3204556, EPI_ISL_3204557, EPI_ISL_3204558, EPI_ISL_3204559, EPI_ISL_3204560 | | | |
| see above | INSA | Instituto Nacional de Saude (INSA) | Borges et al |
| EPI_ISL_3204561, EPI_ISL_3204562, EPI_ISL_3204563, EPI_ISL_3204564, EPI_ISL_3204565, EPI_ISL_3204566, EPI_ISL_3204567, EPI_ISL_3204568, EPI_ISL_3204569, EPI_ISL_3204570 | H Divino Espirito Santo - Ponta Delgada | Instituto Nacional de Saude (INSA) | Borges et al |
| EPI_ISL_3204571, EPI_ISL_3204572, EPI_ISL_3204573, EPI_ISL_3204574, EPI_ISL_3204575, EPI_ISL_3204576, EPI_ISL_3204577, EPI_ISL_3204578, EPI_ISL_3204580, EPI_ISL_3204581, EPI_ISL_3204582, EPI_ISL_3204583, EPI_ISL_3204584, EPI_ISL_3204585, EPI_ISL_3204586, EPI_ISL_3204587, EPI_ISL_3204588, EPI_ISL_3204589, EPI_ISL_3204590, EPI_ISL_3204592, EPI_ISL_3204593, EPI_ISL_3204594, EPI_ISL_3204595, EPI_ISL_3204596, EPI_ISL_3204597, EPI_ISL_3204598, EPI_ISL_3204599, EPI_ISL_3204600, EPI_ISL_3204601, EPI_ISL_3204602, EPI_ISL_3204603, EPI_ISL_3204604, EPI_ISL_3204605, EPI_ISL_3204606, EPI_ISL_3204607, EPI_ISL_3204608, EPI_ISL_3204609, EPI_ISL_3204610, EPI_ISL_3204611, EPI_ISL_3204612, EPI_ISL_3204613, EPI_ISL_3204614, EPI_ISL_3204615, EPI_ISL_3204616, | | | |

| | | | |
|---|---|---|---|
| EPI_ISL_3204617, EPI_ISL_3204618 | | | |
| see above | SESARAM | Instituto Nacional de Saude (INSA) | Borges et al |
| EPI_ISL_3204619, EPI_ISL_3204620, EPI_ISL_3204621, EPI_ISL_3204622, EPI_ISL_3204623, EPI_ISL_3204624, EPI_ISL_3204625, EPI_ISL_3204626, EPI_ISL_3204627, EPI_ISL_3204628, EPI_ISL_3204629, EPI_ISL_3204630, EPI_ISL_3204631, EPI_ISL_3204632, EPI_ISL_3204633, EPI_ISL_3204634, EPI_ISL_3204635, EPI_ISL_3204636, EPI_ISL_3204637, EPI_ISL_3204638, EPI_ISL_3204639, EPI_ISL_3204640, EPI_ISL_3204641, EPI_ISL_3204642, EPI_ISL_3204643, EPI_ISL_3204644, EPI_ISL_3204645, EPI_ISL_3204646, EPI_ISL_3204647, EPI_ISL_3204648, EPI_ISL_3204649, EPI_ISL_3204650 | | | |
| see above | SEEBMO | Instituto Nacional de Saude (INSA) | Borges et al |
| EPI_ISL_3204651, EPI_ISL_3204652, EPI_ISL_3204653, EPI_ISL_3204654, EPI_ISL_3204655, EPI_ISL_3204657, EPI_ISL_3204658, EPI_ISL_3204659, EPI_ISL_3204660, EPI_ISL_3204661, EPI_ISL_3204662, EPI_ISL_3204663, EPI_ISL_3204664, EPI_ISL_3204665 | | | |
| see above | CHEDV | Instituto Nacional de Saude (INSA) | Borges et al |
| EPI_ISL_3204666, EPI_ISL_3204667, EPI_ISL_3204668, EPI_ISL_3204669, EPI_ISL_3204670, EPI_ISL_3204671, EPI_ISL_3204672, EPI_ISL_3204673, EPI_ISL_3204674, EPI_ISL_3204675, EPI_ISL_3204676, EPI_ISL_3204677 | | | |
| see above | CH Tamega e Sousa | Instituto Nacional de Saude (INSA) | Borges et al |
| EPI_ISL_3204678, EPI_ISL_3204679, EPI_ISL_3204680, EPI_ISL_3204681, EPI_ISL_3204682, EPI_ISL_3204683 | ADR Evora | Instituto Nacional de Saude (INSA) | Borges et al |
| EPI_ISL_3204684, EPI_ISL_3204685, EPI_ISL_3204686, EPI_ISL_3204687, EPI_ISL_3204688, EPI_ISL_3204689, EPI_ISL_3204690, EPI_ISL_3204691, EPI_ISL_3204692, EPI_ISL_3204693 | ARS Alentejo | Instituto Nacional de Saude (INSA) | Borges et al |
| EPI_ISL_3204694, EPI_ISL_3204695, EPI_ISL_3204696, EPI_ISL_3204697, EPI_ISL_3204698, EPI_ISL_3204699, EPI_ISL_3204700 | ULS Guarda | Instituto Nacional de Saude (INSA) | Borges et al |
| EPI_ISL_3204701, EPI_ISL_3204702, EPI_ISL_3204703, EPI_ISL_3204704, EPI_ISL_3204705, EPI_ISL_3204706, EPI_ISL_3204707, EPI_ISL_3204708, EPI_ISL_3204709, EPI_ISL_3204710, EPI_ISL_3204711, EPI_ISL_3204712, EPI_ISL_3204713, EPI_ISL_3204714, EPI_ISL_3204715 | | | |
| see above | Avelab | Instituto Nacional de Saude (INSA) | Borges et al |
| EPI_ISL_3204716, EPI_ISL_3204717, EPI_ISL_3204718, EPI_ISL_3204719, EPI_ISL_3204720, EPI_ISL_3204721, EPI_ISL_3204722, EPI_ISL_3204723, EPI_ISL_3204724, EPI_ISL_3204725, EPI_ISL_3204726, EPI_ISL_3204727, EPI_ISL_3204728, EPI_ISL_3204729 | | | |
| see above | H Guimaraes | Instituto Nacional de Saude (INSA) | Borges et al |
| EPI_ISL_3204730, EPI_ISL_3204731, EPI_ISL_3204732, EPI_ISL_3204733, EPI_ISL_3204734, EPI_ISL_3204735, EPI_ISL_3204736, EPI_ISL_3204737, EPI_ISL_3204738, EPI_ISL_3204739, EPI_ISL_3204740, EPI_ISL_3204741, EPI_ISL_3204742, EPI_ISL_3204743, EPI_ISL_3204744, EPI_ISL_3204745, EPI_ISL_3204746, EPI_ISL_3204747, EPI_ISL_3204748, EPI_ISL_3204750, EPI_ISL_3204751, EPI_ISL_3204752, EPI_ISL_3204753, EPI_ISL_3204754, EPI_ISL_3204755, EPI_ISL_3204756, EPI_ISL_3204757, EPI_ISL_3204758, EPI_ISL_3204759, EPI_ISL_3204760, EPI_ISL_3204761, EPI_ISL_3204762, EPI_ISL_3204763, EPI_ISL_3204764, EPI_ISL_3204765, EPI_ISL_3204766, EPI_ISL_3204767, EPI_ISL_3204768, EPI_ISL_3204769, EPI_ISL_3204770, EPI_ISL_3204771, EPI_ISL_3204772, EPI_ISL_3204773, EPI_ISL_3204774, EPI_ISL_3204775, EPI_ISL_3204776, EPI_ISL_3204777, EPI_ISL_3204778, EPI_ISL_3204779, EPI_ISL_3204780, EPI_ISL_3204781, EPI_ISL_3204782, EPI_ISL_3204783, EPI_ISL_3204784, EPI_ISL_3204785, EPI_ISL_3204786, EPI_ISL_3204787, EPI_ISL_3204788, EPI_ISL_3204789, EPI_ISL_3204790, EPI_ISL_3204791, EPI_ISL_3204793, EPI_ISL_3204794, EPI_ISL_3204795, EPI_ISL_3204796, EPI_ISL_3204797, EPI_ISL_3204798, EPI_ISL_3204799, EPI_ISL_3204800, EPI_ISL_3204801, EPI_ISL_3204802, EPI_ISL_3204803, EPI_ISL_3204804, EPI_ISL_3204805, EPI_ISL_3204806, EPI_ISL_3204807, EPI_ISL_3204808, EPI_ISL_3204809, EPI_ISL_3204810, EPI_ISL_3204811, EPI_ISL_3204812, EPI_ISL_3204813, EPI_ISL_3204814, EPI_ISL_3204815, EPI_ISL_3204816, EPI_ISL_3204817, EPI_ISL_3204818, EPI_ISL_3204819, EPI_ISL_3204820, EPI_ISL_3204821, EPI_ISL_3204823, EPI_ISL_3204824, EPI_ISL_3204825, EPI_ISL_3204826 | | | |
| see above | UNILABS | Instituto Nacional de Saude (INSA) | Borges et al |
| EPI_ISL_3204827 | H Braga | Instituto Nacional de Saude (INSA) and Institute of Biomedicine (iBiMed), Universidade de Aveiro | Borges et al |
| EPI_ISL_3204828, EPI_ISL_3204829, EPI_ISL_3204830, EPI_ISL_3204831, EPI_ISL_3204832, EPI_ISL_3204833, EPI_ISL_3204834, EPI_ISL_3204835, EPI_ISL_3204836, EPI_ISL_3204838, EPI_ISL_3204839, EPI_ISL_3204840, EPI_ISL_3204841, EPI_ISL_3204842, EPI_ISL_3204843, EPI_ISL_3204844, EPI_ISL_3204845, EPI_ISL_3204846 | | | |
| see above | CH Tondela Viseu | Instituto Nacional de Saude (INSA) and Institute of Biomedicine (iBiMed), Universidade de Aveiro | Borges et al |
| EPI_ISL_3204847, EPI_ISL_3204848, EPI_ISL_3204849, EPI_ISL_3204850, EPI_ISL_3204851, EPI_ISL_3204852, EPI_ISL_3204853, EPI_ISL_3204854, EPI_ISL_3204855, EPI_ISL_3204856, EPI_ISL_3204857, EPI_ISL_3204858, EPI_ISL_3204859, EPI_ISL_3204860 | | | |
| see above | CHU Sao Joao, Porto | Instituto Nacional de Saude (INSA) and Institute of Biomedicine (iBiMed), Universidade de Aveiro | Borges et al |
| EPI_ISL_3204861, EPI_ISL_3204862, EPI_ISL_3204863, EPI_ISL_3204864, EPI_ISL_3204865, EPI_ISL_3204866, EPI_ISL_3204867, EPI_ISL_3204868, EPI_ISL_3204869, EPI_ISL_3204870, EPI_ISL_3204871, EPI_ISL_3204872, EPI_ISL_3204873, EPI_ISL_3204874, EPI_ISL_3204876, EPI_ISL_3204877, EPI_ISL_3204878, EPI_ISL_3204879, EPI_ISL_3204880 | | | |
| see above | H Divino Espirito Santo - Ponta Delgada | Instituto Nacional de Saude (INSA) and Institute of Biomedicine (iBiMed), Universidade de Aveiro | Borges et al |
| EPI_ISL_3204881, EPI_ISL_3204882, EPI_ISL_3204883, EPI_ISL_3204884, EPI_ISL_3204885, EPI_ISL_3204886, EPI_ISL_3204887, EPI_ISL_3204888, EPI_ISL_3204889, EPI_ISL_3204890, EPI_ISL_3204891, EPI_ISL_3204892, EPI_ISL_3204893, EPI_ISL_3204894, EPI_ISL_3204895 | | | |
| see above | CHTMAD | Instituto Nacional de Saude (INSA) and Institute of Biomedicine (iBiMed), Universidade de Aveiro | Borges et al |
| EPI_ISL_3204896, EPI_ISL_3204897, EPI_ISL_3204898, EPI_ISL_3204899, EPI_ISL_3204900, EPI_ISL_3204901, EPI_ISL_3204902, EPI_ISL_3204903, EPI_ISL_3204904, EPI_ISL_3204905, EPI_ISL_3204906, EPI_ISL_3204907, EPI_ISL_3204908, EPI_ISL_3204909, EPI_ISL_3204910, EPI_ISL_3204911, EPI_ISL_3204912, EPI_ISL_3204913, EPI_ISL_3204914, EPI_ISL_3204915, EPI_ISL_3204916, EPI_ISL_3204917, EPI_ISL_3204918, EPI_ISL_3204919, EPI_ISL_3204920, EPI_ISL_3204921, EPI_ISL_3204922, EPI_ISL_3204923, EPI_ISL_3204924, EPI_ISL_3204925, EPI_ISL_3204926, EPI_ISL_3204927, EPI_ISL_3204928, EPI_ISL_3204929, EPI_ISL_3204930, EPI_ISL_3204931, EPI_ISL_3204932, EPI_ISL_3204933, EPI_ISL_3204934, EPI_ISL_3204935, EPI_ISL_3204936, EPI_ISL_3204937, EPI_ISL_3204938, EPI_ISL_3204939, EPI_ISL_3204940, EPI_ISL_3204941, EPI_ISL_3204942, EPI_ISL_3204943, EPI_ISL_3204944, EPI_ISL_3204945, EPI_ISL_3204946, EPI_ISL_3204947, EPI_ISL_3204948, EPI_ISL_3204949, EPI_ISL_3204950, EPI_ISL_3204951, EPI_ISL_3204952, EPI_ISL_3204953, EPI_ISL_3204954, EPI_ISL_3204955, EPI_ISL_3204956, EPI_ISL_3204957, EPI_ISL_3204958, EPI_ISL_3204959, EPI_ISL_3204960, EPI_ISL_3204961, EPI_ISL_3204962, EPI_ISL_3204963, EPI_ISL_3204964, EPI_ISL_3204965, EPI_ISL_3204966, EPI_ISL_3204967, EPI_ISL_3204968, EPI_ISL_3204969, EPI_ISL_3204970, EPI_ISL_3204971, EPI_ISL_3204972, EPI_ISL_3204973, EPI_ISL_3204974, EPI_ISL_3204975, EPI_ISL_3204976, EPI_ISL_3204977, EPI_ISL_3204978, EPI_ISL_3204979, EPI_ISL_3204980, EPI_ISL_3204981, EPI_ISL_3204982, EPI_ISL_3204983, EPI_ISL_3204984, EPI_ISL_3204985, EPI_ISL_3204986, EPI_ISL_3204987, EPI_ISL_3204988, EPI_ISL_3204989, EPI_ISL_3204990, EPI_ISL_3204991, EPI_ISL_3204992, EPI_ISL_3204993, EPI_ISL_3204994, EPI_ISL_3205007, EPI_ISL_3205008, EPI_ISL_3205013, EPI_ISL_3205024, EPI_ISL_3205031, EPI_ISL_3205036, EPI_ISL_3205037, EPI_ISL_3205038, EPI_ISL_3205039, EPI_ISL_3205040, EPI_ISL_3205041, EPI_ISL_3205042, EPI_ISL_3205046, EPI_ISL_3205047, EPI_ISL_3205048, EPI_ISL_3205049, EPI_ISL_3205050, EPI_ISL_3205051, EPI_ISL_3205052, EPI_ISL_3205053, EPI_ISL_3205054, EPI_ISL_3205055, EPI_ISL_3205056, EPI_ISL_3205057, EPI_ISL_3205058, EPI_ISL_3205059, EPI_ISL_3205060, EPI_ISL_3205061, EPI_ISL_3205062, EPI_ISL_3205063, EPI_ISL_3205064, EPI_ISL_3205065, EPI_ISL_3205066, EPI_ISL_3205067, EPI_ISL_3205068, EPI_ISL_3205069, EPI_ISL_3205070, EPI_ISL_3205071, EPI_ISL_3205072, EPI_ISL_3205073, EPI_ISL_3205074, EPI_ISL_3205075, EPI_ISL_3205076, EPI_ISL_3205077, EPI_ISL_3205078, EPI_ISL_3205079, EPI_ISL_3205080, EPI_ISL_3205081, EPI_ISL_3205082, EPI_ISL_3205083, EPI_ISL_3205084, EPI_ISL_3205085, EPI_ISL_3205086, EPI_ISL_3205087, EPI_ISL_3205088, EPI_ISL_3205089, EPI_ISL_3205090, EPI_ISL_3205091, EPI_ISL_3205092, EPI_ISL_3205093, EPI_ISL_3205094, EPI_ISL_3205095, EPI_ISL_3205096, EPI_ISL_3205097, EPI_ISL_3205099, EPI_ISL_3205100, EPI_ISL_3205101, EPI_ISL_3205102, EPI_ISL_3205103, EPI_ISL_3205104, EPI_ISL_3205105, EPI_ISL_3205106, EPI_ISL_3205107, EPI_ISL_3205108, EPI_ISL_3205109, EPI_ISL_3205110, EPI_ISL_3205111, EPI_ISL_3205112, EPI_ISL_3205113, EPI_ISL_3205114, EPI_ISL_3205115, EPI_ISL_3205116, EPI_ISL_3205117, EPI_ISL_3205118, EPI_ISL_3205119, EPI_ISL_3205121, EPI_ISL_3205122, EPI_ISL_3205123, EPI_ISL_3205124, EPI_ISL_3205125, EPI_ISL_3205126, EPI_ISL_3205127, EPI_ISL_3205128, EPI_ISL_3205129, EPI_ISL_3205130, EPI_ISL_3205131, EPI_ISL_3205132, EPI_ISL_3205133, EPI_ISL_3205134, EPI_ISL_3205135, EPI_ISL_3205136, EPI_ISL_3205137, EPI_ISL_3205138, EPI_ISL_3205139, EPI_ISL_3205140, EPI_ISL_3205141, EPI_ISL_3205142, EPI_ISL_3205143, EPI_ISL_3205144, EPI_ISL_3205145, EPI_ISL_3205146, EPI_ISL_3205148, EPI_ISL_3205150, EPI_ISL_3205159, EPI_ISL_3205160, EPI_ISL_3205161, EPI_ISL_3205162, EPI_ISL_3205163, EPI_ISL_3205164, EPI_ISL_3205165, EPI_ISL_3205166, EPI_ISL_3205167, EPI_ISL_3205168, EPI_ISL_3205170, EPI_ISL_3205171, EPI_ISL_3205177, EPI_ISL_3205178, EPI_ISL_3205179, EPI_ISL_3205180, EPI_ISL_3205181, EPI_ISL_3205182, EPI_ISL_3205183, EPI_ISL_3205184, EPI_ISL_3205185, EPI_ISL_3205186, EPI_ISL_3205187, EPI_ISL_3205188, EPI_ISL_3205189, EPI_ISL_3205190, EPI_ISL_3205191, EPI_ISL_3205192, EPI_ISL_3205193, EPI_ISL_3205194, EPI_ISL_3205195, EPI_ISL_3205196, EPI_ISL_3205197, EPI_ISL_3205198, EPI_ISL_3205199, EPI_ISL_3205200, EPI_ISL_3205201, EPI_ISL_3205202, EPI_ISL_3205203, EPI_ISL_3205204, EPI_ISL_3205205, EPI_ISL_3205206, EPI_ISL_3205207, EPI_ISL_3205208, EPI_ISL_3205209, EPI_ISL_3205210, EPI_ISL_3205211, EPI_ISL_3205212, EPI_ISL_3205213, EPI_ISL_3205214, EPI_ISL_3205215, EPI_ISL_3205216, EPI_ISL_3205217, EPI_ISL_3205218, EPI_ISL_3205219, EPI_ISL_3205220, EPI_ISL_3205221, EPI_ISL_3205222, EPI_ISL_3205223, EPI_ISL_3205224, EPI_ISL_3205225, EPI_ISL_3205226, EPI_ISL_3205227, EPI_ISL_3205228, EPI_ISL_3205229, EPI_ISL_3205230, EPI_ISL_3205231, EPI_ISL_3205232, EPI_ISL_3205233, EPI_ISL_3205234, EPI_ISL_3205235, EPI_ISL_3205236, EPI_ISL_3205237, EPI_ISL_3205238, EPI_ISL_3205239, EPI_ISL_3205240, EPI_ISL_3205241, EPI_ISL_3205242, EPI_ISL_3205243, EPI_ISL_3205244, EPI_ISL_3205245, EPI_ISL_3205246, EPI_ISL_3205247, EPI_ISL_3205248, EPI_ISL_3205249, EPI_ISL_3205250, EPI_ISL_3205251, EPI_ISL_3205252, EPI_ISL_3205253, EPI_ISL_3205254, EPI_ISL_3205255, EPI_ISL_3205256, EPI_ISL_3205257, EPI_ISL_3205258, EPI_ISL_3205259, EPI_ISL_3205260, EPI_ISL_3205261, EPI_ISL_3205262, EPI_ISL_3205263, EPI_ISL_3205264, EPI_ISL_3205265, EPI_ISL_3205266, EPI_ISL_3205267, EPI_ISL_3205268, EPI_ISL_3205269, EPI_ISL_3205270, EPI_ISL_3205271, EPI_ISL_3205272, EPI_ISL_3205273, EPI_ISL_3205274, EPI_ISL_3205275, EPI_ISL_3205276, EPI_ISL_3205277, EPI_ISL_3205278, EPI_ISL_3205279, EPI_ISL_3205280, EPI_ISL_3205281, EPI_ISL_3205282, EPI_ISL_3205283, EPI_ISL_3205284, EPI_ISL_3205285, EPI_ISL_3205286, EPI_ISL_3205287, EPI_ISL_3205288, EPI_ISL_3205289, EPI_ISL_3205290, EPI_ISL_3205291, EPI_ISL_3205292, EPI_ISL_3205293, EPI_ISL_3205294, EPI_ISL_3205295, EPI_ISL_3205296, EPI_ISL_3205297, EPI_ISL_3205298, EPI_ISL_3205299, EPI_ISL_3205300, EPI_ISL_3205301, EPI_ISL_3205302, EPI_ISL_3205303, EPI_ISL_3205304, EPI_ISL_3205305, EPI_ISL_3205306, EPI_ISL_3205307, EPI_ISL_3205308, EPI_ISL_3205309, EPI_ISL_3205310, EPI_ISL_3205311, EPI_ISL_3205312, EPI_ISL_3205313, EPI_ISL_3205314, EPI_ISL_3205315, EPI_ISL_3205316, EPI_ISL_3205317, EPI_ISL_3205318, EPI_ISL_3205319, EPI_ISL_3205320, EPI_ISL_3205321, EPI_ISL_3205322, EPI_ISL_3205323, EPI_ISL_3205324, EPI_ISL_3205325, EPI_ISL_3205326, EPI_ISL_3205327, EPI_ISL_3205328, EPI_ISL_3205329, EPI_ISL_3205330, EPI_ISL_3205331, EPI_ISL_3205332, EPI_ISL_3205333, EPI_ISL_3205334, EPI_ISL_3205335, EPI_ISL_3205336, EPI_ISL_3205337, EPI_ISL_3205338, EPI_ISL_3205339, EPI_ISL_3205340, EPI_ISL_3205342, EPI_ISL_3205343, EPI_ISL_3205344, EPI_ISL_3205345, EPI_ISL_3205346, EPI_ISL_3205348, EPI_ISL_3205349, EPI_ISL_3205350, EPI_ISL_3205351, EPI_ISL_3205352, EPI_ISL_3205353, EPI_ISL_3205354, EPI_ISL_3205355, EPI_ISL_3205356, EPI_ISL_3205357, EPI_ISL_3205358, EPI_ISL_3205359, EPI_ISL_3205360, EPI_ISL_3205361, EPI_ISL_3205362, EPI_ISL_3205363, EPI_ISL_3205364, EPI_ISL_3205365, EPI_ISL_3205366, EPI_ISL_3205367, EPI_ISL_3205368, EPI_ISL_3205369, EPI_ISL_3205370, EPI_ISL_3205371, EPI_ISL_3205372, EPI_ISL_3205373, EPI_ISL_3205374, EPI_ISL_3205375, EPI_ISL_3205376, EPI_ISL_3205377, EPI_ISL_3205378, EPI_ISL_3205379, EPI_ISL_3205380, EPI_ISL_3205381, EPI_ISL_3205382, EPI_ISL_3205383, EPI_ISL_3205384, EPI_ISL_3205385, EPI_ISL_3205386, EPI_ISL_3205387, EPI_ISL_3205388, EPI_ISL_3205389, EPI_ISL_3205390, EPI_ISL_3205391, EPI_ISL_3205392, EPI_ISL_3205393, EPI_ISL_3205394, EPI_ISL_3205395, EPI_ISL_3205396, EPI_ISL_3205397, EPI_ISL_3205398, EPI_ISL_3205399, EPI_ISL_3205400, EPI_ISL_3205401, EPI_ISL_3205403, EPI_ISL_3205404, EPI_ISL_3205405, EPI_ISL_3205406, EPI_ISL_3205407, EPI_ISL_3205408, EPI_ISL_3205409, EPI_ISL_3205410, EPI_ISL_3205411, EPI_ISL_3205412, EPI_ISL_3205413, EPI_ISL_3205414, EPI_ISL_3205415, EPI_ISL_3205416, EPI_ISL_3205417, EPI_ISL_3205418, EPI_ISL_3205419, EPI_ISL_3205420, EPI_ISL_3205421, EPI_ISL_3205422, EPI_ISL_3205423, EPI_ISL_3205424, EPI_ISL_3205425, EPI_ISL_3205426, EPI_ISL_3205427, EPI_ISL_3205428, EPI_ISL_3205429, EPI_ISL_3205430, EPI_ISL_3205431, EPI_ISL_3205432, EPI_ISL_3205433, EPI_ISL_3205434, EPI_ISL_3205435, EPI_ISL_3205436, EPI_ISL_3205437, EPI_ISL_3205438, EPI_ISL_3205439, EPI_ISL_3205440, EPI_ISL_3205441, EPI_ISL_3205442, EPI_ISL_3205443, EPI_ISL_3205444, EPI_ISL_3205445, EPI_ISL_3205446, EPI_ISL_3205447, EPI_ISL_3205448, EPI_ISL_3205449, EPI_ISL_3205450, EPI_ISL_3205451, EPI_ISL_3205452, EPI_ISL_3205453, EPI_ISL_3205454, EPI_ISL_3205455, EPI_ISL_3205456, EPI_ISL_3205457, EPI_ISL_3205458, EPI_ISL_3205459, EPI_ISL_3205460 | | | |
| see above | National Virus Reference Laboratory | National Virus Reference Laboratory | Zoe Yandle, Charlene Bennett, Gabriel Gonzalez, Michael Carr, Jonathan Dean, Cillian F De Gascun |
| EPI_ISL_3205518, EPI_ISL_3205519, EPI_ISL_3205520, EPI_ISL_3205523, EPI_ISL_3205524, EPI_ISL_3205525, EPI_ISL_3205527, EPI_ISL_3205528, EPI_ISL_3205529, EPI_ISL_3205530, EPI_ISL_3205531 | | | |
| see above | LABORATOIRE UNILIANS DECINES | CNR Virus des Infections Respiratoires - France SUD | Antonin Bal, Gregory Destras, Gwendolyne Burfin, Hadrien Regue, Quentin Semanas, Martine Valette, Bruno Lina, Laurence Josset |
| EPI_ISL_3205532, EPI_ISL_3205533, EPI_ISL_3205534 | MIRIALIS CLUSES BECHET | CNR Virus des Infections Respiratoires - France SUD | Antonin Bal, Gregory Destras, Gwendolyne Burfin, Hadrien Regue, Quentin Semanas, Martine Valette, Bruno Lina, Laurence Josset |
| EPI_ISL_3205535 | LABORATOIRE BIOESTEREL | CNR Virus des Infections Respiratoires - France SUD | Antonin Bal, Gregory Destras, Gwendolyne Burfin, Hadrien Regue, Quentin Semanas, Martine Valette, Bruno Lina, Laurence Josset |
| EPI_ISL_3205536 | MIRIALIS CLUSES BECHET | CNR Virus des Infections Respiratoires - France SUD | Antonin Bal, Gregory Destras, Gwendolyne Burfin, Hadrien Regue, Quentin Semanas, Martine Valette, Bruno Lina, Laurence Josset |
| EPI_ISL_3205537, EPI_ISL_3205538, EPI_ISL_3205539 | LABORATOIRE BIOESTEREL | CNR Virus des Infections Respiratoires - France SUD | Antonin Bal, Gregory Destras, Gwendolyne Burfin, Hadrien Regue, Quentin Semanas, Martine Valette, Bruno Lina, Laurence Josset |
| EPI_ISL_3205540 | MIRIALIS CLUSES BECHET | CNR Virus des Infections Respiratoires - France SUD | Antonin Bal, Gregory Destras, Gwendolyne Burfin, Hadrien Regue, Quentin Semanas, Martine Valette, Bruno Lina, Laurence Josset |
| EPI_ISL_3205541, EPI_ISL_3205542, EPI_ISL_3205543, EPI_ISL_3205544 | DYOMEDEA-LABORATOIRE DE LA SAUVEGARDE | CNR Virus des Infections Respiratoires - France SUD | Antonin Bal, Gregory Destras, Gwendolyne Burfin, Hadrien Regue, Quentin Semanas, Martine Valette, Bruno Lina, Laurence Josset |

| | | | |
|---|---|---|---|
| EPI_ISL_3205545 | MIRIALIS CLUSES BECHET | CNR Virus des Infections Respiratoires - France SUD | Antonin Bal, Gregory Destras, Gwendolyne Burfin, Hadrien Regue, Quentin Semanas, Martine Valette, Bruno Lina, Laurence Josset |
| EPI_ISL_3205546, EPI_ISL_3205547 | DYOMEDEA-LABORATOIRE DE LA SAUVEGARDE | CNR Virus des Infections Respiratoires - France SUD | Antonin Bal, Gregory Destras, Gwendolyne Burfin, Hadrien Regue, Quentin Semanas, Martine Valette, Bruno Lina, Laurence Josset |
| EPI_ISL_3205548 | MIRIALIS CLUSES BECHET | CNR Virus des Infections Respiratoires - France SUD | Antonin Bal, Gregory Destras, Gwendolyne Burfin, Hadrien Regue, Quentin Semanas, Martine Valette, Bruno Lina, Laurence Josset |
| EPI_ISL_3205549, EPI_ISL_3205550, EPI_ISL_3205553, EPI_ISL_3205554, EPI_ISL_3205555, EPI_ISL_3205556, EPI_ISL_3205557 | DYOMEDEA-LABORATOIRE DE LA SAUVEGARDE | CNR Virus des Infections Respiratoires - France SUD | Antonin Bal, Gregory Destras, Gwendolyne Burfin, Hadrien Regue, Quentin Semanas, Martine Valette, Bruno Lina, Laurence Josset |
| EPI_ISL_3205558, EPI_ISL_3205559, EPI_ISL_3205560 | MIRIALIS CLUSES BECHET | CNR Virus des Infections Respiratoires - France SUD | Antonin Bal, Gregory Destras, Gwendolyne Burfin, Hadrien Regue, Quentin Semanas, Martine Valette, Bruno Lina, Laurence Josset |
| EPI_ISL_3205561 | LABORATOIRE DES CEDRES | CNR Virus des Infections Respiratoires - France SUD | Antonin Bal, Gregory Destras, Gwendolyne Burfin, Hadrien Regue, Quentin Semanas, Martine Valette, Bruno Lina, Laurence Josset |
| EPI_ISL_3205562, EPI_ISL_3205563, EPI_ISL_3205564 | MIRIALIS CLUSES BECHET | CNR Virus des Infections Respiratoires - France SUD | Antonin Bal, Gregory Destras, Gwendolyne Burfin, Hadrien Regue, Quentin Semanas, Martine Valette, Bruno Lina, Laurence Josset |
| EPI_ISL_3205565, EPI_ISL_3205566, EPI_ISL_3205568, EPI_ISL_3205569, EPI_ISL_3205570 | DYOMEDEA-LABORATOIRE DE LA SAUVEGARDE | CNR Virus des Infections Respiratoires - France SUD | Antonin Bal, Gregory Destras, Gwendolyne Burfin, Hadrien Regue, Quentin Semanas, Martine Valette, Bruno Lina, Laurence Josset |
| EPI_ISL_3205571 | MIRIALIS CLUSES BECHET | CNR Virus des Infections Respiratoires - France SUD | Antonin Bal, Gregory Destras, Gwendolyne Burfin, Hadrien Regue, Quentin Semanas, Martine Valette, Bruno Lina, Laurence Josset |
| EPI_ISL_3205572, EPI_ISL_3205573, EPI_ISL_3205574 | DYOMEDEA-LABORATOIRE DE LA SAUVEGARDE | CNR Virus des Infections Respiratoires - France SUD | Antonin Bal, Gregory Destras, Gwendolyne Burfin, Hadrien Regue, Quentin Semanas, Martine Valette, Bruno Lina, Laurence Josset |
| EPI_ISL_3205575 | MIRIALIS CLUSES BECHET | CNR Virus des Infections Respiratoires - France SUD | Antonin Bal, Gregory Destras, Gwendolyne Burfin, Hadrien Regue, Quentin Semanas, Martine Valette, Bruno Lina, Laurence Josset |
| EPI_ISL_3205576 | LABORATOIRE DES CEDRES | CNR Virus des Infections Respiratoires - France SUD | Antonin Bal, Gregory Destras, Gwendolyne Burfin, Hadrien Regue, Quentin Semanas, Martine Valette, Bruno Lina, Laurence Josset |
| EPI_ISL_3205577, EPI_ISL_3205578 | DYOMEDEA-LABORATOIRE DE LA SAUVEGARDE | CNR Virus des Infections Respiratoires - France SUD | Antonin Bal, Gregory Destras, Gwendolyne Burfin, Hadrien Regue, Quentin Semanas, Martine Valette, Bruno Lina, Laurence Josset |
| EPI_ISL_3205579 | LABORATOIRE BIOESTEREL | CNR Virus des Infections Respiratoires - France SUD | Antonin Bal, Gregory Destras, Gwendolyne Burfin, Hadrien Regue, Quentin Semanas, Martine Valette, Bruno Lina, Laurence Josset |
| EPI_ISL_3205580, EPI_ISL_3205581, EPI_ISL_3205582 | DYOMEDEA-LABORATOIRE DE LA SAUVEGARDE | CNR Virus des Infections Respiratoires - France SUD | Antonin Bal, Gregory Destras, Gwendolyne Burfin, Hadrien Regue, Quentin Semanas, Martine Valette, Bruno Lina, Laurence Josset |
| EPI_ISL_3205583 | MIRIALIS CLUSES BECHET | CNR Virus des Infections Respiratoires - France SUD | Antonin Bal, Gregory Destras, Gwendolyne Burfin, Hadrien Regue, Quentin Semanas, Martine Valette, Bruno Lina, Laurence Josset |
| EPI_ISL_3205584 | DYOMEDEA-LABORATOIRE DE LA SAUVEGARDE | CNR Virus des Infections Respiratoires - France SUD | Antonin Bal, Gregory Destras, Gwendolyne Burfin, Hadrien Regue, Quentin Semanas, Martine Valette, Bruno Lina, Laurence Josset |
| EPI_ISL_3205585 | LABORATOIRE DES CEDRES | CNR Virus des Infections Respiratoires - France SUD | Antonin Bal, Gregory Destras, Gwendolyne Burfin, Hadrien Regue, Quentin Semanas, Martine Valette, Bruno Lina, Laurence Josset |
| EPI_ISL_3205586 | DYOMEDEA-LABORATOIRE DE LA SAUVEGARDE | CNR Virus des Infections Respiratoires - France SUD | Antonin Bal, Gregory Destras, Gwendolyne Burfin, Hadrien Regue, Quentin Semanas, Martine Valette, Bruno Lina, Laurence Josset |
| EPI_ISL_3205587 | MIRIALIS CLUSES BECHET | CNR Virus des Infections Respiratoires - France SUD | Antonin Bal, Gregory Destras, Gwendolyne Burfin, Hadrien Regue, Quentin Semanas, Martine Valette, Bruno Lina, Laurence Josset |
| EPI_ISL_3205588 | LABORATOIRE DES CEDRES | CNR Virus des Infections Respiratoires - France SUD | Antonin Bal, Gregory Destras, Gwendolyne Burfin, Hadrien Regue, Quentin Semanas, Martine Valette, Bruno Lina, Laurence Josset |
| EPI_ISL_3205590, EPI_ISL_3205591, EPI_ISL_3205592, EPI_ISL_3205593, EPI_ISL_3205594 | DYOMEDEA-LABORATOIRE DE LA SAUVEGARDE | CNR Virus des Infections Respiratoires - France SUD | Antonin Bal, Gregory Destras, Gwendolyne Burfin, Hadrien Regue, Quentin Semanas, Martine Valette, Bruno Lina, Laurence Josset |
| EPI_ISL_3205596 | LABORATOIRE BIOESTEREL | CNR Virus des Infections Respiratoires - France SUD | Antonin Bal, Gregory Destras, Gwendolyne Burfin, Hadrien Regue, Quentin Semanas, Martine Valette, Bruno Lina, Laurence Josset |
| EPI_ISL_3205597, EPI_ISL_3205598, EPI_ISL_3205599 | DYOMEDEA-LABORATOIRE DE LA SAUVEGARDE | CNR Virus des Infections Respiratoires - France SUD | Antonin Bal, Gregory Destras, Gwendolyne Burfin, Hadrien Regue, Quentin Semanas, Martine Valette, Bruno Lina, Laurence Josset |
| EPI_ISL_3205600 | LABORATOIRE DES CEDRES | CNR Virus des Infections Respiratoires - France SUD | Antonin Bal, Gregory Destras, Gwendolyne Burfin, Hadrien Regue, Quentin Semanas, Martine Valette, Bruno Lina, Laurence Josset |
| EPI_ISL_3205602, EPI_ISL_3205603, EPI_ISL_3205604, EPI_ISL_3205605 | DYOMEDEA-LABORATOIRE DE LA SAUVEGARDE | CNR Virus des Infections Respiratoires - France SUD | Antonin Bal, Gregory Destras, Gwendolyne Burfin, Hadrien Regue, Quentin Semanas, Martine Valette, Bruno Lina, Laurence Josset |
| EPI_ISL_3205606 | LABORATOIRE DES CEDRES | CNR Virus des Infections Respiratoires - France SUD | Antonin Bal, Gregory Destras, Gwendolyne Burfin, Hadrien Regue, Quentin Semanas, Martine Valette, Bruno Lina, Laurence Josset |
| EPI_ISL_3205607, EPI_ISL_3205608, EPI_ISL_3205609, EPI_ISL_3205610, EPI_ISL_3205611, EPI_ISL_3205612, EPI_ISL_3205613 | DYOMEDEA-LABORATOIRE DE LA SAUVEGARDE | CNR Virus des Infections Respiratoires - France SUD | Antonin Bal, Gregory Destras, Gwendolyne Burfin, Hadrien Regue, Quentin Semanas, Martine Valette, Bruno Lina, Laurence Josset |
| EPI_ISL_3205614 | LABORATOIRE DES CEDRES | CNR Virus des Infections Respiratoires - France SUD | Antonin Bal, Gregory Destras, Gwendolyne Burfin, Hadrien Regue, Quentin Semanas, Martine Valette, Bruno Lina, Laurence Josset |
| EPI_ISL_3205615, EPI_ISL_3205616, EPI_ISL_3205617, EPI_ISL_3205618 | DYOMEDEA-LABORATOIRE DE LA SAUVEGARDE | CNR Virus des Infections Respiratoires - France SUD | Antonin Bal, Gregory Destras, Gwendolyne Burfin, Hadrien Regue, Quentin Semanas, Martine Valette, Bruno Lina, Laurence Josset |
| EPI_ISL_3205619 | LABORATOIRE BIOESTEREL | CNR Virus des Infections Respiratoires - France SUD | Antonin Bal, Gregory Destras, Gwendolyne Burfin, Hadrien Regue, Quentin Semanas, Martine Valette, Bruno Lina, Laurence Josset |
| EPI_ISL_3205620, EPI_ISL_3205621, EPI_ISL_3205622, EPI_ISL_3205623, EPI_ISL_3205624 | LABORATOIRE d'ANALYSES de BIOLOGIE MEDICALES | CNR Virus des Infections Respiratoires - France SUD | Antonin Bal, Gregory Destras, Gwendolyne Burfin, Hadrien Regue, Quentin Semanas, Martine Valette, Bruno Lina, Laurence Josset |
| EPI_ISL_3205625, EPI_ISL_3205626, EPI_ISL_3205627 | LABORATOIRE UNILIANS DECINES | CNR Virus des Infections Respiratoires - France SUD | Antonin Bal, Gregory Destras, Gwendolyne Burfin, Hadrien Regue, Quentin Semanas, Martine Valette, Bruno Lina, Laurence Josset |
| EPI_ISL_3205628 | LABORATOIRE DES CEDRES | CNR Virus des Infections Respiratoires - France SUD | Antonin Bal, Gregory Destras, Gwendolyne Burfin, Hadrien Regue, Quentin Semanas, Martine Valette, Bruno Lina, Laurence Josset |
| EPI_ISL_3205629, EPI_ISL_3205630, EPI_ISL_3205631, EPI_ISL_3205632, EPI_ISL_3205633, EPI_ISL_3205634, EPI_ISL_3205635, EPI_ISL_3205636, EPI_ISL_3205637 | LABORATOIRE UNILIANS DECINES | CNR Virus des Infections Respiratoires - France SUD | Antonin Bal, Gregory Destras, Gwendolyne Burfin, Hadrien Regue, Quentin Semanas, Martine Valette, Bruno Lina, Laurence Josset |
| EPI_ISL_3205638 | LABORATOIRE BOUVIER | CNR Virus des Infections Respiratoires - France SUD | Antonin Bal, Gregory Destras, Gwendolyne Burfin, Hadrien Regue, Quentin Semanas, Martine Valette, Bruno Lina, Laurence Josset |
| EPI_ISL_3205639 | LABORATOIRE UNILIANS DECINES | CNR Virus des Infections Respiratoires - France SUD | Antonin Bal, Gregory Destras, Gwendolyne Burfin, Hadrien Regue, Quentin Semanas, Martine Valette, Bruno Lina, Laurence Josset |
| EPI_ISL_3205640 | LABORATOIRE BOUVIER | CNR Virus des Infections Respiratoires - France SUD | Antonin Bal, Gregory Destras, Gwendolyne Burfin, Hadrien Regue, Quentin Semanas, Martine Valette, Bruno Lina, Laurence Josset |
| EPI_ISL_3205641 | LABORATOIRE BIOESTEREL | CNR Virus des Infections Respiratoires - France SUD | Antonin Bal, Gregory Destras, Gwendolyne Burfin, Hadrien Regue, Quentin Semanas, Martine Valette, Bruno Lina, Laurence Josset |
| EPI_ISL_3205642, EPI_ISL_3205643, | LABORATOIRE BOUVIER | CNR Virus des Infections Respiratoires - France SUD | Antonin Bal, Gregory Destras, Gwendolyne Burfin, Hadrien Regue, Quentin Semanas, Martine Valette, Bruno Lina, Laurence Josset |

| | | | |
|---|---|---|---|
| EPI_ISL_3205644, EPI_ISL_3205645, EPI_ISL_3205646, EPI_ISL_3205647, EPI_ISL_3205648, EPI_ISL_3205649 | | | |
| EPI_ISL_3205650 | LABORATOIRE DES CEDRES | CNR Virus des Infections Respiratoires - France SUD | Antonin Bal, Gregory Destras, Gwendolyne Burfin, Hadrien Regue, Quentin Semanas, Martine Valette, Bruno Lina, Laurence Josset |
| EPI_ISL_3205651 | LABORATOIRE BOUVIER | CNR Virus des Infections Respiratoires - France SUD | Antonin Bal, Gregory Destras, Gwendolyne Burfin, Hadrien Regue, Quentin Semanas, Martine Valette, Bruno Lina, Laurence Josset |
| EPI_ISL_3205652 | LABORATOIRE DES CEDRES | CNR Virus des Infections Respiratoires - France SUD | Antonin Bal, Gregory Destras, Gwendolyne Burfin, Hadrien Regue, Quentin Semanas, Martine Valette, Bruno Lina, Laurence Josset |
| EPI_ISL_3205654 | LABORATOIRE BIOESTEREL | CNR Virus des Infections Respiratoires - France SUD | Antonin Bal, Gregory Destras, Gwendolyne Burfin, Hadrien Regue, Quentin Semanas, Martine Valette, Bruno Lina, Laurence Josset |
| EPI_ISL_3205655, EPI_ISL_3205656, EPI_ISL_3205658, EPI_ISL_3205660, EPI_ISL_3205661 | LABORATOIRE UNILIANS DECINES | CNR Virus des Infections Respiratoires - France SUD | Antonin Bal, Gregory Destras, Gwendolyne Burfin, Hadrien Regue, Quentin Semanas, Martine Valette, Bruno Lina, Laurence Josset |
| EPI_ISL_3205662 | LABORATOIRE BIOESTEREL | CNR Virus des Infections Respiratoires - France SUD | Antonin Bal, Gregory Destras, Gwendolyne Burfin, Hadrien Regue, Quentin Semanas, Martine Valette, Bruno Lina, Laurence Josset |
| EPI_ISL_3205663, EPI_ISL_3205664, EPI_ISL_3205665 | LABORATOIRE UNILIANS DECINES | CNR Virus des Infections Respiratoires - France SUD | Antonin Bal, Gregory Destras, Gwendolyne Burfin, Hadrien Regue, Quentin Semanas, Martine Valette, Bruno Lina, Laurence Josset |
| EPI_ISL_3205666 | LABORATOIRE DES CEDRES | CNR Virus des Infections Respiratoires - France SUD | Antonin Bal, Gregory Destras, Gwendolyne Burfin, Hadrien Regue, Quentin Semanas, Martine Valette, Bruno Lina, Laurence Josset |
| EPI_ISL_3205667 | LABORATOIRE UNILIANS DECINES | CNR Virus des Infections Respiratoires - France SUD | Antonin Bal, Gregory Destras, Gwendolyne Burfin, Hadrien Regue, Quentin Semanas, Martine Valette, Bruno Lina, Laurence Josset |
| EPI_ISL_3205669 | LABORATOIRE BIOESTEREL | CNR Virus des Infections Respiratoires - France SUD | Antonin Bal, Gregory Destras, Gwendolyne Burfin, Hadrien Regue, Quentin Semanas, Martine Valette, Bruno Lina, Laurence Josset |
| EPI_ISL_3205670 | LABORATOIRE ALPIGENE | CNR Virus des Infections Respiratoires - France SUD | Antonin Bal, Gregory Destras, Gwendolyne Burfin, Hadrien Regue, Quentin Semanas, Martine Valette, Bruno Lina, Laurence Josset |
| EPI_ISL_3205671 | LABORATOIRE BIOESTEREL | CNR Virus des Infections Respiratoires - France SUD | Antonin Bal, Gregory Destras, Gwendolyne Burfin, Hadrien Regue, Quentin Semanas, Martine Valette, Bruno Lina, Laurence Josset |
| EPI_ISL_3205672, EPI_ISL_3205674 | LABORATOIRE UNILIANS DECINES | CNR Virus des Infections Respiratoires - France SUD | Antonin Bal, Gregory Destras, Gwendolyne Burfin, Hadrien Regue, Quentin Semanas, Martine Valette, Bruno Lina, Laurence Josset |
| EPI_ISL_3205676 | LABORATOIRE- BIOESTERL | CNR Virus des Infections Respiratoires - France SUD | Antonin Bal, Gregory Destras, Gwendolyne Burfin, Hadrien Regue, Quentin Semanas, Martine Valette, Bruno Lina, Laurence Josset |
| EPI_ISL_3205678 | LABORATOIRE UNILIANS DECINES | CNR Virus des Infections Respiratoires - France SUD | Antonin Bal, Gregory Destras, Gwendolyne Burfin, Hadrien Regue, Quentin Semanas, Martine Valette, Bruno Lina, Laurence Josset |
| EPI_ISL_3205679 | LABORATOIRE BIOESTEREL | CNR Virus des Infections Respiratoires - France SUD | Antonin Bal, Gregory Destras, Gwendolyne Burfin, Hadrien Regue, Quentin Semanas, Martine Valette, Bruno Lina, Laurence Josset |
| EPI_ISL_3205681, EPI_ISL_3205682 | LABORATOIRE- BIOESTERL | CNR Virus des Infections Respiratoires - France SUD | Antonin Bal, Gregory Destras, Gwendolyne Burfin, Hadrien Regue, Quentin Semanas, Martine Valette, Bruno Lina, Laurence Josset |
| EPI_ISL_3205683, EPI_ISL_3205684 | LABORATOIRE BIOESTEREL | CNR Virus des Infections Respiratoires - France SUD | Antonin Bal, Gregory Destras, Gwendolyne Burfin, Hadrien Regue, Quentin Semanas, Martine Valette, Bruno Lina, Laurence Josset |
| EPI_ISL_3205686 | LABORATOIRE UNILIANS DECINES | CNR Virus des Infections Respiratoires - France SUD | Antonin Bal, Gregory Destras, Gwendolyne Burfin, Hadrien Regue, Quentin Semanas, Martine Valette, Bruno Lina, Laurence Josset |
| EPI_ISL_3205688, EPI_ISL_3205689 | LABORATOIRE BIOESTEREL | CNR Virus des Infections Respiratoires - France SUD | Antonin Bal, Gregory Destras, Gwendolyne Burfin, Hadrien Regue, Quentin Semanas, Martine Valette, Bruno Lina, Laurence Josset |
| EPI_ISL_3205690 | LABORATOIRE ALPIGENE | CNR Virus des Infections Respiratoires - France SUD | Antonin Bal, Gregory Destras, Gwendolyne Burfin, Hadrien Regue, Quentin Semanas, Martine Valette, Bruno Lina, Laurence Josset |
| EPI_ISL_3205691, EPI_ISL_3205693, EPI_ISL_3205696 | LABORATOIRE BIOESTEREL | CNR Virus des Infections Respiratoires - France SUD | Antonin Bal, Gregory Destras, Gwendolyne Burfin, Hadrien Regue, Quentin Semanas, Martine Valette, Bruno Lina, Laurence Josset |
| EPI_ISL_3205697 | LABORATOIRE ALPIGENE | CNR Virus des Infections Respiratoires - France SUD | Antonin Bal, Gregory Destras, Gwendolyne Burfin, Hadrien Regue, Quentin Semanas, Martine Valette, Bruno Lina, Laurence Josset |
| EPI_ISL_3205699 | LABORATOIRE BIOESTEREL | CNR Virus des Infections Respiratoires - France SUD | Antonin Bal, Gregory Destras, Gwendolyne Burfin, Hadrien Regue, Quentin Semanas, Martine Valette, Bruno Lina, Laurence Josset |
| EPI_ISL_3205700 | LABORATOIRE ALPIGENE | CNR Virus des Infections Respiratoires - France SUD | Antonin Bal, Gregory Destras, Gwendolyne Burfin, Hadrien Regue, Quentin Semanas, Martine Valette, Bruno Lina, Laurence Josset |
| EPI_ISL_3205702 | LABORATOIRE BIOESTEREL | CNR Virus des Infections Respiratoires - France SUD | Antonin Bal, Gregory Destras, Gwendolyne Burfin, Hadrien Regue, Quentin Semanas, Martine Valette, Bruno Lina, Laurence Josset |
| EPI_ISL_3205706 | LABORATOIRE ALPIGENE | CNR Virus des Infections Respiratoires - France SUD | Antonin Bal, Gregory Destras, Gwendolyne Burfin, Hadrien Regue, Quentin Semanas, Martine Valette, Bruno Lina, Laurence Josset |
| EPI_ISL_3205707 | DYOMEDEA-LABORATOIRE DE LA SAUVEGARDE | CNR Virus des Infections Respiratoires - France SUD | Antonin Bal, Gregory Destras, Gwendolyne Burfin, Hadrien Regue, Quentin Semanas, Martine Valette, Bruno Lina, Laurence Josset |
| EPI_ISL_3205708 | LABORATOIRE ALPIGENE | CNR Virus des Infections Respiratoires - France SUD | Antonin Bal, Gregory Destras, Gwendolyne Burfin, Hadrien Regue, Quentin Semanas, Martine Valette, Bruno Lina, Laurence Josset |
| EPI_ISL_3205710, EPI_ISL_3205711, EPI_ISL_3205712 | DYOMEDEA-LABORATOIRE DE LA SAUVEGARDE | CNR Virus des Infections Respiratoires - France SUD | Antonin Bal, Gregory Destras, Gwendolyne Burfin, Hadrien Regue, Quentin Semanas, Martine Valette, Bruno Lina, Laurence Josset |
| EPI_ISL_3205713 | CENTRE HOSPITALIER DE GUERET | CNR Virus des Infections Respiratoires - France SUD | Antonin Bal, Gregory Destras, Gwendolyne Burfin, Hadrien Regue, Quentin Semanas, Martine Valette, Bruno Lina, Laurence Josset |
| EPI_ISL_3205714 | CENTRE HOSPITALIER DE LA COTE BASQUE | CNR Virus des Infections Respiratoires - France SUD | Antonin Bal, Gregory Destras, Gwendolyne Burfin, Hadrien Regue, Quentin Semanas, Martine Valette, Bruno Lina, Laurence Josset |
| EPI_ISL_3205716 | LABORATOIRE- BIOESTERL | CNR Virus des Infections Respiratoires - France SUD | Antonin Bal, Gregory Destras, Gwendolyne Burfin, Hadrien Regue, Quentin Semanas, Martine Valette, Bruno Lina, Laurence Josset |
| EPI_ISL_3205717, EPI_ISL_3205718 | CENTRE HOSPITALIER DE LA COTE BASQUE | CNR Virus des Infections Respiratoires - France SUD | Antonin Bal, Gregory Destras, Gwendolyne Burfin, Hadrien Regue, Quentin Semanas, Martine Valette, Bruno Lina, Laurence Josset |
| EPI_ISL_3205719 | LABORATOIRE- BIOESTERL | CNR Virus des Infections Respiratoires - France SUD | Antonin Bal, Gregory Destras, Gwendolyne Burfin, Hadrien Regue, Quentin Semanas, Martine Valette, Bruno Lina, Laurence Josset |
| EPI_ISL_3205720 | CNR Virus des Infections Respiratoires - France SUD | CNR Virus des Infections Respiratoires - France SUD | Antonin Bal, Gregory Destras, Gwendolyne Burfin, Hadrien Regue, Quentin Semanas, Martine Valette, Bruno Lina, Laurence Josset |
| EPI_ISL_3205721 | CENTRE HOSPITALIER DE LA COTE BASQUE | CNR Virus des Infections Respiratoires - France SUD | Antonin Bal, Gregory Destras, Gwendolyne Burfin, Hadrien Regue, Quentin Semanas, Martine Valette, Bruno Lina, Laurence Josset |
| EPI_ISL_3205722, EPI_ISL_3205723 | CNR Virus des Infections Respiratoires - France SUD | CNR Virus des Infections Respiratoires - France SUD | Antonin Bal, Gregory Destras, Gwendolyne Burfin, Hadrien Regue, Quentin Semanas, Martine Valette, Bruno Lina, Laurence Josset |
| EPI_ISL_3205724, EPI_ISL_3205726 | LABORATOIRE- BIOESTERL | CNR Virus des Infections Respiratoires - France SUD | Antonin Bal, Gregory Destras, Gwendolyne Burfin, Hadrien Regue, Quentin Semanas, Martine Valette, Bruno Lina, Laurence Josset |
| EPI_ISL_3205727 | CNR Virus des Infections Respiratoires - France SUD | CNR Virus des Infections Respiratoires - France SUD | Antonin Bal, Gregory Destras, Gwendolyne Burfin, Hadrien Regue, Quentin Semanas, Martine Valette, Bruno Lina, Laurence Josset |
| EPI_ISL_3205728 | LAM GEN-BIO GRAVANCHES | CNR Virus des Infections Respiratoires - France SUD | Antonin Bal, Gregory Destras, Gwendolyne Burfin, Hadrien Regue, Quentin Semanas, Martine Valette, Bruno Lina, Laurence Josset |
| EPI_ISL_3205729 | CENTRE HOSPITALIER DE LA COTE BASQUE | CNR Virus des Infections Respiratoires - France SUD | Antonin Bal, Gregory Destras, Gwendolyne Burfin, Hadrien Regue, Quentin Semanas, Martine Valette, Bruno Lina, Laurence Josset |
| EPI_ISL_3205730 | LABORATOIRE- BIOESTERL | CNR Virus des Infections Respiratoires - France SUD | Antonin Bal, Gregory Destras, Gwendolyne Burfin, Hadrien Regue, Quentin Semanas, Martine Valette, Bruno Lina, Laurence Josset |
| EPI_ISL_3205731 | CNR Virus des Infections Respiratoires - France SUD | CNR Virus des Infections Respiratoires - France SUD | Antonin Bal, Gregory Destras, Gwendolyne Burfin, Hadrien Regue, Quentin Semanas, Martine Valette, Bruno Lina, Laurence Josset |
| EPI_ISL_3205732 | LAM GEN-BIO GRAVANCHES | CNR Virus des Infections Respiratoires - France SUD | Antonin Bal, Gregory Destras, Gwendolyne Burfin, Hadrien Regue, Quentin Semanas, Martine Valette, Bruno Lina, Laurence Josset |
| EPI_ISL_3205733 | CNR Virus des Infections Respiratoires - France SUD | CNR Virus des Infections Respiratoires - France SUD | Antonin Bal, Gregory Destras, Gwendolyne Burfin, Hadrien Regue, Quentin Semanas, Martine Valette, Bruno Lina, Laurence Josset |
| EPI_ISL_3205734, EPI_ISL_3205735 | LABORATOIRE- BIOESTERL | CNR Virus des Infections Respiratoires - France SUD | Antonin Bal, Gregory Destras, Gwendolyne Burfin, Hadrien Regue, Quentin Semanas, Martine Valette, Bruno Lina, Laurence Josset |
| EPI_ISL_3205736 | CNR Virus des Infections Respiratoires - France SUD | CNR Virus des Infections Respiratoires - France SUD | Antonin Bal, Gregory Destras, Gwendolyne Burfin, Hadrien Regue, Quentin Semanas, Martine Valette, Bruno Lina, Laurence Josset |
| EPI_ISL_3205737 | AX BIO OCEAN | CNR Virus des Infections Respiratoires - France SUD | Antonin Bal, Gregory Destras, Gwendolyne Burfin, Hadrien Regue, Quentin Semanas, Martine Valette, Bruno Lina, Laurence Josset |
| EPI_ISL_3205738 | CNR Virus des Infections Respiratoires - France SUD | CNR Virus des Infections Respiratoires - France SUD | Antonin Bal, Gregory Destras, Gwendolyne Burfin, Hadrien Regue, Quentin Semanas, Martine Valette, Bruno Lina, Laurence Josset |
| EPI_ISL_3205739 | LABORATOIRE UNILIANS DECINES | CNR Virus des Infections Respiratoires - France SUD | Antonin Bal, Gregory Destras, Gwendolyne Burfin, Hadrien Regue, Quentin Semanas, Martine Valette, Bruno Lina, Laurence Josset |
| EPI_ISL_3205740 | LAM GEN-BIO GRAVANCHES | CNR Virus des Infections Respiratoires - France SUD | Antonin Bal, Gregory Destras, Gwendolyne Burfin, Hadrien Regue, Quentin Semanas, Martine Valette, Bruno Lina, Laurence Josset |
| EPI_ISL_3205741 | AX BIO OCEAN | CNR Virus des Infections Respiratoires - France SUD | Antonin Bal, Gregory Destras, Gwendolyne Burfin, Hadrien Regue, Quentin Semanas, Martine Valette, Bruno Lina, Laurence Josset |
| EPI_ISL_3205742 | LABORATOIRE- BIOESTERL | CNR Virus des Infections Respiratoires - France SUD | Antonin Bal, Gregory Destras, Gwendolyne Burfin, Hadrien Regue, Quentin Semanas, Martine Valette, Bruno Lina, Laurence Josset |
| EPI_ISL_3205743, EPI_ISL_3205745 | CNR Virus des Infections Respiratoires - France SUD | CNR Virus des Infections Respiratoires - France SUD | Antonin Bal, Gregory Destras, Gwendolyne Burfin, Hadrien Regue, Quentin Semanas, Martine Valette, Bruno Lina, Laurence Josset |
| EPI_ISL_3205746 | LABORATOIRE- BIOESTERL | CNR Virus des Infections Respiratoires - France SUD | Antonin Bal, Gregory Destras, Gwendolyne Burfin, Hadrien Regue, Quentin Semanas, Martine Valette, Bruno Lina, Laurence Josset |
| EPI_ISL_3205747 | CNR Virus des Infections Respiratoires - France SUD | CNR Virus des Infections Respiratoires - France SUD | Antonin Bal, Gregory Destras, Gwendolyne Burfin, Hadrien Regue, Quentin Semanas, Martine Valette, Bruno Lina, Laurence Josset |
| EPI_ISL_3205748 | AX BIO OCEAN | CNR Virus des Infections Respiratoires - France SUD | Antonin Bal, Gregory Destras, Gwendolyne Burfin, Hadrien Regue, Quentin Semanas, Martine Valette, Bruno Lina, Laurence Josset |
| EPI_ISL_3205749 | CNR Virus des Infections Respiratoires - France SUD | CNR Virus des Infections Respiratoires - France SUD | Antonin Bal, Gregory Destras, Gwendolyne Burfin, Hadrien Regue, Quentin Semanas, Martine Valette, Bruno Lina, Laurence Josset |

| | | | |
|---|---|---|---|
| EPI_ISL_3205750 | LABORATOIRE- BIOESTERL | CNR Virus des Infections Respiratoires - France SUD | Antonin Bal, Gregory Destras, Gwendolyne Burfin, Hadrien Regue, Quentin Semanas, Martine Valette, Bruno Lina, Laurence Josset |
| EPI_ISL_3205752 | CNR Virus des Infections Respiratoires - France SUD | CNR Virus des Infections Respiratoires - France SUD | Antonin Bal, Gregory Destras, Gwendolyne Burfin, Hadrien Regue, Quentin Semanas, Martine Valette, Bruno Lina, Laurence Josset |
| EPI_ISL_3205753 | LABORATOIRE- BIOESTERL | CNR Virus des Infections Respiratoires - France SUD | Antonin Bal, Gregory Destras, Gwendolyne Burfin, Hadrien Regue, Quentin Semanas, Martine Valette, Bruno Lina, Laurence Josset |
| EPI_ISL_3205754 | AX BIO OCEAN | CNR Virus des Infections Respiratoires - France SUD | Antonin Bal, Gregory Destras, Gwendolyne Burfin, Hadrien Regue, Quentin Semanas, Martine Valette, Bruno Lina, Laurence Josset |
| EPI_ISL_3205755, EPI_ISL_3205757 | CNR Virus des Infections Respiratoires - France SUD | CNR Virus des Infections Respiratoires - France SUD | Antonin Bal, Gregory Destras, Gwendolyne Burfin, Hadrien Regue, Quentin Semanas, Martine Valette, Bruno Lina, Laurence Josset |
| EPI_ISL_3205758 | LABORATOIRE- BIOESTERL | CNR Virus des Infections Respiratoires - France SUD | Antonin Bal, Gregory Destras, Gwendolyne Burfin, Hadrien Regue, Quentin Semanas, Martine Valette, Bruno Lina, Laurence Josset |
| EPI_ISL_3205760 | AX BIO OCEAN | CNR Virus des Infections Respiratoires - France SUD | Antonin Bal, Gregory Destras, Gwendolyne Burfin, Hadrien Regue, Quentin Semanas, Martine Valette, Bruno Lina, Laurence Josset |
| EPI_ISL_3205761 | CNR Virus des Infections Respiratoires - France SUD | CNR Virus des Infections Respiratoires - France SUD | Antonin Bal, Gregory Destras, Gwendolyne Burfin, Hadrien Regue, Quentin Semanas, Martine Valette, Bruno Lina, Laurence Josset |
| EPI_ISL_3205762 | LABORATOIRE- BIOESTERL | CNR Virus des Infections Respiratoires - France SUD | Antonin Bal, Gregory Destras, Gwendolyne Burfin, Hadrien Regue, Quentin Semanas, Martine Valette, Bruno Lina, Laurence Josset |
| EPI_ISL_3205764 | AX BIO OCEAN | CNR Virus des Infections Respiratoires - France SUD | Antonin Bal, Gregory Destras, Gwendolyne Burfin, Hadrien Regue, Quentin Semanas, Martine Valette, Bruno Lina, Laurence Josset |
| EPI_ISL_3205765 | CNR Virus des Infections Respiratoires - France SUD | CNR Virus des Infections Respiratoires - France SUD | Antonin Bal, Gregory Destras, Gwendolyne Burfin, Hadrien Regue, Quentin Semanas, Martine Valette, Bruno Lina, Laurence Josset |
| EPI_ISL_3205767 | LABORATOIRE- BIOESTERL | CNR Virus des Infections Respiratoires - France SUD | Antonin Bal, Gregory Destras, Gwendolyne Burfin, Hadrien Regue, Quentin Semanas, Martine Valette, Bruno Lina, Laurence Josset |
| EPI_ISL_3205768 | CNR Virus des Infections Respiratoires - France SUD | CNR Virus des Infections Respiratoires - France SUD | Antonin Bal, Gregory Destras, Gwendolyne Burfin, Hadrien Regue, Quentin Semanas, Martine Valette, Bruno Lina, Laurence Josset |
| EPI_ISL_3205769 | AX BIO OCEAN | CNR Virus des Infections Respiratoires - France SUD | Antonin Bal, Gregory Destras, Gwendolyne Burfin, Hadrien Regue, Quentin Semanas, Martine Valette, Bruno Lina, Laurence Josset |
| EPI_ISL_3205773 | LABORATOIRE- BIOESTERL | CNR Virus des Infections Respiratoires - France SUD | Antonin Bal, Gregory Destras, Gwendolyne Burfin, Hadrien Regue, Quentin Semanas, Martine Valette, Bruno Lina, Laurence Josset |
| EPI_ISL_3205774, EPI_ISL_3205776, EPI_ISL_3205778 | AX BIO OCEAN | CNR Virus des Infections Respiratoires - France SUD | Antonin Bal, Gregory Destras, Gwendolyne Burfin, Hadrien Regue, Quentin Semanas, Martine Valette, Bruno Lina, Laurence Josset |
| EPI_ISL_3205779, EPI_ISL_3205781, EPI_ISL_3205784 | CNR Virus des Infections Respiratoires - France SUD | CNR Virus des Infections Respiratoires - France SUD | Antonin Bal, Gregory Destras, Gwendolyne Burfin, Hadrien Regue, Quentin Semanas, Martine Valette, Bruno Lina, Laurence Josset |
| EPI_ISL_3205785 | LABORATOIRE- BIOESTERL | CNR Virus des Infections Respiratoires - France SUD | Antonin Bal, Gregory Destras, Gwendolyne Burfin, Hadrien Regue, Quentin Semanas, Martine Valette, Bruno Lina, Laurence Josset |
| EPI_ISL_3205787 | AX BIO OCEAN | CNR Virus des Infections Respiratoires - France SUD | Antonin Bal, Gregory Destras, Gwendolyne Burfin, Hadrien Regue, Quentin Semanas, Martine Valette, Bruno Lina, Laurence Josset |
| EPI_ISL_3205789 | LABORATOIRE- BIOESTERL | CNR Virus des Infections Respiratoires - France SUD | Antonin Bal, Gregory Destras, Gwendolyne Burfin, Hadrien Regue, Quentin Semanas, Martine Valette, Bruno Lina, Laurence Josset |
| EPI_ISL_3205790 | AX BIO OCEAN | CNR Virus des Infections Respiratoires - France SUD | Antonin Bal, Gregory Destras, Gwendolyne Burfin, Hadrien Regue, Quentin Semanas, Martine Valette, Bruno Lina, Laurence Josset |
| EPI_ISL_3205793, EPI_ISL_3205797 | LABORATOIRE- BIOESTERL | CNR Virus des Infections Respiratoires - France SUD | Antonin Bal, Gregory Destras, Gwendolyne Burfin, Hadrien Regue, Quentin Semanas, Martine Valette, Bruno Lina, Laurence Josset |
| EPI_ISL_3205799, EPI_ISL_3205800, EPI_ISL_3205801, EPI_ISL_3205804, EPI_ISL_3205809, EPI_ISL_3205811 | AX BIO OCEAN | CNR Virus des Infections Respiratoires - France SUD | Antonin Bal, Gregory Destras, Gwendolyne Burfin, Hadrien Regue, Quentin Semanas, Martine Valette, Bruno Lina, Laurence Josset |
| EPI_ISL_3205812, EPI_ISL_3205814 | LABORATOIRE- BIOESTERL | CNR Virus des Infections Respiratoires - France SUD | Antonin Bal, Gregory Destras, Gwendolyne Burfin, Hadrien Regue, Quentin Semanas, Martine Valette, Bruno Lina, Laurence Josset |
| EPI_ISL_3205815, EPI_ISL_3205816 | AX BIO OCEAN | CNR Virus des Infections Respiratoires - France SUD | Antonin Bal, Gregory Destras, Gwendolyne Burfin, Hadrien Regue, Quentin Semanas, Martine Valette, Bruno Lina, Laurence Josset |
| EPI_ISL_3205817 | LABORATOIRE- BIOESTERL | CNR Virus des Infections Respiratoires - France SUD | Antonin Bal, Gregory Destras, Gwendolyne Burfin, Hadrien Regue, Quentin Semanas, Martine Valette, Bruno Lina, Laurence Josset |
| EPI_ISL_3205818, EPI_ISL_3205819, EPI_ISL_3205820, EPI_ISL_3205821, EPI_ISL_3205822, EPI_ISL_3205823 | AX BIO OCEAN | CNR Virus des Infections Respiratoires - France SUD | Antonin Bal, Gregory Destras, Gwendolyne Burfin, Hadrien Regue, Quentin Semanas, Martine Valette, Bruno Lina, Laurence Josset |
| EPI_ISL_3205824 | LABORATOIRE- BIOESTERL | CNR Virus des Infections Respiratoires - France SUD | Antonin Bal, Gregory Destras, Gwendolyne Burfin, Hadrien Regue, Quentin Semanas, Martine Valette, Bruno Lina, Laurence Josset |
| EPI_ISL_3205825 | AX BIO OCEAN | CNR Virus des Infections Respiratoires - France SUD | Antonin Bal, Gregory Destras, Gwendolyne Burfin, Hadrien Regue, Quentin Semanas, Martine Valette, Bruno Lina, Laurence Josset |
| EPI_ISL_3205827 | LABORATOIRE- BIOESTERL | CNR Virus des Infections Respiratoires - France SUD | Antonin Bal, Gregory Destras, Gwendolyne Burfin, Hadrien Regue, Quentin Semanas, Martine Valette, Bruno Lina, Laurence Josset |
| EPI_ISL_3205828 | AX BIO OCEAN | CNR Virus des Infections Respiratoires - France SUD | Antonin Bal, Gregory Destras, Gwendolyne Burfin, Hadrien Regue, Quentin Semanas, Martine Valette, Bruno Lina, Laurence Josset |
| EPI_ISL_3205829, EPI_ISL_3205830 | LABORATOIRE- BIOESTERL | CNR Virus des Infections Respiratoires - France SUD | Antonin Bal, Gregory Destras, Gwendolyne Burfin, Hadrien Regue, Quentin Semanas, Martine Valette, Bruno Lina, Laurence Josset |
| EPI_ISL_3205831, EPI_ISL_3205832, EPI_ISL_3205833, EPI_ISL_3205834, EPI_ISL_3205835, EPI_ISL_3205836, EPI_ISL_3205837, EPI_ISL_3205839, EPI_ISL_3205840, EPI_ISL_3205841, EPI_ISL_3205842, EPI_ISL_3205843, EPI_ISL_3205844, EPI_ISL_3205845, EPI_ISL_3205846, EPI_ISL_3205847, EPI_ISL_3205848, EPI_ISL_3205849, EPI_ISL_3205850, EPI_ISL_3205851, EPI_ISL_3205852, EPI_ISL_3205853, EPI_ISL_3205855 | | | |
| see above | AX BIO OCEAN | CNR Virus des Infections Respiratoires - France SUD | Antonin Bal, Gregory Destras, Gwendolyne Burfin, Hadrien Regue, Quentin Semanas, Martine Valette, Bruno Lina, Laurence Josset |
| EPI_ISL_3205856 | CENTRE HOSPITALIER DE BOURG EN BRESSE | CNR Virus des Infections Respiratoires - France SUD | Antonin Bal, Gregory Destras, Gwendolyne Burfin, Hadrien Regue, Quentin Semanas, Martine Valette, Bruno Lina, Laurence Josset |
| EPI_ISL_3205857, EPI_ISL_3205860, EPI_ISL_3205861, EPI_ISL_3205862, EPI_ISL_3205863, EPI_ISL_3205864, EPI_ISL_3205865, EPI_ISL_3205866, EPI_ISL_3205867, EPI_ISL_3205868, EPI_ISL_3205869, EPI_ISL_3205870, EPI_ISL_3205871, EPI_ISL_3205872, EPI_ISL_3205873, EPI_ISL_3205874, EPI_ISL_3205875, EPI_ISL_3205876 | | | |
| see above | AX BIO OCEAN | CNR Virus des Infections Respiratoires - France SUD | Antonin Bal, Gregory Destras, Gwendolyne Burfin, Hadrien Regue, Quentin Semanas, Martine Valette, Bruno Lina, Laurence Josset |
| EPI_ISL_3205877 | LAM GEN-BIO GRAVANCHES | CNR Virus des Infections Respiratoires - France SUD | Antonin Bal, Gregory Destras, Gwendolyne Burfin, Hadrien Regue, Quentin Semanas, Martine Valette, Bruno Lina, Laurence Josset |
| EPI_ISL_3205878 | AX BIO OCEAN | CNR Virus des Infections Respiratoires - France SUD | Antonin Bal, Gregory Destras, Gwendolyne Burfin, Hadrien Regue, Quentin Semanas, Martine Valette, Bruno Lina, Laurence Josset |
| EPI_ISL_3205879 | LAM GEN-BIO GRAVANCHES | CNR Virus des Infections Respiratoires - France SUD | Antonin Bal, Gregory Destras, Gwendolyne Burfin, Hadrien Regue, Quentin Semanas, Martine Valette, Bruno Lina, Laurence Josset |
| EPI_ISL_3205880, EPI_ISL_3205881 | AX BIO OCEAN | CNR Virus des Infections Respiratoires - France SUD | Antonin Bal, Gregory Destras, Gwendolyne Burfin, Hadrien Regue, Quentin Semanas, Martine Valette, Bruno Lina, Laurence Josset |
| EPI_ISL_3205882, EPI_ISL_3205883 | LAM GEN-BIO GRAVANCHES | CNR Virus des Infections Respiratoires - France SUD | Antonin Bal, Gregory Destras, Gwendolyne Burfin, Hadrien Regue, Quentin Semanas, Martine Valette, Bruno Lina, Laurence Josset |
| EPI_ISL_3205884 | AX BIO OCEAN | CNR Virus des Infections Respiratoires - France SUD | Antonin Bal, Gregory Destras, Gwendolyne Burfin, Hadrien Regue, Quentin Semanas, Martine Valette, Bruno Lina, Laurence Josset |
| EPI_ISL_3205885 | LAM GEN-BIO GRAVANCHES | CNR Virus des Infections Respiratoires - France SUD | Antonin Bal, Gregory Destras, Gwendolyne Burfin, Hadrien Regue, Quentin Semanas, Martine Valette, Bruno Lina, Laurence Josset |
| EPI_ISL_3205886, EPI_ISL_3205887 | AX BIO OCEAN | CNR Virus des Infections Respiratoires - France SUD | Antonin Bal, Gregory Destras, Gwendolyne Burfin, Hadrien Regue, Quentin Semanas, Martine Valette, Bruno Lina, Laurence Josset |
| EPI_ISL_3205888, EPI_ISL_3205889, EPI_ISL_3205890 | LAM GEN-BIO GRAVANCHES | CNR Virus des Infections Respiratoires - France SUD | Antonin Bal, Gregory Destras, Gwendolyne Burfin, Hadrien Regue, Quentin Semanas, Martine Valette, Bruno Lina, Laurence Josset |
| EPI_ISL_3205891 | AX BIO OCEAN | CNR Virus des Infections Respiratoires - France SUD | Antonin Bal, Gregory Destras, Gwendolyne Burfin, Hadrien Regue, Quentin Semanas, Martine Valette, Bruno Lina, Laurence Josset |
| EPI_ISL_3205892, EPI_ISL_3205894, EPI_ISL_3205895 | LAM GEN-BIO GRAVANCHES | CNR Virus des Infections Respiratoires - France SUD | Antonin Bal, Gregory Destras, Gwendolyne Burfin, Hadrien Regue, Quentin Semanas, Martine Valette, Bruno Lina, Laurence Josset |
| EPI_ISL_3205897, EPI_ISL_3205898, EPI_ISL_3205899 | LABORATOIRE UNILIANS DECINES | CNR Virus des Infections Respiratoires - France SUD | Antonin Bal, Gregory Destras, Gwendolyne Burfin, Hadrien Regue, Quentin Semanas, Martine Valette, Bruno Lina, Laurence Josset |
| EPI_ISL_3205900 | LAM GEN-BIO GRAVANCHES | CNR Virus des Infections Respiratoires - France SUD | Antonin Bal, Gregory Destras, Gwendolyne Burfin, Hadrien Regue, Quentin Semanas, Martine Valette, Bruno Lina, Laurence Josset |
| EPI_ISL_3205901 | LABORATOIRE UNILIANS DECINES | CNR Virus des Infections Respiratoires - France SUD | Antonin Bal, Gregory Destras, Gwendolyne Burfin, Hadrien Regue, Quentin Semanas, Martine Valette, Bruno Lina, Laurence Josset |
| EPI_ISL_3205902, EPI_ISL_3205903 | LAM GEN-BIO GRAVANCHES | CNR Virus des Infections Respiratoires - France SUD | Antonin Bal, Gregory Destras, Gwendolyne Burfin, Hadrien Regue, Quentin Semanas, Martine Valette, Bruno Lina, Laurence Josset |
| EPI_ISL_3205904 | AX BIO OCEAN | CNR Virus des Infections Respiratoires - France SUD | Antonin Bal, Gregory Destras, Gwendolyne Burfin, Hadrien Regue, Quentin Semanas, Martine Valette, Bruno Lina, Laurence Josset |
| EPI_ISL_3205905, EPI_ISL_3205907 | LABORATOIRE UNILIANS DECINES | CNR Virus des Infections Respiratoires - France SUD | Antonin Bal, Gregory Destras, Gwendolyne Burfin, Hadrien Regue, Quentin Semanas, Martine Valette, Bruno Lina, Laurence Josset |
| EPI_ISL_3205908 | AX BIO OCEAN | CNR Virus des Infections Respiratoires - France SUD | Antonin Bal, Gregory Destras, Gwendolyne Burfin, Hadrien Regue, Quentin Semanas, Martine Valette, Bruno Lina, Laurence Josset |

| | | | |
|---|---|---|---|
| EPI_ISL_3205909, EPI_ISL_3205910, EPI_ISL_3205911, EPI_ISL_3205912, EPI_ISL_3205913, EPI_ISL_3205914, EPI_ISL_3205915, EPI_ISL_3205916, EPI_ISL_3205917, EPI_ISL_3205918 | LABORATOIRE UNILIANS DECINES | CNR Virus des Infections Respiratoires - France SUD | Antonin Bal, Gregory Destras, Gwendolyne Burfin, Hadrien Regue, Quentin Semanas, Martine Valette, Bruno Lina, Laurence Josset |
| EPI_ISL_3205919 | DYOMEDEA-LABORATOIRE DE LA SAUVEGARDE | CNR Virus des Infections Respiratoires - France SUD | Antonin Bal, Gregory Destras, Gwendolyne Burfin, Hadrien Regue, Quentin Semanas, Martine Valette, Bruno Lina, Laurence Josset |
| EPI_ISL_3205920 | INSTITUT BERGONIE | CNR Virus des Infections Respiratoires - France SUD | Antonin Bal, Gregory Destras, Gwendolyne Burfin, Hadrien Regue, Quentin Semanas, Martine Valette, Bruno Lina, Laurence Josset |
| EPI_ISL_3205921 | LAM NOVELAB LVA | CNR Virus des Infections Respiratoires - France SUD | Antonin Bal, Gregory Destras, Gwendolyne Burfin, Hadrien Regue, Quentin Semanas, Martine Valette, Bruno Lina, Laurence Josset |
| EPI_ISL_3205922 | LX BIO | CNR Virus des Infections Respiratoires - France SUD | Antonin Bal, Gregory Destras, Gwendolyne Burfin, Hadrien Regue, Quentin Semanas, Martine Valette, Bruno Lina, Laurence Josset |
| EPI_ISL_3205923, EPI_ISL_3205924 | LABORATOIRE BOUVIER | CNR Virus des Infections Respiratoires - France SUD | Antonin Bal, Gregory Destras, Gwendolyne Burfin, Hadrien Regue, Quentin Semanas, Martine Valette, Bruno Lina, Laurence Josset |
| EPI_ISL_3205925 | LAM NOVELAB LVA | CNR Virus des Infections Respiratoires - France SUD | Antonin Bal, Gregory Destras, Gwendolyne Burfin, Hadrien Regue, Quentin Semanas, Martine Valette, Bruno Lina, Laurence Josset |
| EPI_ISL_3205926, EPI_ISL_3205927 | LABORATOIRE BOUVIER | CNR Virus des Infections Respiratoires - France SUD | Antonin Bal, Gregory Destras, Gwendolyne Burfin, Hadrien Regue, Quentin Semanas, Martine Valette, Bruno Lina, Laurence Josset |
| EPI_ISL_3205928 | LAM NOVELAB LVA | CNR Virus des Infections Respiratoires - France SUD | Antonin Bal, Gregory Destras, Gwendolyne Burfin, Hadrien Regue, Quentin Semanas, Martine Valette, Bruno Lina, Laurence Josset |
| EPI_ISL_3205929 | LABORATOIRE BOUVIER | CNR Virus des Infections Respiratoires - France SUD | Antonin Bal, Gregory Destras, Gwendolyne Burfin, Hadrien Regue, Quentin Semanas, Martine Valette, Bruno Lina, Laurence Josset |
| EPI_ISL_3205930 | LAM NOVELAB LVA | CNR Virus des Infections Respiratoires - France SUD | Antonin Bal, Gregory Destras, Gwendolyne Burfin, Hadrien Regue, Quentin Semanas, Martine Valette, Bruno Lina, Laurence Josset |
| EPI_ISL_3205931 | LAM BIORYLIS | CNR Virus des Infections Respiratoires - France SUD | Antonin Bal, Gregory Destras, Gwendolyne Burfin, Hadrien Regue, Quentin Semanas, Martine Valette, Bruno Lina, Laurence Josset |
| EPI_ISL_3205932, EPI_ISL_3205933, EPI_ISL_3205934 | LAM NOVELAB LVA | CNR Virus des Infections Respiratoires - France SUD | Antonin Bal, Gregory Destras, Gwendolyne Burfin, Hadrien Regue, Quentin Semanas, Martine Valette, Bruno Lina, Laurence Josset |
| EPI_ISL_3205935 | LAM BIORYLIS | CNR Virus des Infections Respiratoires - France SUD | Antonin Bal, Gregory Destras, Gwendolyne Burfin, Hadrien Regue, Quentin Semanas, Martine Valette, Bruno Lina, Laurence Josset |
| EPI_ISL_3205936 | SYNLAB GASCOGNE | CNR Virus des Infections Respiratoires - France SUD | Antonin Bal, Gregory Destras, Gwendolyne Burfin, Hadrien Regue, Quentin Semanas, Martine Valette, Bruno Lina, Laurence Josset |
| EPI_ISL_3205937 | LABORATOIRE MAYMAT | CNR Virus des Infections Respiratoires - France SUD | Antonin Bal, Gregory Destras, Gwendolyne Burfin, Hadrien Regue, Quentin Semanas, Martine Valette, Bruno Lina, Laurence Josset |
| EPI_ISL_3205938 | AX BIO OCEAN | CNR Virus des Infections Respiratoires - France SUD | Antonin Bal, Gregory Destras, Gwendolyne Burfin, Hadrien Regue, Quentin Semanas, Martine Valette, Bruno Lina, Laurence Josset |
| EPI_ISL_3205939, EPI_ISL_3205940 | LABORATOIRE d'ANALYSES de BIOLOGIE MEDICALES | CNR Virus des Infections Respiratoires - France SUD | Antonin Bal, Gregory Destras, Gwendolyne Burfin, Hadrien Regue, Quentin Semanas, Martine Valette, Bruno Lina, Laurence Josset |
| EPI_ISL_3205941 | BIOLITTORAL BIOGROUP PLATEAU TECH | CNR Virus des Infections Respiratoires - France SUD | Antonin Bal, Gregory Destras, Gwendolyne Burfin, Hadrien Regue, Quentin Semanas, Martine Valette, Bruno Lina, Laurence Josset |
| EPI_ISL_3205942 | DYOMEDEA-LABORATOIRE DE LA SAUVEGARDE | CNR Virus des Infections Respiratoires - France SUD | Antonin Bal, Gregory Destras, Gwendolyne Burfin, Hadrien Regue, Quentin Semanas, Martine Valette, Bruno Lina, Laurence Josset |
| EPI_ISL_3205943 | LABORATOIRE BIOGROUP | CNR Virus des Infections Respiratoires - France SUD | Antonin Bal, Gregory Destras, Gwendolyne Burfin, Hadrien Regue, Quentin Semanas, Martine Valette, Bruno Lina, Laurence Josset |
| EPI_ISL_3205944 | LABORATOIRE CERBALLIANCE PLT VILLON | CNR Virus des Infections Respiratoires - France SUD | Antonin Bal, Gregory Destras, Gwendolyne Burfin, Hadrien Regue, Quentin Semanas, Martine Valette, Bruno Lina, Laurence Josset |
| EPI_ISL_3205945, EPI_ISL_3205946, EPI_ISL_3205947, EPI_ISL_3205948 | LABORATOIRE UNILIANS DECINES | CNR Virus des Infections Respiratoires - France SUD | Antonin Bal, Gregory Destras, Gwendolyne Burfin, Hadrien Regue, Quentin Semanas, Martine Valette, Bruno Lina, Laurence Josset |
| EPI_ISL_3205949, EPI_ISL_3205950 | DYOMEDEA-LABORATOIRE DE LA SAUVEGARDE | CNR Virus des Infections Respiratoires - France SUD | Antonin Bal, Gregory Destras, Gwendolyne Burfin, Hadrien Regue, Quentin Semanas, Martine Valette, Bruno Lina, Laurence Josset |
| EPI_ISL_3205951 | LABORATOIRE UNILIANS DECINES | CNR Virus des Infections Respiratoires - France SUD | Antonin Bal, Gregory Destras, Gwendolyne Burfin, Hadrien Regue, Quentin Semanas, Martine Valette, Bruno Lina, Laurence Josset |
| EPI_ISL_3205952 | DYOMEDEA-LABORATOIRE DE LA SAUVEGARDE | CNR Virus des Infections Respiratoires - France SUD | Antonin Bal, Gregory Destras, Gwendolyne Burfin, Hadrien Regue, Quentin Semanas, Martine Valette, Bruno Lina, Laurence Josset |
| EPI_ISL_3205953, EPI_ISL_3205954, EPI_ISL_3205955, EPI_ISL_3205956, EPI_ISL_3205957, EPI_ISL_3205958, EPI_ISL_3205959 | LABORATOIRE UNILIANS DECINES | CNR Virus des Infections Respiratoires - France SUD | Antonin Bal, Gregory Destras, Gwendolyne Burfin, Hadrien Regue, Quentin Semanas, Martine Valette, Bruno Lina, Laurence Josset |
| EPI_ISL_3205960 | LABORATOIRE CERBALLIANCE PLT VILLON | CNR Virus des Infections Respiratoires - France SUD | Antonin Bal, Gregory Destras, Gwendolyne Burfin, Hadrien Regue, Quentin Semanas, Martine Valette, Bruno Lina, Laurence Josset |
| EPI_ISL_3205961, EPI_ISL_3205962, EPI_ISL_3205963, EPI_ISL_3205964, EPI_ISL_3205965, EPI_ISL_3205966, EPI_ISL_3205967, EPI_ISL_3205968, EPI_ISL_3205969, EPI_ISL_3205970 | LABORATOIRE UNILIANS DECINES | CNR Virus des Infections Respiratoires - France SUD | Antonin Bal, Gregory Destras, Gwendolyne Burfin, Hadrien Regue, Quentin Semanas, Martine Valette, Bruno Lina, Laurence Josset |
| EPI_ISL_3205971 | LABORATOIRE CERBALLIANCE PLT VILLON | CNR Virus des Infections Respiratoires - France SUD | Antonin Bal, Gregory Destras, Gwendolyne Burfin, Hadrien Regue, Quentin Semanas, Martine Valette, Bruno Lina, Laurence Josset |
| EPI_ISL_3205972, EPI_ISL_3205973 | LABORATOIRE BOUVIER | CNR Virus des Infections Respiratoires - France SUD | Antonin Bal, Gregory Destras, Gwendolyne Burfin, Hadrien Regue, Quentin Semanas, Martine Valette, Bruno Lina, Laurence Josset |
| EPI_ISL_3205974 | LABORATOIRE UNILIANS DECINES | CNR Virus des Infections Respiratoires - France SUD | Antonin Bal, Gregory Destras, Gwendolyne Burfin, Hadrien Regue, Quentin Semanas, Martine Valette, Bruno Lina, Laurence Josset |
| EPI_ISL_3205975 | LABORATOIRE BOUVIER | CNR Virus des Infections Respiratoires - France SUD | Antonin Bal, Gregory Destras, Gwendolyne Burfin, Hadrien Regue, Quentin Semanas, Martine Valette, Bruno Lina, Laurence Josset |
| EPI_ISL_3205976 | LABORATOIRE UNILIANS DECINES | CNR Virus des Infections Respiratoires - France SUD | Antonin Bal, Gregory Destras, Gwendolyne Burfin, Hadrien Regue, Quentin Semanas, Martine Valette, Bruno Lina, Laurence Josset |
| EPI_ISL_3205977, EPI_ISL_3205978, EPI_ISL_3205979, EPI_ISL_3205980 | LABORATOIRE BOUVIER | CNR Virus des Infections Respiratoires - France SUD | Antonin Bal, Gregory Destras, Gwendolyne Burfin, Hadrien Regue, Quentin Semanas, Martine Valette, Bruno Lina, Laurence Josset |
| EPI_ISL_3205981, EPI_ISL_3205982 | LABORATOIRE CERBALLIANCE PLT VILLON | CNR Virus des Infections Respiratoires - France SUD | Antonin Bal, Gregory Destras, Gwendolyne Burfin, Hadrien Regue, Quentin Semanas, Martine Valette, Bruno Lina, Laurence Josset |
| EPI_ISL_3205983 | LABORATOIRE BOUVIER | CNR Virus des Infections Respiratoires - France SUD | Antonin Bal, Gregory Destras, Gwendolyne Burfin, Hadrien Regue, Quentin Semanas, Martine Valette, Bruno Lina, Laurence Josset |
| EPI_ISL_3205984 | LABORATOIRE CERBALLIANCE PLT VILLON | CNR Virus des Infections Respiratoires - France SUD | Antonin Bal, Gregory Destras, Gwendolyne Burfin, Hadrien Regue, Quentin Semanas, Martine Valette, Bruno Lina, Laurence Josset |
| EPI_ISL_3205985 | LABORATOIRE BOUVIER | CNR Virus des Infections Respiratoires - France SUD | Antonin Bal, Gregory Destras, Gwendolyne Burfin, Hadrien Regue, Quentin Semanas, Martine Valette, Bruno Lina, Laurence Josset |
| EPI_ISL_3205986 | AX BIO OCEAN | CNR Virus des Infections Respiratoires - France SUD | Antonin Bal, Gregory Destras, Gwendolyne Burfin, Hadrien Regue, Quentin Semanas, Martine Valette, Bruno Lina, Laurence Josset |
| EPI_ISL_3205987 | LABORATOIRE BOUVIER | CNR Virus des Infections Respiratoires - France SUD | Antonin Bal, Gregory Destras, Gwendolyne Burfin, Hadrien Regue, Quentin Semanas, Martine Valette, Bruno Lina, Laurence Josset |
| EPI_ISL_3205988, EPI_ISL_3205989 | AX BIO OCEAN | CNR Virus des Infections Respiratoires - France SUD | Antonin Bal, Gregory Destras, Gwendolyne Burfin, Hadrien Regue, Quentin Semanas, Martine Valette, Bruno Lina, Laurence Josset |
| EPI_ISL_3205990 | LABORATOIRE BOUVIER | CNR Virus des Infections Respiratoires - France SUD | Antonin Bal, Gregory Destras, Gwendolyne Burfin, Hadrien Regue, Quentin Semanas, Martine Valette, Bruno Lina, Laurence Josset |
| EPI_ISL_3205991 | AX BIO OCEAN | CNR Virus des Infections Respiratoires - France SUD | Antonin Bal, Gregory Destras, Gwendolyne Burfin, Hadrien Regue, Quentin Semanas, Martine Valette, Bruno Lina, Laurence Josset |
| EPI_ISL_3205992 | LABORATOIRE CERBALLIANCE PLT VILLON | CNR Virus des Infections Respiratoires - France SUD | Antonin Bal, Gregory Destras, Gwendolyne Burfin, Hadrien Regue, Quentin Semanas, Martine Valette, Bruno Lina, Laurence Josset |
| EPI_ISL_3205993 | AX BIO OCEAN | CNR Virus des Infections Respiratoires - France SUD | Antonin Bal, Gregory Destras, Gwendolyne Burfin, Hadrien Regue, Quentin Semanas, Martine Valette, Bruno Lina, Laurence Josset |
| EPI_ISL_3205994 | LABORATOIRE BOUVIER | CNR Virus des Infections Respiratoires - France SUD | Antonin Bal, Gregory Destras, Gwendolyne Burfin, Hadrien Regue, Quentin Semanas, Martine Valette, Bruno Lina, Laurence Josset |
| EPI_ISL_3205995, EPI_ISL_3205996, EPI_ISL_3205997 | AX BIO OCEAN | CNR Virus des Infections Respiratoires - France SUD | Antonin Bal, Gregory Destras, Gwendolyne Burfin, Hadrien Regue, Quentin Semanas, Martine Valette, Bruno Lina, Laurence Josset |

| | | | |
|---|---|---|---|
| EPI_ISL_3205998 | LABORATOIRE BOUVIER | CNR Virus des Infections Respiratoires - France SUD | Antonin Bal, Gregory Destras, Gwendolyne Burfin, Hadrien Regue, Quentin Semanas, Martine Valette, Bruno Lina, Laurence Josset |
| EPI_ISL_3205999, EPI_ISL_3206000 | AX BIO OCEAN | CNR Virus des Infections Respiratoires - France SUD | Antonin Bal, Gregory Destras, Gwendolyne Burfin, Hadrien Regue, Quentin Semanas, Martine Valette, Bruno Lina, Laurence Josset |
| EPI_ISL_3206001 | LABORATOIRE BOUVIER | CNR Virus des Infections Respiratoires - France SUD | Antonin Bal, Gregory Destras, Gwendolyne Burfin, Hadrien Regue, Quentin Semanas, Martine Valette, Bruno Lina, Laurence Josset |
| EPI_ISL_3206002, EPI_ISL_3206003, EPI_ISL_3206004 | AX BIO OCEAN | CNR Virus des Infections Respiratoires - France SUD | Antonin Bal, Gregory Destras, Gwendolyne Burfin, Hadrien Regue, Quentin Semanas, Martine Valette, Bruno Lina, Laurence Josset |
| EPI_ISL_3206005 | LABORATOIRE BOUVIER | CNR Virus des Infections Respiratoires - France SUD | Antonin Bal, Gregory Destras, Gwendolyne Burfin, Hadrien Regue, Quentin Semanas, Martine Valette, Bruno Lina, Laurence Josset |
| EPI_ISL_3206006 | AX BIO OCEAN | CNR Virus des Infections Respiratoires - France SUD | Antonin Bal, Gregory Destras, Gwendolyne Burfin, Hadrien Regue, Quentin Semanas, Martine Valette, Bruno Lina, Laurence Josset |
| EPI_ISL_3206007, EPI_ISL_3206008, EPI_ISL_3206009, EPI_ISL_3206010, EPI_ISL_3206011, EPI_ISL_3206012 | LABORATOIRE BOUVIER | CNR Virus des Infections Respiratoires - France SUD | Antonin Bal, Gregory Destras, Gwendolyne Burfin, Hadrien Regue, Quentin Semanas, Martine Valette, Bruno Lina, Laurence Josset |
| EPI_ISL_3206013, EPI_ISL_3206014, EPI_ISL_3206015 | LABORATOIRE CERBALLIANCE PLT VILLON | CNR Virus des Infections Respiratoires - France SUD | Antonin Bal, Gregory Destras, Gwendolyne Burfin, Hadrien Regue, Quentin Semanas, Martine Valette, Bruno Lina, Laurence Josset |
| EPI_ISL_3206016 | LABORATOIRE d'ANALYSES de BIOLOGIE MEDICALES | CNR Virus des Infections Respiratoires - France SUD | Antonin Bal, Gregory Destras, Gwendolyne Burfin, Hadrien Regue, Quentin Semanas, Martine Valette, Bruno Lina, Laurence Josset |
| EPI_ISL_3206017 | CENTRE HOSPITALIER DE BOURG EN BRESSE | CNR Virus des Infections Respiratoires - France SUD | Antonin Bal, Gregory Destras, Gwendolyne Burfin, Hadrien Regue, Quentin Semanas, Martine Valette, Bruno Lina, Laurence Josset |
| EPI_ISL_3206018 | LABORATOIRE UNILIANS DECINES | CNR Virus des Infections Respiratoires - France SUD | Antonin Bal, Gregory Destras, Gwendolyne Burfin, Hadrien Regue, Quentin Semanas, Martine Valette, Bruno Lina, Laurence Josset |
| EPI_ISL_3206019 | LABORATOIRE CERBALLIANCE PLT VILLON | CNR Virus des Infections Respiratoires - France SUD | Antonin Bal, Gregory Destras, Gwendolyne Burfin, Hadrien Regue, Quentin Semanas, Martine Valette, Bruno Lina, Laurence Josset |
| EPI_ISL_3206020, EPI_ISL_3206021 | LABORATOIRE UNILIANS DECINES | CNR Virus des Infections Respiratoires - France SUD | Antonin Bal, Gregory Destras, Gwendolyne Burfin, Hadrien Regue, Quentin Semanas, Martine Valette, Bruno Lina, Laurence Josset |
| EPI_ISL_3206022 | LABORATOIRE CERBALLIANCE PLT VILLON | CNR Virus des Infections Respiratoires - France SUD | Antonin Bal, Gregory Destras, Gwendolyne Burfin, Hadrien Regue, Quentin Semanas, Martine Valette, Bruno Lina, Laurence Josset |
| EPI_ISL_3206023, EPI_ISL_3206024, EPI_ISL_3206025, EPI_ISL_3206026, EPI_ISL_3206027, EPI_ISL_3206028, EPI_ISL_3206029, EPI_ISL_3206030, EPI_ISL_3206031, EPI_ISL_3206032, EPI_ISL_3206033, EPI_ISL_3206034, EPI_ISL_3206036, EPI_ISL_3206037, EPI_ISL_3206038 | | | |
| see above | AX BIO OCEAN | CNR Virus des Infections Respiratoires - France SUD | Antonin Bal, Gregory Destras, Gwendolyne Burfin, Hadrien Regue, Quentin Semanas, Martine Valette, Bruno Lina, Laurence Josset |
| EPI_ISL_3206039, EPI_ISL_3206040 | GROUPE LCD - UNIBIO LA PLAINE | CNR Virus des Infections Respiratoires - France SUD | Antonin Bal, Gregory Destras, Gwendolyne Burfin, Hadrien Regue, Quentin Semanas, Martine Valette, Bruno Lina, Laurence Josset |
| EPI_ISL_3206048 | LABORATOIRE d'ANALYSES de BIOLOGIE MEDICALES | CNR Virus des Infections Respiratoires - France SUD | Antonin Bal, Gregory Destras, Gwendolyne Burfin, Hadrien Regue, Quentin Semanas, Martine Valette, Bruno Lina, Laurence Josset |
| EPI_ISL_3206050, EPI_ISL_3206051, EPI_ISL_3206052, EPI_ISL_3206054, EPI_ISL_3206055, EPI_ISL_3206056, EPI_ISL_3206057, EPI_ISL_3206058, EPI_ISL_3206059, EPI_ISL_3206060, EPI_ISL_3206061, EPI_ISL_3206062, EPI_ISL_3206063, EPI_ISL_3206064, EPI_ISL_3206065, EPI_ISL_3206066, EPI_ISL_3206067, EPI_ISL_3206068, EPI_ISL_3206069, EPI_ISL_3206070 | | | |
| see above | LABORATOIRE BOUVIER | CNR Virus des Infections Respiratoires - France SUD | Antonin Bal, Gregory Destras, Gwendolyne Burfin, Hadrien Regue, Quentin Semanas, Martine Valette, Bruno Lina, Laurence Josset |
| EPI_ISL_3206071 | DYOMEDEA-LABORATOIRE DE LA SAUVEGARDE | CNR Virus des Infections Respiratoires - France SUD | Antonin Bal, Gregory Destras, Gwendolyne Burfin, Hadrien Regue, Quentin Semanas, Martine Valette, Bruno Lina, Laurence Josset |
| EPI_ISL_3206072, EPI_ISL_3206073, EPI_ISL_3206074, EPI_ISL_3206075, EPI_ISL_3206076, EPI_ISL_3206077 | LABORATOIRE BOUVIER | CNR Virus des Infections Respiratoires - France SUD | Antonin Bal, Gregory Destras, Gwendolyne Burfin, Hadrien Regue, Quentin Semanas, Martine Valette, Bruno Lina, Laurence Josset |
| EPI_ISL_3206078, EPI_ISL_3206079, EPI_ISL_3206080, EPI_ISL_3206081, EPI_ISL_3206082, EPI_ISL_3206083, EPI_ISL_3206084, EPI_ISL_3206085, EPI_ISL_3206086, EPI_ISL_3206087, EPI_ISL_3206088, EPI_ISL_3206089 | | | |
| see above | DYOMEDEA-LABORATOIRE DE LA SAUVEGARDE | CNR Virus des Infections Respiratoires - France SUD | Antonin Bal, Gregory Destras, Gwendolyne Burfin, Hadrien Regue, Quentin Semanas, Martine Valette, Bruno Lina, Laurence Josset |
| EPI_ISL_3206090, EPI_ISL_3206091 | MIRIALIS CLUSES BECHET | CNR Virus des Infections Respiratoires - France SUD | Antonin Bal, Gregory Destras, Gwendolyne Burfin, Hadrien Regue, Quentin Semanas, Martine Valette, Bruno Lina, Laurence Josset |
| EPI_ISL_3206093, EPI_ISL_3206094, EPI_ISL_3206095 | LABORATOIRE BOUVIER | CNR Virus des Infections Respiratoires - France SUD | Antonin Bal, Gregory Destras, Gwendolyne Burfin, Hadrien Regue, Quentin Semanas, Martine Valette, Bruno Lina, Laurence Josset |
| EPI_ISL_3206097 | LBM DYNABIO CARREAU | CNR Virus des Infections Respiratoires - France SUD | Antonin Bal, Gregory Destras, Gwendolyne Burfin, Hadrien Regue, Quentin Semanas, Martine Valette, Bruno Lina, Laurence Josset |
| EPI_ISL_3206099 | DYOMEDEA-LABORATOIRE DE LA SAUVEGARDE | CNR Virus des Infections Respiratoires - France SUD | Antonin Bal, Gregory Destras, Gwendolyne Burfin, Hadrien Regue, Quentin Semanas, Martine Valette, Bruno Lina, Laurence Josset |
| EPI_ISL_3206100, EPI_ISL_3206101, EPI_ISL_3206102, EPI_ISL_3206103, EPI_ISL_3206104, EPI_ISL_3206105, EPI_ISL_3206106, EPI_ISL_3206107, EPI_ISL_3206108, EPI_ISL_3206109, EPI_ISL_3206110 | | | |
| see above | MIRIALIS CLUSES BECHET | CNR Virus des Infections Respiratoires - France SUD | Antonin Bal, Gregory Destras, Gwendolyne Burfin, Hadrien Regue, Quentin Semanas, Martine Valette, Bruno Lina, Laurence Josset |
| EPI_ISL_3206111 | REUNILAB | CNR Virus des Infections Respiratoires - France SUD | Antonin Bal, Gregory Destras, Gwendolyne Burfin, Hadrien Regue, Quentin Semanas, Martine Valette, Bruno Lina, Laurence Josset |
| EPI_ISL_3206120, EPI_ISL_3206121, EPI_ISL_3206122, EPI_ISL_3206123 | AX BIO OCEAN | CNR Virus des Infections Respiratoires - France SUD | Antonin Bal, Gregory Destras, Gwendolyne Burfin, Hadrien Regue, Quentin Semanas, Martine Valette, Bruno Lina, Laurence Josset |
| EPI_ISL_3206124, EPI_ISL_3206125, EPI_ISL_3206126, EPI_ISL_3206127 | MIRIALIS CLUSES BECHET | CNR Virus des Infections Respiratoires - France SUD | Antonin Bal, Gregory Destras, Gwendolyne Burfin, Hadrien Regue, Quentin Semanas, Martine Valette, Bruno Lina, Laurence Josset |
| EPI_ISL_3206128 | LABORATOIRE LABAZUR | CNR Virus des Infections Respiratoires - France SUD | Antonin Bal, Gregory Destras, Gwendolyne Burfin, Hadrien Regue, Quentin Semanas, Martine Valette, Bruno Lina, Laurence Josset |
| EPI_ISL_3206129, EPI_ISL_3206130, EPI_ISL_3206131, EPI_ISL_3206133, EPI_ISL_3206134 | MIRIALIS CLUSES BECHET | CNR Virus des Infections Respiratoires - France SUD | Antonin Bal, Gregory Destras, Gwendolyne Burfin, Hadrien Regue, Quentin Semanas, Martine Valette, Bruno Lina, Laurence Josset |
| EPI_ISL_3206135, EPI_ISL_3206136, EPI_ISL_3206137, EPI_ISL_3206138, EPI_ISL_3206139, EPI_ISL_3206140 | LABORATOIRE BIOESTEREL | CNR Virus des Infections Respiratoires - France SUD | Antonin Bal, Gregory Destras, Gwendolyne Burfin, Hadrien Regue, Quentin Semanas, Martine Valette, Bruno Lina, Laurence Josset |
| EPI_ISL_3206141 | CNR Virus des Infections Respiratoires - France SUD | CNR Virus des Infections Respiratoires - France SUD | Antonin Bal, Gregory Destras, Gwendolyne Burfin, Hadrien Regue, Quentin Semanas, Martine Valette, Bruno Lina, Laurence Josset |
| EPI_ISL_3206142 | LABORATOIRE d'ANALYSES de BIOLOGIE MEDICALES | CNR Virus des Infections Respiratoires - France SUD | Antonin Bal, Gregory Destras, Gwendolyne Burfin, Hadrien Regue, Quentin Semanas, Martine Valette, Bruno Lina, Laurence Josset |
| EPI_ISL_3206143 | BIOLITTORAL BIOGROUP PLATEAU TECH | CNR Virus des Infections Respiratoires - France SUD | Antonin Bal, Gregory Destras, Gwendolyne Burfin, Hadrien Regue, Quentin Semanas, Martine Valette, Bruno Lina, Laurence Josset |
| EPI_ISL_3206144, EPI_ISL_3206145, EPI_ISL_3206146 | LABORATOIRE d'ANALYSES de BIOLOGIE MEDICALES | CNR Virus des Infections Respiratoires - France SUD | Antonin Bal, Gregory Destras, Gwendolyne Burfin, Hadrien Regue, Quentin Semanas, Martine Valette, Bruno Lina, Laurence Josset |
| EPI_ISL_3206147 | LAM NOVELAB LVA | CNR Virus des Infections Respiratoires - France SUD | Antonin Bal, Gregory Destras, Gwendolyne Burfin, Hadrien Regue, Quentin Semanas, Martine Valette, Bruno Lina, Laurence Josset |
| EPI_ISL_3206148 | CENTRE HOSPITALIER DE BOURG EN BRESSE | CNR Virus des Infections Respiratoires - France SUD | Antonin Bal, Gregory Destras, Gwendolyne Burfin, Hadrien Regue, Quentin Semanas, Martine Valette, Bruno Lina, Laurence Josset |
| EPI_ISL_3206150 | LBM UNIBIO VALENTIN | CNR Virus des Infections Respiratoires - France SUD | Antonin Bal, Gregory Destras, Gwendolyne Burfin, Hadrien Regue, Quentin Semanas, Martine Valette, Bruno Lina, Laurence Josset |
| EPI_ISL_3206153, EPI_ISL_3206154, EPI_ISL_3206155, EPI_ISL_3206156, EPI_ISL_3206157, EPI_ISL_3206158, EPI_ISL_3206159, EPI_ISL_3206160, EPI_ISL_3206161, EPI_ISL_3206162, EPI_ISL_3206163, EPI_ISL_3206164, EPI_ISL_3206165, EPI_ISL_3206166, EPI_ISL_3206167, EPI_ISL_3206168, EPI_ISL_3206169, EPI_ISL_3206170, EPI_ISL_3206171, EPI_ISL_3206172, EPI_ISL_3206173 | | | |
| see above | LABORATOIRE LABAZUR | CNR Virus des Infections Respiratoires - France SUD | Antonin Bal, Gregory Destras, Gwendolyne Burfin, Hadrien Regue, Quentin Semanas, Martine Valette, Bruno Lina, Laurence Josset |
| EPI_ISL_3206174, EPI_ISL_3206175, EPI_ISL_3206176 | MIRIALIS CLUSES BECHET | CNR Virus des Infections Respiratoires - France SUD | Antonin Bal, Gregory Destras, Gwendolyne Burfin, Hadrien Regue, Quentin Semanas, Martine Valette, Bruno Lina, Laurence Josset |
| EPI_ISL_3206177, EPI_ISL_3206178, EPI_ISL_3206179, EPI_ISL_3206180, EPI_ISL_3206181, EPI_ISL_3206183, EPI_ISL_3206184, EPI_ISL_3206185, EPI_ISL_3206186, EPI_ISL_3206189, EPI_ISL_3206191, EPI_ISL_3206192, EPI_ISL_3206194, EPI_ISL_3206195, EPI_ISL_3206196, EPI_ISL_3206197, EPI_ISL_3206198, EPI_ISL_3206199, EPI_ISL_3206200, EPI_ISL_3206201, EPI_ISL_3206202, EPI_ISL_3206203, EPI_ISL_3206204, EPI_ISL_3206205, EPI_ISL_3206206, EPI_ISL_3206207, EPI_ISL_3206208, EPI_ISL_3206209, EPI_ISL_3206210, EPI_ISL_3206211, EPI_ISL_3206212, EPI_ISL_3206213, EPI_ISL_3206214, EPI_ISL_3206215, EPI_ISL_3206216, EPI_ISL_3206217, EPI_ISL_3206218, EPI_ISL_3206219, EPI_ISL_3206220, EPI_ISL_3206221, EPI_ISL_3206222 | | | |
| see above | LABORATOIRE UNILIANS DECINES | CNR Virus des Infections Respiratoires - France SUD | Antonin Bal, Gregory Destras, Gwendolyne Burfin, Hadrien Regue, Quentin Semanas, Martine Valette, Bruno Lina, Laurence Josset |

| | | | |
|---|---|---|---|
| EPI_ISL_3206223, EPI_ISL_3206224, EPI_ISL_3206225 | CH GIVORS | CNR Virus des Infections Respiratoires - France SUD | Antonin Bal, Gregory Destras, Gwendolyne Burfin, Hadrien Regue, Quentin Semanas, Martine Valette, Bruno Lina, Laurence Josset |
| EPI_ISL_3206226, EPI_ISL_3206227, EPI_ISL_3206228 | CNR Virus des Infections Respiratoires - France SUD | CNR Virus des Infections Respiratoires - France SUD | Antonin Bal, Gregory Destras, Gwendolyne Burfin, Hadrien Regue, Quentin Semanas, Martine Valette, Bruno Lina, Laurence Josset |
| EPI_ISL_3206229, EPI_ISL_3206230, EPI_ISL_3206231 | CENTRE HOSPITALIER ST JOSEPH ST LUC | CNR Virus des Infections Respiratoires - France SUD | Antonin Bal, Gregory Destras, Gwendolyne Burfin, Hadrien Regue, Quentin Semanas, Martine Valette, Bruno Lina, Laurence Josset |
| EPI_ISL_3206232, EPI_ISL_3206233 | CNR Virus des Infections Respiratoires - France SUD | CNR Virus des Infections Respiratoires - France SUD | Antonin Bal, Gregory Destras, Gwendolyne Burfin, Hadrien Regue, Quentin Semanas, Martine Valette, Bruno Lina, Laurence Josset |
| EPI_ISL_3206234, EPI_ISL_3206235, EPI_ISL_3206236, EPI_ISL_3206237, EPI_ISL_3206238, EPI_ISL_3206239, EPI_ISL_3206240, EPI_ISL_3206241, EPI_ISL_3206242, EPI_ISL_3206243, EPI_ISL_3206244, EPI_ISL_3206245, EPI_ISL_3206246 | | | |
| see above | DYOMEDEA-LABORATOIRE DE LA SAUVEGARDE | CNR Virus des Infections Respiratoires - France SUD | Antonin Bal, Gregory Destras, Gwendolyne Burfin, Hadrien Regue, Quentin Semanas, Martine Valette, Bruno Lina, Laurence Josset |
| EPI_ISL_3206247 | CNR Virus des Infections Respiratoires - France SUD | CNR Virus des Infections Respiratoires - France SUD | Antonin Bal, Gregory Destras, Gwendolyne Burfin, Hadrien Regue, Quentin Semanas, Martine Valette, Bruno Lina, Laurence Josset |
| EPI_ISL_3206248, EPI_ISL_3206249, EPI_ISL_3206250, EPI_ISL_3206251, EPI_ISL_3206252, EPI_ISL_3206253, EPI_ISL_3206254, EPI_ISL_3206255, EPI_ISL_3206256, EPI_ISL_3206257, EPI_ISL_3206258, EPI_ISL_3206259, EPI_ISL_3206260, EPI_ISL_3206261, EPI_ISL_3206262, EPI_ISL_3206263, EPI_ISL_3206264, EPI_ISL_3206265, EPI_ISL_3206266, EPI_ISL_3206267, EPI_ISL_3206268, EPI_ISL_3206269, EPI_ISL_3206270, EPI_ISL_3206271, EPI_ISL_3206272, EPI_ISL_3206273, EPI_ISL_3206274, EPI_ISL_3206275, EPI_ISL_3206276, EPI_ISL_3206277, EPI_ISL_3206278, EPI_ISL_3206279, EPI_ISL_3206280, EPI_ISL_3206281, EPI_ISL_3206282, EPI_ISL_3206283, EPI_ISL_3206284, EPI_ISL_3206285, EPI_ISL_3206286, EPI_ISL_3206287 | | | |
| see above | DYOMEDEA-LABORATOIRE DE LA SAUVEGARDE | CNR Virus des Infections Respiratoires - France SUD | Antonin Bal, Gregory Destras, Gwendolyne Burfin, Hadrien Regue, Quentin Semanas, Martine Valette, Bruno Lina, Laurence Josset |
| EPI_ISL_3206288 | ? (OFFICE SANTE Non Identifiable ) | CNR Virus des Infections Respiratoires - France SUD | Antonin Bal, Gregory Destras, Gwendolyne Burfin, Hadrien Regue, Quentin Semanas, Martine Valette, Bruno Lina, Laurence Josset |
| EPI_ISL_3206289, EPI_ISL_3206290, EPI_ISL_3206291, EPI_ISL_3206292, EPI_ISL_3206293, EPI_ISL_3206294, EPI_ISL_3206295, EPI_ISL_3206296, EPI_ISL_3206297, EPI_ISL_3206298, EPI_ISL_3206299, EPI_ISL_3206300, EPI_ISL_3206301, EPI_ISL_3206302, EPI_ISL_3206303, EPI_ISL_3206304, EPI_ISL_3206305, EPI_ISL_3206306 | | | |
| see above | DYOMEDEA-LABORATOIRE DE LA SAUVEGARDE | CNR Virus des Infections Respiratoires - France SUD | Antonin Bal, Gregory Destras, Gwendolyne Burfin, Hadrien Regue, Quentin Semanas, Martine Valette, Bruno Lina, Laurence Josset |
| EPI_ISL_3206307, EPI_ISL_3206308, EPI_ISL_3206309 | CNR Virus des Infections Respiratoires - France SUD | CNR Virus des Infections Respiratoires - France SUD | Antonin Bal, Gregory Destras, Gwendolyne Burfin, Hadrien Regue, Quentin Semanas, Martine Valette, Bruno Lina, Laurence Josset |
| EPI_ISL_3206310, EPI_ISL_3206311, EPI_ISL_3206312, EPI_ISL_3206313, EPI_ISL_3206314, EPI_ISL_3206315, EPI_ISL_3206316, EPI_ISL_3206317, EPI_ISL_3206318, EPI_ISL_3206319, EPI_ISL_3206320, EPI_ISL_3206321, EPI_ISL_3206322, EPI_ISL_3206323, EPI_ISL_3206324, EPI_ISL_3206325, EPI_ISL_3206326, EPI_ISL_3206327, EPI_ISL_3206328, EPI_ISL_3206329, EPI_ISL_3206330, EPI_ISL_3206331, EPI_ISL_3206332, EPI_ISL_3206333, EPI_ISL_3206334, EPI_ISL_3206335, EPI_ISL_3206336, EPI_ISL_3206337, EPI_ISL_3206338, EPI_ISL_3206339, EPI_ISL_3206340, EPI_ISL_3206341, EPI_ISL_3206342, EPI_ISL_3206343, EPI_ISL_3206344, EPI_ISL_3206345, EPI_ISL_3206346, EPI_ISL_3206347, EPI_ISL_3206348, EPI_ISL_3206349, EPI_ISL_3206350, EPI_ISL_3206351, EPI_ISL_3206352, EPI_ISL_3206353, EPI_ISL_3206355, EPI_ISL_3206356, EPI_ISL_3206357, EPI_ISL_3206358, EPI_ISL_3206359, EPI_ISL_3206360, EPI_ISL_3206361, EPI_ISL_3206362, EPI_ISL_3206363, EPI_ISL_3206364, EPI_ISL_3206365, EPI_ISL_3206366, EPI_ISL_3206367, EPI_ISL_3206368, EPI_ISL_3206369, EPI_ISL_3206370, EPI_ISL_3206371, EPI_ISL_3206372, EPI_ISL_3206373, EPI_ISL_3206374, EPI_ISL_3206375, EPI_ISL_3206376, EPI_ISL_3206377, EPI_ISL_3206378, EPI_ISL_3206379, EPI_ISL_3206380, EPI_ISL_3206381, EPI_ISL_3206382, EPI_ISL_3206383, EPI_ISL_3206384, EPI_ISL_3206385, EPI_ISL_3206386, EPI_ISL_3206387, EPI_ISL_3206388, EPI_ISL_3206389, EPI_ISL_3206390, EPI_ISL_3206391, EPI_ISL_3206392, EPI_ISL_3206393, EPI_ISL_3206394, EPI_ISL_3206395, EPI_ISL_3206396, EPI_ISL_3206397, EPI_ISL_3206398, EPI_ISL_3206399, EPI_ISL_3206400, EPI_ISL_3206401, EPI_ISL_3206402, EPI_ISL_3206403, EPI_ISL_3206404, EPI_ISL_3206405, EPI_ISL_3206406, EPI_ISL_3206407, EPI_ISL_3206408, EPI_ISL_3206410, EPI_ISL_3206411, EPI_ISL_3206412, EPI_ISL_3206413, EPI_ISL_3206414, EPI_ISL_3206415, EPI_ISL_3206416, EPI_ISL_3206417, EPI_ISL_3206418, EPI_ISL_3206419, EPI_ISL_3206420, EPI_ISL_3206421, EPI_ISL_3206422, EPI_ISL_3206423, EPI_ISL_3206424, EPI_ISL_3206425, EPI_ISL_3206426, EPI_ISL_3206427, EPI_ISL_3206428, EPI_ISL_3206429, EPI_ISL_3206430, EPI_ISL_3206431, EPI_ISL_3206432, EPI_ISL_3206433 | | | |
| see above | DYOMEDEA-LABORATOIRE DE LA SAUVEGARDE | CNR Virus des Infections Respiratoires - France SUD | Antonin Bal, Gregory Destras, Gwendolyne Burfin, Hadrien Regue, Quentin Semanas, Martine Valette, Bruno Lina, Laurence Josset |
| EPI_ISL_3206434 | LABORATOIRE CERBALLIANCE PLT VILLON | CNR Virus des Infections Respiratoires - France SUD | Antonin Bal, Gregory Destras, Gwendolyne Burfin, Hadrien Regue, Quentin Semanas, Martine Valette, Bruno Lina, Laurence Josset |
| EPI_ISL_3206435, EPI_ISL_3206436, EPI_ISL_3206437 | DYOMEDEA-LABORATOIRE DE LA SAUVEGARDE | CNR Virus des Infections Respiratoires - France SUD | Antonin Bal, Gregory Destras, Gwendolyne Burfin, Hadrien Regue, Quentin Semanas, Martine Valette, Bruno Lina, Laurence Josset |
| EPI_ISL_3206438 | LABORATOIRE CERBALLIANCE PLT VILLON | CNR Virus des Infections Respiratoires - France SUD | Antonin Bal, Gregory Destras, Gwendolyne Burfin, Hadrien Regue, Quentin Semanas, Martine Valette, Bruno Lina, Laurence Josset |
| EPI_ISL_3206439 | DYOMEDEA-LABORATOIRE DE LA SAUVEGARDE | CNR Virus des Infections Respiratoires - France SUD | Antonin Bal, Gregory Destras, Gwendolyne Burfin, Hadrien Regue, Quentin Semanas, Martine Valette, Bruno Lina, Laurence Josset |
| EPI_ISL_3206440 | LABORATOIRE CERBALLIANCE PLT VILLON | CNR Virus des Infections Respiratoires - France SUD | Antonin Bal, Gregory Destras, Gwendolyne Burfin, Hadrien Regue, Quentin Semanas, Martine Valette, Bruno Lina, Laurence Josset |
| EPI_ISL_3206441, EPI_ISL_3206442 | DYOMEDEA-LABORATOIRE DE LA SAUVEGARDE | CNR Virus des Infections Respiratoires - France SUD | Antonin Bal, Gregory Destras, Gwendolyne Burfin, Hadrien Regue, Quentin Semanas, Martine Valette, Bruno Lina, Laurence Josset |
| EPI_ISL_3206443 | LABORATOIRE CERBALLIANCE PLT VILLON | CNR Virus des Infections Respiratoires - France SUD | Antonin Bal, Gregory Destras, Gwendolyne Burfin, Hadrien Regue, Quentin Semanas, Martine Valette, Bruno Lina, Laurence Josset |
| EPI_ISL_3206444 | DYOMEDEA-LABORATOIRE DE LA SAUVEGARDE | CNR Virus des Infections Respiratoires - France SUD | Antonin Bal, Gregory Destras, Gwendolyne Burfin, Hadrien Regue, Quentin Semanas, Martine Valette, Bruno Lina, Laurence Josset |
| EPI_ISL_3206445, EPI_ISL_3206446, EPI_ISL_3206447 | LABORATOIRE CERBALLIANCE PLT VILLON | CNR Virus des Infections Respiratoires - France SUD | Antonin Bal, Gregory Destras, Gwendolyne Burfin, Hadrien Regue, Quentin Semanas, Martine Valette, Bruno Lina, Laurence Josset |
| EPI_ISL_3206448 | DYOMEDEA-LABORATOIRE DE LA SAUVEGARDE | CNR Virus des Infections Respiratoires - France SUD | Antonin Bal, Gregory Destras, Gwendolyne Burfin, Hadrien Regue, Quentin Semanas, Martine Valette, Bruno Lina, Laurence Josset |
| EPI_ISL_3206449 | LABORATOIRE CERBALLIANCE PLT VILLON | CNR Virus des Infections Respiratoires - France SUD | Antonin Bal, Gregory Destras, Gwendolyne Burfin, Hadrien Regue, Quentin Semanas, Martine Valette, Bruno Lina, Laurence Josset |
| EPI_ISL_3206450 | DYOMEDEA-LABORATOIRE DE LA SAUVEGARDE | CNR Virus des Infections Respiratoires - France SUD | Antonin Bal, Gregory Destras, Gwendolyne Burfin, Hadrien Regue, Quentin Semanas, Martine Valette, Bruno Lina, Laurence Josset |
| EPI_ISL_3206451 | LABORATOIRE CERBALLIANCE PLT VILLON | CNR Virus des Infections Respiratoires - France SUD | Antonin Bal, Gregory Destras, Gwendolyne Burfin, Hadrien Regue, Quentin Semanas, Martine Valette, Bruno Lina, Laurence Josset |
| EPI_ISL_3206452 | DYOMEDEA-LABORATOIRE DE LA SAUVEGARDE | CNR Virus des Infections Respiratoires - France SUD | Antonin Bal, Gregory Destras, Gwendolyne Burfin, Hadrien Regue, Quentin Semanas, Martine Valette, Bruno Lina, Laurence Josset |
| EPI_ISL_3206453 | LABORATOIRE CERBALLIANCE PLT VILLON | CNR Virus des Infections Respiratoires - France SUD | Antonin Bal, Gregory Destras, Gwendolyne Burfin, Hadrien Regue, Quentin Semanas, Martine Valette, Bruno Lina, Laurence Josset |
| EPI_ISL_3206454 | CNR Virus des Infections Respiratoires - France SUD | CNR Virus des Infections Respiratoires - France SUD | Antonin Bal, Gregory Destras, Gwendolyne Burfin, Hadrien Regue, Quentin Semanas, Martine Valette, Bruno Lina, Laurence Josset |
| EPI_ISL_3206455, EPI_ISL_3206456 | LABORATOIRE CERBALLIANCE PLT VILLON | CNR Virus des Infections Respiratoires - France SUD | Antonin Bal, Gregory Destras, Gwendolyne Burfin, Hadrien Regue, Quentin Semanas, Martine Valette, Bruno Lina, Laurence Josset |
| EPI_ISL_3206457 | DYOMEDEA-LABORATOIRE DE LA SAUVEGARDE | CNR Virus des Infections Respiratoires - France SUD | Antonin Bal, Gregory Destras, Gwendolyne Burfin, Hadrien Regue, Quentin Semanas, Martine Valette, Bruno Lina, Laurence Josset |
| EPI_ISL_3206458 | CNR Virus des Infections Respiratoires - France SUD | CNR Virus des Infections Respiratoires - France SUD | Antonin Bal, Gregory Destras, Gwendolyne Burfin, Hadrien Regue, Quentin Semanas, Martine Valette, Bruno Lina, Laurence Josset |
| EPI_ISL_3206459, EPI_ISL_3206460, EPI_ISL_3206461, EPI_ISL_3206462, EPI_ISL_3206463, EPI_ISL_3206464, EPI_ISL_3206465 | DYOMEDEA-LABORATOIRE DE LA SAUVEGARDE | CNR Virus des Infections Respiratoires - France SUD | Antonin Bal, Gregory Destras, Gwendolyne Burfin, Hadrien Regue, Quentin Semanas, Martine Valette, Bruno Lina, Laurence Josset |
| EPI_ISL_3206466 | LABORATOIRE CERBALLIANCE PLT VILLON | CNR Virus des Infections Respiratoires - France SUD | Antonin Bal, Gregory Destras, Gwendolyne Burfin, Hadrien Regue, Quentin Semanas, Martine Valette, Bruno Lina, Laurence Josset |
| EPI_ISL_3206467 | DYOMEDEA-LABORATOIRE DE LA SAUVEGARDE | CNR Virus des Infections Respiratoires - France SUD | Antonin Bal, Gregory Destras, Gwendolyne Burfin, Hadrien Regue, Quentin Semanas, Martine Valette, Bruno Lina, Laurence Josset |
| EPI_ISL_3206468 | LABORATOIRE CERBALLIANCE PLT VILLON | CNR Virus des Infections Respiratoires - France SUD | Antonin Bal, Gregory Destras, Gwendolyne Burfin, Hadrien Regue, Quentin Semanas, Martine Valette, Bruno Lina, Laurence Josset |
| EPI_ISL_3206469, EPI_ISL_3206470, EPI_ISL_3206471 | DYOMEDEA-LABORATOIRE DE LA SAUVEGARDE | CNR Virus des Infections Respiratoires - France SUD | Antonin Bal, Gregory Destras, Gwendolyne Burfin, Hadrien Regue, Quentin Semanas, Martine Valette, Bruno Lina, Laurence Josset |
| EPI_ISL_3206472 | LABORATOIRE CERBALLIANCE PLT VILLON | CNR Virus des Infections Respiratoires - France SUD | Antonin Bal, Gregory Destras, Gwendolyne Burfin, Hadrien Regue, Quentin Semanas, Martine Valette, Bruno Lina, Laurence Josset |
| EPI_ISL_3206473 | DYOMEDEA-LABORATOIRE DE LA SAUVEGARDE | CNR Virus des Infections Respiratoires - France SUD | Antonin Bal, Gregory Destras, Gwendolyne Burfin, Hadrien Regue, Quentin Semanas, Martine Valette, Bruno Lina, Laurence Josset |
| EPI_ISL_3206474, EPI_ISL_3206475, EPI_ISL_3206476, EPI_ISL_3206477, EPI_ISL_3206478 | LABORATOIRE CERBALLIANCE PLT VILLON | CNR Virus des Infections Respiratoires - France SUD | Antonin Bal, Gregory Destras, Gwendolyne Burfin, Hadrien Regue, Quentin Semanas, Martine Valette, Bruno Lina, Laurence Josset |
| EPI_ISL_3206479 | CNR Virus des Infections Respiratoires - France SUD | CNR Virus des Infections Respiratoires - France SUD | Antonin Bal, Gregory Destras, Gwendolyne Burfin, Hadrien Regue, Quentin Semanas, Martine Valette, Bruno Lina, Laurence Josset |
| EPI_ISL_3206480, EPI_ISL_3206481, EPI_ISL_3206482 | LABORATOIRE CERBALLIANCE PLT VILLON | CNR Virus des Infections Respiratoires - France SUD | Antonin Bal, Gregory Destras, Gwendolyne Burfin, Hadrien Regue, Quentin Semanas, Martine Valette, Bruno Lina, Laurence Josset |
| EPI_ISL_3206483 | DYOMEDEA-LABORATOIRE DE LA SAUVEGARDE | CNR Virus des Infections Respiratoires - France SUD | Antonin Bal, Gregory Destras, Gwendolyne Burfin, Hadrien Regue, Quentin Semanas, Martine Valette, Bruno Lina, Laurence Josset |
| EPI_ISL_3206484 | LABORATOIRE CERBALLIANCE PLT VILLON | CNR Virus des Infections Respiratoires - France SUD | Antonin Bal, Gregory Destras, Gwendolyne Burfin, Hadrien Regue, Quentin Semanas, Martine Valette, Bruno Lina, Laurence Josset |
| EPI_ISL_3206485 | DYOMEDEA-LABORATOIRE DE LA SAUVEGARDE | CNR Virus des Infections Respiratoires - France SUD | Antonin Bal, Gregory Destras, Gwendolyne Burfin, Hadrien Regue, Quentin Semanas, Martine Valette, Bruno Lina, Laurence Josset |
| EPI_ISL_3206486, EPI_ISL_3206487 | LABORATOIRE CERBALLIANCE PLT VILLON | CNR Virus des Infections Respiratoires - France SUD | Antonin Bal, Gregory Destras, Gwendolyne Burfin, Hadrien Regue, Quentin Semanas, Martine Valette, Bruno Lina, Laurence Josset |
| EPI_ISL_3206488 | DYOMEDEA-LABORATOIRE DE LA SAUVEGARDE | CNR Virus des Infections Respiratoires - France SUD | Antonin Bal, Gregory Destras, Gwendolyne Burfin, Hadrien Regue, Quentin Semanas, Martine Valette, Bruno Lina, Laurence Josset |
| EPI_ISL_3206489 | LABORATOIRE CERBALLIANCE PLT VILLON | CNR Virus des Infections Respiratoires - France SUD | Antonin Bal, Gregory Destras, Gwendolyne Burfin, Hadrien Regue, Quentin Semanas, Martine Valette, Bruno Lina, Laurence Josset |
| EPI_ISL_3206490, EPI_ISL_3206491 | DYOMEDEA-LABORATOIRE DE LA SAUVEGARDE | CNR Virus des Infections Respiratoires - France SUD | Antonin Bal, Gregory Destras, Gwendolyne Burfin, Hadrien Regue, Quentin Semanas, Martine Valette, Bruno Lina, Laurence Josset |

| | | | |
|---|---|---|---|
| EPI_ISL_3206492, EPI_ISL_3206493, EPI_ISL_3206494, EPI_ISL_3206495, EPI_ISL_3206496, EPI_ISL_3206497, EPI_ISL_3206498, EPI_ISL_3206499, EPI_ISL_3206500, EPI_ISL_3206501, EPI_ISL_3206502, EPI_ISL_3206503, EPI_ISL_3206504, EPI_ISL_3206505, EPI_ISL_3206506, EPI_ISL_3206507, EPI_ISL_3206508, EPI_ISL_3206509, EPI_ISL_3206510, EPI_ISL_3206511, EPI_ISL_3206512, EPI_ISL_3206513, EPI_ISL_3206514, EPI_ISL_3206515, EPI_ISL_3206516, EPI_ISL_3206517, EPI_ISL_3206518, EPI_ISL_3206519, EPI_ISL_3206520, EPI_ISL_3206521, EPI_ISL_3206522 | | | |
| see above | LABORATOIRE CERBALLIANCE PLT VILLON | CNR Virus des Infections Respiratoires - France SUD | Antonin Bal, Gregory Destras, Gwendolyne Burfin, Hadrien Regue, Quentin Semanas, Martine Valette, Bruno Lina, Laurence Josset |
| EPI_ISL_3206523 | CNR Virus des Infections Respiratoires - France SUD | CNR Virus des Infections Respiratoires - France SUD | Antonin Bal, Gregory Destras, Gwendolyne Burfin, Hadrien Regue, Quentin Semanas, Martine Valette, Bruno Lina, Laurence Josset |
| EPI_ISL_3206524, EPI_ISL_3206525, EPI_ISL_3206526, EPI_ISL_3206527, EPI_ISL_3206528, EPI_ISL_3206529, EPI_ISL_3206530, EPI_ISL_3206531, EPI_ISL_3206532, EPI_ISL_3206533, EPI_ISL_3206534, EPI_ISL_3206535, EPI_ISL_3206536, EPI_ISL_3206537, EPI_ISL_3206538, EPI_ISL_3206539, EPI_ISL_3206540, EPI_ISL_3206541, EPI_ISL_3206542, EPI_ISL_3206543, EPI_ISL_3206544, EPI_ISL_3206545, EPI_ISL_3206546, EPI_ISL_3206547, EPI_ISL_3206548, EPI_ISL_3206549, EPI_ISL_3206550, EPI_ISL_3206551, EPI_ISL_3206552, EPI_ISL_3206553, EPI_ISL_3206554, EPI_ISL_3206555, EPI_ISL_3206556, EPI_ISL_3206557, EPI_ISL_3206558 | | | |
| see above | LABORATOIRE CERBALLIANCE PLT VILLON | CNR Virus des Infections Respiratoires - France SUD | Antonin Bal, Gregory Destras, Gwendolyne Burfin, Hadrien Regue, Quentin Semanas, Martine Valette, Bruno Lina, Laurence Josset |
| EPI_ISL_3206559 | CNR Virus des Infections Respiratoires - France SUD | CNR Virus des Infections Respiratoires - France SUD | Antonin Bal, Gregory Destras, Gwendolyne Burfin, Hadrien Regue, Quentin Semanas, Martine Valette, Bruno Lina, Laurence Josset |
| EPI_ISL_3206560, EPI_ISL_3206561, EPI_ISL_3206562, EPI_ISL_3206563, EPI_ISL_3206564, EPI_ISL_3206565 | DYOMEDEA-LABORATOIRE DE LA SAUVEGARDE | CNR Virus des Infections Respiratoires - France SUD | Antonin Bal, Gregory Destras, Gwendolyne Burfin, Hadrien Regue, Quentin Semanas, Martine Valette, Bruno Lina, Laurence Josset |
| EPI_ISL_3206566 | CNR Virus des Infections Respiratoires - France SUD | CNR Virus des Infections Respiratoires - France SUD | Antonin Bal, Gregory Destras, Gwendolyne Burfin, Hadrien Regue, Quentin Semanas, Martine Valette, Bruno Lina, Laurence Josset |
| EPI_ISL_3206567, EPI_ISL_3206568, EPI_ISL_3206569, EPI_ISL_3206570, EPI_ISL_3206571, EPI_ISL_3206572, EPI_ISL_3206574, EPI_ISL_3206575, EPI_ISL_3206576, EPI_ISL_3206577, EPI_ISL_3206578, EPI_ISL_3206579, EPI_ISL_3206580, EPI_ISL_3206581, EPI_ISL_3206582, EPI_ISL_3206583 | | | |
| see above | DYOMEDEA-LABORATOIRE DE LA SAUVEGARDE | CNR Virus des Infections Respiratoires - France SUD | Antonin Bal, Gregory Destras, Gwendolyne Burfin, Hadrien Regue, Quentin Semanas, Martine Valette, Bruno Lina, Laurence Josset |
| EPI_ISL_3206584 | CNR Virus des Infections Respiratoires - France SUD | CNR Virus des Infections Respiratoires - France SUD | Antonin Bal, Gregory Destras, Gwendolyne Burfin, Hadrien Regue, Quentin Semanas, Martine Valette, Bruno Lina, Laurence Josset |
| EPI_ISL_3206585, EPI_ISL_3206586, EPI_ISL_3206587, EPI_ISL_3206588, EPI_ISL_3206589, EPI_ISL_3206590, EPI_ISL_3206591, EPI_ISL_3206592, EPI_ISL_3206593, EPI_ISL_3206594, EPI_ISL_3206595, EPI_ISL_3206596, EPI_ISL_3206597, EPI_ISL_3206598, EPI_ISL_3206599, EPI_ISL_3206600, EPI_ISL_3206601, EPI_ISL_3206602, EPI_ISL_3206603, EPI_ISL_3206604, EPI_ISL_3206605, EPI_ISL_3206606, EPI_ISL_3206607, EPI_ISL_3206608, EPI_ISL_3206609, EPI_ISL_3206610, EPI_ISL_3206611, EPI_ISL_3206612, EPI_ISL_3206613, EPI_ISL_3206614, EPI_ISL_3206615, EPI_ISL_3206616, EPI_ISL_3206617, EPI_ISL_3206618, EPI_ISL_3206619, EPI_ISL_3206620, EPI_ISL_3206621, EPI_ISL_3206622, EPI_ISL_3206623, EPI_ISL_3206624, EPI_ISL_3206625, EPI_ISL_3206626, EPI_ISL_3206627, EPI_ISL_3206628, EPI_ISL_3206629 | | | |
| see above | DYOMEDEA-LABORATOIRE DE LA SAUVEGARDE | CNR Virus des Infections Respiratoires - France SUD | Antonin Bal, Gregory Destras, Gwendolyne Burfin, Hadrien Regue, Quentin Semanas, Martine Valette, Bruno Lina, Laurence Josset |
| EPI_ISL_3206630 | CLINIQUE DE LA SAUVEGARDE | CNR Virus des Infections Respiratoires - France SUD | Antonin Bal, Gregory Destras, Gwendolyne Burfin, Hadrien Regue, Quentin Semanas, Martine Valette, Bruno Lina, Laurence Josset |
| EPI_ISL_3206631, EPI_ISL_3206632 | DYOMEDEA-LABORATOIRE DE LA SAUVEGARDE | CNR Virus des Infections Respiratoires - France SUD | Antonin Bal, Gregory Destras, Gwendolyne Burfin, Hadrien Regue, Quentin Semanas, Martine Valette, Bruno Lina, Laurence Josset |
| EPI_ISL_3206633 | CNR Virus des Infections Respiratoires - France SUD | CNR Virus des Infections Respiratoires - France SUD | Antonin Bal, Gregory Destras, Gwendolyne Burfin, Hadrien Regue, Quentin Semanas, Martine Valette, Bruno Lina, Laurence Josset |
| EPI_ISL_3206634, EPI_ISL_3206635, EPI_ISL_3206636, EPI_ISL_3206637, EPI_ISL_3206638, EPI_ISL_3206639, EPI_ISL_3206640, EPI_ISL_3206641, EPI_ISL_3206642, EPI_ISL_3206643, EPI_ISL_3206644, EPI_ISL_3206645, EPI_ISL_3206646, EPI_ISL_3206647, EPI_ISL_3206648, EPI_ISL_3206649, EPI_ISL_3206650, EPI_ISL_3206651, EPI_ISL_3206652, EPI_ISL_3206653, EPI_ISL_3206654, EPI_ISL_3206655, EPI_ISL_3206656, EPI_ISL_3206657, EPI_ISL_3206658, EPI_ISL_3206659, EPI_ISL_3206660, EPI_ISL_3206661, EPI_ISL_3206662, EPI_ISL_3206663, EPI_ISL_3206664, EPI_ISL_3206665, EPI_ISL_3206666, EPI_ISL_3206667, EPI_ISL_3206668, EPI_ISL_3206669, EPI_ISL_3206670, EPI_ISL_3206671, EPI_ISL_3206672, EPI_ISL_3206673, EPI_ISL_3206674, EPI_ISL_3206675, EPI_ISL_3206676, EPI_ISL_3206677, EPI_ISL_3206678, EPI_ISL_3206679, EPI_ISL_3206680, EPI_ISL_3206681, EPI_ISL_3206682, EPI_ISL_3206683, EPI_ISL_3206684, EPI_ISL_3206685, EPI_ISL_3206686, EPI_ISL_3206687, EPI_ISL_3206688, EPI_ISL_3206689, EPI_ISL_3206690, EPI_ISL_3206691, EPI_ISL_3206692, EPI_ISL_3206693, EPI_ISL_3206694, EPI_ISL_3206695, EPI_ISL_3206696, EPI_ISL_3206697, EPI_ISL_3206698, EPI_ISL_3206699, EPI_ISL_3206700, EPI_ISL_3206701, EPI_ISL_3206702, EPI_ISL_3206703, EPI_ISL_3206704, EPI_ISL_3206705, EPI_ISL_3206706, EPI_ISL_3206707, EPI_ISL_3206708, EPI_ISL_3206709, EPI_ISL_3206710, EPI_ISL_3206711, EPI_ISL_3206712 | | | |
| see above | DYOMEDEA-LABORATOIRE DE LA SAUVEGARDE | CNR Virus des Infections Respiratoires - France SUD | Antonin Bal, Gregory Destras, Gwendolyne Burfin, Hadrien Regue, Quentin Semanas, Martine Valette, Bruno Lina, Laurence Josset |
| EPI_ISL_3206713, EPI_ISL_3206714, EPI_ISL_3206715, EPI_ISL_3206716, EPI_ISL_3206717, EPI_ISL_3206718, EPI_ISL_3206719, EPI_ISL_3206720, EPI_ISL_3206721, EPI_ISL_3206722, EPI_ISL_3206723, EPI_ISL_3206724, EPI_ISL_3206725, EPI_ISL_3206726 | | | |
| see above | LBM DYNABIO CARREAU | CNR Virus des Infections Respiratoires - France SUD | Antonin Bal, Gregory Destras, Gwendolyne Burfin, Hadrien Regue, Quentin Semanas, Martine Valette, Bruno Lina, Laurence Josset |
| EPI_ISL_3206727, EPI_ISL_3206728, EPI_ISL_3206729, EPI_ISL_3206730, EPI_ISL_3206731, EPI_ISL_3206732, EPI_ISL_3206733, EPI_ISL_3206734, EPI_ISL_3206735, EPI_ISL_3206736, EPI_ISL_3206737, EPI_ISL_3206738 | | | |
| see above | LABORATOIRE MAYMAT | CNR Virus des Infections Respiratoires - France SUD | Antonin Bal, Gregory Destras, Gwendolyne Burfin, Hadrien Regue, Quentin Semanas, Martine Valette, Bruno Lina, Laurence Josset |
| EPI_ISL_3206739 | LABORATOIRE BOUVIER | CNR Virus des Infections Respiratoires - France SUD | Antonin Bal, Gregory Destras, Gwendolyne Burfin, Hadrien Regue, Quentin Semanas, Martine Valette, Bruno Lina, Laurence Josset |
| EPI_ISL_3206741 | SCM B12 | CNR Virus des Infections Respiratoires - France SUD | Antonin Bal, Gregory Destras, Gwendolyne Burfin, Hadrien Regue, Quentin Semanas, Martine Valette, Bruno Lina, Laurence Josset |
| EPI_ISL_3206742, EPI_ISL_3206744, EPI_ISL_3206745, EPI_ISL_3206746 | CENTRE HOSPITALIER DE VALENCE | CNR Virus des Infections Respiratoires - France SUD | Antonin Bal, Gregory Destras, Gwendolyne Burfin, Hadrien Regue, Quentin Semanas, Martine Valette, Bruno Lina, Laurence Josset |
| EPI_ISL_3206747 | LABORATOIRE BIOESTEREL | CNR Virus des Infections Respiratoires - France SUD | Antonin Bal, Gregory Destras, Gwendolyne Burfin, Hadrien Regue, Quentin Semanas, Martine Valette, Bruno Lina, Laurence Josset |
| EPI_ISL_3206748 | DYOMEDEA-LABORATOIRE DE LA SAUVEGARDE | CNR Virus des Infections Respiratoires - France SUD | Antonin Bal, Gregory Destras, Gwendolyne Burfin, Hadrien Regue, Quentin Semanas, Martine Valette, Bruno Lina, Laurence Josset |
| EPI_ISL_3206749 | LABORATOIRE d'ANALYSES de BIOLOGIE MEDICALES | CNR Virus des Infections Respiratoires - France SUD | Antonin Bal, Gregory Destras, Gwendolyne Burfin, Hadrien Regue, Quentin Semanas, Martine Valette, Bruno Lina, Laurence Josset |
| EPI_ISL_3206750, EPI_ISL_3206751, EPI_ISL_3206752, EPI_ISL_3206753 | LABORATOIRE CERBALLIANCE PLT VILLON | CNR Virus des Infections Respiratoires - France SUD | Antonin Bal, Gregory Destras, Gwendolyne Burfin, Hadrien Regue, Quentin Semanas, Martine Valette, Bruno Lina, Laurence Josset |
| EPI_ISL_3206754 | LABORATOIRE d'ANALYSES de BIOLOGIE MEDICALES | CNR Virus des Infections Respiratoires - France SUD | Antonin Bal, Gregory Destras, Gwendolyne Burfin, Hadrien Regue, Quentin Semanas, Martine Valette, Bruno Lina, Laurence Josset |
| EPI_ISL_3206755, EPI_ISL_3206756 | LABORATOIRE CERBALLIANCE PLT VILLON | CNR Virus des Infections Respiratoires - France SUD | Antonin Bal, Gregory Destras, Gwendolyne Burfin, Hadrien Regue, Quentin Semanas, Martine Valette, Bruno Lina, Laurence Josset |
| EPI_ISL_3206757 | LABORATOIRE d'ANALYSES de BIOLOGIE MEDICALES | CNR Virus des Infections Respiratoires - France SUD | Antonin Bal, Gregory Destras, Gwendolyne Burfin, Hadrien Regue, Quentin Semanas, Martine Valette, Bruno Lina, Laurence Josset |
| EPI_ISL_3206758 | LABORATOIRE BIOESTEREL | CNR Virus des Infections Respiratoires - France SUD | Antonin Bal, Gregory Destras, Gwendolyne Burfin, Hadrien Regue, Quentin Semanas, Martine Valette, Bruno Lina, Laurence Josset |
| EPI_ISL_3206759 | DYOMEDEA-LABORATOIRE DE LA SAUVEGARDE | CNR Virus des Infections Respiratoires - France SUD | Antonin Bal, Gregory Destras, Gwendolyne Burfin, Hadrien Regue, Quentin Semanas, Martine Valette, Bruno Lina, Laurence Josset |
| EPI_ISL_3206760, EPI_ISL_3206761, EPI_ISL_3206762, EPI_ISL_3206763, EPI_ISL_3206764, EPI_ISL_3206765, EPI_ISL_3206766 | LABORATOIRE BIOESTEREL | CNR Virus des Infections Respiratoires - France SUD | Antonin Bal, Gregory Destras, Gwendolyne Burfin, Hadrien Regue, Quentin Semanas, Martine Valette, Bruno Lina, Laurence Josset |
| EPI_ISL_3206767 | LABORATOIRE d'ANALYSES de BIOLOGIE MEDICALES | CNR Virus des Infections Respiratoires - France SUD | Antonin Bal, Gregory Destras, Gwendolyne Burfin, Hadrien Regue, Quentin Semanas, Martine Valette, Bruno Lina, Laurence Josset |
| EPI_ISL_3206768 | DYOMEDEA-LABORATOIRE DE LA SAUVEGARDE | CNR Virus des Infections Respiratoires - France SUD | Antonin Bal, Gregory Destras, Gwendolyne Burfin, Hadrien Regue, Quentin Semanas, Martine Valette, Bruno Lina, Laurence Josset |
| EPI_ISL_3206769 | LAM NOVELAB LVA | CNR Virus des Infections Respiratoires - France SUD | Antonin Bal, Gregory Destras, Gwendolyne Burfin, Hadrien Regue, Quentin Semanas, Martine Valette, Bruno Lina, Laurence Josset |
| EPI_ISL_3206770 | DYOMEDEA-LABORATOIRE DE LA SAUVEGARDE | CNR Virus des Infections Respiratoires - France SUD | Antonin Bal, Gregory Destras, Gwendolyne Burfin, Hadrien Regue, Quentin Semanas, Martine Valette, Bruno Lina, Laurence Josset |
| EPI_ISL_3206771 | LBM VERMEULIN SITE GUERY CLEMENT | CNR Virus des Infections Respiratoires - France SUD | Antonin Bal, Gregory Destras, Gwendolyne Burfin, Hadrien Regue, Quentin Semanas, Martine Valette, Bruno Lina, Laurence Josset |
| EPI_ISL_3206772, EPI_ISL_3206773, EPI_ISL_3206774, EPI_ISL_3206775, EPI_ISL_3206776, EPI_ISL_3206777, EPI_ISL_3206778, EPI_ISL_3206779, EPI_ISL_3206780 | LAM DU MARAIS | CNR Virus des Infections Respiratoires - France SUD | Antonin Bal, Gregory Destras, Gwendolyne Burfin, Hadrien Regue, Quentin Semanas, Martine Valette, Bruno Lina, Laurence Josset |
| EPI_ISL_3206782, EPI_ISL_3206783, EPI_ISL_3206784, EPI_ISL_3206785, EPI_ISL_3206786, EPI_ISL_3206787, EPI_ISL_3206788, EPI_ISL_3206789, EPI_ISL_3206790, EPI_ISL_3206791, EPI_ISL_3206792, EPI_ISL_3206793, EPI_ISL_3206794, EPI_ISL_3206795, EPI_ISL_3206796, EPI_ISL_3206797, EPI_ISL_3206798, EPI_ISL_3206799, EPI_ISL_3206800, EPI_ISL_3206801, EPI_ISL_3206802, EPI_ISL_3206803, EPI_ISL_3206804, EPI_ISL_3206805, EPI_ISL_3206806, EPI_ISL_3206807, EPI_ISL_3206808, EPI_ISL_3206809, EPI_ISL_3206810, EPI_ISL_3206811, EPI_ISL_3206812, EPI_ISL_3206813, EPI_ISL_3206814, EPI_ISL_3206815, EPI_ISL_3206816, EPI_ISL_3206817, EPI_ISL_3206818, EPI_ISL_3206819, EPI_ISL_3206820, EPI_ISL_3206821, EPI_ISL_3206822, EPI_ISL_3206823, EPI_ISL_3206824, EPI_ISL_3206825, EPI_ISL_3206826, EPI_ISL_3206827, EPI_ISL_3206828, EPI_ISL_3206829, EPI_ISL_3206830, EPI_ISL_3206831, EPI_ISL_3206832, EPI_ISL_3206833, EPI_ISL_3206834, EPI_ISL_3206835, EPI_ISL_3206836, EPI_ISL_3206837, EPI_ISL_3206838 | | | |
| see above | LABORATOIRE BIOGROUP | CNR Virus des Infections Respiratoires - France SUD | Antonin Bal, Gregory Destras, Gwendolyne Burfin, Hadrien Regue, Quentin Semanas, Martine Valette, Bruno Lina, Laurence Josset |
| EPI_ISL_3206839 | LABM DU BOLLWERK | CNR Virus des Infections Respiratoires - France SUD | Antonin Bal, Gregory Destras, Gwendolyne Burfin, Hadrien Regue, Quentin Semanas, Martine Valette, Bruno Lina, Laurence Josset |
| EPI_ISL_3206841, EPI_ISL_3206842, EPI_ISL_3206843, EPI_ISL_3206844, EPI_ISL_3206845, EPI_ISL_3206846, EPI_ISL_3206847, EPI_ISL_3206848, EPI_ISL_3206849, EPI_ISL_3206850, EPI_ISL_3206851, EPI_ISL_3206852 | | | |
| see above | LABORATOIRE BIOESTEREL | CNR Virus des Infections Respiratoires - France SUD | Antonin Bal, Gregory Destras, Gwendolyne Burfin, Hadrien Regue, Quentin Semanas, Martine Valette, Bruno Lina, Laurence Josset |
| EPI_ISL_3206853, EPI_ISL_3206854 | DYOMEDEA-LABORATOIRE DE LA SAUVEGARDE | CNR Virus des Infections Respiratoires - France SUD | Antonin Bal, Gregory Destras, Gwendolyne Burfin, Hadrien Regue, Quentin Semanas, Martine Valette, Bruno Lina, Laurence Josset |
| EPI_ISL_3206855 | LABORATOIRE BIOFUSION | CNR Virus des Infections Respiratoires - France SUD | Antonin Bal, Gregory Destras, Gwendolyne Burfin, Hadrien Regue, Quentin Semanas, Martine Valette, Bruno Lina, Laurence Josset |
| EPI_ISL_3206856 | LABORATOIRE BIOESTEREL | CNR Virus des Infections Respiratoires - France SUD | Antonin Bal, Gregory Destras, Gwendolyne Burfin, Hadrien Regue, Quentin Semanas, Martine Valette, Bruno Lina, Laurence Josset |
| EPI_ISL_3206857 | LABORATOIRE d'ANALYSES de BIOLOGIE MEDICALES | CNR Virus des Infections Respiratoires - France SUD | Antonin Bal, Gregory Destras, Gwendolyne Burfin, Hadrien Regue, Quentin Semanas, Martine Valette, Bruno Lina, Laurence Josset |
| EPI_ISL_3206858 | LAM BIORYLIS | CNR Virus des Infections Respiratoires - France SUD | Antonin Bal, Gregory Destras, Gwendolyne Burfin, Hadrien Regue, Quentin Semanas, Martine Valette, Bruno Lina, Laurence Josset |

| | | | |
|---|---|---|---|
| EPI_ISL_3206859, EPI_ISL_3206860 | LABORATOIRE MAYMAT | CNR Virus des Infections Respiratoires - France SUD | Antonin Bal, Gregory Destras, Gwendolyne Burfin, Hadrien Regue, Quentin Semanas, Martine Valette, Bruno Lina, Laurence Josset |
| EPI_ISL_3206861 | LABORATOIRE d'ANALYSES de BIOLOGIE MEDICALES | CNR Virus des Infections Respiratoires - France SUD | Antonin Bal, Gregory Destras, Gwendolyne Burfin, Hadrien Regue, Quentin Semanas, Martine Valette, Bruno Lina, Laurence Josset |
| EPI_ISL_3206862 | LABORATOIRE CERBALLIANCE PLT VILLON | CNR Virus des Infections Respiratoires - France SUD | Antonin Bal, Gregory Destras, Gwendolyne Burfin, Hadrien Regue, Quentin Semanas, Martine Valette, Bruno Lina, Laurence Josset |
| EPI_ISL_3206863 | LABORATOIRE d'ANALYSES de BIOLOGIE MEDICALES | CNR Virus des Infections Respiratoires - France SUD | Antonin Bal, Gregory Destras, Gwendolyne Burfin, Hadrien Regue, Quentin Semanas, Martine Valette, Bruno Lina, Laurence Josset |
| EPI_ISL_3206864 | LABORATOIRE CERBALLIANCE PLT VILLON | CNR Virus des Infections Respiratoires - France SUD | Antonin Bal, Gregory Destras, Gwendolyne Burfin, Hadrien Regue, Quentin Semanas, Martine Valette, Bruno Lina, Laurence Josset |
| EPI_ISL_3206865, EPI_ISL_3206866 | LABORATOIRE d'ANALYSES de BIOLOGIE MEDICALES | CNR Virus des Infections Respiratoires - France SUD | Antonin Bal, Gregory Destras, Gwendolyne Burfin, Hadrien Regue, Quentin Semanas, Martine Valette, Bruno Lina, Laurence Josset |
| EPI_ISL_3206867, EPI_ISL_3206868, EPI_ISL_3206869, EPI_ISL_3206870 | LABORATOIRE BIOESTEREL | CNR Virus des Infections Respiratoires - France SUD | Antonin Bal, Gregory Destras, Gwendolyne Burfin, Hadrien Regue, Quentin Semanas, Martine Valette, Bruno Lina, Laurence Josset |
| EPI_ISL_3206871 | Microbiology Department, Laboratori Clínic Metropolitana Nord. Hospital Universitari Germans Trias i Pujol | Can Ruti SARS-CoV-2 Sequencing Hub (HUGTiP/IrsiCaixa/IGTP) | Marc Noguera-Julian, Pilar Armengol, Ignacio Blanco, Antoni E Bordoy, Francesc Catala-Moll, Pere-Joan Cardona, Maria Casadellà, Cristina Casañ, Gemma Clara, Bonaventura Clotet, Montserrat Giménez, Mercedes Guerrero, Anna Not, David Panisello, Roger Paredes, Mariona Parera, Alexia París, Verónica Saludes, Alba Sánchez, Laia Soler, Lauro Sumoy, and Elisa Martró on behalf of the Can Ruti SARS-CoV-2 Sequencing Hub |
| EPI_ISL_3206872 | LABORATOIRE BIOESTEREL | CNR Virus des Infections Respiratoires - France SUD | Antonin Bal, Gregory Destras, Gwendolyne Burfin, Hadrien Regue, Quentin Semanas, Martine Valette, Bruno Lina, Laurence Josset |
| EPI_ISL_3206873, EPI_ISL_3206874 | Microbiology Department, Laboratori Clínic Metropolitana Nord. Hospital Universitari Germans Trias i Pujol | Can Ruti SARS-CoV-2 Sequencing Hub (HUGTiP/IrsiCaixa/IGTP) | Marc Noguera-Julian, Pilar Armengol, Ignacio Blanco, Antoni E Bordoy, Francesc Catala-Moll, Pere-Joan Cardona, Maria Casadellà, Cristina Casañ, Gemma Clara, Bonaventura Clotet, Montserrat Giménez, Mercedes Guerrero, Anna Not, David Panisello, Roger Paredes, Mariona Parera, Alexia París, Verónica Saludes, Alba Sánchez, Laia Soler, Lauro Sumoy, and Elisa Martró on behalf of the Can Ruti SARS-CoV-2 Sequencing Hub |
| EPI_ISL_3206875 | LABORATOIRE BIOESTEREL | CNR Virus des Infections Respiratoires - France SUD | Antonin Bal, Gregory Destras, Gwendolyne Burfin, Hadrien Regue, Quentin Semanas, Martine Valette, Bruno Lina, Laurence Josset |
| EPI_ISL_3206876 | Microbiology Department, Laboratori Clínic Metropolitana Nord. Hospital Universitari Germans Trias i Pujol | Can Ruti SARS-CoV-2 Sequencing Hub (HUGTiP/IrsiCaixa/IGTP) | Marc Noguera-Julian, Pilar Armengol, Ignacio Blanco, Antoni E Bordoy, Francesc Catala-Moll, Pere-Joan Cardona, Maria Casadellà, Cristina Casañ, Gemma Clara, Bonaventura Clotet, Montserrat Giménez, Mercedes Guerrero, Anna Not, David Panisello, Roger Paredes, Mariona Parera, Alexia París, Verónica Saludes, Alba Sánchez, Laia Soler, Lauro Sumoy, and Elisa Martró on behalf of the Can Ruti SARS-CoV-2 Sequencing Hub |
| EPI_ISL_3206877 | LABORATOIRE BIOESTEREL | CNR Virus des Infections Respiratoires - France SUD | Antonin Bal, Gregory Destras, Gwendolyne Burfin, Hadrien Regue, Quentin Semanas, Martine Valette, Bruno Lina, Laurence Josset |
| EPI_ISL_3206878 | Microbiology Department, Laboratori Clínic Metropolitana Nord. Hospital Universitari Germans Trias i Pujol | Can Ruti SARS-CoV-2 Sequencing Hub (HUGTiP/IrsiCaixa/IGTP) | Marc Noguera-Julian, Pilar Armengol, Ignacio Blanco, Antoni E Bordoy, Francesc Catala-Moll, Pere-Joan Cardona, Maria Casadellà, Cristina Casañ, Gemma Clara, Bonaventura Clotet, Montserrat Giménez, Mercedes Guerrero, Anna Not, David Panisello, Roger Paredes, Mariona Parera, Alexia París, Verónica Saludes, Alba Sánchez, Laia Soler, Lauro Sumoy, and Elisa Martró on behalf of the Can Ruti SARS-CoV-2 Sequencing Hub |
| EPI_ISL_3206879 | DYOMEDEA-LABORATOIRE DE LA SAUVEGARDE | CNR Virus des Infections Respiratoires - France SUD | Antonin Bal, Gregory Destras, Gwendolyne Burfin, Hadrien Regue, Quentin Semanas, Martine Valette, Bruno Lina, Laurence Josset |
| EPI_ISL_3206880 | Microbiology Department, Laboratori Clínic Metropolitana Nord. Hospital Universitari Germans Trias i Pujol | Can Ruti SARS-CoV-2 Sequencing Hub (HUGTiP/IrsiCaixa/IGTP) | Marc Noguera-Julian, Pilar Armengol, Ignacio Blanco, Antoni E Bordoy, Francesc Catala-Moll, Pere-Joan Cardona, Maria Casadellà, Cristina Casañ, Gemma Clara, Bonaventura Clotet, Montserrat Giménez, Mercedes Guerrero, Anna Not, David Panisello, Roger Paredes, Mariona Parera, Alexia París, Verónica Saludes, Alba Sánchez, Laia Soler, Lauro Sumoy, and Elisa Martró on behalf of the Can Ruti SARS-CoV-2 Sequencing Hub |
| EPI_ISL_3206881 | DYOMEDEA-LABORATOIRE DE LA SAUVEGARDE | CNR Virus des Infections Respiratoires - France SUD | Antonin Bal, Gregory Destras, Gwendolyne Burfin, Hadrien Regue, Quentin Semanas, Martine Valette, Bruno Lina, Laurence Josset |
| EPI_ISL_3206882 | Microbiology Department, Laboratori Clínic Metropolitana Nord. Hospital Universitari Germans Trias i Pujol | Can Ruti SARS-CoV-2 Sequencing Hub (HUGTiP/IrsiCaixa/IGTP) | Marc Noguera-Julian, Pilar Armengol, Ignacio Blanco, Antoni E Bordoy, Francesc Catala-Moll, Pere-Joan Cardona, Maria Casadellà, Cristina Casañ, Gemma Clara, Bonaventura Clotet, Montserrat Giménez, Mercedes Guerrero, Anna Not, David Panisello, Roger Paredes, Mariona Parera, Alexia París, Verónica Saludes, Alba Sánchez, Laia Soler, Lauro Sumoy, and Elisa Martró on behalf of the Can Ruti SARS-CoV-2 Sequencing Hub |
| EPI_ISL_3206883 | LABORATOIRE d'ANALYSES de BIOLOGIE MEDICALES | CNR Virus des Infections Respiratoires - France SUD | Antonin Bal, Gregory Destras, Gwendolyne Burfin, Hadrien Regue, Quentin Semanas, Martine Valette, Bruno Lina, Laurence Josset |
| EPI_ISL_3206884 | Microbiology Department, Laboratori Clínic Metropolitana Nord. Hospital Universitari Germans Trias i Pujol | Can Ruti SARS-CoV-2 Sequencing Hub (HUGTiP/IrsiCaixa/IGTP) | Marc Noguera-Julian, Pilar Armengol, Ignacio Blanco, Antoni E Bordoy, Francesc Catala-Moll, Pere-Joan Cardona, Maria Casadellà, Cristina Casañ, Gemma Clara, Bonaventura Clotet, Montserrat Giménez, Mercedes Guerrero, Anna Not, David Panisello, Roger Paredes, Mariona Parera, Alexia París, Verónica Saludes, Alba Sánchez, Laia Soler, Lauro Sumoy, and Elisa Martró on behalf of the Can Ruti SARS-CoV-2 Sequencing Hub |
| EPI_ISL_3206885 | LABORATOIRE d'ANALYSES de BIOLOGIE MEDICALES | CNR Virus des Infections Respiratoires - France SUD | Antonin Bal, Gregory Destras, Gwendolyne Burfin, Hadrien Regue, Quentin Semanas, Martine Valette, Bruno Lina, Laurence Josset |
| EPI_ISL_3206886 | Microbiology Department, Laboratori Clínic Metropolitana Nord. Hospital Universitari Germans Trias i Pujol | Can Ruti SARS-CoV-2 Sequencing Hub (HUGTiP/IrsiCaixa/IGTP) | Marc Noguera-Julian, Pilar Armengol, Ignacio Blanco, Antoni E Bordoy, Francesc Catala-Moll, Pere-Joan Cardona, Maria Casadellà, Cristina Casañ, Gemma Clara, Bonaventura Clotet, Montserrat Giménez, Mercedes Guerrero, Anna Not, David Panisello, Roger Paredes, Mariona Parera, Alexia París, Verónica Saludes, Alba Sánchez, Laia Soler, Lauro Sumoy, and Elisa Martró on behalf of the Can Ruti SARS-CoV-2 Sequencing Hub |
| EPI_ISL_3206887, EPI_ISL_3206889 | DYOMEDEA-LABORATOIRE DE LA SAUVEGARDE | CNR Virus des Infections Respiratoires - France SUD | Antonin Bal, Gregory Destras, Gwendolyne Burfin, Hadrien Regue, Quentin Semanas, Martine Valette, Bruno Lina, Laurence Josset |
| EPI_ISL_3206891 | LABORATOIRE BIOESTEREL | CNR Virus des Infections Respiratoires - France SUD | Antonin Bal, Gregory Destras, Gwendolyne Burfin, Hadrien Regue, Quentin Semanas, Martine Valette, Bruno Lina, Laurence Josset |
| EPI_ISL_3206892 | Microbiology Department, Laboratori Clínic Metropolitana Nord. Hospital Universitari Germans Trias i Pujol | Can Ruti SARS-CoV-2 Sequencing Hub (HUGTiP/IrsiCaixa/IGTP) | Marc Noguera-Julian, Pilar Armengol, Ignacio Blanco, Antoni E Bordoy, Francesc Catala-Moll, Pere-Joan Cardona, Maria Casadellà, Cristina Casañ, Gemma Clara, Bonaventura Clotet, Montserrat Giménez, Mercedes Guerrero, Anna Not, David Panisello, Roger Paredes, Mariona Parera, Alexia París, Verónica Saludes, Alba Sánchez, Laia Soler, Lauro Sumoy, and Elisa Martró on behalf of the Can Ruti SARS-CoV-2 Sequencing Hub |
| EPI_ISL_3206893 | LABORATOIRE BIOESTEREL | CNR Virus des Infections Respiratoires - France SUD | Antonin Bal, Gregory Destras, Gwendolyne Burfin, Hadrien Regue, Quentin Semanas, Martine Valette, Bruno Lina, Laurence Josset |
| EPI_ISL_3206894 | Microbiology Department, Laboratori Clínic Metropolitana Nord. Hospital Universitari Germans Trias i Pujol | Can Ruti SARS-CoV-2 Sequencing Hub (HUGTiP/IrsiCaixa/IGTP) | Marc Noguera-Julian, Pilar Armengol, Ignacio Blanco, Antoni E Bordoy, Francesc Catala-Moll, Pere-Joan Cardona, Maria Casadellà, Cristina Casañ, Gemma Clara, Bonaventura Clotet, Montserrat Giménez, Mercedes Guerrero, Anna Not, David Panisello, Roger Paredes, Mariona Parera, Alexia París, Verónica Saludes, Alba Sánchez, Laia Soler, Lauro Sumoy, and Elisa Martró on behalf of the Can Ruti SARS-CoV-2 Sequencing Hub |
| EPI_ISL_3206895 | LABORATOIRE BIOESTEREL | CNR Virus des Infections Respiratoires - France SUD | Antonin Bal, Gregory Destras, Gwendolyne Burfin, Hadrien Regue, Quentin Semanas, Martine Valette, Bruno Lina, Laurence Josset |
| EPI_ISL_3206896 | Microbiology Department, Laboratori Clínic Metropolitana Nord. Hospital Universitari Germans Trias i Pujol | Can Ruti SARS-CoV-2 Sequencing Hub (HUGTiP/IrsiCaixa/IGTP) | Marc Noguera-Julian, Pilar Armengol, Ignacio Blanco, Antoni E Bordoy, Francesc Catala-Moll, Pere-Joan Cardona, Maria Casadellà, Cristina Casañ, Gemma Clara, Bonaventura Clotet, Montserrat Giménez, Mercedes Guerrero, Anna Not, David Panisello, Roger Paredes, Mariona Parera, Alexia París, Verónica Saludes, Alba Sánchez, Laia Soler, Lauro Sumoy, and Elisa Martró on behalf of the Can Ruti SARS-CoV-2 Sequencing Hub |
| EPI_ISL_3206897 | LABORATOIRE BIOESTEREL | CNR Virus des Infections Respiratoires - France SUD | Antonin Bal, Gregory Destras, Gwendolyne Burfin, Hadrien Regue, Quentin Semanas, Martine Valette, Bruno Lina, Laurence Josset |
| EPI_ISL_3206898 | Microbiology Department, Laboratori Clínic Metropolitana Nord. Hospital Universitari Germans Trias i Pujol | Can Ruti SARS-CoV-2 Sequencing Hub (HUGTiP/IrsiCaixa/IGTP) | Marc Noguera-Julian, Pilar Armengol, Ignacio Blanco, Antoni E Bordoy, Francesc Catala-Moll, Pere-Joan Cardona, Maria Casadellà, Cristina Casañ, Gemma Clara, Bonaventura Clotet, Montserrat Giménez, Mercedes Guerrero, Anna Not, David Panisello, Roger Paredes, Mariona Parera, Alexia París, Verónica Saludes, Alba Sánchez, Laia Soler, Lauro Sumoy, and Elisa Martró on behalf of the Can Ruti SARS-CoV-2 Sequencing Hub |
| EPI_ISL_3206899 | LABORATOIRE BIOESTEREL | CNR Virus des Infections Respiratoires - France SUD | Antonin Bal, Gregory Destras, Gwendolyne Burfin, Hadrien Regue, Quentin Semanas, Martine Valette, Bruno Lina, Laurence Josset |
| EPI_ISL_3206900 | Microbiology Department, Laboratori Clínic Metropolitana Nord. Hospital Universitari Germans Trias i Pujol | Can Ruti SARS-CoV-2 Sequencing Hub (HUGTiP/IrsiCaixa/IGTP) | Marc Noguera-Julian, Pilar Armengol, Ignacio Blanco, Antoni E Bordoy, Francesc Catala-Moll, Pere-Joan Cardona, Maria Casadellà, Cristina Casañ, Gemma Clara, Bonaventura Clotet, Montserrat Giménez, Mercedes Guerrero, Anna Not, David Panisello, Roger Paredes, Mariona Parera, Alexia París, Verónica Saludes, Alba Sánchez, Laia Soler, Lauro Sumoy, and Elisa Martró on behalf of the Can Ruti SARS-CoV-2 Sequencing Hub |
| EPI_ISL_3206901 | LABORATOIRE BIOESTEREL | CNR Virus des Infections Respiratoires - France SUD | Antonin Bal, Gregory Destras, Gwendolyne Burfin, Hadrien Regue, Quentin Semanas, Martine Valette, Bruno Lina, Laurence Josset |
| EPI_ISL_3206902, EPI_ISL_3206903 | Microbiology Department, Laboratori Clínic Metropolitana Nord. Hospital Universitari Germans Trias i Pujol | Can Ruti SARS-CoV-2 Sequencing Hub (HUGTiP/IrsiCaixa/IGTP) | Marc Noguera-Julian, Pilar Armengol, Ignacio Blanco, Antoni E Bordoy, Francesc Catala-Moll, Pere-Joan Cardona, Maria Casadellà, Cristina Casañ, Gemma Clara, Bonaventura Clotet, Montserrat Giménez, Mercedes Guerrero, Anna Not, David Panisello, Roger Paredes, Mariona Parera, Alexia París, Verónica Saludes, Alba Sánchez, Laia Soler, Lauro Sumoy, and Elisa Martró on behalf of the Can Ruti SARS-CoV-2 Sequencing Hub |
| EPI_ISL_3206904 | LABORATOIRE BIOESTEREL | CNR Virus des Infections Respiratoires - France SUD | Antonin Bal, Gregory Destras, Gwendolyne Burfin, Hadrien Regue, Quentin Semanas, Martine Valette, Bruno Lina, Laurence Josset |
| EPI_ISL_3206905 | Microbiology Department, Laboratori Clínic Metropolitana Nord. Hospital Universitari Germans Trias i Pujol | Can Ruti SARS-CoV-2 Sequencing Hub (HUGTiP/IrsiCaixa/IGTP) | Marc Noguera-Julian, Pilar Armengol, Ignacio Blanco, Antoni E Bordoy, Francesc Catala-Moll, Pere-Joan Cardona, Maria Casadellà, Cristina Casañ, Gemma Clara, Bonaventura Clotet, Montserrat Giménez, Mercedes Guerrero, Anna Not, David Panisello, Roger Paredes, Mariona Parera, Alexia París, Verónica Saludes, Alba Sánchez, Laia Soler, Lauro Sumoy, and Elisa Martró on behalf of the Can Ruti SARS-CoV-2 Sequencing Hub |
| EPI_ISL_3206906 | LABORATOIRE BIOESTEREL | CNR Virus des Infections Respiratoires - France SUD | Antonin Bal, Gregory Destras, Gwendolyne Burfin, Hadrien Regue, Quentin Semanas, Martine Valette, Bruno Lina, Laurence Josset |
| EPI_ISL_3206907 | Microbiology Department, Laboratori Clínic Metropolitana Nord. Hospital Universitari Germans Trias i Pujol | Can Ruti SARS-CoV-2 Sequencing Hub (HUGTiP/IrsiCaixa/IGTP) | Marc Noguera-Julian, Pilar Armengol, Ignacio Blanco, Antoni E Bordoy, Francesc Catala-Moll, Pere-Joan Cardona, Maria Casadellà, Cristina Casañ, Gemma Clara, Bonaventura Clotet, Montserrat Giménez, Mercedes Guerrero, Anna Not, David Panisello, Roger Paredes, Mariona Parera, Alexia París, Verónica Saludes, Alba Sánchez, Laia Soler, Lauro Sumoy, and Elisa Martró on behalf of the Can Ruti SARS-CoV-2 Sequencing Hub |
| EPI_ISL_3206908 | LABORATOIRE BIOESTEREL | CNR Virus des Infections Respiratoires - France SUD | Antonin Bal, Gregory Destras, Gwendolyne Burfin, Hadrien Regue, Quentin Semanas, Martine Valette, Bruno Lina, Laurence Josset |
| EPI_ISL_3206909 | Microbiology Department, Laboratori Clínic Metropolitana Nord. Hospital Universitari Germans Trias i Pujol | Can Ruti SARS-CoV-2 Sequencing Hub (HUGTiP/IrsiCaixa/IGTP) | Marc Noguera-Julian, Pilar Armengol, Ignacio Blanco, Antoni E Bordoy, Francesc Catala-Moll, Pere-Joan Cardona, Maria Casadellà, Cristina Casañ, Gemma Clara, Bonaventura Clotet, Montserrat Giménez, Mercedes Guerrero, Anna Not, David Panisello, Roger Paredes, Mariona Parera, Alexia París, Verónica Saludes, Alba Sánchez, Laia Soler, Lauro Sumoy, and Elisa Martró on behalf of the Can Ruti SARS-CoV-2 Sequencing Hub |
| EPI_ISL_3206910 | DYOMEDEA-LABORATOIRE DE LA SAUVEGARDE | CNR Virus des Infections Respiratoires - France SUD | Antonin Bal, Gregory Destras, Gwendolyne Burfin, Hadrien Regue, Quentin Semanas, Martine Valette, Bruno Lina, Laurence Josset |
| EPI_ISL_3206911 | Microbiology Department, Laboratori Clínic Metropolitana Nord. Hospital Universitari Germans Trias i Pujol | Can Ruti SARS-CoV-2 Sequencing Hub (HUGTiP/IrsiCaixa/IGTP) | Marc Noguera-Julian, Pilar Armengol, Ignacio Blanco, Antoni E Bordoy, Francesc Catala-Moll, Pere-Joan Cardona, Maria Casadellà, Cristina Casañ, Gemma Clara, Bonaventura Clotet, Montserrat Giménez, Mercedes Guerrero, Anna Not, David Panisello, Roger Paredes, Mariona Parera, Alexia París, Verónica Saludes, Alba Sánchez, Laia Soler, Lauro Sumoy, and Elisa Martró on behalf of the Can Ruti SARS-CoV-2 Sequencing Hub |
| EPI_ISL_3206912 | DYOMEDEA-LABORATOIRE DE LA SAUVEGARDE | CNR Virus des Infections Respiratoires - France SUD | Antonin Bal, Gregory Destras, Gwendolyne Burfin, Hadrien Regue, Quentin Semanas, Martine Valette, Bruno Lina, Laurence Josset |
| EPI_ISL_3206913 | Microbiology Department, Laboratori Clínic Metropolitana Nord. Hospital Universitari Germans Trias i Pujol | Can Ruti SARS-CoV-2 Sequencing Hub (HUGTiP/IrsiCaixa/IGTP) | Marc Noguera-Julian, Pilar Armengol, Ignacio Blanco, Antoni E Bordoy, Francesc Catala-Moll, Pere-Joan Cardona, Maria Casadellà, Cristina Casañ, Gemma Clara, Bonaventura Clotet, Montserrat Giménez, Mercedes Guerrero, Anna Not, David Panisello, Roger Paredes, Mariona Parera, Alexia París, Verónica Saludes, Alba Sánchez, Laia Soler, Lauro Sumoy, and Elisa Martró on behalf of the Can Ruti SARS-CoV-2 Sequencing Hub |
| EPI_ISL_3206914 | SYNLAB GASCOGNE | CNR Virus des Infections Respiratoires - France SUD | Antonin Bal, Gregory Destras, Gwendolyne Burfin, Hadrien Regue, Quentin Semanas, Martine Valette, Bruno Lina, Laurence Josset |
| EPI_ISL_3206915 | Microbiology Department, Laboratori Clínic Metropolitana Nord. Hospital Universitari Germans Trias i Pujol | Can Ruti SARS-CoV-2 Sequencing Hub (HUGTiP/IrsiCaixa/IGTP) | Marc Noguera-Julian, Pilar Armengol, Ignacio Blanco, Antoni E Bordoy, Francesc Catala-Moll, Pere-Joan Cardona, Maria Casadellà, Cristina Casañ, Gemma Clara, Bonaventura Clotet, Montserrat Giménez, Mercedes Guerrero, Anna Not, David Panisello, Roger Paredes, Mariona Parera, Alexia París, Verónica Saludes, Alba Sánchez, Laia Soler, Lauro Sumoy, and Elisa Martró on behalf of the Can Ruti SARS-CoV-2 Sequencing Hub |
| EPI_ISL_3206916 | DYOMEDEA-LABORATOIRE DE LA SAUVEGARDE | CNR Virus des Infections Respiratoires - France SUD | Antonin Bal, Gregory Destras, Gwendolyne Burfin, Hadrien Regue, Quentin Semanas, Martine Valette, Bruno Lina, Laurence Josset |
| EPI_ISL_3206917 | Microbiology Department, Laboratori Clínic Metropolitana Nord. Hospital Universitari Germans Trias i Pujol | Can Ruti SARS-CoV-2 Sequencing Hub (HUGTiP/IrsiCaixa/IGTP) | Marc Noguera-Julian, Pilar Armengol, Ignacio Blanco, Antoni E Bordoy, Francesc Catala-Moll, Pere-Joan Cardona, Maria Casadellà, Cristina Casañ, Gemma Clara, Bonaventura Clotet, Montserrat Giménez, Mercedes Guerrero, Anna Not, David Panisello, Roger Paredes, Mariona Parera, Alexia París, Verónica Saludes, Alba Sánchez, Laia Soler, Lauro Sumoy, and Elisa Martró on behalf of the Can Ruti SARS-CoV-2 Sequencing Hub |
| EPI_ISL_3206918 | LABORATOIRE MAYMAT | CNR Virus des Infections Respiratoires - France SUD | Antonin Bal, Gregory Destras, Gwendolyne Burfin, Hadrien Regue, Quentin Semanas, Martine Valette, Bruno Lina, Laurence Josset |
| EPI_ISL_3206919, EPI_ISL_3206920 | Microbiology Department, Laboratori Clínic Metropolitana Nord. Hospital Universitari Germans Trias i Pujol | Can Ruti SARS-CoV-2 Sequencing Hub (HUGTiP/IrsiCaixa/IGTP) | Marc Noguera-Julian, Pilar Armengol, Ignacio Blanco, Antoni E Bordoy, Francesc Catala-Moll, Pere-Joan Cardona, Maria Casadellà, Cristina Casañ, Gemma Clara, Bonaventura Clotet, Montserrat Giménez, Mercedes Guerrero, Anna Not, David Panisello, Roger Paredes, Mariona Parera, Alexia París, Verónica Saludes, Alba Sánchez, Laia Soler, Lauro Sumoy, and Elisa Martró on behalf of the Can Ruti SARS-CoV-2 Sequencing Hub |
| EPI_ISL_3206921 | DYOMEDEA-LABORATOIRE DE LA SAUVEGARDE | CNR Virus des Infections Respiratoires - France SUD | Antonin Bal, Gregory Destras, Gwendolyne Burfin, Hadrien Regue, Quentin Semanas, Martine Valette, Bruno Lina, Laurence Josset |
| EPI_ISL_3206922 | Microbiology Department, Laboratori Clínic Metropolitana Nord. Hospital Universitari Germans Trias i Pujol | Can Ruti SARS-CoV-2 Sequencing Hub (HUGTiP/IrsiCaixa/IGTP) | Marc Noguera-Julian, Pilar Armengol, Ignacio Blanco, Antoni E Bordoy, Francesc Catala-Moll, Pere-Joan Cardona, Maria Casadellà, Cristina Casañ, Gemma Clara, Bonaventura Clotet, Montserrat Giménez, Mercedes Guerrero, Anna Not, David Panisello, Roger Paredes, Mariona Parera, Alexia París, Verónica Saludes, Alba Sánchez, Laia Soler, Lauro Sumoy, and Elisa Martró on behalf of the Can Ruti SARS-CoV-2 Sequencing Hub |
| EPI_ISL_3206923 | DYOMEDEA-LABORATOIRE DE LA SAUVEGARDE | CNR Virus des Infections Respiratoires - France SUD | Antonin Bal, Gregory Destras, Gwendolyne Burfin, Hadrien Regue, Quentin Semanas, Martine Valette, Bruno Lina, Laurence Josset |
| EPI_ISL_3206924 | Microbiology Department, Laboratori Clínic Metropolitana Nord. Hospital Universitari Germans Trias i Pujol | Can Ruti SARS-CoV-2 Sequencing Hub (HUGTiP/IrsiCaixa/IGTP) | Marc Noguera-Julian, Pilar Armengol, Ignacio Blanco, Antoni E Bordoy, Francesc Catala-Moll, Pere-Joan Cardona, Maria Casadellà, Cristina Casañ, Gemma Clara, Bonaventura Clotet, Montserrat Giménez, Mercedes Guerrero, Anna Not, David Panisello, Roger Paredes, Mariona Parera, Alexia París, Verónica Saludes, Alba Sánchez, Laia Soler, Lauro Sumoy, and Elisa Martró on behalf of the Can Ruti SARS-CoV-2 Sequencing Hub |

| | | | |
|---|---|---|---|
| EPI_ISL_3206925 | CENTRE HOSPITALIER BRIVE | CNR Virus des Infections Respiratoires - France SUD | Antonin Bal, Gregory Destras, Gwendolyne Burfin, Hadrien Regue, Quentin Semanas, Martine Valette, Bruno Lina, Laurence Josset |
| EPI_ISL_3206926, EPI_ISL_3206928 | Microbiology Department, Laboratori Clínic Metropolitana Nord. Hospital Universitari Germans Trias i Pujol | Can Ruti SARS-CoV-2 Sequencing Hub (HUGTiP/IrsiCaixa/IGTP) | Marc Noguera-Julian, Pilar Armengol, Ignacio Blanco, Antoni E Bordoy, Francesc Catala-Moll, Pere-Joan Cardona, Maria Casadellà, Cristina Casañ, Gemma Clara, Bonaventura Clotet, Montserrat Giménez, Mercedes Guerrero, Anna Not, David Panisello, Roger Paredes, Mariona Parera, Alexia París, Verónica Saludes, Alba Sánchez, Laia Soler, Lauro Sumoy, and Elisa Martró on behalf of the Can Ruti SARS-CoV-2 Sequencing Hub |
| EPI_ISL_3206929 | CENTRE HOSPITALIER NIORT | CNR Virus des Infections Respiratoires - France SUD | Antonin Bal, Gregory Destras, Gwendolyne Burfin, Hadrien Regue, Quentin Semanas, Martine Valette, Bruno Lina, Laurence Josset |
| EPI_ISL_3206930, EPI_ISL_3206931 | Microbiology Department, Laboratori Clínic Metropolitana Nord. Hospital Universitari Germans Trias i Pujol | Can Ruti SARS-CoV-2 Sequencing Hub (HUGTiP/IrsiCaixa/IGTP) | Marc Noguera-Julian, Pilar Armengol, Ignacio Blanco, Antoni E Bordoy, Francesc Catala-Moll, Pere-Joan Cardona, Maria Casadellà, Cristina Casañ, Gemma Clara, Bonaventura Clotet, Montserrat Giménez, Mercedes Guerrero, Anna Not, David Panisello, Roger Paredes, Mariona Parera, Alexia París, Verónica Saludes, Alba Sánchez, Laia Soler, Lauro Sumoy, and Elisa Martró on behalf of the Can Ruti SARS-CoV-2 Sequencing Hub |
| EPI_ISL_3206932 | CENTRE HOSPITALIER NIORT | CNR Virus des Infections Respiratoires - France SUD | Antonin Bal, Gregory Destras, Gwendolyne Burfin, Hadrien Regue, Quentin Semanas, Martine Valette, Bruno Lina, Laurence Josset |
| EPI_ISL_3206933 | Microbiology Department, Laboratori Clínic Metropolitana Nord. Hospital Universitari Germans Trias i Pujol | Can Ruti SARS-CoV-2 Sequencing Hub (HUGTiP/IrsiCaixa/IGTP) | Marc Noguera-Julian, Pilar Armengol, Ignacio Blanco, Antoni E Bordoy, Francesc Catala-Moll, Pere-Joan Cardona, Maria Casadellà, Cristina Casañ, Gemma Clara, Bonaventura Clotet, Montserrat Giménez, Mercedes Guerrero, Anna Not, David Panisello, Roger Paredes, Mariona Parera, Alexia París, Verónica Saludes, Alba Sánchez, Laia Soler, Lauro Sumoy, and Elisa Martró on behalf of the Can Ruti SARS-CoV-2 Sequencing Hub |
| EPI_ISL_3206934 | LAM BIOPOLE ORTHEZ | CNR Virus des Infections Respiratoires - France SUD | Antonin Bal, Gregory Destras, Gwendolyne Burfin, Hadrien Regue, Quentin Semanas, Martine Valette, Bruno Lina, Laurence Josset |
| EPI_ISL_3206935 | Microbiology Department, Laboratori Clínic Metropolitana Nord. Hospital Universitari Germans Trias i Pujol | Can Ruti SARS-CoV-2 Sequencing Hub (HUGTiP/IrsiCaixa/IGTP) | Marc Noguera-Julian, Pilar Armengol, Ignacio Blanco, Antoni E Bordoy, Francesc Catala-Moll, Pere-Joan Cardona, Maria Casadellà, Cristina Casañ, Gemma Clara, Bonaventura Clotet, Montserrat Giménez, Mercedes Guerrero, Anna Not, David Panisello, Roger Paredes, Mariona Parera, Alexia París, Verónica Saludes, Alba Sánchez, Laia Soler, Lauro Sumoy, and Elisa Martró on behalf of the Can Ruti SARS-CoV-2 Sequencing Hub |
| EPI_ISL_3206936 | LAM BIOPOLE ORTHEZ | CNR Virus des Infections Respiratoires - France SUD | Antonin Bal, Gregory Destras, Gwendolyne Burfin, Hadrien Regue, Quentin Semanas, Martine Valette, Bruno Lina, Laurence Josset |
| EPI_ISL_3206937 | Microbiology Department, Laboratori Clínic Metropolitana Nord. Hospital Universitari Germans Trias i Pujol | Can Ruti SARS-CoV-2 Sequencing Hub (HUGTiP/IrsiCaixa/IGTP) | Marc Noguera-Julian, Pilar Armengol, Ignacio Blanco, Antoni E Bordoy, Francesc Catala-Moll, Pere-Joan Cardona, Maria Casadellà, Cristina Casañ, Gemma Clara, Bonaventura Clotet, Montserrat Giménez, Mercedes Guerrero, Anna Not, David Panisello, Roger Paredes, Mariona Parera, Alexia París, Verónica Saludes, Alba Sánchez, Laia Soler, Lauro Sumoy, and Elisa Martró on behalf of the Can Ruti SARS-CoV-2 Sequencing Hub |
| EPI_ISL_3206938 | LAM BIOPOLE ORTHEZ | CNR Virus des Infections Respiratoires - France SUD | Antonin Bal, Gregory Destras, Gwendolyne Burfin, Hadrien Regue, Quentin Semanas, Martine Valette, Bruno Lina, Laurence Josset |
| EPI_ISL_3206939 | Microbiology Department, Laboratori Clínic Metropolitana Nord. Hospital Universitari Germans Trias i Pujol | Can Ruti SARS-CoV-2 Sequencing Hub (HUGTiP/IrsiCaixa/IGTP) | Marc Noguera-Julian, Pilar Armengol, Ignacio Blanco, Antoni E Bordoy, Francesc Catala-Moll, Pere-Joan Cardona, Maria Casadellà, Cristina Casañ, Gemma Clara, Bonaventura Clotet, Montserrat Giménez, Mercedes Guerrero, Anna Not, David Panisello, Roger Paredes, Mariona Parera, Alexia París, Verónica Saludes, Alba Sánchez, Laia Soler, Lauro Sumoy, and Elisa Martró on behalf of the Can Ruti SARS-CoV-2 Sequencing Hub |
| EPI_ISL_3206940 | LAM BIOPOLE ORTHEZ | CNR Virus des Infections Respiratoires - France SUD | Antonin Bal, Gregory Destras, Gwendolyne Burfin, Hadrien Regue, Quentin Semanas, Martine Valette, Bruno Lina, Laurence Josset |
| EPI_ISL_3206941, EPI_ISL_3206942 | Microbiology Department, Laboratori Clínic Metropolitana Nord. Hospital Universitari Germans Trias i Pujol | Can Ruti SARS-CoV-2 Sequencing Hub (HUGTiP/IrsiCaixa/IGTP) | Marc Noguera-Julian, Pilar Armengol, Ignacio Blanco, Antoni E Bordoy, Francesc Catala-Moll, Pere-Joan Cardona, Maria Casadellà, Cristina Casañ, Gemma Clara, Bonaventura Clotet, Montserrat Giménez, Mercedes Guerrero, Anna Not, David Panisello, Roger Paredes, Mariona Parera, Alexia París, Verónica Saludes, Alba Sánchez, Laia Soler, Lauro Sumoy, and Elisa Martró on behalf of the Can Ruti SARS-CoV-2 Sequencing Hub |
| EPI_ISL_3206943 | LAM BIOPOLE ORTHEZ | CNR Virus des Infections Respiratoires - France SUD | Antonin Bal, Gregory Destras, Gwendolyne Burfin, Hadrien Regue, Quentin Semanas, Martine Valette, Bruno Lina, Laurence Josset |
| EPI_ISL_3206944 | Microbiology Department, Laboratori Clínic Metropolitana Nord. Hospital Universitari Germans Trias i Pujol | Can Ruti SARS-CoV-2 Sequencing Hub (HUGTiP/IrsiCaixa/IGTP) | Marc Noguera-Julian, Pilar Armengol, Ignacio Blanco, Antoni E Bordoy, Francesc Catala-Moll, Pere-Joan Cardona, Maria Casadellà, Cristina Casañ, Gemma Clara, Bonaventura Clotet, Montserrat Giménez, Mercedes Guerrero, Anna Not, David Panisello, Roger Paredes, Mariona Parera, Alexia París, Verónica Saludes, Alba Sánchez, Laia Soler, Lauro Sumoy, and Elisa Martró on behalf of the Can Ruti SARS-CoV-2 Sequencing Hub |
| EPI_ISL_3206945 | LAM BIOPOLE ORTHEZ | CNR Virus des Infections Respiratoires - France SUD | Antonin Bal, Gregory Destras, Gwendolyne Burfin, Hadrien Regue, Quentin Semanas, Martine Valette, Bruno Lina, Laurence Josset |
| EPI_ISL_3206946, EPI_ISL_3206948 | Microbiology Department, Laboratori Clínic Metropolitana Nord. Hospital Universitari Germans Trias i Pujol | Can Ruti SARS-CoV-2 Sequencing Hub (HUGTiP/IrsiCaixa/IGTP) | Marc Noguera-Julian, Pilar Armengol, Ignacio Blanco, Antoni E Bordoy, Francesc Catala-Moll, Pere-Joan Cardona, Maria Casadellà, Cristina Casañ, Gemma Clara, Bonaventura Clotet, Montserrat Giménez, Mercedes Guerrero, Anna Not, David Panisello, Roger Paredes, Mariona Parera, Alexia París, Verónica Saludes, Alba Sánchez, Laia Soler, Lauro Sumoy, and Elisa Martró on behalf of the Can Ruti SARS-CoV-2 Sequencing Hub |
| EPI_ISL_3206949 | LAM BIOPOLE ORTHEZ | CNR Virus des Infections Respiratoires - France SUD | Antonin Bal, Gregory Destras, Gwendolyne Burfin, Hadrien Regue, Quentin Semanas, Martine Valette, Bruno Lina, Laurence Josset |
| EPI_ISL_3206950, EPI_ISL_3206952, EPI_ISL_3206954 | Microbiology Department, Laboratori Clínic Metropolitana Nord. Hospital Universitari Germans Trias i Pujol | Can Ruti SARS-CoV-2 Sequencing Hub (HUGTiP/IrsiCaixa/IGTP) | Marc Noguera-Julian, Pilar Armengol, Ignacio Blanco, Antoni E Bordoy, Francesc Catala-Moll, Pere-Joan Cardona, Maria Casadellà, Cristina Casañ, Gemma Clara, Bonaventura Clotet, Montserrat Giménez, Mercedes Guerrero, Anna Not, David Panisello, Roger Paredes, Mariona Parera, Alexia París, Verónica Saludes, Alba Sánchez, Laia Soler, Lauro Sumoy, and Elisa Martró on behalf of the Can Ruti SARS-CoV-2 Sequencing Hub |
| EPI_ISL_3206955 | LAM BIOPOLE ORTHEZ | CNR Virus des Infections Respiratoires - France SUD | Antonin Bal, Gregory Destras, Gwendolyne Burfin, Hadrien Regue, Quentin Semanas, Martine Valette, Bruno Lina, Laurence Josset |
| EPI_ISL_3206956 | Microbiology Department, Laboratori Clínic Metropolitana Nord. Hospital Universitari Germans Trias i Pujol | Can Ruti SARS-CoV-2 Sequencing Hub (HUGTiP/IrsiCaixa/IGTP) | Marc Noguera-Julian, Pilar Armengol, Ignacio Blanco, Antoni E Bordoy, Francesc Catala-Moll, Pere-Joan Cardona, Maria Casadellà, Cristina Casañ, Gemma Clara, Bonaventura Clotet, Montserrat Giménez, Mercedes Guerrero, Anna Not, David Panisello, Roger Paredes, Mariona Parera, Alexia París, Verónica Saludes, Alba Sánchez, Laia Soler, Lauro Sumoy, and Elisa Martró on behalf of the Can Ruti SARS-CoV-2 Sequencing Hub |
| EPI_ISL_3206957 | LAM BIOPOLE ORTHEZ | CNR Virus des Infections Respiratoires - France SUD | Antonin Bal, Gregory Destras, Gwendolyne Burfin, Hadrien Regue, Quentin Semanas, Martine Valette, Bruno Lina, Laurence Josset |
| EPI_ISL_3206958 | Microbiology Department, Laboratori Clínic Metropolitana Nord. Hospital Universitari Germans Trias i Pujol | Can Ruti SARS-CoV-2 Sequencing Hub (HUGTiP/IrsiCaixa/IGTP) | Marc Noguera-Julian, Pilar Armengol, Ignacio Blanco, Antoni E Bordoy, Francesc Catala-Moll, Pere-Joan Cardona, Maria Casadellà, Cristina Casañ, Gemma Clara, Bonaventura Clotet, Montserrat Giménez, Mercedes Guerrero, Anna Not, David Panisello, Roger Paredes, Mariona Parera, Alexia París, Verónica Saludes, Alba Sánchez, Laia Soler, Lauro Sumoy, and Elisa Martró on behalf of the Can Ruti SARS-CoV-2 Sequencing Hub |
| EPI_ISL_3206959 | LAM BIOPOLE ORTHEZ | CNR Virus des Infections Respiratoires - France SUD | Antonin Bal, Gregory Destras, Gwendolyne Burfin, Hadrien Regue, Quentin Semanas, Martine Valette, Bruno Lina, Laurence Josset |
| EPI_ISL_3206960 | Microbiology Department, Laboratori Clínic Metropolitana Nord. Hospital Universitari Germans Trias i Pujol | Can Ruti SARS-CoV-2 Sequencing Hub (HUGTiP/IrsiCaixa/IGTP) | Marc Noguera-Julian, Pilar Armengol, Ignacio Blanco, Antoni E Bordoy, Francesc Catala-Moll, Pere-Joan Cardona, Maria Casadellà, Cristina Casañ, Gemma Clara, Bonaventura Clotet, Montserrat Giménez, Mercedes Guerrero, Anna Not, David Panisello, Roger Paredes, Mariona Parera, Alexia París, Verónica Saludes, Alba Sánchez, Laia Soler, Lauro Sumoy, and Elisa Martró on behalf of the Can Ruti SARS-CoV-2 Sequencing Hub |
| EPI_ISL_3206961 | LAM BIOPOLE ORTHEZ | CNR Virus des Infections Respiratoires - France SUD | Antonin Bal, Gregory Destras, Gwendolyne Burfin, Hadrien Regue, Quentin Semanas, Martine Valette, Bruno Lina, Laurence Josset |
| EPI_ISL_3206962 | Microbiology Department, Laboratori Clínic Metropolitana Nord. Hospital Universitari Germans Trias i Pujol | Can Ruti SARS-CoV-2 Sequencing Hub (HUGTiP/IrsiCaixa/IGTP) | Marc Noguera-Julian, Pilar Armengol, Ignacio Blanco, Antoni E Bordoy, Francesc Catala-Moll, Pere-Joan Cardona, Maria Casadellà, Cristina Casañ, Gemma Clara, Bonaventura Clotet, Montserrat Giménez, Mercedes Guerrero, Anna Not, David Panisello, Roger Paredes, Mariona Parera, Alexia París, Verónica Saludes, Alba Sánchez, Laia Soler, Lauro Sumoy, and Elisa Martró on behalf of the Can Ruti SARS-CoV-2 Sequencing Hub |
| EPI_ISL_3206963 | LAM BIOPOLE ORTHEZ | CNR Virus des Infections Respiratoires - France SUD | Antonin Bal, Gregory Destras, Gwendolyne Burfin, Hadrien Regue, Quentin Semanas, Martine Valette, Bruno Lina, Laurence Josset |
| EPI_ISL_3206964, EPI_ISL_3206966, EPI_ISL_3206968, EPI_ISL_3206970 | Microbiology Department, Laboratori Clínic Metropolitana Nord. Hospital Universitari Germans Trias i Pujol | Can Ruti SARS-CoV-2 Sequencing Hub (HUGTiP/IrsiCaixa/IGTP) | Marc Noguera-Julian, Pilar Armengol, Ignacio Blanco, Antoni E Bordoy, Francesc Catala-Moll, Pere-Joan Cardona, Maria Casadellà, Cristina Casañ, Gemma Clara, Bonaventura Clotet, Montserrat Giménez, Mercedes Guerrero, Anna Not, David Panisello, Roger Paredes, Mariona Parera, Alexia París, Verónica Saludes, Alba Sánchez, Laia Soler, Lauro Sumoy, and Elisa Martró on behalf of the Can Ruti SARS-CoV-2 Sequencing Hub |
| EPI_ISL_3206971 | LAM BIOPOLE ORTHEZ | CNR Virus des Infections Respiratoires - France SUD | Antonin Bal, Gregory Destras, Gwendolyne Burfin, Hadrien Regue, Quentin Semanas, Martine Valette, Bruno Lina, Laurence Josset |
| EPI_ISL_3206972, EPI_ISL_3206974, EPI_ISL_3206976 | Microbiology Department, Laboratori Clínic Metropolitana Nord. Hospital Universitari Germans Trias i Pujol | Can Ruti SARS-CoV-2 Sequencing Hub (HUGTiP/IrsiCaixa/IGTP) | Marc Noguera-Julian, Pilar Armengol, Ignacio Blanco, Antoni E Bordoy, Francesc Catala-Moll, Pere-Joan Cardona, Maria Casadellà, Cristina Casañ, Gemma Clara, Bonaventura Clotet, Montserrat Giménez, Mercedes Guerrero, Anna Not, David Panisello, Roger Paredes, Mariona Parera, Alexia París, Verónica Saludes, Alba Sánchez, Laia Soler, Lauro Sumoy, and Elisa Martró on behalf of the Can Ruti SARS-CoV-2 Sequencing Hub |
| EPI_ISL_3206977 | LAM BIOPOLE ORTHEZ | CNR Virus des Infections Respiratoires - France SUD | Antonin Bal, Gregory Destras, Gwendolyne Burfin, Hadrien Regue, Quentin Semanas, Martine Valette, Bruno Lina, Laurence Josset |
| EPI_ISL_3206978 | Microbiology Department, Laboratori Clínic Metropolitana Nord. Hospital Universitari Germans Trias i Pujol | Can Ruti SARS-CoV-2 Sequencing Hub (HUGTiP/IrsiCaixa/IGTP) | Marc Noguera-Julian, Pilar Armengol, Ignacio Blanco, Antoni E Bordoy, Francesc Catala-Moll, Pere-Joan Cardona, Maria Casadellà, Cristina Casañ, Gemma Clara, Bonaventura Clotet, Montserrat Giménez, Mercedes Guerrero, Anna Not, David Panisello, Roger Paredes, Mariona Parera, Alexia París, Verónica Saludes, Alba Sánchez, Laia Soler, Lauro Sumoy, and Elisa Martró on behalf of the Can Ruti SARS-CoV-2 Sequencing Hub |
| EPI_ISL_3206979 | LAM BIOPOLE ORTHEZ | CNR Virus des Infections Respiratoires - France SUD | Antonin Bal, Gregory Destras, Gwendolyne Burfin, Hadrien Regue, Quentin Semanas, Martine Valette, Bruno Lina, Laurence Josset |
| EPI_ISL_3206980 | Microbiology Department, Laboratori Clínic Metropolitana Nord. Hospital Universitari Germans Trias i Pujol | Can Ruti SARS-CoV-2 Sequencing Hub (HUGTiP/IrsiCaixa/IGTP) | Marc Noguera-Julian, Pilar Armengol, Ignacio Blanco, Antoni E Bordoy, Francesc Catala-Moll, Pere-Joan Cardona, Maria Casadellà, Cristina Casañ, Gemma Clara, Bonaventura Clotet, Montserrat Giménez, Mercedes Guerrero, Anna Not, David Panisello, Roger Paredes, Mariona Parera, Alexia París, Verónica Saludes, Alba Sánchez, Laia Soler, Lauro Sumoy, and Elisa Martró on behalf of the Can Ruti SARS-CoV-2 Sequencing Hub |
| EPI_ISL_3206981 | LAM BIOPOLE ORTHEZ | CNR Virus des Infections Respiratoires - France SUD | Antonin Bal, Gregory Destras, Gwendolyne Burfin, Hadrien Regue, Quentin Semanas, Martine Valette, Bruno Lina, Laurence Josset |
| EPI_ISL_3206982 | Microbiology Department, Laboratori Clínic Metropolitana Nord. Hospital Universitari Germans Trias i Pujol | Can Ruti SARS-CoV-2 Sequencing Hub (HUGTiP/IrsiCaixa/IGTP) | Marc Noguera-Julian, Pilar Armengol, Ignacio Blanco, Antoni E Bordoy, Francesc Catala-Moll, Pere-Joan Cardona, Maria Casadellà, Cristina Casañ, Gemma Clara, Bonaventura Clotet, Montserrat Giménez, Mercedes Guerrero, Anna Not, David Panisello, Roger Paredes, Mariona Parera, Alexia París, Verónica Saludes, Alba Sánchez, Laia Soler, Lauro Sumoy, and Elisa Martró on behalf of the Can Ruti SARS-CoV-2 Sequencing Hub |
| EPI_ISL_3206983 | LAM BIOPOLE ORTHEZ | CNR Virus des Infections Respiratoires - France SUD | Antonin Bal, Gregory Destras, Gwendolyne Burfin, Hadrien Regue, Quentin Semanas, Martine Valette, Bruno Lina, Laurence Josset |
| EPI_ISL_3206984 | Microbiology Department, Laboratori Clínic Metropolitana Nord. Hospital Universitari Germans Trias i Pujol | Can Ruti SARS-CoV-2 Sequencing Hub (HUGTiP/IrsiCaixa/IGTP) | Marc Noguera-Julian, Pilar Armengol, Ignacio Blanco, Antoni E Bordoy, Francesc Catala-Moll, Pere-Joan Cardona, Maria Casadellà, Cristina Casañ, Gemma Clara, Bonaventura Clotet, Montserrat Giménez, Mercedes Guerrero, Anna Not, David Panisello, Roger Paredes, Mariona Parera, Alexia París, Verónica Saludes, Alba Sánchez, Laia Soler, Lauro Sumoy, and Elisa Martró on behalf of the Can Ruti SARS-CoV-2 Sequencing Hub |
| EPI_ISL_3206986 | LABORATOIRE d'ANALYSES de BIOLOGIE MEDICALES | CNR Virus des Infections Respiratoires - France SUD | Antonin Bal, Gregory Destras, Gwendolyne Burfin, Hadrien Regue, Quentin Semanas, Martine Valette, Bruno Lina, Laurence Josset |
| EPI_ISL_3206987 | Microbiology Department, Laboratori Clínic Metropolitana Nord. Hospital Universitari Germans Trias i Pujol | Can Ruti SARS-CoV-2 Sequencing Hub (HUGTiP/IrsiCaixa/IGTP) | Marc Noguera-Julian, Pilar Armengol, Ignacio Blanco, Antoni E Bordoy, Francesc Catala-Moll, Pere-Joan Cardona, Maria Casadellà, Cristina Casañ, Gemma Clara, Bonaventura Clotet, Montserrat Giménez, Mercedes Guerrero, Anna Not, David Panisello, Roger Paredes, Mariona Parera, Alexia París, Verónica Saludes, Alba Sánchez, Laia Soler, Lauro Sumoy, and Elisa Martró on behalf of the Can Ruti SARS-CoV-2 Sequencing Hub |
| EPI_ISL_3206988 | LABORATOIRE d'ANALYSES de BIOLOGIE MEDICALES | CNR Virus des Infections Respiratoires - France SUD | Antonin Bal, Gregory Destras, Gwendolyne Burfin, Hadrien Regue, Quentin Semanas, Martine Valette, Bruno Lina, Laurence Josset |
| EPI_ISL_3206989 | Microbiology Department, Laboratori Clínic Metropolitana Nord. Hospital Universitari Germans Trias i Pujol | Can Ruti SARS-CoV-2 Sequencing Hub (HUGTiP/IrsiCaixa/IGTP) | Marc Noguera-Julian, Pilar Armengol, Ignacio Blanco, Antoni E Bordoy, Francesc Catala-Moll, Pere-Joan Cardona, Maria Casadellà, Cristina Casañ, Gemma Clara, Bonaventura Clotet, Montserrat Giménez, Mercedes Guerrero, Anna Not, David Panisello, Roger Paredes, Mariona Parera, Alexia París, Verónica Saludes, Alba Sánchez, Laia Soler, Lauro Sumoy, and Elisa Martró on behalf of the Can Ruti SARS-CoV-2 Sequencing Hub |
| EPI_ISL_3206990, EPI_ISL_3206991, EPI_ISL_3206993 | LABORATOIRE d'ANALYSES de BIOLOGIE MEDICALES | CNR Virus des Infections Respiratoires - France SUD | Antonin Bal, Gregory Destras, Gwendolyne Burfin, Hadrien Regue, Quentin Semanas, Martine Valette, Bruno Lina, Laurence Josset |
| EPI_ISL_3206994, EPI_ISL_3206995 | Microbiology Department, Laboratori Clínic Metropolitana Nord. Hospital Universitari Germans Trias i Pujol | Can Ruti SARS-CoV-2 Sequencing Hub (HUGTiP/IrsiCaixa/IGTP) | Marc Noguera-Julian, Pilar Armengol, Ignacio Blanco, Antoni E Bordoy, Francesc Catala-Moll, Pere-Joan Cardona, Maria Casadellà, Cristina Casañ, Gemma Clara, Bonaventura Clotet, Montserrat Giménez, Mercedes Guerrero, Anna Not, David Panisello, Roger Paredes, Mariona Parera, Alexia París, Verónica Saludes, Alba Sánchez, Laia Soler, Lauro Sumoy, and Elisa Martró on behalf of the Can Ruti SARS-CoV-2 Sequencing Hub |
| EPI_ISL_3206996, EPI_ISL_3206997 | LABORATOIRE d'ANALYSES de BIOLOGIE MEDICALES | CNR Virus des Infections Respiratoires - France SUD | Antonin Bal, Gregory Destras, Gwendolyne Burfin, Hadrien Regue, Quentin Semanas, Martine Valette, Bruno Lina, Laurence Josset |
| EPI_ISL_3206998 | Microbiology Department, Laboratori Clínic Metropolitana Nord. Hospital Universitari Germans Trias i Pujol | Can Ruti SARS-CoV-2 Sequencing Hub (HUGTiP/IrsiCaixa/IGTP) | Marc Noguera-Julian, Pilar Armengol, Ignacio Blanco, Antoni E Bordoy, Francesc Catala-Moll, Pere-Joan Cardona, Maria Casadellà, Cristina Casañ, Gemma Clara, Bonaventura Clotet, Montserrat Giménez, Mercedes Guerrero, Anna Not, David Panisello, Roger Paredes, Mariona Parera, Alexia París, Verónica Saludes, Alba Sánchez, Laia Soler, Lauro Sumoy, and Elisa Martró on behalf of the Can Ruti SARS-CoV-2 Sequencing Hub |
| EPI_ISL_3206999 | LABORATOIRE d'ANALYSES de BIOLOGIE MEDICALES | CNR Virus des Infections Respiratoires - France SUD | Antonin Bal, Gregory Destras, Gwendolyne Burfin, Hadrien Regue, Quentin Semanas, Martine Valette, Bruno Lina, Laurence Josset |
| EPI_ISL_3207000 | Microbiology Department, Laboratori Clínic Metropolitana Nord. Hospital Universitari Germans Trias i Pujol | Can Ruti SARS-CoV-2 Sequencing Hub (HUGTiP/IrsiCaixa/IGTP) | Marc Noguera-Julian, Pilar Armengol, Ignacio Blanco, Antoni E Bordoy, Francesc Catala-Moll, Pere-Joan Cardona, Maria Casadellà, Cristina Casañ, Gemma Clara, Bonaventura Clotet, Montserrat Giménez, Mercedes Guerrero, Anna Not, David Panisello, Roger Paredes, Mariona Parera, Alexia París, Verónica Saludes, Alba Sánchez, Laia Soler, Lauro Sumoy, and Elisa Martró on behalf of the Can Ruti SARS-CoV-2 Sequencing Hub |
| EPI_ISL_3207001 | LABORATOIRE d'ANALYSES de BIOLOGIE MEDICALES | CNR Virus des Infections Respiratoires - France SUD | Antonin Bal, Gregory Destras, Gwendolyne Burfin, Hadrien Regue, Quentin Semanas, Martine Valette, Bruno Lina, Laurence Josset |
| EPI_ISL_3207002 | Microbiology Department, Laboratori Clínic Metropolitana Nord. Hospital Universitari Germans Trias i Pujol | Can Ruti SARS-CoV-2 Sequencing Hub (HUGTiP/IrsiCaixa/IGTP) | Marc Noguera-Julian, Pilar Armengol, Ignacio Blanco, Antoni E Bordoy, Francesc Catala-Moll, Pere-Joan Cardona, Maria Casadellà, Cristina Casañ, Gemma Clara, Bonaventura Clotet, Montserrat Giménez, Mercedes Guerrero, Anna Not, David Panisello, Roger Paredes, Mariona Parera, Alexia París, Verónica Saludes, Alba Sánchez, Laia Soler, Lauro Sumoy, and Elisa Martró on behalf of the Can Ruti SARS-CoV-2 Sequencing Hub |
| EPI_ISL_3207003 | LABORATOIRE d'ANALYSES de BIOLOGIE MEDICALES | CNR Virus des Infections Respiratoires - France SUD | Antonin Bal, Gregory Destras, Gwendolyne Burfin, Hadrien Regue, Quentin Semanas, Martine Valette, Bruno Lina, Laurence Josset |

| | | | |
|---|---|---|---|
| EPI_ISL_3207004 | Microbiology Department, Laboratori Clínic Metropolitana Nord. Hospital Universitari Germans Trias i Pujol | Can Ruti SARS-CoV-2 Sequencing Hub (HUGTiP/IrsiCaixa/IGTP) | Marc Noguera-Julian, Pilar Armengol, Ignacio Blanco, Antoni E Bordoy, Francesc Catala-Moll, Pere-Joan Cardona, Maria Casadellà, Cristina Casañ, Gemma Clara, Bonaventura Clotet, Montserrat Giménez, Mercedes Guerrero, Anna Not, David Panisello, Roger Paredes, Mariona Parera, Alexia París, Verónica Saludes, Alba Sánchez, Laia Soler, Lauro Sumoy, and Elisa Martró on behalf of the Can Ruti SARS-CoV-2 Sequencing Hub |
| EPI_ISL_3207005 | LABORATOIRE d'ANALYSES de BIOLOGIE MEDICALES | CNR Virus des Infections Respiratoires - France SUD | Antonin Bal, Gregory Destras, Gwendolyne Burfin, Hadrien Regue, Quentin Semanas, Martine Valette, Bruno Lina, Laurence Josset |
| EPI_ISL_3207006 | Microbiology Department, Laboratori Clínic Metropolitana Nord. Hospital Universitari Germans Trias i Pujol | Can Ruti SARS-CoV-2 Sequencing Hub (HUGTiP/IrsiCaixa/IGTP) | Marc Noguera-Julian, Pilar Armengol, Ignacio Blanco, Antoni E Bordoy, Francesc Catala-Moll, Pere-Joan Cardona, Maria Casadellà, Cristina Casañ, Gemma Clara, Bonaventura Clotet, Montserrat Giménez, Mercedes Guerrero, Anna Not, David Panisello, Roger Paredes, Mariona Parera, Alexia París, Verónica Saludes, Alba Sánchez, Laia Soler, Lauro Sumoy, and Elisa Martró on behalf of the Can Ruti SARS-CoV-2 Sequencing Hub |
| EPI_ISL_3207007 | LABORATOIRE d'ANALYSES de BIOLOGIE MEDICALES | CNR Virus des Infections Respiratoires - France SUD | Antonin Bal, Gregory Destras, Gwendolyne Burfin, Hadrien Regue, Quentin Semanas, Martine Valette, Bruno Lina, Laurence Josset |
| EPI_ISL_3207008 | Microbiology Department, Laboratori Clínic Metropolitana Nord. Hospital Universitari Germans Trias i Pujol | Can Ruti SARS-CoV-2 Sequencing Hub (HUGTiP/IrsiCaixa/IGTP) | Marc Noguera-Julian, Pilar Armengol, Ignacio Blanco, Antoni E Bordoy, Francesc Catala-Moll, Pere-Joan Cardona, Maria Casadellà, Cristina Casañ, Gemma Clara, Bonaventura Clotet, Montserrat Giménez, Mercedes Guerrero, Anna Not, David Panisello, Roger Paredes, Mariona Parera, Alexia París, Verónica Saludes, Alba Sánchez, Laia Soler, Lauro Sumoy, and Elisa Martró on behalf of the Can Ruti SARS-CoV-2 Sequencing Hub |
| EPI_ISL_3207009 | LABORATOIRE d'ANALYSES de BIOLOGIE MEDICALES | CNR Virus des Infections Respiratoires - France SUD | Antonin Bal, Gregory Destras, Gwendolyne Burfin, Hadrien Regue, Quentin Semanas, Martine Valette, Bruno Lina, Laurence Josset |
| EPI_ISL_3207010 | Microbiology Department, Laboratori Clínic Metropolitana Nord. Hospital Universitari Germans Trias i Pujol | Can Ruti SARS-CoV-2 Sequencing Hub (HUGTiP/IrsiCaixa/IGTP) | Marc Noguera-Julian, Pilar Armengol, Ignacio Blanco, Antoni E Bordoy, Francesc Catala-Moll, Pere-Joan Cardona, Maria Casadellà, Cristina Casañ, Gemma Clara, Bonaventura Clotet, Montserrat Giménez, Mercedes Guerrero, Anna Not, David Panisello, Roger Paredes, Mariona Parera, Alexia París, Verónica Saludes, Alba Sánchez, Laia Soler, Lauro Sumoy, and Elisa Martró on behalf of the Can Ruti SARS-CoV-2 Sequencing Hub |
| EPI_ISL_3207011 | LABORATOIRE d'ANALYSES de BIOLOGIE MEDICALES | CNR Virus des Infections Respiratoires - France SUD | Antonin Bal, Gregory Destras, Gwendolyne Burfin, Hadrien Regue, Quentin Semanas, Martine Valette, Bruno Lina, Laurence Josset |
| EPI_ISL_3207012, EPI_ISL_3207013 | Microbiology Department, Laboratori Clínic Metropolitana Nord. Hospital Universitari Germans Trias i Pujol | Can Ruti SARS-CoV-2 Sequencing Hub (HUGTiP/IrsiCaixa/IGTP) | Marc Noguera-Julian, Pilar Armengol, Ignacio Blanco, Antoni E Bordoy, Francesc Catala-Moll, Pere-Joan Cardona, Maria Casadellà, Cristina Casañ, Gemma Clara, Bonaventura Clotet, Montserrat Giménez, Mercedes Guerrero, Anna Not, David Panisello, Roger Paredes, Mariona Parera, Alexia París, Verónica Saludes, Alba Sánchez, Laia Soler, Lauro Sumoy, and Elisa Martró on behalf of the Can Ruti SARS-CoV-2 Sequencing Hub |
| EPI_ISL_3207014 | LABORATOIRE d'ANALYSES de BIOLOGIE MEDICALES | CNR Virus des Infections Respiratoires - France SUD | Antonin Bal, Gregory Destras, Gwendolyne Burfin, Hadrien Regue, Quentin Semanas, Martine Valette, Bruno Lina, Laurence Josset |
| EPI_ISL_3207015 | Microbiology Department, Laboratori Clínic Metropolitana Nord. Hospital Universitari Germans Trias i Pujol | Can Ruti SARS-CoV-2 Sequencing Hub (HUGTiP/IrsiCaixa/IGTP) | Marc Noguera-Julian, Pilar Armengol, Ignacio Blanco, Antoni E Bordoy, Francesc Catala-Moll, Pere-Joan Cardona, Maria Casadellà, Cristina Casañ, Gemma Clara, Bonaventura Clotet, Montserrat Giménez, Mercedes Guerrero, Anna Not, David Panisello, Roger Paredes, Mariona Parera, Alexia París, Verónica Saludes, Alba Sánchez, Laia Soler, Lauro Sumoy, and Elisa Martró on behalf of the Can Ruti SARS-CoV-2 Sequencing Hub |
| EPI_ISL_3207016 | LABORATOIRE d'ANALYSES de BIOLOGIE MEDICALES | CNR Virus des Infections Respiratoires - France SUD | Antonin Bal, Gregory Destras, Gwendolyne Burfin, Hadrien Regue, Quentin Semanas, Martine Valette, Bruno Lina, Laurence Josset |
| EPI_ISL_3207017 | Microbiology Department, Laboratori Clínic Metropolitana Nord. Hospital Universitari Germans Trias i Pujol | Can Ruti SARS-CoV-2 Sequencing Hub (HUGTiP/IrsiCaixa/IGTP) | Marc Noguera-Julian, Pilar Armengol, Ignacio Blanco, Antoni E Bordoy, Francesc Catala-Moll, Pere-Joan Cardona, Maria Casadellà, Cristina Casañ, Gemma Clara, Bonaventura Clotet, Montserrat Giménez, Mercedes Guerrero, Anna Not, David Panisello, Roger Paredes, Mariona Parera, Alexia París, Verónica Saludes, Alba Sánchez, Laia Soler, Lauro Sumoy, and Elisa Martró on behalf of the Can Ruti SARS-CoV-2 Sequencing Hub |
| EPI_ISL_3207018 | LABORATOIRE d'ANALYSES de BIOLOGIE MEDICALES | CNR Virus des Infections Respiratoires - France SUD | Antonin Bal, Gregory Destras, Gwendolyne Burfin, Hadrien Regue, Quentin Semanas, Martine Valette, Bruno Lina, Laurence Josset |
| EPI_ISL_3207019 | Microbiology Department, Laboratori Clínic Metropolitana Nord. Hospital Universitari Germans Trias i Pujol | Can Ruti SARS-CoV-2 Sequencing Hub (HUGTiP/IrsiCaixa/IGTP) | Marc Noguera-Julian, Pilar Armengol, Ignacio Blanco, Antoni E Bordoy, Francesc Catala-Moll, Pere-Joan Cardona, Maria Casadellà, Cristina Casañ, Gemma Clara, Bonaventura Clotet, Montserrat Giménez, Mercedes Guerrero, Anna Not, David Panisello, Roger Paredes, Mariona Parera, Alexia París, Verónica Saludes, Alba Sánchez, Laia Soler, Lauro Sumoy, and Elisa Martró on behalf of the Can Ruti SARS-CoV-2 Sequencing Hub |
| EPI_ISL_3207020 | LABORATOIRE d'ANALYSES de BIOLOGIE MEDICALES | CNR Virus des Infections Respiratoires - France SUD | Antonin Bal, Gregory Destras, Gwendolyne Burfin, Hadrien Regue, Quentin Semanas, Martine Valette, Bruno Lina, Laurence Josset |
| EPI_ISL_3207021 | Microbiology Department, Laboratori Clínic Metropolitana Nord. Hospital Universitari Germans Trias i Pujol | Can Ruti SARS-CoV-2 Sequencing Hub (HUGTiP/IrsiCaixa/IGTP) | Marc Noguera-Julian, Pilar Armengol, Ignacio Blanco, Antoni E Bordoy, Francesc Catala-Moll, Pere-Joan Cardona, Maria Casadellà, Cristina Casañ, Gemma Clara, Bonaventura Clotet, Montserrat Giménez, Mercedes Guerrero, Anna Not, David Panisello, Roger Paredes, Mariona Parera, Alexia París, Verónica Saludes, Alba Sánchez, Laia Soler, Lauro Sumoy, and Elisa Martró on behalf of the Can Ruti SARS-CoV-2 Sequencing Hub |
| EPI_ISL_3207022, EPI_ISL_3207024 | LABORATOIRE d'ANALYSES de BIOLOGIE MEDICALES | CNR Virus des Infections Respiratoires - France SUD | Antonin Bal, Gregory Destras, Gwendolyne Burfin, Hadrien Regue, Quentin Semanas, Martine Valette, Bruno Lina, Laurence Josset |
| EPI_ISL_3207025 | Microbiology Department, Laboratori Clínic Metropolitana Nord. Hospital Universitari Germans Trias i Pujol | Can Ruti SARS-CoV-2 Sequencing Hub (HUGTiP/IrsiCaixa/IGTP) | Marc Noguera-Julian, Pilar Armengol, Ignacio Blanco, Antoni E Bordoy, Francesc Catala-Moll, Pere-Joan Cardona, Maria Casadellà, Cristina Casañ, Gemma Clara, Bonaventura Clotet, Montserrat Giménez, Mercedes Guerrero, Anna Not, David Panisello, Roger Paredes, Mariona Parera, Alexia París, Verónica Saludes, Alba Sánchez, Laia Soler, Lauro Sumoy, and Elisa Martró on behalf of the Can Ruti SARS-CoV-2 Sequencing Hub |
| EPI_ISL_3207026 | LABORATOIRE d'ANALYSES de BIOLOGIE MEDICALES | CNR Virus des Infections Respiratoires - France SUD | Antonin Bal, Gregory Destras, Gwendolyne Burfin, Hadrien Regue, Quentin Semanas, Martine Valette, Bruno Lina, Laurence Josset |
| EPI_ISL_3207027 | Microbiology Department, Laboratori Clínic Metropolitana Nord. Hospital Universitari Germans Trias i Pujol | Can Ruti SARS-CoV-2 Sequencing Hub (HUGTiP/IrsiCaixa/IGTP) | Marc Noguera-Julian, Pilar Armengol, Ignacio Blanco, Antoni E Bordoy, Francesc Catala-Moll, Pere-Joan Cardona, Maria Casadellà, Cristina Casañ, Gemma Clara, Bonaventura Clotet, Montserrat Giménez, Mercedes Guerrero, Anna Not, David Panisello, Roger Paredes, Mariona Parera, Alexia París, Verónica Saludes, Alba Sánchez, Laia Soler, Lauro Sumoy, and Elisa Martró on behalf of the Can Ruti SARS-CoV-2 Sequencing Hub |
| EPI_ISL_3207028 | LABORATOIRE d'ANALYSES de BIOLOGIE MEDICALES | CNR Virus des Infections Respiratoires - France SUD | Antonin Bal, Gregory Destras, Gwendolyne Burfin, Hadrien Regue, Quentin Semanas, Martine Valette, Bruno Lina, Laurence Josset |
| EPI_ISL_3207029 | Microbiology Department, Laboratori Clínic Metropolitana Nord. Hospital Universitari Germans Trias i Pujol | Can Ruti SARS-CoV-2 Sequencing Hub (HUGTiP/IrsiCaixa/IGTP) | Marc Noguera-Julian, Pilar Armengol, Ignacio Blanco, Antoni E Bordoy, Francesc Catala-Moll, Pere-Joan Cardona, Maria Casadellà, Cristina Casañ, Gemma Clara, Bonaventura Clotet, Montserrat Giménez, Mercedes Guerrero, Anna Not, David Panisello, Roger Paredes, Mariona Parera, Alexia París, Verónica Saludes, Alba Sánchez, Laia Soler, Lauro Sumoy, and Elisa Martró on behalf of the Can Ruti SARS-CoV-2 Sequencing Hub |
| EPI_ISL_3207030 | LABORATOIRE d'ANALYSES de BIOLOGIE MEDICALES | CNR Virus des Infections Respiratoires - France SUD | Antonin Bal, Gregory Destras, Gwendolyne Burfin, Hadrien Regue, Quentin Semanas, Martine Valette, Bruno Lina, Laurence Josset |
| EPI_ISL_3207031 | Microbiology Department, Laboratori Clínic Metropolitana Nord. Hospital Universitari Germans Trias i Pujol | Can Ruti SARS-CoV-2 Sequencing Hub (HUGTiP/IrsiCaixa/IGTP) | Marc Noguera-Julian, Pilar Armengol, Ignacio Blanco, Antoni E Bordoy, Francesc Catala-Moll, Pere-Joan Cardona, Maria Casadellà, Cristina Casañ, Gemma Clara, Bonaventura Clotet, Montserrat Giménez, Mercedes Guerrero, Anna Not, David Panisello, Roger Paredes, Mariona Parera, Alexia París, Verónica Saludes, Alba Sánchez, Laia Soler, Lauro Sumoy, and Elisa Martró on behalf of the Can Ruti SARS-CoV-2 Sequencing Hub |
| EPI_ISL_3207032 | LABORATOIRE d'ANALYSES de BIOLOGIE MEDICALES | CNR Virus des Infections Respiratoires - France SUD | Antonin Bal, Gregory Destras, Gwendolyne Burfin, Hadrien Regue, Quentin Semanas, Martine Valette, Bruno Lina, Laurence Josset |
| EPI_ISL_3207033 | Microbiology Department, Laboratori Clínic Metropolitana Nord. Hospital Universitari Germans Trias i Pujol | Can Ruti SARS-CoV-2 Sequencing Hub (HUGTiP/IrsiCaixa/IGTP) | Marc Noguera-Julian, Pilar Armengol, Ignacio Blanco, Antoni E Bordoy, Francesc Catala-Moll, Pere-Joan Cardona, Maria Casadellà, Cristina Casañ, Gemma Clara, Bonaventura Clotet, Montserrat Giménez, Mercedes Guerrero, Anna Not, David Panisello, Roger Paredes, Mariona Parera, Alexia París, Verónica Saludes, Alba Sánchez, Laia Soler, Lauro Sumoy, and Elisa Martró on behalf of the Can Ruti SARS-CoV-2 Sequencing Hub |
| EPI_ISL_3207034 | LABORATOIRE d'ANALYSES de BIOLOGIE MEDICALES | CNR Virus des Infections Respiratoires - France SUD | Antonin Bal, Gregory Destras, Gwendolyne Burfin, Hadrien Regue, Quentin Semanas, Martine Valette, Bruno Lina, Laurence Josset |
| EPI_ISL_3207035 | Microbiology Department, Laboratori Clínic Metropolitana Nord. Hospital Universitari Germans Trias i Pujol | Can Ruti SARS-CoV-2 Sequencing Hub (HUGTiP/IrsiCaixa/IGTP) | Marc Noguera-Julian, Pilar Armengol, Ignacio Blanco, Antoni E Bordoy, Francesc Catala-Moll, Pere-Joan Cardona, Maria Casadellà, Cristina Casañ, Gemma Clara, Bonaventura Clotet, Montserrat Giménez, Mercedes Guerrero, Anna Not, David Panisello, Roger Paredes, Mariona Parera, Alexia París, Verónica Saludes, Alba Sánchez, Laia Soler, Lauro Sumoy, and Elisa Martró on behalf of the Can Ruti SARS-CoV-2 Sequencing Hub |
| EPI_ISL_3207036 | LABORATOIRE d'ANALYSES de BIOLOGIE MEDICALES | CNR Virus des Infections Respiratoires - France SUD | Antonin Bal, Gregory Destras, Gwendolyne Burfin, Hadrien Regue, Quentin Semanas, Martine Valette, Bruno Lina, Laurence Josset |
| EPI_ISL_3207037, EPI_ISL_3207038 | Microbiology Department, Laboratori Clínic Metropolitana Nord. Hospital Universitari Germans Trias i Pujol | Can Ruti SARS-CoV-2 Sequencing Hub (HUGTiP/IrsiCaixa/IGTP) | Marc Noguera-Julian, Pilar Armengol, Ignacio Blanco, Antoni E Bordoy, Francesc Catala-Moll, Pere-Joan Cardona, Maria Casadellà, Cristina Casañ, Gemma Clara, Bonaventura Clotet, Montserrat Giménez, Mercedes Guerrero, Anna Not, David Panisello, Roger Paredes, Mariona Parera, Alexia París, Verónica Saludes, Alba Sánchez, Laia Soler, Lauro Sumoy, and Elisa Martró on behalf of the Can Ruti SARS-CoV-2 Sequencing Hub |
| EPI_ISL_3207039 | LABORATOIRE d'ANALYSES de BIOLOGIE MEDICALES | CNR Virus des Infections Respiratoires - France SUD | Antonin Bal, Gregory Destras, Gwendolyne Burfin, Hadrien Regue, Quentin Semanas, Martine Valette, Bruno Lina, Laurence Josset |
| EPI_ISL_3207040 | Microbiology Department, Laboratori Clínic Metropolitana Nord. Hospital Universitari Germans Trias i Pujol | Can Ruti SARS-CoV-2 Sequencing Hub (HUGTiP/IrsiCaixa/IGTP) | Marc Noguera-Julian, Pilar Armengol, Ignacio Blanco, Antoni E Bordoy, Francesc Catala-Moll, Pere-Joan Cardona, Maria Casadellà, Cristina Casañ, Gemma Clara, Bonaventura Clotet, Montserrat Giménez, Mercedes Guerrero, Anna Not, David Panisello, Roger Paredes, Mariona Parera, Alexia París, Verónica Saludes, Alba Sánchez, Laia Soler, Lauro Sumoy, and Elisa Martró on behalf of the Can Ruti SARS-CoV-2 Sequencing Hub |
| EPI_ISL_3207041 | LABORATOIRE d'ANALYSES de BIOLOGIE MEDICALES | CNR Virus des Infections Respiratoires - France SUD | Antonin Bal, Gregory Destras, Gwendolyne Burfin, Hadrien Regue, Quentin Semanas, Martine Valette, Bruno Lina, Laurence Josset |
| EPI_ISL_3207042 | Microbiology Department, Laboratori Clínic Metropolitana Nord. Hospital Universitari Germans Trias i Pujol | Can Ruti SARS-CoV-2 Sequencing Hub (HUGTiP/IrsiCaixa/IGTP) | Marc Noguera-Julian, Pilar Armengol, Ignacio Blanco, Antoni E Bordoy, Francesc Catala-Moll, Pere-Joan Cardona, Maria Casadellà, Cristina Casañ, Gemma Clara, Bonaventura Clotet, Montserrat Giménez, Mercedes Guerrero, Anna Not, David Panisello, Roger Paredes, Mariona Parera, Alexia París, Verónica Saludes, Alba Sánchez, Laia Soler, Lauro Sumoy, and Elisa Martró on behalf of the Can Ruti SARS-CoV-2 Sequencing Hub |
| EPI_ISL_3207043 | LABORATOIRE d'ANALYSES de BIOLOGIE MEDICALES | CNR Virus des Infections Respiratoires - France SUD | Antonin Bal, Gregory Destras, Gwendolyne Burfin, Hadrien Regue, Quentin Semanas, Martine Valette, Bruno Lina, Laurence Josset |
| EPI_ISL_3207044, EPI_ISL_3207045 | Microbiology Department, Laboratori Clínic Metropolitana Nord. Hospital Universitari Germans Trias i Pujol | Can Ruti SARS-CoV-2 Sequencing Hub (HUGTiP/IrsiCaixa/IGTP) | Marc Noguera-Julian, Pilar Armengol, Ignacio Blanco, Antoni E Bordoy, Francesc Catala-Moll, Pere-Joan Cardona, Maria Casadellà, Cristina Casañ, Gemma Clara, Bonaventura Clotet, Montserrat Giménez, Mercedes Guerrero, Anna Not, David Panisello, Roger Paredes, Mariona Parera, Alexia París, Verónica Saludes, Alba Sánchez, Laia Soler, Lauro Sumoy, and Elisa Martró on behalf of the Can Ruti SARS-CoV-2 Sequencing Hub |
| EPI_ISL_3207046 | LABORATOIRE d'ANALYSES de BIOLOGIE MEDICALES | CNR Virus des Infections Respiratoires - France SUD | Antonin Bal, Gregory Destras, Gwendolyne Burfin, Hadrien Regue, Quentin Semanas, Martine Valette, Bruno Lina, Laurence Josset |
| EPI_ISL_3207047 | Microbiology Department, Laboratori Clínic Metropolitana Nord. Hospital Universitari Germans Trias i Pujol | Can Ruti SARS-CoV-2 Sequencing Hub (HUGTiP/IrsiCaixa/IGTP) | Marc Noguera-Julian, Pilar Armengol, Ignacio Blanco, Antoni E Bordoy, Francesc Catala-Moll, Pere-Joan Cardona, Maria Casadellà, Cristina Casañ, Gemma Clara, Bonaventura Clotet, Montserrat Giménez, Mercedes Guerrero, Anna Not, David Panisello, Roger Paredes, Mariona Parera, Alexia París, Verónica Saludes, Alba Sánchez, Laia Soler, Lauro Sumoy, and Elisa Martró on behalf of the Can Ruti SARS-CoV-2 Sequencing Hub |
| EPI_ISL_3207048 | LABORATOIRE d'ANALYSES de BIOLOGIE MEDICALES | CNR Virus des Infections Respiratoires - France SUD | Antonin Bal, Gregory Destras, Gwendolyne Burfin, Hadrien Regue, Quentin Semanas, Martine Valette, Bruno Lina, Laurence Josset |
| EPI_ISL_3207049 | Microbiology Department, Laboratori Clínic Metropolitana Nord. Hospital Universitari Germans Trias i Pujol | Can Ruti SARS-CoV-2 Sequencing Hub (HUGTiP/IrsiCaixa/IGTP) | Marc Noguera-Julian, Pilar Armengol, Ignacio Blanco, Antoni E Bordoy, Francesc Catala-Moll, Pere-Joan Cardona, Maria Casadellà, Cristina Casañ, Gemma Clara, Bonaventura Clotet, Montserrat Giménez, Mercedes Guerrero, Anna Not, David Panisello, Roger Paredes, Mariona Parera, Alexia París, Verónica Saludes, Alba Sánchez, Laia Soler, Lauro Sumoy, and Elisa Martró on behalf of the Can Ruti SARS-CoV-2 Sequencing Hub |
| EPI_ISL_3207050 | LABORATOIRE d'ANALYSES de BIOLOGIE MEDICALES | CNR Virus des Infections Respiratoires - France SUD | Antonin Bal, Gregory Destras, Gwendolyne Burfin, Hadrien Regue, Quentin Semanas, Martine Valette, Bruno Lina, Laurence Josset |
| EPI_ISL_3207051 | Microbiology Department, Laboratori Clínic Metropolitana Nord. Hospital Universitari Germans Trias i Pujol | Can Ruti SARS-CoV-2 Sequencing Hub (HUGTiP/IrsiCaixa/IGTP) | Marc Noguera-Julian, Pilar Armengol, Ignacio Blanco, Antoni E Bordoy, Francesc Catala-Moll, Pere-Joan Cardona, Maria Casadellà, Cristina Casañ, Gemma Clara, Bonaventura Clotet, Montserrat Giménez, Mercedes Guerrero, Anna Not, David Panisello, Roger Paredes, Mariona Parera, Alexia París, Verónica Saludes, Alba Sánchez, Laia Soler, Lauro Sumoy, and Elisa Martró on behalf of the Can Ruti SARS-CoV-2 Sequencing Hub |
| EPI_ISL_3207052 | LABORATOIRE d'ANALYSES de BIOLOGIE MEDICALES | CNR Virus des Infections Respiratoires - France SUD | Antonin Bal, Gregory Destras, Gwendolyne Burfin, Hadrien Regue, Quentin Semanas, Martine Valette, Bruno Lina, Laurence Josset |
| EPI_ISL_3207053 | Microbiologia CATLAB | Can Ruti SARS-CoV-2 Sequencing Hub (HUGTiP/IrsiCaixa/IGTP) | Marc Noguera-Julian, Pilar Armengol, Ignacio Blanco, Antoni E Bordoy, Francesc Catala-Moll, Pere-Joan Cardona, Maria Casadellà, Cristina Casañ, Gemma Clara, Bonaventura Clotet, Montserrat Giménez, Mercedes Guerrero, Anna Not, David Panisello, Roger Paredes, Mariona Parera, Alexia París, Verónica Saludes, Alba Sánchez, Laia Soler, Lauro Sumoy, and Elisa Martró on behalf of the Can Ruti SARS-CoV-2 Sequencing Hub |
| EPI_ISL_3207054 | LABORATOIRE d'ANALYSES de BIOLOGIE MEDICALES | CNR Virus des Infections Respiratoires - France SUD | Antonin Bal, Gregory Destras, Gwendolyne Burfin, Hadrien Regue, Quentin Semanas, Martine Valette, Bruno Lina, Laurence Josset |
| EPI_ISL_3207055 | Microbiology Department, Laboratori Clínic Metropolitana Nord. Hospital Universitari Germans Trias i Pujol | Can Ruti SARS-CoV-2 Sequencing Hub (HUGTiP/IrsiCaixa/IGTP) | Marc Noguera-Julian, Pilar Armengol, Ignacio Blanco, Antoni E Bordoy, Francesc Catala-Moll, Pere-Joan Cardona, Maria Casadellà, Cristina Casañ, Gemma Clara, Bonaventura Clotet, Montserrat Giménez, Mercedes Guerrero, Anna Not, David Panisello, Roger Paredes, Mariona Parera, Alexia París, Verónica Saludes, Alba Sánchez, Laia Soler, Lauro Sumoy, and Elisa Martró on behalf of the Can Ruti SARS-CoV-2 Sequencing Hub |
| EPI_ISL_3207056 | LABORATOIRE d'ANALYSES de BIOLOGIE MEDICALES | CNR Virus des Infections Respiratoires - France SUD | Antonin Bal, Gregory Destras, Gwendolyne Burfin, Hadrien Regue, Quentin Semanas, Martine Valette, Bruno Lina, Laurence Josset |
| EPI_ISL_3207057 | Microbiology Department, Laboratori Clínic Metropolitana Nord. Hospital Universitari Germans Trias i Pujol | Can Ruti SARS-CoV-2 Sequencing Hub (HUGTiP/IrsiCaixa/IGTP) | Marc Noguera-Julian, Pilar Armengol, Ignacio Blanco, Antoni E Bordoy, Francesc Catala-Moll, Pere-Joan Cardona, Maria Casadellà, Cristina Casañ, Gemma Clara, Bonaventura Clotet, Montserrat Giménez, Mercedes Guerrero, Anna Not, David Panisello, Roger Paredes, Mariona Parera, Alexia París, Verónica Saludes, Alba Sánchez, Laia Soler, Lauro Sumoy, and Elisa Martró on behalf of the Can Ruti SARS-CoV-2 Sequencing Hub |
| EPI_ISL_3207058 | LABORATOIRE d'ANALYSES de BIOLOGIE MEDICALES | CNR Virus des Infections Respiratoires - France SUD | Antonin Bal, Gregory Destras, Gwendolyne Burfin, Hadrien Regue, Quentin Semanas, Martine Valette, Bruno Lina, Laurence Josset |
| EPI_ISL_3207059, EPI_ISL_3207060 | Microbiology Department, Laboratori Clínic Metropolitana Nord. Hospital Universitari Germans Trias i Pujol | Can Ruti SARS-CoV-2 Sequencing Hub (HUGTiP/IrsiCaixa/IGTP) | Marc Noguera-Julian, Pilar Armengol, Ignacio Blanco, Antoni E Bordoy, Francesc Catala-Moll, Pere-Joan Cardona, Maria Casadellà, Cristina Casañ, Gemma Clara, Bonaventura Clotet, Montserrat Giménez, Mercedes Guerrero, Anna Not, David Panisello, Roger Paredes, Mariona Parera, Alexia París, Verónica Saludes, Alba Sánchez, Laia Soler, Lauro Sumoy, and Elisa Martró on behalf of the Can Ruti SARS-CoV-2 Sequencing Hub |
| EPI_ISL_3207061 | LABORATOIRE d'ANALYSES de BIOLOGIE MEDICALES | CNR Virus des Infections Respiratoires - France SUD | Antonin Bal, Gregory Destras, Gwendolyne Burfin, Hadrien Regue, Quentin Semanas, Martine Valette, Bruno Lina, Laurence Josset |
| EPI_ISL_3207062 | Microbiologia CATLAB | Can Ruti SARS-CoV-2 Sequencing Hub (HUGTiP/IrsiCaixa/IGTP) | Marc Noguera-Julian, Pilar Armengol, Ignacio Blanco, Antoni E Bordoy, Francesc Catala-Moll, Pere-Joan Cardona, Maria Casadellà, Cristina Casañ, Gemma Clara, Bonaventura Clotet, Montserrat Giménez, Mercedes Guerrero, Anna Not, David Panisello, Roger Paredes, Mariona Parera, Alexia París, Verónica Saludes, Alba Sánchez, Laia Soler, Lauro Sumoy, and Elisa Martró on behalf of the Can Ruti SARS-CoV-2 Sequencing Hub |
| EPI_ISL_3207063 | LABORATOIRE d'ANALYSES de BIOLOGIE MEDICALES | CNR Virus des Infections Respiratoires - France SUD | Antonin Bal, Gregory Destras, Gwendolyne Burfin, Hadrien Regue, Quentin Semanas, Martine Valette, Bruno Lina, Laurence Josset |
| EPI_ISL_3207064, EPI_ISL_3207065 | Microbiology Department, Laboratori Clínic Metropolitana Nord. Hospital Universitari Germans Trias i Pujol | Can Ruti SARS-CoV-2 Sequencing Hub (HUGTiP/IrsiCaixa/IGTP) | Marc Noguera-Julian, Pilar Armengol, Ignacio Blanco, Antoni E Bordoy, Francesc Catala-Moll, Pere-Joan Cardona, Maria Casadellà, Cristina Casañ, Gemma Clara, Bonaventura Clotet, Montserrat Giménez, Mercedes Guerrero, Anna Not, David Panisello, Roger Paredes, Mariona Parera, Alexia París, Verónica Saludes, Alba Sánchez, Laia Soler, Lauro Sumoy, and Elisa Martró on behalf of the Can Ruti SARS-CoV-2 Sequencing Hub |
| EPI_ISL_3207066 | LABORATOIRE d'ANALYSES de BIOLOGIE MEDICALES | CNR Virus des Infections Respiratoires - France SUD | Antonin Bal, Gregory Destras, Gwendolyne Burfin, Hadrien Regue, Quentin Semanas, Martine Valette, Bruno Lina, Laurence Josset |

| | | | |
|---|---|---|---|
| EPI_ISL_3207067 | Microbiology Department, Laboratori Clínic Metropolitana Nord. Hospital Universitari Germans Trias i Pujol | Can Ruti SARS-CoV-2 Sequencing Hub (HUGTiP/IrsiCaixa/IGTP) | Marc Noguera-Julian, Pilar Armengol, Ignacio Blanco, Antoni E Bordoy, Francesc Catala-Moll, Pere-Joan Cardona, Maria Casadellà, Cristina Casañ, Gemma Clara, Bonaventura Clotet, Montserrat Giménez, Mercedes Guerrero, Anna Not, David Panisello, Roger Paredes, Mariona Parera, Alexia París, Verónica Saludes, Alba Sánchez, Laia Soler, Lauro Sumoy, and Elisa Martró on behalf of the Can Ruti SARS-CoV-2 Sequencing Hub |
| EPI_ISL_3207068 | LABORATOIRE d'ANALYSES de BIOLOGIE MEDICALES | CNR Virus des Infections Respiratoires - France SUD | Antonin Bal, Gregory Destras, Gwendolyne Burfin, Hadrien Regue, Quentin Semanas, Martine Valette, Bruno Lina, Laurence Josset |
| EPI_ISL_3207069 | Microbiology Department, Laboratori Clínic Metropolitana Nord. Hospital Universitari Germans Trias i Pujol | Can Ruti SARS-CoV-2 Sequencing Hub (HUGTiP/IrsiCaixa/IGTP) | Marc Noguera-Julian, Pilar Armengol, Ignacio Blanco, Antoni E Bordoy, Francesc Catala-Moll, Pere-Joan Cardona, Maria Casadellà, Cristina Casañ, Gemma Clara, Bonaventura Clotet, Montserrat Giménez, Mercedes Guerrero, Anna Not, David Panisello, Roger Paredes, Mariona Parera, Alexia París, Verónica Saludes, Alba Sánchez, Laia Soler, Lauro Sumoy, and Elisa Martró on behalf of the Can Ruti SARS-CoV-2 Sequencing Hub |
| EPI_ISL_3207070 | LABORATOIRE d'ANALYSES de BIOLOGIE MEDICALES | CNR Virus des Infections Respiratoires - France SUD | Antonin Bal, Gregory Destras, Gwendolyne Burfin, Hadrien Regue, Quentin Semanas, Martine Valette, Bruno Lina, Laurence Josset |
| EPI_ISL_3207071 | Microbiologia CATLAB | Can Ruti SARS-CoV-2 Sequencing Hub (HUGTiP/IrsiCaixa/IGTP) | Marc Noguera-Julian, Pilar Armengol, Ignacio Blanco, Antoni E Bordoy, Francesc Catala-Moll, Pere-Joan Cardona, Maria Casadellà, Cristina Casañ, Gemma Clara, Bonaventura Clotet, Montserrat Giménez, Mercedes Guerrero, Anna Not, David Panisello, Roger Paredes, Mariona Parera, Alexia París, Verónica Saludes, Alba Sánchez, Laia Soler, Lauro Sumoy, and Elisa Martró on behalf of the Can Ruti SARS-CoV-2 Sequencing Hub |
| EPI_ISL_3207072 | LABORATOIRE d'ANALYSES de BIOLOGIE MEDICALES | CNR Virus des Infections Respiratoires - France SUD | Antonin Bal, Gregory Destras, Gwendolyne Burfin, Hadrien Regue, Quentin Semanas, Martine Valette, Bruno Lina, Laurence Josset |
| EPI_ISL_3207073 | Microbiology Department, Laboratori Clínic Metropolitana Nord. Hospital Universitari Germans Trias i Pujol | Can Ruti SARS-CoV-2 Sequencing Hub (HUGTiP/IrsiCaixa/IGTP) | Marc Noguera-Julian, Pilar Armengol, Ignacio Blanco, Antoni E Bordoy, Francesc Catala-Moll, Pere-Joan Cardona, Maria Casadellà, Cristina Casañ, Gemma Clara, Bonaventura Clotet, Montserrat Giménez, Mercedes Guerrero, Anna Not, David Panisello, Roger Paredes, Mariona Parera, Alexia París, Verónica Saludes, Alba Sánchez, Laia Soler, Lauro Sumoy, and Elisa Martró on behalf of the Can Ruti SARS-CoV-2 Sequencing Hub |
| EPI_ISL_3207074 | LABORATOIRE d'ANALYSES de BIOLOGIE MEDICALES | CNR Virus des Infections Respiratoires - France SUD | Antonin Bal, Gregory Destras, Gwendolyne Burfin, Hadrien Regue, Quentin Semanas, Martine Valette, Bruno Lina, Laurence Josset |
| EPI_ISL_3207075, EPI_ISL_3207076 | Microbiology Department, Laboratori Clínic Metropolitana Nord. Hospital Universitari Germans Trias i Pujol | Can Ruti SARS-CoV-2 Sequencing Hub (HUGTiP/IrsiCaixa/IGTP) | Marc Noguera-Julian, Pilar Armengol, Ignacio Blanco, Antoni E Bordoy, Francesc Catala-Moll, Pere-Joan Cardona, Maria Casadellà, Cristina Casañ, Gemma Clara, Bonaventura Clotet, Montserrat Giménez, Mercedes Guerrero, Anna Not, David Panisello, Roger Paredes, Mariona Parera, Alexia París, Verónica Saludes, Alba Sánchez, Laia Soler, Lauro Sumoy, and Elisa Martró on behalf of the Can Ruti SARS-CoV-2 Sequencing Hub |
| EPI_ISL_3207077 | LABORATOIRE d'ANALYSES de BIOLOGIE MEDICALES | CNR Virus des Infections Respiratoires - France SUD | Antonin Bal, Gregory Destras, Gwendolyne Burfin, Hadrien Regue, Quentin Semanas, Martine Valette, Bruno Lina, Laurence Josset |
| EPI_ISL_3207078 | Microbiology Department, Laboratori Clínic Metropolitana Nord. Hospital Universitari Germans Trias i Pujol | Can Ruti SARS-CoV-2 Sequencing Hub (HUGTiP/IrsiCaixa/IGTP) | Marc Noguera-Julian, Pilar Armengol, Ignacio Blanco, Antoni E Bordoy, Francesc Catala-Moll, Pere-Joan Cardona, Maria Casadellà, Cristina Casañ, Gemma Clara, Bonaventura Clotet, Montserrat Giménez, Mercedes Guerrero, Anna Not, David Panisello, Roger Paredes, Mariona Parera, Alexia París, Verónica Saludes, Alba Sánchez, Laia Soler, Lauro Sumoy, and Elisa Martró on behalf of the Can Ruti SARS-CoV-2 Sequencing Hub |
| EPI_ISL_3207079 | LABORATOIRE d'ANALYSES de BIOLOGIE MEDICALES | CNR Virus des Infections Respiratoires - France SUD | Antonin Bal, Gregory Destras, Gwendolyne Burfin, Hadrien Regue, Quentin Semanas, Martine Valette, Bruno Lina, Laurence Josset |
| EPI_ISL_3207080 | Microbiologia CATLAB | Can Ruti SARS-CoV-2 Sequencing Hub (HUGTiP/IrsiCaixa/IGTP) | Marc Noguera-Julian, Pilar Armengol, Ignacio Blanco, Antoni E Bordoy, Francesc Catala-Moll, Pere-Joan Cardona, Maria Casadellà, Cristina Casañ, Gemma Clara, Bonaventura Clotet, Montserrat Giménez, Mercedes Guerrero, Anna Not, David Panisello, Roger Paredes, Mariona Parera, Alexia París, Verónica Saludes, Alba Sánchez, Laia Soler, Lauro Sumoy, and Elisa Martró on behalf of the Can Ruti SARS-CoV-2 Sequencing Hub |
| EPI_ISL_3207081 | LABORATOIRE d'ANALYSES de BIOLOGIE MEDICALES | CNR Virus des Infections Respiratoires - France SUD | Antonin Bal, Gregory Destras, Gwendolyne Burfin, Hadrien Regue, Quentin Semanas, Martine Valette, Bruno Lina, Laurence Josset |
| EPI_ISL_3207082 | Microbiology Department, Laboratori Clínic Metropolitana Nord. Hospital Universitari Germans Trias i Pujol | Can Ruti SARS-CoV-2 Sequencing Hub (HUGTiP/IrsiCaixa/IGTP) | Marc Noguera-Julian, Pilar Armengol, Ignacio Blanco, Antoni E Bordoy, Francesc Catala-Moll, Pere-Joan Cardona, Maria Casadellà, Cristina Casañ, Gemma Clara, Bonaventura Clotet, Montserrat Giménez, Mercedes Guerrero, Anna Not, David Panisello, Roger Paredes, Mariona Parera, Alexia París, Verónica Saludes, Alba Sánchez, Laia Soler, Lauro Sumoy, and Elisa Martró on behalf of the Can Ruti SARS-CoV-2 Sequencing Hub |
| EPI_ISL_3207083 | LABORATOIRE d'ANALYSES de BIOLOGIE MEDICALES | CNR Virus des Infections Respiratoires - France SUD | Antonin Bal, Gregory Destras, Gwendolyne Burfin, Hadrien Regue, Quentin Semanas, Martine Valette, Bruno Lina, Laurence Josset |
| EPI_ISL_3207084, EPI_ISL_3207085 | Microbiology Department, Laboratori Clínic Metropolitana Nord. Hospital Universitari Germans Trias i Pujol | Can Ruti SARS-CoV-2 Sequencing Hub (HUGTiP/IrsiCaixa/IGTP) | Marc Noguera-Julian, Pilar Armengol, Ignacio Blanco, Antoni E Bordoy, Francesc Catala-Moll, Pere-Joan Cardona, Maria Casadellà, Cristina Casañ, Gemma Clara, Bonaventura Clotet, Montserrat Giménez, Mercedes Guerrero, Anna Not, David Panisello, Roger Paredes, Mariona Parera, Alexia París, Verónica Saludes, Alba Sánchez, Laia Soler, Lauro Sumoy, and Elisa Martró on behalf of the Can Ruti SARS-CoV-2 Sequencing Hub |
| EPI_ISL_3207086, EPI_ISL_3207088 | LABORATOIRE d'ANALYSES de BIOLOGIE MEDICALES | CNR Virus des Infections Respiratoires - France SUD | Antonin Bal, Gregory Destras, Gwendolyne Burfin, Hadrien Regue, Quentin Semanas, Martine Valette, Bruno Lina, Laurence Josset |
| EPI_ISL_3207089 | Microbiologia CATLAB | Can Ruti SARS-CoV-2 Sequencing Hub (HUGTiP/IrsiCaixa/IGTP) | Marc Noguera-Julian, Pilar Armengol, Ignacio Blanco, Antoni E Bordoy, Francesc Catala-Moll, Pere-Joan Cardona, Maria Casadellà, Cristina Casañ, Gemma Clara, Bonaventura Clotet, Montserrat Giménez, Mercedes Guerrero, Anna Not, David Panisello, Roger Paredes, Mariona Parera, Alexia París, Verónica Saludes, Alba Sánchez, Laia Soler, Lauro Sumoy, and Elisa Martró on behalf of the Can Ruti SARS-CoV-2 Sequencing Hub |
| EPI_ISL_3207090 | LABORATOIRE d'ANALYSES de BIOLOGIE MEDICALES | CNR Virus des Infections Respiratoires - France SUD | Antonin Bal, Gregory Destras, Gwendolyne Burfin, Hadrien Regue, Quentin Semanas, Martine Valette, Bruno Lina, Laurence Josset |
| EPI_ISL_3207091, EPI_ISL_3207092 | Microbiology Department, Laboratori Clínic Metropolitana Nord. Hospital Universitari Germans Trias i Pujol | Can Ruti SARS-CoV-2 Sequencing Hub (HUGTiP/IrsiCaixa/IGTP) | Marc Noguera-Julian, Pilar Armengol, Ignacio Blanco, Antoni E Bordoy, Francesc Catala-Moll, Pere-Joan Cardona, Maria Casadellà, Cristina Casañ, Gemma Clara, Bonaventura Clotet, Montserrat Giménez, Mercedes Guerrero, Anna Not, David Panisello, Roger Paredes, Mariona Parera, Alexia París, Verónica Saludes, Alba Sánchez, Laia Soler, Lauro Sumoy, and Elisa Martró on behalf of the Can Ruti SARS-CoV-2 Sequencing Hub |
| EPI_ISL_3207093 | LABORATOIRE d'ANALYSES de BIOLOGIE MEDICALES | CNR Virus des Infections Respiratoires - France SUD | Antonin Bal, Gregory Destras, Gwendolyne Burfin, Hadrien Regue, Quentin Semanas, Martine Valette, Bruno Lina, Laurence Josset |
| EPI_ISL_3207094 | Microbiology Department, Laboratori Clínic Metropolitana Nord. Hospital Universitari Germans Trias i Pujol | Can Ruti SARS-CoV-2 Sequencing Hub (HUGTiP/IrsiCaixa/IGTP) | Marc Noguera-Julian, Pilar Armengol, Ignacio Blanco, Antoni E Bordoy, Francesc Catala-Moll, Pere-Joan Cardona, Maria Casadellà, Cristina Casañ, Gemma Clara, Bonaventura Clotet, Montserrat Giménez, Mercedes Guerrero, Anna Not, David Panisello, Roger Paredes, Mariona Parera, Alexia París, Verónica Saludes, Alba Sánchez, Laia Soler, Lauro Sumoy, and Elisa Martró on behalf of the Can Ruti SARS-CoV-2 Sequencing Hub |
| EPI_ISL_3207095 | LABORATOIRE d'ANALYSES de BIOLOGIE MEDICALES | CNR Virus des Infections Respiratoires - France SUD | Antonin Bal, Gregory Destras, Gwendolyne Burfin, Hadrien Regue, Quentin Semanas, Martine Valette, Bruno Lina, Laurence Josset |
| EPI_ISL_3207096 | Microbiology Department, Laboratori Clínic Metropolitana Nord. Hospital Universitari Germans Trias i Pujol | Can Ruti SARS-CoV-2 Sequencing Hub (HUGTiP/IrsiCaixa/IGTP) | Marc Noguera-Julian, Pilar Armengol, Ignacio Blanco, Antoni E Bordoy, Francesc Catala-Moll, Pere-Joan Cardona, Maria Casadellà, Cristina Casañ, Gemma Clara, Bonaventura Clotet, Montserrat Giménez, Mercedes Guerrero, Anna Not, David Panisello, Roger Paredes, Mariona Parera, Alexia París, Verónica Saludes, Alba Sánchez, Laia Soler, Lauro Sumoy, and Elisa Martró on behalf of the Can Ruti SARS-CoV-2 Sequencing Hub |
| EPI_ISL_3207097 | LABORATOIRE d'ANALYSES de BIOLOGIE MEDICALES | CNR Virus des Infections Respiratoires - France SUD | Antonin Bal, Gregory Destras, Gwendolyne Burfin, Hadrien Regue, Quentin Semanas, Martine Valette, Bruno Lina, Laurence Josset |
| EPI_ISL_3207098, EPI_ISL_3207099 | Microbiology Department, Laboratori Clínic Metropolitana Nord. Hospital Universitari Germans Trias i Pujol | Can Ruti SARS-CoV-2 Sequencing Hub (HUGTiP/IrsiCaixa/IGTP) | Marc Noguera-Julian, Pilar Armengol, Ignacio Blanco, Antoni E Bordoy, Francesc Catala-Moll, Pere-Joan Cardona, Maria Casadellà, Cristina Casañ, Gemma Clara, Bonaventura Clotet, Montserrat Giménez, Mercedes Guerrero, Anna Not, David Panisello, Roger Paredes, Mariona Parera, Alexia París, Verónica Saludes, Alba Sánchez, Laia Soler, Lauro Sumoy, and Elisa Martró on behalf of the Can Ruti SARS-CoV-2 Sequencing Hub |
| EPI_ISL_3207100 | LABORATOIRE d'ANALYSES de BIOLOGIE MEDICALES | CNR Virus des Infections Respiratoires - France SUD | Antonin Bal, Gregory Destras, Gwendolyne Burfin, Hadrien Regue, Quentin Semanas, Martine Valette, Bruno Lina, Laurence Josset |
| EPI_ISL_3207101 | Microbiology Department, Laboratori Clínic Metropolitana Nord. Hospital Universitari Germans Trias i Pujol | Can Ruti SARS-CoV-2 Sequencing Hub (HUGTiP/IrsiCaixa/IGTP) | Marc Noguera-Julian, Pilar Armengol, Ignacio Blanco, Antoni E Bordoy, Francesc Catala-Moll, Pere-Joan Cardona, Maria Casadellà, Cristina Casañ, Gemma Clara, Bonaventura Clotet, Montserrat Giménez, Mercedes Guerrero, Anna Not, David Panisello, Roger Paredes, Mariona Parera, Alexia París, Verónica Saludes, Alba Sánchez, Laia Soler, Lauro Sumoy, and Elisa Martró on behalf of the Can Ruti SARS-CoV-2 Sequencing Hub |
| EPI_ISL_3207102 | LABORATOIRE d'ANALYSES de BIOLOGIE MEDICALES | CNR Virus des Infections Respiratoires - France SUD | Antonin Bal, Gregory Destras, Gwendolyne Burfin, Hadrien Regue, Quentin Semanas, Martine Valette, Bruno Lina, Laurence Josset |
| EPI_ISL_3207103 | Microbiologia CATLAB | Can Ruti SARS-CoV-2 Sequencing Hub (HUGTiP/IrsiCaixa/IGTP) | Marc Noguera-Julian, Pilar Armengol, Ignacio Blanco, Antoni E Bordoy, Francesc Catala-Moll, Pere-Joan Cardona, Maria Casadellà, Cristina Casañ, Gemma Clara, Bonaventura Clotet, Montserrat Giménez, Mercedes Guerrero, Anna Not, David Panisello, Roger Paredes, Mariona Parera, Alexia París, Verónica Saludes, Alba Sánchez, Laia Soler, Lauro Sumoy, and Elisa Martró on behalf of the Can Ruti SARS-CoV-2 Sequencing Hub |
| EPI_ISL_3207104 | Microbiology Department, Laboratori Clínic Metropolitana Nord. Hospital Universitari Germans Trias i Pujol | Can Ruti SARS-CoV-2 Sequencing Hub (HUGTiP/IrsiCaixa/IGTP) | Marc Noguera-Julian, Pilar Armengol, Ignacio Blanco, Antoni E Bordoy, Francesc Catala-Moll, Pere-Joan Cardona, Maria Casadellà, Cristina Casañ, Gemma Clara, Bonaventura Clotet, Montserrat Giménez, Mercedes Guerrero, Anna Not, David Panisello, Roger Paredes, Mariona Parera, Alexia París, Verónica Saludes, Alba Sánchez, Laia Soler, Lauro Sumoy, and Elisa Martró on behalf of the Can Ruti SARS-CoV-2 Sequencing Hub |
| EPI_ISL_3207105, EPI_ISL_3207107 | LABORATOIRE d'ANALYSES de BIOLOGIE MEDICALES | CNR Virus des Infections Respiratoires - France SUD | Antonin Bal, Gregory Destras, Gwendolyne Burfin, Hadrien Regue, Quentin Semanas, Martine Valette, Bruno Lina, Laurence Josset |
| EPI_ISL_3207108 | Microbiology Department, Laboratori Clínic Metropolitana Nord. Hospital Universitari Germans Trias i Pujol | Can Ruti SARS-CoV-2 Sequencing Hub (HUGTiP/IrsiCaixa/IGTP) | Marc Noguera-Julian, Pilar Armengol, Ignacio Blanco, Antoni E Bordoy, Francesc Catala-Moll, Pere-Joan Cardona, Maria Casadellà, Cristina Casañ, Gemma Clara, Bonaventura Clotet, Montserrat Giménez, Mercedes Guerrero, Anna Not, David Panisello, Roger Paredes, Mariona Parera, Alexia París, Verónica Saludes, Alba Sánchez, Laia Soler, Lauro Sumoy, and Elisa Martró on behalf of the Can Ruti SARS-CoV-2 Sequencing Hub |
| EPI_ISL_3207109 | LABORATOIRE d'ANALYSES de BIOLOGIE MEDICALES | CNR Virus des Infections Respiratoires - France SUD | Antonin Bal, Gregory Destras, Gwendolyne Burfin, Hadrien Regue, Quentin Semanas, Martine Valette, Bruno Lina, Laurence Josset |
| EPI_ISL_3207110 | Microbiology Department, Laboratori Clínic Metropolitana Nord. Hospital Universitari Germans Trias i Pujol | Can Ruti SARS-CoV-2 Sequencing Hub (HUGTiP/IrsiCaixa/IGTP) | Marc Noguera-Julian, Pilar Armengol, Ignacio Blanco, Antoni E Bordoy, Francesc Catala-Moll, Pere-Joan Cardona, Maria Casadellà, Cristina Casañ, Gemma Clara, Bonaventura Clotet, Montserrat Giménez, Mercedes Guerrero, Anna Not, David Panisello, Roger Paredes, Mariona Parera, Alexia París, Verónica Saludes, Alba Sánchez, Laia Soler, Lauro Sumoy, and Elisa Martró on behalf of the Can Ruti SARS-CoV-2 Sequencing Hub |
| EPI_ISL_3207111 | LABORATOIRE d'ANALYSES de BIOLOGIE MEDICALES | CNR Virus des Infections Respiratoires - France SUD | Antonin Bal, Gregory Destras, Gwendolyne Burfin, Hadrien Regue, Quentin Semanas, Martine Valette, Bruno Lina, Laurence Josset |
| EPI_ISL_3207112 | Microbiologia CATLAB | Can Ruti SARS-CoV-2 Sequencing Hub (HUGTiP/IrsiCaixa/IGTP) | Marc Noguera-Julian, Pilar Armengol, Ignacio Blanco, Antoni E Bordoy, Francesc Catala-Moll, Pere-Joan Cardona, Maria Casadellà, Cristina Casañ, Gemma Clara, Bonaventura Clotet, Montserrat Giménez, Mercedes Guerrero, Anna Not, David Panisello, Roger Paredes, Mariona Parera, Alexia París, Verónica Saludes, Alba Sánchez, Laia Soler, Lauro Sumoy, and Elisa Martró on behalf of the Can Ruti SARS-CoV-2 Sequencing Hub |
| EPI_ISL_3207113 | Microbiology Department, Laboratori Clínic Metropolitana Nord. Hospital Universitari Germans Trias i Pujol | Can Ruti SARS-CoV-2 Sequencing Hub (HUGTiP/IrsiCaixa/IGTP) | Marc Noguera-Julian, Pilar Armengol, Ignacio Blanco, Antoni E Bordoy, Francesc Catala-Moll, Pere-Joan Cardona, Maria Casadellà, Cristina Casañ, Gemma Clara, Bonaventura Clotet, Montserrat Giménez, Mercedes Guerrero, Anna Not, David Panisello, Roger Paredes, Mariona Parera, Alexia París, Verónica Saludes, Alba Sánchez, Laia Soler, Lauro Sumoy, and Elisa Martró on behalf of the Can Ruti SARS-CoV-2 Sequencing Hub |
| EPI_ISL_3207114 | LABORATOIRE d'ANALYSES de BIOLOGIE MEDICALES | CNR Virus des Infections Respiratoires - France SUD | Antonin Bal, Gregory Destras, Gwendolyne Burfin, Hadrien Regue, Quentin Semanas, Martine Valette, Bruno Lina, Laurence Josset |
| EPI_ISL_3207115 | Microbiology Department, Laboratori Clínic Metropolitana Nord. Hospital Universitari Germans Trias i Pujol | Can Ruti SARS-CoV-2 Sequencing Hub (HUGTiP/IrsiCaixa/IGTP) | Marc Noguera-Julian, Pilar Armengol, Ignacio Blanco, Antoni E Bordoy, Francesc Catala-Moll, Pere-Joan Cardona, Maria Casadellà, Cristina Casañ, Gemma Clara, Bonaventura Clotet, Montserrat Giménez, Mercedes Guerrero, Anna Not, David Panisello, Roger Paredes, Mariona Parera, Alexia París, Verónica Saludes, Alba Sánchez, Laia Soler, Lauro Sumoy, and Elisa Martró on behalf of the Can Ruti SARS-CoV-2 Sequencing Hub |
| EPI_ISL_3207116 | LABORATOIRE d'ANALYSES de BIOLOGIE MEDICALES | CNR Virus des Infections Respiratoires - France SUD | Antonin Bal, Gregory Destras, Gwendolyne Burfin, Hadrien Regue, Quentin Semanas, Martine Valette, Bruno Lina, Laurence Josset |
| EPI_ISL_3207117 | Microbiology Department, Laboratori Clínic Metropolitana Nord. Hospital Universitari Germans Trias i Pujol | Can Ruti SARS-CoV-2 Sequencing Hub (HUGTiP/IrsiCaixa/IGTP) | Marc Noguera-Julian, Pilar Armengol, Ignacio Blanco, Antoni E Bordoy, Francesc Catala-Moll, Pere-Joan Cardona, Maria Casadellà, Cristina Casañ, Gemma Clara, Bonaventura Clotet, Montserrat Giménez, Mercedes Guerrero, Anna Not, David Panisello, Roger Paredes, Mariona Parera, Alexia París, Verónica Saludes, Alba Sánchez, Laia Soler, Lauro Sumoy, and Elisa Martró on behalf of the Can Ruti SARS-CoV-2 Sequencing Hub |
| EPI_ISL_3207118 | LABORATOIRE d'ANALYSES de BIOLOGIE MEDICALES | CNR Virus des Infections Respiratoires - France SUD | Antonin Bal, Gregory Destras, Gwendolyne Burfin, Hadrien Regue, Quentin Semanas, Martine Valette, Bruno Lina, Laurence Josset |
| EPI_ISL_3207119 | Microbiology Department, Laboratori Clínic Metropolitana Nord. Hospital Universitari Germans Trias i Pujol | Can Ruti SARS-CoV-2 Sequencing Hub (HUGTiP/IrsiCaixa/IGTP) | Marc Noguera-Julian, Pilar Armengol, Ignacio Blanco, Antoni E Bordoy, Francesc Catala-Moll, Pere-Joan Cardona, Maria Casadellà, Cristina Casañ, Gemma Clara, Bonaventura Clotet, Montserrat Giménez, Mercedes Guerrero, Anna Not, David Panisello, Roger Paredes, Mariona Parera, Alexia París, Verónica Saludes, Alba Sánchez, Laia Soler, Lauro Sumoy, and Elisa Martró on behalf of the Can Ruti SARS-CoV-2 Sequencing Hub |

EPI_ISL_3207120, EPI_ISL_3207121, EPI_ISL_3207122, EPI_ISL_3207123, EPI_ISL_3207124, EPI_ISL_3207125, EPI_ISL_3207126, EPI_ISL_3207127, EPI_ISL_3207128, EPI_ISL_3207129, EPI_ISL_3207130, EPI_ISL_3207131, EPI_ISL_3207132, EPI_ISL_3207133, EPI_ISL_3207134, EPI_ISL_3207135, EPI_ISL_3207136, EPI_ISL_3207137, EPI_ISL_3207138, EPI_ISL_3207139, EPI_ISL_3207141, EPI_ISL_3207142, EPI_ISL_3207143, EPI_ISL_3207144, EPI_ISL_3207145, EPI_ISL_3207146, EPI_ISL_3207147, EPI_ISL_3207148, EPI_ISL_3207149, EPI_ISL_3207150, EPI_ISL_3207151, EPI_ISL_3207152, EPI_ISL_3207153, EPI_ISL_3207154, EPI_ISL_3207155, EPI_ISL_3207156, EPI_ISL_3207157, EPI_ISL_3207158, EPI_ISL_3207159, EPI_ISL_3207160, EPI_ISL_3207161, EPI_ISL_3207162, EPI_ISL_3207163, EPI_ISL_3207164, EPI_ISL_3207165, EPI_ISL_3207166, EPI_ISL_3207167, EPI_ISL_3207168, EPI_ISL_3207169, EPI_ISL_3207170, EPI_ISL_3207171, EPI_ISL_3207172, EPI_ISL_3207173, EPI_ISL_3207174, EPI_ISL_3207175, EPI_ISL_3207176, EPI_ISL_3207177, EPI_ISL_3207178, EPI_ISL_3207179, EPI_ISL_3207180, EPI_ISL_3207181, EPI_ISL_3207182, EPI_ISL_3207183, EPI_ISL_3207184, EPI_ISL_3207185, EPI_ISL_3207186, EPI_ISL_3207187, EPI_ISL_3207188, EPI_ISL_3207189, EPI_ISL_3207190, EPI_ISL_3207191, EPI_ISL_3207192, EPI_ISL_3207193, EPI_ISL_3207194, EPI_ISL_3207195, EPI_ISL_3207196, EPI_ISL_3207197, EPI_ISL_3207198, EPI_ISL_3207199, EPI_ISL_3207200, EPI_ISL_3207201, EPI_ISL_3207202, EPI_ISL_3207203, EPI_ISL_3207204, EPI_ISL_3207205, EPI_ISL_3207206, EPI_ISL_3207207, EPI_ISL_3207208, EPI_ISL_3207209, EPI_ISL_3207210, EPI_ISL_3207211, EPI_ISL_3207212, EPI_ISL_3207213, EPI_ISL_3207214, EPI_ISL_3207215, EPI_ISL_3207216, EPI_ISL_3207217, EPI_ISL_3207218, EPI_ISL_3207219, EPI_ISL_3207220, EPI_ISL_3207221, EPI_ISL_3207222, EPI_ISL_3207223, EPI_ISL_3207224, EPI_ISL_3207225, EPI_ISL_3207226, EPI_ISL_3207227, EPI_ISL_3207228, EPI_ISL_3207229, EPI_ISL_3207230, EPI_ISL_3207231, EPI_ISL_3207232, EPI_ISL_3207233, EPI_ISL_3207234, EPI_ISL_3207235, EPI_ISL_3207236, EPI_ISL_3207237, EPI_ISL_3207238, EPI_ISL_3207239, EPI_ISL_3207240, EPI_ISL_3207241, EPI_ISL_3207242, EPI_ISL_3207243, EPI_ISL_3207244, EPI_ISL_3207245, EPI_ISL_3207246, EPI_ISL_3207247, EPI_ISL_3207248, EPI_ISL_3207249, EPI_ISL_3207250, EPI_ISL_3207251, EPI_ISL_3207252, EPI_ISL_3207253, EPI_ISL_3207254, EPI_ISL_3207255, EPI_ISL_3207256, EPI_ISL_3207257, EPI_ISL_3207258, EPI_ISL_3207259, EPI_ISL_3207260, EPI_ISL_3207261, EPI_ISL_3207262, EPI_ISL_3207263, EPI_ISL_3207264, EPI_ISL_3207265, EPI_ISL_3207266, EPI_ISL_3207267

| | | | |
|---|---|---|---|
| see above | LABORATOIRE d'ANALYSES de BIOLOGIE MEDICALES | CNR Virus des Infections Respiratoires - France SUD | Antonin Bal, Gregory Destras, Gwendolyne Burfin, Hadrien Regue, Quentin Semanas, Martine Valette, Bruno Lina, Laurence Josset |
| EPI_ISL_3207268, EPI_ISL_3207269, EPI_ISL_3207270, EPI_ISL_3207271, EPI_ISL_3207272, EPI_ISL_3207273, EPI_ISL_3207274, EPI_ISL_3207275 | LABORATOIRE BIOESTEREL | CNR Virus des Infections Respiratoires - France SUD | Antonin Bal, Gregory Destras, Gwendolyne Burfin, Hadrien Regue, Quentin Semanas, Martine Valette, Bruno Lina, Laurence Josset |
| EPI_ISL_3207276 | LABORATOIRE UNILIANS BEYNOST | CNR Virus des Infections Respiratoires - France SUD | Antonin Bal, Gregory Destras, Gwendolyne Burfin, Hadrien Regue, Quentin Semanas, Martine Valette, Bruno Lina, Laurence Josset |

| | | | |
|---|---|---|---|
| EPI_ISL_3207277 | LABORATOIRE MAYMAT | CNR Virus des Infections Respiratoires - France SUD | Antonin Bal, Gregory Destras, Gwendolyne Burfin, Hadrien Regue, Quentin Semanas, Martine Valette, Bruno Lina, Laurence Josset |
| EPI_ISL_3207278 | LAM BIOPOLE ORTHEZ | CNR Virus des Infections Respiratoires - France SUD | Antonin Bal, Gregory Destras, Gwendolyne Burfin, Hadrien Regue, Quentin Semanas, Martine Valette, Bruno Lina, Laurence Josset |
| EPI_ISL_3207279, EPI_ISL_3207280, EPI_ISL_3207282, EPI_ISL_3207283, EPI_ISL_3207284, EPI_ISL_3207285, EPI_ISL_3207286, EPI_ISL_3207287, EPI_ISL_3207288, EPI_ISL_3207289, EPI_ISL_3207290, EPI_ISL_3207291, EPI_ISL_3207292, EPI_ISL_3207293, EPI_ISL_3207294 | | | |
| see above | SYNLAB GASCOGNE | CNR Virus des Infections Respiratoires - France SUD | Antonin Bal, Gregory Destras, Gwendolyne Burfin, Hadrien Regue, Quentin Semanas, Martine Valette, Bruno Lina, Laurence Josset |
| EPI_ISL_3207295, EPI_ISL_3207296, EPI_ISL_3207297, EPI_ISL_3207298, EPI_ISL_3207299, EPI_ISL_3207300, EPI_ISL_3207303, EPI_ISL_3207304, EPI_ISL_3207305, EPI_ISL_3207306, EPI_ISL_3207307, EPI_ISL_3207308, EPI_ISL_3207309, EPI_ISL_3207310, EPI_ISL_3207311, EPI_ISL_3207312, EPI_ISL_3207313, EPI_ISL_3207314, EPI_ISL_3207315, EPI_ISL_3207316, EPI_ISL_3207317, EPI_ISL_3207318, EPI_ISL_3207319, EPI_ISL_3207320, EPI_ISL_3207321, EPI_ISL_3207322, EPI_ISL_3207323, EPI_ISL_3207325, EPI_ISL_3207326, EPI_ISL_3207327, EPI_ISL_3207328, EPI_ISL_3207330, EPI_ISL_3207331, EPI_ISL_3207332, EPI_ISL_3207333, EPI_ISL_3207334, EPI_ISL_3207335, EPI_ISL_3207336, EPI_ISL_3207337, EPI_ISL_3207338, EPI_ISL_3207339, EPI_ISL_3207340, EPI_ISL_3207341, EPI_ISL_3207342, EPI_ISL_3207343, EPI_ISL_3207344, EPI_ISL_3207345 | | | |
| see above | LABORATOIRE DES CEDRES | CNR Virus des Infections Respiratoires - France SUD | Antonin Bal, Gregory Destras, Gwendolyne Burfin, Hadrien Regue, Quentin Semanas, Martine Valette, Bruno Lina, Laurence Josset |
| EPI_ISL_3207346, EPI_ISL_3207347, EPI_ISL_3207348 | LABORATOIRE BIOFUSION | CNR Virus des Infections Respiratoires - France SUD | Antonin Bal, Gregory Destras, Gwendolyne Burfin, Hadrien Regue, Quentin Semanas, Martine Valette, Bruno Lina, Laurence Josset |
| EPI_ISL_3207349, EPI_ISL_3207350, EPI_ISL_3207351, EPI_ISL_3207352, EPI_ISL_3207353, EPI_ISL_3207354, EPI_ISL_3207355, EPI_ISL_3207356 | LABM CH MONTAUBAN | CNR Virus des Infections Respiratoires - France SUD | Antonin Bal, Gregory Destras, Gwendolyne Burfin, Hadrien Regue, Quentin Semanas, Martine Valette, Bruno Lina, Laurence Josset |
| EPI_ISL_3207357, EPI_ISL_3207358, EPI_ISL_3207359, EPI_ISL_3207360, EPI_ISL_3207361, EPI_ISL_3207363, EPI_ISL_3207364, EPI_ISL_3207365, EPI_ISL_3207366, EPI_ISL_3207367 | LX BIO | CNR Virus des Infections Respiratoires - France SUD | Antonin Bal, Gregory Destras, Gwendolyne Burfin, Hadrien Regue, Quentin Semanas, Martine Valette, Bruno Lina, Laurence Josset |
| EPI_ISL_3207368, EPI_ISL_3207369, EPI_ISL_3207370, EPI_ISL_3207371, EPI_ISL_3207372 | LABORATOIRE d'ANALYSES de BIOLOGIE MEDICALES | CNR Virus des Infections Respiratoires - France SUD | Antonin Bal, Gregory Destras, Gwendolyne Burfin, Hadrien Regue, Quentin Semanas, Martine Valette, Bruno Lina, Laurence Josset |
| EPI_ISL_3207373 | LABORATOIRE CERBALLIANCE PLT VILLON | CNR Virus des Infections Respiratoires - France SUD | Antonin Bal, Gregory Destras, Gwendolyne Burfin, Hadrien Regue, Quentin Semanas, Martine Valette, Bruno Lina, Laurence Josset |
| EPI_ISL_3207374, EPI_ISL_3207375 | LABORATOIRE d'ANALYSES de BIOLOGIE MEDICALES | CNR Virus des Infections Respiratoires - France SUD | Antonin Bal, Gregory Destras, Gwendolyne Burfin, Hadrien Regue, Quentin Semanas, Martine Valette, Bruno Lina, Laurence Josset |
| EPI_ISL_3207376, EPI_ISL_3207377, EPI_ISL_3207378, EPI_ISL_3207379, EPI_ISL_3207380, EPI_ISL_3207381, EPI_ISL_3207382, EPI_ISL_3207383, EPI_ISL_3207384 | LAM BIORYLIS | CNR Virus des Infections Respiratoires - France SUD | Antonin Bal, Gregory Destras, Gwendolyne Burfin, Hadrien Regue, Quentin Semanas, Martine Valette, Bruno Lina, Laurence Josset |
| EPI_ISL_3207385 | DYOMEDEA-LABORATOIRE DE LA SAUVEGARDE | CNR Virus des Infections Respiratoires - France SUD | Antonin Bal, Gregory Destras, Gwendolyne Burfin, Hadrien Regue, Quentin Semanas, Martine Valette, Bruno Lina, Laurence Josset |
| EPI_ISL_3207386 | LABORATOIRE d'ANALYSES de BIOLOGIE MEDICALES | CNR Virus des Infections Respiratoires - France SUD | Antonin Bal, Gregory Destras, Gwendolyne Burfin, Hadrien Regue, Quentin Semanas, Martine Valette, Bruno Lina, Laurence Josset |
| EPI_ISL_3207387, EPI_ISL_3207388 | DYOMEDEA-LABORATOIRE DE LA SAUVEGARDE | CNR Virus des Infections Respiratoires - France SUD | Antonin Bal, Gregory Destras, Gwendolyne Burfin, Hadrien Regue, Quentin Semanas, Martine Valette, Bruno Lina, Laurence Josset |
| EPI_ISL_3207389 | LBM DYNABIO CARREAU | CNR Virus des Infections Respiratoires - France SUD | Antonin Bal, Gregory Destras, Gwendolyne Burfin, Hadrien Regue, Quentin Semanas, Martine Valette, Bruno Lina, Laurence Josset |
| EPI_ISL_3207390 | DYOMEDEA-LABORATOIRE DE LA SAUVEGARDE | CNR Virus des Infections Respiratoires - France SUD | Antonin Bal, Gregory Destras, Gwendolyne Burfin, Hadrien Regue, Quentin Semanas, Martine Valette, Bruno Lina, Laurence Josset |
| EPI_ISL_3207391, EPI_ISL_3207392, EPI_ISL_3207393, EPI_ISL_3207394, EPI_ISL_3207395, EPI_ISL_3207396, EPI_ISL_3207397, EPI_ISL_3207398, EPI_ISL_3207399, EPI_ISL_3207400, EPI_ISL_3207401, EPI_ISL_3207402 | | | |
| see above | LABORATOIRE BIOESTEREL | CNR Virus des Infections Respiratoires - France SUD | Antonin Bal, Gregory Destras, Gwendolyne Burfin, Hadrien Regue, Quentin Semanas, Martine Valette, Bruno Lina, Laurence Josset |
| EPI_ISL_3207403, EPI_ISL_3207404, EPI_ISL_3207405, EPI_ISL_3207406, EPI_ISL_3207407 | BIOLITTORAL BIOGROUP PLATEAU TECH | CNR Virus des Infections Respiratoires - France SUD | Antonin Bal, Gregory Destras, Gwendolyne Burfin, Hadrien Regue, Quentin Semanas, Martine Valette, Bruno Lina, Laurence Josset |
| EPI_ISL_3207408 | LABORATOIRE d'ANALYSES de BIOLOGIE MEDICALES | CNR Virus des Infections Respiratoires - France SUD | Antonin Bal, Gregory Destras, Gwendolyne Burfin, Hadrien Regue, Quentin Semanas, Martine Valette, Bruno Lina, Laurence Josset |
| EPI_ISL_3207409, EPI_ISL_3207410, EPI_ISL_3207411, EPI_ISL_3207412, EPI_ISL_3207413, EPI_ISL_3207414, EPI_ISL_3207415, EPI_ISL_3207416, EPI_ISL_3207418, EPI_ISL_3207419, EPI_ISL_3207420, EPI_ISL_3207421, EPI_ISL_3207423, EPI_ISL_3207424, EPI_ISL_3207425, EPI_ISL_3207426, EPI_ISL_3207427, EPI_ISL_3207428, EPI_ISL_3207430, EPI_ISL_3207432, EPI_ISL_3207433, EPI_ISL_3207436, EPI_ISL_3207437, EPI_ISL_3207438, EPI_ISL_3207439, EPI_ISL_3207440, EPI_ISL_3207441, EPI_ISL_3207442, EPI_ISL_3207443, EPI_ISL_3207444, EPI_ISL_3207445, EPI_ISL_3207452, EPI_ISL_3207453, EPI_ISL_3207454, EPI_ISL_3207455, EPI_ISL_3207456, EPI_ISL_3207457, EPI_ISL_3207461, EPI_ISL_3207466, EPI_ISL_3207467 | | | |
| see above | Wyoming Public Health Laboratory | Wyoming Public Health Laboratory | Jim Mildenberger, Taylor Fearing, Channing Weber, Ashley Norberg, Chayse Rowley, Marley Goetz, Brian Dominguez, Elliot Thomasson, Sam Britz, Cari Sloma, Robert Petit, and Rob Christensen |
| EPI_ISL_3207500 | Vredendal Central Clinic wc VDC | NHLS/UCT | Sisonke, Arash Iranzadeh, Deelan Doolabh, Lynn Tyers, Bruna Galvao, Innocent Mudau, Marvin Hsiao, Gert Marais, Diana Hardie, Stephen Korsman, Rageema Joseph, Carolyn Williamson |
| EPI_ISL_3207503 | 2 Military Hospital wc MAA | NHLS/UCT | Sisonke, Arash Iranzadeh, Deelan Doolabh, Lynn Tyers, Bruna Galvao, Innocent Mudau, Marvin Hsiao, Gert Marais, Diana Hardie, Stephen Korsman, Rageema Joseph, Carolyn Williamson |
| EPI_ISL_3207504 | Groote Schuur Hospital wc GSH | NHLS/UCT | Sisonke, Arash Iranzadeh, Deelan Doolabh, Lynn Tyers, Bruna Galvao, Innocent Mudau, Marvin Hsiao, Gert Marais, Diana Hardie, Stephen Korsman, Rageema Joseph, Carolyn Williamson |
| EPI_ISL_3207505, EPI_ISL_3207506, EPI_ISL_3207507, EPI_ISL_3207508, EPI_ISL_3207509 | Lancet lab | NHLS/UCT | Sisonke, Arash Iranzadeh, Deelan Doolabh, Lynn Tyers, Bruna Galvao, Innocent Mudau, Marvin Hsiao, Gert Marais, Diana Hardie, Stephen Korsman, Rageema Joseph, Carolyn Williamson |
| EPI_ISL_3207511 | Pacaltsdorp Clinic wc PAC | NHLS/UCT | Arash Iranzadeh, Deelan Doolabh, Lynn Tyers, Bruna Galvao, Innocent Mudau, Marvin Hsiao, Gert Marais, Diana Hardie, Stephen Korsman, Rageema Joseph, Carolyn Williamson |
| EPI_ISL_3207512 | Conville CDC wc CVC | NHLS/UCT | Arash Iranzadeh, Deelan Doolabh, Lynn Tyers, Bruna Galvao, Innocent Mudau, Marvin Hsiao, Gert Marais, Diana Hardie, Stephen Korsman, Rageema Joseph, Carolyn Williamson |
| EPI_ISL_3207517, EPI_ISL_3207518, EPI_ISL_3207519, EPI_ISL_3207520, EPI_ISL_3207522, EPI_ISL_3207523 | Ampath lab | NHLS/UCT | Arash Iranzadeh, Deelan Doolabh, Lynn Tyers, Bruna Galvao, Innocent Mudau, Marvin Hsiao, Gert Marais, Diana Hardie, Stephen Korsman, Rageema Joseph, Carolyn Williamson |
| EPI_ISL_3207524 | Buffeljagsrivier Clinic wc BBS | NHLS/UCT | Arash Iranzadeh, Deelan Doolabh, Lynn Tyers, Bruna Galvao, Innocent Mudau, Marvin Hsiao, Gert Marais, Diana Hardie, Stephen Korsman, Rageema Joseph, Carolyn Williamson |
| EPI_ISL_3207525 | Swellendam PHC Clinic wc SHC | NHLS/UCT | Arash Iranzadeh, Deelan Doolabh, Lynn Tyers, Bruna Galvao, Innocent Mudau, Marvin Hsiao, Gert Marais, Diana Hardie, Stephen Korsman, Rageema Joseph, Carolyn Williamson |
| EPI_ISL_3207537 | Still Bay Sat Clinic wc SFV | NHLS/UCT | Arash Iranzadeh, Deelan Doolabh, Lynn Tyers, Bruna Galvao, Innocent Mudau, Marvin Hsiao, Gert Marais, Diana Hardie, Stephen Korsman, Rageema Joseph, Carolyn Williamson |
| EPI_ISL_3207538 | Pacaltsdorp Clinic wc PAC | NHLS/UCT | Arash Iranzadeh, Deelan Doolabh, Lynn Tyers, Bruna Galvao, Innocent Mudau, Marvin Hsiao, Gert Marais, Diana Hardie, Stephen Korsman, Rageema Joseph, Carolyn Williamson |
| EPI_ISL_3207539, EPI_ISL_3207540 | Groote Schuur Hospital wc GSH | NHLS/UCT | Arash Iranzadeh, Deelan Doolabh, Lynn Tyers, Bruna Galvao, Innocent Mudau, Marvin Hsiao, Gert Marais, Diana Hardie, Stephen Korsman, Rageema Joseph, Carolyn Williamson |
| EPI_ISL_3207541 | False Bay Hospital wc FBH | NHLS/UCT | Arash Iranzadeh, Deelan Doolabh, Lynn Tyers, Bruna Galvao, Innocent Mudau, Marvin Hsiao, Gert Marais, Diana Hardie, Stephen Korsman, Rageema Joseph, Carolyn Williamson |
| EPI_ISL_3207542 | Groote Schuur Hospital wc GSH | NHLS/UCT | Arash Iranzadeh, Deelan Doolabh, Lynn Tyers, Bruna Galvao, Innocent Mudau, Marvin Hsiao, Gert Marais, Diana Hardie, Stephen Korsman, Rageema Joseph, Carolyn Williamson |
| EPI_ISL_3207543, EPI_ISL_3207544 | Pacaltsdorp Clinic wc PAC | NHLS/UCT | Arash Iranzadeh, Deelan Doolabh, Lynn Tyers, Bruna Galvao, Innocent Mudau, Marvin Hsiao, Gert Marais, Diana Hardie, Stephen Korsman, Rageema Joseph, Carolyn Williamson |
| EPI_ISL_3207545 | D'Almeida Clinic wc DAL | NHLS/UCT | Arash Iranzadeh, Deelan Doolabh, Lynn Tyers, Bruna Galvao, Innocent Mudau, Marvin Hsiao, Gert Marais, Diana Hardie, Stephen Korsman, Rageema Joseph, Carolyn Williamson |
| EPI_ISL_3207546 | Conville CDC wc CVC | NHLS/UCT | Arash Iranzadeh, Deelan Doolabh, Lynn Tyers, Bruna Galvao, Innocent Mudau, Marvin Hsiao, Gert Marais, Diana Hardie, Stephen Korsman, Rageema Joseph, Carolyn Williamson |
| EPI_ISL_3207547, | Groote Schuur Hospital wc GSH | NHLS/UCT | Arash Iranzadeh, Deelan Doolabh, Lynn Tyers, Bruna Galvao, Innocent Mudau, Marvin Hsiao, Gert Marais, Diana Hardie, Stephen Korsman, Rageema Joseph, Carolyn Williamson |

| | | | |
|---|---|---|---|
| EPI_ISL_3207548 | | | |
| EPI_ISL_3207549 | Weltevreden Valley Clinic wc WEL | NHLS/UCT | Arash Iranzadeh, Deelan Doolabh, Lynn Tyers, Bruna Galvao, Innocent Mudau, Marvin Hsiao, Gert Marais, Diana Hardie, Stephen Korsman, Rageema Joseph, Carolyn Williamson |
| EPI_ISL_3207550 | Pacaltsdorp Clinic wc PAC | NHLS/UCT | Arash Iranzadeh, Deelan Doolabh, Lynn Tyers, Bruna Galvao, Innocent Mudau, Marvin Hsiao, Gert Marais, Diana Hardie, Stephen Korsman, Rageema Joseph, Carolyn Williamson |
| EPI_ISL_3207551, EPI_ISL_3207552 | Pacaltsdorp Clinic wc PAC | NHLS/UCT | Sisonke, Arash Iranzadeh, Deelan Doolabh, Lynn Tyers, Bruna Galvao, Innocent Mudau, Marvin Hsiao, Gert Marais, Diana Hardie, Stephen Korsman, Rageema Joseph, Carolyn Williamson |
| EPI_ISL_3207553 | Conville CDC wc CVC | NHLS/UCT | Sisonke, Arash Iranzadeh, Deelan Doolabh, Lynn Tyers, Bruna Galvao, Innocent Mudau, Marvin Hsiao, Gert Marais, Diana Hardie, Stephen Korsman, Rageema Joseph, Carolyn Williamson |
| EPI_ISL_3207554, EPI_ISL_3207555 | Groote Schuur Hospital wc GSH | NHLS/UCT | Sisonke, Arash Iranzadeh, Deelan Doolabh, Lynn Tyers, Bruna Galvao, Innocent Mudau, Marvin Hsiao, Gert Marais, Diana Hardie, Stephen Korsman, Rageema Joseph, Carolyn Williamson |
| EPI_ISL_3207556, EPI_ISL_3207557, EPI_ISL_3207558, EPI_ISL_3207559, EPI_ISL_3207560, EPI_ISL_3207561 | Pacaltsdorp Clinic wc PAC | NHLS/UCT | Sisonke, Arash Iranzadeh, Deelan Doolabh, Lynn Tyers, Bruna Galvao, Innocent Mudau, Marvin Hsiao, Gert Marais, Diana Hardie, Stephen Korsman, Rageema Joseph, Carolyn Williamson |
| EPI_ISL_3207562 | Alma CDC wc AHC | NHLS/UCT | Sisonke, Arash Iranzadeh, Deelan Doolabh, Lynn Tyers, Bruna Galvao, Innocent Mudau, Marvin Hsiao, Gert Marais, Diana Hardie, Stephen Korsman, Rageema Joseph, Carolyn Williamson |
| EPI_ISL_3207563 | Still Bay Sat Clinic wc SFV | NHLS/UCT | Sisonke, Arash Iranzadeh, Deelan Doolabh, Lynn Tyers, Bruna Galvao, Innocent Mudau, Marvin Hsiao, Gert Marais, Diana Hardie, Stephen Korsman, Rageema Joseph, Carolyn Williamson |
| EPI_ISL_3207564 | Pacaltsdorp Clinic wc PAC | NHLS/UCT | Sisonke, Arash Iranzadeh, Deelan Doolabh, Lynn Tyers, Bruna Galvao, Innocent Mudau, Marvin Hsiao, Gert Marais, Diana Hardie, Stephen Korsman, Rageema Joseph, Carolyn Williamson |
| EPI_ISL_3207565 | Groote Schuur Hospital wc GSH | NHLS/UCT | Sisonke, Arash Iranzadeh, Deelan Doolabh, Lynn Tyers, Bruna Galvao, Innocent Mudau, Marvin Hsiao, Gert Marais, Diana Hardie, Stephen Korsman, Rageema Joseph, Carolyn Williamson |
| EPI_ISL_3207566 | Conville CDC wc CVC | NHLS/UCT | Sisonke, Arash Iranzadeh, Deelan Doolabh, Lynn Tyers, Bruna Galvao, Innocent Mudau, Marvin Hsiao, Gert Marais, Diana Hardie, Stephen Korsman, Rageema Joseph, Carolyn Williamson |
| EPI_ISL_3207567, EPI_ISL_3207568, EPI_ISL_3207569 | Pacaltsdorp Clinic wc PAC | NHLS/UCT | Sisonke, Arash Iranzadeh, Deelan Doolabh, Lynn Tyers, Bruna Galvao, Innocent Mudau, Marvin Hsiao, Gert Marais, Diana Hardie, Stephen Korsman, Rageema Joseph, Carolyn Williamson |
| EPI_ISL_3207570 | Red Cross Children's Hospital wc RXH | NHLS/UCT | Sisonke, Arash Iranzadeh, Deelan Doolabh, Lynn Tyers, Bruna Galvao, Innocent Mudau, Marvin Hsiao, Gert Marais, Diana Hardie, Stephen Korsman, Rageema Joseph, Carolyn Williamson |
| EPI_ISL_3207571, EPI_ISL_3207572, EPI_ISL_3207573 | Mitchells Plain Hospital wc MPH | NHLS/UCT | Sisonke, Arash Iranzadeh, Deelan Doolabh, Lynn Tyers, Bruna Galvao, Innocent Mudau, Marvin Hsiao, Gert Marais, Diana Hardie, Stephen Korsman, Rageema Joseph, Carolyn Williamson |
| EPI_ISL_3207574, EPI_ISL_3207575, EPI_ISL_3207577, EPI_ISL_3207578, EPI_ISL_3207579, EPI_ISL_3207580, EPI_ISL_3207582, EPI_ISL_3207583, EPI_ISL_3207584, EPI_ISL_3207585, EPI_ISL_3207586, EPI_ISL_3207587 | | | |
| see above | Lancet lab | NHLS/UCT | Sisonke, Arash Iranzadeh, Deelan Doolabh, Lynn Tyers, Bruna Galvao, Innocent Mudau, Marvin Hsiao, Gert Marais, Diana Hardie, Stephen Korsman, Rageema Joseph, Carolyn Williamson |
| EPI_ISL_3207588 | Hospital Universitari Vall d'Hebron - Vall d'Hebron Institut de Recerca | Hospital Universitari Vall d'Hebron - Vall d'Hebron Institut de Recerca | Cristina Andrés, Maria Piñana, Alejandra González-Sánchez, Damir Garcia-Cehic, Ariadna Rando, Juliana Esperalba, Maria Gema Codina, Carla Castillo, Karen García, Rodrigo Vásquez, Maria Carmen Martin, Tomàs Pumarola, Josep Quer, Andrés Antón |
| EPI_ISL_3207893, EPI_ISL_3207973, EPI_ISL_3208016, EPI_ISL_3208017, EPI_ISL_3208018, EPI_ISL_3208019, EPI_ISL_3208020, EPI_ISL_3208021, EPI_ISL_3208022, EPI_ISL_3208023, EPI_ISL_3208024, EPI_ISL_3208025, EPI_ISL_3208026, EPI_ISL_3208027, EPI_ISL_3208028, EPI_ISL_3208029, EPI_ISL_3208030, EPI_ISL_3208031, EPI_ISL_3208032, EPI_ISL_3208037, EPI_ISL_3208040, EPI_ISL_3208044, EPI_ISL_3208051 | | | |
| see above | Laboratoire de santé publique du Québec | Laboratoire de santé publique du Québec | Sandrine Moreira, Ioannis Ragoussis, Guillaume Bourque, Jesse Shapiro, Mark Lathrop and Michel Roger on behalf of the CoVSeQ research group |
| EPI_ISL_3208058, EPI_ISL_3208059, EPI_ISL_3208062, EPI_ISL_3208063 | Murni Teguh Memorial Hospital | Genomik Solidaritas Indonesia Laboratorium | Vania Gavrila Wikasa, Carissa Sintca Wijaya, Inke Nadia Diniyanti Lubis, Ririe Fachrina Malisie, Lia Kusumawati, Tryna Tania, Mutiara, Reinhart Gabriel, Annisa Muthiah Sukirman, Gracia Felias Enos Korompis, Anuraj Shankar, Meutia Ayuputeri Kumaheri, Ariel Pradipta |
| EPI_ISL_3208067 | Murni Teguh Memorial Hospital | Genomik Solidaritas Indonesia Laboratorium | Carissa Sintca Wijaya, Reinhart Gabriel, Inke Nadia Diniyanti Lubis, Ririe Fachrina Malisie, Lia Kusumawati, Tryna Tania, Mutiara, Vania Gavrila Wikasa, Annisa Muthiah Sukirman, Gracia Felias Enos Korompis, Anuraj Shankar, Meutia Ayuputeri Kumaheri, Ariel Pradipta |
| EPI_ISL_3208075, EPI_ISL_3208076 | LABKESDA / DINAS KESEHATAN DKI JAKARTA | Genomik Solidaritas Indonesia Laboratorium | Reinhart Gabriel, Vania Gavrila Wikasa, Tryna Tania, Carissa Sintca Wijaya, Annisa Muthiah Sukirman, Gracia Felias Enos Korompis, Anuraj Shankar, Meutia Ayuputeri Kumaheri, Ariel Pradipta |
| EPI_ISL_3208077 | RSUD MATRAMAN / DINAS KESEHATAN DKI JAKARTA | Genomik Solidaritas Indonesia Laboratorium | Reinhart Gabriel, Vania Gavrila Wikasa, Tryna Tania, Carissa Sintca Wijaya, Annisa Muthiah Sukirman, Gracia Felias Enos Korompis, Anuraj Shankar, Meutia Ayuputeri Kumaheri, Ariel Pradipta |
| EPI_ISL_3208078 | KIMIA FARMA / DINAS KESEHATAN DKI JAKARTA | Genomik Solidaritas Indonesia Laboratorium | Vania Gavrila Wikasa, Carissa Sintca Wijaya, Latifah Anandari, Reinhart Gabriel, Annisa Muthiah Sukirman, Gracia Felias Enos Korompis, Anuraj Shankar, Meutia Ayuputeri Kumaheri, Ariel Pradipta |
| EPI_ISL_3208079 | KIMIA FARMA / DINAS KESEHATAN DKI JAKARTA | Genomik Solidaritas Indonesia Laboratorium | Carissa Sintca Wijaya, Reinhart Gabriel, Latifah Anandari, Vania Gavrila Wikasa, Annisa Muthiah Sukirman, Gracia Felias Enos Korompis, Anuraj Shankar, Meutia Ayuputeri Kumaheri, Ariel Pradipta |
| EPI_ISL_3208080 | Genomik Solidaritas Indonesia Laboratorium | Genomik Solidaritas Indonesia Laboratorium | Reinhart Gabriel, Vania Gavrila Wikasa, Tryna Tania, Carissa Sintca Wijaya, Annisa Muthiah Sukirman, Gracia Felias Enos Korompis, Anuraj Shankar, Meutia Ayuputeri Kumaheri, Ariel Pradipta |
| EPI_ISL_3208081 | KIMIA FARMA / DINAS KESEHATAN DKI JAKARTA | Genomik Solidaritas Indonesia Laboratorium | Reinhart Gabriel, Vania Gavrila Wikasa, Latifah Anandari, Carissa Sintca Wijaya, Annisa Muthiah Sukirman, Gracia Felias Enos Korompis, Anuraj Shankar, Meutia Ayuputeri Kumaheri, Ariel Pradipta |
| EPI_ISL_3208082, EPI_ISL_3208083 | RSUD KOJA / DINAS KESEHATAN DKI JAKARTA | Genomik Solidaritas Indonesia Laboratorium | Carissa Sintca Wijaya, Reinhart Gabriel,Tryna Tania, Vania Gavrila Wikasa, Annisa Muthiah Sukirman, Gracia Felias Enos Korompis, Anuraj Shankar, Meutia Ayuputeri Kumaheri, Ariel Pradipta |
| EPI_ISL_3208084 | Genomik Solidaritas Indonesia Laboratorium | Genomik Solidaritas Indonesia Laboratorium | Carissa Sintca Wijaya, Reinhart Gabriel,Tryna Tania, Vania Gavrila Wikasa, Annisa Muthiah Sukirman, Gracia Felias Enos Korompis, Anuraj Shankar, Meutia Ayuputeri Kumaheri, Ariel Pradipta |
| EPI_ISL_3208085 | Genomik Solidaritas Indonesia Laboratorium | Genomik Solidaritas Indonesia Laboratorium | Reinhart Gabriel, Vania Gavrila Wikasa, Tryna Tania, Carissa Sintca Wijaya, Annisa Muthiah Sukirman, Gracia Felias Enos Korompis, Anuraj Shankar, Meutia Ayuputeri Kumaheri, Ariel Pradipta |
| EPI_ISL_3208086 | Murni Teguh Memorial Hospital | Genomik Solidaritas Indonesia Laboratorium | Reinhart Gabriel, Vania Gavrila Wikasa, Inke Nadia Diniyanti Lubis, Ririe Fachrina Malisie, Lia Kusumawati, Tryna Tania, Mutiara, Carissa Sintca Wijaya, Annisa Muthiah Sukirman, Gracia Felias Enos Korompis, Anuraj Shankar, Meutia Ayuputeri Kumaheri, Ariel Pradipta |
| EPI_ISL_3208087, EPI_ISL_3208088, EPI_ISL_3208089, EPI_ISL_3208090, EPI_ISL_3208091, EPI_ISL_3208092 | Genomik Solidaritas Indonesia Laboratorium | Genomik Solidaritas Indonesia Laboratorium | Vania Gavrila Wikasa, Carissa Sintca Wijaya, Tryna Tania, Reinhart Gabriel, Annisa Muthiah Sukirman, Gracia Felias Enos Korompis, Anuraj Shankar, Meutia Ayuputeri Kumaheri, Ariel Pradipta |
| EPI_ISL_3208093, EPI_ISL_3208094 | Genomik Solidaritas Indonesia Laboratorium | Genomik Solidaritas Indonesia Laboratorium | Carissa Sintca Wijaya, Reinhart Gabriel,Tryna Tania, Vania Gavrila Wikasa, Annisa Muthiah Sukirman, Gracia Felias Enos Korompis, Anuraj Shankar, Meutia Ayuputeri Kumaheri, Ariel Pradipta |
| EPI_ISL_3208095, EPI_ISL_3208096, EPI_ISL_3208097 | Genomik Solidaritas Indonesia Laboratorium | Genomik Solidaritas Indonesia Laboratorium | Reinhart Gabriel, Vania Gavrila Wikasa, Tryna Tania, Carissa Sintca Wijaya, Annisa Muthiah Sukirman, Gracia Felias Enos Korompis, Anuraj Shankar, Meutia Ayuputeri Kumaheri, Ariel Pradipta |
| EPI_ISL_3208098, EPI_ISL_3208099 | Genomik Solidaritas Indonesia Laboratorium | Genomik Solidaritas Indonesia Laboratorium | Carissa Sintca Wijaya, Reinhart Gabriel,Tryna Tania, Vania Gavrila Wikasa, Annisa Muthiah Sukirman, Gracia Felias Enos Korompis, Anuraj Shankar, Meutia Ayuputeri Kumaheri, Ariel Pradipta |
| EPI_ISL_3208100, EPI_ISL_3208101, EPI_ISL_3208102, EPI_ISL_3208103, EPI_ISL_3208104, EPI_ISL_3208105 | Genomik Solidaritas Indonesia Laboratorium | Genomik Solidaritas Indonesia Laboratorium | Vania Gavrila Wikasa, Reinhart Gabriel, Gracia Felias Enos Korompis, Annisa Muthiah Sukirman, Anuraj Shankar, Meutia Ayuputeri Kumaheri, Ariel Pradipta |
| EPI_ISL_3208106, EPI_ISL_3208107, EPI_ISL_3208108, EPI_ISL_3208109, EPI_ISL_3208110 | Genomik Solidaritas Indonesia Laboratorium | Genomik Solidaritas Indonesia Laboratorium | Reinhart Gabriel, Vania Gavrila Wikasa, Gracia Felias Enos Korompis, Annisa Muthiah Sukirman, Anuraj Shankar, Meutia Ayuputeri Kumaheri, Ariel Pradipta |
| EPI_ISL_3208111 | Genomik Solidaritas Indonesia Laboratorium | Genomik Solidaritas Indonesia Laboratorium | Reinhart Gabriel, Vania Gavrila Wikasa, Carissa Sintca Wijaya, Annisa Muthiah Sukirman, Gracia Felias Enos Korompis, Anuraj Shankar, Meutia Ayuputeri Kumaheri, Ariel Pradipta |
| EPI_ISL_3208112 | Genomik Solidaritas Indonesia Laboratorium | Genomik Solidaritas Indonesia Laboratorium | Vania Gavrila Wikasa, Reinhart Gabriel, Carissa Sintca Wijaya, Annisa Muthiah Sukirman, Gracia Felias Enos Korompis, Anuraj Shankar, Meutia Ayuputeri Kumaheri, Ariel Pradipta |
| EPI_ISL_3208113 | Genomik Solidaritas Indonesia Laboratorium | Genomik Solidaritas Indonesia Laboratorium | Reinhart Gabriel, Vania Gavrila Wikasa, Carissa Sintca Wijaya, Annisa Muthiah Sukirman, Gracia Felias Enos Korompis, Anuraj Shankar, Meutia Ayuputeri Kumaheri, Ariel Pradipta |
| EPI_ISL_3208133 | Department for Virology, Molecular Biology and Genome Research, R. G. Lugar Center for Public Health Research, National Center for Disease Control and Public Health (NCDC) of Georgia. | Department for Virology, Molecular Biology and Genome Research, R. G. Lugar Center for Public Health Research, National Center for Disease Control and Public Health (NCDC) of Georgia. | Meri Pantsulaia, Gvantsa Brachveli, Giorgi Tomashvili, Salome Javashvili, Giorgi Gogoladze, Nino Berishvili, Tata Imnadze, Ana Papkiauri, Gvantsa Chanturia, Ann Machablishvili, Nato Kotaria, Marine Murtskhvaladze, Lela Sabadze, Mari Gavashelidze, Tamar Jashiashvili, Tea Tevdoradze, Ketevan Sidamonidze, Ekaterine Khmaladze, Ekaterine Zhghenti, Roena Sukhiashvili, Mariam Zakalashvili, Lela Urushadze, Magda Dgebuadze, Davit Tsaguria, Ekaterine Zangaladze, Adam Kotorashvili, Maia Alkhazashvili, Irma Burjanadze, Anna Kasradze, Khatuna Zakhashvili, Paata Imnadze, Amiran Gamkrelidze. |
| EPI_ISL_3208138, EPI_ISL_3208139, EPI_ISL_3208140, EPI_ISL_3208141, EPI_ISL_3208142, EPI_ISL_3208143, EPI_ISL_3208144, EPI_ISL_3208145, EPI_ISL_3208146, EPI_ISL_3208147, EPI_ISL_3208148, EPI_ISL_3208149, EPI_ISL_3208150, EPI_ISL_3208151, EPI_ISL_3208152, EPI_ISL_3208154, EPI_ISL_3208155, EPI_ISL_3208158, EPI_ISL_3208159, EPI_ISL_3208160, EPI_ISL_3208161, EPI_ISL_3208162, EPI_ISL_3208163, EPI_ISL_3208164, EPI_ISL_3208166, EPI_ISL_3208167, EPI_ISL_3208168, EPI_ISL_3208169, EPI_ISL_3208170, EPI_ISL_3208172, EPI_ISL_3208173, EPI_ISL_3208174, EPI_ISL_3208175, EPI_ISL_3208176, EPI_ISL_3208177, EPI_ISL_3208178, EPI_ISL_3208179, EPI_ISL_3208180, EPI_ISL_3208181, EPI_ISL_3208182, EPI_ISL_3208183, EPI_ISL_3208184, EPI_ISL_3208185, EPI_ISL_3208186, EPI_ISL_3208187, EPI_ISL_3208188, EPI_ISL_3208189, EPI_ISL_3208190, EPI_ISL_3208191, EPI_ISL_3208192, EPI_ISL_3208193, EPI_ISL_3208194, EPI_ISL_3208195, EPI_ISL_3208197, EPI_ISL_3208198, EPI_ISL_3208199, EPI_ISL_3208200, EPI_ISL_3208201, EPI_ISL_3208202, EPI_ISL_3208203, EPI_ISL_3208205, EPI_ISL_3208206, EPI_ISL_3208207, EPI_ISL_3208208, EPI_ISL_3208209, EPI_ISL_3208210, EPI_ISL_3208211, EPI_ISL_3208212, EPI_ISL_3208214, EPI_ISL_3208215, EPI_ISL_3208216, EPI_ISL_3208217, EPI_ISL_3208218, EPI_ISL_3208219, EPI_ISL_3208220, EPI_ISL_3208221 | | | |
| see above | Fulgent Genetics | Centers for Disease Control and Prevention Division of Viral Diseases, Pathogen Discovery | Dakota Howard, Dhwani Batra, Peter W. Cook, Kara Moser, Adrian Paskey, Jason Caravas, Benjamin Rambo-Martin, Shatavia Morrison, Christopher Gulvick, Scott Sammons, Yvette Unoarumhi, Darlene Wagner, Matthew Schmerer, Harry Gao, Mickey Li, John Gao, Joseph Fierro, Benafsh Sapra, Becky Tsai, Yan Meng, Doreen Ng, James Xie, Clinton R. Paden, Duncan MacCannell |
| EPI_ISL_3208222 | Department for Virology, Molecular Biology and Genome | Department for Virology, Molecular Biology and Genome Research, | Meri Pantsulaia, Gvantsa Brachveli, Giorgi Tomashvili, Salome Javashvili, Giorgi Gogoladze, Nino Berishvili, Tata Imnadze, Ana Papkiauri, Gvantsa Chanturia, Ann Machablishvili, Nato Kotaria, Marine Murtskhvaladze, Lela Sabadze, Mari |

| | | | |
|---|---|---|---|
| | Research, R. G. Lugar Center for Public Health Research, National Center for Disease Control and Public Health (NCDC) of Georgia. | R. G. Lugar Center for Public Health Research, National Center for Disease Control and Public Health (NCDC) of Georgia. | Gavashelidze, Tamar Jashiashvili, Tea Tevdoradze, Ketevan Sidamonidze, Ekaterine Khmaladze, Ekaterine Zhghenti, Roena Sukhiashvili, Mariam Zakalashvili, Lela Urushadze, Magda Dgebuadze, Davit Tsaguria, Ekaterine Zangaladze, Adam Kotorashvili, Maia Alkhazashvili, Irma Burjanadze, Anna Kasradze, Khatuna Zakhashvili, Paata Imnadze, Amiran Gamkrelidze. |
| EPI_ISL_3208223, EPI_ISL_3208224, EPI_ISL_3208225, EPI_ISL_3208226, EPI_ISL_3208227, EPI_ISL_3208228, EPI_ISL_3208229, EPI_ISL_3208231, EPI_ISL_3208232, EPI_ISL_3208233, EPI_ISL_3208234, EPI_ISL_3208235, EPI_ISL_3208236, EPI_ISL_3208237, EPI_ISL_3208238, EPI_ISL_3208239, EPI_ISL_3208240, EPI_ISL_3208241, EPI_ISL_3208242, EPI_ISL_3208244, EPI_ISL_3208245, EPI_ISL_3208246, EPI_ISL_3208247, EPI_ISL_3208248, EPI_ISL_3208250, EPI_ISL_3208251, EPI_ISL_3208252, EPI_ISL_3208253, EPI_ISL_3208254, EPI_ISL_3208255, EPI_ISL_3208257, EPI_ISL_3208258, EPI_ISL_3208259, EPI_ISL_3208260, EPI_ISL_3208261, EPI_ISL_3208263, EPI_ISL_3208264, EPI_ISL_3208265, EPI_ISL_3208267, EPI_ISL_3208268, EPI_ISL_3208270, EPI_ISL_3208271, EPI_ISL_3208272, EPI_ISL_3208273, EPI_ISL_3208275, EPI_ISL_3208276, EPI_ISL_3208277, EPI_ISL_3208278, EPI_ISL_3208279, EPI_ISL_3208281, EPI_ISL_3208282, EPI_ISL_3208283, EPI_ISL_3208288, EPI_ISL_3208289, EPI_ISL_3208291, EPI_ISL_3208292, EPI_ISL_3208293, EPI_ISL_3208295, EPI_ISL_3208296, EPI_ISL_3208298, EPI_ISL_3208300, EPI_ISL_3208301, EPI_ISL_3208302, EPI_ISL_3208303, EPI_ISL_3208305, EPI_ISL_3208306, EPI_ISL_3208307, EPI_ISL_3208308, EPI_ISL_3208309, EPI_ISL_3208311, EPI_ISL_3208312, EPI_ISL_3208313, EPI_ISL_3208314, EPI_ISL_3208315, EPI_ISL_3208316, EPI_ISL_3208317, EPI_ISL_3208318, EPI_ISL_3208319, EPI_ISL_3208321, EPI_ISL_3208322, EPI_ISL_3208323, EPI_ISL_3208324, EPI_ISL_3208325, EPI_ISL_3208326, EPI_ISL_3208327, EPI_ISL_3208328, EPI_ISL_3208329, EPI_ISL_3208330, EPI_ISL_3208331, EPI_ISL_3208334, EPI_ISL_3208335, EPI_ISL_3208336, EPI_ISL_3208337, EPI_ISL_3208338, EPI_ISL_3208339, EPI_ISL_3208340, EPI_ISL_3208342, EPI_ISL_3208343, EPI_ISL_3208344, EPI_ISL_3208346, EPI_ISL_3208347, EPI_ISL_3208350 | | | |
| see above | Fulgent Genetics | Centers for Disease Control and Prevention Division of Viral Diseases, Pathogen Discovery | Dakota Howard, Dhwani Batra, Peter W. Cook, Kara Moser, Adrian Paskey, Jason Caravas, Benjamin Rambo-Martin, Shatavia Morrison, Christopher Gulvick, Scott Sammons, Yvette Unoarumhi, Darlene Wagner, Matthew Schmerer, Harry Gao, Mickey Li, John Gao, Joseph Fierro, Benafsh Sapra, Becky Tsai, Yan Meng, Doreen Ng, James Xie, Clinton R. Paden, Duncan MacCannell |
| EPI_ISL_3208351 | Department for Virology, Molecular Biology and Genome Research, R. G. Lugar Center for Public Health Research, National Center for Disease Control and Public Health (NCDC) of Georgia. | Department for Virology, Molecular Biology and Genome Research, R. G. Lugar Center for Public Health Research, National Center for Disease Control and Public Health (NCDC) of Georgia. | Meri Pantsulaia, Gvantsa Brachveli, Giorgi Tomashvili, Salome Javashvili, Giorgi Gogoladze, Nino Berishvili, Tata Imnadze, Ana Papkiauri, Gvantsa Chanturia, Ann Machablishvili, Nato Kotaria, Marine Murtskhvaladze, Lela Sabadze, Mari Gavashelidze, Tamar Jashiashvili, Tea Tevdoradze, Ketevan Sidamonidze, Ekaterine Khmaladze, Ekaterine Zhghenti, Roena Sukhiashvili, Mariam Zakalashvili, Lela Urushadze, Magda Dgebuadze, Davit Tsaguria, Ekaterine Zangaladze, Adam Kotorashvili, Maia Alkhazashvili, Irma Burjanadze, Anna Kasradze, Khatuna Zakhashvili, Paata Imnadze, Amiran Gamkrelidze. |
| EPI_ISL_3208352, EPI_ISL_3208353, EPI_ISL_3208354, EPI_ISL_3208355, EPI_ISL_3208356, EPI_ISL_3208357, EPI_ISL_3208358, EPI_ISL_3208359, EPI_ISL_3208360, EPI_ISL_3208361, EPI_ISL_3208362, EPI_ISL_3208363, EPI_ISL_3208365, EPI_ISL_3208366, EPI_ISL_3208367, EPI_ISL_3208368, EPI_ISL_3208369, EPI_ISL_3208370, EPI_ISL_3208371, EPI_ISL_3208372, EPI_ISL_3208374, EPI_ISL_3208375, EPI_ISL_3208376, EPI_ISL_3208377, EPI_ISL_3208378, EPI_ISL_3208379, EPI_ISL_3208380, EPI_ISL_3208381, EPI_ISL_3208382, EPI_ISL_3208383, EPI_ISL_3208384, EPI_ISL_3208385, EPI_ISL_3208387, EPI_ISL_3208388, EPI_ISL_3208389, EPI_ISL_3208390, EPI_ISL_3208391, EPI_ISL_3208392, EPI_ISL_3208393, EPI_ISL_3208394, EPI_ISL_3208395, EPI_ISL_3208396, EPI_ISL_3208397, EPI_ISL_3208398, EPI_ISL_3208399, EPI_ISL_3208400, EPI_ISL_3208401, EPI_ISL_3208402, EPI_ISL_3208403, EPI_ISL_3208404, EPI_ISL_3208405, EPI_ISL_3208406, EPI_ISL_3208407, EPI_ISL_3208408, EPI_ISL_3208409, EPI_ISL_3208411, EPI_ISL_3208412, EPI_ISL_3208413, EPI_ISL_3208414, EPI_ISL_3208415, EPI_ISL_3208416, EPI_ISL_3208417, EPI_ISL_3208418, EPI_ISL_3208419, EPI_ISL_3208420, EPI_ISL_3208421, EPI_ISL_3208422, EPI_ISL_3208423, EPI_ISL_3208424, EPI_ISL_3208425, EPI_ISL_3208426, EPI_ISL_3208428, EPI_ISL_3208429, EPI_ISL_3208430, EPI_ISL_3208432, EPI_ISL_3208433, EPI_ISL_3208434, EPI_ISL_3208435, EPI_ISL_3208436, EPI_ISL_3208437, EPI_ISL_3208438, EPI_ISL_3208439, EPI_ISL_3208440, EPI_ISL_3208441, EPI_ISL_3208442, EPI_ISL_3208443, EPI_ISL_3208444 | | | |
| see above | Fulgent Genetics | Centers for Disease Control and Prevention Division of Viral Diseases, Pathogen Discovery | Dakota Howard, Dhwani Batra, Peter W. Cook, Kara Moser, Adrian Paskey, Jason Caravas, Benjamin Rambo-Martin, Shatavia Morrison, Christopher Gulvick, Scott Sammons, Yvette Unoarumhi, Darlene Wagner, Matthew Schmerer, Harry Gao, Mickey Li, John Gao, Joseph Fierro, Benafsh Sapra, Becky Tsai, Yan Meng, Doreen Ng, James Xie, Clinton R. Paden, Duncan MacCannell |
| EPI_ISL_3208445 | Department for Virology, Molecular Biology and Genome Research, R. G. Lugar Center for Public Health Research, National Center for Disease Control and Public Health (NCDC) of Georgia. | Department for Virology, Molecular Biology and Genome Research, R. G. Lugar Center for Public Health Research, National Center for Disease Control and Public Health (NCDC) of Georgia. | Meri Pantsulaia, Gvantsa Brachveli, Giorgi Tomashvili, Salome Javashvili, Giorgi Gogoladze, Nino Berishvili, Tata Imnadze, Ana Papkiauri, Gvantsa Chanturia, Ann Machablishvili, Nato Kotaria, Marine Murtskhvaladze, Lela Sabadze, Mari Gavashelidze, Tamar Jashiashvili, Tea Tevdoradze, Ketevan Sidamonidze, Ekaterine Khmaladze, Ekaterine Zhghenti, Roena Sukhiashvili, Mariam Zakalashvili, Lela Urushadze, Magda Dgebuadze, Davit Tsaguria, Ekaterine Zangaladze, Adam Kotorashvili, Maia Alkhazashvili, Irma Burjanadze, Anna Kasradze, Khatuna Zakhashvili, Paata Imnadze, Amiran Gamkrelidze. |
| EPI_ISL_3208446, EPI_ISL_3208447, EPI_ISL_3208448, EPI_ISL_3208449, EPI_ISL_3208450, EPI_ISL_3208451, EPI_ISL_3208452, EPI_ISL_3208453, EPI_ISL_3208454, EPI_ISL_3208455, EPI_ISL_3208456, EPI_ISL_3208457, EPI_ISL_3208458, EPI_ISL_3208459, EPI_ISL_3208460, EPI_ISL_3208461, EPI_ISL_3208462, EPI_ISL_3208463, EPI_ISL_3208464, EPI_ISL_3208465, EPI_ISL_3208466, EPI_ISL_3208467, EPI_ISL_3208468, EPI_ISL_3208469, EPI_ISL_3208470, EPI_ISL_3208471, EPI_ISL_3208472, EPI_ISL_3208473, EPI_ISL_3208474, EPI_ISL_3208475, EPI_ISL_3208476, EPI_ISL_3208477, EPI_ISL_3208478, EPI_ISL_3208479, EPI_ISL_3208480, EPI_ISL_3208481, EPI_ISL_3208482, EPI_ISL_3208483, EPI_ISL_3208484, EPI_ISL_3208486, EPI_ISL_3208487, EPI_ISL_3208488, EPI_ISL_3208489, EPI_ISL_3208490, EPI_ISL_3208491, EPI_ISL_3208492, EPI_ISL_3208493, EPI_ISL_3208494, EPI_ISL_3208495, EPI_ISL_3208497, EPI_ISL_3208499, EPI_ISL_3208500, EPI_ISL_3208501, EPI_ISL_3208502, EPI_ISL_3208503, EPI_ISL_3208504, EPI_ISL_3208505, EPI_ISL_3208506, EPI_ISL_3208507, EPI_ISL_3208508, EPI_ISL_3208509, EPI_ISL_3208510, EPI_ISL_3208511, EPI_ISL_3208513, EPI_ISL_3208514, EPI_ISL_3208515, EPI_ISL_3208516, EPI_ISL_3208517, EPI_ISL_3208519, EPI_ISL_3208520, EPI_ISL_3208522, EPI_ISL_3208523, EPI_ISL_3208524, EPI_ISL_3208525, EPI_ISL_3208526, EPI_ISL_3208527, EPI_ISL_3208528, EPI_ISL_3208529, EPI_ISL_3208530, EPI_ISL_3208532, EPI_ISL_3208533, EPI_ISL_3208535, EPI_ISL_3208536, EPI_ISL_3208537, EPI_ISL_3208538, EPI_ISL_3208539, EPI_ISL_3208540, EPI_ISL_3208541, EPI_ISL_3208542 | | | |
| see above | Fulgent Genetics | Centers for Disease Control and Prevention Division of Viral Diseases, Pathogen Discovery | Dakota Howard, Dhwani Batra, Peter W. Cook, Kara Moser, Adrian Paskey, Jason Caravas, Benjamin Rambo-Martin, Shatavia Morrison, Christopher Gulvick, Scott Sammons, Yvette Unoarumhi, Darlene Wagner, Matthew Schmerer, Harry Gao, Mickey Li, John Gao, Joseph Fierro, Benafsh Sapra, Becky Tsai, Yan Meng, Doreen Ng, James Xie, Clinton R. Paden, Duncan MacCannell |
| EPI_ISL_3208543 | Department for Virology, Molecular Biology and Genome Research, R. G. Lugar Center for Public Health Research, National Center for Disease Control and Public Health (NCDC) of Georgia. | Department for Virology, Molecular Biology and Genome Research, R. G. Lugar Center for Public Health Research, National Center for Disease Control and Public Health (NCDC) of Georgia. | Meri Pantsulaia, Gvantsa Brachveli, Giorgi Tomashvili, Salome Javashvili, Giorgi Gogoladze, Nino Berishvili, Tata Imnadze, Ana Papkiauri, Gvantsa Chanturia, Ann Machablishvili, Nato Kotaria, Marine Murtskhvaladze, Lela Sabadze, Mari Gavashelidze, Tamar Jashiashvili, Tea Tevdoradze, Ketevan Sidamonidze, Ekaterine Khmaladze, Ekaterine Zhghenti, Roena Sukhiashvili, Mariam Zakalashvili, Lela Urushadze, Magda Dgebuadze, Davit Tsaguria, Ekaterine Zangaladze, Adam Kotorashvili, Maia Alkhazashvili, Irma Burjanadze, Anna Kasradze, Khatuna Zakhashvili, Paata Imnadze, Amiran Gamkrelidze. |
| EPI_ISL_3208544, EPI_ISL_3208545, EPI_ISL_3208546, EPI_ISL_3208547, EPI_ISL_3208548, EPI_ISL_3208549, EPI_ISL_3208550, EPI_ISL_3208551, EPI_ISL_3208552, EPI_ISL_3208553, EPI_ISL_3208554, EPI_ISL_3208555, EPI_ISL_3208556, EPI_ISL_3208557, EPI_ISL_3208558, EPI_ISL_3208559, EPI_ISL_3208560, EPI_ISL_3208561, EPI_ISL_3208562, EPI_ISL_3208563, EPI_ISL_3208564, EPI_ISL_3208565, EPI_ISL_3208566, EPI_ISL_3208567, EPI_ISL_3208568, EPI_ISL_3208569, EPI_ISL_3208570, EPI_ISL_3208571, EPI_ISL_3208572, EPI_ISL_3208573, EPI_ISL_3208575, EPI_ISL_3208578, EPI_ISL_3208579, EPI_ISL_3208580, EPI_ISL_3208581, EPI_ISL_3208582, EPI_ISL_3208583, EPI_ISL_3208584, EPI_ISL_3208585, EPI_ISL_3208586, EPI_ISL_3208587, EPI_ISL_3208589, EPI_ISL_3208590, EPI_ISL_3208591, EPI_ISL_3208592, EPI_ISL_3208593, EPI_ISL_3208594, EPI_ISL_3208595, EPI_ISL_3208596, EPI_ISL_3208597, EPI_ISL_3208598, EPI_ISL_3208599, EPI_ISL_3208600, EPI_ISL_3208601, EPI_ISL_3208602, EPI_ISL_3208603, EPI_ISL_3208606, EPI_ISL_3208607, EPI_ISL_3208608, EPI_ISL_3208609, EPI_ISL_3208611, EPI_ISL_3208612, EPI_ISL_3208613, EPI_ISL_3208614, EPI_ISL_3208615, EPI_ISL_3208616, EPI_ISL_3208617, EPI_ISL_3208619, EPI_ISL_3208620, EPI_ISL_3208621, EPI_ISL_3208622, EPI_ISL_3208623, EPI_ISL_3208625, EPI_ISL_3208626, EPI_ISL_3208627, EPI_ISL_3208628, EPI_ISL_3208629, EPI_ISL_3208630, EPI_ISL_3208631, EPI_ISL_3208632, EPI_ISL_3208633, EPI_ISL_3208634, EPI_ISL_3208636, EPI_ISL_3208637, EPI_ISL_3208638, EPI_ISL_3208639, EPI_ISL_3208640, EPI_ISL_3208642, EPI_ISL_3208644, EPI_ISL_3208646, EPI_ISL_3208647, EPI_ISL_3208648, EPI_ISL_3208649, EPI_ISL_3208650, EPI_ISL_3208651, EPI_ISL_3208652, EPI_ISL_3208653, EPI_ISL_3208654, EPI_ISL_3208655, EPI_ISL_3208656, EPI_ISL_3208657, EPI_ISL_3208658, EPI_ISL_3208660, EPI_ISL_3208661, EPI_ISL_3208662, EPI_ISL_3208663, EPI_ISL_3208664, EPI_ISL_3208665, EPI_ISL_3208668, EPI_ISL_3208669, EPI_ISL_3208670, EPI_ISL_3208671, EPI_ISL_3208672, EPI_ISL_3208673, EPI_ISL_3208674, EPI_ISL_3208675, EPI_ISL_3208676, EPI_ISL_3208677, EPI_ISL_3208678, EPI_ISL_3208679, EPI_ISL_3208680, EPI_ISL_3208681, EPI_ISL_3208682, EPI_ISL_3208683, EPI_ISL_3208684, EPI_ISL_3208686, EPI_ISL_3208687, EPI_ISL_3208688, EPI_ISL_3208690, EPI_ISL_3208691, EPI_ISL_3208692, EPI_ISL_3208693, EPI_ISL_3208694, EPI_ISL_3208695, EPI_ISL_3208696, EPI_ISL_3208697, EPI_ISL_3208698, EPI_ISL_3208699, EPI_ISL_3208700, EPI_ISL_3208701, EPI_ISL_3208702, EPI_ISL_3208703, EPI_ISL_3208704, EPI_ISL_3208705, EPI_ISL_3208707, EPI_ISL_3208708, EPI_ISL_3208709, EPI_ISL_3208710, EPI_ISL_3208711, EPI_ISL_3208712, EPI_ISL_3208713, EPI_ISL_3208714, EPI_ISL_3208715, EPI_ISL_3208716, EPI_ISL_3208717, EPI_ISL_3208720, EPI_ISL_3208721, EPI_ISL_3208722, EPI_ISL_3208723, EPI_ISL_3208724, EPI_ISL_3208725, EPI_ISL_3208726, EPI_ISL_3208727, EPI_ISL_3208728, EPI_ISL_3208729, EPI_ISL_3208730, EPI_ISL_3208731, EPI_ISL_3208732, EPI_ISL_3208733, EPI_ISL_3208734, EPI_ISL_3208735, EPI_ISL_3208736, EPI_ISL_3208737, EPI_ISL_3208738, EPI_ISL_3208739, EPI_ISL_3208740, EPI_ISL_3208741, EPI_ISL_3208742, EPI_ISL_3208744, EPI_ISL_3208745, EPI_ISL_3208746, EPI_ISL_3208747, EPI_ISL_3208748, EPI_ISL_3208749, EPI_ISL_3208750, EPI_ISL_3208751, EPI_ISL_3208752, EPI_ISL_3208753, EPI_ISL_3208754, EPI_ISL_3208755, EPI_ISL_3208756, EPI_ISL_3208757, EPI_ISL_3208758, EPI_ISL_3208759, EPI_ISL_3208760, EPI_ISL_3208761, EPI_ISL_3208763, EPI_ISL_3208764, EPI_ISL_3208765, EPI_ISL_3208766, EPI_ISL_3208767, EPI_ISL_3208768, EPI_ISL_3208769, EPI_ISL_3208770, EPI_ISL_3208771, EPI_ISL_3208772, EPI_ISL_3208773, EPI_ISL_3208774, EPI_ISL_3208775, EPI_ISL_3208776, EPI_ISL_3208777, EPI_ISL_3208778, EPI_ISL_3208779, EPI_ISL_3208780, EPI_ISL_3208782, EPI_ISL_3208783, EPI_ISL_3208784, EPI_ISL_3208785, EPI_ISL_3208786, EPI_ISL_3208787, EPI_ISL_3208788, EPI_ISL_3208789, EPI_ISL_3208790, EPI_ISL_3208791, EPI_ISL_3208792, EPI_ISL_3208793, EPI_ISL_3208794, EPI_ISL_3208795, EPI_ISL_3208796, EPI_ISL_3208797, EPI_ISL_3208798, EPI_ISL_3208799, EPI_ISL_3208800, EPI_ISL_3208801, EPI_ISL_3208802, EPI_ISL_3208803, EPI_ISL_3208804, EPI_ISL_3208805, EPI_ISL_3208807, EPI_ISL_3208808, EPI_ISL_3208810, EPI_ISL_3208811, EPI_ISL_3208812, EPI_ISL_3208813, EPI_ISL_3208814, EPI_ISL_3208816, EPI_ISL_3208817, EPI_ISL_3208818, EPI_ISL_3208820, EPI_ISL_3208821, EPI_ISL_3208822, EPI_ISL_3208823, EPI_ISL_3208824, EPI_ISL_3208825, EPI_ISL_3208826, EPI_ISL_3208828, EPI_ISL_3208829, EPI_ISL_3208830, EPI_ISL_3208831, EPI_ISL_3208832, EPI_ISL_3208833, EPI_ISL_3208835, EPI_ISL_3208836, EPI_ISL_3208838, EPI_ISL_3208842, EPI_ISL_3208843, EPI_ISL_3208846, EPI_ISL_3208847, EPI_ISL_3208848, EPI_ISL_3208849, EPI_ISL_3208850, EPI_ISL_3208851, EPI_ISL_3208853, EPI_ISL_3208854, EPI_ISL_3208855, EPI_ISL_3208856, EPI_ISL_3208859 | | | |
| see above | Fulgent Genetics | Centers for Disease Control and Prevention Division of Viral Diseases, Pathogen Discovery | Dakota Howard, Dhwani Batra, Peter W. Cook, Kara Moser, Adrian Paskey, Jason Caravas, Benjamin Rambo-Martin, Shatavia Morrison, Christopher Gulvick, Scott Sammons, Yvette Unoarumhi, Darlene Wagner, Matthew Schmerer, Harry Gao, Mickey Li, John Gao, Joseph Fierro, Benafsh Sapra, Becky Tsai, Yan Meng, Doreen Ng, James Xie, Clinton R. Paden, Duncan MacCannell |
| EPI_ISL_3208860 | Department for Virology, Molecular Biology and Genome Research, R. G. Lugar Center for Public Health Research, National Center for Disease Control and Public Health (NCDC) of Georgia. | Department for Virology, Molecular Biology and Genome Research, R. G. Lugar Center for Public Health Research, National Center for Disease Control and Public Health (NCDC) of Georgia. | Giorgi Gogoladze, Meri Pantsulaia, Gvantsa Brachveli, Giorgi Tomashvili, Salome Javashvili, Nino Berishvili, Tata Imnadze, Ana Papkiauri, Gvantsa Chanturia, Ann Machablishvili, Nato Kotaria, Marine Murtskhvaladze, Lela Sabadze, Mari Gavashelidze, Tamar Jashiashvili, Tea Tevdoradze, Ketevan Sidamonidze, Ekaterine Khmaladze, Ekaterine Zhghenti, Roena Sukhiashvili, Mariam Zakalashvili, Lela Urushadze, Magda Dgebuadze, Davit Tsaguria, Ekaterine Zangaladze, Adam Kotorashvili, Maia Alkhazashvili, Irma Burjanadze, Anna Kasradze, Khatuna Zakhashvili, Paata Imnadze, Amiran Gamkrelidze. |
| EPI_ISL_3208861, EPI_ISL_3208862, EPI_ISL_3208863, EPI_ISL_3208864, EPI_ISL_3208865, EPI_ISL_3208866, EPI_ISL_3208867, EPI_ISL_3208868, EPI_ISL_3208869, EPI_ISL_3208870, EPI_ISL_3208871, EPI_ISL_3208872, EPI_ISL_3208873, EPI_ISL_3208875, EPI_ISL_3208876, EPI_ISL_3208877, EPI_ISL_3208878, EPI_ISL_3208879, EPI_ISL_3208880, EPI_ISL_3208881, EPI_ISL_3208882, EPI_ISL_3208883, EPI_ISL_3208884, EPI_ISL_3208885, EPI_ISL_3208886, EPI_ISL_3208887, EPI_ISL_3208888, EPI_ISL_3208889, EPI_ISL_3208890, EPI_ISL_3208891, EPI_ISL_3208892, EPI_ISL_3208893 | | | |
| see above | Fulgent Genetics | Centers for Disease Control and Prevention Division of Viral Diseases, Pathogen Discovery | Dakota Howard, Dhwani Batra, Peter W. Cook, Kara Moser, Adrian Paskey, Jason Caravas, Benjamin Rambo-Martin, Shatavia Morrison, Christopher Gulvick, Scott Sammons, Yvette Unoarumhi, Darlene Wagner, Matthew Schmerer, Harry Gao, Mickey Li, John Gao, Joseph Fierro, Benafsh Sapra, Becky Tsai, Yan Meng, Doreen Ng, James Xie, Clinton R. Paden, Duncan MacCannell |
| EPI_ISL_3208894 | Center of Medical Microbiology, Virology, and Hospital Hygiene, University of Duesseldorf | Center of Medical Microbiology, Virology, and Hospital Hygiene, University of Duesseldorf | Maximilian Damagnez;Alexander Dilthey;Torsten Houwaart;Lisanna Hülse;Malte Kohns Vasconcelos;Nadine Lübke;Jessica Nicolai;Klaus Pfeffer;Daniel Strelow;Jörg Timm;Andreas Walker;Tobias Wienemann |
| EPI_ISL_3208895 | ZOTZ KLIMAS MVZ Düsseldorf-Centrum GbR ÜBAG für Labormedizin, Genetik, Zytologie, Pathologie | Center of Medical Microbiology, Virology, and Hospital Hygiene, University of Duesseldorf | Maximilian Damagnez;Alexander Dilthey;Patrick Finzer;Katrin Hoffmann;Torsten Houwaart;Lisanna Hülse;Malte Kohns Vasconcelos;Nadine Lübke;Jessica Nicolai;Klaus Pfeffer;Daniel Strelow;Jörg Timm;Andreas Walker;Tobias Wienemann;Rainer Zotz |
| EPI_ISL_3208896 | Fulgent Genetics | Centers for Disease Control and Prevention Division of Viral Diseases, Pathogen Discovery | Dakota Howard, Dhwani Batra, Peter W. Cook, Kara Moser, Adrian Paskey, Jason Caravas, Benjamin Rambo-Martin, Shatavia Morrison, Christopher Gulvick, Scott Sammons, Yvette Unoarumhi, Darlene Wagner, Matthew Schmerer, Harry Gao, Mickey Li, John Gao, Joseph Fierro, Benafsh Sapra, Becky Tsai, Yan Meng, Doreen Ng, James Xie, Clinton R. Paden, Duncan MacCannell |
| EPI_ISL_3208897, EPI_ISL_3208898 | ZOTZ KLIMAS MVZ Düsseldorf-Centrum GbR ÜBAG für Labormedizin, Genetik, Zytologie, Pathologie | Center of Medical Microbiology, Virology, and Hospital Hygiene, University of Duesseldorf | Maximilian Damagnez;Alexander Dilthey;Patrick Finzer;Katrin Hoffmann;Torsten Houwaart;Lisanna Hülse;Malte Kohns Vasconcelos;Nadine Lübke;Jessica Nicolai;Klaus Pfeffer;Daniel Strelow;Jörg Timm;Andreas Walker;Tobias Wienemann;Rainer Zotz |
| EPI_ISL_3208899 | Fulgent Genetics | Centers for Disease Control and Prevention Division of Viral Diseases, Pathogen Discovery | Dakota Howard, Dhwani Batra, Peter W. Cook, Kara Moser, Adrian Paskey, Jason Caravas, Benjamin Rambo-Martin, Shatavia Morrison, Christopher Gulvick, Scott Sammons, Yvette Unoarumhi, Darlene Wagner, Matthew Schmerer, Harry Gao, Mickey Li, John Gao, Joseph Fierro, Benafsh Sapra, Becky Tsai, Yan Meng, Doreen Ng, James Xie, Clinton R. Paden, Duncan MacCannell |
| EPI_ISL_3208900, EPI_ISL_3208901, EPI_ISL_3208903, EPI_ISL_3208905 | ZOTZ KLIMAS MVZ Düsseldorf-Centrum GbR ÜBAG für Labormedizin, Genetik, Zytologie, Pathologie | Center of Medical Microbiology, Virology, and Hospital Hygiene, University of Duesseldorf | Maximilian Damagnez;Alexander Dilthey;Patrick Finzer;Katrin Hoffmann;Torsten Houwaart;Lisanna Hülse;Malte Kohns Vasconcelos;Nadine Lübke;Jessica Nicolai;Klaus Pfeffer;Daniel Strelow;Jörg Timm;Andreas Walker;Tobias Wienemann;Rainer Zotz |
| EPI_ISL_3208906, EPI_ISL_3208907, EPI_ISL_3208908, EPI_ISL_3208909, EPI_ISL_3208910, EPI_ISL_3208911, EPI_ISL_3208912, EPI_ISL_3208913, EPI_ISL_3208914, EPI_ISL_3208915, EPI_ISL_3208916, EPI_ISL_3208917, EPI_ISL_3208918, EPI_ISL_3208919, EPI_ISL_3208920, EPI_ISL_3208921, EPI_ISL_3208922, EPI_ISL_3208923, EPI_ISL_3208924, EPI_ISL_3208925, EPI_ISL_3208926, EPI_ISL_3208927, EPI_ISL_3208928, EPI_ISL_3208930, EPI_ISL_3208931, EPI_ISL_3208932, EPI_ISL_3208933, EPI_ISL_3208935, EPI_ISL_3208937, EPI_ISL_3208938, EPI_ISL_3208939, EPI_ISL_3208940, EPI_ISL_3208941, EPI_ISL_3208943, EPI_ISL_3208945, EPI_ISL_3208946, EPI_ISL_3208947, EPI_ISL_3208948, EPI_ISL_3208949, EPI_ISL_3208950, EPI_ISL_3208951, EPI_ISL_3208952, EPI_ISL_3208953, EPI_ISL_3208954, EPI_ISL_3208955, EPI_ISL_3208956, EPI_ISL_3208957, EPI_ISL_3208958, EPI_ISL_3208959, EPI_ISL_3208960, EPI_ISL_3208961, EPI_ISL_3208962, EPI_ISL_3208963, EPI_ISL_3208964, EPI_ISL_3208965, EPI_ISL_3208966, EPI_ISL_3208967, EPI_ISL_3208968, EPI_ISL_3208969, EPI_ISL_3208970, EPI_ISL_3208971, EPI_ISL_3208973, EPI_ISL_3208974, EPI_ISL_3208975, EPI_ISL_3208976, EPI_ISL_3208977, EPI_ISL_3208978, EPI_ISL_3208979, EPI_ISL_3208980, EPI_ISL_3208981, EPI_ISL_3208982, EPI_ISL_3208983, EPI_ISL_3208985, EPI_ISL_3208986, EPI_ISL_3208988, EPI_ISL_3208989, EPI_ISL_3208990, EPI_ISL_3208991, EPI_ISL_3208992, EPI_ISL_3208993, EPI_ISL_3208994, EPI_ISL_3208995, EPI_ISL_3208996, EPI_ISL_3208997, EPI_ISL_3208998, EPI_ISL_3208999, EPI_ISL_3209000, EPI_ISL_3209001, EPI_ISL_3209002, EPI_ISL_3209003, EPI_ISL_3209004, EPI_ISL_3209005, EPI_ISL_3209006, EPI_ISL_3209008, EPI_ISL_3209009, EPI_ISL_3209010, EPI_ISL_3209011, EPI_ISL_3209013, EPI_ISL_3209015, EPI_ISL_3209017, EPI_ISL_3209018, EPI_ISL_3209019, EPI_ISL_3209020, EPI_ISL_3209022, EPI_ISL_3209023, EPI_ISL_3209024, EPI_ISL_3209025, EPI_ISL_3209026, EPI_ISL_3209027, EPI_ISL_3209028, EPI_ISL_3209029, EPI_ISL_3209030, EPI_ISL_3209031, EPI_ISL_3209032, EPI_ISL_3209033, EPI_ISL_3209034, EPI_ISL_3209035, EPI_ISL_3209036, EPI_ISL_3209037, EPI_ISL_3209038, EPI_ISL_3209039, EPI_ISL_3209040, EPI_ISL_3209042, EPI_ISL_3209043, EPI_ISL_3209044, EPI_ISL_3209045, EPI_ISL_3209046, EPI_ISL_3209047, EPI_ISL_3209048, EPI_ISL_3209049, EPI_ISL_3209050, EPI_ISL_3209051, EPI_ISL_3209052, EPI_ISL_3209053, EPI_ISL_3209054, EPI_ISL_3209055, EPI_ISL_3209056, EPI_ISL_3209057, EPI_ISL_3209058, EPI_ISL_3209059, EPI_ISL_3209060, EPI_ISL_3209061, EPI_ISL_3209062, EPI_ISL_3209063, EPI_ISL_3209064, EPI_ISL_3209065, EPI_ISL_3209066, EPI_ISL_3209067, EPI_ISL_3209068, EPI_ISL_3209069, EPI_ISL_3209070, EPI_ISL_3209071, EPI_ISL_3209072, EPI_ISL_3209073, EPI_ISL_3209074, EPI_ISL_3209075, EPI_ISL_3209076, EPI_ISL_3209077, EPI_ISL_3209078, EPI_ISL_3209080, EPI_ISL_3209082, EPI_ISL_3209083, EPI_ISL_3209084, EPI_ISL_3209085, EPI_ISL_3209086, EPI_ISL_3209087, EPI_ISL_3209088, EPI_ISL_3209089, EPI_ISL_3209090, EPI_ISL_3209091, EPI_ISL_3209093, EPI_ISL_3209095, EPI_ISL_3209096, EPI_ISL_3209097, EPI_ISL_3209098, EPI_ISL_3209099, EPI_ISL_3209100, EPI_ISL_3209101, EPI_ISL_3209102, EPI_ISL_3209103, EPI_ISL_3209104, EPI_ISL_3209105, EPI_ISL_3209106, EPI_ISL_3209107, EPI_ISL_3209108, EPI_ISL_3209109, EPI_ISL_3209110, EPI_ISL_3209111, EPI_ISL_3209112, EPI_ISL_3209113, EPI_ISL_3209114, EPI_ISL_3209115, EPI_ISL_3209116, EPI_ISL_3209117, EPI_ISL_3209119, EPI_ISL_3209123, EPI_ISL_3209129, EPI_ISL_3209132, EPI_ISL_3209135, EPI_ISL_3209139, EPI_ISL_3209145, EPI_ISL_3209148, EPI_ISL_3209151, EPI_ISL_3209155, EPI_ISL_3209158, EPI_ISL_3209162, EPI_ISL_3209166, EPI_ISL_3209170, EPI_ISL_3209173, EPI_ISL_3209176, EPI_ISL_3209180, EPI_ISL_3209183, EPI_ISL_3209186, EPI_ISL_3209190, EPI_ISL_3209192, EPI_ISL_3209195, EPI_ISL_3209198, EPI_ISL_3209202, EPI_ISL_3209205, EPI_ISL_3209209, EPI_ISL_3209212, EPI_ISL_3209215, EPI_ISL_3209218, EPI_ISL_3209221, EPI_ISL_3209225, EPI_ISL_3209228, EPI_ISL_3209231, EPI_ISL_3209235, EPI_ISL_3209238, EPI_ISL_3209242, EPI_ISL_3209244, EPI_ISL_3209248, EPI_ISL_3209250, EPI_ISL_3209253, EPI_ISL_3209257, EPI_ISL_3209260, EPI_ISL_3209263, EPI_ISL_3209266, EPI_ISL_3209269, EPI_ISL_3209272, EPI_ISL_3209274, EPI_ISL_3209277, EPI_ISL_3209281, EPI_ISL_3209284, EPI_ISL_3209286, EPI_ISL_3209290, EPI_ISL_3209294, EPI_ISL_3209297, EPI_ISL_3209301, EPI_ISL_3209304, EPI_ISL_3209307, EPI_ISL_3209311, EPI_ISL_3209315, EPI_ISL_3209318, EPI_ISL_3209321, EPI_ISL_3209325, EPI_ISL_3209328, EPI_ISL_3209332, EPI_ISL_3209335, EPI_ISL_3209339, EPI_ISL_3209344, EPI_ISL_3209347, EPI_ISL_3209351, EPI_ISL_3209355, EPI_ISL_3209359, EPI_ISL_3209363, EPI_ISL_3209367, EPI_ISL_3209370, EPI_ISL_3209373, EPI_ISL_3209376, EPI_ISL_3209380, EPI_ISL_3209384, EPI_ISL_3209387, EPI_ISL_3209390, EPI_ISL_3209394, EPI_ISL_3209398, EPI_ISL_3209401, EPI_ISL_3209405, EPI_ISL_3209408, EPI_ISL_3209412, EPI_ISL_3209415, EPI_ISL_3209422, EPI_ISL_3209425, EPI_ISL_3209428, EPI_ISL_3209432, EPI_ISL_3209434, EPI_ISL_3209437, EPI_ISL_3209441, EPI_ISL_3209444, EPI_ISL_3209446, EPI_ISL_3209450, EPI_ISL_3209455, EPI_ISL_3209459, EPI_ISL_3209462, EPI_ISL_3209465, EPI_ISL_3209473, EPI_ISL_3209476 | | | |
| see above | Fulgent Genetics | Centers for Disease Control and Prevention Division of Viral Diseases, Pathogen Discovery | Dakota Howard, Dhwani Batra, Peter W. Cook, Kara Moser, Adrian Paskey, Jason Caravas, Benjamin Rambo-Martin, Shatavia Morrison, Christopher Gulvick, Scott Sammons, Yvette Unoarumhi, Darlene Wagner, Matthew Schmerer, Harry Gao, Mickey Li, John Gao, Joseph Fierro, Benafsh Sapra, Becky Tsai, Yan Meng, Doreen Ng, James Xie, Clinton R. Paden, Duncan MacCannell |
| EPI_ISL_3209478 | Department for Virology, Molecular Biology and Genome Research, R. G. Lugar Center for Public Health Research, National Center for Disease Control and Public Health (NCDC) of Georgia. | Department for Virology, Molecular Biology and Genome Research, R. G. Lugar Center for Public Health Research, National Center for Disease Control and Public Health (NCDC) of Georgia. | Giorgi Gogoladze, Meri Pantsulaia, Gvantsa Brachveli, Giorgi Tomashvili, Salome Javashvili, Nino Berishvili, Tata Imnadze, Ana Papkiauri, Gvantsa Chanturia, Ann Machablishvili, Nato Kotaria, Marine Murtskhvaladze, Lela Sabadze, Mari Gavashelidze, Tamar Jashiashvili, Tea Tevdoradze, Ketevan Sidamonidze, Ekaterine Khmaladze, Ekaterine Zhghenti, Roena Sukhiashvili, Mariam Zakalashvili, Lela Urushadze, Magda Dgebuadze, Davit Tsaguria, Ekaterine Zangaladze, Adam Kotorashvili, Maia Alkhazashvili, Irma Burjanadze, Anna Kasradze, Khatuna Zakhashvili, Paata Imnadze, Amiran Gamkrelidze. |
| EPI_ISL_3209480, EPI_ISL_3209483, EPI_ISL_3209486, EPI_ISL_3209489, EPI_ISL_3209493, EPI_ISL_3209496, EPI_ISL_3209500, EPI_ISL_3209519, EPI_ISL_3209522, EPI_ISL_3209525, EPI_ISL_3209528, EPI_ISL_3209531, EPI_ISL_3209538, EPI_ISL_3209542, EPI_ISL_3209546, EPI_ISL_3209549, EPI_ISL_3209553, EPI_ISL_3209556, EPI_ISL_3209563, EPI_ISL_3209566, EPI_ISL_3209568, EPI_ISL_3209572, EPI_ISL_3209575, EPI_ISL_3209579, EPI_ISL_3209581, EPI_ISL_3209582, EPI_ISL_3209583, EPI_ISL_3209584, EPI_ISL_3209586, EPI_ISL_3209587, EPI_ISL_3209588, EPI_ISL_3209589, EPI_ISL_3209590, EPI_ISL_3209591, EPI_ISL_3209592, EPI_ISL_3209593, EPI_ISL_3209594, EPI_ISL_3209595, EPI_ISL_3209596, EPI_ISL_3209597, EPI_ISL_3209598, EPI_ISL_3209599, EPI_ISL_3209600, EPI_ISL_3209601, EPI_ISL_3209602, EPI_ISL_3209603, EPI_ISL_3209604, EPI_ISL_3209605, EPI_ISL_3209607, EPI_ISL_3209608, EPI_ISL_3209609, EPI_ISL_3209610, EPI_ISL_3209611, EPI_ISL_3209612, EPI_ISL_3209614, EPI_ISL_3209615, EPI_ISL_3209616, EPI_ISL_3209617, EPI_ISL_3209618, EPI_ISL_3209620, EPI_ISL_3209622, EPI_ISL_3209623, EPI_ISL_3209624, EPI_ISL_3209627, EPI_ISL_3209628, EPI_ISL_3209629, EPI_ISL_3209630, EPI_ISL_3209631, EPI_ISL_3209632, EPI_ISL_3209633, EPI_ISL_3209634, EPI_ISL_3209635, EPI_ISL_3209636, EPI_ISL_3209637, EPI_ISL_3209638, EPI_ISL_3209639, EPI_ISL_3209640, EPI_ISL_3209641, EPI_ISL_3209642, EPI_ISL_3209643, EPI_ISL_3209644, EPI_ISL_3209645, EPI_ISL_3209646, EPI_ISL_3209647, EPI_ISL_3209648, EPI_ISL_3209649, EPI_ISL_3209650, EPI_ISL_3209651, EPI_ISL_3209652, EPI_ISL_3209653, EPI_ISL_3209654, EPI_ISL_3209655, EPI_ISL_3209656, EPI_ISL_3209657, EPI_ISL_3209658, EPI_ISL_3209659, EPI_ISL_3209660, EPI_ISL_3209661, EPI_ISL_3209662, EPI_ISL_3209663, EPI_ISL_3209664, EPI_ISL_3209665, EPI_ISL_3209666, EPI_ISL_3209667, EPI_ISL_3209669, EPI_ISL_3209670, EPI_ISL_3209671, EPI_ISL_3209672, EPI_ISL_3209673, EPI_ISL_3209674, EPI_ISL_3209675, EPI_ISL_3209676, EPI_ISL_3209677, EPI_ISL_3209678, EPI_ISL_3209679, EPI_ISL_3209680, EPI_ISL_3209681, EPI_ISL_3209682, EPI_ISL_3209684, EPI_ISL_3209685, EPI_ISL_3209686, EPI_ISL_3209688, EPI_ISL_3209689, EPI_ISL_3209690, EPI_ISL_3209691, EPI_ISL_3209692, EPI_ISL_3209693, EPI_ISL_3209694, EPI_ISL_3209695, EPI_ISL_3209696, EPI_ISL_3209697, EPI_ISL_3209698, EPI_ISL_3209699, EPI_ISL_3209700, EPI_ISL_3209701, EPI_ISL_3209702, EPI_ISL_3209703, EPI_ISL_3209704, EPI_ISL_3209705, EPI_ISL_3209706, EPI_ISL_3209707, EPI_ISL_3209708, EPI_ISL_3209709, EPI_ISL_3209710, EPI_ISL_3209711, EPI_ISL_3209712, EPI_ISL_3209713, EPI_ISL_3209714, EPI_ISL_3209715, EPI_ISL_3209716, EPI_ISL_3209717, EPI_ISL_3209718, | | | |

| | | | |
|---|---|---|---|
| EPI_ISL_3209719, EPI_ISL_3209720, EPI_ISL_3209721, EPI_ISL_3209722, EPI_ISL_3209724, EPI_ISL_3209725, EPI_ISL_3209726, EPI_ISL_3209727, EPI_ISL_3209728, EPI_ISL_3209729, EPI_ISL_3209730, EPI_ISL_3209731, EPI_ISL_3209732, EPI_ISL_3209733, EPI_ISL_3209734, EPI_ISL_3209735, EPI_ISL_3209736, EPI_ISL_3209737, EPI_ISL_3209738, EPI_ISL_3209739, EPI_ISL_3209740, EPI_ISL_3209741, EPI_ISL_3209742, EPI_ISL_3209743, EPI_ISL_3209744, EPI_ISL_3209745, EPI_ISL_3209746, EPI_ISL_3209747, EPI_ISL_3209748, EPI_ISL_3209749, EPI_ISL_3209750, EPI_ISL_3209751, EPI_ISL_3209753, EPI_ISL_3209754, EPI_ISL_3209755, EPI_ISL_3209756, EPI_ISL_3209757, EPI_ISL_3209758, EPI_ISL_3209759, EPI_ISL_3209760, EPI_ISL_3209761, EPI_ISL_3209762, EPI_ISL_3209763, EPI_ISL_3209764, EPI_ISL_3209765, EPI_ISL_3209766, EPI_ISL_3209767, EPI_ISL_3209768, EPI_ISL_3209769, EPI_ISL_3209770, EPI_ISL_3209771, EPI_ISL_3209772, EPI_ISL_3209773, EPI_ISL_3209774, EPI_ISL_3209775, EPI_ISL_3209776, EPI_ISL_3209777, EPI_ISL_3209778, EPI_ISL_3209779, EPI_ISL_3209780, EPI_ISL_3209781, EPI_ISL_3209782, EPI_ISL_3209783, EPI_ISL_3209785, EPI_ISL_3209786, EPI_ISL_3209787, EPI_ISL_3209788, EPI_ISL_3209789, EPI_ISL_3209791, EPI_ISL_3209792, EPI_ISL_3209793, EPI_ISL_3209794, EPI_ISL_3209795, EPI_ISL_3209796, EPI_ISL_3209797, EPI_ISL_3209798, EPI_ISL_3209799, EPI_ISL_3209800, EPI_ISL_3209801, EPI_ISL_3209802, EPI_ISL_3209803, EPI_ISL_3209804, EPI_ISL_3209805, EPI_ISL_3209806, EPI_ISL_3209807, EPI_ISL_3209808, EPI_ISL_3209809, EPI_ISL_3209810, EPI_ISL_3209811, EPI_ISL_3209813, EPI_ISL_3209814, EPI_ISL_3209815, EPI_ISL_3209816, EPI_ISL_3209817, EPI_ISL_3209819, EPI_ISL_3209820, EPI_ISL_3209821, EPI_ISL_3209822, EPI_ISL_3209823, EPI_ISL_3209824, EPI_ISL_3209825, EPI_ISL_3209826, EPI_ISL_3209827, EPI_ISL_3209828, EPI_ISL_3209829, EPI_ISL_3209830, EPI_ISL_3209831, EPI_ISL_3209832, EPI_ISL_3209833, EPI_ISL_3209834, EPI_ISL_3209835, EPI_ISL_3209836, EPI_ISL_3209838, EPI_ISL_3209839, EPI_ISL_3209840, EPI_ISL_3209841, EPI_ISL_3209842, EPI_ISL_3209843, EPI_ISL_3209844, EPI_ISL_3209845, EPI_ISL_3209846, EPI_ISL_3209847, EPI_ISL_3209848, EPI_ISL_3209849, EPI_ISL_3209850, EPI_ISL_3209851, EPI_ISL_3209852, EPI_ISL_3209853, EPI_ISL_3209854, EPI_ISL_3209855, EPI_ISL_3209856, EPI_ISL_3209857, EPI_ISL_3209858, EPI_ISL_3209859, EPI_ISL_3209860, EPI_ISL_3209861, EPI_ISL_3209862, EPI_ISL_3209863, EPI_ISL_3209864, EPI_ISL_3209865, EPI_ISL_3209866, EPI_ISL_3209867, EPI_ISL_3209868, EPI_ISL_3209870, EPI_ISL_3209871, EPI_ISL_3209872, EPI_ISL_3209873, EPI_ISL_3209874, EPI_ISL_3209875, EPI_ISL_3209876, EPI_ISL_3209877, EPI_ISL_3209878, EPI_ISL_3209879, EPI_ISL_3209880, EPI_ISL_3209881, EPI_ISL_3209882, EPI_ISL_3209883, EPI_ISL_3209884, EPI_ISL_3209885, EPI_ISL_3209886, EPI_ISL_3209887, EPI_ISL_3209888, EPI_ISL_3209889, EPI_ISL_3209890, EPI_ISL_3209891, EPI_ISL_3209892, EPI_ISL_3209893, EPI_ISL_3209894, EPI_ISL_3209895, EPI_ISL_3209896, EPI_ISL_3209897, EPI_ISL_3209898, EPI_ISL_3209899, EPI_ISL_3209900, EPI_ISL_3209901, EPI_ISL_3209902, EPI_ISL_3209903, EPI_ISL_3209904, EPI_ISL_3209905, EPI_ISL_3209906, EPI_ISL_3209907, EPI_ISL_3209908, EPI_ISL_3209910, EPI_ISL_3209911, EPI_ISL_3209912, EPI_ISL_3209913, EPI_ISL_3209914, EPI_ISL_3209915, EPI_ISL_3209916, EPI_ISL_3209917, EPI_ISL_3209918, EPI_ISL_3209919, EPI_ISL_3209920, EPI_ISL_3209921, EPI_ISL_3209922, EPI_ISL_3209923, EPI_ISL_3209925, EPI_ISL_3209926, EPI_ISL_3209927, EPI_ISL_3209928, EPI_ISL_3209929, EPI_ISL_3209930, EPI_ISL_3209931, EPI_ISL_3209932, EPI_ISL_3209933, EPI_ISL_3209934, EPI_ISL_3209935, EPI_ISL_3209936, EPI_ISL_3209937, EPI_ISL_3209938, EPI_ISL_3209939, EPI_ISL_3209940, EPI_ISL_3209941, EPI_ISL_3209942, EPI_ISL_3209944, EPI_ISL_3209945, EPI_ISL_3209946, EPI_ISL_3209947, EPI_ISL_3209948, EPI_ISL_3209949 | | | |
| see above | Fulgent Genetics | Centers for Disease Control and Prevention Division of Viral Diseases, Pathogen Discovery | Dakota Howard, Dhwani Batra, Peter W. Cook, Kara Moser, Adrian Paskey, Jason Caravas, Benjamin Rambo-Martin, Shatavia Morrison, Christopher Gulvick, Scott Sammons, Yvette Unoarumhi, Darlene Wagner, Matthew Schmerer, Harry Gao, Mickey Li, John Gao, Joseph Fierro, Benafsh Sapra, Becky Tsai, Yan Meng, Doreen Ng, James Xie, Clinton R. Paden, Duncan MacCannell |
| EPI_ISL_3209951 | Department for Virology, Molecular Biology and Genome Research, R. G. Lugar Center for Public Health Research, National Center for Disease Control and Public Health (NCDC) of Georgia. | Department for Virology, Molecular Biology and Genome Research, R. G. Lugar Center for Public Health Research, National Center for Disease Control and Public Health (NCDC) of Georgia. | Giorgi Gogoladze, Meri Pantsulaia, Gvantsa Brachveli, Giorgi Tomashvili, Salome Javashvili, Nino Berishvili, Tata Imnadze, Ana Papkiauri, Gvantsa Chanturia, Ann Machablishvili, Nato Kotaria, Marine Murtskhvaladze, Lela Sabadze, Mari Gavashelidze, Tamar Jashiashvili, Tea Tevdoradze, Ketevan Sidamonidze, Ekaterine Khmaladze, Ekaterine Zhghenti, Roena Sukhiashvili, Mariam Zakalashvili, Lela Urushadze, Magda Dgebuadze, Davit Tsaguria, Ekaterine Zangaladze, Adam Kotorashvili, Maia Alkhazashvili, Irma Burjanadze, Anna Kasradze, Khatuna Zakhashvili, Paata Imnadze, Amiran Gamkrelidze. |
| EPI_ISL_3209952, EPI_ISL_3209953, EPI_ISL_3209955, EPI_ISL_3209956, EPI_ISL_3209957, EPI_ISL_3209959, EPI_ISL_3209960, EPI_ISL_3209961, EPI_ISL_3209962, EPI_ISL_3209963, EPI_ISL_3209964, EPI_ISL_3209966, EPI_ISL_3209967, EPI_ISL_3209968, EPI_ISL_3209969, EPI_ISL_3209971, EPI_ISL_3209972, EPI_ISL_3209973, EPI_ISL_3209974, EPI_ISL_3209975, EPI_ISL_3209976, EPI_ISL_3209977, EPI_ISL_3209978, EPI_ISL_3209979, EPI_ISL_3209980, EPI_ISL_3209981, EPI_ISL_3209982, EPI_ISL_3209983, EPI_ISL_3209984, EPI_ISL_3209985, EPI_ISL_3209986, EPI_ISL_3209988, EPI_ISL_3209989, EPI_ISL_3209990, EPI_ISL_3209991, EPI_ISL_3209992, EPI_ISL_3209993, EPI_ISL_3209994, EPI_ISL_3209995, EPI_ISL_3209996, EPI_ISL_3209997, EPI_ISL_3209998, EPI_ISL_3209999, EPI_ISL_3210000, EPI_ISL_3210001, EPI_ISL_3210002, EPI_ISL_3210003, EPI_ISL_3210004, EPI_ISL_3210005, EPI_ISL_3210006, EPI_ISL_3210007, EPI_ISL_3210008, EPI_ISL_3210009, EPI_ISL_3210010, EPI_ISL_3210011, EPI_ISL_3210012, EPI_ISL_3210013, EPI_ISL_3210014, EPI_ISL_3210015, EPI_ISL_3210016, EPI_ISL_3210017, EPI_ISL_3210018, EPI_ISL_3210019, EPI_ISL_3210020, EPI_ISL_3210021, EPI_ISL_3210022, EPI_ISL_3210023, EPI_ISL_3210024, EPI_ISL_3210025, EPI_ISL_3210026, EPI_ISL_3210027, EPI_ISL_3210028, EPI_ISL_3210029, EPI_ISL_3210030, EPI_ISL_3210031, EPI_ISL_3210032, EPI_ISL_3210033, EPI_ISL_3210034, EPI_ISL_3210035, EPI_ISL_3210036, EPI_ISL_3210037, EPI_ISL_3210038, EPI_ISL_3210039, EPI_ISL_3210040, EPI_ISL_3210041, EPI_ISL_3210042, EPI_ISL_3210044, EPI_ISL_3210045, EPI_ISL_3210046, EPI_ISL_3210047, EPI_ISL_3210048, EPI_ISL_3210049, EPI_ISL_3210051, EPI_ISL_3210052, EPI_ISL_3210053, EPI_ISL_3210054, EPI_ISL_3210055, EPI_ISL_3210056, EPI_ISL_3210057, EPI_ISL_3210058, EPI_ISL_3210059, EPI_ISL_3210060, EPI_ISL_3210061, EPI_ISL_3210062 | | | |
| see above | Fulgent Genetics | Centers for Disease Control and Prevention Division of Viral Diseases, Pathogen Discovery | Dakota Howard, Dhwani Batra, Peter W. Cook, Kara Moser, Adrian Paskey, Jason Caravas, Benjamin Rambo-Martin, Shatavia Morrison, Christopher Gulvick, Scott Sammons, Yvette Unoarumhi, Darlene Wagner, Matthew Schmerer, Harry Gao, Mickey Li, John Gao, Joseph Fierro, Benafsh Sapra, Becky Tsai, Yan Meng, Doreen Ng, James Xie, Clinton R. Paden, Duncan MacCannell |
| EPI_ISL_3210063 | Center of Medical Microbiology, Virology, and Hospital Hygiene, University of Duesseldorf | Center of Medical Microbiology, Virology, and Hospital Hygiene, University of Duesseldorf | Maximilian Damagnez;Alexander Dilthey;Torsten Houwaart;Lisanna Hülse;Malte Kohns Vasconcelos;Nadine Lübke;Jessica Nicolai;Klaus Pfeffer;Daniel Strelow;Jörg Timm;Andreas Walker;Tobias Wienemann |
| EPI_ISL_3210064 | Fulgent Genetics | Centers for Disease Control and Prevention Division of Viral Diseases, Pathogen Discovery | Dakota Howard, Dhwani Batra, Peter W. Cook, Kara Moser, Adrian Paskey, Jason Caravas, Benjamin Rambo-Martin, Shatavia Morrison, Christopher Gulvick, Scott Sammons, Yvette Unoarumhi, Darlene Wagner, Matthew Schmerer, Harry Gao, Mickey Li, John Gao, Joseph Fierro, Benafsh Sapra, Becky Tsai, Yan Meng, Doreen Ng, James Xie, Clinton R. Paden, Duncan MacCannell |
| EPI_ISL_3210065, EPI_ISL_3210066 | ZOTZ KLIMAS MVZ Düsseldorf-Centrum GbR ÜBAG für Labormedizin, Genetik, Zytologie, Pathologie | Center of Medical Microbiology, Virology, and Hospital Hygiene, University of Duesseldorf | Maximilian Damagnez;Alexander Dilthey;Patrick Finzer;Katrin Hoffmann;Torsten Houwaart;Lisanna Hülse;Malte Kohns Vasconcelos;Nadine Lübke;Jessica Nicolai;Klaus Pfeffer;Daniel Strelow;Jörg Timm;Andreas Walker;Tobias Wienemann;Rainer Zotz |
| EPI_ISL_3210067 | Fulgent Genetics | Centers for Disease Control and Prevention Division of Viral Diseases, Pathogen Discovery | Dakota Howard, Dhwani Batra, Peter W. Cook, Kara Moser, Adrian Paskey, Jason Caravas, Benjamin Rambo-Martin, Shatavia Morrison, Christopher Gulvick, Scott Sammons, Yvette Unoarumhi, Darlene Wagner, Matthew Schmerer, Harry Gao, Mickey Li, John Gao, Joseph Fierro, Benafsh Sapra, Becky Tsai, Yan Meng, Doreen Ng, James Xie, Clinton R. Paden, Duncan MacCannell |
| EPI_ISL_3210068, EPI_ISL_3210069 | ZOTZ KLIMAS MVZ Düsseldorf-Centrum GbR ÜBAG für Labormedizin, Genetik, Zytologie, Pathologie | Center of Medical Microbiology, Virology, and Hospital Hygiene, University of Duesseldorf | Maximilian Damagnez;Alexander Dilthey;Patrick Finzer;Katrin Hoffmann;Torsten Houwaart;Lisanna Hülse;Malte Kohns Vasconcelos;Nadine Lübke;Jessica Nicolai;Klaus Pfeffer;Daniel Strelow;Jörg Timm;Andreas Walker;Tobias Wienemann;Rainer Zotz |
| EPI_ISL_3210070 | Fulgent Genetics | Centers for Disease Control and Prevention Division of Viral Diseases, Pathogen Discovery | Dakota Howard, Dhwani Batra, Peter W. Cook, Kara Moser, Adrian Paskey, Jason Caravas, Benjamin Rambo-Martin, Shatavia Morrison, Christopher Gulvick, Scott Sammons, Yvette Unoarumhi, Darlene Wagner, Matthew Schmerer, Harry Gao, Mickey Li, John Gao, Joseph Fierro, Benafsh Sapra, Becky Tsai, Yan Meng, Doreen Ng, James Xie, Clinton R. Paden, Duncan MacCannell |
| EPI_ISL_3210071, EPI_ISL_3210072, EPI_ISL_3210073, EPI_ISL_3210074, EPI_ISL_3210075, EPI_ISL_3210076, EPI_ISL_3210077, EPI_ISL_3210078 | ZOTZ KLIMAS MVZ Düsseldorf-Centrum GbR ÜBAG für Labormedizin, Genetik, Zytologie, Pathologie | Center of Medical Microbiology, Virology, and Hospital Hygiene, University of Duesseldorf | Maximilian Damagnez;Alexander Dilthey;Patrick Finzer;Katrin Hoffmann;Torsten Houwaart;Lisanna Hülse;Malte Kohns Vasconcelos;Nadine Lübke;Jessica Nicolai;Klaus Pfeffer;Daniel Strelow;Jörg Timm;Andreas Walker;Tobias Wienemann;Rainer Zotz |
| EPI_ISL_3210079 | Fulgent Genetics | Centers for Disease Control and Prevention Division of Viral Diseases, Pathogen Discovery | Dakota Howard, Dhwani Batra, Peter W. Cook, Kara Moser, Adrian Paskey, Jason Caravas, Benjamin Rambo-Martin, Shatavia Morrison, Christopher Gulvick, Scott Sammons, Yvette Unoarumhi, Darlene Wagner, Matthew Schmerer, Harry Gao, Mickey Li, John Gao, Joseph Fierro, Benafsh Sapra, Becky Tsai, Yan Meng, Doreen Ng, James Xie, Clinton R. Paden, Duncan MacCannell |
| EPI_ISL_3210080, EPI_ISL_3210081 | ZOTZ KLIMAS MVZ Düsseldorf-Centrum GbR ÜBAG für Labormedizin, Genetik, Zytologie, Pathologie | Center of Medical Microbiology, Virology, and Hospital Hygiene, University of Duesseldorf | Maximilian Damagnez;Alexander Dilthey;Patrick Finzer;Katrin Hoffmann;Torsten Houwaart;Lisanna Hülse;Malte Kohns Vasconcelos;Nadine Lübke;Jessica Nicolai;Klaus Pfeffer;Daniel Strelow;Jörg Timm;Andreas Walker;Tobias Wienemann;Rainer Zotz |
| EPI_ISL_3210082 | Fulgent Genetics | Centers for Disease Control and Prevention Division of Viral Diseases, Pathogen Discovery | Dakota Howard, Dhwani Batra, Peter W. Cook, Kara Moser, Adrian Paskey, Jason Caravas, Benjamin Rambo-Martin, Shatavia Morrison, Christopher Gulvick, Scott Sammons, Yvette Unoarumhi, Darlene Wagner, Matthew Schmerer, Harry Gao, Mickey Li, John Gao, Joseph Fierro, Benafsh Sapra, Becky Tsai, Yan Meng, Doreen Ng, James Xie, Clinton R. Paden, Duncan MacCannell |
| EPI_ISL_3210083, EPI_ISL_3210084 | ZOTZ KLIMAS MVZ Düsseldorf-Centrum GbR ÜBAG für Labormedizin, Genetik, Zytologie, Pathologie | Center of Medical Microbiology, Virology, and Hospital Hygiene, University of Duesseldorf | Maximilian Damagnez;Alexander Dilthey;Patrick Finzer;Katrin Hoffmann;Torsten Houwaart;Lisanna Hülse;Malte Kohns Vasconcelos;Nadine Lübke;Jessica Nicolai;Klaus Pfeffer;Daniel Strelow;Jörg Timm;Andreas Walker;Tobias Wienemann;Rainer Zotz |
| EPI_ISL_3210085 | Fulgent Genetics | Centers for Disease Control and Prevention Division of Viral Diseases, Pathogen Discovery | Dakota Howard, Dhwani Batra, Peter W. Cook, Kara Moser, Adrian Paskey, Jason Caravas, Benjamin Rambo-Martin, Shatavia Morrison, Christopher Gulvick, Scott Sammons, Yvette Unoarumhi, Darlene Wagner, Matthew Schmerer, Harry Gao, Mickey Li, John Gao, Joseph Fierro, Benafsh Sapra, Becky Tsai, Yan Meng, Doreen Ng, James Xie, Clinton R. Paden, Duncan MacCannell |
| EPI_ISL_3210086, EPI_ISL_3210087 | ZOTZ KLIMAS MVZ Düsseldorf-Centrum GbR ÜBAG für Labormedizin, Genetik, Zytologie, Pathologie | Center of Medical Microbiology, Virology, and Hospital Hygiene, University of Duesseldorf | Maximilian Damagnez;Alexander Dilthey;Patrick Finzer;Katrin Hoffmann;Torsten Houwaart;Lisanna Hülse;Malte Kohns Vasconcelos;Nadine Lübke;Jessica Nicolai;Klaus Pfeffer;Daniel Strelow;Jörg Timm;Andreas Walker;Tobias Wienemann;Rainer Zotz |
| EPI_ISL_3210088 | Fulgent Genetics | Centers for Disease Control and Prevention Division of Viral Diseases, Pathogen Discovery | Dakota Howard, Dhwani Batra, Peter W. Cook, Kara Moser, Adrian Paskey, Jason Caravas, Benjamin Rambo-Martin, Shatavia Morrison, Christopher Gulvick, Scott Sammons, Yvette Unoarumhi, Darlene Wagner, Matthew Schmerer, Harry Gao, Mickey Li, John Gao, Joseph Fierro, Benafsh Sapra, Becky Tsai, Yan Meng, Doreen Ng, James Xie, Clinton R. Paden, Duncan MacCannell |
| EPI_ISL_3210089 | ZOTZ KLIMAS MVZ Düsseldorf-Centrum GbR ÜBAG für Labormedizin, Genetik, Zytologie, Pathologie | Center of Medical Microbiology, Virology, and Hospital Hygiene, University of Duesseldorf | Maximilian Damagnez;Alexander Dilthey;Patrick Finzer;Katrin Hoffmann;Torsten Houwaart;Lisanna Hülse;Malte Kohns Vasconcelos;Nadine Lübke;Jessica Nicolai;Klaus Pfeffer;Daniel Strelow;Jörg Timm;Andreas Walker;Tobias Wienemann;Rainer Zotz |
| EPI_ISL_3210090 | Fulgent Genetics | Centers for Disease Control and Prevention Division of Viral Diseases, Pathogen Discovery | Dakota Howard, Dhwani Batra, Peter W. Cook, Kara Moser, Adrian Paskey, Jason Caravas, Benjamin Rambo-Martin, Shatavia Morrison, Christopher Gulvick, Scott Sammons, Yvette Unoarumhi, Darlene Wagner, Matthew Schmerer, Harry Gao, Mickey Li, John Gao, Joseph Fierro, Benafsh Sapra, Becky Tsai, Yan Meng, Doreen Ng, James Xie, Clinton R. Paden, Duncan MacCannell |
| EPI_ISL_3210091, EPI_ISL_3210092 | ZOTZ KLIMAS MVZ Düsseldorf-Centrum GbR ÜBAG für Labormedizin, Genetik, Zytologie, Pathologie | Center of Medical Microbiology, Virology, and Hospital Hygiene, University of Duesseldorf | Maximilian Damagnez;Alexander Dilthey;Patrick Finzer;Katrin Hoffmann;Torsten Houwaart;Lisanna Hülse;Malte Kohns Vasconcelos;Nadine Lübke;Jessica Nicolai;Klaus Pfeffer;Daniel Strelow;Jörg Timm;Andreas Walker;Tobias Wienemann;Rainer Zotz |
| EPI_ISL_3210093 | Fulgent Genetics | Centers for Disease Control and Prevention Division of Viral Diseases, Pathogen Discovery | Dakota Howard, Dhwani Batra, Peter W. Cook, Kara Moser, Adrian Paskey, Jason Caravas, Benjamin Rambo-Martin, Shatavia Morrison, Christopher Gulvick, Scott Sammons, Yvette Unoarumhi, Darlene Wagner, Matthew Schmerer, Harry Gao, Mickey Li, John Gao, Joseph Fierro, Benafsh Sapra, Becky Tsai, Yan Meng, Doreen Ng, James Xie, Clinton R. Paden, Duncan MacCannell |
| EPI_ISL_3210094 | ZOTZ KLIMAS MVZ Düsseldorf-Centrum GbR ÜBAG für Labormedizin, Genetik, Zytologie, Pathologie | Center of Medical Microbiology, Virology, and Hospital Hygiene, University of Duesseldorf | Maximilian Damagnez;Alexander Dilthey;Patrick Finzer;Katrin Hoffmann;Torsten Houwaart;Lisanna Hülse;Malte Kohns Vasconcelos;Nadine Lübke;Jessica Nicolai;Klaus Pfeffer;Daniel Strelow;Jörg Timm;Andreas Walker;Tobias Wienemann;Rainer Zotz |
| EPI_ISL_3210095 | Fulgent Genetics | Centers for Disease Control and Prevention Division of Viral Diseases, Pathogen Discovery | Dakota Howard, Dhwani Batra, Peter W. Cook, Kara Moser, Adrian Paskey, Jason Caravas, Benjamin Rambo-Martin, Shatavia Morrison, Christopher Gulvick, Scott Sammons, Yvette Unoarumhi, Darlene Wagner, Matthew Schmerer, Harry Gao, Mickey Li, John Gao, Joseph Fierro, Benafsh Sapra, Becky Tsai, Yan Meng, Doreen Ng, James Xie, Clinton R. Paden, Duncan MacCannell |
| EPI_ISL_3210096, EPI_ISL_3210097 | ZOTZ KLIMAS MVZ Düsseldorf-Centrum GbR ÜBAG für Labormedizin, Genetik, Zytologie, Pathologie | Center of Medical Microbiology, Virology, and Hospital Hygiene, University of Duesseldorf | Maximilian Damagnez;Alexander Dilthey;Patrick Finzer;Katrin Hoffmann;Torsten Houwaart;Lisanna Hülse;Malte Kohns Vasconcelos;Nadine Lübke;Jessica Nicolai;Klaus Pfeffer;Daniel Strelow;Jörg Timm;Andreas Walker;Tobias Wienemann;Rainer Zotz |
| EPI_ISL_3210098 | Fulgent Genetics | Centers for Disease Control and Prevention Division of Viral Diseases, Pathogen Discovery | Dakota Howard, Dhwani Batra, Peter W. Cook, Kara Moser, Adrian Paskey, Jason Caravas, Benjamin Rambo-Martin, Shatavia Morrison, Christopher Gulvick, Scott Sammons, Yvette Unoarumhi, Darlene Wagner, Matthew Schmerer, Harry Gao, Mickey Li, John Gao, Joseph Fierro, Benafsh Sapra, Becky Tsai, Yan Meng, Doreen Ng, James Xie, Clinton R. Paden, Duncan MacCannell |
| EPI_ISL_3210099, EPI_ISL_3210100 | ZOTZ KLIMAS MVZ Düsseldorf-Centrum GbR ÜBAG für Labormedizin, Genetik, Zytologie, Pathologie | Center of Medical Microbiology, Virology, and Hospital Hygiene, University of Duesseldorf | Maximilian Damagnez;Alexander Dilthey;Patrick Finzer;Katrin Hoffmann;Torsten Houwaart;Lisanna Hülse;Malte Kohns Vasconcelos;Nadine Lübke;Jessica Nicolai;Klaus Pfeffer;Daniel Strelow;Jörg Timm;Andreas Walker;Tobias Wienemann;Rainer Zotz |
| EPI_ISL_3210101 | Fulgent Genetics | Centers for Disease Control and Prevention Division of Viral Diseases, Pathogen Discovery | Dakota Howard, Dhwani Batra, Peter W. Cook, Kara Moser, Adrian Paskey, Jason Caravas, Benjamin Rambo-Martin, Shatavia Morrison, Christopher Gulvick, Scott Sammons, Yvette Unoarumhi, Darlene Wagner, Matthew Schmerer, Harry Gao, Mickey Li, John Gao, Joseph Fierro, Benafsh Sapra, Becky Tsai, Yan Meng, Doreen Ng, James Xie, Clinton R. Paden, Duncan MacCannell |
| EPI_ISL_3210102 | ZOTZ KLIMAS MVZ Düsseldorf-Centrum GbR ÜBAG für Labormedizin, Genetik, Zytologie, Pathologie | Center of Medical Microbiology, Virology, and Hospital Hygiene, University of Duesseldorf | Maximilian Damagnez;Alexander Dilthey;Patrick Finzer;Katrin Hoffmann;Torsten Houwaart;Lisanna Hülse;Malte Kohns Vasconcelos;Nadine Lübke;Jessica Nicolai;Klaus Pfeffer;Daniel Strelow;Jörg Timm;Andreas Walker;Tobias Wienemann;Rainer Zotz |
| EPI_ISL_3210103 | Fulgent Genetics | Centers for Disease Control and Prevention Division of Viral Diseases, Pathogen Discovery | Dakota Howard, Dhwani Batra, Peter W. Cook, Kara Moser, Adrian Paskey, Jason Caravas, Benjamin Rambo-Martin, Shatavia Morrison, Christopher Gulvick, Scott Sammons, Yvette Unoarumhi, Darlene Wagner, Matthew Schmerer, Harry Gao, Mickey Li, John Gao, Joseph Fierro, Benafsh Sapra, Becky Tsai, Yan Meng, Doreen Ng, James Xie, Clinton R. Paden, Duncan MacCannell |
| EPI_ISL_3210104, EPI_ISL_3210105 | ZOTZ KLIMAS MVZ Düsseldorf-Centrum GbR ÜBAG für Labormedizin, Genetik, Zytologie, Pathologie | Center of Medical Microbiology, Virology, and Hospital Hygiene, University of Duesseldorf | Maximilian Damagnez;Alexander Dilthey;Patrick Finzer;Katrin Hoffmann;Torsten Houwaart;Lisanna Hülse;Malte Kohns Vasconcelos;Nadine Lübke;Jessica Nicolai;Klaus Pfeffer;Daniel Strelow;Jörg Timm;Andreas Walker;Tobias Wienemann;Rainer Zotz |
| EPI_ISL_3210106 | Fulgent Genetics | Centers for Disease Control and Prevention Division of Viral Diseases, Pathogen Discovery | Dakota Howard, Dhwani Batra, Peter W. Cook, Kara Moser, Adrian Paskey, Jason Caravas, Benjamin Rambo-Martin, Shatavia Morrison, Christopher Gulvick, Scott Sammons, Yvette Unoarumhi, Darlene Wagner, Matthew Schmerer, Harry Gao, Mickey Li, John Gao, Joseph Fierro, Benafsh Sapra, Becky Tsai, Yan Meng, Doreen Ng, James Xie, Clinton R. Paden, Duncan MacCannell |
| EPI_ISL_3210107 | ZOTZ KLIMAS MVZ Düsseldorf-Centrum GbR ÜBAG für Labormedizin, Genetik, Zytologie, Pathologie | Center of Medical Microbiology, Virology, and Hospital Hygiene, University of Duesseldorf | Maximilian Damagnez;Alexander Dilthey;Patrick Finzer;Katrin Hoffmann;Torsten Houwaart;Lisanna Hülse;Malte Kohns Vasconcelos;Nadine Lübke;Jessica Nicolai;Klaus Pfeffer;Daniel Strelow;Jörg Timm;Andreas Walker;Tobias Wienemann;Rainer Zotz |
| EPI_ISL_3210108 | Fulgent Genetics | Centers for Disease Control and Prevention Division of Viral Diseases, Pathogen Discovery | Dakota Howard, Dhwani Batra, Peter W. Cook, Kara Moser, Adrian Paskey, Jason Caravas, Benjamin Rambo-Martin, Shatavia Morrison, Christopher Gulvick, Scott Sammons, Yvette Unoarumhi, Darlene Wagner, Matthew Schmerer, Harry Gao, Mickey Li, John Gao, Joseph Fierro, Benafsh Sapra, Becky Tsai, Yan Meng, Doreen Ng, James Xie, Clinton R. Paden, Duncan MacCannell |
| EPI_ISL_3210109 | Center of Medical Microbiology, Virology, and Hospital Hygiene, University of Duesseldorf | Center of Medical Microbiology, Virology, and Hospital Hygiene, University of Duesseldorf | Maximilian Damagnez;Alexander Dilthey;Torsten Houwaart;Lisanna Hülse;Malte Kohns Vasconcelos;Nadine Lübke;Jessica Nicolai;Klaus Pfeffer;Daniel Strelow;Jörg Timm;Andreas Walker;Tobias Wienemann |
| EPI_ISL_3210110 | ZOTZ KLIMAS MVZ Düsseldorf-Centrum GbR ÜBAG für Labormedizin, Genetik, Zytologie, Pathologie | Center of Medical Microbiology, Virology, and Hospital Hygiene, University of Duesseldorf | Maximilian Damagnez;Alexander Dilthey;Patrick Finzer;Katrin Hoffmann;Torsten Houwaart;Lisanna Hülse;Malte Kohns Vasconcelos;Nadine Lübke;Jessica Nicolai;Klaus Pfeffer;Daniel Strelow;Jörg Timm;Andreas Walker;Tobias Wienemann;Rainer Zotz |

| | | | |
|---|---|---|---|
| EPI_ISL_3210111 | Fulgent Genetics | Centers for Disease Control and Prevention Division of Viral Diseases, Pathogen Discovery | Dakota Howard, Dhwani Batra, Peter W. Cook, Kara Moser, Adrian Paskey, Jason Caravas, Benjamin Rambo-Martin, Shatavia Morrison, Christopher Gulvick, Scott Sammons, Yvette Unoarumhi, Darlene Wagner, Matthew Schmerer, Harry Gao, Mickey Li, John Gao, Joseph Fierro, Benafsh Sapra, Becky Tsai, Yan Meng, Doreen Ng, James Xie, Clinton R. Paden, Duncan MacCannell |
| EPI_ISL_3210112 | ZOTZ KLIMAS MVZ Düsseldorf-Centrum GbR ÜBAG für Labormedizin, Genetik, Zytologie, Pathologie | Center of Medical Microbiology, Virology, and Hospital Hygiene, University of Duesseldorf | Maximilian Damagnez;Alexander Dilthey;Patrick Finzer;Katrin Hoffmann;Torsten Houwaart;Lisanna Hülse;Malte Kohns Vasconcelos;Nadine Lübke;Jessica Nicolai;Klaus Pfeffer;Daniel Strelow;Jörg Timm;Andreas Walker;Tobias Wienemann;Rainer Zotz |
| EPI_ISL_3210113 | Fulgent Genetics | Centers for Disease Control and Prevention Division of Viral Diseases, Pathogen Discovery | Dakota Howard, Dhwani Batra, Peter W. Cook, Kara Moser, Adrian Paskey, Jason Caravas, Benjamin Rambo-Martin, Shatavia Morrison, Christopher Gulvick, Scott Sammons, Yvette Unoarumhi, Darlene Wagner, Matthew Schmerer, Harry Gao, Mickey Li, John Gao, Joseph Fierro, Benafsh Sapra, Becky Tsai, Yan Meng, Doreen Ng, James Xie, Clinton R. Paden, Duncan MacCannell |
| EPI_ISL_3210114, EPI_ISL_3210115 | ZOTZ KLIMAS MVZ Düsseldorf-Centrum GbR ÜBAG für Labormedizin, Genetik, Zytologie, Pathologie | Center of Medical Microbiology, Virology, and Hospital Hygiene, University of Duesseldorf | Maximilian Damagnez;Alexander Dilthey;Patrick Finzer;Katrin Hoffmann;Torsten Houwaart;Lisanna Hülse;Malte Kohns Vasconcelos;Nadine Lübke;Jessica Nicolai;Klaus Pfeffer;Daniel Strelow;Jörg Timm;Andreas Walker;Tobias Wienemann;Rainer Zotz |
| EPI_ISL_3210116 | Fulgent Genetics | Centers for Disease Control and Prevention Division of Viral Diseases, Pathogen Discovery | Dakota Howard, Dhwani Batra, Peter W. Cook, Kara Moser, Adrian Paskey, Jason Caravas, Benjamin Rambo-Martin, Shatavia Morrison, Christopher Gulvick, Scott Sammons, Yvette Unoarumhi, Darlene Wagner, Matthew Schmerer, Harry Gao, Mickey Li, John Gao, Joseph Fierro, Benafsh Sapra, Becky Tsai, Yan Meng, Doreen Ng, James Xie, Clinton R. Paden, Duncan MacCannell |
| EPI_ISL_3210117, EPI_ISL_3210118 | ZOTZ KLIMAS MVZ Düsseldorf-Centrum GbR ÜBAG für Labormedizin, Genetik, Zytologie, Pathologie | Center of Medical Microbiology, Virology, and Hospital Hygiene, University of Duesseldorf | Maximilian Damagnez;Alexander Dilthey;Patrick Finzer;Katrin Hoffmann;Torsten Houwaart;Lisanna Hülse;Malte Kohns Vasconcelos;Nadine Lübke;Jessica Nicolai;Klaus Pfeffer;Daniel Strelow;Jörg Timm;Andreas Walker;Tobias Wienemann;Rainer Zotz |
| EPI_ISL_3210119 | Fulgent Genetics | Centers for Disease Control and Prevention Division of Viral Diseases, Pathogen Discovery | Dakota Howard, Dhwani Batra, Peter W. Cook, Kara Moser, Adrian Paskey, Jason Caravas, Benjamin Rambo-Martin, Shatavia Morrison, Christopher Gulvick, Scott Sammons, Yvette Unoarumhi, Darlene Wagner, Matthew Schmerer, Harry Gao, Mickey Li, John Gao, Joseph Fierro, Benafsh Sapra, Becky Tsai, Yan Meng, Doreen Ng, James Xie, Clinton R. Paden, Duncan MacCannell |
| EPI_ISL_3210120, EPI_ISL_3210121 | ZOTZ KLIMAS MVZ Düsseldorf-Centrum GbR ÜBAG für Labormedizin, Genetik, Zytologie, Pathologie | Center of Medical Microbiology, Virology, and Hospital Hygiene, University of Duesseldorf | Maximilian Damagnez;Alexander Dilthey;Patrick Finzer;Katrin Hoffmann;Torsten Houwaart;Lisanna Hülse;Malte Kohns Vasconcelos;Nadine Lübke;Jessica Nicolai;Klaus Pfeffer;Daniel Strelow;Jörg Timm;Andreas Walker;Tobias Wienemann;Rainer Zotz |
| EPI_ISL_3210122 | Fulgent Genetics | Centers for Disease Control and Prevention Division of Viral Diseases, Pathogen Discovery | Dakota Howard, Dhwani Batra, Peter W. Cook, Kara Moser, Adrian Paskey, Jason Caravas, Benjamin Rambo-Martin, Shatavia Morrison, Christopher Gulvick, Scott Sammons, Yvette Unoarumhi, Darlene Wagner, Matthew Schmerer, Harry Gao, Mickey Li, John Gao, Joseph Fierro, Benafsh Sapra, Becky Tsai, Yan Meng, Doreen Ng, James Xie, Clinton R. Paden, Duncan MacCannell |
| EPI_ISL_3210123, EPI_ISL_3210124 | ZOTZ KLIMAS MVZ Düsseldorf-Centrum GbR ÜBAG für Labormedizin, Genetik, Zytologie, Pathologie | Center of Medical Microbiology, Virology, and Hospital Hygiene, University of Duesseldorf | Maximilian Damagnez;Alexander Dilthey;Patrick Finzer;Katrin Hoffmann;Torsten Houwaart;Lisanna Hülse;Malte Kohns Vasconcelos;Nadine Lübke;Jessica Nicolai;Klaus Pfeffer;Daniel Strelow;Jörg Timm;Andreas Walker;Tobias Wienemann;Rainer Zotz |
| EPI_ISL_3210125 | Fulgent Genetics | Centers for Disease Control and Prevention Division of Viral Diseases, Pathogen Discovery | Dakota Howard, Dhwani Batra, Peter W. Cook, Kara Moser, Adrian Paskey, Jason Caravas, Benjamin Rambo-Martin, Shatavia Morrison, Christopher Gulvick, Scott Sammons, Yvette Unoarumhi, Darlene Wagner, Matthew Schmerer, Harry Gao, Mickey Li, John Gao, Joseph Fierro, Benafsh Sapra, Becky Tsai, Yan Meng, Doreen Ng, James Xie, Clinton R. Paden, Duncan MacCannell |
| EPI_ISL_3210126 | ZOTZ KLIMAS MVZ Düsseldorf-Centrum GbR ÜBAG für Labormedizin, Genetik, Zytologie, Pathologie | Center of Medical Microbiology, Virology, and Hospital Hygiene, University of Duesseldorf | Maximilian Damagnez;Alexander Dilthey;Patrick Finzer;Katrin Hoffmann;Torsten Houwaart;Lisanna Hülse;Malte Kohns Vasconcelos;Nadine Lübke;Jessica Nicolai;Klaus Pfeffer;Daniel Strelow;Jörg Timm;Andreas Walker;Tobias Wienemann;Rainer Zotz |
| EPI_ISL_3210127, EPI_ISL_3210128, EPI_ISL_3210129, EPI_ISL_3210130, EPI_ISL_3210131, EPI_ISL_3210132, EPI_ISL_3210133, EPI_ISL_3210134, EPI_ISL_3210135, EPI_ISL_3210136, EPI_ISL_3210138, EPI_ISL_3210140, EPI_ISL_3210141, EPI_ISL_3210142, EPI_ISL_3210143, EPI_ISL_3210146, EPI_ISL_3210147, EPI_ISL_3210148, EPI_ISL_3210149, EPI_ISL_3210150, EPI_ISL_3210151, EPI_ISL_3210152, EPI_ISL_3210153, EPI_ISL_3210154, EPI_ISL_3210155, EPI_ISL_3210156, EPI_ISL_3210157, EPI_ISL_3210158, EPI_ISL_3210159, EPI_ISL_3210160, EPI_ISL_3210161, EPI_ISL_3210162, EPI_ISL_3210163, EPI_ISL_3210164, EPI_ISL_3210165, EPI_ISL_3210166, EPI_ISL_3210167, EPI_ISL_3210168, EPI_ISL_3210169, EPI_ISL_3210170, EPI_ISL_3210171, EPI_ISL_3210172, EPI_ISL_3210173, EPI_ISL_3210174, EPI_ISL_3210175, EPI_ISL_3210176, EPI_ISL_3210177, EPI_ISL_3210178, EPI_ISL_3210179, EPI_ISL_3210180, EPI_ISL_3210181, EPI_ISL_3210182, EPI_ISL_3210183, EPI_ISL_3210184, EPI_ISL_3210185, EPI_ISL_3210186, EPI_ISL_3210187, EPI_ISL_3210188, EPI_ISL_3210189, EPI_ISL_3210190, EPI_ISL_3210191, EPI_ISL_3210192, EPI_ISL_3210193, EPI_ISL_3210194, EPI_ISL_3210195, EPI_ISL_3210196, EPI_ISL_3210197, EPI_ISL_3210198, EPI_ISL_3210199, EPI_ISL_3210200, EPI_ISL_3210201, EPI_ISL_3210202, EPI_ISL_3210203, EPI_ISL_3210204, EPI_ISL_3210205, EPI_ISL_3210206, EPI_ISL_3210207, EPI_ISL_3210208, EPI_ISL_3210209, EPI_ISL_3210210, EPI_ISL_3210212, EPI_ISL_3210213, EPI_ISL_3210214, EPI_ISL_3210215, EPI_ISL_3210216, EPI_ISL_3210217, EPI_ISL_3210218, EPI_ISL_3210219, EPI_ISL_3210220, EPI_ISL_3210221, EPI_ISL_3210222, EPI_ISL_3210223, EPI_ISL_3210224, EPI_ISL_3210225, EPI_ISL_3210226, EPI_ISL_3210227, EPI_ISL_3210229, EPI_ISL_3210230, EPI_ISL_3210231, EPI_ISL_3210232, EPI_ISL_3210233, EPI_ISL_3210234, EPI_ISL_3210235, EPI_ISL_3210236, EPI_ISL_3210237, EPI_ISL_3210238, EPI_ISL_3210239, EPI_ISL_3210240, EPI_ISL_3210242, EPI_ISL_3210243, EPI_ISL_3210244, EPI_ISL_3210245, EPI_ISL_3210246, EPI_ISL_3210247, EPI_ISL_3210248, EPI_ISL_3210249, EPI_ISL_3210250, EPI_ISL_3210251, EPI_ISL_3210252, EPI_ISL_3210253, EPI_ISL_3210254, EPI_ISL_3210255, EPI_ISL_3210256, EPI_ISL_3210257, EPI_ISL_3210258, EPI_ISL_3210259, EPI_ISL_3210260, EPI_ISL_3210261, EPI_ISL_3210262, EPI_ISL_3210263, EPI_ISL_3210264, EPI_ISL_3210265, EPI_ISL_3210266, EPI_ISL_3210267, EPI_ISL_3210268, EPI_ISL_3210269, EPI_ISL_3210270, EPI_ISL_3210271, EPI_ISL_3210272, EPI_ISL_3210273, EPI_ISL_3210274, EPI_ISL_3210275, EPI_ISL_3210276, EPI_ISL_3210277, EPI_ISL_3210278, EPI_ISL_3210279, EPI_ISL_3210280, EPI_ISL_3210281, EPI_ISL_3210282, EPI_ISL_3210283, EPI_ISL_3210284, EPI_ISL_3210285, EPI_ISL_3210286, EPI_ISL_3210287, EPI_ISL_3210288, EPI_ISL_3210289, EPI_ISL_3210290, EPI_ISL_3210291, EPI_ISL_3210292, EPI_ISL_3210293, EPI_ISL_3210294, EPI_ISL_3210295, EPI_ISL_3210296, EPI_ISL_3210297, EPI_ISL_3210298, EPI_ISL_3210299, EPI_ISL_3210300, EPI_ISL_3210301, EPI_ISL_3210302, EPI_ISL_3210303, EPI_ISL_3210304, EPI_ISL_3210305, EPI_ISL_3210306, EPI_ISL_3210307, EPI_ISL_3210309, EPI_ISL_3210310, EPI_ISL_3210311, EPI_ISL_3210313, EPI_ISL_3210314, EPI_ISL_3210315, EPI_ISL_3210316, EPI_ISL_3210317, EPI_ISL_3210318, EPI_ISL_3210319, EPI_ISL_3210320, EPI_ISL_3210322, EPI_ISL_3210323, EPI_ISL_3210324, EPI_ISL_3210325, EPI_ISL_3210326, EPI_ISL_3210327, EPI_ISL_3210328, EPI_ISL_3210329, EPI_ISL_3210330, EPI_ISL_3210331, EPI_ISL_3210332, EPI_ISL_3210333, EPI_ISL_3210334, EPI_ISL_3210335, EPI_ISL_3210336, EPI_ISL_3210337, EPI_ISL_3210338, EPI_ISL_3210339, EPI_ISL_3210342, EPI_ISL_3210343, EPI_ISL_3210344, EPI_ISL_3210345, EPI_ISL_3210346, EPI_ISL_3210347, EPI_ISL_3210349, EPI_ISL_3210350, EPI_ISL_3210351, EPI_ISL_3210352, EPI_ISL_3210353, EPI_ISL_3210354, EPI_ISL_3210355, EPI_ISL_3210356, EPI_ISL_3210357, EPI_ISL_3210358, EPI_ISL_3210359, EPI_ISL_3210360, EPI_ISL_3210361, EPI_ISL_3210362, EPI_ISL_3210363, EPI_ISL_3210364, EPI_ISL_3210365, EPI_ISL_3210366, EPI_ISL_3210367, EPI_ISL_3210368, EPI_ISL_3210369, EPI_ISL_3210370, EPI_ISL_3210371, EPI_ISL_3210372, EPI_ISL_3210373, EPI_ISL_3210374, EPI_ISL_3210375, EPI_ISL_3210376, EPI_ISL_3210377, EPI_ISL_3210378, EPI_ISL_3210379, EPI_ISL_3210380, EPI_ISL_3210381, EPI_ISL_3210382, EPI_ISL_3210383, EPI_ISL_3210384, EPI_ISL_3210385, EPI_ISL_3210386, EPI_ISL_3210387, EPI_ISL_3210388, EPI_ISL_3210389, EPI_ISL_3210390, EPI_ISL_3210391, EPI_ISL_3210392, EPI_ISL_3210394, EPI_ISL_3210395, EPI_ISL_3210396, EPI_ISL_3210397, EPI_ISL_3210398, EPI_ISL_3210399, EPI_ISL_3210400, EPI_ISL_3210401, EPI_ISL_3210402, EPI_ISL_3210403, EPI_ISL_3210404, EPI_ISL_3210405, EPI_ISL_3210406, EPI_ISL_3210407, EPI_ISL_3210408, EPI_ISL_3210409, EPI_ISL_3210410, EPI_ISL_3210411, EPI_ISL_3210412, EPI_ISL_3210413, EPI_ISL_3210414, EPI_ISL_3210415, EPI_ISL_3210416, EPI_ISL_3210417, EPI_ISL_3210418, EPI_ISL_3210419, EPI_ISL_3210420, EPI_ISL_3210422, EPI_ISL_3210423, EPI_ISL_3210424, EPI_ISL_3210425, EPI_ISL_3210426, EPI_ISL_3210427, EPI_ISL_3210428, EPI_ISL_3210429, EPI_ISL_3210430, EPI_ISL_3210431, EPI_ISL_3210432, EPI_ISL_3210433, EPI_ISL_3210434, EPI_ISL_3210435, EPI_ISL_3210436, EPI_ISL_3210437, EPI_ISL_3210438, EPI_ISL_3210439, EPI_ISL_3210440, EPI_ISL_3210441, EPI_ISL_3210442, EPI_ISL_3210443, EPI_ISL_3210444, EPI_ISL_3210445, EPI_ISL_3210446, EPI_ISL_3210447, EPI_ISL_3210448, EPI_ISL_3210449, EPI_ISL_3210450, EPI_ISL_3210451, EPI_ISL_3210452, EPI_ISL_3210453, EPI_ISL_3210454, EPI_ISL_3210455, EPI_ISL_3210456, EPI_ISL_3210457, EPI_ISL_3210458, EPI_ISL_3210459, EPI_ISL_3210460, EPI_ISL_3210461, EPI_ISL_3210462, EPI_ISL_3210463, EPI_ISL_3210464, EPI_ISL_3210465, EPI_ISL_3210466, EPI_ISL_3210467, EPI_ISL_3210468, EPI_ISL_3210469, EPI_ISL_3210470, EPI_ISL_3210471, EPI_ISL_3210472, EPI_ISL_3210473, EPI_ISL_3210475, EPI_ISL_3210476, EPI_ISL_3210477, EPI_ISL_3210478, EPI_ISL_3210479, EPI_ISL_3210480, EPI_ISL_3210481, EPI_ISL_3210482, EPI_ISL_3210483, EPI_ISL_3210484, EPI_ISL_3210485, EPI_ISL_3210486, EPI_ISL_3210487, EPI_ISL_3210488, EPI_ISL_3210489, EPI_ISL_3210490, EPI_ISL_3210491, EPI_ISL_3210492, EPI_ISL_3210493, EPI_ISL_3210494, EPI_ISL_3210495, EPI_ISL_3210496, EPI_ISL_3210497, EPI_ISL_3210498, EPI_ISL_3210499, EPI_ISL_3210500, EPI_ISL_3210501, EPI_ISL_3210502, EPI_ISL_3210503, EPI_ISL_3210504, EPI_ISL_3210505, EPI_ISL_3210506, EPI_ISL_3210507, EPI_ISL_3210508, EPI_ISL_3210509, EPI_ISL_3210510, EPI_ISL_3210511, EPI_ISL_3210512, EPI_ISL_3210513, EPI_ISL_3210514, EPI_ISL_3210515, EPI_ISL_3210516, EPI_ISL_3210517, EPI_ISL_3210518, EPI_ISL_3210519, EPI_ISL_3210520, EPI_ISL_3210521, EPI_ISL_3210522, EPI_ISL_3210523, EPI_ISL_3210524, EPI_ISL_3210525, EPI_ISL_3210526, EPI_ISL_3210527, EPI_ISL_3210528, EPI_ISL_3210529, EPI_ISL_3210530, EPI_ISL_3210531, EPI_ISL_3210532, EPI_ISL_3210533, EPI_ISL_3210534, EPI_ISL_3210535, EPI_ISL_3210536, EPI_ISL_3210537, EPI_ISL_3210538, EPI_ISL_3210539, EPI_ISL_3210540, EPI_ISL_3210541, EPI_ISL_3210542, EPI_ISL_3210543, EPI_ISL_3210544, EPI_ISL_3210545, EPI_ISL_3210546, EPI_ISL_3210547, EPI_ISL_3210548, EPI_ISL_3210550, EPI_ISL_3210551, EPI_ISL_3210552, EPI_ISL_3210553, EPI_ISL_3210554, EPI_ISL_3210555, EPI_ISL_3210556, EPI_ISL_3210557, EPI_ISL_3210558, EPI_ISL_3210559, EPI_ISL_3210560, EPI_ISL_3210561, EPI_ISL_3210562, EPI_ISL_3210563, EPI_ISL_3210564, EPI_ISL_3210565, EPI_ISL_3210566, EPI_ISL_3210567, EPI_ISL_3210568, EPI_ISL_3210569, EPI_ISL_3210570, EPI_ISL_3210572, EPI_ISL_3210573, EPI_ISL_3210574, EPI_ISL_3210575, EPI_ISL_3210577, EPI_ISL_3210578, EPI_ISL_3210579, EPI_ISL_3210580, EPI_ISL_3210581, EPI_ISL_3210582, EPI_ISL_3210583, EPI_ISL_3210584, EPI_ISL_3210585, EPI_ISL_3210586, EPI_ISL_3210587, EPI_ISL_3210588, EPI_ISL_3210589, EPI_ISL_3210590, EPI_ISL_3210591, EPI_ISL_3210592, EPI_ISL_3210593, EPI_ISL_3210594, EPI_ISL_3210595, EPI_ISL_3210596, EPI_ISL_3210597, EPI_ISL_3210598, EPI_ISL_3210599, EPI_ISL_3210600, EPI_ISL_3210601, EPI_ISL_3210602, EPI_ISL_3210603, EPI_ISL_3210604, EPI_ISL_3210605, EPI_ISL_3210606, EPI_ISL_3210607, EPI_ISL_3210608, EPI_ISL_3210609, EPI_ISL_3210610, EPI_ISL_3210611, EPI_ISL_3210612, EPI_ISL_3210613, EPI_ISL_3210614, EPI_ISL_3210615, EPI_ISL_3210616, EPI_ISL_3210617, EPI_ISL_3210618, EPI_ISL_3210619, EPI_ISL_3210621, EPI_ISL_3210622, EPI_ISL_3210623, EPI_ISL_3210624, EPI_ISL_3210625, EPI_ISL_3210626, EPI_ISL_3210627, EPI_ISL_3210628, EPI_ISL_3210629, EPI_ISL_3210630, EPI_ISL_3210631, EPI_ISL_3210632, EPI_ISL_3210633, EPI_ISL_3210634, EPI_ISL_3210635, EPI_ISL_3210636, EPI_ISL_3210637, EPI_ISL_3210638, EPI_ISL_3210639, EPI_ISL_3210640, EPI_ISL_3210641, EPI_ISL_3210642, EPI_ISL_3210643, EPI_ISL_3210644, EPI_ISL_3210646, EPI_ISL_3210647, EPI_ISL_3210648, EPI_ISL_3210649, EPI_ISL_3210650, EPI_ISL_3210651, EPI_ISL_3210652, EPI_ISL_3210653, EPI_ISL_3210654, EPI_ISL_3210655, EPI_ISL_3210656, EPI_ISL_3210657, EPI_ISL_3210658, EPI_ISL_3210659, EPI_ISL_3210661, EPI_ISL_3210662, EPI_ISL_3210663, EPI_ISL_3210664, EPI_ISL_3210665, EPI_ISL_3210666, EPI_ISL_3210667, EPI_ISL_3210668, EPI_ISL_3210670, EPI_ISL_3210671, EPI_ISL_3210672, EPI_ISL_3210673, EPI_ISL_3210674, EPI_ISL_3210675, EPI_ISL_3210676, EPI_ISL_3210677, EPI_ISL_3210678, EPI_ISL_3210679, EPI_ISL_3210680, EPI_ISL_3210681, EPI_ISL_3210682, EPI_ISL_3210683, EPI_ISL_3210684, EPI_ISL_3210685, EPI_ISL_3210686, EPI_ISL_3210687, EPI_ISL_3210688, EPI_ISL_3210689, EPI_ISL_3210690, EPI_ISL_3210691, EPI_ISL_3210692, EPI_ISL_3210693, EPI_ISL_3210694, EPI_ISL_3210695, EPI_ISL_3210696, EPI_ISL_3210697, EPI_ISL_3210698, EPI_ISL_3210699, EPI_ISL_3210700, EPI_ISL_3210701, EPI_ISL_3210702, EPI_ISL_3210703, EPI_ISL_3210704, EPI_ISL_3210705, EPI_ISL_3210706, EPI_ISL_3210707, EPI_ISL_3210708, EPI_ISL_3210709, EPI_ISL_3210710, EPI_ISL_3210711, EPI_ISL_3210712, EPI_ISL_3210713, EPI_ISL_3210714, EPI_ISL_3210715, EPI_ISL_3210716, EPI_ISL_3210717, EPI_ISL_3210718, EPI_ISL_3210719, EPI_ISL_3210720, EPI_ISL_3210721, EPI_ISL_3210722, EPI_ISL_3210723, EPI_ISL_3210724, EPI_ISL_3210726, EPI_ISL_3210727, EPI_ISL_3210728, EPI_ISL_3210729, EPI_ISL_3210730, EPI_ISL_3210731, EPI_ISL_3210732, EPI_ISL_3210733, EPI_ISL_3210734, EPI_ISL_3210735, EPI_ISL_3210736, EPI_ISL_3210737, EPI_ISL_3210738, EPI_ISL_3210739, EPI_ISL_3210740, EPI_ISL_3210741, EPI_ISL_3210742, EPI_ISL_3210743, EPI_ISL_3210744, EPI_ISL_3210745, EPI_ISL_3210746, EPI_ISL_3210747, EPI_ISL_3210748, EPI_ISL_3210749, EPI_ISL_3210750, EPI_ISL_3210751, EPI_ISL_3210752, EPI_ISL_3210753, EPI_ISL_3210754, EPI_ISL_3210755, EPI_ISL_3210756, EPI_ISL_3210757, EPI_ISL_3210758, EPI_ISL_3210759, EPI_ISL_3210760, EPI_ISL_3210761, EPI_ISL_3210762, EPI_ISL_3210763, EPI_ISL_3210764, EPI_ISL_3210765, EPI_ISL_3210766, EPI_ISL_3210767, EPI_ISL_3210768, EPI_ISL_3210769, EPI_ISL_3210770, EPI_ISL_3210771, EPI_ISL_3210772, EPI_ISL_3210773, EPI_ISL_3210774, EPI_ISL_3210775, EPI_ISL_3210776, EPI_ISL_3210777, EPI_ISL_3210778, EPI_ISL_3210779, EPI_ISL_3210780, EPI_ISL_3210781, EPI_ISL_3210782, EPI_ISL_3210783, EPI_ISL_3210785, EPI_ISL_3210786, EPI_ISL_3210787, EPI_ISL_3210788, EPI_ISL_3210789, EPI_ISL_3210790, EPI_ISL_3210791, EPI_ISL_3210792, EPI_ISL_3210793, EPI_ISL_3210794, EPI_ISL_3210795, EPI_ISL_3210796, EPI_ISL_3210797, EPI_ISL_3210798, EPI_ISL_3210799, EPI_ISL_3210800, EPI_ISL_3210801, EPI_ISL_3210802, EPI_ISL_3210803, EPI_ISL_3210804, EPI_ISL_3210806, EPI_ISL_3210807, EPI_ISL_3210808, EPI_ISL_3210809, EPI_ISL_3210810, EPI_ISL_3210811, EPI_ISL_3210812, EPI_ISL_3210813, EPI_ISL_3210814, EPI_ISL_3210815, EPI_ISL_3210816, EPI_ISL_3210817, EPI_ISL_3210818, EPI_ISL_3210820, EPI_ISL_3210821, EPI_ISL_3210822, EPI_ISL_3210823, EPI_ISL_3210824, EPI_ISL_3210825, EPI_ISL_3210826, EPI_ISL_3210827, EPI_ISL_3210828, EPI_ISL_3210829, EPI_ISL_3210830, EPI_ISL_3210831, EPI_ISL_3210832, EPI_ISL_3210833, EPI_ISL_3210835, EPI_ISL_3210836, EPI_ISL_3210837, EPI_ISL_3210838, EPI_ISL_3210839, EPI_ISL_3210840, EPI_ISL_3210841, EPI_ISL_3210842, EPI_ISL_3210843, EPI_ISL_3210844, EPI_ISL_3210845, EPI_ISL_3210846, EPI_ISL_3210847, EPI_ISL_3210848, EPI_ISL_3210849, EPI_ISL_3210850, EPI_ISL_3210851, EPI_ISL_3210852, EPI_ISL_3210853, EPI_ISL_3210854, EPI_ISL_3210855, EPI_ISL_3210856, EPI_ISL_3210857, EPI_ISL_3210858, EPI_ISL_3210859, EPI_ISL_3210860, EPI_ISL_3210861, EPI_ISL_3210862, EPI_ISL_3210864, EPI_ISL_3210865, EPI_ISL_3210866, EPI_ISL_3210867, EPI_ISL_3210868, EPI_ISL_3210869, EPI_ISL_3210870, EPI_ISL_3210871, EPI_ISL_3210872, EPI_ISL_3210873, EPI_ISL_3210874, EPI_ISL_3210875, EPI_ISL_3210876, EPI_ISL_3210877, EPI_ISL_3210879, EPI_ISL_3210880, EPI_ISL_3210881, EPI_ISL_3210882, EPI_ISL_3210883, EPI_ISL_3210884, EPI_ISL_3210885, EPI_ISL_3210886, EPI_ISL_3210887, EPI_ISL_3210888, EPI_ISL_3210889, EPI_ISL_3210890, EPI_ISL_3210891, EPI_ISL_3210892, EPI_ISL_3210893, EPI_ISL_3210895, EPI_ISL_3210896, EPI_ISL_3210897, EPI_ISL_3210898, EPI_ISL_3210899, EPI_ISL_3210900, EPI_ISL_3210901, EPI_ISL_3210902, EPI_ISL_3210903, EPI_ISL_3210904, EPI_ISL_3210905, EPI_ISL_3210906, EPI_ISL_3210907, EPI_ISL_3210908, EPI_ISL_3210909, EPI_ISL_3210910, EPI_ISL_3210911, EPI_ISL_3210912, EPI_ISL_3210913, EPI_ISL_3210914, EPI_ISL_3210915, EPI_ISL_3210916, EPI_ISL_3210918, EPI_ISL_3210919, EPI_ISL_3210920, EPI_ISL_3210921, EPI_ISL_3210922, EPI_ISL_3210923, EPI_ISL_3210924, EPI_ISL_3210925, EPI_ISL_3210926, EPI_ISL_3210927, EPI_ISL_3210928 | | | |
| see above | Fulgent Genetics | Centers for Disease Control and Prevention Division of Viral Diseases, Pathogen Discovery | Dakota Howard, Dhwani Batra, Peter W. Cook, Kara Moser, Adrian Paskey, Jason Caravas, Benjamin Rambo-Martin, Shatavia Morrison, Christopher Gulvick, Scott Sammons, Yvette Unoarumhi, Darlene Wagner, Matthew Schmerer, Harry Gao, Mickey Li, John Gao, Joseph Fierro, Benafsh Sapra, Becky Tsai, Yan Meng, Doreen Ng, James Xie, Clinton R. Paden, Duncan MacCannell |
| EPI_ISL_3210929 | ZOTZ KLIMAS MVZ Düsseldorf-Centrum GbR ÜBAG für Labormedizin, Genetik, Zytologie, Pathologie | Center of Medical Microbiology, Virology, and Hospital Hygiene, University of Duesseldorf | Maximilian Damagnez;Alexander Dilthey;Patrick Finzer;Katrin Hoffmann;Torsten Houwaart;Lisanna Hülse;Malte Kohns Vasconcelos;Nadine Lübke;Jessica Nicolai;Klaus Pfeffer;Daniel Strelow;Jörg Timm;Andreas Walker;Tobias Wienemann;Rainer Zotz |
| EPI_ISL_3210930 | Fulgent Genetics | Centers for Disease Control and Prevention Division of Viral Diseases, Pathogen Discovery | Dakota Howard, Dhwani Batra, Peter W. Cook, Kara Moser, Adrian Paskey, Jason Caravas, Benjamin Rambo-Martin, Shatavia Morrison, Christopher Gulvick, Scott Sammons, Yvette Unoarumhi, Darlene Wagner, Matthew Schmerer, Harry Gao, Mickey Li, John Gao, Joseph Fierro, Benafsh Sapra, Becky Tsai, Yan Meng, Doreen Ng, James Xie, Clinton R. Paden, Duncan MacCannell |
| EPI_ISL_3210931, EPI_ISL_3210932 | ZOTZ KLIMAS MVZ Düsseldorf-Centrum GbR ÜBAG für Labormedizin, Genetik, Zytologie, Pathologie | Center of Medical Microbiology, Virology, and Hospital Hygiene, University of Duesseldorf | Maximilian Damagnez;Alexander Dilthey;Patrick Finzer;Katrin Hoffmann;Torsten Houwaart;Lisanna Hülse;Malte Kohns Vasconcelos;Nadine Lübke;Jessica Nicolai;Klaus Pfeffer;Daniel Strelow;Jörg Timm;Andreas Walker;Tobias Wienemann;Rainer Zotz |
| EPI_ISL_3210933 | Fulgent Genetics | Centers for Disease Control and Prevention Division of Viral Diseases, Pathogen Discovery | Dakota Howard, Dhwani Batra, Peter W. Cook, Kara Moser, Adrian Paskey, Jason Caravas, Benjamin Rambo-Martin, Shatavia Morrison, Christopher Gulvick, Scott Sammons, Yvette Unoarumhi, Darlene Wagner, Matthew Schmerer, Harry Gao, Mickey Li, John Gao, Joseph Fierro, Benafsh Sapra, Becky Tsai, Yan Meng, Doreen Ng, James Xie, Clinton R. Paden, Duncan MacCannell |
| EPI_ISL_3210934, EPI_ISL_3210935 | ZOTZ KLIMAS MVZ Düsseldorf-Centrum GbR ÜBAG für Labormedizin, Genetik, Zytologie, Pathologie | Center of Medical Microbiology, Virology, and Hospital Hygiene, University of Duesseldorf | Maximilian Damagnez;Alexander Dilthey;Patrick Finzer;Katrin Hoffmann;Torsten Houwaart;Lisanna Hülse;Malte Kohns Vasconcelos;Nadine Lübke;Jessica Nicolai;Klaus Pfeffer;Daniel Strelow;Jörg Timm;Andreas Walker;Tobias Wienemann;Rainer Zotz |
| EPI_ISL_3210936, EPI_ISL_3210937, EPI_ISL_3210938, EPI_ISL_3210939, EPI_ISL_3210940, EPI_ISL_3210941, EPI_ISL_3210942, EPI_ISL_3210943, EPI_ISL_3210944, EPI_ISL_3210945, EPI_ISL_3210946, EPI_ISL_3210947, EPI_ISL_3210948, EPI_ISL_3210949, EPI_ISL_3210950, EPI_ISL_3210951, EPI_ISL_3210952, EPI_ISL_3210953, EPI_ISL_3210954, EPI_ISL_3210955, EPI_ISL_3210956, EPI_ISL_3210957, EPI_ISL_3210959, EPI_ISL_3210960, EPI_ISL_3210961, EPI_ISL_3210962, EPI_ISL_3210963, EPI_ISL_3210964, EPI_ISL_3210965, EPI_ISL_3210966, EPI_ISL_3210967, EPI_ISL_3210968, EPI_ISL_3210969, EPI_ISL_3210970, EPI_ISL_3210971, EPI_ISL_3210972, EPI_ISL_3210973, EPI_ISL_3210974, EPI_ISL_3210975, EPI_ISL_3210976, EPI_ISL_3210977, EPI_ISL_3210978, EPI_ISL_3210979, EPI_ISL_3210980, EPI_ISL_3210981, EPI_ISL_3210982, EPI_ISL_3210983, EPI_ISL_3210984, EPI_ISL_3210985, EPI_ISL_3210986, EPI_ISL_3210987, EPI_ISL_3210988, EPI_ISL_3210989, EPI_ISL_3210990, EPI_ISL_3210991, EPI_ISL_3210992, EPI_ISL_3210993, EPI_ISL_3210994, EPI_ISL_3210995, EPI_ISL_3210996, EPI_ISL_3210997, EPI_ISL_3210998, EPI_ISL_3210999, EPI_ISL_3211000, EPI_ISL_3211001, EPI_ISL_3211002, EPI_ISL_3211003, EPI_ISL_3211004, EPI_ISL_3211005, EPI_ISL_3211006, EPI_ISL_3211007, EPI_ISL_3211008, EPI_ISL_3211009, EPI_ISL_3211010, EPI_ISL_3211011, EPI_ISL_3211012, EPI_ISL_3211013, EPI_ISL_3211014, EPI_ISL_3211015, EPI_ISL_3211016, EPI_ISL_3211017, EPI_ISL_3211018, EPI_ISL_3211019, EPI_ISL_3211020, EPI_ISL_3211021, EPI_ISL_3211022, EPI_ISL_3211023, EPI_ISL_3211024, EPI_ISL_3211025, EPI_ISL_3211026, EPI_ISL_3211027, EPI_ISL_3211029, EPI_ISL_3211030, EPI_ISL_3211032, EPI_ISL_3211033, EPI_ISL_3211034, EPI_ISL_3211035, EPI_ISL_3211036, EPI_ISL_3211037, EPI_ISL_3211038, EPI_ISL_3211039, EPI_ISL_3211040, EPI_ISL_3211041, EPI_ISL_3211042, EPI_ISL_3211043, EPI_ISL_3211044, EPI_ISL_3211045, EPI_ISL_3211046, EPI_ISL_3211047, EPI_ISL_3211048, EPI_ISL_3211049, EPI_ISL_3211050, EPI_ISL_3211051, EPI_ISL_3211052, EPI_ISL_3211053, EPI_ISL_3211054, EPI_ISL_3211055, EPI_ISL_3211056, EPI_ISL_3211057, EPI_ISL_3211058, EPI_ISL_3211059, EPI_ISL_3211060, EPI_ISL_3211061, EPI_ISL_3211062, EPI_ISL_3211063, EPI_ISL_3211064, EPI_ISL_3211065, EPI_ISL_3211066, EPI_ISL_3211067, EPI_ISL_3211068, EPI_ISL_3211069, EPI_ISL_3211070, EPI_ISL_3211071, EPI_ISL_3211072, EPI_ISL_3211073, EPI_ISL_3211074, EPI_ISL_3211075, EPI_ISL_3211076, EPI_ISL_3211077, EPI_ISL_3211078, EPI_ISL_3211079, EPI_ISL_3211080, EPI_ISL_3211081, EPI_ISL_3211082, EPI_ISL_3211083, EPI_ISL_3211084, EPI_ISL_3211085, EPI_ISL_3211086, EPI_ISL_3211088, EPI_ISL_3211089, EPI_ISL_3211090, EPI_ISL_3211091, EPI_ISL_3211092, EPI_ISL_3211093, EPI_ISL_3211094, EPI_ISL_3211095, EPI_ISL_3211096, EPI_ISL_3211097, EPI_ISL_3211098, EPI_ISL_3211099, EPI_ISL_3211100, EPI_ISL_3211101, EPI_ISL_3211102, EPI_ISL_3211103, EPI_ISL_3211104, EPI_ISL_3211105, EPI_ISL_3211106, EPI_ISL_3211107, EPI_ISL_3211108, EPI_ISL_3211109, EPI_ISL_3211110, EPI_ISL_3211111, EPI_ISL_3211112, EPI_ISL_3211113, EPI_ISL_3211114, EPI_ISL_3211115, EPI_ISL_3211116, EPI_ISL_3211117, EPI_ISL_3211118, EPI_ISL_3211119, EPI_ISL_3211120, EPI_ISL_3211121, EPI_ISL_3211122, EPI_ISL_3211123, EPI_ISL_3211124, EPI_ISL_3211125, EPI_ISL_3211126, EPI_ISL_3211127, EPI_ISL_3211128, EPI_ISL_3211129, EPI_ISL_3211130, EPI_ISL_3211131, EPI_ISL_3211132, EPI_ISL_3211133, EPI_ISL_3211134, EPI_ISL_3211135, EPI_ISL_3211136, EPI_ISL_3211137, EPI_ISL_3211138, EPI_ISL_3211139, EPI_ISL_3211140, EPI_ISL_3211141, EPI_ISL_3211142, EPI_ISL_3211143, EPI_ISL_3211144, EPI_ISL_3211145, EPI_ISL_3211146, EPI_ISL_3211147, EPI_ISL_3211148, EPI_ISL_3211149, EPI_ISL_3211150, EPI_ISL_3211151, EPI_ISL_3211152, EPI_ISL_3211153, EPI_ISL_3211154, EPI_ISL_3211155, EPI_ISL_3211156, EPI_ISL_3211157, EPI_ISL_3211158, EPI_ISL_3211159, EPI_ISL_3211160, EPI_ISL_3211161, EPI_ISL_3211162, EPI_ISL_3211163, EPI_ISL_3211164, EPI_ISL_3211165, EPI_ISL_3211166, EPI_ISL_3211167, EPI_ISL_3211168, EPI_ISL_3211169, EPI_ISL_3211170, EPI_ISL_3211171, EPI_ISL_3211172, EPI_ISL_3211173, EPI_ISL_3211174, EPI_ISL_3211175, EPI_ISL_3211176, EPI_ISL_3211177, EPI_ISL_3211178, EPI_ISL_3211179, EPI_ISL_3211180, EPI_ISL_3211181, EPI_ISL_3211182, EPI_ISL_3211183, EPI_ISL_3211184, EPI_ISL_3211185, EPI_ISL_3211186, EPI_ISL_3211187, EPI_ISL_3211188, EPI_ISL_3211189, EPI_ISL_3211190, EPI_ISL_3211191, EPI_ISL_3211192, EPI_ISL_3211193, EPI_ISL_3211194, EPI_ISL_3211195, EPI_ISL_3211196, EPI_ISL_3211197, EPI_ISL_3211198, EPI_ISL_3211199, EPI_ISL_3211200, EPI_ISL_3211201, EPI_ISL_3211202, EPI_ISL_3211203, EPI_ISL_3211204, EPI_ISL_3211205, EPI_ISL_3211206, EPI_ISL_3211207, EPI_ISL_3211208, EPI_ISL_3211209, EPI_ISL_3211210, EPI_ISL_3211211, EPI_ISL_3211212, EPI_ISL_3211213, EPI_ISL_3211214, EPI_ISL_3211215, EPI_ISL_3211216, EPI_ISL_3211217, EPI_ISL_3211218, EPI_ISL_3211219, EPI_ISL_3211220, EPI_ISL_3211221, EPI_ISL_3211222, EPI_ISL_3211223, EPI_ISL_3211224, EPI_ISL_3211225, EPI_ISL_3211226, EPI_ISL_3211227, EPI_ISL_3211228, EPI_ISL_3211229, EPI_ISL_3211230, EPI_ISL_3211231, EPI_ISL_3211232, EPI_ISL_3211233, EPI_ISL_3211234, EPI_ISL_3211235, EPI_ISL_3211236, EPI_ISL_3211237, EPI_ISL_3211238, EPI_ISL_3211239, EPI_ISL_3211240, EPI_ISL_3211241, EPI_ISL_3211242, EPI_ISL_3211243, EPI_ISL_3211244, EPI_ISL_3211245, EPI_ISL_3211246, EPI_ISL_3211247, EPI_ISL_3211248, EPI_ISL_3211249, EPI_ISL_3211250, EPI_ISL_3211251, EPI_ISL_3211252, EPI_ISL_3211253, EPI_ISL_3211254, EPI_ISL_3211255, EPI_ISL_3211256, EPI_ISL_3211257, EPI_ISL_3211258, EPI_ISL_3211259, EPI_ISL_3211260, EPI_ISL_3211261, EPI_ISL_3211262, EPI_ISL_3211263, EPI_ISL_3211264, EPI_ISL_3211265, EPI_ISL_3211266, EPI_ISL_3211267, EPI_ISL_3211268, EPI_ISL_3211269, EPI_ISL_3211270, EPI_ISL_3211271, EPI_ISL_3211272, EPI_ISL_3211273, EPI_ISL_3211274, EPI_ISL_3211275, EPI_ISL_3211276, EPI_ISL_3211277, EPI_ISL_3211278, EPI_ISL_3211279, EPI_ISL_3211280, EPI_ISL_3211281, EPI_ISL_3211282, EPI_ISL_3211283, EPI_ISL_3211284, EPI_ISL_3211285, EPI_ISL_3211286, EPI_ISL_3211287, EPI_ISL_3211288, EPI_ISL_3211289, EPI_ISL_3211290, EPI_ISL_3211291, EPI_ISL_3211292, EPI_ISL_3211293, EPI_ISL_3211294, EPI_ISL_3211295, EPI_ISL_3211296, EPI_ISL_3211297, EPI_ISL_3211298, EPI_ISL_3211299, EPI_ISL_3211300, EPI_ISL_3211301, EPI_ISL_3211302, EPI_ISL_3211303, EPI_ISL_3211304, EPI_ISL_3211305, EPI_ISL_3211306, EPI_ISL_3211307, EPI_ISL_3211308, EPI_ISL_3211309, EPI_ISL_3211310, EPI_ISL_3211311, EPI_ISL_3211312, EPI_ISL_3211313, EPI_ISL_3211315, EPI_ISL_3211316, EPI_ISL_3211317, EPI_ISL_3211318, EPI_ISL_3211319, EPI_ISL_3211320, EPI_ISL_3211321, EPI_ISL_3211322, EPI_ISL_3211323, EPI_ISL_3211324, EPI_ISL_3211325, EPI_ISL_3211326, EPI_ISL_3211327, EPI_ISL_3211328, EPI_ISL_3211329, EPI_ISL_3211330, EPI_ISL_3211331, EPI_ISL_3211332, EPI_ISL_3211333, EPI_ISL_3211334, EPI_ISL_3211335, EPI_ISL_3211336, EPI_ISL_3211337, EPI_ISL_3211338, EPI_ISL_3211339, EPI_ISL_3211340, EPI_ISL_3211341, EPI_ISL_3211342, EPI_ISL_3211343, EPI_ISL_3211344, EPI_ISL_3211345, EPI_ISL_3211346, EPI_ISL_3211347, EPI_ISL_3211348, EPI_ISL_3211350, EPI_ISL_3211351, EPI_ISL_3211352, EPI_ISL_3211353, EPI_ISL_3211354, EPI_ISL_3211355, EPI_ISL_3211356, EPI_ISL_3211357, EPI_ISL_3211358, EPI_ISL_3211359, EPI_ISL_3211360, EPI_ISL_3211361, EPI_ISL_3211362, EPI_ISL_3211363, EPI_ISL_3211364, EPI_ISL_3211365, EPI_ISL_3211366, EPI_ISL_3211367, EPI_ISL_3211368, EPI_ISL_3211369, EPI_ISL_3211370, EPI_ISL_3211371, EPI_ISL_3211372, EPI_ISL_3211373, EPI_ISL_3211374, EPI_ISL_3211375, EPI_ISL_3211376, EPI_ISL_3211377, EPI_ISL_3211378, EPI_ISL_3211379, EPI_ISL_3211380, EPI_ISL_3211381, EPI_ISL_3211382, EPI_ISL_3211383, EPI_ISL_3211384, EPI_ISL_3211385, EPI_ISL_3211386, EPI_ISL_3211387, EPI_ISL_3211390, EPI_ISL_3211391, EPI_ISL_3211392, EPI_ISL_3211393, EPI_ISL_3211394, EPI_ISL_3211395, EPI_ISL_3211396, EPI_ISL_3211398, EPI_ISL_3211399, EPI_ISL_3211400, EPI_ISL_3211401, EPI_ISL_3211402, EPI_ISL_3211403, EPI_ISL_3211404, EPI_ISL_3211405, EPI_ISL_3211406, EPI_ISL_3211407, EPI_ISL_3211408, EPI_ISL_3211409, EPI_ISL_3211410, EPI_ISL_3211411, EPI_ISL_3211412, EPI_ISL_3211413, EPI_ISL_3211414, EPI_ISL_3211415, EPI_ISL_3211416, EPI_ISL_3211417, EPI_ISL_3211418, EPI_ISL_3211419, EPI_ISL_3211420, EPI_ISL_3211421, EPI_ISL_3211422, EPI_ISL_3211423, EPI_ISL_3211424, EPI_ISL_3211425, EPI_ISL_3211426, EPI_ISL_3211427, EPI_ISL_3211428, EPI_ISL_3211429, EPI_ISL_3211430, EPI_ISL_3211431, EPI_ISL_3211432, EPI_ISL_3211433, EPI_ISL_3211434, EPI_ISL_3211435, EPI_ISL_3211436, EPI_ISL_3211437, EPI_ISL_3211438, EPI_ISL_3211440, EPI_ISL_3211441, EPI_ISL_3211442, EPI_ISL_3211443, EPI_ISL_3211444, EPI_ISL_3211445, EPI_ISL_3211446, EPI_ISL_3211447, EPI_ISL_3211448, EPI_ISL_3211449, EPI_ISL_3211451, EPI_ISL_3211452, EPI_ISL_3211453, EPI_ISL_3211454, EPI_ISL_3211455, EPI_ISL_3211456, EPI_ISL_3211457, EPI_ISL_3211458, EPI_ISL_3211459, EPI_ISL_3211460, EPI_ISL_3211461, EPI_ISL_3211462, EPI_ISL_3211463, EPI_ISL_3211464, EPI_ISL_3211465, EPI_ISL_3211466, EPI_ISL_3211467, EPI_ISL_3211468, EPI_ISL_3211469, EPI_ISL_3211470, EPI_ISL_3211471, EPI_ISL_3211472, EPI_ISL_3211473, EPI_ISL_3211474, EPI_ISL_3211475, EPI_ISL_3211476, EPI_ISL_3211477, EPI_ISL_3211478, EPI_ISL_3211479, EPI_ISL_3211480, EPI_ISL_3211481, EPI_ISL_3211482, EPI_ISL_3211483, EPI_ISL_3211484, EPI_ISL_3211485, EPI_ISL_3211486, EPI_ISL_3211487, EPI_ISL_3211488, EPI_ISL_3211489, EPI_ISL_3211490, EPI_ISL_3211491, EPI_ISL_3211492, EPI_ISL_3211493, EPI_ISL_3211494, EPI_ISL_3211495, EPI_ISL_3211496, EPI_ISL_3211497, EPI_ISL_3211498, EPI_ISL_3211499, EPI_ISL_3211500, EPI_ISL_3211501, EPI_ISL_3211502, EPI_ISL_3211503, EPI_ISL_3211504, EPI_ISL_3211505, EPI_ISL_3211506, EPI_ISL_3211507, EPI_ISL_3211508, EPI_ISL_3211509, EPI_ISL_3211510, EPI_ISL_3211511, EPI_ISL_3211513, EPI_ISL_3211514, EPI_ISL_3211515, EPI_ISL_3211516, EPI_ISL_3211517, EPI_ISL_3211518, EPI_ISL_3211519, EPI_ISL_3211520, EPI_ISL_3211521, EPI_ISL_3211522, EPI_ISL_3211524, EPI_ISL_3211525, EPI_ISL_3211526, EPI_ISL_3211527, EPI_ISL_3211528, EPI_ISL_3211529, EPI_ISL_3211530, EPI_ISL_3211531, EPI_ISL_3211532, EPI_ISL_3211533, EPI_ISL_3211534, EPI_ISL_3211535, EPI_ISL_3211536, EPI_ISL_3211537, EPI_ISL_3211538, EPI_ISL_3211539, EPI_ISL_3211541, EPI_ISL_3211543, EPI_ISL_3211544, EPI_ISL_3211545, EPI_ISL_3211546, EPI_ISL_3211547, EPI_ISL_3211548, EPI_ISL_3211549, EPI_ISL_3211550, EPI_ISL_3211551, EPI_ISL_3211552, EPI_ISL_3211553, EPI_ISL_3211554, EPI_ISL_3211555, EPI_ISL_3211556, EPI_ISL_3211558, EPI_ISL_3211559, EPI_ISL_3211560, EPI_ISL_3211561, EPI_ISL_3211562, EPI_ISL_3211563, EPI_ISL_3211564, EPI_ISL_3211565, EPI_ISL_3211566, EPI_ISL_3211567, EPI_ISL_3211568, EPI_ISL_3211569, EPI_ISL_3211570, EPI_ISL_3211571, EPI_ISL_3211573, EPI_ISL_3211574, EPI_ISL_3211575, EPI_ISL_3211576, EPI_ISL_3211577, EPI_ISL_3211578, EPI_ISL_3211579, EPI_ISL_3211580, EPI_ISL_3211581, EPI_ISL_3211582, EPI_ISL_3211583, EPI_ISL_3211584, EPI_ISL_3211586, EPI_ISL_3211587, EPI_ISL_3211588, EPI_ISL_3211589, EPI_ISL_3211590, EPI_ISL_3211591, EPI_ISL_3211592, EPI_ISL_3211593, EPI_ISL_3211594, EPI_ISL_3211595, EPI_ISL_3211596, EPI_ISL_3211597, EPI_ISL_3211598, EPI_ISL_3211599, EPI_ISL_3211600, EPI_ISL_3211601, EPI_ISL_3211602, EPI_ISL_3211603, EPI_ISL_3211604, EPI_ISL_3211605, EPI_ISL_3211607, EPI_ISL_3211608, EPI_ISL_3211609, EPI_ISL_3211610, EPI_ISL_3211611, EPI_ISL_3211612, EPI_ISL_3211613, EPI_ISL_3211615, | | | |

EPI_ISL_3211616, EPI_ISL_3211617, EPI_ISL_3211618, EPI_ISL_3211619, EPI_ISL_3211620, EPI_ISL_3211621, EPI_ISL_3211622, EPI_ISL_3211623, EPI_ISL_3211624, EPI_ISL_3211625, EPI_ISL_3211626, EPI_ISL_3211627, EPI_ISL_3211628, EPI_ISL_3211629, EPI_ISL_3211630, EPI_ISL_3211631, EPI_ISL_3211632, EPI_ISL_3211633, EPI_ISL_3211634, EPI_ISL_3211635, EPI_ISL_3211636, EPI_ISL_3211637,
EPI_ISL_3211638, EPI_ISL_3211639, EPI_ISL_3211640, EPI_ISL_3211641, EPI_ISL_3211642, EPI_ISL_3211643, EPI_ISL_3211644, EPI_ISL_3211645, EPI_ISL_3211646, EPI_ISL_3211647, EPI_ISL_3211648, EPI_ISL_3211649, EPI_ISL_3211650, EPI_ISL_3211651, EPI_ISL_3211652, EPI_ISL_3211653, EPI_ISL_3211654, EPI_ISL_3211655, EPI_ISL_3211656, EPI_ISL_3211657, EPI_ISL_3211658, EPI_ISL_3211659,
EPI_ISL_3211660, EPI_ISL_3211661, EPI_ISL_3211662, EPI_ISL_3211663, EPI_ISL_3211664, EPI_ISL_3211665, EPI_ISL_3211666, EPI_ISL_3211667, EPI_ISL_3211668, EPI_ISL_3211669, EPI_ISL_3211670, EPI_ISL_3211671, EPI_ISL_3211672, EPI_ISL_3211673, EPI_ISL_3211674, EPI_ISL_3211675, EPI_ISL_3211676, EPI_ISL_3211677, EPI_ISL_3211678, EPI_ISL_3211679, EPI_ISL_3211680, EPI_ISL_3211681,
EPI_ISL_3211682, EPI_ISL_3211683, EPI_ISL_3211684, EPI_ISL_3211685, EPI_ISL_3211687, EPI_ISL_3211688, EPI_ISL_3211690, EPI_ISL_3211691, EPI_ISL_3211692, EPI_ISL_3211693, EPI_ISL_3211694, EPI_ISL_3211695, EPI_ISL_3211696, EPI_ISL_3211697, EPI_ISL_3211698, EPI_ISL_3211699, EPI_ISL_3211700, EPI_ISL_3211701, EPI_ISL_3211702, EPI_ISL_3211703, EPI_ISL_3211704, EPI_ISL_3211705,
EPI_ISL_3211706, EPI_ISL_3211707, EPI_ISL_3211708, EPI_ISL_3211709, EPI_ISL_3211710, EPI_ISL_3211711, EPI_ISL_3211712, EPI_ISL_3211714, EPI_ISL_3211715, EPI_ISL_3211716, EPI_ISL_3211717, EPI_ISL_3211718, EPI_ISL_3211719, EPI_ISL_3211720, EPI_ISL_3211721, EPI_ISL_3211722, EPI_ISL_3211723, EPI_ISL_3211724, EPI_ISL_3211725, EPI_ISL_3211727, EPI_ISL_3211728, EPI_ISL_3211729,
EPI_ISL_3211730, EPI_ISL_3211731, EPI_ISL_3211732, EPI_ISL_3211733, EPI_ISL_3211734, EPI_ISL_3211735, EPI_ISL_3211736, EPI_ISL_3211737, EPI_ISL_3211738, EPI_ISL_3211740, EPI_ISL_3211741, EPI_ISL_3211742, EPI_ISL_3211743, EPI_ISL_3211744, EPI_ISL_3211745, EPI_ISL_3211746, EPI_ISL_3211747, EPI_ISL_3211748, EPI_ISL_3211749, EPI_ISL_3211750, EPI_ISL_3211751, EPI_ISL_3211752,
EPI_ISL_3211753, EPI_ISL_3211754, EPI_ISL_3211755, EPI_ISL_3211756, EPI_ISL_3211757, EPI_ISL_3211758, EPI_ISL_3211759, EPI_ISL_3211760, EPI_ISL_3211761, EPI_ISL_3211762, EPI_ISL_3211763, EPI_ISL_3211764, EPI_ISL_3211765, EPI_ISL_3211766, EPI_ISL_3211767, EPI_ISL_3211768, EPI_ISL_3211769, EPI_ISL_3211770, EPI_ISL_3211772, EPI_ISL_3211773, EPI_ISL_3211774, EPI_ISL_3211775,
EPI_ISL_3211776, EPI_ISL_3211777, EPI_ISL_3211778, EPI_ISL_3211779, EPI_ISL_3211780, EPI_ISL_3211781, EPI_ISL_3211782, EPI_ISL_3211783, EPI_ISL_3211784, EPI_ISL_3211785, EPI_ISL_3211786, EPI_ISL_3211787, EPI_ISL_3211788, EPI_ISL_3211789, EPI_ISL_3211790, EPI_ISL_3211791, EPI_ISL_3211792, EPI_ISL_3211793, EPI_ISL_3211794, EPI_ISL_3211795, EPI_ISL_3211796, EPI_ISL_3211797,
EPI_ISL_3211798, EPI_ISL_3211799, EPI_ISL_3211800, EPI_ISL_3211801, EPI_ISL_3211802, EPI_ISL_3211804, EPI_ISL_3211806, EPI_ISL_3211807, EPI_ISL_3211808, EPI_ISL_3211809, EPI_ISL_3211811, EPI_ISL_3211812, EPI_ISL_3211813, EPI_ISL_3211814, EPI_ISL_3211815, EPI_ISL_3211816, EPI_ISL_3211817, EPI_ISL_3211818, EPI_ISL_3211819, EPI_ISL_3211820, EPI_ISL_3211821, EPI_ISL_3211822,
EPI_ISL_3211823, EPI_ISL_3211824, EPI_ISL_3211825, EPI_ISL_3211826, EPI_ISL_3211827, EPI_ISL_3211828, EPI_ISL_3211829, EPI_ISL_3211830, EPI_ISL_3211831, EPI_ISL_3211832, EPI_ISL_3211833, EPI_ISL_3211835, EPI_ISL_3211836, EPI_ISL_3211837, EPI_ISL_3211838, EPI_ISL_3211839, EPI_ISL_3211840, EPI_ISL_3211841, EPI_ISL_3211842, EPI_ISL_3211843, EPI_ISL_3211844, EPI_ISL_3211845,
EPI_ISL_3211846, EPI_ISL_3211847, EPI_ISL_3211848, EPI_ISL_3211849, EPI_ISL_3211850, EPI_ISL_3211851, EPI_ISL_3211852, EPI_ISL_3211853, EPI_ISL_3211854, EPI_ISL_3211855, EPI_ISL_3211856, EPI_ISL_3211857, EPI_ISL_3211858, EPI_ISL_3211859, EPI_ISL_3211860, EPI_ISL_3211861, EPI_ISL_3211862, EPI_ISL_3211863, EPI_ISL_3211864, EPI_ISL_3211865, EPI_ISL_3211866, EPI_ISL_3211867,
EPI_ISL_3211868, EPI_ISL_3211869, EPI_ISL_3211870, EPI_ISL_3211871, EPI_ISL_3211872, EPI_ISL_3211873, EPI_ISL_3211874, EPI_ISL_3211875, EPI_ISL_3211876, EPI_ISL_3211877, EPI_ISL_3211878, EPI_ISL_3211879, EPI_ISL_3211881, EPI_ISL_3211882, EPI_ISL_3211883, EPI_ISL_3211884, EPI_ISL_3211885, EPI_ISL_3211886, EPI_ISL_3211887, EPI_ISL_3211888, EPI_ISL_3211889, EPI_ISL_3211890,
EPI_ISL_3211891, EPI_ISL_3211892, EPI_ISL_3211893, EPI_ISL_3211894, EPI_ISL_3211895, EPI_ISL_3211896, EPI_ISL_3211897, EPI_ISL_3211898, EPI_ISL_3211899, EPI_ISL_3211900, EPI_ISL_3211901, EPI_ISL_3211902, EPI_ISL_3211903, EPI_ISL_3211904, EPI_ISL_3211905, EPI_ISL_3211906, EPI_ISL_3211907, EPI_ISL_3211908, EPI_ISL_3211909, EPI_ISL_3211910, EPI_ISL_3211911, EPI_ISL_3211912,
EPI_ISL_3211913, EPI_ISL_3211914, EPI_ISL_3211915, EPI_ISL_3211916, EPI_ISL_3211917, EPI_ISL_3211918, EPI_ISL_3211919, EPI_ISL_3211920, EPI_ISL_3211921, EPI_ISL_3211922, EPI_ISL_3211923, EPI_ISL_3211924, EPI_ISL_3211925, EPI_ISL_3211926, EPI_ISL_3211927, EPI_ISL_3211928, EPI_ISL_3211929, EPI_ISL_3211930, EPI_ISL_3211931, EPI_ISL_3211933, EPI_ISL_3211934, EPI_ISL_3211935,
EPI_ISL_3211936, EPI_ISL_3211937, EPI_ISL_3211938, EPI_ISL_3211940, EPI_ISL_3211941, EPI_ISL_3211942, EPI_ISL_3211943, EPI_ISL_3211944, EPI_ISL_3211945, EPI_ISL_3211946, EPI_ISL_3211947, EPI_ISL_3211948, EPI_ISL_3211949, EPI_ISL_3211950, EPI_ISL_3211952, EPI_ISL_3211953, EPI_ISL_3211954, EPI_ISL_3211955, EPI_ISL_3211956, EPI_ISL_3211957, EPI_ISL_3211958, EPI_ISL_3211959,
EPI_ISL_3211960, EPI_ISL_3211961, EPI_ISL_3211963, EPI_ISL_3211964, EPI_ISL_3211965, EPI_ISL_3211966, EPI_ISL_3211968, EPI_ISL_3211969, EPI_ISL_3211970, EPI_ISL_3211971, EPI_ISL_3211972, EPI_ISL_3211973, EPI_ISL_3211974, EPI_ISL_3211975, EPI_ISL_3211977, EPI_ISL_3211978, EPI_ISL_3211979, EPI_ISL_3211980, EPI_ISL_3211981, EPI_ISL_3211982, EPI_ISL_3211983, EPI_ISL_3211984,
EPI_ISL_3211985, EPI_ISL_3211986, EPI_ISL_3211987, EPI_ISL_3211988, EPI_ISL_3211989, EPI_ISL_3211990, EPI_ISL_3211991, EPI_ISL_3211992, EPI_ISL_3211993, EPI_ISL_3211994, EPI_ISL_3211995, EPI_ISL_3211996, EPI_ISL_3211997, EPI_ISL_3211998, EPI_ISL_3211999, EPI_ISL_3212000, EPI_ISL_3212001, EPI_ISL_3212002, EPI_ISL_3212003, EPI_ISL_3212004, EPI_ISL_3212005, EPI_ISL_3212006,
EPI_ISL_3212007, EPI_ISL_3212008, EPI_ISL_3212009, EPI_ISL_3212010, EPI_ISL_3212011, EPI_ISL_3212012, EPI_ISL_3212013, EPI_ISL_3212014, EPI_ISL_3212015, EPI_ISL_3212017, EPI_ISL_3212018, EPI_ISL_3212019, EPI_ISL_3212020, EPI_ISL_3212021, EPI_ISL_3212022, EPI_ISL_3212023, EPI_ISL_3212024, EPI_ISL_3212025, EPI_ISL_3212026, EPI_ISL_3212027, EPI_ISL_3212028, EPI_ISL_3212029,
EPI_ISL_3212030, EPI_ISL_3212031, EPI_ISL_3212032, EPI_ISL_3212033, EPI_ISL_3212034, EPI_ISL_3212035, EPI_ISL_3212036, EPI_ISL_3212037, EPI_ISL_3212038, EPI_ISL_3212039, EPI_ISL_3212040, EPI_ISL_3212041, EPI_ISL_3212042, EPI_ISL_3212043, EPI_ISL_3212044, EPI_ISL_3212045, EPI_ISL_3212046, EPI_ISL_3212047, EPI_ISL_3212048, EPI_ISL_3212049, EPI_ISL_3212050, EPI_ISL_3212051,
EPI_ISL_3212052, EPI_ISL_3212053, EPI_ISL_3212054, EPI_ISL_3212055, EPI_ISL_3212056, EPI_ISL_3212057, EPI_ISL_3212058, EPI_ISL_3212059, EPI_ISL_3212060, EPI_ISL_3212061, EPI_ISL_3212062, EPI_ISL_3212063, EPI_ISL_3212064, EPI_ISL_3212065, EPI_ISL_3212066, EPI_ISL_3212067, EPI_ISL_3212068, EPI_ISL_3212069, EPI_ISL_3212070, EPI_ISL_3212071, EPI_ISL_3212072, EPI_ISL_3212073,
EPI_ISL_3212074, EPI_ISL_3212075, EPI_ISL_3212076, EPI_ISL_3212077, EPI_ISL_3212078, EPI_ISL_3212079, EPI_ISL_3212080, EPI_ISL_3212081, EPI_ISL_3212082, EPI_ISL_3212083, EPI_ISL_3212084, EPI_ISL_3212085, EPI_ISL_3212086, EPI_ISL_3212087, EPI_ISL_3212088, EPI_ISL_3212089, EPI_ISL_3212090, EPI_ISL_3212091, EPI_ISL_3212092, EPI_ISL_3212093, EPI_ISL_3212094, EPI_ISL_3212095,
EPI_ISL_3212096, EPI_ISL_3212097, EPI_ISL_3212098, EPI_ISL_3212099, EPI_ISL_3212100, EPI_ISL_3212101, EPI_ISL_3212102, EPI_ISL_3212103, EPI_ISL_3212104, EPI_ISL_3212105, EPI_ISL_3212106, EPI_ISL_3212107, EPI_ISL_3212108, EPI_ISL_3212109, EPI_ISL_3212110, EPI_ISL_3212111, EPI_ISL_3212112, EPI_ISL_3212114, EPI_ISL_3212115, EPI_ISL_3212116, EPI_ISL_3212117, EPI_ISL_3212118,
EPI_ISL_3212119, EPI_ISL_3212120, EPI_ISL_3212121, EPI_ISL_3212122, EPI_ISL_3212123, EPI_ISL_3212124, EPI_ISL_3212125, EPI_ISL_3212126, EPI_ISL_3212127, EPI_ISL_3212128, EPI_ISL_3212129, EPI_ISL_3212130, EPI_ISL_3212131, EPI_ISL_3212132, EPI_ISL_3212133, EPI_ISL_3212134, EPI_ISL_3212135, EPI_ISL_3212136, EPI_ISL_3212137, EPI_ISL_3212138, EPI_ISL_3212139, EPI_ISL_3212140,
EPI_ISL_3212141, EPI_ISL_3212142, EPI_ISL_3212143, EPI_ISL_3212144, EPI_ISL_3212145, EPI_ISL_3212146, EPI_ISL_3212147, EPI_ISL_3212148, EPI_ISL_3212149, EPI_ISL_3212150, EPI_ISL_3212151, EPI_ISL_3212152, EPI_ISL_3212153, EPI_ISL_3212154, EPI_ISL_3212155, EPI_ISL_3212156, EPI_ISL_3212157, EPI_ISL_3212158, EPI_ISL_3212159, EPI_ISL_3212160, EPI_ISL_3212161, EPI_ISL_3212162,
EPI_ISL_3212163, EPI_ISL_3212164, EPI_ISL_3212165, EPI_ISL_3212166, EPI_ISL_3212167, EPI_ISL_3212168, EPI_ISL_3212169, EPI_ISL_3212170, EPI_ISL_3212171, EPI_ISL_3212172, EPI_ISL_3212173, EPI_ISL_3212174, EPI_ISL_3212176, EPI_ISL_3212177, EPI_ISL_3212178, EPI_ISL_3212179, EPI_ISL_3212180, EPI_ISL_3212181, EPI_ISL_3212182, EPI_ISL_3212183, EPI_ISL_3212184, EPI_ISL_3212185,
EPI_ISL_3212186, EPI_ISL_3212187, EPI_ISL_3212188, EPI_ISL_3212189, EPI_ISL_3212190, EPI_ISL_3212191, EPI_ISL_3212192, EPI_ISL_3212193, EPI_ISL_3212194, EPI_ISL_3212195, EPI_ISL_3212196, EPI_ISL_3212197, EPI_ISL_3212198, EPI_ISL_3212199, EPI_ISL_3212200, EPI_ISL_3212201, EPI_ISL_3212202, EPI_ISL_3212203, EPI_ISL_3212204, EPI_ISL_3212205, EPI_ISL_3212206, EPI_ISL_3212207,
EPI_ISL_3212208, EPI_ISL_3212209, EPI_ISL_3212210, EPI_ISL_3212211, EPI_ISL_3212212, EPI_ISL_3212213, EPI_ISL_3212214, EPI_ISL_3212215, EPI_ISL_3212216, EPI_ISL_3212217, EPI_ISL_3212218, EPI_ISL_3212219, EPI_ISL_3212220, EPI_ISL_3212221, EPI_ISL_3212222, EPI_ISL_3212223, EPI_ISL_3212224, EPI_ISL_3212225, EPI_ISL_3212226, EPI_ISL_3212227, EPI_ISL_3212228, EPI_ISL_3212229,
EPI_ISL_3212230, EPI_ISL_3212231, EPI_ISL_3212232, EPI_ISL_3212233, EPI_ISL_3212234, EPI_ISL_3212235, EPI_ISL_3212236, EPI_ISL_3212237, EPI_ISL_3212238, EPI_ISL_3212239, EPI_ISL_3212240, EPI_ISL_3212241, EPI_ISL_3212242, EPI_ISL_3212243, EPI_ISL_3212244, EPI_ISL_3212245, EPI_ISL_3212246, EPI_ISL_3212247, EPI_ISL_3212248, EPI_ISL_3212250, EPI_ISL_3212251, EPI_ISL_3212252,
EPI_ISL_3212253, EPI_ISL_3212254, EPI_ISL_3212255, EPI_ISL_3212256, EPI_ISL_3212257, EPI_ISL_3212258, EPI_ISL_3212259, EPI_ISL_3212260, EPI_ISL_3212261, EPI_ISL_3212262, EPI_ISL_3212263, EPI_ISL_3212264, EPI_ISL_3212265, EPI_ISL_3212266, EPI_ISL_3212267, EPI_ISL_3212268, EPI_ISL_3212269, EPI_ISL_3212270, EPI_ISL_3212271, EPI_ISL_3212272, EPI_ISL_3212273, EPI_ISL_3212274,
EPI_ISL_3212275, EPI_ISL_3212276, EPI_ISL_3212277, EPI_ISL_3212278, EPI_ISL_3212279, EPI_ISL_3212280, EPI_ISL_3212281, EPI_ISL_3212282, EPI_ISL_3212284, EPI_ISL_3212285, EPI_ISL_3212286, EPI_ISL_3212287, EPI_ISL_3212288, EPI_ISL_3212289, EPI_ISL_3212290, EPI_ISL_3212291, EPI_ISL_3212292, EPI_ISL_3212293, EPI_ISL_3212294, EPI_ISL_3212295, EPI_ISL_3212296, EPI_ISL_3212297,
EPI_ISL_3212298, EPI_ISL_3212299, EPI_ISL_3212300, EPI_ISL_3212301, EPI_ISL_3212302, EPI_ISL_3212303, EPI_ISL_3212304, EPI_ISL_3212305, EPI_ISL_3212306, EPI_ISL_3212307, EPI_ISL_3212308, EPI_ISL_3212309, EPI_ISL_3212310, EPI_ISL_3212311, EPI_ISL_3212312, EPI_ISL_3212313, EPI_ISL_3212314, EPI_ISL_3212315, EPI_ISL_3212316, EPI_ISL_3212317, EPI_ISL_3212318, EPI_ISL_3212319,
EPI_ISL_3212320, EPI_ISL_3212321, EPI_ISL_3212322, EPI_ISL_3212323, EPI_ISL_3212324, EPI_ISL_3212325, EPI_ISL_3212326, EPI_ISL_3212327, EPI_ISL_3212328, EPI_ISL_3212329, EPI_ISL_3212330, EPI_ISL_3212331, EPI_ISL_3212332, EPI_ISL_3212333, EPI_ISL_3212334, EPI_ISL_3212335, EPI_ISL_3212336, EPI_ISL_3212337, EPI_ISL_3212338, EPI_ISL_3212339, EPI_ISL_3212340, EPI_ISL_3212341,
EPI_ISL_3212342, EPI_ISL_3212344, EPI_ISL_3212345, EPI_ISL_3212346, EPI_ISL_3212347, EPI_ISL_3212348, EPI_ISL_3212349, EPI_ISL_3212350, EPI_ISL_3212351, EPI_ISL_3212352, EPI_ISL_3212354, EPI_ISL_3212355, EPI_ISL_3212356, EPI_ISL_3212357, EPI_ISL_3212358, EPI_ISL_3212359, EPI_ISL_3212360, EPI_ISL_3212361, EPI_ISL_3212362, EPI_ISL_3212363, EPI_ISL_3212364, EPI_ISL_3212365,
EPI_ISL_3212366, EPI_ISL_3212367, EPI_ISL_3212368, EPI_ISL_3212369, EPI_ISL_3212370, EPI_ISL_3212371, EPI_ISL_3212372, EPI_ISL_3212373, EPI_ISL_3212374, EPI_ISL_3212375, EPI_ISL_3212376, EPI_ISL_3212378, EPI_ISL_3212379, EPI_ISL_3212380, EPI_ISL_3212381, EPI_ISL_3212382, EPI_ISL_3212383, EPI_ISL_3212384, EPI_ISL_3212385, EPI_ISL_3212386, EPI_ISL_3212387, EPI_ISL_3212388,
EPI_ISL_3212389, EPI_ISL_3212390, EPI_ISL_3212391, EPI_ISL_3212392, EPI_ISL_3212393, EPI_ISL_3212394, EPI_ISL_3212395, EPI_ISL_3212396, EPI_ISL_3212397, EPI_ISL_3212398, EPI_ISL_3212399, EPI_ISL_3212400, EPI_ISL_3212401, EPI_ISL_3212402, EPI_ISL_3212403, EPI_ISL_3212404, EPI_ISL_3212405, EPI_ISL_3212406, EPI_ISL_3212407, EPI_ISL_3212408, EPI_ISL_3212409, EPI_ISL_3212410,
EPI_ISL_3212411, EPI_ISL_3212412, EPI_ISL_3212413, EPI_ISL_3212414, EPI_ISL_3212415, EPI_ISL_3212416, EPI_ISL_3212417, EPI_ISL_3212418, EPI_ISL_3212419, EPI_ISL_3212420, EPI_ISL_3212421, EPI_ISL_3212422, EPI_ISL_3212423, EPI_ISL_3212424, EPI_ISL_3212425, EPI_ISL_3212426, EPI_ISL_3212427, EPI_ISL_3212429, EPI_ISL_3212430, EPI_ISL_3212431, EPI_ISL_3212432, EPI_ISL_3212433,
EPI_ISL_3212434, EPI_ISL_3212435, EPI_ISL_3212436, EPI_ISL_3212437, EPI_ISL_3212438, EPI_ISL_3212439, EPI_ISL_3212440, EPI_ISL_3212441, EPI_ISL_3212442, EPI_ISL_3212443, EPI_ISL_3212444, EPI_ISL_3212445, EPI_ISL_3212446, EPI_ISL_3212447, EPI_ISL_3212448, EPI_ISL_3212449, EPI_ISL_3212450, EPI_ISL_3212451, EPI_ISL_3212452, EPI_ISL_3212453, EPI_ISL_3212454, EPI_ISL_3212455,
EPI_ISL_3212456, EPI_ISL_3212457, EPI_ISL_3212458, EPI_ISL_3212459, EPI_ISL_3212460, EPI_ISL_3212461, EPI_ISL_3212462, EPI_ISL_3212463, EPI_ISL_3212464, EPI_ISL_3212465, EPI_ISL_3212466, EPI_ISL_3212467, EPI_ISL_3212468, EPI_ISL_3212469, EPI_ISL_3212470, EPI_ISL_3212471, EPI_ISL_3212472, EPI_ISL_3212473, EPI_ISL_3212474, EPI_ISL_3212475, EPI_ISL_3212476, EPI_ISL_3212477,
EPI_ISL_3212478, EPI_ISL_3212479, EPI_ISL_3212480, EPI_ISL_3212481, EPI_ISL_3212482, EPI_ISL_3212483, EPI_ISL_3212484, EPI_ISL_3212485, EPI_ISL_3212486, EPI_ISL_3212487, EPI_ISL_3212488, EPI_ISL_3212489, EPI_ISL_3212490, EPI_ISL_3212491, EPI_ISL_3212492, EPI_ISL_3212493, EPI_ISL_3212494, EPI_ISL_3212495, EPI_ISL_3212496, EPI_ISL_3212497, EPI_ISL_3212498, EPI_ISL_3212499,
EPI_ISL_3212500, EPI_ISL_3212501, EPI_ISL_3212502, EPI_ISL_3212503, EPI_ISL_3212504, EPI_ISL_3212505, EPI_ISL_3212507, EPI_ISL_3212508, EPI_ISL_3212509, EPI_ISL_3212510, EPI_ISL_3212511, EPI_ISL_3212512, EPI_ISL_3212513, EPI_ISL_3212514, EPI_ISL_3212515, EPI_ISL_3212516, EPI_ISL_3212517, EPI_ISL_3212518, EPI_ISL_3212519, EPI_ISL_3212520, EPI_ISL_3212521, EPI_ISL_3212522,
EPI_ISL_3212523, EPI_ISL_3212524, EPI_ISL_3212525, EPI_ISL_3212526, EPI_ISL_3212527, EPI_ISL_3212528, EPI_ISL_3212529, EPI_ISL_3212530, EPI_ISL_3212531, EPI_ISL_3212532, EPI_ISL_3212533, EPI_ISL_3212534, EPI_ISL_3212535, EPI_ISL_3212536, EPI_ISL_3212537, EPI_ISL_3212538, EPI_ISL_3212539, EPI_ISL_3212540, EPI_ISL_3212541, EPI_ISL_3212542, EPI_ISL_3212543, EPI_ISL_3212544,
EPI_ISL_3212545, EPI_ISL_3212546, EPI_ISL_3212547, EPI_ISL_3212548, EPI_ISL_3212549, EPI_ISL_3212550, EPI_ISL_3212551, EPI_ISL_3212552, EPI_ISL_3212553, EPI_ISL_3212554, EPI_ISL_3212555, EPI_ISL_3212556, EPI_ISL_3212557, EPI_ISL_3212558, EPI_ISL_3212559, EPI_ISL_3212560, EPI_ISL_3212561, EPI_ISL_3212562, EPI_ISL_3212563, EPI_ISL_3212564, EPI_ISL_3212565, EPI_ISL_3212566,
EPI_ISL_3212567, EPI_ISL_3212568, EPI_ISL_3212569, EPI_ISL_3212570, EPI_ISL_3212571, EPI_ISL_3212572, EPI_ISL_3212573, EPI_ISL_3212574, EPI_ISL_3212575, EPI_ISL_3212576, EPI_ISL_3212577, EPI_ISL_3212579, EPI_ISL_3212580, EPI_ISL_3212581, EPI_ISL_3212582, EPI_ISL_3212584, EPI_ISL_3212585, EPI_ISL_3212586, EPI_ISL_3212587, EPI_ISL_3212588, EPI_ISL_3212589, EPI_ISL_3212590,
EPI_ISL_3212591, EPI_ISL_3212592, EPI_ISL_3212593, EPI_ISL_3212594, EPI_ISL_3212595, EPI_ISL_3212596, EPI_ISL_3212597, EPI_ISL_3212598, EPI_ISL_3212599, EPI_ISL_3212600, EPI_ISL_3212601, EPI_ISL_3212602, EPI_ISL_3212603, EPI_ISL_3212604, EPI_ISL_3212605, EPI_ISL_3212606, EPI_ISL_3212607, EPI_ISL_3212608, EPI_ISL_3212609, EPI_ISL_3212610, EPI_ISL_3212611, EPI_ISL_3212612,
EPI_ISL_3212613, EPI_ISL_3212614, EPI_ISL_3212615, EPI_ISL_3212616, EPI_ISL_3212618, EPI_ISL_3212619, EPI_ISL_3212620, EPI_ISL_3212621, EPI_ISL_3212622, EPI_ISL_3212623, EPI_ISL_3212624, EPI_ISL_3212625, EPI_ISL_3212626, EPI_ISL_3212627, EPI_ISL_3212628, EPI_ISL_3212629, EPI_ISL_3212630, EPI_ISL_3212631, EPI_ISL_3212632, EPI_ISL_3212633, EPI_ISL_3212634, EPI_ISL_3212635,
EPI_ISL_3212636, EPI_ISL_3212637, EPI_ISL_3212638, EPI_ISL_3212639, EPI_ISL_3212640, EPI_ISL_3212641, EPI_ISL_3212642, EPI_ISL_3212643, EPI_ISL_3212644, EPI_ISL_3212645, EPI_ISL_3212646, EPI_ISL_3212647, EPI_ISL_3212648, EPI_ISL_3212649, EPI_ISL_3212650, EPI_ISL_3212651, EPI_ISL_3212652, EPI_ISL_3212653, EPI_ISL_3212654, EPI_ISL_3212655, EPI_ISL_3212656, EPI_ISL_3212657,
EPI_ISL_3212658, EPI_ISL_3212659, EPI_ISL_3212660, EPI_ISL_3212661, EPI_ISL_3212662, EPI_ISL_3212664, EPI_ISL_3212665, EPI_ISL_3212666, EPI_ISL_3212667, EPI_ISL_3212668, EPI_ISL_3212669, EPI_ISL_3212670, EPI_ISL_3212671, EPI_ISL_3212672, EPI_ISL_3212673, EPI_ISL_3212674, EPI_ISL_3212675

| | | | |
|---|---|---|---|
| see above | Fulgent Genetics | Centers for Disease Control and Prevention Division of Viral Diseases, Pathogen Discovery | Dakota Howard, Dhwani Batra, Peter W. Cook, Kara Moser, Adrian Paskey, Jason Caravas, Benjamin Rambo-Martin, Shatavia Morrison, Christopher Gulvick, Scott Sammons, Yvette Unoarumhi, Darlene Wagner, Matthew Schmerer, Harry Gao, Mickey Li, John Gao, Joseph Fierro, Benafsh Sapra, Becky Tsai, Yan Meng, Doreen Ng, James Xie, Clinton R. Paden, Duncan MacCannell |

EPI_ISL_3212676, EPI_ISL_3212677, EPI_ISL_3212678, EPI_ISL_3212679, EPI_ISL_3212680, EPI_ISL_3212681, EPI_ISL_3212682, EPI_ISL_3212683, EPI_ISL_3212684, EPI_ISL_3212685, EPI_ISL_3212686, EPI_ISL_3212687, EPI_ISL_3212688, EPI_ISL_3212689, EPI_ISL_3212690, EPI_ISL_3212691, EPI_ISL_3212692, EPI_ISL_3212693, EPI_ISL_3212694, EPI_ISL_3212695, EPI_ISL_3212696, EPI_ISL_3212697,
EPI_ISL_3212698, EPI_ISL_3212699, EPI_ISL_3212700, EPI_ISL_3212701, EPI_ISL_3212702, EPI_ISL_3212703, EPI_ISL_3212704, EPI_ISL_3212705, EPI_ISL_3212706, EPI_ISL_3212707, EPI_ISL_3212708, EPI_ISL_3212709, EPI_ISL_3212710, EPI_ISL_3212711, EPI_ISL_3212712, EPI_ISL_3212713, EPI_ISL_3212714, EPI_ISL_3212715, EPI_ISL_3212716, EPI_ISL_3212717, EPI_ISL_3212718, EPI_ISL_3212719,
EPI_ISL_3212720, EPI_ISL_3212721, EPI_ISL_3212722, EPI_ISL_3212723, EPI_ISL_3212724, EPI_ISL_3212725, EPI_ISL_3212726, EPI_ISL_3212727, EPI_ISL_3212728, EPI_ISL_3212729, EPI_ISL_3212730, EPI_ISL_3212731, EPI_ISL_3212732, EPI_ISL_3212733, EPI_ISL_3212735, EPI_ISL_3212736, EPI_ISL_3212737, EPI_ISL_3212738, EPI_ISL_3212739, EPI_ISL_3212740, EPI_ISL_3212741, EPI_ISL_3212742,
EPI_ISL_3212744, EPI_ISL_3212745, EPI_ISL_3212747, EPI_ISL_3212748, EPI_ISL_3212749, EPI_ISL_3212750, EPI_ISL_3212751, EPI_ISL_3212752, EPI_ISL_3212754, EPI_ISL_3212755, EPI_ISL_3212756, EPI_ISL_3212758, EPI_ISL_3212759, EPI_ISL_3212760, EPI_ISL_3212761, EPI_ISL_3212762, EPI_ISL_3212763, EPI_ISL_3212764, EPI_ISL_3212765, EPI_ISL_3212766, EPI_ISL_3212767, EPI_ISL_3212769,
EPI_ISL_3212770, EPI_ISL_3212771, EPI_ISL_3212772, EPI_ISL_3212773, EPI_ISL_3212774, EPI_ISL_3212775, EPI_ISL_3212776, EPI_ISL_3212777, EPI_ISL_3212778, EPI_ISL_3212780, EPI_ISL_3212781, EPI_ISL_3212782, EPI_ISL_3212783, EPI_ISL_3212784, EPI_ISL_3212785, EPI_ISL_3212786, EPI_ISL_3212787, EPI_ISL_3212788, EPI_ISL_3212789, EPI_ISL_3212790, EPI_ISL_3212791, EPI_ISL_3212792,
EPI_ISL_3212794, EPI_ISL_3212795, EPI_ISL_3212797, EPI_ISL_3212798, EPI_ISL_3212799, EPI_ISL_3212800, EPI_ISL_3212801, EPI_ISL_3212802, EPI_ISL_3212803, EPI_ISL_3212804, EPI_ISL_3212806, EPI_ISL_3212807, EPI_ISL_3212808, EPI_ISL_3212809, EPI_ISL_3212810, EPI_ISL_3212811, EPI_ISL_3212812, EPI_ISL_3212813, EPI_ISL_3212814, EPI_ISL_3212815, EPI_ISL_3212817, EPI_ISL_3212818,
EPI_ISL_3212819, EPI_ISL_3212820, EPI_ISL_3212821, EPI_ISL_3212822, EPI_ISL_3212823, EPI_ISL_3212824, EPI_ISL_3212826, EPI_ISL_3212827, EPI_ISL_3212828, EPI_ISL_3212829, EPI_ISL_3212830, EPI_ISL_3212831, EPI_ISL_3212832, EPI_ISL_3212833, EPI_ISL_3212835, EPI_ISL_3212836, EPI_ISL_3212837, EPI_ISL_3212838, EPI_ISL_3212839, EPI_ISL_3212840, EPI_ISL_3212841, EPI_ISL_3212842,
EPI_ISL_3212843, EPI_ISL_3212844, EPI_ISL_3212845, EPI_ISL_3212846, EPI_ISL_3212847, EPI_ISL_3212848, EPI_ISL_3212849, EPI_ISL_3212850, EPI_ISL_3212851, EPI_ISL_3212852, EPI_ISL_3212853, EPI_ISL_3212854, EPI_ISL_3212855, EPI_ISL_3212856, EPI_ISL_3212857, EPI_ISL_3212858, EPI_ISL_3212859, EPI_ISL_3212860, EPI_ISL_3212861, EPI_ISL_3212862, EPI_ISL_3212863, EPI_ISL_3212864,
EPI_ISL_3212865, EPI_ISL_3212867, EPI_ISL_3212868, EPI_ISL_3212869, EPI_ISL_3212870, EPI_ISL_3212871, EPI_ISL_3212872, EPI_ISL_3212873, EPI_ISL_3212874, EPI_ISL_3212875, EPI_ISL_3212876, EPI_ISL_3212877, EPI_ISL_3212878, EPI_ISL_3212879, EPI_ISL_3212880, EPI_ISL_3212881, EPI_ISL_3212882, EPI_ISL_3212883, EPI_ISL_3212884, EPI_ISL_3212885, EPI_ISL_3212886, EPI_ISL_3212887,
EPI_ISL_3212888, EPI_ISL_3212889, EPI_ISL_3212890, EPI_ISL_3212891, EPI_ISL_3212892, EPI_ISL_3212893, EPI_ISL_3212894, EPI_ISL_3212895, EPI_ISL_3212897, EPI_ISL_3212898, EPI_ISL_3212899, EPI_ISL_3212900, EPI_ISL_3212902, EPI_ISL_3212903, EPI_ISL_3212904, EPI_ISL_3212905, EPI_ISL_3212906, EPI_ISL_3212907, EPI_ISL_3212908, EPI_ISL_3212909, EPI_ISL_3212910, EPI_ISL_3212911,
EPI_ISL_3212912, EPI_ISL_3212913, EPI_ISL_3212914, EPI_ISL_3212918, EPI_ISL_3212919, EPI_ISL_3212920, EPI_ISL_3212923, EPI_ISL_3212924, EPI_ISL_3212925, EPI_ISL_3212926, EPI_ISL_3212927, EPI_ISL_3212928, EPI_ISL_3212931, EPI_ISL_3212932, EPI_ISL_3212933, EPI_ISL_3212934, EPI_ISL_3212935, EPI_ISL_3212936, EPI_ISL_3212937, EPI_ISL_3212938, EPI_ISL_3212939, EPI_ISL_3212940,
EPI_ISL_3212941, EPI_ISL_3212942, EPI_ISL_3212943, EPI_ISL_3212944, EPI_ISL_3212945, EPI_ISL_3212946, EPI_ISL_3212947, EPI_ISL_3212948, EPI_ISL_3212949, EPI_ISL_3212950, EPI_ISL_3212951, EPI_ISL_3212952, EPI_ISL_3212953, EPI_ISL_3212954, EPI_ISL_3212955, EPI_ISL_3212956, EPI_ISL_3212957, EPI_ISL_3212958, EPI_ISL_3212960, EPI_ISL_3212961, EPI_ISL_3212962, EPI_ISL_3212963,
EPI_ISL_3212964, EPI_ISL_3212965, EPI_ISL_3212966, EPI_ISL_3212967, EPI_ISL_3212968, EPI_ISL_3212969, EPI_ISL_3212970, EPI_ISL_3212972, EPI_ISL_3212973, EPI_ISL_3212974, EPI_ISL_3212975, EPI_ISL_3212977, EPI_ISL_3212978, EPI_ISL_3212979, EPI_ISL_3212981, EPI_ISL_3212982, EPI_ISL_3212983, EPI_ISL_3212984, EPI_ISL_3212986, EPI_ISL_3212987, EPI_ISL_3212988, EPI_ISL_3212990,
EPI_ISL_3212991, EPI_ISL_3212992, EPI_ISL_3212993, EPI_ISL_3212994, EPI_ISL_3212995, EPI_ISL_3212996, EPI_ISL_3212997, EPI_ISL_3212998, EPI_ISL_3212999, EPI_ISL_3213000, EPI_ISL_3213001, EPI_ISL_3213003, EPI_ISL_3213004, EPI_ISL_3213005, EPI_ISL_3213006, EPI_ISL_3213007, EPI_ISL_3213008, EPI_ISL_3213009, EPI_ISL_3213011, EPI_ISL_3213012, EPI_ISL_3213013, EPI_ISL_3213014,
EPI_ISL_3213016, EPI_ISL_3213017, EPI_ISL_3213018, EPI_ISL_3213019, EPI_ISL_3213020, EPI_ISL_3213021, EPI_ISL_3213023, EPI_ISL_3213024, EPI_ISL_3213025, EPI_ISL_3213026, EPI_ISL_3213027, EPI_ISL_3213028, EPI_ISL_3213029, EPI_ISL_3213030, EPI_ISL_3213031, EPI_ISL_3213032, EPI_ISL_3213033, EPI_ISL_3213034, EPI_ISL_3213035, EPI_ISL_3213036, EPI_ISL_3213037, EPI_ISL_3213038,
EPI_ISL_3213039, EPI_ISL_3213041, EPI_ISL_3213045, EPI_ISL_3213046, EPI_ISL_3213048, EPI_ISL_3213051, EPI_ISL_3213052, EPI_ISL_3213053, EPI_ISL_3213055, EPI_ISL_3213056, EPI_ISL_3213057, EPI_ISL_3213058, EPI_ISL_3213059, EPI_ISL_3213060, EPI_ISL_3213061, EPI_ISL_3213062, EPI_ISL_3213063, EPI_ISL_3213064, EPI_ISL_3213065, EPI_ISL_3213066, EPI_ISL_3213067, EPI_ISL_3213068,
EPI_ISL_3213069, EPI_ISL_3213070, EPI_ISL_3213072, EPI_ISL_3213073, EPI_ISL_3213074, EPI_ISL_3213075, EPI_ISL_3213076, EPI_ISL_3213077, EPI_ISL_3213078, EPI_ISL_3213079, EPI_ISL_3213080, EPI_ISL_3213081, EPI_ISL_3213082, EPI_ISL_3213084, EPI_ISL_3213085, EPI_ISL_3213086, EPI_ISL_3213087, EPI_ISL_3213088, EPI_ISL_3213089, EPI_ISL_3213091, EPI_ISL_3213092, EPI_ISL_3213094,
EPI_ISL_3213095, EPI_ISL_3213097, EPI_ISL_3213098, EPI_ISL_3213100, EPI_ISL_3213101, EPI_ISL_3213102, EPI_ISL_3213103, EPI_ISL_3213104, EPI_ISL_3213105, EPI_ISL_3213106, EPI_ISL_3213107, EPI_ISL_3213108, EPI_ISL_3213109, EPI_ISL_3213110, EPI_ISL_3213111, EPI_ISL_3213112, EPI_ISL_3213113, EPI_ISL_3213114, EPI_ISL_3213115, EPI_ISL_3213116, EPI_ISL_3213117, EPI_ISL_3213118,
EPI_ISL_3213119, EPI_ISL_3213120, EPI_ISL_3213121, EPI_ISL_3213122, EPI_ISL_3213123, EPI_ISL_3213124, EPI_ISL_3213125, EPI_ISL_3213126, EPI_ISL_3213127, EPI_ISL_3213128, EPI_ISL_3213131, EPI_ISL_3213132, EPI_ISL_3213133, EPI_ISL_3213134, EPI_ISL_3213136, EPI_ISL_3213137, EPI_ISL_3213138, EPI_ISL_3213140, EPI_ISL_3213141, EPI_ISL_3213143, EPI_ISL_3213146, EPI_ISL_3213147,
EPI_ISL_3213148, EPI_ISL_3213149, EPI_ISL_3213150, EPI_ISL_3213151, EPI_ISL_3213153, EPI_ISL_3213154, EPI_ISL_3213155, EPI_ISL_3213156, EPI_ISL_3213157, EPI_ISL_3213158, EPI_ISL_3213159, EPI_ISL_3213160, EPI_ISL_3213161, EPI_ISL_3213162, EPI_ISL_3213163, EPI_ISL_3213164, EPI_ISL_3213165, EPI_ISL_3213166, EPI_ISL_3213167, EPI_ISL_3213168, EPI_ISL_3213169, EPI_ISL_3213170,
EPI_ISL_3213171, EPI_ISL_3213172, EPI_ISL_3213173, EPI_ISL_3213174, EPI_ISL_3213175, EPI_ISL_3213176, EPI_ISL_3213177, EPI_ISL_3213178, EPI_ISL_3213179, EPI_ISL_3213180, EPI_ISL_3213181, EPI_ISL_3213182, EPI_ISL_3213183, EPI_ISL_3213185, EPI_ISL_3213186, EPI_ISL_3213187, EPI_ISL_3213188, EPI_ISL_3213189, EPI_ISL_3213190, EPI_ISL_3213192, EPI_ISL_3213193, EPI_ISL_3213194,
EPI_ISL_3213195, EPI_ISL_3213196, EPI_ISL_3213197, EPI_ISL_3213198, EPI_ISL_3213199, EPI_ISL_3213200, EPI_ISL_3213201, EPI_ISL_3213202, EPI_ISL_3213203, EPI_ISL_3213204, EPI_ISL_3213205, EPI_ISL_3213206, EPI_ISL_3213207, EPI_ISL_3213208, EPI_ISL_3213209, EPI_ISL_3213210, EPI_ISL_3213211, EPI_ISL_3213212, EPI_ISL_3213213, EPI_ISL_3213214, EPI_ISL_3213215, EPI_ISL_3213216,
EPI_ISL_3213217, EPI_ISL_3213218, EPI_ISL_3213219, EPI_ISL_3213220, EPI_ISL_3213221, EPI_ISL_3213222, EPI_ISL_3213224, EPI_ISL_3213225, EPI_ISL_3213226, EPI_ISL_3213227, EPI_ISL_3213228, EPI_ISL_3213229, EPI_ISL_3213230, EPI_ISL_3213231, EPI_ISL_3213232, EPI_ISL_3213233, EPI_ISL_3213234, EPI_ISL_3213235, EPI_ISL_3213236, EPI_ISL_3213238, EPI_ISL_3213240, EPI_ISL_3213242,
EPI_ISL_3213243, EPI_ISL_3213244, EPI_ISL_3213245, EPI_ISL_3213246, EPI_ISL_3213247, EPI_ISL_3213248, EPI_ISL_3213250, EPI_ISL_3213251, EPI_ISL_3213252, EPI_ISL_3213253, EPI_ISL_3213254, EPI_ISL_3213255, EPI_ISL_3213256, EPI_ISL_3213257, EPI_ISL_3213258, EPI_ISL_3213259, EPI_ISL_3213260, EPI_ISL_3213261, EPI_ISL_3213262, EPI_ISL_3213263, EPI_ISL_3213264, EPI_ISL_3213265,
EPI_ISL_3213266, EPI_ISL_3213267, EPI_ISL_3213268, EPI_ISL_3213269, EPI_ISL_3213270, EPI_ISL_3213272, EPI_ISL_3213273, EPI_ISL_3213274, EPI_ISL_3213275, EPI_ISL_3213276, EPI_ISL_3213277, EPI_ISL_3213278, EPI_ISL_3213279, EPI_ISL_3213280, EPI_ISL_3213281, EPI_ISL_3213282, EPI_ISL_3213283, EPI_ISL_3213284, EPI_ISL_3213285, EPI_ISL_3213286, EPI_ISL_3213287, EPI_ISL_3213288,
EPI_ISL_3213289, EPI_ISL_3213290, EPI_ISL_3213291, EPI_ISL_3213292, EPI_ISL_3213293, EPI_ISL_3213294, EPI_ISL_3213295, EPI_ISL_3213296, EPI_ISL_3213297, EPI_ISL_3213298, EPI_ISL_3213299, EPI_ISL_3213300, EPI_ISL_3213301, EPI_ISL_3213302, EPI_ISL_3213303, EPI_ISL_3213304, EPI_ISL_3213306, EPI_ISL_3213308, EPI_ISL_3213309, EPI_ISL_3213311, EPI_ISL_3213312, EPI_ISL_3213313,
EPI_ISL_3213314, EPI_ISL_3213315, EPI_ISL_3213316, EPI_ISL_3213317, EPI_ISL_3213318, EPI_ISL_3213319, EPI_ISL_3213320, EPI_ISL_3213321, EPI_ISL_3213322, EPI_ISL_3213323, EPI_ISL_3213324, EPI_ISL_3213325, EPI_ISL_3213326, EPI_ISL_3213327, EPI_ISL_3213328, EPI_ISL_3213329, EPI_ISL_3213330, EPI_ISL_3213331, EPI_ISL_3213332, EPI_ISL_3213333, EPI_ISL_3213334, EPI_ISL_3213335,
EPI_ISL_3213336, EPI_ISL_3213337, EPI_ISL_3213338, EPI_ISL_3213339, EPI_ISL_3213340, EPI_ISL_3213341, EPI_ISL_3213342, EPI_ISL_3213343, EPI_ISL_3213344, EPI_ISL_3213345, EPI_ISL_3213346, EPI_ISL_3213347, EPI_ISL_3213348, EPI_ISL_3213349, EPI_ISL_3213350, EPI_ISL_3213351, EPI_ISL_3213352, EPI_ISL_3213353, EPI_ISL_3213354, EPI_ISL_3213355, EPI_ISL_3213356, EPI_ISL_3213357,
EPI_ISL_3213358, EPI_ISL_3213359, EPI_ISL_3213360, EPI_ISL_3213361, EPI_ISL_3213362, EPI_ISL_3213363, EPI_ISL_3213364, EPI_ISL_3213365, EPI_ISL_3213366, EPI_ISL_3213367, EPI_ISL_3213368, EPI_ISL_3213369, EPI_ISL_3213370, EPI_ISL_3213371, EPI_ISL_3213372, EPI_ISL_3213373, EPI_ISL_3213374, EPI_ISL_3213375, EPI_ISL_3213376, EPI_ISL_3213377, EPI_ISL_3213378, EPI_ISL_3213379,
EPI_ISL_3213380, EPI_ISL_3213381, EPI_ISL_3213382, EPI_ISL_3213383, EPI_ISL_3213384, EPI_ISL_3213385, EPI_ISL_3213386, EPI_ISL_3213387, EPI_ISL_3213388, EPI_ISL_3213389, EPI_ISL_3213390, EPI_ISL_3213391, EPI_ISL_3213392, EPI_ISL_3213393, EPI_ISL_3213394, EPI_ISL_3213395, EPI_ISL_3213396, EPI_ISL_3213397, EPI_ISL_3213398, EPI_ISL_3213399, EPI_ISL_3213400, EPI_ISL_3213401,
EPI_ISL_3213402, EPI_ISL_3213403, EPI_ISL_3213404, EPI_ISL_3213405, EPI_ISL_3213406, EPI_ISL_3213407, EPI_ISL_3213408, EPI_ISL_3213409, EPI_ISL_3213410, EPI_ISL_3213411, EPI_ISL_3213412, EPI_ISL_3213413, EPI_ISL_3213414, EPI_ISL_3213415, EPI_ISL_3213416, EPI_ISL_3213417, EPI_ISL_3213418, EPI_ISL_3213419, EPI_ISL_3213420, EPI_ISL_3213421, EPI_ISL_3213422, EPI_ISL_3213423,
EPI_ISL_3213424, EPI_ISL_3213425, EPI_ISL_3213426, EPI_ISL_3213428, EPI_ISL_3213430, EPI_ISL_3213431, EPI_ISL_3213432, EPI_ISL_3213433, EPI_ISL_3213434, EPI_ISL_3213435, EPI_ISL_3213436, EPI_ISL_3213437, EPI_ISL_3213438, EPI_ISL_3213439, EPI_ISL_3213441, EPI_ISL_3213442, EPI_ISL_3213443, EPI_ISL_3213444, EPI_ISL_3213445, EPI_ISL_3213446, EPI_ISL_3213447, EPI_ISL_3213448,
EPI_ISL_3213449, EPI_ISL_3213450, EPI_ISL_3213451, EPI_ISL_3213452, EPI_ISL_3213453, EPI_ISL_3213454, EPI_ISL_3213455, EPI_ISL_3213456, EPI_ISL_3213457, EPI_ISL_3213458, EPI_ISL_3213459, EPI_ISL_3213460, EPI_ISL_3213461, EPI_ISL_3213462, EPI_ISL_3213463, EPI_ISL_3213464, EPI_ISL_3213465, EPI_ISL_3213466, EPI_ISL_3213467, EPI_ISL_3213468, EPI_ISL_3213469, EPI_ISL_3213470,
EPI_ISL_3213471, EPI_ISL_3213472, EPI_ISL_3213474, EPI_ISL_3213475, EPI_ISL_3213476, EPI_ISL_3213477, EPI_ISL_3213478, EPI_ISL_3213479, EPI_ISL_3213480, EPI_ISL_3213481, EPI_ISL_3213483, EPI_ISL_3213484, EPI_ISL_3213485, EPI_ISL_3213486, EPI_ISL_3213487, EPI_ISL_3213488, EPI_ISL_3213490, EPI_ISL_3213491, EPI_ISL_3213492, EPI_ISL_3213493, EPI_ISL_3213494, EPI_ISL_3213495,
EPI_ISL_3213496, EPI_ISL_3213497, EPI_ISL_3213498, EPI_ISL_3213499, EPI_ISL_3213500, EPI_ISL_3213501, EPI_ISL_3213502, EPI_ISL_3213503, EPI_ISL_3213504, EPI_ISL_3213505, EPI_ISL_3213506, EPI_ISL_3213507, EPI_ISL_3213508, EPI_ISL_3213509, EPI_ISL_3213510, EPI_ISL_3213511, EPI_ISL_3213512, EPI_ISL_3213513, EPI_ISL_3213514, EPI_ISL_3213515, EPI_ISL_3213516, EPI_ISL_3213517,
EPI_ISL_3213518, EPI_ISL_3213519, EPI_ISL_3213520, EPI_ISL_3213521, EPI_ISL_3213522, EPI_ISL_3213523, EPI_ISL_3213524, EPI_ISL_3213525, EPI_ISL_3213526, EPI_ISL_3213527, EPI_ISL_3213528, EPI_ISL_3213529, EPI_ISL_3213530, EPI_ISL_3213532, EPI_ISL_3213533, EPI_ISL_3213534, EPI_ISL_3213535, EPI_ISL_3213536, EPI_ISL_3213537, EPI_ISL_3213538, EPI_ISL_3213539, EPI_ISL_3213541,
EPI_ISL_3213542, EPI_ISL_3213543, EPI_ISL_3213544, EPI_ISL_3213545, EPI_ISL_3213546, EPI_ISL_3213547, EPI_ISL_3213548, EPI_ISL_3213549, EPI_ISL_3213550, EPI_ISL_3213551, EPI_ISL_3213552, EPI_ISL_3213553, EPI_ISL_3213554, EPI_ISL_3213555, EPI_ISL_3213556, EPI_ISL_3213557, EPI_ISL_3213558, EPI_ISL_3213559, EPI_ISL_3213560, EPI_ISL_3213561, EPI_ISL_3213562, EPI_ISL_3213564,
EPI_ISL_3213565, EPI_ISL_3213568, EPI_ISL_3213569, EPI_ISL_3213570, EPI_ISL_3213571, EPI_ISL_3213572, EPI_ISL_3213573, EPI_ISL_3213574, EPI_ISL_3213576, EPI_ISL_3213577, EPI_ISL_3213578, EPI_ISL_3213579, EPI_ISL_3213580, EPI_ISL_3213581, EPI_ISL_3213582, EPI_ISL_3213583, EPI_ISL_3213584, EPI_ISL_3213585, EPI_ISL_3213586, EPI_ISL_3213589, EPI_ISL_3213590, EPI_ISL_3213591,
EPI_ISL_3213592, EPI_ISL_3213593, EPI_ISL_3213594, EPI_ISL_3213595, EPI_ISL_3213596, EPI_ISL_3213597, EPI_ISL_3213598, EPI_ISL_3213599, EPI_ISL_3213600, EPI_ISL_3213601, EPI_ISL_3213602, EPI_ISL_3213603, EPI_ISL_3213604, EPI_ISL_3213605, EPI_ISL_3213606, EPI_ISL_3213607, EPI_ISL_3213608, EPI_ISL_3213609, EPI_ISL_3213610, EPI_ISL_3213611, EPI_ISL_3213612, EPI_ISL_3213613,
EPI_ISL_3213614, EPI_ISL_3213615, EPI_ISL_3213616, EPI_ISL_3213617, EPI_ISL_3213618, EPI_ISL_3213619, EPI_ISL_3213620, EPI_ISL_3213621, EPI_ISL_3213622, EPI_ISL_3213623, EPI_ISL_3213624, EPI_ISL_3213625, EPI_ISL_3213626, EPI_ISL_3213627, EPI_ISL_3213628, EPI_ISL_3213629, EPI_ISL_3213630, EPI_ISL_3213631, EPI_ISL_3213633, EPI_ISL_3213635, EPI_ISL_3213636, EPI_ISL_3213637,
EPI_ISL_3213638, EPI_ISL_3213639, EPI_ISL_3213640, EPI_ISL_3213641, EPI_ISL_3213642, EPI_ISL_3213643, EPI_ISL_3213644, EPI_ISL_3213645, EPI_ISL_3213646, EPI_ISL_3213647, EPI_ISL_3213648, EPI_ISL_3213649, EPI_ISL_3213650, EPI_ISL_3213651, EPI_ISL_3213652, EPI_ISL_3213654, EPI_ISL_3213655, EPI_ISL_3213657, EPI_ISL_3213658, EPI_ISL_3213659, EPI_ISL_3213660, EPI_ISL_3213661,
EPI_ISL_3213662, EPI_ISL_3213663, EPI_ISL_3213664, EPI_ISL_3213665, EPI_ISL_3213666, EPI_ISL_3213667, EPI_ISL_3213668, EPI_ISL_3213669, EPI_ISL_3213670, EPI_ISL_3213671, EPI_ISL_3213672, EPI_ISL_3213673, EPI_ISL_3213674, EPI_ISL_3213675, EPI_ISL_3213677, EPI_ISL_3213678, EPI_ISL_3213679, EPI_ISL_3213680, EPI_ISL_3213681, EPI_ISL_3213682, EPI_ISL_3213683, EPI_ISL_3213684,
EPI_ISL_3213685, EPI_ISL_3213686, EPI_ISL_3213687, EPI_ISL_3213688, EPI_ISL_3213689, EPI_ISL_3213690, EPI_ISL_3213691, EPI_ISL_3213692, EPI_ISL_3213693, EPI_ISL_3213694, EPI_ISL_3213695, EPI_ISL_3213696, EPI_ISL_3213697, EPI_ISL_3213698, EPI_ISL_3213699, EPI_ISL_3213700, EPI_ISL_3213701, EPI_ISL_3213702, EPI_ISL_3213705, EPI_ISL_3213706, EPI_ISL_3213707, EPI_ISL_3213708,
EPI_ISL_3213709, EPI_ISL_3213710, EPI_ISL_3213712, EPI_ISL_3213713, EPI_ISL_3213714, EPI_ISL_3213715, EPI_ISL_3213716, EPI_ISL_3213717, EPI_ISL_3213718, EPI_ISL_3213719, EPI_ISL_3213720, EPI_ISL_3213721, EPI_ISL_3213722, EPI_ISL_3213723, EPI_ISL_3213724, EPI_ISL_3213725, EPI_ISL_3213726, EPI_ISL_3213727, EPI_ISL_3213728, EPI_ISL_3213729, EPI_ISL_3213730, EPI_ISL_3213731,
EPI_ISL_3213732, EPI_ISL_3213733, EPI_ISL_3213734, EPI_ISL_3213736, EPI_ISL_3213737, EPI_ISL_3213738, EPI_ISL_3213739, EPI_ISL_3213740, EPI_ISL_3213741, EPI_ISL_3213742, EPI_ISL_3213744, EPI_ISL_3213745, EPI_ISL_3213746, EPI_ISL_3213747, EPI_ISL_3213748, EPI_ISL_3213749, EPI_ISL_3213750, EPI_ISL_3213751, EPI_ISL_3213752, EPI_ISL_3213753, EPI_ISL_3213754, EPI_ISL_3213755,
EPI_ISL_3213756, EPI_ISL_3213758, EPI_ISL_3213760, EPI_ISL_3213761, EPI_ISL_3213762, EPI_ISL_3213763, EPI_ISL_3213764, EPI_ISL_3213765, EPI_ISL_3213767, EPI_ISL_3213768, EPI_ISL_3213769, EPI_ISL_3213770, EPI_ISL_3213771, EPI_ISL_3213772, EPI_ISL_3213774, EPI_ISL_3213775, EPI_ISL_3213776, EPI_ISL_3213777, EPI_ISL_3213778, EPI_ISL_3213779, EPI_ISL_3213780, EPI_ISL_3213781,
EPI_ISL_3213782, EPI_ISL_3213783, EPI_ISL_3213784, EPI_ISL_3213785, EPI_ISL_3213786, EPI_ISL_3213787, EPI_ISL_3213788, EPI_ISL_3213789, EPI_ISL_3213790, EPI_ISL_3213791, EPI_ISL_3213792, EPI_ISL_3213793, EPI_ISL_3213795, EPI_ISL_3213796, EPI_ISL_3213797, EPI_ISL_3213798, EPI_ISL_3213799, EPI_ISL_3213800, EPI_ISL_3213801, EPI_ISL_3213802, EPI_ISL_3213803, EPI_ISL_3213804,
EPI_ISL_3213805, EPI_ISL_3213806, EPI_ISL_3213807, EPI_ISL_3213808, EPI_ISL_3213809, EPI_ISL_3213810, EPI_ISL_3213811, EPI_ISL_3213812, EPI_ISL_3213813, EPI_ISL_3213814, EPI_ISL_3213815, EPI_ISL_3213816, EPI_ISL_3213817, EPI_ISL_3213818, EPI_ISL_3213819, EPI_ISL_3213820, EPI_ISL_3213821, EPI_ISL_3213822, EPI_ISL_3213823, EPI_ISL_3213824, EPI_ISL_3213825, EPI_ISL_3213826,
EPI_ISL_3213827, EPI_ISL_3213828, EPI_ISL_3213830, EPI_ISL_3213831, EPI_ISL_3213832, EPI_ISL_3213833, EPI_ISL_3213834, EPI_ISL_3213836, EPI_ISL_3213837, EPI_ISL_3213838, EPI_ISL_3213839, EPI_ISL_3213840, EPI_ISL_3213841, EPI_ISL_3213842, EPI_ISL_3213843, EPI_ISL_3213844, EPI_ISL_3213845, EPI_ISL_3213846, EPI_ISL_3213848, EPI_ISL_3213850, EPI_ISL_3213851, EPI_ISL_3213853,
EPI_ISL_3213854, EPI_ISL_3213855, EPI_ISL_3213856, EPI_ISL_3213857, EPI_ISL_3213858, EPI_ISL_3213859, EPI_ISL_3213860, EPI_ISL_3213861, EPI_ISL_3213862, EPI_ISL_3213863, EPI_ISL_3213864, EPI_ISL_3213865, EPI_ISL_3213866, EPI_ISL_3213867, EPI_ISL_3213868, EPI_ISL_3213870, EPI_ISL_3213871, EPI_ISL_3213872, EPI_ISL_3213873, EPI_ISL_3213874, EPI_ISL_3213875, EPI_ISL_3213876,
EPI_ISL_3213877, EPI_ISL_3213879, EPI_ISL_3213880, EPI_ISL_3213881, EPI_ISL_3213882, EPI_ISL_3213883, EPI_ISL_3213884, EPI_ISL_3213885, EPI_ISL_3213886, EPI_ISL_3213887, EPI_ISL_3213888, EPI_ISL_3213889, EPI_ISL_3213890, EPI_ISL_3213891, EPI_ISL_3213892, EPI_ISL_3213894, EPI_ISL_3213895, EPI_ISL_3213896, EPI_ISL_3213897, EPI_ISL_3213898, EPI_ISL_3213899, EPI_ISL_3213900,
EPI_ISL_3213901, EPI_ISL_3213902, EPI_ISL_3213903, EPI_ISL_3213904, EPI_ISL_3213905, EPI_ISL_3213906, EPI_ISL_3213907, EPI_ISL_3213908, EPI_ISL_3213909, EPI_ISL_3213910, EPI_ISL_3213913, EPI_ISL_3213914, EPI_ISL_3213915, EPI_ISL_3213916, EPI_ISL_3213917, EPI_ISL_3213918, EPI_ISL_3213919, EPI_ISL_3213920, EPI_ISL_3213921, EPI_ISL_3213922, EPI_ISL_3213923, EPI_ISL_3213924,
EPI_ISL_3213925, EPI_ISL_3213926, EPI_ISL_3213927, EPI_ISL_3213928, EPI_ISL_3213929, EPI_ISL_3213930, EPI_ISL_3213931, EPI_ISL_3213932, EPI_ISL_3213933, EPI_ISL_3213934, EPI_ISL_3213935, EPI_ISL_3213936, EPI_ISL_3213937, EPI_ISL_3213938, EPI_ISL_3213939, EPI_ISL_3213940, EPI_ISL_3213941, EPI_ISL_3213942, EPI_ISL_3213943, EPI_ISL_3213945, EPI_ISL_3213946, EPI_ISL_3213947,
EPI_ISL_3213948, EPI_ISL_3213949, EPI_ISL_3213950, EPI_ISL_3213951, EPI_ISL_3213952, EPI_ISL_3213953, EPI_ISL_3213954, EPI_ISL_3213955, EPI_ISL_3213956, EPI_ISL_3213957, EPI_ISL_3213958, EPI_ISL_3213960, EPI_ISL_3213961, EPI_ISL_3213962, EPI_ISL_3213963, EPI_ISL_3213964, EPI_ISL_3213965, EPI_ISL_3213966, EPI_ISL_3213967, EPI_ISL_3213968, EPI_ISL_3213969, EPI_ISL_3213970,
EPI_ISL_3213971, EPI_ISL_3213972, EPI_ISL_3213973, EPI_ISL_3213974, EPI_ISL_3213975, EPI_ISL_3213977, EPI_ISL_3213978, EPI_ISL_3213979, EPI_ISL_3213980, EPI_ISL_3213981, EPI_ISL_3213982, EPI_ISL_3213983, EPI_ISL_3213984, EPI_ISL_3213985, EPI_ISL_3213986, EPI_ISL_3213987, EPI_ISL_3213988, EPI_ISL_3213989, EPI_ISL_3213990, EPI_ISL_3213991, EPI_ISL_3213992, EPI_ISL_3213993,
EPI_ISL_3213994, EPI_ISL_3213995, EPI_ISL_3213996, EPI_ISL_3213998, EPI_ISL_3213999, EPI_ISL_3214000, EPI_ISL_3214001, EPI_ISL_3214002, EPI_ISL_3214003, EPI_ISL_3214005, EPI_ISL_3214007, EPI_ISL_3214008, EPI_ISL_3214009, EPI_ISL_3214010, EPI_ISL_3214011, EPI_ISL_3214012, EPI_ISL_3214013, EPI_ISL_3214014, EPI_ISL_3214016, EPI_ISL_3214017, EPI_ISL_3214018, EPI_ISL_3214019,
EPI_ISL_3214020, EPI_ISL_3214021, EPI_ISL_3214022, EPI_ISL_3214023, EPI_ISL_3214025, EPI_ISL_3214026, EPI_ISL_3214027, EPI_ISL_3214028, EPI_ISL_3214029, EPI_ISL_3214030, EPI_ISL_3214031, EPI_ISL_3214032, EPI_ISL_3214033, EPI_ISL_3214034, EPI_ISL_3214035, EPI_ISL_3214036, EPI_ISL_3214037, EPI_ISL_3214038, EPI_ISL_3214039, EPI_ISL_3214040, EPI_ISL_3214041, EPI_ISL_3214042,
EPI_ISL_3214043, EPI_ISL_3214044, EPI_ISL_3214045, EPI_ISL_3214046, EPI_ISL_3214048, EPI_ISL_3214050, EPI_ISL_3214051, EPI_ISL_3214052, EPI_ISL_3214053, EPI_ISL_3214054, EPI_ISL_3214055, EPI_ISL_3214056, EPI_ISL_3214057, EPI_ISL_3214058, EPI_ISL_3214059, EPI_ISL_3214060, EPI_ISL_3214061, EPI_ISL_3214062, EPI_ISL_3214063, EPI_ISL_3214064, EPI_ISL_3214065, EPI_ISL_3214066,
EPI_ISL_3214067, EPI_ISL_3214068, EPI_ISL_3214069, EPI_ISL_3214070, EPI_ISL_3214072, EPI_ISL_3214073, EPI_ISL_3214074, EPI_ISL_3214075, EPI_ISL_3214076, EPI_ISL_3214077, EPI_ISL_3214078, EPI_ISL_3214079, EPI_ISL_3214080, EPI_ISL_3214081, EPI_ISL_3214082, EPI_ISL_3214083, EPI_ISL_3214084, EPI_ISL_3214085, EPI_ISL_3214086, EPI_ISL_3214087, EPI_ISL_3214089, EPI_ISL_3214090,
EPI_ISL_3214091, EPI_ISL_3214092, EPI_ISL_3214093, EPI_ISL_3214094, EPI_ISL_3214095, EPI_ISL_3214096, EPI_ISL_3214097, EPI_ISL_3214098, EPI_ISL_3214099, EPI_ISL_3214100, EPI_ISL_3214101, EPI_ISL_3214102, EPI_ISL_3214103, EPI_ISL_3214104, EPI_ISL_3214106, EPI_ISL_3214107, EPI_ISL_3214108, EPI_ISL_3214109, EPI_ISL_3214110, EPI_ISL_3214111, EPI_ISL_3214112, EPI_ISL_3214113,

| | | | |
|---|---|---|---|
| EPI_ISL_3214114, EPI_ISL_3214116, EPI_ISL_3214117, EPI_ISL_3214118, EPI_ISL_3214119, EPI_ISL_3214120, EPI_ISL_3214121, EPI_ISL_3214123, EPI_ISL_3214124, EPI_ISL_3214125, EPI_ISL_3214126, EPI_ISL_3214127, EPI_ISL_3214128, EPI_ISL_3214129, EPI_ISL_3214130, EPI_ISL_3214131, EPI_ISL_3214132, EPI_ISL_3214133, EPI_ISL_3214134, EPI_ISL_3214135, EPI_ISL_3214137, EPI_ISL_3214138, EPI_ISL_3214139, EPI_ISL_3214140, EPI_ISL_3214141, EPI_ISL_3214142, EPI_ISL_3214143, EPI_ISL_3214144, EPI_ISL_3214146, EPI_ISL_3214147, EPI_ISL_3214149, EPI_ISL_3214150, EPI_ISL_3214152, EPI_ISL_3214154, EPI_ISL_3214155, EPI_ISL_3214156, EPI_ISL_3214158, EPI_ISL_3214159, EPI_ISL_3214160, EPI_ISL_3214161, EPI_ISL_3214162, EPI_ISL_3214163, EPI_ISL_3214164, EPI_ISL_3214165, EPI_ISL_3214167, EPI_ISL_3214169, EPI_ISL_3214171, EPI_ISL_3214172, EPI_ISL_3214173, EPI_ISL_3214174, EPI_ISL_3214175, EPI_ISL_3214176, EPI_ISL_3214177, EPI_ISL_3214178, EPI_ISL_3214179, EPI_ISL_3214180, EPI_ISL_3214181, EPI_ISL_3214182, EPI_ISL_3214183, EPI_ISL_3214184, EPI_ISL_3214185, EPI_ISL_3214186, EPI_ISL_3214187, EPI_ISL_3214188, EPI_ISL_3214190, EPI_ISL_3214191, EPI_ISL_3214192, EPI_ISL_3214193, EPI_ISL_3214194, EPI_ISL_3214195, EPI_ISL_3214196, EPI_ISL_3214197, EPI_ISL_3214198, EPI_ISL_3214199, EPI_ISL_3214200, EPI_ISL_3214201, EPI_ISL_3214202, EPI_ISL_3214203, EPI_ISL_3214204, EPI_ISL_3214205, EPI_ISL_3214206, EPI_ISL_3214207, EPI_ISL_3214208, EPI_ISL_3214209, EPI_ISL_3214210, EPI_ISL_3214211, EPI_ISL_3214212, EPI_ISL_3214213, EPI_ISL_3214214, EPI_ISL_3214215, EPI_ISL_3214216, EPI_ISL_3214217, EPI_ISL_3214218, EPI_ISL_3214219, EPI_ISL_3214220, EPI_ISL_3214221, EPI_ISL_3214222, EPI_ISL_3214223, EPI_ISL_3214224, EPI_ISL_3214225, EPI_ISL_3214226, EPI_ISL_3214227, EPI_ISL_3214228, EPI_ISL_3214229, EPI_ISL_3214230, EPI_ISL_3214231, EPI_ISL_3214232, EPI_ISL_3214233, EPI_ISL_3214235, EPI_ISL_3214236, EPI_ISL_3214237, EPI_ISL_3214238, EPI_ISL_3214239, EPI_ISL_3214241, EPI_ISL_3214242, EPI_ISL_3214243, EPI_ISL_3214244, EPI_ISL_3214245, EPI_ISL_3214246, EPI_ISL_3214247, EPI_ISL_3214248, EPI_ISL_3214249, EPI_ISL_3214250, EPI_ISL_3214251, EPI_ISL_3214252, EPI_ISL_3214253, EPI_ISL_3214254, EPI_ISL_3214255, EPI_ISL_3214256, EPI_ISL_3214257, EPI_ISL_3214258, EPI_ISL_3214259, EPI_ISL_3214260, EPI_ISL_3214261, EPI_ISL_3214262, EPI_ISL_3214263, EPI_ISL_3214264, EPI_ISL_3214265, EPI_ISL_3214266, EPI_ISL_3214267, EPI_ISL_3214268, EPI_ISL_3214269, EPI_ISL_3214270, EPI_ISL_3214271, EPI_ISL_3214272, EPI_ISL_3214273, EPI_ISL_3214274, EPI_ISL_3214275, EPI_ISL_3214276, EPI_ISL_3214277, EPI_ISL_3214278, EPI_ISL_3214279, EPI_ISL_3214280, EPI_ISL_3214281, EPI_ISL_3214282, EPI_ISL_3214283, EPI_ISL_3214284, EPI_ISL_3214285, EPI_ISL_3214286, EPI_ISL_3214287, EPI_ISL_3214289, EPI_ISL_3214290, EPI_ISL_3214291, EPI_ISL_3214292, EPI_ISL_3214293, EPI_ISL_3214294, EPI_ISL_3214296, EPI_ISL_3214297, EPI_ISL_3214299, EPI_ISL_3214300, EPI_ISL_3214301, EPI_ISL_3214303, EPI_ISL_3214304, EPI_ISL_3214305, EPI_ISL_3214306, EPI_ISL_3214307, EPI_ISL_3214308, EPI_ISL_3214309, EPI_ISL_3214310, EPI_ISL_3214311, EPI_ISL_3214312, EPI_ISL_3214313, EPI_ISL_3214315, EPI_ISL_3214316, EPI_ISL_3214317, EPI_ISL_3214319, EPI_ISL_3214320, EPI_ISL_3214321, EPI_ISL_3214322, EPI_ISL_3214324, EPI_ISL_3214325, EPI_ISL_3214326, EPI_ISL_3214327, EPI_ISL_3214328, EPI_ISL_3214329, EPI_ISL_3214330, EPI_ISL_3214331, EPI_ISL_3214333, EPI_ISL_3214335, EPI_ISL_3214336, EPI_ISL_3214337, EPI_ISL_3214338, EPI_ISL_3214339, EPI_ISL_3214340, EPI_ISL_3214341, EPI_ISL_3214342, EPI_ISL_3214343, EPI_ISL_3214344, EPI_ISL_3214345, EPI_ISL_3214346, EPI_ISL_3214347, EPI_ISL_3214348, EPI_ISL_3214349, EPI_ISL_3214350, EPI_ISL_3214352, EPI_ISL_3214354, EPI_ISL_3214355, EPI_ISL_3214356, EPI_ISL_3214357, EPI_ISL_3214358, EPI_ISL_3214359, EPI_ISL_3214360, EPI_ISL_3214361, EPI_ISL_3214362, EPI_ISL_3214363, EPI_ISL_3214364, EPI_ISL_3214365, EPI_ISL_3214366, EPI_ISL_3214367, EPI_ISL_3214368, EPI_ISL_3214369, EPI_ISL_3214370, EPI_ISL_3214371, EPI_ISL_3214372, EPI_ISL_3214374, EPI_ISL_3214375, EPI_ISL_3214376, EPI_ISL_3214377, EPI_ISL_3214378, EPI_ISL_3214379, EPI_ISL_3214380, EPI_ISL_3214381, EPI_ISL_3214382, EPI_ISL_3214383, EPI_ISL_3214384, EPI_ISL_3214385, EPI_ISL_3214386 | | | |
| see above | Quest Diagnostics Incorporated | Centers for Disease Control and Prevention Division of Viral Diseases, Pathogen Discovery | Dakota Howard, Dhwani Batra, Peter W. Cook, Kara Moser, Adrian Paskey, Jason Caravas, Benjamin Rambo-Martin, Shatavia Morrison, Christopher Gulvick, Scott Sammons, Yvette Unoarumhi, Darlene Wagner, Matthew Schmerer, S. H. Rosenthal, A. Gerasimova, R. M. Kagan, B. Anderson, M. Hua, Y. Liu, L.E. Bernstein, K.E. Livingston, A. Perez, I. A. Shlyakhter, R. V. Rolando, R. Owen, P. Tanpaiboon, F. Lacbawan, Clinton R. Paden, Duncan MacCannell |
| EPI_ISL_3214388, EPI_ISL_3214393, EPI_ISL_3214394, EPI_ISL_3214395, EPI_ISL_3214396, EPI_ISL_3214399, EPI_ISL_3214400, EPI_ISL_3214401, EPI_ISL_3214402, EPI_ISL_3214403, EPI_ISL_3214404, EPI_ISL_3214407, EPI_ISL_3214408, EPI_ISL_3214409, EPI_ISL_3214410, EPI_ISL_3214411, EPI_ISL_3214412, EPI_ISL_3214413, EPI_ISL_3214414, EPI_ISL_3214415, EPI_ISL_3214416, EPI_ISL_3214417, EPI_ISL_3214418, EPI_ISL_3214419, EPI_ISL_3214420, EPI_ISL_3214421, EPI_ISL_3214422, EPI_ISL_3214423, EPI_ISL_3214424, EPI_ISL_3214425, EPI_ISL_3214427, EPI_ISL_3214428, EPI_ISL_3214429, EPI_ISL_3214431, EPI_ISL_3214432, EPI_ISL_3214433, EPI_ISL_3214435, EPI_ISL_3214436, EPI_ISL_3214437, EPI_ISL_3214438, EPI_ISL_3214439, EPI_ISL_3214440, EPI_ISL_3214442, EPI_ISL_3214443, EPI_ISL_3214444, EPI_ISL_3214445, EPI_ISL_3214446, EPI_ISL_3214447, EPI_ISL_3214448, EPI_ISL_3214450, EPI_ISL_3214452, EPI_ISL_3214454, EPI_ISL_3214455, EPI_ISL_3214456, EPI_ISL_3214457, EPI_ISL_3214458, EPI_ISL_3214459, EPI_ISL_3214460, EPI_ISL_3214461, EPI_ISL_3214463, EPI_ISL_3214464, EPI_ISL_3214465, EPI_ISL_3214466, EPI_ISL_3214467, EPI_ISL_3214468, EPI_ISL_3214469, EPI_ISL_3214470, EPI_ISL_3214471, EPI_ISL_3214472, EPI_ISL_3214473, EPI_ISL_3214474, EPI_ISL_3214476, EPI_ISL_3214477, EPI_ISL_3214478, EPI_ISL_3214479, EPI_ISL_3214480, EPI_ISL_3214481, EPI_ISL_3214482, EPI_ISL_3214483, EPI_ISL_3214484, EPI_ISL_3214485, EPI_ISL_3214486, EPI_ISL_3214487, EPI_ISL_3214489, EPI_ISL_3214490, EPI_ISL_3214491, EPI_ISL_3214492, EPI_ISL_3214493, EPI_ISL_3214494, EPI_ISL_3214495, EPI_ISL_3214496, EPI_ISL_3214497, EPI_ISL_3214498, EPI_ISL_3214499, EPI_ISL_3214500, EPI_ISL_3214501, EPI_ISL_3214502, EPI_ISL_3214503, EPI_ISL_3214504, EPI_ISL_3214505, EPI_ISL_3214506, EPI_ISL_3214507, EPI_ISL_3214508, EPI_ISL_3214510, EPI_ISL_3214511 | | | |
| see above | Lab. Microbiologia e Virologia Cotugno A.O. dei Colli | TIGEM | Antonio Grimaldi Patrizia Annunziata Francesco Panariello Biancamaria Pierri Claudia Tiberio Teresa Giuliano Valentina Bouche Chiara Colantuono Maria Concetta Cuomo Denise Di Concilio Lucio Di Filippo Anna Manfredi Marcello Salvi Antonio Limone Luigi Atripaldi Pellegrino Cerino Andrea Ballabio Davide Cacchiarelli |
| EPI_ISL_3214512 | TXDSHS | TXDSHS | Rashmi Tuladhar, Bonnie Oh, Jenny Zhang, Maliha Rahman, Mayela Pedrueza, Anita Pokharel, Karen Bobier, Lorraine Rodriguez, Myong Koag, Chun Wang, Rachel Lee, Grace Kubin |
| EPI_ISL_3214513, EPI_ISL_3214514 | Lab. Microbiologia e Virologia Cotugno A.O. dei Colli | TIGEM | Antonio Grimaldi Patrizia Annunziata Francesco Panariello Biancamaria Pierri Claudia Tiberio Teresa Giuliano Valentina Bouche Chiara Colantuono Maria Concetta Cuomo Denise Di Concilio Lucio Di Filippo Anna Manfredi Marcello Salvi Antonio Limone Luigi Atripaldi Pellegrino Cerino Andrea Ballabio Davide Cacchiarelli |
| EPI_ISL_3214515 | TXDSHS | TXDSHS | Rashmi Tuladhar, Bonnie Oh, Jenny Zhang, Maliha Rahman, Mayela Pedrueza, Anita Pokharel, Karen Bobier, Lorraine Rodriguez, Myong Koag, Chun Wang, Rachel Lee, Grace Kubin |
| EPI_ISL_3214516, EPI_ISL_3214517 | Lab. Microbiologia e Virologia Cotugno A.O. dei Colli | TIGEM | Antonio Grimaldi Patrizia Annunziata Francesco Panariello Biancamaria Pierri Claudia Tiberio Teresa Giuliano Valentina Bouche Chiara Colantuono Maria Concetta Cuomo Denise Di Concilio Lucio Di Filippo Anna Manfredi Marcello Salvi Antonio Limone Luigi Atripaldi Pellegrino Cerino Andrea Ballabio Davide Cacchiarelli |
| EPI_ISL_3214518 | TXDSHS | TXDSHS | Rashmi Tuladhar, Bonnie Oh, Jenny Zhang, Maliha Rahman, Mayela Pedrueza, Anita Pokharel, Karen Bobier, Lorraine Rodriguez, Myong Koag, Chun Wang, Rachel Lee, Grace Kubin |
| EPI_ISL_3214519 | Lab. Microbiologia e Virologia Cotugno A.O. dei Colli | TIGEM | Antonio Grimaldi Patrizia Annunziata Francesco Panariello Biancamaria Pierri Claudia Tiberio Teresa Giuliano Valentina Bouche Chiara Colantuono Maria Concetta Cuomo Denise Di Concilio Lucio Di Filippo Anna Manfredi Marcello Salvi Antonio Limone Luigi Atripaldi Pellegrino Cerino Andrea Ballabio Davide Cacchiarelli |
| EPI_ISL_3214520 | TXDSHS | TXDSHS | Rashmi Tuladhar, Bonnie Oh, Jenny Zhang, Maliha Rahman, Mayela Pedrueza, Anita Pokharel, Karen Bobier, Lorraine Rodriguez, Myong Koag, Chun Wang, Rachel Lee, Grace Kubin |
| EPI_ISL_3214521, EPI_ISL_3214522 | Lab. Microbiologia e Virologia Cotugno A.O. dei Colli | TIGEM | Antonio Grimaldi Patrizia Annunziata Francesco Panariello Biancamaria Pierri Claudia Tiberio Teresa Giuliano Valentina Bouche Chiara Colantuono Maria Concetta Cuomo Denise Di Concilio Lucio Di Filippo Anna Manfredi Marcello Salvi Antonio Limone Luigi Atripaldi Pellegrino Cerino Andrea Ballabio Davide Cacchiarelli |
| EPI_ISL_3214523, EPI_ISL_3214525 | TXDSHS | TXDSHS | Rashmi Tuladhar, Bonnie Oh, Jenny Zhang, Maliha Rahman, Mayela Pedrueza, Anita Pokharel, Karen Bobier, Lorraine Rodriguez, Myong Koag, Chun Wang, Rachel Lee, Grace Kubin |
| EPI_ISL_3214526, EPI_ISL_3214527 | Lab. Microbiologia e Virologia Cotugno A.O. dei Colli | TIGEM | Antonio Grimaldi Patrizia Annunziata Francesco Panariello Biancamaria Pierri Claudia Tiberio Teresa Giuliano Valentina Bouche Chiara Colantuono Maria Concetta Cuomo Denise Di Concilio Lucio Di Filippo Anna Manfredi Marcello Salvi Antonio Limone Luigi Atripaldi Pellegrino Cerino Andrea Ballabio Davide Cacchiarelli |
| EPI_ISL_3214528 | TXDSHS | TXDSHS | Rashmi Tuladhar, Bonnie Oh, Jenny Zhang, Maliha Rahman, Mayela Pedrueza, Anita Pokharel, Karen Bobier, Lorraine Rodriguez, Myong Koag, Chun Wang, Rachel Lee, Grace Kubin |
| EPI_ISL_3214529, EPI_ISL_3214531 | Lab. Microbiologia e Virologia Cotugno A.O. dei Colli | TIGEM | Antonio Grimaldi Patrizia Annunziata Francesco Panariello Biancamaria Pierri Claudia Tiberio Teresa Giuliano Valentina Bouche Chiara Colantuono Maria Concetta Cuomo Denise Di Concilio Lucio Di Filippo Anna Manfredi Marcello Salvi Antonio Limone Luigi Atripaldi Pellegrino Cerino Andrea Ballabio Davide Cacchiarelli |
| EPI_ISL_3214532 | TXDSHS | TXDSHS | Rashmi Tuladhar, Bonnie Oh, Jenny Zhang, Maliha Rahman, Mayela Pedrueza, Anita Pokharel, Karen Bobier, Lorraine Rodriguez, Myong Koag, Chun Wang, Rachel Lee, Grace Kubin |
| EPI_ISL_3214533 | Lab. Microbiologia e Virologia Cotugno A.O. dei Colli | TIGEM | Antonio Grimaldi Patrizia Annunziata Francesco Panariello Biancamaria Pierri Claudia Tiberio Teresa Giuliano Valentina Bouche Chiara Colantuono Maria Concetta Cuomo Denise Di Concilio Lucio Di Filippo Anna Manfredi Marcello Salvi Antonio Limone Luigi Atripaldi Pellegrino Cerino Andrea Ballabio Davide Cacchiarelli |
| EPI_ISL_3214534 | TXDSHS | TXDSHS | Rashmi Tuladhar, Bonnie Oh, Jenny Zhang, Maliha Rahman, Mayela Pedrueza, Anita Pokharel, Karen Bobier, Lorraine Rodriguez, Myong Koag, Chun Wang, Rachel Lee, Grace Kubin |
| EPI_ISL_3214535, EPI_ISL_3214536 | Lab. Microbiologia e Virologia Cotugno A.O. dei Colli | TIGEM | Antonio Grimaldi Patrizia Annunziata Francesco Panariello Biancamaria Pierri Claudia Tiberio Teresa Giuliano Valentina Bouche Chiara Colantuono Maria Concetta Cuomo Denise Di Concilio Lucio Di Filippo Anna Manfredi Marcello Salvi Antonio Limone Luigi Atripaldi Pellegrino Cerino Andrea Ballabio Davide Cacchiarelli |
| EPI_ISL_3214537 | TXDSHS | TXDSHS | Rashmi Tuladhar, Bonnie Oh, Jenny Zhang, Maliha Rahman, Mayela Pedrueza, Anita Pokharel, Karen Bobier, Lorraine Rodriguez, Myong Koag, Chun Wang, Rachel Lee, Grace Kubin |
| EPI_ISL_3214538 | Lab. Microbiologia e Virologia Cotugno A.O. dei Colli | TIGEM | Antonio Grimaldi Patrizia Annunziata Francesco Panariello Biancamaria Pierri Claudia Tiberio Teresa Giuliano Valentina Bouche Chiara Colantuono Maria Concetta Cuomo Denise Di Concilio Lucio Di Filippo Anna Manfredi Marcello Salvi Antonio Limone Luigi Atripaldi Pellegrino Cerino Andrea Ballabio Davide Cacchiarelli |
| EPI_ISL_3214539 | TXDSHS | TXDSHS | Rashmi Tuladhar, Bonnie Oh, Jenny Zhang, Maliha Rahman, Mayela Pedrueza, Anita Pokharel, Karen Bobier, Lorraine Rodriguez, Myong Koag, Chun Wang, Rachel Lee, Grace Kubin |
| EPI_ISL_3214540, EPI_ISL_3214541 | Lab. Microbiologia e Virologia Cotugno A.O. dei Colli | TIGEM | Antonio Grimaldi Patrizia Annunziata Francesco Panariello Biancamaria Pierri Claudia Tiberio Teresa Giuliano Valentina Bouche Chiara Colantuono Maria Concetta Cuomo Denise Di Concilio Lucio Di Filippo Anna Manfredi Marcello Salvi Antonio Limone Luigi Atripaldi Pellegrino Cerino Andrea Ballabio Davide Cacchiarelli |
| EPI_ISL_3214542 | TXDSHS | TXDSHS | Rashmi Tuladhar, Bonnie Oh, Jenny Zhang, Maliha Rahman, Mayela Pedrueza, Anita Pokharel, Karen Bobier, Lorraine Rodriguez, Myong Koag, Chun Wang, Rachel Lee, Grace Kubin |
| EPI_ISL_3214543, EPI_ISL_3214544 | Lab. Microbiologia e Virologia Cotugno A.O. dei Colli | TIGEM | Antonio Grimaldi Patrizia Annunziata Francesco Panariello Biancamaria Pierri Claudia Tiberio Teresa Giuliano Valentina Bouche Chiara Colantuono Maria Concetta Cuomo Denise Di Concilio Lucio Di Filippo Anna Manfredi Marcello Salvi Antonio Limone Luigi Atripaldi Pellegrino Cerino Andrea Ballabio Davide Cacchiarelli |
| EPI_ISL_3214545 | TXDSHS | TXDSHS | Rashmi Tuladhar, Bonnie Oh, Jenny Zhang, Maliha Rahman, Mayela Pedrueza, Anita Pokharel, Karen Bobier, Lorraine Rodriguez, Myong Koag, Chun Wang, Rachel Lee, Grace Kubin |
| EPI_ISL_3214546 | Lab. Microbiologia e Virologia Cotugno A.O. dei Colli | TIGEM | Antonio Grimaldi Patrizia Annunziata Francesco Panariello Biancamaria Pierri Claudia Tiberio Teresa Giuliano Valentina Bouche Chiara Colantuono Maria Concetta Cuomo Denise Di Concilio Lucio Di Filippo Anna Manfredi Marcello Salvi Antonio Limone Luigi Atripaldi Pellegrino Cerino Andrea Ballabio Davide Cacchiarelli |
| EPI_ISL_3214547 | TXDSHS | TXDSHS | Rashmi Tuladhar, Bonnie Oh, Jenny Zhang, Maliha Rahman, Mayela Pedrueza, Anita Pokharel, Karen Bobier, Lorraine Rodriguez, Myong Koag, Chun Wang, Rachel Lee, Grace Kubin |
| EPI_ISL_3214548, EPI_ISL_3214549 | Lab. Microbiologia e Virologia Cotugno A.O. dei Colli | TIGEM | Antonio Grimaldi Patrizia Annunziata Francesco Panariello Biancamaria Pierri Claudia Tiberio Teresa Giuliano Valentina Bouche Chiara Colantuono Maria Concetta Cuomo Denise Di Concilio Lucio Di Filippo Anna Manfredi Marcello Salvi Antonio Limone Luigi Atripaldi Pellegrino Cerino Andrea Ballabio Davide Cacchiarelli |
| EPI_ISL_3214550 | TXDSHS | TXDSHS | Rashmi Tuladhar, Bonnie Oh, Jenny Zhang, Maliha Rahman, Mayela Pedrueza, Anita Pokharel, Karen Bobier, Lorraine Rodriguez, Myong Koag, Chun Wang, Rachel Lee, Grace Kubin |
| EPI_ISL_3214551 | Lab. Microbiologia e Virologia Cotugno A.O. dei Colli | TIGEM | Antonio Grimaldi Patrizia Annunziata Francesco Panariello Biancamaria Pierri Claudia Tiberio Teresa Giuliano Valentina Bouche Chiara Colantuono Maria Concetta Cuomo Denise Di Concilio Lucio Di Filippo Anna Manfredi Marcello Salvi Antonio Limone Luigi Atripaldi Pellegrino Cerino Andrea Ballabio Davide Cacchiarelli |
| EPI_ISL_3214552 | TXDSHS | TXDSHS | Rashmi Tuladhar, Bonnie Oh, Jenny Zhang, Maliha Rahman, Mayela Pedrueza, Anita Pokharel, Karen Bobier, Lorraine Rodriguez, Myong Koag, Chun Wang, Rachel Lee, Grace Kubin |
| EPI_ISL_3214553, EPI_ISL_3214554 | Lab. Microbiologia e Virologia Cotugno A.O. dei Colli | TIGEM | Antonio Grimaldi Patrizia Annunziata Francesco Panariello Biancamaria Pierri Claudia Tiberio Teresa Giuliano Valentina Bouche Chiara Colantuono Maria Concetta Cuomo Denise Di Concilio Lucio Di Filippo Anna Manfredi Marcello Salvi Antonio Limone Luigi Atripaldi Pellegrino Cerino Andrea Ballabio Davide Cacchiarelli |
| EPI_ISL_3214555 | TXDSHS | TXDSHS | Rashmi Tuladhar, Bonnie Oh, Jenny Zhang, Maliha Rahman, Mayela Pedrueza, Anita Pokharel, Karen Bobier, Lorraine Rodriguez, Myong Koag, Chun Wang, Rachel Lee, Grace Kubin |
| EPI_ISL_3214556 | Lab. Microbiologia e Virologia Cotugno A.O. dei Colli | TIGEM | Antonio Grimaldi Patrizia Annunziata Francesco Panariello Biancamaria Pierri Claudia Tiberio Teresa Giuliano Valentina Bouche Chiara Colantuono Maria Concetta Cuomo Denise Di Concilio Lucio Di Filippo Anna Manfredi Marcello Salvi Antonio Limone Luigi Atripaldi Pellegrino Cerino Andrea Ballabio Davide Cacchiarelli |
| EPI_ISL_3214557 | TXDSHS | TXDSHS | Rashmi Tuladhar, Bonnie Oh, Jenny Zhang, Maliha Rahman, Mayela Pedrueza, Anita Pokharel, Karen Bobier, Lorraine Rodriguez, Myong Koag, Chun Wang, Rachel Lee, Grace Kubin |
| EPI_ISL_3214558, EPI_ISL_3214559 | Lab. Microbiologia e Virologia Cotugno A.O. dei Colli | TIGEM | Antonio Grimaldi Patrizia Annunziata Francesco Panariello Biancamaria Pierri Claudia Tiberio Teresa Giuliano Valentina Bouche Chiara Colantuono Maria Concetta Cuomo Denise Di Concilio Lucio Di Filippo Anna Manfredi Marcello Salvi Antonio Limone Luigi Atripaldi Pellegrino Cerino Andrea Ballabio Davide Cacchiarelli |
| EPI_ISL_3214560 | TXDSHS | TXDSHS | Rashmi Tuladhar, Bonnie Oh, Jenny Zhang, Maliha Rahman, Mayela Pedrueza, Anita Pokharel, Karen Bobier, Lorraine Rodriguez, Myong Koag, Chun Wang, Rachel Lee, Grace Kubin |
| EPI_ISL_3214561 | Lab. Microbiologia e Virologia Cotugno A.O. dei Colli | TIGEM | Antonio Grimaldi Patrizia Annunziata Francesco Panariello Biancamaria Pierri Claudia Tiberio Teresa Giuliano Valentina Bouche Chiara Colantuono Maria Concetta Cuomo Denise Di Concilio Lucio Di Filippo Anna Manfredi Marcello Salvi Antonio Limone Luigi Atripaldi Pellegrino Cerino Andrea Ballabio Davide Cacchiarelli |
| EPI_ISL_3214562 | TXDSHS | TXDSHS | Rashmi Tuladhar, Bonnie Oh, Jenny Zhang, Maliha Rahman, Mayela Pedrueza, Anita Pokharel, Karen Bobier, Lorraine Rodriguez, Myong Koag, Chun Wang, Rachel Lee, Grace Kubin |
| EPI_ISL_3214563, EPI_ISL_3214564, EPI_ISL_3214566, EPI_ISL_3214567, EPI_ISL_3214568, EPI_ISL_3214569, EPI_ISL_3214570, EPI_ISL_3214571, EPI_ISL_3214572, EPI_ISL_3214574, EPI_ISL_3214575, EPI_ISL_3214577, EPI_ISL_3214579, EPI_ISL_3214580, EPI_ISL_3214581, EPI_ISL_3214582, EPI_ISL_3214583, EPI_ISL_3214584, EPI_ISL_3214585, EPI_ISL_3214586, EPI_ISL_3214587, EPI_ISL_3214588, EPI_ISL_3214589, EPI_ISL_3214590, EPI_ISL_3214591, EPI_ISL_3214592, EPI_ISL_3214593, EPI_ISL_3214594, EPI_ISL_3214595, EPI_ISL_3214596, EPI_ISL_3214598, EPI_ISL_3214599, EPI_ISL_3214600, EPI_ISL_3214601, EPI_ISL_3214602, EPI_ISL_3214603, EPI_ISL_3214604, EPI_ISL_3214605, EPI_ISL_3214606, EPI_ISL_3214607, EPI_ISL_3214608, EPI_ISL_3214609, EPI_ISL_3214610, EPI_ISL_3214613, EPI_ISL_3214614, EPI_ISL_3214615, EPI_ISL_3214617, EPI_ISL_3214618, EPI_ISL_3214619, EPI_ISL_3214620, EPI_ISL_3214621, EPI_ISL_3214622, EPI_ISL_3214623, EPI_ISL_3214624, EPI_ISL_3214625, EPI_ISL_3214626, EPI_ISL_3214627, EPI_ISL_3214628, EPI_ISL_3214629, EPI_ISL_3214631, EPI_ISL_3214632, EPI_ISL_3214633, EPI_ISL_3214634, EPI_ISL_3214635, EPI_ISL_3214636, EPI_ISL_3214637, EPI_ISL_3214638, EPI_ISL_3214639, EPI_ISL_3214640, EPI_ISL_3214642 | | | |
| see above | Lab. Microbiologia e Virologia Cotugno A.O. dei Colli | TIGEM | Antonio Grimaldi Patrizia Annunziata Francesco Panariello Biancamaria Pierri Claudia Tiberio Teresa Giuliano Valentina Bouche Chiara Colantuono Maria Concetta Cuomo Denise Di Concilio Lucio Di Filippo Anna Manfredi Marcello Salvi Antonio Limone Luigi Atripaldi Pellegrino Cerino Andrea Ballabio Davide Cacchiarelli |
| EPI_ISL_3214645 | U.O. Microbiologia Laboratorio Unico Centro Servizi - AUSL della Romagna | U.O. Microbiologia, Laboratorio Unico Centro Servizi - AUSL della Romagna | Giorgio Dirani |
| EPI_ISL_3214703, EPI_ISL_3214704, EPI_ISL_3214705, EPI_ISL_3214706, EPI_ISL_3214707, EPI_ISL_3214708, EPI_ISL_3214709, EPI_ISL_3214710, EPI_ISL_3214711, EPI_ISL_3214712, EPI_ISL_3214713, EPI_ISL_3214714, EPI_ISL_3214716, EPI_ISL_3214717, EPI_ISL_3214720, EPI_ISL_3214721, EPI_ISL_3214722, EPI_ISL_3214723, EPI_ISL_3214724, EPI_ISL_3214725, EPI_ISL_3214726, EPI_ISL_3214727, EPI_ISL_3214728, EPI_ISL_3214729, EPI_ISL_3214730, EPI_ISL_3214731, EPI_ISL_3214732, EPI_ISL_3214733, EPI_ISL_3214734, EPI_ISL_3214736, EPI_ISL_3214737, EPI_ISL_3214738, EPI_ISL_3214739 | | | |
| see above | Ochsner Health | Bioinfoexperts | Amy Feehan, Judy Crabtree, Rebecca Rose, Ben Lain, Chris Huston, Samual Moot, David J. Nolan, Tessa LaFleur, Julia-Garcia-Diaz, Lucio Miele, Susanna L. Lamers |
| EPI_ISL_3214740, EPI_ISL_3214741 | Mayo Clinic & Mayo Clinic Laboratories | Minnesota Department of Health, Public Health Laboratory | Alexandra Lorentz, Jacob Garfin, Matt Plumb, and Xiong Wang |

| | | | |
|---|---|---|---|
| EPI_ISL_3214742, EPI_ISL_3214744, EPI_ISL_3214745, EPI_ISL_3214746, EPI_ISL_3214747, EPI_ISL_3214748, EPI_ISL_3214749, EPI_ISL_3214750, EPI_ISL_3214751 | Allina Health Laboratory | Minnesota Department of Health, Public Health Laboratory | Alexandra Lorentz, Jacob Garfin, Matt Plumb, and Xiong Wang |
| EPI_ISL_3214752 | Mayo Clinic & Mayo Clinic Laboratories | Minnesota Department of Health, Public Health Laboratory | Alexandra Lorentz, Jacob Garfin, Matt Plumb, and Xiong Wang |
| EPI_ISL_3214753 | Ridgeview Medical Center | Minnesota Department of Health, Public Health Laboratory | Alexandra Lorentz, Jacob Garfin, Matt Plumb, and Xiong Wang |
| EPI_ISL_3214754 | Centracare Laboratory Services | Minnesota Department of Health, Public Health Laboratory | Alexandra Lorentz, Jacob Garfin, Matt Plumb, and Xiong Wang |
| EPI_ISL_3214756 | Quest Diagnostics | Minnesota Department of Health, Public Health Laboratory | Alexandra Lorentz, Jacob Garfin, Matt Plumb, and Xiong Wang |
| EPI_ISL_3214757, EPI_ISL_3214758 | Allina Health Laboratory | Minnesota Department of Health, Public Health Laboratory | Alexandra Lorentz, Jacob Garfin, Matt Plumb, and Xiong Wang |
| EPI_ISL_3214759 | Minnesota Department of Health, Public Health Laboratory | Minnesota Department of Health, Public Health Laboratory | Alexandra Lorentz, Jacob Garfin, Matt Plumb, and Xiong Wang |
| EPI_ISL_3214760, EPI_ISL_3214761, EPI_ISL_3214762, EPI_ISL_3214763, EPI_ISL_3214764 | Centracare Laboratory Services | Minnesota Department of Health, Public Health Laboratory | Alexandra Lorentz, Jacob Garfin, Matt Plumb, and Xiong Wang |
| EPI_ISL_3214765, EPI_ISL_3214766, EPI_ISL_3214767 | CHI St Gabriels Health- Little Falls | Minnesota Department of Health, Public Health Laboratory | Alexandra Lorentz, Jacob Garfin, Matt Plumb, and Xiong Wang |
| EPI_ISL_3214768, EPI_ISL_3214769, EPI_ISL_3214770, EPI_ISL_3214771, EPI_ISL_3214772, EPI_ISL_3214773, EPI_ISL_3214774, EPI_ISL_3214775, EPI_ISL_3214776, EPI_ISL_3214777, EPI_ISL_3214778, EPI_ISL_3214779, EPI_ISL_3214780, EPI_ISL_3214781, EPI_ISL_3214782 | | | |
| see above | North Memorial | Minnesota Department of Health, Public Health Laboratory | Alexandra Lorentz, Jacob Garfin, Matt Plumb, and Xiong Wang |
| EPI_ISL_3214783 | Minnesota Department of Health, Public Health Laboratory | Minnesota Department of Health, Public Health Laboratory | Alexandra Lorentz, Jacob Garfin, Matt Plumb, and Xiong Wang |
| EPI_ISL_3214784, EPI_ISL_3214785, EPI_ISL_3214786, EPI_ISL_3214787, EPI_ISL_3214788, EPI_ISL_3214789 | North Memorial | Minnesota Department of Health, Public Health Laboratory | Alexandra Lorentz, Jacob Garfin, Matt Plumb, and Xiong Wang |
| EPI_ISL_3214790 | Ridgeview Medical Center | Minnesota Department of Health, Public Health Laboratory | Alexandra Lorentz, Jacob Garfin, Matt Plumb, and Xiong Wang |
| EPI_ISL_3214791, EPI_ISL_3214792 | Regions Hospital | Minnesota Department of Health, Public Health Laboratory | Alexandra Lorentz, Jacob Garfin, Matt Plumb, and Xiong Wang |
| EPI_ISL_3214793, EPI_ISL_3214794 | Park Nicollet Methodist Hospital | Minnesota Department of Health, Public Health Laboratory | Alexandra Lorentz, Jacob Garfin, Matt Plumb, and Xiong Wang |
| EPI_ISL_3214795, EPI_ISL_3214796, EPI_ISL_3214797, EPI_ISL_3214798, EPI_ISL_3214799, EPI_ISL_3214800 | Carris Health Willmar Main Clinic | Minnesota Department of Health, Public Health Laboratory | Alexandra Lorentz, Jacob Garfin, Matt Plumb, and Xiong Wang |
| EPI_ISL_3214801, EPI_ISL_3214802, EPI_ISL_3214803 | Carris Health Rice Memorial Hospital | Minnesota Department of Health, Public Health Laboratory | Alexandra Lorentz, Jacob Garfin, Matt Plumb, and Xiong Wang |
| EPI_ISL_3214804, EPI_ISL_3214805, EPI_ISL_3214806, EPI_ISL_3214807, EPI_ISL_3214809, EPI_ISL_3214810, EPI_ISL_3214811, EPI_ISL_3214812, EPI_ISL_3214813, EPI_ISL_3214814, EPI_ISL_3214815, EPI_ISL_3214816, EPI_ISL_3214817, EPI_ISL_3214818, EPI_ISL_3214819, EPI_ISL_3214820, EPI_ISL_3214821, EPI_ISL_3214822, EPI_ISL_3214823, EPI_ISL_3214824, EPI_ISL_3214825, EPI_ISL_3214826, EPI_ISL_3214827, EPI_ISL_3214828, EPI_ISL_3214829, EPI_ISL_3214830, EPI_ISL_3214831, EPI_ISL_3214832, EPI_ISL_3214833, EPI_ISL_3214834, EPI_ISL_3214836, EPI_ISL_3214837, EPI_ISL_3214838, EPI_ISL_3214839, EPI_ISL_3214840, EPI_ISL_3214841, EPI_ISL_3214842, EPI_ISL_3214843, EPI_ISL_3214844, EPI_ISL_3214845, EPI_ISL_3214846, EPI_ISL_3214847, EPI_ISL_3214848, EPI_ISL_3214849, EPI_ISL_3214850, EPI_ISL_3214851, EPI_ISL_3214852, EPI_ISL_3214853, EPI_ISL_3214854, EPI_ISL_3214855, EPI_ISL_3214856, EPI_ISL_3214857, EPI_ISL_3214858, EPI_ISL_3214859, EPI_ISL_3214860, EPI_ISL_3214861, EPI_ISL_3214862, EPI_ISL_3214863, EPI_ISL_3214864, EPI_ISL_3214865, EPI_ISL_3214866, EPI_ISL_3214867, EPI_ISL_3214868, EPI_ISL_3214869, EPI_ISL_3214870, EPI_ISL_3214871, EPI_ISL_3214872, EPI_ISL_3214873, EPI_ISL_3214874, EPI_ISL_3214875, EPI_ISL_3214876, EPI_ISL_3214877 | | | |
| see above | Baylor Scott & White-Temple | Baylor Scott & White-Temple | Ari Rao, Linden Morales, Kimberly Walker, Marcus Volz, Shelby Johnson |
| EPI_ISL_3214880, EPI_ISL_3214881, EPI_ISL_3214882, EPI_ISL_3214883, EPI_ISL_3214884, EPI_ISL_3214885, EPI_ISL_3214886, EPI_ISL_3214887, EPI_ISL_3214888, EPI_ISL_3214889, EPI_ISL_3214890, EPI_ISL_3214892, EPI_ISL_3214893, EPI_ISL_3214894, EPI_ISL_3214895, EPI_ISL_3214896, EPI_ISL_3214897, EPI_ISL_3214898, EPI_ISL_3214899, EPI_ISL_3214900, EPI_ISL_3214901, EPI_ISL_3214902, EPI_ISL_3214903, EPI_ISL_3214907, EPI_ISL_3214908, EPI_ISL_3214911, EPI_ISL_3214912, EPI_ISL_3214913, EPI_ISL_3214914, EPI_ISL_3214915, EPI_ISL_3214916, EPI_ISL_3214917, EPI_ISL_3214918, EPI_ISL_3214920, EPI_ISL_3214921, EPI_ISL_3214923, EPI_ISL_3214924, EPI_ISL_3214925, EPI_ISL_3214927, EPI_ISL_3214928, EPI_ISL_3214930, EPI_ISL_3214931, EPI_ISL_3214932, EPI_ISL_3214934, EPI_ISL_3214938, EPI_ISL_3214939, EPI_ISL_3214940, EPI_ISL_3214941, EPI_ISL_3214942, EPI_ISL_3214943, EPI_ISL_3214944, EPI_ISL_3214947, EPI_ISL_3214948, EPI_ISL_3214949, EPI_ISL_3214951, EPI_ISL_3214952, EPI_ISL_3214953, EPI_ISL_3214954, EPI_ISL_3214955, EPI_ISL_3214956, EPI_ISL_3214957, EPI_ISL_3214958, EPI_ISL_3214961, EPI_ISL_3214962, EPI_ISL_3214963, EPI_ISL_3214964, EPI_ISL_3214965, EPI_ISL_3214966 | | | |
| see above | Virginia Division of Consolidated Laboratory Services | Virginia Division of Consolidated Laboratory Services | Virginia Division of Consolidated Laboratory Services |
| EPI_ISL_3214970, EPI_ISL_3214971, EPI_ISL_3214972 | UniversitätsSpital Zürich 009 | Institute of Medical Virology | Verena Kufner, Gabriela Ziltener, Maryam Zaheri, Stefan Schmutz, Annette Audigé, Maria Grünberg, Kevin Steiner, Jon Huder, Cyril Shah, Riccarda Capaul, Guido Bloemberg, Jürg Böni, Michael Huber, Alexandra Trkola |
| EPI_ISL_3214973 | Stadtspital Waid | Institute of Medical Virology | Verena Kufner, Gabriela Ziltener, Maryam Zaheri, Stefan Schmutz, Annette Audigé, Maria Grünberg, Kevin Steiner, Jon Huder, Cyril Shah, Riccarda Capaul, Guido Bloemberg, Jürg Böni, Michael Huber, Alexandra Trkola |
| EPI_ISL_3214974, EPI_ISL_3214975 | Kantonsspital Baden AG | Institute of Medical Virology | Verena Kufner, Gabriela Ziltener, Maryam Zaheri, Stefan Schmutz, Annette Audigé, Maria Grünberg, Kevin Steiner, Jon Huder, Cyril Shah, Riccarda Capaul, Guido Bloemberg, Jürg Böni, Michael Huber, Alexandra Trkola |
| EPI_ISL_3214976, EPI_ISL_3214977, EPI_ISL_3214978, EPI_ISL_3214979, EPI_ISL_3214980, EPI_ISL_3214981 | Laborgemeinschaft 1 | Institute of Medical Virology | Verena Kufner, Gabriela Ziltener, Maryam Zaheri, Stefan Schmutz, Annette Audigé, Maria Grünberg, Kevin Steiner, Jon Huder, Cyril Shah, Riccarda Capaul, Guido Bloemberg, Jürg Böni, Michael Huber, Alexandra Trkola |
| EPI_ISL_3214982, EPI_ISL_3214983 | Invenimus AG | Institute of Medical Virology | Verena Kufner, Gabriela Ziltener, Maryam Zaheri, Stefan Schmutz, Annette Audigé, Maria Grünberg, Kevin Steiner, Jon Huder, Cyril Shah, Riccarda Capaul, Guido Bloemberg, Jürg Böni, Michael Huber, Alexandra Trkola |
| EPI_ISL_3214984, EPI_ISL_3214985, EPI_ISL_3214986 | Medica | Institute of Medical Virology | Verena Kufner, Gabriela Ziltener, Maryam Zaheri, Stefan Schmutz, Annette Audigé, Maria Grünberg, Kevin Steiner, Jon Huder, Cyril Shah, Riccarda Capaul, Guido Bloemberg, Jürg Böni, Michael Huber, Alexandra Trkola |
| EPI_ISL_3214987 | Kantonsspital Frauenfeld | Institute of Medical Virology | Verena Kufner, Gabriela Ziltener, Maryam Zaheri, Stefan Schmutz, Annette Audigé, Maria Grünberg, Kevin Steiner, Jon Huder, Cyril Shah, Riccarda Capaul, Guido Bloemberg, Jürg Böni, Michael Huber, Alexandra Trkola |
| EPI_ISL_3214988 | Unilabs | Institute of Medical Virology | Verena Kufner, Gabriela Ziltener, Maryam Zaheri, Stefan Schmutz, Annette Audigé, Maria Grünberg, Kevin Steiner, Jon Huder, Cyril Shah, Riccarda Capaul, Guido Bloemberg, Jürg Böni, Michael Huber, Alexandra Trkola |
| EPI_ISL_3214989, EPI_ISL_3214990, EPI_ISL_3214991, EPI_ISL_3214992, EPI_ISL_3214993, EPI_ISL_3214994, EPI_ISL_3214995 | Analytica | Institute of Medical Virology | Verena Kufner, Gabriela Ziltener, Maryam Zaheri, Stefan Schmutz, Annette Audigé, Maria Grünberg, Kevin Steiner, Jon Huder, Cyril Shah, Riccarda Capaul, Guido Bloemberg, Jürg Böni, Michael Huber, Alexandra Trkola |
| EPI_ISL_3214996 | UniversitätsSpital Zürich | Institute of Medical Virology | Verena Kufner, Gabriela Ziltener, Maryam Zaheri, Stefan Schmutz, Annette Audigé, Maria Grünberg, Kevin Steiner, Jon Huder, Cyril Shah, Riccarda Capaul, Guido Bloemberg, Jürg Böni, Michael Huber, Alexandra Trkola |
| EPI_ISL_3214997, EPI_ISL_3214998, EPI_ISL_3214999 | Zentrum für Labormedizin | Institute of Medical Virology | Verena Kufner, Gabriela Ziltener, Maryam Zaheri, Stefan Schmutz, Annette Audigé, Maria Grünberg, Kevin Steiner, Jon Huder, Cyril Shah, Riccarda Capaul, Guido Bloemberg, Jürg Böni, Michael Huber, Alexandra Trkola |
| EPI_ISL_3215000 | Kantonsspital Aarau | Institute of Medical Virology | Verena Kufner, Gabriela Ziltener, Maryam Zaheri, Stefan Schmutz, Annette Audigé, Maria Grünberg, Kevin Steiner, Jon Huder, Cyril Shah, Riccarda Capaul, Guido Bloemberg, Jürg Böni, Michael Huber, Alexandra Trkola |
| EPI_ISL_3215001 | Invenimus AG | Institute of Medical Virology | Verena Kufner, Gabriela Ziltener, Maryam Zaheri, Stefan Schmutz, Annette Audigé, Maria Grünberg, Kevin Steiner, Jon Huder, Cyril Shah, Riccarda Capaul, Guido Bloemberg, Jürg Böni, Michael Huber, Alexandra Trkola |
| EPI_ISL_3215002 | Medica | Institute of Medical Virology | Verena Kufner, Gabriela Ziltener, Maryam Zaheri, Stefan Schmutz, Annette Audigé, Maria Grünberg, Kevin Steiner, Jon Huder, Cyril Shah, Riccarda Capaul, Guido Bloemberg, Jürg Böni, Michael Huber, Alexandra Trkola |
| EPI_ISL_3215003, EPI_ISL_3215004 | Invenimus AG | Institute of Medical Virology | Verena Kufner, Gabriela Ziltener, Maryam Zaheri, Stefan Schmutz, Annette Audigé, Maria Grünberg, Kevin Steiner, Jon Huder, Cyril Shah, Riccarda Capaul, Guido Bloemberg, Jürg Böni, Michael Huber, Alexandra Trkola |
| EPI_ISL_3215005 | Kantonsspital Aarau | Institute of Medical Virology | Verena Kufner, Gabriela Ziltener, Maryam Zaheri, Stefan Schmutz, Annette Audigé, Maria Grünberg, Kevin Steiner, Jon Huder, Cyril Shah, Riccarda Capaul, Guido Bloemberg, Jürg Böni, Michael Huber, Alexandra Trkola |

| | | | |
|---|---|---|---|
| EPI_ISL_3215006 | Unilabs | Institute of Medical Virology | Verena Kufner, Gabriela Ziltener, Maryam Zaheri, Stefan Schmutz, Annette Audigé, Maria Grünberg, Kevin Steiner, Jon Huder, Cyril Shah, Riccarda Capaul, Guido Bloemberg, Jürg Böni, Michael Huber, Alexandra Trkola |
| EPI_ISL_3215007, EPI_ISL_3215008 | UniversitätsSpital Zürich 009 | Institute of Medical Virology | Verena Kufner, Gabriela Ziltener, Maryam Zaheri, Stefan Schmutz, Annette Audigé, Maria Grünberg, Kevin Steiner, Jon Huder, Cyril Shah, Riccarda Capaul, Guido Bloemberg, Jürg Böni, Michael Huber, Alexandra Trkola |
| EPI_ISL_3215009 | UniversitätsSpital Zürich | Institute of Medical Virology | Verena Kufner, Gabriela Ziltener, Maryam Zaheri, Stefan Schmutz, Annette Audigé, Maria Grünberg, Kevin Steiner, Jon Huder, Cyril Shah, Riccarda Capaul, Guido Bloemberg, Jürg Böni, Michael Huber, Alexandra Trkola |
| EPI_ISL_3215010 | Spital Limmattal | Institute of Medical Virology | Verena Kufner, Gabriela Ziltener, Maryam Zaheri, Stefan Schmutz, Annette Audigé, Maria Grünberg, Kevin Steiner, Jon Huder, Cyril Shah, Riccarda Capaul, Guido Bloemberg, Jürg Böni, Michael Huber, Alexandra Trkola |
| EPI_ISL_3215055, EPI_ISL_3215056, EPI_ISL_3215057 | Diagnostic Genomics Laboratory | Diagnostic Genomics Laboratory | Keely Duff, Carolyn Banister, Phillip Buckhaults |
| EPI_ISL_3215063 | U.O. Microbiologia Laboratorio Unico Centro Servizi - AUSL della Romagna | U.O. Microbiologia, Laboratorio Unico Centro Servizi - AUSL della Romagna | Giorgio Dirani |
| EPI_ISL_3215135, EPI_ISL_3215136, EPI_ISL_3215137, EPI_ISL_3215139, EPI_ISL_3215140, EPI_ISL_3215141, EPI_ISL_3215144, EPI_ISL_3215145, EPI_ISL_3215146, EPI_ISL_3215147, EPI_ISL_3215148, EPI_ISL_3215149, EPI_ISL_3215150, EPI_ISL_3215151, EPI_ISL_3215152, EPI_ISL_3215153, EPI_ISL_3215154, EPI_ISL_3215155, EPI_ISL_3215156, EPI_ISL_3215157, EPI_ISL_3215158, EPI_ISL_3215159, EPI_ISL_3215160, EPI_ISL_3215161, EPI_ISL_3215162 | | | |
| see above | Oregon State Public Health Laboratory | Oregon State Public Health Laboratory | Rafia Razzaque, Eugene Yeboah, Vanda Makris, Laura Tsaknaridis, John Fontana and Shane Sevey |
| EPI_ISL_3215170, EPI_ISL_3215171, EPI_ISL_3215174, EPI_ISL_3215176, EPI_ISL_3215193, EPI_ISL_3215261 | U.O. Microbiologia Laboratorio Unico Centro Servizi - AUSL della Romagna | U.O. Microbiologia, Laboratorio Unico Centro Servizi - AUSL della Romagna | Giorgio Dirani |
| EPI_ISL_3215301, EPI_ISL_3215304, EPI_ISL_3215317, EPI_ISL_3215318, EPI_ISL_3215319, EPI_ISL_3215327, EPI_ISL_3215332, EPI_ISL_3215336, EPI_ISL_3215339, EPI_ISL_3215342, EPI_ISL_3215343, EPI_ISL_3215344, EPI_ISL_3215345, EPI_ISL_3215347, EPI_ISL_3215348, EPI_ISL_3215349, EPI_ISL_3215350, EPI_ISL_3215351, EPI_ISL_3215352, EPI_ISL_3215354, EPI_ISL_3215355 | | | |
| see above | EXCITE Lab | Andersen lab at Scripps Research | David Pride, Sharon Reed, Chip Schooley, Angela Scioscia, Natasha Martin Cheryl Anderson, Sawyer Farmer, Abigail Schnapper, Helena Tubb, Tommy Valles + SEARCH |
| EPI_ISL_3215467 | EXCITE Lab | Andersen lab at Scripps Research | Chip Schooley, Natasha Martin, Cheryl Anderson, Angela Scioscia, Smruthi Karthikeyan, Greg Humphrey, Sawyer Farmer, Abigail Schnapper, Helena Tubb, Tommy Valles + SEARCH |
| EPI_ISL_3215643 | EXCITE Lab | Andersen lab at Scripps Research | David Pride, Sharon Reed, Chip Schooley, Angela Scioscia, Natasha Martin Cheryl Anderson, Sawyer Farmer, Abigail Schnapper, Helena Tubb, Tommy Valles + SEARCH |
| EPI_ISL_3215684 | EXCITE Lab | Andersen lab at Scripps Research | Chip Schooley, Natasha Martin, Cheryl Anderson, Angela Scioscia, Smruthi Karthikeyan, Greg Humphrey, Sawyer Farmer, Abigail Schnapper, Helena Tubb, Tommy Valles + SEARCH |
| EPI_ISL_3215701, EPI_ISL_3215738 | EXCITE Lab | Andersen lab at Scripps Research | David Pride, Sharon Reed, Chip Schooley, Angela Scioscia, Natasha Martin Cheryl Anderson, Sawyer Farmer, Abigail Schnapper, Helena Tubb, Tommy Valles + SEARCH |
| EPI_ISL_3215755, EPI_ISL_3215756, EPI_ISL_3215757, EPI_ISL_3215761, EPI_ISL_3215762, EPI_ISL_3215765, EPI_ISL_3215800 | EXCITE Lab | Andersen lab at Scripps Research | Chip Schooley, Natasha Martin, Cheryl Anderson, Angela Scioscia, Smruthi Karthikeyan, Greg Humphrey, Sawyer Farmer, Abigail Schnapper, Helena Tubb, Tommy Valles + SEARCH |
| EPI_ISL_3215801, EPI_ISL_3215802, EPI_ISL_3215803, EPI_ISL_3215804, EPI_ISL_3215807, EPI_ISL_3215809, EPI_ISL_3215811, EPI_ISL_3215812, EPI_ISL_3215813, EPI_ISL_3215816, EPI_ISL_3215817, EPI_ISL_3215818, EPI_ISL_3215819, EPI_ISL_3215820, EPI_ISL_3215821 | | | |
| see above | EXCITE Lab | Andersen lab at Scripps Research | David Pride, Sharon Reed, Chip Schooley, Angela Scioscia, Natasha Martin Cheryl Anderson, Sawyer Farmer, Abigail Schnapper, Helena Tubb, Tommy Valles + SEARCH |
| EPI_ISL_3215822 | EXCITE Lab | Andersen lab at Scripps Research | Chip Schooley, Natasha Martin, Cheryl Anderson, Angela Scioscia, Smruthi Karthikeyan, Greg Humphrey, Sawyer Farmer, Abigail Schnapper, Helena Tubb, Tommy Valles + SEARCH |
| EPI_ISL_3215824 | EXCITE Lab | Andersen lab at Scripps Research | David Pride, Sharon Reed, Chip Schooley, Angela Scioscia, Natasha Martin Cheryl Anderson, Sawyer Farmer, Abigail Schnapper, Helena Tubb, Tommy Valles + SEARCH |
| EPI_ISL_3215924, EPI_ISL_3215925, EPI_ISL_3215926, EPI_ISL_3215927, EPI_ISL_3215929, EPI_ISL_3215930, EPI_ISL_3215931, EPI_ISL_3215932, EPI_ISL_3215934, EPI_ISL_3215935, EPI_ISL_3215936, EPI_ISL_3215937, EPI_ISL_3215938, EPI_ISL_3215939, EPI_ISL_3215941, EPI_ISL_3215942, EPI_ISL_3215944, EPI_ISL_3215945, EPI_ISL_3215946, EPI_ISL_3215947, EPI_ISL_3215948 | | | |
| see above | Sharp HealthCare Laboratory | Andersen lab at Scripps Research | SEARCH Alliance San Diego with Aaron Harding, Jacquelyn Berumen, Cathy Woerle, Liam McGinnis, Art Mendoza, Omid Bakhtar |
| EPI_ISL_3215952, EPI_ISL_3215953, EPI_ISL_3215954, EPI_ISL_3215955, EPI_ISL_3215956, EPI_ISL_3215959, EPI_ISL_3215960, EPI_ISL_3215961, EPI_ISL_3215962, EPI_ISL_3215965, EPI_ISL_3215966, EPI_ISL_3215967, EPI_ISL_3215968, EPI_ISL_3215970, EPI_ISL_3215971, EPI_ISL_3215972, EPI_ISL_3215973, EPI_ISL_3215974, EPI_ISL_3215975, EPI_ISL_3215977, EPI_ISL_3215978, EPI_ISL_3215979, EPI_ISL_3215980, EPI_ISL_3215981, EPI_ISL_3215983, EPI_ISL_3215984, EPI_ISL_3215985, EPI_ISL_3215986, EPI_ISL_3215987, EPI_ISL_3215990, EPI_ISL_3215991, EPI_ISL_3215992, EPI_ISL_3215993, EPI_ISL_3215994, EPI_ISL_3215995, EPI_ISL_3215996, EPI_ISL_3215997, EPI_ISL_3215998, EPI_ISL_3215999, EPI_ISL_3216001, EPI_ISL_3216002, EPI_ISL_3216003, EPI_ISL_3216004 | | | |
| see above | Genomica Lab Molecular, Mexico | Andersen lab at Scripps Research | SEARCH Alliance San Diego with Jonathan Gonzalez Garcia, Jose Roman Chavez Mendez, Jose Horacio Reyna Verdugo, Martin Gonzalez Ibarra, Luis Alberto Rangel Gonzalez |
| EPI_ISL_3216005, EPI_ISL_3216006, EPI_ISL_3216007, EPI_ISL_3216008, EPI_ISL_3216009, EPI_ISL_3216010, EPI_ISL_3216011, EPI_ISL_3216012, EPI_ISL_3216013, EPI_ISL_3216014, EPI_ISL_3216015, EPI_ISL_3216016, EPI_ISL_3216017, EPI_ISL_3216018, EPI_ISL_3216020, EPI_ISL_3216021, EPI_ISL_3216022, EPI_ISL_3216023, EPI_ISL_3216024, EPI_ISL_3216026, EPI_ISL_3216027, EPI_ISL_3216029, EPI_ISL_3216030, EPI_ISL_3216031, EPI_ISL_3216033, EPI_ISL_3216036, EPI_ISL_3216037, EPI_ISL_3216038, EPI_ISL_3216039, EPI_ISL_3216040, EPI_ISL_3216041, EPI_ISL_3216043, EPI_ISL_3216046, EPI_ISL_3216047, EPI_ISL_3216048, EPI_ISL_3216049, EPI_ISL_3216050, EPI_ISL_3216051, EPI_ISL_3216053, EPI_ISL_3216056, EPI_ISL_3216057, EPI_ISL_3216058, EPI_ISL_3216059, EPI_ISL_3216060, EPI_ISL_3216061, EPI_ISL_3216062, EPI_ISL_3216063, EPI_ISL_3216064, EPI_ISL_3216065, EPI_ISL_3216066, EPI_ISL_3216067, EPI_ISL_3216068, EPI_ISL_3216069, EPI_ISL_3216070, EPI_ISL_3216071, EPI_ISL_3216072, EPI_ISL_3216073, EPI_ISL_3216074, EPI_ISL_3216075, EPI_ISL_3216078, EPI_ISL_3216079, EPI_ISL_3216080, EPI_ISL_3216081, EPI_ISL_3216082, EPI_ISL_3216083, EPI_ISL_3216084, EPI_ISL_3216085, EPI_ISL_3216086, EPI_ISL_3216087, EPI_ISL_3216088, EPI_ISL_3216089, EPI_ISL_3216090, EPI_ISL_3216091, EPI_ISL_3216092, EPI_ISL_3216093, EPI_ISL_3216094, EPI_ISL_3216095, EPI_ISL_3216096, EPI_ISL_3216097, EPI_ISL_3216098, EPI_ISL_3216099, EPI_ISL_3216100, EPI_ISL_3216101, EPI_ISL_3216102, EPI_ISL_3216103, EPI_ISL_3216105, EPI_ISL_3216106, EPI_ISL_3216107 | | | |
| see above | Sharp HealthCare Laboratory | Andersen lab at Scripps Research | SEARCH Alliance San Diego with Aaron Harding, Jacquelyn Berumen, Cathy Woerle, Liam McGinnis, Art Mendoza, Omid Bakhtar |
| EPI_ISL_3216108, EPI_ISL_3216109, EPI_ISL_3216110, EPI_ISL_3216111, EPI_ISL_3216112, EPI_ISL_3216113, EPI_ISL_3216114, EPI_ISL_3216115, EPI_ISL_3216116, EPI_ISL_3216117, EPI_ISL_3216118, EPI_ISL_3216119, EPI_ISL_3216120, EPI_ISL_3216121, EPI_ISL_3216122, EPI_ISL_3216123, EPI_ISL_3216124, EPI_ISL_3216125, EPI_ISL_3216126, EPI_ISL_3216127, EPI_ISL_3216128, EPI_ISL_3216130, EPI_ISL_3216131, EPI_ISL_3216132, EPI_ISL_3216133, EPI_ISL_3216134, EPI_ISL_3216135, EPI_ISL_3216136, EPI_ISL_3216137, EPI_ISL_3216138, EPI_ISL_3216139, EPI_ISL_3216140, EPI_ISL_3216141, EPI_ISL_3216142, EPI_ISL_3216144, EPI_ISL_3216145, EPI_ISL_3216146, EPI_ISL_3216147, EPI_ISL_3216148, EPI_ISL_3216149, EPI_ISL_3216150, EPI_ISL_3216151, EPI_ISL_3216152, EPI_ISL_3216153, EPI_ISL_3216155, EPI_ISL_3216156, EPI_ISL_3216157, EPI_ISL_3216158, EPI_ISL_3216159, EPI_ISL_3216160, EPI_ISL_3216161, EPI_ISL_3216162, EPI_ISL_3216163, EPI_ISL_3216164, EPI_ISL_3216165, EPI_ISL_3216166, EPI_ISL_3216167, EPI_ISL_3216168, EPI_ISL_3216169, EPI_ISL_3216170, EPI_ISL_3216171 | | | |
| see above | Scripps Medical Laboratory | Andersen lab at Scripps Research | SEARCH Alliance San Diego with Michael Quigley, Ellen Stefanski, Ian Mchardy |
| EPI_ISL_3216243, EPI_ISL_3216244, EPI_ISL_3216245, EPI_ISL_3216246, EPI_ISL_3216247, EPI_ISL_3216248, EPI_ISL_3216249, EPI_ISL_3216250, EPI_ISL_3216251, EPI_ISL_3216252, EPI_ISL_3216253, EPI_ISL_3216254, EPI_ISL_3216255, EPI_ISL_3216256, EPI_ISL_3216257, EPI_ISL_3216258, EPI_ISL_3216259, EPI_ISL_3216260, EPI_ISL_3216262, EPI_ISL_3216263, EPI_ISL_3216264, EPI_ISL_3216265, EPI_ISL_3216266, EPI_ISL_3216267, EPI_ISL_3216268, EPI_ISL_3216269, EPI_ISL_3216270, EPI_ISL_3216271 | | | |
| see above | State Hygienic Laboratory at the University of Iowa | State Hygienic Laboratory at the University of Iowa | Valerie Reeb, Erik Twait, Anna Yakos, Mariah Knutson, Kris Eveland, Kristen Zanon, Jeff, Benfer, Davis Rieckenberg, Wes Hottel, Alankar Kampoowale |
| EPI_ISL_3216272, EPI_ISL_3216273, EPI_ISL_3216274, EPI_ISL_3216277, EPI_ISL_3216278, EPI_ISL_3216279, EPI_ISL_3216280, EPI_ISL_3216281, EPI_ISL_3216282, EPI_ISL_3216283, EPI_ISL_3216285, EPI_ISL_3216286, EPI_ISL_3216287, EPI_ISL_3216288, EPI_ISL_3216289, EPI_ISL_3216290, EPI_ISL_3216292, EPI_ISL_3216293, EPI_ISL_3216294, EPI_ISL_3216295, EPI_ISL_3216296, EPI_ISL_3216297, EPI_ISL_3216298, EPI_ISL_3216300, EPI_ISL_3216301, EPI_ISL_3216302, EPI_ISL_3216303, EPI_ISL_3216304, EPI_ISL_3216305, EPI_ISL_3216306, EPI_ISL_3216307, EPI_ISL_3216308, EPI_ISL_3216309, EPI_ISL_3216311, EPI_ISL_3216312, EPI_ISL_3216313, EPI_ISL_3216314, EPI_ISL_3216315, EPI_ISL_3216316, EPI_ISL_3216317, EPI_ISL_3216321, EPI_ISL_3216322, EPI_ISL_3216323, EPI_ISL_3216326, EPI_ISL_3216327, EPI_ISL_3216328, EPI_ISL_3216330, EPI_ISL_3216331, EPI_ISL_3216332, EPI_ISL_3216333, EPI_ISL_3216334 | | | |
| see above | Centro de Investigación Biomédica de La Rioja - Hospital San Pedro Logroño | SeqCOVID-SPAIN consortium/IBV(CSIC) | María de Toro, José Manuel Azcona Gutiérrez, María Pilar Bea Escudero, Miriam Blasco Alberdi and SeqCOVID-SPAIN consortium |
| EPI_ISL_3216335, EPI_ISL_3216336, EPI_ISL_3216337, EPI_ISL_3216338, EPI_ISL_3216339 | County of San Luis Obispo Public Health Laboratory | County of San Luis Obispo Public Health Laboratory | Frances Sidhu, Katrina Erwin |
| EPI_ISL_3216340, EPI_ISL_3216341 | Twin Cities Community Hospital Laboratory | County of San Luis Obispo Public Health Laboratory | Frances Sidhu, Katrina Erwin |
| EPI_ISL_3216342 | County of San Luis Obispo Public Health Laboratory | County of San Luis Obispo Public Health Laboratory | Frances Sidhu, Katrina Erwin |
| EPI_ISL_3216343, EPI_ISL_3216344, EPI_ISL_3216345 | Cal Poly University Health Center | County of San Luis Obispo Public Health Laboratory | Frances Sidhu, Katrina Erwin |
| EPI_ISL_3216346, EPI_ISL_3216347, EPI_ISL_3216348 | County of San Luis Obispo Public Health Laboratory | County of San Luis Obispo Public Health Laboratory | Frances Sidhu, Katrina Erwin |
| EPI_ISL_3216349, EPI_ISL_3216350, EPI_ISL_3216351 | Marian Medical Center Laboratory | County of San Luis Obispo Public Health Laboratory | Frances Sidhu, Katrina Erwin |
| EPI_ISL_3216353, EPI_ISL_3216354, EPI_ISL_3216355, EPI_ISL_3216356, EPI_ISL_3216357, EPI_ISL_3216358, EPI_ISL_3216359, EPI_ISL_3216360, EPI_ISL_3216361, EPI_ISL_3216362, EPI_ISL_3216364, EPI_ISL_3216365, EPI_ISL_3216366, EPI_ISL_3216367, EPI_ISL_3216368, EPI_ISL_3216369, EPI_ISL_3216370, EPI_ISL_3216371, EPI_ISL_3216372, EPI_ISL_3216373, EPI_ISL_3216376, EPI_ISL_3216378, EPI_ISL_3216380, EPI_ISL_3216382, EPI_ISL_3216384, EPI_ISL_3216387, EPI_ISL_3216389, EPI_ISL_3216391 | | | |
| see above | IN State Department of Health Laboratory Services | IN State Department of Health Laboratory Services | Cassandra Campion, Jamie Yeadon, Brian Pope, Lixia Liu, Kyle Brownlee, Melissa Hindenlang, Mark Glazier |
| EPI_ISL_3216399, EPI_ISL_3216400, EPI_ISL_3216405, EPI_ISL_3216410 | Hospital General Universitario de Alicante - Instituto de Investigación Sanitaria y Biomédica de Alicante | SeqCOVID-SPAIN consortium/IBV(CSIC) | Maripaz Ventero Martín, Carmen Molina Pardines and SeqCOVID-SPAIN consortium |
| EPI_ISL_3216417 | DOHMH Jamaica | New York City Public Health Laboratory | Jade Wang, et al. |
| EPI_ISL_3216419, EPI_ISL_3216420 | DOHMH Chelsea | New York City Public Health Laboratory | Jade Wang, et al. |
| EPI_ISL_3216421 | DOHMH Riverside | New York City Public Health Laboratory | Jade Wang, et al. |
| EPI_ISL_3216423 | DOHMH Chelsea | New York City Public Health Laboratory | Jade Wang, et al. |
| EPI_ISL_3216424 | DOHMH Crown Heights | New York City Public Health Laboratory | Jade Wang, et al. |
| EPI_ISL_3216425 | DOHMH Riverside | New York City Public Health Laboratory | Jade Wang, et al. |

| | | | |
|---|---|---|---|
| EPI_ISL_3216426 | DOHMH Fort Greene | New York City Public Health Laboratory | Jade Wang, et al. |
| EPI_ISL_3216427, EPI_ISL_3216428 | DOHMH Jamaica | New York City Public Health Laboratory | Jade Wang, et al. |
| EPI_ISL_3216430 | Quest Diagnostics | New York City Public Health Laboratory | Jade Wang, et al. |
| EPI_ISL_3216431 | Genesis Diagnostics | New York City Public Health Laboratory | Jade Wang, et al. |
| EPI_ISL_3216432 | DOHMH Fort Greene | New York City Public Health Laboratory | Jade Wang, et al. |
| EPI_ISL_3216433 | DOHMH Corona | New York City Public Health Laboratory | Jade Wang, et al. |
| EPI_ISL_3216434, EPI_ISL_3216435 | DOHMH Chelsea | New York City Public Health Laboratory | Jade Wang, et al. |
| EPI_ISL_3216436, EPI_ISL_3216437 | DOHMH Morrisania | New York City Public Health Laboratory | Jade Wang, et al. |
| EPI_ISL_3216438, EPI_ISL_3216439, EPI_ISL_3216440, EPI_ISL_3216441 | DOHMH Crown Heights | New York City Public Health Laboratory | Jade Wang, et al. |
| EPI_ISL_3216442, EPI_ISL_3216443 | DOHMH PHL | New York City Public Health Laboratory | Jade Wang, et al. |
| EPI_ISL_3216444 | DOHMH Riverside | New York City Public Health Laboratory | Jade Wang, et al. |
| EPI_ISL_3216445 | DOHMH Jamaica | New York City Public Health Laboratory | Jade Wang, et al. |
| EPI_ISL_3216446, EPI_ISL_3216447 | DOHMH PHL | New York City Public Health Laboratory | Jade Wang, et al. |
| EPI_ISL_3216448, EPI_ISL_3216449, EPI_ISL_3216450, EPI_ISL_3216451 | Quest Diagnostics | New York City Public Health Laboratory | Jade Wang, et al. |
| EPI_ISL_3216452 | Centers Lab NJ LLC | New York City Public Health Laboratory | Jade Wang, et al. |
| EPI_ISL_3216453, EPI_ISL_3216454, EPI_ISL_3216455 | Quest Diagnostics | New York City Public Health Laboratory | Jade Wang, et al. |
| EPI_ISL_3216458, EPI_ISL_3216459, EPI_ISL_3216463, EPI_ISL_3216466 | LabQ Clinical Diagnostics LLC | New York City Public Health Laboratory | Jade Wang, et al. |
| EPI_ISL_3216467, EPI_ISL_3216468, EPI_ISL_3216469, EPI_ISL_3216470, EPI_ISL_3216471 | Quest Diagnostics | New York City Public Health Laboratory | Jade Wang, et al. |
| EPI_ISL_3216472 | DOHMH Fort Greene | New York City Public Health Laboratory | Jade Wang, et al. |
| EPI_ISL_3216474 | DOHMH Morrisania | New York City Public Health Laboratory | Jade Wang, et al. |
| EPI_ISL_3216475, EPI_ISL_3216476 | DOHMH Fort Greene | New York City Public Health Laboratory | Jade Wang, et al. |
| EPI_ISL_3216477, EPI_ISL_3216478, EPI_ISL_3216479, EPI_ISL_3216480 | DOHMH Crown Heights | New York City Public Health Laboratory | Jade Wang, et al. |
| EPI_ISL_3216481, EPI_ISL_3216482 | DOHMH Riverside | New York City Public Health Laboratory | Jade Wang, et al. |
| EPI_ISL_3216483, EPI_ISL_3216484, EPI_ISL_3216485, EPI_ISL_3216486, EPI_ISL_3216487 | DOHMH Chelsea | New York City Public Health Laboratory | Jade Wang, et al. |
| EPI_ISL_3216488, EPI_ISL_3216489, EPI_ISL_3216490 | DOHMH Corona | New York City Public Health Laboratory | Jade Wang, et al. |
| EPI_ISL_3216491, EPI_ISL_3216492, EPI_ISL_3216493, EPI_ISL_3216494, EPI_ISL_3216495, EPI_ISL_3216496 | DOHMH PHL | New York City Public Health Laboratory | Jade Wang, et al. |
| EPI_ISL_3216497 | DOHMH Crown Heights | New York City Public Health Laboratory | Jade Wang, et al. |
| EPI_ISL_3216498 | DOHMH Riverside | New York City Public Health Laboratory | Jade Wang, et al. |
| EPI_ISL_3216499, EPI_ISL_3216500, EPI_ISL_3216501, EPI_ISL_3216502, EPI_ISL_3216503, EPI_ISL_3216504, EPI_ISL_3216505 | DOHMH PHL | New York City Public Health Laboratory | Jade Wang, et al. |
| EPI_ISL_3216506, EPI_ISL_3216507, EPI_ISL_3216508 | DOHMH Riverside | New York City Public Health Laboratory | Jade Wang, et al. |
| EPI_ISL_3216509 | DOHMH Jamaica | New York City Public Health Laboratory | Jade Wang, et al. |
| EPI_ISL_3216510 | DOHMH Chelsea | New York City Public Health Laboratory | Jade Wang, et al. |
| EPI_ISL_3216511, EPI_ISL_3216512 | DOHMH Crown Heights | New York City Public Health Laboratory | Jade Wang, et al. |
| EPI_ISL_3216513 | DOHMH Chelsea | New York City Public Health Laboratory | Jade Wang, et al. |
| EPI_ISL_3216514 | DOHMH Crown Heights | New York City Public Health Laboratory | Jade Wang, et al. |
| EPI_ISL_3216515 | RSUP FATMAWATI / DINAS KESEHATAN DKI JAKARTA | Genomik Solidaritas Indonesia Laboratorium | Himawan Masyhuri, Neng Hilma Hamidah, Normastuti Adhini Tantyo, Alfin Mohammad Abdillah, Dhahlia Agustina Cahyono, Reinhart Gabriel, Anna Christina Brazia, Vania Gavrila Wikasa, Carissa Sintca Wijaya, Annisa Muthiah Sukirman, Gracia Felias Enos Korompis, Anuraj Shankar, Meutia Ayuputeri Kumaheri, Ariel Pradipta |
| EPI_ISL_3216516 | RSUP FATMAWATI / DINAS KESEHATAN DKI JAKARTA | Genomik Solidaritas Indonesia Laboratorium | Neng Hilma Hamidah, Himawan Masyhuri, Normastuti Adhini Tantyo, Alfin Mohammad Abdillah, Dhahlia Agustina Cahyono, Reinhart Gabriel, Anna Christina Brazia, Vania Gavrila Wikasa, Carissa Sintca Wijaya, Annisa Muthiah Sukirman, Gracia Felias Enos Korompis, Anuraj Shankar, Meutia Ayuputeri Kumaheri, Ariel Pradipta |
| EPI_ISL_3216517 | RSUP FATMAWATI / DINAS KESEHATAN DKI JAKARTA | Genomik Solidaritas Indonesia Laboratorium | Normastuti Adhini Tantyo, Alfin Mohammad Abdillah, Dhahlia Agustina Cahyono, Neng Hilma Hamidah, Himawan Masyhuri, Reinhart Gabriel, Anna Christina Brazia, Vania Gavrila Wikasa, Carissa Sintca Wijaya, Gracia Felias Enos Korompis, Annisa Muthiah Sukirman, Anuraj Shankar, Meutia Ayuputeri Kumaheri, Ariel Pradipta |
| EPI_ISL_3216518 | RSUP FATMAWATI / DINAS KESEHATAN DKI JAKARTA | Genomik Solidaritas Indonesia Laboratorium | Alfin Mohammad Abdillah, Normastuti Adhini Tantyo, Dhahlia Agustina Cahyono, Neng Hilma Hamidah, Himawan Masyhuri, Vania Gavrila Wikasa, Carissa Sintca Wijaya, Anna Christina Brazia, Reinhart Gabriel, Gracia Felias Enos Korompis, Annisa Muthiah Sukirman, Anuraj Shankar, Meutia Ayuputeri Kumaheri, Ariel Pradipta |
| EPI_ISL_3216519 | RSUP FATMAWATI / DINAS KESEHATAN DKI JAKARTA | Genomik Solidaritas Indonesia Laboratorium | Dhahlia Agustina Cahyono, Alfin Mohammad Abdillah, Normastuti Adhini Tantyo, Neng Hilma Hamidah, Himawan Masyhuri, Vania Gavrila Wikasa, Carissa Sintca Wijaya, Anna Christina Brazia, Reinhart Gabriel, Gracia Felias Enos Korompis, Annisa Muthiah Sukirman, Anuraj Shankar, Meutia Ayuputeri Kumaheri, Ariel Pradipta |
| EPI_ISL_3216520 | RSUD CEMPAKA PUTIH / DINAS KESEHATAN DKI JAKARTA | Genomik Solidaritas Indonesia Laboratorium | Himawan Masyhuri, Neng Hilma Hamidah, Normastuti Adhini Tantyo, Alfin Mohammad Abdillah, Dhahlia Agustina Cahyono, Vania Gavrila Wikasa, Carissa Sintca Wijaya, Anna Christina Brazia, Reinhart Gabriel, Annisa Muthiah Sukirman, Gracia Felias Enos Korompis, Anuraj Shankar, Meutia Ayuputeri Kumaheri, Ariel Pradipta |

| | | | |
|---|---|---|---|
| EPI_ISL_3216521 | RSUP FATMAWATI / DINAS KESEHATAN DKI JAKARTA | Genomik Solidaritas Indonesia Laboratorium | Neng Hilma Hamidah, Himawan Masyhuri, Normastuti Adhini Tantyo, Alfin Mohammad Abdillah, Dhahlia Agustina Cahyono, Vania Gavrila Wikasa, Carissa Sintca Wijaya, Anna Christina Brazia, Reinhart Gabriel, Annisa Muthiah Sukirman, Gracia Felias Enos Korompis, Anuraj Shankar, Meutia Ayuputeri Kumaheri, Ariel Pradipta |
| EPI_ISL_3216522 | Genomik Solidaritas Indonesia Laboratorium | Genomik Solidaritas Indonesia Laboratorium | Normastuti Adhini Tantyo, Alfin Mohammad Abdillah, Dhahlia Agustina Cahyono, Neng Hilma Hamidah, Himawan Masyhuri, Vania Gavrila Wikasa, Carissa Sintca Wijaya, Anna Christina Brazia, Reinhart Gabriel, Gracia Felias Enos Korompis, Annisa Muthiah Sukirman, Anuraj Shankar, Meutia Ayuputeri Kumaheri, Ariel Pradipta |
| EPI_ISL_3216523 | Genomik Solidaritas Indonesia Laboratorium | Genomik Solidaritas Indonesia Laboratorium | Dhahlia Agustina Cahyono, Alfin Mohammad Abdillah, Normastuti Adhini Tantyo, Neng Hilma Hamidah, Himawan Masyhuri, Vania Gavrila Wikasa, Carissa Sintca Wijaya, Anna Christina Brazia, Reinhart Gabriel, Gracia Felias Enos Korompis, Annisa Muthiah Sukirman, Anuraj Shankar, Meutia Ayuputeri Kumaheri, Ariel Pradipta |
| EPI_ISL_3216524, EPI_ISL_3216526, EPI_ISL_3216527, EPI_ISL_3216529, EPI_ISL_3216530, EPI_ISL_3216531 | RSUD dr Soehadi Prijonegoro Sragen | National Institute of Health Research and Development | Vivi Setiawaty, Hana Apsari Pawestri, Hartanti Dian Ikawati, Subangkit, Kartika Dewi Puspa, Arie Ardiansyah Nugraha, Fajar Nur Sulistiyohadi, Triyani Soekarso, Krisna Pangesti, Nelly Puspandari |
| EPI_ISL_3216532, EPI_ISL_3216534, EPI_ISL_3216535, EPI_ISL_3216537, EPI_ISL_3216538, EPI_ISL_3216540 | RSUD Arifin Achmad | National Institute of Health Research and Development | Vivi Setiawaty, Hana Apsari Pawestri, Hartanti Dian Ikawati, Subangkit, Kartika Dewi Puspa, Arie Ardiansyah Nugraha, Fajar Nur Sulistiyohadi, Triyani Soekarso, Krisna Pangesti, Fajri Marindra, Nelly Puspandari |
| EPI_ISL_3216541, EPI_ISL_3216543, EPI_ISL_3216544 | Labkesda Kota Tangerang | National Institute of Health Research and Development | Hana Apsari Pawestri, Hartanti Dian Ikawati, Subangkit, Kartika Dewi Puspa, Arie Ardiansyah Nugraha, Fajar Nur Sulistiyohadi, Triyani Soekarso, Krisna Pangesti, Nelly Puspandari, Vivi Setiawaty |
| EPI_ISL_3216546, EPI_ISL_3216547 | RS Jantung dan Pembuluh Darah Harapan Kita | National Institute of Health Research and Development | Hana Apsari Pawestri, Hartanti Dian Ikawati, Subangkit, Kartika Dewi Puspa, Arie Ardiansyah Nugraha, Fajar Nur Sulistiyohadi, Triyani Soekarso, Krisna Pangesti, Nelly Puspandari, Vivi Setiawaty |
| EPI_ISL_3216548, EPI_ISL_3216550, EPI_ISL_3216551, EPI_ISL_3216552, EPI_ISL_3216553, EPI_ISL_3216554 | RSUD Provinsi Nusa Tenggara Barat | National Institute of Health Research and Development | Hana Apsari Pawestri, Hartanti Dian Ikawati, Subangkit, Kartika Dewi Puspa, Arie Ardiansyah Nugraha, Fajar Nur Sulistiyohadi, Triyani Soekarso, Krisna Pangesti, Nelly Puspandari, Laily Indrayanti, Vivi Setiawaty |
| EPI_ISL_3216555, EPI_ISL_3216556 | RSU Siloam, Bekasi | National Institute of Health Research and Development | Hana Apsari Pawestri, Hartanti Dian Ikawati, Subangkit, Kartika Dewi Puspa, Arie Ardiansyah Nugraha, Fajar Nur Sulistiyohadi, Triyani Soekarso, Krisna Pangesti, Nelly Puspandari, Vivi Setiawaty |
| EPI_ISL_3216559, EPI_ISL_3216561 | Lab Genetik Sumbawa Technopark | National Institute of Health Research and Development | Hana Apsari Pawestri, Hartanti Dian Ikawati, Subangkit, Kartika Dewi Puspa, Arie Ardiansyah Nugraha, Fajar Nur Sulistiyohadi, Triyani Soekarso, Krisna Pangesti, Nelly Puspandari, Arief Budi Witarto, Rian Ardha Ardinata, Lalu Hamzi Fikri, Nurhandini Eka Dewi, Laily Indrayanti, Vivi Setiawaty |
| EPI_ISL_3216563, EPI_ISL_3216564, EPI_ISL_3216565, EPI_ISL_3216566, EPI_ISL_3216568, EPI_ISL_3216569 | RSUD Brebes | National Institute of Health Research and Development | Hana Apsari Pawestri, Hartanti Dian Ikawati, Subangkit, Kartika Dewi Puspa, Arie Ardiansyah Nugraha, Fajar Nur Sulistiyohadi, Triyani Soekarso, Krisna Pangesti, Nelly Puspandari, Vivi Setiawaty |
| EPI_ISL_3216571, EPI_ISL_3216572 | Labkesda DKI Jakarta | National Institute of Health Research and Development | Hana Apsari Pawestri, Hartanti Dian Ikawati, Subangkit, Kartika Dewi Puspa, Arie Ardiansyah Nugraha, Fajar Nur Sulistiyohadi, Triyani Soekarso, Krisna Pangesti, Nelly Puspandari, Vivi Setiawaty |
| EPI_ISL_3216576, EPI_ISL_3216577 | Dinkes Provinsi Kalimantan Timur | National Institute of Health Research and Development | Hana Apsari Pawestri, Hartanti Dian Ikawati, Subangkit, Kartika Dewi Puspa, Arie Ardiansyah Nugraha, Fajar Nur Sulistiyohadi, Triyani Soekarso, Krisna Pangesti, Nelly Puspandari, Vivi Setiawaty |
| EPI_ISL_3216578, EPI_ISL_3216580, EPI_ISL_3216581, EPI_ISL_3216583, EPI_ISL_3216584 | Dinkes Provinsi Kalimantan Timur | National Institute of Health Research and Development | Subangkit, Hana Apsari Pawestri, Hartanti Dian Ikawati, Kartika Dewi Puspa, Arie Ardiansyah Nugraha, Fajar Nur Sulistiyohadi, Triyani Soekarso, Krisna Pangesti, Nelly Puspandari, Vivi Setiawaty |
| EPI_ISL_3216585, EPI_ISL_3216586 | Dinkes Provinsi Kalimantan Timur | National Institute of Health Research and Development | Hartanti Dian Ikawati, Hana Apsari Pawestri, Subangkit, Kartika Dewi Puspa, Arie Ardiansyah Nugraha, Fajar Nur Sulistiyohadi, Triyani Soekarso, Krisna Pangesti, Nelly Puspandari, Vivi Setiawaty |
| EPI_ISL_3216587, EPI_ISL_3216588 | RSUD KRMT Wongsonegoro | National Institute of Health Research and Development | Hartanti Dian Ikawati, Hana Apsari Pawestri, Subangkit, Kartika Dewi Puspa, Arie Ardiansyah Nugraha, Fajar Nur Sulistiyohadi, Triyani Soekarso, Krisna Pangesti, Nelly Puspandari, Vivi Setiawaty |
| EPI_ISL_3216589, EPI_ISL_3216590, EPI_ISL_3216591, EPI_ISL_3216592, EPI_ISL_3216593, EPI_ISL_3216594 | RSUD KRMT Wongsonegoro | National Institute of Health Research and Development | Kartika Dewi Puspa, Hana Apsari Pawestri, Subangkit, Hartanti Dian Ikawati, Arie Ardiansyah Nugraha, Fajar Nur Sulistiyohadi, Triyani Soekarso, Krisna Pangesti, Nelly Puspandari, Vivi Setiawaty |
| EPI_ISL_3216596, EPI_ISL_3216597, EPI_ISL_3216598 | RSUD Tarakan Jakarta | National Institute of Health Research and Development | Kartika Dewi Puspa, Hana Apsari Pawestri, Subangkit, Hartanti Dian Ikawati, Arie Ardiansyah Nugraha, Fajar Nur Sulistiyohadi, Triyani Soekarso, Krisna Pangesti, Nelly Puspandari, Vivi Setiawaty |
| EPI_ISL_3216599, EPI_ISL_3216600, EPI_ISL_3216601, EPI_ISL_3216602, EPI_ISL_3216603, EPI_ISL_3216604, EPI_ISL_3216605, EPI_ISL_3216606, EPI_ISL_3216607, EPI_ISL_3216608, EPI_ISL_3216609, EPI_ISL_3216610, EPI_ISL_3216611, EPI_ISL_3216612, EPI_ISL_3216613, EPI_ISL_3216614, EPI_ISL_3216615, EPI_ISL_3216616, EPI_ISL_3216617, EPI_ISL_3216618, EPI_ISL_3216619, EPI_ISL_3216620, EPI_ISL_3216621, EPI_ISL_3216622, EPI_ISL_3216623, EPI_ISL_3216624, EPI_ISL_3216625, EPI_ISL_3216626, EPI_ISL_3216627, EPI_ISL_3216628, EPI_ISL_3216629, EPI_ISL_3216630 | | | |
| see above | Kansas Health and Environmental Lab | Kansas Health and Environmental Lab | Mike Grose, Jonathan Barnell, Ben Olsen, and Phil Adam |
| EPI_ISL_3216631, EPI_ISL_3216632, EPI_ISL_3216634, EPI_ISL_3216635, EPI_ISL_3216636, EPI_ISL_3216637, EPI_ISL_3216638, EPI_ISL_3216639, EPI_ISL_3216640, EPI_ISL_3216641, EPI_ISL_3216642, EPI_ISL_3216643 | | | |
| see above | Minnesota Department of Health, Public Health Laboratory | Minnesota Department of Health, Public Health Laboratory | Alexandra Lorentz, Jacob Garfin, Matt Plumb, and Xiong Wang |
| EPI_ISL_3216644 | Centracare Laboratory Services | Minnesota Department of Health, Public Health Laboratory | Alexandra Lorentz, Jacob Garfin, Matt Plumb, and Xiong Wang |
| EPI_ISL_3216645 | Minnesota Department of Health, Public Health Laboratory | Minnesota Department of Health, Public Health Laboratory | Alexandra Lorentz, Jacob Garfin, Matt Plumb, and Xiong Wang |
| EPI_ISL_3216646, EPI_ISL_3216647 | Allina Health Laboratory | Minnesota Department of Health, Public Health Laboratory | Alexandra Lorentz, Jacob Garfin, Matt Plumb, and Xiong Wang |
| EPI_ISL_3216648, EPI_ISL_3216649, EPI_ISL_3216650, EPI_ISL_3216651, EPI_ISL_3216652, EPI_ISL_3216653, EPI_ISL_3216655, EPI_ISL_3216656, EPI_ISL_3216659, EPI_ISL_3216660 | Minnesota Department of Health, Public Health Laboratory | Minnesota Department of Health, Public Health Laboratory | Alexandra Lorentz, Jacob Garfin, Matt Plumb, and Xiong Wang |
| EPI_ISL_3216661, EPI_ISL_3216662, EPI_ISL_3216663, EPI_ISL_3216664, EPI_ISL_3216665, EPI_ISL_3216666, EPI_ISL_3216667, EPI_ISL_3216668, EPI_ISL_3216669, EPI_ISL_3216670 | Allina Health Laboratory | Minnesota Department of Health, Public Health Laboratory | Alexandra Lorentz, Jacob Garfin, Matt Plumb, and Xiong Wang |
| EPI_ISL_3216671, EPI_ISL_3216672 | Minnesota Department of Health, Public Health Laboratory | Minnesota Department of Health, Public Health Laboratory | Alexandra Lorentz, Jacob Garfin, Matt Plumb, and Xiong Wang |
| EPI_ISL_3216673, EPI_ISL_3216674, EPI_ISL_3216675, EPI_ISL_3216676, EPI_ISL_3216677, | Allina Health Laboratory | Minnesota Department of Health, Public Health Laboratory | Alexandra Lorentz, Jacob Garfin, Matt Plumb, and Xiong Wang |

| | | | |
|---|---|---|---|
| EPI_ISL_3216678, EPI_ISL_3216679 | | | |
| EPI_ISL_3216680, EPI_ISL_3216681, EPI_ISL_3216682, EPI_ISL_3216683, EPI_ISL_3216684, EPI_ISL_3216685, EPI_ISL_3216686, EPI_ISL_3216687, EPI_ISL_3216688, EPI_ISL_3216689, EPI_ISL_3216690, EPI_ISL_3216691, EPI_ISL_3216692, EPI_ISL_3216693, EPI_ISL_3216694 | | | |
| see above | Minnesota Department of Health, Public Health Laboratory | Minnesota Department of Health, Public Health Laboratory | Alexandra Lorentz, Jacob Garfin, Matt Plumb, and Xiong Wang |
| EPI_ISL_3216695, EPI_ISL_3216696, EPI_ISL_3216697, EPI_ISL_3216698, EPI_ISL_3216699, EPI_ISL_3216700 | Allina Health Laboratory | Minnesota Department of Health, Public Health Laboratory | Alexandra Lorentz, Jacob Garfin, Matt Plumb, and Xiong Wang |
| EPI_ISL_3216701, EPI_ISL_3216702, EPI_ISL_3216703, EPI_ISL_3216704, EPI_ISL_3216705, EPI_ISL_3216706, EPI_ISL_3216708 | Minnesota Department of Health, Public Health Laboratory | Minnesota Department of Health, Public Health Laboratory | Alexandra Lorentz, Jacob Garfin, Matt Plumb, and Xiong Wang |
| EPI_ISL_3216709, EPI_ISL_3216710, EPI_ISL_3216711, EPI_ISL_3216712 | Regions Hospital | Minnesota Department of Health, Public Health Laboratory | Alexandra Lorentz, Jacob Garfin, Matt Plumb, and Xiong Wang |
| EPI_ISL_3216713 | Minnesota Department of Health, Public Health Laboratory | Minnesota Department of Health, Public Health Laboratory | Alexandra Lorentz, Jacob Garfin, Matt Plumb, and Xiong Wang |
| EPI_ISL_3216714, EPI_ISL_3216715, EPI_ISL_3216716, EPI_ISL_3216717, EPI_ISL_3216718 | Centracare Laboratory Services | Minnesota Department of Health, Public Health Laboratory | Alexandra Lorentz, Jacob Garfin, Matt Plumb, and Xiong Wang |
| EPI_ISL_3216729, EPI_ISL_3216730, EPI_ISL_3216731, EPI_ISL_3216732, EPI_ISL_3216733, EPI_ISL_3216734, EPI_ISL_3216735, EPI_ISL_3216736, EPI_ISL_3216738, EPI_ISL_3216739, EPI_ISL_3216740, EPI_ISL_3216741, EPI_ISL_3216742, EPI_ISL_3216743, EPI_ISL_3216744, EPI_ISL_3216745, EPI_ISL_3216746, EPI_ISL_3216747, EPI_ISL_3216748, EPI_ISL_3216749, EPI_ISL_3216750, EPI_ISL_3216751, EPI_ISL_3216752, EPI_ISL_3216753, EPI_ISL_3216754 | | | |
| see above | Arizona State Public Health Laboratory | Arizona State Public Health Laboratory | Trung Huynh, Jessica Escobar, Katherine Fullerton, Nobuko Fukushima, Matthew Contursi, Stacy White, Linda Getsinger, Victor Waddell |
| EPI_ISL_3216755, EPI_ISL_3216756, EPI_ISL_3216757, EPI_ISL_3216758, EPI_ISL_3216759, EPI_ISL_3216760, EPI_ISL_3216761, EPI_ISL_3216762, EPI_ISL_3216763, EPI_ISL_3216764, EPI_ISL_3216765, EPI_ISL_3216766, EPI_ISL_3216767 | | | |
| see above | The Jackson Laboratory | The Jackson Laboratory | Uvalic J, Long J, Renzette N, Adams M, Omerza G, Kelly K, Li L |
| EPI_ISL_3216768, EPI_ISL_3216769, EPI_ISL_3216771, EPI_ISL_3216772, EPI_ISL_3216773, EPI_ISL_3216774, EPI_ISL_3216775, EPI_ISL_3216776, EPI_ISL_3216777, EPI_ISL_3216778, EPI_ISL_3216779 | | | |
| see above | Arizona State Public Health Laboratory | Arizona State Public Health Laboratory | Trung Huynh, Jessica Escobar, Katherine Fullerton, Nobuko Fukushima, Matthew Contursi, Stacy White, Linda Getsinger, Victor Waddell |
| EPI_ISL_3216780, EPI_ISL_3216781, EPI_ISL_3216782, EPI_ISL_3216783, EPI_ISL_3216784, EPI_ISL_3216785, EPI_ISL_3216786, EPI_ISL_3216787, EPI_ISL_3216788, EPI_ISL_3216789, EPI_ISL_3216790, EPI_ISL_3216791, EPI_ISL_3216792, EPI_ISL_3216793, EPI_ISL_3216794, EPI_ISL_3216795, EPI_ISL_3216796, EPI_ISL_3216797, EPI_ISL_3216798, EPI_ISL_3216799, EPI_ISL_3216800, EPI_ISL_3216801, EPI_ISL_3216802, EPI_ISL_3216803, EPI_ISL_3216809, EPI_ISL_3216810, EPI_ISL_3216811, EPI_ISL_3216812, EPI_ISL_3216814 | | | |
| see above | Oregon State Public Health Laboratory | Oregon State Public Health Laboratory | Rafia Razzaque, Eugene Yeboah, Vanda Makris, Laura Tsaknaridis, John Fontana and Shane Sevey |
| EPI_ISL_3216815, EPI_ISL_3216817, EPI_ISL_3216818, EPI_ISL_3216819, EPI_ISL_3216821, EPI_ISL_3216822, EPI_ISL_3216823, EPI_ISL_3216824, EPI_ISL_3216825, EPI_ISL_3216826, EPI_ISL_3216827, EPI_ISL_3216828, EPI_ISL_3216829, EPI_ISL_3216830, EPI_ISL_3216831, EPI_ISL_3216832, EPI_ISL_3216833, EPI_ISL_3216834, EPI_ISL_3216835, EPI_ISL_3216837, EPI_ISL_3216838, EPI_ISL_3216839, EPI_ISL_3216840, EPI_ISL_3216841, EPI_ISL_3216842, EPI_ISL_3216843 | | | |
| see above | State Hygienic Laboratory at the University of Iowa | State Hygienic Laboratory at the University of Iowa | Valerie Reeb, Erik Twait, Anna Yakos, Mariah Knutson, Kris Eveland, Kristen Zanon, Jeff, Benfer, Davis Rieckenberg, Wes Hottel, Alankar Kampoowale |
| EPI_ISL_3216844, EPI_ISL_3216845, EPI_ISL_3216846, EPI_ISL_3216847, EPI_ISL_3216848, EPI_ISL_3216849, EPI_ISL_3216850, EPI_ISL_3216851, EPI_ISL_3216852, EPI_ISL_3216853, EPI_ISL_3216855, EPI_ISL_3216856, EPI_ISL_3216857, EPI_ISL_3216858, EPI_ISL_3216859, EPI_ISL_3216860, EPI_ISL_3216861, EPI_ISL_3216862, EPI_ISL_3216863, EPI_ISL_3216864, EPI_ISL_3216865, EPI_ISL_3216866, EPI_ISL_3216867, EPI_ISL_3216868, EPI_ISL_3216869, EPI_ISL_3216870, EPI_ISL_3216871, EPI_ISL_3216872, EPI_ISL_3216874, EPI_ISL_3216875, EPI_ISL_3216876, EPI_ISL_3216877, EPI_ISL_3216878, EPI_ISL_3216879, EPI_ISL_3216880, EPI_ISL_3216881, EPI_ISL_3216882, EPI_ISL_3216883, EPI_ISL_3216884, EPI_ISL_3216885, EPI_ISL_3216886, EPI_ISL_3216887, EPI_ISL_3216888, EPI_ISL_3216889, EPI_ISL_3216890, EPI_ISL_3216891, EPI_ISL_3216892, EPI_ISL_3216893, EPI_ISL_3216894, EPI_ISL_3216895 | | | |
| see above | South Dakota Department Of Health Laboratory | Minnesota Department of Health, Public Health Laboratory | Alexandra Lorentz, Jacob Garfin, Matt Plumb, and Xiong Wang |
| EPI_ISL_3216896, EPI_ISL_3216897, EPI_ISL_3216898, EPI_ISL_3216899, EPI_ISL_3216900, EPI_ISL_3216901, EPI_ISL_3216902 | UCSC Genomics Institute | UCSC Genomics Institute | Mark Akeson, Ikenna Anigbogu, Eric Beraut, Isabel Bjork, Molly Cassatt-Johnstone, Terren Chang, Russell Corbett-Detig, Namrita Dhillon, Maximilian Haeussler, David Haussler, Angie Hinrichs, Miten Jain, Joshua Kapp, A. Marm Kilpatrick, Jakob McBroome, Hugh Olsen, Jeremy Sanford, Beth Shapiro, Michael Stone, Bryan Thornlow, Yatish Turakhia, Ciara Wanket |
| EPI_ISL_3216912, EPI_ISL_3216913, EPI_ISL_3216914, EPI_ISL_3216915, EPI_ISL_3216916, EPI_ISL_3216917, EPI_ISL_3216918, EPI_ISL_3216919, EPI_ISL_3216920, EPI_ISL_3216921, EPI_ISL_3216922, EPI_ISL_3216923, EPI_ISL_3216924, EPI_ISL_3216925, EPI_ISL_3216926, EPI_ISL_3216927, EPI_ISL_3216928, EPI_ISL_3216929, EPI_ISL_3216930, EPI_ISL_3216931, EPI_ISL_3216932, EPI_ISL_3216933, EPI_ISL_3216934, EPI_ISL_3216935, EPI_ISL_3216936, EPI_ISL_3216937, EPI_ISL_3216938, EPI_ISL_3216939, EPI_ISL_3216940, EPI_ISL_3216941, EPI_ISL_3216942, EPI_ISL_3216943 | | | |
| see above | Kansas Health and Environmental Lab | Kansas Health and Environmental Lab | Mike Grose, Jonathan Barnell, Ben Olsen, and Phil Adam |
| EPI_ISL_3216947, EPI_ISL_3216952, EPI_ISL_3216953, EPI_ISL_3216954, EPI_ISL_3216955, EPI_ISL_3216956, EPI_ISL_3216957, EPI_ISL_3216958, EPI_ISL_3216959, EPI_ISL_3216960, EPI_ISL_3216961, EPI_ISL_3216962, EPI_ISL_3216963, EPI_ISL_3216964, EPI_ISL_3216965, EPI_ISL_3216966, EPI_ISL_3216967, EPI_ISL_3216968, EPI_ISL_3216970, EPI_ISL_3216971, EPI_ISL_3216975, EPI_ISL_3216977, EPI_ISL_3216978, EPI_ISL_3216980, EPI_ISL_3216981, EPI_ISL_3216982, EPI_ISL_3216983, EPI_ISL_3216984, EPI_ISL_3216985, EPI_ISL_3216986, EPI_ISL_3216988, EPI_ISL_3216989, EPI_ISL_3216991, EPI_ISL_3216992, EPI_ISL_3216993, EPI_ISL_3216994, EPI_ISL_3216995, EPI_ISL_3216997, EPI_ISL_3216998, EPI_ISL_3216999, EPI_ISL_3217000, EPI_ISL_3217001, EPI_ISL_3217003, EPI_ISL_3217004, EPI_ISL_3217005, EPI_ISL_3217006, EPI_ISL_3217007, EPI_ISL_3217008, EPI_ISL_3217009, EPI_ISL_3217010, EPI_ISL_3217011, EPI_ISL_3217012, EPI_ISL_3217013, EPI_ISL_3217014, EPI_ISL_3217016, EPI_ISL_3217017, EPI_ISL_3217018, EPI_ISL_3217019, EPI_ISL_3217020, EPI_ISL_3217021, EPI_ISL_3217022, EPI_ISL_3217023, EPI_ISL_3217024, EPI_ISL_3217025, EPI_ISL_3217026, EPI_ISL_3217027, EPI_ISL_3217028, EPI_ISL_3217029, EPI_ISL_3217030, EPI_ISL_3217031, EPI_ISL_3217032, EPI_ISL_3217033, EPI_ISL_3217034, EPI_ISL_3217035, EPI_ISL_3217037, EPI_ISL_3217038, EPI_ISL_3217039, EPI_ISL_3217040, EPI_ISL_3217041, EPI_ISL_3217042, EPI_ISL_3217043, EPI_ISL_3217044, EPI_ISL_3217045, EPI_ISL_3217047, EPI_ISL_3217048, EPI_ISL_3217049, EPI_ISL_3217050, EPI_ISL_3217051, EPI_ISL_3217052, EPI_ISL_3217053, EPI_ISL_3217054, EPI_ISL_3217055, EPI_ISL_3217056, EPI_ISL_3217057, EPI_ISL_3217058, EPI_ISL_3217060, EPI_ISL_3217061, EPI_ISL_3217062, EPI_ISL_3217063, EPI_ISL_3217064, EPI_ISL_3217065, EPI_ISL_3217066, EPI_ISL_3217067, EPI_ISL_3217068, EPI_ISL_3217069, EPI_ISL_3217070, EPI_ISL_3217071 | | | |
| see above | Washington State Department of Health Public Health Laboratories | Washington State Department of Health Public Health Laboratories | Drew MacKellar, Philip Dykema, Denny Russell, Joenice Gonzalez, Hannah Gray, Geoff Melly, Vanessa De Los Santos, Darren Lucas, JohnAric Peterson, Avi Singh, Rebecca Cao |
| EPI_ISL_3217072, EPI_ISL_3217073, EPI_ISL_3217074, EPI_ISL_3217075, EPI_ISL_3217077, EPI_ISL_3217078, EPI_ISL_3217079, EPI_ISL_3217080, EPI_ISL_3217081, EPI_ISL_3217082, EPI_ISL_3217083, EPI_ISL_3217084, EPI_ISL_3217086 | | | |
| see above | Tulare County Public Health Laboratory | Tulare County Public Health Laboratory | Denise L Lopez, Joshua Kootstra |
| EPI_ISL_3217097, EPI_ISL_3217098, EPI_ISL_3217099, EPI_ISL_3217100, EPI_ISL_3217101, EPI_ISL_3217102, EPI_ISL_3217103, EPI_ISL_3217105, EPI_ISL_3217107, EPI_ISL_3217108, EPI_ISL_3217109, EPI_ISL_3217110, EPI_ISL_3217111, EPI_ISL_3217113, EPI_ISL_3217114, EPI_ISL_3217115, EPI_ISL_3217116, EPI_ISL_3217117, EPI_ISL_3217118, EPI_ISL_3217119, EPI_ISL_3217120, EPI_ISL_3217121, EPI_ISL_3217122, EPI_ISL_3217123, EPI_ISL_3217124, EPI_ISL_3217125, EPI_ISL_3217126, EPI_ISL_3217127, EPI_ISL_3217128, EPI_ISL_3217129, EPI_ISL_3217131, EPI_ISL_3217132, EPI_ISL_3217133, EPI_ISL_3217134, EPI_ISL_3217135, EPI_ISL_3217136, EPI_ISL_3217137, EPI_ISL_3217138, EPI_ISL_3217139, EPI_ISL_3217140, EPI_ISL_3217141, EPI_ISL_3217142, EPI_ISL_3217145, EPI_ISL_3217146, EPI_ISL_3217147, EPI_ISL_3217149, EPI_ISL_3217150, EPI_ISL_3217151, EPI_ISL_3217152, EPI_ISL_3217153, EPI_ISL_3217154, EPI_ISL_3217155, EPI_ISL_3217156, EPI_ISL_3217157, EPI_ISL_3217158, EPI_ISL_3217159, EPI_ISL_3217160, EPI_ISL_3217161, EPI_ISL_3217162, EPI_ISL_3217163, EPI_ISL_3217164, EPI_ISL_3217165, EPI_ISL_3217166, EPI_ISL_3217167, EPI_ISL_3217168, EPI_ISL_3217169, EPI_ISL_3217170, EPI_ISL_3217171 | | | |
| see above | Arizona State Public Health Laboratory | Arizona State Public Health Laboratory | Trung Huynh, Jessica Escobar, Katherine Fullerton, Nobuko Fukushima, Matthew Contursi, Stacy White, Linda Getsinger, Victor Waddell |
| EPI_ISL_3217175, EPI_ISL_3217176 | South Dakota Department Of Health Laboratory | Minnesota Department of Health, Public Health Laboratory | Alexandra Lorentz, Jacob Garfin, Matt Plumb, and Xiong Wang |
| EPI_ISL_3217270, EPI_ISL_3217271, EPI_ISL_3217272, EPI_ISL_3217273, EPI_ISL_3217274, EPI_ISL_3217275, EPI_ISL_3217276, EPI_ISL_3217277, EPI_ISL_3217278, EPI_ISL_3217279, EPI_ISL_3217280, EPI_ISL_3217283, EPI_ISL_3217284, EPI_ISL_3217288, EPI_ISL_3217289, EPI_ISL_3217290, EPI_ISL_3217291, EPI_ISL_3217292, EPI_ISL_3217293, EPI_ISL_3217295, EPI_ISL_3217296, EPI_ISL_3217297, EPI_ISL_3217298, EPI_ISL_3217300, EPI_ISL_3217301, EPI_ISL_3217302, EPI_ISL_3217303, EPI_ISL_3217304, EPI_ISL_3217305, EPI_ISL_3217306, EPI_ISL_3217307, EPI_ISL_3217308, EPI_ISL_3217309, EPI_ISL_3217310, EPI_ISL_3217311, EPI_ISL_3217312, EPI_ISL_3217313, EPI_ISL_3217314, EPI_ISL_3217315, EPI_ISL_3217316, EPI_ISL_3217318, EPI_ISL_3217319, EPI_ISL_3217320, EPI_ISL_3217322, EPI_ISL_3217323, EPI_ISL_3217325, EPI_ISL_3217326, EPI_ISL_3217328, EPI_ISL_3217329, EPI_ISL_3217330, EPI_ISL_3217331, EPI_ISL_3217332, EPI_ISL_3217333, EPI_ISL_3217334, EPI_ISL_3217335, EPI_ISL_3217338, EPI_ISL_3217339, EPI_ISL_3217340, EPI_ISL_3217341, EPI_ISL_3217343, EPI_ISL_3217344, EPI_ISL_3217345, EPI_ISL_3217347, EPI_ISL_3217348, EPI_ISL_3217349, EPI_ISL_3217350, EPI_ISL_3217351, EPI_ISL_3217352, EPI_ISL_3217353, EPI_ISL_3217354, EPI_ISL_3217356, EPI_ISL_3217357, EPI_ISL_3217358, EPI_ISL_3217359, EPI_ISL_3217360, EPI_ISL_3217361, EPI_ISL_3217362, EPI_ISL_3217363, EPI_ISL_3217364, EPI_ISL_3217366, EPI_ISL_3217367, EPI_ISL_3217368, EPI_ISL_3217369, EPI_ISL_3217370, EPI_ISL_3217371, EPI_ISL_3217372, EPI_ISL_3217373, EPI_ISL_3217374, EPI_ISL_3217375, EPI_ISL_3217376, EPI_ISL_3217377, EPI_ISL_3217378, EPI_ISL_3217379, EPI_ISL_3217380, EPI_ISL_3217381, EPI_ISL_3217384, EPI_ISL_3217385, EPI_ISL_3217386, EPI_ISL_3217387, EPI_ISL_3217388, EPI_ISL_3217389, EPI_ISL_3217390, EPI_ISL_3217392, EPI_ISL_3217394, EPI_ISL_3217396, EPI_ISL_3217397, EPI_ISL_3217398, EPI_ISL_3217399, EPI_ISL_3217400, EPI_ISL_3217401, EPI_ISL_3217402, EPI_ISL_3217403, EPI_ISL_3217404, EPI_ISL_3217405, EPI_ISL_3217407 | | | |
| see above | Aegis Sciences Corporation | Centers for Disease Control and Prevention Division of Viral Diseases, Pathogen Discovery | Dakota Howard, Dhwani Batra, Peter W. Cook, Kara Moser, Adrian Paskey, Jason Caravas, Benjamin Rambo-Martin, Shatavia Morrison, Christopher Gulvick, Scott Sammons, Yvette Unoarumhi, Darlene Wagner, Matthew Schmerer, Cyndi Clark, Patrick Campbell, Rob Case, Vikramsinha Ghorpade, Holly Houdeshell, Ola Kvalvaag, Dillon Nall, Ethan Sanders, Alec Vest, Shaun Westlund, Matthew Hardison, Clinton R. Paden, Duncan MacCannell |
| EPI_ISL_3217408, EPI_ISL_3217410 | Airport Health Laboratory/Central Health Laboratory | Virology Department, Central Health Laboratory ,Victoria Hospital, Candos,Ministry of Health and Wellness, Mauritius | Manraj SS, Sonoo J, Pattoo M, Bahadoor BS, Mathur H, Sujeewon C, Jannoo N, Ramuth M |
| EPI_ISL_3217411 | Aegis Sciences Corporation | Centers for Disease Control and Prevention Division of Viral Diseases, Pathogen Discovery | Dakota Howard, Dhwani Batra, Peter W. Cook, Kara Moser, Adrian Paskey, Jason Caravas, Benjamin Rambo-Martin, Shatavia Morrison, Christopher Gulvick, Scott Sammons, Yvette Unoarumhi, Darlene Wagner, Matthew Schmerer, Cyndi Clark, Patrick Campbell, Rob Case, Vikramsinha Ghorpade, Holly Houdeshell, Ola Kvalvaag, Dillon Nall, Ethan Sanders, Alec Vest, Shaun Westlund, Matthew Hardison, Clinton R. Paden, Duncan MacCannell |
| EPI_ISL_3217412 | Airport Health Laboratory/Central Health Laboratory | Virology Department, Central Health Laboratory ,Victoria Hospital, Candos,Ministry of Health and Wellness, Mauritius | Manraj SS, Sonoo J, Pattoo M, Bahadoor BS, Mathur H, Sujeewon C, Jannoo N, Ramuth M |
| EPI_ISL_3217413 | Aegis Sciences Corporation | Centers for Disease Control and Prevention Division of Viral Diseases, Pathogen Discovery | Dakota Howard, Dhwani Batra, Peter W. Cook, Kara Moser, Adrian Paskey, Jason Caravas, Benjamin Rambo-Martin, Shatavia Morrison, Christopher Gulvick, Scott Sammons, Yvette Unoarumhi, Darlene Wagner, Matthew Schmerer, Cyndi Clark, Patrick Campbell, Rob Case, Vikramsinha Ghorpade, Holly Houdeshell, Ola Kvalvaag, Dillon Nall, Ethan Sanders, Alec Vest, Shaun Westlund, Matthew Hardison, Clinton R. Paden, Duncan MacCannell |
| EPI_ISL_3217414, EPI_ISL_3217415 | Airport Health Laboratory/Central Health Laboratory | Virology Department, Central Health Laboratory ,Victoria Hospital, Candos,Ministry of Health and Wellness, Mauritius | Manraj SS, Sonoo J, Pattoo M, Bahadoor BS, Mathur H, Sujeewon C, Jannoo N, Ramuth M |
| EPI_ISL_3217416 | Aegis Sciences Corporation | Centers for Disease Control and Prevention Division of Viral Diseases, Pathogen Discovery | Dakota Howard, Dhwani Batra, Peter W. Cook, Kara Moser, Adrian Paskey, Jason Caravas, Benjamin Rambo-Martin, Shatavia Morrison, Christopher Gulvick, Scott Sammons, Yvette Unoarumhi, Darlene Wagner, Matthew Schmerer, Cyndi Clark, Patrick Campbell, Rob Case, Vikramsinha Ghorpade, Holly Houdeshell, Ola Kvalvaag, Dillon Nall, Ethan Sanders, Alec Vest, Shaun Westlund, Matthew Hardison, Clinton R. Paden, Duncan MacCannell |
| EPI_ISL_3217417, EPI_ISL_3217418, EPI_ISL_3217420 | Airport Health Laboratory/Central Health Laboratory | Virology Department, Central Health Laboratory ,Victoria Hospital, Candos,Ministry of Health and Wellness, Mauritius | Manraj SS, Sonoo J, Pattoo M, Bahadoor BS, Mathur H, Sujeewon C, Jannoo N, Ramuth M |

| | | | |
|---|---|---|---|
| EPI_ISL_3217421 | Aegis Sciences Corporation | Centers for Disease Control and Prevention Division of Viral Diseases, Pathogen Discovery | Dakota Howard, Dhwani Batra, Peter W. Cook, Kara Moser, Adrian Paskey, Jason Caravas, Benjamin Rambo-Martin, Shatavia Morrison, Christopher Gulvick, Scott Sammons, Yvette Unoarumhi, Darlene Wagner, Matthew Schmerer, Cyndi Clark, Patrick Campbell, Rob Case, Vikramsinha Ghorpade, Holly Houdeshell, Ola Kvalvaag, Dillon Nall, Ethan Sanders, Alec Vest, Shaun Westlund, Matthew Hardison, Clinton R. Paden, Duncan MacCannell |
| EPI_ISL_3217422 | Airport Health Laboratory/Central Health Laboratory | Virology Department, Central Health Laboratory ,Victoria Hospital, Candos,Ministry of Health and Wellness, Mauritius | Manraj SS, Sonoo J, Pattoo M, Bahadoor BS, Mathur H, Sujeewon C, Jannoo N, Ramuth M |
| EPI_ISL_3217423 | Aegis Sciences Corporation | Centers for Disease Control and Prevention Division of Viral Diseases, Pathogen Discovery | Dakota Howard, Dhwani Batra, Peter W. Cook, Kara Moser, Adrian Paskey, Jason Caravas, Benjamin Rambo-Martin, Shatavia Morrison, Christopher Gulvick, Scott Sammons, Yvette Unoarumhi, Darlene Wagner, Matthew Schmerer, Cyndi Clark, Patrick Campbell, Rob Case, Vikramsinha Ghorpade, Holly Houdeshell, Ola Kvalvaag, Dillon Nall, Ethan Sanders, Alec Vest, Shaun Westlund, Matthew Hardison, Clinton R. Paden, Duncan MacCannell |
| EPI_ISL_3217424 | Airport Health Laboratory/Central Health Laboratory | Virology Department, Central Health Laboratory ,Victoria Hospital, Candos,Ministry of Health and Wellness, Mauritius | Manraj SS, Sonoo J, Pattoo M, Bahadoor BS, Mathur H, Sujeewon C, Jannoo N, Ramuth M |
| EPI_ISL_3217426, EPI_ISL_3217428, EPI_ISL_3217430, EPI_ISL_3217433, EPI_ISL_3217438, EPI_ISL_3217441, EPI_ISL_3217443, EPI_ISL_3217446, EPI_ISL_3217448, EPI_ISL_3217449, EPI_ISL_3217450, EPI_ISL_3217451, EPI_ISL_3217452, EPI_ISL_3217453, EPI_ISL_3217454, EPI_ISL_3217457, EPI_ISL_3217458, EPI_ISL_3217459, EPI_ISL_3217460, EPI_ISL_3217461, EPI_ISL_3217462, EPI_ISL_3217465, EPI_ISL_3217466, EPI_ISL_3217468, EPI_ISL_3217470, EPI_ISL_3217471, EPI_ISL_3217472, EPI_ISL_3217473, EPI_ISL_3217475, EPI_ISL_3217476, EPI_ISL_3217477, EPI_ISL_3217478, EPI_ISL_3217479, EPI_ISL_3217480, EPI_ISL_3217481, EPI_ISL_3217482, EPI_ISL_3217483, EPI_ISL_3217484, EPI_ISL_3217485, EPI_ISL_3217486, EPI_ISL_3217487, EPI_ISL_3217489, EPI_ISL_3217492, EPI_ISL_3217493, EPI_ISL_3217494, EPI_ISL_3217496, EPI_ISL_3217497, EPI_ISL_3217498, EPI_ISL_3217500, EPI_ISL_3217501, EPI_ISL_3217502, EPI_ISL_3217504, EPI_ISL_3217505, EPI_ISL_3217506, EPI_ISL_3217507, EPI_ISL_3217508, EPI_ISL_3217509, EPI_ISL_3217511, EPI_ISL_3217513, EPI_ISL_3217514, EPI_ISL_3217515, EPI_ISL_3217516, EPI_ISL_3217517, EPI_ISL_3217518, EPI_ISL_3217519, EPI_ISL_3217521, EPI_ISL_3217523, EPI_ISL_3217524, EPI_ISL_3217525, EPI_ISL_3217526, EPI_ISL_3217527, EPI_ISL_3217529, EPI_ISL_3217531, EPI_ISL_3217532, EPI_ISL_3217533, EPI_ISL_3217534, EPI_ISL_3217536, EPI_ISL_3217537, EPI_ISL_3217538, EPI_ISL_3217540, EPI_ISL_3217541, EPI_ISL_3217543, EPI_ISL_3217544, EPI_ISL_3217545, EPI_ISL_3217546, EPI_ISL_3217547, EPI_ISL_3217549, EPI_ISL_3217550, EPI_ISL_3217551, EPI_ISL_3217552, EPI_ISL_3217553, EPI_ISL_3217554, EPI_ISL_3217555, EPI_ISL_3217556, EPI_ISL_3217557, EPI_ISL_3217558, EPI_ISL_3217559, EPI_ISL_3217560, EPI_ISL_3217561, EPI_ISL_3217562, EPI_ISL_3217563, EPI_ISL_3217564, EPI_ISL_3217565, EPI_ISL_3217566, EPI_ISL_3217567, EPI_ISL_3217568, EPI_ISL_3217569, EPI_ISL_3217570, EPI_ISL_3217571, EPI_ISL_3217573, EPI_ISL_3217574, EPI_ISL_3217575, EPI_ISL_3217577, EPI_ISL_3217578, EPI_ISL_3217579, EPI_ISL_3217580, EPI_ISL_3217582, EPI_ISL_3217583, EPI_ISL_3217584, EPI_ISL_3217585, EPI_ISL_3217586, EPI_ISL_3217587, EPI_ISL_3217588, EPI_ISL_3217589, EPI_ISL_3217590, EPI_ISL_3217592, EPI_ISL_3217596, EPI_ISL_3217597, EPI_ISL_3217598, EPI_ISL_3217600, EPI_ISL_3217601, EPI_ISL_3217602, EPI_ISL_3217603, EPI_ISL_3217604, EPI_ISL_3217605, EPI_ISL_3217606, EPI_ISL_3217608, EPI_ISL_3217610, EPI_ISL_3217611, EPI_ISL_3217612, EPI_ISL_3217616, EPI_ISL_3217617, EPI_ISL_3217618, EPI_ISL_3217620, EPI_ISL_3217621, EPI_ISL_3217622, EPI_ISL_3217623, EPI_ISL_3217624, EPI_ISL_3217625, EPI_ISL_3217627, EPI_ISL_3217628, EPI_ISL_3217629, EPI_ISL_3217630, EPI_ISL_3217632, EPI_ISL_3217633, EPI_ISL_3217634, EPI_ISL_3217636, EPI_ISL_3217640, EPI_ISL_3217641, EPI_ISL_3217644, EPI_ISL_3217645, EPI_ISL_3217647, EPI_ISL_3217648, EPI_ISL_3217649, EPI_ISL_3217650, EPI_ISL_3217651, EPI_ISL_3217652, EPI_ISL_3217653, EPI_ISL_3217654, EPI_ISL_3217655, EPI_ISL_3217657, EPI_ISL_3217658, EPI_ISL_3217660, EPI_ISL_3217661, EPI_ISL_3217662, EPI_ISL_3217665, EPI_ISL_3217666, EPI_ISL_3217667, EPI_ISL_3217668, EPI_ISL_3217669, EPI_ISL_3217670, EPI_ISL_3217671, EPI_ISL_3217672, EPI_ISL_3217673, EPI_ISL_3217677, EPI_ISL_3217680, EPI_ISL_3217681, EPI_ISL_3217683, EPI_ISL_3217685, EPI_ISL_3217686, EPI_ISL_3217687, EPI_ISL_3217688, EPI_ISL_3217689, EPI_ISL_3217691, EPI_ISL_3217692, EPI_ISL_3217693, EPI_ISL_3217695, EPI_ISL_3217696, EPI_ISL_3217698, EPI_ISL_3217699, EPI_ISL_3217700, EPI_ISL_3217701, EPI_ISL_3217702, EPI_ISL_3217703, EPI_ISL_3217704, EPI_ISL_3217706, EPI_ISL_3217707, EPI_ISL_3217708, EPI_ISL_3217709, EPI_ISL_3217710, EPI_ISL_3217711, EPI_ISL_3217712, EPI_ISL_3217713, EPI_ISL_3217714, EPI_ISL_3217716, EPI_ISL_3217718, EPI_ISL_3217719, EPI_ISL_3217720, EPI_ISL_3217721, EPI_ISL_3217722, EPI_ISL_3217724, EPI_ISL_3217725, EPI_ISL_3217726, EPI_ISL_3217727, EPI_ISL_3217728, EPI_ISL_3217729, EPI_ISL_3217730, EPI_ISL_3217733, EPI_ISL_3217734, EPI_ISL_3217735, EPI_ISL_3217736, EPI_ISL_3217737, EPI_ISL_3217738, EPI_ISL_3217739, EPI_ISL_3217740, EPI_ISL_3217744, EPI_ISL_3217745, EPI_ISL_3217746, EPI_ISL_3217747, EPI_ISL_3217750, EPI_ISL_3217751, EPI_ISL_3217752, EPI_ISL_3217753, EPI_ISL_3217755, EPI_ISL_3217756, EPI_ISL_3217758, EPI_ISL_3217759, EPI_ISL_3217761, EPI_ISL_3217762, EPI_ISL_3217763, EPI_ISL_3217764, EPI_ISL_3217765, EPI_ISL_3217766, EPI_ISL_3217768, EPI_ISL_3217769, EPI_ISL_3217770, EPI_ISL_3217771, EPI_ISL_3217773, EPI_ISL_3217775, EPI_ISL_3217776, EPI_ISL_3217778, EPI_ISL_3217779, EPI_ISL_3217780, EPI_ISL_3217782, EPI_ISL_3217783, EPI_ISL_3217784, EPI_ISL_3217785, EPI_ISL_3217787, EPI_ISL_3217788, EPI_ISL_3217789, EPI_ISL_3217790, EPI_ISL_3217791, EPI_ISL_3217793, EPI_ISL_3217794, EPI_ISL_3217795, EPI_ISL_3217796, EPI_ISL_3217797, EPI_ISL_3217798, EPI_ISL_3217799, EPI_ISL_3217800, EPI_ISL_3217801, EPI_ISL_3217802, EPI_ISL_3217805, EPI_ISL_3217806, EPI_ISL_3217807, EPI_ISL_3217808, EPI_ISL_3217809, EPI_ISL_3217810, EPI_ISL_3217811, EPI_ISL_3217812, EPI_ISL_3217813, EPI_ISL_3217815, EPI_ISL_3217816, EPI_ISL_3217817, EPI_ISL_3217818, EPI_ISL_3217819, EPI_ISL_3217820, EPI_ISL_3217821, EPI_ISL_3217823, EPI_ISL_3217824, EPI_ISL_3217828, EPI_ISL_3217830, EPI_ISL_3217833, EPI_ISL_3217834, EPI_ISL_3217835, EPI_ISL_3217836, EPI_ISL_3217838, EPI_ISL_3217839, EPI_ISL_3217840, EPI_ISL_3217841, EPI_ISL_3217842, EPI_ISL_3217843, EPI_ISL_3217845, EPI_ISL_3217846, EPI_ISL_3217847, EPI_ISL_3217848, EPI_ISL_3217849, EPI_ISL_3217850, EPI_ISL_3217851, EPI_ISL_3217852, EPI_ISL_3217853, EPI_ISL_3217855, EPI_ISL_3217856, EPI_ISL_3217857, EPI_ISL_3217858, EPI_ISL_3217860, EPI_ISL_3217861, EPI_ISL_3217863, EPI_ISL_3217864, EPI_ISL_3217865, EPI_ISL_3217866, EPI_ISL_3217869, EPI_ISL_3217870, EPI_ISL_3217871, EPI_ISL_3217872, EPI_ISL_3217873, EPI_ISL_3217874, EPI_ISL_3217876, EPI_ISL_3217877, EPI_ISL_3217878, EPI_ISL_3217879, EPI_ISL_3217880, EPI_ISL_3217881, EPI_ISL_3217882, EPI_ISL_3217883, EPI_ISL_3217884, EPI_ISL_3217885, EPI_ISL_3217886, EPI_ISL_3217889, EPI_ISL_3217890, EPI_ISL_3217891, EPI_ISL_3217893, EPI_ISL_3217894, EPI_ISL_3217895, EPI_ISL_3217896, EPI_ISL_3217897, EPI_ISL_3217898, EPI_ISL_3217899, EPI_ISL_3217900, EPI_ISL_3217901, EPI_ISL_3217902, EPI_ISL_3217904, EPI_ISL_3217905, EPI_ISL_3217906, EPI_ISL_3217907, EPI_ISL_3217908, EPI_ISL_3217909, EPI_ISL_3217910, EPI_ISL_3217911, EPI_ISL_3217913, EPI_ISL_3217914, EPI_ISL_3217915, EPI_ISL_3217916, EPI_ISL_3217917, EPI_ISL_3217918, EPI_ISL_3217919, EPI_ISL_3217920, EPI_ISL_3217921, EPI_ISL_3217922, EPI_ISL_3217923, EPI_ISL_3217924, EPI_ISL_3217925, EPI_ISL_3217926, EPI_ISL_3217927, EPI_ISL_3217928, EPI_ISL_3217929, EPI_ISL_3217931, EPI_ISL_3217932, EPI_ISL_3217933, EPI_ISL_3217934, EPI_ISL_3217935, EPI_ISL_3217936, EPI_ISL_3217937, EPI_ISL_3217938, EPI_ISL_3217939, EPI_ISL_3217940, EPI_ISL_3217941, EPI_ISL_3217942, EPI_ISL_3217943, EPI_ISL_3217945, EPI_ISL_3217946, EPI_ISL_3217947, EPI_ISL_3217948, EPI_ISL_3217949, EPI_ISL_3217950, EPI_ISL_3217952, EPI_ISL_3217953, EPI_ISL_3217954, EPI_ISL_3217955, EPI_ISL_3217957, EPI_ISL_3217959, EPI_ISL_3217960, EPI_ISL_3217961, EPI_ISL_3217962, EPI_ISL_3217963, EPI_ISL_3217964, EPI_ISL_3217968, EPI_ISL_3217971, EPI_ISL_3217973, EPI_ISL_3217974, EPI_ISL_3217975, EPI_ISL_3217976, EPI_ISL_3217977, EPI_ISL_3217978, EPI_ISL_3217979, EPI_ISL_3217980, EPI_ISL_3217981, EPI_ISL_3217982, EPI_ISL_3217983, EPI_ISL_3217984, EPI_ISL_3217985, EPI_ISL_3217986, EPI_ISL_3217988, EPI_ISL_3217989, EPI_ISL_3217990, EPI_ISL_3217991, EPI_ISL_3217992, EPI_ISL_3217993, EPI_ISL_3217994, EPI_ISL_3217995, EPI_ISL_3217996, EPI_ISL_3217997, EPI_ISL_3217998, EPI_ISL_3218000, EPI_ISL_3218001, EPI_ISL_3218002, EPI_ISL_3218003, EPI_ISL_3218004, EPI_ISL_3218005, EPI_ISL_3218006, EPI_ISL_3218008, EPI_ISL_3218010, EPI_ISL_3218011, EPI_ISL_3218013, EPI_ISL_3218014, EPI_ISL_3218015, EPI_ISL_3218016, EPI_ISL_3218017, EPI_ISL_3218018, EPI_ISL_3218019, EPI_ISL_3218020, EPI_ISL_3218021, EPI_ISL_3218022, EPI_ISL_3218023, EPI_ISL_3218024, EPI_ISL_3218025, EPI_ISL_3218026, EPI_ISL_3218028, EPI_ISL_3218029, EPI_ISL_3218030, EPI_ISL_3218031, EPI_ISL_3218032, EPI_ISL_3218033, EPI_ISL_3218035, EPI_ISL_3218036, EPI_ISL_3218037, EPI_ISL_3218038, EPI_ISL_3218039, EPI_ISL_3218041, EPI_ISL_3218042, EPI_ISL_3218044, EPI_ISL_3218045, EPI_ISL_3218047, EPI_ISL_3218048, EPI_ISL_3218050, EPI_ISL_3218051, EPI_ISL_3218052, EPI_ISL_3218054, EPI_ISL_3218055, EPI_ISL_3218057, EPI_ISL_3218058, EPI_ISL_3218059, EPI_ISL_3218060, EPI_ISL_3218061, EPI_ISL_3218062, EPI_ISL_3218064, EPI_ISL_3218067, EPI_ISL_3218069, EPI_ISL_3218070, EPI_ISL_3218071, EPI_ISL_3218072, EPI_ISL_3218073, EPI_ISL_3218074, EPI_ISL_3218075, EPI_ISL_3218076, EPI_ISL_3218077, EPI_ISL_3218078, EPI_ISL_3218079, EPI_ISL_3218080, EPI_ISL_3218081, EPI_ISL_3218082, EPI_ISL_3218085, EPI_ISL_3218086, EPI_ISL_3218087, EPI_ISL_3218088, EPI_ISL_3218089, EPI_ISL_3218090, EPI_ISL_3218091, EPI_ISL_3218092, EPI_ISL_3218093, EPI_ISL_3218094, EPI_ISL_3218096, EPI_ISL_3218097, EPI_ISL_3218098, EPI_ISL_3218099, EPI_ISL_3218100, EPI_ISL_3218102, EPI_ISL_3218103, EPI_ISL_3218104, EPI_ISL_3218105, EPI_ISL_3218106, EPI_ISL_3218107, EPI_ISL_3218108, EPI_ISL_3218109, EPI_ISL_3218112, EPI_ISL_3218113, EPI_ISL_3218115, EPI_ISL_3218118, EPI_ISL_3218120, EPI_ISL_3218122, EPI_ISL_3218123, EPI_ISL_3218128, EPI_ISL_3218129, EPI_ISL_3218132, EPI_ISL_3218133, EPI_ISL_3218134, EPI_ISL_3218135, EPI_ISL_3218136, EPI_ISL_3218137, EPI_ISL_3218138, EPI_ISL_3218140, EPI_ISL_3218141, EPI_ISL_3218142, EPI_ISL_3218143, EPI_ISL_3218144, EPI_ISL_3218146, EPI_ISL_3218147, EPI_ISL_3218150, EPI_ISL_3218151, EPI_ISL_3218152, EPI_ISL_3218153, EPI_ISL_3218154, EPI_ISL_3218155, EPI_ISL_3218156, EPI_ISL_3218157, EPI_ISL_3218158, EPI_ISL_3218159, EPI_ISL_3218160, EPI_ISL_3218161, EPI_ISL_3218162, EPI_ISL_3218163, EPI_ISL_3218164, EPI_ISL_3218165, EPI_ISL_3218166, EPI_ISL_3218167, EPI_ISL_3218168, EPI_ISL_3218169, EPI_ISL_3218171, EPI_ISL_3218172, EPI_ISL_3218173, EPI_ISL_3218174, EPI_ISL_3218175, EPI_ISL_3218176, EPI_ISL_3218177, EPI_ISL_3218178, EPI_ISL_3218180, EPI_ISL_3218181, EPI_ISL_3218182, EPI_ISL_3218183, EPI_ISL_3218184, EPI_ISL_3218186, EPI_ISL_3218187, EPI_ISL_3218188, EPI_ISL_3218189, EPI_ISL_3218190, EPI_ISL_3218191, EPI_ISL_3218192, EPI_ISL_3218193, EPI_ISL_3218194, EPI_ISL_3218195, EPI_ISL_3218196, EPI_ISL_3218198, EPI_ISL_3218199, EPI_ISL_3218201, EPI_ISL_3218203, EPI_ISL_3218204, EPI_ISL_3218205, EPI_ISL_3218206, EPI_ISL_3218207, EPI_ISL_3218209, EPI_ISL_3218211, EPI_ISL_3218214, EPI_ISL_3218218, EPI_ISL_3218223, EPI_ISL_3218225, EPI_ISL_3218227, EPI_ISL_3218230, EPI_ISL_3218234, EPI_ISL_3218236, EPI_ISL_3218248, EPI_ISL_3218252, EPI_ISL_3218255, EPI_ISL_3218257, EPI_ISL_3218262, EPI_ISL_3218267, EPI_ISL_3218272, EPI_ISL_3218274, EPI_ISL_3218276, EPI_ISL_3218279, EPI_ISL_3218281, EPI_ISL_3218282, EPI_ISL_3218284, EPI_ISL_3218286, EPI_ISL_3218287, EPI_ISL_3218288, EPI_ISL_3218289, EPI_ISL_3218290, EPI_ISL_3218291, EPI_ISL_3218292, EPI_ISL_3218293, EPI_ISL_3218294, EPI_ISL_3218295, EPI_ISL_3218296, EPI_ISL_3218297, EPI_ISL_3218301, EPI_ISL_3218302, EPI_ISL_3218303, EPI_ISL_3218304, EPI_ISL_3218305, EPI_ISL_3218306, EPI_ISL_3218307, EPI_ISL_3218308, EPI_ISL_3218309, EPI_ISL_3218310, EPI_ISL_3218311, EPI_ISL_3218312, EPI_ISL_3218313, EPI_ISL_3218314, EPI_ISL_3218315, EPI_ISL_3218316, EPI_ISL_3218317, EPI_ISL_3218318, EPI_ISL_3218321, EPI_ISL_3218322, EPI_ISL_3218323, EPI_ISL_3218324, EPI_ISL_3218325, EPI_ISL_3218326, EPI_ISL_3218327, EPI_ISL_3218328, EPI_ISL_3218329, EPI_ISL_3218330, EPI_ISL_3218331, EPI_ISL_3218333, EPI_ISL_3218334, EPI_ISL_3218335, EPI_ISL_3218336, EPI_ISL_3218337, EPI_ISL_3218339, EPI_ISL_3218340, EPI_ISL_3218341, EPI_ISL_3218343, EPI_ISL_3218344, EPI_ISL_3218345, EPI_ISL_3218346, EPI_ISL_3218347, EPI_ISL_3218348, EPI_ISL_3218349, EPI_ISL_3218350, EPI_ISL_3218351, EPI_ISL_3218352, EPI_ISL_3218353, EPI_ISL_3218355, EPI_ISL_3218356, EPI_ISL_3218358, EPI_ISL_3218360, EPI_ISL_3218361, EPI_ISL_3218362, EPI_ISL_3218363, EPI_ISL_3218364, EPI_ISL_3218365, EPI_ISL_3218366, EPI_ISL_3218367, EPI_ISL_3218368, EPI_ISL_3218369, EPI_ISL_3218370, EPI_ISL_3218371, EPI_ISL_3218372, EPI_ISL_3218373, EPI_ISL_3218374, EPI_ISL_3218375, EPI_ISL_3218376, EPI_ISL_3218378, EPI_ISL_3218379, EPI_ISL_3218380, EPI_ISL_3218381, EPI_ISL_3218384, EPI_ISL_3218386, EPI_ISL_3218387, EPI_ISL_3218388, EPI_ISL_3218389, EPI_ISL_3218390, EPI_ISL_3218391, EPI_ISL_3218392, EPI_ISL_3218397, EPI_ISL_3218398, EPI_ISL_3218399, EPI_ISL_3218400, EPI_ISL_3218401, EPI_ISL_3218403, EPI_ISL_3218404, EPI_ISL_3218405, EPI_ISL_3218406, EPI_ISL_3218407, EPI_ISL_3218408, EPI_ISL_3218409, EPI_ISL_3218410, EPI_ISL_3218411, EPI_ISL_3218413, EPI_ISL_3218414, EPI_ISL_3218416, EPI_ISL_3218417, EPI_ISL_3218418, EPI_ISL_3218419, EPI_ISL_3218420, EPI_ISL_3218421, EPI_ISL_3218422, EPI_ISL_3218424, EPI_ISL_3218425, EPI_ISL_3218426, EPI_ISL_3218429, EPI_ISL_3218430, EPI_ISL_3218431, EPI_ISL_3218432, EPI_ISL_3218433, EPI_ISL_3218434, EPI_ISL_3218435, EPI_ISL_3218436, EPI_ISL_3218437, EPI_ISL_3218438, EPI_ISL_3218439, EPI_ISL_3218440, EPI_ISL_3218441, EPI_ISL_3218442, EPI_ISL_3218443, EPI_ISL_3218445, EPI_ISL_3218447, EPI_ISL_3218449, EPI_ISL_3218450, EPI_ISL_3218451, EPI_ISL_3218452, EPI_ISL_3218453, EPI_ISL_3218454, EPI_ISL_3218455, EPI_ISL_3218456, EPI_ISL_3218457, EPI_ISL_3218458, EPI_ISL_3218459, EPI_ISL_3218460, EPI_ISL_3218462, EPI_ISL_3218464, EPI_ISL_3218465, EPI_ISL_3218467, EPI_ISL_3218468, EPI_ISL_3218471, EPI_ISL_3218472, EPI_ISL_3218473, EPI_ISL_3218474, EPI_ISL_3218475, EPI_ISL_3218476, EPI_ISL_3218477, EPI_ISL_3218478, EPI_ISL_3218479, EPI_ISL_3218480, EPI_ISL_3218481, EPI_ISL_3218482, EPI_ISL_3218483, EPI_ISL_3218484, EPI_ISL_3218485, EPI_ISL_3218487, EPI_ISL_3218488, EPI_ISL_3218489, EPI_ISL_3218490, EPI_ISL_3218491, EPI_ISL_3218492, EPI_ISL_3218493, EPI_ISL_3218494, EPI_ISL_3218495, EPI_ISL_3218496, EPI_ISL_3218497, EPI_ISL_3218498, EPI_ISL_3218499, EPI_ISL_3218500, EPI_ISL_3218503, EPI_ISL_3218505, EPI_ISL_3218506, EPI_ISL_3218507, EPI_ISL_3218508, EPI_ISL_3218509, EPI_ISL_3218510, EPI_ISL_3218516, EPI_ISL_3218517, EPI_ISL_3218518, EPI_ISL_3218519, EPI_ISL_3218521, EPI_ISL_3218522, EPI_ISL_3218523, EPI_ISL_3218524, EPI_ISL_3218525, EPI_ISL_3218527, EPI_ISL_3218528, EPI_ISL_3218529, EPI_ISL_3218530, EPI_ISL_3218531, EPI_ISL_3218532, EPI_ISL_3218533, EPI_ISL_3218534, EPI_ISL_3218535, EPI_ISL_3218536, EPI_ISL_3218537, EPI_ISL_3218538, EPI_ISL_3218539, EPI_ISL_3218540, EPI_ISL_3218541, EPI_ISL_3218542, EPI_ISL_3218543, EPI_ISL_3218544, EPI_ISL_3218545, EPI_ISL_3218546, EPI_ISL_3218547, EPI_ISL_3218550, EPI_ISL_3218551, EPI_ISL_3218552, EPI_ISL_3218553, EPI_ISL_3218554, EPI_ISL_3218556, EPI_ISL_3218558, EPI_ISL_3218559, EPI_ISL_3218560, EPI_ISL_3218561, EPI_ISL_3218562, EPI_ISL_3218563, EPI_ISL_3218564, EPI_ISL_3218565, EPI_ISL_3218566, EPI_ISL_3218567, EPI_ISL_3218568, EPI_ISL_3218569, EPI_ISL_3218570, EPI_ISL_3218571, EPI_ISL_3218572, EPI_ISL_3218573, EPI_ISL_3218574, EPI_ISL_3218575, EPI_ISL_3218576, EPI_ISL_3218577, EPI_ISL_3218578, EPI_ISL_3218579, EPI_ISL_3218580, EPI_ISL_3218581, EPI_ISL_3218582, EPI_ISL_3218583, EPI_ISL_3218584, EPI_ISL_3218585, EPI_ISL_3218587, EPI_ISL_3218588, EPI_ISL_3218589, EPI_ISL_3218590, EPI_ISL_3218591, EPI_ISL_3218592, EPI_ISL_3218593, EPI_ISL_3218594, EPI_ISL_3218595, EPI_ISL_3218596, EPI_ISL_3218597, EPI_ISL_3218600, EPI_ISL_3218603, EPI_ISL_3218604, EPI_ISL_3218605, EPI_ISL_3218607, EPI_ISL_3218608, EPI_ISL_3218609, EPI_ISL_3218610, EPI_ISL_3218611, EPI_ISL_3218612, EPI_ISL_3218613, EPI_ISL_3218614, EPI_ISL_3218616, EPI_ISL_3218617, EPI_ISL_3218618, EPI_ISL_3218619, EPI_ISL_3218621, EPI_ISL_3218622, EPI_ISL_3218624, EPI_ISL_3218625, EPI_ISL_3218626, EPI_ISL_3218627, EPI_ISL_3218628, EPI_ISL_3218629, EPI_ISL_3218630, EPI_ISL_3218631, EPI_ISL_3218632, EPI_ISL_3218633, EPI_ISL_3218634, EPI_ISL_3218635, EPI_ISL_3218636, EPI_ISL_3218637, EPI_ISL_3218638, EPI_ISL_3218639, EPI_ISL_3218640, EPI_ISL_3218642, EPI_ISL_3218643, EPI_ISL_3218644, EPI_ISL_3218645, EPI_ISL_3218646, EPI_ISL_3218647, EPI_ISL_3218648, EPI_ISL_3218649, EPI_ISL_3218650, EPI_ISL_3218652, EPI_ISL_3218653, EPI_ISL_3218654, EPI_ISL_3218655, EPI_ISL_3218657, EPI_ISL_3218658, EPI_ISL_3218659, EPI_ISL_3218661, EPI_ISL_3218662, EPI_ISL_3218663, EPI_ISL_3218664, EPI_ISL_3218665, EPI_ISL_3218666, EPI_ISL_3218667, EPI_ISL_3218668, EPI_ISL_3218670, EPI_ISL_3218672, EPI_ISL_3218673, EPI_ISL_3218674, EPI_ISL_3218675, EPI_ISL_3218677, EPI_ISL_3218679, EPI_ISL_3218680, EPI_ISL_3218681, EPI_ISL_3218682, EPI_ISL_3218683, EPI_ISL_3218684, EPI_ISL_3218685, EPI_ISL_3218686, EPI_ISL_3218687, EPI_ISL_3218688, EPI_ISL_3218689, EPI_ISL_3218690, EPI_ISL_3218692, EPI_ISL_3218693, EPI_ISL_3218694, EPI_ISL_3218695, EPI_ISL_3218696, EPI_ISL_3218697, EPI_ISL_3218698, EPI_ISL_3218699, EPI_ISL_3218700, EPI_ISL_3218702, EPI_ISL_3218703, EPI_ISL_3218704, EPI_ISL_3218705, EPI_ISL_3218706, EPI_ISL_3218707, EPI_ISL_3218709, EPI_ISL_3218710, EPI_ISL_3218711, EPI_ISL_3218712, EPI_ISL_3218713, EPI_ISL_3218714, EPI_ISL_3218715, EPI_ISL_3218716, EPI_ISL_3218717, EPI_ISL_3218718, EPI_ISL_3218719, EPI_ISL_3218720, EPI_ISL_3218721, EPI_ISL_3218722, EPI_ISL_3218724, EPI_ISL_3218726, EPI_ISL_3218727, EPI_ISL_3218728, EPI_ISL_3218729, EPI_ISL_3218730, EPI_ISL_3218731, EPI_ISL_3218732, EPI_ISL_3218733, EPI_ISL_3218735, EPI_ISL_3218737, EPI_ISL_3218739, EPI_ISL_3218740, EPI_ISL_3218741, EPI_ISL_3218742, EPI_ISL_3218743, EPI_ISL_3218744, EPI_ISL_3218745, EPI_ISL_3218746, EPI_ISL_3218747, EPI_ISL_3218748, EPI_ISL_3218749, EPI_ISL_3218750, EPI_ISL_3218751, EPI_ISL_3218752, EPI_ISL_3218753, EPI_ISL_3218754, EPI_ISL_3218755, EPI_ISL_3218756, EPI_ISL_3218758, EPI_ISL_3218761, EPI_ISL_3218762, EPI_ISL_3218763, EPI_ISL_3218764, EPI_ISL_3218765, EPI_ISL_3218766, EPI_ISL_3218767, EPI_ISL_3218768, EPI_ISL_3218769, EPI_ISL_3218770, EPI_ISL_3218771, EPI_ISL_3218772, EPI_ISL_3218773, EPI_ISL_3218774, EPI_ISL_3218775, EPI_ISL_3218777, EPI_ISL_3218778, EPI_ISL_3218779, EPI_ISL_3218780, EPI_ISL_3218781, EPI_ISL_3218782, EPI_ISL_3218783, EPI_ISL_3218785, EPI_ISL_3218786, EPI_ISL_3218787, EPI_ISL_3218788, EPI_ISL_3218789, EPI_ISL_3218790, EPI_ISL_3218791, EPI_ISL_3218792, EPI_ISL_3218793, EPI_ISL_3218794, EPI_ISL_3218795, EPI_ISL_3218796, EPI_ISL_3218797, EPI_ISL_3218798, EPI_ISL_3218799, EPI_ISL_3218800, EPI_ISL_3218801, EPI_ISL_3218802, EPI_ISL_3218803, EPI_ISL_3218804, EPI_ISL_3218805, EPI_ISL_3218807, EPI_ISL_3218808, EPI_ISL_3218809, EPI_ISL_3218810, EPI_ISL_3218811, EPI_ISL_3218812, EPI_ISL_3218813, EPI_ISL_3218815, EPI_ISL_3218816, EPI_ISL_3218818, EPI_ISL_3218819, EPI_ISL_3218822, EPI_ISL_3218823, EPI_ISL_3218824, EPI_ISL_3218825, EPI_ISL_3218826, EPI_ISL_3218827, EPI_ISL_3218831, EPI_ISL_3218832, EPI_ISL_3218833, EPI_ISL_3218835, EPI_ISL_3218836, EPI_ISL_3218838, EPI_ISL_3218840, EPI_ISL_3218841, EPI_ISL_3218842 | | | |
| see above | Aegis Sciences Corporation | Centers for Disease Control and Prevention Division of Viral Diseases, Pathogen Discovery | Dakota Howard, Dhwani Batra, Peter W. Cook, Kara Moser, Adrian Paskey, Jason Caravas, Benjamin Rambo-Martin, Shatavia Morrison, Christopher Gulvick, Scott Sammons, Yvette Unoarumhi, Darlene Wagner, Matthew Schmerer, Cyndi Clark, Patrick Campbell, Rob Case, Vikramsinha Ghorpade, Holly Houdeshell, Ola Kvalvaag, Dillon Nall, Ethan Sanders, Alec Vest, Shaun Westlund, Matthew Hardison, Clinton R. Paden, Duncan MacCannell |
| EPI_ISL_3218846, EPI_ISL_3218847, EPI_ISL_3218848, EPI_ISL_3218849, EPI_ISL_3218850, EPI_ISL_3218852, EPI_ISL_3218853, EPI_ISL_3218854, EPI_ISL_3218855, EPI_ISL_3218856, EPI_ISL_3218857, EPI_ISL_3218858, EPI_ISL_3218859, EPI_ISL_3218860, EPI_ISL_3218861, EPI_ISL_3218862, EPI_ISL_3218863, EPI_ISL_3218864, EPI_ISL_3218865, EPI_ISL_3218866, EPI_ISL_3218868, EPI_ISL_3218869, EPI_ISL_3218870, EPI_ISL_3218871, EPI_ISL_3218872, EPI_ISL_3218873, EPI_ISL_3218874, EPI_ISL_3218875, EPI_ISL_3218876, EPI_ISL_3218877 | | | |
| see above | University of Michigan Clinical Microbiology Laboratory | Lauring Lab, University of Michigan, Department of Microbiology and Immunology | Valesano |
| EPI_ISL_3218878, EPI_ISL_3218880, EPI_ISL_3218881, EPI_ISL_3218882, EPI_ISL_3218883, EPI_ISL_3218884, EPI_ISL_3218886, EPI_ISL_3218887, EPI_ISL_3218890, EPI_ISL_3218891, EPI_ISL_3218893, EPI_ISL_3218894, EPI_ISL_3218897, EPI_ISL_3218899, EPI_ISL_3218901, EPI_ISL_3218905 | | | |
| see above | IVY3 Central Lab, Vanderbilt University Medical Center | Lauring Lab, University of Michigan, Department of Microbiology and Immunology | Valesano |
| EPI_ISL_3218906, EPI_ISL_3218907, EPI_ISL_3218908, EPI_ISL_3218909, EPI_ISL_3218910, EPI_ISL_3218911, EPI_ISL_3218912, EPI_ISL_3218913, EPI_ISL_3218914, EPI_ISL_3218915, EPI_ISL_3218916, EPI_ISL_3218917, EPI_ISL_3218918, EPI_ISL_3218919, EPI_ISL_3218920, EPI_ISL_3218921, EPI_ISL_3218922, EPI_ISL_3218923, EPI_ISL_3218924, EPI_ISL_3218925, EPI_ISL_3218926, EPI_ISL_3218927, EPI_ISL_3218928, EPI_ISL_3218929, EPI_ISL_3218930, EPI_ISL_3218931, EPI_ISL_3218932, EPI_ISL_3218933, EPI_ISL_3218935, EPI_ISL_3218936, EPI_ISL_3218937, EPI_ISL_3218938, EPI_ISL_3218939, EPI_ISL_3218940, EPI_ISL_3218941, EPI_ISL_3218942, EPI_ISL_3218943, EPI_ISL_3218944, EPI_ISL_3218946, EPI_ISL_3218948, EPI_ISL_3218949, EPI_ISL_3218950, EPI_ISL_3218951, EPI_ISL_3218952, EPI_ISL_3218953, EPI_ISL_3218954, EPI_ISL_3218955, EPI_ISL_3218956, EPI_ISL_3218958, EPI_ISL_3218959, EPI_ISL_3218960, EPI_ISL_3218961, EPI_ISL_3218962, EPI_ISL_3218963, EPI_ISL_3218964, EPI_ISL_3218965, EPI_ISL_3218967, EPI_ISL_3218968, EPI_ISL_3218969, EPI_ISL_3218970, EPI_ISL_3218971, EPI_ISL_3218972, EPI_ISL_3218973, EPI_ISL_3218974, EPI_ISL_3218975, EPI_ISL_3218976, EPI_ISL_3218977, EPI_ISL_3218978, EPI_ISL_3218979, EPI_ISL_3218980, EPI_ISL_3218981, EPI_ISL_3218982, EPI_ISL_3218983, EPI_ISL_3218984, EPI_ISL_3218985, EPI_ISL_3218987, EPI_ISL_3218989, EPI_ISL_3218990, EPI_ISL_3218991, EPI_ISL_3218992, EPI_ISL_3218993, EPI_ISL_3218994, EPI_ISL_3218995, EPI_ISL_3218996, EPI_ISL_3218997, EPI_ISL_3218998, EPI_ISL_3218999, EPI_ISL_3219000, EPI_ISL_3219001, EPI_ISL_3219002, EPI_ISL_3219003, EPI_ISL_3219004, EPI_ISL_3219005, EPI_ISL_3219006, EPI_ISL_3219007, EPI_ISL_3219008, EPI_ISL_3219009, EPI_ISL_3219010, EPI_ISL_3219011, EPI_ISL_3219012, EPI_ISL_3219013, EPI_ISL_3219014, EPI_ISL_3219015, EPI_ISL_3219016, EPI_ISL_3219017, EPI_ISL_3219019, EPI_ISL_3219020, EPI_ISL_3219021, EPI_ISL_3219022, EPI_ISL_3219023, EPI_ISL_3219024, EPI_ISL_3219025, EPI_ISL_3219026, EPI_ISL_3219027, EPI_ISL_3219028, EPI_ISL_3219029, EPI_ISL_3219030, EPI_ISL_3219031, EPI_ISL_3219033, EPI_ISL_3219035, EPI_ISL_3219037, EPI_ISL_3219038, EPI_ISL_3219039, EPI_ISL_3219040, EPI_ISL_3219041, EPI_ISL_3219042, EPI_ISL_3219043, EPI_ISL_3219044, EPI_ISL_3219045, EPI_ISL_3219046, EPI_ISL_3219047, EPI_ISL_3219048, EPI_ISL_3219049, EPI_ISL_3219050, EPI_ISL_3219051, EPI_ISL_3219052, EPI_ISL_3219053, EPI_ISL_3219054, EPI_ISL_3219055, EPI_ISL_3219056, EPI_ISL_3219057, EPI_ISL_3219058, EPI_ISL_3219059, EPI_ISL_3219060, EPI_ISL_3219061, EPI_ISL_3219062, EPI_ISL_3219063, EPI_ISL_3219064, EPI_ISL_3219065, EPI_ISL_3219066, EPI_ISL_3219067, EPI_ISL_3219068, EPI_ISL_3219069, EPI_ISL_3219070, EPI_ISL_3219071, EPI_ISL_3219072, EPI_ISL_3219073, EPI_ISL_3219074, EPI_ISL_3219075, EPI_ISL_3219076, EPI_ISL_3219077, EPI_ISL_3219078, EPI_ISL_3219079, EPI_ISL_3219080, EPI_ISL_3219081, EPI_ISL_3219082, EPI_ISL_3219083, EPI_ISL_3219084, EPI_ISL_3219085, EPI_ISL_3219086, EPI_ISL_3219087, EPI_ISL_3219088, EPI_ISL_3219089, EPI_ISL_3219090, EPI_ISL_3219091, EPI_ISL_3219092 | | | |
| see above | Illinois Department of Public Health - Springfield Lab | Illinois Department of Public Health - Springfield Lab | Bryan Sim, Gordon McCall, Marshall Smock |
| EPI_ISL_3219093, EPI_ISL_3219094, EPI_ISL_3219095, EPI_ISL_3219096, EPI_ISL_3219097, EPI_ISL_3219102, EPI_ISL_3219105, EPI_ISL_3219107, EPI_ISL_3219112, EPI_ISL_3219113, EPI_ISL_3219114, EPI_ISL_3219115, EPI_ISL_3219116, EPI_ISL_3219118, EPI_ISL_3219124, EPI_ISL_3219125, EPI_ISL_3219127, EPI_ISL_3219128, EPI_ISL_3219129, EPI_ISL_3219131, EPI_ISL_3219132, EPI_ISL_3219134, EPI_ISL_3219135, EPI_ISL_3219136, EPI_ISL_3219137, EPI_ISL_3219138, EPI_ISL_3219139 | | | |
| see above | Servicio de Microbiologia Hospital Ramon y Cajal | Servicio de Microbiologia Hospital Ramon y Cajal | Ponce M, Galan JC, Martinez L. Abreu M, y Gonzalez-Alba JM |
| EPI_ISL_3219141, EPI_ISL_3219142, EPI_ISL_3219143, EPI_ISL_3219144, EPI_ISL_3219145, EPI_ISL_3219146, EPI_ISL_3219147, EPI_ISL_3219148, EPI_ISL_3219149, EPI_ISL_3219150, EPI_ISL_3219151 | | | |
| see above | The Ohio State University Applied Microbiology Services Laboratory | The Ohio State University Applied Microbiology Services Laboratory | Seth A. Faith PhD |
| EPI_ISL_3219155, EPI_ISL_3219156, EPI_ISL_3219157, EPI_ISL_3219158, EPI_ISL_3219159, EPI_ISL_3219160, EPI_ISL_3219161, EPI_ISL_3219162, EPI_ISL_3219163, EPI_ISL_3219164, EPI_ISL_3219165, EPI_ISL_3219166, EPI_ISL_3219169, EPI_ISL_3219170, EPI_ISL_3219172, EPI_ISL_3219174, EPI_ISL_3219175, EPI_ISL_3219176, EPI_ISL_3219177, EPI_ISL_3219178, EPI_ISL_3219179 | | | |
| see above | Santa Clara Valley Medical Center | Santa Clara County Public Health Laboratory | Santa Clara County Public Health Department |
| EPI_ISL_3219180, EPI_ISL_3219181 | Kaiser San Jose | Santa Clara County Public Health Laboratory | Santa Clara County Public Health Department |
| EPI_ISL_3219182 | El Camino Hospital | Santa Clara County Public Health Laboratory | Santa Clara County Public Health Department |
| EPI_ISL_3219183, EPI_ISL_3219184, EPI_ISL_3219185, EPI_ISL_3219186, EPI_ISL_3219187, EPI_ISL_3219188, EPI_ISL_3219189, EPI_ISL_3219190, EPI_ISL_3219191, EPI_ISL_3219192, EPI_ISL_3219193 | | | |
| see above | Kaiser Santa Clara | Santa Clara County Public Health Laboratory | Santa Clara County Public Health Department |
| EPI_ISL_3219196, EPI_ISL_3219197, EPI_ISL_3219198, EPI_ISL_3219199, EPI_ISL_3219200, EPI_ISL_3219201 | Good Samaritan Hospital | Santa Clara County Public Health Laboratory | Santa Clara County Public Health Department |
| EPI_ISL_3219202, EPI_ISL_3219203, EPI_ISL_3219204, EPI_ISL_3219205, EPI_ISL_3219207, EPI_ISL_3219208, EPI_ISL_3219209, EPI_ISL_3219210, EPI_ISL_3219211, EPI_ISL_3219212, EPI_ISL_3219213, EPI_ISL_3219214, EPI_ISL_3219215, EPI_ISL_3219216, EPI_ISL_3219217, EPI_ISL_3219218 | | | |
| see above | PAMF Palo Alto | Santa Clara County Public Health Laboratory | Santa Clara County Public Health Department |
| EPI_ISL_3219219, | Santa Clara County Public Health Laboratory | Santa Clara County Public Health Laboratory | Santa Clara County Public Health Department |

| | | | |
|---|---|---|---|
| EPI_ISL_3219220 | | | |
| EPI_ISL_3219222, EPI_ISL_3219223, EPI_ISL_3219224, EPI_ISL_3219225, EPI_ISL_3219226, EPI_ISL_3219227, EPI_ISL_3219228, EPI_ISL_3219229, EPI_ISL_3219230, EPI_ISL_3219231, EPI_ISL_3219232, EPI_ISL_3219233, EPI_ISL_3219234, EPI_ISL_3219235, EPI_ISL_3219236, EPI_ISL_3219237, EPI_ISL_3219238, EPI_ISL_3219239, EPI_ISL_3219240, EPI_ISL_3219241, EPI_ISL_3219242, EPI_ISL_3219243, EPI_ISL_3219244, EPI_ISL_3219245, EPI_ISL_3219246, EPI_ISL_3219247, EPI_ISL_3219248, EPI_ISL_3219249, EPI_ISL_3219250, EPI_ISL_3219251, EPI_ISL_3219252, EPI_ISL_3219253, EPI_ISL_3219254, EPI_ISL_3219255, EPI_ISL_3219256, EPI_ISL_3219257, EPI_ISL_3219258, EPI_ISL_3219259, EPI_ISL_3219260, EPI_ISL_3219261, EPI_ISL_3219262, EPI_ISL_3219263, EPI_ISL_3219264, EPI_ISL_3219265, EPI_ISL_3219266, EPI_ISL_3219267, EPI_ISL_3219268, EPI_ISL_3219269, EPI_ISL_3219270, EPI_ISL_3219271, EPI_ISL_3219272, EPI_ISL_3219273, EPI_ISL_3219275, EPI_ISL_3219276, EPI_ISL_3219277, EPI_ISL_3219278, EPI_ISL_3219279, EPI_ISL_3219280, EPI_ISL_3219281, EPI_ISL_3219282, EPI_ISL_3219283, EPI_ISL_3219284, EPI_ISL_3219286, EPI_ISL_3219287, EPI_ISL_3219288, EPI_ISL_3219289, EPI_ISL_3219290, EPI_ISL_3219291, EPI_ISL_3219292, EPI_ISL_3219293, EPI_ISL_3219294, EPI_ISL_3219295, EPI_ISL_3219296, EPI_ISL_3219297, EPI_ISL_3219298, EPI_ISL_3219299, EPI_ISL_3219300, EPI_ISL_3219301, EPI_ISL_3219302, EPI_ISL_3219304, EPI_ISL_3219305, EPI_ISL_3219306, EPI_ISL_3219307, EPI_ISL_3219308, EPI_ISL_3219310, EPI_ISL_3219311, EPI_ISL_3219313, EPI_ISL_3219315, EPI_ISL_3219316, EPI_ISL_3219317, EPI_ISL_3219318, EPI_ISL_3219319, EPI_ISL_3219320, EPI_ISL_3219321, EPI_ISL_3219322 | | | |
| see above | Santa Clara Valley Medical Center | Santa Clara County Public Health Laboratory | Santa Clara County Public Health Department |
| EPI_ISL_3219323 | Santa Clara County Public Health Laboratory | Santa Clara County Public Health Laboratory | Santa Clara County Public Health Department |
| EPI_ISL_3219324, EPI_ISL_3219325, EPI_ISL_3219326, EPI_ISL_3219327, EPI_ISL_3219328, EPI_ISL_3219329, EPI_ISL_3219330, EPI_ISL_3219331, EPI_ISL_3219332, EPI_ISL_3219333, EPI_ISL_3219334, EPI_ISL_3219335, EPI_ISL_3219336, EPI_ISL_3219337, EPI_ISL_3219338, EPI_ISL_3219339, EPI_ISL_3219340, EPI_ISL_3219341, EPI_ISL_3219342, EPI_ISL_3219343, EPI_ISL_3219344, EPI_ISL_3219345, EPI_ISL_3219346, EPI_ISL_3219347, EPI_ISL_3219348, EPI_ISL_3219349, EPI_ISL_3219350, EPI_ISL_3219351, EPI_ISL_3219352, EPI_ISL_3219353, EPI_ISL_3219354, EPI_ISL_3219355, EPI_ISL_3219356 | | | |
| see above | Santa Clara Valley Medical Center | Santa Clara County Public Health Laboratory | Santa Clara County Public Health Department |
| EPI_ISL_3219357, EPI_ISL_3219358, EPI_ISL_3219359, EPI_ISL_3219360, EPI_ISL_3219361, EPI_ISL_3219362 | Kaiser Santa Clara | Santa Clara County Public Health Laboratory | Santa Clara County Public Health Department |
| EPI_ISL_3219363, EPI_ISL_3219364, EPI_ISL_3219365 | Good Samaritan Hospital | Santa Clara County Public Health Laboratory | Santa Clara County Public Health Department |
| EPI_ISL_3219366, EPI_ISL_3219367, EPI_ISL_3219368, EPI_ISL_3219369, EPI_ISL_3219370, EPI_ISL_3219371, EPI_ISL_3219372, EPI_ISL_3219373, EPI_ISL_3219374, EPI_ISL_3219376, EPI_ISL_3219377, EPI_ISL_3219378 | | | |
| see above | El Camino Hospital | Santa Clara County Public Health Laboratory | Santa Clara County Public Health Department |
| EPI_ISL_3219379, EPI_ISL_3219380, EPI_ISL_3219381, EPI_ISL_3219382, EPI_ISL_3219383, EPI_ISL_3219384, EPI_ISL_3219385, EPI_ISL_3219386, EPI_ISL_3219387, EPI_ISL_3219388, EPI_ISL_3219389, EPI_ISL_3219390, EPI_ISL_3219391, EPI_ISL_3219392, EPI_ISL_3219393, EPI_ISL_3219394, EPI_ISL_3219395, EPI_ISL_3219396, EPI_ISL_3219397, EPI_ISL_3219398, EPI_ISL_3219399, EPI_ISL_3219400, EPI_ISL_3219401 | | | |
| see above | PAMF Palo Alto | Santa Clara County Public Health Laboratory | Santa Clara County Public Health Department |
| EPI_ISL_3219403, EPI_ISL_3219404, EPI_ISL_3219405, EPI_ISL_3219406, EPI_ISL_3219407, EPI_ISL_3219408, EPI_ISL_3219409, EPI_ISL_3219410, EPI_ISL_3219411, EPI_ISL_3219412, EPI_ISL_3219413, EPI_ISL_3219414, EPI_ISL_3219415, EPI_ISL_3219416, EPI_ISL_3219417, EPI_ISL_3219418, EPI_ISL_3219419, EPI_ISL_3219420, EPI_ISL_3219421, EPI_ISL_3219422, EPI_ISL_3219423, EPI_ISL_3219424, EPI_ISL_3219425, EPI_ISL_3219427, EPI_ISL_3219428, EPI_ISL_3219429, EPI_ISL_3219430, EPI_ISL_3219431, EPI_ISL_3219432, EPI_ISL_3219433, EPI_ISL_3219434, EPI_ISL_3219435, EPI_ISL_3219436, EPI_ISL_3219438 | | | |
| see above | Santa Clara Valley Medical Center | Santa Clara County Public Health Laboratory | Santa Clara County Public Health Department |
| EPI_ISL_3219439, EPI_ISL_3219440, EPI_ISL_3219441, EPI_ISL_3219442 | Temporary Specimen Collection Centre at the AsiaWorld-Expo | Hong Kong Department of Health | Alan K.L. Tsang, Peter C.W. Yip, Ken H.L. Ng, Edman T.K. Lam, Rickjason C.W. Chan |
| EPI_ISL_3219443 | Quarantine | Hong Kong Department of Health | Alan K.L. Tsang, Peter C.W. Yip, Ken H.L. Ng, Edman T.K. Lam, Rickjason C.W. Chan |
| EPI_ISL_3219444 | Quarantine Camp | Hong Kong Department of Health | Alan K.L. Tsang, Peter C.W. Yip, Ken H.L. Ng, Edman T.K. Lam, Rickjason C.W. Chan |
| EPI_ISL_3219445, EPI_ISL_3219446 | Temporary Specimen Collection Centre at the AsiaWorld-Expo | Hong Kong Department of Health | Alan K.L. Tsang, Peter C.W. Yip, Ken H.L. Ng, Edman T.K. Lam, Rickjason C.W. Chan |
| EPI_ISL_3219453, EPI_ISL_3219454, EPI_ISL_3219459, EPI_ISL_3219460, EPI_ISL_3219461 | Servizo de Microbioloxía. Complexo Hospitalario de Santiago de Compostela | Servizo de Microbioloxía. Complexo Hospitalario de Santiago de Compostela | Xosé Costa, Antonio Aguilera, Gema Barbeito, Amparo Coira, Carlos García_Riestra, Manuela Hernández, José Llovo, Teresa Lopez_Valiño, Laura Millán, Daniel Navarro, Rocío trastoy, Mercedes Treviño, María Luisa Pérez_del_Molino |
| EPI_ISL_3219463, EPI_ISL_3219464, EPI_ISL_3219465, EPI_ISL_3219466, EPI_ISL_3219467, EPI_ISL_3219468, EPI_ISL_3219469, EPI_ISL_3219471, EPI_ISL_3219472, EPI_ISL_3219473, EPI_ISL_3219474, EPI_ISL_3219475, EPI_ISL_3219476, EPI_ISL_3219477, EPI_ISL_3219478, EPI_ISL_3219479, EPI_ISL_3219480, EPI_ISL_3219481, EPI_ISL_3219482, EPI_ISL_3219483, EPI_ISL_3219484, EPI_ISL_3219485, EPI_ISL_3219486, EPI_ISL_3219487, EPI_ISL_3219488, EPI_ISL_3219489, EPI_ISL_3219490, EPI_ISL_3219491, EPI_ISL_3219492, EPI_ISL_3219494, EPI_ISL_3219495, EPI_ISL_3219496, EPI_ISL_3219498, EPI_ISL_3219499, EPI_ISL_3219500, EPI_ISL_3219501, EPI_ISL_3219502, EPI_ISL_3219503, EPI_ISL_3219504, EPI_ISL_3219505, EPI_ISL_3219506, EPI_ISL_3219507, EPI_ISL_3219508, EPI_ISL_3219509 | | | |
| see above | Labo Analyses Med | National Reference Center for Viruses of Respiratory Infections, Institut Pasteur, Paris | Marion Barbet, Sylvie Behillil, Méline Bizard, Angela Brisebarre, Camille Capel, Vincent Enouf, Louise Lefrançois, Frédéric Lemoine, Christophe Malabat, Corinne Maufrais, Hub de Bioinformatique et Biostatistique, Etienne Simon-Lorière, Maud Vanpeene, Sylvie Van der Werf ,Eric Morin |
| EPI_ISL_3219510, EPI_ISL_3219511, EPI_ISL_3219512, EPI_ISL_3219513, EPI_ISL_3219514 | Labo Analyses Med | National Reference Center for Viruses of Respiratory Infections, Institut Pasteur, Paris | Marion Barbet, Sylvie Behillil, Méline Bizard, Angela Brisebarre, Camille Capel, Vincent Enouf, Louise Lefrançois, Frédéric Lemoine, Christophe Malabat, Corinne Maufrais, Hub de Bioinformatique et Biostatistique, Etienne Simon-Lorière, Maud Vanpeene, Sylvie Van der Werf ,AuréLie Dupuis |
| EPI_ISL_3219516, EPI_ISL_3219517 | Labo Analyses Med | National Reference Center for Viruses of Respiratory Infections, Institut Pasteur, Paris | Marion Barbet, Sylvie Behillil, Méline Bizard, Angela Brisebarre, Camille Capel, Vincent Enouf, Louise Lefrançois, Frédéric Lemoine, Christophe Malabat, Corinne Maufrais, Hub de Bioinformatique et Biostatistique, Etienne Simon-Lorière, Maud Vanpeene, Sylvie Van der Werf ,A Feraut |
| EPI_ISL_3219534 | Hospital | National Reference Center for Viruses of Respiratory Infections, Institut Pasteur, Paris | Marion Barbet, Sylvie Behillil, Méline Bizard, Angela Brisebarre, Camille Capel, Vincent Enouf, Louise Lefrançois, Frédéric Lemoine, Christophe Malabat, Corinne Maufrais, Hub de Bioinformatique et Biostatistique, Etienne Simon-Lorière, Maud Vanpeene, Sylvie Van der Werf ,ValéRie Serazin |
| EPI_ISL_3219535 | Hospital | National Reference Center for Viruses of Respiratory Infections, Institut Pasteur, Paris | Marion Barbet, Sylvie Behillil, Méline Bizard, Angela Brisebarre, Camille Capel, Vincent Enouf, Louise Lefrançois, Frédéric Lemoine, Christophe Malabat, Corinne Maufrais, Hub de Bioinformatique et Biostatistique, Etienne Simon-Lorière, Maud Vanpeene, Sylvie Van der Werf ,Marianne Burgard |
| EPI_ISL_3219536, EPI_ISL_3219537 | Hospital | National Reference Center for Viruses of Respiratory Infections, Institut Pasteur, Paris | Marion Barbet, Sylvie Behillil, Méline Bizard, Angela Brisebarre, Camille Capel, Vincent Enouf, Louise Lefrançois, Frédéric Lemoine, Christophe Malabat, Corinne Maufrais, Hub de Bioinformatique et Biostatistique, Etienne Simon-Lorière, Maud Vanpeene, Sylvie Van der Werf ,Pascale Martres |
| EPI_ISL_3219538 | Hospital | National Reference Center for Viruses of Respiratory Infections, Institut Pasteur, Paris | Marion Barbet, Sylvie Behillil, Méline Bizard, Angela Brisebarre, Camille Capel, Vincent Enouf, Louise Lefrançois, Frédéric Lemoine, Christophe Malabat, Corinne Maufrais, Hub de Bioinformatique et Biostatistique, Etienne Simon-Lorière, Maud Vanpeene, Sylvie Van der Werf ,Marianne Burgard |
| EPI_ISL_3219539, EPI_ISL_3219541, EPI_ISL_3219542, EPI_ISL_3219543, EPI_ISL_3219544, EPI_ISL_3219545, EPI_ISL_3219546, EPI_ISL_3219547, EPI_ISL_3219548, EPI_ISL_3219549, EPI_ISL_3219551, EPI_ISL_3219552, EPI_ISL_3219553 | | | |
| see above | Labo Analyses Med | National Reference Center for Viruses of Respiratory Infections, Institut Pasteur, Paris | Marion Barbet, Sylvie Behillil, Méline Bizard, Angela Brisebarre, Camille Capel, Vincent Enouf, Louise Lefrançois, Frédéric Lemoine, Christophe Malabat, Corinne Maufrais, Hub de Bioinformatique et Biostatistique, Etienne Simon-Lorière, Maud Vanpeene, Sylvie Van der Werf ,Jean-Baptiste Foulquier |
| EPI_ISL_3219555, EPI_ISL_3219556, EPI_ISL_3219557, EPI_ISL_3219558, EPI_ISL_3219559 | Labo Analyses Med | National Reference Center for Viruses of Respiratory Infections, Institut Pasteur, Paris | Marion Barbet, Sylvie Behillil, Méline Bizard, Angela Brisebarre, Camille Capel, Vincent Enouf, Louise Lefrançois, Frédéric Lemoine, Christophe Malabat, Corinne Maufrais, Hub de Bioinformatique et Biostatistique, Etienne Simon-Lorière, Maud Vanpeene, Sylvie Van der Werf |
| EPI_ISL_3219560, EPI_ISL_3219561, EPI_ISL_3219562, EPI_ISL_3219563, EPI_ISL_3219564, EPI_ISL_3219565, EPI_ISL_3219566, EPI_ISL_3219567, EPI_ISL_3219568, EPI_ISL_3219569, EPI_ISL_3219570, EPI_ISL_3219571, EPI_ISL_3219572, EPI_ISL_3219573, EPI_ISL_3219574, EPI_ISL_3219575, EPI_ISL_3219576, EPI_ISL_3219577, EPI_ISL_3219578, EPI_ISL_3219579, EPI_ISL_3219580, EPI_ISL_3219581, EPI_ISL_3219582, EPI_ISL_3219583, EPI_ISL_3219584, EPI_ISL_3219585, EPI_ISL_3219586, EPI_ISL_3219587, EPI_ISL_3219588, EPI_ISL_3219589, EPI_ISL_3219590, EPI_ISL_3219591, EPI_ISL_3219592, EPI_ISL_3219593, EPI_ISL_3219594, EPI_ISL_3219595, EPI_ISL_3219596, EPI_ISL_3219597, EPI_ISL_3219598, EPI_ISL_3219599, EPI_ISL_3219600, EPI_ISL_3219601, EPI_ISL_3219602, EPI_ISL_3219603 | | | |
| see above | Labo Analyses Med | National Reference Center for Viruses of Respiratory Infections, Institut Pasteur, Paris | Marion Barbet, Sylvie Behillil, Méline Bizard, Angela Brisebarre, Camille Capel, Vincent Enouf, Louise Lefrançois, Frédéric Lemoine, Christophe Malabat, Corinne Maufrais, Hub de Bioinformatique et Biostatistique, Etienne Simon-Lorière, Maud Vanpeene, Sylvie Van der Werf ,Vanessa Magin |
| EPI_ISL_3219604, EPI_ISL_3219605, EPI_ISL_3219606, EPI_ISL_3219608, EPI_ISL_3219610, EPI_ISL_3219611, EPI_ISL_3219612, EPI_ISL_3219613, EPI_ISL_3219614, EPI_ISL_3219615, EPI_ISL_3219616, EPI_ISL_3219617, EPI_ISL_3219618, EPI_ISL_3219620, EPI_ISL_3219621, EPI_ISL_3219622, EPI_ISL_3219623, EPI_ISL_3219624, EPI_ISL_3219625, EPI_ISL_3219626, EPI_ISL_3219627, EPI_ISL_3219628, EPI_ISL_3219629, EPI_ISL_3219630, EPI_ISL_3219631, EPI_ISL_3219632, EPI_ISL_3219633, EPI_ISL_3219634, EPI_ISL_3219636, EPI_ISL_3219637, EPI_ISL_3219638, EPI_ISL_3219639, EPI_ISL_3219640, EPI_ISL_3219641, EPI_ISL_3219642, EPI_ISL_3219643, EPI_ISL_3219644, EPI_ISL_3219645, EPI_ISL_3219646, EPI_ISL_3219647, EPI_ISL_3219651, EPI_ISL_3219653, EPI_ISL_3219654, EPI_ISL_3219655, EPI_ISL_3219656, EPI_ISL_3219658, EPI_ISL_3219659, EPI_ISL_3219661, EPI_ISL_3219663, EPI_ISL_3219665, EPI_ISL_3219666, EPI_ISL_3219667, EPI_ISL_3219668, EPI_ISL_3219669, EPI_ISL_3219671, EPI_ISL_3219672, EPI_ISL_3219673, EPI_ISL_3219674, EPI_ISL_3219675, EPI_ISL_3219676, EPI_ISL_3219677, EPI_ISL_3219678, EPI_ISL_3219679, EPI_ISL_3219680, EPI_ISL_3219681, EPI_ISL_3219682, EPI_ISL_3219683, EPI_ISL_3219684, EPI_ISL_3219685, EPI_ISL_3219686, EPI_ISL_3219687, EPI_ISL_3219688 | | | |
| see above | Labo Analyses Med | National Reference Center for Viruses of Respiratory Infections, Institut Pasteur, Paris | Marion Barbet, Sylvie Behillil, Méline Bizard, Angela Brisebarre, Camille Capel, Vincent Enouf, Louise Lefrançois, Frédéric Lemoine, Christophe Malabat, Corinne Maufrais, Hub de Bioinformatique et Biostatistique, Etienne Simon-Lorière, Maud Vanpeene, Sylvie Van der Werf ,OphéLie Said-Delattre |
| EPI_ISL_3219689, EPI_ISL_3219690, EPI_ISL_3219692, EPI_ISL_3219693, EPI_ISL_3219694, EPI_ISL_3219695, EPI_ISL_3219696, EPI_ISL_3219698, EPI_ISL_3219699, EPI_ISL_3219700, EPI_ISL_3219701, EPI_ISL_3219702 | | | |
| see above | Hospital | National Reference Center for Viruses of Respiratory Infections, Institut Pasteur, Paris | Marion Barbet, Sylvie Behillil, Méline Bizard, Angela Brisebarre, Camille Capel, Vincent Enouf, Louise Lefrançois, Frédéric Lemoine, Christophe Malabat, Corinne Maufrais, Hub de Bioinformatique et Biostatistique, Etienne Simon-Lorière, Maud Vanpeene, Sylvie Van der Werf ,Nabil Gastli |
| EPI_ISL_3219703 | Hospital | National Reference Center for Viruses of Respiratory Infections, Institut Pasteur, Paris | Marion Barbet, Sylvie Behillil, Méline Bizard, Angela Brisebarre, Camille Capel, Vincent Enouf, Louise Lefrançois, Frédéric Lemoine, Christophe Malabat, Corinne Maufrais, Hub de Bioinformatique et Biostatistique, Etienne Simon-Lorière, Maud Vanpeene, Sylvie Van der Werf ,Salma Ben Hadj Yahia |
| EPI_ISL_3219704, EPI_ISL_3219705, EPI_ISL_3219706, EPI_ISL_3219707, EPI_ISL_3219708 | Labo Analyses Med | National Reference Center for Viruses of Respiratory Infections, Institut Pasteur, Paris | Marion Barbet, Sylvie Behillil, Méline Bizard, Angela Brisebarre, Camille Capel, Vincent Enouf, Louise Lefrançois, Frédéric Lemoine, Christophe Malabat, Corinne Maufrais, Hub de Bioinformatique et Biostatistique, Etienne Simon-Lorière, Maud Vanpeene, Sylvie Van der Werf ,Baptiste Moreira |
| EPI_ISL_3219709, EPI_ISL_3219710, EPI_ISL_3219711, EPI_ISL_3219712, EPI_ISL_3219713, EPI_ISL_3219714, EPI_ISL_3219715, EPI_ISL_3219716, EPI_ISL_3219717, EPI_ISL_3219718, EPI_ISL_3219719, EPI_ISL_3219721 | | | |
| see above | Labo Analyses Med | National Reference Center for Viruses of Respiratory Infections, Institut Pasteur, Paris | Marion Barbet, Sylvie Behillil, Méline Bizard, Angela Brisebarre, Camille Capel, Vincent Enouf, Louise Lefrançois, Frédéric Lemoine, Christophe Malabat, Corinne Maufrais, Hub de Bioinformatique et Biostatistique, Etienne Simon-Lorière, Maud Vanpeene, Sylvie Van der Werf ,Karine Michez |
| EPI_ISL_3219722, EPI_ISL_3219723, | Hospital | National Reference Center for Viruses of Respiratory Infections, Institut Pasteur, Paris | Marion Barbet, Sylvie Behillil, Méline Bizard, Angela Brisebarre, Camille Capel, Vincent Enouf, Louise Lefrançois, Frédéric Lemoine, Christophe Malabat, Corinne Maufrais, Hub de Bioinformatique et Biostatistique, Etienne Simon-Lorière, Maud Vanpeene, Sylvie Van der Werf ,Guillaume Aubin |

| | | | |
|---|---|---|---|
| EPI_ISL_3219724, EPI_ISL_3219725, EPI_ISL_3219726 | | | |
| EPI_ISL_3219727, EPI_ISL_3219728, EPI_ISL_3219729, EPI_ISL_3219730, EPI_ISL_3219731, EPI_ISL_3219732, EPI_ISL_3219735, EPI_ISL_3219736, EPI_ISL_3219738, EPI_ISL_3219739, EPI_ISL_3219740, EPI_ISL_3219741, EPI_ISL_3219742, EPI_ISL_3219743, EPI_ISL_3219744, EPI_ISL_3219745, EPI_ISL_3219746, EPI_ISL_3219747, EPI_ISL_3219748, EPI_ISL_3219749, EPI_ISL_3219750, EPI_ISL_3219751, EPI_ISL_3219752 | | | |
| see above | Labo Analyses Med | National Reference Center for Viruses of Respiratory Infections, Institut Pasteur, Paris | Marion Barbet, Sylvie Behillil, Méline Bizard, Angela Brisebarre, Camille Capel, Vincent Enouf, Louise Lefrançois, Frédéric Lemoine, Christophe Malabat, Corinne Maufrais, Hub de Bioinformatique et Biostatistique, Etienne Simon-Lorière, Maud Vanpeene, Sylvie Van der Werf ,Eric Morin |
| EPI_ISL_3219753, EPI_ISL_3219754, EPI_ISL_3219755, EPI_ISL_3219756, EPI_ISL_3219757, EPI_ISL_3219758, EPI_ISL_3219759 | AZ Zeno Knokke-Blankenberge | AZ Sint Jan | Marijke Reynders, Patrick Descheemaeker, Laurien Hoornaert, Thomas Van Landschoot, Marie Madeleine Chabert-Consen, Jorn Hellemans |
| EPI_ISL_3219760, EPI_ISL_3219761, EPI_ISL_3219762 | AZ Sint-Jan Brugge-Oostende AV | AZ Sint Jan | Marijke Reynders, Patrick Descheemaeker, Laurien Hoornaert, Thomas Van Landschoot, Marie Madeleine Chabert-Consen, Jorn Hellemans |
| EPI_ISL_3219763, EPI_ISL_3219764, EPI_ISL_3219765, EPI_ISL_3219766, EPI_ISL_3219767, EPI_ISL_3219768, EPI_ISL_3219769, EPI_ISL_3219770, EPI_ISL_3219771 | AZ Zeno Knokke-Blankenberge | AZ Sint Jan | Marijke Reynders, Patrick Descheemaeker, Laurien Hoornaert, Thomas Van Landschoot, Marie Madeleine Chabert-Consen, Jorn Hellemans |
| EPI_ISL_3219772, EPI_ISL_3219774, EPI_ISL_3219775, EPI_ISL_3219776, EPI_ISL_3219777, EPI_ISL_3219778, EPI_ISL_3219779, EPI_ISL_3219780, EPI_ISL_3219781, EPI_ISL_3219782, EPI_ISL_3219783, EPI_ISL_3219784, EPI_ISL_3219785, EPI_ISL_3219786, EPI_ISL_3219787, EPI_ISL_3219788, EPI_ISL_3219789, EPI_ISL_3219790, EPI_ISL_3219791, EPI_ISL_3219792, EPI_ISL_3219793, EPI_ISL_3219794, EPI_ISL_3219795, EPI_ISL_3219796, EPI_ISL_3219797, EPI_ISL_3219798, EPI_ISL_3219799, EPI_ISL_3219800, EPI_ISL_3219801, EPI_ISL_3219802, EPI_ISL_3219803, EPI_ISL_3219804, EPI_ISL_3219806, EPI_ISL_3219807, EPI_ISL_3219808, EPI_ISL_3219809, EPI_ISL_3219810, EPI_ISL_3219811, EPI_ISL_3219812, EPI_ISL_3219813, EPI_ISL_3219814, EPI_ISL_3219815, EPI_ISL_3219816, EPI_ISL_3219817, EPI_ISL_3219818, EPI_ISL_3219819, EPI_ISL_3219820, EPI_ISL_3219821, EPI_ISL_3219822, EPI_ISL_3219823, EPI_ISL_3219824, EPI_ISL_3219825, EPI_ISL_3219826, EPI_ISL_3219827, EPI_ISL_3219828, EPI_ISL_3219829, EPI_ISL_3219830, EPI_ISL_3219831, EPI_ISL_3219832, EPI_ISL_3219833, EPI_ISL_3219834, EPI_ISL_3219835, EPI_ISL_3219836, EPI_ISL_3219837, EPI_ISL_3219838, EPI_ISL_3219839, EPI_ISL_3219840, EPI_ISL_3219841, EPI_ISL_3219842, EPI_ISL_3219843, EPI_ISL_3219844, EPI_ISL_3219845, EPI_ISL_3219846, EPI_ISL_3219847, EPI_ISL_3219848, EPI_ISL_3219849, EPI_ISL_3219850, EPI_ISL_3219851, EPI_ISL_3219852, EPI_ISL_3219853, EPI_ISL_3219854, EPI_ISL_3219855, EPI_ISL_3219856, EPI_ISL_3219857, EPI_ISL_3219858, EPI_ISL_3219859, EPI_ISL_3219860, EPI_ISL_3219861, EPI_ISL_3219862, EPI_ISL_3219863, EPI_ISL_3219864, EPI_ISL_3219865, EPI_ISL_3219866, EPI_ISL_3219867, EPI_ISL_3219869, EPI_ISL_3219870, EPI_ISL_3219871, EPI_ISL_3219872, EPI_ISL_3219873, EPI_ISL_3219874, EPI_ISL_3219875, EPI_ISL_3219876, EPI_ISL_3219877, EPI_ISL_3219878, EPI_ISL_3219879, EPI_ISL_3219880, EPI_ISL_3219881, EPI_ISL_3219882, EPI_ISL_3219883, EPI_ISL_3219884, EPI_ISL_3219885, EPI_ISL_3219886, EPI_ISL_3219887, EPI_ISL_3219888, EPI_ISL_3219889, EPI_ISL_3219890, EPI_ISL_3219891, EPI_ISL_3219892, EPI_ISL_3219893, EPI_ISL_3219894, EPI_ISL_3219895, EPI_ISL_3219896, EPI_ISL_3219897, EPI_ISL_3219898, EPI_ISL_3219899, EPI_ISL_3219900, EPI_ISL_3219901, EPI_ISL_3219902, EPI_ISL_3219903, EPI_ISL_3219904, EPI_ISL_3219905, EPI_ISL_3219906, EPI_ISL_3219907, EPI_ISL_3219908, EPI_ISL_3219909, EPI_ISL_3219910, EPI_ISL_3219911, EPI_ISL_3219912, EPI_ISL_3219913, EPI_ISL_3219914, EPI_ISL_3219915, EPI_ISL_3219916, EPI_ISL_3219917, EPI_ISL_3219918, EPI_ISL_3219919, EPI_ISL_3219920, EPI_ISL_3219921 | | | |
| see above | National Institute for Communicable Diseases of the National Health Laboratory Service | National Institute for Communicable Diseases of the National Health Laboratory Service | Amoako DG, Everatt J, Scheepers C, Mohale T, Ntuli N, Mahlangu B, Mnguni A, Ismail A, Bhiman JN |
| EPI_ISL_3219922, EPI_ISL_3219923, EPI_ISL_3219924, EPI_ISL_3219925, EPI_ISL_3219927, EPI_ISL_3219928, EPI_ISL_3219929, EPI_ISL_3219931, EPI_ISL_3219932, EPI_ISL_3219933, EPI_ISL_3219934, EPI_ISL_3219937, EPI_ISL_3219938, EPI_ISL_3219939, EPI_ISL_3219940, EPI_ISL_3219941, EPI_ISL_3219942, EPI_ISL_3219943, EPI_ISL_3219944, EPI_ISL_3219947, EPI_ISL_3219948, EPI_ISL_3219949, EPI_ISL_3219950, EPI_ISL_3219951, EPI_ISL_3219952, EPI_ISL_3219953, EPI_ISL_3219954, EPI_ISL_3219955, EPI_ISL_3219956, EPI_ISL_3219957, EPI_ISL_3219959, EPI_ISL_3219961, EPI_ISL_3219962, EPI_ISL_3219963, EPI_ISL_3219964, EPI_ISL_3219965, EPI_ISL_3219966, EPI_ISL_3219967, EPI_ISL_3219968, EPI_ISL_3219969, EPI_ISL_3219970, EPI_ISL_3219971, EPI_ISL_3219972, EPI_ISL_3219973, EPI_ISL_3219974, EPI_ISL_3219975, EPI_ISL_3219976, EPI_ISL_3219977, EPI_ISL_3219978, EPI_ISL_3219979, EPI_ISL_3219980, EPI_ISL_3219981, EPI_ISL_3219982, EPI_ISL_3219983, EPI_ISL_3219984, EPI_ISL_3219985, EPI_ISL_3219986, EPI_ISL_3219987, EPI_ISL_3219988, EPI_ISL_3219989, EPI_ISL_3219990, EPI_ISL_3219991, EPI_ISL_3219992, EPI_ISL_3219995, EPI_ISL_3219996, EPI_ISL_3219997, EPI_ISL_3219998, EPI_ISL_3219999, EPI_ISL_3220000, EPI_ISL_3220001, EPI_ISL_3220002, EPI_ISL_3220003, EPI_ISL_3220004, EPI_ISL_3220005, EPI_ISL_3220006, EPI_ISL_3220007, EPI_ISL_3220008, EPI_ISL_3220009, EPI_ISL_3220011, EPI_ISL_3220013, EPI_ISL_3220014, EPI_ISL_3220015, EPI_ISL_3220016, EPI_ISL_3220017, EPI_ISL_3220018, EPI_ISL_3220019, EPI_ISL_3220020, EPI_ISL_3220022, EPI_ISL_3220025, EPI_ISL_3220029, EPI_ISL_3220030, EPI_ISL_3220031, EPI_ISL_3220032, EPI_ISL_3220034, EPI_ISL_3220037, EPI_ISL_3220039, EPI_ISL_3220041, EPI_ISL_3220044, EPI_ISL_3220046, EPI_ISL_3220048, EPI_ISL_3220050, EPI_ISL_3220052, EPI_ISL_3220057, EPI_ISL_3220062, EPI_ISL_3220064, EPI_ISL_3220066, EPI_ISL_3220069, EPI_ISL_3220073, EPI_ISL_3220076, EPI_ISL_3220078, EPI_ISL_3220080, EPI_ISL_3220084, EPI_ISL_3220086, EPI_ISL_3220091, EPI_ISL_3220093, EPI_ISL_3220095, EPI_ISL_3220097, EPI_ISL_3220099, EPI_ISL_3220102, EPI_ISL_3220104, EPI_ISL_3220106, EPI_ISL_3220108, EPI_ISL_3220111, EPI_ISL_3220113, EPI_ISL_3220115, EPI_ISL_3220119, EPI_ISL_3220121, EPI_ISL_3220123, EPI_ISL_3220126, EPI_ISL_3220130, EPI_ISL_3220132, EPI_ISL_3220135, EPI_ISL_3220137, EPI_ISL_3220139, EPI_ISL_3220141, EPI_ISL_3220144, EPI_ISL_3220146, EPI_ISL_3220148, EPI_ISL_3220149, EPI_ISL_3220150, EPI_ISL_3220151, EPI_ISL_3220152, EPI_ISL_3220153, EPI_ISL_3220154, EPI_ISL_3220155, EPI_ISL_3220156, EPI_ISL_3220157, EPI_ISL_3220159, EPI_ISL_3220160, EPI_ISL_3220161, EPI_ISL_3220162, EPI_ISL_3220163, EPI_ISL_3220164, EPI_ISL_3220166, EPI_ISL_3220167, EPI_ISL_3220168, EPI_ISL_3220169, EPI_ISL_3220170, EPI_ISL_3220171, EPI_ISL_3220172, EPI_ISL_3220174, EPI_ISL_3220175, EPI_ISL_3220176, EPI_ISL_3220177, EPI_ISL_3220178, EPI_ISL_3220179, EPI_ISL_3220180, EPI_ISL_3220181, EPI_ISL_3220182, EPI_ISL_3220184, EPI_ISL_3220185, EPI_ISL_3220186, EPI_ISL_3220187, EPI_ISL_3220188, EPI_ISL_3220189, EPI_ISL_3220190, EPI_ISL_3220191, EPI_ISL_3220192, EPI_ISL_3220193, EPI_ISL_3220194, EPI_ISL_3220195, EPI_ISL_3220196, EPI_ISL_3220197, EPI_ISL_3220198, EPI_ISL_3220199, EPI_ISL_3220200, EPI_ISL_3220201, EPI_ISL_3220202, EPI_ISL_3220204, EPI_ISL_3220205, EPI_ISL_3220207, EPI_ISL_3220208, EPI_ISL_3220209, EPI_ISL_3220210, EPI_ISL_3220211, EPI_ISL_3220213, EPI_ISL_3220214, EPI_ISL_3220215, EPI_ISL_3220216, EPI_ISL_3220217, EPI_ISL_3220218, EPI_ISL_3220219, EPI_ISL_3220220, EPI_ISL_3220221, EPI_ISL_3220222, EPI_ISL_3220223, EPI_ISL_3220224, EPI_ISL_3220225, EPI_ISL_3220226, EPI_ISL_3220227, EPI_ISL_3220228, EPI_ISL_3220229, EPI_ISL_3220230, EPI_ISL_3220231, EPI_ISL_3220232, EPI_ISL_3220233, EPI_ISL_3220234, EPI_ISL_3220235, EPI_ISL_3220236, EPI_ISL_3220237, EPI_ISL_3220238, EPI_ISL_3220239, EPI_ISL_3220240, EPI_ISL_3220241, EPI_ISL_3220242, EPI_ISL_3220243, EPI_ISL_3220244, EPI_ISL_3220245, EPI_ISL_3220246, EPI_ISL_3220247, EPI_ISL_3220248, EPI_ISL_3220249, EPI_ISL_3220250, EPI_ISL_3220251, EPI_ISL_3220253, EPI_ISL_3220254, EPI_ISL_3220255, EPI_ISL_3220257, EPI_ISL_3220258, EPI_ISL_3220259, EPI_ISL_3220261, EPI_ISL_3220262, EPI_ISL_3220263, EPI_ISL_3220264, EPI_ISL_3220266, EPI_ISL_3220267, EPI_ISL_3220268, EPI_ISL_3220269, EPI_ISL_3220270, EPI_ISL_3220271, EPI_ISL_3220273, EPI_ISL_3220274, EPI_ISL_3220275, EPI_ISL_3220276, EPI_ISL_3220277, EPI_ISL_3220278, EPI_ISL_3220279, EPI_ISL_3220280, EPI_ISL_3220281, EPI_ISL_3220282, EPI_ISL_3220283, EPI_ISL_3220284, EPI_ISL_3220285, EPI_ISL_3220286, EPI_ISL_3220287, EPI_ISL_3220288, EPI_ISL_3220289, EPI_ISL_3220290, EPI_ISL_3220291, EPI_ISL_3220292, EPI_ISL_3220293, EPI_ISL_3220294, EPI_ISL_3220295, EPI_ISL_3220298, EPI_ISL_3220299, EPI_ISL_3220300, EPI_ISL_3220301, EPI_ISL_3220303, EPI_ISL_3220304, EPI_ISL_3220305, EPI_ISL_3220306, EPI_ISL_3220307, EPI_ISL_3220308, EPI_ISL_3220309, EPI_ISL_3220310, EPI_ISL_3220311, EPI_ISL_3220312, EPI_ISL_3220314, EPI_ISL_3220316, EPI_ISL_3220317, EPI_ISL_3220318, EPI_ISL_3220319, EPI_ISL_3220320, EPI_ISL_3220321, EPI_ISL_3220322, EPI_ISL_3220325, EPI_ISL_3220326, EPI_ISL_3220330, EPI_ISL_3220331, EPI_ISL_3220332, EPI_ISL_3220334, EPI_ISL_3220335, EPI_ISL_3220337, EPI_ISL_3220338, EPI_ISL_3220340, EPI_ISL_3220342, EPI_ISL_3220343, EPI_ISL_3220344, EPI_ISL_3220346, EPI_ISL_3220348, EPI_ISL_3220349, EPI_ISL_3220352, EPI_ISL_3220353, EPI_ISL_3220355, EPI_ISL_3220356, EPI_ISL_3220358, EPI_ISL_3220359, EPI_ISL_3220360, EPI_ISL_3220362, EPI_ISL_3220365, EPI_ISL_3220366, EPI_ISL_3220367, EPI_ISL_3220369, EPI_ISL_3220370, EPI_ISL_3220371, EPI_ISL_3220374, EPI_ISL_3220376, EPI_ISL_3220377, EPI_ISL_3220378, EPI_ISL_3220380, EPI_ISL_3220381, EPI_ISL_3220382, EPI_ISL_3220383, EPI_ISL_3220384, EPI_ISL_3220385, EPI_ISL_3220386, EPI_ISL_3220388, EPI_ISL_3220391, EPI_ISL_3220393, EPI_ISL_3220394, EPI_ISL_3220395, EPI_ISL_3220396, EPI_ISL_3220397, EPI_ISL_3220399, EPI_ISL_3220400, EPI_ISL_3220401, EPI_ISL_3220402, EPI_ISL_3220403, EPI_ISL_3220406, EPI_ISL_3220407, EPI_ISL_3220409, EPI_ISL_3220410, EPI_ISL_3220411, EPI_ISL_3220413, EPI_ISL_3220414, EPI_ISL_3220415, EPI_ISL_3220417, EPI_ISL_3220418, EPI_ISL_3220419, EPI_ISL_3220420, EPI_ISL_3220421, EPI_ISL_3220425, EPI_ISL_3220426, EPI_ISL_3220430, EPI_ISL_3220431, EPI_ISL_3220432, EPI_ISL_3220436, EPI_ISL_3220440, EPI_ISL_3220442, EPI_ISL_3220444, EPI_ISL_3220445, EPI_ISL_3220446, EPI_ISL_3220447, EPI_ISL_3220448, EPI_ISL_3220449, EPI_ISL_3220450, EPI_ISL_3220451, EPI_ISL_3220453, EPI_ISL_3220454, EPI_ISL_3220455, EPI_ISL_3220456, EPI_ISL_3220458, EPI_ISL_3220460, EPI_ISL_3220461, EPI_ISL_3220462, EPI_ISL_3220463, EPI_ISL_3220464, EPI_ISL_3220466, EPI_ISL_3220467, EPI_ISL_3220470, EPI_ISL_3220471, EPI_ISL_3220472, EPI_ISL_3220475, EPI_ISL_3220476, EPI_ISL_3220477, EPI_ISL_3220478, EPI_ISL_3220479, EPI_ISL_3220480, EPI_ISL_3220481, EPI_ISL_3220482, EPI_ISL_3220483, EPI_ISL_3220486, EPI_ISL_3220487, EPI_ISL_3220490, EPI_ISL_3220492, EPI_ISL_3220495, EPI_ISL_3220498, EPI_ISL_3220499, EPI_ISL_3220500, EPI_ISL_3220502, EPI_ISL_3220503, EPI_ISL_3220504, EPI_ISL_3220505, EPI_ISL_3220508, EPI_ISL_3220509, EPI_ISL_3220510, EPI_ISL_3220511, EPI_ISL_3220512, EPI_ISL_3220515, EPI_ISL_3220516, EPI_ISL_3220517, EPI_ISL_3220518, EPI_ISL_3220519, EPI_ISL_3220520, EPI_ISL_3220522, EPI_ISL_3220523, EPI_ISL_3220524, EPI_ISL_3220525, EPI_ISL_3220527, EPI_ISL_3220528, EPI_ISL_3220529, EPI_ISL_3220530, EPI_ISL_3220531, EPI_ISL_3220532, EPI_ISL_3220533, EPI_ISL_3220534, EPI_ISL_3220535, EPI_ISL_3220539, EPI_ISL_3220540, EPI_ISL_3220541, EPI_ISL_3220543, EPI_ISL_3220544, EPI_ISL_3220545, EPI_ISL_3220546, EPI_ISL_3220547, EPI_ISL_3220549, EPI_ISL_3220550, EPI_ISL_3220551, EPI_ISL_3220552, EPI_ISL_3220553, EPI_ISL_3220557, EPI_ISL_3220559, EPI_ISL_3220560, EPI_ISL_3220562, EPI_ISL_3220563, EPI_ISL_3220564, EPI_ISL_3220565, EPI_ISL_3220568, EPI_ISL_3220569, EPI_ISL_3220570, EPI_ISL_3220571, EPI_ISL_3220572, EPI_ISL_3220573, EPI_ISL_3220574, EPI_ISL_3220575, EPI_ISL_3220579, EPI_ISL_3220581, EPI_ISL_3220583, EPI_ISL_3220585, EPI_ISL_3220586, EPI_ISL_3220587, EPI_ISL_3220591, EPI_ISL_3220592, EPI_ISL_3220593, EPI_ISL_3220595, EPI_ISL_3220597, EPI_ISL_3220598, EPI_ISL_3220599, EPI_ISL_3220600, EPI_ISL_3220603, EPI_ISL_3220604, EPI_ISL_3220606, EPI_ISL_3220607, EPI_ISL_3220608, EPI_ISL_3220609, EPI_ISL_3220611, EPI_ISL_3220613, EPI_ISL_3220614, EPI_ISL_3220618, EPI_ISL_3220619, EPI_ISL_3220622, EPI_ISL_3220623, EPI_ISL_3220625, EPI_ISL_3220627, EPI_ISL_3220629, EPI_ISL_3220630, EPI_ISL_3220632, EPI_ISL_3220633, EPI_ISL_3220634, EPI_ISL_3220635, EPI_ISL_3220636, EPI_ISL_3220639, EPI_ISL_3220642, EPI_ISL_3220645, EPI_ISL_3220646, EPI_ISL_3220648, EPI_ISL_3220649, EPI_ISL_3220650, EPI_ISL_3220652, EPI_ISL_3220653, EPI_ISL_3220654, EPI_ISL_3220655, EPI_ISL_3220656, EPI_ISL_3220657, EPI_ISL_3220658, EPI_ISL_3220659, EPI_ISL_3220660, EPI_ISL_3220663, EPI_ISL_3220664, EPI_ISL_3220665, EPI_ISL_3220671, EPI_ISL_3220673, EPI_ISL_3220674, EPI_ISL_3220677, EPI_ISL_3220678, EPI_ISL_3220680, EPI_ISL_3220681, EPI_ISL_3220682, EPI_ISL_3220683, EPI_ISL_3220684, EPI_ISL_3220686, EPI_ISL_3220687, EPI_ISL_3220690, EPI_ISL_3220692, EPI_ISL_3220693, EPI_ISL_3220694, EPI_ISL_3220695, EPI_ISL_3220696, EPI_ISL_3220697, EPI_ISL_3220698, EPI_ISL_3220700, EPI_ISL_3220701, EPI_ISL_3220702, EPI_ISL_3220703, EPI_ISL_3220705, EPI_ISL_3220706, EPI_ISL_3220707, EPI_ISL_3220709, EPI_ISL_3220710, EPI_ISL_3220711, EPI_ISL_3220712, EPI_ISL_3220715, EPI_ISL_3220717, EPI_ISL_3220718, EPI_ISL_3220719, EPI_ISL_3220721, EPI_ISL_3220722, EPI_ISL_3220723, EPI_ISL_3220724, EPI_ISL_3220725, EPI_ISL_3220726, EPI_ISL_3220729, EPI_ISL_3220730, EPI_ISL_3220731, EPI_ISL_3220732, EPI_ISL_3220733, EPI_ISL_3220737, EPI_ISL_3220738, EPI_ISL_3220739, EPI_ISL_3220740, EPI_ISL_3220742, EPI_ISL_3220743, EPI_ISL_3220745, EPI_ISL_3220749, EPI_ISL_3220750, EPI_ISL_3220751, EPI_ISL_3220754, EPI_ISL_3220755, EPI_ISL_3220756, EPI_ISL_3220757, EPI_ISL_3220758, EPI_ISL_3220759, EPI_ISL_3220761, EPI_ISL_3220762, EPI_ISL_3220764, EPI_ISL_3220765, EPI_ISL_3220766, EPI_ISL_3220770, EPI_ISL_3220771, EPI_ISL_3220776, EPI_ISL_3220777, EPI_ISL_3220778, EPI_ISL_3220779, EPI_ISL_3220780, EPI_ISL_3220781, EPI_ISL_3220782, EPI_ISL_3220784, EPI_ISL_3220785, EPI_ISL_3220786, EPI_ISL_3220787, EPI_ISL_3220791, EPI_ISL_3220793, EPI_ISL_3220794, EPI_ISL_3220795, EPI_ISL_3220796, EPI_ISL_3220798, EPI_ISL_3220801, EPI_ISL_3220802, EPI_ISL_3220803, EPI_ISL_3220804, EPI_ISL_3220805, EPI_ISL_3220807, EPI_ISL_3220808, EPI_ISL_3220809, EPI_ISL_3220810, EPI_ISL_3220811, EPI_ISL_3220812, EPI_ISL_3220813, EPI_ISL_3220814, EPI_ISL_3220816, EPI_ISL_3220818, EPI_ISL_3220819, EPI_ISL_3220820, EPI_ISL_3220821, EPI_ISL_3220823, EPI_ISL_3220824, EPI_ISL_3220825, EPI_ISL_3220826, EPI_ISL_3220828, EPI_ISL_3220830, EPI_ISL_3220831, EPI_ISL_3220832, EPI_ISL_3220833, EPI_ISL_3220834, EPI_ISL_3220836, EPI_ISL_3220837, EPI_ISL_3220838, EPI_ISL_3220839, EPI_ISL_3220841, EPI_ISL_3220843, EPI_ISL_3220844, EPI_ISL_3220846, EPI_ISL_3220847, EPI_ISL_3220848, EPI_ISL_3220850, EPI_ISL_3220852, EPI_ISL_3220855, EPI_ISL_3220857, EPI_ISL_3220858, EPI_ISL_3220859, EPI_ISL_3220860, EPI_ISL_3220861, EPI_ISL_3220862, EPI_ISL_3220863, EPI_ISL_3220865, EPI_ISL_3220867, EPI_ISL_3220868, EPI_ISL_3220871, EPI_ISL_3220872, EPI_ISL_3220874, EPI_ISL_3220876, EPI_ISL_3220877, EPI_ISL_3220879, EPI_ISL_3220881, EPI_ISL_3220883, EPI_ISL_3220884, EPI_ISL_3220885, EPI_ISL_3220887, EPI_ISL_3220889, EPI_ISL_3220890, EPI_ISL_3220891, EPI_ISL_3220893, EPI_ISL_3220895, EPI_ISL_3220897, EPI_ISL_3220898, EPI_ISL_3220899, EPI_ISL_3220900, EPI_ISL_3220901, EPI_ISL_3220902, EPI_ISL_3220905, EPI_ISL_3220906, EPI_ISL_3220910, EPI_ISL_3220913, EPI_ISL_3220914, EPI_ISL_3220915, EPI_ISL_3220916, EPI_ISL_3220918, EPI_ISL_3220919, EPI_ISL_3220920, EPI_ISL_3220921, EPI_ISL_3220922, EPI_ISL_3220923, EPI_ISL_3220925, EPI_ISL_3220926, EPI_ISL_3220928, EPI_ISL_3220932, EPI_ISL_3220933, EPI_ISL_3220934, EPI_ISL_3220936, EPI_ISL_3220939, EPI_ISL_3220940, EPI_ISL_3220943, EPI_ISL_3220944, EPI_ISL_3220946, EPI_ISL_3220947, EPI_ISL_3220948, EPI_ISL_3220950, EPI_ISL_3220951, EPI_ISL_3220952, EPI_ISL_3220953, EPI_ISL_3220954, EPI_ISL_3220955, EPI_ISL_3220956, EPI_ISL_3220957, EPI_ISL_3220959, EPI_ISL_3220961, EPI_ISL_3220962, EPI_ISL_3220963, EPI_ISL_3220964, EPI_ISL_3220965, EPI_ISL_3220966, EPI_ISL_3220967, EPI_ISL_3220968, EPI_ISL_3220970, EPI_ISL_3220972, EPI_ISL_3220973, EPI_ISL_3220974, EPI_ISL_3220975, EPI_ISL_3220977, EPI_ISL_3220978, EPI_ISL_3220981, EPI_ISL_3220982, EPI_ISL_3220983, EPI_ISL_3220984, EPI_ISL_3220987, EPI_ISL_3220988, EPI_ISL_3220991, EPI_ISL_3220992, EPI_ISL_3220993, EPI_ISL_3220995, EPI_ISL_3220996, EPI_ISL_3220998, EPI_ISL_3220999, EPI_ISL_3221001, EPI_ISL_3221002, EPI_ISL_3221004, EPI_ISL_3221006, EPI_ISL_3221009, EPI_ISL_3221010, EPI_ISL_3221011, EPI_ISL_3221012, EPI_ISL_3221013, EPI_ISL_3221015, EPI_ISL_3221018, EPI_ISL_3221019, EPI_ISL_3221020, EPI_ISL_3221021, EPI_ISL_3221024, EPI_ISL_3221025, EPI_ISL_3221027, EPI_ISL_3221028, EPI_ISL_3221030, EPI_ISL_3221031, EPI_ISL_3221032, EPI_ISL_3221033, EPI_ISL_3221034, EPI_ISL_3221035, EPI_ISL_3221036, EPI_ISL_3221037, EPI_ISL_3221040, EPI_ISL_3221042, EPI_ISL_3221043, EPI_ISL_3221044, EPI_ISL_3221045, EPI_ISL_3221050, EPI_ISL_3221053, EPI_ISL_3221054 | | | |
| see above | Aegis Sciences Corporation | Centers for Disease Control and Prevention Division of Viral Diseases, Pathogen Discovery | Dakota Howard, Dhwani Batra, Peter W. Cook, Kara Moser, Adrian Paskey, Jason Caravas, Benjamin Rambo-Martin, Shatavia Morrison, Christopher Gulvick, Scott Sammons, Yvette Unoarumhi, Darlene Wagner, Matthew Schmerer, Cyndi Clark, Patrick Campbell, Rob Case, Vikramsinha Ghorpade, Holly Houdeshell, Ola Kvalvaag, Dillon Nall, Ethan Sanders, Alec Vest, Shaun Westlund, Matthew Hardison, Clinton R. Paden, Duncan MacCannell |
| EPI_ISL_3221055, EPI_ISL_3221056, EPI_ISL_3221057 | Genome Information Research Center, Research Institute for Microbial Diseases, Osaka University | Genome Information Research Center, Research Institute for Microbial Diseases, Osaka University | Daisuke Motooka, Kentaro Tanaka, Hiroya Oki, and Shota Nakamura |
| EPI_ISL_3221058 | Louisiana Office of Public Heath Laboratory | LSUHS Emerging Viral Threat Laboratory | Gregory L. Ware, Alexander Mijalis, Jeremy P. Kamil, Jennifer L. Carroll, Maarten Van Diest, Rona S. Scott, Andrew D. Yurochko, Christopher G. Kevil, John A. Vanchiere |
| EPI_ISL_3221059 | Green Clinic Tech Care, Lambright Sports and Wellness Center, Louisiana Tech University | LSUHS Emerging Viral Threat Laboratory | Jamie J. Newman, Paul Kim, Nancy B. Adkins |
| EPI_ISL_3221060, EPI_ISL_3221061, EPI_ISL_3221062, EPI_ISL_3221063, EPI_ISL_3221064, EPI_ISL_3221065, EPI_ISL_3221066, EPI_ISL_3221067, EPI_ISL_3221068, EPI_ISL_3221069, EPI_ISL_3221070, EPI_ISL_3221071, EPI_ISL_3221072, EPI_ISL_3221073, EPI_ISL_3221074, EPI_ISL_3221075, EPI_ISL_3221076 | | | |
| see above | Louisiana Office of Public Heath Laboratory | LSUHS Emerging Viral Threat Laboratory | Gregory L. Ware, Alexander Mijalis, Jeremy P. Kamil, Jennifer L. Carroll, Maarten Van Diest, Rona S. Scott, Andrew D. Yurochko, Christopher G. Kevil, John A. Vanchiere |
| EPI_ISL_3221077 | LSUHS Emerging Viral Threat Laboratory | LSUHS Emerging Viral Threat Laboratory | Gregory L. Ware, Alexander Mijalis, Jeremy P. Kamil, Jennifer L. Carroll, Maarten Van Diest, Rona S. Scott, Andrew D. Yurochko, Christopher G. Kevil, John A. Vanchiere |
| EPI_ISL_3221078, EPI_ISL_3221079 | Louisiana Office of Public Heath Laboratory | LSUHS Emerging Viral Threat Laboratory | Gregory L. Ware, Alexander Mijalis, Jeremy P. Kamil, Jennifer L. Carroll, Maarten Van Diest, Rona S. Scott, Andrew D. Yurochko, Christopher G. Kevil, John A. Vanchiere |
| EPI_ISL_3221080 | LSUHS Emerging Viral Threat Laboratory | LSUHS Emerging Viral Threat Laboratory | Gregory L. Ware, Alexander Mijalis, Jeremy P. Kamil, Jennifer L. Carroll, Maarten Van Diest, Rona S. Scott, Andrew D. Yurochko, Christopher G. Kevil, John A. Vanchiere |
| EPI_ISL_3221081, EPI_ISL_3221084, EPI_ISL_3221085, EPI_ISL_3221086 | Louisiana Office of Public Heath Laboratory | LSUHS Emerging Viral Threat Laboratory | Gregory L. Ware, Alexander Mijalis, Jeremy P. Kamil, Jennifer L. Carroll, Maarten Van Diest, Rona S. Scott, Andrew D. Yurochko, Christopher G. Kevil, John A. Vanchiere |
| EPI_ISL_3221087 | Ochsner LSU Health | LSUHS Emerging Viral Threat Laboratory | Gregory L. Ware, Alexander Mijalis, Jeremy P. Kamil, Jennifer L. Carroll, Maarten Van Diest, Rona S. Scott, Andrew D. Yurochko, Christopher G. Kevil, John A. Vanchiere |
| EPI_ISL_3221088, EPI_ISL_3221090, EPI_ISL_3221091 | Louisiana Office of Public Heath Laboratory | LSUHS Emerging Viral Threat Laboratory | Gregory L. Ware, Alexander Mijalis, Jeremy P. Kamil, Jennifer L. Carroll, Maarten Van Diest, Rona S. Scott, Andrew D. Yurochko, Christopher G. Kevil, John A. Vanchiere |
| EPI_ISL_3221092 | Ochsner LSU Health | LSUHS Emerging Viral Threat Laboratory | Gregory L. Ware, Alexander Mijalis, Jeremy P. Kamil, Jennifer L. Carroll, Maarten Van Diest, Rona S. Scott, Andrew D. Yurochko, Christopher G. Kevil, John A. Vanchiere |
| EPI_ISL_3221094 | Louisiana Office of Public Heath Laboratory | LSUHS Emerging Viral Threat Laboratory | Gregory L. Ware, Alexander Mijalis, Jeremy P. Kamil, Jennifer L. Carroll, Maarten Van Diest, Rona S. Scott, Andrew D. Yurochko, Christopher G. Kevil, John A. Vanchiere |
| EPI_ISL_3221095 | LSUHS Emerging Viral Threat Laboratory | LSUHS Emerging Viral Threat Laboratory | Gregory L. Ware, Alexander Mijalis, Jeremy P. Kamil, Jennifer L. Carroll, Maarten Van Diest, Rona S. Scott, Andrew D. Yurochko, Christopher G. Kevil, John A. Vanchiere |
| EPI_ISL_3221096 | Louisiana Office of Public Heath Laboratory | LSUHS Emerging Viral Threat Laboratory | Gregory L. Ware, Alexander Mijalis, Jeremy P. Kamil, Jennifer L. Carroll, Maarten Van Diest, Rona S. Scott, Andrew D. Yurochko, Christopher G. Kevil, John A. Vanchiere |
| EPI_ISL_3221097 | LSUHS Emerging Viral Threat Laboratory | LSUHS Emerging Viral Threat Laboratory | Gregory L. Ware, Alexander Mijalis, Jeremy P. Kamil, Jennifer L. Carroll, Maarten Van Diest, Rona S. Scott, Andrew D. Yurochko, Christopher G. Kevil, John A. Vanchiere |
| EPI_ISL_3221098 | Louisiana Office of Public Heath Laboratory | LSUHS Emerging Viral Threat Laboratory | Gregory L. Ware, Alexander Mijalis, Jeremy P. Kamil, Jennifer L. Carroll, Maarten Van Diest, Rona S. Scott, Andrew D. Yurochko, Christopher G. Kevil, John A. Vanchiere |

| | | | |
|---|---|---|---|
| EPI_ISL_3221099 | Ochsner LSU Health | LSUHS Emerging Viral Threat Laboratory | Gregory L. Ware, Alexander Mijalis, Jeremy P. Kamil, Jennifer L. Carroll, Maarten Van Diest, Rona S. Scott, Andrew D. Yurochko, Christopher G. Kevil, John A. Vanchiere |
| EPI_ISL_3221100, EPI_ISL_3221101, EPI_ISL_3221102, EPI_ISL_3221103, EPI_ISL_3221104 | Louisiana Office of Public Heath Laboratory | LSUHS Emerging Viral Threat Laboratory | Gregory L. Ware, Alexander Mijalis, Jeremy P. Kamil, Jennifer L. Carroll, Maarten Van Diest, Rona S. Scott, Andrew D. Yurochko, Christopher G. Kevil, John A. Vanchiere |
| EPI_ISL_3221105 | LSUHS Emerging Viral Threat Laboratory | LSUHS Emerging Viral Threat Laboratory | Gregory L. Ware, Alexander Mijalis, Jeremy P. Kamil, Jennifer L. Carroll, Maarten Van Diest, Rona S. Scott, Andrew D. Yurochko, Christopher G. Kevil, John A. Vanchiere |
| EPI_ISL_3221106, EPI_ISL_3221107 | Louisiana Office of Public Heath Laboratory | LSUHS Emerging Viral Threat Laboratory | Gregory L. Ware, Alexander Mijalis, Jeremy P. Kamil, Jennifer L. Carroll, Maarten Van Diest, Rona S. Scott, Andrew D. Yurochko, Christopher G. Kevil, John A. Vanchiere |
| EPI_ISL_3221108 | LSUHS Emerging Viral Threat Laboratory | LSUHS Emerging Viral Threat Laboratory | Gregory L. Ware, Alexander Mijalis, Jeremy P. Kamil, Jennifer L. Carroll, Maarten Van Diest, Rona S. Scott, Andrew D. Yurochko, Christopher G. Kevil, John A. Vanchiere |
| EPI_ISL_3221110 | Louisiana Office of Public Heath Laboratory | LSUHS Emerging Viral Threat Laboratory | Gregory L. Ware, Alexander Mijalis, Jeremy P. Kamil, Jennifer L. Carroll, Maarten Van Diest, Rona S. Scott, Andrew D. Yurochko, Christopher G. Kevil, John A. Vanchiere |
| EPI_ISL_3221111, EPI_ISL_3221112 | LSUHS Emerging Viral Threat Laboratory | LSUHS Emerging Viral Threat Laboratory | Gregory L. Ware, Alexander Mijalis, Jeremy P. Kamil, Jennifer L. Carroll, Maarten Van Diest, Rona S. Scott, Andrew D. Yurochko, Christopher G. Kevil, John A. Vanchiere |
| EPI_ISL_3221113, EPI_ISL_3221114, EPI_ISL_3221115, EPI_ISL_3221116, EPI_ISL_3221117, EPI_ISL_3221119, EPI_ISL_3221120, EPI_ISL_3221121, EPI_ISL_3221122, EPI_ISL_3221123 | Louisiana Office of Public Heath Laboratory | LSUHS Emerging Viral Threat Laboratory | Gregory L. Ware, Alexander Mijalis, Jeremy P. Kamil, Jennifer L. Carroll, Maarten Van Diest, Rona S. Scott, Andrew D. Yurochko, Christopher G. Kevil, John A. Vanchiere |
| EPI_ISL_3221124 | LSUHS Emerging Viral Threat Laboratory | LSUHS Emerging Viral Threat Laboratory | Gregory L. Ware, Alexander Mijalis, Jeremy P. Kamil, Jennifer L. Carroll, Maarten Van Diest, Rona S. Scott, Andrew D. Yurochko, Christopher G. Kevil, John A. Vanchiere |
| EPI_ISL_3221125 | Louisiana Office of Public Heath Laboratory | LSUHS Emerging Viral Threat Laboratory | Gregory L. Ware, Alexander Mijalis, Jeremy P. Kamil, Jennifer L. Carroll, Maarten Van Diest, Rona S. Scott, Andrew D. Yurochko, Christopher G. Kevil, John A. Vanchiere |
| EPI_ISL_3221126 | Ochsner LSU Health | LSUHS Emerging Viral Threat Laboratory | Gregory L. Ware, Alexander Mijalis, Jeremy P. Kamil, Jennifer L. Carroll, Maarten Van Diest, Rona S. Scott, Andrew D. Yurochko, Christopher G. Kevil, John A. Vanchiere |
| EPI_ISL_3221127, EPI_ISL_3221128, EPI_ISL_3221129, EPI_ISL_3221130, EPI_ISL_3221131, EPI_ISL_3221132 | Louisiana Office of Public Heath Laboratory | LSUHS Emerging Viral Threat Laboratory | Gregory L. Ware, Alexander Mijalis, Jeremy P. Kamil, Jennifer L. Carroll, Maarten Van Diest, Rona S. Scott, Andrew D. Yurochko, Christopher G. Kevil, John A. Vanchiere |
| EPI_ISL_3221133 | LSUHS Emerging Viral Threat Laboratory | LSUHS Emerging Viral Threat Laboratory | Gregory L. Ware, Alexander Mijalis, Jeremy P. Kamil, Jennifer L. Carroll, Maarten Van Diest, Rona S. Scott, Andrew D. Yurochko, Christopher G. Kevil, John A. Vanchiere |
| EPI_ISL_3221134 | Louisiana Office of Public Heath Laboratory | LSUHS Emerging Viral Threat Laboratory | Gregory L. Ware, Alexander Mijalis, Jeremy P. Kamil, Jennifer L. Carroll, Maarten Van Diest, Rona S. Scott, Andrew D. Yurochko, Christopher G. Kevil, John A. Vanchiere |
| EPI_ISL_3221135 | The Health Hut | LSUHS Emerging Viral Threat Laboratory | Jamie J. Newman, Paul Kim, Jacquelyn K. White |
| EPI_ISL_3221136, EPI_ISL_3221137, EPI_ISL_3221138, EPI_ISL_3221139, EPI_ISL_3221140, EPI_ISL_3221141, EPI_ISL_3221142, EPI_ISL_3221143 | Louisiana Office of Public Heath Laboratory | LSUHS Emerging Viral Threat Laboratory | Gregory L. Ware, Alexander Mijalis, Jeremy P. Kamil, Jennifer L. Carroll, Maarten Van Diest, Rona S. Scott, Andrew D. Yurochko, Christopher G. Kevil, John A. Vanchiere |
| EPI_ISL_3221144 | LSUHS Emerging Viral Threat Laboratory | LSUHS Emerging Viral Threat Laboratory | Gregory L. Ware, Alexander Mijalis, Jeremy P. Kamil, Jennifer L. Carroll, Maarten Van Diest, Rona S. Scott, Andrew D. Yurochko, Christopher G. Kevil, John A. Vanchiere |
| EPI_ISL_3221145, EPI_ISL_3221146 | Louisiana Office of Public Heath Laboratory | LSUHS Emerging Viral Threat Laboratory | Gregory L. Ware, Alexander Mijalis, Jeremy P. Kamil, Jennifer L. Carroll, Maarten Van Diest, Rona S. Scott, Andrew D. Yurochko, Christopher G. Kevil, John A. Vanchiere |
| EPI_ISL_3221147 | LSUHS Emerging Viral Threat Laboratory | LSUHS Emerging Viral Threat Laboratory | Gregory L. Ware, Alexander Mijalis, Jeremy P. Kamil, Jennifer L. Carroll, Maarten Van Diest, Rona S. Scott, Andrew D. Yurochko, Christopher G. Kevil, John A. Vanchiere |
| EPI_ISL_3221148, EPI_ISL_3221149 | Louisiana Office of Public Heath Laboratory | LSUHS Emerging Viral Threat Laboratory | Gregory L. Ware, Alexander Mijalis, Jeremy P. Kamil, Jennifer L. Carroll, Maarten Van Diest, Rona S. Scott, Andrew D. Yurochko, Christopher G. Kevil, John A. Vanchiere |
| EPI_ISL_3221150 | LSUHS Emerging Viral Threat Laboratory | LSUHS Emerging Viral Threat Laboratory | Gregory L. Ware, Alexander Mijalis, Jeremy P. Kamil, Jennifer L. Carroll, Maarten Van Diest, Rona S. Scott, Andrew D. Yurochko, Christopher G. Kevil, John A. Vanchiere |
| EPI_ISL_3221151, EPI_ISL_3221152, EPI_ISL_3221153, EPI_ISL_3221154, EPI_ISL_3221155, EPI_ISL_3221156, EPI_ISL_3221157, EPI_ISL_3221158, EPI_ISL_3221159, EPI_ISL_3221160, EPI_ISL_3221161, EPI_ISL_3221162 | | | |
| see above | Louisiana Office of Public Heath Laboratory | LSUHS Emerging Viral Threat Laboratory | Gregory L. Ware, Alexander Mijalis, Jeremy P. Kamil, Jennifer L. Carroll, Maarten Van Diest, Rona S. Scott, Andrew D. Yurochko, Christopher G. Kevil, John A. Vanchiere |
| EPI_ISL_3221163 | LSUHS Emerging Viral Threat Laboratory | LSUHS Emerging Viral Threat Laboratory | Gregory L. Ware, Alexander Mijalis, Jeremy P. Kamil, Jennifer L. Carroll, Maarten Van Diest, Rona S. Scott, Andrew D. Yurochko, Christopher G. Kevil, John A. Vanchiere |
| EPI_ISL_3221164, EPI_ISL_3221165, EPI_ISL_3221166, EPI_ISL_3221167 | Louisiana Office of Public Heath Laboratory | LSUHS Emerging Viral Threat Laboratory | Gregory L. Ware, Alexander Mijalis, Jeremy P. Kamil, Jennifer L. Carroll, Maarten Van Diest, Rona S. Scott, Andrew D. Yurochko, Christopher G. Kevil, John A. Vanchiere |
| EPI_ISL_3221168, EPI_ISL_3221169 | LSUHS Emerging Viral Threat Laboratory | LSUHS Emerging Viral Threat Laboratory | Gregory L. Ware, Alexander Mijalis, Jeremy P. Kamil, Jennifer L. Carroll, Maarten Van Diest, Rona S. Scott, Andrew D. Yurochko, Christopher G. Kevil, John A. Vanchiere |
| EPI_ISL_3221170, EPI_ISL_3221171 | Louisiana Office of Public Heath Laboratory | LSUHS Emerging Viral Threat Laboratory | Gregory L. Ware, Alexander Mijalis, Jeremy P. Kamil, Jennifer L. Carroll, Maarten Van Diest, Rona S. Scott, Andrew D. Yurochko, Christopher G. Kevil, John A. Vanchiere |
| EPI_ISL_3221172 | LSUHS Emerging Viral Threat Laboratory | LSUHS Emerging Viral Threat Laboratory | Gregory L. Ware, Alexander Mijalis, Jeremy P. Kamil, Jennifer L. Carroll, Maarten Van Diest, Rona S. Scott, Andrew D. Yurochko, Christopher G. Kevil, John A. Vanchiere |
| EPI_ISL_3221173, EPI_ISL_3221174, EPI_ISL_3221175 | Louisiana Office of Public Heath Laboratory | LSUHS Emerging Viral Threat Laboratory | Gregory L. Ware, Alexander Mijalis, Jeremy P. Kamil, Jennifer L. Carroll, Maarten Van Diest, Rona S. Scott, Andrew D. Yurochko, Christopher G. Kevil, John A. Vanchiere |
| EPI_ISL_3221176 | Grambling/Foster Johnson Health Center, Grambling State University | LSUHS Emerging Viral Threat Laboratory | Paul Kim, Jamie J. Newman, Audrey Y. Kim, Patrice Lewis-Outley, Dawn Holmes, Kimbreyana McDuffie, Calandra Moore, Lori Brisco, Sharon Greer, Albertina Riser |
| EPI_ISL_3221177 | LSUHS Emerging Viral Threat Laboratory | LSUHS Emerging Viral Threat Laboratory | Gregory L. Ware, Alexander Mijalis, Jeremy P. Kamil, Jennifer L. Carroll, Maarten Van Diest, Rona S. Scott, Andrew D. Yurochko, Christopher G. Kevil, John A. Vanchiere |
| EPI_ISL_3221178 | Louisiana Office of Public Heath Laboratory | LSUHS Emerging Viral Threat Laboratory | Gregory L. Ware, Alexander Mijalis, Jeremy P. Kamil, Jennifer L. Carroll, Maarten Van Diest, Rona S. Scott, Andrew D. Yurochko, Christopher G. Kevil, John A. Vanchiere |
| EPI_ISL_3221179 | LSUHS Emerging Viral Threat Laboratory | LSUHS Emerging Viral Threat Laboratory | Gregory L. Ware, Alexander Mijalis, Jeremy P. Kamil, Jennifer L. Carroll, Maarten Van Diest, Rona S. Scott, Andrew D. Yurochko, Christopher G. Kevil, John A. Vanchiere |
| EPI_ISL_3221180, EPI_ISL_3221181, EPI_ISL_3221182, EPI_ISL_3221183, EPI_ISL_3221185 | Louisiana Office of Public Heath Laboratory | LSUHS Emerging Viral Threat Laboratory | Gregory L. Ware, Alexander Mijalis, Jeremy P. Kamil, Jennifer L. Carroll, Maarten Van Diest, Rona S. Scott, Andrew D. Yurochko, Christopher G. Kevil, John A. Vanchiere |
| EPI_ISL_3221186 | The Health Hut | LSUHS Emerging Viral Threat Laboratory | Jamie J. Newman, Paul Kim, Jacquelyn K. White |
| EPI_ISL_3221187, EPI_ISL_3221188, EPI_ISL_3221189 | Louisiana Office of Public Heath Laboratory | LSUHS Emerging Viral Threat Laboratory | Gregory L. Ware, Alexander Mijalis, Jeremy P. Kamil, Jennifer L. Carroll, Maarten Van Diest, Rona S. Scott, Andrew D. Yurochko, Christopher G. Kevil, John A. Vanchiere |
| EPI_ISL_3221190 | LSUHS Emerging Viral Threat Laboratory | LSUHS Emerging Viral Threat Laboratory | Gregory L. Ware, Alexander Mijalis, Jeremy P. Kamil, Jennifer L. Carroll, Maarten Van Diest, Rona S. Scott, Andrew D. Yurochko, Christopher G. Kevil, John A. Vanchiere |
| EPI_ISL_3221191, EPI_ISL_3221192 | Louisiana Office of Public Heath Laboratory | LSUHS Emerging Viral Threat Laboratory | Gregory L. Ware, Alexander Mijalis, Jeremy P. Kamil, Jennifer L. Carroll, Maarten Van Diest, Rona S. Scott, Andrew D. Yurochko, Christopher G. Kevil, John A. Vanchiere |
| EPI_ISL_3221193, EPI_ISL_3221194 | LSUHS Emerging Viral Threat Laboratory | LSUHS Emerging Viral Threat Laboratory | Gregory L. Ware, Alexander Mijalis, Jeremy P. Kamil, Jennifer L. Carroll, Maarten Van Diest, Rona S. Scott, Andrew D. Yurochko, Christopher G. Kevil, John A. Vanchiere |
| EPI_ISL_3221195, EPI_ISL_3221196, EPI_ISL_3221198, EPI_ISL_3221199 | Louisiana Office of Public Heath Laboratory | LSUHS Emerging Viral Threat Laboratory | Gregory L. Ware, Alexander Mijalis, Jeremy P. Kamil, Jennifer L. Carroll, Maarten Van Diest, Rona S. Scott, Andrew D. Yurochko, Christopher G. Kevil, John A. Vanchiere |
| EPI_ISL_3221200 | LSUHS Emerging Viral Threat Laboratory | LSUHS Emerging Viral Threat Laboratory | Gregory L. Ware, Alexander Mijalis, Jeremy P. Kamil, Jennifer L. Carroll, Maarten Van Diest, Rona S. Scott, Andrew D. Yurochko, Christopher G. Kevil, John A. Vanchiere |
| EPI_ISL_3221201, EPI_ISL_3221202 | Louisiana Office of Public Heath Laboratory | LSUHS Emerging Viral Threat Laboratory | Gregory L. Ware, Alexander Mijalis, Jeremy P. Kamil, Jennifer L. Carroll, Maarten Van Diest, Rona S. Scott, Andrew D. Yurochko, Christopher G. Kevil, John A. Vanchiere |

| | | | |
|---|---|---|---|
| EPI_ISL_3221203 | Grambling/Foster Johnson Health Center, Grambling State University | LSUHS Emerging Viral Threat Laboratory | Paul Kim, Jamie J. Newman, Audrey Y. Kim, Patrice Lewis-Outley, Dawn Holmes, Kimbreyana McDuffie, Calandra Moore, Lori Brisco, Sharon Greer, Albertina Riser |
| EPI_ISL_3221204 | LSUHS Emerging Viral Threat Laboratory | LSUHS Emerging Viral Threat Laboratory | Gregory L. Ware, Alexander Mijalis, Jeremy P. Kamil, Jennifer L. Carroll, Maarten Van Diest, Rona S. Scott, Andrew D. Yurochko, Christopher G. Kevil, John A. Vanchiere |
| EPI_ISL_3221206, EPI_ISL_3221207, EPI_ISL_3221209, EPI_ISL_3221210, EPI_ISL_3221212 | Louisiana Office of Public Heath Laboratory | LSUHS Emerging Viral Threat Laboratory | Gregory L. Ware, Alexander Mijalis, Jeremy P. Kamil, Jennifer L. Carroll, Maarten Van Diest, Rona S. Scott, Andrew D. Yurochko, Christopher G. Kevil, John A. Vanchiere |
| EPI_ISL_3221213 | LSUHS Emerging Viral Threat Laboratory | LSUHS Emerging Viral Threat Laboratory | Gregory L. Ware, Alexander Mijalis, Jeremy P. Kamil, Jennifer L. Carroll, Maarten Van Diest, Rona S. Scott, Andrew D. Yurochko, Christopher G. Kevil, John A. Vanchiere |
| EPI_ISL_3221214 | The Health Hut | LSUHS Emerging Viral Threat Laboratory | Jamie J. Newman, Paul Kim, Jacquelyn K. White |
| EPI_ISL_3221215 | Louisiana Office of Public Heath Laboratory | LSUHS Emerging Viral Threat Laboratory | Gregory L. Ware, Alexander Mijalis, Jeremy P. Kamil, Jennifer L. Carroll, Maarten Van Diest, Rona S. Scott, Andrew D. Yurochko, Christopher G. Kevil, John A. Vanchiere |
| EPI_ISL_3221216 | The Health Hut | LSUHS Emerging Viral Threat Laboratory | Jamie J. Newman, Paul Kim, Jacquelyn K. White |
| EPI_ISL_3221218 | LSUHS Emerging Viral Threat Laboratory | LSUHS Emerging Viral Threat Laboratory | Gregory L. Ware, Alexander Mijalis, Jeremy P. Kamil, Jennifer L. Carroll, Maarten Van Diest, Rona S. Scott, Andrew D. Yurochko, Christopher G. Kevil, John A. Vanchiere |
| EPI_ISL_3221219 | Willis-Knighton Medical Center Hospital Laboratory | LSUHS Emerging Viral Threat Laboratory | Gregory L. Ware, Alexander Mijalis, Jeremy P. Kamil, Jennifer L. Carroll, Maarten Van Diest, Rona S. Scott, Andrew D. Yurochko, Christopher G. Kevil, John A. Vanchiere |
| EPI_ISL_3221220 | Louisiana Office of Public Heath Laboratory | LSUHS Emerging Viral Threat Laboratory | Gregory L. Ware, Alexander Mijalis, Jeremy P. Kamil, Jennifer L. Carroll, Maarten Van Diest, Rona S. Scott, Andrew D. Yurochko, Christopher G. Kevil, John A. Vanchiere |
| EPI_ISL_3221225, EPI_ISL_3221226, EPI_ISL_3221227, EPI_ISL_3221228, EPI_ISL_3221229, EPI_ISL_3221230, EPI_ISL_3221231, EPI_ISL_3221232, EPI_ISL_3221233, EPI_ISL_3221234, EPI_ISL_3221235 | | | |
| see above | Area of Virology, Serology and Virology Division (SAViD), New South Wales Health Pathology Randwick | Virology Research Laboratory; Area of Virology, Serology and Virology Division (SAViD), New South Wales Health Pathology Randwick | Foster, C.; Au, J.; Ruiz Silva, M.; Deveson, I.; Bull, R.; Van Hal, S.; Rawlinson, W. |
| EPI_ISL_3221240, EPI_ISL_3221241, EPI_ISL_3221243 | HOSPITAL UNIVERSITARIO SON ESPASES | HOSPITAL UNIVERSITARIO SON ESPASES | Dr. Antonio Oliver, Dr. Carla López-Causapé, Dr. Gabriel Cabot, on behalf of Servicio de Microbiología, Hospital Universitario Son Espases |
| EPI_ISL_3221244, EPI_ISL_3221245 | HOSPITAL CAN MISSES | HOSPITAL UNIVERSITARIO SON ESPASES | Dr. Antonio Oliver, Dr. Carla López-Causapé, Dr. Gabriel Cabot, on behalf of Servicio de Microbiología, Hospital Universitario Son Espases |
| EPI_ISL_3221246 | HOSPITAL MATEU ORFILA | HOSPITAL UNIVERSITARIO SON ESPASES | Dr. Antonio Oliver, Dr. Carla López-Causapé, Dr. Gabriel Cabot, on behalf of Servicio de Microbiología, Hospital Universitario Son Espases |
| EPI_ISL_3221247 | HOSPITAL UNIVERSITARIO SON LLATZER | HOSPITAL UNIVERSITARIO SON ESPASES | Dr. Antonio Oliver, Dr. Carla López-Causapé, Dr. Gabriel Cabot, on behalf of Servicio de Microbiología, Hospital Universitario Son Espases |
| EPI_ISL_3221248 | HOSPITAL UNIVERSITARIO SON ESPASES | HOSPITAL UNIVERSITARIO SON ESPASES | Dr. Antonio Oliver, Dr. Carla López-Causapé, Dr. Gabriel Cabot, on behalf of Servicio de Microbiología, Hospital Universitario Son Espases |
| EPI_ISL_3221249 | HOSPITAL MANACOR | HOSPITAL UNIVERSITARIO SON ESPASES | Dr. Antonio Oliver, Dr. Carla López-Causapé, Dr. Gabriel Cabot, on behalf of Servicio de Microbiología, Hospital Universitario Son Espases |
| EPI_ISL_3221252, EPI_ISL_3221253, EPI_ISL_3221254 | HOSPITAL UNIVERSITARIO SON ESPASES | HOSPITAL UNIVERSITARIO SON ESPASES | Dr. Antonio Oliver, Dr. Carla López-Causapé, Dr. Gabriel Cabot, on behalf of Servicio de Microbiología, Hospital Universitario Son Espases |
| EPI_ISL_3221255, EPI_ISL_3221256 | HOSPITAL CAN MISSES | HOSPITAL UNIVERSITARIO SON ESPASES | Dr. Antonio Oliver, Dr. Carla López-Causapé, Dr. Gabriel Cabot, on behalf of Servicio de Microbiología, Hospital Universitario Son Espases |
| EPI_ISL_3221258 | HOSPITAL UNIVERSITARIO SON LLATZER | HOSPITAL UNIVERSITARIO SON ESPASES | Dr. Antonio Oliver, Dr. Carla López-Causapé, Dr. Gabriel Cabot, on behalf of Servicio de Microbiología, Hospital Universitario Son Espases |
| EPI_ISL_3221259 | HOSPITAL UNIVERSITARIO SON ESPASES | HOSPITAL UNIVERSITARIO SON ESPASES | Dr. Antonio Oliver, Dr. Carla López-Causapé, Dr. Gabriel Cabot, on behalf of Servicio de Microbiología, Hospital Universitario Son Espases |
| EPI_ISL_3221260 | HOSPITAL MANACOR | HOSPITAL UNIVERSITARIO SON ESPASES | Dr. Antonio Oliver, Dr. Carla López-Causapé, Dr. Gabriel Cabot, on behalf of Servicio de Microbiología, Hospital Universitario Son Espases |
| EPI_ISL_3221263, EPI_ISL_3221264 | HOSPITAL UNIVERSITARIO SON ESPASES | HOSPITAL UNIVERSITARIO SON ESPASES | Dr. Antonio Oliver, Dr. Carla López-Causapé, Dr. Gabriel Cabot, on behalf of Servicio de Microbiología, Hospital Universitario Son Espases |
| EPI_ISL_3221265, EPI_ISL_3221266, EPI_ISL_3221267 | HOSPITAL CAN MISSES | HOSPITAL UNIVERSITARIO SON ESPASES | Dr. Antonio Oliver, Dr. Carla López-Causapé, Dr. Gabriel Cabot, on behalf of Servicio de Microbiología, Hospital Universitario Son Espases |
| EPI_ISL_3221268 | HOSPITAL UNIVERSITARIO SON LLATZER | HOSPITAL UNIVERSITARIO SON ESPASES | Dr. Antonio Oliver, Dr. Carla López-Causapé, Dr. Gabriel Cabot, on behalf of Servicio de Microbiología, Hospital Universitario Son Espases |
| EPI_ISL_3221269 | HOSPITAL UNIVERSITARIO SON ESPASES | HOSPITAL UNIVERSITARIO SON ESPASES | Dr. Antonio Oliver, Dr. Carla López-Causapé, Dr. Gabriel Cabot, on behalf of Servicio de Microbiología, Hospital Universitario Son Espases |
| EPI_ISL_3221270 | HOSPITAL MANACOR | HOSPITAL UNIVERSITARIO SON ESPASES | Dr. Antonio Oliver, Dr. Carla López-Causapé, Dr. Gabriel Cabot, on behalf of Servicio de Microbiología, Hospital Universitario Son Espases |
| EPI_ISL_3221271, EPI_ISL_3221272, EPI_ISL_3221273, EPI_ISL_3221274 | HOSPITAL UNIVERSITARIO SON ESPASES | HOSPITAL UNIVERSITARIO SON ESPASES | Dr. Antonio Oliver, Dr. Carla López-Causapé, Dr. Gabriel Cabot, on behalf of Servicio de Microbiología, Hospital Universitario Son Espases |
| EPI_ISL_3221275 | HOSPITAL MATEU ORFILA | HOSPITAL UNIVERSITARIO SON ESPASES | Dr. Antonio Oliver, Dr. Carla López-Causapé, Dr. Gabriel Cabot, on behalf of Servicio de Microbiología, Hospital Universitario Son Espases |
| EPI_ISL_3221276 | HOSPITAL CAN MISSES | HOSPITAL UNIVERSITARIO SON ESPASES | Dr. Antonio Oliver, Dr. Carla López-Causapé, Dr. Gabriel Cabot, on behalf of Servicio de Microbiología, Hospital Universitario Son Espases |
| EPI_ISL_3221277, EPI_ISL_3221279 | HOSPITAL UNIVERSITARIO SON ESPASES | HOSPITAL UNIVERSITARIO SON ESPASES | Dr. Antonio Oliver, Dr. Carla López-Causapé, Dr. Gabriel Cabot, on behalf of Servicio de Microbiología, Hospital Universitario Son Espases |
| EPI_ISL_3221280 | HOSPITAL MANACOR | HOSPITAL UNIVERSITARIO SON ESPASES | Dr. Antonio Oliver, Dr. Carla López-Causapé, Dr. Gabriel Cabot, on behalf of Servicio de Microbiología, Hospital Universitario Son Espases |
| EPI_ISL_3221281 | HOSPITAL CAN MISSES | HOSPITAL UNIVERSITARIO SON ESPASES | Dr. Antonio Oliver, Dr. Carla López-Causapé, Dr. Gabriel Cabot, on behalf of Servicio de Microbiología, Hospital Universitario Son Espases |
| EPI_ISL_3221282, EPI_ISL_3221283 | HOSPITAL UNIVERSITARIO SON ESPASES | HOSPITAL UNIVERSITARIO SON ESPASES | Dr. Antonio Oliver, Dr. Carla López-Causapé, Dr. Gabriel Cabot, on behalf of Servicio de Microbiología, Hospital Universitario Son Espases |
| EPI_ISL_3221284, EPI_ISL_3221285 | HOSPITAL MATEU ORFILA | HOSPITAL UNIVERSITARIO SON ESPASES | Dr. Antonio Oliver, Dr. Carla López-Causapé, Dr. Gabriel Cabot, on behalf of Servicio de Microbiología, Hospital Universitario Son Espases |
| EPI_ISL_3221287 | HOSPITAL MANACOR | HOSPITAL UNIVERSITARIO SON ESPASES | Dr. Antonio Oliver, Dr. Carla López-Causapé, Dr. Gabriel Cabot, on behalf of Servicio de Microbiología, Hospital Universitario Son Espases |
| EPI_ISL_3221289, EPI_ISL_3221290, EPI_ISL_3221291, EPI_ISL_3221292 | HOSPITAL UNIVERSITARIO SON ESPASES | HOSPITAL UNIVERSITARIO SON ESPASES | Dr. Antonio Oliver, Dr. Carla López-Causapé, Dr. Gabriel Cabot, on behalf of Servicio de Microbiología, Hospital Universitario Son Espases |
| EPI_ISL_3221293, EPI_ISL_3221294, EPI_ISL_3221295 | HOSPITAL CAN MISSES | HOSPITAL UNIVERSITARIO SON ESPASES | Dr. Antonio Oliver, Dr. Carla López-Causapé, Dr. Gabriel Cabot, on behalf of Servicio de Microbiología, Hospital Universitario Son Espases |
| EPI_ISL_3221296, EPI_ISL_3221297, EPI_ISL_3221298 | HOSPITAL UNIVERSITARIO SON ESPASES | HOSPITAL UNIVERSITARIO SON ESPASES | Dr. Antonio Oliver, Dr. Carla López-Causapé, Dr. Gabriel Cabot, on behalf of Servicio de Microbiología, Hospital Universitario Son Espases |
| EPI_ISL_3221299, EPI_ISL_3221300 | HOSPITAL MANACOR | HOSPITAL UNIVERSITARIO SON ESPASES | Dr. Antonio Oliver, Dr. Carla López-Causapé, Dr. Gabriel Cabot, on behalf of Servicio de Microbiología, Hospital Universitario Son Espases |
| EPI_ISL_3221301, EPI_ISL_3221302, EPI_ISL_3221303 | HOSPITAL UNIVERSITARIO SON ESPASES | HOSPITAL UNIVERSITARIO SON ESPASES | Dr. Antonio Oliver, Dr. Carla López-Causapé, Dr. Gabriel Cabot, on behalf of Servicio de Microbiología, Hospital Universitario Son Espases |
| EPI_ISL_3221305, EPI_ISL_3221306, EPI_ISL_3221307 | HOSPITAL CAN MISSES | HOSPITAL UNIVERSITARIO SON ESPASES | Dr. Antonio Oliver, Dr. Carla López-Causapé, Dr. Gabriel Cabot, on behalf of Servicio de Microbiología, Hospital Universitario Son Espases |
| EPI_ISL_3221308, EPI_ISL_3221309 | HOSPITAL UNIVERSITARIO SON ESPASES | HOSPITAL UNIVERSITARIO SON ESPASES | Dr. Antonio Oliver, Dr. Carla López-Causapé, Dr. Gabriel Cabot, on behalf of Servicio de Microbiología, Hospital Universitario Son Espases |
| EPI_ISL_3221310 | HOSPITAL MANACOR | HOSPITAL UNIVERSITARIO SON ESPASES | Dr. Antonio Oliver, Dr. Carla López-Causapé, Dr. Gabriel Cabot, on behalf of Servicio de Microbiología, Hospital Universitario Son Espases |
| EPI_ISL_3221311, EPI_ISL_3221312, EPI_ISL_3221313 | HOSPITAL UNIVERSITARIO SON ESPASES | HOSPITAL UNIVERSITARIO SON ESPASES | Dr. Antonio Oliver, Dr. Carla López-Causapé, Dr. Gabriel Cabot, on behalf of Servicio de Microbiología, Hospital Universitario Son Espases |
| EPI_ISL_3221314, EPI_ISL_3221315 | HOSPITAL CAN MISSES | HOSPITAL UNIVERSITARIO SON ESPASES | Dr. Antonio Oliver, Dr. Carla López-Causapé, Dr. Gabriel Cabot, on behalf of Servicio de Microbiología, Hospital Universitario Son Espases |
| EPI_ISL_3221317, EPI_ISL_3221318 | HOSPITAL UNIVERSITARIO SON ESPASES | HOSPITAL UNIVERSITARIO SON ESPASES | Dr. Antonio Oliver, Dr. Carla López-Causapé, Dr. Gabriel Cabot, on behalf of Servicio de Microbiología, Hospital Universitario Son Espases |
| EPI_ISL_3221319 | HOSPITAL MANACOR | HOSPITAL UNIVERSITARIO SON ESPASES | Dr. Antonio Oliver, Dr. Carla López-Causapé, Dr. Gabriel Cabot, on behalf of Servicio de Microbiología, Hospital Universitario Son Espases |

| | | | |
|---|---|---|---|
| EPI_ISL_3221320 | HOSPITAL UNIVERSITARIO SON ESPASES | HOSPITAL UNIVERSITARIO SON ESPASES | Dr. Antonio Oliver, Dr. Carla López-Causapé, Dr. Gabriel Cabot, on behalf of Servicio de Microbiología, Hospital Universitario Son Espases |
| EPI_ISL_3221322 | Department of Pathology and Laboratory Medicine, Kulliyyah of Medicine, International Islamic University of Malaysia | Centre for Research in Advanced Tropical Bioscience (Biotropic Centre) | Hajar Fauzan Bin Ahmad |
| EPI_ISL_3221323 | Department of Pathology and Laboratory Medicine, Kulliyyah of Medicine, International Islamic University of Malaysia | Centre for Research in Advanced Tropical Bioscience (Biotropic Centre) | Ummu Afeera Zainulabid, Norhidayah Kamarudin, How Soon Hin and Hajar Fauzan Ahmad |
| EPI_ISL_3221324 | Kulliyyah of Medicine, International Islamic University Malaysia, Bandar Indera Mahkota Campus, Jalan Sultan Ahmad Shah, 25200 Kuantan, Pahang Darul Makmur | Centre for Research in Advanced Tropical Bioscience (Biotropic Centre) | Ummu Afeera Zainulabid, Norhidayah Kamarudin, How Soon Hin and Hajar Fauzan Ahmad |
| EPI_ISL_3221325 | Department of Pathology & Laboratory Medicine | Centre for Research in Advanced Tropical Bioscience (Biotropic Centre) | Ummu Afeera Zainulabid, Norhidayah Kamarudin, How Soon Hin and Hajar Fauzan Ahmad |
| EPI_ISL_3221327 | COVID-19 Detection Lab, Chattogram Veterinary and Animal Sciences University | Genomic Research Lab, BCSIR | Md. Murshed Hasan Sarkar, Abu Sayeed Mohammad Mahmud, Mohammad Samir Uzzaman, Eshrar Osman, Md. Ahasan Habib, Shahina Akter, Tanjina Akhter Banu, Barna Goswami, Iffat Jahan, Mohammad Mohi Uddin, Md. Kamrul Islam, Goutam Buddha Das, Tridip Das, Tanvir Ahmad Nizami, Eaftekhar Ahmed Rana, Md. Sirazul Islam, Pronesh Dutta, Sharmin Chowdhury, Md. Paritosh Kumar Biswas, Md. Salim Khan |
| EPI_ISL_3221329 | Department of Microbiology, The University of Hong Kong | The University of Hong Kong Department of Microbiology | Kelvin K.W. To, Kwok-Yung Yuen |
| EPI_ISL_3221330 | COVID-19 Detection Lab, Chattogram Veterinary and Animal Sciences University | Genomic Research Lab, BCSIR | Md. Murshed Hasan Sarkar, Abu Sayeed Mohammad Mahmud, Mohammad Samir Uzzaman, Eshrar Osman, Md. Ahasan Habib, Shahina Akter, Tanjina Akhter Banu, Barna Goswami, Iffat Jahan, Mohammad Mohi Uddin, Md. Kamrul Islam, Goutam Buddha Das, Tridip Das, Tanvir Ahmad Nizami, Eaftekhar Ahmed Rana, Md. Sirazul Islam, Pronesh Dutta, Sharmin Chowdhury, Md. Paritosh Kumar Biswas, Md. Salim Khan |
| EPI_ISL_3221331 | Lighthouse Laboratory Plymouth | Wellcome Sanger Institute for the COVID-19 Genomics UK (COG-UK) Consortium | Lighthouse Laboratory Plymouth and Alex Alderton, Roberto Amato, Jeffrey Barrett, Sonia Goncalves, Ewan Harrison, David K. Jackson, Ian Johnston, Dominic Kwiatkowski, Cordelia Langford, John Sillitoe on behalf of the Wellcome Sanger Institute COVID-19 Surveillance Team |
| EPI_ISL_3221332, EPI_ISL_3221333, EPI_ISL_3221334, EPI_ISL_3221335 | Lighthouse Lab in Milton Keynes | Wellcome Sanger Institute for the COVID-19 Genomics UK (COG-UK) Consortium | The Lighthouse Lab in Milton Keynes and Alex Alderton, Roberto Amato, Jeffrey Barrett, Sonia Goncalves, Ewan Harrison, David K. Jackson, Ian Johnston, Dominic Kwiatkowski, Cordelia Langford, John Sillitoe on behalf of the Wellcome Sanger Institute COVID-19 Surveillance Team |
| EPI_ISL_3221336 | Lighthouse Laboratory Plymouth | Wellcome Sanger Institute for the COVID-19 Genomics UK (COG-UK) Consortium | Lighthouse Laboratory Plymouth and Alex Alderton, Roberto Amato, Jeffrey Barrett, Sonia Goncalves, Ewan Harrison, David K. Jackson, Ian Johnston, Dominic Kwiatkowski, Cordelia Langford, John Sillitoe on behalf of the Wellcome Sanger Institute COVID-19 Surveillance Team |
| EPI_ISL_3221337 | Lighthouse Lab in Milton Keynes | Wellcome Sanger Institute for the COVID-19 Genomics UK (COG-UK) Consortium | The Lighthouse Lab in Milton Keynes and Alex Alderton, Roberto Amato, Jeffrey Barrett, Sonia Goncalves, Ewan Harrison, David K. Jackson, Ian Johnston, Dominic Kwiatkowski, Cordelia Langford, John Sillitoe on behalf of the Wellcome Sanger Institute COVID-19 Surveillance Team |
| EPI_ISL_3221338 | Lighthouse Laboratory Plymouth | Wellcome Sanger Institute for the COVID-19 Genomics UK (COG-UK) Consortium | Lighthouse Laboratory Plymouth and Alex Alderton, Roberto Amato, Jeffrey Barrett, Sonia Goncalves, Ewan Harrison, David K. Jackson, Ian Johnston, Dominic Kwiatkowski, Cordelia Langford, John Sillitoe on behalf of the Wellcome Sanger Institute COVID-19 Surveillance Team |
| EPI_ISL_3221339 | Lighthouse Lab in Milton Keynes | Wellcome Sanger Institute for the COVID-19 Genomics UK (COG-UK) Consortium | The Lighthouse Lab in Milton Keynes and Alex Alderton, Roberto Amato, Jeffrey Barrett, Sonia Goncalves, Ewan Harrison, David K. Jackson, Ian Johnston, Dominic Kwiatkowski, Cordelia Langford, John Sillitoe on behalf of the Wellcome Sanger Institute COVID-19 Surveillance Team |
| EPI_ISL_3221340 | Lighthouse Laboratory Plymouth | Wellcome Sanger Institute for the COVID-19 Genomics UK (COG-UK) Consortium | Lighthouse Laboratory Plymouth and Alex Alderton, Roberto Amato, Jeffrey Barrett, Sonia Goncalves, Ewan Harrison, David K. Jackson, Ian Johnston, Dominic Kwiatkowski, Cordelia Langford, John Sillitoe on behalf of the Wellcome Sanger Institute COVID-19 Surveillance Team |
| EPI_ISL_3221341, EPI_ISL_3221342 | Lighthouse Lab in Milton Keynes | Wellcome Sanger Institute for the COVID-19 Genomics UK (COG-UK) Consortium | The Lighthouse Lab in Milton Keynes and Alex Alderton, Roberto Amato, Jeffrey Barrett, Sonia Goncalves, Ewan Harrison, David K. Jackson, Ian Johnston, Dominic Kwiatkowski, Cordelia Langford, John Sillitoe on behalf of the Wellcome Sanger Institute COVID-19 Surveillance Team |
| EPI_ISL_3221343 | Lighthouse Laboratory Plymouth | Wellcome Sanger Institute for the COVID-19 Genomics UK (COG-UK) Consortium | Lighthouse Laboratory Plymouth and Alex Alderton, Roberto Amato, Jeffrey Barrett, Sonia Goncalves, Ewan Harrison, David K. Jackson, Ian Johnston, Dominic Kwiatkowski, Cordelia Langford, John Sillitoe on behalf of the Wellcome Sanger Institute COVID-19 Surveillance Team |
| EPI_ISL_3221344, EPI_ISL_3221345, EPI_ISL_3221346 | Lighthouse Lab in Milton Keynes | Wellcome Sanger Institute for the COVID-19 Genomics UK (COG-UK) Consortium | The Lighthouse Lab in Milton Keynes and Alex Alderton, Roberto Amato, Jeffrey Barrett, Sonia Goncalves, Ewan Harrison, David K. Jackson, Ian Johnston, Dominic Kwiatkowski, Cordelia Langford, John Sillitoe on behalf of the Wellcome Sanger Institute COVID-19 Surveillance Team |
| EPI_ISL_3221347, EPI_ISL_3221348, EPI_ISL_3221349 | Lighthouse Laboratory Plymouth | Wellcome Sanger Institute for the COVID-19 Genomics UK (COG-UK) Consortium | Lighthouse Laboratory Plymouth and Alex Alderton, Roberto Amato, Jeffrey Barrett, Sonia Goncalves, Ewan Harrison, David K. Jackson, Ian Johnston, Dominic Kwiatkowski, Cordelia Langford, John Sillitoe on behalf of the Wellcome Sanger Institute COVID-19 Surveillance Team |
| EPI_ISL_3221350, EPI_ISL_3221351 | Lighthouse Lab in Milton Keynes | Wellcome Sanger Institute for the COVID-19 Genomics UK (COG-UK) Consortium | The Lighthouse Lab in Milton Keynes and Alex Alderton, Roberto Amato, Jeffrey Barrett, Sonia Goncalves, Ewan Harrison, David K. Jackson, Ian Johnston, Dominic Kwiatkowski, Cordelia Langford, John Sillitoe on behalf of the Wellcome Sanger Institute COVID-19 Surveillance Team |
| EPI_ISL_3221352 | Lighthouse Laboratory Plymouth | Wellcome Sanger Institute for the COVID-19 Genomics UK (COG-UK) Consortium | Lighthouse Laboratory Plymouth and Alex Alderton, Roberto Amato, Jeffrey Barrett, Sonia Goncalves, Ewan Harrison, David K. Jackson, Ian Johnston, Dominic Kwiatkowski, Cordelia Langford, John Sillitoe on behalf of the Wellcome Sanger Institute COVID-19 Surveillance Team |
| EPI_ISL_3221353 | Lighthouse Lab in Milton Keynes | Wellcome Sanger Institute for the COVID-19 Genomics UK (COG-UK) Consortium | The Lighthouse Lab in Milton Keynes and Alex Alderton, Roberto Amato, Jeffrey Barrett, Sonia Goncalves, Ewan Harrison, David K. Jackson, Ian Johnston, Dominic Kwiatkowski, Cordelia Langford, John Sillitoe on behalf of the Wellcome Sanger Institute COVID-19 Surveillance Team |
| EPI_ISL_3221354 | Lighthouse Laboratory Plymouth | Wellcome Sanger Institute for the COVID-19 Genomics UK (COG-UK) Consortium | Lighthouse Laboratory Plymouth and Alex Alderton, Roberto Amato, Jeffrey Barrett, Sonia Goncalves, Ewan Harrison, David K. Jackson, Ian Johnston, Dominic Kwiatkowski, Cordelia Langford, John Sillitoe on behalf of the Wellcome Sanger Institute COVID-19 Surveillance Team |
| EPI_ISL_3221355 | Lighthouse Lab in Milton Keynes | Wellcome Sanger Institute for the COVID-19 Genomics UK (COG-UK) Consortium | The Lighthouse Lab in Milton Keynes and Alex Alderton, Roberto Amato, Jeffrey Barrett, Sonia Goncalves, Ewan Harrison, David K. Jackson, Ian Johnston, Dominic Kwiatkowski, Cordelia Langford, John Sillitoe on behalf of the Wellcome Sanger Institute COVID-19 Surveillance Team |
| EPI_ISL_3221356 | Lighthouse Laboratory Plymouth | Wellcome Sanger Institute for the COVID-19 Genomics UK (COG-UK) Consortium | Lighthouse Laboratory Plymouth and Alex Alderton, Roberto Amato, Jeffrey Barrett, Sonia Goncalves, Ewan Harrison, David K. Jackson, Ian Johnston, Dominic Kwiatkowski, Cordelia Langford, John Sillitoe on behalf of the Wellcome Sanger Institute COVID-19 Surveillance Team |
| EPI_ISL_3221357, EPI_ISL_3221358 | Lighthouse Lab in Milton Keynes | Wellcome Sanger Institute for the COVID-19 Genomics UK (COG-UK) Consortium | The Lighthouse Lab in Milton Keynes and Alex Alderton, Roberto Amato, Jeffrey Barrett, Sonia Goncalves, Ewan Harrison, David K. Jackson, Ian Johnston, Dominic Kwiatkowski, Cordelia Langford, John Sillitoe on behalf of the Wellcome Sanger Institute COVID-19 Surveillance Team |
| EPI_ISL_3221359, EPI_ISL_3221360 | Lighthouse Laboratory Plymouth | Wellcome Sanger Institute for the COVID-19 Genomics UK (COG-UK) Consortium | Lighthouse Laboratory Plymouth and Alex Alderton, Roberto Amato, Jeffrey Barrett, Sonia Goncalves, Ewan Harrison, David K. Jackson, Ian Johnston, Dominic Kwiatkowski, Cordelia Langford, John Sillitoe on behalf of the Wellcome Sanger Institute COVID-19 Surveillance Team |
| EPI_ISL_3221361 | Lighthouse Lab in Milton Keynes | Wellcome Sanger Institute for the COVID-19 Genomics UK (COG-UK) Consortium | The Lighthouse Lab in Milton Keynes and Alex Alderton, Roberto Amato, Jeffrey Barrett, Sonia Goncalves, Ewan Harrison, David K. Jackson, Ian Johnston, Dominic Kwiatkowski, Cordelia Langford, John Sillitoe on behalf of the Wellcome Sanger Institute COVID-19 Surveillance Team |
| EPI_ISL_3221362, EPI_ISL_3221363, EPI_ISL_3221364 | Lighthouse Laboratory Plymouth | Wellcome Sanger Institute for the COVID-19 Genomics UK (COG-UK) Consortium | Lighthouse Laboratory Plymouth and Alex Alderton, Roberto Amato, Jeffrey Barrett, Sonia Goncalves, Ewan Harrison, David K. Jackson, Ian Johnston, Dominic Kwiatkowski, Cordelia Langford, John Sillitoe on behalf of the Wellcome Sanger Institute COVID-19 Surveillance Team |
| EPI_ISL_3221365 | Lighthouse Lab in Milton Keynes | Wellcome Sanger Institute for the COVID-19 Genomics UK (COG-UK) Consortium | The Lighthouse Lab in Milton Keynes and Alex Alderton, Roberto Amato, Jeffrey Barrett, Sonia Goncalves, Ewan Harrison, David K. Jackson, Ian Johnston, Dominic Kwiatkowski, Cordelia Langford, John Sillitoe on behalf of the Wellcome Sanger Institute COVID-19 Surveillance Team |
| EPI_ISL_3221366 | Lighthouse Laboratory Plymouth | Wellcome Sanger Institute for the COVID-19 Genomics UK (COG-UK) Consortium | Lighthouse Laboratory Plymouth and Alex Alderton, Roberto Amato, Jeffrey Barrett, Sonia Goncalves, Ewan Harrison, David K. Jackson, Ian Johnston, Dominic Kwiatkowski, Cordelia Langford, John Sillitoe on behalf of the Wellcome Sanger Institute COVID-19 Surveillance Team |
| EPI_ISL_3221367 | Lighthouse Lab in Milton Keynes | Wellcome Sanger Institute for the COVID-19 Genomics UK (COG-UK) Consortium | The Lighthouse Lab in Milton Keynes and Alex Alderton, Roberto Amato, Jeffrey Barrett, Sonia Goncalves, Ewan Harrison, David K. Jackson, Ian Johnston, Dominic Kwiatkowski, Cordelia Langford, John Sillitoe on behalf of the Wellcome Sanger Institute COVID-19 Surveillance Team |
| EPI_ISL_3221368 | Lighthouse Laboratory Plymouth | Wellcome Sanger Institute for the COVID-19 Genomics UK (COG-UK) Consortium | Lighthouse Laboratory Plymouth and Alex Alderton, Roberto Amato, Jeffrey Barrett, Sonia Goncalves, Ewan Harrison, David K. Jackson, Ian Johnston, Dominic Kwiatkowski, Cordelia Langford, John Sillitoe on behalf of the Wellcome Sanger Institute COVID-19 Surveillance Team |
| EPI_ISL_3221369, EPI_ISL_3221370, EPI_ISL_3221371 | Lighthouse Lab in Milton Keynes | Wellcome Sanger Institute for the COVID-19 Genomics UK (COG-UK) Consortium | The Lighthouse Lab in Milton Keynes and Alex Alderton, Roberto Amato, Jeffrey Barrett, Sonia Goncalves, Ewan Harrison, David K. Jackson, Ian Johnston, Dominic Kwiatkowski, Cordelia Langford, John Sillitoe on behalf of the Wellcome Sanger Institute COVID-19 Surveillance Team |
| EPI_ISL_3221372 | Lighthouse Laboratory Plymouth | Wellcome Sanger Institute for the COVID-19 Genomics UK (COG-UK) Consortium | Lighthouse Laboratory Plymouth and Alex Alderton, Roberto Amato, Jeffrey Barrett, Sonia Goncalves, Ewan Harrison, David K. Jackson, Ian Johnston, Dominic Kwiatkowski, Cordelia Langford, John Sillitoe on behalf of the Wellcome Sanger Institute COVID-19 Surveillance Team |
| EPI_ISL_3221373, EPI_ISL_3221374 | Lighthouse Lab in Milton Keynes | Wellcome Sanger Institute for the COVID-19 Genomics UK (COG-UK) Consortium | The Lighthouse Lab in Milton Keynes and Alex Alderton, Roberto Amato, Jeffrey Barrett, Sonia Goncalves, Ewan Harrison, David K. Jackson, Ian Johnston, Dominic Kwiatkowski, Cordelia Langford, John Sillitoe on behalf of the Wellcome Sanger Institute COVID-19 Surveillance Team |
| EPI_ISL_3221375 | Lighthouse Laboratory Plymouth | Wellcome Sanger Institute for the COVID-19 Genomics UK (COG-UK) Consortium | Lighthouse Laboratory Plymouth and Alex Alderton, Roberto Amato, Jeffrey Barrett, Sonia Goncalves, Ewan Harrison, David K. Jackson, Ian Johnston, Dominic Kwiatkowski, Cordelia Langford, John Sillitoe on behalf of the Wellcome Sanger Institute COVID-19 Surveillance Team |
| EPI_ISL_3221376 | Lighthouse Lab in Milton Keynes | Wellcome Sanger Institute for the COVID-19 Genomics UK (COG-UK) Consortium | The Lighthouse Lab in Milton Keynes and Alex Alderton, Roberto Amato, Jeffrey Barrett, Sonia Goncalves, Ewan Harrison, David K. Jackson, Ian Johnston, Dominic Kwiatkowski, Cordelia Langford, John Sillitoe on behalf of the Wellcome Sanger Institute COVID-19 Surveillance Team |
| EPI_ISL_3221377 | Lighthouse Laboratory Plymouth | Wellcome Sanger Institute for the COVID-19 Genomics UK (COG-UK) Consortium | Lighthouse Laboratory Plymouth and Alex Alderton, Roberto Amato, Jeffrey Barrett, Sonia Goncalves, Ewan Harrison, David K. Jackson, Ian Johnston, Dominic Kwiatkowski, Cordelia Langford, John Sillitoe on behalf of the Wellcome Sanger Institute COVID-19 Surveillance Team |
| EPI_ISL_3221378 | Lighthouse Lab in Milton Keynes | Wellcome Sanger Institute for the COVID-19 Genomics UK (COG-UK) Consortium | The Lighthouse Lab in Milton Keynes and Alex Alderton, Roberto Amato, Jeffrey Barrett, Sonia Goncalves, Ewan Harrison, David K. Jackson, Ian Johnston, Dominic Kwiatkowski, Cordelia Langford, John Sillitoe on behalf of the Wellcome Sanger Institute COVID-19 Surveillance Team |
| EPI_ISL_3221379, EPI_ISL_3221380 | Lighthouse Laboratory Plymouth | Wellcome Sanger Institute for the COVID-19 Genomics UK (COG-UK) Consortium | Lighthouse Laboratory Plymouth and Alex Alderton, Roberto Amato, Jeffrey Barrett, Sonia Goncalves, Ewan Harrison, David K. Jackson, Ian Johnston, Dominic Kwiatkowski, Cordelia Langford, John Sillitoe on behalf of the Wellcome Sanger Institute COVID-19 Surveillance Team |
| EPI_ISL_3221381 | Lighthouse Lab in Milton Keynes | Wellcome Sanger Institute for the COVID-19 Genomics UK (COG-UK) Consortium | The Lighthouse Lab in Milton Keynes and Alex Alderton, Roberto Amato, Jeffrey Barrett, Sonia Goncalves, Ewan Harrison, David K. Jackson, Ian Johnston, Dominic Kwiatkowski, Cordelia Langford, John Sillitoe on behalf of the Wellcome Sanger Institute COVID-19 Surveillance Team |
| EPI_ISL_3221382, EPI_ISL_3221383 | Lighthouse Laboratory Plymouth | Wellcome Sanger Institute for the COVID-19 Genomics UK (COG-UK) Consortium | Lighthouse Laboratory Plymouth and Alex Alderton, Roberto Amato, Jeffrey Barrett, Sonia Goncalves, Ewan Harrison, David K. Jackson, Ian Johnston, Dominic Kwiatkowski, Cordelia Langford, John Sillitoe on behalf of the Wellcome Sanger Institute COVID-19 Surveillance Team |

| | | | |
|---|---|---|---|
| EPI_ISL_3221384 | Lighthouse Lab in Milton Keynes | Wellcome Sanger Institute for the COVID-19 Genomics UK (COG-UK) Consortium | The Lighthouse Lab in Milton Keynes and Alex Alderton, Roberto Amato, Jeffrey Barrett, Sonia Goncalves, Ewan Harrison, David K. Jackson, Ian Johnston, Dominic Kwiatkowski, Cordelia Langford, John Sillitoe on behalf of the Wellcome Sanger Institute COVID-19 Surveillance Team |
| EPI_ISL_3221385, EPI_ISL_3221386, EPI_ISL_3221387 | Lighthouse Laboratory Plymouth | Wellcome Sanger Institute for the COVID-19 Genomics UK (COG-UK) Consortium | Lighthouse Laboratory Plymouth and Alex Alderton, Roberto Amato, Jeffrey Barrett, Sonia Goncalves, Ewan Harrison, David K. Jackson, Ian Johnston, Dominic Kwiatkowski, Cordelia Langford, John Sillitoe on behalf of the Wellcome Sanger Institute COVID-19 Surveillance Team |
| EPI_ISL_3221388, EPI_ISL_3221389 | Lighthouse Lab in Milton Keynes | Wellcome Sanger Institute for the COVID-19 Genomics UK (COG-UK) Consortium | The Lighthouse Lab in Milton Keynes and Alex Alderton, Roberto Amato, Jeffrey Barrett, Sonia Goncalves, Ewan Harrison, David K. Jackson, Ian Johnston, Dominic Kwiatkowski, Cordelia Langford, John Sillitoe on behalf of the Wellcome Sanger Institute COVID-19 Surveillance Team |
| EPI_ISL_3221390, EPI_ISL_3221391 | Lighthouse Laboratory Plymouth | Wellcome Sanger Institute for the COVID-19 Genomics UK (COG-UK) Consortium | Lighthouse Laboratory Plymouth and Alex Alderton, Roberto Amato, Jeffrey Barrett, Sonia Goncalves, Ewan Harrison, David K. Jackson, Ian Johnston, Dominic Kwiatkowski, Cordelia Langford, John Sillitoe on behalf of the Wellcome Sanger Institute COVID-19 Surveillance Team |
| EPI_ISL_3221392 | Lighthouse Lab in Milton Keynes | Wellcome Sanger Institute for the COVID-19 Genomics UK (COG-UK) Consortium | The Lighthouse Lab in Milton Keynes and Alex Alderton, Roberto Amato, Jeffrey Barrett, Sonia Goncalves, Ewan Harrison, David K. Jackson, Ian Johnston, Dominic Kwiatkowski, Cordelia Langford, John Sillitoe on behalf of the Wellcome Sanger Institute COVID-19 Surveillance Team |
| EPI_ISL_3221393 | Lighthouse Laboratory Plymouth | Wellcome Sanger Institute for the COVID-19 Genomics UK (COG-UK) Consortium | Lighthouse Laboratory Plymouth and Alex Alderton, Roberto Amato, Jeffrey Barrett, Sonia Goncalves, Ewan Harrison, David K. Jackson, Ian Johnston, Dominic Kwiatkowski, Cordelia Langford, John Sillitoe on behalf of the Wellcome Sanger Institute COVID-19 Surveillance Team |
| EPI_ISL_3221394, EPI_ISL_3221395, EPI_ISL_3221396 | Lighthouse Lab in Milton Keynes | Wellcome Sanger Institute for the COVID-19 Genomics UK (COG-UK) Consortium | The Lighthouse Lab in Milton Keynes and Alex Alderton, Roberto Amato, Jeffrey Barrett, Sonia Goncalves, Ewan Harrison, David K. Jackson, Ian Johnston, Dominic Kwiatkowski, Cordelia Langford, John Sillitoe on behalf of the Wellcome Sanger Institute COVID-19 Surveillance Team |
| EPI_ISL_3221397 | Lighthouse Laboratory Plymouth | Wellcome Sanger Institute for the COVID-19 Genomics UK (COG-UK) Consortium | Lighthouse Laboratory Plymouth and Alex Alderton, Roberto Amato, Jeffrey Barrett, Sonia Goncalves, Ewan Harrison, David K. Jackson, Ian Johnston, Dominic Kwiatkowski, Cordelia Langford, John Sillitoe on behalf of the Wellcome Sanger Institute COVID-19 Surveillance Team |
| EPI_ISL_3221398, EPI_ISL_3221399, EPI_ISL_3221400 | Lighthouse Lab in Milton Keynes | Wellcome Sanger Institute for the COVID-19 Genomics UK (COG-UK) Consortium | The Lighthouse Lab in Milton Keynes and Alex Alderton, Roberto Amato, Jeffrey Barrett, Sonia Goncalves, Ewan Harrison, David K. Jackson, Ian Johnston, Dominic Kwiatkowski, Cordelia Langford, John Sillitoe on behalf of the Wellcome Sanger Institute COVID-19 Surveillance Team |
| EPI_ISL_3221401, EPI_ISL_3221402, EPI_ISL_3221403 | Lighthouse Laboratory Plymouth | Wellcome Sanger Institute for the COVID-19 Genomics UK (COG-UK) Consortium | Lighthouse Laboratory Plymouth and Alex Alderton, Roberto Amato, Jeffrey Barrett, Sonia Goncalves, Ewan Harrison, David K. Jackson, Ian Johnston, Dominic Kwiatkowski, Cordelia Langford, John Sillitoe on behalf of the Wellcome Sanger Institute COVID-19 Surveillance Team |
| EPI_ISL_3221404 | Lighthouse Lab in Milton Keynes | Wellcome Sanger Institute for the COVID-19 Genomics UK (COG-UK) Consortium | The Lighthouse Lab in Milton Keynes and Alex Alderton, Roberto Amato, Jeffrey Barrett, Sonia Goncalves, Ewan Harrison, David K. Jackson, Ian Johnston, Dominic Kwiatkowski, Cordelia Langford, John Sillitoe on behalf of the Wellcome Sanger Institute COVID-19 Surveillance Team |
| EPI_ISL_3221405 | Lighthouse Laboratory Plymouth | Wellcome Sanger Institute for the COVID-19 Genomics UK (COG-UK) Consortium | Lighthouse Laboratory Plymouth and Alex Alderton, Roberto Amato, Jeffrey Barrett, Sonia Goncalves, Ewan Harrison, David K. Jackson, Ian Johnston, Dominic Kwiatkowski, Cordelia Langford, John Sillitoe on behalf of the Wellcome Sanger Institute COVID-19 Surveillance Team |
| EPI_ISL_3221406 | Lighthouse Lab in Milton Keynes | Wellcome Sanger Institute for the COVID-19 Genomics UK (COG-UK) Consortium | The Lighthouse Lab in Milton Keynes and Alex Alderton, Roberto Amato, Jeffrey Barrett, Sonia Goncalves, Ewan Harrison, David K. Jackson, Ian Johnston, Dominic Kwiatkowski, Cordelia Langford, John Sillitoe on behalf of the Wellcome Sanger Institute COVID-19 Surveillance Team |
| EPI_ISL_3221407 | Lighthouse Laboratory Plymouth | Wellcome Sanger Institute for the COVID-19 Genomics UK (COG-UK) Consortium | Lighthouse Laboratory Plymouth and Alex Alderton, Roberto Amato, Jeffrey Barrett, Sonia Goncalves, Ewan Harrison, David K. Jackson, Ian Johnston, Dominic Kwiatkowski, Cordelia Langford, John Sillitoe on behalf of the Wellcome Sanger Institute COVID-19 Surveillance Team |
| EPI_ISL_3221408 | Lighthouse Lab in Milton Keynes | Wellcome Sanger Institute for the COVID-19 Genomics UK (COG-UK) Consortium | The Lighthouse Lab in Milton Keynes and Alex Alderton, Roberto Amato, Jeffrey Barrett, Sonia Goncalves, Ewan Harrison, David K. Jackson, Ian Johnston, Dominic Kwiatkowski, Cordelia Langford, John Sillitoe on behalf of the Wellcome Sanger Institute COVID-19 Surveillance Team |
| EPI_ISL_3221409, EPI_ISL_3221410 | Lighthouse Laboratory Plymouth | Wellcome Sanger Institute for the COVID-19 Genomics UK (COG-UK) Consortium | Lighthouse Laboratory Plymouth and Alex Alderton, Roberto Amato, Jeffrey Barrett, Sonia Goncalves, Ewan Harrison, David K. Jackson, Ian Johnston, Dominic Kwiatkowski, Cordelia Langford, John Sillitoe on behalf of the Wellcome Sanger Institute COVID-19 Surveillance Team |
| EPI_ISL_3221411, EPI_ISL_3221412, EPI_ISL_3221413 | Lighthouse Lab in Milton Keynes | Wellcome Sanger Institute for the COVID-19 Genomics UK (COG-UK) Consortium | The Lighthouse Lab in Milton Keynes and Alex Alderton, Roberto Amato, Jeffrey Barrett, Sonia Goncalves, Ewan Harrison, David K. Jackson, Ian Johnston, Dominic Kwiatkowski, Cordelia Langford, John Sillitoe on behalf of the Wellcome Sanger Institute COVID-19 Surveillance Team |
| EPI_ISL_3221414 | Lighthouse Laboratory Plymouth | Wellcome Sanger Institute for the COVID-19 Genomics UK (COG-UK) Consortium | Lighthouse Laboratory Plymouth and Alex Alderton, Roberto Amato, Jeffrey Barrett, Sonia Goncalves, Ewan Harrison, David K. Jackson, Ian Johnston, Dominic Kwiatkowski, Cordelia Langford, John Sillitoe on behalf of the Wellcome Sanger Institute COVID-19 Surveillance Team |
| EPI_ISL_3221415, EPI_ISL_3221416, EPI_ISL_3221417 | Lighthouse Lab in Milton Keynes | Wellcome Sanger Institute for the COVID-19 Genomics UK (COG-UK) Consortium | The Lighthouse Lab in Milton Keynes and Alex Alderton, Roberto Amato, Jeffrey Barrett, Sonia Goncalves, Ewan Harrison, David K. Jackson, Ian Johnston, Dominic Kwiatkowski, Cordelia Langford, John Sillitoe on behalf of the Wellcome Sanger Institute COVID-19 Surveillance Team |
| EPI_ISL_3221418, EPI_ISL_3221419, EPI_ISL_3221420, EPI_ISL_3221421 | Lighthouse Laboratory Plymouth | Wellcome Sanger Institute for the COVID-19 Genomics UK (COG-UK) Consortium | Lighthouse Laboratory Plymouth and Alex Alderton, Roberto Amato, Jeffrey Barrett, Sonia Goncalves, Ewan Harrison, David K. Jackson, Ian Johnston, Dominic Kwiatkowski, Cordelia Langford, John Sillitoe on behalf of the Wellcome Sanger Institute COVID-19 Surveillance Team |
| EPI_ISL_3221422 | Lighthouse Lab in Milton Keynes | Wellcome Sanger Institute for the COVID-19 Genomics UK (COG-UK) Consortium | The Lighthouse Lab in Milton Keynes and Alex Alderton, Roberto Amato, Jeffrey Barrett, Sonia Goncalves, Ewan Harrison, David K. Jackson, Ian Johnston, Dominic Kwiatkowski, Cordelia Langford, John Sillitoe on behalf of the Wellcome Sanger Institute COVID-19 Surveillance Team |
| EPI_ISL_3221423, EPI_ISL_3221425, EPI_ISL_3221426 | Lighthouse Laboratory Plymouth | Wellcome Sanger Institute for the COVID-19 Genomics UK (COG-UK) Consortium | Lighthouse Laboratory Plymouth and Alex Alderton, Roberto Amato, Jeffrey Barrett, Sonia Goncalves, Ewan Harrison, David K. Jackson, Ian Johnston, Dominic Kwiatkowski, Cordelia Langford, John Sillitoe on behalf of the Wellcome Sanger Institute COVID-19 Surveillance Team |
| EPI_ISL_3221427, EPI_ISL_3221428 | Lighthouse Lab in Milton Keynes | Wellcome Sanger Institute for the COVID-19 Genomics UK (COG-UK) Consortium | The Lighthouse Lab in Milton Keynes and Alex Alderton, Roberto Amato, Jeffrey Barrett, Sonia Goncalves, Ewan Harrison, David K. Jackson, Ian Johnston, Dominic Kwiatkowski, Cordelia Langford, John Sillitoe on behalf of the Wellcome Sanger Institute COVID-19 Surveillance Team |
| EPI_ISL_3221429 | Lighthouse Laboratory Plymouth | Wellcome Sanger Institute for the COVID-19 Genomics UK (COG-UK) Consortium | Lighthouse Laboratory Plymouth and Alex Alderton, Roberto Amato, Jeffrey Barrett, Sonia Goncalves, Ewan Harrison, David K. Jackson, Ian Johnston, Dominic Kwiatkowski, Cordelia Langford, John Sillitoe on behalf of the Wellcome Sanger Institute COVID-19 Surveillance Team |
| EPI_ISL_3221430 | Lighthouse Lab in Milton Keynes | Wellcome Sanger Institute for the COVID-19 Genomics UK (COG-UK) Consortium | The Lighthouse Lab in Milton Keynes and Alex Alderton, Roberto Amato, Jeffrey Barrett, Sonia Goncalves, Ewan Harrison, David K. Jackson, Ian Johnston, Dominic Kwiatkowski, Cordelia Langford, John Sillitoe on behalf of the Wellcome Sanger Institute COVID-19 Surveillance Team |
| EPI_ISL_3221431 | Lighthouse Laboratory Plymouth | Wellcome Sanger Institute for the COVID-19 Genomics UK (COG-UK) Consortium | Lighthouse Laboratory Plymouth and Alex Alderton, Roberto Amato, Jeffrey Barrett, Sonia Goncalves, Ewan Harrison, David K. Jackson, Ian Johnston, Dominic Kwiatkowski, Cordelia Langford, John Sillitoe on behalf of the Wellcome Sanger Institute COVID-19 Surveillance Team |
| EPI_ISL_3221432, EPI_ISL_3221433, EPI_ISL_3221434, EPI_ISL_3221435 | Lighthouse Lab in Milton Keynes | Wellcome Sanger Institute for the COVID-19 Genomics UK (COG-UK) Consortium | The Lighthouse Lab in Milton Keynes and Alex Alderton, Roberto Amato, Jeffrey Barrett, Sonia Goncalves, Ewan Harrison, David K. Jackson, Ian Johnston, Dominic Kwiatkowski, Cordelia Langford, John Sillitoe on behalf of the Wellcome Sanger Institute COVID-19 Surveillance Team |
| EPI_ISL_3221436 | Lighthouse Laboratory Plymouth | Wellcome Sanger Institute for the COVID-19 Genomics UK (COG-UK) Consortium | Lighthouse Laboratory Plymouth and Alex Alderton, Roberto Amato, Jeffrey Barrett, Sonia Goncalves, Ewan Harrison, David K. Jackson, Ian Johnston, Dominic Kwiatkowski, Cordelia Langford, John Sillitoe on behalf of the Wellcome Sanger Institute COVID-19 Surveillance Team |
| EPI_ISL_3221437, EPI_ISL_3221438, EPI_ISL_3221439 | Lighthouse Lab in Milton Keynes | Wellcome Sanger Institute for the COVID-19 Genomics UK (COG-UK) Consortium | The Lighthouse Lab in Milton Keynes and Alex Alderton, Roberto Amato, Jeffrey Barrett, Sonia Goncalves, Ewan Harrison, David K. Jackson, Ian Johnston, Dominic Kwiatkowski, Cordelia Langford, John Sillitoe on behalf of the Wellcome Sanger Institute COVID-19 Surveillance Team |
| EPI_ISL_3221440, EPI_ISL_3221441, EPI_ISL_3221442, EPI_ISL_3221443 | Lighthouse Laboratory Plymouth | Wellcome Sanger Institute for the COVID-19 Genomics UK (COG-UK) Consortium | Lighthouse Laboratory Plymouth and Alex Alderton, Roberto Amato, Jeffrey Barrett, Sonia Goncalves, Ewan Harrison, David K. Jackson, Ian Johnston, Dominic Kwiatkowski, Cordelia Langford, John Sillitoe on behalf of the Wellcome Sanger Institute COVID-19 Surveillance Team |
| EPI_ISL_3221444 | Lighthouse Lab in Milton Keynes | Wellcome Sanger Institute for the COVID-19 Genomics UK (COG-UK) Consortium | The Lighthouse Lab in Milton Keynes and Alex Alderton, Roberto Amato, Jeffrey Barrett, Sonia Goncalves, Ewan Harrison, David K. Jackson, Ian Johnston, Dominic Kwiatkowski, Cordelia Langford, John Sillitoe on behalf of the Wellcome Sanger Institute COVID-19 Surveillance Team |
| EPI_ISL_3221446, EPI_ISL_3221447, EPI_ISL_3221448 | Lighthouse Laboratory Plymouth | Wellcome Sanger Institute for the COVID-19 Genomics UK (COG-UK) Consortium | Lighthouse Laboratory Plymouth and Alex Alderton, Roberto Amato, Jeffrey Barrett, Sonia Goncalves, Ewan Harrison, David K. Jackson, Ian Johnston, Dominic Kwiatkowski, Cordelia Langford, John Sillitoe on behalf of the Wellcome Sanger Institute COVID-19 Surveillance Team |
| EPI_ISL_3221449 | Lighthouse Lab in Milton Keynes | Wellcome Sanger Institute for the COVID-19 Genomics UK (COG-UK) Consortium | The Lighthouse Lab in Milton Keynes and Alex Alderton, Roberto Amato, Jeffrey Barrett, Sonia Goncalves, Ewan Harrison, David K. Jackson, Ian Johnston, Dominic Kwiatkowski, Cordelia Langford, John Sillitoe on behalf of the Wellcome Sanger Institute COVID-19 Surveillance Team |
| EPI_ISL_3221450, EPI_ISL_3221451, EPI_ISL_3221452, EPI_ISL_3221453, EPI_ISL_3221454, EPI_ISL_3221455, EPI_ISL_3221456 | Lighthouse Laboratory Plymouth | Wellcome Sanger Institute for the COVID-19 Genomics UK (COG-UK) Consortium | Lighthouse Laboratory Plymouth and Alex Alderton, Roberto Amato, Jeffrey Barrett, Sonia Goncalves, Ewan Harrison, David K. Jackson, Ian Johnston, Dominic Kwiatkowski, Cordelia Langford, John Sillitoe on behalf of the Wellcome Sanger Institute COVID-19 Surveillance Team |
| EPI_ISL_3221457 | Lighthouse Lab in Milton Keynes | Wellcome Sanger Institute for the COVID-19 Genomics UK (COG-UK) Consortium | The Lighthouse Lab in Milton Keynes and Alex Alderton, Roberto Amato, Jeffrey Barrett, Sonia Goncalves, Ewan Harrison, David K. Jackson, Ian Johnston, Dominic Kwiatkowski, Cordelia Langford, John Sillitoe on behalf of the Wellcome Sanger Institute COVID-19 Surveillance Team |
| EPI_ISL_3221458, EPI_ISL_3221459, EPI_ISL_3221460 | Lighthouse Laboratory Plymouth | Wellcome Sanger Institute for the COVID-19 Genomics UK (COG-UK) Consortium | Lighthouse Laboratory Plymouth and Alex Alderton, Roberto Amato, Jeffrey Barrett, Sonia Goncalves, Ewan Harrison, David K. Jackson, Ian Johnston, Dominic Kwiatkowski, Cordelia Langford, John Sillitoe on behalf of the Wellcome Sanger Institute COVID-19 Surveillance Team |
| EPI_ISL_3221461 | Lighthouse Lab in Milton Keynes | Wellcome Sanger Institute for the COVID-19 Genomics UK (COG-UK) | The Lighthouse Lab in Milton Keynes and Alex Alderton, Roberto Amato, Jeffrey Barrett, Sonia Goncalves, Ewan Harrison, David K. Jackson, Ian Johnston, Dominic Kwiatkowski, Cordelia Langford, John Sillitoe on behalf of the Wellcome Sanger |

| | | | |
|---|---|---|---|
| | | Consortium | Institute COVID-19 Surveillance Team |
| EPI_ISL_3221462 | Lighthouse Laboratory Plymouth | Wellcome Sanger Institute for the COVID-19 Genomics UK (COG-UK) Consortium | Lighthouse Laboratory Plymouth and Alex Alderton, Roberto Amato, Jeffrey Barrett, Sonia Goncalves, Ewan Harrison, David K. Jackson, Ian Johnston, Dominic Kwiatkowski, Cordelia Langford, John Sillitoe on behalf of the Wellcome Sanger Institute COVID-19 Surveillance Team |
| EPI_ISL_3221463, EPI_ISL_3221464 | Lighthouse Lab in Milton Keynes | Wellcome Sanger Institute for the COVID-19 Genomics UK (COG-UK) Consortium | The Lighthouse Lab in Milton Keynes and Alex Alderton, Roberto Amato, Jeffrey Barrett, Sonia Goncalves, Ewan Harrison, David K. Jackson, Ian Johnston, Dominic Kwiatkowski, Cordelia Langford, John Sillitoe on behalf of the Wellcome Sanger Institute COVID-19 Surveillance Team |
| EPI_ISL_3221465, EPI_ISL_3221466, EPI_ISL_3221467, EPI_ISL_3221468 | Lighthouse Laboratory Plymouth | Wellcome Sanger Institute for the COVID-19 Genomics UK (COG-UK) Consortium | Lighthouse Laboratory Plymouth and Alex Alderton, Roberto Amato, Jeffrey Barrett, Sonia Goncalves, Ewan Harrison, David K. Jackson, Ian Johnston, Dominic Kwiatkowski, Cordelia Langford, John Sillitoe on behalf of the Wellcome Sanger Institute COVID-19 Surveillance Team |
| EPI_ISL_3221469, EPI_ISL_3221470, EPI_ISL_3221471, EPI_ISL_3221472 | Lighthouse Lab in Milton Keynes | Wellcome Sanger Institute for the COVID-19 Genomics UK (COG-UK) Consortium | The Lighthouse Lab in Milton Keynes and Alex Alderton, Roberto Amato, Jeffrey Barrett, Sonia Goncalves, Ewan Harrison, David K. Jackson, Ian Johnston, Dominic Kwiatkowski, Cordelia Langford, John Sillitoe on behalf of the Wellcome Sanger Institute COVID-19 Surveillance Team |
| EPI_ISL_3221473 | Lighthouse Laboratory Plymouth | Wellcome Sanger Institute for the COVID-19 Genomics UK (COG-UK) Consortium | Lighthouse Laboratory Plymouth and Alex Alderton, Roberto Amato, Jeffrey Barrett, Sonia Goncalves, Ewan Harrison, David K. Jackson, Ian Johnston, Dominic Kwiatkowski, Cordelia Langford, John Sillitoe on behalf of the Wellcome Sanger Institute COVID-19 Surveillance Team |
| EPI_ISL_3221474, EPI_ISL_3221475, EPI_ISL_3221476 | Lighthouse Lab in Milton Keynes | Wellcome Sanger Institute for the COVID-19 Genomics UK (COG-UK) Consortium | The Lighthouse Lab in Milton Keynes and Alex Alderton, Roberto Amato, Jeffrey Barrett, Sonia Goncalves, Ewan Harrison, David K. Jackson, Ian Johnston, Dominic Kwiatkowski, Cordelia Langford, John Sillitoe on behalf of the Wellcome Sanger Institute COVID-19 Surveillance Team |
| EPI_ISL_3221477 | Lighthouse Laboratory Plymouth | Wellcome Sanger Institute for the COVID-19 Genomics UK (COG-UK) Consortium | Lighthouse Laboratory Plymouth and Alex Alderton, Roberto Amato, Jeffrey Barrett, Sonia Goncalves, Ewan Harrison, David K. Jackson, Ian Johnston, Dominic Kwiatkowski, Cordelia Langford, John Sillitoe on behalf of the Wellcome Sanger Institute COVID-19 Surveillance Team |
| EPI_ISL_3221478 | Lighthouse Lab in Milton Keynes | Wellcome Sanger Institute for the COVID-19 Genomics UK (COG-UK) Consortium | The Lighthouse Lab in Milton Keynes and Alex Alderton, Roberto Amato, Jeffrey Barrett, Sonia Goncalves, Ewan Harrison, David K. Jackson, Ian Johnston, Dominic Kwiatkowski, Cordelia Langford, John Sillitoe on behalf of the Wellcome Sanger Institute COVID-19 Surveillance Team |
| EPI_ISL_3221479 | Lighthouse Laboratory Plymouth | Wellcome Sanger Institute for the COVID-19 Genomics UK (COG-UK) Consortium | Lighthouse Laboratory Plymouth and Alex Alderton, Roberto Amato, Jeffrey Barrett, Sonia Goncalves, Ewan Harrison, David K. Jackson, Ian Johnston, Dominic Kwiatkowski, Cordelia Langford, John Sillitoe on behalf of the Wellcome Sanger Institute COVID-19 Surveillance Team |
| EPI_ISL_3221480, EPI_ISL_3221481 | Lighthouse Lab in Milton Keynes | Wellcome Sanger Institute for the COVID-19 Genomics UK (COG-UK) Consortium | The Lighthouse Lab in Milton Keynes and Alex Alderton, Roberto Amato, Jeffrey Barrett, Sonia Goncalves, Ewan Harrison, David K. Jackson, Ian Johnston, Dominic Kwiatkowski, Cordelia Langford, John Sillitoe on behalf of the Wellcome Sanger Institute COVID-19 Surveillance Team |
| EPI_ISL_3221482, EPI_ISL_3221483, EPI_ISL_3221484, EPI_ISL_3221485 | Lighthouse Laboratory Plymouth | Wellcome Sanger Institute for the COVID-19 Genomics UK (COG-UK) Consortium | Lighthouse Laboratory Plymouth and Alex Alderton, Roberto Amato, Jeffrey Barrett, Sonia Goncalves, Ewan Harrison, David K. Jackson, Ian Johnston, Dominic Kwiatkowski, Cordelia Langford, John Sillitoe on behalf of the Wellcome Sanger Institute COVID-19 Surveillance Team |
| EPI_ISL_3221486 | Lighthouse Lab in Milton Keynes | Wellcome Sanger Institute for the COVID-19 Genomics UK (COG-UK) Consortium | The Lighthouse Lab in Milton Keynes and Alex Alderton, Roberto Amato, Jeffrey Barrett, Sonia Goncalves, Ewan Harrison, David K. Jackson, Ian Johnston, Dominic Kwiatkowski, Cordelia Langford, John Sillitoe on behalf of the Wellcome Sanger Institute COVID-19 Surveillance Team |
| EPI_ISL_3221487 | Lighthouse Laboratory Plymouth | Wellcome Sanger Institute for the COVID-19 Genomics UK (COG-UK) Consortium | Lighthouse Laboratory Plymouth and Alex Alderton, Roberto Amato, Jeffrey Barrett, Sonia Goncalves, Ewan Harrison, David K. Jackson, Ian Johnston, Dominic Kwiatkowski, Cordelia Langford, John Sillitoe on behalf of the Wellcome Sanger Institute COVID-19 Surveillance Team |
| EPI_ISL_3221488 | Lighthouse Lab in Milton Keynes | Wellcome Sanger Institute for the COVID-19 Genomics UK (COG-UK) Consortium | The Lighthouse Lab in Milton Keynes and Alex Alderton, Roberto Amato, Jeffrey Barrett, Sonia Goncalves, Ewan Harrison, David K. Jackson, Ian Johnston, Dominic Kwiatkowski, Cordelia Langford, John Sillitoe on behalf of the Wellcome Sanger Institute COVID-19 Surveillance Team |
| EPI_ISL_3221489 | Lighthouse Laboratory Plymouth | Wellcome Sanger Institute for the COVID-19 Genomics UK (COG-UK) Consortium | Lighthouse Laboratory Plymouth and Alex Alderton, Roberto Amato, Jeffrey Barrett, Sonia Goncalves, Ewan Harrison, David K. Jackson, Ian Johnston, Dominic Kwiatkowski, Cordelia Langford, John Sillitoe on behalf of the Wellcome Sanger Institute COVID-19 Surveillance Team |
| EPI_ISL_3221490 | Lighthouse Lab in Milton Keynes | Wellcome Sanger Institute for the COVID-19 Genomics UK (COG-UK) Consortium | The Lighthouse Lab in Milton Keynes and Alex Alderton, Roberto Amato, Jeffrey Barrett, Sonia Goncalves, Ewan Harrison, David K. Jackson, Ian Johnston, Dominic Kwiatkowski, Cordelia Langford, John Sillitoe on behalf of the Wellcome Sanger Institute COVID-19 Surveillance Team |
| EPI_ISL_3221491 | Lighthouse Laboratory Plymouth | Wellcome Sanger Institute for the COVID-19 Genomics UK (COG-UK) Consortium | Lighthouse Laboratory Plymouth and Alex Alderton, Roberto Amato, Jeffrey Barrett, Sonia Goncalves, Ewan Harrison, David K. Jackson, Ian Johnston, Dominic Kwiatkowski, Cordelia Langford, John Sillitoe on behalf of the Wellcome Sanger Institute COVID-19 Surveillance Team |
| EPI_ISL_3221492, EPI_ISL_3221493 | Lighthouse Lab in Milton Keynes | Wellcome Sanger Institute for the COVID-19 Genomics UK (COG-UK) Consortium | The Lighthouse Lab in Milton Keynes and Alex Alderton, Roberto Amato, Jeffrey Barrett, Sonia Goncalves, Ewan Harrison, David K. Jackson, Ian Johnston, Dominic Kwiatkowski, Cordelia Langford, John Sillitoe on behalf of the Wellcome Sanger Institute COVID-19 Surveillance Team |
| EPI_ISL_3221494 | Lighthouse Laboratory Plymouth | Wellcome Sanger Institute for the COVID-19 Genomics UK (COG-UK) Consortium | Lighthouse Laboratory Plymouth and Alex Alderton, Roberto Amato, Jeffrey Barrett, Sonia Goncalves, Ewan Harrison, David K. Jackson, Ian Johnston, Dominic Kwiatkowski, Cordelia Langford, John Sillitoe on behalf of the Wellcome Sanger Institute COVID-19 Surveillance Team |
| EPI_ISL_3221495 | Lighthouse Lab in Milton Keynes | Wellcome Sanger Institute for the COVID-19 Genomics UK (COG-UK) Consortium | The Lighthouse Lab in Milton Keynes and Alex Alderton, Roberto Amato, Jeffrey Barrett, Sonia Goncalves, Ewan Harrison, David K. Jackson, Ian Johnston, Dominic Kwiatkowski, Cordelia Langford, John Sillitoe on behalf of the Wellcome Sanger Institute COVID-19 Surveillance Team |
| EPI_ISL_3221496 | Lighthouse Laboratory Plymouth | Wellcome Sanger Institute for the COVID-19 Genomics UK (COG-UK) Consortium | Lighthouse Laboratory Plymouth and Alex Alderton, Roberto Amato, Jeffrey Barrett, Sonia Goncalves, Ewan Harrison, David K. Jackson, Ian Johnston, Dominic Kwiatkowski, Cordelia Langford, John Sillitoe on behalf of the Wellcome Sanger Institute COVID-19 Surveillance Team |
| EPI_ISL_3221497, EPI_ISL_3221498, EPI_ISL_3221499, EPI_ISL_3221500 | Lighthouse Lab in Milton Keynes | Wellcome Sanger Institute for the COVID-19 Genomics UK (COG-UK) Consortium | The Lighthouse Lab in Milton Keynes and Alex Alderton, Roberto Amato, Jeffrey Barrett, Sonia Goncalves, Ewan Harrison, David K. Jackson, Ian Johnston, Dominic Kwiatkowski, Cordelia Langford, John Sillitoe on behalf of the Wellcome Sanger Institute COVID-19 Surveillance Team |
| EPI_ISL_3221501, EPI_ISL_3221502, EPI_ISL_3221503 | Lighthouse Laboratory Plymouth | Wellcome Sanger Institute for the COVID-19 Genomics UK (COG-UK) Consortium | Lighthouse Laboratory Plymouth and Alex Alderton, Roberto Amato, Jeffrey Barrett, Sonia Goncalves, Ewan Harrison, David K. Jackson, Ian Johnston, Dominic Kwiatkowski, Cordelia Langford, John Sillitoe on behalf of the Wellcome Sanger Institute COVID-19 Surveillance Team |
| EPI_ISL_3221504 | Lighthouse Lab in Milton Keynes | Wellcome Sanger Institute for the COVID-19 Genomics UK (COG-UK) Consortium | The Lighthouse Lab in Milton Keynes and Alex Alderton, Roberto Amato, Jeffrey Barrett, Sonia Goncalves, Ewan Harrison, David K. Jackson, Ian Johnston, Dominic Kwiatkowski, Cordelia Langford, John Sillitoe on behalf of the Wellcome Sanger Institute COVID-19 Surveillance Team |
| EPI_ISL_3221505 | Lighthouse Laboratory Plymouth | Wellcome Sanger Institute for the COVID-19 Genomics UK (COG-UK) Consortium | Lighthouse Laboratory Plymouth and Alex Alderton, Roberto Amato, Jeffrey Barrett, Sonia Goncalves, Ewan Harrison, David K. Jackson, Ian Johnston, Dominic Kwiatkowski, Cordelia Langford, John Sillitoe on behalf of the Wellcome Sanger Institute COVID-19 Surveillance Team |
| EPI_ISL_3221506, EPI_ISL_3221507 | Lighthouse Lab in Milton Keynes | Wellcome Sanger Institute for the COVID-19 Genomics UK (COG-UK) Consortium | The Lighthouse Lab in Milton Keynes and Alex Alderton, Roberto Amato, Jeffrey Barrett, Sonia Goncalves, Ewan Harrison, David K. Jackson, Ian Johnston, Dominic Kwiatkowski, Cordelia Langford, John Sillitoe on behalf of the Wellcome Sanger Institute COVID-19 Surveillance Team |
| EPI_ISL_3221508 | Lighthouse Laboratory Plymouth | Wellcome Sanger Institute for the COVID-19 Genomics UK (COG-UK) Consortium | Lighthouse Laboratory Plymouth and Alex Alderton, Roberto Amato, Jeffrey Barrett, Sonia Goncalves, Ewan Harrison, David K. Jackson, Ian Johnston, Dominic Kwiatkowski, Cordelia Langford, John Sillitoe on behalf of the Wellcome Sanger Institute COVID-19 Surveillance Team |
| EPI_ISL_3221509, EPI_ISL_3221510, EPI_ISL_3221511, EPI_ISL_3221512, EPI_ISL_3221513, EPI_ISL_3221514, EPI_ISL_3221515 | Lighthouse Lab in Milton Keynes | Wellcome Sanger Institute for the COVID-19 Genomics UK (COG-UK) Consortium | The Lighthouse Lab in Milton Keynes and Alex Alderton, Roberto Amato, Jeffrey Barrett, Sonia Goncalves, Ewan Harrison, David K. Jackson, Ian Johnston, Dominic Kwiatkowski, Cordelia Langford, John Sillitoe on behalf of the Wellcome Sanger Institute COVID-19 Surveillance Team |
| EPI_ISL_3221516 | Lighthouse Laboratory Plymouth | Wellcome Sanger Institute for the COVID-19 Genomics UK (COG-UK) Consortium | Lighthouse Laboratory Plymouth and Alex Alderton, Roberto Amato, Jeffrey Barrett, Sonia Goncalves, Ewan Harrison, David K. Jackson, Ian Johnston, Dominic Kwiatkowski, Cordelia Langford, John Sillitoe on behalf of the Wellcome Sanger Institute COVID-19 Surveillance Team |
| EPI_ISL_3221517, EPI_ISL_3221518 | Lighthouse Lab in Milton Keynes | Wellcome Sanger Institute for the COVID-19 Genomics UK (COG-UK) Consortium | The Lighthouse Lab in Milton Keynes and Alex Alderton, Roberto Amato, Jeffrey Barrett, Sonia Goncalves, Ewan Harrison, David K. Jackson, Ian Johnston, Dominic Kwiatkowski, Cordelia Langford, John Sillitoe on behalf of the Wellcome Sanger Institute COVID-19 Surveillance Team |
| EPI_ISL_3221519, EPI_ISL_3221520, EPI_ISL_3221521, EPI_ISL_3221522 | Lighthouse Laboratory Plymouth | Wellcome Sanger Institute for the COVID-19 Genomics UK (COG-UK) Consortium | Lighthouse Laboratory Plymouth and Alex Alderton, Roberto Amato, Jeffrey Barrett, Sonia Goncalves, Ewan Harrison, David K. Jackson, Ian Johnston, Dominic Kwiatkowski, Cordelia Langford, John Sillitoe on behalf of the Wellcome Sanger Institute COVID-19 Surveillance Team |
| EPI_ISL_3221523 | Lighthouse Lab in Milton Keynes | Wellcome Sanger Institute for the COVID-19 Genomics UK (COG-UK) Consortium | The Lighthouse Lab in Milton Keynes and Alex Alderton, Roberto Amato, Jeffrey Barrett, Sonia Goncalves, Ewan Harrison, David K. Jackson, Ian Johnston, Dominic Kwiatkowski, Cordelia Langford, John Sillitoe on behalf of the Wellcome Sanger Institute COVID-19 Surveillance Team |
| EPI_ISL_3221524, EPI_ISL_3221525, EPI_ISL_3221526 | Lighthouse Laboratory Plymouth | Wellcome Sanger Institute for the COVID-19 Genomics UK (COG-UK) Consortium | Lighthouse Laboratory Plymouth and Alex Alderton, Roberto Amato, Jeffrey Barrett, Sonia Goncalves, Ewan Harrison, David K. Jackson, Ian Johnston, Dominic Kwiatkowski, Cordelia Langford, John Sillitoe on behalf of the Wellcome Sanger Institute COVID-19 Surveillance Team |
| EPI_ISL_3221527 | Lighthouse Lab in Milton Keynes | Wellcome Sanger Institute for the COVID-19 Genomics UK (COG-UK) Consortium | The Lighthouse Lab in Milton Keynes and Alex Alderton, Roberto Amato, Jeffrey Barrett, Sonia Goncalves, Ewan Harrison, David K. Jackson, Ian Johnston, Dominic Kwiatkowski, Cordelia Langford, John Sillitoe on behalf of the Wellcome Sanger Institute COVID-19 Surveillance Team |
| EPI_ISL_3221528 | Lighthouse Laboratory Plymouth | Wellcome Sanger Institute for the COVID-19 Genomics UK (COG-UK) Consortium | Lighthouse Laboratory Plymouth and Alex Alderton, Roberto Amato, Jeffrey Barrett, Sonia Goncalves, Ewan Harrison, David K. Jackson, Ian Johnston, Dominic Kwiatkowski, Cordelia Langford, John Sillitoe on behalf of the Wellcome Sanger Institute COVID-19 Surveillance Team |
| EPI_ISL_3221529 | Lighthouse Lab in Milton Keynes | Wellcome Sanger Institute for the COVID-19 Genomics UK (COG-UK) Consortium | The Lighthouse Lab in Milton Keynes and Alex Alderton, Roberto Amato, Jeffrey Barrett, Sonia Goncalves, Ewan Harrison, David K. Jackson, Ian Johnston, Dominic Kwiatkowski, Cordelia Langford, John Sillitoe on behalf of the Wellcome Sanger Institute COVID-19 Surveillance Team |
| EPI_ISL_3221530 | Lighthouse Laboratory Plymouth | Wellcome Sanger Institute for the COVID-19 Genomics UK (COG-UK) Consortium | Lighthouse Laboratory Plymouth and Alex Alderton, Roberto Amato, Jeffrey Barrett, Sonia Goncalves, Ewan Harrison, David K. Jackson, Ian Johnston, Dominic Kwiatkowski, Cordelia Langford, John Sillitoe on behalf of the Wellcome Sanger Institute COVID-19 Surveillance Team |

| | | | |
|---|---|---|---|
| EPI_ISL_3221531 | Lighthouse Lab in Milton Keynes | Wellcome Sanger Institute for the COVID-19 Genomics UK (COG-UK) Consortium | The Lighthouse Lab in Milton Keynes and Alex Alderton, Roberto Amato, Jeffrey Barrett, Sonia Goncalves, Ewan Harrison, David K. Jackson, Ian Johnston, Dominic Kwiatkowski, Cordelia Langford, John Sillitoe on behalf of the Wellcome Sanger Institute COVID-19 Surveillance Team |
| EPI_ISL_3221532 | Lighthouse Laboratory Plymouth | Wellcome Sanger Institute for the COVID-19 Genomics UK (COG-UK) Consortium | Lighthouse Laboratory Plymouth and Alex Alderton, Roberto Amato, Jeffrey Barrett, Sonia Goncalves, Ewan Harrison, David K. Jackson, Ian Johnston, Dominic Kwiatkowski, Cordelia Langford, John Sillitoe on behalf of the Wellcome Sanger Institute COVID-19 Surveillance Team |
| EPI_ISL_3221533, EPI_ISL_3221534 | Lighthouse Lab in Milton Keynes | Wellcome Sanger Institute for the COVID-19 Genomics UK (COG-UK) Consortium | The Lighthouse Lab in Milton Keynes and Alex Alderton, Roberto Amato, Jeffrey Barrett, Sonia Goncalves, Ewan Harrison, David K. Jackson, Ian Johnston, Dominic Kwiatkowski, Cordelia Langford, John Sillitoe on behalf of the Wellcome Sanger Institute COVID-19 Surveillance Team |
| EPI_ISL_3221535, EPI_ISL_3221536 | Lighthouse Laboratory Plymouth | Wellcome Sanger Institute for the COVID-19 Genomics UK (COG-UK) Consortium | Lighthouse Laboratory Plymouth and Alex Alderton, Roberto Amato, Jeffrey Barrett, Sonia Goncalves, Ewan Harrison, David K. Jackson, Ian Johnston, Dominic Kwiatkowski, Cordelia Langford, John Sillitoe on behalf of the Wellcome Sanger Institute COVID-19 Surveillance Team |
| EPI_ISL_3221537 | Lighthouse Lab in Milton Keynes | Wellcome Sanger Institute for the COVID-19 Genomics UK (COG-UK) Consortium | The Lighthouse Lab in Milton Keynes and Alex Alderton, Roberto Amato, Jeffrey Barrett, Sonia Goncalves, Ewan Harrison, David K. Jackson, Ian Johnston, Dominic Kwiatkowski, Cordelia Langford, John Sillitoe on behalf of the Wellcome Sanger Institute COVID-19 Surveillance Team |
| EPI_ISL_3221538 | Lighthouse Laboratory Plymouth | Wellcome Sanger Institute for the COVID-19 Genomics UK (COG-UK) Consortium | Lighthouse Laboratory Plymouth and Alex Alderton, Roberto Amato, Jeffrey Barrett, Sonia Goncalves, Ewan Harrison, David K. Jackson, Ian Johnston, Dominic Kwiatkowski, Cordelia Langford, John Sillitoe on behalf of the Wellcome Sanger Institute COVID-19 Surveillance Team |
| EPI_ISL_3221539 | Lighthouse Lab in Milton Keynes | Wellcome Sanger Institute for the COVID-19 Genomics UK (COG-UK) Consortium | The Lighthouse Lab in Milton Keynes and Alex Alderton, Roberto Amato, Jeffrey Barrett, Sonia Goncalves, Ewan Harrison, David K. Jackson, Ian Johnston, Dominic Kwiatkowski, Cordelia Langford, John Sillitoe on behalf of the Wellcome Sanger Institute COVID-19 Surveillance Team |
| EPI_ISL_3221540 | Lighthouse Laboratory Plymouth | Wellcome Sanger Institute for the COVID-19 Genomics UK (COG-UK) Consortium | Lighthouse Laboratory Plymouth and Alex Alderton, Roberto Amato, Jeffrey Barrett, Sonia Goncalves, Ewan Harrison, David K. Jackson, Ian Johnston, Dominic Kwiatkowski, Cordelia Langford, John Sillitoe on behalf of the Wellcome Sanger Institute COVID-19 Surveillance Team |
| EPI_ISL_3221541 | Lighthouse Lab in Milton Keynes | Wellcome Sanger Institute for the COVID-19 Genomics UK (COG-UK) Consortium | The Lighthouse Lab in Milton Keynes and Alex Alderton, Roberto Amato, Jeffrey Barrett, Sonia Goncalves, Ewan Harrison, David K. Jackson, Ian Johnston, Dominic Kwiatkowski, Cordelia Langford, John Sillitoe on behalf of the Wellcome Sanger Institute COVID-19 Surveillance Team |
| EPI_ISL_3221542, EPI_ISL_3221543, EPI_ISL_3221544 | Lighthouse Laboratory Plymouth | Wellcome Sanger Institute for the COVID-19 Genomics UK (COG-UK) Consortium | Lighthouse Laboratory Plymouth and Alex Alderton, Roberto Amato, Jeffrey Barrett, Sonia Goncalves, Ewan Harrison, David K. Jackson, Ian Johnston, Dominic Kwiatkowski, Cordelia Langford, John Sillitoe on behalf of the Wellcome Sanger Institute COVID-19 Surveillance Team |
| EPI_ISL_3221545 | Lighthouse Lab in Milton Keynes | Wellcome Sanger Institute for the COVID-19 Genomics UK (COG-UK) Consortium | The Lighthouse Lab in Milton Keynes and Alex Alderton, Roberto Amato, Jeffrey Barrett, Sonia Goncalves, Ewan Harrison, David K. Jackson, Ian Johnston, Dominic Kwiatkowski, Cordelia Langford, John Sillitoe on behalf of the Wellcome Sanger Institute COVID-19 Surveillance Team |
| EPI_ISL_3221546 | Lighthouse Laboratory Plymouth | Wellcome Sanger Institute for the COVID-19 Genomics UK (COG-UK) Consortium | Lighthouse Laboratory Plymouth and Alex Alderton, Roberto Amato, Jeffrey Barrett, Sonia Goncalves, Ewan Harrison, David K. Jackson, Ian Johnston, Dominic Kwiatkowski, Cordelia Langford, John Sillitoe on behalf of the Wellcome Sanger Institute COVID-19 Surveillance Team |
| EPI_ISL_3221547, EPI_ISL_3221548 | Lighthouse Lab in Milton Keynes | Wellcome Sanger Institute for the COVID-19 Genomics UK (COG-UK) Consortium | The Lighthouse Lab in Milton Keynes and Alex Alderton, Roberto Amato, Jeffrey Barrett, Sonia Goncalves, Ewan Harrison, David K. Jackson, Ian Johnston, Dominic Kwiatkowski, Cordelia Langford, John Sillitoe on behalf of the Wellcome Sanger Institute COVID-19 Surveillance Team |
| EPI_ISL_3221549, EPI_ISL_3221550 | Lighthouse Laboratory Plymouth | Wellcome Sanger Institute for the COVID-19 Genomics UK (COG-UK) Consortium | Lighthouse Laboratory Plymouth and Alex Alderton, Roberto Amato, Jeffrey Barrett, Sonia Goncalves, Ewan Harrison, David K. Jackson, Ian Johnston, Dominic Kwiatkowski, Cordelia Langford, John Sillitoe on behalf of the Wellcome Sanger Institute COVID-19 Surveillance Team |
| EPI_ISL_3221551 | Lighthouse Lab in Milton Keynes | Wellcome Sanger Institute for the COVID-19 Genomics UK (COG-UK) Consortium | The Lighthouse Lab in Milton Keynes and Alex Alderton, Roberto Amato, Jeffrey Barrett, Sonia Goncalves, Ewan Harrison, David K. Jackson, Ian Johnston, Dominic Kwiatkowski, Cordelia Langford, John Sillitoe on behalf of the Wellcome Sanger Institute COVID-19 Surveillance Team |
| EPI_ISL_3221552 | Lighthouse Laboratory Plymouth | Wellcome Sanger Institute for the COVID-19 Genomics UK (COG-UK) Consortium | Lighthouse Laboratory Plymouth and Alex Alderton, Roberto Amato, Jeffrey Barrett, Sonia Goncalves, Ewan Harrison, David K. Jackson, Ian Johnston, Dominic Kwiatkowski, Cordelia Langford, John Sillitoe on behalf of the Wellcome Sanger Institute COVID-19 Surveillance Team |
| EPI_ISL_3221553 | Lighthouse Lab in Milton Keynes | Wellcome Sanger Institute for the COVID-19 Genomics UK (COG-UK) Consortium | The Lighthouse Lab in Milton Keynes and Alex Alderton, Roberto Amato, Jeffrey Barrett, Sonia Goncalves, Ewan Harrison, David K. Jackson, Ian Johnston, Dominic Kwiatkowski, Cordelia Langford, John Sillitoe on behalf of the Wellcome Sanger Institute COVID-19 Surveillance Team |
| EPI_ISL_3221554 | Lighthouse Laboratory Plymouth | Wellcome Sanger Institute for the COVID-19 Genomics UK (COG-UK) Consortium | Lighthouse Laboratory Plymouth and Alex Alderton, Roberto Amato, Jeffrey Barrett, Sonia Goncalves, Ewan Harrison, David K. Jackson, Ian Johnston, Dominic Kwiatkowski, Cordelia Langford, John Sillitoe on behalf of the Wellcome Sanger Institute COVID-19 Surveillance Team |
| EPI_ISL_3221555 | Lighthouse Lab in Milton Keynes | Wellcome Sanger Institute for the COVID-19 Genomics UK (COG-UK) Consortium | The Lighthouse Lab in Milton Keynes and Alex Alderton, Roberto Amato, Jeffrey Barrett, Sonia Goncalves, Ewan Harrison, David K. Jackson, Ian Johnston, Dominic Kwiatkowski, Cordelia Langford, John Sillitoe on behalf of the Wellcome Sanger Institute COVID-19 Surveillance Team |
| EPI_ISL_3221556 | Lighthouse Laboratory Plymouth | Wellcome Sanger Institute for the COVID-19 Genomics UK (COG-UK) Consortium | Lighthouse Laboratory Plymouth and Alex Alderton, Roberto Amato, Jeffrey Barrett, Sonia Goncalves, Ewan Harrison, David K. Jackson, Ian Johnston, Dominic Kwiatkowski, Cordelia Langford, John Sillitoe on behalf of the Wellcome Sanger Institute COVID-19 Surveillance Team |
| EPI_ISL_3221557, EPI_ISL_3221558 | Lighthouse Lab in Milton Keynes | Wellcome Sanger Institute for the COVID-19 Genomics UK (COG-UK) Consortium | The Lighthouse Lab in Milton Keynes and Alex Alderton, Roberto Amato, Jeffrey Barrett, Sonia Goncalves, Ewan Harrison, David K. Jackson, Ian Johnston, Dominic Kwiatkowski, Cordelia Langford, John Sillitoe on behalf of the Wellcome Sanger Institute COVID-19 Surveillance Team |
| EPI_ISL_3221559 | Lighthouse Laboratory Plymouth | Wellcome Sanger Institute for the COVID-19 Genomics UK (COG-UK) Consortium | Lighthouse Laboratory Plymouth and Alex Alderton, Roberto Amato, Jeffrey Barrett, Sonia Goncalves, Ewan Harrison, David K. Jackson, Ian Johnston, Dominic Kwiatkowski, Cordelia Langford, John Sillitoe on behalf of the Wellcome Sanger Institute COVID-19 Surveillance Team |
| EPI_ISL_3221560, EPI_ISL_3221561 | Lighthouse Lab in Milton Keynes | Wellcome Sanger Institute for the COVID-19 Genomics UK (COG-UK) Consortium | The Lighthouse Lab in Milton Keynes and Alex Alderton, Roberto Amato, Jeffrey Barrett, Sonia Goncalves, Ewan Harrison, David K. Jackson, Ian Johnston, Dominic Kwiatkowski, Cordelia Langford, John Sillitoe on behalf of the Wellcome Sanger Institute COVID-19 Surveillance Team |
| EPI_ISL_3221562, EPI_ISL_3221563 | Lighthouse Laboratory Plymouth | Wellcome Sanger Institute for the COVID-19 Genomics UK (COG-UK) Consortium | Lighthouse Laboratory Plymouth and Alex Alderton, Roberto Amato, Jeffrey Barrett, Sonia Goncalves, Ewan Harrison, David K. Jackson, Ian Johnston, Dominic Kwiatkowski, Cordelia Langford, John Sillitoe on behalf of the Wellcome Sanger Institute COVID-19 Surveillance Team |
| EPI_ISL_3221564, EPI_ISL_3221565, EPI_ISL_3221566, EPI_ISL_3221567 | Lighthouse Lab in Milton Keynes | Wellcome Sanger Institute for the COVID-19 Genomics UK (COG-UK) Consortium | The Lighthouse Lab in Milton Keynes and Alex Alderton, Roberto Amato, Jeffrey Barrett, Sonia Goncalves, Ewan Harrison, David K. Jackson, Ian Johnston, Dominic Kwiatkowski, Cordelia Langford, John Sillitoe on behalf of the Wellcome Sanger Institute COVID-19 Surveillance Team |
| EPI_ISL_3221568, EPI_ISL_3221569 | Lighthouse Laboratory Plymouth | Wellcome Sanger Institute for the COVID-19 Genomics UK (COG-UK) Consortium | Lighthouse Laboratory Plymouth and Alex Alderton, Roberto Amato, Jeffrey Barrett, Sonia Goncalves, Ewan Harrison, David K. Jackson, Ian Johnston, Dominic Kwiatkowski, Cordelia Langford, John Sillitoe on behalf of the Wellcome Sanger Institute COVID-19 Surveillance Team |
| EPI_ISL_3221570 | Lighthouse Lab in Milton Keynes | Wellcome Sanger Institute for the COVID-19 Genomics UK (COG-UK) Consortium | The Lighthouse Lab in Milton Keynes and Alex Alderton, Roberto Amato, Jeffrey Barrett, Sonia Goncalves, Ewan Harrison, David K. Jackson, Ian Johnston, Dominic Kwiatkowski, Cordelia Langford, John Sillitoe on behalf of the Wellcome Sanger Institute COVID-19 Surveillance Team |
| EPI_ISL_3221571, EPI_ISL_3221572, EPI_ISL_3221573 | Lighthouse Laboratory Plymouth | Wellcome Sanger Institute for the COVID-19 Genomics UK (COG-UK) Consortium | Lighthouse Laboratory Plymouth and Alex Alderton, Roberto Amato, Jeffrey Barrett, Sonia Goncalves, Ewan Harrison, David K. Jackson, Ian Johnston, Dominic Kwiatkowski, Cordelia Langford, John Sillitoe on behalf of the Wellcome Sanger Institute COVID-19 Surveillance Team |
| EPI_ISL_3221574, EPI_ISL_3221575 | Lighthouse Lab in Milton Keynes | Wellcome Sanger Institute for the COVID-19 Genomics UK (COG-UK) Consortium | The Lighthouse Lab in Milton Keynes and Alex Alderton, Roberto Amato, Jeffrey Barrett, Sonia Goncalves, Ewan Harrison, David K. Jackson, Ian Johnston, Dominic Kwiatkowski, Cordelia Langford, John Sillitoe on behalf of the Wellcome Sanger Institute COVID-19 Surveillance Team |
| EPI_ISL_3221576 | Lighthouse Laboratory Plymouth | Wellcome Sanger Institute for the COVID-19 Genomics UK (COG-UK) Consortium | Lighthouse Laboratory Plymouth and Alex Alderton, Roberto Amato, Jeffrey Barrett, Sonia Goncalves, Ewan Harrison, David K. Jackson, Ian Johnston, Dominic Kwiatkowski, Cordelia Langford, John Sillitoe on behalf of the Wellcome Sanger Institute COVID-19 Surveillance Team |
| EPI_ISL_3221577, EPI_ISL_3221578 | Lighthouse Lab in Milton Keynes | Wellcome Sanger Institute for the COVID-19 Genomics UK (COG-UK) Consortium | The Lighthouse Lab in Milton Keynes and Alex Alderton, Roberto Amato, Jeffrey Barrett, Sonia Goncalves, Ewan Harrison, David K. Jackson, Ian Johnston, Dominic Kwiatkowski, Cordelia Langford, John Sillitoe on behalf of the Wellcome Sanger Institute COVID-19 Surveillance Team |
| EPI_ISL_3221579 | Lighthouse Laboratory Plymouth | Wellcome Sanger Institute for the COVID-19 Genomics UK (COG-UK) Consortium | Lighthouse Laboratory Plymouth and Alex Alderton, Roberto Amato, Jeffrey Barrett, Sonia Goncalves, Ewan Harrison, David K. Jackson, Ian Johnston, Dominic Kwiatkowski, Cordelia Langford, John Sillitoe on behalf of the Wellcome Sanger Institute COVID-19 Surveillance Team |
| EPI_ISL_3221580 | Lighthouse Lab in Milton Keynes | Wellcome Sanger Institute for the COVID-19 Genomics UK (COG-UK) Consortium | The Lighthouse Lab in Milton Keynes and Alex Alderton, Roberto Amato, Jeffrey Barrett, Sonia Goncalves, Ewan Harrison, David K. Jackson, Ian Johnston, Dominic Kwiatkowski, Cordelia Langford, John Sillitoe on behalf of the Wellcome Sanger Institute COVID-19 Surveillance Team |
| EPI_ISL_3221581, EPI_ISL_3221582, EPI_ISL_3221583, EPI_ISL_3221584 | Lighthouse Laboratory Plymouth | Wellcome Sanger Institute for the COVID-19 Genomics UK (COG-UK) Consortium | Lighthouse Laboratory Plymouth and Alex Alderton, Roberto Amato, Jeffrey Barrett, Sonia Goncalves, Ewan Harrison, David K. Jackson, Ian Johnston, Dominic Kwiatkowski, Cordelia Langford, John Sillitoe on behalf of the Wellcome Sanger Institute COVID-19 Surveillance Team |
| EPI_ISL_3221585, EPI_ISL_3221586, EPI_ISL_3221587 | Lighthouse Lab in Milton Keynes | Wellcome Sanger Institute for the COVID-19 Genomics UK (COG-UK) Consortium | The Lighthouse Lab in Milton Keynes and Alex Alderton, Roberto Amato, Jeffrey Barrett, Sonia Goncalves, Ewan Harrison, David K. Jackson, Ian Johnston, Dominic Kwiatkowski, Cordelia Langford, John Sillitoe on behalf of the Wellcome Sanger Institute COVID-19 Surveillance Team |
| EPI_ISL_3221588, EPI_ISL_3221589, EPI_ISL_3221590, EPI_ISL_3221591 | Lighthouse Laboratory Plymouth | Wellcome Sanger Institute for the COVID-19 Genomics UK (COG-UK) Consortium | Lighthouse Laboratory Plymouth and Alex Alderton, Roberto Amato, Jeffrey Barrett, Sonia Goncalves, Ewan Harrison, David K. Jackson, Ian Johnston, Dominic Kwiatkowski, Cordelia Langford, John Sillitoe on behalf of the Wellcome Sanger Institute COVID-19 Surveillance Team |
| EPI_ISL_3221592, EPI_ISL_3221593 | Lighthouse Lab in Milton Keynes | Wellcome Sanger Institute for the COVID-19 Genomics UK (COG-UK) Consortium | The Lighthouse Lab in Milton Keynes and Alex Alderton, Roberto Amato, Jeffrey Barrett, Sonia Goncalves, Ewan Harrison, David K. Jackson, Ian Johnston, Dominic Kwiatkowski, Cordelia Langford, John Sillitoe on behalf of the Wellcome Sanger Institute COVID-19 Surveillance Team |
| EPI_ISL_3221594, EPI_ISL_3221595 | Lighthouse Laboratory Plymouth | Wellcome Sanger Institute for the COVID-19 Genomics UK (COG-UK) Consortium | Lighthouse Laboratory Plymouth and Alex Alderton, Roberto Amato, Jeffrey Barrett, Sonia Goncalves, Ewan Harrison, David K. Jackson, Ian Johnston, Dominic Kwiatkowski, Cordelia Langford, John Sillitoe on behalf of the Wellcome Sanger Institute COVID-19 Surveillance Team |
| EPI_ISL_3221596 | Lighthouse Lab in Milton Keynes | Wellcome Sanger Institute for the COVID-19 Genomics UK (COG-UK) Consortium | The Lighthouse Lab in Milton Keynes and Alex Alderton, Roberto Amato, Jeffrey Barrett, Sonia Goncalves, Ewan Harrison, David K. Jackson, Ian Johnston, Dominic Kwiatkowski, Cordelia Langford, John Sillitoe on behalf of the Wellcome Sanger Institute COVID-19 Surveillance Team |
| EPI_ISL_3221597 | Lighthouse Laboratory Plymouth | Wellcome Sanger Institute for the COVID-19 Genomics UK (COG-UK) Consortium | Lighthouse Laboratory Plymouth and Alex Alderton, Roberto Amato, Jeffrey Barrett, Sonia Goncalves, Ewan Harrison, David K. Jackson, Ian Johnston, Dominic Kwiatkowski, Cordelia Langford, John Sillitoe on behalf of the Wellcome Sanger Institute COVID-19 Surveillance Team |
| EPI_ISL_3221598, EPI_ISL_3221599, EPI_ISL_3221600, EPI_ISL_3221601, | Lighthouse Lab in Milton Keynes | Wellcome Sanger Institute for the COVID-19 Genomics UK (COG-UK) Consortium | The Lighthouse Lab in Milton Keynes and Alex Alderton, Roberto Amato, Jeffrey Barrett, Sonia Goncalves, Ewan Harrison, David K. Jackson, Ian Johnston, Dominic Kwiatkowski, Cordelia Langford, John Sillitoe on behalf of the Wellcome Sanger Institute COVID-19 Surveillance Team |

| | | | |
|---|---|---|---|
| EPI_ISL_3221602 | | | |
| EPI_ISL_3221603 | Lighthouse Laboratory Plymouth | Wellcome Sanger Institute for the COVID-19 Genomics UK (COG-UK) Consortium | Lighthouse Laboratory Plymouth and Alex Alderton, Roberto Amato, Jeffrey Barrett, Sonia Goncalves, Ewan Harrison, David K. Jackson, Ian Johnston, Dominic Kwiatkowski, Cordelia Langford, John Sillitoe on behalf of the Wellcome Sanger Institute COVID-19 Surveillance Team |
| EPI_ISL_3221604 | Lighthouse Lab in Milton Keynes | Wellcome Sanger Institute for the COVID-19 Genomics UK (COG-UK) Consortium | The Lighthouse Lab in Milton Keynes and Alex Alderton, Roberto Amato, Jeffrey Barrett, Sonia Goncalves, Ewan Harrison, David K. Jackson, Ian Johnston, Dominic Kwiatkowski, Cordelia Langford, John Sillitoe on behalf of the Wellcome Sanger Institute COVID-19 Surveillance Team |
| EPI_ISL_3221605, EPI_ISL_3221606 | Lighthouse Laboratory Plymouth | Wellcome Sanger Institute for the COVID-19 Genomics UK (COG-UK) Consortium | Lighthouse Laboratory Plymouth and Alex Alderton, Roberto Amato, Jeffrey Barrett, Sonia Goncalves, Ewan Harrison, David K. Jackson, Ian Johnston, Dominic Kwiatkowski, Cordelia Langford, John Sillitoe on behalf of the Wellcome Sanger Institute COVID-19 Surveillance Team |
| EPI_ISL_3221607 | Lighthouse Lab in Milton Keynes | Wellcome Sanger Institute for the COVID-19 Genomics UK (COG-UK) Consortium | The Lighthouse Lab in Milton Keynes and Alex Alderton, Roberto Amato, Jeffrey Barrett, Sonia Goncalves, Ewan Harrison, David K. Jackson, Ian Johnston, Dominic Kwiatkowski, Cordelia Langford, John Sillitoe on behalf of the Wellcome Sanger Institute COVID-19 Surveillance Team |
| EPI_ISL_3221608 | Lighthouse Laboratory Plymouth | Wellcome Sanger Institute for the COVID-19 Genomics UK (COG-UK) Consortium | Lighthouse Laboratory Plymouth and Alex Alderton, Roberto Amato, Jeffrey Barrett, Sonia Goncalves, Ewan Harrison, David K. Jackson, Ian Johnston, Dominic Kwiatkowski, Cordelia Langford, John Sillitoe on behalf of the Wellcome Sanger Institute COVID-19 Surveillance Team |
| EPI_ISL_3221609 | Lighthouse Lab in Milton Keynes | Wellcome Sanger Institute for the COVID-19 Genomics UK (COG-UK) Consortium | The Lighthouse Lab in Milton Keynes and Alex Alderton, Roberto Amato, Jeffrey Barrett, Sonia Goncalves, Ewan Harrison, David K. Jackson, Ian Johnston, Dominic Kwiatkowski, Cordelia Langford, John Sillitoe on behalf of the Wellcome Sanger Institute COVID-19 Surveillance Team |
| EPI_ISL_3221610 | Lighthouse Laboratory Plymouth | Wellcome Sanger Institute for the COVID-19 Genomics UK (COG-UK) Consortium | Lighthouse Laboratory Plymouth and Alex Alderton, Roberto Amato, Jeffrey Barrett, Sonia Goncalves, Ewan Harrison, David K. Jackson, Ian Johnston, Dominic Kwiatkowski, Cordelia Langford, John Sillitoe on behalf of the Wellcome Sanger Institute COVID-19 Surveillance Team |
| EPI_ISL_3221611 | Lighthouse Lab in Milton Keynes | Wellcome Sanger Institute for the COVID-19 Genomics UK (COG-UK) Consortium | The Lighthouse Lab in Milton Keynes and Alex Alderton, Roberto Amato, Jeffrey Barrett, Sonia Goncalves, Ewan Harrison, David K. Jackson, Ian Johnston, Dominic Kwiatkowski, Cordelia Langford, John Sillitoe on behalf of the Wellcome Sanger Institute COVID-19 Surveillance Team |
| EPI_ISL_3221612, EPI_ISL_3221613 | Lighthouse Laboratory Plymouth | Wellcome Sanger Institute for the COVID-19 Genomics UK (COG-UK) Consortium | Lighthouse Laboratory Plymouth and Alex Alderton, Roberto Amato, Jeffrey Barrett, Sonia Goncalves, Ewan Harrison, David K. Jackson, Ian Johnston, Dominic Kwiatkowski, Cordelia Langford, John Sillitoe on behalf of the Wellcome Sanger Institute COVID-19 Surveillance Team |
| EPI_ISL_3221614 | Lighthouse Lab in Milton Keynes | Wellcome Sanger Institute for the COVID-19 Genomics UK (COG-UK) Consortium | The Lighthouse Lab in Milton Keynes and Alex Alderton, Roberto Amato, Jeffrey Barrett, Sonia Goncalves, Ewan Harrison, David K. Jackson, Ian Johnston, Dominic Kwiatkowski, Cordelia Langford, John Sillitoe on behalf of the Wellcome Sanger Institute COVID-19 Surveillance Team |
| EPI_ISL_3221615 | Lighthouse Laboratory Plymouth | Wellcome Sanger Institute for the COVID-19 Genomics UK (COG-UK) Consortium | Lighthouse Laboratory Plymouth and Alex Alderton, Roberto Amato, Jeffrey Barrett, Sonia Goncalves, Ewan Harrison, David K. Jackson, Ian Johnston, Dominic Kwiatkowski, Cordelia Langford, John Sillitoe on behalf of the Wellcome Sanger Institute COVID-19 Surveillance Team |
| EPI_ISL_3221616, EPI_ISL_3221617, EPI_ISL_3221618 | Lighthouse Lab in Milton Keynes | Wellcome Sanger Institute for the COVID-19 Genomics UK (COG-UK) Consortium | The Lighthouse Lab in Milton Keynes and Alex Alderton, Roberto Amato, Jeffrey Barrett, Sonia Goncalves, Ewan Harrison, David K. Jackson, Ian Johnston, Dominic Kwiatkowski, Cordelia Langford, John Sillitoe on behalf of the Wellcome Sanger Institute COVID-19 Surveillance Team |
| EPI_ISL_3221619, EPI_ISL_3221620 | Lighthouse Laboratory Plymouth | Wellcome Sanger Institute for the COVID-19 Genomics UK (COG-UK) Consortium | Lighthouse Laboratory Plymouth and Alex Alderton, Roberto Amato, Jeffrey Barrett, Sonia Goncalves, Ewan Harrison, David K. Jackson, Ian Johnston, Dominic Kwiatkowski, Cordelia Langford, John Sillitoe on behalf of the Wellcome Sanger Institute COVID-19 Surveillance Team |
| EPI_ISL_3221621 | Lighthouse Lab in Milton Keynes | Wellcome Sanger Institute for the COVID-19 Genomics UK (COG-UK) Consortium | The Lighthouse Lab in Milton Keynes and Alex Alderton, Roberto Amato, Jeffrey Barrett, Sonia Goncalves, Ewan Harrison, David K. Jackson, Ian Johnston, Dominic Kwiatkowski, Cordelia Langford, John Sillitoe on behalf of the Wellcome Sanger Institute COVID-19 Surveillance Team |
| EPI_ISL_3221622 | Lighthouse Laboratory Plymouth | Wellcome Sanger Institute for the COVID-19 Genomics UK (COG-UK) Consortium | Lighthouse Laboratory Plymouth and Alex Alderton, Roberto Amato, Jeffrey Barrett, Sonia Goncalves, Ewan Harrison, David K. Jackson, Ian Johnston, Dominic Kwiatkowski, Cordelia Langford, John Sillitoe on behalf of the Wellcome Sanger Institute COVID-19 Surveillance Team |
| EPI_ISL_3221623 | Lighthouse Lab in Milton Keynes | Wellcome Sanger Institute for the COVID-19 Genomics UK (COG-UK) Consortium | The Lighthouse Lab in Milton Keynes and Alex Alderton, Roberto Amato, Jeffrey Barrett, Sonia Goncalves, Ewan Harrison, David K. Jackson, Ian Johnston, Dominic Kwiatkowski, Cordelia Langford, John Sillitoe on behalf of the Wellcome Sanger Institute COVID-19 Surveillance Team |
| EPI_ISL_3221624 | Lighthouse Laboratory Plymouth | Wellcome Sanger Institute for the COVID-19 Genomics UK (COG-UK) Consortium | Lighthouse Laboratory Plymouth and Alex Alderton, Roberto Amato, Jeffrey Barrett, Sonia Goncalves, Ewan Harrison, David K. Jackson, Ian Johnston, Dominic Kwiatkowski, Cordelia Langford, John Sillitoe on behalf of the Wellcome Sanger Institute COVID-19 Surveillance Team |
| EPI_ISL_3221625 | Lighthouse Lab in Milton Keynes | Wellcome Sanger Institute for the COVID-19 Genomics UK (COG-UK) Consortium | The Lighthouse Lab in Milton Keynes and Alex Alderton, Roberto Amato, Jeffrey Barrett, Sonia Goncalves, Ewan Harrison, David K. Jackson, Ian Johnston, Dominic Kwiatkowski, Cordelia Langford, John Sillitoe on behalf of the Wellcome Sanger Institute COVID-19 Surveillance Team |
| EPI_ISL_3221626 | Lighthouse Laboratory Plymouth | Wellcome Sanger Institute for the COVID-19 Genomics UK (COG-UK) Consortium | Lighthouse Laboratory Plymouth and Alex Alderton, Roberto Amato, Jeffrey Barrett, Sonia Goncalves, Ewan Harrison, David K. Jackson, Ian Johnston, Dominic Kwiatkowski, Cordelia Langford, John Sillitoe on behalf of the Wellcome Sanger Institute COVID-19 Surveillance Team |
| EPI_ISL_3221627, EPI_ISL_3221628 | Lighthouse Lab in Milton Keynes | Wellcome Sanger Institute for the COVID-19 Genomics UK (COG-UK) Consortium | The Lighthouse Lab in Milton Keynes and Alex Alderton, Roberto Amato, Jeffrey Barrett, Sonia Goncalves, Ewan Harrison, David K. Jackson, Ian Johnston, Dominic Kwiatkowski, Cordelia Langford, John Sillitoe on behalf of the Wellcome Sanger Institute COVID-19 Surveillance Team |
| EPI_ISL_3221629 | Lighthouse Laboratory Plymouth | Wellcome Sanger Institute for the COVID-19 Genomics UK (COG-UK) Consortium | Lighthouse Laboratory Plymouth and Alex Alderton, Roberto Amato, Jeffrey Barrett, Sonia Goncalves, Ewan Harrison, David K. Jackson, Ian Johnston, Dominic Kwiatkowski, Cordelia Langford, John Sillitoe on behalf of the Wellcome Sanger Institute COVID-19 Surveillance Team |
| EPI_ISL_3221630 | Lighthouse Lab in Milton Keynes | Wellcome Sanger Institute for the COVID-19 Genomics UK (COG-UK) Consortium | The Lighthouse Lab in Milton Keynes and Alex Alderton, Roberto Amato, Jeffrey Barrett, Sonia Goncalves, Ewan Harrison, David K. Jackson, Ian Johnston, Dominic Kwiatkowski, Cordelia Langford, John Sillitoe on behalf of the Wellcome Sanger Institute COVID-19 Surveillance Team |
| EPI_ISL_3221631, EPI_ISL_3221632 | Lighthouse Laboratory Plymouth | Wellcome Sanger Institute for the COVID-19 Genomics UK (COG-UK) Consortium | Lighthouse Laboratory Plymouth and Alex Alderton, Roberto Amato, Jeffrey Barrett, Sonia Goncalves, Ewan Harrison, David K. Jackson, Ian Johnston, Dominic Kwiatkowski, Cordelia Langford, John Sillitoe on behalf of the Wellcome Sanger Institute COVID-19 Surveillance Team |
| EPI_ISL_3221633 | Lighthouse Lab in Milton Keynes | Wellcome Sanger Institute for the COVID-19 Genomics UK (COG-UK) Consortium | The Lighthouse Lab in Milton Keynes and Alex Alderton, Roberto Amato, Jeffrey Barrett, Sonia Goncalves, Ewan Harrison, David K. Jackson, Ian Johnston, Dominic Kwiatkowski, Cordelia Langford, John Sillitoe on behalf of the Wellcome Sanger Institute COVID-19 Surveillance Team |
| EPI_ISL_3221634 | Lighthouse Laboratory Plymouth | Wellcome Sanger Institute for the COVID-19 Genomics UK (COG-UK) Consortium | Lighthouse Laboratory Plymouth and Alex Alderton, Roberto Amato, Jeffrey Barrett, Sonia Goncalves, Ewan Harrison, David K. Jackson, Ian Johnston, Dominic Kwiatkowski, Cordelia Langford, John Sillitoe on behalf of the Wellcome Sanger Institute COVID-19 Surveillance Team |
| EPI_ISL_3221635 | Lighthouse Lab in Milton Keynes | Wellcome Sanger Institute for the COVID-19 Genomics UK (COG-UK) Consortium | The Lighthouse Lab in Milton Keynes and Alex Alderton, Roberto Amato, Jeffrey Barrett, Sonia Goncalves, Ewan Harrison, David K. Jackson, Ian Johnston, Dominic Kwiatkowski, Cordelia Langford, John Sillitoe on behalf of the Wellcome Sanger Institute COVID-19 Surveillance Team |
| EPI_ISL_3221636, EPI_ISL_3221637, EPI_ISL_3221638, EPI_ISL_3221639, EPI_ISL_3221640, EPI_ISL_3221641, EPI_ISL_3221642 | Lighthouse Laboratory Plymouth | Wellcome Sanger Institute for the COVID-19 Genomics UK (COG-UK) Consortium | Lighthouse Laboratory Plymouth and Alex Alderton, Roberto Amato, Jeffrey Barrett, Sonia Goncalves, Ewan Harrison, David K. Jackson, Ian Johnston, Dominic Kwiatkowski, Cordelia Langford, John Sillitoe on behalf of the Wellcome Sanger Institute COVID-19 Surveillance Team |
| EPI_ISL_3221643 | Lighthouse Lab in Milton Keynes | Wellcome Sanger Institute for the COVID-19 Genomics UK (COG-UK) Consortium | The Lighthouse Lab in Milton Keynes and Alex Alderton, Roberto Amato, Jeffrey Barrett, Sonia Goncalves, Ewan Harrison, David K. Jackson, Ian Johnston, Dominic Kwiatkowski, Cordelia Langford, John Sillitoe on behalf of the Wellcome Sanger Institute COVID-19 Surveillance Team |
| EPI_ISL_3221644, EPI_ISL_3221645, EPI_ISL_3221646, EPI_ISL_3221647, EPI_ISL_3221648 | Lighthouse Laboratory Plymouth | Wellcome Sanger Institute for the COVID-19 Genomics UK (COG-UK) Consortium | Lighthouse Laboratory Plymouth and Alex Alderton, Roberto Amato, Jeffrey Barrett, Sonia Goncalves, Ewan Harrison, David K. Jackson, Ian Johnston, Dominic Kwiatkowski, Cordelia Langford, John Sillitoe on behalf of the Wellcome Sanger Institute COVID-19 Surveillance Team |
| EPI_ISL_3221649 | Lighthouse Lab in Milton Keynes | Wellcome Sanger Institute for the COVID-19 Genomics UK (COG-UK) Consortium | The Lighthouse Lab in Milton Keynes and Alex Alderton, Roberto Amato, Jeffrey Barrett, Sonia Goncalves, Ewan Harrison, David K. Jackson, Ian Johnston, Dominic Kwiatkowski, Cordelia Langford, John Sillitoe on behalf of the Wellcome Sanger Institute COVID-19 Surveillance Team |
| EPI_ISL_3221650 | Lighthouse Laboratory Plymouth | Wellcome Sanger Institute for the COVID-19 Genomics UK (COG-UK) Consortium | Lighthouse Laboratory Plymouth and Alex Alderton, Roberto Amato, Jeffrey Barrett, Sonia Goncalves, Ewan Harrison, David K. Jackson, Ian Johnston, Dominic Kwiatkowski, Cordelia Langford, John Sillitoe on behalf of the Wellcome Sanger Institute COVID-19 Surveillance Team |
| EPI_ISL_3221651 | Lighthouse Lab in Milton Keynes | Wellcome Sanger Institute for the COVID-19 Genomics UK (COG-UK) Consortium | The Lighthouse Lab in Milton Keynes and Alex Alderton, Roberto Amato, Jeffrey Barrett, Sonia Goncalves, Ewan Harrison, David K. Jackson, Ian Johnston, Dominic Kwiatkowski, Cordelia Langford, John Sillitoe on behalf of the Wellcome Sanger Institute COVID-19 Surveillance Team |
| EPI_ISL_3221652 | Lighthouse Laboratory Plymouth | Wellcome Sanger Institute for the COVID-19 Genomics UK (COG-UK) Consortium | Lighthouse Laboratory Plymouth and Alex Alderton, Roberto Amato, Jeffrey Barrett, Sonia Goncalves, Ewan Harrison, David K. Jackson, Ian Johnston, Dominic Kwiatkowski, Cordelia Langford, John Sillitoe on behalf of the Wellcome Sanger Institute COVID-19 Surveillance Team |
| EPI_ISL_3221653 | Lighthouse Lab in Milton Keynes | Wellcome Sanger Institute for the COVID-19 Genomics UK (COG-UK) Consortium | The Lighthouse Lab in Milton Keynes and Alex Alderton, Roberto Amato, Jeffrey Barrett, Sonia Goncalves, Ewan Harrison, David K. Jackson, Ian Johnston, Dominic Kwiatkowski, Cordelia Langford, John Sillitoe on behalf of the Wellcome Sanger Institute COVID-19 Surveillance Team |
| EPI_ISL_3221654, EPI_ISL_3221655, EPI_ISL_3221656, EPI_ISL_3221657 | Lighthouse Laboratory Plymouth | Wellcome Sanger Institute for the COVID-19 Genomics UK (COG-UK) Consortium | Lighthouse Laboratory Plymouth and Alex Alderton, Roberto Amato, Jeffrey Barrett, Sonia Goncalves, Ewan Harrison, David K. Jackson, Ian Johnston, Dominic Kwiatkowski, Cordelia Langford, John Sillitoe on behalf of the Wellcome Sanger Institute COVID-19 Surveillance Team |
| EPI_ISL_3221658 | Lighthouse Lab in Milton Keynes | Wellcome Sanger Institute for the COVID-19 Genomics UK (COG-UK) Consortium | The Lighthouse Lab in Milton Keynes and Alex Alderton, Roberto Amato, Jeffrey Barrett, Sonia Goncalves, Ewan Harrison, David K. Jackson, Ian Johnston, Dominic Kwiatkowski, Cordelia Langford, John Sillitoe on behalf of the Wellcome Sanger Institute COVID-19 Surveillance Team |
| EPI_ISL_3221659 | Lighthouse Laboratory Plymouth | Wellcome Sanger Institute for the COVID-19 Genomics UK (COG-UK) Consortium | Lighthouse Laboratory Plymouth and Alex Alderton, Roberto Amato, Jeffrey Barrett, Sonia Goncalves, Ewan Harrison, David K. Jackson, Ian Johnston, Dominic Kwiatkowski, Cordelia Langford, John Sillitoe on behalf of the Wellcome Sanger Institute COVID-19 Surveillance Team |
| EPI_ISL_3221660 | Lighthouse Lab in Milton Keynes | Wellcome Sanger Institute for the COVID-19 Genomics UK (COG-UK) Consortium | The Lighthouse Lab in Milton Keynes and Alex Alderton, Roberto Amato, Jeffrey Barrett, Sonia Goncalves, Ewan Harrison, David K. Jackson, Ian Johnston, Dominic Kwiatkowski, Cordelia Langford, John Sillitoe on behalf of the Wellcome Sanger Institute COVID-19 Surveillance Team |
| EPI_ISL_3221661 | Lighthouse Laboratory Plymouth | Wellcome Sanger Institute for the COVID-19 Genomics UK (COG-UK) Consortium | Lighthouse Laboratory Plymouth and Alex Alderton, Roberto Amato, Jeffrey Barrett, Sonia Goncalves, Ewan Harrison, David K. Jackson, Ian Johnston, Dominic Kwiatkowski, Cordelia Langford, John Sillitoe on behalf of the Wellcome Sanger Institute COVID-19 Surveillance Team |
| EPI_ISL_3221662 | Lighthouse Lab in Glasgow | Wellcome Sanger Institute for the COVID-19 Genomics UK (COG-UK) Consortium | Harper VanSteenhouse, Yumi Kasai, David Gray, Carol Clugston, Anna Dominiczak and Alex Alderton, Roberto Amato, Jeffrey Barrett, Sonia Goncalves, Ewan Harrison, David K. Jackson, Ian Johnston, Dominic Kwiatkowski, Cordelia Langford, John Sillitoe on behalf of the Wellcome Sanger Institute COVID-19 Surveillance Team |

| | | | |
|---|---|---|---|
| EPI_ISL_3221663 | Lighthouse Laboratory Plymouth | Wellcome Sanger Institute for the COVID-19 Genomics UK (COG-UK) Consortium | Lighthouse Laboratory Plymouth and Alex Alderton, Roberto Amato, Jeffrey Barrett, Sonia Goncalves, Ewan Harrison, David K. Jackson, Ian Johnston, Dominic Kwiatkowski, Cordelia Langford, John Sillitoe on behalf of the Wellcome Sanger Institute COVID-19 Surveillance Team |
| EPI_ISL_3221664 | Lighthouse Lab in Glasgow | Wellcome Sanger Institute for the COVID-19 Genomics UK (COG-UK) Consortium | Harper VanSteenhouse, Yumi Kasai, David Gray, Carol Clugston, Anna Dominiczak and Alex Alderton, Roberto Amato, Jeffrey Barrett, Sonia Goncalves, Ewan Harrison, David K. Jackson, Ian Johnston, Dominic Kwiatkowski, Cordelia Langford, John Sillitoe on behalf of the Wellcome Sanger Institute COVID-19 Surveillance Team |
| EPI_ISL_3221665 | Lighthouse Lab in Milton Keynes | Wellcome Sanger Institute for the COVID-19 Genomics UK (COG-UK) Consortium | The Lighthouse Lab in Milton Keynes and Alex Alderton, Roberto Amato, Jeffrey Barrett, Sonia Goncalves, Ewan Harrison, David K. Jackson, Ian Johnston, Dominic Kwiatkowski, Cordelia Langford, John Sillitoe on behalf of the Wellcome Sanger Institute COVID-19 Surveillance Team |
| EPI_ISL_3221666, EPI_ISL_3221667, EPI_ISL_3221668 | Lighthouse Lab in Glasgow | Wellcome Sanger Institute for the COVID-19 Genomics UK (COG-UK) Consortium | Harper VanSteenhouse, Yumi Kasai, David Gray, Carol Clugston, Anna Dominiczak and Alex Alderton, Roberto Amato, Jeffrey Barrett, Sonia Goncalves, Ewan Harrison, David K. Jackson, Ian Johnston, Dominic Kwiatkowski, Cordelia Langford, John Sillitoe on behalf of the Wellcome Sanger Institute COVID-19 Surveillance Team |
| EPI_ISL_3221669 | Lighthouse Laboratory Plymouth | Wellcome Sanger Institute for the COVID-19 Genomics UK (COG-UK) Consortium | Lighthouse Laboratory Plymouth and Alex Alderton, Roberto Amato, Jeffrey Barrett, Sonia Goncalves, Ewan Harrison, David K. Jackson, Ian Johnston, Dominic Kwiatkowski, Cordelia Langford, John Sillitoe on behalf of the Wellcome Sanger Institute COVID-19 Surveillance Team |
| EPI_ISL_3221670, EPI_ISL_3221671 | Lighthouse Lab in Milton Keynes | Wellcome Sanger Institute for the COVID-19 Genomics UK (COG-UK) Consortium | The Lighthouse Lab in Milton Keynes and Alex Alderton, Roberto Amato, Jeffrey Barrett, Sonia Goncalves, Ewan Harrison, David K. Jackson, Ian Johnston, Dominic Kwiatkowski, Cordelia Langford, John Sillitoe on behalf of the Wellcome Sanger Institute COVID-19 Surveillance Team |
| EPI_ISL_3221672 | Lighthouse Laboratory Plymouth | Wellcome Sanger Institute for the COVID-19 Genomics UK (COG-UK) Consortium | Lighthouse Laboratory Plymouth and Alex Alderton, Roberto Amato, Jeffrey Barrett, Sonia Goncalves, Ewan Harrison, David K. Jackson, Ian Johnston, Dominic Kwiatkowski, Cordelia Langford, John Sillitoe on behalf of the Wellcome Sanger Institute COVID-19 Surveillance Team |
| EPI_ISL_3221673, EPI_ISL_3221674 | Lighthouse Lab in Glasgow | Wellcome Sanger Institute for the COVID-19 Genomics UK (COG-UK) Consortium | Harper VanSteenhouse, Yumi Kasai, David Gray, Carol Clugston, Anna Dominiczak and Alex Alderton, Roberto Amato, Jeffrey Barrett, Sonia Goncalves, Ewan Harrison, David K. Jackson, Ian Johnston, Dominic Kwiatkowski, Cordelia Langford, John Sillitoe on behalf of the Wellcome Sanger Institute COVID-19 Surveillance Team |
| EPI_ISL_3221675 | Lighthouse Laboratory Plymouth | Wellcome Sanger Institute for the COVID-19 Genomics UK (COG-UK) Consortium | Lighthouse Laboratory Plymouth and Alex Alderton, Roberto Amato, Jeffrey Barrett, Sonia Goncalves, Ewan Harrison, David K. Jackson, Ian Johnston, Dominic Kwiatkowski, Cordelia Langford, John Sillitoe on behalf of the Wellcome Sanger Institute COVID-19 Surveillance Team |
| EPI_ISL_3221676 | Lighthouse Lab in Milton Keynes | Wellcome Sanger Institute for the COVID-19 Genomics UK (COG-UK) Consortium | The Lighthouse Lab in Milton Keynes and Alex Alderton, Roberto Amato, Jeffrey Barrett, Sonia Goncalves, Ewan Harrison, David K. Jackson, Ian Johnston, Dominic Kwiatkowski, Cordelia Langford, John Sillitoe on behalf of the Wellcome Sanger Institute COVID-19 Surveillance Team |
| EPI_ISL_3221677, EPI_ISL_3221678, EPI_ISL_3221679 | Lighthouse Lab in Glasgow | Wellcome Sanger Institute for the COVID-19 Genomics UK (COG-UK) Consortium | Harper VanSteenhouse, Yumi Kasai, David Gray, Carol Clugston, Anna Dominiczak and Alex Alderton, Roberto Amato, Jeffrey Barrett, Sonia Goncalves, Ewan Harrison, David K. Jackson, Ian Johnston, Dominic Kwiatkowski, Cordelia Langford, John Sillitoe on behalf of the Wellcome Sanger Institute COVID-19 Surveillance Team |
| EPI_ISL_3221680, EPI_ISL_3221681, EPI_ISL_3221682 | Lighthouse Lab in Milton Keynes | Wellcome Sanger Institute for the COVID-19 Genomics UK (COG-UK) Consortium | The Lighthouse Lab in Milton Keynes and Alex Alderton, Roberto Amato, Jeffrey Barrett, Sonia Goncalves, Ewan Harrison, David K. Jackson, Ian Johnston, Dominic Kwiatkowski, Cordelia Langford, John Sillitoe on behalf of the Wellcome Sanger Institute COVID-19 Surveillance Team |
| EPI_ISL_3221683, EPI_ISL_3221684 | Lighthouse Lab in Glasgow | Wellcome Sanger Institute for the COVID-19 Genomics UK (COG-UK) Consortium | Harper VanSteenhouse, Yumi Kasai, David Gray, Carol Clugston, Anna Dominiczak and Alex Alderton, Roberto Amato, Jeffrey Barrett, Sonia Goncalves, Ewan Harrison, David K. Jackson, Ian Johnston, Dominic Kwiatkowski, Cordelia Langford, John Sillitoe on behalf of the Wellcome Sanger Institute COVID-19 Surveillance Team |
| EPI_ISL_3221685 | Lighthouse Lab in Milton Keynes | Wellcome Sanger Institute for the COVID-19 Genomics UK (COG-UK) Consortium | The Lighthouse Lab in Milton Keynes and Alex Alderton, Roberto Amato, Jeffrey Barrett, Sonia Goncalves, Ewan Harrison, David K. Jackson, Ian Johnston, Dominic Kwiatkowski, Cordelia Langford, John Sillitoe on behalf of the Wellcome Sanger Institute COVID-19 Surveillance Team |
| EPI_ISL_3221686 | Lighthouse Laboratory Plymouth | Wellcome Sanger Institute for the COVID-19 Genomics UK (COG-UK) Consortium | Lighthouse Laboratory Plymouth and Alex Alderton, Roberto Amato, Jeffrey Barrett, Sonia Goncalves, Ewan Harrison, David K. Jackson, Ian Johnston, Dominic Kwiatkowski, Cordelia Langford, John Sillitoe on behalf of the Wellcome Sanger Institute COVID-19 Surveillance Team |
| EPI_ISL_3221687, EPI_ISL_3221688 | Lighthouse Lab in Milton Keynes | Wellcome Sanger Institute for the COVID-19 Genomics UK (COG-UK) Consortium | The Lighthouse Lab in Milton Keynes and Alex Alderton, Roberto Amato, Jeffrey Barrett, Sonia Goncalves, Ewan Harrison, David K. Jackson, Ian Johnston, Dominic Kwiatkowski, Cordelia Langford, John Sillitoe on behalf of the Wellcome Sanger Institute COVID-19 Surveillance Team |
| EPI_ISL_3221689, EPI_ISL_3221690 | Lighthouse Laboratory Plymouth | Wellcome Sanger Institute for the COVID-19 Genomics UK (COG-UK) Consortium | Lighthouse Laboratory Plymouth and Alex Alderton, Roberto Amato, Jeffrey Barrett, Sonia Goncalves, Ewan Harrison, David K. Jackson, Ian Johnston, Dominic Kwiatkowski, Cordelia Langford, John Sillitoe on behalf of the Wellcome Sanger Institute COVID-19 Surveillance Team |
| EPI_ISL_3221691 | Lighthouse Lab in Milton Keynes | Wellcome Sanger Institute for the COVID-19 Genomics UK (COG-UK) Consortium | The Lighthouse Lab in Milton Keynes and Alex Alderton, Roberto Amato, Jeffrey Barrett, Sonia Goncalves, Ewan Harrison, David K. Jackson, Ian Johnston, Dominic Kwiatkowski, Cordelia Langford, John Sillitoe on behalf of the Wellcome Sanger Institute COVID-19 Surveillance Team |
| EPI_ISL_3221692 | Lighthouse Lab in Glasgow | Wellcome Sanger Institute for the COVID-19 Genomics UK (COG-UK) Consortium | Harper VanSteenhouse, Yumi Kasai, David Gray, Carol Clugston, Anna Dominiczak and Alex Alderton, Roberto Amato, Jeffrey Barrett, Sonia Goncalves, Ewan Harrison, David K. Jackson, Ian Johnston, Dominic Kwiatkowski, Cordelia Langford, John Sillitoe on behalf of the Wellcome Sanger Institute COVID-19 Surveillance Team |
| EPI_ISL_3221693 | Lighthouse Laboratory Plymouth | Wellcome Sanger Institute for the COVID-19 Genomics UK (COG-UK) Consortium | Lighthouse Laboratory Plymouth and Alex Alderton, Roberto Amato, Jeffrey Barrett, Sonia Goncalves, Ewan Harrison, David K. Jackson, Ian Johnston, Dominic Kwiatkowski, Cordelia Langford, John Sillitoe on behalf of the Wellcome Sanger Institute COVID-19 Surveillance Team |
| EPI_ISL_3221694 | Lighthouse Lab in Milton Keynes | Wellcome Sanger Institute for the COVID-19 Genomics UK (COG-UK) Consortium | The Lighthouse Lab in Milton Keynes and Alex Alderton, Roberto Amato, Jeffrey Barrett, Sonia Goncalves, Ewan Harrison, David K. Jackson, Ian Johnston, Dominic Kwiatkowski, Cordelia Langford, John Sillitoe on behalf of the Wellcome Sanger Institute COVID-19 Surveillance Team |
| EPI_ISL_3221695 | Lighthouse Lab in Glasgow | Wellcome Sanger Institute for the COVID-19 Genomics UK (COG-UK) Consortium | Harper VanSteenhouse, Yumi Kasai, David Gray, Carol Clugston, Anna Dominiczak and Alex Alderton, Roberto Amato, Jeffrey Barrett, Sonia Goncalves, Ewan Harrison, David K. Jackson, Ian Johnston, Dominic Kwiatkowski, Cordelia Langford, John Sillitoe on behalf of the Wellcome Sanger Institute COVID-19 Surveillance Team |
| EPI_ISL_3221696 | Lighthouse Laboratory Plymouth | Wellcome Sanger Institute for the COVID-19 Genomics UK (COG-UK) Consortium | Lighthouse Laboratory Plymouth and Alex Alderton, Roberto Amato, Jeffrey Barrett, Sonia Goncalves, Ewan Harrison, David K. Jackson, Ian Johnston, Dominic Kwiatkowski, Cordelia Langford, John Sillitoe on behalf of the Wellcome Sanger Institute COVID-19 Surveillance Team |
| EPI_ISL_3221697 | Lighthouse Lab in Glasgow | Wellcome Sanger Institute for the COVID-19 Genomics UK (COG-UK) Consortium | Harper VanSteenhouse, Yumi Kasai, David Gray, Carol Clugston, Anna Dominiczak and Alex Alderton, Roberto Amato, Jeffrey Barrett, Sonia Goncalves, Ewan Harrison, David K. Jackson, Ian Johnston, Dominic Kwiatkowski, Cordelia Langford, John Sillitoe on behalf of the Wellcome Sanger Institute COVID-19 Surveillance Team |
| EPI_ISL_3221698 | Lighthouse Laboratory Plymouth | Wellcome Sanger Institute for the COVID-19 Genomics UK (COG-UK) Consortium | Lighthouse Laboratory Plymouth and Alex Alderton, Roberto Amato, Jeffrey Barrett, Sonia Goncalves, Ewan Harrison, David K. Jackson, Ian Johnston, Dominic Kwiatkowski, Cordelia Langford, John Sillitoe on behalf of the Wellcome Sanger Institute COVID-19 Surveillance Team |
| EPI_ISL_3221699, EPI_ISL_3221700 | Lighthouse Lab in Milton Keynes | Wellcome Sanger Institute for the COVID-19 Genomics UK (COG-UK) Consortium | The Lighthouse Lab in Milton Keynes and Alex Alderton, Roberto Amato, Jeffrey Barrett, Sonia Goncalves, Ewan Harrison, David K. Jackson, Ian Johnston, Dominic Kwiatkowski, Cordelia Langford, John Sillitoe on behalf of the Wellcome Sanger Institute COVID-19 Surveillance Team |
| EPI_ISL_3221701, EPI_ISL_3221702, EPI_ISL_3221703, EPI_ISL_3221704, EPI_ISL_3221705, EPI_ISL_3221706 | Lighthouse Lab in Glasgow | Wellcome Sanger Institute for the COVID-19 Genomics UK (COG-UK) Consortium | Harper VanSteenhouse, Yumi Kasai, David Gray, Carol Clugston, Anna Dominiczak and Alex Alderton, Roberto Amato, Jeffrey Barrett, Sonia Goncalves, Ewan Harrison, David K. Jackson, Ian Johnston, Dominic Kwiatkowski, Cordelia Langford, John Sillitoe on behalf of the Wellcome Sanger Institute COVID-19 Surveillance Team |
| EPI_ISL_3221707 | Lighthouse Lab in Milton Keynes | Wellcome Sanger Institute for the COVID-19 Genomics UK (COG-UK) Consortium | The Lighthouse Lab in Milton Keynes and Alex Alderton, Roberto Amato, Jeffrey Barrett, Sonia Goncalves, Ewan Harrison, David K. Jackson, Ian Johnston, Dominic Kwiatkowski, Cordelia Langford, John Sillitoe on behalf of the Wellcome Sanger Institute COVID-19 Surveillance Team |
| EPI_ISL_3221708 | Lighthouse Lab in Glasgow | Wellcome Sanger Institute for the COVID-19 Genomics UK (COG-UK) Consortium | Harper VanSteenhouse, Yumi Kasai, David Gray, Carol Clugston, Anna Dominiczak and Alex Alderton, Roberto Amato, Jeffrey Barrett, Sonia Goncalves, Ewan Harrison, David K. Jackson, Ian Johnston, Dominic Kwiatkowski, Cordelia Langford, John Sillitoe on behalf of the Wellcome Sanger Institute COVID-19 Surveillance Team |
| EPI_ISL_3221709 | Lighthouse Laboratory Plymouth | Wellcome Sanger Institute for the COVID-19 Genomics UK (COG-UK) Consortium | Lighthouse Laboratory Plymouth and Alex Alderton, Roberto Amato, Jeffrey Barrett, Sonia Goncalves, Ewan Harrison, David K. Jackson, Ian Johnston, Dominic Kwiatkowski, Cordelia Langford, John Sillitoe on behalf of the Wellcome Sanger Institute COVID-19 Surveillance Team |
| EPI_ISL_3221710 | Lighthouse Lab in Glasgow | Wellcome Sanger Institute for the COVID-19 Genomics UK (COG-UK) Consortium | Harper VanSteenhouse, Yumi Kasai, David Gray, Carol Clugston, Anna Dominiczak and Alex Alderton, Roberto Amato, Jeffrey Barrett, Sonia Goncalves, Ewan Harrison, David K. Jackson, Ian Johnston, Dominic Kwiatkowski, Cordelia Langford, John Sillitoe on behalf of the Wellcome Sanger Institute COVID-19 Surveillance Team |
| EPI_ISL_3221711, EPI_ISL_3221712 | Lighthouse Laboratory Plymouth | Wellcome Sanger Institute for the COVID-19 Genomics UK (COG-UK) Consortium | Lighthouse Laboratory Plymouth and Alex Alderton, Roberto Amato, Jeffrey Barrett, Sonia Goncalves, Ewan Harrison, David K. Jackson, Ian Johnston, Dominic Kwiatkowski, Cordelia Langford, John Sillitoe on behalf of the Wellcome Sanger Institute COVID-19 Surveillance Team |
| EPI_ISL_3221713 | Lighthouse Lab in Glasgow | Wellcome Sanger Institute for the COVID-19 Genomics UK (COG-UK) Consortium | Harper VanSteenhouse, Yumi Kasai, David Gray, Carol Clugston, Anna Dominiczak and Alex Alderton, Roberto Amato, Jeffrey Barrett, Sonia Goncalves, Ewan Harrison, David K. Jackson, Ian Johnston, Dominic Kwiatkowski, Cordelia Langford, John Sillitoe on behalf of the Wellcome Sanger Institute COVID-19 Surveillance Team |
| EPI_ISL_3221714 | Lighthouse Lab in Milton Keynes | Wellcome Sanger Institute for the COVID-19 Genomics UK (COG-UK) Consortium | The Lighthouse Lab in Milton Keynes and Alex Alderton, Roberto Amato, Jeffrey Barrett, Sonia Goncalves, Ewan Harrison, David K. Jackson, Ian Johnston, Dominic Kwiatkowski, Cordelia Langford, John Sillitoe on behalf of the Wellcome Sanger Institute COVID-19 Surveillance Team |
| EPI_ISL_3221715, EPI_ISL_3221716, EPI_ISL_3221717, EPI_ISL_3221718 | Lighthouse Lab in Glasgow | Wellcome Sanger Institute for the COVID-19 Genomics UK (COG-UK) Consortium | Harper VanSteenhouse, Yumi Kasai, David Gray, Carol Clugston, Anna Dominiczak and Alex Alderton, Roberto Amato, Jeffrey Barrett, Sonia Goncalves, Ewan Harrison, David K. Jackson, Ian Johnston, Dominic Kwiatkowski, Cordelia Langford, John Sillitoe on behalf of the Wellcome Sanger Institute COVID-19 Surveillance Team |
| EPI_ISL_3221719 | Lighthouse Laboratory Plymouth | Wellcome Sanger Institute for the COVID-19 Genomics UK (COG-UK) Consortium | Lighthouse Laboratory Plymouth and Alex Alderton, Roberto Amato, Jeffrey Barrett, Sonia Goncalves, Ewan Harrison, David K. Jackson, Ian Johnston, Dominic Kwiatkowski, Cordelia Langford, John Sillitoe on behalf of the Wellcome Sanger Institute COVID-19 Surveillance Team |
| EPI_ISL_3221720, EPI_ISL_3221721, EPI_ISL_3221722, EPI_ISL_3221723 | Lighthouse Lab in Glasgow | Wellcome Sanger Institute for the COVID-19 Genomics UK (COG-UK) Consortium | Harper VanSteenhouse, Yumi Kasai, David Gray, Carol Clugston, Anna Dominiczak and Alex Alderton, Roberto Amato, Jeffrey Barrett, Sonia Goncalves, Ewan Harrison, David K. Jackson, Ian Johnston, Dominic Kwiatkowski, Cordelia Langford, John Sillitoe on behalf of the Wellcome Sanger Institute COVID-19 Surveillance Team |
| EPI_ISL_3221724 | Lighthouse Lab in Milton Keynes | Wellcome Sanger Institute for the COVID-19 Genomics UK (COG-UK) Consortium | The Lighthouse Lab in Milton Keynes and Alex Alderton, Roberto Amato, Jeffrey Barrett, Sonia Goncalves, Ewan Harrison, David K. Jackson, Ian Johnston, Dominic Kwiatkowski, Cordelia Langford, John Sillitoe on behalf of the Wellcome Sanger Institute COVID-19 Surveillance Team |
| EPI_ISL_3221725 | Lighthouse Laboratory Plymouth | Wellcome Sanger Institute for the COVID-19 Genomics UK (COG-UK) Consortium | Lighthouse Laboratory Plymouth and Alex Alderton, Roberto Amato, Jeffrey Barrett, Sonia Goncalves, Ewan Harrison, David K. Jackson, Ian Johnston, Dominic Kwiatkowski, Cordelia Langford, John Sillitoe on behalf of the Wellcome Sanger Institute COVID-19 Surveillance Team |
| EPI_ISL_3221726 | Lighthouse Lab in Glasgow | Wellcome Sanger Institute for the COVID-19 Genomics UK (COG-UK) Consortium | Harper VanSteenhouse, Yumi Kasai, David Gray, Carol Clugston, Anna Dominiczak and Alex Alderton, Roberto Amato, Jeffrey Barrett, Sonia Goncalves, Ewan Harrison, David K. Jackson, Ian Johnston, Dominic Kwiatkowski, Cordelia Langford, John Sillitoe on behalf of the Wellcome Sanger Institute COVID-19 Surveillance Team |
| EPI_ISL_3221727 | Lighthouse Lab in Milton Keynes | Wellcome Sanger Institute for the COVID-19 Genomics UK (COG-UK) | The Lighthouse Lab in Milton Keynes and Alex Alderton, Roberto Amato, Jeffrey Barrett, Sonia Goncalves, Ewan Harrison, David K. Jackson, Ian Johnston, Dominic Kwiatkowski, Cordelia Langford, John Sillitoe on behalf of the Wellcome Sanger |

| | | | |
|---|---|---|---|
| | | Consortium | Institute COVID-19 Surveillance Team |
| EPI_ISL_3221728 | Lighthouse Laboratory Plymouth | Wellcome Sanger Institute for the COVID-19 Genomics UK (COG-UK) Consortium | Lighthouse Laboratory Plymouth and Alex Alderton, Roberto Amato, Jeffrey Barrett, Sonia Goncalves, Ewan Harrison, David K. Jackson, Ian Johnston, Dominic Kwiatkowski, Cordelia Langford, John Sillitoe on behalf of the Wellcome Sanger Institute COVID-19 Surveillance Team |
| EPI_ISL_3221729 | Lighthouse Lab in Glasgow | Wellcome Sanger Institute for the COVID-19 Genomics UK (COG-UK) Consortium | Harper VanSteenhouse, Yumi Kasai, David Gray, Carol Clugston, Anna Dominiczak and Alex Alderton, Roberto Amato, Jeffrey Barrett, Sonia Goncalves, Ewan Harrison, David K. Jackson, Ian Johnston, Dominic Kwiatkowski, Cordelia Langford, John Sillitoe on behalf of the Wellcome Sanger Institute COVID-19 Surveillance Team |
| EPI_ISL_3221730 | Lighthouse Laboratory Plymouth | Wellcome Sanger Institute for the COVID-19 Genomics UK (COG-UK) Consortium | Lighthouse Laboratory Plymouth and Alex Alderton, Roberto Amato, Jeffrey Barrett, Sonia Goncalves, Ewan Harrison, David K. Jackson, Ian Johnston, Dominic Kwiatkowski, Cordelia Langford, John Sillitoe on behalf of the Wellcome Sanger Institute COVID-19 Surveillance Team |
| EPI_ISL_3221731, EPI_ISL_3221732, EPI_ISL_3221733 | Lighthouse Lab in Glasgow | Wellcome Sanger Institute for the COVID-19 Genomics UK (COG-UK) Consortium | Harper VanSteenhouse, Yumi Kasai, David Gray, Carol Clugston, Anna Dominiczak and Alex Alderton, Roberto Amato, Jeffrey Barrett, Sonia Goncalves, Ewan Harrison, David K. Jackson, Ian Johnston, Dominic Kwiatkowski, Cordelia Langford, John Sillitoe on behalf of the Wellcome Sanger Institute COVID-19 Surveillance Team |
| EPI_ISL_3221734, EPI_ISL_3221735 | Lighthouse Laboratory Plymouth | Wellcome Sanger Institute for the COVID-19 Genomics UK (COG-UK) Consortium | Lighthouse Laboratory Plymouth and Alex Alderton, Roberto Amato, Jeffrey Barrett, Sonia Goncalves, Ewan Harrison, David K. Jackson, Ian Johnston, Dominic Kwiatkowski, Cordelia Langford, John Sillitoe on behalf of the Wellcome Sanger Institute COVID-19 Surveillance Team |
| EPI_ISL_3221736 | Lighthouse Lab in Glasgow | Wellcome Sanger Institute for the COVID-19 Genomics UK (COG-UK) Consortium | Harper VanSteenhouse, Yumi Kasai, David Gray, Carol Clugston, Anna Dominiczak and Alex Alderton, Roberto Amato, Jeffrey Barrett, Sonia Goncalves, Ewan Harrison, David K. Jackson, Ian Johnston, Dominic Kwiatkowski, Cordelia Langford, John Sillitoe on behalf of the Wellcome Sanger Institute COVID-19 Surveillance Team |
| EPI_ISL_3221737, EPI_ISL_3221738 | Lighthouse Laboratory Plymouth | Wellcome Sanger Institute for the COVID-19 Genomics UK (COG-UK) Consortium | Lighthouse Laboratory Plymouth and Alex Alderton, Roberto Amato, Jeffrey Barrett, Sonia Goncalves, Ewan Harrison, David K. Jackson, Ian Johnston, Dominic Kwiatkowski, Cordelia Langford, John Sillitoe on behalf of the Wellcome Sanger Institute COVID-19 Surveillance Team |
| EPI_ISL_3221739 | Lighthouse Lab in Milton Keynes | Wellcome Sanger Institute for the COVID-19 Genomics UK (COG-UK) Consortium | The Lighthouse Lab in Milton Keynes and Alex Alderton, Roberto Amato, Jeffrey Barrett, Sonia Goncalves, Ewan Harrison, David K. Jackson, Ian Johnston, Dominic Kwiatkowski, Cordelia Langford, John Sillitoe on behalf of the Wellcome Sanger Institute COVID-19 Surveillance Team |
| EPI_ISL_3221740 | Lighthouse Lab in Glasgow | Wellcome Sanger Institute for the COVID-19 Genomics UK (COG-UK) Consortium | Harper VanSteenhouse, Yumi Kasai, David Gray, Carol Clugston, Anna Dominiczak and Alex Alderton, Roberto Amato, Jeffrey Barrett, Sonia Goncalves, Ewan Harrison, David K. Jackson, Ian Johnston, Dominic Kwiatkowski, Cordelia Langford, John Sillitoe on behalf of the Wellcome Sanger Institute COVID-19 Surveillance Team |
| EPI_ISL_3221741 | Lighthouse Lab in Milton Keynes | Wellcome Sanger Institute for the COVID-19 Genomics UK (COG-UK) Consortium | The Lighthouse Lab in Milton Keynes and Alex Alderton, Roberto Amato, Jeffrey Barrett, Sonia Goncalves, Ewan Harrison, David K. Jackson, Ian Johnston, Dominic Kwiatkowski, Cordelia Langford, John Sillitoe on behalf of the Wellcome Sanger Institute COVID-19 Surveillance Team |
| EPI_ISL_3221742 | Lighthouse Lab in Glasgow | Wellcome Sanger Institute for the COVID-19 Genomics UK (COG-UK) Consortium | Harper VanSteenhouse, Yumi Kasai, David Gray, Carol Clugston, Anna Dominiczak and Alex Alderton, Roberto Amato, Jeffrey Barrett, Sonia Goncalves, Ewan Harrison, David K. Jackson, Ian Johnston, Dominic Kwiatkowski, Cordelia Langford, John Sillitoe on behalf of the Wellcome Sanger Institute COVID-19 Surveillance Team |
| EPI_ISL_3221743, EPI_ISL_3221744 | Lighthouse Lab in Milton Keynes | Wellcome Sanger Institute for the COVID-19 Genomics UK (COG-UK) Consortium | The Lighthouse Lab in Milton Keynes and Alex Alderton, Roberto Amato, Jeffrey Barrett, Sonia Goncalves, Ewan Harrison, David K. Jackson, Ian Johnston, Dominic Kwiatkowski, Cordelia Langford, John Sillitoe on behalf of the Wellcome Sanger Institute COVID-19 Surveillance Team |
| EPI_ISL_3221745, EPI_ISL_3221746 | Lighthouse Laboratory Plymouth | Wellcome Sanger Institute for the COVID-19 Genomics UK (COG-UK) Consortium | Lighthouse Laboratory Plymouth and Alex Alderton, Roberto Amato, Jeffrey Barrett, Sonia Goncalves, Ewan Harrison, David K. Jackson, Ian Johnston, Dominic Kwiatkowski, Cordelia Langford, John Sillitoe on behalf of the Wellcome Sanger Institute COVID-19 Surveillance Team |
| EPI_ISL_3221747, EPI_ISL_3221748, EPI_ISL_3221749 | Lighthouse Lab in Glasgow | Wellcome Sanger Institute for the COVID-19 Genomics UK (COG-UK) Consortium | Harper VanSteenhouse, Yumi Kasai, David Gray, Carol Clugston, Anna Dominiczak and Alex Alderton, Roberto Amato, Jeffrey Barrett, Sonia Goncalves, Ewan Harrison, David K. Jackson, Ian Johnston, Dominic Kwiatkowski, Cordelia Langford, John Sillitoe on behalf of the Wellcome Sanger Institute COVID-19 Surveillance Team |
| EPI_ISL_3221750, EPI_ISL_3221751, EPI_ISL_3221752, EPI_ISL_3221753 | Lighthouse Lab in Milton Keynes | Wellcome Sanger Institute for the COVID-19 Genomics UK (COG-UK) Consortium | The Lighthouse Lab in Milton Keynes and Alex Alderton, Roberto Amato, Jeffrey Barrett, Sonia Goncalves, Ewan Harrison, David K. Jackson, Ian Johnston, Dominic Kwiatkowski, Cordelia Langford, John Sillitoe on behalf of the Wellcome Sanger Institute COVID-19 Surveillance Team |
| EPI_ISL_3221754, EPI_ISL_3221755 | Lighthouse Lab in Glasgow | Wellcome Sanger Institute for the COVID-19 Genomics UK (COG-UK) Consortium | Harper VanSteenhouse, Yumi Kasai, David Gray, Carol Clugston, Anna Dominiczak and Alex Alderton, Roberto Amato, Jeffrey Barrett, Sonia Goncalves, Ewan Harrison, David K. Jackson, Ian Johnston, Dominic Kwiatkowski, Cordelia Langford, John Sillitoe on behalf of the Wellcome Sanger Institute COVID-19 Surveillance Team |
| EPI_ISL_3221756 | Lighthouse Lab in Milton Keynes | Wellcome Sanger Institute for the COVID-19 Genomics UK (COG-UK) Consortium | The Lighthouse Lab in Milton Keynes and Alex Alderton, Roberto Amato, Jeffrey Barrett, Sonia Goncalves, Ewan Harrison, David K. Jackson, Ian Johnston, Dominic Kwiatkowski, Cordelia Langford, John Sillitoe on behalf of the Wellcome Sanger Institute COVID-19 Surveillance Team |
| EPI_ISL_3221757 | Lighthouse Lab in Glasgow | Wellcome Sanger Institute for the COVID-19 Genomics UK (COG-UK) Consortium | Harper VanSteenhouse, Yumi Kasai, David Gray, Carol Clugston, Anna Dominiczak and Alex Alderton, Roberto Amato, Jeffrey Barrett, Sonia Goncalves, Ewan Harrison, David K. Jackson, Ian Johnston, Dominic Kwiatkowski, Cordelia Langford, John Sillitoe on behalf of the Wellcome Sanger Institute COVID-19 Surveillance Team |
| EPI_ISL_3221758, EPI_ISL_3221759 | Lighthouse Laboratory Plymouth | Wellcome Sanger Institute for the COVID-19 Genomics UK (COG-UK) Consortium | Lighthouse Laboratory Plymouth and Alex Alderton, Roberto Amato, Jeffrey Barrett, Sonia Goncalves, Ewan Harrison, David K. Jackson, Ian Johnston, Dominic Kwiatkowski, Cordelia Langford, John Sillitoe on behalf of the Wellcome Sanger Institute COVID-19 Surveillance Team |
| EPI_ISL_3221760 | Lighthouse Lab in Milton Keynes | Wellcome Sanger Institute for the COVID-19 Genomics UK (COG-UK) Consortium | The Lighthouse Lab in Milton Keynes and Alex Alderton, Roberto Amato, Jeffrey Barrett, Sonia Goncalves, Ewan Harrison, David K. Jackson, Ian Johnston, Dominic Kwiatkowski, Cordelia Langford, John Sillitoe on behalf of the Wellcome Sanger Institute COVID-19 Surveillance Team |
| EPI_ISL_3221761 | Lighthouse Laboratory Plymouth | Wellcome Sanger Institute for the COVID-19 Genomics UK (COG-UK) Consortium | Lighthouse Laboratory Plymouth and Alex Alderton, Roberto Amato, Jeffrey Barrett, Sonia Goncalves, Ewan Harrison, David K. Jackson, Ian Johnston, Dominic Kwiatkowski, Cordelia Langford, John Sillitoe on behalf of the Wellcome Sanger Institute COVID-19 Surveillance Team |
| EPI_ISL_3221762, EPI_ISL_3221763 | Lighthouse Lab in Glasgow | Wellcome Sanger Institute for the COVID-19 Genomics UK (COG-UK) Consortium | Harper VanSteenhouse, Yumi Kasai, David Gray, Carol Clugston, Anna Dominiczak and Alex Alderton, Roberto Amato, Jeffrey Barrett, Sonia Goncalves, Ewan Harrison, David K. Jackson, Ian Johnston, Dominic Kwiatkowski, Cordelia Langford, John Sillitoe on behalf of the Wellcome Sanger Institute COVID-19 Surveillance Team |
| EPI_ISL_3221764, EPI_ISL_3221765 | Lighthouse Lab in Milton Keynes | Wellcome Sanger Institute for the COVID-19 Genomics UK (COG-UK) Consortium | The Lighthouse Lab in Milton Keynes and Alex Alderton, Roberto Amato, Jeffrey Barrett, Sonia Goncalves, Ewan Harrison, David K. Jackson, Ian Johnston, Dominic Kwiatkowski, Cordelia Langford, John Sillitoe on behalf of the Wellcome Sanger Institute COVID-19 Surveillance Team |
| EPI_ISL_3221766 | Lighthouse Lab in Glasgow | Wellcome Sanger Institute for the COVID-19 Genomics UK (COG-UK) Consortium | Harper VanSteenhouse, Yumi Kasai, David Gray, Carol Clugston, Anna Dominiczak and Alex Alderton, Roberto Amato, Jeffrey Barrett, Sonia Goncalves, Ewan Harrison, David K. Jackson, Ian Johnston, Dominic Kwiatkowski, Cordelia Langford, John Sillitoe on behalf of the Wellcome Sanger Institute COVID-19 Surveillance Team |
| EPI_ISL_3221767 | Lighthouse Lab in Milton Keynes | Wellcome Sanger Institute for the COVID-19 Genomics UK (COG-UK) Consortium | The Lighthouse Lab in Milton Keynes and Alex Alderton, Roberto Amato, Jeffrey Barrett, Sonia Goncalves, Ewan Harrison, David K. Jackson, Ian Johnston, Dominic Kwiatkowski, Cordelia Langford, John Sillitoe on behalf of the Wellcome Sanger Institute COVID-19 Surveillance Team |
| EPI_ISL_3221768, EPI_ISL_3221769 | Lighthouse Lab in Glasgow | Wellcome Sanger Institute for the COVID-19 Genomics UK (COG-UK) Consortium | Harper VanSteenhouse, Yumi Kasai, David Gray, Carol Clugston, Anna Dominiczak and Alex Alderton, Roberto Amato, Jeffrey Barrett, Sonia Goncalves, Ewan Harrison, David K. Jackson, Ian Johnston, Dominic Kwiatkowski, Cordelia Langford, John Sillitoe on behalf of the Wellcome Sanger Institute COVID-19 Surveillance Team |
| EPI_ISL_3221770 | Lighthouse Laboratory Plymouth | Wellcome Sanger Institute for the COVID-19 Genomics UK (COG-UK) Consortium | Lighthouse Laboratory Plymouth and Alex Alderton, Roberto Amato, Jeffrey Barrett, Sonia Goncalves, Ewan Harrison, David K. Jackson, Ian Johnston, Dominic Kwiatkowski, Cordelia Langford, John Sillitoe on behalf of the Wellcome Sanger Institute COVID-19 Surveillance Team |
| EPI_ISL_3221771 | Lighthouse Lab in Milton Keynes | Wellcome Sanger Institute for the COVID-19 Genomics UK (COG-UK) Consortium | The Lighthouse Lab in Milton Keynes and Alex Alderton, Roberto Amato, Jeffrey Barrett, Sonia Goncalves, Ewan Harrison, David K. Jackson, Ian Johnston, Dominic Kwiatkowski, Cordelia Langford, John Sillitoe on behalf of the Wellcome Sanger Institute COVID-19 Surveillance Team |
| EPI_ISL_3221772 | Lighthouse Lab in Glasgow | Wellcome Sanger Institute for the COVID-19 Genomics UK (COG-UK) Consortium | Harper VanSteenhouse, Yumi Kasai, David Gray, Carol Clugston, Anna Dominiczak and Alex Alderton, Roberto Amato, Jeffrey Barrett, Sonia Goncalves, Ewan Harrison, David K. Jackson, Ian Johnston, Dominic Kwiatkowski, Cordelia Langford, John Sillitoe on behalf of the Wellcome Sanger Institute COVID-19 Surveillance Team |
| EPI_ISL_3221773 | Lighthouse Laboratory Plymouth | Wellcome Sanger Institute for the COVID-19 Genomics UK (COG-UK) Consortium | Lighthouse Laboratory Plymouth and Alex Alderton, Roberto Amato, Jeffrey Barrett, Sonia Goncalves, Ewan Harrison, David K. Jackson, Ian Johnston, Dominic Kwiatkowski, Cordelia Langford, John Sillitoe on behalf of the Wellcome Sanger Institute COVID-19 Surveillance Team |
| EPI_ISL_3221774 | Lighthouse Lab in Milton Keynes | Wellcome Sanger Institute for the COVID-19 Genomics UK (COG-UK) Consortium | The Lighthouse Lab in Milton Keynes and Alex Alderton, Roberto Amato, Jeffrey Barrett, Sonia Goncalves, Ewan Harrison, David K. Jackson, Ian Johnston, Dominic Kwiatkowski, Cordelia Langford, John Sillitoe on behalf of the Wellcome Sanger Institute COVID-19 Surveillance Team |
| EPI_ISL_3221775 | Lighthouse Laboratory Plymouth | Wellcome Sanger Institute for the COVID-19 Genomics UK (COG-UK) Consortium | Lighthouse Laboratory Plymouth and Alex Alderton, Roberto Amato, Jeffrey Barrett, Sonia Goncalves, Ewan Harrison, David K. Jackson, Ian Johnston, Dominic Kwiatkowski, Cordelia Langford, John Sillitoe on behalf of the Wellcome Sanger Institute COVID-19 Surveillance Team |
| EPI_ISL_3221776, EPI_ISL_3221777 | Lighthouse Lab in Glasgow | Wellcome Sanger Institute for the COVID-19 Genomics UK (COG-UK) Consortium | Harper VanSteenhouse, Yumi Kasai, David Gray, Carol Clugston, Anna Dominiczak and Alex Alderton, Roberto Amato, Jeffrey Barrett, Sonia Goncalves, Ewan Harrison, David K. Jackson, Ian Johnston, Dominic Kwiatkowski, Cordelia Langford, John Sillitoe on behalf of the Wellcome Sanger Institute COVID-19 Surveillance Team |
| EPI_ISL_3221778 | Lighthouse Lab in Milton Keynes | Wellcome Sanger Institute for the COVID-19 Genomics UK (COG-UK) Consortium | The Lighthouse Lab in Milton Keynes and Alex Alderton, Roberto Amato, Jeffrey Barrett, Sonia Goncalves, Ewan Harrison, David K. Jackson, Ian Johnston, Dominic Kwiatkowski, Cordelia Langford, John Sillitoe on behalf of the Wellcome Sanger Institute COVID-19 Surveillance Team |
| EPI_ISL_3221779 | Lighthouse Laboratory Plymouth | Wellcome Sanger Institute for the COVID-19 Genomics UK (COG-UK) Consortium | Lighthouse Laboratory Plymouth and Alex Alderton, Roberto Amato, Jeffrey Barrett, Sonia Goncalves, Ewan Harrison, David K. Jackson, Ian Johnston, Dominic Kwiatkowski, Cordelia Langford, John Sillitoe on behalf of the Wellcome Sanger Institute COVID-19 Surveillance Team |
| EPI_ISL_3221780, EPI_ISL_3221781 | Lighthouse Lab in Glasgow | Wellcome Sanger Institute for the COVID-19 Genomics UK (COG-UK) Consortium | Harper VanSteenhouse, Yumi Kasai, David Gray, Carol Clugston, Anna Dominiczak and Alex Alderton, Roberto Amato, Jeffrey Barrett, Sonia Goncalves, Ewan Harrison, David K. Jackson, Ian Johnston, Dominic Kwiatkowski, Cordelia Langford, John Sillitoe on behalf of the Wellcome Sanger Institute COVID-19 Surveillance Team |
| EPI_ISL_3221782, EPI_ISL_3221783, EPI_ISL_3221784 | Lighthouse Lab in Milton Keynes | Wellcome Sanger Institute for the COVID-19 Genomics UK (COG-UK) Consortium | The Lighthouse Lab in Milton Keynes and Alex Alderton, Roberto Amato, Jeffrey Barrett, Sonia Goncalves, Ewan Harrison, David K. Jackson, Ian Johnston, Dominic Kwiatkowski, Cordelia Langford, John Sillitoe on behalf of the Wellcome Sanger Institute COVID-19 Surveillance Team |
| EPI_ISL_3221785, EPI_ISL_3221786, EPI_ISL_3221787, EPI_ISL_3221788 | Lighthouse Lab in Glasgow | Wellcome Sanger Institute for the COVID-19 Genomics UK (COG-UK) Consortium | Harper VanSteenhouse, Yumi Kasai, David Gray, Carol Clugston, Anna Dominiczak and Alex Alderton, Roberto Amato, Jeffrey Barrett, Sonia Goncalves, Ewan Harrison, David K. Jackson, Ian Johnston, Dominic Kwiatkowski, Cordelia Langford, John Sillitoe on behalf of the Wellcome Sanger Institute COVID-19 Surveillance Team |
| EPI_ISL_3221789 | Lighthouse Lab in Milton Keynes | Wellcome Sanger Institute for the COVID-19 Genomics UK (COG-UK) Consortium | The Lighthouse Lab in Milton Keynes and Alex Alderton, Roberto Amato, Jeffrey Barrett, Sonia Goncalves, Ewan Harrison, David K. Jackson, Ian Johnston, Dominic Kwiatkowski, Cordelia Langford, John Sillitoe on behalf of the Wellcome Sanger Institute COVID-19 Surveillance Team |
| EPI_ISL_3221790, EPI_ISL_3221791 | Lighthouse Laboratory Plymouth | Wellcome Sanger Institute for the COVID-19 Genomics UK (COG-UK) Consortium | Lighthouse Laboratory Plymouth and Alex Alderton, Roberto Amato, Jeffrey Barrett, Sonia Goncalves, Ewan Harrison, David K. Jackson, Ian Johnston, Dominic Kwiatkowski, Cordelia Langford, John Sillitoe on behalf of the Wellcome Sanger Institute COVID-19 Surveillance Team |
| EPI_ISL_3221792 | Lighthouse Lab in Milton Keynes | Wellcome Sanger Institute for the COVID-19 Genomics UK (COG-UK) Consortium | The Lighthouse Lab in Milton Keynes and Alex Alderton, Roberto Amato, Jeffrey Barrett, Sonia Goncalves, Ewan Harrison, David K. Jackson, Ian Johnston, Dominic Kwiatkowski, Cordelia Langford, John Sillitoe on behalf of the Wellcome Sanger Institute COVID-19 Surveillance Team |
| EPI_ISL_3221793, EPI_ISL_3221794 | Lighthouse Lab in Glasgow | Wellcome Sanger Institute for the COVID-19 Genomics UK (COG-UK) Consortium | Harper VanSteenhouse, Yumi Kasai, David Gray, Carol Clugston, Anna Dominiczak and Alex Alderton, Roberto Amato, Jeffrey Barrett, Sonia Goncalves, Ewan Harrison, David K. Jackson, Ian Johnston, Dominic Kwiatkowski, Cordelia Langford, John Sillitoe on behalf of the Wellcome Sanger Institute COVID-19 Surveillance Team |

| | | | |
|---|---|---|---|
| EPI_ISL_3221795 | Lighthouse Lab in Milton Keynes | Wellcome Sanger Institute for the COVID-19 Genomics UK (COG-UK) Consortium | The Lighthouse Lab in Milton Keynes and Alex Alderton, Roberto Amato, Jeffrey Barrett, Sonia Goncalves, Ewan Harrison, David K. Jackson, Ian Johnston, Dominic Kwiatkowski, Cordelia Langford, John Sillitoe on behalf of the Wellcome Sanger Institute COVID-19 Surveillance Team |
| EPI_ISL_3221796 | Lighthouse Lab in Glasgow | Wellcome Sanger Institute for the COVID-19 Genomics UK (COG-UK) Consortium | Harper VanSteenhouse, Yumi Kasai, David Gray, Carol Clugston, Anna Dominiczak and Alex Alderton, Roberto Amato, Jeffrey Barrett, Sonia Goncalves, Ewan Harrison, David K. Jackson, Ian Johnston, Dominic Kwiatkowski, Cordelia Langford, John Sillitoe on behalf of the Wellcome Sanger Institute COVID-19 Surveillance Team |
| EPI_ISL_3221797 | Lighthouse Lab in Milton Keynes | Wellcome Sanger Institute for the COVID-19 Genomics UK (COG-UK) Consortium | The Lighthouse Lab in Milton Keynes and Alex Alderton, Roberto Amato, Jeffrey Barrett, Sonia Goncalves, Ewan Harrison, David K. Jackson, Ian Johnston, Dominic Kwiatkowski, Cordelia Langford, John Sillitoe on behalf of the Wellcome Sanger Institute COVID-19 Surveillance Team |
| EPI_ISL_3221798 | Lighthouse Laboratory Plymouth | Wellcome Sanger Institute for the COVID-19 Genomics UK (COG-UK) Consortium | Lighthouse Laboratory Plymouth and Alex Alderton, Roberto Amato, Jeffrey Barrett, Sonia Goncalves, Ewan Harrison, David K. Jackson, Ian Johnston, Dominic Kwiatkowski, Cordelia Langford, John Sillitoe on behalf of the Wellcome Sanger Institute COVID-19 Surveillance Team |
| EPI_ISL_3221799 | Lighthouse Lab in Glasgow | Wellcome Sanger Institute for the COVID-19 Genomics UK (COG-UK) Consortium | Harper VanSteenhouse, Yumi Kasai, David Gray, Carol Clugston, Anna Dominiczak and Alex Alderton, Roberto Amato, Jeffrey Barrett, Sonia Goncalves, Ewan Harrison, David K. Jackson, Ian Johnston, Dominic Kwiatkowski, Cordelia Langford, John Sillitoe on behalf of the Wellcome Sanger Institute COVID-19 Surveillance Team |
| EPI_ISL_3221800 | Lighthouse Lab in Milton Keynes | Wellcome Sanger Institute for the COVID-19 Genomics UK (COG-UK) Consortium | The Lighthouse Lab in Milton Keynes and Alex Alderton, Roberto Amato, Jeffrey Barrett, Sonia Goncalves, Ewan Harrison, David K. Jackson, Ian Johnston, Dominic Kwiatkowski, Cordelia Langford, John Sillitoe on behalf of the Wellcome Sanger Institute COVID-19 Surveillance Team |
| EPI_ISL_3221801, EPI_ISL_3221802 | Lighthouse Lab in Glasgow | Wellcome Sanger Institute for the COVID-19 Genomics UK (COG-UK) Consortium | Harper VanSteenhouse, Yumi Kasai, David Gray, Carol Clugston, Anna Dominiczak and Alex Alderton, Roberto Amato, Jeffrey Barrett, Sonia Goncalves, Ewan Harrison, David K. Jackson, Ian Johnston, Dominic Kwiatkowski, Cordelia Langford, John Sillitoe on behalf of the Wellcome Sanger Institute COVID-19 Surveillance Team |
| EPI_ISL_3221803 | Lighthouse Lab in Milton Keynes | Wellcome Sanger Institute for the COVID-19 Genomics UK (COG-UK) Consortium | The Lighthouse Lab in Milton Keynes and Alex Alderton, Roberto Amato, Jeffrey Barrett, Sonia Goncalves, Ewan Harrison, David K. Jackson, Ian Johnston, Dominic Kwiatkowski, Cordelia Langford, John Sillitoe on behalf of the Wellcome Sanger Institute COVID-19 Surveillance Team |
| EPI_ISL_3221804 | Lighthouse Lab in Glasgow | Wellcome Sanger Institute for the COVID-19 Genomics UK (COG-UK) Consortium | Harper VanSteenhouse, Yumi Kasai, David Gray, Carol Clugston, Anna Dominiczak and Alex Alderton, Roberto Amato, Jeffrey Barrett, Sonia Goncalves, Ewan Harrison, David K. Jackson, Ian Johnston, Dominic Kwiatkowski, Cordelia Langford, John Sillitoe on behalf of the Wellcome Sanger Institute COVID-19 Surveillance Team |
| EPI_ISL_3221805 | Lighthouse Laboratory Plymouth | Wellcome Sanger Institute for the COVID-19 Genomics UK (COG-UK) Consortium | Lighthouse Laboratory Plymouth and Alex Alderton, Roberto Amato, Jeffrey Barrett, Sonia Goncalves, Ewan Harrison, David K. Jackson, Ian Johnston, Dominic Kwiatkowski, Cordelia Langford, John Sillitoe on behalf of the Wellcome Sanger Institute COVID-19 Surveillance Team |
| EPI_ISL_3221806 | Lighthouse Lab in Milton Keynes | Wellcome Sanger Institute for the COVID-19 Genomics UK (COG-UK) Consortium | The Lighthouse Lab in Milton Keynes and Alex Alderton, Roberto Amato, Jeffrey Barrett, Sonia Goncalves, Ewan Harrison, David K. Jackson, Ian Johnston, Dominic Kwiatkowski, Cordelia Langford, John Sillitoe on behalf of the Wellcome Sanger Institute COVID-19 Surveillance Team |
| EPI_ISL_3221807 | Lighthouse Laboratory Plymouth | Wellcome Sanger Institute for the COVID-19 Genomics UK (COG-UK) Consortium | Lighthouse Laboratory Plymouth and Alex Alderton, Roberto Amato, Jeffrey Barrett, Sonia Goncalves, Ewan Harrison, David K. Jackson, Ian Johnston, Dominic Kwiatkowski, Cordelia Langford, John Sillitoe on behalf of the Wellcome Sanger Institute COVID-19 Surveillance Team |
| EPI_ISL_3221808 | Lighthouse Lab in Glasgow | Wellcome Sanger Institute for the COVID-19 Genomics UK (COG-UK) Consortium | Harper VanSteenhouse, Yumi Kasai, David Gray, Carol Clugston, Anna Dominiczak and Alex Alderton, Roberto Amato, Jeffrey Barrett, Sonia Goncalves, Ewan Harrison, David K. Jackson, Ian Johnston, Dominic Kwiatkowski, Cordelia Langford, John Sillitoe on behalf of the Wellcome Sanger Institute COVID-19 Surveillance Team |
| EPI_ISL_3221809 | Lighthouse Laboratory Plymouth | Wellcome Sanger Institute for the COVID-19 Genomics UK (COG-UK) Consortium | Lighthouse Laboratory Plymouth and Alex Alderton, Roberto Amato, Jeffrey Barrett, Sonia Goncalves, Ewan Harrison, David K. Jackson, Ian Johnston, Dominic Kwiatkowski, Cordelia Langford, John Sillitoe on behalf of the Wellcome Sanger Institute COVID-19 Surveillance Team |
| EPI_ISL_3221810 | Lighthouse Lab in Glasgow | Wellcome Sanger Institute for the COVID-19 Genomics UK (COG-UK) Consortium | Harper VanSteenhouse, Yumi Kasai, David Gray, Carol Clugston, Anna Dominiczak and Alex Alderton, Roberto Amato, Jeffrey Barrett, Sonia Goncalves, Ewan Harrison, David K. Jackson, Ian Johnston, Dominic Kwiatkowski, Cordelia Langford, John Sillitoe on behalf of the Wellcome Sanger Institute COVID-19 Surveillance Team |
| EPI_ISL_3221811 | Lighthouse Laboratory Plymouth | Wellcome Sanger Institute for the COVID-19 Genomics UK (COG-UK) Consortium | Lighthouse Laboratory Plymouth and Alex Alderton, Roberto Amato, Jeffrey Barrett, Sonia Goncalves, Ewan Harrison, David K. Jackson, Ian Johnston, Dominic Kwiatkowski, Cordelia Langford, John Sillitoe on behalf of the Wellcome Sanger Institute COVID-19 Surveillance Team |
| EPI_ISL_3221812, EPI_ISL_3221813, EPI_ISL_3221814 | Lighthouse Lab in Glasgow | Wellcome Sanger Institute for the COVID-19 Genomics UK (COG-UK) Consortium | Harper VanSteenhouse, Yumi Kasai, David Gray, Carol Clugston, Anna Dominiczak and Alex Alderton, Roberto Amato, Jeffrey Barrett, Sonia Goncalves, Ewan Harrison, David K. Jackson, Ian Johnston, Dominic Kwiatkowski, Cordelia Langford, John Sillitoe on behalf of the Wellcome Sanger Institute COVID-19 Surveillance Team |
| EPI_ISL_3221815 | Lighthouse Laboratory Plymouth | Wellcome Sanger Institute for the COVID-19 Genomics UK (COG-UK) Consortium | Lighthouse Laboratory Plymouth and Alex Alderton, Roberto Amato, Jeffrey Barrett, Sonia Goncalves, Ewan Harrison, David K. Jackson, Ian Johnston, Dominic Kwiatkowski, Cordelia Langford, John Sillitoe on behalf of the Wellcome Sanger Institute COVID-19 Surveillance Team |
| EPI_ISL_3221816, EPI_ISL_3221817, EPI_ISL_3221818 | Lighthouse Lab in Glasgow | Wellcome Sanger Institute for the COVID-19 Genomics UK (COG-UK) Consortium | Harper VanSteenhouse, Yumi Kasai, David Gray, Carol Clugston, Anna Dominiczak and Alex Alderton, Roberto Amato, Jeffrey Barrett, Sonia Goncalves, Ewan Harrison, David K. Jackson, Ian Johnston, Dominic Kwiatkowski, Cordelia Langford, John Sillitoe on behalf of the Wellcome Sanger Institute COVID-19 Surveillance Team |
| EPI_ISL_3221819 | Lighthouse Lab in Milton Keynes | Wellcome Sanger Institute for the COVID-19 Genomics UK (COG-UK) Consortium | The Lighthouse Lab in Milton Keynes and Alex Alderton, Roberto Amato, Jeffrey Barrett, Sonia Goncalves, Ewan Harrison, David K. Jackson, Ian Johnston, Dominic Kwiatkowski, Cordelia Langford, John Sillitoe on behalf of the Wellcome Sanger Institute COVID-19 Surveillance Team |
| EPI_ISL_3221820 | Lighthouse Laboratory Plymouth | Wellcome Sanger Institute for the COVID-19 Genomics UK (COG-UK) Consortium | Lighthouse Laboratory Plymouth and Alex Alderton, Roberto Amato, Jeffrey Barrett, Sonia Goncalves, Ewan Harrison, David K. Jackson, Ian Johnston, Dominic Kwiatkowski, Cordelia Langford, John Sillitoe on behalf of the Wellcome Sanger Institute COVID-19 Surveillance Team |
| EPI_ISL_3221821 | Lighthouse Lab in Milton Keynes | Wellcome Sanger Institute for the COVID-19 Genomics UK (COG-UK) Consortium | The Lighthouse Lab in Milton Keynes and Alex Alderton, Roberto Amato, Jeffrey Barrett, Sonia Goncalves, Ewan Harrison, David K. Jackson, Ian Johnston, Dominic Kwiatkowski, Cordelia Langford, John Sillitoe on behalf of the Wellcome Sanger Institute COVID-19 Surveillance Team |
| EPI_ISL_3221822, EPI_ISL_3221823 | Lighthouse Lab in Glasgow | Wellcome Sanger Institute for the COVID-19 Genomics UK (COG-UK) Consortium | Harper VanSteenhouse, Yumi Kasai, David Gray, Carol Clugston, Anna Dominiczak and Alex Alderton, Roberto Amato, Jeffrey Barrett, Sonia Goncalves, Ewan Harrison, David K. Jackson, Ian Johnston, Dominic Kwiatkowski, Cordelia Langford, John Sillitoe on behalf of the Wellcome Sanger Institute COVID-19 Surveillance Team |
| EPI_ISL_3221824 | Lighthouse Lab in Milton Keynes | Wellcome Sanger Institute for the COVID-19 Genomics UK (COG-UK) Consortium | The Lighthouse Lab in Milton Keynes and Alex Alderton, Roberto Amato, Jeffrey Barrett, Sonia Goncalves, Ewan Harrison, David K. Jackson, Ian Johnston, Dominic Kwiatkowski, Cordelia Langford, John Sillitoe on behalf of the Wellcome Sanger Institute COVID-19 Surveillance Team |
| EPI_ISL_3221825 | Lighthouse Lab in Glasgow | Wellcome Sanger Institute for the COVID-19 Genomics UK (COG-UK) Consortium | Harper VanSteenhouse, Yumi Kasai, David Gray, Carol Clugston, Anna Dominiczak and Alex Alderton, Roberto Amato, Jeffrey Barrett, Sonia Goncalves, Ewan Harrison, David K. Jackson, Ian Johnston, Dominic Kwiatkowski, Cordelia Langford, John Sillitoe on behalf of the Wellcome Sanger Institute COVID-19 Surveillance Team |
| EPI_ISL_3221826, EPI_ISL_3221827 | Lighthouse Lab in Milton Keynes | Wellcome Sanger Institute for the COVID-19 Genomics UK (COG-UK) Consortium | The Lighthouse Lab in Milton Keynes and Alex Alderton, Roberto Amato, Jeffrey Barrett, Sonia Goncalves, Ewan Harrison, David K. Jackson, Ian Johnston, Dominic Kwiatkowski, Cordelia Langford, John Sillitoe on behalf of the Wellcome Sanger Institute COVID-19 Surveillance Team |
| EPI_ISL_3221828 | Lighthouse Laboratory Plymouth | Wellcome Sanger Institute for the COVID-19 Genomics UK (COG-UK) Consortium | Lighthouse Laboratory Plymouth and Alex Alderton, Roberto Amato, Jeffrey Barrett, Sonia Goncalves, Ewan Harrison, David K. Jackson, Ian Johnston, Dominic Kwiatkowski, Cordelia Langford, John Sillitoe on behalf of the Wellcome Sanger Institute COVID-19 Surveillance Team |
| EPI_ISL_3221829 | Lighthouse Lab in Glasgow | Wellcome Sanger Institute for the COVID-19 Genomics UK (COG-UK) Consortium | Harper VanSteenhouse, Yumi Kasai, David Gray, Carol Clugston, Anna Dominiczak and Alex Alderton, Roberto Amato, Jeffrey Barrett, Sonia Goncalves, Ewan Harrison, David K. Jackson, Ian Johnston, Dominic Kwiatkowski, Cordelia Langford, John Sillitoe on behalf of the Wellcome Sanger Institute COVID-19 Surveillance Team |
| EPI_ISL_3221830, EPI_ISL_3221831, EPI_ISL_3221832 | Lighthouse Laboratory Plymouth | Wellcome Sanger Institute for the COVID-19 Genomics UK (COG-UK) Consortium | Lighthouse Laboratory Plymouth and Alex Alderton, Roberto Amato, Jeffrey Barrett, Sonia Goncalves, Ewan Harrison, David K. Jackson, Ian Johnston, Dominic Kwiatkowski, Cordelia Langford, John Sillitoe on behalf of the Wellcome Sanger Institute COVID-19 Surveillance Team |
| EPI_ISL_3221833 | Lighthouse Lab in Glasgow | Wellcome Sanger Institute for the COVID-19 Genomics UK (COG-UK) Consortium | Harper VanSteenhouse, Yumi Kasai, David Gray, Carol Clugston, Anna Dominiczak and Alex Alderton, Roberto Amato, Jeffrey Barrett, Sonia Goncalves, Ewan Harrison, David K. Jackson, Ian Johnston, Dominic Kwiatkowski, Cordelia Langford, John Sillitoe on behalf of the Wellcome Sanger Institute COVID-19 Surveillance Team |
| EPI_ISL_3221834, EPI_ISL_3221835 | Lighthouse Lab in Milton Keynes | Wellcome Sanger Institute for the COVID-19 Genomics UK (COG-UK) Consortium | The Lighthouse Lab in Milton Keynes and Alex Alderton, Roberto Amato, Jeffrey Barrett, Sonia Goncalves, Ewan Harrison, David K. Jackson, Ian Johnston, Dominic Kwiatkowski, Cordelia Langford, John Sillitoe on behalf of the Wellcome Sanger Institute COVID-19 Surveillance Team |
| EPI_ISL_3221836 | Lighthouse Laboratory Plymouth | Wellcome Sanger Institute for the COVID-19 Genomics UK (COG-UK) Consortium | Lighthouse Laboratory Plymouth and Alex Alderton, Roberto Amato, Jeffrey Barrett, Sonia Goncalves, Ewan Harrison, David K. Jackson, Ian Johnston, Dominic Kwiatkowski, Cordelia Langford, John Sillitoe on behalf of the Wellcome Sanger Institute COVID-19 Surveillance Team |
| EPI_ISL_3221837 | Lighthouse Lab in Milton Keynes | Wellcome Sanger Institute for the COVID-19 Genomics UK (COG-UK) Consortium | The Lighthouse Lab in Milton Keynes and Alex Alderton, Roberto Amato, Jeffrey Barrett, Sonia Goncalves, Ewan Harrison, David K. Jackson, Ian Johnston, Dominic Kwiatkowski, Cordelia Langford, John Sillitoe on behalf of the Wellcome Sanger Institute COVID-19 Surveillance Team |
| EPI_ISL_3221838, EPI_ISL_3221839, EPI_ISL_3221840, EPI_ISL_3221841 | Lighthouse Lab in Glasgow | Wellcome Sanger Institute for the COVID-19 Genomics UK (COG-UK) Consortium | Harper VanSteenhouse, Yumi Kasai, David Gray, Carol Clugston, Anna Dominiczak and Alex Alderton, Roberto Amato, Jeffrey Barrett, Sonia Goncalves, Ewan Harrison, David K. Jackson, Ian Johnston, Dominic Kwiatkowski, Cordelia Langford, John Sillitoe on behalf of the Wellcome Sanger Institute COVID-19 Surveillance Team |
| EPI_ISL_3221842 | Lighthouse Lab in Milton Keynes | Wellcome Sanger Institute for the COVID-19 Genomics UK (COG-UK) Consortium | The Lighthouse Lab in Milton Keynes and Alex Alderton, Roberto Amato, Jeffrey Barrett, Sonia Goncalves, Ewan Harrison, David K. Jackson, Ian Johnston, Dominic Kwiatkowski, Cordelia Langford, John Sillitoe on behalf of the Wellcome Sanger Institute COVID-19 Surveillance Team |
| EPI_ISL_3221843 | Lighthouse Laboratory Plymouth | Wellcome Sanger Institute for the COVID-19 Genomics UK (COG-UK) Consortium | Lighthouse Laboratory Plymouth and Alex Alderton, Roberto Amato, Jeffrey Barrett, Sonia Goncalves, Ewan Harrison, David K. Jackson, Ian Johnston, Dominic Kwiatkowski, Cordelia Langford, John Sillitoe on behalf of the Wellcome Sanger Institute COVID-19 Surveillance Team |
| EPI_ISL_3221844, EPI_ISL_3221845, EPI_ISL_3221846 | Lighthouse Lab in Glasgow | Wellcome Sanger Institute for the COVID-19 Genomics UK (COG-UK) Consortium | Harper VanSteenhouse, Yumi Kasai, David Gray, Carol Clugston, Anna Dominiczak and Alex Alderton, Roberto Amato, Jeffrey Barrett, Sonia Goncalves, Ewan Harrison, David K. Jackson, Ian Johnston, Dominic Kwiatkowski, Cordelia Langford, John Sillitoe on behalf of the Wellcome Sanger Institute COVID-19 Surveillance Team |
| EPI_ISL_3221847 | Lighthouse Lab in Milton Keynes | Wellcome Sanger Institute for the COVID-19 Genomics UK (COG-UK) Consortium | The Lighthouse Lab in Milton Keynes and Alex Alderton, Roberto Amato, Jeffrey Barrett, Sonia Goncalves, Ewan Harrison, David K. Jackson, Ian Johnston, Dominic Kwiatkowski, Cordelia Langford, John Sillitoe on behalf of the Wellcome Sanger Institute COVID-19 Surveillance Team |
| EPI_ISL_3221848 | Lighthouse Laboratory Plymouth | Wellcome Sanger Institute for the COVID-19 Genomics UK (COG-UK) Consortium | Lighthouse Laboratory Plymouth and Alex Alderton, Roberto Amato, Jeffrey Barrett, Sonia Goncalves, Ewan Harrison, David K. Jackson, Ian Johnston, Dominic Kwiatkowski, Cordelia Langford, John Sillitoe on behalf of the Wellcome Sanger Institute COVID-19 Surveillance Team |
| EPI_ISL_3221849, EPI_ISL_3221850 | Lighthouse Lab in Glasgow | Wellcome Sanger Institute for the COVID-19 Genomics UK (COG-UK) Consortium | Harper VanSteenhouse, Yumi Kasai, David Gray, Carol Clugston, Anna Dominiczak and Alex Alderton, Roberto Amato, Jeffrey Barrett, Sonia Goncalves, Ewan Harrison, David K. Jackson, Ian Johnston, Dominic Kwiatkowski, Cordelia Langford, John Sillitoe on behalf of the Wellcome Sanger Institute COVID-19 Surveillance Team |
| EPI_ISL_3221851, EPI_ISL_3221852, EPI_ISL_3221853 | Lighthouse Lab in Milton Keynes | Wellcome Sanger Institute for the COVID-19 Genomics UK (COG-UK) Consortium | The Lighthouse Lab in Milton Keynes and Alex Alderton, Roberto Amato, Jeffrey Barrett, Sonia Goncalves, Ewan Harrison, David K. Jackson, Ian Johnston, Dominic Kwiatkowski, Cordelia Langford, John Sillitoe on behalf of the Wellcome Sanger Institute COVID-19 Surveillance Team |
| EPI_ISL_3221854 | Lighthouse Laboratory Plymouth | Wellcome Sanger Institute for the COVID-19 Genomics UK (COG-UK) Consortium | Lighthouse Laboratory Plymouth and Alex Alderton, Roberto Amato, Jeffrey Barrett, Sonia Goncalves, Ewan Harrison, David K. Jackson, Ian Johnston, Dominic Kwiatkowski, Cordelia Langford, John Sillitoe on behalf of the Wellcome Sanger Institute COVID-19 Surveillance Team |

| | | | |
|---|---|---|---|
| EPI_ISL_3221855 | Lighthouse Lab in Glasgow | Wellcome Sanger Institute for the COVID-19 Genomics UK (COG-UK) Consortium | Harper VanSteenhouse, Yumi Kasai, David Gray, Carol Clugston, Anna Dominiczak and Alex Alderton, Roberto Amato, Jeffrey Barrett, Sonia Goncalves, Ewan Harrison, David K. Jackson, Ian Johnston, Dominic Kwiatkowski, Cordelia Langford, John Sillitoe on behalf of the Wellcome Sanger Institute COVID-19 Surveillance Team |
| EPI_ISL_3221856 | Lighthouse Laboratory Plymouth | Wellcome Sanger Institute for the COVID-19 Genomics UK (COG-UK) Consortium | Lighthouse Laboratory Plymouth and Alex Alderton, Roberto Amato, Jeffrey Barrett, Sonia Goncalves, Ewan Harrison, David K. Jackson, Ian Johnston, Dominic Kwiatkowski, Cordelia Langford, John Sillitoe on behalf of the Wellcome Sanger Institute COVID-19 Surveillance Team |
| EPI_ISL_3221857, EPI_ISL_3221858, EPI_ISL_3221859, EPI_ISL_3221860 | Lighthouse Lab in Glasgow | Wellcome Sanger Institute for the COVID-19 Genomics UK (COG-UK) Consortium | Harper VanSteenhouse, Yumi Kasai, David Gray, Carol Clugston, Anna Dominiczak and Alex Alderton, Roberto Amato, Jeffrey Barrett, Sonia Goncalves, Ewan Harrison, David K. Jackson, Ian Johnston, Dominic Kwiatkowski, Cordelia Langford, John Sillitoe on behalf of the Wellcome Sanger Institute COVID-19 Surveillance Team |
| EPI_ISL_3221861 | Lighthouse Lab in Milton Keynes | Wellcome Sanger Institute for the COVID-19 Genomics UK (COG-UK) Consortium | The Lighthouse Lab in Milton Keynes and Alex Alderton, Roberto Amato, Jeffrey Barrett, Sonia Goncalves, Ewan Harrison, David K. Jackson, Ian Johnston, Dominic Kwiatkowski, Cordelia Langford, John Sillitoe on behalf of the Wellcome Sanger Institute COVID-19 Surveillance Team |
| EPI_ISL_3221862, EPI_ISL_3221863, EPI_ISL_3221864 | Lighthouse Lab in Glasgow | Wellcome Sanger Institute for the COVID-19 Genomics UK (COG-UK) Consortium | Harper VanSteenhouse, Yumi Kasai, David Gray, Carol Clugston, Anna Dominiczak and Alex Alderton, Roberto Amato, Jeffrey Barrett, Sonia Goncalves, Ewan Harrison, David K. Jackson, Ian Johnston, Dominic Kwiatkowski, Cordelia Langford, John Sillitoe on behalf of the Wellcome Sanger Institute COVID-19 Surveillance Team |
| EPI_ISL_3221865 | Lighthouse Laboratory Plymouth | Wellcome Sanger Institute for the COVID-19 Genomics UK (COG-UK) Consortium | Lighthouse Laboratory Plymouth and Alex Alderton, Roberto Amato, Jeffrey Barrett, Sonia Goncalves, Ewan Harrison, David K. Jackson, Ian Johnston, Dominic Kwiatkowski, Cordelia Langford, John Sillitoe on behalf of the Wellcome Sanger Institute COVID-19 Surveillance Team |
| EPI_ISL_3221866, EPI_ISL_3221867, EPI_ISL_3221868 | Lighthouse Lab in Glasgow | Wellcome Sanger Institute for the COVID-19 Genomics UK (COG-UK) Consortium | Harper VanSteenhouse, Yumi Kasai, David Gray, Carol Clugston, Anna Dominiczak and Alex Alderton, Roberto Amato, Jeffrey Barrett, Sonia Goncalves, Ewan Harrison, David K. Jackson, Ian Johnston, Dominic Kwiatkowski, Cordelia Langford, John Sillitoe on behalf of the Wellcome Sanger Institute COVID-19 Surveillance Team |
| EPI_ISL_3221869 | Lighthouse Laboratory Plymouth | Wellcome Sanger Institute for the COVID-19 Genomics UK (COG-UK) Consortium | Lighthouse Laboratory Plymouth and Alex Alderton, Roberto Amato, Jeffrey Barrett, Sonia Goncalves, Ewan Harrison, David K. Jackson, Ian Johnston, Dominic Kwiatkowski, Cordelia Langford, John Sillitoe on behalf of the Wellcome Sanger Institute COVID-19 Surveillance Team |
| EPI_ISL_3221870, EPI_ISL_3221871, EPI_ISL_3221872, EPI_ISL_3221873 | Lighthouse Lab in Glasgow | Wellcome Sanger Institute for the COVID-19 Genomics UK (COG-UK) Consortium | Harper VanSteenhouse, Yumi Kasai, David Gray, Carol Clugston, Anna Dominiczak and Alex Alderton, Roberto Amato, Jeffrey Barrett, Sonia Goncalves, Ewan Harrison, David K. Jackson, Ian Johnston, Dominic Kwiatkowski, Cordelia Langford, John Sillitoe on behalf of the Wellcome Sanger Institute COVID-19 Surveillance Team |
| EPI_ISL_3221874 | Lighthouse Lab in Milton Keynes | Wellcome Sanger Institute for the COVID-19 Genomics UK (COG-UK) Consortium | The Lighthouse Lab in Milton Keynes and Alex Alderton, Roberto Amato, Jeffrey Barrett, Sonia Goncalves, Ewan Harrison, David K. Jackson, Ian Johnston, Dominic Kwiatkowski, Cordelia Langford, John Sillitoe on behalf of the Wellcome Sanger Institute COVID-19 Surveillance Team |
| EPI_ISL_3221875 | Lighthouse Laboratory Plymouth | Wellcome Sanger Institute for the COVID-19 Genomics UK (COG-UK) Consortium | Lighthouse Laboratory Plymouth and Alex Alderton, Roberto Amato, Jeffrey Barrett, Sonia Goncalves, Ewan Harrison, David K. Jackson, Ian Johnston, Dominic Kwiatkowski, Cordelia Langford, John Sillitoe on behalf of the Wellcome Sanger Institute COVID-19 Surveillance Team |
| EPI_ISL_3221876 | Lighthouse Lab in Glasgow | Wellcome Sanger Institute for the COVID-19 Genomics UK (COG-UK) Consortium | Harper VanSteenhouse, Yumi Kasai, David Gray, Carol Clugston, Anna Dominiczak and Alex Alderton, Roberto Amato, Jeffrey Barrett, Sonia Goncalves, Ewan Harrison, David K. Jackson, Ian Johnston, Dominic Kwiatkowski, Cordelia Langford, John Sillitoe on behalf of the Wellcome Sanger Institute COVID-19 Surveillance Team |
| EPI_ISL_3221877 | Lighthouse Laboratory Plymouth | Wellcome Sanger Institute for the COVID-19 Genomics UK (COG-UK) Consortium | Lighthouse Laboratory Plymouth and Alex Alderton, Roberto Amato, Jeffrey Barrett, Sonia Goncalves, Ewan Harrison, David K. Jackson, Ian Johnston, Dominic Kwiatkowski, Cordelia Langford, John Sillitoe on behalf of the Wellcome Sanger Institute COVID-19 Surveillance Team |
| EPI_ISL_3221878, EPI_ISL_3221879, EPI_ISL_3221880 | Lighthouse Lab in Glasgow | Wellcome Sanger Institute for the COVID-19 Genomics UK (COG-UK) Consortium | Harper VanSteenhouse, Yumi Kasai, David Gray, Carol Clugston, Anna Dominiczak and Alex Alderton, Roberto Amato, Jeffrey Barrett, Sonia Goncalves, Ewan Harrison, David K. Jackson, Ian Johnston, Dominic Kwiatkowski, Cordelia Langford, John Sillitoe on behalf of the Wellcome Sanger Institute COVID-19 Surveillance Team |
| EPI_ISL_3221881 | Lighthouse Lab in Milton Keynes | Wellcome Sanger Institute for the COVID-19 Genomics UK (COG-UK) Consortium | The Lighthouse Lab in Milton Keynes and Alex Alderton, Roberto Amato, Jeffrey Barrett, Sonia Goncalves, Ewan Harrison, David K. Jackson, Ian Johnston, Dominic Kwiatkowski, Cordelia Langford, John Sillitoe on behalf of the Wellcome Sanger Institute COVID-19 Surveillance Team |
| EPI_ISL_3221882 | Lighthouse Lab in Glasgow | Wellcome Sanger Institute for the COVID-19 Genomics UK (COG-UK) Consortium | Harper VanSteenhouse, Yumi Kasai, David Gray, Carol Clugston, Anna Dominiczak and Alex Alderton, Roberto Amato, Jeffrey Barrett, Sonia Goncalves, Ewan Harrison, David K. Jackson, Ian Johnston, Dominic Kwiatkowski, Cordelia Langford, John Sillitoe on behalf of the Wellcome Sanger Institute COVID-19 Surveillance Team |
| EPI_ISL_3221883 | Lighthouse Laboratory Plymouth | Wellcome Sanger Institute for the COVID-19 Genomics UK (COG-UK) Consortium | Lighthouse Laboratory Plymouth and Alex Alderton, Roberto Amato, Jeffrey Barrett, Sonia Goncalves, Ewan Harrison, David K. Jackson, Ian Johnston, Dominic Kwiatkowski, Cordelia Langford, John Sillitoe on behalf of the Wellcome Sanger Institute COVID-19 Surveillance Team |
| EPI_ISL_3221884, EPI_ISL_3221885 | Lighthouse Lab in Milton Keynes | Wellcome Sanger Institute for the COVID-19 Genomics UK (COG-UK) Consortium | The Lighthouse Lab in Milton Keynes and Alex Alderton, Roberto Amato, Jeffrey Barrett, Sonia Goncalves, Ewan Harrison, David K. Jackson, Ian Johnston, Dominic Kwiatkowski, Cordelia Langford, John Sillitoe on behalf of the Wellcome Sanger Institute COVID-19 Surveillance Team |
| EPI_ISL_3221886, EPI_ISL_3221887, EPI_ISL_3221888, EPI_ISL_3221889 | Lighthouse Lab in Glasgow | Wellcome Sanger Institute for the COVID-19 Genomics UK (COG-UK) Consortium | Harper VanSteenhouse, Yumi Kasai, David Gray, Carol Clugston, Anna Dominiczak and Alex Alderton, Roberto Amato, Jeffrey Barrett, Sonia Goncalves, Ewan Harrison, David K. Jackson, Ian Johnston, Dominic Kwiatkowski, Cordelia Langford, John Sillitoe on behalf of the Wellcome Sanger Institute COVID-19 Surveillance Team |
| EPI_ISL_3221890 | Lighthouse Lab in Milton Keynes | Wellcome Sanger Institute for the COVID-19 Genomics UK (COG-UK) Consortium | The Lighthouse Lab in Milton Keynes and Alex Alderton, Roberto Amato, Jeffrey Barrett, Sonia Goncalves, Ewan Harrison, David K. Jackson, Ian Johnston, Dominic Kwiatkowski, Cordelia Langford, John Sillitoe on behalf of the Wellcome Sanger Institute COVID-19 Surveillance Team |
| EPI_ISL_3221891 | Lighthouse Lab in Glasgow | Wellcome Sanger Institute for the COVID-19 Genomics UK (COG-UK) Consortium | Harper VanSteenhouse, Yumi Kasai, David Gray, Carol Clugston, Anna Dominiczak and Alex Alderton, Roberto Amato, Jeffrey Barrett, Sonia Goncalves, Ewan Harrison, David K. Jackson, Ian Johnston, Dominic Kwiatkowski, Cordelia Langford, John Sillitoe on behalf of the Wellcome Sanger Institute COVID-19 Surveillance Team |
| EPI_ISL_3221892 | Lighthouse Laboratory Plymouth | Wellcome Sanger Institute for the COVID-19 Genomics UK (COG-UK) Consortium | Lighthouse Laboratory Plymouth and Alex Alderton, Roberto Amato, Jeffrey Barrett, Sonia Goncalves, Ewan Harrison, David K. Jackson, Ian Johnston, Dominic Kwiatkowski, Cordelia Langford, John Sillitoe on behalf of the Wellcome Sanger Institute COVID-19 Surveillance Team |
| EPI_ISL_3221893 | Lighthouse Lab in Milton Keynes | Wellcome Sanger Institute for the COVID-19 Genomics UK (COG-UK) Consortium | The Lighthouse Lab in Milton Keynes and Alex Alderton, Roberto Amato, Jeffrey Barrett, Sonia Goncalves, Ewan Harrison, David K. Jackson, Ian Johnston, Dominic Kwiatkowski, Cordelia Langford, John Sillitoe on behalf of the Wellcome Sanger Institute COVID-19 Surveillance Team |
| EPI_ISL_3221894 | Lighthouse Laboratory Plymouth | Wellcome Sanger Institute for the COVID-19 Genomics UK (COG-UK) Consortium | Lighthouse Laboratory Plymouth and Alex Alderton, Roberto Amato, Jeffrey Barrett, Sonia Goncalves, Ewan Harrison, David K. Jackson, Ian Johnston, Dominic Kwiatkowski, Cordelia Langford, John Sillitoe on behalf of the Wellcome Sanger Institute COVID-19 Surveillance Team |
| EPI_ISL_3221895, EPI_ISL_3221896 | Lighthouse Lab in Glasgow | Wellcome Sanger Institute for the COVID-19 Genomics UK (COG-UK) Consortium | Harper VanSteenhouse, Yumi Kasai, David Gray, Carol Clugston, Anna Dominiczak and Alex Alderton, Roberto Amato, Jeffrey Barrett, Sonia Goncalves, Ewan Harrison, David K. Jackson, Ian Johnston, Dominic Kwiatkowski, Cordelia Langford, John Sillitoe on behalf of the Wellcome Sanger Institute COVID-19 Surveillance Team |
| EPI_ISL_3221897, EPI_ISL_3221898 | Lighthouse Laboratory Plymouth | Wellcome Sanger Institute for the COVID-19 Genomics UK (COG-UK) Consortium | Lighthouse Laboratory Plymouth and Alex Alderton, Roberto Amato, Jeffrey Barrett, Sonia Goncalves, Ewan Harrison, David K. Jackson, Ian Johnston, Dominic Kwiatkowski, Cordelia Langford, John Sillitoe on behalf of the Wellcome Sanger Institute COVID-19 Surveillance Team |
| EPI_ISL_3221900, EPI_ISL_3221901, EPI_ISL_3221902 | Lighthouse Lab in Glasgow | Wellcome Sanger Institute for the COVID-19 Genomics UK (COG-UK) Consortium | Harper VanSteenhouse, Yumi Kasai, David Gray, Carol Clugston, Anna Dominiczak and Alex Alderton, Roberto Amato, Jeffrey Barrett, Sonia Goncalves, Ewan Harrison, David K. Jackson, Ian Johnston, Dominic Kwiatkowski, Cordelia Langford, John Sillitoe on behalf of the Wellcome Sanger Institute COVID-19 Surveillance Team |
| EPI_ISL_3221903 | Lighthouse Lab in Milton Keynes | Wellcome Sanger Institute for the COVID-19 Genomics UK (COG-UK) Consortium | The Lighthouse Lab in Milton Keynes and Alex Alderton, Roberto Amato, Jeffrey Barrett, Sonia Goncalves, Ewan Harrison, David K. Jackson, Ian Johnston, Dominic Kwiatkowski, Cordelia Langford, John Sillitoe on behalf of the Wellcome Sanger Institute COVID-19 Surveillance Team |
| EPI_ISL_3221904, EPI_ISL_3221905 | Lighthouse Lab in Glasgow | Wellcome Sanger Institute for the COVID-19 Genomics UK (COG-UK) Consortium | Harper VanSteenhouse, Yumi Kasai, David Gray, Carol Clugston, Anna Dominiczak and Alex Alderton, Roberto Amato, Jeffrey Barrett, Sonia Goncalves, Ewan Harrison, David K. Jackson, Ian Johnston, Dominic Kwiatkowski, Cordelia Langford, John Sillitoe on behalf of the Wellcome Sanger Institute COVID-19 Surveillance Team |
| EPI_ISL_3221906 | Lighthouse Laboratory Plymouth | Wellcome Sanger Institute for the COVID-19 Genomics UK (COG-UK) Consortium | Lighthouse Laboratory Plymouth and Alex Alderton, Roberto Amato, Jeffrey Barrett, Sonia Goncalves, Ewan Harrison, David K. Jackson, Ian Johnston, Dominic Kwiatkowski, Cordelia Langford, John Sillitoe on behalf of the Wellcome Sanger Institute COVID-19 Surveillance Team |
| EPI_ISL_3221907 | Lighthouse Lab in Glasgow | Wellcome Sanger Institute for the COVID-19 Genomics UK (COG-UK) Consortium | Harper VanSteenhouse, Yumi Kasai, David Gray, Carol Clugston, Anna Dominiczak and Alex Alderton, Roberto Amato, Jeffrey Barrett, Sonia Goncalves, Ewan Harrison, David K. Jackson, Ian Johnston, Dominic Kwiatkowski, Cordelia Langford, John Sillitoe on behalf of the Wellcome Sanger Institute COVID-19 Surveillance Team |
| EPI_ISL_3221908 | Lighthouse Lab in Milton Keynes | Wellcome Sanger Institute for the COVID-19 Genomics UK (COG-UK) Consortium | The Lighthouse Lab in Milton Keynes and Alex Alderton, Roberto Amato, Jeffrey Barrett, Sonia Goncalves, Ewan Harrison, David K. Jackson, Ian Johnston, Dominic Kwiatkowski, Cordelia Langford, John Sillitoe on behalf of the Wellcome Sanger Institute COVID-19 Surveillance Team |
| EPI_ISL_3221909 | Lighthouse Lab in Glasgow | Wellcome Sanger Institute for the COVID-19 Genomics UK (COG-UK) Consortium | Harper VanSteenhouse, Yumi Kasai, David Gray, Carol Clugston, Anna Dominiczak and Alex Alderton, Roberto Amato, Jeffrey Barrett, Sonia Goncalves, Ewan Harrison, David K. Jackson, Ian Johnston, Dominic Kwiatkowski, Cordelia Langford, John Sillitoe on behalf of the Wellcome Sanger Institute COVID-19 Surveillance Team |
| EPI_ISL_3221910 | Lighthouse Lab in Milton Keynes | Wellcome Sanger Institute for the COVID-19 Genomics UK (COG-UK) Consortium | The Lighthouse Lab in Milton Keynes and Alex Alderton, Roberto Amato, Jeffrey Barrett, Sonia Goncalves, Ewan Harrison, David K. Jackson, Ian Johnston, Dominic Kwiatkowski, Cordelia Langford, John Sillitoe on behalf of the Wellcome Sanger Institute COVID-19 Surveillance Team |
| EPI_ISL_3221911 | Lighthouse Lab in Glasgow | Wellcome Sanger Institute for the COVID-19 Genomics UK (COG-UK) Consortium | Harper VanSteenhouse, Yumi Kasai, David Gray, Carol Clugston, Anna Dominiczak and Alex Alderton, Roberto Amato, Jeffrey Barrett, Sonia Goncalves, Ewan Harrison, David K. Jackson, Ian Johnston, Dominic Kwiatkowski, Cordelia Langford, John Sillitoe on behalf of the Wellcome Sanger Institute COVID-19 Surveillance Team |
| EPI_ISL_3221912 | Lighthouse Lab in Milton Keynes | Wellcome Sanger Institute for the COVID-19 Genomics UK (COG-UK) Consortium | The Lighthouse Lab in Milton Keynes and Alex Alderton, Roberto Amato, Jeffrey Barrett, Sonia Goncalves, Ewan Harrison, David K. Jackson, Ian Johnston, Dominic Kwiatkowski, Cordelia Langford, John Sillitoe on behalf of the Wellcome Sanger Institute COVID-19 Surveillance Team |
| EPI_ISL_3221913 | Lighthouse Lab in Glasgow | Wellcome Sanger Institute for the COVID-19 Genomics UK (COG-UK) Consortium | Harper VanSteenhouse, Yumi Kasai, David Gray, Carol Clugston, Anna Dominiczak and Alex Alderton, Roberto Amato, Jeffrey Barrett, Sonia Goncalves, Ewan Harrison, David K. Jackson, Ian Johnston, Dominic Kwiatkowski, Cordelia Langford, John Sillitoe on behalf of the Wellcome Sanger Institute COVID-19 Surveillance Team |
| EPI_ISL_3221914 | Lighthouse Lab in Milton Keynes | Wellcome Sanger Institute for the COVID-19 Genomics UK (COG-UK) Consortium | The Lighthouse Lab in Milton Keynes and Alex Alderton, Roberto Amato, Jeffrey Barrett, Sonia Goncalves, Ewan Harrison, David K. Jackson, Ian Johnston, Dominic Kwiatkowski, Cordelia Langford, John Sillitoe on behalf of the Wellcome Sanger Institute COVID-19 Surveillance Team |
| EPI_ISL_3221915, EPI_ISL_3221916, EPI_ISL_3221917, EPI_ISL_3221918, EPI_ISL_3221919, EPI_ISL_3221920, EPI_ISL_3221921 | Lighthouse Lab in Glasgow | Wellcome Sanger Institute for the COVID-19 Genomics UK (COG-UK) Consortium | Harper VanSteenhouse, Yumi Kasai, David Gray, Carol Clugston, Anna Dominiczak and Alex Alderton, Roberto Amato, Jeffrey Barrett, Sonia Goncalves, Ewan Harrison, David K. Jackson, Ian Johnston, Dominic Kwiatkowski, Cordelia Langford, John Sillitoe on behalf of the Wellcome Sanger Institute COVID-19 Surveillance Team |

| | | | |
|---|---|---|---|
| EPI_ISL_3221922 | Lighthouse Laboratory Plymouth | Wellcome Sanger Institute for the COVID-19 Genomics UK (COG-UK) Consortium | Lighthouse Laboratory Plymouth and Alex Alderton, Roberto Amato, Jeffrey Barrett, Sonia Goncalves, Ewan Harrison, David K. Jackson, Ian Johnston, Dominic Kwiatkowski, Cordelia Langford, John Sillitoe on behalf of the Wellcome Sanger Institute COVID-19 Surveillance Team |
| EPI_ISL_3221923, EPI_ISL_3221924 | Lighthouse Lab in Glasgow | Wellcome Sanger Institute for the COVID-19 Genomics UK (COG-UK) Consortium | Harper VanSteenhouse, Yumi Kasai, David Gray, Carol Clugston, Anna Dominiczak and Alex Alderton, Roberto Amato, Jeffrey Barrett, Sonia Goncalves, Ewan Harrison, David K. Jackson, Ian Johnston, Dominic Kwiatkowski, Cordelia Langford, John Sillitoe on behalf of the Wellcome Sanger Institute COVID-19 Surveillance Team |
| EPI_ISL_3221925 | Lighthouse Laboratory Plymouth | Wellcome Sanger Institute for the COVID-19 Genomics UK (COG-UK) Consortium | Lighthouse Laboratory Plymouth and Alex Alderton, Roberto Amato, Jeffrey Barrett, Sonia Goncalves, Ewan Harrison, David K. Jackson, Ian Johnston, Dominic Kwiatkowski, Cordelia Langford, John Sillitoe on behalf of the Wellcome Sanger Institute COVID-19 Surveillance Team |
| EPI_ISL_3221926 | Lighthouse Lab in Glasgow | Wellcome Sanger Institute for the COVID-19 Genomics UK (COG-UK) Consortium | Harper VanSteenhouse, Yumi Kasai, David Gray, Carol Clugston, Anna Dominiczak and Alex Alderton, Roberto Amato, Jeffrey Barrett, Sonia Goncalves, Ewan Harrison, David K. Jackson, Ian Johnston, Dominic Kwiatkowski, Cordelia Langford, John Sillitoe on behalf of the Wellcome Sanger Institute COVID-19 Surveillance Team |
| EPI_ISL_3221927 | Lighthouse Lab in Milton Keynes | Wellcome Sanger Institute for the COVID-19 Genomics UK (COG-UK) Consortium | The Lighthouse Lab in Milton Keynes and Alex Alderton, Roberto Amato, Jeffrey Barrett, Sonia Goncalves, Ewan Harrison, David K. Jackson, Ian Johnston, Dominic Kwiatkowski, Cordelia Langford, John Sillitoe on behalf of the Wellcome Sanger Institute COVID-19 Surveillance Team |
| EPI_ISL_3221928 | Lighthouse Lab in Glasgow | Wellcome Sanger Institute for the COVID-19 Genomics UK (COG-UK) Consortium | Harper VanSteenhouse, Yumi Kasai, David Gray, Carol Clugston, Anna Dominiczak and Alex Alderton, Roberto Amato, Jeffrey Barrett, Sonia Goncalves, Ewan Harrison, David K. Jackson, Ian Johnston, Dominic Kwiatkowski, Cordelia Langford, John Sillitoe on behalf of the Wellcome Sanger Institute COVID-19 Surveillance Team |
| EPI_ISL_3221929 | Lighthouse Laboratory Plymouth | Wellcome Sanger Institute for the COVID-19 Genomics UK (COG-UK) Consortium | Lighthouse Laboratory Plymouth and Alex Alderton, Roberto Amato, Jeffrey Barrett, Sonia Goncalves, Ewan Harrison, David K. Jackson, Ian Johnston, Dominic Kwiatkowski, Cordelia Langford, John Sillitoe on behalf of the Wellcome Sanger Institute COVID-19 Surveillance Team |
| EPI_ISL_3221930, EPI_ISL_3221931 | Lighthouse Lab in Glasgow | Wellcome Sanger Institute for the COVID-19 Genomics UK (COG-UK) Consortium | Harper VanSteenhouse, Yumi Kasai, David Gray, Carol Clugston, Anna Dominiczak and Alex Alderton, Roberto Amato, Jeffrey Barrett, Sonia Goncalves, Ewan Harrison, David K. Jackson, Ian Johnston, Dominic Kwiatkowski, Cordelia Langford, John Sillitoe on behalf of the Wellcome Sanger Institute COVID-19 Surveillance Team |
| EPI_ISL_3221932 | Lighthouse Laboratory Plymouth | Wellcome Sanger Institute for the COVID-19 Genomics UK (COG-UK) Consortium | Lighthouse Laboratory Plymouth and Alex Alderton, Roberto Amato, Jeffrey Barrett, Sonia Goncalves, Ewan Harrison, David K. Jackson, Ian Johnston, Dominic Kwiatkowski, Cordelia Langford, John Sillitoe on behalf of the Wellcome Sanger Institute COVID-19 Surveillance Team |
| EPI_ISL_3221933 | Lighthouse Lab in Milton Keynes | Wellcome Sanger Institute for the COVID-19 Genomics UK (COG-UK) Consortium | The Lighthouse Lab in Milton Keynes and Alex Alderton, Roberto Amato, Jeffrey Barrett, Sonia Goncalves, Ewan Harrison, David K. Jackson, Ian Johnston, Dominic Kwiatkowski, Cordelia Langford, John Sillitoe on behalf of the Wellcome Sanger Institute COVID-19 Surveillance Team |
| EPI_ISL_3221934 | Lighthouse Lab in Glasgow | Wellcome Sanger Institute for the COVID-19 Genomics UK (COG-UK) Consortium | Harper VanSteenhouse, Yumi Kasai, David Gray, Carol Clugston, Anna Dominiczak and Alex Alderton, Roberto Amato, Jeffrey Barrett, Sonia Goncalves, Ewan Harrison, David K. Jackson, Ian Johnston, Dominic Kwiatkowski, Cordelia Langford, John Sillitoe on behalf of the Wellcome Sanger Institute COVID-19 Surveillance Team |
| EPI_ISL_3221935, EPI_ISL_3221936 | Lighthouse Lab in Milton Keynes | Wellcome Sanger Institute for the COVID-19 Genomics UK (COG-UK) Consortium | The Lighthouse Lab in Milton Keynes and Alex Alderton, Roberto Amato, Jeffrey Barrett, Sonia Goncalves, Ewan Harrison, David K. Jackson, Ian Johnston, Dominic Kwiatkowski, Cordelia Langford, John Sillitoe on behalf of the Wellcome Sanger Institute COVID-19 Surveillance Team |
| EPI_ISL_3221937 | Lighthouse Lab in Glasgow | Wellcome Sanger Institute for the COVID-19 Genomics UK (COG-UK) Consortium | Harper VanSteenhouse, Yumi Kasai, David Gray, Carol Clugston, Anna Dominiczak and Alex Alderton, Roberto Amato, Jeffrey Barrett, Sonia Goncalves, Ewan Harrison, David K. Jackson, Ian Johnston, Dominic Kwiatkowski, Cordelia Langford, John Sillitoe on behalf of the Wellcome Sanger Institute COVID-19 Surveillance Team |
| EPI_ISL_3221938, EPI_ISL_3221939 | Lighthouse Laboratory Plymouth | Wellcome Sanger Institute for the COVID-19 Genomics UK (COG-UK) Consortium | Lighthouse Laboratory Plymouth and Alex Alderton, Roberto Amato, Jeffrey Barrett, Sonia Goncalves, Ewan Harrison, David K. Jackson, Ian Johnston, Dominic Kwiatkowski, Cordelia Langford, John Sillitoe on behalf of the Wellcome Sanger Institute COVID-19 Surveillance Team |
| EPI_ISL_3221940, EPI_ISL_3221941 | Lighthouse Lab in Milton Keynes | Wellcome Sanger Institute for the COVID-19 Genomics UK (COG-UK) Consortium | The Lighthouse Lab in Milton Keynes and Alex Alderton, Roberto Amato, Jeffrey Barrett, Sonia Goncalves, Ewan Harrison, David K. Jackson, Ian Johnston, Dominic Kwiatkowski, Cordelia Langford, John Sillitoe on behalf of the Wellcome Sanger Institute COVID-19 Surveillance Team |
| EPI_ISL_3221942 | Lighthouse Lab in Glasgow | Wellcome Sanger Institute for the COVID-19 Genomics UK (COG-UK) Consortium | Harper VanSteenhouse, Yumi Kasai, David Gray, Carol Clugston, Anna Dominiczak and Alex Alderton, Roberto Amato, Jeffrey Barrett, Sonia Goncalves, Ewan Harrison, David K. Jackson, Ian Johnston, Dominic Kwiatkowski, Cordelia Langford, John Sillitoe on behalf of the Wellcome Sanger Institute COVID-19 Surveillance Team |
| EPI_ISL_3221943 | Lighthouse Lab in Milton Keynes | Wellcome Sanger Institute for the COVID-19 Genomics UK (COG-UK) Consortium | The Lighthouse Lab in Milton Keynes and Alex Alderton, Roberto Amato, Jeffrey Barrett, Sonia Goncalves, Ewan Harrison, David K. Jackson, Ian Johnston, Dominic Kwiatkowski, Cordelia Langford, John Sillitoe on behalf of the Wellcome Sanger Institute COVID-19 Surveillance Team |
| EPI_ISL_3221944 | Lighthouse Lab in Glasgow | Wellcome Sanger Institute for the COVID-19 Genomics UK (COG-UK) Consortium | Harper VanSteenhouse, Yumi Kasai, David Gray, Carol Clugston, Anna Dominiczak and Alex Alderton, Roberto Amato, Jeffrey Barrett, Sonia Goncalves, Ewan Harrison, David K. Jackson, Ian Johnston, Dominic Kwiatkowski, Cordelia Langford, John Sillitoe on behalf of the Wellcome Sanger Institute COVID-19 Surveillance Team |
| EPI_ISL_3221945 | Lighthouse Laboratory Plymouth | Wellcome Sanger Institute for the COVID-19 Genomics UK (COG-UK) Consortium | Lighthouse Laboratory Plymouth and Alex Alderton, Roberto Amato, Jeffrey Barrett, Sonia Goncalves, Ewan Harrison, David K. Jackson, Ian Johnston, Dominic Kwiatkowski, Cordelia Langford, John Sillitoe on behalf of the Wellcome Sanger Institute COVID-19 Surveillance Team |
| EPI_ISL_3221946 | Lighthouse Lab in Milton Keynes | Wellcome Sanger Institute for the COVID-19 Genomics UK (COG-UK) Consortium | The Lighthouse Lab in Milton Keynes and Alex Alderton, Roberto Amato, Jeffrey Barrett, Sonia Goncalves, Ewan Harrison, David K. Jackson, Ian Johnston, Dominic Kwiatkowski, Cordelia Langford, John Sillitoe on behalf of the Wellcome Sanger Institute COVID-19 Surveillance Team |
| EPI_ISL_3221947 | Lighthouse Laboratory Plymouth | Wellcome Sanger Institute for the COVID-19 Genomics UK (COG-UK) Consortium | Lighthouse Laboratory Plymouth and Alex Alderton, Roberto Amato, Jeffrey Barrett, Sonia Goncalves, Ewan Harrison, David K. Jackson, Ian Johnston, Dominic Kwiatkowski, Cordelia Langford, John Sillitoe on behalf of the Wellcome Sanger Institute COVID-19 Surveillance Team |
| EPI_ISL_3221948 | Lighthouse Lab in Glasgow | Wellcome Sanger Institute for the COVID-19 Genomics UK (COG-UK) Consortium | Harper VanSteenhouse, Yumi Kasai, David Gray, Carol Clugston, Anna Dominiczak and Alex Alderton, Roberto Amato, Jeffrey Barrett, Sonia Goncalves, Ewan Harrison, David K. Jackson, Ian Johnston, Dominic Kwiatkowski, Cordelia Langford, John Sillitoe on behalf of the Wellcome Sanger Institute COVID-19 Surveillance Team |
| EPI_ISL_3221949 | Lighthouse Lab in Milton Keynes | Wellcome Sanger Institute for the COVID-19 Genomics UK (COG-UK) Consortium | The Lighthouse Lab in Milton Keynes and Alex Alderton, Roberto Amato, Jeffrey Barrett, Sonia Goncalves, Ewan Harrison, David K. Jackson, Ian Johnston, Dominic Kwiatkowski, Cordelia Langford, John Sillitoe on behalf of the Wellcome Sanger Institute COVID-19 Surveillance Team |
| EPI_ISL_3221950 | Lighthouse Laboratory Plymouth | Wellcome Sanger Institute for the COVID-19 Genomics UK (COG-UK) Consortium | Lighthouse Laboratory Plymouth and Alex Alderton, Roberto Amato, Jeffrey Barrett, Sonia Goncalves, Ewan Harrison, David K. Jackson, Ian Johnston, Dominic Kwiatkowski, Cordelia Langford, John Sillitoe on behalf of the Wellcome Sanger Institute COVID-19 Surveillance Team |
| EPI_ISL_3221951 | Lighthouse Lab in Milton Keynes | Wellcome Sanger Institute for the COVID-19 Genomics UK (COG-UK) Consortium | The Lighthouse Lab in Milton Keynes and Alex Alderton, Roberto Amato, Jeffrey Barrett, Sonia Goncalves, Ewan Harrison, David K. Jackson, Ian Johnston, Dominic Kwiatkowski, Cordelia Langford, John Sillitoe on behalf of the Wellcome Sanger Institute COVID-19 Surveillance Team |
| EPI_ISL_3221952, EPI_ISL_3221953, EPI_ISL_3221954 | Lighthouse Laboratory Plymouth | Wellcome Sanger Institute for the COVID-19 Genomics UK (COG-UK) Consortium | Lighthouse Laboratory Plymouth and Alex Alderton, Roberto Amato, Jeffrey Barrett, Sonia Goncalves, Ewan Harrison, David K. Jackson, Ian Johnston, Dominic Kwiatkowski, Cordelia Langford, John Sillitoe on behalf of the Wellcome Sanger Institute COVID-19 Surveillance Team |
| EPI_ISL_3221955, EPI_ISL_3221956, EPI_ISL_3221957, EPI_ISL_3221958 | Lighthouse Lab in Glasgow | Wellcome Sanger Institute for the COVID-19 Genomics UK (COG-UK) Consortium | Harper VanSteenhouse, Yumi Kasai, David Gray, Carol Clugston, Anna Dominiczak and Alex Alderton, Roberto Amato, Jeffrey Barrett, Sonia Goncalves, Ewan Harrison, David K. Jackson, Ian Johnston, Dominic Kwiatkowski, Cordelia Langford, John Sillitoe on behalf of the Wellcome Sanger Institute COVID-19 Surveillance Team |
| EPI_ISL_3221959 | Lighthouse Laboratory Plymouth | Wellcome Sanger Institute for the COVID-19 Genomics UK (COG-UK) Consortium | Lighthouse Laboratory Plymouth and Alex Alderton, Roberto Amato, Jeffrey Barrett, Sonia Goncalves, Ewan Harrison, David K. Jackson, Ian Johnston, Dominic Kwiatkowski, Cordelia Langford, John Sillitoe on behalf of the Wellcome Sanger Institute COVID-19 Surveillance Team |
| EPI_ISL_3221960 | Lighthouse Lab in Glasgow | Wellcome Sanger Institute for the COVID-19 Genomics UK (COG-UK) Consortium | Harper VanSteenhouse, Yumi Kasai, David Gray, Carol Clugston, Anna Dominiczak and Alex Alderton, Roberto Amato, Jeffrey Barrett, Sonia Goncalves, Ewan Harrison, David K. Jackson, Ian Johnston, Dominic Kwiatkowski, Cordelia Langford, John Sillitoe on behalf of the Wellcome Sanger Institute COVID-19 Surveillance Team |
| EPI_ISL_3221961 | Lighthouse Lab in Milton Keynes | Wellcome Sanger Institute for the COVID-19 Genomics UK (COG-UK) Consortium | The Lighthouse Lab in Milton Keynes and Alex Alderton, Roberto Amato, Jeffrey Barrett, Sonia Goncalves, Ewan Harrison, David K. Jackson, Ian Johnston, Dominic Kwiatkowski, Cordelia Langford, John Sillitoe on behalf of the Wellcome Sanger Institute COVID-19 Surveillance Team |
| EPI_ISL_3221962 | Lighthouse Laboratory Plymouth | Wellcome Sanger Institute for the COVID-19 Genomics UK (COG-UK) Consortium | Lighthouse Laboratory Plymouth and Alex Alderton, Roberto Amato, Jeffrey Barrett, Sonia Goncalves, Ewan Harrison, David K. Jackson, Ian Johnston, Dominic Kwiatkowski, Cordelia Langford, John Sillitoe on behalf of the Wellcome Sanger Institute COVID-19 Surveillance Team |
| EPI_ISL_3221963, EPI_ISL_3221964 | Lighthouse Lab in Glasgow | Wellcome Sanger Institute for the COVID-19 Genomics UK (COG-UK) Consortium | Harper VanSteenhouse, Yumi Kasai, David Gray, Carol Clugston, Anna Dominiczak and Alex Alderton, Roberto Amato, Jeffrey Barrett, Sonia Goncalves, Ewan Harrison, David K. Jackson, Ian Johnston, Dominic Kwiatkowski, Cordelia Langford, John Sillitoe on behalf of the Wellcome Sanger Institute COVID-19 Surveillance Team |
| EPI_ISL_3221965 | Lighthouse Lab in Milton Keynes | Wellcome Sanger Institute for the COVID-19 Genomics UK (COG-UK) Consortium | The Lighthouse Lab in Milton Keynes and Alex Alderton, Roberto Amato, Jeffrey Barrett, Sonia Goncalves, Ewan Harrison, David K. Jackson, Ian Johnston, Dominic Kwiatkowski, Cordelia Langford, John Sillitoe on behalf of the Wellcome Sanger Institute COVID-19 Surveillance Team |
| EPI_ISL_3221966 | Lighthouse Lab in Glasgow | Wellcome Sanger Institute for the COVID-19 Genomics UK (COG-UK) Consortium | Harper VanSteenhouse, Yumi Kasai, David Gray, Carol Clugston, Anna Dominiczak and Alex Alderton, Roberto Amato, Jeffrey Barrett, Sonia Goncalves, Ewan Harrison, David K. Jackson, Ian Johnston, Dominic Kwiatkowski, Cordelia Langford, John Sillitoe on behalf of the Wellcome Sanger Institute COVID-19 Surveillance Team |
| EPI_ISL_3221967 | Lighthouse Laboratory Plymouth | Wellcome Sanger Institute for the COVID-19 Genomics UK (COG-UK) Consortium | Lighthouse Laboratory Plymouth and Alex Alderton, Roberto Amato, Jeffrey Barrett, Sonia Goncalves, Ewan Harrison, David K. Jackson, Ian Johnston, Dominic Kwiatkowski, Cordelia Langford, John Sillitoe on behalf of the Wellcome Sanger Institute COVID-19 Surveillance Team |
| EPI_ISL_3221968 | Lighthouse Lab in Glasgow | Wellcome Sanger Institute for the COVID-19 Genomics UK (COG-UK) Consortium | Harper VanSteenhouse, Yumi Kasai, David Gray, Carol Clugston, Anna Dominiczak and Alex Alderton, Roberto Amato, Jeffrey Barrett, Sonia Goncalves, Ewan Harrison, David K. Jackson, Ian Johnston, Dominic Kwiatkowski, Cordelia Langford, John Sillitoe on behalf of the Wellcome Sanger Institute COVID-19 Surveillance Team |
| EPI_ISL_3221969 | Lighthouse Laboratory Plymouth | Wellcome Sanger Institute for the COVID-19 Genomics UK (COG-UK) Consortium | Lighthouse Laboratory Plymouth and Alex Alderton, Roberto Amato, Jeffrey Barrett, Sonia Goncalves, Ewan Harrison, David K. Jackson, Ian Johnston, Dominic Kwiatkowski, Cordelia Langford, John Sillitoe on behalf of the Wellcome Sanger Institute COVID-19 Surveillance Team |
| EPI_ISL_3221970, EPI_ISL_3221971 | Lighthouse Lab in Milton Keynes | Wellcome Sanger Institute for the COVID-19 Genomics UK (COG-UK) Consortium | The Lighthouse Lab in Milton Keynes and Alex Alderton, Roberto Amato, Jeffrey Barrett, Sonia Goncalves, Ewan Harrison, David K. Jackson, Ian Johnston, Dominic Kwiatkowski, Cordelia Langford, John Sillitoe on behalf of the Wellcome Sanger Institute COVID-19 Surveillance Team |
| EPI_ISL_3221972, EPI_ISL_3221973 | Lighthouse Lab in Glasgow | Wellcome Sanger Institute for the COVID-19 Genomics UK (COG-UK) Consortium | Harper VanSteenhouse, Yumi Kasai, David Gray, Carol Clugston, Anna Dominiczak and Alex Alderton, Roberto Amato, Jeffrey Barrett, Sonia Goncalves, Ewan Harrison, David K. Jackson, Ian Johnston, Dominic Kwiatkowski, Cordelia Langford, John Sillitoe on behalf of the Wellcome Sanger Institute COVID-19 Surveillance Team |
| EPI_ISL_3221974 | Lighthouse Lab in Milton Keynes | Wellcome Sanger Institute for the COVID-19 Genomics UK (COG-UK) Consortium | The Lighthouse Lab in Milton Keynes and Alex Alderton, Roberto Amato, Jeffrey Barrett, Sonia Goncalves, Ewan Harrison, David K. Jackson, Ian Johnston, Dominic Kwiatkowski, Cordelia Langford, John Sillitoe on behalf of the Wellcome Sanger Institute COVID-19 Surveillance Team |
| EPI_ISL_3221975 | Lighthouse Laboratory Plymouth | Wellcome Sanger Institute for the COVID-19 Genomics UK (COG-UK) Consortium | Lighthouse Laboratory Plymouth and Alex Alderton, Roberto Amato, Jeffrey Barrett, Sonia Goncalves, Ewan Harrison, David K. Jackson, Ian Johnston, Dominic Kwiatkowski, Cordelia Langford, John Sillitoe on behalf of the Wellcome Sanger Institute COVID-19 Surveillance Team |
| EPI_ISL_3221976, EPI_ISL_3221977 | Lighthouse Lab in Milton Keynes | Wellcome Sanger Institute for the COVID-19 Genomics UK (COG-UK) Consortium | The Lighthouse Lab in Milton Keynes and Alex Alderton, Roberto Amato, Jeffrey Barrett, Sonia Goncalves, Ewan Harrison, David K. Jackson, Ian Johnston, Dominic Kwiatkowski, Cordelia Langford, John Sillitoe on behalf of the Wellcome Sanger Institute COVID-19 Surveillance Team |
| EPI_ISL_3221978 | Lighthouse Laboratory Plymouth | Wellcome Sanger Institute for the COVID-19 Genomics UK (COG-UK) Consortium | Lighthouse Laboratory Plymouth and Alex Alderton, Roberto Amato, Jeffrey Barrett, Sonia Goncalves, Ewan Harrison, David K. Jackson, Ian Johnston, Dominic Kwiatkowski, Cordelia Langford, John Sillitoe on behalf of the Wellcome Sanger Institute COVID-19 Surveillance Team |
| EPI_ISL_3221979 | Lighthouse Lab in Glasgow | Wellcome Sanger Institute for the COVID-19 Genomics UK (COG-UK) Consortium | Harper VanSteenhouse, Yumi Kasai, David Gray, Carol Clugston, Anna Dominiczak and Alex Alderton, Roberto Amato, Jeffrey Barrett, Sonia Goncalves, Ewan Harrison, David K. Jackson, Ian Johnston, Dominic Kwiatkowski, Cordelia Langford, John Sillitoe on behalf of the Wellcome Sanger Institute COVID-19 Surveillance Team |

| | | | |
|---|---|---|---|
| EPI_ISL_3221980 | Lighthouse Lab in Milton Keynes | Wellcome Sanger Institute for the COVID-19 Genomics UK (COG-UK) Consortium | The Lighthouse Lab in Milton Keynes and Alex Alderton, Roberto Amato, Jeffrey Barrett, Sonia Goncalves, Ewan Harrison, David K. Jackson, Ian Johnston, Dominic Kwiatkowski, Cordelia Langford, John Sillitoe on behalf of the Wellcome Sanger Institute COVID-19 Surveillance Team |
| EPI_ISL_3221981, EPI_ISL_3221982, EPI_ISL_3221983 | Lighthouse Laboratory Plymouth | Wellcome Sanger Institute for the COVID-19 Genomics UK (COG-UK) Consortium | Lighthouse Laboratory Plymouth and Alex Alderton, Roberto Amato, Jeffrey Barrett, Sonia Goncalves, Ewan Harrison, David K. Jackson, Ian Johnston, Dominic Kwiatkowski, Cordelia Langford, John Sillitoe on behalf of the Wellcome Sanger Institute COVID-19 Surveillance Team |
| EPI_ISL_3221984 | Lighthouse Lab in Milton Keynes | Wellcome Sanger Institute for the COVID-19 Genomics UK (COG-UK) Consortium | The Lighthouse Lab in Milton Keynes and Alex Alderton, Roberto Amato, Jeffrey Barrett, Sonia Goncalves, Ewan Harrison, David K. Jackson, Ian Johnston, Dominic Kwiatkowski, Cordelia Langford, John Sillitoe on behalf of the Wellcome Sanger Institute COVID-19 Surveillance Team |
| EPI_ISL_3221985 | Lighthouse Laboratory Plymouth | Wellcome Sanger Institute for the COVID-19 Genomics UK (COG-UK) Consortium | Lighthouse Laboratory Plymouth and Alex Alderton, Roberto Amato, Jeffrey Barrett, Sonia Goncalves, Ewan Harrison, David K. Jackson, Ian Johnston, Dominic Kwiatkowski, Cordelia Langford, John Sillitoe on behalf of the Wellcome Sanger Institute COVID-19 Surveillance Team |
| EPI_ISL_3221986 | Berkshire and Surrey Pathology Services Lighthouse Laboratory | Wellcome Sanger Institute for the COVID-19 Genomics UK (COG-UK) Consortium | Berkshire and Surrey Pathology Services Lighthouse Laboratory and Alex Alderton, Roberto Amato, Jeffrey Barrett, Sonia Goncalves, Ewan Harrison, David K. Jackson, Ian Johnston, Dominic Kwiatkowski, Cordelia Langford, John Sillitoe on behalf of the Wellcome Sanger Institute COVID-19 Surveillance Team |
| EPI_ISL_3221987 | Lighthouse Laboratory Plymouth | Wellcome Sanger Institute for the COVID-19 Genomics UK (COG-UK) Consortium | Lighthouse Laboratory Plymouth and Alex Alderton, Roberto Amato, Jeffrey Barrett, Sonia Goncalves, Ewan Harrison, David K. Jackson, Ian Johnston, Dominic Kwiatkowski, Cordelia Langford, John Sillitoe on behalf of the Wellcome Sanger Institute COVID-19 Surveillance Team |
| EPI_ISL_3221988 | Berkshire and Surrey Pathology Services Lighthouse Laboratory | Wellcome Sanger Institute for the COVID-19 Genomics UK (COG-UK) Consortium | Berkshire and Surrey Pathology Services Lighthouse Laboratory and Alex Alderton, Roberto Amato, Jeffrey Barrett, Sonia Goncalves, Ewan Harrison, David K. Jackson, Ian Johnston, Dominic Kwiatkowski, Cordelia Langford, John Sillitoe on behalf of the Wellcome Sanger Institute COVID-19 Surveillance Team |
| EPI_ISL_3221989, EPI_ISL_3221990 | Lighthouse Laboratory Plymouth | Wellcome Sanger Institute for the COVID-19 Genomics UK (COG-UK) Consortium | Lighthouse Laboratory Plymouth and Alex Alderton, Roberto Amato, Jeffrey Barrett, Sonia Goncalves, Ewan Harrison, David K. Jackson, Ian Johnston, Dominic Kwiatkowski, Cordelia Langford, John Sillitoe on behalf of the Wellcome Sanger Institute COVID-19 Surveillance Team |
| EPI_ISL_3221991 | Lighthouse Lab in Milton Keynes | Wellcome Sanger Institute for the COVID-19 Genomics UK (COG-UK) Consortium | The Lighthouse Lab in Milton Keynes and Alex Alderton, Roberto Amato, Jeffrey Barrett, Sonia Goncalves, Ewan Harrison, David K. Jackson, Ian Johnston, Dominic Kwiatkowski, Cordelia Langford, John Sillitoe on behalf of the Wellcome Sanger Institute COVID-19 Surveillance Team |
| EPI_ISL_3221992, EPI_ISL_3221993 | Lighthouse Laboratory Plymouth | Wellcome Sanger Institute for the COVID-19 Genomics UK (COG-UK) Consortium | Lighthouse Laboratory Plymouth and Alex Alderton, Roberto Amato, Jeffrey Barrett, Sonia Goncalves, Ewan Harrison, David K. Jackson, Ian Johnston, Dominic Kwiatkowski, Cordelia Langford, John Sillitoe on behalf of the Wellcome Sanger Institute COVID-19 Surveillance Team |
| EPI_ISL_3221994 | Lighthouse Lab in Milton Keynes | Wellcome Sanger Institute for the COVID-19 Genomics UK (COG-UK) Consortium | The Lighthouse Lab in Milton Keynes and Alex Alderton, Roberto Amato, Jeffrey Barrett, Sonia Goncalves, Ewan Harrison, David K. Jackson, Ian Johnston, Dominic Kwiatkowski, Cordelia Langford, John Sillitoe on behalf of the Wellcome Sanger Institute COVID-19 Surveillance Team |
| EPI_ISL_3221995 | Lighthouse Laboratory Plymouth | Wellcome Sanger Institute for the COVID-19 Genomics UK (COG-UK) Consortium | Lighthouse Laboratory Plymouth and Alex Alderton, Roberto Amato, Jeffrey Barrett, Sonia Goncalves, Ewan Harrison, David K. Jackson, Ian Johnston, Dominic Kwiatkowski, Cordelia Langford, John Sillitoe on behalf of the Wellcome Sanger Institute COVID-19 Surveillance Team |
| EPI_ISL_3221996 | Lighthouse Lab in Milton Keynes | Wellcome Sanger Institute for the COVID-19 Genomics UK (COG-UK) Consortium | The Lighthouse Lab in Milton Keynes and Alex Alderton, Roberto Amato, Jeffrey Barrett, Sonia Goncalves, Ewan Harrison, David K. Jackson, Ian Johnston, Dominic Kwiatkowski, Cordelia Langford, John Sillitoe on behalf of the Wellcome Sanger Institute COVID-19 Surveillance Team |
| EPI_ISL_3221997 | Lighthouse Laboratory Plymouth | Wellcome Sanger Institute for the COVID-19 Genomics UK (COG-UK) Consortium | Lighthouse Laboratory Plymouth and Alex Alderton, Roberto Amato, Jeffrey Barrett, Sonia Goncalves, Ewan Harrison, David K. Jackson, Ian Johnston, Dominic Kwiatkowski, Cordelia Langford, John Sillitoe on behalf of the Wellcome Sanger Institute COVID-19 Surveillance Team |
| EPI_ISL_3221998 | Lighthouse Lab in Milton Keynes | Wellcome Sanger Institute for the COVID-19 Genomics UK (COG-UK) Consortium | The Lighthouse Lab in Milton Keynes and Alex Alderton, Roberto Amato, Jeffrey Barrett, Sonia Goncalves, Ewan Harrison, David K. Jackson, Ian Johnston, Dominic Kwiatkowski, Cordelia Langford, John Sillitoe on behalf of the Wellcome Sanger Institute COVID-19 Surveillance Team |
| EPI_ISL_3221999 | Lighthouse Laboratory Plymouth | Wellcome Sanger Institute for the COVID-19 Genomics UK (COG-UK) Consortium | Lighthouse Laboratory Plymouth and Alex Alderton, Roberto Amato, Jeffrey Barrett, Sonia Goncalves, Ewan Harrison, David K. Jackson, Ian Johnston, Dominic Kwiatkowski, Cordelia Langford, John Sillitoe on behalf of the Wellcome Sanger Institute COVID-19 Surveillance Team |
| EPI_ISL_3222000 | Lighthouse Lab in Milton Keynes | Wellcome Sanger Institute for the COVID-19 Genomics UK (COG-UK) Consortium | The Lighthouse Lab in Milton Keynes and Alex Alderton, Roberto Amato, Jeffrey Barrett, Sonia Goncalves, Ewan Harrison, David K. Jackson, Ian Johnston, Dominic Kwiatkowski, Cordelia Langford, John Sillitoe on behalf of the Wellcome Sanger Institute COVID-19 Surveillance Team |
| EPI_ISL_3222001, EPI_ISL_3222002 | Lighthouse Laboratory Plymouth | Wellcome Sanger Institute for the COVID-19 Genomics UK (COG-UK) Consortium | Lighthouse Laboratory Plymouth and Alex Alderton, Roberto Amato, Jeffrey Barrett, Sonia Goncalves, Ewan Harrison, David K. Jackson, Ian Johnston, Dominic Kwiatkowski, Cordelia Langford, John Sillitoe on behalf of the Wellcome Sanger Institute COVID-19 Surveillance Team |
| EPI_ISL_3222003 | Berkshire and Surrey Pathology Services Lighthouse Laboratory | Wellcome Sanger Institute for the COVID-19 Genomics UK (COG-UK) Consortium | Berkshire and Surrey Pathology Services Lighthouse Laboratory and Alex Alderton, Roberto Amato, Jeffrey Barrett, Sonia Goncalves, Ewan Harrison, David K. Jackson, Ian Johnston, Dominic Kwiatkowski, Cordelia Langford, John Sillitoe on behalf of the Wellcome Sanger Institute COVID-19 Surveillance Team |
| EPI_ISL_3222004 | Lighthouse Lab in Milton Keynes | Wellcome Sanger Institute for the COVID-19 Genomics UK (COG-UK) Consortium | The Lighthouse Lab in Milton Keynes and Alex Alderton, Roberto Amato, Jeffrey Barrett, Sonia Goncalves, Ewan Harrison, David K. Jackson, Ian Johnston, Dominic Kwiatkowski, Cordelia Langford, John Sillitoe on behalf of the Wellcome Sanger Institute COVID-19 Surveillance Team |
| EPI_ISL_3222005, EPI_ISL_3222006 | Lighthouse Laboratory Plymouth | Wellcome Sanger Institute for the COVID-19 Genomics UK (COG-UK) Consortium | Lighthouse Laboratory Plymouth and Alex Alderton, Roberto Amato, Jeffrey Barrett, Sonia Goncalves, Ewan Harrison, David K. Jackson, Ian Johnston, Dominic Kwiatkowski, Cordelia Langford, John Sillitoe on behalf of the Wellcome Sanger Institute COVID-19 Surveillance Team |
| EPI_ISL_3222007 | Lighthouse Lab in Milton Keynes | Wellcome Sanger Institute for the COVID-19 Genomics UK (COG-UK) Consortium | The Lighthouse Lab in Milton Keynes and Alex Alderton, Roberto Amato, Jeffrey Barrett, Sonia Goncalves, Ewan Harrison, David K. Jackson, Ian Johnston, Dominic Kwiatkowski, Cordelia Langford, John Sillitoe on behalf of the Wellcome Sanger Institute COVID-19 Surveillance Team |
| EPI_ISL_3222008, EPI_ISL_3222009 | Lighthouse Laboratory Plymouth | Wellcome Sanger Institute for the COVID-19 Genomics UK (COG-UK) Consortium | Lighthouse Laboratory Plymouth and Alex Alderton, Roberto Amato, Jeffrey Barrett, Sonia Goncalves, Ewan Harrison, David K. Jackson, Ian Johnston, Dominic Kwiatkowski, Cordelia Langford, John Sillitoe on behalf of the Wellcome Sanger Institute COVID-19 Surveillance Team |
| EPI_ISL_3222010 | Lighthouse Lab in Milton Keynes | Wellcome Sanger Institute for the COVID-19 Genomics UK (COG-UK) Consortium | The Lighthouse Lab in Milton Keynes and Alex Alderton, Roberto Amato, Jeffrey Barrett, Sonia Goncalves, Ewan Harrison, David K. Jackson, Ian Johnston, Dominic Kwiatkowski, Cordelia Langford, John Sillitoe on behalf of the Wellcome Sanger Institute COVID-19 Surveillance Team |
| EPI_ISL_3222011 | Lighthouse Laboratory Plymouth | Wellcome Sanger Institute for the COVID-19 Genomics UK (COG-UK) Consortium | Lighthouse Laboratory Plymouth and Alex Alderton, Roberto Amato, Jeffrey Barrett, Sonia Goncalves, Ewan Harrison, David K. Jackson, Ian Johnston, Dominic Kwiatkowski, Cordelia Langford, John Sillitoe on behalf of the Wellcome Sanger Institute COVID-19 Surveillance Team |
| EPI_ISL_3222012, EPI_ISL_3222013 | Lighthouse Lab in Milton Keynes | Wellcome Sanger Institute for the COVID-19 Genomics UK (COG-UK) Consortium | The Lighthouse Lab in Milton Keynes and Alex Alderton, Roberto Amato, Jeffrey Barrett, Sonia Goncalves, Ewan Harrison, David K. Jackson, Ian Johnston, Dominic Kwiatkowski, Cordelia Langford, John Sillitoe on behalf of the Wellcome Sanger Institute COVID-19 Surveillance Team |
| EPI_ISL_3222014 | Lighthouse Laboratory Plymouth | Wellcome Sanger Institute for the COVID-19 Genomics UK (COG-UK) Consortium | Lighthouse Laboratory Plymouth and Alex Alderton, Roberto Amato, Jeffrey Barrett, Sonia Goncalves, Ewan Harrison, David K. Jackson, Ian Johnston, Dominic Kwiatkowski, Cordelia Langford, John Sillitoe on behalf of the Wellcome Sanger Institute COVID-19 Surveillance Team |
| EPI_ISL_3222015, EPI_ISL_3222016 | Lighthouse Lab in Milton Keynes | Wellcome Sanger Institute for the COVID-19 Genomics UK (COG-UK) Consortium | The Lighthouse Lab in Milton Keynes and Alex Alderton, Roberto Amato, Jeffrey Barrett, Sonia Goncalves, Ewan Harrison, David K. Jackson, Ian Johnston, Dominic Kwiatkowski, Cordelia Langford, John Sillitoe on behalf of the Wellcome Sanger Institute COVID-19 Surveillance Team |
| EPI_ISL_3222017 | Berkshire and Surrey Pathology Services Lighthouse Laboratory | Wellcome Sanger Institute for the COVID-19 Genomics UK (COG-UK) Consortium | Berkshire and Surrey Pathology Services Lighthouse Laboratory and Alex Alderton, Roberto Amato, Jeffrey Barrett, Sonia Goncalves, Ewan Harrison, David K. Jackson, Ian Johnston, Dominic Kwiatkowski, Cordelia Langford, John Sillitoe on behalf of the Wellcome Sanger Institute COVID-19 Surveillance Team |
| EPI_ISL_3222018, EPI_ISL_3222019, EPI_ISL_3222020 | Lighthouse Laboratory Plymouth | Wellcome Sanger Institute for the COVID-19 Genomics UK (COG-UK) Consortium | Lighthouse Laboratory Plymouth and Alex Alderton, Roberto Amato, Jeffrey Barrett, Sonia Goncalves, Ewan Harrison, David K. Jackson, Ian Johnston, Dominic Kwiatkowski, Cordelia Langford, John Sillitoe on behalf of the Wellcome Sanger Institute COVID-19 Surveillance Team |
| EPI_ISL_3222021 | Berkshire and Surrey Pathology Services Lighthouse Laboratory | Wellcome Sanger Institute for the COVID-19 Genomics UK (COG-UK) Consortium | Berkshire and Surrey Pathology Services Lighthouse Laboratory and Alex Alderton, Roberto Amato, Jeffrey Barrett, Sonia Goncalves, Ewan Harrison, David K. Jackson, Ian Johnston, Dominic Kwiatkowski, Cordelia Langford, John Sillitoe on behalf of the Wellcome Sanger Institute COVID-19 Surveillance Team |
| EPI_ISL_3222022, EPI_ISL_3222023, EPI_ISL_3222024, EPI_ISL_3222025, EPI_ISL_3222026 | Lighthouse Laboratory Plymouth | Wellcome Sanger Institute for the COVID-19 Genomics UK (COG-UK) Consortium | Lighthouse Laboratory Plymouth and Alex Alderton, Roberto Amato, Jeffrey Barrett, Sonia Goncalves, Ewan Harrison, David K. Jackson, Ian Johnston, Dominic Kwiatkowski, Cordelia Langford, John Sillitoe on behalf of the Wellcome Sanger Institute COVID-19 Surveillance Team |
| EPI_ISL_3222027 | Lighthouse Lab in Milton Keynes | Wellcome Sanger Institute for the COVID-19 Genomics UK (COG-UK) Consortium | The Lighthouse Lab in Milton Keynes and Alex Alderton, Roberto Amato, Jeffrey Barrett, Sonia Goncalves, Ewan Harrison, David K. Jackson, Ian Johnston, Dominic Kwiatkowski, Cordelia Langford, John Sillitoe on behalf of the Wellcome Sanger Institute COVID-19 Surveillance Team |
| EPI_ISL_3222028, EPI_ISL_3222029 | Lighthouse Laboratory Plymouth | Wellcome Sanger Institute for the COVID-19 Genomics UK (COG-UK) Consortium | Lighthouse Laboratory Plymouth and Alex Alderton, Roberto Amato, Jeffrey Barrett, Sonia Goncalves, Ewan Harrison, David K. Jackson, Ian Johnston, Dominic Kwiatkowski, Cordelia Langford, John Sillitoe on behalf of the Wellcome Sanger Institute COVID-19 Surveillance Team |
| EPI_ISL_3222030 | Lighthouse Lab in Milton Keynes | Wellcome Sanger Institute for the COVID-19 Genomics UK (COG-UK) Consortium | The Lighthouse Lab in Milton Keynes and Alex Alderton, Roberto Amato, Jeffrey Barrett, Sonia Goncalves, Ewan Harrison, David K. Jackson, Ian Johnston, Dominic Kwiatkowski, Cordelia Langford, John Sillitoe on behalf of the Wellcome Sanger Institute COVID-19 Surveillance Team |
| EPI_ISL_3222031, EPI_ISL_3222032 | Lighthouse Laboratory Plymouth | Wellcome Sanger Institute for the COVID-19 Genomics UK (COG-UK) Consortium | Lighthouse Laboratory Plymouth and Alex Alderton, Roberto Amato, Jeffrey Barrett, Sonia Goncalves, Ewan Harrison, David K. Jackson, Ian Johnston, Dominic Kwiatkowski, Cordelia Langford, John Sillitoe on behalf of the Wellcome Sanger Institute COVID-19 Surveillance Team |
| EPI_ISL_3222033 | Lighthouse Lab in Milton Keynes | Wellcome Sanger Institute for the COVID-19 Genomics UK (COG-UK) Consortium | The Lighthouse Lab in Milton Keynes and Alex Alderton, Roberto Amato, Jeffrey Barrett, Sonia Goncalves, Ewan Harrison, David K. Jackson, Ian Johnston, Dominic Kwiatkowski, Cordelia Langford, John Sillitoe on behalf of the Wellcome Sanger Institute COVID-19 Surveillance Team |
| EPI_ISL_3222034 | Lighthouse Laboratory Plymouth | Wellcome Sanger Institute for the COVID-19 Genomics UK (COG-UK) Consortium | Lighthouse Laboratory Plymouth and Alex Alderton, Roberto Amato, Jeffrey Barrett, Sonia Goncalves, Ewan Harrison, David K. Jackson, Ian Johnston, Dominic Kwiatkowski, Cordelia Langford, John Sillitoe on behalf of the Wellcome Sanger Institute COVID-19 Surveillance Team |
| EPI_ISL_3222035 | Lighthouse Lab in Milton Keynes | Wellcome Sanger Institute for the COVID-19 Genomics UK (COG-UK) Consortium | The Lighthouse Lab in Milton Keynes and Alex Alderton, Roberto Amato, Jeffrey Barrett, Sonia Goncalves, Ewan Harrison, David K. Jackson, Ian Johnston, Dominic Kwiatkowski, Cordelia Langford, John Sillitoe on behalf of the Wellcome Sanger Institute COVID-19 Surveillance Team |
| EPI_ISL_3222036 | Lighthouse Laboratory Plymouth | Wellcome Sanger Institute for the COVID-19 Genomics UK (COG-UK) Consortium | Lighthouse Laboratory Plymouth and Alex Alderton, Roberto Amato, Jeffrey Barrett, Sonia Goncalves, Ewan Harrison, David K. Jackson, Ian Johnston, Dominic Kwiatkowski, Cordelia Langford, John Sillitoe on behalf of the Wellcome Sanger Institute COVID-19 Surveillance Team |
| EPI_ISL_3222037 | Lighthouse Lab in Milton Keynes | Wellcome Sanger Institute for the COVID-19 Genomics UK (COG-UK) Consortium | The Lighthouse Lab in Milton Keynes and Alex Alderton, Roberto Amato, Jeffrey Barrett, Sonia Goncalves, Ewan Harrison, David K. Jackson, Ian Johnston, Dominic Kwiatkowski, Cordelia Langford, John Sillitoe on behalf of the Wellcome Sanger Institute COVID-19 Surveillance Team |
| EPI_ISL_3222038, EPI_ISL_3222039, EPI_ISL_3222040, EPI_ISL_3222041, EPI_ISL_3222042 | Lighthouse Laboratory Plymouth | Wellcome Sanger Institute for the COVID-19 Genomics UK (COG-UK) Consortium | Lighthouse Laboratory Plymouth and Alex Alderton, Roberto Amato, Jeffrey Barrett, Sonia Goncalves, Ewan Harrison, David K. Jackson, Ian Johnston, Dominic Kwiatkowski, Cordelia Langford, John Sillitoe on behalf of the Wellcome Sanger Institute COVID-19 Surveillance Team |

| | | | |
|---|---|---|---|
| EPI_ISL_3222043 | Berkshire and Surrey Pathology Services Lighthouse Laboratory | Wellcome Sanger Institute for the COVID-19 Genomics UK (COG-UK) Consortium | Berkshire and Surrey Pathology Services Lighthouse Laboratory and Alex Alderton, Roberto Amato, Jeffrey Barrett, Sonia Goncalves, Ewan Harrison, David K. Jackson, Ian Johnston, Dominic Kwiatkowski, Cordelia Langford, John Sillitoe on behalf of the Wellcome Sanger Institute COVID-19 Surveillance Team |
| EPI_ISL_3222044 | Lighthouse Lab in Milton Keynes | Wellcome Sanger Institute for the COVID-19 Genomics UK (COG-UK) Consortium | The Lighthouse Lab in Milton Keynes and Alex Alderton, Roberto Amato, Jeffrey Barrett, Sonia Goncalves, Ewan Harrison, David K. Jackson, Ian Johnston, Dominic Kwiatkowski, Cordelia Langford, John Sillitoe on behalf of the Wellcome Sanger Institute COVID-19 Surveillance Team |
| EPI_ISL_3222045 | Lighthouse Laboratory Plymouth | Wellcome Sanger Institute for the COVID-19 Genomics UK (COG-UK) Consortium | Lighthouse Laboratory Plymouth and Alex Alderton, Roberto Amato, Jeffrey Barrett, Sonia Goncalves, Ewan Harrison, David K. Jackson, Ian Johnston, Dominic Kwiatkowski, Cordelia Langford, John Sillitoe on behalf of the Wellcome Sanger Institute COVID-19 Surveillance Team |
| EPI_ISL_3222046 | Lighthouse Lab in Milton Keynes | Wellcome Sanger Institute for the COVID-19 Genomics UK (COG-UK) Consortium | The Lighthouse Lab in Milton Keynes and Alex Alderton, Roberto Amato, Jeffrey Barrett, Sonia Goncalves, Ewan Harrison, David K. Jackson, Ian Johnston, Dominic Kwiatkowski, Cordelia Langford, John Sillitoe on behalf of the Wellcome Sanger Institute COVID-19 Surveillance Team |
| EPI_ISL_3222047 | Berkshire and Surrey Pathology Services Lighthouse Laboratory | Wellcome Sanger Institute for the COVID-19 Genomics UK (COG-UK) Consortium | Berkshire and Surrey Pathology Services Lighthouse Laboratory and Alex Alderton, Roberto Amato, Jeffrey Barrett, Sonia Goncalves, Ewan Harrison, David K. Jackson, Ian Johnston, Dominic Kwiatkowski, Cordelia Langford, John Sillitoe on behalf of the Wellcome Sanger Institute COVID-19 Surveillance Team |
| EPI_ISL_3222048 | Lighthouse Laboratory Plymouth | Wellcome Sanger Institute for the COVID-19 Genomics UK (COG-UK) Consortium | Lighthouse Laboratory Plymouth and Alex Alderton, Roberto Amato, Jeffrey Barrett, Sonia Goncalves, Ewan Harrison, David K. Jackson, Ian Johnston, Dominic Kwiatkowski, Cordelia Langford, John Sillitoe on behalf of the Wellcome Sanger Institute COVID-19 Surveillance Team |
| EPI_ISL_3222049, EPI_ISL_3222050, EPI_ISL_3222051, EPI_ISL_3222052 | Lighthouse Lab in Milton Keynes | Wellcome Sanger Institute for the COVID-19 Genomics UK (COG-UK) Consortium | The Lighthouse Lab in Milton Keynes and Alex Alderton, Roberto Amato, Jeffrey Barrett, Sonia Goncalves, Ewan Harrison, David K. Jackson, Ian Johnston, Dominic Kwiatkowski, Cordelia Langford, John Sillitoe on behalf of the Wellcome Sanger Institute COVID-19 Surveillance Team |
| EPI_ISL_3222053, EPI_ISL_3222054 | Berkshire and Surrey Pathology Services Lighthouse Laboratory | Wellcome Sanger Institute for the COVID-19 Genomics UK (COG-UK) Consortium | Berkshire and Surrey Pathology Services Lighthouse Laboratory and Alex Alderton, Roberto Amato, Jeffrey Barrett, Sonia Goncalves, Ewan Harrison, David K. Jackson, Ian Johnston, Dominic Kwiatkowski, Cordelia Langford, John Sillitoe on behalf of the Wellcome Sanger Institute COVID-19 Surveillance Team |
| EPI_ISL_3222055 | Lighthouse Lab in Milton Keynes | Wellcome Sanger Institute for the COVID-19 Genomics UK (COG-UK) Consortium | The Lighthouse Lab in Milton Keynes and Alex Alderton, Roberto Amato, Jeffrey Barrett, Sonia Goncalves, Ewan Harrison, David K. Jackson, Ian Johnston, Dominic Kwiatkowski, Cordelia Langford, John Sillitoe on behalf of the Wellcome Sanger Institute COVID-19 Surveillance Team |
| EPI_ISL_3222056, EPI_ISL_3222057, EPI_ISL_3222058 | Lighthouse Laboratory Plymouth | Wellcome Sanger Institute for the COVID-19 Genomics UK (COG-UK) Consortium | Lighthouse Laboratory Plymouth and Alex Alderton, Roberto Amato, Jeffrey Barrett, Sonia Goncalves, Ewan Harrison, David K. Jackson, Ian Johnston, Dominic Kwiatkowski, Cordelia Langford, John Sillitoe on behalf of the Wellcome Sanger Institute COVID-19 Surveillance Team |
| EPI_ISL_3222059 | Lighthouse Lab in Milton Keynes | Wellcome Sanger Institute for the COVID-19 Genomics UK (COG-UK) Consortium | The Lighthouse Lab in Milton Keynes and Alex Alderton, Roberto Amato, Jeffrey Barrett, Sonia Goncalves, Ewan Harrison, David K. Jackson, Ian Johnston, Dominic Kwiatkowski, Cordelia Langford, John Sillitoe on behalf of the Wellcome Sanger Institute COVID-19 Surveillance Team |
| EPI_ISL_3222060 | Berkshire and Surrey Pathology Services Lighthouse Laboratory | Wellcome Sanger Institute for the COVID-19 Genomics UK (COG-UK) Consortium | Berkshire and Surrey Pathology Services Lighthouse Laboratory and Alex Alderton, Roberto Amato, Jeffrey Barrett, Sonia Goncalves, Ewan Harrison, David K. Jackson, Ian Johnston, Dominic Kwiatkowski, Cordelia Langford, John Sillitoe on behalf of the Wellcome Sanger Institute COVID-19 Surveillance Team |
| EPI_ISL_3222061, EPI_ISL_3222062, EPI_ISL_3222063, EPI_ISL_3222064 | Lighthouse Laboratory Plymouth | Wellcome Sanger Institute for the COVID-19 Genomics UK (COG-UK) Consortium | Lighthouse Laboratory Plymouth and Alex Alderton, Roberto Amato, Jeffrey Barrett, Sonia Goncalves, Ewan Harrison, David K. Jackson, Ian Johnston, Dominic Kwiatkowski, Cordelia Langford, John Sillitoe on behalf of the Wellcome Sanger Institute COVID-19 Surveillance Team |
| EPI_ISL_3222065 | Lighthouse Lab in Milton Keynes | Wellcome Sanger Institute for the COVID-19 Genomics UK (COG-UK) Consortium | The Lighthouse Lab in Milton Keynes and Alex Alderton, Roberto Amato, Jeffrey Barrett, Sonia Goncalves, Ewan Harrison, David K. Jackson, Ian Johnston, Dominic Kwiatkowski, Cordelia Langford, John Sillitoe on behalf of the Wellcome Sanger Institute COVID-19 Surveillance Team |
| EPI_ISL_3222066 | Lighthouse Laboratory Plymouth | Wellcome Sanger Institute for the COVID-19 Genomics UK (COG-UK) Consortium | Lighthouse Laboratory Plymouth and Alex Alderton, Roberto Amato, Jeffrey Barrett, Sonia Goncalves, Ewan Harrison, David K. Jackson, Ian Johnston, Dominic Kwiatkowski, Cordelia Langford, John Sillitoe on behalf of the Wellcome Sanger Institute COVID-19 Surveillance Team |
| EPI_ISL_3222067 | Lighthouse Lab in Milton Keynes | Wellcome Sanger Institute for the COVID-19 Genomics UK (COG-UK) Consortium | The Lighthouse Lab in Milton Keynes and Alex Alderton, Roberto Amato, Jeffrey Barrett, Sonia Goncalves, Ewan Harrison, David K. Jackson, Ian Johnston, Dominic Kwiatkowski, Cordelia Langford, John Sillitoe on behalf of the Wellcome Sanger Institute COVID-19 Surveillance Team |
| EPI_ISL_3222068, EPI_ISL_3222069 | Berkshire and Surrey Pathology Services Lighthouse Laboratory | Wellcome Sanger Institute for the COVID-19 Genomics UK (COG-UK) Consortium | Berkshire and Surrey Pathology Services Lighthouse Laboratory and Alex Alderton, Roberto Amato, Jeffrey Barrett, Sonia Goncalves, Ewan Harrison, David K. Jackson, Ian Johnston, Dominic Kwiatkowski, Cordelia Langford, John Sillitoe on behalf of the Wellcome Sanger Institute COVID-19 Surveillance Team |
| EPI_ISL_3222070, EPI_ISL_3222071 | Lighthouse Laboratory Plymouth | Wellcome Sanger Institute for the COVID-19 Genomics UK (COG-UK) Consortium | Lighthouse Laboratory Plymouth and Alex Alderton, Roberto Amato, Jeffrey Barrett, Sonia Goncalves, Ewan Harrison, David K. Jackson, Ian Johnston, Dominic Kwiatkowski, Cordelia Langford, John Sillitoe on behalf of the Wellcome Sanger Institute COVID-19 Surveillance Team |
| EPI_ISL_3222072 | Lighthouse Lab in Milton Keynes | Wellcome Sanger Institute for the COVID-19 Genomics UK (COG-UK) Consortium | The Lighthouse Lab in Milton Keynes and Alex Alderton, Roberto Amato, Jeffrey Barrett, Sonia Goncalves, Ewan Harrison, David K. Jackson, Ian Johnston, Dominic Kwiatkowski, Cordelia Langford, John Sillitoe on behalf of the Wellcome Sanger Institute COVID-19 Surveillance Team |
| EPI_ISL_3222073, EPI_ISL_3222074 | Lighthouse Laboratory Plymouth | Wellcome Sanger Institute for the COVID-19 Genomics UK (COG-UK) Consortium | Lighthouse Laboratory Plymouth and Alex Alderton, Roberto Amato, Jeffrey Barrett, Sonia Goncalves, Ewan Harrison, David K. Jackson, Ian Johnston, Dominic Kwiatkowski, Cordelia Langford, John Sillitoe on behalf of the Wellcome Sanger Institute COVID-19 Surveillance Team |
| EPI_ISL_3222075 | Lighthouse Lab in Milton Keynes | Wellcome Sanger Institute for the COVID-19 Genomics UK (COG-UK) Consortium | The Lighthouse Lab in Milton Keynes and Alex Alderton, Roberto Amato, Jeffrey Barrett, Sonia Goncalves, Ewan Harrison, David K. Jackson, Ian Johnston, Dominic Kwiatkowski, Cordelia Langford, John Sillitoe on behalf of the Wellcome Sanger Institute COVID-19 Surveillance Team |
| EPI_ISL_3222076, EPI_ISL_3222077 | Lighthouse Laboratory Plymouth | Wellcome Sanger Institute for the COVID-19 Genomics UK (COG-UK) Consortium | Lighthouse Laboratory Plymouth and Alex Alderton, Roberto Amato, Jeffrey Barrett, Sonia Goncalves, Ewan Harrison, David K. Jackson, Ian Johnston, Dominic Kwiatkowski, Cordelia Langford, John Sillitoe on behalf of the Wellcome Sanger Institute COVID-19 Surveillance Team |
| EPI_ISL_3222078 | Berkshire and Surrey Pathology Services Lighthouse Laboratory | Wellcome Sanger Institute for the COVID-19 Genomics UK (COG-UK) Consortium | Berkshire and Surrey Pathology Services Lighthouse Laboratory and Alex Alderton, Roberto Amato, Jeffrey Barrett, Sonia Goncalves, Ewan Harrison, David K. Jackson, Ian Johnston, Dominic Kwiatkowski, Cordelia Langford, John Sillitoe on behalf of the Wellcome Sanger Institute COVID-19 Surveillance Team |
| EPI_ISL_3222079 | Lighthouse Lab in Milton Keynes | Wellcome Sanger Institute for the COVID-19 Genomics UK (COG-UK) Consortium | The Lighthouse Lab in Milton Keynes and Alex Alderton, Roberto Amato, Jeffrey Barrett, Sonia Goncalves, Ewan Harrison, David K. Jackson, Ian Johnston, Dominic Kwiatkowski, Cordelia Langford, John Sillitoe on behalf of the Wellcome Sanger Institute COVID-19 Surveillance Team |
| EPI_ISL_3222080, EPI_ISL_3222081, EPI_ISL_3222082 | Lighthouse Laboratory Plymouth | Wellcome Sanger Institute for the COVID-19 Genomics UK (COG-UK) Consortium | Lighthouse Laboratory Plymouth and Alex Alderton, Roberto Amato, Jeffrey Barrett, Sonia Goncalves, Ewan Harrison, David K. Jackson, Ian Johnston, Dominic Kwiatkowski, Cordelia Langford, John Sillitoe on behalf of the Wellcome Sanger Institute COVID-19 Surveillance Team |
| EPI_ISL_3222083 | Berkshire and Surrey Pathology Services Lighthouse Laboratory | Wellcome Sanger Institute for the COVID-19 Genomics UK (COG-UK) Consortium | Berkshire and Surrey Pathology Services Lighthouse Laboratory and Alex Alderton, Roberto Amato, Jeffrey Barrett, Sonia Goncalves, Ewan Harrison, David K. Jackson, Ian Johnston, Dominic Kwiatkowski, Cordelia Langford, John Sillitoe on behalf of the Wellcome Sanger Institute COVID-19 Surveillance Team |
| EPI_ISL_3222084, EPI_ISL_3222085, EPI_ISL_3222086 | Lighthouse Laboratory Plymouth | Wellcome Sanger Institute for the COVID-19 Genomics UK (COG-UK) Consortium | Lighthouse Laboratory Plymouth and Alex Alderton, Roberto Amato, Jeffrey Barrett, Sonia Goncalves, Ewan Harrison, David K. Jackson, Ian Johnston, Dominic Kwiatkowski, Cordelia Langford, John Sillitoe on behalf of the Wellcome Sanger Institute COVID-19 Surveillance Team |
| EPI_ISL_3222087, EPI_ISL_3222088 | Lighthouse Lab in Milton Keynes | Wellcome Sanger Institute for the COVID-19 Genomics UK (COG-UK) Consortium | The Lighthouse Lab in Milton Keynes and Alex Alderton, Roberto Amato, Jeffrey Barrett, Sonia Goncalves, Ewan Harrison, David K. Jackson, Ian Johnston, Dominic Kwiatkowski, Cordelia Langford, John Sillitoe on behalf of the Wellcome Sanger Institute COVID-19 Surveillance Team |
| EPI_ISL_3222089 | Lighthouse Laboratory Plymouth | Wellcome Sanger Institute for the COVID-19 Genomics UK (COG-UK) Consortium | Lighthouse Laboratory Plymouth and Alex Alderton, Roberto Amato, Jeffrey Barrett, Sonia Goncalves, Ewan Harrison, David K. Jackson, Ian Johnston, Dominic Kwiatkowski, Cordelia Langford, John Sillitoe on behalf of the Wellcome Sanger Institute COVID-19 Surveillance Team |
| EPI_ISL_3222090 | Berkshire and Surrey Pathology Services Lighthouse Laboratory | Wellcome Sanger Institute for the COVID-19 Genomics UK (COG-UK) Consortium | Berkshire and Surrey Pathology Services Lighthouse Laboratory and Alex Alderton, Roberto Amato, Jeffrey Barrett, Sonia Goncalves, Ewan Harrison, David K. Jackson, Ian Johnston, Dominic Kwiatkowski, Cordelia Langford, John Sillitoe on behalf of the Wellcome Sanger Institute COVID-19 Surveillance Team |
| EPI_ISL_3222091, EPI_ISL_3222092, EPI_ISL_3222093 | Lighthouse Lab in Milton Keynes | Wellcome Sanger Institute for the COVID-19 Genomics UK (COG-UK) Consortium | The Lighthouse Lab in Milton Keynes and Alex Alderton, Roberto Amato, Jeffrey Barrett, Sonia Goncalves, Ewan Harrison, David K. Jackson, Ian Johnston, Dominic Kwiatkowski, Cordelia Langford, John Sillitoe on behalf of the Wellcome Sanger Institute COVID-19 Surveillance Team |
| EPI_ISL_3222094 | Lighthouse Laboratory Plymouth | Wellcome Sanger Institute for the COVID-19 Genomics UK (COG-UK) Consortium | Lighthouse Laboratory Plymouth and Alex Alderton, Roberto Amato, Jeffrey Barrett, Sonia Goncalves, Ewan Harrison, David K. Jackson, Ian Johnston, Dominic Kwiatkowski, Cordelia Langford, John Sillitoe on behalf of the Wellcome Sanger Institute COVID-19 Surveillance Team |
| EPI_ISL_3222095 | Lighthouse Lab in Milton Keynes | Wellcome Sanger Institute for the COVID-19 Genomics UK (COG-UK) Consortium | The Lighthouse Lab in Milton Keynes and Alex Alderton, Roberto Amato, Jeffrey Barrett, Sonia Goncalves, Ewan Harrison, David K. Jackson, Ian Johnston, Dominic Kwiatkowski, Cordelia Langford, John Sillitoe on behalf of the Wellcome Sanger Institute COVID-19 Surveillance Team |
| EPI_ISL_3222096 | Berkshire and Surrey Pathology Services Lighthouse Laboratory | Wellcome Sanger Institute for the COVID-19 Genomics UK (COG-UK) Consortium | Berkshire and Surrey Pathology Services Lighthouse Laboratory and Alex Alderton, Roberto Amato, Jeffrey Barrett, Sonia Goncalves, Ewan Harrison, David K. Jackson, Ian Johnston, Dominic Kwiatkowski, Cordelia Langford, John Sillitoe on behalf of the Wellcome Sanger Institute COVID-19 Surveillance Team |
| EPI_ISL_3222097 | Lighthouse Lab in Milton Keynes | Wellcome Sanger Institute for the COVID-19 Genomics UK (COG-UK) Consortium | The Lighthouse Lab in Milton Keynes and Alex Alderton, Roberto Amato, Jeffrey Barrett, Sonia Goncalves, Ewan Harrison, David K. Jackson, Ian Johnston, Dominic Kwiatkowski, Cordelia Langford, John Sillitoe on behalf of the Wellcome Sanger Institute COVID-19 Surveillance Team |
| EPI_ISL_3222098 | Berkshire and Surrey Pathology Services Lighthouse Laboratory | Wellcome Sanger Institute for the COVID-19 Genomics UK (COG-UK) Consortium | Berkshire and Surrey Pathology Services Lighthouse Laboratory and Alex Alderton, Roberto Amato, Jeffrey Barrett, Sonia Goncalves, Ewan Harrison, David K. Jackson, Ian Johnston, Dominic Kwiatkowski, Cordelia Langford, John Sillitoe on behalf of the Wellcome Sanger Institute COVID-19 Surveillance Team |
| EPI_ISL_3222099, EPI_ISL_3222100 | Lighthouse Laboratory Plymouth | Wellcome Sanger Institute for the COVID-19 Genomics UK (COG-UK) Consortium | Lighthouse Laboratory Plymouth and Alex Alderton, Roberto Amato, Jeffrey Barrett, Sonia Goncalves, Ewan Harrison, David K. Jackson, Ian Johnston, Dominic Kwiatkowski, Cordelia Langford, John Sillitoe on behalf of the Wellcome Sanger Institute COVID-19 Surveillance Team |
| EPI_ISL_3222101 | Berkshire and Surrey Pathology Services Lighthouse Laboratory | Wellcome Sanger Institute for the COVID-19 Genomics UK (COG-UK) Consortium | Berkshire and Surrey Pathology Services Lighthouse Laboratory and Alex Alderton, Roberto Amato, Jeffrey Barrett, Sonia Goncalves, Ewan Harrison, David K. Jackson, Ian Johnston, Dominic Kwiatkowski, Cordelia Langford, John Sillitoe on behalf of the Wellcome Sanger Institute COVID-19 Surveillance Team |
| EPI_ISL_3222102 | Lighthouse Laboratory Plymouth | Wellcome Sanger Institute for the COVID-19 Genomics UK (COG-UK) Consortium | Lighthouse Laboratory Plymouth and Alex Alderton, Roberto Amato, Jeffrey Barrett, Sonia Goncalves, Ewan Harrison, David K. Jackson, Ian Johnston, Dominic Kwiatkowski, Cordelia Langford, John Sillitoe on behalf of the Wellcome Sanger Institute COVID-19 Surveillance Team |
| EPI_ISL_3222103 | Berkshire and Surrey Pathology Services Lighthouse Laboratory | Wellcome Sanger Institute for the COVID-19 Genomics UK (COG-UK) Consortium | Berkshire and Surrey Pathology Services Lighthouse Laboratory and Alex Alderton, Roberto Amato, Jeffrey Barrett, Sonia Goncalves, Ewan Harrison, David K. Jackson, Ian Johnston, Dominic Kwiatkowski, Cordelia Langford, John Sillitoe on behalf of the Wellcome Sanger Institute COVID-19 Surveillance Team |
| EPI_ISL_3222104, EPI_ISL_3222105 | Lighthouse Lab in Milton Keynes | Wellcome Sanger Institute for the COVID-19 Genomics UK (COG-UK) Consortium | The Lighthouse Lab in Milton Keynes and Alex Alderton, Roberto Amato, Jeffrey Barrett, Sonia Goncalves, Ewan Harrison, David K. Jackson, Ian Johnston, Dominic Kwiatkowski, Cordelia Langford, John Sillitoe on behalf of the Wellcome Sanger Institute COVID-19 Surveillance Team |
| EPI_ISL_3222106, EPI_ISL_3222107, | Lighthouse Laboratory Plymouth | Wellcome Sanger Institute for the COVID-19 Genomics UK (COG-UK) Consortium | Lighthouse Laboratory Plymouth and Alex Alderton, Roberto Amato, Jeffrey Barrett, Sonia Goncalves, Ewan Harrison, David K. Jackson, Ian Johnston, Dominic Kwiatkowski, Cordelia Langford, John Sillitoe on behalf of the Wellcome Sanger Institute COVID-19 Surveillance Team |

| | | | |
|---|---|---|---|
| EPI_ISL_3222108, EPI_ISL_3222109 | | | |
| EPI_ISL_3222110 | Lighthouse Lab in Milton Keynes | Wellcome Sanger Institute for the COVID-19 Genomics UK (COG-UK) Consortium | The Lighthouse Lab in Milton Keynes and Alex Alderton, Roberto Amato, Jeffrey Barrett, Sonia Goncalves, Ewan Harrison, David K. Jackson, Ian Johnston, Dominic Kwiatkowski, Cordelia Langford, John Sillitoe on behalf of the Wellcome Sanger Institute COVID-19 Surveillance Team |
| EPI_ISL_3222111, EPI_ISL_3222112 | Lighthouse Laboratory Plymouth | Wellcome Sanger Institute for the COVID-19 Genomics UK (COG-UK) Consortium | Lighthouse Laboratory Plymouth and Alex Alderton, Roberto Amato, Jeffrey Barrett, Sonia Goncalves, Ewan Harrison, David K. Jackson, Ian Johnston, Dominic Kwiatkowski, Cordelia Langford, John Sillitoe on behalf of the Wellcome Sanger Institute COVID-19 Surveillance Team |
| EPI_ISL_3222113, EPI_ISL_3222114, EPI_ISL_3222115 | Berkshire and Surrey Pathology Services Lighthouse Laboratory | Wellcome Sanger Institute for the COVID-19 Genomics UK (COG-UK) Consortium | Berkshire and Surrey Pathology Services Lighthouse Laboratory and Alex Alderton, Roberto Amato, Jeffrey Barrett, Sonia Goncalves, Ewan Harrison, David K. Jackson, Ian Johnston, Dominic Kwiatkowski, Cordelia Langford, John Sillitoe on behalf of the Wellcome Sanger Institute COVID-19 Surveillance Team |
| EPI_ISL_3222116 | Lighthouse Lab in Milton Keynes | Wellcome Sanger Institute for the COVID-19 Genomics UK (COG-UK) Consortium | The Lighthouse Lab in Milton Keynes and Alex Alderton, Roberto Amato, Jeffrey Barrett, Sonia Goncalves, Ewan Harrison, David K. Jackson, Ian Johnston, Dominic Kwiatkowski, Cordelia Langford, John Sillitoe on behalf of the Wellcome Sanger Institute COVID-19 Surveillance Team |
| EPI_ISL_3222117, EPI_ISL_3222118 | Lighthouse Laboratory Plymouth | Wellcome Sanger Institute for the COVID-19 Genomics UK (COG-UK) Consortium | Lighthouse Laboratory Plymouth and Alex Alderton, Roberto Amato, Jeffrey Barrett, Sonia Goncalves, Ewan Harrison, David K. Jackson, Ian Johnston, Dominic Kwiatkowski, Cordelia Langford, John Sillitoe on behalf of the Wellcome Sanger Institute COVID-19 Surveillance Team |
| EPI_ISL_3222119, EPI_ISL_3222120 | Lighthouse Lab in Milton Keynes | Wellcome Sanger Institute for the COVID-19 Genomics UK (COG-UK) Consortium | The Lighthouse Lab in Milton Keynes and Alex Alderton, Roberto Amato, Jeffrey Barrett, Sonia Goncalves, Ewan Harrison, David K. Jackson, Ian Johnston, Dominic Kwiatkowski, Cordelia Langford, John Sillitoe on behalf of the Wellcome Sanger Institute COVID-19 Surveillance Team |
| EPI_ISL_3222121, EPI_ISL_3222122, EPI_ISL_3222123, EPI_ISL_3222124, EPI_ISL_3222125 | Lighthouse Laboratory Plymouth | Wellcome Sanger Institute for the COVID-19 Genomics UK (COG-UK) Consortium | Lighthouse Laboratory Plymouth and Alex Alderton, Roberto Amato, Jeffrey Barrett, Sonia Goncalves, Ewan Harrison, David K. Jackson, Ian Johnston, Dominic Kwiatkowski, Cordelia Langford, John Sillitoe on behalf of the Wellcome Sanger Institute COVID-19 Surveillance Team |
| EPI_ISL_3222126 | Lighthouse Lab in Milton Keynes | Wellcome Sanger Institute for the COVID-19 Genomics UK (COG-UK) Consortium | The Lighthouse Lab in Milton Keynes and Alex Alderton, Roberto Amato, Jeffrey Barrett, Sonia Goncalves, Ewan Harrison, David K. Jackson, Ian Johnston, Dominic Kwiatkowski, Cordelia Langford, John Sillitoe on behalf of the Wellcome Sanger Institute COVID-19 Surveillance Team |
| EPI_ISL_3222127 | Lighthouse Laboratory Plymouth | Wellcome Sanger Institute for the COVID-19 Genomics UK (COG-UK) Consortium | Lighthouse Laboratory Plymouth and Alex Alderton, Roberto Amato, Jeffrey Barrett, Sonia Goncalves, Ewan Harrison, David K. Jackson, Ian Johnston, Dominic Kwiatkowski, Cordelia Langford, John Sillitoe on behalf of the Wellcome Sanger Institute COVID-19 Surveillance Team |
| EPI_ISL_3222128 | Lighthouse Lab in Milton Keynes | Wellcome Sanger Institute for the COVID-19 Genomics UK (COG-UK) Consortium | The Lighthouse Lab in Milton Keynes and Alex Alderton, Roberto Amato, Jeffrey Barrett, Sonia Goncalves, Ewan Harrison, David K. Jackson, Ian Johnston, Dominic Kwiatkowski, Cordelia Langford, John Sillitoe on behalf of the Wellcome Sanger Institute COVID-19 Surveillance Team |
| EPI_ISL_3222129 | Berkshire and Surrey Pathology Services Lighthouse Laboratory | Wellcome Sanger Institute for the COVID-19 Genomics UK (COG-UK) Consortium | Berkshire and Surrey Pathology Services Lighthouse Laboratory and Alex Alderton, Roberto Amato, Jeffrey Barrett, Sonia Goncalves, Ewan Harrison, David K. Jackson, Ian Johnston, Dominic Kwiatkowski, Cordelia Langford, John Sillitoe on behalf of the Wellcome Sanger Institute COVID-19 Surveillance Team |
| EPI_ISL_3222130, EPI_ISL_3222131 | Lighthouse Laboratory Plymouth | Wellcome Sanger Institute for the COVID-19 Genomics UK (COG-UK) Consortium | Lighthouse Laboratory Plymouth and Alex Alderton, Roberto Amato, Jeffrey Barrett, Sonia Goncalves, Ewan Harrison, David K. Jackson, Ian Johnston, Dominic Kwiatkowski, Cordelia Langford, John Sillitoe on behalf of the Wellcome Sanger Institute COVID-19 Surveillance Team |
| EPI_ISL_3222132 | Lighthouse Lab in Milton Keynes | Wellcome Sanger Institute for the COVID-19 Genomics UK (COG-UK) Consortium | The Lighthouse Lab in Milton Keynes and Alex Alderton, Roberto Amato, Jeffrey Barrett, Sonia Goncalves, Ewan Harrison, David K. Jackson, Ian Johnston, Dominic Kwiatkowski, Cordelia Langford, John Sillitoe on behalf of the Wellcome Sanger Institute COVID-19 Surveillance Team |
| EPI_ISL_3222133 | Lighthouse Laboratory Plymouth | Wellcome Sanger Institute for the COVID-19 Genomics UK (COG-UK) Consortium | Lighthouse Laboratory Plymouth and Alex Alderton, Roberto Amato, Jeffrey Barrett, Sonia Goncalves, Ewan Harrison, David K. Jackson, Ian Johnston, Dominic Kwiatkowski, Cordelia Langford, John Sillitoe on behalf of the Wellcome Sanger Institute COVID-19 Surveillance Team |
| EPI_ISL_3222134 | Lighthouse Lab in Milton Keynes | Wellcome Sanger Institute for the COVID-19 Genomics UK (COG-UK) Consortium | The Lighthouse Lab in Milton Keynes and Alex Alderton, Roberto Amato, Jeffrey Barrett, Sonia Goncalves, Ewan Harrison, David K. Jackson, Ian Johnston, Dominic Kwiatkowski, Cordelia Langford, John Sillitoe on behalf of the Wellcome Sanger Institute COVID-19 Surveillance Team |
| EPI_ISL_3222135, EPI_ISL_3222136 | Lighthouse Laboratory Plymouth | Wellcome Sanger Institute for the COVID-19 Genomics UK (COG-UK) Consortium | Lighthouse Laboratory Plymouth and Alex Alderton, Roberto Amato, Jeffrey Barrett, Sonia Goncalves, Ewan Harrison, David K. Jackson, Ian Johnston, Dominic Kwiatkowski, Cordelia Langford, John Sillitoe on behalf of the Wellcome Sanger Institute COVID-19 Surveillance Team |
| EPI_ISL_3222137, EPI_ISL_3222138 | Lighthouse Lab in Milton Keynes | Wellcome Sanger Institute for the COVID-19 Genomics UK (COG-UK) Consortium | The Lighthouse Lab in Milton Keynes and Alex Alderton, Roberto Amato, Jeffrey Barrett, Sonia Goncalves, Ewan Harrison, David K. Jackson, Ian Johnston, Dominic Kwiatkowski, Cordelia Langford, John Sillitoe on behalf of the Wellcome Sanger Institute COVID-19 Surveillance Team |
| EPI_ISL_3222139 | Lighthouse Laboratory Plymouth | Wellcome Sanger Institute for the COVID-19 Genomics UK (COG-UK) Consortium | Lighthouse Laboratory Plymouth and Alex Alderton, Roberto Amato, Jeffrey Barrett, Sonia Goncalves, Ewan Harrison, David K. Jackson, Ian Johnston, Dominic Kwiatkowski, Cordelia Langford, John Sillitoe on behalf of the Wellcome Sanger Institute COVID-19 Surveillance Team |
| EPI_ISL_3222140 | Lighthouse Lab in Milton Keynes | Wellcome Sanger Institute for the COVID-19 Genomics UK (COG-UK) Consortium | The Lighthouse Lab in Milton Keynes and Alex Alderton, Roberto Amato, Jeffrey Barrett, Sonia Goncalves, Ewan Harrison, David K. Jackson, Ian Johnston, Dominic Kwiatkowski, Cordelia Langford, John Sillitoe on behalf of the Wellcome Sanger Institute COVID-19 Surveillance Team |
| EPI_ISL_3222141, EPI_ISL_3222142, EPI_ISL_3222143 | Lighthouse Laboratory Plymouth | Wellcome Sanger Institute for the COVID-19 Genomics UK (COG-UK) Consortium | Lighthouse Laboratory Plymouth and Alex Alderton, Roberto Amato, Jeffrey Barrett, Sonia Goncalves, Ewan Harrison, David K. Jackson, Ian Johnston, Dominic Kwiatkowski, Cordelia Langford, John Sillitoe on behalf of the Wellcome Sanger Institute COVID-19 Surveillance Team |
| EPI_ISL_3222144, EPI_ISL_3222145 | Lighthouse Lab in Milton Keynes | Wellcome Sanger Institute for the COVID-19 Genomics UK (COG-UK) Consortium | The Lighthouse Lab in Milton Keynes and Alex Alderton, Roberto Amato, Jeffrey Barrett, Sonia Goncalves, Ewan Harrison, David K. Jackson, Ian Johnston, Dominic Kwiatkowski, Cordelia Langford, John Sillitoe on behalf of the Wellcome Sanger Institute COVID-19 Surveillance Team |
| EPI_ISL_3222146 | Lighthouse Laboratory Plymouth | Wellcome Sanger Institute for the COVID-19 Genomics UK (COG-UK) Consortium | Lighthouse Laboratory Plymouth and Alex Alderton, Roberto Amato, Jeffrey Barrett, Sonia Goncalves, Ewan Harrison, David K. Jackson, Ian Johnston, Dominic Kwiatkowski, Cordelia Langford, John Sillitoe on behalf of the Wellcome Sanger Institute COVID-19 Surveillance Team |
| EPI_ISL_3222147 | Lighthouse Lab in Milton Keynes | Wellcome Sanger Institute for the COVID-19 Genomics UK (COG-UK) Consortium | The Lighthouse Lab in Milton Keynes and Alex Alderton, Roberto Amato, Jeffrey Barrett, Sonia Goncalves, Ewan Harrison, David K. Jackson, Ian Johnston, Dominic Kwiatkowski, Cordelia Langford, John Sillitoe on behalf of the Wellcome Sanger Institute COVID-19 Surveillance Team |
| EPI_ISL_3222148 | Berkshire and Surrey Pathology Services Lighthouse Laboratory | Wellcome Sanger Institute for the COVID-19 Genomics UK (COG-UK) Consortium | Berkshire and Surrey Pathology Services Lighthouse Laboratory and Alex Alderton, Roberto Amato, Jeffrey Barrett, Sonia Goncalves, Ewan Harrison, David K. Jackson, Ian Johnston, Dominic Kwiatkowski, Cordelia Langford, John Sillitoe on behalf of the Wellcome Sanger Institute COVID-19 Surveillance Team |
| EPI_ISL_3222149, EPI_ISL_3222150 | Lighthouse Lab in Milton Keynes | Wellcome Sanger Institute for the COVID-19 Genomics UK (COG-UK) Consortium | The Lighthouse Lab in Milton Keynes and Alex Alderton, Roberto Amato, Jeffrey Barrett, Sonia Goncalves, Ewan Harrison, David K. Jackson, Ian Johnston, Dominic Kwiatkowski, Cordelia Langford, John Sillitoe on behalf of the Wellcome Sanger Institute COVID-19 Surveillance Team |
| EPI_ISL_3222151, EPI_ISL_3222152, EPI_ISL_3222153, EPI_ISL_3222154, EPI_ISL_3222155 | Lighthouse Laboratory Plymouth | Wellcome Sanger Institute for the COVID-19 Genomics UK (COG-UK) Consortium | Lighthouse Laboratory Plymouth and Alex Alderton, Roberto Amato, Jeffrey Barrett, Sonia Goncalves, Ewan Harrison, David K. Jackson, Ian Johnston, Dominic Kwiatkowski, Cordelia Langford, John Sillitoe on behalf of the Wellcome Sanger Institute COVID-19 Surveillance Team |
| EPI_ISL_3222156 | Berkshire and Surrey Pathology Services Lighthouse Laboratory | Wellcome Sanger Institute for the COVID-19 Genomics UK (COG-UK) Consortium | Berkshire and Surrey Pathology Services Lighthouse Laboratory and Alex Alderton, Roberto Amato, Jeffrey Barrett, Sonia Goncalves, Ewan Harrison, David K. Jackson, Ian Johnston, Dominic Kwiatkowski, Cordelia Langford, John Sillitoe on behalf of the Wellcome Sanger Institute COVID-19 Surveillance Team |
| EPI_ISL_3222157 | Lighthouse Lab in Milton Keynes | Wellcome Sanger Institute for the COVID-19 Genomics UK (COG-UK) Consortium | The Lighthouse Lab in Milton Keynes and Alex Alderton, Roberto Amato, Jeffrey Barrett, Sonia Goncalves, Ewan Harrison, David K. Jackson, Ian Johnston, Dominic Kwiatkowski, Cordelia Langford, John Sillitoe on behalf of the Wellcome Sanger Institute COVID-19 Surveillance Team |
| EPI_ISL_3222158, EPI_ISL_3222159 | Lighthouse Laboratory Plymouth | Wellcome Sanger Institute for the COVID-19 Genomics UK (COG-UK) Consortium | Lighthouse Laboratory Plymouth and Alex Alderton, Roberto Amato, Jeffrey Barrett, Sonia Goncalves, Ewan Harrison, David K. Jackson, Ian Johnston, Dominic Kwiatkowski, Cordelia Langford, John Sillitoe on behalf of the Wellcome Sanger Institute COVID-19 Surveillance Team |
| EPI_ISL_3222160, EPI_ISL_3222161, EPI_ISL_3222162 | Berkshire and Surrey Pathology Services Lighthouse Laboratory | Wellcome Sanger Institute for the COVID-19 Genomics UK (COG-UK) Consortium | Berkshire and Surrey Pathology Services Lighthouse Laboratory and Alex Alderton, Roberto Amato, Jeffrey Barrett, Sonia Goncalves, Ewan Harrison, David K. Jackson, Ian Johnston, Dominic Kwiatkowski, Cordelia Langford, John Sillitoe on behalf of the Wellcome Sanger Institute COVID-19 Surveillance Team |
| EPI_ISL_3222163 | Lighthouse Lab in Milton Keynes | Wellcome Sanger Institute for the COVID-19 Genomics UK (COG-UK) Consortium | The Lighthouse Lab in Milton Keynes and Alex Alderton, Roberto Amato, Jeffrey Barrett, Sonia Goncalves, Ewan Harrison, David K. Jackson, Ian Johnston, Dominic Kwiatkowski, Cordelia Langford, John Sillitoe on behalf of the Wellcome Sanger Institute COVID-19 Surveillance Team |
| EPI_ISL_3222164 | Lighthouse Laboratory Plymouth | Wellcome Sanger Institute for the COVID-19 Genomics UK (COG-UK) Consortium | Lighthouse Laboratory Plymouth and Alex Alderton, Roberto Amato, Jeffrey Barrett, Sonia Goncalves, Ewan Harrison, David K. Jackson, Ian Johnston, Dominic Kwiatkowski, Cordelia Langford, John Sillitoe on behalf of the Wellcome Sanger Institute COVID-19 Surveillance Team |
| EPI_ISL_3222165 | Berkshire and Surrey Pathology Services Lighthouse Laboratory | Wellcome Sanger Institute for the COVID-19 Genomics UK (COG-UK) Consortium | Berkshire and Surrey Pathology Services Lighthouse Laboratory and Alex Alderton, Roberto Amato, Jeffrey Barrett, Sonia Goncalves, Ewan Harrison, David K. Jackson, Ian Johnston, Dominic Kwiatkowski, Cordelia Langford, John Sillitoe on behalf of the Wellcome Sanger Institute COVID-19 Surveillance Team |
| EPI_ISL_3222166 | Lighthouse Lab in Milton Keynes | Wellcome Sanger Institute for the COVID-19 Genomics UK (COG-UK) Consortium | The Lighthouse Lab in Milton Keynes and Alex Alderton, Roberto Amato, Jeffrey Barrett, Sonia Goncalves, Ewan Harrison, David K. Jackson, Ian Johnston, Dominic Kwiatkowski, Cordelia Langford, John Sillitoe on behalf of the Wellcome Sanger Institute COVID-19 Surveillance Team |
| EPI_ISL_3222167 | Lighthouse Laboratory Plymouth | Wellcome Sanger Institute for the COVID-19 Genomics UK (COG-UK) Consortium | Lighthouse Laboratory Plymouth and Alex Alderton, Roberto Amato, Jeffrey Barrett, Sonia Goncalves, Ewan Harrison, David K. Jackson, Ian Johnston, Dominic Kwiatkowski, Cordelia Langford, John Sillitoe on behalf of the Wellcome Sanger Institute COVID-19 Surveillance Team |
| EPI_ISL_3222168 | Lighthouse Lab in Milton Keynes | Wellcome Sanger Institute for the COVID-19 Genomics UK (COG-UK) Consortium | The Lighthouse Lab in Milton Keynes and Alex Alderton, Roberto Amato, Jeffrey Barrett, Sonia Goncalves, Ewan Harrison, David K. Jackson, Ian Johnston, Dominic Kwiatkowski, Cordelia Langford, John Sillitoe on behalf of the Wellcome Sanger Institute COVID-19 Surveillance Team |
| EPI_ISL_3222169 | Lighthouse Laboratory Plymouth | Wellcome Sanger Institute for the COVID-19 Genomics UK (COG-UK) Consortium | Lighthouse Laboratory Plymouth and Alex Alderton, Roberto Amato, Jeffrey Barrett, Sonia Goncalves, Ewan Harrison, David K. Jackson, Ian Johnston, Dominic Kwiatkowski, Cordelia Langford, John Sillitoe on behalf of the Wellcome Sanger Institute COVID-19 Surveillance Team |
| EPI_ISL_3222170 | Berkshire and Surrey Pathology Services Lighthouse Laboratory | Wellcome Sanger Institute for the COVID-19 Genomics UK (COG-UK) Consortium | Berkshire and Surrey Pathology Services Lighthouse Laboratory and Alex Alderton, Roberto Amato, Jeffrey Barrett, Sonia Goncalves, Ewan Harrison, David K. Jackson, Ian Johnston, Dominic Kwiatkowski, Cordelia Langford, John Sillitoe on behalf of the Wellcome Sanger Institute COVID-19 Surveillance Team |
| EPI_ISL_3222171, EPI_ISL_3222172, EPI_ISL_3222173 | Lighthouse Laboratory Plymouth | Wellcome Sanger Institute for the COVID-19 Genomics UK (COG-UK) Consortium | Lighthouse Laboratory Plymouth and Alex Alderton, Roberto Amato, Jeffrey Barrett, Sonia Goncalves, Ewan Harrison, David K. Jackson, Ian Johnston, Dominic Kwiatkowski, Cordelia Langford, John Sillitoe on behalf of the Wellcome Sanger Institute COVID-19 Surveillance Team |
| EPI_ISL_3222174, EPI_ISL_3222175, | Lighthouse Lab in Milton Keynes | Wellcome Sanger Institute for the COVID-19 Genomics UK (COG-UK) Consortium | The Lighthouse Lab in Milton Keynes and Alex Alderton, Roberto Amato, Jeffrey Barrett, Sonia Goncalves, Ewan Harrison, David K. Jackson, Ian Johnston, Dominic Kwiatkowski, Cordelia Langford, John Sillitoe on behalf of the Wellcome Sanger Institute COVID-19 Surveillance Team |

| | | | |
|---|---|---|---|
| EPI_ISL_3222176 | | | |
| EPI_ISL_3222177, EPI_ISL_3222178, EPI_ISL_3222179 | Lighthouse Laboratory Plymouth | Wellcome Sanger Institute for the COVID-19 Genomics UK (COG-UK) Consortium | Lighthouse Laboratory Plymouth and Alex Alderton, Roberto Amato, Jeffrey Barrett, Sonia Goncalves, Ewan Harrison, David K. Jackson, Ian Johnston, Dominic Kwiatkowski, Cordelia Langford, John Sillitoe on behalf of the Wellcome Sanger Institute COVID-19 Surveillance Team |
| EPI_ISL_3222180 | Berkshire and Surrey Pathology Services Lighthouse Laboratory | Wellcome Sanger Institute for the COVID-19 Genomics UK (COG-UK) Consortium | Berkshire and Surrey Pathology Services Lighthouse Laboratory and Alex Alderton, Roberto Amato, Jeffrey Barrett, Sonia Goncalves, Ewan Harrison, David K. Jackson, Ian Johnston, Dominic Kwiatkowski, Cordelia Langford, John Sillitoe on behalf of the Wellcome Sanger Institute COVID-19 Surveillance Team |
| EPI_ISL_3222181, EPI_ISL_3222182 | Lighthouse Laboratory Plymouth | Wellcome Sanger Institute for the COVID-19 Genomics UK (COG-UK) Consortium | Lighthouse Laboratory Plymouth and Alex Alderton, Roberto Amato, Jeffrey Barrett, Sonia Goncalves, Ewan Harrison, David K. Jackson, Ian Johnston, Dominic Kwiatkowski, Cordelia Langford, John Sillitoe on behalf of the Wellcome Sanger Institute COVID-19 Surveillance Team |
| EPI_ISL_3222183 | Berkshire and Surrey Pathology Services Lighthouse Laboratory | Wellcome Sanger Institute for the COVID-19 Genomics UK (COG-UK) Consortium | Berkshire and Surrey Pathology Services Lighthouse Laboratory and Alex Alderton, Roberto Amato, Jeffrey Barrett, Sonia Goncalves, Ewan Harrison, David K. Jackson, Ian Johnston, Dominic Kwiatkowski, Cordelia Langford, John Sillitoe on behalf of the Wellcome Sanger Institute COVID-19 Surveillance Team |
| EPI_ISL_3222184 | Lighthouse Laboratory Plymouth | Wellcome Sanger Institute for the COVID-19 Genomics UK (COG-UK) Consortium | Lighthouse Laboratory Plymouth and Alex Alderton, Roberto Amato, Jeffrey Barrett, Sonia Goncalves, Ewan Harrison, David K. Jackson, Ian Johnston, Dominic Kwiatkowski, Cordelia Langford, John Sillitoe on behalf of the Wellcome Sanger Institute COVID-19 Surveillance Team |
| EPI_ISL_3222185 | Berkshire and Surrey Pathology Services Lighthouse Laboratory | Wellcome Sanger Institute for the COVID-19 Genomics UK (COG-UK) Consortium | Berkshire and Surrey Pathology Services Lighthouse Laboratory and Alex Alderton, Roberto Amato, Jeffrey Barrett, Sonia Goncalves, Ewan Harrison, David K. Jackson, Ian Johnston, Dominic Kwiatkowski, Cordelia Langford, John Sillitoe on behalf of the Wellcome Sanger Institute COVID-19 Surveillance Team |
| EPI_ISL_3222186 | Lighthouse Lab in Milton Keynes | Wellcome Sanger Institute for the COVID-19 Genomics UK (COG-UK) Consortium | The Lighthouse Lab in Milton Keynes and Alex Alderton, Roberto Amato, Jeffrey Barrett, Sonia Goncalves, Ewan Harrison, David K. Jackson, Ian Johnston, Dominic Kwiatkowski, Cordelia Langford, John Sillitoe on behalf of the Wellcome Sanger Institute COVID-19 Surveillance Team |
| EPI_ISL_3222187, EPI_ISL_3222188 | Lighthouse Laboratory Plymouth | Wellcome Sanger Institute for the COVID-19 Genomics UK (COG-UK) Consortium | Lighthouse Laboratory Plymouth and Alex Alderton, Roberto Amato, Jeffrey Barrett, Sonia Goncalves, Ewan Harrison, David K. Jackson, Ian Johnston, Dominic Kwiatkowski, Cordelia Langford, John Sillitoe on behalf of the Wellcome Sanger Institute COVID-19 Surveillance Team |
| EPI_ISL_3222189 | Lighthouse Lab in Milton Keynes | Wellcome Sanger Institute for the COVID-19 Genomics UK (COG-UK) Consortium | The Lighthouse Lab in Milton Keynes and Alex Alderton, Roberto Amato, Jeffrey Barrett, Sonia Goncalves, Ewan Harrison, David K. Jackson, Ian Johnston, Dominic Kwiatkowski, Cordelia Langford, John Sillitoe on behalf of the Wellcome Sanger Institute COVID-19 Surveillance Team |
| EPI_ISL_3222190 | Lighthouse Laboratory Plymouth | Wellcome Sanger Institute for the COVID-19 Genomics UK (COG-UK) Consortium | Lighthouse Laboratory Plymouth and Alex Alderton, Roberto Amato, Jeffrey Barrett, Sonia Goncalves, Ewan Harrison, David K. Jackson, Ian Johnston, Dominic Kwiatkowski, Cordelia Langford, John Sillitoe on behalf of the Wellcome Sanger Institute COVID-19 Surveillance Team |
| EPI_ISL_3222191 | Berkshire and Surrey Pathology Services Lighthouse Laboratory | Wellcome Sanger Institute for the COVID-19 Genomics UK (COG-UK) Consortium | Berkshire and Surrey Pathology Services Lighthouse Laboratory and Alex Alderton, Roberto Amato, Jeffrey Barrett, Sonia Goncalves, Ewan Harrison, David K. Jackson, Ian Johnston, Dominic Kwiatkowski, Cordelia Langford, John Sillitoe on behalf of the Wellcome Sanger Institute COVID-19 Surveillance Team |
| EPI_ISL_3222192 | Lighthouse Laboratory Plymouth | Wellcome Sanger Institute for the COVID-19 Genomics UK (COG-UK) Consortium | Lighthouse Laboratory Plymouth and Alex Alderton, Roberto Amato, Jeffrey Barrett, Sonia Goncalves, Ewan Harrison, David K. Jackson, Ian Johnston, Dominic Kwiatkowski, Cordelia Langford, John Sillitoe on behalf of the Wellcome Sanger Institute COVID-19 Surveillance Team |
| EPI_ISL_3222193, EPI_ISL_3222194 | Berkshire and Surrey Pathology Services Lighthouse Laboratory | Wellcome Sanger Institute for the COVID-19 Genomics UK (COG-UK) Consortium | Berkshire and Surrey Pathology Services Lighthouse Laboratory and Alex Alderton, Roberto Amato, Jeffrey Barrett, Sonia Goncalves, Ewan Harrison, David K. Jackson, Ian Johnston, Dominic Kwiatkowski, Cordelia Langford, John Sillitoe on behalf of the Wellcome Sanger Institute COVID-19 Surveillance Team |
| EPI_ISL_3222195 | Lighthouse Lab in Milton Keynes | Wellcome Sanger Institute for the COVID-19 Genomics UK (COG-UK) Consortium | The Lighthouse Lab in Milton Keynes and Alex Alderton, Roberto Amato, Jeffrey Barrett, Sonia Goncalves, Ewan Harrison, David K. Jackson, Ian Johnston, Dominic Kwiatkowski, Cordelia Langford, John Sillitoe on behalf of the Wellcome Sanger Institute COVID-19 Surveillance Team |
| EPI_ISL_3222196 | Berkshire and Surrey Pathology Services Lighthouse Laboratory | Wellcome Sanger Institute for the COVID-19 Genomics UK (COG-UK) Consortium | Berkshire and Surrey Pathology Services Lighthouse Laboratory and Alex Alderton, Roberto Amato, Jeffrey Barrett, Sonia Goncalves, Ewan Harrison, David K. Jackson, Ian Johnston, Dominic Kwiatkowski, Cordelia Langford, John Sillitoe on behalf of the Wellcome Sanger Institute COVID-19 Surveillance Team |
| EPI_ISL_3222197 | Lighthouse Lab in Milton Keynes | Wellcome Sanger Institute for the COVID-19 Genomics UK (COG-UK) Consortium | The Lighthouse Lab in Milton Keynes and Alex Alderton, Roberto Amato, Jeffrey Barrett, Sonia Goncalves, Ewan Harrison, David K. Jackson, Ian Johnston, Dominic Kwiatkowski, Cordelia Langford, John Sillitoe on behalf of the Wellcome Sanger Institute COVID-19 Surveillance Team |
| EPI_ISL_3222198 | Lighthouse Laboratory Plymouth | Wellcome Sanger Institute for the COVID-19 Genomics UK (COG-UK) Consortium | Lighthouse Laboratory Plymouth and Alex Alderton, Roberto Amato, Jeffrey Barrett, Sonia Goncalves, Ewan Harrison, David K. Jackson, Ian Johnston, Dominic Kwiatkowski, Cordelia Langford, John Sillitoe on behalf of the Wellcome Sanger Institute COVID-19 Surveillance Team |
| EPI_ISL_3222199 | Lighthouse Lab in Milton Keynes | Wellcome Sanger Institute for the COVID-19 Genomics UK (COG-UK) Consortium | The Lighthouse Lab in Milton Keynes and Alex Alderton, Roberto Amato, Jeffrey Barrett, Sonia Goncalves, Ewan Harrison, David K. Jackson, Ian Johnston, Dominic Kwiatkowski, Cordelia Langford, John Sillitoe on behalf of the Wellcome Sanger Institute COVID-19 Surveillance Team |
| EPI_ISL_3222200 | Lighthouse Laboratory Plymouth | Wellcome Sanger Institute for the COVID-19 Genomics UK (COG-UK) Consortium | Lighthouse Laboratory Plymouth and Alex Alderton, Roberto Amato, Jeffrey Barrett, Sonia Goncalves, Ewan Harrison, David K. Jackson, Ian Johnston, Dominic Kwiatkowski, Cordelia Langford, John Sillitoe on behalf of the Wellcome Sanger Institute COVID-19 Surveillance Team |
| EPI_ISL_3222201, EPI_ISL_3222202 | Berkshire and Surrey Pathology Services Lighthouse Laboratory | Wellcome Sanger Institute for the COVID-19 Genomics UK (COG-UK) Consortium | Berkshire and Surrey Pathology Services Lighthouse Laboratory and Alex Alderton, Roberto Amato, Jeffrey Barrett, Sonia Goncalves, Ewan Harrison, David K. Jackson, Ian Johnston, Dominic Kwiatkowski, Cordelia Langford, John Sillitoe on behalf of the Wellcome Sanger Institute COVID-19 Surveillance Team |
| EPI_ISL_3222203, EPI_ISL_3222204, EPI_ISL_3222205, EPI_ISL_3222206, EPI_ISL_3222207, EPI_ISL_3222208 | Lighthouse Laboratory Plymouth | Wellcome Sanger Institute for the COVID-19 Genomics UK (COG-UK) Consortium | Lighthouse Laboratory Plymouth and Alex Alderton, Roberto Amato, Jeffrey Barrett, Sonia Goncalves, Ewan Harrison, David K. Jackson, Ian Johnston, Dominic Kwiatkowski, Cordelia Langford, John Sillitoe on behalf of the Wellcome Sanger Institute COVID-19 Surveillance Team |
| EPI_ISL_3222209 | Berkshire and Surrey Pathology Services Lighthouse Laboratory | Wellcome Sanger Institute for the COVID-19 Genomics UK (COG-UK) Consortium | Berkshire and Surrey Pathology Services Lighthouse Laboratory and Alex Alderton, Roberto Amato, Jeffrey Barrett, Sonia Goncalves, Ewan Harrison, David K. Jackson, Ian Johnston, Dominic Kwiatkowski, Cordelia Langford, John Sillitoe on behalf of the Wellcome Sanger Institute COVID-19 Surveillance Team |
| EPI_ISL_3222210, EPI_ISL_3222211, EPI_ISL_3222212, EPI_ISL_3222213, EPI_ISL_3222214, EPI_ISL_3222215, EPI_ISL_3222216 | Lighthouse Laboratory Plymouth | Wellcome Sanger Institute for the COVID-19 Genomics UK (COG-UK) Consortium | Lighthouse Laboratory Plymouth and Alex Alderton, Roberto Amato, Jeffrey Barrett, Sonia Goncalves, Ewan Harrison, David K. Jackson, Ian Johnston, Dominic Kwiatkowski, Cordelia Langford, John Sillitoe on behalf of the Wellcome Sanger Institute COVID-19 Surveillance Team |
| EPI_ISL_3222217 | Lighthouse Lab in Milton Keynes | Wellcome Sanger Institute for the COVID-19 Genomics UK (COG-UK) Consortium | The Lighthouse Lab in Milton Keynes and Alex Alderton, Roberto Amato, Jeffrey Barrett, Sonia Goncalves, Ewan Harrison, David K. Jackson, Ian Johnston, Dominic Kwiatkowski, Cordelia Langford, John Sillitoe on behalf of the Wellcome Sanger Institute COVID-19 Surveillance Team |
| EPI_ISL_3222218 | Lighthouse Laboratory Plymouth | Wellcome Sanger Institute for the COVID-19 Genomics UK (COG-UK) Consortium | Lighthouse Laboratory Plymouth and Alex Alderton, Roberto Amato, Jeffrey Barrett, Sonia Goncalves, Ewan Harrison, David K. Jackson, Ian Johnston, Dominic Kwiatkowski, Cordelia Langford, John Sillitoe on behalf of the Wellcome Sanger Institute COVID-19 Surveillance Team |
| EPI_ISL_3222219 | Berkshire and Surrey Pathology Services Lighthouse Laboratory | Wellcome Sanger Institute for the COVID-19 Genomics UK (COG-UK) Consortium | Berkshire and Surrey Pathology Services Lighthouse Laboratory and Alex Alderton, Roberto Amato, Jeffrey Barrett, Sonia Goncalves, Ewan Harrison, David K. Jackson, Ian Johnston, Dominic Kwiatkowski, Cordelia Langford, John Sillitoe on behalf of the Wellcome Sanger Institute COVID-19 Surveillance Team |
| EPI_ISL_3222220, EPI_ISL_3222221, EPI_ISL_3222222 | Lighthouse Laboratory Plymouth | Wellcome Sanger Institute for the COVID-19 Genomics UK (COG-UK) Consortium | Lighthouse Laboratory Plymouth and Alex Alderton, Roberto Amato, Jeffrey Barrett, Sonia Goncalves, Ewan Harrison, David K. Jackson, Ian Johnston, Dominic Kwiatkowski, Cordelia Langford, John Sillitoe on behalf of the Wellcome Sanger Institute COVID-19 Surveillance Team |
| EPI_ISL_3222223, EPI_ISL_3222224, EPI_ISL_3222225, EPI_ISL_3222226 | Lighthouse Lab in Milton Keynes | Wellcome Sanger Institute for the COVID-19 Genomics UK (COG-UK) Consortium | The Lighthouse Lab in Milton Keynes and Alex Alderton, Roberto Amato, Jeffrey Barrett, Sonia Goncalves, Ewan Harrison, David K. Jackson, Ian Johnston, Dominic Kwiatkowski, Cordelia Langford, John Sillitoe on behalf of the Wellcome Sanger Institute COVID-19 Surveillance Team |
| EPI_ISL_3222227 | Lighthouse Laboratory Plymouth | Wellcome Sanger Institute for the COVID-19 Genomics UK (COG-UK) Consortium | Lighthouse Laboratory Plymouth and Alex Alderton, Roberto Amato, Jeffrey Barrett, Sonia Goncalves, Ewan Harrison, David K. Jackson, Ian Johnston, Dominic Kwiatkowski, Cordelia Langford, John Sillitoe on behalf of the Wellcome Sanger Institute COVID-19 Surveillance Team |
| EPI_ISL_3222228, EPI_ISL_3222229 | Berkshire and Surrey Pathology Services Lighthouse Laboratory | Wellcome Sanger Institute for the COVID-19 Genomics UK (COG-UK) Consortium | Berkshire and Surrey Pathology Services Lighthouse Laboratory and Alex Alderton, Roberto Amato, Jeffrey Barrett, Sonia Goncalves, Ewan Harrison, David K. Jackson, Ian Johnston, Dominic Kwiatkowski, Cordelia Langford, John Sillitoe on behalf of the Wellcome Sanger Institute COVID-19 Surveillance Team |
| EPI_ISL_3222230, EPI_ISL_3222231 | Lighthouse Laboratory Plymouth | Wellcome Sanger Institute for the COVID-19 Genomics UK (COG-UK) Consortium | Lighthouse Laboratory Plymouth and Alex Alderton, Roberto Amato, Jeffrey Barrett, Sonia Goncalves, Ewan Harrison, David K. Jackson, Ian Johnston, Dominic Kwiatkowski, Cordelia Langford, John Sillitoe on behalf of the Wellcome Sanger Institute COVID-19 Surveillance Team |
| EPI_ISL_3222232 | Lighthouse Lab in Milton Keynes | Wellcome Sanger Institute for the COVID-19 Genomics UK (COG-UK) Consortium | The Lighthouse Lab in Milton Keynes and Alex Alderton, Roberto Amato, Jeffrey Barrett, Sonia Goncalves, Ewan Harrison, David K. Jackson, Ian Johnston, Dominic Kwiatkowski, Cordelia Langford, John Sillitoe on behalf of the Wellcome Sanger Institute COVID-19 Surveillance Team |
| EPI_ISL_3222233 | Lighthouse Laboratory Plymouth | Wellcome Sanger Institute for the COVID-19 Genomics UK (COG-UK) Consortium | Lighthouse Laboratory Plymouth and Alex Alderton, Roberto Amato, Jeffrey Barrett, Sonia Goncalves, Ewan Harrison, David K. Jackson, Ian Johnston, Dominic Kwiatkowski, Cordelia Langford, John Sillitoe on behalf of the Wellcome Sanger Institute COVID-19 Surveillance Team |
| EPI_ISL_3222234 | Berkshire and Surrey Pathology Services Lighthouse Laboratory | Wellcome Sanger Institute for the COVID-19 Genomics UK (COG-UK) Consortium | Berkshire and Surrey Pathology Services Lighthouse Laboratory and Alex Alderton, Roberto Amato, Jeffrey Barrett, Sonia Goncalves, Ewan Harrison, David K. Jackson, Ian Johnston, Dominic Kwiatkowski, Cordelia Langford, John Sillitoe on behalf of the Wellcome Sanger Institute COVID-19 Surveillance Team |
| EPI_ISL_3222235 | Lighthouse Lab in Milton Keynes | Wellcome Sanger Institute for the COVID-19 Genomics UK (COG-UK) Consortium | The Lighthouse Lab in Milton Keynes and Alex Alderton, Roberto Amato, Jeffrey Barrett, Sonia Goncalves, Ewan Harrison, David K. Jackson, Ian Johnston, Dominic Kwiatkowski, Cordelia Langford, John Sillitoe on behalf of the Wellcome Sanger Institute COVID-19 Surveillance Team |
| EPI_ISL_3222236 | Berkshire and Surrey Pathology Services Lighthouse Laboratory | Wellcome Sanger Institute for the COVID-19 Genomics UK (COG-UK) Consortium | Berkshire and Surrey Pathology Services Lighthouse Laboratory and Alex Alderton, Roberto Amato, Jeffrey Barrett, Sonia Goncalves, Ewan Harrison, David K. Jackson, Ian Johnston, Dominic Kwiatkowski, Cordelia Langford, John Sillitoe on behalf of the Wellcome Sanger Institute COVID-19 Surveillance Team |
| EPI_ISL_3222237 | Lighthouse Laboratory Plymouth | Wellcome Sanger Institute for the COVID-19 Genomics UK (COG-UK) Consortium | Lighthouse Laboratory Plymouth and Alex Alderton, Roberto Amato, Jeffrey Barrett, Sonia Goncalves, Ewan Harrison, David K. Jackson, Ian Johnston, Dominic Kwiatkowski, Cordelia Langford, John Sillitoe on behalf of the Wellcome Sanger Institute COVID-19 Surveillance Team |
| EPI_ISL_3222238 | Lighthouse Lab in Milton Keynes | Wellcome Sanger Institute for the COVID-19 Genomics UK (COG-UK) Consortium | The Lighthouse Lab in Milton Keynes and Alex Alderton, Roberto Amato, Jeffrey Barrett, Sonia Goncalves, Ewan Harrison, David K. Jackson, Ian Johnston, Dominic Kwiatkowski, Cordelia Langford, John Sillitoe on behalf of the Wellcome Sanger Institute COVID-19 Surveillance Team |
| EPI_ISL_3222239, EPI_ISL_3222240 | Berkshire and Surrey Pathology Services Lighthouse Laboratory | Wellcome Sanger Institute for the COVID-19 Genomics UK (COG-UK) Consortium | Berkshire and Surrey Pathology Services Lighthouse Laboratory and Alex Alderton, Roberto Amato, Jeffrey Barrett, Sonia Goncalves, Ewan Harrison, David K. Jackson, Ian Johnston, Dominic Kwiatkowski, Cordelia Langford, John Sillitoe on behalf of the Wellcome Sanger Institute COVID-19 Surveillance Team |

| | | | |
|---|---|---|---|
| EPI_ISL_3222241, EPI_ISL_3222242 | Lighthouse Laboratory Plymouth | Wellcome Sanger Institute for the COVID-19 Genomics UK (COG-UK) Consortium | Lighthouse Laboratory Plymouth and Alex Alderton, Roberto Amato, Jeffrey Barrett, Sonia Goncalves, Ewan Harrison, David K. Jackson, Ian Johnston, Dominic Kwiatkowski, Cordelia Langford, John Sillitoe on behalf of the Wellcome Sanger Institute COVID-19 Surveillance Team |
| EPI_ISL_3222243, EPI_ISL_3222244 | Lighthouse Lab in Milton Keynes | Wellcome Sanger Institute for the COVID-19 Genomics UK (COG-UK) Consortium | The Lighthouse Lab in Milton Keynes and Alex Alderton, Roberto Amato, Jeffrey Barrett, Sonia Goncalves, Ewan Harrison, David K. Jackson, Ian Johnston, Dominic Kwiatkowski, Cordelia Langford, John Sillitoe on behalf of the Wellcome Sanger Institute COVID-19 Surveillance Team |
| EPI_ISL_3222245 | Berkshire and Surrey Pathology Services Lighthouse Laboratory | Wellcome Sanger Institute for the COVID-19 Genomics UK (COG-UK) Consortium | Berkshire and Surrey Pathology Services Lighthouse Laboratory and Alex Alderton, Roberto Amato, Jeffrey Barrett, Sonia Goncalves, Ewan Harrison, David K. Jackson, Ian Johnston, Dominic Kwiatkowski, Cordelia Langford, John Sillitoe on behalf of the Wellcome Sanger Institute COVID-19 Surveillance Team |
| EPI_ISL_3222246 | Lighthouse Laboratory Plymouth | Wellcome Sanger Institute for the COVID-19 Genomics UK (COG-UK) Consortium | Lighthouse Laboratory Plymouth and Alex Alderton, Roberto Amato, Jeffrey Barrett, Sonia Goncalves, Ewan Harrison, David K. Jackson, Ian Johnston, Dominic Kwiatkowski, Cordelia Langford, John Sillitoe on behalf of the Wellcome Sanger Institute COVID-19 Surveillance Team |
| EPI_ISL_3222247 | Berkshire and Surrey Pathology Services Lighthouse Laboratory | Wellcome Sanger Institute for the COVID-19 Genomics UK (COG-UK) Consortium | Berkshire and Surrey Pathology Services Lighthouse Laboratory and Alex Alderton, Roberto Amato, Jeffrey Barrett, Sonia Goncalves, Ewan Harrison, David K. Jackson, Ian Johnston, Dominic Kwiatkowski, Cordelia Langford, John Sillitoe on behalf of the Wellcome Sanger Institute COVID-19 Surveillance Team |
| EPI_ISL_3222248, EPI_ISL_3222249, EPI_ISL_3222250 | Lighthouse Laboratory Plymouth | Wellcome Sanger Institute for the COVID-19 Genomics UK (COG-UK) Consortium | Lighthouse Laboratory Plymouth and Alex Alderton, Roberto Amato, Jeffrey Barrett, Sonia Goncalves, Ewan Harrison, David K. Jackson, Ian Johnston, Dominic Kwiatkowski, Cordelia Langford, John Sillitoe on behalf of the Wellcome Sanger Institute COVID-19 Surveillance Team |
| EPI_ISL_3222251 | Berkshire and Surrey Pathology Services Lighthouse Laboratory | Wellcome Sanger Institute for the COVID-19 Genomics UK (COG-UK) Consortium | Berkshire and Surrey Pathology Services Lighthouse Laboratory and Alex Alderton, Roberto Amato, Jeffrey Barrett, Sonia Goncalves, Ewan Harrison, David K. Jackson, Ian Johnston, Dominic Kwiatkowski, Cordelia Langford, John Sillitoe on behalf of the Wellcome Sanger Institute COVID-19 Surveillance Team |
| EPI_ISL_3222252 | Lighthouse Laboratory Plymouth | Wellcome Sanger Institute for the COVID-19 Genomics UK (COG-UK) Consortium | Lighthouse Laboratory Plymouth and Alex Alderton, Roberto Amato, Jeffrey Barrett, Sonia Goncalves, Ewan Harrison, David K. Jackson, Ian Johnston, Dominic Kwiatkowski, Cordelia Langford, John Sillitoe on behalf of the Wellcome Sanger Institute COVID-19 Surveillance Team |
| EPI_ISL_3222253 | Lighthouse Lab in Milton Keynes | Wellcome Sanger Institute for the COVID-19 Genomics UK (COG-UK) Consortium | The Lighthouse Lab in Milton Keynes and Alex Alderton, Roberto Amato, Jeffrey Barrett, Sonia Goncalves, Ewan Harrison, David K. Jackson, Ian Johnston, Dominic Kwiatkowski, Cordelia Langford, John Sillitoe on behalf of the Wellcome Sanger Institute COVID-19 Surveillance Team |
| EPI_ISL_3222254 | Lighthouse Laboratory Plymouth | Wellcome Sanger Institute for the COVID-19 Genomics UK (COG-UK) Consortium | Lighthouse Laboratory Plymouth and Alex Alderton, Roberto Amato, Jeffrey Barrett, Sonia Goncalves, Ewan Harrison, David K. Jackson, Ian Johnston, Dominic Kwiatkowski, Cordelia Langford, John Sillitoe on behalf of the Wellcome Sanger Institute COVID-19 Surveillance Team |
| EPI_ISL_3222255 | Berkshire and Surrey Pathology Services Lighthouse Laboratory | Wellcome Sanger Institute for the COVID-19 Genomics UK (COG-UK) Consortium | Berkshire and Surrey Pathology Services Lighthouse Laboratory and Alex Alderton, Roberto Amato, Jeffrey Barrett, Sonia Goncalves, Ewan Harrison, David K. Jackson, Ian Johnston, Dominic Kwiatkowski, Cordelia Langford, John Sillitoe on behalf of the Wellcome Sanger Institute COVID-19 Surveillance Team |
| EPI_ISL_3222256 | Lighthouse Laboratory Plymouth | Wellcome Sanger Institute for the COVID-19 Genomics UK (COG-UK) Consortium | Lighthouse Laboratory Plymouth and Alex Alderton, Roberto Amato, Jeffrey Barrett, Sonia Goncalves, Ewan Harrison, David K. Jackson, Ian Johnston, Dominic Kwiatkowski, Cordelia Langford, John Sillitoe on behalf of the Wellcome Sanger Institute COVID-19 Surveillance Team |
| EPI_ISL_3222257 | Lighthouse Lab in Milton Keynes | Wellcome Sanger Institute for the COVID-19 Genomics UK (COG-UK) Consortium | The Lighthouse Lab in Milton Keynes and Alex Alderton, Roberto Amato, Jeffrey Barrett, Sonia Goncalves, Ewan Harrison, David K. Jackson, Ian Johnston, Dominic Kwiatkowski, Cordelia Langford, John Sillitoe on behalf of the Wellcome Sanger Institute COVID-19 Surveillance Team |
| EPI_ISL_3222258, EPI_ISL_3222259 | Lighthouse Laboratory Plymouth | Wellcome Sanger Institute for the COVID-19 Genomics UK (COG-UK) Consortium | Lighthouse Laboratory Plymouth and Alex Alderton, Roberto Amato, Jeffrey Barrett, Sonia Goncalves, Ewan Harrison, David K. Jackson, Ian Johnston, Dominic Kwiatkowski, Cordelia Langford, John Sillitoe on behalf of the Wellcome Sanger Institute COVID-19 Surveillance Team |
| EPI_ISL_3222260 | Berkshire and Surrey Pathology Services Lighthouse Laboratory | Wellcome Sanger Institute for the COVID-19 Genomics UK (COG-UK) Consortium | Berkshire and Surrey Pathology Services Lighthouse Laboratory and Alex Alderton, Roberto Amato, Jeffrey Barrett, Sonia Goncalves, Ewan Harrison, David K. Jackson, Ian Johnston, Dominic Kwiatkowski, Cordelia Langford, John Sillitoe on behalf of the Wellcome Sanger Institute COVID-19 Surveillance Team |
| EPI_ISL_3222261 | Lighthouse Laboratory Plymouth | Wellcome Sanger Institute for the COVID-19 Genomics UK (COG-UK) Consortium | Lighthouse Laboratory Plymouth and Alex Alderton, Roberto Amato, Jeffrey Barrett, Sonia Goncalves, Ewan Harrison, David K. Jackson, Ian Johnston, Dominic Kwiatkowski, Cordelia Langford, John Sillitoe on behalf of the Wellcome Sanger Institute COVID-19 Surveillance Team |
| EPI_ISL_3222262 | Berkshire and Surrey Pathology Services Lighthouse Laboratory | Wellcome Sanger Institute for the COVID-19 Genomics UK (COG-UK) Consortium | Berkshire and Surrey Pathology Services Lighthouse Laboratory and Alex Alderton, Roberto Amato, Jeffrey Barrett, Sonia Goncalves, Ewan Harrison, David K. Jackson, Ian Johnston, Dominic Kwiatkowski, Cordelia Langford, John Sillitoe on behalf of the Wellcome Sanger Institute COVID-19 Surveillance Team |
| EPI_ISL_3222263, EPI_ISL_3222264, EPI_ISL_3222265, EPI_ISL_3222266 | Lighthouse Laboratory Plymouth | Wellcome Sanger Institute for the COVID-19 Genomics UK (COG-UK) Consortium | Lighthouse Laboratory Plymouth and Alex Alderton, Roberto Amato, Jeffrey Barrett, Sonia Goncalves, Ewan Harrison, David K. Jackson, Ian Johnston, Dominic Kwiatkowski, Cordelia Langford, John Sillitoe on behalf of the Wellcome Sanger Institute COVID-19 Surveillance Team |
| EPI_ISL_3222267 | Berkshire and Surrey Pathology Services Lighthouse Laboratory | Wellcome Sanger Institute for the COVID-19 Genomics UK (COG-UK) Consortium | Berkshire and Surrey Pathology Services Lighthouse Laboratory and Alex Alderton, Roberto Amato, Jeffrey Barrett, Sonia Goncalves, Ewan Harrison, David K. Jackson, Ian Johnston, Dominic Kwiatkowski, Cordelia Langford, John Sillitoe on behalf of the Wellcome Sanger Institute COVID-19 Surveillance Team |
| EPI_ISL_3222268 | Lighthouse Laboratory Plymouth | Wellcome Sanger Institute for the COVID-19 Genomics UK (COG-UK) Consortium | Lighthouse Laboratory Plymouth and Alex Alderton, Roberto Amato, Jeffrey Barrett, Sonia Goncalves, Ewan Harrison, David K. Jackson, Ian Johnston, Dominic Kwiatkowski, Cordelia Langford, John Sillitoe on behalf of the Wellcome Sanger Institute COVID-19 Surveillance Team |
| EPI_ISL_3222269, EPI_ISL_3222270, EPI_ISL_3222271 | Lighthouse Lab in Milton Keynes | Wellcome Sanger Institute for the COVID-19 Genomics UK (COG-UK) Consortium | The Lighthouse Lab in Milton Keynes and Alex Alderton, Roberto Amato, Jeffrey Barrett, Sonia Goncalves, Ewan Harrison, David K. Jackson, Ian Johnston, Dominic Kwiatkowski, Cordelia Langford, John Sillitoe on behalf of the Wellcome Sanger Institute COVID-19 Surveillance Team |
| EPI_ISL_3222272 | Berkshire and Surrey Pathology Services Lighthouse Laboratory | Wellcome Sanger Institute for the COVID-19 Genomics UK (COG-UK) Consortium | Berkshire and Surrey Pathology Services Lighthouse Laboratory and Alex Alderton, Roberto Amato, Jeffrey Barrett, Sonia Goncalves, Ewan Harrison, David K. Jackson, Ian Johnston, Dominic Kwiatkowski, Cordelia Langford, John Sillitoe on behalf of the Wellcome Sanger Institute COVID-19 Surveillance Team |
| EPI_ISL_3222273 | Lighthouse Laboratory Plymouth | Wellcome Sanger Institute for the COVID-19 Genomics UK (COG-UK) Consortium | Lighthouse Laboratory Plymouth and Alex Alderton, Roberto Amato, Jeffrey Barrett, Sonia Goncalves, Ewan Harrison, David K. Jackson, Ian Johnston, Dominic Kwiatkowski, Cordelia Langford, John Sillitoe on behalf of the Wellcome Sanger Institute COVID-19 Surveillance Team |
| EPI_ISL_3222274 | Lighthouse Lab in Milton Keynes | Wellcome Sanger Institute for the COVID-19 Genomics UK (COG-UK) Consortium | The Lighthouse Lab in Milton Keynes and Alex Alderton, Roberto Amato, Jeffrey Barrett, Sonia Goncalves, Ewan Harrison, David K. Jackson, Ian Johnston, Dominic Kwiatkowski, Cordelia Langford, John Sillitoe on behalf of the Wellcome Sanger Institute COVID-19 Surveillance Team |
| EPI_ISL_3222275, EPI_ISL_3222276, EPI_ISL_3222277, EPI_ISL_3222278 | Lighthouse Laboratory Plymouth | Wellcome Sanger Institute for the COVID-19 Genomics UK (COG-UK) Consortium | Lighthouse Laboratory Plymouth and Alex Alderton, Roberto Amato, Jeffrey Barrett, Sonia Goncalves, Ewan Harrison, David K. Jackson, Ian Johnston, Dominic Kwiatkowski, Cordelia Langford, John Sillitoe on behalf of the Wellcome Sanger Institute COVID-19 Surveillance Team |
| EPI_ISL_3222279, EPI_ISL_3222280, EPI_ISL_3222281, EPI_ISL_3222282 | Berkshire and Surrey Pathology Services Lighthouse Laboratory | Wellcome Sanger Institute for the COVID-19 Genomics UK (COG-UK) Consortium | Berkshire and Surrey Pathology Services Lighthouse Laboratory and Alex Alderton, Roberto Amato, Jeffrey Barrett, Sonia Goncalves, Ewan Harrison, David K. Jackson, Ian Johnston, Dominic Kwiatkowski, Cordelia Langford, John Sillitoe on behalf of the Wellcome Sanger Institute COVID-19 Surveillance Team |
| EPI_ISL_3222283, EPI_ISL_3222284 | Lighthouse Laboratory Plymouth | Wellcome Sanger Institute for the COVID-19 Genomics UK (COG-UK) Consortium | Lighthouse Laboratory Plymouth and Alex Alderton, Roberto Amato, Jeffrey Barrett, Sonia Goncalves, Ewan Harrison, David K. Jackson, Ian Johnston, Dominic Kwiatkowski, Cordelia Langford, John Sillitoe on behalf of the Wellcome Sanger Institute COVID-19 Surveillance Team |
| EPI_ISL_3222285 | Lighthouse Lab in Milton Keynes | Wellcome Sanger Institute for the COVID-19 Genomics UK (COG-UK) Consortium | The Lighthouse Lab in Milton Keynes and Alex Alderton, Roberto Amato, Jeffrey Barrett, Sonia Goncalves, Ewan Harrison, David K. Jackson, Ian Johnston, Dominic Kwiatkowski, Cordelia Langford, John Sillitoe on behalf of the Wellcome Sanger Institute COVID-19 Surveillance Team |
| EPI_ISL_3222286 | Lighthouse Laboratory Plymouth | Wellcome Sanger Institute for the COVID-19 Genomics UK (COG-UK) Consortium | Lighthouse Laboratory Plymouth and Alex Alderton, Roberto Amato, Jeffrey Barrett, Sonia Goncalves, Ewan Harrison, David K. Jackson, Ian Johnston, Dominic Kwiatkowski, Cordelia Langford, John Sillitoe on behalf of the Wellcome Sanger Institute COVID-19 Surveillance Team |
| EPI_ISL_3222287 | Lighthouse Lab in Milton Keynes | Wellcome Sanger Institute for the COVID-19 Genomics UK (COG-UK) Consortium | The Lighthouse Lab in Milton Keynes and Alex Alderton, Roberto Amato, Jeffrey Barrett, Sonia Goncalves, Ewan Harrison, David K. Jackson, Ian Johnston, Dominic Kwiatkowski, Cordelia Langford, John Sillitoe on behalf of the Wellcome Sanger Institute COVID-19 Surveillance Team |
| EPI_ISL_3222288, EPI_ISL_3222289 | Berkshire and Surrey Pathology Services Lighthouse Laboratory | Wellcome Sanger Institute for the COVID-19 Genomics UK (COG-UK) Consortium | Berkshire and Surrey Pathology Services Lighthouse Laboratory and Alex Alderton, Roberto Amato, Jeffrey Barrett, Sonia Goncalves, Ewan Harrison, David K. Jackson, Ian Johnston, Dominic Kwiatkowski, Cordelia Langford, John Sillitoe on behalf of the Wellcome Sanger Institute COVID-19 Surveillance Team |
| EPI_ISL_3222290 | Lighthouse Laboratory Plymouth | Wellcome Sanger Institute for the COVID-19 Genomics UK (COG-UK) Consortium | Lighthouse Laboratory Plymouth and Alex Alderton, Roberto Amato, Jeffrey Barrett, Sonia Goncalves, Ewan Harrison, David K. Jackson, Ian Johnston, Dominic Kwiatkowski, Cordelia Langford, John Sillitoe on behalf of the Wellcome Sanger Institute COVID-19 Surveillance Team |
| EPI_ISL_3222291 | Lighthouse Lab in Milton Keynes | Wellcome Sanger Institute for the COVID-19 Genomics UK (COG-UK) Consortium | The Lighthouse Lab in Milton Keynes and Alex Alderton, Roberto Amato, Jeffrey Barrett, Sonia Goncalves, Ewan Harrison, David K. Jackson, Ian Johnston, Dominic Kwiatkowski, Cordelia Langford, John Sillitoe on behalf of the Wellcome Sanger Institute COVID-19 Surveillance Team |
| EPI_ISL_3222292 | Lighthouse Lab in Glasgow | Wellcome Sanger Institute for the COVID-19 Genomics UK (COG-UK) Consortium | Harper VanSteenhouse, Yumi Kasai, David Gray, Carol Clugston, Anna Dominiczak and Alex Alderton, Roberto Amato, Jeffrey Barrett, Sonia Goncalves, Ewan Harrison, David K. Jackson, Ian Johnston, Dominic Kwiatkowski, Cordelia Langford, John Sillitoe on behalf of the Wellcome Sanger Institute COVID-19 Surveillance Team |
| EPI_ISL_3222293 | Lighthouse Laboratory Plymouth | Wellcome Sanger Institute for the COVID-19 Genomics UK (COG-UK) Consortium | Lighthouse Laboratory Plymouth and Alex Alderton, Roberto Amato, Jeffrey Barrett, Sonia Goncalves, Ewan Harrison, David K. Jackson, Ian Johnston, Dominic Kwiatkowski, Cordelia Langford, John Sillitoe on behalf of the Wellcome Sanger Institute COVID-19 Surveillance Team |
| EPI_ISL_3222294 | Lighthouse Lab in Alderley Park | Wellcome Sanger Institute for the COVID-19 Genomics UK (COG-UK) Consortium | Jacquelyn Wynn, Mairead Hyland, The Lighthouse Lab in Alderley Park and Alex Alderton, Roberto Amato, Jeffrey Barrett, Sonia Goncalves, Ewan Harrison, David K. Jackson, Ian Johnston, Dominic Kwiatkowski, Cordelia Langford, John Sillitoe on behalf of the Wellcome Sanger Institute COVID-19 Surveillance Team |
| EPI_ISL_3222295 | Lighthouse Laboratory Plymouth | Wellcome Sanger Institute for the COVID-19 Genomics UK (COG-UK) Consortium | Lighthouse Laboratory Plymouth and Alex Alderton, Roberto Amato, Jeffrey Barrett, Sonia Goncalves, Ewan Harrison, David K. Jackson, Ian Johnston, Dominic Kwiatkowski, Cordelia Langford, John Sillitoe on behalf of the Wellcome Sanger Institute COVID-19 Surveillance Team |
| EPI_ISL_3222296 | Lighthouse Lab in Glasgow | Wellcome Sanger Institute for the COVID-19 Genomics UK (COG-UK) Consortium | Harper VanSteenhouse, Yumi Kasai, David Gray, Carol Clugston, Anna Dominiczak and Alex Alderton, Roberto Amato, Jeffrey Barrett, Sonia Goncalves, Ewan Harrison, David K. Jackson, Ian Johnston, Dominic Kwiatkowski, Cordelia Langford, John Sillitoe on behalf of the Wellcome Sanger Institute COVID-19 Surveillance Team |
| EPI_ISL_3222297 | Lighthouse Lab in Milton Keynes | Wellcome Sanger Institute for the COVID-19 Genomics UK (COG-UK) Consortium | The Lighthouse Lab in Milton Keynes and Alex Alderton, Roberto Amato, Jeffrey Barrett, Sonia Goncalves, Ewan Harrison, David K. Jackson, Ian Johnston, Dominic Kwiatkowski, Cordelia Langford, John Sillitoe on behalf of the Wellcome Sanger Institute COVID-19 Surveillance Team |
| EPI_ISL_3222298 | Lighthouse Lab in Alderley Park | Wellcome Sanger Institute for the COVID-19 Genomics UK (COG-UK) Consortium | Jacquelyn Wynn, Mairead Hyland, The Lighthouse Lab in Alderley Park and Alex Alderton, Roberto Amato, Jeffrey Barrett, Sonia Goncalves, Ewan Harrison, David K. Jackson, Ian Johnston, Dominic Kwiatkowski, Cordelia Langford, John Sillitoe on behalf of the Wellcome Sanger Institute COVID-19 Surveillance Team |
| EPI_ISL_3222299 | Lighthouse Laboratory Plymouth | Wellcome Sanger Institute for the COVID-19 Genomics UK (COG-UK) Consortium | Lighthouse Laboratory Plymouth and Alex Alderton, Roberto Amato, Jeffrey Barrett, Sonia Goncalves, Ewan Harrison, David K. Jackson, Ian Johnston, Dominic Kwiatkowski, Cordelia Langford, John Sillitoe on behalf of the Wellcome Sanger Institute COVID-19 Surveillance Team |
| EPI_ISL_3222300 | Lighthouse Lab in Milton Keynes | Wellcome Sanger Institute for the COVID-19 Genomics UK (COG-UK) Consortium | The Lighthouse Lab in Milton Keynes and Alex Alderton, Roberto Amato, Jeffrey Barrett, Sonia Goncalves, Ewan Harrison, David K. Jackson, Ian Johnston, Dominic Kwiatkowski, Cordelia Langford, John Sillitoe on behalf of the Wellcome Sanger Institute COVID-19 Surveillance Team |

| | | | |
|---|---|---|---|
| EPI_ISL_3222301 | Lighthouse Lab in Alderley Park | Wellcome Sanger Institute for the COVID-19 Genomics UK (COG-UK) Consortium | Jacquelyn Wynn, Mairead Hyland, The Lighthouse Lab in Alderley Park and Alex Alderton, Roberto Amato, Jeffrey Barrett, Sonia Goncalves, Ewan Harrison, David K. Jackson, Ian Johnston, Dominic Kwiatkowski, Cordelia Langford, John Sillitoe on behalf of the Wellcome Sanger Institute COVID-19 Surveillance Team |
| EPI_ISL_3222302, EPI_ISL_3222303 | Lighthouse Lab in Glasgow | Wellcome Sanger Institute for the COVID-19 Genomics UK (COG-UK) Consortium | Harper VanSteenhouse, Yumi Kasai, David Gray, Carol Clugston, Anna Dominiczak and Alex Alderton, Roberto Amato, Jeffrey Barrett, Sonia Goncalves, Ewan Harrison, David K. Jackson, Ian Johnston, Dominic Kwiatkowski, Cordelia Langford, John Sillitoe on behalf of the Wellcome Sanger Institute COVID-19 Surveillance Team |
| EPI_ISL_3222304 | Lighthouse Laboratory Plymouth | Wellcome Sanger Institute for the COVID-19 Genomics UK (COG-UK) Consortium | Lighthouse Laboratory Plymouth and Alex Alderton, Roberto Amato, Jeffrey Barrett, Sonia Goncalves, Ewan Harrison, David K. Jackson, Ian Johnston, Dominic Kwiatkowski, Cordelia Langford, John Sillitoe on behalf of the Wellcome Sanger Institute COVID-19 Surveillance Team |
| EPI_ISL_3222305 | Lighthouse Lab in Milton Keynes | Wellcome Sanger Institute for the COVID-19 Genomics UK (COG-UK) Consortium | The Lighthouse Lab in Milton Keynes and Alex Alderton, Roberto Amato, Jeffrey Barrett, Sonia Goncalves, Ewan Harrison, David K. Jackson, Ian Johnston, Dominic Kwiatkowski, Cordelia Langford, John Sillitoe on behalf of the Wellcome Sanger Institute COVID-19 Surveillance Team |
| EPI_ISL_3222306 | Lighthouse Lab in Alderley Park | Wellcome Sanger Institute for the COVID-19 Genomics UK (COG-UK) Consortium | Jacquelyn Wynn, Mairead Hyland, The Lighthouse Lab in Alderley Park and Alex Alderton, Roberto Amato, Jeffrey Barrett, Sonia Goncalves, Ewan Harrison, David K. Jackson, Ian Johnston, Dominic Kwiatkowski, Cordelia Langford, John Sillitoe on behalf of the Wellcome Sanger Institute COVID-19 Surveillance Team |
| EPI_ISL_3222307 | Lighthouse Laboratory Plymouth | Wellcome Sanger Institute for the COVID-19 Genomics UK (COG-UK) Consortium | Lighthouse Laboratory Plymouth and Alex Alderton, Roberto Amato, Jeffrey Barrett, Sonia Goncalves, Ewan Harrison, David K. Jackson, Ian Johnston, Dominic Kwiatkowski, Cordelia Langford, John Sillitoe on behalf of the Wellcome Sanger Institute COVID-19 Surveillance Team |
| EPI_ISL_3222308 | Lighthouse Lab in Alderley Park | Wellcome Sanger Institute for the COVID-19 Genomics UK (COG-UK) Consortium | Jacquelyn Wynn, Mairead Hyland, The Lighthouse Lab in Alderley Park and Alex Alderton, Roberto Amato, Jeffrey Barrett, Sonia Goncalves, Ewan Harrison, David K. Jackson, Ian Johnston, Dominic Kwiatkowski, Cordelia Langford, John Sillitoe on behalf of the Wellcome Sanger Institute COVID-19 Surveillance Team |
| EPI_ISL_3222309 | Lighthouse Lab in Milton Keynes | Wellcome Sanger Institute for the COVID-19 Genomics UK (COG-UK) Consortium | The Lighthouse Lab in Milton Keynes and Alex Alderton, Roberto Amato, Jeffrey Barrett, Sonia Goncalves, Ewan Harrison, David K. Jackson, Ian Johnston, Dominic Kwiatkowski, Cordelia Langford, John Sillitoe on behalf of the Wellcome Sanger Institute COVID-19 Surveillance Team |
| EPI_ISL_3222310, EPI_ISL_3222311, EPI_ISL_3222312, EPI_ISL_3222313 | Lighthouse Lab in Alderley Park | Wellcome Sanger Institute for the COVID-19 Genomics UK (COG-UK) Consortium | Jacquelyn Wynn, Mairead Hyland, The Lighthouse Lab in Alderley Park and Alex Alderton, Roberto Amato, Jeffrey Barrett, Sonia Goncalves, Ewan Harrison, David K. Jackson, Ian Johnston, Dominic Kwiatkowski, Cordelia Langford, John Sillitoe on behalf of the Wellcome Sanger Institute COVID-19 Surveillance Team |
| EPI_ISL_3222314 | Lighthouse Lab in Glasgow | Wellcome Sanger Institute for the COVID-19 Genomics UK (COG-UK) Consortium | Harper VanSteenhouse, Yumi Kasai, David Gray, Carol Clugston, Anna Dominiczak and Alex Alderton, Roberto Amato, Jeffrey Barrett, Sonia Goncalves, Ewan Harrison, David K. Jackson, Ian Johnston, Dominic Kwiatkowski, Cordelia Langford, John Sillitoe on behalf of the Wellcome Sanger Institute COVID-19 Surveillance Team |
| EPI_ISL_3222315, EPI_ISL_3222316 | Lighthouse Lab in Milton Keynes | Wellcome Sanger Institute for the COVID-19 Genomics UK (COG-UK) Consortium | The Lighthouse Lab in Milton Keynes and Alex Alderton, Roberto Amato, Jeffrey Barrett, Sonia Goncalves, Ewan Harrison, David K. Jackson, Ian Johnston, Dominic Kwiatkowski, Cordelia Langford, John Sillitoe on behalf of the Wellcome Sanger Institute COVID-19 Surveillance Team |
| EPI_ISL_3222317, EPI_ISL_3222318, EPI_ISL_3222319, EPI_ISL_3222320 | Lighthouse Lab in Alderley Park | Wellcome Sanger Institute for the COVID-19 Genomics UK (COG-UK) Consortium | Jacquelyn Wynn, Mairead Hyland, The Lighthouse Lab in Alderley Park and Alex Alderton, Roberto Amato, Jeffrey Barrett, Sonia Goncalves, Ewan Harrison, David K. Jackson, Ian Johnston, Dominic Kwiatkowski, Cordelia Langford, John Sillitoe on behalf of the Wellcome Sanger Institute COVID-19 Surveillance Team |
| EPI_ISL_3222321 | Lighthouse Lab in Glasgow | Wellcome Sanger Institute for the COVID-19 Genomics UK (COG-UK) Consortium | Harper VanSteenhouse, Yumi Kasai, David Gray, Carol Clugston, Anna Dominiczak and Alex Alderton, Roberto Amato, Jeffrey Barrett, Sonia Goncalves, Ewan Harrison, David K. Jackson, Ian Johnston, Dominic Kwiatkowski, Cordelia Langford, John Sillitoe on behalf of the Wellcome Sanger Institute COVID-19 Surveillance Team |
| EPI_ISL_3222322 | Lighthouse Lab in Milton Keynes | Wellcome Sanger Institute for the COVID-19 Genomics UK (COG-UK) Consortium | The Lighthouse Lab in Milton Keynes and Alex Alderton, Roberto Amato, Jeffrey Barrett, Sonia Goncalves, Ewan Harrison, David K. Jackson, Ian Johnston, Dominic Kwiatkowski, Cordelia Langford, John Sillitoe on behalf of the Wellcome Sanger Institute COVID-19 Surveillance Team |
| EPI_ISL_3222323, EPI_ISL_3222324, EPI_ISL_3222325 | Lighthouse Laboratory Plymouth | Wellcome Sanger Institute for the COVID-19 Genomics UK (COG-UK) Consortium | Lighthouse Laboratory Plymouth and Alex Alderton, Roberto Amato, Jeffrey Barrett, Sonia Goncalves, Ewan Harrison, David K. Jackson, Ian Johnston, Dominic Kwiatkowski, Cordelia Langford, John Sillitoe on behalf of the Wellcome Sanger Institute COVID-19 Surveillance Team |
| EPI_ISL_3222326 | Lighthouse Lab in Alderley Park | Wellcome Sanger Institute for the COVID-19 Genomics UK (COG-UK) Consortium | Jacquelyn Wynn, Mairead Hyland, The Lighthouse Lab in Alderley Park and Alex Alderton, Roberto Amato, Jeffrey Barrett, Sonia Goncalves, Ewan Harrison, David K. Jackson, Ian Johnston, Dominic Kwiatkowski, Cordelia Langford, John Sillitoe on behalf of the Wellcome Sanger Institute COVID-19 Surveillance Team |
| EPI_ISL_3222327, EPI_ISL_3222328 | Lighthouse Lab in Milton Keynes | Wellcome Sanger Institute for the COVID-19 Genomics UK (COG-UK) Consortium | The Lighthouse Lab in Milton Keynes and Alex Alderton, Roberto Amato, Jeffrey Barrett, Sonia Goncalves, Ewan Harrison, David K. Jackson, Ian Johnston, Dominic Kwiatkowski, Cordelia Langford, John Sillitoe on behalf of the Wellcome Sanger Institute COVID-19 Surveillance Team |
| EPI_ISL_3222329, EPI_ISL_3222330, EPI_ISL_3222331 | Lighthouse Lab in Alderley Park | Wellcome Sanger Institute for the COVID-19 Genomics UK (COG-UK) Consortium | Jacquelyn Wynn, Mairead Hyland, The Lighthouse Lab in Alderley Park and Alex Alderton, Roberto Amato, Jeffrey Barrett, Sonia Goncalves, Ewan Harrison, David K. Jackson, Ian Johnston, Dominic Kwiatkowski, Cordelia Langford, John Sillitoe on behalf of the Wellcome Sanger Institute COVID-19 Surveillance Team |
| EPI_ISL_3222332 | Lighthouse Lab in Milton Keynes | Wellcome Sanger Institute for the COVID-19 Genomics UK (COG-UK) Consortium | The Lighthouse Lab in Milton Keynes and Alex Alderton, Roberto Amato, Jeffrey Barrett, Sonia Goncalves, Ewan Harrison, David K. Jackson, Ian Johnston, Dominic Kwiatkowski, Cordelia Langford, John Sillitoe on behalf of the Wellcome Sanger Institute COVID-19 Surveillance Team |
| EPI_ISL_3222333 | Lighthouse Lab in Glasgow | Wellcome Sanger Institute for the COVID-19 Genomics UK (COG-UK) Consortium | Harper VanSteenhouse, Yumi Kasai, David Gray, Carol Clugston, Anna Dominiczak and Alex Alderton, Roberto Amato, Jeffrey Barrett, Sonia Goncalves, Ewan Harrison, David K. Jackson, Ian Johnston, Dominic Kwiatkowski, Cordelia Langford, John Sillitoe on behalf of the Wellcome Sanger Institute COVID-19 Surveillance Team |
| EPI_ISL_3222334 | Lighthouse Lab in Alderley Park | Wellcome Sanger Institute for the COVID-19 Genomics UK (COG-UK) Consortium | Jacquelyn Wynn, Mairead Hyland, The Lighthouse Lab in Alderley Park and Alex Alderton, Roberto Amato, Jeffrey Barrett, Sonia Goncalves, Ewan Harrison, David K. Jackson, Ian Johnston, Dominic Kwiatkowski, Cordelia Langford, John Sillitoe on behalf of the Wellcome Sanger Institute COVID-19 Surveillance Team |
| EPI_ISL_3222335 | Lighthouse Lab in Glasgow | Wellcome Sanger Institute for the COVID-19 Genomics UK (COG-UK) Consortium | Harper VanSteenhouse, Yumi Kasai, David Gray, Carol Clugston, Anna Dominiczak and Alex Alderton, Roberto Amato, Jeffrey Barrett, Sonia Goncalves, Ewan Harrison, David K. Jackson, Ian Johnston, Dominic Kwiatkowski, Cordelia Langford, John Sillitoe on behalf of the Wellcome Sanger Institute COVID-19 Surveillance Team |
| EPI_ISL_3222336 | Lighthouse Lab in Milton Keynes | Wellcome Sanger Institute for the COVID-19 Genomics UK (COG-UK) Consortium | The Lighthouse Lab in Milton Keynes and Alex Alderton, Roberto Amato, Jeffrey Barrett, Sonia Goncalves, Ewan Harrison, David K. Jackson, Ian Johnston, Dominic Kwiatkowski, Cordelia Langford, John Sillitoe on behalf of the Wellcome Sanger Institute COVID-19 Surveillance Team |
| EPI_ISL_3222337 | Lighthouse Lab in Alderley Park | Wellcome Sanger Institute for the COVID-19 Genomics UK (COG-UK) Consortium | Jacquelyn Wynn, Mairead Hyland, The Lighthouse Lab in Alderley Park and Alex Alderton, Roberto Amato, Jeffrey Barrett, Sonia Goncalves, Ewan Harrison, David K. Jackson, Ian Johnston, Dominic Kwiatkowski, Cordelia Langford, John Sillitoe on behalf of the Wellcome Sanger Institute COVID-19 Surveillance Team |
| EPI_ISL_3222338, EPI_ISL_3222339, EPI_ISL_3222340 | Lighthouse Lab in Milton Keynes | Wellcome Sanger Institute for the COVID-19 Genomics UK (COG-UK) Consortium | The Lighthouse Lab in Milton Keynes and Alex Alderton, Roberto Amato, Jeffrey Barrett, Sonia Goncalves, Ewan Harrison, David K. Jackson, Ian Johnston, Dominic Kwiatkowski, Cordelia Langford, John Sillitoe on behalf of the Wellcome Sanger Institute COVID-19 Surveillance Team |
| EPI_ISL_3222341 | Lighthouse Lab in Glasgow | Wellcome Sanger Institute for the COVID-19 Genomics UK (COG-UK) Consortium | Harper VanSteenhouse, Yumi Kasai, David Gray, Carol Clugston, Anna Dominiczak and Alex Alderton, Roberto Amato, Jeffrey Barrett, Sonia Goncalves, Ewan Harrison, David K. Jackson, Ian Johnston, Dominic Kwiatkowski, Cordelia Langford, John Sillitoe on behalf of the Wellcome Sanger Institute COVID-19 Surveillance Team |
| EPI_ISL_3222342 | Lighthouse Laboratory Plymouth | Wellcome Sanger Institute for the COVID-19 Genomics UK (COG-UK) Consortium | Lighthouse Laboratory Plymouth and Alex Alderton, Roberto Amato, Jeffrey Barrett, Sonia Goncalves, Ewan Harrison, David K. Jackson, Ian Johnston, Dominic Kwiatkowski, Cordelia Langford, John Sillitoe on behalf of the Wellcome Sanger Institute COVID-19 Surveillance Team |
| EPI_ISL_3222343 | Lighthouse Lab in Alderley Park | Wellcome Sanger Institute for the COVID-19 Genomics UK (COG-UK) Consortium | Jacquelyn Wynn, Mairead Hyland, The Lighthouse Lab in Alderley Park and Alex Alderton, Roberto Amato, Jeffrey Barrett, Sonia Goncalves, Ewan Harrison, David K. Jackson, Ian Johnston, Dominic Kwiatkowski, Cordelia Langford, John Sillitoe on behalf of the Wellcome Sanger Institute COVID-19 Surveillance Team |
| EPI_ISL_3222344 | Lighthouse Lab in Milton Keynes | Wellcome Sanger Institute for the COVID-19 Genomics UK (COG-UK) Consortium | The Lighthouse Lab in Milton Keynes and Alex Alderton, Roberto Amato, Jeffrey Barrett, Sonia Goncalves, Ewan Harrison, David K. Jackson, Ian Johnston, Dominic Kwiatkowski, Cordelia Langford, John Sillitoe on behalf of the Wellcome Sanger Institute COVID-19 Surveillance Team |
| EPI_ISL_3222345 | Lighthouse Laboratory Plymouth | Wellcome Sanger Institute for the COVID-19 Genomics UK (COG-UK) Consortium | Lighthouse Laboratory Plymouth and Alex Alderton, Roberto Amato, Jeffrey Barrett, Sonia Goncalves, Ewan Harrison, David K. Jackson, Ian Johnston, Dominic Kwiatkowski, Cordelia Langford, John Sillitoe on behalf of the Wellcome Sanger Institute COVID-19 Surveillance Team |
| EPI_ISL_3222346, EPI_ISL_3222347 | Lighthouse Lab in Glasgow | Wellcome Sanger Institute for the COVID-19 Genomics UK (COG-UK) Consortium | Harper VanSteenhouse, Yumi Kasai, David Gray, Carol Clugston, Anna Dominiczak and Alex Alderton, Roberto Amato, Jeffrey Barrett, Sonia Goncalves, Ewan Harrison, David K. Jackson, Ian Johnston, Dominic Kwiatkowski, Cordelia Langford, John Sillitoe on behalf of the Wellcome Sanger Institute COVID-19 Surveillance Team |
| EPI_ISL_3222348 | Lighthouse Lab in Alderley Park | Wellcome Sanger Institute for the COVID-19 Genomics UK (COG-UK) Consortium | Jacquelyn Wynn, Mairead Hyland, The Lighthouse Lab in Alderley Park and Alex Alderton, Roberto Amato, Jeffrey Barrett, Sonia Goncalves, Ewan Harrison, David K. Jackson, Ian Johnston, Dominic Kwiatkowski, Cordelia Langford, John Sillitoe on behalf of the Wellcome Sanger Institute COVID-19 Surveillance Team |
| EPI_ISL_3222349 | Lighthouse Lab in Glasgow | Wellcome Sanger Institute for the COVID-19 Genomics UK (COG-UK) Consortium | Harper VanSteenhouse, Yumi Kasai, David Gray, Carol Clugston, Anna Dominiczak and Alex Alderton, Roberto Amato, Jeffrey Barrett, Sonia Goncalves, Ewan Harrison, David K. Jackson, Ian Johnston, Dominic Kwiatkowski, Cordelia Langford, John Sillitoe on behalf of the Wellcome Sanger Institute COVID-19 Surveillance Team |
| EPI_ISL_3222350, EPI_ISL_3222351 | Lighthouse Laboratory Plymouth | Wellcome Sanger Institute for the COVID-19 Genomics UK (COG-UK) Consortium | Lighthouse Laboratory Plymouth and Alex Alderton, Roberto Amato, Jeffrey Barrett, Sonia Goncalves, Ewan Harrison, David K. Jackson, Ian Johnston, Dominic Kwiatkowski, Cordelia Langford, John Sillitoe on behalf of the Wellcome Sanger Institute COVID-19 Surveillance Team |
| EPI_ISL_3222352 | Lighthouse Lab in Glasgow | Wellcome Sanger Institute for the COVID-19 Genomics UK (COG-UK) Consortium | Harper VanSteenhouse, Yumi Kasai, David Gray, Carol Clugston, Anna Dominiczak and Alex Alderton, Roberto Amato, Jeffrey Barrett, Sonia Goncalves, Ewan Harrison, David K. Jackson, Ian Johnston, Dominic Kwiatkowski, Cordelia Langford, John Sillitoe on behalf of the Wellcome Sanger Institute COVID-19 Surveillance Team |
| EPI_ISL_3222353 | Lighthouse Lab in Alderley Park | Wellcome Sanger Institute for the COVID-19 Genomics UK (COG-UK) Consortium | Jacquelyn Wynn, Mairead Hyland, The Lighthouse Lab in Alderley Park and Alex Alderton, Roberto Amato, Jeffrey Barrett, Sonia Goncalves, Ewan Harrison, David K. Jackson, Ian Johnston, Dominic Kwiatkowski, Cordelia Langford, John Sillitoe on behalf of the Wellcome Sanger Institute COVID-19 Surveillance Team |
| EPI_ISL_3222354 | Lighthouse Laboratory Plymouth | Wellcome Sanger Institute for the COVID-19 Genomics UK (COG-UK) Consortium | Lighthouse Laboratory Plymouth and Alex Alderton, Roberto Amato, Jeffrey Barrett, Sonia Goncalves, Ewan Harrison, David K. Jackson, Ian Johnston, Dominic Kwiatkowski, Cordelia Langford, John Sillitoe on behalf of the Wellcome Sanger Institute COVID-19 Surveillance Team |
| EPI_ISL_3222355 | Lighthouse Lab in Milton Keynes | Wellcome Sanger Institute for the COVID-19 Genomics UK (COG-UK) Consortium | The Lighthouse Lab in Milton Keynes and Alex Alderton, Roberto Amato, Jeffrey Barrett, Sonia Goncalves, Ewan Harrison, David K. Jackson, Ian Johnston, Dominic Kwiatkowski, Cordelia Langford, John Sillitoe on behalf of the Wellcome Sanger Institute COVID-19 Surveillance Team |
| EPI_ISL_3222356 | Lighthouse Lab in Alderley Park | Wellcome Sanger Institute for the COVID-19 Genomics UK (COG-UK) Consortium | Jacquelyn Wynn, Mairead Hyland, The Lighthouse Lab in Alderley Park and Alex Alderton, Roberto Amato, Jeffrey Barrett, Sonia Goncalves, Ewan Harrison, David K. Jackson, Ian Johnston, Dominic Kwiatkowski, Cordelia Langford, John Sillitoe on behalf of the Wellcome Sanger Institute COVID-19 Surveillance Team |
| EPI_ISL_3222357 | Lighthouse Laboratory Plymouth | Wellcome Sanger Institute for the COVID-19 Genomics UK (COG-UK) Consortium | Lighthouse Laboratory Plymouth and Alex Alderton, Roberto Amato, Jeffrey Barrett, Sonia Goncalves, Ewan Harrison, David K. Jackson, Ian Johnston, Dominic Kwiatkowski, Cordelia Langford, John Sillitoe on behalf of the Wellcome Sanger Institute COVID-19 Surveillance Team |
| EPI_ISL_3222358 | Lighthouse Lab in Milton Keynes | Wellcome Sanger Institute for the COVID-19 Genomics UK (COG-UK) Consortium | The Lighthouse Lab in Milton Keynes and Alex Alderton, Roberto Amato, Jeffrey Barrett, Sonia Goncalves, Ewan Harrison, David K. Jackson, Ian Johnston, Dominic Kwiatkowski, Cordelia Langford, John Sillitoe on behalf of the Wellcome Sanger Institute COVID-19 Surveillance Team |
| EPI_ISL_3222359 | Lighthouse Lab in Alderley Park | Wellcome Sanger Institute for the COVID-19 Genomics UK (COG-UK) Consortium | Jacquelyn Wynn, Mairead Hyland, The Lighthouse Lab in Alderley Park and Alex Alderton, Roberto Amato, Jeffrey Barrett, Sonia Goncalves, Ewan Harrison, David K. Jackson, Ian Johnston, Dominic Kwiatkowski, Cordelia Langford, John Sillitoe on behalf of the Wellcome Sanger Institute COVID-19 Surveillance Team |

| | | | |
|---|---|---|---|
| EPI_ISL_3222360 | Lighthouse Lab in Glasgow | Wellcome Sanger Institute for the COVID-19 Genomics UK (COG-UK) Consortium | Harper VanSteenhouse, Yumi Kasai, David Gray, Carol Clugston, Anna Dominiczak and Alex Alderton, Roberto Amato, Jeffrey Barrett, Sonia Goncalves, Ewan Harrison, David K. Jackson, Ian Johnston, Dominic Kwiatkowski, Cordelia Langford, John Sillitoe on behalf of the Wellcome Sanger Institute COVID-19 Surveillance Team |
| EPI_ISL_3222361, EPI_ISL_3222362, EPI_ISL_3222363 | Lighthouse Lab in Alderley Park | Wellcome Sanger Institute for the COVID-19 Genomics UK (COG-UK) Consortium | Jacquelyn Wynn, Mairead Hyland, The Lighthouse Lab in Alderley Park and Alex Alderton, Roberto Amato, Jeffrey Barrett, Sonia Goncalves, Ewan Harrison, David K. Jackson, Ian Johnston, Dominic Kwiatkowski, Cordelia Langford, John Sillitoe on behalf of the Wellcome Sanger Institute COVID-19 Surveillance Team |
| EPI_ISL_3222364 | Lighthouse Laboratory Plymouth | Wellcome Sanger Institute for the COVID-19 Genomics UK (COG-UK) Consortium | Lighthouse Laboratory Plymouth and Alex Alderton, Roberto Amato, Jeffrey Barrett, Sonia Goncalves, Ewan Harrison, David K. Jackson, Ian Johnston, Dominic Kwiatkowski, Cordelia Langford, John Sillitoe on behalf of the Wellcome Sanger Institute COVID-19 Surveillance Team |
| EPI_ISL_3222365, EPI_ISL_3222366 | Lighthouse Lab in Glasgow | Wellcome Sanger Institute for the COVID-19 Genomics UK (COG-UK) Consortium | Harper VanSteenhouse, Yumi Kasai, David Gray, Carol Clugston, Anna Dominiczak and Alex Alderton, Roberto Amato, Jeffrey Barrett, Sonia Goncalves, Ewan Harrison, David K. Jackson, Ian Johnston, Dominic Kwiatkowski, Cordelia Langford, John Sillitoe on behalf of the Wellcome Sanger Institute COVID-19 Surveillance Team |
| EPI_ISL_3222367 | Lighthouse Lab in Milton Keynes | Wellcome Sanger Institute for the COVID-19 Genomics UK (COG-UK) Consortium | The Lighthouse Lab in Milton Keynes and Alex Alderton, Roberto Amato, Jeffrey Barrett, Sonia Goncalves, Ewan Harrison, David K. Jackson, Ian Johnston, Dominic Kwiatkowski, Cordelia Langford, John Sillitoe on behalf of the Wellcome Sanger Institute COVID-19 Surveillance Team |
| EPI_ISL_3222368 | Lighthouse Laboratory Plymouth | Wellcome Sanger Institute for the COVID-19 Genomics UK (COG-UK) Consortium | Lighthouse Laboratory Plymouth and Alex Alderton, Roberto Amato, Jeffrey Barrett, Sonia Goncalves, Ewan Harrison, David K. Jackson, Ian Johnston, Dominic Kwiatkowski, Cordelia Langford, John Sillitoe on behalf of the Wellcome Sanger Institute COVID-19 Surveillance Team |
| EPI_ISL_3222369 | Lighthouse Lab in Alderley Park | Wellcome Sanger Institute for the COVID-19 Genomics UK (COG-UK) Consortium | Jacquelyn Wynn, Mairead Hyland, The Lighthouse Lab in Alderley Park and Alex Alderton, Roberto Amato, Jeffrey Barrett, Sonia Goncalves, Ewan Harrison, David K. Jackson, Ian Johnston, Dominic Kwiatkowski, Cordelia Langford, John Sillitoe on behalf of the Wellcome Sanger Institute COVID-19 Surveillance Team |
| EPI_ISL_3222370 | Lighthouse Laboratory Plymouth | Wellcome Sanger Institute for the COVID-19 Genomics UK (COG-UK) Consortium | Lighthouse Laboratory Plymouth and Alex Alderton, Roberto Amato, Jeffrey Barrett, Sonia Goncalves, Ewan Harrison, David K. Jackson, Ian Johnston, Dominic Kwiatkowski, Cordelia Langford, John Sillitoe on behalf of the Wellcome Sanger Institute COVID-19 Surveillance Team |
| EPI_ISL_3222371 | Lighthouse Lab in Milton Keynes | Wellcome Sanger Institute for the COVID-19 Genomics UK (COG-UK) Consortium | The Lighthouse Lab in Milton Keynes and Alex Alderton, Roberto Amato, Jeffrey Barrett, Sonia Goncalves, Ewan Harrison, David K. Jackson, Ian Johnston, Dominic Kwiatkowski, Cordelia Langford, John Sillitoe on behalf of the Wellcome Sanger Institute COVID-19 Surveillance Team |
| EPI_ISL_3222372 | Lighthouse Laboratory Plymouth | Wellcome Sanger Institute for the COVID-19 Genomics UK (COG-UK) Consortium | Lighthouse Laboratory Plymouth and Alex Alderton, Roberto Amato, Jeffrey Barrett, Sonia Goncalves, Ewan Harrison, David K. Jackson, Ian Johnston, Dominic Kwiatkowski, Cordelia Langford, John Sillitoe on behalf of the Wellcome Sanger Institute COVID-19 Surveillance Team |
| EPI_ISL_3222373, EPI_ISL_3222374 | Lighthouse Lab in Milton Keynes | Wellcome Sanger Institute for the COVID-19 Genomics UK (COG-UK) Consortium | The Lighthouse Lab in Milton Keynes and Alex Alderton, Roberto Amato, Jeffrey Barrett, Sonia Goncalves, Ewan Harrison, David K. Jackson, Ian Johnston, Dominic Kwiatkowski, Cordelia Langford, John Sillitoe on behalf of the Wellcome Sanger Institute COVID-19 Surveillance Team |
| EPI_ISL_3222375 | Lighthouse Lab in Glasgow | Wellcome Sanger Institute for the COVID-19 Genomics UK (COG-UK) Consortium | Harper VanSteenhouse, Yumi Kasai, David Gray, Carol Clugston, Anna Dominiczak and Alex Alderton, Roberto Amato, Jeffrey Barrett, Sonia Goncalves, Ewan Harrison, David K. Jackson, Ian Johnston, Dominic Kwiatkowski, Cordelia Langford, John Sillitoe on behalf of the Wellcome Sanger Institute COVID-19 Surveillance Team |
| EPI_ISL_3222376, EPI_ISL_3222377 | Lighthouse Laboratory Plymouth | Wellcome Sanger Institute for the COVID-19 Genomics UK (COG-UK) Consortium | Lighthouse Laboratory Plymouth and Alex Alderton, Roberto Amato, Jeffrey Barrett, Sonia Goncalves, Ewan Harrison, David K. Jackson, Ian Johnston, Dominic Kwiatkowski, Cordelia Langford, John Sillitoe on behalf of the Wellcome Sanger Institute COVID-19 Surveillance Team |
| EPI_ISL_3222378 | Lighthouse Lab in Milton Keynes | Wellcome Sanger Institute for the COVID-19 Genomics UK (COG-UK) Consortium | The Lighthouse Lab in Milton Keynes and Alex Alderton, Roberto Amato, Jeffrey Barrett, Sonia Goncalves, Ewan Harrison, David K. Jackson, Ian Johnston, Dominic Kwiatkowski, Cordelia Langford, John Sillitoe on behalf of the Wellcome Sanger Institute COVID-19 Surveillance Team |
| EPI_ISL_3222379 | Lighthouse Lab in Alderley Park | Wellcome Sanger Institute for the COVID-19 Genomics UK (COG-UK) Consortium | Jacquelyn Wynn, Mairead Hyland, The Lighthouse Lab in Alderley Park and Alex Alderton, Roberto Amato, Jeffrey Barrett, Sonia Goncalves, Ewan Harrison, David K. Jackson, Ian Johnston, Dominic Kwiatkowski, Cordelia Langford, John Sillitoe on behalf of the Wellcome Sanger Institute COVID-19 Surveillance Team |
| EPI_ISL_3222380 | Lighthouse Laboratory Plymouth | Wellcome Sanger Institute for the COVID-19 Genomics UK (COG-UK) Consortium | Lighthouse Laboratory Plymouth and Alex Alderton, Roberto Amato, Jeffrey Barrett, Sonia Goncalves, Ewan Harrison, David K. Jackson, Ian Johnston, Dominic Kwiatkowski, Cordelia Langford, John Sillitoe on behalf of the Wellcome Sanger Institute COVID-19 Surveillance Team |
| EPI_ISL_3222381 | Lighthouse Lab in Glasgow | Wellcome Sanger Institute for the COVID-19 Genomics UK (COG-UK) Consortium | Harper VanSteenhouse, Yumi Kasai, David Gray, Carol Clugston, Anna Dominiczak and Alex Alderton, Roberto Amato, Jeffrey Barrett, Sonia Goncalves, Ewan Harrison, David K. Jackson, Ian Johnston, Dominic Kwiatkowski, Cordelia Langford, John Sillitoe on behalf of the Wellcome Sanger Institute COVID-19 Surveillance Team |
| EPI_ISL_3222382 | Lighthouse Lab in Milton Keynes | Wellcome Sanger Institute for the COVID-19 Genomics UK (COG-UK) Consortium | The Lighthouse Lab in Milton Keynes and Alex Alderton, Roberto Amato, Jeffrey Barrett, Sonia Goncalves, Ewan Harrison, David K. Jackson, Ian Johnston, Dominic Kwiatkowski, Cordelia Langford, John Sillitoe on behalf of the Wellcome Sanger Institute COVID-19 Surveillance Team |
| EPI_ISL_3222383 | Lighthouse Laboratory Plymouth | Wellcome Sanger Institute for the COVID-19 Genomics UK (COG-UK) Consortium | Lighthouse Laboratory Plymouth and Alex Alderton, Roberto Amato, Jeffrey Barrett, Sonia Goncalves, Ewan Harrison, David K. Jackson, Ian Johnston, Dominic Kwiatkowski, Cordelia Langford, John Sillitoe on behalf of the Wellcome Sanger Institute COVID-19 Surveillance Team |
| EPI_ISL_3222384 | Lighthouse Lab in Milton Keynes | Wellcome Sanger Institute for the COVID-19 Genomics UK (COG-UK) Consortium | The Lighthouse Lab in Milton Keynes and Alex Alderton, Roberto Amato, Jeffrey Barrett, Sonia Goncalves, Ewan Harrison, David K. Jackson, Ian Johnston, Dominic Kwiatkowski, Cordelia Langford, John Sillitoe on behalf of the Wellcome Sanger Institute COVID-19 Surveillance Team |
| EPI_ISL_3222385 | Lighthouse Lab in Glasgow | Wellcome Sanger Institute for the COVID-19 Genomics UK (COG-UK) Consortium | Harper VanSteenhouse, Yumi Kasai, David Gray, Carol Clugston, Anna Dominiczak and Alex Alderton, Roberto Amato, Jeffrey Barrett, Sonia Goncalves, Ewan Harrison, David K. Jackson, Ian Johnston, Dominic Kwiatkowski, Cordelia Langford, John Sillitoe on behalf of the Wellcome Sanger Institute COVID-19 Surveillance Team |
| EPI_ISL_3222386 | Lighthouse Lab in Milton Keynes | Wellcome Sanger Institute for the COVID-19 Genomics UK (COG-UK) Consortium | The Lighthouse Lab in Milton Keynes and Alex Alderton, Roberto Amato, Jeffrey Barrett, Sonia Goncalves, Ewan Harrison, David K. Jackson, Ian Johnston, Dominic Kwiatkowski, Cordelia Langford, John Sillitoe on behalf of the Wellcome Sanger Institute COVID-19 Surveillance Team |
| EPI_ISL_3222387 | Lighthouse Lab in Glasgow | Wellcome Sanger Institute for the COVID-19 Genomics UK (COG-UK) Consortium | Harper VanSteenhouse, Yumi Kasai, David Gray, Carol Clugston, Anna Dominiczak and Alex Alderton, Roberto Amato, Jeffrey Barrett, Sonia Goncalves, Ewan Harrison, David K. Jackson, Ian Johnston, Dominic Kwiatkowski, Cordelia Langford, John Sillitoe on behalf of the Wellcome Sanger Institute COVID-19 Surveillance Team |
| EPI_ISL_3222388 | Lighthouse Lab in Alderley Park | Wellcome Sanger Institute for the COVID-19 Genomics UK (COG-UK) Consortium | Jacquelyn Wynn, Mairead Hyland, The Lighthouse Lab in Alderley Park and Alex Alderton, Roberto Amato, Jeffrey Barrett, Sonia Goncalves, Ewan Harrison, David K. Jackson, Ian Johnston, Dominic Kwiatkowski, Cordelia Langford, John Sillitoe on behalf of the Wellcome Sanger Institute COVID-19 Surveillance Team |
| EPI_ISL_3222389 | Lighthouse Lab in Milton Keynes | Wellcome Sanger Institute for the COVID-19 Genomics UK (COG-UK) Consortium | The Lighthouse Lab in Milton Keynes and Alex Alderton, Roberto Amato, Jeffrey Barrett, Sonia Goncalves, Ewan Harrison, David K. Jackson, Ian Johnston, Dominic Kwiatkowski, Cordelia Langford, John Sillitoe on behalf of the Wellcome Sanger Institute COVID-19 Surveillance Team |
| EPI_ISL_3222390 | Lighthouse Lab in Glasgow | Wellcome Sanger Institute for the COVID-19 Genomics UK (COG-UK) Consortium | Harper VanSteenhouse, Yumi Kasai, David Gray, Carol Clugston, Anna Dominiczak and Alex Alderton, Roberto Amato, Jeffrey Barrett, Sonia Goncalves, Ewan Harrison, David K. Jackson, Ian Johnston, Dominic Kwiatkowski, Cordelia Langford, John Sillitoe on behalf of the Wellcome Sanger Institute COVID-19 Surveillance Team |
| EPI_ISL_3222391, EPI_ISL_3222392 | Lighthouse Laboratory Plymouth | Wellcome Sanger Institute for the COVID-19 Genomics UK (COG-UK) Consortium | Lighthouse Laboratory Plymouth and Alex Alderton, Roberto Amato, Jeffrey Barrett, Sonia Goncalves, Ewan Harrison, David K. Jackson, Ian Johnston, Dominic Kwiatkowski, Cordelia Langford, John Sillitoe on behalf of the Wellcome Sanger Institute COVID-19 Surveillance Team |
| EPI_ISL_3222393 | Lighthouse Lab in Alderley Park | Wellcome Sanger Institute for the COVID-19 Genomics UK (COG-UK) Consortium | Jacquelyn Wynn, Mairead Hyland, The Lighthouse Lab in Alderley Park and Alex Alderton, Roberto Amato, Jeffrey Barrett, Sonia Goncalves, Ewan Harrison, David K. Jackson, Ian Johnston, Dominic Kwiatkowski, Cordelia Langford, John Sillitoe on behalf of the Wellcome Sanger Institute COVID-19 Surveillance Team |
| EPI_ISL_3222394 | Lighthouse Lab in Glasgow | Wellcome Sanger Institute for the COVID-19 Genomics UK (COG-UK) Consortium | Harper VanSteenhouse, Yumi Kasai, David Gray, Carol Clugston, Anna Dominiczak and Alex Alderton, Roberto Amato, Jeffrey Barrett, Sonia Goncalves, Ewan Harrison, David K. Jackson, Ian Johnston, Dominic Kwiatkowski, Cordelia Langford, John Sillitoe on behalf of the Wellcome Sanger Institute COVID-19 Surveillance Team |
| EPI_ISL_3222395 | Lighthouse Lab in Alderley Park | Wellcome Sanger Institute for the COVID-19 Genomics UK (COG-UK) Consortium | Jacquelyn Wynn, Mairead Hyland, The Lighthouse Lab in Alderley Park and Alex Alderton, Roberto Amato, Jeffrey Barrett, Sonia Goncalves, Ewan Harrison, David K. Jackson, Ian Johnston, Dominic Kwiatkowski, Cordelia Langford, John Sillitoe on behalf of the Wellcome Sanger Institute COVID-19 Surveillance Team |
| EPI_ISL_3222396, EPI_ISL_3222397 | Lighthouse Lab in Milton Keynes | Wellcome Sanger Institute for the COVID-19 Genomics UK (COG-UK) Consortium | The Lighthouse Lab in Milton Keynes and Alex Alderton, Roberto Amato, Jeffrey Barrett, Sonia Goncalves, Ewan Harrison, David K. Jackson, Ian Johnston, Dominic Kwiatkowski, Cordelia Langford, John Sillitoe on behalf of the Wellcome Sanger Institute COVID-19 Surveillance Team |
| EPI_ISL_3222398 | Lighthouse Lab in Glasgow | Wellcome Sanger Institute for the COVID-19 Genomics UK (COG-UK) Consortium | Harper VanSteenhouse, Yumi Kasai, David Gray, Carol Clugston, Anna Dominiczak and Alex Alderton, Roberto Amato, Jeffrey Barrett, Sonia Goncalves, Ewan Harrison, David K. Jackson, Ian Johnston, Dominic Kwiatkowski, Cordelia Langford, John Sillitoe on behalf of the Wellcome Sanger Institute COVID-19 Surveillance Team |
| EPI_ISL_3222399 | Lighthouse Laboratory Plymouth | Wellcome Sanger Institute for the COVID-19 Genomics UK (COG-UK) Consortium | Lighthouse Laboratory Plymouth and Alex Alderton, Roberto Amato, Jeffrey Barrett, Sonia Goncalves, Ewan Harrison, David K. Jackson, Ian Johnston, Dominic Kwiatkowski, Cordelia Langford, John Sillitoe on behalf of the Wellcome Sanger Institute COVID-19 Surveillance Team |
| EPI_ISL_3222400 | Lighthouse Lab in Alderley Park | Wellcome Sanger Institute for the COVID-19 Genomics UK (COG-UK) Consortium | Jacquelyn Wynn, Mairead Hyland, The Lighthouse Lab in Alderley Park and Alex Alderton, Roberto Amato, Jeffrey Barrett, Sonia Goncalves, Ewan Harrison, David K. Jackson, Ian Johnston, Dominic Kwiatkowski, Cordelia Langford, John Sillitoe on behalf of the Wellcome Sanger Institute COVID-19 Surveillance Team |
| EPI_ISL_3222401, EPI_ISL_3222402 | Lighthouse Lab in Milton Keynes | Wellcome Sanger Institute for the COVID-19 Genomics UK (COG-UK) Consortium | The Lighthouse Lab in Milton Keynes and Alex Alderton, Roberto Amato, Jeffrey Barrett, Sonia Goncalves, Ewan Harrison, David K. Jackson, Ian Johnston, Dominic Kwiatkowski, Cordelia Langford, John Sillitoe on behalf of the Wellcome Sanger Institute COVID-19 Surveillance Team |
| EPI_ISL_3222403 | Lighthouse Laboratory Plymouth | Wellcome Sanger Institute for the COVID-19 Genomics UK (COG-UK) Consortium | Lighthouse Laboratory Plymouth and Alex Alderton, Roberto Amato, Jeffrey Barrett, Sonia Goncalves, Ewan Harrison, David K. Jackson, Ian Johnston, Dominic Kwiatkowski, Cordelia Langford, John Sillitoe on behalf of the Wellcome Sanger Institute COVID-19 Surveillance Team |
| EPI_ISL_3222404, EPI_ISL_3222405 | Lighthouse Lab in Alderley Park | Wellcome Sanger Institute for the COVID-19 Genomics UK (COG-UK) Consortium | Jacquelyn Wynn, Mairead Hyland, The Lighthouse Lab in Alderley Park and Alex Alderton, Roberto Amato, Jeffrey Barrett, Sonia Goncalves, Ewan Harrison, David K. Jackson, Ian Johnston, Dominic Kwiatkowski, Cordelia Langford, John Sillitoe on behalf of the Wellcome Sanger Institute COVID-19 Surveillance Team |
| EPI_ISL_3222406, EPI_ISL_3222407, EPI_ISL_3222408 | Lighthouse Laboratory Plymouth | Wellcome Sanger Institute for the COVID-19 Genomics UK (COG-UK) Consortium | Lighthouse Laboratory Plymouth and Alex Alderton, Roberto Amato, Jeffrey Barrett, Sonia Goncalves, Ewan Harrison, David K. Jackson, Ian Johnston, Dominic Kwiatkowski, Cordelia Langford, John Sillitoe on behalf of the Wellcome Sanger Institute COVID-19 Surveillance Team |
| EPI_ISL_3222409, EPI_ISL_3222410 | Lighthouse Lab in Milton Keynes | Wellcome Sanger Institute for the COVID-19 Genomics UK (COG-UK) Consortium | The Lighthouse Lab in Milton Keynes and Alex Alderton, Roberto Amato, Jeffrey Barrett, Sonia Goncalves, Ewan Harrison, David K. Jackson, Ian Johnston, Dominic Kwiatkowski, Cordelia Langford, John Sillitoe on behalf of the Wellcome Sanger Institute COVID-19 Surveillance Team |
| EPI_ISL_3222411 | Lighthouse Laboratory Plymouth | Wellcome Sanger Institute for the COVID-19 Genomics UK (COG-UK) Consortium | Lighthouse Laboratory Plymouth and Alex Alderton, Roberto Amato, Jeffrey Barrett, Sonia Goncalves, Ewan Harrison, David K. Jackson, Ian Johnston, Dominic Kwiatkowski, Cordelia Langford, John Sillitoe on behalf of the Wellcome Sanger Institute COVID-19 Surveillance Team |
| EPI_ISL_3222412 | Lighthouse Lab in Milton Keynes | Wellcome Sanger Institute for the COVID-19 Genomics UK (COG-UK) Consortium | The Lighthouse Lab in Milton Keynes and Alex Alderton, Roberto Amato, Jeffrey Barrett, Sonia Goncalves, Ewan Harrison, David K. Jackson, Ian Johnston, Dominic Kwiatkowski, Cordelia Langford, John Sillitoe on behalf of the Wellcome Sanger Institute COVID-19 Surveillance Team |
| EPI_ISL_3222413 | Lighthouse Laboratory Plymouth | Wellcome Sanger Institute for the COVID-19 Genomics UK (COG-UK) Consortium | Lighthouse Laboratory Plymouth and Alex Alderton, Roberto Amato, Jeffrey Barrett, Sonia Goncalves, Ewan Harrison, David K. Jackson, Ian Johnston, Dominic Kwiatkowski, Cordelia Langford, John Sillitoe on behalf of the Wellcome Sanger Institute COVID-19 Surveillance Team |
| EPI_ISL_3222414 | Lighthouse Lab in Alderley Park | Wellcome Sanger Institute for the COVID-19 Genomics UK (COG-UK) Consortium | Jacquelyn Wynn, Mairead Hyland, The Lighthouse Lab in Alderley Park and Alex Alderton, Roberto Amato, Jeffrey Barrett, Sonia Goncalves, Ewan Harrison, David K. Jackson, Ian Johnston, Dominic Kwiatkowski, Cordelia Langford, John Sillitoe on behalf of the Wellcome Sanger Institute COVID-19 Surveillance Team |
| EPI_ISL_3222415 | Lighthouse Laboratory Plymouth | Wellcome Sanger Institute for the COVID-19 Genomics UK (COG-UK) Consortium | Lighthouse Laboratory Plymouth and Alex Alderton, Roberto Amato, Jeffrey Barrett, Sonia Goncalves, Ewan Harrison, David K. Jackson, Ian Johnston, Dominic Kwiatkowski, Cordelia Langford, John Sillitoe on behalf of the Wellcome Sanger Institute COVID-19 Surveillance Team |
| EPI_ISL_3222416, | Lighthouse Lab in Milton Keynes | Wellcome Sanger Institute for the COVID-19 Genomics UK (COG-UK) | The Lighthouse Lab in Milton Keynes and Alex Alderton, Roberto Amato, Jeffrey Barrett, Sonia Goncalves, Ewan Harrison, David K. Jackson, Ian Johnston, Dominic Kwiatkowski, Cordelia Langford, John Sillitoe on behalf of the Wellcome Sanger |

| | | | |
|---|---|---|---|
| EPI_ISL_3222417 | | Consortium | Institute COVID-19 Surveillance Team |
| EPI_ISL_3222418 | Lighthouse Laboratory Plymouth | Wellcome Sanger Institute for the COVID-19 Genomics UK (COG-UK) Consortium | Lighthouse Laboratory Plymouth and Alex Alderton, Roberto Amato, Jeffrey Barrett, Sonia Goncalves, Ewan Harrison, David K. Jackson, Ian Johnston, Dominic Kwiatkowski, Cordelia Langford, John Sillitoe on behalf of the Wellcome Sanger Institute COVID-19 Surveillance Team |
| EPI_ISL_3222419 | Lighthouse Lab in Alderley Park | Wellcome Sanger Institute for the COVID-19 Genomics UK (COG-UK) Consortium | Jacquelyn Wynn, Mairead Hyland, The Lighthouse Lab in Alderley Park and Alex Alderton, Roberto Amato, Jeffrey Barrett, Sonia Goncalves, Ewan Harrison, David K. Jackson, Ian Johnston, Dominic Kwiatkowski, Cordelia Langford, John Sillitoe on behalf of the Wellcome Sanger Institute COVID-19 Surveillance Team |
| EPI_ISL_3222420 | Lighthouse Lab in Glasgow | Wellcome Sanger Institute for the COVID-19 Genomics UK (COG-UK) Consortium | Harper VanSteenhouse, Yumi Kasai, David Gray, Carol Clugston, Anna Dominiczak and Alex Alderton, Roberto Amato, Jeffrey Barrett, Sonia Goncalves, Ewan Harrison, David K. Jackson, Ian Johnston, Dominic Kwiatkowski, Cordelia Langford, John Sillitoe on behalf of the Wellcome Sanger Institute COVID-19 Surveillance Team |
| EPI_ISL_3222421 | Lighthouse Laboratory Plymouth | Wellcome Sanger Institute for the COVID-19 Genomics UK (COG-UK) Consortium | Lighthouse Laboratory Plymouth and Alex Alderton, Roberto Amato, Jeffrey Barrett, Sonia Goncalves, Ewan Harrison, David K. Jackson, Ian Johnston, Dominic Kwiatkowski, Cordelia Langford, John Sillitoe on behalf of the Wellcome Sanger Institute COVID-19 Surveillance Team |
| EPI_ISL_3222422 | Lighthouse Lab in Alderley Park | Wellcome Sanger Institute for the COVID-19 Genomics UK (COG-UK) Consortium | Jacquelyn Wynn, Mairead Hyland, The Lighthouse Lab in Alderley Park and Alex Alderton, Roberto Amato, Jeffrey Barrett, Sonia Goncalves, Ewan Harrison, David K. Jackson, Ian Johnston, Dominic Kwiatkowski, Cordelia Langford, John Sillitoe on behalf of the Wellcome Sanger Institute COVID-19 Surveillance Team |
| EPI_ISL_3222423 | Lighthouse Lab in Milton Keynes | Wellcome Sanger Institute for the COVID-19 Genomics UK (COG-UK) Consortium | The Lighthouse Lab in Milton Keynes and Alex Alderton, Roberto Amato, Jeffrey Barrett, Sonia Goncalves, Ewan Harrison, David K. Jackson, Ian Johnston, Dominic Kwiatkowski, Cordelia Langford, John Sillitoe on behalf of the Wellcome Sanger Institute COVID-19 Surveillance Team |
| EPI_ISL_3222424, EPI_ISL_3222425 | Lighthouse Lab in Alderley Park | Wellcome Sanger Institute for the COVID-19 Genomics UK (COG-UK) Consortium | Jacquelyn Wynn, Mairead Hyland, The Lighthouse Lab in Alderley Park and Alex Alderton, Roberto Amato, Jeffrey Barrett, Sonia Goncalves, Ewan Harrison, David K. Jackson, Ian Johnston, Dominic Kwiatkowski, Cordelia Langford, John Sillitoe on behalf of the Wellcome Sanger Institute COVID-19 Surveillance Team |
| EPI_ISL_3222426, EPI_ISL_3222427 | Lighthouse Lab in Milton Keynes | Wellcome Sanger Institute for the COVID-19 Genomics UK (COG-UK) Consortium | The Lighthouse Lab in Milton Keynes and Alex Alderton, Roberto Amato, Jeffrey Barrett, Sonia Goncalves, Ewan Harrison, David K. Jackson, Ian Johnston, Dominic Kwiatkowski, Cordelia Langford, John Sillitoe on behalf of the Wellcome Sanger Institute COVID-19 Surveillance Team |
| EPI_ISL_3222428 | Lighthouse Laboratory Plymouth | Wellcome Sanger Institute for the COVID-19 Genomics UK (COG-UK) Consortium | Lighthouse Laboratory Plymouth and Alex Alderton, Roberto Amato, Jeffrey Barrett, Sonia Goncalves, Ewan Harrison, David K. Jackson, Ian Johnston, Dominic Kwiatkowski, Cordelia Langford, John Sillitoe on behalf of the Wellcome Sanger Institute COVID-19 Surveillance Team |
| EPI_ISL_3222429 | Lighthouse Lab in Alderley Park | Wellcome Sanger Institute for the COVID-19 Genomics UK (COG-UK) Consortium | Jacquelyn Wynn, Mairead Hyland, The Lighthouse Lab in Alderley Park and Alex Alderton, Roberto Amato, Jeffrey Barrett, Sonia Goncalves, Ewan Harrison, David K. Jackson, Ian Johnston, Dominic Kwiatkowski, Cordelia Langford, John Sillitoe on behalf of the Wellcome Sanger Institute COVID-19 Surveillance Team |
| EPI_ISL_3222430 | Lighthouse Lab in Glasgow | Wellcome Sanger Institute for the COVID-19 Genomics UK (COG-UK) Consortium | Harper VanSteenhouse, Yumi Kasai, David Gray, Carol Clugston, Anna Dominiczak and Alex Alderton, Roberto Amato, Jeffrey Barrett, Sonia Goncalves, Ewan Harrison, David K. Jackson, Ian Johnston, Dominic Kwiatkowski, Cordelia Langford, John Sillitoe on behalf of the Wellcome Sanger Institute COVID-19 Surveillance Team |
| EPI_ISL_3222431 | Lighthouse Lab in Milton Keynes | Wellcome Sanger Institute for the COVID-19 Genomics UK (COG-UK) Consortium | The Lighthouse Lab in Milton Keynes and Alex Alderton, Roberto Amato, Jeffrey Barrett, Sonia Goncalves, Ewan Harrison, David K. Jackson, Ian Johnston, Dominic Kwiatkowski, Cordelia Langford, John Sillitoe on behalf of the Wellcome Sanger Institute COVID-19 Surveillance Team |
| EPI_ISL_3222432, EPI_ISL_3222433 | Lighthouse Lab in Glasgow | Wellcome Sanger Institute for the COVID-19 Genomics UK (COG-UK) Consortium | Harper VanSteenhouse, Yumi Kasai, David Gray, Carol Clugston, Anna Dominiczak and Alex Alderton, Roberto Amato, Jeffrey Barrett, Sonia Goncalves, Ewan Harrison, David K. Jackson, Ian Johnston, Dominic Kwiatkowski, Cordelia Langford, John Sillitoe on behalf of the Wellcome Sanger Institute COVID-19 Surveillance Team |
| EPI_ISL_3222434 | Lighthouse Lab in Milton Keynes | Wellcome Sanger Institute for the COVID-19 Genomics UK (COG-UK) Consortium | The Lighthouse Lab in Milton Keynes and Alex Alderton, Roberto Amato, Jeffrey Barrett, Sonia Goncalves, Ewan Harrison, David K. Jackson, Ian Johnston, Dominic Kwiatkowski, Cordelia Langford, John Sillitoe on behalf of the Wellcome Sanger Institute COVID-19 Surveillance Team |
| EPI_ISL_3222435 | Lighthouse Lab in Alderley Park | Wellcome Sanger Institute for the COVID-19 Genomics UK (COG-UK) Consortium | Jacquelyn Wynn, Mairead Hyland, The Lighthouse Lab in Alderley Park and Alex Alderton, Roberto Amato, Jeffrey Barrett, Sonia Goncalves, Ewan Harrison, David K. Jackson, Ian Johnston, Dominic Kwiatkowski, Cordelia Langford, John Sillitoe on behalf of the Wellcome Sanger Institute COVID-19 Surveillance Team |
| EPI_ISL_3222436 | Lighthouse Laboratory Plymouth | Wellcome Sanger Institute for the COVID-19 Genomics UK (COG-UK) Consortium | Lighthouse Laboratory Plymouth and Alex Alderton, Roberto Amato, Jeffrey Barrett, Sonia Goncalves, Ewan Harrison, David K. Jackson, Ian Johnston, Dominic Kwiatkowski, Cordelia Langford, John Sillitoe on behalf of the Wellcome Sanger Institute COVID-19 Surveillance Team |
| EPI_ISL_3222437 | Lighthouse Lab in Glasgow | Wellcome Sanger Institute for the COVID-19 Genomics UK (COG-UK) Consortium | Harper VanSteenhouse, Yumi Kasai, David Gray, Carol Clugston, Anna Dominiczak and Alex Alderton, Roberto Amato, Jeffrey Barrett, Sonia Goncalves, Ewan Harrison, David K. Jackson, Ian Johnston, Dominic Kwiatkowski, Cordelia Langford, John Sillitoe on behalf of the Wellcome Sanger Institute COVID-19 Surveillance Team |
| EPI_ISL_3222438 | Lighthouse Lab in Milton Keynes | Wellcome Sanger Institute for the COVID-19 Genomics UK (COG-UK) Consortium | The Lighthouse Lab in Milton Keynes and Alex Alderton, Roberto Amato, Jeffrey Barrett, Sonia Goncalves, Ewan Harrison, David K. Jackson, Ian Johnston, Dominic Kwiatkowski, Cordelia Langford, John Sillitoe on behalf of the Wellcome Sanger Institute COVID-19 Surveillance Team |
| EPI_ISL_3222439, EPI_ISL_3222440, EPI_ISL_3222441 | Lighthouse Lab in Glasgow | Wellcome Sanger Institute for the COVID-19 Genomics UK (COG-UK) Consortium | Harper VanSteenhouse, Yumi Kasai, David Gray, Carol Clugston, Anna Dominiczak and Alex Alderton, Roberto Amato, Jeffrey Barrett, Sonia Goncalves, Ewan Harrison, David K. Jackson, Ian Johnston, Dominic Kwiatkowski, Cordelia Langford, John Sillitoe on behalf of the Wellcome Sanger Institute COVID-19 Surveillance Team |
| EPI_ISL_3222442 | Lighthouse Laboratory Plymouth | Wellcome Sanger Institute for the COVID-19 Genomics UK (COG-UK) Consortium | Lighthouse Laboratory Plymouth and Alex Alderton, Roberto Amato, Jeffrey Barrett, Sonia Goncalves, Ewan Harrison, David K. Jackson, Ian Johnston, Dominic Kwiatkowski, Cordelia Langford, John Sillitoe on behalf of the Wellcome Sanger Institute COVID-19 Surveillance Team |
| EPI_ISL_3222443 | Lighthouse Lab in Glasgow | Wellcome Sanger Institute for the COVID-19 Genomics UK (COG-UK) Consortium | Harper VanSteenhouse, Yumi Kasai, David Gray, Carol Clugston, Anna Dominiczak and Alex Alderton, Roberto Amato, Jeffrey Barrett, Sonia Goncalves, Ewan Harrison, David K. Jackson, Ian Johnston, Dominic Kwiatkowski, Cordelia Langford, John Sillitoe on behalf of the Wellcome Sanger Institute COVID-19 Surveillance Team |
| EPI_ISL_3222444, EPI_ISL_3222445, EPI_ISL_3222446 | Lighthouse Lab in Alderley Park | Wellcome Sanger Institute for the COVID-19 Genomics UK (COG-UK) Consortium | Jacquelyn Wynn, Mairead Hyland, The Lighthouse Lab in Alderley Park and Alex Alderton, Roberto Amato, Jeffrey Barrett, Sonia Goncalves, Ewan Harrison, David K. Jackson, Ian Johnston, Dominic Kwiatkowski, Cordelia Langford, John Sillitoe on behalf of the Wellcome Sanger Institute COVID-19 Surveillance Team |
| EPI_ISL_3222447 | Lighthouse Laboratory Plymouth | Wellcome Sanger Institute for the COVID-19 Genomics UK (COG-UK) Consortium | Lighthouse Laboratory Plymouth and Alex Alderton, Roberto Amato, Jeffrey Barrett, Sonia Goncalves, Ewan Harrison, David K. Jackson, Ian Johnston, Dominic Kwiatkowski, Cordelia Langford, John Sillitoe on behalf of the Wellcome Sanger Institute COVID-19 Surveillance Team |
| EPI_ISL_3222448 | Lighthouse Lab in Glasgow | Wellcome Sanger Institute for the COVID-19 Genomics UK (COG-UK) Consortium | Harper VanSteenhouse, Yumi Kasai, David Gray, Carol Clugston, Anna Dominiczak and Alex Alderton, Roberto Amato, Jeffrey Barrett, Sonia Goncalves, Ewan Harrison, David K. Jackson, Ian Johnston, Dominic Kwiatkowski, Cordelia Langford, John Sillitoe on behalf of the Wellcome Sanger Institute COVID-19 Surveillance Team |
| EPI_ISL_3222449 | Lighthouse Laboratory Plymouth | Wellcome Sanger Institute for the COVID-19 Genomics UK (COG-UK) Consortium | Lighthouse Laboratory Plymouth and Alex Alderton, Roberto Amato, Jeffrey Barrett, Sonia Goncalves, Ewan Harrison, David K. Jackson, Ian Johnston, Dominic Kwiatkowski, Cordelia Langford, John Sillitoe on behalf of the Wellcome Sanger Institute COVID-19 Surveillance Team |
| EPI_ISL_3222450, EPI_ISL_3222451, EPI_ISL_3222452 | Lighthouse Lab in Alderley Park | Wellcome Sanger Institute for the COVID-19 Genomics UK (COG-UK) Consortium | Jacquelyn Wynn, Mairead Hyland, The Lighthouse Lab in Alderley Park and Alex Alderton, Roberto Amato, Jeffrey Barrett, Sonia Goncalves, Ewan Harrison, David K. Jackson, Ian Johnston, Dominic Kwiatkowski, Cordelia Langford, John Sillitoe on behalf of the Wellcome Sanger Institute COVID-19 Surveillance Team |
| EPI_ISL_3222453 | Lighthouse Laboratory Plymouth | Wellcome Sanger Institute for the COVID-19 Genomics UK (COG-UK) Consortium | Lighthouse Laboratory Plymouth and Alex Alderton, Roberto Amato, Jeffrey Barrett, Sonia Goncalves, Ewan Harrison, David K. Jackson, Ian Johnston, Dominic Kwiatkowski, Cordelia Langford, John Sillitoe on behalf of the Wellcome Sanger Institute COVID-19 Surveillance Team |
| EPI_ISL_3222454 | Lighthouse Lab in Milton Keynes | Wellcome Sanger Institute for the COVID-19 Genomics UK (COG-UK) Consortium | The Lighthouse Lab in Milton Keynes and Alex Alderton, Roberto Amato, Jeffrey Barrett, Sonia Goncalves, Ewan Harrison, David K. Jackson, Ian Johnston, Dominic Kwiatkowski, Cordelia Langford, John Sillitoe on behalf of the Wellcome Sanger Institute COVID-19 Surveillance Team |
| EPI_ISL_3222455 | Lighthouse Lab in Alderley Park | Wellcome Sanger Institute for the COVID-19 Genomics UK (COG-UK) Consortium | Jacquelyn Wynn, Mairead Hyland, The Lighthouse Lab in Alderley Park and Alex Alderton, Roberto Amato, Jeffrey Barrett, Sonia Goncalves, Ewan Harrison, David K. Jackson, Ian Johnston, Dominic Kwiatkowski, Cordelia Langford, John Sillitoe on behalf of the Wellcome Sanger Institute COVID-19 Surveillance Team |
| EPI_ISL_3222456 | Lighthouse Laboratory Plymouth | Wellcome Sanger Institute for the COVID-19 Genomics UK (COG-UK) Consortium | Lighthouse Laboratory Plymouth and Alex Alderton, Roberto Amato, Jeffrey Barrett, Sonia Goncalves, Ewan Harrison, David K. Jackson, Ian Johnston, Dominic Kwiatkowski, Cordelia Langford, John Sillitoe on behalf of the Wellcome Sanger Institute COVID-19 Surveillance Team |
| EPI_ISL_3222457 | Lighthouse Lab in Glasgow | Wellcome Sanger Institute for the COVID-19 Genomics UK (COG-UK) Consortium | Harper VanSteenhouse, Yumi Kasai, David Gray, Carol Clugston, Anna Dominiczak and Alex Alderton, Roberto Amato, Jeffrey Barrett, Sonia Goncalves, Ewan Harrison, David K. Jackson, Ian Johnston, Dominic Kwiatkowski, Cordelia Langford, John Sillitoe on behalf of the Wellcome Sanger Institute COVID-19 Surveillance Team |
| EPI_ISL_3222458 | Lighthouse Lab in Alderley Park | Wellcome Sanger Institute for the COVID-19 Genomics UK (COG-UK) Consortium | Jacquelyn Wynn, Mairead Hyland, The Lighthouse Lab in Alderley Park and Alex Alderton, Roberto Amato, Jeffrey Barrett, Sonia Goncalves, Ewan Harrison, David K. Jackson, Ian Johnston, Dominic Kwiatkowski, Cordelia Langford, John Sillitoe on behalf of the Wellcome Sanger Institute COVID-19 Surveillance Team |
| EPI_ISL_3222459, EPI_ISL_3222460 | Lighthouse Laboratory Plymouth | Wellcome Sanger Institute for the COVID-19 Genomics UK (COG-UK) Consortium | Lighthouse Laboratory Plymouth and Alex Alderton, Roberto Amato, Jeffrey Barrett, Sonia Goncalves, Ewan Harrison, David K. Jackson, Ian Johnston, Dominic Kwiatkowski, Cordelia Langford, John Sillitoe on behalf of the Wellcome Sanger Institute COVID-19 Surveillance Team |
| EPI_ISL_3222461 | Lighthouse Lab in Milton Keynes | Wellcome Sanger Institute for the COVID-19 Genomics UK (COG-UK) Consortium | The Lighthouse Lab in Milton Keynes and Alex Alderton, Roberto Amato, Jeffrey Barrett, Sonia Goncalves, Ewan Harrison, David K. Jackson, Ian Johnston, Dominic Kwiatkowski, Cordelia Langford, John Sillitoe on behalf of the Wellcome Sanger Institute COVID-19 Surveillance Team |
| EPI_ISL_3222462, EPI_ISL_3222463 | Lighthouse Lab in Alderley Park | Wellcome Sanger Institute for the COVID-19 Genomics UK (COG-UK) Consortium | Jacquelyn Wynn, Mairead Hyland, The Lighthouse Lab in Alderley Park and Alex Alderton, Roberto Amato, Jeffrey Barrett, Sonia Goncalves, Ewan Harrison, David K. Jackson, Ian Johnston, Dominic Kwiatkowski, Cordelia Langford, John Sillitoe on behalf of the Wellcome Sanger Institute COVID-19 Surveillance Team |
| EPI_ISL_3222464 | Lighthouse Lab in Milton Keynes | Wellcome Sanger Institute for the COVID-19 Genomics UK (COG-UK) Consortium | The Lighthouse Lab in Milton Keynes and Alex Alderton, Roberto Amato, Jeffrey Barrett, Sonia Goncalves, Ewan Harrison, David K. Jackson, Ian Johnston, Dominic Kwiatkowski, Cordelia Langford, John Sillitoe on behalf of the Wellcome Sanger Institute COVID-19 Surveillance Team |
| EPI_ISL_3222465 | Lighthouse Lab in Alderley Park | Wellcome Sanger Institute for the COVID-19 Genomics UK (COG-UK) Consortium | Jacquelyn Wynn, Mairead Hyland, The Lighthouse Lab in Alderley Park and Alex Alderton, Roberto Amato, Jeffrey Barrett, Sonia Goncalves, Ewan Harrison, David K. Jackson, Ian Johnston, Dominic Kwiatkowski, Cordelia Langford, John Sillitoe on behalf of the Wellcome Sanger Institute COVID-19 Surveillance Team |
| EPI_ISL_3222466, EPI_ISL_3222467, EPI_ISL_3222468 | Lighthouse Laboratory Plymouth | Wellcome Sanger Institute for the COVID-19 Genomics UK (COG-UK) Consortium | Lighthouse Laboratory Plymouth and Alex Alderton, Roberto Amato, Jeffrey Barrett, Sonia Goncalves, Ewan Harrison, David K. Jackson, Ian Johnston, Dominic Kwiatkowski, Cordelia Langford, John Sillitoe on behalf of the Wellcome Sanger Institute COVID-19 Surveillance Team |
| EPI_ISL_3222469 | Lighthouse Lab in Milton Keynes | Wellcome Sanger Institute for the COVID-19 Genomics UK (COG-UK) Consortium | The Lighthouse Lab in Milton Keynes and Alex Alderton, Roberto Amato, Jeffrey Barrett, Sonia Goncalves, Ewan Harrison, David K. Jackson, Ian Johnston, Dominic Kwiatkowski, Cordelia Langford, John Sillitoe on behalf of the Wellcome Sanger Institute COVID-19 Surveillance Team |
| EPI_ISL_3222470 | Lighthouse Lab in Glasgow | Wellcome Sanger Institute for the COVID-19 Genomics UK (COG-UK) Consortium | Harper VanSteenhouse, Yumi Kasai, David Gray, Carol Clugston, Anna Dominiczak and Alex Alderton, Roberto Amato, Jeffrey Barrett, Sonia Goncalves, Ewan Harrison, David K. Jackson, Ian Johnston, Dominic Kwiatkowski, Cordelia Langford, John Sillitoe on behalf of the Wellcome Sanger Institute COVID-19 Surveillance Team |
| EPI_ISL_3222471 | Lighthouse Lab in Milton Keynes | Wellcome Sanger Institute for the COVID-19 Genomics UK (COG-UK) Consortium | The Lighthouse Lab in Milton Keynes and Alex Alderton, Roberto Amato, Jeffrey Barrett, Sonia Goncalves, Ewan Harrison, David K. Jackson, Ian Johnston, Dominic Kwiatkowski, Cordelia Langford, John Sillitoe on behalf of the Wellcome Sanger Institute COVID-19 Surveillance Team |
| EPI_ISL_3222472 | Lighthouse Lab in Alderley Park | Wellcome Sanger Institute for the COVID-19 Genomics UK (COG-UK) Consortium | Jacquelyn Wynn, Mairead Hyland, The Lighthouse Lab in Alderley Park and Alex Alderton, Roberto Amato, Jeffrey Barrett, Sonia Goncalves, Ewan Harrison, David K. Jackson, Ian Johnston, Dominic Kwiatkowski, Cordelia Langford, John Sillitoe on behalf of the Wellcome Sanger Institute COVID-19 Surveillance Team |
| EPI_ISL_3222473 | Lighthouse Lab in Milton Keynes | Wellcome Sanger Institute for the COVID-19 Genomics UK (COG-UK) Consortium | The Lighthouse Lab in Milton Keynes and Alex Alderton, Roberto Amato, Jeffrey Barrett, Sonia Goncalves, Ewan Harrison, David K. Jackson, Ian Johnston, Dominic Kwiatkowski, Cordelia Langford, John Sillitoe on behalf of the Wellcome Sanger Institute COVID-19 Surveillance Team |

| | | | |
|---|---|---|---|
| EPI_ISL_3222474, EPI_ISL_3222475 | Lighthouse Laboratory Plymouth | Wellcome Sanger Institute for the COVID-19 Genomics UK (COG-UK) Consortium | Lighthouse Laboratory Plymouth and Alex Alderton, Roberto Amato, Jeffrey Barrett, Sonia Goncalves, Ewan Harrison, David K. Jackson, Ian Johnston, Dominic Kwiatkowski, Cordelia Langford, John Sillitoe on behalf of the Wellcome Sanger Institute COVID-19 Surveillance Team |
| EPI_ISL_3222476 | Lighthouse Lab in Milton Keynes | Wellcome Sanger Institute for the COVID-19 Genomics UK (COG-UK) Consortium | The Lighthouse Lab in Milton Keynes and Alex Alderton, Roberto Amato, Jeffrey Barrett, Sonia Goncalves, Ewan Harrison, David K. Jackson, Ian Johnston, Dominic Kwiatkowski, Cordelia Langford, John Sillitoe on behalf of the Wellcome Sanger Institute COVID-19 Surveillance Team |
| EPI_ISL_3222477 | Lighthouse Laboratory Plymouth | Wellcome Sanger Institute for the COVID-19 Genomics UK (COG-UK) Consortium | Lighthouse Laboratory Plymouth and Alex Alderton, Roberto Amato, Jeffrey Barrett, Sonia Goncalves, Ewan Harrison, David K. Jackson, Ian Johnston, Dominic Kwiatkowski, Cordelia Langford, John Sillitoe on behalf of the Wellcome Sanger Institute COVID-19 Surveillance Team |
| EPI_ISL_3222478 | Lighthouse Lab in Glasgow | Wellcome Sanger Institute for the COVID-19 Genomics UK (COG-UK) Consortium | Harper VanSteenhouse, Yumi Kasai, David Gray, Carol Clugston, Anna Dominiczak and Alex Alderton, Roberto Amato, Jeffrey Barrett, Sonia Goncalves, Ewan Harrison, David K. Jackson, Ian Johnston, Dominic Kwiatkowski, Cordelia Langford, John Sillitoe on behalf of the Wellcome Sanger Institute COVID-19 Surveillance Team |
| EPI_ISL_3222479 | Lighthouse Lab in Milton Keynes | Wellcome Sanger Institute for the COVID-19 Genomics UK (COG-UK) Consortium | The Lighthouse Lab in Milton Keynes and Alex Alderton, Roberto Amato, Jeffrey Barrett, Sonia Goncalves, Ewan Harrison, David K. Jackson, Ian Johnston, Dominic Kwiatkowski, Cordelia Langford, John Sillitoe on behalf of the Wellcome Sanger Institute COVID-19 Surveillance Team |
| EPI_ISL_3222480 | Lighthouse Lab in Glasgow | Wellcome Sanger Institute for the COVID-19 Genomics UK (COG-UK) Consortium | Harper VanSteenhouse, Yumi Kasai, David Gray, Carol Clugston, Anna Dominiczak and Alex Alderton, Roberto Amato, Jeffrey Barrett, Sonia Goncalves, Ewan Harrison, David K. Jackson, Ian Johnston, Dominic Kwiatkowski, Cordelia Langford, John Sillitoe on behalf of the Wellcome Sanger Institute COVID-19 Surveillance Team |
| EPI_ISL_3222481, EPI_ISL_3222482 | Lighthouse Lab in Milton Keynes | Wellcome Sanger Institute for the COVID-19 Genomics UK (COG-UK) Consortium | The Lighthouse Lab in Milton Keynes and Alex Alderton, Roberto Amato, Jeffrey Barrett, Sonia Goncalves, Ewan Harrison, David K. Jackson, Ian Johnston, Dominic Kwiatkowski, Cordelia Langford, John Sillitoe on behalf of the Wellcome Sanger Institute COVID-19 Surveillance Team |
| EPI_ISL_3222483 | Lighthouse Lab in Alderley Park | Wellcome Sanger Institute for the COVID-19 Genomics UK (COG-UK) Consortium | Jacquelyn Wynn, Mairead Hyland, The Lighthouse Lab in Alderley Park and Alex Alderton, Roberto Amato, Jeffrey Barrett, Sonia Goncalves, Ewan Harrison, David K. Jackson, Ian Johnston, Dominic Kwiatkowski, Cordelia Langford, John Sillitoe on behalf of the Wellcome Sanger Institute COVID-19 Surveillance Team |
| EPI_ISL_3222484 | Lighthouse Lab in Glasgow | Wellcome Sanger Institute for the COVID-19 Genomics UK (COG-UK) Consortium | Harper VanSteenhouse, Yumi Kasai, David Gray, Carol Clugston, Anna Dominiczak and Alex Alderton, Roberto Amato, Jeffrey Barrett, Sonia Goncalves, Ewan Harrison, David K. Jackson, Ian Johnston, Dominic Kwiatkowski, Cordelia Langford, John Sillitoe on behalf of the Wellcome Sanger Institute COVID-19 Surveillance Team |
| EPI_ISL_3222485 | Lighthouse Lab in Milton Keynes | Wellcome Sanger Institute for the COVID-19 Genomics UK (COG-UK) Consortium | The Lighthouse Lab in Milton Keynes and Alex Alderton, Roberto Amato, Jeffrey Barrett, Sonia Goncalves, Ewan Harrison, David K. Jackson, Ian Johnston, Dominic Kwiatkowski, Cordelia Langford, John Sillitoe on behalf of the Wellcome Sanger Institute COVID-19 Surveillance Team |
| EPI_ISL_3222486 | Lighthouse Lab in Glasgow | Wellcome Sanger Institute for the COVID-19 Genomics UK (COG-UK) Consortium | Harper VanSteenhouse, Yumi Kasai, David Gray, Carol Clugston, Anna Dominiczak and Alex Alderton, Roberto Amato, Jeffrey Barrett, Sonia Goncalves, Ewan Harrison, David K. Jackson, Ian Johnston, Dominic Kwiatkowski, Cordelia Langford, John Sillitoe on behalf of the Wellcome Sanger Institute COVID-19 Surveillance Team |
| EPI_ISL_3222487 | Lighthouse Lab in Alderley Park | Wellcome Sanger Institute for the COVID-19 Genomics UK (COG-UK) Consortium | Jacquelyn Wynn, Mairead Hyland, The Lighthouse Lab in Alderley Park and Alex Alderton, Roberto Amato, Jeffrey Barrett, Sonia Goncalves, Ewan Harrison, David K. Jackson, Ian Johnston, Dominic Kwiatkowski, Cordelia Langford, John Sillitoe on behalf of the Wellcome Sanger Institute COVID-19 Surveillance Team |
| EPI_ISL_3222488, EPI_ISL_3222489 | Lighthouse Laboratory Plymouth | Wellcome Sanger Institute for the COVID-19 Genomics UK (COG-UK) Consortium | Lighthouse Laboratory Plymouth and Alex Alderton, Roberto Amato, Jeffrey Barrett, Sonia Goncalves, Ewan Harrison, David K. Jackson, Ian Johnston, Dominic Kwiatkowski, Cordelia Langford, John Sillitoe on behalf of the Wellcome Sanger Institute COVID-19 Surveillance Team |
| EPI_ISL_3222490 | Lighthouse Lab in Alderley Park | Wellcome Sanger Institute for the COVID-19 Genomics UK (COG-UK) Consortium | Jacquelyn Wynn, Mairead Hyland, The Lighthouse Lab in Alderley Park and Alex Alderton, Roberto Amato, Jeffrey Barrett, Sonia Goncalves, Ewan Harrison, David K. Jackson, Ian Johnston, Dominic Kwiatkowski, Cordelia Langford, John Sillitoe on behalf of the Wellcome Sanger Institute COVID-19 Surveillance Team |
| EPI_ISL_3222491, EPI_ISL_3222492 | Lighthouse Lab in Glasgow | Wellcome Sanger Institute for the COVID-19 Genomics UK (COG-UK) Consortium | Harper VanSteenhouse, Yumi Kasai, David Gray, Carol Clugston, Anna Dominiczak and Alex Alderton, Roberto Amato, Jeffrey Barrett, Sonia Goncalves, Ewan Harrison, David K. Jackson, Ian Johnston, Dominic Kwiatkowski, Cordelia Langford, John Sillitoe on behalf of the Wellcome Sanger Institute COVID-19 Surveillance Team |
| EPI_ISL_3222493 | Lighthouse Lab in Milton Keynes | Wellcome Sanger Institute for the COVID-19 Genomics UK (COG-UK) Consortium | The Lighthouse Lab in Milton Keynes and Alex Alderton, Roberto Amato, Jeffrey Barrett, Sonia Goncalves, Ewan Harrison, David K. Jackson, Ian Johnston, Dominic Kwiatkowski, Cordelia Langford, John Sillitoe on behalf of the Wellcome Sanger Institute COVID-19 Surveillance Team |
| EPI_ISL_3222494, EPI_ISL_3222495, EPI_ISL_3222496, EPI_ISL_3222497 | Lighthouse Lab in Alderley Park | Wellcome Sanger Institute for the COVID-19 Genomics UK (COG-UK) Consortium | Jacquelyn Wynn, Mairead Hyland, The Lighthouse Lab in Alderley Park and Alex Alderton, Roberto Amato, Jeffrey Barrett, Sonia Goncalves, Ewan Harrison, David K. Jackson, Ian Johnston, Dominic Kwiatkowski, Cordelia Langford, John Sillitoe on behalf of the Wellcome Sanger Institute COVID-19 Surveillance Team |
| EPI_ISL_3222498, EPI_ISL_3222499, EPI_ISL_3222500, EPI_ISL_3222501 | Lighthouse Lab in Milton Keynes | Wellcome Sanger Institute for the COVID-19 Genomics UK (COG-UK) Consortium | The Lighthouse Lab in Milton Keynes and Alex Alderton, Roberto Amato, Jeffrey Barrett, Sonia Goncalves, Ewan Harrison, David K. Jackson, Ian Johnston, Dominic Kwiatkowski, Cordelia Langford, John Sillitoe on behalf of the Wellcome Sanger Institute COVID-19 Surveillance Team |
| EPI_ISL_3222502 | Lighthouse Lab in Alderley Park | Wellcome Sanger Institute for the COVID-19 Genomics UK (COG-UK) Consortium | Jacquelyn Wynn, Mairead Hyland, The Lighthouse Lab in Alderley Park and Alex Alderton, Roberto Amato, Jeffrey Barrett, Sonia Goncalves, Ewan Harrison, David K. Jackson, Ian Johnston, Dominic Kwiatkowski, Cordelia Langford, John Sillitoe on behalf of the Wellcome Sanger Institute COVID-19 Surveillance Team |
| EPI_ISL_3222503 | Lighthouse Lab in Glasgow | Wellcome Sanger Institute for the COVID-19 Genomics UK (COG-UK) Consortium | Harper VanSteenhouse, Yumi Kasai, David Gray, Carol Clugston, Anna Dominiczak and Alex Alderton, Roberto Amato, Jeffrey Barrett, Sonia Goncalves, Ewan Harrison, David K. Jackson, Ian Johnston, Dominic Kwiatkowski, Cordelia Langford, John Sillitoe on behalf of the Wellcome Sanger Institute COVID-19 Surveillance Team |
| EPI_ISL_3222504 | Lighthouse Lab in Milton Keynes | Wellcome Sanger Institute for the COVID-19 Genomics UK (COG-UK) Consortium | The Lighthouse Lab in Milton Keynes and Alex Alderton, Roberto Amato, Jeffrey Barrett, Sonia Goncalves, Ewan Harrison, David K. Jackson, Ian Johnston, Dominic Kwiatkowski, Cordelia Langford, John Sillitoe on behalf of the Wellcome Sanger Institute COVID-19 Surveillance Team |
| EPI_ISL_3222505, EPI_ISL_3222506, EPI_ISL_3222507 | Lighthouse Laboratory Plymouth | Wellcome Sanger Institute for the COVID-19 Genomics UK (COG-UK) Consortium | Lighthouse Laboratory Plymouth and Alex Alderton, Roberto Amato, Jeffrey Barrett, Sonia Goncalves, Ewan Harrison, David K. Jackson, Ian Johnston, Dominic Kwiatkowski, Cordelia Langford, John Sillitoe on behalf of the Wellcome Sanger Institute COVID-19 Surveillance Team |
| EPI_ISL_3222508, EPI_ISL_3222509 | Lighthouse Lab in Alderley Park | Wellcome Sanger Institute for the COVID-19 Genomics UK (COG-UK) Consortium | Jacquelyn Wynn, Mairead Hyland, The Lighthouse Lab in Alderley Park and Alex Alderton, Roberto Amato, Jeffrey Barrett, Sonia Goncalves, Ewan Harrison, David K. Jackson, Ian Johnston, Dominic Kwiatkowski, Cordelia Langford, John Sillitoe on behalf of the Wellcome Sanger Institute COVID-19 Surveillance Team |
| EPI_ISL_3222510 | Lighthouse Lab in Glasgow | Wellcome Sanger Institute for the COVID-19 Genomics UK (COG-UK) Consortium | Harper VanSteenhouse, Yumi Kasai, David Gray, Carol Clugston, Anna Dominiczak and Alex Alderton, Roberto Amato, Jeffrey Barrett, Sonia Goncalves, Ewan Harrison, David K. Jackson, Ian Johnston, Dominic Kwiatkowski, Cordelia Langford, John Sillitoe on behalf of the Wellcome Sanger Institute COVID-19 Surveillance Team |
| EPI_ISL_3222511 | Lighthouse Laboratory Plymouth | Wellcome Sanger Institute for the COVID-19 Genomics UK (COG-UK) Consortium | Lighthouse Laboratory Plymouth and Alex Alderton, Roberto Amato, Jeffrey Barrett, Sonia Goncalves, Ewan Harrison, David K. Jackson, Ian Johnston, Dominic Kwiatkowski, Cordelia Langford, John Sillitoe on behalf of the Wellcome Sanger Institute COVID-19 Surveillance Team |
| EPI_ISL_3222512, EPI_ISL_3222513 | Lighthouse Lab in Glasgow | Wellcome Sanger Institute for the COVID-19 Genomics UK (COG-UK) Consortium | Harper VanSteenhouse, Yumi Kasai, David Gray, Carol Clugston, Anna Dominiczak and Alex Alderton, Roberto Amato, Jeffrey Barrett, Sonia Goncalves, Ewan Harrison, David K. Jackson, Ian Johnston, Dominic Kwiatkowski, Cordelia Langford, John Sillitoe on behalf of the Wellcome Sanger Institute COVID-19 Surveillance Team |
| EPI_ISL_3222514, EPI_ISL_3222515 | Lighthouse Lab in Milton Keynes | Wellcome Sanger Institute for the COVID-19 Genomics UK (COG-UK) Consortium | The Lighthouse Lab in Milton Keynes and Alex Alderton, Roberto Amato, Jeffrey Barrett, Sonia Goncalves, Ewan Harrison, David K. Jackson, Ian Johnston, Dominic Kwiatkowski, Cordelia Langford, John Sillitoe on behalf of the Wellcome Sanger Institute COVID-19 Surveillance Team |
| EPI_ISL_3222516 | Lighthouse Lab in Glasgow | Wellcome Sanger Institute for the COVID-19 Genomics UK (COG-UK) Consortium | Harper VanSteenhouse, Yumi Kasai, David Gray, Carol Clugston, Anna Dominiczak and Alex Alderton, Roberto Amato, Jeffrey Barrett, Sonia Goncalves, Ewan Harrison, David K. Jackson, Ian Johnston, Dominic Kwiatkowski, Cordelia Langford, John Sillitoe on behalf of the Wellcome Sanger Institute COVID-19 Surveillance Team |
| EPI_ISL_3222517 | Lighthouse Laboratory Plymouth | Wellcome Sanger Institute for the COVID-19 Genomics UK (COG-UK) Consortium | Lighthouse Laboratory Plymouth and Alex Alderton, Roberto Amato, Jeffrey Barrett, Sonia Goncalves, Ewan Harrison, David K. Jackson, Ian Johnston, Dominic Kwiatkowski, Cordelia Langford, John Sillitoe on behalf of the Wellcome Sanger Institute COVID-19 Surveillance Team |
| EPI_ISL_3222518 | Lighthouse Lab in Milton Keynes | Wellcome Sanger Institute for the COVID-19 Genomics UK (COG-UK) Consortium | The Lighthouse Lab in Milton Keynes and Alex Alderton, Roberto Amato, Jeffrey Barrett, Sonia Goncalves, Ewan Harrison, David K. Jackson, Ian Johnston, Dominic Kwiatkowski, Cordelia Langford, John Sillitoe on behalf of the Wellcome Sanger Institute COVID-19 Surveillance Team |
| EPI_ISL_3222519, EPI_ISL_3222520 | Lighthouse Lab in Glasgow | Wellcome Sanger Institute for the COVID-19 Genomics UK (COG-UK) Consortium | Harper VanSteenhouse, Yumi Kasai, David Gray, Carol Clugston, Anna Dominiczak and Alex Alderton, Roberto Amato, Jeffrey Barrett, Sonia Goncalves, Ewan Harrison, David K. Jackson, Ian Johnston, Dominic Kwiatkowski, Cordelia Langford, John Sillitoe on behalf of the Wellcome Sanger Institute COVID-19 Surveillance Team |
| EPI_ISL_3222521, EPI_ISL_3222522 | Lighthouse Lab in Alderley Park | Wellcome Sanger Institute for the COVID-19 Genomics UK (COG-UK) Consortium | Jacquelyn Wynn, Mairead Hyland, The Lighthouse Lab in Alderley Park and Alex Alderton, Roberto Amato, Jeffrey Barrett, Sonia Goncalves, Ewan Harrison, David K. Jackson, Ian Johnston, Dominic Kwiatkowski, Cordelia Langford, John Sillitoe on behalf of the Wellcome Sanger Institute COVID-19 Surveillance Team |
| EPI_ISL_3222523, EPI_ISL_3222524 | Lighthouse Laboratory Plymouth | Wellcome Sanger Institute for the COVID-19 Genomics UK (COG-UK) Consortium | Lighthouse Laboratory Plymouth and Alex Alderton, Roberto Amato, Jeffrey Barrett, Sonia Goncalves, Ewan Harrison, David K. Jackson, Ian Johnston, Dominic Kwiatkowski, Cordelia Langford, John Sillitoe on behalf of the Wellcome Sanger Institute COVID-19 Surveillance Team |
| EPI_ISL_3222525 | Lighthouse Lab in Milton Keynes | Wellcome Sanger Institute for the COVID-19 Genomics UK (COG-UK) Consortium | The Lighthouse Lab in Milton Keynes and Alex Alderton, Roberto Amato, Jeffrey Barrett, Sonia Goncalves, Ewan Harrison, David K. Jackson, Ian Johnston, Dominic Kwiatkowski, Cordelia Langford, John Sillitoe on behalf of the Wellcome Sanger Institute COVID-19 Surveillance Team |
| EPI_ISL_3222526 | Lighthouse Laboratory Plymouth | Wellcome Sanger Institute for the COVID-19 Genomics UK (COG-UK) Consortium | Lighthouse Laboratory Plymouth and Alex Alderton, Roberto Amato, Jeffrey Barrett, Sonia Goncalves, Ewan Harrison, David K. Jackson, Ian Johnston, Dominic Kwiatkowski, Cordelia Langford, John Sillitoe on behalf of the Wellcome Sanger Institute COVID-19 Surveillance Team |
| EPI_ISL_3222527 | Lighthouse Lab in Glasgow | Wellcome Sanger Institute for the COVID-19 Genomics UK (COG-UK) Consortium | Harper VanSteenhouse, Yumi Kasai, David Gray, Carol Clugston, Anna Dominiczak and Alex Alderton, Roberto Amato, Jeffrey Barrett, Sonia Goncalves, Ewan Harrison, David K. Jackson, Ian Johnston, Dominic Kwiatkowski, Cordelia Langford, John Sillitoe on behalf of the Wellcome Sanger Institute COVID-19 Surveillance Team |
| EPI_ISL_3222528 | Lighthouse Lab in Alderley Park | Wellcome Sanger Institute for the COVID-19 Genomics UK (COG-UK) Consortium | Jacquelyn Wynn, Mairead Hyland, The Lighthouse Lab in Alderley Park and Alex Alderton, Roberto Amato, Jeffrey Barrett, Sonia Goncalves, Ewan Harrison, David K. Jackson, Ian Johnston, Dominic Kwiatkowski, Cordelia Langford, John Sillitoe on behalf of the Wellcome Sanger Institute COVID-19 Surveillance Team |
| EPI_ISL_3222529 | Lighthouse Lab in Milton Keynes | Wellcome Sanger Institute for the COVID-19 Genomics UK (COG-UK) Consortium | The Lighthouse Lab in Milton Keynes and Alex Alderton, Roberto Amato, Jeffrey Barrett, Sonia Goncalves, Ewan Harrison, David K. Jackson, Ian Johnston, Dominic Kwiatkowski, Cordelia Langford, John Sillitoe on behalf of the Wellcome Sanger Institute COVID-19 Surveillance Team |
| EPI_ISL_3222530 | Lighthouse Lab in Alderley Park | Wellcome Sanger Institute for the COVID-19 Genomics UK (COG-UK) Consortium | Jacquelyn Wynn, Mairead Hyland, The Lighthouse Lab in Alderley Park and Alex Alderton, Roberto Amato, Jeffrey Barrett, Sonia Goncalves, Ewan Harrison, David K. Jackson, Ian Johnston, Dominic Kwiatkowski, Cordelia Langford, John Sillitoe on behalf of the Wellcome Sanger Institute COVID-19 Surveillance Team |
| EPI_ISL_3222531, EPI_ISL_3222532, EPI_ISL_3222533, EPI_ISL_3222534 | Lighthouse Lab in Milton Keynes | Wellcome Sanger Institute for the COVID-19 Genomics UK (COG-UK) Consortium | The Lighthouse Lab in Milton Keynes and Alex Alderton, Roberto Amato, Jeffrey Barrett, Sonia Goncalves, Ewan Harrison, David K. Jackson, Ian Johnston, Dominic Kwiatkowski, Cordelia Langford, John Sillitoe on behalf of the Wellcome Sanger Institute COVID-19 Surveillance Team |
| EPI_ISL_3222535 | Lighthouse Laboratory Plymouth | Wellcome Sanger Institute for the COVID-19 Genomics UK (COG-UK) Consortium | Lighthouse Laboratory Plymouth and Alex Alderton, Roberto Amato, Jeffrey Barrett, Sonia Goncalves, Ewan Harrison, David K. Jackson, Ian Johnston, Dominic Kwiatkowski, Cordelia Langford, John Sillitoe on behalf of the Wellcome Sanger Institute COVID-19 Surveillance Team |
| EPI_ISL_3222536, EPI_ISL_3222537 | Lighthouse Lab in Milton Keynes | Wellcome Sanger Institute for the COVID-19 Genomics UK (COG-UK) Consortium | The Lighthouse Lab in Milton Keynes and Alex Alderton, Roberto Amato, Jeffrey Barrett, Sonia Goncalves, Ewan Harrison, David K. Jackson, Ian Johnston, Dominic Kwiatkowski, Cordelia Langford, John Sillitoe on behalf of the Wellcome Sanger Institute COVID-19 Surveillance Team |

| | | | |
|---|---|---|---|
| EPI_ISL_3222538 | Lighthouse Lab in Glasgow | Wellcome Sanger Institute for the COVID-19 Genomics UK (COG-UK) Consortium | Harper VanSteenhouse, Yumi Kasai, David Gray, Carol Clugston, Anna Dominiczak and Alex Alderton, Roberto Amato, Jeffrey Barrett, Sonia Goncalves, Ewan Harrison, David K. Jackson, Ian Johnston, Dominic Kwiatkowski, Cordelia Langford, John Sillitoe on behalf of the Wellcome Sanger Institute COVID-19 Surveillance Team |
| EPI_ISL_3222539 | Lighthouse Laboratory Plymouth | Wellcome Sanger Institute for the COVID-19 Genomics UK (COG-UK) Consortium | Lighthouse Laboratory Plymouth and Alex Alderton, Roberto Amato, Jeffrey Barrett, Sonia Goncalves, Ewan Harrison, David K. Jackson, Ian Johnston, Dominic Kwiatkowski, Cordelia Langford, John Sillitoe on behalf of the Wellcome Sanger Institute COVID-19 Surveillance Team |
| EPI_ISL_3222540 | Lighthouse Lab in Glasgow | Wellcome Sanger Institute for the COVID-19 Genomics UK (COG-UK) Consortium | Harper VanSteenhouse, Yumi Kasai, David Gray, Carol Clugston, Anna Dominiczak and Alex Alderton, Roberto Amato, Jeffrey Barrett, Sonia Goncalves, Ewan Harrison, David K. Jackson, Ian Johnston, Dominic Kwiatkowski, Cordelia Langford, John Sillitoe on behalf of the Wellcome Sanger Institute COVID-19 Surveillance Team |
| EPI_ISL_3222541, EPI_ISL_3222542 | Lighthouse Laboratory Plymouth | Wellcome Sanger Institute for the COVID-19 Genomics UK (COG-UK) Consortium | Lighthouse Laboratory Plymouth and Alex Alderton, Roberto Amato, Jeffrey Barrett, Sonia Goncalves, Ewan Harrison, David K. Jackson, Ian Johnston, Dominic Kwiatkowski, Cordelia Langford, John Sillitoe on behalf of the Wellcome Sanger Institute COVID-19 Surveillance Team |
| EPI_ISL_3222543 | Lighthouse Lab in Milton Keynes | Wellcome Sanger Institute for the COVID-19 Genomics UK (COG-UK) Consortium | The Lighthouse Lab in Milton Keynes and Alex Alderton, Roberto Amato, Jeffrey Barrett, Sonia Goncalves, Ewan Harrison, David K. Jackson, Ian Johnston, Dominic Kwiatkowski, Cordelia Langford, John Sillitoe on behalf of the Wellcome Sanger Institute COVID-19 Surveillance Team |
| EPI_ISL_3222544 | Lighthouse Lab in Alderley Park | Wellcome Sanger Institute for the COVID-19 Genomics UK (COG-UK) Consortium | Jacquelyn Wynn, Mairead Hyland, The Lighthouse Lab in Alderley Park and Alex Alderton, Roberto Amato, Jeffrey Barrett, Sonia Goncalves, Ewan Harrison, David K. Jackson, Ian Johnston, Dominic Kwiatkowski, Cordelia Langford, John Sillitoe on behalf of the Wellcome Sanger Institute COVID-19 Surveillance Team |
| EPI_ISL_3222545 | Lighthouse Laboratory Plymouth | Wellcome Sanger Institute for the COVID-19 Genomics UK (COG-UK) Consortium | Lighthouse Laboratory Plymouth and Alex Alderton, Roberto Amato, Jeffrey Barrett, Sonia Goncalves, Ewan Harrison, David K. Jackson, Ian Johnston, Dominic Kwiatkowski, Cordelia Langford, John Sillitoe on behalf of the Wellcome Sanger Institute COVID-19 Surveillance Team |
| EPI_ISL_3222546, EPI_ISL_3222547 | Lighthouse Lab in Glasgow | Wellcome Sanger Institute for the COVID-19 Genomics UK (COG-UK) Consortium | Harper VanSteenhouse, Yumi Kasai, David Gray, Carol Clugston, Anna Dominiczak and Alex Alderton, Roberto Amato, Jeffrey Barrett, Sonia Goncalves, Ewan Harrison, David K. Jackson, Ian Johnston, Dominic Kwiatkowski, Cordelia Langford, John Sillitoe on behalf of the Wellcome Sanger Institute COVID-19 Surveillance Team |
| EPI_ISL_3222548 | Lighthouse Lab in Alderley Park | Wellcome Sanger Institute for the COVID-19 Genomics UK (COG-UK) Consortium | Jacquelyn Wynn, Mairead Hyland, The Lighthouse Lab in Alderley Park and Alex Alderton, Roberto Amato, Jeffrey Barrett, Sonia Goncalves, Ewan Harrison, David K. Jackson, Ian Johnston, Dominic Kwiatkowski, Cordelia Langford, John Sillitoe on behalf of the Wellcome Sanger Institute COVID-19 Surveillance Team |
| EPI_ISL_3222549, EPI_ISL_3222550 | Lighthouse Laboratory Plymouth | Wellcome Sanger Institute for the COVID-19 Genomics UK (COG-UK) Consortium | Lighthouse Laboratory Plymouth and Alex Alderton, Roberto Amato, Jeffrey Barrett, Sonia Goncalves, Ewan Harrison, David K. Jackson, Ian Johnston, Dominic Kwiatkowski, Cordelia Langford, John Sillitoe on behalf of the Wellcome Sanger Institute COVID-19 Surveillance Team |
| EPI_ISL_3222551 | Lighthouse Lab in Alderley Park | Wellcome Sanger Institute for the COVID-19 Genomics UK (COG-UK) Consortium | Jacquelyn Wynn, Mairead Hyland, The Lighthouse Lab in Alderley Park and Alex Alderton, Roberto Amato, Jeffrey Barrett, Sonia Goncalves, Ewan Harrison, David K. Jackson, Ian Johnston, Dominic Kwiatkowski, Cordelia Langford, John Sillitoe on behalf of the Wellcome Sanger Institute COVID-19 Surveillance Team |
| EPI_ISL_3222552, EPI_ISL_3222553 | Lighthouse Lab in Glasgow | Wellcome Sanger Institute for the COVID-19 Genomics UK (COG-UK) Consortium | Harper VanSteenhouse, Yumi Kasai, David Gray, Carol Clugston, Anna Dominiczak and Alex Alderton, Roberto Amato, Jeffrey Barrett, Sonia Goncalves, Ewan Harrison, David K. Jackson, Ian Johnston, Dominic Kwiatkowski, Cordelia Langford, John Sillitoe on behalf of the Wellcome Sanger Institute COVID-19 Surveillance Team |
| EPI_ISL_3222554, EPI_ISL_3222555 | Lighthouse Laboratory Plymouth | Wellcome Sanger Institute for the COVID-19 Genomics UK (COG-UK) Consortium | Lighthouse Laboratory Plymouth and Alex Alderton, Roberto Amato, Jeffrey Barrett, Sonia Goncalves, Ewan Harrison, David K. Jackson, Ian Johnston, Dominic Kwiatkowski, Cordelia Langford, John Sillitoe on behalf of the Wellcome Sanger Institute COVID-19 Surveillance Team |
| EPI_ISL_3222556 | Lighthouse Lab in Alderley Park | Wellcome Sanger Institute for the COVID-19 Genomics UK (COG-UK) Consortium | Jacquelyn Wynn, Mairead Hyland, The Lighthouse Lab in Alderley Park and Alex Alderton, Roberto Amato, Jeffrey Barrett, Sonia Goncalves, Ewan Harrison, David K. Jackson, Ian Johnston, Dominic Kwiatkowski, Cordelia Langford, John Sillitoe on behalf of the Wellcome Sanger Institute COVID-19 Surveillance Team |
| EPI_ISL_3222557 | Lighthouse Laboratory Plymouth | Wellcome Sanger Institute for the COVID-19 Genomics UK (COG-UK) Consortium | Lighthouse Laboratory Plymouth and Alex Alderton, Roberto Amato, Jeffrey Barrett, Sonia Goncalves, Ewan Harrison, David K. Jackson, Ian Johnston, Dominic Kwiatkowski, Cordelia Langford, John Sillitoe on behalf of the Wellcome Sanger Institute COVID-19 Surveillance Team |
| EPI_ISL_3222558 | Lighthouse Lab in Alderley Park | Wellcome Sanger Institute for the COVID-19 Genomics UK (COG-UK) Consortium | Jacquelyn Wynn, Mairead Hyland, The Lighthouse Lab in Alderley Park and Alex Alderton, Roberto Amato, Jeffrey Barrett, Sonia Goncalves, Ewan Harrison, David K. Jackson, Ian Johnston, Dominic Kwiatkowski, Cordelia Langford, John Sillitoe on behalf of the Wellcome Sanger Institute COVID-19 Surveillance Team |
| EPI_ISL_3222559 | Lighthouse Laboratory Plymouth | Wellcome Sanger Institute for the COVID-19 Genomics UK (COG-UK) Consortium | Lighthouse Laboratory Plymouth and Alex Alderton, Roberto Amato, Jeffrey Barrett, Sonia Goncalves, Ewan Harrison, David K. Jackson, Ian Johnston, Dominic Kwiatkowski, Cordelia Langford, John Sillitoe on behalf of the Wellcome Sanger Institute COVID-19 Surveillance Team |
| EPI_ISL_3222560, EPI_ISL_3222561 | Lighthouse Lab in Alderley Park | Wellcome Sanger Institute for the COVID-19 Genomics UK (COG-UK) Consortium | Jacquelyn Wynn, Mairead Hyland, The Lighthouse Lab in Alderley Park and Alex Alderton, Roberto Amato, Jeffrey Barrett, Sonia Goncalves, Ewan Harrison, David K. Jackson, Ian Johnston, Dominic Kwiatkowski, Cordelia Langford, John Sillitoe on behalf of the Wellcome Sanger Institute COVID-19 Surveillance Team |
| EPI_ISL_3222562 | Lighthouse Laboratory Plymouth | Wellcome Sanger Institute for the COVID-19 Genomics UK (COG-UK) Consortium | Lighthouse Laboratory Plymouth and Alex Alderton, Roberto Amato, Jeffrey Barrett, Sonia Goncalves, Ewan Harrison, David K. Jackson, Ian Johnston, Dominic Kwiatkowski, Cordelia Langford, John Sillitoe on behalf of the Wellcome Sanger Institute COVID-19 Surveillance Team |
| EPI_ISL_3222563 | Lighthouse Lab in Alderley Park | Wellcome Sanger Institute for the COVID-19 Genomics UK (COG-UK) Consortium | Jacquelyn Wynn, Mairead Hyland, The Lighthouse Lab in Alderley Park and Alex Alderton, Roberto Amato, Jeffrey Barrett, Sonia Goncalves, Ewan Harrison, David K. Jackson, Ian Johnston, Dominic Kwiatkowski, Cordelia Langford, John Sillitoe on behalf of the Wellcome Sanger Institute COVID-19 Surveillance Team |
| EPI_ISL_3222564 | Lighthouse Laboratory Plymouth | Wellcome Sanger Institute for the COVID-19 Genomics UK (COG-UK) Consortium | Lighthouse Laboratory Plymouth and Alex Alderton, Roberto Amato, Jeffrey Barrett, Sonia Goncalves, Ewan Harrison, David K. Jackson, Ian Johnston, Dominic Kwiatkowski, Cordelia Langford, John Sillitoe on behalf of the Wellcome Sanger Institute COVID-19 Surveillance Team |
| EPI_ISL_3222565, EPI_ISL_3222566 | Lighthouse Lab in Milton Keynes | Wellcome Sanger Institute for the COVID-19 Genomics UK (COG-UK) Consortium | The Lighthouse Lab in Milton Keynes and Alex Alderton, Roberto Amato, Jeffrey Barrett, Sonia Goncalves, Ewan Harrison, David K. Jackson, Ian Johnston, Dominic Kwiatkowski, Cordelia Langford, John Sillitoe on behalf of the Wellcome Sanger Institute COVID-19 Surveillance Team |
| EPI_ISL_3222567, EPI_ISL_3222568, EPI_ISL_3222569 | Lighthouse Lab in Glasgow | Wellcome Sanger Institute for the COVID-19 Genomics UK (COG-UK) Consortium | Harper VanSteenhouse, Yumi Kasai, David Gray, Carol Clugston, Anna Dominiczak and Alex Alderton, Roberto Amato, Jeffrey Barrett, Sonia Goncalves, Ewan Harrison, David K. Jackson, Ian Johnston, Dominic Kwiatkowski, Cordelia Langford, John Sillitoe on behalf of the Wellcome Sanger Institute COVID-19 Surveillance Team |
| EPI_ISL_3222570 | Lighthouse Lab in Alderley Park | Wellcome Sanger Institute for the COVID-19 Genomics UK (COG-UK) Consortium | Jacquelyn Wynn, Mairead Hyland, The Lighthouse Lab in Alderley Park and Alex Alderton, Roberto Amato, Jeffrey Barrett, Sonia Goncalves, Ewan Harrison, David K. Jackson, Ian Johnston, Dominic Kwiatkowski, Cordelia Langford, John Sillitoe on behalf of the Wellcome Sanger Institute COVID-19 Surveillance Team |
| EPI_ISL_3222571, EPI_ISL_3222572 | Lighthouse Laboratory Plymouth | Wellcome Sanger Institute for the COVID-19 Genomics UK (COG-UK) Consortium | Lighthouse Laboratory Plymouth and Alex Alderton, Roberto Amato, Jeffrey Barrett, Sonia Goncalves, Ewan Harrison, David K. Jackson, Ian Johnston, Dominic Kwiatkowski, Cordelia Langford, John Sillitoe on behalf of the Wellcome Sanger Institute COVID-19 Surveillance Team |
| EPI_ISL_3222573 | Lighthouse Lab in Alderley Park | Wellcome Sanger Institute for the COVID-19 Genomics UK (COG-UK) Consortium | Jacquelyn Wynn, Mairead Hyland, The Lighthouse Lab in Alderley Park and Alex Alderton, Roberto Amato, Jeffrey Barrett, Sonia Goncalves, Ewan Harrison, David K. Jackson, Ian Johnston, Dominic Kwiatkowski, Cordelia Langford, John Sillitoe on behalf of the Wellcome Sanger Institute COVID-19 Surveillance Team |
| EPI_ISL_3222574 | Lighthouse Lab in Glasgow | Wellcome Sanger Institute for the COVID-19 Genomics UK (COG-UK) Consortium | Harper VanSteenhouse, Yumi Kasai, David Gray, Carol Clugston, Anna Dominiczak and Alex Alderton, Roberto Amato, Jeffrey Barrett, Sonia Goncalves, Ewan Harrison, David K. Jackson, Ian Johnston, Dominic Kwiatkowski, Cordelia Langford, John Sillitoe on behalf of the Wellcome Sanger Institute COVID-19 Surveillance Team |
| EPI_ISL_3222575 | Lighthouse Laboratory Plymouth | Wellcome Sanger Institute for the COVID-19 Genomics UK (COG-UK) Consortium | Lighthouse Laboratory Plymouth and Alex Alderton, Roberto Amato, Jeffrey Barrett, Sonia Goncalves, Ewan Harrison, David K. Jackson, Ian Johnston, Dominic Kwiatkowski, Cordelia Langford, John Sillitoe on behalf of the Wellcome Sanger Institute COVID-19 Surveillance Team |
| EPI_ISL_3222576 | Lighthouse Lab in Alderley Park | Wellcome Sanger Institute for the COVID-19 Genomics UK (COG-UK) Consortium | Jacquelyn Wynn, Mairead Hyland, The Lighthouse Lab in Alderley Park and Alex Alderton, Roberto Amato, Jeffrey Barrett, Sonia Goncalves, Ewan Harrison, David K. Jackson, Ian Johnston, Dominic Kwiatkowski, Cordelia Langford, John Sillitoe on behalf of the Wellcome Sanger Institute COVID-19 Surveillance Team |
| EPI_ISL_3222577 | Lighthouse Lab in Glasgow | Wellcome Sanger Institute for the COVID-19 Genomics UK (COG-UK) Consortium | Harper VanSteenhouse, Yumi Kasai, David Gray, Carol Clugston, Anna Dominiczak and Alex Alderton, Roberto Amato, Jeffrey Barrett, Sonia Goncalves, Ewan Harrison, David K. Jackson, Ian Johnston, Dominic Kwiatkowski, Cordelia Langford, John Sillitoe on behalf of the Wellcome Sanger Institute COVID-19 Surveillance Team |
| EPI_ISL_3222578 | Lighthouse Lab in Alderley Park | Wellcome Sanger Institute for the COVID-19 Genomics UK (COG-UK) Consortium | Jacquelyn Wynn, Mairead Hyland, The Lighthouse Lab in Alderley Park and Alex Alderton, Roberto Amato, Jeffrey Barrett, Sonia Goncalves, Ewan Harrison, David K. Jackson, Ian Johnston, Dominic Kwiatkowski, Cordelia Langford, John Sillitoe on behalf of the Wellcome Sanger Institute COVID-19 Surveillance Team |
| EPI_ISL_3222579 | Lighthouse Lab in Milton Keynes | Wellcome Sanger Institute for the COVID-19 Genomics UK (COG-UK) Consortium | The Lighthouse Lab in Milton Keynes and Alex Alderton, Roberto Amato, Jeffrey Barrett, Sonia Goncalves, Ewan Harrison, David K. Jackson, Ian Johnston, Dominic Kwiatkowski, Cordelia Langford, John Sillitoe on behalf of the Wellcome Sanger Institute COVID-19 Surveillance Team |
| EPI_ISL_3222580 | Lighthouse Lab in Glasgow | Wellcome Sanger Institute for the COVID-19 Genomics UK (COG-UK) Consortium | Harper VanSteenhouse, Yumi Kasai, David Gray, Carol Clugston, Anna Dominiczak and Alex Alderton, Roberto Amato, Jeffrey Barrett, Sonia Goncalves, Ewan Harrison, David K. Jackson, Ian Johnston, Dominic Kwiatkowski, Cordelia Langford, John Sillitoe on behalf of the Wellcome Sanger Institute COVID-19 Surveillance Team |
| EPI_ISL_3222581, EPI_ISL_3222582 | Lighthouse Lab in Milton Keynes | Wellcome Sanger Institute for the COVID-19 Genomics UK (COG-UK) Consortium | The Lighthouse Lab in Milton Keynes and Alex Alderton, Roberto Amato, Jeffrey Barrett, Sonia Goncalves, Ewan Harrison, David K. Jackson, Ian Johnston, Dominic Kwiatkowski, Cordelia Langford, John Sillitoe on behalf of the Wellcome Sanger Institute COVID-19 Surveillance Team |
| EPI_ISL_3222583, EPI_ISL_3222584 | Lighthouse Lab in Glasgow | Wellcome Sanger Institute for the COVID-19 Genomics UK (COG-UK) Consortium | Harper VanSteenhouse, Yumi Kasai, David Gray, Carol Clugston, Anna Dominiczak and Alex Alderton, Roberto Amato, Jeffrey Barrett, Sonia Goncalves, Ewan Harrison, David K. Jackson, Ian Johnston, Dominic Kwiatkowski, Cordelia Langford, John Sillitoe on behalf of the Wellcome Sanger Institute COVID-19 Surveillance Team |
| EPI_ISL_3222585 | Lighthouse Laboratory Plymouth | Wellcome Sanger Institute for the COVID-19 Genomics UK (COG-UK) Consortium | Lighthouse Laboratory Plymouth and Alex Alderton, Roberto Amato, Jeffrey Barrett, Sonia Goncalves, Ewan Harrison, David K. Jackson, Ian Johnston, Dominic Kwiatkowski, Cordelia Langford, John Sillitoe on behalf of the Wellcome Sanger Institute COVID-19 Surveillance Team |
| EPI_ISL_3222586 | Lighthouse Lab in Milton Keynes | Wellcome Sanger Institute for the COVID-19 Genomics UK (COG-UK) Consortium | The Lighthouse Lab in Milton Keynes and Alex Alderton, Roberto Amato, Jeffrey Barrett, Sonia Goncalves, Ewan Harrison, David K. Jackson, Ian Johnston, Dominic Kwiatkowski, Cordelia Langford, John Sillitoe on behalf of the Wellcome Sanger Institute COVID-19 Surveillance Team |
| EPI_ISL_3222587 | Lighthouse Lab in Alderley Park | Wellcome Sanger Institute for the COVID-19 Genomics UK (COG-UK) Consortium | Jacquelyn Wynn, Mairead Hyland, The Lighthouse Lab in Alderley Park and Alex Alderton, Roberto Amato, Jeffrey Barrett, Sonia Goncalves, Ewan Harrison, David K. Jackson, Ian Johnston, Dominic Kwiatkowski, Cordelia Langford, John Sillitoe on behalf of the Wellcome Sanger Institute COVID-19 Surveillance Team |
| EPI_ISL_3222588, EPI_ISL_3222589 | Lighthouse Lab in Milton Keynes | Wellcome Sanger Institute for the COVID-19 Genomics UK (COG-UK) Consortium | The Lighthouse Lab in Milton Keynes and Alex Alderton, Roberto Amato, Jeffrey Barrett, Sonia Goncalves, Ewan Harrison, David K. Jackson, Ian Johnston, Dominic Kwiatkowski, Cordelia Langford, John Sillitoe on behalf of the Wellcome Sanger Institute COVID-19 Surveillance Team |
| EPI_ISL_3222590 | Lighthouse Lab in Alderley Park | Wellcome Sanger Institute for the COVID-19 Genomics UK (COG-UK) Consortium | Jacquelyn Wynn, Mairead Hyland, The Lighthouse Lab in Alderley Park and Alex Alderton, Roberto Amato, Jeffrey Barrett, Sonia Goncalves, Ewan Harrison, David K. Jackson, Ian Johnston, Dominic Kwiatkowski, Cordelia Langford, John Sillitoe on behalf of the Wellcome Sanger Institute COVID-19 Surveillance Team |
| EPI_ISL_3222591 | Lighthouse Lab in Milton Keynes | Wellcome Sanger Institute for the COVID-19 Genomics UK (COG-UK) Consortium | The Lighthouse Lab in Milton Keynes and Alex Alderton, Roberto Amato, Jeffrey Barrett, Sonia Goncalves, Ewan Harrison, David K. Jackson, Ian Johnston, Dominic Kwiatkowski, Cordelia Langford, John Sillitoe on behalf of the Wellcome Sanger Institute COVID-19 Surveillance Team |
| EPI_ISL_3222592 | Lighthouse Lab in Alderley Park | Wellcome Sanger Institute for the COVID-19 Genomics UK (COG-UK) Consortium | Jacquelyn Wynn, Mairead Hyland, The Lighthouse Lab in Alderley Park and Alex Alderton, Roberto Amato, Jeffrey Barrett, Sonia Goncalves, Ewan Harrison, David K. Jackson, Ian Johnston, Dominic Kwiatkowski, Cordelia Langford, John Sillitoe on behalf of the Wellcome Sanger Institute COVID-19 Surveillance Team |
| EPI_ISL_3222593, EPI_ISL_3222594, EPI_ISL_3222595 | Lighthouse Laboratory Plymouth | Wellcome Sanger Institute for the COVID-19 Genomics UK (COG-UK) Consortium | Lighthouse Laboratory Plymouth and Alex Alderton, Roberto Amato, Jeffrey Barrett, Sonia Goncalves, Ewan Harrison, David K. Jackson, Ian Johnston, Dominic Kwiatkowski, Cordelia Langford, John Sillitoe on behalf of the Wellcome Sanger Institute COVID-19 Surveillance Team |
| EPI_ISL_3222596 | Lighthouse Lab in Milton Keynes | Wellcome Sanger Institute for the COVID-19 Genomics UK (COG-UK) Consortium | The Lighthouse Lab in Milton Keynes and Alex Alderton, Roberto Amato, Jeffrey Barrett, Sonia Goncalves, Ewan Harrison, David K. Jackson, Ian Johnston, Dominic Kwiatkowski, Cordelia Langford, John Sillitoe on behalf of the Wellcome Sanger Institute COVID-19 Surveillance Team |

| | | | |
|---|---|---|---|
| EPI_ISL_3222597, EPI_ISL_3222598 | Lighthouse Lab in Alderley Park | Wellcome Sanger Institute for the COVID-19 Genomics UK (COG-UK) Consortium | Jacquelyn Wynn, Mairead Hyland, The Lighthouse Lab in Alderley Park and Alex Alderton, Roberto Amato, Jeffrey Barrett, Sonia Goncalves, Ewan Harrison, David K. Jackson, Ian Johnston, Dominic Kwiatkowski, Cordelia Langford, John Sillitoe on behalf of the Wellcome Sanger Institute COVID-19 Surveillance Team |
| EPI_ISL_3222599 | Lighthouse Lab in Glasgow | Wellcome Sanger Institute for the COVID-19 Genomics UK (COG-UK) Consortium | Harper VanSteenhouse, Yumi Kasai, David Gray, Carol Clugston, Anna Dominiczak and Alex Alderton, Roberto Amato, Jeffrey Barrett, Sonia Goncalves, Ewan Harrison, David K. Jackson, Ian Johnston, Dominic Kwiatkowski, Cordelia Langford, John Sillitoe on behalf of the Wellcome Sanger Institute COVID-19 Surveillance Team |
| EPI_ISL_3222600 | Lighthouse Lab in Milton Keynes | Wellcome Sanger Institute for the COVID-19 Genomics UK (COG-UK) Consortium | The Lighthouse Lab in Milton Keynes and Alex Alderton, Roberto Amato, Jeffrey Barrett, Sonia Goncalves, Ewan Harrison, David K. Jackson, Ian Johnston, Dominic Kwiatkowski, Cordelia Langford, John Sillitoe on behalf of the Wellcome Sanger Institute COVID-19 Surveillance Team |
| EPI_ISL_3222601 | Lighthouse Lab in Alderley Park | Wellcome Sanger Institute for the COVID-19 Genomics UK (COG-UK) Consortium | Jacquelyn Wynn, Mairead Hyland, The Lighthouse Lab in Alderley Park and Alex Alderton, Roberto Amato, Jeffrey Barrett, Sonia Goncalves, Ewan Harrison, David K. Jackson, Ian Johnston, Dominic Kwiatkowski, Cordelia Langford, John Sillitoe on behalf of the Wellcome Sanger Institute COVID-19 Surveillance Team |
| EPI_ISL_3222602 | Lighthouse Laboratory Plymouth | Wellcome Sanger Institute for the COVID-19 Genomics UK (COG-UK) Consortium | Lighthouse Laboratory Plymouth and Alex Alderton, Roberto Amato, Jeffrey Barrett, Sonia Goncalves, Ewan Harrison, David K. Jackson, Ian Johnston, Dominic Kwiatkowski, Cordelia Langford, John Sillitoe on behalf of the Wellcome Sanger Institute COVID-19 Surveillance Team |
| EPI_ISL_3222603 | Lighthouse Lab in Glasgow | Wellcome Sanger Institute for the COVID-19 Genomics UK (COG-UK) Consortium | Harper VanSteenhouse, Yumi Kasai, David Gray, Carol Clugston, Anna Dominiczak and Alex Alderton, Roberto Amato, Jeffrey Barrett, Sonia Goncalves, Ewan Harrison, David K. Jackson, Ian Johnston, Dominic Kwiatkowski, Cordelia Langford, John Sillitoe on behalf of the Wellcome Sanger Institute COVID-19 Surveillance Team |
| EPI_ISL_3222604, EPI_ISL_3222605 | Lighthouse Lab in Milton Keynes | Wellcome Sanger Institute for the COVID-19 Genomics UK (COG-UK) Consortium | The Lighthouse Lab in Milton Keynes and Alex Alderton, Roberto Amato, Jeffrey Barrett, Sonia Goncalves, Ewan Harrison, David K. Jackson, Ian Johnston, Dominic Kwiatkowski, Cordelia Langford, John Sillitoe on behalf of the Wellcome Sanger Institute COVID-19 Surveillance Team |
| EPI_ISL_3222606 | Lighthouse Laboratory Plymouth | Wellcome Sanger Institute for the COVID-19 Genomics UK (COG-UK) Consortium | Lighthouse Laboratory Plymouth and Alex Alderton, Roberto Amato, Jeffrey Barrett, Sonia Goncalves, Ewan Harrison, David K. Jackson, Ian Johnston, Dominic Kwiatkowski, Cordelia Langford, John Sillitoe on behalf of the Wellcome Sanger Institute COVID-19 Surveillance Team |
| EPI_ISL_3222607, EPI_ISL_3222608 | Lighthouse Lab in Milton Keynes | Wellcome Sanger Institute for the COVID-19 Genomics UK (COG-UK) Consortium | The Lighthouse Lab in Milton Keynes and Alex Alderton, Roberto Amato, Jeffrey Barrett, Sonia Goncalves, Ewan Harrison, David K. Jackson, Ian Johnston, Dominic Kwiatkowski, Cordelia Langford, John Sillitoe on behalf of the Wellcome Sanger Institute COVID-19 Surveillance Team |
| EPI_ISL_3222609 | Lighthouse Laboratory Plymouth | Wellcome Sanger Institute for the COVID-19 Genomics UK (COG-UK) Consortium | Lighthouse Laboratory Plymouth and Alex Alderton, Roberto Amato, Jeffrey Barrett, Sonia Goncalves, Ewan Harrison, David K. Jackson, Ian Johnston, Dominic Kwiatkowski, Cordelia Langford, John Sillitoe on behalf of the Wellcome Sanger Institute COVID-19 Surveillance Team |
| EPI_ISL_3222610 | Lighthouse Lab in Alderley Park | Wellcome Sanger Institute for the COVID-19 Genomics UK (COG-UK) Consortium | Jacquelyn Wynn, Mairead Hyland, The Lighthouse Lab in Alderley Park and Alex Alderton, Roberto Amato, Jeffrey Barrett, Sonia Goncalves, Ewan Harrison, David K. Jackson, Ian Johnston, Dominic Kwiatkowski, Cordelia Langford, John Sillitoe on behalf of the Wellcome Sanger Institute COVID-19 Surveillance Team |
| EPI_ISL_3222611 | Lighthouse Lab in Milton Keynes | Wellcome Sanger Institute for the COVID-19 Genomics UK (COG-UK) Consortium | The Lighthouse Lab in Milton Keynes and Alex Alderton, Roberto Amato, Jeffrey Barrett, Sonia Goncalves, Ewan Harrison, David K. Jackson, Ian Johnston, Dominic Kwiatkowski, Cordelia Langford, John Sillitoe on behalf of the Wellcome Sanger Institute COVID-19 Surveillance Team |
| EPI_ISL_3222612, EPI_ISL_3222613 | Lighthouse Lab in Alderley Park | Wellcome Sanger Institute for the COVID-19 Genomics UK (COG-UK) Consortium | Jacquelyn Wynn, Mairead Hyland, The Lighthouse Lab in Alderley Park and Alex Alderton, Roberto Amato, Jeffrey Barrett, Sonia Goncalves, Ewan Harrison, David K. Jackson, Ian Johnston, Dominic Kwiatkowski, Cordelia Langford, John Sillitoe on behalf of the Wellcome Sanger Institute COVID-19 Surveillance Team |
| EPI_ISL_3222614 | Lighthouse Laboratory Plymouth | Wellcome Sanger Institute for the COVID-19 Genomics UK (COG-UK) Consortium | Lighthouse Laboratory Plymouth and Alex Alderton, Roberto Amato, Jeffrey Barrett, Sonia Goncalves, Ewan Harrison, David K. Jackson, Ian Johnston, Dominic Kwiatkowski, Cordelia Langford, John Sillitoe on behalf of the Wellcome Sanger Institute COVID-19 Surveillance Team |
| EPI_ISL_3222615 | Lighthouse Lab in Glasgow | Wellcome Sanger Institute for the COVID-19 Genomics UK (COG-UK) Consortium | Harper VanSteenhouse, Yumi Kasai, David Gray, Carol Clugston, Anna Dominiczak and Alex Alderton, Roberto Amato, Jeffrey Barrett, Sonia Goncalves, Ewan Harrison, David K. Jackson, Ian Johnston, Dominic Kwiatkowski, Cordelia Langford, John Sillitoe on behalf of the Wellcome Sanger Institute COVID-19 Surveillance Team |
| EPI_ISL_3222616 | Lighthouse Laboratory Plymouth | Wellcome Sanger Institute for the COVID-19 Genomics UK (COG-UK) Consortium | Lighthouse Laboratory Plymouth and Alex Alderton, Roberto Amato, Jeffrey Barrett, Sonia Goncalves, Ewan Harrison, David K. Jackson, Ian Johnston, Dominic Kwiatkowski, Cordelia Langford, John Sillitoe on behalf of the Wellcome Sanger Institute COVID-19 Surveillance Team |
| EPI_ISL_3222617 | Lighthouse Lab in Alderley Park | Wellcome Sanger Institute for the COVID-19 Genomics UK (COG-UK) Consortium | Jacquelyn Wynn, Mairead Hyland, The Lighthouse Lab in Alderley Park and Alex Alderton, Roberto Amato, Jeffrey Barrett, Sonia Goncalves, Ewan Harrison, David K. Jackson, Ian Johnston, Dominic Kwiatkowski, Cordelia Langford, John Sillitoe on behalf of the Wellcome Sanger Institute COVID-19 Surveillance Team |
| EPI_ISL_3222618 | Lighthouse Lab in Milton Keynes | Wellcome Sanger Institute for the COVID-19 Genomics UK (COG-UK) Consortium | The Lighthouse Lab in Milton Keynes and Alex Alderton, Roberto Amato, Jeffrey Barrett, Sonia Goncalves, Ewan Harrison, David K. Jackson, Ian Johnston, Dominic Kwiatkowski, Cordelia Langford, John Sillitoe on behalf of the Wellcome Sanger Institute COVID-19 Surveillance Team |
| EPI_ISL_3222619, EPI_ISL_3222620 | Lighthouse Laboratory Plymouth | Wellcome Sanger Institute for the COVID-19 Genomics UK (COG-UK) Consortium | Lighthouse Laboratory Plymouth and Alex Alderton, Roberto Amato, Jeffrey Barrett, Sonia Goncalves, Ewan Harrison, David K. Jackson, Ian Johnston, Dominic Kwiatkowski, Cordelia Langford, John Sillitoe on behalf of the Wellcome Sanger Institute COVID-19 Surveillance Team |
| EPI_ISL_3222621 | Lighthouse Lab in Milton Keynes | Wellcome Sanger Institute for the COVID-19 Genomics UK (COG-UK) Consortium | The Lighthouse Lab in Milton Keynes and Alex Alderton, Roberto Amato, Jeffrey Barrett, Sonia Goncalves, Ewan Harrison, David K. Jackson, Ian Johnston, Dominic Kwiatkowski, Cordelia Langford, John Sillitoe on behalf of the Wellcome Sanger Institute COVID-19 Surveillance Team |
| EPI_ISL_3222622 | Lighthouse Laboratory Plymouth | Wellcome Sanger Institute for the COVID-19 Genomics UK (COG-UK) Consortium | Lighthouse Laboratory Plymouth and Alex Alderton, Roberto Amato, Jeffrey Barrett, Sonia Goncalves, Ewan Harrison, David K. Jackson, Ian Johnston, Dominic Kwiatkowski, Cordelia Langford, John Sillitoe on behalf of the Wellcome Sanger Institute COVID-19 Surveillance Team |
| EPI_ISL_3222623 | Lighthouse Lab in Alderley Park | Wellcome Sanger Institute for the COVID-19 Genomics UK (COG-UK) Consortium | Jacquelyn Wynn, Mairead Hyland, The Lighthouse Lab in Alderley Park and Alex Alderton, Roberto Amato, Jeffrey Barrett, Sonia Goncalves, Ewan Harrison, David K. Jackson, Ian Johnston, Dominic Kwiatkowski, Cordelia Langford, John Sillitoe on behalf of the Wellcome Sanger Institute COVID-19 Surveillance Team |
| EPI_ISL_3222624, EPI_ISL_3222625 | Lighthouse Lab in Glasgow | Wellcome Sanger Institute for the COVID-19 Genomics UK (COG-UK) Consortium | Harper VanSteenhouse, Yumi Kasai, David Gray, Carol Clugston, Anna Dominiczak and Alex Alderton, Roberto Amato, Jeffrey Barrett, Sonia Goncalves, Ewan Harrison, David K. Jackson, Ian Johnston, Dominic Kwiatkowski, Cordelia Langford, John Sillitoe on behalf of the Wellcome Sanger Institute COVID-19 Surveillance Team |
| EPI_ISL_3222626 | Lighthouse Laboratory Plymouth | Wellcome Sanger Institute for the COVID-19 Genomics UK (COG-UK) Consortium | Lighthouse Laboratory Plymouth and Alex Alderton, Roberto Amato, Jeffrey Barrett, Sonia Goncalves, Ewan Harrison, David K. Jackson, Ian Johnston, Dominic Kwiatkowski, Cordelia Langford, John Sillitoe on behalf of the Wellcome Sanger Institute COVID-19 Surveillance Team |
| EPI_ISL_3222627, EPI_ISL_3222628 | Lighthouse Lab in Alderley Park | Wellcome Sanger Institute for the COVID-19 Genomics UK (COG-UK) Consortium | Jacquelyn Wynn, Mairead Hyland, The Lighthouse Lab in Alderley Park and Alex Alderton, Roberto Amato, Jeffrey Barrett, Sonia Goncalves, Ewan Harrison, David K. Jackson, Ian Johnston, Dominic Kwiatkowski, Cordelia Langford, John Sillitoe on behalf of the Wellcome Sanger Institute COVID-19 Surveillance Team |
| EPI_ISL_3222629 | Lighthouse Laboratory Plymouth | Wellcome Sanger Institute for the COVID-19 Genomics UK (COG-UK) Consortium | Lighthouse Laboratory Plymouth and Alex Alderton, Roberto Amato, Jeffrey Barrett, Sonia Goncalves, Ewan Harrison, David K. Jackson, Ian Johnston, Dominic Kwiatkowski, Cordelia Langford, John Sillitoe on behalf of the Wellcome Sanger Institute COVID-19 Surveillance Team |
| EPI_ISL_3222630 | Lighthouse Lab in Milton Keynes | Wellcome Sanger Institute for the COVID-19 Genomics UK (COG-UK) Consortium | The Lighthouse Lab in Milton Keynes and Alex Alderton, Roberto Amato, Jeffrey Barrett, Sonia Goncalves, Ewan Harrison, David K. Jackson, Ian Johnston, Dominic Kwiatkowski, Cordelia Langford, John Sillitoe on behalf of the Wellcome Sanger Institute COVID-19 Surveillance Team |
| EPI_ISL_3222631 | Lighthouse Laboratory Plymouth | Wellcome Sanger Institute for the COVID-19 Genomics UK (COG-UK) Consortium | Lighthouse Laboratory Plymouth and Alex Alderton, Roberto Amato, Jeffrey Barrett, Sonia Goncalves, Ewan Harrison, David K. Jackson, Ian Johnston, Dominic Kwiatkowski, Cordelia Langford, John Sillitoe on behalf of the Wellcome Sanger Institute COVID-19 Surveillance Team |
| EPI_ISL_3222632 | Lighthouse Lab in Milton Keynes | Wellcome Sanger Institute for the COVID-19 Genomics UK (COG-UK) Consortium | The Lighthouse Lab in Milton Keynes and Alex Alderton, Roberto Amato, Jeffrey Barrett, Sonia Goncalves, Ewan Harrison, David K. Jackson, Ian Johnston, Dominic Kwiatkowski, Cordelia Langford, John Sillitoe on behalf of the Wellcome Sanger Institute COVID-19 Surveillance Team |
| EPI_ISL_3222633 | Lighthouse Lab in Alderley Park | Wellcome Sanger Institute for the COVID-19 Genomics UK (COG-UK) Consortium | Jacquelyn Wynn, Mairead Hyland, The Lighthouse Lab in Alderley Park and Alex Alderton, Roberto Amato, Jeffrey Barrett, Sonia Goncalves, Ewan Harrison, David K. Jackson, Ian Johnston, Dominic Kwiatkowski, Cordelia Langford, John Sillitoe on behalf of the Wellcome Sanger Institute COVID-19 Surveillance Team |
| EPI_ISL_3222634 | Lighthouse Laboratory Plymouth | Wellcome Sanger Institute for the COVID-19 Genomics UK (COG-UK) Consortium | Lighthouse Laboratory Plymouth and Alex Alderton, Roberto Amato, Jeffrey Barrett, Sonia Goncalves, Ewan Harrison, David K. Jackson, Ian Johnston, Dominic Kwiatkowski, Cordelia Langford, John Sillitoe on behalf of the Wellcome Sanger Institute COVID-19 Surveillance Team |
| EPI_ISL_3222635 | Lighthouse Lab in Alderley Park | Wellcome Sanger Institute for the COVID-19 Genomics UK (COG-UK) Consortium | Jacquelyn Wynn, Mairead Hyland, The Lighthouse Lab in Alderley Park and Alex Alderton, Roberto Amato, Jeffrey Barrett, Sonia Goncalves, Ewan Harrison, David K. Jackson, Ian Johnston, Dominic Kwiatkowski, Cordelia Langford, John Sillitoe on behalf of the Wellcome Sanger Institute COVID-19 Surveillance Team |
| EPI_ISL_3222636 | Lighthouse Laboratory Plymouth | Wellcome Sanger Institute for the COVID-19 Genomics UK (COG-UK) Consortium | Lighthouse Laboratory Plymouth and Alex Alderton, Roberto Amato, Jeffrey Barrett, Sonia Goncalves, Ewan Harrison, David K. Jackson, Ian Johnston, Dominic Kwiatkowski, Cordelia Langford, John Sillitoe on behalf of the Wellcome Sanger Institute COVID-19 Surveillance Team |
| EPI_ISL_3222637 | Lighthouse Lab in Alderley Park | Wellcome Sanger Institute for the COVID-19 Genomics UK (COG-UK) Consortium | Jacquelyn Wynn, Mairead Hyland, The Lighthouse Lab in Alderley Park and Alex Alderton, Roberto Amato, Jeffrey Barrett, Sonia Goncalves, Ewan Harrison, David K. Jackson, Ian Johnston, Dominic Kwiatkowski, Cordelia Langford, John Sillitoe on behalf of the Wellcome Sanger Institute COVID-19 Surveillance Team |
| EPI_ISL_3222638 | Lighthouse Laboratory Plymouth | Wellcome Sanger Institute for the COVID-19 Genomics UK (COG-UK) Consortium | Lighthouse Laboratory Plymouth and Alex Alderton, Roberto Amato, Jeffrey Barrett, Sonia Goncalves, Ewan Harrison, David K. Jackson, Ian Johnston, Dominic Kwiatkowski, Cordelia Langford, John Sillitoe on behalf of the Wellcome Sanger Institute COVID-19 Surveillance Team |
| EPI_ISL_3222639, EPI_ISL_3222640 | Lighthouse Lab in Alderley Park | Wellcome Sanger Institute for the COVID-19 Genomics UK (COG-UK) Consortium | Jacquelyn Wynn, Mairead Hyland, The Lighthouse Lab in Alderley Park and Alex Alderton, Roberto Amato, Jeffrey Barrett, Sonia Goncalves, Ewan Harrison, David K. Jackson, Ian Johnston, Dominic Kwiatkowski, Cordelia Langford, John Sillitoe on behalf of the Wellcome Sanger Institute COVID-19 Surveillance Team |
| EPI_ISL_3222641 | Lighthouse Lab in Milton Keynes | Wellcome Sanger Institute for the COVID-19 Genomics UK (COG-UK) Consortium | The Lighthouse Lab in Milton Keynes and Alex Alderton, Roberto Amato, Jeffrey Barrett, Sonia Goncalves, Ewan Harrison, David K. Jackson, Ian Johnston, Dominic Kwiatkowski, Cordelia Langford, John Sillitoe on behalf of the Wellcome Sanger Institute COVID-19 Surveillance Team |
| EPI_ISL_3222642, EPI_ISL_3222643 | Lighthouse Laboratory Plymouth | Wellcome Sanger Institute for the COVID-19 Genomics UK (COG-UK) Consortium | Lighthouse Laboratory Plymouth and Alex Alderton, Roberto Amato, Jeffrey Barrett, Sonia Goncalves, Ewan Harrison, David K. Jackson, Ian Johnston, Dominic Kwiatkowski, Cordelia Langford, John Sillitoe on behalf of the Wellcome Sanger Institute COVID-19 Surveillance Team |
| EPI_ISL_3222644 | Lighthouse Lab in Milton Keynes | Wellcome Sanger Institute for the COVID-19 Genomics UK (COG-UK) Consortium | The Lighthouse Lab in Milton Keynes and Alex Alderton, Roberto Amato, Jeffrey Barrett, Sonia Goncalves, Ewan Harrison, David K. Jackson, Ian Johnston, Dominic Kwiatkowski, Cordelia Langford, John Sillitoe on behalf of the Wellcome Sanger Institute COVID-19 Surveillance Team |
| EPI_ISL_3222645 | Lighthouse Lab in Alderley Park | Wellcome Sanger Institute for the COVID-19 Genomics UK (COG-UK) Consortium | Jacquelyn Wynn, Mairead Hyland, The Lighthouse Lab in Alderley Park and Alex Alderton, Roberto Amato, Jeffrey Barrett, Sonia Goncalves, Ewan Harrison, David K. Jackson, Ian Johnston, Dominic Kwiatkowski, Cordelia Langford, John Sillitoe on behalf of the Wellcome Sanger Institute COVID-19 Surveillance Team |
| EPI_ISL_3222646, EPI_ISL_3222647 | Lighthouse Laboratory Plymouth | Wellcome Sanger Institute for the COVID-19 Genomics UK (COG-UK) Consortium | Lighthouse Laboratory Plymouth and Alex Alderton, Roberto Amato, Jeffrey Barrett, Sonia Goncalves, Ewan Harrison, David K. Jackson, Ian Johnston, Dominic Kwiatkowski, Cordelia Langford, John Sillitoe on behalf of the Wellcome Sanger Institute COVID-19 Surveillance Team |
| EPI_ISL_3222648, EPI_ISL_3222649 | Lighthouse Lab in Milton Keynes | Wellcome Sanger Institute for the COVID-19 Genomics UK (COG-UK) Consortium | The Lighthouse Lab in Milton Keynes and Alex Alderton, Roberto Amato, Jeffrey Barrett, Sonia Goncalves, Ewan Harrison, David K. Jackson, Ian Johnston, Dominic Kwiatkowski, Cordelia Langford, John Sillitoe on behalf of the Wellcome Sanger Institute COVID-19 Surveillance Team |
| EPI_ISL_3222650 | Lighthouse Lab in Alderley Park | Wellcome Sanger Institute for the COVID-19 Genomics UK (COG-UK) Consortium | Jacquelyn Wynn, Mairead Hyland, The Lighthouse Lab in Alderley Park and Alex Alderton, Roberto Amato, Jeffrey Barrett, Sonia Goncalves, Ewan Harrison, David K. Jackson, Ian Johnston, Dominic Kwiatkowski, Cordelia Langford, John Sillitoe on behalf of the Wellcome Sanger Institute COVID-19 Surveillance Team |
| EPI_ISL_3222651 | Lighthouse Lab in Milton Keynes | Wellcome Sanger Institute for the COVID-19 Genomics UK (COG-UK) Consortium | The Lighthouse Lab in Milton Keynes and Alex Alderton, Roberto Amato, Jeffrey Barrett, Sonia Goncalves, Ewan Harrison, David K. Jackson, Ian Johnston, Dominic Kwiatkowski, Cordelia Langford, John Sillitoe on behalf of the Wellcome Sanger Institute COVID-19 Surveillance Team |
| EPI_ISL_3222652 | Lighthouse Laboratory Plymouth | Wellcome Sanger Institute for the COVID-19 Genomics UK (COG-UK) Consortium | Lighthouse Laboratory Plymouth and Alex Alderton, Roberto Amato, Jeffrey Barrett, Sonia Goncalves, Ewan Harrison, David K. Jackson, Ian Johnston, Dominic Kwiatkowski, Cordelia Langford, John Sillitoe on behalf of the Wellcome Sanger Institute COVID-19 Surveillance Team |

| | | | |
|---|---|---|---|
| EPI_ISL_3222653 | Lighthouse Lab in Milton Keynes | Wellcome Sanger Institute for the COVID-19 Genomics UK (COG-UK) Consortium | The Lighthouse Lab in Milton Keynes and Alex Alderton, Roberto Amato, Jeffrey Barrett, Sonia Goncalves, Ewan Harrison, David K. Jackson, Ian Johnston, Dominic Kwiatkowski, Cordelia Langford, John Sillitoe on behalf of the Wellcome Sanger Institute COVID-19 Surveillance Team |
| EPI_ISL_3222654 | Lighthouse Laboratory Plymouth | Wellcome Sanger Institute for the COVID-19 Genomics UK (COG-UK) Consortium | Lighthouse Laboratory Plymouth and Alex Alderton, Roberto Amato, Jeffrey Barrett, Sonia Goncalves, Ewan Harrison, David K. Jackson, Ian Johnston, Dominic Kwiatkowski, Cordelia Langford, John Sillitoe on behalf of the Wellcome Sanger Institute COVID-19 Surveillance Team |
| EPI_ISL_3222655, EPI_ISL_3222656 | Lighthouse Lab in Milton Keynes | Wellcome Sanger Institute for the COVID-19 Genomics UK (COG-UK) Consortium | The Lighthouse Lab in Milton Keynes and Alex Alderton, Roberto Amato, Jeffrey Barrett, Sonia Goncalves, Ewan Harrison, David K. Jackson, Ian Johnston, Dominic Kwiatkowski, Cordelia Langford, John Sillitoe on behalf of the Wellcome Sanger Institute COVID-19 Surveillance Team |
| EPI_ISL_3222657, EPI_ISL_3222658, EPI_ISL_3222659, EPI_ISL_3222660 | Lighthouse Laboratory Plymouth | Wellcome Sanger Institute for the COVID-19 Genomics UK (COG-UK) Consortium | Lighthouse Laboratory Plymouth and Alex Alderton, Roberto Amato, Jeffrey Barrett, Sonia Goncalves, Ewan Harrison, David K. Jackson, Ian Johnston, Dominic Kwiatkowski, Cordelia Langford, John Sillitoe on behalf of the Wellcome Sanger Institute COVID-19 Surveillance Team |
| EPI_ISL_3222661 | Lighthouse Lab in Milton Keynes | Wellcome Sanger Institute for the COVID-19 Genomics UK (COG-UK) Consortium | The Lighthouse Lab in Milton Keynes and Alex Alderton, Roberto Amato, Jeffrey Barrett, Sonia Goncalves, Ewan Harrison, David K. Jackson, Ian Johnston, Dominic Kwiatkowski, Cordelia Langford, John Sillitoe on behalf of the Wellcome Sanger Institute COVID-19 Surveillance Team |
| EPI_ISL_3222662 | Lighthouse Laboratory Plymouth | Wellcome Sanger Institute for the COVID-19 Genomics UK (COG-UK) Consortium | Lighthouse Laboratory Plymouth and Alex Alderton, Roberto Amato, Jeffrey Barrett, Sonia Goncalves, Ewan Harrison, David K. Jackson, Ian Johnston, Dominic Kwiatkowski, Cordelia Langford, John Sillitoe on behalf of the Wellcome Sanger Institute COVID-19 Surveillance Team |
| EPI_ISL_3222663 | Lighthouse Lab in Alderley Park | Wellcome Sanger Institute for the COVID-19 Genomics UK (COG-UK) Consortium | Jacquelyn Wynn, Mairead Hyland, The Lighthouse Lab in Alderley Park and Alex Alderton, Roberto Amato, Jeffrey Barrett, Sonia Goncalves, Ewan Harrison, David K. Jackson, Ian Johnston, Dominic Kwiatkowski, Cordelia Langford, John Sillitoe on behalf of the Wellcome Sanger Institute COVID-19 Surveillance Team |
| EPI_ISL_3222664 | Lighthouse Laboratory Plymouth | Wellcome Sanger Institute for the COVID-19 Genomics UK (COG-UK) Consortium | Lighthouse Laboratory Plymouth and Alex Alderton, Roberto Amato, Jeffrey Barrett, Sonia Goncalves, Ewan Harrison, David K. Jackson, Ian Johnston, Dominic Kwiatkowski, Cordelia Langford, John Sillitoe on behalf of the Wellcome Sanger Institute COVID-19 Surveillance Team |
| EPI_ISL_3222665 | Lighthouse Lab in Milton Keynes | Wellcome Sanger Institute for the COVID-19 Genomics UK (COG-UK) Consortium | The Lighthouse Lab in Milton Keynes and Alex Alderton, Roberto Amato, Jeffrey Barrett, Sonia Goncalves, Ewan Harrison, David K. Jackson, Ian Johnston, Dominic Kwiatkowski, Cordelia Langford, John Sillitoe on behalf of the Wellcome Sanger Institute COVID-19 Surveillance Team |
| EPI_ISL_3222666, EPI_ISL_3222667, EPI_ISL_3222668, EPI_ISL_3222669 | Lighthouse Laboratory Plymouth | Wellcome Sanger Institute for the COVID-19 Genomics UK (COG-UK) Consortium | Lighthouse Laboratory Plymouth and Alex Alderton, Roberto Amato, Jeffrey Barrett, Sonia Goncalves, Ewan Harrison, David K. Jackson, Ian Johnston, Dominic Kwiatkowski, Cordelia Langford, John Sillitoe on behalf of the Wellcome Sanger Institute COVID-19 Surveillance Team |
| EPI_ISL_3222670 | Lighthouse Lab in Alderley Park | Wellcome Sanger Institute for the COVID-19 Genomics UK (COG-UK) Consortium | Jacquelyn Wynn, Mairead Hyland, The Lighthouse Lab in Alderley Park and Alex Alderton, Roberto Amato, Jeffrey Barrett, Sonia Goncalves, Ewan Harrison, David K. Jackson, Ian Johnston, Dominic Kwiatkowski, Cordelia Langford, John Sillitoe on behalf of the Wellcome Sanger Institute COVID-19 Surveillance Team |
| EPI_ISL_3222671, EPI_ISL_3222672 | Lighthouse Lab in Milton Keynes | Wellcome Sanger Institute for the COVID-19 Genomics UK (COG-UK) Consortium | The Lighthouse Lab in Milton Keynes and Alex Alderton, Roberto Amato, Jeffrey Barrett, Sonia Goncalves, Ewan Harrison, David K. Jackson, Ian Johnston, Dominic Kwiatkowski, Cordelia Langford, John Sillitoe on behalf of the Wellcome Sanger Institute COVID-19 Surveillance Team |
| EPI_ISL_3222673 | Lighthouse Laboratory Plymouth | Wellcome Sanger Institute for the COVID-19 Genomics UK (COG-UK) Consortium | Lighthouse Laboratory Plymouth and Alex Alderton, Roberto Amato, Jeffrey Barrett, Sonia Goncalves, Ewan Harrison, David K. Jackson, Ian Johnston, Dominic Kwiatkowski, Cordelia Langford, John Sillitoe on behalf of the Wellcome Sanger Institute COVID-19 Surveillance Team |
| EPI_ISL_3222674 | Lighthouse Lab in Alderley Park | Wellcome Sanger Institute for the COVID-19 Genomics UK (COG-UK) Consortium | Jacquelyn Wynn, Mairead Hyland, The Lighthouse Lab in Alderley Park and Alex Alderton, Roberto Amato, Jeffrey Barrett, Sonia Goncalves, Ewan Harrison, David K. Jackson, Ian Johnston, Dominic Kwiatkowski, Cordelia Langford, John Sillitoe on behalf of the Wellcome Sanger Institute COVID-19 Surveillance Team |
| EPI_ISL_3222675 | Lighthouse Lab in Milton Keynes | Wellcome Sanger Institute for the COVID-19 Genomics UK (COG-UK) Consortium | The Lighthouse Lab in Milton Keynes and Alex Alderton, Roberto Amato, Jeffrey Barrett, Sonia Goncalves, Ewan Harrison, David K. Jackson, Ian Johnston, Dominic Kwiatkowski, Cordelia Langford, John Sillitoe on behalf of the Wellcome Sanger Institute COVID-19 Surveillance Team |
| EPI_ISL_3222676, EPI_ISL_3222677 | Lighthouse Laboratory Plymouth | Wellcome Sanger Institute for the COVID-19 Genomics UK (COG-UK) Consortium | Lighthouse Laboratory Plymouth and Alex Alderton, Roberto Amato, Jeffrey Barrett, Sonia Goncalves, Ewan Harrison, David K. Jackson, Ian Johnston, Dominic Kwiatkowski, Cordelia Langford, John Sillitoe on behalf of the Wellcome Sanger Institute COVID-19 Surveillance Team |
| EPI_ISL_3222678 | Lighthouse Lab in Milton Keynes | Wellcome Sanger Institute for the COVID-19 Genomics UK (COG-UK) Consortium | The Lighthouse Lab in Milton Keynes and Alex Alderton, Roberto Amato, Jeffrey Barrett, Sonia Goncalves, Ewan Harrison, David K. Jackson, Ian Johnston, Dominic Kwiatkowski, Cordelia Langford, John Sillitoe on behalf of the Wellcome Sanger Institute COVID-19 Surveillance Team |
| EPI_ISL_3222679, EPI_ISL_3222680 | Lighthouse Laboratory Plymouth | Wellcome Sanger Institute for the COVID-19 Genomics UK (COG-UK) Consortium | Lighthouse Laboratory Plymouth and Alex Alderton, Roberto Amato, Jeffrey Barrett, Sonia Goncalves, Ewan Harrison, David K. Jackson, Ian Johnston, Dominic Kwiatkowski, Cordelia Langford, John Sillitoe on behalf of the Wellcome Sanger Institute COVID-19 Surveillance Team |
| EPI_ISL_3222681 | Lighthouse Lab in Milton Keynes | Wellcome Sanger Institute for the COVID-19 Genomics UK (COG-UK) Consortium | The Lighthouse Lab in Milton Keynes and Alex Alderton, Roberto Amato, Jeffrey Barrett, Sonia Goncalves, Ewan Harrison, David K. Jackson, Ian Johnston, Dominic Kwiatkowski, Cordelia Langford, John Sillitoe on behalf of the Wellcome Sanger Institute COVID-19 Surveillance Team |
| EPI_ISL_3222682 | Lighthouse Laboratory Plymouth | Wellcome Sanger Institute for the COVID-19 Genomics UK (COG-UK) Consortium | Lighthouse Laboratory Plymouth and Alex Alderton, Roberto Amato, Jeffrey Barrett, Sonia Goncalves, Ewan Harrison, David K. Jackson, Ian Johnston, Dominic Kwiatkowski, Cordelia Langford, John Sillitoe on behalf of the Wellcome Sanger Institute COVID-19 Surveillance Team |
| EPI_ISL_3222683, EPI_ISL_3222684 | Lighthouse Lab in Milton Keynes | Wellcome Sanger Institute for the COVID-19 Genomics UK (COG-UK) Consortium | The Lighthouse Lab in Milton Keynes and Alex Alderton, Roberto Amato, Jeffrey Barrett, Sonia Goncalves, Ewan Harrison, David K. Jackson, Ian Johnston, Dominic Kwiatkowski, Cordelia Langford, John Sillitoe on behalf of the Wellcome Sanger Institute COVID-19 Surveillance Team |
| EPI_ISL_3222685 | Lighthouse Lab in Alderley Park | Wellcome Sanger Institute for the COVID-19 Genomics UK (COG-UK) Consortium | Jacquelyn Wynn, Mairead Hyland, The Lighthouse Lab in Alderley Park and Alex Alderton, Roberto Amato, Jeffrey Barrett, Sonia Goncalves, Ewan Harrison, David K. Jackson, Ian Johnston, Dominic Kwiatkowski, Cordelia Langford, John Sillitoe on behalf of the Wellcome Sanger Institute COVID-19 Surveillance Team |
| EPI_ISL_3222686 | Lighthouse Laboratory Plymouth | Wellcome Sanger Institute for the COVID-19 Genomics UK (COG-UK) Consortium | Lighthouse Laboratory Plymouth and Alex Alderton, Roberto Amato, Jeffrey Barrett, Sonia Goncalves, Ewan Harrison, David K. Jackson, Ian Johnston, Dominic Kwiatkowski, Cordelia Langford, John Sillitoe on behalf of the Wellcome Sanger Institute COVID-19 Surveillance Team |
| EPI_ISL_3222687 | Lighthouse Lab in Alderley Park | Wellcome Sanger Institute for the COVID-19 Genomics UK (COG-UK) Consortium | Jacquelyn Wynn, Mairead Hyland, The Lighthouse Lab in Alderley Park and Alex Alderton, Roberto Amato, Jeffrey Barrett, Sonia Goncalves, Ewan Harrison, David K. Jackson, Ian Johnston, Dominic Kwiatkowski, Cordelia Langford, John Sillitoe on behalf of the Wellcome Sanger Institute COVID-19 Surveillance Team |
| EPI_ISL_3222688 | Lighthouse Laboratory Plymouth | Wellcome Sanger Institute for the COVID-19 Genomics UK (COG-UK) Consortium | Lighthouse Laboratory Plymouth and Alex Alderton, Roberto Amato, Jeffrey Barrett, Sonia Goncalves, Ewan Harrison, David K. Jackson, Ian Johnston, Dominic Kwiatkowski, Cordelia Langford, John Sillitoe on behalf of the Wellcome Sanger Institute COVID-19 Surveillance Team |
| EPI_ISL_3222689 | Lighthouse Lab in Alderley Park | Wellcome Sanger Institute for the COVID-19 Genomics UK (COG-UK) Consortium | Jacquelyn Wynn, Mairead Hyland, The Lighthouse Lab in Alderley Park and Alex Alderton, Roberto Amato, Jeffrey Barrett, Sonia Goncalves, Ewan Harrison, David K. Jackson, Ian Johnston, Dominic Kwiatkowski, Cordelia Langford, John Sillitoe on behalf of the Wellcome Sanger Institute COVID-19 Surveillance Team |
| EPI_ISL_3222690, EPI_ISL_3222691, EPI_ISL_3222692, EPI_ISL_3222693 | Lighthouse Laboratory Plymouth | Wellcome Sanger Institute for the COVID-19 Genomics UK (COG-UK) Consortium | Lighthouse Laboratory Plymouth and Alex Alderton, Roberto Amato, Jeffrey Barrett, Sonia Goncalves, Ewan Harrison, David K. Jackson, Ian Johnston, Dominic Kwiatkowski, Cordelia Langford, John Sillitoe on behalf of the Wellcome Sanger Institute COVID-19 Surveillance Team |
| EPI_ISL_3222694 | Lighthouse Lab in Milton Keynes | Wellcome Sanger Institute for the COVID-19 Genomics UK (COG-UK) Consortium | The Lighthouse Lab in Milton Keynes and Alex Alderton, Roberto Amato, Jeffrey Barrett, Sonia Goncalves, Ewan Harrison, David K. Jackson, Ian Johnston, Dominic Kwiatkowski, Cordelia Langford, John Sillitoe on behalf of the Wellcome Sanger Institute COVID-19 Surveillance Team |
| EPI_ISL_3222695 | Lighthouse Laboratory Plymouth | Wellcome Sanger Institute for the COVID-19 Genomics UK (COG-UK) Consortium | Lighthouse Laboratory Plymouth and Alex Alderton, Roberto Amato, Jeffrey Barrett, Sonia Goncalves, Ewan Harrison, David K. Jackson, Ian Johnston, Dominic Kwiatkowski, Cordelia Langford, John Sillitoe on behalf of the Wellcome Sanger Institute COVID-19 Surveillance Team |
| EPI_ISL_3222696, EPI_ISL_3222697, EPI_ISL_3222698 | Lighthouse Lab in Milton Keynes | Wellcome Sanger Institute for the COVID-19 Genomics UK (COG-UK) Consortium | The Lighthouse Lab in Milton Keynes and Alex Alderton, Roberto Amato, Jeffrey Barrett, Sonia Goncalves, Ewan Harrison, David K. Jackson, Ian Johnston, Dominic Kwiatkowski, Cordelia Langford, John Sillitoe on behalf of the Wellcome Sanger Institute COVID-19 Surveillance Team |
| EPI_ISL_3222699, EPI_ISL_3222700 | Lighthouse Lab in Alderley Park | Wellcome Sanger Institute for the COVID-19 Genomics UK (COG-UK) Consortium | Jacquelyn Wynn, Mairead Hyland, The Lighthouse Lab in Alderley Park and Alex Alderton, Roberto Amato, Jeffrey Barrett, Sonia Goncalves, Ewan Harrison, David K. Jackson, Ian Johnston, Dominic Kwiatkowski, Cordelia Langford, John Sillitoe on behalf of the Wellcome Sanger Institute COVID-19 Surveillance Team |
| EPI_ISL_3222701, EPI_ISL_3222702 | Lighthouse Lab in Milton Keynes | Wellcome Sanger Institute for the COVID-19 Genomics UK (COG-UK) Consortium | The Lighthouse Lab in Milton Keynes and Alex Alderton, Roberto Amato, Jeffrey Barrett, Sonia Goncalves, Ewan Harrison, David K. Jackson, Ian Johnston, Dominic Kwiatkowski, Cordelia Langford, John Sillitoe on behalf of the Wellcome Sanger Institute COVID-19 Surveillance Team |
| EPI_ISL_3222703, EPI_ISL_3222704 | Lighthouse Lab in Alderley Park | Wellcome Sanger Institute for the COVID-19 Genomics UK (COG-UK) Consortium | Jacquelyn Wynn, Mairead Hyland, The Lighthouse Lab in Alderley Park and Alex Alderton, Roberto Amato, Jeffrey Barrett, Sonia Goncalves, Ewan Harrison, David K. Jackson, Ian Johnston, Dominic Kwiatkowski, Cordelia Langford, John Sillitoe on behalf of the Wellcome Sanger Institute COVID-19 Surveillance Team |
| EPI_ISL_3222705, EPI_ISL_3222706, EPI_ISL_3222707, EPI_ISL_3222708 | Lighthouse Laboratory Plymouth | Wellcome Sanger Institute for the COVID-19 Genomics UK (COG-UK) Consortium | Lighthouse Laboratory Plymouth and Alex Alderton, Roberto Amato, Jeffrey Barrett, Sonia Goncalves, Ewan Harrison, David K. Jackson, Ian Johnston, Dominic Kwiatkowski, Cordelia Langford, John Sillitoe on behalf of the Wellcome Sanger Institute COVID-19 Surveillance Team |
| EPI_ISL_3222709 | Lighthouse Lab in Alderley Park | Wellcome Sanger Institute for the COVID-19 Genomics UK (COG-UK) Consortium | Jacquelyn Wynn, Mairead Hyland, The Lighthouse Lab in Alderley Park and Alex Alderton, Roberto Amato, Jeffrey Barrett, Sonia Goncalves, Ewan Harrison, David K. Jackson, Ian Johnston, Dominic Kwiatkowski, Cordelia Langford, John Sillitoe on behalf of the Wellcome Sanger Institute COVID-19 Surveillance Team |
| EPI_ISL_3222710, EPI_ISL_3222711, EPI_ISL_3222712 | Lighthouse Lab in Milton Keynes | Wellcome Sanger Institute for the COVID-19 Genomics UK (COG-UK) Consortium | The Lighthouse Lab in Milton Keynes and Alex Alderton, Roberto Amato, Jeffrey Barrett, Sonia Goncalves, Ewan Harrison, David K. Jackson, Ian Johnston, Dominic Kwiatkowski, Cordelia Langford, John Sillitoe on behalf of the Wellcome Sanger Institute COVID-19 Surveillance Team |
| EPI_ISL_3222713, EPI_ISL_3222714 | Lighthouse Laboratory Plymouth | Wellcome Sanger Institute for the COVID-19 Genomics UK (COG-UK) Consortium | Lighthouse Laboratory Plymouth and Alex Alderton, Roberto Amato, Jeffrey Barrett, Sonia Goncalves, Ewan Harrison, David K. Jackson, Ian Johnston, Dominic Kwiatkowski, Cordelia Langford, John Sillitoe on behalf of the Wellcome Sanger Institute COVID-19 Surveillance Team |
| EPI_ISL_3222715 | Lighthouse Lab in Alderley Park | Wellcome Sanger Institute for the COVID-19 Genomics UK (COG-UK) Consortium | Jacquelyn Wynn, Mairead Hyland, The Lighthouse Lab in Alderley Park and Alex Alderton, Roberto Amato, Jeffrey Barrett, Sonia Goncalves, Ewan Harrison, David K. Jackson, Ian Johnston, Dominic Kwiatkowski, Cordelia Langford, John Sillitoe on behalf of the Wellcome Sanger Institute COVID-19 Surveillance Team |
| EPI_ISL_3222716 | Lighthouse Laboratory Plymouth | Wellcome Sanger Institute for the COVID-19 Genomics UK (COG-UK) Consortium | Lighthouse Laboratory Plymouth and Alex Alderton, Roberto Amato, Jeffrey Barrett, Sonia Goncalves, Ewan Harrison, David K. Jackson, Ian Johnston, Dominic Kwiatkowski, Cordelia Langford, John Sillitoe on behalf of the Wellcome Sanger Institute COVID-19 Surveillance Team |
| EPI_ISL_3222717, EPI_ISL_3222718, EPI_ISL_3222719 | Lighthouse Lab in Alderley Park | Wellcome Sanger Institute for the COVID-19 Genomics UK (COG-UK) Consortium | Jacquelyn Wynn, Mairead Hyland, The Lighthouse Lab in Alderley Park and Alex Alderton, Roberto Amato, Jeffrey Barrett, Sonia Goncalves, Ewan Harrison, David K. Jackson, Ian Johnston, Dominic Kwiatkowski, Cordelia Langford, John Sillitoe on behalf of the Wellcome Sanger Institute COVID-19 Surveillance Team |
| EPI_ISL_3222720 | Lighthouse Lab in Milton Keynes | Wellcome Sanger Institute for the COVID-19 Genomics UK (COG-UK) | The Lighthouse Lab in Milton Keynes and Alex Alderton, Roberto Amato, Jeffrey Barrett, Sonia Goncalves, Ewan Harrison, David K. Jackson, Ian Johnston, Dominic Kwiatkowski, Cordelia Langford, John Sillitoe on behalf of the Wellcome Sanger |

| | | | |
|---|---|---|---|
| | | Consortium | Institute COVID-19 Surveillance Team |
| EPI_ISL_3222721, EPI_ISL_3222722 | Lighthouse Laboratory Plymouth | Wellcome Sanger Institute for the COVID-19 Genomics UK (COG-UK) Consortium | Lighthouse Laboratory Plymouth and Alex Alderton, Roberto Amato, Jeffrey Barrett, Sonia Goncalves, Ewan Harrison, David K. Jackson, Ian Johnston, Dominic Kwiatkowski, Cordelia Langford, John Sillitoe on behalf of the Wellcome Sanger Institute COVID-19 Surveillance Team |
| EPI_ISL_3222723, EPI_ISL_3222724, EPI_ISL_3222725 | Lighthouse Lab in Milton Keynes | Wellcome Sanger Institute for the COVID-19 Genomics UK (COG-UK) Consortium | The Lighthouse Lab in Milton Keynes and Alex Alderton, Roberto Amato, Jeffrey Barrett, Sonia Goncalves, Ewan Harrison, David K. Jackson, Ian Johnston, Dominic Kwiatkowski, Cordelia Langford, John Sillitoe on behalf of the Wellcome Sanger Institute COVID-19 Surveillance Team |
| EPI_ISL_3222726 | Lighthouse Laboratory Plymouth | Wellcome Sanger Institute for the COVID-19 Genomics UK (COG-UK) Consortium | Lighthouse Laboratory Plymouth and Alex Alderton, Roberto Amato, Jeffrey Barrett, Sonia Goncalves, Ewan Harrison, David K. Jackson, Ian Johnston, Dominic Kwiatkowski, Cordelia Langford, John Sillitoe on behalf of the Wellcome Sanger Institute COVID-19 Surveillance Team |
| EPI_ISL_3222727, EPI_ISL_3222728, EPI_ISL_3222729, EPI_ISL_3222730, EPI_ISL_3222731, EPI_ISL_3222732 | Lighthouse Lab in Alderley Park | Wellcome Sanger Institute for the COVID-19 Genomics UK (COG-UK) Consortium | Jacquelyn Wynn, Mairead Hyland, The Lighthouse Lab in Alderley Park and Alex Alderton, Roberto Amato, Jeffrey Barrett, Sonia Goncalves, Ewan Harrison, David K. Jackson, Ian Johnston, Dominic Kwiatkowski, Cordelia Langford, John Sillitoe on behalf of the Wellcome Sanger Institute COVID-19 Surveillance Team |
| EPI_ISL_3222733 | Lighthouse Laboratory Plymouth | Wellcome Sanger Institute for the COVID-19 Genomics UK (COG-UK) Consortium | Lighthouse Laboratory Plymouth and Alex Alderton, Roberto Amato, Jeffrey Barrett, Sonia Goncalves, Ewan Harrison, David K. Jackson, Ian Johnston, Dominic Kwiatkowski, Cordelia Langford, John Sillitoe on behalf of the Wellcome Sanger Institute COVID-19 Surveillance Team |
| EPI_ISL_3222734 | Lighthouse Lab in Alderley Park | Wellcome Sanger Institute for the COVID-19 Genomics UK (COG-UK) Consortium | Jacquelyn Wynn, Mairead Hyland, The Lighthouse Lab in Alderley Park and Alex Alderton, Roberto Amato, Jeffrey Barrett, Sonia Goncalves, Ewan Harrison, David K. Jackson, Ian Johnston, Dominic Kwiatkowski, Cordelia Langford, John Sillitoe on behalf of the Wellcome Sanger Institute COVID-19 Surveillance Team |
| EPI_ISL_3222735 | Lighthouse Lab in Milton Keynes | Wellcome Sanger Institute for the COVID-19 Genomics UK (COG-UK) Consortium | The Lighthouse Lab in Milton Keynes and Alex Alderton, Roberto Amato, Jeffrey Barrett, Sonia Goncalves, Ewan Harrison, David K. Jackson, Ian Johnston, Dominic Kwiatkowski, Cordelia Langford, John Sillitoe on behalf of the Wellcome Sanger Institute COVID-19 Surveillance Team |
| EPI_ISL_3222736, EPI_ISL_3222737 | Lighthouse Laboratory Plymouth | Wellcome Sanger Institute for the COVID-19 Genomics UK (COG-UK) Consortium | Lighthouse Laboratory Plymouth and Alex Alderton, Roberto Amato, Jeffrey Barrett, Sonia Goncalves, Ewan Harrison, David K. Jackson, Ian Johnston, Dominic Kwiatkowski, Cordelia Langford, John Sillitoe on behalf of the Wellcome Sanger Institute COVID-19 Surveillance Team |
| EPI_ISL_3222738 | Lighthouse Lab in Alderley Park | Wellcome Sanger Institute for the COVID-19 Genomics UK (COG-UK) Consortium | Jacquelyn Wynn, Mairead Hyland, The Lighthouse Lab in Alderley Park and Alex Alderton, Roberto Amato, Jeffrey Barrett, Sonia Goncalves, Ewan Harrison, David K. Jackson, Ian Johnston, Dominic Kwiatkowski, Cordelia Langford, John Sillitoe on behalf of the Wellcome Sanger Institute COVID-19 Surveillance Team |
| EPI_ISL_3222739 | Lighthouse Lab in Milton Keynes | Wellcome Sanger Institute for the COVID-19 Genomics UK (COG-UK) Consortium | The Lighthouse Lab in Milton Keynes and Alex Alderton, Roberto Amato, Jeffrey Barrett, Sonia Goncalves, Ewan Harrison, David K. Jackson, Ian Johnston, Dominic Kwiatkowski, Cordelia Langford, John Sillitoe on behalf of the Wellcome Sanger Institute COVID-19 Surveillance Team |
| EPI_ISL_3222740, EPI_ISL_3222741, EPI_ISL_3222742, EPI_ISL_3222743 | Lighthouse Laboratory Plymouth | Wellcome Sanger Institute for the COVID-19 Genomics UK (COG-UK) Consortium | Lighthouse Laboratory Plymouth and Alex Alderton, Roberto Amato, Jeffrey Barrett, Sonia Goncalves, Ewan Harrison, David K. Jackson, Ian Johnston, Dominic Kwiatkowski, Cordelia Langford, John Sillitoe on behalf of the Wellcome Sanger Institute COVID-19 Surveillance Team |
| EPI_ISL_3222744 | Lighthouse Lab in Alderley Park | Wellcome Sanger Institute for the COVID-19 Genomics UK (COG-UK) Consortium | Jacquelyn Wynn, Mairead Hyland, The Lighthouse Lab in Alderley Park and Alex Alderton, Roberto Amato, Jeffrey Barrett, Sonia Goncalves, Ewan Harrison, David K. Jackson, Ian Johnston, Dominic Kwiatkowski, Cordelia Langford, John Sillitoe on behalf of the Wellcome Sanger Institute COVID-19 Surveillance Team |
| EPI_ISL_3222745 | Lighthouse Laboratory Plymouth | Wellcome Sanger Institute for the COVID-19 Genomics UK (COG-UK) Consortium | Lighthouse Laboratory Plymouth and Alex Alderton, Roberto Amato, Jeffrey Barrett, Sonia Goncalves, Ewan Harrison, David K. Jackson, Ian Johnston, Dominic Kwiatkowski, Cordelia Langford, John Sillitoe on behalf of the Wellcome Sanger Institute COVID-19 Surveillance Team |
| EPI_ISL_3222746 | Lighthouse Lab in Milton Keynes | Wellcome Sanger Institute for the COVID-19 Genomics UK (COG-UK) Consortium | The Lighthouse Lab in Milton Keynes and Alex Alderton, Roberto Amato, Jeffrey Barrett, Sonia Goncalves, Ewan Harrison, David K. Jackson, Ian Johnston, Dominic Kwiatkowski, Cordelia Langford, John Sillitoe on behalf of the Wellcome Sanger Institute COVID-19 Surveillance Team |
| EPI_ISL_3222747 | Lighthouse Lab in Alderley Park | Wellcome Sanger Institute for the COVID-19 Genomics UK (COG-UK) Consortium | Jacquelyn Wynn, Mairead Hyland, The Lighthouse Lab in Alderley Park and Alex Alderton, Roberto Amato, Jeffrey Barrett, Sonia Goncalves, Ewan Harrison, David K. Jackson, Ian Johnston, Dominic Kwiatkowski, Cordelia Langford, John Sillitoe on behalf of the Wellcome Sanger Institute COVID-19 Surveillance Team |
| EPI_ISL_3222748 | Lighthouse Lab in Milton Keynes | Wellcome Sanger Institute for the COVID-19 Genomics UK (COG-UK) Consortium | The Lighthouse Lab in Milton Keynes and Alex Alderton, Roberto Amato, Jeffrey Barrett, Sonia Goncalves, Ewan Harrison, David K. Jackson, Ian Johnston, Dominic Kwiatkowski, Cordelia Langford, John Sillitoe on behalf of the Wellcome Sanger Institute COVID-19 Surveillance Team |
| EPI_ISL_3222749, EPI_ISL_3222750, EPI_ISL_3222751, EPI_ISL_3222752 | Lighthouse Laboratory Plymouth | Wellcome Sanger Institute for the COVID-19 Genomics UK (COG-UK) Consortium | Lighthouse Laboratory Plymouth and Alex Alderton, Roberto Amato, Jeffrey Barrett, Sonia Goncalves, Ewan Harrison, David K. Jackson, Ian Johnston, Dominic Kwiatkowski, Cordelia Langford, John Sillitoe on behalf of the Wellcome Sanger Institute COVID-19 Surveillance Team |
| EPI_ISL_3222753 | Lighthouse Lab in Alderley Park | Wellcome Sanger Institute for the COVID-19 Genomics UK (COG-UK) Consortium | Jacquelyn Wynn, Mairead Hyland, The Lighthouse Lab in Alderley Park and Alex Alderton, Roberto Amato, Jeffrey Barrett, Sonia Goncalves, Ewan Harrison, David K. Jackson, Ian Johnston, Dominic Kwiatkowski, Cordelia Langford, John Sillitoe on behalf of the Wellcome Sanger Institute COVID-19 Surveillance Team |
| EPI_ISL_3222754, EPI_ISL_3222755 | Lighthouse Laboratory Plymouth | Wellcome Sanger Institute for the COVID-19 Genomics UK (COG-UK) Consortium | Lighthouse Laboratory Plymouth and Alex Alderton, Roberto Amato, Jeffrey Barrett, Sonia Goncalves, Ewan Harrison, David K. Jackson, Ian Johnston, Dominic Kwiatkowski, Cordelia Langford, John Sillitoe on behalf of the Wellcome Sanger Institute COVID-19 Surveillance Team |
| EPI_ISL_3222756 | Lighthouse Lab in Alderley Park | Wellcome Sanger Institute for the COVID-19 Genomics UK (COG-UK) Consortium | Jacquelyn Wynn, Mairead Hyland, The Lighthouse Lab in Alderley Park and Alex Alderton, Roberto Amato, Jeffrey Barrett, Sonia Goncalves, Ewan Harrison, David K. Jackson, Ian Johnston, Dominic Kwiatkowski, Cordelia Langford, John Sillitoe on behalf of the Wellcome Sanger Institute COVID-19 Surveillance Team |
| EPI_ISL_3222757, EPI_ISL_3222758 | Lighthouse Lab in Milton Keynes | Wellcome Sanger Institute for the COVID-19 Genomics UK (COG-UK) Consortium | The Lighthouse Lab in Milton Keynes and Alex Alderton, Roberto Amato, Jeffrey Barrett, Sonia Goncalves, Ewan Harrison, David K. Jackson, Ian Johnston, Dominic Kwiatkowski, Cordelia Langford, John Sillitoe on behalf of the Wellcome Sanger Institute COVID-19 Surveillance Team |
| EPI_ISL_3222759, EPI_ISL_3222760 | Lighthouse Laboratory Plymouth | Wellcome Sanger Institute for the COVID-19 Genomics UK (COG-UK) Consortium | Lighthouse Laboratory Plymouth and Alex Alderton, Roberto Amato, Jeffrey Barrett, Sonia Goncalves, Ewan Harrison, David K. Jackson, Ian Johnston, Dominic Kwiatkowski, Cordelia Langford, John Sillitoe on behalf of the Wellcome Sanger Institute COVID-19 Surveillance Team |
| EPI_ISL_3222761, EPI_ISL_3222762 | Lighthouse Lab in Milton Keynes | Wellcome Sanger Institute for the COVID-19 Genomics UK (COG-UK) Consortium | The Lighthouse Lab in Milton Keynes and Alex Alderton, Roberto Amato, Jeffrey Barrett, Sonia Goncalves, Ewan Harrison, David K. Jackson, Ian Johnston, Dominic Kwiatkowski, Cordelia Langford, John Sillitoe on behalf of the Wellcome Sanger Institute COVID-19 Surveillance Team |
| EPI_ISL_3222763 | Lighthouse Lab in Alderley Park | Wellcome Sanger Institute for the COVID-19 Genomics UK (COG-UK) Consortium | Jacquelyn Wynn, Mairead Hyland, The Lighthouse Lab in Alderley Park and Alex Alderton, Roberto Amato, Jeffrey Barrett, Sonia Goncalves, Ewan Harrison, David K. Jackson, Ian Johnston, Dominic Kwiatkowski, Cordelia Langford, John Sillitoe on behalf of the Wellcome Sanger Institute COVID-19 Surveillance Team |
| EPI_ISL_3222764, EPI_ISL_3222765 | Lighthouse Lab in Milton Keynes | Wellcome Sanger Institute for the COVID-19 Genomics UK (COG-UK) Consortium | The Lighthouse Lab in Milton Keynes and Alex Alderton, Roberto Amato, Jeffrey Barrett, Sonia Goncalves, Ewan Harrison, David K. Jackson, Ian Johnston, Dominic Kwiatkowski, Cordelia Langford, John Sillitoe on behalf of the Wellcome Sanger Institute COVID-19 Surveillance Team |
| EPI_ISL_3222766, EPI_ISL_3222767 | Lighthouse Laboratory Plymouth | Wellcome Sanger Institute for the COVID-19 Genomics UK (COG-UK) Consortium | Lighthouse Laboratory Plymouth and Alex Alderton, Roberto Amato, Jeffrey Barrett, Sonia Goncalves, Ewan Harrison, David K. Jackson, Ian Johnston, Dominic Kwiatkowski, Cordelia Langford, John Sillitoe on behalf of the Wellcome Sanger Institute COVID-19 Surveillance Team |
| EPI_ISL_3222768 | Lighthouse Lab in Milton Keynes | Wellcome Sanger Institute for the COVID-19 Genomics UK (COG-UK) Consortium | The Lighthouse Lab in Milton Keynes and Alex Alderton, Roberto Amato, Jeffrey Barrett, Sonia Goncalves, Ewan Harrison, David K. Jackson, Ian Johnston, Dominic Kwiatkowski, Cordelia Langford, John Sillitoe on behalf of the Wellcome Sanger Institute COVID-19 Surveillance Team |
| EPI_ISL_3222769, EPI_ISL_3222770 | Lighthouse Laboratory Plymouth | Wellcome Sanger Institute for the COVID-19 Genomics UK (COG-UK) Consortium | Lighthouse Laboratory Plymouth and Alex Alderton, Roberto Amato, Jeffrey Barrett, Sonia Goncalves, Ewan Harrison, David K. Jackson, Ian Johnston, Dominic Kwiatkowski, Cordelia Langford, John Sillitoe on behalf of the Wellcome Sanger Institute COVID-19 Surveillance Team |
| EPI_ISL_3222771 | Lighthouse Lab in Alderley Park | Wellcome Sanger Institute for the COVID-19 Genomics UK (COG-UK) Consortium | Jacquelyn Wynn, Mairead Hyland, The Lighthouse Lab in Alderley Park and Alex Alderton, Roberto Amato, Jeffrey Barrett, Sonia Goncalves, Ewan Harrison, David K. Jackson, Ian Johnston, Dominic Kwiatkowski, Cordelia Langford, John Sillitoe on behalf of the Wellcome Sanger Institute COVID-19 Surveillance Team |
| EPI_ISL_3222772 | Lighthouse Lab in Milton Keynes | Wellcome Sanger Institute for the COVID-19 Genomics UK (COG-UK) Consortium | The Lighthouse Lab in Milton Keynes and Alex Alderton, Roberto Amato, Jeffrey Barrett, Sonia Goncalves, Ewan Harrison, David K. Jackson, Ian Johnston, Dominic Kwiatkowski, Cordelia Langford, John Sillitoe on behalf of the Wellcome Sanger Institute COVID-19 Surveillance Team |
| EPI_ISL_3222773 | Lighthouse Laboratory Plymouth | Wellcome Sanger Institute for the COVID-19 Genomics UK (COG-UK) Consortium | Lighthouse Laboratory Plymouth and Alex Alderton, Roberto Amato, Jeffrey Barrett, Sonia Goncalves, Ewan Harrison, David K. Jackson, Ian Johnston, Dominic Kwiatkowski, Cordelia Langford, John Sillitoe on behalf of the Wellcome Sanger Institute COVID-19 Surveillance Team |
| EPI_ISL_3222774, EPI_ISL_3222775 | Lighthouse Lab in Alderley Park | Wellcome Sanger Institute for the COVID-19 Genomics UK (COG-UK) Consortium | Jacquelyn Wynn, Mairead Hyland, The Lighthouse Lab in Alderley Park and Alex Alderton, Roberto Amato, Jeffrey Barrett, Sonia Goncalves, Ewan Harrison, David K. Jackson, Ian Johnston, Dominic Kwiatkowski, Cordelia Langford, John Sillitoe on behalf of the Wellcome Sanger Institute COVID-19 Surveillance Team |
| EPI_ISL_3222776, EPI_ISL_3222777, EPI_ISL_3222778 | Lighthouse Lab in Milton Keynes | Wellcome Sanger Institute for the COVID-19 Genomics UK (COG-UK) Consortium | The Lighthouse Lab in Milton Keynes and Alex Alderton, Roberto Amato, Jeffrey Barrett, Sonia Goncalves, Ewan Harrison, David K. Jackson, Ian Johnston, Dominic Kwiatkowski, Cordelia Langford, John Sillitoe on behalf of the Wellcome Sanger Institute COVID-19 Surveillance Team |
| EPI_ISL_3222779, EPI_ISL_3222780 | Lighthouse Laboratory Plymouth | Wellcome Sanger Institute for the COVID-19 Genomics UK (COG-UK) Consortium | Lighthouse Laboratory Plymouth and Alex Alderton, Roberto Amato, Jeffrey Barrett, Sonia Goncalves, Ewan Harrison, David K. Jackson, Ian Johnston, Dominic Kwiatkowski, Cordelia Langford, John Sillitoe on behalf of the Wellcome Sanger Institute COVID-19 Surveillance Team |
| EPI_ISL_3222781, EPI_ISL_3222782 | Lighthouse Lab in Alderley Park | Wellcome Sanger Institute for the COVID-19 Genomics UK (COG-UK) Consortium | Jacquelyn Wynn, Mairead Hyland, The Lighthouse Lab in Alderley Park and Alex Alderton, Roberto Amato, Jeffrey Barrett, Sonia Goncalves, Ewan Harrison, David K. Jackson, Ian Johnston, Dominic Kwiatkowski, Cordelia Langford, John Sillitoe on behalf of the Wellcome Sanger Institute COVID-19 Surveillance Team |
| EPI_ISL_3222783 | Lighthouse Laboratory Plymouth | Wellcome Sanger Institute for the COVID-19 Genomics UK (COG-UK) Consortium | Lighthouse Laboratory Plymouth and Alex Alderton, Roberto Amato, Jeffrey Barrett, Sonia Goncalves, Ewan Harrison, David K. Jackson, Ian Johnston, Dominic Kwiatkowski, Cordelia Langford, John Sillitoe on behalf of the Wellcome Sanger Institute COVID-19 Surveillance Team |
| EPI_ISL_3222784 | Lighthouse Lab in Milton Keynes | Wellcome Sanger Institute for the COVID-19 Genomics UK (COG-UK) Consortium | The Lighthouse Lab in Milton Keynes and Alex Alderton, Roberto Amato, Jeffrey Barrett, Sonia Goncalves, Ewan Harrison, David K. Jackson, Ian Johnston, Dominic Kwiatkowski, Cordelia Langford, John Sillitoe on behalf of the Wellcome Sanger Institute COVID-19 Surveillance Team |
| EPI_ISL_3222785, EPI_ISL_3222786 | Lighthouse Laboratory Plymouth | Wellcome Sanger Institute for the COVID-19 Genomics UK (COG-UK) Consortium | Lighthouse Laboratory Plymouth and Alex Alderton, Roberto Amato, Jeffrey Barrett, Sonia Goncalves, Ewan Harrison, David K. Jackson, Ian Johnston, Dominic Kwiatkowski, Cordelia Langford, John Sillitoe on behalf of the Wellcome Sanger Institute COVID-19 Surveillance Team |
| EPI_ISL_3222787, EPI_ISL_3222788, EPI_ISL_3222789 | Lighthouse Lab in Alderley Park | Wellcome Sanger Institute for the COVID-19 Genomics UK (COG-UK) Consortium | Jacquelyn Wynn, Mairead Hyland, The Lighthouse Lab in Alderley Park and Alex Alderton, Roberto Amato, Jeffrey Barrett, Sonia Goncalves, Ewan Harrison, David K. Jackson, Ian Johnston, Dominic Kwiatkowski, Cordelia Langford, John Sillitoe on behalf of the Wellcome Sanger Institute COVID-19 Surveillance Team |
| EPI_ISL_3222790 | Lighthouse Lab in Milton Keynes | Wellcome Sanger Institute for the COVID-19 Genomics UK (COG-UK) Consortium | The Lighthouse Lab in Milton Keynes and Alex Alderton, Roberto Amato, Jeffrey Barrett, Sonia Goncalves, Ewan Harrison, David K. Jackson, Ian Johnston, Dominic Kwiatkowski, Cordelia Langford, John Sillitoe on behalf of the Wellcome Sanger Institute COVID-19 Surveillance Team |

| | | | |
|---|---|---|---|
| EPI_ISL_3222791, EPI_ISL_3222792 | Lighthouse Laboratory Plymouth | Wellcome Sanger Institute for the COVID-19 Genomics UK (COG-UK) Consortium | Lighthouse Laboratory Plymouth and Alex Alderton, Roberto Amato, Jeffrey Barrett, Sonia Goncalves, Ewan Harrison, David K. Jackson, Ian Johnston, Dominic Kwiatkowski, Cordelia Langford, John Sillitoe on behalf of the Wellcome Sanger Institute COVID-19 Surveillance Team |
| EPI_ISL_3222793 | Lighthouse Lab in Milton Keynes | Wellcome Sanger Institute for the COVID-19 Genomics UK (COG-UK) Consortium | The Lighthouse Lab in Milton Keynes and Alex Alderton, Roberto Amato, Jeffrey Barrett, Sonia Goncalves, Ewan Harrison, David K. Jackson, Ian Johnston, Dominic Kwiatkowski, Cordelia Langford, John Sillitoe on behalf of the Wellcome Sanger Institute COVID-19 Surveillance Team |
| EPI_ISL_3222794 | Lighthouse Laboratory Plymouth | Wellcome Sanger Institute for the COVID-19 Genomics UK (COG-UK) Consortium | Lighthouse Laboratory Plymouth and Alex Alderton, Roberto Amato, Jeffrey Barrett, Sonia Goncalves, Ewan Harrison, David K. Jackson, Ian Johnston, Dominic Kwiatkowski, Cordelia Langford, John Sillitoe on behalf of the Wellcome Sanger Institute COVID-19 Surveillance Team |
| EPI_ISL_3222795 | Lighthouse Lab in Alderley Park | Wellcome Sanger Institute for the COVID-19 Genomics UK (COG-UK) Consortium | Jacquelyn Wynn, Mairead Hyland, The Lighthouse Lab in Alderley Park and Alex Alderton, Roberto Amato, Jeffrey Barrett, Sonia Goncalves, Ewan Harrison, David K. Jackson, Ian Johnston, Dominic Kwiatkowski, Cordelia Langford, John Sillitoe on behalf of the Wellcome Sanger Institute COVID-19 Surveillance Team |
| EPI_ISL_3222796 | Lighthouse Laboratory Plymouth | Wellcome Sanger Institute for the COVID-19 Genomics UK (COG-UK) Consortium | Lighthouse Laboratory Plymouth and Alex Alderton, Roberto Amato, Jeffrey Barrett, Sonia Goncalves, Ewan Harrison, David K. Jackson, Ian Johnston, Dominic Kwiatkowski, Cordelia Langford, John Sillitoe on behalf of the Wellcome Sanger Institute COVID-19 Surveillance Team |
| EPI_ISL_3222797 | Lighthouse Lab in Alderley Park | Wellcome Sanger Institute for the COVID-19 Genomics UK (COG-UK) Consortium | Jacquelyn Wynn, Mairead Hyland, The Lighthouse Lab in Alderley Park and Alex Alderton, Roberto Amato, Jeffrey Barrett, Sonia Goncalves, Ewan Harrison, David K. Jackson, Ian Johnston, Dominic Kwiatkowski, Cordelia Langford, John Sillitoe on behalf of the Wellcome Sanger Institute COVID-19 Surveillance Team |
| EPI_ISL_3222798 | Lighthouse Lab in Milton Keynes | Wellcome Sanger Institute for the COVID-19 Genomics UK (COG-UK) Consortium | The Lighthouse Lab in Milton Keynes and Alex Alderton, Roberto Amato, Jeffrey Barrett, Sonia Goncalves, Ewan Harrison, David K. Jackson, Ian Johnston, Dominic Kwiatkowski, Cordelia Langford, John Sillitoe on behalf of the Wellcome Sanger Institute COVID-19 Surveillance Team |
| EPI_ISL_3222799, EPI_ISL_3222800, EPI_ISL_3222801 | Lighthouse Laboratory Plymouth | Wellcome Sanger Institute for the COVID-19 Genomics UK (COG-UK) Consortium | Lighthouse Laboratory Plymouth and Alex Alderton, Roberto Amato, Jeffrey Barrett, Sonia Goncalves, Ewan Harrison, David K. Jackson, Ian Johnston, Dominic Kwiatkowski, Cordelia Langford, John Sillitoe on behalf of the Wellcome Sanger Institute COVID-19 Surveillance Team |
| EPI_ISL_3222802 | Lighthouse Lab in Milton Keynes | Wellcome Sanger Institute for the COVID-19 Genomics UK (COG-UK) Consortium | The Lighthouse Lab in Milton Keynes and Alex Alderton, Roberto Amato, Jeffrey Barrett, Sonia Goncalves, Ewan Harrison, David K. Jackson, Ian Johnston, Dominic Kwiatkowski, Cordelia Langford, John Sillitoe on behalf of the Wellcome Sanger Institute COVID-19 Surveillance Team |
| EPI_ISL_3222803 | Lighthouse Laboratory Plymouth | Wellcome Sanger Institute for the COVID-19 Genomics UK (COG-UK) Consortium | Lighthouse Laboratory Plymouth and Alex Alderton, Roberto Amato, Jeffrey Barrett, Sonia Goncalves, Ewan Harrison, David K. Jackson, Ian Johnston, Dominic Kwiatkowski, Cordelia Langford, John Sillitoe on behalf of the Wellcome Sanger Institute COVID-19 Surveillance Team |
| EPI_ISL_3222804 | Lighthouse Lab in Milton Keynes | Wellcome Sanger Institute for the COVID-19 Genomics UK (COG-UK) Consortium | The Lighthouse Lab in Milton Keynes and Alex Alderton, Roberto Amato, Jeffrey Barrett, Sonia Goncalves, Ewan Harrison, David K. Jackson, Ian Johnston, Dominic Kwiatkowski, Cordelia Langford, John Sillitoe on behalf of the Wellcome Sanger Institute COVID-19 Surveillance Team |
| EPI_ISL_3222805, EPI_ISL_3222806, EPI_ISL_3222807, EPI_ISL_3222808, EPI_ISL_3222809, EPI_ISL_3222810 | Lighthouse Laboratory Plymouth | Wellcome Sanger Institute for the COVID-19 Genomics UK (COG-UK) Consortium | Lighthouse Laboratory Plymouth and Alex Alderton, Roberto Amato, Jeffrey Barrett, Sonia Goncalves, Ewan Harrison, David K. Jackson, Ian Johnston, Dominic Kwiatkowski, Cordelia Langford, John Sillitoe on behalf of the Wellcome Sanger Institute COVID-19 Surveillance Team |
| EPI_ISL_3222811 | Lighthouse Lab in Milton Keynes | Wellcome Sanger Institute for the COVID-19 Genomics UK (COG-UK) Consortium | The Lighthouse Lab in Milton Keynes and Alex Alderton, Roberto Amato, Jeffrey Barrett, Sonia Goncalves, Ewan Harrison, David K. Jackson, Ian Johnston, Dominic Kwiatkowski, Cordelia Langford, John Sillitoe on behalf of the Wellcome Sanger Institute COVID-19 Surveillance Team |
| EPI_ISL_3222812 | Lighthouse Laboratory Plymouth | Wellcome Sanger Institute for the COVID-19 Genomics UK (COG-UK) Consortium | Lighthouse Laboratory Plymouth and Alex Alderton, Roberto Amato, Jeffrey Barrett, Sonia Goncalves, Ewan Harrison, David K. Jackson, Ian Johnston, Dominic Kwiatkowski, Cordelia Langford, John Sillitoe on behalf of the Wellcome Sanger Institute COVID-19 Surveillance Team |
| EPI_ISL_3222813 | Lighthouse Lab in Alderley Park | Wellcome Sanger Institute for the COVID-19 Genomics UK (COG-UK) Consortium | Jacquelyn Wynn, Mairead Hyland, The Lighthouse Lab in Alderley Park and Alex Alderton, Roberto Amato, Jeffrey Barrett, Sonia Goncalves, Ewan Harrison, David K. Jackson, Ian Johnston, Dominic Kwiatkowski, Cordelia Langford, John Sillitoe on behalf of the Wellcome Sanger Institute COVID-19 Surveillance Team |
| EPI_ISL_3222814 | Lighthouse Lab in Milton Keynes | Wellcome Sanger Institute for the COVID-19 Genomics UK (COG-UK) Consortium | The Lighthouse Lab in Milton Keynes and Alex Alderton, Roberto Amato, Jeffrey Barrett, Sonia Goncalves, Ewan Harrison, David K. Jackson, Ian Johnston, Dominic Kwiatkowski, Cordelia Langford, John Sillitoe on behalf of the Wellcome Sanger Institute COVID-19 Surveillance Team |
| EPI_ISL_3222815, EPI_ISL_3222816, EPI_ISL_3222817 | Lighthouse Laboratory Plymouth | Wellcome Sanger Institute for the COVID-19 Genomics UK (COG-UK) Consortium | Lighthouse Laboratory Plymouth and Alex Alderton, Roberto Amato, Jeffrey Barrett, Sonia Goncalves, Ewan Harrison, David K. Jackson, Ian Johnston, Dominic Kwiatkowski, Cordelia Langford, John Sillitoe on behalf of the Wellcome Sanger Institute COVID-19 Surveillance Team |
| EPI_ISL_3222818 | Lighthouse Lab in Alderley Park | Wellcome Sanger Institute for the COVID-19 Genomics UK (COG-UK) Consortium | Jacquelyn Wynn, Mairead Hyland, The Lighthouse Lab in Alderley Park and Alex Alderton, Roberto Amato, Jeffrey Barrett, Sonia Goncalves, Ewan Harrison, David K. Jackson, Ian Johnston, Dominic Kwiatkowski, Cordelia Langford, John Sillitoe on behalf of the Wellcome Sanger Institute COVID-19 Surveillance Team |
| EPI_ISL_3222819 | Lighthouse Lab in Milton Keynes | Wellcome Sanger Institute for the COVID-19 Genomics UK (COG-UK) Consortium | The Lighthouse Lab in Milton Keynes and Alex Alderton, Roberto Amato, Jeffrey Barrett, Sonia Goncalves, Ewan Harrison, David K. Jackson, Ian Johnston, Dominic Kwiatkowski, Cordelia Langford, John Sillitoe on behalf of the Wellcome Sanger Institute COVID-19 Surveillance Team |
| EPI_ISL_3222820 | Lighthouse Laboratory Plymouth | Wellcome Sanger Institute for the COVID-19 Genomics UK (COG-UK) Consortium | Lighthouse Laboratory Plymouth and Alex Alderton, Roberto Amato, Jeffrey Barrett, Sonia Goncalves, Ewan Harrison, David K. Jackson, Ian Johnston, Dominic Kwiatkowski, Cordelia Langford, John Sillitoe on behalf of the Wellcome Sanger Institute COVID-19 Surveillance Team |
| EPI_ISL_3222821, EPI_ISL_3222822, EPI_ISL_3222823 | Lighthouse Lab in Milton Keynes | Wellcome Sanger Institute for the COVID-19 Genomics UK (COG-UK) Consortium | The Lighthouse Lab in Milton Keynes and Alex Alderton, Roberto Amato, Jeffrey Barrett, Sonia Goncalves, Ewan Harrison, David K. Jackson, Ian Johnston, Dominic Kwiatkowski, Cordelia Langford, John Sillitoe on behalf of the Wellcome Sanger Institute COVID-19 Surveillance Team |
| EPI_ISL_3222824, EPI_ISL_3222825 | Lighthouse Laboratory Plymouth | Wellcome Sanger Institute for the COVID-19 Genomics UK (COG-UK) Consortium | Lighthouse Laboratory Plymouth and Alex Alderton, Roberto Amato, Jeffrey Barrett, Sonia Goncalves, Ewan Harrison, David K. Jackson, Ian Johnston, Dominic Kwiatkowski, Cordelia Langford, John Sillitoe on behalf of the Wellcome Sanger Institute COVID-19 Surveillance Team |
| EPI_ISL_3222826 | Lighthouse Lab in Alderley Park | Wellcome Sanger Institute for the COVID-19 Genomics UK (COG-UK) Consortium | Jacquelyn Wynn, Mairead Hyland, The Lighthouse Lab in Alderley Park and Alex Alderton, Roberto Amato, Jeffrey Barrett, Sonia Goncalves, Ewan Harrison, David K. Jackson, Ian Johnston, Dominic Kwiatkowski, Cordelia Langford, John Sillitoe on behalf of the Wellcome Sanger Institute COVID-19 Surveillance Team |
| EPI_ISL_3222827 | Lighthouse Laboratory Plymouth | Wellcome Sanger Institute for the COVID-19 Genomics UK (COG-UK) Consortium | Lighthouse Laboratory Plymouth and Alex Alderton, Roberto Amato, Jeffrey Barrett, Sonia Goncalves, Ewan Harrison, David K. Jackson, Ian Johnston, Dominic Kwiatkowski, Cordelia Langford, John Sillitoe on behalf of the Wellcome Sanger Institute COVID-19 Surveillance Team |
| EPI_ISL_3222828 | Lighthouse Lab in Milton Keynes | Wellcome Sanger Institute for the COVID-19 Genomics UK (COG-UK) Consortium | The Lighthouse Lab in Milton Keynes and Alex Alderton, Roberto Amato, Jeffrey Barrett, Sonia Goncalves, Ewan Harrison, David K. Jackson, Ian Johnston, Dominic Kwiatkowski, Cordelia Langford, John Sillitoe on behalf of the Wellcome Sanger Institute COVID-19 Surveillance Team |
| EPI_ISL_3222829 | Lighthouse Laboratory Plymouth | Wellcome Sanger Institute for the COVID-19 Genomics UK (COG-UK) Consortium | Lighthouse Laboratory Plymouth and Alex Alderton, Roberto Amato, Jeffrey Barrett, Sonia Goncalves, Ewan Harrison, David K. Jackson, Ian Johnston, Dominic Kwiatkowski, Cordelia Langford, John Sillitoe on behalf of the Wellcome Sanger Institute COVID-19 Surveillance Team |
| EPI_ISL_3222830 | Lighthouse Lab in Milton Keynes | Wellcome Sanger Institute for the COVID-19 Genomics UK (COG-UK) Consortium | The Lighthouse Lab in Milton Keynes and Alex Alderton, Roberto Amato, Jeffrey Barrett, Sonia Goncalves, Ewan Harrison, David K. Jackson, Ian Johnston, Dominic Kwiatkowski, Cordelia Langford, John Sillitoe on behalf of the Wellcome Sanger Institute COVID-19 Surveillance Team |
| EPI_ISL_3222831 | Lighthouse Laboratory Plymouth | Wellcome Sanger Institute for the COVID-19 Genomics UK (COG-UK) Consortium | Lighthouse Laboratory Plymouth and Alex Alderton, Roberto Amato, Jeffrey Barrett, Sonia Goncalves, Ewan Harrison, David K. Jackson, Ian Johnston, Dominic Kwiatkowski, Cordelia Langford, John Sillitoe on behalf of the Wellcome Sanger Institute COVID-19 Surveillance Team |
| EPI_ISL_3222832 | Lighthouse Lab in Milton Keynes | Wellcome Sanger Institute for the COVID-19 Genomics UK (COG-UK) Consortium | The Lighthouse Lab in Milton Keynes and Alex Alderton, Roberto Amato, Jeffrey Barrett, Sonia Goncalves, Ewan Harrison, David K. Jackson, Ian Johnston, Dominic Kwiatkowski, Cordelia Langford, John Sillitoe on behalf of the Wellcome Sanger Institute COVID-19 Surveillance Team |
| EPI_ISL_3222833, EPI_ISL_3222834 | Lighthouse Lab in Alderley Park | Wellcome Sanger Institute for the COVID-19 Genomics UK (COG-UK) Consortium | Jacquelyn Wynn, Mairead Hyland, The Lighthouse Lab in Alderley Park and Alex Alderton, Roberto Amato, Jeffrey Barrett, Sonia Goncalves, Ewan Harrison, David K. Jackson, Ian Johnston, Dominic Kwiatkowski, Cordelia Langford, John Sillitoe on behalf of the Wellcome Sanger Institute COVID-19 Surveillance Team |
| EPI_ISL_3222835 | Lighthouse Laboratory Plymouth | Wellcome Sanger Institute for the COVID-19 Genomics UK (COG-UK) Consortium | Lighthouse Laboratory Plymouth and Alex Alderton, Roberto Amato, Jeffrey Barrett, Sonia Goncalves, Ewan Harrison, David K. Jackson, Ian Johnston, Dominic Kwiatkowski, Cordelia Langford, John Sillitoe on behalf of the Wellcome Sanger Institute COVID-19 Surveillance Team |
| EPI_ISL_3222836 | Lighthouse Lab in Milton Keynes | Wellcome Sanger Institute for the COVID-19 Genomics UK (COG-UK) Consortium | The Lighthouse Lab in Milton Keynes and Alex Alderton, Roberto Amato, Jeffrey Barrett, Sonia Goncalves, Ewan Harrison, David K. Jackson, Ian Johnston, Dominic Kwiatkowski, Cordelia Langford, John Sillitoe on behalf of the Wellcome Sanger Institute COVID-19 Surveillance Team |
| EPI_ISL_3222837, EPI_ISL_3222838, EPI_ISL_3222839, EPI_ISL_3222840, EPI_ISL_3222841 | Lighthouse Laboratory Plymouth | Wellcome Sanger Institute for the COVID-19 Genomics UK (COG-UK) Consortium | Lighthouse Laboratory Plymouth and Alex Alderton, Roberto Amato, Jeffrey Barrett, Sonia Goncalves, Ewan Harrison, David K. Jackson, Ian Johnston, Dominic Kwiatkowski, Cordelia Langford, John Sillitoe on behalf of the Wellcome Sanger Institute COVID-19 Surveillance Team |
| EPI_ISL_3222842 | Lighthouse Lab in Alderley Park | Wellcome Sanger Institute for the COVID-19 Genomics UK (COG-UK) Consortium | Jacquelyn Wynn, Mairead Hyland, The Lighthouse Lab in Alderley Park and Alex Alderton, Roberto Amato, Jeffrey Barrett, Sonia Goncalves, Ewan Harrison, David K. Jackson, Ian Johnston, Dominic Kwiatkowski, Cordelia Langford, John Sillitoe on behalf of the Wellcome Sanger Institute COVID-19 Surveillance Team |
| EPI_ISL_3222843, EPI_ISL_3222844 | Lighthouse Laboratory Plymouth | Wellcome Sanger Institute for the COVID-19 Genomics UK (COG-UK) Consortium | Lighthouse Laboratory Plymouth and Alex Alderton, Roberto Amato, Jeffrey Barrett, Sonia Goncalves, Ewan Harrison, David K. Jackson, Ian Johnston, Dominic Kwiatkowski, Cordelia Langford, John Sillitoe on behalf of the Wellcome Sanger Institute COVID-19 Surveillance Team |
| EPI_ISL_3222845 | Lighthouse Lab in Alderley Park | Wellcome Sanger Institute for the COVID-19 Genomics UK (COG-UK) Consortium | Jacquelyn Wynn, Mairead Hyland, The Lighthouse Lab in Alderley Park and Alex Alderton, Roberto Amato, Jeffrey Barrett, Sonia Goncalves, Ewan Harrison, David K. Jackson, Ian Johnston, Dominic Kwiatkowski, Cordelia Langford, John Sillitoe on behalf of the Wellcome Sanger Institute COVID-19 Surveillance Team |
| EPI_ISL_3222846, EPI_ISL_3222847, EPI_ISL_3222848 | Lighthouse Laboratory Plymouth | Wellcome Sanger Institute for the COVID-19 Genomics UK (COG-UK) Consortium | Lighthouse Laboratory Plymouth and Alex Alderton, Roberto Amato, Jeffrey Barrett, Sonia Goncalves, Ewan Harrison, David K. Jackson, Ian Johnston, Dominic Kwiatkowski, Cordelia Langford, John Sillitoe on behalf of the Wellcome Sanger Institute COVID-19 Surveillance Team |
| EPI_ISL_3222849 | Lighthouse Lab in Milton Keynes | Wellcome Sanger Institute for the COVID-19 Genomics UK (COG-UK) Consortium | The Lighthouse Lab in Milton Keynes and Alex Alderton, Roberto Amato, Jeffrey Barrett, Sonia Goncalves, Ewan Harrison, David K. Jackson, Ian Johnston, Dominic Kwiatkowski, Cordelia Langford, John Sillitoe on behalf of the Wellcome Sanger Institute COVID-19 Surveillance Team |
| EPI_ISL_3222850 | Lighthouse Laboratory Plymouth | Wellcome Sanger Institute for the COVID-19 Genomics UK (COG-UK) Consortium | Lighthouse Laboratory Plymouth and Alex Alderton, Roberto Amato, Jeffrey Barrett, Sonia Goncalves, Ewan Harrison, David K. Jackson, Ian Johnston, Dominic Kwiatkowski, Cordelia Langford, John Sillitoe on behalf of the Wellcome Sanger Institute COVID-19 Surveillance Team |
| EPI_ISL_3222851, EPI_ISL_3222852, EPI_ISL_3222853 | Lighthouse Lab in Milton Keynes | Wellcome Sanger Institute for the COVID-19 Genomics UK (COG-UK) Consortium | The Lighthouse Lab in Milton Keynes and Alex Alderton, Roberto Amato, Jeffrey Barrett, Sonia Goncalves, Ewan Harrison, David K. Jackson, Ian Johnston, Dominic Kwiatkowski, Cordelia Langford, John Sillitoe on behalf of the Wellcome Sanger Institute COVID-19 Surveillance Team |
| EPI_ISL_3222854 | Lighthouse Lab in Alderley Park | Wellcome Sanger Institute for the COVID-19 Genomics UK (COG-UK) | Jacquelyn Wynn, Mairead Hyland, The Lighthouse Lab in Alderley Park and Alex Alderton, Roberto Amato, Jeffrey Barrett, Sonia Goncalves, Ewan Harrison, David K. Jackson, Ian Johnston, Dominic Kwiatkowski, Cordelia Langford, John Sillitoe on |

| | | | |
|---|---|---|---|
| | | Consortium | behalf of the Wellcome Sanger Institute COVID-19 Surveillance Team |
| EPI_ISL_3222855 | Lighthouse Laboratory Plymouth | Wellcome Sanger Institute for the COVID-19 Genomics UK (COG-UK) Consortium | Lighthouse Laboratory Plymouth and Alex Alderton, Roberto Amato, Jeffrey Barrett, Sonia Goncalves, Ewan Harrison, David K. Jackson, Ian Johnston, Dominic Kwiatkowski, Cordelia Langford, John Sillitoe on behalf of the Wellcome Sanger Institute COVID-19 Surveillance Team |
| EPI_ISL_3222856 | Lighthouse Lab in Alderley Park | Wellcome Sanger Institute for the COVID-19 Genomics UK (COG-UK) Consortium | Jacquelyn Wynn, Mairead Hyland, The Lighthouse Lab in Alderley Park and Alex Alderton, Roberto Amato, Jeffrey Barrett, Sonia Goncalves, Ewan Harrison, David K. Jackson, Ian Johnston, Dominic Kwiatkowski, Cordelia Langford, John Sillitoe on behalf of the Wellcome Sanger Institute COVID-19 Surveillance Team |
| EPI_ISL_3222857 | Lighthouse Lab in Milton Keynes | Wellcome Sanger Institute for the COVID-19 Genomics UK (COG-UK) Consortium | The Lighthouse Lab in Milton Keynes and Alex Alderton, Roberto Amato, Jeffrey Barrett, Sonia Goncalves, Ewan Harrison, David K. Jackson, Ian Johnston, Dominic Kwiatkowski, Cordelia Langford, John Sillitoe on behalf of the Wellcome Sanger Institute COVID-19 Surveillance Team |
| EPI_ISL_3222858 | Lighthouse Laboratory Plymouth | Wellcome Sanger Institute for the COVID-19 Genomics UK (COG-UK) Consortium | Lighthouse Laboratory Plymouth and Alex Alderton, Roberto Amato, Jeffrey Barrett, Sonia Goncalves, Ewan Harrison, David K. Jackson, Ian Johnston, Dominic Kwiatkowski, Cordelia Langford, John Sillitoe on behalf of the Wellcome Sanger Institute COVID-19 Surveillance Team |
| EPI_ISL_3222859 | Lighthouse Lab in Alderley Park | Wellcome Sanger Institute for the COVID-19 Genomics UK (COG-UK) Consortium | Jacquelyn Wynn, Mairead Hyland, The Lighthouse Lab in Alderley Park and Alex Alderton, Roberto Amato, Jeffrey Barrett, Sonia Goncalves, Ewan Harrison, David K. Jackson, Ian Johnston, Dominic Kwiatkowski, Cordelia Langford, John Sillitoe on behalf of the Wellcome Sanger Institute COVID-19 Surveillance Team |
| EPI_ISL_3222860 | Lighthouse Laboratory Plymouth | Wellcome Sanger Institute for the COVID-19 Genomics UK (COG-UK) Consortium | Lighthouse Laboratory Plymouth and Alex Alderton, Roberto Amato, Jeffrey Barrett, Sonia Goncalves, Ewan Harrison, David K. Jackson, Ian Johnston, Dominic Kwiatkowski, Cordelia Langford, John Sillitoe on behalf of the Wellcome Sanger Institute COVID-19 Surveillance Team |
| EPI_ISL_3222861 | Lighthouse Lab in Alderley Park | Wellcome Sanger Institute for the COVID-19 Genomics UK (COG-UK) Consortium | Jacquelyn Wynn, Mairead Hyland, The Lighthouse Lab in Alderley Park and Alex Alderton, Roberto Amato, Jeffrey Barrett, Sonia Goncalves, Ewan Harrison, David K. Jackson, Ian Johnston, Dominic Kwiatkowski, Cordelia Langford, John Sillitoe on behalf of the Wellcome Sanger Institute COVID-19 Surveillance Team |
| EPI_ISL_3222862, EPI_ISL_3222863 | Lighthouse Laboratory Plymouth | Wellcome Sanger Institute for the COVID-19 Genomics UK (COG-UK) Consortium | Lighthouse Laboratory Plymouth and Alex Alderton, Roberto Amato, Jeffrey Barrett, Sonia Goncalves, Ewan Harrison, David K. Jackson, Ian Johnston, Dominic Kwiatkowski, Cordelia Langford, John Sillitoe on behalf of the Wellcome Sanger Institute COVID-19 Surveillance Team |
| EPI_ISL_3222864 | Lighthouse Lab in Milton Keynes | Wellcome Sanger Institute for the COVID-19 Genomics UK (COG-UK) Consortium | The Lighthouse Lab in Milton Keynes and Alex Alderton, Roberto Amato, Jeffrey Barrett, Sonia Goncalves, Ewan Harrison, David K. Jackson, Ian Johnston, Dominic Kwiatkowski, Cordelia Langford, John Sillitoe on behalf of the Wellcome Sanger Institute COVID-19 Surveillance Team |
| EPI_ISL_3222865 | Lighthouse Lab in Alderley Park | Wellcome Sanger Institute for the COVID-19 Genomics UK (COG-UK) Consortium | Jacquelyn Wynn, Mairead Hyland, The Lighthouse Lab in Alderley Park and Alex Alderton, Roberto Amato, Jeffrey Barrett, Sonia Goncalves, Ewan Harrison, David K. Jackson, Ian Johnston, Dominic Kwiatkowski, Cordelia Langford, John Sillitoe on behalf of the Wellcome Sanger Institute COVID-19 Surveillance Team |
| EPI_ISL_3222866 | Lighthouse Lab in Milton Keynes | Wellcome Sanger Institute for the COVID-19 Genomics UK (COG-UK) Consortium | The Lighthouse Lab in Milton Keynes and Alex Alderton, Roberto Amato, Jeffrey Barrett, Sonia Goncalves, Ewan Harrison, David K. Jackson, Ian Johnston, Dominic Kwiatkowski, Cordelia Langford, John Sillitoe on behalf of the Wellcome Sanger Institute COVID-19 Surveillance Team |
| EPI_ISL_3222867, EPI_ISL_3222868 | Lighthouse Laboratory Plymouth | Wellcome Sanger Institute for the COVID-19 Genomics UK (COG-UK) Consortium | Lighthouse Laboratory Plymouth and Alex Alderton, Roberto Amato, Jeffrey Barrett, Sonia Goncalves, Ewan Harrison, David K. Jackson, Ian Johnston, Dominic Kwiatkowski, Cordelia Langford, John Sillitoe on behalf of the Wellcome Sanger Institute COVID-19 Surveillance Team |
| EPI_ISL_3222869 | Lighthouse Lab in Alderley Park | Wellcome Sanger Institute for the COVID-19 Genomics UK (COG-UK) Consortium | Jacquelyn Wynn, Mairead Hyland, The Lighthouse Lab in Alderley Park and Alex Alderton, Roberto Amato, Jeffrey Barrett, Sonia Goncalves, Ewan Harrison, David K. Jackson, Ian Johnston, Dominic Kwiatkowski, Cordelia Langford, John Sillitoe on behalf of the Wellcome Sanger Institute COVID-19 Surveillance Team |
| EPI_ISL_3222870, EPI_ISL_3222871 | Lighthouse Laboratory Plymouth | Wellcome Sanger Institute for the COVID-19 Genomics UK (COG-UK) Consortium | Lighthouse Laboratory Plymouth and Alex Alderton, Roberto Amato, Jeffrey Barrett, Sonia Goncalves, Ewan Harrison, David K. Jackson, Ian Johnston, Dominic Kwiatkowski, Cordelia Langford, John Sillitoe on behalf of the Wellcome Sanger Institute COVID-19 Surveillance Team |
| EPI_ISL_3222872 | Lighthouse Lab in Alderley Park | Wellcome Sanger Institute for the COVID-19 Genomics UK (COG-UK) Consortium | Jacquelyn Wynn, Mairead Hyland, The Lighthouse Lab in Alderley Park and Alex Alderton, Roberto Amato, Jeffrey Barrett, Sonia Goncalves, Ewan Harrison, David K. Jackson, Ian Johnston, Dominic Kwiatkowski, Cordelia Langford, John Sillitoe on behalf of the Wellcome Sanger Institute COVID-19 Surveillance Team |
| EPI_ISL_3222873 | Lighthouse Laboratory Plymouth | Wellcome Sanger Institute for the COVID-19 Genomics UK (COG-UK) Consortium | Lighthouse Laboratory Plymouth and Alex Alderton, Roberto Amato, Jeffrey Barrett, Sonia Goncalves, Ewan Harrison, David K. Jackson, Ian Johnston, Dominic Kwiatkowski, Cordelia Langford, John Sillitoe on behalf of the Wellcome Sanger Institute COVID-19 Surveillance Team |
| EPI_ISL_3222874, EPI_ISL_3222875 | Lighthouse Lab in Milton Keynes | Wellcome Sanger Institute for the COVID-19 Genomics UK (COG-UK) Consortium | The Lighthouse Lab in Milton Keynes and Alex Alderton, Roberto Amato, Jeffrey Barrett, Sonia Goncalves, Ewan Harrison, David K. Jackson, Ian Johnston, Dominic Kwiatkowski, Cordelia Langford, John Sillitoe on behalf of the Wellcome Sanger Institute COVID-19 Surveillance Team |
| EPI_ISL_3222876, EPI_ISL_3222877 | Lighthouse Laboratory Plymouth | Wellcome Sanger Institute for the COVID-19 Genomics UK (COG-UK) Consortium | Lighthouse Laboratory Plymouth and Alex Alderton, Roberto Amato, Jeffrey Barrett, Sonia Goncalves, Ewan Harrison, David K. Jackson, Ian Johnston, Dominic Kwiatkowski, Cordelia Langford, John Sillitoe on behalf of the Wellcome Sanger Institute COVID-19 Surveillance Team |
| EPI_ISL_3222878 | Lighthouse Lab in Milton Keynes | Wellcome Sanger Institute for the COVID-19 Genomics UK (COG-UK) Consortium | The Lighthouse Lab in Milton Keynes and Alex Alderton, Roberto Amato, Jeffrey Barrett, Sonia Goncalves, Ewan Harrison, David K. Jackson, Ian Johnston, Dominic Kwiatkowski, Cordelia Langford, John Sillitoe on behalf of the Wellcome Sanger Institute COVID-19 Surveillance Team |
| EPI_ISL_3222879, EPI_ISL_3222880, EPI_ISL_3222881 | Lighthouse Laboratory Plymouth | Wellcome Sanger Institute for the COVID-19 Genomics UK (COG-UK) Consortium | Lighthouse Laboratory Plymouth and Alex Alderton, Roberto Amato, Jeffrey Barrett, Sonia Goncalves, Ewan Harrison, David K. Jackson, Ian Johnston, Dominic Kwiatkowski, Cordelia Langford, John Sillitoe on behalf of the Wellcome Sanger Institute COVID-19 Surveillance Team |
| EPI_ISL_3222882 | Lighthouse Lab in Alderley Park | Wellcome Sanger Institute for the COVID-19 Genomics UK (COG-UK) Consortium | Jacquelyn Wynn, Mairead Hyland, The Lighthouse Lab in Alderley Park and Alex Alderton, Roberto Amato, Jeffrey Barrett, Sonia Goncalves, Ewan Harrison, David K. Jackson, Ian Johnston, Dominic Kwiatkowski, Cordelia Langford, John Sillitoe on behalf of the Wellcome Sanger Institute COVID-19 Surveillance Team |
| EPI_ISL_3222883, EPI_ISL_3222884, EPI_ISL_3222885, EPI_ISL_3222886 | Lighthouse Lab in Milton Keynes | Wellcome Sanger Institute for the COVID-19 Genomics UK (COG-UK) Consortium | The Lighthouse Lab in Milton Keynes and Alex Alderton, Roberto Amato, Jeffrey Barrett, Sonia Goncalves, Ewan Harrison, David K. Jackson, Ian Johnston, Dominic Kwiatkowski, Cordelia Langford, John Sillitoe on behalf of the Wellcome Sanger Institute COVID-19 Surveillance Team |
| EPI_ISL_3222887 | Lighthouse Laboratory Plymouth | Wellcome Sanger Institute for the COVID-19 Genomics UK (COG-UK) Consortium | Lighthouse Laboratory Plymouth and Alex Alderton, Roberto Amato, Jeffrey Barrett, Sonia Goncalves, Ewan Harrison, David K. Jackson, Ian Johnston, Dominic Kwiatkowski, Cordelia Langford, John Sillitoe on behalf of the Wellcome Sanger Institute COVID-19 Surveillance Team |
| EPI_ISL_3222888 | Lighthouse Lab in Milton Keynes | Wellcome Sanger Institute for the COVID-19 Genomics UK (COG-UK) Consortium | The Lighthouse Lab in Milton Keynes and Alex Alderton, Roberto Amato, Jeffrey Barrett, Sonia Goncalves, Ewan Harrison, David K. Jackson, Ian Johnston, Dominic Kwiatkowski, Cordelia Langford, John Sillitoe on behalf of the Wellcome Sanger Institute COVID-19 Surveillance Team |
| EPI_ISL_3222889, EPI_ISL_3222890, EPI_ISL_3222891, EPI_ISL_3222892, EPI_ISL_3222893 | Lighthouse Laboratory Plymouth | Wellcome Sanger Institute for the COVID-19 Genomics UK (COG-UK) Consortium | Lighthouse Laboratory Plymouth and Alex Alderton, Roberto Amato, Jeffrey Barrett, Sonia Goncalves, Ewan Harrison, David K. Jackson, Ian Johnston, Dominic Kwiatkowski, Cordelia Langford, John Sillitoe on behalf of the Wellcome Sanger Institute COVID-19 Surveillance Team |
| EPI_ISL_3222894 | Lighthouse Lab in Milton Keynes | Wellcome Sanger Institute for the COVID-19 Genomics UK (COG-UK) Consortium | The Lighthouse Lab in Milton Keynes and Alex Alderton, Roberto Amato, Jeffrey Barrett, Sonia Goncalves, Ewan Harrison, David K. Jackson, Ian Johnston, Dominic Kwiatkowski, Cordelia Langford, John Sillitoe on behalf of the Wellcome Sanger Institute COVID-19 Surveillance Team |
| EPI_ISL_3222895, EPI_ISL_3222896, EPI_ISL_3222897 | Lighthouse Laboratory Plymouth | Wellcome Sanger Institute for the COVID-19 Genomics UK (COG-UK) Consortium | Lighthouse Laboratory Plymouth and Alex Alderton, Roberto Amato, Jeffrey Barrett, Sonia Goncalves, Ewan Harrison, David K. Jackson, Ian Johnston, Dominic Kwiatkowski, Cordelia Langford, John Sillitoe on behalf of the Wellcome Sanger Institute COVID-19 Surveillance Team |
| EPI_ISL_3222898 | Lighthouse Lab in Milton Keynes | Wellcome Sanger Institute for the COVID-19 Genomics UK (COG-UK) Consortium | The Lighthouse Lab in Milton Keynes and Alex Alderton, Roberto Amato, Jeffrey Barrett, Sonia Goncalves, Ewan Harrison, David K. Jackson, Ian Johnston, Dominic Kwiatkowski, Cordelia Langford, John Sillitoe on behalf of the Wellcome Sanger Institute COVID-19 Surveillance Team |
| EPI_ISL_3222899 | Lighthouse Lab in Alderley Park | Wellcome Sanger Institute for the COVID-19 Genomics UK (COG-UK) Consortium | Jacquelyn Wynn, Mairead Hyland, The Lighthouse Lab in Alderley Park and Alex Alderton, Roberto Amato, Jeffrey Barrett, Sonia Goncalves, Ewan Harrison, David K. Jackson, Ian Johnston, Dominic Kwiatkowski, Cordelia Langford, John Sillitoe on behalf of the Wellcome Sanger Institute COVID-19 Surveillance Team |
| EPI_ISL_3222900 | Lighthouse Lab in Milton Keynes | Wellcome Sanger Institute for the COVID-19 Genomics UK (COG-UK) Consortium | The Lighthouse Lab in Milton Keynes and Alex Alderton, Roberto Amato, Jeffrey Barrett, Sonia Goncalves, Ewan Harrison, David K. Jackson, Ian Johnston, Dominic Kwiatkowski, Cordelia Langford, John Sillitoe on behalf of the Wellcome Sanger Institute COVID-19 Surveillance Team |
| EPI_ISL_3222901, EPI_ISL_3222902, EPI_ISL_3222903, EPI_ISL_3222904, EPI_ISL_3222905, EPI_ISL_3222906, EPI_ISL_3222907, EPI_ISL_3222908, EPI_ISL_3222909 | Lighthouse Laboratory Plymouth | Wellcome Sanger Institute for the COVID-19 Genomics UK (COG-UK) Consortium | Lighthouse Laboratory Plymouth and Alex Alderton, Roberto Amato, Jeffrey Barrett, Sonia Goncalves, Ewan Harrison, David K. Jackson, Ian Johnston, Dominic Kwiatkowski, Cordelia Langford, John Sillitoe on behalf of the Wellcome Sanger Institute COVID-19 Surveillance Team |
| EPI_ISL_3222910 | Lighthouse Lab in Milton Keynes | Wellcome Sanger Institute for the COVID-19 Genomics UK (COG-UK) Consortium | The Lighthouse Lab in Milton Keynes and Alex Alderton, Roberto Amato, Jeffrey Barrett, Sonia Goncalves, Ewan Harrison, David K. Jackson, Ian Johnston, Dominic Kwiatkowski, Cordelia Langford, John Sillitoe on behalf of the Wellcome Sanger Institute COVID-19 Surveillance Team |
| EPI_ISL_3222911 | Lighthouse Lab in Alderley Park | Wellcome Sanger Institute for the COVID-19 Genomics UK (COG-UK) Consortium | Jacquelyn Wynn, Mairead Hyland, The Lighthouse Lab in Alderley Park and Alex Alderton, Roberto Amato, Jeffrey Barrett, Sonia Goncalves, Ewan Harrison, David K. Jackson, Ian Johnston, Dominic Kwiatkowski, Cordelia Langford, John Sillitoe on behalf of the Wellcome Sanger Institute COVID-19 Surveillance Team |
| EPI_ISL_3222912 | Lighthouse Lab in Milton Keynes | Wellcome Sanger Institute for the COVID-19 Genomics UK (COG-UK) Consortium | The Lighthouse Lab in Milton Keynes and Alex Alderton, Roberto Amato, Jeffrey Barrett, Sonia Goncalves, Ewan Harrison, David K. Jackson, Ian Johnston, Dominic Kwiatkowski, Cordelia Langford, John Sillitoe on behalf of the Wellcome Sanger Institute COVID-19 Surveillance Team |
| EPI_ISL_3222913, EPI_ISL_3222914, EPI_ISL_3222915 | Lighthouse Laboratory Plymouth | Wellcome Sanger Institute for the COVID-19 Genomics UK (COG-UK) Consortium | Lighthouse Laboratory Plymouth and Alex Alderton, Roberto Amato, Jeffrey Barrett, Sonia Goncalves, Ewan Harrison, David K. Jackson, Ian Johnston, Dominic Kwiatkowski, Cordelia Langford, John Sillitoe on behalf of the Wellcome Sanger Institute COVID-19 Surveillance Team |
| EPI_ISL_3222916 | Lighthouse Lab in Alderley Park | Wellcome Sanger Institute for the COVID-19 Genomics UK (COG-UK) Consortium | Jacquelyn Wynn, Mairead Hyland, The Lighthouse Lab in Alderley Park and Alex Alderton, Roberto Amato, Jeffrey Barrett, Sonia Goncalves, Ewan Harrison, David K. Jackson, Ian Johnston, Dominic Kwiatkowski, Cordelia Langford, John Sillitoe on behalf of the Wellcome Sanger Institute COVID-19 Surveillance Team |
| EPI_ISL_3222917 | Lighthouse Laboratory Plymouth | Wellcome Sanger Institute for the COVID-19 Genomics UK (COG-UK) Consortium | Lighthouse Laboratory Plymouth and Alex Alderton, Roberto Amato, Jeffrey Barrett, Sonia Goncalves, Ewan Harrison, David K. Jackson, Ian Johnston, Dominic Kwiatkowski, Cordelia Langford, John Sillitoe on behalf of the Wellcome Sanger Institute COVID-19 Surveillance Team |
| EPI_ISL_3222918, EPI_ISL_3222919, EPI_ISL_3222920 | Lighthouse Lab in Alderley Park | Wellcome Sanger Institute for the COVID-19 Genomics UK (COG-UK) Consortium | Jacquelyn Wynn, Mairead Hyland, The Lighthouse Lab in Alderley Park and Alex Alderton, Roberto Amato, Jeffrey Barrett, Sonia Goncalves, Ewan Harrison, David K. Jackson, Ian Johnston, Dominic Kwiatkowski, Cordelia Langford, John Sillitoe on behalf of the Wellcome Sanger Institute COVID-19 Surveillance Team |

| | | | |
|---|---|---|---|
| EPI_ISL_3222921 | Lighthouse Laboratory Plymouth | Wellcome Sanger Institute for the COVID-19 Genomics UK (COG-UK) Consortium | Lighthouse Laboratory Plymouth and Alex Alderton, Roberto Amato, Jeffrey Barrett, Sonia Goncalves, Ewan Harrison, David K. Jackson, Ian Johnston, Dominic Kwiatkowski, Cordelia Langford, John Sillitoe on behalf of the Wellcome Sanger Institute COVID-19 Surveillance Team |
| EPI_ISL_3222922, EPI_ISL_3222923 | Lighthouse Lab in Alderley Park | Wellcome Sanger Institute for the COVID-19 Genomics UK (COG-UK) Consortium | Jacquelyn Wynn, Mairead Hyland, The Lighthouse Lab in Alderley Park and Alex Alderton, Roberto Amato, Jeffrey Barrett, Sonia Goncalves, Ewan Harrison, David K. Jackson, Ian Johnston, Dominic Kwiatkowski, Cordelia Langford, John Sillitoe on behalf of the Wellcome Sanger Institute COVID-19 Surveillance Team |
| EPI_ISL_3222924, EPI_ISL_3222925, EPI_ISL_3222926, EPI_ISL_3222927, EPI_ISL_3222928, EPI_ISL_3222929, EPI_ISL_3222930, EPI_ISL_3222931 | Lighthouse Laboratory Plymouth | Wellcome Sanger Institute for the COVID-19 Genomics UK (COG-UK) Consortium | Lighthouse Laboratory Plymouth and Alex Alderton, Roberto Amato, Jeffrey Barrett, Sonia Goncalves, Ewan Harrison, David K. Jackson, Ian Johnston, Dominic Kwiatkowski, Cordelia Langford, John Sillitoe on behalf of the Wellcome Sanger Institute COVID-19 Surveillance Team |
| EPI_ISL_3222932, EPI_ISL_3222933 | Lighthouse Lab in Alderley Park | Wellcome Sanger Institute for the COVID-19 Genomics UK (COG-UK) Consortium | Jacquelyn Wynn, Mairead Hyland, The Lighthouse Lab in Alderley Park and Alex Alderton, Roberto Amato, Jeffrey Barrett, Sonia Goncalves, Ewan Harrison, David K. Jackson, Ian Johnston, Dominic Kwiatkowski, Cordelia Langford, John Sillitoe on behalf of the Wellcome Sanger Institute COVID-19 Surveillance Team |
| EPI_ISL_3222934, EPI_ISL_3222935, EPI_ISL_3222936 | Lighthouse Laboratory Plymouth | Wellcome Sanger Institute for the COVID-19 Genomics UK (COG-UK) Consortium | Lighthouse Laboratory Plymouth and Alex Alderton, Roberto Amato, Jeffrey Barrett, Sonia Goncalves, Ewan Harrison, David K. Jackson, Ian Johnston, Dominic Kwiatkowski, Cordelia Langford, John Sillitoe on behalf of the Wellcome Sanger Institute COVID-19 Surveillance Team |
| EPI_ISL_3222937 | Lighthouse Lab in Milton Keynes | Wellcome Sanger Institute for the COVID-19 Genomics UK (COG-UK) Consortium | The Lighthouse Lab in Milton Keynes and Alex Alderton, Roberto Amato, Jeffrey Barrett, Sonia Goncalves, Ewan Harrison, David K. Jackson, Ian Johnston, Dominic Kwiatkowski, Cordelia Langford, John Sillitoe on behalf of the Wellcome Sanger Institute COVID-19 Surveillance Team |
| EPI_ISL_3222938, EPI_ISL_3222939, EPI_ISL_3222940 | Lighthouse Laboratory Plymouth | Wellcome Sanger Institute for the COVID-19 Genomics UK (COG-UK) Consortium | Lighthouse Laboratory Plymouth and Alex Alderton, Roberto Amato, Jeffrey Barrett, Sonia Goncalves, Ewan Harrison, David K. Jackson, Ian Johnston, Dominic Kwiatkowski, Cordelia Langford, John Sillitoe on behalf of the Wellcome Sanger Institute COVID-19 Surveillance Team |
| EPI_ISL_3222941 | Lighthouse Lab in Alderley Park | Wellcome Sanger Institute for the COVID-19 Genomics UK (COG-UK) Consortium | Jacquelyn Wynn, Mairead Hyland, The Lighthouse Lab in Alderley Park and Alex Alderton, Roberto Amato, Jeffrey Barrett, Sonia Goncalves, Ewan Harrison, David K. Jackson, Ian Johnston, Dominic Kwiatkowski, Cordelia Langford, John Sillitoe on behalf of the Wellcome Sanger Institute COVID-19 Surveillance Team |
| EPI_ISL_3222942, EPI_ISL_3222943, EPI_ISL_3222944, EPI_ISL_3222945, EPI_ISL_3222946 | Lighthouse Laboratory Plymouth | Wellcome Sanger Institute for the COVID-19 Genomics UK (COG-UK) Consortium | Lighthouse Laboratory Plymouth and Alex Alderton, Roberto Amato, Jeffrey Barrett, Sonia Goncalves, Ewan Harrison, David K. Jackson, Ian Johnston, Dominic Kwiatkowski, Cordelia Langford, John Sillitoe on behalf of the Wellcome Sanger Institute COVID-19 Surveillance Team |
| EPI_ISL_3222947 | Lighthouse Lab in Alderley Park | Wellcome Sanger Institute for the COVID-19 Genomics UK (COG-UK) Consortium | Jacquelyn Wynn, Mairead Hyland, The Lighthouse Lab in Alderley Park and Alex Alderton, Roberto Amato, Jeffrey Barrett, Sonia Goncalves, Ewan Harrison, David K. Jackson, Ian Johnston, Dominic Kwiatkowski, Cordelia Langford, John Sillitoe on behalf of the Wellcome Sanger Institute COVID-19 Surveillance Team |
| EPI_ISL_3222948 | Lighthouse Laboratory Plymouth | Wellcome Sanger Institute for the COVID-19 Genomics UK (COG-UK) Consortium | Lighthouse Laboratory Plymouth and Alex Alderton, Roberto Amato, Jeffrey Barrett, Sonia Goncalves, Ewan Harrison, David K. Jackson, Ian Johnston, Dominic Kwiatkowski, Cordelia Langford, John Sillitoe on behalf of the Wellcome Sanger Institute COVID-19 Surveillance Team |
| EPI_ISL_3222949 | Lighthouse Lab in Alderley Park | Wellcome Sanger Institute for the COVID-19 Genomics UK (COG-UK) Consortium | Jacquelyn Wynn, Mairead Hyland, The Lighthouse Lab in Alderley Park and Alex Alderton, Roberto Amato, Jeffrey Barrett, Sonia Goncalves, Ewan Harrison, David K. Jackson, Ian Johnston, Dominic Kwiatkowski, Cordelia Langford, John Sillitoe on behalf of the Wellcome Sanger Institute COVID-19 Surveillance Team |
| EPI_ISL_3222950, EPI_ISL_3222951 | Lighthouse Laboratory Plymouth | Wellcome Sanger Institute for the COVID-19 Genomics UK (COG-UK) Consortium | Lighthouse Laboratory Plymouth and Alex Alderton, Roberto Amato, Jeffrey Barrett, Sonia Goncalves, Ewan Harrison, David K. Jackson, Ian Johnston, Dominic Kwiatkowski, Cordelia Langford, John Sillitoe on behalf of the Wellcome Sanger Institute COVID-19 Surveillance Team |
| EPI_ISL_3222952 | Lighthouse Lab in Milton Keynes | Wellcome Sanger Institute for the COVID-19 Genomics UK (COG-UK) Consortium | The Lighthouse Lab in Milton Keynes and Alex Alderton, Roberto Amato, Jeffrey Barrett, Sonia Goncalves, Ewan Harrison, David K. Jackson, Ian Johnston, Dominic Kwiatkowski, Cordelia Langford, John Sillitoe on behalf of the Wellcome Sanger Institute COVID-19 Surveillance Team |
| EPI_ISL_3222953, EPI_ISL_3222954 | Lighthouse Laboratory Plymouth | Wellcome Sanger Institute for the COVID-19 Genomics UK (COG-UK) Consortium | Lighthouse Laboratory Plymouth and Alex Alderton, Roberto Amato, Jeffrey Barrett, Sonia Goncalves, Ewan Harrison, David K. Jackson, Ian Johnston, Dominic Kwiatkowski, Cordelia Langford, John Sillitoe on behalf of the Wellcome Sanger Institute COVID-19 Surveillance Team |
| EPI_ISL_3222955 | Lighthouse Lab in Milton Keynes | Wellcome Sanger Institute for the COVID-19 Genomics UK (COG-UK) Consortium | The Lighthouse Lab in Milton Keynes and Alex Alderton, Roberto Amato, Jeffrey Barrett, Sonia Goncalves, Ewan Harrison, David K. Jackson, Ian Johnston, Dominic Kwiatkowski, Cordelia Langford, John Sillitoe on behalf of the Wellcome Sanger Institute COVID-19 Surveillance Team |
| EPI_ISL_3222956 | Lighthouse Laboratory Plymouth | Wellcome Sanger Institute for the COVID-19 Genomics UK (COG-UK) Consortium | Lighthouse Laboratory Plymouth and Alex Alderton, Roberto Amato, Jeffrey Barrett, Sonia Goncalves, Ewan Harrison, David K. Jackson, Ian Johnston, Dominic Kwiatkowski, Cordelia Langford, John Sillitoe on behalf of the Wellcome Sanger Institute COVID-19 Surveillance Team |
| EPI_ISL_3222957 | Lighthouse Lab in Alderley Park | Wellcome Sanger Institute for the COVID-19 Genomics UK (COG-UK) Consortium | Jacquelyn Wynn, Mairead Hyland, The Lighthouse Lab in Alderley Park and Alex Alderton, Roberto Amato, Jeffrey Barrett, Sonia Goncalves, Ewan Harrison, David K. Jackson, Ian Johnston, Dominic Kwiatkowski, Cordelia Langford, John Sillitoe on behalf of the Wellcome Sanger Institute COVID-19 Surveillance Team |
| EPI_ISL_3222958 | Lighthouse Lab in Milton Keynes | Wellcome Sanger Institute for the COVID-19 Genomics UK (COG-UK) Consortium | The Lighthouse Lab in Milton Keynes and Alex Alderton, Roberto Amato, Jeffrey Barrett, Sonia Goncalves, Ewan Harrison, David K. Jackson, Ian Johnston, Dominic Kwiatkowski, Cordelia Langford, John Sillitoe on behalf of the Wellcome Sanger Institute COVID-19 Surveillance Team |
| EPI_ISL_3222959 | Lighthouse Laboratory Plymouth | Wellcome Sanger Institute for the COVID-19 Genomics UK (COG-UK) Consortium | Lighthouse Laboratory Plymouth and Alex Alderton, Roberto Amato, Jeffrey Barrett, Sonia Goncalves, Ewan Harrison, David K. Jackson, Ian Johnston, Dominic Kwiatkowski, Cordelia Langford, John Sillitoe on behalf of the Wellcome Sanger Institute COVID-19 Surveillance Team |
| EPI_ISL_3222960, EPI_ISL_3222961 | Lighthouse Lab in Alderley Park | Wellcome Sanger Institute for the COVID-19 Genomics UK (COG-UK) Consortium | Jacquelyn Wynn, Mairead Hyland, The Lighthouse Lab in Alderley Park and Alex Alderton, Roberto Amato, Jeffrey Barrett, Sonia Goncalves, Ewan Harrison, David K. Jackson, Ian Johnston, Dominic Kwiatkowski, Cordelia Langford, John Sillitoe on behalf of the Wellcome Sanger Institute COVID-19 Surveillance Team |
| EPI_ISL_3222962 | Lighthouse Lab in Milton Keynes | Wellcome Sanger Institute for the COVID-19 Genomics UK (COG-UK) Consortium | The Lighthouse Lab in Milton Keynes and Alex Alderton, Roberto Amato, Jeffrey Barrett, Sonia Goncalves, Ewan Harrison, David K. Jackson, Ian Johnston, Dominic Kwiatkowski, Cordelia Langford, John Sillitoe on behalf of the Wellcome Sanger Institute COVID-19 Surveillance Team |
| EPI_ISL_3222963 | Lighthouse Laboratory Plymouth | Wellcome Sanger Institute for the COVID-19 Genomics UK (COG-UK) Consortium | Lighthouse Laboratory Plymouth and Alex Alderton, Roberto Amato, Jeffrey Barrett, Sonia Goncalves, Ewan Harrison, David K. Jackson, Ian Johnston, Dominic Kwiatkowski, Cordelia Langford, John Sillitoe on behalf of the Wellcome Sanger Institute COVID-19 Surveillance Team |
| EPI_ISL_3222964 | Lighthouse Lab in Alderley Park | Wellcome Sanger Institute for the COVID-19 Genomics UK (COG-UK) Consortium | Jacquelyn Wynn, Mairead Hyland, The Lighthouse Lab in Alderley Park and Alex Alderton, Roberto Amato, Jeffrey Barrett, Sonia Goncalves, Ewan Harrison, David K. Jackson, Ian Johnston, Dominic Kwiatkowski, Cordelia Langford, John Sillitoe on behalf of the Wellcome Sanger Institute COVID-19 Surveillance Team |
| EPI_ISL_3222965 | Lighthouse Lab in Milton Keynes | Wellcome Sanger Institute for the COVID-19 Genomics UK (COG-UK) Consortium | The Lighthouse Lab in Milton Keynes and Alex Alderton, Roberto Amato, Jeffrey Barrett, Sonia Goncalves, Ewan Harrison, David K. Jackson, Ian Johnston, Dominic Kwiatkowski, Cordelia Langford, John Sillitoe on behalf of the Wellcome Sanger Institute COVID-19 Surveillance Team |
| EPI_ISL_3222966 | Lighthouse Lab in Alderley Park | Wellcome Sanger Institute for the COVID-19 Genomics UK (COG-UK) Consortium | Jacquelyn Wynn, Mairead Hyland, The Lighthouse Lab in Alderley Park and Alex Alderton, Roberto Amato, Jeffrey Barrett, Sonia Goncalves, Ewan Harrison, David K. Jackson, Ian Johnston, Dominic Kwiatkowski, Cordelia Langford, John Sillitoe on behalf of the Wellcome Sanger Institute COVID-19 Surveillance Team |
| EPI_ISL_3222967, EPI_ISL_3222968 | Lighthouse Lab in Milton Keynes | Wellcome Sanger Institute for the COVID-19 Genomics UK (COG-UK) Consortium | The Lighthouse Lab in Milton Keynes and Alex Alderton, Roberto Amato, Jeffrey Barrett, Sonia Goncalves, Ewan Harrison, David K. Jackson, Ian Johnston, Dominic Kwiatkowski, Cordelia Langford, John Sillitoe on behalf of the Wellcome Sanger Institute COVID-19 Surveillance Team |
| EPI_ISL_3222969, EPI_ISL_3222970, EPI_ISL_3222971 | Lighthouse Laboratory Plymouth | Wellcome Sanger Institute for the COVID-19 Genomics UK (COG-UK) Consortium | Lighthouse Laboratory Plymouth and Alex Alderton, Roberto Amato, Jeffrey Barrett, Sonia Goncalves, Ewan Harrison, David K. Jackson, Ian Johnston, Dominic Kwiatkowski, Cordelia Langford, John Sillitoe on behalf of the Wellcome Sanger Institute COVID-19 Surveillance Team |
| EPI_ISL_3222972 | Lighthouse Lab in Milton Keynes | Wellcome Sanger Institute for the COVID-19 Genomics UK (COG-UK) Consortium | The Lighthouse Lab in Milton Keynes and Alex Alderton, Roberto Amato, Jeffrey Barrett, Sonia Goncalves, Ewan Harrison, David K. Jackson, Ian Johnston, Dominic Kwiatkowski, Cordelia Langford, John Sillitoe on behalf of the Wellcome Sanger Institute COVID-19 Surveillance Team |
| EPI_ISL_3222973 | Lighthouse Laboratory Plymouth | Wellcome Sanger Institute for the COVID-19 Genomics UK (COG-UK) Consortium | Lighthouse Laboratory Plymouth and Alex Alderton, Roberto Amato, Jeffrey Barrett, Sonia Goncalves, Ewan Harrison, David K. Jackson, Ian Johnston, Dominic Kwiatkowski, Cordelia Langford, John Sillitoe on behalf of the Wellcome Sanger Institute COVID-19 Surveillance Team |
| EPI_ISL_3222974 | Lighthouse Lab in Glasgow | Wellcome Sanger Institute for the COVID-19 Genomics UK (COG-UK) Consortium | Harper VanSteenhouse, Yumi Kasai, David Gray, Carol Clugston, Anna Dominiczak and Alex Alderton, Roberto Amato, Jeffrey Barrett, Sonia Goncalves, Ewan Harrison, David K. Jackson, Ian Johnston, Dominic Kwiatkowski, Cordelia Langford, John Sillitoe on behalf of the Wellcome Sanger Institute COVID-19 Surveillance Team |
| EPI_ISL_3222975, EPI_ISL_3222976 | Lighthouse Laboratory Plymouth | Wellcome Sanger Institute for the COVID-19 Genomics UK (COG-UK) Consortium | Lighthouse Laboratory Plymouth and Alex Alderton, Roberto Amato, Jeffrey Barrett, Sonia Goncalves, Ewan Harrison, David K. Jackson, Ian Johnston, Dominic Kwiatkowski, Cordelia Langford, John Sillitoe on behalf of the Wellcome Sanger Institute COVID-19 Surveillance Team |
| EPI_ISL_3222977 | Lighthouse Lab in Milton Keynes | Wellcome Sanger Institute for the COVID-19 Genomics UK (COG-UK) Consortium | The Lighthouse Lab in Milton Keynes and Alex Alderton, Roberto Amato, Jeffrey Barrett, Sonia Goncalves, Ewan Harrison, David K. Jackson, Ian Johnston, Dominic Kwiatkowski, Cordelia Langford, John Sillitoe on behalf of the Wellcome Sanger Institute COVID-19 Surveillance Team |
| EPI_ISL_3222978, EPI_ISL_3222979 | Lighthouse Lab in Alderley Park | Wellcome Sanger Institute for the COVID-19 Genomics UK (COG-UK) Consortium | Jacquelyn Wynn, Mairead Hyland, The Lighthouse Lab in Alderley Park and Alex Alderton, Roberto Amato, Jeffrey Barrett, Sonia Goncalves, Ewan Harrison, David K. Jackson, Ian Johnston, Dominic Kwiatkowski, Cordelia Langford, John Sillitoe on behalf of the Wellcome Sanger Institute COVID-19 Surveillance Team |
| EPI_ISL_3222980 | Lighthouse Lab in Milton Keynes | Wellcome Sanger Institute for the COVID-19 Genomics UK (COG-UK) Consortium | The Lighthouse Lab in Milton Keynes and Alex Alderton, Roberto Amato, Jeffrey Barrett, Sonia Goncalves, Ewan Harrison, David K. Jackson, Ian Johnston, Dominic Kwiatkowski, Cordelia Langford, John Sillitoe on behalf of the Wellcome Sanger Institute COVID-19 Surveillance Team |
| EPI_ISL_3222981 | Lighthouse Lab in Glasgow | Wellcome Sanger Institute for the COVID-19 Genomics UK (COG-UK) Consortium | Harper VanSteenhouse, Yumi Kasai, David Gray, Carol Clugston, Anna Dominiczak and Alex Alderton, Roberto Amato, Jeffrey Barrett, Sonia Goncalves, Ewan Harrison, David K. Jackson, Ian Johnston, Dominic Kwiatkowski, Cordelia Langford, John Sillitoe on behalf of the Wellcome Sanger Institute COVID-19 Surveillance Team |
| EPI_ISL_3222982 | Lighthouse Lab in Alderley Park | Wellcome Sanger Institute for the COVID-19 Genomics UK (COG-UK) Consortium | Jacquelyn Wynn, Mairead Hyland, The Lighthouse Lab in Alderley Park and Alex Alderton, Roberto Amato, Jeffrey Barrett, Sonia Goncalves, Ewan Harrison, David K. Jackson, Ian Johnston, Dominic Kwiatkowski, Cordelia Langford, John Sillitoe on behalf of the Wellcome Sanger Institute COVID-19 Surveillance Team |
| EPI_ISL_3222983 | Lighthouse Laboratory Plymouth | Wellcome Sanger Institute for the COVID-19 Genomics UK (COG-UK) Consortium | Lighthouse Laboratory Plymouth and Alex Alderton, Roberto Amato, Jeffrey Barrett, Sonia Goncalves, Ewan Harrison, David K. Jackson, Ian Johnston, Dominic Kwiatkowski, Cordelia Langford, John Sillitoe on behalf of the Wellcome Sanger Institute COVID-19 Surveillance Team |
| EPI_ISL_3222984, EPI_ISL_3222985 | Lighthouse Lab in Milton Keynes | Wellcome Sanger Institute for the COVID-19 Genomics UK (COG-UK) Consortium | The Lighthouse Lab in Milton Keynes and Alex Alderton, Roberto Amato, Jeffrey Barrett, Sonia Goncalves, Ewan Harrison, David K. Jackson, Ian Johnston, Dominic Kwiatkowski, Cordelia Langford, John Sillitoe on behalf of the Wellcome Sanger Institute COVID-19 Surveillance Team |
| EPI_ISL_3222986 | Lighthouse Lab in Alderley Park | Wellcome Sanger Institute for the COVID-19 Genomics UK (COG-UK) Consortium | Jacquelyn Wynn, Mairead Hyland, The Lighthouse Lab in Alderley Park and Alex Alderton, Roberto Amato, Jeffrey Barrett, Sonia Goncalves, Ewan Harrison, David K. Jackson, Ian Johnston, Dominic Kwiatkowski, Cordelia Langford, John Sillitoe on behalf of the Wellcome Sanger Institute COVID-19 Surveillance Team |

| | | | |
|---|---|---|---|
| EPI_ISL_3222987 | Lighthouse Laboratory Plymouth | Wellcome Sanger Institute for the COVID-19 Genomics UK (COG-UK) Consortium | Lighthouse Laboratory Plymouth and Alex Alderton, Roberto Amato, Jeffrey Barrett, Sonia Goncalves, Ewan Harrison, David K. Jackson, Ian Johnston, Dominic Kwiatkowski, Cordelia Langford, John Sillitoe on behalf of the Wellcome Sanger Institute COVID-19 Surveillance Team |
| EPI_ISL_3222988 | Lighthouse Lab in Milton Keynes | Wellcome Sanger Institute for the COVID-19 Genomics UK (COG-UK) Consortium | The Lighthouse Lab in Milton Keynes and Alex Alderton, Roberto Amato, Jeffrey Barrett, Sonia Goncalves, Ewan Harrison, David K. Jackson, Ian Johnston, Dominic Kwiatkowski, Cordelia Langford, John Sillitoe on behalf of the Wellcome Sanger Institute COVID-19 Surveillance Team |
| EPI_ISL_3222989 | Lighthouse Lab in Glasgow | Wellcome Sanger Institute for the COVID-19 Genomics UK (COG-UK) Consortium | Harper VanSteenhouse, Yumi Kasai, David Gray, Carol Clugston, Anna Dominiczak and Alex Alderton, Roberto Amato, Jeffrey Barrett, Sonia Goncalves, Ewan Harrison, David K. Jackson, Ian Johnston, Dominic Kwiatkowski, Cordelia Langford, John Sillitoe on behalf of the Wellcome Sanger Institute COVID-19 Surveillance Team |
| EPI_ISL_3222990 | Lighthouse Lab in Milton Keynes | Wellcome Sanger Institute for the COVID-19 Genomics UK (COG-UK) Consortium | The Lighthouse Lab in Milton Keynes and Alex Alderton, Roberto Amato, Jeffrey Barrett, Sonia Goncalves, Ewan Harrison, David K. Jackson, Ian Johnston, Dominic Kwiatkowski, Cordelia Langford, John Sillitoe on behalf of the Wellcome Sanger Institute COVID-19 Surveillance Team |
| EPI_ISL_3222991 | Lighthouse Lab in Alderley Park | Wellcome Sanger Institute for the COVID-19 Genomics UK (COG-UK) Consortium | Jacquelyn Wynn, Mairead Hyland, The Lighthouse Lab in Alderley Park and Alex Alderton, Roberto Amato, Jeffrey Barrett, Sonia Goncalves, Ewan Harrison, David K. Jackson, Ian Johnston, Dominic Kwiatkowski, Cordelia Langford, John Sillitoe on behalf of the Wellcome Sanger Institute COVID-19 Surveillance Team |
| EPI_ISL_3222992, EPI_ISL_3222993, EPI_ISL_3222994 | Lighthouse Laboratory Plymouth | Wellcome Sanger Institute for the COVID-19 Genomics UK (COG-UK) Consortium | Lighthouse Laboratory Plymouth and Alex Alderton, Roberto Amato, Jeffrey Barrett, Sonia Goncalves, Ewan Harrison, David K. Jackson, Ian Johnston, Dominic Kwiatkowski, Cordelia Langford, John Sillitoe on behalf of the Wellcome Sanger Institute COVID-19 Surveillance Team |
| EPI_ISL_3222995 | Lighthouse Lab in Milton Keynes | Wellcome Sanger Institute for the COVID-19 Genomics UK (COG-UK) Consortium | The Lighthouse Lab in Milton Keynes and Alex Alderton, Roberto Amato, Jeffrey Barrett, Sonia Goncalves, Ewan Harrison, David K. Jackson, Ian Johnston, Dominic Kwiatkowski, Cordelia Langford, John Sillitoe on behalf of the Wellcome Sanger Institute COVID-19 Surveillance Team |
| EPI_ISL_3222996 | Lighthouse Lab in Alderley Park | Wellcome Sanger Institute for the COVID-19 Genomics UK (COG-UK) Consortium | Jacquelyn Wynn, Mairead Hyland, The Lighthouse Lab in Alderley Park and Alex Alderton, Roberto Amato, Jeffrey Barrett, Sonia Goncalves, Ewan Harrison, David K. Jackson, Ian Johnston, Dominic Kwiatkowski, Cordelia Langford, John Sillitoe on behalf of the Wellcome Sanger Institute COVID-19 Surveillance Team |
| EPI_ISL_3222997 | Lighthouse Lab in Glasgow | Wellcome Sanger Institute for the COVID-19 Genomics UK (COG-UK) Consortium | Harper VanSteenhouse, Yumi Kasai, David Gray, Carol Clugston, Anna Dominiczak and Alex Alderton, Roberto Amato, Jeffrey Barrett, Sonia Goncalves, Ewan Harrison, David K. Jackson, Ian Johnston, Dominic Kwiatkowski, Cordelia Langford, John Sillitoe on behalf of the Wellcome Sanger Institute COVID-19 Surveillance Team |
| EPI_ISL_3222998 | Lighthouse Lab in Milton Keynes | Wellcome Sanger Institute for the COVID-19 Genomics UK (COG-UK) Consortium | The Lighthouse Lab in Milton Keynes and Alex Alderton, Roberto Amato, Jeffrey Barrett, Sonia Goncalves, Ewan Harrison, David K. Jackson, Ian Johnston, Dominic Kwiatkowski, Cordelia Langford, John Sillitoe on behalf of the Wellcome Sanger Institute COVID-19 Surveillance Team |
| EPI_ISL_3222999, EPI_ISL_3223000 | Lighthouse Lab in Alderley Park | Wellcome Sanger Institute for the COVID-19 Genomics UK (COG-UK) Consortium | Jacquelyn Wynn, Mairead Hyland, The Lighthouse Lab in Alderley Park and Alex Alderton, Roberto Amato, Jeffrey Barrett, Sonia Goncalves, Ewan Harrison, David K. Jackson, Ian Johnston, Dominic Kwiatkowski, Cordelia Langford, John Sillitoe on behalf of the Wellcome Sanger Institute COVID-19 Surveillance Team |
| EPI_ISL_3223001 | Lighthouse Laboratory Plymouth | Wellcome Sanger Institute for the COVID-19 Genomics UK (COG-UK) Consortium | Lighthouse Laboratory Plymouth and Alex Alderton, Roberto Amato, Jeffrey Barrett, Sonia Goncalves, Ewan Harrison, David K. Jackson, Ian Johnston, Dominic Kwiatkowski, Cordelia Langford, John Sillitoe on behalf of the Wellcome Sanger Institute COVID-19 Surveillance Team |
| EPI_ISL_3223002, EPI_ISL_3223003 | Lighthouse Lab in Milton Keynes | Wellcome Sanger Institute for the COVID-19 Genomics UK (COG-UK) Consortium | The Lighthouse Lab in Milton Keynes and Alex Alderton, Roberto Amato, Jeffrey Barrett, Sonia Goncalves, Ewan Harrison, David K. Jackson, Ian Johnston, Dominic Kwiatkowski, Cordelia Langford, John Sillitoe on behalf of the Wellcome Sanger Institute COVID-19 Surveillance Team |
| EPI_ISL_3223004 | Lighthouse Lab in Glasgow | Wellcome Sanger Institute for the COVID-19 Genomics UK (COG-UK) Consortium | Harper VanSteenhouse, Yumi Kasai, David Gray, Carol Clugston, Anna Dominiczak and Alex Alderton, Roberto Amato, Jeffrey Barrett, Sonia Goncalves, Ewan Harrison, David K. Jackson, Ian Johnston, Dominic Kwiatkowski, Cordelia Langford, John Sillitoe on behalf of the Wellcome Sanger Institute COVID-19 Surveillance Team |
| EPI_ISL_3223005 | Lighthouse Lab in Milton Keynes | Wellcome Sanger Institute for the COVID-19 Genomics UK (COG-UK) Consortium | The Lighthouse Lab in Milton Keynes and Alex Alderton, Roberto Amato, Jeffrey Barrett, Sonia Goncalves, Ewan Harrison, David K. Jackson, Ian Johnston, Dominic Kwiatkowski, Cordelia Langford, John Sillitoe on behalf of the Wellcome Sanger Institute COVID-19 Surveillance Team |
| EPI_ISL_3223006, EPI_ISL_3223007 | Lighthouse Lab in Alderley Park | Wellcome Sanger Institute for the COVID-19 Genomics UK (COG-UK) Consortium | Jacquelyn Wynn, Mairead Hyland, The Lighthouse Lab in Alderley Park and Alex Alderton, Roberto Amato, Jeffrey Barrett, Sonia Goncalves, Ewan Harrison, David K. Jackson, Ian Johnston, Dominic Kwiatkowski, Cordelia Langford, John Sillitoe on behalf of the Wellcome Sanger Institute COVID-19 Surveillance Team |
| EPI_ISL_3223008 | Lighthouse Laboratory Plymouth | Wellcome Sanger Institute for the COVID-19 Genomics UK (COG-UK) Consortium | Lighthouse Laboratory Plymouth and Alex Alderton, Roberto Amato, Jeffrey Barrett, Sonia Goncalves, Ewan Harrison, David K. Jackson, Ian Johnston, Dominic Kwiatkowski, Cordelia Langford, John Sillitoe on behalf of the Wellcome Sanger Institute COVID-19 Surveillance Team |
| EPI_ISL_3223009, EPI_ISL_3223010, EPI_ISL_3223011, EPI_ISL_3223012 | Lighthouse Lab in Glasgow | Wellcome Sanger Institute for the COVID-19 Genomics UK (COG-UK) Consortium | Harper VanSteenhouse, Yumi Kasai, David Gray, Carol Clugston, Anna Dominiczak and Alex Alderton, Roberto Amato, Jeffrey Barrett, Sonia Goncalves, Ewan Harrison, David K. Jackson, Ian Johnston, Dominic Kwiatkowski, Cordelia Langford, John Sillitoe on behalf of the Wellcome Sanger Institute COVID-19 Surveillance Team |
| EPI_ISL_3223013 | Lighthouse Laboratory Plymouth | Wellcome Sanger Institute for the COVID-19 Genomics UK (COG-UK) Consortium | Lighthouse Laboratory Plymouth and Alex Alderton, Roberto Amato, Jeffrey Barrett, Sonia Goncalves, Ewan Harrison, David K. Jackson, Ian Johnston, Dominic Kwiatkowski, Cordelia Langford, John Sillitoe on behalf of the Wellcome Sanger Institute COVID-19 Surveillance Team |
| EPI_ISL_3223014 | Lighthouse Lab in Milton Keynes | Wellcome Sanger Institute for the COVID-19 Genomics UK (COG-UK) Consortium | The Lighthouse Lab in Milton Keynes and Alex Alderton, Roberto Amato, Jeffrey Barrett, Sonia Goncalves, Ewan Harrison, David K. Jackson, Ian Johnston, Dominic Kwiatkowski, Cordelia Langford, John Sillitoe on behalf of the Wellcome Sanger Institute COVID-19 Surveillance Team |
| EPI_ISL_3223015, EPI_ISL_3223016 | Lighthouse Lab in Glasgow | Wellcome Sanger Institute for the COVID-19 Genomics UK (COG-UK) Consortium | Harper VanSteenhouse, Yumi Kasai, David Gray, Carol Clugston, Anna Dominiczak and Alex Alderton, Roberto Amato, Jeffrey Barrett, Sonia Goncalves, Ewan Harrison, David K. Jackson, Ian Johnston, Dominic Kwiatkowski, Cordelia Langford, John Sillitoe on behalf of the Wellcome Sanger Institute COVID-19 Surveillance Team |
| EPI_ISL_3223017, EPI_ISL_3223018 | Lighthouse Laboratory Plymouth | Wellcome Sanger Institute for the COVID-19 Genomics UK (COG-UK) Consortium | Lighthouse Laboratory Plymouth and Alex Alderton, Roberto Amato, Jeffrey Barrett, Sonia Goncalves, Ewan Harrison, David K. Jackson, Ian Johnston, Dominic Kwiatkowski, Cordelia Langford, John Sillitoe on behalf of the Wellcome Sanger Institute COVID-19 Surveillance Team |
| EPI_ISL_3223019, EPI_ISL_3223020, EPI_ISL_3223021 | Lighthouse Lab in Alderley Park | Wellcome Sanger Institute for the COVID-19 Genomics UK (COG-UK) Consortium | Jacquelyn Wynn, Mairead Hyland, The Lighthouse Lab in Alderley Park and Alex Alderton, Roberto Amato, Jeffrey Barrett, Sonia Goncalves, Ewan Harrison, David K. Jackson, Ian Johnston, Dominic Kwiatkowski, Cordelia Langford, John Sillitoe on behalf of the Wellcome Sanger Institute COVID-19 Surveillance Team |
| EPI_ISL_3223022 | Lighthouse Laboratory Plymouth | Wellcome Sanger Institute for the COVID-19 Genomics UK (COG-UK) Consortium | Lighthouse Laboratory Plymouth and Alex Alderton, Roberto Amato, Jeffrey Barrett, Sonia Goncalves, Ewan Harrison, David K. Jackson, Ian Johnston, Dominic Kwiatkowski, Cordelia Langford, John Sillitoe on behalf of the Wellcome Sanger Institute COVID-19 Surveillance Team |
| EPI_ISL_3223023 | Lighthouse Lab in Alderley Park | Wellcome Sanger Institute for the COVID-19 Genomics UK (COG-UK) Consortium | Jacquelyn Wynn, Mairead Hyland, The Lighthouse Lab in Alderley Park and Alex Alderton, Roberto Amato, Jeffrey Barrett, Sonia Goncalves, Ewan Harrison, David K. Jackson, Ian Johnston, Dominic Kwiatkowski, Cordelia Langford, John Sillitoe on behalf of the Wellcome Sanger Institute COVID-19 Surveillance Team |
| EPI_ISL_3223024 | Lighthouse Lab in Glasgow | Wellcome Sanger Institute for the COVID-19 Genomics UK (COG-UK) Consortium | Harper VanSteenhouse, Yumi Kasai, David Gray, Carol Clugston, Anna Dominiczak and Alex Alderton, Roberto Amato, Jeffrey Barrett, Sonia Goncalves, Ewan Harrison, David K. Jackson, Ian Johnston, Dominic Kwiatkowski, Cordelia Langford, John Sillitoe on behalf of the Wellcome Sanger Institute COVID-19 Surveillance Team |
| EPI_ISL_3223025, EPI_ISL_3223026 | Lighthouse Lab in Milton Keynes | Wellcome Sanger Institute for the COVID-19 Genomics UK (COG-UK) Consortium | The Lighthouse Lab in Milton Keynes and Alex Alderton, Roberto Amato, Jeffrey Barrett, Sonia Goncalves, Ewan Harrison, David K. Jackson, Ian Johnston, Dominic Kwiatkowski, Cordelia Langford, John Sillitoe on behalf of the Wellcome Sanger Institute COVID-19 Surveillance Team |
| EPI_ISL_3223027, EPI_ISL_3223028, EPI_ISL_3223029 | Lighthouse Lab in Alderley Park | Wellcome Sanger Institute for the COVID-19 Genomics UK (COG-UK) Consortium | Jacquelyn Wynn, Mairead Hyland, The Lighthouse Lab in Alderley Park and Alex Alderton, Roberto Amato, Jeffrey Barrett, Sonia Goncalves, Ewan Harrison, David K. Jackson, Ian Johnston, Dominic Kwiatkowski, Cordelia Langford, John Sillitoe on behalf of the Wellcome Sanger Institute COVID-19 Surveillance Team |
| EPI_ISL_3223030 | Lighthouse Laboratory Plymouth | Wellcome Sanger Institute for the COVID-19 Genomics UK (COG-UK) Consortium | Lighthouse Laboratory Plymouth and Alex Alderton, Roberto Amato, Jeffrey Barrett, Sonia Goncalves, Ewan Harrison, David K. Jackson, Ian Johnston, Dominic Kwiatkowski, Cordelia Langford, John Sillitoe on behalf of the Wellcome Sanger Institute COVID-19 Surveillance Team |
| EPI_ISL_3223031 | Lighthouse Lab in Alderley Park | Wellcome Sanger Institute for the COVID-19 Genomics UK (COG-UK) Consortium | Jacquelyn Wynn, Mairead Hyland, The Lighthouse Lab in Alderley Park and Alex Alderton, Roberto Amato, Jeffrey Barrett, Sonia Goncalves, Ewan Harrison, David K. Jackson, Ian Johnston, Dominic Kwiatkowski, Cordelia Langford, John Sillitoe on behalf of the Wellcome Sanger Institute COVID-19 Surveillance Team |
| EPI_ISL_3223032, EPI_ISL_3223033 | Lighthouse Lab in Glasgow | Wellcome Sanger Institute for the COVID-19 Genomics UK (COG-UK) Consortium | Harper VanSteenhouse, Yumi Kasai, David Gray, Carol Clugston, Anna Dominiczak and Alex Alderton, Roberto Amato, Jeffrey Barrett, Sonia Goncalves, Ewan Harrison, David K. Jackson, Ian Johnston, Dominic Kwiatkowski, Cordelia Langford, John Sillitoe on behalf of the Wellcome Sanger Institute COVID-19 Surveillance Team |
| EPI_ISL_3223034 | Lighthouse Lab in Alderley Park | Wellcome Sanger Institute for the COVID-19 Genomics UK (COG-UK) Consortium | Jacquelyn Wynn, Mairead Hyland, The Lighthouse Lab in Alderley Park and Alex Alderton, Roberto Amato, Jeffrey Barrett, Sonia Goncalves, Ewan Harrison, David K. Jackson, Ian Johnston, Dominic Kwiatkowski, Cordelia Langford, John Sillitoe on behalf of the Wellcome Sanger Institute COVID-19 Surveillance Team |
| EPI_ISL_3223035 | Lighthouse Lab in Glasgow | Wellcome Sanger Institute for the COVID-19 Genomics UK (COG-UK) Consortium | Harper VanSteenhouse, Yumi Kasai, David Gray, Carol Clugston, Anna Dominiczak and Alex Alderton, Roberto Amato, Jeffrey Barrett, Sonia Goncalves, Ewan Harrison, David K. Jackson, Ian Johnston, Dominic Kwiatkowski, Cordelia Langford, John Sillitoe on behalf of the Wellcome Sanger Institute COVID-19 Surveillance Team |
| EPI_ISL_3223036 | Lighthouse Lab in Alderley Park | Wellcome Sanger Institute for the COVID-19 Genomics UK (COG-UK) Consortium | Jacquelyn Wynn, Mairead Hyland, The Lighthouse Lab in Alderley Park and Alex Alderton, Roberto Amato, Jeffrey Barrett, Sonia Goncalves, Ewan Harrison, David K. Jackson, Ian Johnston, Dominic Kwiatkowski, Cordelia Langford, John Sillitoe on behalf of the Wellcome Sanger Institute COVID-19 Surveillance Team |
| EPI_ISL_3223037 | Lighthouse Lab in Milton Keynes | Wellcome Sanger Institute for the COVID-19 Genomics UK (COG-UK) Consortium | The Lighthouse Lab in Milton Keynes and Alex Alderton, Roberto Amato, Jeffrey Barrett, Sonia Goncalves, Ewan Harrison, David K. Jackson, Ian Johnston, Dominic Kwiatkowski, Cordelia Langford, John Sillitoe on behalf of the Wellcome Sanger Institute COVID-19 Surveillance Team |
| EPI_ISL_3223038 | Lighthouse Laboratory Plymouth | Wellcome Sanger Institute for the COVID-19 Genomics UK (COG-UK) Consortium | Lighthouse Laboratory Plymouth and Alex Alderton, Roberto Amato, Jeffrey Barrett, Sonia Goncalves, Ewan Harrison, David K. Jackson, Ian Johnston, Dominic Kwiatkowski, Cordelia Langford, John Sillitoe on behalf of the Wellcome Sanger Institute COVID-19 Surveillance Team |
| EPI_ISL_3223039, EPI_ISL_3223040 | Lighthouse Lab in Milton Keynes | Wellcome Sanger Institute for the COVID-19 Genomics UK (COG-UK) Consortium | The Lighthouse Lab in Milton Keynes and Alex Alderton, Roberto Amato, Jeffrey Barrett, Sonia Goncalves, Ewan Harrison, David K. Jackson, Ian Johnston, Dominic Kwiatkowski, Cordelia Langford, John Sillitoe on behalf of the Wellcome Sanger Institute COVID-19 Surveillance Team |
| EPI_ISL_3223041 | Lighthouse Laboratory Plymouth | Wellcome Sanger Institute for the COVID-19 Genomics UK (COG-UK) Consortium | Lighthouse Laboratory Plymouth and Alex Alderton, Roberto Amato, Jeffrey Barrett, Sonia Goncalves, Ewan Harrison, David K. Jackson, Ian Johnston, Dominic Kwiatkowski, Cordelia Langford, John Sillitoe on behalf of the Wellcome Sanger Institute COVID-19 Surveillance Team |
| EPI_ISL_3223042 | Lighthouse Lab in Glasgow | Wellcome Sanger Institute for the COVID-19 Genomics UK (COG-UK) Consortium | Harper VanSteenhouse, Yumi Kasai, David Gray, Carol Clugston, Anna Dominiczak and Alex Alderton, Roberto Amato, Jeffrey Barrett, Sonia Goncalves, Ewan Harrison, David K. Jackson, Ian Johnston, Dominic Kwiatkowski, Cordelia Langford, John Sillitoe on behalf of the Wellcome Sanger Institute COVID-19 Surveillance Team |
| EPI_ISL_3223043, EPI_ISL_3223044 | Lighthouse Lab in Alderley Park | Wellcome Sanger Institute for the COVID-19 Genomics UK (COG-UK) Consortium | Jacquelyn Wynn, Mairead Hyland, The Lighthouse Lab in Alderley Park and Alex Alderton, Roberto Amato, Jeffrey Barrett, Sonia Goncalves, Ewan Harrison, David K. Jackson, Ian Johnston, Dominic Kwiatkowski, Cordelia Langford, John Sillitoe on behalf of the Wellcome Sanger Institute COVID-19 Surveillance Team |
| EPI_ISL_3223045, EPI_ISL_3223046 | Lighthouse Lab in Milton Keynes | Wellcome Sanger Institute for the COVID-19 Genomics UK (COG-UK) Consortium | The Lighthouse Lab in Milton Keynes and Alex Alderton, Roberto Amato, Jeffrey Barrett, Sonia Goncalves, Ewan Harrison, David K. Jackson, Ian Johnston, Dominic Kwiatkowski, Cordelia Langford, John Sillitoe on behalf of the Wellcome Sanger Institute COVID-19 Surveillance Team |
| EPI_ISL_3223047 | Lighthouse Lab in Alderley Park | Wellcome Sanger Institute for the COVID-19 Genomics UK (COG-UK) Consortium | Jacquelyn Wynn, Mairead Hyland, The Lighthouse Lab in Alderley Park and Alex Alderton, Roberto Amato, Jeffrey Barrett, Sonia Goncalves, Ewan Harrison, David K. Jackson, Ian Johnston, Dominic Kwiatkowski, Cordelia Langford, John Sillitoe on behalf of the Wellcome Sanger Institute COVID-19 Surveillance Team |
| EPI_ISL_3223048 | Lighthouse Lab in Milton Keynes | Wellcome Sanger Institute for the COVID-19 Genomics UK (COG-UK) Consortium | The Lighthouse Lab in Milton Keynes and Alex Alderton, Roberto Amato, Jeffrey Barrett, Sonia Goncalves, Ewan Harrison, David K. Jackson, Ian Johnston, Dominic Kwiatkowski, Cordelia Langford, John Sillitoe on behalf of the Wellcome Sanger Institute COVID-19 Surveillance Team |

| | | | |
|---|---|---|---|
| EPI_ISL_3223049, EPI_ISL_3223050 | Lighthouse Lab in Alderley Park | Wellcome Sanger Institute for the COVID-19 Genomics UK (COG-UK) Consortium | Jacquelyn Wynn, Mairead Hyland, The Lighthouse Lab in Alderley Park and Alex Alderton, Roberto Amato, Jeffrey Barrett, Sonia Goncalves, Ewan Harrison, David K. Jackson, Ian Johnston, Dominic Kwiatkowski, Cordelia Langford, John Sillitoe on behalf of the Wellcome Sanger Institute COVID-19 Surveillance Team |
| EPI_ISL_3223051 | Lighthouse Lab in Milton Keynes | Wellcome Sanger Institute for the COVID-19 Genomics UK (COG-UK) Consortium | The Lighthouse Lab in Milton Keynes and Alex Alderton, Roberto Amato, Jeffrey Barrett, Sonia Goncalves, Ewan Harrison, David K. Jackson, Ian Johnston, Dominic Kwiatkowski, Cordelia Langford, John Sillitoe on behalf of the Wellcome Sanger Institute COVID-19 Surveillance Team |
| EPI_ISL_3223052 | Lighthouse Lab in Alderley Park | Wellcome Sanger Institute for the COVID-19 Genomics UK (COG-UK) Consortium | Jacquelyn Wynn, Mairead Hyland, The Lighthouse Lab in Alderley Park and Alex Alderton, Roberto Amato, Jeffrey Barrett, Sonia Goncalves, Ewan Harrison, David K. Jackson, Ian Johnston, Dominic Kwiatkowski, Cordelia Langford, John Sillitoe on behalf of the Wellcome Sanger Institute COVID-19 Surveillance Team |
| EPI_ISL_3223053 | Lighthouse Laboratory Plymouth | Wellcome Sanger Institute for the COVID-19 Genomics UK (COG-UK) Consortium | Lighthouse Laboratory Plymouth and Alex Alderton, Roberto Amato, Jeffrey Barrett, Sonia Goncalves, Ewan Harrison, David K. Jackson, Ian Johnston, Dominic Kwiatkowski, Cordelia Langford, John Sillitoe on behalf of the Wellcome Sanger Institute COVID-19 Surveillance Team |
| EPI_ISL_3223054, EPI_ISL_3223055 | Lighthouse Lab in Glasgow | Wellcome Sanger Institute for the COVID-19 Genomics UK (COG-UK) Consortium | Harper VanSteenhouse, Yumi Kasai, David Gray, Carol Clugston, Anna Dominiczak and Alex Alderton, Roberto Amato, Jeffrey Barrett, Sonia Goncalves, Ewan Harrison, David K. Jackson, Ian Johnston, Dominic Kwiatkowski, Cordelia Langford, John Sillitoe on behalf of the Wellcome Sanger Institute COVID-19 Surveillance Team |
| EPI_ISL_3223056 | Lighthouse Lab in Milton Keynes | Wellcome Sanger Institute for the COVID-19 Genomics UK (COG-UK) Consortium | The Lighthouse Lab in Milton Keynes and Alex Alderton, Roberto Amato, Jeffrey Barrett, Sonia Goncalves, Ewan Harrison, David K. Jackson, Ian Johnston, Dominic Kwiatkowski, Cordelia Langford, John Sillitoe on behalf of the Wellcome Sanger Institute COVID-19 Surveillance Team |
| EPI_ISL_3223057 | Lighthouse Lab in Alderley Park | Wellcome Sanger Institute for the COVID-19 Genomics UK (COG-UK) Consortium | Jacquelyn Wynn, Mairead Hyland, The Lighthouse Lab in Alderley Park and Alex Alderton, Roberto Amato, Jeffrey Barrett, Sonia Goncalves, Ewan Harrison, David K. Jackson, Ian Johnston, Dominic Kwiatkowski, Cordelia Langford, John Sillitoe on behalf of the Wellcome Sanger Institute COVID-19 Surveillance Team |
| EPI_ISL_3223058, EPI_ISL_3223059 | Lighthouse Lab in Milton Keynes | Wellcome Sanger Institute for the COVID-19 Genomics UK (COG-UK) Consortium | The Lighthouse Lab in Milton Keynes and Alex Alderton, Roberto Amato, Jeffrey Barrett, Sonia Goncalves, Ewan Harrison, David K. Jackson, Ian Johnston, Dominic Kwiatkowski, Cordelia Langford, John Sillitoe on behalf of the Wellcome Sanger Institute COVID-19 Surveillance Team |
| EPI_ISL_3223060 | Lighthouse Lab in Glasgow | Wellcome Sanger Institute for the COVID-19 Genomics UK (COG-UK) Consortium | Harper VanSteenhouse, Yumi Kasai, David Gray, Carol Clugston, Anna Dominiczak and Alex Alderton, Roberto Amato, Jeffrey Barrett, Sonia Goncalves, Ewan Harrison, David K. Jackson, Ian Johnston, Dominic Kwiatkowski, Cordelia Langford, John Sillitoe on behalf of the Wellcome Sanger Institute COVID-19 Surveillance Team |
| EPI_ISL_3223061 | Lighthouse Lab in Milton Keynes | Wellcome Sanger Institute for the COVID-19 Genomics UK (COG-UK) Consortium | The Lighthouse Lab in Milton Keynes and Alex Alderton, Roberto Amato, Jeffrey Barrett, Sonia Goncalves, Ewan Harrison, David K. Jackson, Ian Johnston, Dominic Kwiatkowski, Cordelia Langford, John Sillitoe on behalf of the Wellcome Sanger Institute COVID-19 Surveillance Team |
| EPI_ISL_3223062 | Lighthouse Lab in Alderley Park | Wellcome Sanger Institute for the COVID-19 Genomics UK (COG-UK) Consortium | Jacquelyn Wynn, Mairead Hyland, The Lighthouse Lab in Alderley Park and Alex Alderton, Roberto Amato, Jeffrey Barrett, Sonia Goncalves, Ewan Harrison, David K. Jackson, Ian Johnston, Dominic Kwiatkowski, Cordelia Langford, John Sillitoe on behalf of the Wellcome Sanger Institute COVID-19 Surveillance Team |
| EPI_ISL_3223063 | Lighthouse Laboratory Plymouth | Wellcome Sanger Institute for the COVID-19 Genomics UK (COG-UK) Consortium | Lighthouse Laboratory Plymouth and Alex Alderton, Roberto Amato, Jeffrey Barrett, Sonia Goncalves, Ewan Harrison, David K. Jackson, Ian Johnston, Dominic Kwiatkowski, Cordelia Langford, John Sillitoe on behalf of the Wellcome Sanger Institute COVID-19 Surveillance Team |
| EPI_ISL_3223064 | Lighthouse Lab in Glasgow | Wellcome Sanger Institute for the COVID-19 Genomics UK (COG-UK) Consortium | Harper VanSteenhouse, Yumi Kasai, David Gray, Carol Clugston, Anna Dominiczak and Alex Alderton, Roberto Amato, Jeffrey Barrett, Sonia Goncalves, Ewan Harrison, David K. Jackson, Ian Johnston, Dominic Kwiatkowski, Cordelia Langford, John Sillitoe on behalf of the Wellcome Sanger Institute COVID-19 Surveillance Team |
| EPI_ISL_3223065 | Lighthouse Laboratory Plymouth | Wellcome Sanger Institute for the COVID-19 Genomics UK (COG-UK) Consortium | Lighthouse Laboratory Plymouth and Alex Alderton, Roberto Amato, Jeffrey Barrett, Sonia Goncalves, Ewan Harrison, David K. Jackson, Ian Johnston, Dominic Kwiatkowski, Cordelia Langford, John Sillitoe on behalf of the Wellcome Sanger Institute COVID-19 Surveillance Team |
| EPI_ISL_3223066 | Lighthouse Lab in Milton Keynes | Wellcome Sanger Institute for the COVID-19 Genomics UK (COG-UK) Consortium | The Lighthouse Lab in Milton Keynes and Alex Alderton, Roberto Amato, Jeffrey Barrett, Sonia Goncalves, Ewan Harrison, David K. Jackson, Ian Johnston, Dominic Kwiatkowski, Cordelia Langford, John Sillitoe on behalf of the Wellcome Sanger Institute COVID-19 Surveillance Team |
| EPI_ISL_3223067 | Lighthouse Laboratory Plymouth | Wellcome Sanger Institute for the COVID-19 Genomics UK (COG-UK) Consortium | Lighthouse Laboratory Plymouth and Alex Alderton, Roberto Amato, Jeffrey Barrett, Sonia Goncalves, Ewan Harrison, David K. Jackson, Ian Johnston, Dominic Kwiatkowski, Cordelia Langford, John Sillitoe on behalf of the Wellcome Sanger Institute COVID-19 Surveillance Team |
| EPI_ISL_3223068, EPI_ISL_3223069 | Lighthouse Lab in Milton Keynes | Wellcome Sanger Institute for the COVID-19 Genomics UK (COG-UK) Consortium | The Lighthouse Lab in Milton Keynes and Alex Alderton, Roberto Amato, Jeffrey Barrett, Sonia Goncalves, Ewan Harrison, David K. Jackson, Ian Johnston, Dominic Kwiatkowski, Cordelia Langford, John Sillitoe on behalf of the Wellcome Sanger Institute COVID-19 Surveillance Team |
| EPI_ISL_3223070 | Lighthouse Lab in Glasgow | Wellcome Sanger Institute for the COVID-19 Genomics UK (COG-UK) Consortium | Harper VanSteenhouse, Yumi Kasai, David Gray, Carol Clugston, Anna Dominiczak and Alex Alderton, Roberto Amato, Jeffrey Barrett, Sonia Goncalves, Ewan Harrison, David K. Jackson, Ian Johnston, Dominic Kwiatkowski, Cordelia Langford, John Sillitoe on behalf of the Wellcome Sanger Institute COVID-19 Surveillance Team |
| EPI_ISL_3223071 | Lighthouse Laboratory Plymouth | Wellcome Sanger Institute for the COVID-19 Genomics UK (COG-UK) Consortium | Lighthouse Laboratory Plymouth and Alex Alderton, Roberto Amato, Jeffrey Barrett, Sonia Goncalves, Ewan Harrison, David K. Jackson, Ian Johnston, Dominic Kwiatkowski, Cordelia Langford, John Sillitoe on behalf of the Wellcome Sanger Institute COVID-19 Surveillance Team |
| EPI_ISL_3223072, EPI_ISL_3223073 | Lighthouse Lab in Alderley Park | Wellcome Sanger Institute for the COVID-19 Genomics UK (COG-UK) Consortium | Jacquelyn Wynn, Mairead Hyland, The Lighthouse Lab in Alderley Park and Alex Alderton, Roberto Amato, Jeffrey Barrett, Sonia Goncalves, Ewan Harrison, David K. Jackson, Ian Johnston, Dominic Kwiatkowski, Cordelia Langford, John Sillitoe on behalf of the Wellcome Sanger Institute COVID-19 Surveillance Team |
| EPI_ISL_3223074 | Lighthouse Laboratory Plymouth | Wellcome Sanger Institute for the COVID-19 Genomics UK (COG-UK) Consortium | Lighthouse Laboratory Plymouth and Alex Alderton, Roberto Amato, Jeffrey Barrett, Sonia Goncalves, Ewan Harrison, David K. Jackson, Ian Johnston, Dominic Kwiatkowski, Cordelia Langford, John Sillitoe on behalf of the Wellcome Sanger Institute COVID-19 Surveillance Team |
| EPI_ISL_3223075 | Lighthouse Lab in Glasgow | Wellcome Sanger Institute for the COVID-19 Genomics UK (COG-UK) Consortium | Harper VanSteenhouse, Yumi Kasai, David Gray, Carol Clugston, Anna Dominiczak and Alex Alderton, Roberto Amato, Jeffrey Barrett, Sonia Goncalves, Ewan Harrison, David K. Jackson, Ian Johnston, Dominic Kwiatkowski, Cordelia Langford, John Sillitoe on behalf of the Wellcome Sanger Institute COVID-19 Surveillance Team |
| EPI_ISL_3223076, EPI_ISL_3223077, EPI_ISL_3223078, EPI_ISL_3223079 | Lighthouse Lab in Milton Keynes | Wellcome Sanger Institute for the COVID-19 Genomics UK (COG-UK) Consortium | The Lighthouse Lab in Milton Keynes and Alex Alderton, Roberto Amato, Jeffrey Barrett, Sonia Goncalves, Ewan Harrison, David K. Jackson, Ian Johnston, Dominic Kwiatkowski, Cordelia Langford, John Sillitoe on behalf of the Wellcome Sanger Institute COVID-19 Surveillance Team |
| EPI_ISL_3223080, EPI_ISL_3223081, EPI_ISL_3223082 | Lighthouse Lab in Alderley Park | Wellcome Sanger Institute for the COVID-19 Genomics UK (COG-UK) Consortium | Jacquelyn Wynn, Mairead Hyland, The Lighthouse Lab in Alderley Park and Alex Alderton, Roberto Amato, Jeffrey Barrett, Sonia Goncalves, Ewan Harrison, David K. Jackson, Ian Johnston, Dominic Kwiatkowski, Cordelia Langford, John Sillitoe on behalf of the Wellcome Sanger Institute COVID-19 Surveillance Team |
| EPI_ISL_3223083, EPI_ISL_3223084 | Lighthouse Lab in Milton Keynes | Wellcome Sanger Institute for the COVID-19 Genomics UK (COG-UK) Consortium | The Lighthouse Lab in Milton Keynes and Alex Alderton, Roberto Amato, Jeffrey Barrett, Sonia Goncalves, Ewan Harrison, David K. Jackson, Ian Johnston, Dominic Kwiatkowski, Cordelia Langford, John Sillitoe on behalf of the Wellcome Sanger Institute COVID-19 Surveillance Team |
| EPI_ISL_3223085, EPI_ISL_3223086 | Lighthouse Lab in Glasgow | Wellcome Sanger Institute for the COVID-19 Genomics UK (COG-UK) Consortium | Harper VanSteenhouse, Yumi Kasai, David Gray, Carol Clugston, Anna Dominiczak and Alex Alderton, Roberto Amato, Jeffrey Barrett, Sonia Goncalves, Ewan Harrison, David K. Jackson, Ian Johnston, Dominic Kwiatkowski, Cordelia Langford, John Sillitoe on behalf of the Wellcome Sanger Institute COVID-19 Surveillance Team |
| EPI_ISL_3223087 | Lighthouse Laboratory Plymouth | Wellcome Sanger Institute for the COVID-19 Genomics UK (COG-UK) Consortium | Lighthouse Laboratory Plymouth and Alex Alderton, Roberto Amato, Jeffrey Barrett, Sonia Goncalves, Ewan Harrison, David K. Jackson, Ian Johnston, Dominic Kwiatkowski, Cordelia Langford, John Sillitoe on behalf of the Wellcome Sanger Institute COVID-19 Surveillance Team |
| EPI_ISL_3223088 | Lighthouse Lab in Glasgow | Wellcome Sanger Institute for the COVID-19 Genomics UK (COG-UK) Consortium | Harper VanSteenhouse, Yumi Kasai, David Gray, Carol Clugston, Anna Dominiczak and Alex Alderton, Roberto Amato, Jeffrey Barrett, Sonia Goncalves, Ewan Harrison, David K. Jackson, Ian Johnston, Dominic Kwiatkowski, Cordelia Langford, John Sillitoe on behalf of the Wellcome Sanger Institute COVID-19 Surveillance Team |
| EPI_ISL_3223089 | Lighthouse Lab in Alderley Park | Wellcome Sanger Institute for the COVID-19 Genomics UK (COG-UK) Consortium | Jacquelyn Wynn, Mairead Hyland, The Lighthouse Lab in Alderley Park and Alex Alderton, Roberto Amato, Jeffrey Barrett, Sonia Goncalves, Ewan Harrison, David K. Jackson, Ian Johnston, Dominic Kwiatkowski, Cordelia Langford, John Sillitoe on behalf of the Wellcome Sanger Institute COVID-19 Surveillance Team |
| EPI_ISL_3223090 | Lighthouse Laboratory Plymouth | Wellcome Sanger Institute for the COVID-19 Genomics UK (COG-UK) Consortium | Lighthouse Laboratory Plymouth and Alex Alderton, Roberto Amato, Jeffrey Barrett, Sonia Goncalves, Ewan Harrison, David K. Jackson, Ian Johnston, Dominic Kwiatkowski, Cordelia Langford, John Sillitoe on behalf of the Wellcome Sanger Institute COVID-19 Surveillance Team |
| EPI_ISL_3223091 | Lighthouse Lab in Milton Keynes | Wellcome Sanger Institute for the COVID-19 Genomics UK (COG-UK) Consortium | The Lighthouse Lab in Milton Keynes and Alex Alderton, Roberto Amato, Jeffrey Barrett, Sonia Goncalves, Ewan Harrison, David K. Jackson, Ian Johnston, Dominic Kwiatkowski, Cordelia Langford, John Sillitoe on behalf of the Wellcome Sanger Institute COVID-19 Surveillance Team |
| EPI_ISL_3223092 | Lighthouse Lab in Alderley Park | Wellcome Sanger Institute for the COVID-19 Genomics UK (COG-UK) Consortium | Jacquelyn Wynn, Mairead Hyland, The Lighthouse Lab in Alderley Park and Alex Alderton, Roberto Amato, Jeffrey Barrett, Sonia Goncalves, Ewan Harrison, David K. Jackson, Ian Johnston, Dominic Kwiatkowski, Cordelia Langford, John Sillitoe on behalf of the Wellcome Sanger Institute COVID-19 Surveillance Team |
| EPI_ISL_3223093 | Lighthouse Laboratory Plymouth | Wellcome Sanger Institute for the COVID-19 Genomics UK (COG-UK) Consortium | Lighthouse Laboratory Plymouth and Alex Alderton, Roberto Amato, Jeffrey Barrett, Sonia Goncalves, Ewan Harrison, David K. Jackson, Ian Johnston, Dominic Kwiatkowski, Cordelia Langford, John Sillitoe on behalf of the Wellcome Sanger Institute COVID-19 Surveillance Team |
| EPI_ISL_3223094, EPI_ISL_3223095, EPI_ISL_3223096 | Lighthouse Lab in Glasgow | Wellcome Sanger Institute for the COVID-19 Genomics UK (COG-UK) Consortium | Harper VanSteenhouse, Yumi Kasai, David Gray, Carol Clugston, Anna Dominiczak and Alex Alderton, Roberto Amato, Jeffrey Barrett, Sonia Goncalves, Ewan Harrison, David K. Jackson, Ian Johnston, Dominic Kwiatkowski, Cordelia Langford, John Sillitoe on behalf of the Wellcome Sanger Institute COVID-19 Surveillance Team |
| EPI_ISL_3223097 | Lighthouse Lab in Milton Keynes | Wellcome Sanger Institute for the COVID-19 Genomics UK (COG-UK) Consortium | The Lighthouse Lab in Milton Keynes and Alex Alderton, Roberto Amato, Jeffrey Barrett, Sonia Goncalves, Ewan Harrison, David K. Jackson, Ian Johnston, Dominic Kwiatkowski, Cordelia Langford, John Sillitoe on behalf of the Wellcome Sanger Institute COVID-19 Surveillance Team |
| EPI_ISL_3223098 | Lighthouse Laboratory Plymouth | Wellcome Sanger Institute for the COVID-19 Genomics UK (COG-UK) Consortium | Lighthouse Laboratory Plymouth and Alex Alderton, Roberto Amato, Jeffrey Barrett, Sonia Goncalves, Ewan Harrison, David K. Jackson, Ian Johnston, Dominic Kwiatkowski, Cordelia Langford, John Sillitoe on behalf of the Wellcome Sanger Institute COVID-19 Surveillance Team |
| EPI_ISL_3223099, EPI_ISL_3223100, EPI_ISL_3223101 | Lighthouse Lab in Glasgow | Wellcome Sanger Institute for the COVID-19 Genomics UK (COG-UK) Consortium | Harper VanSteenhouse, Yumi Kasai, David Gray, Carol Clugston, Anna Dominiczak and Alex Alderton, Roberto Amato, Jeffrey Barrett, Sonia Goncalves, Ewan Harrison, David K. Jackson, Ian Johnston, Dominic Kwiatkowski, Cordelia Langford, John Sillitoe on behalf of the Wellcome Sanger Institute COVID-19 Surveillance Team |
| EPI_ISL_3223102 | Lighthouse Laboratory Plymouth | Wellcome Sanger Institute for the COVID-19 Genomics UK (COG-UK) Consortium | Lighthouse Laboratory Plymouth and Alex Alderton, Roberto Amato, Jeffrey Barrett, Sonia Goncalves, Ewan Harrison, David K. Jackson, Ian Johnston, Dominic Kwiatkowski, Cordelia Langford, John Sillitoe on behalf of the Wellcome Sanger Institute COVID-19 Surveillance Team |
| EPI_ISL_3223103 | Lighthouse Lab in Alderley Park | Wellcome Sanger Institute for the COVID-19 Genomics UK (COG-UK) Consortium | Jacquelyn Wynn, Mairead Hyland, The Lighthouse Lab in Alderley Park and Alex Alderton, Roberto Amato, Jeffrey Barrett, Sonia Goncalves, Ewan Harrison, David K. Jackson, Ian Johnston, Dominic Kwiatkowski, Cordelia Langford, John Sillitoe on behalf of the Wellcome Sanger Institute COVID-19 Surveillance Team |
| EPI_ISL_3223104, EPI_ISL_3223105 | Lighthouse Laboratory Plymouth | Wellcome Sanger Institute for the COVID-19 Genomics UK (COG-UK) Consortium | Lighthouse Laboratory Plymouth and Alex Alderton, Roberto Amato, Jeffrey Barrett, Sonia Goncalves, Ewan Harrison, David K. Jackson, Ian Johnston, Dominic Kwiatkowski, Cordelia Langford, John Sillitoe on behalf of the Wellcome Sanger Institute COVID-19 Surveillance Team |
| EPI_ISL_3223106 | Lighthouse Lab in Alderley Park | Wellcome Sanger Institute for the COVID-19 Genomics UK (COG-UK) Consortium | Jacquelyn Wynn, Mairead Hyland, The Lighthouse Lab in Alderley Park and Alex Alderton, Roberto Amato, Jeffrey Barrett, Sonia Goncalves, Ewan Harrison, David K. Jackson, Ian Johnston, Dominic Kwiatkowski, Cordelia Langford, John Sillitoe on behalf of the Wellcome Sanger Institute COVID-19 Surveillance Team |
| EPI_ISL_3223107 | Lighthouse Lab in Milton Keynes | Wellcome Sanger Institute for the COVID-19 Genomics UK (COG-UK) Consortium | The Lighthouse Lab in Milton Keynes and Alex Alderton, Roberto Amato, Jeffrey Barrett, Sonia Goncalves, Ewan Harrison, David K. Jackson, Ian Johnston, Dominic Kwiatkowski, Cordelia Langford, John Sillitoe on behalf of the Wellcome Sanger Institute COVID-19 Surveillance Team |
| EPI_ISL_3223108 | Lighthouse Lab in Alderley Park | Wellcome Sanger Institute for the COVID-19 Genomics UK (COG-UK) Consortium | Jacquelyn Wynn, Mairead Hyland, The Lighthouse Lab in Alderley Park and Alex Alderton, Roberto Amato, Jeffrey Barrett, Sonia Goncalves, Ewan Harrison, David K. Jackson, Ian Johnston, Dominic Kwiatkowski, Cordelia Langford, John Sillitoe on behalf of the Wellcome Sanger Institute COVID-19 Surveillance Team |

| | | | |
|---|---|---|---|
| EPI_ISL_3223109, EPI_ISL_3223110 | Lighthouse Lab in Milton Keynes | Wellcome Sanger Institute for the COVID-19 Genomics UK (COG-UK) Consortium | The Lighthouse Lab in Milton Keynes and Alex Alderton, Roberto Amato, Jeffrey Barrett, Sonia Goncalves, Ewan Harrison, David K. Jackson, Ian Johnston, Dominic Kwiatkowski, Cordelia Langford, John Sillitoe on behalf of the Wellcome Sanger Institute COVID-19 Surveillance Team |
| EPI_ISL_3223111 | Lighthouse Lab in Glasgow | Wellcome Sanger Institute for the COVID-19 Genomics UK (COG-UK) Consortium | Harper VanSteenhouse, Yumi Kasai, David Gray, Carol Clugston, Anna Dominiczak and Alex Alderton, Roberto Amato, Jeffrey Barrett, Sonia Goncalves, Ewan Harrison, David K. Jackson, Ian Johnston, Dominic Kwiatkowski, Cordelia Langford, John Sillitoe on behalf of the Wellcome Sanger Institute COVID-19 Surveillance Team |
| EPI_ISL_3223112 | Lighthouse Lab in Milton Keynes | Wellcome Sanger Institute for the COVID-19 Genomics UK (COG-UK) Consortium | The Lighthouse Lab in Milton Keynes and Alex Alderton, Roberto Amato, Jeffrey Barrett, Sonia Goncalves, Ewan Harrison, David K. Jackson, Ian Johnston, Dominic Kwiatkowski, Cordelia Langford, John Sillitoe on behalf of the Wellcome Sanger Institute COVID-19 Surveillance Team |
| EPI_ISL_3223113, EPI_ISL_3223114 | Lighthouse Laboratory Plymouth | Wellcome Sanger Institute for the COVID-19 Genomics UK (COG-UK) Consortium | Lighthouse Laboratory Plymouth and Alex Alderton, Roberto Amato, Jeffrey Barrett, Sonia Goncalves, Ewan Harrison, David K. Jackson, Ian Johnston, Dominic Kwiatkowski, Cordelia Langford, John Sillitoe on behalf of the Wellcome Sanger Institute COVID-19 Surveillance Team |
| EPI_ISL_3223115 | Lighthouse Lab in Alderley Park | Wellcome Sanger Institute for the COVID-19 Genomics UK (COG-UK) Consortium | Jacquelyn Wynn, Mairead Hyland, The Lighthouse Lab in Alderley Park and Alex Alderton, Roberto Amato, Jeffrey Barrett, Sonia Goncalves, Ewan Harrison, David K. Jackson, Ian Johnston, Dominic Kwiatkowski, Cordelia Langford, John Sillitoe on behalf of the Wellcome Sanger Institute COVID-19 Surveillance Team |
| EPI_ISL_3223116 | Lighthouse Lab in Glasgow | Wellcome Sanger Institute for the COVID-19 Genomics UK (COG-UK) Consortium | Harper VanSteenhouse, Yumi Kasai, David Gray, Carol Clugston, Anna Dominiczak and Alex Alderton, Roberto Amato, Jeffrey Barrett, Sonia Goncalves, Ewan Harrison, David K. Jackson, Ian Johnston, Dominic Kwiatkowski, Cordelia Langford, John Sillitoe on behalf of the Wellcome Sanger Institute COVID-19 Surveillance Team |
| EPI_ISL_3223117 | Lighthouse Lab in Milton Keynes | Wellcome Sanger Institute for the COVID-19 Genomics UK (COG-UK) Consortium | The Lighthouse Lab in Milton Keynes and Alex Alderton, Roberto Amato, Jeffrey Barrett, Sonia Goncalves, Ewan Harrison, David K. Jackson, Ian Johnston, Dominic Kwiatkowski, Cordelia Langford, John Sillitoe on behalf of the Wellcome Sanger Institute COVID-19 Surveillance Team |
| EPI_ISL_3223118 | Lighthouse Lab in Alderley Park | Wellcome Sanger Institute for the COVID-19 Genomics UK (COG-UK) Consortium | Jacquelyn Wynn, Mairead Hyland, The Lighthouse Lab in Alderley Park and Alex Alderton, Roberto Amato, Jeffrey Barrett, Sonia Goncalves, Ewan Harrison, David K. Jackson, Ian Johnston, Dominic Kwiatkowski, Cordelia Langford, John Sillitoe on behalf of the Wellcome Sanger Institute COVID-19 Surveillance Team |
| EPI_ISL_3223119 | Lighthouse Lab in Milton Keynes | Wellcome Sanger Institute for the COVID-19 Genomics UK (COG-UK) Consortium | The Lighthouse Lab in Milton Keynes and Alex Alderton, Roberto Amato, Jeffrey Barrett, Sonia Goncalves, Ewan Harrison, David K. Jackson, Ian Johnston, Dominic Kwiatkowski, Cordelia Langford, John Sillitoe on behalf of the Wellcome Sanger Institute COVID-19 Surveillance Team |
| EPI_ISL_3223120, EPI_ISL_3223121, EPI_ISL_3223122, EPI_ISL_3223123 | Lighthouse Lab in Glasgow | Wellcome Sanger Institute for the COVID-19 Genomics UK (COG-UK) Consortium | Harper VanSteenhouse, Yumi Kasai, David Gray, Carol Clugston, Anna Dominiczak and Alex Alderton, Roberto Amato, Jeffrey Barrett, Sonia Goncalves, Ewan Harrison, David K. Jackson, Ian Johnston, Dominic Kwiatkowski, Cordelia Langford, John Sillitoe on behalf of the Wellcome Sanger Institute COVID-19 Surveillance Team |
| EPI_ISL_3223124 | Lighthouse Lab in Milton Keynes | Wellcome Sanger Institute for the COVID-19 Genomics UK (COG-UK) Consortium | The Lighthouse Lab in Milton Keynes and Alex Alderton, Roberto Amato, Jeffrey Barrett, Sonia Goncalves, Ewan Harrison, David K. Jackson, Ian Johnston, Dominic Kwiatkowski, Cordelia Langford, John Sillitoe on behalf of the Wellcome Sanger Institute COVID-19 Surveillance Team |
| EPI_ISL_3223125 | Lighthouse Lab in Alderley Park | Wellcome Sanger Institute for the COVID-19 Genomics UK (COG-UK) Consortium | Jacquelyn Wynn, Mairead Hyland, The Lighthouse Lab in Alderley Park and Alex Alderton, Roberto Amato, Jeffrey Barrett, Sonia Goncalves, Ewan Harrison, David K. Jackson, Ian Johnston, Dominic Kwiatkowski, Cordelia Langford, John Sillitoe on behalf of the Wellcome Sanger Institute COVID-19 Surveillance Team |
| EPI_ISL_3223126 | Lighthouse Lab in Glasgow | Wellcome Sanger Institute for the COVID-19 Genomics UK (COG-UK) Consortium | Harper VanSteenhouse, Yumi Kasai, David Gray, Carol Clugston, Anna Dominiczak and Alex Alderton, Roberto Amato, Jeffrey Barrett, Sonia Goncalves, Ewan Harrison, David K. Jackson, Ian Johnston, Dominic Kwiatkowski, Cordelia Langford, John Sillitoe on behalf of the Wellcome Sanger Institute COVID-19 Surveillance Team |
| EPI_ISL_3223127 | Lighthouse Lab in Alderley Park | Wellcome Sanger Institute for the COVID-19 Genomics UK (COG-UK) Consortium | Jacquelyn Wynn, Mairead Hyland, The Lighthouse Lab in Alderley Park and Alex Alderton, Roberto Amato, Jeffrey Barrett, Sonia Goncalves, Ewan Harrison, David K. Jackson, Ian Johnston, Dominic Kwiatkowski, Cordelia Langford, John Sillitoe on behalf of the Wellcome Sanger Institute COVID-19 Surveillance Team |
| EPI_ISL_3223128 | Lighthouse Lab in Milton Keynes | Wellcome Sanger Institute for the COVID-19 Genomics UK (COG-UK) Consortium | The Lighthouse Lab in Milton Keynes and Alex Alderton, Roberto Amato, Jeffrey Barrett, Sonia Goncalves, Ewan Harrison, David K. Jackson, Ian Johnston, Dominic Kwiatkowski, Cordelia Langford, John Sillitoe on behalf of the Wellcome Sanger Institute COVID-19 Surveillance Team |
| EPI_ISL_3223129, EPI_ISL_3223130 | Lighthouse Lab in Glasgow | Wellcome Sanger Institute for the COVID-19 Genomics UK (COG-UK) Consortium | Harper VanSteenhouse, Yumi Kasai, David Gray, Carol Clugston, Anna Dominiczak and Alex Alderton, Roberto Amato, Jeffrey Barrett, Sonia Goncalves, Ewan Harrison, David K. Jackson, Ian Johnston, Dominic Kwiatkowski, Cordelia Langford, John Sillitoe on behalf of the Wellcome Sanger Institute COVID-19 Surveillance Team |
| EPI_ISL_3223131 | Lighthouse Laboratory Plymouth | Wellcome Sanger Institute for the COVID-19 Genomics UK (COG-UK) Consortium | Lighthouse Laboratory Plymouth and Alex Alderton, Roberto Amato, Jeffrey Barrett, Sonia Goncalves, Ewan Harrison, David K. Jackson, Ian Johnston, Dominic Kwiatkowski, Cordelia Langford, John Sillitoe on behalf of the Wellcome Sanger Institute COVID-19 Surveillance Team |
| EPI_ISL_3223132 | Lighthouse Lab in Glasgow | Wellcome Sanger Institute for the COVID-19 Genomics UK (COG-UK) Consortium | Harper VanSteenhouse, Yumi Kasai, David Gray, Carol Clugston, Anna Dominiczak and Alex Alderton, Roberto Amato, Jeffrey Barrett, Sonia Goncalves, Ewan Harrison, David K. Jackson, Ian Johnston, Dominic Kwiatkowski, Cordelia Langford, John Sillitoe on behalf of the Wellcome Sanger Institute COVID-19 Surveillance Team |
| EPI_ISL_3223133 | Lighthouse Laboratory Plymouth | Wellcome Sanger Institute for the COVID-19 Genomics UK (COG-UK) Consortium | Lighthouse Laboratory Plymouth and Alex Alderton, Roberto Amato, Jeffrey Barrett, Sonia Goncalves, Ewan Harrison, David K. Jackson, Ian Johnston, Dominic Kwiatkowski, Cordelia Langford, John Sillitoe on behalf of the Wellcome Sanger Institute COVID-19 Surveillance Team |
| EPI_ISL_3223134, EPI_ISL_3223135 | Lighthouse Lab in Milton Keynes | Wellcome Sanger Institute for the COVID-19 Genomics UK (COG-UK) Consortium | The Lighthouse Lab in Milton Keynes and Alex Alderton, Roberto Amato, Jeffrey Barrett, Sonia Goncalves, Ewan Harrison, David K. Jackson, Ian Johnston, Dominic Kwiatkowski, Cordelia Langford, John Sillitoe on behalf of the Wellcome Sanger Institute COVID-19 Surveillance Team |
| EPI_ISL_3223136, EPI_ISL_3223137, EPI_ISL_3223138, EPI_ISL_3223139 | Lighthouse Lab in Alderley Park | Wellcome Sanger Institute for the COVID-19 Genomics UK (COG-UK) Consortium | Jacquelyn Wynn, Mairead Hyland, The Lighthouse Lab in Alderley Park and Alex Alderton, Roberto Amato, Jeffrey Barrett, Sonia Goncalves, Ewan Harrison, David K. Jackson, Ian Johnston, Dominic Kwiatkowski, Cordelia Langford, John Sillitoe on behalf of the Wellcome Sanger Institute COVID-19 Surveillance Team |
| EPI_ISL_3223140 | Lighthouse Lab in Milton Keynes | Wellcome Sanger Institute for the COVID-19 Genomics UK (COG-UK) Consortium | The Lighthouse Lab in Milton Keynes and Alex Alderton, Roberto Amato, Jeffrey Barrett, Sonia Goncalves, Ewan Harrison, David K. Jackson, Ian Johnston, Dominic Kwiatkowski, Cordelia Langford, John Sillitoe on behalf of the Wellcome Sanger Institute COVID-19 Surveillance Team |
| EPI_ISL_3223141, EPI_ISL_3223142, EPI_ISL_3223143, EPI_ISL_3223144, EPI_ISL_3223145 | Lighthouse Laboratory Plymouth | Wellcome Sanger Institute for the COVID-19 Genomics UK (COG-UK) Consortium | Lighthouse Laboratory Plymouth and Alex Alderton, Roberto Amato, Jeffrey Barrett, Sonia Goncalves, Ewan Harrison, David K. Jackson, Ian Johnston, Dominic Kwiatkowski, Cordelia Langford, John Sillitoe on behalf of the Wellcome Sanger Institute COVID-19 Surveillance Team |
| EPI_ISL_3223146 | Lighthouse Lab in Alderley Park | Wellcome Sanger Institute for the COVID-19 Genomics UK (COG-UK) Consortium | Jacquelyn Wynn, Mairead Hyland, The Lighthouse Lab in Alderley Park and Alex Alderton, Roberto Amato, Jeffrey Barrett, Sonia Goncalves, Ewan Harrison, David K. Jackson, Ian Johnston, Dominic Kwiatkowski, Cordelia Langford, John Sillitoe on behalf of the Wellcome Sanger Institute COVID-19 Surveillance Team |
| EPI_ISL_3223147 | Lighthouse Laboratory Plymouth | Wellcome Sanger Institute for the COVID-19 Genomics UK (COG-UK) Consortium | Lighthouse Laboratory Plymouth and Alex Alderton, Roberto Amato, Jeffrey Barrett, Sonia Goncalves, Ewan Harrison, David K. Jackson, Ian Johnston, Dominic Kwiatkowski, Cordelia Langford, John Sillitoe on behalf of the Wellcome Sanger Institute COVID-19 Surveillance Team |
| EPI_ISL_3223148, EPI_ISL_3223149 | Lighthouse Lab in Glasgow | Wellcome Sanger Institute for the COVID-19 Genomics UK (COG-UK) Consortium | Harper VanSteenhouse, Yumi Kasai, David Gray, Carol Clugston, Anna Dominiczak and Alex Alderton, Roberto Amato, Jeffrey Barrett, Sonia Goncalves, Ewan Harrison, David K. Jackson, Ian Johnston, Dominic Kwiatkowski, Cordelia Langford, John Sillitoe on behalf of the Wellcome Sanger Institute COVID-19 Surveillance Team |
| EPI_ISL_3223150 | Lighthouse Lab in Milton Keynes | Wellcome Sanger Institute for the COVID-19 Genomics UK (COG-UK) Consortium | The Lighthouse Lab in Milton Keynes and Alex Alderton, Roberto Amato, Jeffrey Barrett, Sonia Goncalves, Ewan Harrison, David K. Jackson, Ian Johnston, Dominic Kwiatkowski, Cordelia Langford, John Sillitoe on behalf of the Wellcome Sanger Institute COVID-19 Surveillance Team |
| EPI_ISL_3223151 | Lighthouse Lab in Alderley Park | Wellcome Sanger Institute for the COVID-19 Genomics UK (COG-UK) Consortium | Jacquelyn Wynn, Mairead Hyland, The Lighthouse Lab in Alderley Park and Alex Alderton, Roberto Amato, Jeffrey Barrett, Sonia Goncalves, Ewan Harrison, David K. Jackson, Ian Johnston, Dominic Kwiatkowski, Cordelia Langford, John Sillitoe on behalf of the Wellcome Sanger Institute COVID-19 Surveillance Team |
| EPI_ISL_3223152 | Lighthouse Lab in Glasgow | Wellcome Sanger Institute for the COVID-19 Genomics UK (COG-UK) Consortium | Harper VanSteenhouse, Yumi Kasai, David Gray, Carol Clugston, Anna Dominiczak and Alex Alderton, Roberto Amato, Jeffrey Barrett, Sonia Goncalves, Ewan Harrison, David K. Jackson, Ian Johnston, Dominic Kwiatkowski, Cordelia Langford, John Sillitoe on behalf of the Wellcome Sanger Institute COVID-19 Surveillance Team |
| EPI_ISL_3223153 | Lighthouse Laboratory Plymouth | Wellcome Sanger Institute for the COVID-19 Genomics UK (COG-UK) Consortium | Lighthouse Laboratory Plymouth and Alex Alderton, Roberto Amato, Jeffrey Barrett, Sonia Goncalves, Ewan Harrison, David K. Jackson, Ian Johnston, Dominic Kwiatkowski, Cordelia Langford, John Sillitoe on behalf of the Wellcome Sanger Institute COVID-19 Surveillance Team |
| EPI_ISL_3223154 | Lighthouse Lab in Alderley Park | Wellcome Sanger Institute for the COVID-19 Genomics UK (COG-UK) Consortium | Jacquelyn Wynn, Mairead Hyland, The Lighthouse Lab in Alderley Park and Alex Alderton, Roberto Amato, Jeffrey Barrett, Sonia Goncalves, Ewan Harrison, David K. Jackson, Ian Johnston, Dominic Kwiatkowski, Cordelia Langford, John Sillitoe on behalf of the Wellcome Sanger Institute COVID-19 Surveillance Team |
| EPI_ISL_3223155 | Lighthouse Lab in Milton Keynes | Wellcome Sanger Institute for the COVID-19 Genomics UK (COG-UK) Consortium | The Lighthouse Lab in Milton Keynes and Alex Alderton, Roberto Amato, Jeffrey Barrett, Sonia Goncalves, Ewan Harrison, David K. Jackson, Ian Johnston, Dominic Kwiatkowski, Cordelia Langford, John Sillitoe on behalf of the Wellcome Sanger Institute COVID-19 Surveillance Team |
| EPI_ISL_3223156 | Lighthouse Lab in Glasgow | Wellcome Sanger Institute for the COVID-19 Genomics UK (COG-UK) Consortium | Harper VanSteenhouse, Yumi Kasai, David Gray, Carol Clugston, Anna Dominiczak and Alex Alderton, Roberto Amato, Jeffrey Barrett, Sonia Goncalves, Ewan Harrison, David K. Jackson, Ian Johnston, Dominic Kwiatkowski, Cordelia Langford, John Sillitoe on behalf of the Wellcome Sanger Institute COVID-19 Surveillance Team |
| EPI_ISL_3223157 | Lighthouse Laboratory Plymouth | Wellcome Sanger Institute for the COVID-19 Genomics UK (COG-UK) Consortium | Lighthouse Laboratory Plymouth and Alex Alderton, Roberto Amato, Jeffrey Barrett, Sonia Goncalves, Ewan Harrison, David K. Jackson, Ian Johnston, Dominic Kwiatkowski, Cordelia Langford, John Sillitoe on behalf of the Wellcome Sanger Institute COVID-19 Surveillance Team |
| EPI_ISL_3223158 | Lighthouse Lab in Milton Keynes | Wellcome Sanger Institute for the COVID-19 Genomics UK (COG-UK) Consortium | The Lighthouse Lab in Milton Keynes and Alex Alderton, Roberto Amato, Jeffrey Barrett, Sonia Goncalves, Ewan Harrison, David K. Jackson, Ian Johnston, Dominic Kwiatkowski, Cordelia Langford, John Sillitoe on behalf of the Wellcome Sanger Institute COVID-19 Surveillance Team |
| EPI_ISL_3223159 | Lighthouse Lab in Glasgow | Wellcome Sanger Institute for the COVID-19 Genomics UK (COG-UK) Consortium | Harper VanSteenhouse, Yumi Kasai, David Gray, Carol Clugston, Anna Dominiczak and Alex Alderton, Roberto Amato, Jeffrey Barrett, Sonia Goncalves, Ewan Harrison, David K. Jackson, Ian Johnston, Dominic Kwiatkowski, Cordelia Langford, John Sillitoe on behalf of the Wellcome Sanger Institute COVID-19 Surveillance Team |
| EPI_ISL_3223160 | Lighthouse Lab in Milton Keynes | Wellcome Sanger Institute for the COVID-19 Genomics UK (COG-UK) Consortium | The Lighthouse Lab in Milton Keynes and Alex Alderton, Roberto Amato, Jeffrey Barrett, Sonia Goncalves, Ewan Harrison, David K. Jackson, Ian Johnston, Dominic Kwiatkowski, Cordelia Langford, John Sillitoe on behalf of the Wellcome Sanger Institute COVID-19 Surveillance Team |
| EPI_ISL_3223161 | Lighthouse Lab in Alderley Park | Wellcome Sanger Institute for the COVID-19 Genomics UK (COG-UK) Consortium | Jacquelyn Wynn, Mairead Hyland, The Lighthouse Lab in Alderley Park and Alex Alderton, Roberto Amato, Jeffrey Barrett, Sonia Goncalves, Ewan Harrison, David K. Jackson, Ian Johnston, Dominic Kwiatkowski, Cordelia Langford, John Sillitoe on behalf of the Wellcome Sanger Institute COVID-19 Surveillance Team |
| EPI_ISL_3223162 | Lighthouse Laboratory Plymouth | Wellcome Sanger Institute for the COVID-19 Genomics UK (COG-UK) Consortium | Lighthouse Laboratory Plymouth and Alex Alderton, Roberto Amato, Jeffrey Barrett, Sonia Goncalves, Ewan Harrison, David K. Jackson, Ian Johnston, Dominic Kwiatkowski, Cordelia Langford, John Sillitoe on behalf of the Wellcome Sanger Institute COVID-19 Surveillance Team |
| EPI_ISL_3223163 | Lighthouse Lab in Alderley Park | Wellcome Sanger Institute for the COVID-19 Genomics UK (COG-UK) Consortium | Jacquelyn Wynn, Mairead Hyland, The Lighthouse Lab in Alderley Park and Alex Alderton, Roberto Amato, Jeffrey Barrett, Sonia Goncalves, Ewan Harrison, David K. Jackson, Ian Johnston, Dominic Kwiatkowski, Cordelia Langford, John Sillitoe on behalf of the Wellcome Sanger Institute COVID-19 Surveillance Team |
| EPI_ISL_3223164, EPI_ISL_3223165 | Lighthouse Lab in Milton Keynes | Wellcome Sanger Institute for the COVID-19 Genomics UK (COG-UK) Consortium | The Lighthouse Lab in Milton Keynes and Alex Alderton, Roberto Amato, Jeffrey Barrett, Sonia Goncalves, Ewan Harrison, David K. Jackson, Ian Johnston, Dominic Kwiatkowski, Cordelia Langford, John Sillitoe on behalf of the Wellcome Sanger Institute COVID-19 Surveillance Team |
| EPI_ISL_3223166 | Lighthouse Laboratory Plymouth | Wellcome Sanger Institute for the COVID-19 Genomics UK (COG-UK) Consortium | Lighthouse Laboratory Plymouth and Alex Alderton, Roberto Amato, Jeffrey Barrett, Sonia Goncalves, Ewan Harrison, David K. Jackson, Ian Johnston, Dominic Kwiatkowski, Cordelia Langford, John Sillitoe on behalf of the Wellcome Sanger Institute COVID-19 Surveillance Team |
| EPI_ISL_3223167 | Lighthouse Lab in Alderley Park | Wellcome Sanger Institute for the COVID-19 Genomics UK (COG-UK) | Jacquelyn Wynn, Mairead Hyland, The Lighthouse Lab in Alderley Park and Alex Alderton, Roberto Amato, Jeffrey Barrett, Sonia Goncalves, Ewan Harrison, David K. Jackson, Ian Johnston, Dominic Kwiatkowski, Cordelia Langford, John Sillitoe on |

| | | | |
|---|---|---|---|
| | | Consortium | behalf of the Wellcome Sanger Institute COVID-19 Surveillance Team |
| EPI_ISL_3223168 | Lighthouse Laboratory Plymouth | Wellcome Sanger Institute for the COVID-19 Genomics UK (COG-UK) Consortium | Lighthouse Laboratory Plymouth and Alex Alderton, Roberto Amato, Jeffrey Barrett, Sonia Goncalves, Ewan Harrison, David K. Jackson, Ian Johnston, Dominic Kwiatkowski, Cordelia Langford, John Sillitoe on behalf of the Wellcome Sanger Institute COVID-19 Surveillance Team |
| EPI_ISL_3223169 | Lighthouse Lab in Glasgow | Wellcome Sanger Institute for the COVID-19 Genomics UK (COG-UK) Consortium | Harper VanSteenhouse, Yumi Kasai, David Gray, Carol Clugston, Anna Dominiczak and Alex Alderton, Roberto Amato, Jeffrey Barrett, Sonia Goncalves, Ewan Harrison, David K. Jackson, Ian Johnston, Dominic Kwiatkowski, Cordelia Langford, John Sillitoe on behalf of the Wellcome Sanger Institute COVID-19 Surveillance Team |
| EPI_ISL_3223170, EPI_ISL_3223171 | Lighthouse Lab in Alderley Park | Wellcome Sanger Institute for the COVID-19 Genomics UK (COG-UK) Consortium | Jacquelyn Wynn, Mairead Hyland, The Lighthouse Lab in Alderley Park and Alex Alderton, Roberto Amato, Jeffrey Barrett, Sonia Goncalves, Ewan Harrison, David K. Jackson, Ian Johnston, Dominic Kwiatkowski, Cordelia Langford, John Sillitoe on behalf of the Wellcome Sanger Institute COVID-19 Surveillance Team |
| EPI_ISL_3223172 | Lighthouse Laboratory Plymouth | Wellcome Sanger Institute for the COVID-19 Genomics UK (COG-UK) Consortium | Lighthouse Laboratory Plymouth and Alex Alderton, Roberto Amato, Jeffrey Barrett, Sonia Goncalves, Ewan Harrison, David K. Jackson, Ian Johnston, Dominic Kwiatkowski, Cordelia Langford, John Sillitoe on behalf of the Wellcome Sanger Institute COVID-19 Surveillance Team |
| EPI_ISL_3223173, EPI_ISL_3223174, EPI_ISL_3223175, EPI_ISL_3223176 | Lighthouse Lab in Alderley Park | Wellcome Sanger Institute for the COVID-19 Genomics UK (COG-UK) Consortium | Jacquelyn Wynn, Mairead Hyland, The Lighthouse Lab in Alderley Park and Alex Alderton, Roberto Amato, Jeffrey Barrett, Sonia Goncalves, Ewan Harrison, David K. Jackson, Ian Johnston, Dominic Kwiatkowski, Cordelia Langford, John Sillitoe on behalf of the Wellcome Sanger Institute COVID-19 Surveillance Team |
| EPI_ISL_3223177 | Lighthouse Laboratory Plymouth | Wellcome Sanger Institute for the COVID-19 Genomics UK (COG-UK) Consortium | Lighthouse Laboratory Plymouth and Alex Alderton, Roberto Amato, Jeffrey Barrett, Sonia Goncalves, Ewan Harrison, David K. Jackson, Ian Johnston, Dominic Kwiatkowski, Cordelia Langford, John Sillitoe on behalf of the Wellcome Sanger Institute COVID-19 Surveillance Team |
| EPI_ISL_3223178 | Lighthouse Lab in Milton Keynes | Wellcome Sanger Institute for the COVID-19 Genomics UK (COG-UK) Consortium | The Lighthouse Lab in Milton Keynes and Alex Alderton, Roberto Amato, Jeffrey Barrett, Sonia Goncalves, Ewan Harrison, David K. Jackson, Ian Johnston, Dominic Kwiatkowski, Cordelia Langford, John Sillitoe on behalf of the Wellcome Sanger Institute COVID-19 Surveillance Team |
| EPI_ISL_3223179 | Lighthouse Lab in Alderley Park | Wellcome Sanger Institute for the COVID-19 Genomics UK (COG-UK) Consortium | Jacquelyn Wynn, Mairead Hyland, The Lighthouse Lab in Alderley Park and Alex Alderton, Roberto Amato, Jeffrey Barrett, Sonia Goncalves, Ewan Harrison, David K. Jackson, Ian Johnston, Dominic Kwiatkowski, Cordelia Langford, John Sillitoe on behalf of the Wellcome Sanger Institute COVID-19 Surveillance Team |
| EPI_ISL_3223180, EPI_ISL_3223181 | Lighthouse Laboratory Plymouth | Wellcome Sanger Institute for the COVID-19 Genomics UK (COG-UK) Consortium | Lighthouse Laboratory Plymouth and Alex Alderton, Roberto Amato, Jeffrey Barrett, Sonia Goncalves, Ewan Harrison, David K. Jackson, Ian Johnston, Dominic Kwiatkowski, Cordelia Langford, John Sillitoe on behalf of the Wellcome Sanger Institute COVID-19 Surveillance Team |
| EPI_ISL_3223182, EPI_ISL_3223183, EPI_ISL_3223184 | Lighthouse Lab in Alderley Park | Wellcome Sanger Institute for the COVID-19 Genomics UK (COG-UK) Consortium | Jacquelyn Wynn, Mairead Hyland, The Lighthouse Lab in Alderley Park and Alex Alderton, Roberto Amato, Jeffrey Barrett, Sonia Goncalves, Ewan Harrison, David K. Jackson, Ian Johnston, Dominic Kwiatkowski, Cordelia Langford, John Sillitoe on behalf of the Wellcome Sanger Institute COVID-19 Surveillance Team |
| EPI_ISL_3223185 | Lighthouse Lab in Milton Keynes | Wellcome Sanger Institute for the COVID-19 Genomics UK (COG-UK) Consortium | The Lighthouse Lab in Milton Keynes and Alex Alderton, Roberto Amato, Jeffrey Barrett, Sonia Goncalves, Ewan Harrison, David K. Jackson, Ian Johnston, Dominic Kwiatkowski, Cordelia Langford, John Sillitoe on behalf of the Wellcome Sanger Institute COVID-19 Surveillance Team |
| EPI_ISL_3223186 | Lighthouse Lab in Alderley Park | Wellcome Sanger Institute for the COVID-19 Genomics UK (COG-UK) Consortium | Jacquelyn Wynn, Mairead Hyland, The Lighthouse Lab in Alderley Park and Alex Alderton, Roberto Amato, Jeffrey Barrett, Sonia Goncalves, Ewan Harrison, David K. Jackson, Ian Johnston, Dominic Kwiatkowski, Cordelia Langford, John Sillitoe on behalf of the Wellcome Sanger Institute COVID-19 Surveillance Team |
| EPI_ISL_3223187, EPI_ISL_3223188, EPI_ISL_3223189, EPI_ISL_3223190 | Lighthouse Lab in Milton Keynes | Wellcome Sanger Institute for the COVID-19 Genomics UK (COG-UK) Consortium | The Lighthouse Lab in Milton Keynes and Alex Alderton, Roberto Amato, Jeffrey Barrett, Sonia Goncalves, Ewan Harrison, David K. Jackson, Ian Johnston, Dominic Kwiatkowski, Cordelia Langford, John Sillitoe on behalf of the Wellcome Sanger Institute COVID-19 Surveillance Team |
| EPI_ISL_3223191 | Lighthouse Lab in Alderley Park | Wellcome Sanger Institute for the COVID-19 Genomics UK (COG-UK) Consortium | Jacquelyn Wynn, Mairead Hyland, The Lighthouse Lab in Alderley Park and Alex Alderton, Roberto Amato, Jeffrey Barrett, Sonia Goncalves, Ewan Harrison, David K. Jackson, Ian Johnston, Dominic Kwiatkowski, Cordelia Langford, John Sillitoe on behalf of the Wellcome Sanger Institute COVID-19 Surveillance Team |
| EPI_ISL_3223192 | Lighthouse Laboratory Plymouth | Wellcome Sanger Institute for the COVID-19 Genomics UK (COG-UK) Consortium | Lighthouse Laboratory Plymouth and Alex Alderton, Roberto Amato, Jeffrey Barrett, Sonia Goncalves, Ewan Harrison, David K. Jackson, Ian Johnston, Dominic Kwiatkowski, Cordelia Langford, John Sillitoe on behalf of the Wellcome Sanger Institute COVID-19 Surveillance Team |
| EPI_ISL_3223193 | Lighthouse Lab in Alderley Park | Wellcome Sanger Institute for the COVID-19 Genomics UK (COG-UK) Consortium | Jacquelyn Wynn, Mairead Hyland, The Lighthouse Lab in Alderley Park and Alex Alderton, Roberto Amato, Jeffrey Barrett, Sonia Goncalves, Ewan Harrison, David K. Jackson, Ian Johnston, Dominic Kwiatkowski, Cordelia Langford, John Sillitoe on behalf of the Wellcome Sanger Institute COVID-19 Surveillance Team |
| EPI_ISL_3223194 | Lighthouse Laboratory Plymouth | Wellcome Sanger Institute for the COVID-19 Genomics UK (COG-UK) Consortium | Lighthouse Laboratory Plymouth and Alex Alderton, Roberto Amato, Jeffrey Barrett, Sonia Goncalves, Ewan Harrison, David K. Jackson, Ian Johnston, Dominic Kwiatkowski, Cordelia Langford, John Sillitoe on behalf of the Wellcome Sanger Institute COVID-19 Surveillance Team |
| EPI_ISL_3223195 | Lighthouse Lab in Glasgow | Wellcome Sanger Institute for the COVID-19 Genomics UK (COG-UK) Consortium | Harper VanSteenhouse, Yumi Kasai, David Gray, Carol Clugston, Anna Dominiczak and Alex Alderton, Roberto Amato, Jeffrey Barrett, Sonia Goncalves, Ewan Harrison, David K. Jackson, Ian Johnston, Dominic Kwiatkowski, Cordelia Langford, John Sillitoe on behalf of the Wellcome Sanger Institute COVID-19 Surveillance Team |
| EPI_ISL_3223196 | Lighthouse Laboratory Plymouth | Wellcome Sanger Institute for the COVID-19 Genomics UK (COG-UK) Consortium | Lighthouse Laboratory Plymouth and Alex Alderton, Roberto Amato, Jeffrey Barrett, Sonia Goncalves, Ewan Harrison, David K. Jackson, Ian Johnston, Dominic Kwiatkowski, Cordelia Langford, John Sillitoe on behalf of the Wellcome Sanger Institute COVID-19 Surveillance Team |
| EPI_ISL_3223197 | Lighthouse Lab in Glasgow | Wellcome Sanger Institute for the COVID-19 Genomics UK (COG-UK) Consortium | Harper VanSteenhouse, Yumi Kasai, David Gray, Carol Clugston, Anna Dominiczak and Alex Alderton, Roberto Amato, Jeffrey Barrett, Sonia Goncalves, Ewan Harrison, David K. Jackson, Ian Johnston, Dominic Kwiatkowski, Cordelia Langford, John Sillitoe on behalf of the Wellcome Sanger Institute COVID-19 Surveillance Team |
| EPI_ISL_3223198, EPI_ISL_3223199 | Lighthouse Laboratory Plymouth | Wellcome Sanger Institute for the COVID-19 Genomics UK (COG-UK) Consortium | Lighthouse Laboratory Plymouth and Alex Alderton, Roberto Amato, Jeffrey Barrett, Sonia Goncalves, Ewan Harrison, David K. Jackson, Ian Johnston, Dominic Kwiatkowski, Cordelia Langford, John Sillitoe on behalf of the Wellcome Sanger Institute COVID-19 Surveillance Team |
| EPI_ISL_3223200 | Lighthouse Lab in Milton Keynes | Wellcome Sanger Institute for the COVID-19 Genomics UK (COG-UK) Consortium | The Lighthouse Lab in Milton Keynes and Alex Alderton, Roberto Amato, Jeffrey Barrett, Sonia Goncalves, Ewan Harrison, David K. Jackson, Ian Johnston, Dominic Kwiatkowski, Cordelia Langford, John Sillitoe on behalf of the Wellcome Sanger Institute COVID-19 Surveillance Team |
| EPI_ISL_3223201 | Lighthouse Laboratory Plymouth | Wellcome Sanger Institute for the COVID-19 Genomics UK (COG-UK) Consortium | Lighthouse Laboratory Plymouth and Alex Alderton, Roberto Amato, Jeffrey Barrett, Sonia Goncalves, Ewan Harrison, David K. Jackson, Ian Johnston, Dominic Kwiatkowski, Cordelia Langford, John Sillitoe on behalf of the Wellcome Sanger Institute COVID-19 Surveillance Team |
| EPI_ISL_3223202 | Lighthouse Lab in Glasgow | Wellcome Sanger Institute for the COVID-19 Genomics UK (COG-UK) Consortium | Harper VanSteenhouse, Yumi Kasai, David Gray, Carol Clugston, Anna Dominiczak and Alex Alderton, Roberto Amato, Jeffrey Barrett, Sonia Goncalves, Ewan Harrison, David K. Jackson, Ian Johnston, Dominic Kwiatkowski, Cordelia Langford, John Sillitoe on behalf of the Wellcome Sanger Institute COVID-19 Surveillance Team |
| EPI_ISL_3223203 | Lighthouse Laboratory Plymouth | Wellcome Sanger Institute for the COVID-19 Genomics UK (COG-UK) Consortium | Lighthouse Laboratory Plymouth and Alex Alderton, Roberto Amato, Jeffrey Barrett, Sonia Goncalves, Ewan Harrison, David K. Jackson, Ian Johnston, Dominic Kwiatkowski, Cordelia Langford, John Sillitoe on behalf of the Wellcome Sanger Institute COVID-19 Surveillance Team |
| EPI_ISL_3223204 | Lighthouse Lab in Alderley Park | Wellcome Sanger Institute for the COVID-19 Genomics UK (COG-UK) Consortium | Jacquelyn Wynn, Mairead Hyland, The Lighthouse Lab in Alderley Park and Alex Alderton, Roberto Amato, Jeffrey Barrett, Sonia Goncalves, Ewan Harrison, David K. Jackson, Ian Johnston, Dominic Kwiatkowski, Cordelia Langford, John Sillitoe on behalf of the Wellcome Sanger Institute COVID-19 Surveillance Team |
| EPI_ISL_3223205 | Lighthouse Lab in Glasgow | Wellcome Sanger Institute for the COVID-19 Genomics UK (COG-UK) Consortium | Harper VanSteenhouse, Yumi Kasai, David Gray, Carol Clugston, Anna Dominiczak and Alex Alderton, Roberto Amato, Jeffrey Barrett, Sonia Goncalves, Ewan Harrison, David K. Jackson, Ian Johnston, Dominic Kwiatkowski, Cordelia Langford, John Sillitoe on behalf of the Wellcome Sanger Institute COVID-19 Surveillance Team |
| EPI_ISL_3223206 | Lighthouse Lab in Alderley Park | Wellcome Sanger Institute for the COVID-19 Genomics UK (COG-UK) Consortium | Jacquelyn Wynn, Mairead Hyland, The Lighthouse Lab in Alderley Park and Alex Alderton, Roberto Amato, Jeffrey Barrett, Sonia Goncalves, Ewan Harrison, David K. Jackson, Ian Johnston, Dominic Kwiatkowski, Cordelia Langford, John Sillitoe on behalf of the Wellcome Sanger Institute COVID-19 Surveillance Team |
| EPI_ISL_3223207 | Lighthouse Laboratory Plymouth | Wellcome Sanger Institute for the COVID-19 Genomics UK (COG-UK) Consortium | Lighthouse Laboratory Plymouth and Alex Alderton, Roberto Amato, Jeffrey Barrett, Sonia Goncalves, Ewan Harrison, David K. Jackson, Ian Johnston, Dominic Kwiatkowski, Cordelia Langford, John Sillitoe on behalf of the Wellcome Sanger Institute COVID-19 Surveillance Team |
| EPI_ISL_3223208, EPI_ISL_3223209 | Lighthouse Lab in Glasgow | Wellcome Sanger Institute for the COVID-19 Genomics UK (COG-UK) Consortium | Harper VanSteenhouse, Yumi Kasai, David Gray, Carol Clugston, Anna Dominiczak and Alex Alderton, Roberto Amato, Jeffrey Barrett, Sonia Goncalves, Ewan Harrison, David K. Jackson, Ian Johnston, Dominic Kwiatkowski, Cordelia Langford, John Sillitoe on behalf of the Wellcome Sanger Institute COVID-19 Surveillance Team |
| EPI_ISL_3223210 | Lighthouse Laboratory Plymouth | Wellcome Sanger Institute for the COVID-19 Genomics UK (COG-UK) Consortium | Lighthouse Laboratory Plymouth and Alex Alderton, Roberto Amato, Jeffrey Barrett, Sonia Goncalves, Ewan Harrison, David K. Jackson, Ian Johnston, Dominic Kwiatkowski, Cordelia Langford, John Sillitoe on behalf of the Wellcome Sanger Institute COVID-19 Surveillance Team |
| EPI_ISL_3223211 | Lighthouse Lab in Milton Keynes | Wellcome Sanger Institute for the COVID-19 Genomics UK (COG-UK) Consortium | The Lighthouse Lab in Milton Keynes and Alex Alderton, Roberto Amato, Jeffrey Barrett, Sonia Goncalves, Ewan Harrison, David K. Jackson, Ian Johnston, Dominic Kwiatkowski, Cordelia Langford, John Sillitoe on behalf of the Wellcome Sanger Institute COVID-19 Surveillance Team |
| EPI_ISL_3223212 | Lighthouse Lab in Glasgow | Wellcome Sanger Institute for the COVID-19 Genomics UK (COG-UK) Consortium | Harper VanSteenhouse, Yumi Kasai, David Gray, Carol Clugston, Anna Dominiczak and Alex Alderton, Roberto Amato, Jeffrey Barrett, Sonia Goncalves, Ewan Harrison, David K. Jackson, Ian Johnston, Dominic Kwiatkowski, Cordelia Langford, John Sillitoe on behalf of the Wellcome Sanger Institute COVID-19 Surveillance Team |
| EPI_ISL_3223213 | Lighthouse Lab in Alderley Park | Wellcome Sanger Institute for the COVID-19 Genomics UK (COG-UK) Consortium | Jacquelyn Wynn, Mairead Hyland, The Lighthouse Lab in Alderley Park and Alex Alderton, Roberto Amato, Jeffrey Barrett, Sonia Goncalves, Ewan Harrison, David K. Jackson, Ian Johnston, Dominic Kwiatkowski, Cordelia Langford, John Sillitoe on behalf of the Wellcome Sanger Institute COVID-19 Surveillance Team |
| EPI_ISL_3223214 | Lighthouse Lab in Milton Keynes | Wellcome Sanger Institute for the COVID-19 Genomics UK (COG-UK) Consortium | The Lighthouse Lab in Milton Keynes and Alex Alderton, Roberto Amato, Jeffrey Barrett, Sonia Goncalves, Ewan Harrison, David K. Jackson, Ian Johnston, Dominic Kwiatkowski, Cordelia Langford, John Sillitoe on behalf of the Wellcome Sanger Institute COVID-19 Surveillance Team |
| EPI_ISL_3223215 | Lighthouse Lab in Glasgow | Wellcome Sanger Institute for the COVID-19 Genomics UK (COG-UK) Consortium | Harper VanSteenhouse, Yumi Kasai, David Gray, Carol Clugston, Anna Dominiczak and Alex Alderton, Roberto Amato, Jeffrey Barrett, Sonia Goncalves, Ewan Harrison, David K. Jackson, Ian Johnston, Dominic Kwiatkowski, Cordelia Langford, John Sillitoe on behalf of the Wellcome Sanger Institute COVID-19 Surveillance Team |
| EPI_ISL_3223216 | Lighthouse Lab in Alderley Park | Wellcome Sanger Institute for the COVID-19 Genomics UK (COG-UK) Consortium | Jacquelyn Wynn, Mairead Hyland, The Lighthouse Lab in Alderley Park and Alex Alderton, Roberto Amato, Jeffrey Barrett, Sonia Goncalves, Ewan Harrison, David K. Jackson, Ian Johnston, Dominic Kwiatkowski, Cordelia Langford, John Sillitoe on behalf of the Wellcome Sanger Institute COVID-19 Surveillance Team |
| EPI_ISL_3223217 | Lighthouse Lab in Milton Keynes | Wellcome Sanger Institute for the COVID-19 Genomics UK (COG-UK) Consortium | The Lighthouse Lab in Milton Keynes and Alex Alderton, Roberto Amato, Jeffrey Barrett, Sonia Goncalves, Ewan Harrison, David K. Jackson, Ian Johnston, Dominic Kwiatkowski, Cordelia Langford, John Sillitoe on behalf of the Wellcome Sanger Institute COVID-19 Surveillance Team |
| EPI_ISL_3223218 | Lighthouse Laboratory Plymouth | Wellcome Sanger Institute for the COVID-19 Genomics UK (COG-UK) Consortium | Lighthouse Laboratory Plymouth and Alex Alderton, Roberto Amato, Jeffrey Barrett, Sonia Goncalves, Ewan Harrison, David K. Jackson, Ian Johnston, Dominic Kwiatkowski, Cordelia Langford, John Sillitoe on behalf of the Wellcome Sanger Institute COVID-19 Surveillance Team |
| EPI_ISL_3223219 | Lighthouse Lab in Milton Keynes | Wellcome Sanger Institute for the COVID-19 Genomics UK (COG-UK) Consortium | The Lighthouse Lab in Milton Keynes and Alex Alderton, Roberto Amato, Jeffrey Barrett, Sonia Goncalves, Ewan Harrison, David K. Jackson, Ian Johnston, Dominic Kwiatkowski, Cordelia Langford, John Sillitoe on behalf of the Wellcome Sanger Institute COVID-19 Surveillance Team |
| EPI_ISL_3223220, EPI_ISL_3223221, EPI_ISL_3223222, EPI_ISL_3223223 | Lighthouse Lab in Glasgow | Wellcome Sanger Institute for the COVID-19 Genomics UK (COG-UK) Consortium | Harper VanSteenhouse, Yumi Kasai, David Gray, Carol Clugston, Anna Dominiczak and Alex Alderton, Roberto Amato, Jeffrey Barrett, Sonia Goncalves, Ewan Harrison, David K. Jackson, Ian Johnston, Dominic Kwiatkowski, Cordelia Langford, John Sillitoe on behalf of the Wellcome Sanger Institute COVID-19 Surveillance Team |
| EPI_ISL_3223224, | Lighthouse Lab in Alderley Park | Wellcome Sanger Institute for the COVID-19 Genomics UK (COG-UK) | Jacquelyn Wynn, Mairead Hyland, The Lighthouse Lab in Alderley Park and Alex Alderton, Roberto Amato, Jeffrey Barrett, Sonia Goncalves, Ewan Harrison, David K. Jackson, Ian Johnston, Dominic Kwiatkowski, Cordelia Langford, John Sillitoe on |

| | | | |
|---|---|---|---|
| EPI_ISL_3223225, EPI_ISL_3223226 | | Consortium | behalf of the Wellcome Sanger Institute COVID-19 Surveillance Team |
| EPI_ISL_3223227 | Lighthouse Lab in Glasgow | Wellcome Sanger Institute for the COVID-19 Genomics UK (COG-UK) Consortium | Harper VanSteenhouse, Yumi Kasai, David Gray, Carol Clugston, Anna Dominiczak and Alex Alderton, Roberto Amato, Jeffrey Barrett, Sonia Goncalves, Ewan Harrison, David K. Jackson, Ian Johnston, Dominic Kwiatkowski, Cordelia Langford, John Sillitoe on behalf of the Wellcome Sanger Institute COVID-19 Surveillance Team |
| EPI_ISL_3223228 | Lighthouse Lab in Alderley Park | Wellcome Sanger Institute for the COVID-19 Genomics UK (COG-UK) Consortium | Jacquelyn Wynn, Mairead Hyland, The Lighthouse Lab in Alderley Park and Alex Alderton, Roberto Amato, Jeffrey Barrett, Sonia Goncalves, Ewan Harrison, David K. Jackson, Ian Johnston, Dominic Kwiatkowski, Cordelia Langford, John Sillitoe on behalf of the Wellcome Sanger Institute COVID-19 Surveillance Team |
| EPI_ISL_3223229 | Lighthouse Lab in Glasgow | Wellcome Sanger Institute for the COVID-19 Genomics UK (COG-UK) Consortium | Harper VanSteenhouse, Yumi Kasai, David Gray, Carol Clugston, Anna Dominiczak and Alex Alderton, Roberto Amato, Jeffrey Barrett, Sonia Goncalves, Ewan Harrison, David K. Jackson, Ian Johnston, Dominic Kwiatkowski, Cordelia Langford, John Sillitoe on behalf of the Wellcome Sanger Institute COVID-19 Surveillance Team |
| EPI_ISL_3223230 | Lighthouse Lab in Milton Keynes | Wellcome Sanger Institute for the COVID-19 Genomics UK (COG-UK) Consortium | The Lighthouse Lab in Milton Keynes and Alex Alderton, Roberto Amato, Jeffrey Barrett, Sonia Goncalves, Ewan Harrison, David K. Jackson, Ian Johnston, Dominic Kwiatkowski, Cordelia Langford, John Sillitoe on behalf of the Wellcome Sanger Institute COVID-19 Surveillance Team |
| EPI_ISL_3223231 | Lighthouse Laboratory Plymouth | Wellcome Sanger Institute for the COVID-19 Genomics UK (COG-UK) Consortium | Lighthouse Laboratory Plymouth and Alex Alderton, Roberto Amato, Jeffrey Barrett, Sonia Goncalves, Ewan Harrison, David K. Jackson, Ian Johnston, Dominic Kwiatkowski, Cordelia Langford, John Sillitoe on behalf of the Wellcome Sanger Institute COVID-19 Surveillance Team |
| EPI_ISL_3223232 | Lighthouse Lab in Alderley Park | Wellcome Sanger Institute for the COVID-19 Genomics UK (COG-UK) Consortium | Jacquelyn Wynn, Mairead Hyland, The Lighthouse Lab in Alderley Park and Alex Alderton, Roberto Amato, Jeffrey Barrett, Sonia Goncalves, Ewan Harrison, David K. Jackson, Ian Johnston, Dominic Kwiatkowski, Cordelia Langford, John Sillitoe on behalf of the Wellcome Sanger Institute COVID-19 Surveillance Team |
| EPI_ISL_3223233 | Lighthouse Lab in Milton Keynes | Wellcome Sanger Institute for the COVID-19 Genomics UK (COG-UK) Consortium | The Lighthouse Lab in Milton Keynes and Alex Alderton, Roberto Amato, Jeffrey Barrett, Sonia Goncalves, Ewan Harrison, David K. Jackson, Ian Johnston, Dominic Kwiatkowski, Cordelia Langford, John Sillitoe on behalf of the Wellcome Sanger Institute COVID-19 Surveillance Team |
| EPI_ISL_3223234 | Lighthouse Laboratory Plymouth | Wellcome Sanger Institute for the COVID-19 Genomics UK (COG-UK) Consortium | Lighthouse Laboratory Plymouth and Alex Alderton, Roberto Amato, Jeffrey Barrett, Sonia Goncalves, Ewan Harrison, David K. Jackson, Ian Johnston, Dominic Kwiatkowski, Cordelia Langford, John Sillitoe on behalf of the Wellcome Sanger Institute COVID-19 Surveillance Team |
| EPI_ISL_3223235 | Lighthouse Lab in Milton Keynes | Wellcome Sanger Institute for the COVID-19 Genomics UK (COG-UK) Consortium | The Lighthouse Lab in Milton Keynes and Alex Alderton, Roberto Amato, Jeffrey Barrett, Sonia Goncalves, Ewan Harrison, David K. Jackson, Ian Johnston, Dominic Kwiatkowski, Cordelia Langford, John Sillitoe on behalf of the Wellcome Sanger Institute COVID-19 Surveillance Team |
| EPI_ISL_3223236 | Lighthouse Laboratory Plymouth | Wellcome Sanger Institute for the COVID-19 Genomics UK (COG-UK) Consortium | Lighthouse Laboratory Plymouth and Alex Alderton, Roberto Amato, Jeffrey Barrett, Sonia Goncalves, Ewan Harrison, David K. Jackson, Ian Johnston, Dominic Kwiatkowski, Cordelia Langford, John Sillitoe on behalf of the Wellcome Sanger Institute COVID-19 Surveillance Team |
| EPI_ISL_3223237 | Lighthouse Lab in Milton Keynes | Wellcome Sanger Institute for the COVID-19 Genomics UK (COG-UK) Consortium | The Lighthouse Lab in Milton Keynes and Alex Alderton, Roberto Amato, Jeffrey Barrett, Sonia Goncalves, Ewan Harrison, David K. Jackson, Ian Johnston, Dominic Kwiatkowski, Cordelia Langford, John Sillitoe on behalf of the Wellcome Sanger Institute COVID-19 Surveillance Team |
| EPI_ISL_3223238 | Lighthouse Laboratory Plymouth | Wellcome Sanger Institute for the COVID-19 Genomics UK (COG-UK) Consortium | Lighthouse Laboratory Plymouth and Alex Alderton, Roberto Amato, Jeffrey Barrett, Sonia Goncalves, Ewan Harrison, David K. Jackson, Ian Johnston, Dominic Kwiatkowski, Cordelia Langford, John Sillitoe on behalf of the Wellcome Sanger Institute COVID-19 Surveillance Team |
| EPI_ISL_3223239 | Lighthouse Lab in Glasgow | Wellcome Sanger Institute for the COVID-19 Genomics UK (COG-UK) Consortium | Harper VanSteenhouse, Yumi Kasai, David Gray, Carol Clugston, Anna Dominiczak and Alex Alderton, Roberto Amato, Jeffrey Barrett, Sonia Goncalves, Ewan Harrison, David K. Jackson, Ian Johnston, Dominic Kwiatkowski, Cordelia Langford, John Sillitoe on behalf of the Wellcome Sanger Institute COVID-19 Surveillance Team |
| EPI_ISL_3223240 | Lighthouse Lab in Milton Keynes | Wellcome Sanger Institute for the COVID-19 Genomics UK (COG-UK) Consortium | The Lighthouse Lab in Milton Keynes and Alex Alderton, Roberto Amato, Jeffrey Barrett, Sonia Goncalves, Ewan Harrison, David K. Jackson, Ian Johnston, Dominic Kwiatkowski, Cordelia Langford, John Sillitoe on behalf of the Wellcome Sanger Institute COVID-19 Surveillance Team |
| EPI_ISL_3223241 | Lighthouse Laboratory Plymouth | Wellcome Sanger Institute for the COVID-19 Genomics UK (COG-UK) Consortium | Lighthouse Laboratory Plymouth and Alex Alderton, Roberto Amato, Jeffrey Barrett, Sonia Goncalves, Ewan Harrison, David K. Jackson, Ian Johnston, Dominic Kwiatkowski, Cordelia Langford, John Sillitoe on behalf of the Wellcome Sanger Institute COVID-19 Surveillance Team |
| EPI_ISL_3223242 | Lighthouse Lab in Milton Keynes | Wellcome Sanger Institute for the COVID-19 Genomics UK (COG-UK) Consortium | The Lighthouse Lab in Milton Keynes and Alex Alderton, Roberto Amato, Jeffrey Barrett, Sonia Goncalves, Ewan Harrison, David K. Jackson, Ian Johnston, Dominic Kwiatkowski, Cordelia Langford, John Sillitoe on behalf of the Wellcome Sanger Institute COVID-19 Surveillance Team |
| EPI_ISL_3223243 | Lighthouse Lab in Glasgow | Wellcome Sanger Institute for the COVID-19 Genomics UK (COG-UK) Consortium | Harper VanSteenhouse, Yumi Kasai, David Gray, Carol Clugston, Anna Dominiczak and Alex Alderton, Roberto Amato, Jeffrey Barrett, Sonia Goncalves, Ewan Harrison, David K. Jackson, Ian Johnston, Dominic Kwiatkowski, Cordelia Langford, John Sillitoe on behalf of the Wellcome Sanger Institute COVID-19 Surveillance Team |
| EPI_ISL_3223244 | Lighthouse Laboratory Plymouth | Wellcome Sanger Institute for the COVID-19 Genomics UK (COG-UK) Consortium | Lighthouse Laboratory Plymouth and Alex Alderton, Roberto Amato, Jeffrey Barrett, Sonia Goncalves, Ewan Harrison, David K. Jackson, Ian Johnston, Dominic Kwiatkowski, Cordelia Langford, John Sillitoe on behalf of the Wellcome Sanger Institute COVID-19 Surveillance Team |
| EPI_ISL_3223245, EPI_ISL_3223246 | Lighthouse Lab in Alderley Park | Wellcome Sanger Institute for the COVID-19 Genomics UK (COG-UK) Consortium | Jacquelyn Wynn, Mairead Hyland, The Lighthouse Lab in Alderley Park and Alex Alderton, Roberto Amato, Jeffrey Barrett, Sonia Goncalves, Ewan Harrison, David K. Jackson, Ian Johnston, Dominic Kwiatkowski, Cordelia Langford, John Sillitoe on behalf of the Wellcome Sanger Institute COVID-19 Surveillance Team |
| EPI_ISL_3223247 | Lighthouse Lab in Glasgow | Wellcome Sanger Institute for the COVID-19 Genomics UK (COG-UK) Consortium | Harper VanSteenhouse, Yumi Kasai, David Gray, Carol Clugston, Anna Dominiczak and Alex Alderton, Roberto Amato, Jeffrey Barrett, Sonia Goncalves, Ewan Harrison, David K. Jackson, Ian Johnston, Dominic Kwiatkowski, Cordelia Langford, John Sillitoe on behalf of the Wellcome Sanger Institute COVID-19 Surveillance Team |
| EPI_ISL_3223248 | Lighthouse Lab in Milton Keynes | Wellcome Sanger Institute for the COVID-19 Genomics UK (COG-UK) Consortium | The Lighthouse Lab in Milton Keynes and Alex Alderton, Roberto Amato, Jeffrey Barrett, Sonia Goncalves, Ewan Harrison, David K. Jackson, Ian Johnston, Dominic Kwiatkowski, Cordelia Langford, John Sillitoe on behalf of the Wellcome Sanger Institute COVID-19 Surveillance Team |
| EPI_ISL_3223249, EPI_ISL_3223250 | Lighthouse Lab in Alderley Park | Wellcome Sanger Institute for the COVID-19 Genomics UK (COG-UK) Consortium | Jacquelyn Wynn, Mairead Hyland, The Lighthouse Lab in Alderley Park and Alex Alderton, Roberto Amato, Jeffrey Barrett, Sonia Goncalves, Ewan Harrison, David K. Jackson, Ian Johnston, Dominic Kwiatkowski, Cordelia Langford, John Sillitoe on behalf of the Wellcome Sanger Institute COVID-19 Surveillance Team |
| EPI_ISL_3223251 | Lighthouse Lab in Glasgow | Wellcome Sanger Institute for the COVID-19 Genomics UK (COG-UK) Consortium | Harper VanSteenhouse, Yumi Kasai, David Gray, Carol Clugston, Anna Dominiczak and Alex Alderton, Roberto Amato, Jeffrey Barrett, Sonia Goncalves, Ewan Harrison, David K. Jackson, Ian Johnston, Dominic Kwiatkowski, Cordelia Langford, John Sillitoe on behalf of the Wellcome Sanger Institute COVID-19 Surveillance Team |
| EPI_ISL_3223252 | Lighthouse Laboratory Plymouth | Wellcome Sanger Institute for the COVID-19 Genomics UK (COG-UK) Consortium | Lighthouse Laboratory Plymouth and Alex Alderton, Roberto Amato, Jeffrey Barrett, Sonia Goncalves, Ewan Harrison, David K. Jackson, Ian Johnston, Dominic Kwiatkowski, Cordelia Langford, John Sillitoe on behalf of the Wellcome Sanger Institute COVID-19 Surveillance Team |
| EPI_ISL_3223253, EPI_ISL_3223254, EPI_ISL_3223255 | Lighthouse Lab in Alderley Park | Wellcome Sanger Institute for the COVID-19 Genomics UK (COG-UK) Consortium | Jacquelyn Wynn, Mairead Hyland, The Lighthouse Lab in Alderley Park and Alex Alderton, Roberto Amato, Jeffrey Barrett, Sonia Goncalves, Ewan Harrison, David K. Jackson, Ian Johnston, Dominic Kwiatkowski, Cordelia Langford, John Sillitoe on behalf of the Wellcome Sanger Institute COVID-19 Surveillance Team |
| EPI_ISL_3223256 | Lighthouse Laboratory Plymouth | Wellcome Sanger Institute for the COVID-19 Genomics UK (COG-UK) Consortium | Lighthouse Laboratory Plymouth and Alex Alderton, Roberto Amato, Jeffrey Barrett, Sonia Goncalves, Ewan Harrison, David K. Jackson, Ian Johnston, Dominic Kwiatkowski, Cordelia Langford, John Sillitoe on behalf of the Wellcome Sanger Institute COVID-19 Surveillance Team |
| EPI_ISL_3223257, EPI_ISL_3223258 | Lighthouse Lab in Glasgow | Wellcome Sanger Institute for the COVID-19 Genomics UK (COG-UK) Consortium | Harper VanSteenhouse, Yumi Kasai, David Gray, Carol Clugston, Anna Dominiczak and Alex Alderton, Roberto Amato, Jeffrey Barrett, Sonia Goncalves, Ewan Harrison, David K. Jackson, Ian Johnston, Dominic Kwiatkowski, Cordelia Langford, John Sillitoe on behalf of the Wellcome Sanger Institute COVID-19 Surveillance Team |
| EPI_ISL_3223259 | Lighthouse Laboratory Plymouth | Wellcome Sanger Institute for the COVID-19 Genomics UK (COG-UK) Consortium | Lighthouse Laboratory Plymouth and Alex Alderton, Roberto Amato, Jeffrey Barrett, Sonia Goncalves, Ewan Harrison, David K. Jackson, Ian Johnston, Dominic Kwiatkowski, Cordelia Langford, John Sillitoe on behalf of the Wellcome Sanger Institute COVID-19 Surveillance Team |
| EPI_ISL_3223260 | Lighthouse Lab in Glasgow | Wellcome Sanger Institute for the COVID-19 Genomics UK (COG-UK) Consortium | Harper VanSteenhouse, Yumi Kasai, David Gray, Carol Clugston, Anna Dominiczak and Alex Alderton, Roberto Amato, Jeffrey Barrett, Sonia Goncalves, Ewan Harrison, David K. Jackson, Ian Johnston, Dominic Kwiatkowski, Cordelia Langford, John Sillitoe on behalf of the Wellcome Sanger Institute COVID-19 Surveillance Team |
| EPI_ISL_3223261 | Lighthouse Lab in Alderley Park | Wellcome Sanger Institute for the COVID-19 Genomics UK (COG-UK) Consortium | Jacquelyn Wynn, Mairead Hyland, The Lighthouse Lab in Alderley Park and Alex Alderton, Roberto Amato, Jeffrey Barrett, Sonia Goncalves, Ewan Harrison, David K. Jackson, Ian Johnston, Dominic Kwiatkowski, Cordelia Langford, John Sillitoe on behalf of the Wellcome Sanger Institute COVID-19 Surveillance Team |
| EPI_ISL_3223262 | Lighthouse Lab in Glasgow | Wellcome Sanger Institute for the COVID-19 Genomics UK (COG-UK) Consortium | Harper VanSteenhouse, Yumi Kasai, David Gray, Carol Clugston, Anna Dominiczak and Alex Alderton, Roberto Amato, Jeffrey Barrett, Sonia Goncalves, Ewan Harrison, David K. Jackson, Ian Johnston, Dominic Kwiatkowski, Cordelia Langford, John Sillitoe on behalf of the Wellcome Sanger Institute COVID-19 Surveillance Team |
| EPI_ISL_3223263, EPI_ISL_3223264, EPI_ISL_3223265 | Lighthouse Lab in Milton Keynes | Wellcome Sanger Institute for the COVID-19 Genomics UK (COG-UK) Consortium | The Lighthouse Lab in Milton Keynes and Alex Alderton, Roberto Amato, Jeffrey Barrett, Sonia Goncalves, Ewan Harrison, David K. Jackson, Ian Johnston, Dominic Kwiatkowski, Cordelia Langford, John Sillitoe on behalf of the Wellcome Sanger Institute COVID-19 Surveillance Team |
| EPI_ISL_3223266 | Lighthouse Laboratory Plymouth | Wellcome Sanger Institute for the COVID-19 Genomics UK (COG-UK) Consortium | Lighthouse Laboratory Plymouth and Alex Alderton, Roberto Amato, Jeffrey Barrett, Sonia Goncalves, Ewan Harrison, David K. Jackson, Ian Johnston, Dominic Kwiatkowski, Cordelia Langford, John Sillitoe on behalf of the Wellcome Sanger Institute COVID-19 Surveillance Team |
| EPI_ISL_3223267 | Lighthouse Lab in Glasgow | Wellcome Sanger Institute for the COVID-19 Genomics UK (COG-UK) Consortium | Harper VanSteenhouse, Yumi Kasai, David Gray, Carol Clugston, Anna Dominiczak and Alex Alderton, Roberto Amato, Jeffrey Barrett, Sonia Goncalves, Ewan Harrison, David K. Jackson, Ian Johnston, Dominic Kwiatkowski, Cordelia Langford, John Sillitoe on behalf of the Wellcome Sanger Institute COVID-19 Surveillance Team |
| EPI_ISL_3223268 | Lighthouse Lab in Alderley Park | Wellcome Sanger Institute for the COVID-19 Genomics UK (COG-UK) Consortium | Jacquelyn Wynn, Mairead Hyland, The Lighthouse Lab in Alderley Park and Alex Alderton, Roberto Amato, Jeffrey Barrett, Sonia Goncalves, Ewan Harrison, David K. Jackson, Ian Johnston, Dominic Kwiatkowski, Cordelia Langford, John Sillitoe on behalf of the Wellcome Sanger Institute COVID-19 Surveillance Team |
| EPI_ISL_3223269 | Lighthouse Laboratory Plymouth | Wellcome Sanger Institute for the COVID-19 Genomics UK (COG-UK) Consortium | Lighthouse Laboratory Plymouth and Alex Alderton, Roberto Amato, Jeffrey Barrett, Sonia Goncalves, Ewan Harrison, David K. Jackson, Ian Johnston, Dominic Kwiatkowski, Cordelia Langford, John Sillitoe on behalf of the Wellcome Sanger Institute COVID-19 Surveillance Team |
| EPI_ISL_3223270 | Lighthouse Lab in Alderley Park | Wellcome Sanger Institute for the COVID-19 Genomics UK (COG-UK) Consortium | Jacquelyn Wynn, Mairead Hyland, The Lighthouse Lab in Alderley Park and Alex Alderton, Roberto Amato, Jeffrey Barrett, Sonia Goncalves, Ewan Harrison, David K. Jackson, Ian Johnston, Dominic Kwiatkowski, Cordelia Langford, John Sillitoe on behalf of the Wellcome Sanger Institute COVID-19 Surveillance Team |
| EPI_ISL_3223271 | Lighthouse Laboratory Plymouth | Wellcome Sanger Institute for the COVID-19 Genomics UK (COG-UK) Consortium | Lighthouse Laboratory Plymouth and Alex Alderton, Roberto Amato, Jeffrey Barrett, Sonia Goncalves, Ewan Harrison, David K. Jackson, Ian Johnston, Dominic Kwiatkowski, Cordelia Langford, John Sillitoe on behalf of the Wellcome Sanger Institute COVID-19 Surveillance Team |
| EPI_ISL_3223272, EPI_ISL_3223273 | Lighthouse Lab in Milton Keynes | Wellcome Sanger Institute for the COVID-19 Genomics UK (COG-UK) Consortium | The Lighthouse Lab in Milton Keynes and Alex Alderton, Roberto Amato, Jeffrey Barrett, Sonia Goncalves, Ewan Harrison, David K. Jackson, Ian Johnston, Dominic Kwiatkowski, Cordelia Langford, John Sillitoe on behalf of the Wellcome Sanger Institute COVID-19 Surveillance Team |
| EPI_ISL_3223274 | Lighthouse Lab in Alderley Park | Wellcome Sanger Institute for the COVID-19 Genomics UK (COG-UK) Consortium | Jacquelyn Wynn, Mairead Hyland, The Lighthouse Lab in Alderley Park and Alex Alderton, Roberto Amato, Jeffrey Barrett, Sonia Goncalves, Ewan Harrison, David K. Jackson, Ian Johnston, Dominic Kwiatkowski, Cordelia Langford, John Sillitoe on behalf of the Wellcome Sanger Institute COVID-19 Surveillance Team |
| EPI_ISL_3223275, EPI_ISL_3223276 | Lighthouse Laboratory Plymouth | Wellcome Sanger Institute for the COVID-19 Genomics UK (COG-UK) Consortium | Lighthouse Laboratory Plymouth and Alex Alderton, Roberto Amato, Jeffrey Barrett, Sonia Goncalves, Ewan Harrison, David K. Jackson, Ian Johnston, Dominic Kwiatkowski, Cordelia Langford, John Sillitoe on behalf of the Wellcome Sanger Institute COVID-19 Surveillance Team |
| EPI_ISL_3223277 | Lighthouse Lab in Alderley Park | Wellcome Sanger Institute for the COVID-19 Genomics UK (COG-UK) Consortium | Jacquelyn Wynn, Mairead Hyland, The Lighthouse Lab in Alderley Park and Alex Alderton, Roberto Amato, Jeffrey Barrett, Sonia Goncalves, Ewan Harrison, David K. Jackson, Ian Johnston, Dominic Kwiatkowski, Cordelia Langford, John Sillitoe on behalf of the Wellcome Sanger Institute COVID-19 Surveillance Team |
| EPI_ISL_3223278, EPI_ISL_3223279, EPI_ISL_3223280, | Lighthouse Lab in Milton Keynes | Wellcome Sanger Institute for the COVID-19 Genomics UK (COG-UK) Consortium | The Lighthouse Lab in Milton Keynes and Alex Alderton, Roberto Amato, Jeffrey Barrett, Sonia Goncalves, Ewan Harrison, David K. Jackson, Ian Johnston, Dominic Kwiatkowski, Cordelia Langford, John Sillitoe on behalf of the Wellcome Sanger Institute COVID-19 Surveillance Team |

| | | | |
|---|---|---|---|
| EPI_ISL_3223281 | | | |
| EPI_ISL_3223282, EPI_ISL_3223283 | Lighthouse Lab in Alderley Park | Wellcome Sanger Institute for the COVID-19 Genomics UK (COG-UK) Consortium | Jacquelyn Wynn, Mairead Hyland, The Lighthouse Lab in Alderley Park and Alex Alderton, Roberto Amato, Jeffrey Barrett, Sonia Goncalves, Ewan Harrison, David K. Jackson, Ian Johnston, Dominic Kwiatkowski, Cordelia Langford, John Sillitoe on behalf of the Wellcome Sanger Institute COVID-19 Surveillance Team |
| EPI_ISL_3223284, EPI_ISL_3223285 | Lighthouse Lab in Glasgow | Wellcome Sanger Institute for the COVID-19 Genomics UK (COG-UK) Consortium | Harper VanSteenhouse, Yumi Kasai, David Gray, Carol Clugston, Anna Dominiczak and Alex Alderton, Roberto Amato, Jeffrey Barrett, Sonia Goncalves, Ewan Harrison, David K. Jackson, Ian Johnston, Dominic Kwiatkowski, Cordelia Langford, John Sillitoe on behalf of the Wellcome Sanger Institute COVID-19 Surveillance Team |
| EPI_ISL_3223286 | Lighthouse Lab in Milton Keynes | Wellcome Sanger Institute for the COVID-19 Genomics UK (COG-UK) Consortium | The Lighthouse Lab in Milton Keynes and Alex Alderton, Roberto Amato, Jeffrey Barrett, Sonia Goncalves, Ewan Harrison, David K. Jackson, Ian Johnston, Dominic Kwiatkowski, Cordelia Langford, John Sillitoe on behalf of the Wellcome Sanger Institute COVID-19 Surveillance Team |
| EPI_ISL_3223287, EPI_ISL_3223288 | Lighthouse Laboratory Plymouth | Wellcome Sanger Institute for the COVID-19 Genomics UK (COG-UK) Consortium | Lighthouse Laboratory Plymouth and Alex Alderton, Roberto Amato, Jeffrey Barrett, Sonia Goncalves, Ewan Harrison, David K. Jackson, Ian Johnston, Dominic Kwiatkowski, Cordelia Langford, John Sillitoe on behalf of the Wellcome Sanger Institute COVID-19 Surveillance Team |
| EPI_ISL_3223289 | Lighthouse Lab in Alderley Park | Wellcome Sanger Institute for the COVID-19 Genomics UK (COG-UK) Consortium | Jacquelyn Wynn, Mairead Hyland, The Lighthouse Lab in Alderley Park and Alex Alderton, Roberto Amato, Jeffrey Barrett, Sonia Goncalves, Ewan Harrison, David K. Jackson, Ian Johnston, Dominic Kwiatkowski, Cordelia Langford, John Sillitoe on behalf of the Wellcome Sanger Institute COVID-19 Surveillance Team |
| EPI_ISL_3223290 | Lighthouse Lab in Glasgow | Wellcome Sanger Institute for the COVID-19 Genomics UK (COG-UK) Consortium | Harper VanSteenhouse, Yumi Kasai, David Gray, Carol Clugston, Anna Dominiczak and Alex Alderton, Roberto Amato, Jeffrey Barrett, Sonia Goncalves, Ewan Harrison, David K. Jackson, Ian Johnston, Dominic Kwiatkowski, Cordelia Langford, John Sillitoe on behalf of the Wellcome Sanger Institute COVID-19 Surveillance Team |
| EPI_ISL_3223291 | Lighthouse Lab in Milton Keynes | Wellcome Sanger Institute for the COVID-19 Genomics UK (COG-UK) Consortium | The Lighthouse Lab in Milton Keynes and Alex Alderton, Roberto Amato, Jeffrey Barrett, Sonia Goncalves, Ewan Harrison, David K. Jackson, Ian Johnston, Dominic Kwiatkowski, Cordelia Langford, John Sillitoe on behalf of the Wellcome Sanger Institute COVID-19 Surveillance Team |
| EPI_ISL_3223292 | Lighthouse Lab in Glasgow | Wellcome Sanger Institute for the COVID-19 Genomics UK (COG-UK) Consortium | Harper VanSteenhouse, Yumi Kasai, David Gray, Carol Clugston, Anna Dominiczak and Alex Alderton, Roberto Amato, Jeffrey Barrett, Sonia Goncalves, Ewan Harrison, David K. Jackson, Ian Johnston, Dominic Kwiatkowski, Cordelia Langford, John Sillitoe on behalf of the Wellcome Sanger Institute COVID-19 Surveillance Team |
| EPI_ISL_3223293, EPI_ISL_3223294, EPI_ISL_3223295 | Lighthouse Laboratory Plymouth | Wellcome Sanger Institute for the COVID-19 Genomics UK (COG-UK) Consortium | Lighthouse Laboratory Plymouth and Alex Alderton, Roberto Amato, Jeffrey Barrett, Sonia Goncalves, Ewan Harrison, David K. Jackson, Ian Johnston, Dominic Kwiatkowski, Cordelia Langford, John Sillitoe on behalf of the Wellcome Sanger Institute COVID-19 Surveillance Team |
| EPI_ISL_3223296 | Lighthouse Lab in Glasgow | Wellcome Sanger Institute for the COVID-19 Genomics UK (COG-UK) Consortium | Harper VanSteenhouse, Yumi Kasai, David Gray, Carol Clugston, Anna Dominiczak and Alex Alderton, Roberto Amato, Jeffrey Barrett, Sonia Goncalves, Ewan Harrison, David K. Jackson, Ian Johnston, Dominic Kwiatkowski, Cordelia Langford, John Sillitoe on behalf of the Wellcome Sanger Institute COVID-19 Surveillance Team |
| EPI_ISL_3223297, EPI_ISL_3223298 | Lighthouse Lab in Milton Keynes | Wellcome Sanger Institute for the COVID-19 Genomics UK (COG-UK) Consortium | The Lighthouse Lab in Milton Keynes and Alex Alderton, Roberto Amato, Jeffrey Barrett, Sonia Goncalves, Ewan Harrison, David K. Jackson, Ian Johnston, Dominic Kwiatkowski, Cordelia Langford, John Sillitoe on behalf of the Wellcome Sanger Institute COVID-19 Surveillance Team |
| EPI_ISL_3223299 | Lighthouse Lab in Glasgow | Wellcome Sanger Institute for the COVID-19 Genomics UK (COG-UK) Consortium | Harper VanSteenhouse, Yumi Kasai, David Gray, Carol Clugston, Anna Dominiczak and Alex Alderton, Roberto Amato, Jeffrey Barrett, Sonia Goncalves, Ewan Harrison, David K. Jackson, Ian Johnston, Dominic Kwiatkowski, Cordelia Langford, John Sillitoe on behalf of the Wellcome Sanger Institute COVID-19 Surveillance Team |
| EPI_ISL_3223300 | Lighthouse Lab in Milton Keynes | Wellcome Sanger Institute for the COVID-19 Genomics UK (COG-UK) Consortium | The Lighthouse Lab in Milton Keynes and Alex Alderton, Roberto Amato, Jeffrey Barrett, Sonia Goncalves, Ewan Harrison, David K. Jackson, Ian Johnston, Dominic Kwiatkowski, Cordelia Langford, John Sillitoe on behalf of the Wellcome Sanger Institute COVID-19 Surveillance Team |
| EPI_ISL_3223301 | Lighthouse Lab in Glasgow | Wellcome Sanger Institute for the COVID-19 Genomics UK (COG-UK) Consortium | Harper VanSteenhouse, Yumi Kasai, David Gray, Carol Clugston, Anna Dominiczak and Alex Alderton, Roberto Amato, Jeffrey Barrett, Sonia Goncalves, Ewan Harrison, David K. Jackson, Ian Johnston, Dominic Kwiatkowski, Cordelia Langford, John Sillitoe on behalf of the Wellcome Sanger Institute COVID-19 Surveillance Team |
| EPI_ISL_3223302, EPI_ISL_3223303 | Lighthouse Laboratory Plymouth | Wellcome Sanger Institute for the COVID-19 Genomics UK (COG-UK) Consortium | Lighthouse Laboratory Plymouth and Alex Alderton, Roberto Amato, Jeffrey Barrett, Sonia Goncalves, Ewan Harrison, David K. Jackson, Ian Johnston, Dominic Kwiatkowski, Cordelia Langford, John Sillitoe on behalf of the Wellcome Sanger Institute COVID-19 Surveillance Team |
| EPI_ISL_3223304 | Lighthouse Lab in Milton Keynes | Wellcome Sanger Institute for the COVID-19 Genomics UK (COG-UK) Consortium | The Lighthouse Lab in Milton Keynes and Alex Alderton, Roberto Amato, Jeffrey Barrett, Sonia Goncalves, Ewan Harrison, David K. Jackson, Ian Johnston, Dominic Kwiatkowski, Cordelia Langford, John Sillitoe on behalf of the Wellcome Sanger Institute COVID-19 Surveillance Team |
| EPI_ISL_3223305 | Lighthouse Lab in Glasgow | Wellcome Sanger Institute for the COVID-19 Genomics UK (COG-UK) Consortium | Harper VanSteenhouse, Yumi Kasai, David Gray, Carol Clugston, Anna Dominiczak and Alex Alderton, Roberto Amato, Jeffrey Barrett, Sonia Goncalves, Ewan Harrison, David K. Jackson, Ian Johnston, Dominic Kwiatkowski, Cordelia Langford, John Sillitoe on behalf of the Wellcome Sanger Institute COVID-19 Surveillance Team |
| EPI_ISL_3223306, EPI_ISL_3223307 | Lighthouse Laboratory Plymouth | Wellcome Sanger Institute for the COVID-19 Genomics UK (COG-UK) Consortium | Lighthouse Laboratory Plymouth and Alex Alderton, Roberto Amato, Jeffrey Barrett, Sonia Goncalves, Ewan Harrison, David K. Jackson, Ian Johnston, Dominic Kwiatkowski, Cordelia Langford, John Sillitoe on behalf of the Wellcome Sanger Institute COVID-19 Surveillance Team |
| EPI_ISL_3223308 | Lighthouse Lab in Glasgow | Wellcome Sanger Institute for the COVID-19 Genomics UK (COG-UK) Consortium | Harper VanSteenhouse, Yumi Kasai, David Gray, Carol Clugston, Anna Dominiczak and Alex Alderton, Roberto Amato, Jeffrey Barrett, Sonia Goncalves, Ewan Harrison, David K. Jackson, Ian Johnston, Dominic Kwiatkowski, Cordelia Langford, John Sillitoe on behalf of the Wellcome Sanger Institute COVID-19 Surveillance Team |
| EPI_ISL_3223309 | Lighthouse Laboratory Plymouth | Wellcome Sanger Institute for the COVID-19 Genomics UK (COG-UK) Consortium | Lighthouse Laboratory Plymouth and Alex Alderton, Roberto Amato, Jeffrey Barrett, Sonia Goncalves, Ewan Harrison, David K. Jackson, Ian Johnston, Dominic Kwiatkowski, Cordelia Langford, John Sillitoe on behalf of the Wellcome Sanger Institute COVID-19 Surveillance Team |
| EPI_ISL_3223310, EPI_ISL_3223311 | Lighthouse Lab in Milton Keynes | Wellcome Sanger Institute for the COVID-19 Genomics UK (COG-UK) Consortium | The Lighthouse Lab in Milton Keynes and Alex Alderton, Roberto Amato, Jeffrey Barrett, Sonia Goncalves, Ewan Harrison, David K. Jackson, Ian Johnston, Dominic Kwiatkowski, Cordelia Langford, John Sillitoe on behalf of the Wellcome Sanger Institute COVID-19 Surveillance Team |
| EPI_ISL_3223312, EPI_ISL_3223313 | Lighthouse Laboratory Plymouth | Wellcome Sanger Institute for the COVID-19 Genomics UK (COG-UK) Consortium | Lighthouse Laboratory Plymouth and Alex Alderton, Roberto Amato, Jeffrey Barrett, Sonia Goncalves, Ewan Harrison, David K. Jackson, Ian Johnston, Dominic Kwiatkowski, Cordelia Langford, John Sillitoe on behalf of the Wellcome Sanger Institute COVID-19 Surveillance Team |
| EPI_ISL_3223314 | Lighthouse Lab in Milton Keynes | Wellcome Sanger Institute for the COVID-19 Genomics UK (COG-UK) Consortium | The Lighthouse Lab in Milton Keynes and Alex Alderton, Roberto Amato, Jeffrey Barrett, Sonia Goncalves, Ewan Harrison, David K. Jackson, Ian Johnston, Dominic Kwiatkowski, Cordelia Langford, John Sillitoe on behalf of the Wellcome Sanger Institute COVID-19 Surveillance Team |
| EPI_ISL_3223315, EPI_ISL_3223316 | Lighthouse Laboratory Plymouth | Wellcome Sanger Institute for the COVID-19 Genomics UK (COG-UK) Consortium | Lighthouse Laboratory Plymouth and Alex Alderton, Roberto Amato, Jeffrey Barrett, Sonia Goncalves, Ewan Harrison, David K. Jackson, Ian Johnston, Dominic Kwiatkowski, Cordelia Langford, John Sillitoe on behalf of the Wellcome Sanger Institute COVID-19 Surveillance Team |
| EPI_ISL_3223317, EPI_ISL_3223318 | Lighthouse Lab in Milton Keynes | Wellcome Sanger Institute for the COVID-19 Genomics UK (COG-UK) Consortium | The Lighthouse Lab in Milton Keynes and Alex Alderton, Roberto Amato, Jeffrey Barrett, Sonia Goncalves, Ewan Harrison, David K. Jackson, Ian Johnston, Dominic Kwiatkowski, Cordelia Langford, John Sillitoe on behalf of the Wellcome Sanger Institute COVID-19 Surveillance Team |
| EPI_ISL_3223319, EPI_ISL_3223320, EPI_ISL_3223321, EPI_ISL_3223322 | Lighthouse Laboratory Plymouth | Wellcome Sanger Institute for the COVID-19 Genomics UK (COG-UK) Consortium | Lighthouse Laboratory Plymouth and Alex Alderton, Roberto Amato, Jeffrey Barrett, Sonia Goncalves, Ewan Harrison, David K. Jackson, Ian Johnston, Dominic Kwiatkowski, Cordelia Langford, John Sillitoe on behalf of the Wellcome Sanger Institute COVID-19 Surveillance Team |
| EPI_ISL_3223323, EPI_ISL_3223324, EPI_ISL_3223325 | Lighthouse Lab in Glasgow | Wellcome Sanger Institute for the COVID-19 Genomics UK (COG-UK) Consortium | Harper VanSteenhouse, Yumi Kasai, David Gray, Carol Clugston, Anna Dominiczak and Alex Alderton, Roberto Amato, Jeffrey Barrett, Sonia Goncalves, Ewan Harrison, David K. Jackson, Ian Johnston, Dominic Kwiatkowski, Cordelia Langford, John Sillitoe on behalf of the Wellcome Sanger Institute COVID-19 Surveillance Team |
| EPI_ISL_3223326 | Lighthouse Laboratory Plymouth | Wellcome Sanger Institute for the COVID-19 Genomics UK (COG-UK) Consortium | Lighthouse Laboratory Plymouth and Alex Alderton, Roberto Amato, Jeffrey Barrett, Sonia Goncalves, Ewan Harrison, David K. Jackson, Ian Johnston, Dominic Kwiatkowski, Cordelia Langford, John Sillitoe on behalf of the Wellcome Sanger Institute COVID-19 Surveillance Team |
| EPI_ISL_3223327 | Lighthouse Lab in Milton Keynes | Wellcome Sanger Institute for the COVID-19 Genomics UK (COG-UK) Consortium | The Lighthouse Lab in Milton Keynes and Alex Alderton, Roberto Amato, Jeffrey Barrett, Sonia Goncalves, Ewan Harrison, David K. Jackson, Ian Johnston, Dominic Kwiatkowski, Cordelia Langford, John Sillitoe on behalf of the Wellcome Sanger Institute COVID-19 Surveillance Team |
| EPI_ISL_3223328 | Lighthouse Lab in Glasgow | Wellcome Sanger Institute for the COVID-19 Genomics UK (COG-UK) Consortium | Harper VanSteenhouse, Yumi Kasai, David Gray, Carol Clugston, Anna Dominiczak and Alex Alderton, Roberto Amato, Jeffrey Barrett, Sonia Goncalves, Ewan Harrison, David K. Jackson, Ian Johnston, Dominic Kwiatkowski, Cordelia Langford, John Sillitoe on behalf of the Wellcome Sanger Institute COVID-19 Surveillance Team |
| EPI_ISL_3223329 | Lighthouse Laboratory Plymouth | Wellcome Sanger Institute for the COVID-19 Genomics UK (COG-UK) Consortium | Lighthouse Laboratory Plymouth and Alex Alderton, Roberto Amato, Jeffrey Barrett, Sonia Goncalves, Ewan Harrison, David K. Jackson, Ian Johnston, Dominic Kwiatkowski, Cordelia Langford, John Sillitoe on behalf of the Wellcome Sanger Institute COVID-19 Surveillance Team |
| EPI_ISL_3223330, EPI_ISL_3223331 | Lighthouse Lab in Milton Keynes | Wellcome Sanger Institute for the COVID-19 Genomics UK (COG-UK) Consortium | The Lighthouse Lab in Milton Keynes and Alex Alderton, Roberto Amato, Jeffrey Barrett, Sonia Goncalves, Ewan Harrison, David K. Jackson, Ian Johnston, Dominic Kwiatkowski, Cordelia Langford, John Sillitoe on behalf of the Wellcome Sanger Institute COVID-19 Surveillance Team |
| EPI_ISL_3223332, EPI_ISL_3223333 | Lighthouse Laboratory Plymouth | Wellcome Sanger Institute for the COVID-19 Genomics UK (COG-UK) Consortium | Lighthouse Laboratory Plymouth and Alex Alderton, Roberto Amato, Jeffrey Barrett, Sonia Goncalves, Ewan Harrison, David K. Jackson, Ian Johnston, Dominic Kwiatkowski, Cordelia Langford, John Sillitoe on behalf of the Wellcome Sanger Institute COVID-19 Surveillance Team |
| EPI_ISL_3223334 | Lighthouse Lab in Glasgow | Wellcome Sanger Institute for the COVID-19 Genomics UK (COG-UK) Consortium | Harper VanSteenhouse, Yumi Kasai, David Gray, Carol Clugston, Anna Dominiczak and Alex Alderton, Roberto Amato, Jeffrey Barrett, Sonia Goncalves, Ewan Harrison, David K. Jackson, Ian Johnston, Dominic Kwiatkowski, Cordelia Langford, John Sillitoe on behalf of the Wellcome Sanger Institute COVID-19 Surveillance Team |
| EPI_ISL_3223335, EPI_ISL_3223336, EPI_ISL_3223337 | Lighthouse Laboratory Plymouth | Wellcome Sanger Institute for the COVID-19 Genomics UK (COG-UK) Consortium | Lighthouse Laboratory Plymouth and Alex Alderton, Roberto Amato, Jeffrey Barrett, Sonia Goncalves, Ewan Harrison, David K. Jackson, Ian Johnston, Dominic Kwiatkowski, Cordelia Langford, John Sillitoe on behalf of the Wellcome Sanger Institute COVID-19 Surveillance Team |
| EPI_ISL_3223338 | Lighthouse Lab in Milton Keynes | Wellcome Sanger Institute for the COVID-19 Genomics UK (COG-UK) Consortium | The Lighthouse Lab in Milton Keynes and Alex Alderton, Roberto Amato, Jeffrey Barrett, Sonia Goncalves, Ewan Harrison, David K. Jackson, Ian Johnston, Dominic Kwiatkowski, Cordelia Langford, John Sillitoe on behalf of the Wellcome Sanger Institute COVID-19 Surveillance Team |
| EPI_ISL_3223339, EPI_ISL_3223340 | Lighthouse Lab in Glasgow | Wellcome Sanger Institute for the COVID-19 Genomics UK (COG-UK) Consortium | Harper VanSteenhouse, Yumi Kasai, David Gray, Carol Clugston, Anna Dominiczak and Alex Alderton, Roberto Amato, Jeffrey Barrett, Sonia Goncalves, Ewan Harrison, David K. Jackson, Ian Johnston, Dominic Kwiatkowski, Cordelia Langford, John Sillitoe on behalf of the Wellcome Sanger Institute COVID-19 Surveillance Team |
| EPI_ISL_3223341, EPI_ISL_3223342 | Lighthouse Lab in Milton Keynes | Wellcome Sanger Institute for the COVID-19 Genomics UK (COG-UK) Consortium | The Lighthouse Lab in Milton Keynes and Alex Alderton, Roberto Amato, Jeffrey Barrett, Sonia Goncalves, Ewan Harrison, David K. Jackson, Ian Johnston, Dominic Kwiatkowski, Cordelia Langford, John Sillitoe on behalf of the Wellcome Sanger Institute COVID-19 Surveillance Team |
| EPI_ISL_3223343 | Lighthouse Lab in Glasgow | Wellcome Sanger Institute for the COVID-19 Genomics UK (COG-UK) Consortium | Harper VanSteenhouse, Yumi Kasai, David Gray, Carol Clugston, Anna Dominiczak and Alex Alderton, Roberto Amato, Jeffrey Barrett, Sonia Goncalves, Ewan Harrison, David K. Jackson, Ian Johnston, Dominic Kwiatkowski, Cordelia Langford, John Sillitoe on behalf of the Wellcome Sanger Institute COVID-19 Surveillance Team |
| EPI_ISL_3223344, EPI_ISL_3223345 | Lighthouse Lab in Milton Keynes | Wellcome Sanger Institute for the COVID-19 Genomics UK (COG-UK) Consortium | The Lighthouse Lab in Milton Keynes and Alex Alderton, Roberto Amato, Jeffrey Barrett, Sonia Goncalves, Ewan Harrison, David K. Jackson, Ian Johnston, Dominic Kwiatkowski, Cordelia Langford, John Sillitoe on behalf of the Wellcome Sanger Institute COVID-19 Surveillance Team |
| EPI_ISL_3223346, EPI_ISL_3223347, EPI_ISL_3223348, EPI_ISL_3223349, EPI_ISL_3223350, EPI_ISL_3223351, | Lighthouse Laboratory Plymouth | Wellcome Sanger Institute for the COVID-19 Genomics UK (COG-UK) Consortium | Lighthouse Laboratory Plymouth and Alex Alderton, Roberto Amato, Jeffrey Barrett, Sonia Goncalves, Ewan Harrison, David K. Jackson, Ian Johnston, Dominic Kwiatkowski, Cordelia Langford, John Sillitoe on behalf of the Wellcome Sanger Institute COVID-19 Surveillance Team |

| | | | |
|---|---|---|---|
| EPI_ISL_3223352, EPI_ISL_3223353, EPI_ISL_3223354, EPI_ISL_3223355 | | | |
| EPI_ISL_3223356 | Lighthouse Lab in Glasgow | Wellcome Sanger Institute for the COVID-19 Genomics UK (COG-UK) Consortium | Harper VanSteenhouse, Yumi Kasai, David Gray, Carol Clugston, Anna Dominiczak and Alex Alderton, Roberto Amato, Jeffrey Barrett, Sonia Goncalves, Ewan Harrison, David K. Jackson, Ian Johnston, Dominic Kwiatkowski, Cordelia Langford, John Sillitoe on behalf of the Wellcome Sanger Institute COVID-19 Surveillance Team |
| EPI_ISL_3223357, EPI_ISL_3223358 | Lighthouse Laboratory Plymouth | Wellcome Sanger Institute for the COVID-19 Genomics UK (COG-UK) Consortium | Lighthouse Laboratory Plymouth and Alex Alderton, Roberto Amato, Jeffrey Barrett, Sonia Goncalves, Ewan Harrison, David K. Jackson, Ian Johnston, Dominic Kwiatkowski, Cordelia Langford, John Sillitoe on behalf of the Wellcome Sanger Institute COVID-19 Surveillance Team |
| EPI_ISL_3223359 | Lighthouse Lab in Glasgow | Wellcome Sanger Institute for the COVID-19 Genomics UK (COG-UK) Consortium | Harper VanSteenhouse, Yumi Kasai, David Gray, Carol Clugston, Anna Dominiczak and Alex Alderton, Roberto Amato, Jeffrey Barrett, Sonia Goncalves, Ewan Harrison, David K. Jackson, Ian Johnston, Dominic Kwiatkowski, Cordelia Langford, John Sillitoe on behalf of the Wellcome Sanger Institute COVID-19 Surveillance Team |
| EPI_ISL_3223360, EPI_ISL_3223361 | Lighthouse Laboratory Plymouth | Wellcome Sanger Institute for the COVID-19 Genomics UK (COG-UK) Consortium | Lighthouse Laboratory Plymouth and Alex Alderton, Roberto Amato, Jeffrey Barrett, Sonia Goncalves, Ewan Harrison, David K. Jackson, Ian Johnston, Dominic Kwiatkowski, Cordelia Langford, John Sillitoe on behalf of the Wellcome Sanger Institute COVID-19 Surveillance Team |
| EPI_ISL_3223362 | Lighthouse Lab in Milton Keynes | Wellcome Sanger Institute for the COVID-19 Genomics UK (COG-UK) Consortium | The Lighthouse Lab in Milton Keynes and Alex Alderton, Roberto Amato, Jeffrey Barrett, Sonia Goncalves, Ewan Harrison, David K. Jackson, Ian Johnston, Dominic Kwiatkowski, Cordelia Langford, John Sillitoe on behalf of the Wellcome Sanger Institute COVID-19 Surveillance Team |
| EPI_ISL_3223363, EPI_ISL_3223364 | Lighthouse Laboratory Plymouth | Wellcome Sanger Institute for the COVID-19 Genomics UK (COG-UK) Consortium | Lighthouse Laboratory Plymouth and Alex Alderton, Roberto Amato, Jeffrey Barrett, Sonia Goncalves, Ewan Harrison, David K. Jackson, Ian Johnston, Dominic Kwiatkowski, Cordelia Langford, John Sillitoe on behalf of the Wellcome Sanger Institute COVID-19 Surveillance Team |
| EPI_ISL_3223365 | Lighthouse Lab in Glasgow | Wellcome Sanger Institute for the COVID-19 Genomics UK (COG-UK) Consortium | Harper VanSteenhouse, Yumi Kasai, David Gray, Carol Clugston, Anna Dominiczak and Alex Alderton, Roberto Amato, Jeffrey Barrett, Sonia Goncalves, Ewan Harrison, David K. Jackson, Ian Johnston, Dominic Kwiatkowski, Cordelia Langford, John Sillitoe on behalf of the Wellcome Sanger Institute COVID-19 Surveillance Team |
| EPI_ISL_3223366, EPI_ISL_3223367, EPI_ISL_3223368 | Lighthouse Laboratory Plymouth | Wellcome Sanger Institute for the COVID-19 Genomics UK (COG-UK) Consortium | Lighthouse Laboratory Plymouth and Alex Alderton, Roberto Amato, Jeffrey Barrett, Sonia Goncalves, Ewan Harrison, David K. Jackson, Ian Johnston, Dominic Kwiatkowski, Cordelia Langford, John Sillitoe on behalf of the Wellcome Sanger Institute COVID-19 Surveillance Team |
| EPI_ISL_3223369 | Lighthouse Lab in Milton Keynes | Wellcome Sanger Institute for the COVID-19 Genomics UK (COG-UK) Consortium | The Lighthouse Lab in Milton Keynes and Alex Alderton, Roberto Amato, Jeffrey Barrett, Sonia Goncalves, Ewan Harrison, David K. Jackson, Ian Johnston, Dominic Kwiatkowski, Cordelia Langford, John Sillitoe on behalf of the Wellcome Sanger Institute COVID-19 Surveillance Team |
| EPI_ISL_3223370 | Lighthouse Lab in Glasgow | Wellcome Sanger Institute for the COVID-19 Genomics UK (COG-UK) Consortium | Harper VanSteenhouse, Yumi Kasai, David Gray, Carol Clugston, Anna Dominiczak and Alex Alderton, Roberto Amato, Jeffrey Barrett, Sonia Goncalves, Ewan Harrison, David K. Jackson, Ian Johnston, Dominic Kwiatkowski, Cordelia Langford, John Sillitoe on behalf of the Wellcome Sanger Institute COVID-19 Surveillance Team |
| EPI_ISL_3223371 | Lighthouse Laboratory Plymouth | Wellcome Sanger Institute for the COVID-19 Genomics UK (COG-UK) Consortium | Lighthouse Laboratory Plymouth and Alex Alderton, Roberto Amato, Jeffrey Barrett, Sonia Goncalves, Ewan Harrison, David K. Jackson, Ian Johnston, Dominic Kwiatkowski, Cordelia Langford, John Sillitoe on behalf of the Wellcome Sanger Institute COVID-19 Surveillance Team |
| EPI_ISL_3223372 | Lighthouse Lab in Milton Keynes | Wellcome Sanger Institute for the COVID-19 Genomics UK (COG-UK) Consortium | The Lighthouse Lab in Milton Keynes and Alex Alderton, Roberto Amato, Jeffrey Barrett, Sonia Goncalves, Ewan Harrison, David K. Jackson, Ian Johnston, Dominic Kwiatkowski, Cordelia Langford, John Sillitoe on behalf of the Wellcome Sanger Institute COVID-19 Surveillance Team |
| EPI_ISL_3223373 | Lighthouse Laboratory Plymouth | Wellcome Sanger Institute for the COVID-19 Genomics UK (COG-UK) Consortium | Lighthouse Laboratory Plymouth and Alex Alderton, Roberto Amato, Jeffrey Barrett, Sonia Goncalves, Ewan Harrison, David K. Jackson, Ian Johnston, Dominic Kwiatkowski, Cordelia Langford, John Sillitoe on behalf of the Wellcome Sanger Institute COVID-19 Surveillance Team |
| EPI_ISL_3223374 | Lighthouse Lab in Milton Keynes | Wellcome Sanger Institute for the COVID-19 Genomics UK (COG-UK) Consortium | The Lighthouse Lab in Milton Keynes and Alex Alderton, Roberto Amato, Jeffrey Barrett, Sonia Goncalves, Ewan Harrison, David K. Jackson, Ian Johnston, Dominic Kwiatkowski, Cordelia Langford, John Sillitoe on behalf of the Wellcome Sanger Institute COVID-19 Surveillance Team |
| EPI_ISL_3223375, EPI_ISL_3223376 | Lighthouse Laboratory Plymouth | Wellcome Sanger Institute for the COVID-19 Genomics UK (COG-UK) Consortium | Lighthouse Laboratory Plymouth and Alex Alderton, Roberto Amato, Jeffrey Barrett, Sonia Goncalves, Ewan Harrison, David K. Jackson, Ian Johnston, Dominic Kwiatkowski, Cordelia Langford, John Sillitoe on behalf of the Wellcome Sanger Institute COVID-19 Surveillance Team |
| EPI_ISL_3223377 | Lighthouse Lab in Glasgow | Wellcome Sanger Institute for the COVID-19 Genomics UK (COG-UK) Consortium | Harper VanSteenhouse, Yumi Kasai, David Gray, Carol Clugston, Anna Dominiczak and Alex Alderton, Roberto Amato, Jeffrey Barrett, Sonia Goncalves, Ewan Harrison, David K. Jackson, Ian Johnston, Dominic Kwiatkowski, Cordelia Langford, John Sillitoe on behalf of the Wellcome Sanger Institute COVID-19 Surveillance Team |
| EPI_ISL_3223378 | Lighthouse Laboratory Plymouth | Wellcome Sanger Institute for the COVID-19 Genomics UK (COG-UK) Consortium | Lighthouse Laboratory Plymouth and Alex Alderton, Roberto Amato, Jeffrey Barrett, Sonia Goncalves, Ewan Harrison, David K. Jackson, Ian Johnston, Dominic Kwiatkowski, Cordelia Langford, John Sillitoe on behalf of the Wellcome Sanger Institute COVID-19 Surveillance Team |
| EPI_ISL_3223379 | Lighthouse Lab in Glasgow | Wellcome Sanger Institute for the COVID-19 Genomics UK (COG-UK) Consortium | Harper VanSteenhouse, Yumi Kasai, David Gray, Carol Clugston, Anna Dominiczak and Alex Alderton, Roberto Amato, Jeffrey Barrett, Sonia Goncalves, Ewan Harrison, David K. Jackson, Ian Johnston, Dominic Kwiatkowski, Cordelia Langford, John Sillitoe on behalf of the Wellcome Sanger Institute COVID-19 Surveillance Team |
| EPI_ISL_3223380, EPI_ISL_3223381 | Lighthouse Laboratory Plymouth | Wellcome Sanger Institute for the COVID-19 Genomics UK (COG-UK) Consortium | Lighthouse Laboratory Plymouth and Alex Alderton, Roberto Amato, Jeffrey Barrett, Sonia Goncalves, Ewan Harrison, David K. Jackson, Ian Johnston, Dominic Kwiatkowski, Cordelia Langford, John Sillitoe on behalf of the Wellcome Sanger Institute COVID-19 Surveillance Team |
| EPI_ISL_3223382 | Lighthouse Lab in Milton Keynes | Wellcome Sanger Institute for the COVID-19 Genomics UK (COG-UK) Consortium | The Lighthouse Lab in Milton Keynes and Alex Alderton, Roberto Amato, Jeffrey Barrett, Sonia Goncalves, Ewan Harrison, David K. Jackson, Ian Johnston, Dominic Kwiatkowski, Cordelia Langford, John Sillitoe on behalf of the Wellcome Sanger Institute COVID-19 Surveillance Team |
| EPI_ISL_3223383 | Lighthouse Lab in Glasgow | Wellcome Sanger Institute for the COVID-19 Genomics UK (COG-UK) Consortium | Harper VanSteenhouse, Yumi Kasai, David Gray, Carol Clugston, Anna Dominiczak and Alex Alderton, Roberto Amato, Jeffrey Barrett, Sonia Goncalves, Ewan Harrison, David K. Jackson, Ian Johnston, Dominic Kwiatkowski, Cordelia Langford, John Sillitoe on behalf of the Wellcome Sanger Institute COVID-19 Surveillance Team |
| EPI_ISL_3223384 | Lighthouse Laboratory Plymouth | Wellcome Sanger Institute for the COVID-19 Genomics UK (COG-UK) Consortium | Lighthouse Laboratory Plymouth and Alex Alderton, Roberto Amato, Jeffrey Barrett, Sonia Goncalves, Ewan Harrison, David K. Jackson, Ian Johnston, Dominic Kwiatkowski, Cordelia Langford, John Sillitoe on behalf of the Wellcome Sanger Institute COVID-19 Surveillance Team |
| EPI_ISL_3223385, EPI_ISL_3223386 | Lighthouse Lab in Milton Keynes | Wellcome Sanger Institute for the COVID-19 Genomics UK (COG-UK) Consortium | The Lighthouse Lab in Milton Keynes and Alex Alderton, Roberto Amato, Jeffrey Barrett, Sonia Goncalves, Ewan Harrison, David K. Jackson, Ian Johnston, Dominic Kwiatkowski, Cordelia Langford, John Sillitoe on behalf of the Wellcome Sanger Institute COVID-19 Surveillance Team |
| EPI_ISL_3223387, EPI_ISL_3223388 | Lighthouse Laboratory Plymouth | Wellcome Sanger Institute for the COVID-19 Genomics UK (COG-UK) Consortium | Lighthouse Laboratory Plymouth and Alex Alderton, Roberto Amato, Jeffrey Barrett, Sonia Goncalves, Ewan Harrison, David K. Jackson, Ian Johnston, Dominic Kwiatkowski, Cordelia Langford, John Sillitoe on behalf of the Wellcome Sanger Institute COVID-19 Surveillance Team |
| EPI_ISL_3223389 | Lighthouse Lab in Milton Keynes | Wellcome Sanger Institute for the COVID-19 Genomics UK (COG-UK) Consortium | The Lighthouse Lab in Milton Keynes and Alex Alderton, Roberto Amato, Jeffrey Barrett, Sonia Goncalves, Ewan Harrison, David K. Jackson, Ian Johnston, Dominic Kwiatkowski, Cordelia Langford, John Sillitoe on behalf of the Wellcome Sanger Institute COVID-19 Surveillance Team |
| EPI_ISL_3223390, EPI_ISL_3223391 | Lighthouse Lab in Glasgow | Wellcome Sanger Institute for the COVID-19 Genomics UK (COG-UK) Consortium | Harper VanSteenhouse, Yumi Kasai, David Gray, Carol Clugston, Anna Dominiczak and Alex Alderton, Roberto Amato, Jeffrey Barrett, Sonia Goncalves, Ewan Harrison, David K. Jackson, Ian Johnston, Dominic Kwiatkowski, Cordelia Langford, John Sillitoe on behalf of the Wellcome Sanger Institute COVID-19 Surveillance Team |
| EPI_ISL_3223392 | Lighthouse Laboratory Plymouth | Wellcome Sanger Institute for the COVID-19 Genomics UK (COG-UK) Consortium | Lighthouse Laboratory Plymouth and Alex Alderton, Roberto Amato, Jeffrey Barrett, Sonia Goncalves, Ewan Harrison, David K. Jackson, Ian Johnston, Dominic Kwiatkowski, Cordelia Langford, John Sillitoe on behalf of the Wellcome Sanger Institute COVID-19 Surveillance Team |
| EPI_ISL_3223393, EPI_ISL_3223394 | Lighthouse Lab in Glasgow | Wellcome Sanger Institute for the COVID-19 Genomics UK (COG-UK) Consortium | Harper VanSteenhouse, Yumi Kasai, David Gray, Carol Clugston, Anna Dominiczak and Alex Alderton, Roberto Amato, Jeffrey Barrett, Sonia Goncalves, Ewan Harrison, David K. Jackson, Ian Johnston, Dominic Kwiatkowski, Cordelia Langford, John Sillitoe on behalf of the Wellcome Sanger Institute COVID-19 Surveillance Team |
| EPI_ISL_3223395 | Lighthouse Lab in Milton Keynes | Wellcome Sanger Institute for the COVID-19 Genomics UK (COG-UK) Consortium | The Lighthouse Lab in Milton Keynes and Alex Alderton, Roberto Amato, Jeffrey Barrett, Sonia Goncalves, Ewan Harrison, David K. Jackson, Ian Johnston, Dominic Kwiatkowski, Cordelia Langford, John Sillitoe on behalf of the Wellcome Sanger Institute COVID-19 Surveillance Team |
| EPI_ISL_3223396 | Lighthouse Lab in Glasgow | Wellcome Sanger Institute for the COVID-19 Genomics UK (COG-UK) Consortium | Harper VanSteenhouse, Yumi Kasai, David Gray, Carol Clugston, Anna Dominiczak and Alex Alderton, Roberto Amato, Jeffrey Barrett, Sonia Goncalves, Ewan Harrison, David K. Jackson, Ian Johnston, Dominic Kwiatkowski, Cordelia Langford, John Sillitoe on behalf of the Wellcome Sanger Institute COVID-19 Surveillance Team |
| EPI_ISL_3223397 | Lighthouse Lab in Milton Keynes | Wellcome Sanger Institute for the COVID-19 Genomics UK (COG-UK) Consortium | The Lighthouse Lab in Milton Keynes and Alex Alderton, Roberto Amato, Jeffrey Barrett, Sonia Goncalves, Ewan Harrison, David K. Jackson, Ian Johnston, Dominic Kwiatkowski, Cordelia Langford, John Sillitoe on behalf of the Wellcome Sanger Institute COVID-19 Surveillance Team |
| EPI_ISL_3223398, EPI_ISL_3223399, EPI_ISL_3223400, EPI_ISL_3223401 | Lighthouse Laboratory Plymouth | Wellcome Sanger Institute for the COVID-19 Genomics UK (COG-UK) Consortium | Lighthouse Laboratory Plymouth and Alex Alderton, Roberto Amato, Jeffrey Barrett, Sonia Goncalves, Ewan Harrison, David K. Jackson, Ian Johnston, Dominic Kwiatkowski, Cordelia Langford, John Sillitoe on behalf of the Wellcome Sanger Institute COVID-19 Surveillance Team |
| EPI_ISL_3223402, EPI_ISL_3223403 | Lighthouse Lab in Glasgow | Wellcome Sanger Institute for the COVID-19 Genomics UK (COG-UK) Consortium | Harper VanSteenhouse, Yumi Kasai, David Gray, Carol Clugston, Anna Dominiczak and Alex Alderton, Roberto Amato, Jeffrey Barrett, Sonia Goncalves, Ewan Harrison, David K. Jackson, Ian Johnston, Dominic Kwiatkowski, Cordelia Langford, John Sillitoe on behalf of the Wellcome Sanger Institute COVID-19 Surveillance Team |
| EPI_ISL_3223404, EPI_ISL_3223405 | Lighthouse Laboratory Plymouth | Wellcome Sanger Institute for the COVID-19 Genomics UK (COG-UK) Consortium | Lighthouse Laboratory Plymouth and Alex Alderton, Roberto Amato, Jeffrey Barrett, Sonia Goncalves, Ewan Harrison, David K. Jackson, Ian Johnston, Dominic Kwiatkowski, Cordelia Langford, John Sillitoe on behalf of the Wellcome Sanger Institute COVID-19 Surveillance Team |
| EPI_ISL_3223406, EPI_ISL_3223407, EPI_ISL_3223408, EPI_ISL_3223409, EPI_ISL_3223410, EPI_ISL_3223411 | Lighthouse Lab in Milton Keynes | Wellcome Sanger Institute for the COVID-19 Genomics UK (COG-UK) Consortium | The Lighthouse Lab in Milton Keynes and Alex Alderton, Roberto Amato, Jeffrey Barrett, Sonia Goncalves, Ewan Harrison, David K. Jackson, Ian Johnston, Dominic Kwiatkowski, Cordelia Langford, John Sillitoe on behalf of the Wellcome Sanger Institute COVID-19 Surveillance Team |
| EPI_ISL_3223412 | Lighthouse Laboratory Plymouth | Wellcome Sanger Institute for the COVID-19 Genomics UK (COG-UK) Consortium | Lighthouse Laboratory Plymouth and Alex Alderton, Roberto Amato, Jeffrey Barrett, Sonia Goncalves, Ewan Harrison, David K. Jackson, Ian Johnston, Dominic Kwiatkowski, Cordelia Langford, John Sillitoe on behalf of the Wellcome Sanger Institute COVID-19 Surveillance Team |
| EPI_ISL_3223413 | Lighthouse Lab in Milton Keynes | Wellcome Sanger Institute for the COVID-19 Genomics UK (COG-UK) Consortium | The Lighthouse Lab in Milton Keynes and Alex Alderton, Roberto Amato, Jeffrey Barrett, Sonia Goncalves, Ewan Harrison, David K. Jackson, Ian Johnston, Dominic Kwiatkowski, Cordelia Langford, John Sillitoe on behalf of the Wellcome Sanger Institute COVID-19 Surveillance Team |
| EPI_ISL_3223414 | Lighthouse Lab in Glasgow | Wellcome Sanger Institute for the COVID-19 Genomics UK (COG-UK) Consortium | Harper VanSteenhouse, Yumi Kasai, David Gray, Carol Clugston, Anna Dominiczak and Alex Alderton, Roberto Amato, Jeffrey Barrett, Sonia Goncalves, Ewan Harrison, David K. Jackson, Ian Johnston, Dominic Kwiatkowski, Cordelia Langford, John Sillitoe on behalf of the Wellcome Sanger Institute COVID-19 Surveillance Team |
| EPI_ISL_3223415, EPI_ISL_3223416, EPI_ISL_3223417 | Lighthouse Laboratory Plymouth | Wellcome Sanger Institute for the COVID-19 Genomics UK (COG-UK) Consortium | Lighthouse Laboratory Plymouth and Alex Alderton, Roberto Amato, Jeffrey Barrett, Sonia Goncalves, Ewan Harrison, David K. Jackson, Ian Johnston, Dominic Kwiatkowski, Cordelia Langford, John Sillitoe on behalf of the Wellcome Sanger Institute COVID-19 Surveillance Team |
| EPI_ISL_3223418, EPI_ISL_3223419, EPI_ISL_3223420 | Lighthouse Lab in Milton Keynes | Wellcome Sanger Institute for the COVID-19 Genomics UK (COG-UK) Consortium | The Lighthouse Lab in Milton Keynes and Alex Alderton, Roberto Amato, Jeffrey Barrett, Sonia Goncalves, Ewan Harrison, David K. Jackson, Ian Johnston, Dominic Kwiatkowski, Cordelia Langford, John Sillitoe on behalf of the Wellcome Sanger Institute COVID-19 Surveillance Team |

| | | | |
|---|---|---|---|
| EPI_ISL_3223421, EPI_ISL_3223422 | Lighthouse Laboratory Plymouth | Wellcome Sanger Institute for the COVID-19 Genomics UK (COG-UK) Consortium | Lighthouse Laboratory Plymouth and Alex Alderton, Roberto Amato, Jeffrey Barrett, Sonia Goncalves, Ewan Harrison, David K. Jackson, Ian Johnston, Dominic Kwiatkowski, Cordelia Langford, John Sillitoe on behalf of the Wellcome Sanger Institute COVID-19 Surveillance Team |
| EPI_ISL_3223423 | Lighthouse Lab in Milton Keynes | Wellcome Sanger Institute for the COVID-19 Genomics UK (COG-UK) Consortium | The Lighthouse Lab in Milton Keynes and Alex Alderton, Roberto Amato, Jeffrey Barrett, Sonia Goncalves, Ewan Harrison, David K. Jackson, Ian Johnston, Dominic Kwiatkowski, Cordelia Langford, John Sillitoe on behalf of the Wellcome Sanger Institute COVID-19 Surveillance Team |
| EPI_ISL_3223424, EPI_ISL_3223425, EPI_ISL_3223426 | Lighthouse Laboratory Plymouth | Wellcome Sanger Institute for the COVID-19 Genomics UK (COG-UK) Consortium | Lighthouse Laboratory Plymouth and Alex Alderton, Roberto Amato, Jeffrey Barrett, Sonia Goncalves, Ewan Harrison, David K. Jackson, Ian Johnston, Dominic Kwiatkowski, Cordelia Langford, John Sillitoe on behalf of the Wellcome Sanger Institute COVID-19 Surveillance Team |
| EPI_ISL_3223427 | Lighthouse Lab in Glasgow | Wellcome Sanger Institute for the COVID-19 Genomics UK (COG-UK) Consortium | Harper VanSteenhouse, Yumi Kasai, David Gray, Carol Clugston, Anna Dominiczak and Alex Alderton, Roberto Amato, Jeffrey Barrett, Sonia Goncalves, Ewan Harrison, David K. Jackson, Ian Johnston, Dominic Kwiatkowski, Cordelia Langford, John Sillitoe on behalf of the Wellcome Sanger Institute COVID-19 Surveillance Team |
| EPI_ISL_3223428, EPI_ISL_3223429 | Lighthouse Lab in Milton Keynes | Wellcome Sanger Institute for the COVID-19 Genomics UK (COG-UK) Consortium | The Lighthouse Lab in Milton Keynes and Alex Alderton, Roberto Amato, Jeffrey Barrett, Sonia Goncalves, Ewan Harrison, David K. Jackson, Ian Johnston, Dominic Kwiatkowski, Cordelia Langford, John Sillitoe on behalf of the Wellcome Sanger Institute COVID-19 Surveillance Team |
| EPI_ISL_3223430, EPI_ISL_3223431, EPI_ISL_3223432, EPI_ISL_3223433 | Lighthouse Laboratory Plymouth | Wellcome Sanger Institute for the COVID-19 Genomics UK (COG-UK) Consortium | Lighthouse Laboratory Plymouth and Alex Alderton, Roberto Amato, Jeffrey Barrett, Sonia Goncalves, Ewan Harrison, David K. Jackson, Ian Johnston, Dominic Kwiatkowski, Cordelia Langford, John Sillitoe on behalf of the Wellcome Sanger Institute COVID-19 Surveillance Team |
| EPI_ISL_3223434 | Lighthouse Lab in Milton Keynes | Wellcome Sanger Institute for the COVID-19 Genomics UK (COG-UK) Consortium | The Lighthouse Lab in Milton Keynes and Alex Alderton, Roberto Amato, Jeffrey Barrett, Sonia Goncalves, Ewan Harrison, David K. Jackson, Ian Johnston, Dominic Kwiatkowski, Cordelia Langford, John Sillitoe on behalf of the Wellcome Sanger Institute COVID-19 Surveillance Team |
| EPI_ISL_3223435 | Lighthouse Laboratory Plymouth | Wellcome Sanger Institute for the COVID-19 Genomics UK (COG-UK) Consortium | Lighthouse Laboratory Plymouth and Alex Alderton, Roberto Amato, Jeffrey Barrett, Sonia Goncalves, Ewan Harrison, David K. Jackson, Ian Johnston, Dominic Kwiatkowski, Cordelia Langford, John Sillitoe on behalf of the Wellcome Sanger Institute COVID-19 Surveillance Team |
| EPI_ISL_3223436, EPI_ISL_3223437, EPI_ISL_3223438 | Lighthouse Lab in Milton Keynes | Wellcome Sanger Institute for the COVID-19 Genomics UK (COG-UK) Consortium | The Lighthouse Lab in Milton Keynes and Alex Alderton, Roberto Amato, Jeffrey Barrett, Sonia Goncalves, Ewan Harrison, David K. Jackson, Ian Johnston, Dominic Kwiatkowski, Cordelia Langford, John Sillitoe on behalf of the Wellcome Sanger Institute COVID-19 Surveillance Team |
| EPI_ISL_3223439, EPI_ISL_3223440 | Lighthouse Lab in Glasgow | Wellcome Sanger Institute for the COVID-19 Genomics UK (COG-UK) Consortium | Harper VanSteenhouse, Yumi Kasai, David Gray, Carol Clugston, Anna Dominiczak and Alex Alderton, Roberto Amato, Jeffrey Barrett, Sonia Goncalves, Ewan Harrison, David K. Jackson, Ian Johnston, Dominic Kwiatkowski, Cordelia Langford, John Sillitoe on behalf of the Wellcome Sanger Institute COVID-19 Surveillance Team |
| EPI_ISL_3223441, EPI_ISL_3223442, EPI_ISL_3223443 | Lighthouse Lab in Milton Keynes | Wellcome Sanger Institute for the COVID-19 Genomics UK (COG-UK) Consortium | The Lighthouse Lab in Milton Keynes and Alex Alderton, Roberto Amato, Jeffrey Barrett, Sonia Goncalves, Ewan Harrison, David K. Jackson, Ian Johnston, Dominic Kwiatkowski, Cordelia Langford, John Sillitoe on behalf of the Wellcome Sanger Institute COVID-19 Surveillance Team |
| EPI_ISL_3223444, EPI_ISL_3223445, EPI_ISL_3223446, EPI_ISL_3223447 | Lighthouse Laboratory Plymouth | Wellcome Sanger Institute for the COVID-19 Genomics UK (COG-UK) Consortium | Lighthouse Laboratory Plymouth and Alex Alderton, Roberto Amato, Jeffrey Barrett, Sonia Goncalves, Ewan Harrison, David K. Jackson, Ian Johnston, Dominic Kwiatkowski, Cordelia Langford, John Sillitoe on behalf of the Wellcome Sanger Institute COVID-19 Surveillance Team |
| EPI_ISL_3223448, EPI_ISL_3223449 | Lighthouse Lab in Milton Keynes | Wellcome Sanger Institute for the COVID-19 Genomics UK (COG-UK) Consortium | The Lighthouse Lab in Milton Keynes and Alex Alderton, Roberto Amato, Jeffrey Barrett, Sonia Goncalves, Ewan Harrison, David K. Jackson, Ian Johnston, Dominic Kwiatkowski, Cordelia Langford, John Sillitoe on behalf of the Wellcome Sanger Institute COVID-19 Surveillance Team |
| EPI_ISL_3223450, EPI_ISL_3223451 | Lighthouse Laboratory Plymouth | Wellcome Sanger Institute for the COVID-19 Genomics UK (COG-UK) Consortium | Lighthouse Laboratory Plymouth and Alex Alderton, Roberto Amato, Jeffrey Barrett, Sonia Goncalves, Ewan Harrison, David K. Jackson, Ian Johnston, Dominic Kwiatkowski, Cordelia Langford, John Sillitoe on behalf of the Wellcome Sanger Institute COVID-19 Surveillance Team |
| EPI_ISL_3223452 | Lighthouse Lab in Milton Keynes | Wellcome Sanger Institute for the COVID-19 Genomics UK (COG-UK) Consortium | The Lighthouse Lab in Milton Keynes and Alex Alderton, Roberto Amato, Jeffrey Barrett, Sonia Goncalves, Ewan Harrison, David K. Jackson, Ian Johnston, Dominic Kwiatkowski, Cordelia Langford, John Sillitoe on behalf of the Wellcome Sanger Institute COVID-19 Surveillance Team |
| EPI_ISL_3223453 | Lighthouse Laboratory Plymouth | Wellcome Sanger Institute for the COVID-19 Genomics UK (COG-UK) Consortium | Lighthouse Laboratory Plymouth and Alex Alderton, Roberto Amato, Jeffrey Barrett, Sonia Goncalves, Ewan Harrison, David K. Jackson, Ian Johnston, Dominic Kwiatkowski, Cordelia Langford, John Sillitoe on behalf of the Wellcome Sanger Institute COVID-19 Surveillance Team |
| EPI_ISL_3223454 | Lighthouse Lab in Glasgow | Wellcome Sanger Institute for the COVID-19 Genomics UK (COG-UK) Consortium | Harper VanSteenhouse, Yumi Kasai, David Gray, Carol Clugston, Anna Dominiczak and Alex Alderton, Roberto Amato, Jeffrey Barrett, Sonia Goncalves, Ewan Harrison, David K. Jackson, Ian Johnston, Dominic Kwiatkowski, Cordelia Langford, John Sillitoe on behalf of the Wellcome Sanger Institute COVID-19 Surveillance Team |
| EPI_ISL_3223455 | Lighthouse Laboratory Plymouth | Wellcome Sanger Institute for the COVID-19 Genomics UK (COG-UK) Consortium | Lighthouse Laboratory Plymouth and Alex Alderton, Roberto Amato, Jeffrey Barrett, Sonia Goncalves, Ewan Harrison, David K. Jackson, Ian Johnston, Dominic Kwiatkowski, Cordelia Langford, John Sillitoe on behalf of the Wellcome Sanger Institute COVID-19 Surveillance Team |
| EPI_ISL_3223456 | Lighthouse Lab in Milton Keynes | Wellcome Sanger Institute for the COVID-19 Genomics UK (COG-UK) Consortium | The Lighthouse Lab in Milton Keynes and Alex Alderton, Roberto Amato, Jeffrey Barrett, Sonia Goncalves, Ewan Harrison, David K. Jackson, Ian Johnston, Dominic Kwiatkowski, Cordelia Langford, John Sillitoe on behalf of the Wellcome Sanger Institute COVID-19 Surveillance Team |
| EPI_ISL_3223457 | Lighthouse Laboratory Plymouth | Wellcome Sanger Institute for the COVID-19 Genomics UK (COG-UK) Consortium | Lighthouse Laboratory Plymouth and Alex Alderton, Roberto Amato, Jeffrey Barrett, Sonia Goncalves, Ewan Harrison, David K. Jackson, Ian Johnston, Dominic Kwiatkowski, Cordelia Langford, John Sillitoe on behalf of the Wellcome Sanger Institute COVID-19 Surveillance Team |
| EPI_ISL_3223458 | Lighthouse Lab in Milton Keynes | Wellcome Sanger Institute for the COVID-19 Genomics UK (COG-UK) Consortium | The Lighthouse Lab in Milton Keynes and Alex Alderton, Roberto Amato, Jeffrey Barrett, Sonia Goncalves, Ewan Harrison, David K. Jackson, Ian Johnston, Dominic Kwiatkowski, Cordelia Langford, John Sillitoe on behalf of the Wellcome Sanger Institute COVID-19 Surveillance Team |
| EPI_ISL_3223459, EPI_ISL_3223460, EPI_ISL_3223461, EPI_ISL_3223462, EPI_ISL_3223463 | Lighthouse Laboratory Plymouth | Wellcome Sanger Institute for the COVID-19 Genomics UK (COG-UK) Consortium | Lighthouse Laboratory Plymouth and Alex Alderton, Roberto Amato, Jeffrey Barrett, Sonia Goncalves, Ewan Harrison, David K. Jackson, Ian Johnston, Dominic Kwiatkowski, Cordelia Langford, John Sillitoe on behalf of the Wellcome Sanger Institute COVID-19 Surveillance Team |
| EPI_ISL_3223464 | Lighthouse Lab in Milton Keynes | Wellcome Sanger Institute for the COVID-19 Genomics UK (COG-UK) Consortium | The Lighthouse Lab in Milton Keynes and Alex Alderton, Roberto Amato, Jeffrey Barrett, Sonia Goncalves, Ewan Harrison, David K. Jackson, Ian Johnston, Dominic Kwiatkowski, Cordelia Langford, John Sillitoe on behalf of the Wellcome Sanger Institute COVID-19 Surveillance Team |
| EPI_ISL_3223465 | Lighthouse Lab in Glasgow | Wellcome Sanger Institute for the COVID-19 Genomics UK (COG-UK) Consortium | Harper VanSteenhouse, Yumi Kasai, David Gray, Carol Clugston, Anna Dominiczak and Alex Alderton, Roberto Amato, Jeffrey Barrett, Sonia Goncalves, Ewan Harrison, David K. Jackson, Ian Johnston, Dominic Kwiatkowski, Cordelia Langford, John Sillitoe on behalf of the Wellcome Sanger Institute COVID-19 Surveillance Team |
| EPI_ISL_3223466 | Lighthouse Lab in Milton Keynes | Wellcome Sanger Institute for the COVID-19 Genomics UK (COG-UK) Consortium | The Lighthouse Lab in Milton Keynes and Alex Alderton, Roberto Amato, Jeffrey Barrett, Sonia Goncalves, Ewan Harrison, David K. Jackson, Ian Johnston, Dominic Kwiatkowski, Cordelia Langford, John Sillitoe on behalf of the Wellcome Sanger Institute COVID-19 Surveillance Team |
| EPI_ISL_3223467, EPI_ISL_3223468 | Lighthouse Lab in Glasgow | Wellcome Sanger Institute for the COVID-19 Genomics UK (COG-UK) Consortium | Harper VanSteenhouse, Yumi Kasai, David Gray, Carol Clugston, Anna Dominiczak and Alex Alderton, Roberto Amato, Jeffrey Barrett, Sonia Goncalves, Ewan Harrison, David K. Jackson, Ian Johnston, Dominic Kwiatkowski, Cordelia Langford, John Sillitoe on behalf of the Wellcome Sanger Institute COVID-19 Surveillance Team |
| EPI_ISL_3223469 | Lighthouse Laboratory Plymouth | Wellcome Sanger Institute for the COVID-19 Genomics UK (COG-UK) Consortium | Lighthouse Laboratory Plymouth and Alex Alderton, Roberto Amato, Jeffrey Barrett, Sonia Goncalves, Ewan Harrison, David K. Jackson, Ian Johnston, Dominic Kwiatkowski, Cordelia Langford, John Sillitoe on behalf of the Wellcome Sanger Institute COVID-19 Surveillance Team |
| EPI_ISL_3223470 | Lighthouse Lab in Milton Keynes | Wellcome Sanger Institute for the COVID-19 Genomics UK (COG-UK) Consortium | The Lighthouse Lab in Milton Keynes and Alex Alderton, Roberto Amato, Jeffrey Barrett, Sonia Goncalves, Ewan Harrison, David K. Jackson, Ian Johnston, Dominic Kwiatkowski, Cordelia Langford, John Sillitoe on behalf of the Wellcome Sanger Institute COVID-19 Surveillance Team |
| EPI_ISL_3223471, EPI_ISL_3223472 | Lighthouse Laboratory Plymouth | Wellcome Sanger Institute for the COVID-19 Genomics UK (COG-UK) Consortium | Lighthouse Laboratory Plymouth and Alex Alderton, Roberto Amato, Jeffrey Barrett, Sonia Goncalves, Ewan Harrison, David K. Jackson, Ian Johnston, Dominic Kwiatkowski, Cordelia Langford, John Sillitoe on behalf of the Wellcome Sanger Institute COVID-19 Surveillance Team |
| EPI_ISL_3223473 | Lighthouse Lab in Glasgow | Wellcome Sanger Institute for the COVID-19 Genomics UK (COG-UK) Consortium | Harper VanSteenhouse, Yumi Kasai, David Gray, Carol Clugston, Anna Dominiczak and Alex Alderton, Roberto Amato, Jeffrey Barrett, Sonia Goncalves, Ewan Harrison, David K. Jackson, Ian Johnston, Dominic Kwiatkowski, Cordelia Langford, John Sillitoe on behalf of the Wellcome Sanger Institute COVID-19 Surveillance Team |
| EPI_ISL_3223474 | Lighthouse Laboratory Plymouth | Wellcome Sanger Institute for the COVID-19 Genomics UK (COG-UK) Consortium | Lighthouse Laboratory Plymouth and Alex Alderton, Roberto Amato, Jeffrey Barrett, Sonia Goncalves, Ewan Harrison, David K. Jackson, Ian Johnston, Dominic Kwiatkowski, Cordelia Langford, John Sillitoe on behalf of the Wellcome Sanger Institute COVID-19 Surveillance Team |
| EPI_ISL_3223475 | Lighthouse Lab in Glasgow | Wellcome Sanger Institute for the COVID-19 Genomics UK (COG-UK) Consortium | Harper VanSteenhouse, Yumi Kasai, David Gray, Carol Clugston, Anna Dominiczak and Alex Alderton, Roberto Amato, Jeffrey Barrett, Sonia Goncalves, Ewan Harrison, David K. Jackson, Ian Johnston, Dominic Kwiatkowski, Cordelia Langford, John Sillitoe on behalf of the Wellcome Sanger Institute COVID-19 Surveillance Team |
| EPI_ISL_3223476 | Lighthouse Laboratory Plymouth | Wellcome Sanger Institute for the COVID-19 Genomics UK (COG-UK) Consortium | Lighthouse Laboratory Plymouth and Alex Alderton, Roberto Amato, Jeffrey Barrett, Sonia Goncalves, Ewan Harrison, David K. Jackson, Ian Johnston, Dominic Kwiatkowski, Cordelia Langford, John Sillitoe on behalf of the Wellcome Sanger Institute COVID-19 Surveillance Team |
| EPI_ISL_3223477 | Lighthouse Lab in Milton Keynes | Wellcome Sanger Institute for the COVID-19 Genomics UK (COG-UK) Consortium | The Lighthouse Lab in Milton Keynes and Alex Alderton, Roberto Amato, Jeffrey Barrett, Sonia Goncalves, Ewan Harrison, David K. Jackson, Ian Johnston, Dominic Kwiatkowski, Cordelia Langford, John Sillitoe on behalf of the Wellcome Sanger Institute COVID-19 Surveillance Team |
| EPI_ISL_3223478, EPI_ISL_3223479, EPI_ISL_3223480, EPI_ISL_3223481, EPI_ISL_3223482 | Lighthouse Laboratory Plymouth | Wellcome Sanger Institute for the COVID-19 Genomics UK (COG-UK) Consortium | Lighthouse Laboratory Plymouth and Alex Alderton, Roberto Amato, Jeffrey Barrett, Sonia Goncalves, Ewan Harrison, David K. Jackson, Ian Johnston, Dominic Kwiatkowski, Cordelia Langford, John Sillitoe on behalf of the Wellcome Sanger Institute COVID-19 Surveillance Team |
| EPI_ISL_3223483 | Lighthouse Lab in Glasgow | Wellcome Sanger Institute for the COVID-19 Genomics UK (COG-UK) Consortium | Harper VanSteenhouse, Yumi Kasai, David Gray, Carol Clugston, Anna Dominiczak and Alex Alderton, Roberto Amato, Jeffrey Barrett, Sonia Goncalves, Ewan Harrison, David K. Jackson, Ian Johnston, Dominic Kwiatkowski, Cordelia Langford, John Sillitoe on behalf of the Wellcome Sanger Institute COVID-19 Surveillance Team |
| EPI_ISL_3223484, EPI_ISL_3223485, EPI_ISL_3223486, EPI_ISL_3223487, EPI_ISL_3223488 | Lighthouse Laboratory Plymouth | Wellcome Sanger Institute for the COVID-19 Genomics UK (COG-UK) Consortium | Lighthouse Laboratory Plymouth and Alex Alderton, Roberto Amato, Jeffrey Barrett, Sonia Goncalves, Ewan Harrison, David K. Jackson, Ian Johnston, Dominic Kwiatkowski, Cordelia Langford, John Sillitoe on behalf of the Wellcome Sanger Institute COVID-19 Surveillance Team |
| EPI_ISL_3223489 | Lighthouse Lab in Milton Keynes | Wellcome Sanger Institute for the COVID-19 Genomics UK (COG-UK) Consortium | The Lighthouse Lab in Milton Keynes and Alex Alderton, Roberto Amato, Jeffrey Barrett, Sonia Goncalves, Ewan Harrison, David K. Jackson, Ian Johnston, Dominic Kwiatkowski, Cordelia Langford, John Sillitoe on behalf of the Wellcome Sanger Institute COVID-19 Surveillance Team |

| | | | |
|---|---|---|---|
| EPI_ISL_3223490, EPI_ISL_3223491, EPI_ISL_3223492, EPI_ISL_3223493 | Lighthouse Laboratory Plymouth | Wellcome Sanger Institute for the COVID-19 Genomics UK (COG-UK) Consortium | Lighthouse Laboratory Plymouth and Alex Alderton, Roberto Amato, Jeffrey Barrett, Sonia Goncalves, Ewan Harrison, David K. Jackson, Ian Johnston, Dominic Kwiatkowski, Cordelia Langford, John Sillitoe on behalf of the Wellcome Sanger Institute COVID-19 Surveillance Team |
| EPI_ISL_3223494 | Lighthouse Lab in Glasgow | Wellcome Sanger Institute for the COVID-19 Genomics UK (COG-UK) Consortium | Harper VanSteenhouse, Yumi Kasai, David Gray, Carol Clugston, Anna Dominiczak and Alex Alderton, Roberto Amato, Jeffrey Barrett, Sonia Goncalves, Ewan Harrison, David K. Jackson, Ian Johnston, Dominic Kwiatkowski, Cordelia Langford, John Sillitoe on behalf of the Wellcome Sanger Institute COVID-19 Surveillance Team |
| EPI_ISL_3223495, EPI_ISL_3223496 | Lighthouse Lab in Milton Keynes | Wellcome Sanger Institute for the COVID-19 Genomics UK (COG-UK) Consortium | The Lighthouse Lab in Milton Keynes and Alex Alderton, Roberto Amato, Jeffrey Barrett, Sonia Goncalves, Ewan Harrison, David K. Jackson, Ian Johnston, Dominic Kwiatkowski, Cordelia Langford, John Sillitoe on behalf of the Wellcome Sanger Institute COVID-19 Surveillance Team |
| EPI_ISL_3223497 | Lighthouse Lab in Glasgow | Wellcome Sanger Institute for the COVID-19 Genomics UK (COG-UK) Consortium | Harper VanSteenhouse, Yumi Kasai, David Gray, Carol Clugston, Anna Dominiczak and Alex Alderton, Roberto Amato, Jeffrey Barrett, Sonia Goncalves, Ewan Harrison, David K. Jackson, Ian Johnston, Dominic Kwiatkowski, Cordelia Langford, John Sillitoe on behalf of the Wellcome Sanger Institute COVID-19 Surveillance Team |
| EPI_ISL_3223498 | Lighthouse Laboratory Plymouth | Wellcome Sanger Institute for the COVID-19 Genomics UK (COG-UK) Consortium | Lighthouse Laboratory Plymouth and Alex Alderton, Roberto Amato, Jeffrey Barrett, Sonia Goncalves, Ewan Harrison, David K. Jackson, Ian Johnston, Dominic Kwiatkowski, Cordelia Langford, John Sillitoe on behalf of the Wellcome Sanger Institute COVID-19 Surveillance Team |
| EPI_ISL_3223499, EPI_ISL_3223500, EPI_ISL_3223501, EPI_ISL_3223502 | Lighthouse Lab in Milton Keynes | Wellcome Sanger Institute for the COVID-19 Genomics UK (COG-UK) Consortium | The Lighthouse Lab in Milton Keynes and Alex Alderton, Roberto Amato, Jeffrey Barrett, Sonia Goncalves, Ewan Harrison, David K. Jackson, Ian Johnston, Dominic Kwiatkowski, Cordelia Langford, John Sillitoe on behalf of the Wellcome Sanger Institute COVID-19 Surveillance Team |
| EPI_ISL_3223503 | Lighthouse Laboratory Plymouth | Wellcome Sanger Institute for the COVID-19 Genomics UK (COG-UK) Consortium | Lighthouse Laboratory Plymouth and Alex Alderton, Roberto Amato, Jeffrey Barrett, Sonia Goncalves, Ewan Harrison, David K. Jackson, Ian Johnston, Dominic Kwiatkowski, Cordelia Langford, John Sillitoe on behalf of the Wellcome Sanger Institute COVID-19 Surveillance Team |
| EPI_ISL_3223504 | Lighthouse Lab in Glasgow | Wellcome Sanger Institute for the COVID-19 Genomics UK (COG-UK) Consortium | Harper VanSteenhouse, Yumi Kasai, David Gray, Carol Clugston, Anna Dominiczak and Alex Alderton, Roberto Amato, Jeffrey Barrett, Sonia Goncalves, Ewan Harrison, David K. Jackson, Ian Johnston, Dominic Kwiatkowski, Cordelia Langford, John Sillitoe on behalf of the Wellcome Sanger Institute COVID-19 Surveillance Team |
| EPI_ISL_3223505 | Lighthouse Laboratory Plymouth | Wellcome Sanger Institute for the COVID-19 Genomics UK (COG-UK) Consortium | Lighthouse Laboratory Plymouth and Alex Alderton, Roberto Amato, Jeffrey Barrett, Sonia Goncalves, Ewan Harrison, David K. Jackson, Ian Johnston, Dominic Kwiatkowski, Cordelia Langford, John Sillitoe on behalf of the Wellcome Sanger Institute COVID-19 Surveillance Team |
| EPI_ISL_3223506 | Lighthouse Lab in Glasgow | Wellcome Sanger Institute for the COVID-19 Genomics UK (COG-UK) Consortium | Harper VanSteenhouse, Yumi Kasai, David Gray, Carol Clugston, Anna Dominiczak and Alex Alderton, Roberto Amato, Jeffrey Barrett, Sonia Goncalves, Ewan Harrison, David K. Jackson, Ian Johnston, Dominic Kwiatkowski, Cordelia Langford, John Sillitoe on behalf of the Wellcome Sanger Institute COVID-19 Surveillance Team |
| EPI_ISL_3223507, EPI_ISL_3223508 | Lighthouse Laboratory Plymouth | Wellcome Sanger Institute for the COVID-19 Genomics UK (COG-UK) Consortium | Lighthouse Laboratory Plymouth and Alex Alderton, Roberto Amato, Jeffrey Barrett, Sonia Goncalves, Ewan Harrison, David K. Jackson, Ian Johnston, Dominic Kwiatkowski, Cordelia Langford, John Sillitoe on behalf of the Wellcome Sanger Institute COVID-19 Surveillance Team |
| EPI_ISL_3223509 | Lighthouse Lab in Milton Keynes | Wellcome Sanger Institute for the COVID-19 Genomics UK (COG-UK) Consortium | The Lighthouse Lab in Milton Keynes and Alex Alderton, Roberto Amato, Jeffrey Barrett, Sonia Goncalves, Ewan Harrison, David K. Jackson, Ian Johnston, Dominic Kwiatkowski, Cordelia Langford, John Sillitoe on behalf of the Wellcome Sanger Institute COVID-19 Surveillance Team |
| EPI_ISL_3223510, EPI_ISL_3223511, EPI_ISL_3223512 | Lighthouse Laboratory Plymouth | Wellcome Sanger Institute for the COVID-19 Genomics UK (COG-UK) Consortium | Lighthouse Laboratory Plymouth and Alex Alderton, Roberto Amato, Jeffrey Barrett, Sonia Goncalves, Ewan Harrison, David K. Jackson, Ian Johnston, Dominic Kwiatkowski, Cordelia Langford, John Sillitoe on behalf of the Wellcome Sanger Institute COVID-19 Surveillance Team |
| EPI_ISL_3223513 | Lighthouse Lab in Glasgow | Wellcome Sanger Institute for the COVID-19 Genomics UK (COG-UK) Consortium | Harper VanSteenhouse, Yumi Kasai, David Gray, Carol Clugston, Anna Dominiczak and Alex Alderton, Roberto Amato, Jeffrey Barrett, Sonia Goncalves, Ewan Harrison, David K. Jackson, Ian Johnston, Dominic Kwiatkowski, Cordelia Langford, John Sillitoe on behalf of the Wellcome Sanger Institute COVID-19 Surveillance Team |
| EPI_ISL_3223514 | Lighthouse Lab in Milton Keynes | Wellcome Sanger Institute for the COVID-19 Genomics UK (COG-UK) Consortium | The Lighthouse Lab in Milton Keynes and Alex Alderton, Roberto Amato, Jeffrey Barrett, Sonia Goncalves, Ewan Harrison, David K. Jackson, Ian Johnston, Dominic Kwiatkowski, Cordelia Langford, John Sillitoe on behalf of the Wellcome Sanger Institute COVID-19 Surveillance Team |
| EPI_ISL_3223515 | Lighthouse Lab in Glasgow | Wellcome Sanger Institute for the COVID-19 Genomics UK (COG-UK) Consortium | Harper VanSteenhouse, Yumi Kasai, David Gray, Carol Clugston, Anna Dominiczak and Alex Alderton, Roberto Amato, Jeffrey Barrett, Sonia Goncalves, Ewan Harrison, David K. Jackson, Ian Johnston, Dominic Kwiatkowski, Cordelia Langford, John Sillitoe on behalf of the Wellcome Sanger Institute COVID-19 Surveillance Team |
| EPI_ISL_3223516, EPI_ISL_3223517 | Lighthouse Laboratory Plymouth | Wellcome Sanger Institute for the COVID-19 Genomics UK (COG-UK) Consortium | Lighthouse Laboratory Plymouth and Alex Alderton, Roberto Amato, Jeffrey Barrett, Sonia Goncalves, Ewan Harrison, David K. Jackson, Ian Johnston, Dominic Kwiatkowski, Cordelia Langford, John Sillitoe on behalf of the Wellcome Sanger Institute COVID-19 Surveillance Team |
| EPI_ISL_3223518 | Lighthouse Lab in Milton Keynes | Wellcome Sanger Institute for the COVID-19 Genomics UK (COG-UK) Consortium | The Lighthouse Lab in Milton Keynes and Alex Alderton, Roberto Amato, Jeffrey Barrett, Sonia Goncalves, Ewan Harrison, David K. Jackson, Ian Johnston, Dominic Kwiatkowski, Cordelia Langford, John Sillitoe on behalf of the Wellcome Sanger Institute COVID-19 Surveillance Team |
| EPI_ISL_3223519 | Lighthouse Laboratory Plymouth | Wellcome Sanger Institute for the COVID-19 Genomics UK (COG-UK) Consortium | Lighthouse Laboratory Plymouth and Alex Alderton, Roberto Amato, Jeffrey Barrett, Sonia Goncalves, Ewan Harrison, David K. Jackson, Ian Johnston, Dominic Kwiatkowski, Cordelia Langford, John Sillitoe on behalf of the Wellcome Sanger Institute COVID-19 Surveillance Team |
| EPI_ISL_3223520 | Lighthouse Lab in Milton Keynes | Wellcome Sanger Institute for the COVID-19 Genomics UK (COG-UK) Consortium | The Lighthouse Lab in Milton Keynes and Alex Alderton, Roberto Amato, Jeffrey Barrett, Sonia Goncalves, Ewan Harrison, David K. Jackson, Ian Johnston, Dominic Kwiatkowski, Cordelia Langford, John Sillitoe on behalf of the Wellcome Sanger Institute COVID-19 Surveillance Team |
| EPI_ISL_3223521 | Lighthouse Laboratory Plymouth | Wellcome Sanger Institute for the COVID-19 Genomics UK (COG-UK) Consortium | Lighthouse Laboratory Plymouth and Alex Alderton, Roberto Amato, Jeffrey Barrett, Sonia Goncalves, Ewan Harrison, David K. Jackson, Ian Johnston, Dominic Kwiatkowski, Cordelia Langford, John Sillitoe on behalf of the Wellcome Sanger Institute COVID-19 Surveillance Team |
| EPI_ISL_3223522, EPI_ISL_3223523 | Lighthouse Lab in Glasgow | Wellcome Sanger Institute for the COVID-19 Genomics UK (COG-UK) Consortium | Harper VanSteenhouse, Yumi Kasai, David Gray, Carol Clugston, Anna Dominiczak and Alex Alderton, Roberto Amato, Jeffrey Barrett, Sonia Goncalves, Ewan Harrison, David K. Jackson, Ian Johnston, Dominic Kwiatkowski, Cordelia Langford, John Sillitoe on behalf of the Wellcome Sanger Institute COVID-19 Surveillance Team |
| EPI_ISL_3223524 | Lighthouse Lab in Milton Keynes | Wellcome Sanger Institute for the COVID-19 Genomics UK (COG-UK) Consortium | The Lighthouse Lab in Milton Keynes and Alex Alderton, Roberto Amato, Jeffrey Barrett, Sonia Goncalves, Ewan Harrison, David K. Jackson, Ian Johnston, Dominic Kwiatkowski, Cordelia Langford, John Sillitoe on behalf of the Wellcome Sanger Institute COVID-19 Surveillance Team |
| EPI_ISL_3223525 | Lighthouse Laboratory Plymouth | Wellcome Sanger Institute for the COVID-19 Genomics UK (COG-UK) Consortium | Lighthouse Laboratory Plymouth and Alex Alderton, Roberto Amato, Jeffrey Barrett, Sonia Goncalves, Ewan Harrison, David K. Jackson, Ian Johnston, Dominic Kwiatkowski, Cordelia Langford, John Sillitoe on behalf of the Wellcome Sanger Institute COVID-19 Surveillance Team |
| EPI_ISL_3223526 | Lighthouse Lab in Glasgow | Wellcome Sanger Institute for the COVID-19 Genomics UK (COG-UK) Consortium | Harper VanSteenhouse, Yumi Kasai, David Gray, Carol Clugston, Anna Dominiczak and Alex Alderton, Roberto Amato, Jeffrey Barrett, Sonia Goncalves, Ewan Harrison, David K. Jackson, Ian Johnston, Dominic Kwiatkowski, Cordelia Langford, John Sillitoe on behalf of the Wellcome Sanger Institute COVID-19 Surveillance Team |
| EPI_ISL_3223527 | Lighthouse Lab in Milton Keynes | Wellcome Sanger Institute for the COVID-19 Genomics UK (COG-UK) Consortium | The Lighthouse Lab in Milton Keynes and Alex Alderton, Roberto Amato, Jeffrey Barrett, Sonia Goncalves, Ewan Harrison, David K. Jackson, Ian Johnston, Dominic Kwiatkowski, Cordelia Langford, John Sillitoe on behalf of the Wellcome Sanger Institute COVID-19 Surveillance Team |
| EPI_ISL_3223528 | Lighthouse Laboratory Plymouth | Wellcome Sanger Institute for the COVID-19 Genomics UK (COG-UK) Consortium | Lighthouse Laboratory Plymouth and Alex Alderton, Roberto Amato, Jeffrey Barrett, Sonia Goncalves, Ewan Harrison, David K. Jackson, Ian Johnston, Dominic Kwiatkowski, Cordelia Langford, John Sillitoe on behalf of the Wellcome Sanger Institute COVID-19 Surveillance Team |
| EPI_ISL_3223529 | Lighthouse Lab in Milton Keynes | Wellcome Sanger Institute for the COVID-19 Genomics UK (COG-UK) Consortium | The Lighthouse Lab in Milton Keynes and Alex Alderton, Roberto Amato, Jeffrey Barrett, Sonia Goncalves, Ewan Harrison, David K. Jackson, Ian Johnston, Dominic Kwiatkowski, Cordelia Langford, John Sillitoe on behalf of the Wellcome Sanger Institute COVID-19 Surveillance Team |
| EPI_ISL_3223530, EPI_ISL_3223531, EPI_ISL_3223532 | Lighthouse Laboratory Plymouth | Wellcome Sanger Institute for the COVID-19 Genomics UK (COG-UK) Consortium | Lighthouse Laboratory Plymouth and Alex Alderton, Roberto Amato, Jeffrey Barrett, Sonia Goncalves, Ewan Harrison, David K. Jackson, Ian Johnston, Dominic Kwiatkowski, Cordelia Langford, John Sillitoe on behalf of the Wellcome Sanger Institute COVID-19 Surveillance Team |
| EPI_ISL_3223533 | Lighthouse Lab in Glasgow | Wellcome Sanger Institute for the COVID-19 Genomics UK (COG-UK) Consortium | Harper VanSteenhouse, Yumi Kasai, David Gray, Carol Clugston, Anna Dominiczak and Alex Alderton, Roberto Amato, Jeffrey Barrett, Sonia Goncalves, Ewan Harrison, David K. Jackson, Ian Johnston, Dominic Kwiatkowski, Cordelia Langford, John Sillitoe on behalf of the Wellcome Sanger Institute COVID-19 Surveillance Team |
| EPI_ISL_3223534 | Lighthouse Lab in Milton Keynes | Wellcome Sanger Institute for the COVID-19 Genomics UK (COG-UK) Consortium | The Lighthouse Lab in Milton Keynes and Alex Alderton, Roberto Amato, Jeffrey Barrett, Sonia Goncalves, Ewan Harrison, David K. Jackson, Ian Johnston, Dominic Kwiatkowski, Cordelia Langford, John Sillitoe on behalf of the Wellcome Sanger Institute COVID-19 Surveillance Team |
| EPI_ISL_3223535 | Lighthouse Laboratory Plymouth | Wellcome Sanger Institute for the COVID-19 Genomics UK (COG-UK) Consortium | Lighthouse Laboratory Plymouth and Alex Alderton, Roberto Amato, Jeffrey Barrett, Sonia Goncalves, Ewan Harrison, David K. Jackson, Ian Johnston, Dominic Kwiatkowski, Cordelia Langford, John Sillitoe on behalf of the Wellcome Sanger Institute COVID-19 Surveillance Team |
| EPI_ISL_3223536, EPI_ISL_3223537 | Lighthouse Lab in Milton Keynes | Wellcome Sanger Institute for the COVID-19 Genomics UK (COG-UK) Consortium | The Lighthouse Lab in Milton Keynes and Alex Alderton, Roberto Amato, Jeffrey Barrett, Sonia Goncalves, Ewan Harrison, David K. Jackson, Ian Johnston, Dominic Kwiatkowski, Cordelia Langford, John Sillitoe on behalf of the Wellcome Sanger Institute COVID-19 Surveillance Team |
| EPI_ISL_3223538, EPI_ISL_3223539, EPI_ISL_3223540, EPI_ISL_3223541 | Lighthouse Laboratory Plymouth | Wellcome Sanger Institute for the COVID-19 Genomics UK (COG-UK) Consortium | Lighthouse Laboratory Plymouth and Alex Alderton, Roberto Amato, Jeffrey Barrett, Sonia Goncalves, Ewan Harrison, David K. Jackson, Ian Johnston, Dominic Kwiatkowski, Cordelia Langford, John Sillitoe on behalf of the Wellcome Sanger Institute COVID-19 Surveillance Team |
| EPI_ISL_3223542 | Lighthouse Lab in Milton Keynes | Wellcome Sanger Institute for the COVID-19 Genomics UK (COG-UK) Consortium | The Lighthouse Lab in Milton Keynes and Alex Alderton, Roberto Amato, Jeffrey Barrett, Sonia Goncalves, Ewan Harrison, David K. Jackson, Ian Johnston, Dominic Kwiatkowski, Cordelia Langford, John Sillitoe on behalf of the Wellcome Sanger Institute COVID-19 Surveillance Team |
| EPI_ISL_3223543 | Lighthouse Laboratory Plymouth | Wellcome Sanger Institute for the COVID-19 Genomics UK (COG-UK) Consortium | Lighthouse Laboratory Plymouth and Alex Alderton, Roberto Amato, Jeffrey Barrett, Sonia Goncalves, Ewan Harrison, David K. Jackson, Ian Johnston, Dominic Kwiatkowski, Cordelia Langford, John Sillitoe on behalf of the Wellcome Sanger Institute COVID-19 Surveillance Team |
| EPI_ISL_3223544 | Lighthouse Lab in Milton Keynes | Wellcome Sanger Institute for the COVID-19 Genomics UK (COG-UK) Consortium | The Lighthouse Lab in Milton Keynes and Alex Alderton, Roberto Amato, Jeffrey Barrett, Sonia Goncalves, Ewan Harrison, David K. Jackson, Ian Johnston, Dominic Kwiatkowski, Cordelia Langford, John Sillitoe on behalf of the Wellcome Sanger Institute COVID-19 Surveillance Team |
| EPI_ISL_3223545, EPI_ISL_3223546, EPI_ISL_3223547 | Lighthouse Laboratory Plymouth | Wellcome Sanger Institute for the COVID-19 Genomics UK (COG-UK) Consortium | Lighthouse Laboratory Plymouth and Alex Alderton, Roberto Amato, Jeffrey Barrett, Sonia Goncalves, Ewan Harrison, David K. Jackson, Ian Johnston, Dominic Kwiatkowski, Cordelia Langford, John Sillitoe on behalf of the Wellcome Sanger Institute COVID-19 Surveillance Team |
| EPI_ISL_3223548, EPI_ISL_3223549 | Lighthouse Lab in Glasgow | Wellcome Sanger Institute for the COVID-19 Genomics UK (COG-UK) Consortium | Harper VanSteenhouse, Yumi Kasai, David Gray, Carol Clugston, Anna Dominiczak and Alex Alderton, Roberto Amato, Jeffrey Barrett, Sonia Goncalves, Ewan Harrison, David K. Jackson, Ian Johnston, Dominic Kwiatkowski, Cordelia Langford, John Sillitoe on behalf of the Wellcome Sanger Institute COVID-19 Surveillance Team |
| EPI_ISL_3223550 | Lighthouse Laboratory Plymouth | Wellcome Sanger Institute for the COVID-19 Genomics UK (COG-UK) Consortium | Lighthouse Laboratory Plymouth and Alex Alderton, Roberto Amato, Jeffrey Barrett, Sonia Goncalves, Ewan Harrison, David K. Jackson, Ian Johnston, Dominic Kwiatkowski, Cordelia Langford, John Sillitoe on behalf of the Wellcome Sanger Institute COVID-19 Surveillance Team |
| EPI_ISL_3223551 | Lighthouse Lab in Milton Keynes | Wellcome Sanger Institute for the COVID-19 Genomics UK (COG-UK) Consortium | The Lighthouse Lab in Milton Keynes and Alex Alderton, Roberto Amato, Jeffrey Barrett, Sonia Goncalves, Ewan Harrison, David K. Jackson, Ian Johnston, Dominic Kwiatkowski, Cordelia Langford, John Sillitoe on behalf of the Wellcome Sanger Institute COVID-19 Surveillance Team |
| EPI_ISL_3223552 | Lighthouse Lab in Glasgow | Wellcome Sanger Institute for the COVID-19 Genomics UK (COG-UK) | Harper VanSteenhouse, Yumi Kasai, David Gray, Carol Clugston, Anna Dominiczak and Alex Alderton, Roberto Amato, Jeffrey Barrett, Sonia Goncalves, Ewan Harrison, David K. Jackson, Ian Johnston, Dominic Kwiatkowski, Cordelia Langford, |

| | | | |
|---|---|---|---|
| | | Consortium | John Sillitoe on behalf of the Wellcome Sanger Institute COVID-19 Surveillance Team |
| EPI_ISL_3223553 | Lighthouse Lab in Milton Keynes | Wellcome Sanger Institute for the COVID-19 Genomics UK (COG-UK) Consortium | The Lighthouse Lab in Milton Keynes and Alex Alderton, Roberto Amato, Jeffrey Barrett, Sonia Goncalves, Ewan Harrison, David K. Jackson, Ian Johnston, Dominic Kwiatkowski, Cordelia Langford, John Sillitoe on behalf of the Wellcome Sanger Institute COVID-19 Surveillance Team |
| EPI_ISL_3223554 | Lighthouse Laboratory Plymouth | Wellcome Sanger Institute for the COVID-19 Genomics UK (COG-UK) Consortium | Lighthouse Laboratory Plymouth and Alex Alderton, Roberto Amato, Jeffrey Barrett, Sonia Goncalves, Ewan Harrison, David K. Jackson, Ian Johnston, Dominic Kwiatkowski, Cordelia Langford, John Sillitoe on behalf of the Wellcome Sanger Institute COVID-19 Surveillance Team |
| EPI_ISL_3223555 | Lighthouse Lab in Glasgow | Wellcome Sanger Institute for the COVID-19 Genomics UK (COG-UK) Consortium | Harper VanSteenhouse, Yumi Kasai, David Gray, Carol Clugston, Anna Dominiczak and Alex Alderton, Roberto Amato, Jeffrey Barrett, Sonia Goncalves, Ewan Harrison, David K. Jackson, Ian Johnston, Dominic Kwiatkowski, Cordelia Langford, John Sillitoe on behalf of the Wellcome Sanger Institute COVID-19 Surveillance Team |
| EPI_ISL_3223556, EPI_ISL_3223557, EPI_ISL_3223558, EPI_ISL_3223559 | Lighthouse Laboratory Plymouth | Wellcome Sanger Institute for the COVID-19 Genomics UK (COG-UK) Consortium | Lighthouse Laboratory Plymouth and Alex Alderton, Roberto Amato, Jeffrey Barrett, Sonia Goncalves, Ewan Harrison, David K. Jackson, Ian Johnston, Dominic Kwiatkowski, Cordelia Langford, John Sillitoe on behalf of the Wellcome Sanger Institute COVID-19 Surveillance Team |
| EPI_ISL_3223560 | Lighthouse Lab in Milton Keynes | Wellcome Sanger Institute for the COVID-19 Genomics UK (COG-UK) Consortium | The Lighthouse Lab in Milton Keynes and Alex Alderton, Roberto Amato, Jeffrey Barrett, Sonia Goncalves, Ewan Harrison, David K. Jackson, Ian Johnston, Dominic Kwiatkowski, Cordelia Langford, John Sillitoe on behalf of the Wellcome Sanger Institute COVID-19 Surveillance Team |
| EPI_ISL_3223561 | Lighthouse Lab in Glasgow | Wellcome Sanger Institute for the COVID-19 Genomics UK (COG-UK) Consortium | Harper VanSteenhouse, Yumi Kasai, David Gray, Carol Clugston, Anna Dominiczak and Alex Alderton, Roberto Amato, Jeffrey Barrett, Sonia Goncalves, Ewan Harrison, David K. Jackson, Ian Johnston, Dominic Kwiatkowski, Cordelia Langford, John Sillitoe on behalf of the Wellcome Sanger Institute COVID-19 Surveillance Team |
| EPI_ISL_3223562, EPI_ISL_3223563 | Lighthouse Laboratory Plymouth | Wellcome Sanger Institute for the COVID-19 Genomics UK (COG-UK) Consortium | Lighthouse Laboratory Plymouth and Alex Alderton, Roberto Amato, Jeffrey Barrett, Sonia Goncalves, Ewan Harrison, David K. Jackson, Ian Johnston, Dominic Kwiatkowski, Cordelia Langford, John Sillitoe on behalf of the Wellcome Sanger Institute COVID-19 Surveillance Team |
| EPI_ISL_3223564, EPI_ISL_3223565 | Lighthouse Lab in Glasgow | Wellcome Sanger Institute for the COVID-19 Genomics UK (COG-UK) Consortium | Harper VanSteenhouse, Yumi Kasai, David Gray, Carol Clugston, Anna Dominiczak and Alex Alderton, Roberto Amato, Jeffrey Barrett, Sonia Goncalves, Ewan Harrison, David K. Jackson, Ian Johnston, Dominic Kwiatkowski, Cordelia Langford, John Sillitoe on behalf of the Wellcome Sanger Institute COVID-19 Surveillance Team |
| EPI_ISL_3223566, EPI_ISL_3223567 | Lighthouse Laboratory Plymouth | Wellcome Sanger Institute for the COVID-19 Genomics UK (COG-UK) Consortium | Lighthouse Laboratory Plymouth and Alex Alderton, Roberto Amato, Jeffrey Barrett, Sonia Goncalves, Ewan Harrison, David K. Jackson, Ian Johnston, Dominic Kwiatkowski, Cordelia Langford, John Sillitoe on behalf of the Wellcome Sanger Institute COVID-19 Surveillance Team |
| EPI_ISL_3223568, EPI_ISL_3223569 | Lighthouse Lab in Milton Keynes | Wellcome Sanger Institute for the COVID-19 Genomics UK (COG-UK) Consortium | The Lighthouse Lab in Milton Keynes and Alex Alderton, Roberto Amato, Jeffrey Barrett, Sonia Goncalves, Ewan Harrison, David K. Jackson, Ian Johnston, Dominic Kwiatkowski, Cordelia Langford, John Sillitoe on behalf of the Wellcome Sanger Institute COVID-19 Surveillance Team |
| EPI_ISL_3223570, EPI_ISL_3223571 | Lighthouse Laboratory Plymouth | Wellcome Sanger Institute for the COVID-19 Genomics UK (COG-UK) Consortium | Lighthouse Laboratory Plymouth and Alex Alderton, Roberto Amato, Jeffrey Barrett, Sonia Goncalves, Ewan Harrison, David K. Jackson, Ian Johnston, Dominic Kwiatkowski, Cordelia Langford, John Sillitoe on behalf of the Wellcome Sanger Institute COVID-19 Surveillance Team |
| EPI_ISL_3223572 | Lighthouse Lab in Glasgow | Wellcome Sanger Institute for the COVID-19 Genomics UK (COG-UK) Consortium | Harper VanSteenhouse, Yumi Kasai, David Gray, Carol Clugston, Anna Dominiczak and Alex Alderton, Roberto Amato, Jeffrey Barrett, Sonia Goncalves, Ewan Harrison, David K. Jackson, Ian Johnston, Dominic Kwiatkowski, Cordelia Langford, John Sillitoe on behalf of the Wellcome Sanger Institute COVID-19 Surveillance Team |
| EPI_ISL_3223573 | Lighthouse Laboratory Plymouth | Wellcome Sanger Institute for the COVID-19 Genomics UK (COG-UK) Consortium | Lighthouse Laboratory Plymouth and Alex Alderton, Roberto Amato, Jeffrey Barrett, Sonia Goncalves, Ewan Harrison, David K. Jackson, Ian Johnston, Dominic Kwiatkowski, Cordelia Langford, John Sillitoe on behalf of the Wellcome Sanger Institute COVID-19 Surveillance Team |
| EPI_ISL_3223574 | Lighthouse Lab in Glasgow | Wellcome Sanger Institute for the COVID-19 Genomics UK (COG-UK) Consortium | Harper VanSteenhouse, Yumi Kasai, David Gray, Carol Clugston, Anna Dominiczak and Alex Alderton, Roberto Amato, Jeffrey Barrett, Sonia Goncalves, Ewan Harrison, David K. Jackson, Ian Johnston, Dominic Kwiatkowski, Cordelia Langford, John Sillitoe on behalf of the Wellcome Sanger Institute COVID-19 Surveillance Team |
| EPI_ISL_3223575, EPI_ISL_3223576 | Lighthouse Laboratory Plymouth | Wellcome Sanger Institute for the COVID-19 Genomics UK (COG-UK) Consortium | Lighthouse Laboratory Plymouth and Alex Alderton, Roberto Amato, Jeffrey Barrett, Sonia Goncalves, Ewan Harrison, David K. Jackson, Ian Johnston, Dominic Kwiatkowski, Cordelia Langford, John Sillitoe on behalf of the Wellcome Sanger Institute COVID-19 Surveillance Team |
| EPI_ISL_3223577 | Lighthouse Lab in Glasgow | Wellcome Sanger Institute for the COVID-19 Genomics UK (COG-UK) Consortium | Harper VanSteenhouse, Yumi Kasai, David Gray, Carol Clugston, Anna Dominiczak and Alex Alderton, Roberto Amato, Jeffrey Barrett, Sonia Goncalves, Ewan Harrison, David K. Jackson, Ian Johnston, Dominic Kwiatkowski, Cordelia Langford, John Sillitoe on behalf of the Wellcome Sanger Institute COVID-19 Surveillance Team |
| EPI_ISL_3223578, EPI_ISL_3223579, EPI_ISL_3223580, EPI_ISL_3223581, EPI_ISL_3223582 | Lighthouse Laboratory Plymouth | Wellcome Sanger Institute for the COVID-19 Genomics UK (COG-UK) Consortium | Lighthouse Laboratory Plymouth and Alex Alderton, Roberto Amato, Jeffrey Barrett, Sonia Goncalves, Ewan Harrison, David K. Jackson, Ian Johnston, Dominic Kwiatkowski, Cordelia Langford, John Sillitoe on behalf of the Wellcome Sanger Institute COVID-19 Surveillance Team |
| EPI_ISL_3223583, EPI_ISL_3223584 | Lighthouse Lab in Glasgow | Wellcome Sanger Institute for the COVID-19 Genomics UK (COG-UK) Consortium | Harper VanSteenhouse, Yumi Kasai, David Gray, Carol Clugston, Anna Dominiczak and Alex Alderton, Roberto Amato, Jeffrey Barrett, Sonia Goncalves, Ewan Harrison, David K. Jackson, Ian Johnston, Dominic Kwiatkowski, Cordelia Langford, John Sillitoe on behalf of the Wellcome Sanger Institute COVID-19 Surveillance Team |
| EPI_ISL_3223585, EPI_ISL_3223586 | Lighthouse Lab in Milton Keynes | Wellcome Sanger Institute for the COVID-19 Genomics UK (COG-UK) Consortium | The Lighthouse Lab in Milton Keynes and Alex Alderton, Roberto Amato, Jeffrey Barrett, Sonia Goncalves, Ewan Harrison, David K. Jackson, Ian Johnston, Dominic Kwiatkowski, Cordelia Langford, John Sillitoe on behalf of the Wellcome Sanger Institute COVID-19 Surveillance Team |
| EPI_ISL_3223587, EPI_ISL_3223588, EPI_ISL_3223589 | Lighthouse Laboratory Plymouth | Wellcome Sanger Institute for the COVID-19 Genomics UK (COG-UK) Consortium | Lighthouse Laboratory Plymouth and Alex Alderton, Roberto Amato, Jeffrey Barrett, Sonia Goncalves, Ewan Harrison, David K. Jackson, Ian Johnston, Dominic Kwiatkowski, Cordelia Langford, John Sillitoe on behalf of the Wellcome Sanger Institute COVID-19 Surveillance Team |
| EPI_ISL_3223590 | Lighthouse Lab in Milton Keynes | Wellcome Sanger Institute for the COVID-19 Genomics UK (COG-UK) Consortium | The Lighthouse Lab in Milton Keynes and Alex Alderton, Roberto Amato, Jeffrey Barrett, Sonia Goncalves, Ewan Harrison, David K. Jackson, Ian Johnston, Dominic Kwiatkowski, Cordelia Langford, John Sillitoe on behalf of the Wellcome Sanger Institute COVID-19 Surveillance Team |
| EPI_ISL_3223591 | Lighthouse Lab in Glasgow | Wellcome Sanger Institute for the COVID-19 Genomics UK (COG-UK) Consortium | Harper VanSteenhouse, Yumi Kasai, David Gray, Carol Clugston, Anna Dominiczak and Alex Alderton, Roberto Amato, Jeffrey Barrett, Sonia Goncalves, Ewan Harrison, David K. Jackson, Ian Johnston, Dominic Kwiatkowski, Cordelia Langford, John Sillitoe on behalf of the Wellcome Sanger Institute COVID-19 Surveillance Team |
| EPI_ISL_3223592 | Lighthouse Lab in Milton Keynes | Wellcome Sanger Institute for the COVID-19 Genomics UK (COG-UK) Consortium | The Lighthouse Lab in Milton Keynes and Alex Alderton, Roberto Amato, Jeffrey Barrett, Sonia Goncalves, Ewan Harrison, David K. Jackson, Ian Johnston, Dominic Kwiatkowski, Cordelia Langford, John Sillitoe on behalf of the Wellcome Sanger Institute COVID-19 Surveillance Team |
| EPI_ISL_3223593 | Lighthouse Laboratory Plymouth | Wellcome Sanger Institute for the COVID-19 Genomics UK (COG-UK) Consortium | Lighthouse Laboratory Plymouth and Alex Alderton, Roberto Amato, Jeffrey Barrett, Sonia Goncalves, Ewan Harrison, David K. Jackson, Ian Johnston, Dominic Kwiatkowski, Cordelia Langford, John Sillitoe on behalf of the Wellcome Sanger Institute COVID-19 Surveillance Team |
| EPI_ISL_3223594 | Lighthouse Lab in Glasgow | Wellcome Sanger Institute for the COVID-19 Genomics UK (COG-UK) Consortium | Harper VanSteenhouse, Yumi Kasai, David Gray, Carol Clugston, Anna Dominiczak and Alex Alderton, Roberto Amato, Jeffrey Barrett, Sonia Goncalves, Ewan Harrison, David K. Jackson, Ian Johnston, Dominic Kwiatkowski, Cordelia Langford, John Sillitoe on behalf of the Wellcome Sanger Institute COVID-19 Surveillance Team |
| EPI_ISL_3223595, EPI_ISL_3223596, EPI_ISL_3223597, EPI_ISL_3223598 | Lighthouse Laboratory Plymouth | Wellcome Sanger Institute for the COVID-19 Genomics UK (COG-UK) Consortium | Lighthouse Laboratory Plymouth and Alex Alderton, Roberto Amato, Jeffrey Barrett, Sonia Goncalves, Ewan Harrison, David K. Jackson, Ian Johnston, Dominic Kwiatkowski, Cordelia Langford, John Sillitoe on behalf of the Wellcome Sanger Institute COVID-19 Surveillance Team |
| EPI_ISL_3223599, EPI_ISL_3223600 | Lighthouse Lab in Glasgow | Wellcome Sanger Institute for the COVID-19 Genomics UK (COG-UK) Consortium | Harper VanSteenhouse, Yumi Kasai, David Gray, Carol Clugston, Anna Dominiczak and Alex Alderton, Roberto Amato, Jeffrey Barrett, Sonia Goncalves, Ewan Harrison, David K. Jackson, Ian Johnston, Dominic Kwiatkowski, Cordelia Langford, John Sillitoe on behalf of the Wellcome Sanger Institute COVID-19 Surveillance Team |
| EPI_ISL_3223601 | Lighthouse Lab in Milton Keynes | Wellcome Sanger Institute for the COVID-19 Genomics UK (COG-UK) Consortium | The Lighthouse Lab in Milton Keynes and Alex Alderton, Roberto Amato, Jeffrey Barrett, Sonia Goncalves, Ewan Harrison, David K. Jackson, Ian Johnston, Dominic Kwiatkowski, Cordelia Langford, John Sillitoe on behalf of the Wellcome Sanger Institute COVID-19 Surveillance Team |
| EPI_ISL_3223602, EPI_ISL_3223603 | Lighthouse Laboratory Plymouth | Wellcome Sanger Institute for the COVID-19 Genomics UK (COG-UK) Consortium | Lighthouse Laboratory Plymouth and Alex Alderton, Roberto Amato, Jeffrey Barrett, Sonia Goncalves, Ewan Harrison, David K. Jackson, Ian Johnston, Dominic Kwiatkowski, Cordelia Langford, John Sillitoe on behalf of the Wellcome Sanger Institute COVID-19 Surveillance Team |
| EPI_ISL_3223604 | Lighthouse Lab in Glasgow | Wellcome Sanger Institute for the COVID-19 Genomics UK (COG-UK) Consortium | Harper VanSteenhouse, Yumi Kasai, David Gray, Carol Clugston, Anna Dominiczak and Alex Alderton, Roberto Amato, Jeffrey Barrett, Sonia Goncalves, Ewan Harrison, David K. Jackson, Ian Johnston, Dominic Kwiatkowski, Cordelia Langford, John Sillitoe on behalf of the Wellcome Sanger Institute COVID-19 Surveillance Team |
| EPI_ISL_3223605 | Lighthouse Laboratory Plymouth | Wellcome Sanger Institute for the COVID-19 Genomics UK (COG-UK) Consortium | Lighthouse Laboratory Plymouth and Alex Alderton, Roberto Amato, Jeffrey Barrett, Sonia Goncalves, Ewan Harrison, David K. Jackson, Ian Johnston, Dominic Kwiatkowski, Cordelia Langford, John Sillitoe on behalf of the Wellcome Sanger Institute COVID-19 Surveillance Team |
| EPI_ISL_3223606 | Lighthouse Lab in Glasgow | Wellcome Sanger Institute for the COVID-19 Genomics UK (COG-UK) Consortium | Harper VanSteenhouse, Yumi Kasai, David Gray, Carol Clugston, Anna Dominiczak and Alex Alderton, Roberto Amato, Jeffrey Barrett, Sonia Goncalves, Ewan Harrison, David K. Jackson, Ian Johnston, Dominic Kwiatkowski, Cordelia Langford, John Sillitoe on behalf of the Wellcome Sanger Institute COVID-19 Surveillance Team |
| EPI_ISL_3223607, EPI_ISL_3223608 | Lighthouse Lab in Milton Keynes | Wellcome Sanger Institute for the COVID-19 Genomics UK (COG-UK) Consortium | The Lighthouse Lab in Milton Keynes and Alex Alderton, Roberto Amato, Jeffrey Barrett, Sonia Goncalves, Ewan Harrison, David K. Jackson, Ian Johnston, Dominic Kwiatkowski, Cordelia Langford, John Sillitoe on behalf of the Wellcome Sanger Institute COVID-19 Surveillance Team |
| EPI_ISL_3223609 | Lighthouse Lab in Glasgow | Wellcome Sanger Institute for the COVID-19 Genomics UK (COG-UK) Consortium | Harper VanSteenhouse, Yumi Kasai, David Gray, Carol Clugston, Anna Dominiczak and Alex Alderton, Roberto Amato, Jeffrey Barrett, Sonia Goncalves, Ewan Harrison, David K. Jackson, Ian Johnston, Dominic Kwiatkowski, Cordelia Langford, John Sillitoe on behalf of the Wellcome Sanger Institute COVID-19 Surveillance Team |
| EPI_ISL_3223610, EPI_ISL_3223611, EPI_ISL_3223612 | Lighthouse Laboratory Plymouth | Wellcome Sanger Institute for the COVID-19 Genomics UK (COG-UK) Consortium | Lighthouse Laboratory Plymouth and Alex Alderton, Roberto Amato, Jeffrey Barrett, Sonia Goncalves, Ewan Harrison, David K. Jackson, Ian Johnston, Dominic Kwiatkowski, Cordelia Langford, John Sillitoe on behalf of the Wellcome Sanger Institute COVID-19 Surveillance Team |
| EPI_ISL_3223613, EPI_ISL_3223614 | Lighthouse Lab in Alderley Park | Wellcome Sanger Institute for the COVID-19 Genomics UK (COG-UK) Consortium | Jacquelyn Wynn, Mairead Hyland, The Lighthouse Lab in Alderley Park and Alex Alderton, Roberto Amato, Jeffrey Barrett, Sonia Goncalves, Ewan Harrison, David K. Jackson, Ian Johnston, Dominic Kwiatkowski, Cordelia Langford, John Sillitoe on behalf of the Wellcome Sanger Institute COVID-19 Surveillance Team |
| EPI_ISL_3223615 | Lighthouse Lab in Glasgow | Wellcome Sanger Institute for the COVID-19 Genomics UK (COG-UK) Consortium | Harper VanSteenhouse, Yumi Kasai, David Gray, Carol Clugston, Anna Dominiczak and Alex Alderton, Roberto Amato, Jeffrey Barrett, Sonia Goncalves, Ewan Harrison, David K. Jackson, Ian Johnston, Dominic Kwiatkowski, Cordelia Langford, John Sillitoe on behalf of the Wellcome Sanger Institute COVID-19 Surveillance Team |
| EPI_ISL_3223616 | Lighthouse Lab in Alderley Park | Wellcome Sanger Institute for the COVID-19 Genomics UK (COG-UK) Consortium | Jacquelyn Wynn, Mairead Hyland, The Lighthouse Lab in Alderley Park and Alex Alderton, Roberto Amato, Jeffrey Barrett, Sonia Goncalves, Ewan Harrison, David K. Jackson, Ian Johnston, Dominic Kwiatkowski, Cordelia Langford, John Sillitoe on behalf of the Wellcome Sanger Institute COVID-19 Surveillance Team |
| EPI_ISL_3223617 | Lighthouse Lab in Milton Keynes | Wellcome Sanger Institute for the COVID-19 Genomics UK (COG-UK) Consortium | The Lighthouse Lab in Milton Keynes and Alex Alderton, Roberto Amato, Jeffrey Barrett, Sonia Goncalves, Ewan Harrison, David K. Jackson, Ian Johnston, Dominic Kwiatkowski, Cordelia Langford, John Sillitoe on behalf of the Wellcome Sanger Institute COVID-19 Surveillance Team |
| EPI_ISL_3223618 | Lighthouse Laboratory Plymouth | Wellcome Sanger Institute for the COVID-19 Genomics UK (COG-UK) Consortium | Lighthouse Laboratory Plymouth and Alex Alderton, Roberto Amato, Jeffrey Barrett, Sonia Goncalves, Ewan Harrison, David K. Jackson, Ian Johnston, Dominic Kwiatkowski, Cordelia Langford, John Sillitoe on behalf of the Wellcome Sanger Institute COVID-19 Surveillance Team |
| EPI_ISL_3223619, EPI_ISL_3223620 | Lighthouse Lab in Glasgow | Wellcome Sanger Institute for the COVID-19 Genomics UK (COG-UK) Consortium | Harper VanSteenhouse, Yumi Kasai, David Gray, Carol Clugston, Anna Dominiczak and Alex Alderton, Roberto Amato, Jeffrey Barrett, Sonia Goncalves, Ewan Harrison, David K. Jackson, Ian Johnston, Dominic Kwiatkowski, Cordelia Langford, John Sillitoe on behalf of the Wellcome Sanger Institute COVID-19 Surveillance Team |

| | | | |
|---|---|---|---|
| EPI_ISL_3223621 | Lighthouse Lab in Alderley Park | Wellcome Sanger Institute for the COVID-19 Genomics UK (COG-UK) Consortium | Jacquelyn Wynn, Mairead Hyland, The Lighthouse Lab in Alderley Park and Alex Alderton, Roberto Amato, Jeffrey Barrett, Sonia Goncalves, Ewan Harrison, David K. Jackson, Ian Johnston, Dominic Kwiatkowski, Cordelia Langford, John Sillitoe on behalf of the Wellcome Sanger Institute COVID-19 Surveillance Team |
| EPI_ISL_3223622 | Lighthouse Laboratory Plymouth | Wellcome Sanger Institute for the COVID-19 Genomics UK (COG-UK) Consortium | Lighthouse Laboratory Plymouth and Alex Alderton, Roberto Amato, Jeffrey Barrett, Sonia Goncalves, Ewan Harrison, David K. Jackson, Ian Johnston, Dominic Kwiatkowski, Cordelia Langford, John Sillitoe on behalf of the Wellcome Sanger Institute COVID-19 Surveillance Team |
| EPI_ISL_3223623 | Lighthouse Lab in Alderley Park | Wellcome Sanger Institute for the COVID-19 Genomics UK (COG-UK) Consortium | Jacquelyn Wynn, Mairead Hyland, The Lighthouse Lab in Alderley Park and Alex Alderton, Roberto Amato, Jeffrey Barrett, Sonia Goncalves, Ewan Harrison, David K. Jackson, Ian Johnston, Dominic Kwiatkowski, Cordelia Langford, John Sillitoe on behalf of the Wellcome Sanger Institute COVID-19 Surveillance Team |
| EPI_ISL_3223624 | Lighthouse Laboratory Plymouth | Wellcome Sanger Institute for the COVID-19 Genomics UK (COG-UK) Consortium | Lighthouse Laboratory Plymouth and Alex Alderton, Roberto Amato, Jeffrey Barrett, Sonia Goncalves, Ewan Harrison, David K. Jackson, Ian Johnston, Dominic Kwiatkowski, Cordelia Langford, John Sillitoe on behalf of the Wellcome Sanger Institute COVID-19 Surveillance Team |
| EPI_ISL_3223625, EPI_ISL_3223626 | Lighthouse Lab in Alderley Park | Wellcome Sanger Institute for the COVID-19 Genomics UK (COG-UK) Consortium | Jacquelyn Wynn, Mairead Hyland, The Lighthouse Lab in Alderley Park and Alex Alderton, Roberto Amato, Jeffrey Barrett, Sonia Goncalves, Ewan Harrison, David K. Jackson, Ian Johnston, Dominic Kwiatkowski, Cordelia Langford, John Sillitoe on behalf of the Wellcome Sanger Institute COVID-19 Surveillance Team |
| EPI_ISL_3223627 | Lighthouse Laboratory Plymouth | Wellcome Sanger Institute for the COVID-19 Genomics UK (COG-UK) Consortium | Lighthouse Laboratory Plymouth and Alex Alderton, Roberto Amato, Jeffrey Barrett, Sonia Goncalves, Ewan Harrison, David K. Jackson, Ian Johnston, Dominic Kwiatkowski, Cordelia Langford, John Sillitoe on behalf of the Wellcome Sanger Institute COVID-19 Surveillance Team |
| EPI_ISL_3223628 | Lighthouse Lab in Glasgow | Wellcome Sanger Institute for the COVID-19 Genomics UK (COG-UK) Consortium | Harper VanSteenhouse, Yumi Kasai, David Gray, Carol Clugston, Anna Dominiczak and Alex Alderton, Roberto Amato, Jeffrey Barrett, Sonia Goncalves, Ewan Harrison, David K. Jackson, Ian Johnston, Dominic Kwiatkowski, Cordelia Langford, John Sillitoe on behalf of the Wellcome Sanger Institute COVID-19 Surveillance Team |
| EPI_ISL_3223629 | Lighthouse Laboratory Plymouth | Wellcome Sanger Institute for the COVID-19 Genomics UK (COG-UK) Consortium | Lighthouse Laboratory Plymouth and Alex Alderton, Roberto Amato, Jeffrey Barrett, Sonia Goncalves, Ewan Harrison, David K. Jackson, Ian Johnston, Dominic Kwiatkowski, Cordelia Langford, John Sillitoe on behalf of the Wellcome Sanger Institute COVID-19 Surveillance Team |
| EPI_ISL_3223630 | Lighthouse Lab in Glasgow | Wellcome Sanger Institute for the COVID-19 Genomics UK (COG-UK) Consortium | Harper VanSteenhouse, Yumi Kasai, David Gray, Carol Clugston, Anna Dominiczak and Alex Alderton, Roberto Amato, Jeffrey Barrett, Sonia Goncalves, Ewan Harrison, David K. Jackson, Ian Johnston, Dominic Kwiatkowski, Cordelia Langford, John Sillitoe on behalf of the Wellcome Sanger Institute COVID-19 Surveillance Team |
| EPI_ISL_3223631 | Lighthouse Laboratory Plymouth | Wellcome Sanger Institute for the COVID-19 Genomics UK (COG-UK) Consortium | Lighthouse Laboratory Plymouth and Alex Alderton, Roberto Amato, Jeffrey Barrett, Sonia Goncalves, Ewan Harrison, David K. Jackson, Ian Johnston, Dominic Kwiatkowski, Cordelia Langford, John Sillitoe on behalf of the Wellcome Sanger Institute COVID-19 Surveillance Team |
| EPI_ISL_3223632, EPI_ISL_3223633 | Lighthouse Lab in Glasgow | Wellcome Sanger Institute for the COVID-19 Genomics UK (COG-UK) Consortium | Harper VanSteenhouse, Yumi Kasai, David Gray, Carol Clugston, Anna Dominiczak and Alex Alderton, Roberto Amato, Jeffrey Barrett, Sonia Goncalves, Ewan Harrison, David K. Jackson, Ian Johnston, Dominic Kwiatkowski, Cordelia Langford, John Sillitoe on behalf of the Wellcome Sanger Institute COVID-19 Surveillance Team |
| EPI_ISL_3223634 | Lighthouse Lab in Milton Keynes | Wellcome Sanger Institute for the COVID-19 Genomics UK (COG-UK) Consortium | The Lighthouse Lab in Milton Keynes and Alex Alderton, Roberto Amato, Jeffrey Barrett, Sonia Goncalves, Ewan Harrison, David K. Jackson, Ian Johnston, Dominic Kwiatkowski, Cordelia Langford, John Sillitoe on behalf of the Wellcome Sanger Institute COVID-19 Surveillance Team |
| EPI_ISL_3223635 | Lighthouse Laboratory Plymouth | Wellcome Sanger Institute for the COVID-19 Genomics UK (COG-UK) Consortium | Lighthouse Laboratory Plymouth and Alex Alderton, Roberto Amato, Jeffrey Barrett, Sonia Goncalves, Ewan Harrison, David K. Jackson, Ian Johnston, Dominic Kwiatkowski, Cordelia Langford, John Sillitoe on behalf of the Wellcome Sanger Institute COVID-19 Surveillance Team |
| EPI_ISL_3223636 | Lighthouse Lab in Alderley Park | Wellcome Sanger Institute for the COVID-19 Genomics UK (COG-UK) Consortium | Jacquelyn Wynn, Mairead Hyland, The Lighthouse Lab in Alderley Park and Alex Alderton, Roberto Amato, Jeffrey Barrett, Sonia Goncalves, Ewan Harrison, David K. Jackson, Ian Johnston, Dominic Kwiatkowski, Cordelia Langford, John Sillitoe on behalf of the Wellcome Sanger Institute COVID-19 Surveillance Team |
| EPI_ISL_3223637, EPI_ISL_3223638 | Lighthouse Lab in Milton Keynes | Wellcome Sanger Institute for the COVID-19 Genomics UK (COG-UK) Consortium | The Lighthouse Lab in Milton Keynes and Alex Alderton, Roberto Amato, Jeffrey Barrett, Sonia Goncalves, Ewan Harrison, David K. Jackson, Ian Johnston, Dominic Kwiatkowski, Cordelia Langford, John Sillitoe on behalf of the Wellcome Sanger Institute COVID-19 Surveillance Team |
| EPI_ISL_3223639 | Lighthouse Laboratory Plymouth | Wellcome Sanger Institute for the COVID-19 Genomics UK (COG-UK) Consortium | Lighthouse Laboratory Plymouth and Alex Alderton, Roberto Amato, Jeffrey Barrett, Sonia Goncalves, Ewan Harrison, David K. Jackson, Ian Johnston, Dominic Kwiatkowski, Cordelia Langford, John Sillitoe on behalf of the Wellcome Sanger Institute COVID-19 Surveillance Team |
| EPI_ISL_3223640 | Lighthouse Lab in Alderley Park | Wellcome Sanger Institute for the COVID-19 Genomics UK (COG-UK) Consortium | Jacquelyn Wynn, Mairead Hyland, The Lighthouse Lab in Alderley Park and Alex Alderton, Roberto Amato, Jeffrey Barrett, Sonia Goncalves, Ewan Harrison, David K. Jackson, Ian Johnston, Dominic Kwiatkowski, Cordelia Langford, John Sillitoe on behalf of the Wellcome Sanger Institute COVID-19 Surveillance Team |
| EPI_ISL_3223641 | Lighthouse Lab in Glasgow | Wellcome Sanger Institute for the COVID-19 Genomics UK (COG-UK) Consortium | Harper VanSteenhouse, Yumi Kasai, David Gray, Carol Clugston, Anna Dominiczak and Alex Alderton, Roberto Amato, Jeffrey Barrett, Sonia Goncalves, Ewan Harrison, David K. Jackson, Ian Johnston, Dominic Kwiatkowski, Cordelia Langford, John Sillitoe on behalf of the Wellcome Sanger Institute COVID-19 Surveillance Team |
| EPI_ISL_3223642 | Lighthouse Lab in Alderley Park | Wellcome Sanger Institute for the COVID-19 Genomics UK (COG-UK) Consortium | Jacquelyn Wynn, Mairead Hyland, The Lighthouse Lab in Alderley Park and Alex Alderton, Roberto Amato, Jeffrey Barrett, Sonia Goncalves, Ewan Harrison, David K. Jackson, Ian Johnston, Dominic Kwiatkowski, Cordelia Langford, John Sillitoe on behalf of the Wellcome Sanger Institute COVID-19 Surveillance Team |
| EPI_ISL_3223643 | Lighthouse Lab in Milton Keynes | Wellcome Sanger Institute for the COVID-19 Genomics UK (COG-UK) Consortium | The Lighthouse Lab in Milton Keynes and Alex Alderton, Roberto Amato, Jeffrey Barrett, Sonia Goncalves, Ewan Harrison, David K. Jackson, Ian Johnston, Dominic Kwiatkowski, Cordelia Langford, John Sillitoe on behalf of the Wellcome Sanger Institute COVID-19 Surveillance Team |
| EPI_ISL_3223644 | Lighthouse Lab in Alderley Park | Wellcome Sanger Institute for the COVID-19 Genomics UK (COG-UK) Consortium | Jacquelyn Wynn, Mairead Hyland, The Lighthouse Lab in Alderley Park and Alex Alderton, Roberto Amato, Jeffrey Barrett, Sonia Goncalves, Ewan Harrison, David K. Jackson, Ian Johnston, Dominic Kwiatkowski, Cordelia Langford, John Sillitoe on behalf of the Wellcome Sanger Institute COVID-19 Surveillance Team |
| EPI_ISL_3223645 | Lighthouse Lab in Glasgow | Wellcome Sanger Institute for the COVID-19 Genomics UK (COG-UK) Consortium | Harper VanSteenhouse, Yumi Kasai, David Gray, Carol Clugston, Anna Dominiczak and Alex Alderton, Roberto Amato, Jeffrey Barrett, Sonia Goncalves, Ewan Harrison, David K. Jackson, Ian Johnston, Dominic Kwiatkowski, Cordelia Langford, John Sillitoe on behalf of the Wellcome Sanger Institute COVID-19 Surveillance Team |
| EPI_ISL_3223646 | Lighthouse Lab in Alderley Park | Wellcome Sanger Institute for the COVID-19 Genomics UK (COG-UK) Consortium | Jacquelyn Wynn, Mairead Hyland, The Lighthouse Lab in Alderley Park and Alex Alderton, Roberto Amato, Jeffrey Barrett, Sonia Goncalves, Ewan Harrison, David K. Jackson, Ian Johnston, Dominic Kwiatkowski, Cordelia Langford, John Sillitoe on behalf of the Wellcome Sanger Institute COVID-19 Surveillance Team |
| EPI_ISL_3223647 | Lighthouse Lab in Milton Keynes | Wellcome Sanger Institute for the COVID-19 Genomics UK (COG-UK) Consortium | The Lighthouse Lab in Milton Keynes and Alex Alderton, Roberto Amato, Jeffrey Barrett, Sonia Goncalves, Ewan Harrison, David K. Jackson, Ian Johnston, Dominic Kwiatkowski, Cordelia Langford, John Sillitoe on behalf of the Wellcome Sanger Institute COVID-19 Surveillance Team |
| EPI_ISL_3223648 | Lighthouse Laboratory Plymouth | Wellcome Sanger Institute for the COVID-19 Genomics UK (COG-UK) Consortium | Lighthouse Laboratory Plymouth and Alex Alderton, Roberto Amato, Jeffrey Barrett, Sonia Goncalves, Ewan Harrison, David K. Jackson, Ian Johnston, Dominic Kwiatkowski, Cordelia Langford, John Sillitoe on behalf of the Wellcome Sanger Institute COVID-19 Surveillance Team |
| EPI_ISL_3223649, EPI_ISL_3223650 | Lighthouse Lab in Alderley Park | Wellcome Sanger Institute for the COVID-19 Genomics UK (COG-UK) Consortium | Jacquelyn Wynn, Mairead Hyland, The Lighthouse Lab in Alderley Park and Alex Alderton, Roberto Amato, Jeffrey Barrett, Sonia Goncalves, Ewan Harrison, David K. Jackson, Ian Johnston, Dominic Kwiatkowski, Cordelia Langford, John Sillitoe on behalf of the Wellcome Sanger Institute COVID-19 Surveillance Team |
| EPI_ISL_3223651 | Lighthouse Lab in Milton Keynes | Wellcome Sanger Institute for the COVID-19 Genomics UK (COG-UK) Consortium | The Lighthouse Lab in Milton Keynes and Alex Alderton, Roberto Amato, Jeffrey Barrett, Sonia Goncalves, Ewan Harrison, David K. Jackson, Ian Johnston, Dominic Kwiatkowski, Cordelia Langford, John Sillitoe on behalf of the Wellcome Sanger Institute COVID-19 Surveillance Team |
| EPI_ISL_3223652 | Lighthouse Lab in Glasgow | Wellcome Sanger Institute for the COVID-19 Genomics UK (COG-UK) Consortium | Harper VanSteenhouse, Yumi Kasai, David Gray, Carol Clugston, Anna Dominiczak and Alex Alderton, Roberto Amato, Jeffrey Barrett, Sonia Goncalves, Ewan Harrison, David K. Jackson, Ian Johnston, Dominic Kwiatkowski, Cordelia Langford, John Sillitoe on behalf of the Wellcome Sanger Institute COVID-19 Surveillance Team |
| EPI_ISL_3223653, EPI_ISL_3223654, EPI_ISL_3223655 | Lighthouse Lab in Alderley Park | Wellcome Sanger Institute for the COVID-19 Genomics UK (COG-UK) Consortium | Jacquelyn Wynn, Mairead Hyland, The Lighthouse Lab in Alderley Park and Alex Alderton, Roberto Amato, Jeffrey Barrett, Sonia Goncalves, Ewan Harrison, David K. Jackson, Ian Johnston, Dominic Kwiatkowski, Cordelia Langford, John Sillitoe on behalf of the Wellcome Sanger Institute COVID-19 Surveillance Team |
| EPI_ISL_3223656, EPI_ISL_3223657 | Lighthouse Lab in Glasgow | Wellcome Sanger Institute for the COVID-19 Genomics UK (COG-UK) Consortium | Harper VanSteenhouse, Yumi Kasai, David Gray, Carol Clugston, Anna Dominiczak and Alex Alderton, Roberto Amato, Jeffrey Barrett, Sonia Goncalves, Ewan Harrison, David K. Jackson, Ian Johnston, Dominic Kwiatkowski, Cordelia Langford, John Sillitoe on behalf of the Wellcome Sanger Institute COVID-19 Surveillance Team |
| EPI_ISL_3223658, EPI_ISL_3223659 | Lighthouse Laboratory Plymouth | Wellcome Sanger Institute for the COVID-19 Genomics UK (COG-UK) Consortium | Lighthouse Laboratory Plymouth and Alex Alderton, Roberto Amato, Jeffrey Barrett, Sonia Goncalves, Ewan Harrison, David K. Jackson, Ian Johnston, Dominic Kwiatkowski, Cordelia Langford, John Sillitoe on behalf of the Wellcome Sanger Institute COVID-19 Surveillance Team |
| EPI_ISL_3223660 | Lighthouse Lab in Alderley Park | Wellcome Sanger Institute for the COVID-19 Genomics UK (COG-UK) Consortium | Jacquelyn Wynn, Mairead Hyland, The Lighthouse Lab in Alderley Park and Alex Alderton, Roberto Amato, Jeffrey Barrett, Sonia Goncalves, Ewan Harrison, David K. Jackson, Ian Johnston, Dominic Kwiatkowski, Cordelia Langford, John Sillitoe on behalf of the Wellcome Sanger Institute COVID-19 Surveillance Team |
| EPI_ISL_3223661 | Lighthouse Lab in Glasgow | Wellcome Sanger Institute for the COVID-19 Genomics UK (COG-UK) Consortium | Harper VanSteenhouse, Yumi Kasai, David Gray, Carol Clugston, Anna Dominiczak and Alex Alderton, Roberto Amato, Jeffrey Barrett, Sonia Goncalves, Ewan Harrison, David K. Jackson, Ian Johnston, Dominic Kwiatkowski, Cordelia Langford, John Sillitoe on behalf of the Wellcome Sanger Institute COVID-19 Surveillance Team |
| EPI_ISL_3223662 | Lighthouse Laboratory Plymouth | Wellcome Sanger Institute for the COVID-19 Genomics UK (COG-UK) Consortium | Lighthouse Laboratory Plymouth and Alex Alderton, Roberto Amato, Jeffrey Barrett, Sonia Goncalves, Ewan Harrison, David K. Jackson, Ian Johnston, Dominic Kwiatkowski, Cordelia Langford, John Sillitoe on behalf of the Wellcome Sanger Institute COVID-19 Surveillance Team |
| EPI_ISL_3223663 | Lighthouse Lab in Alderley Park | Wellcome Sanger Institute for the COVID-19 Genomics UK (COG-UK) Consortium | Jacquelyn Wynn, Mairead Hyland, The Lighthouse Lab in Alderley Park and Alex Alderton, Roberto Amato, Jeffrey Barrett, Sonia Goncalves, Ewan Harrison, David K. Jackson, Ian Johnston, Dominic Kwiatkowski, Cordelia Langford, John Sillitoe on behalf of the Wellcome Sanger Institute COVID-19 Surveillance Team |
| EPI_ISL_3223664, EPI_ISL_3223665 | Lighthouse Lab in Milton Keynes | Wellcome Sanger Institute for the COVID-19 Genomics UK (COG-UK) Consortium | The Lighthouse Lab in Milton Keynes and Alex Alderton, Roberto Amato, Jeffrey Barrett, Sonia Goncalves, Ewan Harrison, David K. Jackson, Ian Johnston, Dominic Kwiatkowski, Cordelia Langford, John Sillitoe on behalf of the Wellcome Sanger Institute COVID-19 Surveillance Team |
| EPI_ISL_3223666 | Lighthouse Laboratory Plymouth | Wellcome Sanger Institute for the COVID-19 Genomics UK (COG-UK) Consortium | Lighthouse Laboratory Plymouth and Alex Alderton, Roberto Amato, Jeffrey Barrett, Sonia Goncalves, Ewan Harrison, David K. Jackson, Ian Johnston, Dominic Kwiatkowski, Cordelia Langford, John Sillitoe on behalf of the Wellcome Sanger Institute COVID-19 Surveillance Team |
| EPI_ISL_3223667 | Lighthouse Lab in Alderley Park | Wellcome Sanger Institute for the COVID-19 Genomics UK (COG-UK) Consortium | Jacquelyn Wynn, Mairead Hyland, The Lighthouse Lab in Alderley Park and Alex Alderton, Roberto Amato, Jeffrey Barrett, Sonia Goncalves, Ewan Harrison, David K. Jackson, Ian Johnston, Dominic Kwiatkowski, Cordelia Langford, John Sillitoe on behalf of the Wellcome Sanger Institute COVID-19 Surveillance Team |
| EPI_ISL_3223668 | Lighthouse Lab in Glasgow | Wellcome Sanger Institute for the COVID-19 Genomics UK (COG-UK) Consortium | Harper VanSteenhouse, Yumi Kasai, David Gray, Carol Clugston, Anna Dominiczak and Alex Alderton, Roberto Amato, Jeffrey Barrett, Sonia Goncalves, Ewan Harrison, David K. Jackson, Ian Johnston, Dominic Kwiatkowski, Cordelia Langford, John Sillitoe on behalf of the Wellcome Sanger Institute COVID-19 Surveillance Team |
| EPI_ISL_3223669 | Lighthouse Lab in Alderley Park | Wellcome Sanger Institute for the COVID-19 Genomics UK (COG-UK) Consortium | Jacquelyn Wynn, Mairead Hyland, The Lighthouse Lab in Alderley Park and Alex Alderton, Roberto Amato, Jeffrey Barrett, Sonia Goncalves, Ewan Harrison, David K. Jackson, Ian Johnston, Dominic Kwiatkowski, Cordelia Langford, John Sillitoe on behalf of the Wellcome Sanger Institute COVID-19 Surveillance Team |
| EPI_ISL_3223670, EPI_ISL_3223671 | Lighthouse Lab in Glasgow | Wellcome Sanger Institute for the COVID-19 Genomics UK (COG-UK) Consortium | Harper VanSteenhouse, Yumi Kasai, David Gray, Carol Clugston, Anna Dominiczak and Alex Alderton, Roberto Amato, Jeffrey Barrett, Sonia Goncalves, Ewan Harrison, David K. Jackson, Ian Johnston, Dominic Kwiatkowski, Cordelia Langford, John Sillitoe on behalf of the Wellcome Sanger Institute COVID-19 Surveillance Team |
| EPI_ISL_3223672, EPI_ISL_3223673 | Lighthouse Laboratory Plymouth | Wellcome Sanger Institute for the COVID-19 Genomics UK (COG-UK) Consortium | Lighthouse Laboratory Plymouth and Alex Alderton, Roberto Amato, Jeffrey Barrett, Sonia Goncalves, Ewan Harrison, David K. Jackson, Ian Johnston, Dominic Kwiatkowski, Cordelia Langford, John Sillitoe on behalf of the Wellcome Sanger Institute COVID-19 Surveillance Team |
| EPI_ISL_3223674, EPI_ISL_3223675 | Lighthouse Lab in Glasgow | Wellcome Sanger Institute for the COVID-19 Genomics UK (COG-UK) Consortium | Harper VanSteenhouse, Yumi Kasai, David Gray, Carol Clugston, Anna Dominiczak and Alex Alderton, Roberto Amato, Jeffrey Barrett, Sonia Goncalves, Ewan Harrison, David K. Jackson, Ian Johnston, Dominic Kwiatkowski, Cordelia Langford, John Sillitoe on behalf of the Wellcome Sanger Institute COVID-19 Surveillance Team |
| EPI_ISL_3223676 | Lighthouse Lab in Alderley Park | Wellcome Sanger Institute for the COVID-19 Genomics UK (COG-UK) Consortium | Jacquelyn Wynn, Mairead Hyland, The Lighthouse Lab in Alderley Park and Alex Alderton, Roberto Amato, Jeffrey Barrett, Sonia Goncalves, Ewan Harrison, David K. Jackson, Ian Johnston, Dominic Kwiatkowski, Cordelia Langford, John Sillitoe on behalf of the Wellcome Sanger Institute COVID-19 Surveillance Team |
| EPI_ISL_3223677 | Lighthouse Lab in Milton Keynes | Wellcome Sanger Institute for the COVID-19 Genomics UK (COG-UK) | The Lighthouse Lab in Milton Keynes and Alex Alderton, Roberto Amato, Jeffrey Barrett, Sonia Goncalves, Ewan Harrison, David K. Jackson, Ian Johnston, Dominic Kwiatkowski, Cordelia Langford, John Sillitoe on behalf of the Wellcome Sanger |

| | | | |
|---|---|---|---|
| | | Consortium | Institute COVID-19 Surveillance Team |
| EPI_ISL_3223678, EPI_ISL_3223679 | Lighthouse Lab in Glasgow | Wellcome Sanger Institute for the COVID-19 Genomics UK (COG-UK) Consortium | Harper VanSteenhouse, Yumi Kasai, David Gray, Carol Clugston, Anna Dominiczak and Alex Alderton, Roberto Amato, Jeffrey Barrett, Sonia Goncalves, Ewan Harrison, David K. Jackson, Ian Johnston, Dominic Kwiatkowski, Cordelia Langford, John Sillitoe on behalf of the Wellcome Sanger Institute COVID-19 Surveillance Team |
| EPI_ISL_3223680 | Lighthouse Laboratory Plymouth | Wellcome Sanger Institute for the COVID-19 Genomics UK (COG-UK) Consortium | Lighthouse Laboratory Plymouth and Alex Alderton, Roberto Amato, Jeffrey Barrett, Sonia Goncalves, Ewan Harrison, David K. Jackson, Ian Johnston, Dominic Kwiatkowski, Cordelia Langford, John Sillitoe on behalf of the Wellcome Sanger Institute COVID-19 Surveillance Team |
| EPI_ISL_3223681, EPI_ISL_3223682 | Lighthouse Lab in Glasgow | Wellcome Sanger Institute for the COVID-19 Genomics UK (COG-UK) Consortium | Harper VanSteenhouse, Yumi Kasai, David Gray, Carol Clugston, Anna Dominiczak and Alex Alderton, Roberto Amato, Jeffrey Barrett, Sonia Goncalves, Ewan Harrison, David K. Jackson, Ian Johnston, Dominic Kwiatkowski, Cordelia Langford, John Sillitoe on behalf of the Wellcome Sanger Institute COVID-19 Surveillance Team |
| EPI_ISL_3223683 | Lighthouse Lab in Milton Keynes | Wellcome Sanger Institute for the COVID-19 Genomics UK (COG-UK) Consortium | The Lighthouse Lab in Milton Keynes and Alex Alderton, Roberto Amato, Jeffrey Barrett, Sonia Goncalves, Ewan Harrison, David K. Jackson, Ian Johnston, Dominic Kwiatkowski, Cordelia Langford, John Sillitoe on behalf of the Wellcome Sanger Institute COVID-19 Surveillance Team |
| EPI_ISL_3223684, EPI_ISL_3223685 | Lighthouse Lab in Alderley Park | Wellcome Sanger Institute for the COVID-19 Genomics UK (COG-UK) Consortium | Jacquelyn Wynn, Mairead Hyland, The Lighthouse Lab in Alderley Park and Alex Alderton, Roberto Amato, Jeffrey Barrett, Sonia Goncalves, Ewan Harrison, David K. Jackson, Ian Johnston, Dominic Kwiatkowski, Cordelia Langford, John Sillitoe on behalf of the Wellcome Sanger Institute COVID-19 Surveillance Team |
| EPI_ISL_3223686, EPI_ISL_3223687 | Lighthouse Laboratory Plymouth | Wellcome Sanger Institute for the COVID-19 Genomics UK (COG-UK) Consortium | Lighthouse Laboratory Plymouth and Alex Alderton, Roberto Amato, Jeffrey Barrett, Sonia Goncalves, Ewan Harrison, David K. Jackson, Ian Johnston, Dominic Kwiatkowski, Cordelia Langford, John Sillitoe on behalf of the Wellcome Sanger Institute COVID-19 Surveillance Team |
| EPI_ISL_3223688 | Lighthouse Lab in Glasgow | Wellcome Sanger Institute for the COVID-19 Genomics UK (COG-UK) Consortium | Harper VanSteenhouse, Yumi Kasai, David Gray, Carol Clugston, Anna Dominiczak and Alex Alderton, Roberto Amato, Jeffrey Barrett, Sonia Goncalves, Ewan Harrison, David K. Jackson, Ian Johnston, Dominic Kwiatkowski, Cordelia Langford, John Sillitoe on behalf of the Wellcome Sanger Institute COVID-19 Surveillance Team |
| EPI_ISL_3223689 | Lighthouse Lab in Alderley Park | Wellcome Sanger Institute for the COVID-19 Genomics UK (COG-UK) Consortium | Jacquelyn Wynn, Mairead Hyland, The Lighthouse Lab in Alderley Park and Alex Alderton, Roberto Amato, Jeffrey Barrett, Sonia Goncalves, Ewan Harrison, David K. Jackson, Ian Johnston, Dominic Kwiatkowski, Cordelia Langford, John Sillitoe on behalf of the Wellcome Sanger Institute COVID-19 Surveillance Team |
| EPI_ISL_3223690 | Lighthouse Lab in Glasgow | Wellcome Sanger Institute for the COVID-19 Genomics UK (COG-UK) Consortium | Harper VanSteenhouse, Yumi Kasai, David Gray, Carol Clugston, Anna Dominiczak and Alex Alderton, Roberto Amato, Jeffrey Barrett, Sonia Goncalves, Ewan Harrison, David K. Jackson, Ian Johnston, Dominic Kwiatkowski, Cordelia Langford, John Sillitoe on behalf of the Wellcome Sanger Institute COVID-19 Surveillance Team |
| EPI_ISL_3223691 | Lighthouse Laboratory Plymouth | Wellcome Sanger Institute for the COVID-19 Genomics UK (COG-UK) Consortium | Lighthouse Laboratory Plymouth and Alex Alderton, Roberto Amato, Jeffrey Barrett, Sonia Goncalves, Ewan Harrison, David K. Jackson, Ian Johnston, Dominic Kwiatkowski, Cordelia Langford, John Sillitoe on behalf of the Wellcome Sanger Institute COVID-19 Surveillance Team |
| EPI_ISL_3223692 | Lighthouse Lab in Glasgow | Wellcome Sanger Institute for the COVID-19 Genomics UK (COG-UK) Consortium | Harper VanSteenhouse, Yumi Kasai, David Gray, Carol Clugston, Anna Dominiczak and Alex Alderton, Roberto Amato, Jeffrey Barrett, Sonia Goncalves, Ewan Harrison, David K. Jackson, Ian Johnston, Dominic Kwiatkowski, Cordelia Langford, John Sillitoe on behalf of the Wellcome Sanger Institute COVID-19 Surveillance Team |
| EPI_ISL_3223693 | Lighthouse Laboratory Plymouth | Wellcome Sanger Institute for the COVID-19 Genomics UK (COG-UK) Consortium | Lighthouse Laboratory Plymouth and Alex Alderton, Roberto Amato, Jeffrey Barrett, Sonia Goncalves, Ewan Harrison, David K. Jackson, Ian Johnston, Dominic Kwiatkowski, Cordelia Langford, John Sillitoe on behalf of the Wellcome Sanger Institute COVID-19 Surveillance Team |
| EPI_ISL_3223694 | Lighthouse Lab in Alderley Park | Wellcome Sanger Institute for the COVID-19 Genomics UK (COG-UK) Consortium | Jacquelyn Wynn, Mairead Hyland, The Lighthouse Lab in Alderley Park and Alex Alderton, Roberto Amato, Jeffrey Barrett, Sonia Goncalves, Ewan Harrison, David K. Jackson, Ian Johnston, Dominic Kwiatkowski, Cordelia Langford, John Sillitoe on behalf of the Wellcome Sanger Institute COVID-19 Surveillance Team |
| EPI_ISL_3223695 | Lighthouse Lab in Glasgow | Wellcome Sanger Institute for the COVID-19 Genomics UK (COG-UK) Consortium | Harper VanSteenhouse, Yumi Kasai, David Gray, Carol Clugston, Anna Dominiczak and Alex Alderton, Roberto Amato, Jeffrey Barrett, Sonia Goncalves, Ewan Harrison, David K. Jackson, Ian Johnston, Dominic Kwiatkowski, Cordelia Langford, John Sillitoe on behalf of the Wellcome Sanger Institute COVID-19 Surveillance Team |
| EPI_ISL_3223696 | Lighthouse Lab in Milton Keynes | Wellcome Sanger Institute for the COVID-19 Genomics UK (COG-UK) Consortium | The Lighthouse Lab in Milton Keynes and Alex Alderton, Roberto Amato, Jeffrey Barrett, Sonia Goncalves, Ewan Harrison, David K. Jackson, Ian Johnston, Dominic Kwiatkowski, Cordelia Langford, John Sillitoe on behalf of the Wellcome Sanger Institute COVID-19 Surveillance Team |
| EPI_ISL_3223697 | Lighthouse Lab in Alderley Park | Wellcome Sanger Institute for the COVID-19 Genomics UK (COG-UK) Consortium | Jacquelyn Wynn, Mairead Hyland, The Lighthouse Lab in Alderley Park and Alex Alderton, Roberto Amato, Jeffrey Barrett, Sonia Goncalves, Ewan Harrison, David K. Jackson, Ian Johnston, Dominic Kwiatkowski, Cordelia Langford, John Sillitoe on behalf of the Wellcome Sanger Institute COVID-19 Surveillance Team |
| EPI_ISL_3223698, EPI_ISL_3223699 | Lighthouse Lab in Glasgow | Wellcome Sanger Institute for the COVID-19 Genomics UK (COG-UK) Consortium | Harper VanSteenhouse, Yumi Kasai, David Gray, Carol Clugston, Anna Dominiczak and Alex Alderton, Roberto Amato, Jeffrey Barrett, Sonia Goncalves, Ewan Harrison, David K. Jackson, Ian Johnston, Dominic Kwiatkowski, Cordelia Langford, John Sillitoe on behalf of the Wellcome Sanger Institute COVID-19 Surveillance Team |
| EPI_ISL_3223700, EPI_ISL_3223701 | Lighthouse Lab in Milton Keynes | Wellcome Sanger Institute for the COVID-19 Genomics UK (COG-UK) Consortium | The Lighthouse Lab in Milton Keynes and Alex Alderton, Roberto Amato, Jeffrey Barrett, Sonia Goncalves, Ewan Harrison, David K. Jackson, Ian Johnston, Dominic Kwiatkowski, Cordelia Langford, John Sillitoe on behalf of the Wellcome Sanger Institute COVID-19 Surveillance Team |
| EPI_ISL_3223702 | Lighthouse Laboratory Plymouth | Wellcome Sanger Institute for the COVID-19 Genomics UK (COG-UK) Consortium | Lighthouse Laboratory Plymouth and Alex Alderton, Roberto Amato, Jeffrey Barrett, Sonia Goncalves, Ewan Harrison, David K. Jackson, Ian Johnston, Dominic Kwiatkowski, Cordelia Langford, John Sillitoe on behalf of the Wellcome Sanger Institute COVID-19 Surveillance Team |
| EPI_ISL_3223703 | Lighthouse Lab in Glasgow | Wellcome Sanger Institute for the COVID-19 Genomics UK (COG-UK) Consortium | Harper VanSteenhouse, Yumi Kasai, David Gray, Carol Clugston, Anna Dominiczak and Alex Alderton, Roberto Amato, Jeffrey Barrett, Sonia Goncalves, Ewan Harrison, David K. Jackson, Ian Johnston, Dominic Kwiatkowski, Cordelia Langford, John Sillitoe on behalf of the Wellcome Sanger Institute COVID-19 Surveillance Team |
| EPI_ISL_3223704 | Lighthouse Lab in Alderley Park | Wellcome Sanger Institute for the COVID-19 Genomics UK (COG-UK) Consortium | Jacquelyn Wynn, Mairead Hyland, The Lighthouse Lab in Alderley Park and Alex Alderton, Roberto Amato, Jeffrey Barrett, Sonia Goncalves, Ewan Harrison, David K. Jackson, Ian Johnston, Dominic Kwiatkowski, Cordelia Langford, John Sillitoe on behalf of the Wellcome Sanger Institute COVID-19 Surveillance Team |
| EPI_ISL_3223705, EPI_ISL_3223706 | Lighthouse Laboratory Plymouth | Wellcome Sanger Institute for the COVID-19 Genomics UK (COG-UK) Consortium | Lighthouse Laboratory Plymouth and Alex Alderton, Roberto Amato, Jeffrey Barrett, Sonia Goncalves, Ewan Harrison, David K. Jackson, Ian Johnston, Dominic Kwiatkowski, Cordelia Langford, John Sillitoe on behalf of the Wellcome Sanger Institute COVID-19 Surveillance Team |
| EPI_ISL_3223707 | Lighthouse Lab in Milton Keynes | Wellcome Sanger Institute for the COVID-19 Genomics UK (COG-UK) Consortium | The Lighthouse Lab in Milton Keynes and Alex Alderton, Roberto Amato, Jeffrey Barrett, Sonia Goncalves, Ewan Harrison, David K. Jackson, Ian Johnston, Dominic Kwiatkowski, Cordelia Langford, John Sillitoe on behalf of the Wellcome Sanger Institute COVID-19 Surveillance Team |
| EPI_ISL_3223708 | Lighthouse Lab in Alderley Park | Wellcome Sanger Institute for the COVID-19 Genomics UK (COG-UK) Consortium | Jacquelyn Wynn, Mairead Hyland, The Lighthouse Lab in Alderley Park and Alex Alderton, Roberto Amato, Jeffrey Barrett, Sonia Goncalves, Ewan Harrison, David K. Jackson, Ian Johnston, Dominic Kwiatkowski, Cordelia Langford, John Sillitoe on behalf of the Wellcome Sanger Institute COVID-19 Surveillance Team |
| EPI_ISL_3223709 | Lighthouse Laboratory Plymouth | Wellcome Sanger Institute for the COVID-19 Genomics UK (COG-UK) Consortium | Lighthouse Laboratory Plymouth and Alex Alderton, Roberto Amato, Jeffrey Barrett, Sonia Goncalves, Ewan Harrison, David K. Jackson, Ian Johnston, Dominic Kwiatkowski, Cordelia Langford, John Sillitoe on behalf of the Wellcome Sanger Institute COVID-19 Surveillance Team |
| EPI_ISL_3223710 | Lighthouse Lab in Glasgow | Wellcome Sanger Institute for the COVID-19 Genomics UK (COG-UK) Consortium | Harper VanSteenhouse, Yumi Kasai, David Gray, Carol Clugston, Anna Dominiczak and Alex Alderton, Roberto Amato, Jeffrey Barrett, Sonia Goncalves, Ewan Harrison, David K. Jackson, Ian Johnston, Dominic Kwiatkowski, Cordelia Langford, John Sillitoe on behalf of the Wellcome Sanger Institute COVID-19 Surveillance Team |
| EPI_ISL_3223711 | Lighthouse Lab in Alderley Park | Wellcome Sanger Institute for the COVID-19 Genomics UK (COG-UK) Consortium | Jacquelyn Wynn, Mairead Hyland, The Lighthouse Lab in Alderley Park and Alex Alderton, Roberto Amato, Jeffrey Barrett, Sonia Goncalves, Ewan Harrison, David K. Jackson, Ian Johnston, Dominic Kwiatkowski, Cordelia Langford, John Sillitoe on behalf of the Wellcome Sanger Institute COVID-19 Surveillance Team |
| EPI_ISL_3223712, EPI_ISL_3223713 | Lighthouse Lab in Milton Keynes | Wellcome Sanger Institute for the COVID-19 Genomics UK (COG-UK) Consortium | The Lighthouse Lab in Milton Keynes and Alex Alderton, Roberto Amato, Jeffrey Barrett, Sonia Goncalves, Ewan Harrison, David K. Jackson, Ian Johnston, Dominic Kwiatkowski, Cordelia Langford, John Sillitoe on behalf of the Wellcome Sanger Institute COVID-19 Surveillance Team |
| EPI_ISL_3223714 | Lighthouse Lab in Alderley Park | Wellcome Sanger Institute for the COVID-19 Genomics UK (COG-UK) Consortium | Jacquelyn Wynn, Mairead Hyland, The Lighthouse Lab in Alderley Park and Alex Alderton, Roberto Amato, Jeffrey Barrett, Sonia Goncalves, Ewan Harrison, David K. Jackson, Ian Johnston, Dominic Kwiatkowski, Cordelia Langford, John Sillitoe on behalf of the Wellcome Sanger Institute COVID-19 Surveillance Team |
| EPI_ISL_3223715 | Lighthouse Laboratory Plymouth | Wellcome Sanger Institute for the COVID-19 Genomics UK (COG-UK) Consortium | Lighthouse Laboratory Plymouth and Alex Alderton, Roberto Amato, Jeffrey Barrett, Sonia Goncalves, Ewan Harrison, David K. Jackson, Ian Johnston, Dominic Kwiatkowski, Cordelia Langford, John Sillitoe on behalf of the Wellcome Sanger Institute COVID-19 Surveillance Team |
| EPI_ISL_3223716 | Lighthouse Lab in Milton Keynes | Wellcome Sanger Institute for the COVID-19 Genomics UK (COG-UK) Consortium | The Lighthouse Lab in Milton Keynes and Alex Alderton, Roberto Amato, Jeffrey Barrett, Sonia Goncalves, Ewan Harrison, David K. Jackson, Ian Johnston, Dominic Kwiatkowski, Cordelia Langford, John Sillitoe on behalf of the Wellcome Sanger Institute COVID-19 Surveillance Team |
| EPI_ISL_3223717, EPI_ISL_3223718, EPI_ISL_3223719 | Lighthouse Lab in Alderley Park | Wellcome Sanger Institute for the COVID-19 Genomics UK (COG-UK) Consortium | Jacquelyn Wynn, Mairead Hyland, The Lighthouse Lab in Alderley Park and Alex Alderton, Roberto Amato, Jeffrey Barrett, Sonia Goncalves, Ewan Harrison, David K. Jackson, Ian Johnston, Dominic Kwiatkowski, Cordelia Langford, John Sillitoe on behalf of the Wellcome Sanger Institute COVID-19 Surveillance Team |
| EPI_ISL_3223720 | Lighthouse Lab in Glasgow | Wellcome Sanger Institute for the COVID-19 Genomics UK (COG-UK) Consortium | Harper VanSteenhouse, Yumi Kasai, David Gray, Carol Clugston, Anna Dominiczak and Alex Alderton, Roberto Amato, Jeffrey Barrett, Sonia Goncalves, Ewan Harrison, David K. Jackson, Ian Johnston, Dominic Kwiatkowski, Cordelia Langford, John Sillitoe on behalf of the Wellcome Sanger Institute COVID-19 Surveillance Team |
| EPI_ISL_3223721 | Lighthouse Laboratory Plymouth | Wellcome Sanger Institute for the COVID-19 Genomics UK (COG-UK) Consortium | Lighthouse Laboratory Plymouth and Alex Alderton, Roberto Amato, Jeffrey Barrett, Sonia Goncalves, Ewan Harrison, David K. Jackson, Ian Johnston, Dominic Kwiatkowski, Cordelia Langford, John Sillitoe on behalf of the Wellcome Sanger Institute COVID-19 Surveillance Team |
| EPI_ISL_3223722 | Lighthouse Lab in Alderley Park | Wellcome Sanger Institute for the COVID-19 Genomics UK (COG-UK) Consortium | Jacquelyn Wynn, Mairead Hyland, The Lighthouse Lab in Alderley Park and Alex Alderton, Roberto Amato, Jeffrey Barrett, Sonia Goncalves, Ewan Harrison, David K. Jackson, Ian Johnston, Dominic Kwiatkowski, Cordelia Langford, John Sillitoe on behalf of the Wellcome Sanger Institute COVID-19 Surveillance Team |
| EPI_ISL_3223723, EPI_ISL_3223724 | Lighthouse Lab in Glasgow | Wellcome Sanger Institute for the COVID-19 Genomics UK (COG-UK) Consortium | Harper VanSteenhouse, Yumi Kasai, David Gray, Carol Clugston, Anna Dominiczak and Alex Alderton, Roberto Amato, Jeffrey Barrett, Sonia Goncalves, Ewan Harrison, David K. Jackson, Ian Johnston, Dominic Kwiatkowski, Cordelia Langford, John Sillitoe on behalf of the Wellcome Sanger Institute COVID-19 Surveillance Team |
| EPI_ISL_3223725 | Lighthouse Lab in Milton Keynes | Wellcome Sanger Institute for the COVID-19 Genomics UK (COG-UK) Consortium | The Lighthouse Lab in Milton Keynes and Alex Alderton, Roberto Amato, Jeffrey Barrett, Sonia Goncalves, Ewan Harrison, David K. Jackson, Ian Johnston, Dominic Kwiatkowski, Cordelia Langford, John Sillitoe on behalf of the Wellcome Sanger Institute COVID-19 Surveillance Team |
| EPI_ISL_3223726 | Lighthouse Lab in Glasgow | Wellcome Sanger Institute for the COVID-19 Genomics UK (COG-UK) Consortium | Harper VanSteenhouse, Yumi Kasai, David Gray, Carol Clugston, Anna Dominiczak and Alex Alderton, Roberto Amato, Jeffrey Barrett, Sonia Goncalves, Ewan Harrison, David K. Jackson, Ian Johnston, Dominic Kwiatkowski, Cordelia Langford, John Sillitoe on behalf of the Wellcome Sanger Institute COVID-19 Surveillance Team |
| EPI_ISL_3223727 | Lighthouse Lab in Milton Keynes | Wellcome Sanger Institute for the COVID-19 Genomics UK (COG-UK) Consortium | The Lighthouse Lab in Milton Keynes and Alex Alderton, Roberto Amato, Jeffrey Barrett, Sonia Goncalves, Ewan Harrison, David K. Jackson, Ian Johnston, Dominic Kwiatkowski, Cordelia Langford, John Sillitoe on behalf of the Wellcome Sanger Institute COVID-19 Surveillance Team |
| EPI_ISL_3223728 | Lighthouse Lab in Glasgow | Wellcome Sanger Institute for the COVID-19 Genomics UK (COG-UK) Consortium | Harper VanSteenhouse, Yumi Kasai, David Gray, Carol Clugston, Anna Dominiczak and Alex Alderton, Roberto Amato, Jeffrey Barrett, Sonia Goncalves, Ewan Harrison, David K. Jackson, Ian Johnston, Dominic Kwiatkowski, Cordelia Langford, John Sillitoe on behalf of the Wellcome Sanger Institute COVID-19 Surveillance Team |
| EPI_ISL_3223729 | Lighthouse Laboratory Plymouth | Wellcome Sanger Institute for the COVID-19 Genomics UK (COG-UK) Consortium | Lighthouse Laboratory Plymouth and Alex Alderton, Roberto Amato, Jeffrey Barrett, Sonia Goncalves, Ewan Harrison, David K. Jackson, Ian Johnston, Dominic Kwiatkowski, Cordelia Langford, John Sillitoe on behalf of the Wellcome Sanger Institute COVID-19 Surveillance Team |
| EPI_ISL_3223730, EPI_ISL_3223731, EPI_ISL_3223732, EPI_ISL_3223733 | Lighthouse Lab in Alderley Park | Wellcome Sanger Institute for the COVID-19 Genomics UK (COG-UK) Consortium | Jacquelyn Wynn, Mairead Hyland, The Lighthouse Lab in Alderley Park and Alex Alderton, Roberto Amato, Jeffrey Barrett, Sonia Goncalves, Ewan Harrison, David K. Jackson, Ian Johnston, Dominic Kwiatkowski, Cordelia Langford, John Sillitoe on behalf of the Wellcome Sanger Institute COVID-19 Surveillance Team |
| EPI_ISL_3223734 | Lighthouse Laboratory Plymouth | Wellcome Sanger Institute for the COVID-19 Genomics UK (COG-UK) Consortium | Lighthouse Laboratory Plymouth and Alex Alderton, Roberto Amato, Jeffrey Barrett, Sonia Goncalves, Ewan Harrison, David K. Jackson, Ian Johnston, Dominic Kwiatkowski, Cordelia Langford, John Sillitoe on behalf of the Wellcome Sanger Institute COVID-19 Surveillance Team |

| | | | |
|---|---|---|---|
| EPI_ISL_3223735 | Lighthouse Lab in Alderley Park | Wellcome Sanger Institute for the COVID-19 Genomics UK (COG-UK) Consortium | Jacquelyn Wynn, Mairead Hyland, The Lighthouse Lab in Alderley Park and Alex Alderton, Roberto Amato, Jeffrey Barrett, Sonia Goncalves, Ewan Harrison, David K. Jackson, Ian Johnston, Dominic Kwiatkowski, Cordelia Langford, John Sillitoe on behalf of the Wellcome Sanger Institute COVID-19 Surveillance Team |
| EPI_ISL_3223736 | Lighthouse Laboratory Plymouth | Wellcome Sanger Institute for the COVID-19 Genomics UK (COG-UK) Consortium | Lighthouse Laboratory Plymouth and Alex Alderton, Roberto Amato, Jeffrey Barrett, Sonia Goncalves, Ewan Harrison, David K. Jackson, Ian Johnston, Dominic Kwiatkowski, Cordelia Langford, John Sillitoe on behalf of the Wellcome Sanger Institute COVID-19 Surveillance Team |
| EPI_ISL_3223737 | Lighthouse Lab in Glasgow | Wellcome Sanger Institute for the COVID-19 Genomics UK (COG-UK) Consortium | Harper VanSteenhouse, Yumi Kasai, David Gray, Carol Clugston, Anna Dominiczak and Alex Alderton, Roberto Amato, Jeffrey Barrett, Sonia Goncalves, Ewan Harrison, David K. Jackson, Ian Johnston, Dominic Kwiatkowski, Cordelia Langford, John Sillitoe on behalf of the Wellcome Sanger Institute COVID-19 Surveillance Team |
| EPI_ISL_3223738, EPI_ISL_3223739, EPI_ISL_3223740 | Lighthouse Laboratory Plymouth | Wellcome Sanger Institute for the COVID-19 Genomics UK (COG-UK) Consortium | Lighthouse Laboratory Plymouth and Alex Alderton, Roberto Amato, Jeffrey Barrett, Sonia Goncalves, Ewan Harrison, David K. Jackson, Ian Johnston, Dominic Kwiatkowski, Cordelia Langford, John Sillitoe on behalf of the Wellcome Sanger Institute COVID-19 Surveillance Team |
| EPI_ISL_3223741 | Lighthouse Lab in Glasgow | Wellcome Sanger Institute for the COVID-19 Genomics UK (COG-UK) Consortium | Harper VanSteenhouse, Yumi Kasai, David Gray, Carol Clugston, Anna Dominiczak and Alex Alderton, Roberto Amato, Jeffrey Barrett, Sonia Goncalves, Ewan Harrison, David K. Jackson, Ian Johnston, Dominic Kwiatkowski, Cordelia Langford, John Sillitoe on behalf of the Wellcome Sanger Institute COVID-19 Surveillance Team |
| EPI_ISL_3223742 | Lighthouse Lab in Alderley Park | Wellcome Sanger Institute for the COVID-19 Genomics UK (COG-UK) Consortium | Jacquelyn Wynn, Mairead Hyland, The Lighthouse Lab in Alderley Park and Alex Alderton, Roberto Amato, Jeffrey Barrett, Sonia Goncalves, Ewan Harrison, David K. Jackson, Ian Johnston, Dominic Kwiatkowski, Cordelia Langford, John Sillitoe on behalf of the Wellcome Sanger Institute COVID-19 Surveillance Team |
| EPI_ISL_3223743, EPI_ISL_3223744, EPI_ISL_3223745 | Lighthouse Laboratory Plymouth | Wellcome Sanger Institute for the COVID-19 Genomics UK (COG-UK) Consortium | Lighthouse Laboratory Plymouth and Alex Alderton, Roberto Amato, Jeffrey Barrett, Sonia Goncalves, Ewan Harrison, David K. Jackson, Ian Johnston, Dominic Kwiatkowski, Cordelia Langford, John Sillitoe on behalf of the Wellcome Sanger Institute COVID-19 Surveillance Team |
| EPI_ISL_3223746 | Lighthouse Lab in Milton Keynes | Wellcome Sanger Institute for the COVID-19 Genomics UK (COG-UK) Consortium | The Lighthouse Lab in Milton Keynes and Alex Alderton, Roberto Amato, Jeffrey Barrett, Sonia Goncalves, Ewan Harrison, David K. Jackson, Ian Johnston, Dominic Kwiatkowski, Cordelia Langford, John Sillitoe on behalf of the Wellcome Sanger Institute COVID-19 Surveillance Team |
| EPI_ISL_3223747 | Lighthouse Lab in Glasgow | Wellcome Sanger Institute for the COVID-19 Genomics UK (COG-UK) Consortium | Harper VanSteenhouse, Yumi Kasai, David Gray, Carol Clugston, Anna Dominiczak and Alex Alderton, Roberto Amato, Jeffrey Barrett, Sonia Goncalves, Ewan Harrison, David K. Jackson, Ian Johnston, Dominic Kwiatkowski, Cordelia Langford, John Sillitoe on behalf of the Wellcome Sanger Institute COVID-19 Surveillance Team |
| EPI_ISL_3223748 | Lighthouse Lab in Milton Keynes | Wellcome Sanger Institute for the COVID-19 Genomics UK (COG-UK) Consortium | The Lighthouse Lab in Milton Keynes and Alex Alderton, Roberto Amato, Jeffrey Barrett, Sonia Goncalves, Ewan Harrison, David K. Jackson, Ian Johnston, Dominic Kwiatkowski, Cordelia Langford, John Sillitoe on behalf of the Wellcome Sanger Institute COVID-19 Surveillance Team |
| EPI_ISL_3223749 | Lighthouse Lab in Glasgow | Wellcome Sanger Institute for the COVID-19 Genomics UK (COG-UK) Consortium | Harper VanSteenhouse, Yumi Kasai, David Gray, Carol Clugston, Anna Dominiczak and Alex Alderton, Roberto Amato, Jeffrey Barrett, Sonia Goncalves, Ewan Harrison, David K. Jackson, Ian Johnston, Dominic Kwiatkowski, Cordelia Langford, John Sillitoe on behalf of the Wellcome Sanger Institute COVID-19 Surveillance Team |
| EPI_ISL_3223750, EPI_ISL_3223751 | Lighthouse Laboratory Plymouth | Wellcome Sanger Institute for the COVID-19 Genomics UK (COG-UK) Consortium | Lighthouse Laboratory Plymouth and Alex Alderton, Roberto Amato, Jeffrey Barrett, Sonia Goncalves, Ewan Harrison, David K. Jackson, Ian Johnston, Dominic Kwiatkowski, Cordelia Langford, John Sillitoe on behalf of the Wellcome Sanger Institute COVID-19 Surveillance Team |
| EPI_ISL_3223752 | Lighthouse Lab in Milton Keynes | Wellcome Sanger Institute for the COVID-19 Genomics UK (COG-UK) Consortium | The Lighthouse Lab in Milton Keynes and Alex Alderton, Roberto Amato, Jeffrey Barrett, Sonia Goncalves, Ewan Harrison, David K. Jackson, Ian Johnston, Dominic Kwiatkowski, Cordelia Langford, John Sillitoe on behalf of the Wellcome Sanger Institute COVID-19 Surveillance Team |
| EPI_ISL_3223753, EPI_ISL_3223754, EPI_ISL_3223755 | Lighthouse Lab in Glasgow | Wellcome Sanger Institute for the COVID-19 Genomics UK (COG-UK) Consortium | Harper VanSteenhouse, Yumi Kasai, David Gray, Carol Clugston, Anna Dominiczak and Alex Alderton, Roberto Amato, Jeffrey Barrett, Sonia Goncalves, Ewan Harrison, David K. Jackson, Ian Johnston, Dominic Kwiatkowski, Cordelia Langford, John Sillitoe on behalf of the Wellcome Sanger Institute COVID-19 Surveillance Team |
| EPI_ISL_3223756 | Lighthouse Lab in Milton Keynes | Wellcome Sanger Institute for the COVID-19 Genomics UK (COG-UK) Consortium | The Lighthouse Lab in Milton Keynes and Alex Alderton, Roberto Amato, Jeffrey Barrett, Sonia Goncalves, Ewan Harrison, David K. Jackson, Ian Johnston, Dominic Kwiatkowski, Cordelia Langford, John Sillitoe on behalf of the Wellcome Sanger Institute COVID-19 Surveillance Team |
| EPI_ISL_3223757 | Lighthouse Lab in Alderley Park | Wellcome Sanger Institute for the COVID-19 Genomics UK (COG-UK) Consortium | Jacquelyn Wynn, Mairead Hyland, The Lighthouse Lab in Alderley Park and Alex Alderton, Roberto Amato, Jeffrey Barrett, Sonia Goncalves, Ewan Harrison, David K. Jackson, Ian Johnston, Dominic Kwiatkowski, Cordelia Langford, John Sillitoe on behalf of the Wellcome Sanger Institute COVID-19 Surveillance Team |
| EPI_ISL_3223758 | Lighthouse Laboratory Plymouth | Wellcome Sanger Institute for the COVID-19 Genomics UK (COG-UK) Consortium | Lighthouse Laboratory Plymouth and Alex Alderton, Roberto Amato, Jeffrey Barrett, Sonia Goncalves, Ewan Harrison, David K. Jackson, Ian Johnston, Dominic Kwiatkowski, Cordelia Langford, John Sillitoe on behalf of the Wellcome Sanger Institute COVID-19 Surveillance Team |
| EPI_ISL_3223759 | Lighthouse Lab in Glasgow | Wellcome Sanger Institute for the COVID-19 Genomics UK (COG-UK) Consortium | Harper VanSteenhouse, Yumi Kasai, David Gray, Carol Clugston, Anna Dominiczak and Alex Alderton, Roberto Amato, Jeffrey Barrett, Sonia Goncalves, Ewan Harrison, David K. Jackson, Ian Johnston, Dominic Kwiatkowski, Cordelia Langford, John Sillitoe on behalf of the Wellcome Sanger Institute COVID-19 Surveillance Team |
| EPI_ISL_3223760 | Lighthouse Laboratory Plymouth | Wellcome Sanger Institute for the COVID-19 Genomics UK (COG-UK) Consortium | Lighthouse Laboratory Plymouth and Alex Alderton, Roberto Amato, Jeffrey Barrett, Sonia Goncalves, Ewan Harrison, David K. Jackson, Ian Johnston, Dominic Kwiatkowski, Cordelia Langford, John Sillitoe on behalf of the Wellcome Sanger Institute COVID-19 Surveillance Team |
| EPI_ISL_3223761, EPI_ISL_3223762 | Lighthouse Lab in Glasgow | Wellcome Sanger Institute for the COVID-19 Genomics UK (COG-UK) Consortium | Harper VanSteenhouse, Yumi Kasai, David Gray, Carol Clugston, Anna Dominiczak and Alex Alderton, Roberto Amato, Jeffrey Barrett, Sonia Goncalves, Ewan Harrison, David K. Jackson, Ian Johnston, Dominic Kwiatkowski, Cordelia Langford, John Sillitoe on behalf of the Wellcome Sanger Institute COVID-19 Surveillance Team |
| EPI_ISL_3223763 | Lighthouse Lab in Milton Keynes | Wellcome Sanger Institute for the COVID-19 Genomics UK (COG-UK) Consortium | The Lighthouse Lab in Milton Keynes and Alex Alderton, Roberto Amato, Jeffrey Barrett, Sonia Goncalves, Ewan Harrison, David K. Jackson, Ian Johnston, Dominic Kwiatkowski, Cordelia Langford, John Sillitoe on behalf of the Wellcome Sanger Institute COVID-19 Surveillance Team |
| EPI_ISL_3223764 | Lighthouse Lab in Glasgow | Wellcome Sanger Institute for the COVID-19 Genomics UK (COG-UK) Consortium | Harper VanSteenhouse, Yumi Kasai, David Gray, Carol Clugston, Anna Dominiczak and Alex Alderton, Roberto Amato, Jeffrey Barrett, Sonia Goncalves, Ewan Harrison, David K. Jackson, Ian Johnston, Dominic Kwiatkowski, Cordelia Langford, John Sillitoe on behalf of the Wellcome Sanger Institute COVID-19 Surveillance Team |
| EPI_ISL_3223765 | Lighthouse Lab in Alderley Park | Wellcome Sanger Institute for the COVID-19 Genomics UK (COG-UK) Consortium | Jacquelyn Wynn, Mairead Hyland, The Lighthouse Lab in Alderley Park and Alex Alderton, Roberto Amato, Jeffrey Barrett, Sonia Goncalves, Ewan Harrison, David K. Jackson, Ian Johnston, Dominic Kwiatkowski, Cordelia Langford, John Sillitoe on behalf of the Wellcome Sanger Institute COVID-19 Surveillance Team |
| EPI_ISL_3223766 | Lighthouse Lab in Milton Keynes | Wellcome Sanger Institute for the COVID-19 Genomics UK (COG-UK) Consortium | The Lighthouse Lab in Milton Keynes and Alex Alderton, Roberto Amato, Jeffrey Barrett, Sonia Goncalves, Ewan Harrison, David K. Jackson, Ian Johnston, Dominic Kwiatkowski, Cordelia Langford, John Sillitoe on behalf of the Wellcome Sanger Institute COVID-19 Surveillance Team |
| EPI_ISL_3223767 | Lighthouse Lab in Alderley Park | Wellcome Sanger Institute for the COVID-19 Genomics UK (COG-UK) Consortium | Jacquelyn Wynn, Mairead Hyland, The Lighthouse Lab in Alderley Park and Alex Alderton, Roberto Amato, Jeffrey Barrett, Sonia Goncalves, Ewan Harrison, David K. Jackson, Ian Johnston, Dominic Kwiatkowski, Cordelia Langford, John Sillitoe on behalf of the Wellcome Sanger Institute COVID-19 Surveillance Team |
| EPI_ISL_3223768 | Lighthouse Lab in Glasgow | Wellcome Sanger Institute for the COVID-19 Genomics UK (COG-UK) Consortium | Harper VanSteenhouse, Yumi Kasai, David Gray, Carol Clugston, Anna Dominiczak and Alex Alderton, Roberto Amato, Jeffrey Barrett, Sonia Goncalves, Ewan Harrison, David K. Jackson, Ian Johnston, Dominic Kwiatkowski, Cordelia Langford, John Sillitoe on behalf of the Wellcome Sanger Institute COVID-19 Surveillance Team |
| EPI_ISL_3223769, EPI_ISL_3223770, EPI_ISL_3223771 | Lighthouse Lab in Milton Keynes | Wellcome Sanger Institute for the COVID-19 Genomics UK (COG-UK) Consortium | The Lighthouse Lab in Milton Keynes and Alex Alderton, Roberto Amato, Jeffrey Barrett, Sonia Goncalves, Ewan Harrison, David K. Jackson, Ian Johnston, Dominic Kwiatkowski, Cordelia Langford, John Sillitoe on behalf of the Wellcome Sanger Institute COVID-19 Surveillance Team |
| EPI_ISL_3223772 | Lighthouse Lab in Glasgow | Wellcome Sanger Institute for the COVID-19 Genomics UK (COG-UK) Consortium | Harper VanSteenhouse, Yumi Kasai, David Gray, Carol Clugston, Anna Dominiczak and Alex Alderton, Roberto Amato, Jeffrey Barrett, Sonia Goncalves, Ewan Harrison, David K. Jackson, Ian Johnston, Dominic Kwiatkowski, Cordelia Langford, John Sillitoe on behalf of the Wellcome Sanger Institute COVID-19 Surveillance Team |
| EPI_ISL_3223773 | Lighthouse Lab in Milton Keynes | Wellcome Sanger Institute for the COVID-19 Genomics UK (COG-UK) Consortium | The Lighthouse Lab in Milton Keynes and Alex Alderton, Roberto Amato, Jeffrey Barrett, Sonia Goncalves, Ewan Harrison, David K. Jackson, Ian Johnston, Dominic Kwiatkowski, Cordelia Langford, John Sillitoe on behalf of the Wellcome Sanger Institute COVID-19 Surveillance Team |
| EPI_ISL_3223774, EPI_ISL_3223775 | Lighthouse Lab in Alderley Park | Wellcome Sanger Institute for the COVID-19 Genomics UK (COG-UK) Consortium | Jacquelyn Wynn, Mairead Hyland, The Lighthouse Lab in Alderley Park and Alex Alderton, Roberto Amato, Jeffrey Barrett, Sonia Goncalves, Ewan Harrison, David K. Jackson, Ian Johnston, Dominic Kwiatkowski, Cordelia Langford, John Sillitoe on behalf of the Wellcome Sanger Institute COVID-19 Surveillance Team |
| EPI_ISL_3223776 | Lighthouse Lab in Glasgow | Wellcome Sanger Institute for the COVID-19 Genomics UK (COG-UK) Consortium | Harper VanSteenhouse, Yumi Kasai, David Gray, Carol Clugston, Anna Dominiczak and Alex Alderton, Roberto Amato, Jeffrey Barrett, Sonia Goncalves, Ewan Harrison, David K. Jackson, Ian Johnston, Dominic Kwiatkowski, Cordelia Langford, John Sillitoe on behalf of the Wellcome Sanger Institute COVID-19 Surveillance Team |
| EPI_ISL_3223777, EPI_ISL_3223778 | Lighthouse Lab in Alderley Park | Wellcome Sanger Institute for the COVID-19 Genomics UK (COG-UK) Consortium | Jacquelyn Wynn, Mairead Hyland, The Lighthouse Lab in Alderley Park and Alex Alderton, Roberto Amato, Jeffrey Barrett, Sonia Goncalves, Ewan Harrison, David K. Jackson, Ian Johnston, Dominic Kwiatkowski, Cordelia Langford, John Sillitoe on behalf of the Wellcome Sanger Institute COVID-19 Surveillance Team |
| EPI_ISL_3223779, EPI_ISL_3223780 | Lighthouse Laboratory Plymouth | Wellcome Sanger Institute for the COVID-19 Genomics UK (COG-UK) Consortium | Lighthouse Laboratory Plymouth and Alex Alderton, Roberto Amato, Jeffrey Barrett, Sonia Goncalves, Ewan Harrison, David K. Jackson, Ian Johnston, Dominic Kwiatkowski, Cordelia Langford, John Sillitoe on behalf of the Wellcome Sanger Institute COVID-19 Surveillance Team |
| EPI_ISL_3223781, EPI_ISL_3223782 | Lighthouse Lab in Glasgow | Wellcome Sanger Institute for the COVID-19 Genomics UK (COG-UK) Consortium | Harper VanSteenhouse, Yumi Kasai, David Gray, Carol Clugston, Anna Dominiczak and Alex Alderton, Roberto Amato, Jeffrey Barrett, Sonia Goncalves, Ewan Harrison, David K. Jackson, Ian Johnston, Dominic Kwiatkowski, Cordelia Langford, John Sillitoe on behalf of the Wellcome Sanger Institute COVID-19 Surveillance Team |
| EPI_ISL_3223783 | Lighthouse Laboratory Plymouth | Wellcome Sanger Institute for the COVID-19 Genomics UK (COG-UK) Consortium | Lighthouse Laboratory Plymouth and Alex Alderton, Roberto Amato, Jeffrey Barrett, Sonia Goncalves, Ewan Harrison, David K. Jackson, Ian Johnston, Dominic Kwiatkowski, Cordelia Langford, John Sillitoe on behalf of the Wellcome Sanger Institute COVID-19 Surveillance Team |
| EPI_ISL_3223784, EPI_ISL_3223785 | Lighthouse Lab in Alderley Park | Wellcome Sanger Institute for the COVID-19 Genomics UK (COG-UK) Consortium | Jacquelyn Wynn, Mairead Hyland, The Lighthouse Lab in Alderley Park and Alex Alderton, Roberto Amato, Jeffrey Barrett, Sonia Goncalves, Ewan Harrison, David K. Jackson, Ian Johnston, Dominic Kwiatkowski, Cordelia Langford, John Sillitoe on behalf of the Wellcome Sanger Institute COVID-19 Surveillance Team |
| EPI_ISL_3223786 | Lighthouse Lab in Milton Keynes | Wellcome Sanger Institute for the COVID-19 Genomics UK (COG-UK) Consortium | The Lighthouse Lab in Milton Keynes and Alex Alderton, Roberto Amato, Jeffrey Barrett, Sonia Goncalves, Ewan Harrison, David K. Jackson, Ian Johnston, Dominic Kwiatkowski, Cordelia Langford, John Sillitoe on behalf of the Wellcome Sanger Institute COVID-19 Surveillance Team |
| EPI_ISL_3223787 | Lighthouse Laboratory Plymouth | Wellcome Sanger Institute for the COVID-19 Genomics UK (COG-UK) Consortium | Lighthouse Laboratory Plymouth and Alex Alderton, Roberto Amato, Jeffrey Barrett, Sonia Goncalves, Ewan Harrison, David K. Jackson, Ian Johnston, Dominic Kwiatkowski, Cordelia Langford, John Sillitoe on behalf of the Wellcome Sanger Institute COVID-19 Surveillance Team |
| EPI_ISL_3223788 | Lighthouse Lab in Glasgow | Wellcome Sanger Institute for the COVID-19 Genomics UK (COG-UK) Consortium | Harper VanSteenhouse, Yumi Kasai, David Gray, Carol Clugston, Anna Dominiczak and Alex Alderton, Roberto Amato, Jeffrey Barrett, Sonia Goncalves, Ewan Harrison, David K. Jackson, Ian Johnston, Dominic Kwiatkowski, Cordelia Langford, John Sillitoe on behalf of the Wellcome Sanger Institute COVID-19 Surveillance Team |
| EPI_ISL_3223789 | Lighthouse Lab in Alderley Park | Wellcome Sanger Institute for the COVID-19 Genomics UK (COG-UK) Consortium | Jacquelyn Wynn, Mairead Hyland, The Lighthouse Lab in Alderley Park and Alex Alderton, Roberto Amato, Jeffrey Barrett, Sonia Goncalves, Ewan Harrison, David K. Jackson, Ian Johnston, Dominic Kwiatkowski, Cordelia Langford, John Sillitoe on behalf of the Wellcome Sanger Institute COVID-19 Surveillance Team |
| EPI_ISL_3223790 | Lighthouse Laboratory Plymouth | Wellcome Sanger Institute for the COVID-19 Genomics UK (COG-UK) Consortium | Lighthouse Laboratory Plymouth and Alex Alderton, Roberto Amato, Jeffrey Barrett, Sonia Goncalves, Ewan Harrison, David K. Jackson, Ian Johnston, Dominic Kwiatkowski, Cordelia Langford, John Sillitoe on behalf of the Wellcome Sanger Institute COVID-19 Surveillance Team |
| EPI_ISL_3223791, EPI_ISL_3223792 | Lighthouse Lab in Milton Keynes | Wellcome Sanger Institute for the COVID-19 Genomics UK (COG-UK) Consortium | The Lighthouse Lab in Milton Keynes and Alex Alderton, Roberto Amato, Jeffrey Barrett, Sonia Goncalves, Ewan Harrison, David K. Jackson, Ian Johnston, Dominic Kwiatkowski, Cordelia Langford, John Sillitoe on behalf of the Wellcome Sanger Institute COVID-19 Surveillance Team |
| EPI_ISL_3223793, EPI_ISL_3223794 | Lighthouse Laboratory Plymouth | Wellcome Sanger Institute for the COVID-19 Genomics UK (COG-UK) Consortium | Lighthouse Laboratory Plymouth and Alex Alderton, Roberto Amato, Jeffrey Barrett, Sonia Goncalves, Ewan Harrison, David K. Jackson, Ian Johnston, Dominic Kwiatkowski, Cordelia Langford, John Sillitoe on behalf of the Wellcome Sanger Institute COVID-19 Surveillance Team |
| EPI_ISL_3223795 | Lighthouse Lab in Glasgow | Wellcome Sanger Institute for the COVID-19 Genomics UK (COG-UK) | Harper VanSteenhouse, Yumi Kasai, David Gray, Carol Clugston, Anna Dominiczak and Alex Alderton, Roberto Amato, Jeffrey Barrett, Sonia Goncalves, Ewan Harrison, David K. Jackson, Ian Johnston, Dominic Kwiatkowski, Cordelia Langford, |

| | | | |
|---|---|---|---|
| | | Consortium | John Sillitoe on behalf of the Wellcome Sanger Institute COVID-19 Surveillance Team |
| EPI_ISL_3223796, EPI_ISL_3223797 | Lighthouse Lab in Alderley Park | Wellcome Sanger Institute for the COVID-19 Genomics UK (COG-UK) Consortium | Jacquelyn Wynn, Mairead Hyland, The Lighthouse Lab in Alderley Park and Alex Alderton, Roberto Amato, Jeffrey Barrett, Sonia Goncalves, Ewan Harrison, David K. Jackson, Ian Johnston, Dominic Kwiatkowski, Cordelia Langford, John Sillitoe on behalf of the Wellcome Sanger Institute COVID-19 Surveillance Team |
| EPI_ISL_3223798 | Lighthouse Laboratory Plymouth | Wellcome Sanger Institute for the COVID-19 Genomics UK (COG-UK) Consortium | Lighthouse Laboratory Plymouth and Alex Alderton, Roberto Amato, Jeffrey Barrett, Sonia Goncalves, Ewan Harrison, David K. Jackson, Ian Johnston, Dominic Kwiatkowski, Cordelia Langford, John Sillitoe on behalf of the Wellcome Sanger Institute COVID-19 Surveillance Team |
| EPI_ISL_3223799 | Lighthouse Lab in Glasgow | Wellcome Sanger Institute for the COVID-19 Genomics UK (COG-UK) Consortium | Harper VanSteenhouse, Yumi Kasai, David Gray, Carol Clugston, Anna Dominiczak and Alex Alderton, Roberto Amato, Jeffrey Barrett, Sonia Goncalves, Ewan Harrison, David K. Jackson, Ian Johnston, Dominic Kwiatkowski, Cordelia Langford, John Sillitoe on behalf of the Wellcome Sanger Institute COVID-19 Surveillance Team |
| EPI_ISL_3223800 | Lighthouse Lab in Milton Keynes | Wellcome Sanger Institute for the COVID-19 Genomics UK (COG-UK) Consortium | The Lighthouse Lab in Milton Keynes and Alex Alderton, Roberto Amato, Jeffrey Barrett, Sonia Goncalves, Ewan Harrison, David K. Jackson, Ian Johnston, Dominic Kwiatkowski, Cordelia Langford, John Sillitoe on behalf of the Wellcome Sanger Institute COVID-19 Surveillance Team |
| EPI_ISL_3223801 | Lighthouse Lab in Alderley Park | Wellcome Sanger Institute for the COVID-19 Genomics UK (COG-UK) Consortium | Jacquelyn Wynn, Mairead Hyland, The Lighthouse Lab in Alderley Park and Alex Alderton, Roberto Amato, Jeffrey Barrett, Sonia Goncalves, Ewan Harrison, David K. Jackson, Ian Johnston, Dominic Kwiatkowski, Cordelia Langford, John Sillitoe on behalf of the Wellcome Sanger Institute COVID-19 Surveillance Team |
| EPI_ISL_3223802 | Lighthouse Lab in Glasgow | Wellcome Sanger Institute for the COVID-19 Genomics UK (COG-UK) Consortium | Harper VanSteenhouse, Yumi Kasai, David Gray, Carol Clugston, Anna Dominiczak and Alex Alderton, Roberto Amato, Jeffrey Barrett, Sonia Goncalves, Ewan Harrison, David K. Jackson, Ian Johnston, Dominic Kwiatkowski, Cordelia Langford, John Sillitoe on behalf of the Wellcome Sanger Institute COVID-19 Surveillance Team |
| EPI_ISL_3223803 | Lighthouse Lab in Alderley Park | Wellcome Sanger Institute for the COVID-19 Genomics UK (COG-UK) Consortium | Jacquelyn Wynn, Mairead Hyland, The Lighthouse Lab in Alderley Park and Alex Alderton, Roberto Amato, Jeffrey Barrett, Sonia Goncalves, Ewan Harrison, David K. Jackson, Ian Johnston, Dominic Kwiatkowski, Cordelia Langford, John Sillitoe on behalf of the Wellcome Sanger Institute COVID-19 Surveillance Team |
| EPI_ISL_3223804 | Lighthouse Lab in Glasgow | Wellcome Sanger Institute for the COVID-19 Genomics UK (COG-UK) Consortium | Harper VanSteenhouse, Yumi Kasai, David Gray, Carol Clugston, Anna Dominiczak and Alex Alderton, Roberto Amato, Jeffrey Barrett, Sonia Goncalves, Ewan Harrison, David K. Jackson, Ian Johnston, Dominic Kwiatkowski, Cordelia Langford, John Sillitoe on behalf of the Wellcome Sanger Institute COVID-19 Surveillance Team |
| EPI_ISL_3223805 | Lighthouse Laboratory Plymouth | Wellcome Sanger Institute for the COVID-19 Genomics UK (COG-UK) Consortium | Lighthouse Laboratory Plymouth and Alex Alderton, Roberto Amato, Jeffrey Barrett, Sonia Goncalves, Ewan Harrison, David K. Jackson, Ian Johnston, Dominic Kwiatkowski, Cordelia Langford, John Sillitoe on behalf of the Wellcome Sanger Institute COVID-19 Surveillance Team |
| EPI_ISL_3223806 | Lighthouse Lab in Glasgow | Wellcome Sanger Institute for the COVID-19 Genomics UK (COG-UK) Consortium | Harper VanSteenhouse, Yumi Kasai, David Gray, Carol Clugston, Anna Dominiczak and Alex Alderton, Roberto Amato, Jeffrey Barrett, Sonia Goncalves, Ewan Harrison, David K. Jackson, Ian Johnston, Dominic Kwiatkowski, Cordelia Langford, John Sillitoe on behalf of the Wellcome Sanger Institute COVID-19 Surveillance Team |
| EPI_ISL_3223807 | Lighthouse Laboratory Plymouth | Wellcome Sanger Institute for the COVID-19 Genomics UK (COG-UK) Consortium | Lighthouse Laboratory Plymouth and Alex Alderton, Roberto Amato, Jeffrey Barrett, Sonia Goncalves, Ewan Harrison, David K. Jackson, Ian Johnston, Dominic Kwiatkowski, Cordelia Langford, John Sillitoe on behalf of the Wellcome Sanger Institute COVID-19 Surveillance Team |
| EPI_ISL_3223808, EPI_ISL_3223809 | Lighthouse Lab in Milton Keynes | Wellcome Sanger Institute for the COVID-19 Genomics UK (COG-UK) Consortium | The Lighthouse Lab in Milton Keynes and Alex Alderton, Roberto Amato, Jeffrey Barrett, Sonia Goncalves, Ewan Harrison, David K. Jackson, Ian Johnston, Dominic Kwiatkowski, Cordelia Langford, John Sillitoe on behalf of the Wellcome Sanger Institute COVID-19 Surveillance Team |
| EPI_ISL_3223810 | Lighthouse Lab in Glasgow | Wellcome Sanger Institute for the COVID-19 Genomics UK (COG-UK) Consortium | Harper VanSteenhouse, Yumi Kasai, David Gray, Carol Clugston, Anna Dominiczak and Alex Alderton, Roberto Amato, Jeffrey Barrett, Sonia Goncalves, Ewan Harrison, David K. Jackson, Ian Johnston, Dominic Kwiatkowski, Cordelia Langford, John Sillitoe on behalf of the Wellcome Sanger Institute COVID-19 Surveillance Team |
| EPI_ISL_3223811, EPI_ISL_3223812, EPI_ISL_3223813 | Lighthouse Laboratory Plymouth | Wellcome Sanger Institute for the COVID-19 Genomics UK (COG-UK) Consortium | Lighthouse Laboratory Plymouth and Alex Alderton, Roberto Amato, Jeffrey Barrett, Sonia Goncalves, Ewan Harrison, David K. Jackson, Ian Johnston, Dominic Kwiatkowski, Cordelia Langford, John Sillitoe on behalf of the Wellcome Sanger Institute COVID-19 Surveillance Team |
| EPI_ISL_3223814 | Lighthouse Lab in Glasgow | Wellcome Sanger Institute for the COVID-19 Genomics UK (COG-UK) Consortium | Harper VanSteenhouse, Yumi Kasai, David Gray, Carol Clugston, Anna Dominiczak and Alex Alderton, Roberto Amato, Jeffrey Barrett, Sonia Goncalves, Ewan Harrison, David K. Jackson, Ian Johnston, Dominic Kwiatkowski, Cordelia Langford, John Sillitoe on behalf of the Wellcome Sanger Institute COVID-19 Surveillance Team |
| EPI_ISL_3223815 | Lighthouse Laboratory Plymouth | Wellcome Sanger Institute for the COVID-19 Genomics UK (COG-UK) Consortium | Lighthouse Laboratory Plymouth and Alex Alderton, Roberto Amato, Jeffrey Barrett, Sonia Goncalves, Ewan Harrison, David K. Jackson, Ian Johnston, Dominic Kwiatkowski, Cordelia Langford, John Sillitoe on behalf of the Wellcome Sanger Institute COVID-19 Surveillance Team |
| EPI_ISL_3223816, EPI_ISL_3223817 | Lighthouse Lab in Glasgow | Wellcome Sanger Institute for the COVID-19 Genomics UK (COG-UK) Consortium | Harper VanSteenhouse, Yumi Kasai, David Gray, Carol Clugston, Anna Dominiczak and Alex Alderton, Roberto Amato, Jeffrey Barrett, Sonia Goncalves, Ewan Harrison, David K. Jackson, Ian Johnston, Dominic Kwiatkowski, Cordelia Langford, John Sillitoe on behalf of the Wellcome Sanger Institute COVID-19 Surveillance Team |
| EPI_ISL_3223818, EPI_ISL_3223819, EPI_ISL_3223820, EPI_ISL_3223821 | Lighthouse Lab in Alderley Park | Wellcome Sanger Institute for the COVID-19 Genomics UK (COG-UK) Consortium | Jacquelyn Wynn, Mairead Hyland, The Lighthouse Lab in Alderley Park and Alex Alderton, Roberto Amato, Jeffrey Barrett, Sonia Goncalves, Ewan Harrison, David K. Jackson, Ian Johnston, Dominic Kwiatkowski, Cordelia Langford, John Sillitoe on behalf of the Wellcome Sanger Institute COVID-19 Surveillance Team |
| EPI_ISL_3223822 | Lighthouse Laboratory Plymouth | Wellcome Sanger Institute for the COVID-19 Genomics UK (COG-UK) Consortium | Lighthouse Laboratory Plymouth and Alex Alderton, Roberto Amato, Jeffrey Barrett, Sonia Goncalves, Ewan Harrison, David K. Jackson, Ian Johnston, Dominic Kwiatkowski, Cordelia Langford, John Sillitoe on behalf of the Wellcome Sanger Institute COVID-19 Surveillance Team |
| EPI_ISL_3223823 | Lighthouse Lab in Alderley Park | Wellcome Sanger Institute for the COVID-19 Genomics UK (COG-UK) Consortium | Jacquelyn Wynn, Mairead Hyland, The Lighthouse Lab in Alderley Park and Alex Alderton, Roberto Amato, Jeffrey Barrett, Sonia Goncalves, Ewan Harrison, David K. Jackson, Ian Johnston, Dominic Kwiatkowski, Cordelia Langford, John Sillitoe on behalf of the Wellcome Sanger Institute COVID-19 Surveillance Team |
| EPI_ISL_3223824, EPI_ISL_3223825, EPI_ISL_3223826 | Lighthouse Lab in Milton Keynes | Wellcome Sanger Institute for the COVID-19 Genomics UK (COG-UK) Consortium | The Lighthouse Lab in Milton Keynes and Alex Alderton, Roberto Amato, Jeffrey Barrett, Sonia Goncalves, Ewan Harrison, David K. Jackson, Ian Johnston, Dominic Kwiatkowski, Cordelia Langford, John Sillitoe on behalf of the Wellcome Sanger Institute COVID-19 Surveillance Team |
| EPI_ISL_3223827, EPI_ISL_3223828, EPI_ISL_3223829 | Lighthouse Lab in Glasgow | Wellcome Sanger Institute for the COVID-19 Genomics UK (COG-UK) Consortium | Harper VanSteenhouse, Yumi Kasai, David Gray, Carol Clugston, Anna Dominiczak and Alex Alderton, Roberto Amato, Jeffrey Barrett, Sonia Goncalves, Ewan Harrison, David K. Jackson, Ian Johnston, Dominic Kwiatkowski, Cordelia Langford, John Sillitoe on behalf of the Wellcome Sanger Institute COVID-19 Surveillance Team |
| EPI_ISL_3223830 | Lighthouse Laboratory Plymouth | Wellcome Sanger Institute for the COVID-19 Genomics UK (COG-UK) Consortium | Lighthouse Laboratory Plymouth and Alex Alderton, Roberto Amato, Jeffrey Barrett, Sonia Goncalves, Ewan Harrison, David K. Jackson, Ian Johnston, Dominic Kwiatkowski, Cordelia Langford, John Sillitoe on behalf of the Wellcome Sanger Institute COVID-19 Surveillance Team |
| EPI_ISL_3223831, EPI_ISL_3223832 | Lighthouse Lab in Glasgow | Wellcome Sanger Institute for the COVID-19 Genomics UK (COG-UK) Consortium | Harper VanSteenhouse, Yumi Kasai, David Gray, Carol Clugston, Anna Dominiczak and Alex Alderton, Roberto Amato, Jeffrey Barrett, Sonia Goncalves, Ewan Harrison, David K. Jackson, Ian Johnston, Dominic Kwiatkowski, Cordelia Langford, John Sillitoe on behalf of the Wellcome Sanger Institute COVID-19 Surveillance Team |
| EPI_ISL_3223833 | Lighthouse Laboratory Plymouth | Wellcome Sanger Institute for the COVID-19 Genomics UK (COG-UK) Consortium | Lighthouse Laboratory Plymouth and Alex Alderton, Roberto Amato, Jeffrey Barrett, Sonia Goncalves, Ewan Harrison, David K. Jackson, Ian Johnston, Dominic Kwiatkowski, Cordelia Langford, John Sillitoe on behalf of the Wellcome Sanger Institute COVID-19 Surveillance Team |
| EPI_ISL_3223834 | Lighthouse Lab in Milton Keynes | Wellcome Sanger Institute for the COVID-19 Genomics UK (COG-UK) Consortium | The Lighthouse Lab in Milton Keynes and Alex Alderton, Roberto Amato, Jeffrey Barrett, Sonia Goncalves, Ewan Harrison, David K. Jackson, Ian Johnston, Dominic Kwiatkowski, Cordelia Langford, John Sillitoe on behalf of the Wellcome Sanger Institute COVID-19 Surveillance Team |
| EPI_ISL_3223835, EPI_ISL_3223836 | Lighthouse Lab in Alderley Park | Wellcome Sanger Institute for the COVID-19 Genomics UK (COG-UK) Consortium | Jacquelyn Wynn, Mairead Hyland, The Lighthouse Lab in Alderley Park and Alex Alderton, Roberto Amato, Jeffrey Barrett, Sonia Goncalves, Ewan Harrison, David K. Jackson, Ian Johnston, Dominic Kwiatkowski, Cordelia Langford, John Sillitoe on behalf of the Wellcome Sanger Institute COVID-19 Surveillance Team |
| EPI_ISL_3223837 | Lighthouse Lab in Glasgow | Wellcome Sanger Institute for the COVID-19 Genomics UK (COG-UK) Consortium | Harper VanSteenhouse, Yumi Kasai, David Gray, Carol Clugston, Anna Dominiczak and Alex Alderton, Roberto Amato, Jeffrey Barrett, Sonia Goncalves, Ewan Harrison, David K. Jackson, Ian Johnston, Dominic Kwiatkowski, Cordelia Langford, John Sillitoe on behalf of the Wellcome Sanger Institute COVID-19 Surveillance Team |
| EPI_ISL_3223838 | Lighthouse Lab in Milton Keynes | Wellcome Sanger Institute for the COVID-19 Genomics UK (COG-UK) Consortium | The Lighthouse Lab in Milton Keynes and Alex Alderton, Roberto Amato, Jeffrey Barrett, Sonia Goncalves, Ewan Harrison, David K. Jackson, Ian Johnston, Dominic Kwiatkowski, Cordelia Langford, John Sillitoe on behalf of the Wellcome Sanger Institute COVID-19 Surveillance Team |
| EPI_ISL_3223839 | Lighthouse Laboratory Plymouth | Wellcome Sanger Institute for the COVID-19 Genomics UK (COG-UK) Consortium | Lighthouse Laboratory Plymouth and Alex Alderton, Roberto Amato, Jeffrey Barrett, Sonia Goncalves, Ewan Harrison, David K. Jackson, Ian Johnston, Dominic Kwiatkowski, Cordelia Langford, John Sillitoe on behalf of the Wellcome Sanger Institute COVID-19 Surveillance Team |
| EPI_ISL_3223840 | Lighthouse Lab in Glasgow | Wellcome Sanger Institute for the COVID-19 Genomics UK (COG-UK) Consortium | Harper VanSteenhouse, Yumi Kasai, David Gray, Carol Clugston, Anna Dominiczak and Alex Alderton, Roberto Amato, Jeffrey Barrett, Sonia Goncalves, Ewan Harrison, David K. Jackson, Ian Johnston, Dominic Kwiatkowski, Cordelia Langford, John Sillitoe on behalf of the Wellcome Sanger Institute COVID-19 Surveillance Team |
| EPI_ISL_3223841 | Lighthouse Laboratory Plymouth | Wellcome Sanger Institute for the COVID-19 Genomics UK (COG-UK) Consortium | Lighthouse Laboratory Plymouth and Alex Alderton, Roberto Amato, Jeffrey Barrett, Sonia Goncalves, Ewan Harrison, David K. Jackson, Ian Johnston, Dominic Kwiatkowski, Cordelia Langford, John Sillitoe on behalf of the Wellcome Sanger Institute COVID-19 Surveillance Team |
| EPI_ISL_3223842 | Lighthouse Lab in Glasgow | Wellcome Sanger Institute for the COVID-19 Genomics UK (COG-UK) Consortium | Harper VanSteenhouse, Yumi Kasai, David Gray, Carol Clugston, Anna Dominiczak and Alex Alderton, Roberto Amato, Jeffrey Barrett, Sonia Goncalves, Ewan Harrison, David K. Jackson, Ian Johnston, Dominic Kwiatkowski, Cordelia Langford, John Sillitoe on behalf of the Wellcome Sanger Institute COVID-19 Surveillance Team |
| EPI_ISL_3223843 | Lighthouse Lab in Milton Keynes | Wellcome Sanger Institute for the COVID-19 Genomics UK (COG-UK) Consortium | The Lighthouse Lab in Milton Keynes and Alex Alderton, Roberto Amato, Jeffrey Barrett, Sonia Goncalves, Ewan Harrison, David K. Jackson, Ian Johnston, Dominic Kwiatkowski, Cordelia Langford, John Sillitoe on behalf of the Wellcome Sanger Institute COVID-19 Surveillance Team |
| EPI_ISL_3223844 | Lighthouse Lab in Alderley Park | Wellcome Sanger Institute for the COVID-19 Genomics UK (COG-UK) Consortium | Jacquelyn Wynn, Mairead Hyland, The Lighthouse Lab in Alderley Park and Alex Alderton, Roberto Amato, Jeffrey Barrett, Sonia Goncalves, Ewan Harrison, David K. Jackson, Ian Johnston, Dominic Kwiatkowski, Cordelia Langford, John Sillitoe on behalf of the Wellcome Sanger Institute COVID-19 Surveillance Team |
| EPI_ISL_3223845 | Lighthouse Lab in Glasgow | Wellcome Sanger Institute for the COVID-19 Genomics UK (COG-UK) Consortium | Harper VanSteenhouse, Yumi Kasai, David Gray, Carol Clugston, Anna Dominiczak and Alex Alderton, Roberto Amato, Jeffrey Barrett, Sonia Goncalves, Ewan Harrison, David K. Jackson, Ian Johnston, Dominic Kwiatkowski, Cordelia Langford, John Sillitoe on behalf of the Wellcome Sanger Institute COVID-19 Surveillance Team |
| EPI_ISL_3223846, EPI_ISL_3223847 | Lighthouse Laboratory Plymouth | Wellcome Sanger Institute for the COVID-19 Genomics UK (COG-UK) Consortium | Lighthouse Laboratory Plymouth and Alex Alderton, Roberto Amato, Jeffrey Barrett, Sonia Goncalves, Ewan Harrison, David K. Jackson, Ian Johnston, Dominic Kwiatkowski, Cordelia Langford, John Sillitoe on behalf of the Wellcome Sanger Institute COVID-19 Surveillance Team |
| EPI_ISL_3223848 | Lighthouse Lab in Glasgow | Wellcome Sanger Institute for the COVID-19 Genomics UK (COG-UK) Consortium | Harper VanSteenhouse, Yumi Kasai, David Gray, Carol Clugston, Anna Dominiczak and Alex Alderton, Roberto Amato, Jeffrey Barrett, Sonia Goncalves, Ewan Harrison, David K. Jackson, Ian Johnston, Dominic Kwiatkowski, Cordelia Langford, John Sillitoe on behalf of the Wellcome Sanger Institute COVID-19 Surveillance Team |
| EPI_ISL_3223849 | Lighthouse Lab in Milton Keynes | Wellcome Sanger Institute for the COVID-19 Genomics UK (COG-UK) Consortium | The Lighthouse Lab in Milton Keynes and Alex Alderton, Roberto Amato, Jeffrey Barrett, Sonia Goncalves, Ewan Harrison, David K. Jackson, Ian Johnston, Dominic Kwiatkowski, Cordelia Langford, John Sillitoe on behalf of the Wellcome Sanger Institute COVID-19 Surveillance Team |
| EPI_ISL_3223850 | Lighthouse Lab in Glasgow | Wellcome Sanger Institute for the COVID-19 Genomics UK (COG-UK) Consortium | Harper VanSteenhouse, Yumi Kasai, David Gray, Carol Clugston, Anna Dominiczak and Alex Alderton, Roberto Amato, Jeffrey Barrett, Sonia Goncalves, Ewan Harrison, David K. Jackson, Ian Johnston, Dominic Kwiatkowski, Cordelia Langford, John Sillitoe on behalf of the Wellcome Sanger Institute COVID-19 Surveillance Team |
| EPI_ISL_3223851 | Lighthouse Lab in Alderley Park | Wellcome Sanger Institute for the COVID-19 Genomics UK (COG-UK) Consortium | Jacquelyn Wynn, Mairead Hyland, The Lighthouse Lab in Alderley Park and Alex Alderton, Roberto Amato, Jeffrey Barrett, Sonia Goncalves, Ewan Harrison, David K. Jackson, Ian Johnston, Dominic Kwiatkowski, Cordelia Langford, John Sillitoe on behalf of the Wellcome Sanger Institute COVID-19 Surveillance Team |
| EPI_ISL_3223852 | Lighthouse Lab in Milton Keynes | Wellcome Sanger Institute for the COVID-19 Genomics UK (COG-UK) Consortium | The Lighthouse Lab in Milton Keynes and Alex Alderton, Roberto Amato, Jeffrey Barrett, Sonia Goncalves, Ewan Harrison, David K. Jackson, Ian Johnston, Dominic Kwiatkowski, Cordelia Langford, John Sillitoe on behalf of the Wellcome Sanger Institute COVID-19 Surveillance Team |
| EPI_ISL_3223853 | Lighthouse Laboratory Plymouth | Wellcome Sanger Institute for the COVID-19 Genomics UK (COG-UK) | Lighthouse Laboratory Plymouth and Alex Alderton, Roberto Amato, Jeffrey Barrett, Sonia Goncalves, Ewan Harrison, David K. Jackson, Ian Johnston, Dominic Kwiatkowski, Cordelia Langford, John Sillitoe on behalf of the Wellcome Sanger |

| | | | |
|---|---|---|---|
| | | Consortium | Institute COVID-19 Surveillance Team |
| EPI_ISL_3223854 | Lighthouse Lab in Milton Keynes | Wellcome Sanger Institute for the COVID-19 Genomics UK (COG-UK) Consortium | The Lighthouse Lab in Milton Keynes and Alex Alderton, Roberto Amato, Jeffrey Barrett, Sonia Goncalves, Ewan Harrison, David K. Jackson, Ian Johnston, Dominic Kwiatkowski, Cordelia Langford, John Sillitoe on behalf of the Wellcome Sanger Institute COVID-19 Surveillance Team |
| EPI_ISL_3223855 | Lighthouse Lab in Glasgow | Wellcome Sanger Institute for the COVID-19 Genomics UK (COG-UK) Consortium | Harper VanSteenhouse, Yumi Kasai, David Gray, Carol Clugston, Anna Dominiczak and Alex Alderton, Roberto Amato, Jeffrey Barrett, Sonia Goncalves, Ewan Harrison, David K. Jackson, Ian Johnston, Dominic Kwiatkowski, Cordelia Langford, John Sillitoe on behalf of the Wellcome Sanger Institute COVID-19 Surveillance Team |
| EPI_ISL_3223856 | Lighthouse Lab in Alderley Park | Wellcome Sanger Institute for the COVID-19 Genomics UK (COG-UK) Consortium | Jacquelyn Wynn, Mairead Hyland, The Lighthouse Lab in Alderley Park and Alex Alderton, Roberto Amato, Jeffrey Barrett, Sonia Goncalves, Ewan Harrison, David K. Jackson, Ian Johnston, Dominic Kwiatkowski, Cordelia Langford, John Sillitoe on behalf of the Wellcome Sanger Institute COVID-19 Surveillance Team |
| EPI_ISL_3223857, EPI_ISL_3223858, EPI_ISL_3223859 | Lighthouse Lab in Glasgow | Wellcome Sanger Institute for the COVID-19 Genomics UK (COG-UK) Consortium | Harper VanSteenhouse, Yumi Kasai, David Gray, Carol Clugston, Anna Dominiczak and Alex Alderton, Roberto Amato, Jeffrey Barrett, Sonia Goncalves, Ewan Harrison, David K. Jackson, Ian Johnston, Dominic Kwiatkowski, Cordelia Langford, John Sillitoe on behalf of the Wellcome Sanger Institute COVID-19 Surveillance Team |
| EPI_ISL_3223860 | Lighthouse Lab in Alderley Park | Wellcome Sanger Institute for the COVID-19 Genomics UK (COG-UK) Consortium | Jacquelyn Wynn, Mairead Hyland, The Lighthouse Lab in Alderley Park and Alex Alderton, Roberto Amato, Jeffrey Barrett, Sonia Goncalves, Ewan Harrison, David K. Jackson, Ian Johnston, Dominic Kwiatkowski, Cordelia Langford, John Sillitoe on behalf of the Wellcome Sanger Institute COVID-19 Surveillance Team |
| EPI_ISL_3223861 | Lighthouse Lab in Milton Keynes | Wellcome Sanger Institute for the COVID-19 Genomics UK (COG-UK) Consortium | The Lighthouse Lab in Milton Keynes and Alex Alderton, Roberto Amato, Jeffrey Barrett, Sonia Goncalves, Ewan Harrison, David K. Jackson, Ian Johnston, Dominic Kwiatkowski, Cordelia Langford, John Sillitoe on behalf of the Wellcome Sanger Institute COVID-19 Surveillance Team |
| EPI_ISL_3223862, EPI_ISL_3223863 | Lighthouse Lab in Glasgow | Wellcome Sanger Institute for the COVID-19 Genomics UK (COG-UK) Consortium | Harper VanSteenhouse, Yumi Kasai, David Gray, Carol Clugston, Anna Dominiczak and Alex Alderton, Roberto Amato, Jeffrey Barrett, Sonia Goncalves, Ewan Harrison, David K. Jackson, Ian Johnston, Dominic Kwiatkowski, Cordelia Langford, John Sillitoe on behalf of the Wellcome Sanger Institute COVID-19 Surveillance Team |
| EPI_ISL_3223864 | Lighthouse Laboratory Plymouth | Wellcome Sanger Institute for the COVID-19 Genomics UK (COG-UK) Consortium | Lighthouse Laboratory Plymouth and Alex Alderton, Roberto Amato, Jeffrey Barrett, Sonia Goncalves, Ewan Harrison, David K. Jackson, Ian Johnston, Dominic Kwiatkowski, Cordelia Langford, John Sillitoe on behalf of the Wellcome Sanger Institute COVID-19 Surveillance Team |
| EPI_ISL_3223865 | Lighthouse Lab in Milton Keynes | Wellcome Sanger Institute for the COVID-19 Genomics UK (COG-UK) Consortium | The Lighthouse Lab in Milton Keynes and Alex Alderton, Roberto Amato, Jeffrey Barrett, Sonia Goncalves, Ewan Harrison, David K. Jackson, Ian Johnston, Dominic Kwiatkowski, Cordelia Langford, John Sillitoe on behalf of the Wellcome Sanger Institute COVID-19 Surveillance Team |
| EPI_ISL_3223866, EPI_ISL_3223867 | Lighthouse Laboratory Plymouth | Wellcome Sanger Institute for the COVID-19 Genomics UK (COG-UK) Consortium | Lighthouse Laboratory Plymouth and Alex Alderton, Roberto Amato, Jeffrey Barrett, Sonia Goncalves, Ewan Harrison, David K. Jackson, Ian Johnston, Dominic Kwiatkowski, Cordelia Langford, John Sillitoe on behalf of the Wellcome Sanger Institute COVID-19 Surveillance Team |
| EPI_ISL_3223868 | Lighthouse Lab in Glasgow | Wellcome Sanger Institute for the COVID-19 Genomics UK (COG-UK) Consortium | Harper VanSteenhouse, Yumi Kasai, David Gray, Carol Clugston, Anna Dominiczak and Alex Alderton, Roberto Amato, Jeffrey Barrett, Sonia Goncalves, Ewan Harrison, David K. Jackson, Ian Johnston, Dominic Kwiatkowski, Cordelia Langford, John Sillitoe on behalf of the Wellcome Sanger Institute COVID-19 Surveillance Team |
| EPI_ISL_3223869 | Lighthouse Lab in Alderley Park | Wellcome Sanger Institute for the COVID-19 Genomics UK (COG-UK) Consortium | Jacquelyn Wynn, Mairead Hyland, The Lighthouse Lab in Alderley Park and Alex Alderton, Roberto Amato, Jeffrey Barrett, Sonia Goncalves, Ewan Harrison, David K. Jackson, Ian Johnston, Dominic Kwiatkowski, Cordelia Langford, John Sillitoe on behalf of the Wellcome Sanger Institute COVID-19 Surveillance Team |
| EPI_ISL_3223870 | Lighthouse Laboratory Plymouth | Wellcome Sanger Institute for the COVID-19 Genomics UK (COG-UK) Consortium | Lighthouse Laboratory Plymouth and Alex Alderton, Roberto Amato, Jeffrey Barrett, Sonia Goncalves, Ewan Harrison, David K. Jackson, Ian Johnston, Dominic Kwiatkowski, Cordelia Langford, John Sillitoe on behalf of the Wellcome Sanger Institute COVID-19 Surveillance Team |
| EPI_ISL_3223871 | Lighthouse Lab in Milton Keynes | Wellcome Sanger Institute for the COVID-19 Genomics UK (COG-UK) Consortium | The Lighthouse Lab in Milton Keynes and Alex Alderton, Roberto Amato, Jeffrey Barrett, Sonia Goncalves, Ewan Harrison, David K. Jackson, Ian Johnston, Dominic Kwiatkowski, Cordelia Langford, John Sillitoe on behalf of the Wellcome Sanger Institute COVID-19 Surveillance Team |
| EPI_ISL_3223872 | Lighthouse Lab in Glasgow | Wellcome Sanger Institute for the COVID-19 Genomics UK (COG-UK) Consortium | Harper VanSteenhouse, Yumi Kasai, David Gray, Carol Clugston, Anna Dominiczak and Alex Alderton, Roberto Amato, Jeffrey Barrett, Sonia Goncalves, Ewan Harrison, David K. Jackson, Ian Johnston, Dominic Kwiatkowski, Cordelia Langford, John Sillitoe on behalf of the Wellcome Sanger Institute COVID-19 Surveillance Team |
| EPI_ISL_3223873, EPI_ISL_3223874 | Lighthouse Lab in Milton Keynes | Wellcome Sanger Institute for the COVID-19 Genomics UK (COG-UK) Consortium | The Lighthouse Lab in Milton Keynes and Alex Alderton, Roberto Amato, Jeffrey Barrett, Sonia Goncalves, Ewan Harrison, David K. Jackson, Ian Johnston, Dominic Kwiatkowski, Cordelia Langford, John Sillitoe on behalf of the Wellcome Sanger Institute COVID-19 Surveillance Team |
| EPI_ISL_3223875 | Lighthouse Laboratory Plymouth | Wellcome Sanger Institute for the COVID-19 Genomics UK (COG-UK) Consortium | Lighthouse Laboratory Plymouth and Alex Alderton, Roberto Amato, Jeffrey Barrett, Sonia Goncalves, Ewan Harrison, David K. Jackson, Ian Johnston, Dominic Kwiatkowski, Cordelia Langford, John Sillitoe on behalf of the Wellcome Sanger Institute COVID-19 Surveillance Team |
| EPI_ISL_3223876, EPI_ISL_3223877 | Lighthouse Lab in Milton Keynes | Wellcome Sanger Institute for the COVID-19 Genomics UK (COG-UK) Consortium | The Lighthouse Lab in Milton Keynes and Alex Alderton, Roberto Amato, Jeffrey Barrett, Sonia Goncalves, Ewan Harrison, David K. Jackson, Ian Johnston, Dominic Kwiatkowski, Cordelia Langford, John Sillitoe on behalf of the Wellcome Sanger Institute COVID-19 Surveillance Team |
| EPI_ISL_3223878 | Lighthouse Lab in Alderley Park | Wellcome Sanger Institute for the COVID-19 Genomics UK (COG-UK) Consortium | Jacquelyn Wynn, Mairead Hyland, The Lighthouse Lab in Alderley Park and Alex Alderton, Roberto Amato, Jeffrey Barrett, Sonia Goncalves, Ewan Harrison, David K. Jackson, Ian Johnston, Dominic Kwiatkowski, Cordelia Langford, John Sillitoe on behalf of the Wellcome Sanger Institute COVID-19 Surveillance Team |
| EPI_ISL_3223879 | Lighthouse Laboratory Plymouth | Wellcome Sanger Institute for the COVID-19 Genomics UK (COG-UK) Consortium | Lighthouse Laboratory Plymouth and Alex Alderton, Roberto Amato, Jeffrey Barrett, Sonia Goncalves, Ewan Harrison, David K. Jackson, Ian Johnston, Dominic Kwiatkowski, Cordelia Langford, John Sillitoe on behalf of the Wellcome Sanger Institute COVID-19 Surveillance Team |
| EPI_ISL_3223880 | Lighthouse Lab in Milton Keynes | Wellcome Sanger Institute for the COVID-19 Genomics UK (COG-UK) Consortium | The Lighthouse Lab in Milton Keynes and Alex Alderton, Roberto Amato, Jeffrey Barrett, Sonia Goncalves, Ewan Harrison, David K. Jackson, Ian Johnston, Dominic Kwiatkowski, Cordelia Langford, John Sillitoe on behalf of the Wellcome Sanger Institute COVID-19 Surveillance Team |
| EPI_ISL_3223881 | Lighthouse Laboratory Plymouth | Wellcome Sanger Institute for the COVID-19 Genomics UK (COG-UK) Consortium | Lighthouse Laboratory Plymouth and Alex Alderton, Roberto Amato, Jeffrey Barrett, Sonia Goncalves, Ewan Harrison, David K. Jackson, Ian Johnston, Dominic Kwiatkowski, Cordelia Langford, John Sillitoe on behalf of the Wellcome Sanger Institute COVID-19 Surveillance Team |
| EPI_ISL_3223882 | Lighthouse Lab in Glasgow | Wellcome Sanger Institute for the COVID-19 Genomics UK (COG-UK) Consortium | Harper VanSteenhouse, Yumi Kasai, David Gray, Carol Clugston, Anna Dominiczak and Alex Alderton, Roberto Amato, Jeffrey Barrett, Sonia Goncalves, Ewan Harrison, David K. Jackson, Ian Johnston, Dominic Kwiatkowski, Cordelia Langford, John Sillitoe on behalf of the Wellcome Sanger Institute COVID-19 Surveillance Team |
| EPI_ISL_3223883, EPI_ISL_3223884 | Lighthouse Lab in Alderley Park | Wellcome Sanger Institute for the COVID-19 Genomics UK (COG-UK) Consortium | Jacquelyn Wynn, Mairead Hyland, The Lighthouse Lab in Alderley Park and Alex Alderton, Roberto Amato, Jeffrey Barrett, Sonia Goncalves, Ewan Harrison, David K. Jackson, Ian Johnston, Dominic Kwiatkowski, Cordelia Langford, John Sillitoe on behalf of the Wellcome Sanger Institute COVID-19 Surveillance Team |
| EPI_ISL_3223885 | Lighthouse Lab in Glasgow | Wellcome Sanger Institute for the COVID-19 Genomics UK (COG-UK) Consortium | Harper VanSteenhouse, Yumi Kasai, David Gray, Carol Clugston, Anna Dominiczak and Alex Alderton, Roberto Amato, Jeffrey Barrett, Sonia Goncalves, Ewan Harrison, David K. Jackson, Ian Johnston, Dominic Kwiatkowski, Cordelia Langford, John Sillitoe on behalf of the Wellcome Sanger Institute COVID-19 Surveillance Team |
| EPI_ISL_3223886 | Lighthouse Lab in Alderley Park | Wellcome Sanger Institute for the COVID-19 Genomics UK (COG-UK) Consortium | Jacquelyn Wynn, Mairead Hyland, The Lighthouse Lab in Alderley Park and Alex Alderton, Roberto Amato, Jeffrey Barrett, Sonia Goncalves, Ewan Harrison, David K. Jackson, Ian Johnston, Dominic Kwiatkowski, Cordelia Langford, John Sillitoe on behalf of the Wellcome Sanger Institute COVID-19 Surveillance Team |
| EPI_ISL_3223887, EPI_ISL_3223888 | Lighthouse Laboratory Plymouth | Wellcome Sanger Institute for the COVID-19 Genomics UK (COG-UK) Consortium | Lighthouse Laboratory Plymouth and Alex Alderton, Roberto Amato, Jeffrey Barrett, Sonia Goncalves, Ewan Harrison, David K. Jackson, Ian Johnston, Dominic Kwiatkowski, Cordelia Langford, John Sillitoe on behalf of the Wellcome Sanger Institute COVID-19 Surveillance Team |
| EPI_ISL_3223889 | Lighthouse Lab in Glasgow | Wellcome Sanger Institute for the COVID-19 Genomics UK (COG-UK) Consortium | Harper VanSteenhouse, Yumi Kasai, David Gray, Carol Clugston, Anna Dominiczak and Alex Alderton, Roberto Amato, Jeffrey Barrett, Sonia Goncalves, Ewan Harrison, David K. Jackson, Ian Johnston, Dominic Kwiatkowski, Cordelia Langford, John Sillitoe on behalf of the Wellcome Sanger Institute COVID-19 Surveillance Team |
| EPI_ISL_3223890, EPI_ISL_3223891, EPI_ISL_3223892, EPI_ISL_3223893 | Lighthouse Lab in Alderley Park | Wellcome Sanger Institute for the COVID-19 Genomics UK (COG-UK) Consortium | Jacquelyn Wynn, Mairead Hyland, The Lighthouse Lab in Alderley Park and Alex Alderton, Roberto Amato, Jeffrey Barrett, Sonia Goncalves, Ewan Harrison, David K. Jackson, Ian Johnston, Dominic Kwiatkowski, Cordelia Langford, John Sillitoe on behalf of the Wellcome Sanger Institute COVID-19 Surveillance Team |
| EPI_ISL_3223894 | Lighthouse Lab in Glasgow | Wellcome Sanger Institute for the COVID-19 Genomics UK (COG-UK) Consortium | Harper VanSteenhouse, Yumi Kasai, David Gray, Carol Clugston, Anna Dominiczak and Alex Alderton, Roberto Amato, Jeffrey Barrett, Sonia Goncalves, Ewan Harrison, David K. Jackson, Ian Johnston, Dominic Kwiatkowski, Cordelia Langford, John Sillitoe on behalf of the Wellcome Sanger Institute COVID-19 Surveillance Team |
| EPI_ISL_3223895, EPI_ISL_3223896 | Lighthouse Lab in Milton Keynes | Wellcome Sanger Institute for the COVID-19 Genomics UK (COG-UK) Consortium | The Lighthouse Lab in Milton Keynes and Alex Alderton, Roberto Amato, Jeffrey Barrett, Sonia Goncalves, Ewan Harrison, David K. Jackson, Ian Johnston, Dominic Kwiatkowski, Cordelia Langford, John Sillitoe on behalf of the Wellcome Sanger Institute COVID-19 Surveillance Team |
| EPI_ISL_3223897 | Lighthouse Laboratory Plymouth | Wellcome Sanger Institute for the COVID-19 Genomics UK (COG-UK) Consortium | Lighthouse Laboratory Plymouth and Alex Alderton, Roberto Amato, Jeffrey Barrett, Sonia Goncalves, Ewan Harrison, David K. Jackson, Ian Johnston, Dominic Kwiatkowski, Cordelia Langford, John Sillitoe on behalf of the Wellcome Sanger Institute COVID-19 Surveillance Team |
| EPI_ISL_3223898, EPI_ISL_3223899 | Lighthouse Lab in Alderley Park | Wellcome Sanger Institute for the COVID-19 Genomics UK (COG-UK) Consortium | Jacquelyn Wynn, Mairead Hyland, The Lighthouse Lab in Alderley Park and Alex Alderton, Roberto Amato, Jeffrey Barrett, Sonia Goncalves, Ewan Harrison, David K. Jackson, Ian Johnston, Dominic Kwiatkowski, Cordelia Langford, John Sillitoe on behalf of the Wellcome Sanger Institute COVID-19 Surveillance Team |
| EPI_ISL_3223900 | Lighthouse Laboratory Plymouth | Wellcome Sanger Institute for the COVID-19 Genomics UK (COG-UK) Consortium | Lighthouse Laboratory Plymouth and Alex Alderton, Roberto Amato, Jeffrey Barrett, Sonia Goncalves, Ewan Harrison, David K. Jackson, Ian Johnston, Dominic Kwiatkowski, Cordelia Langford, John Sillitoe on behalf of the Wellcome Sanger Institute COVID-19 Surveillance Team |
| EPI_ISL_3223901 | Lighthouse Lab in Milton Keynes | Wellcome Sanger Institute for the COVID-19 Genomics UK (COG-UK) Consortium | The Lighthouse Lab in Milton Keynes and Alex Alderton, Roberto Amato, Jeffrey Barrett, Sonia Goncalves, Ewan Harrison, David K. Jackson, Ian Johnston, Dominic Kwiatkowski, Cordelia Langford, John Sillitoe on behalf of the Wellcome Sanger Institute COVID-19 Surveillance Team |
| EPI_ISL_3223902, EPI_ISL_3223903, EPI_ISL_3223904 | Lighthouse Lab in Alderley Park | Wellcome Sanger Institute for the COVID-19 Genomics UK (COG-UK) Consortium | Jacquelyn Wynn, Mairead Hyland, The Lighthouse Lab in Alderley Park and Alex Alderton, Roberto Amato, Jeffrey Barrett, Sonia Goncalves, Ewan Harrison, David K. Jackson, Ian Johnston, Dominic Kwiatkowski, Cordelia Langford, John Sillitoe on behalf of the Wellcome Sanger Institute COVID-19 Surveillance Team |
| EPI_ISL_3223905, EPI_ISL_3223906 | Lighthouse Laboratory Plymouth | Wellcome Sanger Institute for the COVID-19 Genomics UK (COG-UK) Consortium | Lighthouse Laboratory Plymouth and Alex Alderton, Roberto Amato, Jeffrey Barrett, Sonia Goncalves, Ewan Harrison, David K. Jackson, Ian Johnston, Dominic Kwiatkowski, Cordelia Langford, John Sillitoe on behalf of the Wellcome Sanger Institute COVID-19 Surveillance Team |
| EPI_ISL_3223907 | Lighthouse Lab in Alderley Park | Wellcome Sanger Institute for the COVID-19 Genomics UK (COG-UK) Consortium | Jacquelyn Wynn, Mairead Hyland, The Lighthouse Lab in Alderley Park and Alex Alderton, Roberto Amato, Jeffrey Barrett, Sonia Goncalves, Ewan Harrison, David K. Jackson, Ian Johnston, Dominic Kwiatkowski, Cordelia Langford, John Sillitoe on behalf of the Wellcome Sanger Institute COVID-19 Surveillance Team |
| EPI_ISL_3223908, EPI_ISL_3223909, EPI_ISL_3223910, EPI_ISL_3223911, EPI_ISL_3223912 | Lighthouse Lab in Milton Keynes | Wellcome Sanger Institute for the COVID-19 Genomics UK (COG-UK) Consortium | The Lighthouse Lab in Milton Keynes and Alex Alderton, Roberto Amato, Jeffrey Barrett, Sonia Goncalves, Ewan Harrison, David K. Jackson, Ian Johnston, Dominic Kwiatkowski, Cordelia Langford, John Sillitoe on behalf of the Wellcome Sanger Institute COVID-19 Surveillance Team |
| EPI_ISL_3223913, EPI_ISL_3223914 | Lighthouse Lab in Alderley Park | Wellcome Sanger Institute for the COVID-19 Genomics UK (COG-UK) Consortium | Jacquelyn Wynn, Mairead Hyland, The Lighthouse Lab in Alderley Park and Alex Alderton, Roberto Amato, Jeffrey Barrett, Sonia Goncalves, Ewan Harrison, David K. Jackson, Ian Johnston, Dominic Kwiatkowski, Cordelia Langford, John Sillitoe on behalf of the Wellcome Sanger Institute COVID-19 Surveillance Team |
| EPI_ISL_3223915 | Lighthouse Lab in Milton Keynes | Wellcome Sanger Institute for the COVID-19 Genomics UK (COG-UK) Consortium | The Lighthouse Lab in Milton Keynes and Alex Alderton, Roberto Amato, Jeffrey Barrett, Sonia Goncalves, Ewan Harrison, David K. Jackson, Ian Johnston, Dominic Kwiatkowski, Cordelia Langford, John Sillitoe on behalf of the Wellcome Sanger Institute COVID-19 Surveillance Team |
| EPI_ISL_3223916, EPI_ISL_3223917, | Lighthouse Lab in Glasgow | Wellcome Sanger Institute for the COVID-19 Genomics UK (COG-UK) Consortium | Harper VanSteenhouse, Yumi Kasai, David Gray, Carol Clugston, Anna Dominiczak and Alex Alderton, Roberto Amato, Jeffrey Barrett, Sonia Goncalves, Ewan Harrison, David K. Jackson, Ian Johnston, Dominic Kwiatkowski, Cordelia Langford, John Sillitoe on behalf of the Wellcome Sanger Institute COVID-19 Surveillance Team |

| | | | |
|---|---|---|---|
| EPI_ISL_3223918 | | | |
| EPI_ISL_3223919 | Lighthouse Lab in Alderley Park | Wellcome Sanger Institute for the COVID-19 Genomics UK (COG-UK) Consortium | Jacquelyn Wynn, Mairead Hyland, The Lighthouse Lab in Alderley Park and Alex Alderton, Roberto Amato, Jeffrey Barrett, Sonia Goncalves, Ewan Harrison, David K. Jackson, Ian Johnston, Dominic Kwiatkowski, Cordelia Langford, John Sillitoe on behalf of the Wellcome Sanger Institute COVID-19 Surveillance Team |
| EPI_ISL_3223920 | Lighthouse Lab in Milton Keynes | Wellcome Sanger Institute for the COVID-19 Genomics UK (COG-UK) Consortium | The Lighthouse Lab in Milton Keynes and Alex Alderton, Roberto Amato, Jeffrey Barrett, Sonia Goncalves, Ewan Harrison, David K. Jackson, Ian Johnston, Dominic Kwiatkowski, Cordelia Langford, John Sillitoe on behalf of the Wellcome Sanger Institute COVID-19 Surveillance Team |
| EPI_ISL_3223921, EPI_ISL_3223922 | Lighthouse Laboratory Plymouth | Wellcome Sanger Institute for the COVID-19 Genomics UK (COG-UK) Consortium | Lighthouse Laboratory Plymouth and Alex Alderton, Roberto Amato, Jeffrey Barrett, Sonia Goncalves, Ewan Harrison, David K. Jackson, Ian Johnston, Dominic Kwiatkowski, Cordelia Langford, John Sillitoe on behalf of the Wellcome Sanger Institute COVID-19 Surveillance Team |
| EPI_ISL_3223923 | Lighthouse Lab in Glasgow | Wellcome Sanger Institute for the COVID-19 Genomics UK (COG-UK) Consortium | Harper VanSteenhouse, Yumi Kasai, David Gray, Carol Clugston, Anna Dominiczak and Alex Alderton, Roberto Amato, Jeffrey Barrett, Sonia Goncalves, Ewan Harrison, David K. Jackson, Ian Johnston, Dominic Kwiatkowski, Cordelia Langford, John Sillitoe on behalf of the Wellcome Sanger Institute COVID-19 Surveillance Team |
| EPI_ISL_3223924 | Lighthouse Lab in Milton Keynes | Wellcome Sanger Institute for the COVID-19 Genomics UK (COG-UK) Consortium | The Lighthouse Lab in Milton Keynes and Alex Alderton, Roberto Amato, Jeffrey Barrett, Sonia Goncalves, Ewan Harrison, David K. Jackson, Ian Johnston, Dominic Kwiatkowski, Cordelia Langford, John Sillitoe on behalf of the Wellcome Sanger Institute COVID-19 Surveillance Team |
| EPI_ISL_3223925, EPI_ISL_3223926 | Lighthouse Lab in Alderley Park | Wellcome Sanger Institute for the COVID-19 Genomics UK (COG-UK) Consortium | Jacquelyn Wynn, Mairead Hyland, The Lighthouse Lab in Alderley Park and Alex Alderton, Roberto Amato, Jeffrey Barrett, Sonia Goncalves, Ewan Harrison, David K. Jackson, Ian Johnston, Dominic Kwiatkowski, Cordelia Langford, John Sillitoe on behalf of the Wellcome Sanger Institute COVID-19 Surveillance Team |
| EPI_ISL_3223927 | Lighthouse Lab in Glasgow | Wellcome Sanger Institute for the COVID-19 Genomics UK (COG-UK) Consortium | Harper VanSteenhouse, Yumi Kasai, David Gray, Carol Clugston, Anna Dominiczak and Alex Alderton, Roberto Amato, Jeffrey Barrett, Sonia Goncalves, Ewan Harrison, David K. Jackson, Ian Johnston, Dominic Kwiatkowski, Cordelia Langford, John Sillitoe on behalf of the Wellcome Sanger Institute COVID-19 Surveillance Team |
| EPI_ISL_3223928 | Lighthouse Laboratory Plymouth | Wellcome Sanger Institute for the COVID-19 Genomics UK (COG-UK) Consortium | Lighthouse Laboratory Plymouth and Alex Alderton, Roberto Amato, Jeffrey Barrett, Sonia Goncalves, Ewan Harrison, David K. Jackson, Ian Johnston, Dominic Kwiatkowski, Cordelia Langford, John Sillitoe on behalf of the Wellcome Sanger Institute COVID-19 Surveillance Team |
| EPI_ISL_3223929 | Lighthouse Lab in Glasgow | Wellcome Sanger Institute for the COVID-19 Genomics UK (COG-UK) Consortium | Harper VanSteenhouse, Yumi Kasai, David Gray, Carol Clugston, Anna Dominiczak and Alex Alderton, Roberto Amato, Jeffrey Barrett, Sonia Goncalves, Ewan Harrison, David K. Jackson, Ian Johnston, Dominic Kwiatkowski, Cordelia Langford, John Sillitoe on behalf of the Wellcome Sanger Institute COVID-19 Surveillance Team |
| EPI_ISL_3223930 | Lighthouse Laboratory Plymouth | Wellcome Sanger Institute for the COVID-19 Genomics UK (COG-UK) Consortium | Lighthouse Laboratory Plymouth and Alex Alderton, Roberto Amato, Jeffrey Barrett, Sonia Goncalves, Ewan Harrison, David K. Jackson, Ian Johnston, Dominic Kwiatkowski, Cordelia Langford, John Sillitoe on behalf of the Wellcome Sanger Institute COVID-19 Surveillance Team |
| EPI_ISL_3223931 | Lighthouse Lab in Alderley Park | Wellcome Sanger Institute for the COVID-19 Genomics UK (COG-UK) Consortium | Jacquelyn Wynn, Mairead Hyland, The Lighthouse Lab in Alderley Park and Alex Alderton, Roberto Amato, Jeffrey Barrett, Sonia Goncalves, Ewan Harrison, David K. Jackson, Ian Johnston, Dominic Kwiatkowski, Cordelia Langford, John Sillitoe on behalf of the Wellcome Sanger Institute COVID-19 Surveillance Team |
| EPI_ISL_3223932 | Lighthouse Lab in Milton Keynes | Wellcome Sanger Institute for the COVID-19 Genomics UK (COG-UK) Consortium | The Lighthouse Lab in Milton Keynes and Alex Alderton, Roberto Amato, Jeffrey Barrett, Sonia Goncalves, Ewan Harrison, David K. Jackson, Ian Johnston, Dominic Kwiatkowski, Cordelia Langford, John Sillitoe on behalf of the Wellcome Sanger Institute COVID-19 Surveillance Team |
| EPI_ISL_3223933 | Lighthouse Lab in Glasgow | Wellcome Sanger Institute for the COVID-19 Genomics UK (COG-UK) Consortium | Harper VanSteenhouse, Yumi Kasai, David Gray, Carol Clugston, Anna Dominiczak and Alex Alderton, Roberto Amato, Jeffrey Barrett, Sonia Goncalves, Ewan Harrison, David K. Jackson, Ian Johnston, Dominic Kwiatkowski, Cordelia Langford, John Sillitoe on behalf of the Wellcome Sanger Institute COVID-19 Surveillance Team |
| EPI_ISL_3223934 | Lighthouse Lab in Alderley Park | Wellcome Sanger Institute for the COVID-19 Genomics UK (COG-UK) Consortium | Jacquelyn Wynn, Mairead Hyland, The Lighthouse Lab in Alderley Park and Alex Alderton, Roberto Amato, Jeffrey Barrett, Sonia Goncalves, Ewan Harrison, David K. Jackson, Ian Johnston, Dominic Kwiatkowski, Cordelia Langford, John Sillitoe on behalf of the Wellcome Sanger Institute COVID-19 Surveillance Team |
| EPI_ISL_3223935 | Lighthouse Lab in Milton Keynes | Wellcome Sanger Institute for the COVID-19 Genomics UK (COG-UK) Consortium | The Lighthouse Lab in Milton Keynes and Alex Alderton, Roberto Amato, Jeffrey Barrett, Sonia Goncalves, Ewan Harrison, David K. Jackson, Ian Johnston, Dominic Kwiatkowski, Cordelia Langford, John Sillitoe on behalf of the Wellcome Sanger Institute COVID-19 Surveillance Team |
| EPI_ISL_3223936 | Lighthouse Lab in Glasgow | Wellcome Sanger Institute for the COVID-19 Genomics UK (COG-UK) Consortium | Harper VanSteenhouse, Yumi Kasai, David Gray, Carol Clugston, Anna Dominiczak and Alex Alderton, Roberto Amato, Jeffrey Barrett, Sonia Goncalves, Ewan Harrison, David K. Jackson, Ian Johnston, Dominic Kwiatkowski, Cordelia Langford, John Sillitoe on behalf of the Wellcome Sanger Institute COVID-19 Surveillance Team |
| EPI_ISL_3223937 | Lighthouse Laboratory Plymouth | Wellcome Sanger Institute for the COVID-19 Genomics UK (COG-UK) Consortium | Lighthouse Laboratory Plymouth and Alex Alderton, Roberto Amato, Jeffrey Barrett, Sonia Goncalves, Ewan Harrison, David K. Jackson, Ian Johnston, Dominic Kwiatkowski, Cordelia Langford, John Sillitoe on behalf of the Wellcome Sanger Institute COVID-19 Surveillance Team |
| EPI_ISL_3223938, EPI_ISL_3223939 | Lighthouse Lab in Glasgow | Wellcome Sanger Institute for the COVID-19 Genomics UK (COG-UK) Consortium | Harper VanSteenhouse, Yumi Kasai, David Gray, Carol Clugston, Anna Dominiczak and Alex Alderton, Roberto Amato, Jeffrey Barrett, Sonia Goncalves, Ewan Harrison, David K. Jackson, Ian Johnston, Dominic Kwiatkowski, Cordelia Langford, John Sillitoe on behalf of the Wellcome Sanger Institute COVID-19 Surveillance Team |
| EPI_ISL_3223940, EPI_ISL_3223941, EPI_ISL_3223942 | Lighthouse Laboratory Plymouth | Wellcome Sanger Institute for the COVID-19 Genomics UK (COG-UK) Consortium | Lighthouse Laboratory Plymouth and Alex Alderton, Roberto Amato, Jeffrey Barrett, Sonia Goncalves, Ewan Harrison, David K. Jackson, Ian Johnston, Dominic Kwiatkowski, Cordelia Langford, John Sillitoe on behalf of the Wellcome Sanger Institute COVID-19 Surveillance Team |
| EPI_ISL_3223943 | Lighthouse Lab in Glasgow | Wellcome Sanger Institute for the COVID-19 Genomics UK (COG-UK) Consortium | Harper VanSteenhouse, Yumi Kasai, David Gray, Carol Clugston, Anna Dominiczak and Alex Alderton, Roberto Amato, Jeffrey Barrett, Sonia Goncalves, Ewan Harrison, David K. Jackson, Ian Johnston, Dominic Kwiatkowski, Cordelia Langford, John Sillitoe on behalf of the Wellcome Sanger Institute COVID-19 Surveillance Team |
| EPI_ISL_3223944 | Lighthouse Lab in Milton Keynes | Wellcome Sanger Institute for the COVID-19 Genomics UK (COG-UK) Consortium | The Lighthouse Lab in Milton Keynes and Alex Alderton, Roberto Amato, Jeffrey Barrett, Sonia Goncalves, Ewan Harrison, David K. Jackson, Ian Johnston, Dominic Kwiatkowski, Cordelia Langford, John Sillitoe on behalf of the Wellcome Sanger Institute COVID-19 Surveillance Team |
| EPI_ISL_3223945 | Lighthouse Lab in Alderley Park | Wellcome Sanger Institute for the COVID-19 Genomics UK (COG-UK) Consortium | Jacquelyn Wynn, Mairead Hyland, The Lighthouse Lab in Alderley Park and Alex Alderton, Roberto Amato, Jeffrey Barrett, Sonia Goncalves, Ewan Harrison, David K. Jackson, Ian Johnston, Dominic Kwiatkowski, Cordelia Langford, John Sillitoe on behalf of the Wellcome Sanger Institute COVID-19 Surveillance Team |
| EPI_ISL_3223946 | Lighthouse Lab in Glasgow | Wellcome Sanger Institute for the COVID-19 Genomics UK (COG-UK) Consortium | Harper VanSteenhouse, Yumi Kasai, David Gray, Carol Clugston, Anna Dominiczak and Alex Alderton, Roberto Amato, Jeffrey Barrett, Sonia Goncalves, Ewan Harrison, David K. Jackson, Ian Johnston, Dominic Kwiatkowski, Cordelia Langford, John Sillitoe on behalf of the Wellcome Sanger Institute COVID-19 Surveillance Team |
| EPI_ISL_3223947 | Lighthouse Laboratory Plymouth | Wellcome Sanger Institute for the COVID-19 Genomics UK (COG-UK) Consortium | Lighthouse Laboratory Plymouth and Alex Alderton, Roberto Amato, Jeffrey Barrett, Sonia Goncalves, Ewan Harrison, David K. Jackson, Ian Johnston, Dominic Kwiatkowski, Cordelia Langford, John Sillitoe on behalf of the Wellcome Sanger Institute COVID-19 Surveillance Team |
| EPI_ISL_3223948 | Lighthouse Lab in Milton Keynes | Wellcome Sanger Institute for the COVID-19 Genomics UK (COG-UK) Consortium | The Lighthouse Lab in Milton Keynes and Alex Alderton, Roberto Amato, Jeffrey Barrett, Sonia Goncalves, Ewan Harrison, David K. Jackson, Ian Johnston, Dominic Kwiatkowski, Cordelia Langford, John Sillitoe on behalf of the Wellcome Sanger Institute COVID-19 Surveillance Team |
| EPI_ISL_3223949 | Lighthouse Lab in Glasgow | Wellcome Sanger Institute for the COVID-19 Genomics UK (COG-UK) Consortium | Harper VanSteenhouse, Yumi Kasai, David Gray, Carol Clugston, Anna Dominiczak and Alex Alderton, Roberto Amato, Jeffrey Barrett, Sonia Goncalves, Ewan Harrison, David K. Jackson, Ian Johnston, Dominic Kwiatkowski, Cordelia Langford, John Sillitoe on behalf of the Wellcome Sanger Institute COVID-19 Surveillance Team |
| EPI_ISL_3223950 | Lighthouse Lab in Alderley Park | Wellcome Sanger Institute for the COVID-19 Genomics UK (COG-UK) Consortium | Jacquelyn Wynn, Mairead Hyland, The Lighthouse Lab in Alderley Park and Alex Alderton, Roberto Amato, Jeffrey Barrett, Sonia Goncalves, Ewan Harrison, David K. Jackson, Ian Johnston, Dominic Kwiatkowski, Cordelia Langford, John Sillitoe on behalf of the Wellcome Sanger Institute COVID-19 Surveillance Team |
| EPI_ISL_3223951 | Lighthouse Lab in Glasgow | Wellcome Sanger Institute for the COVID-19 Genomics UK (COG-UK) Consortium | Harper VanSteenhouse, Yumi Kasai, David Gray, Carol Clugston, Anna Dominiczak and Alex Alderton, Roberto Amato, Jeffrey Barrett, Sonia Goncalves, Ewan Harrison, David K. Jackson, Ian Johnston, Dominic Kwiatkowski, Cordelia Langford, John Sillitoe on behalf of the Wellcome Sanger Institute COVID-19 Surveillance Team |
| EPI_ISL_3223952 | Lighthouse Lab in Alderley Park | Wellcome Sanger Institute for the COVID-19 Genomics UK (COG-UK) Consortium | Jacquelyn Wynn, Mairead Hyland, The Lighthouse Lab in Alderley Park and Alex Alderton, Roberto Amato, Jeffrey Barrett, Sonia Goncalves, Ewan Harrison, David K. Jackson, Ian Johnston, Dominic Kwiatkowski, Cordelia Langford, John Sillitoe on behalf of the Wellcome Sanger Institute COVID-19 Surveillance Team |
| EPI_ISL_3223953, EPI_ISL_3223954, EPI_ISL_3223955 | Lighthouse Laboratory Plymouth | Wellcome Sanger Institute for the COVID-19 Genomics UK (COG-UK) Consortium | Lighthouse Laboratory Plymouth and Alex Alderton, Roberto Amato, Jeffrey Barrett, Sonia Goncalves, Ewan Harrison, David K. Jackson, Ian Johnston, Dominic Kwiatkowski, Cordelia Langford, John Sillitoe on behalf of the Wellcome Sanger Institute COVID-19 Surveillance Team |
| EPI_ISL_3223956 | Lighthouse Lab in Glasgow | Wellcome Sanger Institute for the COVID-19 Genomics UK (COG-UK) Consortium | Harper VanSteenhouse, Yumi Kasai, David Gray, Carol Clugston, Anna Dominiczak and Alex Alderton, Roberto Amato, Jeffrey Barrett, Sonia Goncalves, Ewan Harrison, David K. Jackson, Ian Johnston, Dominic Kwiatkowski, Cordelia Langford, John Sillitoe on behalf of the Wellcome Sanger Institute COVID-19 Surveillance Team |
| EPI_ISL_3223957 | Lighthouse Laboratory Plymouth | Wellcome Sanger Institute for the COVID-19 Genomics UK (COG-UK) Consortium | Lighthouse Laboratory Plymouth and Alex Alderton, Roberto Amato, Jeffrey Barrett, Sonia Goncalves, Ewan Harrison, David K. Jackson, Ian Johnston, Dominic Kwiatkowski, Cordelia Langford, John Sillitoe on behalf of the Wellcome Sanger Institute COVID-19 Surveillance Team |
| EPI_ISL_3223958 | Lighthouse Lab in Milton Keynes | Wellcome Sanger Institute for the COVID-19 Genomics UK (COG-UK) Consortium | The Lighthouse Lab in Milton Keynes and Alex Alderton, Roberto Amato, Jeffrey Barrett, Sonia Goncalves, Ewan Harrison, David K. Jackson, Ian Johnston, Dominic Kwiatkowski, Cordelia Langford, John Sillitoe on behalf of the Wellcome Sanger Institute COVID-19 Surveillance Team |
| EPI_ISL_3223959 | Lighthouse Lab in Alderley Park | Wellcome Sanger Institute for the COVID-19 Genomics UK (COG-UK) Consortium | Jacquelyn Wynn, Mairead Hyland, The Lighthouse Lab in Alderley Park and Alex Alderton, Roberto Amato, Jeffrey Barrett, Sonia Goncalves, Ewan Harrison, David K. Jackson, Ian Johnston, Dominic Kwiatkowski, Cordelia Langford, John Sillitoe on behalf of the Wellcome Sanger Institute COVID-19 Surveillance Team |
| EPI_ISL_3223960 | Lighthouse Lab in Milton Keynes | Wellcome Sanger Institute for the COVID-19 Genomics UK (COG-UK) Consortium | The Lighthouse Lab in Milton Keynes and Alex Alderton, Roberto Amato, Jeffrey Barrett, Sonia Goncalves, Ewan Harrison, David K. Jackson, Ian Johnston, Dominic Kwiatkowski, Cordelia Langford, John Sillitoe on behalf of the Wellcome Sanger Institute COVID-19 Surveillance Team |
| EPI_ISL_3223961, EPI_ISL_3223962 | Lighthouse Lab in Glasgow | Wellcome Sanger Institute for the COVID-19 Genomics UK (COG-UK) Consortium | Harper VanSteenhouse, Yumi Kasai, David Gray, Carol Clugston, Anna Dominiczak and Alex Alderton, Roberto Amato, Jeffrey Barrett, Sonia Goncalves, Ewan Harrison, David K. Jackson, Ian Johnston, Dominic Kwiatkowski, Cordelia Langford, John Sillitoe on behalf of the Wellcome Sanger Institute COVID-19 Surveillance Team |
| EPI_ISL_3223963, EPI_ISL_3223964 | Lighthouse Lab in Milton Keynes | Wellcome Sanger Institute for the COVID-19 Genomics UK (COG-UK) Consortium | The Lighthouse Lab in Milton Keynes and Alex Alderton, Roberto Amato, Jeffrey Barrett, Sonia Goncalves, Ewan Harrison, David K. Jackson, Ian Johnston, Dominic Kwiatkowski, Cordelia Langford, John Sillitoe on behalf of the Wellcome Sanger Institute COVID-19 Surveillance Team |
| EPI_ISL_3223965 | Lighthouse Lab in Glasgow | Wellcome Sanger Institute for the COVID-19 Genomics UK (COG-UK) Consortium | Harper VanSteenhouse, Yumi Kasai, David Gray, Carol Clugston, Anna Dominiczak and Alex Alderton, Roberto Amato, Jeffrey Barrett, Sonia Goncalves, Ewan Harrison, David K. Jackson, Ian Johnston, Dominic Kwiatkowski, Cordelia Langford, John Sillitoe on behalf of the Wellcome Sanger Institute COVID-19 Surveillance Team |
| EPI_ISL_3223966, EPI_ISL_3223967, EPI_ISL_3223968, EPI_ISL_3223969 | Lighthouse Laboratory Plymouth | Wellcome Sanger Institute for the COVID-19 Genomics UK (COG-UK) Consortium | Lighthouse Laboratory Plymouth and Alex Alderton, Roberto Amato, Jeffrey Barrett, Sonia Goncalves, Ewan Harrison, David K. Jackson, Ian Johnston, Dominic Kwiatkowski, Cordelia Langford, John Sillitoe on behalf of the Wellcome Sanger Institute COVID-19 Surveillance Team |
| EPI_ISL_3223970 | Lighthouse Lab in Alderley Park | Wellcome Sanger Institute for the COVID-19 Genomics UK (COG-UK) Consortium | Jacquelyn Wynn, Mairead Hyland, The Lighthouse Lab in Alderley Park and Alex Alderton, Roberto Amato, Jeffrey Barrett, Sonia Goncalves, Ewan Harrison, David K. Jackson, Ian Johnston, Dominic Kwiatkowski, Cordelia Langford, John Sillitoe on behalf of the Wellcome Sanger Institute COVID-19 Surveillance Team |
| EPI_ISL_3223971, EPI_ISL_3223972 | Lighthouse Lab in Glasgow | Wellcome Sanger Institute for the COVID-19 Genomics UK (COG-UK) Consortium | Harper VanSteenhouse, Yumi Kasai, David Gray, Carol Clugston, Anna Dominiczak and Alex Alderton, Roberto Amato, Jeffrey Barrett, Sonia Goncalves, Ewan Harrison, David K. Jackson, Ian Johnston, Dominic Kwiatkowski, Cordelia Langford, John Sillitoe on behalf of the Wellcome Sanger Institute COVID-19 Surveillance Team |
| EPI_ISL_3223973, EPI_ISL_3223974, EPI_ISL_3223975 | Lighthouse Lab in Milton Keynes | Wellcome Sanger Institute for the COVID-19 Genomics UK (COG-UK) Consortium | The Lighthouse Lab in Milton Keynes and Alex Alderton, Roberto Amato, Jeffrey Barrett, Sonia Goncalves, Ewan Harrison, David K. Jackson, Ian Johnston, Dominic Kwiatkowski, Cordelia Langford, John Sillitoe on behalf of the Wellcome Sanger Institute COVID-19 Surveillance Team |

| | | | |
|---|---|---|---|
| EPI_ISL_3223976 | Lighthouse Laboratory Plymouth | Wellcome Sanger Institute for the COVID-19 Genomics UK (COG-UK) Consortium | Lighthouse Laboratory Plymouth and Alex Alderton, Roberto Amato, Jeffrey Barrett, Sonia Goncalves, Ewan Harrison, David K. Jackson, Ian Johnston, Dominic Kwiatkowski, Cordelia Langford, John Sillitoe on behalf of the Wellcome Sanger Institute COVID-19 Surveillance Team |
| EPI_ISL_3223977 | Lighthouse Lab in Glasgow | Wellcome Sanger Institute for the COVID-19 Genomics UK (COG-UK) Consortium | Harper VanSteenhouse, Yumi Kasai, David Gray, Carol Clugston, Anna Dominiczak and Alex Alderton, Roberto Amato, Jeffrey Barrett, Sonia Goncalves, Ewan Harrison, David K. Jackson, Ian Johnston, Dominic Kwiatkowski, Cordelia Langford, John Sillitoe on behalf of the Wellcome Sanger Institute COVID-19 Surveillance Team |
| EPI_ISL_3223978 | Lighthouse Lab in Milton Keynes | Wellcome Sanger Institute for the COVID-19 Genomics UK (COG-UK) Consortium | The Lighthouse Lab in Milton Keynes and Alex Alderton, Roberto Amato, Jeffrey Barrett, Sonia Goncalves, Ewan Harrison, David K. Jackson, Ian Johnston, Dominic Kwiatkowski, Cordelia Langford, John Sillitoe on behalf of the Wellcome Sanger Institute COVID-19 Surveillance Team |
| EPI_ISL_3223979 | Lighthouse Lab in Glasgow | Wellcome Sanger Institute for the COVID-19 Genomics UK (COG-UK) Consortium | Harper VanSteenhouse, Yumi Kasai, David Gray, Carol Clugston, Anna Dominiczak and Alex Alderton, Roberto Amato, Jeffrey Barrett, Sonia Goncalves, Ewan Harrison, David K. Jackson, Ian Johnston, Dominic Kwiatkowski, Cordelia Langford, John Sillitoe on behalf of the Wellcome Sanger Institute COVID-19 Surveillance Team |
| EPI_ISL_3223980 | Lighthouse Lab in Alderley Park | Wellcome Sanger Institute for the COVID-19 Genomics UK (COG-UK) Consortium | Jacquelyn Wynn, Mairead Hyland, The Lighthouse Lab in Alderley Park and Alex Alderton, Roberto Amato, Jeffrey Barrett, Sonia Goncalves, Ewan Harrison, David K. Jackson, Ian Johnston, Dominic Kwiatkowski, Cordelia Langford, John Sillitoe on behalf of the Wellcome Sanger Institute COVID-19 Surveillance Team |
| EPI_ISL_3223981, EPI_ISL_3223982 | Lighthouse Lab in Milton Keynes | Wellcome Sanger Institute for the COVID-19 Genomics UK (COG-UK) Consortium | The Lighthouse Lab in Milton Keynes and Alex Alderton, Roberto Amato, Jeffrey Barrett, Sonia Goncalves, Ewan Harrison, David K. Jackson, Ian Johnston, Dominic Kwiatkowski, Cordelia Langford, John Sillitoe on behalf of the Wellcome Sanger Institute COVID-19 Surveillance Team |
| EPI_ISL_3223983 | Lighthouse Laboratory Plymouth | Wellcome Sanger Institute for the COVID-19 Genomics UK (COG-UK) Consortium | Lighthouse Laboratory Plymouth and Alex Alderton, Roberto Amato, Jeffrey Barrett, Sonia Goncalves, Ewan Harrison, David K. Jackson, Ian Johnston, Dominic Kwiatkowski, Cordelia Langford, John Sillitoe on behalf of the Wellcome Sanger Institute COVID-19 Surveillance Team |
| EPI_ISL_3223984, EPI_ISL_3223985, EPI_ISL_3223986, EPI_ISL_3223987 | Lighthouse Lab in Milton Keynes | Wellcome Sanger Institute for the COVID-19 Genomics UK (COG-UK) Consortium | The Lighthouse Lab in Milton Keynes and Alex Alderton, Roberto Amato, Jeffrey Barrett, Sonia Goncalves, Ewan Harrison, David K. Jackson, Ian Johnston, Dominic Kwiatkowski, Cordelia Langford, John Sillitoe on behalf of the Wellcome Sanger Institute COVID-19 Surveillance Team |
| EPI_ISL_3223988 | Lighthouse Laboratory Plymouth | Wellcome Sanger Institute for the COVID-19 Genomics UK (COG-UK) Consortium | Lighthouse Laboratory Plymouth and Alex Alderton, Roberto Amato, Jeffrey Barrett, Sonia Goncalves, Ewan Harrison, David K. Jackson, Ian Johnston, Dominic Kwiatkowski, Cordelia Langford, John Sillitoe on behalf of the Wellcome Sanger Institute COVID-19 Surveillance Team |
| EPI_ISL_3223989 | Lighthouse Lab in Glasgow | Wellcome Sanger Institute for the COVID-19 Genomics UK (COG-UK) Consortium | Harper VanSteenhouse, Yumi Kasai, David Gray, Carol Clugston, Anna Dominiczak and Alex Alderton, Roberto Amato, Jeffrey Barrett, Sonia Goncalves, Ewan Harrison, David K. Jackson, Ian Johnston, Dominic Kwiatkowski, Cordelia Langford, John Sillitoe on behalf of the Wellcome Sanger Institute COVID-19 Surveillance Team |
| EPI_ISL_3223990 | Lighthouse Lab in Alderley Park | Wellcome Sanger Institute for the COVID-19 Genomics UK (COG-UK) Consortium | Jacquelyn Wynn, Mairead Hyland, The Lighthouse Lab in Alderley Park and Alex Alderton, Roberto Amato, Jeffrey Barrett, Sonia Goncalves, Ewan Harrison, David K. Jackson, Ian Johnston, Dominic Kwiatkowski, Cordelia Langford, John Sillitoe on behalf of the Wellcome Sanger Institute COVID-19 Surveillance Team |
| EPI_ISL_3223991, EPI_ISL_3223992 | Lighthouse Lab in Glasgow | Wellcome Sanger Institute for the COVID-19 Genomics UK (COG-UK) Consortium | Harper VanSteenhouse, Yumi Kasai, David Gray, Carol Clugston, Anna Dominiczak and Alex Alderton, Roberto Amato, Jeffrey Barrett, Sonia Goncalves, Ewan Harrison, David K. Jackson, Ian Johnston, Dominic Kwiatkowski, Cordelia Langford, John Sillitoe on behalf of the Wellcome Sanger Institute COVID-19 Surveillance Team |
| EPI_ISL_3223993 | Lighthouse Lab in Milton Keynes | Wellcome Sanger Institute for the COVID-19 Genomics UK (COG-UK) Consortium | The Lighthouse Lab in Milton Keynes and Alex Alderton, Roberto Amato, Jeffrey Barrett, Sonia Goncalves, Ewan Harrison, David K. Jackson, Ian Johnston, Dominic Kwiatkowski, Cordelia Langford, John Sillitoe on behalf of the Wellcome Sanger Institute COVID-19 Surveillance Team |
| EPI_ISL_3223994 | Lighthouse Laboratory Plymouth | Wellcome Sanger Institute for the COVID-19 Genomics UK (COG-UK) Consortium | Lighthouse Laboratory Plymouth and Alex Alderton, Roberto Amato, Jeffrey Barrett, Sonia Goncalves, Ewan Harrison, David K. Jackson, Ian Johnston, Dominic Kwiatkowski, Cordelia Langford, John Sillitoe on behalf of the Wellcome Sanger Institute COVID-19 Surveillance Team |
| EPI_ISL_3223995 | Lighthouse Lab in Milton Keynes | Wellcome Sanger Institute for the COVID-19 Genomics UK (COG-UK) Consortium | The Lighthouse Lab in Milton Keynes and Alex Alderton, Roberto Amato, Jeffrey Barrett, Sonia Goncalves, Ewan Harrison, David K. Jackson, Ian Johnston, Dominic Kwiatkowski, Cordelia Langford, John Sillitoe on behalf of the Wellcome Sanger Institute COVID-19 Surveillance Team |
| EPI_ISL_3223996 | Lighthouse Lab in Alderley Park | Wellcome Sanger Institute for the COVID-19 Genomics UK (COG-UK) Consortium | Jacquelyn Wynn, Mairead Hyland, The Lighthouse Lab in Alderley Park and Alex Alderton, Roberto Amato, Jeffrey Barrett, Sonia Goncalves, Ewan Harrison, David K. Jackson, Ian Johnston, Dominic Kwiatkowski, Cordelia Langford, John Sillitoe on behalf of the Wellcome Sanger Institute COVID-19 Surveillance Team |
| EPI_ISL_3223997, EPI_ISL_3223998 | Lighthouse Lab in Glasgow | Wellcome Sanger Institute for the COVID-19 Genomics UK (COG-UK) Consortium | Harper VanSteenhouse, Yumi Kasai, David Gray, Carol Clugston, Anna Dominiczak and Alex Alderton, Roberto Amato, Jeffrey Barrett, Sonia Goncalves, Ewan Harrison, David K. Jackson, Ian Johnston, Dominic Kwiatkowski, Cordelia Langford, John Sillitoe on behalf of the Wellcome Sanger Institute COVID-19 Surveillance Team |
| EPI_ISL_3223999, EPI_ISL_3224000 | Lighthouse Laboratory Plymouth | Wellcome Sanger Institute for the COVID-19 Genomics UK (COG-UK) Consortium | Lighthouse Laboratory Plymouth and Alex Alderton, Roberto Amato, Jeffrey Barrett, Sonia Goncalves, Ewan Harrison, David K. Jackson, Ian Johnston, Dominic Kwiatkowski, Cordelia Langford, John Sillitoe on behalf of the Wellcome Sanger Institute COVID-19 Surveillance Team |
| EPI_ISL_3224001 | Lighthouse Lab in Milton Keynes | Wellcome Sanger Institute for the COVID-19 Genomics UK (COG-UK) Consortium | The Lighthouse Lab in Milton Keynes and Alex Alderton, Roberto Amato, Jeffrey Barrett, Sonia Goncalves, Ewan Harrison, David K. Jackson, Ian Johnston, Dominic Kwiatkowski, Cordelia Langford, John Sillitoe on behalf of the Wellcome Sanger Institute COVID-19 Surveillance Team |
| EPI_ISL_3224002 | Lighthouse Lab in Glasgow | Wellcome Sanger Institute for the COVID-19 Genomics UK (COG-UK) Consortium | Harper VanSteenhouse, Yumi Kasai, David Gray, Carol Clugston, Anna Dominiczak and Alex Alderton, Roberto Amato, Jeffrey Barrett, Sonia Goncalves, Ewan Harrison, David K. Jackson, Ian Johnston, Dominic Kwiatkowski, Cordelia Langford, John Sillitoe on behalf of the Wellcome Sanger Institute COVID-19 Surveillance Team |
| EPI_ISL_3224003 | Lighthouse Lab in Milton Keynes | Wellcome Sanger Institute for the COVID-19 Genomics UK (COG-UK) Consortium | The Lighthouse Lab in Milton Keynes and Alex Alderton, Roberto Amato, Jeffrey Barrett, Sonia Goncalves, Ewan Harrison, David K. Jackson, Ian Johnston, Dominic Kwiatkowski, Cordelia Langford, John Sillitoe on behalf of the Wellcome Sanger Institute COVID-19 Surveillance Team |
| EPI_ISL_3224004 | Lighthouse Laboratory Plymouth | Wellcome Sanger Institute for the COVID-19 Genomics UK (COG-UK) Consortium | Lighthouse Laboratory Plymouth and Alex Alderton, Roberto Amato, Jeffrey Barrett, Sonia Goncalves, Ewan Harrison, David K. Jackson, Ian Johnston, Dominic Kwiatkowski, Cordelia Langford, John Sillitoe on behalf of the Wellcome Sanger Institute COVID-19 Surveillance Team |
| EPI_ISL_3224005 | Lighthouse Lab in Glasgow | Wellcome Sanger Institute for the COVID-19 Genomics UK (COG-UK) Consortium | Harper VanSteenhouse, Yumi Kasai, David Gray, Carol Clugston, Anna Dominiczak and Alex Alderton, Roberto Amato, Jeffrey Barrett, Sonia Goncalves, Ewan Harrison, David K. Jackson, Ian Johnston, Dominic Kwiatkowski, Cordelia Langford, John Sillitoe on behalf of the Wellcome Sanger Institute COVID-19 Surveillance Team |
| EPI_ISL_3224006 | Lighthouse Laboratory Plymouth | Wellcome Sanger Institute for the COVID-19 Genomics UK (COG-UK) Consortium | Lighthouse Laboratory Plymouth and Alex Alderton, Roberto Amato, Jeffrey Barrett, Sonia Goncalves, Ewan Harrison, David K. Jackson, Ian Johnston, Dominic Kwiatkowski, Cordelia Langford, John Sillitoe on behalf of the Wellcome Sanger Institute COVID-19 Surveillance Team |
| EPI_ISL_3224007 | Lighthouse Lab in Alderley Park | Wellcome Sanger Institute for the COVID-19 Genomics UK (COG-UK) Consortium | Jacquelyn Wynn, Mairead Hyland, The Lighthouse Lab in Alderley Park and Alex Alderton, Roberto Amato, Jeffrey Barrett, Sonia Goncalves, Ewan Harrison, David K. Jackson, Ian Johnston, Dominic Kwiatkowski, Cordelia Langford, John Sillitoe on behalf of the Wellcome Sanger Institute COVID-19 Surveillance Team |
| EPI_ISL_3224008 | Lighthouse Lab in Milton Keynes | Wellcome Sanger Institute for the COVID-19 Genomics UK (COG-UK) Consortium | The Lighthouse Lab in Milton Keynes and Alex Alderton, Roberto Amato, Jeffrey Barrett, Sonia Goncalves, Ewan Harrison, David K. Jackson, Ian Johnston, Dominic Kwiatkowski, Cordelia Langford, John Sillitoe on behalf of the Wellcome Sanger Institute COVID-19 Surveillance Team |
| EPI_ISL_3224009, EPI_ISL_3224010 | Lighthouse Lab in Glasgow | Wellcome Sanger Institute for the COVID-19 Genomics UK (COG-UK) Consortium | Harper VanSteenhouse, Yumi Kasai, David Gray, Carol Clugston, Anna Dominiczak and Alex Alderton, Roberto Amato, Jeffrey Barrett, Sonia Goncalves, Ewan Harrison, David K. Jackson, Ian Johnston, Dominic Kwiatkowski, Cordelia Langford, John Sillitoe on behalf of the Wellcome Sanger Institute COVID-19 Surveillance Team |
| EPI_ISL_3224011 | Lighthouse Lab in Alderley Park | Wellcome Sanger Institute for the COVID-19 Genomics UK (COG-UK) Consortium | Jacquelyn Wynn, Mairead Hyland, The Lighthouse Lab in Alderley Park and Alex Alderton, Roberto Amato, Jeffrey Barrett, Sonia Goncalves, Ewan Harrison, David K. Jackson, Ian Johnston, Dominic Kwiatkowski, Cordelia Langford, John Sillitoe on behalf of the Wellcome Sanger Institute COVID-19 Surveillance Team |
| EPI_ISL_3224012, EPI_ISL_3224013 | Lighthouse Lab in Milton Keynes | Wellcome Sanger Institute for the COVID-19 Genomics UK (COG-UK) Consortium | The Lighthouse Lab in Milton Keynes and Alex Alderton, Roberto Amato, Jeffrey Barrett, Sonia Goncalves, Ewan Harrison, David K. Jackson, Ian Johnston, Dominic Kwiatkowski, Cordelia Langford, John Sillitoe on behalf of the Wellcome Sanger Institute COVID-19 Surveillance Team |
| EPI_ISL_3224014 | Lighthouse Lab in Glasgow | Wellcome Sanger Institute for the COVID-19 Genomics UK (COG-UK) Consortium | Harper VanSteenhouse, Yumi Kasai, David Gray, Carol Clugston, Anna Dominiczak and Alex Alderton, Roberto Amato, Jeffrey Barrett, Sonia Goncalves, Ewan Harrison, David K. Jackson, Ian Johnston, Dominic Kwiatkowski, Cordelia Langford, John Sillitoe on behalf of the Wellcome Sanger Institute COVID-19 Surveillance Team |
| EPI_ISL_3224015 | Lighthouse Lab in Milton Keynes | Wellcome Sanger Institute for the COVID-19 Genomics UK (COG-UK) Consortium | The Lighthouse Lab in Milton Keynes and Alex Alderton, Roberto Amato, Jeffrey Barrett, Sonia Goncalves, Ewan Harrison, David K. Jackson, Ian Johnston, Dominic Kwiatkowski, Cordelia Langford, John Sillitoe on behalf of the Wellcome Sanger Institute COVID-19 Surveillance Team |
| EPI_ISL_3224016 | Lighthouse Laboratory Plymouth | Wellcome Sanger Institute for the COVID-19 Genomics UK (COG-UK) Consortium | Lighthouse Laboratory Plymouth and Alex Alderton, Roberto Amato, Jeffrey Barrett, Sonia Goncalves, Ewan Harrison, David K. Jackson, Ian Johnston, Dominic Kwiatkowski, Cordelia Langford, John Sillitoe on behalf of the Wellcome Sanger Institute COVID-19 Surveillance Team |
| EPI_ISL_3224017 | Lighthouse Lab in Glasgow | Wellcome Sanger Institute for the COVID-19 Genomics UK (COG-UK) Consortium | Harper VanSteenhouse, Yumi Kasai, David Gray, Carol Clugston, Anna Dominiczak and Alex Alderton, Roberto Amato, Jeffrey Barrett, Sonia Goncalves, Ewan Harrison, David K. Jackson, Ian Johnston, Dominic Kwiatkowski, Cordelia Langford, John Sillitoe on behalf of the Wellcome Sanger Institute COVID-19 Surveillance Team |
| EPI_ISL_3224018 | Lighthouse Lab in Milton Keynes | Wellcome Sanger Institute for the COVID-19 Genomics UK (COG-UK) Consortium | The Lighthouse Lab in Milton Keynes and Alex Alderton, Roberto Amato, Jeffrey Barrett, Sonia Goncalves, Ewan Harrison, David K. Jackson, Ian Johnston, Dominic Kwiatkowski, Cordelia Langford, John Sillitoe on behalf of the Wellcome Sanger Institute COVID-19 Surveillance Team |
| EPI_ISL_3224019 | Lighthouse Lab in Alderley Park | Wellcome Sanger Institute for the COVID-19 Genomics UK (COG-UK) Consortium | Jacquelyn Wynn, Mairead Hyland, The Lighthouse Lab in Alderley Park and Alex Alderton, Roberto Amato, Jeffrey Barrett, Sonia Goncalves, Ewan Harrison, David K. Jackson, Ian Johnston, Dominic Kwiatkowski, Cordelia Langford, John Sillitoe on behalf of the Wellcome Sanger Institute COVID-19 Surveillance Team |
| EPI_ISL_3224020, EPI_ISL_3224021, EPI_ISL_3224022 | Lighthouse Lab in Milton Keynes | Wellcome Sanger Institute for the COVID-19 Genomics UK (COG-UK) Consortium | The Lighthouse Lab in Milton Keynes and Alex Alderton, Roberto Amato, Jeffrey Barrett, Sonia Goncalves, Ewan Harrison, David K. Jackson, Ian Johnston, Dominic Kwiatkowski, Cordelia Langford, John Sillitoe on behalf of the Wellcome Sanger Institute COVID-19 Surveillance Team |
| EPI_ISL_3224023 | Lighthouse Laboratory Plymouth | Wellcome Sanger Institute for the COVID-19 Genomics UK (COG-UK) Consortium | Lighthouse Laboratory Plymouth and Alex Alderton, Roberto Amato, Jeffrey Barrett, Sonia Goncalves, Ewan Harrison, David K. Jackson, Ian Johnston, Dominic Kwiatkowski, Cordelia Langford, John Sillitoe on behalf of the Wellcome Sanger Institute COVID-19 Surveillance Team |
| EPI_ISL_3224024 | Lighthouse Lab in Glasgow | Wellcome Sanger Institute for the COVID-19 Genomics UK (COG-UK) Consortium | Harper VanSteenhouse, Yumi Kasai, David Gray, Carol Clugston, Anna Dominiczak and Alex Alderton, Roberto Amato, Jeffrey Barrett, Sonia Goncalves, Ewan Harrison, David K. Jackson, Ian Johnston, Dominic Kwiatkowski, Cordelia Langford, John Sillitoe on behalf of the Wellcome Sanger Institute COVID-19 Surveillance Team |
| EPI_ISL_3224025 | Lighthouse Lab in Alderley Park | Wellcome Sanger Institute for the COVID-19 Genomics UK (COG-UK) Consortium | Jacquelyn Wynn, Mairead Hyland, The Lighthouse Lab in Alderley Park and Alex Alderton, Roberto Amato, Jeffrey Barrett, Sonia Goncalves, Ewan Harrison, David K. Jackson, Ian Johnston, Dominic Kwiatkowski, Cordelia Langford, John Sillitoe on behalf of the Wellcome Sanger Institute COVID-19 Surveillance Team |
| EPI_ISL_3224026 | Lighthouse Laboratory Plymouth | Wellcome Sanger Institute for the COVID-19 Genomics UK (COG-UK) Consortium | Lighthouse Laboratory Plymouth and Alex Alderton, Roberto Amato, Jeffrey Barrett, Sonia Goncalves, Ewan Harrison, David K. Jackson, Ian Johnston, Dominic Kwiatkowski, Cordelia Langford, John Sillitoe on behalf of the Wellcome Sanger Institute COVID-19 Surveillance Team |
| EPI_ISL_3224027 | Lighthouse Lab in Glasgow | Wellcome Sanger Institute for the COVID-19 Genomics UK (COG-UK) Consortium | Harper VanSteenhouse, Yumi Kasai, David Gray, Carol Clugston, Anna Dominiczak and Alex Alderton, Roberto Amato, Jeffrey Barrett, Sonia Goncalves, Ewan Harrison, David K. Jackson, Ian Johnston, Dominic Kwiatkowski, Cordelia Langford, John Sillitoe on behalf of the Wellcome Sanger Institute COVID-19 Surveillance Team |
| EPI_ISL_3224028 | Lighthouse Lab in Alderley Park | Wellcome Sanger Institute for the COVID-19 Genomics UK (COG-UK) Consortium | Jacquelyn Wynn, Mairead Hyland, The Lighthouse Lab in Alderley Park and Alex Alderton, Roberto Amato, Jeffrey Barrett, Sonia Goncalves, Ewan Harrison, David K. Jackson, Ian Johnston, Dominic Kwiatkowski, Cordelia Langford, John Sillitoe on behalf of the Wellcome Sanger Institute COVID-19 Surveillance Team |
| EPI_ISL_3224029, EPI_ISL_3224030 | Lighthouse Lab in Glasgow | Wellcome Sanger Institute for the COVID-19 Genomics UK (COG-UK) Consortium | Harper VanSteenhouse, Yumi Kasai, David Gray, Carol Clugston, Anna Dominiczak and Alex Alderton, Roberto Amato, Jeffrey Barrett, Sonia Goncalves, Ewan Harrison, David K. Jackson, Ian Johnston, Dominic Kwiatkowski, Cordelia Langford, John Sillitoe on behalf of the Wellcome Sanger Institute COVID-19 Surveillance Team |
| EPI_ISL_3224031 | Lighthouse Laboratory Plymouth | Wellcome Sanger Institute for the COVID-19 Genomics UK (COG-UK) Consortium | Lighthouse Laboratory Plymouth and Alex Alderton, Roberto Amato, Jeffrey Barrett, Sonia Goncalves, Ewan Harrison, David K. Jackson, Ian Johnston, Dominic Kwiatkowski, Cordelia Langford, John Sillitoe on behalf of the Wellcome Sanger Institute COVID-19 Surveillance Team |

| | | | |
|---|---|---|---|
| EPI_ISL_3224032 | Lighthouse Lab in Milton Keynes | Wellcome Sanger Institute for the COVID-19 Genomics UK (COG-UK) Consortium | The Lighthouse Lab in Milton Keynes and Alex Alderton, Roberto Amato, Jeffrey Barrett, Sonia Goncalves, Ewan Harrison, David K. Jackson, Ian Johnston, Dominic Kwiatkowski, Cordelia Langford, John Sillitoe on behalf of the Wellcome Sanger Institute COVID-19 Surveillance Team |
| EPI_ISL_3224033 | Lighthouse Lab in Glasgow | Wellcome Sanger Institute for the COVID-19 Genomics UK (COG-UK) Consortium | Harper VanSteenhouse, Yumi Kasai, David Gray, Carol Clugston, Anna Dominiczak and Alex Alderton, Roberto Amato, Jeffrey Barrett, Sonia Goncalves, Ewan Harrison, David K. Jackson, Ian Johnston, Dominic Kwiatkowski, Cordelia Langford, John Sillitoe on behalf of the Wellcome Sanger Institute COVID-19 Surveillance Team |
| EPI_ISL_3224034 | Lighthouse Lab in Alderley Park | Wellcome Sanger Institute for the COVID-19 Genomics UK (COG-UK) Consortium | Jacquelyn Wynn, Mairead Hyland, The Lighthouse Lab in Alderley Park and Alex Alderton, Roberto Amato, Jeffrey Barrett, Sonia Goncalves, Ewan Harrison, David K. Jackson, Ian Johnston, Dominic Kwiatkowski, Cordelia Langford, John Sillitoe on behalf of the Wellcome Sanger Institute COVID-19 Surveillance Team |
| EPI_ISL_3224035 | Lighthouse Laboratory Plymouth | Wellcome Sanger Institute for the COVID-19 Genomics UK (COG-UK) Consortium | Lighthouse Laboratory Plymouth and Alex Alderton, Roberto Amato, Jeffrey Barrett, Sonia Goncalves, Ewan Harrison, David K. Jackson, Ian Johnston, Dominic Kwiatkowski, Cordelia Langford, John Sillitoe on behalf of the Wellcome Sanger Institute COVID-19 Surveillance Team |
| EPI_ISL_3224036 | Lighthouse Lab in Milton Keynes | Wellcome Sanger Institute for the COVID-19 Genomics UK (COG-UK) Consortium | The Lighthouse Lab in Milton Keynes and Alex Alderton, Roberto Amato, Jeffrey Barrett, Sonia Goncalves, Ewan Harrison, David K. Jackson, Ian Johnston, Dominic Kwiatkowski, Cordelia Langford, John Sillitoe on behalf of the Wellcome Sanger Institute COVID-19 Surveillance Team |
| EPI_ISL_3224037, EPI_ISL_3224038 | Lighthouse Lab in Alderley Park | Wellcome Sanger Institute for the COVID-19 Genomics UK (COG-UK) Consortium | Jacquelyn Wynn, Mairead Hyland, The Lighthouse Lab in Alderley Park and Alex Alderton, Roberto Amato, Jeffrey Barrett, Sonia Goncalves, Ewan Harrison, David K. Jackson, Ian Johnston, Dominic Kwiatkowski, Cordelia Langford, John Sillitoe on behalf of the Wellcome Sanger Institute COVID-19 Surveillance Team |
| EPI_ISL_3224039, EPI_ISL_3224040, EPI_ISL_3224041 | Lighthouse Lab in Milton Keynes | Wellcome Sanger Institute for the COVID-19 Genomics UK (COG-UK) Consortium | The Lighthouse Lab in Milton Keynes and Alex Alderton, Roberto Amato, Jeffrey Barrett, Sonia Goncalves, Ewan Harrison, David K. Jackson, Ian Johnston, Dominic Kwiatkowski, Cordelia Langford, John Sillitoe on behalf of the Wellcome Sanger Institute COVID-19 Surveillance Team |
| EPI_ISL_3224042 | Lighthouse Laboratory Plymouth | Wellcome Sanger Institute for the COVID-19 Genomics UK (COG-UK) Consortium | Lighthouse Laboratory Plymouth and Alex Alderton, Roberto Amato, Jeffrey Barrett, Sonia Goncalves, Ewan Harrison, David K. Jackson, Ian Johnston, Dominic Kwiatkowski, Cordelia Langford, John Sillitoe on behalf of the Wellcome Sanger Institute COVID-19 Surveillance Team |
| EPI_ISL_3224043 | Lighthouse Lab in Glasgow | Wellcome Sanger Institute for the COVID-19 Genomics UK (COG-UK) Consortium | Harper VanSteenhouse, Yumi Kasai, David Gray, Carol Clugston, Anna Dominiczak and Alex Alderton, Roberto Amato, Jeffrey Barrett, Sonia Goncalves, Ewan Harrison, David K. Jackson, Ian Johnston, Dominic Kwiatkowski, Cordelia Langford, John Sillitoe on behalf of the Wellcome Sanger Institute COVID-19 Surveillance Team |
| EPI_ISL_3224044 | Lighthouse Laboratory Plymouth | Wellcome Sanger Institute for the COVID-19 Genomics UK (COG-UK) Consortium | Lighthouse Laboratory Plymouth and Alex Alderton, Roberto Amato, Jeffrey Barrett, Sonia Goncalves, Ewan Harrison, David K. Jackson, Ian Johnston, Dominic Kwiatkowski, Cordelia Langford, John Sillitoe on behalf of the Wellcome Sanger Institute COVID-19 Surveillance Team |
| EPI_ISL_3224045 | Lighthouse Lab in Alderley Park | Wellcome Sanger Institute for the COVID-19 Genomics UK (COG-UK) Consortium | Jacquelyn Wynn, Mairead Hyland, The Lighthouse Lab in Alderley Park and Alex Alderton, Roberto Amato, Jeffrey Barrett, Sonia Goncalves, Ewan Harrison, David K. Jackson, Ian Johnston, Dominic Kwiatkowski, Cordelia Langford, John Sillitoe on behalf of the Wellcome Sanger Institute COVID-19 Surveillance Team |
| EPI_ISL_3224046, EPI_ISL_3224047 | Lighthouse Laboratory Plymouth | Wellcome Sanger Institute for the COVID-19 Genomics UK (COG-UK) Consortium | Lighthouse Laboratory Plymouth and Alex Alderton, Roberto Amato, Jeffrey Barrett, Sonia Goncalves, Ewan Harrison, David K. Jackson, Ian Johnston, Dominic Kwiatkowski, Cordelia Langford, John Sillitoe on behalf of the Wellcome Sanger Institute COVID-19 Surveillance Team |
| EPI_ISL_3224048 | Lighthouse Lab in Glasgow | Wellcome Sanger Institute for the COVID-19 Genomics UK (COG-UK) Consortium | Harper VanSteenhouse, Yumi Kasai, David Gray, Carol Clugston, Anna Dominiczak and Alex Alderton, Roberto Amato, Jeffrey Barrett, Sonia Goncalves, Ewan Harrison, David K. Jackson, Ian Johnston, Dominic Kwiatkowski, Cordelia Langford, John Sillitoe on behalf of the Wellcome Sanger Institute COVID-19 Surveillance Team |
| EPI_ISL_3224049 | Lighthouse Lab in Alderley Park | Wellcome Sanger Institute for the COVID-19 Genomics UK (COG-UK) Consortium | Jacquelyn Wynn, Mairead Hyland, The Lighthouse Lab in Alderley Park and Alex Alderton, Roberto Amato, Jeffrey Barrett, Sonia Goncalves, Ewan Harrison, David K. Jackson, Ian Johnston, Dominic Kwiatkowski, Cordelia Langford, John Sillitoe on behalf of the Wellcome Sanger Institute COVID-19 Surveillance Team |
| EPI_ISL_3224050 | Lighthouse Lab in Glasgow | Wellcome Sanger Institute for the COVID-19 Genomics UK (COG-UK) Consortium | Harper VanSteenhouse, Yumi Kasai, David Gray, Carol Clugston, Anna Dominiczak and Alex Alderton, Roberto Amato, Jeffrey Barrett, Sonia Goncalves, Ewan Harrison, David K. Jackson, Ian Johnston, Dominic Kwiatkowski, Cordelia Langford, John Sillitoe on behalf of the Wellcome Sanger Institute COVID-19 Surveillance Team |
| EPI_ISL_3224051, EPI_ISL_3224052 | Lighthouse Lab in Milton Keynes | Wellcome Sanger Institute for the COVID-19 Genomics UK (COG-UK) Consortium | The Lighthouse Lab in Milton Keynes and Alex Alderton, Roberto Amato, Jeffrey Barrett, Sonia Goncalves, Ewan Harrison, David K. Jackson, Ian Johnston, Dominic Kwiatkowski, Cordelia Langford, John Sillitoe on behalf of the Wellcome Sanger Institute COVID-19 Surveillance Team |
| EPI_ISL_3224053 | Lighthouse Laboratory Plymouth | Wellcome Sanger Institute for the COVID-19 Genomics UK (COG-UK) Consortium | Lighthouse Laboratory Plymouth and Alex Alderton, Roberto Amato, Jeffrey Barrett, Sonia Goncalves, Ewan Harrison, David K. Jackson, Ian Johnston, Dominic Kwiatkowski, Cordelia Langford, John Sillitoe on behalf of the Wellcome Sanger Institute COVID-19 Surveillance Team |
| EPI_ISL_3224054 | Lighthouse Lab in Glasgow | Wellcome Sanger Institute for the COVID-19 Genomics UK (COG-UK) Consortium | Harper VanSteenhouse, Yumi Kasai, David Gray, Carol Clugston, Anna Dominiczak and Alex Alderton, Roberto Amato, Jeffrey Barrett, Sonia Goncalves, Ewan Harrison, David K. Jackson, Ian Johnston, Dominic Kwiatkowski, Cordelia Langford, John Sillitoe on behalf of the Wellcome Sanger Institute COVID-19 Surveillance Team |
| EPI_ISL_3224055 | Lighthouse Laboratory Plymouth | Wellcome Sanger Institute for the COVID-19 Genomics UK (COG-UK) Consortium | Lighthouse Laboratory Plymouth and Alex Alderton, Roberto Amato, Jeffrey Barrett, Sonia Goncalves, Ewan Harrison, David K. Jackson, Ian Johnston, Dominic Kwiatkowski, Cordelia Langford, John Sillitoe on behalf of the Wellcome Sanger Institute COVID-19 Surveillance Team |
| EPI_ISL_3224056 | Lighthouse Lab in Milton Keynes | Wellcome Sanger Institute for the COVID-19 Genomics UK (COG-UK) Consortium | The Lighthouse Lab in Milton Keynes and Alex Alderton, Roberto Amato, Jeffrey Barrett, Sonia Goncalves, Ewan Harrison, David K. Jackson, Ian Johnston, Dominic Kwiatkowski, Cordelia Langford, John Sillitoe on behalf of the Wellcome Sanger Institute COVID-19 Surveillance Team |
| EPI_ISL_3224057, EPI_ISL_3224058, EPI_ISL_3224059, EPI_ISL_3224060 | Lighthouse Lab in Alderley Park | Wellcome Sanger Institute for the COVID-19 Genomics UK (COG-UK) Consortium | Jacquelyn Wynn, Mairead Hyland, The Lighthouse Lab in Alderley Park and Alex Alderton, Roberto Amato, Jeffrey Barrett, Sonia Goncalves, Ewan Harrison, David K. Jackson, Ian Johnston, Dominic Kwiatkowski, Cordelia Langford, John Sillitoe on behalf of the Wellcome Sanger Institute COVID-19 Surveillance Team |
| EPI_ISL_3224061 | Lighthouse Lab in Glasgow | Wellcome Sanger Institute for the COVID-19 Genomics UK (COG-UK) Consortium | Harper VanSteenhouse, Yumi Kasai, David Gray, Carol Clugston, Anna Dominiczak and Alex Alderton, Roberto Amato, Jeffrey Barrett, Sonia Goncalves, Ewan Harrison, David K. Jackson, Ian Johnston, Dominic Kwiatkowski, Cordelia Langford, John Sillitoe on behalf of the Wellcome Sanger Institute COVID-19 Surveillance Team |
| EPI_ISL_3224062 | Lighthouse Laboratory Plymouth | Wellcome Sanger Institute for the COVID-19 Genomics UK (COG-UK) Consortium | Lighthouse Laboratory Plymouth and Alex Alderton, Roberto Amato, Jeffrey Barrett, Sonia Goncalves, Ewan Harrison, David K. Jackson, Ian Johnston, Dominic Kwiatkowski, Cordelia Langford, John Sillitoe on behalf of the Wellcome Sanger Institute COVID-19 Surveillance Team |
| EPI_ISL_3224063, EPI_ISL_3224064 | Lighthouse Lab in Alderley Park | Wellcome Sanger Institute for the COVID-19 Genomics UK (COG-UK) Consortium | Jacquelyn Wynn, Mairead Hyland, The Lighthouse Lab in Alderley Park and Alex Alderton, Roberto Amato, Jeffrey Barrett, Sonia Goncalves, Ewan Harrison, David K. Jackson, Ian Johnston, Dominic Kwiatkowski, Cordelia Langford, John Sillitoe on behalf of the Wellcome Sanger Institute COVID-19 Surveillance Team |
| EPI_ISL_3224065 | Lighthouse Lab in Milton Keynes | Wellcome Sanger Institute for the COVID-19 Genomics UK (COG-UK) Consortium | The Lighthouse Lab in Milton Keynes and Alex Alderton, Roberto Amato, Jeffrey Barrett, Sonia Goncalves, Ewan Harrison, David K. Jackson, Ian Johnston, Dominic Kwiatkowski, Cordelia Langford, John Sillitoe on behalf of the Wellcome Sanger Institute COVID-19 Surveillance Team |
| EPI_ISL_3224066, EPI_ISL_3224067 | Lighthouse Lab in Alderley Park | Wellcome Sanger Institute for the COVID-19 Genomics UK (COG-UK) Consortium | Jacquelyn Wynn, Mairead Hyland, The Lighthouse Lab in Alderley Park and Alex Alderton, Roberto Amato, Jeffrey Barrett, Sonia Goncalves, Ewan Harrison, David K. Jackson, Ian Johnston, Dominic Kwiatkowski, Cordelia Langford, John Sillitoe on behalf of the Wellcome Sanger Institute COVID-19 Surveillance Team |
| EPI_ISL_3224068, EPI_ISL_3224069 | Lighthouse Lab in Milton Keynes | Wellcome Sanger Institute for the COVID-19 Genomics UK (COG-UK) Consortium | The Lighthouse Lab in Milton Keynes and Alex Alderton, Roberto Amato, Jeffrey Barrett, Sonia Goncalves, Ewan Harrison, David K. Jackson, Ian Johnston, Dominic Kwiatkowski, Cordelia Langford, John Sillitoe on behalf of the Wellcome Sanger Institute COVID-19 Surveillance Team |
| EPI_ISL_3224070, EPI_ISL_3224071 | Lighthouse Lab in Alderley Park | Wellcome Sanger Institute for the COVID-19 Genomics UK (COG-UK) Consortium | Jacquelyn Wynn, Mairead Hyland, The Lighthouse Lab in Alderley Park and Alex Alderton, Roberto Amato, Jeffrey Barrett, Sonia Goncalves, Ewan Harrison, David K. Jackson, Ian Johnston, Dominic Kwiatkowski, Cordelia Langford, John Sillitoe on behalf of the Wellcome Sanger Institute COVID-19 Surveillance Team |
| EPI_ISL_3224072 | Lighthouse Laboratory Plymouth | Wellcome Sanger Institute for the COVID-19 Genomics UK (COG-UK) Consortium | Lighthouse Laboratory Plymouth and Alex Alderton, Roberto Amato, Jeffrey Barrett, Sonia Goncalves, Ewan Harrison, David K. Jackson, Ian Johnston, Dominic Kwiatkowski, Cordelia Langford, John Sillitoe on behalf of the Wellcome Sanger Institute COVID-19 Surveillance Team |
| EPI_ISL_3224073, EPI_ISL_3224074, EPI_ISL_3224075 | Lighthouse Lab in Glasgow | Wellcome Sanger Institute for the COVID-19 Genomics UK (COG-UK) Consortium | Harper VanSteenhouse, Yumi Kasai, David Gray, Carol Clugston, Anna Dominiczak and Alex Alderton, Roberto Amato, Jeffrey Barrett, Sonia Goncalves, Ewan Harrison, David K. Jackson, Ian Johnston, Dominic Kwiatkowski, Cordelia Langford, John Sillitoe on behalf of the Wellcome Sanger Institute COVID-19 Surveillance Team |
| EPI_ISL_3224076 | Lighthouse Lab in Milton Keynes | Wellcome Sanger Institute for the COVID-19 Genomics UK (COG-UK) Consortium | The Lighthouse Lab in Milton Keynes and Alex Alderton, Roberto Amato, Jeffrey Barrett, Sonia Goncalves, Ewan Harrison, David K. Jackson, Ian Johnston, Dominic Kwiatkowski, Cordelia Langford, John Sillitoe on behalf of the Wellcome Sanger Institute COVID-19 Surveillance Team |
| EPI_ISL_3224077 | Lighthouse Lab in Alderley Park | Wellcome Sanger Institute for the COVID-19 Genomics UK (COG-UK) Consortium | Jacquelyn Wynn, Mairead Hyland, The Lighthouse Lab in Alderley Park and Alex Alderton, Roberto Amato, Jeffrey Barrett, Sonia Goncalves, Ewan Harrison, David K. Jackson, Ian Johnston, Dominic Kwiatkowski, Cordelia Langford, John Sillitoe on behalf of the Wellcome Sanger Institute COVID-19 Surveillance Team |
| EPI_ISL_3224078 | Lighthouse Lab in Glasgow | Wellcome Sanger Institute for the COVID-19 Genomics UK (COG-UK) Consortium | Harper VanSteenhouse, Yumi Kasai, David Gray, Carol Clugston, Anna Dominiczak and Alex Alderton, Roberto Amato, Jeffrey Barrett, Sonia Goncalves, Ewan Harrison, David K. Jackson, Ian Johnston, Dominic Kwiatkowski, Cordelia Langford, John Sillitoe on behalf of the Wellcome Sanger Institute COVID-19 Surveillance Team |
| EPI_ISL_3224079 | Lighthouse Lab in Alderley Park | Wellcome Sanger Institute for the COVID-19 Genomics UK (COG-UK) Consortium | Jacquelyn Wynn, Mairead Hyland, The Lighthouse Lab in Alderley Park and Alex Alderton, Roberto Amato, Jeffrey Barrett, Sonia Goncalves, Ewan Harrison, David K. Jackson, Ian Johnston, Dominic Kwiatkowski, Cordelia Langford, John Sillitoe on behalf of the Wellcome Sanger Institute COVID-19 Surveillance Team |
| EPI_ISL_3224080, EPI_ISL_3224081 | Lighthouse Lab in Milton Keynes | Wellcome Sanger Institute for the COVID-19 Genomics UK (COG-UK) Consortium | The Lighthouse Lab in Milton Keynes and Alex Alderton, Roberto Amato, Jeffrey Barrett, Sonia Goncalves, Ewan Harrison, David K. Jackson, Ian Johnston, Dominic Kwiatkowski, Cordelia Langford, John Sillitoe on behalf of the Wellcome Sanger Institute COVID-19 Surveillance Team |
| EPI_ISL_3224082 | Lighthouse Lab in Alderley Park | Wellcome Sanger Institute for the COVID-19 Genomics UK (COG-UK) Consortium | Jacquelyn Wynn, Mairead Hyland, The Lighthouse Lab in Alderley Park and Alex Alderton, Roberto Amato, Jeffrey Barrett, Sonia Goncalves, Ewan Harrison, David K. Jackson, Ian Johnston, Dominic Kwiatkowski, Cordelia Langford, John Sillitoe on behalf of the Wellcome Sanger Institute COVID-19 Surveillance Team |
| EPI_ISL_3224083 | Lighthouse Lab in Glasgow | Wellcome Sanger Institute for the COVID-19 Genomics UK (COG-UK) Consortium | Harper VanSteenhouse, Yumi Kasai, David Gray, Carol Clugston, Anna Dominiczak and Alex Alderton, Roberto Amato, Jeffrey Barrett, Sonia Goncalves, Ewan Harrison, David K. Jackson, Ian Johnston, Dominic Kwiatkowski, Cordelia Langford, John Sillitoe on behalf of the Wellcome Sanger Institute COVID-19 Surveillance Team |
| EPI_ISL_3224084, EPI_ISL_3224085 | Lighthouse Lab in Milton Keynes | Wellcome Sanger Institute for the COVID-19 Genomics UK (COG-UK) Consortium | The Lighthouse Lab in Milton Keynes and Alex Alderton, Roberto Amato, Jeffrey Barrett, Sonia Goncalves, Ewan Harrison, David K. Jackson, Ian Johnston, Dominic Kwiatkowski, Cordelia Langford, John Sillitoe on behalf of the Wellcome Sanger Institute COVID-19 Surveillance Team |
| EPI_ISL_3224086 | Lighthouse Laboratory Plymouth | Wellcome Sanger Institute for the COVID-19 Genomics UK (COG-UK) Consortium | Lighthouse Laboratory Plymouth and Alex Alderton, Roberto Amato, Jeffrey Barrett, Sonia Goncalves, Ewan Harrison, David K. Jackson, Ian Johnston, Dominic Kwiatkowski, Cordelia Langford, John Sillitoe on behalf of the Wellcome Sanger Institute COVID-19 Surveillance Team |
| EPI_ISL_3224087 | Lighthouse Lab in Alderley Park | Wellcome Sanger Institute for the COVID-19 Genomics UK (COG-UK) Consortium | Jacquelyn Wynn, Mairead Hyland, The Lighthouse Lab in Alderley Park and Alex Alderton, Roberto Amato, Jeffrey Barrett, Sonia Goncalves, Ewan Harrison, David K. Jackson, Ian Johnston, Dominic Kwiatkowski, Cordelia Langford, John Sillitoe on behalf of the Wellcome Sanger Institute COVID-19 Surveillance Team |
| EPI_ISL_3224088 | Lighthouse Laboratory Plymouth | Wellcome Sanger Institute for the COVID-19 Genomics UK (COG-UK) Consortium | Lighthouse Laboratory Plymouth and Alex Alderton, Roberto Amato, Jeffrey Barrett, Sonia Goncalves, Ewan Harrison, David K. Jackson, Ian Johnston, Dominic Kwiatkowski, Cordelia Langford, John Sillitoe on behalf of the Wellcome Sanger Institute COVID-19 Surveillance Team |
| EPI_ISL_3224089 | Lighthouse Lab in Milton Keynes | Wellcome Sanger Institute for the COVID-19 Genomics UK (COG-UK) Consortium | The Lighthouse Lab in Milton Keynes and Alex Alderton, Roberto Amato, Jeffrey Barrett, Sonia Goncalves, Ewan Harrison, David K. Jackson, Ian Johnston, Dominic Kwiatkowski, Cordelia Langford, John Sillitoe on behalf of the Wellcome Sanger Institute COVID-19 Surveillance Team |
| EPI_ISL_3224090, EPI_ISL_3224091 | Lighthouse Laboratory Plymouth | Wellcome Sanger Institute for the COVID-19 Genomics UK (COG-UK) Consortium | Lighthouse Laboratory Plymouth and Alex Alderton, Roberto Amato, Jeffrey Barrett, Sonia Goncalves, Ewan Harrison, David K. Jackson, Ian Johnston, Dominic Kwiatkowski, Cordelia Langford, John Sillitoe on behalf of the Wellcome Sanger Institute COVID-19 Surveillance Team |
| EPI_ISL_3224092, | Lighthouse Lab in Alderley Park | Wellcome Sanger Institute for the COVID-19 Genomics UK (COG-UK) | Jacquelyn Wynn, Mairead Hyland, The Lighthouse Lab in Alderley Park and Alex Alderton, Roberto Amato, Jeffrey Barrett, Sonia Goncalves, Ewan Harrison, David K. Jackson, Ian Johnston, Dominic Kwiatkowski, Cordelia Langford, John Sillitoe on |

| | | | |
|---|---|---|---|
| EPI_ISL_3224093 | | Consortium | behalf of the Wellcome Sanger Institute COVID-19 Surveillance Team |
| EPI_ISL_3224094 | Lighthouse Laboratory Plymouth | Wellcome Sanger Institute for the COVID-19 Genomics UK (COG-UK) Consortium | Lighthouse Laboratory Plymouth and Alex Alderton, Roberto Amato, Jeffrey Barrett, Sonia Goncalves, Ewan Harrison, David K. Jackson, Ian Johnston, Dominic Kwiatkowski, Cordelia Langford, John Sillitoe on behalf of the Wellcome Sanger Institute COVID-19 Surveillance Team |
| EPI_ISL_3224095 | Lighthouse Lab in Glasgow | Wellcome Sanger Institute for the COVID-19 Genomics UK (COG-UK) Consortium | Harper VanSteenhouse, Yumi Kasai, David Gray, Carol Clugston, Anna Dominiczak and Alex Alderton, Roberto Amato, Jeffrey Barrett, Sonia Goncalves, Ewan Harrison, David K. Jackson, Ian Johnston, Dominic Kwiatkowski, Cordelia Langford, John Sillitoe on behalf of the Wellcome Sanger Institute COVID-19 Surveillance Team |
| EPI_ISL_3224096 | Lighthouse Laboratory Plymouth | Wellcome Sanger Institute for the COVID-19 Genomics UK (COG-UK) Consortium | Lighthouse Laboratory Plymouth and Alex Alderton, Roberto Amato, Jeffrey Barrett, Sonia Goncalves, Ewan Harrison, David K. Jackson, Ian Johnston, Dominic Kwiatkowski, Cordelia Langford, John Sillitoe on behalf of the Wellcome Sanger Institute COVID-19 Surveillance Team |
| EPI_ISL_3224097, EPI_ISL_3224098, EPI_ISL_3224099, EPI_ISL_3224100 | Lighthouse Lab in Alderley Park | Wellcome Sanger Institute for the COVID-19 Genomics UK (COG-UK) Consortium | Jacquelyn Wynn, Mairead Hyland, The Lighthouse Lab in Alderley Park and Alex Alderton, Roberto Amato, Jeffrey Barrett, Sonia Goncalves, Ewan Harrison, David K. Jackson, Ian Johnston, Dominic Kwiatkowski, Cordelia Langford, John Sillitoe on behalf of the Wellcome Sanger Institute COVID-19 Surveillance Team |
| EPI_ISL_3224101 | Lighthouse Laboratory Plymouth | Wellcome Sanger Institute for the COVID-19 Genomics UK (COG-UK) Consortium | Lighthouse Laboratory Plymouth and Alex Alderton, Roberto Amato, Jeffrey Barrett, Sonia Goncalves, Ewan Harrison, David K. Jackson, Ian Johnston, Dominic Kwiatkowski, Cordelia Langford, John Sillitoe on behalf of the Wellcome Sanger Institute COVID-19 Surveillance Team |
| EPI_ISL_3224102 | Lighthouse Lab in Alderley Park | Wellcome Sanger Institute for the COVID-19 Genomics UK (COG-UK) Consortium | Jacquelyn Wynn, Mairead Hyland, The Lighthouse Lab in Alderley Park and Alex Alderton, Roberto Amato, Jeffrey Barrett, Sonia Goncalves, Ewan Harrison, David K. Jackson, Ian Johnston, Dominic Kwiatkowski, Cordelia Langford, John Sillitoe on behalf of the Wellcome Sanger Institute COVID-19 Surveillance Team |
| EPI_ISL_3224103, EPI_ISL_3224104 | Lighthouse Laboratory Plymouth | Wellcome Sanger Institute for the COVID-19 Genomics UK (COG-UK) Consortium | Lighthouse Laboratory Plymouth and Alex Alderton, Roberto Amato, Jeffrey Barrett, Sonia Goncalves, Ewan Harrison, David K. Jackson, Ian Johnston, Dominic Kwiatkowski, Cordelia Langford, John Sillitoe on behalf of the Wellcome Sanger Institute COVID-19 Surveillance Team |
| EPI_ISL_3224105 | Lighthouse Lab in Milton Keynes | Wellcome Sanger Institute for the COVID-19 Genomics UK (COG-UK) Consortium | The Lighthouse Lab in Milton Keynes and Alex Alderton, Roberto Amato, Jeffrey Barrett, Sonia Goncalves, Ewan Harrison, David K. Jackson, Ian Johnston, Dominic Kwiatkowski, Cordelia Langford, John Sillitoe on behalf of the Wellcome Sanger Institute COVID-19 Surveillance Team |
| EPI_ISL_3224106, EPI_ISL_3224107 | Lighthouse Laboratory Plymouth | Wellcome Sanger Institute for the COVID-19 Genomics UK (COG-UK) Consortium | Lighthouse Laboratory Plymouth and Alex Alderton, Roberto Amato, Jeffrey Barrett, Sonia Goncalves, Ewan Harrison, David K. Jackson, Ian Johnston, Dominic Kwiatkowski, Cordelia Langford, John Sillitoe on behalf of the Wellcome Sanger Institute COVID-19 Surveillance Team |
| EPI_ISL_3224108, EPI_ISL_3224109 | Lighthouse Lab in Glasgow | Wellcome Sanger Institute for the COVID-19 Genomics UK (COG-UK) Consortium | Harper VanSteenhouse, Yumi Kasai, David Gray, Carol Clugston, Anna Dominiczak and Alex Alderton, Roberto Amato, Jeffrey Barrett, Sonia Goncalves, Ewan Harrison, David K. Jackson, Ian Johnston, Dominic Kwiatkowski, Cordelia Langford, John Sillitoe on behalf of the Wellcome Sanger Institute COVID-19 Surveillance Team |
| EPI_ISL_3224110 | Lighthouse Laboratory Plymouth | Wellcome Sanger Institute for the COVID-19 Genomics UK (COG-UK) Consortium | Lighthouse Laboratory Plymouth and Alex Alderton, Roberto Amato, Jeffrey Barrett, Sonia Goncalves, Ewan Harrison, David K. Jackson, Ian Johnston, Dominic Kwiatkowski, Cordelia Langford, John Sillitoe on behalf of the Wellcome Sanger Institute COVID-19 Surveillance Team |
| EPI_ISL_3224111 | Lighthouse Lab in Milton Keynes | Wellcome Sanger Institute for the COVID-19 Genomics UK (COG-UK) Consortium | The Lighthouse Lab in Milton Keynes and Alex Alderton, Roberto Amato, Jeffrey Barrett, Sonia Goncalves, Ewan Harrison, David K. Jackson, Ian Johnston, Dominic Kwiatkowski, Cordelia Langford, John Sillitoe on behalf of the Wellcome Sanger Institute COVID-19 Surveillance Team |
| EPI_ISL_3224112 | Lighthouse Lab in Alderley Park | Wellcome Sanger Institute for the COVID-19 Genomics UK (COG-UK) Consortium | Jacquelyn Wynn, Mairead Hyland, The Lighthouse Lab in Alderley Park and Alex Alderton, Roberto Amato, Jeffrey Barrett, Sonia Goncalves, Ewan Harrison, David K. Jackson, Ian Johnston, Dominic Kwiatkowski, Cordelia Langford, John Sillitoe on behalf of the Wellcome Sanger Institute COVID-19 Surveillance Team |
| EPI_ISL_3224113 | Lighthouse Lab in Milton Keynes | Wellcome Sanger Institute for the COVID-19 Genomics UK (COG-UK) Consortium | The Lighthouse Lab in Milton Keynes and Alex Alderton, Roberto Amato, Jeffrey Barrett, Sonia Goncalves, Ewan Harrison, David K. Jackson, Ian Johnston, Dominic Kwiatkowski, Cordelia Langford, John Sillitoe on behalf of the Wellcome Sanger Institute COVID-19 Surveillance Team |
| EPI_ISL_3224114 | Lighthouse Lab in Glasgow | Wellcome Sanger Institute for the COVID-19 Genomics UK (COG-UK) Consortium | Harper VanSteenhouse, Yumi Kasai, David Gray, Carol Clugston, Anna Dominiczak and Alex Alderton, Roberto Amato, Jeffrey Barrett, Sonia Goncalves, Ewan Harrison, David K. Jackson, Ian Johnston, Dominic Kwiatkowski, Cordelia Langford, John Sillitoe on behalf of the Wellcome Sanger Institute COVID-19 Surveillance Team |
| EPI_ISL_3224115, EPI_ISL_3224116 | Lighthouse Lab in Alderley Park | Wellcome Sanger Institute for the COVID-19 Genomics UK (COG-UK) Consortium | Jacquelyn Wynn, Mairead Hyland, The Lighthouse Lab in Alderley Park and Alex Alderton, Roberto Amato, Jeffrey Barrett, Sonia Goncalves, Ewan Harrison, David K. Jackson, Ian Johnston, Dominic Kwiatkowski, Cordelia Langford, John Sillitoe on behalf of the Wellcome Sanger Institute COVID-19 Surveillance Team |
| EPI_ISL_3224117 | Lighthouse Laboratory Plymouth | Wellcome Sanger Institute for the COVID-19 Genomics UK (COG-UK) Consortium | Lighthouse Laboratory Plymouth and Alex Alderton, Roberto Amato, Jeffrey Barrett, Sonia Goncalves, Ewan Harrison, David K. Jackson, Ian Johnston, Dominic Kwiatkowski, Cordelia Langford, John Sillitoe on behalf of the Wellcome Sanger Institute COVID-19 Surveillance Team |
| EPI_ISL_3224118, EPI_ISL_3224119 | Lighthouse Lab in Alderley Park | Wellcome Sanger Institute for the COVID-19 Genomics UK (COG-UK) Consortium | Jacquelyn Wynn, Mairead Hyland, The Lighthouse Lab in Alderley Park and Alex Alderton, Roberto Amato, Jeffrey Barrett, Sonia Goncalves, Ewan Harrison, David K. Jackson, Ian Johnston, Dominic Kwiatkowski, Cordelia Langford, John Sillitoe on behalf of the Wellcome Sanger Institute COVID-19 Surveillance Team |
| EPI_ISL_3224120, EPI_ISL_3224121 | Lighthouse Lab in Glasgow | Wellcome Sanger Institute for the COVID-19 Genomics UK (COG-UK) Consortium | Harper VanSteenhouse, Yumi Kasai, David Gray, Carol Clugston, Anna Dominiczak and Alex Alderton, Roberto Amato, Jeffrey Barrett, Sonia Goncalves, Ewan Harrison, David K. Jackson, Ian Johnston, Dominic Kwiatkowski, Cordelia Langford, John Sillitoe on behalf of the Wellcome Sanger Institute COVID-19 Surveillance Team |
| EPI_ISL_3224122 | Lighthouse Lab in Alderley Park | Wellcome Sanger Institute for the COVID-19 Genomics UK (COG-UK) Consortium | Jacquelyn Wynn, Mairead Hyland, The Lighthouse Lab in Alderley Park and Alex Alderton, Roberto Amato, Jeffrey Barrett, Sonia Goncalves, Ewan Harrison, David K. Jackson, Ian Johnston, Dominic Kwiatkowski, Cordelia Langford, John Sillitoe on behalf of the Wellcome Sanger Institute COVID-19 Surveillance Team |
| EPI_ISL_3224123 | Lighthouse Laboratory Plymouth | Wellcome Sanger Institute for the COVID-19 Genomics UK (COG-UK) Consortium | Lighthouse Laboratory Plymouth and Alex Alderton, Roberto Amato, Jeffrey Barrett, Sonia Goncalves, Ewan Harrison, David K. Jackson, Ian Johnston, Dominic Kwiatkowski, Cordelia Langford, John Sillitoe on behalf of the Wellcome Sanger Institute COVID-19 Surveillance Team |
| EPI_ISL_3224124, EPI_ISL_3224125, EPI_ISL_3224126 | Lighthouse Lab in Glasgow | Wellcome Sanger Institute for the COVID-19 Genomics UK (COG-UK) Consortium | Harper VanSteenhouse, Yumi Kasai, David Gray, Carol Clugston, Anna Dominiczak and Alex Alderton, Roberto Amato, Jeffrey Barrett, Sonia Goncalves, Ewan Harrison, David K. Jackson, Ian Johnston, Dominic Kwiatkowski, Cordelia Langford, John Sillitoe on behalf of the Wellcome Sanger Institute COVID-19 Surveillance Team |
| EPI_ISL_3224127, EPI_ISL_3224128 | Lighthouse Laboratory Plymouth | Wellcome Sanger Institute for the COVID-19 Genomics UK (COG-UK) Consortium | Lighthouse Laboratory Plymouth and Alex Alderton, Roberto Amato, Jeffrey Barrett, Sonia Goncalves, Ewan Harrison, David K. Jackson, Ian Johnston, Dominic Kwiatkowski, Cordelia Langford, John Sillitoe on behalf of the Wellcome Sanger Institute COVID-19 Surveillance Team |
| EPI_ISL_3224129, EPI_ISL_3224130 | Lighthouse Lab in Glasgow | Wellcome Sanger Institute for the COVID-19 Genomics UK (COG-UK) Consortium | Harper VanSteenhouse, Yumi Kasai, David Gray, Carol Clugston, Anna Dominiczak and Alex Alderton, Roberto Amato, Jeffrey Barrett, Sonia Goncalves, Ewan Harrison, David K. Jackson, Ian Johnston, Dominic Kwiatkowski, Cordelia Langford, John Sillitoe on behalf of the Wellcome Sanger Institute COVID-19 Surveillance Team |
| EPI_ISL_3224131 | Lighthouse Lab in Alderley Park | Wellcome Sanger Institute for the COVID-19 Genomics UK (COG-UK) Consortium | Jacquelyn Wynn, Mairead Hyland, The Lighthouse Lab in Alderley Park and Alex Alderton, Roberto Amato, Jeffrey Barrett, Sonia Goncalves, Ewan Harrison, David K. Jackson, Ian Johnston, Dominic Kwiatkowski, Cordelia Langford, John Sillitoe on behalf of the Wellcome Sanger Institute COVID-19 Surveillance Team |
| EPI_ISL_3224132 | Lighthouse Laboratory Plymouth | Wellcome Sanger Institute for the COVID-19 Genomics UK (COG-UK) Consortium | Lighthouse Laboratory Plymouth and Alex Alderton, Roberto Amato, Jeffrey Barrett, Sonia Goncalves, Ewan Harrison, David K. Jackson, Ian Johnston, Dominic Kwiatkowski, Cordelia Langford, John Sillitoe on behalf of the Wellcome Sanger Institute COVID-19 Surveillance Team |
| EPI_ISL_3224133 | Lighthouse Lab in Glasgow | Wellcome Sanger Institute for the COVID-19 Genomics UK (COG-UK) Consortium | Harper VanSteenhouse, Yumi Kasai, David Gray, Carol Clugston, Anna Dominiczak and Alex Alderton, Roberto Amato, Jeffrey Barrett, Sonia Goncalves, Ewan Harrison, David K. Jackson, Ian Johnston, Dominic Kwiatkowski, Cordelia Langford, John Sillitoe on behalf of the Wellcome Sanger Institute COVID-19 Surveillance Team |
| EPI_ISL_3224134, EPI_ISL_3224135 | Lighthouse Laboratory Plymouth | Wellcome Sanger Institute for the COVID-19 Genomics UK (COG-UK) Consortium | Lighthouse Laboratory Plymouth and Alex Alderton, Roberto Amato, Jeffrey Barrett, Sonia Goncalves, Ewan Harrison, David K. Jackson, Ian Johnston, Dominic Kwiatkowski, Cordelia Langford, John Sillitoe on behalf of the Wellcome Sanger Institute COVID-19 Surveillance Team |
| EPI_ISL_3224136 | Lighthouse Lab in Milton Keynes | Wellcome Sanger Institute for the COVID-19 Genomics UK (COG-UK) Consortium | The Lighthouse Lab in Milton Keynes and Alex Alderton, Roberto Amato, Jeffrey Barrett, Sonia Goncalves, Ewan Harrison, David K. Jackson, Ian Johnston, Dominic Kwiatkowski, Cordelia Langford, John Sillitoe on behalf of the Wellcome Sanger Institute COVID-19 Surveillance Team |
| EPI_ISL_3224137 | Lighthouse Lab in Glasgow | Wellcome Sanger Institute for the COVID-19 Genomics UK (COG-UK) Consortium | Harper VanSteenhouse, Yumi Kasai, David Gray, Carol Clugston, Anna Dominiczak and Alex Alderton, Roberto Amato, Jeffrey Barrett, Sonia Goncalves, Ewan Harrison, David K. Jackson, Ian Johnston, Dominic Kwiatkowski, Cordelia Langford, John Sillitoe on behalf of the Wellcome Sanger Institute COVID-19 Surveillance Team |
| EPI_ISL_3224138 | Lighthouse Lab in Alderley Park | Wellcome Sanger Institute for the COVID-19 Genomics UK (COG-UK) Consortium | Jacquelyn Wynn, Mairead Hyland, The Lighthouse Lab in Alderley Park and Alex Alderton, Roberto Amato, Jeffrey Barrett, Sonia Goncalves, Ewan Harrison, David K. Jackson, Ian Johnston, Dominic Kwiatkowski, Cordelia Langford, John Sillitoe on behalf of the Wellcome Sanger Institute COVID-19 Surveillance Team |
| EPI_ISL_3224139 | Lighthouse Lab in Glasgow | Wellcome Sanger Institute for the COVID-19 Genomics UK (COG-UK) Consortium | Harper VanSteenhouse, Yumi Kasai, David Gray, Carol Clugston, Anna Dominiczak and Alex Alderton, Roberto Amato, Jeffrey Barrett, Sonia Goncalves, Ewan Harrison, David K. Jackson, Ian Johnston, Dominic Kwiatkowski, Cordelia Langford, John Sillitoe on behalf of the Wellcome Sanger Institute COVID-19 Surveillance Team |
| EPI_ISL_3224140 | Lighthouse Lab in Milton Keynes | Wellcome Sanger Institute for the COVID-19 Genomics UK (COG-UK) Consortium | The Lighthouse Lab in Milton Keynes and Alex Alderton, Roberto Amato, Jeffrey Barrett, Sonia Goncalves, Ewan Harrison, David K. Jackson, Ian Johnston, Dominic Kwiatkowski, Cordelia Langford, John Sillitoe on behalf of the Wellcome Sanger Institute COVID-19 Surveillance Team |
| EPI_ISL_3224141, EPI_ISL_3224142 | Lighthouse Lab in Glasgow | Wellcome Sanger Institute for the COVID-19 Genomics UK (COG-UK) Consortium | Harper VanSteenhouse, Yumi Kasai, David Gray, Carol Clugston, Anna Dominiczak and Alex Alderton, Roberto Amato, Jeffrey Barrett, Sonia Goncalves, Ewan Harrison, David K. Jackson, Ian Johnston, Dominic Kwiatkowski, Cordelia Langford, John Sillitoe on behalf of the Wellcome Sanger Institute COVID-19 Surveillance Team |
| EPI_ISL_3224143 | Lighthouse Lab in Milton Keynes | Wellcome Sanger Institute for the COVID-19 Genomics UK (COG-UK) Consortium | The Lighthouse Lab in Milton Keynes and Alex Alderton, Roberto Amato, Jeffrey Barrett, Sonia Goncalves, Ewan Harrison, David K. Jackson, Ian Johnston, Dominic Kwiatkowski, Cordelia Langford, John Sillitoe on behalf of the Wellcome Sanger Institute COVID-19 Surveillance Team |
| EPI_ISL_3224144 | Lighthouse Lab in Glasgow | Wellcome Sanger Institute for the COVID-19 Genomics UK (COG-UK) Consortium | Harper VanSteenhouse, Yumi Kasai, David Gray, Carol Clugston, Anna Dominiczak and Alex Alderton, Roberto Amato, Jeffrey Barrett, Sonia Goncalves, Ewan Harrison, David K. Jackson, Ian Johnston, Dominic Kwiatkowski, Cordelia Langford, John Sillitoe on behalf of the Wellcome Sanger Institute COVID-19 Surveillance Team |
| EPI_ISL_3224145, EPI_ISL_3224146 | Lighthouse Lab in Alderley Park | Wellcome Sanger Institute for the COVID-19 Genomics UK (COG-UK) Consortium | Jacquelyn Wynn, Mairead Hyland, The Lighthouse Lab in Alderley Park and Alex Alderton, Roberto Amato, Jeffrey Barrett, Sonia Goncalves, Ewan Harrison, David K. Jackson, Ian Johnston, Dominic Kwiatkowski, Cordelia Langford, John Sillitoe on behalf of the Wellcome Sanger Institute COVID-19 Surveillance Team |
| EPI_ISL_3224147 | Lighthouse Lab in Milton Keynes | Wellcome Sanger Institute for the COVID-19 Genomics UK (COG-UK) Consortium | The Lighthouse Lab in Milton Keynes and Alex Alderton, Roberto Amato, Jeffrey Barrett, Sonia Goncalves, Ewan Harrison, David K. Jackson, Ian Johnston, Dominic Kwiatkowski, Cordelia Langford, John Sillitoe on behalf of the Wellcome Sanger Institute COVID-19 Surveillance Team |
| EPI_ISL_3224148 | Lighthouse Lab in Alderley Park | Wellcome Sanger Institute for the COVID-19 Genomics UK (COG-UK) Consortium | Jacquelyn Wynn, Mairead Hyland, The Lighthouse Lab in Alderley Park and Alex Alderton, Roberto Amato, Jeffrey Barrett, Sonia Goncalves, Ewan Harrison, David K. Jackson, Ian Johnston, Dominic Kwiatkowski, Cordelia Langford, John Sillitoe on behalf of the Wellcome Sanger Institute COVID-19 Surveillance Team |
| EPI_ISL_3224149 | Lighthouse Lab in Glasgow | Wellcome Sanger Institute for the COVID-19 Genomics UK (COG-UK) Consortium | Harper VanSteenhouse, Yumi Kasai, David Gray, Carol Clugston, Anna Dominiczak and Alex Alderton, Roberto Amato, Jeffrey Barrett, Sonia Goncalves, Ewan Harrison, David K. Jackson, Ian Johnston, Dominic Kwiatkowski, Cordelia Langford, John Sillitoe on behalf of the Wellcome Sanger Institute COVID-19 Surveillance Team |
| EPI_ISL_3224150 | Lighthouse Lab in Alderley Park | Wellcome Sanger Institute for the COVID-19 Genomics UK (COG-UK) Consortium | Jacquelyn Wynn, Mairead Hyland, The Lighthouse Lab in Alderley Park and Alex Alderton, Roberto Amato, Jeffrey Barrett, Sonia Goncalves, Ewan Harrison, David K. Jackson, Ian Johnston, Dominic Kwiatkowski, Cordelia Langford, John Sillitoe on behalf of the Wellcome Sanger Institute COVID-19 Surveillance Team |
| EPI_ISL_3224151 | Lighthouse Lab in Glasgow | Wellcome Sanger Institute for the COVID-19 Genomics UK (COG-UK) Consortium | Harper VanSteenhouse, Yumi Kasai, David Gray, Carol Clugston, Anna Dominiczak and Alex Alderton, Roberto Amato, Jeffrey Barrett, Sonia Goncalves, Ewan Harrison, David K. Jackson, Ian Johnston, Dominic Kwiatkowski, Cordelia Langford, John Sillitoe on behalf of the Wellcome Sanger Institute COVID-19 Surveillance Team |
| EPI_ISL_3224152 | Lighthouse Lab in Alderley Park | Wellcome Sanger Institute for the COVID-19 Genomics UK (COG-UK) Consortium | Jacquelyn Wynn, Mairead Hyland, The Lighthouse Lab in Alderley Park and Alex Alderton, Roberto Amato, Jeffrey Barrett, Sonia Goncalves, Ewan Harrison, David K. Jackson, Ian Johnston, Dominic Kwiatkowski, Cordelia Langford, John Sillitoe on behalf of the Wellcome Sanger Institute COVID-19 Surveillance Team |

| | | | |
|---|---|---|---|
| EPI_ISL_3224153 | Lighthouse Lab in Glasgow | Wellcome Sanger Institute for the COVID-19 Genomics UK (COG-UK) Consortium | Harper VanSteenhouse, Yumi Kasai, David Gray, Carol Clugston, Anna Dominiczak and Alex Alderton, Roberto Amato, Jeffrey Barrett, Sonia Goncalves, Ewan Harrison, David K. Jackson, Ian Johnston, Dominic Kwiatkowski, Cordelia Langford, John Sillitoe on behalf of the Wellcome Sanger Institute COVID-19 Surveillance Team |
| EPI_ISL_3224154, EPI_ISL_3224155 | Lighthouse Laboratory Plymouth | Wellcome Sanger Institute for the COVID-19 Genomics UK (COG-UK) Consortium | Lighthouse Laboratory Plymouth and Alex Alderton, Roberto Amato, Jeffrey Barrett, Sonia Goncalves, Ewan Harrison, David K. Jackson, Ian Johnston, Dominic Kwiatkowski, Cordelia Langford, John Sillitoe on behalf of the Wellcome Sanger Institute COVID-19 Surveillance Team |
| EPI_ISL_3224156 | Lighthouse Lab in Milton Keynes | Wellcome Sanger Institute for the COVID-19 Genomics UK (COG-UK) Consortium | The Lighthouse Lab in Milton Keynes and Alex Alderton, Roberto Amato, Jeffrey Barrett, Sonia Goncalves, Ewan Harrison, David K. Jackson, Ian Johnston, Dominic Kwiatkowski, Cordelia Langford, John Sillitoe on behalf of the Wellcome Sanger Institute COVID-19 Surveillance Team |
| EPI_ISL_3224157 | Lighthouse Lab in Glasgow | Wellcome Sanger Institute for the COVID-19 Genomics UK (COG-UK) Consortium | Harper VanSteenhouse, Yumi Kasai, David Gray, Carol Clugston, Anna Dominiczak and Alex Alderton, Roberto Amato, Jeffrey Barrett, Sonia Goncalves, Ewan Harrison, David K. Jackson, Ian Johnston, Dominic Kwiatkowski, Cordelia Langford, John Sillitoe on behalf of the Wellcome Sanger Institute COVID-19 Surveillance Team |
| EPI_ISL_3224158 | Lighthouse Lab in Alderley Park | Wellcome Sanger Institute for the COVID-19 Genomics UK (COG-UK) Consortium | Jacquelyn Wynn, Mairead Hyland, The Lighthouse Lab in Alderley Park and Alex Alderton, Roberto Amato, Jeffrey Barrett, Sonia Goncalves, Ewan Harrison, David K. Jackson, Ian Johnston, Dominic Kwiatkowski, Cordelia Langford, John Sillitoe on behalf of the Wellcome Sanger Institute COVID-19 Surveillance Team |
| EPI_ISL_3224159, EPI_ISL_3224160 | Lighthouse Laboratory Plymouth | Wellcome Sanger Institute for the COVID-19 Genomics UK (COG-UK) Consortium | Lighthouse Laboratory Plymouth and Alex Alderton, Roberto Amato, Jeffrey Barrett, Sonia Goncalves, Ewan Harrison, David K. Jackson, Ian Johnston, Dominic Kwiatkowski, Cordelia Langford, John Sillitoe on behalf of the Wellcome Sanger Institute COVID-19 Surveillance Team |
| EPI_ISL_3224161 | Lighthouse Lab in Milton Keynes | Wellcome Sanger Institute for the COVID-19 Genomics UK (COG-UK) Consortium | The Lighthouse Lab in Milton Keynes and Alex Alderton, Roberto Amato, Jeffrey Barrett, Sonia Goncalves, Ewan Harrison, David K. Jackson, Ian Johnston, Dominic Kwiatkowski, Cordelia Langford, John Sillitoe on behalf of the Wellcome Sanger Institute COVID-19 Surveillance Team |
| EPI_ISL_3224162 | Lighthouse Lab in Alderley Park | Wellcome Sanger Institute for the COVID-19 Genomics UK (COG-UK) Consortium | Jacquelyn Wynn, Mairead Hyland, The Lighthouse Lab in Alderley Park and Alex Alderton, Roberto Amato, Jeffrey Barrett, Sonia Goncalves, Ewan Harrison, David K. Jackson, Ian Johnston, Dominic Kwiatkowski, Cordelia Langford, John Sillitoe on behalf of the Wellcome Sanger Institute COVID-19 Surveillance Team |
| EPI_ISL_3224163 | Lighthouse Laboratory Plymouth | Wellcome Sanger Institute for the COVID-19 Genomics UK (COG-UK) Consortium | Lighthouse Laboratory Plymouth and Alex Alderton, Roberto Amato, Jeffrey Barrett, Sonia Goncalves, Ewan Harrison, David K. Jackson, Ian Johnston, Dominic Kwiatkowski, Cordelia Langford, John Sillitoe on behalf of the Wellcome Sanger Institute COVID-19 Surveillance Team |
| EPI_ISL_3224164, EPI_ISL_3224165, EPI_ISL_3224166, EPI_ISL_3224167 | Lighthouse Lab in Milton Keynes | Wellcome Sanger Institute for the COVID-19 Genomics UK (COG-UK) Consortium | The Lighthouse Lab in Milton Keynes and Alex Alderton, Roberto Amato, Jeffrey Barrett, Sonia Goncalves, Ewan Harrison, David K. Jackson, Ian Johnston, Dominic Kwiatkowski, Cordelia Langford, John Sillitoe on behalf of the Wellcome Sanger Institute COVID-19 Surveillance Team |
| EPI_ISL_3224168 | Lighthouse Laboratory Plymouth | Wellcome Sanger Institute for the COVID-19 Genomics UK (COG-UK) Consortium | Lighthouse Laboratory Plymouth and Alex Alderton, Roberto Amato, Jeffrey Barrett, Sonia Goncalves, Ewan Harrison, David K. Jackson, Ian Johnston, Dominic Kwiatkowski, Cordelia Langford, John Sillitoe on behalf of the Wellcome Sanger Institute COVID-19 Surveillance Team |
| EPI_ISL_3224169 | Lighthouse Lab in Glasgow | Wellcome Sanger Institute for the COVID-19 Genomics UK (COG-UK) Consortium | Harper VanSteenhouse, Yumi Kasai, David Gray, Carol Clugston, Anna Dominiczak and Alex Alderton, Roberto Amato, Jeffrey Barrett, Sonia Goncalves, Ewan Harrison, David K. Jackson, Ian Johnston, Dominic Kwiatkowski, Cordelia Langford, John Sillitoe on behalf of the Wellcome Sanger Institute COVID-19 Surveillance Team |
| EPI_ISL_3224170 | Lighthouse Laboratory Plymouth | Wellcome Sanger Institute for the COVID-19 Genomics UK (COG-UK) Consortium | Lighthouse Laboratory Plymouth and Alex Alderton, Roberto Amato, Jeffrey Barrett, Sonia Goncalves, Ewan Harrison, David K. Jackson, Ian Johnston, Dominic Kwiatkowski, Cordelia Langford, John Sillitoe on behalf of the Wellcome Sanger Institute COVID-19 Surveillance Team |
| EPI_ISL_3224171 | Lighthouse Lab in Glasgow | Wellcome Sanger Institute for the COVID-19 Genomics UK (COG-UK) Consortium | Harper VanSteenhouse, Yumi Kasai, David Gray, Carol Clugston, Anna Dominiczak and Alex Alderton, Roberto Amato, Jeffrey Barrett, Sonia Goncalves, Ewan Harrison, David K. Jackson, Ian Johnston, Dominic Kwiatkowski, Cordelia Langford, John Sillitoe on behalf of the Wellcome Sanger Institute COVID-19 Surveillance Team |
| EPI_ISL_3224172, EPI_ISL_3224173 | Lighthouse Lab in Alderley Park | Wellcome Sanger Institute for the COVID-19 Genomics UK (COG-UK) Consortium | Jacquelyn Wynn, Mairead Hyland, The Lighthouse Lab in Alderley Park and Alex Alderton, Roberto Amato, Jeffrey Barrett, Sonia Goncalves, Ewan Harrison, David K. Jackson, Ian Johnston, Dominic Kwiatkowski, Cordelia Langford, John Sillitoe on behalf of the Wellcome Sanger Institute COVID-19 Surveillance Team |
| EPI_ISL_3224174, EPI_ISL_3224175 | Lighthouse Lab in Milton Keynes | Wellcome Sanger Institute for the COVID-19 Genomics UK (COG-UK) Consortium | The Lighthouse Lab in Milton Keynes and Alex Alderton, Roberto Amato, Jeffrey Barrett, Sonia Goncalves, Ewan Harrison, David K. Jackson, Ian Johnston, Dominic Kwiatkowski, Cordelia Langford, John Sillitoe on behalf of the Wellcome Sanger Institute COVID-19 Surveillance Team |
| EPI_ISL_3224176 | Lighthouse Lab in Alderley Park | Wellcome Sanger Institute for the COVID-19 Genomics UK (COG-UK) Consortium | Jacquelyn Wynn, Mairead Hyland, The Lighthouse Lab in Alderley Park and Alex Alderton, Roberto Amato, Jeffrey Barrett, Sonia Goncalves, Ewan Harrison, David K. Jackson, Ian Johnston, Dominic Kwiatkowski, Cordelia Langford, John Sillitoe on behalf of the Wellcome Sanger Institute COVID-19 Surveillance Team |
| EPI_ISL_3224177, EPI_ISL_3224178, EPI_ISL_3224179 | Lighthouse Laboratory Plymouth | Wellcome Sanger Institute for the COVID-19 Genomics UK (COG-UK) Consortium | Lighthouse Laboratory Plymouth and Alex Alderton, Roberto Amato, Jeffrey Barrett, Sonia Goncalves, Ewan Harrison, David K. Jackson, Ian Johnston, Dominic Kwiatkowski, Cordelia Langford, John Sillitoe on behalf of the Wellcome Sanger Institute COVID-19 Surveillance Team |
| EPI_ISL_3224180, EPI_ISL_3224181, EPI_ISL_3224182, EPI_ISL_3224183 | Lighthouse Lab in Milton Keynes | Wellcome Sanger Institute for the COVID-19 Genomics UK (COG-UK) Consortium | The Lighthouse Lab in Milton Keynes and Alex Alderton, Roberto Amato, Jeffrey Barrett, Sonia Goncalves, Ewan Harrison, David K. Jackson, Ian Johnston, Dominic Kwiatkowski, Cordelia Langford, John Sillitoe on behalf of the Wellcome Sanger Institute COVID-19 Surveillance Team |
| EPI_ISL_3224184 | Lighthouse Laboratory Plymouth | Wellcome Sanger Institute for the COVID-19 Genomics UK (COG-UK) Consortium | Lighthouse Laboratory Plymouth and Alex Alderton, Roberto Amato, Jeffrey Barrett, Sonia Goncalves, Ewan Harrison, David K. Jackson, Ian Johnston, Dominic Kwiatkowski, Cordelia Langford, John Sillitoe on behalf of the Wellcome Sanger Institute COVID-19 Surveillance Team |
| EPI_ISL_3224185 | Lighthouse Lab in Glasgow | Wellcome Sanger Institute for the COVID-19 Genomics UK (COG-UK) Consortium | Harper VanSteenhouse, Yumi Kasai, David Gray, Carol Clugston, Anna Dominiczak and Alex Alderton, Roberto Amato, Jeffrey Barrett, Sonia Goncalves, Ewan Harrison, David K. Jackson, Ian Johnston, Dominic Kwiatkowski, Cordelia Langford, John Sillitoe on behalf of the Wellcome Sanger Institute COVID-19 Surveillance Team |
| EPI_ISL_3224186 | Lighthouse Lab in Milton Keynes | Wellcome Sanger Institute for the COVID-19 Genomics UK (COG-UK) Consortium | The Lighthouse Lab in Milton Keynes and Alex Alderton, Roberto Amato, Jeffrey Barrett, Sonia Goncalves, Ewan Harrison, David K. Jackson, Ian Johnston, Dominic Kwiatkowski, Cordelia Langford, John Sillitoe on behalf of the Wellcome Sanger Institute COVID-19 Surveillance Team |
| EPI_ISL_3224187 | Lighthouse Lab in Alderley Park | Wellcome Sanger Institute for the COVID-19 Genomics UK (COG-UK) Consortium | Jacquelyn Wynn, Mairead Hyland, The Lighthouse Lab in Alderley Park and Alex Alderton, Roberto Amato, Jeffrey Barrett, Sonia Goncalves, Ewan Harrison, David K. Jackson, Ian Johnston, Dominic Kwiatkowski, Cordelia Langford, John Sillitoe on behalf of the Wellcome Sanger Institute COVID-19 Surveillance Team |
| EPI_ISL_3224188 | Lighthouse Lab in Glasgow | Wellcome Sanger Institute for the COVID-19 Genomics UK (COG-UK) Consortium | Harper VanSteenhouse, Yumi Kasai, David Gray, Carol Clugston, Anna Dominiczak and Alex Alderton, Roberto Amato, Jeffrey Barrett, Sonia Goncalves, Ewan Harrison, David K. Jackson, Ian Johnston, Dominic Kwiatkowski, Cordelia Langford, John Sillitoe on behalf of the Wellcome Sanger Institute COVID-19 Surveillance Team |
| EPI_ISL_3224189, EPI_ISL_3224190 | Lighthouse Laboratory Plymouth | Wellcome Sanger Institute for the COVID-19 Genomics UK (COG-UK) Consortium | Lighthouse Laboratory Plymouth and Alex Alderton, Roberto Amato, Jeffrey Barrett, Sonia Goncalves, Ewan Harrison, David K. Jackson, Ian Johnston, Dominic Kwiatkowski, Cordelia Langford, John Sillitoe on behalf of the Wellcome Sanger Institute COVID-19 Surveillance Team |
| EPI_ISL_3224191 | Lighthouse Lab in Milton Keynes | Wellcome Sanger Institute for the COVID-19 Genomics UK (COG-UK) Consortium | The Lighthouse Lab in Milton Keynes and Alex Alderton, Roberto Amato, Jeffrey Barrett, Sonia Goncalves, Ewan Harrison, David K. Jackson, Ian Johnston, Dominic Kwiatkowski, Cordelia Langford, John Sillitoe on behalf of the Wellcome Sanger Institute COVID-19 Surveillance Team |
| EPI_ISL_3224192, EPI_ISL_3224193 | Lighthouse Laboratory Plymouth | Wellcome Sanger Institute for the COVID-19 Genomics UK (COG-UK) Consortium | Lighthouse Laboratory Plymouth and Alex Alderton, Roberto Amato, Jeffrey Barrett, Sonia Goncalves, Ewan Harrison, David K. Jackson, Ian Johnston, Dominic Kwiatkowski, Cordelia Langford, John Sillitoe on behalf of the Wellcome Sanger Institute COVID-19 Surveillance Team |
| EPI_ISL_3224194, EPI_ISL_3224195 | Lighthouse Lab in Glasgow | Wellcome Sanger Institute for the COVID-19 Genomics UK (COG-UK) Consortium | Harper VanSteenhouse, Yumi Kasai, David Gray, Carol Clugston, Anna Dominiczak and Alex Alderton, Roberto Amato, Jeffrey Barrett, Sonia Goncalves, Ewan Harrison, David K. Jackson, Ian Johnston, Dominic Kwiatkowski, Cordelia Langford, John Sillitoe on behalf of the Wellcome Sanger Institute COVID-19 Surveillance Team |
| EPI_ISL_3224196 | Lighthouse Lab in Milton Keynes | Wellcome Sanger Institute for the COVID-19 Genomics UK (COG-UK) Consortium | The Lighthouse Lab in Milton Keynes and Alex Alderton, Roberto Amato, Jeffrey Barrett, Sonia Goncalves, Ewan Harrison, David K. Jackson, Ian Johnston, Dominic Kwiatkowski, Cordelia Langford, John Sillitoe on behalf of the Wellcome Sanger Institute COVID-19 Surveillance Team |
| EPI_ISL_3224197 | Lighthouse Laboratory Plymouth | Wellcome Sanger Institute for the COVID-19 Genomics UK (COG-UK) Consortium | Lighthouse Laboratory Plymouth and Alex Alderton, Roberto Amato, Jeffrey Barrett, Sonia Goncalves, Ewan Harrison, David K. Jackson, Ian Johnston, Dominic Kwiatkowski, Cordelia Langford, John Sillitoe on behalf of the Wellcome Sanger Institute COVID-19 Surveillance Team |
| EPI_ISL_3224198 | Lighthouse Lab in Glasgow | Wellcome Sanger Institute for the COVID-19 Genomics UK (COG-UK) Consortium | Harper VanSteenhouse, Yumi Kasai, David Gray, Carol Clugston, Anna Dominiczak and Alex Alderton, Roberto Amato, Jeffrey Barrett, Sonia Goncalves, Ewan Harrison, David K. Jackson, Ian Johnston, Dominic Kwiatkowski, Cordelia Langford, John Sillitoe on behalf of the Wellcome Sanger Institute COVID-19 Surveillance Team |
| EPI_ISL_3224199, EPI_ISL_3224200 | Lighthouse Lab in Milton Keynes | Wellcome Sanger Institute for the COVID-19 Genomics UK (COG-UK) Consortium | The Lighthouse Lab in Milton Keynes and Alex Alderton, Roberto Amato, Jeffrey Barrett, Sonia Goncalves, Ewan Harrison, David K. Jackson, Ian Johnston, Dominic Kwiatkowski, Cordelia Langford, John Sillitoe on behalf of the Wellcome Sanger Institute COVID-19 Surveillance Team |
| EPI_ISL_3224201 | Lighthouse Laboratory Plymouth | Wellcome Sanger Institute for the COVID-19 Genomics UK (COG-UK) Consortium | Lighthouse Laboratory Plymouth and Alex Alderton, Roberto Amato, Jeffrey Barrett, Sonia Goncalves, Ewan Harrison, David K. Jackson, Ian Johnston, Dominic Kwiatkowski, Cordelia Langford, John Sillitoe on behalf of the Wellcome Sanger Institute COVID-19 Surveillance Team |
| EPI_ISL_3224202 | Lighthouse Lab in Alderley Park | Wellcome Sanger Institute for the COVID-19 Genomics UK (COG-UK) Consortium | Jacquelyn Wynn, Mairead Hyland, The Lighthouse Lab in Alderley Park and Alex Alderton, Roberto Amato, Jeffrey Barrett, Sonia Goncalves, Ewan Harrison, David K. Jackson, Ian Johnston, Dominic Kwiatkowski, Cordelia Langford, John Sillitoe on behalf of the Wellcome Sanger Institute COVID-19 Surveillance Team |
| EPI_ISL_3224203 | Lighthouse Lab in Milton Keynes | Wellcome Sanger Institute for the COVID-19 Genomics UK (COG-UK) Consortium | The Lighthouse Lab in Milton Keynes and Alex Alderton, Roberto Amato, Jeffrey Barrett, Sonia Goncalves, Ewan Harrison, David K. Jackson, Ian Johnston, Dominic Kwiatkowski, Cordelia Langford, John Sillitoe on behalf of the Wellcome Sanger Institute COVID-19 Surveillance Team |
| EPI_ISL_3224204 | Lighthouse Lab in Glasgow | Wellcome Sanger Institute for the COVID-19 Genomics UK (COG-UK) Consortium | Harper VanSteenhouse, Yumi Kasai, David Gray, Carol Clugston, Anna Dominiczak and Alex Alderton, Roberto Amato, Jeffrey Barrett, Sonia Goncalves, Ewan Harrison, David K. Jackson, Ian Johnston, Dominic Kwiatkowski, Cordelia Langford, John Sillitoe on behalf of the Wellcome Sanger Institute COVID-19 Surveillance Team |
| EPI_ISL_3224205 | Lighthouse Laboratory Plymouth | Wellcome Sanger Institute for the COVID-19 Genomics UK (COG-UK) Consortium | Lighthouse Laboratory Plymouth and Alex Alderton, Roberto Amato, Jeffrey Barrett, Sonia Goncalves, Ewan Harrison, David K. Jackson, Ian Johnston, Dominic Kwiatkowski, Cordelia Langford, John Sillitoe on behalf of the Wellcome Sanger Institute COVID-19 Surveillance Team |
| EPI_ISL_3224206 | Lighthouse Lab in Glasgow | Wellcome Sanger Institute for the COVID-19 Genomics UK (COG-UK) Consortium | Harper VanSteenhouse, Yumi Kasai, David Gray, Carol Clugston, Anna Dominiczak and Alex Alderton, Roberto Amato, Jeffrey Barrett, Sonia Goncalves, Ewan Harrison, David K. Jackson, Ian Johnston, Dominic Kwiatkowski, Cordelia Langford, John Sillitoe on behalf of the Wellcome Sanger Institute COVID-19 Surveillance Team |
| EPI_ISL_3224207 | Lighthouse Lab in Alderley Park | Wellcome Sanger Institute for the COVID-19 Genomics UK (COG-UK) Consortium | Jacquelyn Wynn, Mairead Hyland, The Lighthouse Lab in Alderley Park and Alex Alderton, Roberto Amato, Jeffrey Barrett, Sonia Goncalves, Ewan Harrison, David K. Jackson, Ian Johnston, Dominic Kwiatkowski, Cordelia Langford, John Sillitoe on behalf of the Wellcome Sanger Institute COVID-19 Surveillance Team |
| EPI_ISL_3224208 | Lighthouse Laboratory Plymouth | Wellcome Sanger Institute for the COVID-19 Genomics UK (COG-UK) Consortium | Lighthouse Laboratory Plymouth and Alex Alderton, Roberto Amato, Jeffrey Barrett, Sonia Goncalves, Ewan Harrison, David K. Jackson, Ian Johnston, Dominic Kwiatkowski, Cordelia Langford, John Sillitoe on behalf of the Wellcome Sanger Institute COVID-19 Surveillance Team |
| EPI_ISL_3224209 | Lighthouse Lab in Alderley Park | Wellcome Sanger Institute for the COVID-19 Genomics UK (COG-UK) Consortium | Jacquelyn Wynn, Mairead Hyland, The Lighthouse Lab in Alderley Park and Alex Alderton, Roberto Amato, Jeffrey Barrett, Sonia Goncalves, Ewan Harrison, David K. Jackson, Ian Johnston, Dominic Kwiatkowski, Cordelia Langford, John Sillitoe on behalf of the Wellcome Sanger Institute COVID-19 Surveillance Team |
| EPI_ISL_3224210 | Lighthouse Lab in Milton Keynes | Wellcome Sanger Institute for the COVID-19 Genomics UK (COG-UK) Consortium | The Lighthouse Lab in Milton Keynes and Alex Alderton, Roberto Amato, Jeffrey Barrett, Sonia Goncalves, Ewan Harrison, David K. Jackson, Ian Johnston, Dominic Kwiatkowski, Cordelia Langford, John Sillitoe on behalf of the Wellcome Sanger Institute COVID-19 Surveillance Team |
| EPI_ISL_3224211 | Lighthouse Lab in Glasgow | Wellcome Sanger Institute for the COVID-19 Genomics UK (COG-UK) Consortium | Harper VanSteenhouse, Yumi Kasai, David Gray, Carol Clugston, Anna Dominiczak and Alex Alderton, Roberto Amato, Jeffrey Barrett, Sonia Goncalves, Ewan Harrison, David K. Jackson, Ian Johnston, Dominic Kwiatkowski, Cordelia Langford, John Sillitoe on behalf of the Wellcome Sanger Institute COVID-19 Surveillance Team |

| | | | |
|---|---|---|---|
| EPI_ISL_3224212 | Lighthouse Lab in Milton Keynes | Wellcome Sanger Institute for the COVID-19 Genomics UK (COG-UK) Consortium | The Lighthouse Lab in Milton Keynes and Alex Alderton, Roberto Amato, Jeffrey Barrett, Sonia Goncalves, Ewan Harrison, David K. Jackson, Ian Johnston, Dominic Kwiatkowski, Cordelia Langford, John Sillitoe on behalf of the Wellcome Sanger Institute COVID-19 Surveillance Team |
| EPI_ISL_3224213 | Lighthouse Laboratory Plymouth | Wellcome Sanger Institute for the COVID-19 Genomics UK (COG-UK) Consortium | Lighthouse Laboratory Plymouth and Alex Alderton, Roberto Amato, Jeffrey Barrett, Sonia Goncalves, Ewan Harrison, David K. Jackson, Ian Johnston, Dominic Kwiatkowski, Cordelia Langford, John Sillitoe on behalf of the Wellcome Sanger Institute COVID-19 Surveillance Team |
| EPI_ISL_3224214 | Lighthouse Lab in Alderley Park | Wellcome Sanger Institute for the COVID-19 Genomics UK (COG-UK) Consortium | Jacquelyn Wynn, Mairead Hyland, The Lighthouse Lab in Alderley Park and Alex Alderton, Roberto Amato, Jeffrey Barrett, Sonia Goncalves, Ewan Harrison, David K. Jackson, Ian Johnston, Dominic Kwiatkowski, Cordelia Langford, John Sillitoe on behalf of the Wellcome Sanger Institute COVID-19 Surveillance Team |
| EPI_ISL_3224215, EPI_ISL_3224216, EPI_ISL_3224217 | Lighthouse Laboratory Plymouth | Wellcome Sanger Institute for the COVID-19 Genomics UK (COG-UK) Consortium | Lighthouse Laboratory Plymouth and Alex Alderton, Roberto Amato, Jeffrey Barrett, Sonia Goncalves, Ewan Harrison, David K. Jackson, Ian Johnston, Dominic Kwiatkowski, Cordelia Langford, John Sillitoe on behalf of the Wellcome Sanger Institute COVID-19 Surveillance Team |
| EPI_ISL_3224218 | Lighthouse Lab in Milton Keynes | Wellcome Sanger Institute for the COVID-19 Genomics UK (COG-UK) Consortium | The Lighthouse Lab in Milton Keynes and Alex Alderton, Roberto Amato, Jeffrey Barrett, Sonia Goncalves, Ewan Harrison, David K. Jackson, Ian Johnston, Dominic Kwiatkowski, Cordelia Langford, John Sillitoe on behalf of the Wellcome Sanger Institute COVID-19 Surveillance Team |
| EPI_ISL_3224219 | Lighthouse Lab in Alderley Park | Wellcome Sanger Institute for the COVID-19 Genomics UK (COG-UK) Consortium | Jacquelyn Wynn, Mairead Hyland, The Lighthouse Lab in Alderley Park and Alex Alderton, Roberto Amato, Jeffrey Barrett, Sonia Goncalves, Ewan Harrison, David K. Jackson, Ian Johnston, Dominic Kwiatkowski, Cordelia Langford, John Sillitoe on behalf of the Wellcome Sanger Institute COVID-19 Surveillance Team |
| EPI_ISL_3224220, EPI_ISL_3224221 | Lighthouse Lab in Glasgow | Wellcome Sanger Institute for the COVID-19 Genomics UK (COG-UK) Consortium | Harper VanSteenhouse, Yumi Kasai, David Gray, Carol Clugston, Anna Dominiczak and Alex Alderton, Roberto Amato, Jeffrey Barrett, Sonia Goncalves, Ewan Harrison, David K. Jackson, Ian Johnston, Dominic Kwiatkowski, Cordelia Langford, John Sillitoe on behalf of the Wellcome Sanger Institute COVID-19 Surveillance Team |
| EPI_ISL_3224222, EPI_ISL_3224223 | Lighthouse Laboratory Plymouth | Wellcome Sanger Institute for the COVID-19 Genomics UK (COG-UK) Consortium | Lighthouse Laboratory Plymouth and Alex Alderton, Roberto Amato, Jeffrey Barrett, Sonia Goncalves, Ewan Harrison, David K. Jackson, Ian Johnston, Dominic Kwiatkowski, Cordelia Langford, John Sillitoe on behalf of the Wellcome Sanger Institute COVID-19 Surveillance Team |
| EPI_ISL_3224224 | Lighthouse Lab in Glasgow | Wellcome Sanger Institute for the COVID-19 Genomics UK (COG-UK) Consortium | Harper VanSteenhouse, Yumi Kasai, David Gray, Carol Clugston, Anna Dominiczak and Alex Alderton, Roberto Amato, Jeffrey Barrett, Sonia Goncalves, Ewan Harrison, David K. Jackson, Ian Johnston, Dominic Kwiatkowski, Cordelia Langford, John Sillitoe on behalf of the Wellcome Sanger Institute COVID-19 Surveillance Team |
| EPI_ISL_3224225 | Lighthouse Lab in Milton Keynes | Wellcome Sanger Institute for the COVID-19 Genomics UK (COG-UK) Consortium | The Lighthouse Lab in Milton Keynes and Alex Alderton, Roberto Amato, Jeffrey Barrett, Sonia Goncalves, Ewan Harrison, David K. Jackson, Ian Johnston, Dominic Kwiatkowski, Cordelia Langford, John Sillitoe on behalf of the Wellcome Sanger Institute COVID-19 Surveillance Team |
| EPI_ISL_3224226 | Lighthouse Laboratory Plymouth | Wellcome Sanger Institute for the COVID-19 Genomics UK (COG-UK) Consortium | Lighthouse Laboratory Plymouth and Alex Alderton, Roberto Amato, Jeffrey Barrett, Sonia Goncalves, Ewan Harrison, David K. Jackson, Ian Johnston, Dominic Kwiatkowski, Cordelia Langford, John Sillitoe on behalf of the Wellcome Sanger Institute COVID-19 Surveillance Team |
| EPI_ISL_3224227 | Lighthouse Lab in Milton Keynes | Wellcome Sanger Institute for the COVID-19 Genomics UK (COG-UK) Consortium | The Lighthouse Lab in Milton Keynes and Alex Alderton, Roberto Amato, Jeffrey Barrett, Sonia Goncalves, Ewan Harrison, David K. Jackson, Ian Johnston, Dominic Kwiatkowski, Cordelia Langford, John Sillitoe on behalf of the Wellcome Sanger Institute COVID-19 Surveillance Team |
| EPI_ISL_3224228 | Lighthouse Lab in Glasgow | Wellcome Sanger Institute for the COVID-19 Genomics UK (COG-UK) Consortium | Harper VanSteenhouse, Yumi Kasai, David Gray, Carol Clugston, Anna Dominiczak and Alex Alderton, Roberto Amato, Jeffrey Barrett, Sonia Goncalves, Ewan Harrison, David K. Jackson, Ian Johnston, Dominic Kwiatkowski, Cordelia Langford, John Sillitoe on behalf of the Wellcome Sanger Institute COVID-19 Surveillance Team |
| EPI_ISL_3224229, EPI_ISL_3224230, EPI_ISL_3224231 | Lighthouse Lab in Alderley Park | Wellcome Sanger Institute for the COVID-19 Genomics UK (COG-UK) Consortium | Jacquelyn Wynn, Mairead Hyland, The Lighthouse Lab in Alderley Park and Alex Alderton, Roberto Amato, Jeffrey Barrett, Sonia Goncalves, Ewan Harrison, David K. Jackson, Ian Johnston, Dominic Kwiatkowski, Cordelia Langford, John Sillitoe on behalf of the Wellcome Sanger Institute COVID-19 Surveillance Team |
| EPI_ISL_3224232 | Lighthouse Laboratory Plymouth | Wellcome Sanger Institute for the COVID-19 Genomics UK (COG-UK) Consortium | Lighthouse Laboratory Plymouth and Alex Alderton, Roberto Amato, Jeffrey Barrett, Sonia Goncalves, Ewan Harrison, David K. Jackson, Ian Johnston, Dominic Kwiatkowski, Cordelia Langford, John Sillitoe on behalf of the Wellcome Sanger Institute COVID-19 Surveillance Team |
| EPI_ISL_3224233, EPI_ISL_3224234 | Lighthouse Lab in Alderley Park | Wellcome Sanger Institute for the COVID-19 Genomics UK (COG-UK) Consortium | Jacquelyn Wynn, Mairead Hyland, The Lighthouse Lab in Alderley Park and Alex Alderton, Roberto Amato, Jeffrey Barrett, Sonia Goncalves, Ewan Harrison, David K. Jackson, Ian Johnston, Dominic Kwiatkowski, Cordelia Langford, John Sillitoe on behalf of the Wellcome Sanger Institute COVID-19 Surveillance Team |
| EPI_ISL_3224235 | Lighthouse Lab in Glasgow | Wellcome Sanger Institute for the COVID-19 Genomics UK (COG-UK) Consortium | Harper VanSteenhouse, Yumi Kasai, David Gray, Carol Clugston, Anna Dominiczak and Alex Alderton, Roberto Amato, Jeffrey Barrett, Sonia Goncalves, Ewan Harrison, David K. Jackson, Ian Johnston, Dominic Kwiatkowski, Cordelia Langford, John Sillitoe on behalf of the Wellcome Sanger Institute COVID-19 Surveillance Team |
| EPI_ISL_3224236 | Lighthouse Laboratory Plymouth | Wellcome Sanger Institute for the COVID-19 Genomics UK (COG-UK) Consortium | Lighthouse Laboratory Plymouth and Alex Alderton, Roberto Amato, Jeffrey Barrett, Sonia Goncalves, Ewan Harrison, David K. Jackson, Ian Johnston, Dominic Kwiatkowski, Cordelia Langford, John Sillitoe on behalf of the Wellcome Sanger Institute COVID-19 Surveillance Team |
| EPI_ISL_3224237, EPI_ISL_3224238 | Lighthouse Lab in Glasgow | Wellcome Sanger Institute for the COVID-19 Genomics UK (COG-UK) Consortium | Harper VanSteenhouse, Yumi Kasai, David Gray, Carol Clugston, Anna Dominiczak and Alex Alderton, Roberto Amato, Jeffrey Barrett, Sonia Goncalves, Ewan Harrison, David K. Jackson, Ian Johnston, Dominic Kwiatkowski, Cordelia Langford, John Sillitoe on behalf of the Wellcome Sanger Institute COVID-19 Surveillance Team |
| EPI_ISL_3224239 | Lighthouse Lab in Alderley Park | Wellcome Sanger Institute for the COVID-19 Genomics UK (COG-UK) Consortium | Jacquelyn Wynn, Mairead Hyland, The Lighthouse Lab in Alderley Park and Alex Alderton, Roberto Amato, Jeffrey Barrett, Sonia Goncalves, Ewan Harrison, David K. Jackson, Ian Johnston, Dominic Kwiatkowski, Cordelia Langford, John Sillitoe on behalf of the Wellcome Sanger Institute COVID-19 Surveillance Team |
| EPI_ISL_3224240 | Lighthouse Lab in Glasgow | Wellcome Sanger Institute for the COVID-19 Genomics UK (COG-UK) Consortium | Harper VanSteenhouse, Yumi Kasai, David Gray, Carol Clugston, Anna Dominiczak and Alex Alderton, Roberto Amato, Jeffrey Barrett, Sonia Goncalves, Ewan Harrison, David K. Jackson, Ian Johnston, Dominic Kwiatkowski, Cordelia Langford, John Sillitoe on behalf of the Wellcome Sanger Institute COVID-19 Surveillance Team |
| EPI_ISL_3224241 | Lighthouse Laboratory Plymouth | Wellcome Sanger Institute for the COVID-19 Genomics UK (COG-UK) Consortium | Lighthouse Laboratory Plymouth and Alex Alderton, Roberto Amato, Jeffrey Barrett, Sonia Goncalves, Ewan Harrison, David K. Jackson, Ian Johnston, Dominic Kwiatkowski, Cordelia Langford, John Sillitoe on behalf of the Wellcome Sanger Institute COVID-19 Surveillance Team |
| EPI_ISL_3224242 | Lighthouse Lab in Alderley Park | Wellcome Sanger Institute for the COVID-19 Genomics UK (COG-UK) Consortium | Jacquelyn Wynn, Mairead Hyland, The Lighthouse Lab in Alderley Park and Alex Alderton, Roberto Amato, Jeffrey Barrett, Sonia Goncalves, Ewan Harrison, David K. Jackson, Ian Johnston, Dominic Kwiatkowski, Cordelia Langford, John Sillitoe on behalf of the Wellcome Sanger Institute COVID-19 Surveillance Team |
| EPI_ISL_3224243 | Lighthouse Lab in Glasgow | Wellcome Sanger Institute for the COVID-19 Genomics UK (COG-UK) Consortium | Harper VanSteenhouse, Yumi Kasai, David Gray, Carol Clugston, Anna Dominiczak and Alex Alderton, Roberto Amato, Jeffrey Barrett, Sonia Goncalves, Ewan Harrison, David K. Jackson, Ian Johnston, Dominic Kwiatkowski, Cordelia Langford, John Sillitoe on behalf of the Wellcome Sanger Institute COVID-19 Surveillance Team |
| EPI_ISL_3224244 | Lighthouse Lab in Alderley Park | Wellcome Sanger Institute for the COVID-19 Genomics UK (COG-UK) Consortium | Jacquelyn Wynn, Mairead Hyland, The Lighthouse Lab in Alderley Park and Alex Alderton, Roberto Amato, Jeffrey Barrett, Sonia Goncalves, Ewan Harrison, David K. Jackson, Ian Johnston, Dominic Kwiatkowski, Cordelia Langford, John Sillitoe on behalf of the Wellcome Sanger Institute COVID-19 Surveillance Team |
| EPI_ISL_3224245, EPI_ISL_3224246 | Lighthouse Lab in Milton Keynes | Wellcome Sanger Institute for the COVID-19 Genomics UK (COG-UK) Consortium | The Lighthouse Lab in Milton Keynes and Alex Alderton, Roberto Amato, Jeffrey Barrett, Sonia Goncalves, Ewan Harrison, David K. Jackson, Ian Johnston, Dominic Kwiatkowski, Cordelia Langford, John Sillitoe on behalf of the Wellcome Sanger Institute COVID-19 Surveillance Team |
| EPI_ISL_3224247, EPI_ISL_3224248 | Lighthouse Lab in Glasgow | Wellcome Sanger Institute for the COVID-19 Genomics UK (COG-UK) Consortium | Harper VanSteenhouse, Yumi Kasai, David Gray, Carol Clugston, Anna Dominiczak and Alex Alderton, Roberto Amato, Jeffrey Barrett, Sonia Goncalves, Ewan Harrison, David K. Jackson, Ian Johnston, Dominic Kwiatkowski, Cordelia Langford, John Sillitoe on behalf of the Wellcome Sanger Institute COVID-19 Surveillance Team |
| EPI_ISL_3224249 | Lighthouse Lab in Milton Keynes | Wellcome Sanger Institute for the COVID-19 Genomics UK (COG-UK) Consortium | The Lighthouse Lab in Milton Keynes and Alex Alderton, Roberto Amato, Jeffrey Barrett, Sonia Goncalves, Ewan Harrison, David K. Jackson, Ian Johnston, Dominic Kwiatkowski, Cordelia Langford, John Sillitoe on behalf of the Wellcome Sanger Institute COVID-19 Surveillance Team |
| EPI_ISL_3224250 | Lighthouse Lab in Glasgow | Wellcome Sanger Institute for the COVID-19 Genomics UK (COG-UK) Consortium | Harper VanSteenhouse, Yumi Kasai, David Gray, Carol Clugston, Anna Dominiczak and Alex Alderton, Roberto Amato, Jeffrey Barrett, Sonia Goncalves, Ewan Harrison, David K. Jackson, Ian Johnston, Dominic Kwiatkowski, Cordelia Langford, John Sillitoe on behalf of the Wellcome Sanger Institute COVID-19 Surveillance Team |
| EPI_ISL_3224251 | Lighthouse Laboratory Plymouth | Wellcome Sanger Institute for the COVID-19 Genomics UK (COG-UK) Consortium | Lighthouse Laboratory Plymouth and Alex Alderton, Roberto Amato, Jeffrey Barrett, Sonia Goncalves, Ewan Harrison, David K. Jackson, Ian Johnston, Dominic Kwiatkowski, Cordelia Langford, John Sillitoe on behalf of the Wellcome Sanger Institute COVID-19 Surveillance Team |
| EPI_ISL_3224252, EPI_ISL_3224253, EPI_ISL_3224254, EPI_ISL_3224255 | Lighthouse Lab in Alderley Park | Wellcome Sanger Institute for the COVID-19 Genomics UK (COG-UK) Consortium | Jacquelyn Wynn, Mairead Hyland, The Lighthouse Lab in Alderley Park and Alex Alderton, Roberto Amato, Jeffrey Barrett, Sonia Goncalves, Ewan Harrison, David K. Jackson, Ian Johnston, Dominic Kwiatkowski, Cordelia Langford, John Sillitoe on behalf of the Wellcome Sanger Institute COVID-19 Surveillance Team |
| EPI_ISL_3224256 | Lighthouse Lab in Milton Keynes | Wellcome Sanger Institute for the COVID-19 Genomics UK (COG-UK) Consortium | The Lighthouse Lab in Milton Keynes and Alex Alderton, Roberto Amato, Jeffrey Barrett, Sonia Goncalves, Ewan Harrison, David K. Jackson, Ian Johnston, Dominic Kwiatkowski, Cordelia Langford, John Sillitoe on behalf of the Wellcome Sanger Institute COVID-19 Surveillance Team |
| EPI_ISL_3224257, EPI_ISL_3224258 | Lighthouse Laboratory Plymouth | Wellcome Sanger Institute for the COVID-19 Genomics UK (COG-UK) Consortium | Lighthouse Laboratory Plymouth and Alex Alderton, Roberto Amato, Jeffrey Barrett, Sonia Goncalves, Ewan Harrison, David K. Jackson, Ian Johnston, Dominic Kwiatkowski, Cordelia Langford, John Sillitoe on behalf of the Wellcome Sanger Institute COVID-19 Surveillance Team |
| EPI_ISL_3224259 | Lighthouse Lab in Glasgow | Wellcome Sanger Institute for the COVID-19 Genomics UK (COG-UK) Consortium | Harper VanSteenhouse, Yumi Kasai, David Gray, Carol Clugston, Anna Dominiczak and Alex Alderton, Roberto Amato, Jeffrey Barrett, Sonia Goncalves, Ewan Harrison, David K. Jackson, Ian Johnston, Dominic Kwiatkowski, Cordelia Langford, John Sillitoe on behalf of the Wellcome Sanger Institute COVID-19 Surveillance Team |
| EPI_ISL_3224260, EPI_ISL_3224261 | Lighthouse Lab in Alderley Park | Wellcome Sanger Institute for the COVID-19 Genomics UK (COG-UK) Consortium | Jacquelyn Wynn, Mairead Hyland, The Lighthouse Lab in Alderley Park and Alex Alderton, Roberto Amato, Jeffrey Barrett, Sonia Goncalves, Ewan Harrison, David K. Jackson, Ian Johnston, Dominic Kwiatkowski, Cordelia Langford, John Sillitoe on behalf of the Wellcome Sanger Institute COVID-19 Surveillance Team |
| EPI_ISL_3224262 | Lighthouse Lab in Milton Keynes | Wellcome Sanger Institute for the COVID-19 Genomics UK (COG-UK) Consortium | The Lighthouse Lab in Milton Keynes and Alex Alderton, Roberto Amato, Jeffrey Barrett, Sonia Goncalves, Ewan Harrison, David K. Jackson, Ian Johnston, Dominic Kwiatkowski, Cordelia Langford, John Sillitoe on behalf of the Wellcome Sanger Institute COVID-19 Surveillance Team |
| EPI_ISL_3224263 | Lighthouse Laboratory Plymouth | Wellcome Sanger Institute for the COVID-19 Genomics UK (COG-UK) Consortium | Lighthouse Laboratory Plymouth and Alex Alderton, Roberto Amato, Jeffrey Barrett, Sonia Goncalves, Ewan Harrison, David K. Jackson, Ian Johnston, Dominic Kwiatkowski, Cordelia Langford, John Sillitoe on behalf of the Wellcome Sanger Institute COVID-19 Surveillance Team |
| EPI_ISL_3224264 | Lighthouse Lab in Glasgow | Wellcome Sanger Institute for the COVID-19 Genomics UK (COG-UK) Consortium | Harper VanSteenhouse, Yumi Kasai, David Gray, Carol Clugston, Anna Dominiczak and Alex Alderton, Roberto Amato, Jeffrey Barrett, Sonia Goncalves, Ewan Harrison, David K. Jackson, Ian Johnston, Dominic Kwiatkowski, Cordelia Langford, John Sillitoe on behalf of the Wellcome Sanger Institute COVID-19 Surveillance Team |
| EPI_ISL_3224265 | Lighthouse Lab in Milton Keynes | Wellcome Sanger Institute for the COVID-19 Genomics UK (COG-UK) Consortium | The Lighthouse Lab in Milton Keynes and Alex Alderton, Roberto Amato, Jeffrey Barrett, Sonia Goncalves, Ewan Harrison, David K. Jackson, Ian Johnston, Dominic Kwiatkowski, Cordelia Langford, John Sillitoe on behalf of the Wellcome Sanger Institute COVID-19 Surveillance Team |
| EPI_ISL_3224266 | Lighthouse Lab in Glasgow | Wellcome Sanger Institute for the COVID-19 Genomics UK (COG-UK) Consortium | Harper VanSteenhouse, Yumi Kasai, David Gray, Carol Clugston, Anna Dominiczak and Alex Alderton, Roberto Amato, Jeffrey Barrett, Sonia Goncalves, Ewan Harrison, David K. Jackson, Ian Johnston, Dominic Kwiatkowski, Cordelia Langford, John Sillitoe on behalf of the Wellcome Sanger Institute COVID-19 Surveillance Team |
| EPI_ISL_3224267 | Lighthouse Lab in Milton Keynes | Wellcome Sanger Institute for the COVID-19 Genomics UK (COG-UK) Consortium | The Lighthouse Lab in Milton Keynes and Alex Alderton, Roberto Amato, Jeffrey Barrett, Sonia Goncalves, Ewan Harrison, David K. Jackson, Ian Johnston, Dominic Kwiatkowski, Cordelia Langford, John Sillitoe on behalf of the Wellcome Sanger Institute COVID-19 Surveillance Team |
| EPI_ISL_3224268 | Lighthouse Lab in Glasgow | Wellcome Sanger Institute for the COVID-19 Genomics UK (COG-UK) Consortium | Harper VanSteenhouse, Yumi Kasai, David Gray, Carol Clugston, Anna Dominiczak and Alex Alderton, Roberto Amato, Jeffrey Barrett, Sonia Goncalves, Ewan Harrison, David K. Jackson, Ian Johnston, Dominic Kwiatkowski, Cordelia Langford, John Sillitoe on behalf of the Wellcome Sanger Institute COVID-19 Surveillance Team |
| EPI_ISL_3224269 | Lighthouse Lab in Alderley Park | Wellcome Sanger Institute for the COVID-19 Genomics UK (COG-UK) Consortium | Jacquelyn Wynn, Mairead Hyland, The Lighthouse Lab in Alderley Park and Alex Alderton, Roberto Amato, Jeffrey Barrett, Sonia Goncalves, Ewan Harrison, David K. Jackson, Ian Johnston, Dominic Kwiatkowski, Cordelia Langford, John Sillitoe on behalf of the Wellcome Sanger Institute COVID-19 Surveillance Team |
| EPI_ISL_3224270, | Lighthouse Lab in Milton Keynes | Wellcome Sanger Institute for the COVID-19 Genomics UK (COG-UK) | The Lighthouse Lab in Milton Keynes and Alex Alderton, Roberto Amato, Jeffrey Barrett, Sonia Goncalves, Ewan Harrison, David K. Jackson, Ian Johnston, Dominic Kwiatkowski, Cordelia Langford, John Sillitoe on behalf of the Wellcome Sanger |

| | | | |
|---|---|---|---|
| EPI_ISL_3224271 | | Consortium | Institute COVID-19 Surveillance Team |
| EPI_ISL_3224272, EPI_ISL_3224273 | Lighthouse Lab in Alderley Park | Wellcome Sanger Institute for the COVID-19 Genomics UK (COG-UK) Consortium | Jacquelyn Wynn, Mairead Hyland, The Lighthouse Lab in Alderley Park and Alex Alderton, Roberto Amato, Jeffrey Barrett, Sonia Goncalves, Ewan Harrison, David K. Jackson, Ian Johnston, Dominic Kwiatkowski, Cordelia Langford, John Sillitoe on behalf of the Wellcome Sanger Institute COVID-19 Surveillance Team |
| EPI_ISL_3224274, EPI_ISL_3224275 | Lighthouse Lab in Milton Keynes | Wellcome Sanger Institute for the COVID-19 Genomics UK (COG-UK) Consortium | The Lighthouse Lab in Milton Keynes and Alex Alderton, Roberto Amato, Jeffrey Barrett, Sonia Goncalves, Ewan Harrison, David K. Jackson, Ian Johnston, Dominic Kwiatkowski, Cordelia Langford, John Sillitoe on behalf of the Wellcome Sanger Institute COVID-19 Surveillance Team |
| EPI_ISL_3224276 | Lighthouse Lab in Glasgow | Wellcome Sanger Institute for the COVID-19 Genomics UK (COG-UK) Consortium | Harper VanSteenhouse, Yumi Kasai, David Gray, Carol Clugston, Anna Dominiczak and Alex Alderton, Roberto Amato, Jeffrey Barrett, Sonia Goncalves, Ewan Harrison, David K. Jackson, Ian Johnston, Dominic Kwiatkowski, Cordelia Langford, John Sillitoe on behalf of the Wellcome Sanger Institute COVID-19 Surveillance Team |
| EPI_ISL_3224277 | Lighthouse Lab in Milton Keynes | Wellcome Sanger Institute for the COVID-19 Genomics UK (COG-UK) Consortium | The Lighthouse Lab in Milton Keynes and Alex Alderton, Roberto Amato, Jeffrey Barrett, Sonia Goncalves, Ewan Harrison, David K. Jackson, Ian Johnston, Dominic Kwiatkowski, Cordelia Langford, John Sillitoe on behalf of the Wellcome Sanger Institute COVID-19 Surveillance Team |
| EPI_ISL_3224278 | Lighthouse Lab in Alderley Park | Wellcome Sanger Institute for the COVID-19 Genomics UK (COG-UK) Consortium | Jacquelyn Wynn, Mairead Hyland, The Lighthouse Lab in Alderley Park and Alex Alderton, Roberto Amato, Jeffrey Barrett, Sonia Goncalves, Ewan Harrison, David K. Jackson, Ian Johnston, Dominic Kwiatkowski, Cordelia Langford, John Sillitoe on behalf of the Wellcome Sanger Institute COVID-19 Surveillance Team |
| EPI_ISL_3224279, EPI_ISL_3224280, EPI_ISL_3224281, EPI_ISL_3224282 | Lighthouse Lab in Milton Keynes | Wellcome Sanger Institute for the COVID-19 Genomics UK (COG-UK) Consortium | The Lighthouse Lab in Milton Keynes and Alex Alderton, Roberto Amato, Jeffrey Barrett, Sonia Goncalves, Ewan Harrison, David K. Jackson, Ian Johnston, Dominic Kwiatkowski, Cordelia Langford, John Sillitoe on behalf of the Wellcome Sanger Institute COVID-19 Surveillance Team |
| EPI_ISL_3224283 | Lighthouse Lab in Glasgow | Wellcome Sanger Institute for the COVID-19 Genomics UK (COG-UK) Consortium | Harper VanSteenhouse, Yumi Kasai, David Gray, Carol Clugston, Anna Dominiczak and Alex Alderton, Roberto Amato, Jeffrey Barrett, Sonia Goncalves, Ewan Harrison, David K. Jackson, Ian Johnston, Dominic Kwiatkowski, Cordelia Langford, John Sillitoe on behalf of the Wellcome Sanger Institute COVID-19 Surveillance Team |
| EPI_ISL_3224284 | Lighthouse Lab in Alderley Park | Wellcome Sanger Institute for the COVID-19 Genomics UK (COG-UK) Consortium | Jacquelyn Wynn, Mairead Hyland, The Lighthouse Lab in Alderley Park and Alex Alderton, Roberto Amato, Jeffrey Barrett, Sonia Goncalves, Ewan Harrison, David K. Jackson, Ian Johnston, Dominic Kwiatkowski, Cordelia Langford, John Sillitoe on behalf of the Wellcome Sanger Institute COVID-19 Surveillance Team |
| EPI_ISL_3224285, EPI_ISL_3224286, EPI_ISL_3224287, EPI_ISL_3224288 | Lighthouse Lab in Milton Keynes | Wellcome Sanger Institute for the COVID-19 Genomics UK (COG-UK) Consortium | The Lighthouse Lab in Milton Keynes and Alex Alderton, Roberto Amato, Jeffrey Barrett, Sonia Goncalves, Ewan Harrison, David K. Jackson, Ian Johnston, Dominic Kwiatkowski, Cordelia Langford, John Sillitoe on behalf of the Wellcome Sanger Institute COVID-19 Surveillance Team |
| EPI_ISL_3224289 | Lighthouse Lab in Glasgow | Wellcome Sanger Institute for the COVID-19 Genomics UK (COG-UK) Consortium | Harper VanSteenhouse, Yumi Kasai, David Gray, Carol Clugston, Anna Dominiczak and Alex Alderton, Roberto Amato, Jeffrey Barrett, Sonia Goncalves, Ewan Harrison, David K. Jackson, Ian Johnston, Dominic Kwiatkowski, Cordelia Langford, John Sillitoe on behalf of the Wellcome Sanger Institute COVID-19 Surveillance Team |
| EPI_ISL_3224290, EPI_ISL_3224291, EPI_ISL_3224292 | Lighthouse Lab in Milton Keynes | Wellcome Sanger Institute for the COVID-19 Genomics UK (COG-UK) Consortium | The Lighthouse Lab in Milton Keynes and Alex Alderton, Roberto Amato, Jeffrey Barrett, Sonia Goncalves, Ewan Harrison, David K. Jackson, Ian Johnston, Dominic Kwiatkowski, Cordelia Langford, John Sillitoe on behalf of the Wellcome Sanger Institute COVID-19 Surveillance Team |
| EPI_ISL_3224293 | Lighthouse Lab in Glasgow | Wellcome Sanger Institute for the COVID-19 Genomics UK (COG-UK) Consortium | Harper VanSteenhouse, Yumi Kasai, David Gray, Carol Clugston, Anna Dominiczak and Alex Alderton, Roberto Amato, Jeffrey Barrett, Sonia Goncalves, Ewan Harrison, David K. Jackson, Ian Johnston, Dominic Kwiatkowski, Cordelia Langford, John Sillitoe on behalf of the Wellcome Sanger Institute COVID-19 Surveillance Team |
| EPI_ISL_3224294 | Lighthouse Lab in Alderley Park | Wellcome Sanger Institute for the COVID-19 Genomics UK (COG-UK) Consortium | Jacquelyn Wynn, Mairead Hyland, The Lighthouse Lab in Alderley Park and Alex Alderton, Roberto Amato, Jeffrey Barrett, Sonia Goncalves, Ewan Harrison, David K. Jackson, Ian Johnston, Dominic Kwiatkowski, Cordelia Langford, John Sillitoe on behalf of the Wellcome Sanger Institute COVID-19 Surveillance Team |
| EPI_ISL_3224295 | Lighthouse Lab in Glasgow | Wellcome Sanger Institute for the COVID-19 Genomics UK (COG-UK) Consortium | Harper VanSteenhouse, Yumi Kasai, David Gray, Carol Clugston, Anna Dominiczak and Alex Alderton, Roberto Amato, Jeffrey Barrett, Sonia Goncalves, Ewan Harrison, David K. Jackson, Ian Johnston, Dominic Kwiatkowski, Cordelia Langford, John Sillitoe on behalf of the Wellcome Sanger Institute COVID-19 Surveillance Team |
| EPI_ISL_3224296 | Lighthouse Lab in Alderley Park | Wellcome Sanger Institute for the COVID-19 Genomics UK (COG-UK) Consortium | Jacquelyn Wynn, Mairead Hyland, The Lighthouse Lab in Alderley Park and Alex Alderton, Roberto Amato, Jeffrey Barrett, Sonia Goncalves, Ewan Harrison, David K. Jackson, Ian Johnston, Dominic Kwiatkowski, Cordelia Langford, John Sillitoe on behalf of the Wellcome Sanger Institute COVID-19 Surveillance Team |
| EPI_ISL_3224297 | Lighthouse Lab in Glasgow | Wellcome Sanger Institute for the COVID-19 Genomics UK (COG-UK) Consortium | Harper VanSteenhouse, Yumi Kasai, David Gray, Carol Clugston, Anna Dominiczak and Alex Alderton, Roberto Amato, Jeffrey Barrett, Sonia Goncalves, Ewan Harrison, David K. Jackson, Ian Johnston, Dominic Kwiatkowski, Cordelia Langford, John Sillitoe on behalf of the Wellcome Sanger Institute COVID-19 Surveillance Team |
| EPI_ISL_3224298, EPI_ISL_3224299, EPI_ISL_3224300, EPI_ISL_3224301 | Lighthouse Lab in Milton Keynes | Wellcome Sanger Institute for the COVID-19 Genomics UK (COG-UK) Consortium | The Lighthouse Lab in Milton Keynes and Alex Alderton, Roberto Amato, Jeffrey Barrett, Sonia Goncalves, Ewan Harrison, David K. Jackson, Ian Johnston, Dominic Kwiatkowski, Cordelia Langford, John Sillitoe on behalf of the Wellcome Sanger Institute COVID-19 Surveillance Team |
| EPI_ISL_3224302 | Lighthouse Lab in Alderley Park | Wellcome Sanger Institute for the COVID-19 Genomics UK (COG-UK) Consortium | Jacquelyn Wynn, Mairead Hyland, The Lighthouse Lab in Alderley Park and Alex Alderton, Roberto Amato, Jeffrey Barrett, Sonia Goncalves, Ewan Harrison, David K. Jackson, Ian Johnston, Dominic Kwiatkowski, Cordelia Langford, John Sillitoe on behalf of the Wellcome Sanger Institute COVID-19 Surveillance Team |
| EPI_ISL_3224303, EPI_ISL_3224304 | Lighthouse Lab in Glasgow | Wellcome Sanger Institute for the COVID-19 Genomics UK (COG-UK) Consortium | Harper VanSteenhouse, Yumi Kasai, David Gray, Carol Clugston, Anna Dominiczak and Alex Alderton, Roberto Amato, Jeffrey Barrett, Sonia Goncalves, Ewan Harrison, David K. Jackson, Ian Johnston, Dominic Kwiatkowski, Cordelia Langford, John Sillitoe on behalf of the Wellcome Sanger Institute COVID-19 Surveillance Team |
| EPI_ISL_3224305, EPI_ISL_3224306 | Lighthouse Lab in Alderley Park | Wellcome Sanger Institute for the COVID-19 Genomics UK (COG-UK) Consortium | Jacquelyn Wynn, Mairead Hyland, The Lighthouse Lab in Alderley Park and Alex Alderton, Roberto Amato, Jeffrey Barrett, Sonia Goncalves, Ewan Harrison, David K. Jackson, Ian Johnston, Dominic Kwiatkowski, Cordelia Langford, John Sillitoe on behalf of the Wellcome Sanger Institute COVID-19 Surveillance Team |
| EPI_ISL_3224307 | Lighthouse Lab in Milton Keynes | Wellcome Sanger Institute for the COVID-19 Genomics UK (COG-UK) Consortium | The Lighthouse Lab in Milton Keynes and Alex Alderton, Roberto Amato, Jeffrey Barrett, Sonia Goncalves, Ewan Harrison, David K. Jackson, Ian Johnston, Dominic Kwiatkowski, Cordelia Langford, John Sillitoe on behalf of the Wellcome Sanger Institute COVID-19 Surveillance Team |
| EPI_ISL_3224308, EPI_ISL_3224309 | Lighthouse Lab in Alderley Park | Wellcome Sanger Institute for the COVID-19 Genomics UK (COG-UK) Consortium | Jacquelyn Wynn, Mairead Hyland, The Lighthouse Lab in Alderley Park and Alex Alderton, Roberto Amato, Jeffrey Barrett, Sonia Goncalves, Ewan Harrison, David K. Jackson, Ian Johnston, Dominic Kwiatkowski, Cordelia Langford, John Sillitoe on behalf of the Wellcome Sanger Institute COVID-19 Surveillance Team |
| EPI_ISL_3224310 | Lighthouse Lab in Milton Keynes | Wellcome Sanger Institute for the COVID-19 Genomics UK (COG-UK) Consortium | The Lighthouse Lab in Milton Keynes and Alex Alderton, Roberto Amato, Jeffrey Barrett, Sonia Goncalves, Ewan Harrison, David K. Jackson, Ian Johnston, Dominic Kwiatkowski, Cordelia Langford, John Sillitoe on behalf of the Wellcome Sanger Institute COVID-19 Surveillance Team |
| EPI_ISL_3224311, EPI_ISL_3224312 | Lighthouse Lab in Alderley Park | Wellcome Sanger Institute for the COVID-19 Genomics UK (COG-UK) Consortium | Jacquelyn Wynn, Mairead Hyland, The Lighthouse Lab in Alderley Park and Alex Alderton, Roberto Amato, Jeffrey Barrett, Sonia Goncalves, Ewan Harrison, David K. Jackson, Ian Johnston, Dominic Kwiatkowski, Cordelia Langford, John Sillitoe on behalf of the Wellcome Sanger Institute COVID-19 Surveillance Team |
| EPI_ISL_3224313, EPI_ISL_3224314, EPI_ISL_3224315 | Lighthouse Lab in Milton Keynes | Wellcome Sanger Institute for the COVID-19 Genomics UK (COG-UK) Consortium | The Lighthouse Lab in Milton Keynes and Alex Alderton, Roberto Amato, Jeffrey Barrett, Sonia Goncalves, Ewan Harrison, David K. Jackson, Ian Johnston, Dominic Kwiatkowski, Cordelia Langford, John Sillitoe on behalf of the Wellcome Sanger Institute COVID-19 Surveillance Team |
| EPI_ISL_3224316 | Lighthouse Lab in Alderley Park | Wellcome Sanger Institute for the COVID-19 Genomics UK (COG-UK) Consortium | Jacquelyn Wynn, Mairead Hyland, The Lighthouse Lab in Alderley Park and Alex Alderton, Roberto Amato, Jeffrey Barrett, Sonia Goncalves, Ewan Harrison, David K. Jackson, Ian Johnston, Dominic Kwiatkowski, Cordelia Langford, John Sillitoe on behalf of the Wellcome Sanger Institute COVID-19 Surveillance Team |
| EPI_ISL_3224317 | Lighthouse Lab in Glasgow | Wellcome Sanger Institute for the COVID-19 Genomics UK (COG-UK) Consortium | Harper VanSteenhouse, Yumi Kasai, David Gray, Carol Clugston, Anna Dominiczak and Alex Alderton, Roberto Amato, Jeffrey Barrett, Sonia Goncalves, Ewan Harrison, David K. Jackson, Ian Johnston, Dominic Kwiatkowski, Cordelia Langford, John Sillitoe on behalf of the Wellcome Sanger Institute COVID-19 Surveillance Team |
| EPI_ISL_3224318 | Lighthouse Lab in Milton Keynes | Wellcome Sanger Institute for the COVID-19 Genomics UK (COG-UK) Consortium | The Lighthouse Lab in Milton Keynes and Alex Alderton, Roberto Amato, Jeffrey Barrett, Sonia Goncalves, Ewan Harrison, David K. Jackson, Ian Johnston, Dominic Kwiatkowski, Cordelia Langford, John Sillitoe on behalf of the Wellcome Sanger Institute COVID-19 Surveillance Team |
| EPI_ISL_3224319 | Lighthouse Lab in Alderley Park | Wellcome Sanger Institute for the COVID-19 Genomics UK (COG-UK) Consortium | Jacquelyn Wynn, Mairead Hyland, The Lighthouse Lab in Alderley Park and Alex Alderton, Roberto Amato, Jeffrey Barrett, Sonia Goncalves, Ewan Harrison, David K. Jackson, Ian Johnston, Dominic Kwiatkowski, Cordelia Langford, John Sillitoe on behalf of the Wellcome Sanger Institute COVID-19 Surveillance Team |
| EPI_ISL_3224320 | Lighthouse Lab in Milton Keynes | Wellcome Sanger Institute for the COVID-19 Genomics UK (COG-UK) Consortium | The Lighthouse Lab in Milton Keynes and Alex Alderton, Roberto Amato, Jeffrey Barrett, Sonia Goncalves, Ewan Harrison, David K. Jackson, Ian Johnston, Dominic Kwiatkowski, Cordelia Langford, John Sillitoe on behalf of the Wellcome Sanger Institute COVID-19 Surveillance Team |
| EPI_ISL_3224321 | Lighthouse Lab in Glasgow | Wellcome Sanger Institute for the COVID-19 Genomics UK (COG-UK) Consortium | Harper VanSteenhouse, Yumi Kasai, David Gray, Carol Clugston, Anna Dominiczak and Alex Alderton, Roberto Amato, Jeffrey Barrett, Sonia Goncalves, Ewan Harrison, David K. Jackson, Ian Johnston, Dominic Kwiatkowski, Cordelia Langford, John Sillitoe on behalf of the Wellcome Sanger Institute COVID-19 Surveillance Team |
| EPI_ISL_3224322 | Lighthouse Lab in Milton Keynes | Wellcome Sanger Institute for the COVID-19 Genomics UK (COG-UK) Consortium | The Lighthouse Lab in Milton Keynes and Alex Alderton, Roberto Amato, Jeffrey Barrett, Sonia Goncalves, Ewan Harrison, David K. Jackson, Ian Johnston, Dominic Kwiatkowski, Cordelia Langford, John Sillitoe on behalf of the Wellcome Sanger Institute COVID-19 Surveillance Team |
| EPI_ISL_3224323 | Lighthouse Lab in Alderley Park | Wellcome Sanger Institute for the COVID-19 Genomics UK (COG-UK) Consortium | Jacquelyn Wynn, Mairead Hyland, The Lighthouse Lab in Alderley Park and Alex Alderton, Roberto Amato, Jeffrey Barrett, Sonia Goncalves, Ewan Harrison, David K. Jackson, Ian Johnston, Dominic Kwiatkowski, Cordelia Langford, John Sillitoe on behalf of the Wellcome Sanger Institute COVID-19 Surveillance Team |
| EPI_ISL_3224324, EPI_ISL_3224325 | Lighthouse Lab in Milton Keynes | Wellcome Sanger Institute for the COVID-19 Genomics UK (COG-UK) Consortium | The Lighthouse Lab in Milton Keynes and Alex Alderton, Roberto Amato, Jeffrey Barrett, Sonia Goncalves, Ewan Harrison, David K. Jackson, Ian Johnston, Dominic Kwiatkowski, Cordelia Langford, John Sillitoe on behalf of the Wellcome Sanger Institute COVID-19 Surveillance Team |
| EPI_ISL_3224326 | Lighthouse Lab in Glasgow | Wellcome Sanger Institute for the COVID-19 Genomics UK (COG-UK) Consortium | Harper VanSteenhouse, Yumi Kasai, David Gray, Carol Clugston, Anna Dominiczak and Alex Alderton, Roberto Amato, Jeffrey Barrett, Sonia Goncalves, Ewan Harrison, David K. Jackson, Ian Johnston, Dominic Kwiatkowski, Cordelia Langford, John Sillitoe on behalf of the Wellcome Sanger Institute COVID-19 Surveillance Team |
| EPI_ISL_3224327, EPI_ISL_3224328 | Lighthouse Lab in Milton Keynes | Wellcome Sanger Institute for the COVID-19 Genomics UK (COG-UK) Consortium | The Lighthouse Lab in Milton Keynes and Alex Alderton, Roberto Amato, Jeffrey Barrett, Sonia Goncalves, Ewan Harrison, David K. Jackson, Ian Johnston, Dominic Kwiatkowski, Cordelia Langford, John Sillitoe on behalf of the Wellcome Sanger Institute COVID-19 Surveillance Team |
| EPI_ISL_3224329 | Lighthouse Lab in Glasgow | Wellcome Sanger Institute for the COVID-19 Genomics UK (COG-UK) Consortium | Harper VanSteenhouse, Yumi Kasai, David Gray, Carol Clugston, Anna Dominiczak and Alex Alderton, Roberto Amato, Jeffrey Barrett, Sonia Goncalves, Ewan Harrison, David K. Jackson, Ian Johnston, Dominic Kwiatkowski, Cordelia Langford, John Sillitoe on behalf of the Wellcome Sanger Institute COVID-19 Surveillance Team |
| EPI_ISL_3224330, EPI_ISL_3224331, EPI_ISL_3224332, EPI_ISL_3224333, EPI_ISL_3224334, EPI_ISL_3224335 | Lighthouse Lab in Milton Keynes | Wellcome Sanger Institute for the COVID-19 Genomics UK (COG-UK) Consortium | The Lighthouse Lab in Milton Keynes and Alex Alderton, Roberto Amato, Jeffrey Barrett, Sonia Goncalves, Ewan Harrison, David K. Jackson, Ian Johnston, Dominic Kwiatkowski, Cordelia Langford, John Sillitoe on behalf of the Wellcome Sanger Institute COVID-19 Surveillance Team |
| EPI_ISL_3224336 | Lighthouse Lab in Glasgow | Wellcome Sanger Institute for the COVID-19 Genomics UK (COG-UK) Consortium | Harper VanSteenhouse, Yumi Kasai, David Gray, Carol Clugston, Anna Dominiczak and Alex Alderton, Roberto Amato, Jeffrey Barrett, Sonia Goncalves, Ewan Harrison, David K. Jackson, Ian Johnston, Dominic Kwiatkowski, Cordelia Langford, John Sillitoe on behalf of the Wellcome Sanger Institute COVID-19 Surveillance Team |
| EPI_ISL_3224337 | Lighthouse Lab in Milton Keynes | Wellcome Sanger Institute for the COVID-19 Genomics UK (COG-UK) Consortium | The Lighthouse Lab in Milton Keynes and Alex Alderton, Roberto Amato, Jeffrey Barrett, Sonia Goncalves, Ewan Harrison, David K. Jackson, Ian Johnston, Dominic Kwiatkowski, Cordelia Langford, John Sillitoe on behalf of the Wellcome Sanger Institute COVID-19 Surveillance Team |

| | | | |
|---|---|---|---|
| EPI_ISL_3224338 | Lighthouse Lab in Alderley Park | Wellcome Sanger Institute for the COVID-19 Genomics UK (COG-UK) Consortium | Jacquelyn Wynn, Mairead Hyland, The Lighthouse Lab in Alderley Park and Alex Alderton, Roberto Amato, Jeffrey Barrett, Sonia Goncalves, Ewan Harrison, David K. Jackson, Ian Johnston, Dominic Kwiatkowski, Cordelia Langford, John Sillitoe on behalf of the Wellcome Sanger Institute COVID-19 Surveillance Team |
| EPI_ISL_3224339 | Lighthouse Lab in Milton Keynes | Wellcome Sanger Institute for the COVID-19 Genomics UK (COG-UK) Consortium | The Lighthouse Lab in Milton Keynes and Alex Alderton, Roberto Amato, Jeffrey Barrett, Sonia Goncalves, Ewan Harrison, David K. Jackson, Ian Johnston, Dominic Kwiatkowski, Cordelia Langford, John Sillitoe on behalf of the Wellcome Sanger Institute COVID-19 Surveillance Team |
| EPI_ISL_3224340 | Lighthouse Lab in Alderley Park | Wellcome Sanger Institute for the COVID-19 Genomics UK (COG-UK) Consortium | Jacquelyn Wynn, Mairead Hyland, The Lighthouse Lab in Alderley Park and Alex Alderton, Roberto Amato, Jeffrey Barrett, Sonia Goncalves, Ewan Harrison, David K. Jackson, Ian Johnston, Dominic Kwiatkowski, Cordelia Langford, John Sillitoe on behalf of the Wellcome Sanger Institute COVID-19 Surveillance Team |
| EPI_ISL_3224341 | Lighthouse Lab in Milton Keynes | Wellcome Sanger Institute for the COVID-19 Genomics UK (COG-UK) Consortium | The Lighthouse Lab in Milton Keynes and Alex Alderton, Roberto Amato, Jeffrey Barrett, Sonia Goncalves, Ewan Harrison, David K. Jackson, Ian Johnston, Dominic Kwiatkowski, Cordelia Langford, John Sillitoe on behalf of the Wellcome Sanger Institute COVID-19 Surveillance Team |
| EPI_ISL_3224342 | Lighthouse Lab in Alderley Park | Wellcome Sanger Institute for the COVID-19 Genomics UK (COG-UK) Consortium | Jacquelyn Wynn, Mairead Hyland, The Lighthouse Lab in Alderley Park and Alex Alderton, Roberto Amato, Jeffrey Barrett, Sonia Goncalves, Ewan Harrison, David K. Jackson, Ian Johnston, Dominic Kwiatkowski, Cordelia Langford, John Sillitoe on behalf of the Wellcome Sanger Institute COVID-19 Surveillance Team |
| EPI_ISL_3224343, EPI_ISL_3224344 | Lighthouse Lab in Milton Keynes | Wellcome Sanger Institute for the COVID-19 Genomics UK (COG-UK) Consortium | The Lighthouse Lab in Milton Keynes and Alex Alderton, Roberto Amato, Jeffrey Barrett, Sonia Goncalves, Ewan Harrison, David K. Jackson, Ian Johnston, Dominic Kwiatkowski, Cordelia Langford, John Sillitoe on behalf of the Wellcome Sanger Institute COVID-19 Surveillance Team |
| EPI_ISL_3224345 | Lighthouse Lab in Glasgow | Wellcome Sanger Institute for the COVID-19 Genomics UK (COG-UK) Consortium | Harper VanSteenhouse, Yumi Kasai, David Gray, Carol Clugston, Anna Dominiczak and Alex Alderton, Roberto Amato, Jeffrey Barrett, Sonia Goncalves, Ewan Harrison, David K. Jackson, Ian Johnston, Dominic Kwiatkowski, Cordelia Langford, John Sillitoe on behalf of the Wellcome Sanger Institute COVID-19 Surveillance Team |
| EPI_ISL_3224346 | Lighthouse Lab in Alderley Park | Wellcome Sanger Institute for the COVID-19 Genomics UK (COG-UK) Consortium | Jacquelyn Wynn, Mairead Hyland, The Lighthouse Lab in Alderley Park and Alex Alderton, Roberto Amato, Jeffrey Barrett, Sonia Goncalves, Ewan Harrison, David K. Jackson, Ian Johnston, Dominic Kwiatkowski, Cordelia Langford, John Sillitoe on behalf of the Wellcome Sanger Institute COVID-19 Surveillance Team |
| EPI_ISL_3224347 | Lighthouse Lab in Glasgow | Wellcome Sanger Institute for the COVID-19 Genomics UK (COG-UK) Consortium | Harper VanSteenhouse, Yumi Kasai, David Gray, Carol Clugston, Anna Dominiczak and Alex Alderton, Roberto Amato, Jeffrey Barrett, Sonia Goncalves, Ewan Harrison, David K. Jackson, Ian Johnston, Dominic Kwiatkowski, Cordelia Langford, John Sillitoe on behalf of the Wellcome Sanger Institute COVID-19 Surveillance Team |
| EPI_ISL_3224348 | Lighthouse Lab in Alderley Park | Wellcome Sanger Institute for the COVID-19 Genomics UK (COG-UK) Consortium | Jacquelyn Wynn, Mairead Hyland, The Lighthouse Lab in Alderley Park and Alex Alderton, Roberto Amato, Jeffrey Barrett, Sonia Goncalves, Ewan Harrison, David K. Jackson, Ian Johnston, Dominic Kwiatkowski, Cordelia Langford, John Sillitoe on behalf of the Wellcome Sanger Institute COVID-19 Surveillance Team |
| EPI_ISL_3224349, EPI_ISL_3224350, EPI_ISL_3224351, EPI_ISL_3224352 | Lighthouse Lab in Milton Keynes | Wellcome Sanger Institute for the COVID-19 Genomics UK (COG-UK) Consortium | The Lighthouse Lab in Milton Keynes and Alex Alderton, Roberto Amato, Jeffrey Barrett, Sonia Goncalves, Ewan Harrison, David K. Jackson, Ian Johnston, Dominic Kwiatkowski, Cordelia Langford, John Sillitoe on behalf of the Wellcome Sanger Institute COVID-19 Surveillance Team |
| EPI_ISL_3224353, EPI_ISL_3224354 | Lighthouse Lab in Alderley Park | Wellcome Sanger Institute for the COVID-19 Genomics UK (COG-UK) Consortium | Jacquelyn Wynn, Mairead Hyland, The Lighthouse Lab in Alderley Park and Alex Alderton, Roberto Amato, Jeffrey Barrett, Sonia Goncalves, Ewan Harrison, David K. Jackson, Ian Johnston, Dominic Kwiatkowski, Cordelia Langford, John Sillitoe on behalf of the Wellcome Sanger Institute COVID-19 Surveillance Team |
| EPI_ISL_3224355, EPI_ISL_3224356, EPI_ISL_3224357, EPI_ISL_3224358, EPI_ISL_3224359, EPI_ISL_3224360 | Lighthouse Lab in Milton Keynes | Wellcome Sanger Institute for the COVID-19 Genomics UK (COG-UK) Consortium | The Lighthouse Lab in Milton Keynes and Alex Alderton, Roberto Amato, Jeffrey Barrett, Sonia Goncalves, Ewan Harrison, David K. Jackson, Ian Johnston, Dominic Kwiatkowski, Cordelia Langford, John Sillitoe on behalf of the Wellcome Sanger Institute COVID-19 Surveillance Team |
| EPI_ISL_3224361 | Lighthouse Lab in Glasgow | Wellcome Sanger Institute for the COVID-19 Genomics UK (COG-UK) Consortium | Harper VanSteenhouse, Yumi Kasai, David Gray, Carol Clugston, Anna Dominiczak and Alex Alderton, Roberto Amato, Jeffrey Barrett, Sonia Goncalves, Ewan Harrison, David K. Jackson, Ian Johnston, Dominic Kwiatkowski, Cordelia Langford, John Sillitoe on behalf of the Wellcome Sanger Institute COVID-19 Surveillance Team |
| EPI_ISL_3224362 | Lighthouse Lab in Milton Keynes | Wellcome Sanger Institute for the COVID-19 Genomics UK (COG-UK) Consortium | The Lighthouse Lab in Milton Keynes and Alex Alderton, Roberto Amato, Jeffrey Barrett, Sonia Goncalves, Ewan Harrison, David K. Jackson, Ian Johnston, Dominic Kwiatkowski, Cordelia Langford, John Sillitoe on behalf of the Wellcome Sanger Institute COVID-19 Surveillance Team |
| EPI_ISL_3224363 | Lighthouse Lab in Alderley Park | Wellcome Sanger Institute for the COVID-19 Genomics UK (COG-UK) Consortium | Jacquelyn Wynn, Mairead Hyland, The Lighthouse Lab in Alderley Park and Alex Alderton, Roberto Amato, Jeffrey Barrett, Sonia Goncalves, Ewan Harrison, David K. Jackson, Ian Johnston, Dominic Kwiatkowski, Cordelia Langford, John Sillitoe on behalf of the Wellcome Sanger Institute COVID-19 Surveillance Team |
| EPI_ISL_3224364 | Lighthouse Lab in Milton Keynes | Wellcome Sanger Institute for the COVID-19 Genomics UK (COG-UK) Consortium | The Lighthouse Lab in Milton Keynes and Alex Alderton, Roberto Amato, Jeffrey Barrett, Sonia Goncalves, Ewan Harrison, David K. Jackson, Ian Johnston, Dominic Kwiatkowski, Cordelia Langford, John Sillitoe on behalf of the Wellcome Sanger Institute COVID-19 Surveillance Team |
| EPI_ISL_3224365, EPI_ISL_3224366 | Lighthouse Lab in Alderley Park | Wellcome Sanger Institute for the COVID-19 Genomics UK (COG-UK) Consortium | Jacquelyn Wynn, Mairead Hyland, The Lighthouse Lab in Alderley Park and Alex Alderton, Roberto Amato, Jeffrey Barrett, Sonia Goncalves, Ewan Harrison, David K. Jackson, Ian Johnston, Dominic Kwiatkowski, Cordelia Langford, John Sillitoe on behalf of the Wellcome Sanger Institute COVID-19 Surveillance Team |
| EPI_ISL_3224367 | Lighthouse Lab in Glasgow | Wellcome Sanger Institute for the COVID-19 Genomics UK (COG-UK) Consortium | Harper VanSteenhouse, Yumi Kasai, David Gray, Carol Clugston, Anna Dominiczak and Alex Alderton, Roberto Amato, Jeffrey Barrett, Sonia Goncalves, Ewan Harrison, David K. Jackson, Ian Johnston, Dominic Kwiatkowski, Cordelia Langford, John Sillitoe on behalf of the Wellcome Sanger Institute COVID-19 Surveillance Team |
| EPI_ISL_3224368 | Lighthouse Lab in Milton Keynes | Wellcome Sanger Institute for the COVID-19 Genomics UK (COG-UK) Consortium | The Lighthouse Lab in Milton Keynes and Alex Alderton, Roberto Amato, Jeffrey Barrett, Sonia Goncalves, Ewan Harrison, David K. Jackson, Ian Johnston, Dominic Kwiatkowski, Cordelia Langford, John Sillitoe on behalf of the Wellcome Sanger Institute COVID-19 Surveillance Team |
| EPI_ISL_3224369 | Lighthouse Lab in Alderley Park | Wellcome Sanger Institute for the COVID-19 Genomics UK (COG-UK) Consortium | Jacquelyn Wynn, Mairead Hyland, The Lighthouse Lab in Alderley Park and Alex Alderton, Roberto Amato, Jeffrey Barrett, Sonia Goncalves, Ewan Harrison, David K. Jackson, Ian Johnston, Dominic Kwiatkowski, Cordelia Langford, John Sillitoe on behalf of the Wellcome Sanger Institute COVID-19 Surveillance Team |
| EPI_ISL_3224370 | Lighthouse Lab in Glasgow | Wellcome Sanger Institute for the COVID-19 Genomics UK (COG-UK) Consortium | Harper VanSteenhouse, Yumi Kasai, David Gray, Carol Clugston, Anna Dominiczak and Alex Alderton, Roberto Amato, Jeffrey Barrett, Sonia Goncalves, Ewan Harrison, David K. Jackson, Ian Johnston, Dominic Kwiatkowski, Cordelia Langford, John Sillitoe on behalf of the Wellcome Sanger Institute COVID-19 Surveillance Team |
| EPI_ISL_3224371 | Lighthouse Lab in Milton Keynes | Wellcome Sanger Institute for the COVID-19 Genomics UK (COG-UK) Consortium | The Lighthouse Lab in Milton Keynes and Alex Alderton, Roberto Amato, Jeffrey Barrett, Sonia Goncalves, Ewan Harrison, David K. Jackson, Ian Johnston, Dominic Kwiatkowski, Cordelia Langford, John Sillitoe on behalf of the Wellcome Sanger Institute COVID-19 Surveillance Team |
| EPI_ISL_3224372 | Lighthouse Lab in Glasgow | Wellcome Sanger Institute for the COVID-19 Genomics UK (COG-UK) Consortium | Harper VanSteenhouse, Yumi Kasai, David Gray, Carol Clugston, Anna Dominiczak and Alex Alderton, Roberto Amato, Jeffrey Barrett, Sonia Goncalves, Ewan Harrison, David K. Jackson, Ian Johnston, Dominic Kwiatkowski, Cordelia Langford, John Sillitoe on behalf of the Wellcome Sanger Institute COVID-19 Surveillance Team |
| EPI_ISL_3224373, EPI_ISL_3224374 | Lighthouse Lab in Alderley Park | Wellcome Sanger Institute for the COVID-19 Genomics UK (COG-UK) Consortium | Jacquelyn Wynn, Mairead Hyland, The Lighthouse Lab in Alderley Park and Alex Alderton, Roberto Amato, Jeffrey Barrett, Sonia Goncalves, Ewan Harrison, David K. Jackson, Ian Johnston, Dominic Kwiatkowski, Cordelia Langford, John Sillitoe on behalf of the Wellcome Sanger Institute COVID-19 Surveillance Team |
| EPI_ISL_3224375 | Lighthouse Lab in Milton Keynes | Wellcome Sanger Institute for the COVID-19 Genomics UK (COG-UK) Consortium | The Lighthouse Lab in Milton Keynes and Alex Alderton, Roberto Amato, Jeffrey Barrett, Sonia Goncalves, Ewan Harrison, David K. Jackson, Ian Johnston, Dominic Kwiatkowski, Cordelia Langford, John Sillitoe on behalf of the Wellcome Sanger Institute COVID-19 Surveillance Team |
| EPI_ISL_3224376 | Lighthouse Lab in Alderley Park | Wellcome Sanger Institute for the COVID-19 Genomics UK (COG-UK) Consortium | Jacquelyn Wynn, Mairead Hyland, The Lighthouse Lab in Alderley Park and Alex Alderton, Roberto Amato, Jeffrey Barrett, Sonia Goncalves, Ewan Harrison, David K. Jackson, Ian Johnston, Dominic Kwiatkowski, Cordelia Langford, John Sillitoe on behalf of the Wellcome Sanger Institute COVID-19 Surveillance Team |
| EPI_ISL_3224377 | Lighthouse Lab in Glasgow | Wellcome Sanger Institute for the COVID-19 Genomics UK (COG-UK) Consortium | Harper VanSteenhouse, Yumi Kasai, David Gray, Carol Clugston, Anna Dominiczak and Alex Alderton, Roberto Amato, Jeffrey Barrett, Sonia Goncalves, Ewan Harrison, David K. Jackson, Ian Johnston, Dominic Kwiatkowski, Cordelia Langford, John Sillitoe on behalf of the Wellcome Sanger Institute COVID-19 Surveillance Team |
| EPI_ISL_3224378 | Lighthouse Lab in Milton Keynes | Wellcome Sanger Institute for the COVID-19 Genomics UK (COG-UK) Consortium | The Lighthouse Lab in Milton Keynes and Alex Alderton, Roberto Amato, Jeffrey Barrett, Sonia Goncalves, Ewan Harrison, David K. Jackson, Ian Johnston, Dominic Kwiatkowski, Cordelia Langford, John Sillitoe on behalf of the Wellcome Sanger Institute COVID-19 Surveillance Team |
| EPI_ISL_3224379 | Lighthouse Lab in Glasgow | Wellcome Sanger Institute for the COVID-19 Genomics UK (COG-UK) Consortium | Harper VanSteenhouse, Yumi Kasai, David Gray, Carol Clugston, Anna Dominiczak and Alex Alderton, Roberto Amato, Jeffrey Barrett, Sonia Goncalves, Ewan Harrison, David K. Jackson, Ian Johnston, Dominic Kwiatkowski, Cordelia Langford, John Sillitoe on behalf of the Wellcome Sanger Institute COVID-19 Surveillance Team |
| EPI_ISL_3224380, EPI_ISL_3224381, EPI_ISL_3224382, EPI_ISL_3224383 | Lighthouse Lab in Alderley Park | Wellcome Sanger Institute for the COVID-19 Genomics UK (COG-UK) Consortium | Jacquelyn Wynn, Mairead Hyland, The Lighthouse Lab in Alderley Park and Alex Alderton, Roberto Amato, Jeffrey Barrett, Sonia Goncalves, Ewan Harrison, David K. Jackson, Ian Johnston, Dominic Kwiatkowski, Cordelia Langford, John Sillitoe on behalf of the Wellcome Sanger Institute COVID-19 Surveillance Team |
| EPI_ISL_3224384 | Lighthouse Lab in Milton Keynes | Wellcome Sanger Institute for the COVID-19 Genomics UK (COG-UK) Consortium | The Lighthouse Lab in Milton Keynes and Alex Alderton, Roberto Amato, Jeffrey Barrett, Sonia Goncalves, Ewan Harrison, David K. Jackson, Ian Johnston, Dominic Kwiatkowski, Cordelia Langford, John Sillitoe on behalf of the Wellcome Sanger Institute COVID-19 Surveillance Team |
| EPI_ISL_3224385 | Lighthouse Lab in Glasgow | Wellcome Sanger Institute for the COVID-19 Genomics UK (COG-UK) Consortium | Harper VanSteenhouse, Yumi Kasai, David Gray, Carol Clugston, Anna Dominiczak and Alex Alderton, Roberto Amato, Jeffrey Barrett, Sonia Goncalves, Ewan Harrison, David K. Jackson, Ian Johnston, Dominic Kwiatkowski, Cordelia Langford, John Sillitoe on behalf of the Wellcome Sanger Institute COVID-19 Surveillance Team |
| EPI_ISL_3224386, EPI_ISL_3224387 | Lighthouse Lab in Milton Keynes | Wellcome Sanger Institute for the COVID-19 Genomics UK (COG-UK) Consortium | The Lighthouse Lab in Milton Keynes and Alex Alderton, Roberto Amato, Jeffrey Barrett, Sonia Goncalves, Ewan Harrison, David K. Jackson, Ian Johnston, Dominic Kwiatkowski, Cordelia Langford, John Sillitoe on behalf of the Wellcome Sanger Institute COVID-19 Surveillance Team |
| EPI_ISL_3224388 | Lighthouse Lab in Glasgow | Wellcome Sanger Institute for the COVID-19 Genomics UK (COG-UK) Consortium | Harper VanSteenhouse, Yumi Kasai, David Gray, Carol Clugston, Anna Dominiczak and Alex Alderton, Roberto Amato, Jeffrey Barrett, Sonia Goncalves, Ewan Harrison, David K. Jackson, Ian Johnston, Dominic Kwiatkowski, Cordelia Langford, John Sillitoe on behalf of the Wellcome Sanger Institute COVID-19 Surveillance Team |
| EPI_ISL_3224389 | Lighthouse Lab in Alderley Park | Wellcome Sanger Institute for the COVID-19 Genomics UK (COG-UK) Consortium | Jacquelyn Wynn, Mairead Hyland, The Lighthouse Lab in Alderley Park and Alex Alderton, Roberto Amato, Jeffrey Barrett, Sonia Goncalves, Ewan Harrison, David K. Jackson, Ian Johnston, Dominic Kwiatkowski, Cordelia Langford, John Sillitoe on behalf of the Wellcome Sanger Institute COVID-19 Surveillance Team |
| EPI_ISL_3224390 | Lighthouse Lab in Milton Keynes | Wellcome Sanger Institute for the COVID-19 Genomics UK (COG-UK) Consortium | The Lighthouse Lab in Milton Keynes and Alex Alderton, Roberto Amato, Jeffrey Barrett, Sonia Goncalves, Ewan Harrison, David K. Jackson, Ian Johnston, Dominic Kwiatkowski, Cordelia Langford, John Sillitoe on behalf of the Wellcome Sanger Institute COVID-19 Surveillance Team |
| EPI_ISL_3224391, EPI_ISL_3224392 | Lighthouse Lab in Glasgow | Wellcome Sanger Institute for the COVID-19 Genomics UK (COG-UK) Consortium | Harper VanSteenhouse, Yumi Kasai, David Gray, Carol Clugston, Anna Dominiczak and Alex Alderton, Roberto Amato, Jeffrey Barrett, Sonia Goncalves, Ewan Harrison, David K. Jackson, Ian Johnston, Dominic Kwiatkowski, Cordelia Langford, John Sillitoe on behalf of the Wellcome Sanger Institute COVID-19 Surveillance Team |
| EPI_ISL_3224393 | Lighthouse Lab in Alderley Park | Wellcome Sanger Institute for the COVID-19 Genomics UK (COG-UK) Consortium | Jacquelyn Wynn, Mairead Hyland, The Lighthouse Lab in Alderley Park and Alex Alderton, Roberto Amato, Jeffrey Barrett, Sonia Goncalves, Ewan Harrison, David K. Jackson, Ian Johnston, Dominic Kwiatkowski, Cordelia Langford, John Sillitoe on behalf of the Wellcome Sanger Institute COVID-19 Surveillance Team |
| EPI_ISL_3224394, EPI_ISL_3224395 | Lighthouse Lab in Milton Keynes | Wellcome Sanger Institute for the COVID-19 Genomics UK (COG-UK) Consortium | The Lighthouse Lab in Milton Keynes and Alex Alderton, Roberto Amato, Jeffrey Barrett, Sonia Goncalves, Ewan Harrison, David K. Jackson, Ian Johnston, Dominic Kwiatkowski, Cordelia Langford, John Sillitoe on behalf of the Wellcome Sanger Institute COVID-19 Surveillance Team |
| EPI_ISL_3224396 | Lighthouse Lab in Glasgow | Wellcome Sanger Institute for the COVID-19 Genomics UK (COG-UK) Consortium | Harper VanSteenhouse, Yumi Kasai, David Gray, Carol Clugston, Anna Dominiczak and Alex Alderton, Roberto Amato, Jeffrey Barrett, Sonia Goncalves, Ewan Harrison, David K. Jackson, Ian Johnston, Dominic Kwiatkowski, Cordelia Langford, John Sillitoe on behalf of the Wellcome Sanger Institute COVID-19 Surveillance Team |
| EPI_ISL_3224397, | Lighthouse Lab in Alderley Park | Wellcome Sanger Institute for the COVID-19 Genomics UK (COG-UK) | Jacquelyn Wynn, Mairead Hyland, The Lighthouse Lab in Alderley Park and Alex Alderton, Roberto Amato, Jeffrey Barrett, Sonia Goncalves, Ewan Harrison, David K. Jackson, Ian Johnston, Dominic Kwiatkowski, Cordelia Langford, John Sillitoe on |

| | | | |
|---|---|---|---|
| EPI_ISL_3224398 | | Consortium | behalf of the Wellcome Sanger Institute COVID-19 Surveillance Team |
| EPI_ISL_3224399 | Lighthouse Lab in Milton Keynes | Wellcome Sanger Institute for the COVID-19 Genomics UK (COG-UK) Consortium | The Lighthouse Lab in Milton Keynes and Alex Alderton, Roberto Amato, Jeffrey Barrett, Sonia Goncalves, Ewan Harrison, David K. Jackson, Ian Johnston, Dominic Kwiatkowski, Cordelia Langford, John Sillitoe on behalf of the Wellcome Sanger Institute COVID-19 Surveillance Team |
| EPI_ISL_3224400 | Lighthouse Lab in Glasgow | Wellcome Sanger Institute for the COVID-19 Genomics UK (COG-UK) Consortium | Harper VanSteenhouse, Yumi Kasai, David Gray, Carol Clugston, Anna Dominiczak and Alex Alderton, Roberto Amato, Jeffrey Barrett, Sonia Goncalves, Ewan Harrison, David K. Jackson, Ian Johnston, Dominic Kwiatkowski, Cordelia Langford, John Sillitoe on behalf of the Wellcome Sanger Institute COVID-19 Surveillance Team |
| EPI_ISL_3224401, EPI_ISL_3224402 | Lighthouse Lab in Milton Keynes | Wellcome Sanger Institute for the COVID-19 Genomics UK (COG-UK) Consortium | The Lighthouse Lab in Milton Keynes and Alex Alderton, Roberto Amato, Jeffrey Barrett, Sonia Goncalves, Ewan Harrison, David K. Jackson, Ian Johnston, Dominic Kwiatkowski, Cordelia Langford, John Sillitoe on behalf of the Wellcome Sanger Institute COVID-19 Surveillance Team |
| EPI_ISL_3224403 | Lighthouse Lab in Glasgow | Wellcome Sanger Institute for the COVID-19 Genomics UK (COG-UK) Consortium | Harper VanSteenhouse, Yumi Kasai, David Gray, Carol Clugston, Anna Dominiczak and Alex Alderton, Roberto Amato, Jeffrey Barrett, Sonia Goncalves, Ewan Harrison, David K. Jackson, Ian Johnston, Dominic Kwiatkowski, Cordelia Langford, John Sillitoe on behalf of the Wellcome Sanger Institute COVID-19 Surveillance Team |
| EPI_ISL_3224404 | Lighthouse Lab in Milton Keynes | Wellcome Sanger Institute for the COVID-19 Genomics UK (COG-UK) Consortium | The Lighthouse Lab in Milton Keynes and Alex Alderton, Roberto Amato, Jeffrey Barrett, Sonia Goncalves, Ewan Harrison, David K. Jackson, Ian Johnston, Dominic Kwiatkowski, Cordelia Langford, John Sillitoe on behalf of the Wellcome Sanger Institute COVID-19 Surveillance Team |
| EPI_ISL_3224405 | Lighthouse Lab in Alderley Park | Wellcome Sanger Institute for the COVID-19 Genomics UK (COG-UK) Consortium | Jacquelyn Wynn, Mairead Hyland, The Lighthouse Lab in Alderley Park and Alex Alderton, Roberto Amato, Jeffrey Barrett, Sonia Goncalves, Ewan Harrison, David K. Jackson, Ian Johnston, Dominic Kwiatkowski, Cordelia Langford, John Sillitoe on behalf of the Wellcome Sanger Institute COVID-19 Surveillance Team |
| EPI_ISL_3224406 | Lighthouse Lab in Glasgow | Wellcome Sanger Institute for the COVID-19 Genomics UK (COG-UK) Consortium | Harper VanSteenhouse, Yumi Kasai, David Gray, Carol Clugston, Anna Dominiczak and Alex Alderton, Roberto Amato, Jeffrey Barrett, Sonia Goncalves, Ewan Harrison, David K. Jackson, Ian Johnston, Dominic Kwiatkowski, Cordelia Langford, John Sillitoe on behalf of the Wellcome Sanger Institute COVID-19 Surveillance Team |
| EPI_ISL_3224407 | Lighthouse Lab in Alderley Park | Wellcome Sanger Institute for the COVID-19 Genomics UK (COG-UK) Consortium | Jacquelyn Wynn, Mairead Hyland, The Lighthouse Lab in Alderley Park and Alex Alderton, Roberto Amato, Jeffrey Barrett, Sonia Goncalves, Ewan Harrison, David K. Jackson, Ian Johnston, Dominic Kwiatkowski, Cordelia Langford, John Sillitoe on behalf of the Wellcome Sanger Institute COVID-19 Surveillance Team |
| EPI_ISL_3224408, EPI_ISL_3224409 | Lighthouse Lab in Milton Keynes | Wellcome Sanger Institute for the COVID-19 Genomics UK (COG-UK) Consortium | The Lighthouse Lab in Milton Keynes and Alex Alderton, Roberto Amato, Jeffrey Barrett, Sonia Goncalves, Ewan Harrison, David K. Jackson, Ian Johnston, Dominic Kwiatkowski, Cordelia Langford, John Sillitoe on behalf of the Wellcome Sanger Institute COVID-19 Surveillance Team |
| EPI_ISL_3224410 | Lighthouse Lab in Glasgow | Wellcome Sanger Institute for the COVID-19 Genomics UK (COG-UK) Consortium | Harper VanSteenhouse, Yumi Kasai, David Gray, Carol Clugston, Anna Dominiczak and Alex Alderton, Roberto Amato, Jeffrey Barrett, Sonia Goncalves, Ewan Harrison, David K. Jackson, Ian Johnston, Dominic Kwiatkowski, Cordelia Langford, John Sillitoe on behalf of the Wellcome Sanger Institute COVID-19 Surveillance Team |
| EPI_ISL_3224411 | Lighthouse Lab in Milton Keynes | Wellcome Sanger Institute for the COVID-19 Genomics UK (COG-UK) Consortium | The Lighthouse Lab in Milton Keynes and Alex Alderton, Roberto Amato, Jeffrey Barrett, Sonia Goncalves, Ewan Harrison, David K. Jackson, Ian Johnston, Dominic Kwiatkowski, Cordelia Langford, John Sillitoe on behalf of the Wellcome Sanger Institute COVID-19 Surveillance Team |
| EPI_ISL_3224412 | Lighthouse Lab in Glasgow | Wellcome Sanger Institute for the COVID-19 Genomics UK (COG-UK) Consortium | Harper VanSteenhouse, Yumi Kasai, David Gray, Carol Clugston, Anna Dominiczak and Alex Alderton, Roberto Amato, Jeffrey Barrett, Sonia Goncalves, Ewan Harrison, David K. Jackson, Ian Johnston, Dominic Kwiatkowski, Cordelia Langford, John Sillitoe on behalf of the Wellcome Sanger Institute COVID-19 Surveillance Team |
| EPI_ISL_3224413, EPI_ISL_3224414 | Lighthouse Lab in Milton Keynes | Wellcome Sanger Institute for the COVID-19 Genomics UK (COG-UK) Consortium | The Lighthouse Lab in Milton Keynes and Alex Alderton, Roberto Amato, Jeffrey Barrett, Sonia Goncalves, Ewan Harrison, David K. Jackson, Ian Johnston, Dominic Kwiatkowski, Cordelia Langford, John Sillitoe on behalf of the Wellcome Sanger Institute COVID-19 Surveillance Team |
| EPI_ISL_3224415, EPI_ISL_3224416 | Lighthouse Lab in Glasgow | Wellcome Sanger Institute for the COVID-19 Genomics UK (COG-UK) Consortium | Harper VanSteenhouse, Yumi Kasai, David Gray, Carol Clugston, Anna Dominiczak and Alex Alderton, Roberto Amato, Jeffrey Barrett, Sonia Goncalves, Ewan Harrison, David K. Jackson, Ian Johnston, Dominic Kwiatkowski, Cordelia Langford, John Sillitoe on behalf of the Wellcome Sanger Institute COVID-19 Surveillance Team |
| EPI_ISL_3224417 | Lighthouse Lab in Milton Keynes | Wellcome Sanger Institute for the COVID-19 Genomics UK (COG-UK) Consortium | The Lighthouse Lab in Milton Keynes and Alex Alderton, Roberto Amato, Jeffrey Barrett, Sonia Goncalves, Ewan Harrison, David K. Jackson, Ian Johnston, Dominic Kwiatkowski, Cordelia Langford, John Sillitoe on behalf of the Wellcome Sanger Institute COVID-19 Surveillance Team |
| EPI_ISL_3224418, EPI_ISL_3224419 | Lighthouse Lab in Alderley Park | Wellcome Sanger Institute for the COVID-19 Genomics UK (COG-UK) Consortium | Jacquelyn Wynn, Mairead Hyland, The Lighthouse Lab in Alderley Park and Alex Alderton, Roberto Amato, Jeffrey Barrett, Sonia Goncalves, Ewan Harrison, David K. Jackson, Ian Johnston, Dominic Kwiatkowski, Cordelia Langford, John Sillitoe on behalf of the Wellcome Sanger Institute COVID-19 Surveillance Team |
| EPI_ISL_3224420, EPI_ISL_3224421 | Lighthouse Lab in Milton Keynes | Wellcome Sanger Institute for the COVID-19 Genomics UK (COG-UK) Consortium | The Lighthouse Lab in Milton Keynes and Alex Alderton, Roberto Amato, Jeffrey Barrett, Sonia Goncalves, Ewan Harrison, David K. Jackson, Ian Johnston, Dominic Kwiatkowski, Cordelia Langford, John Sillitoe on behalf of the Wellcome Sanger Institute COVID-19 Surveillance Team |
| EPI_ISL_3224422 | Lighthouse Lab in Glasgow | Wellcome Sanger Institute for the COVID-19 Genomics UK (COG-UK) Consortium | Harper VanSteenhouse, Yumi Kasai, David Gray, Carol Clugston, Anna Dominiczak and Alex Alderton, Roberto Amato, Jeffrey Barrett, Sonia Goncalves, Ewan Harrison, David K. Jackson, Ian Johnston, Dominic Kwiatkowski, Cordelia Langford, John Sillitoe on behalf of the Wellcome Sanger Institute COVID-19 Surveillance Team |
| EPI_ISL_3224423, EPI_ISL_3224424, EPI_ISL_3224425 | Lighthouse Lab in Milton Keynes | Wellcome Sanger Institute for the COVID-19 Genomics UK (COG-UK) Consortium | The Lighthouse Lab in Milton Keynes and Alex Alderton, Roberto Amato, Jeffrey Barrett, Sonia Goncalves, Ewan Harrison, David K. Jackson, Ian Johnston, Dominic Kwiatkowski, Cordelia Langford, John Sillitoe on behalf of the Wellcome Sanger Institute COVID-19 Surveillance Team |
| EPI_ISL_3224426 | Lighthouse Lab in Glasgow | Wellcome Sanger Institute for the COVID-19 Genomics UK (COG-UK) Consortium | Harper VanSteenhouse, Yumi Kasai, David Gray, Carol Clugston, Anna Dominiczak and Alex Alderton, Roberto Amato, Jeffrey Barrett, Sonia Goncalves, Ewan Harrison, David K. Jackson, Ian Johnston, Dominic Kwiatkowski, Cordelia Langford, John Sillitoe on behalf of the Wellcome Sanger Institute COVID-19 Surveillance Team |
| EPI_ISL_3224427 | Lighthouse Lab in Alderley Park | Wellcome Sanger Institute for the COVID-19 Genomics UK (COG-UK) Consortium | Jacquelyn Wynn, Mairead Hyland, The Lighthouse Lab in Alderley Park and Alex Alderton, Roberto Amato, Jeffrey Barrett, Sonia Goncalves, Ewan Harrison, David K. Jackson, Ian Johnston, Dominic Kwiatkowski, Cordelia Langford, John Sillitoe on behalf of the Wellcome Sanger Institute COVID-19 Surveillance Team |
| EPI_ISL_3224428, EPI_ISL_3224429 | Lighthouse Lab in Milton Keynes | Wellcome Sanger Institute for the COVID-19 Genomics UK (COG-UK) Consortium | The Lighthouse Lab in Milton Keynes and Alex Alderton, Roberto Amato, Jeffrey Barrett, Sonia Goncalves, Ewan Harrison, David K. Jackson, Ian Johnston, Dominic Kwiatkowski, Cordelia Langford, John Sillitoe on behalf of the Wellcome Sanger Institute COVID-19 Surveillance Team |
| EPI_ISL_3224430 | Lighthouse Lab in Alderley Park | Wellcome Sanger Institute for the COVID-19 Genomics UK (COG-UK) Consortium | Jacquelyn Wynn, Mairead Hyland, The Lighthouse Lab in Alderley Park and Alex Alderton, Roberto Amato, Jeffrey Barrett, Sonia Goncalves, Ewan Harrison, David K. Jackson, Ian Johnston, Dominic Kwiatkowski, Cordelia Langford, John Sillitoe on behalf of the Wellcome Sanger Institute COVID-19 Surveillance Team |
| EPI_ISL_3224431 | Lighthouse Lab in Glasgow | Wellcome Sanger Institute for the COVID-19 Genomics UK (COG-UK) Consortium | Harper VanSteenhouse, Yumi Kasai, David Gray, Carol Clugston, Anna Dominiczak and Alex Alderton, Roberto Amato, Jeffrey Barrett, Sonia Goncalves, Ewan Harrison, David K. Jackson, Ian Johnston, Dominic Kwiatkowski, Cordelia Langford, John Sillitoe on behalf of the Wellcome Sanger Institute COVID-19 Surveillance Team |
| EPI_ISL_3224432 | Lighthouse Lab in Alderley Park | Wellcome Sanger Institute for the COVID-19 Genomics UK (COG-UK) Consortium | Jacquelyn Wynn, Mairead Hyland, The Lighthouse Lab in Alderley Park and Alex Alderton, Roberto Amato, Jeffrey Barrett, Sonia Goncalves, Ewan Harrison, David K. Jackson, Ian Johnston, Dominic Kwiatkowski, Cordelia Langford, John Sillitoe on behalf of the Wellcome Sanger Institute COVID-19 Surveillance Team |
| EPI_ISL_3224433, EPI_ISL_3224434, EPI_ISL_3224435 | Lighthouse Lab in Milton Keynes | Wellcome Sanger Institute for the COVID-19 Genomics UK (COG-UK) Consortium | The Lighthouse Lab in Milton Keynes and Alex Alderton, Roberto Amato, Jeffrey Barrett, Sonia Goncalves, Ewan Harrison, David K. Jackson, Ian Johnston, Dominic Kwiatkowski, Cordelia Langford, John Sillitoe on behalf of the Wellcome Sanger Institute COVID-19 Surveillance Team |
| EPI_ISL_3224436 | Lighthouse Lab in Alderley Park | Wellcome Sanger Institute for the COVID-19 Genomics UK (COG-UK) Consortium | Jacquelyn Wynn, Mairead Hyland, The Lighthouse Lab in Alderley Park and Alex Alderton, Roberto Amato, Jeffrey Barrett, Sonia Goncalves, Ewan Harrison, David K. Jackson, Ian Johnston, Dominic Kwiatkowski, Cordelia Langford, John Sillitoe on behalf of the Wellcome Sanger Institute COVID-19 Surveillance Team |
| EPI_ISL_3224437 | Lighthouse Lab in Milton Keynes | Wellcome Sanger Institute for the COVID-19 Genomics UK (COG-UK) Consortium | The Lighthouse Lab in Milton Keynes and Alex Alderton, Roberto Amato, Jeffrey Barrett, Sonia Goncalves, Ewan Harrison, David K. Jackson, Ian Johnston, Dominic Kwiatkowski, Cordelia Langford, John Sillitoe on behalf of the Wellcome Sanger Institute COVID-19 Surveillance Team |
| EPI_ISL_3224438 | Lighthouse Lab in Glasgow | Wellcome Sanger Institute for the COVID-19 Genomics UK (COG-UK) Consortium | Harper VanSteenhouse, Yumi Kasai, David Gray, Carol Clugston, Anna Dominiczak and Alex Alderton, Roberto Amato, Jeffrey Barrett, Sonia Goncalves, Ewan Harrison, David K. Jackson, Ian Johnston, Dominic Kwiatkowski, Cordelia Langford, John Sillitoe on behalf of the Wellcome Sanger Institute COVID-19 Surveillance Team |
| EPI_ISL_3224439 | Lighthouse Lab in Alderley Park | Wellcome Sanger Institute for the COVID-19 Genomics UK (COG-UK) Consortium | Jacquelyn Wynn, Mairead Hyland, The Lighthouse Lab in Alderley Park and Alex Alderton, Roberto Amato, Jeffrey Barrett, Sonia Goncalves, Ewan Harrison, David K. Jackson, Ian Johnston, Dominic Kwiatkowski, Cordelia Langford, John Sillitoe on behalf of the Wellcome Sanger Institute COVID-19 Surveillance Team |
| EPI_ISL_3224440, EPI_ISL_3224441, EPI_ISL_3224442, EPI_ISL_3224443 | Lighthouse Lab in Milton Keynes | Wellcome Sanger Institute for the COVID-19 Genomics UK (COG-UK) Consortium | The Lighthouse Lab in Milton Keynes and Alex Alderton, Roberto Amato, Jeffrey Barrett, Sonia Goncalves, Ewan Harrison, David K. Jackson, Ian Johnston, Dominic Kwiatkowski, Cordelia Langford, John Sillitoe on behalf of the Wellcome Sanger Institute COVID-19 Surveillance Team |
| EPI_ISL_3224444 | Lighthouse Lab in Glasgow | Wellcome Sanger Institute for the COVID-19 Genomics UK (COG-UK) Consortium | Harper VanSteenhouse, Yumi Kasai, David Gray, Carol Clugston, Anna Dominiczak and Alex Alderton, Roberto Amato, Jeffrey Barrett, Sonia Goncalves, Ewan Harrison, David K. Jackson, Ian Johnston, Dominic Kwiatkowski, Cordelia Langford, John Sillitoe on behalf of the Wellcome Sanger Institute COVID-19 Surveillance Team |
| EPI_ISL_3224445, EPI_ISL_3224446, EPI_ISL_3224447, EPI_ISL_3224448, EPI_ISL_3224449 | Lighthouse Lab in Milton Keynes | Wellcome Sanger Institute for the COVID-19 Genomics UK (COG-UK) Consortium | The Lighthouse Lab in Milton Keynes and Alex Alderton, Roberto Amato, Jeffrey Barrett, Sonia Goncalves, Ewan Harrison, David K. Jackson, Ian Johnston, Dominic Kwiatkowski, Cordelia Langford, John Sillitoe on behalf of the Wellcome Sanger Institute COVID-19 Surveillance Team |
| EPI_ISL_3224450, EPI_ISL_3224451 | Lighthouse Lab in Alderley Park | Wellcome Sanger Institute for the COVID-19 Genomics UK (COG-UK) Consortium | Jacquelyn Wynn, Mairead Hyland, The Lighthouse Lab in Alderley Park and Alex Alderton, Roberto Amato, Jeffrey Barrett, Sonia Goncalves, Ewan Harrison, David K. Jackson, Ian Johnston, Dominic Kwiatkowski, Cordelia Langford, John Sillitoe on behalf of the Wellcome Sanger Institute COVID-19 Surveillance Team |
| EPI_ISL_3224452 | Lighthouse Lab in Glasgow | Wellcome Sanger Institute for the COVID-19 Genomics UK (COG-UK) Consortium | Harper VanSteenhouse, Yumi Kasai, David Gray, Carol Clugston, Anna Dominiczak and Alex Alderton, Roberto Amato, Jeffrey Barrett, Sonia Goncalves, Ewan Harrison, David K. Jackson, Ian Johnston, Dominic Kwiatkowski, Cordelia Langford, John Sillitoe on behalf of the Wellcome Sanger Institute COVID-19 Surveillance Team |
| EPI_ISL_3224453, EPI_ISL_3224454 | Lighthouse Lab in Milton Keynes | Wellcome Sanger Institute for the COVID-19 Genomics UK (COG-UK) Consortium | The Lighthouse Lab in Milton Keynes and Alex Alderton, Roberto Amato, Jeffrey Barrett, Sonia Goncalves, Ewan Harrison, David K. Jackson, Ian Johnston, Dominic Kwiatkowski, Cordelia Langford, John Sillitoe on behalf of the Wellcome Sanger Institute COVID-19 Surveillance Team |
| EPI_ISL_3224455 | Lighthouse Lab in Alderley Park | Wellcome Sanger Institute for the COVID-19 Genomics UK (COG-UK) Consortium | Jacquelyn Wynn, Mairead Hyland, The Lighthouse Lab in Alderley Park and Alex Alderton, Roberto Amato, Jeffrey Barrett, Sonia Goncalves, Ewan Harrison, David K. Jackson, Ian Johnston, Dominic Kwiatkowski, Cordelia Langford, John Sillitoe on behalf of the Wellcome Sanger Institute COVID-19 Surveillance Team |
| EPI_ISL_3224456 | Lighthouse Lab in Glasgow | Wellcome Sanger Institute for the COVID-19 Genomics UK (COG-UK) Consortium | Harper VanSteenhouse, Yumi Kasai, David Gray, Carol Clugston, Anna Dominiczak and Alex Alderton, Roberto Amato, Jeffrey Barrett, Sonia Goncalves, Ewan Harrison, David K. Jackson, Ian Johnston, Dominic Kwiatkowski, Cordelia Langford, John Sillitoe on behalf of the Wellcome Sanger Institute COVID-19 Surveillance Team |
| EPI_ISL_3224457 | Lighthouse Lab in Milton Keynes | Wellcome Sanger Institute for the COVID-19 Genomics UK (COG-UK) Consortium | The Lighthouse Lab in Milton Keynes and Alex Alderton, Roberto Amato, Jeffrey Barrett, Sonia Goncalves, Ewan Harrison, David K. Jackson, Ian Johnston, Dominic Kwiatkowski, Cordelia Langford, John Sillitoe on behalf of the Wellcome Sanger Institute COVID-19 Surveillance Team |
| EPI_ISL_3224458, EPI_ISL_3224459 | Lighthouse Lab in Glasgow | Wellcome Sanger Institute for the COVID-19 Genomics UK (COG-UK) Consortium | Harper VanSteenhouse, Yumi Kasai, David Gray, Carol Clugston, Anna Dominiczak and Alex Alderton, Roberto Amato, Jeffrey Barrett, Sonia Goncalves, Ewan Harrison, David K. Jackson, Ian Johnston, Dominic Kwiatkowski, Cordelia Langford, John Sillitoe on behalf of the Wellcome Sanger Institute COVID-19 Surveillance Team |
| EPI_ISL_3224460, EPI_ISL_3224461, EPI_ISL_3224462, | Lighthouse Lab in Milton Keynes | Wellcome Sanger Institute for the COVID-19 Genomics UK (COG-UK) Consortium | The Lighthouse Lab in Milton Keynes and Alex Alderton, Roberto Amato, Jeffrey Barrett, Sonia Goncalves, Ewan Harrison, David K. Jackson, Ian Johnston, Dominic Kwiatkowski, Cordelia Langford, John Sillitoe on behalf of the Wellcome Sanger Institute COVID-19 Surveillance Team |

| | | | |
|---|---|---|---|
| EPI_ISL_3224463, EPI_ISL_3224464 | | | |
| EPI_ISL_3224465 | Lighthouse Lab in Glasgow | Wellcome Sanger Institute for the COVID-19 Genomics UK (COG-UK) Consortium | Harper VanSteenhouse, Yumi Kasai, David Gray, Carol Clugston, Anna Dominiczak and Alex Alderton, Roberto Amato, Jeffrey Barrett, Sonia Goncalves, Ewan Harrison, David K. Jackson, Ian Johnston, Dominic Kwiatkowski, Cordelia Langford, John Sillitoe on behalf of the Wellcome Sanger Institute COVID-19 Surveillance Team |
| EPI_ISL_3224466, EPI_ISL_3224467 | Lighthouse Lab in Milton Keynes | Wellcome Sanger Institute for the COVID-19 Genomics UK (COG-UK) Consortium | The Lighthouse Lab in Milton Keynes and Alex Alderton, Roberto Amato, Jeffrey Barrett, Sonia Goncalves, Ewan Harrison, David K. Jackson, Ian Johnston, Dominic Kwiatkowski, Cordelia Langford, John Sillitoe on behalf of the Wellcome Sanger Institute COVID-19 Surveillance Team |
| EPI_ISL_3224468, EPI_ISL_3224469 | Lighthouse Lab in Alderley Park | Wellcome Sanger Institute for the COVID-19 Genomics UK (COG-UK) Consortium | Jacquelyn Wynn, Mairead Hyland, The Lighthouse Lab in Alderley Park and Alex Alderton, Roberto Amato, Jeffrey Barrett, Sonia Goncalves, Ewan Harrison, David K. Jackson, Ian Johnston, Dominic Kwiatkowski, Cordelia Langford, John Sillitoe on behalf of the Wellcome Sanger Institute COVID-19 Surveillance Team |
| EPI_ISL_3224470, EPI_ISL_3224471 | Lighthouse Lab in Milton Keynes | Wellcome Sanger Institute for the COVID-19 Genomics UK (COG-UK) Consortium | The Lighthouse Lab in Milton Keynes and Alex Alderton, Roberto Amato, Jeffrey Barrett, Sonia Goncalves, Ewan Harrison, David K. Jackson, Ian Johnston, Dominic Kwiatkowski, Cordelia Langford, John Sillitoe on behalf of the Wellcome Sanger Institute COVID-19 Surveillance Team |
| EPI_ISL_3224472 | Lighthouse Lab in Alderley Park | Wellcome Sanger Institute for the COVID-19 Genomics UK (COG-UK) Consortium | Jacquelyn Wynn, Mairead Hyland, The Lighthouse Lab in Alderley Park and Alex Alderton, Roberto Amato, Jeffrey Barrett, Sonia Goncalves, Ewan Harrison, David K. Jackson, Ian Johnston, Dominic Kwiatkowski, Cordelia Langford, John Sillitoe on behalf of the Wellcome Sanger Institute COVID-19 Surveillance Team |
| EPI_ISL_3224473 | Lighthouse Lab in Glasgow | Wellcome Sanger Institute for the COVID-19 Genomics UK (COG-UK) Consortium | Harper VanSteenhouse, Yumi Kasai, David Gray, Carol Clugston, Anna Dominiczak and Alex Alderton, Roberto Amato, Jeffrey Barrett, Sonia Goncalves, Ewan Harrison, David K. Jackson, Ian Johnston, Dominic Kwiatkowski, Cordelia Langford, John Sillitoe on behalf of the Wellcome Sanger Institute COVID-19 Surveillance Team |
| EPI_ISL_3224474 | Lighthouse Lab in Milton Keynes | Wellcome Sanger Institute for the COVID-19 Genomics UK (COG-UK) Consortium | The Lighthouse Lab in Milton Keynes and Alex Alderton, Roberto Amato, Jeffrey Barrett, Sonia Goncalves, Ewan Harrison, David K. Jackson, Ian Johnston, Dominic Kwiatkowski, Cordelia Langford, John Sillitoe on behalf of the Wellcome Sanger Institute COVID-19 Surveillance Team |
| EPI_ISL_3224475, EPI_ISL_3224476 | Lighthouse Lab in Alderley Park | Wellcome Sanger Institute for the COVID-19 Genomics UK (COG-UK) Consortium | Jacquelyn Wynn, Mairead Hyland, The Lighthouse Lab in Alderley Park and Alex Alderton, Roberto Amato, Jeffrey Barrett, Sonia Goncalves, Ewan Harrison, David K. Jackson, Ian Johnston, Dominic Kwiatkowski, Cordelia Langford, John Sillitoe on behalf of the Wellcome Sanger Institute COVID-19 Surveillance Team |
| EPI_ISL_3224477, EPI_ISL_3224478 | Lighthouse Lab in Milton Keynes | Wellcome Sanger Institute for the COVID-19 Genomics UK (COG-UK) Consortium | The Lighthouse Lab in Milton Keynes and Alex Alderton, Roberto Amato, Jeffrey Barrett, Sonia Goncalves, Ewan Harrison, David K. Jackson, Ian Johnston, Dominic Kwiatkowski, Cordelia Langford, John Sillitoe on behalf of the Wellcome Sanger Institute COVID-19 Surveillance Team |
| EPI_ISL_3224479 | Lighthouse Lab in Glasgow | Wellcome Sanger Institute for the COVID-19 Genomics UK (COG-UK) Consortium | Harper VanSteenhouse, Yumi Kasai, David Gray, Carol Clugston, Anna Dominiczak and Alex Alderton, Roberto Amato, Jeffrey Barrett, Sonia Goncalves, Ewan Harrison, David K. Jackson, Ian Johnston, Dominic Kwiatkowski, Cordelia Langford, John Sillitoe on behalf of the Wellcome Sanger Institute COVID-19 Surveillance Team |
| EPI_ISL_3224480 | Lighthouse Lab in Alderley Park | Wellcome Sanger Institute for the COVID-19 Genomics UK (COG-UK) Consortium | Jacquelyn Wynn, Mairead Hyland, The Lighthouse Lab in Alderley Park and Alex Alderton, Roberto Amato, Jeffrey Barrett, Sonia Goncalves, Ewan Harrison, David K. Jackson, Ian Johnston, Dominic Kwiatkowski, Cordelia Langford, John Sillitoe on behalf of the Wellcome Sanger Institute COVID-19 Surveillance Team |
| EPI_ISL_3224481, EPI_ISL_3224482 | Lighthouse Lab in Milton Keynes | Wellcome Sanger Institute for the COVID-19 Genomics UK (COG-UK) Consortium | The Lighthouse Lab in Milton Keynes and Alex Alderton, Roberto Amato, Jeffrey Barrett, Sonia Goncalves, Ewan Harrison, David K. Jackson, Ian Johnston, Dominic Kwiatkowski, Cordelia Langford, John Sillitoe on behalf of the Wellcome Sanger Institute COVID-19 Surveillance Team |
| EPI_ISL_3224483, EPI_ISL_3224484 | Lighthouse Lab in Glasgow | Wellcome Sanger Institute for the COVID-19 Genomics UK (COG-UK) Consortium | Harper VanSteenhouse, Yumi Kasai, David Gray, Carol Clugston, Anna Dominiczak and Alex Alderton, Roberto Amato, Jeffrey Barrett, Sonia Goncalves, Ewan Harrison, David K. Jackson, Ian Johnston, Dominic Kwiatkowski, Cordelia Langford, John Sillitoe on behalf of the Wellcome Sanger Institute COVID-19 Surveillance Team |
| EPI_ISL_3224485, EPI_ISL_3224486, EPI_ISL_3224487, EPI_ISL_3224488, EPI_ISL_3224489, EPI_ISL_3224490, EPI_ISL_3224491, EPI_ISL_3224492, EPI_ISL_3224493, EPI_ISL_3224494 | Lighthouse Lab in Milton Keynes | Wellcome Sanger Institute for the COVID-19 Genomics UK (COG-UK) Consortium | The Lighthouse Lab in Milton Keynes and Alex Alderton, Roberto Amato, Jeffrey Barrett, Sonia Goncalves, Ewan Harrison, David K. Jackson, Ian Johnston, Dominic Kwiatkowski, Cordelia Langford, John Sillitoe on behalf of the Wellcome Sanger Institute COVID-19 Surveillance Team |
| EPI_ISL_3224495, EPI_ISL_3224496 | Lighthouse Lab in Alderley Park | Wellcome Sanger Institute for the COVID-19 Genomics UK (COG-UK) Consortium | Jacquelyn Wynn, Mairead Hyland, The Lighthouse Lab in Alderley Park and Alex Alderton, Roberto Amato, Jeffrey Barrett, Sonia Goncalves, Ewan Harrison, David K. Jackson, Ian Johnston, Dominic Kwiatkowski, Cordelia Langford, John Sillitoe on behalf of the Wellcome Sanger Institute COVID-19 Surveillance Team |
| EPI_ISL_3224497, EPI_ISL_3224498 | Lighthouse Lab in Glasgow | Wellcome Sanger Institute for the COVID-19 Genomics UK (COG-UK) Consortium | Harper VanSteenhouse, Yumi Kasai, David Gray, Carol Clugston, Anna Dominiczak and Alex Alderton, Roberto Amato, Jeffrey Barrett, Sonia Goncalves, Ewan Harrison, David K. Jackson, Ian Johnston, Dominic Kwiatkowski, Cordelia Langford, John Sillitoe on behalf of the Wellcome Sanger Institute COVID-19 Surveillance Team |
| EPI_ISL_3224499 | Lighthouse Lab in Alderley Park | Wellcome Sanger Institute for the COVID-19 Genomics UK (COG-UK) Consortium | Jacquelyn Wynn, Mairead Hyland, The Lighthouse Lab in Alderley Park and Alex Alderton, Roberto Amato, Jeffrey Barrett, Sonia Goncalves, Ewan Harrison, David K. Jackson, Ian Johnston, Dominic Kwiatkowski, Cordelia Langford, John Sillitoe on behalf of the Wellcome Sanger Institute COVID-19 Surveillance Team |
| EPI_ISL_3224500, EPI_ISL_3224501 | Lighthouse Lab in Glasgow | Wellcome Sanger Institute for the COVID-19 Genomics UK (COG-UK) Consortium | Harper VanSteenhouse, Yumi Kasai, David Gray, Carol Clugston, Anna Dominiczak and Alex Alderton, Roberto Amato, Jeffrey Barrett, Sonia Goncalves, Ewan Harrison, David K. Jackson, Ian Johnston, Dominic Kwiatkowski, Cordelia Langford, John Sillitoe on behalf of the Wellcome Sanger Institute COVID-19 Surveillance Team |
| EPI_ISL_3224502, EPI_ISL_3224503, EPI_ISL_3224504 | Lighthouse Lab in Milton Keynes | Wellcome Sanger Institute for the COVID-19 Genomics UK (COG-UK) Consortium | The Lighthouse Lab in Milton Keynes and Alex Alderton, Roberto Amato, Jeffrey Barrett, Sonia Goncalves, Ewan Harrison, David K. Jackson, Ian Johnston, Dominic Kwiatkowski, Cordelia Langford, John Sillitoe on behalf of the Wellcome Sanger Institute COVID-19 Surveillance Team |
| EPI_ISL_3224505 | Lighthouse Lab in Glasgow | Wellcome Sanger Institute for the COVID-19 Genomics UK (COG-UK) Consortium | Harper VanSteenhouse, Yumi Kasai, David Gray, Carol Clugston, Anna Dominiczak and Alex Alderton, Roberto Amato, Jeffrey Barrett, Sonia Goncalves, Ewan Harrison, David K. Jackson, Ian Johnston, Dominic Kwiatkowski, Cordelia Langford, John Sillitoe on behalf of the Wellcome Sanger Institute COVID-19 Surveillance Team |
| EPI_ISL_3224506 | Lighthouse Lab in Milton Keynes | Wellcome Sanger Institute for the COVID-19 Genomics UK (COG-UK) Consortium | The Lighthouse Lab in Milton Keynes and Alex Alderton, Roberto Amato, Jeffrey Barrett, Sonia Goncalves, Ewan Harrison, David K. Jackson, Ian Johnston, Dominic Kwiatkowski, Cordelia Langford, John Sillitoe on behalf of the Wellcome Sanger Institute COVID-19 Surveillance Team |
| EPI_ISL_3224507, EPI_ISL_3224508 | Lighthouse Lab in Alderley Park | Wellcome Sanger Institute for the COVID-19 Genomics UK (COG-UK) Consortium | Jacquelyn Wynn, Mairead Hyland, The Lighthouse Lab in Alderley Park and Alex Alderton, Roberto Amato, Jeffrey Barrett, Sonia Goncalves, Ewan Harrison, David K. Jackson, Ian Johnston, Dominic Kwiatkowski, Cordelia Langford, John Sillitoe on behalf of the Wellcome Sanger Institute COVID-19 Surveillance Team |
| EPI_ISL_3224509 | Lighthouse Lab in Milton Keynes | Wellcome Sanger Institute for the COVID-19 Genomics UK (COG-UK) Consortium | The Lighthouse Lab in Milton Keynes and Alex Alderton, Roberto Amato, Jeffrey Barrett, Sonia Goncalves, Ewan Harrison, David K. Jackson, Ian Johnston, Dominic Kwiatkowski, Cordelia Langford, John Sillitoe on behalf of the Wellcome Sanger Institute COVID-19 Surveillance Team |
| EPI_ISL_3224510, EPI_ISL_3224511 | Lighthouse Lab in Alderley Park | Wellcome Sanger Institute for the COVID-19 Genomics UK (COG-UK) Consortium | Jacquelyn Wynn, Mairead Hyland, The Lighthouse Lab in Alderley Park and Alex Alderton, Roberto Amato, Jeffrey Barrett, Sonia Goncalves, Ewan Harrison, David K. Jackson, Ian Johnston, Dominic Kwiatkowski, Cordelia Langford, John Sillitoe on behalf of the Wellcome Sanger Institute COVID-19 Surveillance Team |
| EPI_ISL_3224512 | Lighthouse Lab in Glasgow | Wellcome Sanger Institute for the COVID-19 Genomics UK (COG-UK) Consortium | Harper VanSteenhouse, Yumi Kasai, David Gray, Carol Clugston, Anna Dominiczak and Alex Alderton, Roberto Amato, Jeffrey Barrett, Sonia Goncalves, Ewan Harrison, David K. Jackson, Ian Johnston, Dominic Kwiatkowski, Cordelia Langford, John Sillitoe on behalf of the Wellcome Sanger Institute COVID-19 Surveillance Team |
| EPI_ISL_3224513 | Lighthouse Lab in Alderley Park | Wellcome Sanger Institute for the COVID-19 Genomics UK (COG-UK) Consortium | Jacquelyn Wynn, Mairead Hyland, The Lighthouse Lab in Alderley Park and Alex Alderton, Roberto Amato, Jeffrey Barrett, Sonia Goncalves, Ewan Harrison, David K. Jackson, Ian Johnston, Dominic Kwiatkowski, Cordelia Langford, John Sillitoe on behalf of the Wellcome Sanger Institute COVID-19 Surveillance Team |
| EPI_ISL_3224514, EPI_ISL_3224515 | Lighthouse Lab in Glasgow | Wellcome Sanger Institute for the COVID-19 Genomics UK (COG-UK) Consortium | Harper VanSteenhouse, Yumi Kasai, David Gray, Carol Clugston, Anna Dominiczak and Alex Alderton, Roberto Amato, Jeffrey Barrett, Sonia Goncalves, Ewan Harrison, David K. Jackson, Ian Johnston, Dominic Kwiatkowski, Cordelia Langford, John Sillitoe on behalf of the Wellcome Sanger Institute COVID-19 Surveillance Team |
| EPI_ISL_3224516 | Lighthouse Lab in Milton Keynes | Wellcome Sanger Institute for the COVID-19 Genomics UK (COG-UK) Consortium | The Lighthouse Lab in Milton Keynes and Alex Alderton, Roberto Amato, Jeffrey Barrett, Sonia Goncalves, Ewan Harrison, David K. Jackson, Ian Johnston, Dominic Kwiatkowski, Cordelia Langford, John Sillitoe on behalf of the Wellcome Sanger Institute COVID-19 Surveillance Team |
| EPI_ISL_3224517 | Lighthouse Lab in Alderley Park | Wellcome Sanger Institute for the COVID-19 Genomics UK (COG-UK) Consortium | Jacquelyn Wynn, Mairead Hyland, The Lighthouse Lab in Alderley Park and Alex Alderton, Roberto Amato, Jeffrey Barrett, Sonia Goncalves, Ewan Harrison, David K. Jackson, Ian Johnston, Dominic Kwiatkowski, Cordelia Langford, John Sillitoe on behalf of the Wellcome Sanger Institute COVID-19 Surveillance Team |
| EPI_ISL_3224518 | Lighthouse Lab in Milton Keynes | Wellcome Sanger Institute for the COVID-19 Genomics UK (COG-UK) Consortium | The Lighthouse Lab in Milton Keynes and Alex Alderton, Roberto Amato, Jeffrey Barrett, Sonia Goncalves, Ewan Harrison, David K. Jackson, Ian Johnston, Dominic Kwiatkowski, Cordelia Langford, John Sillitoe on behalf of the Wellcome Sanger Institute COVID-19 Surveillance Team |
| EPI_ISL_3224519 | Lighthouse Lab in Alderley Park | Wellcome Sanger Institute for the COVID-19 Genomics UK (COG-UK) Consortium | Jacquelyn Wynn, Mairead Hyland, The Lighthouse Lab in Alderley Park and Alex Alderton, Roberto Amato, Jeffrey Barrett, Sonia Goncalves, Ewan Harrison, David K. Jackson, Ian Johnston, Dominic Kwiatkowski, Cordelia Langford, John Sillitoe on behalf of the Wellcome Sanger Institute COVID-19 Surveillance Team |
| EPI_ISL_3224520, EPI_ISL_3224521, EPI_ISL_3224522, EPI_ISL_3224523 | Lighthouse Lab in Milton Keynes | Wellcome Sanger Institute for the COVID-19 Genomics UK (COG-UK) Consortium | The Lighthouse Lab in Milton Keynes and Alex Alderton, Roberto Amato, Jeffrey Barrett, Sonia Goncalves, Ewan Harrison, David K. Jackson, Ian Johnston, Dominic Kwiatkowski, Cordelia Langford, John Sillitoe on behalf of the Wellcome Sanger Institute COVID-19 Surveillance Team |
| EPI_ISL_3224524 | Lighthouse Lab in Alderley Park | Wellcome Sanger Institute for the COVID-19 Genomics UK (COG-UK) Consortium | Jacquelyn Wynn, Mairead Hyland, The Lighthouse Lab in Alderley Park and Alex Alderton, Roberto Amato, Jeffrey Barrett, Sonia Goncalves, Ewan Harrison, David K. Jackson, Ian Johnston, Dominic Kwiatkowski, Cordelia Langford, John Sillitoe on behalf of the Wellcome Sanger Institute COVID-19 Surveillance Team |
| EPI_ISL_3224525 | Lighthouse Lab in Milton Keynes | Wellcome Sanger Institute for the COVID-19 Genomics UK (COG-UK) Consortium | The Lighthouse Lab in Milton Keynes and Alex Alderton, Roberto Amato, Jeffrey Barrett, Sonia Goncalves, Ewan Harrison, David K. Jackson, Ian Johnston, Dominic Kwiatkowski, Cordelia Langford, John Sillitoe on behalf of the Wellcome Sanger Institute COVID-19 Surveillance Team |
| EPI_ISL_3224526 | Lighthouse Lab in Alderley Park | Wellcome Sanger Institute for the COVID-19 Genomics UK (COG-UK) Consortium | Jacquelyn Wynn, Mairead Hyland, The Lighthouse Lab in Alderley Park and Alex Alderton, Roberto Amato, Jeffrey Barrett, Sonia Goncalves, Ewan Harrison, David K. Jackson, Ian Johnston, Dominic Kwiatkowski, Cordelia Langford, John Sillitoe on behalf of the Wellcome Sanger Institute COVID-19 Surveillance Team |
| EPI_ISL_3224527 | Lighthouse Lab in Glasgow | Wellcome Sanger Institute for the COVID-19 Genomics UK (COG-UK) Consortium | Harper VanSteenhouse, Yumi Kasai, David Gray, Carol Clugston, Anna Dominiczak and Alex Alderton, Roberto Amato, Jeffrey Barrett, Sonia Goncalves, Ewan Harrison, David K. Jackson, Ian Johnston, Dominic Kwiatkowski, Cordelia Langford, John Sillitoe on behalf of the Wellcome Sanger Institute COVID-19 Surveillance Team |
| EPI_ISL_3224528 | Lighthouse Lab in Alderley Park | Wellcome Sanger Institute for the COVID-19 Genomics UK (COG-UK) Consortium | Jacquelyn Wynn, Mairead Hyland, The Lighthouse Lab in Alderley Park and Alex Alderton, Roberto Amato, Jeffrey Barrett, Sonia Goncalves, Ewan Harrison, David K. Jackson, Ian Johnston, Dominic Kwiatkowski, Cordelia Langford, John Sillitoe on behalf of the Wellcome Sanger Institute COVID-19 Surveillance Team |
| EPI_ISL_3224529 | Lighthouse Lab in Milton Keynes | Wellcome Sanger Institute for the COVID-19 Genomics UK (COG-UK) Consortium | The Lighthouse Lab in Milton Keynes and Alex Alderton, Roberto Amato, Jeffrey Barrett, Sonia Goncalves, Ewan Harrison, David K. Jackson, Ian Johnston, Dominic Kwiatkowski, Cordelia Langford, John Sillitoe on behalf of the Wellcome Sanger Institute COVID-19 Surveillance Team |
| EPI_ISL_3224530 | Lighthouse Lab in Glasgow | Wellcome Sanger Institute for the COVID-19 Genomics UK (COG-UK) Consortium | Harper VanSteenhouse, Yumi Kasai, David Gray, Carol Clugston, Anna Dominiczak and Alex Alderton, Roberto Amato, Jeffrey Barrett, Sonia Goncalves, Ewan Harrison, David K. Jackson, Ian Johnston, Dominic Kwiatkowski, Cordelia Langford, John Sillitoe on behalf of the Wellcome Sanger Institute COVID-19 Surveillance Team |
| EPI_ISL_3224531, EPI_ISL_3224532 | Lighthouse Lab in Alderley Park | Wellcome Sanger Institute for the COVID-19 Genomics UK (COG-UK) Consortium | Jacquelyn Wynn, Mairead Hyland, The Lighthouse Lab in Alderley Park and Alex Alderton, Roberto Amato, Jeffrey Barrett, Sonia Goncalves, Ewan Harrison, David K. Jackson, Ian Johnston, Dominic Kwiatkowski, Cordelia Langford, John Sillitoe on behalf of the Wellcome Sanger Institute COVID-19 Surveillance Team |

| | | | |
|---|---|---|---|
| EPI_ISL_3224533, EPI_ISL_3224534, EPI_ISL_3224535 | Lighthouse Lab in Glasgow | Wellcome Sanger Institute for the COVID-19 Genomics UK (COG-UK) Consortium | Harper VanSteenhouse, Yumi Kasai, David Gray, Carol Clugston, Anna Dominiczak and Alex Alderton, Roberto Amato, Jeffrey Barrett, Sonia Goncalves, Ewan Harrison, David K. Jackson, Ian Johnston, Dominic Kwiatkowski, Cordelia Langford, John Sillitoe on behalf of the Wellcome Sanger Institute COVID-19 Surveillance Team |
| EPI_ISL_3224536 | Lighthouse Lab in Milton Keynes | Wellcome Sanger Institute for the COVID-19 Genomics UK (COG-UK) Consortium | The Lighthouse Lab in Milton Keynes and Alex Alderton, Roberto Amato, Jeffrey Barrett, Sonia Goncalves, Ewan Harrison, David K. Jackson, Ian Johnston, Dominic Kwiatkowski, Cordelia Langford, John Sillitoe on behalf of the Wellcome Sanger Institute COVID-19 Surveillance Team |
| EPI_ISL_3224537 | Lighthouse Lab in Alderley Park | Wellcome Sanger Institute for the COVID-19 Genomics UK (COG-UK) Consortium | Jacquelyn Wynn, Mairead Hyland, The Lighthouse Lab in Alderley Park and Alex Alderton, Roberto Amato, Jeffrey Barrett, Sonia Goncalves, Ewan Harrison, David K. Jackson, Ian Johnston, Dominic Kwiatkowski, Cordelia Langford, John Sillitoe on behalf of the Wellcome Sanger Institute COVID-19 Surveillance Team |
| EPI_ISL_3224538 | Lighthouse Lab in Glasgow | Wellcome Sanger Institute for the COVID-19 Genomics UK (COG-UK) Consortium | Harper VanSteenhouse, Yumi Kasai, David Gray, Carol Clugston, Anna Dominiczak and Alex Alderton, Roberto Amato, Jeffrey Barrett, Sonia Goncalves, Ewan Harrison, David K. Jackson, Ian Johnston, Dominic Kwiatkowski, Cordelia Langford, John Sillitoe on behalf of the Wellcome Sanger Institute COVID-19 Surveillance Team |
| EPI_ISL_3224539, EPI_ISL_3224540 | Lighthouse Lab in Alderley Park | Wellcome Sanger Institute for the COVID-19 Genomics UK (COG-UK) Consortium | Jacquelyn Wynn, Mairead Hyland, The Lighthouse Lab in Alderley Park and Alex Alderton, Roberto Amato, Jeffrey Barrett, Sonia Goncalves, Ewan Harrison, David K. Jackson, Ian Johnston, Dominic Kwiatkowski, Cordelia Langford, John Sillitoe on behalf of the Wellcome Sanger Institute COVID-19 Surveillance Team |
| EPI_ISL_3224541, EPI_ISL_3224542 | Lighthouse Lab in Milton Keynes | Wellcome Sanger Institute for the COVID-19 Genomics UK (COG-UK) Consortium | The Lighthouse Lab in Milton Keynes and Alex Alderton, Roberto Amato, Jeffrey Barrett, Sonia Goncalves, Ewan Harrison, David K. Jackson, Ian Johnston, Dominic Kwiatkowski, Cordelia Langford, John Sillitoe on behalf of the Wellcome Sanger Institute COVID-19 Surveillance Team |
| EPI_ISL_3224543 | Lighthouse Lab in Alderley Park | Wellcome Sanger Institute for the COVID-19 Genomics UK (COG-UK) Consortium | Jacquelyn Wynn, Mairead Hyland, The Lighthouse Lab in Alderley Park and Alex Alderton, Roberto Amato, Jeffrey Barrett, Sonia Goncalves, Ewan Harrison, David K. Jackson, Ian Johnston, Dominic Kwiatkowski, Cordelia Langford, John Sillitoe on behalf of the Wellcome Sanger Institute COVID-19 Surveillance Team |
| EPI_ISL_3224544 | Lighthouse Lab in Glasgow | Wellcome Sanger Institute for the COVID-19 Genomics UK (COG-UK) Consortium | Harper VanSteenhouse, Yumi Kasai, David Gray, Carol Clugston, Anna Dominiczak and Alex Alderton, Roberto Amato, Jeffrey Barrett, Sonia Goncalves, Ewan Harrison, David K. Jackson, Ian Johnston, Dominic Kwiatkowski, Cordelia Langford, John Sillitoe on behalf of the Wellcome Sanger Institute COVID-19 Surveillance Team |
| EPI_ISL_3224545 | Lighthouse Lab in Alderley Park | Wellcome Sanger Institute for the COVID-19 Genomics UK (COG-UK) Consortium | Jacquelyn Wynn, Mairead Hyland, The Lighthouse Lab in Alderley Park and Alex Alderton, Roberto Amato, Jeffrey Barrett, Sonia Goncalves, Ewan Harrison, David K. Jackson, Ian Johnston, Dominic Kwiatkowski, Cordelia Langford, John Sillitoe on behalf of the Wellcome Sanger Institute COVID-19 Surveillance Team |
| EPI_ISL_3224546, EPI_ISL_3224547 | Lighthouse Lab in Milton Keynes | Wellcome Sanger Institute for the COVID-19 Genomics UK (COG-UK) Consortium | The Lighthouse Lab in Milton Keynes and Alex Alderton, Roberto Amato, Jeffrey Barrett, Sonia Goncalves, Ewan Harrison, David K. Jackson, Ian Johnston, Dominic Kwiatkowski, Cordelia Langford, John Sillitoe on behalf of the Wellcome Sanger Institute COVID-19 Surveillance Team |
| EPI_ISL_3224548 | Lighthouse Lab in Glasgow | Wellcome Sanger Institute for the COVID-19 Genomics UK (COG-UK) Consortium | Harper VanSteenhouse, Yumi Kasai, David Gray, Carol Clugston, Anna Dominiczak and Alex Alderton, Roberto Amato, Jeffrey Barrett, Sonia Goncalves, Ewan Harrison, David K. Jackson, Ian Johnston, Dominic Kwiatkowski, Cordelia Langford, John Sillitoe on behalf of the Wellcome Sanger Institute COVID-19 Surveillance Team |
| EPI_ISL_3224549 | Lighthouse Lab in Alderley Park | Wellcome Sanger Institute for the COVID-19 Genomics UK (COG-UK) Consortium | Jacquelyn Wynn, Mairead Hyland, The Lighthouse Lab in Alderley Park and Alex Alderton, Roberto Amato, Jeffrey Barrett, Sonia Goncalves, Ewan Harrison, David K. Jackson, Ian Johnston, Dominic Kwiatkowski, Cordelia Langford, John Sillitoe on behalf of the Wellcome Sanger Institute COVID-19 Surveillance Team |
| EPI_ISL_3224550, EPI_ISL_3224551, EPI_ISL_3224552, EPI_ISL_3224553, EPI_ISL_3224554, EPI_ISL_3224555 | Lighthouse Lab in Milton Keynes | Wellcome Sanger Institute for the COVID-19 Genomics UK (COG-UK) Consortium | The Lighthouse Lab in Milton Keynes and Alex Alderton, Roberto Amato, Jeffrey Barrett, Sonia Goncalves, Ewan Harrison, David K. Jackson, Ian Johnston, Dominic Kwiatkowski, Cordelia Langford, John Sillitoe on behalf of the Wellcome Sanger Institute COVID-19 Surveillance Team |
| EPI_ISL_3224556, EPI_ISL_3224557 | Lighthouse Lab in Glasgow | Wellcome Sanger Institute for the COVID-19 Genomics UK (COG-UK) Consortium | Harper VanSteenhouse, Yumi Kasai, David Gray, Carol Clugston, Anna Dominiczak and Alex Alderton, Roberto Amato, Jeffrey Barrett, Sonia Goncalves, Ewan Harrison, David K. Jackson, Ian Johnston, Dominic Kwiatkowski, Cordelia Langford, John Sillitoe on behalf of the Wellcome Sanger Institute COVID-19 Surveillance Team |
| EPI_ISL_3224558, EPI_ISL_3224559, EPI_ISL_3224560 | Lighthouse Lab in Milton Keynes | Wellcome Sanger Institute for the COVID-19 Genomics UK (COG-UK) Consortium | The Lighthouse Lab in Milton Keynes and Alex Alderton, Roberto Amato, Jeffrey Barrett, Sonia Goncalves, Ewan Harrison, David K. Jackson, Ian Johnston, Dominic Kwiatkowski, Cordelia Langford, John Sillitoe on behalf of the Wellcome Sanger Institute COVID-19 Surveillance Team |
| EPI_ISL_3224561 | Lighthouse Lab in Alderley Park | Wellcome Sanger Institute for the COVID-19 Genomics UK (COG-UK) Consortium | Jacquelyn Wynn, Mairead Hyland, The Lighthouse Lab in Alderley Park and Alex Alderton, Roberto Amato, Jeffrey Barrett, Sonia Goncalves, Ewan Harrison, David K. Jackson, Ian Johnston, Dominic Kwiatkowski, Cordelia Langford, John Sillitoe on behalf of the Wellcome Sanger Institute COVID-19 Surveillance Team |
| EPI_ISL_3224562 | Lighthouse Lab in Milton Keynes | Wellcome Sanger Institute for the COVID-19 Genomics UK (COG-UK) Consortium | The Lighthouse Lab in Milton Keynes and Alex Alderton, Roberto Amato, Jeffrey Barrett, Sonia Goncalves, Ewan Harrison, David K. Jackson, Ian Johnston, Dominic Kwiatkowski, Cordelia Langford, John Sillitoe on behalf of the Wellcome Sanger Institute COVID-19 Surveillance Team |
| EPI_ISL_3224563 | Lighthouse Lab in Alderley Park | Wellcome Sanger Institute for the COVID-19 Genomics UK (COG-UK) Consortium | Jacquelyn Wynn, Mairead Hyland, The Lighthouse Lab in Alderley Park and Alex Alderton, Roberto Amato, Jeffrey Barrett, Sonia Goncalves, Ewan Harrison, David K. Jackson, Ian Johnston, Dominic Kwiatkowski, Cordelia Langford, John Sillitoe on behalf of the Wellcome Sanger Institute COVID-19 Surveillance Team |
| EPI_ISL_3224564 | Lighthouse Lab in Glasgow | Wellcome Sanger Institute for the COVID-19 Genomics UK (COG-UK) Consortium | Harper VanSteenhouse, Yumi Kasai, David Gray, Carol Clugston, Anna Dominiczak and Alex Alderton, Roberto Amato, Jeffrey Barrett, Sonia Goncalves, Ewan Harrison, David K. Jackson, Ian Johnston, Dominic Kwiatkowski, Cordelia Langford, John Sillitoe on behalf of the Wellcome Sanger Institute COVID-19 Surveillance Team |
| EPI_ISL_3224565, EPI_ISL_3224566 | Lighthouse Lab in Milton Keynes | Wellcome Sanger Institute for the COVID-19 Genomics UK (COG-UK) Consortium | The Lighthouse Lab in Milton Keynes and Alex Alderton, Roberto Amato, Jeffrey Barrett, Sonia Goncalves, Ewan Harrison, David K. Jackson, Ian Johnston, Dominic Kwiatkowski, Cordelia Langford, John Sillitoe on behalf of the Wellcome Sanger Institute COVID-19 Surveillance Team |
| EPI_ISL_3224567 | Lighthouse Lab in Glasgow | Wellcome Sanger Institute for the COVID-19 Genomics UK (COG-UK) Consortium | Harper VanSteenhouse, Yumi Kasai, David Gray, Carol Clugston, Anna Dominiczak and Alex Alderton, Roberto Amato, Jeffrey Barrett, Sonia Goncalves, Ewan Harrison, David K. Jackson, Ian Johnston, Dominic Kwiatkowski, Cordelia Langford, John Sillitoe on behalf of the Wellcome Sanger Institute COVID-19 Surveillance Team |
| EPI_ISL_3224568, EPI_ISL_3224569 | Lighthouse Lab in Alderley Park | Wellcome Sanger Institute for the COVID-19 Genomics UK (COG-UK) Consortium | Jacquelyn Wynn, Mairead Hyland, The Lighthouse Lab in Alderley Park and Alex Alderton, Roberto Amato, Jeffrey Barrett, Sonia Goncalves, Ewan Harrison, David K. Jackson, Ian Johnston, Dominic Kwiatkowski, Cordelia Langford, John Sillitoe on behalf of the Wellcome Sanger Institute COVID-19 Surveillance Team |
| EPI_ISL_3224570, EPI_ISL_3224571, EPI_ISL_3224572 | Lighthouse Lab in Milton Keynes | Wellcome Sanger Institute for the COVID-19 Genomics UK (COG-UK) Consortium | The Lighthouse Lab in Milton Keynes and Alex Alderton, Roberto Amato, Jeffrey Barrett, Sonia Goncalves, Ewan Harrison, David K. Jackson, Ian Johnston, Dominic Kwiatkowski, Cordelia Langford, John Sillitoe on behalf of the Wellcome Sanger Institute COVID-19 Surveillance Team |
| EPI_ISL_3224573 | Lighthouse Lab in Alderley Park | Wellcome Sanger Institute for the COVID-19 Genomics UK (COG-UK) Consortium | Jacquelyn Wynn, Mairead Hyland, The Lighthouse Lab in Alderley Park and Alex Alderton, Roberto Amato, Jeffrey Barrett, Sonia Goncalves, Ewan Harrison, David K. Jackson, Ian Johnston, Dominic Kwiatkowski, Cordelia Langford, John Sillitoe on behalf of the Wellcome Sanger Institute COVID-19 Surveillance Team |
| EPI_ISL_3224574 | Lighthouse Lab in Milton Keynes | Wellcome Sanger Institute for the COVID-19 Genomics UK (COG-UK) Consortium | The Lighthouse Lab in Milton Keynes and Alex Alderton, Roberto Amato, Jeffrey Barrett, Sonia Goncalves, Ewan Harrison, David K. Jackson, Ian Johnston, Dominic Kwiatkowski, Cordelia Langford, John Sillitoe on behalf of the Wellcome Sanger Institute COVID-19 Surveillance Team |
| EPI_ISL_3224575 | Lighthouse Lab in Alderley Park | Wellcome Sanger Institute for the COVID-19 Genomics UK (COG-UK) Consortium | Jacquelyn Wynn, Mairead Hyland, The Lighthouse Lab in Alderley Park and Alex Alderton, Roberto Amato, Jeffrey Barrett, Sonia Goncalves, Ewan Harrison, David K. Jackson, Ian Johnston, Dominic Kwiatkowski, Cordelia Langford, John Sillitoe on behalf of the Wellcome Sanger Institute COVID-19 Surveillance Team |
| EPI_ISL_3224576 | Lighthouse Lab in Milton Keynes | Wellcome Sanger Institute for the COVID-19 Genomics UK (COG-UK) Consortium | The Lighthouse Lab in Milton Keynes and Alex Alderton, Roberto Amato, Jeffrey Barrett, Sonia Goncalves, Ewan Harrison, David K. Jackson, Ian Johnston, Dominic Kwiatkowski, Cordelia Langford, John Sillitoe on behalf of the Wellcome Sanger Institute COVID-19 Surveillance Team |
| EPI_ISL_3224577 | Lighthouse Lab in Glasgow | Wellcome Sanger Institute for the COVID-19 Genomics UK (COG-UK) Consortium | Harper VanSteenhouse, Yumi Kasai, David Gray, Carol Clugston, Anna Dominiczak and Alex Alderton, Roberto Amato, Jeffrey Barrett, Sonia Goncalves, Ewan Harrison, David K. Jackson, Ian Johnston, Dominic Kwiatkowski, Cordelia Langford, John Sillitoe on behalf of the Wellcome Sanger Institute COVID-19 Surveillance Team |
| EPI_ISL_3224578, EPI_ISL_3224579 | Lighthouse Lab in Milton Keynes | Wellcome Sanger Institute for the COVID-19 Genomics UK (COG-UK) Consortium | The Lighthouse Lab in Milton Keynes and Alex Alderton, Roberto Amato, Jeffrey Barrett, Sonia Goncalves, Ewan Harrison, David K. Jackson, Ian Johnston, Dominic Kwiatkowski, Cordelia Langford, John Sillitoe on behalf of the Wellcome Sanger Institute COVID-19 Surveillance Team |
| EPI_ISL_3224580 | Lighthouse Lab in Glasgow | Wellcome Sanger Institute for the COVID-19 Genomics UK (COG-UK) Consortium | Harper VanSteenhouse, Yumi Kasai, David Gray, Carol Clugston, Anna Dominiczak and Alex Alderton, Roberto Amato, Jeffrey Barrett, Sonia Goncalves, Ewan Harrison, David K. Jackson, Ian Johnston, Dominic Kwiatkowski, Cordelia Langford, John Sillitoe on behalf of the Wellcome Sanger Institute COVID-19 Surveillance Team |
| EPI_ISL_3224581, EPI_ISL_3224582, EPI_ISL_3224583 | Lighthouse Lab in Milton Keynes | Wellcome Sanger Institute for the COVID-19 Genomics UK (COG-UK) Consortium | The Lighthouse Lab in Milton Keynes and Alex Alderton, Roberto Amato, Jeffrey Barrett, Sonia Goncalves, Ewan Harrison, David K. Jackson, Ian Johnston, Dominic Kwiatkowski, Cordelia Langford, John Sillitoe on behalf of the Wellcome Sanger Institute COVID-19 Surveillance Team |
| EPI_ISL_3224584 | Lighthouse Lab in Alderley Park | Wellcome Sanger Institute for the COVID-19 Genomics UK (COG-UK) Consortium | Jacquelyn Wynn, Mairead Hyland, The Lighthouse Lab in Alderley Park and Alex Alderton, Roberto Amato, Jeffrey Barrett, Sonia Goncalves, Ewan Harrison, David K. Jackson, Ian Johnston, Dominic Kwiatkowski, Cordelia Langford, John Sillitoe on behalf of the Wellcome Sanger Institute COVID-19 Surveillance Team |
| EPI_ISL_3224585, EPI_ISL_3224586, EPI_ISL_3224587, EPI_ISL_3224588, EPI_ISL_3224589, EPI_ISL_3224590, EPI_ISL_3224591, EPI_ISL_3224592, EPI_ISL_3224593, EPI_ISL_3224594 | Lighthouse Lab in Milton Keynes | Wellcome Sanger Institute for the COVID-19 Genomics UK (COG-UK) Consortium | The Lighthouse Lab in Milton Keynes and Alex Alderton, Roberto Amato, Jeffrey Barrett, Sonia Goncalves, Ewan Harrison, David K. Jackson, Ian Johnston, Dominic Kwiatkowski, Cordelia Langford, John Sillitoe on behalf of the Wellcome Sanger Institute COVID-19 Surveillance Team |
| EPI_ISL_3224595 | Berkshire and Surrey Pathology Services Lighthouse Laboratory | Wellcome Sanger Institute for the COVID-19 Genomics UK (COG-UK) Consortium | Berkshire and Surrey Pathology Services Lighthouse Laboratory and Alex Alderton, Roberto Amato, Jeffrey Barrett, Sonia Goncalves, Ewan Harrison, David K. Jackson, Ian Johnston, Dominic Kwiatkowski, Cordelia Langford, John Sillitoe on behalf of the Wellcome Sanger Institute COVID-19 Surveillance Team |
| EPI_ISL_3224596 | Lighthouse Lab in Glasgow | Wellcome Sanger Institute for the COVID-19 Genomics UK (COG-UK) Consortium | Harper VanSteenhouse, Yumi Kasai, David Gray, Carol Clugston, Anna Dominiczak and Alex Alderton, Roberto Amato, Jeffrey Barrett, Sonia Goncalves, Ewan Harrison, David K. Jackson, Ian Johnston, Dominic Kwiatkowski, Cordelia Langford, John Sillitoe on behalf of the Wellcome Sanger Institute COVID-19 Surveillance Team |
| EPI_ISL_3224597 | Berkshire and Surrey Pathology Services Lighthouse Laboratory | Wellcome Sanger Institute for the COVID-19 Genomics UK (COG-UK) Consortium | Berkshire and Surrey Pathology Services Lighthouse Laboratory and Alex Alderton, Roberto Amato, Jeffrey Barrett, Sonia Goncalves, Ewan Harrison, David K. Jackson, Ian Johnston, Dominic Kwiatkowski, Cordelia Langford, John Sillitoe on behalf of the Wellcome Sanger Institute COVID-19 Surveillance Team |
| EPI_ISL_3224598, EPI_ISL_3224599, EPI_ISL_3224600 | Lighthouse Lab in Glasgow | Wellcome Sanger Institute for the COVID-19 Genomics UK (COG-UK) Consortium | Harper VanSteenhouse, Yumi Kasai, David Gray, Carol Clugston, Anna Dominiczak and Alex Alderton, Roberto Amato, Jeffrey Barrett, Sonia Goncalves, Ewan Harrison, David K. Jackson, Ian Johnston, Dominic Kwiatkowski, Cordelia Langford, John Sillitoe on behalf of the Wellcome Sanger Institute COVID-19 Surveillance Team |
| EPI_ISL_3224601 | Lighthouse Lab in Milton Keynes | Wellcome Sanger Institute for the COVID-19 Genomics UK (COG-UK) Consortium | The Lighthouse Lab in Milton Keynes and Alex Alderton, Roberto Amato, Jeffrey Barrett, Sonia Goncalves, Ewan Harrison, David K. Jackson, Ian Johnston, Dominic Kwiatkowski, Cordelia Langford, John Sillitoe on behalf of the Wellcome Sanger Institute COVID-19 Surveillance Team |

| | | | |
|---|---|---|---|
| EPI_ISL_3224602 | Lighthouse Lab in Glasgow | Wellcome Sanger Institute for the COVID-19 Genomics UK (COG-UK) Consortium | Harper VanSteenhouse, Yumi Kasai, David Gray, Carol Clugston, Anna Dominiczak and Alex Alderton, Roberto Amato, Jeffrey Barrett, Sonia Goncalves, Ewan Harrison, David K. Jackson, Ian Johnston, Dominic Kwiatkowski, Cordelia Langford, John Sillitoe on behalf of the Wellcome Sanger Institute COVID-19 Surveillance Team |
| EPI_ISL_3224603 | Lighthouse Laboratory Plymouth | Wellcome Sanger Institute for the COVID-19 Genomics UK (COG-UK) Consortium | Lighthouse Laboratory Plymouth and Alex Alderton, Roberto Amato, Jeffrey Barrett, Sonia Goncalves, Ewan Harrison, David K. Jackson, Ian Johnston, Dominic Kwiatkowski, Cordelia Langford, John Sillitoe on behalf of the Wellcome Sanger Institute COVID-19 Surveillance Team |
| EPI_ISL_3224604, EPI_ISL_3224605, EPI_ISL_3224606 | Lighthouse Lab in Milton Keynes | Wellcome Sanger Institute for the COVID-19 Genomics UK (COG-UK) Consortium | The Lighthouse Lab in Milton Keynes and Alex Alderton, Roberto Amato, Jeffrey Barrett, Sonia Goncalves, Ewan Harrison, David K. Jackson, Ian Johnston, Dominic Kwiatkowski, Cordelia Langford, John Sillitoe on behalf of the Wellcome Sanger Institute COVID-19 Surveillance Team |
| EPI_ISL_3224607 | Lighthouse Lab in Glasgow | Wellcome Sanger Institute for the COVID-19 Genomics UK (COG-UK) Consortium | Harper VanSteenhouse, Yumi Kasai, David Gray, Carol Clugston, Anna Dominiczak and Alex Alderton, Roberto Amato, Jeffrey Barrett, Sonia Goncalves, Ewan Harrison, David K. Jackson, Ian Johnston, Dominic Kwiatkowski, Cordelia Langford, John Sillitoe on behalf of the Wellcome Sanger Institute COVID-19 Surveillance Team |
| EPI_ISL_3224608 | Berkshire and Surrey Pathology Services Lighthouse Laboratory | Wellcome Sanger Institute for the COVID-19 Genomics UK (COG-UK) Consortium | Berkshire and Surrey Pathology Services Lighthouse Laboratory and Alex Alderton, Roberto Amato, Jeffrey Barrett, Sonia Goncalves, Ewan Harrison, David K. Jackson, Ian Johnston, Dominic Kwiatkowski, Cordelia Langford, John Sillitoe on behalf of the Wellcome Sanger Institute COVID-19 Surveillance Team |
| EPI_ISL_3224609 | Lighthouse Lab in Glasgow | Wellcome Sanger Institute for the COVID-19 Genomics UK (COG-UK) Consortium | Harper VanSteenhouse, Yumi Kasai, David Gray, Carol Clugston, Anna Dominiczak and Alex Alderton, Roberto Amato, Jeffrey Barrett, Sonia Goncalves, Ewan Harrison, David K. Jackson, Ian Johnston, Dominic Kwiatkowski, Cordelia Langford, John Sillitoe on behalf of the Wellcome Sanger Institute COVID-19 Surveillance Team |
| EPI_ISL_3224610 | Lighthouse Laboratory Plymouth | Wellcome Sanger Institute for the COVID-19 Genomics UK (COG-UK) Consortium | Lighthouse Laboratory Plymouth and Alex Alderton, Roberto Amato, Jeffrey Barrett, Sonia Goncalves, Ewan Harrison, David K. Jackson, Ian Johnston, Dominic Kwiatkowski, Cordelia Langford, John Sillitoe on behalf of the Wellcome Sanger Institute COVID-19 Surveillance Team |
| EPI_ISL_3224611 | Lighthouse Lab in Glasgow | Wellcome Sanger Institute for the COVID-19 Genomics UK (COG-UK) Consortium | Harper VanSteenhouse, Yumi Kasai, David Gray, Carol Clugston, Anna Dominiczak and Alex Alderton, Roberto Amato, Jeffrey Barrett, Sonia Goncalves, Ewan Harrison, David K. Jackson, Ian Johnston, Dominic Kwiatkowski, Cordelia Langford, John Sillitoe on behalf of the Wellcome Sanger Institute COVID-19 Surveillance Team |
| EPI_ISL_3224612, EPI_ISL_3224613 | Lighthouse Lab in Milton Keynes | Wellcome Sanger Institute for the COVID-19 Genomics UK (COG-UK) Consortium | The Lighthouse Lab in Milton Keynes and Alex Alderton, Roberto Amato, Jeffrey Barrett, Sonia Goncalves, Ewan Harrison, David K. Jackson, Ian Johnston, Dominic Kwiatkowski, Cordelia Langford, John Sillitoe on behalf of the Wellcome Sanger Institute COVID-19 Surveillance Team |
| EPI_ISL_3224614, EPI_ISL_3224615 | Lighthouse Laboratory Plymouth | Wellcome Sanger Institute for the COVID-19 Genomics UK (COG-UK) Consortium | Lighthouse Laboratory Plymouth and Alex Alderton, Roberto Amato, Jeffrey Barrett, Sonia Goncalves, Ewan Harrison, David K. Jackson, Ian Johnston, Dominic Kwiatkowski, Cordelia Langford, John Sillitoe on behalf of the Wellcome Sanger Institute COVID-19 Surveillance Team |
| EPI_ISL_3224616 | Lighthouse Lab in Glasgow | Wellcome Sanger Institute for the COVID-19 Genomics UK (COG-UK) Consortium | Harper VanSteenhouse, Yumi Kasai, David Gray, Carol Clugston, Anna Dominiczak and Alex Alderton, Roberto Amato, Jeffrey Barrett, Sonia Goncalves, Ewan Harrison, David K. Jackson, Ian Johnston, Dominic Kwiatkowski, Cordelia Langford, John Sillitoe on behalf of the Wellcome Sanger Institute COVID-19 Surveillance Team |
| EPI_ISL_3224617 | Lighthouse Lab in Milton Keynes | Wellcome Sanger Institute for the COVID-19 Genomics UK (COG-UK) Consortium | The Lighthouse Lab in Milton Keynes and Alex Alderton, Roberto Amato, Jeffrey Barrett, Sonia Goncalves, Ewan Harrison, David K. Jackson, Ian Johnston, Dominic Kwiatkowski, Cordelia Langford, John Sillitoe on behalf of the Wellcome Sanger Institute COVID-19 Surveillance Team |
| EPI_ISL_3224618, EPI_ISL_3224619, EPI_ISL_3224620 | Lighthouse Laboratory Plymouth | Wellcome Sanger Institute for the COVID-19 Genomics UK (COG-UK) Consortium | Lighthouse Laboratory Plymouth and Alex Alderton, Roberto Amato, Jeffrey Barrett, Sonia Goncalves, Ewan Harrison, David K. Jackson, Ian Johnston, Dominic Kwiatkowski, Cordelia Langford, John Sillitoe on behalf of the Wellcome Sanger Institute COVID-19 Surveillance Team |
| EPI_ISL_3224621 | Lighthouse Lab in Glasgow | Wellcome Sanger Institute for the COVID-19 Genomics UK (COG-UK) Consortium | Harper VanSteenhouse, Yumi Kasai, David Gray, Carol Clugston, Anna Dominiczak and Alex Alderton, Roberto Amato, Jeffrey Barrett, Sonia Goncalves, Ewan Harrison, David K. Jackson, Ian Johnston, Dominic Kwiatkowski, Cordelia Langford, John Sillitoe on behalf of the Wellcome Sanger Institute COVID-19 Surveillance Team |
| EPI_ISL_3224622, EPI_ISL_3224623, EPI_ISL_3224624, EPI_ISL_3224625, EPI_ISL_3224626 | Lighthouse Laboratory Plymouth | Wellcome Sanger Institute for the COVID-19 Genomics UK (COG-UK) Consortium | Lighthouse Laboratory Plymouth and Alex Alderton, Roberto Amato, Jeffrey Barrett, Sonia Goncalves, Ewan Harrison, David K. Jackson, Ian Johnston, Dominic Kwiatkowski, Cordelia Langford, John Sillitoe on behalf of the Wellcome Sanger Institute COVID-19 Surveillance Team |
| EPI_ISL_3224627 | Lighthouse Lab in Glasgow | Wellcome Sanger Institute for the COVID-19 Genomics UK (COG-UK) Consortium | Harper VanSteenhouse, Yumi Kasai, David Gray, Carol Clugston, Anna Dominiczak and Alex Alderton, Roberto Amato, Jeffrey Barrett, Sonia Goncalves, Ewan Harrison, David K. Jackson, Ian Johnston, Dominic Kwiatkowski, Cordelia Langford, John Sillitoe on behalf of the Wellcome Sanger Institute COVID-19 Surveillance Team |
| EPI_ISL_3224628 | Lighthouse Laboratory Plymouth | Wellcome Sanger Institute for the COVID-19 Genomics UK (COG-UK) Consortium | Lighthouse Laboratory Plymouth and Alex Alderton, Roberto Amato, Jeffrey Barrett, Sonia Goncalves, Ewan Harrison, David K. Jackson, Ian Johnston, Dominic Kwiatkowski, Cordelia Langford, John Sillitoe on behalf of the Wellcome Sanger Institute COVID-19 Surveillance Team |
| EPI_ISL_3224629 | Lighthouse Lab in Glasgow | Wellcome Sanger Institute for the COVID-19 Genomics UK (COG-UK) Consortium | Harper VanSteenhouse, Yumi Kasai, David Gray, Carol Clugston, Anna Dominiczak and Alex Alderton, Roberto Amato, Jeffrey Barrett, Sonia Goncalves, Ewan Harrison, David K. Jackson, Ian Johnston, Dominic Kwiatkowski, Cordelia Langford, John Sillitoe on behalf of the Wellcome Sanger Institute COVID-19 Surveillance Team |
| EPI_ISL_3224630, EPI_ISL_3224631 | Berkshire and Surrey Pathology Services Lighthouse Laboratory | Wellcome Sanger Institute for the COVID-19 Genomics UK (COG-UK) Consortium | Berkshire and Surrey Pathology Services Lighthouse Laboratory and Alex Alderton, Roberto Amato, Jeffrey Barrett, Sonia Goncalves, Ewan Harrison, David K. Jackson, Ian Johnston, Dominic Kwiatkowski, Cordelia Langford, John Sillitoe on behalf of the Wellcome Sanger Institute COVID-19 Surveillance Team |
| EPI_ISL_3224632 | Lighthouse Lab in Milton Keynes | Wellcome Sanger Institute for the COVID-19 Genomics UK (COG-UK) Consortium | The Lighthouse Lab in Milton Keynes and Alex Alderton, Roberto Amato, Jeffrey Barrett, Sonia Goncalves, Ewan Harrison, David K. Jackson, Ian Johnston, Dominic Kwiatkowski, Cordelia Langford, John Sillitoe on behalf of the Wellcome Sanger Institute COVID-19 Surveillance Team |
| EPI_ISL_3224633 | Lighthouse Lab in Glasgow | Wellcome Sanger Institute for the COVID-19 Genomics UK (COG-UK) Consortium | Harper VanSteenhouse, Yumi Kasai, David Gray, Carol Clugston, Anna Dominiczak and Alex Alderton, Roberto Amato, Jeffrey Barrett, Sonia Goncalves, Ewan Harrison, David K. Jackson, Ian Johnston, Dominic Kwiatkowski, Cordelia Langford, John Sillitoe on behalf of the Wellcome Sanger Institute COVID-19 Surveillance Team |
| EPI_ISL_3224634, EPI_ISL_3224635 | Lighthouse Laboratory Plymouth | Wellcome Sanger Institute for the COVID-19 Genomics UK (COG-UK) Consortium | Lighthouse Laboratory Plymouth and Alex Alderton, Roberto Amato, Jeffrey Barrett, Sonia Goncalves, Ewan Harrison, David K. Jackson, Ian Johnston, Dominic Kwiatkowski, Cordelia Langford, John Sillitoe on behalf of the Wellcome Sanger Institute COVID-19 Surveillance Team |
| EPI_ISL_3224636, EPI_ISL_3224637 | Berkshire and Surrey Pathology Services Lighthouse Laboratory | Wellcome Sanger Institute for the COVID-19 Genomics UK (COG-UK) Consortium | Berkshire and Surrey Pathology Services Lighthouse Laboratory and Alex Alderton, Roberto Amato, Jeffrey Barrett, Sonia Goncalves, Ewan Harrison, David K. Jackson, Ian Johnston, Dominic Kwiatkowski, Cordelia Langford, John Sillitoe on behalf of the Wellcome Sanger Institute COVID-19 Surveillance Team |
| EPI_ISL_3224638 | Lighthouse Lab in Glasgow | Wellcome Sanger Institute for the COVID-19 Genomics UK (COG-UK) Consortium | Harper VanSteenhouse, Yumi Kasai, David Gray, Carol Clugston, Anna Dominiczak and Alex Alderton, Roberto Amato, Jeffrey Barrett, Sonia Goncalves, Ewan Harrison, David K. Jackson, Ian Johnston, Dominic Kwiatkowski, Cordelia Langford, John Sillitoe on behalf of the Wellcome Sanger Institute COVID-19 Surveillance Team |
| EPI_ISL_3224639 | Lighthouse Lab in Milton Keynes | Wellcome Sanger Institute for the COVID-19 Genomics UK (COG-UK) Consortium | The Lighthouse Lab in Milton Keynes and Alex Alderton, Roberto Amato, Jeffrey Barrett, Sonia Goncalves, Ewan Harrison, David K. Jackson, Ian Johnston, Dominic Kwiatkowski, Cordelia Langford, John Sillitoe on behalf of the Wellcome Sanger Institute COVID-19 Surveillance Team |
| EPI_ISL_3224640 | Berkshire and Surrey Pathology Services Lighthouse Laboratory | Wellcome Sanger Institute for the COVID-19 Genomics UK (COG-UK) Consortium | Berkshire and Surrey Pathology Services Lighthouse Laboratory and Alex Alderton, Roberto Amato, Jeffrey Barrett, Sonia Goncalves, Ewan Harrison, David K. Jackson, Ian Johnston, Dominic Kwiatkowski, Cordelia Langford, John Sillitoe on behalf of the Wellcome Sanger Institute COVID-19 Surveillance Team |
| EPI_ISL_3224641 | Lighthouse Lab in Glasgow | Wellcome Sanger Institute for the COVID-19 Genomics UK (COG-UK) Consortium | Harper VanSteenhouse, Yumi Kasai, David Gray, Carol Clugston, Anna Dominiczak and Alex Alderton, Roberto Amato, Jeffrey Barrett, Sonia Goncalves, Ewan Harrison, David K. Jackson, Ian Johnston, Dominic Kwiatkowski, Cordelia Langford, John Sillitoe on behalf of the Wellcome Sanger Institute COVID-19 Surveillance Team |
| EPI_ISL_3224642 | Lighthouse Lab in Milton Keynes | Wellcome Sanger Institute for the COVID-19 Genomics UK (COG-UK) Consortium | The Lighthouse Lab in Milton Keynes and Alex Alderton, Roberto Amato, Jeffrey Barrett, Sonia Goncalves, Ewan Harrison, David K. Jackson, Ian Johnston, Dominic Kwiatkowski, Cordelia Langford, John Sillitoe on behalf of the Wellcome Sanger Institute COVID-19 Surveillance Team |
| EPI_ISL_3224643, EPI_ISL_3224644 | Berkshire and Surrey Pathology Services Lighthouse Laboratory | Wellcome Sanger Institute for the COVID-19 Genomics UK (COG-UK) Consortium | Berkshire and Surrey Pathology Services Lighthouse Laboratory and Alex Alderton, Roberto Amato, Jeffrey Barrett, Sonia Goncalves, Ewan Harrison, David K. Jackson, Ian Johnston, Dominic Kwiatkowski, Cordelia Langford, John Sillitoe on behalf of the Wellcome Sanger Institute COVID-19 Surveillance Team |
| EPI_ISL_3224645, EPI_ISL_3224646 | Lighthouse Lab in Milton Keynes | Wellcome Sanger Institute for the COVID-19 Genomics UK (COG-UK) Consortium | The Lighthouse Lab in Milton Keynes and Alex Alderton, Roberto Amato, Jeffrey Barrett, Sonia Goncalves, Ewan Harrison, David K. Jackson, Ian Johnston, Dominic Kwiatkowski, Cordelia Langford, John Sillitoe on behalf of the Wellcome Sanger Institute COVID-19 Surveillance Team |
| EPI_ISL_3224647 | Lighthouse Laboratory Plymouth | Wellcome Sanger Institute for the COVID-19 Genomics UK (COG-UK) Consortium | Lighthouse Laboratory Plymouth and Alex Alderton, Roberto Amato, Jeffrey Barrett, Sonia Goncalves, Ewan Harrison, David K. Jackson, Ian Johnston, Dominic Kwiatkowski, Cordelia Langford, John Sillitoe on behalf of the Wellcome Sanger Institute COVID-19 Surveillance Team |
| EPI_ISL_3224648 | Lighthouse Lab in Milton Keynes | Wellcome Sanger Institute for the COVID-19 Genomics UK (COG-UK) Consortium | The Lighthouse Lab in Milton Keynes and Alex Alderton, Roberto Amato, Jeffrey Barrett, Sonia Goncalves, Ewan Harrison, David K. Jackson, Ian Johnston, Dominic Kwiatkowski, Cordelia Langford, John Sillitoe on behalf of the Wellcome Sanger Institute COVID-19 Surveillance Team |
| EPI_ISL_3224649 | Lighthouse Lab in Glasgow | Wellcome Sanger Institute for the COVID-19 Genomics UK (COG-UK) Consortium | Harper VanSteenhouse, Yumi Kasai, David Gray, Carol Clugston, Anna Dominiczak and Alex Alderton, Roberto Amato, Jeffrey Barrett, Sonia Goncalves, Ewan Harrison, David K. Jackson, Ian Johnston, Dominic Kwiatkowski, Cordelia Langford, John Sillitoe on behalf of the Wellcome Sanger Institute COVID-19 Surveillance Team |
| EPI_ISL_3224650, EPI_ISL_3224651, EPI_ISL_3224652 | Lighthouse Laboratory Plymouth | Wellcome Sanger Institute for the COVID-19 Genomics UK (COG-UK) Consortium | Lighthouse Laboratory Plymouth and Alex Alderton, Roberto Amato, Jeffrey Barrett, Sonia Goncalves, Ewan Harrison, David K. Jackson, Ian Johnston, Dominic Kwiatkowski, Cordelia Langford, John Sillitoe on behalf of the Wellcome Sanger Institute COVID-19 Surveillance Team |
| EPI_ISL_3224653 | Lighthouse Lab in Milton Keynes | Wellcome Sanger Institute for the COVID-19 Genomics UK (COG-UK) Consortium | The Lighthouse Lab in Milton Keynes and Alex Alderton, Roberto Amato, Jeffrey Barrett, Sonia Goncalves, Ewan Harrison, David K. Jackson, Ian Johnston, Dominic Kwiatkowski, Cordelia Langford, John Sillitoe on behalf of the Wellcome Sanger Institute COVID-19 Surveillance Team |
| EPI_ISL_3224654 | Lighthouse Laboratory Plymouth | Wellcome Sanger Institute for the COVID-19 Genomics UK (COG-UK) Consortium | Lighthouse Laboratory Plymouth and Alex Alderton, Roberto Amato, Jeffrey Barrett, Sonia Goncalves, Ewan Harrison, David K. Jackson, Ian Johnston, Dominic Kwiatkowski, Cordelia Langford, John Sillitoe on behalf of the Wellcome Sanger Institute COVID-19 Surveillance Team |
| EPI_ISL_3224655 | Lighthouse Lab in Glasgow | Wellcome Sanger Institute for the COVID-19 Genomics UK (COG-UK) Consortium | Harper VanSteenhouse, Yumi Kasai, David Gray, Carol Clugston, Anna Dominiczak and Alex Alderton, Roberto Amato, Jeffrey Barrett, Sonia Goncalves, Ewan Harrison, David K. Jackson, Ian Johnston, Dominic Kwiatkowski, Cordelia Langford, John Sillitoe on behalf of the Wellcome Sanger Institute COVID-19 Surveillance Team |
| EPI_ISL_3224656, EPI_ISL_3224657 | Lighthouse Laboratory Plymouth | Wellcome Sanger Institute for the COVID-19 Genomics UK (COG-UK) Consortium | Lighthouse Laboratory Plymouth and Alex Alderton, Roberto Amato, Jeffrey Barrett, Sonia Goncalves, Ewan Harrison, David K. Jackson, Ian Johnston, Dominic Kwiatkowski, Cordelia Langford, John Sillitoe on behalf of the Wellcome Sanger Institute COVID-19 Surveillance Team |
| EPI_ISL_3224658 | Lighthouse Lab in Glasgow | Wellcome Sanger Institute for the COVID-19 Genomics UK (COG-UK) Consortium | Harper VanSteenhouse, Yumi Kasai, David Gray, Carol Clugston, Anna Dominiczak and Alex Alderton, Roberto Amato, Jeffrey Barrett, Sonia Goncalves, Ewan Harrison, David K. Jackson, Ian Johnston, Dominic Kwiatkowski, Cordelia Langford, John Sillitoe on behalf of the Wellcome Sanger Institute COVID-19 Surveillance Team |
| EPI_ISL_3224659, EPI_ISL_3224660 | Berkshire and Surrey Pathology Services Lighthouse Laboratory | Wellcome Sanger Institute for the COVID-19 Genomics UK (COG-UK) Consortium | Berkshire and Surrey Pathology Services Lighthouse Laboratory and Alex Alderton, Roberto Amato, Jeffrey Barrett, Sonia Goncalves, Ewan Harrison, David K. Jackson, Ian Johnston, Dominic Kwiatkowski, Cordelia Langford, John Sillitoe on behalf of the Wellcome Sanger Institute COVID-19 Surveillance Team |
| EPI_ISL_3224661, EPI_ISL_3224662 | Lighthouse Lab in Glasgow | Wellcome Sanger Institute for the COVID-19 Genomics UK (COG-UK) Consortium | Harper VanSteenhouse, Yumi Kasai, David Gray, Carol Clugston, Anna Dominiczak and Alex Alderton, Roberto Amato, Jeffrey Barrett, Sonia Goncalves, Ewan Harrison, David K. Jackson, Ian Johnston, Dominic Kwiatkowski, Cordelia Langford, John Sillitoe on behalf of the Wellcome Sanger Institute COVID-19 Surveillance Team |
| EPI_ISL_3224663 | Lighthouse Lab in Milton Keynes | Wellcome Sanger Institute for the COVID-19 Genomics UK (COG-UK) Consortium | The Lighthouse Lab in Milton Keynes and Alex Alderton, Roberto Amato, Jeffrey Barrett, Sonia Goncalves, Ewan Harrison, David K. Jackson, Ian Johnston, Dominic Kwiatkowski, Cordelia Langford, John Sillitoe on behalf of the Wellcome Sanger Institute COVID-19 Surveillance Team |
| EPI_ISL_3224664 | Lighthouse Laboratory Plymouth | Wellcome Sanger Institute for the COVID-19 Genomics UK (COG-UK) | Lighthouse Laboratory Plymouth and Alex Alderton, Roberto Amato, Jeffrey Barrett, Sonia Goncalves, Ewan Harrison, David K. Jackson, Ian Johnston, Dominic Kwiatkowski, Cordelia Langford, John Sillitoe on behalf of the Wellcome Sanger |

| | | | |
|---|---|---|---|
| | | Consortium | Institute COVID-19 Surveillance Team |
| EPI_ISL_3224665 | Lighthouse Lab in Glasgow | Wellcome Sanger Institute for the COVID-19 Genomics UK (COG-UK) Consortium | Harper VanSteenhouse, Yumi Kasai, David Gray, Carol Clugston, Anna Dominiczak and Alex Alderton, Roberto Amato, Jeffrey Barrett, Sonia Goncalves, Ewan Harrison, David K. Jackson, Ian Johnston, Dominic Kwiatkowski, Cordelia Langford, John Sillitoe on behalf of the Wellcome Sanger Institute COVID-19 Surveillance Team |
| EPI_ISL_3224666 | Lighthouse Lab in Milton Keynes | Wellcome Sanger Institute for the COVID-19 Genomics UK (COG-UK) Consortium | The Lighthouse Lab in Milton Keynes and Alex Alderton, Roberto Amato, Jeffrey Barrett, Sonia Goncalves, Ewan Harrison, David K. Jackson, Ian Johnston, Dominic Kwiatkowski, Cordelia Langford, John Sillitoe on behalf of the Wellcome Sanger Institute COVID-19 Surveillance Team |
| EPI_ISL_3224667 | Lighthouse Laboratory Plymouth | Wellcome Sanger Institute for the COVID-19 Genomics UK (COG-UK) Consortium | Lighthouse Laboratory Plymouth and Alex Alderton, Roberto Amato, Jeffrey Barrett, Sonia Goncalves, Ewan Harrison, David K. Jackson, Ian Johnston, Dominic Kwiatkowski, Cordelia Langford, John Sillitoe on behalf of the Wellcome Sanger Institute COVID-19 Surveillance Team |
| EPI_ISL_3224668, EPI_ISL_3224669 | Lighthouse Lab in Milton Keynes | Wellcome Sanger Institute for the COVID-19 Genomics UK (COG-UK) Consortium | The Lighthouse Lab in Milton Keynes and Alex Alderton, Roberto Amato, Jeffrey Barrett, Sonia Goncalves, Ewan Harrison, David K. Jackson, Ian Johnston, Dominic Kwiatkowski, Cordelia Langford, John Sillitoe on behalf of the Wellcome Sanger Institute COVID-19 Surveillance Team |
| EPI_ISL_3224670 | Lighthouse Laboratory Plymouth | Wellcome Sanger Institute for the COVID-19 Genomics UK (COG-UK) Consortium | Lighthouse Laboratory Plymouth and Alex Alderton, Roberto Amato, Jeffrey Barrett, Sonia Goncalves, Ewan Harrison, David K. Jackson, Ian Johnston, Dominic Kwiatkowski, Cordelia Langford, John Sillitoe on behalf of the Wellcome Sanger Institute COVID-19 Surveillance Team |
| EPI_ISL_3224671, EPI_ISL_3224672, EPI_ISL_3224673, EPI_ISL_3224674 | Lighthouse Lab in Glasgow | Wellcome Sanger Institute for the COVID-19 Genomics UK (COG-UK) Consortium | Harper VanSteenhouse, Yumi Kasai, David Gray, Carol Clugston, Anna Dominiczak and Alex Alderton, Roberto Amato, Jeffrey Barrett, Sonia Goncalves, Ewan Harrison, David K. Jackson, Ian Johnston, Dominic Kwiatkowski, Cordelia Langford, John Sillitoe on behalf of the Wellcome Sanger Institute COVID-19 Surveillance Team |
| EPI_ISL_3224675 | Lighthouse Lab in Milton Keynes | Wellcome Sanger Institute for the COVID-19 Genomics UK (COG-UK) Consortium | The Lighthouse Lab in Milton Keynes and Alex Alderton, Roberto Amato, Jeffrey Barrett, Sonia Goncalves, Ewan Harrison, David K. Jackson, Ian Johnston, Dominic Kwiatkowski, Cordelia Langford, John Sillitoe on behalf of the Wellcome Sanger Institute COVID-19 Surveillance Team |
| EPI_ISL_3224676, EPI_ISL_3224677, EPI_ISL_3224678, EPI_ISL_3224679 | Lighthouse Laboratory Plymouth | Wellcome Sanger Institute for the COVID-19 Genomics UK (COG-UK) Consortium | Lighthouse Laboratory Plymouth and Alex Alderton, Roberto Amato, Jeffrey Barrett, Sonia Goncalves, Ewan Harrison, David K. Jackson, Ian Johnston, Dominic Kwiatkowski, Cordelia Langford, John Sillitoe on behalf of the Wellcome Sanger Institute COVID-19 Surveillance Team |
| EPI_ISL_3224680 | Lighthouse Lab in Glasgow | Wellcome Sanger Institute for the COVID-19 Genomics UK (COG-UK) Consortium | Harper VanSteenhouse, Yumi Kasai, David Gray, Carol Clugston, Anna Dominiczak and Alex Alderton, Roberto Amato, Jeffrey Barrett, Sonia Goncalves, Ewan Harrison, David K. Jackson, Ian Johnston, Dominic Kwiatkowski, Cordelia Langford, John Sillitoe on behalf of the Wellcome Sanger Institute COVID-19 Surveillance Team |
| EPI_ISL_3224681, EPI_ISL_3224682 | Lighthouse Lab in Milton Keynes | Wellcome Sanger Institute for the COVID-19 Genomics UK (COG-UK) Consortium | The Lighthouse Lab in Milton Keynes and Alex Alderton, Roberto Amato, Jeffrey Barrett, Sonia Goncalves, Ewan Harrison, David K. Jackson, Ian Johnston, Dominic Kwiatkowski, Cordelia Langford, John Sillitoe on behalf of the Wellcome Sanger Institute COVID-19 Surveillance Team |
| EPI_ISL_3224683 | Lighthouse Laboratory Plymouth | Wellcome Sanger Institute for the COVID-19 Genomics UK (COG-UK) Consortium | Lighthouse Laboratory Plymouth and Alex Alderton, Roberto Amato, Jeffrey Barrett, Sonia Goncalves, Ewan Harrison, David K. Jackson, Ian Johnston, Dominic Kwiatkowski, Cordelia Langford, John Sillitoe on behalf of the Wellcome Sanger Institute COVID-19 Surveillance Team |
| EPI_ISL_3224684 | Lighthouse Lab in Milton Keynes | Wellcome Sanger Institute for the COVID-19 Genomics UK (COG-UK) Consortium | The Lighthouse Lab in Milton Keynes and Alex Alderton, Roberto Amato, Jeffrey Barrett, Sonia Goncalves, Ewan Harrison, David K. Jackson, Ian Johnston, Dominic Kwiatkowski, Cordelia Langford, John Sillitoe on behalf of the Wellcome Sanger Institute COVID-19 Surveillance Team |
| EPI_ISL_3224685 | Berkshire and Surrey Pathology Services Lighthouse Laboratory | Wellcome Sanger Institute for the COVID-19 Genomics UK (COG-UK) Consortium | Berkshire and Surrey Pathology Services Lighthouse Laboratory and Alex Alderton, Roberto Amato, Jeffrey Barrett, Sonia Goncalves, Ewan Harrison, David K. Jackson, Ian Johnston, Dominic Kwiatkowski, Cordelia Langford, John Sillitoe on behalf of the Wellcome Sanger Institute COVID-19 Surveillance Team |
| EPI_ISL_3224686 | Lighthouse Laboratory Plymouth | Wellcome Sanger Institute for the COVID-19 Genomics UK (COG-UK) Consortium | Lighthouse Laboratory Plymouth and Alex Alderton, Roberto Amato, Jeffrey Barrett, Sonia Goncalves, Ewan Harrison, David K. Jackson, Ian Johnston, Dominic Kwiatkowski, Cordelia Langford, John Sillitoe on behalf of the Wellcome Sanger Institute COVID-19 Surveillance Team |
| EPI_ISL_3224687 | Lighthouse Lab in Glasgow | Wellcome Sanger Institute for the COVID-19 Genomics UK (COG-UK) Consortium | Harper VanSteenhouse, Yumi Kasai, David Gray, Carol Clugston, Anna Dominiczak and Alex Alderton, Roberto Amato, Jeffrey Barrett, Sonia Goncalves, Ewan Harrison, David K. Jackson, Ian Johnston, Dominic Kwiatkowski, Cordelia Langford, John Sillitoe on behalf of the Wellcome Sanger Institute COVID-19 Surveillance Team |
| EPI_ISL_3224688 | Lighthouse Laboratory Plymouth | Wellcome Sanger Institute for the COVID-19 Genomics UK (COG-UK) Consortium | Lighthouse Laboratory Plymouth and Alex Alderton, Roberto Amato, Jeffrey Barrett, Sonia Goncalves, Ewan Harrison, David K. Jackson, Ian Johnston, Dominic Kwiatkowski, Cordelia Langford, John Sillitoe on behalf of the Wellcome Sanger Institute COVID-19 Surveillance Team |
| EPI_ISL_3224689, EPI_ISL_3224690 | Lighthouse Lab in Glasgow | Wellcome Sanger Institute for the COVID-19 Genomics UK (COG-UK) Consortium | Harper VanSteenhouse, Yumi Kasai, David Gray, Carol Clugston, Anna Dominiczak and Alex Alderton, Roberto Amato, Jeffrey Barrett, Sonia Goncalves, Ewan Harrison, David K. Jackson, Ian Johnston, Dominic Kwiatkowski, Cordelia Langford, John Sillitoe on behalf of the Wellcome Sanger Institute COVID-19 Surveillance Team |
| EPI_ISL_3224691 | Lighthouse Lab in Milton Keynes | Wellcome Sanger Institute for the COVID-19 Genomics UK (COG-UK) Consortium | The Lighthouse Lab in Milton Keynes and Alex Alderton, Roberto Amato, Jeffrey Barrett, Sonia Goncalves, Ewan Harrison, David K. Jackson, Ian Johnston, Dominic Kwiatkowski, Cordelia Langford, John Sillitoe on behalf of the Wellcome Sanger Institute COVID-19 Surveillance Team |
| EPI_ISL_3224692 | Berkshire and Surrey Pathology Services Lighthouse Laboratory | Wellcome Sanger Institute for the COVID-19 Genomics UK (COG-UK) Consortium | Berkshire and Surrey Pathology Services Lighthouse Laboratory and Alex Alderton, Roberto Amato, Jeffrey Barrett, Sonia Goncalves, Ewan Harrison, David K. Jackson, Ian Johnston, Dominic Kwiatkowski, Cordelia Langford, John Sillitoe on behalf of the Wellcome Sanger Institute COVID-19 Surveillance Team |
| EPI_ISL_3224693 | Lighthouse Lab in Glasgow | Wellcome Sanger Institute for the COVID-19 Genomics UK (COG-UK) Consortium | Harper VanSteenhouse, Yumi Kasai, David Gray, Carol Clugston, Anna Dominiczak and Alex Alderton, Roberto Amato, Jeffrey Barrett, Sonia Goncalves, Ewan Harrison, David K. Jackson, Ian Johnston, Dominic Kwiatkowski, Cordelia Langford, John Sillitoe on behalf of the Wellcome Sanger Institute COVID-19 Surveillance Team |
| EPI_ISL_3224694 | Lighthouse Lab in Milton Keynes | Wellcome Sanger Institute for the COVID-19 Genomics UK (COG-UK) Consortium | The Lighthouse Lab in Milton Keynes and Alex Alderton, Roberto Amato, Jeffrey Barrett, Sonia Goncalves, Ewan Harrison, David K. Jackson, Ian Johnston, Dominic Kwiatkowski, Cordelia Langford, John Sillitoe on behalf of the Wellcome Sanger Institute COVID-19 Surveillance Team |
| EPI_ISL_3224695 | Lighthouse Laboratory Plymouth | Wellcome Sanger Institute for the COVID-19 Genomics UK (COG-UK) Consortium | Lighthouse Laboratory Plymouth and Alex Alderton, Roberto Amato, Jeffrey Barrett, Sonia Goncalves, Ewan Harrison, David K. Jackson, Ian Johnston, Dominic Kwiatkowski, Cordelia Langford, John Sillitoe on behalf of the Wellcome Sanger Institute COVID-19 Surveillance Team |
| EPI_ISL_3224696 | Berkshire and Surrey Pathology Services Lighthouse Laboratory | Wellcome Sanger Institute for the COVID-19 Genomics UK (COG-UK) Consortium | Berkshire and Surrey Pathology Services Lighthouse Laboratory and Alex Alderton, Roberto Amato, Jeffrey Barrett, Sonia Goncalves, Ewan Harrison, David K. Jackson, Ian Johnston, Dominic Kwiatkowski, Cordelia Langford, John Sillitoe on behalf of the Wellcome Sanger Institute COVID-19 Surveillance Team |
| EPI_ISL_3224697 | Lighthouse Lab in Glasgow | Wellcome Sanger Institute for the COVID-19 Genomics UK (COG-UK) Consortium | Harper VanSteenhouse, Yumi Kasai, David Gray, Carol Clugston, Anna Dominiczak and Alex Alderton, Roberto Amato, Jeffrey Barrett, Sonia Goncalves, Ewan Harrison, David K. Jackson, Ian Johnston, Dominic Kwiatkowski, Cordelia Langford, John Sillitoe on behalf of the Wellcome Sanger Institute COVID-19 Surveillance Team |
| EPI_ISL_3224698 | Lighthouse Lab in Milton Keynes | Wellcome Sanger Institute for the COVID-19 Genomics UK (COG-UK) Consortium | The Lighthouse Lab in Milton Keynes and Alex Alderton, Roberto Amato, Jeffrey Barrett, Sonia Goncalves, Ewan Harrison, David K. Jackson, Ian Johnston, Dominic Kwiatkowski, Cordelia Langford, John Sillitoe on behalf of the Wellcome Sanger Institute COVID-19 Surveillance Team |
| EPI_ISL_3224699 | Berkshire and Surrey Pathology Services Lighthouse Laboratory | Wellcome Sanger Institute for the COVID-19 Genomics UK (COG-UK) Consortium | Berkshire and Surrey Pathology Services Lighthouse Laboratory and Alex Alderton, Roberto Amato, Jeffrey Barrett, Sonia Goncalves, Ewan Harrison, David K. Jackson, Ian Johnston, Dominic Kwiatkowski, Cordelia Langford, John Sillitoe on behalf of the Wellcome Sanger Institute COVID-19 Surveillance Team |
| EPI_ISL_3224700 | Lighthouse Lab in Glasgow | Wellcome Sanger Institute for the COVID-19 Genomics UK (COG-UK) Consortium | Harper VanSteenhouse, Yumi Kasai, David Gray, Carol Clugston, Anna Dominiczak and Alex Alderton, Roberto Amato, Jeffrey Barrett, Sonia Goncalves, Ewan Harrison, David K. Jackson, Ian Johnston, Dominic Kwiatkowski, Cordelia Langford, John Sillitoe on behalf of the Wellcome Sanger Institute COVID-19 Surveillance Team |
| EPI_ISL_3224701, EPI_ISL_3224702 | Lighthouse Lab in Milton Keynes | Wellcome Sanger Institute for the COVID-19 Genomics UK (COG-UK) Consortium | The Lighthouse Lab in Milton Keynes and Alex Alderton, Roberto Amato, Jeffrey Barrett, Sonia Goncalves, Ewan Harrison, David K. Jackson, Ian Johnston, Dominic Kwiatkowski, Cordelia Langford, John Sillitoe on behalf of the Wellcome Sanger Institute COVID-19 Surveillance Team |
| EPI_ISL_3224703 | Lighthouse Lab in Glasgow | Wellcome Sanger Institute for the COVID-19 Genomics UK (COG-UK) Consortium | Harper VanSteenhouse, Yumi Kasai, David Gray, Carol Clugston, Anna Dominiczak and Alex Alderton, Roberto Amato, Jeffrey Barrett, Sonia Goncalves, Ewan Harrison, David K. Jackson, Ian Johnston, Dominic Kwiatkowski, Cordelia Langford, John Sillitoe on behalf of the Wellcome Sanger Institute COVID-19 Surveillance Team |
| EPI_ISL_3224704, EPI_ISL_3224705, EPI_ISL_3224706 | Berkshire and Surrey Pathology Services Lighthouse Laboratory | Wellcome Sanger Institute for the COVID-19 Genomics UK (COG-UK) Consortium | Berkshire and Surrey Pathology Services Lighthouse Laboratory and Alex Alderton, Roberto Amato, Jeffrey Barrett, Sonia Goncalves, Ewan Harrison, David K. Jackson, Ian Johnston, Dominic Kwiatkowski, Cordelia Langford, John Sillitoe on behalf of the Wellcome Sanger Institute COVID-19 Surveillance Team |
| EPI_ISL_3224707, EPI_ISL_3224708 | Lighthouse Lab in Milton Keynes | Wellcome Sanger Institute for the COVID-19 Genomics UK (COG-UK) Consortium | The Lighthouse Lab in Milton Keynes and Alex Alderton, Roberto Amato, Jeffrey Barrett, Sonia Goncalves, Ewan Harrison, David K. Jackson, Ian Johnston, Dominic Kwiatkowski, Cordelia Langford, John Sillitoe on behalf of the Wellcome Sanger Institute COVID-19 Surveillance Team |
| EPI_ISL_3224709 | Lighthouse Laboratory Plymouth | Wellcome Sanger Institute for the COVID-19 Genomics UK (COG-UK) Consortium | Lighthouse Laboratory Plymouth and Alex Alderton, Roberto Amato, Jeffrey Barrett, Sonia Goncalves, Ewan Harrison, David K. Jackson, Ian Johnston, Dominic Kwiatkowski, Cordelia Langford, John Sillitoe on behalf of the Wellcome Sanger Institute COVID-19 Surveillance Team |
| EPI_ISL_3224710 | Lighthouse Lab in Glasgow | Wellcome Sanger Institute for the COVID-19 Genomics UK (COG-UK) Consortium | Harper VanSteenhouse, Yumi Kasai, David Gray, Carol Clugston, Anna Dominiczak and Alex Alderton, Roberto Amato, Jeffrey Barrett, Sonia Goncalves, Ewan Harrison, David K. Jackson, Ian Johnston, Dominic Kwiatkowski, Cordelia Langford, John Sillitoe on behalf of the Wellcome Sanger Institute COVID-19 Surveillance Team |
| EPI_ISL_3224711 | Lighthouse Lab in Milton Keynes | Wellcome Sanger Institute for the COVID-19 Genomics UK (COG-UK) Consortium | The Lighthouse Lab in Milton Keynes and Alex Alderton, Roberto Amato, Jeffrey Barrett, Sonia Goncalves, Ewan Harrison, David K. Jackson, Ian Johnston, Dominic Kwiatkowski, Cordelia Langford, John Sillitoe on behalf of the Wellcome Sanger Institute COVID-19 Surveillance Team |
| EPI_ISL_3224712 | Berkshire and Surrey Pathology Services Lighthouse Laboratory | Wellcome Sanger Institute for the COVID-19 Genomics UK (COG-UK) Consortium | Berkshire and Surrey Pathology Services Lighthouse Laboratory and Alex Alderton, Roberto Amato, Jeffrey Barrett, Sonia Goncalves, Ewan Harrison, David K. Jackson, Ian Johnston, Dominic Kwiatkowski, Cordelia Langford, John Sillitoe on behalf of the Wellcome Sanger Institute COVID-19 Surveillance Team |
| EPI_ISL_3224713 | Lighthouse Laboratory Plymouth | Wellcome Sanger Institute for the COVID-19 Genomics UK (COG-UK) Consortium | Lighthouse Laboratory Plymouth and Alex Alderton, Roberto Amato, Jeffrey Barrett, Sonia Goncalves, Ewan Harrison, David K. Jackson, Ian Johnston, Dominic Kwiatkowski, Cordelia Langford, John Sillitoe on behalf of the Wellcome Sanger Institute COVID-19 Surveillance Team |
| EPI_ISL_3224714 | Berkshire and Surrey Pathology Services Lighthouse Laboratory | Wellcome Sanger Institute for the COVID-19 Genomics UK (COG-UK) Consortium | Berkshire and Surrey Pathology Services Lighthouse Laboratory and Alex Alderton, Roberto Amato, Jeffrey Barrett, Sonia Goncalves, Ewan Harrison, David K. Jackson, Ian Johnston, Dominic Kwiatkowski, Cordelia Langford, John Sillitoe on behalf of the Wellcome Sanger Institute COVID-19 Surveillance Team |
| EPI_ISL_3224715, EPI_ISL_3224716 | Lighthouse Laboratory Plymouth | Wellcome Sanger Institute for the COVID-19 Genomics UK (COG-UK) Consortium | Lighthouse Laboratory Plymouth and Alex Alderton, Roberto Amato, Jeffrey Barrett, Sonia Goncalves, Ewan Harrison, David K. Jackson, Ian Johnston, Dominic Kwiatkowski, Cordelia Langford, John Sillitoe on behalf of the Wellcome Sanger Institute COVID-19 Surveillance Team |
| EPI_ISL_3224717 | Lighthouse Lab in Glasgow | Wellcome Sanger Institute for the COVID-19 Genomics UK (COG-UK) Consortium | Harper VanSteenhouse, Yumi Kasai, David Gray, Carol Clugston, Anna Dominiczak and Alex Alderton, Roberto Amato, Jeffrey Barrett, Sonia Goncalves, Ewan Harrison, David K. Jackson, Ian Johnston, Dominic Kwiatkowski, Cordelia Langford, John Sillitoe on behalf of the Wellcome Sanger Institute COVID-19 Surveillance Team |
| EPI_ISL_3224718 | Berkshire and Surrey Pathology Services Lighthouse Laboratory | Wellcome Sanger Institute for the COVID-19 Genomics UK (COG-UK) Consortium | Berkshire and Surrey Pathology Services Lighthouse Laboratory and Alex Alderton, Roberto Amato, Jeffrey Barrett, Sonia Goncalves, Ewan Harrison, David K. Jackson, Ian Johnston, Dominic Kwiatkowski, Cordelia Langford, John Sillitoe on behalf of the Wellcome Sanger Institute COVID-19 Surveillance Team |
| EPI_ISL_3224719 | Lighthouse Lab in Milton Keynes | Wellcome Sanger Institute for the COVID-19 Genomics UK (COG-UK) Consortium | The Lighthouse Lab in Milton Keynes and Alex Alderton, Roberto Amato, Jeffrey Barrett, Sonia Goncalves, Ewan Harrison, David K. Jackson, Ian Johnston, Dominic Kwiatkowski, Cordelia Langford, John Sillitoe on behalf of the Wellcome Sanger Institute COVID-19 Surveillance Team |
| EPI_ISL_3224720 | Berkshire and Surrey Pathology Services Lighthouse Laboratory | Wellcome Sanger Institute for the COVID-19 Genomics UK (COG-UK) Consortium | Berkshire and Surrey Pathology Services Lighthouse Laboratory and Alex Alderton, Roberto Amato, Jeffrey Barrett, Sonia Goncalves, Ewan Harrison, David K. Jackson, Ian Johnston, Dominic Kwiatkowski, Cordelia Langford, John Sillitoe on behalf of the Wellcome Sanger Institute COVID-19 Surveillance Team |
| EPI_ISL_3224721 | Lighthouse Lab in Milton Keynes | Wellcome Sanger Institute for the COVID-19 Genomics UK (COG-UK) | The Lighthouse Lab in Milton Keynes and Alex Alderton, Roberto Amato, Jeffrey Barrett, Sonia Goncalves, Ewan Harrison, David K. Jackson, Ian Johnston, Dominic Kwiatkowski, Cordelia Langford, John Sillitoe on behalf of the Wellcome Sanger |

| | | | |
|---|---|---|---|
| | | Consortium | Institute COVID-19 Surveillance Team |
| EPI_ISL_3224722 | Berkshire and Surrey Pathology Services Lighthouse Laboratory | Wellcome Sanger Institute for the COVID-19 Genomics UK (COG-UK) Consortium | Berkshire and Surrey Pathology Services Lighthouse Laboratory and Alex Alderton, Roberto Amato, Jeffrey Barrett, Sonia Goncalves, Ewan Harrison, David K. Jackson, Ian Johnston, Dominic Kwiatkowski, Cordelia Langford, John Sillitoe on behalf of the Wellcome Sanger Institute COVID-19 Surveillance Team |
| EPI_ISL_3224723 | Lighthouse Laboratory Plymouth | Wellcome Sanger Institute for the COVID-19 Genomics UK (COG-UK) Consortium | Lighthouse Laboratory Plymouth and Alex Alderton, Roberto Amato, Jeffrey Barrett, Sonia Goncalves, Ewan Harrison, David K. Jackson, Ian Johnston, Dominic Kwiatkowski, Cordelia Langford, John Sillitoe on behalf of the Wellcome Sanger Institute COVID-19 Surveillance Team |
| EPI_ISL_3224724, EPI_ISL_3224725, EPI_ISL_3224726, EPI_ISL_3224727 | Lighthouse Lab in Milton Keynes | Wellcome Sanger Institute for the COVID-19 Genomics UK (COG-UK) Consortium | The Lighthouse Lab in Milton Keynes and Alex Alderton, Roberto Amato, Jeffrey Barrett, Sonia Goncalves, Ewan Harrison, David K. Jackson, Ian Johnston, Dominic Kwiatkowski, Cordelia Langford, John Sillitoe on behalf of the Wellcome Sanger Institute COVID-19 Surveillance Team |
| EPI_ISL_3224728 | Lighthouse Laboratory Plymouth | Wellcome Sanger Institute for the COVID-19 Genomics UK (COG-UK) Consortium | Lighthouse Laboratory Plymouth and Alex Alderton, Roberto Amato, Jeffrey Barrett, Sonia Goncalves, Ewan Harrison, David K. Jackson, Ian Johnston, Dominic Kwiatkowski, Cordelia Langford, John Sillitoe on behalf of the Wellcome Sanger Institute COVID-19 Surveillance Team |
| EPI_ISL_3224729 | Lighthouse Lab in Milton Keynes | Wellcome Sanger Institute for the COVID-19 Genomics UK (COG-UK) Consortium | The Lighthouse Lab in Milton Keynes and Alex Alderton, Roberto Amato, Jeffrey Barrett, Sonia Goncalves, Ewan Harrison, David K. Jackson, Ian Johnston, Dominic Kwiatkowski, Cordelia Langford, John Sillitoe on behalf of the Wellcome Sanger Institute COVID-19 Surveillance Team |
| EPI_ISL_3224730, EPI_ISL_3224731, EPI_ISL_3224732, EPI_ISL_3224733, EPI_ISL_3224734, EPI_ISL_3224735 | Lighthouse Lab in Glasgow | Wellcome Sanger Institute for the COVID-19 Genomics UK (COG-UK) Consortium | Harper VanSteenhouse, Yumi Kasai, David Gray, Carol Clugston, Anna Dominiczak and Alex Alderton, Roberto Amato, Jeffrey Barrett, Sonia Goncalves, Ewan Harrison, David K. Jackson, Ian Johnston, Dominic Kwiatkowski, Cordelia Langford, John Sillitoe on behalf of the Wellcome Sanger Institute COVID-19 Surveillance Team |
| EPI_ISL_3224736 | Lighthouse Lab in Milton Keynes | Wellcome Sanger Institute for the COVID-19 Genomics UK (COG-UK) Consortium | The Lighthouse Lab in Milton Keynes and Alex Alderton, Roberto Amato, Jeffrey Barrett, Sonia Goncalves, Ewan Harrison, David K. Jackson, Ian Johnston, Dominic Kwiatkowski, Cordelia Langford, John Sillitoe on behalf of the Wellcome Sanger Institute COVID-19 Surveillance Team |
| EPI_ISL_3224737 | Lighthouse Laboratory Plymouth | Wellcome Sanger Institute for the COVID-19 Genomics UK (COG-UK) Consortium | Lighthouse Laboratory Plymouth and Alex Alderton, Roberto Amato, Jeffrey Barrett, Sonia Goncalves, Ewan Harrison, David K. Jackson, Ian Johnston, Dominic Kwiatkowski, Cordelia Langford, John Sillitoe on behalf of the Wellcome Sanger Institute COVID-19 Surveillance Team |
| EPI_ISL_3224738 | Lighthouse Lab in Glasgow | Wellcome Sanger Institute for the COVID-19 Genomics UK (COG-UK) Consortium | Harper VanSteenhouse, Yumi Kasai, David Gray, Carol Clugston, Anna Dominiczak and Alex Alderton, Roberto Amato, Jeffrey Barrett, Sonia Goncalves, Ewan Harrison, David K. Jackson, Ian Johnston, Dominic Kwiatkowski, Cordelia Langford, John Sillitoe on behalf of the Wellcome Sanger Institute COVID-19 Surveillance Team |
| EPI_ISL_3224739 | Lighthouse Lab in Milton Keynes | Wellcome Sanger Institute for the COVID-19 Genomics UK (COG-UK) Consortium | The Lighthouse Lab in Milton Keynes and Alex Alderton, Roberto Amato, Jeffrey Barrett, Sonia Goncalves, Ewan Harrison, David K. Jackson, Ian Johnston, Dominic Kwiatkowski, Cordelia Langford, John Sillitoe on behalf of the Wellcome Sanger Institute COVID-19 Surveillance Team |
| EPI_ISL_3224740 | Lighthouse Laboratory Plymouth | Wellcome Sanger Institute for the COVID-19 Genomics UK (COG-UK) Consortium | Lighthouse Laboratory Plymouth and Alex Alderton, Roberto Amato, Jeffrey Barrett, Sonia Goncalves, Ewan Harrison, David K. Jackson, Ian Johnston, Dominic Kwiatkowski, Cordelia Langford, John Sillitoe on behalf of the Wellcome Sanger Institute COVID-19 Surveillance Team |
| EPI_ISL_3224741 | Berkshire and Surrey Pathology Services Lighthouse Laboratory | Wellcome Sanger Institute for the COVID-19 Genomics UK (COG-UK) Consortium | Berkshire and Surrey Pathology Services Lighthouse Laboratory and Alex Alderton, Roberto Amato, Jeffrey Barrett, Sonia Goncalves, Ewan Harrison, David K. Jackson, Ian Johnston, Dominic Kwiatkowski, Cordelia Langford, John Sillitoe on behalf of the Wellcome Sanger Institute COVID-19 Surveillance Team |
| EPI_ISL_3224742 | Lighthouse Lab in Milton Keynes | Wellcome Sanger Institute for the COVID-19 Genomics UK (COG-UK) Consortium | The Lighthouse Lab in Milton Keynes and Alex Alderton, Roberto Amato, Jeffrey Barrett, Sonia Goncalves, Ewan Harrison, David K. Jackson, Ian Johnston, Dominic Kwiatkowski, Cordelia Langford, John Sillitoe on behalf of the Wellcome Sanger Institute COVID-19 Surveillance Team |
| EPI_ISL_3224743 | Lighthouse Lab in Glasgow | Wellcome Sanger Institute for the COVID-19 Genomics UK (COG-UK) Consortium | Harper VanSteenhouse, Yumi Kasai, David Gray, Carol Clugston, Anna Dominiczak and Alex Alderton, Roberto Amato, Jeffrey Barrett, Sonia Goncalves, Ewan Harrison, David K. Jackson, Ian Johnston, Dominic Kwiatkowski, Cordelia Langford, John Sillitoe on behalf of the Wellcome Sanger Institute COVID-19 Surveillance Team |
| EPI_ISL_3224744 | Lighthouse Laboratory Plymouth | Wellcome Sanger Institute for the COVID-19 Genomics UK (COG-UK) Consortium | Lighthouse Laboratory Plymouth and Alex Alderton, Roberto Amato, Jeffrey Barrett, Sonia Goncalves, Ewan Harrison, David K. Jackson, Ian Johnston, Dominic Kwiatkowski, Cordelia Langford, John Sillitoe on behalf of the Wellcome Sanger Institute COVID-19 Surveillance Team |
| EPI_ISL_3224745 | Berkshire and Surrey Pathology Services Lighthouse Laboratory | Wellcome Sanger Institute for the COVID-19 Genomics UK (COG-UK) Consortium | Berkshire and Surrey Pathology Services Lighthouse Laboratory and Alex Alderton, Roberto Amato, Jeffrey Barrett, Sonia Goncalves, Ewan Harrison, David K. Jackson, Ian Johnston, Dominic Kwiatkowski, Cordelia Langford, John Sillitoe on behalf of the Wellcome Sanger Institute COVID-19 Surveillance Team |
| EPI_ISL_3224746 | Lighthouse Lab in Milton Keynes | Wellcome Sanger Institute for the COVID-19 Genomics UK (COG-UK) Consortium | The Lighthouse Lab in Milton Keynes and Alex Alderton, Roberto Amato, Jeffrey Barrett, Sonia Goncalves, Ewan Harrison, David K. Jackson, Ian Johnston, Dominic Kwiatkowski, Cordelia Langford, John Sillitoe on behalf of the Wellcome Sanger Institute COVID-19 Surveillance Team |
| EPI_ISL_3224747 | Lighthouse Laboratory Plymouth | Wellcome Sanger Institute for the COVID-19 Genomics UK (COG-UK) Consortium | Lighthouse Laboratory Plymouth and Alex Alderton, Roberto Amato, Jeffrey Barrett, Sonia Goncalves, Ewan Harrison, David K. Jackson, Ian Johnston, Dominic Kwiatkowski, Cordelia Langford, John Sillitoe on behalf of the Wellcome Sanger Institute COVID-19 Surveillance Team |
| EPI_ISL_3224748 | Lighthouse Lab in Glasgow | Wellcome Sanger Institute for the COVID-19 Genomics UK (COG-UK) Consortium | Harper VanSteenhouse, Yumi Kasai, David Gray, Carol Clugston, Anna Dominiczak and Alex Alderton, Roberto Amato, Jeffrey Barrett, Sonia Goncalves, Ewan Harrison, David K. Jackson, Ian Johnston, Dominic Kwiatkowski, Cordelia Langford, John Sillitoe on behalf of the Wellcome Sanger Institute COVID-19 Surveillance Team |
| EPI_ISL_3224749 | Lighthouse Lab in Milton Keynes | Wellcome Sanger Institute for the COVID-19 Genomics UK (COG-UK) Consortium | The Lighthouse Lab in Milton Keynes and Alex Alderton, Roberto Amato, Jeffrey Barrett, Sonia Goncalves, Ewan Harrison, David K. Jackson, Ian Johnston, Dominic Kwiatkowski, Cordelia Langford, John Sillitoe on behalf of the Wellcome Sanger Institute COVID-19 Surveillance Team |
| EPI_ISL_3224750 | Lighthouse Lab in Glasgow | Wellcome Sanger Institute for the COVID-19 Genomics UK (COG-UK) Consortium | Harper VanSteenhouse, Yumi Kasai, David Gray, Carol Clugston, Anna Dominiczak and Alex Alderton, Roberto Amato, Jeffrey Barrett, Sonia Goncalves, Ewan Harrison, David K. Jackson, Ian Johnston, Dominic Kwiatkowski, Cordelia Langford, John Sillitoe on behalf of the Wellcome Sanger Institute COVID-19 Surveillance Team |
| EPI_ISL_3224751 | Berkshire and Surrey Pathology Services Lighthouse Laboratory | Wellcome Sanger Institute for the COVID-19 Genomics UK (COG-UK) Consortium | Berkshire and Surrey Pathology Services Lighthouse Laboratory and Alex Alderton, Roberto Amato, Jeffrey Barrett, Sonia Goncalves, Ewan Harrison, David K. Jackson, Ian Johnston, Dominic Kwiatkowski, Cordelia Langford, John Sillitoe on behalf of the Wellcome Sanger Institute COVID-19 Surveillance Team |
| EPI_ISL_3224752 | Lighthouse Laboratory Plymouth | Wellcome Sanger Institute for the COVID-19 Genomics UK (COG-UK) Consortium | Lighthouse Laboratory Plymouth and Alex Alderton, Roberto Amato, Jeffrey Barrett, Sonia Goncalves, Ewan Harrison, David K. Jackson, Ian Johnston, Dominic Kwiatkowski, Cordelia Langford, John Sillitoe on behalf of the Wellcome Sanger Institute COVID-19 Surveillance Team |
| EPI_ISL_3224753 | Berkshire and Surrey Pathology Services Lighthouse Laboratory | Wellcome Sanger Institute for the COVID-19 Genomics UK (COG-UK) Consortium | Berkshire and Surrey Pathology Services Lighthouse Laboratory and Alex Alderton, Roberto Amato, Jeffrey Barrett, Sonia Goncalves, Ewan Harrison, David K. Jackson, Ian Johnston, Dominic Kwiatkowski, Cordelia Langford, John Sillitoe on behalf of the Wellcome Sanger Institute COVID-19 Surveillance Team |
| EPI_ISL_3224754 | Lighthouse Lab in Milton Keynes | Wellcome Sanger Institute for the COVID-19 Genomics UK (COG-UK) Consortium | The Lighthouse Lab in Milton Keynes and Alex Alderton, Roberto Amato, Jeffrey Barrett, Sonia Goncalves, Ewan Harrison, David K. Jackson, Ian Johnston, Dominic Kwiatkowski, Cordelia Langford, John Sillitoe on behalf of the Wellcome Sanger Institute COVID-19 Surveillance Team |
| EPI_ISL_3224755 | Berkshire and Surrey Pathology Services Lighthouse Laboratory | Wellcome Sanger Institute for the COVID-19 Genomics UK (COG-UK) Consortium | Berkshire and Surrey Pathology Services Lighthouse Laboratory and Alex Alderton, Roberto Amato, Jeffrey Barrett, Sonia Goncalves, Ewan Harrison, David K. Jackson, Ian Johnston, Dominic Kwiatkowski, Cordelia Langford, John Sillitoe on behalf of the Wellcome Sanger Institute COVID-19 Surveillance Team |
| EPI_ISL_3224756 | Lighthouse Lab in Glasgow | Wellcome Sanger Institute for the COVID-19 Genomics UK (COG-UK) Consortium | Harper VanSteenhouse, Yumi Kasai, David Gray, Carol Clugston, Anna Dominiczak and Alex Alderton, Roberto Amato, Jeffrey Barrett, Sonia Goncalves, Ewan Harrison, David K. Jackson, Ian Johnston, Dominic Kwiatkowski, Cordelia Langford, John Sillitoe on behalf of the Wellcome Sanger Institute COVID-19 Surveillance Team |
| EPI_ISL_3224757 | Lighthouse Laboratory Plymouth | Wellcome Sanger Institute for the COVID-19 Genomics UK (COG-UK) Consortium | Lighthouse Laboratory Plymouth and Alex Alderton, Roberto Amato, Jeffrey Barrett, Sonia Goncalves, Ewan Harrison, David K. Jackson, Ian Johnston, Dominic Kwiatkowski, Cordelia Langford, John Sillitoe on behalf of the Wellcome Sanger Institute COVID-19 Surveillance Team |
| EPI_ISL_3224758 | Lighthouse Lab in Milton Keynes | Wellcome Sanger Institute for the COVID-19 Genomics UK (COG-UK) Consortium | The Lighthouse Lab in Milton Keynes and Alex Alderton, Roberto Amato, Jeffrey Barrett, Sonia Goncalves, Ewan Harrison, David K. Jackson, Ian Johnston, Dominic Kwiatkowski, Cordelia Langford, John Sillitoe on behalf of the Wellcome Sanger Institute COVID-19 Surveillance Team |
| EPI_ISL_3224759 | Lighthouse Lab in Glasgow | Wellcome Sanger Institute for the COVID-19 Genomics UK (COG-UK) Consortium | Harper VanSteenhouse, Yumi Kasai, David Gray, Carol Clugston, Anna Dominiczak and Alex Alderton, Roberto Amato, Jeffrey Barrett, Sonia Goncalves, Ewan Harrison, David K. Jackson, Ian Johnston, Dominic Kwiatkowski, Cordelia Langford, John Sillitoe on behalf of the Wellcome Sanger Institute COVID-19 Surveillance Team |
| EPI_ISL_3224760, EPI_ISL_3224761 | Lighthouse Lab in Milton Keynes | Wellcome Sanger Institute for the COVID-19 Genomics UK (COG-UK) Consortium | The Lighthouse Lab in Milton Keynes and Alex Alderton, Roberto Amato, Jeffrey Barrett, Sonia Goncalves, Ewan Harrison, David K. Jackson, Ian Johnston, Dominic Kwiatkowski, Cordelia Langford, John Sillitoe on behalf of the Wellcome Sanger Institute COVID-19 Surveillance Team |
| EPI_ISL_3224762 | Lighthouse Laboratory Plymouth | Wellcome Sanger Institute for the COVID-19 Genomics UK (COG-UK) Consortium | Lighthouse Laboratory Plymouth and Alex Alderton, Roberto Amato, Jeffrey Barrett, Sonia Goncalves, Ewan Harrison, David K. Jackson, Ian Johnston, Dominic Kwiatkowski, Cordelia Langford, John Sillitoe on behalf of the Wellcome Sanger Institute COVID-19 Surveillance Team |
| EPI_ISL_3224763 | Lighthouse Lab in Milton Keynes | Wellcome Sanger Institute for the COVID-19 Genomics UK (COG-UK) Consortium | The Lighthouse Lab in Milton Keynes and Alex Alderton, Roberto Amato, Jeffrey Barrett, Sonia Goncalves, Ewan Harrison, David K. Jackson, Ian Johnston, Dominic Kwiatkowski, Cordelia Langford, John Sillitoe on behalf of the Wellcome Sanger Institute COVID-19 Surveillance Team |
| EPI_ISL_3224764 | Berkshire and Surrey Pathology Services Lighthouse Laboratory | Wellcome Sanger Institute for the COVID-19 Genomics UK (COG-UK) Consortium | Berkshire and Surrey Pathology Services Lighthouse Laboratory and Alex Alderton, Roberto Amato, Jeffrey Barrett, Sonia Goncalves, Ewan Harrison, David K. Jackson, Ian Johnston, Dominic Kwiatkowski, Cordelia Langford, John Sillitoe on behalf of the Wellcome Sanger Institute COVID-19 Surveillance Team |
| EPI_ISL_3224765 | Lighthouse Laboratory Plymouth | Wellcome Sanger Institute for the COVID-19 Genomics UK (COG-UK) Consortium | Lighthouse Laboratory Plymouth and Alex Alderton, Roberto Amato, Jeffrey Barrett, Sonia Goncalves, Ewan Harrison, David K. Jackson, Ian Johnston, Dominic Kwiatkowski, Cordelia Langford, John Sillitoe on behalf of the Wellcome Sanger Institute COVID-19 Surveillance Team |
| EPI_ISL_3224766 | Lighthouse Lab in Milton Keynes | Wellcome Sanger Institute for the COVID-19 Genomics UK (COG-UK) Consortium | The Lighthouse Lab in Milton Keynes and Alex Alderton, Roberto Amato, Jeffrey Barrett, Sonia Goncalves, Ewan Harrison, David K. Jackson, Ian Johnston, Dominic Kwiatkowski, Cordelia Langford, John Sillitoe on behalf of the Wellcome Sanger Institute COVID-19 Surveillance Team |
| EPI_ISL_3224767, EPI_ISL_3224768 | Berkshire and Surrey Pathology Services Lighthouse Laboratory | Wellcome Sanger Institute for the COVID-19 Genomics UK (COG-UK) Consortium | Berkshire and Surrey Pathology Services Lighthouse Laboratory and Alex Alderton, Roberto Amato, Jeffrey Barrett, Sonia Goncalves, Ewan Harrison, David K. Jackson, Ian Johnston, Dominic Kwiatkowski, Cordelia Langford, John Sillitoe on behalf of the Wellcome Sanger Institute COVID-19 Surveillance Team |
| EPI_ISL_3224769 | Lighthouse Lab in Milton Keynes | Wellcome Sanger Institute for the COVID-19 Genomics UK (COG-UK) Consortium | The Lighthouse Lab in Milton Keynes and Alex Alderton, Roberto Amato, Jeffrey Barrett, Sonia Goncalves, Ewan Harrison, David K. Jackson, Ian Johnston, Dominic Kwiatkowski, Cordelia Langford, John Sillitoe on behalf of the Wellcome Sanger Institute COVID-19 Surveillance Team |
| EPI_ISL_3224770 | Lighthouse Lab in Glasgow | Wellcome Sanger Institute for the COVID-19 Genomics UK (COG-UK) Consortium | Harper VanSteenhouse, Yumi Kasai, David Gray, Carol Clugston, Anna Dominiczak and Alex Alderton, Roberto Amato, Jeffrey Barrett, Sonia Goncalves, Ewan Harrison, David K. Jackson, Ian Johnston, Dominic Kwiatkowski, Cordelia Langford, John Sillitoe on behalf of the Wellcome Sanger Institute COVID-19 Surveillance Team |
| EPI_ISL_3224771 | Lighthouse Lab in Milton Keynes | Wellcome Sanger Institute for the COVID-19 Genomics UK (COG-UK) Consortium | The Lighthouse Lab in Milton Keynes and Alex Alderton, Roberto Amato, Jeffrey Barrett, Sonia Goncalves, Ewan Harrison, David K. Jackson, Ian Johnston, Dominic Kwiatkowski, Cordelia Langford, John Sillitoe on behalf of the Wellcome Sanger Institute COVID-19 Surveillance Team |
| EPI_ISL_3224772 | Lighthouse Lab in Glasgow | Wellcome Sanger Institute for the COVID-19 Genomics UK (COG-UK) Consortium | Harper VanSteenhouse, Yumi Kasai, David Gray, Carol Clugston, Anna Dominiczak and Alex Alderton, Roberto Amato, Jeffrey Barrett, Sonia Goncalves, Ewan Harrison, David K. Jackson, Ian Johnston, Dominic Kwiatkowski, Cordelia Langford, John Sillitoe on behalf of the Wellcome Sanger Institute COVID-19 Surveillance Team |
| EPI_ISL_3224773 | Lighthouse Lab in Milton Keynes | Wellcome Sanger Institute for the COVID-19 Genomics UK (COG-UK) Consortium | The Lighthouse Lab in Milton Keynes and Alex Alderton, Roberto Amato, Jeffrey Barrett, Sonia Goncalves, Ewan Harrison, David K. Jackson, Ian Johnston, Dominic Kwiatkowski, Cordelia Langford, John Sillitoe on behalf of the Wellcome Sanger Institute COVID-19 Surveillance Team |

| | | | |
|---|---|---|---|
| EPI_ISL_3224774 | Lighthouse Laboratory Plymouth | Wellcome Sanger Institute for the COVID-19 Genomics UK (COG-UK) Consortium | Lighthouse Laboratory Plymouth and Alex Alderton, Roberto Amato, Jeffrey Barrett, Sonia Goncalves, Ewan Harrison, David K. Jackson, Ian Johnston, Dominic Kwiatkowski, Cordelia Langford, John Sillitoe on behalf of the Wellcome Sanger Institute COVID-19 Surveillance Team |
| EPI_ISL_3224775 | Lighthouse Lab in Glasgow | Wellcome Sanger Institute for the COVID-19 Genomics UK (COG-UK) Consortium | Harper VanSteenhouse, Yumi Kasai, David Gray, Carol Clugston, Anna Dominiczak and Alex Alderton, Roberto Amato, Jeffrey Barrett, Sonia Goncalves, Ewan Harrison, David K. Jackson, Ian Johnston, Dominic Kwiatkowski, Cordelia Langford, John Sillitoe on behalf of the Wellcome Sanger Institute COVID-19 Surveillance Team |
| EPI_ISL_3224776 | Lighthouse Laboratory Plymouth | Wellcome Sanger Institute for the COVID-19 Genomics UK (COG-UK) Consortium | Lighthouse Laboratory Plymouth and Alex Alderton, Roberto Amato, Jeffrey Barrett, Sonia Goncalves, Ewan Harrison, David K. Jackson, Ian Johnston, Dominic Kwiatkowski, Cordelia Langford, John Sillitoe on behalf of the Wellcome Sanger Institute COVID-19 Surveillance Team |
| EPI_ISL_3224777 | Lighthouse Lab in Glasgow | Wellcome Sanger Institute for the COVID-19 Genomics UK (COG-UK) Consortium | Harper VanSteenhouse, Yumi Kasai, David Gray, Carol Clugston, Anna Dominiczak and Alex Alderton, Roberto Amato, Jeffrey Barrett, Sonia Goncalves, Ewan Harrison, David K. Jackson, Ian Johnston, Dominic Kwiatkowski, Cordelia Langford, John Sillitoe on behalf of the Wellcome Sanger Institute COVID-19 Surveillance Team |
| EPI_ISL_3224778, EPI_ISL_3224779 | Berkshire and Surrey Pathology Services Lighthouse Laboratory | Wellcome Sanger Institute for the COVID-19 Genomics UK (COG-UK) Consortium | Berkshire and Surrey Pathology Services Lighthouse Laboratory and Alex Alderton, Roberto Amato, Jeffrey Barrett, Sonia Goncalves, Ewan Harrison, David K. Jackson, Ian Johnston, Dominic Kwiatkowski, Cordelia Langford, John Sillitoe on behalf of the Wellcome Sanger Institute COVID-19 Surveillance Team |
| EPI_ISL_3224780, EPI_ISL_3224781, EPI_ISL_3224782 | Lighthouse Lab in Milton Keynes | Wellcome Sanger Institute for the COVID-19 Genomics UK (COG-UK) Consortium | The Lighthouse Lab in Milton Keynes and Alex Alderton, Roberto Amato, Jeffrey Barrett, Sonia Goncalves, Ewan Harrison, David K. Jackson, Ian Johnston, Dominic Kwiatkowski, Cordelia Langford, John Sillitoe on behalf of the Wellcome Sanger Institute COVID-19 Surveillance Team |
| EPI_ISL_3224783, EPI_ISL_3224784 | Lighthouse Laboratory Plymouth | Wellcome Sanger Institute for the COVID-19 Genomics UK (COG-UK) Consortium | Lighthouse Laboratory Plymouth and Alex Alderton, Roberto Amato, Jeffrey Barrett, Sonia Goncalves, Ewan Harrison, David K. Jackson, Ian Johnston, Dominic Kwiatkowski, Cordelia Langford, John Sillitoe on behalf of the Wellcome Sanger Institute COVID-19 Surveillance Team |
| EPI_ISL_3224785, EPI_ISL_3224786 | Berkshire and Surrey Pathology Services Lighthouse Laboratory | Wellcome Sanger Institute for the COVID-19 Genomics UK (COG-UK) Consortium | Berkshire and Surrey Pathology Services Lighthouse Laboratory and Alex Alderton, Roberto Amato, Jeffrey Barrett, Sonia Goncalves, Ewan Harrison, David K. Jackson, Ian Johnston, Dominic Kwiatkowski, Cordelia Langford, John Sillitoe on behalf of the Wellcome Sanger Institute COVID-19 Surveillance Team |
| EPI_ISL_3224787 | Lighthouse Lab in Glasgow | Wellcome Sanger Institute for the COVID-19 Genomics UK (COG-UK) Consortium | Harper VanSteenhouse, Yumi Kasai, David Gray, Carol Clugston, Anna Dominiczak and Alex Alderton, Roberto Amato, Jeffrey Barrett, Sonia Goncalves, Ewan Harrison, David K. Jackson, Ian Johnston, Dominic Kwiatkowski, Cordelia Langford, John Sillitoe on behalf of the Wellcome Sanger Institute COVID-19 Surveillance Team |
| EPI_ISL_3224788 | Berkshire and Surrey Pathology Services Lighthouse Laboratory | Wellcome Sanger Institute for the COVID-19 Genomics UK (COG-UK) Consortium | Berkshire and Surrey Pathology Services Lighthouse Laboratory and Alex Alderton, Roberto Amato, Jeffrey Barrett, Sonia Goncalves, Ewan Harrison, David K. Jackson, Ian Johnston, Dominic Kwiatkowski, Cordelia Langford, John Sillitoe on behalf of the Wellcome Sanger Institute COVID-19 Surveillance Team |
| EPI_ISL_3224789 | Lighthouse Lab in Glasgow | Wellcome Sanger Institute for the COVID-19 Genomics UK (COG-UK) Consortium | Harper VanSteenhouse, Yumi Kasai, David Gray, Carol Clugston, Anna Dominiczak and Alex Alderton, Roberto Amato, Jeffrey Barrett, Sonia Goncalves, Ewan Harrison, David K. Jackson, Ian Johnston, Dominic Kwiatkowski, Cordelia Langford, John Sillitoe on behalf of the Wellcome Sanger Institute COVID-19 Surveillance Team |
| EPI_ISL_3224790 | Lighthouse Lab in Milton Keynes | Wellcome Sanger Institute for the COVID-19 Genomics UK (COG-UK) Consortium | The Lighthouse Lab in Milton Keynes and Alex Alderton, Roberto Amato, Jeffrey Barrett, Sonia Goncalves, Ewan Harrison, David K. Jackson, Ian Johnston, Dominic Kwiatkowski, Cordelia Langford, John Sillitoe on behalf of the Wellcome Sanger Institute COVID-19 Surveillance Team |
| EPI_ISL_3224791 | Berkshire and Surrey Pathology Services Lighthouse Laboratory | Wellcome Sanger Institute for the COVID-19 Genomics UK (COG-UK) Consortium | Berkshire and Surrey Pathology Services Lighthouse Laboratory and Alex Alderton, Roberto Amato, Jeffrey Barrett, Sonia Goncalves, Ewan Harrison, David K. Jackson, Ian Johnston, Dominic Kwiatkowski, Cordelia Langford, John Sillitoe on behalf of the Wellcome Sanger Institute COVID-19 Surveillance Team |
| EPI_ISL_3224792 | Lighthouse Lab in Milton Keynes | Wellcome Sanger Institute for the COVID-19 Genomics UK (COG-UK) Consortium | The Lighthouse Lab in Milton Keynes and Alex Alderton, Roberto Amato, Jeffrey Barrett, Sonia Goncalves, Ewan Harrison, David K. Jackson, Ian Johnston, Dominic Kwiatkowski, Cordelia Langford, John Sillitoe on behalf of the Wellcome Sanger Institute COVID-19 Surveillance Team |
| EPI_ISL_3224793, EPI_ISL_3224794, EPI_ISL_3224795 | Berkshire and Surrey Pathology Services Lighthouse Laboratory | Wellcome Sanger Institute for the COVID-19 Genomics UK (COG-UK) Consortium | Berkshire and Surrey Pathology Services Lighthouse Laboratory and Alex Alderton, Roberto Amato, Jeffrey Barrett, Sonia Goncalves, Ewan Harrison, David K. Jackson, Ian Johnston, Dominic Kwiatkowski, Cordelia Langford, John Sillitoe on behalf of the Wellcome Sanger Institute COVID-19 Surveillance Team |
| EPI_ISL_3224796 | Lighthouse Lab in Milton Keynes | Wellcome Sanger Institute for the COVID-19 Genomics UK (COG-UK) Consortium | The Lighthouse Lab in Milton Keynes and Alex Alderton, Roberto Amato, Jeffrey Barrett, Sonia Goncalves, Ewan Harrison, David K. Jackson, Ian Johnston, Dominic Kwiatkowski, Cordelia Langford, John Sillitoe on behalf of the Wellcome Sanger Institute COVID-19 Surveillance Team |
| EPI_ISL_3224797 | Berkshire and Surrey Pathology Services Lighthouse Laboratory | Wellcome Sanger Institute for the COVID-19 Genomics UK (COG-UK) Consortium | Berkshire and Surrey Pathology Services Lighthouse Laboratory and Alex Alderton, Roberto Amato, Jeffrey Barrett, Sonia Goncalves, Ewan Harrison, David K. Jackson, Ian Johnston, Dominic Kwiatkowski, Cordelia Langford, John Sillitoe on behalf of the Wellcome Sanger Institute COVID-19 Surveillance Team |
| EPI_ISL_3224798 | Lighthouse Lab in Glasgow | Wellcome Sanger Institute for the COVID-19 Genomics UK (COG-UK) Consortium | Harper VanSteenhouse, Yumi Kasai, David Gray, Carol Clugston, Anna Dominiczak and Alex Alderton, Roberto Amato, Jeffrey Barrett, Sonia Goncalves, Ewan Harrison, David K. Jackson, Ian Johnston, Dominic Kwiatkowski, Cordelia Langford, John Sillitoe on behalf of the Wellcome Sanger Institute COVID-19 Surveillance Team |
| EPI_ISL_3224799 | Lighthouse Laboratory Plymouth | Wellcome Sanger Institute for the COVID-19 Genomics UK (COG-UK) Consortium | Lighthouse Laboratory Plymouth and Alex Alderton, Roberto Amato, Jeffrey Barrett, Sonia Goncalves, Ewan Harrison, David K. Jackson, Ian Johnston, Dominic Kwiatkowski, Cordelia Langford, John Sillitoe on behalf of the Wellcome Sanger Institute COVID-19 Surveillance Team |
| EPI_ISL_3224800 | Lighthouse Lab in Milton Keynes | Wellcome Sanger Institute for the COVID-19 Genomics UK (COG-UK) Consortium | The Lighthouse Lab in Milton Keynes and Alex Alderton, Roberto Amato, Jeffrey Barrett, Sonia Goncalves, Ewan Harrison, David K. Jackson, Ian Johnston, Dominic Kwiatkowski, Cordelia Langford, John Sillitoe on behalf of the Wellcome Sanger Institute COVID-19 Surveillance Team |
| EPI_ISL_3224801 | Lighthouse Lab in Glasgow | Wellcome Sanger Institute for the COVID-19 Genomics UK (COG-UK) Consortium | Harper VanSteenhouse, Yumi Kasai, David Gray, Carol Clugston, Anna Dominiczak and Alex Alderton, Roberto Amato, Jeffrey Barrett, Sonia Goncalves, Ewan Harrison, David K. Jackson, Ian Johnston, Dominic Kwiatkowski, Cordelia Langford, John Sillitoe on behalf of the Wellcome Sanger Institute COVID-19 Surveillance Team |
| EPI_ISL_3224802 | Lighthouse Lab in Milton Keynes | Wellcome Sanger Institute for the COVID-19 Genomics UK (COG-UK) Consortium | The Lighthouse Lab in Milton Keynes and Alex Alderton, Roberto Amato, Jeffrey Barrett, Sonia Goncalves, Ewan Harrison, David K. Jackson, Ian Johnston, Dominic Kwiatkowski, Cordelia Langford, John Sillitoe on behalf of the Wellcome Sanger Institute COVID-19 Surveillance Team |
| EPI_ISL_3224803 | Lighthouse Lab in Glasgow | Wellcome Sanger Institute for the COVID-19 Genomics UK (COG-UK) Consortium | Harper VanSteenhouse, Yumi Kasai, David Gray, Carol Clugston, Anna Dominiczak and Alex Alderton, Roberto Amato, Jeffrey Barrett, Sonia Goncalves, Ewan Harrison, David K. Jackson, Ian Johnston, Dominic Kwiatkowski, Cordelia Langford, John Sillitoe on behalf of the Wellcome Sanger Institute COVID-19 Surveillance Team |
| EPI_ISL_3224804 | Berkshire and Surrey Pathology Services Lighthouse Laboratory | Wellcome Sanger Institute for the COVID-19 Genomics UK (COG-UK) Consortium | Berkshire and Surrey Pathology Services Lighthouse Laboratory and Alex Alderton, Roberto Amato, Jeffrey Barrett, Sonia Goncalves, Ewan Harrison, David K. Jackson, Ian Johnston, Dominic Kwiatkowski, Cordelia Langford, John Sillitoe on behalf of the Wellcome Sanger Institute COVID-19 Surveillance Team |
| EPI_ISL_3224805 | Lighthouse Laboratory Plymouth | Wellcome Sanger Institute for the COVID-19 Genomics UK (COG-UK) Consortium | Lighthouse Laboratory Plymouth and Alex Alderton, Roberto Amato, Jeffrey Barrett, Sonia Goncalves, Ewan Harrison, David K. Jackson, Ian Johnston, Dominic Kwiatkowski, Cordelia Langford, John Sillitoe on behalf of the Wellcome Sanger Institute COVID-19 Surveillance Team |
| EPI_ISL_3224806 | Lighthouse Lab in Glasgow | Wellcome Sanger Institute for the COVID-19 Genomics UK (COG-UK) Consortium | Harper VanSteenhouse, Yumi Kasai, David Gray, Carol Clugston, Anna Dominiczak and Alex Alderton, Roberto Amato, Jeffrey Barrett, Sonia Goncalves, Ewan Harrison, David K. Jackson, Ian Johnston, Dominic Kwiatkowski, Cordelia Langford, John Sillitoe on behalf of the Wellcome Sanger Institute COVID-19 Surveillance Team |
| EPI_ISL_3224807 | Lighthouse Lab in Milton Keynes | Wellcome Sanger Institute for the COVID-19 Genomics UK (COG-UK) Consortium | The Lighthouse Lab in Milton Keynes and Alex Alderton, Roberto Amato, Jeffrey Barrett, Sonia Goncalves, Ewan Harrison, David K. Jackson, Ian Johnston, Dominic Kwiatkowski, Cordelia Langford, John Sillitoe on behalf of the Wellcome Sanger Institute COVID-19 Surveillance Team |
| EPI_ISL_3224808 | Lighthouse Lab in Glasgow | Wellcome Sanger Institute for the COVID-19 Genomics UK (COG-UK) Consortium | Harper VanSteenhouse, Yumi Kasai, David Gray, Carol Clugston, Anna Dominiczak and Alex Alderton, Roberto Amato, Jeffrey Barrett, Sonia Goncalves, Ewan Harrison, David K. Jackson, Ian Johnston, Dominic Kwiatkowski, Cordelia Langford, John Sillitoe on behalf of the Wellcome Sanger Institute COVID-19 Surveillance Team |
| EPI_ISL_3224809 | Lighthouse Laboratory Plymouth | Wellcome Sanger Institute for the COVID-19 Genomics UK (COG-UK) Consortium | Lighthouse Laboratory Plymouth and Alex Alderton, Roberto Amato, Jeffrey Barrett, Sonia Goncalves, Ewan Harrison, David K. Jackson, Ian Johnston, Dominic Kwiatkowski, Cordelia Langford, John Sillitoe on behalf of the Wellcome Sanger Institute COVID-19 Surveillance Team |
| EPI_ISL_3224810 | Berkshire and Surrey Pathology Services Lighthouse Laboratory | Wellcome Sanger Institute for the COVID-19 Genomics UK (COG-UK) Consortium | Berkshire and Surrey Pathology Services Lighthouse Laboratory and Alex Alderton, Roberto Amato, Jeffrey Barrett, Sonia Goncalves, Ewan Harrison, David K. Jackson, Ian Johnston, Dominic Kwiatkowski, Cordelia Langford, John Sillitoe on behalf of the Wellcome Sanger Institute COVID-19 Surveillance Team |
| EPI_ISL_3224811 | Lighthouse Laboratory Plymouth | Wellcome Sanger Institute for the COVID-19 Genomics UK (COG-UK) Consortium | Lighthouse Laboratory Plymouth and Alex Alderton, Roberto Amato, Jeffrey Barrett, Sonia Goncalves, Ewan Harrison, David K. Jackson, Ian Johnston, Dominic Kwiatkowski, Cordelia Langford, John Sillitoe on behalf of the Wellcome Sanger Institute COVID-19 Surveillance Team |
| EPI_ISL_3224812 | Berkshire and Surrey Pathology Services Lighthouse Laboratory | Wellcome Sanger Institute for the COVID-19 Genomics UK (COG-UK) Consortium | Berkshire and Surrey Pathology Services Lighthouse Laboratory and Alex Alderton, Roberto Amato, Jeffrey Barrett, Sonia Goncalves, Ewan Harrison, David K. Jackson, Ian Johnston, Dominic Kwiatkowski, Cordelia Langford, John Sillitoe on behalf of the Wellcome Sanger Institute COVID-19 Surveillance Team |
| EPI_ISL_3224813, EPI_ISL_3224814 | Lighthouse Laboratory Plymouth | Wellcome Sanger Institute for the COVID-19 Genomics UK (COG-UK) Consortium | Lighthouse Laboratory Plymouth and Alex Alderton, Roberto Amato, Jeffrey Barrett, Sonia Goncalves, Ewan Harrison, David K. Jackson, Ian Johnston, Dominic Kwiatkowski, Cordelia Langford, John Sillitoe on behalf of the Wellcome Sanger Institute COVID-19 Surveillance Team |
| EPI_ISL_3224815, EPI_ISL_3224816 | Lighthouse Lab in Milton Keynes | Wellcome Sanger Institute for the COVID-19 Genomics UK (COG-UK) Consortium | The Lighthouse Lab in Milton Keynes and Alex Alderton, Roberto Amato, Jeffrey Barrett, Sonia Goncalves, Ewan Harrison, David K. Jackson, Ian Johnston, Dominic Kwiatkowski, Cordelia Langford, John Sillitoe on behalf of the Wellcome Sanger Institute COVID-19 Surveillance Team |
| EPI_ISL_3224817 | Lighthouse Lab in Glasgow | Wellcome Sanger Institute for the COVID-19 Genomics UK (COG-UK) Consortium | Harper VanSteenhouse, Yumi Kasai, David Gray, Carol Clugston, Anna Dominiczak and Alex Alderton, Roberto Amato, Jeffrey Barrett, Sonia Goncalves, Ewan Harrison, David K. Jackson, Ian Johnston, Dominic Kwiatkowski, Cordelia Langford, John Sillitoe on behalf of the Wellcome Sanger Institute COVID-19 Surveillance Team |
| EPI_ISL_3224818 | Lighthouse Lab in Milton Keynes | Wellcome Sanger Institute for the COVID-19 Genomics UK (COG-UK) Consortium | The Lighthouse Lab in Milton Keynes and Alex Alderton, Roberto Amato, Jeffrey Barrett, Sonia Goncalves, Ewan Harrison, David K. Jackson, Ian Johnston, Dominic Kwiatkowski, Cordelia Langford, John Sillitoe on behalf of the Wellcome Sanger Institute COVID-19 Surveillance Team |
| EPI_ISL_3224819, EPI_ISL_3224820, EPI_ISL_3224821 | Lighthouse Laboratory Plymouth | Wellcome Sanger Institute for the COVID-19 Genomics UK (COG-UK) Consortium | Lighthouse Laboratory Plymouth and Alex Alderton, Roberto Amato, Jeffrey Barrett, Sonia Goncalves, Ewan Harrison, David K. Jackson, Ian Johnston, Dominic Kwiatkowski, Cordelia Langford, John Sillitoe on behalf of the Wellcome Sanger Institute COVID-19 Surveillance Team |
| EPI_ISL_3224822 | Berkshire and Surrey Pathology Services Lighthouse Laboratory | Wellcome Sanger Institute for the COVID-19 Genomics UK (COG-UK) Consortium | Berkshire and Surrey Pathology Services Lighthouse Laboratory and Alex Alderton, Roberto Amato, Jeffrey Barrett, Sonia Goncalves, Ewan Harrison, David K. Jackson, Ian Johnston, Dominic Kwiatkowski, Cordelia Langford, John Sillitoe on behalf of the Wellcome Sanger Institute COVID-19 Surveillance Team |
| EPI_ISL_3224823 | Lighthouse Lab in Milton Keynes | Wellcome Sanger Institute for the COVID-19 Genomics UK (COG-UK) Consortium | The Lighthouse Lab in Milton Keynes and Alex Alderton, Roberto Amato, Jeffrey Barrett, Sonia Goncalves, Ewan Harrison, David K. Jackson, Ian Johnston, Dominic Kwiatkowski, Cordelia Langford, John Sillitoe on behalf of the Wellcome Sanger Institute COVID-19 Surveillance Team |
| EPI_ISL_3224824 | Berkshire and Surrey Pathology Services Lighthouse Laboratory | Wellcome Sanger Institute for the COVID-19 Genomics UK (COG-UK) Consortium | Berkshire and Surrey Pathology Services Lighthouse Laboratory and Alex Alderton, Roberto Amato, Jeffrey Barrett, Sonia Goncalves, Ewan Harrison, David K. Jackson, Ian Johnston, Dominic Kwiatkowski, Cordelia Langford, John Sillitoe on behalf of the Wellcome Sanger Institute COVID-19 Surveillance Team |
| EPI_ISL_3224825 | Lighthouse Laboratory Plymouth | Wellcome Sanger Institute for the COVID-19 Genomics UK (COG-UK) Consortium | Lighthouse Laboratory Plymouth and Alex Alderton, Roberto Amato, Jeffrey Barrett, Sonia Goncalves, Ewan Harrison, David K. Jackson, Ian Johnston, Dominic Kwiatkowski, Cordelia Langford, John Sillitoe on behalf of the Wellcome Sanger Institute COVID-19 Surveillance Team |
| EPI_ISL_3224826 | Berkshire and Surrey Pathology Services Lighthouse Laboratory | Wellcome Sanger Institute for the COVID-19 Genomics UK (COG-UK) Consortium | Berkshire and Surrey Pathology Services Lighthouse Laboratory and Alex Alderton, Roberto Amato, Jeffrey Barrett, Sonia Goncalves, Ewan Harrison, David K. Jackson, Ian Johnston, Dominic Kwiatkowski, Cordelia Langford, John Sillitoe on behalf of the Wellcome Sanger Institute COVID-19 Surveillance Team |
| EPI_ISL_3224827 | Lighthouse Lab in Milton Keynes | Wellcome Sanger Institute for the COVID-19 Genomics UK (COG-UK) Consortium | The Lighthouse Lab in Milton Keynes and Alex Alderton, Roberto Amato, Jeffrey Barrett, Sonia Goncalves, Ewan Harrison, David K. Jackson, Ian Johnston, Dominic Kwiatkowski, Cordelia Langford, John Sillitoe on behalf of the Wellcome Sanger Institute COVID-19 Surveillance Team |
| EPI_ISL_3224828, | Berkshire and Surrey Pathology Services Lighthouse Laboratory | Wellcome Sanger Institute for the COVID-19 Genomics UK (COG-UK) | Berkshire and Surrey Pathology Services Lighthouse Laboratory and Alex Alderton, Roberto Amato, Jeffrey Barrett, Sonia Goncalves, Ewan Harrison, David K. Jackson, Ian Johnston, Dominic Kwiatkowski, Cordelia Langford, John Sillitoe on behalf |

| | | | |
|---|---|---|---|
| EPI_ISL_3224829 | | Consortium | of the Wellcome Sanger Institute COVID-19 Surveillance Team |
| EPI_ISL_3224830, EPI_ISL_3224831, EPI_ISL_3224832 | Lighthouse Laboratory Plymouth | Wellcome Sanger Institute for the COVID-19 Genomics UK (COG-UK) Consortium | Lighthouse Laboratory Plymouth and Alex Alderton, Roberto Amato, Jeffrey Barrett, Sonia Goncalves, Ewan Harrison, David K. Jackson, Ian Johnston, Dominic Kwiatkowski, Cordelia Langford, John Sillitoe on behalf of the Wellcome Sanger Institute COVID-19 Surveillance Team |
| EPI_ISL_3224833 | Lighthouse Lab in Milton Keynes | Wellcome Sanger Institute for the COVID-19 Genomics UK (COG-UK) Consortium | The Lighthouse Lab in Milton Keynes and Alex Alderton, Roberto Amato, Jeffrey Barrett, Sonia Goncalves, Ewan Harrison, David K. Jackson, Ian Johnston, Dominic Kwiatkowski, Cordelia Langford, John Sillitoe on behalf of the Wellcome Sanger Institute COVID-19 Surveillance Team |
| EPI_ISL_3224834 | Lighthouse Laboratory Plymouth | Wellcome Sanger Institute for the COVID-19 Genomics UK (COG-UK) Consortium | Lighthouse Laboratory Plymouth and Alex Alderton, Roberto Amato, Jeffrey Barrett, Sonia Goncalves, Ewan Harrison, David K. Jackson, Ian Johnston, Dominic Kwiatkowski, Cordelia Langford, John Sillitoe on behalf of the Wellcome Sanger Institute COVID-19 Surveillance Team |
| EPI_ISL_3224835, EPI_ISL_3224836, EPI_ISL_3224837 | Berkshire and Surrey Pathology Services Lighthouse Laboratory | Wellcome Sanger Institute for the COVID-19 Genomics UK (COG-UK) Consortium | Berkshire and Surrey Pathology Services Lighthouse Laboratory and Alex Alderton, Roberto Amato, Jeffrey Barrett, Sonia Goncalves, Ewan Harrison, David K. Jackson, Ian Johnston, Dominic Kwiatkowski, Cordelia Langford, John Sillitoe on behalf of the Wellcome Sanger Institute COVID-19 Surveillance Team |
| EPI_ISL_3224838 | Lighthouse Laboratory Plymouth | Wellcome Sanger Institute for the COVID-19 Genomics UK (COG-UK) Consortium | Lighthouse Laboratory Plymouth and Alex Alderton, Roberto Amato, Jeffrey Barrett, Sonia Goncalves, Ewan Harrison, David K. Jackson, Ian Johnston, Dominic Kwiatkowski, Cordelia Langford, John Sillitoe on behalf of the Wellcome Sanger Institute COVID-19 Surveillance Team |
| EPI_ISL_3224839 | Lighthouse Lab in Glasgow | Wellcome Sanger Institute for the COVID-19 Genomics UK (COG-UK) Consortium | Harper VanSteenhouse, Yumi Kasai, David Gray, Carol Clugston, Anna Dominiczak and Alex Alderton, Roberto Amato, Jeffrey Barrett, Sonia Goncalves, Ewan Harrison, David K. Jackson, Ian Johnston, Dominic Kwiatkowski, Cordelia Langford, John Sillitoe on behalf of the Wellcome Sanger Institute COVID-19 Surveillance Team |
| EPI_ISL_3224840 | Berkshire and Surrey Pathology Services Lighthouse Laboratory | Wellcome Sanger Institute for the COVID-19 Genomics UK (COG-UK) Consortium | Berkshire and Surrey Pathology Services Lighthouse Laboratory and Alex Alderton, Roberto Amato, Jeffrey Barrett, Sonia Goncalves, Ewan Harrison, David K. Jackson, Ian Johnston, Dominic Kwiatkowski, Cordelia Langford, John Sillitoe on behalf of the Wellcome Sanger Institute COVID-19 Surveillance Team |
| EPI_ISL_3224841 | Lighthouse Laboratory Plymouth | Wellcome Sanger Institute for the COVID-19 Genomics UK (COG-UK) Consortium | Lighthouse Laboratory Plymouth and Alex Alderton, Roberto Amato, Jeffrey Barrett, Sonia Goncalves, Ewan Harrison, David K. Jackson, Ian Johnston, Dominic Kwiatkowski, Cordelia Langford, John Sillitoe on behalf of the Wellcome Sanger Institute COVID-19 Surveillance Team |
| EPI_ISL_3224842, EPI_ISL_3224843 | Lighthouse Lab in Glasgow | Wellcome Sanger Institute for the COVID-19 Genomics UK (COG-UK) Consortium | Harper VanSteenhouse, Yumi Kasai, David Gray, Carol Clugston, Anna Dominiczak and Alex Alderton, Roberto Amato, Jeffrey Barrett, Sonia Goncalves, Ewan Harrison, David K. Jackson, Ian Johnston, Dominic Kwiatkowski, Cordelia Langford, John Sillitoe on behalf of the Wellcome Sanger Institute COVID-19 Surveillance Team |
| EPI_ISL_3224844 | Lighthouse Lab in Milton Keynes | Wellcome Sanger Institute for the COVID-19 Genomics UK (COG-UK) Consortium | The Lighthouse Lab in Milton Keynes and Alex Alderton, Roberto Amato, Jeffrey Barrett, Sonia Goncalves, Ewan Harrison, David K. Jackson, Ian Johnston, Dominic Kwiatkowski, Cordelia Langford, John Sillitoe on behalf of the Wellcome Sanger Institute COVID-19 Surveillance Team |
| EPI_ISL_3224845 | Lighthouse Laboratory Plymouth | Wellcome Sanger Institute for the COVID-19 Genomics UK (COG-UK) Consortium | Lighthouse Laboratory Plymouth and Alex Alderton, Roberto Amato, Jeffrey Barrett, Sonia Goncalves, Ewan Harrison, David K. Jackson, Ian Johnston, Dominic Kwiatkowski, Cordelia Langford, John Sillitoe on behalf of the Wellcome Sanger Institute COVID-19 Surveillance Team |
| EPI_ISL_3224846 | Berkshire and Surrey Pathology Services Lighthouse Laboratory | Wellcome Sanger Institute for the COVID-19 Genomics UK (COG-UK) Consortium | Berkshire and Surrey Pathology Services Lighthouse Laboratory and Alex Alderton, Roberto Amato, Jeffrey Barrett, Sonia Goncalves, Ewan Harrison, David K. Jackson, Ian Johnston, Dominic Kwiatkowski, Cordelia Langford, John Sillitoe on behalf of the Wellcome Sanger Institute COVID-19 Surveillance Team |
| EPI_ISL_3224847, EPI_ISL_3224848 | Lighthouse Lab in Glasgow | Wellcome Sanger Institute for the COVID-19 Genomics UK (COG-UK) Consortium | Harper VanSteenhouse, Yumi Kasai, David Gray, Carol Clugston, Anna Dominiczak and Alex Alderton, Roberto Amato, Jeffrey Barrett, Sonia Goncalves, Ewan Harrison, David K. Jackson, Ian Johnston, Dominic Kwiatkowski, Cordelia Langford, John Sillitoe on behalf of the Wellcome Sanger Institute COVID-19 Surveillance Team |
| EPI_ISL_3224849, EPI_ISL_3224850 | Lighthouse Laboratory Plymouth | Wellcome Sanger Institute for the COVID-19 Genomics UK (COG-UK) Consortium | Lighthouse Laboratory Plymouth and Alex Alderton, Roberto Amato, Jeffrey Barrett, Sonia Goncalves, Ewan Harrison, David K. Jackson, Ian Johnston, Dominic Kwiatkowski, Cordelia Langford, John Sillitoe on behalf of the Wellcome Sanger Institute COVID-19 Surveillance Team |
| EPI_ISL_3224851 | Lighthouse Lab in Milton Keynes | Wellcome Sanger Institute for the COVID-19 Genomics UK (COG-UK) Consortium | The Lighthouse Lab in Milton Keynes and Alex Alderton, Roberto Amato, Jeffrey Barrett, Sonia Goncalves, Ewan Harrison, David K. Jackson, Ian Johnston, Dominic Kwiatkowski, Cordelia Langford, John Sillitoe on behalf of the Wellcome Sanger Institute COVID-19 Surveillance Team |
| EPI_ISL_3224852 | Lighthouse Lab in Glasgow | Wellcome Sanger Institute for the COVID-19 Genomics UK (COG-UK) Consortium | Harper VanSteenhouse, Yumi Kasai, David Gray, Carol Clugston, Anna Dominiczak and Alex Alderton, Roberto Amato, Jeffrey Barrett, Sonia Goncalves, Ewan Harrison, David K. Jackson, Ian Johnston, Dominic Kwiatkowski, Cordelia Langford, John Sillitoe on behalf of the Wellcome Sanger Institute COVID-19 Surveillance Team |
| EPI_ISL_3224853, EPI_ISL_3224854 | Berkshire and Surrey Pathology Services Lighthouse Laboratory | Wellcome Sanger Institute for the COVID-19 Genomics UK (COG-UK) Consortium | Berkshire and Surrey Pathology Services Lighthouse Laboratory and Alex Alderton, Roberto Amato, Jeffrey Barrett, Sonia Goncalves, Ewan Harrison, David K. Jackson, Ian Johnston, Dominic Kwiatkowski, Cordelia Langford, John Sillitoe on behalf of the Wellcome Sanger Institute COVID-19 Surveillance Team |
| EPI_ISL_3224855, EPI_ISL_3224856 | Lighthouse Laboratory Plymouth | Wellcome Sanger Institute for the COVID-19 Genomics UK (COG-UK) Consortium | Lighthouse Laboratory Plymouth and Alex Alderton, Roberto Amato, Jeffrey Barrett, Sonia Goncalves, Ewan Harrison, David K. Jackson, Ian Johnston, Dominic Kwiatkowski, Cordelia Langford, John Sillitoe on behalf of the Wellcome Sanger Institute COVID-19 Surveillance Team |
| EPI_ISL_3224857 | Lighthouse Lab in Milton Keynes | Wellcome Sanger Institute for the COVID-19 Genomics UK (COG-UK) Consortium | The Lighthouse Lab in Milton Keynes and Alex Alderton, Roberto Amato, Jeffrey Barrett, Sonia Goncalves, Ewan Harrison, David K. Jackson, Ian Johnston, Dominic Kwiatkowski, Cordelia Langford, John Sillitoe on behalf of the Wellcome Sanger Institute COVID-19 Surveillance Team |
| EPI_ISL_3224858 | Lighthouse Lab in Glasgow | Wellcome Sanger Institute for the COVID-19 Genomics UK (COG-UK) Consortium | Harper VanSteenhouse, Yumi Kasai, David Gray, Carol Clugston, Anna Dominiczak and Alex Alderton, Roberto Amato, Jeffrey Barrett, Sonia Goncalves, Ewan Harrison, David K. Jackson, Ian Johnston, Dominic Kwiatkowski, Cordelia Langford, John Sillitoe on behalf of the Wellcome Sanger Institute COVID-19 Surveillance Team |
| EPI_ISL_3224859 | Berkshire and Surrey Pathology Services Lighthouse Laboratory | Wellcome Sanger Institute for the COVID-19 Genomics UK (COG-UK) Consortium | Berkshire and Surrey Pathology Services Lighthouse Laboratory and Alex Alderton, Roberto Amato, Jeffrey Barrett, Sonia Goncalves, Ewan Harrison, David K. Jackson, Ian Johnston, Dominic Kwiatkowski, Cordelia Langford, John Sillitoe on behalf of the Wellcome Sanger Institute COVID-19 Surveillance Team |
| EPI_ISL_3224860 | Lighthouse Lab in Glasgow | Wellcome Sanger Institute for the COVID-19 Genomics UK (COG-UK) Consortium | Harper VanSteenhouse, Yumi Kasai, David Gray, Carol Clugston, Anna Dominiczak and Alex Alderton, Roberto Amato, Jeffrey Barrett, Sonia Goncalves, Ewan Harrison, David K. Jackson, Ian Johnston, Dominic Kwiatkowski, Cordelia Langford, John Sillitoe on behalf of the Wellcome Sanger Institute COVID-19 Surveillance Team |
| EPI_ISL_3224861 | Berkshire and Surrey Pathology Services Lighthouse Laboratory | Wellcome Sanger Institute for the COVID-19 Genomics UK (COG-UK) Consortium | Berkshire and Surrey Pathology Services Lighthouse Laboratory and Alex Alderton, Roberto Amato, Jeffrey Barrett, Sonia Goncalves, Ewan Harrison, David K. Jackson, Ian Johnston, Dominic Kwiatkowski, Cordelia Langford, John Sillitoe on behalf of the Wellcome Sanger Institute COVID-19 Surveillance Team |
| EPI_ISL_3224862 | Lighthouse Lab in Glasgow | Wellcome Sanger Institute for the COVID-19 Genomics UK (COG-UK) Consortium | Harper VanSteenhouse, Yumi Kasai, David Gray, Carol Clugston, Anna Dominiczak and Alex Alderton, Roberto Amato, Jeffrey Barrett, Sonia Goncalves, Ewan Harrison, David K. Jackson, Ian Johnston, Dominic Kwiatkowski, Cordelia Langford, John Sillitoe on behalf of the Wellcome Sanger Institute COVID-19 Surveillance Team |
| EPI_ISL_3224863, EPI_ISL_3224864 | Lighthouse Lab in Milton Keynes | Wellcome Sanger Institute for the COVID-19 Genomics UK (COG-UK) Consortium | The Lighthouse Lab in Milton Keynes and Alex Alderton, Roberto Amato, Jeffrey Barrett, Sonia Goncalves, Ewan Harrison, David K. Jackson, Ian Johnston, Dominic Kwiatkowski, Cordelia Langford, John Sillitoe on behalf of the Wellcome Sanger Institute COVID-19 Surveillance Team |
| EPI_ISL_3224865 | Lighthouse Lab in Glasgow | Wellcome Sanger Institute for the COVID-19 Genomics UK (COG-UK) Consortium | Harper VanSteenhouse, Yumi Kasai, David Gray, Carol Clugston, Anna Dominiczak and Alex Alderton, Roberto Amato, Jeffrey Barrett, Sonia Goncalves, Ewan Harrison, David K. Jackson, Ian Johnston, Dominic Kwiatkowski, Cordelia Langford, John Sillitoe on behalf of the Wellcome Sanger Institute COVID-19 Surveillance Team |
| EPI_ISL_3224866 | Lighthouse Lab in Milton Keynes | Wellcome Sanger Institute for the COVID-19 Genomics UK (COG-UK) Consortium | The Lighthouse Lab in Milton Keynes and Alex Alderton, Roberto Amato, Jeffrey Barrett, Sonia Goncalves, Ewan Harrison, David K. Jackson, Ian Johnston, Dominic Kwiatkowski, Cordelia Langford, John Sillitoe on behalf of the Wellcome Sanger Institute COVID-19 Surveillance Team |
| EPI_ISL_3224867 | Lighthouse Lab in Glasgow | Wellcome Sanger Institute for the COVID-19 Genomics UK (COG-UK) Consortium | Harper VanSteenhouse, Yumi Kasai, David Gray, Carol Clugston, Anna Dominiczak and Alex Alderton, Roberto Amato, Jeffrey Barrett, Sonia Goncalves, Ewan Harrison, David K. Jackson, Ian Johnston, Dominic Kwiatkowski, Cordelia Langford, John Sillitoe on behalf of the Wellcome Sanger Institute COVID-19 Surveillance Team |
| EPI_ISL_3224868 | Lighthouse Lab in Milton Keynes | Wellcome Sanger Institute for the COVID-19 Genomics UK (COG-UK) Consortium | The Lighthouse Lab in Milton Keynes and Alex Alderton, Roberto Amato, Jeffrey Barrett, Sonia Goncalves, Ewan Harrison, David K. Jackson, Ian Johnston, Dominic Kwiatkowski, Cordelia Langford, John Sillitoe on behalf of the Wellcome Sanger Institute COVID-19 Surveillance Team |
| EPI_ISL_3224869 | Berkshire and Surrey Pathology Services Lighthouse Laboratory | Wellcome Sanger Institute for the COVID-19 Genomics UK (COG-UK) Consortium | Berkshire and Surrey Pathology Services Lighthouse Laboratory and Alex Alderton, Roberto Amato, Jeffrey Barrett, Sonia Goncalves, Ewan Harrison, David K. Jackson, Ian Johnston, Dominic Kwiatkowski, Cordelia Langford, John Sillitoe on behalf of the Wellcome Sanger Institute COVID-19 Surveillance Team |
| EPI_ISL_3224870, EPI_ISL_3224871 | Lighthouse Lab in Milton Keynes | Wellcome Sanger Institute for the COVID-19 Genomics UK (COG-UK) Consortium | The Lighthouse Lab in Milton Keynes and Alex Alderton, Roberto Amato, Jeffrey Barrett, Sonia Goncalves, Ewan Harrison, David K. Jackson, Ian Johnston, Dominic Kwiatkowski, Cordelia Langford, John Sillitoe on behalf of the Wellcome Sanger Institute COVID-19 Surveillance Team |
| EPI_ISL_3224872, EPI_ISL_3224873 | Berkshire and Surrey Pathology Services Lighthouse Laboratory | Wellcome Sanger Institute for the COVID-19 Genomics UK (COG-UK) Consortium | Berkshire and Surrey Pathology Services Lighthouse Laboratory and Alex Alderton, Roberto Amato, Jeffrey Barrett, Sonia Goncalves, Ewan Harrison, David K. Jackson, Ian Johnston, Dominic Kwiatkowski, Cordelia Langford, John Sillitoe on behalf of the Wellcome Sanger Institute COVID-19 Surveillance Team |
| EPI_ISL_3224874 | Lighthouse Lab in Glasgow | Wellcome Sanger Institute for the COVID-19 Genomics UK (COG-UK) Consortium | Harper VanSteenhouse, Yumi Kasai, David Gray, Carol Clugston, Anna Dominiczak and Alex Alderton, Roberto Amato, Jeffrey Barrett, Sonia Goncalves, Ewan Harrison, David K. Jackson, Ian Johnston, Dominic Kwiatkowski, Cordelia Langford, John Sillitoe on behalf of the Wellcome Sanger Institute COVID-19 Surveillance Team |
| EPI_ISL_3224875, EPI_ISL_3224876, EPI_ISL_3224877, EPI_ISL_3224878 | Lighthouse Laboratory Plymouth | Wellcome Sanger Institute for the COVID-19 Genomics UK (COG-UK) Consortium | Lighthouse Laboratory Plymouth and Alex Alderton, Roberto Amato, Jeffrey Barrett, Sonia Goncalves, Ewan Harrison, David K. Jackson, Ian Johnston, Dominic Kwiatkowski, Cordelia Langford, John Sillitoe on behalf of the Wellcome Sanger Institute COVID-19 Surveillance Team |
| EPI_ISL_3224879 | Lighthouse Lab in Glasgow | Wellcome Sanger Institute for the COVID-19 Genomics UK (COG-UK) Consortium | Harper VanSteenhouse, Yumi Kasai, David Gray, Carol Clugston, Anna Dominiczak and Alex Alderton, Roberto Amato, Jeffrey Barrett, Sonia Goncalves, Ewan Harrison, David K. Jackson, Ian Johnston, Dominic Kwiatkowski, Cordelia Langford, John Sillitoe on behalf of the Wellcome Sanger Institute COVID-19 Surveillance Team |
| EPI_ISL_3224880 | Lighthouse Laboratory Plymouth | Wellcome Sanger Institute for the COVID-19 Genomics UK (COG-UK) Consortium | Lighthouse Laboratory Plymouth and Alex Alderton, Roberto Amato, Jeffrey Barrett, Sonia Goncalves, Ewan Harrison, David K. Jackson, Ian Johnston, Dominic Kwiatkowski, Cordelia Langford, John Sillitoe on behalf of the Wellcome Sanger Institute COVID-19 Surveillance Team |
| EPI_ISL_3224881 | Berkshire and Surrey Pathology Services Lighthouse Laboratory | Wellcome Sanger Institute for the COVID-19 Genomics UK (COG-UK) Consortium | Berkshire and Surrey Pathology Services Lighthouse Laboratory and Alex Alderton, Roberto Amato, Jeffrey Barrett, Sonia Goncalves, Ewan Harrison, David K. Jackson, Ian Johnston, Dominic Kwiatkowski, Cordelia Langford, John Sillitoe on behalf of the Wellcome Sanger Institute COVID-19 Surveillance Team |
| EPI_ISL_3224882, EPI_ISL_3224883 | Lighthouse Lab in Glasgow | Wellcome Sanger Institute for the COVID-19 Genomics UK (COG-UK) Consortium | Harper VanSteenhouse, Yumi Kasai, David Gray, Carol Clugston, Anna Dominiczak and Alex Alderton, Roberto Amato, Jeffrey Barrett, Sonia Goncalves, Ewan Harrison, David K. Jackson, Ian Johnston, Dominic Kwiatkowski, Cordelia Langford, John Sillitoe on behalf of the Wellcome Sanger Institute COVID-19 Surveillance Team |
| EPI_ISL_3224884, EPI_ISL_3224885 | Lighthouse Lab in Milton Keynes | Wellcome Sanger Institute for the COVID-19 Genomics UK (COG-UK) Consortium | The Lighthouse Lab in Milton Keynes and Alex Alderton, Roberto Amato, Jeffrey Barrett, Sonia Goncalves, Ewan Harrison, David K. Jackson, Ian Johnston, Dominic Kwiatkowski, Cordelia Langford, John Sillitoe on behalf of the Wellcome Sanger Institute COVID-19 Surveillance Team |
| EPI_ISL_3224886 | Lighthouse Lab in Glasgow | Wellcome Sanger Institute for the COVID-19 Genomics UK (COG-UK) Consortium | Harper VanSteenhouse, Yumi Kasai, David Gray, Carol Clugston, Anna Dominiczak and Alex Alderton, Roberto Amato, Jeffrey Barrett, Sonia Goncalves, Ewan Harrison, David K. Jackson, Ian Johnston, Dominic Kwiatkowski, Cordelia Langford, John Sillitoe on behalf of the Wellcome Sanger Institute COVID-19 Surveillance Team |
| EPI_ISL_3224887 | Lighthouse Laboratory Plymouth | Wellcome Sanger Institute for the COVID-19 Genomics UK (COG-UK) Consortium | Lighthouse Laboratory Plymouth and Alex Alderton, Roberto Amato, Jeffrey Barrett, Sonia Goncalves, Ewan Harrison, David K. Jackson, Ian Johnston, Dominic Kwiatkowski, Cordelia Langford, John Sillitoe on behalf of the Wellcome Sanger Institute COVID-19 Surveillance Team |
| EPI_ISL_3224888 | Lighthouse Lab in Milton Keynes | Wellcome Sanger Institute for the COVID-19 Genomics UK (COG-UK) Consortium | The Lighthouse Lab in Milton Keynes and Alex Alderton, Roberto Amato, Jeffrey Barrett, Sonia Goncalves, Ewan Harrison, David K. Jackson, Ian Johnston, Dominic Kwiatkowski, Cordelia Langford, John Sillitoe on behalf of the Wellcome Sanger Institute COVID-19 Surveillance Team |
| EPI_ISL_3224889 | Lighthouse Lab in Glasgow | Wellcome Sanger Institute for the COVID-19 Genomics UK (COG-UK) | Harper VanSteenhouse, Yumi Kasai, David Gray, Carol Clugston, Anna Dominiczak and Alex Alderton, Roberto Amato, Jeffrey Barrett, Sonia Goncalves, Ewan Harrison, David K. Jackson, Ian Johnston, Dominic Kwiatkowski, Cordelia Langford, |

| | | | |
|---|---|---|---|
| | | Consortium | John Sillitoe on behalf of the Wellcome Sanger Institute COVID-19 Surveillance Team |
| EPI_ISL_3224890 | Lighthouse Laboratory Plymouth | Wellcome Sanger Institute for the COVID-19 Genomics UK (COG-UK) Consortium | Lighthouse Laboratory Plymouth and Alex Alderton, Roberto Amato, Jeffrey Barrett, Sonia Goncalves, Ewan Harrison, David K. Jackson, Ian Johnston, Dominic Kwiatkowski, Cordelia Langford, John Sillitoe on behalf of the Wellcome Sanger Institute COVID-19 Surveillance Team |
| EPI_ISL_3224891 | Lighthouse Lab in Glasgow | Wellcome Sanger Institute for the COVID-19 Genomics UK (COG-UK) Consortium | Harper VanSteenhouse, Yumi Kasai, David Gray, Carol Clugston, Anna Dominiczak and Alex Alderton, Roberto Amato, Jeffrey Barrett, Sonia Goncalves, Ewan Harrison, David K. Jackson, Ian Johnston, Dominic Kwiatkowski, Cordelia Langford, John Sillitoe on behalf of the Wellcome Sanger Institute COVID-19 Surveillance Team |
| EPI_ISL_3224892, EPI_ISL_3224893 | Lighthouse Lab in Milton Keynes | Wellcome Sanger Institute for the COVID-19 Genomics UK (COG-UK) Consortium | The Lighthouse Lab in Milton Keynes and Alex Alderton, Roberto Amato, Jeffrey Barrett, Sonia Goncalves, Ewan Harrison, David K. Jackson, Ian Johnston, Dominic Kwiatkowski, Cordelia Langford, John Sillitoe on behalf of the Wellcome Sanger Institute COVID-19 Surveillance Team |
| EPI_ISL_3224894 | Berkshire and Surrey Pathology Services Lighthouse Laboratory | Wellcome Sanger Institute for the COVID-19 Genomics UK (COG-UK) Consortium | Berkshire and Surrey Pathology Services Lighthouse Laboratory and Alex Alderton, Roberto Amato, Jeffrey Barrett, Sonia Goncalves, Ewan Harrison, David K. Jackson, Ian Johnston, Dominic Kwiatkowski, Cordelia Langford, John Sillitoe on behalf of the Wellcome Sanger Institute COVID-19 Surveillance Team |
| EPI_ISL_3224895, EPI_ISL_3224896 | Lighthouse Lab in Glasgow | Wellcome Sanger Institute for the COVID-19 Genomics UK (COG-UK) Consortium | Harper VanSteenhouse, Yumi Kasai, David Gray, Carol Clugston, Anna Dominiczak and Alex Alderton, Roberto Amato, Jeffrey Barrett, Sonia Goncalves, Ewan Harrison, David K. Jackson, Ian Johnston, Dominic Kwiatkowski, Cordelia Langford, John Sillitoe on behalf of the Wellcome Sanger Institute COVID-19 Surveillance Team |
| EPI_ISL_3224897 | Berkshire and Surrey Pathology Services Lighthouse Laboratory | Wellcome Sanger Institute for the COVID-19 Genomics UK (COG-UK) Consortium | Berkshire and Surrey Pathology Services Lighthouse Laboratory and Alex Alderton, Roberto Amato, Jeffrey Barrett, Sonia Goncalves, Ewan Harrison, David K. Jackson, Ian Johnston, Dominic Kwiatkowski, Cordelia Langford, John Sillitoe on behalf of the Wellcome Sanger Institute COVID-19 Surveillance Team |
| EPI_ISL_3224898 | Lighthouse Lab in Milton Keynes | Wellcome Sanger Institute for the COVID-19 Genomics UK (COG-UK) Consortium | The Lighthouse Lab in Milton Keynes and Alex Alderton, Roberto Amato, Jeffrey Barrett, Sonia Goncalves, Ewan Harrison, David K. Jackson, Ian Johnston, Dominic Kwiatkowski, Cordelia Langford, John Sillitoe on behalf of the Wellcome Sanger Institute COVID-19 Surveillance Team |
| EPI_ISL_3224899 | Lighthouse Lab in Glasgow | Wellcome Sanger Institute for the COVID-19 Genomics UK (COG-UK) Consortium | Harper VanSteenhouse, Yumi Kasai, David Gray, Carol Clugston, Anna Dominiczak and Alex Alderton, Roberto Amato, Jeffrey Barrett, Sonia Goncalves, Ewan Harrison, David K. Jackson, Ian Johnston, Dominic Kwiatkowski, Cordelia Langford, John Sillitoe on behalf of the Wellcome Sanger Institute COVID-19 Surveillance Team |
| EPI_ISL_3224900, EPI_ISL_3224901 | Lighthouse Laboratory Plymouth | Wellcome Sanger Institute for the COVID-19 Genomics UK (COG-UK) Consortium | Lighthouse Laboratory Plymouth and Alex Alderton, Roberto Amato, Jeffrey Barrett, Sonia Goncalves, Ewan Harrison, David K. Jackson, Ian Johnston, Dominic Kwiatkowski, Cordelia Langford, John Sillitoe on behalf of the Wellcome Sanger Institute COVID-19 Surveillance Team |
| EPI_ISL_3224902 | Lighthouse Lab in Glasgow | Wellcome Sanger Institute for the COVID-19 Genomics UK (COG-UK) Consortium | Harper VanSteenhouse, Yumi Kasai, David Gray, Carol Clugston, Anna Dominiczak and Alex Alderton, Roberto Amato, Jeffrey Barrett, Sonia Goncalves, Ewan Harrison, David K. Jackson, Ian Johnston, Dominic Kwiatkowski, Cordelia Langford, John Sillitoe on behalf of the Wellcome Sanger Institute COVID-19 Surveillance Team |
| EPI_ISL_3224903 | Lighthouse Lab in Milton Keynes | Wellcome Sanger Institute for the COVID-19 Genomics UK (COG-UK) Consortium | The Lighthouse Lab in Milton Keynes and Alex Alderton, Roberto Amato, Jeffrey Barrett, Sonia Goncalves, Ewan Harrison, David K. Jackson, Ian Johnston, Dominic Kwiatkowski, Cordelia Langford, John Sillitoe on behalf of the Wellcome Sanger Institute COVID-19 Surveillance Team |
| EPI_ISL_3224904 | Lighthouse Laboratory Plymouth | Wellcome Sanger Institute for the COVID-19 Genomics UK (COG-UK) Consortium | Lighthouse Laboratory Plymouth and Alex Alderton, Roberto Amato, Jeffrey Barrett, Sonia Goncalves, Ewan Harrison, David K. Jackson, Ian Johnston, Dominic Kwiatkowski, Cordelia Langford, John Sillitoe on behalf of the Wellcome Sanger Institute COVID-19 Surveillance Team |
| EPI_ISL_3224905 | Lighthouse Lab in Glasgow | Wellcome Sanger Institute for the COVID-19 Genomics UK (COG-UK) Consortium | Harper VanSteenhouse, Yumi Kasai, David Gray, Carol Clugston, Anna Dominiczak and Alex Alderton, Roberto Amato, Jeffrey Barrett, Sonia Goncalves, Ewan Harrison, David K. Jackson, Ian Johnston, Dominic Kwiatkowski, Cordelia Langford, John Sillitoe on behalf of the Wellcome Sanger Institute COVID-19 Surveillance Team |
| EPI_ISL_3224906 | Lighthouse Lab in Milton Keynes | Wellcome Sanger Institute for the COVID-19 Genomics UK (COG-UK) Consortium | The Lighthouse Lab in Milton Keynes and Alex Alderton, Roberto Amato, Jeffrey Barrett, Sonia Goncalves, Ewan Harrison, David K. Jackson, Ian Johnston, Dominic Kwiatkowski, Cordelia Langford, John Sillitoe on behalf of the Wellcome Sanger Institute COVID-19 Surveillance Team |
| EPI_ISL_3224907 | Lighthouse Laboratory Plymouth | Wellcome Sanger Institute for the COVID-19 Genomics UK (COG-UK) Consortium | Lighthouse Laboratory Plymouth and Alex Alderton, Roberto Amato, Jeffrey Barrett, Sonia Goncalves, Ewan Harrison, David K. Jackson, Ian Johnston, Dominic Kwiatkowski, Cordelia Langford, John Sillitoe on behalf of the Wellcome Sanger Institute COVID-19 Surveillance Team |
| EPI_ISL_3224908 | Lighthouse Lab in Glasgow | Wellcome Sanger Institute for the COVID-19 Genomics UK (COG-UK) Consortium | Harper VanSteenhouse, Yumi Kasai, David Gray, Carol Clugston, Anna Dominiczak and Alex Alderton, Roberto Amato, Jeffrey Barrett, Sonia Goncalves, Ewan Harrison, David K. Jackson, Ian Johnston, Dominic Kwiatkowski, Cordelia Langford, John Sillitoe on behalf of the Wellcome Sanger Institute COVID-19 Surveillance Team |
| EPI_ISL_3224909 | Lighthouse Laboratory Plymouth | Wellcome Sanger Institute for the COVID-19 Genomics UK (COG-UK) Consortium | Lighthouse Laboratory Plymouth and Alex Alderton, Roberto Amato, Jeffrey Barrett, Sonia Goncalves, Ewan Harrison, David K. Jackson, Ian Johnston, Dominic Kwiatkowski, Cordelia Langford, John Sillitoe on behalf of the Wellcome Sanger Institute COVID-19 Surveillance Team |
| EPI_ISL_3224910 | Berkshire and Surrey Pathology Services Lighthouse Laboratory | Wellcome Sanger Institute for the COVID-19 Genomics UK (COG-UK) Consortium | Berkshire and Surrey Pathology Services Lighthouse Laboratory and Alex Alderton, Roberto Amato, Jeffrey Barrett, Sonia Goncalves, Ewan Harrison, David K. Jackson, Ian Johnston, Dominic Kwiatkowski, Cordelia Langford, John Sillitoe on behalf of the Wellcome Sanger Institute COVID-19 Surveillance Team |
| EPI_ISL_3224911 | Lighthouse Lab in Milton Keynes | Wellcome Sanger Institute for the COVID-19 Genomics UK (COG-UK) Consortium | The Lighthouse Lab in Milton Keynes and Alex Alderton, Roberto Amato, Jeffrey Barrett, Sonia Goncalves, Ewan Harrison, David K. Jackson, Ian Johnston, Dominic Kwiatkowski, Cordelia Langford, John Sillitoe on behalf of the Wellcome Sanger Institute COVID-19 Surveillance Team |
| EPI_ISL_3224912, EPI_ISL_3224913, EPI_ISL_3224914, EPI_ISL_3224915, EPI_ISL_3224916, EPI_ISL_3224917 | Lighthouse Lab in Glasgow | Wellcome Sanger Institute for the COVID-19 Genomics UK (COG-UK) Consortium | Harper VanSteenhouse, Yumi Kasai, David Gray, Carol Clugston, Anna Dominiczak and Alex Alderton, Roberto Amato, Jeffrey Barrett, Sonia Goncalves, Ewan Harrison, David K. Jackson, Ian Johnston, Dominic Kwiatkowski, Cordelia Langford, John Sillitoe on behalf of the Wellcome Sanger Institute COVID-19 Surveillance Team |
| EPI_ISL_3224918, EPI_ISL_3224919 | Lighthouse Lab in Alderley Park | Wellcome Sanger Institute for the COVID-19 Genomics UK (COG-UK) Consortium | Jacquelyn Wynn, Mairead Hyland, The Lighthouse Lab in Alderley Park and Alex Alderton, Roberto Amato, Jeffrey Barrett, Sonia Goncalves, Ewan Harrison, David K. Jackson, Ian Johnston, Dominic Kwiatkowski, Cordelia Langford, John Sillitoe on behalf of the Wellcome Sanger Institute COVID-19 Surveillance Team |
| EPI_ISL_3224920 | Lighthouse Laboratory Plymouth | Wellcome Sanger Institute for the COVID-19 Genomics UK (COG-UK) Consortium | Lighthouse Laboratory Plymouth and Alex Alderton, Roberto Amato, Jeffrey Barrett, Sonia Goncalves, Ewan Harrison, David K. Jackson, Ian Johnston, Dominic Kwiatkowski, Cordelia Langford, John Sillitoe on behalf of the Wellcome Sanger Institute COVID-19 Surveillance Team |
| EPI_ISL_3224921 | Lighthouse Lab in Glasgow | Wellcome Sanger Institute for the COVID-19 Genomics UK (COG-UK) Consortium | Harper VanSteenhouse, Yumi Kasai, David Gray, Carol Clugston, Anna Dominiczak and Alex Alderton, Roberto Amato, Jeffrey Barrett, Sonia Goncalves, Ewan Harrison, David K. Jackson, Ian Johnston, Dominic Kwiatkowski, Cordelia Langford, John Sillitoe on behalf of the Wellcome Sanger Institute COVID-19 Surveillance Team |
| EPI_ISL_3224922 | Berkshire and Surrey Pathology Services Lighthouse Laboratory | Wellcome Sanger Institute for the COVID-19 Genomics UK (COG-UK) Consortium | Berkshire and Surrey Pathology Services Lighthouse Laboratory and Alex Alderton, Roberto Amato, Jeffrey Barrett, Sonia Goncalves, Ewan Harrison, David K. Jackson, Ian Johnston, Dominic Kwiatkowski, Cordelia Langford, John Sillitoe on behalf of the Wellcome Sanger Institute COVID-19 Surveillance Team |
| EPI_ISL_3224923 | Lighthouse Laboratory Plymouth | Wellcome Sanger Institute for the COVID-19 Genomics UK (COG-UK) Consortium | Lighthouse Laboratory Plymouth and Alex Alderton, Roberto Amato, Jeffrey Barrett, Sonia Goncalves, Ewan Harrison, David K. Jackson, Ian Johnston, Dominic Kwiatkowski, Cordelia Langford, John Sillitoe on behalf of the Wellcome Sanger Institute COVID-19 Surveillance Team |
| EPI_ISL_3224924 | Lighthouse Lab in Glasgow | Wellcome Sanger Institute for the COVID-19 Genomics UK (COG-UK) Consortium | Harper VanSteenhouse, Yumi Kasai, David Gray, Carol Clugston, Anna Dominiczak and Alex Alderton, Roberto Amato, Jeffrey Barrett, Sonia Goncalves, Ewan Harrison, David K. Jackson, Ian Johnston, Dominic Kwiatkowski, Cordelia Langford, John Sillitoe on behalf of the Wellcome Sanger Institute COVID-19 Surveillance Team |
| EPI_ISL_3224925, EPI_ISL_3224926 | Berkshire and Surrey Pathology Services Lighthouse Laboratory | Wellcome Sanger Institute for the COVID-19 Genomics UK (COG-UK) Consortium | Berkshire and Surrey Pathology Services Lighthouse Laboratory and Alex Alderton, Roberto Amato, Jeffrey Barrett, Sonia Goncalves, Ewan Harrison, David K. Jackson, Ian Johnston, Dominic Kwiatkowski, Cordelia Langford, John Sillitoe on behalf of the Wellcome Sanger Institute COVID-19 Surveillance Team |
| EPI_ISL_3224927 | Lighthouse Lab in Glasgow | Wellcome Sanger Institute for the COVID-19 Genomics UK (COG-UK) Consortium | Harper VanSteenhouse, Yumi Kasai, David Gray, Carol Clugston, Anna Dominiczak and Alex Alderton, Roberto Amato, Jeffrey Barrett, Sonia Goncalves, Ewan Harrison, David K. Jackson, Ian Johnston, Dominic Kwiatkowski, Cordelia Langford, John Sillitoe on behalf of the Wellcome Sanger Institute COVID-19 Surveillance Team |
| EPI_ISL_3224928, EPI_ISL_3224929 | Berkshire and Surrey Pathology Services Lighthouse Laboratory | Wellcome Sanger Institute for the COVID-19 Genomics UK (COG-UK) Consortium | Berkshire and Surrey Pathology Services Lighthouse Laboratory and Alex Alderton, Roberto Amato, Jeffrey Barrett, Sonia Goncalves, Ewan Harrison, David K. Jackson, Ian Johnston, Dominic Kwiatkowski, Cordelia Langford, John Sillitoe on behalf of the Wellcome Sanger Institute COVID-19 Surveillance Team |
| EPI_ISL_3224930 | Lighthouse Lab in Glasgow | Wellcome Sanger Institute for the COVID-19 Genomics UK (COG-UK) Consortium | Harper VanSteenhouse, Yumi Kasai, David Gray, Carol Clugston, Anna Dominiczak and Alex Alderton, Roberto Amato, Jeffrey Barrett, Sonia Goncalves, Ewan Harrison, David K. Jackson, Ian Johnston, Dominic Kwiatkowski, Cordelia Langford, John Sillitoe on behalf of the Wellcome Sanger Institute COVID-19 Surveillance Team |
| EPI_ISL_3224931, EPI_ISL_3224932 | Lighthouse Lab in Alderley Park | Wellcome Sanger Institute for the COVID-19 Genomics UK (COG-UK) Consortium | Jacquelyn Wynn, Mairead Hyland, The Lighthouse Lab in Alderley Park and Alex Alderton, Roberto Amato, Jeffrey Barrett, Sonia Goncalves, Ewan Harrison, David K. Jackson, Ian Johnston, Dominic Kwiatkowski, Cordelia Langford, John Sillitoe on behalf of the Wellcome Sanger Institute COVID-19 Surveillance Team |
| EPI_ISL_3224933 | Lighthouse Lab in Glasgow | Wellcome Sanger Institute for the COVID-19 Genomics UK (COG-UK) Consortium | Harper VanSteenhouse, Yumi Kasai, David Gray, Carol Clugston, Anna Dominiczak and Alex Alderton, Roberto Amato, Jeffrey Barrett, Sonia Goncalves, Ewan Harrison, David K. Jackson, Ian Johnston, Dominic Kwiatkowski, Cordelia Langford, John Sillitoe on behalf of the Wellcome Sanger Institute COVID-19 Surveillance Team |
| EPI_ISL_3224934 | Lighthouse Lab in Alderley Park | Wellcome Sanger Institute for the COVID-19 Genomics UK (COG-UK) Consortium | Jacquelyn Wynn, Mairead Hyland, The Lighthouse Lab in Alderley Park and Alex Alderton, Roberto Amato, Jeffrey Barrett, Sonia Goncalves, Ewan Harrison, David K. Jackson, Ian Johnston, Dominic Kwiatkowski, Cordelia Langford, John Sillitoe on behalf of the Wellcome Sanger Institute COVID-19 Surveillance Team |
| EPI_ISL_3224935 | Berkshire and Surrey Pathology Services Lighthouse Laboratory | Wellcome Sanger Institute for the COVID-19 Genomics UK (COG-UK) Consortium | Berkshire and Surrey Pathology Services Lighthouse Laboratory and Alex Alderton, Roberto Amato, Jeffrey Barrett, Sonia Goncalves, Ewan Harrison, David K. Jackson, Ian Johnston, Dominic Kwiatkowski, Cordelia Langford, John Sillitoe on behalf of the Wellcome Sanger Institute COVID-19 Surveillance Team |
| EPI_ISL_3224936 | Lighthouse Lab in Alderley Park | Wellcome Sanger Institute for the COVID-19 Genomics UK (COG-UK) Consortium | Jacquelyn Wynn, Mairead Hyland, The Lighthouse Lab in Alderley Park and Alex Alderton, Roberto Amato, Jeffrey Barrett, Sonia Goncalves, Ewan Harrison, David K. Jackson, Ian Johnston, Dominic Kwiatkowski, Cordelia Langford, John Sillitoe on behalf of the Wellcome Sanger Institute COVID-19 Surveillance Team |
| EPI_ISL_3224937, EPI_ISL_3224938, EPI_ISL_3224939 | Lighthouse Lab in Glasgow | Wellcome Sanger Institute for the COVID-19 Genomics UK (COG-UK) Consortium | Harper VanSteenhouse, Yumi Kasai, David Gray, Carol Clugston, Anna Dominiczak and Alex Alderton, Roberto Amato, Jeffrey Barrett, Sonia Goncalves, Ewan Harrison, David K. Jackson, Ian Johnston, Dominic Kwiatkowski, Cordelia Langford, John Sillitoe on behalf of the Wellcome Sanger Institute COVID-19 Surveillance Team |
| EPI_ISL_3224940 | Lighthouse Laboratory Plymouth | Wellcome Sanger Institute for the COVID-19 Genomics UK (COG-UK) Consortium | Lighthouse Laboratory Plymouth and Alex Alderton, Roberto Amato, Jeffrey Barrett, Sonia Goncalves, Ewan Harrison, David K. Jackson, Ian Johnston, Dominic Kwiatkowski, Cordelia Langford, John Sillitoe on behalf of the Wellcome Sanger Institute COVID-19 Surveillance Team |
| EPI_ISL_3224941 | Lighthouse Lab in Alderley Park | Wellcome Sanger Institute for the COVID-19 Genomics UK (COG-UK) Consortium | Jacquelyn Wynn, Mairead Hyland, The Lighthouse Lab in Alderley Park and Alex Alderton, Roberto Amato, Jeffrey Barrett, Sonia Goncalves, Ewan Harrison, David K. Jackson, Ian Johnston, Dominic Kwiatkowski, Cordelia Langford, John Sillitoe on behalf of the Wellcome Sanger Institute COVID-19 Surveillance Team |
| EPI_ISL_3224942, EPI_ISL_3224943, EPI_ISL_3224944 | Lighthouse Laboratory Plymouth | Wellcome Sanger Institute for the COVID-19 Genomics UK (COG-UK) Consortium | Lighthouse Laboratory Plymouth and Alex Alderton, Roberto Amato, Jeffrey Barrett, Sonia Goncalves, Ewan Harrison, David K. Jackson, Ian Johnston, Dominic Kwiatkowski, Cordelia Langford, John Sillitoe on behalf of the Wellcome Sanger Institute COVID-19 Surveillance Team |
| EPI_ISL_3224945, EPI_ISL_3224946 | Lighthouse Lab in Alderley Park | Wellcome Sanger Institute for the COVID-19 Genomics UK (COG-UK) Consortium | Jacquelyn Wynn, Mairead Hyland, The Lighthouse Lab in Alderley Park and Alex Alderton, Roberto Amato, Jeffrey Barrett, Sonia Goncalves, Ewan Harrison, David K. Jackson, Ian Johnston, Dominic Kwiatkowski, Cordelia Langford, John Sillitoe on behalf of the Wellcome Sanger Institute COVID-19 Surveillance Team |
| EPI_ISL_3224947 | Lighthouse Laboratory Plymouth | Wellcome Sanger Institute for the COVID-19 Genomics UK (COG-UK) Consortium | Lighthouse Laboratory Plymouth and Alex Alderton, Roberto Amato, Jeffrey Barrett, Sonia Goncalves, Ewan Harrison, David K. Jackson, Ian Johnston, Dominic Kwiatkowski, Cordelia Langford, John Sillitoe on behalf of the Wellcome Sanger Institute COVID-19 Surveillance Team |
| EPI_ISL_3224948 | Lighthouse Lab in Glasgow | Wellcome Sanger Institute for the COVID-19 Genomics UK (COG-UK) Consortium | Harper VanSteenhouse, Yumi Kasai, David Gray, Carol Clugston, Anna Dominiczak and Alex Alderton, Roberto Amato, Jeffrey Barrett, Sonia Goncalves, Ewan Harrison, David K. Jackson, Ian Johnston, Dominic Kwiatkowski, Cordelia Langford, John Sillitoe on behalf of the Wellcome Sanger Institute COVID-19 Surveillance Team |

| | | | |
|---|---|---|---|
| EPI_ISL_3224949 | Lighthouse Lab in Alderley Park | Wellcome Sanger Institute for the COVID-19 Genomics UK (COG-UK) Consortium | Jacquelyn Wynn, Mairead Hyland, The Lighthouse Lab in Alderley Park and Alex Alderton, Roberto Amato, Jeffrey Barrett, Sonia Goncalves, Ewan Harrison, David K. Jackson, Ian Johnston, Dominic Kwiatkowski, Cordelia Langford, John Sillitoe on behalf of the Wellcome Sanger Institute COVID-19 Surveillance Team |
| EPI_ISL_3224950 | Lighthouse Lab in Glasgow | Wellcome Sanger Institute for the COVID-19 Genomics UK (COG-UK) Consortium | Harper VanSteenhouse, Yumi Kasai, David Gray, Carol Clugston, Anna Dominiczak and Alex Alderton, Roberto Amato, Jeffrey Barrett, Sonia Goncalves, Ewan Harrison, David K. Jackson, Ian Johnston, Dominic Kwiatkowski, Cordelia Langford, John Sillitoe on behalf of the Wellcome Sanger Institute COVID-19 Surveillance Team |
| EPI_ISL_3224951 | Lighthouse Laboratory Plymouth | Wellcome Sanger Institute for the COVID-19 Genomics UK (COG-UK) Consortium | Lighthouse Laboratory Plymouth and Alex Alderton, Roberto Amato, Jeffrey Barrett, Sonia Goncalves, Ewan Harrison, David K. Jackson, Ian Johnston, Dominic Kwiatkowski, Cordelia Langford, John Sillitoe on behalf of the Wellcome Sanger Institute COVID-19 Surveillance Team |
| EPI_ISL_3224952 | Lighthouse Lab in Glasgow | Wellcome Sanger Institute for the COVID-19 Genomics UK (COG-UK) Consortium | Harper VanSteenhouse, Yumi Kasai, David Gray, Carol Clugston, Anna Dominiczak and Alex Alderton, Roberto Amato, Jeffrey Barrett, Sonia Goncalves, Ewan Harrison, David K. Jackson, Ian Johnston, Dominic Kwiatkowski, Cordelia Langford, John Sillitoe on behalf of the Wellcome Sanger Institute COVID-19 Surveillance Team |
| EPI_ISL_3224953, EPI_ISL_3224954, EPI_ISL_3224955 | Lighthouse Laboratory Plymouth | Wellcome Sanger Institute for the COVID-19 Genomics UK (COG-UK) Consortium | Lighthouse Laboratory Plymouth and Alex Alderton, Roberto Amato, Jeffrey Barrett, Sonia Goncalves, Ewan Harrison, David K. Jackson, Ian Johnston, Dominic Kwiatkowski, Cordelia Langford, John Sillitoe on behalf of the Wellcome Sanger Institute COVID-19 Surveillance Team |
| EPI_ISL_3224956 | Lighthouse Lab in Alderley Park | Wellcome Sanger Institute for the COVID-19 Genomics UK (COG-UK) Consortium | Jacquelyn Wynn, Mairead Hyland, The Lighthouse Lab in Alderley Park and Alex Alderton, Roberto Amato, Jeffrey Barrett, Sonia Goncalves, Ewan Harrison, David K. Jackson, Ian Johnston, Dominic Kwiatkowski, Cordelia Langford, John Sillitoe on behalf of the Wellcome Sanger Institute COVID-19 Surveillance Team |
| EPI_ISL_3224957 | Berkshire and Surrey Pathology Services Lighthouse Laboratory | Wellcome Sanger Institute for the COVID-19 Genomics UK (COG-UK) Consortium | Berkshire and Surrey Pathology Services Lighthouse Laboratory and Alex Alderton, Roberto Amato, Jeffrey Barrett, Sonia Goncalves, Ewan Harrison, David K. Jackson, Ian Johnston, Dominic Kwiatkowski, Cordelia Langford, John Sillitoe on behalf of the Wellcome Sanger Institute COVID-19 Surveillance Team |
| EPI_ISL_3224958 | Lighthouse Lab in Glasgow | Wellcome Sanger Institute for the COVID-19 Genomics UK (COG-UK) Consortium | Harper VanSteenhouse, Yumi Kasai, David Gray, Carol Clugston, Anna Dominiczak and Alex Alderton, Roberto Amato, Jeffrey Barrett, Sonia Goncalves, Ewan Harrison, David K. Jackson, Ian Johnston, Dominic Kwiatkowski, Cordelia Langford, John Sillitoe on behalf of the Wellcome Sanger Institute COVID-19 Surveillance Team |
| EPI_ISL_3224959 | Lighthouse Lab in Alderley Park | Wellcome Sanger Institute for the COVID-19 Genomics UK (COG-UK) Consortium | Jacquelyn Wynn, Mairead Hyland, The Lighthouse Lab in Alderley Park and Alex Alderton, Roberto Amato, Jeffrey Barrett, Sonia Goncalves, Ewan Harrison, David K. Jackson, Ian Johnston, Dominic Kwiatkowski, Cordelia Langford, John Sillitoe on behalf of the Wellcome Sanger Institute COVID-19 Surveillance Team |
| EPI_ISL_3224960 | Berkshire and Surrey Pathology Services Lighthouse Laboratory | Wellcome Sanger Institute for the COVID-19 Genomics UK (COG-UK) Consortium | Berkshire and Surrey Pathology Services Lighthouse Laboratory and Alex Alderton, Roberto Amato, Jeffrey Barrett, Sonia Goncalves, Ewan Harrison, David K. Jackson, Ian Johnston, Dominic Kwiatkowski, Cordelia Langford, John Sillitoe on behalf of the Wellcome Sanger Institute COVID-19 Surveillance Team |
| EPI_ISL_3224961 | Lighthouse Lab in Glasgow | Wellcome Sanger Institute for the COVID-19 Genomics UK (COG-UK) Consortium | Harper VanSteenhouse, Yumi Kasai, David Gray, Carol Clugston, Anna Dominiczak and Alex Alderton, Roberto Amato, Jeffrey Barrett, Sonia Goncalves, Ewan Harrison, David K. Jackson, Ian Johnston, Dominic Kwiatkowski, Cordelia Langford, John Sillitoe on behalf of the Wellcome Sanger Institute COVID-19 Surveillance Team |
| EPI_ISL_3224962 | Lighthouse Lab in Alderley Park | Wellcome Sanger Institute for the COVID-19 Genomics UK (COG-UK) Consortium | Jacquelyn Wynn, Mairead Hyland, The Lighthouse Lab in Alderley Park and Alex Alderton, Roberto Amato, Jeffrey Barrett, Sonia Goncalves, Ewan Harrison, David K. Jackson, Ian Johnston, Dominic Kwiatkowski, Cordelia Langford, John Sillitoe on behalf of the Wellcome Sanger Institute COVID-19 Surveillance Team |
| EPI_ISL_3224963 | Lighthouse Laboratory Plymouth | Wellcome Sanger Institute for the COVID-19 Genomics UK (COG-UK) Consortium | Lighthouse Laboratory Plymouth and Alex Alderton, Roberto Amato, Jeffrey Barrett, Sonia Goncalves, Ewan Harrison, David K. Jackson, Ian Johnston, Dominic Kwiatkowski, Cordelia Langford, John Sillitoe on behalf of the Wellcome Sanger Institute COVID-19 Surveillance Team |
| EPI_ISL_3224964, EPI_ISL_3224965, EPI_ISL_3224966, EPI_ISL_3224967 | Lighthouse Lab in Alderley Park | Wellcome Sanger Institute for the COVID-19 Genomics UK (COG-UK) Consortium | Jacquelyn Wynn, Mairead Hyland, The Lighthouse Lab in Alderley Park and Alex Alderton, Roberto Amato, Jeffrey Barrett, Sonia Goncalves, Ewan Harrison, David K. Jackson, Ian Johnston, Dominic Kwiatkowski, Cordelia Langford, John Sillitoe on behalf of the Wellcome Sanger Institute COVID-19 Surveillance Team |
| EPI_ISL_3224968 | Lighthouse Lab in Glasgow | Wellcome Sanger Institute for the COVID-19 Genomics UK (COG-UK) Consortium | Harper VanSteenhouse, Yumi Kasai, David Gray, Carol Clugston, Anna Dominiczak and Alex Alderton, Roberto Amato, Jeffrey Barrett, Sonia Goncalves, Ewan Harrison, David K. Jackson, Ian Johnston, Dominic Kwiatkowski, Cordelia Langford, John Sillitoe on behalf of the Wellcome Sanger Institute COVID-19 Surveillance Team |
| EPI_ISL_3224969 | Lighthouse Laboratory Plymouth | Wellcome Sanger Institute for the COVID-19 Genomics UK (COG-UK) Consortium | Lighthouse Laboratory Plymouth and Alex Alderton, Roberto Amato, Jeffrey Barrett, Sonia Goncalves, Ewan Harrison, David K. Jackson, Ian Johnston, Dominic Kwiatkowski, Cordelia Langford, John Sillitoe on behalf of the Wellcome Sanger Institute COVID-19 Surveillance Team |
| EPI_ISL_3224970 | Lighthouse Lab in Alderley Park | Wellcome Sanger Institute for the COVID-19 Genomics UK (COG-UK) Consortium | Jacquelyn Wynn, Mairead Hyland, The Lighthouse Lab in Alderley Park and Alex Alderton, Roberto Amato, Jeffrey Barrett, Sonia Goncalves, Ewan Harrison, David K. Jackson, Ian Johnston, Dominic Kwiatkowski, Cordelia Langford, John Sillitoe on behalf of the Wellcome Sanger Institute COVID-19 Surveillance Team |
| EPI_ISL_3224971 | Berkshire and Surrey Pathology Services Lighthouse Laboratory | Wellcome Sanger Institute for the COVID-19 Genomics UK (COG-UK) Consortium | Berkshire and Surrey Pathology Services Lighthouse Laboratory and Alex Alderton, Roberto Amato, Jeffrey Barrett, Sonia Goncalves, Ewan Harrison, David K. Jackson, Ian Johnston, Dominic Kwiatkowski, Cordelia Langford, John Sillitoe on behalf of the Wellcome Sanger Institute COVID-19 Surveillance Team |
| EPI_ISL_3224972 | Health Services Laboratories | Wellcome Sanger Institute for the COVID-19 Genomics UK (COG-UK) Consortium | Health Services Laboratories and Alex Alderton, Roberto Amato, Jeffrey Barrett, Sonia Goncalves, Ewan Harrison, David K. Jackson, Ian Johnston, Dominic Kwiatkowski, Cordelia Langford, John Sillitoe on behalf of the Wellcome Sanger Institute COVID-19 Surveillance Team |
| EPI_ISL_3224973, EPI_ISL_3224974 | Lighthouse Lab in Alderley Park | Wellcome Sanger Institute for the COVID-19 Genomics UK (COG-UK) Consortium | Jacquelyn Wynn, Mairead Hyland, The Lighthouse Lab in Alderley Park and Alex Alderton, Roberto Amato, Jeffrey Barrett, Sonia Goncalves, Ewan Harrison, David K. Jackson, Ian Johnston, Dominic Kwiatkowski, Cordelia Langford, John Sillitoe on behalf of the Wellcome Sanger Institute COVID-19 Surveillance Team |
| EPI_ISL_3224975 | Lighthouse Laboratory Plymouth | Wellcome Sanger Institute for the COVID-19 Genomics UK (COG-UK) Consortium | Lighthouse Laboratory Plymouth and Alex Alderton, Roberto Amato, Jeffrey Barrett, Sonia Goncalves, Ewan Harrison, David K. Jackson, Ian Johnston, Dominic Kwiatkowski, Cordelia Langford, John Sillitoe on behalf of the Wellcome Sanger Institute COVID-19 Surveillance Team |
| EPI_ISL_3224976, EPI_ISL_3224977, EPI_ISL_3224978 | Lighthouse Lab in Alderley Park | Wellcome Sanger Institute for the COVID-19 Genomics UK (COG-UK) Consortium | Jacquelyn Wynn, Mairead Hyland, The Lighthouse Lab in Alderley Park and Alex Alderton, Roberto Amato, Jeffrey Barrett, Sonia Goncalves, Ewan Harrison, David K. Jackson, Ian Johnston, Dominic Kwiatkowski, Cordelia Langford, John Sillitoe on behalf of the Wellcome Sanger Institute COVID-19 Surveillance Team |
| EPI_ISL_3224979 | Lighthouse Lab in Glasgow | Wellcome Sanger Institute for the COVID-19 Genomics UK (COG-UK) Consortium | Harper VanSteenhouse, Yumi Kasai, David Gray, Carol Clugston, Anna Dominiczak and Alex Alderton, Roberto Amato, Jeffrey Barrett, Sonia Goncalves, Ewan Harrison, David K. Jackson, Ian Johnston, Dominic Kwiatkowski, Cordelia Langford, John Sillitoe on behalf of the Wellcome Sanger Institute COVID-19 Surveillance Team |
| EPI_ISL_3224980 | Lighthouse Lab in Alderley Park | Wellcome Sanger Institute for the COVID-19 Genomics UK (COG-UK) Consortium | Jacquelyn Wynn, Mairead Hyland, The Lighthouse Lab in Alderley Park and Alex Alderton, Roberto Amato, Jeffrey Barrett, Sonia Goncalves, Ewan Harrison, David K. Jackson, Ian Johnston, Dominic Kwiatkowski, Cordelia Langford, John Sillitoe on behalf of the Wellcome Sanger Institute COVID-19 Surveillance Team |
| EPI_ISL_3224981, EPI_ISL_3224982 | Berkshire and Surrey Pathology Services Lighthouse Laboratory | Wellcome Sanger Institute for the COVID-19 Genomics UK (COG-UK) Consortium | Berkshire and Surrey Pathology Services Lighthouse Laboratory and Alex Alderton, Roberto Amato, Jeffrey Barrett, Sonia Goncalves, Ewan Harrison, David K. Jackson, Ian Johnston, Dominic Kwiatkowski, Cordelia Langford, John Sillitoe on behalf of the Wellcome Sanger Institute COVID-19 Surveillance Team |
| EPI_ISL_3224983 | Lighthouse Lab in Glasgow | Wellcome Sanger Institute for the COVID-19 Genomics UK (COG-UK) Consortium | Harper VanSteenhouse, Yumi Kasai, David Gray, Carol Clugston, Anna Dominiczak and Alex Alderton, Roberto Amato, Jeffrey Barrett, Sonia Goncalves, Ewan Harrison, David K. Jackson, Ian Johnston, Dominic Kwiatkowski, Cordelia Langford, John Sillitoe on behalf of the Wellcome Sanger Institute COVID-19 Surveillance Team |
| EPI_ISL_3224984, EPI_ISL_3224985 | Lighthouse Lab in Alderley Park | Wellcome Sanger Institute for the COVID-19 Genomics UK (COG-UK) Consortium | Jacquelyn Wynn, Mairead Hyland, The Lighthouse Lab in Alderley Park and Alex Alderton, Roberto Amato, Jeffrey Barrett, Sonia Goncalves, Ewan Harrison, David K. Jackson, Ian Johnston, Dominic Kwiatkowski, Cordelia Langford, John Sillitoe on behalf of the Wellcome Sanger Institute COVID-19 Surveillance Team |
| EPI_ISL_3224986 | Lighthouse Lab in Glasgow | Wellcome Sanger Institute for the COVID-19 Genomics UK (COG-UK) Consortium | Harper VanSteenhouse, Yumi Kasai, David Gray, Carol Clugston, Anna Dominiczak and Alex Alderton, Roberto Amato, Jeffrey Barrett, Sonia Goncalves, Ewan Harrison, David K. Jackson, Ian Johnston, Dominic Kwiatkowski, Cordelia Langford, John Sillitoe on behalf of the Wellcome Sanger Institute COVID-19 Surveillance Team |
| EPI_ISL_3224987 | Lighthouse Lab in Alderley Park | Wellcome Sanger Institute for the COVID-19 Genomics UK (COG-UK) Consortium | Jacquelyn Wynn, Mairead Hyland, The Lighthouse Lab in Alderley Park and Alex Alderton, Roberto Amato, Jeffrey Barrett, Sonia Goncalves, Ewan Harrison, David K. Jackson, Ian Johnston, Dominic Kwiatkowski, Cordelia Langford, John Sillitoe on behalf of the Wellcome Sanger Institute COVID-19 Surveillance Team |
| EPI_ISL_3224988 | Berkshire and Surrey Pathology Services Lighthouse Laboratory | Wellcome Sanger Institute for the COVID-19 Genomics UK (COG-UK) Consortium | Berkshire and Surrey Pathology Services Lighthouse Laboratory and Alex Alderton, Roberto Amato, Jeffrey Barrett, Sonia Goncalves, Ewan Harrison, David K. Jackson, Ian Johnston, Dominic Kwiatkowski, Cordelia Langford, John Sillitoe on behalf of the Wellcome Sanger Institute COVID-19 Surveillance Team |
| EPI_ISL_3224989, EPI_ISL_3224990 | Lighthouse Laboratory Plymouth | Wellcome Sanger Institute for the COVID-19 Genomics UK (COG-UK) Consortium | Lighthouse Laboratory Plymouth and Alex Alderton, Roberto Amato, Jeffrey Barrett, Sonia Goncalves, Ewan Harrison, David K. Jackson, Ian Johnston, Dominic Kwiatkowski, Cordelia Langford, John Sillitoe on behalf of the Wellcome Sanger Institute COVID-19 Surveillance Team |
| EPI_ISL_3224991, EPI_ISL_3224992 | Lighthouse Lab in Alderley Park | Wellcome Sanger Institute for the COVID-19 Genomics UK (COG-UK) Consortium | Jacquelyn Wynn, Mairead Hyland, The Lighthouse Lab in Alderley Park and Alex Alderton, Roberto Amato, Jeffrey Barrett, Sonia Goncalves, Ewan Harrison, David K. Jackson, Ian Johnston, Dominic Kwiatkowski, Cordelia Langford, John Sillitoe on behalf of the Wellcome Sanger Institute COVID-19 Surveillance Team |
| EPI_ISL_3224993 | Lighthouse Laboratory Plymouth | Wellcome Sanger Institute for the COVID-19 Genomics UK (COG-UK) Consortium | Lighthouse Laboratory Plymouth and Alex Alderton, Roberto Amato, Jeffrey Barrett, Sonia Goncalves, Ewan Harrison, David K. Jackson, Ian Johnston, Dominic Kwiatkowski, Cordelia Langford, John Sillitoe on behalf of the Wellcome Sanger Institute COVID-19 Surveillance Team |
| EPI_ISL_3224994, EPI_ISL_3224995, EPI_ISL_3224996 | Lighthouse Lab in Alderley Park | Wellcome Sanger Institute for the COVID-19 Genomics UK (COG-UK) Consortium | Jacquelyn Wynn, Mairead Hyland, The Lighthouse Lab in Alderley Park and Alex Alderton, Roberto Amato, Jeffrey Barrett, Sonia Goncalves, Ewan Harrison, David K. Jackson, Ian Johnston, Dominic Kwiatkowski, Cordelia Langford, John Sillitoe on behalf of the Wellcome Sanger Institute COVID-19 Surveillance Team |
| EPI_ISL_3224997 | Lighthouse Lab in Glasgow | Wellcome Sanger Institute for the COVID-19 Genomics UK (COG-UK) Consortium | Harper VanSteenhouse, Yumi Kasai, David Gray, Carol Clugston, Anna Dominiczak and Alex Alderton, Roberto Amato, Jeffrey Barrett, Sonia Goncalves, Ewan Harrison, David K. Jackson, Ian Johnston, Dominic Kwiatkowski, Cordelia Langford, John Sillitoe on behalf of the Wellcome Sanger Institute COVID-19 Surveillance Team |
| EPI_ISL_3224998 | Lighthouse Lab in Alderley Park | Wellcome Sanger Institute for the COVID-19 Genomics UK (COG-UK) Consortium | Jacquelyn Wynn, Mairead Hyland, The Lighthouse Lab in Alderley Park and Alex Alderton, Roberto Amato, Jeffrey Barrett, Sonia Goncalves, Ewan Harrison, David K. Jackson, Ian Johnston, Dominic Kwiatkowski, Cordelia Langford, John Sillitoe on behalf of the Wellcome Sanger Institute COVID-19 Surveillance Team |
| EPI_ISL_3224999 | Berkshire and Surrey Pathology Services Lighthouse Laboratory | Wellcome Sanger Institute for the COVID-19 Genomics UK (COG-UK) Consortium | Berkshire and Surrey Pathology Services Lighthouse Laboratory and Alex Alderton, Roberto Amato, Jeffrey Barrett, Sonia Goncalves, Ewan Harrison, David K. Jackson, Ian Johnston, Dominic Kwiatkowski, Cordelia Langford, John Sillitoe on behalf of the Wellcome Sanger Institute COVID-19 Surveillance Team |
| EPI_ISL_3225000 | Lighthouse Laboratory Plymouth | Wellcome Sanger Institute for the COVID-19 Genomics UK (COG-UK) Consortium | Lighthouse Laboratory Plymouth and Alex Alderton, Roberto Amato, Jeffrey Barrett, Sonia Goncalves, Ewan Harrison, David K. Jackson, Ian Johnston, Dominic Kwiatkowski, Cordelia Langford, John Sillitoe on behalf of the Wellcome Sanger Institute COVID-19 Surveillance Team |
| EPI_ISL_3225001, EPI_ISL_3225002, EPI_ISL_3225003 | Lighthouse Lab in Glasgow | Wellcome Sanger Institute for the COVID-19 Genomics UK (COG-UK) Consortium | Harper VanSteenhouse, Yumi Kasai, David Gray, Carol Clugston, Anna Dominiczak and Alex Alderton, Roberto Amato, Jeffrey Barrett, Sonia Goncalves, Ewan Harrison, David K. Jackson, Ian Johnston, Dominic Kwiatkowski, Cordelia Langford, John Sillitoe on behalf of the Wellcome Sanger Institute COVID-19 Surveillance Team |
| EPI_ISL_3225004, EPI_ISL_3225005 | Lighthouse Laboratory Plymouth | Wellcome Sanger Institute for the COVID-19 Genomics UK (COG-UK) Consortium | Lighthouse Laboratory Plymouth and Alex Alderton, Roberto Amato, Jeffrey Barrett, Sonia Goncalves, Ewan Harrison, David K. Jackson, Ian Johnston, Dominic Kwiatkowski, Cordelia Langford, John Sillitoe on behalf of the Wellcome Sanger Institute COVID-19 Surveillance Team |
| EPI_ISL_3225006 | Berkshire and Surrey Pathology Services Lighthouse Laboratory | Wellcome Sanger Institute for the COVID-19 Genomics UK (COG-UK) Consortium | Berkshire and Surrey Pathology Services Lighthouse Laboratory and Alex Alderton, Roberto Amato, Jeffrey Barrett, Sonia Goncalves, Ewan Harrison, David K. Jackson, Ian Johnston, Dominic Kwiatkowski, Cordelia Langford, John Sillitoe on behalf of the Wellcome Sanger Institute COVID-19 Surveillance Team |
| EPI_ISL_3225007 | Lighthouse Laboratory Plymouth | Wellcome Sanger Institute for the COVID-19 Genomics UK (COG-UK) Consortium | Lighthouse Laboratory Plymouth and Alex Alderton, Roberto Amato, Jeffrey Barrett, Sonia Goncalves, Ewan Harrison, David K. Jackson, Ian Johnston, Dominic Kwiatkowski, Cordelia Langford, John Sillitoe on behalf of the Wellcome Sanger Institute COVID-19 Surveillance Team |
| EPI_ISL_3225008 | Lighthouse Lab in Alderley Park | Wellcome Sanger Institute for the COVID-19 Genomics UK (COG-UK) | Jacquelyn Wynn, Mairead Hyland, The Lighthouse Lab in Alderley Park and Alex Alderton, Roberto Amato, Jeffrey Barrett, Sonia Goncalves, Ewan Harrison, David K. Jackson, Ian Johnston, Dominic Kwiatkowski, Cordelia Langford, John Sillitoe on |

| | | | |
|---|---|---|---|
| | | Consortium | behalf of the Wellcome Sanger Institute COVID-19 Surveillance Team |
| EPI_ISL_3225009 | Lighthouse Lab in Glasgow | Wellcome Sanger Institute for the COVID-19 Genomics UK (COG-UK) Consortium | Harper VanSteenhouse, Yumi Kasai, David Gray, Carol Clugston, Anna Dominiczak and Alex Alderton, Roberto Amato, Jeffrey Barrett, Sonia Goncalves, Ewan Harrison, David K. Jackson, Ian Johnston, Dominic Kwiatkowski, Cordelia Langford, John Sillitoe on behalf of the Wellcome Sanger Institute COVID-19 Surveillance Team |
| EPI_ISL_3225010 | Lighthouse Laboratory Plymouth | Wellcome Sanger Institute for the COVID-19 Genomics UK (COG-UK) Consortium | Lighthouse Laboratory Plymouth and Alex Alderton, Roberto Amato, Jeffrey Barrett, Sonia Goncalves, Ewan Harrison, David K. Jackson, Ian Johnston, Dominic Kwiatkowski, Cordelia Langford, John Sillitoe on behalf of the Wellcome Sanger Institute COVID-19 Surveillance Team |
| EPI_ISL_3225011 | Lighthouse Lab in Glasgow | Wellcome Sanger Institute for the COVID-19 Genomics UK (COG-UK) Consortium | Harper VanSteenhouse, Yumi Kasai, David Gray, Carol Clugston, Anna Dominiczak and Alex Alderton, Roberto Amato, Jeffrey Barrett, Sonia Goncalves, Ewan Harrison, David K. Jackson, Ian Johnston, Dominic Kwiatkowski, Cordelia Langford, John Sillitoe on behalf of the Wellcome Sanger Institute COVID-19 Surveillance Team |
| EPI_ISL_3225012, EPI_ISL_3225013, EPI_ISL_3225014 | Lighthouse Laboratory Plymouth | Wellcome Sanger Institute for the COVID-19 Genomics UK (COG-UK) Consortium | Lighthouse Laboratory Plymouth and Alex Alderton, Roberto Amato, Jeffrey Barrett, Sonia Goncalves, Ewan Harrison, David K. Jackson, Ian Johnston, Dominic Kwiatkowski, Cordelia Langford, John Sillitoe on behalf of the Wellcome Sanger Institute COVID-19 Surveillance Team |
| EPI_ISL_3225015, EPI_ISL_3225016 | Berkshire and Surrey Pathology Services Lighthouse Laboratory | Wellcome Sanger Institute for the COVID-19 Genomics UK (COG-UK) Consortium | Berkshire and Surrey Pathology Services Lighthouse Laboratory and Alex Alderton, Roberto Amato, Jeffrey Barrett, Sonia Goncalves, Ewan Harrison, David K. Jackson, Ian Johnston, Dominic Kwiatkowski, Cordelia Langford, John Sillitoe on behalf of the Wellcome Sanger Institute COVID-19 Surveillance Team |
| EPI_ISL_3225017 | Lighthouse Lab in Alderley Park | Wellcome Sanger Institute for the COVID-19 Genomics UK (COG-UK) Consortium | Jacquelyn Wynn, Mairead Hyland, The Lighthouse Lab in Alderley Park and Alex Alderton, Roberto Amato, Jeffrey Barrett, Sonia Goncalves, Ewan Harrison, David K. Jackson, Ian Johnston, Dominic Kwiatkowski, Cordelia Langford, John Sillitoe on behalf of the Wellcome Sanger Institute COVID-19 Surveillance Team |
| EPI_ISL_3225018 | Lighthouse Lab in Glasgow | Wellcome Sanger Institute for the COVID-19 Genomics UK (COG-UK) Consortium | Harper VanSteenhouse, Yumi Kasai, David Gray, Carol Clugston, Anna Dominiczak and Alex Alderton, Roberto Amato, Jeffrey Barrett, Sonia Goncalves, Ewan Harrison, David K. Jackson, Ian Johnston, Dominic Kwiatkowski, Cordelia Langford, John Sillitoe on behalf of the Wellcome Sanger Institute COVID-19 Surveillance Team |
| EPI_ISL_3225019, EPI_ISL_3225020 | Berkshire and Surrey Pathology Services Lighthouse Laboratory | Wellcome Sanger Institute for the COVID-19 Genomics UK (COG-UK) Consortium | Berkshire and Surrey Pathology Services Lighthouse Laboratory and Alex Alderton, Roberto Amato, Jeffrey Barrett, Sonia Goncalves, Ewan Harrison, David K. Jackson, Ian Johnston, Dominic Kwiatkowski, Cordelia Langford, John Sillitoe on behalf of the Wellcome Sanger Institute COVID-19 Surveillance Team |
| EPI_ISL_3225021 | Lighthouse Lab in Glasgow | Wellcome Sanger Institute for the COVID-19 Genomics UK (COG-UK) Consortium | Harper VanSteenhouse, Yumi Kasai, David Gray, Carol Clugston, Anna Dominiczak and Alex Alderton, Roberto Amato, Jeffrey Barrett, Sonia Goncalves, Ewan Harrison, David K. Jackson, Ian Johnston, Dominic Kwiatkowski, Cordelia Langford, John Sillitoe on behalf of the Wellcome Sanger Institute COVID-19 Surveillance Team |
| EPI_ISL_3225022 | Lighthouse Laboratory Plymouth | Wellcome Sanger Institute for the COVID-19 Genomics UK (COG-UK) Consortium | Lighthouse Laboratory Plymouth and Alex Alderton, Roberto Amato, Jeffrey Barrett, Sonia Goncalves, Ewan Harrison, David K. Jackson, Ian Johnston, Dominic Kwiatkowski, Cordelia Langford, John Sillitoe on behalf of the Wellcome Sanger Institute COVID-19 Surveillance Team |
| EPI_ISL_3225023 | Berkshire and Surrey Pathology Services Lighthouse Laboratory | Wellcome Sanger Institute for the COVID-19 Genomics UK (COG-UK) Consortium | Berkshire and Surrey Pathology Services Lighthouse Laboratory and Alex Alderton, Roberto Amato, Jeffrey Barrett, Sonia Goncalves, Ewan Harrison, David K. Jackson, Ian Johnston, Dominic Kwiatkowski, Cordelia Langford, John Sillitoe on behalf of the Wellcome Sanger Institute COVID-19 Surveillance Team |
| EPI_ISL_3225024, EPI_ISL_3225025 | Lighthouse Laboratory Plymouth | Wellcome Sanger Institute for the COVID-19 Genomics UK (COG-UK) Consortium | Lighthouse Laboratory Plymouth and Alex Alderton, Roberto Amato, Jeffrey Barrett, Sonia Goncalves, Ewan Harrison, David K. Jackson, Ian Johnston, Dominic Kwiatkowski, Cordelia Langford, John Sillitoe on behalf of the Wellcome Sanger Institute COVID-19 Surveillance Team |
| EPI_ISL_3225026 | Berkshire and Surrey Pathology Services Lighthouse Laboratory | Wellcome Sanger Institute for the COVID-19 Genomics UK (COG-UK) Consortium | Berkshire and Surrey Pathology Services Lighthouse Laboratory and Alex Alderton, Roberto Amato, Jeffrey Barrett, Sonia Goncalves, Ewan Harrison, David K. Jackson, Ian Johnston, Dominic Kwiatkowski, Cordelia Langford, John Sillitoe on behalf of the Wellcome Sanger Institute COVID-19 Surveillance Team |
| EPI_ISL_3225027 | Lighthouse Lab in Glasgow | Wellcome Sanger Institute for the COVID-19 Genomics UK (COG-UK) Consortium | Harper VanSteenhouse, Yumi Kasai, David Gray, Carol Clugston, Anna Dominiczak and Alex Alderton, Roberto Amato, Jeffrey Barrett, Sonia Goncalves, Ewan Harrison, David K. Jackson, Ian Johnston, Dominic Kwiatkowski, Cordelia Langford, John Sillitoe on behalf of the Wellcome Sanger Institute COVID-19 Surveillance Team |
| EPI_ISL_3225028 | Berkshire and Surrey Pathology Services Lighthouse Laboratory | Wellcome Sanger Institute for the COVID-19 Genomics UK (COG-UK) Consortium | Berkshire and Surrey Pathology Services Lighthouse Laboratory and Alex Alderton, Roberto Amato, Jeffrey Barrett, Sonia Goncalves, Ewan Harrison, David K. Jackson, Ian Johnston, Dominic Kwiatkowski, Cordelia Langford, John Sillitoe on behalf of the Wellcome Sanger Institute COVID-19 Surveillance Team |
| EPI_ISL_3225029, EPI_ISL_3225030, EPI_ISL_3225031 | Lighthouse Lab in Alderley Park | Wellcome Sanger Institute for the COVID-19 Genomics UK (COG-UK) Consortium | Jacquelyn Wynn, Mairead Hyland, The Lighthouse Lab in Alderley Park and Alex Alderton, Roberto Amato, Jeffrey Barrett, Sonia Goncalves, Ewan Harrison, David K. Jackson, Ian Johnston, Dominic Kwiatkowski, Cordelia Langford, John Sillitoe on behalf of the Wellcome Sanger Institute COVID-19 Surveillance Team |
| EPI_ISL_3225032 | Berkshire and Surrey Pathology Services Lighthouse Laboratory | Wellcome Sanger Institute for the COVID-19 Genomics UK (COG-UK) Consortium | Berkshire and Surrey Pathology Services Lighthouse Laboratory and Alex Alderton, Roberto Amato, Jeffrey Barrett, Sonia Goncalves, Ewan Harrison, David K. Jackson, Ian Johnston, Dominic Kwiatkowski, Cordelia Langford, John Sillitoe on behalf of the Wellcome Sanger Institute COVID-19 Surveillance Team |
| EPI_ISL_3225033 | Lighthouse Lab in Alderley Park | Wellcome Sanger Institute for the COVID-19 Genomics UK (COG-UK) Consortium | Jacquelyn Wynn, Mairead Hyland, The Lighthouse Lab in Alderley Park and Alex Alderton, Roberto Amato, Jeffrey Barrett, Sonia Goncalves, Ewan Harrison, David K. Jackson, Ian Johnston, Dominic Kwiatkowski, Cordelia Langford, John Sillitoe on behalf of the Wellcome Sanger Institute COVID-19 Surveillance Team |
| EPI_ISL_3225034 | Lighthouse Laboratory Plymouth | Wellcome Sanger Institute for the COVID-19 Genomics UK (COG-UK) Consortium | Lighthouse Laboratory Plymouth and Alex Alderton, Roberto Amato, Jeffrey Barrett, Sonia Goncalves, Ewan Harrison, David K. Jackson, Ian Johnston, Dominic Kwiatkowski, Cordelia Langford, John Sillitoe on behalf of the Wellcome Sanger Institute COVID-19 Surveillance Team |
| EPI_ISL_3225035 | Lighthouse Lab in Glasgow | Wellcome Sanger Institute for the COVID-19 Genomics UK (COG-UK) Consortium | Harper VanSteenhouse, Yumi Kasai, David Gray, Carol Clugston, Anna Dominiczak and Alex Alderton, Roberto Amato, Jeffrey Barrett, Sonia Goncalves, Ewan Harrison, David K. Jackson, Ian Johnston, Dominic Kwiatkowski, Cordelia Langford, John Sillitoe on behalf of the Wellcome Sanger Institute COVID-19 Surveillance Team |
| EPI_ISL_3225036, EPI_ISL_3225037, EPI_ISL_3225038, EPI_ISL_3225039 | Lighthouse Laboratory Plymouth | Wellcome Sanger Institute for the COVID-19 Genomics UK (COG-UK) Consortium | Lighthouse Laboratory Plymouth and Alex Alderton, Roberto Amato, Jeffrey Barrett, Sonia Goncalves, Ewan Harrison, David K. Jackson, Ian Johnston, Dominic Kwiatkowski, Cordelia Langford, John Sillitoe on behalf of the Wellcome Sanger Institute COVID-19 Surveillance Team |
| EPI_ISL_3225040, EPI_ISL_3225041, EPI_ISL_3225042 | Lighthouse Lab in Glasgow | Wellcome Sanger Institute for the COVID-19 Genomics UK (COG-UK) Consortium | Harper VanSteenhouse, Yumi Kasai, David Gray, Carol Clugston, Anna Dominiczak and Alex Alderton, Roberto Amato, Jeffrey Barrett, Sonia Goncalves, Ewan Harrison, David K. Jackson, Ian Johnston, Dominic Kwiatkowski, Cordelia Langford, John Sillitoe on behalf of the Wellcome Sanger Institute COVID-19 Surveillance Team |
| EPI_ISL_3225043 | Berkshire and Surrey Pathology Services Lighthouse Laboratory | Wellcome Sanger Institute for the COVID-19 Genomics UK (COG-UK) Consortium | Berkshire and Surrey Pathology Services Lighthouse Laboratory and Alex Alderton, Roberto Amato, Jeffrey Barrett, Sonia Goncalves, Ewan Harrison, David K. Jackson, Ian Johnston, Dominic Kwiatkowski, Cordelia Langford, John Sillitoe on behalf of the Wellcome Sanger Institute COVID-19 Surveillance Team |
| EPI_ISL_3225044, EPI_ISL_3225045, EPI_ISL_3225046 | Lighthouse Lab in Glasgow | Wellcome Sanger Institute for the COVID-19 Genomics UK (COG-UK) Consortium | Harper VanSteenhouse, Yumi Kasai, David Gray, Carol Clugston, Anna Dominiczak and Alex Alderton, Roberto Amato, Jeffrey Barrett, Sonia Goncalves, Ewan Harrison, David K. Jackson, Ian Johnston, Dominic Kwiatkowski, Cordelia Langford, John Sillitoe on behalf of the Wellcome Sanger Institute COVID-19 Surveillance Team |
| EPI_ISL_3225047 | Lighthouse Lab in Alderley Park | Wellcome Sanger Institute for the COVID-19 Genomics UK (COG-UK) Consortium | Jacquelyn Wynn, Mairead Hyland, The Lighthouse Lab in Alderley Park and Alex Alderton, Roberto Amato, Jeffrey Barrett, Sonia Goncalves, Ewan Harrison, David K. Jackson, Ian Johnston, Dominic Kwiatkowski, Cordelia Langford, John Sillitoe on behalf of the Wellcome Sanger Institute COVID-19 Surveillance Team |
| EPI_ISL_3225048, EPI_ISL_3225049 | Lighthouse Lab in Glasgow | Wellcome Sanger Institute for the COVID-19 Genomics UK (COG-UK) Consortium | Harper VanSteenhouse, Yumi Kasai, David Gray, Carol Clugston, Anna Dominiczak and Alex Alderton, Roberto Amato, Jeffrey Barrett, Sonia Goncalves, Ewan Harrison, David K. Jackson, Ian Johnston, Dominic Kwiatkowski, Cordelia Langford, John Sillitoe on behalf of the Wellcome Sanger Institute COVID-19 Surveillance Team |
| EPI_ISL_3225050 | Lighthouse Laboratory Plymouth | Wellcome Sanger Institute for the COVID-19 Genomics UK (COG-UK) Consortium | Lighthouse Laboratory Plymouth and Alex Alderton, Roberto Amato, Jeffrey Barrett, Sonia Goncalves, Ewan Harrison, David K. Jackson, Ian Johnston, Dominic Kwiatkowski, Cordelia Langford, John Sillitoe on behalf of the Wellcome Sanger Institute COVID-19 Surveillance Team |
| EPI_ISL_3225051 | Berkshire and Surrey Pathology Services Lighthouse Laboratory | Wellcome Sanger Institute for the COVID-19 Genomics UK (COG-UK) Consortium | Berkshire and Surrey Pathology Services Lighthouse Laboratory and Alex Alderton, Roberto Amato, Jeffrey Barrett, Sonia Goncalves, Ewan Harrison, David K. Jackson, Ian Johnston, Dominic Kwiatkowski, Cordelia Langford, John Sillitoe on behalf of the Wellcome Sanger Institute COVID-19 Surveillance Team |
| EPI_ISL_3225052 | Lighthouse Lab in Glasgow | Wellcome Sanger Institute for the COVID-19 Genomics UK (COG-UK) Consortium | Harper VanSteenhouse, Yumi Kasai, David Gray, Carol Clugston, Anna Dominiczak and Alex Alderton, Roberto Amato, Jeffrey Barrett, Sonia Goncalves, Ewan Harrison, David K. Jackson, Ian Johnston, Dominic Kwiatkowski, Cordelia Langford, John Sillitoe on behalf of the Wellcome Sanger Institute COVID-19 Surveillance Team |
| EPI_ISL_3225053 | Lighthouse Laboratory Plymouth | Wellcome Sanger Institute for the COVID-19 Genomics UK (COG-UK) Consortium | Lighthouse Laboratory Plymouth and Alex Alderton, Roberto Amato, Jeffrey Barrett, Sonia Goncalves, Ewan Harrison, David K. Jackson, Ian Johnston, Dominic Kwiatkowski, Cordelia Langford, John Sillitoe on behalf of the Wellcome Sanger Institute COVID-19 Surveillance Team |
| EPI_ISL_3225054 | Lighthouse Lab in Alderley Park | Wellcome Sanger Institute for the COVID-19 Genomics UK (COG-UK) Consortium | Jacquelyn Wynn, Mairead Hyland, The Lighthouse Lab in Alderley Park and Alex Alderton, Roberto Amato, Jeffrey Barrett, Sonia Goncalves, Ewan Harrison, David K. Jackson, Ian Johnston, Dominic Kwiatkowski, Cordelia Langford, John Sillitoe on behalf of the Wellcome Sanger Institute COVID-19 Surveillance Team |
| EPI_ISL_3225055 | Berkshire and Surrey Pathology Services Lighthouse Laboratory | Wellcome Sanger Institute for the COVID-19 Genomics UK (COG-UK) Consortium | Berkshire and Surrey Pathology Services Lighthouse Laboratory and Alex Alderton, Roberto Amato, Jeffrey Barrett, Sonia Goncalves, Ewan Harrison, David K. Jackson, Ian Johnston, Dominic Kwiatkowski, Cordelia Langford, John Sillitoe on behalf of the Wellcome Sanger Institute COVID-19 Surveillance Team |
| EPI_ISL_3225056 | Lighthouse Laboratory Plymouth | Wellcome Sanger Institute for the COVID-19 Genomics UK (COG-UK) Consortium | Lighthouse Laboratory Plymouth and Alex Alderton, Roberto Amato, Jeffrey Barrett, Sonia Goncalves, Ewan Harrison, David K. Jackson, Ian Johnston, Dominic Kwiatkowski, Cordelia Langford, John Sillitoe on behalf of the Wellcome Sanger Institute COVID-19 Surveillance Team |
| EPI_ISL_3225057 | Lighthouse Lab in Glasgow | Wellcome Sanger Institute for the COVID-19 Genomics UK (COG-UK) Consortium | Harper VanSteenhouse, Yumi Kasai, David Gray, Carol Clugston, Anna Dominiczak and Alex Alderton, Roberto Amato, Jeffrey Barrett, Sonia Goncalves, Ewan Harrison, David K. Jackson, Ian Johnston, Dominic Kwiatkowski, Cordelia Langford, John Sillitoe on behalf of the Wellcome Sanger Institute COVID-19 Surveillance Team |
| EPI_ISL_3225058 | Lighthouse Lab in Alderley Park | Wellcome Sanger Institute for the COVID-19 Genomics UK (COG-UK) Consortium | Jacquelyn Wynn, Mairead Hyland, The Lighthouse Lab in Alderley Park and Alex Alderton, Roberto Amato, Jeffrey Barrett, Sonia Goncalves, Ewan Harrison, David K. Jackson, Ian Johnston, Dominic Kwiatkowski, Cordelia Langford, John Sillitoe on behalf of the Wellcome Sanger Institute COVID-19 Surveillance Team |
| EPI_ISL_3225059 | Berkshire and Surrey Pathology Services Lighthouse Laboratory | Wellcome Sanger Institute for the COVID-19 Genomics UK (COG-UK) Consortium | Berkshire and Surrey Pathology Services Lighthouse Laboratory and Alex Alderton, Roberto Amato, Jeffrey Barrett, Sonia Goncalves, Ewan Harrison, David K. Jackson, Ian Johnston, Dominic Kwiatkowski, Cordelia Langford, John Sillitoe on behalf of the Wellcome Sanger Institute COVID-19 Surveillance Team |
| EPI_ISL_3225060, EPI_ISL_3225061, EPI_ISL_3225062, EPI_ISL_3225063 | Lighthouse Laboratory Plymouth | Wellcome Sanger Institute for the COVID-19 Genomics UK (COG-UK) Consortium | Lighthouse Laboratory Plymouth and Alex Alderton, Roberto Amato, Jeffrey Barrett, Sonia Goncalves, Ewan Harrison, David K. Jackson, Ian Johnston, Dominic Kwiatkowski, Cordelia Langford, John Sillitoe on behalf of the Wellcome Sanger Institute COVID-19 Surveillance Team |
| EPI_ISL_3225064 | Lighthouse Lab in Alderley Park | Wellcome Sanger Institute for the COVID-19 Genomics UK (COG-UK) Consortium | Jacquelyn Wynn, Mairead Hyland, The Lighthouse Lab in Alderley Park and Alex Alderton, Roberto Amato, Jeffrey Barrett, Sonia Goncalves, Ewan Harrison, David K. Jackson, Ian Johnston, Dominic Kwiatkowski, Cordelia Langford, John Sillitoe on behalf of the Wellcome Sanger Institute COVID-19 Surveillance Team |
| EPI_ISL_3225065, EPI_ISL_3225066 | Lighthouse Lab in Glasgow | Wellcome Sanger Institute for the COVID-19 Genomics UK (COG-UK) Consortium | Harper VanSteenhouse, Yumi Kasai, David Gray, Carol Clugston, Anna Dominiczak and Alex Alderton, Roberto Amato, Jeffrey Barrett, Sonia Goncalves, Ewan Harrison, David K. Jackson, Ian Johnston, Dominic Kwiatkowski, Cordelia Langford, John Sillitoe on behalf of the Wellcome Sanger Institute COVID-19 Surveillance Team |
| EPI_ISL_3225067 | Lighthouse Lab in Alderley Park | Wellcome Sanger Institute for the COVID-19 Genomics UK (COG-UK) Consortium | Jacquelyn Wynn, Mairead Hyland, The Lighthouse Lab in Alderley Park and Alex Alderton, Roberto Amato, Jeffrey Barrett, Sonia Goncalves, Ewan Harrison, David K. Jackson, Ian Johnston, Dominic Kwiatkowski, Cordelia Langford, John Sillitoe on behalf of the Wellcome Sanger Institute COVID-19 Surveillance Team |
| EPI_ISL_3225068 | Lighthouse Laboratory Plymouth | Wellcome Sanger Institute for the COVID-19 Genomics UK (COG-UK) Consortium | Lighthouse Laboratory Plymouth and Alex Alderton, Roberto Amato, Jeffrey Barrett, Sonia Goncalves, Ewan Harrison, David K. Jackson, Ian Johnston, Dominic Kwiatkowski, Cordelia Langford, John Sillitoe on behalf of the Wellcome Sanger Institute COVID-19 Surveillance Team |
| EPI_ISL_3225069 | Berkshire and Surrey Pathology Services Lighthouse Laboratory | Wellcome Sanger Institute for the COVID-19 Genomics UK (COG-UK) | Berkshire and Surrey Pathology Services Lighthouse Laboratory and Alex Alderton, Roberto Amato, Jeffrey Barrett, Sonia Goncalves, Ewan Harrison, David K. Jackson, Ian Johnston, Dominic Kwiatkowski, Cordelia Langford, John Sillitoe on behalf |

| | | | |
|---|---|---|---|
| | | Consortium | of the Wellcome Sanger Institute COVID-19 Surveillance Team |
| EPI_ISL_3225070, EPI_ISL_3225071 | Lighthouse Laboratory Plymouth | Wellcome Sanger Institute for the COVID-19 Genomics UK (COG-UK) Consortium | Lighthouse Laboratory Plymouth and Alex Alderton, Roberto Amato, Jeffrey Barrett, Sonia Goncalves, Ewan Harrison, David K. Jackson, Ian Johnston, Dominic Kwiatkowski, Cordelia Langford, John Sillitoe on behalf of the Wellcome Sanger Institute COVID-19 Surveillance Team |
| EPI_ISL_3225072 | Lighthouse Lab in Glasgow | Wellcome Sanger Institute for the COVID-19 Genomics UK (COG-UK) Consortium | Harper VanSteenhouse, Yumi Kasai, David Gray, Carol Clugston, Anna Dominiczak and Alex Alderton, Roberto Amato, Jeffrey Barrett, Sonia Goncalves, Ewan Harrison, David K. Jackson, Ian Johnston, Dominic Kwiatkowski, Cordelia Langford, John Sillitoe on behalf of the Wellcome Sanger Institute COVID-19 Surveillance Team |
| EPI_ISL_3225073 | Lighthouse Lab in Alderley Park | Wellcome Sanger Institute for the COVID-19 Genomics UK (COG-UK) Consortium | Jacquelyn Wynn, Mairead Hyland, The Lighthouse Lab in Alderley Park and Alex Alderton, Roberto Amato, Jeffrey Barrett, Sonia Goncalves, Ewan Harrison, David K. Jackson, Ian Johnston, Dominic Kwiatkowski, Cordelia Langford, John Sillitoe on behalf of the Wellcome Sanger Institute COVID-19 Surveillance Team |
| EPI_ISL_3225074 | Lighthouse Lab in Glasgow | Wellcome Sanger Institute for the COVID-19 Genomics UK (COG-UK) Consortium | Harper VanSteenhouse, Yumi Kasai, David Gray, Carol Clugston, Anna Dominiczak and Alex Alderton, Roberto Amato, Jeffrey Barrett, Sonia Goncalves, Ewan Harrison, David K. Jackson, Ian Johnston, Dominic Kwiatkowski, Cordelia Langford, John Sillitoe on behalf of the Wellcome Sanger Institute COVID-19 Surveillance Team |
| EPI_ISL_3225075 | Health Services Laboratories | Wellcome Sanger Institute for the COVID-19 Genomics UK (COG-UK) Consortium | Health Services Laboratories and Alex Alderton, Roberto Amato, Jeffrey Barrett, Sonia Goncalves, Ewan Harrison, David K. Jackson, Ian Johnston, Dominic Kwiatkowski, Cordelia Langford, John Sillitoe on behalf of the Wellcome Sanger Institute COVID-19 Surveillance Team |
| EPI_ISL_3225076 | Lighthouse Lab in Glasgow | Wellcome Sanger Institute for the COVID-19 Genomics UK (COG-UK) Consortium | Harper VanSteenhouse, Yumi Kasai, David Gray, Carol Clugston, Anna Dominiczak and Alex Alderton, Roberto Amato, Jeffrey Barrett, Sonia Goncalves, Ewan Harrison, David K. Jackson, Ian Johnston, Dominic Kwiatkowski, Cordelia Langford, John Sillitoe on behalf of the Wellcome Sanger Institute COVID-19 Surveillance Team |
| EPI_ISL_3225077, EPI_ISL_3225078, EPI_ISL_3225079 | Lighthouse Laboratory Plymouth | Wellcome Sanger Institute for the COVID-19 Genomics UK (COG-UK) Consortium | Lighthouse Laboratory Plymouth and Alex Alderton, Roberto Amato, Jeffrey Barrett, Sonia Goncalves, Ewan Harrison, David K. Jackson, Ian Johnston, Dominic Kwiatkowski, Cordelia Langford, John Sillitoe on behalf of the Wellcome Sanger Institute COVID-19 Surveillance Team |
| EPI_ISL_3225080 | Lighthouse Lab in Glasgow | Wellcome Sanger Institute for the COVID-19 Genomics UK (COG-UK) Consortium | Harper VanSteenhouse, Yumi Kasai, David Gray, Carol Clugston, Anna Dominiczak and Alex Alderton, Roberto Amato, Jeffrey Barrett, Sonia Goncalves, Ewan Harrison, David K. Jackson, Ian Johnston, Dominic Kwiatkowski, Cordelia Langford, John Sillitoe on behalf of the Wellcome Sanger Institute COVID-19 Surveillance Team |
| EPI_ISL_3225081, EPI_ISL_3225082 | Lighthouse Laboratory Plymouth | Wellcome Sanger Institute for the COVID-19 Genomics UK (COG-UK) Consortium | Lighthouse Laboratory Plymouth and Alex Alderton, Roberto Amato, Jeffrey Barrett, Sonia Goncalves, Ewan Harrison, David K. Jackson, Ian Johnston, Dominic Kwiatkowski, Cordelia Langford, John Sillitoe on behalf of the Wellcome Sanger Institute COVID-19 Surveillance Team |
| EPI_ISL_3225083 | Berkshire and Surrey Pathology Services Lighthouse Laboratory | Wellcome Sanger Institute for the COVID-19 Genomics UK (COG-UK) Consortium | Berkshire and Surrey Pathology Services Lighthouse Laboratory and Alex Alderton, Roberto Amato, Jeffrey Barrett, Sonia Goncalves, Ewan Harrison, David K. Jackson, Ian Johnston, Dominic Kwiatkowski, Cordelia Langford, John Sillitoe on behalf of the Wellcome Sanger Institute COVID-19 Surveillance Team |
| EPI_ISL_3225084 | Lighthouse Lab in Glasgow | Wellcome Sanger Institute for the COVID-19 Genomics UK (COG-UK) Consortium | Harper VanSteenhouse, Yumi Kasai, David Gray, Carol Clugston, Anna Dominiczak and Alex Alderton, Roberto Amato, Jeffrey Barrett, Sonia Goncalves, Ewan Harrison, David K. Jackson, Ian Johnston, Dominic Kwiatkowski, Cordelia Langford, John Sillitoe on behalf of the Wellcome Sanger Institute COVID-19 Surveillance Team |
| EPI_ISL_3225085 | Berkshire and Surrey Pathology Services Lighthouse Laboratory | Wellcome Sanger Institute for the COVID-19 Genomics UK (COG-UK) Consortium | Berkshire and Surrey Pathology Services Lighthouse Laboratory and Alex Alderton, Roberto Amato, Jeffrey Barrett, Sonia Goncalves, Ewan Harrison, David K. Jackson, Ian Johnston, Dominic Kwiatkowski, Cordelia Langford, John Sillitoe on behalf of the Wellcome Sanger Institute COVID-19 Surveillance Team |
| EPI_ISL_3225086 | Lighthouse Lab in Glasgow | Wellcome Sanger Institute for the COVID-19 Genomics UK (COG-UK) Consortium | Harper VanSteenhouse, Yumi Kasai, David Gray, Carol Clugston, Anna Dominiczak and Alex Alderton, Roberto Amato, Jeffrey Barrett, Sonia Goncalves, Ewan Harrison, David K. Jackson, Ian Johnston, Dominic Kwiatkowski, Cordelia Langford, John Sillitoe on behalf of the Wellcome Sanger Institute COVID-19 Surveillance Team |
| EPI_ISL_3225087, EPI_ISL_3225088 | Lighthouse Lab in Alderley Park | Wellcome Sanger Institute for the COVID-19 Genomics UK (COG-UK) Consortium | Jacquelyn Wynn, Mairead Hyland, The Lighthouse Lab in Alderley Park and Alex Alderton, Roberto Amato, Jeffrey Barrett, Sonia Goncalves, Ewan Harrison, David K. Jackson, Ian Johnston, Dominic Kwiatkowski, Cordelia Langford, John Sillitoe on behalf of the Wellcome Sanger Institute COVID-19 Surveillance Team |
| EPI_ISL_3225089 | Lighthouse Laboratory Plymouth | Wellcome Sanger Institute for the COVID-19 Genomics UK (COG-UK) Consortium | Lighthouse Laboratory Plymouth and Alex Alderton, Roberto Amato, Jeffrey Barrett, Sonia Goncalves, Ewan Harrison, David K. Jackson, Ian Johnston, Dominic Kwiatkowski, Cordelia Langford, John Sillitoe on behalf of the Wellcome Sanger Institute COVID-19 Surveillance Team |
| EPI_ISL_3225090 | Berkshire and Surrey Pathology Services Lighthouse Laboratory | Wellcome Sanger Institute for the COVID-19 Genomics UK (COG-UK) Consortium | Berkshire and Surrey Pathology Services Lighthouse Laboratory and Alex Alderton, Roberto Amato, Jeffrey Barrett, Sonia Goncalves, Ewan Harrison, David K. Jackson, Ian Johnston, Dominic Kwiatkowski, Cordelia Langford, John Sillitoe on behalf of the Wellcome Sanger Institute COVID-19 Surveillance Team |
| EPI_ISL_3225091 | Lighthouse Lab in Glasgow | Wellcome Sanger Institute for the COVID-19 Genomics UK (COG-UK) Consortium | Harper VanSteenhouse, Yumi Kasai, David Gray, Carol Clugston, Anna Dominiczak and Alex Alderton, Roberto Amato, Jeffrey Barrett, Sonia Goncalves, Ewan Harrison, David K. Jackson, Ian Johnston, Dominic Kwiatkowski, Cordelia Langford, John Sillitoe on behalf of the Wellcome Sanger Institute COVID-19 Surveillance Team |
| EPI_ISL_3225092 | Lighthouse Lab in Alderley Park | Wellcome Sanger Institute for the COVID-19 Genomics UK (COG-UK) Consortium | Jacquelyn Wynn, Mairead Hyland, The Lighthouse Lab in Alderley Park and Alex Alderton, Roberto Amato, Jeffrey Barrett, Sonia Goncalves, Ewan Harrison, David K. Jackson, Ian Johnston, Dominic Kwiatkowski, Cordelia Langford, John Sillitoe on behalf of the Wellcome Sanger Institute COVID-19 Surveillance Team |
| EPI_ISL_3225093 | Berkshire and Surrey Pathology Services Lighthouse Laboratory | Wellcome Sanger Institute for the COVID-19 Genomics UK (COG-UK) Consortium | Berkshire and Surrey Pathology Services Lighthouse Laboratory and Alex Alderton, Roberto Amato, Jeffrey Barrett, Sonia Goncalves, Ewan Harrison, David K. Jackson, Ian Johnston, Dominic Kwiatkowski, Cordelia Langford, John Sillitoe on behalf of the Wellcome Sanger Institute COVID-19 Surveillance Team |
| EPI_ISL_3225094 | Lighthouse Laboratory Plymouth | Wellcome Sanger Institute for the COVID-19 Genomics UK (COG-UK) Consortium | Lighthouse Laboratory Plymouth and Alex Alderton, Roberto Amato, Jeffrey Barrett, Sonia Goncalves, Ewan Harrison, David K. Jackson, Ian Johnston, Dominic Kwiatkowski, Cordelia Langford, John Sillitoe on behalf of the Wellcome Sanger Institute COVID-19 Surveillance Team |
| EPI_ISL_3225095 | Lighthouse Lab in Alderley Park | Wellcome Sanger Institute for the COVID-19 Genomics UK (COG-UK) Consortium | Jacquelyn Wynn, Mairead Hyland, The Lighthouse Lab in Alderley Park and Alex Alderton, Roberto Amato, Jeffrey Barrett, Sonia Goncalves, Ewan Harrison, David K. Jackson, Ian Johnston, Dominic Kwiatkowski, Cordelia Langford, John Sillitoe on behalf of the Wellcome Sanger Institute COVID-19 Surveillance Team |
| EPI_ISL_3225096 | Berkshire and Surrey Pathology Services Lighthouse Laboratory | Wellcome Sanger Institute for the COVID-19 Genomics UK (COG-UK) Consortium | Berkshire and Surrey Pathology Services Lighthouse Laboratory and Alex Alderton, Roberto Amato, Jeffrey Barrett, Sonia Goncalves, Ewan Harrison, David K. Jackson, Ian Johnston, Dominic Kwiatkowski, Cordelia Langford, John Sillitoe on behalf of the Wellcome Sanger Institute COVID-19 Surveillance Team |
| EPI_ISL_3225097 | Lighthouse Lab in Alderley Park | Wellcome Sanger Institute for the COVID-19 Genomics UK (COG-UK) Consortium | Jacquelyn Wynn, Mairead Hyland, The Lighthouse Lab in Alderley Park and Alex Alderton, Roberto Amato, Jeffrey Barrett, Sonia Goncalves, Ewan Harrison, David K. Jackson, Ian Johnston, Dominic Kwiatkowski, Cordelia Langford, John Sillitoe on behalf of the Wellcome Sanger Institute COVID-19 Surveillance Team |
| EPI_ISL_3225098 | Berkshire and Surrey Pathology Services Lighthouse Laboratory | Wellcome Sanger Institute for the COVID-19 Genomics UK (COG-UK) Consortium | Berkshire and Surrey Pathology Services Lighthouse Laboratory and Alex Alderton, Roberto Amato, Jeffrey Barrett, Sonia Goncalves, Ewan Harrison, David K. Jackson, Ian Johnston, Dominic Kwiatkowski, Cordelia Langford, John Sillitoe on behalf of the Wellcome Sanger Institute COVID-19 Surveillance Team |
| EPI_ISL_3225099 | Lighthouse Lab in Glasgow | Wellcome Sanger Institute for the COVID-19 Genomics UK (COG-UK) Consortium | Harper VanSteenhouse, Yumi Kasai, David Gray, Carol Clugston, Anna Dominiczak and Alex Alderton, Roberto Amato, Jeffrey Barrett, Sonia Goncalves, Ewan Harrison, David K. Jackson, Ian Johnston, Dominic Kwiatkowski, Cordelia Langford, John Sillitoe on behalf of the Wellcome Sanger Institute COVID-19 Surveillance Team |
| EPI_ISL_3225100 | Berkshire and Surrey Pathology Services Lighthouse Laboratory | Wellcome Sanger Institute for the COVID-19 Genomics UK (COG-UK) Consortium | Berkshire and Surrey Pathology Services Lighthouse Laboratory and Alex Alderton, Roberto Amato, Jeffrey Barrett, Sonia Goncalves, Ewan Harrison, David K. Jackson, Ian Johnston, Dominic Kwiatkowski, Cordelia Langford, John Sillitoe on behalf of the Wellcome Sanger Institute COVID-19 Surveillance Team |
| EPI_ISL_3225101 | Lighthouse Lab in Glasgow | Wellcome Sanger Institute for the COVID-19 Genomics UK (COG-UK) Consortium | Harper VanSteenhouse, Yumi Kasai, David Gray, Carol Clugston, Anna Dominiczak and Alex Alderton, Roberto Amato, Jeffrey Barrett, Sonia Goncalves, Ewan Harrison, David K. Jackson, Ian Johnston, Dominic Kwiatkowski, Cordelia Langford, John Sillitoe on behalf of the Wellcome Sanger Institute COVID-19 Surveillance Team |
| EPI_ISL_3225102 | Lighthouse Laboratory Plymouth | Wellcome Sanger Institute for the COVID-19 Genomics UK (COG-UK) Consortium | Lighthouse Laboratory Plymouth and Alex Alderton, Roberto Amato, Jeffrey Barrett, Sonia Goncalves, Ewan Harrison, David K. Jackson, Ian Johnston, Dominic Kwiatkowski, Cordelia Langford, John Sillitoe on behalf of the Wellcome Sanger Institute COVID-19 Surveillance Team |
| EPI_ISL_3225103 | Lighthouse Lab in Glasgow | Wellcome Sanger Institute for the COVID-19 Genomics UK (COG-UK) Consortium | Harper VanSteenhouse, Yumi Kasai, David Gray, Carol Clugston, Anna Dominiczak and Alex Alderton, Roberto Amato, Jeffrey Barrett, Sonia Goncalves, Ewan Harrison, David K. Jackson, Ian Johnston, Dominic Kwiatkowski, Cordelia Langford, John Sillitoe on behalf of the Wellcome Sanger Institute COVID-19 Surveillance Team |
| EPI_ISL_3225104 | Lighthouse Laboratory Plymouth | Wellcome Sanger Institute for the COVID-19 Genomics UK (COG-UK) Consortium | Lighthouse Laboratory Plymouth and Alex Alderton, Roberto Amato, Jeffrey Barrett, Sonia Goncalves, Ewan Harrison, David K. Jackson, Ian Johnston, Dominic Kwiatkowski, Cordelia Langford, John Sillitoe on behalf of the Wellcome Sanger Institute COVID-19 Surveillance Team |
| EPI_ISL_3225105 | Berkshire and Surrey Pathology Services Lighthouse Laboratory | Wellcome Sanger Institute for the COVID-19 Genomics UK (COG-UK) Consortium | Berkshire and Surrey Pathology Services Lighthouse Laboratory and Alex Alderton, Roberto Amato, Jeffrey Barrett, Sonia Goncalves, Ewan Harrison, David K. Jackson, Ian Johnston, Dominic Kwiatkowski, Cordelia Langford, John Sillitoe on behalf of the Wellcome Sanger Institute COVID-19 Surveillance Team |
| EPI_ISL_3225106, EPI_ISL_3225107, EPI_ISL_3225108 | Lighthouse Lab in Alderley Park | Wellcome Sanger Institute for the COVID-19 Genomics UK (COG-UK) Consortium | Jacquelyn Wynn, Mairead Hyland, The Lighthouse Lab in Alderley Park and Alex Alderton, Roberto Amato, Jeffrey Barrett, Sonia Goncalves, Ewan Harrison, David K. Jackson, Ian Johnston, Dominic Kwiatkowski, Cordelia Langford, John Sillitoe on behalf of the Wellcome Sanger Institute COVID-19 Surveillance Team |
| EPI_ISL_3225109, EPI_ISL_3225110 | Berkshire and Surrey Pathology Services Lighthouse Laboratory | Wellcome Sanger Institute for the COVID-19 Genomics UK (COG-UK) Consortium | Berkshire and Surrey Pathology Services Lighthouse Laboratory and Alex Alderton, Roberto Amato, Jeffrey Barrett, Sonia Goncalves, Ewan Harrison, David K. Jackson, Ian Johnston, Dominic Kwiatkowski, Cordelia Langford, John Sillitoe on behalf of the Wellcome Sanger Institute COVID-19 Surveillance Team |
| EPI_ISL_3225111 | Lighthouse Laboratory Plymouth | Wellcome Sanger Institute for the COVID-19 Genomics UK (COG-UK) Consortium | Lighthouse Laboratory Plymouth and Alex Alderton, Roberto Amato, Jeffrey Barrett, Sonia Goncalves, Ewan Harrison, David K. Jackson, Ian Johnston, Dominic Kwiatkowski, Cordelia Langford, John Sillitoe on behalf of the Wellcome Sanger Institute COVID-19 Surveillance Team |
| EPI_ISL_3225112, EPI_ISL_3225113 | Lighthouse Lab in Alderley Park | Wellcome Sanger Institute for the COVID-19 Genomics UK (COG-UK) Consortium | Jacquelyn Wynn, Mairead Hyland, The Lighthouse Lab in Alderley Park and Alex Alderton, Roberto Amato, Jeffrey Barrett, Sonia Goncalves, Ewan Harrison, David K. Jackson, Ian Johnston, Dominic Kwiatkowski, Cordelia Langford, John Sillitoe on behalf of the Wellcome Sanger Institute COVID-19 Surveillance Team |
| EPI_ISL_3225114 | Berkshire and Surrey Pathology Services Lighthouse Laboratory | Wellcome Sanger Institute for the COVID-19 Genomics UK (COG-UK) Consortium | Berkshire and Surrey Pathology Services Lighthouse Laboratory and Alex Alderton, Roberto Amato, Jeffrey Barrett, Sonia Goncalves, Ewan Harrison, David K. Jackson, Ian Johnston, Dominic Kwiatkowski, Cordelia Langford, John Sillitoe on behalf of the Wellcome Sanger Institute COVID-19 Surveillance Team |
| EPI_ISL_3225115 | Health Services Laboratories | Wellcome Sanger Institute for the COVID-19 Genomics UK (COG-UK) Consortium | Health Services Laboratories and Alex Alderton, Roberto Amato, Jeffrey Barrett, Sonia Goncalves, Ewan Harrison, David K. Jackson, Ian Johnston, Dominic Kwiatkowski, Cordelia Langford, John Sillitoe on behalf of the Wellcome Sanger Institute COVID-19 Surveillance Team |
| EPI_ISL_3225116 | Berkshire and Surrey Pathology Services Lighthouse Laboratory | Wellcome Sanger Institute for the COVID-19 Genomics UK (COG-UK) Consortium | Berkshire and Surrey Pathology Services Lighthouse Laboratory and Alex Alderton, Roberto Amato, Jeffrey Barrett, Sonia Goncalves, Ewan Harrison, David K. Jackson, Ian Johnston, Dominic Kwiatkowski, Cordelia Langford, John Sillitoe on behalf of the Wellcome Sanger Institute COVID-19 Surveillance Team |
| EPI_ISL_3225117 | Lighthouse Lab in Alderley Park | Wellcome Sanger Institute for the COVID-19 Genomics UK (COG-UK) Consortium | Jacquelyn Wynn, Mairead Hyland, The Lighthouse Lab in Alderley Park and Alex Alderton, Roberto Amato, Jeffrey Barrett, Sonia Goncalves, Ewan Harrison, David K. Jackson, Ian Johnston, Dominic Kwiatkowski, Cordelia Langford, John Sillitoe on behalf of the Wellcome Sanger Institute COVID-19 Surveillance Team |
| EPI_ISL_3225118, EPI_ISL_3225119 | Lighthouse Laboratory Plymouth | Wellcome Sanger Institute for the COVID-19 Genomics UK (COG-UK) Consortium | Lighthouse Laboratory Plymouth and Alex Alderton, Roberto Amato, Jeffrey Barrett, Sonia Goncalves, Ewan Harrison, David K. Jackson, Ian Johnston, Dominic Kwiatkowski, Cordelia Langford, John Sillitoe on behalf of the Wellcome Sanger Institute COVID-19 Surveillance Team |
| EPI_ISL_3225120 | Lighthouse Lab in Glasgow | Wellcome Sanger Institute for the COVID-19 Genomics UK (COG-UK) Consortium | Harper VanSteenhouse, Yumi Kasai, David Gray, Carol Clugston, Anna Dominiczak and Alex Alderton, Roberto Amato, Jeffrey Barrett, Sonia Goncalves, Ewan Harrison, David K. Jackson, Ian Johnston, Dominic Kwiatkowski, Cordelia Langford, John Sillitoe on behalf of the Wellcome Sanger Institute COVID-19 Surveillance Team |
| EPI_ISL_3225121 | Lighthouse Laboratory Plymouth | Wellcome Sanger Institute for the COVID-19 Genomics UK (COG-UK) Consortium | Lighthouse Laboratory Plymouth and Alex Alderton, Roberto Amato, Jeffrey Barrett, Sonia Goncalves, Ewan Harrison, David K. Jackson, Ian Johnston, Dominic Kwiatkowski, Cordelia Langford, John Sillitoe on behalf of the Wellcome Sanger Institute COVID-19 Surveillance Team |
| EPI_ISL_3225122, EPI_ISL_3225123 | Lighthouse Lab in Alderley Park | Wellcome Sanger Institute for the COVID-19 Genomics UK (COG-UK) Consortium | Jacquelyn Wynn, Mairead Hyland, The Lighthouse Lab in Alderley Park and Alex Alderton, Roberto Amato, Jeffrey Barrett, Sonia Goncalves, Ewan Harrison, David K. Jackson, Ian Johnston, Dominic Kwiatkowski, Cordelia Langford, John Sillitoe on behalf of the Wellcome Sanger Institute COVID-19 Surveillance Team |
| EPI_ISL_3225124 | Lighthouse Laboratory Plymouth | Wellcome Sanger Institute for the COVID-19 Genomics UK (COG-UK) Consortium | Lighthouse Laboratory Plymouth and Alex Alderton, Roberto Amato, Jeffrey Barrett, Sonia Goncalves, Ewan Harrison, David K. Jackson, Ian Johnston, Dominic Kwiatkowski, Cordelia Langford, John Sillitoe on behalf of the Wellcome Sanger Institute COVID-19 Surveillance Team |

| | | | |
|---|---|---|---|
| EPI_ISL_3225125, EPI_ISL_3225126, EPI_ISL_3225127 | Lighthouse Lab in Alderley Park | Wellcome Sanger Institute for the COVID-19 Genomics UK (COG-UK) Consortium | Jacquelyn Wynn, Mairead Hyland, The Lighthouse Lab in Alderley Park and Alex Alderton, Roberto Amato, Jeffrey Barrett, Sonia Goncalves, Ewan Harrison, David K. Jackson, Ian Johnston, Dominic Kwiatkowski, Cordelia Langford, John Sillitoe on behalf of the Wellcome Sanger Institute COVID-19 Surveillance Team |
| EPI_ISL_3225128 | Lighthouse Lab in Glasgow | Wellcome Sanger Institute for the COVID-19 Genomics UK (COG-UK) Consortium | Harper VanSteenhouse, Yumi Kasai, David Gray, Carol Clugston, Anna Dominiczak and Alex Alderton, Roberto Amato, Jeffrey Barrett, Sonia Goncalves, Ewan Harrison, David K. Jackson, Ian Johnston, Dominic Kwiatkowski, Cordelia Langford, John Sillitoe on behalf of the Wellcome Sanger Institute COVID-19 Surveillance Team |
| EPI_ISL_3225129 | Berkshire and Surrey Pathology Services Lighthouse Laboratory | Wellcome Sanger Institute for the COVID-19 Genomics UK (COG-UK) Consortium | Berkshire and Surrey Pathology Services Lighthouse Laboratory and Alex Alderton, Roberto Amato, Jeffrey Barrett, Sonia Goncalves, Ewan Harrison, David K. Jackson, Ian Johnston, Dominic Kwiatkowski, Cordelia Langford, John Sillitoe on behalf of the Wellcome Sanger Institute COVID-19 Surveillance Team |
| EPI_ISL_3225130 | Lighthouse Laboratory Plymouth | Wellcome Sanger Institute for the COVID-19 Genomics UK (COG-UK) Consortium | Lighthouse Laboratory Plymouth and Alex Alderton, Roberto Amato, Jeffrey Barrett, Sonia Goncalves, Ewan Harrison, David K. Jackson, Ian Johnston, Dominic Kwiatkowski, Cordelia Langford, John Sillitoe on behalf of the Wellcome Sanger Institute COVID-19 Surveillance Team |
| EPI_ISL_3225131 | Lighthouse Lab in Glasgow | Wellcome Sanger Institute for the COVID-19 Genomics UK (COG-UK) Consortium | Harper VanSteenhouse, Yumi Kasai, David Gray, Carol Clugston, Anna Dominiczak and Alex Alderton, Roberto Amato, Jeffrey Barrett, Sonia Goncalves, Ewan Harrison, David K. Jackson, Ian Johnston, Dominic Kwiatkowski, Cordelia Langford, John Sillitoe on behalf of the Wellcome Sanger Institute COVID-19 Surveillance Team |
| EPI_ISL_3225132 | Health Services Laboratories | Wellcome Sanger Institute for the COVID-19 Genomics UK (COG-UK) Consortium | Health Services Laboratories and Alex Alderton, Roberto Amato, Jeffrey Barrett, Sonia Goncalves, Ewan Harrison, David K. Jackson, Ian Johnston, Dominic Kwiatkowski, Cordelia Langford, John Sillitoe on behalf of the Wellcome Sanger Institute COVID-19 Surveillance Team |
| EPI_ISL_3225133 | Lighthouse Laboratory Plymouth | Wellcome Sanger Institute for the COVID-19 Genomics UK (COG-UK) Consortium | Lighthouse Laboratory Plymouth and Alex Alderton, Roberto Amato, Jeffrey Barrett, Sonia Goncalves, Ewan Harrison, David K. Jackson, Ian Johnston, Dominic Kwiatkowski, Cordelia Langford, John Sillitoe on behalf of the Wellcome Sanger Institute COVID-19 Surveillance Team |
| EPI_ISL_3225134, EPI_ISL_3225135 | Lighthouse Lab in Glasgow | Wellcome Sanger Institute for the COVID-19 Genomics UK (COG-UK) Consortium | Harper VanSteenhouse, Yumi Kasai, David Gray, Carol Clugston, Anna Dominiczak and Alex Alderton, Roberto Amato, Jeffrey Barrett, Sonia Goncalves, Ewan Harrison, David K. Jackson, Ian Johnston, Dominic Kwiatkowski, Cordelia Langford, John Sillitoe on behalf of the Wellcome Sanger Institute COVID-19 Surveillance Team |
| EPI_ISL_3225136 | Lighthouse Laboratory Plymouth | Wellcome Sanger Institute for the COVID-19 Genomics UK (COG-UK) Consortium | Lighthouse Laboratory Plymouth and Alex Alderton, Roberto Amato, Jeffrey Barrett, Sonia Goncalves, Ewan Harrison, David K. Jackson, Ian Johnston, Dominic Kwiatkowski, Cordelia Langford, John Sillitoe on behalf of the Wellcome Sanger Institute COVID-19 Surveillance Team |
| EPI_ISL_3225137 | Berkshire and Surrey Pathology Services Lighthouse Laboratory | Wellcome Sanger Institute for the COVID-19 Genomics UK (COG-UK) Consortium | Berkshire and Surrey Pathology Services Lighthouse Laboratory and Alex Alderton, Roberto Amato, Jeffrey Barrett, Sonia Goncalves, Ewan Harrison, David K. Jackson, Ian Johnston, Dominic Kwiatkowski, Cordelia Langford, John Sillitoe on behalf of the Wellcome Sanger Institute COVID-19 Surveillance Team |
| EPI_ISL_3225138 | Lighthouse Lab in Alderley Park | Wellcome Sanger Institute for the COVID-19 Genomics UK (COG-UK) Consortium | Jacquelyn Wynn, Mairead Hyland, The Lighthouse Lab in Alderley Park and Alex Alderton, Roberto Amato, Jeffrey Barrett, Sonia Goncalves, Ewan Harrison, David K. Jackson, Ian Johnston, Dominic Kwiatkowski, Cordelia Langford, John Sillitoe on behalf of the Wellcome Sanger Institute COVID-19 Surveillance Team |
| EPI_ISL_3225139 | Lighthouse Lab in Glasgow | Wellcome Sanger Institute for the COVID-19 Genomics UK (COG-UK) Consortium | Harper VanSteenhouse, Yumi Kasai, David Gray, Carol Clugston, Anna Dominiczak and Alex Alderton, Roberto Amato, Jeffrey Barrett, Sonia Goncalves, Ewan Harrison, David K. Jackson, Ian Johnston, Dominic Kwiatkowski, Cordelia Langford, John Sillitoe on behalf of the Wellcome Sanger Institute COVID-19 Surveillance Team |
| EPI_ISL_3225140 | Lighthouse Lab in Alderley Park | Wellcome Sanger Institute for the COVID-19 Genomics UK (COG-UK) Consortium | Jacquelyn Wynn, Mairead Hyland, The Lighthouse Lab in Alderley Park and Alex Alderton, Roberto Amato, Jeffrey Barrett, Sonia Goncalves, Ewan Harrison, David K. Jackson, Ian Johnston, Dominic Kwiatkowski, Cordelia Langford, John Sillitoe on behalf of the Wellcome Sanger Institute COVID-19 Surveillance Team |
| EPI_ISL_3225141 | Berkshire and Surrey Pathology Services Lighthouse Laboratory | Wellcome Sanger Institute for the COVID-19 Genomics UK (COG-UK) Consortium | Berkshire and Surrey Pathology Services Lighthouse Laboratory and Alex Alderton, Roberto Amato, Jeffrey Barrett, Sonia Goncalves, Ewan Harrison, David K. Jackson, Ian Johnston, Dominic Kwiatkowski, Cordelia Langford, John Sillitoe on behalf of the Wellcome Sanger Institute COVID-19 Surveillance Team |
| EPI_ISL_3225142 | Lighthouse Laboratory Plymouth | Wellcome Sanger Institute for the COVID-19 Genomics UK (COG-UK) Consortium | Lighthouse Laboratory Plymouth and Alex Alderton, Roberto Amato, Jeffrey Barrett, Sonia Goncalves, Ewan Harrison, David K. Jackson, Ian Johnston, Dominic Kwiatkowski, Cordelia Langford, John Sillitoe on behalf of the Wellcome Sanger Institute COVID-19 Surveillance Team |
| EPI_ISL_3225143 | Lighthouse Lab in Glasgow | Wellcome Sanger Institute for the COVID-19 Genomics UK (COG-UK) Consortium | Harper VanSteenhouse, Yumi Kasai, David Gray, Carol Clugston, Anna Dominiczak and Alex Alderton, Roberto Amato, Jeffrey Barrett, Sonia Goncalves, Ewan Harrison, David K. Jackson, Ian Johnston, Dominic Kwiatkowski, Cordelia Langford, John Sillitoe on behalf of the Wellcome Sanger Institute COVID-19 Surveillance Team |
| EPI_ISL_3225144 | Lighthouse Laboratory Plymouth | Wellcome Sanger Institute for the COVID-19 Genomics UK (COG-UK) Consortium | Lighthouse Laboratory Plymouth and Alex Alderton, Roberto Amato, Jeffrey Barrett, Sonia Goncalves, Ewan Harrison, David K. Jackson, Ian Johnston, Dominic Kwiatkowski, Cordelia Langford, John Sillitoe on behalf of the Wellcome Sanger Institute COVID-19 Surveillance Team |
| EPI_ISL_3225145 | Berkshire and Surrey Pathology Services Lighthouse Laboratory | Wellcome Sanger Institute for the COVID-19 Genomics UK (COG-UK) Consortium | Berkshire and Surrey Pathology Services Lighthouse Laboratory and Alex Alderton, Roberto Amato, Jeffrey Barrett, Sonia Goncalves, Ewan Harrison, David K. Jackson, Ian Johnston, Dominic Kwiatkowski, Cordelia Langford, John Sillitoe on behalf of the Wellcome Sanger Institute COVID-19 Surveillance Team |
| EPI_ISL_3225146, EPI_ISL_3225147 | Lighthouse Lab in Glasgow | Wellcome Sanger Institute for the COVID-19 Genomics UK (COG-UK) Consortium | Harper VanSteenhouse, Yumi Kasai, David Gray, Carol Clugston, Anna Dominiczak and Alex Alderton, Roberto Amato, Jeffrey Barrett, Sonia Goncalves, Ewan Harrison, David K. Jackson, Ian Johnston, Dominic Kwiatkowski, Cordelia Langford, John Sillitoe on behalf of the Wellcome Sanger Institute COVID-19 Surveillance Team |
| EPI_ISL_3225148, EPI_ISL_3225149 | Lighthouse Laboratory Plymouth | Wellcome Sanger Institute for the COVID-19 Genomics UK (COG-UK) Consortium | Lighthouse Laboratory Plymouth and Alex Alderton, Roberto Amato, Jeffrey Barrett, Sonia Goncalves, Ewan Harrison, David K. Jackson, Ian Johnston, Dominic Kwiatkowski, Cordelia Langford, John Sillitoe on behalf of the Wellcome Sanger Institute COVID-19 Surveillance Team |
| EPI_ISL_3225150 | Berkshire and Surrey Pathology Services Lighthouse Laboratory | Wellcome Sanger Institute for the COVID-19 Genomics UK (COG-UK) Consortium | Berkshire and Surrey Pathology Services Lighthouse Laboratory and Alex Alderton, Roberto Amato, Jeffrey Barrett, Sonia Goncalves, Ewan Harrison, David K. Jackson, Ian Johnston, Dominic Kwiatkowski, Cordelia Langford, John Sillitoe on behalf of the Wellcome Sanger Institute COVID-19 Surveillance Team |
| EPI_ISL_3225151 | Health Services Laboratories | Wellcome Sanger Institute for the COVID-19 Genomics UK (COG-UK) Consortium | Health Services Laboratories and Alex Alderton, Roberto Amato, Jeffrey Barrett, Sonia Goncalves, Ewan Harrison, David K. Jackson, Ian Johnston, Dominic Kwiatkowski, Cordelia Langford, John Sillitoe on behalf of the Wellcome Sanger Institute COVID-19 Surveillance Team |
| EPI_ISL_3225152 | Lighthouse Lab in Glasgow | Wellcome Sanger Institute for the COVID-19 Genomics UK (COG-UK) Consortium | Harper VanSteenhouse, Yumi Kasai, David Gray, Carol Clugston, Anna Dominiczak and Alex Alderton, Roberto Amato, Jeffrey Barrett, Sonia Goncalves, Ewan Harrison, David K. Jackson, Ian Johnston, Dominic Kwiatkowski, Cordelia Langford, John Sillitoe on behalf of the Wellcome Sanger Institute COVID-19 Surveillance Team |
| EPI_ISL_3225153 | Lighthouse Laboratory Plymouth | Wellcome Sanger Institute for the COVID-19 Genomics UK (COG-UK) Consortium | Lighthouse Laboratory Plymouth and Alex Alderton, Roberto Amato, Jeffrey Barrett, Sonia Goncalves, Ewan Harrison, David K. Jackson, Ian Johnston, Dominic Kwiatkowski, Cordelia Langford, John Sillitoe on behalf of the Wellcome Sanger Institute COVID-19 Surveillance Team |
| EPI_ISL_3225154 | Lighthouse Lab in Alderley Park | Wellcome Sanger Institute for the COVID-19 Genomics UK (COG-UK) Consortium | Jacquelyn Wynn, Mairead Hyland, The Lighthouse Lab in Alderley Park and Alex Alderton, Roberto Amato, Jeffrey Barrett, Sonia Goncalves, Ewan Harrison, David K. Jackson, Ian Johnston, Dominic Kwiatkowski, Cordelia Langford, John Sillitoe on behalf of the Wellcome Sanger Institute COVID-19 Surveillance Team |
| EPI_ISL_3225155 | Lighthouse Laboratory Plymouth | Wellcome Sanger Institute for the COVID-19 Genomics UK (COG-UK) Consortium | Lighthouse Laboratory Plymouth and Alex Alderton, Roberto Amato, Jeffrey Barrett, Sonia Goncalves, Ewan Harrison, David K. Jackson, Ian Johnston, Dominic Kwiatkowski, Cordelia Langford, John Sillitoe on behalf of the Wellcome Sanger Institute COVID-19 Surveillance Team |
| EPI_ISL_3225156, EPI_ISL_3225157 | Lighthouse Lab in Glasgow | Wellcome Sanger Institute for the COVID-19 Genomics UK (COG-UK) Consortium | Harper VanSteenhouse, Yumi Kasai, David Gray, Carol Clugston, Anna Dominiczak and Alex Alderton, Roberto Amato, Jeffrey Barrett, Sonia Goncalves, Ewan Harrison, David K. Jackson, Ian Johnston, Dominic Kwiatkowski, Cordelia Langford, John Sillitoe on behalf of the Wellcome Sanger Institute COVID-19 Surveillance Team |
| EPI_ISL_3225158 | Lighthouse Laboratory Plymouth | Wellcome Sanger Institute for the COVID-19 Genomics UK (COG-UK) Consortium | Lighthouse Laboratory Plymouth and Alex Alderton, Roberto Amato, Jeffrey Barrett, Sonia Goncalves, Ewan Harrison, David K. Jackson, Ian Johnston, Dominic Kwiatkowski, Cordelia Langford, John Sillitoe on behalf of the Wellcome Sanger Institute COVID-19 Surveillance Team |
| EPI_ISL_3225159 | Berkshire and Surrey Pathology Services Lighthouse Laboratory | Wellcome Sanger Institute for the COVID-19 Genomics UK (COG-UK) Consortium | Berkshire and Surrey Pathology Services Lighthouse Laboratory and Alex Alderton, Roberto Amato, Jeffrey Barrett, Sonia Goncalves, Ewan Harrison, David K. Jackson, Ian Johnston, Dominic Kwiatkowski, Cordelia Langford, John Sillitoe on behalf of the Wellcome Sanger Institute COVID-19 Surveillance Team |
| EPI_ISL_3225160 | Lighthouse Lab in Glasgow | Wellcome Sanger Institute for the COVID-19 Genomics UK (COG-UK) Consortium | Harper VanSteenhouse, Yumi Kasai, David Gray, Carol Clugston, Anna Dominiczak and Alex Alderton, Roberto Amato, Jeffrey Barrett, Sonia Goncalves, Ewan Harrison, David K. Jackson, Ian Johnston, Dominic Kwiatkowski, Cordelia Langford, John Sillitoe on behalf of the Wellcome Sanger Institute COVID-19 Surveillance Team |
| EPI_ISL_3225161 | Berkshire and Surrey Pathology Services Lighthouse Laboratory | Wellcome Sanger Institute for the COVID-19 Genomics UK (COG-UK) Consortium | Berkshire and Surrey Pathology Services Lighthouse Laboratory and Alex Alderton, Roberto Amato, Jeffrey Barrett, Sonia Goncalves, Ewan Harrison, David K. Jackson, Ian Johnston, Dominic Kwiatkowski, Cordelia Langford, John Sillitoe on behalf of the Wellcome Sanger Institute COVID-19 Surveillance Team |
| EPI_ISL_3225162 | Lighthouse Lab in Glasgow | Wellcome Sanger Institute for the COVID-19 Genomics UK (COG-UK) Consortium | Harper VanSteenhouse, Yumi Kasai, David Gray, Carol Clugston, Anna Dominiczak and Alex Alderton, Roberto Amato, Jeffrey Barrett, Sonia Goncalves, Ewan Harrison, David K. Jackson, Ian Johnston, Dominic Kwiatkowski, Cordelia Langford, John Sillitoe on behalf of the Wellcome Sanger Institute COVID-19 Surveillance Team |
| EPI_ISL_3225163 | Berkshire and Surrey Pathology Services Lighthouse Laboratory | Wellcome Sanger Institute for the COVID-19 Genomics UK (COG-UK) Consortium | Berkshire and Surrey Pathology Services Lighthouse Laboratory and Alex Alderton, Roberto Amato, Jeffrey Barrett, Sonia Goncalves, Ewan Harrison, David K. Jackson, Ian Johnston, Dominic Kwiatkowski, Cordelia Langford, John Sillitoe on behalf of the Wellcome Sanger Institute COVID-19 Surveillance Team |
| EPI_ISL_3225164 | Lighthouse Lab in Glasgow | Wellcome Sanger Institute for the COVID-19 Genomics UK (COG-UK) Consortium | Harper VanSteenhouse, Yumi Kasai, David Gray, Carol Clugston, Anna Dominiczak and Alex Alderton, Roberto Amato, Jeffrey Barrett, Sonia Goncalves, Ewan Harrison, David K. Jackson, Ian Johnston, Dominic Kwiatkowski, Cordelia Langford, John Sillitoe on behalf of the Wellcome Sanger Institute COVID-19 Surveillance Team |
| EPI_ISL_3225165 | Berkshire and Surrey Pathology Services Lighthouse Laboratory | Wellcome Sanger Institute for the COVID-19 Genomics UK (COG-UK) Consortium | Berkshire and Surrey Pathology Services Lighthouse Laboratory and Alex Alderton, Roberto Amato, Jeffrey Barrett, Sonia Goncalves, Ewan Harrison, David K. Jackson, Ian Johnston, Dominic Kwiatkowski, Cordelia Langford, John Sillitoe on behalf of the Wellcome Sanger Institute COVID-19 Surveillance Team |
| EPI_ISL_3225166 | Lighthouse Laboratory Plymouth | Wellcome Sanger Institute for the COVID-19 Genomics UK (COG-UK) Consortium | Lighthouse Laboratory Plymouth and Alex Alderton, Roberto Amato, Jeffrey Barrett, Sonia Goncalves, Ewan Harrison, David K. Jackson, Ian Johnston, Dominic Kwiatkowski, Cordelia Langford, John Sillitoe on behalf of the Wellcome Sanger Institute COVID-19 Surveillance Team |
| EPI_ISL_3225167 | Berkshire and Surrey Pathology Services Lighthouse Laboratory | Wellcome Sanger Institute for the COVID-19 Genomics UK (COG-UK) Consortium | Berkshire and Surrey Pathology Services Lighthouse Laboratory and Alex Alderton, Roberto Amato, Jeffrey Barrett, Sonia Goncalves, Ewan Harrison, David K. Jackson, Ian Johnston, Dominic Kwiatkowski, Cordelia Langford, John Sillitoe on behalf of the Wellcome Sanger Institute COVID-19 Surveillance Team |
| EPI_ISL_3225168, EPI_ISL_3225169, EPI_ISL_3225170 | Lighthouse Lab in Glasgow | Wellcome Sanger Institute for the COVID-19 Genomics UK (COG-UK) Consortium | Harper VanSteenhouse, Yumi Kasai, David Gray, Carol Clugston, Anna Dominiczak and Alex Alderton, Roberto Amato, Jeffrey Barrett, Sonia Goncalves, Ewan Harrison, David K. Jackson, Ian Johnston, Dominic Kwiatkowski, Cordelia Langford, John Sillitoe on behalf of the Wellcome Sanger Institute COVID-19 Surveillance Team |
| EPI_ISL_3225171, EPI_ISL_3225172 | Lighthouse Laboratory Plymouth | Wellcome Sanger Institute for the COVID-19 Genomics UK (COG-UK) Consortium | Lighthouse Laboratory Plymouth and Alex Alderton, Roberto Amato, Jeffrey Barrett, Sonia Goncalves, Ewan Harrison, David K. Jackson, Ian Johnston, Dominic Kwiatkowski, Cordelia Langford, John Sillitoe on behalf of the Wellcome Sanger Institute COVID-19 Surveillance Team |
| EPI_ISL_3225173 | Lighthouse Lab in Alderley Park | Wellcome Sanger Institute for the COVID-19 Genomics UK (COG-UK) Consortium | Jacquelyn Wynn, Mairead Hyland, The Lighthouse Lab in Alderley Park and Alex Alderton, Roberto Amato, Jeffrey Barrett, Sonia Goncalves, Ewan Harrison, David K. Jackson, Ian Johnston, Dominic Kwiatkowski, Cordelia Langford, John Sillitoe on behalf of the Wellcome Sanger Institute COVID-19 Surveillance Team |
| EPI_ISL_3225174 | Berkshire and Surrey Pathology Services Lighthouse Laboratory | Wellcome Sanger Institute for the COVID-19 Genomics UK (COG-UK) Consortium | Berkshire and Surrey Pathology Services Lighthouse Laboratory and Alex Alderton, Roberto Amato, Jeffrey Barrett, Sonia Goncalves, Ewan Harrison, David K. Jackson, Ian Johnston, Dominic Kwiatkowski, Cordelia Langford, John Sillitoe on behalf of the Wellcome Sanger Institute COVID-19 Surveillance Team |
| EPI_ISL_3225175 | Lighthouse Lab in Alderley Park | Wellcome Sanger Institute for the COVID-19 Genomics UK (COG-UK) Consortium | Jacquelyn Wynn, Mairead Hyland, The Lighthouse Lab in Alderley Park and Alex Alderton, Roberto Amato, Jeffrey Barrett, Sonia Goncalves, Ewan Harrison, David K. Jackson, Ian Johnston, Dominic Kwiatkowski, Cordelia Langford, John Sillitoe on behalf of the Wellcome Sanger Institute COVID-19 Surveillance Team |
| EPI_ISL_3225176 | Lighthouse Laboratory Plymouth | Wellcome Sanger Institute for the COVID-19 Genomics UK (COG-UK) Consortium | Lighthouse Laboratory Plymouth and Alex Alderton, Roberto Amato, Jeffrey Barrett, Sonia Goncalves, Ewan Harrison, David K. Jackson, Ian Johnston, Dominic Kwiatkowski, Cordelia Langford, John Sillitoe on behalf of the Wellcome Sanger Institute COVID-19 Surveillance Team |
| EPI_ISL_3225177 | Berkshire and Surrey Pathology Services Lighthouse Laboratory | Wellcome Sanger Institute for the COVID-19 Genomics UK (COG-UK) Consortium | Berkshire and Surrey Pathology Services Lighthouse Laboratory and Alex Alderton, Roberto Amato, Jeffrey Barrett, Sonia Goncalves, Ewan Harrison, David K. Jackson, Ian Johnston, Dominic Kwiatkowski, Cordelia Langford, John Sillitoe on behalf of the Wellcome Sanger Institute COVID-19 Surveillance Team |

| | | | |
|---|---|---|---|
| EPI_ISL_3225178 | Lighthouse Lab in Glasgow | Wellcome Sanger Institute for the COVID-19 Genomics UK (COG-UK) Consortium | Harper VanSteenhouse, Yumi Kasai, David Gray, Carol Clugston, Anna Dominiczak and Alex Alderton, Roberto Amato, Jeffrey Barrett, Sonia Goncalves, Ewan Harrison, David K. Jackson, Ian Johnston, Dominic Kwiatkowski, Cordelia Langford, John Sillitoe on behalf of the Wellcome Sanger Institute COVID-19 Surveillance Team |
| EPI_ISL_3225179 | Berkshire and Surrey Pathology Services Lighthouse Laboratory | Wellcome Sanger Institute for the COVID-19 Genomics UK (COG-UK) Consortium | Berkshire and Surrey Pathology Services Lighthouse Laboratory and Alex Alderton, Roberto Amato, Jeffrey Barrett, Sonia Goncalves, Ewan Harrison, David K. Jackson, Ian Johnston, Dominic Kwiatkowski, Cordelia Langford, John Sillitoe on behalf of the Wellcome Sanger Institute COVID-19 Surveillance Team |
| EPI_ISL_3225180 | Lighthouse Laboratory Plymouth | Wellcome Sanger Institute for the COVID-19 Genomics UK (COG-UK) Consortium | Lighthouse Laboratory Plymouth and Alex Alderton, Roberto Amato, Jeffrey Barrett, Sonia Goncalves, Ewan Harrison, David K. Jackson, Ian Johnston, Dominic Kwiatkowski, Cordelia Langford, John Sillitoe on behalf of the Wellcome Sanger Institute COVID-19 Surveillance Team |
| EPI_ISL_3225181 | Lighthouse Lab in Glasgow | Wellcome Sanger Institute for the COVID-19 Genomics UK (COG-UK) Consortium | Harper VanSteenhouse, Yumi Kasai, David Gray, Carol Clugston, Anna Dominiczak and Alex Alderton, Roberto Amato, Jeffrey Barrett, Sonia Goncalves, Ewan Harrison, David K. Jackson, Ian Johnston, Dominic Kwiatkowski, Cordelia Langford, John Sillitoe on behalf of the Wellcome Sanger Institute COVID-19 Surveillance Team |
| EPI_ISL_3225182 | Berkshire and Surrey Pathology Services Lighthouse Laboratory | Wellcome Sanger Institute for the COVID-19 Genomics UK (COG-UK) Consortium | Berkshire and Surrey Pathology Services Lighthouse Laboratory and Alex Alderton, Roberto Amato, Jeffrey Barrett, Sonia Goncalves, Ewan Harrison, David K. Jackson, Ian Johnston, Dominic Kwiatkowski, Cordelia Langford, John Sillitoe on behalf of the Wellcome Sanger Institute COVID-19 Surveillance Team |
| EPI_ISL_3225183 | Lighthouse Lab in Glasgow | Wellcome Sanger Institute for the COVID-19 Genomics UK (COG-UK) Consortium | Harper VanSteenhouse, Yumi Kasai, David Gray, Carol Clugston, Anna Dominiczak and Alex Alderton, Roberto Amato, Jeffrey Barrett, Sonia Goncalves, Ewan Harrison, David K. Jackson, Ian Johnston, Dominic Kwiatkowski, Cordelia Langford, John Sillitoe on behalf of the Wellcome Sanger Institute COVID-19 Surveillance Team |
| EPI_ISL_3225184 | Lighthouse Lab in Alderley Park | Wellcome Sanger Institute for the COVID-19 Genomics UK (COG-UK) Consortium | Jacquelyn Wynn, Mairead Hyland, The Lighthouse Lab in Alderley Park and Alex Alderton, Roberto Amato, Jeffrey Barrett, Sonia Goncalves, Ewan Harrison, David K. Jackson, Ian Johnston, Dominic Kwiatkowski, Cordelia Langford, John Sillitoe on behalf of the Wellcome Sanger Institute COVID-19 Surveillance Team |
| EPI_ISL_3225185, EPI_ISL_3225186 | Lighthouse Laboratory Plymouth | Wellcome Sanger Institute for the COVID-19 Genomics UK (COG-UK) Consortium | Lighthouse Laboratory Plymouth and Alex Alderton, Roberto Amato, Jeffrey Barrett, Sonia Goncalves, Ewan Harrison, David K. Jackson, Ian Johnston, Dominic Kwiatkowski, Cordelia Langford, John Sillitoe on behalf of the Wellcome Sanger Institute COVID-19 Surveillance Team |
| EPI_ISL_3225187, EPI_ISL_3225188 | Lighthouse Lab in Alderley Park | Wellcome Sanger Institute for the COVID-19 Genomics UK (COG-UK) Consortium | Jacquelyn Wynn, Mairead Hyland, The Lighthouse Lab in Alderley Park and Alex Alderton, Roberto Amato, Jeffrey Barrett, Sonia Goncalves, Ewan Harrison, David K. Jackson, Ian Johnston, Dominic Kwiatkowski, Cordelia Langford, John Sillitoe on behalf of the Wellcome Sanger Institute COVID-19 Surveillance Team |
| EPI_ISL_3225189, EPI_ISL_3225190 | Lighthouse Laboratory Plymouth | Wellcome Sanger Institute for the COVID-19 Genomics UK (COG-UK) Consortium | Lighthouse Laboratory Plymouth and Alex Alderton, Roberto Amato, Jeffrey Barrett, Sonia Goncalves, Ewan Harrison, David K. Jackson, Ian Johnston, Dominic Kwiatkowski, Cordelia Langford, John Sillitoe on behalf of the Wellcome Sanger Institute COVID-19 Surveillance Team |
| EPI_ISL_3225191 | Lighthouse Lab in Glasgow | Wellcome Sanger Institute for the COVID-19 Genomics UK (COG-UK) Consortium | Harper VanSteenhouse, Yumi Kasai, David Gray, Carol Clugston, Anna Dominiczak and Alex Alderton, Roberto Amato, Jeffrey Barrett, Sonia Goncalves, Ewan Harrison, David K. Jackson, Ian Johnston, Dominic Kwiatkowski, Cordelia Langford, John Sillitoe on behalf of the Wellcome Sanger Institute COVID-19 Surveillance Team |
| EPI_ISL_3225192 | Lighthouse Laboratory Plymouth | Wellcome Sanger Institute for the COVID-19 Genomics UK (COG-UK) Consortium | Lighthouse Laboratory Plymouth and Alex Alderton, Roberto Amato, Jeffrey Barrett, Sonia Goncalves, Ewan Harrison, David K. Jackson, Ian Johnston, Dominic Kwiatkowski, Cordelia Langford, John Sillitoe on behalf of the Wellcome Sanger Institute COVID-19 Surveillance Team |
| EPI_ISL_3225193 | Lighthouse Lab in Glasgow | Wellcome Sanger Institute for the COVID-19 Genomics UK (COG-UK) Consortium | Harper VanSteenhouse, Yumi Kasai, David Gray, Carol Clugston, Anna Dominiczak and Alex Alderton, Roberto Amato, Jeffrey Barrett, Sonia Goncalves, Ewan Harrison, David K. Jackson, Ian Johnston, Dominic Kwiatkowski, Cordelia Langford, John Sillitoe on behalf of the Wellcome Sanger Institute COVID-19 Surveillance Team |
| EPI_ISL_3225194 | Health Services Laboratories | Wellcome Sanger Institute for the COVID-19 Genomics UK (COG-UK) Consortium | Health Services Laboratories and Alex Alderton, Roberto Amato, Jeffrey Barrett, Sonia Goncalves, Ewan Harrison, David K. Jackson, Ian Johnston, Dominic Kwiatkowski, Cordelia Langford, John Sillitoe on behalf of the Wellcome Sanger Institute COVID-19 Surveillance Team |
| EPI_ISL_3225195, EPI_ISL_3225196, EPI_ISL_3225197 | Lighthouse Lab in Glasgow | Wellcome Sanger Institute for the COVID-19 Genomics UK (COG-UK) Consortium | Harper VanSteenhouse, Yumi Kasai, David Gray, Carol Clugston, Anna Dominiczak and Alex Alderton, Roberto Amato, Jeffrey Barrett, Sonia Goncalves, Ewan Harrison, David K. Jackson, Ian Johnston, Dominic Kwiatkowski, Cordelia Langford, John Sillitoe on behalf of the Wellcome Sanger Institute COVID-19 Surveillance Team |
| EPI_ISL_3225198 | Lighthouse Laboratory Plymouth | Wellcome Sanger Institute for the COVID-19 Genomics UK (COG-UK) Consortium | Lighthouse Laboratory Plymouth and Alex Alderton, Roberto Amato, Jeffrey Barrett, Sonia Goncalves, Ewan Harrison, David K. Jackson, Ian Johnston, Dominic Kwiatkowski, Cordelia Langford, John Sillitoe on behalf of the Wellcome Sanger Institute COVID-19 Surveillance Team |
| EPI_ISL_3225199 | Lighthouse Lab in Alderley Park | Wellcome Sanger Institute for the COVID-19 Genomics UK (COG-UK) Consortium | Jacquelyn Wynn, Mairead Hyland, The Lighthouse Lab in Alderley Park and Alex Alderton, Roberto Amato, Jeffrey Barrett, Sonia Goncalves, Ewan Harrison, David K. Jackson, Ian Johnston, Dominic Kwiatkowski, Cordelia Langford, John Sillitoe on behalf of the Wellcome Sanger Institute COVID-19 Surveillance Team |
| EPI_ISL_3225200 | Lighthouse Lab in Glasgow | Wellcome Sanger Institute for the COVID-19 Genomics UK (COG-UK) Consortium | Harper VanSteenhouse, Yumi Kasai, David Gray, Carol Clugston, Anna Dominiczak and Alex Alderton, Roberto Amato, Jeffrey Barrett, Sonia Goncalves, Ewan Harrison, David K. Jackson, Ian Johnston, Dominic Kwiatkowski, Cordelia Langford, John Sillitoe on behalf of the Wellcome Sanger Institute COVID-19 Surveillance Team |
| EPI_ISL_3225201 | Lighthouse Lab in Alderley Park | Wellcome Sanger Institute for the COVID-19 Genomics UK (COG-UK) Consortium | Jacquelyn Wynn, Mairead Hyland, The Lighthouse Lab in Alderley Park and Alex Alderton, Roberto Amato, Jeffrey Barrett, Sonia Goncalves, Ewan Harrison, David K. Jackson, Ian Johnston, Dominic Kwiatkowski, Cordelia Langford, John Sillitoe on behalf of the Wellcome Sanger Institute COVID-19 Surveillance Team |
| EPI_ISL_3225202 | Berkshire and Surrey Pathology Services Lighthouse Laboratory | Wellcome Sanger Institute for the COVID-19 Genomics UK (COG-UK) Consortium | Berkshire and Surrey Pathology Services Lighthouse Laboratory and Alex Alderton, Roberto Amato, Jeffrey Barrett, Sonia Goncalves, Ewan Harrison, David K. Jackson, Ian Johnston, Dominic Kwiatkowski, Cordelia Langford, John Sillitoe on behalf of the Wellcome Sanger Institute COVID-19 Surveillance Team |
| EPI_ISL_3225203 | Lighthouse Laboratory Plymouth | Wellcome Sanger Institute for the COVID-19 Genomics UK (COG-UK) Consortium | Lighthouse Laboratory Plymouth and Alex Alderton, Roberto Amato, Jeffrey Barrett, Sonia Goncalves, Ewan Harrison, David K. Jackson, Ian Johnston, Dominic Kwiatkowski, Cordelia Langford, John Sillitoe on behalf of the Wellcome Sanger Institute COVID-19 Surveillance Team |
| EPI_ISL_3225204 | Lighthouse Lab in Alderley Park | Wellcome Sanger Institute for the COVID-19 Genomics UK (COG-UK) Consortium | Jacquelyn Wynn, Mairead Hyland, The Lighthouse Lab in Alderley Park and Alex Alderton, Roberto Amato, Jeffrey Barrett, Sonia Goncalves, Ewan Harrison, David K. Jackson, Ian Johnston, Dominic Kwiatkowski, Cordelia Langford, John Sillitoe on behalf of the Wellcome Sanger Institute COVID-19 Surveillance Team |
| EPI_ISL_3225205 | Health Services Laboratories | Wellcome Sanger Institute for the COVID-19 Genomics UK (COG-UK) Consortium | Health Services Laboratories and Alex Alderton, Roberto Amato, Jeffrey Barrett, Sonia Goncalves, Ewan Harrison, David K. Jackson, Ian Johnston, Dominic Kwiatkowski, Cordelia Langford, John Sillitoe on behalf of the Wellcome Sanger Institute COVID-19 Surveillance Team |
| EPI_ISL_3225206 | Lighthouse Laboratory Plymouth | Wellcome Sanger Institute for the COVID-19 Genomics UK (COG-UK) Consortium | Lighthouse Laboratory Plymouth and Alex Alderton, Roberto Amato, Jeffrey Barrett, Sonia Goncalves, Ewan Harrison, David K. Jackson, Ian Johnston, Dominic Kwiatkowski, Cordelia Langford, John Sillitoe on behalf of the Wellcome Sanger Institute COVID-19 Surveillance Team |
| EPI_ISL_3225207 | Lighthouse Lab in Alderley Park | Wellcome Sanger Institute for the COVID-19 Genomics UK (COG-UK) Consortium | Jacquelyn Wynn, Mairead Hyland, The Lighthouse Lab in Alderley Park and Alex Alderton, Roberto Amato, Jeffrey Barrett, Sonia Goncalves, Ewan Harrison, David K. Jackson, Ian Johnston, Dominic Kwiatkowski, Cordelia Langford, John Sillitoe on behalf of the Wellcome Sanger Institute COVID-19 Surveillance Team |
| EPI_ISL_3225208, EPI_ISL_3225209 | Lighthouse Lab in Glasgow | Wellcome Sanger Institute for the COVID-19 Genomics UK (COG-UK) Consortium | Harper VanSteenhouse, Yumi Kasai, David Gray, Carol Clugston, Anna Dominiczak and Alex Alderton, Roberto Amato, Jeffrey Barrett, Sonia Goncalves, Ewan Harrison, David K. Jackson, Ian Johnston, Dominic Kwiatkowski, Cordelia Langford, John Sillitoe on behalf of the Wellcome Sanger Institute COVID-19 Surveillance Team |
| EPI_ISL_3225210 | Lighthouse Lab in Alderley Park | Wellcome Sanger Institute for the COVID-19 Genomics UK (COG-UK) Consortium | Jacquelyn Wynn, Mairead Hyland, The Lighthouse Lab in Alderley Park and Alex Alderton, Roberto Amato, Jeffrey Barrett, Sonia Goncalves, Ewan Harrison, David K. Jackson, Ian Johnston, Dominic Kwiatkowski, Cordelia Langford, John Sillitoe on behalf of the Wellcome Sanger Institute COVID-19 Surveillance Team |
| EPI_ISL_3225211 | Berkshire and Surrey Pathology Services Lighthouse Laboratory | Wellcome Sanger Institute for the COVID-19 Genomics UK (COG-UK) Consortium | Berkshire and Surrey Pathology Services Lighthouse Laboratory and Alex Alderton, Roberto Amato, Jeffrey Barrett, Sonia Goncalves, Ewan Harrison, David K. Jackson, Ian Johnston, Dominic Kwiatkowski, Cordelia Langford, John Sillitoe on behalf of the Wellcome Sanger Institute COVID-19 Surveillance Team |
| EPI_ISL_3225212 | Lighthouse Lab in Alderley Park | Wellcome Sanger Institute for the COVID-19 Genomics UK (COG-UK) Consortium | Jacquelyn Wynn, Mairead Hyland, The Lighthouse Lab in Alderley Park and Alex Alderton, Roberto Amato, Jeffrey Barrett, Sonia Goncalves, Ewan Harrison, David K. Jackson, Ian Johnston, Dominic Kwiatkowski, Cordelia Langford, John Sillitoe on behalf of the Wellcome Sanger Institute COVID-19 Surveillance Team |
| EPI_ISL_3225213, EPI_ISL_3225214, EPI_ISL_3225215, EPI_ISL_3225216 | Berkshire and Surrey Pathology Services Lighthouse Laboratory | Wellcome Sanger Institute for the COVID-19 Genomics UK (COG-UK) Consortium | Berkshire and Surrey Pathology Services Lighthouse Laboratory and Alex Alderton, Roberto Amato, Jeffrey Barrett, Sonia Goncalves, Ewan Harrison, David K. Jackson, Ian Johnston, Dominic Kwiatkowski, Cordelia Langford, John Sillitoe on behalf of the Wellcome Sanger Institute COVID-19 Surveillance Team |
| EPI_ISL_3225217 | Lighthouse Lab in Glasgow | Wellcome Sanger Institute for the COVID-19 Genomics UK (COG-UK) Consortium | Harper VanSteenhouse, Yumi Kasai, David Gray, Carol Clugston, Anna Dominiczak and Alex Alderton, Roberto Amato, Jeffrey Barrett, Sonia Goncalves, Ewan Harrison, David K. Jackson, Ian Johnston, Dominic Kwiatkowski, Cordelia Langford, John Sillitoe on behalf of the Wellcome Sanger Institute COVID-19 Surveillance Team |
| EPI_ISL_3225219 | Health Services Laboratories | Wellcome Sanger Institute for the COVID-19 Genomics UK (COG-UK) Consortium | Health Services Laboratories and Alex Alderton, Roberto Amato, Jeffrey Barrett, Sonia Goncalves, Ewan Harrison, David K. Jackson, Ian Johnston, Dominic Kwiatkowski, Cordelia Langford, John Sillitoe on behalf of the Wellcome Sanger Institute COVID-19 Surveillance Team |
| EPI_ISL_3225220 | Lighthouse Lab in Alderley Park | Wellcome Sanger Institute for the COVID-19 Genomics UK (COG-UK) Consortium | Jacquelyn Wynn, Mairead Hyland, The Lighthouse Lab in Alderley Park and Alex Alderton, Roberto Amato, Jeffrey Barrett, Sonia Goncalves, Ewan Harrison, David K. Jackson, Ian Johnston, Dominic Kwiatkowski, Cordelia Langford, John Sillitoe on behalf of the Wellcome Sanger Institute COVID-19 Surveillance Team |
| EPI_ISL_3225221 | Lighthouse Laboratory Plymouth | Wellcome Sanger Institute for the COVID-19 Genomics UK (COG-UK) Consortium | Lighthouse Laboratory Plymouth and Alex Alderton, Roberto Amato, Jeffrey Barrett, Sonia Goncalves, Ewan Harrison, David K. Jackson, Ian Johnston, Dominic Kwiatkowski, Cordelia Langford, John Sillitoe on behalf of the Wellcome Sanger Institute COVID-19 Surveillance Team |
| EPI_ISL_3225222 | Lighthouse Lab in Glasgow | Wellcome Sanger Institute for the COVID-19 Genomics UK (COG-UK) Consortium | Harper VanSteenhouse, Yumi Kasai, David Gray, Carol Clugston, Anna Dominiczak and Alex Alderton, Roberto Amato, Jeffrey Barrett, Sonia Goncalves, Ewan Harrison, David K. Jackson, Ian Johnston, Dominic Kwiatkowski, Cordelia Langford, John Sillitoe on behalf of the Wellcome Sanger Institute COVID-19 Surveillance Team |
| EPI_ISL_3225223, EPI_ISL_3225224 | Lighthouse Laboratory Plymouth | Wellcome Sanger Institute for the COVID-19 Genomics UK (COG-UK) Consortium | Lighthouse Laboratory Plymouth and Alex Alderton, Roberto Amato, Jeffrey Barrett, Sonia Goncalves, Ewan Harrison, David K. Jackson, Ian Johnston, Dominic Kwiatkowski, Cordelia Langford, John Sillitoe on behalf of the Wellcome Sanger Institute COVID-19 Surveillance Team |
| EPI_ISL_3225225 | Lighthouse Lab in Alderley Park | Wellcome Sanger Institute for the COVID-19 Genomics UK (COG-UK) Consortium | Jacquelyn Wynn, Mairead Hyland, The Lighthouse Lab in Alderley Park and Alex Alderton, Roberto Amato, Jeffrey Barrett, Sonia Goncalves, Ewan Harrison, David K. Jackson, Ian Johnston, Dominic Kwiatkowski, Cordelia Langford, John Sillitoe on behalf of the Wellcome Sanger Institute COVID-19 Surveillance Team |
| EPI_ISL_3225226, EPI_ISL_3225227, EPI_ISL_3225228 | Lighthouse Lab in Glasgow | Wellcome Sanger Institute for the COVID-19 Genomics UK (COG-UK) Consortium | Harper VanSteenhouse, Yumi Kasai, David Gray, Carol Clugston, Anna Dominiczak and Alex Alderton, Roberto Amato, Jeffrey Barrett, Sonia Goncalves, Ewan Harrison, David K. Jackson, Ian Johnston, Dominic Kwiatkowski, Cordelia Langford, John Sillitoe on behalf of the Wellcome Sanger Institute COVID-19 Surveillance Team |
| EPI_ISL_3225229 | Lighthouse Laboratory Plymouth | Wellcome Sanger Institute for the COVID-19 Genomics UK (COG-UK) Consortium | Lighthouse Laboratory Plymouth and Alex Alderton, Roberto Amato, Jeffrey Barrett, Sonia Goncalves, Ewan Harrison, David K. Jackson, Ian Johnston, Dominic Kwiatkowski, Cordelia Langford, John Sillitoe on behalf of the Wellcome Sanger Institute COVID-19 Surveillance Team |
| EPI_ISL_3225230, EPI_ISL_3225231 | Lighthouse Lab in Alderley Park | Wellcome Sanger Institute for the COVID-19 Genomics UK (COG-UK) Consortium | Jacquelyn Wynn, Mairead Hyland, The Lighthouse Lab in Alderley Park and Alex Alderton, Roberto Amato, Jeffrey Barrett, Sonia Goncalves, Ewan Harrison, David K. Jackson, Ian Johnston, Dominic Kwiatkowski, Cordelia Langford, John Sillitoe on behalf of the Wellcome Sanger Institute COVID-19 Surveillance Team |
| EPI_ISL_3225232, EPI_ISL_3225233 | Lighthouse Laboratory Plymouth | Wellcome Sanger Institute for the COVID-19 Genomics UK (COG-UK) Consortium | Lighthouse Laboratory Plymouth and Alex Alderton, Roberto Amato, Jeffrey Barrett, Sonia Goncalves, Ewan Harrison, David K. Jackson, Ian Johnston, Dominic Kwiatkowski, Cordelia Langford, John Sillitoe on behalf of the Wellcome Sanger Institute COVID-19 Surveillance Team |
| EPI_ISL_3225234 | Lighthouse Lab in Glasgow | Wellcome Sanger Institute for the COVID-19 Genomics UK (COG-UK) Consortium | Harper VanSteenhouse, Yumi Kasai, David Gray, Carol Clugston, Anna Dominiczak and Alex Alderton, Roberto Amato, Jeffrey Barrett, Sonia Goncalves, Ewan Harrison, David K. Jackson, Ian Johnston, Dominic Kwiatkowski, Cordelia Langford, John Sillitoe on behalf of the Wellcome Sanger Institute COVID-19 Surveillance Team |
| EPI_ISL_3225235 | Lighthouse Laboratory Plymouth | Wellcome Sanger Institute for the COVID-19 Genomics UK (COG-UK) Consortium | Lighthouse Laboratory Plymouth and Alex Alderton, Roberto Amato, Jeffrey Barrett, Sonia Goncalves, Ewan Harrison, David K. Jackson, Ian Johnston, Dominic Kwiatkowski, Cordelia Langford, John Sillitoe on behalf of the Wellcome Sanger Institute COVID-19 Surveillance Team |

| | | | |
|---|---|---|---|
| EPI_ISL_3225236, EPI_ISL_3225237 | Lighthouse Lab in Alderley Park | Wellcome Sanger Institute for the COVID-19 Genomics UK (COG-UK) Consortium | Jacquelyn Wynn, Mairead Hyland, The Lighthouse Lab in Alderley Park and Alex Alderton, Roberto Amato, Jeffrey Barrett, Sonia Goncalves, Ewan Harrison, David K. Jackson, Ian Johnston, Dominic Kwiatkowski, Cordelia Langford, John Sillitoe on behalf of the Wellcome Sanger Institute COVID-19 Surveillance Team |
| EPI_ISL_3225238 | Lighthouse Lab in Glasgow | Wellcome Sanger Institute for the COVID-19 Genomics UK (COG-UK) Consortium | Harper VanSteenhouse, Yumi Kasai, David Gray, Carol Clugston, Anna Dominiczak and Alex Alderton, Roberto Amato, Jeffrey Barrett, Sonia Goncalves, Ewan Harrison, David K. Jackson, Ian Johnston, Dominic Kwiatkowski, Cordelia Langford, John Sillitoe on behalf of the Wellcome Sanger Institute COVID-19 Surveillance Team |
| EPI_ISL_3225239 | Lighthouse Lab in Alderley Park | Wellcome Sanger Institute for the COVID-19 Genomics UK (COG-UK) Consortium | Jacquelyn Wynn, Mairead Hyland, The Lighthouse Lab in Alderley Park and Alex Alderton, Roberto Amato, Jeffrey Barrett, Sonia Goncalves, Ewan Harrison, David K. Jackson, Ian Johnston, Dominic Kwiatkowski, Cordelia Langford, John Sillitoe on behalf of the Wellcome Sanger Institute COVID-19 Surveillance Team |
| EPI_ISL_3225240, EPI_ISL_3225241 | Lighthouse Lab in Glasgow | Wellcome Sanger Institute for the COVID-19 Genomics UK (COG-UK) Consortium | Harper VanSteenhouse, Yumi Kasai, David Gray, Carol Clugston, Anna Dominiczak and Alex Alderton, Roberto Amato, Jeffrey Barrett, Sonia Goncalves, Ewan Harrison, David K. Jackson, Ian Johnston, Dominic Kwiatkowski, Cordelia Langford, John Sillitoe on behalf of the Wellcome Sanger Institute COVID-19 Surveillance Team |
| EPI_ISL_3225242, EPI_ISL_3225243 | Lighthouse Lab in Alderley Park | Wellcome Sanger Institute for the COVID-19 Genomics UK (COG-UK) Consortium | Jacquelyn Wynn, Mairead Hyland, The Lighthouse Lab in Alderley Park and Alex Alderton, Roberto Amato, Jeffrey Barrett, Sonia Goncalves, Ewan Harrison, David K. Jackson, Ian Johnston, Dominic Kwiatkowski, Cordelia Langford, John Sillitoe on behalf of the Wellcome Sanger Institute COVID-19 Surveillance Team |
| EPI_ISL_3225244 | Lighthouse Lab in Glasgow | Wellcome Sanger Institute for the COVID-19 Genomics UK (COG-UK) Consortium | Harper VanSteenhouse, Yumi Kasai, David Gray, Carol Clugston, Anna Dominiczak and Alex Alderton, Roberto Amato, Jeffrey Barrett, Sonia Goncalves, Ewan Harrison, David K. Jackson, Ian Johnston, Dominic Kwiatkowski, Cordelia Langford, John Sillitoe on behalf of the Wellcome Sanger Institute COVID-19 Surveillance Team |
| EPI_ISL_3225245 | Lighthouse Laboratory Plymouth | Wellcome Sanger Institute for the COVID-19 Genomics UK (COG-UK) Consortium | Lighthouse Laboratory Plymouth and Alex Alderton, Roberto Amato, Jeffrey Barrett, Sonia Goncalves, Ewan Harrison, David K. Jackson, Ian Johnston, Dominic Kwiatkowski, Cordelia Langford, John Sillitoe on behalf of the Wellcome Sanger Institute COVID-19 Surveillance Team |
| EPI_ISL_3225246 | Lighthouse Lab in Alderley Park | Wellcome Sanger Institute for the COVID-19 Genomics UK (COG-UK) Consortium | Jacquelyn Wynn, Mairead Hyland, The Lighthouse Lab in Alderley Park and Alex Alderton, Roberto Amato, Jeffrey Barrett, Sonia Goncalves, Ewan Harrison, David K. Jackson, Ian Johnston, Dominic Kwiatkowski, Cordelia Langford, John Sillitoe on behalf of the Wellcome Sanger Institute COVID-19 Surveillance Team |
| EPI_ISL_3225247 | Lighthouse Lab in Glasgow | Wellcome Sanger Institute for the COVID-19 Genomics UK (COG-UK) Consortium | Harper VanSteenhouse, Yumi Kasai, David Gray, Carol Clugston, Anna Dominiczak and Alex Alderton, Roberto Amato, Jeffrey Barrett, Sonia Goncalves, Ewan Harrison, David K. Jackson, Ian Johnston, Dominic Kwiatkowski, Cordelia Langford, John Sillitoe on behalf of the Wellcome Sanger Institute COVID-19 Surveillance Team |
| EPI_ISL_3225248, EPI_ISL_3225249 | Lighthouse Laboratory Plymouth | Wellcome Sanger Institute for the COVID-19 Genomics UK (COG-UK) Consortium | Lighthouse Laboratory Plymouth and Alex Alderton, Roberto Amato, Jeffrey Barrett, Sonia Goncalves, Ewan Harrison, David K. Jackson, Ian Johnston, Dominic Kwiatkowski, Cordelia Langford, John Sillitoe on behalf of the Wellcome Sanger Institute COVID-19 Surveillance Team |
| EPI_ISL_3225250, EPI_ISL_3225251 | Lighthouse Lab in Glasgow | Wellcome Sanger Institute for the COVID-19 Genomics UK (COG-UK) Consortium | Harper VanSteenhouse, Yumi Kasai, David Gray, Carol Clugston, Anna Dominiczak and Alex Alderton, Roberto Amato, Jeffrey Barrett, Sonia Goncalves, Ewan Harrison, David K. Jackson, Ian Johnston, Dominic Kwiatkowski, Cordelia Langford, John Sillitoe on behalf of the Wellcome Sanger Institute COVID-19 Surveillance Team |
| EPI_ISL_3225252, EPI_ISL_3225253 | Lighthouse Lab in Alderley Park | Wellcome Sanger Institute for the COVID-19 Genomics UK (COG-UK) Consortium | Jacquelyn Wynn, Mairead Hyland, The Lighthouse Lab in Alderley Park and Alex Alderton, Roberto Amato, Jeffrey Barrett, Sonia Goncalves, Ewan Harrison, David K. Jackson, Ian Johnston, Dominic Kwiatkowski, Cordelia Langford, John Sillitoe on behalf of the Wellcome Sanger Institute COVID-19 Surveillance Team |
| EPI_ISL_3225254 | Lighthouse Lab in Glasgow | Wellcome Sanger Institute for the COVID-19 Genomics UK (COG-UK) Consortium | Harper VanSteenhouse, Yumi Kasai, David Gray, Carol Clugston, Anna Dominiczak and Alex Alderton, Roberto Amato, Jeffrey Barrett, Sonia Goncalves, Ewan Harrison, David K. Jackson, Ian Johnston, Dominic Kwiatkowski, Cordelia Langford, John Sillitoe on behalf of the Wellcome Sanger Institute COVID-19 Surveillance Team |
| EPI_ISL_3225255 | Lighthouse Lab in Alderley Park | Wellcome Sanger Institute for the COVID-19 Genomics UK (COG-UK) Consortium | Jacquelyn Wynn, Mairead Hyland, The Lighthouse Lab in Alderley Park and Alex Alderton, Roberto Amato, Jeffrey Barrett, Sonia Goncalves, Ewan Harrison, David K. Jackson, Ian Johnston, Dominic Kwiatkowski, Cordelia Langford, John Sillitoe on behalf of the Wellcome Sanger Institute COVID-19 Surveillance Team |
| EPI_ISL_3225256 | Lighthouse Lab in Glasgow | Wellcome Sanger Institute for the COVID-19 Genomics UK (COG-UK) Consortium | Harper VanSteenhouse, Yumi Kasai, David Gray, Carol Clugston, Anna Dominiczak and Alex Alderton, Roberto Amato, Jeffrey Barrett, Sonia Goncalves, Ewan Harrison, David K. Jackson, Ian Johnston, Dominic Kwiatkowski, Cordelia Langford, John Sillitoe on behalf of the Wellcome Sanger Institute COVID-19 Surveillance Team |
| EPI_ISL_3225257, EPI_ISL_3225258 | Lighthouse Lab in Alderley Park | Wellcome Sanger Institute for the COVID-19 Genomics UK (COG-UK) Consortium | Jacquelyn Wynn, Mairead Hyland, The Lighthouse Lab in Alderley Park and Alex Alderton, Roberto Amato, Jeffrey Barrett, Sonia Goncalves, Ewan Harrison, David K. Jackson, Ian Johnston, Dominic Kwiatkowski, Cordelia Langford, John Sillitoe on behalf of the Wellcome Sanger Institute COVID-19 Surveillance Team |
| EPI_ISL_3225259, EPI_ISL_3225260 | Lighthouse Laboratory Plymouth | Wellcome Sanger Institute for the COVID-19 Genomics UK (COG-UK) Consortium | Lighthouse Laboratory Plymouth and Alex Alderton, Roberto Amato, Jeffrey Barrett, Sonia Goncalves, Ewan Harrison, David K. Jackson, Ian Johnston, Dominic Kwiatkowski, Cordelia Langford, John Sillitoe on behalf of the Wellcome Sanger Institute COVID-19 Surveillance Team |
| EPI_ISL_3225261 | Lighthouse Lab in Glasgow | Wellcome Sanger Institute for the COVID-19 Genomics UK (COG-UK) Consortium | Harper VanSteenhouse, Yumi Kasai, David Gray, Carol Clugston, Anna Dominiczak and Alex Alderton, Roberto Amato, Jeffrey Barrett, Sonia Goncalves, Ewan Harrison, David K. Jackson, Ian Johnston, Dominic Kwiatkowski, Cordelia Langford, John Sillitoe on behalf of the Wellcome Sanger Institute COVID-19 Surveillance Team |
| EPI_ISL_3225262 | Lighthouse Lab in Alderley Park | Wellcome Sanger Institute for the COVID-19 Genomics UK (COG-UK) Consortium | Jacquelyn Wynn, Mairead Hyland, The Lighthouse Lab in Alderley Park and Alex Alderton, Roberto Amato, Jeffrey Barrett, Sonia Goncalves, Ewan Harrison, David K. Jackson, Ian Johnston, Dominic Kwiatkowski, Cordelia Langford, John Sillitoe on behalf of the Wellcome Sanger Institute COVID-19 Surveillance Team |
| EPI_ISL_3225263 | Lighthouse Lab in Glasgow | Wellcome Sanger Institute for the COVID-19 Genomics UK (COG-UK) Consortium | Harper VanSteenhouse, Yumi Kasai, David Gray, Carol Clugston, Anna Dominiczak and Alex Alderton, Roberto Amato, Jeffrey Barrett, Sonia Goncalves, Ewan Harrison, David K. Jackson, Ian Johnston, Dominic Kwiatkowski, Cordelia Langford, John Sillitoe on behalf of the Wellcome Sanger Institute COVID-19 Surveillance Team |
| EPI_ISL_3225264 | Lighthouse Lab in Alderley Park | Wellcome Sanger Institute for the COVID-19 Genomics UK (COG-UK) Consortium | Jacquelyn Wynn, Mairead Hyland, The Lighthouse Lab in Alderley Park and Alex Alderton, Roberto Amato, Jeffrey Barrett, Sonia Goncalves, Ewan Harrison, David K. Jackson, Ian Johnston, Dominic Kwiatkowski, Cordelia Langford, John Sillitoe on behalf of the Wellcome Sanger Institute COVID-19 Surveillance Team |
| EPI_ISL_3225265 | Lighthouse Lab in Glasgow | Wellcome Sanger Institute for the COVID-19 Genomics UK (COG-UK) Consortium | Harper VanSteenhouse, Yumi Kasai, David Gray, Carol Clugston, Anna Dominiczak and Alex Alderton, Roberto Amato, Jeffrey Barrett, Sonia Goncalves, Ewan Harrison, David K. Jackson, Ian Johnston, Dominic Kwiatkowski, Cordelia Langford, John Sillitoe on behalf of the Wellcome Sanger Institute COVID-19 Surveillance Team |
| EPI_ISL_3225266 | Lighthouse Laboratory Plymouth | Wellcome Sanger Institute for the COVID-19 Genomics UK (COG-UK) Consortium | Lighthouse Laboratory Plymouth and Alex Alderton, Roberto Amato, Jeffrey Barrett, Sonia Goncalves, Ewan Harrison, David K. Jackson, Ian Johnston, Dominic Kwiatkowski, Cordelia Langford, John Sillitoe on behalf of the Wellcome Sanger Institute COVID-19 Surveillance Team |
| EPI_ISL_3225267, EPI_ISL_3225268 | Lighthouse Lab in Alderley Park | Wellcome Sanger Institute for the COVID-19 Genomics UK (COG-UK) Consortium | Jacquelyn Wynn, Mairead Hyland, The Lighthouse Lab in Alderley Park and Alex Alderton, Roberto Amato, Jeffrey Barrett, Sonia Goncalves, Ewan Harrison, David K. Jackson, Ian Johnston, Dominic Kwiatkowski, Cordelia Langford, John Sillitoe on behalf of the Wellcome Sanger Institute COVID-19 Surveillance Team |
| EPI_ISL_3225269 | Lighthouse Laboratory Plymouth | Wellcome Sanger Institute for the COVID-19 Genomics UK (COG-UK) Consortium | Lighthouse Laboratory Plymouth and Alex Alderton, Roberto Amato, Jeffrey Barrett, Sonia Goncalves, Ewan Harrison, David K. Jackson, Ian Johnston, Dominic Kwiatkowski, Cordelia Langford, John Sillitoe on behalf of the Wellcome Sanger Institute COVID-19 Surveillance Team |
| EPI_ISL_3225270 | Lighthouse Lab in Alderley Park | Wellcome Sanger Institute for the COVID-19 Genomics UK (COG-UK) Consortium | Jacquelyn Wynn, Mairead Hyland, The Lighthouse Lab in Alderley Park and Alex Alderton, Roberto Amato, Jeffrey Barrett, Sonia Goncalves, Ewan Harrison, David K. Jackson, Ian Johnston, Dominic Kwiatkowski, Cordelia Langford, John Sillitoe on behalf of the Wellcome Sanger Institute COVID-19 Surveillance Team |
| EPI_ISL_3225271 | Lighthouse Laboratory Plymouth | Wellcome Sanger Institute for the COVID-19 Genomics UK (COG-UK) Consortium | Lighthouse Laboratory Plymouth and Alex Alderton, Roberto Amato, Jeffrey Barrett, Sonia Goncalves, Ewan Harrison, David K. Jackson, Ian Johnston, Dominic Kwiatkowski, Cordelia Langford, John Sillitoe on behalf of the Wellcome Sanger Institute COVID-19 Surveillance Team |
| EPI_ISL_3225272 | Lighthouse Lab in Alderley Park | Wellcome Sanger Institute for the COVID-19 Genomics UK (COG-UK) Consortium | Jacquelyn Wynn, Mairead Hyland, The Lighthouse Lab in Alderley Park and Alex Alderton, Roberto Amato, Jeffrey Barrett, Sonia Goncalves, Ewan Harrison, David K. Jackson, Ian Johnston, Dominic Kwiatkowski, Cordelia Langford, John Sillitoe on behalf of the Wellcome Sanger Institute COVID-19 Surveillance Team |
| EPI_ISL_3225273 | Lighthouse Laboratory Plymouth | Wellcome Sanger Institute for the COVID-19 Genomics UK (COG-UK) Consortium | Lighthouse Laboratory Plymouth and Alex Alderton, Roberto Amato, Jeffrey Barrett, Sonia Goncalves, Ewan Harrison, David K. Jackson, Ian Johnston, Dominic Kwiatkowski, Cordelia Langford, John Sillitoe on behalf of the Wellcome Sanger Institute COVID-19 Surveillance Team |
| EPI_ISL_3225274 | Lighthouse Lab in Alderley Park | Wellcome Sanger Institute for the COVID-19 Genomics UK (COG-UK) Consortium | Jacquelyn Wynn, Mairead Hyland, The Lighthouse Lab in Alderley Park and Alex Alderton, Roberto Amato, Jeffrey Barrett, Sonia Goncalves, Ewan Harrison, David K. Jackson, Ian Johnston, Dominic Kwiatkowski, Cordelia Langford, John Sillitoe on behalf of the Wellcome Sanger Institute COVID-19 Surveillance Team |
| EPI_ISL_3225275 | Lighthouse Lab in Glasgow | Wellcome Sanger Institute for the COVID-19 Genomics UK (COG-UK) Consortium | Harper VanSteenhouse, Yumi Kasai, David Gray, Carol Clugston, Anna Dominiczak and Alex Alderton, Roberto Amato, Jeffrey Barrett, Sonia Goncalves, Ewan Harrison, David K. Jackson, Ian Johnston, Dominic Kwiatkowski, Cordelia Langford, John Sillitoe on behalf of the Wellcome Sanger Institute COVID-19 Surveillance Team |
| EPI_ISL_3225276, EPI_ISL_3225277, EPI_ISL_3225278, EPI_ISL_3225279, EPI_ISL_3225280, EPI_ISL_3225281 | Lighthouse Lab in Alderley Park | Wellcome Sanger Institute for the COVID-19 Genomics UK (COG-UK) Consortium | Jacquelyn Wynn, Mairead Hyland, The Lighthouse Lab in Alderley Park and Alex Alderton, Roberto Amato, Jeffrey Barrett, Sonia Goncalves, Ewan Harrison, David K. Jackson, Ian Johnston, Dominic Kwiatkowski, Cordelia Langford, John Sillitoe on behalf of the Wellcome Sanger Institute COVID-19 Surveillance Team |
| EPI_ISL_3225282 | Lighthouse Laboratory Plymouth | Wellcome Sanger Institute for the COVID-19 Genomics UK (COG-UK) Consortium | Lighthouse Laboratory Plymouth and Alex Alderton, Roberto Amato, Jeffrey Barrett, Sonia Goncalves, Ewan Harrison, David K. Jackson, Ian Johnston, Dominic Kwiatkowski, Cordelia Langford, John Sillitoe on behalf of the Wellcome Sanger Institute COVID-19 Surveillance Team |
| EPI_ISL_3225283 | Lighthouse Lab in Glasgow | Wellcome Sanger Institute for the COVID-19 Genomics UK (COG-UK) Consortium | Harper VanSteenhouse, Yumi Kasai, David Gray, Carol Clugston, Anna Dominiczak and Alex Alderton, Roberto Amato, Jeffrey Barrett, Sonia Goncalves, Ewan Harrison, David K. Jackson, Ian Johnston, Dominic Kwiatkowski, Cordelia Langford, John Sillitoe on behalf of the Wellcome Sanger Institute COVID-19 Surveillance Team |
| EPI_ISL_3225284 | Lighthouse Lab in Alderley Park | Wellcome Sanger Institute for the COVID-19 Genomics UK (COG-UK) Consortium | Jacquelyn Wynn, Mairead Hyland, The Lighthouse Lab in Alderley Park and Alex Alderton, Roberto Amato, Jeffrey Barrett, Sonia Goncalves, Ewan Harrison, David K. Jackson, Ian Johnston, Dominic Kwiatkowski, Cordelia Langford, John Sillitoe on behalf of the Wellcome Sanger Institute COVID-19 Surveillance Team |
| EPI_ISL_3225285 | Lighthouse Laboratory Plymouth | Wellcome Sanger Institute for the COVID-19 Genomics UK (COG-UK) Consortium | Lighthouse Laboratory Plymouth and Alex Alderton, Roberto Amato, Jeffrey Barrett, Sonia Goncalves, Ewan Harrison, David K. Jackson, Ian Johnston, Dominic Kwiatkowski, Cordelia Langford, John Sillitoe on behalf of the Wellcome Sanger Institute COVID-19 Surveillance Team |
| EPI_ISL_3225286, EPI_ISL_3225287, EPI_ISL_3225288 | Lighthouse Lab in Alderley Park | Wellcome Sanger Institute for the COVID-19 Genomics UK (COG-UK) Consortium | Jacquelyn Wynn, Mairead Hyland, The Lighthouse Lab in Alderley Park and Alex Alderton, Roberto Amato, Jeffrey Barrett, Sonia Goncalves, Ewan Harrison, David K. Jackson, Ian Johnston, Dominic Kwiatkowski, Cordelia Langford, John Sillitoe on behalf of the Wellcome Sanger Institute COVID-19 Surveillance Team |
| EPI_ISL_3225289 | Lighthouse Lab in Glasgow | Wellcome Sanger Institute for the COVID-19 Genomics UK (COG-UK) Consortium | Harper VanSteenhouse, Yumi Kasai, David Gray, Carol Clugston, Anna Dominiczak and Alex Alderton, Roberto Amato, Jeffrey Barrett, Sonia Goncalves, Ewan Harrison, David K. Jackson, Ian Johnston, Dominic Kwiatkowski, Cordelia Langford, John Sillitoe on behalf of the Wellcome Sanger Institute COVID-19 Surveillance Team |
| EPI_ISL_3225290, EPI_ISL_3225291 | Lighthouse Laboratory Plymouth | Wellcome Sanger Institute for the COVID-19 Genomics UK (COG-UK) Consortium | Lighthouse Laboratory Plymouth and Alex Alderton, Roberto Amato, Jeffrey Barrett, Sonia Goncalves, Ewan Harrison, David K. Jackson, Ian Johnston, Dominic Kwiatkowski, Cordelia Langford, John Sillitoe on behalf of the Wellcome Sanger Institute COVID-19 Surveillance Team |
| EPI_ISL_3225292, EPI_ISL_3225293 | Lighthouse Lab in Glasgow | Wellcome Sanger Institute for the COVID-19 Genomics UK (COG-UK) Consortium | Harper VanSteenhouse, Yumi Kasai, David Gray, Carol Clugston, Anna Dominiczak and Alex Alderton, Roberto Amato, Jeffrey Barrett, Sonia Goncalves, Ewan Harrison, David K. Jackson, Ian Johnston, Dominic Kwiatkowski, Cordelia Langford, John Sillitoe on behalf of the Wellcome Sanger Institute COVID-19 Surveillance Team |
| EPI_ISL_3225294, EPI_ISL_3225295, EPI_ISL_3225296, EPI_ISL_3225297, EPI_ISL_3225298 | Lighthouse Lab in Alderley Park | Wellcome Sanger Institute for the COVID-19 Genomics UK (COG-UK) Consortium | Jacquelyn Wynn, Mairead Hyland, The Lighthouse Lab in Alderley Park and Alex Alderton, Roberto Amato, Jeffrey Barrett, Sonia Goncalves, Ewan Harrison, David K. Jackson, Ian Johnston, Dominic Kwiatkowski, Cordelia Langford, John Sillitoe on behalf of the Wellcome Sanger Institute COVID-19 Surveillance Team |
| EPI_ISL_3225299 | Lighthouse Laboratory Plymouth | Wellcome Sanger Institute for the COVID-19 Genomics UK (COG-UK) | Lighthouse Laboratory Plymouth and Alex Alderton, Roberto Amato, Jeffrey Barrett, Sonia Goncalves, Ewan Harrison, David K. Jackson, Ian Johnston, Dominic Kwiatkowski, Cordelia Langford, John Sillitoe on behalf of the Wellcome Sanger |

| | | | |
|---|---|---|---|
| | | Consortium | Institute COVID-19 Surveillance Team |
| EPI_ISL_3225300 | Lighthouse Lab in Glasgow | Wellcome Sanger Institute for the COVID-19 Genomics UK (COG-UK) Consortium | Harper VanSteenhouse, Yumi Kasai, David Gray, Carol Clugston, Anna Dominiczak and Alex Alderton, Roberto Amato, Jeffrey Barrett, Sonia Goncalves, Ewan Harrison, David K. Jackson, Ian Johnston, Dominic Kwiatkowski, Cordelia Langford, John Sillitoe on behalf of the Wellcome Sanger Institute COVID-19 Surveillance Team |
| EPI_ISL_3225301 | Lighthouse Laboratory Plymouth | Wellcome Sanger Institute for the COVID-19 Genomics UK (COG-UK) Consortium | Lighthouse Laboratory Plymouth and Alex Alderton, Roberto Amato, Jeffrey Barrett, Sonia Goncalves, Ewan Harrison, David K. Jackson, Ian Johnston, Dominic Kwiatkowski, Cordelia Langford, John Sillitoe on behalf of the Wellcome Sanger Institute COVID-19 Surveillance Team |
| EPI_ISL_3225302 | Lighthouse Lab in Glasgow | Wellcome Sanger Institute for the COVID-19 Genomics UK (COG-UK) Consortium | Harper VanSteenhouse, Yumi Kasai, David Gray, Carol Clugston, Anna Dominiczak and Alex Alderton, Roberto Amato, Jeffrey Barrett, Sonia Goncalves, Ewan Harrison, David K. Jackson, Ian Johnston, Dominic Kwiatkowski, Cordelia Langford, John Sillitoe on behalf of the Wellcome Sanger Institute COVID-19 Surveillance Team |
| EPI_ISL_3225303, EPI_ISL_3225304 | Lighthouse Laboratory Plymouth | Wellcome Sanger Institute for the COVID-19 Genomics UK (COG-UK) Consortium | Lighthouse Laboratory Plymouth and Alex Alderton, Roberto Amato, Jeffrey Barrett, Sonia Goncalves, Ewan Harrison, David K. Jackson, Ian Johnston, Dominic Kwiatkowski, Cordelia Langford, John Sillitoe on behalf of the Wellcome Sanger Institute COVID-19 Surveillance Team |
| EPI_ISL_3225305, EPI_ISL_3225306, EPI_ISL_3225307, EPI_ISL_3225308 | Lighthouse Lab in Alderley Park | Wellcome Sanger Institute for the COVID-19 Genomics UK (COG-UK) Consortium | Jacquelyn Wynn, Mairead Hyland, The Lighthouse Lab in Alderley Park and Alex Alderton, Roberto Amato, Jeffrey Barrett, Sonia Goncalves, Ewan Harrison, David K. Jackson, Ian Johnston, Dominic Kwiatkowski, Cordelia Langford, John Sillitoe on behalf of the Wellcome Sanger Institute COVID-19 Surveillance Team |
| EPI_ISL_3225309 | Lighthouse Lab in Glasgow | Wellcome Sanger Institute for the COVID-19 Genomics UK (COG-UK) Consortium | Harper VanSteenhouse, Yumi Kasai, David Gray, Carol Clugston, Anna Dominiczak and Alex Alderton, Roberto Amato, Jeffrey Barrett, Sonia Goncalves, Ewan Harrison, David K. Jackson, Ian Johnston, Dominic Kwiatkowski, Cordelia Langford, John Sillitoe on behalf of the Wellcome Sanger Institute COVID-19 Surveillance Team |
| EPI_ISL_3225310, EPI_ISL_3225311, EPI_ISL_3225312, EPI_ISL_3225313, EPI_ISL_3225314, EPI_ISL_3225315, EPI_ISL_3225316 | Lighthouse Lab in Alderley Park | Wellcome Sanger Institute for the COVID-19 Genomics UK (COG-UK) Consortium | Jacquelyn Wynn, Mairead Hyland, The Lighthouse Lab in Alderley Park and Alex Alderton, Roberto Amato, Jeffrey Barrett, Sonia Goncalves, Ewan Harrison, David K. Jackson, Ian Johnston, Dominic Kwiatkowski, Cordelia Langford, John Sillitoe on behalf of the Wellcome Sanger Institute COVID-19 Surveillance Team |
| EPI_ISL_3225317 | Lighthouse Lab in Glasgow | Wellcome Sanger Institute for the COVID-19 Genomics UK (COG-UK) Consortium | Harper VanSteenhouse, Yumi Kasai, David Gray, Carol Clugston, Anna Dominiczak and Alex Alderton, Roberto Amato, Jeffrey Barrett, Sonia Goncalves, Ewan Harrison, David K. Jackson, Ian Johnston, Dominic Kwiatkowski, Cordelia Langford, John Sillitoe on behalf of the Wellcome Sanger Institute COVID-19 Surveillance Team |
| EPI_ISL_3225318, EPI_ISL_3225319 | Lighthouse Lab in Alderley Park | Wellcome Sanger Institute for the COVID-19 Genomics UK (COG-UK) Consortium | Jacquelyn Wynn, Mairead Hyland, The Lighthouse Lab in Alderley Park and Alex Alderton, Roberto Amato, Jeffrey Barrett, Sonia Goncalves, Ewan Harrison, David K. Jackson, Ian Johnston, Dominic Kwiatkowski, Cordelia Langford, John Sillitoe on behalf of the Wellcome Sanger Institute COVID-19 Surveillance Team |
| EPI_ISL_3225320 | Lighthouse Lab in Glasgow | Wellcome Sanger Institute for the COVID-19 Genomics UK (COG-UK) Consortium | Harper VanSteenhouse, Yumi Kasai, David Gray, Carol Clugston, Anna Dominiczak and Alex Alderton, Roberto Amato, Jeffrey Barrett, Sonia Goncalves, Ewan Harrison, David K. Jackson, Ian Johnston, Dominic Kwiatkowski, Cordelia Langford, John Sillitoe on behalf of the Wellcome Sanger Institute COVID-19 Surveillance Team |
| EPI_ISL_3225321 | Lighthouse Laboratory Plymouth | Wellcome Sanger Institute for the COVID-19 Genomics UK (COG-UK) Consortium | Lighthouse Laboratory Plymouth and Alex Alderton, Roberto Amato, Jeffrey Barrett, Sonia Goncalves, Ewan Harrison, David K. Jackson, Ian Johnston, Dominic Kwiatkowski, Cordelia Langford, John Sillitoe on behalf of the Wellcome Sanger Institute COVID-19 Surveillance Team |
| EPI_ISL_3225322 | Lighthouse Lab in Alderley Park | Wellcome Sanger Institute for the COVID-19 Genomics UK (COG-UK) Consortium | Jacquelyn Wynn, Mairead Hyland, The Lighthouse Lab in Alderley Park and Alex Alderton, Roberto Amato, Jeffrey Barrett, Sonia Goncalves, Ewan Harrison, David K. Jackson, Ian Johnston, Dominic Kwiatkowski, Cordelia Langford, John Sillitoe on behalf of the Wellcome Sanger Institute COVID-19 Surveillance Team |
| EPI_ISL_3225323 | Lighthouse Lab in Glasgow | Wellcome Sanger Institute for the COVID-19 Genomics UK (COG-UK) Consortium | Harper VanSteenhouse, Yumi Kasai, David Gray, Carol Clugston, Anna Dominiczak and Alex Alderton, Roberto Amato, Jeffrey Barrett, Sonia Goncalves, Ewan Harrison, David K. Jackson, Ian Johnston, Dominic Kwiatkowski, Cordelia Langford, John Sillitoe on behalf of the Wellcome Sanger Institute COVID-19 Surveillance Team |
| EPI_ISL_3225324 | Lighthouse Laboratory Plymouth | Wellcome Sanger Institute for the COVID-19 Genomics UK (COG-UK) Consortium | Lighthouse Laboratory Plymouth and Alex Alderton, Roberto Amato, Jeffrey Barrett, Sonia Goncalves, Ewan Harrison, David K. Jackson, Ian Johnston, Dominic Kwiatkowski, Cordelia Langford, John Sillitoe on behalf of the Wellcome Sanger Institute COVID-19 Surveillance Team |
| EPI_ISL_3225325 | Lighthouse Lab in Alderley Park | Wellcome Sanger Institute for the COVID-19 Genomics UK (COG-UK) Consortium | Jacquelyn Wynn, Mairead Hyland, The Lighthouse Lab in Alderley Park and Alex Alderton, Roberto Amato, Jeffrey Barrett, Sonia Goncalves, Ewan Harrison, David K. Jackson, Ian Johnston, Dominic Kwiatkowski, Cordelia Langford, John Sillitoe on behalf of the Wellcome Sanger Institute COVID-19 Surveillance Team |
| EPI_ISL_3225326 | Lighthouse Lab in Glasgow | Wellcome Sanger Institute for the COVID-19 Genomics UK (COG-UK) Consortium | Harper VanSteenhouse, Yumi Kasai, David Gray, Carol Clugston, Anna Dominiczak and Alex Alderton, Roberto Amato, Jeffrey Barrett, Sonia Goncalves, Ewan Harrison, David K. Jackson, Ian Johnston, Dominic Kwiatkowski, Cordelia Langford, John Sillitoe on behalf of the Wellcome Sanger Institute COVID-19 Surveillance Team |
| EPI_ISL_3225327, EPI_ISL_3225328, EPI_ISL_3225329 | Lighthouse Lab in Alderley Park | Wellcome Sanger Institute for the COVID-19 Genomics UK (COG-UK) Consortium | Jacquelyn Wynn, Mairead Hyland, The Lighthouse Lab in Alderley Park and Alex Alderton, Roberto Amato, Jeffrey Barrett, Sonia Goncalves, Ewan Harrison, David K. Jackson, Ian Johnston, Dominic Kwiatkowski, Cordelia Langford, John Sillitoe on behalf of the Wellcome Sanger Institute COVID-19 Surveillance Team |
| EPI_ISL_3225330, EPI_ISL_3225331, EPI_ISL_3225332 | Lighthouse Lab in Glasgow | Wellcome Sanger Institute for the COVID-19 Genomics UK (COG-UK) Consortium | Harper VanSteenhouse, Yumi Kasai, David Gray, Carol Clugston, Anna Dominiczak and Alex Alderton, Roberto Amato, Jeffrey Barrett, Sonia Goncalves, Ewan Harrison, David K. Jackson, Ian Johnston, Dominic Kwiatkowski, Cordelia Langford, John Sillitoe on behalf of the Wellcome Sanger Institute COVID-19 Surveillance Team |
| EPI_ISL_3225333 | Lighthouse Laboratory Plymouth | Wellcome Sanger Institute for the COVID-19 Genomics UK (COG-UK) Consortium | Lighthouse Laboratory Plymouth and Alex Alderton, Roberto Amato, Jeffrey Barrett, Sonia Goncalves, Ewan Harrison, David K. Jackson, Ian Johnston, Dominic Kwiatkowski, Cordelia Langford, John Sillitoe on behalf of the Wellcome Sanger Institute COVID-19 Surveillance Team |
| EPI_ISL_3225334, EPI_ISL_3225335, EPI_ISL_3225336, EPI_ISL_3225337 | Lighthouse Lab in Alderley Park | Wellcome Sanger Institute for the COVID-19 Genomics UK (COG-UK) Consortium | Jacquelyn Wynn, Mairead Hyland, The Lighthouse Lab in Alderley Park and Alex Alderton, Roberto Amato, Jeffrey Barrett, Sonia Goncalves, Ewan Harrison, David K. Jackson, Ian Johnston, Dominic Kwiatkowski, Cordelia Langford, John Sillitoe on behalf of the Wellcome Sanger Institute COVID-19 Surveillance Team |
| EPI_ISL_3225338 | Lighthouse Lab in Glasgow | Wellcome Sanger Institute for the COVID-19 Genomics UK (COG-UK) Consortium | Harper VanSteenhouse, Yumi Kasai, David Gray, Carol Clugston, Anna Dominiczak and Alex Alderton, Roberto Amato, Jeffrey Barrett, Sonia Goncalves, Ewan Harrison, David K. Jackson, Ian Johnston, Dominic Kwiatkowski, Cordelia Langford, John Sillitoe on behalf of the Wellcome Sanger Institute COVID-19 Surveillance Team |
| EPI_ISL_3225339, EPI_ISL_3225340 | Lighthouse Lab in Alderley Park | Wellcome Sanger Institute for the COVID-19 Genomics UK (COG-UK) Consortium | Jacquelyn Wynn, Mairead Hyland, The Lighthouse Lab in Alderley Park and Alex Alderton, Roberto Amato, Jeffrey Barrett, Sonia Goncalves, Ewan Harrison, David K. Jackson, Ian Johnston, Dominic Kwiatkowski, Cordelia Langford, John Sillitoe on behalf of the Wellcome Sanger Institute COVID-19 Surveillance Team |
| EPI_ISL_3225341 | Lighthouse Laboratory Plymouth | Wellcome Sanger Institute for the COVID-19 Genomics UK (COG-UK) Consortium | Lighthouse Laboratory Plymouth and Alex Alderton, Roberto Amato, Jeffrey Barrett, Sonia Goncalves, Ewan Harrison, David K. Jackson, Ian Johnston, Dominic Kwiatkowski, Cordelia Langford, John Sillitoe on behalf of the Wellcome Sanger Institute COVID-19 Surveillance Team |
| EPI_ISL_3225342 | Lighthouse Lab in Alderley Park | Wellcome Sanger Institute for the COVID-19 Genomics UK (COG-UK) Consortium | Jacquelyn Wynn, Mairead Hyland, The Lighthouse Lab in Alderley Park and Alex Alderton, Roberto Amato, Jeffrey Barrett, Sonia Goncalves, Ewan Harrison, David K. Jackson, Ian Johnston, Dominic Kwiatkowski, Cordelia Langford, John Sillitoe on behalf of the Wellcome Sanger Institute COVID-19 Surveillance Team |
| EPI_ISL_3225343 | Lighthouse Laboratory Plymouth | Wellcome Sanger Institute for the COVID-19 Genomics UK (COG-UK) Consortium | Lighthouse Laboratory Plymouth and Alex Alderton, Roberto Amato, Jeffrey Barrett, Sonia Goncalves, Ewan Harrison, David K. Jackson, Ian Johnston, Dominic Kwiatkowski, Cordelia Langford, John Sillitoe on behalf of the Wellcome Sanger Institute COVID-19 Surveillance Team |
| EPI_ISL_3225344, EPI_ISL_3225345, EPI_ISL_3225346, EPI_ISL_3225347, EPI_ISL_3225348, EPI_ISL_3225349 | Lighthouse Lab in Alderley Park | Wellcome Sanger Institute for the COVID-19 Genomics UK (COG-UK) Consortium | Jacquelyn Wynn, Mairead Hyland, The Lighthouse Lab in Alderley Park and Alex Alderton, Roberto Amato, Jeffrey Barrett, Sonia Goncalves, Ewan Harrison, David K. Jackson, Ian Johnston, Dominic Kwiatkowski, Cordelia Langford, John Sillitoe on behalf of the Wellcome Sanger Institute COVID-19 Surveillance Team |
| EPI_ISL_3225350 | Lighthouse Lab in Glasgow | Wellcome Sanger Institute for the COVID-19 Genomics UK (COG-UK) Consortium | Harper VanSteenhouse, Yumi Kasai, David Gray, Carol Clugston, Anna Dominiczak and Alex Alderton, Roberto Amato, Jeffrey Barrett, Sonia Goncalves, Ewan Harrison, David K. Jackson, Ian Johnston, Dominic Kwiatkowski, Cordelia Langford, John Sillitoe on behalf of the Wellcome Sanger Institute COVID-19 Surveillance Team |
| EPI_ISL_3225351, EPI_ISL_3225352, EPI_ISL_3225353 | Lighthouse Lab in Alderley Park | Wellcome Sanger Institute for the COVID-19 Genomics UK (COG-UK) Consortium | Jacquelyn Wynn, Mairead Hyland, The Lighthouse Lab in Alderley Park and Alex Alderton, Roberto Amato, Jeffrey Barrett, Sonia Goncalves, Ewan Harrison, David K. Jackson, Ian Johnston, Dominic Kwiatkowski, Cordelia Langford, John Sillitoe on behalf of the Wellcome Sanger Institute COVID-19 Surveillance Team |
| EPI_ISL_3225354 | Lighthouse Lab in Glasgow | Wellcome Sanger Institute for the COVID-19 Genomics UK (COG-UK) Consortium | Harper VanSteenhouse, Yumi Kasai, David Gray, Carol Clugston, Anna Dominiczak and Alex Alderton, Roberto Amato, Jeffrey Barrett, Sonia Goncalves, Ewan Harrison, David K. Jackson, Ian Johnston, Dominic Kwiatkowski, Cordelia Langford, John Sillitoe on behalf of the Wellcome Sanger Institute COVID-19 Surveillance Team |
| EPI_ISL_3225355, EPI_ISL_3225356, EPI_ISL_3225357 | Lighthouse Lab in Alderley Park | Wellcome Sanger Institute for the COVID-19 Genomics UK (COG-UK) Consortium | Jacquelyn Wynn, Mairead Hyland, The Lighthouse Lab in Alderley Park and Alex Alderton, Roberto Amato, Jeffrey Barrett, Sonia Goncalves, Ewan Harrison, David K. Jackson, Ian Johnston, Dominic Kwiatkowski, Cordelia Langford, John Sillitoe on behalf of the Wellcome Sanger Institute COVID-19 Surveillance Team |
| EPI_ISL_3225358, EPI_ISL_3225359 | Lighthouse Lab in Glasgow | Wellcome Sanger Institute for the COVID-19 Genomics UK (COG-UK) Consortium | Harper VanSteenhouse, Yumi Kasai, David Gray, Carol Clugston, Anna Dominiczak and Alex Alderton, Roberto Amato, Jeffrey Barrett, Sonia Goncalves, Ewan Harrison, David K. Jackson, Ian Johnston, Dominic Kwiatkowski, Cordelia Langford, John Sillitoe on behalf of the Wellcome Sanger Institute COVID-19 Surveillance Team |
| EPI_ISL_3225360, EPI_ISL_3225361 | Lighthouse Laboratory Plymouth | Wellcome Sanger Institute for the COVID-19 Genomics UK (COG-UK) Consortium | Lighthouse Laboratory Plymouth and Alex Alderton, Roberto Amato, Jeffrey Barrett, Sonia Goncalves, Ewan Harrison, David K. Jackson, Ian Johnston, Dominic Kwiatkowski, Cordelia Langford, John Sillitoe on behalf of the Wellcome Sanger Institute COVID-19 Surveillance Team |
| EPI_ISL_3225362 | Lighthouse Lab in Glasgow | Wellcome Sanger Institute for the COVID-19 Genomics UK (COG-UK) Consortium | Harper VanSteenhouse, Yumi Kasai, David Gray, Carol Clugston, Anna Dominiczak and Alex Alderton, Roberto Amato, Jeffrey Barrett, Sonia Goncalves, Ewan Harrison, David K. Jackson, Ian Johnston, Dominic Kwiatkowski, Cordelia Langford, John Sillitoe on behalf of the Wellcome Sanger Institute COVID-19 Surveillance Team |
| EPI_ISL_3225363 | Lighthouse Laboratory Plymouth | Wellcome Sanger Institute for the COVID-19 Genomics UK (COG-UK) Consortium | Lighthouse Laboratory Plymouth and Alex Alderton, Roberto Amato, Jeffrey Barrett, Sonia Goncalves, Ewan Harrison, David K. Jackson, Ian Johnston, Dominic Kwiatkowski, Cordelia Langford, John Sillitoe on behalf of the Wellcome Sanger Institute COVID-19 Surveillance Team |
| EPI_ISL_3225364, EPI_ISL_3225365 | Lighthouse Lab in Alderley Park | Wellcome Sanger Institute for the COVID-19 Genomics UK (COG-UK) Consortium | Jacquelyn Wynn, Mairead Hyland, The Lighthouse Lab in Alderley Park and Alex Alderton, Roberto Amato, Jeffrey Barrett, Sonia Goncalves, Ewan Harrison, David K. Jackson, Ian Johnston, Dominic Kwiatkowski, Cordelia Langford, John Sillitoe on behalf of the Wellcome Sanger Institute COVID-19 Surveillance Team |
| EPI_ISL_3225366 | Lighthouse Laboratory Plymouth | Wellcome Sanger Institute for the COVID-19 Genomics UK (COG-UK) Consortium | Lighthouse Laboratory Plymouth and Alex Alderton, Roberto Amato, Jeffrey Barrett, Sonia Goncalves, Ewan Harrison, David K. Jackson, Ian Johnston, Dominic Kwiatkowski, Cordelia Langford, John Sillitoe on behalf of the Wellcome Sanger Institute COVID-19 Surveillance Team |
| EPI_ISL_3225367, EPI_ISL_3225368, EPI_ISL_3225369 | Lighthouse Lab in Alderley Park | Wellcome Sanger Institute for the COVID-19 Genomics UK (COG-UK) Consortium | Jacquelyn Wynn, Mairead Hyland, The Lighthouse Lab in Alderley Park and Alex Alderton, Roberto Amato, Jeffrey Barrett, Sonia Goncalves, Ewan Harrison, David K. Jackson, Ian Johnston, Dominic Kwiatkowski, Cordelia Langford, John Sillitoe on behalf of the Wellcome Sanger Institute COVID-19 Surveillance Team |

| | | | |
|---|---|---|---|
| EPI_ISL_3225370 | Lighthouse Lab in Glasgow | Wellcome Sanger Institute for the COVID-19 Genomics UK (COG-UK) Consortium | Harper VanSteenhouse, Yumi Kasai, David Gray, Carol Clugston, Anna Dominiczak and Alex Alderton, Roberto Amato, Jeffrey Barrett, Sonia Goncalves, Ewan Harrison, David K. Jackson, Ian Johnston, Dominic Kwiatkowski, Cordelia Langford, John Sillitoe on behalf of the Wellcome Sanger Institute COVID-19 Surveillance Team |
| EPI_ISL_3225371, EPI_ISL_3225372 | Lighthouse Lab in Alderley Park | Wellcome Sanger Institute for the COVID-19 Genomics UK (COG-UK) Consortium | Jacquelyn Wynn, Mairead Hyland, The Lighthouse Lab in Alderley Park and Alex Alderton, Roberto Amato, Jeffrey Barrett, Sonia Goncalves, Ewan Harrison, David K. Jackson, Ian Johnston, Dominic Kwiatkowski, Cordelia Langford, John Sillitoe on behalf of the Wellcome Sanger Institute COVID-19 Surveillance Team |
| EPI_ISL_3225373, EPI_ISL_3225374 | Lighthouse Laboratory Plymouth | Wellcome Sanger Institute for the COVID-19 Genomics UK (COG-UK) Consortium | Lighthouse Laboratory Plymouth and Alex Alderton, Roberto Amato, Jeffrey Barrett, Sonia Goncalves, Ewan Harrison, David K. Jackson, Ian Johnston, Dominic Kwiatkowski, Cordelia Langford, John Sillitoe on behalf of the Wellcome Sanger Institute COVID-19 Surveillance Team |
| EPI_ISL_3225375 | Lighthouse Lab in Glasgow | Wellcome Sanger Institute for the COVID-19 Genomics UK (COG-UK) Consortium | Harper VanSteenhouse, Yumi Kasai, David Gray, Carol Clugston, Anna Dominiczak and Alex Alderton, Roberto Amato, Jeffrey Barrett, Sonia Goncalves, Ewan Harrison, David K. Jackson, Ian Johnston, Dominic Kwiatkowski, Cordelia Langford, John Sillitoe on behalf of the Wellcome Sanger Institute COVID-19 Surveillance Team |
| EPI_ISL_3225376 | Lighthouse Laboratory Plymouth | Wellcome Sanger Institute for the COVID-19 Genomics UK (COG-UK) Consortium | Lighthouse Laboratory Plymouth and Alex Alderton, Roberto Amato, Jeffrey Barrett, Sonia Goncalves, Ewan Harrison, David K. Jackson, Ian Johnston, Dominic Kwiatkowski, Cordelia Langford, John Sillitoe on behalf of the Wellcome Sanger Institute COVID-19 Surveillance Team |
| EPI_ISL_3225377, EPI_ISL_3225378 | Lighthouse Lab in Alderley Park | Wellcome Sanger Institute for the COVID-19 Genomics UK (COG-UK) Consortium | Jacquelyn Wynn, Mairead Hyland, The Lighthouse Lab in Alderley Park and Alex Alderton, Roberto Amato, Jeffrey Barrett, Sonia Goncalves, Ewan Harrison, David K. Jackson, Ian Johnston, Dominic Kwiatkowski, Cordelia Langford, John Sillitoe on behalf of the Wellcome Sanger Institute COVID-19 Surveillance Team |
| EPI_ISL_3225379 | Lighthouse Laboratory Plymouth | Wellcome Sanger Institute for the COVID-19 Genomics UK (COG-UK) Consortium | Lighthouse Laboratory Plymouth and Alex Alderton, Roberto Amato, Jeffrey Barrett, Sonia Goncalves, Ewan Harrison, David K. Jackson, Ian Johnston, Dominic Kwiatkowski, Cordelia Langford, John Sillitoe on behalf of the Wellcome Sanger Institute COVID-19 Surveillance Team |
| EPI_ISL_3225380 | Lighthouse Lab in Alderley Park | Wellcome Sanger Institute for the COVID-19 Genomics UK (COG-UK) Consortium | Jacquelyn Wynn, Mairead Hyland, The Lighthouse Lab in Alderley Park and Alex Alderton, Roberto Amato, Jeffrey Barrett, Sonia Goncalves, Ewan Harrison, David K. Jackson, Ian Johnston, Dominic Kwiatkowski, Cordelia Langford, John Sillitoe on behalf of the Wellcome Sanger Institute COVID-19 Surveillance Team |
| EPI_ISL_3225381, EPI_ISL_3225382 | Lighthouse Laboratory Plymouth | Wellcome Sanger Institute for the COVID-19 Genomics UK (COG-UK) Consortium | Lighthouse Laboratory Plymouth and Alex Alderton, Roberto Amato, Jeffrey Barrett, Sonia Goncalves, Ewan Harrison, David K. Jackson, Ian Johnston, Dominic Kwiatkowski, Cordelia Langford, John Sillitoe on behalf of the Wellcome Sanger Institute COVID-19 Surveillance Team |
| EPI_ISL_3225383 | Lighthouse Lab in Alderley Park | Wellcome Sanger Institute for the COVID-19 Genomics UK (COG-UK) Consortium | Jacquelyn Wynn, Mairead Hyland, The Lighthouse Lab in Alderley Park and Alex Alderton, Roberto Amato, Jeffrey Barrett, Sonia Goncalves, Ewan Harrison, David K. Jackson, Ian Johnston, Dominic Kwiatkowski, Cordelia Langford, John Sillitoe on behalf of the Wellcome Sanger Institute COVID-19 Surveillance Team |
| EPI_ISL_3225384 | Lighthouse Laboratory Plymouth | Wellcome Sanger Institute for the COVID-19 Genomics UK (COG-UK) Consortium | Lighthouse Laboratory Plymouth and Alex Alderton, Roberto Amato, Jeffrey Barrett, Sonia Goncalves, Ewan Harrison, David K. Jackson, Ian Johnston, Dominic Kwiatkowski, Cordelia Langford, John Sillitoe on behalf of the Wellcome Sanger Institute COVID-19 Surveillance Team |
| EPI_ISL_3225385 | Lighthouse Lab in Alderley Park | Wellcome Sanger Institute for the COVID-19 Genomics UK (COG-UK) Consortium | Jacquelyn Wynn, Mairead Hyland, The Lighthouse Lab in Alderley Park and Alex Alderton, Roberto Amato, Jeffrey Barrett, Sonia Goncalves, Ewan Harrison, David K. Jackson, Ian Johnston, Dominic Kwiatkowski, Cordelia Langford, John Sillitoe on behalf of the Wellcome Sanger Institute COVID-19 Surveillance Team |
| EPI_ISL_3225386 | Lighthouse Laboratory Plymouth | Wellcome Sanger Institute for the COVID-19 Genomics UK (COG-UK) Consortium | Lighthouse Laboratory Plymouth and Alex Alderton, Roberto Amato, Jeffrey Barrett, Sonia Goncalves, Ewan Harrison, David K. Jackson, Ian Johnston, Dominic Kwiatkowski, Cordelia Langford, John Sillitoe on behalf of the Wellcome Sanger Institute COVID-19 Surveillance Team |
| EPI_ISL_3225387 | Lighthouse Lab in Alderley Park | Wellcome Sanger Institute for the COVID-19 Genomics UK (COG-UK) Consortium | Jacquelyn Wynn, Mairead Hyland, The Lighthouse Lab in Alderley Park and Alex Alderton, Roberto Amato, Jeffrey Barrett, Sonia Goncalves, Ewan Harrison, David K. Jackson, Ian Johnston, Dominic Kwiatkowski, Cordelia Langford, John Sillitoe on behalf of the Wellcome Sanger Institute COVID-19 Surveillance Team |
| EPI_ISL_3225388 | Lighthouse Laboratory Plymouth | Wellcome Sanger Institute for the COVID-19 Genomics UK (COG-UK) Consortium | Lighthouse Laboratory Plymouth and Alex Alderton, Roberto Amato, Jeffrey Barrett, Sonia Goncalves, Ewan Harrison, David K. Jackson, Ian Johnston, Dominic Kwiatkowski, Cordelia Langford, John Sillitoe on behalf of the Wellcome Sanger Institute COVID-19 Surveillance Team |
| EPI_ISL_3225389 | Lighthouse Lab in Glasgow | Wellcome Sanger Institute for the COVID-19 Genomics UK (COG-UK) Consortium | Harper VanSteenhouse, Yumi Kasai, David Gray, Carol Clugston, Anna Dominiczak and Alex Alderton, Roberto Amato, Jeffrey Barrett, Sonia Goncalves, Ewan Harrison, David K. Jackson, Ian Johnston, Dominic Kwiatkowski, Cordelia Langford, John Sillitoe on behalf of the Wellcome Sanger Institute COVID-19 Surveillance Team |
| EPI_ISL_3225390, EPI_ISL_3225391, EPI_ISL_3225392 | Lighthouse Lab in Alderley Park | Wellcome Sanger Institute for the COVID-19 Genomics UK (COG-UK) Consortium | Jacquelyn Wynn, Mairead Hyland, The Lighthouse Lab in Alderley Park and Alex Alderton, Roberto Amato, Jeffrey Barrett, Sonia Goncalves, Ewan Harrison, David K. Jackson, Ian Johnston, Dominic Kwiatkowski, Cordelia Langford, John Sillitoe on behalf of the Wellcome Sanger Institute COVID-19 Surveillance Team |
| EPI_ISL_3225393 | Lighthouse Lab in Glasgow | Wellcome Sanger Institute for the COVID-19 Genomics UK (COG-UK) Consortium | Harper VanSteenhouse, Yumi Kasai, David Gray, Carol Clugston, Anna Dominiczak and Alex Alderton, Roberto Amato, Jeffrey Barrett, Sonia Goncalves, Ewan Harrison, David K. Jackson, Ian Johnston, Dominic Kwiatkowski, Cordelia Langford, John Sillitoe on behalf of the Wellcome Sanger Institute COVID-19 Surveillance Team |
| EPI_ISL_3225394, EPI_ISL_3225395 | Lighthouse Lab in Alderley Park | Wellcome Sanger Institute for the COVID-19 Genomics UK (COG-UK) Consortium | Jacquelyn Wynn, Mairead Hyland, The Lighthouse Lab in Alderley Park and Alex Alderton, Roberto Amato, Jeffrey Barrett, Sonia Goncalves, Ewan Harrison, David K. Jackson, Ian Johnston, Dominic Kwiatkowski, Cordelia Langford, John Sillitoe on behalf of the Wellcome Sanger Institute COVID-19 Surveillance Team |
| EPI_ISL_3225396 | Lighthouse Lab in Glasgow | Wellcome Sanger Institute for the COVID-19 Genomics UK (COG-UK) Consortium | Harper VanSteenhouse, Yumi Kasai, David Gray, Carol Clugston, Anna Dominiczak and Alex Alderton, Roberto Amato, Jeffrey Barrett, Sonia Goncalves, Ewan Harrison, David K. Jackson, Ian Johnston, Dominic Kwiatkowski, Cordelia Langford, John Sillitoe on behalf of the Wellcome Sanger Institute COVID-19 Surveillance Team |
| EPI_ISL_3225397 | Lighthouse Laboratory Plymouth | Wellcome Sanger Institute for the COVID-19 Genomics UK (COG-UK) Consortium | Lighthouse Laboratory Plymouth and Alex Alderton, Roberto Amato, Jeffrey Barrett, Sonia Goncalves, Ewan Harrison, David K. Jackson, Ian Johnston, Dominic Kwiatkowski, Cordelia Langford, John Sillitoe on behalf of the Wellcome Sanger Institute COVID-19 Surveillance Team |
| EPI_ISL_3225398, EPI_ISL_3225399, EPI_ISL_3225400 | Lighthouse Lab in Alderley Park | Wellcome Sanger Institute for the COVID-19 Genomics UK (COG-UK) Consortium | Jacquelyn Wynn, Mairead Hyland, The Lighthouse Lab in Alderley Park and Alex Alderton, Roberto Amato, Jeffrey Barrett, Sonia Goncalves, Ewan Harrison, David K. Jackson, Ian Johnston, Dominic Kwiatkowski, Cordelia Langford, John Sillitoe on behalf of the Wellcome Sanger Institute COVID-19 Surveillance Team |
| EPI_ISL_3225401 | Lighthouse Lab in Glasgow | Wellcome Sanger Institute for the COVID-19 Genomics UK (COG-UK) Consortium | Harper VanSteenhouse, Yumi Kasai, David Gray, Carol Clugston, Anna Dominiczak and Alex Alderton, Roberto Amato, Jeffrey Barrett, Sonia Goncalves, Ewan Harrison, David K. Jackson, Ian Johnston, Dominic Kwiatkowski, Cordelia Langford, John Sillitoe on behalf of the Wellcome Sanger Institute COVID-19 Surveillance Team |
| EPI_ISL_3225402 | Lighthouse Laboratory Plymouth | Wellcome Sanger Institute for the COVID-19 Genomics UK (COG-UK) Consortium | Lighthouse Laboratory Plymouth and Alex Alderton, Roberto Amato, Jeffrey Barrett, Sonia Goncalves, Ewan Harrison, David K. Jackson, Ian Johnston, Dominic Kwiatkowski, Cordelia Langford, John Sillitoe on behalf of the Wellcome Sanger Institute COVID-19 Surveillance Team |
| EPI_ISL_3225403 | Lighthouse Lab in Alderley Park | Wellcome Sanger Institute for the COVID-19 Genomics UK (COG-UK) Consortium | Jacquelyn Wynn, Mairead Hyland, The Lighthouse Lab in Alderley Park and Alex Alderton, Roberto Amato, Jeffrey Barrett, Sonia Goncalves, Ewan Harrison, David K. Jackson, Ian Johnston, Dominic Kwiatkowski, Cordelia Langford, John Sillitoe on behalf of the Wellcome Sanger Institute COVID-19 Surveillance Team |
| EPI_ISL_3225404, EPI_ISL_3225405 | Lighthouse Lab in Glasgow | Wellcome Sanger Institute for the COVID-19 Genomics UK (COG-UK) Consortium | Harper VanSteenhouse, Yumi Kasai, David Gray, Carol Clugston, Anna Dominiczak and Alex Alderton, Roberto Amato, Jeffrey Barrett, Sonia Goncalves, Ewan Harrison, David K. Jackson, Ian Johnston, Dominic Kwiatkowski, Cordelia Langford, John Sillitoe on behalf of the Wellcome Sanger Institute COVID-19 Surveillance Team |
| EPI_ISL_3225406 | Lighthouse Laboratory Plymouth | Wellcome Sanger Institute for the COVID-19 Genomics UK (COG-UK) Consortium | Lighthouse Laboratory Plymouth and Alex Alderton, Roberto Amato, Jeffrey Barrett, Sonia Goncalves, Ewan Harrison, David K. Jackson, Ian Johnston, Dominic Kwiatkowski, Cordelia Langford, John Sillitoe on behalf of the Wellcome Sanger Institute COVID-19 Surveillance Team |
| EPI_ISL_3225407 | Lighthouse Lab in Glasgow | Wellcome Sanger Institute for the COVID-19 Genomics UK (COG-UK) Consortium | Harper VanSteenhouse, Yumi Kasai, David Gray, Carol Clugston, Anna Dominiczak and Alex Alderton, Roberto Amato, Jeffrey Barrett, Sonia Goncalves, Ewan Harrison, David K. Jackson, Ian Johnston, Dominic Kwiatkowski, Cordelia Langford, John Sillitoe on behalf of the Wellcome Sanger Institute COVID-19 Surveillance Team |
| EPI_ISL_3225408, EPI_ISL_3225409, EPI_ISL_3225410 | Lighthouse Laboratory Plymouth | Wellcome Sanger Institute for the COVID-19 Genomics UK (COG-UK) Consortium | Lighthouse Laboratory Plymouth and Alex Alderton, Roberto Amato, Jeffrey Barrett, Sonia Goncalves, Ewan Harrison, David K. Jackson, Ian Johnston, Dominic Kwiatkowski, Cordelia Langford, John Sillitoe on behalf of the Wellcome Sanger Institute COVID-19 Surveillance Team |
| EPI_ISL_3225411, EPI_ISL_3225412 | Lighthouse Lab in Alderley Park | Wellcome Sanger Institute for the COVID-19 Genomics UK (COG-UK) Consortium | Jacquelyn Wynn, Mairead Hyland, The Lighthouse Lab in Alderley Park and Alex Alderton, Roberto Amato, Jeffrey Barrett, Sonia Goncalves, Ewan Harrison, David K. Jackson, Ian Johnston, Dominic Kwiatkowski, Cordelia Langford, John Sillitoe on behalf of the Wellcome Sanger Institute COVID-19 Surveillance Team |
| EPI_ISL_3225413 | Lighthouse Lab in Glasgow | Wellcome Sanger Institute for the COVID-19 Genomics UK (COG-UK) Consortium | Harper VanSteenhouse, Yumi Kasai, David Gray, Carol Clugston, Anna Dominiczak and Alex Alderton, Roberto Amato, Jeffrey Barrett, Sonia Goncalves, Ewan Harrison, David K. Jackson, Ian Johnston, Dominic Kwiatkowski, Cordelia Langford, John Sillitoe on behalf of the Wellcome Sanger Institute COVID-19 Surveillance Team |
| EPI_ISL_3225414, EPI_ISL_3225415, EPI_ISL_3225416 | Lighthouse Lab in Alderley Park | Wellcome Sanger Institute for the COVID-19 Genomics UK (COG-UK) Consortium | Jacquelyn Wynn, Mairead Hyland, The Lighthouse Lab in Alderley Park and Alex Alderton, Roberto Amato, Jeffrey Barrett, Sonia Goncalves, Ewan Harrison, David K. Jackson, Ian Johnston, Dominic Kwiatkowski, Cordelia Langford, John Sillitoe on behalf of the Wellcome Sanger Institute COVID-19 Surveillance Team |
| EPI_ISL_3225417 | Lighthouse Laboratory Plymouth | Wellcome Sanger Institute for the COVID-19 Genomics UK (COG-UK) Consortium | Lighthouse Laboratory Plymouth and Alex Alderton, Roberto Amato, Jeffrey Barrett, Sonia Goncalves, Ewan Harrison, David K. Jackson, Ian Johnston, Dominic Kwiatkowski, Cordelia Langford, John Sillitoe on behalf of the Wellcome Sanger Institute COVID-19 Surveillance Team |
| EPI_ISL_3225418 | Lighthouse Lab in Alderley Park | Wellcome Sanger Institute for the COVID-19 Genomics UK (COG-UK) Consortium | Jacquelyn Wynn, Mairead Hyland, The Lighthouse Lab in Alderley Park and Alex Alderton, Roberto Amato, Jeffrey Barrett, Sonia Goncalves, Ewan Harrison, David K. Jackson, Ian Johnston, Dominic Kwiatkowski, Cordelia Langford, John Sillitoe on behalf of the Wellcome Sanger Institute COVID-19 Surveillance Team |
| EPI_ISL_3225419, EPI_ISL_3225420 | Lighthouse Laboratory Plymouth | Wellcome Sanger Institute for the COVID-19 Genomics UK (COG-UK) Consortium | Lighthouse Laboratory Plymouth and Alex Alderton, Roberto Amato, Jeffrey Barrett, Sonia Goncalves, Ewan Harrison, David K. Jackson, Ian Johnston, Dominic Kwiatkowski, Cordelia Langford, John Sillitoe on behalf of the Wellcome Sanger Institute COVID-19 Surveillance Team |
| EPI_ISL_3225421, EPI_ISL_3225422, EPI_ISL_3225423 | Lighthouse Lab in Alderley Park | Wellcome Sanger Institute for the COVID-19 Genomics UK (COG-UK) Consortium | Jacquelyn Wynn, Mairead Hyland, The Lighthouse Lab in Alderley Park and Alex Alderton, Roberto Amato, Jeffrey Barrett, Sonia Goncalves, Ewan Harrison, David K. Jackson, Ian Johnston, Dominic Kwiatkowski, Cordelia Langford, John Sillitoe on behalf of the Wellcome Sanger Institute COVID-19 Surveillance Team |
| EPI_ISL_3225424 | Lighthouse Laboratory Plymouth | Wellcome Sanger Institute for the COVID-19 Genomics UK (COG-UK) Consortium | Lighthouse Laboratory Plymouth and Alex Alderton, Roberto Amato, Jeffrey Barrett, Sonia Goncalves, Ewan Harrison, David K. Jackson, Ian Johnston, Dominic Kwiatkowski, Cordelia Langford, John Sillitoe on behalf of the Wellcome Sanger Institute COVID-19 Surveillance Team |
| EPI_ISL_3225425, EPI_ISL_3225426 | Lighthouse Lab in Alderley Park | Wellcome Sanger Institute for the COVID-19 Genomics UK (COG-UK) Consortium | Jacquelyn Wynn, Mairead Hyland, The Lighthouse Lab in Alderley Park and Alex Alderton, Roberto Amato, Jeffrey Barrett, Sonia Goncalves, Ewan Harrison, David K. Jackson, Ian Johnston, Dominic Kwiatkowski, Cordelia Langford, John Sillitoe on behalf of the Wellcome Sanger Institute COVID-19 Surveillance Team |
| EPI_ISL_3225427, EPI_ISL_3225428 | Lighthouse Laboratory Plymouth | Wellcome Sanger Institute for the COVID-19 Genomics UK (COG-UK) Consortium | Lighthouse Laboratory Plymouth and Alex Alderton, Roberto Amato, Jeffrey Barrett, Sonia Goncalves, Ewan Harrison, David K. Jackson, Ian Johnston, Dominic Kwiatkowski, Cordelia Langford, John Sillitoe on behalf of the Wellcome Sanger Institute COVID-19 Surveillance Team |
| EPI_ISL_3225429, EPI_ISL_3225430, EPI_ISL_3225431 | Lighthouse Lab in Alderley Park | Wellcome Sanger Institute for the COVID-19 Genomics UK (COG-UK) Consortium | Jacquelyn Wynn, Mairead Hyland, The Lighthouse Lab in Alderley Park and Alex Alderton, Roberto Amato, Jeffrey Barrett, Sonia Goncalves, Ewan Harrison, David K. Jackson, Ian Johnston, Dominic Kwiatkowski, Cordelia Langford, John Sillitoe on behalf of the Wellcome Sanger Institute COVID-19 Surveillance Team |
| EPI_ISL_3225432 | Lighthouse Laboratory Plymouth | Wellcome Sanger Institute for the COVID-19 Genomics UK (COG-UK) Consortium | Lighthouse Laboratory Plymouth and Alex Alderton, Roberto Amato, Jeffrey Barrett, Sonia Goncalves, Ewan Harrison, David K. Jackson, Ian Johnston, Dominic Kwiatkowski, Cordelia Langford, John Sillitoe on behalf of the Wellcome Sanger Institute COVID-19 Surveillance Team |
| EPI_ISL_3225433 | Lighthouse Lab in Glasgow | Wellcome Sanger Institute for the COVID-19 Genomics UK (COG-UK) Consortium | Harper VanSteenhouse, Yumi Kasai, David Gray, Carol Clugston, Anna Dominiczak and Alex Alderton, Roberto Amato, Jeffrey Barrett, Sonia Goncalves, Ewan Harrison, David K. Jackson, Ian Johnston, Dominic Kwiatkowski, Cordelia Langford, John Sillitoe on behalf of the Wellcome Sanger Institute COVID-19 Surveillance Team |
| EPI_ISL_3225434 | Lighthouse Lab in Alderley Park | Wellcome Sanger Institute for the COVID-19 Genomics UK (COG-UK) Consortium | Jacquelyn Wynn, Mairead Hyland, The Lighthouse Lab in Alderley Park and Alex Alderton, Roberto Amato, Jeffrey Barrett, Sonia Goncalves, Ewan Harrison, David K. Jackson, Ian Johnston, Dominic Kwiatkowski, Cordelia Langford, John Sillitoe on behalf of the Wellcome Sanger Institute COVID-19 Surveillance Team |

| | | | |
|---|---|---|---|
| EPI_ISL_3225435, EPI_ISL_3225436 | Lighthouse Lab in Glasgow | Wellcome Sanger Institute for the COVID-19 Genomics UK (COG-UK) Consortium | Harper VanSteenhouse, Yumi Kasai, David Gray, Carol Clugston, Anna Dominiczak and Alex Alderton, Roberto Amato, Jeffrey Barrett, Sonia Goncalves, Ewan Harrison, David K. Jackson, Ian Johnston, Dominic Kwiatkowski, Cordelia Langford, John Sillitoe on behalf of the Wellcome Sanger Institute COVID-19 Surveillance Team |
| EPI_ISL_3225437 | Lighthouse Lab in Alderley Park | Wellcome Sanger Institute for the COVID-19 Genomics UK (COG-UK) Consortium | Jacquelyn Wynn, Mairead Hyland, The Lighthouse Lab in Alderley Park and Alex Alderton, Roberto Amato, Jeffrey Barrett, Sonia Goncalves, Ewan Harrison, David K. Jackson, Ian Johnston, Dominic Kwiatkowski, Cordelia Langford, John Sillitoe on behalf of the Wellcome Sanger Institute COVID-19 Surveillance Team |
| EPI_ISL_3225438 | Lighthouse Laboratory Plymouth | Wellcome Sanger Institute for the COVID-19 Genomics UK (COG-UK) Consortium | Lighthouse Laboratory Plymouth and Alex Alderton, Roberto Amato, Jeffrey Barrett, Sonia Goncalves, Ewan Harrison, David K. Jackson, Ian Johnston, Dominic Kwiatkowski, Cordelia Langford, John Sillitoe on behalf of the Wellcome Sanger Institute COVID-19 Surveillance Team |
| EPI_ISL_3225439, EPI_ISL_3225440 | Lighthouse Lab in Glasgow | Wellcome Sanger Institute for the COVID-19 Genomics UK (COG-UK) Consortium | Harper VanSteenhouse, Yumi Kasai, David Gray, Carol Clugston, Anna Dominiczak and Alex Alderton, Roberto Amato, Jeffrey Barrett, Sonia Goncalves, Ewan Harrison, David K. Jackson, Ian Johnston, Dominic Kwiatkowski, Cordelia Langford, John Sillitoe on behalf of the Wellcome Sanger Institute COVID-19 Surveillance Team |
| EPI_ISL_3225441, EPI_ISL_3225442, EPI_ISL_3225443 | Lighthouse Lab in Alderley Park | Wellcome Sanger Institute for the COVID-19 Genomics UK (COG-UK) Consortium | Jacquelyn Wynn, Mairead Hyland, The Lighthouse Lab in Alderley Park and Alex Alderton, Roberto Amato, Jeffrey Barrett, Sonia Goncalves, Ewan Harrison, David K. Jackson, Ian Johnston, Dominic Kwiatkowski, Cordelia Langford, John Sillitoe on behalf of the Wellcome Sanger Institute COVID-19 Surveillance Team |
| EPI_ISL_3225444, EPI_ISL_3225445 | Lighthouse Lab in Glasgow | Wellcome Sanger Institute for the COVID-19 Genomics UK (COG-UK) Consortium | Harper VanSteenhouse, Yumi Kasai, David Gray, Carol Clugston, Anna Dominiczak and Alex Alderton, Roberto Amato, Jeffrey Barrett, Sonia Goncalves, Ewan Harrison, David K. Jackson, Ian Johnston, Dominic Kwiatkowski, Cordelia Langford, John Sillitoe on behalf of the Wellcome Sanger Institute COVID-19 Surveillance Team |
| EPI_ISL_3225446 | Lighthouse Laboratory Plymouth | Wellcome Sanger Institute for the COVID-19 Genomics UK (COG-UK) Consortium | Lighthouse Laboratory Plymouth and Alex Alderton, Roberto Amato, Jeffrey Barrett, Sonia Goncalves, Ewan Harrison, David K. Jackson, Ian Johnston, Dominic Kwiatkowski, Cordelia Langford, John Sillitoe on behalf of the Wellcome Sanger Institute COVID-19 Surveillance Team |
| EPI_ISL_3225447, EPI_ISL_3225448 | Lighthouse Lab in Alderley Park | Wellcome Sanger Institute for the COVID-19 Genomics UK (COG-UK) Consortium | Jacquelyn Wynn, Mairead Hyland, The Lighthouse Lab in Alderley Park and Alex Alderton, Roberto Amato, Jeffrey Barrett, Sonia Goncalves, Ewan Harrison, David K. Jackson, Ian Johnston, Dominic Kwiatkowski, Cordelia Langford, John Sillitoe on behalf of the Wellcome Sanger Institute COVID-19 Surveillance Team |
| EPI_ISL_3225449, EPI_ISL_3225450 | Lighthouse Lab in Glasgow | Wellcome Sanger Institute for the COVID-19 Genomics UK (COG-UK) Consortium | Harper VanSteenhouse, Yumi Kasai, David Gray, Carol Clugston, Anna Dominiczak and Alex Alderton, Roberto Amato, Jeffrey Barrett, Sonia Goncalves, Ewan Harrison, David K. Jackson, Ian Johnston, Dominic Kwiatkowski, Cordelia Langford, John Sillitoe on behalf of the Wellcome Sanger Institute COVID-19 Surveillance Team |
| EPI_ISL_3225451 | Lighthouse Lab in Alderley Park | Wellcome Sanger Institute for the COVID-19 Genomics UK (COG-UK) Consortium | Jacquelyn Wynn, Mairead Hyland, The Lighthouse Lab in Alderley Park and Alex Alderton, Roberto Amato, Jeffrey Barrett, Sonia Goncalves, Ewan Harrison, David K. Jackson, Ian Johnston, Dominic Kwiatkowski, Cordelia Langford, John Sillitoe on behalf of the Wellcome Sanger Institute COVID-19 Surveillance Team |
| EPI_ISL_3225452 | Lighthouse Lab in Glasgow | Wellcome Sanger Institute for the COVID-19 Genomics UK (COG-UK) Consortium | Harper VanSteenhouse, Yumi Kasai, David Gray, Carol Clugston, Anna Dominiczak and Alex Alderton, Roberto Amato, Jeffrey Barrett, Sonia Goncalves, Ewan Harrison, David K. Jackson, Ian Johnston, Dominic Kwiatkowski, Cordelia Langford, John Sillitoe on behalf of the Wellcome Sanger Institute COVID-19 Surveillance Team |
| EPI_ISL_3225453 | Lighthouse Laboratory Plymouth | Wellcome Sanger Institute for the COVID-19 Genomics UK (COG-UK) Consortium | Lighthouse Laboratory Plymouth and Alex Alderton, Roberto Amato, Jeffrey Barrett, Sonia Goncalves, Ewan Harrison, David K. Jackson, Ian Johnston, Dominic Kwiatkowski, Cordelia Langford, John Sillitoe on behalf of the Wellcome Sanger Institute COVID-19 Surveillance Team |
| EPI_ISL_3225454, EPI_ISL_3225455, EPI_ISL_3225456, EPI_ISL_3225457, EPI_ISL_3225458 | Lighthouse Lab in Alderley Park | Wellcome Sanger Institute for the COVID-19 Genomics UK (COG-UK) Consortium | Jacquelyn Wynn, Mairead Hyland, The Lighthouse Lab in Alderley Park and Alex Alderton, Roberto Amato, Jeffrey Barrett, Sonia Goncalves, Ewan Harrison, David K. Jackson, Ian Johnston, Dominic Kwiatkowski, Cordelia Langford, John Sillitoe on behalf of the Wellcome Sanger Institute COVID-19 Surveillance Team |
| EPI_ISL_3225459 | Lighthouse Laboratory Plymouth | Wellcome Sanger Institute for the COVID-19 Genomics UK (COG-UK) Consortium | Lighthouse Laboratory Plymouth and Alex Alderton, Roberto Amato, Jeffrey Barrett, Sonia Goncalves, Ewan Harrison, David K. Jackson, Ian Johnston, Dominic Kwiatkowski, Cordelia Langford, John Sillitoe on behalf of the Wellcome Sanger Institute COVID-19 Surveillance Team |
| EPI_ISL_3225460, EPI_ISL_3225461 | Lighthouse Lab in Alderley Park | Wellcome Sanger Institute for the COVID-19 Genomics UK (COG-UK) Consortium | Jacquelyn Wynn, Mairead Hyland, The Lighthouse Lab in Alderley Park and Alex Alderton, Roberto Amato, Jeffrey Barrett, Sonia Goncalves, Ewan Harrison, David K. Jackson, Ian Johnston, Dominic Kwiatkowski, Cordelia Langford, John Sillitoe on behalf of the Wellcome Sanger Institute COVID-19 Surveillance Team |
| EPI_ISL_3225462 | Lighthouse Laboratory Plymouth | Wellcome Sanger Institute for the COVID-19 Genomics UK (COG-UK) Consortium | Lighthouse Laboratory Plymouth and Alex Alderton, Roberto Amato, Jeffrey Barrett, Sonia Goncalves, Ewan Harrison, David K. Jackson, Ian Johnston, Dominic Kwiatkowski, Cordelia Langford, John Sillitoe on behalf of the Wellcome Sanger Institute COVID-19 Surveillance Team |
| EPI_ISL_3225463 | Lighthouse Lab in Alderley Park | Wellcome Sanger Institute for the COVID-19 Genomics UK (COG-UK) Consortium | Jacquelyn Wynn, Mairead Hyland, The Lighthouse Lab in Alderley Park and Alex Alderton, Roberto Amato, Jeffrey Barrett, Sonia Goncalves, Ewan Harrison, David K. Jackson, Ian Johnston, Dominic Kwiatkowski, Cordelia Langford, John Sillitoe on behalf of the Wellcome Sanger Institute COVID-19 Surveillance Team |
| EPI_ISL_3225464 | Lighthouse Lab in Glasgow | Wellcome Sanger Institute for the COVID-19 Genomics UK (COG-UK) Consortium | Harper VanSteenhouse, Yumi Kasai, David Gray, Carol Clugston, Anna Dominiczak and Alex Alderton, Roberto Amato, Jeffrey Barrett, Sonia Goncalves, Ewan Harrison, David K. Jackson, Ian Johnston, Dominic Kwiatkowski, Cordelia Langford, John Sillitoe on behalf of the Wellcome Sanger Institute COVID-19 Surveillance Team |
| EPI_ISL_3225465, EPI_ISL_3225466 | Lighthouse Laboratory Plymouth | Wellcome Sanger Institute for the COVID-19 Genomics UK (COG-UK) Consortium | Lighthouse Laboratory Plymouth and Alex Alderton, Roberto Amato, Jeffrey Barrett, Sonia Goncalves, Ewan Harrison, David K. Jackson, Ian Johnston, Dominic Kwiatkowski, Cordelia Langford, John Sillitoe on behalf of the Wellcome Sanger Institute COVID-19 Surveillance Team |
| EPI_ISL_3225467 | Lighthouse Lab in Glasgow | Wellcome Sanger Institute for the COVID-19 Genomics UK (COG-UK) Consortium | Harper VanSteenhouse, Yumi Kasai, David Gray, Carol Clugston, Anna Dominiczak and Alex Alderton, Roberto Amato, Jeffrey Barrett, Sonia Goncalves, Ewan Harrison, David K. Jackson, Ian Johnston, Dominic Kwiatkowski, Cordelia Langford, John Sillitoe on behalf of the Wellcome Sanger Institute COVID-19 Surveillance Team |
| EPI_ISL_3225468 | Lighthouse Lab in Alderley Park | Wellcome Sanger Institute for the COVID-19 Genomics UK (COG-UK) Consortium | Jacquelyn Wynn, Mairead Hyland, The Lighthouse Lab in Alderley Park and Alex Alderton, Roberto Amato, Jeffrey Barrett, Sonia Goncalves, Ewan Harrison, David K. Jackson, Ian Johnston, Dominic Kwiatkowski, Cordelia Langford, John Sillitoe on behalf of the Wellcome Sanger Institute COVID-19 Surveillance Team |
| EPI_ISL_3225469 | Lighthouse Lab in Glasgow | Wellcome Sanger Institute for the COVID-19 Genomics UK (COG-UK) Consortium | Harper VanSteenhouse, Yumi Kasai, David Gray, Carol Clugston, Anna Dominiczak and Alex Alderton, Roberto Amato, Jeffrey Barrett, Sonia Goncalves, Ewan Harrison, David K. Jackson, Ian Johnston, Dominic Kwiatkowski, Cordelia Langford, John Sillitoe on behalf of the Wellcome Sanger Institute COVID-19 Surveillance Team |
| EPI_ISL_3225470 | Lighthouse Lab in Alderley Park | Wellcome Sanger Institute for the COVID-19 Genomics UK (COG-UK) Consortium | Jacquelyn Wynn, Mairead Hyland, The Lighthouse Lab in Alderley Park and Alex Alderton, Roberto Amato, Jeffrey Barrett, Sonia Goncalves, Ewan Harrison, David K. Jackson, Ian Johnston, Dominic Kwiatkowski, Cordelia Langford, John Sillitoe on behalf of the Wellcome Sanger Institute COVID-19 Surveillance Team |
| EPI_ISL_3225471 | Lighthouse Lab in Glasgow | Wellcome Sanger Institute for the COVID-19 Genomics UK (COG-UK) Consortium | Harper VanSteenhouse, Yumi Kasai, David Gray, Carol Clugston, Anna Dominiczak and Alex Alderton, Roberto Amato, Jeffrey Barrett, Sonia Goncalves, Ewan Harrison, David K. Jackson, Ian Johnston, Dominic Kwiatkowski, Cordelia Langford, John Sillitoe on behalf of the Wellcome Sanger Institute COVID-19 Surveillance Team |
| EPI_ISL_3225472 | Lighthouse Laboratory Plymouth | Wellcome Sanger Institute for the COVID-19 Genomics UK (COG-UK) Consortium | Lighthouse Laboratory Plymouth and Alex Alderton, Roberto Amato, Jeffrey Barrett, Sonia Goncalves, Ewan Harrison, David K. Jackson, Ian Johnston, Dominic Kwiatkowski, Cordelia Langford, John Sillitoe on behalf of the Wellcome Sanger Institute COVID-19 Surveillance Team |
| EPI_ISL_3225473, EPI_ISL_3225474 | Lighthouse Lab in Glasgow | Wellcome Sanger Institute for the COVID-19 Genomics UK (COG-UK) Consortium | Harper VanSteenhouse, Yumi Kasai, David Gray, Carol Clugston, Anna Dominiczak and Alex Alderton, Roberto Amato, Jeffrey Barrett, Sonia Goncalves, Ewan Harrison, David K. Jackson, Ian Johnston, Dominic Kwiatkowski, Cordelia Langford, John Sillitoe on behalf of the Wellcome Sanger Institute COVID-19 Surveillance Team |
| EPI_ISL_3225475 | Lighthouse Lab in Alderley Park | Wellcome Sanger Institute for the COVID-19 Genomics UK (COG-UK) Consortium | Jacquelyn Wynn, Mairead Hyland, The Lighthouse Lab in Alderley Park and Alex Alderton, Roberto Amato, Jeffrey Barrett, Sonia Goncalves, Ewan Harrison, David K. Jackson, Ian Johnston, Dominic Kwiatkowski, Cordelia Langford, John Sillitoe on behalf of the Wellcome Sanger Institute COVID-19 Surveillance Team |
| EPI_ISL_3225476 | Lighthouse Laboratory Plymouth | Wellcome Sanger Institute for the COVID-19 Genomics UK (COG-UK) Consortium | Lighthouse Laboratory Plymouth and Alex Alderton, Roberto Amato, Jeffrey Barrett, Sonia Goncalves, Ewan Harrison, David K. Jackson, Ian Johnston, Dominic Kwiatkowski, Cordelia Langford, John Sillitoe on behalf of the Wellcome Sanger Institute COVID-19 Surveillance Team |
| EPI_ISL_3225477, EPI_ISL_3225478 | Lighthouse Lab in Alderley Park | Wellcome Sanger Institute for the COVID-19 Genomics UK (COG-UK) Consortium | Jacquelyn Wynn, Mairead Hyland, The Lighthouse Lab in Alderley Park and Alex Alderton, Roberto Amato, Jeffrey Barrett, Sonia Goncalves, Ewan Harrison, David K. Jackson, Ian Johnston, Dominic Kwiatkowski, Cordelia Langford, John Sillitoe on behalf of the Wellcome Sanger Institute COVID-19 Surveillance Team |
| EPI_ISL_3225479 | Lighthouse Laboratory Plymouth | Wellcome Sanger Institute for the COVID-19 Genomics UK (COG-UK) Consortium | Lighthouse Laboratory Plymouth and Alex Alderton, Roberto Amato, Jeffrey Barrett, Sonia Goncalves, Ewan Harrison, David K. Jackson, Ian Johnston, Dominic Kwiatkowski, Cordelia Langford, John Sillitoe on behalf of the Wellcome Sanger Institute COVID-19 Surveillance Team |
| EPI_ISL_3225480, EPI_ISL_3225481 | Lighthouse Lab in Alderley Park | Wellcome Sanger Institute for the COVID-19 Genomics UK (COG-UK) Consortium | Jacquelyn Wynn, Mairead Hyland, The Lighthouse Lab in Alderley Park and Alex Alderton, Roberto Amato, Jeffrey Barrett, Sonia Goncalves, Ewan Harrison, David K. Jackson, Ian Johnston, Dominic Kwiatkowski, Cordelia Langford, John Sillitoe on behalf of the Wellcome Sanger Institute COVID-19 Surveillance Team |
| EPI_ISL_3225482 | Lighthouse Lab in Glasgow | Wellcome Sanger Institute for the COVID-19 Genomics UK (COG-UK) Consortium | Harper VanSteenhouse, Yumi Kasai, David Gray, Carol Clugston, Anna Dominiczak and Alex Alderton, Roberto Amato, Jeffrey Barrett, Sonia Goncalves, Ewan Harrison, David K. Jackson, Ian Johnston, Dominic Kwiatkowski, Cordelia Langford, John Sillitoe on behalf of the Wellcome Sanger Institute COVID-19 Surveillance Team |
| EPI_ISL_3225483, EPI_ISL_3225484, EPI_ISL_3225485, EPI_ISL_3225486 | Lighthouse Lab in Alderley Park | Wellcome Sanger Institute for the COVID-19 Genomics UK (COG-UK) Consortium | Jacquelyn Wynn, Mairead Hyland, The Lighthouse Lab in Alderley Park and Alex Alderton, Roberto Amato, Jeffrey Barrett, Sonia Goncalves, Ewan Harrison, David K. Jackson, Ian Johnston, Dominic Kwiatkowski, Cordelia Langford, John Sillitoe on behalf of the Wellcome Sanger Institute COVID-19 Surveillance Team |
| EPI_ISL_3225487, EPI_ISL_3225488 | Lighthouse Lab in Glasgow | Wellcome Sanger Institute for the COVID-19 Genomics UK (COG-UK) Consortium | Harper VanSteenhouse, Yumi Kasai, David Gray, Carol Clugston, Anna Dominiczak and Alex Alderton, Roberto Amato, Jeffrey Barrett, Sonia Goncalves, Ewan Harrison, David K. Jackson, Ian Johnston, Dominic Kwiatkowski, Cordelia Langford, John Sillitoe on behalf of the Wellcome Sanger Institute COVID-19 Surveillance Team |
| EPI_ISL_3225489, EPI_ISL_3225490, EPI_ISL_3225491 | Lighthouse Lab in Alderley Park | Wellcome Sanger Institute for the COVID-19 Genomics UK (COG-UK) Consortium | Jacquelyn Wynn, Mairead Hyland, The Lighthouse Lab in Alderley Park and Alex Alderton, Roberto Amato, Jeffrey Barrett, Sonia Goncalves, Ewan Harrison, David K. Jackson, Ian Johnston, Dominic Kwiatkowski, Cordelia Langford, John Sillitoe on behalf of the Wellcome Sanger Institute COVID-19 Surveillance Team |
| EPI_ISL_3225492, EPI_ISL_3225493 | Lighthouse Laboratory Plymouth | Wellcome Sanger Institute for the COVID-19 Genomics UK (COG-UK) Consortium | Lighthouse Laboratory Plymouth and Alex Alderton, Roberto Amato, Jeffrey Barrett, Sonia Goncalves, Ewan Harrison, David K. Jackson, Ian Johnston, Dominic Kwiatkowski, Cordelia Langford, John Sillitoe on behalf of the Wellcome Sanger Institute COVID-19 Surveillance Team |
| EPI_ISL_3225494, EPI_ISL_3225495, EPI_ISL_3225496 | Lighthouse Lab in Alderley Park | Wellcome Sanger Institute for the COVID-19 Genomics UK (COG-UK) Consortium | Jacquelyn Wynn, Mairead Hyland, The Lighthouse Lab in Alderley Park and Alex Alderton, Roberto Amato, Jeffrey Barrett, Sonia Goncalves, Ewan Harrison, David K. Jackson, Ian Johnston, Dominic Kwiatkowski, Cordelia Langford, John Sillitoe on behalf of the Wellcome Sanger Institute COVID-19 Surveillance Team |
| EPI_ISL_3225497 | Lighthouse Laboratory Plymouth | Wellcome Sanger Institute for the COVID-19 Genomics UK (COG-UK) Consortium | Lighthouse Laboratory Plymouth and Alex Alderton, Roberto Amato, Jeffrey Barrett, Sonia Goncalves, Ewan Harrison, David K. Jackson, Ian Johnston, Dominic Kwiatkowski, Cordelia Langford, John Sillitoe on behalf of the Wellcome Sanger Institute COVID-19 Surveillance Team |
| EPI_ISL_3225498, EPI_ISL_3225499, EPI_ISL_3225500 | Lighthouse Lab in Glasgow | Wellcome Sanger Institute for the COVID-19 Genomics UK (COG-UK) Consortium | Harper VanSteenhouse, Yumi Kasai, David Gray, Carol Clugston, Anna Dominiczak and Alex Alderton, Roberto Amato, Jeffrey Barrett, Sonia Goncalves, Ewan Harrison, David K. Jackson, Ian Johnston, Dominic Kwiatkowski, Cordelia Langford, John Sillitoe on behalf of the Wellcome Sanger Institute COVID-19 Surveillance Team |
| EPI_ISL_3225501, EPI_ISL_3225502 | Lighthouse Lab in Alderley Park | Wellcome Sanger Institute for the COVID-19 Genomics UK (COG-UK) Consortium | Jacquelyn Wynn, Mairead Hyland, The Lighthouse Lab in Alderley Park and Alex Alderton, Roberto Amato, Jeffrey Barrett, Sonia Goncalves, Ewan Harrison, David K. Jackson, Ian Johnston, Dominic Kwiatkowski, Cordelia Langford, John Sillitoe on behalf of the Wellcome Sanger Institute COVID-19 Surveillance Team |
| EPI_ISL_3225503 | Lighthouse Lab in Glasgow | Wellcome Sanger Institute for the COVID-19 Genomics UK (COG-UK) Consortium | Harper VanSteenhouse, Yumi Kasai, David Gray, Carol Clugston, Anna Dominiczak and Alex Alderton, Roberto Amato, Jeffrey Barrett, Sonia Goncalves, Ewan Harrison, David K. Jackson, Ian Johnston, Dominic Kwiatkowski, Cordelia Langford, John Sillitoe on behalf of the Wellcome Sanger Institute COVID-19 Surveillance Team |

| | | | |
|---|---|---|---|
| EPI_ISL_3225504 | Lighthouse Lab in Alderley Park | Wellcome Sanger Institute for the COVID-19 Genomics UK (COG-UK) Consortium | Jacquelyn Wynn, Mairead Hyland, The Lighthouse Lab in Alderley Park and Alex Alderton, Roberto Amato, Jeffrey Barrett, Sonia Goncalves, Ewan Harrison, David K. Jackson, Ian Johnston, Dominic Kwiatkowski, Cordelia Langford, John Sillitoe on behalf of the Wellcome Sanger Institute COVID-19 Surveillance Team |
| EPI_ISL_3225505 | Lighthouse Lab in Glasgow | Wellcome Sanger Institute for the COVID-19 Genomics UK (COG-UK) Consortium | Harper VanSteenhouse, Yumi Kasai, David Gray, Carol Clugston, Anna Dominiczak and Alex Alderton, Roberto Amato, Jeffrey Barrett, Sonia Goncalves, Ewan Harrison, David K. Jackson, Ian Johnston, Dominic Kwiatkowski, Cordelia Langford, John Sillitoe on behalf of the Wellcome Sanger Institute COVID-19 Surveillance Team |
| EPI_ISL_3225506 | Lighthouse Laboratory Plymouth | Wellcome Sanger Institute for the COVID-19 Genomics UK (COG-UK) Consortium | Lighthouse Laboratory Plymouth and Alex Alderton, Roberto Amato, Jeffrey Barrett, Sonia Goncalves, Ewan Harrison, David K. Jackson, Ian Johnston, Dominic Kwiatkowski, Cordelia Langford, John Sillitoe on behalf of the Wellcome Sanger Institute COVID-19 Surveillance Team |
| EPI_ISL_3225507 | Lighthouse Lab in Alderley Park | Wellcome Sanger Institute for the COVID-19 Genomics UK (COG-UK) Consortium | Jacquelyn Wynn, Mairead Hyland, The Lighthouse Lab in Alderley Park and Alex Alderton, Roberto Amato, Jeffrey Barrett, Sonia Goncalves, Ewan Harrison, David K. Jackson, Ian Johnston, Dominic Kwiatkowski, Cordelia Langford, John Sillitoe on behalf of the Wellcome Sanger Institute COVID-19 Surveillance Team |
| EPI_ISL_3225508 | Lighthouse Lab in Glasgow | Wellcome Sanger Institute for the COVID-19 Genomics UK (COG-UK) Consortium | Harper VanSteenhouse, Yumi Kasai, David Gray, Carol Clugston, Anna Dominiczak and Alex Alderton, Roberto Amato, Jeffrey Barrett, Sonia Goncalves, Ewan Harrison, David K. Jackson, Ian Johnston, Dominic Kwiatkowski, Cordelia Langford, John Sillitoe on behalf of the Wellcome Sanger Institute COVID-19 Surveillance Team |
| EPI_ISL_3225509 | Lighthouse Lab in Alderley Park | Wellcome Sanger Institute for the COVID-19 Genomics UK (COG-UK) Consortium | Jacquelyn Wynn, Mairead Hyland, The Lighthouse Lab in Alderley Park and Alex Alderton, Roberto Amato, Jeffrey Barrett, Sonia Goncalves, Ewan Harrison, David K. Jackson, Ian Johnston, Dominic Kwiatkowski, Cordelia Langford, John Sillitoe on behalf of the Wellcome Sanger Institute COVID-19 Surveillance Team |
| EPI_ISL_3225510 | Lighthouse Laboratory Plymouth | Wellcome Sanger Institute for the COVID-19 Genomics UK (COG-UK) Consortium | Lighthouse Laboratory Plymouth and Alex Alderton, Roberto Amato, Jeffrey Barrett, Sonia Goncalves, Ewan Harrison, David K. Jackson, Ian Johnston, Dominic Kwiatkowski, Cordelia Langford, John Sillitoe on behalf of the Wellcome Sanger Institute COVID-19 Surveillance Team |
| EPI_ISL_3225511, EPI_ISL_3225512, EPI_ISL_3225513 | Lighthouse Lab in Alderley Park | Wellcome Sanger Institute for the COVID-19 Genomics UK (COG-UK) Consortium | Jacquelyn Wynn, Mairead Hyland, The Lighthouse Lab in Alderley Park and Alex Alderton, Roberto Amato, Jeffrey Barrett, Sonia Goncalves, Ewan Harrison, David K. Jackson, Ian Johnston, Dominic Kwiatkowski, Cordelia Langford, John Sillitoe on behalf of the Wellcome Sanger Institute COVID-19 Surveillance Team |
| EPI_ISL_3225514 | Lighthouse Laboratory Plymouth | Wellcome Sanger Institute for the COVID-19 Genomics UK (COG-UK) Consortium | Lighthouse Laboratory Plymouth and Alex Alderton, Roberto Amato, Jeffrey Barrett, Sonia Goncalves, Ewan Harrison, David K. Jackson, Ian Johnston, Dominic Kwiatkowski, Cordelia Langford, John Sillitoe on behalf of the Wellcome Sanger Institute COVID-19 Surveillance Team |
| EPI_ISL_3225515, EPI_ISL_3225516 | Lighthouse Lab in Glasgow | Wellcome Sanger Institute for the COVID-19 Genomics UK (COG-UK) Consortium | Harper VanSteenhouse, Yumi Kasai, David Gray, Carol Clugston, Anna Dominiczak and Alex Alderton, Roberto Amato, Jeffrey Barrett, Sonia Goncalves, Ewan Harrison, David K. Jackson, Ian Johnston, Dominic Kwiatkowski, Cordelia Langford, John Sillitoe on behalf of the Wellcome Sanger Institute COVID-19 Surveillance Team |
| EPI_ISL_3225517 | Lighthouse Lab in Alderley Park | Wellcome Sanger Institute for the COVID-19 Genomics UK (COG-UK) Consortium | Jacquelyn Wynn, Mairead Hyland, The Lighthouse Lab in Alderley Park and Alex Alderton, Roberto Amato, Jeffrey Barrett, Sonia Goncalves, Ewan Harrison, David K. Jackson, Ian Johnston, Dominic Kwiatkowski, Cordelia Langford, John Sillitoe on behalf of the Wellcome Sanger Institute COVID-19 Surveillance Team |
| EPI_ISL_3225518 | Lighthouse Lab in Glasgow | Wellcome Sanger Institute for the COVID-19 Genomics UK (COG-UK) Consortium | Harper VanSteenhouse, Yumi Kasai, David Gray, Carol Clugston, Anna Dominiczak and Alex Alderton, Roberto Amato, Jeffrey Barrett, Sonia Goncalves, Ewan Harrison, David K. Jackson, Ian Johnston, Dominic Kwiatkowski, Cordelia Langford, John Sillitoe on behalf of the Wellcome Sanger Institute COVID-19 Surveillance Team |
| EPI_ISL_3225519 | Lighthouse Laboratory Plymouth | Wellcome Sanger Institute for the COVID-19 Genomics UK (COG-UK) Consortium | Lighthouse Laboratory Plymouth and Alex Alderton, Roberto Amato, Jeffrey Barrett, Sonia Goncalves, Ewan Harrison, David K. Jackson, Ian Johnston, Dominic Kwiatkowski, Cordelia Langford, John Sillitoe on behalf of the Wellcome Sanger Institute COVID-19 Surveillance Team |
| EPI_ISL_3225520 | Lighthouse Lab in Alderley Park | Wellcome Sanger Institute for the COVID-19 Genomics UK (COG-UK) Consortium | Jacquelyn Wynn, Mairead Hyland, The Lighthouse Lab in Alderley Park and Alex Alderton, Roberto Amato, Jeffrey Barrett, Sonia Goncalves, Ewan Harrison, David K. Jackson, Ian Johnston, Dominic Kwiatkowski, Cordelia Langford, John Sillitoe on behalf of the Wellcome Sanger Institute COVID-19 Surveillance Team |
| EPI_ISL_3225521, EPI_ISL_3225522 | Lighthouse Laboratory Plymouth | Wellcome Sanger Institute for the COVID-19 Genomics UK (COG-UK) Consortium | Lighthouse Laboratory Plymouth and Alex Alderton, Roberto Amato, Jeffrey Barrett, Sonia Goncalves, Ewan Harrison, David K. Jackson, Ian Johnston, Dominic Kwiatkowski, Cordelia Langford, John Sillitoe on behalf of the Wellcome Sanger Institute COVID-19 Surveillance Team |
| EPI_ISL_3225523 | Lighthouse Lab in Glasgow | Wellcome Sanger Institute for the COVID-19 Genomics UK (COG-UK) Consortium | Harper VanSteenhouse, Yumi Kasai, David Gray, Carol Clugston, Anna Dominiczak and Alex Alderton, Roberto Amato, Jeffrey Barrett, Sonia Goncalves, Ewan Harrison, David K. Jackson, Ian Johnston, Dominic Kwiatkowski, Cordelia Langford, John Sillitoe on behalf of the Wellcome Sanger Institute COVID-19 Surveillance Team |
| EPI_ISL_3225524 | Lighthouse Lab in Alderley Park | Wellcome Sanger Institute for the COVID-19 Genomics UK (COG-UK) Consortium | Jacquelyn Wynn, Mairead Hyland, The Lighthouse Lab in Alderley Park and Alex Alderton, Roberto Amato, Jeffrey Barrett, Sonia Goncalves, Ewan Harrison, David K. Jackson, Ian Johnston, Dominic Kwiatkowski, Cordelia Langford, John Sillitoe on behalf of the Wellcome Sanger Institute COVID-19 Surveillance Team |
| EPI_ISL_3225525 | Lighthouse Lab in Glasgow | Wellcome Sanger Institute for the COVID-19 Genomics UK (COG-UK) Consortium | Harper VanSteenhouse, Yumi Kasai, David Gray, Carol Clugston, Anna Dominiczak and Alex Alderton, Roberto Amato, Jeffrey Barrett, Sonia Goncalves, Ewan Harrison, David K. Jackson, Ian Johnston, Dominic Kwiatkowski, Cordelia Langford, John Sillitoe on behalf of the Wellcome Sanger Institute COVID-19 Surveillance Team |
| EPI_ISL_3225526 | Lighthouse Lab in Alderley Park | Wellcome Sanger Institute for the COVID-19 Genomics UK (COG-UK) Consortium | Jacquelyn Wynn, Mairead Hyland, The Lighthouse Lab in Alderley Park and Alex Alderton, Roberto Amato, Jeffrey Barrett, Sonia Goncalves, Ewan Harrison, David K. Jackson, Ian Johnston, Dominic Kwiatkowski, Cordelia Langford, John Sillitoe on behalf of the Wellcome Sanger Institute COVID-19 Surveillance Team |
| EPI_ISL_3225527 | Lighthouse Lab in Glasgow | Wellcome Sanger Institute for the COVID-19 Genomics UK (COG-UK) Consortium | Harper VanSteenhouse, Yumi Kasai, David Gray, Carol Clugston, Anna Dominiczak and Alex Alderton, Roberto Amato, Jeffrey Barrett, Sonia Goncalves, Ewan Harrison, David K. Jackson, Ian Johnston, Dominic Kwiatkowski, Cordelia Langford, John Sillitoe on behalf of the Wellcome Sanger Institute COVID-19 Surveillance Team |
| EPI_ISL_3225528 | Lighthouse Lab in Alderley Park | Wellcome Sanger Institute for the COVID-19 Genomics UK (COG-UK) Consortium | Jacquelyn Wynn, Mairead Hyland, The Lighthouse Lab in Alderley Park and Alex Alderton, Roberto Amato, Jeffrey Barrett, Sonia Goncalves, Ewan Harrison, David K. Jackson, Ian Johnston, Dominic Kwiatkowski, Cordelia Langford, John Sillitoe on behalf of the Wellcome Sanger Institute COVID-19 Surveillance Team |
| EPI_ISL_3225529 | Lighthouse Lab in Glasgow | Wellcome Sanger Institute for the COVID-19 Genomics UK (COG-UK) Consortium | Harper VanSteenhouse, Yumi Kasai, David Gray, Carol Clugston, Anna Dominiczak and Alex Alderton, Roberto Amato, Jeffrey Barrett, Sonia Goncalves, Ewan Harrison, David K. Jackson, Ian Johnston, Dominic Kwiatkowski, Cordelia Langford, John Sillitoe on behalf of the Wellcome Sanger Institute COVID-19 Surveillance Team |
| EPI_ISL_3225530, EPI_ISL_3225531, EPI_ISL_3225532 | Lighthouse Lab in Alderley Park | Wellcome Sanger Institute for the COVID-19 Genomics UK (COG-UK) Consortium | Jacquelyn Wynn, Mairead Hyland, The Lighthouse Lab in Alderley Park and Alex Alderton, Roberto Amato, Jeffrey Barrett, Sonia Goncalves, Ewan Harrison, David K. Jackson, Ian Johnston, Dominic Kwiatkowski, Cordelia Langford, John Sillitoe on behalf of the Wellcome Sanger Institute COVID-19 Surveillance Team |
| EPI_ISL_3225533, EPI_ISL_3225534 | Lighthouse Laboratory Plymouth | Wellcome Sanger Institute for the COVID-19 Genomics UK (COG-UK) Consortium | Lighthouse Laboratory Plymouth and Alex Alderton, Roberto Amato, Jeffrey Barrett, Sonia Goncalves, Ewan Harrison, David K. Jackson, Ian Johnston, Dominic Kwiatkowski, Cordelia Langford, John Sillitoe on behalf of the Wellcome Sanger Institute COVID-19 Surveillance Team |
| EPI_ISL_3225535, EPI_ISL_3225536, EPI_ISL_3225537 | Lighthouse Lab in Alderley Park | Wellcome Sanger Institute for the COVID-19 Genomics UK (COG-UK) Consortium | Jacquelyn Wynn, Mairead Hyland, The Lighthouse Lab in Alderley Park and Alex Alderton, Roberto Amato, Jeffrey Barrett, Sonia Goncalves, Ewan Harrison, David K. Jackson, Ian Johnston, Dominic Kwiatkowski, Cordelia Langford, John Sillitoe on behalf of the Wellcome Sanger Institute COVID-19 Surveillance Team |
| EPI_ISL_3225538 | Lighthouse Lab in Glasgow | Wellcome Sanger Institute for the COVID-19 Genomics UK (COG-UK) Consortium | Harper VanSteenhouse, Yumi Kasai, David Gray, Carol Clugston, Anna Dominiczak and Alex Alderton, Roberto Amato, Jeffrey Barrett, Sonia Goncalves, Ewan Harrison, David K. Jackson, Ian Johnston, Dominic Kwiatkowski, Cordelia Langford, John Sillitoe on behalf of the Wellcome Sanger Institute COVID-19 Surveillance Team |
| EPI_ISL_3225539 | Lighthouse Lab in Alderley Park | Wellcome Sanger Institute for the COVID-19 Genomics UK (COG-UK) Consortium | Jacquelyn Wynn, Mairead Hyland, The Lighthouse Lab in Alderley Park and Alex Alderton, Roberto Amato, Jeffrey Barrett, Sonia Goncalves, Ewan Harrison, David K. Jackson, Ian Johnston, Dominic Kwiatkowski, Cordelia Langford, John Sillitoe on behalf of the Wellcome Sanger Institute COVID-19 Surveillance Team |
| EPI_ISL_3225540, EPI_ISL_3225541, EPI_ISL_3225542 | Lighthouse Laboratory Plymouth | Wellcome Sanger Institute for the COVID-19 Genomics UK (COG-UK) Consortium | Lighthouse Laboratory Plymouth and Alex Alderton, Roberto Amato, Jeffrey Barrett, Sonia Goncalves, Ewan Harrison, David K. Jackson, Ian Johnston, Dominic Kwiatkowski, Cordelia Langford, John Sillitoe on behalf of the Wellcome Sanger Institute COVID-19 Surveillance Team |
| EPI_ISL_3225543 | Lighthouse Lab in Alderley Park | Wellcome Sanger Institute for the COVID-19 Genomics UK (COG-UK) Consortium | Jacquelyn Wynn, Mairead Hyland, The Lighthouse Lab in Alderley Park and Alex Alderton, Roberto Amato, Jeffrey Barrett, Sonia Goncalves, Ewan Harrison, David K. Jackson, Ian Johnston, Dominic Kwiatkowski, Cordelia Langford, John Sillitoe on behalf of the Wellcome Sanger Institute COVID-19 Surveillance Team |
| EPI_ISL_3225544, EPI_ISL_3225545, EPI_ISL_3225546 | Lighthouse Lab in Glasgow | Wellcome Sanger Institute for the COVID-19 Genomics UK (COG-UK) Consortium | Harper VanSteenhouse, Yumi Kasai, David Gray, Carol Clugston, Anna Dominiczak and Alex Alderton, Roberto Amato, Jeffrey Barrett, Sonia Goncalves, Ewan Harrison, David K. Jackson, Ian Johnston, Dominic Kwiatkowski, Cordelia Langford, John Sillitoe on behalf of the Wellcome Sanger Institute COVID-19 Surveillance Team |
| EPI_ISL_3225547 | Lighthouse Laboratory Plymouth | Wellcome Sanger Institute for the COVID-19 Genomics UK (COG-UK) Consortium | Lighthouse Laboratory Plymouth and Alex Alderton, Roberto Amato, Jeffrey Barrett, Sonia Goncalves, Ewan Harrison, David K. Jackson, Ian Johnston, Dominic Kwiatkowski, Cordelia Langford, John Sillitoe on behalf of the Wellcome Sanger Institute COVID-19 Surveillance Team |
| EPI_ISL_3225548 | Lighthouse Lab in Glasgow | Wellcome Sanger Institute for the COVID-19 Genomics UK (COG-UK) Consortium | Harper VanSteenhouse, Yumi Kasai, David Gray, Carol Clugston, Anna Dominiczak and Alex Alderton, Roberto Amato, Jeffrey Barrett, Sonia Goncalves, Ewan Harrison, David K. Jackson, Ian Johnston, Dominic Kwiatkowski, Cordelia Langford, John Sillitoe on behalf of the Wellcome Sanger Institute COVID-19 Surveillance Team |
| EPI_ISL_3225549, EPI_ISL_3225550, EPI_ISL_3225551, EPI_ISL_3225552, EPI_ISL_3225553 | Lighthouse Lab in Alderley Park | Wellcome Sanger Institute for the COVID-19 Genomics UK (COG-UK) Consortium | Jacquelyn Wynn, Mairead Hyland, The Lighthouse Lab in Alderley Park and Alex Alderton, Roberto Amato, Jeffrey Barrett, Sonia Goncalves, Ewan Harrison, David K. Jackson, Ian Johnston, Dominic Kwiatkowski, Cordelia Langford, John Sillitoe on behalf of the Wellcome Sanger Institute COVID-19 Surveillance Team |
| EPI_ISL_3225554 | Lighthouse Lab in Glasgow | Wellcome Sanger Institute for the COVID-19 Genomics UK (COG-UK) Consortium | Harper VanSteenhouse, Yumi Kasai, David Gray, Carol Clugston, Anna Dominiczak and Alex Alderton, Roberto Amato, Jeffrey Barrett, Sonia Goncalves, Ewan Harrison, David K. Jackson, Ian Johnston, Dominic Kwiatkowski, Cordelia Langford, John Sillitoe on behalf of the Wellcome Sanger Institute COVID-19 Surveillance Team |
| EPI_ISL_3225555 | Lighthouse Lab in Alderley Park | Wellcome Sanger Institute for the COVID-19 Genomics UK (COG-UK) Consortium | Jacquelyn Wynn, Mairead Hyland, The Lighthouse Lab in Alderley Park and Alex Alderton, Roberto Amato, Jeffrey Barrett, Sonia Goncalves, Ewan Harrison, David K. Jackson, Ian Johnston, Dominic Kwiatkowski, Cordelia Langford, John Sillitoe on behalf of the Wellcome Sanger Institute COVID-19 Surveillance Team |
| EPI_ISL_3225556, EPI_ISL_3225557 | Lighthouse Laboratory Plymouth | Wellcome Sanger Institute for the COVID-19 Genomics UK (COG-UK) Consortium | Lighthouse Laboratory Plymouth and Alex Alderton, Roberto Amato, Jeffrey Barrett, Sonia Goncalves, Ewan Harrison, David K. Jackson, Ian Johnston, Dominic Kwiatkowski, Cordelia Langford, John Sillitoe on behalf of the Wellcome Sanger Institute COVID-19 Surveillance Team |
| EPI_ISL_3225558 | Lighthouse Lab in Glasgow | Wellcome Sanger Institute for the COVID-19 Genomics UK (COG-UK) Consortium | Harper VanSteenhouse, Yumi Kasai, David Gray, Carol Clugston, Anna Dominiczak and Alex Alderton, Roberto Amato, Jeffrey Barrett, Sonia Goncalves, Ewan Harrison, David K. Jackson, Ian Johnston, Dominic Kwiatkowski, Cordelia Langford, John Sillitoe on behalf of the Wellcome Sanger Institute COVID-19 Surveillance Team |
| EPI_ISL_3225559 | Lighthouse Lab in Alderley Park | Wellcome Sanger Institute for the COVID-19 Genomics UK (COG-UK) Consortium | Jacquelyn Wynn, Mairead Hyland, The Lighthouse Lab in Alderley Park and Alex Alderton, Roberto Amato, Jeffrey Barrett, Sonia Goncalves, Ewan Harrison, David K. Jackson, Ian Johnston, Dominic Kwiatkowski, Cordelia Langford, John Sillitoe on behalf of the Wellcome Sanger Institute COVID-19 Surveillance Team |
| EPI_ISL_3225560 | Lighthouse Laboratory Plymouth | Wellcome Sanger Institute for the COVID-19 Genomics UK (COG-UK) Consortium | Lighthouse Laboratory Plymouth and Alex Alderton, Roberto Amato, Jeffrey Barrett, Sonia Goncalves, Ewan Harrison, David K. Jackson, Ian Johnston, Dominic Kwiatkowski, Cordelia Langford, John Sillitoe on behalf of the Wellcome Sanger Institute COVID-19 Surveillance Team |
| EPI_ISL_3225561, EPI_ISL_3225562 | Lighthouse Lab in Milton Keynes | Wellcome Sanger Institute for the COVID-19 Genomics UK (COG-UK) Consortium | The Lighthouse Lab in Milton Keynes and Alex Alderton, Roberto Amato, Jeffrey Barrett, Sonia Goncalves, Ewan Harrison, David K. Jackson, Ian Johnston, Dominic Kwiatkowski, Cordelia Langford, John Sillitoe on behalf of the Wellcome Sanger Institute COVID-19 Surveillance Team |
| EPI_ISL_3225563, EPI_ISL_3225564, EPI_ISL_3225565 | Lighthouse Laboratory Plymouth | Wellcome Sanger Institute for the COVID-19 Genomics UK (COG-UK) Consortium | Lighthouse Laboratory Plymouth and Alex Alderton, Roberto Amato, Jeffrey Barrett, Sonia Goncalves, Ewan Harrison, David K. Jackson, Ian Johnston, Dominic Kwiatkowski, Cordelia Langford, John Sillitoe on behalf of the Wellcome Sanger Institute COVID-19 Surveillance Team |

| | | | |
|---|---|---|---|
| EPI_ISL_3225566 | Lighthouse Lab in Milton Keynes | Wellcome Sanger Institute for the COVID-19 Genomics UK (COG-UK) Consortium | The Lighthouse Lab in Milton Keynes and Alex Alderton, Roberto Amato, Jeffrey Barrett, Sonia Goncalves, Ewan Harrison, David K. Jackson, Ian Johnston, Dominic Kwiatkowski, Cordelia Langford, John Sillitoe on behalf of the Wellcome Sanger Institute COVID-19 Surveillance Team |
| EPI_ISL_3225567 | Lighthouse Laboratory Plymouth | Wellcome Sanger Institute for the COVID-19 Genomics UK (COG-UK) Consortium | Lighthouse Laboratory Plymouth and Alex Alderton, Roberto Amato, Jeffrey Barrett, Sonia Goncalves, Ewan Harrison, David K. Jackson, Ian Johnston, Dominic Kwiatkowski, Cordelia Langford, John Sillitoe on behalf of the Wellcome Sanger Institute COVID-19 Surveillance Team |
| EPI_ISL_3225568 | Lighthouse Lab in Milton Keynes | Wellcome Sanger Institute for the COVID-19 Genomics UK (COG-UK) Consortium | The Lighthouse Lab in Milton Keynes and Alex Alderton, Roberto Amato, Jeffrey Barrett, Sonia Goncalves, Ewan Harrison, David K. Jackson, Ian Johnston, Dominic Kwiatkowski, Cordelia Langford, John Sillitoe on behalf of the Wellcome Sanger Institute COVID-19 Surveillance Team |
| EPI_ISL_3225569 | Lighthouse Lab in Glasgow | Wellcome Sanger Institute for the COVID-19 Genomics UK (COG-UK) Consortium | Harper VanSteenhouse, Yumi Kasai, David Gray, Carol Clugston, Anna Dominiczak and Alex Alderton, Roberto Amato, Jeffrey Barrett, Sonia Goncalves, Ewan Harrison, David K. Jackson, Ian Johnston, Dominic Kwiatkowski, Cordelia Langford, John Sillitoe on behalf of the Wellcome Sanger Institute COVID-19 Surveillance Team |
| EPI_ISL_3225570, EPI_ISL_3225571 | Lighthouse Lab in Milton Keynes | Wellcome Sanger Institute for the COVID-19 Genomics UK (COG-UK) Consortium | The Lighthouse Lab in Milton Keynes and Alex Alderton, Roberto Amato, Jeffrey Barrett, Sonia Goncalves, Ewan Harrison, David K. Jackson, Ian Johnston, Dominic Kwiatkowski, Cordelia Langford, John Sillitoe on behalf of the Wellcome Sanger Institute COVID-19 Surveillance Team |
| EPI_ISL_3225572 | Lighthouse Laboratory Plymouth | Wellcome Sanger Institute for the COVID-19 Genomics UK (COG-UK) Consortium | Lighthouse Laboratory Plymouth and Alex Alderton, Roberto Amato, Jeffrey Barrett, Sonia Goncalves, Ewan Harrison, David K. Jackson, Ian Johnston, Dominic Kwiatkowski, Cordelia Langford, John Sillitoe on behalf of the Wellcome Sanger Institute COVID-19 Surveillance Team |
| EPI_ISL_3225573, EPI_ISL_3225574, EPI_ISL_3225575, EPI_ISL_3225576 | Lighthouse Lab in Milton Keynes | Wellcome Sanger Institute for the COVID-19 Genomics UK (COG-UK) Consortium | The Lighthouse Lab in Milton Keynes and Alex Alderton, Roberto Amato, Jeffrey Barrett, Sonia Goncalves, Ewan Harrison, David K. Jackson, Ian Johnston, Dominic Kwiatkowski, Cordelia Langford, John Sillitoe on behalf of the Wellcome Sanger Institute COVID-19 Surveillance Team |
| EPI_ISL_3225577 | Lighthouse Lab in Glasgow | Wellcome Sanger Institute for the COVID-19 Genomics UK (COG-UK) Consortium | Harper VanSteenhouse, Yumi Kasai, David Gray, Carol Clugston, Anna Dominiczak and Alex Alderton, Roberto Amato, Jeffrey Barrett, Sonia Goncalves, Ewan Harrison, David K. Jackson, Ian Johnston, Dominic Kwiatkowski, Cordelia Langford, John Sillitoe on behalf of the Wellcome Sanger Institute COVID-19 Surveillance Team |
| EPI_ISL_3225578 | Lighthouse Laboratory Plymouth | Wellcome Sanger Institute for the COVID-19 Genomics UK (COG-UK) Consortium | Lighthouse Laboratory Plymouth and Alex Alderton, Roberto Amato, Jeffrey Barrett, Sonia Goncalves, Ewan Harrison, David K. Jackson, Ian Johnston, Dominic Kwiatkowski, Cordelia Langford, John Sillitoe on behalf of the Wellcome Sanger Institute COVID-19 Surveillance Team |
| EPI_ISL_3225579 | Lighthouse Lab in Milton Keynes | Wellcome Sanger Institute for the COVID-19 Genomics UK (COG-UK) Consortium | The Lighthouse Lab in Milton Keynes and Alex Alderton, Roberto Amato, Jeffrey Barrett, Sonia Goncalves, Ewan Harrison, David K. Jackson, Ian Johnston, Dominic Kwiatkowski, Cordelia Langford, John Sillitoe on behalf of the Wellcome Sanger Institute COVID-19 Surveillance Team |
| EPI_ISL_3225580 | Lighthouse Laboratory Plymouth | Wellcome Sanger Institute for the COVID-19 Genomics UK (COG-UK) Consortium | Lighthouse Laboratory Plymouth and Alex Alderton, Roberto Amato, Jeffrey Barrett, Sonia Goncalves, Ewan Harrison, David K. Jackson, Ian Johnston, Dominic Kwiatkowski, Cordelia Langford, John Sillitoe on behalf of the Wellcome Sanger Institute COVID-19 Surveillance Team |
| EPI_ISL_3225581, EPI_ISL_3225582, EPI_ISL_3225583 | Lighthouse Lab in Glasgow | Wellcome Sanger Institute for the COVID-19 Genomics UK (COG-UK) Consortium | Harper VanSteenhouse, Yumi Kasai, David Gray, Carol Clugston, Anna Dominiczak and Alex Alderton, Roberto Amato, Jeffrey Barrett, Sonia Goncalves, Ewan Harrison, David K. Jackson, Ian Johnston, Dominic Kwiatkowski, Cordelia Langford, John Sillitoe on behalf of the Wellcome Sanger Institute COVID-19 Surveillance Team |
| EPI_ISL_3225584 | Lighthouse Lab in Milton Keynes | Wellcome Sanger Institute for the COVID-19 Genomics UK (COG-UK) Consortium | The Lighthouse Lab in Milton Keynes and Alex Alderton, Roberto Amato, Jeffrey Barrett, Sonia Goncalves, Ewan Harrison, David K. Jackson, Ian Johnston, Dominic Kwiatkowski, Cordelia Langford, John Sillitoe on behalf of the Wellcome Sanger Institute COVID-19 Surveillance Team |
| EPI_ISL_3225585, EPI_ISL_3225586, EPI_ISL_3225587 | Lighthouse Lab in Glasgow | Wellcome Sanger Institute for the COVID-19 Genomics UK (COG-UK) Consortium | Harper VanSteenhouse, Yumi Kasai, David Gray, Carol Clugston, Anna Dominiczak and Alex Alderton, Roberto Amato, Jeffrey Barrett, Sonia Goncalves, Ewan Harrison, David K. Jackson, Ian Johnston, Dominic Kwiatkowski, Cordelia Langford, John Sillitoe on behalf of the Wellcome Sanger Institute COVID-19 Surveillance Team |
| EPI_ISL_3225588, EPI_ISL_3225589, EPI_ISL_3225590 | Lighthouse Lab in Milton Keynes | Wellcome Sanger Institute for the COVID-19 Genomics UK (COG-UK) Consortium | The Lighthouse Lab in Milton Keynes and Alex Alderton, Roberto Amato, Jeffrey Barrett, Sonia Goncalves, Ewan Harrison, David K. Jackson, Ian Johnston, Dominic Kwiatkowski, Cordelia Langford, John Sillitoe on behalf of the Wellcome Sanger Institute COVID-19 Surveillance Team |
| EPI_ISL_3225591 | Lighthouse Lab in Glasgow | Wellcome Sanger Institute for the COVID-19 Genomics UK (COG-UK) Consortium | Harper VanSteenhouse, Yumi Kasai, David Gray, Carol Clugston, Anna Dominiczak and Alex Alderton, Roberto Amato, Jeffrey Barrett, Sonia Goncalves, Ewan Harrison, David K. Jackson, Ian Johnston, Dominic Kwiatkowski, Cordelia Langford, John Sillitoe on behalf of the Wellcome Sanger Institute COVID-19 Surveillance Team |
| EPI_ISL_3225592, EPI_ISL_3225593 | Lighthouse Lab in Milton Keynes | Wellcome Sanger Institute for the COVID-19 Genomics UK (COG-UK) Consortium | The Lighthouse Lab in Milton Keynes and Alex Alderton, Roberto Amato, Jeffrey Barrett, Sonia Goncalves, Ewan Harrison, David K. Jackson, Ian Johnston, Dominic Kwiatkowski, Cordelia Langford, John Sillitoe on behalf of the Wellcome Sanger Institute COVID-19 Surveillance Team |
| EPI_ISL_3225594 | Lighthouse Lab in Glasgow | Wellcome Sanger Institute for the COVID-19 Genomics UK (COG-UK) Consortium | Harper VanSteenhouse, Yumi Kasai, David Gray, Carol Clugston, Anna Dominiczak and Alex Alderton, Roberto Amato, Jeffrey Barrett, Sonia Goncalves, Ewan Harrison, David K. Jackson, Ian Johnston, Dominic Kwiatkowski, Cordelia Langford, John Sillitoe on behalf of the Wellcome Sanger Institute COVID-19 Surveillance Team |
| EPI_ISL_3225595 | Lighthouse Lab in Milton Keynes | Wellcome Sanger Institute for the COVID-19 Genomics UK (COG-UK) Consortium | The Lighthouse Lab in Milton Keynes and Alex Alderton, Roberto Amato, Jeffrey Barrett, Sonia Goncalves, Ewan Harrison, David K. Jackson, Ian Johnston, Dominic Kwiatkowski, Cordelia Langford, John Sillitoe on behalf of the Wellcome Sanger Institute COVID-19 Surveillance Team |
| EPI_ISL_3225596, EPI_ISL_3225597 | Lighthouse Laboratory Plymouth | Wellcome Sanger Institute for the COVID-19 Genomics UK (COG-UK) Consortium | Lighthouse Laboratory Plymouth and Alex Alderton, Roberto Amato, Jeffrey Barrett, Sonia Goncalves, Ewan Harrison, David K. Jackson, Ian Johnston, Dominic Kwiatkowski, Cordelia Langford, John Sillitoe on behalf of the Wellcome Sanger Institute COVID-19 Surveillance Team |
| EPI_ISL_3225598 | Lighthouse Lab in Milton Keynes | Wellcome Sanger Institute for the COVID-19 Genomics UK (COG-UK) Consortium | The Lighthouse Lab in Milton Keynes and Alex Alderton, Roberto Amato, Jeffrey Barrett, Sonia Goncalves, Ewan Harrison, David K. Jackson, Ian Johnston, Dominic Kwiatkowski, Cordelia Langford, John Sillitoe on behalf of the Wellcome Sanger Institute COVID-19 Surveillance Team |
| EPI_ISL_3225599 | Lighthouse Lab in Glasgow | Wellcome Sanger Institute for the COVID-19 Genomics UK (COG-UK) Consortium | Harper VanSteenhouse, Yumi Kasai, David Gray, Carol Clugston, Anna Dominiczak and Alex Alderton, Roberto Amato, Jeffrey Barrett, Sonia Goncalves, Ewan Harrison, David K. Jackson, Ian Johnston, Dominic Kwiatkowski, Cordelia Langford, John Sillitoe on behalf of the Wellcome Sanger Institute COVID-19 Surveillance Team |
| EPI_ISL_3225600 | Lighthouse Laboratory Plymouth | Wellcome Sanger Institute for the COVID-19 Genomics UK (COG-UK) Consortium | Lighthouse Laboratory Plymouth and Alex Alderton, Roberto Amato, Jeffrey Barrett, Sonia Goncalves, Ewan Harrison, David K. Jackson, Ian Johnston, Dominic Kwiatkowski, Cordelia Langford, John Sillitoe on behalf of the Wellcome Sanger Institute COVID-19 Surveillance Team |
| EPI_ISL_3225601 | Lighthouse Lab in Milton Keynes | Wellcome Sanger Institute for the COVID-19 Genomics UK (COG-UK) Consortium | The Lighthouse Lab in Milton Keynes and Alex Alderton, Roberto Amato, Jeffrey Barrett, Sonia Goncalves, Ewan Harrison, David K. Jackson, Ian Johnston, Dominic Kwiatkowski, Cordelia Langford, John Sillitoe on behalf of the Wellcome Sanger Institute COVID-19 Surveillance Team |
| EPI_ISL_3225602, EPI_ISL_3225603 | Lighthouse Laboratory Plymouth | Wellcome Sanger Institute for the COVID-19 Genomics UK (COG-UK) Consortium | Lighthouse Laboratory Plymouth and Alex Alderton, Roberto Amato, Jeffrey Barrett, Sonia Goncalves, Ewan Harrison, David K. Jackson, Ian Johnston, Dominic Kwiatkowski, Cordelia Langford, John Sillitoe on behalf of the Wellcome Sanger Institute COVID-19 Surveillance Team |
| EPI_ISL_3225604, EPI_ISL_3225605 | Lighthouse Lab in Glasgow | Wellcome Sanger Institute for the COVID-19 Genomics UK (COG-UK) Consortium | Harper VanSteenhouse, Yumi Kasai, David Gray, Carol Clugston, Anna Dominiczak and Alex Alderton, Roberto Amato, Jeffrey Barrett, Sonia Goncalves, Ewan Harrison, David K. Jackson, Ian Johnston, Dominic Kwiatkowski, Cordelia Langford, John Sillitoe on behalf of the Wellcome Sanger Institute COVID-19 Surveillance Team |
| EPI_ISL_3225606 | Lighthouse Lab in Milton Keynes | Wellcome Sanger Institute for the COVID-19 Genomics UK (COG-UK) Consortium | The Lighthouse Lab in Milton Keynes and Alex Alderton, Roberto Amato, Jeffrey Barrett, Sonia Goncalves, Ewan Harrison, David K. Jackson, Ian Johnston, Dominic Kwiatkowski, Cordelia Langford, John Sillitoe on behalf of the Wellcome Sanger Institute COVID-19 Surveillance Team |
| EPI_ISL_3225607 | Lighthouse Laboratory Plymouth | Wellcome Sanger Institute for the COVID-19 Genomics UK (COG-UK) Consortium | Lighthouse Laboratory Plymouth and Alex Alderton, Roberto Amato, Jeffrey Barrett, Sonia Goncalves, Ewan Harrison, David K. Jackson, Ian Johnston, Dominic Kwiatkowski, Cordelia Langford, John Sillitoe on behalf of the Wellcome Sanger Institute COVID-19 Surveillance Team |
| EPI_ISL_3225608, EPI_ISL_3225609 | Lighthouse Lab in Glasgow | Wellcome Sanger Institute for the COVID-19 Genomics UK (COG-UK) Consortium | Harper VanSteenhouse, Yumi Kasai, David Gray, Carol Clugston, Anna Dominiczak and Alex Alderton, Roberto Amato, Jeffrey Barrett, Sonia Goncalves, Ewan Harrison, David K. Jackson, Ian Johnston, Dominic Kwiatkowski, Cordelia Langford, John Sillitoe on behalf of the Wellcome Sanger Institute COVID-19 Surveillance Team |
| EPI_ISL_3225610 | Lighthouse Laboratory Plymouth | Wellcome Sanger Institute for the COVID-19 Genomics UK (COG-UK) Consortium | Lighthouse Laboratory Plymouth and Alex Alderton, Roberto Amato, Jeffrey Barrett, Sonia Goncalves, Ewan Harrison, David K. Jackson, Ian Johnston, Dominic Kwiatkowski, Cordelia Langford, John Sillitoe on behalf of the Wellcome Sanger Institute COVID-19 Surveillance Team |
| EPI_ISL_3225611, EPI_ISL_3225612 | Lighthouse Lab in Milton Keynes | Wellcome Sanger Institute for the COVID-19 Genomics UK (COG-UK) Consortium | The Lighthouse Lab in Milton Keynes and Alex Alderton, Roberto Amato, Jeffrey Barrett, Sonia Goncalves, Ewan Harrison, David K. Jackson, Ian Johnston, Dominic Kwiatkowski, Cordelia Langford, John Sillitoe on behalf of the Wellcome Sanger Institute COVID-19 Surveillance Team |
| EPI_ISL_3225613 | Lighthouse Lab in Glasgow | Wellcome Sanger Institute for the COVID-19 Genomics UK (COG-UK) Consortium | Harper VanSteenhouse, Yumi Kasai, David Gray, Carol Clugston, Anna Dominiczak and Alex Alderton, Roberto Amato, Jeffrey Barrett, Sonia Goncalves, Ewan Harrison, David K. Jackson, Ian Johnston, Dominic Kwiatkowski, Cordelia Langford, John Sillitoe on behalf of the Wellcome Sanger Institute COVID-19 Surveillance Team |
| EPI_ISL_3225614, EPI_ISL_3225615 | Lighthouse Lab in Milton Keynes | Wellcome Sanger Institute for the COVID-19 Genomics UK (COG-UK) Consortium | The Lighthouse Lab in Milton Keynes and Alex Alderton, Roberto Amato, Jeffrey Barrett, Sonia Goncalves, Ewan Harrison, David K. Jackson, Ian Johnston, Dominic Kwiatkowski, Cordelia Langford, John Sillitoe on behalf of the Wellcome Sanger Institute COVID-19 Surveillance Team |
| EPI_ISL_3225616, EPI_ISL_3225617, EPI_ISL_3225618 | Lighthouse Laboratory Plymouth | Wellcome Sanger Institute for the COVID-19 Genomics UK (COG-UK) Consortium | Lighthouse Laboratory Plymouth and Alex Alderton, Roberto Amato, Jeffrey Barrett, Sonia Goncalves, Ewan Harrison, David K. Jackson, Ian Johnston, Dominic Kwiatkowski, Cordelia Langford, John Sillitoe on behalf of the Wellcome Sanger Institute COVID-19 Surveillance Team |
| EPI_ISL_3225619 | Lighthouse Lab in Glasgow | Wellcome Sanger Institute for the COVID-19 Genomics UK (COG-UK) Consortium | Harper VanSteenhouse, Yumi Kasai, David Gray, Carol Clugston, Anna Dominiczak and Alex Alderton, Roberto Amato, Jeffrey Barrett, Sonia Goncalves, Ewan Harrison, David K. Jackson, Ian Johnston, Dominic Kwiatkowski, Cordelia Langford, John Sillitoe on behalf of the Wellcome Sanger Institute COVID-19 Surveillance Team |
| EPI_ISL_3225620, EPI_ISL_3225621, EPI_ISL_3225622 | Lighthouse Lab in Milton Keynes | Wellcome Sanger Institute for the COVID-19 Genomics UK (COG-UK) Consortium | The Lighthouse Lab in Milton Keynes and Alex Alderton, Roberto Amato, Jeffrey Barrett, Sonia Goncalves, Ewan Harrison, David K. Jackson, Ian Johnston, Dominic Kwiatkowski, Cordelia Langford, John Sillitoe on behalf of the Wellcome Sanger Institute COVID-19 Surveillance Team |
| EPI_ISL_3225623 | Lighthouse Lab in Glasgow | Wellcome Sanger Institute for the COVID-19 Genomics UK (COG-UK) Consortium | Harper VanSteenhouse, Yumi Kasai, David Gray, Carol Clugston, Anna Dominiczak and Alex Alderton, Roberto Amato, Jeffrey Barrett, Sonia Goncalves, Ewan Harrison, David K. Jackson, Ian Johnston, Dominic Kwiatkowski, Cordelia Langford, John Sillitoe on behalf of the Wellcome Sanger Institute COVID-19 Surveillance Team |
| EPI_ISL_3225624 | Lighthouse Lab in Milton Keynes | Wellcome Sanger Institute for the COVID-19 Genomics UK (COG-UK) Consortium | The Lighthouse Lab in Milton Keynes and Alex Alderton, Roberto Amato, Jeffrey Barrett, Sonia Goncalves, Ewan Harrison, David K. Jackson, Ian Johnston, Dominic Kwiatkowski, Cordelia Langford, John Sillitoe on behalf of the Wellcome Sanger Institute COVID-19 Surveillance Team |
| EPI_ISL_3225625 | Lighthouse Lab in Glasgow | Wellcome Sanger Institute for the COVID-19 Genomics UK (COG-UK) Consortium | Harper VanSteenhouse, Yumi Kasai, David Gray, Carol Clugston, Anna Dominiczak and Alex Alderton, Roberto Amato, Jeffrey Barrett, Sonia Goncalves, Ewan Harrison, David K. Jackson, Ian Johnston, Dominic Kwiatkowski, Cordelia Langford, John Sillitoe on behalf of the Wellcome Sanger Institute COVID-19 Surveillance Team |
| EPI_ISL_3225626 | Lighthouse Laboratory Plymouth | Wellcome Sanger Institute for the COVID-19 Genomics UK (COG-UK) Consortium | Lighthouse Laboratory Plymouth and Alex Alderton, Roberto Amato, Jeffrey Barrett, Sonia Goncalves, Ewan Harrison, David K. Jackson, Ian Johnston, Dominic Kwiatkowski, Cordelia Langford, John Sillitoe on behalf of the Wellcome Sanger Institute COVID-19 Surveillance Team |
| EPI_ISL_3225627 | Lighthouse Lab in Milton Keynes | Wellcome Sanger Institute for the COVID-19 Genomics UK (COG-UK) Consortium | The Lighthouse Lab in Milton Keynes and Alex Alderton, Roberto Amato, Jeffrey Barrett, Sonia Goncalves, Ewan Harrison, David K. Jackson, Ian Johnston, Dominic Kwiatkowski, Cordelia Langford, John Sillitoe on behalf of the Wellcome Sanger Institute COVID-19 Surveillance Team |
| EPI_ISL_3225628 | Lighthouse Lab in Glasgow | Wellcome Sanger Institute for the COVID-19 Genomics UK (COG-UK) Consortium | Harper VanSteenhouse, Yumi Kasai, David Gray, Carol Clugston, Anna Dominiczak and Alex Alderton, Roberto Amato, Jeffrey Barrett, Sonia Goncalves, Ewan Harrison, David K. Jackson, Ian Johnston, Dominic Kwiatkowski, Cordelia Langford, John Sillitoe on behalf of the Wellcome Sanger Institute COVID-19 Surveillance Team |

| | | | |
|---|---|---|---|
| EPI_ISL_3225629 | Lighthouse Lab in Milton Keynes | Wellcome Sanger Institute for the COVID-19 Genomics UK (COG-UK) Consortium | The Lighthouse Lab in Milton Keynes and Alex Alderton, Roberto Amato, Jeffrey Barrett, Sonia Goncalves, Ewan Harrison, David K. Jackson, Ian Johnston, Dominic Kwiatkowski, Cordelia Langford, John Sillitoe on behalf of the Wellcome Sanger Institute COVID-19 Surveillance Team |
| EPI_ISL_3225630 | Lighthouse Laboratory Plymouth | Wellcome Sanger Institute for the COVID-19 Genomics UK (COG-UK) Consortium | Lighthouse Laboratory Plymouth and Alex Alderton, Roberto Amato, Jeffrey Barrett, Sonia Goncalves, Ewan Harrison, David K. Jackson, Ian Johnston, Dominic Kwiatkowski, Cordelia Langford, John Sillitoe on behalf of the Wellcome Sanger Institute COVID-19 Surveillance Team |
| EPI_ISL_3225631 | Lighthouse Lab in Milton Keynes | Wellcome Sanger Institute for the COVID-19 Genomics UK (COG-UK) Consortium | The Lighthouse Lab in Milton Keynes and Alex Alderton, Roberto Amato, Jeffrey Barrett, Sonia Goncalves, Ewan Harrison, David K. Jackson, Ian Johnston, Dominic Kwiatkowski, Cordelia Langford, John Sillitoe on behalf of the Wellcome Sanger Institute COVID-19 Surveillance Team |
| EPI_ISL_3225632 | Lighthouse Laboratory Plymouth | Wellcome Sanger Institute for the COVID-19 Genomics UK (COG-UK) Consortium | Lighthouse Laboratory Plymouth and Alex Alderton, Roberto Amato, Jeffrey Barrett, Sonia Goncalves, Ewan Harrison, David K. Jackson, Ian Johnston, Dominic Kwiatkowski, Cordelia Langford, John Sillitoe on behalf of the Wellcome Sanger Institute COVID-19 Surveillance Team |
| EPI_ISL_3225633, EPI_ISL_3225634, EPI_ISL_3225635, EPI_ISL_3225636 | Lighthouse Lab in Milton Keynes | Wellcome Sanger Institute for the COVID-19 Genomics UK (COG-UK) Consortium | The Lighthouse Lab in Milton Keynes and Alex Alderton, Roberto Amato, Jeffrey Barrett, Sonia Goncalves, Ewan Harrison, David K. Jackson, Ian Johnston, Dominic Kwiatkowski, Cordelia Langford, John Sillitoe on behalf of the Wellcome Sanger Institute COVID-19 Surveillance Team |
| EPI_ISL_3225637 | Lighthouse Laboratory Plymouth | Wellcome Sanger Institute for the COVID-19 Genomics UK (COG-UK) Consortium | Lighthouse Laboratory Plymouth and Alex Alderton, Roberto Amato, Jeffrey Barrett, Sonia Goncalves, Ewan Harrison, David K. Jackson, Ian Johnston, Dominic Kwiatkowski, Cordelia Langford, John Sillitoe on behalf of the Wellcome Sanger Institute COVID-19 Surveillance Team |
| EPI_ISL_3225638, EPI_ISL_3225639 | Lighthouse Lab in Milton Keynes | Wellcome Sanger Institute for the COVID-19 Genomics UK (COG-UK) Consortium | The Lighthouse Lab in Milton Keynes and Alex Alderton, Roberto Amato, Jeffrey Barrett, Sonia Goncalves, Ewan Harrison, David K. Jackson, Ian Johnston, Dominic Kwiatkowski, Cordelia Langford, John Sillitoe on behalf of the Wellcome Sanger Institute COVID-19 Surveillance Team |
| EPI_ISL_3225640 | Lighthouse Laboratory Plymouth | Wellcome Sanger Institute for the COVID-19 Genomics UK (COG-UK) Consortium | Lighthouse Laboratory Plymouth and Alex Alderton, Roberto Amato, Jeffrey Barrett, Sonia Goncalves, Ewan Harrison, David K. Jackson, Ian Johnston, Dominic Kwiatkowski, Cordelia Langford, John Sillitoe on behalf of the Wellcome Sanger Institute COVID-19 Surveillance Team |
| EPI_ISL_3225641 | Lighthouse Lab in Glasgow | Wellcome Sanger Institute for the COVID-19 Genomics UK (COG-UK) Consortium | Harper VanSteenhouse, Yumi Kasai, David Gray, Carol Clugston, Anna Dominiczak and Alex Alderton, Roberto Amato, Jeffrey Barrett, Sonia Goncalves, Ewan Harrison, David K. Jackson, Ian Johnston, Dominic Kwiatkowski, Cordelia Langford, John Sillitoe on behalf of the Wellcome Sanger Institute COVID-19 Surveillance Team |
| EPI_ISL_3225642, EPI_ISL_3225643 | Lighthouse Lab in Milton Keynes | Wellcome Sanger Institute for the COVID-19 Genomics UK (COG-UK) Consortium | The Lighthouse Lab in Milton Keynes and Alex Alderton, Roberto Amato, Jeffrey Barrett, Sonia Goncalves, Ewan Harrison, David K. Jackson, Ian Johnston, Dominic Kwiatkowski, Cordelia Langford, John Sillitoe on behalf of the Wellcome Sanger Institute COVID-19 Surveillance Team |
| EPI_ISL_3225644, EPI_ISL_3225645 | Lighthouse Lab in Glasgow | Wellcome Sanger Institute for the COVID-19 Genomics UK (COG-UK) Consortium | Harper VanSteenhouse, Yumi Kasai, David Gray, Carol Clugston, Anna Dominiczak and Alex Alderton, Roberto Amato, Jeffrey Barrett, Sonia Goncalves, Ewan Harrison, David K. Jackson, Ian Johnston, Dominic Kwiatkowski, Cordelia Langford, John Sillitoe on behalf of the Wellcome Sanger Institute COVID-19 Surveillance Team |
| EPI_ISL_3225646 | Lighthouse Lab in Milton Keynes | Wellcome Sanger Institute for the COVID-19 Genomics UK (COG-UK) Consortium | The Lighthouse Lab in Milton Keynes and Alex Alderton, Roberto Amato, Jeffrey Barrett, Sonia Goncalves, Ewan Harrison, David K. Jackson, Ian Johnston, Dominic Kwiatkowski, Cordelia Langford, John Sillitoe on behalf of the Wellcome Sanger Institute COVID-19 Surveillance Team |
| EPI_ISL_3225647 | Lighthouse Laboratory Plymouth | Wellcome Sanger Institute for the COVID-19 Genomics UK (COG-UK) Consortium | Lighthouse Laboratory Plymouth and Alex Alderton, Roberto Amato, Jeffrey Barrett, Sonia Goncalves, Ewan Harrison, David K. Jackson, Ian Johnston, Dominic Kwiatkowski, Cordelia Langford, John Sillitoe on behalf of the Wellcome Sanger Institute COVID-19 Surveillance Team |
| EPI_ISL_3225648 | Lighthouse Lab in Milton Keynes | Wellcome Sanger Institute for the COVID-19 Genomics UK (COG-UK) Consortium | The Lighthouse Lab in Milton Keynes and Alex Alderton, Roberto Amato, Jeffrey Barrett, Sonia Goncalves, Ewan Harrison, David K. Jackson, Ian Johnston, Dominic Kwiatkowski, Cordelia Langford, John Sillitoe on behalf of the Wellcome Sanger Institute COVID-19 Surveillance Team |
| EPI_ISL_3225649 | Lighthouse Laboratory Plymouth | Wellcome Sanger Institute for the COVID-19 Genomics UK (COG-UK) Consortium | Lighthouse Laboratory Plymouth and Alex Alderton, Roberto Amato, Jeffrey Barrett, Sonia Goncalves, Ewan Harrison, David K. Jackson, Ian Johnston, Dominic Kwiatkowski, Cordelia Langford, John Sillitoe on behalf of the Wellcome Sanger Institute COVID-19 Surveillance Team |
| EPI_ISL_3225650, EPI_ISL_3225651, EPI_ISL_3225652, EPI_ISL_3225653 | Lighthouse Lab in Milton Keynes | Wellcome Sanger Institute for the COVID-19 Genomics UK (COG-UK) Consortium | The Lighthouse Lab in Milton Keynes and Alex Alderton, Roberto Amato, Jeffrey Barrett, Sonia Goncalves, Ewan Harrison, David K. Jackson, Ian Johnston, Dominic Kwiatkowski, Cordelia Langford, John Sillitoe on behalf of the Wellcome Sanger Institute COVID-19 Surveillance Team |
| EPI_ISL_3225654 | Lighthouse Lab in Glasgow | Wellcome Sanger Institute for the COVID-19 Genomics UK (COG-UK) Consortium | Harper VanSteenhouse, Yumi Kasai, David Gray, Carol Clugston, Anna Dominiczak and Alex Alderton, Roberto Amato, Jeffrey Barrett, Sonia Goncalves, Ewan Harrison, David K. Jackson, Ian Johnston, Dominic Kwiatkowski, Cordelia Langford, John Sillitoe on behalf of the Wellcome Sanger Institute COVID-19 Surveillance Team |
| EPI_ISL_3225655, EPI_ISL_3225656 | Lighthouse Lab in Milton Keynes | Wellcome Sanger Institute for the COVID-19 Genomics UK (COG-UK) Consortium | The Lighthouse Lab in Milton Keynes and Alex Alderton, Roberto Amato, Jeffrey Barrett, Sonia Goncalves, Ewan Harrison, David K. Jackson, Ian Johnston, Dominic Kwiatkowski, Cordelia Langford, John Sillitoe on behalf of the Wellcome Sanger Institute COVID-19 Surveillance Team |
| EPI_ISL_3225657 | Lighthouse Lab in Glasgow | Wellcome Sanger Institute for the COVID-19 Genomics UK (COG-UK) Consortium | Harper VanSteenhouse, Yumi Kasai, David Gray, Carol Clugston, Anna Dominiczak and Alex Alderton, Roberto Amato, Jeffrey Barrett, Sonia Goncalves, Ewan Harrison, David K. Jackson, Ian Johnston, Dominic Kwiatkowski, Cordelia Langford, John Sillitoe on behalf of the Wellcome Sanger Institute COVID-19 Surveillance Team |
| EPI_ISL_3225658 | Lighthouse Lab in Milton Keynes | Wellcome Sanger Institute for the COVID-19 Genomics UK (COG-UK) Consortium | The Lighthouse Lab in Milton Keynes and Alex Alderton, Roberto Amato, Jeffrey Barrett, Sonia Goncalves, Ewan Harrison, David K. Jackson, Ian Johnston, Dominic Kwiatkowski, Cordelia Langford, John Sillitoe on behalf of the Wellcome Sanger Institute COVID-19 Surveillance Team |
| EPI_ISL_3225659 | Lighthouse Laboratory Plymouth | Wellcome Sanger Institute for the COVID-19 Genomics UK (COG-UK) Consortium | Lighthouse Laboratory Plymouth and Alex Alderton, Roberto Amato, Jeffrey Barrett, Sonia Goncalves, Ewan Harrison, David K. Jackson, Ian Johnston, Dominic Kwiatkowski, Cordelia Langford, John Sillitoe on behalf of the Wellcome Sanger Institute COVID-19 Surveillance Team |
| EPI_ISL_3225660, EPI_ISL_3225661, EPI_ISL_3225662 | Lighthouse Lab in Milton Keynes | Wellcome Sanger Institute for the COVID-19 Genomics UK (COG-UK) Consortium | The Lighthouse Lab in Milton Keynes and Alex Alderton, Roberto Amato, Jeffrey Barrett, Sonia Goncalves, Ewan Harrison, David K. Jackson, Ian Johnston, Dominic Kwiatkowski, Cordelia Langford, John Sillitoe on behalf of the Wellcome Sanger Institute COVID-19 Surveillance Team |
| EPI_ISL_3225663 | Lighthouse Laboratory Plymouth | Wellcome Sanger Institute for the COVID-19 Genomics UK (COG-UK) Consortium | Lighthouse Laboratory Plymouth and Alex Alderton, Roberto Amato, Jeffrey Barrett, Sonia Goncalves, Ewan Harrison, David K. Jackson, Ian Johnston, Dominic Kwiatkowski, Cordelia Langford, John Sillitoe on behalf of the Wellcome Sanger Institute COVID-19 Surveillance Team |
| EPI_ISL_3225664, EPI_ISL_3225665 | Lighthouse Lab in Milton Keynes | Wellcome Sanger Institute for the COVID-19 Genomics UK (COG-UK) Consortium | The Lighthouse Lab in Milton Keynes and Alex Alderton, Roberto Amato, Jeffrey Barrett, Sonia Goncalves, Ewan Harrison, David K. Jackson, Ian Johnston, Dominic Kwiatkowski, Cordelia Langford, John Sillitoe on behalf of the Wellcome Sanger Institute COVID-19 Surveillance Team |
| EPI_ISL_3225666, EPI_ISL_3225667 | Lighthouse Laboratory Plymouth | Wellcome Sanger Institute for the COVID-19 Genomics UK (COG-UK) Consortium | Lighthouse Laboratory Plymouth and Alex Alderton, Roberto Amato, Jeffrey Barrett, Sonia Goncalves, Ewan Harrison, David K. Jackson, Ian Johnston, Dominic Kwiatkowski, Cordelia Langford, John Sillitoe on behalf of the Wellcome Sanger Institute COVID-19 Surveillance Team |
| EPI_ISL_3225668 | Lighthouse Lab in Milton Keynes | Wellcome Sanger Institute for the COVID-19 Genomics UK (COG-UK) Consortium | The Lighthouse Lab in Milton Keynes and Alex Alderton, Roberto Amato, Jeffrey Barrett, Sonia Goncalves, Ewan Harrison, David K. Jackson, Ian Johnston, Dominic Kwiatkowski, Cordelia Langford, John Sillitoe on behalf of the Wellcome Sanger Institute COVID-19 Surveillance Team |
| EPI_ISL_3225669 | Lighthouse Lab in Glasgow | Wellcome Sanger Institute for the COVID-19 Genomics UK (COG-UK) Consortium | Harper VanSteenhouse, Yumi Kasai, David Gray, Carol Clugston, Anna Dominiczak and Alex Alderton, Roberto Amato, Jeffrey Barrett, Sonia Goncalves, Ewan Harrison, David K. Jackson, Ian Johnston, Dominic Kwiatkowski, Cordelia Langford, John Sillitoe on behalf of the Wellcome Sanger Institute COVID-19 Surveillance Team |
| EPI_ISL_3225670, EPI_ISL_3225671, EPI_ISL_3225672 | Lighthouse Laboratory Plymouth | Wellcome Sanger Institute for the COVID-19 Genomics UK (COG-UK) Consortium | Lighthouse Laboratory Plymouth and Alex Alderton, Roberto Amato, Jeffrey Barrett, Sonia Goncalves, Ewan Harrison, David K. Jackson, Ian Johnston, Dominic Kwiatkowski, Cordelia Langford, John Sillitoe on behalf of the Wellcome Sanger Institute COVID-19 Surveillance Team |
| EPI_ISL_3225673, EPI_ISL_3225674, EPI_ISL_3225675, EPI_ISL_3225676, EPI_ISL_3225677, EPI_ISL_3225678 | Lighthouse Lab in Milton Keynes | Wellcome Sanger Institute for the COVID-19 Genomics UK (COG-UK) Consortium | The Lighthouse Lab in Milton Keynes and Alex Alderton, Roberto Amato, Jeffrey Barrett, Sonia Goncalves, Ewan Harrison, David K. Jackson, Ian Johnston, Dominic Kwiatkowski, Cordelia Langford, John Sillitoe on behalf of the Wellcome Sanger Institute COVID-19 Surveillance Team |
| EPI_ISL_3225679 | Lighthouse Laboratory Plymouth | Wellcome Sanger Institute for the COVID-19 Genomics UK (COG-UK) Consortium | Lighthouse Laboratory Plymouth and Alex Alderton, Roberto Amato, Jeffrey Barrett, Sonia Goncalves, Ewan Harrison, David K. Jackson, Ian Johnston, Dominic Kwiatkowski, Cordelia Langford, John Sillitoe on behalf of the Wellcome Sanger Institute COVID-19 Surveillance Team |
| EPI_ISL_3225680 | Lighthouse Lab in Glasgow | Wellcome Sanger Institute for the COVID-19 Genomics UK (COG-UK) Consortium | Harper VanSteenhouse, Yumi Kasai, David Gray, Carol Clugston, Anna Dominiczak and Alex Alderton, Roberto Amato, Jeffrey Barrett, Sonia Goncalves, Ewan Harrison, David K. Jackson, Ian Johnston, Dominic Kwiatkowski, Cordelia Langford, John Sillitoe on behalf of the Wellcome Sanger Institute COVID-19 Surveillance Team |
| EPI_ISL_3225681 | Lighthouse Laboratory Plymouth | Wellcome Sanger Institute for the COVID-19 Genomics UK (COG-UK) Consortium | Lighthouse Laboratory Plymouth and Alex Alderton, Roberto Amato, Jeffrey Barrett, Sonia Goncalves, Ewan Harrison, David K. Jackson, Ian Johnston, Dominic Kwiatkowski, Cordelia Langford, John Sillitoe on behalf of the Wellcome Sanger Institute COVID-19 Surveillance Team |
| EPI_ISL_3225682 | Lighthouse Lab in Glasgow | Wellcome Sanger Institute for the COVID-19 Genomics UK (COG-UK) Consortium | Harper VanSteenhouse, Yumi Kasai, David Gray, Carol Clugston, Anna Dominiczak and Alex Alderton, Roberto Amato, Jeffrey Barrett, Sonia Goncalves, Ewan Harrison, David K. Jackson, Ian Johnston, Dominic Kwiatkowski, Cordelia Langford, John Sillitoe on behalf of the Wellcome Sanger Institute COVID-19 Surveillance Team |
| EPI_ISL_3225683 | Lighthouse Laboratory Plymouth | Wellcome Sanger Institute for the COVID-19 Genomics UK (COG-UK) Consortium | Lighthouse Laboratory Plymouth and Alex Alderton, Roberto Amato, Jeffrey Barrett, Sonia Goncalves, Ewan Harrison, David K. Jackson, Ian Johnston, Dominic Kwiatkowski, Cordelia Langford, John Sillitoe on behalf of the Wellcome Sanger Institute COVID-19 Surveillance Team |
| EPI_ISL_3225684 | Lighthouse Lab in Milton Keynes | Wellcome Sanger Institute for the COVID-19 Genomics UK (COG-UK) Consortium | The Lighthouse Lab in Milton Keynes and Alex Alderton, Roberto Amato, Jeffrey Barrett, Sonia Goncalves, Ewan Harrison, David K. Jackson, Ian Johnston, Dominic Kwiatkowski, Cordelia Langford, John Sillitoe on behalf of the Wellcome Sanger Institute COVID-19 Surveillance Team |
| EPI_ISL_3225685 | Lighthouse Laboratory Plymouth | Wellcome Sanger Institute for the COVID-19 Genomics UK (COG-UK) Consortium | Lighthouse Laboratory Plymouth and Alex Alderton, Roberto Amato, Jeffrey Barrett, Sonia Goncalves, Ewan Harrison, David K. Jackson, Ian Johnston, Dominic Kwiatkowski, Cordelia Langford, John Sillitoe on behalf of the Wellcome Sanger Institute COVID-19 Surveillance Team |
| EPI_ISL_3225686 | Lighthouse Lab in Milton Keynes | Wellcome Sanger Institute for the COVID-19 Genomics UK (COG-UK) Consortium | The Lighthouse Lab in Milton Keynes and Alex Alderton, Roberto Amato, Jeffrey Barrett, Sonia Goncalves, Ewan Harrison, David K. Jackson, Ian Johnston, Dominic Kwiatkowski, Cordelia Langford, John Sillitoe on behalf of the Wellcome Sanger Institute COVID-19 Surveillance Team |
| EPI_ISL_3225687 | Lighthouse Laboratory Plymouth | Wellcome Sanger Institute for the COVID-19 Genomics UK (COG-UK) Consortium | Lighthouse Laboratory Plymouth and Alex Alderton, Roberto Amato, Jeffrey Barrett, Sonia Goncalves, Ewan Harrison, David K. Jackson, Ian Johnston, Dominic Kwiatkowski, Cordelia Langford, John Sillitoe on behalf of the Wellcome Sanger Institute COVID-19 Surveillance Team |
| EPI_ISL_3225688, EPI_ISL_3225689 | Lighthouse Lab in Milton Keynes | Wellcome Sanger Institute for the COVID-19 Genomics UK (COG-UK) Consortium | The Lighthouse Lab in Milton Keynes and Alex Alderton, Roberto Amato, Jeffrey Barrett, Sonia Goncalves, Ewan Harrison, David K. Jackson, Ian Johnston, Dominic Kwiatkowski, Cordelia Langford, John Sillitoe on behalf of the Wellcome Sanger Institute COVID-19 Surveillance Team |
| EPI_ISL_3225690 | Lighthouse Laboratory Plymouth | Wellcome Sanger Institute for the COVID-19 Genomics UK (COG-UK) Consortium | Lighthouse Laboratory Plymouth and Alex Alderton, Roberto Amato, Jeffrey Barrett, Sonia Goncalves, Ewan Harrison, David K. Jackson, Ian Johnston, Dominic Kwiatkowski, Cordelia Langford, John Sillitoe on behalf of the Wellcome Sanger Institute COVID-19 Surveillance Team |
| EPI_ISL_3225691, EPI_ISL_3225692, | Lighthouse Lab in Milton Keynes | Wellcome Sanger Institute for the COVID-19 Genomics UK (COG-UK) Consortium | The Lighthouse Lab in Milton Keynes and Alex Alderton, Roberto Amato, Jeffrey Barrett, Sonia Goncalves, Ewan Harrison, David K. Jackson, Ian Johnston, Dominic Kwiatkowski, Cordelia Langford, John Sillitoe on behalf of the Wellcome Sanger Institute COVID-19 Surveillance Team |

| | | | |
|---|---|---|---|
| EPI_ISL_3225693, EPI_ISL_3225694 | | | |
| EPI_ISL_3225695 | Lighthouse Lab in Glasgow | Wellcome Sanger Institute for the COVID-19 Genomics UK (COG-UK) Consortium | Harper VanSteenhouse, Yumi Kasai, David Gray, Carol Clugston, Anna Dominiczak and Alex Alderton, Roberto Amato, Jeffrey Barrett, Sonia Goncalves, Ewan Harrison, David K. Jackson, Ian Johnston, Dominic Kwiatkowski, Cordelia Langford, John Sillitoe on behalf of the Wellcome Sanger Institute COVID-19 Surveillance Team |
| EPI_ISL_3225696 | Lighthouse Lab in Milton Keynes | Wellcome Sanger Institute for the COVID-19 Genomics UK (COG-UK) Consortium | The Lighthouse Lab in Milton Keynes and Alex Alderton, Roberto Amato, Jeffrey Barrett, Sonia Goncalves, Ewan Harrison, David K. Jackson, Ian Johnston, Dominic Kwiatkowski, Cordelia Langford, John Sillitoe on behalf of the Wellcome Sanger Institute COVID-19 Surveillance Team |
| EPI_ISL_3225697 | Lighthouse Laboratory Plymouth | Wellcome Sanger Institute for the COVID-19 Genomics UK (COG-UK) Consortium | Lighthouse Laboratory Plymouth and Alex Alderton, Roberto Amato, Jeffrey Barrett, Sonia Goncalves, Ewan Harrison, David K. Jackson, Ian Johnston, Dominic Kwiatkowski, Cordelia Langford, John Sillitoe on behalf of the Wellcome Sanger Institute COVID-19 Surveillance Team |
| EPI_ISL_3225698 | Lighthouse Lab in Milton Keynes | Wellcome Sanger Institute for the COVID-19 Genomics UK (COG-UK) Consortium | The Lighthouse Lab in Milton Keynes and Alex Alderton, Roberto Amato, Jeffrey Barrett, Sonia Goncalves, Ewan Harrison, David K. Jackson, Ian Johnston, Dominic Kwiatkowski, Cordelia Langford, John Sillitoe on behalf of the Wellcome Sanger Institute COVID-19 Surveillance Team |
| EPI_ISL_3225699, EPI_ISL_3225700 | Lighthouse Lab in Glasgow | Wellcome Sanger Institute for the COVID-19 Genomics UK (COG-UK) Consortium | Harper VanSteenhouse, Yumi Kasai, David Gray, Carol Clugston, Anna Dominiczak and Alex Alderton, Roberto Amato, Jeffrey Barrett, Sonia Goncalves, Ewan Harrison, David K. Jackson, Ian Johnston, Dominic Kwiatkowski, Cordelia Langford, John Sillitoe on behalf of the Wellcome Sanger Institute COVID-19 Surveillance Team |
| EPI_ISL_3225701 | Lighthouse Laboratory Plymouth | Wellcome Sanger Institute for the COVID-19 Genomics UK (COG-UK) Consortium | Lighthouse Laboratory Plymouth and Alex Alderton, Roberto Amato, Jeffrey Barrett, Sonia Goncalves, Ewan Harrison, David K. Jackson, Ian Johnston, Dominic Kwiatkowski, Cordelia Langford, John Sillitoe on behalf of the Wellcome Sanger Institute COVID-19 Surveillance Team |
| EPI_ISL_3225702, EPI_ISL_3225703, EPI_ISL_3225704 | Lighthouse Lab in Glasgow | Wellcome Sanger Institute for the COVID-19 Genomics UK (COG-UK) Consortium | Harper VanSteenhouse, Yumi Kasai, David Gray, Carol Clugston, Anna Dominiczak and Alex Alderton, Roberto Amato, Jeffrey Barrett, Sonia Goncalves, Ewan Harrison, David K. Jackson, Ian Johnston, Dominic Kwiatkowski, Cordelia Langford, John Sillitoe on behalf of the Wellcome Sanger Institute COVID-19 Surveillance Team |
| EPI_ISL_3225705 | Lighthouse Laboratory Plymouth | Wellcome Sanger Institute for the COVID-19 Genomics UK (COG-UK) Consortium | Lighthouse Laboratory Plymouth and Alex Alderton, Roberto Amato, Jeffrey Barrett, Sonia Goncalves, Ewan Harrison, David K. Jackson, Ian Johnston, Dominic Kwiatkowski, Cordelia Langford, John Sillitoe on behalf of the Wellcome Sanger Institute COVID-19 Surveillance Team |
| EPI_ISL_3225706, EPI_ISL_3225707, EPI_ISL_3225708, EPI_ISL_3225709, EPI_ISL_3225710 | Lighthouse Lab in Milton Keynes | Wellcome Sanger Institute for the COVID-19 Genomics UK (COG-UK) Consortium | The Lighthouse Lab in Milton Keynes and Alex Alderton, Roberto Amato, Jeffrey Barrett, Sonia Goncalves, Ewan Harrison, David K. Jackson, Ian Johnston, Dominic Kwiatkowski, Cordelia Langford, John Sillitoe on behalf of the Wellcome Sanger Institute COVID-19 Surveillance Team |
| EPI_ISL_3225711, EPI_ISL_3225712, EPI_ISL_3225713 | Lighthouse Laboratory Plymouth | Wellcome Sanger Institute for the COVID-19 Genomics UK (COG-UK) Consortium | Lighthouse Laboratory Plymouth and Alex Alderton, Roberto Amato, Jeffrey Barrett, Sonia Goncalves, Ewan Harrison, David K. Jackson, Ian Johnston, Dominic Kwiatkowski, Cordelia Langford, John Sillitoe on behalf of the Wellcome Sanger Institute COVID-19 Surveillance Team |
| EPI_ISL_3225714, EPI_ISL_3225715 | Lighthouse Lab in Milton Keynes | Wellcome Sanger Institute for the COVID-19 Genomics UK (COG-UK) Consortium | The Lighthouse Lab in Milton Keynes and Alex Alderton, Roberto Amato, Jeffrey Barrett, Sonia Goncalves, Ewan Harrison, David K. Jackson, Ian Johnston, Dominic Kwiatkowski, Cordelia Langford, John Sillitoe on behalf of the Wellcome Sanger Institute COVID-19 Surveillance Team |
| EPI_ISL_3225716 | Lighthouse Lab in Glasgow | Wellcome Sanger Institute for the COVID-19 Genomics UK (COG-UK) Consortium | Harper VanSteenhouse, Yumi Kasai, David Gray, Carol Clugston, Anna Dominiczak and Alex Alderton, Roberto Amato, Jeffrey Barrett, Sonia Goncalves, Ewan Harrison, David K. Jackson, Ian Johnston, Dominic Kwiatkowski, Cordelia Langford, John Sillitoe on behalf of the Wellcome Sanger Institute COVID-19 Surveillance Team |
| EPI_ISL_3225717, EPI_ISL_3225718 | Lighthouse Lab in Milton Keynes | Wellcome Sanger Institute for the COVID-19 Genomics UK (COG-UK) Consortium | The Lighthouse Lab in Milton Keynes and Alex Alderton, Roberto Amato, Jeffrey Barrett, Sonia Goncalves, Ewan Harrison, David K. Jackson, Ian Johnston, Dominic Kwiatkowski, Cordelia Langford, John Sillitoe on behalf of the Wellcome Sanger Institute COVID-19 Surveillance Team |
| EPI_ISL_3225719, EPI_ISL_3225720, EPI_ISL_3225721, EPI_ISL_3225722 | Lighthouse Lab in Glasgow | Wellcome Sanger Institute for the COVID-19 Genomics UK (COG-UK) Consortium | Harper VanSteenhouse, Yumi Kasai, David Gray, Carol Clugston, Anna Dominiczak and Alex Alderton, Roberto Amato, Jeffrey Barrett, Sonia Goncalves, Ewan Harrison, David K. Jackson, Ian Johnston, Dominic Kwiatkowski, Cordelia Langford, John Sillitoe on behalf of the Wellcome Sanger Institute COVID-19 Surveillance Team |
| EPI_ISL_3225723 | Lighthouse Lab in Milton Keynes | Wellcome Sanger Institute for the COVID-19 Genomics UK (COG-UK) Consortium | The Lighthouse Lab in Milton Keynes and Alex Alderton, Roberto Amato, Jeffrey Barrett, Sonia Goncalves, Ewan Harrison, David K. Jackson, Ian Johnston, Dominic Kwiatkowski, Cordelia Langford, John Sillitoe on behalf of the Wellcome Sanger Institute COVID-19 Surveillance Team |
| EPI_ISL_3225724 | Lighthouse Lab in Glasgow | Wellcome Sanger Institute for the COVID-19 Genomics UK (COG-UK) Consortium | Harper VanSteenhouse, Yumi Kasai, David Gray, Carol Clugston, Anna Dominiczak and Alex Alderton, Roberto Amato, Jeffrey Barrett, Sonia Goncalves, Ewan Harrison, David K. Jackson, Ian Johnston, Dominic Kwiatkowski, Cordelia Langford, John Sillitoe on behalf of the Wellcome Sanger Institute COVID-19 Surveillance Team |
| EPI_ISL_3225725 | Lighthouse Laboratory Plymouth | Wellcome Sanger Institute for the COVID-19 Genomics UK (COG-UK) Consortium | Lighthouse Laboratory Plymouth and Alex Alderton, Roberto Amato, Jeffrey Barrett, Sonia Goncalves, Ewan Harrison, David K. Jackson, Ian Johnston, Dominic Kwiatkowski, Cordelia Langford, John Sillitoe on behalf of the Wellcome Sanger Institute COVID-19 Surveillance Team |
| EPI_ISL_3225726, EPI_ISL_3225727, EPI_ISL_3225728 | Lighthouse Lab in Milton Keynes | Wellcome Sanger Institute for the COVID-19 Genomics UK (COG-UK) Consortium | The Lighthouse Lab in Milton Keynes and Alex Alderton, Roberto Amato, Jeffrey Barrett, Sonia Goncalves, Ewan Harrison, David K. Jackson, Ian Johnston, Dominic Kwiatkowski, Cordelia Langford, John Sillitoe on behalf of the Wellcome Sanger Institute COVID-19 Surveillance Team |
| EPI_ISL_3225729 | Lighthouse Lab in Glasgow | Wellcome Sanger Institute for the COVID-19 Genomics UK (COG-UK) Consortium | Harper VanSteenhouse, Yumi Kasai, David Gray, Carol Clugston, Anna Dominiczak and Alex Alderton, Roberto Amato, Jeffrey Barrett, Sonia Goncalves, Ewan Harrison, David K. Jackson, Ian Johnston, Dominic Kwiatkowski, Cordelia Langford, John Sillitoe on behalf of the Wellcome Sanger Institute COVID-19 Surveillance Team |
| EPI_ISL_3225730 | Lighthouse Lab in Milton Keynes | Wellcome Sanger Institute for the COVID-19 Genomics UK (COG-UK) Consortium | The Lighthouse Lab in Milton Keynes and Alex Alderton, Roberto Amato, Jeffrey Barrett, Sonia Goncalves, Ewan Harrison, David K. Jackson, Ian Johnston, Dominic Kwiatkowski, Cordelia Langford, John Sillitoe on behalf of the Wellcome Sanger Institute COVID-19 Surveillance Team |
| EPI_ISL_3225731 | Lighthouse Lab in Glasgow | Wellcome Sanger Institute for the COVID-19 Genomics UK (COG-UK) Consortium | Harper VanSteenhouse, Yumi Kasai, David Gray, Carol Clugston, Anna Dominiczak and Alex Alderton, Roberto Amato, Jeffrey Barrett, Sonia Goncalves, Ewan Harrison, David K. Jackson, Ian Johnston, Dominic Kwiatkowski, Cordelia Langford, John Sillitoe on behalf of the Wellcome Sanger Institute COVID-19 Surveillance Team |
| EPI_ISL_3225732, EPI_ISL_3225733, EPI_ISL_3225734 | Lighthouse Lab in Milton Keynes | Wellcome Sanger Institute for the COVID-19 Genomics UK (COG-UK) Consortium | The Lighthouse Lab in Milton Keynes and Alex Alderton, Roberto Amato, Jeffrey Barrett, Sonia Goncalves, Ewan Harrison, David K. Jackson, Ian Johnston, Dominic Kwiatkowski, Cordelia Langford, John Sillitoe on behalf of the Wellcome Sanger Institute COVID-19 Surveillance Team |
| EPI_ISL_3225735 | Lighthouse Laboratory Plymouth | Wellcome Sanger Institute for the COVID-19 Genomics UK (COG-UK) Consortium | Lighthouse Laboratory Plymouth and Alex Alderton, Roberto Amato, Jeffrey Barrett, Sonia Goncalves, Ewan Harrison, David K. Jackson, Ian Johnston, Dominic Kwiatkowski, Cordelia Langford, John Sillitoe on behalf of the Wellcome Sanger Institute COVID-19 Surveillance Team |
| EPI_ISL_3225736, EPI_ISL_3225737, EPI_ISL_3225738, EPI_ISL_3225739, EPI_ISL_3225740 | Lighthouse Lab in Milton Keynes | Wellcome Sanger Institute for the COVID-19 Genomics UK (COG-UK) Consortium | The Lighthouse Lab in Milton Keynes and Alex Alderton, Roberto Amato, Jeffrey Barrett, Sonia Goncalves, Ewan Harrison, David K. Jackson, Ian Johnston, Dominic Kwiatkowski, Cordelia Langford, John Sillitoe on behalf of the Wellcome Sanger Institute COVID-19 Surveillance Team |
| EPI_ISL_3225741 | Lighthouse Laboratory Plymouth | Wellcome Sanger Institute for the COVID-19 Genomics UK (COG-UK) Consortium | Lighthouse Laboratory Plymouth and Alex Alderton, Roberto Amato, Jeffrey Barrett, Sonia Goncalves, Ewan Harrison, David K. Jackson, Ian Johnston, Dominic Kwiatkowski, Cordelia Langford, John Sillitoe on behalf of the Wellcome Sanger Institute COVID-19 Surveillance Team |
| EPI_ISL_3225742, EPI_ISL_3225743 | Lighthouse Lab in Glasgow | Wellcome Sanger Institute for the COVID-19 Genomics UK (COG-UK) Consortium | Harper VanSteenhouse, Yumi Kasai, David Gray, Carol Clugston, Anna Dominiczak and Alex Alderton, Roberto Amato, Jeffrey Barrett, Sonia Goncalves, Ewan Harrison, David K. Jackson, Ian Johnston, Dominic Kwiatkowski, Cordelia Langford, John Sillitoe on behalf of the Wellcome Sanger Institute COVID-19 Surveillance Team |
| EPI_ISL_3225744 | Lighthouse Lab in Milton Keynes | Wellcome Sanger Institute for the COVID-19 Genomics UK (COG-UK) Consortium | The Lighthouse Lab in Milton Keynes and Alex Alderton, Roberto Amato, Jeffrey Barrett, Sonia Goncalves, Ewan Harrison, David K. Jackson, Ian Johnston, Dominic Kwiatkowski, Cordelia Langford, John Sillitoe on behalf of the Wellcome Sanger Institute COVID-19 Surveillance Team |
| EPI_ISL_3225745, EPI_ISL_3225746 | Lighthouse Laboratory Plymouth | Wellcome Sanger Institute for the COVID-19 Genomics UK (COG-UK) Consortium | Lighthouse Laboratory Plymouth and Alex Alderton, Roberto Amato, Jeffrey Barrett, Sonia Goncalves, Ewan Harrison, David K. Jackson, Ian Johnston, Dominic Kwiatkowski, Cordelia Langford, John Sillitoe on behalf of the Wellcome Sanger Institute COVID-19 Surveillance Team |
| EPI_ISL_3225747 | Lighthouse Lab in Milton Keynes | Wellcome Sanger Institute for the COVID-19 Genomics UK (COG-UK) Consortium | The Lighthouse Lab in Milton Keynes and Alex Alderton, Roberto Amato, Jeffrey Barrett, Sonia Goncalves, Ewan Harrison, David K. Jackson, Ian Johnston, Dominic Kwiatkowski, Cordelia Langford, John Sillitoe on behalf of the Wellcome Sanger Institute COVID-19 Surveillance Team |
| EPI_ISL_3225748 | Lighthouse Lab in Glasgow | Wellcome Sanger Institute for the COVID-19 Genomics UK (COG-UK) Consortium | Harper VanSteenhouse, Yumi Kasai, David Gray, Carol Clugston, Anna Dominiczak and Alex Alderton, Roberto Amato, Jeffrey Barrett, Sonia Goncalves, Ewan Harrison, David K. Jackson, Ian Johnston, Dominic Kwiatkowski, Cordelia Langford, John Sillitoe on behalf of the Wellcome Sanger Institute COVID-19 Surveillance Team |
| EPI_ISL_3225749 | Lighthouse Lab in Milton Keynes | Wellcome Sanger Institute for the COVID-19 Genomics UK (COG-UK) Consortium | The Lighthouse Lab in Milton Keynes and Alex Alderton, Roberto Amato, Jeffrey Barrett, Sonia Goncalves, Ewan Harrison, David K. Jackson, Ian Johnston, Dominic Kwiatkowski, Cordelia Langford, John Sillitoe on behalf of the Wellcome Sanger Institute COVID-19 Surveillance Team |
| EPI_ISL_3225750 | Lighthouse Laboratory Plymouth | Wellcome Sanger Institute for the COVID-19 Genomics UK (COG-UK) Consortium | Lighthouse Laboratory Plymouth and Alex Alderton, Roberto Amato, Jeffrey Barrett, Sonia Goncalves, Ewan Harrison, David K. Jackson, Ian Johnston, Dominic Kwiatkowski, Cordelia Langford, John Sillitoe on behalf of the Wellcome Sanger Institute COVID-19 Surveillance Team |
| EPI_ISL_3225751 | Lighthouse Lab in Milton Keynes | Wellcome Sanger Institute for the COVID-19 Genomics UK (COG-UK) Consortium | The Lighthouse Lab in Milton Keynes and Alex Alderton, Roberto Amato, Jeffrey Barrett, Sonia Goncalves, Ewan Harrison, David K. Jackson, Ian Johnston, Dominic Kwiatkowski, Cordelia Langford, John Sillitoe on behalf of the Wellcome Sanger Institute COVID-19 Surveillance Team |
| EPI_ISL_3225752 | Lighthouse Laboratory Plymouth | Wellcome Sanger Institute for the COVID-19 Genomics UK (COG-UK) Consortium | Lighthouse Laboratory Plymouth and Alex Alderton, Roberto Amato, Jeffrey Barrett, Sonia Goncalves, Ewan Harrison, David K. Jackson, Ian Johnston, Dominic Kwiatkowski, Cordelia Langford, John Sillitoe on behalf of the Wellcome Sanger Institute COVID-19 Surveillance Team |
| EPI_ISL_3225753 | Lighthouse Lab in Milton Keynes | Wellcome Sanger Institute for the COVID-19 Genomics UK (COG-UK) Consortium | The Lighthouse Lab in Milton Keynes and Alex Alderton, Roberto Amato, Jeffrey Barrett, Sonia Goncalves, Ewan Harrison, David K. Jackson, Ian Johnston, Dominic Kwiatkowski, Cordelia Langford, John Sillitoe on behalf of the Wellcome Sanger Institute COVID-19 Surveillance Team |
| EPI_ISL_3225754 | Lighthouse Lab in Glasgow | Wellcome Sanger Institute for the COVID-19 Genomics UK (COG-UK) Consortium | Harper VanSteenhouse, Yumi Kasai, David Gray, Carol Clugston, Anna Dominiczak and Alex Alderton, Roberto Amato, Jeffrey Barrett, Sonia Goncalves, Ewan Harrison, David K. Jackson, Ian Johnston, Dominic Kwiatkowski, Cordelia Langford, John Sillitoe on behalf of the Wellcome Sanger Institute COVID-19 Surveillance Team |
| EPI_ISL_3225755, EPI_ISL_3225756 | Lighthouse Lab in Milton Keynes | Wellcome Sanger Institute for the COVID-19 Genomics UK (COG-UK) Consortium | The Lighthouse Lab in Milton Keynes and Alex Alderton, Roberto Amato, Jeffrey Barrett, Sonia Goncalves, Ewan Harrison, David K. Jackson, Ian Johnston, Dominic Kwiatkowski, Cordelia Langford, John Sillitoe on behalf of the Wellcome Sanger Institute COVID-19 Surveillance Team |
| EPI_ISL_3225757, EPI_ISL_3225758 | Lighthouse Lab in Glasgow | Wellcome Sanger Institute for the COVID-19 Genomics UK (COG-UK) Consortium | Harper VanSteenhouse, Yumi Kasai, David Gray, Carol Clugston, Anna Dominiczak and Alex Alderton, Roberto Amato, Jeffrey Barrett, Sonia Goncalves, Ewan Harrison, David K. Jackson, Ian Johnston, Dominic Kwiatkowski, Cordelia Langford, John Sillitoe on behalf of the Wellcome Sanger Institute COVID-19 Surveillance Team |
| EPI_ISL_3225759, EPI_ISL_3225760, EPI_ISL_3225761 | Lighthouse Lab in Milton Keynes | Wellcome Sanger Institute for the COVID-19 Genomics UK (COG-UK) Consortium | The Lighthouse Lab in Milton Keynes and Alex Alderton, Roberto Amato, Jeffrey Barrett, Sonia Goncalves, Ewan Harrison, David K. Jackson, Ian Johnston, Dominic Kwiatkowski, Cordelia Langford, John Sillitoe on behalf of the Wellcome Sanger Institute COVID-19 Surveillance Team |

| | | | |
|---|---|---|---|
| EPI_ISL_3225762 | Lighthouse Lab in Glasgow | Wellcome Sanger Institute for the COVID-19 Genomics UK (COG-UK) Consortium | Harper VanSteenhouse, Yumi Kasai, David Gray, Carol Clugston, Anna Dominiczak and Alex Alderton, Roberto Amato, Jeffrey Barrett, Sonia Goncalves, Ewan Harrison, David K. Jackson, Ian Johnston, Dominic Kwiatkowski, Cordelia Langford, John Sillitoe on behalf of the Wellcome Sanger Institute COVID-19 Surveillance Team |
| EPI_ISL_3225763 | Lighthouse Lab in Milton Keynes | Wellcome Sanger Institute for the COVID-19 Genomics UK (COG-UK) Consortium | The Lighthouse Lab in Milton Keynes and Alex Alderton, Roberto Amato, Jeffrey Barrett, Sonia Goncalves, Ewan Harrison, David K. Jackson, Ian Johnston, Dominic Kwiatkowski, Cordelia Langford, John Sillitoe on behalf of the Wellcome Sanger Institute COVID-19 Surveillance Team |
| EPI_ISL_3225764 | Lighthouse Lab in Glasgow | Wellcome Sanger Institute for the COVID-19 Genomics UK (COG-UK) Consortium | Harper VanSteenhouse, Yumi Kasai, David Gray, Carol Clugston, Anna Dominiczak and Alex Alderton, Roberto Amato, Jeffrey Barrett, Sonia Goncalves, Ewan Harrison, David K. Jackson, Ian Johnston, Dominic Kwiatkowski, Cordelia Langford, John Sillitoe on behalf of the Wellcome Sanger Institute COVID-19 Surveillance Team |
| EPI_ISL_3225765 | Lighthouse Lab in Milton Keynes | Wellcome Sanger Institute for the COVID-19 Genomics UK (COG-UK) Consortium | The Lighthouse Lab in Milton Keynes and Alex Alderton, Roberto Amato, Jeffrey Barrett, Sonia Goncalves, Ewan Harrison, David K. Jackson, Ian Johnston, Dominic Kwiatkowski, Cordelia Langford, John Sillitoe on behalf of the Wellcome Sanger Institute COVID-19 Surveillance Team |
| EPI_ISL_3225766 | Lighthouse Lab in Glasgow | Wellcome Sanger Institute for the COVID-19 Genomics UK (COG-UK) Consortium | Harper VanSteenhouse, Yumi Kasai, David Gray, Carol Clugston, Anna Dominiczak and Alex Alderton, Roberto Amato, Jeffrey Barrett, Sonia Goncalves, Ewan Harrison, David K. Jackson, Ian Johnston, Dominic Kwiatkowski, Cordelia Langford, John Sillitoe on behalf of the Wellcome Sanger Institute COVID-19 Surveillance Team |
| EPI_ISL_3225767 | Lighthouse Laboratory Plymouth | Wellcome Sanger Institute for the COVID-19 Genomics UK (COG-UK) Consortium | Lighthouse Laboratory Plymouth and Alex Alderton, Roberto Amato, Jeffrey Barrett, Sonia Goncalves, Ewan Harrison, David K. Jackson, Ian Johnston, Dominic Kwiatkowski, Cordelia Langford, John Sillitoe on behalf of the Wellcome Sanger Institute COVID-19 Surveillance Team |
| EPI_ISL_3225768 | Lighthouse Lab in Milton Keynes | Wellcome Sanger Institute for the COVID-19 Genomics UK (COG-UK) Consortium | The Lighthouse Lab in Milton Keynes and Alex Alderton, Roberto Amato, Jeffrey Barrett, Sonia Goncalves, Ewan Harrison, David K. Jackson, Ian Johnston, Dominic Kwiatkowski, Cordelia Langford, John Sillitoe on behalf of the Wellcome Sanger Institute COVID-19 Surveillance Team |
| EPI_ISL_3225769 | Lighthouse Lab in Glasgow | Wellcome Sanger Institute for the COVID-19 Genomics UK (COG-UK) Consortium | Harper VanSteenhouse, Yumi Kasai, David Gray, Carol Clugston, Anna Dominiczak and Alex Alderton, Roberto Amato, Jeffrey Barrett, Sonia Goncalves, Ewan Harrison, David K. Jackson, Ian Johnston, Dominic Kwiatkowski, Cordelia Langford, John Sillitoe on behalf of the Wellcome Sanger Institute COVID-19 Surveillance Team |
| EPI_ISL_3225770 | Lighthouse Lab in Milton Keynes | Wellcome Sanger Institute for the COVID-19 Genomics UK (COG-UK) Consortium | The Lighthouse Lab in Milton Keynes and Alex Alderton, Roberto Amato, Jeffrey Barrett, Sonia Goncalves, Ewan Harrison, David K. Jackson, Ian Johnston, Dominic Kwiatkowski, Cordelia Langford, John Sillitoe on behalf of the Wellcome Sanger Institute COVID-19 Surveillance Team |
| EPI_ISL_3225771 | Lighthouse Laboratory Plymouth | Wellcome Sanger Institute for the COVID-19 Genomics UK (COG-UK) Consortium | Lighthouse Laboratory Plymouth and Alex Alderton, Roberto Amato, Jeffrey Barrett, Sonia Goncalves, Ewan Harrison, David K. Jackson, Ian Johnston, Dominic Kwiatkowski, Cordelia Langford, John Sillitoe on behalf of the Wellcome Sanger Institute COVID-19 Surveillance Team |
| EPI_ISL_3225772, EPI_ISL_3225773, EPI_ISL_3225774 | Lighthouse Lab in Glasgow | Wellcome Sanger Institute for the COVID-19 Genomics UK (COG-UK) Consortium | Harper VanSteenhouse, Yumi Kasai, David Gray, Carol Clugston, Anna Dominiczak and Alex Alderton, Roberto Amato, Jeffrey Barrett, Sonia Goncalves, Ewan Harrison, David K. Jackson, Ian Johnston, Dominic Kwiatkowski, Cordelia Langford, John Sillitoe on behalf of the Wellcome Sanger Institute COVID-19 Surveillance Team |
| EPI_ISL_3225775, EPI_ISL_3225776, EPI_ISL_3225777, EPI_ISL_3225778 | Lighthouse Lab in Milton Keynes | Wellcome Sanger Institute for the COVID-19 Genomics UK (COG-UK) Consortium | The Lighthouse Lab in Milton Keynes and Alex Alderton, Roberto Amato, Jeffrey Barrett, Sonia Goncalves, Ewan Harrison, David K. Jackson, Ian Johnston, Dominic Kwiatkowski, Cordelia Langford, John Sillitoe on behalf of the Wellcome Sanger Institute COVID-19 Surveillance Team |
| EPI_ISL_3225779, EPI_ISL_3225780 | Lighthouse Laboratory Plymouth | Wellcome Sanger Institute for the COVID-19 Genomics UK (COG-UK) Consortium | Lighthouse Laboratory Plymouth and Alex Alderton, Roberto Amato, Jeffrey Barrett, Sonia Goncalves, Ewan Harrison, David K. Jackson, Ian Johnston, Dominic Kwiatkowski, Cordelia Langford, John Sillitoe on behalf of the Wellcome Sanger Institute COVID-19 Surveillance Team |
| EPI_ISL_3225781 | Lighthouse Lab in Glasgow | Wellcome Sanger Institute for the COVID-19 Genomics UK (COG-UK) Consortium | Harper VanSteenhouse, Yumi Kasai, David Gray, Carol Clugston, Anna Dominiczak and Alex Alderton, Roberto Amato, Jeffrey Barrett, Sonia Goncalves, Ewan Harrison, David K. Jackson, Ian Johnston, Dominic Kwiatkowski, Cordelia Langford, John Sillitoe on behalf of the Wellcome Sanger Institute COVID-19 Surveillance Team |
| EPI_ISL_3225782, EPI_ISL_3225783 | Lighthouse Lab in Milton Keynes | Wellcome Sanger Institute for the COVID-19 Genomics UK (COG-UK) Consortium | The Lighthouse Lab in Milton Keynes and Alex Alderton, Roberto Amato, Jeffrey Barrett, Sonia Goncalves, Ewan Harrison, David K. Jackson, Ian Johnston, Dominic Kwiatkowski, Cordelia Langford, John Sillitoe on behalf of the Wellcome Sanger Institute COVID-19 Surveillance Team |
| EPI_ISL_3225784 | Lighthouse Laboratory Plymouth | Wellcome Sanger Institute for the COVID-19 Genomics UK (COG-UK) Consortium | Lighthouse Laboratory Plymouth and Alex Alderton, Roberto Amato, Jeffrey Barrett, Sonia Goncalves, Ewan Harrison, David K. Jackson, Ian Johnston, Dominic Kwiatkowski, Cordelia Langford, John Sillitoe on behalf of the Wellcome Sanger Institute COVID-19 Surveillance Team |
| EPI_ISL_3225785 | Lighthouse Lab in Glasgow | Wellcome Sanger Institute for the COVID-19 Genomics UK (COG-UK) Consortium | Harper VanSteenhouse, Yumi Kasai, David Gray, Carol Clugston, Anna Dominiczak and Alex Alderton, Roberto Amato, Jeffrey Barrett, Sonia Goncalves, Ewan Harrison, David K. Jackson, Ian Johnston, Dominic Kwiatkowski, Cordelia Langford, John Sillitoe on behalf of the Wellcome Sanger Institute COVID-19 Surveillance Team |
| EPI_ISL_3225786, EPI_ISL_3225787 | Lighthouse Laboratory Plymouth | Wellcome Sanger Institute for the COVID-19 Genomics UK (COG-UK) Consortium | Lighthouse Laboratory Plymouth and Alex Alderton, Roberto Amato, Jeffrey Barrett, Sonia Goncalves, Ewan Harrison, David K. Jackson, Ian Johnston, Dominic Kwiatkowski, Cordelia Langford, John Sillitoe on behalf of the Wellcome Sanger Institute COVID-19 Surveillance Team |
| EPI_ISL_3225788, EPI_ISL_3225789 | Lighthouse Lab in Glasgow | Wellcome Sanger Institute for the COVID-19 Genomics UK (COG-UK) Consortium | Harper VanSteenhouse, Yumi Kasai, David Gray, Carol Clugston, Anna Dominiczak and Alex Alderton, Roberto Amato, Jeffrey Barrett, Sonia Goncalves, Ewan Harrison, David K. Jackson, Ian Johnston, Dominic Kwiatkowski, Cordelia Langford, John Sillitoe on behalf of the Wellcome Sanger Institute COVID-19 Surveillance Team |
| EPI_ISL_3225790 | Lighthouse Lab in Milton Keynes | Wellcome Sanger Institute for the COVID-19 Genomics UK (COG-UK) Consortium | The Lighthouse Lab in Milton Keynes and Alex Alderton, Roberto Amato, Jeffrey Barrett, Sonia Goncalves, Ewan Harrison, David K. Jackson, Ian Johnston, Dominic Kwiatkowski, Cordelia Langford, John Sillitoe on behalf of the Wellcome Sanger Institute COVID-19 Surveillance Team |
| EPI_ISL_3225791, EPI_ISL_3225792 | Lighthouse Lab in Glasgow | Wellcome Sanger Institute for the COVID-19 Genomics UK (COG-UK) Consortium | Harper VanSteenhouse, Yumi Kasai, David Gray, Carol Clugston, Anna Dominiczak and Alex Alderton, Roberto Amato, Jeffrey Barrett, Sonia Goncalves, Ewan Harrison, David K. Jackson, Ian Johnston, Dominic Kwiatkowski, Cordelia Langford, John Sillitoe on behalf of the Wellcome Sanger Institute COVID-19 Surveillance Team |
| EPI_ISL_3225793 | Lighthouse Laboratory Plymouth | Wellcome Sanger Institute for the COVID-19 Genomics UK (COG-UK) Consortium | Lighthouse Laboratory Plymouth and Alex Alderton, Roberto Amato, Jeffrey Barrett, Sonia Goncalves, Ewan Harrison, David K. Jackson, Ian Johnston, Dominic Kwiatkowski, Cordelia Langford, John Sillitoe on behalf of the Wellcome Sanger Institute COVID-19 Surveillance Team |
| EPI_ISL_3225794 | Lighthouse Lab in Glasgow | Wellcome Sanger Institute for the COVID-19 Genomics UK (COG-UK) Consortium | Harper VanSteenhouse, Yumi Kasai, David Gray, Carol Clugston, Anna Dominiczak and Alex Alderton, Roberto Amato, Jeffrey Barrett, Sonia Goncalves, Ewan Harrison, David K. Jackson, Ian Johnston, Dominic Kwiatkowski, Cordelia Langford, John Sillitoe on behalf of the Wellcome Sanger Institute COVID-19 Surveillance Team |
| EPI_ISL_3225795 | Lighthouse Laboratory Plymouth | Wellcome Sanger Institute for the COVID-19 Genomics UK (COG-UK) Consortium | Lighthouse Laboratory Plymouth and Alex Alderton, Roberto Amato, Jeffrey Barrett, Sonia Goncalves, Ewan Harrison, David K. Jackson, Ian Johnston, Dominic Kwiatkowski, Cordelia Langford, John Sillitoe on behalf of the Wellcome Sanger Institute COVID-19 Surveillance Team |
| EPI_ISL_3225796 | Lighthouse Lab in Milton Keynes | Wellcome Sanger Institute for the COVID-19 Genomics UK (COG-UK) Consortium | The Lighthouse Lab in Milton Keynes and Alex Alderton, Roberto Amato, Jeffrey Barrett, Sonia Goncalves, Ewan Harrison, David K. Jackson, Ian Johnston, Dominic Kwiatkowski, Cordelia Langford, John Sillitoe on behalf of the Wellcome Sanger Institute COVID-19 Surveillance Team |
| EPI_ISL_3225797 | Lighthouse Lab in Glasgow | Wellcome Sanger Institute for the COVID-19 Genomics UK (COG-UK) Consortium | Harper VanSteenhouse, Yumi Kasai, David Gray, Carol Clugston, Anna Dominiczak and Alex Alderton, Roberto Amato, Jeffrey Barrett, Sonia Goncalves, Ewan Harrison, David K. Jackson, Ian Johnston, Dominic Kwiatkowski, Cordelia Langford, John Sillitoe on behalf of the Wellcome Sanger Institute COVID-19 Surveillance Team |
| EPI_ISL_3225798 | Lighthouse Lab in Milton Keynes | Wellcome Sanger Institute for the COVID-19 Genomics UK (COG-UK) Consortium | The Lighthouse Lab in Milton Keynes and Alex Alderton, Roberto Amato, Jeffrey Barrett, Sonia Goncalves, Ewan Harrison, David K. Jackson, Ian Johnston, Dominic Kwiatkowski, Cordelia Langford, John Sillitoe on behalf of the Wellcome Sanger Institute COVID-19 Surveillance Team |
| EPI_ISL_3225799 | Lighthouse Lab in Glasgow | Wellcome Sanger Institute for the COVID-19 Genomics UK (COG-UK) Consortium | Harper VanSteenhouse, Yumi Kasai, David Gray, Carol Clugston, Anna Dominiczak and Alex Alderton, Roberto Amato, Jeffrey Barrett, Sonia Goncalves, Ewan Harrison, David K. Jackson, Ian Johnston, Dominic Kwiatkowski, Cordelia Langford, John Sillitoe on behalf of the Wellcome Sanger Institute COVID-19 Surveillance Team |
| EPI_ISL_3225800, EPI_ISL_3225801 | Lighthouse Lab in Milton Keynes | Wellcome Sanger Institute for the COVID-19 Genomics UK (COG-UK) Consortium | The Lighthouse Lab in Milton Keynes and Alex Alderton, Roberto Amato, Jeffrey Barrett, Sonia Goncalves, Ewan Harrison, David K. Jackson, Ian Johnston, Dominic Kwiatkowski, Cordelia Langford, John Sillitoe on behalf of the Wellcome Sanger Institute COVID-19 Surveillance Team |
| EPI_ISL_3225802, EPI_ISL_3225803, EPI_ISL_3225804, EPI_ISL_3225805 | Lighthouse Lab in Glasgow | Wellcome Sanger Institute for the COVID-19 Genomics UK (COG-UK) Consortium | Harper VanSteenhouse, Yumi Kasai, David Gray, Carol Clugston, Anna Dominiczak and Alex Alderton, Roberto Amato, Jeffrey Barrett, Sonia Goncalves, Ewan Harrison, David K. Jackson, Ian Johnston, Dominic Kwiatkowski, Cordelia Langford, John Sillitoe on behalf of the Wellcome Sanger Institute COVID-19 Surveillance Team |
| EPI_ISL_3225806 | Lighthouse Laboratory Plymouth | Wellcome Sanger Institute for the COVID-19 Genomics UK (COG-UK) Consortium | Lighthouse Laboratory Plymouth and Alex Alderton, Roberto Amato, Jeffrey Barrett, Sonia Goncalves, Ewan Harrison, David K. Jackson, Ian Johnston, Dominic Kwiatkowski, Cordelia Langford, John Sillitoe on behalf of the Wellcome Sanger Institute COVID-19 Surveillance Team |
| EPI_ISL_3225807, EPI_ISL_3225808, EPI_ISL_3225809 | Lighthouse Lab in Milton Keynes | Wellcome Sanger Institute for the COVID-19 Genomics UK (COG-UK) Consortium | The Lighthouse Lab in Milton Keynes and Alex Alderton, Roberto Amato, Jeffrey Barrett, Sonia Goncalves, Ewan Harrison, David K. Jackson, Ian Johnston, Dominic Kwiatkowski, Cordelia Langford, John Sillitoe on behalf of the Wellcome Sanger Institute COVID-19 Surveillance Team |
| EPI_ISL_3225810 | Lighthouse Laboratory Plymouth | Wellcome Sanger Institute for the COVID-19 Genomics UK (COG-UK) Consortium | Lighthouse Laboratory Plymouth and Alex Alderton, Roberto Amato, Jeffrey Barrett, Sonia Goncalves, Ewan Harrison, David K. Jackson, Ian Johnston, Dominic Kwiatkowski, Cordelia Langford, John Sillitoe on behalf of the Wellcome Sanger Institute COVID-19 Surveillance Team |
| EPI_ISL_3225811 | Lighthouse Lab in Milton Keynes | Wellcome Sanger Institute for the COVID-19 Genomics UK (COG-UK) Consortium | The Lighthouse Lab in Milton Keynes and Alex Alderton, Roberto Amato, Jeffrey Barrett, Sonia Goncalves, Ewan Harrison, David K. Jackson, Ian Johnston, Dominic Kwiatkowski, Cordelia Langford, John Sillitoe on behalf of the Wellcome Sanger Institute COVID-19 Surveillance Team |
| EPI_ISL_3225812 | Lighthouse Lab in Glasgow | Wellcome Sanger Institute for the COVID-19 Genomics UK (COG-UK) Consortium | Harper VanSteenhouse, Yumi Kasai, David Gray, Carol Clugston, Anna Dominiczak and Alex Alderton, Roberto Amato, Jeffrey Barrett, Sonia Goncalves, Ewan Harrison, David K. Jackson, Ian Johnston, Dominic Kwiatkowski, Cordelia Langford, John Sillitoe on behalf of the Wellcome Sanger Institute COVID-19 Surveillance Team |
| EPI_ISL_3225813 | Lighthouse Lab in Milton Keynes | Wellcome Sanger Institute for the COVID-19 Genomics UK (COG-UK) Consortium | The Lighthouse Lab in Milton Keynes and Alex Alderton, Roberto Amato, Jeffrey Barrett, Sonia Goncalves, Ewan Harrison, David K. Jackson, Ian Johnston, Dominic Kwiatkowski, Cordelia Langford, John Sillitoe on behalf of the Wellcome Sanger Institute COVID-19 Surveillance Team |
| EPI_ISL_3225814 | Lighthouse Laboratory Plymouth | Wellcome Sanger Institute for the COVID-19 Genomics UK (COG-UK) Consortium | Lighthouse Laboratory Plymouth and Alex Alderton, Roberto Amato, Jeffrey Barrett, Sonia Goncalves, Ewan Harrison, David K. Jackson, Ian Johnston, Dominic Kwiatkowski, Cordelia Langford, John Sillitoe on behalf of the Wellcome Sanger Institute COVID-19 Surveillance Team |
| EPI_ISL_3225815 | Lighthouse Lab in Milton Keynes | Wellcome Sanger Institute for the COVID-19 Genomics UK (COG-UK) Consortium | The Lighthouse Lab in Milton Keynes and Alex Alderton, Roberto Amato, Jeffrey Barrett, Sonia Goncalves, Ewan Harrison, David K. Jackson, Ian Johnston, Dominic Kwiatkowski, Cordelia Langford, John Sillitoe on behalf of the Wellcome Sanger Institute COVID-19 Surveillance Team |
| EPI_ISL_3225816 | Lighthouse Lab in Glasgow | Wellcome Sanger Institute for the COVID-19 Genomics UK (COG-UK) Consortium | Harper VanSteenhouse, Yumi Kasai, David Gray, Carol Clugston, Anna Dominiczak and Alex Alderton, Roberto Amato, Jeffrey Barrett, Sonia Goncalves, Ewan Harrison, David K. Jackson, Ian Johnston, Dominic Kwiatkowski, Cordelia Langford, John Sillitoe on behalf of the Wellcome Sanger Institute COVID-19 Surveillance Team |
| EPI_ISL_3225817 | Lighthouse Laboratory Plymouth | Wellcome Sanger Institute for the COVID-19 Genomics UK (COG-UK) Consortium | Lighthouse Laboratory Plymouth and Alex Alderton, Roberto Amato, Jeffrey Barrett, Sonia Goncalves, Ewan Harrison, David K. Jackson, Ian Johnston, Dominic Kwiatkowski, Cordelia Langford, John Sillitoe on behalf of the Wellcome Sanger Institute COVID-19 Surveillance Team |
| EPI_ISL_3225818, EPI_ISL_3225819, EPI_ISL_3225820 | Lighthouse Lab in Milton Keynes | Wellcome Sanger Institute for the COVID-19 Genomics UK (COG-UK) Consortium | The Lighthouse Lab in Milton Keynes and Alex Alderton, Roberto Amato, Jeffrey Barrett, Sonia Goncalves, Ewan Harrison, David K. Jackson, Ian Johnston, Dominic Kwiatkowski, Cordelia Langford, John Sillitoe on behalf of the Wellcome Sanger Institute COVID-19 Surveillance Team |
| EPI_ISL_3225821 | Lighthouse Laboratory Plymouth | Wellcome Sanger Institute for the COVID-19 Genomics UK (COG-UK) Consortium | Lighthouse Laboratory Plymouth and Alex Alderton, Roberto Amato, Jeffrey Barrett, Sonia Goncalves, Ewan Harrison, David K. Jackson, Ian Johnston, Dominic Kwiatkowski, Cordelia Langford, John Sillitoe on behalf of the Wellcome Sanger Institute COVID-19 Surveillance Team |

| | | | |
|---|---|---|---|
| EPI_ISL_3225822 | Lighthouse Lab in Milton Keynes | Wellcome Sanger Institute for the COVID-19 Genomics UK (COG-UK) Consortium | The Lighthouse Lab in Milton Keynes and Alex Alderton, Roberto Amato, Jeffrey Barrett, Sonia Goncalves, Ewan Harrison, David K. Jackson, Ian Johnston, Dominic Kwiatkowski, Cordelia Langford, John Sillitoe on behalf of the Wellcome Sanger Institute COVID-19 Surveillance Team |
| EPI_ISL_3225823 | Lighthouse Lab in Glasgow | Wellcome Sanger Institute for the COVID-19 Genomics UK (COG-UK) Consortium | Harper VanSteenhouse, Yumi Kasai, David Gray, Carol Clugston, Anna Dominiczak and Alex Alderton, Roberto Amato, Jeffrey Barrett, Sonia Goncalves, Ewan Harrison, David K. Jackson, Ian Johnston, Dominic Kwiatkowski, Cordelia Langford, John Sillitoe on behalf of the Wellcome Sanger Institute COVID-19 Surveillance Team |
| EPI_ISL_3225824 | Lighthouse Laboratory Plymouth | Wellcome Sanger Institute for the COVID-19 Genomics UK (COG-UK) Consortium | Lighthouse Laboratory Plymouth and Alex Alderton, Roberto Amato, Jeffrey Barrett, Sonia Goncalves, Ewan Harrison, David K. Jackson, Ian Johnston, Dominic Kwiatkowski, Cordelia Langford, John Sillitoe on behalf of the Wellcome Sanger Institute COVID-19 Surveillance Team |
| EPI_ISL_3225825, EPI_ISL_3225826, EPI_ISL_3225827 | Lighthouse Lab in Milton Keynes | Wellcome Sanger Institute for the COVID-19 Genomics UK (COG-UK) Consortium | The Lighthouse Lab in Milton Keynes and Alex Alderton, Roberto Amato, Jeffrey Barrett, Sonia Goncalves, Ewan Harrison, David K. Jackson, Ian Johnston, Dominic Kwiatkowski, Cordelia Langford, John Sillitoe on behalf of the Wellcome Sanger Institute COVID-19 Surveillance Team |
| EPI_ISL_3225828 | Lighthouse Laboratory Plymouth | Wellcome Sanger Institute for the COVID-19 Genomics UK (COG-UK) Consortium | Lighthouse Laboratory Plymouth and Alex Alderton, Roberto Amato, Jeffrey Barrett, Sonia Goncalves, Ewan Harrison, David K. Jackson, Ian Johnston, Dominic Kwiatkowski, Cordelia Langford, John Sillitoe on behalf of the Wellcome Sanger Institute COVID-19 Surveillance Team |
| EPI_ISL_3225829 | Lighthouse Lab in Milton Keynes | Wellcome Sanger Institute for the COVID-19 Genomics UK (COG-UK) Consortium | The Lighthouse Lab in Milton Keynes and Alex Alderton, Roberto Amato, Jeffrey Barrett, Sonia Goncalves, Ewan Harrison, David K. Jackson, Ian Johnston, Dominic Kwiatkowski, Cordelia Langford, John Sillitoe on behalf of the Wellcome Sanger Institute COVID-19 Surveillance Team |
| EPI_ISL_3225830 | Lighthouse Laboratory Plymouth | Wellcome Sanger Institute for the COVID-19 Genomics UK (COG-UK) Consortium | Lighthouse Laboratory Plymouth and Alex Alderton, Roberto Amato, Jeffrey Barrett, Sonia Goncalves, Ewan Harrison, David K. Jackson, Ian Johnston, Dominic Kwiatkowski, Cordelia Langford, John Sillitoe on behalf of the Wellcome Sanger Institute COVID-19 Surveillance Team |
| EPI_ISL_3225831 | Lighthouse Lab in Milton Keynes | Wellcome Sanger Institute for the COVID-19 Genomics UK (COG-UK) Consortium | The Lighthouse Lab in Milton Keynes and Alex Alderton, Roberto Amato, Jeffrey Barrett, Sonia Goncalves, Ewan Harrison, David K. Jackson, Ian Johnston, Dominic Kwiatkowski, Cordelia Langford, John Sillitoe on behalf of the Wellcome Sanger Institute COVID-19 Surveillance Team |
| EPI_ISL_3225832 | Lighthouse Laboratory Plymouth | Wellcome Sanger Institute for the COVID-19 Genomics UK (COG-UK) Consortium | Lighthouse Laboratory Plymouth and Alex Alderton, Roberto Amato, Jeffrey Barrett, Sonia Goncalves, Ewan Harrison, David K. Jackson, Ian Johnston, Dominic Kwiatkowski, Cordelia Langford, John Sillitoe on behalf of the Wellcome Sanger Institute COVID-19 Surveillance Team |
| EPI_ISL_3225833 | Lighthouse Lab in Milton Keynes | Wellcome Sanger Institute for the COVID-19 Genomics UK (COG-UK) Consortium | The Lighthouse Lab in Milton Keynes and Alex Alderton, Roberto Amato, Jeffrey Barrett, Sonia Goncalves, Ewan Harrison, David K. Jackson, Ian Johnston, Dominic Kwiatkowski, Cordelia Langford, John Sillitoe on behalf of the Wellcome Sanger Institute COVID-19 Surveillance Team |
| EPI_ISL_3225834 | Lighthouse Laboratory Plymouth | Wellcome Sanger Institute for the COVID-19 Genomics UK (COG-UK) Consortium | Lighthouse Laboratory Plymouth and Alex Alderton, Roberto Amato, Jeffrey Barrett, Sonia Goncalves, Ewan Harrison, David K. Jackson, Ian Johnston, Dominic Kwiatkowski, Cordelia Langford, John Sillitoe on behalf of the Wellcome Sanger Institute COVID-19 Surveillance Team |
| EPI_ISL_3225835 | Lighthouse Lab in Milton Keynes | Wellcome Sanger Institute for the COVID-19 Genomics UK (COG-UK) Consortium | The Lighthouse Lab in Milton Keynes and Alex Alderton, Roberto Amato, Jeffrey Barrett, Sonia Goncalves, Ewan Harrison, David K. Jackson, Ian Johnston, Dominic Kwiatkowski, Cordelia Langford, John Sillitoe on behalf of the Wellcome Sanger Institute COVID-19 Surveillance Team |
| EPI_ISL_3225836 | Lighthouse Laboratory Plymouth | Wellcome Sanger Institute for the COVID-19 Genomics UK (COG-UK) Consortium | Lighthouse Laboratory Plymouth and Alex Alderton, Roberto Amato, Jeffrey Barrett, Sonia Goncalves, Ewan Harrison, David K. Jackson, Ian Johnston, Dominic Kwiatkowski, Cordelia Langford, John Sillitoe on behalf of the Wellcome Sanger Institute COVID-19 Surveillance Team |
| EPI_ISL_3225837, EPI_ISL_3225838, EPI_ISL_3225839, EPI_ISL_3225840, EPI_ISL_3225841, EPI_ISL_3225842, EPI_ISL_3225843 | Lighthouse Lab in Milton Keynes | Wellcome Sanger Institute for the COVID-19 Genomics UK (COG-UK) Consortium | The Lighthouse Lab in Milton Keynes and Alex Alderton, Roberto Amato, Jeffrey Barrett, Sonia Goncalves, Ewan Harrison, David K. Jackson, Ian Johnston, Dominic Kwiatkowski, Cordelia Langford, John Sillitoe on behalf of the Wellcome Sanger Institute COVID-19 Surveillance Team |
| EPI_ISL_3225844, EPI_ISL_3225845, EPI_ISL_3225846 | Lighthouse Laboratory Plymouth | Wellcome Sanger Institute for the COVID-19 Genomics UK (COG-UK) Consortium | Lighthouse Laboratory Plymouth and Alex Alderton, Roberto Amato, Jeffrey Barrett, Sonia Goncalves, Ewan Harrison, David K. Jackson, Ian Johnston, Dominic Kwiatkowski, Cordelia Langford, John Sillitoe on behalf of the Wellcome Sanger Institute COVID-19 Surveillance Team |
| EPI_ISL_3225847, EPI_ISL_3225848, EPI_ISL_3225849 | Lighthouse Lab in Milton Keynes | Wellcome Sanger Institute for the COVID-19 Genomics UK (COG-UK) Consortium | The Lighthouse Lab in Milton Keynes and Alex Alderton, Roberto Amato, Jeffrey Barrett, Sonia Goncalves, Ewan Harrison, David K. Jackson, Ian Johnston, Dominic Kwiatkowski, Cordelia Langford, John Sillitoe on behalf of the Wellcome Sanger Institute COVID-19 Surveillance Team |
| EPI_ISL_3225850, EPI_ISL_3225851, EPI_ISL_3225852, EPI_ISL_3225853 | Lighthouse Laboratory Plymouth | Wellcome Sanger Institute for the COVID-19 Genomics UK (COG-UK) Consortium | Lighthouse Laboratory Plymouth and Alex Alderton, Roberto Amato, Jeffrey Barrett, Sonia Goncalves, Ewan Harrison, David K. Jackson, Ian Johnston, Dominic Kwiatkowski, Cordelia Langford, John Sillitoe on behalf of the Wellcome Sanger Institute COVID-19 Surveillance Team |
| EPI_ISL_3225854 | Lighthouse Lab in Milton Keynes | Wellcome Sanger Institute for the COVID-19 Genomics UK (COG-UK) Consortium | The Lighthouse Lab in Milton Keynes and Alex Alderton, Roberto Amato, Jeffrey Barrett, Sonia Goncalves, Ewan Harrison, David K. Jackson, Ian Johnston, Dominic Kwiatkowski, Cordelia Langford, John Sillitoe on behalf of the Wellcome Sanger Institute COVID-19 Surveillance Team |
| EPI_ISL_3225855 | Lighthouse Laboratory Plymouth | Wellcome Sanger Institute for the COVID-19 Genomics UK (COG-UK) Consortium | Lighthouse Laboratory Plymouth and Alex Alderton, Roberto Amato, Jeffrey Barrett, Sonia Goncalves, Ewan Harrison, David K. Jackson, Ian Johnston, Dominic Kwiatkowski, Cordelia Langford, John Sillitoe on behalf of the Wellcome Sanger Institute COVID-19 Surveillance Team |
| EPI_ISL_3225856 | Lighthouse Lab in Milton Keynes | Wellcome Sanger Institute for the COVID-19 Genomics UK (COG-UK) Consortium | The Lighthouse Lab in Milton Keynes and Alex Alderton, Roberto Amato, Jeffrey Barrett, Sonia Goncalves, Ewan Harrison, David K. Jackson, Ian Johnston, Dominic Kwiatkowski, Cordelia Langford, John Sillitoe on behalf of the Wellcome Sanger Institute COVID-19 Surveillance Team |
| EPI_ISL_3225857, EPI_ISL_3225858 | Lighthouse Laboratory Plymouth | Wellcome Sanger Institute for the COVID-19 Genomics UK (COG-UK) Consortium | Lighthouse Laboratory Plymouth and Alex Alderton, Roberto Amato, Jeffrey Barrett, Sonia Goncalves, Ewan Harrison, David K. Jackson, Ian Johnston, Dominic Kwiatkowski, Cordelia Langford, John Sillitoe on behalf of the Wellcome Sanger Institute COVID-19 Surveillance Team |
| EPI_ISL_3225859 | Lighthouse Lab in Milton Keynes | Wellcome Sanger Institute for the COVID-19 Genomics UK (COG-UK) Consortium | The Lighthouse Lab in Milton Keynes and Alex Alderton, Roberto Amato, Jeffrey Barrett, Sonia Goncalves, Ewan Harrison, David K. Jackson, Ian Johnston, Dominic Kwiatkowski, Cordelia Langford, John Sillitoe on behalf of the Wellcome Sanger Institute COVID-19 Surveillance Team |
| EPI_ISL_3225860 | Lighthouse Laboratory Plymouth | Wellcome Sanger Institute for the COVID-19 Genomics UK (COG-UK) Consortium | Lighthouse Laboratory Plymouth and Alex Alderton, Roberto Amato, Jeffrey Barrett, Sonia Goncalves, Ewan Harrison, David K. Jackson, Ian Johnston, Dominic Kwiatkowski, Cordelia Langford, John Sillitoe on behalf of the Wellcome Sanger Institute COVID-19 Surveillance Team |
| EPI_ISL_3225861, EPI_ISL_3225862, EPI_ISL_3225863, EPI_ISL_3225864 | Lighthouse Lab in Milton Keynes | Wellcome Sanger Institute for the COVID-19 Genomics UK (COG-UK) Consortium | The Lighthouse Lab in Milton Keynes and Alex Alderton, Roberto Amato, Jeffrey Barrett, Sonia Goncalves, Ewan Harrison, David K. Jackson, Ian Johnston, Dominic Kwiatkowski, Cordelia Langford, John Sillitoe on behalf of the Wellcome Sanger Institute COVID-19 Surveillance Team |
| EPI_ISL_3225865 | Lighthouse Lab in Glasgow | Wellcome Sanger Institute for the COVID-19 Genomics UK (COG-UK) Consortium | Harper VanSteenhouse, Yumi Kasai, David Gray, Carol Clugston, Anna Dominiczak and Alex Alderton, Roberto Amato, Jeffrey Barrett, Sonia Goncalves, Ewan Harrison, David K. Jackson, Ian Johnston, Dominic Kwiatkowski, Cordelia Langford, John Sillitoe on behalf of the Wellcome Sanger Institute COVID-19 Surveillance Team |
| EPI_ISL_3225866 | Lighthouse Lab in Milton Keynes | Wellcome Sanger Institute for the COVID-19 Genomics UK (COG-UK) Consortium | The Lighthouse Lab in Milton Keynes and Alex Alderton, Roberto Amato, Jeffrey Barrett, Sonia Goncalves, Ewan Harrison, David K. Jackson, Ian Johnston, Dominic Kwiatkowski, Cordelia Langford, John Sillitoe on behalf of the Wellcome Sanger Institute COVID-19 Surveillance Team |
| EPI_ISL_3225867 | Lighthouse Lab in Glasgow | Wellcome Sanger Institute for the COVID-19 Genomics UK (COG-UK) Consortium | Harper VanSteenhouse, Yumi Kasai, David Gray, Carol Clugston, Anna Dominiczak and Alex Alderton, Roberto Amato, Jeffrey Barrett, Sonia Goncalves, Ewan Harrison, David K. Jackson, Ian Johnston, Dominic Kwiatkowski, Cordelia Langford, John Sillitoe on behalf of the Wellcome Sanger Institute COVID-19 Surveillance Team |
| EPI_ISL_3225868 | Lighthouse Laboratory Plymouth | Wellcome Sanger Institute for the COVID-19 Genomics UK (COG-UK) Consortium | Lighthouse Laboratory Plymouth and Alex Alderton, Roberto Amato, Jeffrey Barrett, Sonia Goncalves, Ewan Harrison, David K. Jackson, Ian Johnston, Dominic Kwiatkowski, Cordelia Langford, John Sillitoe on behalf of the Wellcome Sanger Institute COVID-19 Surveillance Team |
| EPI_ISL_3225869, EPI_ISL_3225870 | Lighthouse Lab in Glasgow | Wellcome Sanger Institute for the COVID-19 Genomics UK (COG-UK) Consortium | Harper VanSteenhouse, Yumi Kasai, David Gray, Carol Clugston, Anna Dominiczak and Alex Alderton, Roberto Amato, Jeffrey Barrett, Sonia Goncalves, Ewan Harrison, David K. Jackson, Ian Johnston, Dominic Kwiatkowski, Cordelia Langford, John Sillitoe on behalf of the Wellcome Sanger Institute COVID-19 Surveillance Team |
| EPI_ISL_3225871, EPI_ISL_3225872 | Lighthouse Lab in Milton Keynes | Wellcome Sanger Institute for the COVID-19 Genomics UK (COG-UK) Consortium | The Lighthouse Lab in Milton Keynes and Alex Alderton, Roberto Amato, Jeffrey Barrett, Sonia Goncalves, Ewan Harrison, David K. Jackson, Ian Johnston, Dominic Kwiatkowski, Cordelia Langford, John Sillitoe on behalf of the Wellcome Sanger Institute COVID-19 Surveillance Team |
| EPI_ISL_3225873 | Lighthouse Laboratory Plymouth | Wellcome Sanger Institute for the COVID-19 Genomics UK (COG-UK) Consortium | Lighthouse Laboratory Plymouth and Alex Alderton, Roberto Amato, Jeffrey Barrett, Sonia Goncalves, Ewan Harrison, David K. Jackson, Ian Johnston, Dominic Kwiatkowski, Cordelia Langford, John Sillitoe on behalf of the Wellcome Sanger Institute COVID-19 Surveillance Team |
| EPI_ISL_3225874 | Lighthouse Lab in Glasgow | Wellcome Sanger Institute for the COVID-19 Genomics UK (COG-UK) Consortium | Harper VanSteenhouse, Yumi Kasai, David Gray, Carol Clugston, Anna Dominiczak and Alex Alderton, Roberto Amato, Jeffrey Barrett, Sonia Goncalves, Ewan Harrison, David K. Jackson, Ian Johnston, Dominic Kwiatkowski, Cordelia Langford, John Sillitoe on behalf of the Wellcome Sanger Institute COVID-19 Surveillance Team |
| EPI_ISL_3225875, EPI_ISL_3225876, EPI_ISL_3225877 | Lighthouse Lab in Milton Keynes | Wellcome Sanger Institute for the COVID-19 Genomics UK (COG-UK) Consortium | The Lighthouse Lab in Milton Keynes and Alex Alderton, Roberto Amato, Jeffrey Barrett, Sonia Goncalves, Ewan Harrison, David K. Jackson, Ian Johnston, Dominic Kwiatkowski, Cordelia Langford, John Sillitoe on behalf of the Wellcome Sanger Institute COVID-19 Surveillance Team |
| EPI_ISL_3225878 | Lighthouse Lab in Glasgow | Wellcome Sanger Institute for the COVID-19 Genomics UK (COG-UK) Consortium | Harper VanSteenhouse, Yumi Kasai, David Gray, Carol Clugston, Anna Dominiczak and Alex Alderton, Roberto Amato, Jeffrey Barrett, Sonia Goncalves, Ewan Harrison, David K. Jackson, Ian Johnston, Dominic Kwiatkowski, Cordelia Langford, John Sillitoe on behalf of the Wellcome Sanger Institute COVID-19 Surveillance Team |
| EPI_ISL_3225879 | Lighthouse Lab in Milton Keynes | Wellcome Sanger Institute for the COVID-19 Genomics UK (COG-UK) Consortium | The Lighthouse Lab in Milton Keynes and Alex Alderton, Roberto Amato, Jeffrey Barrett, Sonia Goncalves, Ewan Harrison, David K. Jackson, Ian Johnston, Dominic Kwiatkowski, Cordelia Langford, John Sillitoe on behalf of the Wellcome Sanger Institute COVID-19 Surveillance Team |
| EPI_ISL_3225880 | Lighthouse Laboratory Plymouth | Wellcome Sanger Institute for the COVID-19 Genomics UK (COG-UK) Consortium | Lighthouse Laboratory Plymouth and Alex Alderton, Roberto Amato, Jeffrey Barrett, Sonia Goncalves, Ewan Harrison, David K. Jackson, Ian Johnston, Dominic Kwiatkowski, Cordelia Langford, John Sillitoe on behalf of the Wellcome Sanger Institute COVID-19 Surveillance Team |
| EPI_ISL_3225881, EPI_ISL_3225883 | Lighthouse Lab in Glasgow | Wellcome Sanger Institute for the COVID-19 Genomics UK (COG-UK) Consortium | Harper VanSteenhouse, Yumi Kasai, David Gray, Carol Clugston, Anna Dominiczak and Alex Alderton, Roberto Amato, Jeffrey Barrett, Sonia Goncalves, Ewan Harrison, David K. Jackson, Ian Johnston, Dominic Kwiatkowski, Cordelia Langford, John Sillitoe on behalf of the Wellcome Sanger Institute COVID-19 Surveillance Team |
| EPI_ISL_3225884, EPI_ISL_3225885, EPI_ISL_3225886 | Lighthouse Lab in Milton Keynes | Wellcome Sanger Institute for the COVID-19 Genomics UK (COG-UK) Consortium | The Lighthouse Lab in Milton Keynes and Alex Alderton, Roberto Amato, Jeffrey Barrett, Sonia Goncalves, Ewan Harrison, David K. Jackson, Ian Johnston, Dominic Kwiatkowski, Cordelia Langford, John Sillitoe on behalf of the Wellcome Sanger Institute COVID-19 Surveillance Team |
| EPI_ISL_3225887 | Lighthouse Laboratory Plymouth | Wellcome Sanger Institute for the COVID-19 Genomics UK (COG-UK) Consortium | Lighthouse Laboratory Plymouth and Alex Alderton, Roberto Amato, Jeffrey Barrett, Sonia Goncalves, Ewan Harrison, David K. Jackson, Ian Johnston, Dominic Kwiatkowski, Cordelia Langford, John Sillitoe on behalf of the Wellcome Sanger Institute COVID-19 Surveillance Team |
| EPI_ISL_3225888, | Lighthouse Lab in Milton Keynes | Wellcome Sanger Institute for the COVID-19 Genomics UK (COG-UK) | The Lighthouse Lab in Milton Keynes and Alex Alderton, Roberto Amato, Jeffrey Barrett, Sonia Goncalves, Ewan Harrison, David K. Jackson, Ian Johnston, Dominic Kwiatkowski, Cordelia Langford, John Sillitoe on behalf of the Wellcome Sanger |

| | | | |
|---|---|---|---|
| EPI_ISL_3225889 | | Consortium | Institute COVID-19 Surveillance Team |
| EPI_ISL_3225890, EPI_ISL_3225891 | Lighthouse Lab in Alderley Park | Wellcome Sanger Institute for the COVID-19 Genomics UK (COG-UK) Consortium | Jacquelyn Wynn, Mairead Hyland, The Lighthouse Lab in Alderley Park and Alex Alderton, Roberto Amato, Jeffrey Barrett, Sonia Goncalves, Ewan Harrison, David K. Jackson, Ian Johnston, Dominic Kwiatkowski, Cordelia Langford, John Sillitoe on behalf of the Wellcome Sanger Institute COVID-19 Surveillance Team |
| EPI_ISL_3225892 | Lighthouse Laboratory Plymouth | Wellcome Sanger Institute for the COVID-19 Genomics UK (COG-UK) Consortium | Lighthouse Laboratory Plymouth and Alex Alderton, Roberto Amato, Jeffrey Barrett, Sonia Goncalves, Ewan Harrison, David K. Jackson, Ian Johnston, Dominic Kwiatkowski, Cordelia Langford, John Sillitoe on behalf of the Wellcome Sanger Institute COVID-19 Surveillance Team |
| EPI_ISL_3225893, EPI_ISL_3225894 | Lighthouse Lab in Milton Keynes | Wellcome Sanger Institute for the COVID-19 Genomics UK (COG-UK) Consortium | The Lighthouse Lab in Milton Keynes and Alex Alderton, Roberto Amato, Jeffrey Barrett, Sonia Goncalves, Ewan Harrison, David K. Jackson, Ian Johnston, Dominic Kwiatkowski, Cordelia Langford, John Sillitoe on behalf of the Wellcome Sanger Institute COVID-19 Surveillance Team |
| EPI_ISL_3225895 | Lighthouse Lab in Alderley Park | Wellcome Sanger Institute for the COVID-19 Genomics UK (COG-UK) Consortium | Jacquelyn Wynn, Mairead Hyland, The Lighthouse Lab in Alderley Park and Alex Alderton, Roberto Amato, Jeffrey Barrett, Sonia Goncalves, Ewan Harrison, David K. Jackson, Ian Johnston, Dominic Kwiatkowski, Cordelia Langford, John Sillitoe on behalf of the Wellcome Sanger Institute COVID-19 Surveillance Team |
| EPI_ISL_3225896, EPI_ISL_3225897 | Lighthouse Laboratory Plymouth | Wellcome Sanger Institute for the COVID-19 Genomics UK (COG-UK) Consortium | Lighthouse Laboratory Plymouth and Alex Alderton, Roberto Amato, Jeffrey Barrett, Sonia Goncalves, Ewan Harrison, David K. Jackson, Ian Johnston, Dominic Kwiatkowski, Cordelia Langford, John Sillitoe on behalf of the Wellcome Sanger Institute COVID-19 Surveillance Team |
| EPI_ISL_3225898 | Lighthouse Lab in Glasgow | Wellcome Sanger Institute for the COVID-19 Genomics UK (COG-UK) Consortium | Harper VanSteenhouse, Yumi Kasai, David Gray, Carol Clugston, Anna Dominiczak and Alex Alderton, Roberto Amato, Jeffrey Barrett, Sonia Goncalves, Ewan Harrison, David K. Jackson, Ian Johnston, Dominic Kwiatkowski, Cordelia Langford, John Sillitoe on behalf of the Wellcome Sanger Institute COVID-19 Surveillance Team |
| EPI_ISL_3225899 | Lighthouse Lab in Alderley Park | Wellcome Sanger Institute for the COVID-19 Genomics UK (COG-UK) Consortium | Jacquelyn Wynn, Mairead Hyland, The Lighthouse Lab in Alderley Park and Alex Alderton, Roberto Amato, Jeffrey Barrett, Sonia Goncalves, Ewan Harrison, David K. Jackson, Ian Johnston, Dominic Kwiatkowski, Cordelia Langford, John Sillitoe on behalf of the Wellcome Sanger Institute COVID-19 Surveillance Team |
| EPI_ISL_3225900, EPI_ISL_3225901 | Lighthouse Lab in Glasgow | Wellcome Sanger Institute for the COVID-19 Genomics UK (COG-UK) Consortium | Harper VanSteenhouse, Yumi Kasai, David Gray, Carol Clugston, Anna Dominiczak and Alex Alderton, Roberto Amato, Jeffrey Barrett, Sonia Goncalves, Ewan Harrison, David K. Jackson, Ian Johnston, Dominic Kwiatkowski, Cordelia Langford, John Sillitoe on behalf of the Wellcome Sanger Institute COVID-19 Surveillance Team |
| EPI_ISL_3225902 | Lighthouse Laboratory Plymouth | Wellcome Sanger Institute for the COVID-19 Genomics UK (COG-UK) Consortium | Lighthouse Laboratory Plymouth and Alex Alderton, Roberto Amato, Jeffrey Barrett, Sonia Goncalves, Ewan Harrison, David K. Jackson, Ian Johnston, Dominic Kwiatkowski, Cordelia Langford, John Sillitoe on behalf of the Wellcome Sanger Institute COVID-19 Surveillance Team |
| EPI_ISL_3225903 | Lighthouse Lab in Alderley Park | Wellcome Sanger Institute for the COVID-19 Genomics UK (COG-UK) Consortium | Jacquelyn Wynn, Mairead Hyland, The Lighthouse Lab in Alderley Park and Alex Alderton, Roberto Amato, Jeffrey Barrett, Sonia Goncalves, Ewan Harrison, David K. Jackson, Ian Johnston, Dominic Kwiatkowski, Cordelia Langford, John Sillitoe on behalf of the Wellcome Sanger Institute COVID-19 Surveillance Team |
| EPI_ISL_3225904 | Lighthouse Lab in Milton Keynes | Wellcome Sanger Institute for the COVID-19 Genomics UK (COG-UK) Consortium | The Lighthouse Lab in Milton Keynes and Alex Alderton, Roberto Amato, Jeffrey Barrett, Sonia Goncalves, Ewan Harrison, David K. Jackson, Ian Johnston, Dominic Kwiatkowski, Cordelia Langford, John Sillitoe on behalf of the Wellcome Sanger Institute COVID-19 Surveillance Team |
| EPI_ISL_3225905, EPI_ISL_3225906, EPI_ISL_3225907 | Lighthouse Laboratory Plymouth | Wellcome Sanger Institute for the COVID-19 Genomics UK (COG-UK) Consortium | Lighthouse Laboratory Plymouth and Alex Alderton, Roberto Amato, Jeffrey Barrett, Sonia Goncalves, Ewan Harrison, David K. Jackson, Ian Johnston, Dominic Kwiatkowski, Cordelia Langford, John Sillitoe on behalf of the Wellcome Sanger Institute COVID-19 Surveillance Team |
| EPI_ISL_3225908, EPI_ISL_3225909 | Lighthouse Lab in Glasgow | Wellcome Sanger Institute for the COVID-19 Genomics UK (COG-UK) Consortium | Harper VanSteenhouse, Yumi Kasai, David Gray, Carol Clugston, Anna Dominiczak and Alex Alderton, Roberto Amato, Jeffrey Barrett, Sonia Goncalves, Ewan Harrison, David K. Jackson, Ian Johnston, Dominic Kwiatkowski, Cordelia Langford, John Sillitoe on behalf of the Wellcome Sanger Institute COVID-19 Surveillance Team |
| EPI_ISL_3225910 | Lighthouse Lab in Alderley Park | Wellcome Sanger Institute for the COVID-19 Genomics UK (COG-UK) Consortium | Jacquelyn Wynn, Mairead Hyland, The Lighthouse Lab in Alderley Park and Alex Alderton, Roberto Amato, Jeffrey Barrett, Sonia Goncalves, Ewan Harrison, David K. Jackson, Ian Johnston, Dominic Kwiatkowski, Cordelia Langford, John Sillitoe on behalf of the Wellcome Sanger Institute COVID-19 Surveillance Team |
| EPI_ISL_3225911 | Lighthouse Lab in Milton Keynes | Wellcome Sanger Institute for the COVID-19 Genomics UK (COG-UK) Consortium | The Lighthouse Lab in Milton Keynes and Alex Alderton, Roberto Amato, Jeffrey Barrett, Sonia Goncalves, Ewan Harrison, David K. Jackson, Ian Johnston, Dominic Kwiatkowski, Cordelia Langford, John Sillitoe on behalf of the Wellcome Sanger Institute COVID-19 Surveillance Team |
| EPI_ISL_3225912, EPI_ISL_3225913, EPI_ISL_3225914 | Lighthouse Laboratory Plymouth | Wellcome Sanger Institute for the COVID-19 Genomics UK (COG-UK) Consortium | Lighthouse Laboratory Plymouth and Alex Alderton, Roberto Amato, Jeffrey Barrett, Sonia Goncalves, Ewan Harrison, David K. Jackson, Ian Johnston, Dominic Kwiatkowski, Cordelia Langford, John Sillitoe on behalf of the Wellcome Sanger Institute COVID-19 Surveillance Team |
| EPI_ISL_3225915 | Lighthouse Lab in Glasgow | Wellcome Sanger Institute for the COVID-19 Genomics UK (COG-UK) Consortium | Harper VanSteenhouse, Yumi Kasai, David Gray, Carol Clugston, Anna Dominiczak and Alex Alderton, Roberto Amato, Jeffrey Barrett, Sonia Goncalves, Ewan Harrison, David K. Jackson, Ian Johnston, Dominic Kwiatkowski, Cordelia Langford, John Sillitoe on behalf of the Wellcome Sanger Institute COVID-19 Surveillance Team |
| EPI_ISL_3225916, EPI_ISL_3225917, EPI_ISL_3225918 | Lighthouse Laboratory Plymouth | Wellcome Sanger Institute for the COVID-19 Genomics UK (COG-UK) Consortium | Lighthouse Laboratory Plymouth and Alex Alderton, Roberto Amato, Jeffrey Barrett, Sonia Goncalves, Ewan Harrison, David K. Jackson, Ian Johnston, Dominic Kwiatkowski, Cordelia Langford, John Sillitoe on behalf of the Wellcome Sanger Institute COVID-19 Surveillance Team |
| EPI_ISL_3225919 | Lighthouse Lab in Alderley Park | Wellcome Sanger Institute for the COVID-19 Genomics UK (COG-UK) Consortium | Jacquelyn Wynn, Mairead Hyland, The Lighthouse Lab in Alderley Park and Alex Alderton, Roberto Amato, Jeffrey Barrett, Sonia Goncalves, Ewan Harrison, David K. Jackson, Ian Johnston, Dominic Kwiatkowski, Cordelia Langford, John Sillitoe on behalf of the Wellcome Sanger Institute COVID-19 Surveillance Team |
| EPI_ISL_3225920 | Lighthouse Lab in Milton Keynes | Wellcome Sanger Institute for the COVID-19 Genomics UK (COG-UK) Consortium | The Lighthouse Lab in Milton Keynes and Alex Alderton, Roberto Amato, Jeffrey Barrett, Sonia Goncalves, Ewan Harrison, David K. Jackson, Ian Johnston, Dominic Kwiatkowski, Cordelia Langford, John Sillitoe on behalf of the Wellcome Sanger Institute COVID-19 Surveillance Team |
| EPI_ISL_3225921 | Lighthouse Lab in Glasgow | Wellcome Sanger Institute for the COVID-19 Genomics UK (COG-UK) Consortium | Harper VanSteenhouse, Yumi Kasai, David Gray, Carol Clugston, Anna Dominiczak and Alex Alderton, Roberto Amato, Jeffrey Barrett, Sonia Goncalves, Ewan Harrison, David K. Jackson, Ian Johnston, Dominic Kwiatkowski, Cordelia Langford, John Sillitoe on behalf of the Wellcome Sanger Institute COVID-19 Surveillance Team |
| EPI_ISL_3225922 | Lighthouse Lab in Alderley Park | Wellcome Sanger Institute for the COVID-19 Genomics UK (COG-UK) Consortium | Jacquelyn Wynn, Mairead Hyland, The Lighthouse Lab in Alderley Park and Alex Alderton, Roberto Amato, Jeffrey Barrett, Sonia Goncalves, Ewan Harrison, David K. Jackson, Ian Johnston, Dominic Kwiatkowski, Cordelia Langford, John Sillitoe on behalf of the Wellcome Sanger Institute COVID-19 Surveillance Team |
| EPI_ISL_3225923 | Lighthouse Laboratory Plymouth | Wellcome Sanger Institute for the COVID-19 Genomics UK (COG-UK) Consortium | Lighthouse Laboratory Plymouth and Alex Alderton, Roberto Amato, Jeffrey Barrett, Sonia Goncalves, Ewan Harrison, David K. Jackson, Ian Johnston, Dominic Kwiatkowski, Cordelia Langford, John Sillitoe on behalf of the Wellcome Sanger Institute COVID-19 Surveillance Team |
| EPI_ISL_3225924 | Lighthouse Lab in Milton Keynes | Wellcome Sanger Institute for the COVID-19 Genomics UK (COG-UK) Consortium | The Lighthouse Lab in Milton Keynes and Alex Alderton, Roberto Amato, Jeffrey Barrett, Sonia Goncalves, Ewan Harrison, David K. Jackson, Ian Johnston, Dominic Kwiatkowski, Cordelia Langford, John Sillitoe on behalf of the Wellcome Sanger Institute COVID-19 Surveillance Team |
| EPI_ISL_3225925 | Lighthouse Lab in Alderley Park | Wellcome Sanger Institute for the COVID-19 Genomics UK (COG-UK) Consortium | Jacquelyn Wynn, Mairead Hyland, The Lighthouse Lab in Alderley Park and Alex Alderton, Roberto Amato, Jeffrey Barrett, Sonia Goncalves, Ewan Harrison, David K. Jackson, Ian Johnston, Dominic Kwiatkowski, Cordelia Langford, John Sillitoe on behalf of the Wellcome Sanger Institute COVID-19 Surveillance Team |
| EPI_ISL_3225926 | Lighthouse Laboratory Plymouth | Wellcome Sanger Institute for the COVID-19 Genomics UK (COG-UK) Consortium | Lighthouse Laboratory Plymouth and Alex Alderton, Roberto Amato, Jeffrey Barrett, Sonia Goncalves, Ewan Harrison, David K. Jackson, Ian Johnston, Dominic Kwiatkowski, Cordelia Langford, John Sillitoe on behalf of the Wellcome Sanger Institute COVID-19 Surveillance Team |
| EPI_ISL_3225927 | Lighthouse Lab in Glasgow | Wellcome Sanger Institute for the COVID-19 Genomics UK (COG-UK) Consortium | Harper VanSteenhouse, Yumi Kasai, David Gray, Carol Clugston, Anna Dominiczak and Alex Alderton, Roberto Amato, Jeffrey Barrett, Sonia Goncalves, Ewan Harrison, David K. Jackson, Ian Johnston, Dominic Kwiatkowski, Cordelia Langford, John Sillitoe on behalf of the Wellcome Sanger Institute COVID-19 Surveillance Team |
| EPI_ISL_3225928 | Lighthouse Lab in Alderley Park | Wellcome Sanger Institute for the COVID-19 Genomics UK (COG-UK) Consortium | Jacquelyn Wynn, Mairead Hyland, The Lighthouse Lab in Alderley Park and Alex Alderton, Roberto Amato, Jeffrey Barrett, Sonia Goncalves, Ewan Harrison, David K. Jackson, Ian Johnston, Dominic Kwiatkowski, Cordelia Langford, John Sillitoe on behalf of the Wellcome Sanger Institute COVID-19 Surveillance Team |
| EPI_ISL_3225929 | Lighthouse Lab in Milton Keynes | Wellcome Sanger Institute for the COVID-19 Genomics UK (COG-UK) Consortium | The Lighthouse Lab in Milton Keynes and Alex Alderton, Roberto Amato, Jeffrey Barrett, Sonia Goncalves, Ewan Harrison, David K. Jackson, Ian Johnston, Dominic Kwiatkowski, Cordelia Langford, John Sillitoe on behalf of the Wellcome Sanger Institute COVID-19 Surveillance Team |
| EPI_ISL_3225930 | Lighthouse Lab in Alderley Park | Wellcome Sanger Institute for the COVID-19 Genomics UK (COG-UK) Consortium | Jacquelyn Wynn, Mairead Hyland, The Lighthouse Lab in Alderley Park and Alex Alderton, Roberto Amato, Jeffrey Barrett, Sonia Goncalves, Ewan Harrison, David K. Jackson, Ian Johnston, Dominic Kwiatkowski, Cordelia Langford, John Sillitoe on behalf of the Wellcome Sanger Institute COVID-19 Surveillance Team |
| EPI_ISL_3225931, EPI_ISL_3225932 | Lighthouse Lab in Milton Keynes | Wellcome Sanger Institute for the COVID-19 Genomics UK (COG-UK) Consortium | The Lighthouse Lab in Milton Keynes and Alex Alderton, Roberto Amato, Jeffrey Barrett, Sonia Goncalves, Ewan Harrison, David K. Jackson, Ian Johnston, Dominic Kwiatkowski, Cordelia Langford, John Sillitoe on behalf of the Wellcome Sanger Institute COVID-19 Surveillance Team |
| EPI_ISL_3225933 | Lighthouse Lab in Glasgow | Wellcome Sanger Institute for the COVID-19 Genomics UK (COG-UK) Consortium | Harper VanSteenhouse, Yumi Kasai, David Gray, Carol Clugston, Anna Dominiczak and Alex Alderton, Roberto Amato, Jeffrey Barrett, Sonia Goncalves, Ewan Harrison, David K. Jackson, Ian Johnston, Dominic Kwiatkowski, Cordelia Langford, John Sillitoe on behalf of the Wellcome Sanger Institute COVID-19 Surveillance Team |
| EPI_ISL_3225934 | Lighthouse Lab in Alderley Park | Wellcome Sanger Institute for the COVID-19 Genomics UK (COG-UK) Consortium | Jacquelyn Wynn, Mairead Hyland, The Lighthouse Lab in Alderley Park and Alex Alderton, Roberto Amato, Jeffrey Barrett, Sonia Goncalves, Ewan Harrison, David K. Jackson, Ian Johnston, Dominic Kwiatkowski, Cordelia Langford, John Sillitoe on behalf of the Wellcome Sanger Institute COVID-19 Surveillance Team |
| EPI_ISL_3225935 | Lighthouse Lab in Milton Keynes | Wellcome Sanger Institute for the COVID-19 Genomics UK (COG-UK) Consortium | The Lighthouse Lab in Milton Keynes and Alex Alderton, Roberto Amato, Jeffrey Barrett, Sonia Goncalves, Ewan Harrison, David K. Jackson, Ian Johnston, Dominic Kwiatkowski, Cordelia Langford, John Sillitoe on behalf of the Wellcome Sanger Institute COVID-19 Surveillance Team |
| EPI_ISL_3225936 | Lighthouse Lab in Glasgow | Wellcome Sanger Institute for the COVID-19 Genomics UK (COG-UK) Consortium | Harper VanSteenhouse, Yumi Kasai, David Gray, Carol Clugston, Anna Dominiczak and Alex Alderton, Roberto Amato, Jeffrey Barrett, Sonia Goncalves, Ewan Harrison, David K. Jackson, Ian Johnston, Dominic Kwiatkowski, Cordelia Langford, John Sillitoe on behalf of the Wellcome Sanger Institute COVID-19 Surveillance Team |
| EPI_ISL_3225937 | Lighthouse Lab in Milton Keynes | Wellcome Sanger Institute for the COVID-19 Genomics UK (COG-UK) Consortium | The Lighthouse Lab in Milton Keynes and Alex Alderton, Roberto Amato, Jeffrey Barrett, Sonia Goncalves, Ewan Harrison, David K. Jackson, Ian Johnston, Dominic Kwiatkowski, Cordelia Langford, John Sillitoe on behalf of the Wellcome Sanger Institute COVID-19 Surveillance Team |
| EPI_ISL_3225938 | Lighthouse Laboratory Plymouth | Wellcome Sanger Institute for the COVID-19 Genomics UK (COG-UK) Consortium | Lighthouse Laboratory Plymouth and Alex Alderton, Roberto Amato, Jeffrey Barrett, Sonia Goncalves, Ewan Harrison, David K. Jackson, Ian Johnston, Dominic Kwiatkowski, Cordelia Langford, John Sillitoe on behalf of the Wellcome Sanger Institute COVID-19 Surveillance Team |
| EPI_ISL_3225939 | Lighthouse Lab in Milton Keynes | Wellcome Sanger Institute for the COVID-19 Genomics UK (COG-UK) Consortium | The Lighthouse Lab in Milton Keynes and Alex Alderton, Roberto Amato, Jeffrey Barrett, Sonia Goncalves, Ewan Harrison, David K. Jackson, Ian Johnston, Dominic Kwiatkowski, Cordelia Langford, John Sillitoe on behalf of the Wellcome Sanger Institute COVID-19 Surveillance Team |
| EPI_ISL_3225940, EPI_ISL_3225941, EPI_ISL_3225942 | Lighthouse Laboratory Plymouth | Wellcome Sanger Institute for the COVID-19 Genomics UK (COG-UK) Consortium | Lighthouse Laboratory Plymouth and Alex Alderton, Roberto Amato, Jeffrey Barrett, Sonia Goncalves, Ewan Harrison, David K. Jackson, Ian Johnston, Dominic Kwiatkowski, Cordelia Langford, John Sillitoe on behalf of the Wellcome Sanger Institute COVID-19 Surveillance Team |
| EPI_ISL_3225943 | Lighthouse Lab in Milton Keynes | Wellcome Sanger Institute for the COVID-19 Genomics UK (COG-UK) Consortium | The Lighthouse Lab in Milton Keynes and Alex Alderton, Roberto Amato, Jeffrey Barrett, Sonia Goncalves, Ewan Harrison, David K. Jackson, Ian Johnston, Dominic Kwiatkowski, Cordelia Langford, John Sillitoe on behalf of the Wellcome Sanger Institute COVID-19 Surveillance Team |
| EPI_ISL_3225944 | Lighthouse Laboratory Plymouth | Wellcome Sanger Institute for the COVID-19 Genomics UK (COG-UK) Consortium | Lighthouse Laboratory Plymouth and Alex Alderton, Roberto Amato, Jeffrey Barrett, Sonia Goncalves, Ewan Harrison, David K. Jackson, Ian Johnston, Dominic Kwiatkowski, Cordelia Langford, John Sillitoe on behalf of the Wellcome Sanger Institute COVID-19 Surveillance Team |
| EPI_ISL_3225945, EPI_ISL_3225946, EPI_ISL_3225947, EPI_ISL_3225948 | Lighthouse Lab in Milton Keynes | Wellcome Sanger Institute for the COVID-19 Genomics UK (COG-UK) Consortium | The Lighthouse Lab in Milton Keynes and Alex Alderton, Roberto Amato, Jeffrey Barrett, Sonia Goncalves, Ewan Harrison, David K. Jackson, Ian Johnston, Dominic Kwiatkowski, Cordelia Langford, John Sillitoe on behalf of the Wellcome Sanger Institute COVID-19 Surveillance Team |

| | | | |
|---|---|---|---|
| EPI_ISL_3225949 | Lighthouse Lab in Alderley Park | Wellcome Sanger Institute for the COVID-19 Genomics UK (COG-UK) Consortium | Jacquelyn Wynn, Mairead Hyland, The Lighthouse Lab in Alderley Park and Alex Alderton, Roberto Amato, Jeffrey Barrett, Sonia Goncalves, Ewan Harrison, David K. Jackson, Ian Johnston, Dominic Kwiatkowski, Cordelia Langford, John Sillitoe on behalf of the Wellcome Sanger Institute COVID-19 Surveillance Team |
| EPI_ISL_3225950, EPI_ISL_3225951 | Lighthouse Lab in Glasgow | Wellcome Sanger Institute for the COVID-19 Genomics UK (COG-UK) Consortium | Harper VanSteenhouse, Yumi Kasai, David Gray, Carol Clugston, Anna Dominiczak and Alex Alderton, Roberto Amato, Jeffrey Barrett, Sonia Goncalves, Ewan Harrison, David K. Jackson, Ian Johnston, Dominic Kwiatkowski, Cordelia Langford, John Sillitoe on behalf of the Wellcome Sanger Institute COVID-19 Surveillance Team |
| EPI_ISL_3225952 | Lighthouse Laboratory Plymouth | Wellcome Sanger Institute for the COVID-19 Genomics UK (COG-UK) Consortium | Lighthouse Laboratory Plymouth and Alex Alderton, Roberto Amato, Jeffrey Barrett, Sonia Goncalves, Ewan Harrison, David K. Jackson, Ian Johnston, Dominic Kwiatkowski, Cordelia Langford, John Sillitoe on behalf of the Wellcome Sanger Institute COVID-19 Surveillance Team |
| EPI_ISL_3225953, EPI_ISL_3225954, EPI_ISL_3225955 | Lighthouse Lab in Milton Keynes | Wellcome Sanger Institute for the COVID-19 Genomics UK (COG-UK) Consortium | The Lighthouse Lab in Milton Keynes and Alex Alderton, Roberto Amato, Jeffrey Barrett, Sonia Goncalves, Ewan Harrison, David K. Jackson, Ian Johnston, Dominic Kwiatkowski, Cordelia Langford, John Sillitoe on behalf of the Wellcome Sanger Institute COVID-19 Surveillance Team |
| EPI_ISL_3225956, EPI_ISL_3225957 | Lighthouse Laboratory Plymouth | Wellcome Sanger Institute for the COVID-19 Genomics UK (COG-UK) Consortium | Lighthouse Laboratory Plymouth and Alex Alderton, Roberto Amato, Jeffrey Barrett, Sonia Goncalves, Ewan Harrison, David K. Jackson, Ian Johnston, Dominic Kwiatkowski, Cordelia Langford, John Sillitoe on behalf of the Wellcome Sanger Institute COVID-19 Surveillance Team |
| EPI_ISL_3225958, EPI_ISL_3225959 | Lighthouse Lab in Alderley Park | Wellcome Sanger Institute for the COVID-19 Genomics UK (COG-UK) Consortium | Jacquelyn Wynn, Mairead Hyland, The Lighthouse Lab in Alderley Park and Alex Alderton, Roberto Amato, Jeffrey Barrett, Sonia Goncalves, Ewan Harrison, David K. Jackson, Ian Johnston, Dominic Kwiatkowski, Cordelia Langford, John Sillitoe on behalf of the Wellcome Sanger Institute COVID-19 Surveillance Team |
| EPI_ISL_3225960 | Lighthouse Lab in Milton Keynes | Wellcome Sanger Institute for the COVID-19 Genomics UK (COG-UK) Consortium | The Lighthouse Lab in Milton Keynes and Alex Alderton, Roberto Amato, Jeffrey Barrett, Sonia Goncalves, Ewan Harrison, David K. Jackson, Ian Johnston, Dominic Kwiatkowski, Cordelia Langford, John Sillitoe on behalf of the Wellcome Sanger Institute COVID-19 Surveillance Team |
| EPI_ISL_3225961 | Lighthouse Lab in Glasgow | Wellcome Sanger Institute for the COVID-19 Genomics UK (COG-UK) Consortium | Harper VanSteenhouse, Yumi Kasai, David Gray, Carol Clugston, Anna Dominiczak and Alex Alderton, Roberto Amato, Jeffrey Barrett, Sonia Goncalves, Ewan Harrison, David K. Jackson, Ian Johnston, Dominic Kwiatkowski, Cordelia Langford, John Sillitoe on behalf of the Wellcome Sanger Institute COVID-19 Surveillance Team |
| EPI_ISL_3225962 | Lighthouse Lab in Milton Keynes | Wellcome Sanger Institute for the COVID-19 Genomics UK (COG-UK) Consortium | The Lighthouse Lab in Milton Keynes and Alex Alderton, Roberto Amato, Jeffrey Barrett, Sonia Goncalves, Ewan Harrison, David K. Jackson, Ian Johnston, Dominic Kwiatkowski, Cordelia Langford, John Sillitoe on behalf of the Wellcome Sanger Institute COVID-19 Surveillance Team |
| EPI_ISL_3225963 | Lighthouse Lab in Alderley Park | Wellcome Sanger Institute for the COVID-19 Genomics UK (COG-UK) Consortium | Jacquelyn Wynn, Mairead Hyland, The Lighthouse Lab in Alderley Park and Alex Alderton, Roberto Amato, Jeffrey Barrett, Sonia Goncalves, Ewan Harrison, David K. Jackson, Ian Johnston, Dominic Kwiatkowski, Cordelia Langford, John Sillitoe on behalf of the Wellcome Sanger Institute COVID-19 Surveillance Team |
| EPI_ISL_3225964 | Lighthouse Lab in Milton Keynes | Wellcome Sanger Institute for the COVID-19 Genomics UK (COG-UK) Consortium | The Lighthouse Lab in Milton Keynes and Alex Alderton, Roberto Amato, Jeffrey Barrett, Sonia Goncalves, Ewan Harrison, David K. Jackson, Ian Johnston, Dominic Kwiatkowski, Cordelia Langford, John Sillitoe on behalf of the Wellcome Sanger Institute COVID-19 Surveillance Team |
| EPI_ISL_3225965 | Lighthouse Lab in Glasgow | Wellcome Sanger Institute for the COVID-19 Genomics UK (COG-UK) Consortium | Harper VanSteenhouse, Yumi Kasai, David Gray, Carol Clugston, Anna Dominiczak and Alex Alderton, Roberto Amato, Jeffrey Barrett, Sonia Goncalves, Ewan Harrison, David K. Jackson, Ian Johnston, Dominic Kwiatkowski, Cordelia Langford, John Sillitoe on behalf of the Wellcome Sanger Institute COVID-19 Surveillance Team |
| EPI_ISL_3225966, EPI_ISL_3225967, EPI_ISL_3225968 | Lighthouse Laboratory Plymouth | Wellcome Sanger Institute for the COVID-19 Genomics UK (COG-UK) Consortium | Lighthouse Laboratory Plymouth and Alex Alderton, Roberto Amato, Jeffrey Barrett, Sonia Goncalves, Ewan Harrison, David K. Jackson, Ian Johnston, Dominic Kwiatkowski, Cordelia Langford, John Sillitoe on behalf of the Wellcome Sanger Institute COVID-19 Surveillance Team |
| EPI_ISL_3225969 | Lighthouse Lab in Alderley Park | Wellcome Sanger Institute for the COVID-19 Genomics UK (COG-UK) Consortium | Jacquelyn Wynn, Mairead Hyland, The Lighthouse Lab in Alderley Park and Alex Alderton, Roberto Amato, Jeffrey Barrett, Sonia Goncalves, Ewan Harrison, David K. Jackson, Ian Johnston, Dominic Kwiatkowski, Cordelia Langford, John Sillitoe on behalf of the Wellcome Sanger Institute COVID-19 Surveillance Team |
| EPI_ISL_3225970 | Lighthouse Lab in Milton Keynes | Wellcome Sanger Institute for the COVID-19 Genomics UK (COG-UK) Consortium | The Lighthouse Lab in Milton Keynes and Alex Alderton, Roberto Amato, Jeffrey Barrett, Sonia Goncalves, Ewan Harrison, David K. Jackson, Ian Johnston, Dominic Kwiatkowski, Cordelia Langford, John Sillitoe on behalf of the Wellcome Sanger Institute COVID-19 Surveillance Team |
| EPI_ISL_3225971 | Lighthouse Lab in Alderley Park | Wellcome Sanger Institute for the COVID-19 Genomics UK (COG-UK) Consortium | Jacquelyn Wynn, Mairead Hyland, The Lighthouse Lab in Alderley Park and Alex Alderton, Roberto Amato, Jeffrey Barrett, Sonia Goncalves, Ewan Harrison, David K. Jackson, Ian Johnston, Dominic Kwiatkowski, Cordelia Langford, John Sillitoe on behalf of the Wellcome Sanger Institute COVID-19 Surveillance Team |
| EPI_ISL_3225972, EPI_ISL_3225973 | Lighthouse Lab in Glasgow | Wellcome Sanger Institute for the COVID-19 Genomics UK (COG-UK) Consortium | Harper VanSteenhouse, Yumi Kasai, David Gray, Carol Clugston, Anna Dominiczak and Alex Alderton, Roberto Amato, Jeffrey Barrett, Sonia Goncalves, Ewan Harrison, David K. Jackson, Ian Johnston, Dominic Kwiatkowski, Cordelia Langford, John Sillitoe on behalf of the Wellcome Sanger Institute COVID-19 Surveillance Team |
| EPI_ISL_3225974 | Lighthouse Lab in Alderley Park | Wellcome Sanger Institute for the COVID-19 Genomics UK (COG-UK) Consortium | Jacquelyn Wynn, Mairead Hyland, The Lighthouse Lab in Alderley Park and Alex Alderton, Roberto Amato, Jeffrey Barrett, Sonia Goncalves, Ewan Harrison, David K. Jackson, Ian Johnston, Dominic Kwiatkowski, Cordelia Langford, John Sillitoe on behalf of the Wellcome Sanger Institute COVID-19 Surveillance Team |
| EPI_ISL_3225975, EPI_ISL_3225976, EPI_ISL_3225977 | Lighthouse Lab in Milton Keynes | Wellcome Sanger Institute for the COVID-19 Genomics UK (COG-UK) Consortium | The Lighthouse Lab in Milton Keynes and Alex Alderton, Roberto Amato, Jeffrey Barrett, Sonia Goncalves, Ewan Harrison, David K. Jackson, Ian Johnston, Dominic Kwiatkowski, Cordelia Langford, John Sillitoe on behalf of the Wellcome Sanger Institute COVID-19 Surveillance Team |
| EPI_ISL_3225978 | Lighthouse Lab in Alderley Park | Wellcome Sanger Institute for the COVID-19 Genomics UK (COG-UK) Consortium | Jacquelyn Wynn, Mairead Hyland, The Lighthouse Lab in Alderley Park and Alex Alderton, Roberto Amato, Jeffrey Barrett, Sonia Goncalves, Ewan Harrison, David K. Jackson, Ian Johnston, Dominic Kwiatkowski, Cordelia Langford, John Sillitoe on behalf of the Wellcome Sanger Institute COVID-19 Surveillance Team |
| EPI_ISL_3225979 | Lighthouse Lab in Milton Keynes | Wellcome Sanger Institute for the COVID-19 Genomics UK (COG-UK) Consortium | The Lighthouse Lab in Milton Keynes and Alex Alderton, Roberto Amato, Jeffrey Barrett, Sonia Goncalves, Ewan Harrison, David K. Jackson, Ian Johnston, Dominic Kwiatkowski, Cordelia Langford, John Sillitoe on behalf of the Wellcome Sanger Institute COVID-19 Surveillance Team |
| EPI_ISL_3225980 | Lighthouse Lab in Alderley Park | Wellcome Sanger Institute for the COVID-19 Genomics UK (COG-UK) Consortium | Jacquelyn Wynn, Mairead Hyland, The Lighthouse Lab in Alderley Park and Alex Alderton, Roberto Amato, Jeffrey Barrett, Sonia Goncalves, Ewan Harrison, David K. Jackson, Ian Johnston, Dominic Kwiatkowski, Cordelia Langford, John Sillitoe on behalf of the Wellcome Sanger Institute COVID-19 Surveillance Team |
| EPI_ISL_3225981, EPI_ISL_3225982 | Lighthouse Lab in Glasgow | Wellcome Sanger Institute for the COVID-19 Genomics UK (COG-UK) Consortium | Harper VanSteenhouse, Yumi Kasai, David Gray, Carol Clugston, Anna Dominiczak and Alex Alderton, Roberto Amato, Jeffrey Barrett, Sonia Goncalves, Ewan Harrison, David K. Jackson, Ian Johnston, Dominic Kwiatkowski, Cordelia Langford, John Sillitoe on behalf of the Wellcome Sanger Institute COVID-19 Surveillance Team |
| EPI_ISL_3225983 | Lighthouse Lab in Alderley Park | Wellcome Sanger Institute for the COVID-19 Genomics UK (COG-UK) Consortium | Jacquelyn Wynn, Mairead Hyland, The Lighthouse Lab in Alderley Park and Alex Alderton, Roberto Amato, Jeffrey Barrett, Sonia Goncalves, Ewan Harrison, David K. Jackson, Ian Johnston, Dominic Kwiatkowski, Cordelia Langford, John Sillitoe on behalf of the Wellcome Sanger Institute COVID-19 Surveillance Team |
| EPI_ISL_3225984 | Lighthouse Lab in Glasgow | Wellcome Sanger Institute for the COVID-19 Genomics UK (COG-UK) Consortium | Harper VanSteenhouse, Yumi Kasai, David Gray, Carol Clugston, Anna Dominiczak and Alex Alderton, Roberto Amato, Jeffrey Barrett, Sonia Goncalves, Ewan Harrison, David K. Jackson, Ian Johnston, Dominic Kwiatkowski, Cordelia Langford, John Sillitoe on behalf of the Wellcome Sanger Institute COVID-19 Surveillance Team |
| EPI_ISL_3225985 | Lighthouse Lab in Alderley Park | Wellcome Sanger Institute for the COVID-19 Genomics UK (COG-UK) Consortium | Jacquelyn Wynn, Mairead Hyland, The Lighthouse Lab in Alderley Park and Alex Alderton, Roberto Amato, Jeffrey Barrett, Sonia Goncalves, Ewan Harrison, David K. Jackson, Ian Johnston, Dominic Kwiatkowski, Cordelia Langford, John Sillitoe on behalf of the Wellcome Sanger Institute COVID-19 Surveillance Team |
| EPI_ISL_3225986, EPI_ISL_3225987, EPI_ISL_3225988 | Lighthouse Laboratory Plymouth | Wellcome Sanger Institute for the COVID-19 Genomics UK (COG-UK) Consortium | Lighthouse Laboratory Plymouth and Alex Alderton, Roberto Amato, Jeffrey Barrett, Sonia Goncalves, Ewan Harrison, David K. Jackson, Ian Johnston, Dominic Kwiatkowski, Cordelia Langford, John Sillitoe on behalf of the Wellcome Sanger Institute COVID-19 Surveillance Team |
| EPI_ISL_3225989 | Lighthouse Lab in Alderley Park | Wellcome Sanger Institute for the COVID-19 Genomics UK (COG-UK) Consortium | Jacquelyn Wynn, Mairead Hyland, The Lighthouse Lab in Alderley Park and Alex Alderton, Roberto Amato, Jeffrey Barrett, Sonia Goncalves, Ewan Harrison, David K. Jackson, Ian Johnston, Dominic Kwiatkowski, Cordelia Langford, John Sillitoe on behalf of the Wellcome Sanger Institute COVID-19 Surveillance Team |
| EPI_ISL_3225990 | Lighthouse Lab in Milton Keynes | Wellcome Sanger Institute for the COVID-19 Genomics UK (COG-UK) Consortium | The Lighthouse Lab in Milton Keynes and Alex Alderton, Roberto Amato, Jeffrey Barrett, Sonia Goncalves, Ewan Harrison, David K. Jackson, Ian Johnston, Dominic Kwiatkowski, Cordelia Langford, John Sillitoe on behalf of the Wellcome Sanger Institute COVID-19 Surveillance Team |
| EPI_ISL_3225991, EPI_ISL_3225992 | Lighthouse Laboratory Plymouth | Wellcome Sanger Institute for the COVID-19 Genomics UK (COG-UK) Consortium | Lighthouse Laboratory Plymouth and Alex Alderton, Roberto Amato, Jeffrey Barrett, Sonia Goncalves, Ewan Harrison, David K. Jackson, Ian Johnston, Dominic Kwiatkowski, Cordelia Langford, John Sillitoe on behalf of the Wellcome Sanger Institute COVID-19 Surveillance Team |
| EPI_ISL_3225993 | Lighthouse Lab in Glasgow | Wellcome Sanger Institute for the COVID-19 Genomics UK (COG-UK) Consortium | Harper VanSteenhouse, Yumi Kasai, David Gray, Carol Clugston, Anna Dominiczak and Alex Alderton, Roberto Amato, Jeffrey Barrett, Sonia Goncalves, Ewan Harrison, David K. Jackson, Ian Johnston, Dominic Kwiatkowski, Cordelia Langford, John Sillitoe on behalf of the Wellcome Sanger Institute COVID-19 Surveillance Team |
| EPI_ISL_3225994 | Lighthouse Lab in Milton Keynes | Wellcome Sanger Institute for the COVID-19 Genomics UK (COG-UK) Consortium | The Lighthouse Lab in Milton Keynes and Alex Alderton, Roberto Amato, Jeffrey Barrett, Sonia Goncalves, Ewan Harrison, David K. Jackson, Ian Johnston, Dominic Kwiatkowski, Cordelia Langford, John Sillitoe on behalf of the Wellcome Sanger Institute COVID-19 Surveillance Team |
| EPI_ISL_3225995 | Lighthouse Laboratory Plymouth | Wellcome Sanger Institute for the COVID-19 Genomics UK (COG-UK) Consortium | Lighthouse Laboratory Plymouth and Alex Alderton, Roberto Amato, Jeffrey Barrett, Sonia Goncalves, Ewan Harrison, David K. Jackson, Ian Johnston, Dominic Kwiatkowski, Cordelia Langford, John Sillitoe on behalf of the Wellcome Sanger Institute COVID-19 Surveillance Team |
| EPI_ISL_3225996 | Lighthouse Lab in Alderley Park | Wellcome Sanger Institute for the COVID-19 Genomics UK (COG-UK) Consortium | Jacquelyn Wynn, Mairead Hyland, The Lighthouse Lab in Alderley Park and Alex Alderton, Roberto Amato, Jeffrey Barrett, Sonia Goncalves, Ewan Harrison, David K. Jackson, Ian Johnston, Dominic Kwiatkowski, Cordelia Langford, John Sillitoe on behalf of the Wellcome Sanger Institute COVID-19 Surveillance Team |
| EPI_ISL_3225997 | Lighthouse Laboratory Plymouth | Wellcome Sanger Institute for the COVID-19 Genomics UK (COG-UK) Consortium | Lighthouse Laboratory Plymouth and Alex Alderton, Roberto Amato, Jeffrey Barrett, Sonia Goncalves, Ewan Harrison, David K. Jackson, Ian Johnston, Dominic Kwiatkowski, Cordelia Langford, John Sillitoe on behalf of the Wellcome Sanger Institute COVID-19 Surveillance Team |
| EPI_ISL_3225998, EPI_ISL_3225999 | Lighthouse Lab in Glasgow | Wellcome Sanger Institute for the COVID-19 Genomics UK (COG-UK) Consortium | Harper VanSteenhouse, Yumi Kasai, David Gray, Carol Clugston, Anna Dominiczak and Alex Alderton, Roberto Amato, Jeffrey Barrett, Sonia Goncalves, Ewan Harrison, David K. Jackson, Ian Johnston, Dominic Kwiatkowski, Cordelia Langford, John Sillitoe on behalf of the Wellcome Sanger Institute COVID-19 Surveillance Team |
| EPI_ISL_3226000 | Lighthouse Lab in Alderley Park | Wellcome Sanger Institute for the COVID-19 Genomics UK (COG-UK) Consortium | Jacquelyn Wynn, Mairead Hyland, The Lighthouse Lab in Alderley Park and Alex Alderton, Roberto Amato, Jeffrey Barrett, Sonia Goncalves, Ewan Harrison, David K. Jackson, Ian Johnston, Dominic Kwiatkowski, Cordelia Langford, John Sillitoe on behalf of the Wellcome Sanger Institute COVID-19 Surveillance Team |
| EPI_ISL_3226001 | Lighthouse Lab in Milton Keynes | Wellcome Sanger Institute for the COVID-19 Genomics UK (COG-UK) Consortium | The Lighthouse Lab in Milton Keynes and Alex Alderton, Roberto Amato, Jeffrey Barrett, Sonia Goncalves, Ewan Harrison, David K. Jackson, Ian Johnston, Dominic Kwiatkowski, Cordelia Langford, John Sillitoe on behalf of the Wellcome Sanger Institute COVID-19 Surveillance Team |
| EPI_ISL_3226002 | Lighthouse Lab in Alderley Park | Wellcome Sanger Institute for the COVID-19 Genomics UK (COG-UK) Consortium | Jacquelyn Wynn, Mairead Hyland, The Lighthouse Lab in Alderley Park and Alex Alderton, Roberto Amato, Jeffrey Barrett, Sonia Goncalves, Ewan Harrison, David K. Jackson, Ian Johnston, Dominic Kwiatkowski, Cordelia Langford, John Sillitoe on behalf of the Wellcome Sanger Institute COVID-19 Surveillance Team |
| EPI_ISL_3226003 | Lighthouse Lab in Milton Keynes | Wellcome Sanger Institute for the COVID-19 Genomics UK (COG-UK) Consortium | The Lighthouse Lab in Milton Keynes and Alex Alderton, Roberto Amato, Jeffrey Barrett, Sonia Goncalves, Ewan Harrison, David K. Jackson, Ian Johnston, Dominic Kwiatkowski, Cordelia Langford, John Sillitoe on behalf of the Wellcome Sanger Institute COVID-19 Surveillance Team |
| EPI_ISL_3226004 | Lighthouse Lab in Glasgow | Wellcome Sanger Institute for the COVID-19 Genomics UK (COG-UK) Consortium | Harper VanSteenhouse, Yumi Kasai, David Gray, Carol Clugston, Anna Dominiczak and Alex Alderton, Roberto Amato, Jeffrey Barrett, Sonia Goncalves, Ewan Harrison, David K. Jackson, Ian Johnston, Dominic Kwiatkowski, Cordelia Langford, John Sillitoe on behalf of the Wellcome Sanger Institute COVID-19 Surveillance Team |
| EPI_ISL_3226005 | Lighthouse Laboratory Plymouth | Wellcome Sanger Institute for the COVID-19 Genomics UK (COG-UK) Consortium | Lighthouse Laboratory Plymouth and Alex Alderton, Roberto Amato, Jeffrey Barrett, Sonia Goncalves, Ewan Harrison, David K. Jackson, Ian Johnston, Dominic Kwiatkowski, Cordelia Langford, John Sillitoe on behalf of the Wellcome Sanger Institute COVID-19 Surveillance Team |
| EPI_ISL_3226006 | Lighthouse Lab in Alderley Park | Wellcome Sanger Institute for the COVID-19 Genomics UK (COG-UK) Consortium | Jacquelyn Wynn, Mairead Hyland, The Lighthouse Lab in Alderley Park and Alex Alderton, Roberto Amato, Jeffrey Barrett, Sonia Goncalves, Ewan Harrison, David K. Jackson, Ian Johnston, Dominic Kwiatkowski, Cordelia Langford, John Sillitoe on behalf of the Wellcome Sanger Institute COVID-19 Surveillance Team |
| EPI_ISL_3226007, | Lighthouse Laboratory Plymouth | Wellcome Sanger Institute for the COVID-19 Genomics UK (COG-UK) | Lighthouse Laboratory Plymouth and Alex Alderton, Roberto Amato, Jeffrey Barrett, Sonia Goncalves, Ewan Harrison, David K. Jackson, Ian Johnston, Dominic Kwiatkowski, Cordelia Langford, John Sillitoe on behalf of the Wellcome Sanger |

| | | | |
|---|---|---|---|
| EPI_ISL_3226008 | | Consortium | Institute COVID-19 Surveillance Team |
| EPI_ISL_3226009 | Lighthouse Lab in Milton Keynes | Wellcome Sanger Institute for the COVID-19 Genomics UK (COG-UK) Consortium | The Lighthouse Lab in Milton Keynes and Alex Alderton, Roberto Amato, Jeffrey Barrett, Sonia Goncalves, Ewan Harrison, David K. Jackson, Ian Johnston, Dominic Kwiatkowski, Cordelia Langford, John Sillitoe on behalf of the Wellcome Sanger Institute COVID-19 Surveillance Team |
| EPI_ISL_3226010 | Lighthouse Lab in Alderley Park | Wellcome Sanger Institute for the COVID-19 Genomics UK (COG-UK) Consortium | Jacquelyn Wynn, Mairead Hyland, The Lighthouse Lab in Alderley Park and Alex Alderton, Roberto Amato, Jeffrey Barrett, Sonia Goncalves, Ewan Harrison, David K. Jackson, Ian Johnston, Dominic Kwiatkowski, Cordelia Langford, John Sillitoe on behalf of the Wellcome Sanger Institute COVID-19 Surveillance Team |
| EPI_ISL_3226011 | Lighthouse Lab in Glasgow | Wellcome Sanger Institute for the COVID-19 Genomics UK (COG-UK) Consortium | Harper VanSteenhouse, Yumi Kasai, David Gray, Carol Clugston, Anna Dominiczak and Alex Alderton, Roberto Amato, Jeffrey Barrett, Sonia Goncalves, Ewan Harrison, David K. Jackson, Ian Johnston, Dominic Kwiatkowski, Cordelia Langford, John Sillitoe on behalf of the Wellcome Sanger Institute COVID-19 Surveillance Team |
| EPI_ISL_3226012 | Lighthouse Lab in Alderley Park | Wellcome Sanger Institute for the COVID-19 Genomics UK (COG-UK) Consortium | Jacquelyn Wynn, Mairead Hyland, The Lighthouse Lab in Alderley Park and Alex Alderton, Roberto Amato, Jeffrey Barrett, Sonia Goncalves, Ewan Harrison, David K. Jackson, Ian Johnston, Dominic Kwiatkowski, Cordelia Langford, John Sillitoe on behalf of the Wellcome Sanger Institute COVID-19 Surveillance Team |
| EPI_ISL_3226013 | Lighthouse Lab in Milton Keynes | Wellcome Sanger Institute for the COVID-19 Genomics UK (COG-UK) Consortium | The Lighthouse Lab in Milton Keynes and Alex Alderton, Roberto Amato, Jeffrey Barrett, Sonia Goncalves, Ewan Harrison, David K. Jackson, Ian Johnston, Dominic Kwiatkowski, Cordelia Langford, John Sillitoe on behalf of the Wellcome Sanger Institute COVID-19 Surveillance Team |
| EPI_ISL_3226014 | Lighthouse Laboratory Plymouth | Wellcome Sanger Institute for the COVID-19 Genomics UK (COG-UK) Consortium | Lighthouse Laboratory Plymouth and Alex Alderton, Roberto Amato, Jeffrey Barrett, Sonia Goncalves, Ewan Harrison, David K. Jackson, Ian Johnston, Dominic Kwiatkowski, Cordelia Langford, John Sillitoe on behalf of the Wellcome Sanger Institute COVID-19 Surveillance Team |
| EPI_ISL_3226015 | Lighthouse Lab in Glasgow | Wellcome Sanger Institute for the COVID-19 Genomics UK (COG-UK) Consortium | Harper VanSteenhouse, Yumi Kasai, David Gray, Carol Clugston, Anna Dominiczak and Alex Alderton, Roberto Amato, Jeffrey Barrett, Sonia Goncalves, Ewan Harrison, David K. Jackson, Ian Johnston, Dominic Kwiatkowski, Cordelia Langford, John Sillitoe on behalf of the Wellcome Sanger Institute COVID-19 Surveillance Team |
| EPI_ISL_3226016 | Lighthouse Lab in Alderley Park | Wellcome Sanger Institute for the COVID-19 Genomics UK (COG-UK) Consortium | Jacquelyn Wynn, Mairead Hyland, The Lighthouse Lab in Alderley Park and Alex Alderton, Roberto Amato, Jeffrey Barrett, Sonia Goncalves, Ewan Harrison, David K. Jackson, Ian Johnston, Dominic Kwiatkowski, Cordelia Langford, John Sillitoe on behalf of the Wellcome Sanger Institute COVID-19 Surveillance Team |
| EPI_ISL_3226017 | Lighthouse Laboratory Plymouth | Wellcome Sanger Institute for the COVID-19 Genomics UK (COG-UK) Consortium | Lighthouse Laboratory Plymouth and Alex Alderton, Roberto Amato, Jeffrey Barrett, Sonia Goncalves, Ewan Harrison, David K. Jackson, Ian Johnston, Dominic Kwiatkowski, Cordelia Langford, John Sillitoe on behalf of the Wellcome Sanger Institute COVID-19 Surveillance Team |
| EPI_ISL_3226018, EPI_ISL_3226019 | Lighthouse Lab in Alderley Park | Wellcome Sanger Institute for the COVID-19 Genomics UK (COG-UK) Consortium | Jacquelyn Wynn, Mairead Hyland, The Lighthouse Lab in Alderley Park and Alex Alderton, Roberto Amato, Jeffrey Barrett, Sonia Goncalves, Ewan Harrison, David K. Jackson, Ian Johnston, Dominic Kwiatkowski, Cordelia Langford, John Sillitoe on behalf of the Wellcome Sanger Institute COVID-19 Surveillance Team |
| EPI_ISL_3226020 | Lighthouse Laboratory Plymouth | Wellcome Sanger Institute for the COVID-19 Genomics UK (COG-UK) Consortium | Lighthouse Laboratory Plymouth and Alex Alderton, Roberto Amato, Jeffrey Barrett, Sonia Goncalves, Ewan Harrison, David K. Jackson, Ian Johnston, Dominic Kwiatkowski, Cordelia Langford, John Sillitoe on behalf of the Wellcome Sanger Institute COVID-19 Surveillance Team |
| EPI_ISL_3226021 | Lighthouse Lab in Milton Keynes | Wellcome Sanger Institute for the COVID-19 Genomics UK (COG-UK) Consortium | The Lighthouse Lab in Milton Keynes and Alex Alderton, Roberto Amato, Jeffrey Barrett, Sonia Goncalves, Ewan Harrison, David K. Jackson, Ian Johnston, Dominic Kwiatkowski, Cordelia Langford, John Sillitoe on behalf of the Wellcome Sanger Institute COVID-19 Surveillance Team |
| EPI_ISL_3226022 | Lighthouse Lab in Alderley Park | Wellcome Sanger Institute for the COVID-19 Genomics UK (COG-UK) Consortium | Jacquelyn Wynn, Mairead Hyland, The Lighthouse Lab in Alderley Park and Alex Alderton, Roberto Amato, Jeffrey Barrett, Sonia Goncalves, Ewan Harrison, David K. Jackson, Ian Johnston, Dominic Kwiatkowski, Cordelia Langford, John Sillitoe on behalf of the Wellcome Sanger Institute COVID-19 Surveillance Team |
| EPI_ISL_3226023, EPI_ISL_3226024 | Lighthouse Lab in Glasgow | Wellcome Sanger Institute for the COVID-19 Genomics UK (COG-UK) Consortium | Harper VanSteenhouse, Yumi Kasai, David Gray, Carol Clugston, Anna Dominiczak and Alex Alderton, Roberto Amato, Jeffrey Barrett, Sonia Goncalves, Ewan Harrison, David K. Jackson, Ian Johnston, Dominic Kwiatkowski, Cordelia Langford, John Sillitoe on behalf of the Wellcome Sanger Institute COVID-19 Surveillance Team |
| EPI_ISL_3226025 | Lighthouse Laboratory Plymouth | Wellcome Sanger Institute for the COVID-19 Genomics UK (COG-UK) Consortium | Lighthouse Laboratory Plymouth and Alex Alderton, Roberto Amato, Jeffrey Barrett, Sonia Goncalves, Ewan Harrison, David K. Jackson, Ian Johnston, Dominic Kwiatkowski, Cordelia Langford, John Sillitoe on behalf of the Wellcome Sanger Institute COVID-19 Surveillance Team |
| EPI_ISL_3226026 | Lighthouse Lab in Glasgow | Wellcome Sanger Institute for the COVID-19 Genomics UK (COG-UK) Consortium | Harper VanSteenhouse, Yumi Kasai, David Gray, Carol Clugston, Anna Dominiczak and Alex Alderton, Roberto Amato, Jeffrey Barrett, Sonia Goncalves, Ewan Harrison, David K. Jackson, Ian Johnston, Dominic Kwiatkowski, Cordelia Langford, John Sillitoe on behalf of the Wellcome Sanger Institute COVID-19 Surveillance Team |
| EPI_ISL_3226027 | Lighthouse Lab in Milton Keynes | Wellcome Sanger Institute for the COVID-19 Genomics UK (COG-UK) Consortium | The Lighthouse Lab in Milton Keynes and Alex Alderton, Roberto Amato, Jeffrey Barrett, Sonia Goncalves, Ewan Harrison, David K. Jackson, Ian Johnston, Dominic Kwiatkowski, Cordelia Langford, John Sillitoe on behalf of the Wellcome Sanger Institute COVID-19 Surveillance Team |
| EPI_ISL_3226028 | Lighthouse Lab in Alderley Park | Wellcome Sanger Institute for the COVID-19 Genomics UK (COG-UK) Consortium | Jacquelyn Wynn, Mairead Hyland, The Lighthouse Lab in Alderley Park and Alex Alderton, Roberto Amato, Jeffrey Barrett, Sonia Goncalves, Ewan Harrison, David K. Jackson, Ian Johnston, Dominic Kwiatkowski, Cordelia Langford, John Sillitoe on behalf of the Wellcome Sanger Institute COVID-19 Surveillance Team |
| EPI_ISL_3226029, EPI_ISL_3226030 | Lighthouse Lab in Glasgow | Wellcome Sanger Institute for the COVID-19 Genomics UK (COG-UK) Consortium | Harper VanSteenhouse, Yumi Kasai, David Gray, Carol Clugston, Anna Dominiczak and Alex Alderton, Roberto Amato, Jeffrey Barrett, Sonia Goncalves, Ewan Harrison, David K. Jackson, Ian Johnston, Dominic Kwiatkowski, Cordelia Langford, John Sillitoe on behalf of the Wellcome Sanger Institute COVID-19 Surveillance Team |
| EPI_ISL_3226031, EPI_ISL_3226032 | Lighthouse Laboratory Plymouth | Wellcome Sanger Institute for the COVID-19 Genomics UK (COG-UK) Consortium | Lighthouse Laboratory Plymouth and Alex Alderton, Roberto Amato, Jeffrey Barrett, Sonia Goncalves, Ewan Harrison, David K. Jackson, Ian Johnston, Dominic Kwiatkowski, Cordelia Langford, John Sillitoe on behalf of the Wellcome Sanger Institute COVID-19 Surveillance Team |
| EPI_ISL_3226033, EPI_ISL_3226034 | Lighthouse Lab in Milton Keynes | Wellcome Sanger Institute for the COVID-19 Genomics UK (COG-UK) Consortium | The Lighthouse Lab in Milton Keynes and Alex Alderton, Roberto Amato, Jeffrey Barrett, Sonia Goncalves, Ewan Harrison, David K. Jackson, Ian Johnston, Dominic Kwiatkowski, Cordelia Langford, John Sillitoe on behalf of the Wellcome Sanger Institute COVID-19 Surveillance Team |
| EPI_ISL_3226035 | Lighthouse Lab in Glasgow | Wellcome Sanger Institute for the COVID-19 Genomics UK (COG-UK) Consortium | Harper VanSteenhouse, Yumi Kasai, David Gray, Carol Clugston, Anna Dominiczak and Alex Alderton, Roberto Amato, Jeffrey Barrett, Sonia Goncalves, Ewan Harrison, David K. Jackson, Ian Johnston, Dominic Kwiatkowski, Cordelia Langford, John Sillitoe on behalf of the Wellcome Sanger Institute COVID-19 Surveillance Team |
| EPI_ISL_3226036, EPI_ISL_3226037 | Lighthouse Lab in Alderley Park | Wellcome Sanger Institute for the COVID-19 Genomics UK (COG-UK) Consortium | Jacquelyn Wynn, Mairead Hyland, The Lighthouse Lab in Alderley Park and Alex Alderton, Roberto Amato, Jeffrey Barrett, Sonia Goncalves, Ewan Harrison, David K. Jackson, Ian Johnston, Dominic Kwiatkowski, Cordelia Langford, John Sillitoe on behalf of the Wellcome Sanger Institute COVID-19 Surveillance Team |
| EPI_ISL_3226038 | Lighthouse Lab in Milton Keynes | Wellcome Sanger Institute for the COVID-19 Genomics UK (COG-UK) Consortium | The Lighthouse Lab in Milton Keynes and Alex Alderton, Roberto Amato, Jeffrey Barrett, Sonia Goncalves, Ewan Harrison, David K. Jackson, Ian Johnston, Dominic Kwiatkowski, Cordelia Langford, John Sillitoe on behalf of the Wellcome Sanger Institute COVID-19 Surveillance Team |
| EPI_ISL_3226039 | Lighthouse Laboratory Plymouth | Wellcome Sanger Institute for the COVID-19 Genomics UK (COG-UK) Consortium | Lighthouse Laboratory Plymouth and Alex Alderton, Roberto Amato, Jeffrey Barrett, Sonia Goncalves, Ewan Harrison, David K. Jackson, Ian Johnston, Dominic Kwiatkowski, Cordelia Langford, John Sillitoe on behalf of the Wellcome Sanger Institute COVID-19 Surveillance Team |
| EPI_ISL_3226040 | Lighthouse Lab in Alderley Park | Wellcome Sanger Institute for the COVID-19 Genomics UK (COG-UK) Consortium | Jacquelyn Wynn, Mairead Hyland, The Lighthouse Lab in Alderley Park and Alex Alderton, Roberto Amato, Jeffrey Barrett, Sonia Goncalves, Ewan Harrison, David K. Jackson, Ian Johnston, Dominic Kwiatkowski, Cordelia Langford, John Sillitoe on behalf of the Wellcome Sanger Institute COVID-19 Surveillance Team |
| EPI_ISL_3226041 | Lighthouse Lab in Glasgow | Wellcome Sanger Institute for the COVID-19 Genomics UK (COG-UK) Consortium | Harper VanSteenhouse, Yumi Kasai, David Gray, Carol Clugston, Anna Dominiczak and Alex Alderton, Roberto Amato, Jeffrey Barrett, Sonia Goncalves, Ewan Harrison, David K. Jackson, Ian Johnston, Dominic Kwiatkowski, Cordelia Langford, John Sillitoe on behalf of the Wellcome Sanger Institute COVID-19 Surveillance Team |
| EPI_ISL_3226042, EPI_ISL_3226043 | Lighthouse Lab in Milton Keynes | Wellcome Sanger Institute for the COVID-19 Genomics UK (COG-UK) Consortium | The Lighthouse Lab in Milton Keynes and Alex Alderton, Roberto Amato, Jeffrey Barrett, Sonia Goncalves, Ewan Harrison, David K. Jackson, Ian Johnston, Dominic Kwiatkowski, Cordelia Langford, John Sillitoe on behalf of the Wellcome Sanger Institute COVID-19 Surveillance Team |
| EPI_ISL_3226044 | Lighthouse Lab in Alderley Park | Wellcome Sanger Institute for the COVID-19 Genomics UK (COG-UK) Consortium | Jacquelyn Wynn, Mairead Hyland, The Lighthouse Lab in Alderley Park and Alex Alderton, Roberto Amato, Jeffrey Barrett, Sonia Goncalves, Ewan Harrison, David K. Jackson, Ian Johnston, Dominic Kwiatkowski, Cordelia Langford, John Sillitoe on behalf of the Wellcome Sanger Institute COVID-19 Surveillance Team |
| EPI_ISL_3226045 | Lighthouse Lab in Glasgow | Wellcome Sanger Institute for the COVID-19 Genomics UK (COG-UK) Consortium | Harper VanSteenhouse, Yumi Kasai, David Gray, Carol Clugston, Anna Dominiczak and Alex Alderton, Roberto Amato, Jeffrey Barrett, Sonia Goncalves, Ewan Harrison, David K. Jackson, Ian Johnston, Dominic Kwiatkowski, Cordelia Langford, John Sillitoe on behalf of the Wellcome Sanger Institute COVID-19 Surveillance Team |
| EPI_ISL_3226046 | Lighthouse Lab in Milton Keynes | Wellcome Sanger Institute for the COVID-19 Genomics UK (COG-UK) Consortium | The Lighthouse Lab in Milton Keynes and Alex Alderton, Roberto Amato, Jeffrey Barrett, Sonia Goncalves, Ewan Harrison, David K. Jackson, Ian Johnston, Dominic Kwiatkowski, Cordelia Langford, John Sillitoe on behalf of the Wellcome Sanger Institute COVID-19 Surveillance Team |
| EPI_ISL_3226047 | Lighthouse Laboratory Plymouth | Wellcome Sanger Institute for the COVID-19 Genomics UK (COG-UK) Consortium | Lighthouse Laboratory Plymouth and Alex Alderton, Roberto Amato, Jeffrey Barrett, Sonia Goncalves, Ewan Harrison, David K. Jackson, Ian Johnston, Dominic Kwiatkowski, Cordelia Langford, John Sillitoe on behalf of the Wellcome Sanger Institute COVID-19 Surveillance Team |
| EPI_ISL_3226048, EPI_ISL_3226049 | Lighthouse Lab in Milton Keynes | Wellcome Sanger Institute for the COVID-19 Genomics UK (COG-UK) Consortium | The Lighthouse Lab in Milton Keynes and Alex Alderton, Roberto Amato, Jeffrey Barrett, Sonia Goncalves, Ewan Harrison, David K. Jackson, Ian Johnston, Dominic Kwiatkowski, Cordelia Langford, John Sillitoe on behalf of the Wellcome Sanger Institute COVID-19 Surveillance Team |
| EPI_ISL_3226050 | Lighthouse Lab in Glasgow | Wellcome Sanger Institute for the COVID-19 Genomics UK (COG-UK) Consortium | Harper VanSteenhouse, Yumi Kasai, David Gray, Carol Clugston, Anna Dominiczak and Alex Alderton, Roberto Amato, Jeffrey Barrett, Sonia Goncalves, Ewan Harrison, David K. Jackson, Ian Johnston, Dominic Kwiatkowski, Cordelia Langford, John Sillitoe on behalf of the Wellcome Sanger Institute COVID-19 Surveillance Team |
| EPI_ISL_3226051 | Lighthouse Lab in Alderley Park | Wellcome Sanger Institute for the COVID-19 Genomics UK (COG-UK) Consortium | Jacquelyn Wynn, Mairead Hyland, The Lighthouse Lab in Alderley Park and Alex Alderton, Roberto Amato, Jeffrey Barrett, Sonia Goncalves, Ewan Harrison, David K. Jackson, Ian Johnston, Dominic Kwiatkowski, Cordelia Langford, John Sillitoe on behalf of the Wellcome Sanger Institute COVID-19 Surveillance Team |
| EPI_ISL_3226052 | Lighthouse Lab in Glasgow | Wellcome Sanger Institute for the COVID-19 Genomics UK (COG-UK) Consortium | Harper VanSteenhouse, Yumi Kasai, David Gray, Carol Clugston, Anna Dominiczak and Alex Alderton, Roberto Amato, Jeffrey Barrett, Sonia Goncalves, Ewan Harrison, David K. Jackson, Ian Johnston, Dominic Kwiatkowski, Cordelia Langford, John Sillitoe on behalf of the Wellcome Sanger Institute COVID-19 Surveillance Team |
| EPI_ISL_3226053, EPI_ISL_3226054, EPI_ISL_3226055 | Lighthouse Lab in Alderley Park | Wellcome Sanger Institute for the COVID-19 Genomics UK (COG-UK) Consortium | Jacquelyn Wynn, Mairead Hyland, The Lighthouse Lab in Alderley Park and Alex Alderton, Roberto Amato, Jeffrey Barrett, Sonia Goncalves, Ewan Harrison, David K. Jackson, Ian Johnston, Dominic Kwiatkowski, Cordelia Langford, John Sillitoe on behalf of the Wellcome Sanger Institute COVID-19 Surveillance Team |
| EPI_ISL_3226056 | Lighthouse Lab in Milton Keynes | Wellcome Sanger Institute for the COVID-19 Genomics UK (COG-UK) Consortium | The Lighthouse Lab in Milton Keynes and Alex Alderton, Roberto Amato, Jeffrey Barrett, Sonia Goncalves, Ewan Harrison, David K. Jackson, Ian Johnston, Dominic Kwiatkowski, Cordelia Langford, John Sillitoe on behalf of the Wellcome Sanger Institute COVID-19 Surveillance Team |
| EPI_ISL_3226057 | Lighthouse Laboratory Plymouth | Wellcome Sanger Institute for the COVID-19 Genomics UK (COG-UK) Consortium | Lighthouse Laboratory Plymouth and Alex Alderton, Roberto Amato, Jeffrey Barrett, Sonia Goncalves, Ewan Harrison, David K. Jackson, Ian Johnston, Dominic Kwiatkowski, Cordelia Langford, John Sillitoe on behalf of the Wellcome Sanger Institute COVID-19 Surveillance Team |
| EPI_ISL_3226058 | Lighthouse Lab in Glasgow | Wellcome Sanger Institute for the COVID-19 Genomics UK (COG-UK) Consortium | Harper VanSteenhouse, Yumi Kasai, David Gray, Carol Clugston, Anna Dominiczak and Alex Alderton, Roberto Amato, Jeffrey Barrett, Sonia Goncalves, Ewan Harrison, David K. Jackson, Ian Johnston, Dominic Kwiatkowski, Cordelia Langford, John Sillitoe on behalf of the Wellcome Sanger Institute COVID-19 Surveillance Team |
| EPI_ISL_3226059, EPI_ISL_3226060 | Lighthouse Laboratory Plymouth | Wellcome Sanger Institute for the COVID-19 Genomics UK (COG-UK) Consortium | Lighthouse Laboratory Plymouth and Alex Alderton, Roberto Amato, Jeffrey Barrett, Sonia Goncalves, Ewan Harrison, David K. Jackson, Ian Johnston, Dominic Kwiatkowski, Cordelia Langford, John Sillitoe on behalf of the Wellcome Sanger Institute COVID-19 Surveillance Team |
| EPI_ISL_3226061, EPI_ISL_3226062 | Lighthouse Lab in Glasgow | Wellcome Sanger Institute for the COVID-19 Genomics UK (COG-UK) Consortium | Harper VanSteenhouse, Yumi Kasai, David Gray, Carol Clugston, Anna Dominiczak and Alex Alderton, Roberto Amato, Jeffrey Barrett, Sonia Goncalves, Ewan Harrison, David K. Jackson, Ian Johnston, Dominic Kwiatkowski, Cordelia Langford, John Sillitoe on behalf of the Wellcome Sanger Institute COVID-19 Surveillance Team |
| EPI_ISL_3226063, EPI_ISL_3226064 | Lighthouse Lab in Milton Keynes | Wellcome Sanger Institute for the COVID-19 Genomics UK (COG-UK) Consortium | The Lighthouse Lab in Milton Keynes and Alex Alderton, Roberto Amato, Jeffrey Barrett, Sonia Goncalves, Ewan Harrison, David K. Jackson, Ian Johnston, Dominic Kwiatkowski, Cordelia Langford, John Sillitoe on behalf of the Wellcome Sanger Institute COVID-19 Surveillance Team |
| EPI_ISL_3226065, EPI_ISL_3226066, EPI_ISL_3226067 | Lighthouse Lab in Alderley Park | Wellcome Sanger Institute for the COVID-19 Genomics UK (COG-UK) Consortium | Jacquelyn Wynn, Mairead Hyland, The Lighthouse Lab in Alderley Park and Alex Alderton, Roberto Amato, Jeffrey Barrett, Sonia Goncalves, Ewan Harrison, David K. Jackson, Ian Johnston, Dominic Kwiatkowski, Cordelia Langford, John Sillitoe on behalf of the Wellcome Sanger Institute COVID-19 Surveillance Team |

| | | | |
|---|---|---|---|
| EPI_ISL_3226068 | Lighthouse Laboratory Plymouth | Wellcome Sanger Institute for the COVID-19 Genomics UK (COG-UK) Consortium | Lighthouse Laboratory Plymouth and Alex Alderton, Roberto Amato, Jeffrey Barrett, Sonia Goncalves, Ewan Harrison, David K. Jackson, Ian Johnston, Dominic Kwiatkowski, Cordelia Langford, John Sillitoe on behalf of the Wellcome Sanger Institute COVID-19 Surveillance Team |
| EPI_ISL_3226069 | Lighthouse Lab in Alderley Park | Wellcome Sanger Institute for the COVID-19 Genomics UK (COG-UK) Consortium | Jacquelyn Wynn, Mairead Hyland, The Lighthouse Lab in Alderley Park and Alex Alderton, Roberto Amato, Jeffrey Barrett, Sonia Goncalves, Ewan Harrison, David K. Jackson, Ian Johnston, Dominic Kwiatkowski, Cordelia Langford, John Sillitoe on behalf of the Wellcome Sanger Institute COVID-19 Surveillance Team |
| EPI_ISL_3226070 | Lighthouse Lab in Milton Keynes | Wellcome Sanger Institute for the COVID-19 Genomics UK (COG-UK) Consortium | The Lighthouse Lab in Milton Keynes and Alex Alderton, Roberto Amato, Jeffrey Barrett, Sonia Goncalves, Ewan Harrison, David K. Jackson, Ian Johnston, Dominic Kwiatkowski, Cordelia Langford, John Sillitoe on behalf of the Wellcome Sanger Institute COVID-19 Surveillance Team |
| EPI_ISL_3226071, EPI_ISL_3226072 | Lighthouse Lab in Glasgow | Wellcome Sanger Institute for the COVID-19 Genomics UK (COG-UK) Consortium | Harper VanSteenhouse, Yumi Kasai, David Gray, Carol Clugston, Anna Dominiczak and Alex Alderton, Roberto Amato, Jeffrey Barrett, Sonia Goncalves, Ewan Harrison, David K. Jackson, Ian Johnston, Dominic Kwiatkowski, Cordelia Langford, John Sillitoe on behalf of the Wellcome Sanger Institute COVID-19 Surveillance Team |
| EPI_ISL_3226073 | Lighthouse Lab in Milton Keynes | Wellcome Sanger Institute for the COVID-19 Genomics UK (COG-UK) Consortium | The Lighthouse Lab in Milton Keynes and Alex Alderton, Roberto Amato, Jeffrey Barrett, Sonia Goncalves, Ewan Harrison, David K. Jackson, Ian Johnston, Dominic Kwiatkowski, Cordelia Langford, John Sillitoe on behalf of the Wellcome Sanger Institute COVID-19 Surveillance Team |
| EPI_ISL_3226074 | Lighthouse Laboratory Plymouth | Wellcome Sanger Institute for the COVID-19 Genomics UK (COG-UK) Consortium | Lighthouse Laboratory Plymouth and Alex Alderton, Roberto Amato, Jeffrey Barrett, Sonia Goncalves, Ewan Harrison, David K. Jackson, Ian Johnston, Dominic Kwiatkowski, Cordelia Langford, John Sillitoe on behalf of the Wellcome Sanger Institute COVID-19 Surveillance Team |
| EPI_ISL_3226075, EPI_ISL_3226076, EPI_ISL_3226077 | Lighthouse Lab in Glasgow | Wellcome Sanger Institute for the COVID-19 Genomics UK (COG-UK) Consortium | Harper VanSteenhouse, Yumi Kasai, David Gray, Carol Clugston, Anna Dominiczak and Alex Alderton, Roberto Amato, Jeffrey Barrett, Sonia Goncalves, Ewan Harrison, David K. Jackson, Ian Johnston, Dominic Kwiatkowski, Cordelia Langford, John Sillitoe on behalf of the Wellcome Sanger Institute COVID-19 Surveillance Team |
| EPI_ISL_3226078 | Lighthouse Lab in Milton Keynes | Wellcome Sanger Institute for the COVID-19 Genomics UK (COG-UK) Consortium | The Lighthouse Lab in Milton Keynes and Alex Alderton, Roberto Amato, Jeffrey Barrett, Sonia Goncalves, Ewan Harrison, David K. Jackson, Ian Johnston, Dominic Kwiatkowski, Cordelia Langford, John Sillitoe on behalf of the Wellcome Sanger Institute COVID-19 Surveillance Team |
| EPI_ISL_3226079 | Lighthouse Lab in Glasgow | Wellcome Sanger Institute for the COVID-19 Genomics UK (COG-UK) Consortium | Harper VanSteenhouse, Yumi Kasai, David Gray, Carol Clugston, Anna Dominiczak and Alex Alderton, Roberto Amato, Jeffrey Barrett, Sonia Goncalves, Ewan Harrison, David K. Jackson, Ian Johnston, Dominic Kwiatkowski, Cordelia Langford, John Sillitoe on behalf of the Wellcome Sanger Institute COVID-19 Surveillance Team |
| EPI_ISL_3226080 | Lighthouse Lab in Alderley Park | Wellcome Sanger Institute for the COVID-19 Genomics UK (COG-UK) Consortium | Jacquelyn Wynn, Mairead Hyland, The Lighthouse Lab in Alderley Park and Alex Alderton, Roberto Amato, Jeffrey Barrett, Sonia Goncalves, Ewan Harrison, David K. Jackson, Ian Johnston, Dominic Kwiatkowski, Cordelia Langford, John Sillitoe on behalf of the Wellcome Sanger Institute COVID-19 Surveillance Team |
| EPI_ISL_3226081 | Lighthouse Lab in Milton Keynes | Wellcome Sanger Institute for the COVID-19 Genomics UK (COG-UK) Consortium | The Lighthouse Lab in Milton Keynes and Alex Alderton, Roberto Amato, Jeffrey Barrett, Sonia Goncalves, Ewan Harrison, David K. Jackson, Ian Johnston, Dominic Kwiatkowski, Cordelia Langford, John Sillitoe on behalf of the Wellcome Sanger Institute COVID-19 Surveillance Team |
| EPI_ISL_3226082 | Lighthouse Lab in Alderley Park | Wellcome Sanger Institute for the COVID-19 Genomics UK (COG-UK) Consortium | Jacquelyn Wynn, Mairead Hyland, The Lighthouse Lab in Alderley Park and Alex Alderton, Roberto Amato, Jeffrey Barrett, Sonia Goncalves, Ewan Harrison, David K. Jackson, Ian Johnston, Dominic Kwiatkowski, Cordelia Langford, John Sillitoe on behalf of the Wellcome Sanger Institute COVID-19 Surveillance Team |
| EPI_ISL_3226083 | Lighthouse Lab in Glasgow | Wellcome Sanger Institute for the COVID-19 Genomics UK (COG-UK) Consortium | Harper VanSteenhouse, Yumi Kasai, David Gray, Carol Clugston, Anna Dominiczak and Alex Alderton, Roberto Amato, Jeffrey Barrett, Sonia Goncalves, Ewan Harrison, David K. Jackson, Ian Johnston, Dominic Kwiatkowski, Cordelia Langford, John Sillitoe on behalf of the Wellcome Sanger Institute COVID-19 Surveillance Team |
| EPI_ISL_3226084 | Lighthouse Laboratory Plymouth | Wellcome Sanger Institute for the COVID-19 Genomics UK (COG-UK) Consortium | Lighthouse Laboratory Plymouth and Alex Alderton, Roberto Amato, Jeffrey Barrett, Sonia Goncalves, Ewan Harrison, David K. Jackson, Ian Johnston, Dominic Kwiatkowski, Cordelia Langford, John Sillitoe on behalf of the Wellcome Sanger Institute COVID-19 Surveillance Team |
| EPI_ISL_3226085 | Lighthouse Lab in Alderley Park | Wellcome Sanger Institute for the COVID-19 Genomics UK (COG-UK) Consortium | Jacquelyn Wynn, Mairead Hyland, The Lighthouse Lab in Alderley Park and Alex Alderton, Roberto Amato, Jeffrey Barrett, Sonia Goncalves, Ewan Harrison, David K. Jackson, Ian Johnston, Dominic Kwiatkowski, Cordelia Langford, John Sillitoe on behalf of the Wellcome Sanger Institute COVID-19 Surveillance Team |
| EPI_ISL_3226086 | Lighthouse Lab in Glasgow | Wellcome Sanger Institute for the COVID-19 Genomics UK (COG-UK) Consortium | Harper VanSteenhouse, Yumi Kasai, David Gray, Carol Clugston, Anna Dominiczak and Alex Alderton, Roberto Amato, Jeffrey Barrett, Sonia Goncalves, Ewan Harrison, David K. Jackson, Ian Johnston, Dominic Kwiatkowski, Cordelia Langford, John Sillitoe on behalf of the Wellcome Sanger Institute COVID-19 Surveillance Team |
| EPI_ISL_3226087 | Lighthouse Lab in Milton Keynes | Wellcome Sanger Institute for the COVID-19 Genomics UK (COG-UK) Consortium | The Lighthouse Lab in Milton Keynes and Alex Alderton, Roberto Amato, Jeffrey Barrett, Sonia Goncalves, Ewan Harrison, David K. Jackson, Ian Johnston, Dominic Kwiatkowski, Cordelia Langford, John Sillitoe on behalf of the Wellcome Sanger Institute COVID-19 Surveillance Team |
| EPI_ISL_3226088 | Lighthouse Laboratory Plymouth | Wellcome Sanger Institute for the COVID-19 Genomics UK (COG-UK) Consortium | Lighthouse Laboratory Plymouth and Alex Alderton, Roberto Amato, Jeffrey Barrett, Sonia Goncalves, Ewan Harrison, David K. Jackson, Ian Johnston, Dominic Kwiatkowski, Cordelia Langford, John Sillitoe on behalf of the Wellcome Sanger Institute COVID-19 Surveillance Team |
| EPI_ISL_3226089 | Lighthouse Lab in Milton Keynes | Wellcome Sanger Institute for the COVID-19 Genomics UK (COG-UK) Consortium | The Lighthouse Lab in Milton Keynes and Alex Alderton, Roberto Amato, Jeffrey Barrett, Sonia Goncalves, Ewan Harrison, David K. Jackson, Ian Johnston, Dominic Kwiatkowski, Cordelia Langford, John Sillitoe on behalf of the Wellcome Sanger Institute COVID-19 Surveillance Team |
| EPI_ISL_3226090 | Lighthouse Lab in Alderley Park | Wellcome Sanger Institute for the COVID-19 Genomics UK (COG-UK) Consortium | Jacquelyn Wynn, Mairead Hyland, The Lighthouse Lab in Alderley Park and Alex Alderton, Roberto Amato, Jeffrey Barrett, Sonia Goncalves, Ewan Harrison, David K. Jackson, Ian Johnston, Dominic Kwiatkowski, Cordelia Langford, John Sillitoe on behalf of the Wellcome Sanger Institute COVID-19 Surveillance Team |
| EPI_ISL_3226091, EPI_ISL_3226092 | Lighthouse Lab in Milton Keynes | Wellcome Sanger Institute for the COVID-19 Genomics UK (COG-UK) Consortium | The Lighthouse Lab in Milton Keynes and Alex Alderton, Roberto Amato, Jeffrey Barrett, Sonia Goncalves, Ewan Harrison, David K. Jackson, Ian Johnston, Dominic Kwiatkowski, Cordelia Langford, John Sillitoe on behalf of the Wellcome Sanger Institute COVID-19 Surveillance Team |
| EPI_ISL_3226093, EPI_ISL_3226094 | Lighthouse Lab in Glasgow | Wellcome Sanger Institute for the COVID-19 Genomics UK (COG-UK) Consortium | Harper VanSteenhouse, Yumi Kasai, David Gray, Carol Clugston, Anna Dominiczak and Alex Alderton, Roberto Amato, Jeffrey Barrett, Sonia Goncalves, Ewan Harrison, David K. Jackson, Ian Johnston, Dominic Kwiatkowski, Cordelia Langford, John Sillitoe on behalf of the Wellcome Sanger Institute COVID-19 Surveillance Team |
| EPI_ISL_3226095, EPI_ISL_3226096 | Lighthouse Lab in Milton Keynes | Wellcome Sanger Institute for the COVID-19 Genomics UK (COG-UK) Consortium | The Lighthouse Lab in Milton Keynes and Alex Alderton, Roberto Amato, Jeffrey Barrett, Sonia Goncalves, Ewan Harrison, David K. Jackson, Ian Johnston, Dominic Kwiatkowski, Cordelia Langford, John Sillitoe on behalf of the Wellcome Sanger Institute COVID-19 Surveillance Team |
| EPI_ISL_3226097 | Lighthouse Lab in Glasgow | Wellcome Sanger Institute for the COVID-19 Genomics UK (COG-UK) Consortium | Harper VanSteenhouse, Yumi Kasai, David Gray, Carol Clugston, Anna Dominiczak and Alex Alderton, Roberto Amato, Jeffrey Barrett, Sonia Goncalves, Ewan Harrison, David K. Jackson, Ian Johnston, Dominic Kwiatkowski, Cordelia Langford, John Sillitoe on behalf of the Wellcome Sanger Institute COVID-19 Surveillance Team |
| EPI_ISL_3226098, EPI_ISL_3226099 | Lighthouse Laboratory Plymouth | Wellcome Sanger Institute for the COVID-19 Genomics UK (COG-UK) Consortium | Lighthouse Laboratory Plymouth and Alex Alderton, Roberto Amato, Jeffrey Barrett, Sonia Goncalves, Ewan Harrison, David K. Jackson, Ian Johnston, Dominic Kwiatkowski, Cordelia Langford, John Sillitoe on behalf of the Wellcome Sanger Institute COVID-19 Surveillance Team |
| EPI_ISL_3226100 | Lighthouse Lab in Alderley Park | Wellcome Sanger Institute for the COVID-19 Genomics UK (COG-UK) Consortium | Jacquelyn Wynn, Mairead Hyland, The Lighthouse Lab in Alderley Park and Alex Alderton, Roberto Amato, Jeffrey Barrett, Sonia Goncalves, Ewan Harrison, David K. Jackson, Ian Johnston, Dominic Kwiatkowski, Cordelia Langford, John Sillitoe on behalf of the Wellcome Sanger Institute COVID-19 Surveillance Team |
| EPI_ISL_3226101 | Lighthouse Lab in Milton Keynes | Wellcome Sanger Institute for the COVID-19 Genomics UK (COG-UK) Consortium | The Lighthouse Lab in Milton Keynes and Alex Alderton, Roberto Amato, Jeffrey Barrett, Sonia Goncalves, Ewan Harrison, David K. Jackson, Ian Johnston, Dominic Kwiatkowski, Cordelia Langford, John Sillitoe on behalf of the Wellcome Sanger Institute COVID-19 Surveillance Team |
| EPI_ISL_3226102 | Lighthouse Lab in Alderley Park | Wellcome Sanger Institute for the COVID-19 Genomics UK (COG-UK) Consortium | Jacquelyn Wynn, Mairead Hyland, The Lighthouse Lab in Alderley Park and Alex Alderton, Roberto Amato, Jeffrey Barrett, Sonia Goncalves, Ewan Harrison, David K. Jackson, Ian Johnston, Dominic Kwiatkowski, Cordelia Langford, John Sillitoe on behalf of the Wellcome Sanger Institute COVID-19 Surveillance Team |
| EPI_ISL_3226103 | Lighthouse Laboratory Plymouth | Wellcome Sanger Institute for the COVID-19 Genomics UK (COG-UK) Consortium | Lighthouse Laboratory Plymouth and Alex Alderton, Roberto Amato, Jeffrey Barrett, Sonia Goncalves, Ewan Harrison, David K. Jackson, Ian Johnston, Dominic Kwiatkowski, Cordelia Langford, John Sillitoe on behalf of the Wellcome Sanger Institute COVID-19 Surveillance Team |
| EPI_ISL_3226104 | Lighthouse Lab in Alderley Park | Wellcome Sanger Institute for the COVID-19 Genomics UK (COG-UK) Consortium | Jacquelyn Wynn, Mairead Hyland, The Lighthouse Lab in Alderley Park and Alex Alderton, Roberto Amato, Jeffrey Barrett, Sonia Goncalves, Ewan Harrison, David K. Jackson, Ian Johnston, Dominic Kwiatkowski, Cordelia Langford, John Sillitoe on behalf of the Wellcome Sanger Institute COVID-19 Surveillance Team |
| EPI_ISL_3226105, EPI_ISL_3226106 | Lighthouse Lab in Glasgow | Wellcome Sanger Institute for the COVID-19 Genomics UK (COG-UK) Consortium | Harper VanSteenhouse, Yumi Kasai, David Gray, Carol Clugston, Anna Dominiczak and Alex Alderton, Roberto Amato, Jeffrey Barrett, Sonia Goncalves, Ewan Harrison, David K. Jackson, Ian Johnston, Dominic Kwiatkowski, Cordelia Langford, John Sillitoe on behalf of the Wellcome Sanger Institute COVID-19 Surveillance Team |
| EPI_ISL_3226107, EPI_ISL_3226108 | Lighthouse Laboratory Plymouth | Wellcome Sanger Institute for the COVID-19 Genomics UK (COG-UK) Consortium | Lighthouse Laboratory Plymouth and Alex Alderton, Roberto Amato, Jeffrey Barrett, Sonia Goncalves, Ewan Harrison, David K. Jackson, Ian Johnston, Dominic Kwiatkowski, Cordelia Langford, John Sillitoe on behalf of the Wellcome Sanger Institute COVID-19 Surveillance Team |
| EPI_ISL_3226109 | Lighthouse Lab in Milton Keynes | Wellcome Sanger Institute for the COVID-19 Genomics UK (COG-UK) Consortium | The Lighthouse Lab in Milton Keynes and Alex Alderton, Roberto Amato, Jeffrey Barrett, Sonia Goncalves, Ewan Harrison, David K. Jackson, Ian Johnston, Dominic Kwiatkowski, Cordelia Langford, John Sillitoe on behalf of the Wellcome Sanger Institute COVID-19 Surveillance Team |
| EPI_ISL_3226110, EPI_ISL_3226111 | Lighthouse Lab in Glasgow | Wellcome Sanger Institute for the COVID-19 Genomics UK (COG-UK) Consortium | Harper VanSteenhouse, Yumi Kasai, David Gray, Carol Clugston, Anna Dominiczak and Alex Alderton, Roberto Amato, Jeffrey Barrett, Sonia Goncalves, Ewan Harrison, David K. Jackson, Ian Johnston, Dominic Kwiatkowski, Cordelia Langford, John Sillitoe on behalf of the Wellcome Sanger Institute COVID-19 Surveillance Team |
| EPI_ISL_3226112 | Lighthouse Lab in Alderley Park | Wellcome Sanger Institute for the COVID-19 Genomics UK (COG-UK) Consortium | Jacquelyn Wynn, Mairead Hyland, The Lighthouse Lab in Alderley Park and Alex Alderton, Roberto Amato, Jeffrey Barrett, Sonia Goncalves, Ewan Harrison, David K. Jackson, Ian Johnston, Dominic Kwiatkowski, Cordelia Langford, John Sillitoe on behalf of the Wellcome Sanger Institute COVID-19 Surveillance Team |
| EPI_ISL_3226113 | Lighthouse Lab in Glasgow | Wellcome Sanger Institute for the COVID-19 Genomics UK (COG-UK) Consortium | Harper VanSteenhouse, Yumi Kasai, David Gray, Carol Clugston, Anna Dominiczak and Alex Alderton, Roberto Amato, Jeffrey Barrett, Sonia Goncalves, Ewan Harrison, David K. Jackson, Ian Johnston, Dominic Kwiatkowski, Cordelia Langford, John Sillitoe on behalf of the Wellcome Sanger Institute COVID-19 Surveillance Team |
| EPI_ISL_3226114 | Lighthouse Lab in Milton Keynes | Wellcome Sanger Institute for the COVID-19 Genomics UK (COG-UK) Consortium | The Lighthouse Lab in Milton Keynes and Alex Alderton, Roberto Amato, Jeffrey Barrett, Sonia Goncalves, Ewan Harrison, David K. Jackson, Ian Johnston, Dominic Kwiatkowski, Cordelia Langford, John Sillitoe on behalf of the Wellcome Sanger Institute COVID-19 Surveillance Team |
| EPI_ISL_3226115 | Lighthouse Lab in Alderley Park | Wellcome Sanger Institute for the COVID-19 Genomics UK (COG-UK) Consortium | Jacquelyn Wynn, Mairead Hyland, The Lighthouse Lab in Alderley Park and Alex Alderton, Roberto Amato, Jeffrey Barrett, Sonia Goncalves, Ewan Harrison, David K. Jackson, Ian Johnston, Dominic Kwiatkowski, Cordelia Langford, John Sillitoe on behalf of the Wellcome Sanger Institute COVID-19 Surveillance Team |
| EPI_ISL_3226116 | Lighthouse Lab in Milton Keynes | Wellcome Sanger Institute for the COVID-19 Genomics UK (COG-UK) Consortium | The Lighthouse Lab in Milton Keynes and Alex Alderton, Roberto Amato, Jeffrey Barrett, Sonia Goncalves, Ewan Harrison, David K. Jackson, Ian Johnston, Dominic Kwiatkowski, Cordelia Langford, John Sillitoe on behalf of the Wellcome Sanger Institute COVID-19 Surveillance Team |
| EPI_ISL_3226117 | Lighthouse Laboratory Plymouth | Wellcome Sanger Institute for the COVID-19 Genomics UK (COG-UK) Consortium | Lighthouse Laboratory Plymouth and Alex Alderton, Roberto Amato, Jeffrey Barrett, Sonia Goncalves, Ewan Harrison, David K. Jackson, Ian Johnston, Dominic Kwiatkowski, Cordelia Langford, John Sillitoe on behalf of the Wellcome Sanger Institute COVID-19 Surveillance Team |
| EPI_ISL_3226118 | Lighthouse Lab in Milton Keynes | Wellcome Sanger Institute for the COVID-19 Genomics UK (COG-UK) Consortium | The Lighthouse Lab in Milton Keynes and Alex Alderton, Roberto Amato, Jeffrey Barrett, Sonia Goncalves, Ewan Harrison, David K. Jackson, Ian Johnston, Dominic Kwiatkowski, Cordelia Langford, John Sillitoe on behalf of the Wellcome Sanger Institute COVID-19 Surveillance Team |
| EPI_ISL_3226119 | Lighthouse Laboratory Plymouth | Wellcome Sanger Institute for the COVID-19 Genomics UK (COG-UK) Consortium | Lighthouse Laboratory Plymouth and Alex Alderton, Roberto Amato, Jeffrey Barrett, Sonia Goncalves, Ewan Harrison, David K. Jackson, Ian Johnston, Dominic Kwiatkowski, Cordelia Langford, John Sillitoe on behalf of the Wellcome Sanger Institute COVID-19 Surveillance Team |
| EPI_ISL_3226120 | Lighthouse Lab in Alderley Park | Wellcome Sanger Institute for the COVID-19 Genomics UK (COG-UK) Consortium | Jacquelyn Wynn, Mairead Hyland, The Lighthouse Lab in Alderley Park and Alex Alderton, Roberto Amato, Jeffrey Barrett, Sonia Goncalves, Ewan Harrison, David K. Jackson, Ian Johnston, Dominic Kwiatkowski, Cordelia Langford, John Sillitoe on behalf of the Wellcome Sanger Institute COVID-19 Surveillance Team |
| EPI_ISL_3226121 | Lighthouse Laboratory Plymouth | Wellcome Sanger Institute for the COVID-19 Genomics UK (COG-UK) Consortium | Lighthouse Laboratory Plymouth and Alex Alderton, Roberto Amato, Jeffrey Barrett, Sonia Goncalves, Ewan Harrison, David K. Jackson, Ian Johnston, Dominic Kwiatkowski, Cordelia Langford, John Sillitoe on behalf of the Wellcome Sanger Institute COVID-19 Surveillance Team |
| EPI_ISL_3226122 | Lighthouse Lab in Alderley Park | Wellcome Sanger Institute for the COVID-19 Genomics UK (COG-UK) | Jacquelyn Wynn, Mairead Hyland, The Lighthouse Lab in Alderley Park and Alex Alderton, Roberto Amato, Jeffrey Barrett, Sonia Goncalves, Ewan Harrison, David K. Jackson, Ian Johnston, Dominic Kwiatkowski, Cordelia Langford, John Sillitoe on |

| | | | |
|---|---|---|---|
| | | Consortium | behalf of the Wellcome Sanger Institute COVID-19 Surveillance Team |
| EPI_ISL_3226123 | Lighthouse Lab in Milton Keynes | Wellcome Sanger Institute for the COVID-19 Genomics UK (COG-UK) Consortium | The Lighthouse Lab in Milton Keynes and Alex Alderton, Roberto Amato, Jeffrey Barrett, Sonia Goncalves, Ewan Harrison, David K. Jackson, Ian Johnston, Dominic Kwiatkowski, Cordelia Langford, John Sillitoe on behalf of the Wellcome Sanger Institute COVID-19 Surveillance Team |
| EPI_ISL_3226124 | Lighthouse Lab in Glasgow | Wellcome Sanger Institute for the COVID-19 Genomics UK (COG-UK) Consortium | Harper VanSteenhouse, Yumi Kasai, David Gray, Carol Clugston, Anna Dominiczak and Alex Alderton, Roberto Amato, Jeffrey Barrett, Sonia Goncalves, Ewan Harrison, David K. Jackson, Ian Johnston, Dominic Kwiatkowski, Cordelia Langford, John Sillitoe on behalf of the Wellcome Sanger Institute COVID-19 Surveillance Team |
| EPI_ISL_3226125 | Lighthouse Lab in Milton Keynes | Wellcome Sanger Institute for the COVID-19 Genomics UK (COG-UK) Consortium | The Lighthouse Lab in Milton Keynes and Alex Alderton, Roberto Amato, Jeffrey Barrett, Sonia Goncalves, Ewan Harrison, David K. Jackson, Ian Johnston, Dominic Kwiatkowski, Cordelia Langford, John Sillitoe on behalf of the Wellcome Sanger Institute COVID-19 Surveillance Team |
| EPI_ISL_3226126, EPI_ISL_3226127 | Lighthouse Lab in Glasgow | Wellcome Sanger Institute for the COVID-19 Genomics UK (COG-UK) Consortium | Harper VanSteenhouse, Yumi Kasai, David Gray, Carol Clugston, Anna Dominiczak and Alex Alderton, Roberto Amato, Jeffrey Barrett, Sonia Goncalves, Ewan Harrison, David K. Jackson, Ian Johnston, Dominic Kwiatkowski, Cordelia Langford, John Sillitoe on behalf of the Wellcome Sanger Institute COVID-19 Surveillance Team |
| EPI_ISL_3226128 | Lighthouse Lab in Milton Keynes | Wellcome Sanger Institute for the COVID-19 Genomics UK (COG-UK) Consortium | The Lighthouse Lab in Milton Keynes and Alex Alderton, Roberto Amato, Jeffrey Barrett, Sonia Goncalves, Ewan Harrison, David K. Jackson, Ian Johnston, Dominic Kwiatkowski, Cordelia Langford, John Sillitoe on behalf of the Wellcome Sanger Institute COVID-19 Surveillance Team |
| EPI_ISL_3226129 | Lighthouse Laboratory Plymouth | Wellcome Sanger Institute for the COVID-19 Genomics UK (COG-UK) Consortium | Lighthouse Laboratory Plymouth and Alex Alderton, Roberto Amato, Jeffrey Barrett, Sonia Goncalves, Ewan Harrison, David K. Jackson, Ian Johnston, Dominic Kwiatkowski, Cordelia Langford, John Sillitoe on behalf of the Wellcome Sanger Institute COVID-19 Surveillance Team |
| EPI_ISL_3226130 | Lighthouse Lab in Glasgow | Wellcome Sanger Institute for the COVID-19 Genomics UK (COG-UK) Consortium | Harper VanSteenhouse, Yumi Kasai, David Gray, Carol Clugston, Anna Dominiczak and Alex Alderton, Roberto Amato, Jeffrey Barrett, Sonia Goncalves, Ewan Harrison, David K. Jackson, Ian Johnston, Dominic Kwiatkowski, Cordelia Langford, John Sillitoe on behalf of the Wellcome Sanger Institute COVID-19 Surveillance Team |
| EPI_ISL_3226131, EPI_ISL_3226132, EPI_ISL_3226133 | Lighthouse Lab in Alderley Park | Wellcome Sanger Institute for the COVID-19 Genomics UK (COG-UK) Consortium | Jacquelyn Wynn, Mairead Hyland, The Lighthouse Lab in Alderley Park and Alex Alderton, Roberto Amato, Jeffrey Barrett, Sonia Goncalves, Ewan Harrison, David K. Jackson, Ian Johnston, Dominic Kwiatkowski, Cordelia Langford, John Sillitoe on behalf of the Wellcome Sanger Institute COVID-19 Surveillance Team |
| EPI_ISL_3226134, EPI_ISL_3226135 | Lighthouse Lab in Milton Keynes | Wellcome Sanger Institute for the COVID-19 Genomics UK (COG-UK) Consortium | The Lighthouse Lab in Milton Keynes and Alex Alderton, Roberto Amato, Jeffrey Barrett, Sonia Goncalves, Ewan Harrison, David K. Jackson, Ian Johnston, Dominic Kwiatkowski, Cordelia Langford, John Sillitoe on behalf of the Wellcome Sanger Institute COVID-19 Surveillance Team |
| EPI_ISL_3226136, EPI_ISL_3226137 | Lighthouse Lab in Alderley Park | Wellcome Sanger Institute for the COVID-19 Genomics UK (COG-UK) Consortium | Jacquelyn Wynn, Mairead Hyland, The Lighthouse Lab in Alderley Park and Alex Alderton, Roberto Amato, Jeffrey Barrett, Sonia Goncalves, Ewan Harrison, David K. Jackson, Ian Johnston, Dominic Kwiatkowski, Cordelia Langford, John Sillitoe on behalf of the Wellcome Sanger Institute COVID-19 Surveillance Team |
| EPI_ISL_3226138 | Lighthouse Lab in Milton Keynes | Wellcome Sanger Institute for the COVID-19 Genomics UK (COG-UK) Consortium | The Lighthouse Lab in Milton Keynes and Alex Alderton, Roberto Amato, Jeffrey Barrett, Sonia Goncalves, Ewan Harrison, David K. Jackson, Ian Johnston, Dominic Kwiatkowski, Cordelia Langford, John Sillitoe on behalf of the Wellcome Sanger Institute COVID-19 Surveillance Team |
| EPI_ISL_3226139 | Lighthouse Lab in Alderley Park | Wellcome Sanger Institute for the COVID-19 Genomics UK (COG-UK) Consortium | Jacquelyn Wynn, Mairead Hyland, The Lighthouse Lab in Alderley Park and Alex Alderton, Roberto Amato, Jeffrey Barrett, Sonia Goncalves, Ewan Harrison, David K. Jackson, Ian Johnston, Dominic Kwiatkowski, Cordelia Langford, John Sillitoe on behalf of the Wellcome Sanger Institute COVID-19 Surveillance Team |
| EPI_ISL_3226140 | Lighthouse Laboratory Plymouth | Wellcome Sanger Institute for the COVID-19 Genomics UK (COG-UK) Consortium | Lighthouse Laboratory Plymouth and Alex Alderton, Roberto Amato, Jeffrey Barrett, Sonia Goncalves, Ewan Harrison, David K. Jackson, Ian Johnston, Dominic Kwiatkowski, Cordelia Langford, John Sillitoe on behalf of the Wellcome Sanger Institute COVID-19 Surveillance Team |
| EPI_ISL_3226141, EPI_ISL_3226142, EPI_ISL_3226143 | Lighthouse Lab in Glasgow | Wellcome Sanger Institute for the COVID-19 Genomics UK (COG-UK) Consortium | Harper VanSteenhouse, Yumi Kasai, David Gray, Carol Clugston, Anna Dominiczak and Alex Alderton, Roberto Amato, Jeffrey Barrett, Sonia Goncalves, Ewan Harrison, David K. Jackson, Ian Johnston, Dominic Kwiatkowski, Cordelia Langford, John Sillitoe on behalf of the Wellcome Sanger Institute COVID-19 Surveillance Team |
| EPI_ISL_3226144, EPI_ISL_3226145 | Lighthouse Laboratory Plymouth | Wellcome Sanger Institute for the COVID-19 Genomics UK (COG-UK) Consortium | Lighthouse Laboratory Plymouth and Alex Alderton, Roberto Amato, Jeffrey Barrett, Sonia Goncalves, Ewan Harrison, David K. Jackson, Ian Johnston, Dominic Kwiatkowski, Cordelia Langford, John Sillitoe on behalf of the Wellcome Sanger Institute COVID-19 Surveillance Team |
| EPI_ISL_3226146 | Lighthouse Lab in Milton Keynes | Wellcome Sanger Institute for the COVID-19 Genomics UK (COG-UK) Consortium | The Lighthouse Lab in Milton Keynes and Alex Alderton, Roberto Amato, Jeffrey Barrett, Sonia Goncalves, Ewan Harrison, David K. Jackson, Ian Johnston, Dominic Kwiatkowski, Cordelia Langford, John Sillitoe on behalf of the Wellcome Sanger Institute COVID-19 Surveillance Team |
| EPI_ISL_3226147 | Lighthouse Laboratory Plymouth | Wellcome Sanger Institute for the COVID-19 Genomics UK (COG-UK) Consortium | Lighthouse Laboratory Plymouth and Alex Alderton, Roberto Amato, Jeffrey Barrett, Sonia Goncalves, Ewan Harrison, David K. Jackson, Ian Johnston, Dominic Kwiatkowski, Cordelia Langford, John Sillitoe on behalf of the Wellcome Sanger Institute COVID-19 Surveillance Team |
| EPI_ISL_3226148 | Lighthouse Lab in Alderley Park | Wellcome Sanger Institute for the COVID-19 Genomics UK (COG-UK) Consortium | Jacquelyn Wynn, Mairead Hyland, The Lighthouse Lab in Alderley Park and Alex Alderton, Roberto Amato, Jeffrey Barrett, Sonia Goncalves, Ewan Harrison, David K. Jackson, Ian Johnston, Dominic Kwiatkowski, Cordelia Langford, John Sillitoe on behalf of the Wellcome Sanger Institute COVID-19 Surveillance Team |
| EPI_ISL_3226149 | Lighthouse Laboratory Plymouth | Wellcome Sanger Institute for the COVID-19 Genomics UK (COG-UK) Consortium | Lighthouse Laboratory Plymouth and Alex Alderton, Roberto Amato, Jeffrey Barrett, Sonia Goncalves, Ewan Harrison, David K. Jackson, Ian Johnston, Dominic Kwiatkowski, Cordelia Langford, John Sillitoe on behalf of the Wellcome Sanger Institute COVID-19 Surveillance Team |
| EPI_ISL_3226150 | Lighthouse Lab in Alderley Park | Wellcome Sanger Institute for the COVID-19 Genomics UK (COG-UK) Consortium | Jacquelyn Wynn, Mairead Hyland, The Lighthouse Lab in Alderley Park and Alex Alderton, Roberto Amato, Jeffrey Barrett, Sonia Goncalves, Ewan Harrison, David K. Jackson, Ian Johnston, Dominic Kwiatkowski, Cordelia Langford, John Sillitoe on behalf of the Wellcome Sanger Institute COVID-19 Surveillance Team |
| EPI_ISL_3226151, EPI_ISL_3226152 | Lighthouse Lab in Milton Keynes | Wellcome Sanger Institute for the COVID-19 Genomics UK (COG-UK) Consortium | The Lighthouse Lab in Milton Keynes and Alex Alderton, Roberto Amato, Jeffrey Barrett, Sonia Goncalves, Ewan Harrison, David K. Jackson, Ian Johnston, Dominic Kwiatkowski, Cordelia Langford, John Sillitoe on behalf of the Wellcome Sanger Institute COVID-19 Surveillance Team |
| EPI_ISL_3226153 | Lighthouse Lab in Alderley Park | Wellcome Sanger Institute for the COVID-19 Genomics UK (COG-UK) Consortium | Jacquelyn Wynn, Mairead Hyland, The Lighthouse Lab in Alderley Park and Alex Alderton, Roberto Amato, Jeffrey Barrett, Sonia Goncalves, Ewan Harrison, David K. Jackson, Ian Johnston, Dominic Kwiatkowski, Cordelia Langford, John Sillitoe on behalf of the Wellcome Sanger Institute COVID-19 Surveillance Team |
| EPI_ISL_3226154 | Lighthouse Laboratory Plymouth | Wellcome Sanger Institute for the COVID-19 Genomics UK (COG-UK) Consortium | Lighthouse Laboratory Plymouth and Alex Alderton, Roberto Amato, Jeffrey Barrett, Sonia Goncalves, Ewan Harrison, David K. Jackson, Ian Johnston, Dominic Kwiatkowski, Cordelia Langford, John Sillitoe on behalf of the Wellcome Sanger Institute COVID-19 Surveillance Team |
| EPI_ISL_3226155 | Lighthouse Lab in Glasgow | Wellcome Sanger Institute for the COVID-19 Genomics UK (COG-UK) Consortium | Harper VanSteenhouse, Yumi Kasai, David Gray, Carol Clugston, Anna Dominiczak and Alex Alderton, Roberto Amato, Jeffrey Barrett, Sonia Goncalves, Ewan Harrison, David K. Jackson, Ian Johnston, Dominic Kwiatkowski, Cordelia Langford, John Sillitoe on behalf of the Wellcome Sanger Institute COVID-19 Surveillance Team |
| EPI_ISL_3226156, EPI_ISL_3226157, EPI_ISL_3226158 | Lighthouse Laboratory Plymouth | Wellcome Sanger Institute for the COVID-19 Genomics UK (COG-UK) Consortium | Lighthouse Laboratory Plymouth and Alex Alderton, Roberto Amato, Jeffrey Barrett, Sonia Goncalves, Ewan Harrison, David K. Jackson, Ian Johnston, Dominic Kwiatkowski, Cordelia Langford, John Sillitoe on behalf of the Wellcome Sanger Institute COVID-19 Surveillance Team |
| EPI_ISL_3226159 | Lighthouse Lab in Glasgow | Wellcome Sanger Institute for the COVID-19 Genomics UK (COG-UK) Consortium | Harper VanSteenhouse, Yumi Kasai, David Gray, Carol Clugston, Anna Dominiczak and Alex Alderton, Roberto Amato, Jeffrey Barrett, Sonia Goncalves, Ewan Harrison, David K. Jackson, Ian Johnston, Dominic Kwiatkowski, Cordelia Langford, John Sillitoe on behalf of the Wellcome Sanger Institute COVID-19 Surveillance Team |
| EPI_ISL_3226160 | Lighthouse Lab in Alderley Park | Wellcome Sanger Institute for the COVID-19 Genomics UK (COG-UK) Consortium | Jacquelyn Wynn, Mairead Hyland, The Lighthouse Lab in Alderley Park and Alex Alderton, Roberto Amato, Jeffrey Barrett, Sonia Goncalves, Ewan Harrison, David K. Jackson, Ian Johnston, Dominic Kwiatkowski, Cordelia Langford, John Sillitoe on behalf of the Wellcome Sanger Institute COVID-19 Surveillance Team |
| EPI_ISL_3226161, EPI_ISL_3226162 | Lighthouse Laboratory Plymouth | Wellcome Sanger Institute for the COVID-19 Genomics UK (COG-UK) Consortium | Lighthouse Laboratory Plymouth and Alex Alderton, Roberto Amato, Jeffrey Barrett, Sonia Goncalves, Ewan Harrison, David K. Jackson, Ian Johnston, Dominic Kwiatkowski, Cordelia Langford, John Sillitoe on behalf of the Wellcome Sanger Institute COVID-19 Surveillance Team |
| EPI_ISL_3226163 | Lighthouse Lab in Alderley Park | Wellcome Sanger Institute for the COVID-19 Genomics UK (COG-UK) Consortium | Jacquelyn Wynn, Mairead Hyland, The Lighthouse Lab in Alderley Park and Alex Alderton, Roberto Amato, Jeffrey Barrett, Sonia Goncalves, Ewan Harrison, David K. Jackson, Ian Johnston, Dominic Kwiatkowski, Cordelia Langford, John Sillitoe on behalf of the Wellcome Sanger Institute COVID-19 Surveillance Team |
| EPI_ISL_3226164 | Lighthouse Lab in Milton Keynes | Wellcome Sanger Institute for the COVID-19 Genomics UK (COG-UK) Consortium | The Lighthouse Lab in Milton Keynes and Alex Alderton, Roberto Amato, Jeffrey Barrett, Sonia Goncalves, Ewan Harrison, David K. Jackson, Ian Johnston, Dominic Kwiatkowski, Cordelia Langford, John Sillitoe on behalf of the Wellcome Sanger Institute COVID-19 Surveillance Team |
| EPI_ISL_3226165 | Lighthouse Lab in Glasgow | Wellcome Sanger Institute for the COVID-19 Genomics UK (COG-UK) Consortium | Harper VanSteenhouse, Yumi Kasai, David Gray, Carol Clugston, Anna Dominiczak and Alex Alderton, Roberto Amato, Jeffrey Barrett, Sonia Goncalves, Ewan Harrison, David K. Jackson, Ian Johnston, Dominic Kwiatkowski, Cordelia Langford, John Sillitoe on behalf of the Wellcome Sanger Institute COVID-19 Surveillance Team |
| EPI_ISL_3226166 | Lighthouse Lab in Alderley Park | Wellcome Sanger Institute for the COVID-19 Genomics UK (COG-UK) Consortium | Jacquelyn Wynn, Mairead Hyland, The Lighthouse Lab in Alderley Park and Alex Alderton, Roberto Amato, Jeffrey Barrett, Sonia Goncalves, Ewan Harrison, David K. Jackson, Ian Johnston, Dominic Kwiatkowski, Cordelia Langford, John Sillitoe on behalf of the Wellcome Sanger Institute COVID-19 Surveillance Team |
| EPI_ISL_3226167 | Lighthouse Lab in Glasgow | Wellcome Sanger Institute for the COVID-19 Genomics UK (COG-UK) Consortium | Harper VanSteenhouse, Yumi Kasai, David Gray, Carol Clugston, Anna Dominiczak and Alex Alderton, Roberto Amato, Jeffrey Barrett, Sonia Goncalves, Ewan Harrison, David K. Jackson, Ian Johnston, Dominic Kwiatkowski, Cordelia Langford, John Sillitoe on behalf of the Wellcome Sanger Institute COVID-19 Surveillance Team |
| EPI_ISL_3226168 | Lighthouse Lab in Milton Keynes | Wellcome Sanger Institute for the COVID-19 Genomics UK (COG-UK) Consortium | The Lighthouse Lab in Milton Keynes and Alex Alderton, Roberto Amato, Jeffrey Barrett, Sonia Goncalves, Ewan Harrison, David K. Jackson, Ian Johnston, Dominic Kwiatkowski, Cordelia Langford, John Sillitoe on behalf of the Wellcome Sanger Institute COVID-19 Surveillance Team |
| EPI_ISL_3226169, EPI_ISL_3226170 | Lighthouse Lab in Glasgow | Wellcome Sanger Institute for the COVID-19 Genomics UK (COG-UK) Consortium | Harper VanSteenhouse, Yumi Kasai, David Gray, Carol Clugston, Anna Dominiczak and Alex Alderton, Roberto Amato, Jeffrey Barrett, Sonia Goncalves, Ewan Harrison, David K. Jackson, Ian Johnston, Dominic Kwiatkowski, Cordelia Langford, John Sillitoe on behalf of the Wellcome Sanger Institute COVID-19 Surveillance Team |
| EPI_ISL_3226171 | Lighthouse Lab in Milton Keynes | Wellcome Sanger Institute for the COVID-19 Genomics UK (COG-UK) Consortium | The Lighthouse Lab in Milton Keynes and Alex Alderton, Roberto Amato, Jeffrey Barrett, Sonia Goncalves, Ewan Harrison, David K. Jackson, Ian Johnston, Dominic Kwiatkowski, Cordelia Langford, John Sillitoe on behalf of the Wellcome Sanger Institute COVID-19 Surveillance Team |
| EPI_ISL_3226172 | Lighthouse Lab in Alderley Park | Wellcome Sanger Institute for the COVID-19 Genomics UK (COG-UK) Consortium | Jacquelyn Wynn, Mairead Hyland, The Lighthouse Lab in Alderley Park and Alex Alderton, Roberto Amato, Jeffrey Barrett, Sonia Goncalves, Ewan Harrison, David K. Jackson, Ian Johnston, Dominic Kwiatkowski, Cordelia Langford, John Sillitoe on behalf of the Wellcome Sanger Institute COVID-19 Surveillance Team |
| EPI_ISL_3226173, EPI_ISL_3226174 | Lighthouse Laboratory Plymouth | Wellcome Sanger Institute for the COVID-19 Genomics UK (COG-UK) Consortium | Lighthouse Laboratory Plymouth and Alex Alderton, Roberto Amato, Jeffrey Barrett, Sonia Goncalves, Ewan Harrison, David K. Jackson, Ian Johnston, Dominic Kwiatkowski, Cordelia Langford, John Sillitoe on behalf of the Wellcome Sanger Institute COVID-19 Surveillance Team |
| EPI_ISL_3226175 | Lighthouse Lab in Milton Keynes | Wellcome Sanger Institute for the COVID-19 Genomics UK (COG-UK) Consortium | The Lighthouse Lab in Milton Keynes and Alex Alderton, Roberto Amato, Jeffrey Barrett, Sonia Goncalves, Ewan Harrison, David K. Jackson, Ian Johnston, Dominic Kwiatkowski, Cordelia Langford, John Sillitoe on behalf of the Wellcome Sanger Institute COVID-19 Surveillance Team |
| EPI_ISL_3226176 | Lighthouse Laboratory Plymouth | Wellcome Sanger Institute for the COVID-19 Genomics UK (COG-UK) Consortium | Lighthouse Laboratory Plymouth and Alex Alderton, Roberto Amato, Jeffrey Barrett, Sonia Goncalves, Ewan Harrison, David K. Jackson, Ian Johnston, Dominic Kwiatkowski, Cordelia Langford, John Sillitoe on behalf of the Wellcome Sanger Institute COVID-19 Surveillance Team |
| EPI_ISL_3226177 | Lighthouse Lab in Alderley Park | Wellcome Sanger Institute for the COVID-19 Genomics UK (COG-UK) Consortium | Jacquelyn Wynn, Mairead Hyland, The Lighthouse Lab in Alderley Park and Alex Alderton, Roberto Amato, Jeffrey Barrett, Sonia Goncalves, Ewan Harrison, David K. Jackson, Ian Johnston, Dominic Kwiatkowski, Cordelia Langford, John Sillitoe on behalf of the Wellcome Sanger Institute COVID-19 Surveillance Team |
| EPI_ISL_3226178 | Lighthouse Lab in Glasgow | Wellcome Sanger Institute for the COVID-19 Genomics UK (COG-UK) Consortium | Harper VanSteenhouse, Yumi Kasai, David Gray, Carol Clugston, Anna Dominiczak and Alex Alderton, Roberto Amato, Jeffrey Barrett, Sonia Goncalves, Ewan Harrison, David K. Jackson, Ian Johnston, Dominic Kwiatkowski, Cordelia Langford, John Sillitoe on behalf of the Wellcome Sanger Institute COVID-19 Surveillance Team |
| EPI_ISL_3226179 | Lighthouse Lab in Milton Keynes | Wellcome Sanger Institute for the COVID-19 Genomics UK (COG-UK) Consortium | The Lighthouse Lab in Milton Keynes and Alex Alderton, Roberto Amato, Jeffrey Barrett, Sonia Goncalves, Ewan Harrison, David K. Jackson, Ian Johnston, Dominic Kwiatkowski, Cordelia Langford, John Sillitoe on behalf of the Wellcome Sanger Institute COVID-19 Surveillance Team |

| | | | |
|---|---|---|---|
| EPI_ISL_3226180 | Lighthouse Lab in Glasgow | Wellcome Sanger Institute for the COVID-19 Genomics UK (COG-UK) Consortium | Harper VanSteenhouse, Yumi Kasai, David Gray, Carol Clugston, Anna Dominiczak and Alex Alderton, Roberto Amato, Jeffrey Barrett, Sonia Goncalves, Ewan Harrison, David K. Jackson, Ian Johnston, Dominic Kwiatkowski, Cordelia Langford, John Sillitoe on behalf of the Wellcome Sanger Institute COVID-19 Surveillance Team |
| EPI_ISL_3226181, EPI_ISL_3226182 | Lighthouse Laboratory Plymouth | Wellcome Sanger Institute for the COVID-19 Genomics UK (COG-UK) Consortium | Lighthouse Laboratory Plymouth and Alex Alderton, Roberto Amato, Jeffrey Barrett, Sonia Goncalves, Ewan Harrison, David K. Jackson, Ian Johnston, Dominic Kwiatkowski, Cordelia Langford, John Sillitoe on behalf of the Wellcome Sanger Institute COVID-19 Surveillance Team |
| EPI_ISL_3226183 | Lighthouse Lab in Glasgow | Wellcome Sanger Institute for the COVID-19 Genomics UK (COG-UK) Consortium | Harper VanSteenhouse, Yumi Kasai, David Gray, Carol Clugston, Anna Dominiczak and Alex Alderton, Roberto Amato, Jeffrey Barrett, Sonia Goncalves, Ewan Harrison, David K. Jackson, Ian Johnston, Dominic Kwiatkowski, Cordelia Langford, John Sillitoe on behalf of the Wellcome Sanger Institute COVID-19 Surveillance Team |
| EPI_ISL_3226184 | Lighthouse Lab in Alderley Park | Wellcome Sanger Institute for the COVID-19 Genomics UK (COG-UK) Consortium | Jacquelyn Wynn, Mairead Hyland, The Lighthouse Lab in Alderley Park and Alex Alderton, Roberto Amato, Jeffrey Barrett, Sonia Goncalves, Ewan Harrison, David K. Jackson, Ian Johnston, Dominic Kwiatkowski, Cordelia Langford, John Sillitoe on behalf of the Wellcome Sanger Institute COVID-19 Surveillance Team |
| EPI_ISL_3226185 | Lighthouse Lab in Milton Keynes | Wellcome Sanger Institute for the COVID-19 Genomics UK (COG-UK) Consortium | The Lighthouse Lab in Milton Keynes and Alex Alderton, Roberto Amato, Jeffrey Barrett, Sonia Goncalves, Ewan Harrison, David K. Jackson, Ian Johnston, Dominic Kwiatkowski, Cordelia Langford, John Sillitoe on behalf of the Wellcome Sanger Institute COVID-19 Surveillance Team |
| EPI_ISL_3226186 | Lighthouse Lab in Alderley Park | Wellcome Sanger Institute for the COVID-19 Genomics UK (COG-UK) Consortium | Jacquelyn Wynn, Mairead Hyland, The Lighthouse Lab in Alderley Park and Alex Alderton, Roberto Amato, Jeffrey Barrett, Sonia Goncalves, Ewan Harrison, David K. Jackson, Ian Johnston, Dominic Kwiatkowski, Cordelia Langford, John Sillitoe on behalf of the Wellcome Sanger Institute COVID-19 Surveillance Team |
| EPI_ISL_3226187 | Lighthouse Lab in Milton Keynes | Wellcome Sanger Institute for the COVID-19 Genomics UK (COG-UK) Consortium | The Lighthouse Lab in Milton Keynes and Alex Alderton, Roberto Amato, Jeffrey Barrett, Sonia Goncalves, Ewan Harrison, David K. Jackson, Ian Johnston, Dominic Kwiatkowski, Cordelia Langford, John Sillitoe on behalf of the Wellcome Sanger Institute COVID-19 Surveillance Team |
| EPI_ISL_3226188 | Lighthouse Laboratory Plymouth | Wellcome Sanger Institute for the COVID-19 Genomics UK (COG-UK) Consortium | Lighthouse Laboratory Plymouth and Alex Alderton, Roberto Amato, Jeffrey Barrett, Sonia Goncalves, Ewan Harrison, David K. Jackson, Ian Johnston, Dominic Kwiatkowski, Cordelia Langford, John Sillitoe on behalf of the Wellcome Sanger Institute COVID-19 Surveillance Team |
| EPI_ISL_3226189, EPI_ISL_3226190 | Lighthouse Lab in Milton Keynes | Wellcome Sanger Institute for the COVID-19 Genomics UK (COG-UK) Consortium | The Lighthouse Lab in Milton Keynes and Alex Alderton, Roberto Amato, Jeffrey Barrett, Sonia Goncalves, Ewan Harrison, David K. Jackson, Ian Johnston, Dominic Kwiatkowski, Cordelia Langford, John Sillitoe on behalf of the Wellcome Sanger Institute COVID-19 Surveillance Team |
| EPI_ISL_3226191, EPI_ISL_3226192 | Lighthouse Lab in Glasgow | Wellcome Sanger Institute for the COVID-19 Genomics UK (COG-UK) Consortium | Harper VanSteenhouse, Yumi Kasai, David Gray, Carol Clugston, Anna Dominiczak and Alex Alderton, Roberto Amato, Jeffrey Barrett, Sonia Goncalves, Ewan Harrison, David K. Jackson, Ian Johnston, Dominic Kwiatkowski, Cordelia Langford, John Sillitoe on behalf of the Wellcome Sanger Institute COVID-19 Surveillance Team |
| EPI_ISL_3226193 | Lighthouse Laboratory Plymouth | Wellcome Sanger Institute for the COVID-19 Genomics UK (COG-UK) Consortium | Lighthouse Laboratory Plymouth and Alex Alderton, Roberto Amato, Jeffrey Barrett, Sonia Goncalves, Ewan Harrison, David K. Jackson, Ian Johnston, Dominic Kwiatkowski, Cordelia Langford, John Sillitoe on behalf of the Wellcome Sanger Institute COVID-19 Surveillance Team |
| EPI_ISL_3226194 | Lighthouse Lab in Alderley Park | Wellcome Sanger Institute for the COVID-19 Genomics UK (COG-UK) Consortium | Jacquelyn Wynn, Mairead Hyland, The Lighthouse Lab in Alderley Park and Alex Alderton, Roberto Amato, Jeffrey Barrett, Sonia Goncalves, Ewan Harrison, David K. Jackson, Ian Johnston, Dominic Kwiatkowski, Cordelia Langford, John Sillitoe on behalf of the Wellcome Sanger Institute COVID-19 Surveillance Team |
| EPI_ISL_3226195 | Lighthouse Lab in Glasgow | Wellcome Sanger Institute for the COVID-19 Genomics UK (COG-UK) Consortium | Harper VanSteenhouse, Yumi Kasai, David Gray, Carol Clugston, Anna Dominiczak and Alex Alderton, Roberto Amato, Jeffrey Barrett, Sonia Goncalves, Ewan Harrison, David K. Jackson, Ian Johnston, Dominic Kwiatkowski, Cordelia Langford, John Sillitoe on behalf of the Wellcome Sanger Institute COVID-19 Surveillance Team |
| EPI_ISL_3226196 | Lighthouse Lab in Milton Keynes | Wellcome Sanger Institute for the COVID-19 Genomics UK (COG-UK) Consortium | The Lighthouse Lab in Milton Keynes and Alex Alderton, Roberto Amato, Jeffrey Barrett, Sonia Goncalves, Ewan Harrison, David K. Jackson, Ian Johnston, Dominic Kwiatkowski, Cordelia Langford, John Sillitoe on behalf of the Wellcome Sanger Institute COVID-19 Surveillance Team |
| EPI_ISL_3226197 | Lighthouse Lab in Glasgow | Wellcome Sanger Institute for the COVID-19 Genomics UK (COG-UK) Consortium | Harper VanSteenhouse, Yumi Kasai, David Gray, Carol Clugston, Anna Dominiczak and Alex Alderton, Roberto Amato, Jeffrey Barrett, Sonia Goncalves, Ewan Harrison, David K. Jackson, Ian Johnston, Dominic Kwiatkowski, Cordelia Langford, John Sillitoe on behalf of the Wellcome Sanger Institute COVID-19 Surveillance Team |
| EPI_ISL_3226198 | Lighthouse Lab in Alderley Park | Wellcome Sanger Institute for the COVID-19 Genomics UK (COG-UK) Consortium | Jacquelyn Wynn, Mairead Hyland, The Lighthouse Lab in Alderley Park and Alex Alderton, Roberto Amato, Jeffrey Barrett, Sonia Goncalves, Ewan Harrison, David K. Jackson, Ian Johnston, Dominic Kwiatkowski, Cordelia Langford, John Sillitoe on behalf of the Wellcome Sanger Institute COVID-19 Surveillance Team |
| EPI_ISL_3226199, EPI_ISL_3226200 | Lighthouse Lab in Milton Keynes | Wellcome Sanger Institute for the COVID-19 Genomics UK (COG-UK) Consortium | The Lighthouse Lab in Milton Keynes and Alex Alderton, Roberto Amato, Jeffrey Barrett, Sonia Goncalves, Ewan Harrison, David K. Jackson, Ian Johnston, Dominic Kwiatkowski, Cordelia Langford, John Sillitoe on behalf of the Wellcome Sanger Institute COVID-19 Surveillance Team |
| EPI_ISL_3226201, EPI_ISL_3226202, EPI_ISL_3226203, EPI_ISL_3226204 | Lighthouse Lab in Glasgow | Wellcome Sanger Institute for the COVID-19 Genomics UK (COG-UK) Consortium | Harper VanSteenhouse, Yumi Kasai, David Gray, Carol Clugston, Anna Dominiczak and Alex Alderton, Roberto Amato, Jeffrey Barrett, Sonia Goncalves, Ewan Harrison, David K. Jackson, Ian Johnston, Dominic Kwiatkowski, Cordelia Langford, John Sillitoe on behalf of the Wellcome Sanger Institute COVID-19 Surveillance Team |
| EPI_ISL_3226205 | Lighthouse Lab in Milton Keynes | Wellcome Sanger Institute for the COVID-19 Genomics UK (COG-UK) Consortium | The Lighthouse Lab in Milton Keynes and Alex Alderton, Roberto Amato, Jeffrey Barrett, Sonia Goncalves, Ewan Harrison, David K. Jackson, Ian Johnston, Dominic Kwiatkowski, Cordelia Langford, John Sillitoe on behalf of the Wellcome Sanger Institute COVID-19 Surveillance Team |
| EPI_ISL_3226206 | Lighthouse Laboratory Plymouth | Wellcome Sanger Institute for the COVID-19 Genomics UK (COG-UK) Consortium | Lighthouse Laboratory Plymouth and Alex Alderton, Roberto Amato, Jeffrey Barrett, Sonia Goncalves, Ewan Harrison, David K. Jackson, Ian Johnston, Dominic Kwiatkowski, Cordelia Langford, John Sillitoe on behalf of the Wellcome Sanger Institute COVID-19 Surveillance Team |
| EPI_ISL_3226207 | Lighthouse Lab in Alderley Park | Wellcome Sanger Institute for the COVID-19 Genomics UK (COG-UK) Consortium | Jacquelyn Wynn, Mairead Hyland, The Lighthouse Lab in Alderley Park and Alex Alderton, Roberto Amato, Jeffrey Barrett, Sonia Goncalves, Ewan Harrison, David K. Jackson, Ian Johnston, Dominic Kwiatkowski, Cordelia Langford, John Sillitoe on behalf of the Wellcome Sanger Institute COVID-19 Surveillance Team |
| EPI_ISL_3226208 | Lighthouse Laboratory Plymouth | Wellcome Sanger Institute for the COVID-19 Genomics UK (COG-UK) Consortium | Lighthouse Laboratory Plymouth and Alex Alderton, Roberto Amato, Jeffrey Barrett, Sonia Goncalves, Ewan Harrison, David K. Jackson, Ian Johnston, Dominic Kwiatkowski, Cordelia Langford, John Sillitoe on behalf of the Wellcome Sanger Institute COVID-19 Surveillance Team |
| EPI_ISL_3226209 | Lighthouse Lab in Glasgow | Wellcome Sanger Institute for the COVID-19 Genomics UK (COG-UK) Consortium | Harper VanSteenhouse, Yumi Kasai, David Gray, Carol Clugston, Anna Dominiczak and Alex Alderton, Roberto Amato, Jeffrey Barrett, Sonia Goncalves, Ewan Harrison, David K. Jackson, Ian Johnston, Dominic Kwiatkowski, Cordelia Langford, John Sillitoe on behalf of the Wellcome Sanger Institute COVID-19 Surveillance Team |
| EPI_ISL_3226210 | Lighthouse Laboratory Plymouth | Wellcome Sanger Institute for the COVID-19 Genomics UK (COG-UK) Consortium | Lighthouse Laboratory Plymouth and Alex Alderton, Roberto Amato, Jeffrey Barrett, Sonia Goncalves, Ewan Harrison, David K. Jackson, Ian Johnston, Dominic Kwiatkowski, Cordelia Langford, John Sillitoe on behalf of the Wellcome Sanger Institute COVID-19 Surveillance Team |
| EPI_ISL_3226211, EPI_ISL_3226212 | Lighthouse Lab in Alderley Park | Wellcome Sanger Institute for the COVID-19 Genomics UK (COG-UK) Consortium | Jacquelyn Wynn, Mairead Hyland, The Lighthouse Lab in Alderley Park and Alex Alderton, Roberto Amato, Jeffrey Barrett, Sonia Goncalves, Ewan Harrison, David K. Jackson, Ian Johnston, Dominic Kwiatkowski, Cordelia Langford, John Sillitoe on behalf of the Wellcome Sanger Institute COVID-19 Surveillance Team |
| EPI_ISL_3226213 | Lighthouse Lab in Milton Keynes | Wellcome Sanger Institute for the COVID-19 Genomics UK (COG-UK) Consortium | The Lighthouse Lab in Milton Keynes and Alex Alderton, Roberto Amato, Jeffrey Barrett, Sonia Goncalves, Ewan Harrison, David K. Jackson, Ian Johnston, Dominic Kwiatkowski, Cordelia Langford, John Sillitoe on behalf of the Wellcome Sanger Institute COVID-19 Surveillance Team |
| EPI_ISL_3226214 | Lighthouse Laboratory Plymouth | Wellcome Sanger Institute for the COVID-19 Genomics UK (COG-UK) Consortium | Lighthouse Laboratory Plymouth and Alex Alderton, Roberto Amato, Jeffrey Barrett, Sonia Goncalves, Ewan Harrison, David K. Jackson, Ian Johnston, Dominic Kwiatkowski, Cordelia Langford, John Sillitoe on behalf of the Wellcome Sanger Institute COVID-19 Surveillance Team |
| EPI_ISL_3226215, EPI_ISL_3226216 | Lighthouse Lab in Glasgow | Wellcome Sanger Institute for the COVID-19 Genomics UK (COG-UK) Consortium | Harper VanSteenhouse, Yumi Kasai, David Gray, Carol Clugston, Anna Dominiczak and Alex Alderton, Roberto Amato, Jeffrey Barrett, Sonia Goncalves, Ewan Harrison, David K. Jackson, Ian Johnston, Dominic Kwiatkowski, Cordelia Langford, John Sillitoe on behalf of the Wellcome Sanger Institute COVID-19 Surveillance Team |
| EPI_ISL_3226217 | Lighthouse Laboratory Plymouth | Wellcome Sanger Institute for the COVID-19 Genomics UK (COG-UK) Consortium | Lighthouse Laboratory Plymouth and Alex Alderton, Roberto Amato, Jeffrey Barrett, Sonia Goncalves, Ewan Harrison, David K. Jackson, Ian Johnston, Dominic Kwiatkowski, Cordelia Langford, John Sillitoe on behalf of the Wellcome Sanger Institute COVID-19 Surveillance Team |
| EPI_ISL_3226218 | Lighthouse Lab in Glasgow | Wellcome Sanger Institute for the COVID-19 Genomics UK (COG-UK) Consortium | Harper VanSteenhouse, Yumi Kasai, David Gray, Carol Clugston, Anna Dominiczak and Alex Alderton, Roberto Amato, Jeffrey Barrett, Sonia Goncalves, Ewan Harrison, David K. Jackson, Ian Johnston, Dominic Kwiatkowski, Cordelia Langford, John Sillitoe on behalf of the Wellcome Sanger Institute COVID-19 Surveillance Team |
| EPI_ISL_3226219 | Lighthouse Laboratory Plymouth | Wellcome Sanger Institute for the COVID-19 Genomics UK (COG-UK) Consortium | Lighthouse Laboratory Plymouth and Alex Alderton, Roberto Amato, Jeffrey Barrett, Sonia Goncalves, Ewan Harrison, David K. Jackson, Ian Johnston, Dominic Kwiatkowski, Cordelia Langford, John Sillitoe on behalf of the Wellcome Sanger Institute COVID-19 Surveillance Team |
| EPI_ISL_3226220 | Lighthouse Lab in Alderley Park | Wellcome Sanger Institute for the COVID-19 Genomics UK (COG-UK) Consortium | Jacquelyn Wynn, Mairead Hyland, The Lighthouse Lab in Alderley Park and Alex Alderton, Roberto Amato, Jeffrey Barrett, Sonia Goncalves, Ewan Harrison, David K. Jackson, Ian Johnston, Dominic Kwiatkowski, Cordelia Langford, John Sillitoe on behalf of the Wellcome Sanger Institute COVID-19 Surveillance Team |
| EPI_ISL_3226221 | Lighthouse Laboratory Plymouth | Wellcome Sanger Institute for the COVID-19 Genomics UK (COG-UK) Consortium | Lighthouse Laboratory Plymouth and Alex Alderton, Roberto Amato, Jeffrey Barrett, Sonia Goncalves, Ewan Harrison, David K. Jackson, Ian Johnston, Dominic Kwiatkowski, Cordelia Langford, John Sillitoe on behalf of the Wellcome Sanger Institute COVID-19 Surveillance Team |
| EPI_ISL_3226222 | Lighthouse Lab in Milton Keynes | Wellcome Sanger Institute for the COVID-19 Genomics UK (COG-UK) Consortium | The Lighthouse Lab in Milton Keynes and Alex Alderton, Roberto Amato, Jeffrey Barrett, Sonia Goncalves, Ewan Harrison, David K. Jackson, Ian Johnston, Dominic Kwiatkowski, Cordelia Langford, John Sillitoe on behalf of the Wellcome Sanger Institute COVID-19 Surveillance Team |
| EPI_ISL_3226223 | Lighthouse Lab in Glasgow | Wellcome Sanger Institute for the COVID-19 Genomics UK (COG-UK) Consortium | Harper VanSteenhouse, Yumi Kasai, David Gray, Carol Clugston, Anna Dominiczak and Alex Alderton, Roberto Amato, Jeffrey Barrett, Sonia Goncalves, Ewan Harrison, David K. Jackson, Ian Johnston, Dominic Kwiatkowski, Cordelia Langford, John Sillitoe on behalf of the Wellcome Sanger Institute COVID-19 Surveillance Team |
| EPI_ISL_3226224 | Lighthouse Lab in Alderley Park | Wellcome Sanger Institute for the COVID-19 Genomics UK (COG-UK) Consortium | Jacquelyn Wynn, Mairead Hyland, The Lighthouse Lab in Alderley Park and Alex Alderton, Roberto Amato, Jeffrey Barrett, Sonia Goncalves, Ewan Harrison, David K. Jackson, Ian Johnston, Dominic Kwiatkowski, Cordelia Langford, John Sillitoe on behalf of the Wellcome Sanger Institute COVID-19 Surveillance Team |
| EPI_ISL_3226225 | Lighthouse Lab in Glasgow | Wellcome Sanger Institute for the COVID-19 Genomics UK (COG-UK) Consortium | Harper VanSteenhouse, Yumi Kasai, David Gray, Carol Clugston, Anna Dominiczak and Alex Alderton, Roberto Amato, Jeffrey Barrett, Sonia Goncalves, Ewan Harrison, David K. Jackson, Ian Johnston, Dominic Kwiatkowski, Cordelia Langford, John Sillitoe on behalf of the Wellcome Sanger Institute COVID-19 Surveillance Team |
| EPI_ISL_3226226 | Lighthouse Laboratory Plymouth | Wellcome Sanger Institute for the COVID-19 Genomics UK (COG-UK) Consortium | Lighthouse Laboratory Plymouth and Alex Alderton, Roberto Amato, Jeffrey Barrett, Sonia Goncalves, Ewan Harrison, David K. Jackson, Ian Johnston, Dominic Kwiatkowski, Cordelia Langford, John Sillitoe on behalf of the Wellcome Sanger Institute COVID-19 Surveillance Team |
| EPI_ISL_3226227 | Lighthouse Lab in Milton Keynes | Wellcome Sanger Institute for the COVID-19 Genomics UK (COG-UK) Consortium | The Lighthouse Lab in Milton Keynes and Alex Alderton, Roberto Amato, Jeffrey Barrett, Sonia Goncalves, Ewan Harrison, David K. Jackson, Ian Johnston, Dominic Kwiatkowski, Cordelia Langford, John Sillitoe on behalf of the Wellcome Sanger Institute COVID-19 Surveillance Team |
| EPI_ISL_3226228 | Lighthouse Lab in Alderley Park | Wellcome Sanger Institute for the COVID-19 Genomics UK (COG-UK) Consortium | Jacquelyn Wynn, Mairead Hyland, The Lighthouse Lab in Alderley Park and Alex Alderton, Roberto Amato, Jeffrey Barrett, Sonia Goncalves, Ewan Harrison, David K. Jackson, Ian Johnston, Dominic Kwiatkowski, Cordelia Langford, John Sillitoe on behalf of the Wellcome Sanger Institute COVID-19 Surveillance Team |
| EPI_ISL_3226229 | Lighthouse Lab in Milton Keynes | Wellcome Sanger Institute for the COVID-19 Genomics UK (COG-UK) Consortium | The Lighthouse Lab in Milton Keynes and Alex Alderton, Roberto Amato, Jeffrey Barrett, Sonia Goncalves, Ewan Harrison, David K. Jackson, Ian Johnston, Dominic Kwiatkowski, Cordelia Langford, John Sillitoe on behalf of the Wellcome Sanger Institute COVID-19 Surveillance Team |
| EPI_ISL_3226230 | Lighthouse Lab in Alderley Park | Wellcome Sanger Institute for the COVID-19 Genomics UK (COG-UK) Consortium | Jacquelyn Wynn, Mairead Hyland, The Lighthouse Lab in Alderley Park and Alex Alderton, Roberto Amato, Jeffrey Barrett, Sonia Goncalves, Ewan Harrison, David K. Jackson, Ian Johnston, Dominic Kwiatkowski, Cordelia Langford, John Sillitoe on behalf of the Wellcome Sanger Institute COVID-19 Surveillance Team |
| EPI_ISL_3226231 | Lighthouse Lab in Milton Keynes | Wellcome Sanger Institute for the COVID-19 Genomics UK (COG-UK) Consortium | The Lighthouse Lab in Milton Keynes and Alex Alderton, Roberto Amato, Jeffrey Barrett, Sonia Goncalves, Ewan Harrison, David K. Jackson, Ian Johnston, Dominic Kwiatkowski, Cordelia Langford, John Sillitoe on behalf of the Wellcome Sanger Institute COVID-19 Surveillance Team |
| EPI_ISL_3226232, EPI_ISL_3226233 | Lighthouse Laboratory Plymouth | Wellcome Sanger Institute for the COVID-19 Genomics UK (COG-UK) Consortium | Lighthouse Laboratory Plymouth and Alex Alderton, Roberto Amato, Jeffrey Barrett, Sonia Goncalves, Ewan Harrison, David K. Jackson, Ian Johnston, Dominic Kwiatkowski, Cordelia Langford, John Sillitoe on behalf of the Wellcome Sanger Institute COVID-19 Surveillance Team |

| | | | |
|---|---|---|---|
| EPI_ISL_3226234 | Lighthouse Lab in Alderley Park | Wellcome Sanger Institute for the COVID-19 Genomics UK (COG-UK) Consortium | Jacquelyn Wynn, Mairead Hyland, The Lighthouse Lab in Alderley Park and Alex Alderton, Roberto Amato, Jeffrey Barrett, Sonia Goncalves, Ewan Harrison, David K. Jackson, Ian Johnston, Dominic Kwiatkowski, Cordelia Langford, John Sillitoe on behalf of the Wellcome Sanger Institute COVID-19 Surveillance Team |
| EPI_ISL_3226235 | Lighthouse Laboratory Plymouth | Wellcome Sanger Institute for the COVID-19 Genomics UK (COG-UK) Consortium | Lighthouse Laboratory Plymouth and Alex Alderton, Roberto Amato, Jeffrey Barrett, Sonia Goncalves, Ewan Harrison, David K. Jackson, Ian Johnston, Dominic Kwiatkowski, Cordelia Langford, John Sillitoe on behalf of the Wellcome Sanger Institute COVID-19 Surveillance Team |
| EPI_ISL_3226236, EPI_ISL_3226237 | Lighthouse Lab in Milton Keynes | Wellcome Sanger Institute for the COVID-19 Genomics UK (COG-UK) Consortium | The Lighthouse Lab in Milton Keynes and Alex Alderton, Roberto Amato, Jeffrey Barrett, Sonia Goncalves, Ewan Harrison, David K. Jackson, Ian Johnston, Dominic Kwiatkowski, Cordelia Langford, John Sillitoe on behalf of the Wellcome Sanger Institute COVID-19 Surveillance Team |
| EPI_ISL_3226238 | Lighthouse Lab in Alderley Park | Wellcome Sanger Institute for the COVID-19 Genomics UK (COG-UK) Consortium | Jacquelyn Wynn, Mairead Hyland, The Lighthouse Lab in Alderley Park and Alex Alderton, Roberto Amato, Jeffrey Barrett, Sonia Goncalves, Ewan Harrison, David K. Jackson, Ian Johnston, Dominic Kwiatkowski, Cordelia Langford, John Sillitoe on behalf of the Wellcome Sanger Institute COVID-19 Surveillance Team |
| EPI_ISL_3226239, EPI_ISL_3226240 | Lighthouse Lab in Milton Keynes | Wellcome Sanger Institute for the COVID-19 Genomics UK (COG-UK) Consortium | The Lighthouse Lab in Milton Keynes and Alex Alderton, Roberto Amato, Jeffrey Barrett, Sonia Goncalves, Ewan Harrison, David K. Jackson, Ian Johnston, Dominic Kwiatkowski, Cordelia Langford, John Sillitoe on behalf of the Wellcome Sanger Institute COVID-19 Surveillance Team |
| EPI_ISL_3226241 | Lighthouse Lab in Alderley Park | Wellcome Sanger Institute for the COVID-19 Genomics UK (COG-UK) Consortium | Jacquelyn Wynn, Mairead Hyland, The Lighthouse Lab in Alderley Park and Alex Alderton, Roberto Amato, Jeffrey Barrett, Sonia Goncalves, Ewan Harrison, David K. Jackson, Ian Johnston, Dominic Kwiatkowski, Cordelia Langford, John Sillitoe on behalf of the Wellcome Sanger Institute COVID-19 Surveillance Team |
| EPI_ISL_3226242 | Lighthouse Lab in Glasgow | Wellcome Sanger Institute for the COVID-19 Genomics UK (COG-UK) Consortium | Harper VanSteenhouse, Yumi Kasai, David Gray, Carol Clugston, Anna Dominiczak and Alex Alderton, Roberto Amato, Jeffrey Barrett, Sonia Goncalves, Ewan Harrison, David K. Jackson, Ian Johnston, Dominic Kwiatkowski, Cordelia Langford, John Sillitoe on behalf of the Wellcome Sanger Institute COVID-19 Surveillance Team |
| EPI_ISL_3226243 | Lighthouse Lab in Milton Keynes | Wellcome Sanger Institute for the COVID-19 Genomics UK (COG-UK) Consortium | The Lighthouse Lab in Milton Keynes and Alex Alderton, Roberto Amato, Jeffrey Barrett, Sonia Goncalves, Ewan Harrison, David K. Jackson, Ian Johnston, Dominic Kwiatkowski, Cordelia Langford, John Sillitoe on behalf of the Wellcome Sanger Institute COVID-19 Surveillance Team |
| EPI_ISL_3226244 | Lighthouse Laboratory Plymouth | Wellcome Sanger Institute for the COVID-19 Genomics UK (COG-UK) Consortium | Lighthouse Laboratory Plymouth and Alex Alderton, Roberto Amato, Jeffrey Barrett, Sonia Goncalves, Ewan Harrison, David K. Jackson, Ian Johnston, Dominic Kwiatkowski, Cordelia Langford, John Sillitoe on behalf of the Wellcome Sanger Institute COVID-19 Surveillance Team |
| EPI_ISL_3226245, EPI_ISL_3226246 | Lighthouse Lab in Alderley Park | Wellcome Sanger Institute for the COVID-19 Genomics UK (COG-UK) Consortium | Jacquelyn Wynn, Mairead Hyland, The Lighthouse Lab in Alderley Park and Alex Alderton, Roberto Amato, Jeffrey Barrett, Sonia Goncalves, Ewan Harrison, David K. Jackson, Ian Johnston, Dominic Kwiatkowski, Cordelia Langford, John Sillitoe on behalf of the Wellcome Sanger Institute COVID-19 Surveillance Team |
| EPI_ISL_3226247 | Lighthouse Lab in Glasgow | Wellcome Sanger Institute for the COVID-19 Genomics UK (COG-UK) Consortium | Harper VanSteenhouse, Yumi Kasai, David Gray, Carol Clugston, Anna Dominiczak and Alex Alderton, Roberto Amato, Jeffrey Barrett, Sonia Goncalves, Ewan Harrison, David K. Jackson, Ian Johnston, Dominic Kwiatkowski, Cordelia Langford, John Sillitoe on behalf of the Wellcome Sanger Institute COVID-19 Surveillance Team |
| EPI_ISL_3226248, EPI_ISL_3226249, EPI_ISL_3226250 | Lighthouse Lab in Milton Keynes | Wellcome Sanger Institute for the COVID-19 Genomics UK (COG-UK) Consortium | The Lighthouse Lab in Milton Keynes and Alex Alderton, Roberto Amato, Jeffrey Barrett, Sonia Goncalves, Ewan Harrison, David K. Jackson, Ian Johnston, Dominic Kwiatkowski, Cordelia Langford, John Sillitoe on behalf of the Wellcome Sanger Institute COVID-19 Surveillance Team |
| EPI_ISL_3226251 | Lighthouse Lab in Alderley Park | Wellcome Sanger Institute for the COVID-19 Genomics UK (COG-UK) Consortium | Jacquelyn Wynn, Mairead Hyland, The Lighthouse Lab in Alderley Park and Alex Alderton, Roberto Amato, Jeffrey Barrett, Sonia Goncalves, Ewan Harrison, David K. Jackson, Ian Johnston, Dominic Kwiatkowski, Cordelia Langford, John Sillitoe on behalf of the Wellcome Sanger Institute COVID-19 Surveillance Team |
| EPI_ISL_3226252 | Lighthouse Lab in Milton Keynes | Wellcome Sanger Institute for the COVID-19 Genomics UK (COG-UK) Consortium | The Lighthouse Lab in Milton Keynes and Alex Alderton, Roberto Amato, Jeffrey Barrett, Sonia Goncalves, Ewan Harrison, David K. Jackson, Ian Johnston, Dominic Kwiatkowski, Cordelia Langford, John Sillitoe on behalf of the Wellcome Sanger Institute COVID-19 Surveillance Team |
| EPI_ISL_3226253 | Lighthouse Laboratory Plymouth | Wellcome Sanger Institute for the COVID-19 Genomics UK (COG-UK) Consortium | Lighthouse Laboratory Plymouth and Alex Alderton, Roberto Amato, Jeffrey Barrett, Sonia Goncalves, Ewan Harrison, David K. Jackson, Ian Johnston, Dominic Kwiatkowski, Cordelia Langford, John Sillitoe on behalf of the Wellcome Sanger Institute COVID-19 Surveillance Team |
| EPI_ISL_3226254 | Lighthouse Lab in Milton Keynes | Wellcome Sanger Institute for the COVID-19 Genomics UK (COG-UK) Consortium | The Lighthouse Lab in Milton Keynes and Alex Alderton, Roberto Amato, Jeffrey Barrett, Sonia Goncalves, Ewan Harrison, David K. Jackson, Ian Johnston, Dominic Kwiatkowski, Cordelia Langford, John Sillitoe on behalf of the Wellcome Sanger Institute COVID-19 Surveillance Team |
| EPI_ISL_3226255, EPI_ISL_3226256 | Lighthouse Lab in Alderley Park | Wellcome Sanger Institute for the COVID-19 Genomics UK (COG-UK) Consortium | Jacquelyn Wynn, Mairead Hyland, The Lighthouse Lab in Alderley Park and Alex Alderton, Roberto Amato, Jeffrey Barrett, Sonia Goncalves, Ewan Harrison, David K. Jackson, Ian Johnston, Dominic Kwiatkowski, Cordelia Langford, John Sillitoe on behalf of the Wellcome Sanger Institute COVID-19 Surveillance Team |
| EPI_ISL_3226257 | Lighthouse Laboratory Plymouth | Wellcome Sanger Institute for the COVID-19 Genomics UK (COG-UK) Consortium | Lighthouse Laboratory Plymouth and Alex Alderton, Roberto Amato, Jeffrey Barrett, Sonia Goncalves, Ewan Harrison, David K. Jackson, Ian Johnston, Dominic Kwiatkowski, Cordelia Langford, John Sillitoe on behalf of the Wellcome Sanger Institute COVID-19 Surveillance Team |
| EPI_ISL_3226258 | Lighthouse Lab in Glasgow | Wellcome Sanger Institute for the COVID-19 Genomics UK (COG-UK) Consortium | Harper VanSteenhouse, Yumi Kasai, David Gray, Carol Clugston, Anna Dominiczak and Alex Alderton, Roberto Amato, Jeffrey Barrett, Sonia Goncalves, Ewan Harrison, David K. Jackson, Ian Johnston, Dominic Kwiatkowski, Cordelia Langford, John Sillitoe on behalf of the Wellcome Sanger Institute COVID-19 Surveillance Team |
| EPI_ISL_3226259 | Lighthouse Lab in Alderley Park | Wellcome Sanger Institute for the COVID-19 Genomics UK (COG-UK) Consortium | Jacquelyn Wynn, Mairead Hyland, The Lighthouse Lab in Alderley Park and Alex Alderton, Roberto Amato, Jeffrey Barrett, Sonia Goncalves, Ewan Harrison, David K. Jackson, Ian Johnston, Dominic Kwiatkowski, Cordelia Langford, John Sillitoe on behalf of the Wellcome Sanger Institute COVID-19 Surveillance Team |
| EPI_ISL_3226260 | Lighthouse Laboratory Plymouth | Wellcome Sanger Institute for the COVID-19 Genomics UK (COG-UK) Consortium | Lighthouse Laboratory Plymouth and Alex Alderton, Roberto Amato, Jeffrey Barrett, Sonia Goncalves, Ewan Harrison, David K. Jackson, Ian Johnston, Dominic Kwiatkowski, Cordelia Langford, John Sillitoe on behalf of the Wellcome Sanger Institute COVID-19 Surveillance Team |
| EPI_ISL_3226261 | Lighthouse Lab in Milton Keynes | Wellcome Sanger Institute for the COVID-19 Genomics UK (COG-UK) Consortium | The Lighthouse Lab in Milton Keynes and Alex Alderton, Roberto Amato, Jeffrey Barrett, Sonia Goncalves, Ewan Harrison, David K. Jackson, Ian Johnston, Dominic Kwiatkowski, Cordelia Langford, John Sillitoe on behalf of the Wellcome Sanger Institute COVID-19 Surveillance Team |
| EPI_ISL_3226262 | Lighthouse Lab in Glasgow | Wellcome Sanger Institute for the COVID-19 Genomics UK (COG-UK) Consortium | Harper VanSteenhouse, Yumi Kasai, David Gray, Carol Clugston, Anna Dominiczak and Alex Alderton, Roberto Amato, Jeffrey Barrett, Sonia Goncalves, Ewan Harrison, David K. Jackson, Ian Johnston, Dominic Kwiatkowski, Cordelia Langford, John Sillitoe on behalf of the Wellcome Sanger Institute COVID-19 Surveillance Team |
| EPI_ISL_3226263 | Lighthouse Laboratory Plymouth | Wellcome Sanger Institute for the COVID-19 Genomics UK (COG-UK) Consortium | Lighthouse Laboratory Plymouth and Alex Alderton, Roberto Amato, Jeffrey Barrett, Sonia Goncalves, Ewan Harrison, David K. Jackson, Ian Johnston, Dominic Kwiatkowski, Cordelia Langford, John Sillitoe on behalf of the Wellcome Sanger Institute COVID-19 Surveillance Team |
| EPI_ISL_3226264 | Lighthouse Lab in Milton Keynes | Wellcome Sanger Institute for the COVID-19 Genomics UK (COG-UK) Consortium | The Lighthouse Lab in Milton Keynes and Alex Alderton, Roberto Amato, Jeffrey Barrett, Sonia Goncalves, Ewan Harrison, David K. Jackson, Ian Johnston, Dominic Kwiatkowski, Cordelia Langford, John Sillitoe on behalf of the Wellcome Sanger Institute COVID-19 Surveillance Team |
| EPI_ISL_3226265 | Lighthouse Lab in Alderley Park | Wellcome Sanger Institute for the COVID-19 Genomics UK (COG-UK) Consortium | Jacquelyn Wynn, Mairead Hyland, The Lighthouse Lab in Alderley Park and Alex Alderton, Roberto Amato, Jeffrey Barrett, Sonia Goncalves, Ewan Harrison, David K. Jackson, Ian Johnston, Dominic Kwiatkowski, Cordelia Langford, John Sillitoe on behalf of the Wellcome Sanger Institute COVID-19 Surveillance Team |
| EPI_ISL_3226266 | Lighthouse Laboratory Plymouth | Wellcome Sanger Institute for the COVID-19 Genomics UK (COG-UK) Consortium | Lighthouse Laboratory Plymouth and Alex Alderton, Roberto Amato, Jeffrey Barrett, Sonia Goncalves, Ewan Harrison, David K. Jackson, Ian Johnston, Dominic Kwiatkowski, Cordelia Langford, John Sillitoe on behalf of the Wellcome Sanger Institute COVID-19 Surveillance Team |
| EPI_ISL_3226267, EPI_ISL_3226268 | Lighthouse Lab in Milton Keynes | Wellcome Sanger Institute for the COVID-19 Genomics UK (COG-UK) Consortium | The Lighthouse Lab in Milton Keynes and Alex Alderton, Roberto Amato, Jeffrey Barrett, Sonia Goncalves, Ewan Harrison, David K. Jackson, Ian Johnston, Dominic Kwiatkowski, Cordelia Langford, John Sillitoe on behalf of the Wellcome Sanger Institute COVID-19 Surveillance Team |
| EPI_ISL_3226269 | Lighthouse Lab in Glasgow | Wellcome Sanger Institute for the COVID-19 Genomics UK (COG-UK) Consortium | Harper VanSteenhouse, Yumi Kasai, David Gray, Carol Clugston, Anna Dominiczak and Alex Alderton, Roberto Amato, Jeffrey Barrett, Sonia Goncalves, Ewan Harrison, David K. Jackson, Ian Johnston, Dominic Kwiatkowski, Cordelia Langford, John Sillitoe on behalf of the Wellcome Sanger Institute COVID-19 Surveillance Team |
| EPI_ISL_3226270 | Lighthouse Laboratory Plymouth | Wellcome Sanger Institute for the COVID-19 Genomics UK (COG-UK) Consortium | Lighthouse Laboratory Plymouth and Alex Alderton, Roberto Amato, Jeffrey Barrett, Sonia Goncalves, Ewan Harrison, David K. Jackson, Ian Johnston, Dominic Kwiatkowski, Cordelia Langford, John Sillitoe on behalf of the Wellcome Sanger Institute COVID-19 Surveillance Team |
| EPI_ISL_3226271 | Lighthouse Lab in Alderley Park | Wellcome Sanger Institute for the COVID-19 Genomics UK (COG-UK) Consortium | Jacquelyn Wynn, Mairead Hyland, The Lighthouse Lab in Alderley Park and Alex Alderton, Roberto Amato, Jeffrey Barrett, Sonia Goncalves, Ewan Harrison, David K. Jackson, Ian Johnston, Dominic Kwiatkowski, Cordelia Langford, John Sillitoe on behalf of the Wellcome Sanger Institute COVID-19 Surveillance Team |
| EPI_ISL_3226272, EPI_ISL_3226273, EPI_ISL_3226274, EPI_ISL_3226275 | Lighthouse Lab in Glasgow | Wellcome Sanger Institute for the COVID-19 Genomics UK (COG-UK) Consortium | Harper VanSteenhouse, Yumi Kasai, David Gray, Carol Clugston, Anna Dominiczak and Alex Alderton, Roberto Amato, Jeffrey Barrett, Sonia Goncalves, Ewan Harrison, David K. Jackson, Ian Johnston, Dominic Kwiatkowski, Cordelia Langford, John Sillitoe on behalf of the Wellcome Sanger Institute COVID-19 Surveillance Team |
| EPI_ISL_3226276 | Lighthouse Laboratory Plymouth | Wellcome Sanger Institute for the COVID-19 Genomics UK (COG-UK) Consortium | Lighthouse Laboratory Plymouth and Alex Alderton, Roberto Amato, Jeffrey Barrett, Sonia Goncalves, Ewan Harrison, David K. Jackson, Ian Johnston, Dominic Kwiatkowski, Cordelia Langford, John Sillitoe on behalf of the Wellcome Sanger Institute COVID-19 Surveillance Team |
| EPI_ISL_3226277, EPI_ISL_3226278 | Lighthouse Lab in Glasgow | Wellcome Sanger Institute for the COVID-19 Genomics UK (COG-UK) Consortium | Harper VanSteenhouse, Yumi Kasai, David Gray, Carol Clugston, Anna Dominiczak and Alex Alderton, Roberto Amato, Jeffrey Barrett, Sonia Goncalves, Ewan Harrison, David K. Jackson, Ian Johnston, Dominic Kwiatkowski, Cordelia Langford, John Sillitoe on behalf of the Wellcome Sanger Institute COVID-19 Surveillance Team |
| EPI_ISL_3226279 | Lighthouse Lab in Alderley Park | Wellcome Sanger Institute for the COVID-19 Genomics UK (COG-UK) Consortium | Jacquelyn Wynn, Mairead Hyland, The Lighthouse Lab in Alderley Park and Alex Alderton, Roberto Amato, Jeffrey Barrett, Sonia Goncalves, Ewan Harrison, David K. Jackson, Ian Johnston, Dominic Kwiatkowski, Cordelia Langford, John Sillitoe on behalf of the Wellcome Sanger Institute COVID-19 Surveillance Team |
| EPI_ISL_3226280 | Lighthouse Lab in Milton Keynes | Wellcome Sanger Institute for the COVID-19 Genomics UK (COG-UK) Consortium | The Lighthouse Lab in Milton Keynes and Alex Alderton, Roberto Amato, Jeffrey Barrett, Sonia Goncalves, Ewan Harrison, David K. Jackson, Ian Johnston, Dominic Kwiatkowski, Cordelia Langford, John Sillitoe on behalf of the Wellcome Sanger Institute COVID-19 Surveillance Team |
| EPI_ISL_3226281 | Lighthouse Lab in Alderley Park | Wellcome Sanger Institute for the COVID-19 Genomics UK (COG-UK) Consortium | Jacquelyn Wynn, Mairead Hyland, The Lighthouse Lab in Alderley Park and Alex Alderton, Roberto Amato, Jeffrey Barrett, Sonia Goncalves, Ewan Harrison, David K. Jackson, Ian Johnston, Dominic Kwiatkowski, Cordelia Langford, John Sillitoe on behalf of the Wellcome Sanger Institute COVID-19 Surveillance Team |
| EPI_ISL_3226282, EPI_ISL_3226283 | Lighthouse Laboratory Plymouth | Wellcome Sanger Institute for the COVID-19 Genomics UK (COG-UK) Consortium | Lighthouse Laboratory Plymouth and Alex Alderton, Roberto Amato, Jeffrey Barrett, Sonia Goncalves, Ewan Harrison, David K. Jackson, Ian Johnston, Dominic Kwiatkowski, Cordelia Langford, John Sillitoe on behalf of the Wellcome Sanger Institute COVID-19 Surveillance Team |
| EPI_ISL_3226284 | Lighthouse Lab in Milton Keynes | Wellcome Sanger Institute for the COVID-19 Genomics UK (COG-UK) Consortium | The Lighthouse Lab in Milton Keynes and Alex Alderton, Roberto Amato, Jeffrey Barrett, Sonia Goncalves, Ewan Harrison, David K. Jackson, Ian Johnston, Dominic Kwiatkowski, Cordelia Langford, John Sillitoe on behalf of the Wellcome Sanger Institute COVID-19 Surveillance Team |
| EPI_ISL_3226285 | Lighthouse Lab in Alderley Park | Wellcome Sanger Institute for the COVID-19 Genomics UK (COG-UK) Consortium | Jacquelyn Wynn, Mairead Hyland, The Lighthouse Lab in Alderley Park and Alex Alderton, Roberto Amato, Jeffrey Barrett, Sonia Goncalves, Ewan Harrison, David K. Jackson, Ian Johnston, Dominic Kwiatkowski, Cordelia Langford, John Sillitoe on behalf of the Wellcome Sanger Institute COVID-19 Surveillance Team |
| EPI_ISL_3226286, EPI_ISL_3226287 | Lighthouse Lab in Glasgow | Wellcome Sanger Institute for the COVID-19 Genomics UK (COG-UK) Consortium | Harper VanSteenhouse, Yumi Kasai, David Gray, Carol Clugston, Anna Dominiczak and Alex Alderton, Roberto Amato, Jeffrey Barrett, Sonia Goncalves, Ewan Harrison, David K. Jackson, Ian Johnston, Dominic Kwiatkowski, Cordelia Langford, John Sillitoe on behalf of the Wellcome Sanger Institute COVID-19 Surveillance Team |
| EPI_ISL_3226288 | Lighthouse Lab in Milton Keynes | Wellcome Sanger Institute for the COVID-19 Genomics UK (COG-UK) Consortium | The Lighthouse Lab in Milton Keynes and Alex Alderton, Roberto Amato, Jeffrey Barrett, Sonia Goncalves, Ewan Harrison, David K. Jackson, Ian Johnston, Dominic Kwiatkowski, Cordelia Langford, John Sillitoe on behalf of the Wellcome Sanger Institute COVID-19 Surveillance Team |
| EPI_ISL_3226289 | Lighthouse Laboratory Plymouth | Wellcome Sanger Institute for the COVID-19 Genomics UK (COG-UK) Consortium | Lighthouse Laboratory Plymouth and Alex Alderton, Roberto Amato, Jeffrey Barrett, Sonia Goncalves, Ewan Harrison, David K. Jackson, Ian Johnston, Dominic Kwiatkowski, Cordelia Langford, John Sillitoe on behalf of the Wellcome Sanger Institute COVID-19 Surveillance Team |
| EPI_ISL_3226290 | Lighthouse Lab in Milton Keynes | Wellcome Sanger Institute for the COVID-19 Genomics UK (COG-UK) Consortium | The Lighthouse Lab in Milton Keynes and Alex Alderton, Roberto Amato, Jeffrey Barrett, Sonia Goncalves, Ewan Harrison, David K. Jackson, Ian Johnston, Dominic Kwiatkowski, Cordelia Langford, John Sillitoe on behalf of the Wellcome Sanger Institute COVID-19 Surveillance Team |

| | | | |
|---|---|---|---|
| EPI_ISL_3226291 | Lighthouse Laboratory Plymouth | Wellcome Sanger Institute for the COVID-19 Genomics UK (COG-UK) Consortium | Lighthouse Laboratory Plymouth and Alex Alderton, Roberto Amato, Jeffrey Barrett, Sonia Goncalves, Ewan Harrison, David K. Jackson, Ian Johnston, Dominic Kwiatkowski, Cordelia Langford, John Sillitoe on behalf of the Wellcome Sanger Institute COVID-19 Surveillance Team |
| EPI_ISL_3226292 | Lighthouse Lab in Alderley Park | Wellcome Sanger Institute for the COVID-19 Genomics UK (COG-UK) Consortium | Jacquelyn Wynn, Mairead Hyland, The Lighthouse Lab in Alderley Park and Alex Alderton, Roberto Amato, Jeffrey Barrett, Sonia Goncalves, Ewan Harrison, David K. Jackson, Ian Johnston, Dominic Kwiatkowski, Cordelia Langford, John Sillitoe on behalf of the Wellcome Sanger Institute COVID-19 Surveillance Team |
| EPI_ISL_3226293 | Lighthouse Lab in Glasgow | Wellcome Sanger Institute for the COVID-19 Genomics UK (COG-UK) Consortium | Harper VanSteenhouse, Yumi Kasai, David Gray, Carol Clugston, Anna Dominiczak and Alex Alderton, Roberto Amato, Jeffrey Barrett, Sonia Goncalves, Ewan Harrison, David K. Jackson, Ian Johnston, Dominic Kwiatkowski, Cordelia Langford, John Sillitoe on behalf of the Wellcome Sanger Institute COVID-19 Surveillance Team |
| EPI_ISL_3226294 | Lighthouse Laboratory Plymouth | Wellcome Sanger Institute for the COVID-19 Genomics UK (COG-UK) Consortium | Lighthouse Laboratory Plymouth and Alex Alderton, Roberto Amato, Jeffrey Barrett, Sonia Goncalves, Ewan Harrison, David K. Jackson, Ian Johnston, Dominic Kwiatkowski, Cordelia Langford, John Sillitoe on behalf of the Wellcome Sanger Institute COVID-19 Surveillance Team |
| EPI_ISL_3226295, EPI_ISL_3226296 | Lighthouse Lab in Milton Keynes | Wellcome Sanger Institute for the COVID-19 Genomics UK (COG-UK) Consortium | The Lighthouse Lab in Milton Keynes and Alex Alderton, Roberto Amato, Jeffrey Barrett, Sonia Goncalves, Ewan Harrison, David K. Jackson, Ian Johnston, Dominic Kwiatkowski, Cordelia Langford, John Sillitoe on behalf of the Wellcome Sanger Institute COVID-19 Surveillance Team |
| EPI_ISL_3226297 | Lighthouse Lab in Glasgow | Wellcome Sanger Institute for the COVID-19 Genomics UK (COG-UK) Consortium | Harper VanSteenhouse, Yumi Kasai, David Gray, Carol Clugston, Anna Dominiczak and Alex Alderton, Roberto Amato, Jeffrey Barrett, Sonia Goncalves, Ewan Harrison, David K. Jackson, Ian Johnston, Dominic Kwiatkowski, Cordelia Langford, John Sillitoe on behalf of the Wellcome Sanger Institute COVID-19 Surveillance Team |
| EPI_ISL_3226298 | Lighthouse Lab in Alderley Park | Wellcome Sanger Institute for the COVID-19 Genomics UK (COG-UK) Consortium | Jacquelyn Wynn, Mairead Hyland, The Lighthouse Lab in Alderley Park and Alex Alderton, Roberto Amato, Jeffrey Barrett, Sonia Goncalves, Ewan Harrison, David K. Jackson, Ian Johnston, Dominic Kwiatkowski, Cordelia Langford, John Sillitoe on behalf of the Wellcome Sanger Institute COVID-19 Surveillance Team |
| EPI_ISL_3226299, EPI_ISL_3226300 | Lighthouse Laboratory Plymouth | Wellcome Sanger Institute for the COVID-19 Genomics UK (COG-UK) Consortium | Lighthouse Laboratory Plymouth and Alex Alderton, Roberto Amato, Jeffrey Barrett, Sonia Goncalves, Ewan Harrison, David K. Jackson, Ian Johnston, Dominic Kwiatkowski, Cordelia Langford, John Sillitoe on behalf of the Wellcome Sanger Institute COVID-19 Surveillance Team |
| EPI_ISL_3226301, EPI_ISL_3226302 | Lighthouse Lab in Glasgow | Wellcome Sanger Institute for the COVID-19 Genomics UK (COG-UK) Consortium | Harper VanSteenhouse, Yumi Kasai, David Gray, Carol Clugston, Anna Dominiczak and Alex Alderton, Roberto Amato, Jeffrey Barrett, Sonia Goncalves, Ewan Harrison, David K. Jackson, Ian Johnston, Dominic Kwiatkowski, Cordelia Langford, John Sillitoe on behalf of the Wellcome Sanger Institute COVID-19 Surveillance Team |
| EPI_ISL_3226303 | Lighthouse Laboratory Plymouth | Wellcome Sanger Institute for the COVID-19 Genomics UK (COG-UK) Consortium | Lighthouse Laboratory Plymouth and Alex Alderton, Roberto Amato, Jeffrey Barrett, Sonia Goncalves, Ewan Harrison, David K. Jackson, Ian Johnston, Dominic Kwiatkowski, Cordelia Langford, John Sillitoe on behalf of the Wellcome Sanger Institute COVID-19 Surveillance Team |
| EPI_ISL_3226304 | Lighthouse Lab in Milton Keynes | Wellcome Sanger Institute for the COVID-19 Genomics UK (COG-UK) Consortium | The Lighthouse Lab in Milton Keynes and Alex Alderton, Roberto Amato, Jeffrey Barrett, Sonia Goncalves, Ewan Harrison, David K. Jackson, Ian Johnston, Dominic Kwiatkowski, Cordelia Langford, John Sillitoe on behalf of the Wellcome Sanger Institute COVID-19 Surveillance Team |
| EPI_ISL_3226305, EPI_ISL_3226306 | Lighthouse Lab in Glasgow | Wellcome Sanger Institute for the COVID-19 Genomics UK (COG-UK) Consortium | Harper VanSteenhouse, Yumi Kasai, David Gray, Carol Clugston, Anna Dominiczak and Alex Alderton, Roberto Amato, Jeffrey Barrett, Sonia Goncalves, Ewan Harrison, David K. Jackson, Ian Johnston, Dominic Kwiatkowski, Cordelia Langford, John Sillitoe on behalf of the Wellcome Sanger Institute COVID-19 Surveillance Team |
| EPI_ISL_3226307 | Lighthouse Lab in Milton Keynes | Wellcome Sanger Institute for the COVID-19 Genomics UK (COG-UK) Consortium | The Lighthouse Lab in Milton Keynes and Alex Alderton, Roberto Amato, Jeffrey Barrett, Sonia Goncalves, Ewan Harrison, David K. Jackson, Ian Johnston, Dominic Kwiatkowski, Cordelia Langford, John Sillitoe on behalf of the Wellcome Sanger Institute COVID-19 Surveillance Team |
| EPI_ISL_3226308, EPI_ISL_3226309 | Lighthouse Lab in Alderley Park | Wellcome Sanger Institute for the COVID-19 Genomics UK (COG-UK) Consortium | Jacquelyn Wynn, Mairead Hyland, The Lighthouse Lab in Alderley Park and Alex Alderton, Roberto Amato, Jeffrey Barrett, Sonia Goncalves, Ewan Harrison, David K. Jackson, Ian Johnston, Dominic Kwiatkowski, Cordelia Langford, John Sillitoe on behalf of the Wellcome Sanger Institute COVID-19 Surveillance Team |
| EPI_ISL_3226310 | Lighthouse Laboratory Plymouth | Wellcome Sanger Institute for the COVID-19 Genomics UK (COG-UK) Consortium | Lighthouse Laboratory Plymouth and Alex Alderton, Roberto Amato, Jeffrey Barrett, Sonia Goncalves, Ewan Harrison, David K. Jackson, Ian Johnston, Dominic Kwiatkowski, Cordelia Langford, John Sillitoe on behalf of the Wellcome Sanger Institute COVID-19 Surveillance Team |
| EPI_ISL_3226311, EPI_ISL_3226312, EPI_ISL_3226313 | Lighthouse Lab in Glasgow | Wellcome Sanger Institute for the COVID-19 Genomics UK (COG-UK) Consortium | Harper VanSteenhouse, Yumi Kasai, David Gray, Carol Clugston, Anna Dominiczak and Alex Alderton, Roberto Amato, Jeffrey Barrett, Sonia Goncalves, Ewan Harrison, David K. Jackson, Ian Johnston, Dominic Kwiatkowski, Cordelia Langford, John Sillitoe on behalf of the Wellcome Sanger Institute COVID-19 Surveillance Team |
| EPI_ISL_3226314 | Lighthouse Lab in Alderley Park | Wellcome Sanger Institute for the COVID-19 Genomics UK (COG-UK) Consortium | Jacquelyn Wynn, Mairead Hyland, The Lighthouse Lab in Alderley Park and Alex Alderton, Roberto Amato, Jeffrey Barrett, Sonia Goncalves, Ewan Harrison, David K. Jackson, Ian Johnston, Dominic Kwiatkowski, Cordelia Langford, John Sillitoe on behalf of the Wellcome Sanger Institute COVID-19 Surveillance Team |
| EPI_ISL_3226315, EPI_ISL_3226316 | Lighthouse Lab in Milton Keynes | Wellcome Sanger Institute for the COVID-19 Genomics UK (COG-UK) Consortium | The Lighthouse Lab in Milton Keynes and Alex Alderton, Roberto Amato, Jeffrey Barrett, Sonia Goncalves, Ewan Harrison, David K. Jackson, Ian Johnston, Dominic Kwiatkowski, Cordelia Langford, John Sillitoe on behalf of the Wellcome Sanger Institute COVID-19 Surveillance Team |
| EPI_ISL_3226317 | Lighthouse Laboratory Plymouth | Wellcome Sanger Institute for the COVID-19 Genomics UK (COG-UK) Consortium | Lighthouse Laboratory Plymouth and Alex Alderton, Roberto Amato, Jeffrey Barrett, Sonia Goncalves, Ewan Harrison, David K. Jackson, Ian Johnston, Dominic Kwiatkowski, Cordelia Langford, John Sillitoe on behalf of the Wellcome Sanger Institute COVID-19 Surveillance Team |
| EPI_ISL_3226318, EPI_ISL_3226319 | Lighthouse Lab in Glasgow | Wellcome Sanger Institute for the COVID-19 Genomics UK (COG-UK) Consortium | Harper VanSteenhouse, Yumi Kasai, David Gray, Carol Clugston, Anna Dominiczak and Alex Alderton, Roberto Amato, Jeffrey Barrett, Sonia Goncalves, Ewan Harrison, David K. Jackson, Ian Johnston, Dominic Kwiatkowski, Cordelia Langford, John Sillitoe on behalf of the Wellcome Sanger Institute COVID-19 Surveillance Team |
| EPI_ISL_3226320, EPI_ISL_3226321 | Lighthouse Lab in Alderley Park | Wellcome Sanger Institute for the COVID-19 Genomics UK (COG-UK) Consortium | Jacquelyn Wynn, Mairead Hyland, The Lighthouse Lab in Alderley Park and Alex Alderton, Roberto Amato, Jeffrey Barrett, Sonia Goncalves, Ewan Harrison, David K. Jackson, Ian Johnston, Dominic Kwiatkowski, Cordelia Langford, John Sillitoe on behalf of the Wellcome Sanger Institute COVID-19 Surveillance Team |
| EPI_ISL_3226322 | Lighthouse Lab in Milton Keynes | Wellcome Sanger Institute for the COVID-19 Genomics UK (COG-UK) Consortium | The Lighthouse Lab in Milton Keynes and Alex Alderton, Roberto Amato, Jeffrey Barrett, Sonia Goncalves, Ewan Harrison, David K. Jackson, Ian Johnston, Dominic Kwiatkowski, Cordelia Langford, John Sillitoe on behalf of the Wellcome Sanger Institute COVID-19 Surveillance Team |
| EPI_ISL_3226323 | Lighthouse Lab in Alderley Park | Wellcome Sanger Institute for the COVID-19 Genomics UK (COG-UK) Consortium | Jacquelyn Wynn, Mairead Hyland, The Lighthouse Lab in Alderley Park and Alex Alderton, Roberto Amato, Jeffrey Barrett, Sonia Goncalves, Ewan Harrison, David K. Jackson, Ian Johnston, Dominic Kwiatkowski, Cordelia Langford, John Sillitoe on behalf of the Wellcome Sanger Institute COVID-19 Surveillance Team |
| EPI_ISL_3226324 | Lighthouse Lab in Glasgow | Wellcome Sanger Institute for the COVID-19 Genomics UK (COG-UK) Consortium | Harper VanSteenhouse, Yumi Kasai, David Gray, Carol Clugston, Anna Dominiczak and Alex Alderton, Roberto Amato, Jeffrey Barrett, Sonia Goncalves, Ewan Harrison, David K. Jackson, Ian Johnston, Dominic Kwiatkowski, Cordelia Langford, John Sillitoe on behalf of the Wellcome Sanger Institute COVID-19 Surveillance Team |
| EPI_ISL_3226325 | Lighthouse Lab in Alderley Park | Wellcome Sanger Institute for the COVID-19 Genomics UK (COG-UK) Consortium | Jacquelyn Wynn, Mairead Hyland, The Lighthouse Lab in Alderley Park and Alex Alderton, Roberto Amato, Jeffrey Barrett, Sonia Goncalves, Ewan Harrison, David K. Jackson, Ian Johnston, Dominic Kwiatkowski, Cordelia Langford, John Sillitoe on behalf of the Wellcome Sanger Institute COVID-19 Surveillance Team |
| EPI_ISL_3226326, EPI_ISL_3226327, EPI_ISL_3226328 | Lighthouse Laboratory Plymouth | Wellcome Sanger Institute for the COVID-19 Genomics UK (COG-UK) Consortium | Lighthouse Laboratory Plymouth and Alex Alderton, Roberto Amato, Jeffrey Barrett, Sonia Goncalves, Ewan Harrison, David K. Jackson, Ian Johnston, Dominic Kwiatkowski, Cordelia Langford, John Sillitoe on behalf of the Wellcome Sanger Institute COVID-19 Surveillance Team |
| EPI_ISL_3226329 | Lighthouse Lab in Alderley Park | Wellcome Sanger Institute for the COVID-19 Genomics UK (COG-UK) Consortium | Jacquelyn Wynn, Mairead Hyland, The Lighthouse Lab in Alderley Park and Alex Alderton, Roberto Amato, Jeffrey Barrett, Sonia Goncalves, Ewan Harrison, David K. Jackson, Ian Johnston, Dominic Kwiatkowski, Cordelia Langford, John Sillitoe on behalf of the Wellcome Sanger Institute COVID-19 Surveillance Team |
| EPI_ISL_3226330, EPI_ISL_3226332 | Lighthouse Lab in Glasgow | Wellcome Sanger Institute for the COVID-19 Genomics UK (COG-UK) Consortium | Harper VanSteenhouse, Yumi Kasai, David Gray, Carol Clugston, Anna Dominiczak and Alex Alderton, Roberto Amato, Jeffrey Barrett, Sonia Goncalves, Ewan Harrison, David K. Jackson, Ian Johnston, Dominic Kwiatkowski, Cordelia Langford, John Sillitoe on behalf of the Wellcome Sanger Institute COVID-19 Surveillance Team |
| EPI_ISL_3226333 | Lighthouse Laboratory Plymouth | Wellcome Sanger Institute for the COVID-19 Genomics UK (COG-UK) Consortium | Lighthouse Laboratory Plymouth and Alex Alderton, Roberto Amato, Jeffrey Barrett, Sonia Goncalves, Ewan Harrison, David K. Jackson, Ian Johnston, Dominic Kwiatkowski, Cordelia Langford, John Sillitoe on behalf of the Wellcome Sanger Institute COVID-19 Surveillance Team |
| EPI_ISL_3226334, EPI_ISL_3226335 | Lighthouse Lab in Glasgow | Wellcome Sanger Institute for the COVID-19 Genomics UK (COG-UK) Consortium | Harper VanSteenhouse, Yumi Kasai, David Gray, Carol Clugston, Anna Dominiczak and Alex Alderton, Roberto Amato, Jeffrey Barrett, Sonia Goncalves, Ewan Harrison, David K. Jackson, Ian Johnston, Dominic Kwiatkowski, Cordelia Langford, John Sillitoe on behalf of the Wellcome Sanger Institute COVID-19 Surveillance Team |
| EPI_ISL_3226336 | Lighthouse Laboratory Plymouth | Wellcome Sanger Institute for the COVID-19 Genomics UK (COG-UK) Consortium | Lighthouse Laboratory Plymouth and Alex Alderton, Roberto Amato, Jeffrey Barrett, Sonia Goncalves, Ewan Harrison, David K. Jackson, Ian Johnston, Dominic Kwiatkowski, Cordelia Langford, John Sillitoe on behalf of the Wellcome Sanger Institute COVID-19 Surveillance Team |
| EPI_ISL_3226337 | Lighthouse Lab in Alderley Park | Wellcome Sanger Institute for the COVID-19 Genomics UK (COG-UK) Consortium | Jacquelyn Wynn, Mairead Hyland, The Lighthouse Lab in Alderley Park and Alex Alderton, Roberto Amato, Jeffrey Barrett, Sonia Goncalves, Ewan Harrison, David K. Jackson, Ian Johnston, Dominic Kwiatkowski, Cordelia Langford, John Sillitoe on behalf of the Wellcome Sanger Institute COVID-19 Surveillance Team |
| EPI_ISL_3226338 | Lighthouse Lab in Glasgow | Wellcome Sanger Institute for the COVID-19 Genomics UK (COG-UK) Consortium | Harper VanSteenhouse, Yumi Kasai, David Gray, Carol Clugston, Anna Dominiczak and Alex Alderton, Roberto Amato, Jeffrey Barrett, Sonia Goncalves, Ewan Harrison, David K. Jackson, Ian Johnston, Dominic Kwiatkowski, Cordelia Langford, John Sillitoe on behalf of the Wellcome Sanger Institute COVID-19 Surveillance Team |
| EPI_ISL_3226339, EPI_ISL_3226340, EPI_ISL_3226341 | Lighthouse Lab in Alderley Park | Wellcome Sanger Institute for the COVID-19 Genomics UK (COG-UK) Consortium | Jacquelyn Wynn, Mairead Hyland, The Lighthouse Lab in Alderley Park and Alex Alderton, Roberto Amato, Jeffrey Barrett, Sonia Goncalves, Ewan Harrison, David K. Jackson, Ian Johnston, Dominic Kwiatkowski, Cordelia Langford, John Sillitoe on behalf of the Wellcome Sanger Institute COVID-19 Surveillance Team |
| EPI_ISL_3226342 | Lighthouse Laboratory Plymouth | Wellcome Sanger Institute for the COVID-19 Genomics UK (COG-UK) Consortium | Lighthouse Laboratory Plymouth and Alex Alderton, Roberto Amato, Jeffrey Barrett, Sonia Goncalves, Ewan Harrison, David K. Jackson, Ian Johnston, Dominic Kwiatkowski, Cordelia Langford, John Sillitoe on behalf of the Wellcome Sanger Institute COVID-19 Surveillance Team |
| EPI_ISL_3226343 | Lighthouse Lab in Milton Keynes | Wellcome Sanger Institute for the COVID-19 Genomics UK (COG-UK) Consortium | The Lighthouse Lab in Milton Keynes and Alex Alderton, Roberto Amato, Jeffrey Barrett, Sonia Goncalves, Ewan Harrison, David K. Jackson, Ian Johnston, Dominic Kwiatkowski, Cordelia Langford, John Sillitoe on behalf of the Wellcome Sanger Institute COVID-19 Surveillance Team |
| EPI_ISL_3226344 | Lighthouse Lab in Glasgow | Wellcome Sanger Institute for the COVID-19 Genomics UK (COG-UK) Consortium | Harper VanSteenhouse, Yumi Kasai, David Gray, Carol Clugston, Anna Dominiczak and Alex Alderton, Roberto Amato, Jeffrey Barrett, Sonia Goncalves, Ewan Harrison, David K. Jackson, Ian Johnston, Dominic Kwiatkowski, Cordelia Langford, John Sillitoe on behalf of the Wellcome Sanger Institute COVID-19 Surveillance Team |
| EPI_ISL_3226345 | Lighthouse Laboratory Plymouth | Wellcome Sanger Institute for the COVID-19 Genomics UK (COG-UK) Consortium | Lighthouse Laboratory Plymouth and Alex Alderton, Roberto Amato, Jeffrey Barrett, Sonia Goncalves, Ewan Harrison, David K. Jackson, Ian Johnston, Dominic Kwiatkowski, Cordelia Langford, John Sillitoe on behalf of the Wellcome Sanger Institute COVID-19 Surveillance Team |
| EPI_ISL_3226346, EPI_ISL_3226347 | Lighthouse Lab in Glasgow | Wellcome Sanger Institute for the COVID-19 Genomics UK (COG-UK) Consortium | Harper VanSteenhouse, Yumi Kasai, David Gray, Carol Clugston, Anna Dominiczak and Alex Alderton, Roberto Amato, Jeffrey Barrett, Sonia Goncalves, Ewan Harrison, David K. Jackson, Ian Johnston, Dominic Kwiatkowski, Cordelia Langford, John Sillitoe on behalf of the Wellcome Sanger Institute COVID-19 Surveillance Team |
| EPI_ISL_3226348 | Lighthouse Lab in Milton Keynes | Wellcome Sanger Institute for the COVID-19 Genomics UK (COG-UK) Consortium | The Lighthouse Lab in Milton Keynes and Alex Alderton, Roberto Amato, Jeffrey Barrett, Sonia Goncalves, Ewan Harrison, David K. Jackson, Ian Johnston, Dominic Kwiatkowski, Cordelia Langford, John Sillitoe on behalf of the Wellcome Sanger Institute COVID-19 Surveillance Team |
| EPI_ISL_3226349 | Lighthouse Lab in Alderley Park | Wellcome Sanger Institute for the COVID-19 Genomics UK (COG-UK) Consortium | Jacquelyn Wynn, Mairead Hyland, The Lighthouse Lab in Alderley Park and Alex Alderton, Roberto Amato, Jeffrey Barrett, Sonia Goncalves, Ewan Harrison, David K. Jackson, Ian Johnston, Dominic Kwiatkowski, Cordelia Langford, John Sillitoe on behalf of the Wellcome Sanger Institute COVID-19 Surveillance Team |
| EPI_ISL_3226350 | Lighthouse Laboratory Plymouth | Wellcome Sanger Institute for the COVID-19 Genomics UK (COG-UK) Consortium | Lighthouse Laboratory Plymouth and Alex Alderton, Roberto Amato, Jeffrey Barrett, Sonia Goncalves, Ewan Harrison, David K. Jackson, Ian Johnston, Dominic Kwiatkowski, Cordelia Langford, John Sillitoe on behalf of the Wellcome Sanger Institute COVID-19 Surveillance Team |
| EPI_ISL_3226351, EPI_ISL_3226352 | Lighthouse Lab in Milton Keynes | Wellcome Sanger Institute for the COVID-19 Genomics UK (COG-UK) Consortium | The Lighthouse Lab in Milton Keynes and Alex Alderton, Roberto Amato, Jeffrey Barrett, Sonia Goncalves, Ewan Harrison, David K. Jackson, Ian Johnston, Dominic Kwiatkowski, Cordelia Langford, John Sillitoe on behalf of the Wellcome Sanger Institute COVID-19 Surveillance Team |
| EPI_ISL_3226353 | Lighthouse Lab in Alderley Park | Wellcome Sanger Institute for the COVID-19 Genomics UK (COG-UK) Consortium | Jacquelyn Wynn, Mairead Hyland, The Lighthouse Lab in Alderley Park and Alex Alderton, Roberto Amato, Jeffrey Barrett, Sonia Goncalves, Ewan Harrison, David K. Jackson, Ian Johnston, Dominic Kwiatkowski, Cordelia Langford, John Sillitoe on behalf of the Wellcome Sanger Institute COVID-19 Surveillance Team |

| | | | |
|---|---|---|---|
| EPI_ISL_3226354 | Lighthouse Lab in Milton Keynes | Wellcome Sanger Institute for the COVID-19 Genomics UK (COG-UK) Consortium | The Lighthouse Lab in Milton Keynes and Alex Alderton, Roberto Amato, Jeffrey Barrett, Sonia Goncalves, Ewan Harrison, David K. Jackson, Ian Johnston, Dominic Kwiatkowski, Cordelia Langford, John Sillitoe on behalf of the Wellcome Sanger Institute COVID-19 Surveillance Team |
| EPI_ISL_3226355 | Lighthouse Lab in Alderley Park | Wellcome Sanger Institute for the COVID-19 Genomics UK (COG-UK) Consortium | Jacquelyn Wynn, Mairead Hyland, The Lighthouse Lab in Alderley Park and Alex Alderton, Roberto Amato, Jeffrey Barrett, Sonia Goncalves, Ewan Harrison, David K. Jackson, Ian Johnston, Dominic Kwiatkowski, Cordelia Langford, John Sillitoe on behalf of the Wellcome Sanger Institute COVID-19 Surveillance Team |
| EPI_ISL_3226356 | Lighthouse Lab in Glasgow | Wellcome Sanger Institute for the COVID-19 Genomics UK (COG-UK) Consortium | Harper VanSteenhouse, Yumi Kasai, David Gray, Carol Clugston, Anna Dominiczak and Alex Alderton, Roberto Amato, Jeffrey Barrett, Sonia Goncalves, Ewan Harrison, David K. Jackson, Ian Johnston, Dominic Kwiatkowski, Cordelia Langford, John Sillitoe on behalf of the Wellcome Sanger Institute COVID-19 Surveillance Team |
| EPI_ISL_3226357 | Lighthouse Lab in Alderley Park | Wellcome Sanger Institute for the COVID-19 Genomics UK (COG-UK) Consortium | Jacquelyn Wynn, Mairead Hyland, The Lighthouse Lab in Alderley Park and Alex Alderton, Roberto Amato, Jeffrey Barrett, Sonia Goncalves, Ewan Harrison, David K. Jackson, Ian Johnston, Dominic Kwiatkowski, Cordelia Langford, John Sillitoe on behalf of the Wellcome Sanger Institute COVID-19 Surveillance Team |
| EPI_ISL_3226358, EPI_ISL_3226359 | Lighthouse Laboratory Plymouth | Wellcome Sanger Institute for the COVID-19 Genomics UK (COG-UK) Consortium | Lighthouse Laboratory Plymouth and Alex Alderton, Roberto Amato, Jeffrey Barrett, Sonia Goncalves, Ewan Harrison, David K. Jackson, Ian Johnston, Dominic Kwiatkowski, Cordelia Langford, John Sillitoe on behalf of the Wellcome Sanger Institute COVID-19 Surveillance Team |
| EPI_ISL_3226360, EPI_ISL_3226361 | Lighthouse Lab in Milton Keynes | Wellcome Sanger Institute for the COVID-19 Genomics UK (COG-UK) Consortium | The Lighthouse Lab in Milton Keynes and Alex Alderton, Roberto Amato, Jeffrey Barrett, Sonia Goncalves, Ewan Harrison, David K. Jackson, Ian Johnston, Dominic Kwiatkowski, Cordelia Langford, John Sillitoe on behalf of the Wellcome Sanger Institute COVID-19 Surveillance Team |
| EPI_ISL_3226362 | Lighthouse Lab in Alderley Park | Wellcome Sanger Institute for the COVID-19 Genomics UK (COG-UK) Consortium | Jacquelyn Wynn, Mairead Hyland, The Lighthouse Lab in Alderley Park and Alex Alderton, Roberto Amato, Jeffrey Barrett, Sonia Goncalves, Ewan Harrison, David K. Jackson, Ian Johnston, Dominic Kwiatkowski, Cordelia Langford, John Sillitoe on behalf of the Wellcome Sanger Institute COVID-19 Surveillance Team |
| EPI_ISL_3226363 | Lighthouse Lab in Glasgow | Wellcome Sanger Institute for the COVID-19 Genomics UK (COG-UK) Consortium | Harper VanSteenhouse, Yumi Kasai, David Gray, Carol Clugston, Anna Dominiczak and Alex Alderton, Roberto Amato, Jeffrey Barrett, Sonia Goncalves, Ewan Harrison, David K. Jackson, Ian Johnston, Dominic Kwiatkowski, Cordelia Langford, John Sillitoe on behalf of the Wellcome Sanger Institute COVID-19 Surveillance Team |
| EPI_ISL_3226364 | Lighthouse Laboratory Plymouth | Wellcome Sanger Institute for the COVID-19 Genomics UK (COG-UK) Consortium | Lighthouse Laboratory Plymouth and Alex Alderton, Roberto Amato, Jeffrey Barrett, Sonia Goncalves, Ewan Harrison, David K. Jackson, Ian Johnston, Dominic Kwiatkowski, Cordelia Langford, John Sillitoe on behalf of the Wellcome Sanger Institute COVID-19 Surveillance Team |
| EPI_ISL_3226365, EPI_ISL_3226366 | Lighthouse Lab in Milton Keynes | Wellcome Sanger Institute for the COVID-19 Genomics UK (COG-UK) Consortium | The Lighthouse Lab in Milton Keynes and Alex Alderton, Roberto Amato, Jeffrey Barrett, Sonia Goncalves, Ewan Harrison, David K. Jackson, Ian Johnston, Dominic Kwiatkowski, Cordelia Langford, John Sillitoe on behalf of the Wellcome Sanger Institute COVID-19 Surveillance Team |
| EPI_ISL_3226367, EPI_ISL_3226368 | Lighthouse Lab in Alderley Park | Wellcome Sanger Institute for the COVID-19 Genomics UK (COG-UK) Consortium | Jacquelyn Wynn, Mairead Hyland, The Lighthouse Lab in Alderley Park and Alex Alderton, Roberto Amato, Jeffrey Barrett, Sonia Goncalves, Ewan Harrison, David K. Jackson, Ian Johnston, Dominic Kwiatkowski, Cordelia Langford, John Sillitoe on behalf of the Wellcome Sanger Institute COVID-19 Surveillance Team |
| EPI_ISL_3226369 | Lighthouse Lab in Glasgow | Wellcome Sanger Institute for the COVID-19 Genomics UK (COG-UK) Consortium | Harper VanSteenhouse, Yumi Kasai, David Gray, Carol Clugston, Anna Dominiczak and Alex Alderton, Roberto Amato, Jeffrey Barrett, Sonia Goncalves, Ewan Harrison, David K. Jackson, Ian Johnston, Dominic Kwiatkowski, Cordelia Langford, John Sillitoe on behalf of the Wellcome Sanger Institute COVID-19 Surveillance Team |
| EPI_ISL_3226370 | Lighthouse Lab in Alderley Park | Wellcome Sanger Institute for the COVID-19 Genomics UK (COG-UK) Consortium | Jacquelyn Wynn, Mairead Hyland, The Lighthouse Lab in Alderley Park and Alex Alderton, Roberto Amato, Jeffrey Barrett, Sonia Goncalves, Ewan Harrison, David K. Jackson, Ian Johnston, Dominic Kwiatkowski, Cordelia Langford, John Sillitoe on behalf of the Wellcome Sanger Institute COVID-19 Surveillance Team |
| EPI_ISL_3226371, EPI_ISL_3226372 | Lighthouse Lab in Glasgow | Wellcome Sanger Institute for the COVID-19 Genomics UK (COG-UK) Consortium | Harper VanSteenhouse, Yumi Kasai, David Gray, Carol Clugston, Anna Dominiczak and Alex Alderton, Roberto Amato, Jeffrey Barrett, Sonia Goncalves, Ewan Harrison, David K. Jackson, Ian Johnston, Dominic Kwiatkowski, Cordelia Langford, John Sillitoe on behalf of the Wellcome Sanger Institute COVID-19 Surveillance Team |
| EPI_ISL_3226373 | Lighthouse Lab in Alderley Park | Wellcome Sanger Institute for the COVID-19 Genomics UK (COG-UK) Consortium | Jacquelyn Wynn, Mairead Hyland, The Lighthouse Lab in Alderley Park and Alex Alderton, Roberto Amato, Jeffrey Barrett, Sonia Goncalves, Ewan Harrison, David K. Jackson, Ian Johnston, Dominic Kwiatkowski, Cordelia Langford, John Sillitoe on behalf of the Wellcome Sanger Institute COVID-19 Surveillance Team |
| EPI_ISL_3226374 | Lighthouse Laboratory Plymouth | Wellcome Sanger Institute for the COVID-19 Genomics UK (COG-UK) Consortium | Lighthouse Laboratory Plymouth and Alex Alderton, Roberto Amato, Jeffrey Barrett, Sonia Goncalves, Ewan Harrison, David K. Jackson, Ian Johnston, Dominic Kwiatkowski, Cordelia Langford, John Sillitoe on behalf of the Wellcome Sanger Institute COVID-19 Surveillance Team |
| EPI_ISL_3226375, EPI_ISL_3226376 | Lighthouse Lab in Glasgow | Wellcome Sanger Institute for the COVID-19 Genomics UK (COG-UK) Consortium | Harper VanSteenhouse, Yumi Kasai, David Gray, Carol Clugston, Anna Dominiczak and Alex Alderton, Roberto Amato, Jeffrey Barrett, Sonia Goncalves, Ewan Harrison, David K. Jackson, Ian Johnston, Dominic Kwiatkowski, Cordelia Langford, John Sillitoe on behalf of the Wellcome Sanger Institute COVID-19 Surveillance Team |
| EPI_ISL_3226377 | Lighthouse Laboratory Plymouth | Wellcome Sanger Institute for the COVID-19 Genomics UK (COG-UK) Consortium | Lighthouse Laboratory Plymouth and Alex Alderton, Roberto Amato, Jeffrey Barrett, Sonia Goncalves, Ewan Harrison, David K. Jackson, Ian Johnston, Dominic Kwiatkowski, Cordelia Langford, John Sillitoe on behalf of the Wellcome Sanger Institute COVID-19 Surveillance Team |
| EPI_ISL_3226378 | Lighthouse Lab in Glasgow | Wellcome Sanger Institute for the COVID-19 Genomics UK (COG-UK) Consortium | Harper VanSteenhouse, Yumi Kasai, David Gray, Carol Clugston, Anna Dominiczak and Alex Alderton, Roberto Amato, Jeffrey Barrett, Sonia Goncalves, Ewan Harrison, David K. Jackson, Ian Johnston, Dominic Kwiatkowski, Cordelia Langford, John Sillitoe on behalf of the Wellcome Sanger Institute COVID-19 Surveillance Team |
| EPI_ISL_3226379 | Lighthouse Lab in Alderley Park | Wellcome Sanger Institute for the COVID-19 Genomics UK (COG-UK) Consortium | Jacquelyn Wynn, Mairead Hyland, The Lighthouse Lab in Alderley Park and Alex Alderton, Roberto Amato, Jeffrey Barrett, Sonia Goncalves, Ewan Harrison, David K. Jackson, Ian Johnston, Dominic Kwiatkowski, Cordelia Langford, John Sillitoe on behalf of the Wellcome Sanger Institute COVID-19 Surveillance Team |
| EPI_ISL_3226380 | Lighthouse Lab in Milton Keynes | Wellcome Sanger Institute for the COVID-19 Genomics UK (COG-UK) Consortium | The Lighthouse Lab in Milton Keynes and Alex Alderton, Roberto Amato, Jeffrey Barrett, Sonia Goncalves, Ewan Harrison, David K. Jackson, Ian Johnston, Dominic Kwiatkowski, Cordelia Langford, John Sillitoe on behalf of the Wellcome Sanger Institute COVID-19 Surveillance Team |
| EPI_ISL_3226381 | Lighthouse Lab in Glasgow | Wellcome Sanger Institute for the COVID-19 Genomics UK (COG-UK) Consortium | Harper VanSteenhouse, Yumi Kasai, David Gray, Carol Clugston, Anna Dominiczak and Alex Alderton, Roberto Amato, Jeffrey Barrett, Sonia Goncalves, Ewan Harrison, David K. Jackson, Ian Johnston, Dominic Kwiatkowski, Cordelia Langford, John Sillitoe on behalf of the Wellcome Sanger Institute COVID-19 Surveillance Team |
| EPI_ISL_3226382 | Lighthouse Laboratory Plymouth | Wellcome Sanger Institute for the COVID-19 Genomics UK (COG-UK) Consortium | Lighthouse Laboratory Plymouth and Alex Alderton, Roberto Amato, Jeffrey Barrett, Sonia Goncalves, Ewan Harrison, David K. Jackson, Ian Johnston, Dominic Kwiatkowski, Cordelia Langford, John Sillitoe on behalf of the Wellcome Sanger Institute COVID-19 Surveillance Team |
| EPI_ISL_3226383 | Lighthouse Lab in Glasgow | Wellcome Sanger Institute for the COVID-19 Genomics UK (COG-UK) Consortium | Harper VanSteenhouse, Yumi Kasai, David Gray, Carol Clugston, Anna Dominiczak and Alex Alderton, Roberto Amato, Jeffrey Barrett, Sonia Goncalves, Ewan Harrison, David K. Jackson, Ian Johnston, Dominic Kwiatkowski, Cordelia Langford, John Sillitoe on behalf of the Wellcome Sanger Institute COVID-19 Surveillance Team |
| EPI_ISL_3226384, EPI_ISL_3226385 | Lighthouse Lab in Alderley Park | Wellcome Sanger Institute for the COVID-19 Genomics UK (COG-UK) Consortium | Jacquelyn Wynn, Mairead Hyland, The Lighthouse Lab in Alderley Park and Alex Alderton, Roberto Amato, Jeffrey Barrett, Sonia Goncalves, Ewan Harrison, David K. Jackson, Ian Johnston, Dominic Kwiatkowski, Cordelia Langford, John Sillitoe on behalf of the Wellcome Sanger Institute COVID-19 Surveillance Team |
| EPI_ISL_3226386 | Lighthouse Laboratory Plymouth | Wellcome Sanger Institute for the COVID-19 Genomics UK (COG-UK) Consortium | Lighthouse Laboratory Plymouth and Alex Alderton, Roberto Amato, Jeffrey Barrett, Sonia Goncalves, Ewan Harrison, David K. Jackson, Ian Johnston, Dominic Kwiatkowski, Cordelia Langford, John Sillitoe on behalf of the Wellcome Sanger Institute COVID-19 Surveillance Team |
| EPI_ISL_3226387, EPI_ISL_3226388 | Lighthouse Lab in Milton Keynes | Wellcome Sanger Institute for the COVID-19 Genomics UK (COG-UK) Consortium | The Lighthouse Lab in Milton Keynes and Alex Alderton, Roberto Amato, Jeffrey Barrett, Sonia Goncalves, Ewan Harrison, David K. Jackson, Ian Johnston, Dominic Kwiatkowski, Cordelia Langford, John Sillitoe on behalf of the Wellcome Sanger Institute COVID-19 Surveillance Team |
| EPI_ISL_3226389 | Lighthouse Lab in Glasgow | Wellcome Sanger Institute for the COVID-19 Genomics UK (COG-UK) Consortium | Harper VanSteenhouse, Yumi Kasai, David Gray, Carol Clugston, Anna Dominiczak and Alex Alderton, Roberto Amato, Jeffrey Barrett, Sonia Goncalves, Ewan Harrison, David K. Jackson, Ian Johnston, Dominic Kwiatkowski, Cordelia Langford, John Sillitoe on behalf of the Wellcome Sanger Institute COVID-19 Surveillance Team |
| EPI_ISL_3226390, EPI_ISL_3226391 | Lighthouse Laboratory Plymouth | Wellcome Sanger Institute for the COVID-19 Genomics UK (COG-UK) Consortium | Lighthouse Laboratory Plymouth and Alex Alderton, Roberto Amato, Jeffrey Barrett, Sonia Goncalves, Ewan Harrison, David K. Jackson, Ian Johnston, Dominic Kwiatkowski, Cordelia Langford, John Sillitoe on behalf of the Wellcome Sanger Institute COVID-19 Surveillance Team |
| EPI_ISL_3226392 | Lighthouse Lab in Milton Keynes | Wellcome Sanger Institute for the COVID-19 Genomics UK (COG-UK) Consortium | The Lighthouse Lab in Milton Keynes and Alex Alderton, Roberto Amato, Jeffrey Barrett, Sonia Goncalves, Ewan Harrison, David K. Jackson, Ian Johnston, Dominic Kwiatkowski, Cordelia Langford, John Sillitoe on behalf of the Wellcome Sanger Institute COVID-19 Surveillance Team |
| EPI_ISL_3226393, EPI_ISL_3226394 | Lighthouse Lab in Glasgow | Wellcome Sanger Institute for the COVID-19 Genomics UK (COG-UK) Consortium | Harper VanSteenhouse, Yumi Kasai, David Gray, Carol Clugston, Anna Dominiczak and Alex Alderton, Roberto Amato, Jeffrey Barrett, Sonia Goncalves, Ewan Harrison, David K. Jackson, Ian Johnston, Dominic Kwiatkowski, Cordelia Langford, John Sillitoe on behalf of the Wellcome Sanger Institute COVID-19 Surveillance Team |
| EPI_ISL_3226395, EPI_ISL_3226396 | Lighthouse Laboratory Plymouth | Wellcome Sanger Institute for the COVID-19 Genomics UK (COG-UK) Consortium | Lighthouse Laboratory Plymouth and Alex Alderton, Roberto Amato, Jeffrey Barrett, Sonia Goncalves, Ewan Harrison, David K. Jackson, Ian Johnston, Dominic Kwiatkowski, Cordelia Langford, John Sillitoe on behalf of the Wellcome Sanger Institute COVID-19 Surveillance Team |
| EPI_ISL_3226397, EPI_ISL_3226398 | Lighthouse Lab in Alderley Park | Wellcome Sanger Institute for the COVID-19 Genomics UK (COG-UK) Consortium | Jacquelyn Wynn, Mairead Hyland, The Lighthouse Lab in Alderley Park and Alex Alderton, Roberto Amato, Jeffrey Barrett, Sonia Goncalves, Ewan Harrison, David K. Jackson, Ian Johnston, Dominic Kwiatkowski, Cordelia Langford, John Sillitoe on behalf of the Wellcome Sanger Institute COVID-19 Surveillance Team |
| EPI_ISL_3226399 | Lighthouse Lab in Milton Keynes | Wellcome Sanger Institute for the COVID-19 Genomics UK (COG-UK) Consortium | The Lighthouse Lab in Milton Keynes and Alex Alderton, Roberto Amato, Jeffrey Barrett, Sonia Goncalves, Ewan Harrison, David K. Jackson, Ian Johnston, Dominic Kwiatkowski, Cordelia Langford, John Sillitoe on behalf of the Wellcome Sanger Institute COVID-19 Surveillance Team |
| EPI_ISL_3226400 | Lighthouse Laboratory Plymouth | Wellcome Sanger Institute for the COVID-19 Genomics UK (COG-UK) Consortium | Lighthouse Laboratory Plymouth and Alex Alderton, Roberto Amato, Jeffrey Barrett, Sonia Goncalves, Ewan Harrison, David K. Jackson, Ian Johnston, Dominic Kwiatkowski, Cordelia Langford, John Sillitoe on behalf of the Wellcome Sanger Institute COVID-19 Surveillance Team |
| EPI_ISL_3226401 | Lighthouse Lab in Glasgow | Wellcome Sanger Institute for the COVID-19 Genomics UK (COG-UK) Consortium | Harper VanSteenhouse, Yumi Kasai, David Gray, Carol Clugston, Anna Dominiczak and Alex Alderton, Roberto Amato, Jeffrey Barrett, Sonia Goncalves, Ewan Harrison, David K. Jackson, Ian Johnston, Dominic Kwiatkowski, Cordelia Langford, John Sillitoe on behalf of the Wellcome Sanger Institute COVID-19 Surveillance Team |
| EPI_ISL_3226402, EPI_ISL_3226403, EPI_ISL_3226404 | Lighthouse Laboratory Plymouth | Wellcome Sanger Institute for the COVID-19 Genomics UK (COG-UK) Consortium | Lighthouse Laboratory Plymouth and Alex Alderton, Roberto Amato, Jeffrey Barrett, Sonia Goncalves, Ewan Harrison, David K. Jackson, Ian Johnston, Dominic Kwiatkowski, Cordelia Langford, John Sillitoe on behalf of the Wellcome Sanger Institute COVID-19 Surveillance Team |
| EPI_ISL_3226405 | Lighthouse Lab in Alderley Park | Wellcome Sanger Institute for the COVID-19 Genomics UK (COG-UK) Consortium | Jacquelyn Wynn, Mairead Hyland, The Lighthouse Lab in Alderley Park and Alex Alderton, Roberto Amato, Jeffrey Barrett, Sonia Goncalves, Ewan Harrison, David K. Jackson, Ian Johnston, Dominic Kwiatkowski, Cordelia Langford, John Sillitoe on behalf of the Wellcome Sanger Institute COVID-19 Surveillance Team |
| EPI_ISL_3226406 | Lighthouse Lab in Milton Keynes | Wellcome Sanger Institute for the COVID-19 Genomics UK (COG-UK) Consortium | The Lighthouse Lab in Milton Keynes and Alex Alderton, Roberto Amato, Jeffrey Barrett, Sonia Goncalves, Ewan Harrison, David K. Jackson, Ian Johnston, Dominic Kwiatkowski, Cordelia Langford, John Sillitoe on behalf of the Wellcome Sanger Institute COVID-19 Surveillance Team |
| EPI_ISL_3226407 | Lighthouse Lab in Alderley Park | Wellcome Sanger Institute for the COVID-19 Genomics UK (COG-UK) Consortium | Jacquelyn Wynn, Mairead Hyland, The Lighthouse Lab in Alderley Park and Alex Alderton, Roberto Amato, Jeffrey Barrett, Sonia Goncalves, Ewan Harrison, David K. Jackson, Ian Johnston, Dominic Kwiatkowski, Cordelia Langford, John Sillitoe on behalf of the Wellcome Sanger Institute COVID-19 Surveillance Team |
| EPI_ISL_3226408 | Lighthouse Lab in Milton Keynes | Wellcome Sanger Institute for the COVID-19 Genomics UK (COG-UK) Consortium | The Lighthouse Lab in Milton Keynes and Alex Alderton, Roberto Amato, Jeffrey Barrett, Sonia Goncalves, Ewan Harrison, David K. Jackson, Ian Johnston, Dominic Kwiatkowski, Cordelia Langford, John Sillitoe on behalf of the Wellcome Sanger Institute COVID-19 Surveillance Team |
| EPI_ISL_3226409 | Lighthouse Lab in Alderley Park | Wellcome Sanger Institute for the COVID-19 Genomics UK (COG-UK) Consortium | Jacquelyn Wynn, Mairead Hyland, The Lighthouse Lab in Alderley Park and Alex Alderton, Roberto Amato, Jeffrey Barrett, Sonia Goncalves, Ewan Harrison, David K. Jackson, Ian Johnston, Dominic Kwiatkowski, Cordelia Langford, John Sillitoe on behalf of the Wellcome Sanger Institute COVID-19 Surveillance Team |
| EPI_ISL_3226410 | Lighthouse Lab in Milton Keynes | Wellcome Sanger Institute for the COVID-19 Genomics UK (COG-UK) Consortium | The Lighthouse Lab in Milton Keynes and Alex Alderton, Roberto Amato, Jeffrey Barrett, Sonia Goncalves, Ewan Harrison, David K. Jackson, Ian Johnston, Dominic Kwiatkowski, Cordelia Langford, John Sillitoe on behalf of the Wellcome Sanger Institute COVID-19 Surveillance Team |
| EPI_ISL_3226411 | Lighthouse Lab in Alderley Park | Wellcome Sanger Institute for the COVID-19 Genomics UK (COG-UK) Consortium | Jacquelyn Wynn, Mairead Hyland, The Lighthouse Lab in Alderley Park and Alex Alderton, Roberto Amato, Jeffrey Barrett, Sonia Goncalves, Ewan Harrison, David K. Jackson, Ian Johnston, Dominic Kwiatkowski, Cordelia Langford, John Sillitoe on behalf of the Wellcome Sanger Institute COVID-19 Surveillance Team |
| EPI_ISL_3226412, | Lighthouse Lab in Milton Keynes | Wellcome Sanger Institute for the COVID-19 Genomics UK (COG-UK) | The Lighthouse Lab in Milton Keynes and Alex Alderton, Roberto Amato, Jeffrey Barrett, Sonia Goncalves, Ewan Harrison, David K. Jackson, Ian Johnston, Dominic Kwiatkowski, Cordelia Langford, John Sillitoe on behalf of the Wellcome Sanger |

| | | | |
|---|---|---|---|
| EPI_ISL_3226413, EPI_ISL_3226414 | | Consortium | Institute COVID-19 Surveillance Team |
| EPI_ISL_3226415 | Lighthouse Laboratory Plymouth | Wellcome Sanger Institute for the COVID-19 Genomics UK (COG-UK) Consortium | Lighthouse Laboratory Plymouth and Alex Alderton, Roberto Amato, Jeffrey Barrett, Sonia Goncalves, Ewan Harrison, David K. Jackson, Ian Johnston, Dominic Kwiatkowski, Cordelia Langford, John Sillitoe on behalf of the Wellcome Sanger Institute COVID-19 Surveillance Team |
| EPI_ISL_3226416 | Lighthouse Lab in Milton Keynes | Wellcome Sanger Institute for the COVID-19 Genomics UK (COG-UK) Consortium | The Lighthouse Lab in Milton Keynes and Alex Alderton, Roberto Amato, Jeffrey Barrett, Sonia Goncalves, Ewan Harrison, David K. Jackson, Ian Johnston, Dominic Kwiatkowski, Cordelia Langford, John Sillitoe on behalf of the Wellcome Sanger Institute COVID-19 Surveillance Team |
| EPI_ISL_3226417, EPI_ISL_3226418, EPI_ISL_3226419, EPI_ISL_3226420 | Lighthouse Lab in Glasgow | Wellcome Sanger Institute for the COVID-19 Genomics UK (COG-UK) Consortium | Harper VanSteenhouse, Yumi Kasai, David Gray, Carol Clugston, Anna Dominiczak and Alex Alderton, Roberto Amato, Jeffrey Barrett, Sonia Goncalves, Ewan Harrison, David K. Jackson, Ian Johnston, Dominic Kwiatkowski, Cordelia Langford, John Sillitoe on behalf of the Wellcome Sanger Institute COVID-19 Surveillance Team |
| EPI_ISL_3226421, EPI_ISL_3226422 | Lighthouse Lab in Alderley Park | Wellcome Sanger Institute for the COVID-19 Genomics UK (COG-UK) Consortium | Jacquelyn Wynn, Mairead Hyland, The Lighthouse Lab in Alderley Park and Alex Alderton, Roberto Amato, Jeffrey Barrett, Sonia Goncalves, Ewan Harrison, David K. Jackson, Ian Johnston, Dominic Kwiatkowski, Cordelia Langford, John Sillitoe on behalf of the Wellcome Sanger Institute COVID-19 Surveillance Team |
| EPI_ISL_3226423 | Lighthouse Laboratory Plymouth | Wellcome Sanger Institute for the COVID-19 Genomics UK (COG-UK) Consortium | Lighthouse Laboratory Plymouth and Alex Alderton, Roberto Amato, Jeffrey Barrett, Sonia Goncalves, Ewan Harrison, David K. Jackson, Ian Johnston, Dominic Kwiatkowski, Cordelia Langford, John Sillitoe on behalf of the Wellcome Sanger Institute COVID-19 Surveillance Team |
| EPI_ISL_3226424 | Lighthouse Lab in Milton Keynes | Wellcome Sanger Institute for the COVID-19 Genomics UK (COG-UK) Consortium | The Lighthouse Lab in Milton Keynes and Alex Alderton, Roberto Amato, Jeffrey Barrett, Sonia Goncalves, Ewan Harrison, David K. Jackson, Ian Johnston, Dominic Kwiatkowski, Cordelia Langford, John Sillitoe on behalf of the Wellcome Sanger Institute COVID-19 Surveillance Team |
| EPI_ISL_3226425 | Lighthouse Lab in Glasgow | Wellcome Sanger Institute for the COVID-19 Genomics UK (COG-UK) Consortium | Harper VanSteenhouse, Yumi Kasai, David Gray, Carol Clugston, Anna Dominiczak and Alex Alderton, Roberto Amato, Jeffrey Barrett, Sonia Goncalves, Ewan Harrison, David K. Jackson, Ian Johnston, Dominic Kwiatkowski, Cordelia Langford, John Sillitoe on behalf of the Wellcome Sanger Institute COVID-19 Surveillance Team |
| EPI_ISL_3226426 | Lighthouse Laboratory Plymouth | Wellcome Sanger Institute for the COVID-19 Genomics UK (COG-UK) Consortium | Lighthouse Laboratory Plymouth and Alex Alderton, Roberto Amato, Jeffrey Barrett, Sonia Goncalves, Ewan Harrison, David K. Jackson, Ian Johnston, Dominic Kwiatkowski, Cordelia Langford, John Sillitoe on behalf of the Wellcome Sanger Institute COVID-19 Surveillance Team |
| EPI_ISL_3226427 | Lighthouse Lab in Alderley Park | Wellcome Sanger Institute for the COVID-19 Genomics UK (COG-UK) Consortium | Jacquelyn Wynn, Mairead Hyland, The Lighthouse Lab in Alderley Park and Alex Alderton, Roberto Amato, Jeffrey Barrett, Sonia Goncalves, Ewan Harrison, David K. Jackson, Ian Johnston, Dominic Kwiatkowski, Cordelia Langford, John Sillitoe on behalf of the Wellcome Sanger Institute COVID-19 Surveillance Team |
| EPI_ISL_3226428, EPI_ISL_3226429 | Lighthouse Lab in Milton Keynes | Wellcome Sanger Institute for the COVID-19 Genomics UK (COG-UK) Consortium | The Lighthouse Lab in Milton Keynes and Alex Alderton, Roberto Amato, Jeffrey Barrett, Sonia Goncalves, Ewan Harrison, David K. Jackson, Ian Johnston, Dominic Kwiatkowski, Cordelia Langford, John Sillitoe on behalf of the Wellcome Sanger Institute COVID-19 Surveillance Team |
| EPI_ISL_3226430, EPI_ISL_3226431 | Lighthouse Lab in Alderley Park | Wellcome Sanger Institute for the COVID-19 Genomics UK (COG-UK) Consortium | Jacquelyn Wynn, Mairead Hyland, The Lighthouse Lab in Alderley Park and Alex Alderton, Roberto Amato, Jeffrey Barrett, Sonia Goncalves, Ewan Harrison, David K. Jackson, Ian Johnston, Dominic Kwiatkowski, Cordelia Langford, John Sillitoe on behalf of the Wellcome Sanger Institute COVID-19 Surveillance Team |
| EPI_ISL_3226432, EPI_ISL_3226433 | Lighthouse Lab in Milton Keynes | Wellcome Sanger Institute for the COVID-19 Genomics UK (COG-UK) Consortium | The Lighthouse Lab in Milton Keynes and Alex Alderton, Roberto Amato, Jeffrey Barrett, Sonia Goncalves, Ewan Harrison, David K. Jackson, Ian Johnston, Dominic Kwiatkowski, Cordelia Langford, John Sillitoe on behalf of the Wellcome Sanger Institute COVID-19 Surveillance Team |
| EPI_ISL_3226434 | Lighthouse Lab in Alderley Park | Wellcome Sanger Institute for the COVID-19 Genomics UK (COG-UK) Consortium | Jacquelyn Wynn, Mairead Hyland, The Lighthouse Lab in Alderley Park and Alex Alderton, Roberto Amato, Jeffrey Barrett, Sonia Goncalves, Ewan Harrison, David K. Jackson, Ian Johnston, Dominic Kwiatkowski, Cordelia Langford, John Sillitoe on behalf of the Wellcome Sanger Institute COVID-19 Surveillance Team |
| EPI_ISL_3226435 | Lighthouse Lab in Milton Keynes | Wellcome Sanger Institute for the COVID-19 Genomics UK (COG-UK) Consortium | The Lighthouse Lab in Milton Keynes and Alex Alderton, Roberto Amato, Jeffrey Barrett, Sonia Goncalves, Ewan Harrison, David K. Jackson, Ian Johnston, Dominic Kwiatkowski, Cordelia Langford, John Sillitoe on behalf of the Wellcome Sanger Institute COVID-19 Surveillance Team |
| EPI_ISL_3226436 | Lighthouse Lab in Glasgow | Wellcome Sanger Institute for the COVID-19 Genomics UK (COG-UK) Consortium | Harper VanSteenhouse, Yumi Kasai, David Gray, Carol Clugston, Anna Dominiczak and Alex Alderton, Roberto Amato, Jeffrey Barrett, Sonia Goncalves, Ewan Harrison, David K. Jackson, Ian Johnston, Dominic Kwiatkowski, Cordelia Langford, John Sillitoe on behalf of the Wellcome Sanger Institute COVID-19 Surveillance Team |
| EPI_ISL_3226437 | Lighthouse Laboratory Plymouth | Wellcome Sanger Institute for the COVID-19 Genomics UK (COG-UK) Consortium | Lighthouse Laboratory Plymouth and Alex Alderton, Roberto Amato, Jeffrey Barrett, Sonia Goncalves, Ewan Harrison, David K. Jackson, Ian Johnston, Dominic Kwiatkowski, Cordelia Langford, John Sillitoe on behalf of the Wellcome Sanger Institute COVID-19 Surveillance Team |
| EPI_ISL_3226438 | Lighthouse Lab in Alderley Park | Wellcome Sanger Institute for the COVID-19 Genomics UK (COG-UK) Consortium | Jacquelyn Wynn, Mairead Hyland, The Lighthouse Lab in Alderley Park and Alex Alderton, Roberto Amato, Jeffrey Barrett, Sonia Goncalves, Ewan Harrison, David K. Jackson, Ian Johnston, Dominic Kwiatkowski, Cordelia Langford, John Sillitoe on behalf of the Wellcome Sanger Institute COVID-19 Surveillance Team |
| EPI_ISL_3226439 | Lighthouse Laboratory Plymouth | Wellcome Sanger Institute for the COVID-19 Genomics UK (COG-UK) Consortium | Lighthouse Laboratory Plymouth and Alex Alderton, Roberto Amato, Jeffrey Barrett, Sonia Goncalves, Ewan Harrison, David K. Jackson, Ian Johnston, Dominic Kwiatkowski, Cordelia Langford, John Sillitoe on behalf of the Wellcome Sanger Institute COVID-19 Surveillance Team |
| EPI_ISL_3226440 | Lighthouse Lab in Glasgow | Wellcome Sanger Institute for the COVID-19 Genomics UK (COG-UK) Consortium | Harper VanSteenhouse, Yumi Kasai, David Gray, Carol Clugston, Anna Dominiczak and Alex Alderton, Roberto Amato, Jeffrey Barrett, Sonia Goncalves, Ewan Harrison, David K. Jackson, Ian Johnston, Dominic Kwiatkowski, Cordelia Langford, John Sillitoe on behalf of the Wellcome Sanger Institute COVID-19 Surveillance Team |
| EPI_ISL_3226441 | Lighthouse Laboratory Plymouth | Wellcome Sanger Institute for the COVID-19 Genomics UK (COG-UK) Consortium | Lighthouse Laboratory Plymouth and Alex Alderton, Roberto Amato, Jeffrey Barrett, Sonia Goncalves, Ewan Harrison, David K. Jackson, Ian Johnston, Dominic Kwiatkowski, Cordelia Langford, John Sillitoe on behalf of the Wellcome Sanger Institute COVID-19 Surveillance Team |
| EPI_ISL_3226442, EPI_ISL_3226443 | Lighthouse Lab in Alderley Park | Wellcome Sanger Institute for the COVID-19 Genomics UK (COG-UK) Consortium | Jacquelyn Wynn, Mairead Hyland, The Lighthouse Lab in Alderley Park and Alex Alderton, Roberto Amato, Jeffrey Barrett, Sonia Goncalves, Ewan Harrison, David K. Jackson, Ian Johnston, Dominic Kwiatkowski, Cordelia Langford, John Sillitoe on behalf of the Wellcome Sanger Institute COVID-19 Surveillance Team |
| EPI_ISL_3226444 | Lighthouse Laboratory Plymouth | Wellcome Sanger Institute for the COVID-19 Genomics UK (COG-UK) Consortium | Lighthouse Laboratory Plymouth and Alex Alderton, Roberto Amato, Jeffrey Barrett, Sonia Goncalves, Ewan Harrison, David K. Jackson, Ian Johnston, Dominic Kwiatkowski, Cordelia Langford, John Sillitoe on behalf of the Wellcome Sanger Institute COVID-19 Surveillance Team |
| EPI_ISL_3226445, EPI_ISL_3226446 | Lighthouse Lab in Milton Keynes | Wellcome Sanger Institute for the COVID-19 Genomics UK (COG-UK) Consortium | The Lighthouse Lab in Milton Keynes and Alex Alderton, Roberto Amato, Jeffrey Barrett, Sonia Goncalves, Ewan Harrison, David K. Jackson, Ian Johnston, Dominic Kwiatkowski, Cordelia Langford, John Sillitoe on behalf of the Wellcome Sanger Institute COVID-19 Surveillance Team |
| EPI_ISL_3226447 | Lighthouse Laboratory Plymouth | Wellcome Sanger Institute for the COVID-19 Genomics UK (COG-UK) Consortium | Lighthouse Laboratory Plymouth and Alex Alderton, Roberto Amato, Jeffrey Barrett, Sonia Goncalves, Ewan Harrison, David K. Jackson, Ian Johnston, Dominic Kwiatkowski, Cordelia Langford, John Sillitoe on behalf of the Wellcome Sanger Institute COVID-19 Surveillance Team |
| EPI_ISL_3226448 | Lighthouse Lab in Milton Keynes | Wellcome Sanger Institute for the COVID-19 Genomics UK (COG-UK) Consortium | The Lighthouse Lab in Milton Keynes and Alex Alderton, Roberto Amato, Jeffrey Barrett, Sonia Goncalves, Ewan Harrison, David K. Jackson, Ian Johnston, Dominic Kwiatkowski, Cordelia Langford, John Sillitoe on behalf of the Wellcome Sanger Institute COVID-19 Surveillance Team |
| EPI_ISL_3226449 | Lighthouse Lab in Alderley Park | Wellcome Sanger Institute for the COVID-19 Genomics UK (COG-UK) Consortium | Jacquelyn Wynn, Mairead Hyland, The Lighthouse Lab in Alderley Park and Alex Alderton, Roberto Amato, Jeffrey Barrett, Sonia Goncalves, Ewan Harrison, David K. Jackson, Ian Johnston, Dominic Kwiatkowski, Cordelia Langford, John Sillitoe on behalf of the Wellcome Sanger Institute COVID-19 Surveillance Team |
| EPI_ISL_3226450 | Lighthouse Laboratory Plymouth | Wellcome Sanger Institute for the COVID-19 Genomics UK (COG-UK) Consortium | Lighthouse Laboratory Plymouth and Alex Alderton, Roberto Amato, Jeffrey Barrett, Sonia Goncalves, Ewan Harrison, David K. Jackson, Ian Johnston, Dominic Kwiatkowski, Cordelia Langford, John Sillitoe on behalf of the Wellcome Sanger Institute COVID-19 Surveillance Team |
| EPI_ISL_3226451, EPI_ISL_3226452 | Lighthouse Lab in Alderley Park | Wellcome Sanger Institute for the COVID-19 Genomics UK (COG-UK) Consortium | Jacquelyn Wynn, Mairead Hyland, The Lighthouse Lab in Alderley Park and Alex Alderton, Roberto Amato, Jeffrey Barrett, Sonia Goncalves, Ewan Harrison, David K. Jackson, Ian Johnston, Dominic Kwiatkowski, Cordelia Langford, John Sillitoe on behalf of the Wellcome Sanger Institute COVID-19 Surveillance Team |
| EPI_ISL_3226453, EPI_ISL_3226454 | Lighthouse Lab in Milton Keynes | Wellcome Sanger Institute for the COVID-19 Genomics UK (COG-UK) Consortium | The Lighthouse Lab in Milton Keynes and Alex Alderton, Roberto Amato, Jeffrey Barrett, Sonia Goncalves, Ewan Harrison, David K. Jackson, Ian Johnston, Dominic Kwiatkowski, Cordelia Langford, John Sillitoe on behalf of the Wellcome Sanger Institute COVID-19 Surveillance Team |
| EPI_ISL_3226455, EPI_ISL_3226456 | Lighthouse Laboratory Plymouth | Wellcome Sanger Institute for the COVID-19 Genomics UK (COG-UK) Consortium | Lighthouse Laboratory Plymouth and Alex Alderton, Roberto Amato, Jeffrey Barrett, Sonia Goncalves, Ewan Harrison, David K. Jackson, Ian Johnston, Dominic Kwiatkowski, Cordelia Langford, John Sillitoe on behalf of the Wellcome Sanger Institute COVID-19 Surveillance Team |
| EPI_ISL_3226457 | Lighthouse Lab in Alderley Park | Wellcome Sanger Institute for the COVID-19 Genomics UK (COG-UK) Consortium | Jacquelyn Wynn, Mairead Hyland, The Lighthouse Lab in Alderley Park and Alex Alderton, Roberto Amato, Jeffrey Barrett, Sonia Goncalves, Ewan Harrison, David K. Jackson, Ian Johnston, Dominic Kwiatkowski, Cordelia Langford, John Sillitoe on behalf of the Wellcome Sanger Institute COVID-19 Surveillance Team |
| EPI_ISL_3226458 | Lighthouse Lab in Glasgow | Wellcome Sanger Institute for the COVID-19 Genomics UK (COG-UK) Consortium | Harper VanSteenhouse, Yumi Kasai, David Gray, Carol Clugston, Anna Dominiczak and Alex Alderton, Roberto Amato, Jeffrey Barrett, Sonia Goncalves, Ewan Harrison, David K. Jackson, Ian Johnston, Dominic Kwiatkowski, Cordelia Langford, John Sillitoe on behalf of the Wellcome Sanger Institute COVID-19 Surveillance Team |
| EPI_ISL_3226459 | Lighthouse Laboratory Plymouth | Wellcome Sanger Institute for the COVID-19 Genomics UK (COG-UK) Consortium | Lighthouse Laboratory Plymouth and Alex Alderton, Roberto Amato, Jeffrey Barrett, Sonia Goncalves, Ewan Harrison, David K. Jackson, Ian Johnston, Dominic Kwiatkowski, Cordelia Langford, John Sillitoe on behalf of the Wellcome Sanger Institute COVID-19 Surveillance Team |
| EPI_ISL_3226460, EPI_ISL_3226461, EPI_ISL_3226462 | Lighthouse Lab in Glasgow | Wellcome Sanger Institute for the COVID-19 Genomics UK (COG-UK) Consortium | Harper VanSteenhouse, Yumi Kasai, David Gray, Carol Clugston, Anna Dominiczak and Alex Alderton, Roberto Amato, Jeffrey Barrett, Sonia Goncalves, Ewan Harrison, David K. Jackson, Ian Johnston, Dominic Kwiatkowski, Cordelia Langford, John Sillitoe on behalf of the Wellcome Sanger Institute COVID-19 Surveillance Team |
| EPI_ISL_3226463, EPI_ISL_3226464 | Lighthouse Lab in Alderley Park | Wellcome Sanger Institute for the COVID-19 Genomics UK (COG-UK) Consortium | Jacquelyn Wynn, Mairead Hyland, The Lighthouse Lab in Alderley Park and Alex Alderton, Roberto Amato, Jeffrey Barrett, Sonia Goncalves, Ewan Harrison, David K. Jackson, Ian Johnston, Dominic Kwiatkowski, Cordelia Langford, John Sillitoe on behalf of the Wellcome Sanger Institute COVID-19 Surveillance Team |
| EPI_ISL_3226465 | Lighthouse Lab in Glasgow | Wellcome Sanger Institute for the COVID-19 Genomics UK (COG-UK) Consortium | Harper VanSteenhouse, Yumi Kasai, David Gray, Carol Clugston, Anna Dominiczak and Alex Alderton, Roberto Amato, Jeffrey Barrett, Sonia Goncalves, Ewan Harrison, David K. Jackson, Ian Johnston, Dominic Kwiatkowski, Cordelia Langford, John Sillitoe on behalf of the Wellcome Sanger Institute COVID-19 Surveillance Team |
| EPI_ISL_3226466 | Lighthouse Laboratory Plymouth | Wellcome Sanger Institute for the COVID-19 Genomics UK (COG-UK) Consortium | Lighthouse Laboratory Plymouth and Alex Alderton, Roberto Amato, Jeffrey Barrett, Sonia Goncalves, Ewan Harrison, David K. Jackson, Ian Johnston, Dominic Kwiatkowski, Cordelia Langford, John Sillitoe on behalf of the Wellcome Sanger Institute COVID-19 Surveillance Team |
| EPI_ISL_3226467 | Lighthouse Lab in Alderley Park | Wellcome Sanger Institute for the COVID-19 Genomics UK (COG-UK) Consortium | Jacquelyn Wynn, Mairead Hyland, The Lighthouse Lab in Alderley Park and Alex Alderton, Roberto Amato, Jeffrey Barrett, Sonia Goncalves, Ewan Harrison, David K. Jackson, Ian Johnston, Dominic Kwiatkowski, Cordelia Langford, John Sillitoe on behalf of the Wellcome Sanger Institute COVID-19 Surveillance Team |
| EPI_ISL_3226468 | Lighthouse Lab in Milton Keynes | Wellcome Sanger Institute for the COVID-19 Genomics UK (COG-UK) Consortium | The Lighthouse Lab in Milton Keynes and Alex Alderton, Roberto Amato, Jeffrey Barrett, Sonia Goncalves, Ewan Harrison, David K. Jackson, Ian Johnston, Dominic Kwiatkowski, Cordelia Langford, John Sillitoe on behalf of the Wellcome Sanger Institute COVID-19 Surveillance Team |
| EPI_ISL_3226469, EPI_ISL_3226470 | Lighthouse Laboratory Plymouth | Wellcome Sanger Institute for the COVID-19 Genomics UK (COG-UK) Consortium | Lighthouse Laboratory Plymouth and Alex Alderton, Roberto Amato, Jeffrey Barrett, Sonia Goncalves, Ewan Harrison, David K. Jackson, Ian Johnston, Dominic Kwiatkowski, Cordelia Langford, John Sillitoe on behalf of the Wellcome Sanger Institute COVID-19 Surveillance Team |
| EPI_ISL_3226471 | Lighthouse Lab in Milton Keynes | Wellcome Sanger Institute for the COVID-19 Genomics UK (COG-UK) Consortium | The Lighthouse Lab in Milton Keynes and Alex Alderton, Roberto Amato, Jeffrey Barrett, Sonia Goncalves, Ewan Harrison, David K. Jackson, Ian Johnston, Dominic Kwiatkowski, Cordelia Langford, John Sillitoe on behalf of the Wellcome Sanger Institute COVID-19 Surveillance Team |
| EPI_ISL_3226472 | Lighthouse Lab in Glasgow | Wellcome Sanger Institute for the COVID-19 Genomics UK (COG-UK) Consortium | Harper VanSteenhouse, Yumi Kasai, David Gray, Carol Clugston, Anna Dominiczak and Alex Alderton, Roberto Amato, Jeffrey Barrett, Sonia Goncalves, Ewan Harrison, David K. Jackson, Ian Johnston, Dominic Kwiatkowski, Cordelia Langford, John Sillitoe on behalf of the Wellcome Sanger Institute COVID-19 Surveillance Team |
| EPI_ISL_3226473 | Lighthouse Laboratory Plymouth | Wellcome Sanger Institute for the COVID-19 Genomics UK (COG-UK) Consortium | Lighthouse Laboratory Plymouth and Alex Alderton, Roberto Amato, Jeffrey Barrett, Sonia Goncalves, Ewan Harrison, David K. Jackson, Ian Johnston, Dominic Kwiatkowski, Cordelia Langford, John Sillitoe on behalf of the Wellcome Sanger Institute COVID-19 Surveillance Team |

| | | | |
|---|---|---|---|
| EPI_ISL_3226474 | Lighthouse Lab in Milton Keynes | Wellcome Sanger Institute for the COVID-19 Genomics UK (COG-UK) Consortium | The Lighthouse Lab in Milton Keynes and Alex Alderton, Roberto Amato, Jeffrey Barrett, Sonia Goncalves, Ewan Harrison, David K. Jackson, Ian Johnston, Dominic Kwiatkowski, Cordelia Langford, John Sillitoe on behalf of the Wellcome Sanger Institute COVID-19 Surveillance Team |
| EPI_ISL_3226475 | Lighthouse Lab in Alderley Park | Wellcome Sanger Institute for the COVID-19 Genomics UK (COG-UK) Consortium | Jacquelyn Wynn, Mairead Hyland, The Lighthouse Lab in Alderley Park and Alex Alderton, Roberto Amato, Jeffrey Barrett, Sonia Goncalves, Ewan Harrison, David K. Jackson, Ian Johnston, Dominic Kwiatkowski, Cordelia Langford, John Sillitoe on behalf of the Wellcome Sanger Institute COVID-19 Surveillance Team |
| EPI_ISL_3226476, EPI_ISL_3226477 | Lighthouse Lab in Milton Keynes | Wellcome Sanger Institute for the COVID-19 Genomics UK (COG-UK) Consortium | The Lighthouse Lab in Milton Keynes and Alex Alderton, Roberto Amato, Jeffrey Barrett, Sonia Goncalves, Ewan Harrison, David K. Jackson, Ian Johnston, Dominic Kwiatkowski, Cordelia Langford, John Sillitoe on behalf of the Wellcome Sanger Institute COVID-19 Surveillance Team |
| EPI_ISL_3226478 | Lighthouse Lab in Glasgow | Wellcome Sanger Institute for the COVID-19 Genomics UK (COG-UK) Consortium | Harper VanSteenhouse, Yumi Kasai, David Gray, Carol Clugston, Anna Dominiczak and Alex Alderton, Roberto Amato, Jeffrey Barrett, Sonia Goncalves, Ewan Harrison, David K. Jackson, Ian Johnston, Dominic Kwiatkowski, Cordelia Langford, John Sillitoe on behalf of the Wellcome Sanger Institute COVID-19 Surveillance Team |
| EPI_ISL_3226479, EPI_ISL_3226480 | Lighthouse Laboratory Plymouth | Wellcome Sanger Institute for the COVID-19 Genomics UK (COG-UK) Consortium | Lighthouse Laboratory Plymouth and Alex Alderton, Roberto Amato, Jeffrey Barrett, Sonia Goncalves, Ewan Harrison, David K. Jackson, Ian Johnston, Dominic Kwiatkowski, Cordelia Langford, John Sillitoe on behalf of the Wellcome Sanger Institute COVID-19 Surveillance Team |
| EPI_ISL_3226481, EPI_ISL_3226482 | Lighthouse Lab in Alderley Park | Wellcome Sanger Institute for the COVID-19 Genomics UK (COG-UK) Consortium | Jacquelyn Wynn, Mairead Hyland, The Lighthouse Lab in Alderley Park and Alex Alderton, Roberto Amato, Jeffrey Barrett, Sonia Goncalves, Ewan Harrison, David K. Jackson, Ian Johnston, Dominic Kwiatkowski, Cordelia Langford, John Sillitoe on behalf of the Wellcome Sanger Institute COVID-19 Surveillance Team |
| EPI_ISL_3226483 | Lighthouse Laboratory Plymouth | Wellcome Sanger Institute for the COVID-19 Genomics UK (COG-UK) Consortium | Lighthouse Laboratory Plymouth and Alex Alderton, Roberto Amato, Jeffrey Barrett, Sonia Goncalves, Ewan Harrison, David K. Jackson, Ian Johnston, Dominic Kwiatkowski, Cordelia Langford, John Sillitoe on behalf of the Wellcome Sanger Institute COVID-19 Surveillance Team |
| EPI_ISL_3226484 | Lighthouse Lab in Alderley Park | Wellcome Sanger Institute for the COVID-19 Genomics UK (COG-UK) Consortium | Jacquelyn Wynn, Mairead Hyland, The Lighthouse Lab in Alderley Park and Alex Alderton, Roberto Amato, Jeffrey Barrett, Sonia Goncalves, Ewan Harrison, David K. Jackson, Ian Johnston, Dominic Kwiatkowski, Cordelia Langford, John Sillitoe on behalf of the Wellcome Sanger Institute COVID-19 Surveillance Team |
| EPI_ISL_3226485 | Lighthouse Lab in Glasgow | Wellcome Sanger Institute for the COVID-19 Genomics UK (COG-UK) Consortium | Harper VanSteenhouse, Yumi Kasai, David Gray, Carol Clugston, Anna Dominiczak and Alex Alderton, Roberto Amato, Jeffrey Barrett, Sonia Goncalves, Ewan Harrison, David K. Jackson, Ian Johnston, Dominic Kwiatkowski, Cordelia Langford, John Sillitoe on behalf of the Wellcome Sanger Institute COVID-19 Surveillance Team |
| EPI_ISL_3226486, EPI_ISL_3226487 | Lighthouse Lab in Milton Keynes | Wellcome Sanger Institute for the COVID-19 Genomics UK (COG-UK) Consortium | The Lighthouse Lab in Milton Keynes and Alex Alderton, Roberto Amato, Jeffrey Barrett, Sonia Goncalves, Ewan Harrison, David K. Jackson, Ian Johnston, Dominic Kwiatkowski, Cordelia Langford, John Sillitoe on behalf of the Wellcome Sanger Institute COVID-19 Surveillance Team |
| EPI_ISL_3226488, EPI_ISL_3226489 | Lighthouse Lab in Glasgow | Wellcome Sanger Institute for the COVID-19 Genomics UK (COG-UK) Consortium | Harper VanSteenhouse, Yumi Kasai, David Gray, Carol Clugston, Anna Dominiczak and Alex Alderton, Roberto Amato, Jeffrey Barrett, Sonia Goncalves, Ewan Harrison, David K. Jackson, Ian Johnston, Dominic Kwiatkowski, Cordelia Langford, John Sillitoe on behalf of the Wellcome Sanger Institute COVID-19 Surveillance Team |
| EPI_ISL_3226490 | Lighthouse Lab in Alderley Park | Wellcome Sanger Institute for the COVID-19 Genomics UK (COG-UK) Consortium | Jacquelyn Wynn, Mairead Hyland, The Lighthouse Lab in Alderley Park and Alex Alderton, Roberto Amato, Jeffrey Barrett, Sonia Goncalves, Ewan Harrison, David K. Jackson, Ian Johnston, Dominic Kwiatkowski, Cordelia Langford, John Sillitoe on behalf of the Wellcome Sanger Institute COVID-19 Surveillance Team |
| EPI_ISL_3226491 | Lighthouse Lab in Glasgow | Wellcome Sanger Institute for the COVID-19 Genomics UK (COG-UK) Consortium | Harper VanSteenhouse, Yumi Kasai, David Gray, Carol Clugston, Anna Dominiczak and Alex Alderton, Roberto Amato, Jeffrey Barrett, Sonia Goncalves, Ewan Harrison, David K. Jackson, Ian Johnston, Dominic Kwiatkowski, Cordelia Langford, John Sillitoe on behalf of the Wellcome Sanger Institute COVID-19 Surveillance Team |
| EPI_ISL_3226492, EPI_ISL_3226493 | Lighthouse Lab in Milton Keynes | Wellcome Sanger Institute for the COVID-19 Genomics UK (COG-UK) Consortium | The Lighthouse Lab in Milton Keynes and Alex Alderton, Roberto Amato, Jeffrey Barrett, Sonia Goncalves, Ewan Harrison, David K. Jackson, Ian Johnston, Dominic Kwiatkowski, Cordelia Langford, John Sillitoe on behalf of the Wellcome Sanger Institute COVID-19 Surveillance Team |
| EPI_ISL_3226494 | Lighthouse Lab in Alderley Park | Wellcome Sanger Institute for the COVID-19 Genomics UK (COG-UK) Consortium | Jacquelyn Wynn, Mairead Hyland, The Lighthouse Lab in Alderley Park and Alex Alderton, Roberto Amato, Jeffrey Barrett, Sonia Goncalves, Ewan Harrison, David K. Jackson, Ian Johnston, Dominic Kwiatkowski, Cordelia Langford, John Sillitoe on behalf of the Wellcome Sanger Institute COVID-19 Surveillance Team |
| EPI_ISL_3226495, EPI_ISL_3226496 | Lighthouse Laboratory Plymouth | Wellcome Sanger Institute for the COVID-19 Genomics UK (COG-UK) Consortium | Lighthouse Laboratory Plymouth and Alex Alderton, Roberto Amato, Jeffrey Barrett, Sonia Goncalves, Ewan Harrison, David K. Jackson, Ian Johnston, Dominic Kwiatkowski, Cordelia Langford, John Sillitoe on behalf of the Wellcome Sanger Institute COVID-19 Surveillance Team |
| EPI_ISL_3226497 | Lighthouse Lab in Glasgow | Wellcome Sanger Institute for the COVID-19 Genomics UK (COG-UK) Consortium | Harper VanSteenhouse, Yumi Kasai, David Gray, Carol Clugston, Anna Dominiczak and Alex Alderton, Roberto Amato, Jeffrey Barrett, Sonia Goncalves, Ewan Harrison, David K. Jackson, Ian Johnston, Dominic Kwiatkowski, Cordelia Langford, John Sillitoe on behalf of the Wellcome Sanger Institute COVID-19 Surveillance Team |
| EPI_ISL_3226498 | Lighthouse Laboratory Plymouth | Wellcome Sanger Institute for the COVID-19 Genomics UK (COG-UK) Consortium | Lighthouse Laboratory Plymouth and Alex Alderton, Roberto Amato, Jeffrey Barrett, Sonia Goncalves, Ewan Harrison, David K. Jackson, Ian Johnston, Dominic Kwiatkowski, Cordelia Langford, John Sillitoe on behalf of the Wellcome Sanger Institute COVID-19 Surveillance Team |
| EPI_ISL_3226499, EPI_ISL_3226500 | Lighthouse Lab in Alderley Park | Wellcome Sanger Institute for the COVID-19 Genomics UK (COG-UK) Consortium | Jacquelyn Wynn, Mairead Hyland, The Lighthouse Lab in Alderley Park and Alex Alderton, Roberto Amato, Jeffrey Barrett, Sonia Goncalves, Ewan Harrison, David K. Jackson, Ian Johnston, Dominic Kwiatkowski, Cordelia Langford, John Sillitoe on behalf of the Wellcome Sanger Institute COVID-19 Surveillance Team |
| EPI_ISL_3226501, EPI_ISL_3226502 | Lighthouse Lab in Milton Keynes | Wellcome Sanger Institute for the COVID-19 Genomics UK (COG-UK) Consortium | The Lighthouse Lab in Milton Keynes and Alex Alderton, Roberto Amato, Jeffrey Barrett, Sonia Goncalves, Ewan Harrison, David K. Jackson, Ian Johnston, Dominic Kwiatkowski, Cordelia Langford, John Sillitoe on behalf of the Wellcome Sanger Institute COVID-19 Surveillance Team |
| EPI_ISL_3226503 | Lighthouse Laboratory Plymouth | Wellcome Sanger Institute for the COVID-19 Genomics UK (COG-UK) Consortium | Lighthouse Laboratory Plymouth and Alex Alderton, Roberto Amato, Jeffrey Barrett, Sonia Goncalves, Ewan Harrison, David K. Jackson, Ian Johnston, Dominic Kwiatkowski, Cordelia Langford, John Sillitoe on behalf of the Wellcome Sanger Institute COVID-19 Surveillance Team |
| EPI_ISL_3226504 | Lighthouse Lab in Glasgow | Wellcome Sanger Institute for the COVID-19 Genomics UK (COG-UK) Consortium | Harper VanSteenhouse, Yumi Kasai, David Gray, Carol Clugston, Anna Dominiczak and Alex Alderton, Roberto Amato, Jeffrey Barrett, Sonia Goncalves, Ewan Harrison, David K. Jackson, Ian Johnston, Dominic Kwiatkowski, Cordelia Langford, John Sillitoe on behalf of the Wellcome Sanger Institute COVID-19 Surveillance Team |
| EPI_ISL_3226505 | Lighthouse Laboratory Plymouth | Wellcome Sanger Institute for the COVID-19 Genomics UK (COG-UK) Consortium | Lighthouse Laboratory Plymouth and Alex Alderton, Roberto Amato, Jeffrey Barrett, Sonia Goncalves, Ewan Harrison, David K. Jackson, Ian Johnston, Dominic Kwiatkowski, Cordelia Langford, John Sillitoe on behalf of the Wellcome Sanger Institute COVID-19 Surveillance Team |
| EPI_ISL_3226506, EPI_ISL_3226507 | Lighthouse Lab in Glasgow | Wellcome Sanger Institute for the COVID-19 Genomics UK (COG-UK) Consortium | Harper VanSteenhouse, Yumi Kasai, David Gray, Carol Clugston, Anna Dominiczak and Alex Alderton, Roberto Amato, Jeffrey Barrett, Sonia Goncalves, Ewan Harrison, David K. Jackson, Ian Johnston, Dominic Kwiatkowski, Cordelia Langford, John Sillitoe on behalf of the Wellcome Sanger Institute COVID-19 Surveillance Team |
| EPI_ISL_3226508 | Lighthouse Lab in Alderley Park | Wellcome Sanger Institute for the COVID-19 Genomics UK (COG-UK) Consortium | Jacquelyn Wynn, Mairead Hyland, The Lighthouse Lab in Alderley Park and Alex Alderton, Roberto Amato, Jeffrey Barrett, Sonia Goncalves, Ewan Harrison, David K. Jackson, Ian Johnston, Dominic Kwiatkowski, Cordelia Langford, John Sillitoe on behalf of the Wellcome Sanger Institute COVID-19 Surveillance Team |
| EPI_ISL_3226509, EPI_ISL_3226510 | Lighthouse Laboratory Plymouth | Wellcome Sanger Institute for the COVID-19 Genomics UK (COG-UK) Consortium | Lighthouse Laboratory Plymouth and Alex Alderton, Roberto Amato, Jeffrey Barrett, Sonia Goncalves, Ewan Harrison, David K. Jackson, Ian Johnston, Dominic Kwiatkowski, Cordelia Langford, John Sillitoe on behalf of the Wellcome Sanger Institute COVID-19 Surveillance Team |
| EPI_ISL_3226511 | Lighthouse Lab in Glasgow | Wellcome Sanger Institute for the COVID-19 Genomics UK (COG-UK) Consortium | Harper VanSteenhouse, Yumi Kasai, David Gray, Carol Clugston, Anna Dominiczak and Alex Alderton, Roberto Amato, Jeffrey Barrett, Sonia Goncalves, Ewan Harrison, David K. Jackson, Ian Johnston, Dominic Kwiatkowski, Cordelia Langford, John Sillitoe on behalf of the Wellcome Sanger Institute COVID-19 Surveillance Team |
| EPI_ISL_3226512 | Lighthouse Lab in Alderley Park | Wellcome Sanger Institute for the COVID-19 Genomics UK (COG-UK) Consortium | Jacquelyn Wynn, Mairead Hyland, The Lighthouse Lab in Alderley Park and Alex Alderton, Roberto Amato, Jeffrey Barrett, Sonia Goncalves, Ewan Harrison, David K. Jackson, Ian Johnston, Dominic Kwiatkowski, Cordelia Langford, John Sillitoe on behalf of the Wellcome Sanger Institute COVID-19 Surveillance Team |
| EPI_ISL_3226513 | Lighthouse Lab in Milton Keynes | Wellcome Sanger Institute for the COVID-19 Genomics UK (COG-UK) Consortium | The Lighthouse Lab in Milton Keynes and Alex Alderton, Roberto Amato, Jeffrey Barrett, Sonia Goncalves, Ewan Harrison, David K. Jackson, Ian Johnston, Dominic Kwiatkowski, Cordelia Langford, John Sillitoe on behalf of the Wellcome Sanger Institute COVID-19 Surveillance Team |
| EPI_ISL_3226514 | Lighthouse Lab in Alderley Park | Wellcome Sanger Institute for the COVID-19 Genomics UK (COG-UK) Consortium | Jacquelyn Wynn, Mairead Hyland, The Lighthouse Lab in Alderley Park and Alex Alderton, Roberto Amato, Jeffrey Barrett, Sonia Goncalves, Ewan Harrison, David K. Jackson, Ian Johnston, Dominic Kwiatkowski, Cordelia Langford, John Sillitoe on behalf of the Wellcome Sanger Institute COVID-19 Surveillance Team |
| EPI_ISL_3226515, EPI_ISL_3226516 | Lighthouse Lab in Glasgow | Wellcome Sanger Institute for the COVID-19 Genomics UK (COG-UK) Consortium | Harper VanSteenhouse, Yumi Kasai, David Gray, Carol Clugston, Anna Dominiczak and Alex Alderton, Roberto Amato, Jeffrey Barrett, Sonia Goncalves, Ewan Harrison, David K. Jackson, Ian Johnston, Dominic Kwiatkowski, Cordelia Langford, John Sillitoe on behalf of the Wellcome Sanger Institute COVID-19 Surveillance Team |
| EPI_ISL_3226517 | Lighthouse Laboratory Plymouth | Wellcome Sanger Institute for the COVID-19 Genomics UK (COG-UK) Consortium | Lighthouse Laboratory Plymouth and Alex Alderton, Roberto Amato, Jeffrey Barrett, Sonia Goncalves, Ewan Harrison, David K. Jackson, Ian Johnston, Dominic Kwiatkowski, Cordelia Langford, John Sillitoe on behalf of the Wellcome Sanger Institute COVID-19 Surveillance Team |
| EPI_ISL_3226518 | Lighthouse Lab in Milton Keynes | Wellcome Sanger Institute for the COVID-19 Genomics UK (COG-UK) Consortium | The Lighthouse Lab in Milton Keynes and Alex Alderton, Roberto Amato, Jeffrey Barrett, Sonia Goncalves, Ewan Harrison, David K. Jackson, Ian Johnston, Dominic Kwiatkowski, Cordelia Langford, John Sillitoe on behalf of the Wellcome Sanger Institute COVID-19 Surveillance Team |
| EPI_ISL_3226519 | Lighthouse Lab in Glasgow | Wellcome Sanger Institute for the COVID-19 Genomics UK (COG-UK) Consortium | Harper VanSteenhouse, Yumi Kasai, David Gray, Carol Clugston, Anna Dominiczak and Alex Alderton, Roberto Amato, Jeffrey Barrett, Sonia Goncalves, Ewan Harrison, David K. Jackson, Ian Johnston, Dominic Kwiatkowski, Cordelia Langford, John Sillitoe on behalf of the Wellcome Sanger Institute COVID-19 Surveillance Team |
| EPI_ISL_3226520 | Lighthouse Lab in Alderley Park | Wellcome Sanger Institute for the COVID-19 Genomics UK (COG-UK) Consortium | Jacquelyn Wynn, Mairead Hyland, The Lighthouse Lab in Alderley Park and Alex Alderton, Roberto Amato, Jeffrey Barrett, Sonia Goncalves, Ewan Harrison, David K. Jackson, Ian Johnston, Dominic Kwiatkowski, Cordelia Langford, John Sillitoe on behalf of the Wellcome Sanger Institute COVID-19 Surveillance Team |
| EPI_ISL_3226521 | Lighthouse Lab in Glasgow | Wellcome Sanger Institute for the COVID-19 Genomics UK (COG-UK) Consortium | Harper VanSteenhouse, Yumi Kasai, David Gray, Carol Clugston, Anna Dominiczak and Alex Alderton, Roberto Amato, Jeffrey Barrett, Sonia Goncalves, Ewan Harrison, David K. Jackson, Ian Johnston, Dominic Kwiatkowski, Cordelia Langford, John Sillitoe on behalf of the Wellcome Sanger Institute COVID-19 Surveillance Team |
| EPI_ISL_3226522 | Lighthouse Lab in Alderley Park | Wellcome Sanger Institute for the COVID-19 Genomics UK (COG-UK) Consortium | Jacquelyn Wynn, Mairead Hyland, The Lighthouse Lab in Alderley Park and Alex Alderton, Roberto Amato, Jeffrey Barrett, Sonia Goncalves, Ewan Harrison, David K. Jackson, Ian Johnston, Dominic Kwiatkowski, Cordelia Langford, John Sillitoe on behalf of the Wellcome Sanger Institute COVID-19 Surveillance Team |
| EPI_ISL_3226523 | Lighthouse Lab in Glasgow | Wellcome Sanger Institute for the COVID-19 Genomics UK (COG-UK) Consortium | Harper VanSteenhouse, Yumi Kasai, David Gray, Carol Clugston, Anna Dominiczak and Alex Alderton, Roberto Amato, Jeffrey Barrett, Sonia Goncalves, Ewan Harrison, David K. Jackson, Ian Johnston, Dominic Kwiatkowski, Cordelia Langford, John Sillitoe on behalf of the Wellcome Sanger Institute COVID-19 Surveillance Team |
| EPI_ISL_3226524 | Lighthouse Lab in Alderley Park | Wellcome Sanger Institute for the COVID-19 Genomics UK (COG-UK) Consortium | Jacquelyn Wynn, Mairead Hyland, The Lighthouse Lab in Alderley Park and Alex Alderton, Roberto Amato, Jeffrey Barrett, Sonia Goncalves, Ewan Harrison, David K. Jackson, Ian Johnston, Dominic Kwiatkowski, Cordelia Langford, John Sillitoe on behalf of the Wellcome Sanger Institute COVID-19 Surveillance Team |
| EPI_ISL_3226525, EPI_ISL_3226526 | Lighthouse Lab in Milton Keynes | Wellcome Sanger Institute for the COVID-19 Genomics UK (COG-UK) Consortium | The Lighthouse Lab in Milton Keynes and Alex Alderton, Roberto Amato, Jeffrey Barrett, Sonia Goncalves, Ewan Harrison, David K. Jackson, Ian Johnston, Dominic Kwiatkowski, Cordelia Langford, John Sillitoe on behalf of the Wellcome Sanger Institute COVID-19 Surveillance Team |
| EPI_ISL_3226527 | Lighthouse Lab in Alderley Park | Wellcome Sanger Institute for the COVID-19 Genomics UK (COG-UK) Consortium | Jacquelyn Wynn, Mairead Hyland, The Lighthouse Lab in Alderley Park and Alex Alderton, Roberto Amato, Jeffrey Barrett, Sonia Goncalves, Ewan Harrison, David K. Jackson, Ian Johnston, Dominic Kwiatkowski, Cordelia Langford, John Sillitoe on behalf of the Wellcome Sanger Institute COVID-19 Surveillance Team |
| EPI_ISL_3226528, EPI_ISL_3226529, EPI_ISL_3226530, EPI_ISL_3226531, EPI_ISL_3226532 | Lighthouse Laboratory Plymouth | Wellcome Sanger Institute for the COVID-19 Genomics UK (COG-UK) Consortium | Lighthouse Laboratory Plymouth and Alex Alderton, Roberto Amato, Jeffrey Barrett, Sonia Goncalves, Ewan Harrison, David K. Jackson, Ian Johnston, Dominic Kwiatkowski, Cordelia Langford, John Sillitoe on behalf of the Wellcome Sanger Institute COVID-19 Surveillance Team |
| EPI_ISL_3226533, EPI_ISL_3226534 | Lighthouse Lab in Alderley Park | Wellcome Sanger Institute for the COVID-19 Genomics UK (COG-UK) Consortium | Jacquelyn Wynn, Mairead Hyland, The Lighthouse Lab in Alderley Park and Alex Alderton, Roberto Amato, Jeffrey Barrett, Sonia Goncalves, Ewan Harrison, David K. Jackson, Ian Johnston, Dominic Kwiatkowski, Cordelia Langford, John Sillitoe on behalf of the Wellcome Sanger Institute COVID-19 Surveillance Team |
| EPI_ISL_3226535, EPI_ISL_3226536 | Lighthouse Laboratory Plymouth | Wellcome Sanger Institute for the COVID-19 Genomics UK (COG-UK) Consortium | Lighthouse Laboratory Plymouth and Alex Alderton, Roberto Amato, Jeffrey Barrett, Sonia Goncalves, Ewan Harrison, David K. Jackson, Ian Johnston, Dominic Kwiatkowski, Cordelia Langford, John Sillitoe on behalf of the Wellcome Sanger Institute COVID-19 Surveillance Team |

| | | | |
|---|---|---|---|
| EPI_ISL_3226537 | Lighthouse Lab in Glasgow | Wellcome Sanger Institute for the COVID-19 Genomics UK (COG-UK) Consortium | Harper VanSteenhouse, Yumi Kasai, David Gray, Carol Clugston, Anna Dominiczak and Alex Alderton, Roberto Amato, Jeffrey Barrett, Sonia Goncalves, Ewan Harrison, David K. Jackson, Ian Johnston, Dominic Kwiatkowski, Cordelia Langford, John Sillitoe on behalf of the Wellcome Sanger Institute COVID-19 Surveillance Team |
| EPI_ISL_3226538, EPI_ISL_3226539 | Lighthouse Lab in Milton Keynes | Wellcome Sanger Institute for the COVID-19 Genomics UK (COG-UK) Consortium | The Lighthouse Lab in Milton Keynes and Alex Alderton, Roberto Amato, Jeffrey Barrett, Sonia Goncalves, Ewan Harrison, David K. Jackson, Ian Johnston, Dominic Kwiatkowski, Cordelia Langford, John Sillitoe on behalf of the Wellcome Sanger Institute COVID-19 Surveillance Team |
| EPI_ISL_3226540 | Lighthouse Lab in Alderley Park | Wellcome Sanger Institute for the COVID-19 Genomics UK (COG-UK) Consortium | Jacquelyn Wynn, Mairead Hyland, The Lighthouse Lab in Alderley Park and Alex Alderton, Roberto Amato, Jeffrey Barrett, Sonia Goncalves, Ewan Harrison, David K. Jackson, Ian Johnston, Dominic Kwiatkowski, Cordelia Langford, John Sillitoe on behalf of the Wellcome Sanger Institute COVID-19 Surveillance Team |
| EPI_ISL_3226541 | Lighthouse Lab in Milton Keynes | Wellcome Sanger Institute for the COVID-19 Genomics UK (COG-UK) Consortium | The Lighthouse Lab in Milton Keynes and Alex Alderton, Roberto Amato, Jeffrey Barrett, Sonia Goncalves, Ewan Harrison, David K. Jackson, Ian Johnston, Dominic Kwiatkowski, Cordelia Langford, John Sillitoe on behalf of the Wellcome Sanger Institute COVID-19 Surveillance Team |
| EPI_ISL_3226542 | Lighthouse Lab in Alderley Park | Wellcome Sanger Institute for the COVID-19 Genomics UK (COG-UK) Consortium | Jacquelyn Wynn, Mairead Hyland, The Lighthouse Lab in Alderley Park and Alex Alderton, Roberto Amato, Jeffrey Barrett, Sonia Goncalves, Ewan Harrison, David K. Jackson, Ian Johnston, Dominic Kwiatkowski, Cordelia Langford, John Sillitoe on behalf of the Wellcome Sanger Institute COVID-19 Surveillance Team |
| EPI_ISL_3226543, EPI_ISL_3226544 | Lighthouse Lab in Milton Keynes | Wellcome Sanger Institute for the COVID-19 Genomics UK (COG-UK) Consortium | The Lighthouse Lab in Milton Keynes and Alex Alderton, Roberto Amato, Jeffrey Barrett, Sonia Goncalves, Ewan Harrison, David K. Jackson, Ian Johnston, Dominic Kwiatkowski, Cordelia Langford, John Sillitoe on behalf of the Wellcome Sanger Institute COVID-19 Surveillance Team |
| EPI_ISL_3226545 | Lighthouse Laboratory Plymouth | Wellcome Sanger Institute for the COVID-19 Genomics UK (COG-UK) Consortium | Lighthouse Laboratory Plymouth and Alex Alderton, Roberto Amato, Jeffrey Barrett, Sonia Goncalves, Ewan Harrison, David K. Jackson, Ian Johnston, Dominic Kwiatkowski, Cordelia Langford, John Sillitoe on behalf of the Wellcome Sanger Institute COVID-19 Surveillance Team |
| EPI_ISL_3226546 | Lighthouse Lab in Alderley Park | Wellcome Sanger Institute for the COVID-19 Genomics UK (COG-UK) Consortium | Jacquelyn Wynn, Mairead Hyland, The Lighthouse Lab in Alderley Park and Alex Alderton, Roberto Amato, Jeffrey Barrett, Sonia Goncalves, Ewan Harrison, David K. Jackson, Ian Johnston, Dominic Kwiatkowski, Cordelia Langford, John Sillitoe on behalf of the Wellcome Sanger Institute COVID-19 Surveillance Team |
| EPI_ISL_3226547, EPI_ISL_3226548, EPI_ISL_3226549 | Lighthouse Laboratory Plymouth | Wellcome Sanger Institute for the COVID-19 Genomics UK (COG-UK) Consortium | Lighthouse Laboratory Plymouth and Alex Alderton, Roberto Amato, Jeffrey Barrett, Sonia Goncalves, Ewan Harrison, David K. Jackson, Ian Johnston, Dominic Kwiatkowski, Cordelia Langford, John Sillitoe on behalf of the Wellcome Sanger Institute COVID-19 Surveillance Team |
| EPI_ISL_3226550 | Lighthouse Lab in Alderley Park | Wellcome Sanger Institute for the COVID-19 Genomics UK (COG-UK) Consortium | Jacquelyn Wynn, Mairead Hyland, The Lighthouse Lab in Alderley Park and Alex Alderton, Roberto Amato, Jeffrey Barrett, Sonia Goncalves, Ewan Harrison, David K. Jackson, Ian Johnston, Dominic Kwiatkowski, Cordelia Langford, John Sillitoe on behalf of the Wellcome Sanger Institute COVID-19 Surveillance Team |
| EPI_ISL_3226551 | Lighthouse Lab in Milton Keynes | Wellcome Sanger Institute for the COVID-19 Genomics UK (COG-UK) Consortium | The Lighthouse Lab in Milton Keynes and Alex Alderton, Roberto Amato, Jeffrey Barrett, Sonia Goncalves, Ewan Harrison, David K. Jackson, Ian Johnston, Dominic Kwiatkowski, Cordelia Langford, John Sillitoe on behalf of the Wellcome Sanger Institute COVID-19 Surveillance Team |
| EPI_ISL_3226552 | Lighthouse Lab in Alderley Park | Wellcome Sanger Institute for the COVID-19 Genomics UK (COG-UK) Consortium | Jacquelyn Wynn, Mairead Hyland, The Lighthouse Lab in Alderley Park and Alex Alderton, Roberto Amato, Jeffrey Barrett, Sonia Goncalves, Ewan Harrison, David K. Jackson, Ian Johnston, Dominic Kwiatkowski, Cordelia Langford, John Sillitoe on behalf of the Wellcome Sanger Institute COVID-19 Surveillance Team |
| EPI_ISL_3226553 | Lighthouse Lab in Milton Keynes | Wellcome Sanger Institute for the COVID-19 Genomics UK (COG-UK) Consortium | The Lighthouse Lab in Milton Keynes and Alex Alderton, Roberto Amato, Jeffrey Barrett, Sonia Goncalves, Ewan Harrison, David K. Jackson, Ian Johnston, Dominic Kwiatkowski, Cordelia Langford, John Sillitoe on behalf of the Wellcome Sanger Institute COVID-19 Surveillance Team |
| EPI_ISL_3226554, EPI_ISL_3226555 | Lighthouse Lab in Alderley Park | Wellcome Sanger Institute for the COVID-19 Genomics UK (COG-UK) Consortium | Jacquelyn Wynn, Mairead Hyland, The Lighthouse Lab in Alderley Park and Alex Alderton, Roberto Amato, Jeffrey Barrett, Sonia Goncalves, Ewan Harrison, David K. Jackson, Ian Johnston, Dominic Kwiatkowski, Cordelia Langford, John Sillitoe on behalf of the Wellcome Sanger Institute COVID-19 Surveillance Team |
| EPI_ISL_3226556 | Lighthouse Lab in Glasgow | Wellcome Sanger Institute for the COVID-19 Genomics UK (COG-UK) Consortium | Harper VanSteenhouse, Yumi Kasai, David Gray, Carol Clugston, Anna Dominiczak and Alex Alderton, Roberto Amato, Jeffrey Barrett, Sonia Goncalves, Ewan Harrison, David K. Jackson, Ian Johnston, Dominic Kwiatkowski, Cordelia Langford, John Sillitoe on behalf of the Wellcome Sanger Institute COVID-19 Surveillance Team |
| EPI_ISL_3226557, EPI_ISL_3226558 | Lighthouse Lab in Alderley Park | Wellcome Sanger Institute for the COVID-19 Genomics UK (COG-UK) Consortium | Jacquelyn Wynn, Mairead Hyland, The Lighthouse Lab in Alderley Park and Alex Alderton, Roberto Amato, Jeffrey Barrett, Sonia Goncalves, Ewan Harrison, David K. Jackson, Ian Johnston, Dominic Kwiatkowski, Cordelia Langford, John Sillitoe on behalf of the Wellcome Sanger Institute COVID-19 Surveillance Team |
| EPI_ISL_3226559, EPI_ISL_3226560 | Lighthouse Lab in Milton Keynes | Wellcome Sanger Institute for the COVID-19 Genomics UK (COG-UK) Consortium | The Lighthouse Lab in Milton Keynes and Alex Alderton, Roberto Amato, Jeffrey Barrett, Sonia Goncalves, Ewan Harrison, David K. Jackson, Ian Johnston, Dominic Kwiatkowski, Cordelia Langford, John Sillitoe on behalf of the Wellcome Sanger Institute COVID-19 Surveillance Team |
| EPI_ISL_3226561, EPI_ISL_3226562 | Lighthouse Lab in Alderley Park | Wellcome Sanger Institute for the COVID-19 Genomics UK (COG-UK) Consortium | Jacquelyn Wynn, Mairead Hyland, The Lighthouse Lab in Alderley Park and Alex Alderton, Roberto Amato, Jeffrey Barrett, Sonia Goncalves, Ewan Harrison, David K. Jackson, Ian Johnston, Dominic Kwiatkowski, Cordelia Langford, John Sillitoe on behalf of the Wellcome Sanger Institute COVID-19 Surveillance Team |
| EPI_ISL_3226563 | Lighthouse Lab in Milton Keynes | Wellcome Sanger Institute for the COVID-19 Genomics UK (COG-UK) Consortium | The Lighthouse Lab in Milton Keynes and Alex Alderton, Roberto Amato, Jeffrey Barrett, Sonia Goncalves, Ewan Harrison, David K. Jackson, Ian Johnston, Dominic Kwiatkowski, Cordelia Langford, John Sillitoe on behalf of the Wellcome Sanger Institute COVID-19 Surveillance Team |
| EPI_ISL_3226564, EPI_ISL_3226565, EPI_ISL_3226566 | Lighthouse Lab in Alderley Park | Wellcome Sanger Institute for the COVID-19 Genomics UK (COG-UK) Consortium | Jacquelyn Wynn, Mairead Hyland, The Lighthouse Lab in Alderley Park and Alex Alderton, Roberto Amato, Jeffrey Barrett, Sonia Goncalves, Ewan Harrison, David K. Jackson, Ian Johnston, Dominic Kwiatkowski, Cordelia Langford, John Sillitoe on behalf of the Wellcome Sanger Institute COVID-19 Surveillance Team |
| EPI_ISL_3226567, EPI_ISL_3226568 | Lighthouse Lab in Milton Keynes | Wellcome Sanger Institute for the COVID-19 Genomics UK (COG-UK) Consortium | The Lighthouse Lab in Milton Keynes and Alex Alderton, Roberto Amato, Jeffrey Barrett, Sonia Goncalves, Ewan Harrison, David K. Jackson, Ian Johnston, Dominic Kwiatkowski, Cordelia Langford, John Sillitoe on behalf of the Wellcome Sanger Institute COVID-19 Surveillance Team |
| EPI_ISL_3226569, EPI_ISL_3226570 | Lighthouse Lab in Alderley Park | Wellcome Sanger Institute for the COVID-19 Genomics UK (COG-UK) Consortium | Jacquelyn Wynn, Mairead Hyland, The Lighthouse Lab in Alderley Park and Alex Alderton, Roberto Amato, Jeffrey Barrett, Sonia Goncalves, Ewan Harrison, David K. Jackson, Ian Johnston, Dominic Kwiatkowski, Cordelia Langford, John Sillitoe on behalf of the Wellcome Sanger Institute COVID-19 Surveillance Team |
| EPI_ISL_3226571 | Lighthouse Lab in Glasgow | Wellcome Sanger Institute for the COVID-19 Genomics UK (COG-UK) Consortium | Harper VanSteenhouse, Yumi Kasai, David Gray, Carol Clugston, Anna Dominiczak and Alex Alderton, Roberto Amato, Jeffrey Barrett, Sonia Goncalves, Ewan Harrison, David K. Jackson, Ian Johnston, Dominic Kwiatkowski, Cordelia Langford, John Sillitoe on behalf of the Wellcome Sanger Institute COVID-19 Surveillance Team |
| EPI_ISL_3226572 | Lighthouse Lab in Milton Keynes | Wellcome Sanger Institute for the COVID-19 Genomics UK (COG-UK) Consortium | The Lighthouse Lab in Milton Keynes and Alex Alderton, Roberto Amato, Jeffrey Barrett, Sonia Goncalves, Ewan Harrison, David K. Jackson, Ian Johnston, Dominic Kwiatkowski, Cordelia Langford, John Sillitoe on behalf of the Wellcome Sanger Institute COVID-19 Surveillance Team |
| EPI_ISL_3226573, EPI_ISL_3226574 | Lighthouse Lab in Glasgow | Wellcome Sanger Institute for the COVID-19 Genomics UK (COG-UK) Consortium | Harper VanSteenhouse, Yumi Kasai, David Gray, Carol Clugston, Anna Dominiczak and Alex Alderton, Roberto Amato, Jeffrey Barrett, Sonia Goncalves, Ewan Harrison, David K. Jackson, Ian Johnston, Dominic Kwiatkowski, Cordelia Langford, John Sillitoe on behalf of the Wellcome Sanger Institute COVID-19 Surveillance Team |
| EPI_ISL_3226575, EPI_ISL_3226576, EPI_ISL_3226577 | Lighthouse Lab in Alderley Park | Wellcome Sanger Institute for the COVID-19 Genomics UK (COG-UK) Consortium | Jacquelyn Wynn, Mairead Hyland, The Lighthouse Lab in Alderley Park and Alex Alderton, Roberto Amato, Jeffrey Barrett, Sonia Goncalves, Ewan Harrison, David K. Jackson, Ian Johnston, Dominic Kwiatkowski, Cordelia Langford, John Sillitoe on behalf of the Wellcome Sanger Institute COVID-19 Surveillance Team |
| EPI_ISL_3226578, EPI_ISL_3226579 | Lighthouse Lab in Glasgow | Wellcome Sanger Institute for the COVID-19 Genomics UK (COG-UK) Consortium | Harper VanSteenhouse, Yumi Kasai, David Gray, Carol Clugston, Anna Dominiczak and Alex Alderton, Roberto Amato, Jeffrey Barrett, Sonia Goncalves, Ewan Harrison, David K. Jackson, Ian Johnston, Dominic Kwiatkowski, Cordelia Langford, John Sillitoe on behalf of the Wellcome Sanger Institute COVID-19 Surveillance Team |
| EPI_ISL_3226580 | Lighthouse Lab in Milton Keynes | Wellcome Sanger Institute for the COVID-19 Genomics UK (COG-UK) Consortium | The Lighthouse Lab in Milton Keynes and Alex Alderton, Roberto Amato, Jeffrey Barrett, Sonia Goncalves, Ewan Harrison, David K. Jackson, Ian Johnston, Dominic Kwiatkowski, Cordelia Langford, John Sillitoe on behalf of the Wellcome Sanger Institute COVID-19 Surveillance Team |
| EPI_ISL_3226581 | Lighthouse Lab in Alderley Park | Wellcome Sanger Institute for the COVID-19 Genomics UK (COG-UK) Consortium | Jacquelyn Wynn, Mairead Hyland, The Lighthouse Lab in Alderley Park and Alex Alderton, Roberto Amato, Jeffrey Barrett, Sonia Goncalves, Ewan Harrison, David K. Jackson, Ian Johnston, Dominic Kwiatkowski, Cordelia Langford, John Sillitoe on behalf of the Wellcome Sanger Institute COVID-19 Surveillance Team |
| EPI_ISL_3226582 | Lighthouse Lab in Milton Keynes | Wellcome Sanger Institute for the COVID-19 Genomics UK (COG-UK) Consortium | The Lighthouse Lab in Milton Keynes and Alex Alderton, Roberto Amato, Jeffrey Barrett, Sonia Goncalves, Ewan Harrison, David K. Jackson, Ian Johnston, Dominic Kwiatkowski, Cordelia Langford, John Sillitoe on behalf of the Wellcome Sanger Institute COVID-19 Surveillance Team |
| EPI_ISL_3226583, EPI_ISL_3226584 | Lighthouse Lab in Glasgow | Wellcome Sanger Institute for the COVID-19 Genomics UK (COG-UK) Consortium | Harper VanSteenhouse, Yumi Kasai, David Gray, Carol Clugston, Anna Dominiczak and Alex Alderton, Roberto Amato, Jeffrey Barrett, Sonia Goncalves, Ewan Harrison, David K. Jackson, Ian Johnston, Dominic Kwiatkowski, Cordelia Langford, John Sillitoe on behalf of the Wellcome Sanger Institute COVID-19 Surveillance Team |
| EPI_ISL_3226585 | Lighthouse Lab in Alderley Park | Wellcome Sanger Institute for the COVID-19 Genomics UK (COG-UK) Consortium | Jacquelyn Wynn, Mairead Hyland, The Lighthouse Lab in Alderley Park and Alex Alderton, Roberto Amato, Jeffrey Barrett, Sonia Goncalves, Ewan Harrison, David K. Jackson, Ian Johnston, Dominic Kwiatkowski, Cordelia Langford, John Sillitoe on behalf of the Wellcome Sanger Institute COVID-19 Surveillance Team |
| EPI_ISL_3226586 | Lighthouse Lab in Glasgow | Wellcome Sanger Institute for the COVID-19 Genomics UK (COG-UK) Consortium | Harper VanSteenhouse, Yumi Kasai, David Gray, Carol Clugston, Anna Dominiczak and Alex Alderton, Roberto Amato, Jeffrey Barrett, Sonia Goncalves, Ewan Harrison, David K. Jackson, Ian Johnston, Dominic Kwiatkowski, Cordelia Langford, John Sillitoe on behalf of the Wellcome Sanger Institute COVID-19 Surveillance Team |
| EPI_ISL_3226587, EPI_ISL_3226588 | Lighthouse Lab in Alderley Park | Wellcome Sanger Institute for the COVID-19 Genomics UK (COG-UK) Consortium | Jacquelyn Wynn, Mairead Hyland, The Lighthouse Lab in Alderley Park and Alex Alderton, Roberto Amato, Jeffrey Barrett, Sonia Goncalves, Ewan Harrison, David K. Jackson, Ian Johnston, Dominic Kwiatkowski, Cordelia Langford, John Sillitoe on behalf of the Wellcome Sanger Institute COVID-19 Surveillance Team |
| EPI_ISL_3226589 | Lighthouse Lab in Milton Keynes | Wellcome Sanger Institute for the COVID-19 Genomics UK (COG-UK) Consortium | The Lighthouse Lab in Milton Keynes and Alex Alderton, Roberto Amato, Jeffrey Barrett, Sonia Goncalves, Ewan Harrison, David K. Jackson, Ian Johnston, Dominic Kwiatkowski, Cordelia Langford, John Sillitoe on behalf of the Wellcome Sanger Institute COVID-19 Surveillance Team |
| EPI_ISL_3226590 | Lighthouse Lab in Glasgow | Wellcome Sanger Institute for the COVID-19 Genomics UK (COG-UK) Consortium | Harper VanSteenhouse, Yumi Kasai, David Gray, Carol Clugston, Anna Dominiczak and Alex Alderton, Roberto Amato, Jeffrey Barrett, Sonia Goncalves, Ewan Harrison, David K. Jackson, Ian Johnston, Dominic Kwiatkowski, Cordelia Langford, John Sillitoe on behalf of the Wellcome Sanger Institute COVID-19 Surveillance Team |
| EPI_ISL_3226591, EPI_ISL_3226592 | Lighthouse Lab in Milton Keynes | Wellcome Sanger Institute for the COVID-19 Genomics UK (COG-UK) Consortium | The Lighthouse Lab in Milton Keynes and Alex Alderton, Roberto Amato, Jeffrey Barrett, Sonia Goncalves, Ewan Harrison, David K. Jackson, Ian Johnston, Dominic Kwiatkowski, Cordelia Langford, John Sillitoe on behalf of the Wellcome Sanger Institute COVID-19 Surveillance Team |
| EPI_ISL_3226593 | Lighthouse Lab in Glasgow | Wellcome Sanger Institute for the COVID-19 Genomics UK (COG-UK) Consortium | Harper VanSteenhouse, Yumi Kasai, David Gray, Carol Clugston, Anna Dominiczak and Alex Alderton, Roberto Amato, Jeffrey Barrett, Sonia Goncalves, Ewan Harrison, David K. Jackson, Ian Johnston, Dominic Kwiatkowski, Cordelia Langford, John Sillitoe on behalf of the Wellcome Sanger Institute COVID-19 Surveillance Team |
| EPI_ISL_3226594 | Lighthouse Lab in Alderley Park | Wellcome Sanger Institute for the COVID-19 Genomics UK (COG-UK) Consortium | Jacquelyn Wynn, Mairead Hyland, The Lighthouse Lab in Alderley Park and Alex Alderton, Roberto Amato, Jeffrey Barrett, Sonia Goncalves, Ewan Harrison, David K. Jackson, Ian Johnston, Dominic Kwiatkowski, Cordelia Langford, John Sillitoe on behalf of the Wellcome Sanger Institute COVID-19 Surveillance Team |
| EPI_ISL_3226595 | Lighthouse Lab in Glasgow | Wellcome Sanger Institute for the COVID-19 Genomics UK (COG-UK) Consortium | Harper VanSteenhouse, Yumi Kasai, David Gray, Carol Clugston, Anna Dominiczak and Alex Alderton, Roberto Amato, Jeffrey Barrett, Sonia Goncalves, Ewan Harrison, David K. Jackson, Ian Johnston, Dominic Kwiatkowski, Cordelia Langford, John Sillitoe on behalf of the Wellcome Sanger Institute COVID-19 Surveillance Team |
| EPI_ISL_3226596 | Lighthouse Lab in Milton Keynes | Wellcome Sanger Institute for the COVID-19 Genomics UK (COG-UK) Consortium | The Lighthouse Lab in Milton Keynes and Alex Alderton, Roberto Amato, Jeffrey Barrett, Sonia Goncalves, Ewan Harrison, David K. Jackson, Ian Johnston, Dominic Kwiatkowski, Cordelia Langford, John Sillitoe on behalf of the Wellcome Sanger Institute COVID-19 Surveillance Team |
| EPI_ISL_3226597, EPI_ISL_3226598, EPI_ISL_3226599, EPI_ISL_3226600 | Lighthouse Lab in Alderley Park | Wellcome Sanger Institute for the COVID-19 Genomics UK (COG-UK) Consortium | Jacquelyn Wynn, Mairead Hyland, The Lighthouse Lab in Alderley Park and Alex Alderton, Roberto Amato, Jeffrey Barrett, Sonia Goncalves, Ewan Harrison, David K. Jackson, Ian Johnston, Dominic Kwiatkowski, Cordelia Langford, John Sillitoe on behalf of the Wellcome Sanger Institute COVID-19 Surveillance Team |
| EPI_ISL_3226601, | Lighthouse Lab in Milton Keynes | Wellcome Sanger Institute for the COVID-19 Genomics UK (COG-UK) | The Lighthouse Lab in Milton Keynes and Alex Alderton, Roberto Amato, Jeffrey Barrett, Sonia Goncalves, Ewan Harrison, David K. Jackson, Ian Johnston, Dominic Kwiatkowski, Cordelia Langford, John Sillitoe on behalf of the Wellcome Sanger |

| | | | |
|---|---|---|---|
| EPI_ISL_3226602 | | Consortium | Institute COVID-19 Surveillance Team |
| EPI_ISL_3226603 | Lighthouse Lab in Alderley Park | Wellcome Sanger Institute for the COVID-19 Genomics UK (COG-UK) Consortium | Jacquelyn Wynn, Mairead Hyland, The Lighthouse Lab in Alderley Park and Alex Alderton, Roberto Amato, Jeffrey Barrett, Sonia Goncalves, Ewan Harrison, David K. Jackson, Ian Johnston, Dominic Kwiatkowski, Cordelia Langford, John Sillitoe on behalf of the Wellcome Sanger Institute COVID-19 Surveillance Team |
| EPI_ISL_3226604 | Lighthouse Lab in Glasgow | Wellcome Sanger Institute for the COVID-19 Genomics UK (COG-UK) Consortium | Harper VanSteenhouse, Yumi Kasai, David Gray, Carol Clugston, Anna Dominiczak and Alex Alderton, Roberto Amato, Jeffrey Barrett, Sonia Goncalves, Ewan Harrison, David K. Jackson, Ian Johnston, Dominic Kwiatkowski, Cordelia Langford, John Sillitoe on behalf of the Wellcome Sanger Institute COVID-19 Surveillance Team |
| EPI_ISL_3226605, EPI_ISL_3226606 | Lighthouse Lab in Alderley Park | Wellcome Sanger Institute for the COVID-19 Genomics UK (COG-UK) Consortium | Jacquelyn Wynn, Mairead Hyland, The Lighthouse Lab in Alderley Park and Alex Alderton, Roberto Amato, Jeffrey Barrett, Sonia Goncalves, Ewan Harrison, David K. Jackson, Ian Johnston, Dominic Kwiatkowski, Cordelia Langford, John Sillitoe on behalf of the Wellcome Sanger Institute COVID-19 Surveillance Team |
| EPI_ISL_3226607 | Lighthouse Lab in Glasgow | Wellcome Sanger Institute for the COVID-19 Genomics UK (COG-UK) Consortium | Harper VanSteenhouse, Yumi Kasai, David Gray, Carol Clugston, Anna Dominiczak and Alex Alderton, Roberto Amato, Jeffrey Barrett, Sonia Goncalves, Ewan Harrison, David K. Jackson, Ian Johnston, Dominic Kwiatkowski, Cordelia Langford, John Sillitoe on behalf of the Wellcome Sanger Institute COVID-19 Surveillance Team |
| EPI_ISL_3226608 | Lighthouse Lab in Alderley Park | Wellcome Sanger Institute for the COVID-19 Genomics UK (COG-UK) Consortium | Jacquelyn Wynn, Mairead Hyland, The Lighthouse Lab in Alderley Park and Alex Alderton, Roberto Amato, Jeffrey Barrett, Sonia Goncalves, Ewan Harrison, David K. Jackson, Ian Johnston, Dominic Kwiatkowski, Cordelia Langford, John Sillitoe on behalf of the Wellcome Sanger Institute COVID-19 Surveillance Team |
| EPI_ISL_3226609, EPI_ISL_3226610 | Lighthouse Lab in Glasgow | Wellcome Sanger Institute for the COVID-19 Genomics UK (COG-UK) Consortium | Harper VanSteenhouse, Yumi Kasai, David Gray, Carol Clugston, Anna Dominiczak and Alex Alderton, Roberto Amato, Jeffrey Barrett, Sonia Goncalves, Ewan Harrison, David K. Jackson, Ian Johnston, Dominic Kwiatkowski, Cordelia Langford, John Sillitoe on behalf of the Wellcome Sanger Institute COVID-19 Surveillance Team |
| EPI_ISL_3226611 | Lighthouse Lab in Milton Keynes | Wellcome Sanger Institute for the COVID-19 Genomics UK (COG-UK) Consortium | The Lighthouse Lab in Milton Keynes and Alex Alderton, Roberto Amato, Jeffrey Barrett, Sonia Goncalves, Ewan Harrison, David K. Jackson, Ian Johnston, Dominic Kwiatkowski, Cordelia Langford, John Sillitoe on behalf of the Wellcome Sanger Institute COVID-19 Surveillance Team |
| EPI_ISL_3226612, EPI_ISL_3226613 | Lighthouse Lab in Alderley Park | Wellcome Sanger Institute for the COVID-19 Genomics UK (COG-UK) Consortium | Jacquelyn Wynn, Mairead Hyland, The Lighthouse Lab in Alderley Park and Alex Alderton, Roberto Amato, Jeffrey Barrett, Sonia Goncalves, Ewan Harrison, David K. Jackson, Ian Johnston, Dominic Kwiatkowski, Cordelia Langford, John Sillitoe on behalf of the Wellcome Sanger Institute COVID-19 Surveillance Team |
| EPI_ISL_3226614 | Lighthouse Lab in Milton Keynes | Wellcome Sanger Institute for the COVID-19 Genomics UK (COG-UK) Consortium | The Lighthouse Lab in Milton Keynes and Alex Alderton, Roberto Amato, Jeffrey Barrett, Sonia Goncalves, Ewan Harrison, David K. Jackson, Ian Johnston, Dominic Kwiatkowski, Cordelia Langford, John Sillitoe on behalf of the Wellcome Sanger Institute COVID-19 Surveillance Team |
| EPI_ISL_3226615, EPI_ISL_3226616 | Lighthouse Lab in Alderley Park | Wellcome Sanger Institute for the COVID-19 Genomics UK (COG-UK) Consortium | Jacquelyn Wynn, Mairead Hyland, The Lighthouse Lab in Alderley Park and Alex Alderton, Roberto Amato, Jeffrey Barrett, Sonia Goncalves, Ewan Harrison, David K. Jackson, Ian Johnston, Dominic Kwiatkowski, Cordelia Langford, John Sillitoe on behalf of the Wellcome Sanger Institute COVID-19 Surveillance Team |
| EPI_ISL_3226617 | Lighthouse Lab in Milton Keynes | Wellcome Sanger Institute for the COVID-19 Genomics UK (COG-UK) Consortium | The Lighthouse Lab in Milton Keynes and Alex Alderton, Roberto Amato, Jeffrey Barrett, Sonia Goncalves, Ewan Harrison, David K. Jackson, Ian Johnston, Dominic Kwiatkowski, Cordelia Langford, John Sillitoe on behalf of the Wellcome Sanger Institute COVID-19 Surveillance Team |
| EPI_ISL_3226618 | Lighthouse Lab in Alderley Park | Wellcome Sanger Institute for the COVID-19 Genomics UK (COG-UK) Consortium | Jacquelyn Wynn, Mairead Hyland, The Lighthouse Lab in Alderley Park and Alex Alderton, Roberto Amato, Jeffrey Barrett, Sonia Goncalves, Ewan Harrison, David K. Jackson, Ian Johnston, Dominic Kwiatkowski, Cordelia Langford, John Sillitoe on behalf of the Wellcome Sanger Institute COVID-19 Surveillance Team |
| EPI_ISL_3226619 | Lighthouse Lab in Milton Keynes | Wellcome Sanger Institute for the COVID-19 Genomics UK (COG-UK) Consortium | The Lighthouse Lab in Milton Keynes and Alex Alderton, Roberto Amato, Jeffrey Barrett, Sonia Goncalves, Ewan Harrison, David K. Jackson, Ian Johnston, Dominic Kwiatkowski, Cordelia Langford, John Sillitoe on behalf of the Wellcome Sanger Institute COVID-19 Surveillance Team |
| EPI_ISL_3226620 | Lighthouse Lab in Alderley Park | Wellcome Sanger Institute for the COVID-19 Genomics UK (COG-UK) Consortium | Jacquelyn Wynn, Mairead Hyland, The Lighthouse Lab in Alderley Park and Alex Alderton, Roberto Amato, Jeffrey Barrett, Sonia Goncalves, Ewan Harrison, David K. Jackson, Ian Johnston, Dominic Kwiatkowski, Cordelia Langford, John Sillitoe on behalf of the Wellcome Sanger Institute COVID-19 Surveillance Team |
| EPI_ISL_3226621, EPI_ISL_3226622 | Lighthouse Lab in Milton Keynes | Wellcome Sanger Institute for the COVID-19 Genomics UK (COG-UK) Consortium | The Lighthouse Lab in Milton Keynes and Alex Alderton, Roberto Amato, Jeffrey Barrett, Sonia Goncalves, Ewan Harrison, David K. Jackson, Ian Johnston, Dominic Kwiatkowski, Cordelia Langford, John Sillitoe on behalf of the Wellcome Sanger Institute COVID-19 Surveillance Team |
| EPI_ISL_3226623 | Lighthouse Lab in Glasgow | Wellcome Sanger Institute for the COVID-19 Genomics UK (COG-UK) Consortium | Harper VanSteenhouse, Yumi Kasai, David Gray, Carol Clugston, Anna Dominiczak and Alex Alderton, Roberto Amato, Jeffrey Barrett, Sonia Goncalves, Ewan Harrison, David K. Jackson, Ian Johnston, Dominic Kwiatkowski, Cordelia Langford, John Sillitoe on behalf of the Wellcome Sanger Institute COVID-19 Surveillance Team |
| EPI_ISL_3226624 | Lighthouse Lab in Milton Keynes | Wellcome Sanger Institute for the COVID-19 Genomics UK (COG-UK) Consortium | The Lighthouse Lab in Milton Keynes and Alex Alderton, Roberto Amato, Jeffrey Barrett, Sonia Goncalves, Ewan Harrison, David K. Jackson, Ian Johnston, Dominic Kwiatkowski, Cordelia Langford, John Sillitoe on behalf of the Wellcome Sanger Institute COVID-19 Surveillance Team |
| EPI_ISL_3226625 | Lighthouse Lab in Alderley Park | Wellcome Sanger Institute for the COVID-19 Genomics UK (COG-UK) Consortium | Jacquelyn Wynn, Mairead Hyland, The Lighthouse Lab in Alderley Park and Alex Alderton, Roberto Amato, Jeffrey Barrett, Sonia Goncalves, Ewan Harrison, David K. Jackson, Ian Johnston, Dominic Kwiatkowski, Cordelia Langford, John Sillitoe on behalf of the Wellcome Sanger Institute COVID-19 Surveillance Team |
| EPI_ISL_3226626 | Lighthouse Lab in Milton Keynes | Wellcome Sanger Institute for the COVID-19 Genomics UK (COG-UK) Consortium | The Lighthouse Lab in Milton Keynes and Alex Alderton, Roberto Amato, Jeffrey Barrett, Sonia Goncalves, Ewan Harrison, David K. Jackson, Ian Johnston, Dominic Kwiatkowski, Cordelia Langford, John Sillitoe on behalf of the Wellcome Sanger Institute COVID-19 Surveillance Team |
| EPI_ISL_3226627 | Lighthouse Lab in Alderley Park | Wellcome Sanger Institute for the COVID-19 Genomics UK (COG-UK) Consortium | Jacquelyn Wynn, Mairead Hyland, The Lighthouse Lab in Alderley Park and Alex Alderton, Roberto Amato, Jeffrey Barrett, Sonia Goncalves, Ewan Harrison, David K. Jackson, Ian Johnston, Dominic Kwiatkowski, Cordelia Langford, John Sillitoe on behalf of the Wellcome Sanger Institute COVID-19 Surveillance Team |
| EPI_ISL_3226628 | Lighthouse Lab in Milton Keynes | Wellcome Sanger Institute for the COVID-19 Genomics UK (COG-UK) Consortium | The Lighthouse Lab in Milton Keynes and Alex Alderton, Roberto Amato, Jeffrey Barrett, Sonia Goncalves, Ewan Harrison, David K. Jackson, Ian Johnston, Dominic Kwiatkowski, Cordelia Langford, John Sillitoe on behalf of the Wellcome Sanger Institute COVID-19 Surveillance Team |
| EPI_ISL_3226629 | Lighthouse Lab in Alderley Park | Wellcome Sanger Institute for the COVID-19 Genomics UK (COG-UK) Consortium | Jacquelyn Wynn, Mairead Hyland, The Lighthouse Lab in Alderley Park and Alex Alderton, Roberto Amato, Jeffrey Barrett, Sonia Goncalves, Ewan Harrison, David K. Jackson, Ian Johnston, Dominic Kwiatkowski, Cordelia Langford, John Sillitoe on behalf of the Wellcome Sanger Institute COVID-19 Surveillance Team |
| EPI_ISL_3226630 | Lighthouse Lab in Milton Keynes | Wellcome Sanger Institute for the COVID-19 Genomics UK (COG-UK) Consortium | The Lighthouse Lab in Milton Keynes and Alex Alderton, Roberto Amato, Jeffrey Barrett, Sonia Goncalves, Ewan Harrison, David K. Jackson, Ian Johnston, Dominic Kwiatkowski, Cordelia Langford, John Sillitoe on behalf of the Wellcome Sanger Institute COVID-19 Surveillance Team |
| EPI_ISL_3226631 | Lighthouse Lab in Glasgow | Wellcome Sanger Institute for the COVID-19 Genomics UK (COG-UK) Consortium | Harper VanSteenhouse, Yumi Kasai, David Gray, Carol Clugston, Anna Dominiczak and Alex Alderton, Roberto Amato, Jeffrey Barrett, Sonia Goncalves, Ewan Harrison, David K. Jackson, Ian Johnston, Dominic Kwiatkowski, Cordelia Langford, John Sillitoe on behalf of the Wellcome Sanger Institute COVID-19 Surveillance Team |
| EPI_ISL_3226632, EPI_ISL_3226633, EPI_ISL_3226634 | Lighthouse Lab in Alderley Park | Wellcome Sanger Institute for the COVID-19 Genomics UK (COG-UK) Consortium | Jacquelyn Wynn, Mairead Hyland, The Lighthouse Lab in Alderley Park and Alex Alderton, Roberto Amato, Jeffrey Barrett, Sonia Goncalves, Ewan Harrison, David K. Jackson, Ian Johnston, Dominic Kwiatkowski, Cordelia Langford, John Sillitoe on behalf of the Wellcome Sanger Institute COVID-19 Surveillance Team |
| EPI_ISL_3226635, EPI_ISL_3226636, EPI_ISL_3226637, EPI_ISL_3226638 | Lighthouse Lab in Milton Keynes | Wellcome Sanger Institute for the COVID-19 Genomics UK (COG-UK) Consortium | The Lighthouse Lab in Milton Keynes and Alex Alderton, Roberto Amato, Jeffrey Barrett, Sonia Goncalves, Ewan Harrison, David K. Jackson, Ian Johnston, Dominic Kwiatkowski, Cordelia Langford, John Sillitoe on behalf of the Wellcome Sanger Institute COVID-19 Surveillance Team |
| EPI_ISL_3226639, EPI_ISL_3226640, EPI_ISL_3226641, EPI_ISL_3226642 | Lighthouse Lab in Alderley Park | Wellcome Sanger Institute for the COVID-19 Genomics UK (COG-UK) Consortium | Jacquelyn Wynn, Mairead Hyland, The Lighthouse Lab in Alderley Park and Alex Alderton, Roberto Amato, Jeffrey Barrett, Sonia Goncalves, Ewan Harrison, David K. Jackson, Ian Johnston, Dominic Kwiatkowski, Cordelia Langford, John Sillitoe on behalf of the Wellcome Sanger Institute COVID-19 Surveillance Team |
| EPI_ISL_3226643 | Lighthouse Lab in Milton Keynes | Wellcome Sanger Institute for the COVID-19 Genomics UK (COG-UK) Consortium | The Lighthouse Lab in Milton Keynes and Alex Alderton, Roberto Amato, Jeffrey Barrett, Sonia Goncalves, Ewan Harrison, David K. Jackson, Ian Johnston, Dominic Kwiatkowski, Cordelia Langford, John Sillitoe on behalf of the Wellcome Sanger Institute COVID-19 Surveillance Team |
| EPI_ISL_3226644 | Lighthouse Lab in Alderley Park | Wellcome Sanger Institute for the COVID-19 Genomics UK (COG-UK) Consortium | Jacquelyn Wynn, Mairead Hyland, The Lighthouse Lab in Alderley Park and Alex Alderton, Roberto Amato, Jeffrey Barrett, Sonia Goncalves, Ewan Harrison, David K. Jackson, Ian Johnston, Dominic Kwiatkowski, Cordelia Langford, John Sillitoe on behalf of the Wellcome Sanger Institute COVID-19 Surveillance Team |
| EPI_ISL_3226645 | Lighthouse Lab in Glasgow | Wellcome Sanger Institute for the COVID-19 Genomics UK (COG-UK) Consortium | Harper VanSteenhouse, Yumi Kasai, David Gray, Carol Clugston, Anna Dominiczak and Alex Alderton, Roberto Amato, Jeffrey Barrett, Sonia Goncalves, Ewan Harrison, David K. Jackson, Ian Johnston, Dominic Kwiatkowski, Cordelia Langford, John Sillitoe on behalf of the Wellcome Sanger Institute COVID-19 Surveillance Team |
| EPI_ISL_3226646, EPI_ISL_3226647, EPI_ISL_3226648 | Lighthouse Lab in Alderley Park | Wellcome Sanger Institute for the COVID-19 Genomics UK (COG-UK) Consortium | Jacquelyn Wynn, Mairead Hyland, The Lighthouse Lab in Alderley Park and Alex Alderton, Roberto Amato, Jeffrey Barrett, Sonia Goncalves, Ewan Harrison, David K. Jackson, Ian Johnston, Dominic Kwiatkowski, Cordelia Langford, John Sillitoe on behalf of the Wellcome Sanger Institute COVID-19 Surveillance Team |
| EPI_ISL_3226649, EPI_ISL_3226650 | Lighthouse Lab in Glasgow | Wellcome Sanger Institute for the COVID-19 Genomics UK (COG-UK) Consortium | Harper VanSteenhouse, Yumi Kasai, David Gray, Carol Clugston, Anna Dominiczak and Alex Alderton, Roberto Amato, Jeffrey Barrett, Sonia Goncalves, Ewan Harrison, David K. Jackson, Ian Johnston, Dominic Kwiatkowski, Cordelia Langford, John Sillitoe on behalf of the Wellcome Sanger Institute COVID-19 Surveillance Team |
| EPI_ISL_3226651 | Lighthouse Lab in Milton Keynes | Wellcome Sanger Institute for the COVID-19 Genomics UK (COG-UK) Consortium | The Lighthouse Lab in Milton Keynes and Alex Alderton, Roberto Amato, Jeffrey Barrett, Sonia Goncalves, Ewan Harrison, David K. Jackson, Ian Johnston, Dominic Kwiatkowski, Cordelia Langford, John Sillitoe on behalf of the Wellcome Sanger Institute COVID-19 Surveillance Team |
| EPI_ISL_3226652 | Lighthouse Lab in Glasgow | Wellcome Sanger Institute for the COVID-19 Genomics UK (COG-UK) Consortium | Harper VanSteenhouse, Yumi Kasai, David Gray, Carol Clugston, Anna Dominiczak and Alex Alderton, Roberto Amato, Jeffrey Barrett, Sonia Goncalves, Ewan Harrison, David K. Jackson, Ian Johnston, Dominic Kwiatkowski, Cordelia Langford, John Sillitoe on behalf of the Wellcome Sanger Institute COVID-19 Surveillance Team |
| EPI_ISL_3226653 | Lighthouse Lab in Milton Keynes | Wellcome Sanger Institute for the COVID-19 Genomics UK (COG-UK) Consortium | The Lighthouse Lab in Milton Keynes and Alex Alderton, Roberto Amato, Jeffrey Barrett, Sonia Goncalves, Ewan Harrison, David K. Jackson, Ian Johnston, Dominic Kwiatkowski, Cordelia Langford, John Sillitoe on behalf of the Wellcome Sanger Institute COVID-19 Surveillance Team |
| EPI_ISL_3226654, EPI_ISL_3226655 | Lighthouse Lab in Alderley Park | Wellcome Sanger Institute for the COVID-19 Genomics UK (COG-UK) Consortium | Jacquelyn Wynn, Mairead Hyland, The Lighthouse Lab in Alderley Park and Alex Alderton, Roberto Amato, Jeffrey Barrett, Sonia Goncalves, Ewan Harrison, David K. Jackson, Ian Johnston, Dominic Kwiatkowski, Cordelia Langford, John Sillitoe on behalf of the Wellcome Sanger Institute COVID-19 Surveillance Team |
| EPI_ISL_3226656 | Lighthouse Lab in Glasgow | Wellcome Sanger Institute for the COVID-19 Genomics UK (COG-UK) Consortium | Harper VanSteenhouse, Yumi Kasai, David Gray, Carol Clugston, Anna Dominiczak and Alex Alderton, Roberto Amato, Jeffrey Barrett, Sonia Goncalves, Ewan Harrison, David K. Jackson, Ian Johnston, Dominic Kwiatkowski, Cordelia Langford, John Sillitoe on behalf of the Wellcome Sanger Institute COVID-19 Surveillance Team |
| EPI_ISL_3226657, EPI_ISL_3226658, EPI_ISL_3226659, EPI_ISL_3226660 | Lighthouse Lab in Alderley Park | Wellcome Sanger Institute for the COVID-19 Genomics UK (COG-UK) Consortium | Jacquelyn Wynn, Mairead Hyland, The Lighthouse Lab in Alderley Park and Alex Alderton, Roberto Amato, Jeffrey Barrett, Sonia Goncalves, Ewan Harrison, David K. Jackson, Ian Johnston, Dominic Kwiatkowski, Cordelia Langford, John Sillitoe on behalf of the Wellcome Sanger Institute COVID-19 Surveillance Team |
| EPI_ISL_3226661, EPI_ISL_3226662 | Lighthouse Lab in Milton Keynes | Wellcome Sanger Institute for the COVID-19 Genomics UK (COG-UK) Consortium | The Lighthouse Lab in Milton Keynes and Alex Alderton, Roberto Amato, Jeffrey Barrett, Sonia Goncalves, Ewan Harrison, David K. Jackson, Ian Johnston, Dominic Kwiatkowski, Cordelia Langford, John Sillitoe on behalf of the Wellcome Sanger Institute COVID-19 Surveillance Team |
| EPI_ISL_3226663 | Lighthouse Lab in Alderley Park | Wellcome Sanger Institute for the COVID-19 Genomics UK (COG-UK) Consortium | Jacquelyn Wynn, Mairead Hyland, The Lighthouse Lab in Alderley Park and Alex Alderton, Roberto Amato, Jeffrey Barrett, Sonia Goncalves, Ewan Harrison, David K. Jackson, Ian Johnston, Dominic Kwiatkowski, Cordelia Langford, John Sillitoe on behalf of the Wellcome Sanger Institute COVID-19 Surveillance Team |
| EPI_ISL_3226664, EPI_ISL_3226665 | Lighthouse Lab in Glasgow | Wellcome Sanger Institute for the COVID-19 Genomics UK (COG-UK) Consortium | Harper VanSteenhouse, Yumi Kasai, David Gray, Carol Clugston, Anna Dominiczak and Alex Alderton, Roberto Amato, Jeffrey Barrett, Sonia Goncalves, Ewan Harrison, David K. Jackson, Ian Johnston, Dominic Kwiatkowski, Cordelia Langford, John Sillitoe on behalf of the Wellcome Sanger Institute COVID-19 Surveillance Team |

| | | | |
|---|---|---|---|
| EPI_ISL_3226666 | Lighthouse Lab in Milton Keynes | Wellcome Sanger Institute for the COVID-19 Genomics UK (COG-UK) Consortium | The Lighthouse Lab in Milton Keynes and Alex Alderton, Roberto Amato, Jeffrey Barrett, Sonia Goncalves, Ewan Harrison, David K. Jackson, Ian Johnston, Dominic Kwiatkowski, Cordelia Langford, John Sillitoe on behalf of the Wellcome Sanger Institute COVID-19 Surveillance Team |
| EPI_ISL_3226667 | Lighthouse Lab in Alderley Park | Wellcome Sanger Institute for the COVID-19 Genomics UK (COG-UK) Consortium | Jacquelyn Wynn, Mairead Hyland, The Lighthouse Lab in Alderley Park and Alex Alderton, Roberto Amato, Jeffrey Barrett, Sonia Goncalves, Ewan Harrison, David K. Jackson, Ian Johnston, Dominic Kwiatkowski, Cordelia Langford, John Sillitoe on behalf of the Wellcome Sanger Institute COVID-19 Surveillance Team |
| EPI_ISL_3226668 | Lighthouse Lab in Milton Keynes | Wellcome Sanger Institute for the COVID-19 Genomics UK (COG-UK) Consortium | The Lighthouse Lab in Milton Keynes and Alex Alderton, Roberto Amato, Jeffrey Barrett, Sonia Goncalves, Ewan Harrison, David K. Jackson, Ian Johnston, Dominic Kwiatkowski, Cordelia Langford, John Sillitoe on behalf of the Wellcome Sanger Institute COVID-19 Surveillance Team |
| EPI_ISL_3226669, EPI_ISL_3226670, EPI_ISL_3226671 | Lighthouse Lab in Alderley Park | Wellcome Sanger Institute for the COVID-19 Genomics UK (COG-UK) Consortium | Jacquelyn Wynn, Mairead Hyland, The Lighthouse Lab in Alderley Park and Alex Alderton, Roberto Amato, Jeffrey Barrett, Sonia Goncalves, Ewan Harrison, David K. Jackson, Ian Johnston, Dominic Kwiatkowski, Cordelia Langford, John Sillitoe on behalf of the Wellcome Sanger Institute COVID-19 Surveillance Team |
| EPI_ISL_3226672 | Lighthouse Lab in Milton Keynes | Wellcome Sanger Institute for the COVID-19 Genomics UK (COG-UK) Consortium | The Lighthouse Lab in Milton Keynes and Alex Alderton, Roberto Amato, Jeffrey Barrett, Sonia Goncalves, Ewan Harrison, David K. Jackson, Ian Johnston, Dominic Kwiatkowski, Cordelia Langford, John Sillitoe on behalf of the Wellcome Sanger Institute COVID-19 Surveillance Team |
| EPI_ISL_3226673 | Lighthouse Lab in Glasgow | Wellcome Sanger Institute for the COVID-19 Genomics UK (COG-UK) Consortium | Harper VanSteenhouse, Yumi Kasai, David Gray, Carol Clugston, Anna Dominiczak and Alex Alderton, Roberto Amato, Jeffrey Barrett, Sonia Goncalves, Ewan Harrison, David K. Jackson, Ian Johnston, Dominic Kwiatkowski, Cordelia Langford, John Sillitoe on behalf of the Wellcome Sanger Institute COVID-19 Surveillance Team |
| EPI_ISL_3226674 | Lighthouse Lab in Alderley Park | Wellcome Sanger Institute for the COVID-19 Genomics UK (COG-UK) Consortium | Jacquelyn Wynn, Mairead Hyland, The Lighthouse Lab in Alderley Park and Alex Alderton, Roberto Amato, Jeffrey Barrett, Sonia Goncalves, Ewan Harrison, David K. Jackson, Ian Johnston, Dominic Kwiatkowski, Cordelia Langford, John Sillitoe on behalf of the Wellcome Sanger Institute COVID-19 Surveillance Team |
| EPI_ISL_3226675, EPI_ISL_3226676 | Lighthouse Lab in Glasgow | Wellcome Sanger Institute for the COVID-19 Genomics UK (COG-UK) Consortium | Harper VanSteenhouse, Yumi Kasai, David Gray, Carol Clugston, Anna Dominiczak and Alex Alderton, Roberto Amato, Jeffrey Barrett, Sonia Goncalves, Ewan Harrison, David K. Jackson, Ian Johnston, Dominic Kwiatkowski, Cordelia Langford, John Sillitoe on behalf of the Wellcome Sanger Institute COVID-19 Surveillance Team |
| EPI_ISL_3226677, EPI_ISL_3226678, EPI_ISL_3226679 | Lighthouse Lab in Alderley Park | Wellcome Sanger Institute for the COVID-19 Genomics UK (COG-UK) Consortium | Jacquelyn Wynn, Mairead Hyland, The Lighthouse Lab in Alderley Park and Alex Alderton, Roberto Amato, Jeffrey Barrett, Sonia Goncalves, Ewan Harrison, David K. Jackson, Ian Johnston, Dominic Kwiatkowski, Cordelia Langford, John Sillitoe on behalf of the Wellcome Sanger Institute COVID-19 Surveillance Team |
| EPI_ISL_3226680 | Lighthouse Lab in Milton Keynes | Wellcome Sanger Institute for the COVID-19 Genomics UK (COG-UK) Consortium | The Lighthouse Lab in Milton Keynes and Alex Alderton, Roberto Amato, Jeffrey Barrett, Sonia Goncalves, Ewan Harrison, David K. Jackson, Ian Johnston, Dominic Kwiatkowski, Cordelia Langford, John Sillitoe on behalf of the Wellcome Sanger Institute COVID-19 Surveillance Team |
| EPI_ISL_3226681, EPI_ISL_3226682 | Lighthouse Lab in Alderley Park | Wellcome Sanger Institute for the COVID-19 Genomics UK (COG-UK) Consortium | Jacquelyn Wynn, Mairead Hyland, The Lighthouse Lab in Alderley Park and Alex Alderton, Roberto Amato, Jeffrey Barrett, Sonia Goncalves, Ewan Harrison, David K. Jackson, Ian Johnston, Dominic Kwiatkowski, Cordelia Langford, John Sillitoe on behalf of the Wellcome Sanger Institute COVID-19 Surveillance Team |
| EPI_ISL_3226683 | Lighthouse Lab in Milton Keynes | Wellcome Sanger Institute for the COVID-19 Genomics UK (COG-UK) Consortium | The Lighthouse Lab in Milton Keynes and Alex Alderton, Roberto Amato, Jeffrey Barrett, Sonia Goncalves, Ewan Harrison, David K. Jackson, Ian Johnston, Dominic Kwiatkowski, Cordelia Langford, John Sillitoe on behalf of the Wellcome Sanger Institute COVID-19 Surveillance Team |
| EPI_ISL_3226684 | Lighthouse Lab in Glasgow | Wellcome Sanger Institute for the COVID-19 Genomics UK (COG-UK) Consortium | Harper VanSteenhouse, Yumi Kasai, David Gray, Carol Clugston, Anna Dominiczak and Alex Alderton, Roberto Amato, Jeffrey Barrett, Sonia Goncalves, Ewan Harrison, David K. Jackson, Ian Johnston, Dominic Kwiatkowski, Cordelia Langford, John Sillitoe on behalf of the Wellcome Sanger Institute COVID-19 Surveillance Team |
| EPI_ISL_3226685 | Lighthouse Lab in Alderley Park | Wellcome Sanger Institute for the COVID-19 Genomics UK (COG-UK) Consortium | Jacquelyn Wynn, Mairead Hyland, The Lighthouse Lab in Alderley Park and Alex Alderton, Roberto Amato, Jeffrey Barrett, Sonia Goncalves, Ewan Harrison, David K. Jackson, Ian Johnston, Dominic Kwiatkowski, Cordelia Langford, John Sillitoe on behalf of the Wellcome Sanger Institute COVID-19 Surveillance Team |
| EPI_ISL_3226686, EPI_ISL_3226687 | Lighthouse Lab in Glasgow | Wellcome Sanger Institute for the COVID-19 Genomics UK (COG-UK) Consortium | Harper VanSteenhouse, Yumi Kasai, David Gray, Carol Clugston, Anna Dominiczak and Alex Alderton, Roberto Amato, Jeffrey Barrett, Sonia Goncalves, Ewan Harrison, David K. Jackson, Ian Johnston, Dominic Kwiatkowski, Cordelia Langford, John Sillitoe on behalf of the Wellcome Sanger Institute COVID-19 Surveillance Team |
| EPI_ISL_3226688 | Lighthouse Lab in Milton Keynes | Wellcome Sanger Institute for the COVID-19 Genomics UK (COG-UK) Consortium | The Lighthouse Lab in Milton Keynes and Alex Alderton, Roberto Amato, Jeffrey Barrett, Sonia Goncalves, Ewan Harrison, David K. Jackson, Ian Johnston, Dominic Kwiatkowski, Cordelia Langford, John Sillitoe on behalf of the Wellcome Sanger Institute COVID-19 Surveillance Team |
| EPI_ISL_3226689 | Lighthouse Lab in Glasgow | Wellcome Sanger Institute for the COVID-19 Genomics UK (COG-UK) Consortium | Harper VanSteenhouse, Yumi Kasai, David Gray, Carol Clugston, Anna Dominiczak and Alex Alderton, Roberto Amato, Jeffrey Barrett, Sonia Goncalves, Ewan Harrison, David K. Jackson, Ian Johnston, Dominic Kwiatkowski, Cordelia Langford, John Sillitoe on behalf of the Wellcome Sanger Institute COVID-19 Surveillance Team |
| EPI_ISL_3226690 | Lighthouse Lab in Milton Keynes | Wellcome Sanger Institute for the COVID-19 Genomics UK (COG-UK) Consortium | The Lighthouse Lab in Milton Keynes and Alex Alderton, Roberto Amato, Jeffrey Barrett, Sonia Goncalves, Ewan Harrison, David K. Jackson, Ian Johnston, Dominic Kwiatkowski, Cordelia Langford, John Sillitoe on behalf of the Wellcome Sanger Institute COVID-19 Surveillance Team |
| EPI_ISL_3226691, EPI_ISL_3226692 | Lighthouse Lab in Alderley Park | Wellcome Sanger Institute for the COVID-19 Genomics UK (COG-UK) Consortium | Jacquelyn Wynn, Mairead Hyland, The Lighthouse Lab in Alderley Park and Alex Alderton, Roberto Amato, Jeffrey Barrett, Sonia Goncalves, Ewan Harrison, David K. Jackson, Ian Johnston, Dominic Kwiatkowski, Cordelia Langford, John Sillitoe on behalf of the Wellcome Sanger Institute COVID-19 Surveillance Team |
| EPI_ISL_3226693 | Lighthouse Lab in Glasgow | Wellcome Sanger Institute for the COVID-19 Genomics UK (COG-UK) Consortium | Harper VanSteenhouse, Yumi Kasai, David Gray, Carol Clugston, Anna Dominiczak and Alex Alderton, Roberto Amato, Jeffrey Barrett, Sonia Goncalves, Ewan Harrison, David K. Jackson, Ian Johnston, Dominic Kwiatkowski, Cordelia Langford, John Sillitoe on behalf of the Wellcome Sanger Institute COVID-19 Surveillance Team |
| EPI_ISL_3226694, EPI_ISL_3226695, EPI_ISL_3226696, EPI_ISL_3226697, EPI_ISL_3226698 | Lighthouse Lab in Alderley Park | Wellcome Sanger Institute for the COVID-19 Genomics UK (COG-UK) Consortium | Jacquelyn Wynn, Mairead Hyland, The Lighthouse Lab in Alderley Park and Alex Alderton, Roberto Amato, Jeffrey Barrett, Sonia Goncalves, Ewan Harrison, David K. Jackson, Ian Johnston, Dominic Kwiatkowski, Cordelia Langford, John Sillitoe on behalf of the Wellcome Sanger Institute COVID-19 Surveillance Team |
| EPI_ISL_3226699 | Lighthouse Lab in Glasgow | Wellcome Sanger Institute for the COVID-19 Genomics UK (COG-UK) Consortium | Harper VanSteenhouse, Yumi Kasai, David Gray, Carol Clugston, Anna Dominiczak and Alex Alderton, Roberto Amato, Jeffrey Barrett, Sonia Goncalves, Ewan Harrison, David K. Jackson, Ian Johnston, Dominic Kwiatkowski, Cordelia Langford, John Sillitoe on behalf of the Wellcome Sanger Institute COVID-19 Surveillance Team |
| EPI_ISL_3226700, EPI_ISL_3226701, EPI_ISL_3226702, EPI_ISL_3226703 | Lighthouse Lab in Alderley Park | Wellcome Sanger Institute for the COVID-19 Genomics UK (COG-UK) Consortium | Jacquelyn Wynn, Mairead Hyland, The Lighthouse Lab in Alderley Park and Alex Alderton, Roberto Amato, Jeffrey Barrett, Sonia Goncalves, Ewan Harrison, David K. Jackson, Ian Johnston, Dominic Kwiatkowski, Cordelia Langford, John Sillitoe on behalf of the Wellcome Sanger Institute COVID-19 Surveillance Team |
| EPI_ISL_3226704 | Lighthouse Lab in Milton Keynes | Wellcome Sanger Institute for the COVID-19 Genomics UK (COG-UK) Consortium | The Lighthouse Lab in Milton Keynes and Alex Alderton, Roberto Amato, Jeffrey Barrett, Sonia Goncalves, Ewan Harrison, David K. Jackson, Ian Johnston, Dominic Kwiatkowski, Cordelia Langford, John Sillitoe on behalf of the Wellcome Sanger Institute COVID-19 Surveillance Team |
| EPI_ISL_3226705, EPI_ISL_3226706, EPI_ISL_3226707, EPI_ISL_3226708 | Lighthouse Lab in Alderley Park | Wellcome Sanger Institute for the COVID-19 Genomics UK (COG-UK) Consortium | Jacquelyn Wynn, Mairead Hyland, The Lighthouse Lab in Alderley Park and Alex Alderton, Roberto Amato, Jeffrey Barrett, Sonia Goncalves, Ewan Harrison, David K. Jackson, Ian Johnston, Dominic Kwiatkowski, Cordelia Langford, John Sillitoe on behalf of the Wellcome Sanger Institute COVID-19 Surveillance Team |
| EPI_ISL_3226709 | Lighthouse Lab in Milton Keynes | Wellcome Sanger Institute for the COVID-19 Genomics UK (COG-UK) Consortium | The Lighthouse Lab in Milton Keynes and Alex Alderton, Roberto Amato, Jeffrey Barrett, Sonia Goncalves, Ewan Harrison, David K. Jackson, Ian Johnston, Dominic Kwiatkowski, Cordelia Langford, John Sillitoe on behalf of the Wellcome Sanger Institute COVID-19 Surveillance Team |
| EPI_ISL_3226710, EPI_ISL_3226711 | Lighthouse Lab in Glasgow | Wellcome Sanger Institute for the COVID-19 Genomics UK (COG-UK) Consortium | Harper VanSteenhouse, Yumi Kasai, David Gray, Carol Clugston, Anna Dominiczak and Alex Alderton, Roberto Amato, Jeffrey Barrett, Sonia Goncalves, Ewan Harrison, David K. Jackson, Ian Johnston, Dominic Kwiatkowski, Cordelia Langford, John Sillitoe on behalf of the Wellcome Sanger Institute COVID-19 Surveillance Team |
| EPI_ISL_3226712 | Lighthouse Lab in Milton Keynes | Wellcome Sanger Institute for the COVID-19 Genomics UK (COG-UK) Consortium | The Lighthouse Lab in Milton Keynes and Alex Alderton, Roberto Amato, Jeffrey Barrett, Sonia Goncalves, Ewan Harrison, David K. Jackson, Ian Johnston, Dominic Kwiatkowski, Cordelia Langford, John Sillitoe on behalf of the Wellcome Sanger Institute COVID-19 Surveillance Team |
| EPI_ISL_3226713, EPI_ISL_3226714, EPI_ISL_3226715, EPI_ISL_3226716 | Lighthouse Lab in Alderley Park | Wellcome Sanger Institute for the COVID-19 Genomics UK (COG-UK) Consortium | Jacquelyn Wynn, Mairead Hyland, The Lighthouse Lab in Alderley Park and Alex Alderton, Roberto Amato, Jeffrey Barrett, Sonia Goncalves, Ewan Harrison, David K. Jackson, Ian Johnston, Dominic Kwiatkowski, Cordelia Langford, John Sillitoe on behalf of the Wellcome Sanger Institute COVID-19 Surveillance Team |
| EPI_ISL_3226717 | Lighthouse Lab in Glasgow | Wellcome Sanger Institute for the COVID-19 Genomics UK (COG-UK) Consortium | Harper VanSteenhouse, Yumi Kasai, David Gray, Carol Clugston, Anna Dominiczak and Alex Alderton, Roberto Amato, Jeffrey Barrett, Sonia Goncalves, Ewan Harrison, David K. Jackson, Ian Johnston, Dominic Kwiatkowski, Cordelia Langford, John Sillitoe on behalf of the Wellcome Sanger Institute COVID-19 Surveillance Team |
| EPI_ISL_3226718 | Lighthouse Lab in Alderley Park | Wellcome Sanger Institute for the COVID-19 Genomics UK (COG-UK) Consortium | Jacquelyn Wynn, Mairead Hyland, The Lighthouse Lab in Alderley Park and Alex Alderton, Roberto Amato, Jeffrey Barrett, Sonia Goncalves, Ewan Harrison, David K. Jackson, Ian Johnston, Dominic Kwiatkowski, Cordelia Langford, John Sillitoe on behalf of the Wellcome Sanger Institute COVID-19 Surveillance Team |
| EPI_ISL_3226719 | Lighthouse Lab in Glasgow | Wellcome Sanger Institute for the COVID-19 Genomics UK (COG-UK) Consortium | Harper VanSteenhouse, Yumi Kasai, David Gray, Carol Clugston, Anna Dominiczak and Alex Alderton, Roberto Amato, Jeffrey Barrett, Sonia Goncalves, Ewan Harrison, David K. Jackson, Ian Johnston, Dominic Kwiatkowski, Cordelia Langford, John Sillitoe on behalf of the Wellcome Sanger Institute COVID-19 Surveillance Team |
| EPI_ISL_3226720 | Lighthouse Lab in Alderley Park | Wellcome Sanger Institute for the COVID-19 Genomics UK (COG-UK) Consortium | Jacquelyn Wynn, Mairead Hyland, The Lighthouse Lab in Alderley Park and Alex Alderton, Roberto Amato, Jeffrey Barrett, Sonia Goncalves, Ewan Harrison, David K. Jackson, Ian Johnston, Dominic Kwiatkowski, Cordelia Langford, John Sillitoe on behalf of the Wellcome Sanger Institute COVID-19 Surveillance Team |
| EPI_ISL_3226721, EPI_ISL_3226722 | Lighthouse Lab in Glasgow | Wellcome Sanger Institute for the COVID-19 Genomics UK (COG-UK) Consortium | Harper VanSteenhouse, Yumi Kasai, David Gray, Carol Clugston, Anna Dominiczak and Alex Alderton, Roberto Amato, Jeffrey Barrett, Sonia Goncalves, Ewan Harrison, David K. Jackson, Ian Johnston, Dominic Kwiatkowski, Cordelia Langford, John Sillitoe on behalf of the Wellcome Sanger Institute COVID-19 Surveillance Team |
| EPI_ISL_3226723 | Lighthouse Lab in Milton Keynes | Wellcome Sanger Institute for the COVID-19 Genomics UK (COG-UK) Consortium | The Lighthouse Lab in Milton Keynes and Alex Alderton, Roberto Amato, Jeffrey Barrett, Sonia Goncalves, Ewan Harrison, David K. Jackson, Ian Johnston, Dominic Kwiatkowski, Cordelia Langford, John Sillitoe on behalf of the Wellcome Sanger Institute COVID-19 Surveillance Team |
| EPI_ISL_3226724 | Lighthouse Lab in Alderley Park | Wellcome Sanger Institute for the COVID-19 Genomics UK (COG-UK) Consortium | Jacquelyn Wynn, Mairead Hyland, The Lighthouse Lab in Alderley Park and Alex Alderton, Roberto Amato, Jeffrey Barrett, Sonia Goncalves, Ewan Harrison, David K. Jackson, Ian Johnston, Dominic Kwiatkowski, Cordelia Langford, John Sillitoe on behalf of the Wellcome Sanger Institute COVID-19 Surveillance Team |
| EPI_ISL_3226725, EPI_ISL_3226726 | Lighthouse Lab in Milton Keynes | Wellcome Sanger Institute for the COVID-19 Genomics UK (COG-UK) Consortium | The Lighthouse Lab in Milton Keynes and Alex Alderton, Roberto Amato, Jeffrey Barrett, Sonia Goncalves, Ewan Harrison, David K. Jackson, Ian Johnston, Dominic Kwiatkowski, Cordelia Langford, John Sillitoe on behalf of the Wellcome Sanger Institute COVID-19 Surveillance Team |
| EPI_ISL_3226727, EPI_ISL_3226728 | Lighthouse Lab in Alderley Park | Wellcome Sanger Institute for the COVID-19 Genomics UK (COG-UK) Consortium | Jacquelyn Wynn, Mairead Hyland, The Lighthouse Lab in Alderley Park and Alex Alderton, Roberto Amato, Jeffrey Barrett, Sonia Goncalves, Ewan Harrison, David K. Jackson, Ian Johnston, Dominic Kwiatkowski, Cordelia Langford, John Sillitoe on behalf of the Wellcome Sanger Institute COVID-19 Surveillance Team |
| EPI_ISL_3226729 | Lighthouse Lab in Milton Keynes | Wellcome Sanger Institute for the COVID-19 Genomics UK (COG-UK) Consortium | The Lighthouse Lab in Milton Keynes and Alex Alderton, Roberto Amato, Jeffrey Barrett, Sonia Goncalves, Ewan Harrison, David K. Jackson, Ian Johnston, Dominic Kwiatkowski, Cordelia Langford, John Sillitoe on behalf of the Wellcome Sanger Institute COVID-19 Surveillance Team |
| EPI_ISL_3226730 | Lighthouse Lab in Glasgow | Wellcome Sanger Institute for the COVID-19 Genomics UK (COG-UK) Consortium | Harper VanSteenhouse, Yumi Kasai, David Gray, Carol Clugston, Anna Dominiczak and Alex Alderton, Roberto Amato, Jeffrey Barrett, Sonia Goncalves, Ewan Harrison, David K. Jackson, Ian Johnston, Dominic Kwiatkowski, Cordelia Langford, John Sillitoe on behalf of the Wellcome Sanger Institute COVID-19 Surveillance Team |
| EPI_ISL_3226731 | Lighthouse Lab in Alderley Park | Wellcome Sanger Institute for the COVID-19 Genomics UK (COG-UK) Consortium | Jacquelyn Wynn, Mairead Hyland, The Lighthouse Lab in Alderley Park and Alex Alderton, Roberto Amato, Jeffrey Barrett, Sonia Goncalves, Ewan Harrison, David K. Jackson, Ian Johnston, Dominic Kwiatkowski, Cordelia Langford, John Sillitoe on behalf of the Wellcome Sanger Institute COVID-19 Surveillance Team |

| | | | |
|---|---|---|---|
| EPI_ISL_3226732 | Lighthouse Lab in Milton Keynes | Wellcome Sanger Institute for the COVID-19 Genomics UK (COG-UK) Consortium | The Lighthouse Lab in Milton Keynes and Alex Alderton, Roberto Amato, Jeffrey Barrett, Sonia Goncalves, Ewan Harrison, David K. Jackson, Ian Johnston, Dominic Kwiatkowski, Cordelia Langford, John Sillitoe on behalf of the Wellcome Sanger Institute COVID-19 Surveillance Team |
| EPI_ISL_3226733, EPI_ISL_3226734, EPI_ISL_3226735 | Lighthouse Lab in Alderley Park | Wellcome Sanger Institute for the COVID-19 Genomics UK (COG-UK) Consortium | Jacquelyn Wynn, Mairead Hyland, The Lighthouse Lab in Alderley Park and Alex Alderton, Roberto Amato, Jeffrey Barrett, Sonia Goncalves, Ewan Harrison, David K. Jackson, Ian Johnston, Dominic Kwiatkowski, Cordelia Langford, John Sillitoe on behalf of the Wellcome Sanger Institute COVID-19 Surveillance Team |
| EPI_ISL_3226736 | Lighthouse Lab in Milton Keynes | Wellcome Sanger Institute for the COVID-19 Genomics UK (COG-UK) Consortium | The Lighthouse Lab in Milton Keynes and Alex Alderton, Roberto Amato, Jeffrey Barrett, Sonia Goncalves, Ewan Harrison, David K. Jackson, Ian Johnston, Dominic Kwiatkowski, Cordelia Langford, John Sillitoe on behalf of the Wellcome Sanger Institute COVID-19 Surveillance Team |
| EPI_ISL_3226737 | Lighthouse Lab in Alderley Park | Wellcome Sanger Institute for the COVID-19 Genomics UK (COG-UK) Consortium | Jacquelyn Wynn, Mairead Hyland, The Lighthouse Lab in Alderley Park and Alex Alderton, Roberto Amato, Jeffrey Barrett, Sonia Goncalves, Ewan Harrison, David K. Jackson, Ian Johnston, Dominic Kwiatkowski, Cordelia Langford, John Sillitoe on behalf of the Wellcome Sanger Institute COVID-19 Surveillance Team |
| EPI_ISL_3226738 | Lighthouse Lab in Glasgow | Wellcome Sanger Institute for the COVID-19 Genomics UK (COG-UK) Consortium | Harper VanSteenhouse, Yumi Kasai, David Gray, Carol Clugston, Anna Dominiczak and Alex Alderton, Roberto Amato, Jeffrey Barrett, Sonia Goncalves, Ewan Harrison, David K. Jackson, Ian Johnston, Dominic Kwiatkowski, Cordelia Langford, John Sillitoe on behalf of the Wellcome Sanger Institute COVID-19 Surveillance Team |
| EPI_ISL_3226739, EPI_ISL_3226740 | Lighthouse Lab in Milton Keynes | Wellcome Sanger Institute for the COVID-19 Genomics UK (COG-UK) Consortium | The Lighthouse Lab in Milton Keynes and Alex Alderton, Roberto Amato, Jeffrey Barrett, Sonia Goncalves, Ewan Harrison, David K. Jackson, Ian Johnston, Dominic Kwiatkowski, Cordelia Langford, John Sillitoe on behalf of the Wellcome Sanger Institute COVID-19 Surveillance Team |
| EPI_ISL_3226741 | Lighthouse Lab in Glasgow | Wellcome Sanger Institute for the COVID-19 Genomics UK (COG-UK) Consortium | Harper VanSteenhouse, Yumi Kasai, David Gray, Carol Clugston, Anna Dominiczak and Alex Alderton, Roberto Amato, Jeffrey Barrett, Sonia Goncalves, Ewan Harrison, David K. Jackson, Ian Johnston, Dominic Kwiatkowski, Cordelia Langford, John Sillitoe on behalf of the Wellcome Sanger Institute COVID-19 Surveillance Team |
| EPI_ISL_3226742, EPI_ISL_3226743 | Lighthouse Lab in Alderley Park | Wellcome Sanger Institute for the COVID-19 Genomics UK (COG-UK) Consortium | Jacquelyn Wynn, Mairead Hyland, The Lighthouse Lab in Alderley Park and Alex Alderton, Roberto Amato, Jeffrey Barrett, Sonia Goncalves, Ewan Harrison, David K. Jackson, Ian Johnston, Dominic Kwiatkowski, Cordelia Langford, John Sillitoe on behalf of the Wellcome Sanger Institute COVID-19 Surveillance Team |
| EPI_ISL_3226744 | Lighthouse Lab in Milton Keynes | Wellcome Sanger Institute for the COVID-19 Genomics UK (COG-UK) Consortium | The Lighthouse Lab in Milton Keynes and Alex Alderton, Roberto Amato, Jeffrey Barrett, Sonia Goncalves, Ewan Harrison, David K. Jackson, Ian Johnston, Dominic Kwiatkowski, Cordelia Langford, John Sillitoe on behalf of the Wellcome Sanger Institute COVID-19 Surveillance Team |
| EPI_ISL_3226745, EPI_ISL_3226746, EPI_ISL_3226747 | Lighthouse Lab in Alderley Park | Wellcome Sanger Institute for the COVID-19 Genomics UK (COG-UK) Consortium | Jacquelyn Wynn, Mairead Hyland, The Lighthouse Lab in Alderley Park and Alex Alderton, Roberto Amato, Jeffrey Barrett, Sonia Goncalves, Ewan Harrison, David K. Jackson, Ian Johnston, Dominic Kwiatkowski, Cordelia Langford, John Sillitoe on behalf of the Wellcome Sanger Institute COVID-19 Surveillance Team |
| EPI_ISL_3226748 | Lighthouse Lab in Glasgow | Wellcome Sanger Institute for the COVID-19 Genomics UK (COG-UK) Consortium | Harper VanSteenhouse, Yumi Kasai, David Gray, Carol Clugston, Anna Dominiczak and Alex Alderton, Roberto Amato, Jeffrey Barrett, Sonia Goncalves, Ewan Harrison, David K. Jackson, Ian Johnston, Dominic Kwiatkowski, Cordelia Langford, John Sillitoe on behalf of the Wellcome Sanger Institute COVID-19 Surveillance Team |
| EPI_ISL_3226749, EPI_ISL_3226750 | Lighthouse Lab in Alderley Park | Wellcome Sanger Institute for the COVID-19 Genomics UK (COG-UK) Consortium | Jacquelyn Wynn, Mairead Hyland, The Lighthouse Lab in Alderley Park and Alex Alderton, Roberto Amato, Jeffrey Barrett, Sonia Goncalves, Ewan Harrison, David K. Jackson, Ian Johnston, Dominic Kwiatkowski, Cordelia Langford, John Sillitoe on behalf of the Wellcome Sanger Institute COVID-19 Surveillance Team |
| EPI_ISL_3226751 | Lighthouse Lab in Milton Keynes | Wellcome Sanger Institute for the COVID-19 Genomics UK (COG-UK) Consortium | The Lighthouse Lab in Milton Keynes and Alex Alderton, Roberto Amato, Jeffrey Barrett, Sonia Goncalves, Ewan Harrison, David K. Jackson, Ian Johnston, Dominic Kwiatkowski, Cordelia Langford, John Sillitoe on behalf of the Wellcome Sanger Institute COVID-19 Surveillance Team |
| EPI_ISL_3226752, EPI_ISL_3226753 | Lighthouse Lab in Alderley Park | Wellcome Sanger Institute for the COVID-19 Genomics UK (COG-UK) Consortium | Jacquelyn Wynn, Mairead Hyland, The Lighthouse Lab in Alderley Park and Alex Alderton, Roberto Amato, Jeffrey Barrett, Sonia Goncalves, Ewan Harrison, David K. Jackson, Ian Johnston, Dominic Kwiatkowski, Cordelia Langford, John Sillitoe on behalf of the Wellcome Sanger Institute COVID-19 Surveillance Team |
| EPI_ISL_3226754, EPI_ISL_3226755 | Lighthouse Lab in Glasgow | Wellcome Sanger Institute for the COVID-19 Genomics UK (COG-UK) Consortium | Harper VanSteenhouse, Yumi Kasai, David Gray, Carol Clugston, Anna Dominiczak and Alex Alderton, Roberto Amato, Jeffrey Barrett, Sonia Goncalves, Ewan Harrison, David K. Jackson, Ian Johnston, Dominic Kwiatkowski, Cordelia Langford, John Sillitoe on behalf of the Wellcome Sanger Institute COVID-19 Surveillance Team |
| EPI_ISL_3226756, EPI_ISL_3226757, EPI_ISL_3226758, EPI_ISL_3226759, EPI_ISL_3226760 | Lighthouse Lab in Alderley Park | Wellcome Sanger Institute for the COVID-19 Genomics UK (COG-UK) Consortium | Jacquelyn Wynn, Mairead Hyland, The Lighthouse Lab in Alderley Park and Alex Alderton, Roberto Amato, Jeffrey Barrett, Sonia Goncalves, Ewan Harrison, David K. Jackson, Ian Johnston, Dominic Kwiatkowski, Cordelia Langford, John Sillitoe on behalf of the Wellcome Sanger Institute COVID-19 Surveillance Team |
| EPI_ISL_3226761 | Lighthouse Lab in Glasgow | Wellcome Sanger Institute for the COVID-19 Genomics UK (COG-UK) Consortium | Harper VanSteenhouse, Yumi Kasai, David Gray, Carol Clugston, Anna Dominiczak and Alex Alderton, Roberto Amato, Jeffrey Barrett, Sonia Goncalves, Ewan Harrison, David K. Jackson, Ian Johnston, Dominic Kwiatkowski, Cordelia Langford, John Sillitoe on behalf of the Wellcome Sanger Institute COVID-19 Surveillance Team |
| EPI_ISL_3226762 | Lighthouse Lab in Alderley Park | Wellcome Sanger Institute for the COVID-19 Genomics UK (COG-UK) Consortium | Jacquelyn Wynn, Mairead Hyland, The Lighthouse Lab in Alderley Park and Alex Alderton, Roberto Amato, Jeffrey Barrett, Sonia Goncalves, Ewan Harrison, David K. Jackson, Ian Johnston, Dominic Kwiatkowski, Cordelia Langford, John Sillitoe on behalf of the Wellcome Sanger Institute COVID-19 Surveillance Team |
| EPI_ISL_3226763, EPI_ISL_3226764 | Lighthouse Lab in Glasgow | Wellcome Sanger Institute for the COVID-19 Genomics UK (COG-UK) Consortium | Harper VanSteenhouse, Yumi Kasai, David Gray, Carol Clugston, Anna Dominiczak and Alex Alderton, Roberto Amato, Jeffrey Barrett, Sonia Goncalves, Ewan Harrison, David K. Jackson, Ian Johnston, Dominic Kwiatkowski, Cordelia Langford, John Sillitoe on behalf of the Wellcome Sanger Institute COVID-19 Surveillance Team |
| EPI_ISL_3226765, EPI_ISL_3226766 | Lighthouse Lab in Alderley Park | Wellcome Sanger Institute for the COVID-19 Genomics UK (COG-UK) Consortium | Jacquelyn Wynn, Mairead Hyland, The Lighthouse Lab in Alderley Park and Alex Alderton, Roberto Amato, Jeffrey Barrett, Sonia Goncalves, Ewan Harrison, David K. Jackson, Ian Johnston, Dominic Kwiatkowski, Cordelia Langford, John Sillitoe on behalf of the Wellcome Sanger Institute COVID-19 Surveillance Team |
| EPI_ISL_3226767 | Lighthouse Lab in Glasgow | Wellcome Sanger Institute for the COVID-19 Genomics UK (COG-UK) Consortium | Harper VanSteenhouse, Yumi Kasai, David Gray, Carol Clugston, Anna Dominiczak and Alex Alderton, Roberto Amato, Jeffrey Barrett, Sonia Goncalves, Ewan Harrison, David K. Jackson, Ian Johnston, Dominic Kwiatkowski, Cordelia Langford, John Sillitoe on behalf of the Wellcome Sanger Institute COVID-19 Surveillance Team |
| EPI_ISL_3226768 | Lighthouse Lab in Alderley Park | Wellcome Sanger Institute for the COVID-19 Genomics UK (COG-UK) Consortium | Jacquelyn Wynn, Mairead Hyland, The Lighthouse Lab in Alderley Park and Alex Alderton, Roberto Amato, Jeffrey Barrett, Sonia Goncalves, Ewan Harrison, David K. Jackson, Ian Johnston, Dominic Kwiatkowski, Cordelia Langford, John Sillitoe on behalf of the Wellcome Sanger Institute COVID-19 Surveillance Team |
| EPI_ISL_3226769 | Lighthouse Lab in Milton Keynes | Wellcome Sanger Institute for the COVID-19 Genomics UK (COG-UK) Consortium | The Lighthouse Lab in Milton Keynes and Alex Alderton, Roberto Amato, Jeffrey Barrett, Sonia Goncalves, Ewan Harrison, David K. Jackson, Ian Johnston, Dominic Kwiatkowski, Cordelia Langford, John Sillitoe on behalf of the Wellcome Sanger Institute COVID-19 Surveillance Team |
| EPI_ISL_3226770 | Lighthouse Lab in Alderley Park | Wellcome Sanger Institute for the COVID-19 Genomics UK (COG-UK) Consortium | Jacquelyn Wynn, Mairead Hyland, The Lighthouse Lab in Alderley Park and Alex Alderton, Roberto Amato, Jeffrey Barrett, Sonia Goncalves, Ewan Harrison, David K. Jackson, Ian Johnston, Dominic Kwiatkowski, Cordelia Langford, John Sillitoe on behalf of the Wellcome Sanger Institute COVID-19 Surveillance Team |
| EPI_ISL_3226771 | Lighthouse Lab in Milton Keynes | Wellcome Sanger Institute for the COVID-19 Genomics UK (COG-UK) Consortium | The Lighthouse Lab in Milton Keynes and Alex Alderton, Roberto Amato, Jeffrey Barrett, Sonia Goncalves, Ewan Harrison, David K. Jackson, Ian Johnston, Dominic Kwiatkowski, Cordelia Langford, John Sillitoe on behalf of the Wellcome Sanger Institute COVID-19 Surveillance Team |
| EPI_ISL_3226772, EPI_ISL_3226773, EPI_ISL_3226774, EPI_ISL_3226775, EPI_ISL_3226776, EPI_ISL_3226777 | Lighthouse Lab in Alderley Park | Wellcome Sanger Institute for the COVID-19 Genomics UK (COG-UK) Consortium | Jacquelyn Wynn, Mairead Hyland, The Lighthouse Lab in Alderley Park and Alex Alderton, Roberto Amato, Jeffrey Barrett, Sonia Goncalves, Ewan Harrison, David K. Jackson, Ian Johnston, Dominic Kwiatkowski, Cordelia Langford, John Sillitoe on behalf of the Wellcome Sanger Institute COVID-19 Surveillance Team |
| EPI_ISL_3226778 | Lighthouse Lab in Milton Keynes | Wellcome Sanger Institute for the COVID-19 Genomics UK (COG-UK) Consortium | The Lighthouse Lab in Milton Keynes and Alex Alderton, Roberto Amato, Jeffrey Barrett, Sonia Goncalves, Ewan Harrison, David K. Jackson, Ian Johnston, Dominic Kwiatkowski, Cordelia Langford, John Sillitoe on behalf of the Wellcome Sanger Institute COVID-19 Surveillance Team |
| EPI_ISL_3226779, EPI_ISL_3226780, EPI_ISL_3226781 | Lighthouse Lab in Glasgow | Wellcome Sanger Institute for the COVID-19 Genomics UK (COG-UK) Consortium | Harper VanSteenhouse, Yumi Kasai, David Gray, Carol Clugston, Anna Dominiczak and Alex Alderton, Roberto Amato, Jeffrey Barrett, Sonia Goncalves, Ewan Harrison, David K. Jackson, Ian Johnston, Dominic Kwiatkowski, Cordelia Langford, John Sillitoe on behalf of the Wellcome Sanger Institute COVID-19 Surveillance Team |
| EPI_ISL_3226782 | Lighthouse Lab in Milton Keynes | Wellcome Sanger Institute for the COVID-19 Genomics UK (COG-UK) Consortium | The Lighthouse Lab in Milton Keynes and Alex Alderton, Roberto Amato, Jeffrey Barrett, Sonia Goncalves, Ewan Harrison, David K. Jackson, Ian Johnston, Dominic Kwiatkowski, Cordelia Langford, John Sillitoe on behalf of the Wellcome Sanger Institute COVID-19 Surveillance Team |
| EPI_ISL_3226783 | Lighthouse Lab in Alderley Park | Wellcome Sanger Institute for the COVID-19 Genomics UK (COG-UK) Consortium | Jacquelyn Wynn, Mairead Hyland, The Lighthouse Lab in Alderley Park and Alex Alderton, Roberto Amato, Jeffrey Barrett, Sonia Goncalves, Ewan Harrison, David K. Jackson, Ian Johnston, Dominic Kwiatkowski, Cordelia Langford, John Sillitoe on behalf of the Wellcome Sanger Institute COVID-19 Surveillance Team |
| EPI_ISL_3226784 | Lighthouse Lab in Glasgow | Wellcome Sanger Institute for the COVID-19 Genomics UK (COG-UK) Consortium | Harper VanSteenhouse, Yumi Kasai, David Gray, Carol Clugston, Anna Dominiczak and Alex Alderton, Roberto Amato, Jeffrey Barrett, Sonia Goncalves, Ewan Harrison, David K. Jackson, Ian Johnston, Dominic Kwiatkowski, Cordelia Langford, John Sillitoe on behalf of the Wellcome Sanger Institute COVID-19 Surveillance Team |
| EPI_ISL_3226785 | Lighthouse Lab in Alderley Park | Wellcome Sanger Institute for the COVID-19 Genomics UK (COG-UK) Consortium | Jacquelyn Wynn, Mairead Hyland, The Lighthouse Lab in Alderley Park and Alex Alderton, Roberto Amato, Jeffrey Barrett, Sonia Goncalves, Ewan Harrison, David K. Jackson, Ian Johnston, Dominic Kwiatkowski, Cordelia Langford, John Sillitoe on behalf of the Wellcome Sanger Institute COVID-19 Surveillance Team |
| EPI_ISL_3226786, EPI_ISL_3226787 | Lighthouse Lab in Glasgow | Wellcome Sanger Institute for the COVID-19 Genomics UK (COG-UK) Consortium | Harper VanSteenhouse, Yumi Kasai, David Gray, Carol Clugston, Anna Dominiczak and Alex Alderton, Roberto Amato, Jeffrey Barrett, Sonia Goncalves, Ewan Harrison, David K. Jackson, Ian Johnston, Dominic Kwiatkowski, Cordelia Langford, John Sillitoe on behalf of the Wellcome Sanger Institute COVID-19 Surveillance Team |
| EPI_ISL_3226788 | Lighthouse Lab in Alderley Park | Wellcome Sanger Institute for the COVID-19 Genomics UK (COG-UK) Consortium | Jacquelyn Wynn, Mairead Hyland, The Lighthouse Lab in Alderley Park and Alex Alderton, Roberto Amato, Jeffrey Barrett, Sonia Goncalves, Ewan Harrison, David K. Jackson, Ian Johnston, Dominic Kwiatkowski, Cordelia Langford, John Sillitoe on behalf of the Wellcome Sanger Institute COVID-19 Surveillance Team |
| EPI_ISL_3226789 | Lighthouse Lab in Glasgow | Wellcome Sanger Institute for the COVID-19 Genomics UK (COG-UK) Consortium | Harper VanSteenhouse, Yumi Kasai, David Gray, Carol Clugston, Anna Dominiczak and Alex Alderton, Roberto Amato, Jeffrey Barrett, Sonia Goncalves, Ewan Harrison, David K. Jackson, Ian Johnston, Dominic Kwiatkowski, Cordelia Langford, John Sillitoe on behalf of the Wellcome Sanger Institute COVID-19 Surveillance Team |
| EPI_ISL_3226790 | Lighthouse Lab in Alderley Park | Wellcome Sanger Institute for the COVID-19 Genomics UK (COG-UK) Consortium | Jacquelyn Wynn, Mairead Hyland, The Lighthouse Lab in Alderley Park and Alex Alderton, Roberto Amato, Jeffrey Barrett, Sonia Goncalves, Ewan Harrison, David K. Jackson, Ian Johnston, Dominic Kwiatkowski, Cordelia Langford, John Sillitoe on behalf of the Wellcome Sanger Institute COVID-19 Surveillance Team |
| EPI_ISL_3226791, EPI_ISL_3226792 | Lighthouse Lab in Milton Keynes | Wellcome Sanger Institute for the COVID-19 Genomics UK (COG-UK) Consortium | The Lighthouse Lab in Milton Keynes and Alex Alderton, Roberto Amato, Jeffrey Barrett, Sonia Goncalves, Ewan Harrison, David K. Jackson, Ian Johnston, Dominic Kwiatkowski, Cordelia Langford, John Sillitoe on behalf of the Wellcome Sanger Institute COVID-19 Surveillance Team |
| EPI_ISL_3226793 | Lighthouse Lab in Alderley Park | Wellcome Sanger Institute for the COVID-19 Genomics UK (COG-UK) Consortium | Jacquelyn Wynn, Mairead Hyland, The Lighthouse Lab in Alderley Park and Alex Alderton, Roberto Amato, Jeffrey Barrett, Sonia Goncalves, Ewan Harrison, David K. Jackson, Ian Johnston, Dominic Kwiatkowski, Cordelia Langford, John Sillitoe on behalf of the Wellcome Sanger Institute COVID-19 Surveillance Team |
| EPI_ISL_3226794 | Lighthouse Lab in Glasgow | Wellcome Sanger Institute for the COVID-19 Genomics UK (COG-UK) Consortium | Harper VanSteenhouse, Yumi Kasai, David Gray, Carol Clugston, Anna Dominiczak and Alex Alderton, Roberto Amato, Jeffrey Barrett, Sonia Goncalves, Ewan Harrison, David K. Jackson, Ian Johnston, Dominic Kwiatkowski, Cordelia Langford, John Sillitoe on behalf of the Wellcome Sanger Institute COVID-19 Surveillance Team |
| EPI_ISL_3226795 | Lighthouse Lab in Alderley Park | Wellcome Sanger Institute for the COVID-19 Genomics UK (COG-UK) Consortium | Jacquelyn Wynn, Mairead Hyland, The Lighthouse Lab in Alderley Park and Alex Alderton, Roberto Amato, Jeffrey Barrett, Sonia Goncalves, Ewan Harrison, David K. Jackson, Ian Johnston, Dominic Kwiatkowski, Cordelia Langford, John Sillitoe on behalf of the Wellcome Sanger Institute COVID-19 Surveillance Team |
| EPI_ISL_3226796 | Lighthouse Lab in Milton Keynes | Wellcome Sanger Institute for the COVID-19 Genomics UK (COG-UK) | The Lighthouse Lab in Milton Keynes and Alex Alderton, Roberto Amato, Jeffrey Barrett, Sonia Goncalves, Ewan Harrison, David K. Jackson, Ian Johnston, Dominic Kwiatkowski, Cordelia Langford, John Sillitoe on behalf of the Wellcome Sanger |

| | | | |
|---|---|---|---|
| | | Consortium | Institute COVID-19 Surveillance Team |
| EPI_ISL_3226797 | Lighthouse Lab in Alderley Park | Wellcome Sanger Institute for the COVID-19 Genomics UK (COG-UK) Consortium | Jacquelyn Wynn, Mairead Hyland, The Lighthouse Lab in Alderley Park and Alex Alderton, Roberto Amato, Jeffrey Barrett, Sonia Goncalves, Ewan Harrison, David K. Jackson, Ian Johnston, Dominic Kwiatkowski, Cordelia Langford, John Sillitoe on behalf of the Wellcome Sanger Institute COVID-19 Surveillance Team |
| EPI_ISL_3226798 | Lighthouse Lab in Milton Keynes | Wellcome Sanger Institute for the COVID-19 Genomics UK (COG-UK) Consortium | The Lighthouse Lab in Milton Keynes and Alex Alderton, Roberto Amato, Jeffrey Barrett, Sonia Goncalves, Ewan Harrison, David K. Jackson, Ian Johnston, Dominic Kwiatkowski, Cordelia Langford, John Sillitoe on behalf of the Wellcome Sanger Institute COVID-19 Surveillance Team |
| EPI_ISL_3226799 | Lighthouse Lab in Alderley Park | Wellcome Sanger Institute for the COVID-19 Genomics UK (COG-UK) Consortium | Jacquelyn Wynn, Mairead Hyland, The Lighthouse Lab in Alderley Park and Alex Alderton, Roberto Amato, Jeffrey Barrett, Sonia Goncalves, Ewan Harrison, David K. Jackson, Ian Johnston, Dominic Kwiatkowski, Cordelia Langford, John Sillitoe on behalf of the Wellcome Sanger Institute COVID-19 Surveillance Team |
| EPI_ISL_3226800 | Lighthouse Lab in Milton Keynes | Wellcome Sanger Institute for the COVID-19 Genomics UK (COG-UK) Consortium | The Lighthouse Lab in Milton Keynes and Alex Alderton, Roberto Amato, Jeffrey Barrett, Sonia Goncalves, Ewan Harrison, David K. Jackson, Ian Johnston, Dominic Kwiatkowski, Cordelia Langford, John Sillitoe on behalf of the Wellcome Sanger Institute COVID-19 Surveillance Team |
| EPI_ISL_3226801 | Lighthouse Lab in Alderley Park | Wellcome Sanger Institute for the COVID-19 Genomics UK (COG-UK) Consortium | Jacquelyn Wynn, Mairead Hyland, The Lighthouse Lab in Alderley Park and Alex Alderton, Roberto Amato, Jeffrey Barrett, Sonia Goncalves, Ewan Harrison, David K. Jackson, Ian Johnston, Dominic Kwiatkowski, Cordelia Langford, John Sillitoe on behalf of the Wellcome Sanger Institute COVID-19 Surveillance Team |
| EPI_ISL_3226802 | Lighthouse Lab in Milton Keynes | Wellcome Sanger Institute for the COVID-19 Genomics UK (COG-UK) Consortium | The Lighthouse Lab in Milton Keynes and Alex Alderton, Roberto Amato, Jeffrey Barrett, Sonia Goncalves, Ewan Harrison, David K. Jackson, Ian Johnston, Dominic Kwiatkowski, Cordelia Langford, John Sillitoe on behalf of the Wellcome Sanger Institute COVID-19 Surveillance Team |
| EPI_ISL_3226803 | Lighthouse Lab in Alderley Park | Wellcome Sanger Institute for the COVID-19 Genomics UK (COG-UK) Consortium | Jacquelyn Wynn, Mairead Hyland, The Lighthouse Lab in Alderley Park and Alex Alderton, Roberto Amato, Jeffrey Barrett, Sonia Goncalves, Ewan Harrison, David K. Jackson, Ian Johnston, Dominic Kwiatkowski, Cordelia Langford, John Sillitoe on behalf of the Wellcome Sanger Institute COVID-19 Surveillance Team |
| EPI_ISL_3226804 | Lighthouse Lab in Glasgow | Wellcome Sanger Institute for the COVID-19 Genomics UK (COG-UK) Consortium | Harper VanSteenhouse, Yumi Kasai, David Gray, Carol Clugston, Anna Dominiczak and Alex Alderton, Roberto Amato, Jeffrey Barrett, Sonia Goncalves, Ewan Harrison, David K. Jackson, Ian Johnston, Dominic Kwiatkowski, Cordelia Langford, John Sillitoe on behalf of the Wellcome Sanger Institute COVID-19 Surveillance Team |
| EPI_ISL_3226805 | Lighthouse Lab in Milton Keynes | Wellcome Sanger Institute for the COVID-19 Genomics UK (COG-UK) Consortium | The Lighthouse Lab in Milton Keynes and Alex Alderton, Roberto Amato, Jeffrey Barrett, Sonia Goncalves, Ewan Harrison, David K. Jackson, Ian Johnston, Dominic Kwiatkowski, Cordelia Langford, John Sillitoe on behalf of the Wellcome Sanger Institute COVID-19 Surveillance Team |
| EPI_ISL_3226806 | Lighthouse Lab in Glasgow | Wellcome Sanger Institute for the COVID-19 Genomics UK (COG-UK) Consortium | Harper VanSteenhouse, Yumi Kasai, David Gray, Carol Clugston, Anna Dominiczak and Alex Alderton, Roberto Amato, Jeffrey Barrett, Sonia Goncalves, Ewan Harrison, David K. Jackson, Ian Johnston, Dominic Kwiatkowski, Cordelia Langford, John Sillitoe on behalf of the Wellcome Sanger Institute COVID-19 Surveillance Team |
| EPI_ISL_3226807 | Lighthouse Lab in Alderley Park | Wellcome Sanger Institute for the COVID-19 Genomics UK (COG-UK) Consortium | Jacquelyn Wynn, Mairead Hyland, The Lighthouse Lab in Alderley Park and Alex Alderton, Roberto Amato, Jeffrey Barrett, Sonia Goncalves, Ewan Harrison, David K. Jackson, Ian Johnston, Dominic Kwiatkowski, Cordelia Langford, John Sillitoe on behalf of the Wellcome Sanger Institute COVID-19 Surveillance Team |
| EPI_ISL_3226808 | Lighthouse Lab in Glasgow | Wellcome Sanger Institute for the COVID-19 Genomics UK (COG-UK) Consortium | Harper VanSteenhouse, Yumi Kasai, David Gray, Carol Clugston, Anna Dominiczak and Alex Alderton, Roberto Amato, Jeffrey Barrett, Sonia Goncalves, Ewan Harrison, David K. Jackson, Ian Johnston, Dominic Kwiatkowski, Cordelia Langford, John Sillitoe on behalf of the Wellcome Sanger Institute COVID-19 Surveillance Team |
| EPI_ISL_3226809, EPI_ISL_3226810, EPI_ISL_3226811 | Lighthouse Lab in Alderley Park | Wellcome Sanger Institute for the COVID-19 Genomics UK (COG-UK) Consortium | Jacquelyn Wynn, Mairead Hyland, The Lighthouse Lab in Alderley Park and Alex Alderton, Roberto Amato, Jeffrey Barrett, Sonia Goncalves, Ewan Harrison, David K. Jackson, Ian Johnston, Dominic Kwiatkowski, Cordelia Langford, John Sillitoe on behalf of the Wellcome Sanger Institute COVID-19 Surveillance Team |
| EPI_ISL_3226812 | Lighthouse Lab in Glasgow | Wellcome Sanger Institute for the COVID-19 Genomics UK (COG-UK) Consortium | Harper VanSteenhouse, Yumi Kasai, David Gray, Carol Clugston, Anna Dominiczak and Alex Alderton, Roberto Amato, Jeffrey Barrett, Sonia Goncalves, Ewan Harrison, David K. Jackson, Ian Johnston, Dominic Kwiatkowski, Cordelia Langford, John Sillitoe on behalf of the Wellcome Sanger Institute COVID-19 Surveillance Team |
| EPI_ISL_3226813 | Lighthouse Lab in Alderley Park | Wellcome Sanger Institute for the COVID-19 Genomics UK (COG-UK) Consortium | Jacquelyn Wynn, Mairead Hyland, The Lighthouse Lab in Alderley Park and Alex Alderton, Roberto Amato, Jeffrey Barrett, Sonia Goncalves, Ewan Harrison, David K. Jackson, Ian Johnston, Dominic Kwiatkowski, Cordelia Langford, John Sillitoe on behalf of the Wellcome Sanger Institute COVID-19 Surveillance Team |
| EPI_ISL_3226814 | Lighthouse Lab in Milton Keynes | Wellcome Sanger Institute for the COVID-19 Genomics UK (COG-UK) Consortium | The Lighthouse Lab in Milton Keynes and Alex Alderton, Roberto Amato, Jeffrey Barrett, Sonia Goncalves, Ewan Harrison, David K. Jackson, Ian Johnston, Dominic Kwiatkowski, Cordelia Langford, John Sillitoe on behalf of the Wellcome Sanger Institute COVID-19 Surveillance Team |
| EPI_ISL_3226815, EPI_ISL_3226816 | Lighthouse Lab in Alderley Park | Wellcome Sanger Institute for the COVID-19 Genomics UK (COG-UK) Consortium | Jacquelyn Wynn, Mairead Hyland, The Lighthouse Lab in Alderley Park and Alex Alderton, Roberto Amato, Jeffrey Barrett, Sonia Goncalves, Ewan Harrison, David K. Jackson, Ian Johnston, Dominic Kwiatkowski, Cordelia Langford, John Sillitoe on behalf of the Wellcome Sanger Institute COVID-19 Surveillance Team |
| EPI_ISL_3226817 | Lighthouse Lab in Glasgow | Wellcome Sanger Institute for the COVID-19 Genomics UK (COG-UK) Consortium | Harper VanSteenhouse, Yumi Kasai, David Gray, Carol Clugston, Anna Dominiczak and Alex Alderton, Roberto Amato, Jeffrey Barrett, Sonia Goncalves, Ewan Harrison, David K. Jackson, Ian Johnston, Dominic Kwiatkowski, Cordelia Langford, John Sillitoe on behalf of the Wellcome Sanger Institute COVID-19 Surveillance Team |
| EPI_ISL_3226818 | Lighthouse Lab in Milton Keynes | Wellcome Sanger Institute for the COVID-19 Genomics UK (COG-UK) Consortium | The Lighthouse Lab in Milton Keynes and Alex Alderton, Roberto Amato, Jeffrey Barrett, Sonia Goncalves, Ewan Harrison, David K. Jackson, Ian Johnston, Dominic Kwiatkowski, Cordelia Langford, John Sillitoe on behalf of the Wellcome Sanger Institute COVID-19 Surveillance Team |
| EPI_ISL_3226819 | Lighthouse Lab in Alderley Park | Wellcome Sanger Institute for the COVID-19 Genomics UK (COG-UK) Consortium | Jacquelyn Wynn, Mairead Hyland, The Lighthouse Lab in Alderley Park and Alex Alderton, Roberto Amato, Jeffrey Barrett, Sonia Goncalves, Ewan Harrison, David K. Jackson, Ian Johnston, Dominic Kwiatkowski, Cordelia Langford, John Sillitoe on behalf of the Wellcome Sanger Institute COVID-19 Surveillance Team |
| EPI_ISL_3226820 | Lighthouse Lab in Glasgow | Wellcome Sanger Institute for the COVID-19 Genomics UK (COG-UK) Consortium | Harper VanSteenhouse, Yumi Kasai, David Gray, Carol Clugston, Anna Dominiczak and Alex Alderton, Roberto Amato, Jeffrey Barrett, Sonia Goncalves, Ewan Harrison, David K. Jackson, Ian Johnston, Dominic Kwiatkowski, Cordelia Langford, John Sillitoe on behalf of the Wellcome Sanger Institute COVID-19 Surveillance Team |
| EPI_ISL_3226821 | Lighthouse Lab in Alderley Park | Wellcome Sanger Institute for the COVID-19 Genomics UK (COG-UK) Consortium | Jacquelyn Wynn, Mairead Hyland, The Lighthouse Lab in Alderley Park and Alex Alderton, Roberto Amato, Jeffrey Barrett, Sonia Goncalves, Ewan Harrison, David K. Jackson, Ian Johnston, Dominic Kwiatkowski, Cordelia Langford, John Sillitoe on behalf of the Wellcome Sanger Institute COVID-19 Surveillance Team |
| EPI_ISL_3226822 | Lighthouse Lab in Glasgow | Wellcome Sanger Institute for the COVID-19 Genomics UK (COG-UK) Consortium | Harper VanSteenhouse, Yumi Kasai, David Gray, Carol Clugston, Anna Dominiczak and Alex Alderton, Roberto Amato, Jeffrey Barrett, Sonia Goncalves, Ewan Harrison, David K. Jackson, Ian Johnston, Dominic Kwiatkowski, Cordelia Langford, John Sillitoe on behalf of the Wellcome Sanger Institute COVID-19 Surveillance Team |
| EPI_ISL_3226823 | Lighthouse Lab in Milton Keynes | Wellcome Sanger Institute for the COVID-19 Genomics UK (COG-UK) Consortium | The Lighthouse Lab in Milton Keynes and Alex Alderton, Roberto Amato, Jeffrey Barrett, Sonia Goncalves, Ewan Harrison, David K. Jackson, Ian Johnston, Dominic Kwiatkowski, Cordelia Langford, John Sillitoe on behalf of the Wellcome Sanger Institute COVID-19 Surveillance Team |
| EPI_ISL_3226824, EPI_ISL_3226825 | Lighthouse Lab in Alderley Park | Wellcome Sanger Institute for the COVID-19 Genomics UK (COG-UK) Consortium | Jacquelyn Wynn, Mairead Hyland, The Lighthouse Lab in Alderley Park and Alex Alderton, Roberto Amato, Jeffrey Barrett, Sonia Goncalves, Ewan Harrison, David K. Jackson, Ian Johnston, Dominic Kwiatkowski, Cordelia Langford, John Sillitoe on behalf of the Wellcome Sanger Institute COVID-19 Surveillance Team |
| EPI_ISL_3226826 | Lighthouse Lab in Milton Keynes | Wellcome Sanger Institute for the COVID-19 Genomics UK (COG-UK) Consortium | The Lighthouse Lab in Milton Keynes and Alex Alderton, Roberto Amato, Jeffrey Barrett, Sonia Goncalves, Ewan Harrison, David K. Jackson, Ian Johnston, Dominic Kwiatkowski, Cordelia Langford, John Sillitoe on behalf of the Wellcome Sanger Institute COVID-19 Surveillance Team |
| EPI_ISL_3226827, EPI_ISL_3226828 | Lighthouse Lab in Alderley Park | Wellcome Sanger Institute for the COVID-19 Genomics UK (COG-UK) Consortium | Jacquelyn Wynn, Mairead Hyland, The Lighthouse Lab in Alderley Park and Alex Alderton, Roberto Amato, Jeffrey Barrett, Sonia Goncalves, Ewan Harrison, David K. Jackson, Ian Johnston, Dominic Kwiatkowski, Cordelia Langford, John Sillitoe on behalf of the Wellcome Sanger Institute COVID-19 Surveillance Team |
| EPI_ISL_3226829, EPI_ISL_3226830, EPI_ISL_3226831 | Lighthouse Lab in Milton Keynes | Wellcome Sanger Institute for the COVID-19 Genomics UK (COG-UK) Consortium | The Lighthouse Lab in Milton Keynes and Alex Alderton, Roberto Amato, Jeffrey Barrett, Sonia Goncalves, Ewan Harrison, David K. Jackson, Ian Johnston, Dominic Kwiatkowski, Cordelia Langford, John Sillitoe on behalf of the Wellcome Sanger Institute COVID-19 Surveillance Team |
| EPI_ISL_3226832 | Lighthouse Lab in Alderley Park | Wellcome Sanger Institute for the COVID-19 Genomics UK (COG-UK) Consortium | Jacquelyn Wynn, Mairead Hyland, The Lighthouse Lab in Alderley Park and Alex Alderton, Roberto Amato, Jeffrey Barrett, Sonia Goncalves, Ewan Harrison, David K. Jackson, Ian Johnston, Dominic Kwiatkowski, Cordelia Langford, John Sillitoe on behalf of the Wellcome Sanger Institute COVID-19 Surveillance Team |
| EPI_ISL_3226833 | Lighthouse Lab in Glasgow | Wellcome Sanger Institute for the COVID-19 Genomics UK (COG-UK) Consortium | Harper VanSteenhouse, Yumi Kasai, David Gray, Carol Clugston, Anna Dominiczak and Alex Alderton, Roberto Amato, Jeffrey Barrett, Sonia Goncalves, Ewan Harrison, David K. Jackson, Ian Johnston, Dominic Kwiatkowski, Cordelia Langford, John Sillitoe on behalf of the Wellcome Sanger Institute COVID-19 Surveillance Team |
| EPI_ISL_3226834 | Lighthouse Lab in Milton Keynes | Wellcome Sanger Institute for the COVID-19 Genomics UK (COG-UK) Consortium | The Lighthouse Lab in Milton Keynes and Alex Alderton, Roberto Amato, Jeffrey Barrett, Sonia Goncalves, Ewan Harrison, David K. Jackson, Ian Johnston, Dominic Kwiatkowski, Cordelia Langford, John Sillitoe on behalf of the Wellcome Sanger Institute COVID-19 Surveillance Team |
| EPI_ISL_3226835, EPI_ISL_3226836, EPI_ISL_3226837, EPI_ISL_3226838 | Lighthouse Lab in Alderley Park | Wellcome Sanger Institute for the COVID-19 Genomics UK (COG-UK) Consortium | Jacquelyn Wynn, Mairead Hyland, The Lighthouse Lab in Alderley Park and Alex Alderton, Roberto Amato, Jeffrey Barrett, Sonia Goncalves, Ewan Harrison, David K. Jackson, Ian Johnston, Dominic Kwiatkowski, Cordelia Langford, John Sillitoe on behalf of the Wellcome Sanger Institute COVID-19 Surveillance Team |
| EPI_ISL_3226839 | Lighthouse Lab in Glasgow | Wellcome Sanger Institute for the COVID-19 Genomics UK (COG-UK) Consortium | Harper VanSteenhouse, Yumi Kasai, David Gray, Carol Clugston, Anna Dominiczak and Alex Alderton, Roberto Amato, Jeffrey Barrett, Sonia Goncalves, Ewan Harrison, David K. Jackson, Ian Johnston, Dominic Kwiatkowski, Cordelia Langford, John Sillitoe on behalf of the Wellcome Sanger Institute COVID-19 Surveillance Team |
| EPI_ISL_3226840 | Lighthouse Lab in Alderley Park | Wellcome Sanger Institute for the COVID-19 Genomics UK (COG-UK) Consortium | Jacquelyn Wynn, Mairead Hyland, The Lighthouse Lab in Alderley Park and Alex Alderton, Roberto Amato, Jeffrey Barrett, Sonia Goncalves, Ewan Harrison, David K. Jackson, Ian Johnston, Dominic Kwiatkowski, Cordelia Langford, John Sillitoe on behalf of the Wellcome Sanger Institute COVID-19 Surveillance Team |
| EPI_ISL_3226841 | Lighthouse Lab in Milton Keynes | Wellcome Sanger Institute for the COVID-19 Genomics UK (COG-UK) Consortium | The Lighthouse Lab in Milton Keynes and Alex Alderton, Roberto Amato, Jeffrey Barrett, Sonia Goncalves, Ewan Harrison, David K. Jackson, Ian Johnston, Dominic Kwiatkowski, Cordelia Langford, John Sillitoe on behalf of the Wellcome Sanger Institute COVID-19 Surveillance Team |
| EPI_ISL_3226842, EPI_ISL_3226843 | Lighthouse Lab in Glasgow | Wellcome Sanger Institute for the COVID-19 Genomics UK (COG-UK) Consortium | Harper VanSteenhouse, Yumi Kasai, David Gray, Carol Clugston, Anna Dominiczak and Alex Alderton, Roberto Amato, Jeffrey Barrett, Sonia Goncalves, Ewan Harrison, David K. Jackson, Ian Johnston, Dominic Kwiatkowski, Cordelia Langford, John Sillitoe on behalf of the Wellcome Sanger Institute COVID-19 Surveillance Team |
| EPI_ISL_3226844 | Lighthouse Lab in Alderley Park | Wellcome Sanger Institute for the COVID-19 Genomics UK (COG-UK) Consortium | Jacquelyn Wynn, Mairead Hyland, The Lighthouse Lab in Alderley Park and Alex Alderton, Roberto Amato, Jeffrey Barrett, Sonia Goncalves, Ewan Harrison, David K. Jackson, Ian Johnston, Dominic Kwiatkowski, Cordelia Langford, John Sillitoe on behalf of the Wellcome Sanger Institute COVID-19 Surveillance Team |
| EPI_ISL_3226845 | Lighthouse Lab in Glasgow | Wellcome Sanger Institute for the COVID-19 Genomics UK (COG-UK) Consortium | Harper VanSteenhouse, Yumi Kasai, David Gray, Carol Clugston, Anna Dominiczak and Alex Alderton, Roberto Amato, Jeffrey Barrett, Sonia Goncalves, Ewan Harrison, David K. Jackson, Ian Johnston, Dominic Kwiatkowski, Cordelia Langford, John Sillitoe on behalf of the Wellcome Sanger Institute COVID-19 Surveillance Team |
| EPI_ISL_3226846, EPI_ISL_3226847 | Lighthouse Lab in Alderley Park | Wellcome Sanger Institute for the COVID-19 Genomics UK (COG-UK) Consortium | Jacquelyn Wynn, Mairead Hyland, The Lighthouse Lab in Alderley Park and Alex Alderton, Roberto Amato, Jeffrey Barrett, Sonia Goncalves, Ewan Harrison, David K. Jackson, Ian Johnston, Dominic Kwiatkowski, Cordelia Langford, John Sillitoe on behalf of the Wellcome Sanger Institute COVID-19 Surveillance Team |
| EPI_ISL_3226848, EPI_ISL_3226849 | Lighthouse Lab in Milton Keynes | Wellcome Sanger Institute for the COVID-19 Genomics UK (COG-UK) Consortium | The Lighthouse Lab in Milton Keynes and Alex Alderton, Roberto Amato, Jeffrey Barrett, Sonia Goncalves, Ewan Harrison, David K. Jackson, Ian Johnston, Dominic Kwiatkowski, Cordelia Langford, John Sillitoe on behalf of the Wellcome Sanger Institute COVID-19 Surveillance Team |
| EPI_ISL_3226850, EPI_ISL_3226851, EPI_ISL_3226852 | Lighthouse Lab in Alderley Park | Wellcome Sanger Institute for the COVID-19 Genomics UK (COG-UK) Consortium | Jacquelyn Wynn, Mairead Hyland, The Lighthouse Lab in Alderley Park and Alex Alderton, Roberto Amato, Jeffrey Barrett, Sonia Goncalves, Ewan Harrison, David K. Jackson, Ian Johnston, Dominic Kwiatkowski, Cordelia Langford, John Sillitoe on behalf of the Wellcome Sanger Institute COVID-19 Surveillance Team |
| EPI_ISL_3226853 | Lighthouse Lab in Glasgow | Wellcome Sanger Institute for the COVID-19 Genomics UK (COG-UK) Consortium | Harper VanSteenhouse, Yumi Kasai, David Gray, Carol Clugston, Anna Dominiczak and Alex Alderton, Roberto Amato, Jeffrey Barrett, Sonia Goncalves, Ewan Harrison, David K. Jackson, Ian Johnston, Dominic Kwiatkowski, Cordelia Langford, John Sillitoe on behalf of the Wellcome Sanger Institute COVID-19 Surveillance Team |
| EPI_ISL_3226854 | Lighthouse Lab in Alderley Park | Wellcome Sanger Institute for the COVID-19 Genomics UK (COG-UK) | Jacquelyn Wynn, Mairead Hyland, The Lighthouse Lab in Alderley Park and Alex Alderton, Roberto Amato, Jeffrey Barrett, Sonia Goncalves, Ewan Harrison, David K. Jackson, Ian Johnston, Dominic Kwiatkowski, Cordelia Langford, John Sillitoe on |

| | | | |
|---|---|---|---|
| | | Consortium | behalf of the Wellcome Sanger Institute COVID-19 Surveillance Team |
| EPI_ISL_3226855 | Lighthouse Lab in Glasgow | Wellcome Sanger Institute for the COVID-19 Genomics UK (COG-UK) Consortium | Harper VanSteenhouse, Yumi Kasai, David Gray, Carol Clugston, Anna Dominiczak and Alex Alderton, Roberto Amato, Jeffrey Barrett, Sonia Goncalves, Ewan Harrison, David K. Jackson, Ian Johnston, Dominic Kwiatkowski, Cordelia Langford, John Sillitoe on behalf of the Wellcome Sanger Institute COVID-19 Surveillance Team |
| EPI_ISL_3226856 | Lighthouse Lab in Milton Keynes | Wellcome Sanger Institute for the COVID-19 Genomics UK (COG-UK) Consortium | The Lighthouse Lab in Milton Keynes and Alex Alderton, Roberto Amato, Jeffrey Barrett, Sonia Goncalves, Ewan Harrison, David K. Jackson, Ian Johnston, Dominic Kwiatkowski, Cordelia Langford, John Sillitoe on behalf of the Wellcome Sanger Institute COVID-19 Surveillance Team |
| EPI_ISL_3226857, EPI_ISL_3226858 | Lighthouse Lab in Alderley Park | Wellcome Sanger Institute for the COVID-19 Genomics UK (COG-UK) Consortium | Jacquelyn Wynn, Mairead Hyland, The Lighthouse Lab in Alderley Park and Alex Alderton, Roberto Amato, Jeffrey Barrett, Sonia Goncalves, Ewan Harrison, David K. Jackson, Ian Johnston, Dominic Kwiatkowski, Cordelia Langford, John Sillitoe on behalf of the Wellcome Sanger Institute COVID-19 Surveillance Team |
| EPI_ISL_3226859 | Lighthouse Lab in Glasgow | Wellcome Sanger Institute for the COVID-19 Genomics UK (COG-UK) Consortium | Harper VanSteenhouse, Yumi Kasai, David Gray, Carol Clugston, Anna Dominiczak and Alex Alderton, Roberto Amato, Jeffrey Barrett, Sonia Goncalves, Ewan Harrison, David K. Jackson, Ian Johnston, Dominic Kwiatkowski, Cordelia Langford, John Sillitoe on behalf of the Wellcome Sanger Institute COVID-19 Surveillance Team |
| EPI_ISL_3226860 | Lighthouse Lab in Milton Keynes | Wellcome Sanger Institute for the COVID-19 Genomics UK (COG-UK) Consortium | The Lighthouse Lab in Milton Keynes and Alex Alderton, Roberto Amato, Jeffrey Barrett, Sonia Goncalves, Ewan Harrison, David K. Jackson, Ian Johnston, Dominic Kwiatkowski, Cordelia Langford, John Sillitoe on behalf of the Wellcome Sanger Institute COVID-19 Surveillance Team |
| EPI_ISL_3226861 | Lighthouse Lab in Glasgow | Wellcome Sanger Institute for the COVID-19 Genomics UK (COG-UK) Consortium | Harper VanSteenhouse, Yumi Kasai, David Gray, Carol Clugston, Anna Dominiczak and Alex Alderton, Roberto Amato, Jeffrey Barrett, Sonia Goncalves, Ewan Harrison, David K. Jackson, Ian Johnston, Dominic Kwiatkowski, Cordelia Langford, John Sillitoe on behalf of the Wellcome Sanger Institute COVID-19 Surveillance Team |
| EPI_ISL_3226862, EPI_ISL_3226863 | Lighthouse Lab in Alderley Park | Wellcome Sanger Institute for the COVID-19 Genomics UK (COG-UK) Consortium | Jacquelyn Wynn, Mairead Hyland, The Lighthouse Lab in Alderley Park and Alex Alderton, Roberto Amato, Jeffrey Barrett, Sonia Goncalves, Ewan Harrison, David K. Jackson, Ian Johnston, Dominic Kwiatkowski, Cordelia Langford, John Sillitoe on behalf of the Wellcome Sanger Institute COVID-19 Surveillance Team |
| EPI_ISL_3226864 | Lighthouse Lab in Glasgow | Wellcome Sanger Institute for the COVID-19 Genomics UK (COG-UK) Consortium | Harper VanSteenhouse, Yumi Kasai, David Gray, Carol Clugston, Anna Dominiczak and Alex Alderton, Roberto Amato, Jeffrey Barrett, Sonia Goncalves, Ewan Harrison, David K. Jackson, Ian Johnston, Dominic Kwiatkowski, Cordelia Langford, John Sillitoe on behalf of the Wellcome Sanger Institute COVID-19 Surveillance Team |
| EPI_ISL_3226865 | Lighthouse Lab in Alderley Park | Wellcome Sanger Institute for the COVID-19 Genomics UK (COG-UK) Consortium | Jacquelyn Wynn, Mairead Hyland, The Lighthouse Lab in Alderley Park and Alex Alderton, Roberto Amato, Jeffrey Barrett, Sonia Goncalves, Ewan Harrison, David K. Jackson, Ian Johnston, Dominic Kwiatkowski, Cordelia Langford, John Sillitoe on behalf of the Wellcome Sanger Institute COVID-19 Surveillance Team |
| EPI_ISL_3226866 | Lighthouse Lab in Milton Keynes | Wellcome Sanger Institute for the COVID-19 Genomics UK (COG-UK) Consortium | The Lighthouse Lab in Milton Keynes and Alex Alderton, Roberto Amato, Jeffrey Barrett, Sonia Goncalves, Ewan Harrison, David K. Jackson, Ian Johnston, Dominic Kwiatkowski, Cordelia Langford, John Sillitoe on behalf of the Wellcome Sanger Institute COVID-19 Surveillance Team |
| EPI_ISL_3226867 | Lighthouse Lab in Alderley Park | Wellcome Sanger Institute for the COVID-19 Genomics UK (COG-UK) Consortium | Jacquelyn Wynn, Mairead Hyland, The Lighthouse Lab in Alderley Park and Alex Alderton, Roberto Amato, Jeffrey Barrett, Sonia Goncalves, Ewan Harrison, David K. Jackson, Ian Johnston, Dominic Kwiatkowski, Cordelia Langford, John Sillitoe on behalf of the Wellcome Sanger Institute COVID-19 Surveillance Team |
| EPI_ISL_3226868 | Lighthouse Lab in Milton Keynes | Wellcome Sanger Institute for the COVID-19 Genomics UK (COG-UK) Consortium | The Lighthouse Lab in Milton Keynes and Alex Alderton, Roberto Amato, Jeffrey Barrett, Sonia Goncalves, Ewan Harrison, David K. Jackson, Ian Johnston, Dominic Kwiatkowski, Cordelia Langford, John Sillitoe on behalf of the Wellcome Sanger Institute COVID-19 Surveillance Team |
| EPI_ISL_3226869 | Lighthouse Lab in Glasgow | Wellcome Sanger Institute for the COVID-19 Genomics UK (COG-UK) Consortium | Harper VanSteenhouse, Yumi Kasai, David Gray, Carol Clugston, Anna Dominiczak and Alex Alderton, Roberto Amato, Jeffrey Barrett, Sonia Goncalves, Ewan Harrison, David K. Jackson, Ian Johnston, Dominic Kwiatkowski, Cordelia Langford, John Sillitoe on behalf of the Wellcome Sanger Institute COVID-19 Surveillance Team |
| EPI_ISL_3226870, EPI_ISL_3226871 | Lighthouse Lab in Milton Keynes | Wellcome Sanger Institute for the COVID-19 Genomics UK (COG-UK) Consortium | The Lighthouse Lab in Milton Keynes and Alex Alderton, Roberto Amato, Jeffrey Barrett, Sonia Goncalves, Ewan Harrison, David K. Jackson, Ian Johnston, Dominic Kwiatkowski, Cordelia Langford, John Sillitoe on behalf of the Wellcome Sanger Institute COVID-19 Surveillance Team |
| EPI_ISL_3226872, EPI_ISL_3226873 | Lighthouse Lab in Alderley Park | Wellcome Sanger Institute for the COVID-19 Genomics UK (COG-UK) Consortium | Jacquelyn Wynn, Mairead Hyland, The Lighthouse Lab in Alderley Park and Alex Alderton, Roberto Amato, Jeffrey Barrett, Sonia Goncalves, Ewan Harrison, David K. Jackson, Ian Johnston, Dominic Kwiatkowski, Cordelia Langford, John Sillitoe on behalf of the Wellcome Sanger Institute COVID-19 Surveillance Team |
| EPI_ISL_3226874 | Lighthouse Lab in Glasgow | Wellcome Sanger Institute for the COVID-19 Genomics UK (COG-UK) Consortium | Harper VanSteenhouse, Yumi Kasai, David Gray, Carol Clugston, Anna Dominiczak and Alex Alderton, Roberto Amato, Jeffrey Barrett, Sonia Goncalves, Ewan Harrison, David K. Jackson, Ian Johnston, Dominic Kwiatkowski, Cordelia Langford, John Sillitoe on behalf of the Wellcome Sanger Institute COVID-19 Surveillance Team |
| EPI_ISL_3226875, EPI_ISL_3226876, EPI_ISL_3226877 | Lighthouse Lab in Milton Keynes | Wellcome Sanger Institute for the COVID-19 Genomics UK (COG-UK) Consortium | The Lighthouse Lab in Milton Keynes and Alex Alderton, Roberto Amato, Jeffrey Barrett, Sonia Goncalves, Ewan Harrison, David K. Jackson, Ian Johnston, Dominic Kwiatkowski, Cordelia Langford, John Sillitoe on behalf of the Wellcome Sanger Institute COVID-19 Surveillance Team |
| EPI_ISL_3226878, EPI_ISL_3226879 | Lighthouse Lab in Alderley Park | Wellcome Sanger Institute for the COVID-19 Genomics UK (COG-UK) Consortium | Jacquelyn Wynn, Mairead Hyland, The Lighthouse Lab in Alderley Park and Alex Alderton, Roberto Amato, Jeffrey Barrett, Sonia Goncalves, Ewan Harrison, David K. Jackson, Ian Johnston, Dominic Kwiatkowski, Cordelia Langford, John Sillitoe on behalf of the Wellcome Sanger Institute COVID-19 Surveillance Team |
| EPI_ISL_3226880, EPI_ISL_3226881 | Lighthouse Lab in Milton Keynes | Wellcome Sanger Institute for the COVID-19 Genomics UK (COG-UK) Consortium | The Lighthouse Lab in Milton Keynes and Alex Alderton, Roberto Amato, Jeffrey Barrett, Sonia Goncalves, Ewan Harrison, David K. Jackson, Ian Johnston, Dominic Kwiatkowski, Cordelia Langford, John Sillitoe on behalf of the Wellcome Sanger Institute COVID-19 Surveillance Team |
| EPI_ISL_3226882 | Lighthouse Lab in Alderley Park | Wellcome Sanger Institute for the COVID-19 Genomics UK (COG-UK) Consortium | Jacquelyn Wynn, Mairead Hyland, The Lighthouse Lab in Alderley Park and Alex Alderton, Roberto Amato, Jeffrey Barrett, Sonia Goncalves, Ewan Harrison, David K. Jackson, Ian Johnston, Dominic Kwiatkowski, Cordelia Langford, John Sillitoe on behalf of the Wellcome Sanger Institute COVID-19 Surveillance Team |
| EPI_ISL_3226883, EPI_ISL_3226884 | Lighthouse Lab in Glasgow | Wellcome Sanger Institute for the COVID-19 Genomics UK (COG-UK) Consortium | Harper VanSteenhouse, Yumi Kasai, David Gray, Carol Clugston, Anna Dominiczak and Alex Alderton, Roberto Amato, Jeffrey Barrett, Sonia Goncalves, Ewan Harrison, David K. Jackson, Ian Johnston, Dominic Kwiatkowski, Cordelia Langford, John Sillitoe on behalf of the Wellcome Sanger Institute COVID-19 Surveillance Team |
| EPI_ISL_3226885 | Lighthouse Lab in Alderley Park | Wellcome Sanger Institute for the COVID-19 Genomics UK (COG-UK) Consortium | Jacquelyn Wynn, Mairead Hyland, The Lighthouse Lab in Alderley Park and Alex Alderton, Roberto Amato, Jeffrey Barrett, Sonia Goncalves, Ewan Harrison, David K. Jackson, Ian Johnston, Dominic Kwiatkowski, Cordelia Langford, John Sillitoe on behalf of the Wellcome Sanger Institute COVID-19 Surveillance Team |
| EPI_ISL_3226886 | Lighthouse Lab in Milton Keynes | Wellcome Sanger Institute for the COVID-19 Genomics UK (COG-UK) Consortium | The Lighthouse Lab in Milton Keynes and Alex Alderton, Roberto Amato, Jeffrey Barrett, Sonia Goncalves, Ewan Harrison, David K. Jackson, Ian Johnston, Dominic Kwiatkowski, Cordelia Langford, John Sillitoe on behalf of the Wellcome Sanger Institute COVID-19 Surveillance Team |
| EPI_ISL_3226887, EPI_ISL_3226888 | Lighthouse Lab in Glasgow | Wellcome Sanger Institute for the COVID-19 Genomics UK (COG-UK) Consortium | Harper VanSteenhouse, Yumi Kasai, David Gray, Carol Clugston, Anna Dominiczak and Alex Alderton, Roberto Amato, Jeffrey Barrett, Sonia Goncalves, Ewan Harrison, David K. Jackson, Ian Johnston, Dominic Kwiatkowski, Cordelia Langford, John Sillitoe on behalf of the Wellcome Sanger Institute COVID-19 Surveillance Team |
| EPI_ISL_3226889 | Lighthouse Lab in Alderley Park | Wellcome Sanger Institute for the COVID-19 Genomics UK (COG-UK) Consortium | Jacquelyn Wynn, Mairead Hyland, The Lighthouse Lab in Alderley Park and Alex Alderton, Roberto Amato, Jeffrey Barrett, Sonia Goncalves, Ewan Harrison, David K. Jackson, Ian Johnston, Dominic Kwiatkowski, Cordelia Langford, John Sillitoe on behalf of the Wellcome Sanger Institute COVID-19 Surveillance Team |
| EPI_ISL_3226890 | Lighthouse Lab in Glasgow | Wellcome Sanger Institute for the COVID-19 Genomics UK (COG-UK) Consortium | Harper VanSteenhouse, Yumi Kasai, David Gray, Carol Clugston, Anna Dominiczak and Alex Alderton, Roberto Amato, Jeffrey Barrett, Sonia Goncalves, Ewan Harrison, David K. Jackson, Ian Johnston, Dominic Kwiatkowski, Cordelia Langford, John Sillitoe on behalf of the Wellcome Sanger Institute COVID-19 Surveillance Team |
| EPI_ISL_3226891, EPI_ISL_3226892, EPI_ISL_3226893, EPI_ISL_3226894, EPI_ISL_3226895 | Lighthouse Lab in Milton Keynes | Wellcome Sanger Institute for the COVID-19 Genomics UK (COG-UK) Consortium | The Lighthouse Lab in Milton Keynes and Alex Alderton, Roberto Amato, Jeffrey Barrett, Sonia Goncalves, Ewan Harrison, David K. Jackson, Ian Johnston, Dominic Kwiatkowski, Cordelia Langford, John Sillitoe on behalf of the Wellcome Sanger Institute COVID-19 Surveillance Team |
| EPI_ISL_3226896 | Lighthouse Lab in Alderley Park | Wellcome Sanger Institute for the COVID-19 Genomics UK (COG-UK) Consortium | Jacquelyn Wynn, Mairead Hyland, The Lighthouse Lab in Alderley Park and Alex Alderton, Roberto Amato, Jeffrey Barrett, Sonia Goncalves, Ewan Harrison, David K. Jackson, Ian Johnston, Dominic Kwiatkowski, Cordelia Langford, John Sillitoe on behalf of the Wellcome Sanger Institute COVID-19 Surveillance Team |
| EPI_ISL_3226897 | Lighthouse Lab in Milton Keynes | Wellcome Sanger Institute for the COVID-19 Genomics UK (COG-UK) Consortium | The Lighthouse Lab in Milton Keynes and Alex Alderton, Roberto Amato, Jeffrey Barrett, Sonia Goncalves, Ewan Harrison, David K. Jackson, Ian Johnston, Dominic Kwiatkowski, Cordelia Langford, John Sillitoe on behalf of the Wellcome Sanger Institute COVID-19 Surveillance Team |
| EPI_ISL_3226898 | Lighthouse Lab in Alderley Park | Wellcome Sanger Institute for the COVID-19 Genomics UK (COG-UK) Consortium | Jacquelyn Wynn, Mairead Hyland, The Lighthouse Lab in Alderley Park and Alex Alderton, Roberto Amato, Jeffrey Barrett, Sonia Goncalves, Ewan Harrison, David K. Jackson, Ian Johnston, Dominic Kwiatkowski, Cordelia Langford, John Sillitoe on behalf of the Wellcome Sanger Institute COVID-19 Surveillance Team |
| EPI_ISL_3226899 | Lighthouse Lab in Milton Keynes | Wellcome Sanger Institute for the COVID-19 Genomics UK (COG-UK) Consortium | The Lighthouse Lab in Milton Keynes and Alex Alderton, Roberto Amato, Jeffrey Barrett, Sonia Goncalves, Ewan Harrison, David K. Jackson, Ian Johnston, Dominic Kwiatkowski, Cordelia Langford, John Sillitoe on behalf of the Wellcome Sanger Institute COVID-19 Surveillance Team |
| EPI_ISL_3226900, EPI_ISL_3226901 | Lighthouse Lab in Glasgow | Wellcome Sanger Institute for the COVID-19 Genomics UK (COG-UK) Consortium | Harper VanSteenhouse, Yumi Kasai, David Gray, Carol Clugston, Anna Dominiczak and Alex Alderton, Roberto Amato, Jeffrey Barrett, Sonia Goncalves, Ewan Harrison, David K. Jackson, Ian Johnston, Dominic Kwiatkowski, Cordelia Langford, John Sillitoe on behalf of the Wellcome Sanger Institute COVID-19 Surveillance Team |
| EPI_ISL_3226902 | Lighthouse Lab in Alderley Park | Wellcome Sanger Institute for the COVID-19 Genomics UK (COG-UK) Consortium | Jacquelyn Wynn, Mairead Hyland, The Lighthouse Lab in Alderley Park and Alex Alderton, Roberto Amato, Jeffrey Barrett, Sonia Goncalves, Ewan Harrison, David K. Jackson, Ian Johnston, Dominic Kwiatkowski, Cordelia Langford, John Sillitoe on behalf of the Wellcome Sanger Institute COVID-19 Surveillance Team |
| EPI_ISL_3226903 | Lighthouse Lab in Milton Keynes | Wellcome Sanger Institute for the COVID-19 Genomics UK (COG-UK) Consortium | The Lighthouse Lab in Milton Keynes and Alex Alderton, Roberto Amato, Jeffrey Barrett, Sonia Goncalves, Ewan Harrison, David K. Jackson, Ian Johnston, Dominic Kwiatkowski, Cordelia Langford, John Sillitoe on behalf of the Wellcome Sanger Institute COVID-19 Surveillance Team |
| EPI_ISL_3226904, EPI_ISL_3226905, EPI_ISL_3226906 | Lighthouse Lab in Alderley Park | Wellcome Sanger Institute for the COVID-19 Genomics UK (COG-UK) Consortium | Jacquelyn Wynn, Mairead Hyland, The Lighthouse Lab in Alderley Park and Alex Alderton, Roberto Amato, Jeffrey Barrett, Sonia Goncalves, Ewan Harrison, David K. Jackson, Ian Johnston, Dominic Kwiatkowski, Cordelia Langford, John Sillitoe on behalf of the Wellcome Sanger Institute COVID-19 Surveillance Team |
| EPI_ISL_3226907, EPI_ISL_3226908 | Lighthouse Lab in Milton Keynes | Wellcome Sanger Institute for the COVID-19 Genomics UK (COG-UK) Consortium | The Lighthouse Lab in Milton Keynes and Alex Alderton, Roberto Amato, Jeffrey Barrett, Sonia Goncalves, Ewan Harrison, David K. Jackson, Ian Johnston, Dominic Kwiatkowski, Cordelia Langford, John Sillitoe on behalf of the Wellcome Sanger Institute COVID-19 Surveillance Team |
| EPI_ISL_3226909 | Lighthouse Lab in Glasgow | Wellcome Sanger Institute for the COVID-19 Genomics UK (COG-UK) Consortium | Harper VanSteenhouse, Yumi Kasai, David Gray, Carol Clugston, Anna Dominiczak and Alex Alderton, Roberto Amato, Jeffrey Barrett, Sonia Goncalves, Ewan Harrison, David K. Jackson, Ian Johnston, Dominic Kwiatkowski, Cordelia Langford, John Sillitoe on behalf of the Wellcome Sanger Institute COVID-19 Surveillance Team |
| EPI_ISL_3226910, EPI_ISL_3226911 | Lighthouse Lab in Alderley Park | Wellcome Sanger Institute for the COVID-19 Genomics UK (COG-UK) Consortium | Jacquelyn Wynn, Mairead Hyland, The Lighthouse Lab in Alderley Park and Alex Alderton, Roberto Amato, Jeffrey Barrett, Sonia Goncalves, Ewan Harrison, David K. Jackson, Ian Johnston, Dominic Kwiatkowski, Cordelia Langford, John Sillitoe on behalf of the Wellcome Sanger Institute COVID-19 Surveillance Team |
| EPI_ISL_3226912 | Lighthouse Lab in Milton Keynes | Wellcome Sanger Institute for the COVID-19 Genomics UK (COG-UK) Consortium | The Lighthouse Lab in Milton Keynes and Alex Alderton, Roberto Amato, Jeffrey Barrett, Sonia Goncalves, Ewan Harrison, David K. Jackson, Ian Johnston, Dominic Kwiatkowski, Cordelia Langford, John Sillitoe on behalf of the Wellcome Sanger Institute COVID-19 Surveillance Team |
| EPI_ISL_3226913, EPI_ISL_3226914, EPI_ISL_3226915 | Lighthouse Lab in Alderley Park | Wellcome Sanger Institute for the COVID-19 Genomics UK (COG-UK) Consortium | Jacquelyn Wynn, Mairead Hyland, The Lighthouse Lab in Alderley Park and Alex Alderton, Roberto Amato, Jeffrey Barrett, Sonia Goncalves, Ewan Harrison, David K. Jackson, Ian Johnston, Dominic Kwiatkowski, Cordelia Langford, John Sillitoe on behalf of the Wellcome Sanger Institute COVID-19 Surveillance Team |
| EPI_ISL_3226916 | Lighthouse Lab in Milton Keynes | Wellcome Sanger Institute for the COVID-19 Genomics UK (COG-UK) Consortium | The Lighthouse Lab in Milton Keynes and Alex Alderton, Roberto Amato, Jeffrey Barrett, Sonia Goncalves, Ewan Harrison, David K. Jackson, Ian Johnston, Dominic Kwiatkowski, Cordelia Langford, John Sillitoe on behalf of the Wellcome Sanger Institute COVID-19 Surveillance Team |
| EPI_ISL_3226917, EPI_ISL_3226918 | Lighthouse Lab in Alderley Park | Wellcome Sanger Institute for the COVID-19 Genomics UK (COG-UK) Consortium | Jacquelyn Wynn, Mairead Hyland, The Lighthouse Lab in Alderley Park and Alex Alderton, Roberto Amato, Jeffrey Barrett, Sonia Goncalves, Ewan Harrison, David K. Jackson, Ian Johnston, Dominic Kwiatkowski, Cordelia Langford, John Sillitoe on behalf of the Wellcome Sanger Institute COVID-19 Surveillance Team |

| | | | |
|---|---|---|---|
| EPI_ISL_3226919 | Lighthouse Lab in Milton Keynes | Wellcome Sanger Institute for the COVID-19 Genomics UK (COG-UK) Consortium | The Lighthouse Lab in Milton Keynes and Alex Alderton, Roberto Amato, Jeffrey Barrett, Sonia Goncalves, Ewan Harrison, David K. Jackson, Ian Johnston, Dominic Kwiatkowski, Cordelia Langford, John Sillitoe on behalf of the Wellcome Sanger Institute COVID-19 Surveillance Team |
| EPI_ISL_3226920, EPI_ISL_3226921 | Lighthouse Lab in Alderley Park | Wellcome Sanger Institute for the COVID-19 Genomics UK (COG-UK) Consortium | Jacquelyn Wynn, Mairead Hyland, The Lighthouse Lab in Alderley Park and Alex Alderton, Roberto Amato, Jeffrey Barrett, Sonia Goncalves, Ewan Harrison, David K. Jackson, Ian Johnston, Dominic Kwiatkowski, Cordelia Langford, John Sillitoe on behalf of the Wellcome Sanger Institute COVID-19 Surveillance Team |
| EPI_ISL_3226922 | Lighthouse Lab in Milton Keynes | Wellcome Sanger Institute for the COVID-19 Genomics UK (COG-UK) Consortium | The Lighthouse Lab in Milton Keynes and Alex Alderton, Roberto Amato, Jeffrey Barrett, Sonia Goncalves, Ewan Harrison, David K. Jackson, Ian Johnston, Dominic Kwiatkowski, Cordelia Langford, John Sillitoe on behalf of the Wellcome Sanger Institute COVID-19 Surveillance Team |
| EPI_ISL_3226923 | Lighthouse Lab in Alderley Park | Wellcome Sanger Institute for the COVID-19 Genomics UK (COG-UK) Consortium | Jacquelyn Wynn, Mairead Hyland, The Lighthouse Lab in Alderley Park and Alex Alderton, Roberto Amato, Jeffrey Barrett, Sonia Goncalves, Ewan Harrison, David K. Jackson, Ian Johnston, Dominic Kwiatkowski, Cordelia Langford, John Sillitoe on behalf of the Wellcome Sanger Institute COVID-19 Surveillance Team |
| EPI_ISL_3226924 | Lighthouse Lab in Glasgow | Wellcome Sanger Institute for the COVID-19 Genomics UK (COG-UK) Consortium | Harper VanSteenhouse, Yumi Kasai, David Gray, Carol Clugston, Anna Dominiczak and Alex Alderton, Roberto Amato, Jeffrey Barrett, Sonia Goncalves, Ewan Harrison, David K. Jackson, Ian Johnston, Dominic Kwiatkowski, Cordelia Langford, John Sillitoe on behalf of the Wellcome Sanger Institute COVID-19 Surveillance Team |
| EPI_ISL_3226925, EPI_ISL_3226926 | Lighthouse Lab in Alderley Park | Wellcome Sanger Institute for the COVID-19 Genomics UK (COG-UK) Consortium | Jacquelyn Wynn, Mairead Hyland, The Lighthouse Lab in Alderley Park and Alex Alderton, Roberto Amato, Jeffrey Barrett, Sonia Goncalves, Ewan Harrison, David K. Jackson, Ian Johnston, Dominic Kwiatkowski, Cordelia Langford, John Sillitoe on behalf of the Wellcome Sanger Institute COVID-19 Surveillance Team |
| EPI_ISL_3226927 | Lighthouse Lab in Glasgow | Wellcome Sanger Institute for the COVID-19 Genomics UK (COG-UK) Consortium | Harper VanSteenhouse, Yumi Kasai, David Gray, Carol Clugston, Anna Dominiczak and Alex Alderton, Roberto Amato, Jeffrey Barrett, Sonia Goncalves, Ewan Harrison, David K. Jackson, Ian Johnston, Dominic Kwiatkowski, Cordelia Langford, John Sillitoe on behalf of the Wellcome Sanger Institute COVID-19 Surveillance Team |
| EPI_ISL_3226928, EPI_ISL_3226929 | Lighthouse Lab in Alderley Park | Wellcome Sanger Institute for the COVID-19 Genomics UK (COG-UK) Consortium | Jacquelyn Wynn, Mairead Hyland, The Lighthouse Lab in Alderley Park and Alex Alderton, Roberto Amato, Jeffrey Barrett, Sonia Goncalves, Ewan Harrison, David K. Jackson, Ian Johnston, Dominic Kwiatkowski, Cordelia Langford, John Sillitoe on behalf of the Wellcome Sanger Institute COVID-19 Surveillance Team |
| EPI_ISL_3226930 | Lighthouse Lab in Glasgow | Wellcome Sanger Institute for the COVID-19 Genomics UK (COG-UK) Consortium | Harper VanSteenhouse, Yumi Kasai, David Gray, Carol Clugston, Anna Dominiczak and Alex Alderton, Roberto Amato, Jeffrey Barrett, Sonia Goncalves, Ewan Harrison, David K. Jackson, Ian Johnston, Dominic Kwiatkowski, Cordelia Langford, John Sillitoe on behalf of the Wellcome Sanger Institute COVID-19 Surveillance Team |
| EPI_ISL_3226931 | Lighthouse Lab in Alderley Park | Wellcome Sanger Institute for the COVID-19 Genomics UK (COG-UK) Consortium | Jacquelyn Wynn, Mairead Hyland, The Lighthouse Lab in Alderley Park and Alex Alderton, Roberto Amato, Jeffrey Barrett, Sonia Goncalves, Ewan Harrison, David K. Jackson, Ian Johnston, Dominic Kwiatkowski, Cordelia Langford, John Sillitoe on behalf of the Wellcome Sanger Institute COVID-19 Surveillance Team |
| EPI_ISL_3226932 | Lighthouse Lab in Glasgow | Wellcome Sanger Institute for the COVID-19 Genomics UK (COG-UK) Consortium | Harper VanSteenhouse, Yumi Kasai, David Gray, Carol Clugston, Anna Dominiczak and Alex Alderton, Roberto Amato, Jeffrey Barrett, Sonia Goncalves, Ewan Harrison, David K. Jackson, Ian Johnston, Dominic Kwiatkowski, Cordelia Langford, John Sillitoe on behalf of the Wellcome Sanger Institute COVID-19 Surveillance Team |
| EPI_ISL_3226933, EPI_ISL_3226934 | Lighthouse Lab in Alderley Park | Wellcome Sanger Institute for the COVID-19 Genomics UK (COG-UK) Consortium | Jacquelyn Wynn, Mairead Hyland, The Lighthouse Lab in Alderley Park and Alex Alderton, Roberto Amato, Jeffrey Barrett, Sonia Goncalves, Ewan Harrison, David K. Jackson, Ian Johnston, Dominic Kwiatkowski, Cordelia Langford, John Sillitoe on behalf of the Wellcome Sanger Institute COVID-19 Surveillance Team |
| EPI_ISL_3226935, EPI_ISL_3226936 | Lighthouse Lab in Glasgow | Wellcome Sanger Institute for the COVID-19 Genomics UK (COG-UK) Consortium | Harper VanSteenhouse, Yumi Kasai, David Gray, Carol Clugston, Anna Dominiczak and Alex Alderton, Roberto Amato, Jeffrey Barrett, Sonia Goncalves, Ewan Harrison, David K. Jackson, Ian Johnston, Dominic Kwiatkowski, Cordelia Langford, John Sillitoe on behalf of the Wellcome Sanger Institute COVID-19 Surveillance Team |
| EPI_ISL_3226937 | Lighthouse Lab in Alderley Park | Wellcome Sanger Institute for the COVID-19 Genomics UK (COG-UK) Consortium | Jacquelyn Wynn, Mairead Hyland, The Lighthouse Lab in Alderley Park and Alex Alderton, Roberto Amato, Jeffrey Barrett, Sonia Goncalves, Ewan Harrison, David K. Jackson, Ian Johnston, Dominic Kwiatkowski, Cordelia Langford, John Sillitoe on behalf of the Wellcome Sanger Institute COVID-19 Surveillance Team |
| EPI_ISL_3226938 | Lighthouse Lab in Glasgow | Wellcome Sanger Institute for the COVID-19 Genomics UK (COG-UK) Consortium | Harper VanSteenhouse, Yumi Kasai, David Gray, Carol Clugston, Anna Dominiczak and Alex Alderton, Roberto Amato, Jeffrey Barrett, Sonia Goncalves, Ewan Harrison, David K. Jackson, Ian Johnston, Dominic Kwiatkowski, Cordelia Langford, John Sillitoe on behalf of the Wellcome Sanger Institute COVID-19 Surveillance Team |
| EPI_ISL_3226939, EPI_ISL_3226940, EPI_ISL_3226941 | Lighthouse Lab in Alderley Park | Wellcome Sanger Institute for the COVID-19 Genomics UK (COG-UK) Consortium | Jacquelyn Wynn, Mairead Hyland, The Lighthouse Lab in Alderley Park and Alex Alderton, Roberto Amato, Jeffrey Barrett, Sonia Goncalves, Ewan Harrison, David K. Jackson, Ian Johnston, Dominic Kwiatkowski, Cordelia Langford, John Sillitoe on behalf of the Wellcome Sanger Institute COVID-19 Surveillance Team |
| EPI_ISL_3226942 | Lighthouse Lab in Milton Keynes | Wellcome Sanger Institute for the COVID-19 Genomics UK (COG-UK) Consortium | The Lighthouse Lab in Milton Keynes and Alex Alderton, Roberto Amato, Jeffrey Barrett, Sonia Goncalves, Ewan Harrison, David K. Jackson, Ian Johnston, Dominic Kwiatkowski, Cordelia Langford, John Sillitoe on behalf of the Wellcome Sanger Institute COVID-19 Surveillance Team |
| EPI_ISL_3226943 | Lighthouse Lab in Glasgow | Wellcome Sanger Institute for the COVID-19 Genomics UK (COG-UK) Consortium | Harper VanSteenhouse, Yumi Kasai, David Gray, Carol Clugston, Anna Dominiczak and Alex Alderton, Roberto Amato, Jeffrey Barrett, Sonia Goncalves, Ewan Harrison, David K. Jackson, Ian Johnston, Dominic Kwiatkowski, Cordelia Langford, John Sillitoe on behalf of the Wellcome Sanger Institute COVID-19 Surveillance Team |
| EPI_ISL_3226944, EPI_ISL_3226945 | Lighthouse Lab in Milton Keynes | Wellcome Sanger Institute for the COVID-19 Genomics UK (COG-UK) Consortium | The Lighthouse Lab in Milton Keynes and Alex Alderton, Roberto Amato, Jeffrey Barrett, Sonia Goncalves, Ewan Harrison, David K. Jackson, Ian Johnston, Dominic Kwiatkowski, Cordelia Langford, John Sillitoe on behalf of the Wellcome Sanger Institute COVID-19 Surveillance Team |
| EPI_ISL_3226946 | Lighthouse Lab in Glasgow | Wellcome Sanger Institute for the COVID-19 Genomics UK (COG-UK) Consortium | Harper VanSteenhouse, Yumi Kasai, David Gray, Carol Clugston, Anna Dominiczak and Alex Alderton, Roberto Amato, Jeffrey Barrett, Sonia Goncalves, Ewan Harrison, David K. Jackson, Ian Johnston, Dominic Kwiatkowski, Cordelia Langford, John Sillitoe on behalf of the Wellcome Sanger Institute COVID-19 Surveillance Team |
| EPI_ISL_3226947, EPI_ISL_3226948, EPI_ISL_3226949 | Lighthouse Lab in Alderley Park | Wellcome Sanger Institute for the COVID-19 Genomics UK (COG-UK) Consortium | Jacquelyn Wynn, Mairead Hyland, The Lighthouse Lab in Alderley Park and Alex Alderton, Roberto Amato, Jeffrey Barrett, Sonia Goncalves, Ewan Harrison, David K. Jackson, Ian Johnston, Dominic Kwiatkowski, Cordelia Langford, John Sillitoe on behalf of the Wellcome Sanger Institute COVID-19 Surveillance Team |
| EPI_ISL_3226950, EPI_ISL_3226951 | Lighthouse Lab in Glasgow | Wellcome Sanger Institute for the COVID-19 Genomics UK (COG-UK) Consortium | Harper VanSteenhouse, Yumi Kasai, David Gray, Carol Clugston, Anna Dominiczak and Alex Alderton, Roberto Amato, Jeffrey Barrett, Sonia Goncalves, Ewan Harrison, David K. Jackson, Ian Johnston, Dominic Kwiatkowski, Cordelia Langford, John Sillitoe on behalf of the Wellcome Sanger Institute COVID-19 Surveillance Team |
| EPI_ISL_3226952, EPI_ISL_3226953 | Lighthouse Lab in Alderley Park | Wellcome Sanger Institute for the COVID-19 Genomics UK (COG-UK) Consortium | Jacquelyn Wynn, Mairead Hyland, The Lighthouse Lab in Alderley Park and Alex Alderton, Roberto Amato, Jeffrey Barrett, Sonia Goncalves, Ewan Harrison, David K. Jackson, Ian Johnston, Dominic Kwiatkowski, Cordelia Langford, John Sillitoe on behalf of the Wellcome Sanger Institute COVID-19 Surveillance Team |
| EPI_ISL_3226954, EPI_ISL_3226955 | Lighthouse Lab in Milton Keynes | Wellcome Sanger Institute for the COVID-19 Genomics UK (COG-UK) Consortium | The Lighthouse Lab in Milton Keynes and Alex Alderton, Roberto Amato, Jeffrey Barrett, Sonia Goncalves, Ewan Harrison, David K. Jackson, Ian Johnston, Dominic Kwiatkowski, Cordelia Langford, John Sillitoe on behalf of the Wellcome Sanger Institute COVID-19 Surveillance Team |
| EPI_ISL_3226956 | Lighthouse Lab in Glasgow | Wellcome Sanger Institute for the COVID-19 Genomics UK (COG-UK) Consortium | Harper VanSteenhouse, Yumi Kasai, David Gray, Carol Clugston, Anna Dominiczak and Alex Alderton, Roberto Amato, Jeffrey Barrett, Sonia Goncalves, Ewan Harrison, David K. Jackson, Ian Johnston, Dominic Kwiatkowski, Cordelia Langford, John Sillitoe on behalf of the Wellcome Sanger Institute COVID-19 Surveillance Team |
| EPI_ISL_3226957 | Lighthouse Lab in Milton Keynes | Wellcome Sanger Institute for the COVID-19 Genomics UK (COG-UK) Consortium | The Lighthouse Lab in Milton Keynes and Alex Alderton, Roberto Amato, Jeffrey Barrett, Sonia Goncalves, Ewan Harrison, David K. Jackson, Ian Johnston, Dominic Kwiatkowski, Cordelia Langford, John Sillitoe on behalf of the Wellcome Sanger Institute COVID-19 Surveillance Team |
| EPI_ISL_3226958, EPI_ISL_3226959, EPI_ISL_3226960, EPI_ISL_3226961 | Lighthouse Lab in Alderley Park | Wellcome Sanger Institute for the COVID-19 Genomics UK (COG-UK) Consortium | Jacquelyn Wynn, Mairead Hyland, The Lighthouse Lab in Alderley Park and Alex Alderton, Roberto Amato, Jeffrey Barrett, Sonia Goncalves, Ewan Harrison, David K. Jackson, Ian Johnston, Dominic Kwiatkowski, Cordelia Langford, John Sillitoe on behalf of the Wellcome Sanger Institute COVID-19 Surveillance Team |
| EPI_ISL_3226962 | Lighthouse Lab in Glasgow | Wellcome Sanger Institute for the COVID-19 Genomics UK (COG-UK) Consortium | Harper VanSteenhouse, Yumi Kasai, David Gray, Carol Clugston, Anna Dominiczak and Alex Alderton, Roberto Amato, Jeffrey Barrett, Sonia Goncalves, Ewan Harrison, David K. Jackson, Ian Johnston, Dominic Kwiatkowski, Cordelia Langford, John Sillitoe on behalf of the Wellcome Sanger Institute COVID-19 Surveillance Team |
| EPI_ISL_3226963 | Lighthouse Lab in Alderley Park | Wellcome Sanger Institute for the COVID-19 Genomics UK (COG-UK) Consortium | Jacquelyn Wynn, Mairead Hyland, The Lighthouse Lab in Alderley Park and Alex Alderton, Roberto Amato, Jeffrey Barrett, Sonia Goncalves, Ewan Harrison, David K. Jackson, Ian Johnston, Dominic Kwiatkowski, Cordelia Langford, John Sillitoe on behalf of the Wellcome Sanger Institute COVID-19 Surveillance Team |
| EPI_ISL_3226964 | Lighthouse Lab in Milton Keynes | Wellcome Sanger Institute for the COVID-19 Genomics UK (COG-UK) Consortium | The Lighthouse Lab in Milton Keynes and Alex Alderton, Roberto Amato, Jeffrey Barrett, Sonia Goncalves, Ewan Harrison, David K. Jackson, Ian Johnston, Dominic Kwiatkowski, Cordelia Langford, John Sillitoe on behalf of the Wellcome Sanger Institute COVID-19 Surveillance Team |
| EPI_ISL_3226965, EPI_ISL_3226966, EPI_ISL_3226967 | Lighthouse Lab in Alderley Park | Wellcome Sanger Institute for the COVID-19 Genomics UK (COG-UK) Consortium | Jacquelyn Wynn, Mairead Hyland, The Lighthouse Lab in Alderley Park and Alex Alderton, Roberto Amato, Jeffrey Barrett, Sonia Goncalves, Ewan Harrison, David K. Jackson, Ian Johnston, Dominic Kwiatkowski, Cordelia Langford, John Sillitoe on behalf of the Wellcome Sanger Institute COVID-19 Surveillance Team |
| EPI_ISL_3226968, EPI_ISL_3226969 | Lighthouse Lab in Glasgow | Wellcome Sanger Institute for the COVID-19 Genomics UK (COG-UK) Consortium | Harper VanSteenhouse, Yumi Kasai, David Gray, Carol Clugston, Anna Dominiczak and Alex Alderton, Roberto Amato, Jeffrey Barrett, Sonia Goncalves, Ewan Harrison, David K. Jackson, Ian Johnston, Dominic Kwiatkowski, Cordelia Langford, John Sillitoe on behalf of the Wellcome Sanger Institute COVID-19 Surveillance Team |
| EPI_ISL_3226970 | Lighthouse Lab in Milton Keynes | Wellcome Sanger Institute for the COVID-19 Genomics UK (COG-UK) Consortium | The Lighthouse Lab in Milton Keynes and Alex Alderton, Roberto Amato, Jeffrey Barrett, Sonia Goncalves, Ewan Harrison, David K. Jackson, Ian Johnston, Dominic Kwiatkowski, Cordelia Langford, John Sillitoe on behalf of the Wellcome Sanger Institute COVID-19 Surveillance Team |
| EPI_ISL_3226971, EPI_ISL_3226972, EPI_ISL_3226973 | Lighthouse Lab in Alderley Park | Wellcome Sanger Institute for the COVID-19 Genomics UK (COG-UK) Consortium | Jacquelyn Wynn, Mairead Hyland, The Lighthouse Lab in Alderley Park and Alex Alderton, Roberto Amato, Jeffrey Barrett, Sonia Goncalves, Ewan Harrison, David K. Jackson, Ian Johnston, Dominic Kwiatkowski, Cordelia Langford, John Sillitoe on behalf of the Wellcome Sanger Institute COVID-19 Surveillance Team |
| EPI_ISL_3226974 | Lighthouse Lab in Milton Keynes | Wellcome Sanger Institute for the COVID-19 Genomics UK (COG-UK) Consortium | The Lighthouse Lab in Milton Keynes and Alex Alderton, Roberto Amato, Jeffrey Barrett, Sonia Goncalves, Ewan Harrison, David K. Jackson, Ian Johnston, Dominic Kwiatkowski, Cordelia Langford, John Sillitoe on behalf of the Wellcome Sanger Institute COVID-19 Surveillance Team |
| EPI_ISL_3226975 | Lighthouse Lab in Alderley Park | Wellcome Sanger Institute for the COVID-19 Genomics UK (COG-UK) Consortium | Jacquelyn Wynn, Mairead Hyland, The Lighthouse Lab in Alderley Park and Alex Alderton, Roberto Amato, Jeffrey Barrett, Sonia Goncalves, Ewan Harrison, David K. Jackson, Ian Johnston, Dominic Kwiatkowski, Cordelia Langford, John Sillitoe on behalf of the Wellcome Sanger Institute COVID-19 Surveillance Team |
| EPI_ISL_3226976 | Lighthouse Lab in Milton Keynes | Wellcome Sanger Institute for the COVID-19 Genomics UK (COG-UK) Consortium | The Lighthouse Lab in Milton Keynes and Alex Alderton, Roberto Amato, Jeffrey Barrett, Sonia Goncalves, Ewan Harrison, David K. Jackson, Ian Johnston, Dominic Kwiatkowski, Cordelia Langford, John Sillitoe on behalf of the Wellcome Sanger Institute COVID-19 Surveillance Team |
| EPI_ISL_3226977, EPI_ISL_3226978, EPI_ISL_3226979, EPI_ISL_3226980 | Lighthouse Lab in Alderley Park | Wellcome Sanger Institute for the COVID-19 Genomics UK (COG-UK) Consortium | Jacquelyn Wynn, Mairead Hyland, The Lighthouse Lab in Alderley Park and Alex Alderton, Roberto Amato, Jeffrey Barrett, Sonia Goncalves, Ewan Harrison, David K. Jackson, Ian Johnston, Dominic Kwiatkowski, Cordelia Langford, John Sillitoe on behalf of the Wellcome Sanger Institute COVID-19 Surveillance Team |
| EPI_ISL_3226981 | Lighthouse Lab in Glasgow | Wellcome Sanger Institute for the COVID-19 Genomics UK (COG-UK) Consortium | Harper VanSteenhouse, Yumi Kasai, David Gray, Carol Clugston, Anna Dominiczak and Alex Alderton, Roberto Amato, Jeffrey Barrett, Sonia Goncalves, Ewan Harrison, David K. Jackson, Ian Johnston, Dominic Kwiatkowski, Cordelia Langford, John Sillitoe on behalf of the Wellcome Sanger Institute COVID-19 Surveillance Team |
| EPI_ISL_3226982 | Lighthouse Lab in Milton Keynes | Wellcome Sanger Institute for the COVID-19 Genomics UK (COG-UK) Consortium | The Lighthouse Lab in Milton Keynes and Alex Alderton, Roberto Amato, Jeffrey Barrett, Sonia Goncalves, Ewan Harrison, David K. Jackson, Ian Johnston, Dominic Kwiatkowski, Cordelia Langford, John Sillitoe on behalf of the Wellcome Sanger Institute COVID-19 Surveillance Team |
| EPI_ISL_3226983 | Lighthouse Lab in Alderley Park | Wellcome Sanger Institute for the COVID-19 Genomics UK (COG-UK) Consortium | Jacquelyn Wynn, Mairead Hyland, The Lighthouse Lab in Alderley Park and Alex Alderton, Roberto Amato, Jeffrey Barrett, Sonia Goncalves, Ewan Harrison, David K. Jackson, Ian Johnston, Dominic Kwiatkowski, Cordelia Langford, John Sillitoe on behalf of the Wellcome Sanger Institute COVID-19 Surveillance Team |
| EPI_ISL_3226984 | Lighthouse Lab in Milton Keynes | Wellcome Sanger Institute for the COVID-19 Genomics UK (COG-UK) Consortium | The Lighthouse Lab in Milton Keynes and Alex Alderton, Roberto Amato, Jeffrey Barrett, Sonia Goncalves, Ewan Harrison, David K. Jackson, Ian Johnston, Dominic Kwiatkowski, Cordelia Langford, John Sillitoe on behalf of the Wellcome Sanger Institute COVID-19 Surveillance Team |

| | | | |
|---|---|---|---|
| EPI_ISL_3226985, EPI_ISL_3226986 | Lighthouse Lab in Glasgow | Wellcome Sanger Institute for the COVID-19 Genomics UK (COG-UK) Consortium | Harper VanSteenhouse, Yumi Kasai, David Gray, Carol Clugston, Anna Dominiczak and Alex Alderton, Roberto Amato, Jeffrey Barrett, Sonia Goncalves, Ewan Harrison, David K. Jackson, Ian Johnston, Dominic Kwiatkowski, Cordelia Langford, John Sillitoe on behalf of the Wellcome Sanger Institute COVID-19 Surveillance Team |
| EPI_ISL_3226987 | Lighthouse Lab in Alderley Park | Wellcome Sanger Institute for the COVID-19 Genomics UK (COG-UK) Consortium | Jacquelyn Wynn, Mairead Hyland, The Lighthouse Lab in Alderley Park and Alex Alderton, Roberto Amato, Jeffrey Barrett, Sonia Goncalves, Ewan Harrison, David K. Jackson, Ian Johnston, Dominic Kwiatkowski, Cordelia Langford, John Sillitoe on behalf of the Wellcome Sanger Institute COVID-19 Surveillance Team |
| EPI_ISL_3226988 | Lighthouse Lab in Milton Keynes | Wellcome Sanger Institute for the COVID-19 Genomics UK (COG-UK) Consortium | The Lighthouse Lab in Milton Keynes and Alex Alderton, Roberto Amato, Jeffrey Barrett, Sonia Goncalves, Ewan Harrison, David K. Jackson, Ian Johnston, Dominic Kwiatkowski, Cordelia Langford, John Sillitoe on behalf of the Wellcome Sanger Institute COVID-19 Surveillance Team |
| EPI_ISL_3226989 | Lighthouse Lab in Alderley Park | Wellcome Sanger Institute for the COVID-19 Genomics UK (COG-UK) Consortium | Jacquelyn Wynn, Mairead Hyland, The Lighthouse Lab in Alderley Park and Alex Alderton, Roberto Amato, Jeffrey Barrett, Sonia Goncalves, Ewan Harrison, David K. Jackson, Ian Johnston, Dominic Kwiatkowski, Cordelia Langford, John Sillitoe on behalf of the Wellcome Sanger Institute COVID-19 Surveillance Team |
| EPI_ISL_3226990 | Lighthouse Lab in Glasgow | Wellcome Sanger Institute for the COVID-19 Genomics UK (COG-UK) Consortium | Harper VanSteenhouse, Yumi Kasai, David Gray, Carol Clugston, Anna Dominiczak and Alex Alderton, Roberto Amato, Jeffrey Barrett, Sonia Goncalves, Ewan Harrison, David K. Jackson, Ian Johnston, Dominic Kwiatkowski, Cordelia Langford, John Sillitoe on behalf of the Wellcome Sanger Institute COVID-19 Surveillance Team |
| EPI_ISL_3226991, EPI_ISL_3226992, EPI_ISL_3226993, EPI_ISL_3226994, EPI_ISL_3226995, EPI_ISL_3226996, EPI_ISL_3226997, EPI_ISL_3226998, EPI_ISL_3226999 | Lighthouse Lab in Alderley Park | Wellcome Sanger Institute for the COVID-19 Genomics UK (COG-UK) Consortium | Jacquelyn Wynn, Mairead Hyland, The Lighthouse Lab in Alderley Park and Alex Alderton, Roberto Amato, Jeffrey Barrett, Sonia Goncalves, Ewan Harrison, David K. Jackson, Ian Johnston, Dominic Kwiatkowski, Cordelia Langford, John Sillitoe on behalf of the Wellcome Sanger Institute COVID-19 Surveillance Team |
| EPI_ISL_3227000 | Lighthouse Lab in Milton Keynes | Wellcome Sanger Institute for the COVID-19 Genomics UK (COG-UK) Consortium | The Lighthouse Lab in Milton Keynes and Alex Alderton, Roberto Amato, Jeffrey Barrett, Sonia Goncalves, Ewan Harrison, David K. Jackson, Ian Johnston, Dominic Kwiatkowski, Cordelia Langford, John Sillitoe on behalf of the Wellcome Sanger Institute COVID-19 Surveillance Team |
| EPI_ISL_3227001, EPI_ISL_3227002, EPI_ISL_3227003 | Lighthouse Lab in Alderley Park | Wellcome Sanger Institute for the COVID-19 Genomics UK (COG-UK) Consortium | Jacquelyn Wynn, Mairead Hyland, The Lighthouse Lab in Alderley Park and Alex Alderton, Roberto Amato, Jeffrey Barrett, Sonia Goncalves, Ewan Harrison, David K. Jackson, Ian Johnston, Dominic Kwiatkowski, Cordelia Langford, John Sillitoe on behalf of the Wellcome Sanger Institute COVID-19 Surveillance Team |
| EPI_ISL_3227004 | Lighthouse Lab in Glasgow | Wellcome Sanger Institute for the COVID-19 Genomics UK (COG-UK) Consortium | Harper VanSteenhouse, Yumi Kasai, David Gray, Carol Clugston, Anna Dominiczak and Alex Alderton, Roberto Amato, Jeffrey Barrett, Sonia Goncalves, Ewan Harrison, David K. Jackson, Ian Johnston, Dominic Kwiatkowski, Cordelia Langford, John Sillitoe on behalf of the Wellcome Sanger Institute COVID-19 Surveillance Team |
| EPI_ISL_3227005 | Lighthouse Lab in Milton Keynes | Wellcome Sanger Institute for the COVID-19 Genomics UK (COG-UK) Consortium | The Lighthouse Lab in Milton Keynes and Alex Alderton, Roberto Amato, Jeffrey Barrett, Sonia Goncalves, Ewan Harrison, David K. Jackson, Ian Johnston, Dominic Kwiatkowski, Cordelia Langford, John Sillitoe on behalf of the Wellcome Sanger Institute COVID-19 Surveillance Team |
| EPI_ISL_3227006 | Lighthouse Lab in Alderley Park | Wellcome Sanger Institute for the COVID-19 Genomics UK (COG-UK) Consortium | Jacquelyn Wynn, Mairead Hyland, The Lighthouse Lab in Alderley Park and Alex Alderton, Roberto Amato, Jeffrey Barrett, Sonia Goncalves, Ewan Harrison, David K. Jackson, Ian Johnston, Dominic Kwiatkowski, Cordelia Langford, John Sillitoe on behalf of the Wellcome Sanger Institute COVID-19 Surveillance Team |
| EPI_ISL_3227007 | Lighthouse Lab in Milton Keynes | Wellcome Sanger Institute for the COVID-19 Genomics UK (COG-UK) Consortium | The Lighthouse Lab in Milton Keynes and Alex Alderton, Roberto Amato, Jeffrey Barrett, Sonia Goncalves, Ewan Harrison, David K. Jackson, Ian Johnston, Dominic Kwiatkowski, Cordelia Langford, John Sillitoe on behalf of the Wellcome Sanger Institute COVID-19 Surveillance Team |
| EPI_ISL_3227008 | Lighthouse Lab in Glasgow | Wellcome Sanger Institute for the COVID-19 Genomics UK (COG-UK) Consortium | Harper VanSteenhouse, Yumi Kasai, David Gray, Carol Clugston, Anna Dominiczak and Alex Alderton, Roberto Amato, Jeffrey Barrett, Sonia Goncalves, Ewan Harrison, David K. Jackson, Ian Johnston, Dominic Kwiatkowski, Cordelia Langford, John Sillitoe on behalf of the Wellcome Sanger Institute COVID-19 Surveillance Team |
| EPI_ISL_3227009, EPI_ISL_3227010 | Lighthouse Lab in Alderley Park | Wellcome Sanger Institute for the COVID-19 Genomics UK (COG-UK) Consortium | Jacquelyn Wynn, Mairead Hyland, The Lighthouse Lab in Alderley Park and Alex Alderton, Roberto Amato, Jeffrey Barrett, Sonia Goncalves, Ewan Harrison, David K. Jackson, Ian Johnston, Dominic Kwiatkowski, Cordelia Langford, John Sillitoe on behalf of the Wellcome Sanger Institute COVID-19 Surveillance Team |
| EPI_ISL_3227011, EPI_ISL_3227012 | Lighthouse Lab in Milton Keynes | Wellcome Sanger Institute for the COVID-19 Genomics UK (COG-UK) Consortium | The Lighthouse Lab in Milton Keynes and Alex Alderton, Roberto Amato, Jeffrey Barrett, Sonia Goncalves, Ewan Harrison, David K. Jackson, Ian Johnston, Dominic Kwiatkowski, Cordelia Langford, John Sillitoe on behalf of the Wellcome Sanger Institute COVID-19 Surveillance Team |
| EPI_ISL_3227013 | Lighthouse Lab in Glasgow | Wellcome Sanger Institute for the COVID-19 Genomics UK (COG-UK) Consortium | Harper VanSteenhouse, Yumi Kasai, David Gray, Carol Clugston, Anna Dominiczak and Alex Alderton, Roberto Amato, Jeffrey Barrett, Sonia Goncalves, Ewan Harrison, David K. Jackson, Ian Johnston, Dominic Kwiatkowski, Cordelia Langford, John Sillitoe on behalf of the Wellcome Sanger Institute COVID-19 Surveillance Team |
| EPI_ISL_3227014, EPI_ISL_3227015 | Lighthouse Lab in Alderley Park | Wellcome Sanger Institute for the COVID-19 Genomics UK (COG-UK) Consortium | Jacquelyn Wynn, Mairead Hyland, The Lighthouse Lab in Alderley Park and Alex Alderton, Roberto Amato, Jeffrey Barrett, Sonia Goncalves, Ewan Harrison, David K. Jackson, Ian Johnston, Dominic Kwiatkowski, Cordelia Langford, John Sillitoe on behalf of the Wellcome Sanger Institute COVID-19 Surveillance Team |
| EPI_ISL_3227016, EPI_ISL_3227017, EPI_ISL_3227018, EPI_ISL_3227019, EPI_ISL_3227020 | Lighthouse Lab in Milton Keynes | Wellcome Sanger Institute for the COVID-19 Genomics UK (COG-UK) Consortium | The Lighthouse Lab in Milton Keynes and Alex Alderton, Roberto Amato, Jeffrey Barrett, Sonia Goncalves, Ewan Harrison, David K. Jackson, Ian Johnston, Dominic Kwiatkowski, Cordelia Langford, John Sillitoe on behalf of the Wellcome Sanger Institute COVID-19 Surveillance Team |
| EPI_ISL_3227021 | Lighthouse Lab in Glasgow | Wellcome Sanger Institute for the COVID-19 Genomics UK (COG-UK) Consortium | Harper VanSteenhouse, Yumi Kasai, David Gray, Carol Clugston, Anna Dominiczak and Alex Alderton, Roberto Amato, Jeffrey Barrett, Sonia Goncalves, Ewan Harrison, David K. Jackson, Ian Johnston, Dominic Kwiatkowski, Cordelia Langford, John Sillitoe on behalf of the Wellcome Sanger Institute COVID-19 Surveillance Team |
| EPI_ISL_3227022, EPI_ISL_3227023, EPI_ISL_3227024, EPI_ISL_3227025 | Lighthouse Lab in Alderley Park | Wellcome Sanger Institute for the COVID-19 Genomics UK (COG-UK) Consortium | Jacquelyn Wynn, Mairead Hyland, The Lighthouse Lab in Alderley Park and Alex Alderton, Roberto Amato, Jeffrey Barrett, Sonia Goncalves, Ewan Harrison, David K. Jackson, Ian Johnston, Dominic Kwiatkowski, Cordelia Langford, John Sillitoe on behalf of the Wellcome Sanger Institute COVID-19 Surveillance Team |
| EPI_ISL_3227026, EPI_ISL_3227027 | Lighthouse Lab in Milton Keynes | Wellcome Sanger Institute for the COVID-19 Genomics UK (COG-UK) Consortium | The Lighthouse Lab in Milton Keynes and Alex Alderton, Roberto Amato, Jeffrey Barrett, Sonia Goncalves, Ewan Harrison, David K. Jackson, Ian Johnston, Dominic Kwiatkowski, Cordelia Langford, John Sillitoe on behalf of the Wellcome Sanger Institute COVID-19 Surveillance Team |
| EPI_ISL_3227028 | Lighthouse Lab in Alderley Park | Wellcome Sanger Institute for the COVID-19 Genomics UK (COG-UK) Consortium | Jacquelyn Wynn, Mairead Hyland, The Lighthouse Lab in Alderley Park and Alex Alderton, Roberto Amato, Jeffrey Barrett, Sonia Goncalves, Ewan Harrison, David K. Jackson, Ian Johnston, Dominic Kwiatkowski, Cordelia Langford, John Sillitoe on behalf of the Wellcome Sanger Institute COVID-19 Surveillance Team |
| EPI_ISL_3227029 | Lighthouse Lab in Glasgow | Wellcome Sanger Institute for the COVID-19 Genomics UK (COG-UK) Consortium | Harper VanSteenhouse, Yumi Kasai, David Gray, Carol Clugston, Anna Dominiczak and Alex Alderton, Roberto Amato, Jeffrey Barrett, Sonia Goncalves, Ewan Harrison, David K. Jackson, Ian Johnston, Dominic Kwiatkowski, Cordelia Langford, John Sillitoe on behalf of the Wellcome Sanger Institute COVID-19 Surveillance Team |
| EPI_ISL_3227030, EPI_ISL_3227031, EPI_ISL_3227032, EPI_ISL_3227033 | Lighthouse Lab in Alderley Park | Wellcome Sanger Institute for the COVID-19 Genomics UK (COG-UK) Consortium | Jacquelyn Wynn, Mairead Hyland, The Lighthouse Lab in Alderley Park and Alex Alderton, Roberto Amato, Jeffrey Barrett, Sonia Goncalves, Ewan Harrison, David K. Jackson, Ian Johnston, Dominic Kwiatkowski, Cordelia Langford, John Sillitoe on behalf of the Wellcome Sanger Institute COVID-19 Surveillance Team |
| EPI_ISL_3227034 | Lighthouse Lab in Milton Keynes | Wellcome Sanger Institute for the COVID-19 Genomics UK (COG-UK) Consortium | The Lighthouse Lab in Milton Keynes and Alex Alderton, Roberto Amato, Jeffrey Barrett, Sonia Goncalves, Ewan Harrison, David K. Jackson, Ian Johnston, Dominic Kwiatkowski, Cordelia Langford, John Sillitoe on behalf of the Wellcome Sanger Institute COVID-19 Surveillance Team |
| EPI_ISL_3227035 | Lighthouse Lab in Alderley Park | Wellcome Sanger Institute for the COVID-19 Genomics UK (COG-UK) Consortium | Jacquelyn Wynn, Mairead Hyland, The Lighthouse Lab in Alderley Park and Alex Alderton, Roberto Amato, Jeffrey Barrett, Sonia Goncalves, Ewan Harrison, David K. Jackson, Ian Johnston, Dominic Kwiatkowski, Cordelia Langford, John Sillitoe on behalf of the Wellcome Sanger Institute COVID-19 Surveillance Team |
| EPI_ISL_3227036, EPI_ISL_3227037 | Lighthouse Lab in Glasgow | Wellcome Sanger Institute for the COVID-19 Genomics UK (COG-UK) Consortium | Harper VanSteenhouse, Yumi Kasai, David Gray, Carol Clugston, Anna Dominiczak and Alex Alderton, Roberto Amato, Jeffrey Barrett, Sonia Goncalves, Ewan Harrison, David K. Jackson, Ian Johnston, Dominic Kwiatkowski, Cordelia Langford, John Sillitoe on behalf of the Wellcome Sanger Institute COVID-19 Surveillance Team |
| EPI_ISL_3227038 | Lighthouse Lab in Milton Keynes | Wellcome Sanger Institute for the COVID-19 Genomics UK (COG-UK) Consortium | The Lighthouse Lab in Milton Keynes and Alex Alderton, Roberto Amato, Jeffrey Barrett, Sonia Goncalves, Ewan Harrison, David K. Jackson, Ian Johnston, Dominic Kwiatkowski, Cordelia Langford, John Sillitoe on behalf of the Wellcome Sanger Institute COVID-19 Surveillance Team |
| EPI_ISL_3227039, EPI_ISL_3227040, EPI_ISL_3227041 | Lighthouse Lab in Glasgow | Wellcome Sanger Institute for the COVID-19 Genomics UK (COG-UK) Consortium | Harper VanSteenhouse, Yumi Kasai, David Gray, Carol Clugston, Anna Dominiczak and Alex Alderton, Roberto Amato, Jeffrey Barrett, Sonia Goncalves, Ewan Harrison, David K. Jackson, Ian Johnston, Dominic Kwiatkowski, Cordelia Langford, John Sillitoe on behalf of the Wellcome Sanger Institute COVID-19 Surveillance Team |
| EPI_ISL_3227042 | Lighthouse Lab in Alderley Park | Wellcome Sanger Institute for the COVID-19 Genomics UK (COG-UK) Consortium | Jacquelyn Wynn, Mairead Hyland, The Lighthouse Lab in Alderley Park and Alex Alderton, Roberto Amato, Jeffrey Barrett, Sonia Goncalves, Ewan Harrison, David K. Jackson, Ian Johnston, Dominic Kwiatkowski, Cordelia Langford, John Sillitoe on behalf of the Wellcome Sanger Institute COVID-19 Surveillance Team |
| EPI_ISL_3227043 | Lighthouse Lab in Glasgow | Wellcome Sanger Institute for the COVID-19 Genomics UK (COG-UK) Consortium | Harper VanSteenhouse, Yumi Kasai, David Gray, Carol Clugston, Anna Dominiczak and Alex Alderton, Roberto Amato, Jeffrey Barrett, Sonia Goncalves, Ewan Harrison, David K. Jackson, Ian Johnston, Dominic Kwiatkowski, Cordelia Langford, John Sillitoe on behalf of the Wellcome Sanger Institute COVID-19 Surveillance Team |
| EPI_ISL_3227044, EPI_ISL_3227045, EPI_ISL_3227046 | Lighthouse Lab in Milton Keynes | Wellcome Sanger Institute for the COVID-19 Genomics UK (COG-UK) Consortium | The Lighthouse Lab in Milton Keynes and Alex Alderton, Roberto Amato, Jeffrey Barrett, Sonia Goncalves, Ewan Harrison, David K. Jackson, Ian Johnston, Dominic Kwiatkowski, Cordelia Langford, John Sillitoe on behalf of the Wellcome Sanger Institute COVID-19 Surveillance Team |
| EPI_ISL_3227047 | Lighthouse Lab in Glasgow | Wellcome Sanger Institute for the COVID-19 Genomics UK (COG-UK) Consortium | Harper VanSteenhouse, Yumi Kasai, David Gray, Carol Clugston, Anna Dominiczak and Alex Alderton, Roberto Amato, Jeffrey Barrett, Sonia Goncalves, Ewan Harrison, David K. Jackson, Ian Johnston, Dominic Kwiatkowski, Cordelia Langford, John Sillitoe on behalf of the Wellcome Sanger Institute COVID-19 Surveillance Team |
| EPI_ISL_3227048, EPI_ISL_3227049, EPI_ISL_3227050, EPI_ISL_3227051 | Lighthouse Lab in Alderley Park | Wellcome Sanger Institute for the COVID-19 Genomics UK (COG-UK) Consortium | Jacquelyn Wynn, Mairead Hyland, The Lighthouse Lab in Alderley Park and Alex Alderton, Roberto Amato, Jeffrey Barrett, Sonia Goncalves, Ewan Harrison, David K. Jackson, Ian Johnston, Dominic Kwiatkowski, Cordelia Langford, John Sillitoe on behalf of the Wellcome Sanger Institute COVID-19 Surveillance Team |
| EPI_ISL_3227052 | Lighthouse Lab in Milton Keynes | Wellcome Sanger Institute for the COVID-19 Genomics UK (COG-UK) Consortium | The Lighthouse Lab in Milton Keynes and Alex Alderton, Roberto Amato, Jeffrey Barrett, Sonia Goncalves, Ewan Harrison, David K. Jackson, Ian Johnston, Dominic Kwiatkowski, Cordelia Langford, John Sillitoe on behalf of the Wellcome Sanger Institute COVID-19 Surveillance Team |
| EPI_ISL_3227053 | Lighthouse Lab in Alderley Park | Wellcome Sanger Institute for the COVID-19 Genomics UK (COG-UK) Consortium | Jacquelyn Wynn, Mairead Hyland, The Lighthouse Lab in Alderley Park and Alex Alderton, Roberto Amato, Jeffrey Barrett, Sonia Goncalves, Ewan Harrison, David K. Jackson, Ian Johnston, Dominic Kwiatkowski, Cordelia Langford, John Sillitoe on behalf of the Wellcome Sanger Institute COVID-19 Surveillance Team |
| EPI_ISL_3227054 | Lighthouse Lab in Milton Keynes | Wellcome Sanger Institute for the COVID-19 Genomics UK (COG-UK) Consortium | The Lighthouse Lab in Milton Keynes and Alex Alderton, Roberto Amato, Jeffrey Barrett, Sonia Goncalves, Ewan Harrison, David K. Jackson, Ian Johnston, Dominic Kwiatkowski, Cordelia Langford, John Sillitoe on behalf of the Wellcome Sanger Institute COVID-19 Surveillance Team |
| EPI_ISL_3227055 | Lighthouse Lab in Glasgow | Wellcome Sanger Institute for the COVID-19 Genomics UK (COG-UK) | Harper VanSteenhouse, Yumi Kasai, David Gray, Carol Clugston, Anna Dominiczak and Alex Alderton, Roberto Amato, Jeffrey Barrett, Sonia Goncalves, Ewan Harrison, David K. Jackson, Ian Johnston, Dominic Kwiatkowski, Cordelia Langford, |

| | | | |
|---|---|---|---|
| | | Consortium | John Sillitoe on behalf of the Wellcome Sanger Institute COVID-19 Surveillance Team |
| EPI_ISL_3227056 | Lighthouse Lab in Milton Keynes | Wellcome Sanger Institute for the COVID-19 Genomics UK (COG-UK) Consortium | The Lighthouse Lab in Milton Keynes and Alex Alderton, Roberto Amato, Jeffrey Barrett, Sonia Goncalves, Ewan Harrison, David K. Jackson, Ian Johnston, Dominic Kwiatkowski, Cordelia Langford, John Sillitoe on behalf of the Wellcome Sanger Institute COVID-19 Surveillance Team |
| EPI_ISL_3227057 | Lighthouse Lab in Glasgow | Wellcome Sanger Institute for the COVID-19 Genomics UK (COG-UK) Consortium | Harper VanSteenhouse, Yumi Kasai, David Gray, Carol Clugston, Anna Dominiczak and Alex Alderton, Roberto Amato, Jeffrey Barrett, Sonia Goncalves, Ewan Harrison, David K. Jackson, Ian Johnston, Dominic Kwiatkowski, Cordelia Langford, John Sillitoe on behalf of the Wellcome Sanger Institute COVID-19 Surveillance Team |
| EPI_ISL_3227058 | Lighthouse Lab in Milton Keynes | Wellcome Sanger Institute for the COVID-19 Genomics UK (COG-UK) Consortium | The Lighthouse Lab in Milton Keynes and Alex Alderton, Roberto Amato, Jeffrey Barrett, Sonia Goncalves, Ewan Harrison, David K. Jackson, Ian Johnston, Dominic Kwiatkowski, Cordelia Langford, John Sillitoe on behalf of the Wellcome Sanger Institute COVID-19 Surveillance Team |
| EPI_ISL_3227059, EPI_ISL_3227060, EPI_ISL_3227061, EPI_ISL_3227062, EPI_ISL_3227063 | Lighthouse Lab in Alderley Park | Wellcome Sanger Institute for the COVID-19 Genomics UK (COG-UK) Consortium | Jacquelyn Wynn, Mairead Hyland, The Lighthouse Lab in Alderley Park and Alex Alderton, Roberto Amato, Jeffrey Barrett, Sonia Goncalves, Ewan Harrison, David K. Jackson, Ian Johnston, Dominic Kwiatkowski, Cordelia Langford, John Sillitoe on behalf of the Wellcome Sanger Institute COVID-19 Surveillance Team |
| EPI_ISL_3227064, EPI_ISL_3227065 | Lighthouse Lab in Milton Keynes | Wellcome Sanger Institute for the COVID-19 Genomics UK (COG-UK) Consortium | The Lighthouse Lab in Milton Keynes and Alex Alderton, Roberto Amato, Jeffrey Barrett, Sonia Goncalves, Ewan Harrison, David K. Jackson, Ian Johnston, Dominic Kwiatkowski, Cordelia Langford, John Sillitoe on behalf of the Wellcome Sanger Institute COVID-19 Surveillance Team |
| EPI_ISL_3227066, EPI_ISL_3227067 | Lighthouse Lab in Alderley Park | Wellcome Sanger Institute for the COVID-19 Genomics UK (COG-UK) Consortium | Jacquelyn Wynn, Mairead Hyland, The Lighthouse Lab in Alderley Park and Alex Alderton, Roberto Amato, Jeffrey Barrett, Sonia Goncalves, Ewan Harrison, David K. Jackson, Ian Johnston, Dominic Kwiatkowski, Cordelia Langford, John Sillitoe on behalf of the Wellcome Sanger Institute COVID-19 Surveillance Team |
| EPI_ISL_3227068 | Lighthouse Lab in Milton Keynes | Wellcome Sanger Institute for the COVID-19 Genomics UK (COG-UK) Consortium | The Lighthouse Lab in Milton Keynes and Alex Alderton, Roberto Amato, Jeffrey Barrett, Sonia Goncalves, Ewan Harrison, David K. Jackson, Ian Johnston, Dominic Kwiatkowski, Cordelia Langford, John Sillitoe on behalf of the Wellcome Sanger Institute COVID-19 Surveillance Team |
| EPI_ISL_3227069 | Lighthouse Lab in Glasgow | Wellcome Sanger Institute for the COVID-19 Genomics UK (COG-UK) Consortium | Harper VanSteenhouse, Yumi Kasai, David Gray, Carol Clugston, Anna Dominiczak and Alex Alderton, Roberto Amato, Jeffrey Barrett, Sonia Goncalves, Ewan Harrison, David K. Jackson, Ian Johnston, Dominic Kwiatkowski, Cordelia Langford, John Sillitoe on behalf of the Wellcome Sanger Institute COVID-19 Surveillance Team |
| EPI_ISL_3227070, EPI_ISL_3227071, EPI_ISL_3227072, EPI_ISL_3227073 | Lighthouse Lab in Alderley Park | Wellcome Sanger Institute for the COVID-19 Genomics UK (COG-UK) Consortium | Jacquelyn Wynn, Mairead Hyland, The Lighthouse Lab in Alderley Park and Alex Alderton, Roberto Amato, Jeffrey Barrett, Sonia Goncalves, Ewan Harrison, David K. Jackson, Ian Johnston, Dominic Kwiatkowski, Cordelia Langford, John Sillitoe on behalf of the Wellcome Sanger Institute COVID-19 Surveillance Team |
| EPI_ISL_3227074 | Lighthouse Lab in Glasgow | Wellcome Sanger Institute for the COVID-19 Genomics UK (COG-UK) Consortium | Harper VanSteenhouse, Yumi Kasai, David Gray, Carol Clugston, Anna Dominiczak and Alex Alderton, Roberto Amato, Jeffrey Barrett, Sonia Goncalves, Ewan Harrison, David K. Jackson, Ian Johnston, Dominic Kwiatkowski, Cordelia Langford, John Sillitoe on behalf of the Wellcome Sanger Institute COVID-19 Surveillance Team |
| EPI_ISL_3227075 | Lighthouse Lab in Milton Keynes | Wellcome Sanger Institute for the COVID-19 Genomics UK (COG-UK) Consortium | The Lighthouse Lab in Milton Keynes and Alex Alderton, Roberto Amato, Jeffrey Barrett, Sonia Goncalves, Ewan Harrison, David K. Jackson, Ian Johnston, Dominic Kwiatkowski, Cordelia Langford, John Sillitoe on behalf of the Wellcome Sanger Institute COVID-19 Surveillance Team |
| EPI_ISL_3227076 | Lighthouse Lab in Alderley Park | Wellcome Sanger Institute for the COVID-19 Genomics UK (COG-UK) Consortium | Jacquelyn Wynn, Mairead Hyland, The Lighthouse Lab in Alderley Park and Alex Alderton, Roberto Amato, Jeffrey Barrett, Sonia Goncalves, Ewan Harrison, David K. Jackson, Ian Johnston, Dominic Kwiatkowski, Cordelia Langford, John Sillitoe on behalf of the Wellcome Sanger Institute COVID-19 Surveillance Team |
| EPI_ISL_3227077 | Lighthouse Lab in Glasgow | Wellcome Sanger Institute for the COVID-19 Genomics UK (COG-UK) Consortium | Harper VanSteenhouse, Yumi Kasai, David Gray, Carol Clugston, Anna Dominiczak and Alex Alderton, Roberto Amato, Jeffrey Barrett, Sonia Goncalves, Ewan Harrison, David K. Jackson, Ian Johnston, Dominic Kwiatkowski, Cordelia Langford, John Sillitoe on behalf of the Wellcome Sanger Institute COVID-19 Surveillance Team |
| EPI_ISL_3227078, EPI_ISL_3227079, EPI_ISL_3227080 | Lighthouse Lab in Alderley Park | Wellcome Sanger Institute for the COVID-19 Genomics UK (COG-UK) Consortium | Jacquelyn Wynn, Mairead Hyland, The Lighthouse Lab in Alderley Park and Alex Alderton, Roberto Amato, Jeffrey Barrett, Sonia Goncalves, Ewan Harrison, David K. Jackson, Ian Johnston, Dominic Kwiatkowski, Cordelia Langford, John Sillitoe on behalf of the Wellcome Sanger Institute COVID-19 Surveillance Team |
| EPI_ISL_3227081 | Lighthouse Lab in Glasgow | Wellcome Sanger Institute for the COVID-19 Genomics UK (COG-UK) Consortium | Harper VanSteenhouse, Yumi Kasai, David Gray, Carol Clugston, Anna Dominiczak and Alex Alderton, Roberto Amato, Jeffrey Barrett, Sonia Goncalves, Ewan Harrison, David K. Jackson, Ian Johnston, Dominic Kwiatkowski, Cordelia Langford, John Sillitoe on behalf of the Wellcome Sanger Institute COVID-19 Surveillance Team |
| EPI_ISL_3227082, EPI_ISL_3227083, EPI_ISL_3227084, EPI_ISL_3227085, EPI_ISL_3227086, EPI_ISL_3227087 | Lighthouse Lab in Alderley Park | Wellcome Sanger Institute for the COVID-19 Genomics UK (COG-UK) Consortium | Jacquelyn Wynn, Mairead Hyland, The Lighthouse Lab in Alderley Park and Alex Alderton, Roberto Amato, Jeffrey Barrett, Sonia Goncalves, Ewan Harrison, David K. Jackson, Ian Johnston, Dominic Kwiatkowski, Cordelia Langford, John Sillitoe on behalf of the Wellcome Sanger Institute COVID-19 Surveillance Team |
| EPI_ISL_3227088, EPI_ISL_3227089, EPI_ISL_3227090, EPI_ISL_3227091, EPI_ISL_3227092 | Lighthouse Lab in Glasgow | Wellcome Sanger Institute for the COVID-19 Genomics UK (COG-UK) Consortium | Harper VanSteenhouse, Yumi Kasai, David Gray, Carol Clugston, Anna Dominiczak and Alex Alderton, Roberto Amato, Jeffrey Barrett, Sonia Goncalves, Ewan Harrison, David K. Jackson, Ian Johnston, Dominic Kwiatkowski, Cordelia Langford, John Sillitoe on behalf of the Wellcome Sanger Institute COVID-19 Surveillance Team |
| EPI_ISL_3227093 | Lighthouse Lab in Milton Keynes | Wellcome Sanger Institute for the COVID-19 Genomics UK (COG-UK) Consortium | The Lighthouse Lab in Milton Keynes and Alex Alderton, Roberto Amato, Jeffrey Barrett, Sonia Goncalves, Ewan Harrison, David K. Jackson, Ian Johnston, Dominic Kwiatkowski, Cordelia Langford, John Sillitoe on behalf of the Wellcome Sanger Institute COVID-19 Surveillance Team |
| EPI_ISL_3227094 | Lighthouse Lab in Alderley Park | Wellcome Sanger Institute for the COVID-19 Genomics UK (COG-UK) Consortium | Jacquelyn Wynn, Mairead Hyland, The Lighthouse Lab in Alderley Park and Alex Alderton, Roberto Amato, Jeffrey Barrett, Sonia Goncalves, Ewan Harrison, David K. Jackson, Ian Johnston, Dominic Kwiatkowski, Cordelia Langford, John Sillitoe on behalf of the Wellcome Sanger Institute COVID-19 Surveillance Team |
| EPI_ISL_3227095 | Lighthouse Lab in Milton Keynes | Wellcome Sanger Institute for the COVID-19 Genomics UK (COG-UK) Consortium | The Lighthouse Lab in Milton Keynes and Alex Alderton, Roberto Amato, Jeffrey Barrett, Sonia Goncalves, Ewan Harrison, David K. Jackson, Ian Johnston, Dominic Kwiatkowski, Cordelia Langford, John Sillitoe on behalf of the Wellcome Sanger Institute COVID-19 Surveillance Team |
| EPI_ISL_3227096, EPI_ISL_3227097, EPI_ISL_3227098, EPI_ISL_3227099 | Lighthouse Lab in Alderley Park | Wellcome Sanger Institute for the COVID-19 Genomics UK (COG-UK) Consortium | Jacquelyn Wynn, Mairead Hyland, The Lighthouse Lab in Alderley Park and Alex Alderton, Roberto Amato, Jeffrey Barrett, Sonia Goncalves, Ewan Harrison, David K. Jackson, Ian Johnston, Dominic Kwiatkowski, Cordelia Langford, John Sillitoe on behalf of the Wellcome Sanger Institute COVID-19 Surveillance Team |
| EPI_ISL_3227100, EPI_ISL_3227101 | Lighthouse Lab in Glasgow | Wellcome Sanger Institute for the COVID-19 Genomics UK (COG-UK) Consortium | Harper VanSteenhouse, Yumi Kasai, David Gray, Carol Clugston, Anna Dominiczak and Alex Alderton, Roberto Amato, Jeffrey Barrett, Sonia Goncalves, Ewan Harrison, David K. Jackson, Ian Johnston, Dominic Kwiatkowski, Cordelia Langford, John Sillitoe on behalf of the Wellcome Sanger Institute COVID-19 Surveillance Team |
| EPI_ISL_3227102 | Lighthouse Lab in Milton Keynes | Wellcome Sanger Institute for the COVID-19 Genomics UK (COG-UK) Consortium | The Lighthouse Lab in Milton Keynes and Alex Alderton, Roberto Amato, Jeffrey Barrett, Sonia Goncalves, Ewan Harrison, David K. Jackson, Ian Johnston, Dominic Kwiatkowski, Cordelia Langford, John Sillitoe on behalf of the Wellcome Sanger Institute COVID-19 Surveillance Team |
| EPI_ISL_3227103 | Lighthouse Lab in Glasgow | Wellcome Sanger Institute for the COVID-19 Genomics UK (COG-UK) Consortium | Harper VanSteenhouse, Yumi Kasai, David Gray, Carol Clugston, Anna Dominiczak and Alex Alderton, Roberto Amato, Jeffrey Barrett, Sonia Goncalves, Ewan Harrison, David K. Jackson, Ian Johnston, Dominic Kwiatkowski, Cordelia Langford, John Sillitoe on behalf of the Wellcome Sanger Institute COVID-19 Surveillance Team |
| EPI_ISL_3227104, EPI_ISL_3227105 | Lighthouse Lab in Alderley Park | Wellcome Sanger Institute for the COVID-19 Genomics UK (COG-UK) Consortium | Jacquelyn Wynn, Mairead Hyland, The Lighthouse Lab in Alderley Park and Alex Alderton, Roberto Amato, Jeffrey Barrett, Sonia Goncalves, Ewan Harrison, David K. Jackson, Ian Johnston, Dominic Kwiatkowski, Cordelia Langford, John Sillitoe on behalf of the Wellcome Sanger Institute COVID-19 Surveillance Team |
| EPI_ISL_3227106 | Lighthouse Lab in Glasgow | Wellcome Sanger Institute for the COVID-19 Genomics UK (COG-UK) Consortium | Harper VanSteenhouse, Yumi Kasai, David Gray, Carol Clugston, Anna Dominiczak and Alex Alderton, Roberto Amato, Jeffrey Barrett, Sonia Goncalves, Ewan Harrison, David K. Jackson, Ian Johnston, Dominic Kwiatkowski, Cordelia Langford, John Sillitoe on behalf of the Wellcome Sanger Institute COVID-19 Surveillance Team |
| EPI_ISL_3227107 | Lighthouse Lab in Alderley Park | Wellcome Sanger Institute for the COVID-19 Genomics UK (COG-UK) Consortium | Jacquelyn Wynn, Mairead Hyland, The Lighthouse Lab in Alderley Park and Alex Alderton, Roberto Amato, Jeffrey Barrett, Sonia Goncalves, Ewan Harrison, David K. Jackson, Ian Johnston, Dominic Kwiatkowski, Cordelia Langford, John Sillitoe on behalf of the Wellcome Sanger Institute COVID-19 Surveillance Team |
| EPI_ISL_3227108 | Lighthouse Lab in Milton Keynes | Wellcome Sanger Institute for the COVID-19 Genomics UK (COG-UK) Consortium | The Lighthouse Lab in Milton Keynes and Alex Alderton, Roberto Amato, Jeffrey Barrett, Sonia Goncalves, Ewan Harrison, David K. Jackson, Ian Johnston, Dominic Kwiatkowski, Cordelia Langford, John Sillitoe on behalf of the Wellcome Sanger Institute COVID-19 Surveillance Team |
| EPI_ISL_3227109, EPI_ISL_3227110 | Lighthouse Lab in Glasgow | Wellcome Sanger Institute for the COVID-19 Genomics UK (COG-UK) Consortium | Harper VanSteenhouse, Yumi Kasai, David Gray, Carol Clugston, Anna Dominiczak and Alex Alderton, Roberto Amato, Jeffrey Barrett, Sonia Goncalves, Ewan Harrison, David K. Jackson, Ian Johnston, Dominic Kwiatkowski, Cordelia Langford, John Sillitoe on behalf of the Wellcome Sanger Institute COVID-19 Surveillance Team |
| EPI_ISL_3227111, EPI_ISL_3227112, EPI_ISL_3227113, EPI_ISL_3227114, EPI_ISL_3227115 | Lighthouse Lab in Alderley Park | Wellcome Sanger Institute for the COVID-19 Genomics UK (COG-UK) Consortium | Jacquelyn Wynn, Mairead Hyland, The Lighthouse Lab in Alderley Park and Alex Alderton, Roberto Amato, Jeffrey Barrett, Sonia Goncalves, Ewan Harrison, David K. Jackson, Ian Johnston, Dominic Kwiatkowski, Cordelia Langford, John Sillitoe on behalf of the Wellcome Sanger Institute COVID-19 Surveillance Team |
| EPI_ISL_3227116 | Lighthouse Lab in Glasgow | Wellcome Sanger Institute for the COVID-19 Genomics UK (COG-UK) Consortium | Harper VanSteenhouse, Yumi Kasai, David Gray, Carol Clugston, Anna Dominiczak and Alex Alderton, Roberto Amato, Jeffrey Barrett, Sonia Goncalves, Ewan Harrison, David K. Jackson, Ian Johnston, Dominic Kwiatkowski, Cordelia Langford, John Sillitoe on behalf of the Wellcome Sanger Institute COVID-19 Surveillance Team |
| EPI_ISL_3227117 | Lighthouse Lab in Milton Keynes | Wellcome Sanger Institute for the COVID-19 Genomics UK (COG-UK) Consortium | The Lighthouse Lab in Milton Keynes and Alex Alderton, Roberto Amato, Jeffrey Barrett, Sonia Goncalves, Ewan Harrison, David K. Jackson, Ian Johnston, Dominic Kwiatkowski, Cordelia Langford, John Sillitoe on behalf of the Wellcome Sanger Institute COVID-19 Surveillance Team |
| EPI_ISL_3227118 | Lighthouse Lab in Alderley Park | Wellcome Sanger Institute for the COVID-19 Genomics UK (COG-UK) Consortium | Jacquelyn Wynn, Mairead Hyland, The Lighthouse Lab in Alderley Park and Alex Alderton, Roberto Amato, Jeffrey Barrett, Sonia Goncalves, Ewan Harrison, David K. Jackson, Ian Johnston, Dominic Kwiatkowski, Cordelia Langford, John Sillitoe on behalf of the Wellcome Sanger Institute COVID-19 Surveillance Team |
| EPI_ISL_3227119 | Lighthouse Lab in Milton Keynes | Wellcome Sanger Institute for the COVID-19 Genomics UK (COG-UK) Consortium | The Lighthouse Lab in Milton Keynes and Alex Alderton, Roberto Amato, Jeffrey Barrett, Sonia Goncalves, Ewan Harrison, David K. Jackson, Ian Johnston, Dominic Kwiatkowski, Cordelia Langford, John Sillitoe on behalf of the Wellcome Sanger Institute COVID-19 Surveillance Team |
| EPI_ISL_3227120, EPI_ISL_3227121 | Lighthouse Lab in Glasgow | Wellcome Sanger Institute for the COVID-19 Genomics UK (COG-UK) Consortium | Harper VanSteenhouse, Yumi Kasai, David Gray, Carol Clugston, Anna Dominiczak and Alex Alderton, Roberto Amato, Jeffrey Barrett, Sonia Goncalves, Ewan Harrison, David K. Jackson, Ian Johnston, Dominic Kwiatkowski, Cordelia Langford, John Sillitoe on behalf of the Wellcome Sanger Institute COVID-19 Surveillance Team |
| EPI_ISL_3227122 | Lighthouse Lab in Alderley Park | Wellcome Sanger Institute for the COVID-19 Genomics UK (COG-UK) Consortium | Jacquelyn Wynn, Mairead Hyland, The Lighthouse Lab in Alderley Park and Alex Alderton, Roberto Amato, Jeffrey Barrett, Sonia Goncalves, Ewan Harrison, David K. Jackson, Ian Johnston, Dominic Kwiatkowski, Cordelia Langford, John Sillitoe on behalf of the Wellcome Sanger Institute COVID-19 Surveillance Team |
| EPI_ISL_3227123, EPI_ISL_3227124 | Lighthouse Lab in Milton Keynes | Wellcome Sanger Institute for the COVID-19 Genomics UK (COG-UK) Consortium | The Lighthouse Lab in Milton Keynes and Alex Alderton, Roberto Amato, Jeffrey Barrett, Sonia Goncalves, Ewan Harrison, David K. Jackson, Ian Johnston, Dominic Kwiatkowski, Cordelia Langford, John Sillitoe on behalf of the Wellcome Sanger Institute COVID-19 Surveillance Team |

| | | | |
|---|---|---|---|
| EPI_ISL_3227125 | Lighthouse Lab in Alderley Park | Wellcome Sanger Institute for the COVID-19 Genomics UK (COG-UK) Consortium | Jacquelyn Wynn, Mairead Hyland, The Lighthouse Lab in Alderley Park and Alex Alderton, Roberto Amato, Jeffrey Barrett, Sonia Goncalves, Ewan Harrison, David K. Jackson, Ian Johnston, Dominic Kwiatkowski, Cordelia Langford, John Sillitoe on behalf of the Wellcome Sanger Institute COVID-19 Surveillance Team |
| EPI_ISL_3227126, EPI_ISL_3227127 | Lighthouse Lab in Milton Keynes | Wellcome Sanger Institute for the COVID-19 Genomics UK (COG-UK) Consortium | The Lighthouse Lab in Milton Keynes and Alex Alderton, Roberto Amato, Jeffrey Barrett, Sonia Goncalves, Ewan Harrison, David K. Jackson, Ian Johnston, Dominic Kwiatkowski, Cordelia Langford, John Sillitoe on behalf of the Wellcome Sanger Institute COVID-19 Surveillance Team |
| EPI_ISL_3227128 | Lighthouse Lab in Alderley Park | Wellcome Sanger Institute for the COVID-19 Genomics UK (COG-UK) Consortium | Jacquelyn Wynn, Mairead Hyland, The Lighthouse Lab in Alderley Park and Alex Alderton, Roberto Amato, Jeffrey Barrett, Sonia Goncalves, Ewan Harrison, David K. Jackson, Ian Johnston, Dominic Kwiatkowski, Cordelia Langford, John Sillitoe on behalf of the Wellcome Sanger Institute COVID-19 Surveillance Team |
| EPI_ISL_3227129 | Lighthouse Lab in Glasgow | Wellcome Sanger Institute for the COVID-19 Genomics UK (COG-UK) Consortium | Harper VanSteenhouse, Yumi Kasai, David Gray, Carol Clugston, Anna Dominiczak and Alex Alderton, Roberto Amato, Jeffrey Barrett, Sonia Goncalves, Ewan Harrison, David K. Jackson, Ian Johnston, Dominic Kwiatkowski, Cordelia Langford, John Sillitoe on behalf of the Wellcome Sanger Institute COVID-19 Surveillance Team |
| EPI_ISL_3227130, EPI_ISL_3227131 | Lighthouse Lab in Alderley Park | Wellcome Sanger Institute for the COVID-19 Genomics UK (COG-UK) Consortium | Jacquelyn Wynn, Mairead Hyland, The Lighthouse Lab in Alderley Park and Alex Alderton, Roberto Amato, Jeffrey Barrett, Sonia Goncalves, Ewan Harrison, David K. Jackson, Ian Johnston, Dominic Kwiatkowski, Cordelia Langford, John Sillitoe on behalf of the Wellcome Sanger Institute COVID-19 Surveillance Team |
| EPI_ISL_3227132, EPI_ISL_3227133 | Lighthouse Lab in Milton Keynes | Wellcome Sanger Institute for the COVID-19 Genomics UK (COG-UK) Consortium | The Lighthouse Lab in Milton Keynes and Alex Alderton, Roberto Amato, Jeffrey Barrett, Sonia Goncalves, Ewan Harrison, David K. Jackson, Ian Johnston, Dominic Kwiatkowski, Cordelia Langford, John Sillitoe on behalf of the Wellcome Sanger Institute COVID-19 Surveillance Team |
| EPI_ISL_3227134 | Lighthouse Lab in Alderley Park | Wellcome Sanger Institute for the COVID-19 Genomics UK (COG-UK) Consortium | Jacquelyn Wynn, Mairead Hyland, The Lighthouse Lab in Alderley Park and Alex Alderton, Roberto Amato, Jeffrey Barrett, Sonia Goncalves, Ewan Harrison, David K. Jackson, Ian Johnston, Dominic Kwiatkowski, Cordelia Langford, John Sillitoe on behalf of the Wellcome Sanger Institute COVID-19 Surveillance Team |
| EPI_ISL_3227135 | Lighthouse Lab in Glasgow | Wellcome Sanger Institute for the COVID-19 Genomics UK (COG-UK) Consortium | Harper VanSteenhouse, Yumi Kasai, David Gray, Carol Clugston, Anna Dominiczak and Alex Alderton, Roberto Amato, Jeffrey Barrett, Sonia Goncalves, Ewan Harrison, David K. Jackson, Ian Johnston, Dominic Kwiatkowski, Cordelia Langford, John Sillitoe on behalf of the Wellcome Sanger Institute COVID-19 Surveillance Team |
| EPI_ISL_3227136, EPI_ISL_3227137 | Lighthouse Lab in Alderley Park | Wellcome Sanger Institute for the COVID-19 Genomics UK (COG-UK) Consortium | Jacquelyn Wynn, Mairead Hyland, The Lighthouse Lab in Alderley Park and Alex Alderton, Roberto Amato, Jeffrey Barrett, Sonia Goncalves, Ewan Harrison, David K. Jackson, Ian Johnston, Dominic Kwiatkowski, Cordelia Langford, John Sillitoe on behalf of the Wellcome Sanger Institute COVID-19 Surveillance Team |
| EPI_ISL_3227138 | Lighthouse Lab in Milton Keynes | Wellcome Sanger Institute for the COVID-19 Genomics UK (COG-UK) Consortium | The Lighthouse Lab in Milton Keynes and Alex Alderton, Roberto Amato, Jeffrey Barrett, Sonia Goncalves, Ewan Harrison, David K. Jackson, Ian Johnston, Dominic Kwiatkowski, Cordelia Langford, John Sillitoe on behalf of the Wellcome Sanger Institute COVID-19 Surveillance Team |
| EPI_ISL_3227139, EPI_ISL_3227140 | Lighthouse Lab in Alderley Park | Wellcome Sanger Institute for the COVID-19 Genomics UK (COG-UK) Consortium | Jacquelyn Wynn, Mairead Hyland, The Lighthouse Lab in Alderley Park and Alex Alderton, Roberto Amato, Jeffrey Barrett, Sonia Goncalves, Ewan Harrison, David K. Jackson, Ian Johnston, Dominic Kwiatkowski, Cordelia Langford, John Sillitoe on behalf of the Wellcome Sanger Institute COVID-19 Surveillance Team |
| EPI_ISL_3227141 | Lighthouse Lab in Milton Keynes | Wellcome Sanger Institute for the COVID-19 Genomics UK (COG-UK) Consortium | The Lighthouse Lab in Milton Keynes and Alex Alderton, Roberto Amato, Jeffrey Barrett, Sonia Goncalves, Ewan Harrison, David K. Jackson, Ian Johnston, Dominic Kwiatkowski, Cordelia Langford, John Sillitoe on behalf of the Wellcome Sanger Institute COVID-19 Surveillance Team |
| EPI_ISL_3227142 | Lighthouse Lab in Glasgow | Wellcome Sanger Institute for the COVID-19 Genomics UK (COG-UK) Consortium | Harper VanSteenhouse, Yumi Kasai, David Gray, Carol Clugston, Anna Dominiczak and Alex Alderton, Roberto Amato, Jeffrey Barrett, Sonia Goncalves, Ewan Harrison, David K. Jackson, Ian Johnston, Dominic Kwiatkowski, Cordelia Langford, John Sillitoe on behalf of the Wellcome Sanger Institute COVID-19 Surveillance Team |
| EPI_ISL_3227143, EPI_ISL_3227144, EPI_ISL_3227145 | Lighthouse Lab in Alderley Park | Wellcome Sanger Institute for the COVID-19 Genomics UK (COG-UK) Consortium | Jacquelyn Wynn, Mairead Hyland, The Lighthouse Lab in Alderley Park and Alex Alderton, Roberto Amato, Jeffrey Barrett, Sonia Goncalves, Ewan Harrison, David K. Jackson, Ian Johnston, Dominic Kwiatkowski, Cordelia Langford, John Sillitoe on behalf of the Wellcome Sanger Institute COVID-19 Surveillance Team |
| EPI_ISL_3227146 | Lighthouse Lab in Milton Keynes | Wellcome Sanger Institute for the COVID-19 Genomics UK (COG-UK) Consortium | The Lighthouse Lab in Milton Keynes and Alex Alderton, Roberto Amato, Jeffrey Barrett, Sonia Goncalves, Ewan Harrison, David K. Jackson, Ian Johnston, Dominic Kwiatkowski, Cordelia Langford, John Sillitoe on behalf of the Wellcome Sanger Institute COVID-19 Surveillance Team |
| EPI_ISL_3227147 | Lighthouse Lab in Alderley Park | Wellcome Sanger Institute for the COVID-19 Genomics UK (COG-UK) Consortium | Jacquelyn Wynn, Mairead Hyland, The Lighthouse Lab in Alderley Park and Alex Alderton, Roberto Amato, Jeffrey Barrett, Sonia Goncalves, Ewan Harrison, David K. Jackson, Ian Johnston, Dominic Kwiatkowski, Cordelia Langford, John Sillitoe on behalf of the Wellcome Sanger Institute COVID-19 Surveillance Team |
| EPI_ISL_3227148 | Lighthouse Lab in Glasgow | Wellcome Sanger Institute for the COVID-19 Genomics UK (COG-UK) Consortium | Harper VanSteenhouse, Yumi Kasai, David Gray, Carol Clugston, Anna Dominiczak and Alex Alderton, Roberto Amato, Jeffrey Barrett, Sonia Goncalves, Ewan Harrison, David K. Jackson, Ian Johnston, Dominic Kwiatkowski, Cordelia Langford, John Sillitoe on behalf of the Wellcome Sanger Institute COVID-19 Surveillance Team |
| EPI_ISL_3227149, EPI_ISL_3227150, EPI_ISL_3227151 | Lighthouse Lab in Milton Keynes | Wellcome Sanger Institute for the COVID-19 Genomics UK (COG-UK) Consortium | The Lighthouse Lab in Milton Keynes and Alex Alderton, Roberto Amato, Jeffrey Barrett, Sonia Goncalves, Ewan Harrison, David K. Jackson, Ian Johnston, Dominic Kwiatkowski, Cordelia Langford, John Sillitoe on behalf of the Wellcome Sanger Institute COVID-19 Surveillance Team |
| EPI_ISL_3227152 | Lighthouse Lab in Alderley Park | Wellcome Sanger Institute for the COVID-19 Genomics UK (COG-UK) Consortium | Jacquelyn Wynn, Mairead Hyland, The Lighthouse Lab in Alderley Park and Alex Alderton, Roberto Amato, Jeffrey Barrett, Sonia Goncalves, Ewan Harrison, David K. Jackson, Ian Johnston, Dominic Kwiatkowski, Cordelia Langford, John Sillitoe on behalf of the Wellcome Sanger Institute COVID-19 Surveillance Team |
| EPI_ISL_3227153, EPI_ISL_3227154 | Lighthouse Lab in Glasgow | Wellcome Sanger Institute for the COVID-19 Genomics UK (COG-UK) Consortium | Harper VanSteenhouse, Yumi Kasai, David Gray, Carol Clugston, Anna Dominiczak and Alex Alderton, Roberto Amato, Jeffrey Barrett, Sonia Goncalves, Ewan Harrison, David K. Jackson, Ian Johnston, Dominic Kwiatkowski, Cordelia Langford, John Sillitoe on behalf of the Wellcome Sanger Institute COVID-19 Surveillance Team |
| EPI_ISL_3227155 | Lighthouse Lab in Alderley Park | Wellcome Sanger Institute for the COVID-19 Genomics UK (COG-UK) Consortium | Jacquelyn Wynn, Mairead Hyland, The Lighthouse Lab in Alderley Park and Alex Alderton, Roberto Amato, Jeffrey Barrett, Sonia Goncalves, Ewan Harrison, David K. Jackson, Ian Johnston, Dominic Kwiatkowski, Cordelia Langford, John Sillitoe on behalf of the Wellcome Sanger Institute COVID-19 Surveillance Team |
| EPI_ISL_3227156 | Lighthouse Lab in Glasgow | Wellcome Sanger Institute for the COVID-19 Genomics UK (COG-UK) Consortium | Harper VanSteenhouse, Yumi Kasai, David Gray, Carol Clugston, Anna Dominiczak and Alex Alderton, Roberto Amato, Jeffrey Barrett, Sonia Goncalves, Ewan Harrison, David K. Jackson, Ian Johnston, Dominic Kwiatkowski, Cordelia Langford, John Sillitoe on behalf of the Wellcome Sanger Institute COVID-19 Surveillance Team |
| EPI_ISL_3227157 | Lighthouse Lab in Milton Keynes | Wellcome Sanger Institute for the COVID-19 Genomics UK (COG-UK) Consortium | The Lighthouse Lab in Milton Keynes and Alex Alderton, Roberto Amato, Jeffrey Barrett, Sonia Goncalves, Ewan Harrison, David K. Jackson, Ian Johnston, Dominic Kwiatkowski, Cordelia Langford, John Sillitoe on behalf of the Wellcome Sanger Institute COVID-19 Surveillance Team |
| EPI_ISL_3227158 | Lighthouse Lab in Alderley Park | Wellcome Sanger Institute for the COVID-19 Genomics UK (COG-UK) Consortium | Jacquelyn Wynn, Mairead Hyland, The Lighthouse Lab in Alderley Park and Alex Alderton, Roberto Amato, Jeffrey Barrett, Sonia Goncalves, Ewan Harrison, David K. Jackson, Ian Johnston, Dominic Kwiatkowski, Cordelia Langford, John Sillitoe on behalf of the Wellcome Sanger Institute COVID-19 Surveillance Team |
| EPI_ISL_3227159 | Lighthouse Lab in Glasgow | Wellcome Sanger Institute for the COVID-19 Genomics UK (COG-UK) Consortium | Harper VanSteenhouse, Yumi Kasai, David Gray, Carol Clugston, Anna Dominiczak and Alex Alderton, Roberto Amato, Jeffrey Barrett, Sonia Goncalves, Ewan Harrison, David K. Jackson, Ian Johnston, Dominic Kwiatkowski, Cordelia Langford, John Sillitoe on behalf of the Wellcome Sanger Institute COVID-19 Surveillance Team |
| EPI_ISL_3227160, EPI_ISL_3227161, EPI_ISL_3227162 | Lighthouse Lab in Alderley Park | Wellcome Sanger Institute for the COVID-19 Genomics UK (COG-UK) Consortium | Jacquelyn Wynn, Mairead Hyland, The Lighthouse Lab in Alderley Park and Alex Alderton, Roberto Amato, Jeffrey Barrett, Sonia Goncalves, Ewan Harrison, David K. Jackson, Ian Johnston, Dominic Kwiatkowski, Cordelia Langford, John Sillitoe on behalf of the Wellcome Sanger Institute COVID-19 Surveillance Team |
| EPI_ISL_3227163, EPI_ISL_3227164 | Lighthouse Lab in Milton Keynes | Wellcome Sanger Institute for the COVID-19 Genomics UK (COG-UK) Consortium | The Lighthouse Lab in Milton Keynes and Alex Alderton, Roberto Amato, Jeffrey Barrett, Sonia Goncalves, Ewan Harrison, David K. Jackson, Ian Johnston, Dominic Kwiatkowski, Cordelia Langford, John Sillitoe on behalf of the Wellcome Sanger Institute COVID-19 Surveillance Team |
| EPI_ISL_3227165, EPI_ISL_3227166 | Lighthouse Lab in Alderley Park | Wellcome Sanger Institute for the COVID-19 Genomics UK (COG-UK) Consortium | Jacquelyn Wynn, Mairead Hyland, The Lighthouse Lab in Alderley Park and Alex Alderton, Roberto Amato, Jeffrey Barrett, Sonia Goncalves, Ewan Harrison, David K. Jackson, Ian Johnston, Dominic Kwiatkowski, Cordelia Langford, John Sillitoe on behalf of the Wellcome Sanger Institute COVID-19 Surveillance Team |
| EPI_ISL_3227167, EPI_ISL_3227168 | Lighthouse Lab in Glasgow | Wellcome Sanger Institute for the COVID-19 Genomics UK (COG-UK) Consortium | Harper VanSteenhouse, Yumi Kasai, David Gray, Carol Clugston, Anna Dominiczak and Alex Alderton, Roberto Amato, Jeffrey Barrett, Sonia Goncalves, Ewan Harrison, David K. Jackson, Ian Johnston, Dominic Kwiatkowski, Cordelia Langford, John Sillitoe on behalf of the Wellcome Sanger Institute COVID-19 Surveillance Team |
| EPI_ISL_3227169, EPI_ISL_3227170 | Lighthouse Lab in Alderley Park | Wellcome Sanger Institute for the COVID-19 Genomics UK (COG-UK) Consortium | Jacquelyn Wynn, Mairead Hyland, The Lighthouse Lab in Alderley Park and Alex Alderton, Roberto Amato, Jeffrey Barrett, Sonia Goncalves, Ewan Harrison, David K. Jackson, Ian Johnston, Dominic Kwiatkowski, Cordelia Langford, John Sillitoe on behalf of the Wellcome Sanger Institute COVID-19 Surveillance Team |
| EPI_ISL_3227171 | Lighthouse Lab in Milton Keynes | Wellcome Sanger Institute for the COVID-19 Genomics UK (COG-UK) Consortium | The Lighthouse Lab in Milton Keynes and Alex Alderton, Roberto Amato, Jeffrey Barrett, Sonia Goncalves, Ewan Harrison, David K. Jackson, Ian Johnston, Dominic Kwiatkowski, Cordelia Langford, John Sillitoe on behalf of the Wellcome Sanger Institute COVID-19 Surveillance Team |
| EPI_ISL_3227172, EPI_ISL_3227173 | Lighthouse Lab in Alderley Park | Wellcome Sanger Institute for the COVID-19 Genomics UK (COG-UK) Consortium | Jacquelyn Wynn, Mairead Hyland, The Lighthouse Lab in Alderley Park and Alex Alderton, Roberto Amato, Jeffrey Barrett, Sonia Goncalves, Ewan Harrison, David K. Jackson, Ian Johnston, Dominic Kwiatkowski, Cordelia Langford, John Sillitoe on behalf of the Wellcome Sanger Institute COVID-19 Surveillance Team |
| EPI_ISL_3227174 | Lighthouse Lab in Milton Keynes | Wellcome Sanger Institute for the COVID-19 Genomics UK (COG-UK) Consortium | The Lighthouse Lab in Milton Keynes and Alex Alderton, Roberto Amato, Jeffrey Barrett, Sonia Goncalves, Ewan Harrison, David K. Jackson, Ian Johnston, Dominic Kwiatkowski, Cordelia Langford, John Sillitoe on behalf of the Wellcome Sanger Institute COVID-19 Surveillance Team |
| EPI_ISL_3227175 | Lighthouse Lab in Alderley Park | Wellcome Sanger Institute for the COVID-19 Genomics UK (COG-UK) Consortium | Jacquelyn Wynn, Mairead Hyland, The Lighthouse Lab in Alderley Park and Alex Alderton, Roberto Amato, Jeffrey Barrett, Sonia Goncalves, Ewan Harrison, David K. Jackson, Ian Johnston, Dominic Kwiatkowski, Cordelia Langford, John Sillitoe on behalf of the Wellcome Sanger Institute COVID-19 Surveillance Team |
| EPI_ISL_3227176, EPI_ISL_3227177 | Lighthouse Lab in Glasgow | Wellcome Sanger Institute for the COVID-19 Genomics UK (COG-UK) Consortium | Harper VanSteenhouse, Yumi Kasai, David Gray, Carol Clugston, Anna Dominiczak and Alex Alderton, Roberto Amato, Jeffrey Barrett, Sonia Goncalves, Ewan Harrison, David K. Jackson, Ian Johnston, Dominic Kwiatkowski, Cordelia Langford, John Sillitoe on behalf of the Wellcome Sanger Institute COVID-19 Surveillance Team |
| EPI_ISL_3227178 | Lighthouse Lab in Milton Keynes | Wellcome Sanger Institute for the COVID-19 Genomics UK (COG-UK) Consortium | The Lighthouse Lab in Milton Keynes and Alex Alderton, Roberto Amato, Jeffrey Barrett, Sonia Goncalves, Ewan Harrison, David K. Jackson, Ian Johnston, Dominic Kwiatkowski, Cordelia Langford, John Sillitoe on behalf of the Wellcome Sanger Institute COVID-19 Surveillance Team |
| EPI_ISL_3227179, EPI_ISL_3227180 | Lighthouse Lab in Alderley Park | Wellcome Sanger Institute for the COVID-19 Genomics UK (COG-UK) Consortium | Jacquelyn Wynn, Mairead Hyland, The Lighthouse Lab in Alderley Park and Alex Alderton, Roberto Amato, Jeffrey Barrett, Sonia Goncalves, Ewan Harrison, David K. Jackson, Ian Johnston, Dominic Kwiatkowski, Cordelia Langford, John Sillitoe on behalf of the Wellcome Sanger Institute COVID-19 Surveillance Team |
| EPI_ISL_3227181 | Lighthouse Lab in Glasgow | Wellcome Sanger Institute for the COVID-19 Genomics UK (COG-UK) Consortium | Harper VanSteenhouse, Yumi Kasai, David Gray, Carol Clugston, Anna Dominiczak and Alex Alderton, Roberto Amato, Jeffrey Barrett, Sonia Goncalves, Ewan Harrison, David K. Jackson, Ian Johnston, Dominic Kwiatkowski, Cordelia Langford, John Sillitoe on behalf of the Wellcome Sanger Institute COVID-19 Surveillance Team |
| EPI_ISL_3227182, EPI_ISL_3227183 | Lighthouse Lab in Milton Keynes | Wellcome Sanger Institute for the COVID-19 Genomics UK (COG-UK) Consortium | The Lighthouse Lab in Milton Keynes and Alex Alderton, Roberto Amato, Jeffrey Barrett, Sonia Goncalves, Ewan Harrison, David K. Jackson, Ian Johnston, Dominic Kwiatkowski, Cordelia Langford, John Sillitoe on behalf of the Wellcome Sanger Institute COVID-19 Surveillance Team |
| EPI_ISL_3227184, EPI_ISL_3227185, EPI_ISL_3227186 | Lighthouse Lab in Alderley Park | Wellcome Sanger Institute for the COVID-19 Genomics UK (COG-UK) Consortium | Jacquelyn Wynn, Mairead Hyland, The Lighthouse Lab in Alderley Park and Alex Alderton, Roberto Amato, Jeffrey Barrett, Sonia Goncalves, Ewan Harrison, David K. Jackson, Ian Johnston, Dominic Kwiatkowski, Cordelia Langford, John Sillitoe on behalf of the Wellcome Sanger Institute COVID-19 Surveillance Team |
| EPI_ISL_3227187 | Lighthouse Lab in Milton Keynes | Wellcome Sanger Institute for the COVID-19 Genomics UK (COG-UK) Consortium | The Lighthouse Lab in Milton Keynes and Alex Alderton, Roberto Amato, Jeffrey Barrett, Sonia Goncalves, Ewan Harrison, David K. Jackson, Ian Johnston, Dominic Kwiatkowski, Cordelia Langford, John Sillitoe on behalf of the Wellcome Sanger Institute COVID-19 Surveillance Team |
| EPI_ISL_3227188, EPI_ISL_3227189 | Lighthouse Lab in Alderley Park | Wellcome Sanger Institute for the COVID-19 Genomics UK (COG-UK) Consortium | Jacquelyn Wynn, Mairead Hyland, The Lighthouse Lab in Alderley Park and Alex Alderton, Roberto Amato, Jeffrey Barrett, Sonia Goncalves, Ewan Harrison, David K. Jackson, Ian Johnston, Dominic Kwiatkowski, Cordelia Langford, John Sillitoe on behalf of the Wellcome Sanger Institute COVID-19 Surveillance Team |
| EPI_ISL_3227190 | Lighthouse Lab in Milton Keynes | Wellcome Sanger Institute for the COVID-19 Genomics UK (COG-UK) Consortium | The Lighthouse Lab in Milton Keynes and Alex Alderton, Roberto Amato, Jeffrey Barrett, Sonia Goncalves, Ewan Harrison, David K. Jackson, Ian Johnston, Dominic Kwiatkowski, Cordelia Langford, John Sillitoe on behalf of the Wellcome Sanger Institute COVID-19 Surveillance Team |
| EPI_ISL_3227191 | Lighthouse Lab in Alderley Park | Wellcome Sanger Institute for the COVID-19 Genomics UK (COG-UK) | Jacquelyn Wynn, Mairead Hyland, The Lighthouse Lab in Alderley Park and Alex Alderton, Roberto Amato, Jeffrey Barrett, Sonia Goncalves, Ewan Harrison, David K. Jackson, Ian Johnston, Dominic Kwiatkowski, Cordelia Langford, John Sillitoe on |

| | | | |
|---|---|---|---|
| | | Consortium | behalf of the Wellcome Sanger Institute COVID-19 Surveillance Team |
| EPI_ISL_3227192 | Lighthouse Lab in Glasgow | Wellcome Sanger Institute for the COVID-19 Genomics UK (COG-UK) Consortium | Harper VanSteenhouse, Yumi Kasai, David Gray, Carol Clugston, Anna Dominiczak and Alex Alderton, Roberto Amato, Jeffrey Barrett, Sonia Goncalves, Ewan Harrison, David K. Jackson, Ian Johnston, Dominic Kwiatkowski, Cordelia Langford, John Sillitoe on behalf of the Wellcome Sanger Institute COVID-19 Surveillance Team |
| EPI_ISL_3227193, EPI_ISL_3227194 | Lighthouse Lab in Milton Keynes | Wellcome Sanger Institute for the COVID-19 Genomics UK (COG-UK) Consortium | The Lighthouse Lab in Milton Keynes and Alex Alderton, Roberto Amato, Jeffrey Barrett, Sonia Goncalves, Ewan Harrison, David K. Jackson, Ian Johnston, Dominic Kwiatkowski, Cordelia Langford, John Sillitoe on behalf of the Wellcome Sanger Institute COVID-19 Surveillance Team |
| EPI_ISL_3227195 | Lighthouse Lab in Alderley Park | Wellcome Sanger Institute for the COVID-19 Genomics UK (COG-UK) Consortium | Jacquelyn Wynn, Mairead Hyland, The Lighthouse Lab in Alderley Park and Alex Alderton, Roberto Amato, Jeffrey Barrett, Sonia Goncalves, Ewan Harrison, David K. Jackson, Ian Johnston, Dominic Kwiatkowski, Cordelia Langford, John Sillitoe on behalf of the Wellcome Sanger Institute COVID-19 Surveillance Team |
| EPI_ISL_3227196 | Lighthouse Lab in Milton Keynes | Wellcome Sanger Institute for the COVID-19 Genomics UK (COG-UK) Consortium | The Lighthouse Lab in Milton Keynes and Alex Alderton, Roberto Amato, Jeffrey Barrett, Sonia Goncalves, Ewan Harrison, David K. Jackson, Ian Johnston, Dominic Kwiatkowski, Cordelia Langford, John Sillitoe on behalf of the Wellcome Sanger Institute COVID-19 Surveillance Team |
| EPI_ISL_3227197 | Lighthouse Lab in Glasgow | Wellcome Sanger Institute for the COVID-19 Genomics UK (COG-UK) Consortium | Harper VanSteenhouse, Yumi Kasai, David Gray, Carol Clugston, Anna Dominiczak and Alex Alderton, Roberto Amato, Jeffrey Barrett, Sonia Goncalves, Ewan Harrison, David K. Jackson, Ian Johnston, Dominic Kwiatkowski, Cordelia Langford, John Sillitoe on behalf of the Wellcome Sanger Institute COVID-19 Surveillance Team |
| EPI_ISL_3227198, EPI_ISL_3227199 | Lighthouse Lab in Alderley Park | Wellcome Sanger Institute for the COVID-19 Genomics UK (COG-UK) Consortium | Jacquelyn Wynn, Mairead Hyland, The Lighthouse Lab in Alderley Park and Alex Alderton, Roberto Amato, Jeffrey Barrett, Sonia Goncalves, Ewan Harrison, David K. Jackson, Ian Johnston, Dominic Kwiatkowski, Cordelia Langford, John Sillitoe on behalf of the Wellcome Sanger Institute COVID-19 Surveillance Team |
| EPI_ISL_3227200 | Lighthouse Lab in Milton Keynes | Wellcome Sanger Institute for the COVID-19 Genomics UK (COG-UK) Consortium | The Lighthouse Lab in Milton Keynes and Alex Alderton, Roberto Amato, Jeffrey Barrett, Sonia Goncalves, Ewan Harrison, David K. Jackson, Ian Johnston, Dominic Kwiatkowski, Cordelia Langford, John Sillitoe on behalf of the Wellcome Sanger Institute COVID-19 Surveillance Team |
| EPI_ISL_3227201 | Lighthouse Lab in Alderley Park | Wellcome Sanger Institute for the COVID-19 Genomics UK (COG-UK) Consortium | Jacquelyn Wynn, Mairead Hyland, The Lighthouse Lab in Alderley Park and Alex Alderton, Roberto Amato, Jeffrey Barrett, Sonia Goncalves, Ewan Harrison, David K. Jackson, Ian Johnston, Dominic Kwiatkowski, Cordelia Langford, John Sillitoe on behalf of the Wellcome Sanger Institute COVID-19 Surveillance Team |
| EPI_ISL_3227202 | Lighthouse Lab in Glasgow | Wellcome Sanger Institute for the COVID-19 Genomics UK (COG-UK) Consortium | Harper VanSteenhouse, Yumi Kasai, David Gray, Carol Clugston, Anna Dominiczak and Alex Alderton, Roberto Amato, Jeffrey Barrett, Sonia Goncalves, Ewan Harrison, David K. Jackson, Ian Johnston, Dominic Kwiatkowski, Cordelia Langford, John Sillitoe on behalf of the Wellcome Sanger Institute COVID-19 Surveillance Team |
| EPI_ISL_3227203 | Lighthouse Lab in Alderley Park | Wellcome Sanger Institute for the COVID-19 Genomics UK (COG-UK) Consortium | Jacquelyn Wynn, Mairead Hyland, The Lighthouse Lab in Alderley Park and Alex Alderton, Roberto Amato, Jeffrey Barrett, Sonia Goncalves, Ewan Harrison, David K. Jackson, Ian Johnston, Dominic Kwiatkowski, Cordelia Langford, John Sillitoe on behalf of the Wellcome Sanger Institute COVID-19 Surveillance Team |
| EPI_ISL_3227204 | Lighthouse Lab in Glasgow | Wellcome Sanger Institute for the COVID-19 Genomics UK (COG-UK) Consortium | Harper VanSteenhouse, Yumi Kasai, David Gray, Carol Clugston, Anna Dominiczak and Alex Alderton, Roberto Amato, Jeffrey Barrett, Sonia Goncalves, Ewan Harrison, David K. Jackson, Ian Johnston, Dominic Kwiatkowski, Cordelia Langford, John Sillitoe on behalf of the Wellcome Sanger Institute COVID-19 Surveillance Team |
| EPI_ISL_3227205 | Lighthouse Lab in Alderley Park | Wellcome Sanger Institute for the COVID-19 Genomics UK (COG-UK) Consortium | Jacquelyn Wynn, Mairead Hyland, The Lighthouse Lab in Alderley Park and Alex Alderton, Roberto Amato, Jeffrey Barrett, Sonia Goncalves, Ewan Harrison, David K. Jackson, Ian Johnston, Dominic Kwiatkowski, Cordelia Langford, John Sillitoe on behalf of the Wellcome Sanger Institute COVID-19 Surveillance Team |
| EPI_ISL_3227206 | Lighthouse Lab in Milton Keynes | Wellcome Sanger Institute for the COVID-19 Genomics UK (COG-UK) Consortium | The Lighthouse Lab in Milton Keynes and Alex Alderton, Roberto Amato, Jeffrey Barrett, Sonia Goncalves, Ewan Harrison, David K. Jackson, Ian Johnston, Dominic Kwiatkowski, Cordelia Langford, John Sillitoe on behalf of the Wellcome Sanger Institute COVID-19 Surveillance Team |
| EPI_ISL_3227207 | Lighthouse Lab in Glasgow | Wellcome Sanger Institute for the COVID-19 Genomics UK (COG-UK) Consortium | Harper VanSteenhouse, Yumi Kasai, David Gray, Carol Clugston, Anna Dominiczak and Alex Alderton, Roberto Amato, Jeffrey Barrett, Sonia Goncalves, Ewan Harrison, David K. Jackson, Ian Johnston, Dominic Kwiatkowski, Cordelia Langford, John Sillitoe on behalf of the Wellcome Sanger Institute COVID-19 Surveillance Team |
| EPI_ISL_3227208, EPI_ISL_3227209, EPI_ISL_3227210, EPI_ISL_3227211, EPI_ISL_3227212 | Lighthouse Lab in Alderley Park | Wellcome Sanger Institute for the COVID-19 Genomics UK (COG-UK) Consortium | Jacquelyn Wynn, Mairead Hyland, The Lighthouse Lab in Alderley Park and Alex Alderton, Roberto Amato, Jeffrey Barrett, Sonia Goncalves, Ewan Harrison, David K. Jackson, Ian Johnston, Dominic Kwiatkowski, Cordelia Langford, John Sillitoe on behalf of the Wellcome Sanger Institute COVID-19 Surveillance Team |
| EPI_ISL_3227213 | Lighthouse Lab in Milton Keynes | Wellcome Sanger Institute for the COVID-19 Genomics UK (COG-UK) Consortium | The Lighthouse Lab in Milton Keynes and Alex Alderton, Roberto Amato, Jeffrey Barrett, Sonia Goncalves, Ewan Harrison, David K. Jackson, Ian Johnston, Dominic Kwiatkowski, Cordelia Langford, John Sillitoe on behalf of the Wellcome Sanger Institute COVID-19 Surveillance Team |
| EPI_ISL_3227214 | Lighthouse Lab in Glasgow | Wellcome Sanger Institute for the COVID-19 Genomics UK (COG-UK) Consortium | Harper VanSteenhouse, Yumi Kasai, David Gray, Carol Clugston, Anna Dominiczak and Alex Alderton, Roberto Amato, Jeffrey Barrett, Sonia Goncalves, Ewan Harrison, David K. Jackson, Ian Johnston, Dominic Kwiatkowski, Cordelia Langford, John Sillitoe on behalf of the Wellcome Sanger Institute COVID-19 Surveillance Team |
| EPI_ISL_3227215 | Lighthouse Lab in Milton Keynes | Wellcome Sanger Institute for the COVID-19 Genomics UK (COG-UK) Consortium | The Lighthouse Lab in Milton Keynes and Alex Alderton, Roberto Amato, Jeffrey Barrett, Sonia Goncalves, Ewan Harrison, David K. Jackson, Ian Johnston, Dominic Kwiatkowski, Cordelia Langford, John Sillitoe on behalf of the Wellcome Sanger Institute COVID-19 Surveillance Team |
| EPI_ISL_3227216, EPI_ISL_3227217 | Lighthouse Lab in Glasgow | Wellcome Sanger Institute for the COVID-19 Genomics UK (COG-UK) Consortium | Harper VanSteenhouse, Yumi Kasai, David Gray, Carol Clugston, Anna Dominiczak and Alex Alderton, Roberto Amato, Jeffrey Barrett, Sonia Goncalves, Ewan Harrison, David K. Jackson, Ian Johnston, Dominic Kwiatkowski, Cordelia Langford, John Sillitoe on behalf of the Wellcome Sanger Institute COVID-19 Surveillance Team |
| EPI_ISL_3227218, EPI_ISL_3227219 | Lighthouse Lab in Alderley Park | Wellcome Sanger Institute for the COVID-19 Genomics UK (COG-UK) Consortium | Jacquelyn Wynn, Mairead Hyland, The Lighthouse Lab in Alderley Park and Alex Alderton, Roberto Amato, Jeffrey Barrett, Sonia Goncalves, Ewan Harrison, David K. Jackson, Ian Johnston, Dominic Kwiatkowski, Cordelia Langford, John Sillitoe on behalf of the Wellcome Sanger Institute COVID-19 Surveillance Team |
| EPI_ISL_3227220 | Lighthouse Lab in Milton Keynes | Wellcome Sanger Institute for the COVID-19 Genomics UK (COG-UK) Consortium | The Lighthouse Lab in Milton Keynes and Alex Alderton, Roberto Amato, Jeffrey Barrett, Sonia Goncalves, Ewan Harrison, David K. Jackson, Ian Johnston, Dominic Kwiatkowski, Cordelia Langford, John Sillitoe on behalf of the Wellcome Sanger Institute COVID-19 Surveillance Team |
| EPI_ISL_3227221 | Lighthouse Lab in Glasgow | Wellcome Sanger Institute for the COVID-19 Genomics UK (COG-UK) Consortium | Harper VanSteenhouse, Yumi Kasai, David Gray, Carol Clugston, Anna Dominiczak and Alex Alderton, Roberto Amato, Jeffrey Barrett, Sonia Goncalves, Ewan Harrison, David K. Jackson, Ian Johnston, Dominic Kwiatkowski, Cordelia Langford, John Sillitoe on behalf of the Wellcome Sanger Institute COVID-19 Surveillance Team |
| EPI_ISL_3227222, EPI_ISL_3227223 | Lighthouse Laboratory Plymouth | Wellcome Sanger Institute for the COVID-19 Genomics UK (COG-UK) Consortium | Lighthouse Laboratory Plymouth and Alex Alderton, Roberto Amato, Jeffrey Barrett, Sonia Goncalves, Ewan Harrison, David K. Jackson, Ian Johnston, Dominic Kwiatkowski, Cordelia Langford, John Sillitoe on behalf of the Wellcome Sanger Institute COVID-19 Surveillance Team |
| EPI_ISL_3227224 | Lighthouse Lab in Milton Keynes | Wellcome Sanger Institute for the COVID-19 Genomics UK (COG-UK) Consortium | The Lighthouse Lab in Milton Keynes and Alex Alderton, Roberto Amato, Jeffrey Barrett, Sonia Goncalves, Ewan Harrison, David K. Jackson, Ian Johnston, Dominic Kwiatkowski, Cordelia Langford, John Sillitoe on behalf of the Wellcome Sanger Institute COVID-19 Surveillance Team |
| EPI_ISL_3227225, EPI_ISL_3227226, EPI_ISL_3227227 | Lighthouse Laboratory Plymouth | Wellcome Sanger Institute for the COVID-19 Genomics UK (COG-UK) Consortium | Lighthouse Laboratory Plymouth and Alex Alderton, Roberto Amato, Jeffrey Barrett, Sonia Goncalves, Ewan Harrison, David K. Jackson, Ian Johnston, Dominic Kwiatkowski, Cordelia Langford, John Sillitoe on behalf of the Wellcome Sanger Institute COVID-19 Surveillance Team |
| EPI_ISL_3227228 | Lighthouse Lab in Milton Keynes | Wellcome Sanger Institute for the COVID-19 Genomics UK (COG-UK) Consortium | The Lighthouse Lab in Milton Keynes and Alex Alderton, Roberto Amato, Jeffrey Barrett, Sonia Goncalves, Ewan Harrison, David K. Jackson, Ian Johnston, Dominic Kwiatkowski, Cordelia Langford, John Sillitoe on behalf of the Wellcome Sanger Institute COVID-19 Surveillance Team |
| EPI_ISL_3227229, EPI_ISL_3227230 | Lighthouse Laboratory Plymouth | Wellcome Sanger Institute for the COVID-19 Genomics UK (COG-UK) Consortium | Lighthouse Laboratory Plymouth and Alex Alderton, Roberto Amato, Jeffrey Barrett, Sonia Goncalves, Ewan Harrison, David K. Jackson, Ian Johnston, Dominic Kwiatkowski, Cordelia Langford, John Sillitoe on behalf of the Wellcome Sanger Institute COVID-19 Surveillance Team |
| EPI_ISL_3227231 | Lighthouse Lab in Milton Keynes | Wellcome Sanger Institute for the COVID-19 Genomics UK (COG-UK) Consortium | The Lighthouse Lab in Milton Keynes and Alex Alderton, Roberto Amato, Jeffrey Barrett, Sonia Goncalves, Ewan Harrison, David K. Jackson, Ian Johnston, Dominic Kwiatkowski, Cordelia Langford, John Sillitoe on behalf of the Wellcome Sanger Institute COVID-19 Surveillance Team |
| EPI_ISL_3227232 | Lighthouse Laboratory Plymouth | Wellcome Sanger Institute for the COVID-19 Genomics UK (COG-UK) Consortium | Lighthouse Laboratory Plymouth and Alex Alderton, Roberto Amato, Jeffrey Barrett, Sonia Goncalves, Ewan Harrison, David K. Jackson, Ian Johnston, Dominic Kwiatkowski, Cordelia Langford, John Sillitoe on behalf of the Wellcome Sanger Institute COVID-19 Surveillance Team |
| EPI_ISL_3227233, EPI_ISL_3227234 | Lighthouse Lab in Milton Keynes | Wellcome Sanger Institute for the COVID-19 Genomics UK (COG-UK) Consortium | The Lighthouse Lab in Milton Keynes and Alex Alderton, Roberto Amato, Jeffrey Barrett, Sonia Goncalves, Ewan Harrison, David K. Jackson, Ian Johnston, Dominic Kwiatkowski, Cordelia Langford, John Sillitoe on behalf of the Wellcome Sanger Institute COVID-19 Surveillance Team |
| EPI_ISL_3227235 | Lighthouse Laboratory Plymouth | Wellcome Sanger Institute for the COVID-19 Genomics UK (COG-UK) Consortium | Lighthouse Laboratory Plymouth and Alex Alderton, Roberto Amato, Jeffrey Barrett, Sonia Goncalves, Ewan Harrison, David K. Jackson, Ian Johnston, Dominic Kwiatkowski, Cordelia Langford, John Sillitoe on behalf of the Wellcome Sanger Institute COVID-19 Surveillance Team |
| EPI_ISL_3227236, EPI_ISL_3227237, EPI_ISL_3227238, EPI_ISL_3227239, EPI_ISL_3227240, EPI_ISL_3227241, EPI_ISL_3227242 | Lighthouse Lab in Milton Keynes | Wellcome Sanger Institute for the COVID-19 Genomics UK (COG-UK) Consortium | The Lighthouse Lab in Milton Keynes and Alex Alderton, Roberto Amato, Jeffrey Barrett, Sonia Goncalves, Ewan Harrison, David K. Jackson, Ian Johnston, Dominic Kwiatkowski, Cordelia Langford, John Sillitoe on behalf of the Wellcome Sanger Institute COVID-19 Surveillance Team |
| EPI_ISL_3227243, EPI_ISL_3227244 | Lighthouse Laboratory Plymouth | Wellcome Sanger Institute for the COVID-19 Genomics UK (COG-UK) Consortium | Lighthouse Laboratory Plymouth and Alex Alderton, Roberto Amato, Jeffrey Barrett, Sonia Goncalves, Ewan Harrison, David K. Jackson, Ian Johnston, Dominic Kwiatkowski, Cordelia Langford, John Sillitoe on behalf of the Wellcome Sanger Institute COVID-19 Surveillance Team |
| EPI_ISL_3227245, EPI_ISL_3227246, EPI_ISL_3227247, EPI_ISL_3227248, EPI_ISL_3227249, EPI_ISL_3227250 | Lighthouse Lab in Milton Keynes | Wellcome Sanger Institute for the COVID-19 Genomics UK (COG-UK) Consortium | The Lighthouse Lab in Milton Keynes and Alex Alderton, Roberto Amato, Jeffrey Barrett, Sonia Goncalves, Ewan Harrison, David K. Jackson, Ian Johnston, Dominic Kwiatkowski, Cordelia Langford, John Sillitoe on behalf of the Wellcome Sanger Institute COVID-19 Surveillance Team |
| EPI_ISL_3227251, EPI_ISL_3227252, EPI_ISL_3227253 | Lighthouse Laboratory Plymouth | Wellcome Sanger Institute for the COVID-19 Genomics UK (COG-UK) Consortium | Lighthouse Laboratory Plymouth and Alex Alderton, Roberto Amato, Jeffrey Barrett, Sonia Goncalves, Ewan Harrison, David K. Jackson, Ian Johnston, Dominic Kwiatkowski, Cordelia Langford, John Sillitoe on behalf of the Wellcome Sanger Institute COVID-19 Surveillance Team |
| EPI_ISL_3227254, EPI_ISL_3227255 | Lighthouse Lab in Milton Keynes | Wellcome Sanger Institute for the COVID-19 Genomics UK (COG-UK) Consortium | The Lighthouse Lab in Milton Keynes and Alex Alderton, Roberto Amato, Jeffrey Barrett, Sonia Goncalves, Ewan Harrison, David K. Jackson, Ian Johnston, Dominic Kwiatkowski, Cordelia Langford, John Sillitoe on behalf of the Wellcome Sanger Institute COVID-19 Surveillance Team |
| EPI_ISL_3227256, EPI_ISL_3227257 | Lighthouse Laboratory Plymouth | Wellcome Sanger Institute for the COVID-19 Genomics UK (COG-UK) Consortium | Lighthouse Laboratory Plymouth and Alex Alderton, Roberto Amato, Jeffrey Barrett, Sonia Goncalves, Ewan Harrison, David K. Jackson, Ian Johnston, Dominic Kwiatkowski, Cordelia Langford, John Sillitoe on behalf of the Wellcome Sanger Institute COVID-19 Surveillance Team |
| EPI_ISL_3227258 | Lighthouse Lab in Milton Keynes | Wellcome Sanger Institute for the COVID-19 Genomics UK (COG-UK) Consortium | The Lighthouse Lab in Milton Keynes and Alex Alderton, Roberto Amato, Jeffrey Barrett, Sonia Goncalves, Ewan Harrison, David K. Jackson, Ian Johnston, Dominic Kwiatkowski, Cordelia Langford, John Sillitoe on behalf of the Wellcome Sanger Institute COVID-19 Surveillance Team |
| EPI_ISL_3227259 | Lighthouse Laboratory Plymouth | Wellcome Sanger Institute for the COVID-19 Genomics UK (COG-UK) Consortium | Lighthouse Laboratory Plymouth and Alex Alderton, Roberto Amato, Jeffrey Barrett, Sonia Goncalves, Ewan Harrison, David K. Jackson, Ian Johnston, Dominic Kwiatkowski, Cordelia Langford, John Sillitoe on behalf of the Wellcome Sanger Institute COVID-19 Surveillance Team |

| | | | |
|---|---|---|---|
| EPI_ISL_3227260, EPI_ISL_3227261 | Lighthouse Lab in Milton Keynes | Wellcome Sanger Institute for the COVID-19 Genomics UK (COG-UK) Consortium | The Lighthouse Lab in Milton Keynes and Alex Alderton, Roberto Amato, Jeffrey Barrett, Sonia Goncalves, Ewan Harrison, David K. Jackson, Ian Johnston, Dominic Kwiatkowski, Cordelia Langford, John Sillitoe on behalf of the Wellcome Sanger Institute COVID-19 Surveillance Team |
| EPI_ISL_3227262 | Lighthouse Laboratory Plymouth | Wellcome Sanger Institute for the COVID-19 Genomics UK (COG-UK) Consortium | Lighthouse Laboratory Plymouth and Alex Alderton, Roberto Amato, Jeffrey Barrett, Sonia Goncalves, Ewan Harrison, David K. Jackson, Ian Johnston, Dominic Kwiatkowski, Cordelia Langford, John Sillitoe on behalf of the Wellcome Sanger Institute COVID-19 Surveillance Team |
| EPI_ISL_3227263 | Lighthouse Lab in Milton Keynes | Wellcome Sanger Institute for the COVID-19 Genomics UK (COG-UK) Consortium | The Lighthouse Lab in Milton Keynes and Alex Alderton, Roberto Amato, Jeffrey Barrett, Sonia Goncalves, Ewan Harrison, David K. Jackson, Ian Johnston, Dominic Kwiatkowski, Cordelia Langford, John Sillitoe on behalf of the Wellcome Sanger Institute COVID-19 Surveillance Team |
| EPI_ISL_3227264, EPI_ISL_3227265 | Lighthouse Laboratory Plymouth | Wellcome Sanger Institute for the COVID-19 Genomics UK (COG-UK) Consortium | Lighthouse Laboratory Plymouth and Alex Alderton, Roberto Amato, Jeffrey Barrett, Sonia Goncalves, Ewan Harrison, David K. Jackson, Ian Johnston, Dominic Kwiatkowski, Cordelia Langford, John Sillitoe on behalf of the Wellcome Sanger Institute COVID-19 Surveillance Team |
| EPI_ISL_3227266, EPI_ISL_3227267, EPI_ISL_3227268 | Lighthouse Lab in Milton Keynes | Wellcome Sanger Institute for the COVID-19 Genomics UK (COG-UK) Consortium | The Lighthouse Lab in Milton Keynes and Alex Alderton, Roberto Amato, Jeffrey Barrett, Sonia Goncalves, Ewan Harrison, David K. Jackson, Ian Johnston, Dominic Kwiatkowski, Cordelia Langford, John Sillitoe on behalf of the Wellcome Sanger Institute COVID-19 Surveillance Team |
| EPI_ISL_3227269 | Lighthouse Laboratory Plymouth | Wellcome Sanger Institute for the COVID-19 Genomics UK (COG-UK) Consortium | Lighthouse Laboratory Plymouth and Alex Alderton, Roberto Amato, Jeffrey Barrett, Sonia Goncalves, Ewan Harrison, David K. Jackson, Ian Johnston, Dominic Kwiatkowski, Cordelia Langford, John Sillitoe on behalf of the Wellcome Sanger Institute COVID-19 Surveillance Team |
| EPI_ISL_3227270, EPI_ISL_3227271, EPI_ISL_3227272 | Lighthouse Lab in Milton Keynes | Wellcome Sanger Institute for the COVID-19 Genomics UK (COG-UK) Consortium | The Lighthouse Lab in Milton Keynes and Alex Alderton, Roberto Amato, Jeffrey Barrett, Sonia Goncalves, Ewan Harrison, David K. Jackson, Ian Johnston, Dominic Kwiatkowski, Cordelia Langford, John Sillitoe on behalf of the Wellcome Sanger Institute COVID-19 Surveillance Team |
| EPI_ISL_3227273, EPI_ISL_3227274 | Lighthouse Laboratory Plymouth | Wellcome Sanger Institute for the COVID-19 Genomics UK (COG-UK) Consortium | Lighthouse Laboratory Plymouth and Alex Alderton, Roberto Amato, Jeffrey Barrett, Sonia Goncalves, Ewan Harrison, David K. Jackson, Ian Johnston, Dominic Kwiatkowski, Cordelia Langford, John Sillitoe on behalf of the Wellcome Sanger Institute COVID-19 Surveillance Team |
| EPI_ISL_3227275, EPI_ISL_3227276, EPI_ISL_3227277 | Lighthouse Lab in Milton Keynes | Wellcome Sanger Institute for the COVID-19 Genomics UK (COG-UK) Consortium | The Lighthouse Lab in Milton Keynes and Alex Alderton, Roberto Amato, Jeffrey Barrett, Sonia Goncalves, Ewan Harrison, David K. Jackson, Ian Johnston, Dominic Kwiatkowski, Cordelia Langford, John Sillitoe on behalf of the Wellcome Sanger Institute COVID-19 Surveillance Team |
| EPI_ISL_3227278 | Lighthouse Laboratory Plymouth | Wellcome Sanger Institute for the COVID-19 Genomics UK (COG-UK) Consortium | Lighthouse Laboratory Plymouth and Alex Alderton, Roberto Amato, Jeffrey Barrett, Sonia Goncalves, Ewan Harrison, David K. Jackson, Ian Johnston, Dominic Kwiatkowski, Cordelia Langford, John Sillitoe on behalf of the Wellcome Sanger Institute COVID-19 Surveillance Team |
| EPI_ISL_3227279 | Lighthouse Lab in Milton Keynes | Wellcome Sanger Institute for the COVID-19 Genomics UK (COG-UK) Consortium | The Lighthouse Lab in Milton Keynes and Alex Alderton, Roberto Amato, Jeffrey Barrett, Sonia Goncalves, Ewan Harrison, David K. Jackson, Ian Johnston, Dominic Kwiatkowski, Cordelia Langford, John Sillitoe on behalf of the Wellcome Sanger Institute COVID-19 Surveillance Team |
| EPI_ISL_3227280, EPI_ISL_3227281 | Lighthouse Laboratory Plymouth | Wellcome Sanger Institute for the COVID-19 Genomics UK (COG-UK) Consortium | Lighthouse Laboratory Plymouth and Alex Alderton, Roberto Amato, Jeffrey Barrett, Sonia Goncalves, Ewan Harrison, David K. Jackson, Ian Johnston, Dominic Kwiatkowski, Cordelia Langford, John Sillitoe on behalf of the Wellcome Sanger Institute COVID-19 Surveillance Team |
| EPI_ISL_3227282 | Lighthouse Lab in Milton Keynes | Wellcome Sanger Institute for the COVID-19 Genomics UK (COG-UK) Consortium | The Lighthouse Lab in Milton Keynes and Alex Alderton, Roberto Amato, Jeffrey Barrett, Sonia Goncalves, Ewan Harrison, David K. Jackson, Ian Johnston, Dominic Kwiatkowski, Cordelia Langford, John Sillitoe on behalf of the Wellcome Sanger Institute COVID-19 Surveillance Team |
| EPI_ISL_3227283 | Lighthouse Laboratory Plymouth | Wellcome Sanger Institute for the COVID-19 Genomics UK (COG-UK) Consortium | Lighthouse Laboratory Plymouth and Alex Alderton, Roberto Amato, Jeffrey Barrett, Sonia Goncalves, Ewan Harrison, David K. Jackson, Ian Johnston, Dominic Kwiatkowski, Cordelia Langford, John Sillitoe on behalf of the Wellcome Sanger Institute COVID-19 Surveillance Team |
| EPI_ISL_3227284 | Lighthouse Lab in Milton Keynes | Wellcome Sanger Institute for the COVID-19 Genomics UK (COG-UK) Consortium | The Lighthouse Lab in Milton Keynes and Alex Alderton, Roberto Amato, Jeffrey Barrett, Sonia Goncalves, Ewan Harrison, David K. Jackson, Ian Johnston, Dominic Kwiatkowski, Cordelia Langford, John Sillitoe on behalf of the Wellcome Sanger Institute COVID-19 Surveillance Team |
| EPI_ISL_3227285, EPI_ISL_3227286 | Lighthouse Laboratory Plymouth | Wellcome Sanger Institute for the COVID-19 Genomics UK (COG-UK) Consortium | Lighthouse Laboratory Plymouth and Alex Alderton, Roberto Amato, Jeffrey Barrett, Sonia Goncalves, Ewan Harrison, David K. Jackson, Ian Johnston, Dominic Kwiatkowski, Cordelia Langford, John Sillitoe on behalf of the Wellcome Sanger Institute COVID-19 Surveillance Team |
| EPI_ISL_3227287 | Lighthouse Lab in Milton Keynes | Wellcome Sanger Institute for the COVID-19 Genomics UK (COG-UK) Consortium | The Lighthouse Lab in Milton Keynes and Alex Alderton, Roberto Amato, Jeffrey Barrett, Sonia Goncalves, Ewan Harrison, David K. Jackson, Ian Johnston, Dominic Kwiatkowski, Cordelia Langford, John Sillitoe on behalf of the Wellcome Sanger Institute COVID-19 Surveillance Team |
| EPI_ISL_3227288 | Lighthouse Laboratory Plymouth | Wellcome Sanger Institute for the COVID-19 Genomics UK (COG-UK) Consortium | Lighthouse Laboratory Plymouth and Alex Alderton, Roberto Amato, Jeffrey Barrett, Sonia Goncalves, Ewan Harrison, David K. Jackson, Ian Johnston, Dominic Kwiatkowski, Cordelia Langford, John Sillitoe on behalf of the Wellcome Sanger Institute COVID-19 Surveillance Team |
| EPI_ISL_3227289 | Lighthouse Lab in Milton Keynes | Wellcome Sanger Institute for the COVID-19 Genomics UK (COG-UK) Consortium | The Lighthouse Lab in Milton Keynes and Alex Alderton, Roberto Amato, Jeffrey Barrett, Sonia Goncalves, Ewan Harrison, David K. Jackson, Ian Johnston, Dominic Kwiatkowski, Cordelia Langford, John Sillitoe on behalf of the Wellcome Sanger Institute COVID-19 Surveillance Team |
| EPI_ISL_3227290, EPI_ISL_3227291, EPI_ISL_3227292, EPI_ISL_3227293 | Lighthouse Laboratory Plymouth | Wellcome Sanger Institute for the COVID-19 Genomics UK (COG-UK) Consortium | Lighthouse Laboratory Plymouth and Alex Alderton, Roberto Amato, Jeffrey Barrett, Sonia Goncalves, Ewan Harrison, David K. Jackson, Ian Johnston, Dominic Kwiatkowski, Cordelia Langford, John Sillitoe on behalf of the Wellcome Sanger Institute COVID-19 Surveillance Team |
| EPI_ISL_3227294 | Lighthouse Lab in Milton Keynes | Wellcome Sanger Institute for the COVID-19 Genomics UK (COG-UK) Consortium | The Lighthouse Lab in Milton Keynes and Alex Alderton, Roberto Amato, Jeffrey Barrett, Sonia Goncalves, Ewan Harrison, David K. Jackson, Ian Johnston, Dominic Kwiatkowski, Cordelia Langford, John Sillitoe on behalf of the Wellcome Sanger Institute COVID-19 Surveillance Team |
| EPI_ISL_3227295 | Lighthouse Laboratory Plymouth | Wellcome Sanger Institute for the COVID-19 Genomics UK (COG-UK) Consortium | Lighthouse Laboratory Plymouth and Alex Alderton, Roberto Amato, Jeffrey Barrett, Sonia Goncalves, Ewan Harrison, David K. Jackson, Ian Johnston, Dominic Kwiatkowski, Cordelia Langford, John Sillitoe on behalf of the Wellcome Sanger Institute COVID-19 Surveillance Team |
| EPI_ISL_3227296, EPI_ISL_3227297, EPI_ISL_3227298, EPI_ISL_3227299, EPI_ISL_3227300, EPI_ISL_3227301, EPI_ISL_3227302, EPI_ISL_3227303 | Lighthouse Lab in Milton Keynes | Wellcome Sanger Institute for the COVID-19 Genomics UK (COG-UK) Consortium | The Lighthouse Lab in Milton Keynes and Alex Alderton, Roberto Amato, Jeffrey Barrett, Sonia Goncalves, Ewan Harrison, David K. Jackson, Ian Johnston, Dominic Kwiatkowski, Cordelia Langford, John Sillitoe on behalf of the Wellcome Sanger Institute COVID-19 Surveillance Team |
| EPI_ISL_3227304 | Lighthouse Laboratory Plymouth | Wellcome Sanger Institute for the COVID-19 Genomics UK (COG-UK) Consortium | Lighthouse Laboratory Plymouth and Alex Alderton, Roberto Amato, Jeffrey Barrett, Sonia Goncalves, Ewan Harrison, David K. Jackson, Ian Johnston, Dominic Kwiatkowski, Cordelia Langford, John Sillitoe on behalf of the Wellcome Sanger Institute COVID-19 Surveillance Team |
| EPI_ISL_3227305, EPI_ISL_3227306 | Lighthouse Lab in Milton Keynes | Wellcome Sanger Institute for the COVID-19 Genomics UK (COG-UK) Consortium | The Lighthouse Lab in Milton Keynes and Alex Alderton, Roberto Amato, Jeffrey Barrett, Sonia Goncalves, Ewan Harrison, David K. Jackson, Ian Johnston, Dominic Kwiatkowski, Cordelia Langford, John Sillitoe on behalf of the Wellcome Sanger Institute COVID-19 Surveillance Team |
| EPI_ISL_3227307, EPI_ISL_3227308, EPI_ISL_3227309 | Lighthouse Laboratory Plymouth | Wellcome Sanger Institute for the COVID-19 Genomics UK (COG-UK) Consortium | Lighthouse Laboratory Plymouth and Alex Alderton, Roberto Amato, Jeffrey Barrett, Sonia Goncalves, Ewan Harrison, David K. Jackson, Ian Johnston, Dominic Kwiatkowski, Cordelia Langford, John Sillitoe on behalf of the Wellcome Sanger Institute COVID-19 Surveillance Team |
| EPI_ISL_3227310 | Lighthouse Lab in Milton Keynes | Wellcome Sanger Institute for the COVID-19 Genomics UK (COG-UK) Consortium | The Lighthouse Lab in Milton Keynes and Alex Alderton, Roberto Amato, Jeffrey Barrett, Sonia Goncalves, Ewan Harrison, David K. Jackson, Ian Johnston, Dominic Kwiatkowski, Cordelia Langford, John Sillitoe on behalf of the Wellcome Sanger Institute COVID-19 Surveillance Team |
| EPI_ISL_3227311, EPI_ISL_3227312 | Lighthouse Laboratory Plymouth | Wellcome Sanger Institute for the COVID-19 Genomics UK (COG-UK) Consortium | Lighthouse Laboratory Plymouth and Alex Alderton, Roberto Amato, Jeffrey Barrett, Sonia Goncalves, Ewan Harrison, David K. Jackson, Ian Johnston, Dominic Kwiatkowski, Cordelia Langford, John Sillitoe on behalf of the Wellcome Sanger Institute COVID-19 Surveillance Team |
| EPI_ISL_3227313, EPI_ISL_3227314 | Lighthouse Lab in Milton Keynes | Wellcome Sanger Institute for the COVID-19 Genomics UK (COG-UK) Consortium | The Lighthouse Lab in Milton Keynes and Alex Alderton, Roberto Amato, Jeffrey Barrett, Sonia Goncalves, Ewan Harrison, David K. Jackson, Ian Johnston, Dominic Kwiatkowski, Cordelia Langford, John Sillitoe on behalf of the Wellcome Sanger Institute COVID-19 Surveillance Team |
| EPI_ISL_3227315, EPI_ISL_3227316 | Lighthouse Laboratory Plymouth | Wellcome Sanger Institute for the COVID-19 Genomics UK (COG-UK) Consortium | Lighthouse Laboratory Plymouth and Alex Alderton, Roberto Amato, Jeffrey Barrett, Sonia Goncalves, Ewan Harrison, David K. Jackson, Ian Johnston, Dominic Kwiatkowski, Cordelia Langford, John Sillitoe on behalf of the Wellcome Sanger Institute COVID-19 Surveillance Team |
| EPI_ISL_3227317 | Lighthouse Lab in Milton Keynes | Wellcome Sanger Institute for the COVID-19 Genomics UK (COG-UK) Consortium | The Lighthouse Lab in Milton Keynes and Alex Alderton, Roberto Amato, Jeffrey Barrett, Sonia Goncalves, Ewan Harrison, David K. Jackson, Ian Johnston, Dominic Kwiatkowski, Cordelia Langford, John Sillitoe on behalf of the Wellcome Sanger Institute COVID-19 Surveillance Team |
| EPI_ISL_3227318 | Lighthouse Laboratory Plymouth | Wellcome Sanger Institute for the COVID-19 Genomics UK (COG-UK) Consortium | Lighthouse Laboratory Plymouth and Alex Alderton, Roberto Amato, Jeffrey Barrett, Sonia Goncalves, Ewan Harrison, David K. Jackson, Ian Johnston, Dominic Kwiatkowski, Cordelia Langford, John Sillitoe on behalf of the Wellcome Sanger Institute COVID-19 Surveillance Team |
| EPI_ISL_3227319, EPI_ISL_3227320, EPI_ISL_3227321, EPI_ISL_3227322, EPI_ISL_3227323 | Lighthouse Lab in Milton Keynes | Wellcome Sanger Institute for the COVID-19 Genomics UK (COG-UK) Consortium | The Lighthouse Lab in Milton Keynes and Alex Alderton, Roberto Amato, Jeffrey Barrett, Sonia Goncalves, Ewan Harrison, David K. Jackson, Ian Johnston, Dominic Kwiatkowski, Cordelia Langford, John Sillitoe on behalf of the Wellcome Sanger Institute COVID-19 Surveillance Team |
| EPI_ISL_3227324 | Lighthouse Laboratory Plymouth | Wellcome Sanger Institute for the COVID-19 Genomics UK (COG-UK) Consortium | Lighthouse Laboratory Plymouth and Alex Alderton, Roberto Amato, Jeffrey Barrett, Sonia Goncalves, Ewan Harrison, David K. Jackson, Ian Johnston, Dominic Kwiatkowski, Cordelia Langford, John Sillitoe on behalf of the Wellcome Sanger Institute COVID-19 Surveillance Team |
| EPI_ISL_3227325, EPI_ISL_3227326 | Lighthouse Lab in Milton Keynes | Wellcome Sanger Institute for the COVID-19 Genomics UK (COG-UK) Consortium | The Lighthouse Lab in Milton Keynes and Alex Alderton, Roberto Amato, Jeffrey Barrett, Sonia Goncalves, Ewan Harrison, David K. Jackson, Ian Johnston, Dominic Kwiatkowski, Cordelia Langford, John Sillitoe on behalf of the Wellcome Sanger Institute COVID-19 Surveillance Team |
| EPI_ISL_3227327, EPI_ISL_3227328 | Lighthouse Laboratory Plymouth | Wellcome Sanger Institute for the COVID-19 Genomics UK (COG-UK) Consortium | Lighthouse Laboratory Plymouth and Alex Alderton, Roberto Amato, Jeffrey Barrett, Sonia Goncalves, Ewan Harrison, David K. Jackson, Ian Johnston, Dominic Kwiatkowski, Cordelia Langford, John Sillitoe on behalf of the Wellcome Sanger Institute COVID-19 Surveillance Team |
| EPI_ISL_3227329 | Lighthouse Lab in Milton Keynes | Wellcome Sanger Institute for the COVID-19 Genomics UK (COG-UK) Consortium | The Lighthouse Lab in Milton Keynes and Alex Alderton, Roberto Amato, Jeffrey Barrett, Sonia Goncalves, Ewan Harrison, David K. Jackson, Ian Johnston, Dominic Kwiatkowski, Cordelia Langford, John Sillitoe on behalf of the Wellcome Sanger Institute COVID-19 Surveillance Team |
| EPI_ISL_3227330, EPI_ISL_3227331 | Lighthouse Laboratory Plymouth | Wellcome Sanger Institute for the COVID-19 Genomics UK (COG-UK) Consortium | Lighthouse Laboratory Plymouth and Alex Alderton, Roberto Amato, Jeffrey Barrett, Sonia Goncalves, Ewan Harrison, David K. Jackson, Ian Johnston, Dominic Kwiatkowski, Cordelia Langford, John Sillitoe on behalf of the Wellcome Sanger Institute COVID-19 Surveillance Team |
| EPI_ISL_3227332 | Lighthouse Lab in Milton Keynes | Wellcome Sanger Institute for the COVID-19 Genomics UK (COG-UK) Consortium | The Lighthouse Lab in Milton Keynes and Alex Alderton, Roberto Amato, Jeffrey Barrett, Sonia Goncalves, Ewan Harrison, David K. Jackson, Ian Johnston, Dominic Kwiatkowski, Cordelia Langford, John Sillitoe on behalf of the Wellcome Sanger Institute COVID-19 Surveillance Team |

| | | | |
|---|---|---|---|
| EPI_ISL_3227333 | Lighthouse Laboratory Plymouth | Wellcome Sanger Institute for the COVID-19 Genomics UK (COG-UK) Consortium | Lighthouse Laboratory Plymouth and Alex Alderton, Roberto Amato, Jeffrey Barrett, Sonia Goncalves, Ewan Harrison, David K. Jackson, Ian Johnston, Dominic Kwiatkowski, Cordelia Langford, John Sillitoe on behalf of the Wellcome Sanger Institute COVID-19 Surveillance Team |
| EPI_ISL_3227334 | Lighthouse Lab in Milton Keynes | Wellcome Sanger Institute for the COVID-19 Genomics UK (COG-UK) Consortium | The Lighthouse Lab in Milton Keynes and Alex Alderton, Roberto Amato, Jeffrey Barrett, Sonia Goncalves, Ewan Harrison, David K. Jackson, Ian Johnston, Dominic Kwiatkowski, Cordelia Langford, John Sillitoe on behalf of the Wellcome Sanger Institute COVID-19 Surveillance Team |
| EPI_ISL_3227335 | Lighthouse Laboratory Plymouth | Wellcome Sanger Institute for the COVID-19 Genomics UK (COG-UK) Consortium | Lighthouse Laboratory Plymouth and Alex Alderton, Roberto Amato, Jeffrey Barrett, Sonia Goncalves, Ewan Harrison, David K. Jackson, Ian Johnston, Dominic Kwiatkowski, Cordelia Langford, John Sillitoe on behalf of the Wellcome Sanger Institute COVID-19 Surveillance Team |
| EPI_ISL_3227336 | Lighthouse Lab in Milton Keynes | Wellcome Sanger Institute for the COVID-19 Genomics UK (COG-UK) Consortium | The Lighthouse Lab in Milton Keynes and Alex Alderton, Roberto Amato, Jeffrey Barrett, Sonia Goncalves, Ewan Harrison, David K. Jackson, Ian Johnston, Dominic Kwiatkowski, Cordelia Langford, John Sillitoe on behalf of the Wellcome Sanger Institute COVID-19 Surveillance Team |
| EPI_ISL_3227337 | Lighthouse Laboratory Plymouth | Wellcome Sanger Institute for the COVID-19 Genomics UK (COG-UK) Consortium | Lighthouse Laboratory Plymouth and Alex Alderton, Roberto Amato, Jeffrey Barrett, Sonia Goncalves, Ewan Harrison, David K. Jackson, Ian Johnston, Dominic Kwiatkowski, Cordelia Langford, John Sillitoe on behalf of the Wellcome Sanger Institute COVID-19 Surveillance Team |
| EPI_ISL_3227338 | Lighthouse Lab in Milton Keynes | Wellcome Sanger Institute for the COVID-19 Genomics UK (COG-UK) Consortium | The Lighthouse Lab in Milton Keynes and Alex Alderton, Roberto Amato, Jeffrey Barrett, Sonia Goncalves, Ewan Harrison, David K. Jackson, Ian Johnston, Dominic Kwiatkowski, Cordelia Langford, John Sillitoe on behalf of the Wellcome Sanger Institute COVID-19 Surveillance Team |
| EPI_ISL_3227339, EPI_ISL_3227340, EPI_ISL_3227341, EPI_ISL_3227342, EPI_ISL_3227343, EPI_ISL_3227344, EPI_ISL_3227345, EPI_ISL_3227346, EPI_ISL_3227347 | Lighthouse Laboratory Plymouth | Wellcome Sanger Institute for the COVID-19 Genomics UK (COG-UK) Consortium | Lighthouse Laboratory Plymouth and Alex Alderton, Roberto Amato, Jeffrey Barrett, Sonia Goncalves, Ewan Harrison, David K. Jackson, Ian Johnston, Dominic Kwiatkowski, Cordelia Langford, John Sillitoe on behalf of the Wellcome Sanger Institute COVID-19 Surveillance Team |
| EPI_ISL_3227348, EPI_ISL_3227349 | Lighthouse Lab in Milton Keynes | Wellcome Sanger Institute for the COVID-19 Genomics UK (COG-UK) Consortium | The Lighthouse Lab in Milton Keynes and Alex Alderton, Roberto Amato, Jeffrey Barrett, Sonia Goncalves, Ewan Harrison, David K. Jackson, Ian Johnston, Dominic Kwiatkowski, Cordelia Langford, John Sillitoe on behalf of the Wellcome Sanger Institute COVID-19 Surveillance Team |
| EPI_ISL_3227350, EPI_ISL_3227351, EPI_ISL_3227352 | Lighthouse Laboratory Plymouth | Wellcome Sanger Institute for the COVID-19 Genomics UK (COG-UK) Consortium | Lighthouse Laboratory Plymouth and Alex Alderton, Roberto Amato, Jeffrey Barrett, Sonia Goncalves, Ewan Harrison, David K. Jackson, Ian Johnston, Dominic Kwiatkowski, Cordelia Langford, John Sillitoe on behalf of the Wellcome Sanger Institute COVID-19 Surveillance Team |
| EPI_ISL_3227353, EPI_ISL_3227354 | Lighthouse Lab in Milton Keynes | Wellcome Sanger Institute for the COVID-19 Genomics UK (COG-UK) Consortium | The Lighthouse Lab in Milton Keynes and Alex Alderton, Roberto Amato, Jeffrey Barrett, Sonia Goncalves, Ewan Harrison, David K. Jackson, Ian Johnston, Dominic Kwiatkowski, Cordelia Langford, John Sillitoe on behalf of the Wellcome Sanger Institute COVID-19 Surveillance Team |
| EPI_ISL_3227355, EPI_ISL_3227356, EPI_ISL_3227357 | Lighthouse Laboratory Plymouth | Wellcome Sanger Institute for the COVID-19 Genomics UK (COG-UK) Consortium | Lighthouse Laboratory Plymouth and Alex Alderton, Roberto Amato, Jeffrey Barrett, Sonia Goncalves, Ewan Harrison, David K. Jackson, Ian Johnston, Dominic Kwiatkowski, Cordelia Langford, John Sillitoe on behalf of the Wellcome Sanger Institute COVID-19 Surveillance Team |
| EPI_ISL_3227358 | Lighthouse Lab in Milton Keynes | Wellcome Sanger Institute for the COVID-19 Genomics UK (COG-UK) Consortium | The Lighthouse Lab in Milton Keynes and Alex Alderton, Roberto Amato, Jeffrey Barrett, Sonia Goncalves, Ewan Harrison, David K. Jackson, Ian Johnston, Dominic Kwiatkowski, Cordelia Langford, John Sillitoe on behalf of the Wellcome Sanger Institute COVID-19 Surveillance Team |
| EPI_ISL_3227359 | Lighthouse Laboratory Plymouth | Wellcome Sanger Institute for the COVID-19 Genomics UK (COG-UK) Consortium | Lighthouse Laboratory Plymouth and Alex Alderton, Roberto Amato, Jeffrey Barrett, Sonia Goncalves, Ewan Harrison, David K. Jackson, Ian Johnston, Dominic Kwiatkowski, Cordelia Langford, John Sillitoe on behalf of the Wellcome Sanger Institute COVID-19 Surveillance Team |
| EPI_ISL_3227360, EPI_ISL_3227361, EPI_ISL_3227362 | Lighthouse Lab in Milton Keynes | Wellcome Sanger Institute for the COVID-19 Genomics UK (COG-UK) Consortium | The Lighthouse Lab in Milton Keynes and Alex Alderton, Roberto Amato, Jeffrey Barrett, Sonia Goncalves, Ewan Harrison, David K. Jackson, Ian Johnston, Dominic Kwiatkowski, Cordelia Langford, John Sillitoe on behalf of the Wellcome Sanger Institute COVID-19 Surveillance Team |
| EPI_ISL_3227363, EPI_ISL_3227364, EPI_ISL_3227365, EPI_ISL_3227366, EPI_ISL_3227367 | Lighthouse Laboratory Plymouth | Wellcome Sanger Institute for the COVID-19 Genomics UK (COG-UK) Consortium | Lighthouse Laboratory Plymouth and Alex Alderton, Roberto Amato, Jeffrey Barrett, Sonia Goncalves, Ewan Harrison, David K. Jackson, Ian Johnston, Dominic Kwiatkowski, Cordelia Langford, John Sillitoe on behalf of the Wellcome Sanger Institute COVID-19 Surveillance Team |
| EPI_ISL_3227368, EPI_ISL_3227369, EPI_ISL_3227370 | Lighthouse Lab in Milton Keynes | Wellcome Sanger Institute for the COVID-19 Genomics UK (COG-UK) Consortium | The Lighthouse Lab in Milton Keynes and Alex Alderton, Roberto Amato, Jeffrey Barrett, Sonia Goncalves, Ewan Harrison, David K. Jackson, Ian Johnston, Dominic Kwiatkowski, Cordelia Langford, John Sillitoe on behalf of the Wellcome Sanger Institute COVID-19 Surveillance Team |
| EPI_ISL_3227371, EPI_ISL_3227372, EPI_ISL_3227373, EPI_ISL_3227374 | Lighthouse Laboratory Plymouth | Wellcome Sanger Institute for the COVID-19 Genomics UK (COG-UK) Consortium | Lighthouse Laboratory Plymouth and Alex Alderton, Roberto Amato, Jeffrey Barrett, Sonia Goncalves, Ewan Harrison, David K. Jackson, Ian Johnston, Dominic Kwiatkowski, Cordelia Langford, John Sillitoe on behalf of the Wellcome Sanger Institute COVID-19 Surveillance Team |
| EPI_ISL_3227375, EPI_ISL_3227376, EPI_ISL_3227377 | Lighthouse Lab in Milton Keynes | Wellcome Sanger Institute for the COVID-19 Genomics UK (COG-UK) Consortium | The Lighthouse Lab in Milton Keynes and Alex Alderton, Roberto Amato, Jeffrey Barrett, Sonia Goncalves, Ewan Harrison, David K. Jackson, Ian Johnston, Dominic Kwiatkowski, Cordelia Langford, John Sillitoe on behalf of the Wellcome Sanger Institute COVID-19 Surveillance Team |
| EPI_ISL_3227378 | Lighthouse Laboratory Plymouth | Wellcome Sanger Institute for the COVID-19 Genomics UK (COG-UK) Consortium | Lighthouse Laboratory Plymouth and Alex Alderton, Roberto Amato, Jeffrey Barrett, Sonia Goncalves, Ewan Harrison, David K. Jackson, Ian Johnston, Dominic Kwiatkowski, Cordelia Langford, John Sillitoe on behalf of the Wellcome Sanger Institute COVID-19 Surveillance Team |
| EPI_ISL_3227379 | Lighthouse Lab in Milton Keynes | Wellcome Sanger Institute for the COVID-19 Genomics UK (COG-UK) Consortium | The Lighthouse Lab in Milton Keynes and Alex Alderton, Roberto Amato, Jeffrey Barrett, Sonia Goncalves, Ewan Harrison, David K. Jackson, Ian Johnston, Dominic Kwiatkowski, Cordelia Langford, John Sillitoe on behalf of the Wellcome Sanger Institute COVID-19 Surveillance Team |
| EPI_ISL_3227380, EPI_ISL_3227381, EPI_ISL_3227382 | Lighthouse Laboratory Plymouth | Wellcome Sanger Institute for the COVID-19 Genomics UK (COG-UK) Consortium | Lighthouse Laboratory Plymouth and Alex Alderton, Roberto Amato, Jeffrey Barrett, Sonia Goncalves, Ewan Harrison, David K. Jackson, Ian Johnston, Dominic Kwiatkowski, Cordelia Langford, John Sillitoe on behalf of the Wellcome Sanger Institute COVID-19 Surveillance Team |
| EPI_ISL_3227383 | Lighthouse Lab in Milton Keynes | Wellcome Sanger Institute for the COVID-19 Genomics UK (COG-UK) Consortium | The Lighthouse Lab in Milton Keynes and Alex Alderton, Roberto Amato, Jeffrey Barrett, Sonia Goncalves, Ewan Harrison, David K. Jackson, Ian Johnston, Dominic Kwiatkowski, Cordelia Langford, John Sillitoe on behalf of the Wellcome Sanger Institute COVID-19 Surveillance Team |
| EPI_ISL_3227384 | Lighthouse Laboratory Plymouth | Wellcome Sanger Institute for the COVID-19 Genomics UK (COG-UK) Consortium | Lighthouse Laboratory Plymouth and Alex Alderton, Roberto Amato, Jeffrey Barrett, Sonia Goncalves, Ewan Harrison, David K. Jackson, Ian Johnston, Dominic Kwiatkowski, Cordelia Langford, John Sillitoe on behalf of the Wellcome Sanger Institute COVID-19 Surveillance Team |
| EPI_ISL_3227385, EPI_ISL_3227386 | Lighthouse Lab in Milton Keynes | Wellcome Sanger Institute for the COVID-19 Genomics UK (COG-UK) Consortium | The Lighthouse Lab in Milton Keynes and Alex Alderton, Roberto Amato, Jeffrey Barrett, Sonia Goncalves, Ewan Harrison, David K. Jackson, Ian Johnston, Dominic Kwiatkowski, Cordelia Langford, John Sillitoe on behalf of the Wellcome Sanger Institute COVID-19 Surveillance Team |
| EPI_ISL_3227387 | Lighthouse Laboratory Plymouth | Wellcome Sanger Institute for the COVID-19 Genomics UK (COG-UK) Consortium | Lighthouse Laboratory Plymouth and Alex Alderton, Roberto Amato, Jeffrey Barrett, Sonia Goncalves, Ewan Harrison, David K. Jackson, Ian Johnston, Dominic Kwiatkowski, Cordelia Langford, John Sillitoe on behalf of the Wellcome Sanger Institute COVID-19 Surveillance Team |
| EPI_ISL_3227388, EPI_ISL_3227389, EPI_ISL_3227390 | Lighthouse Lab in Milton Keynes | Wellcome Sanger Institute for the COVID-19 Genomics UK (COG-UK) Consortium | The Lighthouse Lab in Milton Keynes and Alex Alderton, Roberto Amato, Jeffrey Barrett, Sonia Goncalves, Ewan Harrison, David K. Jackson, Ian Johnston, Dominic Kwiatkowski, Cordelia Langford, John Sillitoe on behalf of the Wellcome Sanger Institute COVID-19 Surveillance Team |
| EPI_ISL_3227391 | Lighthouse Laboratory Plymouth | Wellcome Sanger Institute for the COVID-19 Genomics UK (COG-UK) Consortium | Lighthouse Laboratory Plymouth and Alex Alderton, Roberto Amato, Jeffrey Barrett, Sonia Goncalves, Ewan Harrison, David K. Jackson, Ian Johnston, Dominic Kwiatkowski, Cordelia Langford, John Sillitoe on behalf of the Wellcome Sanger Institute COVID-19 Surveillance Team |
| EPI_ISL_3227392 | Lighthouse Lab in Milton Keynes | Wellcome Sanger Institute for the COVID-19 Genomics UK (COG-UK) Consortium | The Lighthouse Lab in Milton Keynes and Alex Alderton, Roberto Amato, Jeffrey Barrett, Sonia Goncalves, Ewan Harrison, David K. Jackson, Ian Johnston, Dominic Kwiatkowski, Cordelia Langford, John Sillitoe on behalf of the Wellcome Sanger Institute COVID-19 Surveillance Team |
| EPI_ISL_3227393, EPI_ISL_3227394, EPI_ISL_3227395 | Lighthouse Laboratory Plymouth | Wellcome Sanger Institute for the COVID-19 Genomics UK (COG-UK) Consortium | Lighthouse Laboratory Plymouth and Alex Alderton, Roberto Amato, Jeffrey Barrett, Sonia Goncalves, Ewan Harrison, David K. Jackson, Ian Johnston, Dominic Kwiatkowski, Cordelia Langford, John Sillitoe on behalf of the Wellcome Sanger Institute COVID-19 Surveillance Team |
| EPI_ISL_3227396 | Lighthouse Lab in Milton Keynes | Wellcome Sanger Institute for the COVID-19 Genomics UK (COG-UK) Consortium | The Lighthouse Lab in Milton Keynes and Alex Alderton, Roberto Amato, Jeffrey Barrett, Sonia Goncalves, Ewan Harrison, David K. Jackson, Ian Johnston, Dominic Kwiatkowski, Cordelia Langford, John Sillitoe on behalf of the Wellcome Sanger Institute COVID-19 Surveillance Team |
| EPI_ISL_3227397 | Lighthouse Laboratory Plymouth | Wellcome Sanger Institute for the COVID-19 Genomics UK (COG-UK) Consortium | Lighthouse Laboratory Plymouth and Alex Alderton, Roberto Amato, Jeffrey Barrett, Sonia Goncalves, Ewan Harrison, David K. Jackson, Ian Johnston, Dominic Kwiatkowski, Cordelia Langford, John Sillitoe on behalf of the Wellcome Sanger Institute COVID-19 Surveillance Team |
| EPI_ISL_3227398, EPI_ISL_3227399 | Lighthouse Lab in Milton Keynes | Wellcome Sanger Institute for the COVID-19 Genomics UK (COG-UK) Consortium | The Lighthouse Lab in Milton Keynes and Alex Alderton, Roberto Amato, Jeffrey Barrett, Sonia Goncalves, Ewan Harrison, David K. Jackson, Ian Johnston, Dominic Kwiatkowski, Cordelia Langford, John Sillitoe on behalf of the Wellcome Sanger Institute COVID-19 Surveillance Team |
| EPI_ISL_3227400, EPI_ISL_3227401, EPI_ISL_3227402 | Lighthouse Laboratory Plymouth | Wellcome Sanger Institute for the COVID-19 Genomics UK (COG-UK) Consortium | Lighthouse Laboratory Plymouth and Alex Alderton, Roberto Amato, Jeffrey Barrett, Sonia Goncalves, Ewan Harrison, David K. Jackson, Ian Johnston, Dominic Kwiatkowski, Cordelia Langford, John Sillitoe on behalf of the Wellcome Sanger Institute COVID-19 Surveillance Team |
| EPI_ISL_3227403, EPI_ISL_3227404, EPI_ISL_3227405 | Lighthouse Lab in Milton Keynes | Wellcome Sanger Institute for the COVID-19 Genomics UK (COG-UK) Consortium | The Lighthouse Lab in Milton Keynes and Alex Alderton, Roberto Amato, Jeffrey Barrett, Sonia Goncalves, Ewan Harrison, David K. Jackson, Ian Johnston, Dominic Kwiatkowski, Cordelia Langford, John Sillitoe on behalf of the Wellcome Sanger Institute COVID-19 Surveillance Team |
| EPI_ISL_3227406 | Lighthouse Laboratory Plymouth | Wellcome Sanger Institute for the COVID-19 Genomics UK (COG-UK) Consortium | Lighthouse Laboratory Plymouth and Alex Alderton, Roberto Amato, Jeffrey Barrett, Sonia Goncalves, Ewan Harrison, David K. Jackson, Ian Johnston, Dominic Kwiatkowski, Cordelia Langford, John Sillitoe on behalf of the Wellcome Sanger Institute COVID-19 Surveillance Team |
| EPI_ISL_3227407, EPI_ISL_3227408 | Lighthouse Lab in Milton Keynes | Wellcome Sanger Institute for the COVID-19 Genomics UK (COG-UK) Consortium | The Lighthouse Lab in Milton Keynes and Alex Alderton, Roberto Amato, Jeffrey Barrett, Sonia Goncalves, Ewan Harrison, David K. Jackson, Ian Johnston, Dominic Kwiatkowski, Cordelia Langford, John Sillitoe on behalf of the Wellcome Sanger Institute COVID-19 Surveillance Team |

| | | | |
|---|---|---|---|
| EPI_ISL_3227409 | Lighthouse Laboratory Plymouth | Wellcome Sanger Institute for the COVID-19 Genomics UK (COG-UK) Consortium | Lighthouse Laboratory Plymouth and Alex Alderton, Roberto Amato, Jeffrey Barrett, Sonia Goncalves, Ewan Harrison, David K. Jackson, Ian Johnston, Dominic Kwiatkowski, Cordelia Langford, John Sillitoe on behalf of the Wellcome Sanger Institute COVID-19 Surveillance Team |
| EPI_ISL_3227410 | Lighthouse Lab in Milton Keynes | Wellcome Sanger Institute for the COVID-19 Genomics UK (COG-UK) Consortium | The Lighthouse Lab in Milton Keynes and Alex Alderton, Roberto Amato, Jeffrey Barrett, Sonia Goncalves, Ewan Harrison, David K. Jackson, Ian Johnston, Dominic Kwiatkowski, Cordelia Langford, John Sillitoe on behalf of the Wellcome Sanger Institute COVID-19 Surveillance Team |
| EPI_ISL_3227411, EPI_ISL_3227412, EPI_ISL_3227413 | Lighthouse Laboratory Plymouth | Wellcome Sanger Institute for the COVID-19 Genomics UK (COG-UK) Consortium | Lighthouse Laboratory Plymouth and Alex Alderton, Roberto Amato, Jeffrey Barrett, Sonia Goncalves, Ewan Harrison, David K. Jackson, Ian Johnston, Dominic Kwiatkowski, Cordelia Langford, John Sillitoe on behalf of the Wellcome Sanger Institute COVID-19 Surveillance Team |
| EPI_ISL_3227414 | Lighthouse Lab in Milton Keynes | Wellcome Sanger Institute for the COVID-19 Genomics UK (COG-UK) Consortium | The Lighthouse Lab in Milton Keynes and Alex Alderton, Roberto Amato, Jeffrey Barrett, Sonia Goncalves, Ewan Harrison, David K. Jackson, Ian Johnston, Dominic Kwiatkowski, Cordelia Langford, John Sillitoe on behalf of the Wellcome Sanger Institute COVID-19 Surveillance Team |
| EPI_ISL_3227415 | Lighthouse Laboratory Plymouth | Wellcome Sanger Institute for the COVID-19 Genomics UK (COG-UK) Consortium | Lighthouse Laboratory Plymouth and Alex Alderton, Roberto Amato, Jeffrey Barrett, Sonia Goncalves, Ewan Harrison, David K. Jackson, Ian Johnston, Dominic Kwiatkowski, Cordelia Langford, John Sillitoe on behalf of the Wellcome Sanger Institute COVID-19 Surveillance Team |
| EPI_ISL_3227416, EPI_ISL_3227417 | Lighthouse Lab in Milton Keynes | Wellcome Sanger Institute for the COVID-19 Genomics UK (COG-UK) Consortium | The Lighthouse Lab in Milton Keynes and Alex Alderton, Roberto Amato, Jeffrey Barrett, Sonia Goncalves, Ewan Harrison, David K. Jackson, Ian Johnston, Dominic Kwiatkowski, Cordelia Langford, John Sillitoe on behalf of the Wellcome Sanger Institute COVID-19 Surveillance Team |
| EPI_ISL_3227418 | Lighthouse Laboratory Plymouth | Wellcome Sanger Institute for the COVID-19 Genomics UK (COG-UK) Consortium | Lighthouse Laboratory Plymouth and Alex Alderton, Roberto Amato, Jeffrey Barrett, Sonia Goncalves, Ewan Harrison, David K. Jackson, Ian Johnston, Dominic Kwiatkowski, Cordelia Langford, John Sillitoe on behalf of the Wellcome Sanger Institute COVID-19 Surveillance Team |
| EPI_ISL_3227419, EPI_ISL_3227420 | Lighthouse Lab in Milton Keynes | Wellcome Sanger Institute for the COVID-19 Genomics UK (COG-UK) Consortium | The Lighthouse Lab in Milton Keynes and Alex Alderton, Roberto Amato, Jeffrey Barrett, Sonia Goncalves, Ewan Harrison, David K. Jackson, Ian Johnston, Dominic Kwiatkowski, Cordelia Langford, John Sillitoe on behalf of the Wellcome Sanger Institute COVID-19 Surveillance Team |
| EPI_ISL_3227421, EPI_ISL_3227422 | Lighthouse Laboratory Plymouth | Wellcome Sanger Institute for the COVID-19 Genomics UK (COG-UK) Consortium | Lighthouse Laboratory Plymouth and Alex Alderton, Roberto Amato, Jeffrey Barrett, Sonia Goncalves, Ewan Harrison, David K. Jackson, Ian Johnston, Dominic Kwiatkowski, Cordelia Langford, John Sillitoe on behalf of the Wellcome Sanger Institute COVID-19 Surveillance Team |
| EPI_ISL_3227423 | Lighthouse Lab in Milton Keynes | Wellcome Sanger Institute for the COVID-19 Genomics UK (COG-UK) Consortium | The Lighthouse Lab in Milton Keynes and Alex Alderton, Roberto Amato, Jeffrey Barrett, Sonia Goncalves, Ewan Harrison, David K. Jackson, Ian Johnston, Dominic Kwiatkowski, Cordelia Langford, John Sillitoe on behalf of the Wellcome Sanger Institute COVID-19 Surveillance Team |
| EPI_ISL_3227424, EPI_ISL_3227425 | Lighthouse Laboratory Plymouth | Wellcome Sanger Institute for the COVID-19 Genomics UK (COG-UK) Consortium | Lighthouse Laboratory Plymouth and Alex Alderton, Roberto Amato, Jeffrey Barrett, Sonia Goncalves, Ewan Harrison, David K. Jackson, Ian Johnston, Dominic Kwiatkowski, Cordelia Langford, John Sillitoe on behalf of the Wellcome Sanger Institute COVID-19 Surveillance Team |
| EPI_ISL_3227426 | Lighthouse Lab in Milton Keynes | Wellcome Sanger Institute for the COVID-19 Genomics UK (COG-UK) Consortium | The Lighthouse Lab in Milton Keynes and Alex Alderton, Roberto Amato, Jeffrey Barrett, Sonia Goncalves, Ewan Harrison, David K. Jackson, Ian Johnston, Dominic Kwiatkowski, Cordelia Langford, John Sillitoe on behalf of the Wellcome Sanger Institute COVID-19 Surveillance Team |
| EPI_ISL_3227427 | Lighthouse Laboratory Plymouth | Wellcome Sanger Institute for the COVID-19 Genomics UK (COG-UK) Consortium | Lighthouse Laboratory Plymouth and Alex Alderton, Roberto Amato, Jeffrey Barrett, Sonia Goncalves, Ewan Harrison, David K. Jackson, Ian Johnston, Dominic Kwiatkowski, Cordelia Langford, John Sillitoe on behalf of the Wellcome Sanger Institute COVID-19 Surveillance Team |
| EPI_ISL_3227428 | Lighthouse Lab in Milton Keynes | Wellcome Sanger Institute for the COVID-19 Genomics UK (COG-UK) Consortium | The Lighthouse Lab in Milton Keynes and Alex Alderton, Roberto Amato, Jeffrey Barrett, Sonia Goncalves, Ewan Harrison, David K. Jackson, Ian Johnston, Dominic Kwiatkowski, Cordelia Langford, John Sillitoe on behalf of the Wellcome Sanger Institute COVID-19 Surveillance Team |
| EPI_ISL_3227429, EPI_ISL_3227430 | Lighthouse Laboratory Plymouth | Wellcome Sanger Institute for the COVID-19 Genomics UK (COG-UK) Consortium | Lighthouse Laboratory Plymouth and Alex Alderton, Roberto Amato, Jeffrey Barrett, Sonia Goncalves, Ewan Harrison, David K. Jackson, Ian Johnston, Dominic Kwiatkowski, Cordelia Langford, John Sillitoe on behalf of the Wellcome Sanger Institute COVID-19 Surveillance Team |
| EPI_ISL_3227431, EPI_ISL_3227432, EPI_ISL_3227433, EPI_ISL_3227434 | Lighthouse Lab in Milton Keynes | Wellcome Sanger Institute for the COVID-19 Genomics UK (COG-UK) Consortium | The Lighthouse Lab in Milton Keynes and Alex Alderton, Roberto Amato, Jeffrey Barrett, Sonia Goncalves, Ewan Harrison, David K. Jackson, Ian Johnston, Dominic Kwiatkowski, Cordelia Langford, John Sillitoe on behalf of the Wellcome Sanger Institute COVID-19 Surveillance Team |
| EPI_ISL_3227435, EPI_ISL_3227436, EPI_ISL_3227437 | Lighthouse Laboratory Plymouth | Wellcome Sanger Institute for the COVID-19 Genomics UK (COG-UK) Consortium | Lighthouse Laboratory Plymouth and Alex Alderton, Roberto Amato, Jeffrey Barrett, Sonia Goncalves, Ewan Harrison, David K. Jackson, Ian Johnston, Dominic Kwiatkowski, Cordelia Langford, John Sillitoe on behalf of the Wellcome Sanger Institute COVID-19 Surveillance Team |
| EPI_ISL_3227438 | Lighthouse Lab in Milton Keynes | Wellcome Sanger Institute for the COVID-19 Genomics UK (COG-UK) Consortium | The Lighthouse Lab in Milton Keynes and Alex Alderton, Roberto Amato, Jeffrey Barrett, Sonia Goncalves, Ewan Harrison, David K. Jackson, Ian Johnston, Dominic Kwiatkowski, Cordelia Langford, John Sillitoe on behalf of the Wellcome Sanger Institute COVID-19 Surveillance Team |
| EPI_ISL_3227439, EPI_ISL_3227440 | Lighthouse Laboratory Plymouth | Wellcome Sanger Institute for the COVID-19 Genomics UK (COG-UK) Consortium | Lighthouse Laboratory Plymouth and Alex Alderton, Roberto Amato, Jeffrey Barrett, Sonia Goncalves, Ewan Harrison, David K. Jackson, Ian Johnston, Dominic Kwiatkowski, Cordelia Langford, John Sillitoe on behalf of the Wellcome Sanger Institute COVID-19 Surveillance Team |
| EPI_ISL_3227441 | Lighthouse Lab in Milton Keynes | Wellcome Sanger Institute for the COVID-19 Genomics UK (COG-UK) Consortium | The Lighthouse Lab in Milton Keynes and Alex Alderton, Roberto Amato, Jeffrey Barrett, Sonia Goncalves, Ewan Harrison, David K. Jackson, Ian Johnston, Dominic Kwiatkowski, Cordelia Langford, John Sillitoe on behalf of the Wellcome Sanger Institute COVID-19 Surveillance Team |
| EPI_ISL_3227442, EPI_ISL_3227443 | Lighthouse Laboratory Plymouth | Wellcome Sanger Institute for the COVID-19 Genomics UK (COG-UK) Consortium | Lighthouse Laboratory Plymouth and Alex Alderton, Roberto Amato, Jeffrey Barrett, Sonia Goncalves, Ewan Harrison, David K. Jackson, Ian Johnston, Dominic Kwiatkowski, Cordelia Langford, John Sillitoe on behalf of the Wellcome Sanger Institute COVID-19 Surveillance Team |
| EPI_ISL_3227444 | Lighthouse Lab in Milton Keynes | Wellcome Sanger Institute for the COVID-19 Genomics UK (COG-UK) Consortium | The Lighthouse Lab in Milton Keynes and Alex Alderton, Roberto Amato, Jeffrey Barrett, Sonia Goncalves, Ewan Harrison, David K. Jackson, Ian Johnston, Dominic Kwiatkowski, Cordelia Langford, John Sillitoe on behalf of the Wellcome Sanger Institute COVID-19 Surveillance Team |
| EPI_ISL_3227445, EPI_ISL_3227446, EPI_ISL_3227447, EPI_ISL_3227448 | Lighthouse Laboratory Plymouth | Wellcome Sanger Institute for the COVID-19 Genomics UK (COG-UK) Consortium | Lighthouse Laboratory Plymouth and Alex Alderton, Roberto Amato, Jeffrey Barrett, Sonia Goncalves, Ewan Harrison, David K. Jackson, Ian Johnston, Dominic Kwiatkowski, Cordelia Langford, John Sillitoe on behalf of the Wellcome Sanger Institute COVID-19 Surveillance Team |
| EPI_ISL_3227449 | Lighthouse Lab in Milton Keynes | Wellcome Sanger Institute for the COVID-19 Genomics UK (COG-UK) Consortium | The Lighthouse Lab in Milton Keynes and Alex Alderton, Roberto Amato, Jeffrey Barrett, Sonia Goncalves, Ewan Harrison, David K. Jackson, Ian Johnston, Dominic Kwiatkowski, Cordelia Langford, John Sillitoe on behalf of the Wellcome Sanger Institute COVID-19 Surveillance Team |
| EPI_ISL_3227450, EPI_ISL_3227451, EPI_ISL_3227452 | Lighthouse Laboratory Plymouth | Wellcome Sanger Institute for the COVID-19 Genomics UK (COG-UK) Consortium | Lighthouse Laboratory Plymouth and Alex Alderton, Roberto Amato, Jeffrey Barrett, Sonia Goncalves, Ewan Harrison, David K. Jackson, Ian Johnston, Dominic Kwiatkowski, Cordelia Langford, John Sillitoe on behalf of the Wellcome Sanger Institute COVID-19 Surveillance Team |
| EPI_ISL_3227453 | Lighthouse Lab in Milton Keynes | Wellcome Sanger Institute for the COVID-19 Genomics UK (COG-UK) Consortium | The Lighthouse Lab in Milton Keynes and Alex Alderton, Roberto Amato, Jeffrey Barrett, Sonia Goncalves, Ewan Harrison, David K. Jackson, Ian Johnston, Dominic Kwiatkowski, Cordelia Langford, John Sillitoe on behalf of the Wellcome Sanger Institute COVID-19 Surveillance Team |
| EPI_ISL_3227454 | Lighthouse Laboratory Plymouth | Wellcome Sanger Institute for the COVID-19 Genomics UK (COG-UK) Consortium | Lighthouse Laboratory Plymouth and Alex Alderton, Roberto Amato, Jeffrey Barrett, Sonia Goncalves, Ewan Harrison, David K. Jackson, Ian Johnston, Dominic Kwiatkowski, Cordelia Langford, John Sillitoe on behalf of the Wellcome Sanger Institute COVID-19 Surveillance Team |
| EPI_ISL_3227455 | Lighthouse Lab in Milton Keynes | Wellcome Sanger Institute for the COVID-19 Genomics UK (COG-UK) Consortium | The Lighthouse Lab in Milton Keynes and Alex Alderton, Roberto Amato, Jeffrey Barrett, Sonia Goncalves, Ewan Harrison, David K. Jackson, Ian Johnston, Dominic Kwiatkowski, Cordelia Langford, John Sillitoe on behalf of the Wellcome Sanger Institute COVID-19 Surveillance Team |
| EPI_ISL_3227456 | Lighthouse Laboratory Plymouth | Wellcome Sanger Institute for the COVID-19 Genomics UK (COG-UK) Consortium | Lighthouse Laboratory Plymouth and Alex Alderton, Roberto Amato, Jeffrey Barrett, Sonia Goncalves, Ewan Harrison, David K. Jackson, Ian Johnston, Dominic Kwiatkowski, Cordelia Langford, John Sillitoe on behalf of the Wellcome Sanger Institute COVID-19 Surveillance Team |
| EPI_ISL_3227457 | Lighthouse Lab in Milton Keynes | Wellcome Sanger Institute for the COVID-19 Genomics UK (COG-UK) Consortium | The Lighthouse Lab in Milton Keynes and Alex Alderton, Roberto Amato, Jeffrey Barrett, Sonia Goncalves, Ewan Harrison, David K. Jackson, Ian Johnston, Dominic Kwiatkowski, Cordelia Langford, John Sillitoe on behalf of the Wellcome Sanger Institute COVID-19 Surveillance Team |
| EPI_ISL_3227458, EPI_ISL_3227459, EPI_ISL_3227460 | Lighthouse Laboratory Plymouth | Wellcome Sanger Institute for the COVID-19 Genomics UK (COG-UK) Consortium | Lighthouse Laboratory Plymouth and Alex Alderton, Roberto Amato, Jeffrey Barrett, Sonia Goncalves, Ewan Harrison, David K. Jackson, Ian Johnston, Dominic Kwiatkowski, Cordelia Langford, John Sillitoe on behalf of the Wellcome Sanger Institute COVID-19 Surveillance Team |
| EPI_ISL_3227461, EPI_ISL_3227462, EPI_ISL_3227463, EPI_ISL_3227464, EPI_ISL_3227465 | Lighthouse Lab in Milton Keynes | Wellcome Sanger Institute for the COVID-19 Genomics UK (COG-UK) Consortium | The Lighthouse Lab in Milton Keynes and Alex Alderton, Roberto Amato, Jeffrey Barrett, Sonia Goncalves, Ewan Harrison, David K. Jackson, Ian Johnston, Dominic Kwiatkowski, Cordelia Langford, John Sillitoe on behalf of the Wellcome Sanger Institute COVID-19 Surveillance Team |
| EPI_ISL_3227466 | Lighthouse Laboratory Plymouth | Wellcome Sanger Institute for the COVID-19 Genomics UK (COG-UK) Consortium | Lighthouse Laboratory Plymouth and Alex Alderton, Roberto Amato, Jeffrey Barrett, Sonia Goncalves, Ewan Harrison, David K. Jackson, Ian Johnston, Dominic Kwiatkowski, Cordelia Langford, John Sillitoe on behalf of the Wellcome Sanger Institute COVID-19 Surveillance Team |
| EPI_ISL_3227467 | Lighthouse Lab in Milton Keynes | Wellcome Sanger Institute for the COVID-19 Genomics UK (COG-UK) Consortium | The Lighthouse Lab in Milton Keynes and Alex Alderton, Roberto Amato, Jeffrey Barrett, Sonia Goncalves, Ewan Harrison, David K. Jackson, Ian Johnston, Dominic Kwiatkowski, Cordelia Langford, John Sillitoe on behalf of the Wellcome Sanger Institute COVID-19 Surveillance Team |
| EPI_ISL_3227468, EPI_ISL_3227469 | Lighthouse Laboratory Plymouth | Wellcome Sanger Institute for the COVID-19 Genomics UK (COG-UK) Consortium | Lighthouse Laboratory Plymouth and Alex Alderton, Roberto Amato, Jeffrey Barrett, Sonia Goncalves, Ewan Harrison, David K. Jackson, Ian Johnston, Dominic Kwiatkowski, Cordelia Langford, John Sillitoe on behalf of the Wellcome Sanger Institute COVID-19 Surveillance Team |
| EPI_ISL_3227470, EPI_ISL_3227471, EPI_ISL_3227472 | Lighthouse Lab in Milton Keynes | Wellcome Sanger Institute for the COVID-19 Genomics UK (COG-UK) Consortium | The Lighthouse Lab in Milton Keynes and Alex Alderton, Roberto Amato, Jeffrey Barrett, Sonia Goncalves, Ewan Harrison, David K. Jackson, Ian Johnston, Dominic Kwiatkowski, Cordelia Langford, John Sillitoe on behalf of the Wellcome Sanger Institute COVID-19 Surveillance Team |
| EPI_ISL_3227473, EPI_ISL_3227474, EPI_ISL_3227475 | Lighthouse Laboratory Plymouth | Wellcome Sanger Institute for the COVID-19 Genomics UK (COG-UK) Consortium | Lighthouse Laboratory Plymouth and Alex Alderton, Roberto Amato, Jeffrey Barrett, Sonia Goncalves, Ewan Harrison, David K. Jackson, Ian Johnston, Dominic Kwiatkowski, Cordelia Langford, John Sillitoe on behalf of the Wellcome Sanger Institute COVID-19 Surveillance Team |
| EPI_ISL_3227476, EPI_ISL_3227477, EPI_ISL_3227478 | Lighthouse Lab in Milton Keynes | Wellcome Sanger Institute for the COVID-19 Genomics UK (COG-UK) Consortium | The Lighthouse Lab in Milton Keynes and Alex Alderton, Roberto Amato, Jeffrey Barrett, Sonia Goncalves, Ewan Harrison, David K. Jackson, Ian Johnston, Dominic Kwiatkowski, Cordelia Langford, John Sillitoe on behalf of the Wellcome Sanger Institute COVID-19 Surveillance Team |
| EPI_ISL_3227479 | Lighthouse Laboratory Plymouth | Wellcome Sanger Institute for the COVID-19 Genomics UK (COG-UK) Consortium | Lighthouse Laboratory Plymouth and Alex Alderton, Roberto Amato, Jeffrey Barrett, Sonia Goncalves, Ewan Harrison, David K. Jackson, Ian Johnston, Dominic Kwiatkowski, Cordelia Langford, John Sillitoe on behalf of the Wellcome Sanger Institute COVID-19 Surveillance Team |
| EPI_ISL_3227480, | Lighthouse Lab in Milton Keynes | Wellcome Sanger Institute for the COVID-19 Genomics UK (COG-UK) | The Lighthouse Lab in Milton Keynes and Alex Alderton, Roberto Amato, Jeffrey Barrett, Sonia Goncalves, Ewan Harrison, David K. Jackson, Ian Johnston, Dominic Kwiatkowski, Cordelia Langford, John Sillitoe on behalf of the Wellcome Sanger |

| | | | |
|---|---|---|---|
| EPI_ISL_3227481 | | Consortium | Institute COVID-19 Surveillance Team |
| EPI_ISL_3227482 | Lighthouse Laboratory Plymouth | Wellcome Sanger Institute for the COVID-19 Genomics UK (COG-UK) Consortium | Lighthouse Laboratory Plymouth and Alex Alderton, Roberto Amato, Jeffrey Barrett, Sonia Goncalves, Ewan Harrison, David K. Jackson, Ian Johnston, Dominic Kwiatkowski, Cordelia Langford, John Sillitoe on behalf of the Wellcome Sanger Institute COVID-19 Surveillance Team |
| EPI_ISL_3227483 | Lighthouse Lab in Milton Keynes | Wellcome Sanger Institute for the COVID-19 Genomics UK (COG-UK) Consortium | The Lighthouse Lab in Milton Keynes and Alex Alderton, Roberto Amato, Jeffrey Barrett, Sonia Goncalves, Ewan Harrison, David K. Jackson, Ian Johnston, Dominic Kwiatkowski, Cordelia Langford, John Sillitoe on behalf of the Wellcome Sanger Institute COVID-19 Surveillance Team |
| EPI_ISL_3227484, EPI_ISL_3227485, EPI_ISL_3227486 | Lighthouse Laboratory Plymouth | Wellcome Sanger Institute for the COVID-19 Genomics UK (COG-UK) Consortium | Lighthouse Laboratory Plymouth and Alex Alderton, Roberto Amato, Jeffrey Barrett, Sonia Goncalves, Ewan Harrison, David K. Jackson, Ian Johnston, Dominic Kwiatkowski, Cordelia Langford, John Sillitoe on behalf of the Wellcome Sanger Institute COVID-19 Surveillance Team |
| EPI_ISL_3227487, EPI_ISL_3227488 | Lighthouse Lab in Milton Keynes | Wellcome Sanger Institute for the COVID-19 Genomics UK (COG-UK) Consortium | The Lighthouse Lab in Milton Keynes and Alex Alderton, Roberto Amato, Jeffrey Barrett, Sonia Goncalves, Ewan Harrison, David K. Jackson, Ian Johnston, Dominic Kwiatkowski, Cordelia Langford, John Sillitoe on behalf of the Wellcome Sanger Institute COVID-19 Surveillance Team |
| EPI_ISL_3227489, EPI_ISL_3227490, EPI_ISL_3227491 | Lighthouse Laboratory Plymouth | Wellcome Sanger Institute for the COVID-19 Genomics UK (COG-UK) Consortium | Lighthouse Laboratory Plymouth and Alex Alderton, Roberto Amato, Jeffrey Barrett, Sonia Goncalves, Ewan Harrison, David K. Jackson, Ian Johnston, Dominic Kwiatkowski, Cordelia Langford, John Sillitoe on behalf of the Wellcome Sanger Institute COVID-19 Surveillance Team |
| EPI_ISL_3227492 | Lighthouse Lab in Milton Keynes | Wellcome Sanger Institute for the COVID-19 Genomics UK (COG-UK) Consortium | The Lighthouse Lab in Milton Keynes and Alex Alderton, Roberto Amato, Jeffrey Barrett, Sonia Goncalves, Ewan Harrison, David K. Jackson, Ian Johnston, Dominic Kwiatkowski, Cordelia Langford, John Sillitoe on behalf of the Wellcome Sanger Institute COVID-19 Surveillance Team |
| EPI_ISL_3227493 | Lighthouse Laboratory Plymouth | Wellcome Sanger Institute for the COVID-19 Genomics UK (COG-UK) Consortium | Lighthouse Laboratory Plymouth and Alex Alderton, Roberto Amato, Jeffrey Barrett, Sonia Goncalves, Ewan Harrison, David K. Jackson, Ian Johnston, Dominic Kwiatkowski, Cordelia Langford, John Sillitoe on behalf of the Wellcome Sanger Institute COVID-19 Surveillance Team |
| EPI_ISL_3227494, EPI_ISL_3227495 | Lighthouse Lab in Milton Keynes | Wellcome Sanger Institute for the COVID-19 Genomics UK (COG-UK) Consortium | The Lighthouse Lab in Milton Keynes and Alex Alderton, Roberto Amato, Jeffrey Barrett, Sonia Goncalves, Ewan Harrison, David K. Jackson, Ian Johnston, Dominic Kwiatkowski, Cordelia Langford, John Sillitoe on behalf of the Wellcome Sanger Institute COVID-19 Surveillance Team |
| EPI_ISL_3227496 | Lighthouse Laboratory Plymouth | Wellcome Sanger Institute for the COVID-19 Genomics UK (COG-UK) Consortium | Lighthouse Laboratory Plymouth and Alex Alderton, Roberto Amato, Jeffrey Barrett, Sonia Goncalves, Ewan Harrison, David K. Jackson, Ian Johnston, Dominic Kwiatkowski, Cordelia Langford, John Sillitoe on behalf of the Wellcome Sanger Institute COVID-19 Surveillance Team |
| EPI_ISL_3227497 | Lighthouse Lab in Milton Keynes | Wellcome Sanger Institute for the COVID-19 Genomics UK (COG-UK) Consortium | The Lighthouse Lab in Milton Keynes and Alex Alderton, Roberto Amato, Jeffrey Barrett, Sonia Goncalves, Ewan Harrison, David K. Jackson, Ian Johnston, Dominic Kwiatkowski, Cordelia Langford, John Sillitoe on behalf of the Wellcome Sanger Institute COVID-19 Surveillance Team |
| EPI_ISL_3227498 | Lighthouse Laboratory Plymouth | Wellcome Sanger Institute for the COVID-19 Genomics UK (COG-UK) Consortium | Lighthouse Laboratory Plymouth and Alex Alderton, Roberto Amato, Jeffrey Barrett, Sonia Goncalves, Ewan Harrison, David K. Jackson, Ian Johnston, Dominic Kwiatkowski, Cordelia Langford, John Sillitoe on behalf of the Wellcome Sanger Institute COVID-19 Surveillance Team |
| EPI_ISL_3227499, EPI_ISL_3227500, EPI_ISL_3227501, EPI_ISL_3227502, EPI_ISL_3227503 | Lighthouse Lab in Milton Keynes | Wellcome Sanger Institute for the COVID-19 Genomics UK (COG-UK) Consortium | The Lighthouse Lab in Milton Keynes and Alex Alderton, Roberto Amato, Jeffrey Barrett, Sonia Goncalves, Ewan Harrison, David K. Jackson, Ian Johnston, Dominic Kwiatkowski, Cordelia Langford, John Sillitoe on behalf of the Wellcome Sanger Institute COVID-19 Surveillance Team |
| EPI_ISL_3227504, EPI_ISL_3227505 | Lighthouse Laboratory Plymouth | Wellcome Sanger Institute for the COVID-19 Genomics UK (COG-UK) Consortium | Lighthouse Laboratory Plymouth and Alex Alderton, Roberto Amato, Jeffrey Barrett, Sonia Goncalves, Ewan Harrison, David K. Jackson, Ian Johnston, Dominic Kwiatkowski, Cordelia Langford, John Sillitoe on behalf of the Wellcome Sanger Institute COVID-19 Surveillance Team |
| EPI_ISL_3227506, EPI_ISL_3227507 | Lighthouse Lab in Milton Keynes | Wellcome Sanger Institute for the COVID-19 Genomics UK (COG-UK) Consortium | The Lighthouse Lab in Milton Keynes and Alex Alderton, Roberto Amato, Jeffrey Barrett, Sonia Goncalves, Ewan Harrison, David K. Jackson, Ian Johnston, Dominic Kwiatkowski, Cordelia Langford, John Sillitoe on behalf of the Wellcome Sanger Institute COVID-19 Surveillance Team |
| EPI_ISL_3227508 | Lighthouse Laboratory Plymouth | Wellcome Sanger Institute for the COVID-19 Genomics UK (COG-UK) Consortium | Lighthouse Laboratory Plymouth and Alex Alderton, Roberto Amato, Jeffrey Barrett, Sonia Goncalves, Ewan Harrison, David K. Jackson, Ian Johnston, Dominic Kwiatkowski, Cordelia Langford, John Sillitoe on behalf of the Wellcome Sanger Institute COVID-19 Surveillance Team |
| EPI_ISL_3227509, EPI_ISL_3227510 | Lighthouse Lab in Milton Keynes | Wellcome Sanger Institute for the COVID-19 Genomics UK (COG-UK) Consortium | The Lighthouse Lab in Milton Keynes and Alex Alderton, Roberto Amato, Jeffrey Barrett, Sonia Goncalves, Ewan Harrison, David K. Jackson, Ian Johnston, Dominic Kwiatkowski, Cordelia Langford, John Sillitoe on behalf of the Wellcome Sanger Institute COVID-19 Surveillance Team |
| EPI_ISL_3227511, EPI_ISL_3227512 | Lighthouse Laboratory Plymouth | Wellcome Sanger Institute for the COVID-19 Genomics UK (COG-UK) Consortium | Lighthouse Laboratory Plymouth and Alex Alderton, Roberto Amato, Jeffrey Barrett, Sonia Goncalves, Ewan Harrison, David K. Jackson, Ian Johnston, Dominic Kwiatkowski, Cordelia Langford, John Sillitoe on behalf of the Wellcome Sanger Institute COVID-19 Surveillance Team |
| EPI_ISL_3227513, EPI_ISL_3227514 | Lighthouse Lab in Milton Keynes | Wellcome Sanger Institute for the COVID-19 Genomics UK (COG-UK) Consortium | The Lighthouse Lab in Milton Keynes and Alex Alderton, Roberto Amato, Jeffrey Barrett, Sonia Goncalves, Ewan Harrison, David K. Jackson, Ian Johnston, Dominic Kwiatkowski, Cordelia Langford, John Sillitoe on behalf of the Wellcome Sanger Institute COVID-19 Surveillance Team |
| EPI_ISL_3227515 | Lighthouse Laboratory Plymouth | Wellcome Sanger Institute for the COVID-19 Genomics UK (COG-UK) Consortium | Lighthouse Laboratory Plymouth and Alex Alderton, Roberto Amato, Jeffrey Barrett, Sonia Goncalves, Ewan Harrison, David K. Jackson, Ian Johnston, Dominic Kwiatkowski, Cordelia Langford, John Sillitoe on behalf of the Wellcome Sanger Institute COVID-19 Surveillance Team |
| EPI_ISL_3227516, EPI_ISL_3227517, EPI_ISL_3227518 | Lighthouse Lab in Milton Keynes | Wellcome Sanger Institute for the COVID-19 Genomics UK (COG-UK) Consortium | The Lighthouse Lab in Milton Keynes and Alex Alderton, Roberto Amato, Jeffrey Barrett, Sonia Goncalves, Ewan Harrison, David K. Jackson, Ian Johnston, Dominic Kwiatkowski, Cordelia Langford, John Sillitoe on behalf of the Wellcome Sanger Institute COVID-19 Surveillance Team |
| EPI_ISL_3227519, EPI_ISL_3227520 | Lighthouse Laboratory Plymouth | Wellcome Sanger Institute for the COVID-19 Genomics UK (COG-UK) Consortium | Lighthouse Laboratory Plymouth and Alex Alderton, Roberto Amato, Jeffrey Barrett, Sonia Goncalves, Ewan Harrison, David K. Jackson, Ian Johnston, Dominic Kwiatkowski, Cordelia Langford, John Sillitoe on behalf of the Wellcome Sanger Institute COVID-19 Surveillance Team |
| EPI_ISL_3227521, EPI_ISL_3227522, EPI_ISL_3227523, EPI_ISL_3227524 | Lighthouse Lab in Milton Keynes | Wellcome Sanger Institute for the COVID-19 Genomics UK (COG-UK) Consortium | The Lighthouse Lab in Milton Keynes and Alex Alderton, Roberto Amato, Jeffrey Barrett, Sonia Goncalves, Ewan Harrison, David K. Jackson, Ian Johnston, Dominic Kwiatkowski, Cordelia Langford, John Sillitoe on behalf of the Wellcome Sanger Institute COVID-19 Surveillance Team |
| EPI_ISL_3227525 | Lighthouse Laboratory Plymouth | Wellcome Sanger Institute for the COVID-19 Genomics UK (COG-UK) Consortium | Lighthouse Laboratory Plymouth and Alex Alderton, Roberto Amato, Jeffrey Barrett, Sonia Goncalves, Ewan Harrison, David K. Jackson, Ian Johnston, Dominic Kwiatkowski, Cordelia Langford, John Sillitoe on behalf of the Wellcome Sanger Institute COVID-19 Surveillance Team |
| EPI_ISL_3227526 | Lighthouse Lab in Milton Keynes | Wellcome Sanger Institute for the COVID-19 Genomics UK (COG-UK) Consortium | The Lighthouse Lab in Milton Keynes and Alex Alderton, Roberto Amato, Jeffrey Barrett, Sonia Goncalves, Ewan Harrison, David K. Jackson, Ian Johnston, Dominic Kwiatkowski, Cordelia Langford, John Sillitoe on behalf of the Wellcome Sanger Institute COVID-19 Surveillance Team |
| EPI_ISL_3227527 | Lighthouse Laboratory Plymouth | Wellcome Sanger Institute for the COVID-19 Genomics UK (COG-UK) Consortium | Lighthouse Laboratory Plymouth and Alex Alderton, Roberto Amato, Jeffrey Barrett, Sonia Goncalves, Ewan Harrison, David K. Jackson, Ian Johnston, Dominic Kwiatkowski, Cordelia Langford, John Sillitoe on behalf of the Wellcome Sanger Institute COVID-19 Surveillance Team |
| EPI_ISL_3227528, EPI_ISL_3227529, EPI_ISL_3227530, EPI_ISL_3227531 | Lighthouse Lab in Milton Keynes | Wellcome Sanger Institute for the COVID-19 Genomics UK (COG-UK) Consortium | The Lighthouse Lab in Milton Keynes and Alex Alderton, Roberto Amato, Jeffrey Barrett, Sonia Goncalves, Ewan Harrison, David K. Jackson, Ian Johnston, Dominic Kwiatkowski, Cordelia Langford, John Sillitoe on behalf of the Wellcome Sanger Institute COVID-19 Surveillance Team |
| EPI_ISL_3227532 | Lighthouse Laboratory Plymouth | Wellcome Sanger Institute for the COVID-19 Genomics UK (COG-UK) Consortium | Lighthouse Laboratory Plymouth and Alex Alderton, Roberto Amato, Jeffrey Barrett, Sonia Goncalves, Ewan Harrison, David K. Jackson, Ian Johnston, Dominic Kwiatkowski, Cordelia Langford, John Sillitoe on behalf of the Wellcome Sanger Institute COVID-19 Surveillance Team |
| EPI_ISL_3227533, EPI_ISL_3227534, EPI_ISL_3227535 | Lighthouse Lab in Milton Keynes | Wellcome Sanger Institute for the COVID-19 Genomics UK (COG-UK) Consortium | The Lighthouse Lab in Milton Keynes and Alex Alderton, Roberto Amato, Jeffrey Barrett, Sonia Goncalves, Ewan Harrison, David K. Jackson, Ian Johnston, Dominic Kwiatkowski, Cordelia Langford, John Sillitoe on behalf of the Wellcome Sanger Institute COVID-19 Surveillance Team |
| EPI_ISL_3227536 | Lighthouse Laboratory Plymouth | Wellcome Sanger Institute for the COVID-19 Genomics UK (COG-UK) Consortium | Lighthouse Laboratory Plymouth and Alex Alderton, Roberto Amato, Jeffrey Barrett, Sonia Goncalves, Ewan Harrison, David K. Jackson, Ian Johnston, Dominic Kwiatkowski, Cordelia Langford, John Sillitoe on behalf of the Wellcome Sanger Institute COVID-19 Surveillance Team |
| EPI_ISL_3227537, EPI_ISL_3227538 | Lighthouse Lab in Milton Keynes | Wellcome Sanger Institute for the COVID-19 Genomics UK (COG-UK) Consortium | The Lighthouse Lab in Milton Keynes and Alex Alderton, Roberto Amato, Jeffrey Barrett, Sonia Goncalves, Ewan Harrison, David K. Jackson, Ian Johnston, Dominic Kwiatkowski, Cordelia Langford, John Sillitoe on behalf of the Wellcome Sanger Institute COVID-19 Surveillance Team |
| EPI_ISL_3227539, EPI_ISL_3227540 | Lighthouse Laboratory Plymouth | Wellcome Sanger Institute for the COVID-19 Genomics UK (COG-UK) Consortium | Lighthouse Laboratory Plymouth and Alex Alderton, Roberto Amato, Jeffrey Barrett, Sonia Goncalves, Ewan Harrison, David K. Jackson, Ian Johnston, Dominic Kwiatkowski, Cordelia Langford, John Sillitoe on behalf of the Wellcome Sanger Institute COVID-19 Surveillance Team |
| EPI_ISL_3227541, EPI_ISL_3227542 | Lighthouse Lab in Milton Keynes | Wellcome Sanger Institute for the COVID-19 Genomics UK (COG-UK) Consortium | The Lighthouse Lab in Milton Keynes and Alex Alderton, Roberto Amato, Jeffrey Barrett, Sonia Goncalves, Ewan Harrison, David K. Jackson, Ian Johnston, Dominic Kwiatkowski, Cordelia Langford, John Sillitoe on behalf of the Wellcome Sanger Institute COVID-19 Surveillance Team |
| EPI_ISL_3227543, EPI_ISL_3227544, EPI_ISL_3227545, EPI_ISL_3227546 | Lighthouse Laboratory Plymouth | Wellcome Sanger Institute for the COVID-19 Genomics UK (COG-UK) Consortium | Lighthouse Laboratory Plymouth and Alex Alderton, Roberto Amato, Jeffrey Barrett, Sonia Goncalves, Ewan Harrison, David K. Jackson, Ian Johnston, Dominic Kwiatkowski, Cordelia Langford, John Sillitoe on behalf of the Wellcome Sanger Institute COVID-19 Surveillance Team |
| EPI_ISL_3227547, EPI_ISL_3227548, EPI_ISL_3227549, EPI_ISL_3227550 | Lighthouse Lab in Milton Keynes | Wellcome Sanger Institute for the COVID-19 Genomics UK (COG-UK) Consortium | The Lighthouse Lab in Milton Keynes and Alex Alderton, Roberto Amato, Jeffrey Barrett, Sonia Goncalves, Ewan Harrison, David K. Jackson, Ian Johnston, Dominic Kwiatkowski, Cordelia Langford, John Sillitoe on behalf of the Wellcome Sanger Institute COVID-19 Surveillance Team |
| EPI_ISL_3227551, EPI_ISL_3227552 | Lighthouse Laboratory Plymouth | Wellcome Sanger Institute for the COVID-19 Genomics UK (COG-UK) Consortium | Lighthouse Laboratory Plymouth and Alex Alderton, Roberto Amato, Jeffrey Barrett, Sonia Goncalves, Ewan Harrison, David K. Jackson, Ian Johnston, Dominic Kwiatkowski, Cordelia Langford, John Sillitoe on behalf of the Wellcome Sanger Institute COVID-19 Surveillance Team |
| EPI_ISL_3227553 | Lighthouse Lab in Milton Keynes | Wellcome Sanger Institute for the COVID-19 Genomics UK (COG-UK) Consortium | The Lighthouse Lab in Milton Keynes and Alex Alderton, Roberto Amato, Jeffrey Barrett, Sonia Goncalves, Ewan Harrison, David K. Jackson, Ian Johnston, Dominic Kwiatkowski, Cordelia Langford, John Sillitoe on behalf of the Wellcome Sanger Institute COVID-19 Surveillance Team |
| EPI_ISL_3227554 | Lighthouse Laboratory Plymouth | Wellcome Sanger Institute for the COVID-19 Genomics UK (COG-UK) Consortium | Lighthouse Laboratory Plymouth and Alex Alderton, Roberto Amato, Jeffrey Barrett, Sonia Goncalves, Ewan Harrison, David K. Jackson, Ian Johnston, Dominic Kwiatkowski, Cordelia Langford, John Sillitoe on behalf of the Wellcome Sanger Institute COVID-19 Surveillance Team |
| EPI_ISL_3227555 | Lighthouse Lab in Milton Keynes | Wellcome Sanger Institute for the COVID-19 Genomics UK (COG-UK) Consortium | The Lighthouse Lab in Milton Keynes and Alex Alderton, Roberto Amato, Jeffrey Barrett, Sonia Goncalves, Ewan Harrison, David K. Jackson, Ian Johnston, Dominic Kwiatkowski, Cordelia Langford, John Sillitoe on behalf of the Wellcome Sanger Institute COVID-19 Surveillance Team |

| | | | |
|---|---|---|---|
| EPI_ISL_3227556, EPI_ISL_3227557, EPI_ISL_3227558, EPI_ISL_3227559 | Lighthouse Laboratory Plymouth | Wellcome Sanger Institute for the COVID-19 Genomics UK (COG-UK) Consortium | Lighthouse Laboratory Plymouth and Alex Alderton, Roberto Amato, Jeffrey Barrett, Sonia Goncalves, Ewan Harrison, David K. Jackson, Ian Johnston, Dominic Kwiatkowski, Cordelia Langford, John Sillitoe on behalf of the Wellcome Sanger Institute COVID-19 Surveillance Team |
| EPI_ISL_3227560, EPI_ISL_3227561 | Lighthouse Lab in Milton Keynes | Wellcome Sanger Institute for the COVID-19 Genomics UK (COG-UK) Consortium | The Lighthouse Lab in Milton Keynes and Alex Alderton, Roberto Amato, Jeffrey Barrett, Sonia Goncalves, Ewan Harrison, David K. Jackson, Ian Johnston, Dominic Kwiatkowski, Cordelia Langford, John Sillitoe on behalf of the Wellcome Sanger Institute COVID-19 Surveillance Team |
| EPI_ISL_3227562 | Lighthouse Laboratory Plymouth | Wellcome Sanger Institute for the COVID-19 Genomics UK (COG-UK) Consortium | Lighthouse Laboratory Plymouth and Alex Alderton, Roberto Amato, Jeffrey Barrett, Sonia Goncalves, Ewan Harrison, David K. Jackson, Ian Johnston, Dominic Kwiatkowski, Cordelia Langford, John Sillitoe on behalf of the Wellcome Sanger Institute COVID-19 Surveillance Team |
| EPI_ISL_3227563 | Lighthouse Lab in Milton Keynes | Wellcome Sanger Institute for the COVID-19 Genomics UK (COG-UK) Consortium | The Lighthouse Lab in Milton Keynes and Alex Alderton, Roberto Amato, Jeffrey Barrett, Sonia Goncalves, Ewan Harrison, David K. Jackson, Ian Johnston, Dominic Kwiatkowski, Cordelia Langford, John Sillitoe on behalf of the Wellcome Sanger Institute COVID-19 Surveillance Team |
| EPI_ISL_3227564, EPI_ISL_3227565, EPI_ISL_3227566 | Lighthouse Laboratory Plymouth | Wellcome Sanger Institute for the COVID-19 Genomics UK (COG-UK) Consortium | Lighthouse Laboratory Plymouth and Alex Alderton, Roberto Amato, Jeffrey Barrett, Sonia Goncalves, Ewan Harrison, David K. Jackson, Ian Johnston, Dominic Kwiatkowski, Cordelia Langford, John Sillitoe on behalf of the Wellcome Sanger Institute COVID-19 Surveillance Team |
| EPI_ISL_3227567, EPI_ISL_3227568 | Lighthouse Lab in Milton Keynes | Wellcome Sanger Institute for the COVID-19 Genomics UK (COG-UK) Consortium | The Lighthouse Lab in Milton Keynes and Alex Alderton, Roberto Amato, Jeffrey Barrett, Sonia Goncalves, Ewan Harrison, David K. Jackson, Ian Johnston, Dominic Kwiatkowski, Cordelia Langford, John Sillitoe on behalf of the Wellcome Sanger Institute COVID-19 Surveillance Team |
| EPI_ISL_3227569 | Lighthouse Laboratory Plymouth | Wellcome Sanger Institute for the COVID-19 Genomics UK (COG-UK) Consortium | Lighthouse Laboratory Plymouth and Alex Alderton, Roberto Amato, Jeffrey Barrett, Sonia Goncalves, Ewan Harrison, David K. Jackson, Ian Johnston, Dominic Kwiatkowski, Cordelia Langford, John Sillitoe on behalf of the Wellcome Sanger Institute COVID-19 Surveillance Team |
| EPI_ISL_3227570, EPI_ISL_3227571 | Lighthouse Lab in Milton Keynes | Wellcome Sanger Institute for the COVID-19 Genomics UK (COG-UK) Consortium | The Lighthouse Lab in Milton Keynes and Alex Alderton, Roberto Amato, Jeffrey Barrett, Sonia Goncalves, Ewan Harrison, David K. Jackson, Ian Johnston, Dominic Kwiatkowski, Cordelia Langford, John Sillitoe on behalf of the Wellcome Sanger Institute COVID-19 Surveillance Team |
| EPI_ISL_3227572 | Lighthouse Laboratory Plymouth | Wellcome Sanger Institute for the COVID-19 Genomics UK (COG-UK) Consortium | Lighthouse Laboratory Plymouth and Alex Alderton, Roberto Amato, Jeffrey Barrett, Sonia Goncalves, Ewan Harrison, David K. Jackson, Ian Johnston, Dominic Kwiatkowski, Cordelia Langford, John Sillitoe on behalf of the Wellcome Sanger Institute COVID-19 Surveillance Team |
| EPI_ISL_3227573 | Lighthouse Lab in Milton Keynes | Wellcome Sanger Institute for the COVID-19 Genomics UK (COG-UK) Consortium | The Lighthouse Lab in Milton Keynes and Alex Alderton, Roberto Amato, Jeffrey Barrett, Sonia Goncalves, Ewan Harrison, David K. Jackson, Ian Johnston, Dominic Kwiatkowski, Cordelia Langford, John Sillitoe on behalf of the Wellcome Sanger Institute COVID-19 Surveillance Team |
| EPI_ISL_3227574, EPI_ISL_3227575, EPI_ISL_3227576, EPI_ISL_3227577, EPI_ISL_3227578, EPI_ISL_3227579, EPI_ISL_3227580, EPI_ISL_3227581, EPI_ISL_3227582 | Lighthouse Lab in Alderley Park | Wellcome Sanger Institute for the COVID-19 Genomics UK (COG-UK) Consortium | Jacquelyn Wynn, Mairead Hyland, The Lighthouse Lab in Alderley Park and Alex Alderton, Roberto Amato, Jeffrey Barrett, Sonia Goncalves, Ewan Harrison, David K. Jackson, Ian Johnston, Dominic Kwiatkowski, Cordelia Langford, John Sillitoe on behalf of the Wellcome Sanger Institute COVID-19 Surveillance Team |
| EPI_ISL_3227583 | Lighthouse Laboratory Plymouth | Wellcome Sanger Institute for the COVID-19 Genomics UK (COG-UK) Consortium | Lighthouse Laboratory Plymouth and Alex Alderton, Roberto Amato, Jeffrey Barrett, Sonia Goncalves, Ewan Harrison, David K. Jackson, Ian Johnston, Dominic Kwiatkowski, Cordelia Langford, John Sillitoe on behalf of the Wellcome Sanger Institute COVID-19 Surveillance Team |
| EPI_ISL_3227584, EPI_ISL_3227585, EPI_ISL_3227586, EPI_ISL_3227587, EPI_ISL_3227588, EPI_ISL_3227589, EPI_ISL_3227590, EPI_ISL_3227591, EPI_ISL_3227592, EPI_ISL_3227593 | Lighthouse Lab in Alderley Park | Wellcome Sanger Institute for the COVID-19 Genomics UK (COG-UK) Consortium | Jacquelyn Wynn, Mairead Hyland, The Lighthouse Lab in Alderley Park and Alex Alderton, Roberto Amato, Jeffrey Barrett, Sonia Goncalves, Ewan Harrison, David K. Jackson, Ian Johnston, Dominic Kwiatkowski, Cordelia Langford, John Sillitoe on behalf of the Wellcome Sanger Institute COVID-19 Surveillance Team |
| EPI_ISL_3227594 | Lighthouse Laboratory Plymouth | Wellcome Sanger Institute for the COVID-19 Genomics UK (COG-UK) Consortium | Lighthouse Laboratory Plymouth and Alex Alderton, Roberto Amato, Jeffrey Barrett, Sonia Goncalves, Ewan Harrison, David K. Jackson, Ian Johnston, Dominic Kwiatkowski, Cordelia Langford, John Sillitoe on behalf of the Wellcome Sanger Institute COVID-19 Surveillance Team |
| EPI_ISL_3227595, EPI_ISL_3227596 | Lighthouse Lab in Alderley Park | Wellcome Sanger Institute for the COVID-19 Genomics UK (COG-UK) Consortium | Jacquelyn Wynn, Mairead Hyland, The Lighthouse Lab in Alderley Park and Alex Alderton, Roberto Amato, Jeffrey Barrett, Sonia Goncalves, Ewan Harrison, David K. Jackson, Ian Johnston, Dominic Kwiatkowski, Cordelia Langford, John Sillitoe on behalf of the Wellcome Sanger Institute COVID-19 Surveillance Team |
| EPI_ISL_3227597 | Lighthouse Laboratory Plymouth | Wellcome Sanger Institute for the COVID-19 Genomics UK (COG-UK) Consortium | Lighthouse Laboratory Plymouth and Alex Alderton, Roberto Amato, Jeffrey Barrett, Sonia Goncalves, Ewan Harrison, David K. Jackson, Ian Johnston, Dominic Kwiatkowski, Cordelia Langford, John Sillitoe on behalf of the Wellcome Sanger Institute COVID-19 Surveillance Team |
| EPI_ISL_3227598 | Lighthouse Lab in Alderley Park | Wellcome Sanger Institute for the COVID-19 Genomics UK (COG-UK) Consortium | Jacquelyn Wynn, Mairead Hyland, The Lighthouse Lab in Alderley Park and Alex Alderton, Roberto Amato, Jeffrey Barrett, Sonia Goncalves, Ewan Harrison, David K. Jackson, Ian Johnston, Dominic Kwiatkowski, Cordelia Langford, John Sillitoe on behalf of the Wellcome Sanger Institute COVID-19 Surveillance Team |
| EPI_ISL_3227599 | Lighthouse Laboratory Plymouth | Wellcome Sanger Institute for the COVID-19 Genomics UK (COG-UK) Consortium | Lighthouse Laboratory Plymouth and Alex Alderton, Roberto Amato, Jeffrey Barrett, Sonia Goncalves, Ewan Harrison, David K. Jackson, Ian Johnston, Dominic Kwiatkowski, Cordelia Langford, John Sillitoe on behalf of the Wellcome Sanger Institute COVID-19 Surveillance Team |
| EPI_ISL_3227600, EPI_ISL_3227601, EPI_ISL_3227602, EPI_ISL_3227603 | Lighthouse Lab in Alderley Park | Wellcome Sanger Institute for the COVID-19 Genomics UK (COG-UK) Consortium | Jacquelyn Wynn, Mairead Hyland, The Lighthouse Lab in Alderley Park and Alex Alderton, Roberto Amato, Jeffrey Barrett, Sonia Goncalves, Ewan Harrison, David K. Jackson, Ian Johnston, Dominic Kwiatkowski, Cordelia Langford, John Sillitoe on behalf of the Wellcome Sanger Institute COVID-19 Surveillance Team |
| EPI_ISL_3227604 | Lighthouse Laboratory Plymouth | Wellcome Sanger Institute for the COVID-19 Genomics UK (COG-UK) Consortium | Lighthouse Laboratory Plymouth and Alex Alderton, Roberto Amato, Jeffrey Barrett, Sonia Goncalves, Ewan Harrison, David K. Jackson, Ian Johnston, Dominic Kwiatkowski, Cordelia Langford, John Sillitoe on behalf of the Wellcome Sanger Institute COVID-19 Surveillance Team |
| EPI_ISL_3227605 | Lighthouse Lab in Alderley Park | Wellcome Sanger Institute for the COVID-19 Genomics UK (COG-UK) Consortium | Jacquelyn Wynn, Mairead Hyland, The Lighthouse Lab in Alderley Park and Alex Alderton, Roberto Amato, Jeffrey Barrett, Sonia Goncalves, Ewan Harrison, David K. Jackson, Ian Johnston, Dominic Kwiatkowski, Cordelia Langford, John Sillitoe on behalf of the Wellcome Sanger Institute COVID-19 Surveillance Team |
| EPI_ISL_3227606 | Lighthouse Laboratory Plymouth | Wellcome Sanger Institute for the COVID-19 Genomics UK (COG-UK) Consortium | Lighthouse Laboratory Plymouth and Alex Alderton, Roberto Amato, Jeffrey Barrett, Sonia Goncalves, Ewan Harrison, David K. Jackson, Ian Johnston, Dominic Kwiatkowski, Cordelia Langford, John Sillitoe on behalf of the Wellcome Sanger Institute COVID-19 Surveillance Team |
| EPI_ISL_3227607, EPI_ISL_3227608, EPI_ISL_3227609, EPI_ISL_3227610, EPI_ISL_3227611 | Lighthouse Lab in Alderley Park | Wellcome Sanger Institute for the COVID-19 Genomics UK (COG-UK) Consortium | Jacquelyn Wynn, Mairead Hyland, The Lighthouse Lab in Alderley Park and Alex Alderton, Roberto Amato, Jeffrey Barrett, Sonia Goncalves, Ewan Harrison, David K. Jackson, Ian Johnston, Dominic Kwiatkowski, Cordelia Langford, John Sillitoe on behalf of the Wellcome Sanger Institute COVID-19 Surveillance Team |
| EPI_ISL_3227612, EPI_ISL_3227613, EPI_ISL_3227614 | Lighthouse Laboratory Plymouth | Wellcome Sanger Institute for the COVID-19 Genomics UK (COG-UK) Consortium | Lighthouse Laboratory Plymouth and Alex Alderton, Roberto Amato, Jeffrey Barrett, Sonia Goncalves, Ewan Harrison, David K. Jackson, Ian Johnston, Dominic Kwiatkowski, Cordelia Langford, John Sillitoe on behalf of the Wellcome Sanger Institute COVID-19 Surveillance Team |
| EPI_ISL_3227615, EPI_ISL_3227616 | Lighthouse Lab in Alderley Park | Wellcome Sanger Institute for the COVID-19 Genomics UK (COG-UK) Consortium | Jacquelyn Wynn, Mairead Hyland, The Lighthouse Lab in Alderley Park and Alex Alderton, Roberto Amato, Jeffrey Barrett, Sonia Goncalves, Ewan Harrison, David K. Jackson, Ian Johnston, Dominic Kwiatkowski, Cordelia Langford, John Sillitoe on behalf of the Wellcome Sanger Institute COVID-19 Surveillance Team |
| EPI_ISL_3227617 | Lighthouse Laboratory Plymouth | Wellcome Sanger Institute for the COVID-19 Genomics UK (COG-UK) Consortium | Lighthouse Laboratory Plymouth and Alex Alderton, Roberto Amato, Jeffrey Barrett, Sonia Goncalves, Ewan Harrison, David K. Jackson, Ian Johnston, Dominic Kwiatkowski, Cordelia Langford, John Sillitoe on behalf of the Wellcome Sanger Institute COVID-19 Surveillance Team |
| EPI_ISL_3227618, EPI_ISL_3227619 | Lighthouse Lab in Alderley Park | Wellcome Sanger Institute for the COVID-19 Genomics UK (COG-UK) Consortium | Jacquelyn Wynn, Mairead Hyland, The Lighthouse Lab in Alderley Park and Alex Alderton, Roberto Amato, Jeffrey Barrett, Sonia Goncalves, Ewan Harrison, David K. Jackson, Ian Johnston, Dominic Kwiatkowski, Cordelia Langford, John Sillitoe on behalf of the Wellcome Sanger Institute COVID-19 Surveillance Team |
| EPI_ISL_3227620 | Lighthouse Laboratory Plymouth | Wellcome Sanger Institute for the COVID-19 Genomics UK (COG-UK) Consortium | Lighthouse Laboratory Plymouth and Alex Alderton, Roberto Amato, Jeffrey Barrett, Sonia Goncalves, Ewan Harrison, David K. Jackson, Ian Johnston, Dominic Kwiatkowski, Cordelia Langford, John Sillitoe on behalf of the Wellcome Sanger Institute COVID-19 Surveillance Team |
| EPI_ISL_3227621, EPI_ISL_3227622, EPI_ISL_3227623, EPI_ISL_3227624, EPI_ISL_3227625, EPI_ISL_3227626, EPI_ISL_3227627, EPI_ISL_3227628, EPI_ISL_3227629, EPI_ISL_3227630 | Lighthouse Lab in Alderley Park | Wellcome Sanger Institute for the COVID-19 Genomics UK (COG-UK) Consortium | Jacquelyn Wynn, Mairead Hyland, The Lighthouse Lab in Alderley Park and Alex Alderton, Roberto Amato, Jeffrey Barrett, Sonia Goncalves, Ewan Harrison, David K. Jackson, Ian Johnston, Dominic Kwiatkowski, Cordelia Langford, John Sillitoe on behalf of the Wellcome Sanger Institute COVID-19 Surveillance Team |
| EPI_ISL_3227631 | Lighthouse Laboratory Plymouth | Wellcome Sanger Institute for the COVID-19 Genomics UK (COG-UK) Consortium | Lighthouse Laboratory Plymouth and Alex Alderton, Roberto Amato, Jeffrey Barrett, Sonia Goncalves, Ewan Harrison, David K. Jackson, Ian Johnston, Dominic Kwiatkowski, Cordelia Langford, John Sillitoe on behalf of the Wellcome Sanger Institute COVID-19 Surveillance Team |
| EPI_ISL_3227632 | Lighthouse Lab in Alderley Park | Wellcome Sanger Institute for the COVID-19 Genomics UK (COG-UK) Consortium | Jacquelyn Wynn, Mairead Hyland, The Lighthouse Lab in Alderley Park and Alex Alderton, Roberto Amato, Jeffrey Barrett, Sonia Goncalves, Ewan Harrison, David K. Jackson, Ian Johnston, Dominic Kwiatkowski, Cordelia Langford, John Sillitoe on behalf of the Wellcome Sanger Institute COVID-19 Surveillance Team |
| EPI_ISL_3227633, EPI_ISL_3227634 | Lighthouse Laboratory Plymouth | Wellcome Sanger Institute for the COVID-19 Genomics UK (COG-UK) Consortium | Lighthouse Laboratory Plymouth and Alex Alderton, Roberto Amato, Jeffrey Barrett, Sonia Goncalves, Ewan Harrison, David K. Jackson, Ian Johnston, Dominic Kwiatkowski, Cordelia Langford, John Sillitoe on behalf of the Wellcome Sanger Institute COVID-19 Surveillance Team |

| | | | |
|---|---|---|---|
| EPI_ISL_3227635 | Lighthouse Lab in Alderley Park | Wellcome Sanger Institute for the COVID-19 Genomics UK (COG-UK) Consortium | Jacquelyn Wynn, Mairead Hyland, The Lighthouse Lab in Alderley Park and Alex Alderton, Roberto Amato, Jeffrey Barrett, Sonia Goncalves, Ewan Harrison, David K. Jackson, Ian Johnston, Dominic Kwiatkowski, Cordelia Langford, John Sillitoe on behalf of the Wellcome Sanger Institute COVID-19 Surveillance Team |
| EPI_ISL_3227636, EPI_ISL_3227637 | Lighthouse Laboratory Plymouth | Wellcome Sanger Institute for the COVID-19 Genomics UK (COG-UK) Consortium | Lighthouse Laboratory Plymouth and Alex Alderton, Roberto Amato, Jeffrey Barrett, Sonia Goncalves, Ewan Harrison, David K. Jackson, Ian Johnston, Dominic Kwiatkowski, Cordelia Langford, John Sillitoe on behalf of the Wellcome Sanger Institute COVID-19 Surveillance Team |
| EPI_ISL_3227638, EPI_ISL_3227639, EPI_ISL_3227640, EPI_ISL_3227641 | Lighthouse Lab in Alderley Park | Wellcome Sanger Institute for the COVID-19 Genomics UK (COG-UK) Consortium | Jacquelyn Wynn, Mairead Hyland, The Lighthouse Lab in Alderley Park and Alex Alderton, Roberto Amato, Jeffrey Barrett, Sonia Goncalves, Ewan Harrison, David K. Jackson, Ian Johnston, Dominic Kwiatkowski, Cordelia Langford, John Sillitoe on behalf of the Wellcome Sanger Institute COVID-19 Surveillance Team |
| EPI_ISL_3227642 | Lighthouse Laboratory Plymouth | Wellcome Sanger Institute for the COVID-19 Genomics UK (COG-UK) Consortium | Lighthouse Laboratory Plymouth and Alex Alderton, Roberto Amato, Jeffrey Barrett, Sonia Goncalves, Ewan Harrison, David K. Jackson, Ian Johnston, Dominic Kwiatkowski, Cordelia Langford, John Sillitoe on behalf of the Wellcome Sanger Institute COVID-19 Surveillance Team |
| EPI_ISL_3227643, EPI_ISL_3227644, EPI_ISL_3227645 | Lighthouse Lab in Alderley Park | Wellcome Sanger Institute for the COVID-19 Genomics UK (COG-UK) Consortium | Jacquelyn Wynn, Mairead Hyland, The Lighthouse Lab in Alderley Park and Alex Alderton, Roberto Amato, Jeffrey Barrett, Sonia Goncalves, Ewan Harrison, David K. Jackson, Ian Johnston, Dominic Kwiatkowski, Cordelia Langford, John Sillitoe on behalf of the Wellcome Sanger Institute COVID-19 Surveillance Team |
| EPI_ISL_3227646, EPI_ISL_3227647 | Lighthouse Laboratory Plymouth | Wellcome Sanger Institute for the COVID-19 Genomics UK (COG-UK) Consortium | Lighthouse Laboratory Plymouth and Alex Alderton, Roberto Amato, Jeffrey Barrett, Sonia Goncalves, Ewan Harrison, David K. Jackson, Ian Johnston, Dominic Kwiatkowski, Cordelia Langford, John Sillitoe on behalf of the Wellcome Sanger Institute COVID-19 Surveillance Team |
| EPI_ISL_3227648, EPI_ISL_3227649 | Lighthouse Lab in Alderley Park | Wellcome Sanger Institute for the COVID-19 Genomics UK (COG-UK) Consortium | Jacquelyn Wynn, Mairead Hyland, The Lighthouse Lab in Alderley Park and Alex Alderton, Roberto Amato, Jeffrey Barrett, Sonia Goncalves, Ewan Harrison, David K. Jackson, Ian Johnston, Dominic Kwiatkowski, Cordelia Langford, John Sillitoe on behalf of the Wellcome Sanger Institute COVID-19 Surveillance Team |
| EPI_ISL_3227650 | Lighthouse Laboratory Plymouth | Wellcome Sanger Institute for the COVID-19 Genomics UK (COG-UK) Consortium | Lighthouse Laboratory Plymouth and Alex Alderton, Roberto Amato, Jeffrey Barrett, Sonia Goncalves, Ewan Harrison, David K. Jackson, Ian Johnston, Dominic Kwiatkowski, Cordelia Langford, John Sillitoe on behalf of the Wellcome Sanger Institute COVID-19 Surveillance Team |
| EPI_ISL_3227651, EPI_ISL_3227652 | Lighthouse Lab in Alderley Park | Wellcome Sanger Institute for the COVID-19 Genomics UK (COG-UK) Consortium | Jacquelyn Wynn, Mairead Hyland, The Lighthouse Lab in Alderley Park and Alex Alderton, Roberto Amato, Jeffrey Barrett, Sonia Goncalves, Ewan Harrison, David K. Jackson, Ian Johnston, Dominic Kwiatkowski, Cordelia Langford, John Sillitoe on behalf of the Wellcome Sanger Institute COVID-19 Surveillance Team |
| EPI_ISL_3227653 | Lighthouse Laboratory Plymouth | Wellcome Sanger Institute for the COVID-19 Genomics UK (COG-UK) Consortium | Lighthouse Laboratory Plymouth and Alex Alderton, Roberto Amato, Jeffrey Barrett, Sonia Goncalves, Ewan Harrison, David K. Jackson, Ian Johnston, Dominic Kwiatkowski, Cordelia Langford, John Sillitoe on behalf of the Wellcome Sanger Institute COVID-19 Surveillance Team |
| EPI_ISL_3227654 | Lighthouse Lab in Alderley Park | Wellcome Sanger Institute for the COVID-19 Genomics UK (COG-UK) Consortium | Jacquelyn Wynn, Mairead Hyland, The Lighthouse Lab in Alderley Park and Alex Alderton, Roberto Amato, Jeffrey Barrett, Sonia Goncalves, Ewan Harrison, David K. Jackson, Ian Johnston, Dominic Kwiatkowski, Cordelia Langford, John Sillitoe on behalf of the Wellcome Sanger Institute COVID-19 Surveillance Team |
| EPI_ISL_3227655, EPI_ISL_3227656, EPI_ISL_3227657, EPI_ISL_3227658 | Lighthouse Laboratory Plymouth | Wellcome Sanger Institute for the COVID-19 Genomics UK (COG-UK) Consortium | Lighthouse Laboratory Plymouth and Alex Alderton, Roberto Amato, Jeffrey Barrett, Sonia Goncalves, Ewan Harrison, David K. Jackson, Ian Johnston, Dominic Kwiatkowski, Cordelia Langford, John Sillitoe on behalf of the Wellcome Sanger Institute COVID-19 Surveillance Team |
| EPI_ISL_3227659 | Lighthouse Lab in Alderley Park | Wellcome Sanger Institute for the COVID-19 Genomics UK (COG-UK) Consortium | Jacquelyn Wynn, Mairead Hyland, The Lighthouse Lab in Alderley Park and Alex Alderton, Roberto Amato, Jeffrey Barrett, Sonia Goncalves, Ewan Harrison, David K. Jackson, Ian Johnston, Dominic Kwiatkowski, Cordelia Langford, John Sillitoe on behalf of the Wellcome Sanger Institute COVID-19 Surveillance Team |
| EPI_ISL_3227660 | Lighthouse Laboratory Plymouth | Wellcome Sanger Institute for the COVID-19 Genomics UK (COG-UK) Consortium | Lighthouse Laboratory Plymouth and Alex Alderton, Roberto Amato, Jeffrey Barrett, Sonia Goncalves, Ewan Harrison, David K. Jackson, Ian Johnston, Dominic Kwiatkowski, Cordelia Langford, John Sillitoe on behalf of the Wellcome Sanger Institute COVID-19 Surveillance Team |
| EPI_ISL_3227661, EPI_ISL_3227662, EPI_ISL_3227663 | Lighthouse Lab in Alderley Park | Wellcome Sanger Institute for the COVID-19 Genomics UK (COG-UK) Consortium | Jacquelyn Wynn, Mairead Hyland, The Lighthouse Lab in Alderley Park and Alex Alderton, Roberto Amato, Jeffrey Barrett, Sonia Goncalves, Ewan Harrison, David K. Jackson, Ian Johnston, Dominic Kwiatkowski, Cordelia Langford, John Sillitoe on behalf of the Wellcome Sanger Institute COVID-19 Surveillance Team |
| EPI_ISL_3227664 | Lighthouse Laboratory Plymouth | Wellcome Sanger Institute for the COVID-19 Genomics UK (COG-UK) Consortium | Lighthouse Laboratory Plymouth and Alex Alderton, Roberto Amato, Jeffrey Barrett, Sonia Goncalves, Ewan Harrison, David K. Jackson, Ian Johnston, Dominic Kwiatkowski, Cordelia Langford, John Sillitoe on behalf of the Wellcome Sanger Institute COVID-19 Surveillance Team |
| EPI_ISL_3227665, EPI_ISL_3227666, EPI_ISL_3227667 | Lighthouse Lab in Alderley Park | Wellcome Sanger Institute for the COVID-19 Genomics UK (COG-UK) Consortium | Jacquelyn Wynn, Mairead Hyland, The Lighthouse Lab in Alderley Park and Alex Alderton, Roberto Amato, Jeffrey Barrett, Sonia Goncalves, Ewan Harrison, David K. Jackson, Ian Johnston, Dominic Kwiatkowski, Cordelia Langford, John Sillitoe on behalf of the Wellcome Sanger Institute COVID-19 Surveillance Team |
| EPI_ISL_3227668 | Lighthouse Laboratory Plymouth | Wellcome Sanger Institute for the COVID-19 Genomics UK (COG-UK) Consortium | Lighthouse Laboratory Plymouth and Alex Alderton, Roberto Amato, Jeffrey Barrett, Sonia Goncalves, Ewan Harrison, David K. Jackson, Ian Johnston, Dominic Kwiatkowski, Cordelia Langford, John Sillitoe on behalf of the Wellcome Sanger Institute COVID-19 Surveillance Team |
| EPI_ISL_3227669, EPI_ISL_3227670, EPI_ISL_3227671, EPI_ISL_3227672, EPI_ISL_3227673, EPI_ISL_3227674, EPI_ISL_3227675, EPI_ISL_3227676, EPI_ISL_3227677, EPI_ISL_3227678, EPI_ISL_3227679, EPI_ISL_3227680, EPI_ISL_3227681, EPI_ISL_3227682, EPI_ISL_3227683 | | | |
| see above | Lighthouse Lab in Alderley Park | Wellcome Sanger Institute for the COVID-19 Genomics UK (COG-UK) Consortium | Jacquelyn Wynn, Mairead Hyland, The Lighthouse Lab in Alderley Park and Alex Alderton, Roberto Amato, Jeffrey Barrett, Sonia Goncalves, Ewan Harrison, David K. Jackson, Ian Johnston, Dominic Kwiatkowski, Cordelia Langford, John Sillitoe on behalf of the Wellcome Sanger Institute COVID-19 Surveillance Team |
| EPI_ISL_3227684 | Lighthouse Laboratory Plymouth | Wellcome Sanger Institute for the COVID-19 Genomics UK (COG-UK) Consortium | Lighthouse Laboratory Plymouth and Alex Alderton, Roberto Amato, Jeffrey Barrett, Sonia Goncalves, Ewan Harrison, David K. Jackson, Ian Johnston, Dominic Kwiatkowski, Cordelia Langford, John Sillitoe on behalf of the Wellcome Sanger Institute COVID-19 Surveillance Team |
| EPI_ISL_3227685 | Lighthouse Lab in Alderley Park | Wellcome Sanger Institute for the COVID-19 Genomics UK (COG-UK) Consortium | Jacquelyn Wynn, Mairead Hyland, The Lighthouse Lab in Alderley Park and Alex Alderton, Roberto Amato, Jeffrey Barrett, Sonia Goncalves, Ewan Harrison, David K. Jackson, Ian Johnston, Dominic Kwiatkowski, Cordelia Langford, John Sillitoe on behalf of the Wellcome Sanger Institute COVID-19 Surveillance Team |
| EPI_ISL_3227686, EPI_ISL_3227687 | Lighthouse Laboratory Plymouth | Wellcome Sanger Institute for the COVID-19 Genomics UK (COG-UK) Consortium | Lighthouse Laboratory Plymouth and Alex Alderton, Roberto Amato, Jeffrey Barrett, Sonia Goncalves, Ewan Harrison, David K. Jackson, Ian Johnston, Dominic Kwiatkowski, Cordelia Langford, John Sillitoe on behalf of the Wellcome Sanger Institute COVID-19 Surveillance Team |
| EPI_ISL_3227688, EPI_ISL_3227689 | Lighthouse Lab in Alderley Park | Wellcome Sanger Institute for the COVID-19 Genomics UK (COG-UK) Consortium | Jacquelyn Wynn, Mairead Hyland, The Lighthouse Lab in Alderley Park and Alex Alderton, Roberto Amato, Jeffrey Barrett, Sonia Goncalves, Ewan Harrison, David K. Jackson, Ian Johnston, Dominic Kwiatkowski, Cordelia Langford, John Sillitoe on behalf of the Wellcome Sanger Institute COVID-19 Surveillance Team |
| EPI_ISL_3227690 | Lighthouse Laboratory Plymouth | Wellcome Sanger Institute for the COVID-19 Genomics UK (COG-UK) Consortium | Lighthouse Laboratory Plymouth and Alex Alderton, Roberto Amato, Jeffrey Barrett, Sonia Goncalves, Ewan Harrison, David K. Jackson, Ian Johnston, Dominic Kwiatkowski, Cordelia Langford, John Sillitoe on behalf of the Wellcome Sanger Institute COVID-19 Surveillance Team |
| EPI_ISL_3227691, EPI_ISL_3227692, EPI_ISL_3227693, EPI_ISL_3227694, EPI_ISL_3227695, EPI_ISL_3227696, EPI_ISL_3227697, EPI_ISL_3227698 | Lighthouse Lab in Alderley Park | Wellcome Sanger Institute for the COVID-19 Genomics UK (COG-UK) Consortium | Jacquelyn Wynn, Mairead Hyland, The Lighthouse Lab in Alderley Park and Alex Alderton, Roberto Amato, Jeffrey Barrett, Sonia Goncalves, Ewan Harrison, David K. Jackson, Ian Johnston, Dominic Kwiatkowski, Cordelia Langford, John Sillitoe on behalf of the Wellcome Sanger Institute COVID-19 Surveillance Team |
| EPI_ISL_3227699 | Lighthouse Laboratory Plymouth | Wellcome Sanger Institute for the COVID-19 Genomics UK (COG-UK) Consortium | Lighthouse Laboratory Plymouth and Alex Alderton, Roberto Amato, Jeffrey Barrett, Sonia Goncalves, Ewan Harrison, David K. Jackson, Ian Johnston, Dominic Kwiatkowski, Cordelia Langford, John Sillitoe on behalf of the Wellcome Sanger Institute COVID-19 Surveillance Team |
| EPI_ISL_3227700, EPI_ISL_3227701 | Lighthouse Lab in Alderley Park | Wellcome Sanger Institute for the COVID-19 Genomics UK (COG-UK) Consortium | Jacquelyn Wynn, Mairead Hyland, The Lighthouse Lab in Alderley Park and Alex Alderton, Roberto Amato, Jeffrey Barrett, Sonia Goncalves, Ewan Harrison, David K. Jackson, Ian Johnston, Dominic Kwiatkowski, Cordelia Langford, John Sillitoe on behalf of the Wellcome Sanger Institute COVID-19 Surveillance Team |
| EPI_ISL_3227702 | Lighthouse Laboratory Plymouth | Wellcome Sanger Institute for the COVID-19 Genomics UK (COG-UK) Consortium | Lighthouse Laboratory Plymouth and Alex Alderton, Roberto Amato, Jeffrey Barrett, Sonia Goncalves, Ewan Harrison, David K. Jackson, Ian Johnston, Dominic Kwiatkowski, Cordelia Langford, John Sillitoe on behalf of the Wellcome Sanger Institute COVID-19 Surveillance Team |
| EPI_ISL_3227703, EPI_ISL_3227704, EPI_ISL_3227705, EPI_ISL_3227706, EPI_ISL_3227707, EPI_ISL_3227708, EPI_ISL_3227709, EPI_ISL_3227710, EPI_ISL_3227711, EPI_ISL_3227712, EPI_ISL_3227713, EPI_ISL_3227714, EPI_ISL_3227715, EPI_ISL_3227716, EPI_ISL_3227717, EPI_ISL_3227718, EPI_ISL_3227719, EPI_ISL_3227720 | | | |
| see above | Lighthouse Lab in Alderley Park | Wellcome Sanger Institute for the COVID-19 Genomics UK (COG-UK) Consortium | Jacquelyn Wynn, Mairead Hyland, The Lighthouse Lab in Alderley Park and Alex Alderton, Roberto Amato, Jeffrey Barrett, Sonia Goncalves, Ewan Harrison, David K. Jackson, Ian Johnston, Dominic Kwiatkowski, Cordelia Langford, John Sillitoe on behalf of the Wellcome Sanger Institute COVID-19 Surveillance Team |
| EPI_ISL_3227721 | Lighthouse Laboratory Plymouth | Wellcome Sanger Institute for the COVID-19 Genomics UK (COG-UK) Consortium | Lighthouse Laboratory Plymouth and Alex Alderton, Roberto Amato, Jeffrey Barrett, Sonia Goncalves, Ewan Harrison, David K. Jackson, Ian Johnston, Dominic Kwiatkowski, Cordelia Langford, John Sillitoe on behalf of the Wellcome Sanger Institute COVID-19 Surveillance Team |
| EPI_ISL_3227722, EPI_ISL_3227723 | Lighthouse Lab in Alderley Park | Wellcome Sanger Institute for the COVID-19 Genomics UK (COG-UK) Consortium | Jacquelyn Wynn, Mairead Hyland, The Lighthouse Lab in Alderley Park and Alex Alderton, Roberto Amato, Jeffrey Barrett, Sonia Goncalves, Ewan Harrison, David K. Jackson, Ian Johnston, Dominic Kwiatkowski, Cordelia Langford, John Sillitoe on behalf of the Wellcome Sanger Institute COVID-19 Surveillance Team |
| EPI_ISL_3227724 | Lighthouse Laboratory Plymouth | Wellcome Sanger Institute for the COVID-19 Genomics UK (COG-UK) Consortium | Lighthouse Laboratory Plymouth and Alex Alderton, Roberto Amato, Jeffrey Barrett, Sonia Goncalves, Ewan Harrison, David K. Jackson, Ian Johnston, Dominic Kwiatkowski, Cordelia Langford, John Sillitoe on behalf of the Wellcome Sanger Institute COVID-19 Surveillance Team |
| EPI_ISL_3227725, EPI_ISL_3227726, EPI_ISL_3227727, EPI_ISL_3227728, EPI_ISL_3227729, EPI_ISL_3227730 | Lighthouse Lab in Alderley Park | Wellcome Sanger Institute for the COVID-19 Genomics UK (COG-UK) Consortium | Jacquelyn Wynn, Mairead Hyland, The Lighthouse Lab in Alderley Park and Alex Alderton, Roberto Amato, Jeffrey Barrett, Sonia Goncalves, Ewan Harrison, David K. Jackson, Ian Johnston, Dominic Kwiatkowski, Cordelia Langford, John Sillitoe on behalf of the Wellcome Sanger Institute COVID-19 Surveillance Team |
| EPI_ISL_3227731, EPI_ISL_3227732 | Lighthouse Laboratory Plymouth | Wellcome Sanger Institute for the COVID-19 Genomics UK (COG-UK) Consortium | Lighthouse Laboratory Plymouth and Alex Alderton, Roberto Amato, Jeffrey Barrett, Sonia Goncalves, Ewan Harrison, David K. Jackson, Ian Johnston, Dominic Kwiatkowski, Cordelia Langford, John Sillitoe on behalf of the Wellcome Sanger Institute COVID-19 Surveillance Team |
| EPI_ISL_3227733 | Lighthouse Lab in Alderley Park | Wellcome Sanger Institute for the COVID-19 Genomics UK (COG-UK) Consortium | Jacquelyn Wynn, Mairead Hyland, The Lighthouse Lab in Alderley Park and Alex Alderton, Roberto Amato, Jeffrey Barrett, Sonia Goncalves, Ewan Harrison, David K. Jackson, Ian Johnston, Dominic Kwiatkowski, Cordelia Langford, John Sillitoe on behalf of the Wellcome Sanger Institute COVID-19 Surveillance Team |
| EPI_ISL_3227734, EPI_ISL_3227735, EPI_ISL_3227736 | Lighthouse Laboratory Plymouth | Wellcome Sanger Institute for the COVID-19 Genomics UK (COG-UK) Consortium | Lighthouse Laboratory Plymouth and Alex Alderton, Roberto Amato, Jeffrey Barrett, Sonia Goncalves, Ewan Harrison, David K. Jackson, Ian Johnston, Dominic Kwiatkowski, Cordelia Langford, John Sillitoe on behalf of the Wellcome Sanger Institute COVID-19 Surveillance Team |
| EPI_ISL_3227737, | Lighthouse Lab in Alderley Park | Wellcome Sanger Institute for the COVID-19 Genomics UK (COG-UK) | Jacquelyn Wynn, Mairead Hyland, The Lighthouse Lab in Alderley Park and Alex Alderton, Roberto Amato, Jeffrey Barrett, Sonia Goncalves, Ewan Harrison, David K. Jackson, Ian Johnston, Dominic Kwiatkowski, Cordelia Langford, John Sillitoe on |

| | | | |
|---|---|---|---|
| EPI_ISL_3227738, EPI_ISL_3227739, EPI_ISL_3227740, EPI_ISL_3227741, EPI_ISL_3227742, EPI_ISL_3227743, EPI_ISL_3227744, EPI_ISL_3227745 | | Consortium | behalf of the Wellcome Sanger Institute COVID-19 Surveillance Team |
| EPI_ISL_3227746 | Lighthouse Laboratory Plymouth | Wellcome Sanger Institute for the COVID-19 Genomics UK (COG-UK) Consortium | Lighthouse Laboratory Plymouth and Alex Alderton, Roberto Amato, Jeffrey Barrett, Sonia Goncalves, Ewan Harrison, David K. Jackson, Ian Johnston, Dominic Kwiatkowski, Cordelia Langford, John Sillitoe on behalf of the Wellcome Sanger Institute COVID-19 Surveillance Team |
| EPI_ISL_3227747, EPI_ISL_3227748, EPI_ISL_3227749, EPI_ISL_3227750, EPI_ISL_3227751, EPI_ISL_3227752, EPI_ISL_3227753 | Lighthouse Lab in Alderley Park | Wellcome Sanger Institute for the COVID-19 Genomics UK (COG-UK) Consortium | Jacquelyn Wynn, Mairead Hyland, The Lighthouse Lab in Alderley Park and Alex Alderton, Roberto Amato, Jeffrey Barrett, Sonia Goncalves, Ewan Harrison, David K. Jackson, Ian Johnston, Dominic Kwiatkowski, Cordelia Langford, John Sillitoe on behalf of the Wellcome Sanger Institute COVID-19 Surveillance Team |
| EPI_ISL_3227754 | Lighthouse Laboratory Plymouth | Wellcome Sanger Institute for the COVID-19 Genomics UK (COG-UK) Consortium | Lighthouse Laboratory Plymouth and Alex Alderton, Roberto Amato, Jeffrey Barrett, Sonia Goncalves, Ewan Harrison, David K. Jackson, Ian Johnston, Dominic Kwiatkowski, Cordelia Langford, John Sillitoe on behalf of the Wellcome Sanger Institute COVID-19 Surveillance Team |
| EPI_ISL_3227755, EPI_ISL_3227756, EPI_ISL_3227757, EPI_ISL_3227758, EPI_ISL_3227759, EPI_ISL_3227760, EPI_ISL_3227761, EPI_ISL_3227762, EPI_ISL_3227763 | Lighthouse Lab in Alderley Park | Wellcome Sanger Institute for the COVID-19 Genomics UK (COG-UK) Consortium | Jacquelyn Wynn, Mairead Hyland, The Lighthouse Lab in Alderley Park and Alex Alderton, Roberto Amato, Jeffrey Barrett, Sonia Goncalves, Ewan Harrison, David K. Jackson, Ian Johnston, Dominic Kwiatkowski, Cordelia Langford, John Sillitoe on behalf of the Wellcome Sanger Institute COVID-19 Surveillance Team |
| EPI_ISL_3227764 | Lighthouse Laboratory Plymouth | Wellcome Sanger Institute for the COVID-19 Genomics UK (COG-UK) Consortium | Lighthouse Laboratory Plymouth and Alex Alderton, Roberto Amato, Jeffrey Barrett, Sonia Goncalves, Ewan Harrison, David K. Jackson, Ian Johnston, Dominic Kwiatkowski, Cordelia Langford, John Sillitoe on behalf of the Wellcome Sanger Institute COVID-19 Surveillance Team |
| EPI_ISL_3227765 | Lighthouse Lab in Alderley Park | Wellcome Sanger Institute for the COVID-19 Genomics UK (COG-UK) Consortium | Jacquelyn Wynn, Mairead Hyland, The Lighthouse Lab in Alderley Park and Alex Alderton, Roberto Amato, Jeffrey Barrett, Sonia Goncalves, Ewan Harrison, David K. Jackson, Ian Johnston, Dominic Kwiatkowski, Cordelia Langford, John Sillitoe on behalf of the Wellcome Sanger Institute COVID-19 Surveillance Team |
| EPI_ISL_3227766 | Lighthouse Laboratory Plymouth | Wellcome Sanger Institute for the COVID-19 Genomics UK (COG-UK) Consortium | Lighthouse Laboratory Plymouth and Alex Alderton, Roberto Amato, Jeffrey Barrett, Sonia Goncalves, Ewan Harrison, David K. Jackson, Ian Johnston, Dominic Kwiatkowski, Cordelia Langford, John Sillitoe on behalf of the Wellcome Sanger Institute COVID-19 Surveillance Team |
| EPI_ISL_3227767, EPI_ISL_3227768 | Lighthouse Lab in Alderley Park | Wellcome Sanger Institute for the COVID-19 Genomics UK (COG-UK) Consortium | Jacquelyn Wynn, Mairead Hyland, The Lighthouse Lab in Alderley Park and Alex Alderton, Roberto Amato, Jeffrey Barrett, Sonia Goncalves, Ewan Harrison, David K. Jackson, Ian Johnston, Dominic Kwiatkowski, Cordelia Langford, John Sillitoe on behalf of the Wellcome Sanger Institute COVID-19 Surveillance Team |
| EPI_ISL_3227769 | Lighthouse Laboratory Plymouth | Wellcome Sanger Institute for the COVID-19 Genomics UK (COG-UK) Consortium | Lighthouse Laboratory Plymouth and Alex Alderton, Roberto Amato, Jeffrey Barrett, Sonia Goncalves, Ewan Harrison, David K. Jackson, Ian Johnston, Dominic Kwiatkowski, Cordelia Langford, John Sillitoe on behalf of the Wellcome Sanger Institute COVID-19 Surveillance Team |
| EPI_ISL_3227770, EPI_ISL_3227771, EPI_ISL_3227772, EPI_ISL_3227773, EPI_ISL_3227774, EPI_ISL_3227775 | Lighthouse Lab in Alderley Park | Wellcome Sanger Institute for the COVID-19 Genomics UK (COG-UK) Consortium | Jacquelyn Wynn, Mairead Hyland, The Lighthouse Lab in Alderley Park and Alex Alderton, Roberto Amato, Jeffrey Barrett, Sonia Goncalves, Ewan Harrison, David K. Jackson, Ian Johnston, Dominic Kwiatkowski, Cordelia Langford, John Sillitoe on behalf of the Wellcome Sanger Institute COVID-19 Surveillance Team |
| EPI_ISL_3227776 | Lighthouse Laboratory Plymouth | Wellcome Sanger Institute for the COVID-19 Genomics UK (COG-UK) Consortium | Lighthouse Laboratory Plymouth and Alex Alderton, Roberto Amato, Jeffrey Barrett, Sonia Goncalves, Ewan Harrison, David K. Jackson, Ian Johnston, Dominic Kwiatkowski, Cordelia Langford, John Sillitoe on behalf of the Wellcome Sanger Institute COVID-19 Surveillance Team |
| EPI_ISL_3227777 | Lighthouse Lab in Alderley Park | Wellcome Sanger Institute for the COVID-19 Genomics UK (COG-UK) Consortium | Jacquelyn Wynn, Mairead Hyland, The Lighthouse Lab in Alderley Park and Alex Alderton, Roberto Amato, Jeffrey Barrett, Sonia Goncalves, Ewan Harrison, David K. Jackson, Ian Johnston, Dominic Kwiatkowski, Cordelia Langford, John Sillitoe on behalf of the Wellcome Sanger Institute COVID-19 Surveillance Team |
| EPI_ISL_3227778 | Lighthouse Laboratory Plymouth | Wellcome Sanger Institute for the COVID-19 Genomics UK (COG-UK) Consortium | Lighthouse Laboratory Plymouth and Alex Alderton, Roberto Amato, Jeffrey Barrett, Sonia Goncalves, Ewan Harrison, David K. Jackson, Ian Johnston, Dominic Kwiatkowski, Cordelia Langford, John Sillitoe on behalf of the Wellcome Sanger Institute COVID-19 Surveillance Team |
| EPI_ISL_3227779, EPI_ISL_3227780, EPI_ISL_3227781, EPI_ISL_3227782 | Lighthouse Lab in Alderley Park | Wellcome Sanger Institute for the COVID-19 Genomics UK (COG-UK) Consortium | Jacquelyn Wynn, Mairead Hyland, The Lighthouse Lab in Alderley Park and Alex Alderton, Roberto Amato, Jeffrey Barrett, Sonia Goncalves, Ewan Harrison, David K. Jackson, Ian Johnston, Dominic Kwiatkowski, Cordelia Langford, John Sillitoe on behalf of the Wellcome Sanger Institute COVID-19 Surveillance Team |
| EPI_ISL_3227783 | Lighthouse Laboratory Plymouth | Wellcome Sanger Institute for the COVID-19 Genomics UK (COG-UK) Consortium | Lighthouse Laboratory Plymouth and Alex Alderton, Roberto Amato, Jeffrey Barrett, Sonia Goncalves, Ewan Harrison, David K. Jackson, Ian Johnston, Dominic Kwiatkowski, Cordelia Langford, John Sillitoe on behalf of the Wellcome Sanger Institute COVID-19 Surveillance Team |
| EPI_ISL_3227784, EPI_ISL_3227785 | Lighthouse Lab in Alderley Park | Wellcome Sanger Institute for the COVID-19 Genomics UK (COG-UK) Consortium | Jacquelyn Wynn, Mairead Hyland, The Lighthouse Lab in Alderley Park and Alex Alderton, Roberto Amato, Jeffrey Barrett, Sonia Goncalves, Ewan Harrison, David K. Jackson, Ian Johnston, Dominic Kwiatkowski, Cordelia Langford, John Sillitoe on behalf of the Wellcome Sanger Institute COVID-19 Surveillance Team |
| EPI_ISL_3227786 | Lighthouse Laboratory Plymouth | Wellcome Sanger Institute for the COVID-19 Genomics UK (COG-UK) Consortium | Lighthouse Laboratory Plymouth and Alex Alderton, Roberto Amato, Jeffrey Barrett, Sonia Goncalves, Ewan Harrison, David K. Jackson, Ian Johnston, Dominic Kwiatkowski, Cordelia Langford, John Sillitoe on behalf of the Wellcome Sanger Institute COVID-19 Surveillance Team |
| EPI_ISL_3227787, EPI_ISL_3227788 | Lighthouse Lab in Alderley Park | Wellcome Sanger Institute for the COVID-19 Genomics UK (COG-UK) Consortium | Jacquelyn Wynn, Mairead Hyland, The Lighthouse Lab in Alderley Park and Alex Alderton, Roberto Amato, Jeffrey Barrett, Sonia Goncalves, Ewan Harrison, David K. Jackson, Ian Johnston, Dominic Kwiatkowski, Cordelia Langford, John Sillitoe on behalf of the Wellcome Sanger Institute COVID-19 Surveillance Team |
| EPI_ISL_3227789, EPI_ISL_3227790 | Lighthouse Laboratory Plymouth | Wellcome Sanger Institute for the COVID-19 Genomics UK (COG-UK) Consortium | Lighthouse Laboratory Plymouth and Alex Alderton, Roberto Amato, Jeffrey Barrett, Sonia Goncalves, Ewan Harrison, David K. Jackson, Ian Johnston, Dominic Kwiatkowski, Cordelia Langford, John Sillitoe on behalf of the Wellcome Sanger Institute COVID-19 Surveillance Team |
| EPI_ISL_3227791, EPI_ISL_3227792, EPI_ISL_3227793, EPI_ISL_3227794, EPI_ISL_3227795 | Lighthouse Lab in Alderley Park | Wellcome Sanger Institute for the COVID-19 Genomics UK (COG-UK) Consortium | Jacquelyn Wynn, Mairead Hyland, The Lighthouse Lab in Alderley Park and Alex Alderton, Roberto Amato, Jeffrey Barrett, Sonia Goncalves, Ewan Harrison, David K. Jackson, Ian Johnston, Dominic Kwiatkowski, Cordelia Langford, John Sillitoe on behalf of the Wellcome Sanger Institute COVID-19 Surveillance Team |
| EPI_ISL_3227796 | Lighthouse Laboratory Plymouth | Wellcome Sanger Institute for the COVID-19 Genomics UK (COG-UK) Consortium | Lighthouse Laboratory Plymouth and Alex Alderton, Roberto Amato, Jeffrey Barrett, Sonia Goncalves, Ewan Harrison, David K. Jackson, Ian Johnston, Dominic Kwiatkowski, Cordelia Langford, John Sillitoe on behalf of the Wellcome Sanger Institute COVID-19 Surveillance Team |
| EPI_ISL_3227797, EPI_ISL_3227798, EPI_ISL_3227799, EPI_ISL_3227800, EPI_ISL_3227801, EPI_ISL_3227802, EPI_ISL_3227803 | Lighthouse Lab in Alderley Park | Wellcome Sanger Institute for the COVID-19 Genomics UK (COG-UK) Consortium | Jacquelyn Wynn, Mairead Hyland, The Lighthouse Lab in Alderley Park and Alex Alderton, Roberto Amato, Jeffrey Barrett, Sonia Goncalves, Ewan Harrison, David K. Jackson, Ian Johnston, Dominic Kwiatkowski, Cordelia Langford, John Sillitoe on behalf of the Wellcome Sanger Institute COVID-19 Surveillance Team |
| EPI_ISL_3227804, EPI_ISL_3227805 | Lighthouse Laboratory Plymouth | Wellcome Sanger Institute for the COVID-19 Genomics UK (COG-UK) Consortium | Lighthouse Laboratory Plymouth and Alex Alderton, Roberto Amato, Jeffrey Barrett, Sonia Goncalves, Ewan Harrison, David K. Jackson, Ian Johnston, Dominic Kwiatkowski, Cordelia Langford, John Sillitoe on behalf of the Wellcome Sanger Institute COVID-19 Surveillance Team |
| EPI_ISL_3227806, EPI_ISL_3227807, EPI_ISL_3227808 | Lighthouse Lab in Alderley Park | Wellcome Sanger Institute for the COVID-19 Genomics UK (COG-UK) Consortium | Jacquelyn Wynn, Mairead Hyland, The Lighthouse Lab in Alderley Park and Alex Alderton, Roberto Amato, Jeffrey Barrett, Sonia Goncalves, Ewan Harrison, David K. Jackson, Ian Johnston, Dominic Kwiatkowski, Cordelia Langford, John Sillitoe on behalf of the Wellcome Sanger Institute COVID-19 Surveillance Team |
| EPI_ISL_3227809 | Lighthouse Laboratory Plymouth | Wellcome Sanger Institute for the COVID-19 Genomics UK (COG-UK) Consortium | Lighthouse Laboratory Plymouth and Alex Alderton, Roberto Amato, Jeffrey Barrett, Sonia Goncalves, Ewan Harrison, David K. Jackson, Ian Johnston, Dominic Kwiatkowski, Cordelia Langford, John Sillitoe on behalf of the Wellcome Sanger Institute COVID-19 Surveillance Team |
| EPI_ISL_3227810, EPI_ISL_3227811, EPI_ISL_3227812, EPI_ISL_3227813 | Lighthouse Lab in Alderley Park | Wellcome Sanger Institute for the COVID-19 Genomics UK (COG-UK) Consortium | Jacquelyn Wynn, Mairead Hyland, The Lighthouse Lab in Alderley Park and Alex Alderton, Roberto Amato, Jeffrey Barrett, Sonia Goncalves, Ewan Harrison, David K. Jackson, Ian Johnston, Dominic Kwiatkowski, Cordelia Langford, John Sillitoe on behalf of the Wellcome Sanger Institute COVID-19 Surveillance Team |
| EPI_ISL_3227814 | Lighthouse Laboratory Plymouth | Wellcome Sanger Institute for the COVID-19 Genomics UK (COG-UK) Consortium | Lighthouse Laboratory Plymouth and Alex Alderton, Roberto Amato, Jeffrey Barrett, Sonia Goncalves, Ewan Harrison, David K. Jackson, Ian Johnston, Dominic Kwiatkowski, Cordelia Langford, John Sillitoe on behalf of the Wellcome Sanger Institute COVID-19 Surveillance Team |
| EPI_ISL_3227815, EPI_ISL_3227816, EPI_ISL_3227817, EPI_ISL_3227818 | Lighthouse Lab in Alderley Park | Wellcome Sanger Institute for the COVID-19 Genomics UK (COG-UK) Consortium | Jacquelyn Wynn, Mairead Hyland, The Lighthouse Lab in Alderley Park and Alex Alderton, Roberto Amato, Jeffrey Barrett, Sonia Goncalves, Ewan Harrison, David K. Jackson, Ian Johnston, Dominic Kwiatkowski, Cordelia Langford, John Sillitoe on behalf of the Wellcome Sanger Institute COVID-19 Surveillance Team |
| EPI_ISL_3227819 | Lighthouse Laboratory Plymouth | Wellcome Sanger Institute for the COVID-19 Genomics UK (COG-UK) Consortium | Lighthouse Laboratory Plymouth and Alex Alderton, Roberto Amato, Jeffrey Barrett, Sonia Goncalves, Ewan Harrison, David K. Jackson, Ian Johnston, Dominic Kwiatkowski, Cordelia Langford, John Sillitoe on behalf of the Wellcome Sanger Institute COVID-19 Surveillance Team |

| | | | |
|---|---|---|---|
| EPI_ISL_3227820 | Lighthouse Lab in Alderley Park | Wellcome Sanger Institute for the COVID-19 Genomics UK (COG-UK) Consortium | Jacquelyn Wynn, Mairead Hyland, The Lighthouse Lab in Alderley Park and Alex Alderton, Roberto Amato, Jeffrey Barrett, Sonia Goncalves, Ewan Harrison, David K. Jackson, Ian Johnston, Dominic Kwiatkowski, Cordelia Langford, John Sillitoe on behalf of the Wellcome Sanger Institute COVID-19 Surveillance Team |
| EPI_ISL_3227821 | Lighthouse Laboratory Plymouth | Wellcome Sanger Institute for the COVID-19 Genomics UK (COG-UK) Consortium | Lighthouse Laboratory Plymouth and Alex Alderton, Roberto Amato, Jeffrey Barrett, Sonia Goncalves, Ewan Harrison, David K. Jackson, Ian Johnston, Dominic Kwiatkowski, Cordelia Langford, John Sillitoe on behalf of the Wellcome Sanger Institute COVID-19 Surveillance Team |
| EPI_ISL_3227822, EPI_ISL_3227823 | Lighthouse Lab in Alderley Park | Wellcome Sanger Institute for the COVID-19 Genomics UK (COG-UK) Consortium | Jacquelyn Wynn, Mairead Hyland, The Lighthouse Lab in Alderley Park and Alex Alderton, Roberto Amato, Jeffrey Barrett, Sonia Goncalves, Ewan Harrison, David K. Jackson, Ian Johnston, Dominic Kwiatkowski, Cordelia Langford, John Sillitoe on behalf of the Wellcome Sanger Institute COVID-19 Surveillance Team |
| EPI_ISL_3227824 | Lighthouse Laboratory Plymouth | Wellcome Sanger Institute for the COVID-19 Genomics UK (COG-UK) Consortium | Lighthouse Laboratory Plymouth and Alex Alderton, Roberto Amato, Jeffrey Barrett, Sonia Goncalves, Ewan Harrison, David K. Jackson, Ian Johnston, Dominic Kwiatkowski, Cordelia Langford, John Sillitoe on behalf of the Wellcome Sanger Institute COVID-19 Surveillance Team |
| EPI_ISL_3227825, EPI_ISL_3227826 | Lighthouse Lab in Alderley Park | Wellcome Sanger Institute for the COVID-19 Genomics UK (COG-UK) Consortium | Jacquelyn Wynn, Mairead Hyland, The Lighthouse Lab in Alderley Park and Alex Alderton, Roberto Amato, Jeffrey Barrett, Sonia Goncalves, Ewan Harrison, David K. Jackson, Ian Johnston, Dominic Kwiatkowski, Cordelia Langford, John Sillitoe on behalf of the Wellcome Sanger Institute COVID-19 Surveillance Team |
| EPI_ISL_3227827, EPI_ISL_3227828, EPI_ISL_3227829 | Lighthouse Laboratory Plymouth | Wellcome Sanger Institute for the COVID-19 Genomics UK (COG-UK) Consortium | Lighthouse Laboratory Plymouth and Alex Alderton, Roberto Amato, Jeffrey Barrett, Sonia Goncalves, Ewan Harrison, David K. Jackson, Ian Johnston, Dominic Kwiatkowski, Cordelia Langford, John Sillitoe on behalf of the Wellcome Sanger Institute COVID-19 Surveillance Team |
| EPI_ISL_3227830, EPI_ISL_3227831, EPI_ISL_3227832, EPI_ISL_3227833, EPI_ISL_3227834, EPI_ISL_3227835 | Lighthouse Lab in Alderley Park | Wellcome Sanger Institute for the COVID-19 Genomics UK (COG-UK) Consortium | Jacquelyn Wynn, Mairead Hyland, The Lighthouse Lab in Alderley Park and Alex Alderton, Roberto Amato, Jeffrey Barrett, Sonia Goncalves, Ewan Harrison, David K. Jackson, Ian Johnston, Dominic Kwiatkowski, Cordelia Langford, John Sillitoe on behalf of the Wellcome Sanger Institute COVID-19 Surveillance Team |
| EPI_ISL_3227836 | Lighthouse Laboratory Plymouth | Wellcome Sanger Institute for the COVID-19 Genomics UK (COG-UK) Consortium | Lighthouse Laboratory Plymouth and Alex Alderton, Roberto Amato, Jeffrey Barrett, Sonia Goncalves, Ewan Harrison, David K. Jackson, Ian Johnston, Dominic Kwiatkowski, Cordelia Langford, John Sillitoe on behalf of the Wellcome Sanger Institute COVID-19 Surveillance Team |
| EPI_ISL_3227837, EPI_ISL_3227838, EPI_ISL_3227839 | Lighthouse Lab in Alderley Park | Wellcome Sanger Institute for the COVID-19 Genomics UK (COG-UK) Consortium | Jacquelyn Wynn, Mairead Hyland, The Lighthouse Lab in Alderley Park and Alex Alderton, Roberto Amato, Jeffrey Barrett, Sonia Goncalves, Ewan Harrison, David K. Jackson, Ian Johnston, Dominic Kwiatkowski, Cordelia Langford, John Sillitoe on behalf of the Wellcome Sanger Institute COVID-19 Surveillance Team |
| EPI_ISL_3227840 | Lighthouse Laboratory Plymouth | Wellcome Sanger Institute for the COVID-19 Genomics UK (COG-UK) Consortium | Lighthouse Laboratory Plymouth and Alex Alderton, Roberto Amato, Jeffrey Barrett, Sonia Goncalves, Ewan Harrison, David K. Jackson, Ian Johnston, Dominic Kwiatkowski, Cordelia Langford, John Sillitoe on behalf of the Wellcome Sanger Institute COVID-19 Surveillance Team |
| EPI_ISL_3227841, EPI_ISL_3227842 | Lighthouse Lab in Alderley Park | Wellcome Sanger Institute for the COVID-19 Genomics UK (COG-UK) Consortium | Jacquelyn Wynn, Mairead Hyland, The Lighthouse Lab in Alderley Park and Alex Alderton, Roberto Amato, Jeffrey Barrett, Sonia Goncalves, Ewan Harrison, David K. Jackson, Ian Johnston, Dominic Kwiatkowski, Cordelia Langford, John Sillitoe on behalf of the Wellcome Sanger Institute COVID-19 Surveillance Team |
| EPI_ISL_3227843 | Lighthouse Laboratory Plymouth | Wellcome Sanger Institute for the COVID-19 Genomics UK (COG-UK) Consortium | Lighthouse Laboratory Plymouth and Alex Alderton, Roberto Amato, Jeffrey Barrett, Sonia Goncalves, Ewan Harrison, David K. Jackson, Ian Johnston, Dominic Kwiatkowski, Cordelia Langford, John Sillitoe on behalf of the Wellcome Sanger Institute COVID-19 Surveillance Team |
| EPI_ISL_3227844, EPI_ISL_3227845, EPI_ISL_3227846 | Lighthouse Lab in Alderley Park | Wellcome Sanger Institute for the COVID-19 Genomics UK (COG-UK) Consortium | Jacquelyn Wynn, Mairead Hyland, The Lighthouse Lab in Alderley Park and Alex Alderton, Roberto Amato, Jeffrey Barrett, Sonia Goncalves, Ewan Harrison, David K. Jackson, Ian Johnston, Dominic Kwiatkowski, Cordelia Langford, John Sillitoe on behalf of the Wellcome Sanger Institute COVID-19 Surveillance Team |
| EPI_ISL_3227847 | Lighthouse Laboratory Plymouth | Wellcome Sanger Institute for the COVID-19 Genomics UK (COG-UK) Consortium | Lighthouse Laboratory Plymouth and Alex Alderton, Roberto Amato, Jeffrey Barrett, Sonia Goncalves, Ewan Harrison, David K. Jackson, Ian Johnston, Dominic Kwiatkowski, Cordelia Langford, John Sillitoe on behalf of the Wellcome Sanger Institute COVID-19 Surveillance Team |
| EPI_ISL_3227848, EPI_ISL_3227849, EPI_ISL_3227850, EPI_ISL_3227851, EPI_ISL_3227852 | Lighthouse Lab in Alderley Park | Wellcome Sanger Institute for the COVID-19 Genomics UK (COG-UK) Consortium | Jacquelyn Wynn, Mairead Hyland, The Lighthouse Lab in Alderley Park and Alex Alderton, Roberto Amato, Jeffrey Barrett, Sonia Goncalves, Ewan Harrison, David K. Jackson, Ian Johnston, Dominic Kwiatkowski, Cordelia Langford, John Sillitoe on behalf of the Wellcome Sanger Institute COVID-19 Surveillance Team |
| EPI_ISL_3227853 | Lighthouse Laboratory Plymouth | Wellcome Sanger Institute for the COVID-19 Genomics UK (COG-UK) Consortium | Lighthouse Laboratory Plymouth and Alex Alderton, Roberto Amato, Jeffrey Barrett, Sonia Goncalves, Ewan Harrison, David K. Jackson, Ian Johnston, Dominic Kwiatkowski, Cordelia Langford, John Sillitoe on behalf of the Wellcome Sanger Institute COVID-19 Surveillance Team |
| EPI_ISL_3227854, EPI_ISL_3227855, EPI_ISL_3227856, EPI_ISL_3227857 | Lighthouse Lab in Alderley Park | Wellcome Sanger Institute for the COVID-19 Genomics UK (COG-UK) Consortium | Jacquelyn Wynn, Mairead Hyland, The Lighthouse Lab in Alderley Park and Alex Alderton, Roberto Amato, Jeffrey Barrett, Sonia Goncalves, Ewan Harrison, David K. Jackson, Ian Johnston, Dominic Kwiatkowski, Cordelia Langford, John Sillitoe on behalf of the Wellcome Sanger Institute COVID-19 Surveillance Team |
| EPI_ISL_3227858 | Lighthouse Laboratory Plymouth | Wellcome Sanger Institute for the COVID-19 Genomics UK (COG-UK) Consortium | Lighthouse Laboratory Plymouth and Alex Alderton, Roberto Amato, Jeffrey Barrett, Sonia Goncalves, Ewan Harrison, David K. Jackson, Ian Johnston, Dominic Kwiatkowski, Cordelia Langford, John Sillitoe on behalf of the Wellcome Sanger Institute COVID-19 Surveillance Team |
| EPI_ISL_3227859, EPI_ISL_3227860, EPI_ISL_3227861, EPI_ISL_3227862 | Lighthouse Lab in Alderley Park | Wellcome Sanger Institute for the COVID-19 Genomics UK (COG-UK) Consortium | Jacquelyn Wynn, Mairead Hyland, The Lighthouse Lab in Alderley Park and Alex Alderton, Roberto Amato, Jeffrey Barrett, Sonia Goncalves, Ewan Harrison, David K. Jackson, Ian Johnston, Dominic Kwiatkowski, Cordelia Langford, John Sillitoe on behalf of the Wellcome Sanger Institute COVID-19 Surveillance Team |
| EPI_ISL_3227863, EPI_ISL_3227864, EPI_ISL_3227865 | Lighthouse Laboratory Plymouth | Wellcome Sanger Institute for the COVID-19 Genomics UK (COG-UK) Consortium | Lighthouse Laboratory Plymouth and Alex Alderton, Roberto Amato, Jeffrey Barrett, Sonia Goncalves, Ewan Harrison, David K. Jackson, Ian Johnston, Dominic Kwiatkowski, Cordelia Langford, John Sillitoe on behalf of the Wellcome Sanger Institute COVID-19 Surveillance Team |
| EPI_ISL_3227866, EPI_ISL_3227867, EPI_ISL_3227868, EPI_ISL_3227869, EPI_ISL_3227870 | Lighthouse Lab in Alderley Park | Wellcome Sanger Institute for the COVID-19 Genomics UK (COG-UK) Consortium | Jacquelyn Wynn, Mairead Hyland, The Lighthouse Lab in Alderley Park and Alex Alderton, Roberto Amato, Jeffrey Barrett, Sonia Goncalves, Ewan Harrison, David K. Jackson, Ian Johnston, Dominic Kwiatkowski, Cordelia Langford, John Sillitoe on behalf of the Wellcome Sanger Institute COVID-19 Surveillance Team |
| EPI_ISL_3227871 | Lighthouse Laboratory Plymouth | Wellcome Sanger Institute for the COVID-19 Genomics UK (COG-UK) Consortium | Lighthouse Laboratory Plymouth and Alex Alderton, Roberto Amato, Jeffrey Barrett, Sonia Goncalves, Ewan Harrison, David K. Jackson, Ian Johnston, Dominic Kwiatkowski, Cordelia Langford, John Sillitoe on behalf of the Wellcome Sanger Institute COVID-19 Surveillance Team |
| EPI_ISL_3227872 | Lighthouse Lab in Alderley Park | Wellcome Sanger Institute for the COVID-19 Genomics UK (COG-UK) Consortium | Jacquelyn Wynn, Mairead Hyland, The Lighthouse Lab in Alderley Park and Alex Alderton, Roberto Amato, Jeffrey Barrett, Sonia Goncalves, Ewan Harrison, David K. Jackson, Ian Johnston, Dominic Kwiatkowski, Cordelia Langford, John Sillitoe on behalf of the Wellcome Sanger Institute COVID-19 Surveillance Team |
| EPI_ISL_3227873 | Lighthouse Laboratory Plymouth | Wellcome Sanger Institute for the COVID-19 Genomics UK (COG-UK) Consortium | Lighthouse Laboratory Plymouth and Alex Alderton, Roberto Amato, Jeffrey Barrett, Sonia Goncalves, Ewan Harrison, David K. Jackson, Ian Johnston, Dominic Kwiatkowski, Cordelia Langford, John Sillitoe on behalf of the Wellcome Sanger Institute COVID-19 Surveillance Team |
| EPI_ISL_3227874 | Lighthouse Lab in Alderley Park | Wellcome Sanger Institute for the COVID-19 Genomics UK (COG-UK) Consortium | Jacquelyn Wynn, Mairead Hyland, The Lighthouse Lab in Alderley Park and Alex Alderton, Roberto Amato, Jeffrey Barrett, Sonia Goncalves, Ewan Harrison, David K. Jackson, Ian Johnston, Dominic Kwiatkowski, Cordelia Langford, John Sillitoe on behalf of the Wellcome Sanger Institute COVID-19 Surveillance Team |
| EPI_ISL_3227875 | Berkshire and Surrey Pathology Services Lighthouse Laboratory | Wellcome Sanger Institute for the COVID-19 Genomics UK (COG-UK) Consortium | Berkshire and Surrey Pathology Services Lighthouse Laboratory and Alex Alderton, Roberto Amato, Jeffrey Barrett, Sonia Goncalves, Ewan Harrison, David K. Jackson, Ian Johnston, Dominic Kwiatkowski, Cordelia Langford, John Sillitoe on behalf of the Wellcome Sanger Institute COVID-19 Surveillance Team |
| EPI_ISL_3227876 | Lighthouse Lab in Alderley Park | Wellcome Sanger Institute for the COVID-19 Genomics UK (COG-UK) Consortium | Jacquelyn Wynn, Mairead Hyland, The Lighthouse Lab in Alderley Park and Alex Alderton, Roberto Amato, Jeffrey Barrett, Sonia Goncalves, Ewan Harrison, David K. Jackson, Ian Johnston, Dominic Kwiatkowski, Cordelia Langford, John Sillitoe on behalf of the Wellcome Sanger Institute COVID-19 Surveillance Team |
| EPI_ISL_3227877 | Berkshire and Surrey Pathology Services Lighthouse Laboratory | Wellcome Sanger Institute for the COVID-19 Genomics UK (COG-UK) Consortium | Berkshire and Surrey Pathology Services Lighthouse Laboratory and Alex Alderton, Roberto Amato, Jeffrey Barrett, Sonia Goncalves, Ewan Harrison, David K. Jackson, Ian Johnston, Dominic Kwiatkowski, Cordelia Langford, John Sillitoe on behalf of the Wellcome Sanger Institute COVID-19 Surveillance Team |
| EPI_ISL_3227878 | Lighthouse Lab in Alderley Park | Wellcome Sanger Institute for the COVID-19 Genomics UK (COG-UK) Consortium | Jacquelyn Wynn, Mairead Hyland, The Lighthouse Lab in Alderley Park and Alex Alderton, Roberto Amato, Jeffrey Barrett, Sonia Goncalves, Ewan Harrison, David K. Jackson, Ian Johnston, Dominic Kwiatkowski, Cordelia Langford, John Sillitoe on behalf of the Wellcome Sanger Institute COVID-19 Surveillance Team |
| EPI_ISL_3227879, EPI_ISL_3227880, EPI_ISL_3227881 | Berkshire and Surrey Pathology Services Lighthouse Laboratory | Wellcome Sanger Institute for the COVID-19 Genomics UK (COG-UK) Consortium | Berkshire and Surrey Pathology Services Lighthouse Laboratory and Alex Alderton, Roberto Amato, Jeffrey Barrett, Sonia Goncalves, Ewan Harrison, David K. Jackson, Ian Johnston, Dominic Kwiatkowski, Cordelia Langford, John Sillitoe on behalf of the Wellcome Sanger Institute COVID-19 Surveillance Team |
| EPI_ISL_3227882, EPI_ISL_3227883 | Lighthouse Lab in Alderley Park | Wellcome Sanger Institute for the COVID-19 Genomics UK (COG-UK) Consortium | Jacquelyn Wynn, Mairead Hyland, The Lighthouse Lab in Alderley Park and Alex Alderton, Roberto Amato, Jeffrey Barrett, Sonia Goncalves, Ewan Harrison, David K. Jackson, Ian Johnston, Dominic Kwiatkowski, Cordelia Langford, John Sillitoe on behalf of the Wellcome Sanger Institute COVID-19 Surveillance Team |
| EPI_ISL_3227884, EPI_ISL_3227885, EPI_ISL_3227886, EPI_ISL_3227887, EPI_ISL_3227888 | Berkshire and Surrey Pathology Services Lighthouse Laboratory | Wellcome Sanger Institute for the COVID-19 Genomics UK (COG-UK) Consortium | Berkshire and Surrey Pathology Services Lighthouse Laboratory and Alex Alderton, Roberto Amato, Jeffrey Barrett, Sonia Goncalves, Ewan Harrison, David K. Jackson, Ian Johnston, Dominic Kwiatkowski, Cordelia Langford, John Sillitoe on behalf of the Wellcome Sanger Institute COVID-19 Surveillance Team |
| EPI_ISL_3227889, EPI_ISL_3227890, EPI_ISL_3227891 | Lighthouse Lab in Alderley Park | Wellcome Sanger Institute for the COVID-19 Genomics UK (COG-UK) Consortium | Jacquelyn Wynn, Mairead Hyland, The Lighthouse Lab in Alderley Park and Alex Alderton, Roberto Amato, Jeffrey Barrett, Sonia Goncalves, Ewan Harrison, David K. Jackson, Ian Johnston, Dominic Kwiatkowski, Cordelia Langford, John Sillitoe on behalf of the Wellcome Sanger Institute COVID-19 Surveillance Team |
| EPI_ISL_3227892 | Berkshire and Surrey Pathology Services Lighthouse Laboratory | Wellcome Sanger Institute for the COVID-19 Genomics UK (COG-UK) Consortium | Berkshire and Surrey Pathology Services Lighthouse Laboratory and Alex Alderton, Roberto Amato, Jeffrey Barrett, Sonia Goncalves, Ewan Harrison, David K. Jackson, Ian Johnston, Dominic Kwiatkowski, Cordelia Langford, John Sillitoe on behalf of the Wellcome Sanger Institute COVID-19 Surveillance Team |
| EPI_ISL_3227893 | Lighthouse Lab in Alderley Park | Wellcome Sanger Institute for the COVID-19 Genomics UK (COG-UK) Consortium | Jacquelyn Wynn, Mairead Hyland, The Lighthouse Lab in Alderley Park and Alex Alderton, Roberto Amato, Jeffrey Barrett, Sonia Goncalves, Ewan Harrison, David K. Jackson, Ian Johnston, Dominic Kwiatkowski, Cordelia Langford, John Sillitoe on behalf of the Wellcome Sanger Institute COVID-19 Surveillance Team |
| EPI_ISL_3227894, | Berkshire and Surrey Pathology Services Lighthouse Laboratory | Wellcome Sanger Institute for the COVID-19 Genomics UK (COG-UK) | Berkshire and Surrey Pathology Services Lighthouse Laboratory and Alex Alderton, Roberto Amato, Jeffrey Barrett, Sonia Goncalves, Ewan Harrison, David K. Jackson, Ian Johnston, Dominic Kwiatkowski, Cordelia Langford, John Sillitoe on behalf |

| | | | |
|---|---|---|---|
| EPI_ISL_3227895 | | Consortium | of the Wellcome Sanger Institute COVID-19 Surveillance Team |
| EPI_ISL_3227896 | Lighthouse Lab in Alderley Park | Wellcome Sanger Institute for the COVID-19 Genomics UK (COG-UK) Consortium | Jacquelyn Wynn, Mairead Hyland, The Lighthouse Lab in Alderley Park and Alex Alderton, Roberto Amato, Jeffrey Barrett, Sonia Goncalves, Ewan Harrison, David K. Jackson, Ian Johnston, Dominic Kwiatkowski, Cordelia Langford, John Sillitoe on behalf of the Wellcome Sanger Institute COVID-19 Surveillance Team |
| EPI_ISL_3227897, EPI_ISL_3227898 | Berkshire and Surrey Pathology Services Lighthouse Laboratory | Wellcome Sanger Institute for the COVID-19 Genomics UK (COG-UK) Consortium | Berkshire and Surrey Pathology Services Lighthouse Laboratory and Alex Alderton, Roberto Amato, Jeffrey Barrett, Sonia Goncalves, Ewan Harrison, David K. Jackson, Ian Johnston, Dominic Kwiatkowski, Cordelia Langford, John Sillitoe on behalf of the Wellcome Sanger Institute COVID-19 Surveillance Team |
| EPI_ISL_3227899, EPI_ISL_3227900, EPI_ISL_3227901, EPI_ISL_3227902 | Lighthouse Lab in Alderley Park | Wellcome Sanger Institute for the COVID-19 Genomics UK (COG-UK) Consortium | Jacquelyn Wynn, Mairead Hyland, The Lighthouse Lab in Alderley Park and Alex Alderton, Roberto Amato, Jeffrey Barrett, Sonia Goncalves, Ewan Harrison, David K. Jackson, Ian Johnston, Dominic Kwiatkowski, Cordelia Langford, John Sillitoe on behalf of the Wellcome Sanger Institute COVID-19 Surveillance Team |
| EPI_ISL_3227903 | Berkshire and Surrey Pathology Services Lighthouse Laboratory | Wellcome Sanger Institute for the COVID-19 Genomics UK (COG-UK) Consortium | Berkshire and Surrey Pathology Services Lighthouse Laboratory and Alex Alderton, Roberto Amato, Jeffrey Barrett, Sonia Goncalves, Ewan Harrison, David K. Jackson, Ian Johnston, Dominic Kwiatkowski, Cordelia Langford, John Sillitoe on behalf of the Wellcome Sanger Institute COVID-19 Surveillance Team |
| EPI_ISL_3227904, EPI_ISL_3227905, EPI_ISL_3227906 | Lighthouse Lab in Alderley Park | Wellcome Sanger Institute for the COVID-19 Genomics UK (COG-UK) Consortium | Jacquelyn Wynn, Mairead Hyland, The Lighthouse Lab in Alderley Park and Alex Alderton, Roberto Amato, Jeffrey Barrett, Sonia Goncalves, Ewan Harrison, David K. Jackson, Ian Johnston, Dominic Kwiatkowski, Cordelia Langford, John Sillitoe on behalf of the Wellcome Sanger Institute COVID-19 Surveillance Team |
| EPI_ISL_3227907, EPI_ISL_3227908 | Berkshire and Surrey Pathology Services Lighthouse Laboratory | Wellcome Sanger Institute for the COVID-19 Genomics UK (COG-UK) Consortium | Berkshire and Surrey Pathology Services Lighthouse Laboratory and Alex Alderton, Roberto Amato, Jeffrey Barrett, Sonia Goncalves, Ewan Harrison, David K. Jackson, Ian Johnston, Dominic Kwiatkowski, Cordelia Langford, John Sillitoe on behalf of the Wellcome Sanger Institute COVID-19 Surveillance Team |
| EPI_ISL_3227909 | Lighthouse Lab in Alderley Park | Wellcome Sanger Institute for the COVID-19 Genomics UK (COG-UK) Consortium | Jacquelyn Wynn, Mairead Hyland, The Lighthouse Lab in Alderley Park and Alex Alderton, Roberto Amato, Jeffrey Barrett, Sonia Goncalves, Ewan Harrison, David K. Jackson, Ian Johnston, Dominic Kwiatkowski, Cordelia Langford, John Sillitoe on behalf of the Wellcome Sanger Institute COVID-19 Surveillance Team |
| EPI_ISL_3227910 | Berkshire and Surrey Pathology Services Lighthouse Laboratory | Wellcome Sanger Institute for the COVID-19 Genomics UK (COG-UK) Consortium | Berkshire and Surrey Pathology Services Lighthouse Laboratory and Alex Alderton, Roberto Amato, Jeffrey Barrett, Sonia Goncalves, Ewan Harrison, David K. Jackson, Ian Johnston, Dominic Kwiatkowski, Cordelia Langford, John Sillitoe on behalf of the Wellcome Sanger Institute COVID-19 Surveillance Team |
| EPI_ISL_3227911, EPI_ISL_3227912 | Lighthouse Lab in Alderley Park | Wellcome Sanger Institute for the COVID-19 Genomics UK (COG-UK) Consortium | Jacquelyn Wynn, Mairead Hyland, The Lighthouse Lab in Alderley Park and Alex Alderton, Roberto Amato, Jeffrey Barrett, Sonia Goncalves, Ewan Harrison, David K. Jackson, Ian Johnston, Dominic Kwiatkowski, Cordelia Langford, John Sillitoe on behalf of the Wellcome Sanger Institute COVID-19 Surveillance Team |
| EPI_ISL_3227913 | Berkshire and Surrey Pathology Services Lighthouse Laboratory | Wellcome Sanger Institute for the COVID-19 Genomics UK (COG-UK) Consortium | Berkshire and Surrey Pathology Services Lighthouse Laboratory and Alex Alderton, Roberto Amato, Jeffrey Barrett, Sonia Goncalves, Ewan Harrison, David K. Jackson, Ian Johnston, Dominic Kwiatkowski, Cordelia Langford, John Sillitoe on behalf of the Wellcome Sanger Institute COVID-19 Surveillance Team |
| EPI_ISL_3227914 | Lighthouse Lab in Alderley Park | Wellcome Sanger Institute for the COVID-19 Genomics UK (COG-UK) Consortium | Jacquelyn Wynn, Mairead Hyland, The Lighthouse Lab in Alderley Park and Alex Alderton, Roberto Amato, Jeffrey Barrett, Sonia Goncalves, Ewan Harrison, David K. Jackson, Ian Johnston, Dominic Kwiatkowski, Cordelia Langford, John Sillitoe on behalf of the Wellcome Sanger Institute COVID-19 Surveillance Team |
| EPI_ISL_3227915 | Berkshire and Surrey Pathology Services Lighthouse Laboratory | Wellcome Sanger Institute for the COVID-19 Genomics UK (COG-UK) Consortium | Berkshire and Surrey Pathology Services Lighthouse Laboratory and Alex Alderton, Roberto Amato, Jeffrey Barrett, Sonia Goncalves, Ewan Harrison, David K. Jackson, Ian Johnston, Dominic Kwiatkowski, Cordelia Langford, John Sillitoe on behalf of the Wellcome Sanger Institute COVID-19 Surveillance Team |
| EPI_ISL_3227916 | Lighthouse Lab in Alderley Park | Wellcome Sanger Institute for the COVID-19 Genomics UK (COG-UK) Consortium | Jacquelyn Wynn, Mairead Hyland, The Lighthouse Lab in Alderley Park and Alex Alderton, Roberto Amato, Jeffrey Barrett, Sonia Goncalves, Ewan Harrison, David K. Jackson, Ian Johnston, Dominic Kwiatkowski, Cordelia Langford, John Sillitoe on behalf of the Wellcome Sanger Institute COVID-19 Surveillance Team |
| EPI_ISL_3227917, EPI_ISL_3227918 | Berkshire and Surrey Pathology Services Lighthouse Laboratory | Wellcome Sanger Institute for the COVID-19 Genomics UK (COG-UK) Consortium | Berkshire and Surrey Pathology Services Lighthouse Laboratory and Alex Alderton, Roberto Amato, Jeffrey Barrett, Sonia Goncalves, Ewan Harrison, David K. Jackson, Ian Johnston, Dominic Kwiatkowski, Cordelia Langford, John Sillitoe on behalf of the Wellcome Sanger Institute COVID-19 Surveillance Team |
| EPI_ISL_3227919 | Lighthouse Lab in Alderley Park | Wellcome Sanger Institute for the COVID-19 Genomics UK (COG-UK) Consortium | Jacquelyn Wynn, Mairead Hyland, The Lighthouse Lab in Alderley Park and Alex Alderton, Roberto Amato, Jeffrey Barrett, Sonia Goncalves, Ewan Harrison, David K. Jackson, Ian Johnston, Dominic Kwiatkowski, Cordelia Langford, John Sillitoe on behalf of the Wellcome Sanger Institute COVID-19 Surveillance Team |
| EPI_ISL_3227920 | Berkshire and Surrey Pathology Services Lighthouse Laboratory | Wellcome Sanger Institute for the COVID-19 Genomics UK (COG-UK) Consortium | Berkshire and Surrey Pathology Services Lighthouse Laboratory and Alex Alderton, Roberto Amato, Jeffrey Barrett, Sonia Goncalves, Ewan Harrison, David K. Jackson, Ian Johnston, Dominic Kwiatkowski, Cordelia Langford, John Sillitoe on behalf of the Wellcome Sanger Institute COVID-19 Surveillance Team |
| EPI_ISL_3227921 | Lighthouse Lab in Alderley Park | Wellcome Sanger Institute for the COVID-19 Genomics UK (COG-UK) Consortium | Jacquelyn Wynn, Mairead Hyland, The Lighthouse Lab in Alderley Park and Alex Alderton, Roberto Amato, Jeffrey Barrett, Sonia Goncalves, Ewan Harrison, David K. Jackson, Ian Johnston, Dominic Kwiatkowski, Cordelia Langford, John Sillitoe on behalf of the Wellcome Sanger Institute COVID-19 Surveillance Team |
| EPI_ISL_3227922, EPI_ISL_3227923 | Berkshire and Surrey Pathology Services Lighthouse Laboratory | Wellcome Sanger Institute for the COVID-19 Genomics UK (COG-UK) Consortium | Berkshire and Surrey Pathology Services Lighthouse Laboratory and Alex Alderton, Roberto Amato, Jeffrey Barrett, Sonia Goncalves, Ewan Harrison, David K. Jackson, Ian Johnston, Dominic Kwiatkowski, Cordelia Langford, John Sillitoe on behalf of the Wellcome Sanger Institute COVID-19 Surveillance Team |
| EPI_ISL_3227924, EPI_ISL_3227925, EPI_ISL_3227926, EPI_ISL_3227927, EPI_ISL_3227928, EPI_ISL_3227929, EPI_ISL_3227930, EPI_ISL_3227931 | Lighthouse Lab in Alderley Park | Wellcome Sanger Institute for the COVID-19 Genomics UK (COG-UK) Consortium | Jacquelyn Wynn, Mairead Hyland, The Lighthouse Lab in Alderley Park and Alex Alderton, Roberto Amato, Jeffrey Barrett, Sonia Goncalves, Ewan Harrison, David K. Jackson, Ian Johnston, Dominic Kwiatkowski, Cordelia Langford, John Sillitoe on behalf of the Wellcome Sanger Institute COVID-19 Surveillance Team |
| EPI_ISL_3227932 | Berkshire and Surrey Pathology Services Lighthouse Laboratory | Wellcome Sanger Institute for the COVID-19 Genomics UK (COG-UK) Consortium | Berkshire and Surrey Pathology Services Lighthouse Laboratory and Alex Alderton, Roberto Amato, Jeffrey Barrett, Sonia Goncalves, Ewan Harrison, David K. Jackson, Ian Johnston, Dominic Kwiatkowski, Cordelia Langford, John Sillitoe on behalf of the Wellcome Sanger Institute COVID-19 Surveillance Team |
| EPI_ISL_3227933, EPI_ISL_3227934, EPI_ISL_3227935 | Lighthouse Lab in Alderley Park | Wellcome Sanger Institute for the COVID-19 Genomics UK (COG-UK) Consortium | Jacquelyn Wynn, Mairead Hyland, The Lighthouse Lab in Alderley Park and Alex Alderton, Roberto Amato, Jeffrey Barrett, Sonia Goncalves, Ewan Harrison, David K. Jackson, Ian Johnston, Dominic Kwiatkowski, Cordelia Langford, John Sillitoe on behalf of the Wellcome Sanger Institute COVID-19 Surveillance Team |
| EPI_ISL_3227936 | Berkshire and Surrey Pathology Services Lighthouse Laboratory | Wellcome Sanger Institute for the COVID-19 Genomics UK (COG-UK) Consortium | Berkshire and Surrey Pathology Services Lighthouse Laboratory and Alex Alderton, Roberto Amato, Jeffrey Barrett, Sonia Goncalves, Ewan Harrison, David K. Jackson, Ian Johnston, Dominic Kwiatkowski, Cordelia Langford, John Sillitoe on behalf of the Wellcome Sanger Institute COVID-19 Surveillance Team |
| EPI_ISL_3227937 | Lighthouse Lab in Alderley Park | Wellcome Sanger Institute for the COVID-19 Genomics UK (COG-UK) Consortium | Jacquelyn Wynn, Mairead Hyland, The Lighthouse Lab in Alderley Park and Alex Alderton, Roberto Amato, Jeffrey Barrett, Sonia Goncalves, Ewan Harrison, David K. Jackson, Ian Johnston, Dominic Kwiatkowski, Cordelia Langford, John Sillitoe on behalf of the Wellcome Sanger Institute COVID-19 Surveillance Team |
| EPI_ISL_3227938, EPI_ISL_3227939 | Berkshire and Surrey Pathology Services Lighthouse Laboratory | Wellcome Sanger Institute for the COVID-19 Genomics UK (COG-UK) Consortium | Berkshire and Surrey Pathology Services Lighthouse Laboratory and Alex Alderton, Roberto Amato, Jeffrey Barrett, Sonia Goncalves, Ewan Harrison, David K. Jackson, Ian Johnston, Dominic Kwiatkowski, Cordelia Langford, John Sillitoe on behalf of the Wellcome Sanger Institute COVID-19 Surveillance Team |
| EPI_ISL_3227940, EPI_ISL_3227941 | Lighthouse Lab in Alderley Park | Wellcome Sanger Institute for the COVID-19 Genomics UK (COG-UK) Consortium | Jacquelyn Wynn, Mairead Hyland, The Lighthouse Lab in Alderley Park and Alex Alderton, Roberto Amato, Jeffrey Barrett, Sonia Goncalves, Ewan Harrison, David K. Jackson, Ian Johnston, Dominic Kwiatkowski, Cordelia Langford, John Sillitoe on behalf of the Wellcome Sanger Institute COVID-19 Surveillance Team |
| EPI_ISL_3227942, EPI_ISL_3227943, EPI_ISL_3227944 | Berkshire and Surrey Pathology Services Lighthouse Laboratory | Wellcome Sanger Institute for the COVID-19 Genomics UK (COG-UK) Consortium | Berkshire and Surrey Pathology Services Lighthouse Laboratory and Alex Alderton, Roberto Amato, Jeffrey Barrett, Sonia Goncalves, Ewan Harrison, David K. Jackson, Ian Johnston, Dominic Kwiatkowski, Cordelia Langford, John Sillitoe on behalf of the Wellcome Sanger Institute COVID-19 Surveillance Team |
| EPI_ISL_3227945, EPI_ISL_3227946, EPI_ISL_3227947 | Lighthouse Lab in Alderley Park | Wellcome Sanger Institute for the COVID-19 Genomics UK (COG-UK) Consortium | Jacquelyn Wynn, Mairead Hyland, The Lighthouse Lab in Alderley Park and Alex Alderton, Roberto Amato, Jeffrey Barrett, Sonia Goncalves, Ewan Harrison, David K. Jackson, Ian Johnston, Dominic Kwiatkowski, Cordelia Langford, John Sillitoe on behalf of the Wellcome Sanger Institute COVID-19 Surveillance Team |
| EPI_ISL_3227948, EPI_ISL_3227949 | Berkshire and Surrey Pathology Services Lighthouse Laboratory | Wellcome Sanger Institute for the COVID-19 Genomics UK (COG-UK) Consortium | Berkshire and Surrey Pathology Services Lighthouse Laboratory and Alex Alderton, Roberto Amato, Jeffrey Barrett, Sonia Goncalves, Ewan Harrison, David K. Jackson, Ian Johnston, Dominic Kwiatkowski, Cordelia Langford, John Sillitoe on behalf of the Wellcome Sanger Institute COVID-19 Surveillance Team |
| EPI_ISL_3227950 | Lighthouse Lab in Alderley Park | Wellcome Sanger Institute for the COVID-19 Genomics UK (COG-UK) Consortium | Jacquelyn Wynn, Mairead Hyland, The Lighthouse Lab in Alderley Park and Alex Alderton, Roberto Amato, Jeffrey Barrett, Sonia Goncalves, Ewan Harrison, David K. Jackson, Ian Johnston, Dominic Kwiatkowski, Cordelia Langford, John Sillitoe on behalf of the Wellcome Sanger Institute COVID-19 Surveillance Team |
| EPI_ISL_3227951 | Berkshire and Surrey Pathology Services Lighthouse Laboratory | Wellcome Sanger Institute for the COVID-19 Genomics UK (COG-UK) Consortium | Berkshire and Surrey Pathology Services Lighthouse Laboratory and Alex Alderton, Roberto Amato, Jeffrey Barrett, Sonia Goncalves, Ewan Harrison, David K. Jackson, Ian Johnston, Dominic Kwiatkowski, Cordelia Langford, John Sillitoe on behalf of the Wellcome Sanger Institute COVID-19 Surveillance Team |
| EPI_ISL_3227952 | Lighthouse Lab in Alderley Park | Wellcome Sanger Institute for the COVID-19 Genomics UK (COG-UK) Consortium | Jacquelyn Wynn, Mairead Hyland, The Lighthouse Lab in Alderley Park and Alex Alderton, Roberto Amato, Jeffrey Barrett, Sonia Goncalves, Ewan Harrison, David K. Jackson, Ian Johnston, Dominic Kwiatkowski, Cordelia Langford, John Sillitoe on behalf of the Wellcome Sanger Institute COVID-19 Surveillance Team |
| EPI_ISL_3227953 | Berkshire and Surrey Pathology Services Lighthouse Laboratory | Wellcome Sanger Institute for the COVID-19 Genomics UK (COG-UK) Consortium | Berkshire and Surrey Pathology Services Lighthouse Laboratory and Alex Alderton, Roberto Amato, Jeffrey Barrett, Sonia Goncalves, Ewan Harrison, David K. Jackson, Ian Johnston, Dominic Kwiatkowski, Cordelia Langford, John Sillitoe on behalf of the Wellcome Sanger Institute COVID-19 Surveillance Team |
| EPI_ISL_3227954, EPI_ISL_3227955, EPI_ISL_3227956 | Lighthouse Lab in Alderley Park | Wellcome Sanger Institute for the COVID-19 Genomics UK (COG-UK) Consortium | Jacquelyn Wynn, Mairead Hyland, The Lighthouse Lab in Alderley Park and Alex Alderton, Roberto Amato, Jeffrey Barrett, Sonia Goncalves, Ewan Harrison, David K. Jackson, Ian Johnston, Dominic Kwiatkowski, Cordelia Langford, John Sillitoe on behalf of the Wellcome Sanger Institute COVID-19 Surveillance Team |
| EPI_ISL_3227957 | Berkshire and Surrey Pathology Services Lighthouse Laboratory | Wellcome Sanger Institute for the COVID-19 Genomics UK (COG-UK) Consortium | Berkshire and Surrey Pathology Services Lighthouse Laboratory and Alex Alderton, Roberto Amato, Jeffrey Barrett, Sonia Goncalves, Ewan Harrison, David K. Jackson, Ian Johnston, Dominic Kwiatkowski, Cordelia Langford, John Sillitoe on behalf of the Wellcome Sanger Institute COVID-19 Surveillance Team |
| EPI_ISL_3227958, EPI_ISL_3227959 | Lighthouse Lab in Alderley Park | Wellcome Sanger Institute for the COVID-19 Genomics UK (COG-UK) Consortium | Jacquelyn Wynn, Mairead Hyland, The Lighthouse Lab in Alderley Park and Alex Alderton, Roberto Amato, Jeffrey Barrett, Sonia Goncalves, Ewan Harrison, David K. Jackson, Ian Johnston, Dominic Kwiatkowski, Cordelia Langford, John Sillitoe on behalf of the Wellcome Sanger Institute COVID-19 Surveillance Team |
| EPI_ISL_3227960 | Berkshire and Surrey Pathology Services Lighthouse Laboratory | Wellcome Sanger Institute for the COVID-19 Genomics UK (COG-UK) Consortium | Berkshire and Surrey Pathology Services Lighthouse Laboratory and Alex Alderton, Roberto Amato, Jeffrey Barrett, Sonia Goncalves, Ewan Harrison, David K. Jackson, Ian Johnston, Dominic Kwiatkowski, Cordelia Langford, John Sillitoe on behalf of the Wellcome Sanger Institute COVID-19 Surveillance Team |
| EPI_ISL_3227961, EPI_ISL_3227962 | Lighthouse Lab in Alderley Park | Wellcome Sanger Institute for the COVID-19 Genomics UK (COG-UK) Consortium | Jacquelyn Wynn, Mairead Hyland, The Lighthouse Lab in Alderley Park and Alex Alderton, Roberto Amato, Jeffrey Barrett, Sonia Goncalves, Ewan Harrison, David K. Jackson, Ian Johnston, Dominic Kwiatkowski, Cordelia Langford, John Sillitoe on behalf of the Wellcome Sanger Institute COVID-19 Surveillance Team |
| EPI_ISL_3227963 | Berkshire and Surrey Pathology Services Lighthouse Laboratory | Wellcome Sanger Institute for the COVID-19 Genomics UK (COG-UK) Consortium | Berkshire and Surrey Pathology Services Lighthouse Laboratory and Alex Alderton, Roberto Amato, Jeffrey Barrett, Sonia Goncalves, Ewan Harrison, David K. Jackson, Ian Johnston, Dominic Kwiatkowski, Cordelia Langford, John Sillitoe on behalf of the Wellcome Sanger Institute COVID-19 Surveillance Team |
| EPI_ISL_3227964, EPI_ISL_3227965 | Lighthouse Lab in Alderley Park | Wellcome Sanger Institute for the COVID-19 Genomics UK (COG-UK) Consortium | Jacquelyn Wynn, Mairead Hyland, The Lighthouse Lab in Alderley Park and Alex Alderton, Roberto Amato, Jeffrey Barrett, Sonia Goncalves, Ewan Harrison, David K. Jackson, Ian Johnston, Dominic Kwiatkowski, Cordelia Langford, John Sillitoe on behalf of the Wellcome Sanger Institute COVID-19 Surveillance Team |
| EPI_ISL_3227966, | Berkshire and Surrey Pathology Services Lighthouse Laboratory | Wellcome Sanger Institute for the COVID-19 Genomics UK (COG-UK) | Berkshire and Surrey Pathology Services Lighthouse Laboratory and Alex Alderton, Roberto Amato, Jeffrey Barrett, Sonia Goncalves, Ewan Harrison, David K. Jackson, Ian Johnston, Dominic Kwiatkowski, Cordelia Langford, John Sillitoe on behalf |

| | | | |
|---|---|---|---|
| EPI_ISL_3227967 | | Consortium | of the Wellcome Sanger Institute COVID-19 Surveillance Team |
| EPI_ISL_3227968, EPI_ISL_3227969, EPI_ISL_3227970 | Lighthouse Lab in Alderley Park | Wellcome Sanger Institute for the COVID-19 Genomics UK (COG-UK) Consortium | Jacquelyn Wynn, Mairead Hyland, The Lighthouse Lab in Alderley Park and Alex Alderton, Roberto Amato, Jeffrey Barrett, Sonia Goncalves, Ewan Harrison, David K. Jackson, Ian Johnston, Dominic Kwiatkowski, Cordelia Langford, John Sillitoe on behalf of the Wellcome Sanger Institute COVID-19 Surveillance Team |
| EPI_ISL_3227971 | Berkshire and Surrey Pathology Services Lighthouse Laboratory | Wellcome Sanger Institute for the COVID-19 Genomics UK (COG-UK) Consortium | Berkshire and Surrey Pathology Services Lighthouse Laboratory and Alex Alderton, Roberto Amato, Jeffrey Barrett, Sonia Goncalves, Ewan Harrison, David K. Jackson, Ian Johnston, Dominic Kwiatkowski, Cordelia Langford, John Sillitoe on behalf of the Wellcome Sanger Institute COVID-19 Surveillance Team |
| EPI_ISL_3227972 | Lighthouse Lab in Alderley Park | Wellcome Sanger Institute for the COVID-19 Genomics UK (COG-UK) Consortium | Jacquelyn Wynn, Mairead Hyland, The Lighthouse Lab in Alderley Park and Alex Alderton, Roberto Amato, Jeffrey Barrett, Sonia Goncalves, Ewan Harrison, David K. Jackson, Ian Johnston, Dominic Kwiatkowski, Cordelia Langford, John Sillitoe on behalf of the Wellcome Sanger Institute COVID-19 Surveillance Team |
| EPI_ISL_3227973, EPI_ISL_3227974 | Berkshire and Surrey Pathology Services Lighthouse Laboratory | Wellcome Sanger Institute for the COVID-19 Genomics UK (COG-UK) Consortium | Berkshire and Surrey Pathology Services Lighthouse Laboratory and Alex Alderton, Roberto Amato, Jeffrey Barrett, Sonia Goncalves, Ewan Harrison, David K. Jackson, Ian Johnston, Dominic Kwiatkowski, Cordelia Langford, John Sillitoe on behalf of the Wellcome Sanger Institute COVID-19 Surveillance Team |
| EPI_ISL_3227975, EPI_ISL_3227976, EPI_ISL_3227977 | Lighthouse Lab in Alderley Park | Wellcome Sanger Institute for the COVID-19 Genomics UK (COG-UK) Consortium | Jacquelyn Wynn, Mairead Hyland, The Lighthouse Lab in Alderley Park and Alex Alderton, Roberto Amato, Jeffrey Barrett, Sonia Goncalves, Ewan Harrison, David K. Jackson, Ian Johnston, Dominic Kwiatkowski, Cordelia Langford, John Sillitoe on behalf of the Wellcome Sanger Institute COVID-19 Surveillance Team |
| EPI_ISL_3227978 | Berkshire and Surrey Pathology Services Lighthouse Laboratory | Wellcome Sanger Institute for the COVID-19 Genomics UK (COG-UK) Consortium | Berkshire and Surrey Pathology Services Lighthouse Laboratory and Alex Alderton, Roberto Amato, Jeffrey Barrett, Sonia Goncalves, Ewan Harrison, David K. Jackson, Ian Johnston, Dominic Kwiatkowski, Cordelia Langford, John Sillitoe on behalf of the Wellcome Sanger Institute COVID-19 Surveillance Team |
| EPI_ISL_3227979, EPI_ISL_3227980 | Lighthouse Lab in Alderley Park | Wellcome Sanger Institute for the COVID-19 Genomics UK (COG-UK) Consortium | Jacquelyn Wynn, Mairead Hyland, The Lighthouse Lab in Alderley Park and Alex Alderton, Roberto Amato, Jeffrey Barrett, Sonia Goncalves, Ewan Harrison, David K. Jackson, Ian Johnston, Dominic Kwiatkowski, Cordelia Langford, John Sillitoe on behalf of the Wellcome Sanger Institute COVID-19 Surveillance Team |
| EPI_ISL_3227981, EPI_ISL_3227982, EPI_ISL_3227983 | Berkshire and Surrey Pathology Services Lighthouse Laboratory | Wellcome Sanger Institute for the COVID-19 Genomics UK (COG-UK) Consortium | Berkshire and Surrey Pathology Services Lighthouse Laboratory and Alex Alderton, Roberto Amato, Jeffrey Barrett, Sonia Goncalves, Ewan Harrison, David K. Jackson, Ian Johnston, Dominic Kwiatkowski, Cordelia Langford, John Sillitoe on behalf of the Wellcome Sanger Institute COVID-19 Surveillance Team |
| EPI_ISL_3227984, EPI_ISL_3227985 | Lighthouse Lab in Alderley Park | Wellcome Sanger Institute for the COVID-19 Genomics UK (COG-UK) Consortium | Jacquelyn Wynn, Mairead Hyland, The Lighthouse Lab in Alderley Park and Alex Alderton, Roberto Amato, Jeffrey Barrett, Sonia Goncalves, Ewan Harrison, David K. Jackson, Ian Johnston, Dominic Kwiatkowski, Cordelia Langford, John Sillitoe on behalf of the Wellcome Sanger Institute COVID-19 Surveillance Team |
| EPI_ISL_3227986 | Berkshire and Surrey Pathology Services Lighthouse Laboratory | Wellcome Sanger Institute for the COVID-19 Genomics UK (COG-UK) Consortium | Berkshire and Surrey Pathology Services Lighthouse Laboratory and Alex Alderton, Roberto Amato, Jeffrey Barrett, Sonia Goncalves, Ewan Harrison, David K. Jackson, Ian Johnston, Dominic Kwiatkowski, Cordelia Langford, John Sillitoe on behalf of the Wellcome Sanger Institute COVID-19 Surveillance Team |
| EPI_ISL_3227987, EPI_ISL_3227988, EPI_ISL_3227989, EPI_ISL_3227990 | Lighthouse Lab in Alderley Park | Wellcome Sanger Institute for the COVID-19 Genomics UK (COG-UK) Consortium | Jacquelyn Wynn, Mairead Hyland, The Lighthouse Lab in Alderley Park and Alex Alderton, Roberto Amato, Jeffrey Barrett, Sonia Goncalves, Ewan Harrison, David K. Jackson, Ian Johnston, Dominic Kwiatkowski, Cordelia Langford, John Sillitoe on behalf of the Wellcome Sanger Institute COVID-19 Surveillance Team |
| EPI_ISL_3227991, EPI_ISL_3227992 | Berkshire and Surrey Pathology Services Lighthouse Laboratory | Wellcome Sanger Institute for the COVID-19 Genomics UK (COG-UK) Consortium | Berkshire and Surrey Pathology Services Lighthouse Laboratory and Alex Alderton, Roberto Amato, Jeffrey Barrett, Sonia Goncalves, Ewan Harrison, David K. Jackson, Ian Johnston, Dominic Kwiatkowski, Cordelia Langford, John Sillitoe on behalf of the Wellcome Sanger Institute COVID-19 Surveillance Team |
| EPI_ISL_3227993, EPI_ISL_3227994, EPI_ISL_3227995, EPI_ISL_3227996, EPI_ISL_3227997 | Lighthouse Lab in Alderley Park | Wellcome Sanger Institute for the COVID-19 Genomics UK (COG-UK) Consortium | Jacquelyn Wynn, Mairead Hyland, The Lighthouse Lab in Alderley Park and Alex Alderton, Roberto Amato, Jeffrey Barrett, Sonia Goncalves, Ewan Harrison, David K. Jackson, Ian Johnston, Dominic Kwiatkowski, Cordelia Langford, John Sillitoe on behalf of the Wellcome Sanger Institute COVID-19 Surveillance Team |
| EPI_ISL_3227998, EPI_ISL_3227999 | Berkshire and Surrey Pathology Services Lighthouse Laboratory | Wellcome Sanger Institute for the COVID-19 Genomics UK (COG-UK) Consortium | Berkshire and Surrey Pathology Services Lighthouse Laboratory and Alex Alderton, Roberto Amato, Jeffrey Barrett, Sonia Goncalves, Ewan Harrison, David K. Jackson, Ian Johnston, Dominic Kwiatkowski, Cordelia Langford, John Sillitoe on behalf of the Wellcome Sanger Institute COVID-19 Surveillance Team |
| EPI_ISL_3228000 | Lighthouse Lab in Alderley Park | Wellcome Sanger Institute for the COVID-19 Genomics UK (COG-UK) Consortium | Jacquelyn Wynn, Mairead Hyland, The Lighthouse Lab in Alderley Park and Alex Alderton, Roberto Amato, Jeffrey Barrett, Sonia Goncalves, Ewan Harrison, David K. Jackson, Ian Johnston, Dominic Kwiatkowski, Cordelia Langford, John Sillitoe on behalf of the Wellcome Sanger Institute COVID-19 Surveillance Team |
| EPI_ISL_3228001, EPI_ISL_3228002, EPI_ISL_3228003, EPI_ISL_3228004 | Berkshire and Surrey Pathology Services Lighthouse Laboratory | Wellcome Sanger Institute for the COVID-19 Genomics UK (COG-UK) Consortium | Berkshire and Surrey Pathology Services Lighthouse Laboratory and Alex Alderton, Roberto Amato, Jeffrey Barrett, Sonia Goncalves, Ewan Harrison, David K. Jackson, Ian Johnston, Dominic Kwiatkowski, Cordelia Langford, John Sillitoe on behalf of the Wellcome Sanger Institute COVID-19 Surveillance Team |
| EPI_ISL_3228005 | Lighthouse Lab in Alderley Park | Wellcome Sanger Institute for the COVID-19 Genomics UK (COG-UK) Consortium | Jacquelyn Wynn, Mairead Hyland, The Lighthouse Lab in Alderley Park and Alex Alderton, Roberto Amato, Jeffrey Barrett, Sonia Goncalves, Ewan Harrison, David K. Jackson, Ian Johnston, Dominic Kwiatkowski, Cordelia Langford, John Sillitoe on behalf of the Wellcome Sanger Institute COVID-19 Surveillance Team |
| EPI_ISL_3228006 | Berkshire and Surrey Pathology Services Lighthouse Laboratory | Wellcome Sanger Institute for the COVID-19 Genomics UK (COG-UK) Consortium | Berkshire and Surrey Pathology Services Lighthouse Laboratory and Alex Alderton, Roberto Amato, Jeffrey Barrett, Sonia Goncalves, Ewan Harrison, David K. Jackson, Ian Johnston, Dominic Kwiatkowski, Cordelia Langford, John Sillitoe on behalf of the Wellcome Sanger Institute COVID-19 Surveillance Team |
| EPI_ISL_3228007, EPI_ISL_3228008, EPI_ISL_3228009 | Lighthouse Lab in Alderley Park | Wellcome Sanger Institute for the COVID-19 Genomics UK (COG-UK) Consortium | Jacquelyn Wynn, Mairead Hyland, The Lighthouse Lab in Alderley Park and Alex Alderton, Roberto Amato, Jeffrey Barrett, Sonia Goncalves, Ewan Harrison, David K. Jackson, Ian Johnston, Dominic Kwiatkowski, Cordelia Langford, John Sillitoe on behalf of the Wellcome Sanger Institute COVID-19 Surveillance Team |
| EPI_ISL_3228010 | Berkshire and Surrey Pathology Services Lighthouse Laboratory | Wellcome Sanger Institute for the COVID-19 Genomics UK (COG-UK) Consortium | Berkshire and Surrey Pathology Services Lighthouse Laboratory and Alex Alderton, Roberto Amato, Jeffrey Barrett, Sonia Goncalves, Ewan Harrison, David K. Jackson, Ian Johnston, Dominic Kwiatkowski, Cordelia Langford, John Sillitoe on behalf of the Wellcome Sanger Institute COVID-19 Surveillance Team |
| EPI_ISL_3228011, EPI_ISL_3228012 | Lighthouse Lab in Alderley Park | Wellcome Sanger Institute for the COVID-19 Genomics UK (COG-UK) Consortium | Jacquelyn Wynn, Mairead Hyland, The Lighthouse Lab in Alderley Park and Alex Alderton, Roberto Amato, Jeffrey Barrett, Sonia Goncalves, Ewan Harrison, David K. Jackson, Ian Johnston, Dominic Kwiatkowski, Cordelia Langford, John Sillitoe on behalf of the Wellcome Sanger Institute COVID-19 Surveillance Team |
| EPI_ISL_3228013 | Berkshire and Surrey Pathology Services Lighthouse Laboratory | Wellcome Sanger Institute for the COVID-19 Genomics UK (COG-UK) Consortium | Berkshire and Surrey Pathology Services Lighthouse Laboratory and Alex Alderton, Roberto Amato, Jeffrey Barrett, Sonia Goncalves, Ewan Harrison, David K. Jackson, Ian Johnston, Dominic Kwiatkowski, Cordelia Langford, John Sillitoe on behalf of the Wellcome Sanger Institute COVID-19 Surveillance Team |
| EPI_ISL_3228014, EPI_ISL_3228015 | Lighthouse Lab in Alderley Park | Wellcome Sanger Institute for the COVID-19 Genomics UK (COG-UK) Consortium | Jacquelyn Wynn, Mairead Hyland, The Lighthouse Lab in Alderley Park and Alex Alderton, Roberto Amato, Jeffrey Barrett, Sonia Goncalves, Ewan Harrison, David K. Jackson, Ian Johnston, Dominic Kwiatkowski, Cordelia Langford, John Sillitoe on behalf of the Wellcome Sanger Institute COVID-19 Surveillance Team |
| EPI_ISL_3228016 | Berkshire and Surrey Pathology Services Lighthouse Laboratory | Wellcome Sanger Institute for the COVID-19 Genomics UK (COG-UK) Consortium | Berkshire and Surrey Pathology Services Lighthouse Laboratory and Alex Alderton, Roberto Amato, Jeffrey Barrett, Sonia Goncalves, Ewan Harrison, David K. Jackson, Ian Johnston, Dominic Kwiatkowski, Cordelia Langford, John Sillitoe on behalf of the Wellcome Sanger Institute COVID-19 Surveillance Team |
| EPI_ISL_3228017, EPI_ISL_3228018, EPI_ISL_3228019 | Lighthouse Lab in Alderley Park | Wellcome Sanger Institute for the COVID-19 Genomics UK (COG-UK) Consortium | Jacquelyn Wynn, Mairead Hyland, The Lighthouse Lab in Alderley Park and Alex Alderton, Roberto Amato, Jeffrey Barrett, Sonia Goncalves, Ewan Harrison, David K. Jackson, Ian Johnston, Dominic Kwiatkowski, Cordelia Langford, John Sillitoe on behalf of the Wellcome Sanger Institute COVID-19 Surveillance Team |
| EPI_ISL_3228020 | Berkshire and Surrey Pathology Services Lighthouse Laboratory | Wellcome Sanger Institute for the COVID-19 Genomics UK (COG-UK) Consortium | Berkshire and Surrey Pathology Services Lighthouse Laboratory and Alex Alderton, Roberto Amato, Jeffrey Barrett, Sonia Goncalves, Ewan Harrison, David K. Jackson, Ian Johnston, Dominic Kwiatkowski, Cordelia Langford, John Sillitoe on behalf of the Wellcome Sanger Institute COVID-19 Surveillance Team |
| EPI_ISL_3228021 | Lighthouse Lab in Alderley Park | Wellcome Sanger Institute for the COVID-19 Genomics UK (COG-UK) Consortium | Jacquelyn Wynn, Mairead Hyland, The Lighthouse Lab in Alderley Park and Alex Alderton, Roberto Amato, Jeffrey Barrett, Sonia Goncalves, Ewan Harrison, David K. Jackson, Ian Johnston, Dominic Kwiatkowski, Cordelia Langford, John Sillitoe on behalf of the Wellcome Sanger Institute COVID-19 Surveillance Team |
| EPI_ISL_3228022 | Berkshire and Surrey Pathology Services Lighthouse Laboratory | Wellcome Sanger Institute for the COVID-19 Genomics UK (COG-UK) Consortium | Berkshire and Surrey Pathology Services Lighthouse Laboratory and Alex Alderton, Roberto Amato, Jeffrey Barrett, Sonia Goncalves, Ewan Harrison, David K. Jackson, Ian Johnston, Dominic Kwiatkowski, Cordelia Langford, John Sillitoe on behalf of the Wellcome Sanger Institute COVID-19 Surveillance Team |
| EPI_ISL_3228023 | Lighthouse Lab in Alderley Park | Wellcome Sanger Institute for the COVID-19 Genomics UK (COG-UK) Consortium | Jacquelyn Wynn, Mairead Hyland, The Lighthouse Lab in Alderley Park and Alex Alderton, Roberto Amato, Jeffrey Barrett, Sonia Goncalves, Ewan Harrison, David K. Jackson, Ian Johnston, Dominic Kwiatkowski, Cordelia Langford, John Sillitoe on behalf of the Wellcome Sanger Institute COVID-19 Surveillance Team |
| EPI_ISL_3228024 | Berkshire and Surrey Pathology Services Lighthouse Laboratory | Wellcome Sanger Institute for the COVID-19 Genomics UK (COG-UK) Consortium | Berkshire and Surrey Pathology Services Lighthouse Laboratory and Alex Alderton, Roberto Amato, Jeffrey Barrett, Sonia Goncalves, Ewan Harrison, David K. Jackson, Ian Johnston, Dominic Kwiatkowski, Cordelia Langford, John Sillitoe on behalf of the Wellcome Sanger Institute COVID-19 Surveillance Team |
| EPI_ISL_3228025 | Lighthouse Lab in Alderley Park | Wellcome Sanger Institute for the COVID-19 Genomics UK (COG-UK) Consortium | Jacquelyn Wynn, Mairead Hyland, The Lighthouse Lab in Alderley Park and Alex Alderton, Roberto Amato, Jeffrey Barrett, Sonia Goncalves, Ewan Harrison, David K. Jackson, Ian Johnston, Dominic Kwiatkowski, Cordelia Langford, John Sillitoe on behalf of the Wellcome Sanger Institute COVID-19 Surveillance Team |
| EPI_ISL_3228026 | Berkshire and Surrey Pathology Services Lighthouse Laboratory | Wellcome Sanger Institute for the COVID-19 Genomics UK (COG-UK) Consortium | Berkshire and Surrey Pathology Services Lighthouse Laboratory and Alex Alderton, Roberto Amato, Jeffrey Barrett, Sonia Goncalves, Ewan Harrison, David K. Jackson, Ian Johnston, Dominic Kwiatkowski, Cordelia Langford, John Sillitoe on behalf of the Wellcome Sanger Institute COVID-19 Surveillance Team |
| EPI_ISL_3228027, EPI_ISL_3228028, EPI_ISL_3228029, EPI_ISL_3228030 | Lighthouse Lab in Alderley Park | Wellcome Sanger Institute for the COVID-19 Genomics UK (COG-UK) Consortium | Jacquelyn Wynn, Mairead Hyland, The Lighthouse Lab in Alderley Park and Alex Alderton, Roberto Amato, Jeffrey Barrett, Sonia Goncalves, Ewan Harrison, David K. Jackson, Ian Johnston, Dominic Kwiatkowski, Cordelia Langford, John Sillitoe on behalf of the Wellcome Sanger Institute COVID-19 Surveillance Team |
| EPI_ISL_3228031 | Berkshire and Surrey Pathology Services Lighthouse Laboratory | Wellcome Sanger Institute for the COVID-19 Genomics UK (COG-UK) Consortium | Berkshire and Surrey Pathology Services Lighthouse Laboratory and Alex Alderton, Roberto Amato, Jeffrey Barrett, Sonia Goncalves, Ewan Harrison, David K. Jackson, Ian Johnston, Dominic Kwiatkowski, Cordelia Langford, John Sillitoe on behalf of the Wellcome Sanger Institute COVID-19 Surveillance Team |
| EPI_ISL_3228032 | Lighthouse Lab in Alderley Park | Wellcome Sanger Institute for the COVID-19 Genomics UK (COG-UK) Consortium | Jacquelyn Wynn, Mairead Hyland, The Lighthouse Lab in Alderley Park and Alex Alderton, Roberto Amato, Jeffrey Barrett, Sonia Goncalves, Ewan Harrison, David K. Jackson, Ian Johnston, Dominic Kwiatkowski, Cordelia Langford, John Sillitoe on behalf of the Wellcome Sanger Institute COVID-19 Surveillance Team |
| EPI_ISL_3228033, EPI_ISL_3228034, EPI_ISL_3228035 | Berkshire and Surrey Pathology Services Lighthouse Laboratory | Wellcome Sanger Institute for the COVID-19 Genomics UK (COG-UK) Consortium | Berkshire and Surrey Pathology Services Lighthouse Laboratory and Alex Alderton, Roberto Amato, Jeffrey Barrett, Sonia Goncalves, Ewan Harrison, David K. Jackson, Ian Johnston, Dominic Kwiatkowski, Cordelia Langford, John Sillitoe on behalf of the Wellcome Sanger Institute COVID-19 Surveillance Team |
| EPI_ISL_3228036 | Lighthouse Lab in Alderley Park | Wellcome Sanger Institute for the COVID-19 Genomics UK (COG-UK) Consortium | Jacquelyn Wynn, Mairead Hyland, The Lighthouse Lab in Alderley Park and Alex Alderton, Roberto Amato, Jeffrey Barrett, Sonia Goncalves, Ewan Harrison, David K. Jackson, Ian Johnston, Dominic Kwiatkowski, Cordelia Langford, John Sillitoe on behalf of the Wellcome Sanger Institute COVID-19 Surveillance Team |
| EPI_ISL_3228037 | Berkshire and Surrey Pathology Services Lighthouse Laboratory | Wellcome Sanger Institute for the COVID-19 Genomics UK (COG-UK) Consortium | Berkshire and Surrey Pathology Services Lighthouse Laboratory and Alex Alderton, Roberto Amato, Jeffrey Barrett, Sonia Goncalves, Ewan Harrison, David K. Jackson, Ian Johnston, Dominic Kwiatkowski, Cordelia Langford, John Sillitoe on behalf of the Wellcome Sanger Institute COVID-19 Surveillance Team |
| EPI_ISL_3228038 | Lighthouse Lab in Alderley Park | Wellcome Sanger Institute for the COVID-19 Genomics UK (COG-UK) | Jacquelyn Wynn, Mairead Hyland, The Lighthouse Lab in Alderley Park and Alex Alderton, Roberto Amato, Jeffrey Barrett, Sonia Goncalves, Ewan Harrison, David K. Jackson, Ian Johnston, Dominic Kwiatkowski, Cordelia Langford, John Sillitoe on |

| | | | |
|---|---|---|---|
| | | Consortium | behalf of the Wellcome Sanger Institute COVID-19 Surveillance Team |
| EPI_ISL_3228039 | Berkshire and Surrey Pathology Services Lighthouse Laboratory | Wellcome Sanger Institute for the COVID-19 Genomics UK (COG-UK) Consortium | Berkshire and Surrey Pathology Services Lighthouse Laboratory and Alex Alderton, Roberto Amato, Jeffrey Barrett, Sonia Goncalves, Ewan Harrison, David K. Jackson, Ian Johnston, Dominic Kwiatkowski, Cordelia Langford, John Sillitoe on behalf of the Wellcome Sanger Institute COVID-19 Surveillance Team |
| EPI_ISL_3228040, EPI_ISL_3228041 | Lighthouse Lab in Alderley Park | Wellcome Sanger Institute for the COVID-19 Genomics UK (COG-UK) Consortium | Jacquelyn Wynn, Mairead Hyland, The Lighthouse Lab in Alderley Park and Alex Alderton, Roberto Amato, Jeffrey Barrett, Sonia Goncalves, Ewan Harrison, David K. Jackson, Ian Johnston, Dominic Kwiatkowski, Cordelia Langford, John Sillitoe on behalf of the Wellcome Sanger Institute COVID-19 Surveillance Team |
| EPI_ISL_3228042 | Berkshire and Surrey Pathology Services Lighthouse Laboratory | Wellcome Sanger Institute for the COVID-19 Genomics UK (COG-UK) Consortium | Berkshire and Surrey Pathology Services Lighthouse Laboratory and Alex Alderton, Roberto Amato, Jeffrey Barrett, Sonia Goncalves, Ewan Harrison, David K. Jackson, Ian Johnston, Dominic Kwiatkowski, Cordelia Langford, John Sillitoe on behalf of the Wellcome Sanger Institute COVID-19 Surveillance Team |
| EPI_ISL_3228043 | Lighthouse Lab in Alderley Park | Wellcome Sanger Institute for the COVID-19 Genomics UK (COG-UK) Consortium | Jacquelyn Wynn, Mairead Hyland, The Lighthouse Lab in Alderley Park and Alex Alderton, Roberto Amato, Jeffrey Barrett, Sonia Goncalves, Ewan Harrison, David K. Jackson, Ian Johnston, Dominic Kwiatkowski, Cordelia Langford, John Sillitoe on behalf of the Wellcome Sanger Institute COVID-19 Surveillance Team |
| EPI_ISL_3228044, EPI_ISL_3228045, EPI_ISL_3228046 | Berkshire and Surrey Pathology Services Lighthouse Laboratory | Wellcome Sanger Institute for the COVID-19 Genomics UK (COG-UK) Consortium | Berkshire and Surrey Pathology Services Lighthouse Laboratory and Alex Alderton, Roberto Amato, Jeffrey Barrett, Sonia Goncalves, Ewan Harrison, David K. Jackson, Ian Johnston, Dominic Kwiatkowski, Cordelia Langford, John Sillitoe on behalf of the Wellcome Sanger Institute COVID-19 Surveillance Team |
| EPI_ISL_3228047, EPI_ISL_3228048 | Lighthouse Lab in Alderley Park | Wellcome Sanger Institute for the COVID-19 Genomics UK (COG-UK) Consortium | Jacquelyn Wynn, Mairead Hyland, The Lighthouse Lab in Alderley Park and Alex Alderton, Roberto Amato, Jeffrey Barrett, Sonia Goncalves, Ewan Harrison, David K. Jackson, Ian Johnston, Dominic Kwiatkowski, Cordelia Langford, John Sillitoe on behalf of the Wellcome Sanger Institute COVID-19 Surveillance Team |
| EPI_ISL_3228049 | Berkshire and Surrey Pathology Services Lighthouse Laboratory | Wellcome Sanger Institute for the COVID-19 Genomics UK (COG-UK) Consortium | Berkshire and Surrey Pathology Services Lighthouse Laboratory and Alex Alderton, Roberto Amato, Jeffrey Barrett, Sonia Goncalves, Ewan Harrison, David K. Jackson, Ian Johnston, Dominic Kwiatkowski, Cordelia Langford, John Sillitoe on behalf of the Wellcome Sanger Institute COVID-19 Surveillance Team |
| EPI_ISL_3228050, EPI_ISL_3228051 | Lighthouse Lab in Alderley Park | Wellcome Sanger Institute for the COVID-19 Genomics UK (COG-UK) Consortium | Jacquelyn Wynn, Mairead Hyland, The Lighthouse Lab in Alderley Park and Alex Alderton, Roberto Amato, Jeffrey Barrett, Sonia Goncalves, Ewan Harrison, David K. Jackson, Ian Johnston, Dominic Kwiatkowski, Cordelia Langford, John Sillitoe on behalf of the Wellcome Sanger Institute COVID-19 Surveillance Team |
| EPI_ISL_3228052, EPI_ISL_3228053 | Berkshire and Surrey Pathology Services Lighthouse Laboratory | Wellcome Sanger Institute for the COVID-19 Genomics UK (COG-UK) Consortium | Berkshire and Surrey Pathology Services Lighthouse Laboratory and Alex Alderton, Roberto Amato, Jeffrey Barrett, Sonia Goncalves, Ewan Harrison, David K. Jackson, Ian Johnston, Dominic Kwiatkowski, Cordelia Langford, John Sillitoe on behalf of the Wellcome Sanger Institute COVID-19 Surveillance Team |
| EPI_ISL_3228054 | Lighthouse Lab in Alderley Park | Wellcome Sanger Institute for the COVID-19 Genomics UK (COG-UK) Consortium | Jacquelyn Wynn, Mairead Hyland, The Lighthouse Lab in Alderley Park and Alex Alderton, Roberto Amato, Jeffrey Barrett, Sonia Goncalves, Ewan Harrison, David K. Jackson, Ian Johnston, Dominic Kwiatkowski, Cordelia Langford, John Sillitoe on behalf of the Wellcome Sanger Institute COVID-19 Surveillance Team |
| EPI_ISL_3228055 | Berkshire and Surrey Pathology Services Lighthouse Laboratory | Wellcome Sanger Institute for the COVID-19 Genomics UK (COG-UK) Consortium | Berkshire and Surrey Pathology Services Lighthouse Laboratory and Alex Alderton, Roberto Amato, Jeffrey Barrett, Sonia Goncalves, Ewan Harrison, David K. Jackson, Ian Johnston, Dominic Kwiatkowski, Cordelia Langford, John Sillitoe on behalf of the Wellcome Sanger Institute COVID-19 Surveillance Team |
| EPI_ISL_3228056, EPI_ISL_3228057, EPI_ISL_3228058, EPI_ISL_3228059 | Lighthouse Lab in Alderley Park | Wellcome Sanger Institute for the COVID-19 Genomics UK (COG-UK) Consortium | Jacquelyn Wynn, Mairead Hyland, The Lighthouse Lab in Alderley Park and Alex Alderton, Roberto Amato, Jeffrey Barrett, Sonia Goncalves, Ewan Harrison, David K. Jackson, Ian Johnston, Dominic Kwiatkowski, Cordelia Langford, John Sillitoe on behalf of the Wellcome Sanger Institute COVID-19 Surveillance Team |
| EPI_ISL_3228060, EPI_ISL_3228061 | Berkshire and Surrey Pathology Services Lighthouse Laboratory | Wellcome Sanger Institute for the COVID-19 Genomics UK (COG-UK) Consortium | Berkshire and Surrey Pathology Services Lighthouse Laboratory and Alex Alderton, Roberto Amato, Jeffrey Barrett, Sonia Goncalves, Ewan Harrison, David K. Jackson, Ian Johnston, Dominic Kwiatkowski, Cordelia Langford, John Sillitoe on behalf of the Wellcome Sanger Institute COVID-19 Surveillance Team |
| EPI_ISL_3228062, EPI_ISL_3228063, EPI_ISL_3228064 | Lighthouse Lab in Alderley Park | Wellcome Sanger Institute for the COVID-19 Genomics UK (COG-UK) Consortium | Jacquelyn Wynn, Mairead Hyland, The Lighthouse Lab in Alderley Park and Alex Alderton, Roberto Amato, Jeffrey Barrett, Sonia Goncalves, Ewan Harrison, David K. Jackson, Ian Johnston, Dominic Kwiatkowski, Cordelia Langford, John Sillitoe on behalf of the Wellcome Sanger Institute COVID-19 Surveillance Team |
| EPI_ISL_3228065, EPI_ISL_3228066 | Berkshire and Surrey Pathology Services Lighthouse Laboratory | Wellcome Sanger Institute for the COVID-19 Genomics UK (COG-UK) Consortium | Berkshire and Surrey Pathology Services Lighthouse Laboratory and Alex Alderton, Roberto Amato, Jeffrey Barrett, Sonia Goncalves, Ewan Harrison, David K. Jackson, Ian Johnston, Dominic Kwiatkowski, Cordelia Langford, John Sillitoe on behalf of the Wellcome Sanger Institute COVID-19 Surveillance Team |
| EPI_ISL_3228067 | Lighthouse Lab in Alderley Park | Wellcome Sanger Institute for the COVID-19 Genomics UK (COG-UK) Consortium | Jacquelyn Wynn, Mairead Hyland, The Lighthouse Lab in Alderley Park and Alex Alderton, Roberto Amato, Jeffrey Barrett, Sonia Goncalves, Ewan Harrison, David K. Jackson, Ian Johnston, Dominic Kwiatkowski, Cordelia Langford, John Sillitoe on behalf of the Wellcome Sanger Institute COVID-19 Surveillance Team |
| EPI_ISL_3228068, EPI_ISL_3228069, EPI_ISL_3228070 | Berkshire and Surrey Pathology Services Lighthouse Laboratory | Wellcome Sanger Institute for the COVID-19 Genomics UK (COG-UK) Consortium | Berkshire and Surrey Pathology Services Lighthouse Laboratory and Alex Alderton, Roberto Amato, Jeffrey Barrett, Sonia Goncalves, Ewan Harrison, David K. Jackson, Ian Johnston, Dominic Kwiatkowski, Cordelia Langford, John Sillitoe on behalf of the Wellcome Sanger Institute COVID-19 Surveillance Team |
| EPI_ISL_3228071, EPI_ISL_3228072, EPI_ISL_3228073, EPI_ISL_3228074 | Lighthouse Lab in Alderley Park | Wellcome Sanger Institute for the COVID-19 Genomics UK (COG-UK) Consortium | Jacquelyn Wynn, Mairead Hyland, The Lighthouse Lab in Alderley Park and Alex Alderton, Roberto Amato, Jeffrey Barrett, Sonia Goncalves, Ewan Harrison, David K. Jackson, Ian Johnston, Dominic Kwiatkowski, Cordelia Langford, John Sillitoe on behalf of the Wellcome Sanger Institute COVID-19 Surveillance Team |
| EPI_ISL_3228075 | Berkshire and Surrey Pathology Services Lighthouse Laboratory | Wellcome Sanger Institute for the COVID-19 Genomics UK (COG-UK) Consortium | Berkshire and Surrey Pathology Services Lighthouse Laboratory and Alex Alderton, Roberto Amato, Jeffrey Barrett, Sonia Goncalves, Ewan Harrison, David K. Jackson, Ian Johnston, Dominic Kwiatkowski, Cordelia Langford, John Sillitoe on behalf of the Wellcome Sanger Institute COVID-19 Surveillance Team |
| EPI_ISL_3228076 | Lighthouse Lab in Alderley Park | Wellcome Sanger Institute for the COVID-19 Genomics UK (COG-UK) Consortium | Jacquelyn Wynn, Mairead Hyland, The Lighthouse Lab in Alderley Park and Alex Alderton, Roberto Amato, Jeffrey Barrett, Sonia Goncalves, Ewan Harrison, David K. Jackson, Ian Johnston, Dominic Kwiatkowski, Cordelia Langford, John Sillitoe on behalf of the Wellcome Sanger Institute COVID-19 Surveillance Team |
| EPI_ISL_3228077 | COVID-19 Detection Lab, Chattogram Veterinary and Animal Sciences University | Genomic Research Lab, BCSIR | Md. Murshed Hasan Sarkar, Abu Sayeed Mohammad Mahmud, Mohammad Samir Uzzaman, Eshrar Osman, Md. Ahasan Habib, Shahina Akter, Tanjina Akhter Banu, Barna Goswami, Iffat Jahan, Mohammad Mohi Uddin, Md. Kamrul Islam, Goutam Buddha Das, Tridip Das, Tanvir Ahmad Nizami, Eaftekhar Ahmed Rana, Md. Sirazul Islam, Pronesh Dutta, Sharmin Chowdhury, Md. Paritosh Kumar Biswas, Md. Salim Khan |
| EPI_ISL_3228078 | Berkshire and Surrey Pathology Services Lighthouse Laboratory | Wellcome Sanger Institute for the COVID-19 Genomics UK (COG-UK) Consortium | Berkshire and Surrey Pathology Services Lighthouse Laboratory and Alex Alderton, Roberto Amato, Jeffrey Barrett, Sonia Goncalves, Ewan Harrison, David K. Jackson, Ian Johnston, Dominic Kwiatkowski, Cordelia Langford, John Sillitoe on behalf of the Wellcome Sanger Institute COVID-19 Surveillance Team |
| EPI_ISL_3228079, EPI_ISL_3228080 | Lighthouse Lab in Alderley Park | Wellcome Sanger Institute for the COVID-19 Genomics UK (COG-UK) Consortium | Jacquelyn Wynn, Mairead Hyland, The Lighthouse Lab in Alderley Park and Alex Alderton, Roberto Amato, Jeffrey Barrett, Sonia Goncalves, Ewan Harrison, David K. Jackson, Ian Johnston, Dominic Kwiatkowski, Cordelia Langford, John Sillitoe on behalf of the Wellcome Sanger Institute COVID-19 Surveillance Team |
| EPI_ISL_3228081, EPI_ISL_3228082 | Berkshire and Surrey Pathology Services Lighthouse Laboratory | Wellcome Sanger Institute for the COVID-19 Genomics UK (COG-UK) Consortium | Berkshire and Surrey Pathology Services Lighthouse Laboratory and Alex Alderton, Roberto Amato, Jeffrey Barrett, Sonia Goncalves, Ewan Harrison, David K. Jackson, Ian Johnston, Dominic Kwiatkowski, Cordelia Langford, John Sillitoe on behalf of the Wellcome Sanger Institute COVID-19 Surveillance Team |
| EPI_ISL_3228083 | Lighthouse Lab in Alderley Park | Wellcome Sanger Institute for the COVID-19 Genomics UK (COG-UK) Consortium | Jacquelyn Wynn, Mairead Hyland, The Lighthouse Lab in Alderley Park and Alex Alderton, Roberto Amato, Jeffrey Barrett, Sonia Goncalves, Ewan Harrison, David K. Jackson, Ian Johnston, Dominic Kwiatkowski, Cordelia Langford, John Sillitoe on behalf of the Wellcome Sanger Institute COVID-19 Surveillance Team |
| EPI_ISL_3228084 | Berkshire and Surrey Pathology Services Lighthouse Laboratory | Wellcome Sanger Institute for the COVID-19 Genomics UK (COG-UK) Consortium | Berkshire and Surrey Pathology Services Lighthouse Laboratory and Alex Alderton, Roberto Amato, Jeffrey Barrett, Sonia Goncalves, Ewan Harrison, David K. Jackson, Ian Johnston, Dominic Kwiatkowski, Cordelia Langford, John Sillitoe on behalf of the Wellcome Sanger Institute COVID-19 Surveillance Team |
| EPI_ISL_3228085, EPI_ISL_3228086 | Lighthouse Lab in Alderley Park | Wellcome Sanger Institute for the COVID-19 Genomics UK (COG-UK) Consortium | Jacquelyn Wynn, Mairead Hyland, The Lighthouse Lab in Alderley Park and Alex Alderton, Roberto Amato, Jeffrey Barrett, Sonia Goncalves, Ewan Harrison, David K. Jackson, Ian Johnston, Dominic Kwiatkowski, Cordelia Langford, John Sillitoe on behalf of the Wellcome Sanger Institute COVID-19 Surveillance Team |
| EPI_ISL_3228087, EPI_ISL_3228088 | Berkshire and Surrey Pathology Services Lighthouse Laboratory | Wellcome Sanger Institute for the COVID-19 Genomics UK (COG-UK) Consortium | Berkshire and Surrey Pathology Services Lighthouse Laboratory and Alex Alderton, Roberto Amato, Jeffrey Barrett, Sonia Goncalves, Ewan Harrison, David K. Jackson, Ian Johnston, Dominic Kwiatkowski, Cordelia Langford, John Sillitoe on behalf of the Wellcome Sanger Institute COVID-19 Surveillance Team |
| EPI_ISL_3228089, EPI_ISL_3228090 | Lighthouse Lab in Alderley Park | Wellcome Sanger Institute for the COVID-19 Genomics UK (COG-UK) Consortium | Jacquelyn Wynn, Mairead Hyland, The Lighthouse Lab in Alderley Park and Alex Alderton, Roberto Amato, Jeffrey Barrett, Sonia Goncalves, Ewan Harrison, David K. Jackson, Ian Johnston, Dominic Kwiatkowski, Cordelia Langford, John Sillitoe on behalf of the Wellcome Sanger Institute COVID-19 Surveillance Team |
| EPI_ISL_3228091 | Berkshire and Surrey Pathology Services Lighthouse Laboratory | Wellcome Sanger Institute for the COVID-19 Genomics UK (COG-UK) Consortium | Berkshire and Surrey Pathology Services Lighthouse Laboratory and Alex Alderton, Roberto Amato, Jeffrey Barrett, Sonia Goncalves, Ewan Harrison, David K. Jackson, Ian Johnston, Dominic Kwiatkowski, Cordelia Langford, John Sillitoe on behalf of the Wellcome Sanger Institute COVID-19 Surveillance Team |
| EPI_ISL_3228092 | Lighthouse Lab in Alderley Park | Wellcome Sanger Institute for the COVID-19 Genomics UK (COG-UK) Consortium | Jacquelyn Wynn, Mairead Hyland, The Lighthouse Lab in Alderley Park and Alex Alderton, Roberto Amato, Jeffrey Barrett, Sonia Goncalves, Ewan Harrison, David K. Jackson, Ian Johnston, Dominic Kwiatkowski, Cordelia Langford, John Sillitoe on behalf of the Wellcome Sanger Institute COVID-19 Surveillance Team |
| EPI_ISL_3228093 | Berkshire and Surrey Pathology Services Lighthouse Laboratory | Wellcome Sanger Institute for the COVID-19 Genomics UK (COG-UK) Consortium | Berkshire and Surrey Pathology Services Lighthouse Laboratory and Alex Alderton, Roberto Amato, Jeffrey Barrett, Sonia Goncalves, Ewan Harrison, David K. Jackson, Ian Johnston, Dominic Kwiatkowski, Cordelia Langford, John Sillitoe on behalf of the Wellcome Sanger Institute COVID-19 Surveillance Team |
| EPI_ISL_3228094 | Lighthouse Lab in Alderley Park | Wellcome Sanger Institute for the COVID-19 Genomics UK (COG-UK) Consortium | Jacquelyn Wynn, Mairead Hyland, The Lighthouse Lab in Alderley Park and Alex Alderton, Roberto Amato, Jeffrey Barrett, Sonia Goncalves, Ewan Harrison, David K. Jackson, Ian Johnston, Dominic Kwiatkowski, Cordelia Langford, John Sillitoe on behalf of the Wellcome Sanger Institute COVID-19 Surveillance Team |
| EPI_ISL_3228095, EPI_ISL_3228096, EPI_ISL_3228097, EPI_ISL_3228098 | Berkshire and Surrey Pathology Services Lighthouse Laboratory | Wellcome Sanger Institute for the COVID-19 Genomics UK (COG-UK) Consortium | Berkshire and Surrey Pathology Services Lighthouse Laboratory and Alex Alderton, Roberto Amato, Jeffrey Barrett, Sonia Goncalves, Ewan Harrison, David K. Jackson, Ian Johnston, Dominic Kwiatkowski, Cordelia Langford, John Sillitoe on behalf of the Wellcome Sanger Institute COVID-19 Surveillance Team |
| EPI_ISL_3228099 | Lighthouse Lab in Alderley Park | Wellcome Sanger Institute for the COVID-19 Genomics UK (COG-UK) Consortium | Jacquelyn Wynn, Mairead Hyland, The Lighthouse Lab in Alderley Park and Alex Alderton, Roberto Amato, Jeffrey Barrett, Sonia Goncalves, Ewan Harrison, David K. Jackson, Ian Johnston, Dominic Kwiatkowski, Cordelia Langford, John Sillitoe on behalf of the Wellcome Sanger Institute COVID-19 Surveillance Team |
| EPI_ISL_3228100 | Berkshire and Surrey Pathology Services Lighthouse Laboratory | Wellcome Sanger Institute for the COVID-19 Genomics UK (COG-UK) Consortium | Berkshire and Surrey Pathology Services Lighthouse Laboratory and Alex Alderton, Roberto Amato, Jeffrey Barrett, Sonia Goncalves, Ewan Harrison, David K. Jackson, Ian Johnston, Dominic Kwiatkowski, Cordelia Langford, John Sillitoe on behalf of the Wellcome Sanger Institute COVID-19 Surveillance Team |
| EPI_ISL_3228101, EPI_ISL_3228102, EPI_ISL_3228103, EPI_ISL_3228104 | Lighthouse Lab in Alderley Park | Wellcome Sanger Institute for the COVID-19 Genomics UK (COG-UK) Consortium | Jacquelyn Wynn, Mairead Hyland, The Lighthouse Lab in Alderley Park and Alex Alderton, Roberto Amato, Jeffrey Barrett, Sonia Goncalves, Ewan Harrison, David K. Jackson, Ian Johnston, Dominic Kwiatkowski, Cordelia Langford, John Sillitoe on behalf of the Wellcome Sanger Institute COVID-19 Surveillance Team |
| EPI_ISL_3228105, EPI_ISL_3228106, EPI_ISL_3228107, EPI_ISL_3228108, EPI_ISL_3228109, EPI_ISL_3228110 | Berkshire and Surrey Pathology Services Lighthouse Laboratory | Wellcome Sanger Institute for the COVID-19 Genomics UK (COG-UK) Consortium | Berkshire and Surrey Pathology Services Lighthouse Laboratory and Alex Alderton, Roberto Amato, Jeffrey Barrett, Sonia Goncalves, Ewan Harrison, David K. Jackson, Ian Johnston, Dominic Kwiatkowski, Cordelia Langford, John Sillitoe on behalf of the Wellcome Sanger Institute COVID-19 Surveillance Team |

| | | | |
|---|---|---|---|
| EPI_ISL_3228111, EPI_ISL_3228112 | Lighthouse Lab in Alderley Park | Wellcome Sanger Institute for the COVID-19 Genomics UK (COG-UK) Consortium | Jacquelyn Wynn, Mairead Hyland, The Lighthouse Lab in Alderley Park and Alex Alderton, Roberto Amato, Jeffrey Barrett, Sonia Goncalves, Ewan Harrison, David K. Jackson, Ian Johnston, Dominic Kwiatkowski, Cordelia Langford, John Sillitoe on behalf of the Wellcome Sanger Institute COVID-19 Surveillance Team |
| EPI_ISL_3228113 | Berkshire and Surrey Pathology Services Lighthouse Laboratory | Wellcome Sanger Institute for the COVID-19 Genomics UK (COG-UK) Consortium | Berkshire and Surrey Pathology Services Lighthouse Laboratory and Alex Alderton, Roberto Amato, Jeffrey Barrett, Sonia Goncalves, Ewan Harrison, David K. Jackson, Ian Johnston, Dominic Kwiatkowski, Cordelia Langford, John Sillitoe on behalf of the Wellcome Sanger Institute COVID-19 Surveillance Team |
| EPI_ISL_3228114, EPI_ISL_3228115 | Lighthouse Lab in Alderley Park | Wellcome Sanger Institute for the COVID-19 Genomics UK (COG-UK) Consortium | Jacquelyn Wynn, Mairead Hyland, The Lighthouse Lab in Alderley Park and Alex Alderton, Roberto Amato, Jeffrey Barrett, Sonia Goncalves, Ewan Harrison, David K. Jackson, Ian Johnston, Dominic Kwiatkowski, Cordelia Langford, John Sillitoe on behalf of the Wellcome Sanger Institute COVID-19 Surveillance Team |
| EPI_ISL_3228116 | Berkshire and Surrey Pathology Services Lighthouse Laboratory | Wellcome Sanger Institute for the COVID-19 Genomics UK (COG-UK) Consortium | Berkshire and Surrey Pathology Services Lighthouse Laboratory and Alex Alderton, Roberto Amato, Jeffrey Barrett, Sonia Goncalves, Ewan Harrison, David K. Jackson, Ian Johnston, Dominic Kwiatkowski, Cordelia Langford, John Sillitoe on behalf of the Wellcome Sanger Institute COVID-19 Surveillance Team |
| EPI_ISL_3228117, EPI_ISL_3228118, EPI_ISL_3228119 | Lighthouse Lab in Alderley Park | Wellcome Sanger Institute for the COVID-19 Genomics UK (COG-UK) Consortium | Jacquelyn Wynn, Mairead Hyland, The Lighthouse Lab in Alderley Park and Alex Alderton, Roberto Amato, Jeffrey Barrett, Sonia Goncalves, Ewan Harrison, David K. Jackson, Ian Johnston, Dominic Kwiatkowski, Cordelia Langford, John Sillitoe on behalf of the Wellcome Sanger Institute COVID-19 Surveillance Team |
| EPI_ISL_3228120 | Berkshire and Surrey Pathology Services Lighthouse Laboratory | Wellcome Sanger Institute for the COVID-19 Genomics UK (COG-UK) Consortium | Berkshire and Surrey Pathology Services Lighthouse Laboratory and Alex Alderton, Roberto Amato, Jeffrey Barrett, Sonia Goncalves, Ewan Harrison, David K. Jackson, Ian Johnston, Dominic Kwiatkowski, Cordelia Langford, John Sillitoe on behalf of the Wellcome Sanger Institute COVID-19 Surveillance Team |
| EPI_ISL_3228121 | Lighthouse Lab in Alderley Park | Wellcome Sanger Institute for the COVID-19 Genomics UK (COG-UK) Consortium | Jacquelyn Wynn, Mairead Hyland, The Lighthouse Lab in Alderley Park and Alex Alderton, Roberto Amato, Jeffrey Barrett, Sonia Goncalves, Ewan Harrison, David K. Jackson, Ian Johnston, Dominic Kwiatkowski, Cordelia Langford, John Sillitoe on behalf of the Wellcome Sanger Institute COVID-19 Surveillance Team |
| EPI_ISL_3228122, EPI_ISL_3228123 | Berkshire and Surrey Pathology Services Lighthouse Laboratory | Wellcome Sanger Institute for the COVID-19 Genomics UK (COG-UK) Consortium | Berkshire and Surrey Pathology Services Lighthouse Laboratory and Alex Alderton, Roberto Amato, Jeffrey Barrett, Sonia Goncalves, Ewan Harrison, David K. Jackson, Ian Johnston, Dominic Kwiatkowski, Cordelia Langford, John Sillitoe on behalf of the Wellcome Sanger Institute COVID-19 Surveillance Team |
| EPI_ISL_3228124 | Lighthouse Lab in Alderley Park | Wellcome Sanger Institute for the COVID-19 Genomics UK (COG-UK) Consortium | Jacquelyn Wynn, Mairead Hyland, The Lighthouse Lab in Alderley Park and Alex Alderton, Roberto Amato, Jeffrey Barrett, Sonia Goncalves, Ewan Harrison, David K. Jackson, Ian Johnston, Dominic Kwiatkowski, Cordelia Langford, John Sillitoe on behalf of the Wellcome Sanger Institute COVID-19 Surveillance Team |
| EPI_ISL_3228125 | Berkshire and Surrey Pathology Services Lighthouse Laboratory | Wellcome Sanger Institute for the COVID-19 Genomics UK (COG-UK) Consortium | Berkshire and Surrey Pathology Services Lighthouse Laboratory and Alex Alderton, Roberto Amato, Jeffrey Barrett, Sonia Goncalves, Ewan Harrison, David K. Jackson, Ian Johnston, Dominic Kwiatkowski, Cordelia Langford, John Sillitoe on behalf of the Wellcome Sanger Institute COVID-19 Surveillance Team |
| EPI_ISL_3228126 | Lighthouse Lab in Alderley Park | Wellcome Sanger Institute for the COVID-19 Genomics UK (COG-UK) Consortium | Jacquelyn Wynn, Mairead Hyland, The Lighthouse Lab in Alderley Park and Alex Alderton, Roberto Amato, Jeffrey Barrett, Sonia Goncalves, Ewan Harrison, David K. Jackson, Ian Johnston, Dominic Kwiatkowski, Cordelia Langford, John Sillitoe on behalf of the Wellcome Sanger Institute COVID-19 Surveillance Team |
| EPI_ISL_3228127, EPI_ISL_3228128 | Berkshire and Surrey Pathology Services Lighthouse Laboratory | Wellcome Sanger Institute for the COVID-19 Genomics UK (COG-UK) Consortium | Berkshire and Surrey Pathology Services Lighthouse Laboratory and Alex Alderton, Roberto Amato, Jeffrey Barrett, Sonia Goncalves, Ewan Harrison, David K. Jackson, Ian Johnston, Dominic Kwiatkowski, Cordelia Langford, John Sillitoe on behalf of the Wellcome Sanger Institute COVID-19 Surveillance Team |
| EPI_ISL_3228129, EPI_ISL_3228130, EPI_ISL_3228131 | Lighthouse Lab in Alderley Park | Wellcome Sanger Institute for the COVID-19 Genomics UK (COG-UK) Consortium | Jacquelyn Wynn, Mairead Hyland, The Lighthouse Lab in Alderley Park and Alex Alderton, Roberto Amato, Jeffrey Barrett, Sonia Goncalves, Ewan Harrison, David K. Jackson, Ian Johnston, Dominic Kwiatkowski, Cordelia Langford, John Sillitoe on behalf of the Wellcome Sanger Institute COVID-19 Surveillance Team |
| EPI_ISL_3228132 | Berkshire and Surrey Pathology Services Lighthouse Laboratory | Wellcome Sanger Institute for the COVID-19 Genomics UK (COG-UK) Consortium | Berkshire and Surrey Pathology Services Lighthouse Laboratory and Alex Alderton, Roberto Amato, Jeffrey Barrett, Sonia Goncalves, Ewan Harrison, David K. Jackson, Ian Johnston, Dominic Kwiatkowski, Cordelia Langford, John Sillitoe on behalf of the Wellcome Sanger Institute COVID-19 Surveillance Team |
| EPI_ISL_3228133 | Lighthouse Lab in Alderley Park | Wellcome Sanger Institute for the COVID-19 Genomics UK (COG-UK) Consortium | Jacquelyn Wynn, Mairead Hyland, The Lighthouse Lab in Alderley Park and Alex Alderton, Roberto Amato, Jeffrey Barrett, Sonia Goncalves, Ewan Harrison, David K. Jackson, Ian Johnston, Dominic Kwiatkowski, Cordelia Langford, John Sillitoe on behalf of the Wellcome Sanger Institute COVID-19 Surveillance Team |
| EPI_ISL_3228134 | Berkshire and Surrey Pathology Services Lighthouse Laboratory | Wellcome Sanger Institute for the COVID-19 Genomics UK (COG-UK) Consortium | Berkshire and Surrey Pathology Services Lighthouse Laboratory and Alex Alderton, Roberto Amato, Jeffrey Barrett, Sonia Goncalves, Ewan Harrison, David K. Jackson, Ian Johnston, Dominic Kwiatkowski, Cordelia Langford, John Sillitoe on behalf of the Wellcome Sanger Institute COVID-19 Surveillance Team |
| EPI_ISL_3228135 | Lighthouse Lab in Alderley Park | Wellcome Sanger Institute for the COVID-19 Genomics UK (COG-UK) Consortium | Jacquelyn Wynn, Mairead Hyland, The Lighthouse Lab in Alderley Park and Alex Alderton, Roberto Amato, Jeffrey Barrett, Sonia Goncalves, Ewan Harrison, David K. Jackson, Ian Johnston, Dominic Kwiatkowski, Cordelia Langford, John Sillitoe on behalf of the Wellcome Sanger Institute COVID-19 Surveillance Team |
| EPI_ISL_3228136, EPI_ISL_3228137, EPI_ISL_3228138 | Berkshire and Surrey Pathology Services Lighthouse Laboratory | Wellcome Sanger Institute for the COVID-19 Genomics UK (COG-UK) Consortium | Berkshire and Surrey Pathology Services Lighthouse Laboratory and Alex Alderton, Roberto Amato, Jeffrey Barrett, Sonia Goncalves, Ewan Harrison, David K. Jackson, Ian Johnston, Dominic Kwiatkowski, Cordelia Langford, John Sillitoe on behalf of the Wellcome Sanger Institute COVID-19 Surveillance Team |
| EPI_ISL_3228139 | Lighthouse Lab in Alderley Park | Wellcome Sanger Institute for the COVID-19 Genomics UK (COG-UK) Consortium | Jacquelyn Wynn, Mairead Hyland, The Lighthouse Lab in Alderley Park and Alex Alderton, Roberto Amato, Jeffrey Barrett, Sonia Goncalves, Ewan Harrison, David K. Jackson, Ian Johnston, Dominic Kwiatkowski, Cordelia Langford, John Sillitoe on behalf of the Wellcome Sanger Institute COVID-19 Surveillance Team |
| EPI_ISL_3228140, EPI_ISL_3228141 | Berkshire and Surrey Pathology Services Lighthouse Laboratory | Wellcome Sanger Institute for the COVID-19 Genomics UK (COG-UK) Consortium | Berkshire and Surrey Pathology Services Lighthouse Laboratory and Alex Alderton, Roberto Amato, Jeffrey Barrett, Sonia Goncalves, Ewan Harrison, David K. Jackson, Ian Johnston, Dominic Kwiatkowski, Cordelia Langford, John Sillitoe on behalf of the Wellcome Sanger Institute COVID-19 Surveillance Team |
| EPI_ISL_3228142 | Lighthouse Lab in Alderley Park | Wellcome Sanger Institute for the COVID-19 Genomics UK (COG-UK) Consortium | Jacquelyn Wynn, Mairead Hyland, The Lighthouse Lab in Alderley Park and Alex Alderton, Roberto Amato, Jeffrey Barrett, Sonia Goncalves, Ewan Harrison, David K. Jackson, Ian Johnston, Dominic Kwiatkowski, Cordelia Langford, John Sillitoe on behalf of the Wellcome Sanger Institute COVID-19 Surveillance Team |
| EPI_ISL_3228143 | Berkshire and Surrey Pathology Services Lighthouse Laboratory | Wellcome Sanger Institute for the COVID-19 Genomics UK (COG-UK) Consortium | Berkshire and Surrey Pathology Services Lighthouse Laboratory and Alex Alderton, Roberto Amato, Jeffrey Barrett, Sonia Goncalves, Ewan Harrison, David K. Jackson, Ian Johnston, Dominic Kwiatkowski, Cordelia Langford, John Sillitoe on behalf of the Wellcome Sanger Institute COVID-19 Surveillance Team |
| EPI_ISL_3228144 | Lighthouse Lab in Alderley Park | Wellcome Sanger Institute for the COVID-19 Genomics UK (COG-UK) Consortium | Jacquelyn Wynn, Mairead Hyland, The Lighthouse Lab in Alderley Park and Alex Alderton, Roberto Amato, Jeffrey Barrett, Sonia Goncalves, Ewan Harrison, David K. Jackson, Ian Johnston, Dominic Kwiatkowski, Cordelia Langford, John Sillitoe on behalf of the Wellcome Sanger Institute COVID-19 Surveillance Team |
| EPI_ISL_3228145, EPI_ISL_3228146, EPI_ISL_3228147 | Berkshire and Surrey Pathology Services Lighthouse Laboratory | Wellcome Sanger Institute for the COVID-19 Genomics UK (COG-UK) Consortium | Berkshire and Surrey Pathology Services Lighthouse Laboratory and Alex Alderton, Roberto Amato, Jeffrey Barrett, Sonia Goncalves, Ewan Harrison, David K. Jackson, Ian Johnston, Dominic Kwiatkowski, Cordelia Langford, John Sillitoe on behalf of the Wellcome Sanger Institute COVID-19 Surveillance Team |
| EPI_ISL_3228148, EPI_ISL_3228149, EPI_ISL_3228150, EPI_ISL_3228151, EPI_ISL_3228152, EPI_ISL_3228153, EPI_ISL_3228154, EPI_ISL_3228155, EPI_ISL_3228156, EPI_ISL_3228157, EPI_ISL_3228158, EPI_ISL_3228159, EPI_ISL_3228160, EPI_ISL_3228161, EPI_ISL_3228162, EPI_ISL_3228163, EPI_ISL_3228164, EPI_ISL_3228165, EPI_ISL_3228166, EPI_ISL_3228167, EPI_ISL_3228168, EPI_ISL_3228169, EPI_ISL_3228170, EPI_ISL_3228171, EPI_ISL_3228172, EPI_ISL_3228173, EPI_ISL_3228174, EPI_ISL_3228175, EPI_ISL_3228176, EPI_ISL_3228177, EPI_ISL_3228178, EPI_ISL_3228179, EPI_ISL_3228180, EPI_ISL_3228181, EPI_ISL_3228182, EPI_ISL_3228183, EPI_ISL_3228184, EPI_ISL_3228185, EPI_ISL_3228186, EPI_ISL_3228187, EPI_ISL_3228188, EPI_ISL_3228189, EPI_ISL_3228190, EPI_ISL_3228191, EPI_ISL_3228192, EPI_ISL_3228193, EPI_ISL_3228194, EPI_ISL_3228195, EPI_ISL_3228196, EPI_ISL_3228197, EPI_ISL_3228198, EPI_ISL_3228199, EPI_ISL_3228200, EPI_ISL_3228201, EPI_ISL_3228202, EPI_ISL_3228203, EPI_ISL_3228204, EPI_ISL_3228205, EPI_ISL_3228206, EPI_ISL_3228207, EPI_ISL_3228208, EPI_ISL_3228209, EPI_ISL_3228210, EPI_ISL_3228211, EPI_ISL_3228212, EPI_ISL_3228213, EPI_ISL_3228214, EPI_ISL_3228215, EPI_ISL_3228216, EPI_ISL_3228217, EPI_ISL_3228218, EPI_ISL_3228219 | | | |
| see above | Lighthouse Lab in Glasgow | Wellcome Sanger Institute for the COVID-19 Genomics UK (COG-UK) Consortium | Harper VanSteenhouse, Yumi Kasai, David Gray, Carol Clugston, Anna Dominiczak and Alex Alderton, Roberto Amato, Jeffrey Barrett, Sonia Goncalves, Ewan Harrison, David K. Jackson, Ian Johnston, Dominic Kwiatkowski, Cordelia Langford, John Sillitoe on behalf of the Wellcome Sanger Institute COVID-19 Surveillance Team |
| EPI_ISL_3228220 | Lighthouse Lab in Milton Keynes | Wellcome Sanger Institute for the COVID-19 Genomics UK (COG-UK) Consortium | The Lighthouse Lab in Milton Keynes and Alex Alderton, Roberto Amato, Jeffrey Barrett, Sonia Goncalves, Ewan Harrison, David K. Jackson, Ian Johnston, Dominic Kwiatkowski, Cordelia Langford, John Sillitoe on behalf of the Wellcome Sanger Institute COVID-19 Surveillance Team |
| EPI_ISL_3228221, EPI_ISL_3228222, EPI_ISL_3228223, EPI_ISL_3228224, EPI_ISL_3228225, EPI_ISL_3228226, EPI_ISL_3228227, EPI_ISL_3228228 | Lighthouse Lab in Glasgow | Wellcome Sanger Institute for the COVID-19 Genomics UK (COG-UK) Consortium | Harper VanSteenhouse, Yumi Kasai, David Gray, Carol Clugston, Anna Dominiczak and Alex Alderton, Roberto Amato, Jeffrey Barrett, Sonia Goncalves, Ewan Harrison, David K. Jackson, Ian Johnston, Dominic Kwiatkowski, Cordelia Langford, John Sillitoe on behalf of the Wellcome Sanger Institute COVID-19 Surveillance Team |
| EPI_ISL_3228229 | Lighthouse Lab in Milton Keynes | Wellcome Sanger Institute for the COVID-19 Genomics UK (COG-UK) Consortium | The Lighthouse Lab in Milton Keynes and Alex Alderton, Roberto Amato, Jeffrey Barrett, Sonia Goncalves, Ewan Harrison, David K. Jackson, Ian Johnston, Dominic Kwiatkowski, Cordelia Langford, John Sillitoe on behalf of the Wellcome Sanger Institute COVID-19 Surveillance Team |
| EPI_ISL_3228230, EPI_ISL_3228231, EPI_ISL_3228232, EPI_ISL_3228233, EPI_ISL_3228234, EPI_ISL_3228235, EPI_ISL_3228236, EPI_ISL_3228237, EPI_ISL_3228238, EPI_ISL_3228239, EPI_ISL_3228240 | | | |
| see above | Lighthouse Lab in Glasgow | Wellcome Sanger Institute for the COVID-19 Genomics UK (COG-UK) Consortium | Harper VanSteenhouse, Yumi Kasai, David Gray, Carol Clugston, Anna Dominiczak and Alex Alderton, Roberto Amato, Jeffrey Barrett, Sonia Goncalves, Ewan Harrison, David K. Jackson, Ian Johnston, Dominic Kwiatkowski, Cordelia Langford, John Sillitoe on behalf of the Wellcome Sanger Institute COVID-19 Surveillance Team |
| EPI_ISL_3228242, EPI_ISL_3228243, EPI_ISL_3228244, EPI_ISL_3228245 | Lighthouse Laboratory Plymouth | Wellcome Sanger Institute for the COVID-19 Genomics UK (COG-UK) Consortium | Lighthouse Laboratory Plymouth and Alex Alderton, Roberto Amato, Jeffrey Barrett, Sonia Goncalves, Ewan Harrison, David K. Jackson, Ian Johnston, Dominic Kwiatkowski, Cordelia Langford, John Sillitoe on behalf of the Wellcome Sanger Institute COVID-19 Surveillance Team |
| EPI_ISL_3228246, EPI_ISL_3228247 | Lighthouse Lab in Alderley Park | Wellcome Sanger Institute for the COVID-19 Genomics UK (COG-UK) Consortium | Jacquelyn Wynn, Mairead Hyland, The Lighthouse Lab in Alderley Park and Alex Alderton, Roberto Amato, Jeffrey Barrett, Sonia Goncalves, Ewan Harrison, David K. Jackson, Ian Johnston, Dominic Kwiatkowski, Cordelia Langford, John Sillitoe on behalf of the Wellcome Sanger Institute COVID-19 Surveillance Team |
| EPI_ISL_3228248 | Berkshire and Surrey Pathology Services Lighthouse Laboratory | Wellcome Sanger Institute for the COVID-19 Genomics UK (COG-UK) Consortium | Berkshire and Surrey Pathology Services Lighthouse Laboratory and Alex Alderton, Roberto Amato, Jeffrey Barrett, Sonia Goncalves, Ewan Harrison, David K. Jackson, Ian Johnston, Dominic Kwiatkowski, Cordelia Langford, John Sillitoe on behalf of the Wellcome Sanger Institute COVID-19 Surveillance Team |
| EPI_ISL_3228249 | Lighthouse Lab in Alderley Park | Wellcome Sanger Institute for the COVID-19 Genomics UK (COG-UK) Consortium | Jacquelyn Wynn, Mairead Hyland, The Lighthouse Lab in Alderley Park and Alex Alderton, Roberto Amato, Jeffrey Barrett, Sonia Goncalves, Ewan Harrison, David K. Jackson, Ian Johnston, Dominic Kwiatkowski, Cordelia Langford, John Sillitoe on behalf of the Wellcome Sanger Institute COVID-19 Surveillance Team |
| EPI_ISL_3228250 | Health Services Laboratories | Wellcome Sanger Institute for the COVID-19 Genomics UK (COG-UK) Consortium | Health Services Laboratories and Alex Alderton, Roberto Amato, Jeffrey Barrett, Sonia Goncalves, Ewan Harrison, David K. Jackson, Ian Johnston, Dominic Kwiatkowski, Cordelia Langford, John Sillitoe on behalf of the Wellcome Sanger Institute COVID-19 Surveillance Team |
| EPI_ISL_3228251 | Lighthouse Lab in Alderley Park | Wellcome Sanger Institute for the COVID-19 Genomics UK (COG-UK) Consortium | Jacquelyn Wynn, Mairead Hyland, The Lighthouse Lab in Alderley Park and Alex Alderton, Roberto Amato, Jeffrey Barrett, Sonia Goncalves, Ewan Harrison, David K. Jackson, Ian Johnston, Dominic Kwiatkowski, Cordelia Langford, John Sillitoe on behalf of the Wellcome Sanger Institute COVID-19 Surveillance Team |
| EPI_ISL_3228252 | Health Services Laboratories | Wellcome Sanger Institute for the COVID-19 Genomics UK (COG-UK) Consortium | Health Services Laboratories and Alex Alderton, Roberto Amato, Jeffrey Barrett, Sonia Goncalves, Ewan Harrison, David K. Jackson, Ian Johnston, Dominic Kwiatkowski, Cordelia Langford, John Sillitoe on behalf of the Wellcome Sanger Institute COVID-19 Surveillance Team |
| EPI_ISL_3228253 | Lighthouse Lab in Glasgow | Wellcome Sanger Institute for the COVID-19 Genomics UK (COG-UK) Consortium | Harper VanSteenhouse, Yumi Kasai, David Gray, Carol Clugston, Anna Dominiczak and Alex Alderton, Roberto Amato, Jeffrey Barrett, Sonia Goncalves, Ewan Harrison, David K. Jackson, Ian Johnston, Dominic Kwiatkowski, Cordelia Langford, John Sillitoe on behalf of the Wellcome Sanger Institute COVID-19 Surveillance Team |
| EPI_ISL_3228254 | Lighthouse Lab in Alderley Park | Wellcome Sanger Institute for the COVID-19 Genomics UK (COG-UK) | Jacquelyn Wynn, Mairead Hyland, The Lighthouse Lab in Alderley Park and Alex Alderton, Roberto Amato, Jeffrey Barrett, Sonia Goncalves, Ewan Harrison, David K. Jackson, Ian Johnston, Dominic Kwiatkowski, Cordelia Langford, John Sillitoe on |

| | | | |
|---|---|---|---|
| | | Consortium | behalf of the Wellcome Sanger Institute COVID-19 Surveillance Team |
| EPI_ISL_3228255 | Berkshire and Surrey Pathology Services Lighthouse Laboratory | Wellcome Sanger Institute for the COVID-19 Genomics UK (COG-UK) Consortium | Berkshire and Surrey Pathology Services Lighthouse Laboratory and Alex Alderton, Roberto Amato, Jeffrey Barrett, Sonia Goncalves, Ewan Harrison, David K. Jackson, Ian Johnston, Dominic Kwiatkowski, Cordelia Langford, John Sillitoe on behalf of the Wellcome Sanger Institute COVID-19 Surveillance Team |
| EPI_ISL_3228256 | Lighthouse Lab in Glasgow | Wellcome Sanger Institute for the COVID-19 Genomics UK (COG-UK) Consortium | Harper VanSteenhouse, Yumi Kasai, David Gray, Carol Clugston, Anna Dominiczak and Alex Alderton, Roberto Amato, Jeffrey Barrett, Sonia Goncalves, Ewan Harrison, David K. Jackson, Ian Johnston, Dominic Kwiatkowski, Cordelia Langford, John Sillitoe on behalf of the Wellcome Sanger Institute COVID-19 Surveillance Team |
| EPI_ISL_3228257, EPI_ISL_3228258 | Health Services Laboratories | Wellcome Sanger Institute for the COVID-19 Genomics UK (COG-UK) Consortium | Health Services Laboratories and Alex Alderton, Roberto Amato, Jeffrey Barrett, Sonia Goncalves, Ewan Harrison, David K. Jackson, Ian Johnston, Dominic Kwiatkowski, Cordelia Langford, John Sillitoe on behalf of the Wellcome Sanger Institute COVID-19 Surveillance Team |
| EPI_ISL_3228259 | Lighthouse Lab in Alderley Park | Wellcome Sanger Institute for the COVID-19 Genomics UK (COG-UK) Consortium | Jacquelyn Wynn, Mairead Hyland, The Lighthouse Lab in Alderley Park and Alex Alderton, Roberto Amato, Jeffrey Barrett, Sonia Goncalves, Ewan Harrison, David K. Jackson, Ian Johnston, Dominic Kwiatkowski, Cordelia Langford, John Sillitoe on behalf of the Wellcome Sanger Institute COVID-19 Surveillance Team |
| EPI_ISL_3228260 | Berkshire and Surrey Pathology Services Lighthouse Laboratory | Wellcome Sanger Institute for the COVID-19 Genomics UK (COG-UK) Consortium | Berkshire and Surrey Pathology Services Lighthouse Laboratory and Alex Alderton, Roberto Amato, Jeffrey Barrett, Sonia Goncalves, Ewan Harrison, David K. Jackson, Ian Johnston, Dominic Kwiatkowski, Cordelia Langford, John Sillitoe on behalf of the Wellcome Sanger Institute COVID-19 Surveillance Team |
| EPI_ISL_3228261, EPI_ISL_3228262, EPI_ISL_3228263 | Lighthouse Lab in Alderley Park | Wellcome Sanger Institute for the COVID-19 Genomics UK (COG-UK) Consortium | Jacquelyn Wynn, Mairead Hyland, The Lighthouse Lab in Alderley Park and Alex Alderton, Roberto Amato, Jeffrey Barrett, Sonia Goncalves, Ewan Harrison, David K. Jackson, Ian Johnston, Dominic Kwiatkowski, Cordelia Langford, John Sillitoe on behalf of the Wellcome Sanger Institute COVID-19 Surveillance Team |
| EPI_ISL_3228264 | Berkshire and Surrey Pathology Services Lighthouse Laboratory | Wellcome Sanger Institute for the COVID-19 Genomics UK (COG-UK) Consortium | Berkshire and Surrey Pathology Services Lighthouse Laboratory and Alex Alderton, Roberto Amato, Jeffrey Barrett, Sonia Goncalves, Ewan Harrison, David K. Jackson, Ian Johnston, Dominic Kwiatkowski, Cordelia Langford, John Sillitoe on behalf of the Wellcome Sanger Institute COVID-19 Surveillance Team |
| EPI_ISL_3228265 | Lighthouse Lab in Glasgow | Wellcome Sanger Institute for the COVID-19 Genomics UK (COG-UK) Consortium | Harper VanSteenhouse, Yumi Kasai, David Gray, Carol Clugston, Anna Dominiczak and Alex Alderton, Roberto Amato, Jeffrey Barrett, Sonia Goncalves, Ewan Harrison, David K. Jackson, Ian Johnston, Dominic Kwiatkowski, Cordelia Langford, John Sillitoe on behalf of the Wellcome Sanger Institute COVID-19 Surveillance Team |
| EPI_ISL_3228266, EPI_ISL_3228267 | Berkshire and Surrey Pathology Services Lighthouse Laboratory | Wellcome Sanger Institute for the COVID-19 Genomics UK (COG-UK) Consortium | Berkshire and Surrey Pathology Services Lighthouse Laboratory and Alex Alderton, Roberto Amato, Jeffrey Barrett, Sonia Goncalves, Ewan Harrison, David K. Jackson, Ian Johnston, Dominic Kwiatkowski, Cordelia Langford, John Sillitoe on behalf of the Wellcome Sanger Institute COVID-19 Surveillance Team |
| EPI_ISL_3228268 | Health Services Laboratories | Wellcome Sanger Institute for the COVID-19 Genomics UK (COG-UK) Consortium | Health Services Laboratories and Alex Alderton, Roberto Amato, Jeffrey Barrett, Sonia Goncalves, Ewan Harrison, David K. Jackson, Ian Johnston, Dominic Kwiatkowski, Cordelia Langford, John Sillitoe on behalf of the Wellcome Sanger Institute COVID-19 Surveillance Team |
| EPI_ISL_3228269, EPI_ISL_3228270 | Lighthouse Lab in Alderley Park | Wellcome Sanger Institute for the COVID-19 Genomics UK (COG-UK) Consortium | Jacquelyn Wynn, Mairead Hyland, The Lighthouse Lab in Alderley Park and Alex Alderton, Roberto Amato, Jeffrey Barrett, Sonia Goncalves, Ewan Harrison, David K. Jackson, Ian Johnston, Dominic Kwiatkowski, Cordelia Langford, John Sillitoe on behalf of the Wellcome Sanger Institute COVID-19 Surveillance Team |
| EPI_ISL_3228271, EPI_ISL_3228272 | Lighthouse Lab in Glasgow | Wellcome Sanger Institute for the COVID-19 Genomics UK (COG-UK) Consortium | Harper VanSteenhouse, Yumi Kasai, David Gray, Carol Clugston, Anna Dominiczak and Alex Alderton, Roberto Amato, Jeffrey Barrett, Sonia Goncalves, Ewan Harrison, David K. Jackson, Ian Johnston, Dominic Kwiatkowski, Cordelia Langford, John Sillitoe on behalf of the Wellcome Sanger Institute COVID-19 Surveillance Team |
| EPI_ISL_3228273 | Lighthouse Laboratory Plymouth | Wellcome Sanger Institute for the COVID-19 Genomics UK (COG-UK) Consortium | Lighthouse Laboratory Plymouth and Alex Alderton, Roberto Amato, Jeffrey Barrett, Sonia Goncalves, Ewan Harrison, David K. Jackson, Ian Johnston, Dominic Kwiatkowski, Cordelia Langford, John Sillitoe on behalf of the Wellcome Sanger Institute COVID-19 Surveillance Team |
| EPI_ISL_3228274, EPI_ISL_3228275, EPI_ISL_3228276, EPI_ISL_3228277, EPI_ISL_3228278 | Lighthouse Lab in Glasgow | Wellcome Sanger Institute for the COVID-19 Genomics UK (COG-UK) Consortium | Harper VanSteenhouse, Yumi Kasai, David Gray, Carol Clugston, Anna Dominiczak and Alex Alderton, Roberto Amato, Jeffrey Barrett, Sonia Goncalves, Ewan Harrison, David K. Jackson, Ian Johnston, Dominic Kwiatkowski, Cordelia Langford, John Sillitoe on behalf of the Wellcome Sanger Institute COVID-19 Surveillance Team |
| EPI_ISL_3228279 | Lighthouse Lab in Alderley Park | Wellcome Sanger Institute for the COVID-19 Genomics UK (COG-UK) Consortium | Jacquelyn Wynn, Mairead Hyland, The Lighthouse Lab in Alderley Park and Alex Alderton, Roberto Amato, Jeffrey Barrett, Sonia Goncalves, Ewan Harrison, David K. Jackson, Ian Johnston, Dominic Kwiatkowski, Cordelia Langford, John Sillitoe on behalf of the Wellcome Sanger Institute COVID-19 Surveillance Team |
| EPI_ISL_3228280 | Lighthouse Lab in Glasgow | Wellcome Sanger Institute for the COVID-19 Genomics UK (COG-UK) Consortium | Harper VanSteenhouse, Yumi Kasai, David Gray, Carol Clugston, Anna Dominiczak and Alex Alderton, Roberto Amato, Jeffrey Barrett, Sonia Goncalves, Ewan Harrison, David K. Jackson, Ian Johnston, Dominic Kwiatkowski, Cordelia Langford, John Sillitoe on behalf of the Wellcome Sanger Institute COVID-19 Surveillance Team |
| EPI_ISL_3228281 | Lighthouse Laboratory Plymouth | Wellcome Sanger Institute for the COVID-19 Genomics UK (COG-UK) Consortium | Lighthouse Laboratory Plymouth and Alex Alderton, Roberto Amato, Jeffrey Barrett, Sonia Goncalves, Ewan Harrison, David K. Jackson, Ian Johnston, Dominic Kwiatkowski, Cordelia Langford, John Sillitoe on behalf of the Wellcome Sanger Institute COVID-19 Surveillance Team |
| EPI_ISL_3228282 | Lighthouse Lab in Glasgow | Wellcome Sanger Institute for the COVID-19 Genomics UK (COG-UK) Consortium | Harper VanSteenhouse, Yumi Kasai, David Gray, Carol Clugston, Anna Dominiczak and Alex Alderton, Roberto Amato, Jeffrey Barrett, Sonia Goncalves, Ewan Harrison, David K. Jackson, Ian Johnston, Dominic Kwiatkowski, Cordelia Langford, John Sillitoe on behalf of the Wellcome Sanger Institute COVID-19 Surveillance Team |
| EPI_ISL_3228283 | Lighthouse Lab in Alderley Park | Wellcome Sanger Institute for the COVID-19 Genomics UK (COG-UK) Consortium | Jacquelyn Wynn, Mairead Hyland, The Lighthouse Lab in Alderley Park and Alex Alderton, Roberto Amato, Jeffrey Barrett, Sonia Goncalves, Ewan Harrison, David K. Jackson, Ian Johnston, Dominic Kwiatkowski, Cordelia Langford, John Sillitoe on behalf of the Wellcome Sanger Institute COVID-19 Surveillance Team |
| EPI_ISL_3228284, EPI_ISL_3228285, EPI_ISL_3228286, EPI_ISL_3228287, EPI_ISL_3228288, EPI_ISL_3228289, EPI_ISL_3228290, EPI_ISL_3228291, EPI_ISL_3228292, EPI_ISL_3228293, EPI_ISL_3228294, EPI_ISL_3228295, EPI_ISL_3228296, EPI_ISL_3228297, EPI_ISL_3228298, EPI_ISL_3228299, EPI_ISL_3228300, EPI_ISL_3228301, EPI_ISL_3228302, EPI_ISL_3228303, EPI_ISL_3228304, EPI_ISL_3228305, EPI_ISL_3228306 | | | |
| see above | Lighthouse Lab in Glasgow | Wellcome Sanger Institute for the COVID-19 Genomics UK (COG-UK) Consortium | Harper VanSteenhouse, Yumi Kasai, David Gray, Carol Clugston, Anna Dominiczak and Alex Alderton, Roberto Amato, Jeffrey Barrett, Sonia Goncalves, Ewan Harrison, David K. Jackson, Ian Johnston, Dominic Kwiatkowski, Cordelia Langford, John Sillitoe on behalf of the Wellcome Sanger Institute COVID-19 Surveillance Team |
| EPI_ISL_3228307 | Lighthouse Laboratory Plymouth | Wellcome Sanger Institute for the COVID-19 Genomics UK (COG-UK) Consortium | Lighthouse Laboratory Plymouth and Alex Alderton, Roberto Amato, Jeffrey Barrett, Sonia Goncalves, Ewan Harrison, David K. Jackson, Ian Johnston, Dominic Kwiatkowski, Cordelia Langford, John Sillitoe on behalf of the Wellcome Sanger Institute COVID-19 Surveillance Team |
| EPI_ISL_3228308 | Lighthouse Lab in Glasgow | Wellcome Sanger Institute for the COVID-19 Genomics UK (COG-UK) Consortium | Harper VanSteenhouse, Yumi Kasai, David Gray, Carol Clugston, Anna Dominiczak and Alex Alderton, Roberto Amato, Jeffrey Barrett, Sonia Goncalves, Ewan Harrison, David K. Jackson, Ian Johnston, Dominic Kwiatkowski, Cordelia Langford, John Sillitoe on behalf of the Wellcome Sanger Institute COVID-19 Surveillance Team |
| EPI_ISL_3228309, EPI_ISL_3228311 | COVID-19 Detection Lab, Chattogram Veterinary and Animal Sciences University | Genomic Research Lab, BCSIR | Md. Murshed Hasan Sarkar, Abu Sayeed Mohammad Mahmud, Mohammad Samir Uzzaman, Eshrar Osman, Md. Ahasan Habib, Shahina Akter, Tanjina Akhter Banu, Barna Goswami, Iffat Jahan, Mohammad Mohi Uddin, Md. Kamrul Islam, Goutam Buddha Das, Tridip Das, Tanvir Ahmad Nizami, Eaftekhar Ahmed Rana, Md. Sirazul Islam, Pronesh Dutta, Sharmin Chowdhury, Md. Paritosh Kumar Biswas, Md. Salim Khan |
| EPI_ISL_3228320, EPI_ISL_3228321, EPI_ISL_3228322, EPI_ISL_3228323, EPI_ISL_3228324, EPI_ISL_3228325, EPI_ISL_3228326, EPI_ISL_3228327, EPI_ISL_3228328, EPI_ISL_3228329, EPI_ISL_3228330, EPI_ISL_3228331, EPI_ISL_3228332, EPI_ISL_3228333, EPI_ISL_3228334, EPI_ISL_3228335, EPI_ISL_3228336, EPI_ISL_3228337, EPI_ISL_3228338, EPI_ISL_3228339, EPI_ISL_3228340, EPI_ISL_3228341, EPI_ISL_3228342, EPI_ISL_3228343, EPI_ISL_3228344, EPI_ISL_3228345, EPI_ISL_3228346, EPI_ISL_3228347, EPI_ISL_3228348, EPI_ISL_3228349, EPI_ISL_3228350, EPI_ISL_3228351, EPI_ISL_3228352, EPI_ISL_3228353, EPI_ISL_3228354, EPI_ISL_3228355, EPI_ISL_3228356, EPI_ISL_3228357, EPI_ISL_3228358, EPI_ISL_3228359, EPI_ISL_3228360, EPI_ISL_3228361, EPI_ISL_3228362, EPI_ISL_3228363, EPI_ISL_3228364, EPI_ISL_3228365, EPI_ISL_3228366, EPI_ISL_3228367, EPI_ISL_3228368, EPI_ISL_3228369, EPI_ISL_3228370, EPI_ISL_3228371, EPI_ISL_3228372, EPI_ISL_3228373, EPI_ISL_3228375, EPI_ISL_3228376, EPI_ISL_3228377, EPI_ISL_3228378, EPI_ISL_3228379, EPI_ISL_3228380, EPI_ISL_3228381, EPI_ISL_3228382, EPI_ISL_3228383, EPI_ISL_3228384, EPI_ISL_3228385, EPI_ISL_3228386, EPI_ISL_3228387, EPI_ISL_3228388, EPI_ISL_3228389, EPI_ISL_3228390, EPI_ISL_3228391, EPI_ISL_3228392, EPI_ISL_3228393, EPI_ISL_3228394, EPI_ISL_3228395, EPI_ISL_3228396, EPI_ISL_3228397, EPI_ISL_3228398, EPI_ISL_3228399, EPI_ISL_3228400, EPI_ISL_3228401, EPI_ISL_3228402, EPI_ISL_3228403, EPI_ISL_3228404, EPI_ISL_3228405, EPI_ISL_3228406, EPI_ISL_3228407, EPI_ISL_3228408, EPI_ISL_3228409, EPI_ISL_3228410, EPI_ISL_3228411, EPI_ISL_3228412, EPI_ISL_3228413, EPI_ISL_3228414, EPI_ISL_3228415, EPI_ISL_3228416, EPI_ISL_3228417, EPI_ISL_3228418, EPI_ISL_3228419, EPI_ISL_3228420, EPI_ISL_3228421, EPI_ISL_3228422, EPI_ISL_3228423, EPI_ISL_3228424, EPI_ISL_3228425, EPI_ISL_3228426, EPI_ISL_3228427, EPI_ISL_3228428, EPI_ISL_3228429, EPI_ISL_3228430, EPI_ISL_3228431, EPI_ISL_3228432, EPI_ISL_3228433, EPI_ISL_3228434, EPI_ISL_3228435, EPI_ISL_3228436, EPI_ISL_3228437, EPI_ISL_3228438, EPI_ISL_3228439, EPI_ISL_3228440, EPI_ISL_3228441, EPI_ISL_3228442, EPI_ISL_3228443, EPI_ISL_3228444, EPI_ISL_3228445, EPI_ISL_3228446, EPI_ISL_3228447, EPI_ISL_3228448, EPI_ISL_3228449, EPI_ISL_3228450, EPI_ISL_3228451, EPI_ISL_3228452, EPI_ISL_3228453, EPI_ISL_3228454, EPI_ISL_3228455, EPI_ISL_3228456, EPI_ISL_3228457, EPI_ISL_3228458, EPI_ISL_3228459, EPI_ISL_3228460, EPI_ISL_3228461, EPI_ISL_3228462, EPI_ISL_3228463, EPI_ISL_3228464, EPI_ISL_3228465, EPI_ISL_3228466, EPI_ISL_3228467, EPI_ISL_3228468, EPI_ISL_3228469, EPI_ISL_3228470, EPI_ISL_3228471, EPI_ISL_3228472, EPI_ISL_3228473, EPI_ISL_3228474, EPI_ISL_3228475, EPI_ISL_3228476, EPI_ISL_3228477, EPI_ISL_3228478, EPI_ISL_3228479, EPI_ISL_3228480, EPI_ISL_3228481, EPI_ISL_3228482, EPI_ISL_3228483, EPI_ISL_3228484, EPI_ISL_3228485, EPI_ISL_3228486, EPI_ISL_3228487, EPI_ISL_3228488, EPI_ISL_3228489, EPI_ISL_3228490, EPI_ISL_3228491, EPI_ISL_3228492, EPI_ISL_3228493, EPI_ISL_3228494, EPI_ISL_3228495, EPI_ISL_3228496, EPI_ISL_3228497, EPI_ISL_3228498, EPI_ISL_3228499, EPI_ISL_3228500, EPI_ISL_3228501, EPI_ISL_3228502, EPI_ISL_3228503, EPI_ISL_3228504, EPI_ISL_3228505, EPI_ISL_3228506, EPI_ISL_3228507, EPI_ISL_3228509, EPI_ISL_3228510, EPI_ISL_3228511, EPI_ISL_3228512, EPI_ISL_3228513, EPI_ISL_3228514, EPI_ISL_3228515, EPI_ISL_3228516, EPI_ISL_3228517, EPI_ISL_3228518, EPI_ISL_3228519, EPI_ISL_3228520, EPI_ISL_3228521, EPI_ISL_3228522, EPI_ISL_3228523, EPI_ISL_3228524, EPI_ISL_3228525, EPI_ISL_3228526, EPI_ISL_3228527, EPI_ISL_3228528, EPI_ISL_3228529, EPI_ISL_3228530, EPI_ISL_3228531, EPI_ISL_3228532, EPI_ISL_3228533, EPI_ISL_3228534, EPI_ISL_3228535, EPI_ISL_3228536, EPI_ISL_3228537, EPI_ISL_3228538, EPI_ISL_3228539, EPI_ISL_3228540, EPI_ISL_3228541, EPI_ISL_3228542, EPI_ISL_3228543, EPI_ISL_3228544, EPI_ISL_3228545, EPI_ISL_3228546, EPI_ISL_3228547, EPI_ISL_3228548, EPI_ISL_3228549, EPI_ISL_3228550, EPI_ISL_3228551, EPI_ISL_3228552, EPI_ISL_3228553, EPI_ISL_3228554, EPI_ISL_3228555, EPI_ISL_3228556, EPI_ISL_3228557 | | | |
| see above | Department of Bacteria, Parasites and Fungi, Statens Serum Institut, Copenhagen, Denmark | Statens Serum Institut Bioinformatics and Microbial Genomics | Danish Covid-19 Genome Consortium |
| EPI_ISL_3228558 | Molekylær Medicinsk Afdeling, Aarhus University Hospital, Aarhus, Denmark | Statens Serum Institut Bioinformatics and Microbial Genomics | Danish Covid-19 Genome Consortium |
| EPI_ISL_3228559, EPI_ISL_3228560, EPI_ISL_3228561, EPI_ISL_3228562, EPI_ISL_3228563, EPI_ISL_3228564, EPI_ISL_3228565, EPI_ISL_3228566, EPI_ISL_3228567, EPI_ISL_3228568, EPI_ISL_3228569, EPI_ISL_3228570, EPI_ISL_3228571, EPI_ISL_3228572, EPI_ISL_3228573, EPI_ISL_3228574, EPI_ISL_3228575, EPI_ISL_3228576, EPI_ISL_3228577, EPI_ISL_3228578, EPI_ISL_3228579, EPI_ISL_3228580, EPI_ISL_3228581, EPI_ISL_3228582, EPI_ISL_3228583, EPI_ISL_3228584, EPI_ISL_3228585, EPI_ISL_3228586, EPI_ISL_3228587, EPI_ISL_3228588, EPI_ISL_3228589, EPI_ISL_3228590, EPI_ISL_3228591, EPI_ISL_3228592, EPI_ISL_3228593, EPI_ISL_3228594, EPI_ISL_3228595, EPI_ISL_3228596, EPI_ISL_3228597, EPI_ISL_3228598, EPI_ISL_3228599 | | | |
| see above | Department of Bacteria, Parasites and Fungi, Statens Serum Institut, Copenhagen, Denmark | Statens Serum Institut Bioinformatics and Microbial Genomics | Danish Covid-19 Genome Consortium |
| EPI_ISL_3228600 | Molekylær Medicinsk Afdeling, Aarhus University Hospital, Aarhus, Denmark | Statens Serum Institut Bioinformatics and Microbial Genomics | Danish Covid-19 Genome Consortium |
| EPI_ISL_3228601, EPI_ISL_3228602, EPI_ISL_3228603, EPI_ISL_3228604, EPI_ISL_3228605, EPI_ISL_3228606, EPI_ISL_3228607, EPI_ISL_3228608, EPI_ISL_3228609, EPI_ISL_3228610, EPI_ISL_3228611, EPI_ISL_3228612, EPI_ISL_3228613, EPI_ISL_3228614, EPI_ISL_3228615, EPI_ISL_3228616, EPI_ISL_3228617, EPI_ISL_3228618, EPI_ISL_3228619, EPI_ISL_3228620, EPI_ISL_3228621, EPI_ISL_3228622, EPI_ISL_3228623, EPI_ISL_3228624, EPI_ISL_3228625, EPI_ISL_3228626, EPI_ISL_3228627, EPI_ISL_3228628, EPI_ISL_3228629, EPI_ISL_3228630, EPI_ISL_3228631, EPI_ISL_3228632, EPI_ISL_3228633, EPI_ISL_3228634, EPI_ISL_3228635, EPI_ISL_3228636, EPI_ISL_3228637, EPI_ISL_3228638, EPI_ISL_3228639, EPI_ISL_3228640, EPI_ISL_3228641, EPI_ISL_3228642, EPI_ISL_3228643, EPI_ISL_3228644, EPI_ISL_3228645, EPI_ISL_3228646, EPI_ISL_3228647, EPI_ISL_3228648, EPI_ISL_3228649, EPI_ISL_3228650, EPI_ISL_3228651, EPI_ISL_3228652, EPI_ISL_3228653, EPI_ISL_3228654, EPI_ISL_3228655, EPI_ISL_3228656, EPI_ISL_3228657, EPI_ISL_3228658, EPI_ISL_3228659, EPI_ISL_3228660, EPI_ISL_3228661, EPI_ISL_3228662, EPI_ISL_3228663, EPI_ISL_3228664, EPI_ISL_3228665, EPI_ISL_3228666, EPI_ISL_3228667, EPI_ISL_3228668, EPI_ISL_3228669, EPI_ISL_3228670, EPI_ISL_3228671, EPI_ISL_3228672, EPI_ISL_3228673, EPI_ISL_3228674, EPI_ISL_3228675, EPI_ISL_3228676, EPI_ISL_3228677, EPI_ISL_3228678, EPI_ISL_3228679, EPI_ISL_3228680, EPI_ISL_3228681, EPI_ISL_3228682, EPI_ISL_3228683, EPI_ISL_3228684, EPI_ISL_3228685, EPI_ISL_3228686, EPI_ISL_3228687, EPI_ISL_3228688, EPI_ISL_3228689, EPI_ISL_3228690, EPI_ISL_3228691, EPI_ISL_3228692, EPI_ISL_3228693, EPI_ISL_3228694, EPI_ISL_3228695, EPI_ISL_3228696, EPI_ISL_3228697, EPI_ISL_3228698, EPI_ISL_3228699, EPI_ISL_3228700, EPI_ISL_3228701, EPI_ISL_3228702, EPI_ISL_3228703, EPI_ISL_3228704, EPI_ISL_3228705, EPI_ISL_3228706, EPI_ISL_3228707, EPI_ISL_3228708, EPI_ISL_3228709, EPI_ISL_3228710, EPI_ISL_3228711, EPI_ISL_3228712, EPI_ISL_3228713, EPI_ISL_3228714, EPI_ISL_3228715, EPI_ISL_3228716, EPI_ISL_3228717, EPI_ISL_3228718, EPI_ISL_3228719, EPI_ISL_3228720, EPI_ISL_3228721, EPI_ISL_3228722, EPI_ISL_3228723, EPI_ISL_3228724, EPI_ISL_3228725, EPI_ISL_3228726, EPI_ISL_3228727, EPI_ISL_3228728, EPI_ISL_3228729, EPI_ISL_3228730, EPI_ISL_3228731, EPI_ISL_3228732, EPI_ISL_3228733, EPI_ISL_3228734, EPI_ISL_3228735, EPI_ISL_3228736, EPI_ISL_3228737, EPI_ISL_3228738, EPI_ISL_3228739, EPI_ISL_3228740, EPI_ISL_3228741, EPI_ISL_3228742, EPI_ISL_3228743, EPI_ISL_3228744, EPI_ISL_3228745, EPI_ISL_3228746, EPI_ISL_3228747, EPI_ISL_3228748, EPI_ISL_3228749, EPI_ISL_3228750, EPI_ISL_3228751, EPI_ISL_3228752, EPI_ISL_3228753, EPI_ISL_3228754, EPI_ISL_3228755, EPI_ISL_3228756, EPI_ISL_3228757, EPI_ISL_3228758, EPI_ISL_3228759, EPI_ISL_3228760, EPI_ISL_3228761, EPI_ISL_3228762, EPI_ISL_3228763, EPI_ISL_3228764, EPI_ISL_3228765, EPI_ISL_3228766 | | | |
| see above | Department of Bacteria, Parasites and Fungi, Statens Serum Institut, Copenhagen, Denmark | Statens Serum Institut Bioinformatics and Microbial Genomics | Danish Covid-19 Genome Consortium |
| EPI_ISL_3228767 | Molekylær Medicinsk Afdeling, Aarhus University Hospital, Aarhus, Denmark | Statens Serum Institut Bioinformatics and Microbial Genomics | Danish Covid-19 Genome Consortium |
| EPI_ISL_3228768, EPI_ISL_3228769, EPI_ISL_3228770, EPI_ISL_3228771, EPI_ISL_3228772, EPI_ISL_3228773, EPI_ISL_3228775, EPI_ISL_3228776, EPI_ISL_3228777, EPI_ISL_3228778, EPI_ISL_3228779, EPI_ISL_3228780, EPI_ISL_3228781, EPI_ISL_3228783, EPI_ISL_3228784, EPI_ISL_3228785, EPI_ISL_3228786, EPI_ISL_3228787, EPI_ISL_3228788, EPI_ISL_3228789, EPI_ISL_3228790, EPI_ISL_3228791, EPI_ISL_3228792, EPI_ISL_3228793, EPI_ISL_3228794, EPI_ISL_3228795, EPI_ISL_3228796, EPI_ISL_3228797, EPI_ISL_3228798, EPI_ISL_3228799, EPI_ISL_3228800, EPI_ISL_3228801, EPI_ISL_3228802, EPI_ISL_3228803, EPI_ISL_3228804, EPI_ISL_3228805, EPI_ISL_3228806, EPI_ISL_3228807, EPI_ISL_3228808, EPI_ISL_3228809, EPI_ISL_3228810, EPI_ISL_3228811, EPI_ISL_3228812, EPI_ISL_3228813, EPI_ISL_3228814, EPI_ISL_3228815, EPI_ISL_3228816, EPI_ISL_3228817, EPI_ISL_3228818, EPI_ISL_3228819, EPI_ISL_3228820, EPI_ISL_3228821, EPI_ISL_3228822, EPI_ISL_3228823, EPI_ISL_3228824, EPI_ISL_3228825 | | | |
| see above | Department of Bacteria, Parasites and Fungi, Statens Serum Institut, Copenhagen, Denmark | Statens Serum Institut Bioinformatics and Microbial Genomics | Danish Covid-19 Genome Consortium |
| EPI_ISL_3228826 | Molekylær Medicinsk Afdeling, Aarhus University Hospital, Aarhus, Denmark | Statens Serum Institut Bioinformatics and Microbial Genomics | Danish Covid-19 Genome Consortium |
| EPI_ISL_3228827, EPI_ISL_3228828, EPI_ISL_3228829, EPI_ISL_3228830, EPI_ISL_3228831, EPI_ISL_3228832, EPI_ISL_3228833, EPI_ISL_3228834, EPI_ISL_3228835, EPI_ISL_3228836, EPI_ISL_3228837, EPI_ISL_3228838, EPI_ISL_3228839, EPI_ISL_3228840, EPI_ISL_3228841, EPI_ISL_3228842, EPI_ISL_3228843, EPI_ISL_3228844, EPI_ISL_3228845, EPI_ISL_3228846, EPI_ISL_3228847, EPI_ISL_3228848, | | | |

| | | | |
|---|---|---|---|
| EPI_ISL_3228849, EPI_ISL_3228850, EPI_ISL_3228851, EPI_ISL_3228852, EPI_ISL_3228853, EPI_ISL_3228854, EPI_ISL_3228855, EPI_ISL_3228856, EPI_ISL_3228857, EPI_ISL_3228858, EPI_ISL_3228859, EPI_ISL_3228860, EPI_ISL_3228862, EPI_ISL_3228863, EPI_ISL_3228864, EPI_ISL_3228865, EPI_ISL_3228866, EPI_ISL_3228867, EPI_ISL_3228868, EPI_ISL_3228869, EPI_ISL_3228870, EPI_ISL_3228871, EPI_ISL_3228872, EPI_ISL_3228873, EPI_ISL_3228874, EPI_ISL_3228875, EPI_ISL_3228876, EPI_ISL_3228877, EPI_ISL_3228878, EPI_ISL_3228879, EPI_ISL_3228880, EPI_ISL_3228881, EPI_ISL_3228882, EPI_ISL_3228883, EPI_ISL_3228884, EPI_ISL_3228885, EPI_ISL_3228886, EPI_ISL_3228887, EPI_ISL_3228888, EPI_ISL_3228889, EPI_ISL_3228890, EPI_ISL_3228891, EPI_ISL_3228892 | | | |
| see above | Department of Bacteria, Parasites and Fungi, Statens Serum Institut, Copenhagen, Denmark | Statens Serum Institut Bioinformatics and Microbial Genomics | Danish Covid-19 Genome Consortium |
| EPI_ISL_3228893 | Molekylær Medicinsk Afdeling, Aarhus University Hospital, Aarhus, Denmark | Statens Serum Institut Bioinformatics and Microbial Genomics | Danish Covid-19 Genome Consortium |
| EPI_ISL_3228895, EPI_ISL_3228896, EPI_ISL_3228897, EPI_ISL_3228898, EPI_ISL_3228899, EPI_ISL_3228900, EPI_ISL_3228901, EPI_ISL_3228903, EPI_ISL_3228904, EPI_ISL_3228905, EPI_ISL_3228906, EPI_ISL_3228907, EPI_ISL_3228908, EPI_ISL_3228909, EPI_ISL_3228910, EPI_ISL_3228911, EPI_ISL_3228912, EPI_ISL_3228913, EPI_ISL_3228914, EPI_ISL_3228915, EPI_ISL_3228916, EPI_ISL_3228917, EPI_ISL_3228918, EPI_ISL_3228919, EPI_ISL_3228920, EPI_ISL_3228921, EPI_ISL_3228922, EPI_ISL_3228923, EPI_ISL_3228924, EPI_ISL_3228925, EPI_ISL_3228926, EPI_ISL_3228927, EPI_ISL_3228928, EPI_ISL_3228929, EPI_ISL_3228930, EPI_ISL_3228931, EPI_ISL_3228932, EPI_ISL_3228933, EPI_ISL_3228934, EPI_ISL_3228935, EPI_ISL_3228936, EPI_ISL_3228937, EPI_ISL_3228938, EPI_ISL_3228939, EPI_ISL_3228940, EPI_ISL_3228941, EPI_ISL_3228942, EPI_ISL_3228943, EPI_ISL_3228944, EPI_ISL_3228945, EPI_ISL_3228946, EPI_ISL_3228947, EPI_ISL_3228948, EPI_ISL_3228949, EPI_ISL_3228950, EPI_ISL_3228951, EPI_ISL_3228952, EPI_ISL_3228953, EPI_ISL_3228954, EPI_ISL_3228955, EPI_ISL_3228956, EPI_ISL_3228957, EPI_ISL_3228958, EPI_ISL_3228959, EPI_ISL_3228960, EPI_ISL_3228961, EPI_ISL_3228962, EPI_ISL_3228963, EPI_ISL_3228964, EPI_ISL_3228965, EPI_ISL_3228966, EPI_ISL_3228967, EPI_ISL_3228968, EPI_ISL_3228969, EPI_ISL_3228970, EPI_ISL_3228971, EPI_ISL_3228972, EPI_ISL_3228973, EPI_ISL_3228974, EPI_ISL_3228975, EPI_ISL_3228976, EPI_ISL_3228978, EPI_ISL_3228979, EPI_ISL_3228980, EPI_ISL_3228981, EPI_ISL_3228982, EPI_ISL_3228983, EPI_ISL_3228984, EPI_ISL_3228985, EPI_ISL_3228986, EPI_ISL_3228987, EPI_ISL_3228988, EPI_ISL_3228989, EPI_ISL_3228990, EPI_ISL_3228991, EPI_ISL_3228992, EPI_ISL_3228993 | | | |
| see above | Department of Bacteria, Parasites and Fungi, Statens Serum Institut, Copenhagen, Denmark | Statens Serum Institut Bioinformatics and Microbial Genomics | Danish Covid-19 Genome Consortium |
| EPI_ISL_3228994 | Molekylær Medicinsk Afdeling, Aarhus University Hospital, Aarhus, Denmark | Statens Serum Institut Bioinformatics and Microbial Genomics | Danish Covid-19 Genome Consortium |
| EPI_ISL_3228995, EPI_ISL_3228996, EPI_ISL_3228997, EPI_ISL_3228998, EPI_ISL_3228999, EPI_ISL_3229000, EPI_ISL_3229001, EPI_ISL_3229002, EPI_ISL_3229003, EPI_ISL_3229004, EPI_ISL_3229005, EPI_ISL_3229006, EPI_ISL_3229007, EPI_ISL_3229008, EPI_ISL_3229009, EPI_ISL_3229010, EPI_ISL_3229011, EPI_ISL_3229013, EPI_ISL_3229014, EPI_ISL_3229015, EPI_ISL_3229016, EPI_ISL_3229017, EPI_ISL_3229018, EPI_ISL_3229019, EPI_ISL_3229020, EPI_ISL_3229021, EPI_ISL_3229022, EPI_ISL_3229023, EPI_ISL_3229024, EPI_ISL_3229025, EPI_ISL_3229026, EPI_ISL_3229027, EPI_ISL_3229028, EPI_ISL_3229029, EPI_ISL_3229030, EPI_ISL_3229031, EPI_ISL_3229032 | | | |
| see above | Department of Bacteria, Parasites and Fungi, Statens Serum Institut, Copenhagen, Denmark | Statens Serum Institut Bioinformatics and Microbial Genomics | Danish Covid-19 Genome Consortium |
| EPI_ISL_3229033 | Molekylær Medicinsk Afdeling, Aarhus University Hospital, Aarhus, Denmark | Statens Serum Institut Bioinformatics and Microbial Genomics | Danish Covid-19 Genome Consortium |
| EPI_ISL_3229034, EPI_ISL_3229035, EPI_ISL_3229036, EPI_ISL_3229037, EPI_ISL_3229038, EPI_ISL_3229039, EPI_ISL_3229040, EPI_ISL_3229041, EPI_ISL_3229042, EPI_ISL_3229043, EPI_ISL_3229044, EPI_ISL_3229045, EPI_ISL_3229046, EPI_ISL_3229047, EPI_ISL_3229048, EPI_ISL_3229049, EPI_ISL_3229050, EPI_ISL_3229051, EPI_ISL_3229052, EPI_ISL_3229053, EPI_ISL_3229054, EPI_ISL_3229055, EPI_ISL_3229056, EPI_ISL_3229057, EPI_ISL_3229058, EPI_ISL_3229059, EPI_ISL_3229060, EPI_ISL_3229061, EPI_ISL_3229062, EPI_ISL_3229063, EPI_ISL_3229064, EPI_ISL_3229065, EPI_ISL_3229066, EPI_ISL_3229067, EPI_ISL_3229068, EPI_ISL_3229069, EPI_ISL_3229070, EPI_ISL_3229071, EPI_ISL_3229072, EPI_ISL_3229073, EPI_ISL_3229074, EPI_ISL_3229075, EPI_ISL_3229076, EPI_ISL_3229077, EPI_ISL_3229078, EPI_ISL_3229079, EPI_ISL_3229080, EPI_ISL_3229081, EPI_ISL_3229082, EPI_ISL_3229083, EPI_ISL_3229084, EPI_ISL_3229085, EPI_ISL_3229086, EPI_ISL_3229087, EPI_ISL_3229088, EPI_ISL_3229089, EPI_ISL_3229090, EPI_ISL_3229091, EPI_ISL_3229092, EPI_ISL_3229093, EPI_ISL_3229094, EPI_ISL_3229095, EPI_ISL_3229096, EPI_ISL_3229097, EPI_ISL_3229098, EPI_ISL_3229099, EPI_ISL_3229100, EPI_ISL_3229101, EPI_ISL_3229102, EPI_ISL_3229103, EPI_ISL_3229104, EPI_ISL_3229105, EPI_ISL_3229106, EPI_ISL_3229107, EPI_ISL_3229108, EPI_ISL_3229109, EPI_ISL_3229110, EPI_ISL_3229111, EPI_ISL_3229112, EPI_ISL_3229113, EPI_ISL_3229114, EPI_ISL_3229115 | | | |
| see above | Department of Bacteria, Parasites and Fungi, Statens Serum Institut, Copenhagen, Denmark | Statens Serum Institut Bioinformatics and Microbial Genomics | Danish Covid-19 Genome Consortium |
| EPI_ISL_3229116 | Molekylær Medicinsk Afdeling, Aarhus University Hospital, Aarhus, Denmark | Statens Serum Institut Bioinformatics and Microbial Genomics | Danish Covid-19 Genome Consortium |
| EPI_ISL_3229117, EPI_ISL_3229118, EPI_ISL_3229119, EPI_ISL_3229120, EPI_ISL_3229121, EPI_ISL_3229122, EPI_ISL_3229123, EPI_ISL_3229124, EPI_ISL_3229125, EPI_ISL_3229126, EPI_ISL_3229127, EPI_ISL_3229128, EPI_ISL_3229129, EPI_ISL_3229131, EPI_ISL_3229132, EPI_ISL_3229133, EPI_ISL_3229134, EPI_ISL_3229135, EPI_ISL_3229136 | | | |
| see above | Department of Bacteria, Parasites and Fungi, Statens Serum Institut, Copenhagen, Denmark | Statens Serum Institut Bioinformatics and Microbial Genomics | Danish Covid-19 Genome Consortium |
| EPI_ISL_3229137 | Molekylær Medicinsk Afdeling, Aarhus University Hospital, Aarhus, Denmark | Statens Serum Institut Bioinformatics and Microbial Genomics | Danish Covid-19 Genome Consortium |
| EPI_ISL_3229138, EPI_ISL_3229139, EPI_ISL_3229140, EPI_ISL_3229141, EPI_ISL_3229142, EPI_ISL_3229143 | Department of Bacteria, Parasites and Fungi, Statens Serum Institut, Copenhagen, Denmark | Statens Serum Institut Bioinformatics and Microbial Genomics | Danish Covid-19 Genome Consortium |
| EPI_ISL_3229144 | Molekylær Medicinsk Afdeling, Aarhus University Hospital, Aarhus, Denmark | Statens Serum Institut Bioinformatics and Microbial Genomics | Danish Covid-19 Genome Consortium |
| EPI_ISL_3229145, EPI_ISL_3229146, EPI_ISL_3229147, EPI_ISL_3229148, EPI_ISL_3229149, EPI_ISL_3229150, EPI_ISL_3229151, EPI_ISL_3229152, EPI_ISL_3229153, EPI_ISL_3229154, EPI_ISL_3229155, EPI_ISL_3229156, EPI_ISL_3229157, EPI_ISL_3229158, EPI_ISL_3229159, EPI_ISL_3229160, EPI_ISL_3229161, EPI_ISL_3229162, EPI_ISL_3229163, EPI_ISL_3229164, EPI_ISL_3229165, EPI_ISL_3229166, EPI_ISL_3229167, EPI_ISL_3229168, EPI_ISL_3229169, EPI_ISL_3229170, EPI_ISL_3229171, EPI_ISL_3229172, EPI_ISL_3229173, EPI_ISL_3229174, EPI_ISL_3229175, EPI_ISL_3229176, EPI_ISL_3229177, EPI_ISL_3229178, EPI_ISL_3229179 | | | |
| see above | Department of Bacteria, Parasites and Fungi, Statens Serum Institut, Copenhagen, Denmark | Statens Serum Institut Bioinformatics and Microbial Genomics | Danish Covid-19 Genome Consortium |
| EPI_ISL_3229189, EPI_ISL_3229190, EPI_ISL_3229193, EPI_ISL_3229194, EPI_ISL_3229195, EPI_ISL_3229196, EPI_ISL_3229197, EPI_ISL_3229198, EPI_ISL_3229199, EPI_ISL_3229200, EPI_ISL_3229201, EPI_ISL_3229202, EPI_ISL_3229203, EPI_ISL_3229204, EPI_ISL_3229205, EPI_ISL_3229206, EPI_ISL_3229207, EPI_ISL_3229208, EPI_ISL_3229209, EPI_ISL_3229210, EPI_ISL_3229211 | | | |
| see above | Clinical Virology | Clinical Bacteriology | Tim Roloff, Fanny Wegner, Helena MB Seth-Smith, Alfredo Mari, Karoline Leuzinger, Julia Bielicki, Manuel Battegay, Hans Hirsch, Adrian Egli |
| EPI_ISL_3229212 | Institut für Rechtsmedizin Basel | Clinical Bacteriology | Tim Roloff, Fanny Wegner, Helena MB Seth-Smith, Alfredo Mari, Karoline Leuzinger, Julia Bielicki, Manuel Battegay, Hans Hirsch, Adrian Egli |
| EPI_ISL_3229213, EPI_ISL_3229214, EPI_ISL_3229215, EPI_ISL_3229216, EPI_ISL_3229217 | Clinical Virology | Clinical Bacteriology | Tim Roloff, Fanny Wegner, Helena MB Seth-Smith, Alfredo Mari, Karoline Leuzinger, Julia Bielicki, Manuel Battegay, Hans Hirsch, Adrian Egli |
| EPI_ISL_3229218, EPI_ISL_3229219 | Rothen Medizinische Laboratorien AG | Clinical Bacteriology | Tim Roloff, Fanny Wegner, Helena MB Seth-Smith, Alfredo Mari, Karoline Leuzinger, Julia Bielicki, Ingrid Steffen, Manuel Battegay, Hans Hirsch, Adrian Egli |
| EPI_ISL_3229220, EPI_ISL_3229221, EPI_ISL_3229222, EPI_ISL_3229223 | Clinical Virology | Clinical Bacteriology | Tim Roloff, Fanny Wegner, Helena MB Seth-Smith, Alfredo Mari, Karoline Leuzinger, Julia Bielicki, Manuel Battegay, Hans Hirsch, Adrian Egli |
| EPI_ISL_3229224 | Rothen Medizinische Laboratorien AG | Clinical Bacteriology | Tim Roloff, Fanny Wegner, Helena MB Seth-Smith, Alfredo Mari, Karoline Leuzinger, Julia Bielicki, Ingrid Steffen, Manuel Battegay, Hans Hirsch, Adrian Egli |
| EPI_ISL_3229225, EPI_ISL_3229226, EPI_ISL_3229227 | Clinical Virology | Clinical Bacteriology | Tim Roloff, Fanny Wegner, Helena MB Seth-Smith, Alfredo Mari, Karoline Leuzinger, Julia Bielicki, Manuel Battegay, Hans Hirsch, Adrian Egli |
| EPI_ISL_3229228 | Synlab Suisse SA | Clinical Bacteriology | Tim Roloff, Fanny Wegner, Helena MB Seth-Smith, Alfredo Mari, Karoline Leuzinger, Julia Bielicki, Manuel Battegay, Hans Hirsch, Adrian Egli |
| EPI_ISL_3229230, EPI_ISL_3229231, EPI_ISL_3229232 | Clinical Virology | Clinical Bacteriology | Tim Roloff, Fanny Wegner, Helena MB Seth-Smith, Alfredo Mari, Karoline Leuzinger, Julia Bielicki, Manuel Battegay, Hans Hirsch, Adrian Egli |
| EPI_ISL_3229233, EPI_ISL_3229234, EPI_ISL_3229235, EPI_ISL_3229236, EPI_ISL_3229237, EPI_ISL_3229238, EPI_ISL_3229239, EPI_ISL_3229240 | Medics Labor AG | Clinical Bacteriology | Tim Roloff, Fanny Wegner, Helena MB Seth-Smith, Alfredo Mari, Karoline Leuzinger, Julia Bielicki, Manuel Battegay, Hans Hirsch, Adrian Egli |
| EPI_ISL_3229242, EPI_ISL_3229243, EPI_ISL_3229244 | Synlab Suisse SA | Clinical Bacteriology | Tim Roloff, Fanny Wegner, Helena MB Seth-Smith, Alfredo Mari, Karoline Leuzinger, Julia Bielicki, Manuel Battegay, Hans Hirsch, Adrian Egli |
| EPI_ISL_3229245, EPI_ISL_3229246, EPI_ISL_3229247, EPI_ISL_3229248, EPI_ISL_3229249, EPI_ISL_3229250, EPI_ISL_3229251, EPI_ISL_3229252, EPI_ISL_3229253, EPI_ISL_3229254, EPI_ISL_3229255 | | | |
| see above | Clinical Virology | Clinical Bacteriology | Tim Roloff, Fanny Wegner, Helena MB Seth-Smith, Alfredo Mari, Karoline Leuzinger, Julia Bielicki, Manuel Battegay, Hans Hirsch, Adrian Egli |
| EPI_ISL_3229256 | Medics Labor AG | Clinical Bacteriology | Tim Roloff, Fanny Wegner, Helena MB Seth-Smith, Alfredo Mari, Karoline Leuzinger, Julia Bielicki, Manuel Battegay, Hans Hirsch, Adrian Egli |
| EPI_ISL_3229257, EPI_ISL_3229258, EPI_ISL_3229259 | Clinical Virology | Clinical Bacteriology | Tim Roloff, Fanny Wegner, Helena MB Seth-Smith, Alfredo Mari, Karoline Leuzinger, Julia Bielicki, Manuel Battegay, Hans Hirsch, Adrian Egli |
| EPI_ISL_3229262, EPI_ISL_3229263 | New Mexico Department of Health Scientific Laboratory | New Mexico Department of Health Scientific Laboratory | Ellie Johnson, D'eldra Malone, Jennifer Benoit, Ratheesh Rajan, T. Mark Willmon, Linda Salazar Anastacia Griego-Fisher |
| EPI_ISL_3229264, EPI_ISL_3229267, EPI_ISL_3229268, EPI_ISL_3229269, EPI_ISL_3229270, EPI_ISL_3229271, EPI_ISL_3229273, EPI_ISL_3229274, EPI_ISL_3229275, EPI_ISL_3229277, EPI_ISL_3229278, EPI_ISL_3229279, EPI_ISL_3229281, EPI_ISL_3229282, EPI_ISL_3229283, EPI_ISL_3229284, EPI_ISL_3229286, EPI_ISL_3229287, EPI_ISL_3229288, EPI_ISL_3229289, EPI_ISL_3229290, EPI_ISL_3229291, EPI_ISL_3229292 | | | |
| see above | Curative Labs | New Mexico Department of Health Scientific Laboratory | Ellie Johnson, D'eldra Malone, Jennifer Benoit, Ratheesh Rajan, T. Mark Willmon, Linda Salazar Anastacia Griego-Fisher |
| EPI_ISL_3229293, EPI_ISL_3229294, EPI_ISL_3229295, EPI_ISL_3229296, EPI_ISL_3229297, EPI_ISL_3229298, EPI_ISL_3229299, EPI_ISL_3229300, EPI_ISL_3229301, EPI_ISL_3229302, EPI_ISL_3229303, EPI_ISL_3229304, EPI_ISL_3229305, EPI_ISL_3229306, EPI_ISL_3229307, EPI_ISL_3229308, EPI_ISL_3229309, EPI_ISL_3229310, EPI_ISL_3229311, EPI_ISL_3229312, EPI_ISL_3229313, EPI_ISL_3229314, EPI_ISL_3229315, EPI_ISL_3229316, EPI_ISL_3229317, EPI_ISL_3229318, EPI_ISL_3229319, EPI_ISL_3229320, EPI_ISL_3229321, EPI_ISL_3229322, EPI_ISL_3229323, EPI_ISL_3229324, EPI_ISL_3229325, EPI_ISL_3229326, EPI_ISL_3229327, EPI_ISL_3229328, EPI_ISL_3229329, EPI_ISL_3229330, EPI_ISL_3229331, EPI_ISL_3229332, EPI_ISL_3229333, EPI_ISL_3229334 | | | |
| see above | New Mexico Department of Health Scientific Laboratory | New Mexico Department of Health Scientific Laboratory | Ellie Johnson, D'eldra Malone, Jennifer Benoit, Ratheesh Rajan, T. Mark Willmon, Linda Salazar Anastacia Griego-Fisher |

| | | | |
|---|---|---|---|
| EPI_ISL_3229335 | Public Health Authority of the Slovak Republic | Laboratory of Genomics and Bioinformatics, Comenius University Science Park | Tatiana Sedláčková, Diana Rusňáková, Miroslav Böhmer, Anna Gičová, Jaroslav Budiš, Tomáš Szemes, Jakub Styk |
| EPI_ISL_3229336 | New Mexico Department of Health Scientific Laboratory | New Mexico Department of Health Scientific Laboratory | Ellie Johnson, D'eldra Malone, Jennifer Benoit, Ratheesh Rajan, T. Mark Willmon, Linda Salazar Anastacia Griego-Fisher |
| EPI_ISL_3229337, EPI_ISL_3229339, EPI_ISL_3229340 | Public Health Authority of the Slovak Republic | Laboratory of Genomics and Bioinformatics, Comenius University Science Park | Tatiana Sedláčková, Diana Rusňáková, Miroslav Böhmer, Anna Gičová, Jaroslav Budiš, Tomáš Szemes, Jakub Styk |
| EPI_ISL_3229341 | New Mexico Department of Health Scientific Laboratory | New Mexico Department of Health Scientific Laboratory | Ellie Johnson, D'eldra Malone, Jennifer Benoit, Ratheesh Rajan, T. Mark Willmon, Linda Salazar Anastacia Griego-Fisher |
| EPI_ISL_3229342, EPI_ISL_3229343 | Public Health Authority of the Slovak Republic | Laboratory of Genomics and Bioinformatics, Comenius University Science Park | Tatiana Sedláčková, Diana Rusňáková, Miroslav Böhmer, Anna Gičová, Jaroslav Budiš, Tomáš Szemes, Jakub Styk |
| EPI_ISL_3229344 | New Mexico Department of Health Scientific Laboratory | New Mexico Department of Health Scientific Laboratory | Ellie Johnson, D'eldra Malone, Jennifer Benoit, Ratheesh Rajan, T. Mark Willmon, Linda Salazar Anastacia Griego-Fisher |
| EPI_ISL_3229345, EPI_ISL_3229346 | Public Health Authority of the Slovak Republic | Laboratory of Genomics and Bioinformatics, Comenius University Science Park | Tatiana Sedláčková, Diana Rusňáková, Miroslav Böhmer, Anna Gičová, Jaroslav Budiš, Tomáš Szemes, Jakub Styk |
| EPI_ISL_3229347 | New Mexico Department of Health Scientific Laboratory | New Mexico Department of Health Scientific Laboratory | Ellie Johnson, D'eldra Malone, Jennifer Benoit, Ratheesh Rajan, T. Mark Willmon, Linda Salazar Anastacia Griego-Fisher |
| EPI_ISL_3229348, EPI_ISL_3229349 | Public Health Authority of the Slovak Republic | Laboratory of Genomics and Bioinformatics, Comenius University Science Park | Tatiana Sedláčková, Diana Rusňáková, Miroslav Böhmer, Anna Gičová, Jaroslav Budiš, Tomáš Szemes, Jakub Styk |
| EPI_ISL_3229350 | New Mexico Department of Health Scientific Laboratory | New Mexico Department of Health Scientific Laboratory | Ellie Johnson, D'eldra Malone, Jennifer Benoit, Ratheesh Rajan, T. Mark Willmon, Linda Salazar Anastacia Griego-Fisher |
| EPI_ISL_3229352 | Public Health Authority of the Slovak Republic | Laboratory of Genomics and Bioinformatics, Comenius University Science Park | Tatiana Sedláčková, Diana Rusňáková, Miroslav Böhmer, Anna Gičová, Jaroslav Budiš, Tomáš Szemes, Jakub Styk |
| EPI_ISL_3229353 | New Mexico Department of Health Scientific Laboratory | New Mexico Department of Health Scientific Laboratory | Ellie Johnson, D'eldra Malone, Jennifer Benoit, Ratheesh Rajan, T. Mark Willmon, Linda Salazar Anastacia Griego-Fisher |
| EPI_ISL_3229354, EPI_ISL_3229355 | Public Health Authority of the Slovak Republic | Laboratory of Genomics and Bioinformatics, Comenius University Science Park | Tatiana Sedláčková, Diana Rusňáková, Miroslav Böhmer, Anna Gičová, Jaroslav Budiš, Tomáš Szemes, Jakub Styk |
| EPI_ISL_3229356 | New Mexico Department of Health Scientific Laboratory | New Mexico Department of Health Scientific Laboratory | Ellie Johnson, D'eldra Malone, Jennifer Benoit, Ratheesh Rajan, T. Mark Willmon, Linda Salazar Anastacia Griego-Fisher |
| EPI_ISL_3229357, EPI_ISL_3229358 | Public Health Authority of the Slovak Republic | Laboratory of Genomics and Bioinformatics, Comenius University Science Park | Tatiana Sedláčková, Diana Rusňáková, Miroslav Böhmer, Anna Gičová, Jaroslav Budiš, Tomáš Szemes, Jakub Styk |
| EPI_ISL_3229359 | New Mexico Department of Health Scientific Laboratory | New Mexico Department of Health Scientific Laboratory | Ellie Johnson, D'eldra Malone, Jennifer Benoit, Ratheesh Rajan, T. Mark Willmon, Linda Salazar Anastacia Griego-Fisher |
| EPI_ISL_3229360, EPI_ISL_3229361 | Public Health Authority of the Slovak Republic | Laboratory of Genomics and Bioinformatics, Comenius University Science Park | Tatiana Sedláčková, Diana Rusňáková, Miroslav Böhmer, Anna Gičová, Jaroslav Budiš, Tomáš Szemes, Jakub Styk |
| EPI_ISL_3229362 | New Mexico Department of Health Scientific Laboratory | New Mexico Department of Health Scientific Laboratory | Ellie Johnson, D'eldra Malone, Jennifer Benoit, Ratheesh Rajan, T. Mark Willmon, Linda Salazar Anastacia Griego-Fisher |
| EPI_ISL_3229363 | Public Health Authority of the Slovak Republic | Laboratory of Genomics and Bioinformatics, Comenius University Science Park | Tatiana Sedláčková, Diana Rusňáková, Miroslav Böhmer, Anna Gičová, Jaroslav Budiš, Tomáš Szemes, Jakub Styk |
| EPI_ISL_3229364 | New Mexico Department of Health Scientific Laboratory | New Mexico Department of Health Scientific Laboratory | Ellie Johnson, D'eldra Malone, Jennifer Benoit, Ratheesh Rajan, T. Mark Willmon, Linda Salazar Anastacia Griego-Fisher |
| EPI_ISL_3229365, EPI_ISL_3229366 | Public Health Authority of the Slovak Republic | Laboratory of Genomics and Bioinformatics, Comenius University Science Park | Tatiana Sedláčková, Diana Rusňáková, Miroslav Böhmer, Anna Gičová, Jaroslav Budiš, Tomáš Szemes, Jakub Styk |
| EPI_ISL_3229367 | New Mexico Department of Health Scientific Laboratory | New Mexico Department of Health Scientific Laboratory | Ellie Johnson, D'eldra Malone, Jennifer Benoit, Ratheesh Rajan, T. Mark Willmon, Linda Salazar Anastacia Griego-Fisher |
| EPI_ISL_3229368, EPI_ISL_3229369 | Public Health Authority of the Slovak Republic | Laboratory of Genomics and Bioinformatics, Comenius University Science Park | Tatiana Sedláčková, Diana Rusňáková, Miroslav Böhmer, Anna Gičová, Jaroslav Budiš, Tomáš Szemes, Jakub Styk |
| EPI_ISL_3229370 | New Mexico Department of Health Scientific Laboratory | New Mexico Department of Health Scientific Laboratory | Ellie Johnson, D'eldra Malone, Jennifer Benoit, Ratheesh Rajan, T. Mark Willmon, Linda Salazar Anastacia Griego-Fisher |
| EPI_ISL_3229371 | Public Health Authority of the Slovak Republic | Laboratory of Genomics and Bioinformatics, Comenius University Science Park | Tatiana Sedláčková, Diana Rusňáková, Miroslav Böhmer, Anna Gičová, Jaroslav Budiš, Tomáš Szemes, Jakub Styk |
| EPI_ISL_3229372 | New Mexico Department of Health Scientific Laboratory | New Mexico Department of Health Scientific Laboratory | Ellie Johnson, D'eldra Malone, Jennifer Benoit, Ratheesh Rajan, T. Mark Willmon, Linda Salazar Anastacia Griego-Fisher |
| EPI_ISL_3229373, EPI_ISL_3229374 | Public Health Authority of the Slovak Republic | Laboratory of Genomics and Bioinformatics, Comenius University Science Park | Tatiana Sedláčková, Diana Rusňáková, Miroslav Böhmer, Anna Gičová, Jaroslav Budiš, Tomáš Szemes, Jakub Styk |
| EPI_ISL_3229375 | New Mexico Department of Health Scientific Laboratory | New Mexico Department of Health Scientific Laboratory | Ellie Johnson, D'eldra Malone, Jennifer Benoit, Ratheesh Rajan, T. Mark Willmon, Linda Salazar Anastacia Griego-Fisher |
| EPI_ISL_3229376, EPI_ISL_3229377 | Public Health Authority of the Slovak Republic | Laboratory of Genomics and Bioinformatics, Comenius University Science Park | Tatiana Sedláčková, Diana Rusňáková, Miroslav Böhmer, Anna Gičová, Jaroslav Budiš, Tomáš Szemes, Jakub Styk |
| EPI_ISL_3229378 | New Mexico Department of Health Scientific Laboratory | New Mexico Department of Health Scientific Laboratory | Ellie Johnson, D'eldra Malone, Jennifer Benoit, Ratheesh Rajan, T. Mark Willmon, Linda Salazar Anastacia Griego-Fisher |
| EPI_ISL_3229379, EPI_ISL_3229380 | Public Health Authority of the Slovak Republic | Laboratory of Genomics and Bioinformatics, Comenius University Science Park | Tatiana Sedláčková, Diana Rusňáková, Miroslav Böhmer, Anna Gičová, Jaroslav Budiš, Tomáš Szemes, Jakub Styk |
| EPI_ISL_3229381 | New Mexico Department of Health Scientific Laboratory | New Mexico Department of Health Scientific Laboratory | Ellie Johnson, D'eldra Malone, Jennifer Benoit, Ratheesh Rajan, T. Mark Willmon, Linda Salazar Anastacia Griego-Fisher |
| EPI_ISL_3229382 | Public Health Authority of the Slovak Republic | Laboratory of Genomics and Bioinformatics, Comenius University Science Park | Tatiana Sedláčková, Diana Rusňáková, Miroslav Böhmer, Anna Gičová, Jaroslav Budiš, Tomáš Szemes, Jakub Styk |
| EPI_ISL_3229383 | New Mexico Department of Health Scientific Laboratory | New Mexico Department of Health Scientific Laboratory | Ellie Johnson, D'eldra Malone, Jennifer Benoit, Ratheesh Rajan, T. Mark Willmon, Linda Salazar Anastacia Griego-Fisher |
| EPI_ISL_3229384, EPI_ISL_3229385 | Public Health Authority of the Slovak Republic | Laboratory of Genomics and Bioinformatics, Comenius University Science Park | Tatiana Sedláčková, Diana Rusňáková, Miroslav Böhmer, Anna Gičová, Jaroslav Budiš, Tomáš Szemes, Jakub Styk |
| EPI_ISL_3229386 | New Mexico Department of Health Scientific Laboratory | New Mexico Department of Health Scientific Laboratory | Ellie Johnson, D'eldra Malone, Jennifer Benoit, Ratheesh Rajan, T. Mark Willmon, Linda Salazar Anastacia Griego-Fisher |
| EPI_ISL_3229387, EPI_ISL_3229388, EPI_ISL_3229389, EPI_ISL_3229390, EPI_ISL_3229391, EPI_ISL_3229392, EPI_ISL_3229393, EPI_ISL_3229394, EPI_ISL_3229397, EPI_ISL_3229398, EPI_ISL_3229399, EPI_ISL_3229400 | | | |
| see above | Public Health Authority of the Slovak Republic | Laboratory of Genomics and Bioinformatics, Comenius University Science Park | Tatiana Sedláčková, Diana Rusňáková, Miroslav Böhmer, Anna Gičová, Jaroslav Budiš, Tomáš Szemes, Jakub Styk |
| EPI_ISL_3229402 | Servicio de Microbiología. Hospital General Universitario de Castellón | SeqCOVID-SPAIN consortium/IBV(CSIC) | Rosario Moreno Muñoz, María Dolores Tirado Balaguer and SeqCOVID-SPAIN consortium |
| EPI_ISL_3229403, EPI_ISL_3229404 | Servicio de Microbiología. Hospital Arnau de Vilanova | SeqCOVID-SPAIN consortium/IBV(CSIC) | Victoria Domínguez, Rocío Falcón, Amparo Farga and SeqCOVID-SPAIN consortium |
| EPI_ISL_3229405, EPI_ISL_3229407, EPI_ISL_3229408, EPI_ISL_3229409, EPI_ISL_3229410, EPI_ISL_3229411, EPI_ISL_3229412, EPI_ISL_3229413, EPI_ISL_3229414, EPI_ISL_3229415, EPI_ISL_3229416, EPI_ISL_3229417, EPI_ISL_3229418, EPI_ISL_3229419, EPI_ISL_3229420, EPI_ISL_3229421, EPI_ISL_3229423, EPI_ISL_3229424, EPI_ISL_3229425, EPI_ISL_3229426, EPI_ISL_3229427, EPI_ISL_3229428, EPI_ISL_3229431, EPI_ISL_3229432, EPI_ISL_3229433, EPI_ISL_3229434, EPI_ISL_3229435, EPI_ISL_3229436, EPI_ISL_3229437, EPI_ISL_3229438, EPI_ISL_3229439, EPI_ISL_3229440, EPI_ISL_3229441, EPI_ISL_3229442, EPI_ISL_3229443, EPI_ISL_3229445, EPI_ISL_3229447, EPI_ISL_3229448, EPI_ISL_3229449, EPI_ISL_3229451, EPI_ISL_3229454, EPI_ISL_3229455, EPI_ISL_3229456, EPI_ISL_3229457, EPI_ISL_3229458, EPI_ISL_3229459, EPI_ISL_3229460, EPI_ISL_3229461, EPI_ISL_3229462, EPI_ISL_3229463, EPI_ISL_3229464, EPI_ISL_3229465, EPI_ISL_3229466, EPI_ISL_3229467, EPI_ISL_3229468, EPI_ISL_3229469, EPI_ISL_3229470, EPI_ISL_3229471, EPI_ISL_3229472, EPI_ISL_3229473, EPI_ISL_3229474, EPI_ISL_3229475, EPI_ISL_3229476, EPI_ISL_3229477, EPI_ISL_3229478, EPI_ISL_3229479, EPI_ISL_3229480, EPI_ISL_3229481, EPI_ISL_3229482, EPI_ISL_3229483, EPI_ISL_3229485, EPI_ISL_3229486, EPI_ISL_3229487, EPI_ISL_3229488, EPI_ISL_3229489, EPI_ISL_3229490, EPI_ISL_3229491, EPI_ISL_3229492, EPI_ISL_3229493, EPI_ISL_3229494 | | | |
| see above | Hospital Universitari i Politècnic La Fe de València | SeqCOVID-SPAIN consortium/IBV(CSIC) | María Dolores Gómez Ruiz, Eva González Barberá, Ana Gil Brusola, Salvador Giner Almaraz, José Luis López Hontangas and SeqCOVID-SPAIN consortium |
| EPI_ISL_3229495, EPI_ISL_3229496 | Platform BIS UZA/UAntwerpen | Labo Klinische Biologie, UZA | Marie Le Mercier, Jasmine Coppens, Kathleen Holemans, Ines Verbesselt, Basil Britto Xavier, Christine Lammens, Veerle Matheeussen, Herman Goossens |
| EPI_ISL_3229497 | Labo Klinische Biologie, UZA | Labo Klinische Biologie, UZA | Marie Le Mercier, Jasmine Coppens, Kathleen Holemans, Ines Verbesselt, Basil Britto Xavier, Christine Lammens, Veerle Matheeussen, Herman Goossens |
| EPI_ISL_3229498, EPI_ISL_3229499 | Platform BIS UZA/UAntwerpen | Labo Klinische Biologie, UZA | Marie Le Mercier, Jasmine Coppens, Kathleen Holemans, Ines Verbesselt, Basil Britto Xavier, Christine Lammens, Veerle Matheeussen, Herman Goossens |
| EPI_ISL_3229500 | Instituut Tropische Geneeskunde | Labo Klinische Biologie, UZA | Marie Le Mercier, Jasmine Coppens, Kathleen Holemans, Ines Verbesselt, Basil Britto Xavier, Christine Lammens, Veerle Matheeussen, Herman Goossens |
| EPI_ISL_3229503, EPI_ISL_3229504, EPI_ISL_3229505, EPI_ISL_3229506, EPI_ISL_3229507, EPI_ISL_3229508, EPI_ISL_3229509, EPI_ISL_3229510, EPI_ISL_3229511, EPI_ISL_3229512, EPI_ISL_3229513, EPI_ISL_3229514, EPI_ISL_3229515, EPI_ISL_3229516, EPI_ISL_3229517 | | | |
| see above | Platform BIS UZA/UAntwerpen | Labo Klinische Biologie, UZA | Marie Le Mercier, Jasmine Coppens, Kathleen Holemans, Ines Verbesselt, Basil Britto Xavier, Christine Lammens, Veerle Matheeussen, Herman Goossens |
| EPI_ISL_3229518 | Algemeen Klinisch Labo | Labo Klinische Biologie, UZA | Marie Le Mercier, Jasmine Coppens, Kathleen Holemans, Ines Verbesselt, Basil Britto Xavier, Christine Lammens, Veerle Matheeussen, Herman Goossens |
| EPI_ISL_3229519, EPI_ISL_3229520, EPI_ISL_3229521, EPI_ISL_3229522, EPI_ISL_3229523, EPI_ISL_3229525, EPI_ISL_3229526, EPI_ISL_3229527, EPI_ISL_3229528, EPI_ISL_3229529 | Centrum voor Medische Analyse | Labo Klinische Biologie, UZA | Marie Le Mercier, Jasmine Coppens, Kathleen Holemans, Ines Verbesselt, Basil Britto Xavier, Christine Lammens, Veerle Matheeussen, Herman Goossens |
| EPI_ISL_3229530, EPI_ISL_3229531, EPI_ISL_3229534, EPI_ISL_3229535, EPI_ISL_3229536, EPI_ISL_3229537, EPI_ISL_3229538, EPI_ISL_3229539, EPI_ISL_3229540, EPI_ISL_3229541, EPI_ISL_3229542, EPI_ISL_3229543, EPI_ISL_3229544, EPI_ISL_3229545, EPI_ISL_3229546, EPI_ISL_3229547, EPI_ISL_3229548, EPI_ISL_3229549, EPI_ISL_3229550, EPI_ISL_3229551, EPI_ISL_3229552, EPI_ISL_3229553 | | | |
| see above | Platform BIS UZA/UAntwerpen | Labo Klinische Biologie, UZA | Marie Le Mercier, Jasmine Coppens, Kathleen Holemans, Ines Verbesselt, Basil Britto Xavier, Christine Lammens, Veerle Matheeussen, Herman Goossens |

| | | | |
|---|---|---|---|
| EPI_ISL_3229554, EPI_ISL_3229555, EPI_ISL_3229556, EPI_ISL_3229557, EPI_ISL_3229558, EPI_ISL_3229559, EPI_ISL_3229560, EPI_ISL_3229561, EPI_ISL_3229562, EPI_ISL_3229563, EPI_ISL_3229564, EPI_ISL_3229565, EPI_ISL_3229566, EPI_ISL_3229567, EPI_ISL_3229568, EPI_ISL_3229569, EPI_ISL_3229570, EPI_ISL_3229571, EPI_ISL_3229572, EPI_ISL_3229573 | | | |
| see above | Algemeen Medisch Labo | Labo Klinische Biologie, UZA | Marie Le Mercier, Jasmine Coppens, Kathleen Holemans, Ines Verbesselt, Basil Britto Xavier, Christine Lammens, Veerle Matheeussen, Herman Goossens |
| EPI_ISL_3229574 | INT Fondazione Pascale | INT Fondazione Pascale | N. Normanno INT Fondazione Pascale |
| EPI_ISL_3229576, EPI_ISL_3229577, EPI_ISL_3229578, EPI_ISL_3229579, EPI_ISL_3229580, EPI_ISL_3229581, EPI_ISL_3229582, EPI_ISL_3229583, EPI_ISL_3229585, EPI_ISL_3229586, EPI_ISL_3229587, EPI_ISL_3229588, EPI_ISL_3229589 | | | |
| see above | University of Liège COVID-19 testing center | GIGA Medical Genomics | Keith Durkin, Maria Artesi, Bouchra Boujemla, Nathalie Renotte, Nadine Cambisano, Sébastien Bontems, Cécile Meex, Claire Gourzonès, Olivier Ek, Laurent Gillet, Marie-Pierre Hayette, Vincent Bours |
| EPI_ISL_3229590, EPI_ISL_3229591 | Department of Clinical Microbiology | GIGA Medical Genomics | Keith Durkin, Maria Artesi, Bouchra Boujemla, Nathalie Renotte, Nadine Cambisano, Sébastien Bontems, Cécile Meex, Claire Gourzonès, Olivier Ek, Laurent Gillet, Marie-Pierre Hayette, Vincent Bours |
| EPI_ISL_3229592, EPI_ISL_3229593, EPI_ISL_3229594, EPI_ISL_3229595, EPI_ISL_3229596, EPI_ISL_3229597, EPI_ISL_3229598, EPI_ISL_3229599, EPI_ISL_3229600, EPI_ISL_3229601, EPI_ISL_3229602, EPI_ISL_3229603, EPI_ISL_3229604, EPI_ISL_3229605, EPI_ISL_3229606, EPI_ISL_3229607, EPI_ISL_3229608, EPI_ISL_3229609, EPI_ISL_3229610, EPI_ISL_3229611, EPI_ISL_3229612 | | | |
| see above | SYNLAB | GIGA Medical Genomics | Keith Durkin, Maria Artesi, Bouchra Boujemla, Nathalie Renotte, Nadine Cambisano, Sébastien Bontems, Cécile Meex, Claire Gourzonès, Olivier Ek, Laurent Gillet, Marie-Pierre Hayette, Vincent Bours |
| EPI_ISL_3229613 | University of Liège COVID-19 testing center | GIGA Medical Genomics | Keith Durkin, Maria Artesi, Bouchra Boujemla, Nathalie Renotte, Nadine Cambisano, Sébastien Bontems, Cécile Meex, Claire Gourzonès, Olivier Ek, Laurent Gillet, Marie-Pierre Hayette, Vincent Bours |
| EPI_ISL_3229614 | SYNLAB | GIGA Medical Genomics | Keith Durkin, Maria Artesi, Bouchra Boujemla, Nathalie Renotte, Nadine Cambisano, Sébastien Bontems, Cécile Meex, Claire Gourzonès, Olivier Ek, Laurent Gillet, Marie-Pierre Hayette, Vincent Bours |
| EPI_ISL_3229615 | University of Liège COVID-19 testing center | GIGA Medical Genomics | Keith Durkin, Maria Artesi, Bouchra Boujemla, Nathalie Renotte, Nadine Cambisano, Sébastien Bontems, Cécile Meex, Claire Gourzonès, Olivier Ek, Laurent Gillet, Marie-Pierre Hayette, Vincent Bours |
| EPI_ISL_3229616 | Department of Clinical Microbiology | GIGA Medical Genomics | Keith Durkin, Maria Artesi, Bouchra Boujemla, Nathalie Renotte, Nadine Cambisano, Sébastien Bontems, Cécile Meex, Claire Gourzonès, Olivier Ek, Laurent Gillet, Marie-Pierre Hayette, Vincent Bours |
| EPI_ISL_3229617 | SYNLAB | GIGA Medical Genomics | Keith Durkin, Maria Artesi, Bouchra Boujemla, Nathalie Renotte, Nadine Cambisano, Sébastien Bontems, Cécile Meex, Claire Gourzonès, Olivier Ek, Laurent Gillet, Marie-Pierre Hayette, Vincent Bours |
| EPI_ISL_3229620 | University of Liège COVID-19 testing center | GIGA Medical Genomics | Keith Durkin, Maria Artesi, Bouchra Boujemla, Nathalie Renotte, Nadine Cambisano, Sébastien Bontems, Cécile Meex, Claire Gourzonès, Olivier Ek, Laurent Gillet, Marie-Pierre Hayette, Vincent Bours |
| EPI_ISL_3229621, EPI_ISL_3229622, EPI_ISL_3229623 | SYNLAB | GIGA Medical Genomics | Keith Durkin, Maria Artesi, Bouchra Boujemla, Nathalie Renotte, Nadine Cambisano, Sébastien Bontems, Cécile Meex, Claire Gourzonès, Olivier Ek, Laurent Gillet, Marie-Pierre Hayette, Vincent Bours |
| EPI_ISL_3229624, EPI_ISL_3229625, EPI_ISL_3229626, EPI_ISL_3229627 | University of Liège COVID-19 testing center | GIGA Medical Genomics | Keith Durkin, Maria Artesi, Bouchra Boujemla, Nathalie Renotte, Nadine Cambisano, Sébastien Bontems, Cécile Meex, Claire Gourzonès, Olivier Ek, Laurent Gillet, Marie-Pierre Hayette, Vincent Bours |
| EPI_ISL_3229628, EPI_ISL_3229629 | SYNLAB | GIGA Medical Genomics | Keith Durkin, Maria Artesi, Bouchra Boujemla, Nathalie Renotte, Nadine Cambisano, Sébastien Bontems, Cécile Meex, Claire Gourzonès, Olivier Ek, Laurent Gillet, Marie-Pierre Hayette, Vincent Bours |
| EPI_ISL_3229630, EPI_ISL_3229631 | University of Liège COVID-19 testing center | GIGA Medical Genomics | Keith Durkin, Maria Artesi, Bouchra Boujemla, Nathalie Renotte, Nadine Cambisano, Sébastien Bontems, Cécile Meex, Claire Gourzonès, Olivier Ek, Laurent Gillet, Marie-Pierre Hayette, Vincent Bours |
| EPI_ISL_3229632, EPI_ISL_3229633 | Department of Clinical Microbiology | GIGA Medical Genomics | Keith Durkin, Maria Artesi, Bouchra Boujemla, Nathalie Renotte, Nadine Cambisano, Sébastien Bontems, Cécile Meex, Claire Gourzonès, Olivier Ek, Laurent Gillet, Marie-Pierre Hayette, Vincent Bours |
| EPI_ISL_3229634, EPI_ISL_3229635 | SYNLAB | GIGA Medical Genomics | Keith Durkin, Maria Artesi, Bouchra Boujemla, Nathalie Renotte, Nadine Cambisano, Sébastien Bontems, Cécile Meex, Claire Gourzonès, Olivier Ek, Laurent Gillet, Marie-Pierre Hayette, Vincent Bours |
| EPI_ISL_3229639 | University of Liège COVID-19 testing center | GIGA Medical Genomics | Keith Durkin, Maria Artesi, Bouchra Boujemla, Nathalie Renotte, Nadine Cambisano, Sébastien Bontems, Cécile Meex, Claire Gourzonès, Olivier Ek, Laurent Gillet, Marie-Pierre Hayette, Vincent Bours |
| EPI_ISL_3229640, EPI_ISL_3229641 | SYNLAB | GIGA Medical Genomics | Keith Durkin, Maria Artesi, Bouchra Boujemla, Nathalie Renotte, Nadine Cambisano, Sébastien Bontems, Cécile Meex, Claire Gourzonès, Olivier Ek, Laurent Gillet, Marie-Pierre Hayette, Vincent Bours |
| EPI_ISL_3229642 | INT Fondazione Pascale | INT Fondazione Pascale | N. Normanno INT Fondazione Pascale |
| EPI_ISL_3229648 | Diagnostic Genomics Laboratory | Diagnostic Genomics Laboratory | Keely Duff, Carolyn Banister, Phillip Buckhaults |
| EPI_ISL_3229649, EPI_ISL_3229650, EPI_ISL_3229651, EPI_ISL_3229652, EPI_ISL_3229653, EPI_ISL_3229654, EPI_ISL_3229655, EPI_ISL_3229657, EPI_ISL_3229658, EPI_ISL_3229660, EPI_ISL_3229662, EPI_ISL_3229664, EPI_ISL_3229665, EPI_ISL_3229668, EPI_ISL_3229669, EPI_ISL_3229670, EPI_ISL_3229671, EPI_ISL_3229672, EPI_ISL_3229673, EPI_ISL_3229674, EPI_ISL_3229675, EPI_ISL_3229676, EPI_ISL_3229677, EPI_ISL_3229678, EPI_ISL_3229679, EPI_ISL_3229680, EPI_ISL_3229681, EPI_ISL_3229682, EPI_ISL_3229683, EPI_ISL_3229684, EPI_ISL_3229685 | | | |
| see above | National Institute of Public Health | National Institute of Public Health | Helena Jirincova, Jaromira Vecerova, Timotej Suri, Dusan Trnka, Alexander Nagy |
| EPI_ISL_3229687 | HOPITAL DE VICHY | CHU Clermont-Ferrand, service de virologie | Bisseux Maxime, Mirand Audrey, Combes Patricia, Henquell Cécile |
| EPI_ISL_3229688 | Centre Hospitalier Universitaire Clermont-Ferrand | CHU Clermont-Ferrand, service de virologie | Bisseux Maxime, Mirand Audrey, Combes Patricia, Henquell Cécile |
| EPI_ISL_3229689 | HOPITAL DE VICHY | CHU Clermont-Ferrand, service de virologie | Bisseux Maxime, Mirand Audrey, Combes Patricia, Henquell Cécile |
| EPI_ISL_3229690 | Centre hospitalier D'AURILLAC | CHU Clermont-Ferrand, service de virologie | Bisseux Maxime, Mirand Audrey, Combes Patricia, Henquell Cécile |
| EPI_ISL_3229691 | Centre Hospitalier Universitaire Clermont-Ferrand | CHU Clermont-Ferrand, service de virologie | Bisseux Maxime, Mirand Audrey, Combes Patricia, Henquell Cécile |
| EPI_ISL_3229692 | Centre Hospitalier Paul Ardier | CHU Clermont-Ferrand, service de virologie | Bisseux Maxime, Mirand Audrey, Combes Patricia, Henquell Cécile |
| EPI_ISL_3229693, EPI_ISL_3229694, EPI_ISL_3229696 | Centre Hospitalier Universitaire Clermont-Ferrand | CHU Clermont-Ferrand, service de virologie | Bisseux Maxime, Mirand Audrey, Combes Patricia, Henquell Cécile |
| EPI_ISL_3229697, EPI_ISL_3229698 | Centre hospitalier D'AURILLAC | CHU Clermont-Ferrand, service de virologie | Bisseux Maxime, Mirand Audrey, Combes Patricia, Henquell Cécile |
| EPI_ISL_3229699 | HOPITAL DE VICHY | CHU Clermont-Ferrand, service de virologie | Bisseux Maxime, Mirand Audrey, Combes Patricia, Henquell Cécile |
| EPI_ISL_3229701 | HOPITAL DE MONTLUCON | CHU Clermont-Ferrand, service de virologie | Bisseux Maxime, Mirand Audrey, Combes Patricia, Henquell Cécile |
| EPI_ISL_3229703 | Centre Hospitalier Universitaire Clermont-Ferrand | CHU Clermont-Ferrand, service de virologie | Bisseux Maxime, Mirand Audrey, Combes Patricia, Henquell Cécile |
| EPI_ISL_3229704, EPI_ISL_3229705, EPI_ISL_3229706, EPI_ISL_3229707, EPI_ISL_3229708 | HOPITAL DU MONT DORE | CHU Clermont-Ferrand, service de virologie | Bisseux Maxime, Mirand Audrey, Combes Patricia, Henquell Cécile |
| EPI_ISL_3229709 | Centre De Prelevement COVID RIOM | CHU Clermont-Ferrand, service de virologie | Bisseux Maxime, Mirand Audrey, Combes Patricia, Henquell Cécile |
| EPI_ISL_3229710, EPI_ISL_3229711, EPI_ISL_3229712, EPI_ISL_3229713, EPI_ISL_3229714 | Centre Hospitalier Universitaire Clermont-Ferrand | CHU Clermont-Ferrand, service de virologie | Bisseux Maxime, Mirand Audrey, Combes Patricia, Henquell Cécile |
| EPI_ISL_3229715 | Diagnostic Genomics Laboratory | Diagnostic Genomics Laboratory | Keely Duff, Carolyn Banister, Phillip Buckhaults |
| EPI_ISL_3229716, EPI_ISL_3229718 | NCSLPH | NCSLPH | Chase K, Miller MC, Greene S, Glover W |
| EPI_ISL_3229719 | University Campus Bio-Medico of Rome (UCBM) | University Campus Bio-Medico of Rome (UCBM) | Riva E., De Florio L., Fogolari M., Francesconi M., Lintas C., Veralli R., Angeletti S. |
| EPI_ISL_3229721 | NCSLPH | NCSLPH | Chase K, Miller MC, Greene S, Glover W |
| EPI_ISL_3229744, EPI_ISL_3229745, EPI_ISL_3229746, EPI_ISL_3229747, EPI_ISL_3229748, EPI_ISL_3229749, EPI_ISL_3229750, EPI_ISL_3229751, EPI_ISL_3229752, EPI_ISL_3229753, EPI_ISL_3229754, EPI_ISL_3229755, EPI_ISL_3229756, EPI_ISL_3229757, EPI_ISL_3229758, EPI_ISL_3229759, EPI_ISL_3229760 | | | |
| see above | Platform BIS UZA/UAntwerpen | Labo Klinische Biologie, UZA | Marie Le Mercier, Jasmine Coppens, Kathleen Holemans, Ines Verbesselt, Basil Britto Xavier, Christine Lammens, Veerle Matheeussen, Herman Goossens |
| EPI_ISL_3229761, EPI_ISL_3229762 | Labo Klinische Biologie, UZA | Labo Klinische Biologie, UZA | Marie Le Mercier, Jasmine Coppens, Kathleen Holemans, Ines Verbesselt, Basil Britto Xavier, Christine Lammens, Veerle Matheeussen, Herman Goossens |
| EPI_ISL_3229763, EPI_ISL_3229764, EPI_ISL_3229765, EPI_ISL_3229766, EPI_ISL_3229767, EPI_ISL_3229768, EPI_ISL_3229769, EPI_ISL_3229770, EPI_ISL_3229771, EPI_ISL_3229772, EPI_ISL_3229773, EPI_ISL_3229774, EPI_ISL_3229775, EPI_ISL_3229776, EPI_ISL_3229777 | | | |
| see above | Platform BIS UZA/UAntwerpen | Labo Klinische Biologie, UZA | Marie Le Mercier, Jasmine Coppens, Kathleen Holemans, Ines Verbesselt, Basil Britto Xavier, Christine Lammens, Veerle Matheeussen, Herman Goossens |
| EPI_ISL_3229778 | Centrum voor Medische Analyse | Labo Klinische Biologie, UZA | Marie Le Mercier, Jasmine Coppens, Kathleen Holemans, Ines Verbesselt, Basil Britto Xavier, Christine Lammens, Veerle Matheeussen, Herman Goossens |
| EPI_ISL_3229779, EPI_ISL_3229780, EPI_ISL_3229781, EPI_ISL_3229782, EPI_ISL_3229783, EPI_ISL_3229784, EPI_ISL_3229785 | Platform BIS UZA/UAntwerpen | Labo Klinische Biologie, UZA | Marie Le Mercier, Jasmine Coppens, Kathleen Holemans, Ines Verbesselt, Basil Britto Xavier, Christine Lammens, Veerle Matheeussen, Herman Goossens |
| EPI_ISL_3229786, EPI_ISL_3229787, EPI_ISL_3229788 | Centrum voor Medische Analyse | Labo Klinische Biologie, UZA | Marie Le Mercier, Jasmine Coppens, Kathleen Holemans, Ines Verbesselt, Basil Britto Xavier, Christine Lammens, Veerle Matheeussen, Herman Goossens |
| EPI_ISL_3229789, EPI_ISL_3229790, | Algemeen Medisch Labo | Labo Klinische Biologie, UZA | Marie Le Mercier, Jasmine Coppens, Kathleen Holemans, Ines Verbesselt, Basil Britto Xavier, Christine Lammens, Veerle Matheeussen, Herman Goossens |

| | | | |
|---|---|---|---|
| EPI_ISL_3229791, EPI_ISL_3229792 | | | |
| EPI_ISL_3229793, EPI_ISL_3229794, EPI_ISL_3229795, EPI_ISL_3229796, EPI_ISL_3229797, EPI_ISL_3229798, EPI_ISL_3229799, EPI_ISL_3229800, EPI_ISL_3229801, EPI_ISL_3229802, EPI_ISL_3229803, EPI_ISL_3229804, EPI_ISL_3229805, EPI_ISL_3229806, EPI_ISL_3229807, EPI_ISL_3229808 | | | |
| see above | Platform BIS UZA/UAntwerpen | Labo Klinische Biologie, UZA | Marie Le Mercier, Jasmine Coppens, Kathleen Holemans, Ines Verbesselt, Basil Britto Xavier, Christine Lammens, Veerle Matheeussen, Herman Goossens |
| EPI_ISL_3229809, EPI_ISL_3229810, EPI_ISL_3229811, EPI_ISL_3229812, EPI_ISL_3229813, EPI_ISL_3229814 | Centrum voor Medische Analyse | Labo Klinische Biologie, UZA | Marie Le Mercier, Jasmine Coppens, Kathleen Holemans, Ines Verbesselt, Basil Britto Xavier, Christine Lammens, Veerle Matheeussen, Herman Goossens |
| EPI_ISL_3229815, EPI_ISL_3229816, EPI_ISL_3229817, EPI_ISL_3229818, EPI_ISL_3229819, EPI_ISL_3229820, EPI_ISL_3229821, EPI_ISL_3229822 | Platform BIS UZA/UAntwerpen | Labo Klinische Biologie, UZA | Marie Le Mercier, Jasmine Coppens, Kathleen Holemans, Ines Verbesselt, Basil Britto Xavier, Christine Lammens, Veerle Matheeussen, Herman Goossens |
| EPI_ISL_3229823 | Centrum voor Medische Analyse | Labo Klinische Biologie, UZA | Marie Le Mercier, Jasmine Coppens, Kathleen Holemans, Ines Verbesselt, Basil Britto Xavier, Christine Lammens, Veerle Matheeussen, Herman Goossens |
| EPI_ISL_3229824, EPI_ISL_3229825, EPI_ISL_3229826, EPI_ISL_3229827, EPI_ISL_3229828, EPI_ISL_3229829, EPI_ISL_3229830, EPI_ISL_3229831, EPI_ISL_3229832 | Platform BIS UZA/UAntwerpen | Labo Klinische Biologie, UZA | Marie Le Mercier, Jasmine Coppens, Kathleen Holemans, Ines Verbesselt, Basil Britto Xavier, Christine Lammens, Veerle Matheeussen, Herman Goossens |
| EPI_ISL_3229834, EPI_ISL_3229835, EPI_ISL_3229836, EPI_ISL_3229837, EPI_ISL_3229838, EPI_ISL_3229839, EPI_ISL_3229842 | University Campus Bio-Medico of Rome (UCBM) | University Campus Bio-Medico of Rome (UCBM) | Riva E., De Florio L., Fogolari M., Francesconi M., Lintas C., Veralli R., Angeletti S. |
| EPI_ISL_3229899 | Medical Microbiology Unit, Department for Laboratory Medicine, Drammen Hospital, Vestre Viken Health Trust, | Norwegian Institute of Public Health, Department of Virology | Kathrine Stene-Johansen, Kamilla Heddeland Instefjord, Hilde Elshaug, Garcia Llorente Ignacio, Jon Bråte, Engebretsen Serina Beate,Pedersen Benedikte Nevjen, Line Victoria Moen, Debech Nadia, Atiya R Ali,Marie Paulsen Madsen, Rasmus Riis Kopperud, Hilde Vollan, Karoline Bragstad, Olav Hungnes |
| EPI_ISL_3229900 | Ostfold Hospital Trust - Kalnes, Centre for Laboratory Medicine, Section for gene technology and infection serology | Norwegian Institute of Public Health, Department of Virology | Kathrine Stene-Johansen, Kamilla Heddeland Instefjord, Hilde Elshaug, Garcia Llorente Ignacio, Jon Bråte, Engebretsen Serina Beate,Pedersen Benedikte Nevjen, Line Victoria Moen, Debech Nadia, Atiya R Ali,Marie Paulsen Madsen, Rasmus Riis Kopperud, Hilde Vollan, Karoline Bragstad, Olav Hungnes |
| EPI_ISL_3229901 | Medical Microbiology Unit, Department for Laboratory Medicine, Drammen Hospital, Vestre Viken Health Trust, | Norwegian Institute of Public Health, Department of Virology | Kathrine Stene-Johansen, Kamilla Heddeland Instefjord, Hilde Elshaug, Garcia Llorente Ignacio, Jon Bråte, Engebretsen Serina Beate,Pedersen Benedikte Nevjen, Line Victoria Moen, Debech Nadia, Atiya R Ali,Marie Paulsen Madsen, Rasmus Riis Kopperud, Hilde Vollan, Karoline Bragstad, Olav Hungnes |
| EPI_ISL_3229902 | Hospital of Southern Norway - Kristiansand, Department of Medical Microbiology | Norwegian Institute of Public Health, Department of Virology | Kathrine Stene-Johansen, Kamilla Heddeland Instefjord, Hilde Elshaug, Garcia Llorente Ignacio, Jon Bråte, Engebretsen Serina Beate,Pedersen Benedikte Nevjen, Line Victoria Moen, Debech Nadia, Atiya R Ali,Marie Paulsen Madsen, Rasmus Riis Kopperud, Hilde Vollan, Karoline Bragstad, Olav Hungnes |
| EPI_ISL_3229903, EPI_ISL_3229904 | Medical Microbiology Unit, Department for Laboratory Medicine, Drammen Hospital, Vestre Viken Health Trust, | Norwegian Institute of Public Health, Department of Virology | Kathrine Stene-Johansen, Kamilla Heddeland Instefjord, Hilde Elshaug, Garcia Llorente Ignacio, Jon Bråte, Engebretsen Serina Beate,Pedersen Benedikte Nevjen, Line Victoria Moen, Debech Nadia, Atiya R Ali,Marie Paulsen Madsen, Rasmus Riis Kopperud, Hilde Vollan, Karoline Bragstad, Olav Hungnes |
| EPI_ISL_3229905 | Hospital of Southern Norway - Kristiansand, Department of Medical Microbiology | Norwegian Institute of Public Health, Department of Virology | Kathrine Stene-Johansen, Kamilla Heddeland Instefjord, Hilde Elshaug, Garcia Llorente Ignacio, Jon Bråte, Engebretsen Serina Beate,Pedersen Benedikte Nevjen, Line Victoria Moen, Debech Nadia, Atiya R Ali,Marie Paulsen Madsen, Rasmus Riis Kopperud, Hilde Vollan, Karoline Bragstad, Olav Hungnes |
| EPI_ISL_3229914 | Haukeland University Hospital, Dept. of Microbiology | Norwegian Institute of Public Health, Department of Virology | Kathrine Stene-Johansen, Kamilla Heddeland Instefjord, Hilde Elshaug, Garcia Llorente Ignacio, Jon Bråte, Engebretsen Serina Beate,Pedersen Benedikte Nevjen, Line Victoria Moen, Debech Nadia, Atiya R Ali,Marie Paulsen Madsen, Rasmus Riis Kopperud, Hilde Vollan, Karoline Bragstad, Olav Hungnes |
| EPI_ISL_3229915, EPI_ISL_3229916, EPI_ISL_3229917, EPI_ISL_3229918, EPI_ISL_3229919 | Medical Microbiology Unit, Department for Laboratory Medicine, Drammen Hospital, Vestre Viken Health Trust, | Norwegian Institute of Public Health, Department of Virology | Kathrine Stene-Johansen, Kamilla Heddeland Instefjord, Hilde Elshaug, Garcia Llorente Ignacio, Jon Bråte, Engebretsen Serina Beate,Pedersen Benedikte Nevjen, Line Victoria Moen, Debech Nadia, Atiya R Ali,Marie Paulsen Madsen, Rasmus Riis Kopperud, Hilde Vollan, Karoline Bragstad, Olav Hungnes |
| EPI_ISL_3229921 | Nordland Hospital - Bodo, Laboratory Department, Molecular Biology Unit | Norwegian Institute of Public Health, Department of Virology | Kathrine Stene-Johansen, Kamilla Heddeland Instefjord, Hilde Elshaug, Garcia Llorente Ignacio, Jon Bråte, Engebretsen Serina Beate,Pedersen Benedikte Nevjen, Line Victoria Moen, Debech Nadia, Atiya R Ali,Marie Paulsen Madsen, Rasmus Riis Kopperud, Hilde Vollan, Karoline Bragstad, Olav Hungnes |
| EPI_ISL_3229928 | Furst Medical Laboratory | Norwegian Institute of Public Health, Department of Virology | Kathrine Stene-Johansen, Kamilla Heddeland Instefjord, Hilde Elshaug, Garcia Llorente Ignacio, Jon Bråte, Engebretsen Serina Beate,Pedersen Benedikte Nevjen, Line Victoria Moen, Debech Nadia, Atiya R Ali,Marie Paulsen Madsen, Rasmus Riis Kopperud, Hilde Vollan, Karoline Bragstad, Olav Hungnes |
| EPI_ISL_3229929 | Department of Medical Microbiology, St. Olavs hospital | Norwegian Institute of Public Health, Department of Virology | Kathrine Stene-Johansen, Kamilla Heddeland Instefjord, Hilde Elshaug, Garcia Llorente Ignacio, Jon Bråte, Engebretsen Serina Beate,Pedersen Benedikte Nevjen, Line Victoria Moen, Debech Nadia, Atiya R Ali,Marie Paulsen Madsen, Rasmus Riis Kopperud, Hilde Vollan, Karoline Bragstad, Olav Hungnes |
| EPI_ISL_3229930 | Haukeland University Hospital, Dept. of Microbiology | Norwegian Institute of Public Health, Department of Virology | Kathrine Stene-Johansen, Kamilla Heddeland Instefjord, Hilde Elshaug, Garcia Llorente Ignacio, Jon Bråte, Engebretsen Serina Beate,Pedersen Benedikte Nevjen, Line Victoria Moen, Debech Nadia, Atiya R Ali,Marie Paulsen Madsen, Rasmus Riis Kopperud, Hilde Vollan, Karoline Bragstad, Olav Hungnes |
| EPI_ISL_3229931, EPI_ISL_3229932 | Furst Medical Laboratory | Norwegian Institute of Public Health, Department of Virology | Kathrine Stene-Johansen, Kamilla Heddeland Instefjord, Hilde Elshaug, Garcia Llorente Ignacio, Jon Bråte, Engebretsen Serina Beate,Pedersen Benedikte Nevjen, Line Victoria Moen, Debech Nadia, Atiya R Ali,Marie Paulsen Madsen, Rasmus Riis Kopperud, Hilde Vollan, Karoline Bragstad, Olav Hungnes |
| EPI_ISL_3229933 | Department of Medical Microbiology, St. Olavs hospital | Norwegian Institute of Public Health, Department of Virology | Kathrine Stene-Johansen, Kamilla Heddeland Instefjord, Hilde Elshaug, Garcia Llorente Ignacio, Jon Bråte, Engebretsen Serina Beate,Pedersen Benedikte Nevjen, Line Victoria Moen, Debech Nadia, Atiya R Ali,Marie Paulsen Madsen, Rasmus Riis Kopperud, Hilde Vollan, Karoline Bragstad, Olav Hungnes |
| EPI_ISL_3229934 | Haukeland University Hospital, Dept. of Microbiology | Norwegian Institute of Public Health, Department of Virology | Kathrine Stene-Johansen, Kamilla Heddeland Instefjord, Hilde Elshaug, Garcia Llorente Ignacio, Jon Bråte, Engebretsen Serina Beate,Pedersen Benedikte Nevjen, Line Victoria Moen, Debech Nadia, Atiya R Ali,Marie Paulsen Madsen, Rasmus Riis Kopperud, Hilde Vollan, Karoline Bragstad, Olav Hungnes |
| EPI_ISL_3229935 | Furst Medical Laboratory | Norwegian Institute of Public Health, Department of Virology | Kathrine Stene-Johansen, Kamilla Heddeland Instefjord, Hilde Elshaug, Garcia Llorente Ignacio, Jon Bråte, Engebretsen Serina Beate,Pedersen Benedikte Nevjen, Line Victoria Moen, Debech Nadia, Atiya R Ali,Marie Paulsen Madsen, Rasmus Riis Kopperud, Hilde Vollan, Karoline Bragstad, Olav Hungnes |
| EPI_ISL_3229936, EPI_ISL_3229939 | Haukeland University Hospital, Dept. of Microbiology | Norwegian Institute of Public Health, Department of Virology | Kathrine Stene-Johansen, Kamilla Heddeland Instefjord, Hilde Elshaug, Garcia Llorente Ignacio, Jon Bråte, Engebretsen Serina Beate,Pedersen Benedikte Nevjen, Line Victoria Moen, Debech Nadia, Atiya R Ali,Marie Paulsen Madsen, Rasmus Riis Kopperud, Hilde Vollan, Karoline Bragstad, Olav Hungnes |
| EPI_ISL_3229940 | Furst Medical Laboratory | Norwegian Institute of Public Health, Department of Virology | Kathrine Stene-Johansen, Kamilla Heddeland Instefjord, Hilde Elshaug, Garcia Llorente Ignacio, Jon Bråte, Engebretsen Serina Beate,Pedersen Benedikte Nevjen, Line Victoria Moen, Debech Nadia, Atiya R Ali,Marie Paulsen Madsen, Rasmus Riis Kopperud, Hilde Vollan, Karoline Bragstad, Olav Hungnes |
| EPI_ISL_3229942, EPI_ISL_3229943 | Haukeland University Hospital, Dept. of Microbiology | Norwegian Institute of Public Health, Department of Virology | Kathrine Stene-Johansen, Kamilla Heddeland Instefjord, Hilde Elshaug, Garcia Llorente Ignacio, Jon Bråte, Engebretsen Serina Beate,Pedersen Benedikte Nevjen, Line Victoria Moen, Debech Nadia, Atiya R Ali,Marie Paulsen Madsen, Rasmus Riis Kopperud, Hilde Vollan, Karoline Bragstad, Olav Hungnes |
| EPI_ISL_3229944 | Furst Medical Laboratory | Norwegian Institute of Public Health, Department of Virology | Kathrine Stene-Johansen, Kamilla Heddeland Instefjord, Hilde Elshaug, Garcia Llorente Ignacio, Jon Bråte, Engebretsen Serina Beate,Pedersen Benedikte Nevjen, Line Victoria Moen, Debech Nadia, Atiya R Ali,Marie Paulsen Madsen, Rasmus Riis Kopperud, Hilde Vollan, Karoline Bragstad, Olav Hungnes |
| EPI_ISL_3229946 | Haukeland University Hospital, Dept. of Microbiology | Norwegian Institute of Public Health, Department of Virology | Kathrine Stene-Johansen, Kamilla Heddeland Instefjord, Hilde Elshaug, Garcia Llorente Ignacio, Jon Bråte, Engebretsen Serina Beate,Pedersen Benedikte Nevjen, Line Victoria Moen, Debech Nadia, Atiya R Ali,Marie Paulsen Madsen, Rasmus Riis Kopperud, Hilde Vollan, Karoline Bragstad, Olav Hungnes |
| EPI_ISL_3229947 | Furst Medical Laboratory | Norwegian Institute of Public Health, Department of Virology | Kathrine Stene-Johansen, Kamilla Heddeland Instefjord, Hilde Elshaug, Garcia Llorente Ignacio, Jon Bråte, Engebretsen Serina Beate,Pedersen Benedikte Nevjen, Line Victoria Moen, Debech Nadia, Atiya R Ali,Marie Paulsen Madsen, Rasmus Riis Kopperud, Hilde Vollan, Karoline Bragstad, Olav Hungnes |
| EPI_ISL_3229949 | Hospital of Southern Norway - Kristiansand, Department of Medical Microbiology | Norwegian Institute of Public Health, Department of Virology | Kathrine Stene-Johansen, Kamilla Heddeland Instefjord, Hilde Elshaug, Garcia Llorente Ignacio, Jon Bråte, Engebretsen Serina Beate,Pedersen Benedikte Nevjen, Line Victoria Moen, Debech Nadia, Atiya R Ali,Marie Paulsen Madsen, Rasmus Riis Kopperud, Hilde Vollan, Karoline Bragstad, Olav Hungnes |
| EPI_ISL_3229950 | University Campus Bio-Medico of Rome (UCBM) | University Campus Bio-Medico of Rome (UCBM) | Riva E., De Florio L., Fogolari M., Francesconi M., Lintas C., Veralli R., Angeletti S. |
| EPI_ISL_3229951 | Haukeland University Hospital, Dept. of Microbiology | Norwegian Institute of Public Health, Department of Virology | Kathrine Stene-Johansen, Kamilla Heddeland Instefjord, Hilde Elshaug, Garcia Llorente Ignacio, Jon Bråte, Engebretsen Serina Beate,Pedersen Benedikte Nevjen, Line Victoria Moen, Debech Nadia, Atiya R Ali,Marie Paulsen Madsen, Rasmus Riis Kopperud, Hilde Vollan, Karoline Bragstad, Olav Hungnes |
| EPI_ISL_3229952 | Hospital of Southern Norway - Kristiansand, Department of Medical Microbiology | Norwegian Institute of Public Health, Department of Virology | Kathrine Stene-Johansen, Kamilla Heddeland Instefjord, Hilde Elshaug, Garcia Llorente Ignacio, Jon Bråte, Engebretsen Serina Beate,Pedersen Benedikte Nevjen, Line Victoria Moen, Debech Nadia, Atiya R Ali,Marie Paulsen Madsen, Rasmus Riis Kopperud, Hilde Vollan, Karoline Bragstad, Olav Hungnes |
| EPI_ISL_3229953 | Haukeland University Hospital, Dept. of Microbiology | Norwegian Institute of Public Health, Department of Virology | Kathrine Stene-Johansen, Kamilla Heddeland Instefjord, Hilde Elshaug, Garcia Llorente Ignacio, Jon Bråte, Engebretsen Serina Beate,Pedersen Benedikte Nevjen, Line Victoria Moen, Debech Nadia, Atiya R Ali,Marie Paulsen Madsen, Rasmus Riis Kopperud, Hilde Vollan, Karoline Bragstad, Olav Hungnes |
| EPI_ISL_3229954 | Hospital of Southern Norway - Kristiansand, Department of Medical Microbiology | Norwegian Institute of Public Health, Department of Virology | Kathrine Stene-Johansen, Kamilla Heddeland Instefjord, Hilde Elshaug, Garcia Llorente Ignacio, Jon Bråte, Engebretsen Serina Beate,Pedersen Benedikte Nevjen, Line Victoria Moen, Debech Nadia, Atiya R Ali,Marie Paulsen Madsen, Rasmus Riis Kopperud, Hilde Vollan, Karoline Bragstad, Olav Hungnes |
| EPI_ISL_3229955, | Haukeland University Hospital, Dept. of Microbiology | Norwegian Institute of Public Health, Department of Virology | Kathrine Stene-Johansen, Kamilla Heddeland Instefjord, Hilde Elshaug, Garcia Llorente Ignacio, Jon Bråte, Engebretsen Serina Beate,Pedersen Benedikte Nevjen, Line Victoria Moen, Debech Nadia, Atiya R Ali,Marie Paulsen Madsen, Rasmus Riis |

| | | | |
|---|---|---|---|
| EPI_ISL_3229956, EPI_ISL_3229957 | | | Kopperud, Hilde Vollan, Karoline Bragstad, Olav Hungnes |
| EPI_ISL_3229958 | Hospital of Southern Norway - Kristiansand, Department of Medical Microbiology | Norwegian Institute of Public Health, Department of Virology | Kathrine Stene-Johansen, Kamilla Heddeland Instefjord, Hilde Elshaug, Garcia Llorente Ignacio, Jon Bråte, Engebretsen Serina Beate,Pedersen Benedikte Nevjen, Line Victoria Moen, Debech Nadia, Atiya R Ali,Marie Paulsen Madsen, Rasmus Riis Kopperud, Hilde Vollan, Karoline Bragstad, Olav Hungnes |
| EPI_ISL_3229959, EPI_ISL_3229960, EPI_ISL_3229962 | Haukeland University Hospital, Dept. of Microbiology | Norwegian Institute of Public Health, Department of Virology | Kathrine Stene-Johansen, Kamilla Heddeland Instefjord, Hilde Elshaug, Garcia Llorente Ignacio, Jon Bråte, Engebretsen Serina Beate,Pedersen Benedikte Nevjen, Line Victoria Moen, Debech Nadia, Atiya R Ali,Marie Paulsen Madsen, Rasmus Riis Kopperud, Hilde Vollan, Karoline Bragstad, Olav Hungnes |
| EPI_ISL_3229963 | Hospital of Southern Norway - Kristiansand, Department of Medical Microbiology | Norwegian Institute of Public Health, Department of Virology | Kathrine Stene-Johansen, Kamilla Heddeland Instefjord, Hilde Elshaug, Garcia Llorente Ignacio, Jon Bråte, Engebretsen Serina Beate,Pedersen Benedikte Nevjen, Line Victoria Moen, Debech Nadia, Atiya R Ali,Marie Paulsen Madsen, Rasmus Riis Kopperud, Hilde Vollan, Karoline Bragstad, Olav Hungnes |
| EPI_ISL_3229964, EPI_ISL_3229965, EPI_ISL_3229966, EPI_ISL_3229967 | Haukeland University Hospital, Dept. of Microbiology | Norwegian Institute of Public Health, Department of Virology | Kathrine Stene-Johansen, Kamilla Heddeland Instefjord, Hilde Elshaug, Garcia Llorente Ignacio, Jon Bråte, Engebretsen Serina Beate,Pedersen Benedikte Nevjen, Line Victoria Moen, Debech Nadia, Atiya R Ali,Marie Paulsen Madsen, Rasmus Riis Kopperud, Hilde Vollan, Karoline Bragstad, Olav Hungnes |
| EPI_ISL_3229968, EPI_ISL_3229969, EPI_ISL_3229970 | Furst Medical Laboratory | Norwegian Institute of Public Health, Department of Virology | Kathrine Stene-Johansen, Kamilla Heddeland Instefjord, Hilde Elshaug, Garcia Llorente Ignacio, Jon Bråte, Engebretsen Serina Beate,Pedersen Benedikte Nevjen, Line Victoria Moen, Debech Nadia, Atiya R Ali,Marie Paulsen Madsen, Rasmus Riis Kopperud, Hilde Vollan, Karoline Bragstad, Olav Hungnes |
| EPI_ISL_3229972 | Haukeland University Hospital, Dept. of Microbiology | Norwegian Institute of Public Health, Department of Virology | Kathrine Stene-Johansen, Kamilla Heddeland Instefjord, Hilde Elshaug, Garcia Llorente Ignacio, Jon Bråte, Engebretsen Serina Beate,Pedersen Benedikte Nevjen, Line Victoria Moen, Debech Nadia, Atiya R Ali,Marie Paulsen Madsen, Rasmus Riis Kopperud, Hilde Vollan, Karoline Bragstad, Olav Hungnes |
| EPI_ISL_3229973 | Furst Medical Laboratory | Norwegian Institute of Public Health, Department of Virology | Kathrine Stene-Johansen, Kamilla Heddeland Instefjord, Hilde Elshaug, Garcia Llorente Ignacio, Jon Bråte, Engebretsen Serina Beate,Pedersen Benedikte Nevjen, Line Victoria Moen, Debech Nadia, Atiya R Ali,Marie Paulsen Madsen, Rasmus Riis Kopperud, Hilde Vollan, Karoline Bragstad, Olav Hungnes |
| EPI_ISL_3229974 | Haukeland University Hospital, Dept. of Microbiology | Norwegian Institute of Public Health, Department of Virology | Kathrine Stene-Johansen, Kamilla Heddeland Instefjord, Hilde Elshaug, Garcia Llorente Ignacio, Jon Bråte, Engebretsen Serina Beate,Pedersen Benedikte Nevjen, Line Victoria Moen, Debech Nadia, Atiya R Ali,Marie Paulsen Madsen, Rasmus Riis Kopperud, Hilde Vollan, Karoline Bragstad, Olav Hungnes |
| EPI_ISL_3229975, EPI_ISL_3229977, EPI_ISL_3229978, EPI_ISL_3229979 | Furst Medical Laboratory | Norwegian Institute of Public Health, Department of Virology | Kathrine Stene-Johansen, Kamilla Heddeland Instefjord, Hilde Elshaug, Garcia Llorente Ignacio, Jon Bråte, Engebretsen Serina Beate,Pedersen Benedikte Nevjen, Line Victoria Moen, Debech Nadia, Atiya R Ali,Marie Paulsen Madsen, Rasmus Riis Kopperud, Hilde Vollan, Karoline Bragstad, Olav Hungnes |
| EPI_ISL_3229980 | University Campus Bio-Medico of Rome (UCBM) | University Campus Bio-Medico of Rome (UCBM) | Riva E., De Florio L., Fogolari M., Francesconi M., Lintas C., Veralli R., Angeletti S. |
| EPI_ISL_3229981, EPI_ISL_3229982, EPI_ISL_3229983, EPI_ISL_3229984, EPI_ISL_3229985, EPI_ISL_3229986, EPI_ISL_3229987, EPI_ISL_3229988, EPI_ISL_3229989, EPI_ISL_3229990, EPI_ISL_3229991 | | | |
| see above | Furst Medical Laboratory | Norwegian Institute of Public Health, Department of Virology | Kathrine Stene-Johansen, Kamilla Heddeland Instefjord, Hilde Elshaug, Garcia Llorente Ignacio, Jon Bråte, Engebretsen Serina Beate,Pedersen Benedikte Nevjen, Line Victoria Moen, Debech Nadia, Atiya R Ali,Marie Paulsen Madsen, Rasmus Riis Kopperud, Hilde Vollan, Karoline Bragstad, Olav Hungnes |
| EPI_ISL_3229992 | Innlandet Hospital Trust, Division Lillehammer, Department for Medical Microbiology | Norwegian Institute of Public Health, Department of Virology | Kathrine Stene-Johansen, Kamilla Heddeland Instefjord, Hilde Elshaug, Garcia Llorente Ignacio, Jon Bråte, Engebretsen Serina Beate,Pedersen Benedikte Nevjen, Line Victoria Moen, Debech Nadia, Atiya R Ali,Marie Paulsen Madsen, Rasmus Riis Kopperud, Hilde Vollan, Karoline Bragstad, Olav Hungnes |
| EPI_ISL_3229994 | Haukeland University Hospital, Dept. of Microbiology | Norwegian Institute of Public Health, Department of Virology | Kathrine Stene-Johansen, Kamilla Heddeland Instefjord, Hilde Elshaug, Garcia Llorente Ignacio, Jon Bråte, Engebretsen Serina Beate,Pedersen Benedikte Nevjen, Line Victoria Moen, Debech Nadia, Atiya R Ali,Marie Paulsen Madsen, Rasmus Riis Kopperud, Hilde Vollan, Karoline Bragstad, Olav Hungnes |
| EPI_ISL_3229995 | Department of Medical Microbiology - section Molde, Molde Hospital | Norwegian Institute of Public Health, Department of Virology | Kathrine Stene-Johansen, Kamilla Heddeland Instefjord, Hilde Elshaug, Garcia Llorente Ignacio, Jon Bråte, Engebretsen Serina Beate,Pedersen Benedikte Nevjen, Line Victoria Moen, Debech Nadia, Atiya R Ali,Marie Paulsen Madsen, Rasmus Riis Kopperud, Hilde Vollan, Karoline Bragstad, Olav Hungnes |
| EPI_ISL_3229997, EPI_ISL_3229998 | Haukeland University Hospital, Dept. of Microbiology | Norwegian Institute of Public Health, Department of Virology | Kathrine Stene-Johansen, Kamilla Heddeland Instefjord, Hilde Elshaug, Garcia Llorente Ignacio, Jon Bråte, Engebretsen Serina Beate,Pedersen Benedikte Nevjen, Line Victoria Moen, Debech Nadia, Atiya R Ali,Marie Paulsen Madsen, Rasmus Riis Kopperud, Hilde Vollan, Karoline Bragstad, Olav Hungnes |
| EPI_ISL_3230002 | Department of Medical Microbiology - section Molde, Molde Hospital | Norwegian Institute of Public Health, Department of Virology | Kathrine Stene-Johansen, Kamilla Heddeland Instefjord, Hilde Elshaug, Garcia Llorente Ignacio, Jon Bråte, Engebretsen Serina Beate,Pedersen Benedikte Nevjen, Line Victoria Moen, Debech Nadia, Atiya R Ali,Marie Paulsen Madsen, Rasmus Riis Kopperud, Hilde Vollan, Karoline Bragstad, Olav Hungnes |
| EPI_ISL_3230003 | Unilabs Laboratory Medicine | Norwegian Institute of Public Health, Department of Virology | Kathrine Stene-Johansen, Kamilla Heddeland Instefjord, Hilde Elshaug, Garcia Llorente Ignacio, Jon Bråte, Engebretsen Serina Beate,Pedersen Benedikte Nevjen, Line Victoria Moen, Debech Nadia, Atiya R Ali,Marie Paulsen Madsen, Rasmus Riis Kopperud, Hilde Vollan, Karoline Bragstad, Olav Hungnes |
| EPI_ISL_3230004 | Department of Medical Microbiology - section Molde, Molde Hospital | Norwegian Institute of Public Health, Department of Virology | Kathrine Stene-Johansen, Kamilla Heddeland Instefjord, Hilde Elshaug, Garcia Llorente Ignacio, Jon Bråte, Engebretsen Serina Beate,Pedersen Benedikte Nevjen, Line Victoria Moen, Debech Nadia, Atiya R Ali,Marie Paulsen Madsen, Rasmus Riis Kopperud, Hilde Vollan, Karoline Bragstad, Olav Hungnes |
| EPI_ISL_3230006, EPI_ISL_3230007, EPI_ISL_3230008, EPI_ISL_3230009, EPI_ISL_3230010, EPI_ISL_3230011, EPI_ISL_3230012 | Haukeland University Hospital, Dept. of Microbiology | Norwegian Institute of Public Health, Department of Virology | Kathrine Stene-Johansen, Kamilla Heddeland Instefjord, Hilde Elshaug, Garcia Llorente Ignacio, Jon Bråte, Engebretsen Serina Beate,Pedersen Benedikte Nevjen, Line Victoria Moen, Debech Nadia, Atiya R Ali,Marie Paulsen Madsen, Rasmus Riis Kopperud, Hilde Vollan, Karoline Bragstad, Olav Hungnes |
| EPI_ISL_3230013, EPI_ISL_3230014 | University Campus Bio-Medico of Rome (UCBM) | University Campus Bio-Medico of Rome (UCBM) | Riva E., De Florio L., Fogolari M., Francesconi M., Lintas C., Veralli R., Angeletti S. |
| EPI_ISL_3230016, EPI_ISL_3230017, EPI_ISL_3230018 | Laboratorio Central, Ministerio de Salud Córdoba | Instituto de Patologia Vegetal (CIAP-INTA) on behalf of 'Proyecto Argentino Interinstitucional de genomica de SARS-CoV-2' (PAIS Consortium) | Fernández, FD; Marquez, N.; Debat, HJ.; M; Re, V.; Pisano, M.B.; Castro, G.; Barbas, G. |
| EPI_ISL_3230028, EPI_ISL_3230029 | University Campus Bio-Medico of Rome (UCBM) | University Campus Bio-Medico of Rome (UCBM) | Riva E., De Florio L., Fogolari M., Francesconi M., Lintas C., Veralli R., Angeletti S. |
| EPI_ISL_3230031, EPI_ISL_3230033, EPI_ISL_3230035, EPI_ISL_3230036, EPI_ISL_3230037, EPI_ISL_3230039, EPI_ISL_3230045, EPI_ISL_3230046, EPI_ISL_3230050, EPI_ISL_3230051, EPI_ISL_3230052 | | | |
| see above | CHUGA-IBP-laboratoire de Virologie | IBP-laboratoire de virologie | Sylvie Larrat; Patrice Morand; Pascal Poignard; Raphaële Germi; Julien Lupo; Anne Signori -Schmuck; Anne-Karen Faure; Benjamin Nemoz; Aurélie Truffot; Léa Ponderand; Julien Andréani; Alban Caporossi; Hugo Jardin |
| EPI_ISL_3230053 | UCC (Universidad Central del Caribe) | Grubaugh Lab - Yale School of Public Health | Joseph Fauver, Mallery Breban, Isabel Ott, Tara Alpert, Mary Petrone, Anderson Brito, Chantal Vogels, Annie Watkins, Chaney Kalinich, Jessica Rothman, Kendall Billig, Tobias Koch, Fabiola Cruz-Lopez, David Rivera Aponte, Fabiola Fontanet, Natalie Machargo, Andrea Firpo, Vianca Aponte, Ernesto Vazquez, Patricia Serrano, Tania Mitwalli, Karolane Gonzalez, Nathan Grubaugh |
| EPI_ISL_3230054, EPI_ISL_3230055, EPI_ISL_3230056, EPI_ISL_3230057, EPI_ISL_3230058, EPI_ISL_3230059, EPI_ISL_3230060, EPI_ISL_3230061 | University Campus Bio-Medico of Rome (UCBM) | University Campus Bio-Medico of Rome (UCBM) | Riva E., De Florio L., Fogolari M., Francesconi M., Lintas C., Veralli R., Angeletti S. |
| EPI_ISL_3230064 | COVID-19 Detection Lab, Chattogram Veterinary and Animal Sciences University | Genomic Research Lab, Bangladesh Council of Scientific and Industrial Research | Goutam Buddha Das, Tridip Das, Tanvir Ahmad Nizami, Md. Sirazul Islam, Pronesh Dutta, Eaftekhar Ahmed Rana, Md. Murshed Hasan Sarkar, Md. Ahasan Habib, Shahina Akter, Tanjina Akhter Banu, Barna Goswami, Iffat Jahan, Md. Kamrul Islam, Sharmin Chowdhury, Md. Salim Khan, Paritosh Kumar Biswas |
| EPI_ISL_3230078, EPI_ISL_3230110, EPI_ISL_3230111 | University Campus Bio-Medico of Rome (UCBM) | University Campus Bio-Medico of Rome (UCBM) | Riva E., De Florio L., Fogolari M., Francesconi M., Lintas C., Veralli R., Angeletti S. |
| EPI_ISL_3230112 | COVID-19 Detection Lab, Chattogram Veterinary and Animal Sciences University | Genomic Research Lab, Bangladesh Council of Scientific and Industrial Research | Goutam Buddha Das, Tridip Das, Tanvir Ahmad Nizami, Md. Sirazul Islam, Pronesh Dutta, Eaftekhar Ahmed Rana, Md. Murshed Hasan Sarkar, Md. Ahasan Habib, Shahina Akter, Tanjina Akhter Banu, Barna Goswami, Iffat Jahan, Md. Kamrul Islam, Sharmin Chowdhury, Md. Salim Khan, Paritosh Kumar Biswas |
| EPI_ISL_3230113, EPI_ISL_3230114, EPI_ISL_3230115, EPI_ISL_3230116, EPI_ISL_3230117, EPI_ISL_3230118, EPI_ISL_3230119, EPI_ISL_3230120, EPI_ISL_3230121, EPI_ISL_3230122 | Michigan Department of Health and Human Services, Bureau of Laboratories | Michigan Department of Health and Human Services, Bureau of Laboratories | Blankenship HM, Riner D, Soehnlen MK |
| EPI_ISL_3230123 | University Campus Bio-Medico of Rome (UCBM) | University Campus Bio-Medico of Rome (UCBM) | Riva E., De Florio L., Fogolari M., Francesconi M., Lintas C., Veralli R., Angeletti S. |
| EPI_ISL_3230124, EPI_ISL_3230125, EPI_ISL_3230126, EPI_ISL_3230127, EPI_ISL_3230128, EPI_ISL_3230129, EPI_ISL_3230130, EPI_ISL_3230131, EPI_ISL_3230133, EPI_ISL_3230134, EPI_ISL_3230135, EPI_ISL_3230136, EPI_ISL_3230137, EPI_ISL_3230138, EPI_ISL_3230139, EPI_ISL_3230141, EPI_ISL_3230142, EPI_ISL_3230143, EPI_ISL_3230144, EPI_ISL_3230145, EPI_ISL_3230146, EPI_ISL_3230147, EPI_ISL_3230148, EPI_ISL_3230149, EPI_ISL_3230150, EPI_ISL_3230151, EPI_ISL_3230152, EPI_ISL_3230153, EPI_ISL_3230154, EPI_ISL_3230155, EPI_ISL_3230156, EPI_ISL_3230157, EPI_ISL_3230158, EPI_ISL_3230159, EPI_ISL_3230160, EPI_ISL_3230161 | | | |
| see above | Michigan Department of Health and Human Services, Bureau of Laboratories | Michigan Department of Health and Human Services, Bureau of Laboratories | Blankenship HM, Riner D, Soehnlen MK |
| EPI_ISL_3230162 | University Campus Bio-Medico of Rome (UCBM) | University Campus Bio-Medico of Rome (UCBM) | Riva E., De Florio L., Fogolari M., Francesconi M., Lintas C., Veralli R., Angeletti S. |
| EPI_ISL_3230163, | Michigan Department of Health and Human Services, Bureau of | Michigan Department of Health and Human Services, Bureau of | Blankenship HM, Riner D, Soehnlen MK |

| | | | |
|---|---|---|---|
| EPI_ISL_3230164, EPI_ISL_3230165, EPI_ISL_3230167, EPI_ISL_3230168, EPI_ISL_3230169, EPI_ISL_3230170, EPI_ISL_3230171, EPI_ISL_3230172, EPI_ISL_3230173 | Laboratories | Laboratories | |
| EPI_ISL_3230174, EPI_ISL_3230175 | University Campus Bio-Medico of Rome (UCBM) | University Campus Bio-Medico of Rome (UCBM) | Riva E., De Florio L., Fogolari M., Francesconi M., Lintas C., Veralli R., Angeletti S. |
| EPI_ISL_3230176, EPI_ISL_3230177 | RSUD KEBAYORAN BARU | Genomik Solidaritas Indonesia Laboratorium | Vania Gavrila Wikasa, Carissa Sintca Wijaya, Anna Christina Brazia, Annisa Muthiah Sukirman, Gracia Felias Enos Korompis, Anuraj Shankar, Louisa Markus, Meutia Ayuputeri Kumaheri, Ariel Pradipta |
| EPI_ISL_3230178, EPI_ISL_3230179, EPI_ISL_3230180, EPI_ISL_3230181, EPI_ISL_3230182 | Genomik Solidaritas Indonesia Laboratorium | Genomik Solidaritas Indonesia Laboratorium | Vania Gavrila Wikasa, Carissa Sintca Wijaya, Anna Christina Brazia, Annisa Muthiah Sukirman, Gracia Felias Enos Korompis, Anuraj Shankar, Louisa Markus, Meutia Ayuputeri Kumaheri, Ariel Pradipta |
| EPI_ISL_3230183, EPI_ISL_3230184, EPI_ISL_3230185, EPI_ISL_3230186, EPI_ISL_3230187, EPI_ISL_3230188, EPI_ISL_3230189, EPI_ISL_3230190 | Genomik Solidaritas Indonesia Laboratorium | Genomik Solidaritas Indonesia Laboratorium | Carissa Sintca Wijaya, Vania Gavrila Wikasa, Anna Christina Brazia, Annisa Muthiah Sukirman, Gracia Felias Enos Korompis, Anuraj Shankar, Louisa Markus, Meutia Ayuputeri Kumaheri, Ariel Pradipta |
| EPI_ISL_3230191, EPI_ISL_3230192, EPI_ISL_3230193, EPI_ISL_3230194, EPI_ISL_3230195, EPI_ISL_3230196, EPI_ISL_3230197 | Genomik Solidaritas Indonesia Laboratorium | Genomik Solidaritas Indonesia Laboratorium | Anna Christina Brazia, Vania Gavrila Wikasa, Carissa Sintca Wijaya, Annisa Muthiah Sukirman, Gracia Felias Enos Korompis, Anuraj Shankar, Louisa Markus, Meutia Ayuputeri Kumaheri, Ariel Pradipta |
| EPI_ISL_3230198 | Genomik Solidaritas Indonesia Laboratorium | Genomik Solidaritas Indonesia Laboratorium | Vania Gavrila Wikasa, Carissa Sintca Wijaya, Anna Christina Brazia, Annisa Muthiah Sukirman, Gracia Felias Enos Korompis, Anuraj Shankar, Louisa Markus, Meutia Ayuputeri Kumaheri, Ariel Pradipta |
| EPI_ISL_3230199 | University Campus Bio-Medico of Rome (UCBM) | University Campus Bio-Medico of Rome (UCBM) | Riva E., De Florio L., Fogolari M., Francesconi M., Lintas C., Veralli R., Angeletti S. |
| EPI_ISL_3230200 | COVID-19 Detection Lab, Chattogram Veterinary and Animal Sciences University | Genomic Research Lab, Bangladesh Council of Scientific and Industrial Research | Goutam Buddha Das, Tridip Das, Tanvir Ahmad Nizami, Md. Sirazul Islam, Pronesh Dutta, Eaftekhar Ahmed Rana, Md. Murshed Hasan Sarkar, Md. Ahasan Habib, Shahina Akter, Tanjina Akhter Banu, Barna Goswami, Iffat Jahan, Md. Kamrul Islam, Sharmin Chowdhury, Md. Salim Khan, Paritosh Kumar Biswas |
| EPI_ISL_3230201, EPI_ISL_3230202, EPI_ISL_3230203, EPI_ISL_3230204, EPI_ISL_3230205, EPI_ISL_3230206 | University Campus Bio-Medico of Rome (UCBM) | University Campus Bio-Medico of Rome (UCBM) | Riva E., De Florio L., Fogolari M., Francesconi M., Lintas C., Veralli R., Angeletti S. |
| EPI_ISL_3230211 | INMI Lazzaro Spallanzani IRCCS | INMI Lazzaro Spallanzani IRCCS | O Butera, CEM Gruber, F Messina, F Santini, M Rueca, G Bonfiglio, S Meschi, F Colavita, B Bartolini, E Giombini, , A Di Caro, MR Capobianchi |
| EPI_ISL_3230212, EPI_ISL_3230213 | University Campus Bio-Medico of Rome (UCBM) | University Campus Bio-Medico of Rome (UCBM) | Riva E., De Florio L., Fogolari M., Francesconi M., Lintas C., Veralli R., Angeletti S. |
| EPI_ISL_3230220, EPI_ISL_3230226, EPI_ISL_3230227, EPI_ISL_3230228, EPI_ISL_3230229, EPI_ISL_3230230, EPI_ISL_3230231, EPI_ISL_3230232 | Servizo de Microbioloxía. Complexo Hospitalario de Santiago de Compostela | Servizo de Microbioloxía. Complexo Hospitalario de Santiago de Compostela | Xosé Costa, Antonio Aguilera, Gema Barbeito, Amparo Coira, Carlos García Riestra, Manuela Hernández, José Llovo, Teresa Lopez Valiño, Laura Millán, Daniel Navarro, Rocío trastoy, Mercedes Treviño, María Luisa Pérez del Molino |
| EPI_ISL_3230233 | Genomik Solidaritas Indonesia Laboratorium | Genomik Solidaritas Indonesia Laboratorium | Reinhart Gabriel, Vania Gavrila Wikasa, Carissa Sintca Wijaya, Gracia Felias Enos Korompis, Annisa Muthiah Sukirman, Anuraj Shankar, Meutia Ayuputeri Kumaheri, Ariel Pradipta |
| EPI_ISL_3230236 | Rumah Sakit Pantai Indah Kapuk / DINAS KESEHATAN DKI JAKARTA | Genomik Solidaritas Indonesia Laboratorium | Carissa Sintca Wijaya, Vania Gavrila Wikasa, Leonardo H Simanjuntak, Hartini Cahyadi, Irwan Idris, Reinhart Gabriel, Gracia Felias Enos Korompis, Annisa Muthiah Sukirman, Anuraj Shankar, Meutia Ayuputeri Kumaheri, Ariel Pradipta |
| EPI_ISL_3230238 | RSUP FATMAWATI / DINAS KESEHATAN DKI JAKARTA | Genomik Solidaritas Indonesia Laboratorium | Carissa Sintca Wijaya, Vania Gavrila Wikasa, Jerry, Lidya Utami, Loli Simajuntak, Reinhart Gabriel, Gracia Felias Enos Korompis, Annisa Muthiah Sukirman, Anuraj Shankar, Meutia Ayuputeri Kumaheri, Ariel Pradipta |
| EPI_ISL_3230239 | RSUP FATMAWATI / DINAS KESEHATAN DKI JAKARTA | Genomik Solidaritas Indonesia Laboratorium | Carissa Sintca Wijaya, Vania Gavrila Wikasa, Jerry, Lidya Utami, Loli Simajuntak, Reinhart Gabriel, Gracia Felias Enos Korompis, Annisa Muthiah Sukirman, Anuraj Shankar, Meutia Ayuputeri Kumaheri, Ariel Pradipta |
| EPI_ISL_3230240 | RSUP FATMAWATI / DINAS KESEHATAN DKI JAKARTA | Genomik Solidaritas Indonesia Laboratorium | Reinhart Gabriel, Vania Gavrila Wikasa, Jerry, Lidya Utami, Loli Simajuntak, Carissa Sintca Wijaya, Gracia Felias Enos Korompis, Annisa Muthiah Sukirman, Anuraj Shankar, Meutia Ayuputeri Kumaheri, Ariel Pradipta |
| EPI_ISL_3230241 | RSUP FATMAWATI / DINAS KESEHATAN DKI JAKARTA | Genomik Solidaritas Indonesia Laboratorium | Vania Gavrila Wikasa, Carissa Sintca Wijaya, Jerry, Lidya Utami, Loli Simajuntak, Reinhart Gabriel, Gracia Felias Enos Korompis, Annisa Muthiah Sukirman, Anuraj Shankar, Meutia Ayuputeri Kumaheri, Ariel Pradipta |
| EPI_ISL_3230242 | RSUP FATMAWATI / DINAS KESEHATAN DKI JAKARTA | Genomik Solidaritas Indonesia Laboratorium | Reinhart Gabriel, Vania Gavrila Wikasa, Jerry, Lidya Utami, Loli Simajuntak, Carissa Sintca Wijaya, Gracia Felias Enos Korompis, Annisa Muthiah Sukirman, Anuraj Shankar, Meutia Ayuputeri Kumaheri, Ariel Pradipta |
| EPI_ISL_3230243 | RSUP FATMAWATI / DINAS KESEHATAN DKI JAKARTA | Genomik Solidaritas Indonesia Laboratorium | Carissa Sintca Wijaya, Vania Gavrila Wikasa, Leonardo H Simanjuntak, Hartini Cahyadi, Irwan Idris, Reinhart Gabriel, Gracia Felias Enos Korompis, Annisa Muthiah Sukirman, Anuraj Shankar, Meutia Ayuputeri Kumaheri, Ariel Pradipta |
| EPI_ISL_3230244 | Genomik Solidaritas Indonesia Laboratorium | Genomik Solidaritas Indonesia Laboratorium | Reinhart Gabriel, Vania Gavrila Wikasa, Carissa Sintca Wijaya, Gracia Felias Enos Korompis, Annisa Muthiah Sukirman, Anuraj Shankar, Meutia Ayuputeri Kumaheri, Ariel Pradipta |
| EPI_ISL_3230245, EPI_ISL_3230246 | Genomik Solidaritas Indonesia Laboratorium | Genomik Solidaritas Indonesia Laboratorium | Vania Gavrila Wikasa, Carissa Sintca Wijaya, Reinhart Gabriel, Gracia Felias Enos Korompis, Annisa Muthiah Sukirman, Anuraj Shankar, Meutia Ayuputeri Kumaheri, Ariel Pradipta |
| EPI_ISL_3230247 | RSUP FATMAWATI / DINAS KESEHATAN DKI JAKARTA | Genomik Solidaritas Indonesia Laboratorium | Vania Gavrila Wikasa, Carissa Sintca Wijaya, Jerry, Lidya Utami, Loli Simajuntak, Reinhart Gabriel, Gracia Felias Enos Korompis, Annisa Muthiah Sukirman, Anuraj Shankar, Meutia Ayuputeri Kumaheri, Ariel Pradipta |
| EPI_ISL_3230248 | Genomik Solidaritas Indonesia Laboratorium | Genomik Solidaritas Indonesia Laboratorium | Reinhart Gabriel, Vania Gavrila Wikasa, Carissa Sintca Wijaya, Gracia Felias Enos Korompis, Annisa Muthiah Sukirman, Anuraj Shankar, Meutia Ayuputeri Kumaheri, Ariel Pradipta |
| EPI_ISL_3230249 | GOOD DOCTOR / DINAS KESEHATAN DKI JAKARTA | Genomik Solidaritas Indonesia Laboratorium | Vania Gavrila Wikasa, Carissa Sintca Wijaya, Venny Beauty, Reinhart Gabriel, Gracia Felias Enos Korompis, Annisa Muthiah Sukirman, Anuraj Shankar, Meutia Ayuputeri Kumaheri, Ariel Pradipta |
| EPI_ISL_3230250 | COVID-19 Detection Lab, Chattogram Veterinary and Animal Sciences University | Genomic Research Lab, Bangladesh Council of Scientific and Industrial Research | Goutam Buddha Das, Tridip Das, Tanvir Ahmad Nizami, Md. Sirazul Islam, Pronesh Dutta, Eaftekhar Ahmed Rana, Md. Murshed Hasan Sarkar, Md. Ahasan Habib, Shahina Akter, Tanjina Akhter Banu, Barna Goswami, Iffat Jahan, Md. Kamrul Islam, Sharmin Chowdhury, Md. Salim Khan, Paritosh Kumar Biswas |
| EPI_ISL_3230251, EPI_ISL_3230252 | University Campus Bio-Medico of Rome (UCBM) | University Campus Bio-Medico of Rome (UCBM) | Riva E., De Florio L., Fogolari M., Francesconi M., Lintas C., Veralli R., Angeletti S. |
| EPI_ISL_3230253, EPI_ISL_3230254 | COVID-19 Detection Lab, Chattogram Veterinary and Animal Sciences University | Genomic Research Lab, Bangladesh Council of Scientific and Industrial Research | Goutam Buddha Das, Tridip Das, Tanvir Ahmad Nizami, Md. Sirazul Islam, Pronesh Dutta, Eaftekhar Ahmed Rana, Md. Murshed Hasan Sarkar, Md. Ahasan Habib, Shahina Akter, Tanjina Akhter Banu, Barna Goswami, Iffat Jahan, Md. Kamrul Islam, Sharmin Chowdhury, Md. Salim Khan, Paritosh Kumar Biswas |
| EPI_ISL_3230255 | General Hospital "Abdulah Nakas" | Alela Genetic Centre | Rijad Konjhodzic, Dino Pecar, Lana Salihefendic, Ivana Ceko, Adis Kandic, Enis Kandic |
| EPI_ISL_3230262, EPI_ISL_3230263, EPI_ISL_3230267 | University Campus Bio-Medico of Rome (UCBM) | University Campus Bio-Medico of Rome (UCBM) | Riva E., De Florio L., Fogolari M., Francesconi M., Lintas C., Veralli R., Angeletti S. |
| EPI_ISL_3230272, EPI_ISL_3230273 | TYKS, Kliininen mikrobiologia | Expert Microbiology, National Institute for Health and Welfare | Soile Blomqvist, Jani Halkilahti, Kirsi Liitsola, Haider al-Hello, Päivi Laurila, Erika Lindh, Teemu Smura, Ravi Kant, Phuoc Truong, Olli Vapalahti, Sari Hannula, Pekka Ellonen, Niina Ikonen, Carita Savolainen-Kopra |
| EPI_ISL_3230278 | Islab, Pohjois-Savon aluelaboratorio | Expert Microbiology, National Institute for Health and Welfare | Soile Blomqvist, Jani Halkilahti, Kirsi Liitsola, Haider al-Hello, Päivi Laurila, Erika Lindh, Teemu Smura, Ravi Kant, Phuoc Truong, Olli Vapalahti, Sari Hannula, Pekka Ellonen, Niina Ikonen, Carita Savolainen-Kopra |
| EPI_ISL_3230279 | SYNLAB Suomi | Expert Microbiology, National Institute for Health and Welfare | Soile Blomqvist, Jani Halkilahti, Kirsi Liitsola, Haider al-Hello, Päivi Laurila, Erika Lindh, Teemu Smura, Ravi Kant, Phuoc Truong, Olli Vapalahti, Sari Hannula, Pekka Ellonen, Niina Ikonen, Carita Savolainen-Kopra |
| EPI_ISL_3230280 | Islab, Pohjois-Savon aluelaboratorio | Expert Microbiology, National Institute for Health and Welfare | Soile Blomqvist, Jani Halkilahti, Kirsi Liitsola, Haider al-Hello, Päivi Laurila, Erika Lindh, Teemu Smura, Ravi Kant, Phuoc Truong, Olli Vapalahti, Sari Hannula, Pekka Ellonen, Niina Ikonen, Carita Savolainen-Kopra |
| EPI_ISL_3230302 | Vita Laboratoriot Oy | Expert Microbiology, National Institute for Health and Welfare | Soile Blomqvist, Jani Halkilahti, Kirsi Liitsola, Haider al-Hello, Päivi Laurila, Erika Lindh, Teemu Smura, Ravi Kant, Phuoc Truong, Olli Vapalahti, Sari Hannula, Pekka Ellonen, Niina Ikonen, Carita Savolainen-Kopra |
| EPI_ISL_3230304 | Islab, Pohjois-Savon aluelaboratorio | Expert Microbiology, National Institute for Health and Welfare | Soile Blomqvist, Jani Halkilahti, Kirsi Liitsola, Haider al-Hello, Päivi Laurila, Erika Lindh, Teemu Smura, Ravi Kant, Phuoc Truong, Olli Vapalahti, Sari Hannula, Pekka Ellonen, Niina Ikonen, Carita Savolainen-Kopra |
| EPI_ISL_3230307 | TYKS, Kliininen mikrobiologia | Expert Microbiology, National Institute for Health and Welfare | Soile Blomqvist, Jani Halkilahti, Kirsi Liitsola, Haider al-Hello, Päivi Laurila, Erika Lindh, Teemu Smura, Ravi Kant, Phuoc Truong, Olli Vapalahti, Sari Hannula, Pekka Ellonen, Niina Ikonen, Carita Savolainen-Kopra |
| EPI_ISL_3230308 | Fimlab Laboratoriot Oy Tampere | Expert Microbiology, National Institute for Health and Welfare | Soile Blomqvist, Jani Halkilahti, Kirsi Liitsola, Haider al-Hello, Päivi Laurila, Erika Lindh, Teemu Smura, Ravi Kant, Phuoc Truong, Olli Vapalahti, Sari Hannula, Pekka Ellonen, Niina Ikonen, Carita Savolainen-Kopra |
| EPI_ISL_3230309 | Islab, Pohjois-Savon aluelaboratorio | Expert Microbiology, National Institute for Health and Welfare | Soile Blomqvist, Jani Halkilahti, Kirsi Liitsola, Haider al-Hello, Päivi Laurila, Erika Lindh, Teemu Smura, Ravi Kant, Phuoc Truong, Olli Vapalahti, Sari Hannula, Pekka Ellonen, Niina Ikonen, Carita Savolainen-Kopra |
| EPI_ISL_3230310 | TYKS, Kliininen mikrobiologia | Expert Microbiology, National Institute for Health and Welfare | Soile Blomqvist, Jani Halkilahti, Kirsi Liitsola, Haider al-Hello, Päivi Laurila, Erika Lindh, Teemu Smura, Ravi Kant, Phuoc Truong, Olli Vapalahti, Sari Hannula, Pekka Ellonen, Niina Ikonen, Carita Savolainen-Kopra |
| EPI_ISL_3230312, EPI_ISL_3230313, | Islab, Pohjois-Savon aluelaboratorio | Expert Microbiology, National Institute for Health and Welfare | Soile Blomqvist, Jani Halkilahti, Kirsi Liitsola, Haider al-Hello, Päivi Laurila, Erika Lindh, Teemu Smura, Ravi Kant, Phuoc Truong, Olli Vapalahti, Sari Hannula, Pekka Ellonen, Niina Ikonen, Carita Savolainen-Kopra |

| | | | |
|---|---|---|---|
| EPI_ISL_3230314, EPI_ISL_3230315, EPI_ISL_3230316 | | | |
| EPI_ISL_3230317 | NordLab Kajaani | Expert Microbiology, National Institute for Health and Welfare | Soile Blomqvist, Jani Halkilahti, Kirsi Liitsola, Haider al-Hello, Päivi Laurila, Erika Lindh, Teemu Smura, Ravi Kant, Phuoc Truong, Olli Vapalahti, Sari Hannula, Pekka Ellonen, Niina Ikonen, Carita Savolainen-Kopra |
| EPI_ISL_3230318, EPI_ISL_3230319, EPI_ISL_3230320 | Islab, Pohjois-Karjalan aluelaboratorio | Expert Microbiology, National Institute for Health and Welfare | Soile Blomqvist, Jani Halkilahti, Kirsi Liitsola, Haider al-Hello, Päivi Laurila, Erika Lindh, Teemu Smura, Ravi Kant, Phuoc Truong, Olli Vapalahti, Sari Hannula, Pekka Ellonen, Niina Ikonen, Carita Savolainen-Kopra |
| EPI_ISL_3230321, EPI_ISL_3230322 | NordLab Kajaani | Expert Microbiology, National Institute for Health and Welfare | Soile Blomqvist, Jani Halkilahti, Kirsi Liitsola, Haider al-Hello, Päivi Laurila, Erika Lindh, Teemu Smura, Ravi Kant, Phuoc Truong, Olli Vapalahti, Sari Hannula, Pekka Ellonen, Niina Ikonen, Carita Savolainen-Kopra |
| EPI_ISL_3230323 | Islab, Pohjois-Karjalan aluelaboratorio | Expert Microbiology, National Institute for Health and Welfare | Soile Blomqvist, Jani Halkilahti, Kirsi Liitsola, Haider al-Hello, Päivi Laurila, Erika Lindh, Teemu Smura, Ravi Kant, Phuoc Truong, Olli Vapalahti, Sari Hannula, Pekka Ellonen, Niina Ikonen, Carita Savolainen-Kopra |
| EPI_ISL_3230324, EPI_ISL_3230325, EPI_ISL_3230326, EPI_ISL_3230327, EPI_ISL_3230328, EPI_ISL_3230329 | Islab, Pohjois-Savon aluelaboratorio | Expert Microbiology, National Institute for Health and Welfare | Soile Blomqvist, Jani Halkilahti, Kirsi Liitsola, Haider al-Hello, Päivi Laurila, Erika Lindh, Teemu Smura, Ravi Kant, Phuoc Truong, Olli Vapalahti, Sari Hannula, Pekka Ellonen, Niina Ikonen, Carita Savolainen-Kopra |
| EPI_ISL_3230330 | University Campus Bio-Medico of Rome (UCBM) | University Campus Bio-Medico of Rome (UCBM) | Riva E., De Florio L., Fogolari M., Francesconi M., Lintas C., Veralli R., Angeletti S. |
| EPI_ISL_3230331, EPI_ISL_3230332, EPI_ISL_3230333, EPI_ISL_3230334, EPI_ISL_3230335, EPI_ISL_3230336, EPI_ISL_3230337, EPI_ISL_3230338 | Islab, Pohjois-Savon aluelaboratorio | Expert Microbiology, National Institute for Health and Welfare | Soile Blomqvist, Jani Halkilahti, Kirsi Liitsola, Haider al-Hello, Päivi Laurila, Erika Lindh, Teemu Smura, Ravi Kant, Phuoc Truong, Olli Vapalahti, Sari Hannula, Pekka Ellonen, Niina Ikonen, Carita Savolainen-Kopra |
| EPI_ISL_3230339, EPI_ISL_3230340, EPI_ISL_3230341, EPI_ISL_3230342, EPI_ISL_3230343, EPI_ISL_3230344, EPI_ISL_3230345, EPI_ISL_3230346, EPI_ISL_3230347, EPI_ISL_3230348, EPI_ISL_3230349, EPI_ISL_3230350, EPI_ISL_3230351 | | | |
| see above | Fimlab Laboratoriot Oy Tampere | Expert Microbiology, National Institute for Health and Welfare | Soile Blomqvist, Jani Halkilahti, Kirsi Liitsola, Haider al-Hello, Päivi Laurila, Erika Lindh, Teemu Smura, Ravi Kant, Phuoc Truong, Olli Vapalahti, Sari Hannula, Pekka Ellonen, Niina Ikonen, Carita Savolainen-Kopra |
| EPI_ISL_3230352 | Bimelix Ab | Expert Microbiology, National Institute for Health and Welfare | Soile Blomqvist, Jani Halkilahti, Kirsi Liitsola, Haider al-Hello, Päivi Laurila, Erika Lindh, Teemu Smura, Ravi Kant, Phuoc Truong, Olli Vapalahti, Sari Hannula, Pekka Ellonen, Niina Ikonen, Carita Savolainen-Kopra |
| EPI_ISL_3230353 | SataDiag | Expert Microbiology, National Institute for Health and Welfare | Soile Blomqvist, Jani Halkilahti, Kirsi Liitsola, Haider al-Hello, Päivi Laurila, Erika Lindh, Teemu Smura, Ravi Kant, Phuoc Truong, Olli Vapalahti, Sari Hannula, Pekka Ellonen, Niina Ikonen, Carita Savolainen-Kopra |
| EPI_ISL_3230354 | Bimelix Ab | Expert Microbiology, National Institute for Health and Welfare | Soile Blomqvist, Jani Halkilahti, Kirsi Liitsola, Haider al-Hello, Päivi Laurila, Erika Lindh, Teemu Smura, Ravi Kant, Phuoc Truong, Olli Vapalahti, Sari Hannula, Pekka Ellonen, Niina Ikonen, Carita Savolainen-Kopra |
| EPI_ISL_3230355, EPI_ISL_3230356, EPI_ISL_3230357, EPI_ISL_3230358 | Vita Laboratoriot Oy | Expert Microbiology, National Institute for Health and Welfare | Soile Blomqvist, Jani Halkilahti, Kirsi Liitsola, Haider al-Hello, Päivi Laurila, Erika Lindh, Teemu Smura, Ravi Kant, Phuoc Truong, Olli Vapalahti, Sari Hannula, Pekka Ellonen, Niina Ikonen, Carita Savolainen-Kopra |
| EPI_ISL_3230359, EPI_ISL_3230360, EPI_ISL_3230361, EPI_ISL_3230362, EPI_ISL_3230363, EPI_ISL_3230364, EPI_ISL_3230365, EPI_ISL_3230366, EPI_ISL_3230367, EPI_ISL_3230368, EPI_ISL_3230369, EPI_ISL_3230370, EPI_ISL_3230371, EPI_ISL_3230372, EPI_ISL_3230373, EPI_ISL_3230374, EPI_ISL_3230375, EPI_ISL_3230376, EPI_ISL_3230377, EPI_ISL_3230378, EPI_ISL_3230379, EPI_ISL_3230380, EPI_ISL_3230381, EPI_ISL_3230382, EPI_ISL_3230383, EPI_ISL_3230384, EPI_ISL_3230385, EPI_ISL_3230386, EPI_ISL_3230387, EPI_ISL_3230388, EPI_ISL_3230389 | | | |
| see above | TYKS, Kliininen mikrobiologia | Expert Microbiology, National Institute for Health and Welfare | Soile Blomqvist, Jani Halkilahti, Kirsi Liitsola, Haider al-Hello, Päivi Laurila, Erika Lindh, Teemu Smura, Ravi Kant, Phuoc Truong, Olli Vapalahti, Sari Hannula, Pekka Ellonen, Niina Ikonen, Carita Savolainen-Kopra |
| EPI_ISL_3230390, EPI_ISL_3230391, EPI_ISL_3230392, EPI_ISL_3230393, EPI_ISL_3230394, EPI_ISL_3230395, EPI_ISL_3230396, EPI_ISL_3230397, EPI_ISL_3230398, EPI_ISL_3230399 | Fimlab Laboratoriot Oy Tampere | Expert Microbiology, National Institute for Health and Welfare | Soile Blomqvist, Jani Halkilahti, Kirsi Liitsola, Haider al-Hello, Päivi Laurila, Erika Lindh, Teemu Smura, Ravi Kant, Phuoc Truong, Olli Vapalahti, Sari Hannula, Pekka Ellonen, Niina Ikonen, Carita Savolainen-Kopra |
| EPI_ISL_3230400 | NordLab Kajaani | Expert Microbiology, National Institute for Health and Welfare | Soile Blomqvist, Jani Halkilahti, Kirsi Liitsola, Haider al-Hello, Päivi Laurila, Erika Lindh, Teemu Smura, Ravi Kant, Phuoc Truong, Olli Vapalahti, Sari Hannula, Pekka Ellonen, Niina Ikonen, Carita Savolainen-Kopra |
| EPI_ISL_3230401 | Fimlab Laboratoriot Oy Tampere | Expert Microbiology, National Institute for Health and Welfare | Soile Blomqvist, Jani Halkilahti, Kirsi Liitsola, Haider al-Hello, Päivi Laurila, Erika Lindh, Teemu Smura, Ravi Kant, Phuoc Truong, Olli Vapalahti, Sari Hannula, Pekka Ellonen, Niina Ikonen, Carita Savolainen-Kopra |
| EPI_ISL_3230402 | TYKS, Kliininen mikrobiologia | Expert Microbiology, National Institute for Health and Welfare | Soile Blomqvist, Jani Halkilahti, Kirsi Liitsola, Haider al-Hello, Päivi Laurila, Erika Lindh, Teemu Smura, Ravi Kant, Phuoc Truong, Olli Vapalahti, Sari Hannula, Pekka Ellonen, Niina Ikonen, Carita Savolainen-Kopra |
| EPI_ISL_3230403 | NordLab Kajaani | Expert Microbiology, National Institute for Health and Welfare | Soile Blomqvist, Jani Halkilahti, Kirsi Liitsola, Haider al-Hello, Päivi Laurila, Erika Lindh, Teemu Smura, Ravi Kant, Phuoc Truong, Olli Vapalahti, Sari Hannula, Pekka Ellonen, Niina Ikonen, Carita Savolainen-Kopra |
| EPI_ISL_3230404 | TYKS, Kliininen mikrobiologia | Expert Microbiology, National Institute for Health and Welfare | Soile Blomqvist, Jani Halkilahti, Kirsi Liitsola, Haider al-Hello, Päivi Laurila, Erika Lindh, Teemu Smura, Ravi Kant, Phuoc Truong, Olli Vapalahti, Sari Hannula, Pekka Ellonen, Niina Ikonen, Carita Savolainen-Kopra |
| EPI_ISL_3230405 | Islab, Pohjois-Karjalan aluelaboratorio | Expert Microbiology, National Institute for Health and Welfare | Soile Blomqvist, Jani Halkilahti, Kirsi Liitsola, Haider al-Hello, Päivi Laurila, Erika Lindh, Teemu Smura, Ravi Kant, Phuoc Truong, Olli Vapalahti, Sari Hannula, Pekka Ellonen, Niina Ikonen, Carita Savolainen-Kopra |
| EPI_ISL_3230406 | Vita Laboratoriot Oy | Expert Microbiology, National Institute for Health and Welfare | Soile Blomqvist, Jani Halkilahti, Kirsi Liitsola, Haider al-Hello, Päivi Laurila, Erika Lindh, Teemu Smura, Ravi Kant, Phuoc Truong, Olli Vapalahti, Sari Hannula, Pekka Ellonen, Niina Ikonen, Carita Savolainen-Kopra |
| EPI_ISL_3230407, EPI_ISL_3230408, EPI_ISL_3230409 | Fimlab Laboratoriot Oy Tampere | Expert Microbiology, National Institute for Health and Welfare | Soile Blomqvist, Jani Halkilahti, Kirsi Liitsola, Haider al-Hello, Päivi Laurila, Erika Lindh, Teemu Smura, Ravi Kant, Phuoc Truong, Olli Vapalahti, Sari Hannula, Pekka Ellonen, Niina Ikonen, Carita Savolainen-Kopra |
| EPI_ISL_3230410, EPI_ISL_3230411, EPI_ISL_3230412, EPI_ISL_3230413, EPI_ISL_3230414 | TYKS, Kliininen mikrobiologia | Expert Microbiology, National Institute for Health and Welfare | Soile Blomqvist, Jani Halkilahti, Kirsi Liitsola, Haider al-Hello, Päivi Laurila, Erika Lindh, Teemu Smura, Ravi Kant, Phuoc Truong, Olli Vapalahti, Sari Hannula, Pekka Ellonen, Niina Ikonen, Carita Savolainen-Kopra |
| EPI_ISL_3230415, EPI_ISL_3230416, EPI_ISL_3230417 | Fimlab Laboratoriot Oy Tampere | Expert Microbiology, National Institute for Health and Welfare | Soile Blomqvist, Jani Halkilahti, Kirsi Liitsola, Haider al-Hello, Päivi Laurila, Erika Lindh, Teemu Smura, Ravi Kant, Phuoc Truong, Olli Vapalahti, Sari Hannula, Pekka Ellonen, Niina Ikonen, Carita Savolainen-Kopra |
| EPI_ISL_3230418 | TYKS, Kliininen mikrobiologia | Expert Microbiology, National Institute for Health and Welfare | Soile Blomqvist, Jani Halkilahti, Kirsi Liitsola, Haider al-Hello, Päivi Laurila, Erika Lindh, Teemu Smura, Ravi Kant, Phuoc Truong, Olli Vapalahti, Sari Hannula, Pekka Ellonen, Niina Ikonen, Carita Savolainen-Kopra |
| EPI_ISL_3230419 | Islab, Pohjois-Savon aluelaboratorio | Expert Microbiology, National Institute for Health and Welfare | Soile Blomqvist, Jani Halkilahti, Kirsi Liitsola, Haider al-Hello, Päivi Laurila, Erika Lindh, Teemu Smura, Ravi Kant, Phuoc Truong, Olli Vapalahti, Sari Hannula, Pekka Ellonen, Niina Ikonen, Carita Savolainen-Kopra |
| EPI_ISL_3230420 | Fimlab Laboratoriot Oy Tampere | Expert Microbiology, National Institute for Health and Welfare | Soile Blomqvist, Jani Halkilahti, Kirsi Liitsola, Haider al-Hello, Päivi Laurila, Erika Lindh, Teemu Smura, Ravi Kant, Phuoc Truong, Olli Vapalahti, Sari Hannula, Pekka Ellonen, Niina Ikonen, Carita Savolainen-Kopra |
| EPI_ISL_3230421 | NordLab Kajaani | Expert Microbiology, National Institute for Health and Welfare | Soile Blomqvist, Jani Halkilahti, Kirsi Liitsola, Haider al-Hello, Päivi Laurila, Erika Lindh, Teemu Smura, Ravi Kant, Phuoc Truong, Olli Vapalahti, Sari Hannula, Pekka Ellonen, Niina Ikonen, Carita Savolainen-Kopra |
| EPI_ISL_3230422 | NordLab Oulu | Expert Microbiology, National Institute for Health and Welfare | Soile Blomqvist, Jani Halkilahti, Kirsi Liitsola, Haider al-Hello, Päivi Laurila, Erika Lindh, Teemu Smura, Ravi Kant, Phuoc Truong, Olli Vapalahti, Sari Hannula, Pekka Ellonen, Niina Ikonen, Carita Savolainen-Kopra |
| EPI_ISL_3230424 | Fimlab Laboratoriot Oy Tampere | Expert Microbiology, National Institute for Health and Welfare | Soile Blomqvist, Jani Halkilahti, Kirsi Liitsola, Haider al-Hello, Päivi Laurila, Erika Lindh, Teemu Smura, Ravi Kant, Phuoc Truong, Olli Vapalahti, Sari Hannula, Pekka Ellonen, Niina Ikonen, Carita Savolainen-Kopra |
| EPI_ISL_3230425 | TYKS, Kliininen mikrobiologia | Expert Microbiology, National Institute for Health and Welfare | Soile Blomqvist, Jani Halkilahti, Kirsi Liitsola, Haider al-Hello, Päivi Laurila, Erika Lindh, Teemu Smura, Ravi Kant, Phuoc Truong, Olli Vapalahti, Sari Hannula, Pekka Ellonen, Niina Ikonen, Carita Savolainen-Kopra |
| EPI_ISL_3230426 | Vita Laboratoriot Oy | Expert Microbiology, National Institute for Health and Welfare | Soile Blomqvist, Jani Halkilahti, Kirsi Liitsola, Haider al-Hello, Päivi Laurila, Erika Lindh, Teemu Smura, Ravi Kant, Phuoc Truong, Olli Vapalahti, Sari Hannula, Pekka Ellonen, Niina Ikonen, Carita Savolainen-Kopra |
| EPI_ISL_3230427 | Fimlab Laboratoriot Oy Tampere | Expert Microbiology, National Institute for Health and Welfare | Soile Blomqvist, Jani Halkilahti, Kirsi Liitsola, Haider al-Hello, Päivi Laurila, Erika Lindh, Teemu Smura, Ravi Kant, Phuoc Truong, Olli Vapalahti, Sari Hannula, Pekka Ellonen, Niina Ikonen, Carita Savolainen-Kopra |
| EPI_ISL_3230428 | Islab, Pohjois-Savon aluelaboratorio | Expert Microbiology, National Institute for Health and Welfare | Soile Blomqvist, Jani Halkilahti, Kirsi Liitsola, Haider al-Hello, Päivi Laurila, Erika Lindh, Teemu Smura, Ravi Kant, Phuoc Truong, Olli Vapalahti, Sari Hannula, Pekka Ellonen, Niina Ikonen, Carita Savolainen-Kopra |
| EPI_ISL_3230429 | Fimlab Laboratoriot Oy Tampere | Expert Microbiology, National Institute for Health and Welfare | Soile Blomqvist, Jani Halkilahti, Kirsi Liitsola, Haider al-Hello, Päivi Laurila, Erika Lindh, Teemu Smura, Ravi Kant, Phuoc Truong, Olli Vapalahti, Sari Hannula, Pekka Ellonen, Niina Ikonen, Carita Savolainen-Kopra |
| EPI_ISL_3230431 | Islab, Etelä-Savon aluelaboratorio | Expert Microbiology, National Institute for Health and Welfare | Soile Blomqvist, Jani Halkilahti, Kirsi Liitsola, Haider al-Hello, Päivi Laurila, Erika Lindh, Teemu Smura, Ravi Kant, Phuoc Truong, Olli Vapalahti, Sari Hannula, Pekka Ellonen, Niina Ikonen, Carita Savolainen-Kopra |
| EPI_ISL_3230432 | TYKS, Kliininen mikrobiologia | Expert Microbiology, National Institute for Health and Welfare | Soile Blomqvist, Jani Halkilahti, Kirsi Liitsola, Haider al-Hello, Päivi Laurila, Erika Lindh, Teemu Smura, Ravi Kant, Phuoc Truong, Olli Vapalahti, Sari Hannula, Pekka Ellonen, Niina Ikonen, Carita Savolainen-Kopra |
| EPI_ISL_3230433 | NordLab Oulu | Expert Microbiology, National Institute for Health and Welfare | Soile Blomqvist, Jani Halkilahti, Kirsi Liitsola, Haider al-Hello, Päivi Laurila, Erika Lindh, Teemu Smura, Ravi Kant, Phuoc Truong, Olli Vapalahti, Sari Hannula, Pekka Ellonen, Niina Ikonen, Carita Savolainen-Kopra |
| EPI_ISL_3230434, EPI_ISL_3230435, EPI_ISL_3230436 | SataDiag | Expert Microbiology, National Institute for Health and Welfare | Soile Blomqvist, Jani Halkilahti, Kirsi Liitsola, Haider al-Hello, Päivi Laurila, Erika Lindh, Teemu Smura, Ravi Kant, Phuoc Truong, Olli Vapalahti, Sari Hannula, Pekka Ellonen, Niina Ikonen, Carita Savolainen-Kopra |
| EPI_ISL_3230437 | Helsinki University Hospital, Huslab | Expert Microbiology, National Institute for Health and Welfare | Soile Blomqvist, Jani Halkilahti, Kirsi Liitsola, Haider al-Hello, Päivi Laurila, Erika Lindh, Teemu Smura, Ravi Kant, Phuoc Truong, Olli Vapalahti, Sari Hannula, Pekka Ellonen, Niina Ikonen, Carita Savolainen-Kopra |
| EPI_ISL_3230438 | Bimelix Ab | Expert Microbiology, National Institute for Health and Welfare | Soile Blomqvist, Jani Halkilahti, Kirsi Liitsola, Haider al-Hello, Päivi Laurila, Erika Lindh, Teemu Smura, Ravi Kant, Phuoc Truong, Olli Vapalahti, Sari Hannula, Pekka Ellonen, Niina Ikonen, Carita Savolainen-Kopra |
| EPI_ISL_3230439, EPI_ISL_3230440 | SataDiag | Expert Microbiology, National Institute for Health and Welfare | Soile Blomqvist, Jani Halkilahti, Kirsi Liitsola, Haider al-Hello, Päivi Laurila, Erika Lindh, Teemu Smura, Ravi Kant, Phuoc Truong, Olli Vapalahti, Sari Hannula, Pekka Ellonen, Niina Ikonen, Carita Savolainen-Kopra |
| EPI_ISL_3230442, EPI_ISL_3230443 | Islab, Pohjois-Karjalan aluelaboratorio | Expert Microbiology, National Institute for Health and Welfare | Soile Blomqvist, Jani Halkilahti, Kirsi Liitsola, Haider al-Hello, Päivi Laurila, Erika Lindh, Teemu Smura, Ravi Kant, Phuoc Truong, Olli Vapalahti, Sari Hannula, Pekka Ellonen, Niina Ikonen, Carita Savolainen-Kopra |
| EPI_ISL_3230447, | NordLab Oulu | Expert Microbiology, National Institute for Health and Welfare | Soile Blomqvist, Jani Halkilahti, Kirsi Liitsola, Haider al-Hello, Päivi Laurila, Erika Lindh, Teemu Smura, Ravi Kant, Phuoc Truong, Olli Vapalahti, Sari Hannula, Pekka Ellonen, Niina Ikonen, Carita Savolainen-Kopra |

| | | | |
|---|---|---|---|
| EPI_ISL_3230448, EPI_ISL_3230450, EPI_ISL_3230451, EPI_ISL_3230452 | | | |
| EPI_ISL_3230453 | Vita Laboratoriot Oy | Expert Microbiology, National Institute for Health and Welfare | Soile Blomqvist, Jani Halkilahti, Kirsi Liitsola, Haider al-Hello, Päivi Laurila, Erika Lindh, Teemu Smura, Ravi Kant, Phuoc Truong, Olli Vapalahti, Sari Hannula, Pekka Ellonen, Niina Ikonen, Carita Savolainen-Kopra |
| EPI_ISL_3230454 | COVID-19 Detection Lab, Chattogram Veterinary and Animal Sciences University | Genomic Research Lab, Bangladesh Council of Scientific and Industrial Research | Goutam Buddha Das, Tridip Das, Tanvir Ahmad Nizami, Md. Sirazul Islam, Pronesh Dutta, Eaftekhar Ahmed Rana, Md. Murshed Hasan Sarkar, Md. Ahasan Habib, Shahina Akter, Tanjina Akhter Banu, Barna Goswami, Iffat Jahan, Md. Kamrul Islam, Sharmin Chowdhury, Md. Salim Khan, Paritosh Kumar Biswas |
| EPI_ISL_3230457, EPI_ISL_3230458, EPI_ISL_3230459 | Islab, Pohjois-Savon aluelaboratorio | Expert Microbiology, National Institute for Health and Welfare | Soile Blomqvist, Jani Halkilahti, Kirsi Liitsola, Haider al-Hello, Päivi Laurila, Erika Lindh, Teemu Smura, Ravi Kant, Phuoc Truong, Olli Vapalahti, Sari Hannula, Pekka Ellonen, Niina Ikonen, Carita Savolainen-Kopra |
| EPI_ISL_3230461 | Vita Laboratoriot Oy | Expert Microbiology, National Institute for Health and Welfare | Soile Blomqvist, Jani Halkilahti, Kirsi Liitsola, Haider al-Hello, Päivi Laurila, Erika Lindh, Teemu Smura, Ravi Kant, Phuoc Truong, Olli Vapalahti, Sari Hannula, Pekka Ellonen, Niina Ikonen, Carita Savolainen-Kopra |
| EPI_ISL_3230462 | Bimelix Ab | Expert Microbiology, National Institute for Health and Welfare | Soile Blomqvist, Jani Halkilahti, Kirsi Liitsola, Haider al-Hello, Päivi Laurila, Erika Lindh, Teemu Smura, Ravi Kant, Phuoc Truong, Olli Vapalahti, Sari Hannula, Pekka Ellonen, Niina Ikonen, Carita Savolainen-Kopra |
| EPI_ISL_3230463, EPI_ISL_3230465 | NordLab Kajaani | Expert Microbiology, National Institute for Health and Welfare | Soile Blomqvist, Jani Halkilahti, Kirsi Liitsola, Haider al-Hello, Päivi Laurila, Erika Lindh, Teemu Smura, Ravi Kant, Phuoc Truong, Olli Vapalahti, Sari Hannula, Pekka Ellonen, Niina Ikonen, Carita Savolainen-Kopra |
| EPI_ISL_3230466, EPI_ISL_3230467, EPI_ISL_3230468, EPI_ISL_3230469, EPI_ISL_3230470 | Fimlab Laboratoriot Oy Tampere | Expert Microbiology, National Institute for Health and Welfare | Soile Blomqvist, Jani Halkilahti, Kirsi Liitsola, Haider al-Hello, Päivi Laurila, Erika Lindh, Teemu Smura, Ravi Kant, Phuoc Truong, Olli Vapalahti, Sari Hannula, Pekka Ellonen, Niina Ikonen, Carita Savolainen-Kopra |
| EPI_ISL_3230471 | Poliklinika Vitalis | Alea Genetic Centre | Dino Pecar, Lana Salihefendic, Ivana Ceko, Enis Kandic, Adis Kandic, Rijad Konjhodzic |
| EPI_ISL_3230473, EPI_ISL_3230474, EPI_ISL_3230476, EPI_ISL_3230477, EPI_ISL_3230478, EPI_ISL_3230479, EPI_ISL_3230480, EPI_ISL_3230481, EPI_ISL_3230482, EPI_ISL_3230483, EPI_ISL_3230484, EPI_ISL_3230485, EPI_ISL_3230486, EPI_ISL_3230488, EPI_ISL_3230489, EPI_ISL_3230490, EPI_ISL_3230492, EPI_ISL_3230493, EPI_ISL_3230494 | | | |
| see above | Fimlab Laboratoriot Oy Tampere | Expert Microbiology, National Institute for Health and Welfare | Soile Blomqvist, Jani Halkilahti, Kirsi Liitsola, Haider al-Hello, Päivi Laurila, Erika Lindh, Teemu Smura, Ravi Kant, Phuoc Truong, Olli Vapalahti, Sari Hannula, Pekka Ellonen, Niina Ikonen, Carita Savolainen-Kopra |
| EPI_ISL_3230495, EPI_ISL_3230496, EPI_ISL_3230497, EPI_ISL_3230498, EPI_ISL_3230499, EPI_ISL_3230500, EPI_ISL_3230501, EPI_ISL_3230502, EPI_ISL_3230503, EPI_ISL_3230504, EPI_ISL_3230505, EPI_ISL_3230506, EPI_ISL_3230507, EPI_ISL_3230508 | | | |
| see above | SYNLAB Suomi | Expert Microbiology, National Institute for Health and Welfare | Soile Blomqvist, Jani Halkilahti, Kirsi Liitsola, Haider al-Hello, Päivi Laurila, Erika Lindh, Teemu Smura, Ravi Kant, Phuoc Truong, Olli Vapalahti, Sari Hannula, Pekka Ellonen, Niina Ikonen, Carita Savolainen-Kopra |
| EPI_ISL_3230509, EPI_ISL_3230511, EPI_ISL_3230512, EPI_ISL_3230513 | Vita Laboratoriot Oy | Expert Microbiology, National Institute for Health and Welfare | Soile Blomqvist, Jani Halkilahti, Kirsi Liitsola, Haider al-Hello, Päivi Laurila, Erika Lindh, Teemu Smura, Ravi Kant, Phuoc Truong, Olli Vapalahti, Sari Hannula, Pekka Ellonen, Niina Ikonen, Carita Savolainen-Kopra |
| EPI_ISL_3230514, EPI_ISL_3230515, EPI_ISL_3230516, EPI_ISL_3230517, EPI_ISL_3230518 | SYNLAB Suomi | Expert Microbiology, National Institute for Health and Welfare | Soile Blomqvist, Jani Halkilahti, Kirsi Liitsola, Haider al-Hello, Päivi Laurila, Erika Lindh, Teemu Smura, Ravi Kant, Phuoc Truong, Olli Vapalahti, Sari Hannula, Pekka Ellonen, Niina Ikonen, Carita Savolainen-Kopra |
| EPI_ISL_3230519 | SataDiag | Expert Microbiology, National Institute for Health and Welfare | Soile Blomqvist, Jani Halkilahti, Kirsi Liitsola, Haider al-Hello, Päivi Laurila, Erika Lindh, Teemu Smura, Ravi Kant, Phuoc Truong, Olli Vapalahti, Sari Hannula, Pekka Ellonen, Niina Ikonen, Carita Savolainen-Kopra |
| EPI_ISL_3230521, EPI_ISL_3230522 | NordLab Kajaani | Expert Microbiology, National Institute for Health and Welfare | Soile Blomqvist, Jani Halkilahti, Kirsi Liitsola, Haider al-Hello, Päivi Laurila, Erika Lindh, Teemu Smura, Ravi Kant, Phuoc Truong, Olli Vapalahti, Sari Hannula, Pekka Ellonen, Niina Ikonen, Carita Savolainen-Kopra |
| EPI_ISL_3230523, EPI_ISL_3230524 | Islab, Pohjois-Karjalan aluelaboratorio | Expert Microbiology, National Institute for Health and Welfare | Soile Blomqvist, Jani Halkilahti, Kirsi Liitsola, Haider al-Hello, Päivi Laurila, Erika Lindh, Teemu Smura, Ravi Kant, Phuoc Truong, Olli Vapalahti, Sari Hannula, Pekka Ellonen, Niina Ikonen, Carita Savolainen-Kopra |
| EPI_ISL_3230527, EPI_ISL_3230528, EPI_ISL_3230529, EPI_ISL_3230532, EPI_ISL_3230533, EPI_ISL_3230534, EPI_ISL_3230535, EPI_ISL_3230536 | Seinäjoen keskussairaala | Expert Microbiology, National Institute for Health and Welfare | Soile Blomqvist, Jani Halkilahti, Kirsi Liitsola, Haider al-Hello, Päivi Laurila, Erika Lindh, Teemu Smura, Ravi Kant, Phuoc Truong, Olli Vapalahti, Sari Hannula, Pekka Ellonen, Niina Ikonen, Carita Savolainen-Kopra |
| EPI_ISL_3230537, EPI_ISL_3230538, EPI_ISL_3230539, EPI_ISL_3230540, EPI_ISL_3230541 | NordLab Oulu | Expert Microbiology, National Institute for Health and Welfare | Soile Blomqvist, Jani Halkilahti, Kirsi Liitsola, Haider al-Hello, Päivi Laurila, Erika Lindh, Teemu Smura, Ravi Kant, Phuoc Truong, Olli Vapalahti, Sari Hannula, Pekka Ellonen, Niina Ikonen, Carita Savolainen-Kopra |
| EPI_ISL_3230542, EPI_ISL_3230545, EPI_ISL_3230546, EPI_ISL_3230547, EPI_ISL_3230548, EPI_ISL_3230549, EPI_ISL_3230550, EPI_ISL_3230551, EPI_ISL_3230552, EPI_ISL_3230553, EPI_ISL_3230554, EPI_ISL_3230555, EPI_ISL_3230557, EPI_ISL_3230559, EPI_ISL_3230560, EPI_ISL_3230561, EPI_ISL_3230562, EPI_ISL_3230563, EPI_ISL_3230564, EPI_ISL_3230565, EPI_ISL_3230566, EPI_ISL_3230567, EPI_ISL_3230568, EPI_ISL_3230569, EPI_ISL_3230570, EPI_ISL_3230571, EPI_ISL_3230572, EPI_ISL_3230573, EPI_ISL_3230574, EPI_ISL_3230575, EPI_ISL_3230576, EPI_ISL_3230577, EPI_ISL_3230578, EPI_ISL_3230579, EPI_ISL_3230580, EPI_ISL_3230581, EPI_ISL_3230583, EPI_ISL_3230585, EPI_ISL_3230586, EPI_ISL_3230587, EPI_ISL_3230588, EPI_ISL_3230589, EPI_ISL_3230591, EPI_ISL_3230592, EPI_ISL_3230594, EPI_ISL_3230595, EPI_ISL_3230596, EPI_ISL_3230597, EPI_ISL_3230598, EPI_ISL_3230599, EPI_ISL_3230600 | | | |
| see above | Fimlab Laboratoriot Oy Tampere | Expert Microbiology, National Institute for Health and Welfare | Soile Blomqvist, Jani Halkilahti, Kirsi Liitsola, Haider al-Hello, Päivi Laurila, Erika Lindh, Teemu Smura, Ravi Kant, Phuoc Truong, Olli Vapalahti, Sari Hannula, Pekka Ellonen, Niina Ikonen, Carita Savolainen-Kopra |
| EPI_ISL_3230601, EPI_ISL_3230602, EPI_ISL_3230603, EPI_ISL_3230604, EPI_ISL_3230605, EPI_ISL_3230606 | Islab, Etelä-Savon aluelaboratorio | Expert Microbiology, National Institute for Health and Welfare | Soile Blomqvist, Jani Halkilahti, Kirsi Liitsola, Haider al-Hello, Päivi Laurila, Erika Lindh, Teemu Smura, Ravi Kant, Phuoc Truong, Olli Vapalahti, Sari Hannula, Pekka Ellonen, Niina Ikonen, Carita Savolainen-Kopra |
| EPI_ISL_3230607, EPI_ISL_3230608, EPI_ISL_3230610, EPI_ISL_3230611, EPI_ISL_3230612, EPI_ISL_3230613, EPI_ISL_3230616, EPI_ISL_3230617, EPI_ISL_3230618, EPI_ISL_3230619, EPI_ISL_3230620, EPI_ISL_3230621, EPI_ISL_3230622 | | | |
| see above | Islab, Pohjois-Savon aluelaboratorio | Expert Microbiology, National Institute for Health and Welfare | Soile Blomqvist, Jani Halkilahti, Kirsi Liitsola, Haider al-Hello, Päivi Laurila, Erika Lindh, Teemu Smura, Ravi Kant, Phuoc Truong, Olli Vapalahti, Sari Hannula, Pekka Ellonen, Niina Ikonen, Carita Savolainen-Kopra |
| EPI_ISL_3230624, EPI_ISL_3230625, EPI_ISL_3230626, EPI_ISL_3230627, EPI_ISL_3230628 | Vita Laboratoriot Oy | Expert Microbiology, National Institute for Health and Welfare | Soile Blomqvist, Jani Halkilahti, Kirsi Liitsola, Haider al-Hello, Päivi Laurila, Erika Lindh, Teemu Smura, Ravi Kant, Phuoc Truong, Olli Vapalahti, Sari Hannula, Pekka Ellonen, Niina Ikonen, Carita Savolainen-Kopra |
| EPI_ISL_3230629, EPI_ISL_3230630, EPI_ISL_3230632, EPI_ISL_3230633, EPI_ISL_3230634 | SYNLAB Suomi | Expert Microbiology, National Institute for Health and Welfare | Soile Blomqvist, Jani Halkilahti, Kirsi Liitsola, Haider al-Hello, Päivi Laurila, Erika Lindh, Teemu Smura, Ravi Kant, Phuoc Truong, Olli Vapalahti, Sari Hannula, Pekka Ellonen, Niina Ikonen, Carita Savolainen-Kopra |
| EPI_ISL_3230635, EPI_ISL_3230636, EPI_ISL_3230637, EPI_ISL_3230638, EPI_ISL_3230639 | NordLab Kokkola | Expert Microbiology, National Institute for Health and Welfare | Soile Blomqvist, Jani Halkilahti, Kirsi Liitsola, Haider al-Hello, Päivi Laurila, Erika Lindh, Teemu Smura, Ravi Kant, Phuoc Truong, Olli Vapalahti, Sari Hannula, Pekka Ellonen, Niina Ikonen, Carita Savolainen-Kopra |
| EPI_ISL_3230640, EPI_ISL_3230641, EPI_ISL_3230642, EPI_ISL_3230643, EPI_ISL_3230644 | NordLab Rovaniemi | Expert Microbiology, National Institute for Health and Welfare | Soile Blomqvist, Jani Halkilahti, Kirsi Liitsola, Haider al-Hello, Päivi Laurila, Erika Lindh, Teemu Smura, Ravi Kant, Phuoc Truong, Olli Vapalahti, Sari Hannula, Pekka Ellonen, Niina Ikonen, Carita Savolainen-Kopra |
| EPI_ISL_3230645, EPI_ISL_3230646, EPI_ISL_3230647, EPI_ISL_3230648, EPI_ISL_3230649, EPI_ISL_3230650, EPI_ISL_3230651, EPI_ISL_3230652, EPI_ISL_3230653, EPI_ISL_3230654, EPI_ISL_3230655, EPI_ISL_3230656, EPI_ISL_3230657 | | | |
| see above | Islab, Pohjois-Savon aluelaboratorio | Expert Microbiology, National Institute for Health and Welfare | Soile Blomqvist, Jani Halkilahti, Kirsi Liitsola, Haider al-Hello, Päivi Laurila, Erika Lindh, Teemu Smura, Ravi Kant, Phuoc Truong, Olli Vapalahti, Sari Hannula, Pekka Ellonen, Niina Ikonen, Carita Savolainen-Kopra |
| EPI_ISL_3230658, EPI_ISL_3230659 | NordLab Kajaani | Expert Microbiology, National Institute for Health and Welfare | Soile Blomqvist, Jani Halkilahti, Kirsi Liitsola, Haider al-Hello, Päivi Laurila, Erika Lindh, Teemu Smura, Ravi Kant, Phuoc Truong, Olli Vapalahti, Sari Hannula, Pekka Ellonen, Niina Ikonen, Carita Savolainen-Kopra |
| EPI_ISL_3230660, EPI_ISL_3230661, EPI_ISL_3230662, EPI_ISL_3230663 | Seinäjoen keskussairaala | Expert Microbiology, National Institute for Health and Welfare | Soile Blomqvist, Jani Halkilahti, Kirsi Liitsola, Haider al-Hello, Päivi Laurila, Erika Lindh, Teemu Smura, Ravi Kant, Phuoc Truong, Olli Vapalahti, Sari Hannula, Pekka Ellonen, Niina Ikonen, Carita Savolainen-Kopra |
| EPI_ISL_3230664, EPI_ISL_3230665, EPI_ISL_3230666, EPI_ISL_3230667, | NordLab Oulu | Expert Microbiology, National Institute for Health and Welfare | Soile Blomqvist, Jani Halkilahti, Kirsi Liitsola, Haider al-Hello, Päivi Laurila, Erika Lindh, Teemu Smura, Ravi Kant, Phuoc Truong, Olli Vapalahti, Sari Hannula, Pekka Ellonen, Niina Ikonen, Carita Savolainen-Kopra |

| | | | |
|---|---|---|---|
| EPI_ISL_3230668, EPI_ISL_3230669, EPI_ISL_3230670, EPI_ISL_3230671 | | | |
| EPI_ISL_3230672, EPI_ISL_3230673, EPI_ISL_3230674, EPI_ISL_3230675, EPI_ISL_3230676, EPI_ISL_3230677, EPI_ISL_3230678, EPI_ISL_3230679, EPI_ISL_3230680, EPI_ISL_3230681, EPI_ISL_3230682, EPI_ISL_3230683, EPI_ISL_3230684, EPI_ISL_3230685, EPI_ISL_3230686, EPI_ISL_3230687, EPI_ISL_3230688, EPI_ISL_3230689, EPI_ISL_3230690, EPI_ISL_3230691, EPI_ISL_3230692, EPI_ISL_3230693, EPI_ISL_3230694, EPI_ISL_3230695, EPI_ISL_3230696 | | | |
| see above | Fimlab Laboratoriot Oy Tampere | Expert Microbiology, National Institute for Health and Welfare | Soile Blomqvist, Jani Halkilahti, Kirsi Liitsola, Haider al-Hello, Päivi Laurila, Erika Lindh, Teemu Smura, Ravi Kant, Phuoc Truong, Olli Vapalahti, Sari Hannula, Pekka Ellonen, Niina Ikonen, Carita Savolainen-Kopra |
| EPI_ISL_3230697, EPI_ISL_3230698, EPI_ISL_3230699, EPI_ISL_3230700, EPI_ISL_3230701, EPI_ISL_3230702, EPI_ISL_3230703, EPI_ISL_3230704, EPI_ISL_3230705, EPI_ISL_3230706, EPI_ISL_3230707, EPI_ISL_3230708, EPI_ISL_3230709, EPI_ISL_3230710, EPI_ISL_3230711, EPI_ISL_3230712, EPI_ISL_3230713, EPI_ISL_3230714, EPI_ISL_3230715, EPI_ISL_3230716, EPI_ISL_3230717, EPI_ISL_3230718, EPI_ISL_3230719, EPI_ISL_3230720, EPI_ISL_3230721, EPI_ISL_3230722, EPI_ISL_3230723, EPI_ISL_3230724, EPI_ISL_3230725, EPI_ISL_3230726, EPI_ISL_3230727, EPI_ISL_3230728, EPI_ISL_3230729, EPI_ISL_3230730, EPI_ISL_3230731, EPI_ISL_3230732, EPI_ISL_3230733, EPI_ISL_3230734, EPI_ISL_3230735, EPI_ISL_3230736, EPI_ISL_3230737 | | | |
| see above | TYKS, Kliininen mikrobiologia | Expert Microbiology, National Institute for Health and Welfare | Soile Blomqvist, Jani Halkilahti, Kirsi Liitsola, Haider al-Hello, Päivi Laurila, Erika Lindh, Teemu Smura, Ravi Kant, Phuoc Truong, Olli Vapalahti, Sari Hannula, Pekka Ellonen, Niina Ikonen, Carita Savolainen-Kopra |
| EPI_ISL_3230738 | COVID-19 Detection Lab, Chattogram Veterinary and Animal Sciences University | Genomic Research Lab, Bangladesh Council of Scientific and Industrial Research | Goutam Buddha Das, Tridip Das, Tanvir Ahmad Nizami, Md. Sirazul Islam, Pronesh Dutta, Eaftekhar Ahmed Rana, Md. Murshed Hasan Sarkar, Md. Ahasan Habib, Shahina Akter, Tanjina Akhter Banu, Barna Goswami, Iffat Jahan, Md. Kamrul Islam, Sharmin Chowdhury, Md. Salim Khan, Paritosh Kumar Biswas |
| EPI_ISL_3230739 | MedLab Čapljina | Alea Genetic Centre | Lana Salihefendic, Dino Pecar, Ivana Ceko, Rijad Konjhodzic |
| EPI_ISL_3230740, EPI_ISL_3230741, EPI_ISL_3230742, EPI_ISL_3230743, EPI_ISL_3230744, EPI_ISL_3230745, EPI_ISL_3230747, EPI_ISL_3230748, EPI_ISL_3230749, EPI_ISL_3230750, EPI_ISL_3230751 | | | |
| see above | COVID-19 Detection Lab, Chattogram Veterinary and Animal Sciences University | Genomic Research Lab, Bangladesh Council of Scientific and Industrial Research | Goutam Buddha Das, Tridip Das, Tanvir Ahmad Nizami, Md. Sirazul Islam, Pronesh Dutta, Eaftekhar Ahmed Rana, Md. Murshed Hasan Sarkar, Md. Ahasan Habib, Shahina Akter, Tanjina Akhter Banu, Barna Goswami, Iffat Jahan, Md. Kamrul Islam, Sharmin Chowdhury, Md. Salim Khan, Paritosh Kumar Biswas |
| EPI_ISL_3230758 | CMSI | CHU Poitiers | Caroline MICHAUD, Maxime PICHON, Manon PRAT, Valentin BON-BARET, Birama N'DIAYE, Magali GARCIA, Agnes BEBY-DEFAUX, Nicolas LEVEQUE |
| EPI_ISL_3230759 | CH NIORT | CHU Poitiers | Caroline MICHAUD, Maxime PICHON, Manon PRAT, Valentin BON-BARET, Birama N'DIAYE, Magali GARCIA, Agnes BEBY-DEFAUX, Nicolas LEVEQUE |
| EPI_ISL_3230760 | MEDILAB | CHU Poitiers | Caroline MICHAUD, Maxime PICHON, Manon PRAT, Valentin BON-BARET, Birama N'DIAYE, Magali GARCIA, Agnes BEBY-DEFAUX, Nicolas LEVEQUE |
| EPI_ISL_3230765 | CMSI | CHU Poitiers | Caroline MICHAUD, Maxime PICHON, Manon PRAT, Valentin BON-BARET, Birama N'DIAYE, Magali GARCIA, Agnes BEBY-DEFAUX, Nicolas LEVEQUE |
| EPI_ISL_3230768, EPI_ISL_3230769, EPI_ISL_3230770, EPI_ISL_3230771, EPI_ISL_3230772, EPI_ISL_3230773, EPI_ISL_3230774, EPI_ISL_3230775, EPI_ISL_3230776, EPI_ISL_3230777, EPI_ISL_3230778, EPI_ISL_3230779, EPI_ISL_3230780, EPI_ISL_3230781, EPI_ISL_3230782, EPI_ISL_3230783, EPI_ISL_3230784, EPI_ISL_3230785, EPI_ISL_3230786, EPI_ISL_3230787, EPI_ISL_3230788, EPI_ISL_3230789, EPI_ISL_3230790 | | | |
| see above | FORTEBIO - UNILABS | CHU Poitiers | Caroline MICHAUD, Maxime PICHON, Manon PRAT, Valentin BON-BARET, Birama N'DIAYE, Magali GARCIA, Agnes BEBY-DEFAUX, Nicolas LEVEQUE |
| EPI_ISL_3230791 | CH ANGOULEME | CHU Poitiers | Caroline MICHAUD, Maxime PICHON, Manon PRAT, Valentin BON-BARET, Birama N'DIAYE, Magali GARCIA, Agnes BEBY-DEFAUX, Nicolas LEVEQUE |
| EPI_ISL_3230792, EPI_ISL_3230793 | CH NIORT | CHU Poitiers | Caroline MICHAUD, Maxime PICHON, Manon PRAT, Valentin BON-BARET, Birama N'DIAYE, Magali GARCIA, Agnes BEBY-DEFAUX, Nicolas LEVEQUE |
| EPI_ISL_3230794 | CH SAINT JEAN D'ANGELY | CHU Poitiers | Caroline MICHAUD, Maxime PICHON, Manon PRAT, Valentin BON-BARET, Birama N'DIAYE, Magali GARCIA, Agnes BEBY-DEFAUX, Nicolas LEVEQUE |
| EPI_ISL_3230795 | CH LA ROCHELLE | CHU Poitiers | Caroline MICHAUD, Maxime PICHON, Manon PRAT, Valentin BON-BARET, Birama N'DIAYE, Magali GARCIA, Agnes BEBY-DEFAUX, Nicolas LEVEQUE |
| EPI_ISL_3230796 | CH NIORT | CHU Poitiers | Caroline MICHAUD, Maxime PICHON, Manon PRAT, Valentin BON-BARET, Birama N'DIAYE, Magali GARCIA, Agnes BEBY-DEFAUX, Nicolas LEVEQUE |
| EPI_ISL_3230797 | CH ANGOULEME | CHU Poitiers | Caroline MICHAUD, Maxime PICHON, Manon PRAT, Valentin BON-BARET, Birama N'DIAYE, Magali GARCIA, Agnes BEBY-DEFAUX, Nicolas LEVEQUE |
| EPI_ISL_3230798 | LABOFFICE | CHU Poitiers | Caroline MICHAUD, Maxime PICHON, Manon PRAT, Valentin BON-BARET, Birama N'DIAYE, Magali GARCIA, Agnes BEBY-DEFAUX, Nicolas LEVEQUE |
| EPI_ISL_3230799 | MEDILAB | CHU Poitiers | Caroline MICHAUD, Maxime PICHON, Manon PRAT, Valentin BON-BARET, Birama N'DIAYE, Magali GARCIA, Agnes BEBY-DEFAUX, Nicolas LEVEQUE |
| EPI_ISL_3230800, EPI_ISL_3230801, EPI_ISL_3230802 | LABOFFICE | CHU Poitiers | Caroline MICHAUD, Maxime PICHON, Manon PRAT, Valentin BON-BARET, Birama N'DIAYE, Magali GARCIA, Agnes BEBY-DEFAUX, Nicolas LEVEQUE |
| EPI_ISL_3230803, EPI_ISL_3230804, EPI_ISL_3230805 | CH NIORT | CHU Poitiers | Caroline MICHAUD, Maxime PICHON, Manon PRAT, Valentin BON-BARET, Birama N'DIAYE, Magali GARCIA, Agnes BEBY-DEFAUX, Nicolas LEVEQUE |
| EPI_ISL_3230807, EPI_ISL_3230808 | CH LA ROCHELLE | CHU Poitiers | Caroline MICHAUD, Maxime PICHON, Manon PRAT, Valentin BON-BARET, Birama N'DIAYE, Magali GARCIA, Agnes BEBY-DEFAUX, Nicolas LEVEQUE |
| EPI_ISL_3230809, EPI_ISL_3230810 | CH NIORT | CHU Poitiers | Caroline MICHAUD, Maxime PICHON, Manon PRAT, Valentin BON-BARET, Birama N'DIAYE, Magali GARCIA, Agnes BEBY-DEFAUX, Nicolas LEVEQUE |
| EPI_ISL_3230811 | CHU POITIERS | CHU Poitiers | Caroline MICHAUD, Maxime PICHON, Manon PRAT, Valentin BON-BARET, Birama N'DIAYE, Magali GARCIA, Agnes BEBY-DEFAUX, Nicolas LEVEQUE |
| EPI_ISL_3230812, EPI_ISL_3230813, EPI_ISL_3230814, EPI_ISL_3230816, EPI_ISL_3230817, EPI_ISL_3230818, EPI_ISL_3230819, EPI_ISL_3230821, EPI_ISL_3230822, EPI_ISL_3230823, EPI_ISL_3230824 | | | |
| see above | LABOFFICE | CHU Poitiers | Caroline MICHAUD, Maxime PICHON, Manon PRAT, Valentin BON-BARET, Birama N'DIAYE, Magali GARCIA, Agnes BEBY-DEFAUX, Nicolas LEVEQUE |
| EPI_ISL_3230825, EPI_ISL_3230826, EPI_ISL_3230827, EPI_ISL_3230828, EPI_ISL_3230829, EPI_ISL_3230830, EPI_ISL_3230831, EPI_ISL_3230832, EPI_ISL_3230833 | MEDILAB | CHU Poitiers | Caroline MICHAUD, Maxime PICHON, Manon PRAT, Valentin BON-BARET, Birama N'DIAYE, Magali GARCIA, Agnes BEBY-DEFAUX, Nicolas LEVEQUE |
| EPI_ISL_3230834 | CH ANGOULEME | CHU Poitiers | Caroline MICHAUD, Maxime PICHON, Manon PRAT, Valentin BON-BARET, Birama N'DIAYE, Magali GARCIA, Agnes BEBY-DEFAUX, Nicolas LEVEQUE |
| EPI_ISL_3230835 | LABORATOIRE CHNDS | CHU Poitiers | Caroline MICHAUD, Maxime PICHON, Manon PRAT, Valentin BON-BARET, Birama N'DIAYE, Magali GARCIA, Agnes BEBY-DEFAUX, Nicolas LEVEQUE |
| EPI_ISL_3230837 | Universität Innsbruck, Institut für Mikrobiologie | Bergthaler laboratory, CeMM Research Center for Molecular Medicine of the Austrian Academy of Sciences | Lukas Endler, Anna Schedl, Fabian Amman, Petr Triska, Matthew Thornton, Thomas Penz, Benedikt Agerer, Maelle Le Moing, Michael Schuster, Bekir Erguner, Jan Laine, Martin Senekowitsch, Christoph Bock, Andreas Bergthaler |
| EPI_ISL_3230838, EPI_ISL_3230839, EPI_ISL_3230840, EPI_ISL_3230841, EPI_ISL_3230842, EPI_ISL_3230843 | Austrian Agency for Health and Food Safety (AGES) | Bergthaler laboratory, CeMM Research Center for Molecular Medicine of the Austrian Academy of Sciences | Lukas Endler, Anna Schedl, Fabian Amman, Petr Triska, Matthew Thornton, Thomas Penz, Benedikt Agerer, Maelle Le Moing, Michael Schuster, Bekir Erguner, Jan Laine, Martin Senekowitsch, Christoph Bock, Andreas Bergthaler |
| EPI_ISL_3230844 | Universität Innsbruck, Institut für Mikrobiologie | Bergthaler laboratory, CeMM Research Center for Molecular Medicine of the Austrian Academy of Sciences | Lukas Endler, Anna Schedl, Fabian Amman, Petr Triska, Matthew Thornton, Thomas Penz, Benedikt Agerer, Maelle Le Moing, Michael Schuster, Bekir Erguner, Jan Laine, Martin Senekowitsch, Christoph Bock, Andreas Bergthaler |
| EPI_ISL_3230845, EPI_ISL_3230846, EPI_ISL_3230847, EPI_ISL_3230848, EPI_ISL_3230849, EPI_ISL_3230850, EPI_ISL_3230851, EPI_ISL_3230852, EPI_ISL_3230853, EPI_ISL_3230854, EPI_ISL_3230855, EPI_ISL_3230856, EPI_ISL_3230857, EPI_ISL_3230858, EPI_ISL_3230859, EPI_ISL_3230860, EPI_ISL_3230861, EPI_ISL_3230862, EPI_ISL_3230863, EPI_ISL_3230864, EPI_ISL_3230865, EPI_ISL_3230866, EPI_ISL_3230867, EPI_ISL_3230868, EPI_ISL_3230869, EPI_ISL_3230870, EPI_ISL_3230871 | | | |
| see above | Austrian Agency for Health and Food Safety (AGES) | Bergthaler laboratory, CeMM Research Center for Molecular Medicine of the Austrian Academy of Sciences | Lukas Endler, Anna Schedl, Fabian Amman, Petr Triska, Matthew Thornton, Thomas Penz, Benedikt Agerer, Maelle Le Moing, Michael Schuster, Bekir Erguner, Jan Laine, Martin Senekowitsch, Christoph Bock, Andreas Bergthaler |
| EPI_ISL_3230872, EPI_ISL_3230873, EPI_ISL_3230874, EPI_ISL_3230875 | Klinikum Wels-Grieskirchen | Bergthaler laboratory, CeMM Research Center for Molecular Medicine of the Austrian Academy of Sciences | Lukas Endler, Anna Schedl, Fabian Amman, Petr Triska, Matthew Thornton, Thomas Penz, Benedikt Agerer, Maelle Le Moing, Michael Schuster, Bekir Erguner, Jan Laine, Martin Senekowitsch, Christoph Bock, Andreas Bergthaler |
| EPI_ISL_3230876, EPI_ISL_3230877, EPI_ISL_3230878 | Austrian Agency for Health and Food Safety (AGES) | Bergthaler laboratory, CeMM Research Center for Molecular Medicine of the Austrian Academy of Sciences | Lukas Endler, Anna Schedl, Fabian Amman, Petr Triska, Matthew Thornton, Thomas Penz, Benedikt Agerer, Maelle Le Moing, Michael Schuster, Bekir Erguner, Jan Laine, Martin Senekowitsch, Christoph Bock, Andreas Bergthaler |
| EPI_ISL_3230879, EPI_ISL_3230880, EPI_ISL_3230881, EPI_ISL_3230882, EPI_ISL_3230883, EPI_ISL_3230884, EPI_ISL_3230885, EPI_ISL_3230886, EPI_ISL_3230887, EPI_ISL_3230888, EPI_ISL_3230889 | | | |
| see above | Klinikum Wels-Grieskirchen | Bergthaler laboratory, CeMM Research Center for Molecular Medicine of the Austrian Academy of Sciences | Lukas Endler, Anna Schedl, Fabian Amman, Petr Triska, Matthew Thornton, Thomas Penz, Benedikt Agerer, Maelle Le Moing, Michael Schuster, Bekir Erguner, Jan Laine, Martin Senekowitsch, Christoph Bock, Andreas Bergthaler |
| EPI_ISL_3230890, EPI_ISL_3230891, EPI_ISL_3230892, EPI_ISL_3230893, EPI_ISL_3230894, EPI_ISL_3230895, EPI_ISL_3230896, EPI_ISL_3230897, EPI_ISL_3230898, EPI_ISL_3230899, EPI_ISL_3230900, EPI_ISL_3230901, EPI_ISL_3230902, EPI_ISL_3230903, EPI_ISL_3230904, EPI_ISL_3230905 | | | |
| see above | Austrian Agency for Health and Food Safety (AGES) | Bergthaler laboratory, CeMM Research Center for Molecular Medicine of the Austrian Academy of Sciences | Lukas Endler, Anna Schedl, Fabian Amman, Petr Triska, Matthew Thornton, Thomas Penz, Benedikt Agerer, Maelle Le Moing, Michael Schuster, Bekir Erguner, Jan Laine, Martin Senekowitsch, Christoph Bock, Andreas Bergthaler |
| EPI_ISL_3230906 | Klinikum Wels-Grieskirchen | Bergthaler laboratory, CeMM Research Center for Molecular Medicine of the Austrian Academy of Sciences | Lukas Endler, Anna Schedl, Fabian Amman, Petr Triska, Matthew Thornton, Thomas Penz, Benedikt Agerer, Maelle Le Moing, Michael Schuster, Bekir Erguner, Jan Laine, Martin Senekowitsch, Christoph Bock, Andreas Bergthaler |
| EPI_ISL_3230907, EPI_ISL_3230908, EPI_ISL_3230909 | Austrian Agency for Health and Food Safety (AGES) | Bergthaler laboratory, CeMM Research Center for Molecular Medicine of the Austrian Academy of Sciences | Lukas Endler, Anna Schedl, Fabian Amman, Petr Triska, Matthew Thornton, Thomas Penz, Benedikt Agerer, Maelle Le Moing, Michael Schuster, Bekir Erguner, Jan Laine, Martin Senekowitsch, Christoph Bock, Andreas Bergthaler |
| EPI_ISL_3230910 | Klinikum Wels-Grieskirchen | Bergthaler laboratory, CeMM Research Center for Molecular | Lukas Endler, Anna Schedl, Fabian Amman, Petr Triska, Matthew Thornton, Thomas Penz, Benedikt Agerer, Maelle Le Moing, Michael Schuster, Bekir Erguner, Jan Laine, Martin Senekowitsch, Christoph Bock, Andreas Bergthaler |

| | | | |
|---|---|---|---|
| | | Medicine of the Austrian Academy of Sciences | |
| EPI_ISL_3230911, EPI_ISL_3230912, EPI_ISL_3230913 | Austrian Agency for Health and Food Safety (AGES) | Bergthaler laboratory, CeMM Research Center for Molecular Medicine of the Austrian Academy of Sciences | Lukas Endler, Anna Schedl, Fabian Amman, Petr Triska, Matthew Thornton, Thomas Penz, Benedikt Agerer, Maelle Le Moing, Michael Schuster, Bekir Erguner, Jan Laine, Martin Senekowitsch, Christoph Bock, Andreas Bergthaler |
| EPI_ISL_3230914 | Klinikum Wels-Grieskirchen | Bergthaler laboratory, CeMM Research Center for Molecular Medicine of the Austrian Academy of Sciences | Lukas Endler, Anna Schedl, Fabian Amman, Petr Triska, Matthew Thornton, Thomas Penz, Benedikt Agerer, Maelle Le Moing, Michael Schuster, Bekir Erguner, Jan Laine, Martin Senekowitsch, Christoph Bock, Andreas Bergthaler |
| EPI_ISL_3230915 | Austrian Agency for Health and Food Safety (AGES) | Bergthaler laboratory, CeMM Research Center for Molecular Medicine of the Austrian Academy of Sciences | Lukas Endler, Anna Schedl, Fabian Amman, Petr Triska, Matthew Thornton, Thomas Penz, Benedikt Agerer, Maelle Le Moing, Michael Schuster, Bekir Erguner, Jan Laine, Martin Senekowitsch, Christoph Bock, Andreas Bergthaler |
| EPI_ISL_3230916 | Klinikum Wels-Grieskirchen | Bergthaler laboratory, CeMM Research Center for Molecular Medicine of the Austrian Academy of Sciences | Lukas Endler, Anna Schedl, Fabian Amman, Petr Triska, Matthew Thornton, Thomas Penz, Benedikt Agerer, Maelle Le Moing, Michael Schuster, Bekir Erguner, Jan Laine, Martin Senekowitsch, Christoph Bock, Andreas Bergthaler |
| EPI_ISL_3230917, EPI_ISL_3230918, EPI_ISL_3230919 | Austrian Agency for Health and Food Safety (AGES) | Bergthaler laboratory, CeMM Research Center for Molecular Medicine of the Austrian Academy of Sciences | Lukas Endler, Anna Schedl, Fabian Amman, Petr Triska, Matthew Thornton, Thomas Penz, Benedikt Agerer, Maelle Le Moing, Michael Schuster, Bekir Erguner, Jan Laine, Martin Senekowitsch, Christoph Bock, Andreas Bergthaler |
| EPI_ISL_3230920, EPI_ISL_3230921, EPI_ISL_3230922 | Klinikum Wels-Grieskirchen | Bergthaler laboratory, CeMM Research Center for Molecular Medicine of the Austrian Academy of Sciences | Lukas Endler, Anna Schedl, Fabian Amman, Petr Triska, Matthew Thornton, Thomas Penz, Benedikt Agerer, Maelle Le Moing, Michael Schuster, Bekir Erguner, Jan Laine, Martin Senekowitsch, Christoph Bock, Andreas Bergthaler |
| EPI_ISL_3230923, EPI_ISL_3230930, EPI_ISL_3230931, EPI_ISL_3230932, EPI_ISL_3230933 | Austrian Agency for Health and Food Safety (AGES) | Bergthaler laboratory, CeMM Research Center for Molecular Medicine of the Austrian Academy of Sciences | Lukas Endler, Anna Schedl, Fabian Amman, Petr Triska, Matthew Thornton, Thomas Penz, Benedikt Agerer, Maelle Le Moing, Michael Schuster, Bekir Erguner, Jan Laine, Martin Senekowitsch, Christoph Bock, Andreas Bergthaler |
| EPI_ISL_3230934 | Klinikum Wels-Grieskirchen | Bergthaler laboratory, CeMM Research Center for Molecular Medicine of the Austrian Academy of Sciences | Lukas Endler, Anna Schedl, Fabian Amman, Petr Triska, Matthew Thornton, Thomas Penz, Benedikt Agerer, Maelle Le Moing, Michael Schuster, Bekir Erguner, Jan Laine, Martin Senekowitsch, Christoph Bock, Andreas Bergthaler |
| EPI_ISL_3230935 | Austrian Agency for Health and Food Safety (AGES) | Bergthaler laboratory, CeMM Research Center for Molecular Medicine of the Austrian Academy of Sciences | Lukas Endler, Anna Schedl, Fabian Amman, Petr Triska, Matthew Thornton, Thomas Penz, Benedikt Agerer, Maelle Le Moing, Michael Schuster, Bekir Erguner, Jan Laine, Martin Senekowitsch, Christoph Bock, Andreas Bergthaler |
| EPI_ISL_3230936, EPI_ISL_3230937, EPI_ISL_3230943, EPI_ISL_3230944, EPI_ISL_3230945 | Universität Innsbruck, Institut für Mikrobiologie | Bergthaler laboratory, CeMM Research Center for Molecular Medicine of the Austrian Academy of Sciences | Lukas Endler, Anna Schedl, Fabian Amman, Petr Triska, Matthew Thornton, Thomas Penz, Benedikt Agerer, Maelle Le Moing, Michael Schuster, Bekir Erguner, Jan Laine, Martin Senekowitsch, Christoph Bock, Andreas Bergthaler |
| EPI_ISL_3230946 | Austrian Agency for Health and Food Safety (AGES) | Bergthaler laboratory, CeMM Research Center for Molecular Medicine of the Austrian Academy of Sciences | Lukas Endler, Anna Schedl, Fabian Amman, Petr Triska, Matthew Thornton, Thomas Penz, Benedikt Agerer, Maelle Le Moing, Michael Schuster, Bekir Erguner, Jan Laine, Martin Senekowitsch, Christoph Bock, Andreas Bergthaler |
| EPI_ISL_3230947 | Institute of Legal Medicine, Medical University of Innsbruck | Bergthaler laboratory, CeMM Research Center for Molecular Medicine of the Austrian Academy of Sciences | Lukas Endler, Anna Schedl, Fabian Amman, Petr Triska, Matthew Thornton, Thomas Penz, Benedikt Agerer, Maelle Le Moing, Michael Schuster, Bekir Erguner, Jan Laine, Martin Senekowitsch, Christoph Bock, Andreas Bergthaler |
| EPI_ISL_3230948 | Austrian Agency for Health and Food Safety (AGES) | Bergthaler laboratory, CeMM Research Center for Molecular Medicine of the Austrian Academy of Sciences | Lukas Endler, Anna Schedl, Fabian Amman, Petr Triska, Matthew Thornton, Thomas Penz, Benedikt Agerer, Maelle Le Moing, Michael Schuster, Bekir Erguner, Jan Laine, Martin Senekowitsch, Christoph Bock, Andreas Bergthaler |
| EPI_ISL_3230949, EPI_ISL_3230951 | Universität Innsbruck, Institut für Mikrobiologie | Bergthaler laboratory, CeMM Research Center for Molecular Medicine of the Austrian Academy of Sciences | Lukas Endler, Anna Schedl, Fabian Amman, Petr Triska, Matthew Thornton, Thomas Penz, Benedikt Agerer, Maelle Le Moing, Michael Schuster, Bekir Erguner, Jan Laine, Martin Senekowitsch, Christoph Bock, Andreas Bergthaler |
| EPI_ISL_3230952, EPI_ISL_3230953, EPI_ISL_3230954, EPI_ISL_3230955, EPI_ISL_3230956, EPI_ISL_3230957, EPI_ISL_3230958, EPI_ISL_3230959, EPI_ISL_3230960, EPI_ISL_3230961, EPI_ISL_3230962, EPI_ISL_3230963, EPI_ISL_3230964, EPI_ISL_3230965, EPI_ISL_3230966, EPI_ISL_3230967, EPI_ISL_3230968, EPI_ISL_3230969, EPI_ISL_3230970, EPI_ISL_3230971, EPI_ISL_3230972, EPI_ISL_3230973, EPI_ISL_3230974, EPI_ISL_3230975, EPI_ISL_3230976 | | | |
| see above | Austrian Agency for Health and Food Safety (AGES) | Bergthaler laboratory, CeMM Research Center for Molecular Medicine of the Austrian Academy of Sciences | Lukas Endler, Anna Schedl, Fabian Amman, Petr Triska, Matthew Thornton, Thomas Penz, Benedikt Agerer, Maelle Le Moing, Michael Schuster, Bekir Erguner, Jan Laine, Martin Senekowitsch, Christoph Bock, Andreas Bergthaler |
| EPI_ISL_3230977 | Klinikum Wels-Grieskirchen | Bergthaler laboratory, CeMM Research Center for Molecular Medicine of the Austrian Academy of Sciences | Lukas Endler, Anna Schedl, Fabian Amman, Petr Triska, Matthew Thornton, Thomas Penz, Benedikt Agerer, Maelle Le Moing, Michael Schuster, Bekir Erguner, Jan Laine, Martin Senekowitsch, Christoph Bock, Andreas Bergthaler |
| EPI_ISL_3230978, EPI_ISL_3230979, EPI_ISL_3230980, EPI_ISL_3230981, EPI_ISL_3230982, EPI_ISL_3230983, EPI_ISL_3230984, EPI_ISL_3230985, EPI_ISL_3230986, EPI_ISL_3230987, EPI_ISL_3230988, EPI_ISL_3230989, EPI_ISL_3230990, EPI_ISL_3230991, EPI_ISL_3230992, EPI_ISL_3230993, EPI_ISL_3230994, EPI_ISL_3230995, EPI_ISL_3230996, EPI_ISL_3230997, EPI_ISL_3230998, EPI_ISL_3230999, EPI_ISL_3231000, EPI_ISL_3231001, EPI_ISL_3231002, EPI_ISL_3231003, EPI_ISL_3231004, EPI_ISL_3231005, EPI_ISL_3231006, EPI_ISL_3231007, EPI_ISL_3231008, EPI_ISL_3231009, EPI_ISL_3231010, EPI_ISL_3231011, EPI_ISL_3231012, EPI_ISL_3231013, EPI_ISL_3231014, EPI_ISL_3231015, EPI_ISL_3231016, EPI_ISL_3231017 | | | |
| see above | Austrian Agency for Health and Food Safety (AGES) | Bergthaler laboratory, CeMM Research Center for Molecular Medicine of the Austrian Academy of Sciences | Lukas Endler, Anna Schedl, Fabian Amman, Petr Triska, Matthew Thornton, Thomas Penz, Benedikt Agerer, Maelle Le Moing, Michael Schuster, Bekir Erguner, Jan Laine, Martin Senekowitsch, Christoph Bock, Andreas Bergthaler |
| EPI_ISL_3231018, EPI_ISL_3231019, EPI_ISL_3231020, EPI_ISL_3231021, EPI_ISL_3231022, EPI_ISL_3231023, EPI_ISL_3231024, EPI_ISL_3231025, EPI_ISL_3231026, EPI_ISL_3231027, EPI_ISL_3231028, EPI_ISL_3231029, EPI_ISL_3231030, EPI_ISL_3231031, EPI_ISL_3231032, EPI_ISL_3231033, EPI_ISL_3231034, EPI_ISL_3231035, EPI_ISL_3231036, EPI_ISL_3231037, EPI_ISL_3231038, EPI_ISL_3231039, EPI_ISL_3231040, EPI_ISL_3231041, EPI_ISL_3231042, EPI_ISL_3231043 | | | |
| see above | Klinikum Wels-Grieskirchen | Bergthaler laboratory, CeMM Research Center for Molecular Medicine of the Austrian Academy of Sciences | Lukas Endler, Anna Schedl, Fabian Amman, Petr Triska, Matthew Thornton, Thomas Penz, Benedikt Agerer, Maelle Le Moing, Michael Schuster, Bekir Erguner, Jan Laine, Martin Senekowitsch, Christoph Bock, Andreas Bergthaler |
| EPI_ISL_3231044, EPI_ISL_3231045, EPI_ISL_3231046, EPI_ISL_3231047, EPI_ISL_3231048, EPI_ISL_3231049, EPI_ISL_3231050, EPI_ISL_3231051, EPI_ISL_3231052, EPI_ISL_3231053, EPI_ISL_3231054, EPI_ISL_3231055, EPI_ISL_3231056, EPI_ISL_3231057, EPI_ISL_3231058, EPI_ISL_3231059, EPI_ISL_3231060, EPI_ISL_3231061, EPI_ISL_3231062, EPI_ISL_3231063, EPI_ISL_3231064, EPI_ISL_3231065, EPI_ISL_3231066, EPI_ISL_3231067, EPI_ISL_3231068, EPI_ISL_3231069, EPI_ISL_3231070, EPI_ISL_3231071, EPI_ISL_3231072, EPI_ISL_3231073, EPI_ISL_3231074, EPI_ISL_3231075, EPI_ISL_3231076, EPI_ISL_3231077, EPI_ISL_3231078, EPI_ISL_3231079, EPI_ISL_3231080, EPI_ISL_3231081, EPI_ISL_3231082, EPI_ISL_3231083, EPI_ISL_3231084, EPI_ISL_3231085, EPI_ISL_3231086, EPI_ISL_3231087, EPI_ISL_3231088, EPI_ISL_3231089, EPI_ISL_3231090, EPI_ISL_3231091, EPI_ISL_3231092, EPI_ISL_3231093, EPI_ISL_3231094, EPI_ISL_3231095, EPI_ISL_3231096, EPI_ISL_3231097, EPI_ISL_3231098, EPI_ISL_3231099, EPI_ISL_3231100, EPI_ISL_3231101, EPI_ISL_3231102, EPI_ISL_3231103, EPI_ISL_3231104, EPI_ISL_3231105, EPI_ISL_3231106, EPI_ISL_3231107, EPI_ISL_3231108, EPI_ISL_3231109, EPI_ISL_3231110, EPI_ISL_3231111, EPI_ISL_3231112, EPI_ISL_3231113, EPI_ISL_3231114, EPI_ISL_3231115, EPI_ISL_3231116, EPI_ISL_3231117, EPI_ISL_3231118, EPI_ISL_3231119, EPI_ISL_3231120, EPI_ISL_3231121, EPI_ISL_3231122, EPI_ISL_3231123, EPI_ISL_3231124, EPI_ISL_3231125, EPI_ISL_3231126, EPI_ISL_3231127, EPI_ISL_3231128, EPI_ISL_3231129, EPI_ISL_3231130, EPI_ISL_3231131, EPI_ISL_3231132, EPI_ISL_3231133, EPI_ISL_3231134, EPI_ISL_3231135, EPI_ISL_3231136, EPI_ISL_3231137, EPI_ISL_3231139, EPI_ISL_3231140, EPI_ISL_3231141, EPI_ISL_3231142, EPI_ISL_3231143, EPI_ISL_3231144, EPI_ISL_3231145, EPI_ISL_3231146, EPI_ISL_3231147, EPI_ISL_3231148, EPI_ISL_3231149, EPI_ISL_3231150, EPI_ISL_3231151, EPI_ISL_3231152, EPI_ISL_3231153, EPI_ISL_3231154, EPI_ISL_3231155, EPI_ISL_3231156, EPI_ISL_3231157, EPI_ISL_3231158, EPI_ISL_3231159, EPI_ISL_3231160, EPI_ISL_3231161, EPI_ISL_3231162, EPI_ISL_3231163, EPI_ISL_3231164, EPI_ISL_3231165, EPI_ISL_3231166, EPI_ISL_3231167, EPI_ISL_3231168, EPI_ISL_3231169, EPI_ISL_3231170, EPI_ISL_3231171, EPI_ISL_3231172, EPI_ISL_3231173, EPI_ISL_3231174, EPI_ISL_3231175 | | | |
| see above | Austrian Agency for Health and Food Safety (AGES) | Bergthaler laboratory, CeMM Research Center for Molecular Medicine of the Austrian Academy of Sciences | Lukas Endler, Anna Schedl, Fabian Amman, Petr Triska, Matthew Thornton, Thomas Penz, Benedikt Agerer, Maelle Le Moing, Michael Schuster, Bekir Erguner, Jan Laine, Martin Senekowitsch, Christoph Bock, Andreas Bergthaler |
| EPI_ISL_3231189, EPI_ISL_3231197, EPI_ISL_3231198, EPI_ISL_3231199, EPI_ISL_3231200, EPI_ISL_3231201, EPI_ISL_3231202, EPI_ISL_3231203 | Università degli Studi di Perugia | Istituto Zooprofilattico Sperimentale dell'Abruzzo e Molise "G. Caporale" | Mencacci A, Camilloni B, Biagetti M, Giammarioli M, Lorusso A, Marcacci M, Di Domenico M, Ancora M, Curini V, Mangone I, Rinaldi A, Delli Compagni E, Scialabba S, Di Pasquale A, Cammà C, Puglia I, Calistri P, Savini G |
| EPI_ISL_3231204 | Azienda Ospedaliera Terni | Istituto Zooprofilattico Sperimentale dell'Abruzzo e Molise "G. Caporale" | Palumbo M, Scaccetti A, Lorusso A, Marcacci M, Di Domenico M, Ancora M, Curini V, Mangone I, Rinaldi A, Delli Compagni E, Scialabba S, Di Pasquale A, Cammà C, Puglia I, Calistri P, Savini G |
| EPI_ISL_3231205, EPI_ISL_3231206, EPI_ISL_3231207, EPI_ISL_3231208, EPI_ISL_3231209, EPI_ISL_3231210, EPI_ISL_3231211, EPI_ISL_3231212, EPI_ISL_3231213, EPI_ISL_3231214, EPI_ISL_3231215, EPI_ISL_3231216, EPI_ISL_3231217, EPI_ISL_3231218, EPI_ISL_3231219, EPI_ISL_3231220, EPI_ISL_3231221, EPI_ISL_3231222, EPI_ISL_3231223, EPI_ISL_3231224 | | | |
| see above | Laboratorio Analisi Osp. Città di Castello - Azienda USL Umbria1 | Istituto Zooprofilattico Sperimentale dell'Abruzzo e Molise "G. Caporale" | Malagigi V, Tacconi P,Lorusso A, Marcacci M, Di Domenico M, Ancora M, Curini V, Mangone I, Rinaldi A, Delli Compagni E, Scialabba S, Di Pasquale A, Cammà C, Puglia I, Calistri P, Savini G |
| EPI_ISL_3231225, EPI_ISL_3231226, EPI_ISL_3231227, EPI_ISL_3231228, EPI_ISL_3231229, EPI_ISL_3231230, EPI_ISL_3231231, EPI_ISL_3231232, EPI_ISL_3231233, EPI_ISL_3231234, EPI_ISL_3231235, EPI_ISL_3231236, EPI_ISL_3231237 | | | |
| see above | Università degli Studi di Perugia | Istituto Zooprofilattico Sperimentale dell'Abruzzo e Molise "G. Caporale" | Mencacci A, Camilloni B, Biagetti M, Giammarioli M, Lorusso A, Marcacci M, Di Domenico M, Ancora M, Curini V, Mangone I, Rinaldi A, Delli Compagni E, Scialabba S, Di Pasquale A, Cammà C, Puglia I, Calistri P, Savini G |
| EPI_ISL_3231238, EPI_ISL_3231239, EPI_ISL_3231240, EPI_ISL_3231241, EPI_ISL_3231242, EPI_ISL_3231243, EPI_ISL_3231244, EPI_ISL_3231245, EPI_ISL_3231246, EPI_ISL_3231247, EPI_ISL_3231248, EPI_ISL_3231249, EPI_ISL_3231250, EPI_ISL_3231251, EPI_ISL_3231252, EPI_ISL_3231253, EPI_ISL_3231254, EPI_ISL_3231255, EPI_ISL_3231256, EPI_ISL_3231257, EPI_ISL_3231258, EPI_ISL_3231259, EPI_ISL_3231260, EPI_ISL_3231261, EPI_ISL_3231262 | | | |
| see above | Azienda USL Umbria 2 | Istituto Zooprofilattico Sperimentale dell'Abruzzo e Molise "G. Caporale" | Proietti A, Pistoni E, Lorusso A, Marcacci M, Di Domenico M, Ancora M, Curini V, Mangone I, Rinaldi A, Delli Compagni E, Scialabba S, Di Pasquale A, Cammà C, Puglia I, Calistri P, Savini G |
| EPI_ISL_3231263 | Azienda Ospedaliera Terni | Istituto Zooprofilattico Sperimentale dell'Abruzzo e Molise "G. Caporale" | Palumbo M, Scaccetti A, Lorusso A, Marcacci M, Di Domenico M, Ancora M, Curini V, Mangone I, Rinaldi A, Delli Compagni E, Scialabba S, Di Pasquale A, Cammà C, Puglia I, Calistri P, Savini G |
| EPI_ISL_3231264, EPI_ISL_3231265, EPI_ISL_3231266, EPI_ISL_3231267, EPI_ISL_3231268, EPI_ISL_3231269, EPI_ISL_3231270 | Lafene Health Center | Kansas State Veterinary Diagnostic Laboratory | Tyler Doerksen, Andreea Lu, Lance Noll, Elizabeth Porter, Emily Cox, Jianfa Bai, Jamie Henningson, Rachel Palinski |
| EPI_ISL_3231271, EPI_ISL_3231272, EPI_ISL_3231273, EPI_ISL_3231274, EPI_ISL_3231284, EPI_ISL_3231285, EPI_ISL_3231286, EPI_ISL_3231287, EPI_ISL_3231288, EPI_ISL_3231289, EPI_ISL_3231290, EPI_ISL_3231291, EPI_ISL_3231292, EPI_ISL_3231293, EPI_ISL_3231294, EPI_ISL_3231295, EPI_ISL_3231296, EPI_ISL_3231297, EPI_ISL_3231298, EPI_ISL_3231299, EPI_ISL_3231300, EPI_ISL_3231301, EPI_ISL_3231302, EPI_ISL_3231303, EPI_ISL_3231304, EPI_ISL_3231305, EPI_ISL_3231306, EPI_ISL_3231307, EPI_ISL_3231308, EPI_ISL_3231309, EPI_ISL_3231310, EPI_ISL_3231311, EPI_ISL_3231312, EPI_ISL_3231313, EPI_ISL_3231314, EPI_ISL_3231315, EPI_ISL_3231316, EPI_ISL_3231317, EPI_ISL_3231318, EPI_ISL_3231319, EPI_ISL_3231320, EPI_ISL_3231321, EPI_ISL_3231322, EPI_ISL_3231323, EPI_ISL_3231324, EPI_ISL_3231325, EPI_ISL_3231326, EPI_ISL_3231327, EPI_ISL_3231328, EPI_ISL_3231329, EPI_ISL_3231330, EPI_ISL_3231331, EPI_ISL_3231332, EPI_ISL_3231333, EPI_ISL_3231334, EPI_ISL_3231335, EPI_ISL_3231336, EPI_ISL_3231337, EPI_ISL_3231338, EPI_ISL_3231339, EPI_ISL_3231340, EPI_ISL_3231341, EPI_ISL_3231342, EPI_ISL_3231343, EPI_ISL_3231344, EPI_ISL_3231345 | | | |
| see above | Public Health Authority of the Slovak Republic | Laboratory of Genomics and Bioinformatics, Comenius University Science Park | Tatiana Sedláčková, Diana Rusňáková, Miroslav Böhmer, Anna Gičová, Jaroslav Budiš, Tomáš Szemes, Jakub Styk |

| | | | |
|---|---|---|---|
| EPI_ISL_3231346 | COVID-19 Detection Lab, Chattogram Veterinary and Animal Sciences University | Genomic Research Lab, BCSIR | Md. Murshed Hasan Sarkar, Abu Sayeed Mohammad Mahmud, Mohammad Samir Uzzaman, Eshrar Osman, Md. Ahasan Habib, Shahina Akter, Tanjina Akhter Banu, Barna Goswami, Iffat Jahan, Mohammad Mohi Uddin, Md. Kamrul Islam, Goutam Buddha Das, Tridip Das, Tanvir Ahmad Nizami, Eaftekhar Ahmed Rana, Md. Sirazul Islam, Pronesh Dutta, Sharmin Chowdhury, Md. Paritosh Kumar Biswas, Md. Salim Khan |
| EPI_ISL_3231348, EPI_ISL_3231349, EPI_ISL_3231350, EPI_ISL_3231351, EPI_ISL_3231352, EPI_ISL_3231353, EPI_ISL_3231354, EPI_ISL_3231355, EPI_ISL_3231356, EPI_ISL_3231361, EPI_ISL_3231362, EPI_ISL_3231363, EPI_ISL_3231364, EPI_ISL_3231365, EPI_ISL_3231366, EPI_ISL_3231367, EPI_ISL_3231368, EPI_ISL_3231369, EPI_ISL_3231371, EPI_ISL_3231372, EPI_ISL_3231373, EPI_ISL_3231374, EPI_ISL_3231375, EPI_ISL_3231376, EPI_ISL_3231377, EPI_ISL_3231378, EPI_ISL_3231379 | | | |
| see above | Azarian Lab at Burnett School of Biomedical Sciences | Azarian Lab at Burnett School of Biomedical Sciences | Catherine Johnston, Eleonora Cella, Taj Azarian |
| EPI_ISL_3231410 | COVID-19 Detection Lab, Chattogram Veterinary and Animal Sciences University | Genomic Research Lab, BCSIR | Md. Murshed Hasan Sarkar, Abu Sayeed Mohammad Mahmud, Mohammad Samir Uzzaman, Eshrar Osman, Md. Ahasan Habib, Shahina Akter, Tanjina Akhter Banu, Barna Goswami, Iffat Jahan, Mohammad Mohi Uddin, Md. Kamrul Islam, Goutam Buddha Das, Tridip Das, Tanvir Ahmad Nizami, Eaftekhar Ahmed Rana, Md. Sirazul Islam, Pronesh Dutta, Sharmin Chowdhury, Md. Paritosh Kumar Biswas, Md. Salim Khan |
| EPI_ISL_3231411, EPI_ISL_3231413, EPI_ISL_3231414, EPI_ISL_3231415, EPI_ISL_3231416, EPI_ISL_3231417, EPI_ISL_3231418, EPI_ISL_3231419, EPI_ISL_3231420, EPI_ISL_3231421, EPI_ISL_3231422, EPI_ISL_3231423, EPI_ISL_3231424, EPI_ISL_3231425, EPI_ISL_3231426, EPI_ISL_3231427, EPI_ISL_3231428, EPI_ISL_3231429, EPI_ISL_3231430, EPI_ISL_3231431, EPI_ISL_3231432, EPI_ISL_3231433, EPI_ISL_3231434, EPI_ISL_3231435, EPI_ISL_3231436, EPI_ISL_3231437, EPI_ISL_3231438, EPI_ISL_3231439, EPI_ISL_3231441, EPI_ISL_3231442, EPI_ISL_3231443, EPI_ISL_3231444, EPI_ISL_3231445 | | | |
| see above | Alaska State Virology Laboratory | Alaska State Virology Laboratory | Stephanie DeRonde, Elva House, Jacob Zidek, Lisa Smith, Ph.D., Jack Chen, Ph.D. |
| EPI_ISL_3231446, EPI_ISL_3231447, EPI_ISL_3231448, EPI_ISL_3231449, EPI_ISL_3231451, EPI_ISL_3231452, EPI_ISL_3231453, EPI_ISL_3231454, EPI_ISL_3231455, EPI_ISL_3231458, EPI_ISL_3231459, EPI_ISL_3231460, EPI_ISL_3231462, EPI_ISL_3231463, EPI_ISL_3231464, EPI_ISL_3231466, EPI_ISL_3231467, EPI_ISL_3231469, EPI_ISL_3231470, EPI_ISL_3231471, EPI_ISL_3231472, EPI_ISL_3231474, EPI_ISL_3231475, EPI_ISL_3231476, EPI_ISL_3231477, EPI_ISL_3231478, EPI_ISL_3231479, EPI_ISL_3231480, EPI_ISL_3231481, EPI_ISL_3231482, EPI_ISL_3231483, EPI_ISL_3231485, EPI_ISL_3231487, EPI_ISL_3231488, EPI_ISL_3231489, EPI_ISL_3231490, EPI_ISL_3231493, EPI_ISL_3231498, EPI_ISL_3231499, EPI_ISL_3231501, EPI_ISL_3231502, EPI_ISL_3231503, EPI_ISL_3231505, EPI_ISL_3231506, EPI_ISL_3231507, EPI_ISL_3231508, EPI_ISL_3231509, EPI_ISL_3231510, EPI_ISL_3231511, EPI_ISL_3231512 | | | |
| see above | Clinical Molecular Microbiology Laboratory, UNC Hospitals | Jeremy Wang | Jeremy Wang, Shawn Hawken, Corbin Jones, Melissa Miller |
| EPI_ISL_3231514, EPI_ISL_3231515, EPI_ISL_3231516, EPI_ISL_3231517, EPI_ISL_3231518 | Stadtspital Triemli | Institute of Medical Virology | Daniel Ehrsam, Isabel Stürmer, Catharine Aquino, Joel Wirz, Weihong Qi, Hubert Rehrauer, Verena Kufner, Gabriela Ziltener, Maryam Zaheri, Stefan Schmutz, Annette Audigé, Maria Grünberg, Kevin Steiner, Jon Huder, Cyril Shah, Riccarda Capaul, Guido Bloemberg, Jürg Böni, Michael Huber, Alexandra Trkola |
| EPI_ISL_3231519, EPI_ISL_3231520 | Stadtspital Waid | Institute of Medical Virology | Daniel Ehrsam, Isabel Stürmer, Catharine Aquino, Joel Wirz, Weihong Qi, Hubert Rehrauer, Verena Kufner, Gabriela Ziltener, Maryam Zaheri, Stefan Schmutz, Annette Audigé, Maria Grünberg, Kevin Steiner, Jon Huder, Cyril Shah, Riccarda Capaul, Guido Bloemberg, Jürg Böni, Michael Huber, Alexandra Trkola |
| EPI_ISL_3231521 | Kantonsspital Münsterlingen | Institute of Medical Virology | Daniel Ehrsam, Isabel Stürmer, Catharine Aquino, Joel Wirz, Weihong Qi, Hubert Rehrauer, Verena Kufner, Gabriela Ziltener, Maryam Zaheri, Stefan Schmutz, Annette Audigé, Maria Grünberg, Kevin Steiner, Jon Huder, Cyril Shah, Riccarda Capaul, Guido Bloemberg, Jürg Böni, Michael Huber, Alexandra Trkola |
| EPI_ISL_3231522, EPI_ISL_3231523, EPI_ISL_3231524 | UniversitätsSpital Zürich | Institute of Medical Virology | Daniel Ehrsam, Isabel Stürmer, Catharine Aquino, Joel Wirz, Weihong Qi, Hubert Rehrauer, Verena Kufner, Gabriela Ziltener, Maryam Zaheri, Stefan Schmutz, Annette Audigé, Maria Grünberg, Kevin Steiner, Jon Huder, Cyril Shah, Riccarda Capaul, Guido Bloemberg, Jürg Böni, Michael Huber, Alexandra Trkola |
| EPI_ISL_3231525, EPI_ISL_3231526, EPI_ISL_3231527, EPI_ISL_3231528, EPI_ISL_3231529, EPI_ISL_3231530, EPI_ISL_3231531, EPI_ISL_3231532, EPI_ISL_3231533, EPI_ISL_3231534, EPI_ISL_3231535, EPI_ISL_3231536, EPI_ISL_3231537, EPI_ISL_3231538, EPI_ISL_3231539, EPI_ISL_3231540, EPI_ISL_3231541, EPI_ISL_3231542, EPI_ISL_3231543, EPI_ISL_3231544, EPI_ISL_3231545, EPI_ISL_3231546, EPI_ISL_3231547, EPI_ISL_3231548, EPI_ISL_3231549, EPI_ISL_3231550, EPI_ISL_3231551, EPI_ISL_3231552 | | | |
| see above | Kantonsspital Winterthur | Institute of Medical Virology | Daniel Ehrsam, Isabel Stürmer, Catharine Aquino, Joel Wirz, Weihong Qi, Hubert Rehrauer, Verena Kufner, Gabriela Ziltener, Maryam Zaheri, Stefan Schmutz, Annette Audigé, Maria Grünberg, Kevin Steiner, Jon Huder, Cyril Shah, Riccarda Capaul, Guido Bloemberg, Jürg Böni, Michael Huber, Alexandra Trkola |
| EPI_ISL_3231553, EPI_ISL_3231554 | Spital Limmattal | Institute of Medical Virology | Daniel Ehrsam, Isabel Stürmer, Catharine Aquino, Joel Wirz, Weihong Qi, Hubert Rehrauer, Verena Kufner, Gabriela Ziltener, Maryam Zaheri, Stefan Schmutz, Annette Audigé, Maria Grünberg, Kevin Steiner, Jon Huder, Cyril Shah, Riccarda Capaul, Guido Bloemberg, Jürg Böni, Michael Huber, Alexandra Trkola |
| EPI_ISL_3231555 | Stadtspital Triemli | Institute of Medical Virology | Daniel Ehrsam, Isabel Stürmer, Catharine Aquino, Joel Wirz, Weihong Qi, Hubert Rehrauer, Verena Kufner, Gabriela Ziltener, Maryam Zaheri, Stefan Schmutz, Annette Audigé, Maria Grünberg, Kevin Steiner, Jon Huder, Cyril Shah, Riccarda Capaul, Guido Bloemberg, Jürg Böni, Michael Huber, Alexandra Trkola |
| EPI_ISL_3231556 | Spital Limmattal | Institute of Medical Virology | Daniel Ehrsam, Isabel Stürmer, Catharine Aquino, Joel Wirz, Weihong Qi, Hubert Rehrauer, Verena Kufner, Gabriela Ziltener, Maryam Zaheri, Stefan Schmutz, Annette Audigé, Maria Grünberg, Kevin Steiner, Jon Huder, Cyril Shah, Riccarda Capaul, Guido Bloemberg, Jürg Böni, Michael Huber, Alexandra Trkola |
| EPI_ISL_3231557 | Spital Männedorf AG | Institute of Medical Virology | Daniel Ehrsam, Isabel Stürmer, Catharine Aquino, Joel Wirz, Weihong Qi, Hubert Rehrauer, Verena Kufner, Gabriela Ziltener, Maryam Zaheri, Stefan Schmutz, Annette Audigé, Maria Grünberg, Kevin Steiner, Jon Huder, Cyril Shah, Riccarda Capaul, Guido Bloemberg, Jürg Böni, Michael Huber, Alexandra Trkola |
| EPI_ISL_3231559, EPI_ISL_3231560, EPI_ISL_3231562, EPI_ISL_3231563, EPI_ISL_3231564, EPI_ISL_3231565, EPI_ISL_3231566, EPI_ISL_3231567, EPI_ISL_3231568, EPI_ISL_3231569, EPI_ISL_3231570, EPI_ISL_3231571, EPI_ISL_3231572 | | | |
| see above | Stadtspital Triemli | Institute of Medical Virology | Daniel Ehrsam, Isabel Stürmer, Catharine Aquino, Joel Wirz, Weihong Qi, Hubert Rehrauer, Verena Kufner, Gabriela Ziltener, Maryam Zaheri, Stefan Schmutz, Annette Audigé, Maria Grünberg, Kevin Steiner, Jon Huder, Cyril Shah, Riccarda Capaul, Guido Bloemberg, Jürg Böni, Michael Huber, Alexandra Trkola |
| EPI_ISL_3231573 | Spital Männedorf AG | Institute of Medical Virology | Daniel Ehrsam, Isabel Stürmer, Catharine Aquino, Joel Wirz, Weihong Qi, Hubert Rehrauer, Verena Kufner, Gabriela Ziltener, Maryam Zaheri, Stefan Schmutz, Annette Audigé, Maria Grünberg, Kevin Steiner, Jon Huder, Cyril Shah, Riccarda Capaul, Guido Bloemberg, Jürg Böni, Michael Huber, Alexandra Trkola |
| EPI_ISL_3231574 | UniversitätsSpital Zürich | Institute of Medical Virology | Daniel Ehrsam, Isabel Stürmer, Catharine Aquino, Joel Wirz, Weihong Qi, Hubert Rehrauer, Verena Kufner, Gabriela Ziltener, Maryam Zaheri, Stefan Schmutz, Annette Audigé, Maria Grünberg, Kevin Steiner, Jon Huder, Cyril Shah, Riccarda Capaul, Guido Bloemberg, Jürg Böni, Michael Huber, Alexandra Trkola |
| EPI_ISL_3231575 | Zentrallabor Zürich | Institute of Medical Virology | Daniel Ehrsam, Isabel Stürmer, Catharine Aquino, Joel Wirz, Weihong Qi, Hubert Rehrauer, Verena Kufner, Gabriela Ziltener, Maryam Zaheri, Stefan Schmutz, Annette Audigé, Maria Grünberg, Kevin Steiner, Jon Huder, Cyril Shah, Riccarda Capaul, Guido Bloemberg, Jürg Böni, Michael Huber, Alexandra Trkola |
| EPI_ISL_3231576, EPI_ISL_3231577, EPI_ISL_3231578, EPI_ISL_3231579, EPI_ISL_3231580, EPI_ISL_3231581 | Stadtspital Triemli | Institute of Medical Virology | Daniel Ehrsam, Isabel Stürmer, Catharine Aquino, Joel Wirz, Weihong Qi, Hubert Rehrauer, Verena Kufner, Gabriela Ziltener, Maryam Zaheri, Stefan Schmutz, Annette Audigé, Maria Grünberg, Kevin Steiner, Jon Huder, Cyril Shah, Riccarda Capaul, Guido Bloemberg, Jürg Böni, Michael Huber, Alexandra Trkola |
| EPI_ISL_3231582, EPI_ISL_3231583, EPI_ISL_3231584, EPI_ISL_3231585, EPI_ISL_3231586, EPI_ISL_3231587, EPI_ISL_3231588, EPI_ISL_3231589, EPI_ISL_3231590, EPI_ISL_3231591 | UniversitätsSpital Zürich | Institute of Medical Virology | Daniel Ehrsam, Isabel Stürmer, Catharine Aquino, Joel Wirz, Weihong Qi, Hubert Rehrauer, Verena Kufner, Gabriela Ziltener, Maryam Zaheri, Stefan Schmutz, Annette Audigé, Maria Grünberg, Kevin Steiner, Jon Huder, Cyril Shah, Riccarda Capaul, Guido Bloemberg, Jürg Böni, Michael Huber, Alexandra Trkola |
| EPI_ISL_3231592, EPI_ISL_3231593 | Stadtspital Triemli | Institute of Medical Virology | Daniel Ehrsam, Isabel Stürmer, Catharine Aquino, Joel Wirz, Weihong Qi, Hubert Rehrauer, Verena Kufner, Gabriela Ziltener, Maryam Zaheri, Stefan Schmutz, Annette Audigé, Maria Grünberg, Kevin Steiner, Jon Huder, Cyril Shah, Riccarda Capaul, Guido Bloemberg, Jürg Böni, Michael Huber, Alexandra Trkola |
| EPI_ISL_3231735, EPI_ISL_3231737, EPI_ISL_3231738, EPI_ISL_3231741, EPI_ISL_3231742 | EXCITE Lab | Andersen lab at Scripps Research | Chip Schooley, Natasha Martin, Cheryl Anderson, Angela Scioscia, Smruthi Karthikeyan, Greg Humphrey, Sawyer Farmer, Abigail Schnapper, Helena Tubb, Tommy Valles + SEARCH |
| EPI_ISL_3231767, EPI_ISL_3231768, EPI_ISL_3231769, EPI_ISL_3231770, EPI_ISL_3231771, EPI_ISL_3231772, EPI_ISL_3231773, EPI_ISL_3231774, EPI_ISL_3231775, EPI_ISL_3231776, EPI_ISL_3231777, EPI_ISL_3231778, EPI_ISL_3231779, EPI_ISL_3231780, EPI_ISL_3231781, EPI_ISL_3231782, EPI_ISL_3231783, EPI_ISL_3231784, EPI_ISL_3231785, EPI_ISL_3231786, EPI_ISL_3231787, EPI_ISL_3231788, EPI_ISL_3231789, EPI_ISL_3231790, EPI_ISL_3231791, EPI_ISL_3231792, EPI_ISL_3231793, EPI_ISL_3231794, EPI_ISL_3231795, EPI_ISL_3231796, EPI_ISL_3231797, EPI_ISL_3231798, EPI_ISL_3231799, EPI_ISL_3231800, EPI_ISL_3231801, EPI_ISL_3231802, EPI_ISL_3231803, EPI_ISL_3231804, EPI_ISL_3231805, EPI_ISL_3231806, EPI_ISL_3231807, EPI_ISL_3231808, EPI_ISL_3231809, EPI_ISL_3231810, EPI_ISL_3231811, EPI_ISL_3231812, EPI_ISL_3231813, EPI_ISL_3231814, EPI_ISL_3231815, EPI_ISL_3231816, EPI_ISL_3231817, EPI_ISL_3231818, EPI_ISL_3231819, EPI_ISL_3231820, EPI_ISL_3231821, EPI_ISL_3231822, EPI_ISL_3231823, EPI_ISL_3231824, EPI_ISL_3231825, EPI_ISL_3231826, EPI_ISL_3231827, EPI_ISL_3231828, EPI_ISL_3231829, EPI_ISL_3231830, EPI_ISL_3231831, EPI_ISL_3231832, EPI_ISL_3231833, EPI_ISL_3231834, EPI_ISL_3231835, EPI_ISL_3231836, EPI_ISL_3231837, EPI_ISL_3231838, EPI_ISL_3231839, EPI_ISL_3231840, EPI_ISL_3231841, EPI_ISL_3231842, EPI_ISL_3231843, EPI_ISL_3231844, EPI_ISL_3231845, EPI_ISL_3231846, EPI_ISL_3231847, EPI_ISL_3231848, EPI_ISL_3231849, EPI_ISL_3231850, EPI_ISL_3231851, EPI_ISL_3231852, EPI_ISL_3231853, EPI_ISL_3231854, EPI_ISL_3231855, EPI_ISL_3231856, EPI_ISL_3231857, EPI_ISL_3231858, EPI_ISL_3231859, EPI_ISL_3231860, EPI_ISL_3231861, EPI_ISL_3231862, EPI_ISL_3231863, EPI_ISL_3231864, EPI_ISL_3231865, EPI_ISL_3231866, EPI_ISL_3231867, EPI_ISL_3231868, EPI_ISL_3231869, EPI_ISL_3231870, EPI_ISL_3231871, EPI_ISL_3231872, EPI_ISL_3231873, EPI_ISL_3231874, EPI_ISL_3231875 | | | |
| see above | GA Department of Public Health | GA Department of Public Health | Stacy Reeves, Sharmila Talekar, Jonathan Edwards, Cynthia Dixey, Tonia Parrott, Aliyah Fields, Taylor Smith |
| EPI_ISL_3231887, EPI_ISL_3231889, EPI_ISL_3231890, EPI_ISL_3231892, EPI_ISL_3231893, EPI_ISL_3231894, EPI_ISL_3231895, EPI_ISL_3231897, EPI_ISL_3231898, EPI_ISL_3231899, EPI_ISL_3231901, EPI_ISL_3231902, EPI_ISL_3231903, EPI_ISL_3231906, EPI_ISL_3231907, EPI_ISL_3231909, EPI_ISL_3231910, EPI_ISL_3231912, EPI_ISL_3231913, EPI_ISL_3231916, EPI_ISL_3231917, EPI_ISL_3231921, EPI_ISL_3231922, EPI_ISL_3231923, EPI_ISL_3231926, EPI_ISL_3231927, EPI_ISL_3231928, EPI_ISL_3231930, EPI_ISL_3231931, EPI_ISL_3231932, EPI_ISL_3231933, EPI_ISL_3231934, EPI_ISL_3231935, EPI_ISL_3231936, EPI_ISL_3231938, EPI_ISL_3231939, EPI_ISL_3231941, EPI_ISL_3231947, EPI_ISL_3231948, EPI_ISL_3231949, EPI_ISL_3231950, EPI_ISL_3231951, EPI_ISL_3231952, EPI_ISL_3231953, EPI_ISL_3231954, EPI_ISL_3231955, EPI_ISL_3231956, EPI_ISL_3231958, EPI_ISL_3231959, EPI_ISL_3231960, EPI_ISL_3231961, EPI_ISL_3231962, EPI_ISL_3231963, EPI_ISL_3231964, EPI_ISL_3231965, EPI_ISL_3231966, EPI_ISL_3231967, EPI_ISL_3231968, EPI_ISL_3231969, EPI_ISL_3231970, EPI_ISL_3231989, EPI_ISL_3231990, EPI_ISL_3231991, EPI_ISL_3231992, EPI_ISL_3231993, EPI_ISL_3231995, EPI_ISL_3231997, EPI_ISL_3231998, EPI_ISL_3232000, EPI_ISL_3232005, EPI_ISL_3232023, EPI_ISL_3232024, EPI_ISL_3232026, EPI_ISL_3232027, EPI_ISL_3232029, EPI_ISL_3232030, EPI_ISL_3232032, EPI_ISL_3232033, EPI_ISL_3232035, EPI_ISL_3232036, EPI_ISL_3232037, EPI_ISL_3232038, EPI_ISL_3232039, EPI_ISL_3232040, EPI_ISL_3232041, EPI_ISL_3232042, EPI_ISL_3232043, EPI_ISL_3232044, EPI_ISL_3232045, EPI_ISL_3232046, EPI_ISL_3232047, EPI_ISL_3232048, EPI_ISL_3232049, EPI_ISL_3232050, EPI_ISL_3232051, EPI_ISL_3232052, EPI_ISL_3232053, EPI_ISL_3232054, EPI_ISL_3232055, EPI_ISL_3232056, EPI_ISL_3232057, EPI_ISL_3232058, EPI_ISL_3232059, EPI_ISL_3232060, EPI_ISL_3232061, EPI_ISL_3232062, EPI_ISL_3232063, EPI_ISL_3232064, EPI_ISL_3232065, EPI_ISL_3232066, EPI_ISL_3232067, EPI_ISL_3232068, EPI_ISL_3232069, EPI_ISL_3232070, EPI_ISL_3232071, EPI_ISL_3232072, EPI_ISL_3232073, EPI_ISL_3232074, EPI_ISL_3232075, EPI_ISL_3232077, EPI_ISL_3232078 | | | |
| see above | Illinois Department of Public Health | Illinois Department of Public Health - Chicago Lab | Vineet K. Dhiman, Ira Heimler, Joel Price |
| EPI_ISL_3232079 | LSUHS Emerging Viral Threat Lab | LSUHS Emerging Viral Threat Laboratory | Gregory L. Ware, Alexander Mijalis, Jeremy P. Kamil, Jennifer L. Carroll, Maarten Van Diest, Rona S. Scott, Andrew D. Yurochko, Christopher G. Kevil, John A. Vanchiere |
| EPI_ISL_3232080, EPI_ISL_3232081, EPI_ISL_3232083, EPI_ISL_3232084, EPI_ISL_3232085, EPI_ISL_3232086, EPI_ISL_3232087, EPI_ISL_3232088, EPI_ISL_3232089, EPI_ISL_3232090, EPI_ISL_3232091, EPI_ISL_3232092, EPI_ISL_3232093, EPI_ISL_3232094, EPI_ISL_3232095, EPI_ISL_3232096, EPI_ISL_3232097, EPI_ISL_3232098, EPI_ISL_3232099, EPI_ISL_3232100, EPI_ISL_3232101, EPI_ISL_3232102, EPI_ISL_3232103, EPI_ISL_3232104, EPI_ISL_3232105, EPI_ISL_3232106, EPI_ISL_3232107, EPI_ISL_3232108, EPI_ISL_3232109, EPI_ISL_3232110, EPI_ISL_3232111, EPI_ISL_3232112, EPI_ISL_3232113, EPI_ISL_3232114, EPI_ISL_3232115, EPI_ISL_3232116, EPI_ISL_3232117, EPI_ISL_3232118, EPI_ISL_3232119, EPI_ISL_3232120, EPI_ISL_3232121, EPI_ISL_3232122, EPI_ISL_3232123, EPI_ISL_3232124, EPI_ISL_3232125, EPI_ISL_3232126, EPI_ISL_3232127, EPI_ISL_3232128, EPI_ISL_3232129, EPI_ISL_3232130, EPI_ISL_3232131, EPI_ISL_3232132, EPI_ISL_3232133, EPI_ISL_3232134, EPI_ISL_3232135, EPI_ISL_3232136, EPI_ISL_3232137, EPI_ISL_3232138, EPI_ISL_3232139, EPI_ISL_3232140, EPI_ISL_3232141, EPI_ISL_3232142, EPI_ISL_3232143 | | | |
| see above | Illinois Department of Public Health | Illinois Department of Public Health - Chicago Lab | Vineet K. Dhiman, Ira Heimler, Joel Price |
| EPI_ISL_3232146, EPI_ISL_3232148, EPI_ISL_3232149, EPI_ISL_3232150, EPI_ISL_3232152, EPI_ISL_3232153, EPI_ISL_3232154, EPI_ISL_3232155, EPI_ISL_3232156, EPI_ISL_3232166, EPI_ISL_3232168, EPI_ISL_3232169, EPI_ISL_3232170, EPI_ISL_3232171, EPI_ISL_3232174, EPI_ISL_3232175, EPI_ISL_3232181, EPI_ISL_3232182, EPI_ISL_3232183, EPI_ISL_3232184, EPI_ISL_3232185, EPI_ISL_3232186, EPI_ISL_3232187, EPI_ISL_3232188, EPI_ISL_3232189, EPI_ISL_3232190, EPI_ISL_3232191, EPI_ISL_3232192, EPI_ISL_3232193, EPI_ISL_3232194, EPI_ISL_3232196, EPI_ISL_3232197, EPI_ISL_3232198, EPI_ISL_3232199, EPI_ISL_3232200, EPI_ISL_3232201, EPI_ISL_3232202, EPI_ISL_3232203, EPI_ISL_3232204, EPI_ISL_3232205, EPI_ISL_3232206, EPI_ISL_3232207, EPI_ISL_3232208, EPI_ISL_3232209, EPI_ISL_3232210, EPI_ISL_3232211, EPI_ISL_3232212, EPI_ISL_3232213, EPI_ISL_3232214, EPI_ISL_3232215, EPI_ISL_3232216, EPI_ISL_3232217, EPI_ISL_3232218, EPI_ISL_3232219, EPI_ISL_3232220, EPI_ISL_3232221 | | | |
| see above | NJDOH, Public Health and Environmental Laboratories | NJ_PHEL | Lindsey Bodnar, Shiv K. Verma, Ryan Pachucki, Chelsea San Filippo, Jacquelyn Deverell, Dana Woell, Allison Roder, Byeong Jeong |
| EPI_ISL_3232267, EPI_ISL_3232283, EPI_ISL_3232304, EPI_ISL_3232334, EPI_ISL_3232336, EPI_ISL_3232340, EPI_ISL_3232394, EPI_ISL_3232395, EPI_ISL_3232407, EPI_ISL_3232408, EPI_ISL_3232409, EPI_ISL_3232410, EPI_ISL_3232411, EPI_ISL_3232412, EPI_ISL_3232413, EPI_ISL_3232414, EPI_ISL_3232415, EPI_ISL_3232416, EPI_ISL_3232417, EPI_ISL_3232418, EPI_ISL_3232419 | | | |
| see above | Clinical Microbiology Laboratory, NewYork Presbyterian | Uhlemann Laboratory, Columbia University Irving Medical Center | Medini K. Annavajhala, Anne Kelley, Maya Tagliavia, Anne-Catrin Uhlemann |

| | | | |
|---|---|---|---|
| | Hospital/Columbia University Irving Medical Center | | |
| EPI_ISL_3232432 | Laboratory of Microbiology, ASST Ospedale di Circolo, Varese | Laboratory of Microbiology, ASST Ospedale di Circolo, Varese | Federica Novazzi, Francesca Dragoferrante, Angelo Genoni, Daniela Dalla Gasperina, Daniele Focosi, Andreina Baj, Fabrizio Maggi |
| EPI_ISL_3232471, EPI_ISL_3232472, EPI_ISL_3232473, EPI_ISL_3232474, EPI_ISL_3232475, EPI_ISL_3232476, EPI_ISL_3232477, EPI_ISL_3232478, EPI_ISL_3232479, EPI_ISL_3232480, EPI_ISL_3232481, EPI_ISL_3232482, EPI_ISL_3232483, EPI_ISL_3232484, EPI_ISL_3232485, EPI_ISL_3232486, EPI_ISL_3232487, EPI_ISL_3232488, EPI_ISL_3232489, EPI_ISL_3232490, EPI_ISL_3232492, EPI_ISL_3232493, EPI_ISL_3232494, EPI_ISL_3232495, EPI_ISL_3232496, EPI_ISL_3232497, EPI_ISL_3232498, EPI_ISL_3232499, EPI_ISL_3232500, EPI_ISL_3232501, EPI_ISL_3232502, EPI_ISL_3232503, EPI_ISL_3232504, EPI_ISL_3232505, EPI_ISL_3232506 | | | |
| see above | Alaska State Virology Laboratory | Alaska State Virology Laboratory | Stephanie DeRonde, Elva House, Jacob Zidek, Lisa Smith, Ph.D., Jack Chen, Ph.D. |
| EPI_ISL_3232507 | Laboratory of Microbiology, Varese | Laboratory of Microbiology, Varese | Federica Novazzi, Francesca Dragoferrante, Angelo Genoni, Daniela Dalla Gasperina, Daniele Focosi, Andreina Baj, Fabrizio Maggi Submitter information |
| EPI_ISL_3232508 | Laboratory of Microbiology, Varese | Laboratory of Microbiology, Varese | Federica Novazzi, Francesca Dragoferrante, Angelo Genoni, Daniele Focosi, Andreina Baj, Fabrizio Maggi |
| EPI_ISL_3232509, EPI_ISL_3232510, EPI_ISL_3232511, EPI_ISL_3232512, EPI_ISL_3232513 | Genome Information Research Center, Research Institute for Microbial Diseases, Osaka University | Genome Information Research Center, Research Institute for Microbial Diseases, Osaka University | Daisuke Motooka, Kentaro Tanaka, Hiroya Oki, and Shota Nakamura |
| EPI_ISL_3232514, EPI_ISL_3232515, EPI_ISL_3232516, EPI_ISL_3232517, EPI_ISL_3232518, EPI_ISL_3232519, EPI_ISL_3232520, EPI_ISL_3232521, EPI_ISL_3232522 | CHWAPI - SITE NOTRE DAME | Institut de Pathologie et Genetique (IPG) | Pascale Hilbert, Jérémie Gras |
| EPI_ISL_3232523, EPI_ISL_3232524, EPI_ISL_3232525, EPI_ISL_3232526, EPI_ISL_3232527, EPI_ISL_3232529, EPI_ISL_3232530, EPI_ISL_3232531, EPI_ISL_3232532, EPI_ISL_3232533, EPI_ISL_3232534, EPI_ISL_3232535 | | | |
| see above | CLINIQUE N-D DE GRACE | Institut de Pathologie et Genetique (IPG) | Pascale Hilbert, Jérémie Gras |
| EPI_ISL_3232536, EPI_ISL_3232538, EPI_ISL_3232542, EPI_ISL_3232543 | Juntendo University Hospital | Juntendo University Hospital | Koji Tsuchiya, Yoshie Hosaka,Yoko Tabe |
| EPI_ISL_3232553 | Unilabs Laboratory Medicine | Norwegian Institute of Public Health, Department of Virology | Kathrine Stene-Johansen, Kamilla Heddeland Instefjord, Hilde Elshaug, Garcia Llorente Ignacio, Jon Bråte, Engebretsen Serina Beate,Pedersen Benedikte Nevjen, Line Victoria Moen, Debech Nadia, Atiya R Ali,Marie Paulsen Madsen, Rasmus Riis Kopperud, Hilde Vollan, Karoline Bragstad, Olav Hungnes |
| EPI_ISL_3232554 | Haukeland University Hospital, Dept. of Microbiology | Norwegian Institute of Public Health, Department of Virology | Kathrine Stene-Johansen, Kamilla Heddeland Instefjord, Hilde Elshaug, Garcia Llorente Ignacio, Jon Bråte, Engebretsen Serina Beate,Pedersen Benedikte Nevjen, Line Victoria Moen, Debech Nadia, Atiya R Ali,Marie Paulsen Madsen, Rasmus Riis Kopperud, Hilde Vollan, Karoline Bragstad, Olav Hungnes |
| EPI_ISL_3232555 | Hospital of Southern Norway - Kristiansand, Department of Medical Microbiology | Norwegian Institute of Public Health, Department of Virology | Kathrine Stene-Johansen, Kamilla Heddeland Instefjord, Hilde Elshaug, Garcia Llorente Ignacio, Jon Bråte, Engebretsen Serina Beate,Pedersen Benedikte Nevjen, Line Victoria Moen, Debech Nadia, Atiya R Ali,Marie Paulsen Madsen, Rasmus Riis Kopperud, Hilde Vollan, Karoline Bragstad, Olav Hungnes |
| EPI_ISL_3232556 | Haukeland University Hospital, Dept. of Microbiology | Norwegian Institute of Public Health, Department of Virology | Kathrine Stene-Johansen, Kamilla Heddeland Instefjord, Hilde Elshaug, Garcia Llorente Ignacio, Jon Bråte, Engebretsen Serina Beate,Pedersen Benedikte Nevjen, Line Victoria Moen, Debech Nadia, Atiya R Ali,Marie Paulsen Madsen, Rasmus Riis Kopperud, Hilde Vollan, Karoline Bragstad, Olav Hungnes |
| EPI_ISL_3232559 | Department of Medical Microbiology - section Molde, Molde Hospital | Norwegian Institute of Public Health, Department of Virology | Kathrine Stene-Johansen, Kamilla Heddeland Instefjord, Hilde Elshaug, Garcia Llorente Ignacio, Jon Bråte, Engebretsen Serina Beate,Pedersen Benedikte Nevjen, Line Victoria Moen, Debech Nadia, Atiya R Ali,Marie Paulsen Madsen, Rasmus Riis Kopperud, Hilde Vollan, Karoline Bragstad, Olav Hungnes |
| EPI_ISL_3232560 | Department of Medical Microbiology, St. Olavs hospital | Norwegian Institute of Public Health, Department of Virology | Kathrine Stene-Johansen, Kamilla Heddeland Instefjord, Hilde Elshaug, Garcia Llorente Ignacio, Jon Bråte, Engebretsen Serina Beate,Pedersen Benedikte Nevjen, Line Victoria Moen, Debech Nadia, Atiya R Ali,Marie Paulsen Madsen, Rasmus Riis Kopperud, Hilde Vollan, Karoline Bragstad, Olav Hungnes |
| EPI_ISL_3232561 | Department of Medical Microbiology - section Molde, Molde Hospital | Norwegian Institute of Public Health, Department of Virology | Kathrine Stene-Johansen, Kamilla Heddeland Instefjord, Hilde Elshaug, Garcia Llorente Ignacio, Jon Bråte, Engebretsen Serina Beate,Pedersen Benedikte Nevjen, Line Victoria Moen, Debech Nadia, Atiya R Ali,Marie Paulsen Madsen, Rasmus Riis Kopperud, Hilde Vollan, Karoline Bragstad, Olav Hungnes |
| EPI_ISL_3232562 | Haukeland University Hospital, Dept. of Microbiology | Norwegian Institute of Public Health, Department of Virology | Kathrine Stene-Johansen, Kamilla Heddeland Instefjord, Hilde Elshaug, Garcia Llorente Ignacio, Jon Bråte, Engebretsen Serina Beate,Pedersen Benedikte Nevjen, Line Victoria Moen, Debech Nadia, Atiya R Ali,Marie Paulsen Madsen, Rasmus Riis Kopperud, Hilde Vollan, Karoline Bragstad, Olav Hungnes |
| EPI_ISL_3232563, EPI_ISL_3232564 | Department of Medical Microbiology - section Molde, Molde Hospital | Norwegian Institute of Public Health, Department of Virology | Kathrine Stene-Johansen, Kamilla Heddeland Instefjord, Hilde Elshaug, Garcia Llorente Ignacio, Jon Bråte, Engebretsen Serina Beate,Pedersen Benedikte Nevjen, Line Victoria Moen, Debech Nadia, Atiya R Ali,Marie Paulsen Madsen, Rasmus Riis Kopperud, Hilde Vollan, Karoline Bragstad, Olav Hungnes |
| EPI_ISL_3232567 | Department of Medical Microbiology, St. Olavs hospital | Norwegian Institute of Public Health, Department of Virology | Kathrine Stene-Johansen, Kamilla Heddeland Instefjord, Hilde Elshaug, Garcia Llorente Ignacio, Jon Bråte, Engebretsen Serina Beate,Pedersen Benedikte Nevjen, Line Victoria Moen, Debech Nadia, Atiya R Ali,Marie Paulsen Madsen, Rasmus Riis Kopperud, Hilde Vollan, Karoline Bragstad, Olav Hungnes |
| EPI_ISL_3232569 | Department of Medical Microbiology - section Molde, Molde Hospital | Norwegian Institute of Public Health, Department of Virology | Kathrine Stene-Johansen, Kamilla Heddeland Instefjord, Hilde Elshaug, Garcia Llorente Ignacio, Jon Bråte, Engebretsen Serina Beate,Pedersen Benedikte Nevjen, Line Victoria Moen, Debech Nadia, Atiya R Ali,Marie Paulsen Madsen, Rasmus Riis Kopperud, Hilde Vollan, Karoline Bragstad, Olav Hungnes |
| EPI_ISL_3232570 | Unilabs Laboratory Medicine | Norwegian Institute of Public Health, Department of Virology | Kathrine Stene-Johansen, Kamilla Heddeland Instefjord, Hilde Elshaug, Garcia Llorente Ignacio, Jon Bråte, Engebretsen Serina Beate,Pedersen Benedikte Nevjen, Line Victoria Moen, Debech Nadia, Atiya R Ali,Marie Paulsen Madsen, Rasmus Riis Kopperud, Hilde Vollan, Karoline Bragstad, Olav Hungnes |
| EPI_ISL_3232571, EPI_ISL_3232573 | Haukeland University Hospital, Dept. of Microbiology | Norwegian Institute of Public Health, Department of Virology | Kathrine Stene-Johansen, Kamilla Heddeland Instefjord, Hilde Elshaug, Garcia Llorente Ignacio, Jon Bråte, Engebretsen Serina Beate,Pedersen Benedikte Nevjen, Line Victoria Moen, Debech Nadia, Atiya R Ali,Marie Paulsen Madsen, Rasmus Riis Kopperud, Hilde Vollan, Karoline Bragstad, Olav Hungnes |
| EPI_ISL_3232574 | Hospital of Southern Norway - Kristiansand, Department of Medical Microbiology | Norwegian Institute of Public Health, Department of Virology | Kathrine Stene-Johansen, Kamilla Heddeland Instefjord, Hilde Elshaug, Garcia Llorente Ignacio, Jon Bråte, Engebretsen Serina Beate,Pedersen Benedikte Nevjen, Line Victoria Moen, Debech Nadia, Atiya R Ali,Marie Paulsen Madsen, Rasmus Riis Kopperud, Hilde Vollan, Karoline Bragstad, Olav Hungnes |
| EPI_ISL_3232575 | Haukeland University Hospital, Dept. of Microbiology | Norwegian Institute of Public Health, Department of Virology | Kathrine Stene-Johansen, Kamilla Heddeland Instefjord, Hilde Elshaug, Garcia Llorente Ignacio, Jon Bråte, Engebretsen Serina Beate,Pedersen Benedikte Nevjen, Line Victoria Moen, Debech Nadia, Atiya R Ali,Marie Paulsen Madsen, Rasmus Riis Kopperud, Hilde Vollan, Karoline Bragstad, Olav Hungnes |
| EPI_ISL_3232576 | Unilabs Laboratory Medicine | Norwegian Institute of Public Health, Department of Virology | Kathrine Stene-Johansen, Kamilla Heddeland Instefjord, Hilde Elshaug, Garcia Llorente Ignacio, Jon Bråte, Engebretsen Serina Beate,Pedersen Benedikte Nevjen, Line Victoria Moen, Debech Nadia, Atiya R Ali,Marie Paulsen Madsen, Rasmus Riis Kopperud, Hilde Vollan, Karoline Bragstad, Olav Hungnes |
| EPI_ISL_3232577, EPI_ISL_3232580, EPI_ISL_3232581, EPI_ISL_3232582, EPI_ISL_3232583, EPI_ISL_3232584, EPI_ISL_3232585 | Haukeland University Hospital, Dept. of Microbiology | Norwegian Institute of Public Health, Department of Virology | Kathrine Stene-Johansen, Kamilla Heddeland Instefjord, Hilde Elshaug, Garcia Llorente Ignacio, Jon Bråte, Engebretsen Serina Beate,Pedersen Benedikte Nevjen, Line Victoria Moen, Debech Nadia, Atiya R Ali,Marie Paulsen Madsen, Rasmus Riis Kopperud, Hilde Vollan, Karoline Bragstad, Olav Hungnes |
| EPI_ISL_3232586 | Hospital of Southern Norway - Kristiansand, Department of Medical Microbiology | Norwegian Institute of Public Health, Department of Virology | Kathrine Stene-Johansen, Kamilla Heddeland Instefjord, Hilde Elshaug, Garcia Llorente Ignacio, Jon Bråte, Engebretsen Serina Beate,Pedersen Benedikte Nevjen, Line Victoria Moen, Debech Nadia, Atiya R Ali,Marie Paulsen Madsen, Rasmus Riis Kopperud, Hilde Vollan, Karoline Bragstad, Olav Hungnes |
| EPI_ISL_3232587, EPI_ISL_3232588 | Haukeland University Hospital, Dept. of Microbiology | Norwegian Institute of Public Health, Department of Virology | Kathrine Stene-Johansen, Kamilla Heddeland Instefjord, Hilde Elshaug, Garcia Llorente Ignacio, Jon Bråte, Engebretsen Serina Beate,Pedersen Benedikte Nevjen, Line Victoria Moen, Debech Nadia, Atiya R Ali,Marie Paulsen Madsen, Rasmus Riis Kopperud, Hilde Vollan, Karoline Bragstad, Olav Hungnes |
| EPI_ISL_3232590 | Department of Medical Microbiology, St. Olavs hospital | Norwegian Institute of Public Health, Department of Virology | Kathrine Stene-Johansen, Kamilla Heddeland Instefjord, Hilde Elshaug, Garcia Llorente Ignacio, Jon Bråte, Engebretsen Serina Beate,Pedersen Benedikte Nevjen, Line Victoria Moen, Debech Nadia, Atiya R Ali,Marie Paulsen Madsen, Rasmus Riis Kopperud, Hilde Vollan, Karoline Bragstad, Olav Hungnes |
| EPI_ISL_3232592 | Haukeland University Hospital, Dept. of Microbiology | Norwegian Institute of Public Health, Department of Virology | Kathrine Stene-Johansen, Kamilla Heddeland Instefjord, Hilde Elshaug, Garcia Llorente Ignacio, Jon Bråte, Engebretsen Serina Beate,Pedersen Benedikte Nevjen, Line Victoria Moen, Debech Nadia, Atiya R Ali,Marie Paulsen Madsen, Rasmus Riis Kopperud, Hilde Vollan, Karoline Bragstad, Olav Hungnes |
| EPI_ISL_3232593 | Department of Medical Microbiology, St. Olavs hospital | Norwegian Institute of Public Health, Department of Virology | Kathrine Stene-Johansen, Kamilla Heddeland Instefjord, Hilde Elshaug, Garcia Llorente Ignacio, Jon Bråte, Engebretsen Serina Beate,Pedersen Benedikte Nevjen, Line Victoria Moen, Debech Nadia, Atiya R Ali,Marie Paulsen Madsen, Rasmus Riis Kopperud, Hilde Vollan, Karoline Bragstad, Olav Hungnes |
| EPI_ISL_3232594 | Haukeland University Hospital, Dept. of Microbiology | Norwegian Institute of Public Health, Department of Virology | Kathrine Stene-Johansen, Kamilla Heddeland Instefjord, Hilde Elshaug, Garcia Llorente Ignacio, Jon Bråte, Engebretsen Serina Beate,Pedersen Benedikte Nevjen, Line Victoria Moen, Debech Nadia, Atiya R Ali,Marie Paulsen Madsen, Rasmus Riis Kopperud, Hilde Vollan, Karoline Bragstad, Olav Hungnes |
| EPI_ISL_3232598 | Department of Medical Microbiology - section Molde, Molde Hospital | Norwegian Institute of Public Health, Department of Virology | Kathrine Stene-Johansen, Kamilla Heddeland Instefjord, Hilde Elshaug, Garcia Llorente Ignacio, Jon Bråte, Engebretsen Serina Beate,Pedersen Benedikte Nevjen, Line Victoria Moen, Debech Nadia, Atiya R Ali,Marie Paulsen Madsen, Rasmus Riis Kopperud, Hilde Vollan, Karoline Bragstad, Olav Hungnes |
| EPI_ISL_3232600, EPI_ISL_3232601 | Haukeland University Hospital, Dept. of Microbiology | Norwegian Institute of Public Health, Department of Virology | Kathrine Stene-Johansen, Kamilla Heddeland Instefjord, Hilde Elshaug, Garcia Llorente Ignacio, Jon Bråte, Engebretsen Serina Beate,Pedersen Benedikte Nevjen, Line Victoria Moen, Debech Nadia, Atiya R Ali,Marie Paulsen Madsen, Rasmus Riis Kopperud, Hilde Vollan, Karoline Bragstad, Olav Hungnes |
| EPI_ISL_3232603, EPI_ISL_3232604 | Hospital of Southern Norway - Kristiansand, Department of Medical Microbiology | Norwegian Institute of Public Health, Department of Virology | Kathrine Stene-Johansen, Kamilla Heddeland Instefjord, Hilde Elshaug, Garcia Llorente Ignacio, Jon Bråte, Engebretsen Serina Beate,Pedersen Benedikte Nevjen, Line Victoria Moen, Debech Nadia, Atiya R Ali,Marie Paulsen Madsen, Rasmus Riis Kopperud, Hilde Vollan, Karoline Bragstad, Olav Hungnes |
| EPI_ISL_3232605, EPI_ISL_3232606 | Haukeland University Hospital, Dept. of Microbiology | Norwegian Institute of Public Health, Department of Virology | Kathrine Stene-Johansen, Kamilla Heddeland Instefjord, Hilde Elshaug, Garcia Llorente Ignacio, Jon Bråte, Engebretsen Serina Beate,Pedersen Benedikte Nevjen, Line Victoria Moen, Debech Nadia, Atiya R Ali,Marie Paulsen Madsen, Rasmus Riis Kopperud, Hilde Vollan, Karoline Bragstad, Olav Hungnes |
| EPI_ISL_3232607 | Hospital of Southern Norway - Kristiansand, Department of Medical Microbiology | Norwegian Institute of Public Health, Department of Virology | Kathrine Stene-Johansen, Kamilla Heddeland Instefjord, Hilde Elshaug, Garcia Llorente Ignacio, Jon Bråte, Engebretsen Serina Beate,Pedersen Benedikte Nevjen, Line Victoria Moen, Debech Nadia, Atiya R Ali,Marie Paulsen Madsen, Rasmus Riis Kopperud, Hilde Vollan, Karoline Bragstad, Olav Hungnes |
| EPI_ISL_3232624, EPI_ISL_3232625 | University Hospital Brno, OKMI | University Hospital Brno, CMBG | Matej Bezdicek, Jan Svaton, Martina Lengerova, Kristyna Dufkova |
| EPI_ISL_3232627, EPI_ISL_3232628, EPI_ISL_3232632, EPI_ISL_3232634, EPI_ISL_3232635, EPI_ISL_3232636, EPI_ISL_3232637, EPI_ISL_3232640, EPI_ISL_3232641, EPI_ISL_3232643, EPI_ISL_3232644, EPI_ISL_3232645, EPI_ISL_3232646, EPI_ISL_3232648, EPI_ISL_3232649, EPI_ISL_3232650, EPI_ISL_3232651, EPI_ISL_3232658, EPI_ISL_3232659, EPI_ISL_3232662, EPI_ISL_3232663, EPI_ISL_3232664, EPI_ISL_3232665, EPI_ISL_3232666, EPI_ISL_3232670, EPI_ISL_3232673, EPI_ISL_3232674, EPI_ISL_3232675, EPI_ISL_3232676, EPI_ISL_3232677, EPI_ISL_3232678, EPI_ISL_3232679, EPI_ISL_3232680, EPI_ISL_3232681, EPI_ISL_3232682, EPI_ISL_3232685, EPI_ISL_3232686, EPI_ISL_3232687, EPI_ISL_3232688, EPI_ISL_3232689, EPI_ISL_3232690, EPI_ISL_3232691, EPI_ISL_3232692, EPI_ISL_3232693, | | | |

| | | | |
|---|---|---|---|
| EPI_ISL_3232694, EPI_ISL_3232696, EPI_ISL_3232697, EPI_ISL_3232701, EPI_ISL_3232702, EPI_ISL_3232703, EPI_ISL_3232704, EPI_ISL_3232705, EPI_ISL_3232706, EPI_ISL_3232708, EPI_ISL_3232709, EPI_ISL_3232710, EPI_ISL_3232711 | | | |
| see above | National Center of Infectious and Parasitic Diseases | National Center of Infectious and Parasitic Diseases | Alexiev et al |
| EPI_ISL_3232751, EPI_ISL_3232752, EPI_ISL_3232753, EPI_ISL_3232754, EPI_ISL_3232756, EPI_ISL_3232757, EPI_ISL_3232758 | VRDL-GIMS | INSACOG-KA, NIMHANS | Chitra Pattabiraman, Anson Kunjumon George,Darshan Sreenivas, Ellango Ramasamy, Sony Sharma, Mahesh Kumar.C.S, Ananthapadmanabha Kotambail, Chetan G K, Gautham Arunachal Udupi, Anita S Desai, V Ravi |
| EPI_ISL_3232759, EPI_ISL_3232762, EPI_ISL_3232763, EPI_ISL_3232764, EPI_ISL_3232765, EPI_ISL_3232766, EPI_ISL_3232767 | VRDL-VIJAYAPURA | INSACOG-KA, NIMHANS | Chitra Pattabiraman, Anson Kunjumon George,Darshan Sreenivas, Ellango Ramasamy, Sony Sharma, Mahesh Kumar.C.S, Ananthapadmanabha Kotambail, Chetan G K, Gautham Arunachal Udupi, Anita S Desai, V Ravi |
| EPI_ISL_3232769 | DSU- CHIKKABALLAPURA | INSACOG-KA, NIMHANS | Chitra Pattabiraman, Anson Kunjumon George,Darshan Sreenivas, Ellango Ramasamy, Sony Sharma, Mahesh Kumar.C.S, Ananthapadmanabha Kotambail, Chetan G K, Gautham Arunachal Udupi, Anita S Desai, V Ravi |
| EPI_ISL_3232770 | AURIGA | INSACOG-KA, NIMHANS | Chitra Pattabiraman, Anson Kunjumon George,Darshan Sreenivas, Ellango Ramasamy, Sony Sharma, Mahesh Kumar.C.S, Ananthapadmanabha Kotambail, Chetan G K, Gautham Arunachal Udupi, Anita S Desai, V Ravi |
| EPI_ISL_3232771, EPI_ISL_3232772, EPI_ISL_3232773, EPI_ISL_3232774, EPI_ISL_3232775, EPI_ISL_3232776, EPI_ISL_3232777, EPI_ISL_3232778, EPI_ISL_3232779, EPI_ISL_3232781, EPI_ISL_3232782 | | | |
| see above | MMCRI | INSACOG-KA, NIMHANS | Chitra Pattabiraman, Anson Kunjumon George,Darshan Sreenivas, Ellango Ramasamy, Sony Sharma, Mahesh Kumar.C.S, Ananthapadmanabha Kotambail, Chetan G K, Gautham Arunachal Udupi, Anita S Desai, V Ravi |
| EPI_ISL_3232783 | AURIGA LAB | INSACOG-KA, NIMHANS | Chitra Pattabiraman, Anson Kunjumon George,Darshan Sreenivas, Ellango Ramasamy, Sony Sharma, Mahesh Kumar.C.S, Ananthapadmanabha Kotambail, Chetan G K, Gautham Arunachal Udupi, Anita S Desai, V Ravi |
| EPI_ISL_3232784, EPI_ISL_3232785, EPI_ISL_3232786, EPI_ISL_3232787, EPI_ISL_3232788, EPI_ISL_3232789, EPI_ISL_3232790, EPI_ISL_3232791 | NIMHANS | INSACOG-KA, NIMHANS | Chitra Pattabiraman, Anson Kunjumon George,Darshan Sreenivas, Ellango Ramasamy, Sony Sharma, Mahesh Kumar.C.S, Ananthapadmanabha Kotambail, Chetan G K, Gautham Arunachal Udupi, Anita S Desai, V Ravi |
| EPI_ISL_3232824, EPI_ISL_3232827 | VRDL-GIMS | INSACOG-KA, NIMHANS | Chitra Pattabiraman, Anson Kunjumon George,Darshan Sreenivas, Ellango Ramasamy, Sony Sharma, Mahesh Kumar.C.S, Ananthapadmanabha Kotambail, Chetan G K, Gautham Arunachal Udupi, Anita S Desai, V Ravi |
| EPI_ISL_3232828 | ASHOK PURAM DISPENSARY UPHC | INSACOG-KA, NIMHANS | Chitra Pattabiraman, Anson Kunjumon George,Darshan Sreenivas, Ellango Ramasamy, Sony Sharma, Mahesh Kumar.C.S, Ananthapadmanabha Kotambail, Chetan G K, Gautham Arunachal Udupi, Anita S Desai, V Ravi |
| EPI_ISL_3232829 | NIMHANS | INSACOG-KA, NIMHANS | Chitra Pattabiraman, Anson Kunjumon George,Darshan Sreenivas, Ellango Ramasamy, Sony Sharma, Mahesh Kumar.C.S, Ananthapadmanabha Kotambail, Chetan G K, Gautham Arunachal Udupi, Anita S Desai, V Ravi |
| EPI_ISL_3232830 | UPHC SHANKARNAGAR | INSACOG-KA, NIMHANS | Chitra Pattabiraman, Anson Kunjumon George,Darshan Sreenivas, Ellango Ramasamy, Sony Sharma, Mahesh Kumar.C.S, Ananthapadmanabha Kotambail, Chetan G K, Gautham Arunachal Udupi, Anita S Desai, V Ravi |
| EPI_ISL_3232832 | RAILWAY HOSPITAL | INSACOG-KA, NIMHANS | Chitra Pattabiraman, Anson Kunjumon George,Darshan Sreenivas, Ellango Ramasamy, Sony Sharma, Mahesh Kumar.C.S, Ananthapadmanabha Kotambail, Chetan G K, Gautham Arunachal Udupi, Anita S Desai, V Ravi |
| EPI_ISL_3232833 | NIMHANS | INSACOG-KA, NIMHANS | Chitra Pattabiraman, Anson Kunjumon George,Darshan Sreenivas, Ellango Ramasamy, Sony Sharma, Mahesh Kumar.C.S, Ananthapadmanabha Kotambail, Chetan G K, Gautham Arunachal Udupi, Anita S Desai, V Ravi |
| EPI_ISL_3232834 | UPHC SHANKARNAGAR | INSACOG-KA, NIMHANS | Chitra Pattabiraman, Anson Kunjumon George,Darshan Sreenivas, Ellango Ramasamy, Sony Sharma, Mahesh Kumar.C.S, Ananthapadmanabha Kotambail, Chetan G K, Gautham Arunachal Udupi, Anita S Desai, V Ravi |
| EPI_ISL_3232835, EPI_ISL_3232836, EPI_ISL_3232837, EPI_ISL_3232838, EPI_ISL_3232840, EPI_ISL_3232841, EPI_ISL_3232842 | NIMHANS | INSACOG-KA, NIMHANS | Chitra Pattabiraman, Anson Kunjumon George,Darshan Sreenivas, Ellango Ramasamy, Sony Sharma, Mahesh Kumar.C.S, Ananthapadmanabha Kotambail, Chetan G K, Gautham Arunachal Udupi, Anita S Desai, V Ravi |
| EPI_ISL_3232843 | JAYADEVA H | INSACOG-KA, NIMHANS | Chitra Pattabiraman, Anson Kunjumon George,Darshan Sreenivas, Ellango Ramasamy, Sony Sharma, Mahesh Kumar.C.S, Ananthapadmanabha Kotambail, Chetan G K, Gautham Arunachal Udupi, Anita S Desai, V Ravi |
| EPI_ISL_3232844 | RAILWAY HOSPITAL | INSACOG-KA, NIMHANS | Chitra Pattabiraman, Anson Kunjumon George,Darshan Sreenivas, Ellango Ramasamy, Sony Sharma, Mahesh Kumar.C.S, Ananthapadmanabha Kotambail, Chetan G K, Gautham Arunachal Udupi, Anita S Desai, V Ravi |
| EPI_ISL_3232845, EPI_ISL_3232846, EPI_ISL_3232847, EPI_ISL_3232848, EPI_ISL_3232849, EPI_ISL_3232850, EPI_ISL_3232851, EPI_ISL_3232852 | NIMHANS | INSACOG-KA, NIMHANS | Chitra Pattabiraman, Anson Kunjumon George,Darshan Sreenivas, Ellango Ramasamy, Sony Sharma, Mahesh Kumar.C.S, Ananthapadmanabha Kotambail, Chetan G K, Gautham Arunachal Udupi, Anita S Desai, V Ravi |
| EPI_ISL_3232853 | JAYADEEVA H | INSACOG-KA, NIMHANS | Chitra Pattabiraman, Anson Kunjumon George,Darshan Sreenivas, Ellango Ramasamy, Sony Sharma, Mahesh Kumar.C.S, Ananthapadmanabha Kotambail, Chetan G K, Gautham Arunachal Udupi, Anita S Desai, V Ravi |
| EPI_ISL_3232854, EPI_ISL_3232855 | NIMHANS | INSACOG-KA, NIMHANS | Chitra Pattabiraman, Anson Kunjumon George,Darshan Sreenivas, Ellango Ramasamy, Sony Sharma, Mahesh Kumar.C.S, Ananthapadmanabha Kotambail, Chetan G K, Gautham Arunachal Udupi, Anita S Desai, V Ravi |
| EPI_ISL_3232857 | SANJAY GANDHI INSTITUTE OF TRAUMA AND ORTHOPAEDICS | INSACOG-KA, NIMHANS | Chitra Pattabiraman, Anson Kunjumon George,Darshan Sreenivas, Ellango Ramasamy, Sony Sharma, Mahesh Kumar.C.S, Ananthapadmanabha Kotambail, Chetan G K, Gautham Arunachal Udupi, Anita S Desai, V Ravi |
| EPI_ISL_3232858, EPI_ISL_3232859, EPI_ISL_3232860, EPI_ISL_3232861, EPI_ISL_3232862, EPI_ISL_3232863, EPI_ISL_3232864, EPI_ISL_3232865, EPI_ISL_3232866, EPI_ISL_3232867, EPI_ISL_3232868, EPI_ISL_3232869, EPI_ISL_3232870, EPI_ISL_3232871 | | | |
| see above | NIMHANS | INSACOG-KA, NIMHANS | Chitra Pattabiraman, Anson Kunjumon George,Darshan Sreenivas, Ellango Ramasamy, Sony Sharma, Mahesh Kumar.C.S, Ananthapadmanabha Kotambail, Chetan G K, Gautham Arunachal Udupi, Anita S Desai, V Ravi |
| EPI_ISL_3232872 | AURIGA | INSACOG-KA, NIMHANS | Chitra Pattabiraman, Anson Kunjumon George,Darshan Sreenivas, Ellango Ramasamy, Sony Sharma, Mahesh Kumar.C.S, Ananthapadmanabha Kotambail, Chetan G K, Gautham Arunachal Udupi, Anita S Desai, V Ravi |
| EPI_ISL_3232873, EPI_ISL_3232874, EPI_ISL_3232875, EPI_ISL_3232876, EPI_ISL_3232877, EPI_ISL_3232878, EPI_ISL_3232879, EPI_ISL_3232880, EPI_ISL_3232881, EPI_ISL_3232882, EPI_ISL_3232883, EPI_ISL_3232884, EPI_ISL_3232885, EPI_ISL_3232886, EPI_ISL_3232887, EPI_ISL_3232888, EPI_ISL_3232889, EPI_ISL_3232890, EPI_ISL_3232891, EPI_ISL_3232892, EPI_ISL_3232893, EPI_ISL_3232894, EPI_ISL_3232895, EPI_ISL_3232896, EPI_ISL_3232897, EPI_ISL_3232898, EPI_ISL_3232899, EPI_ISL_3232900, EPI_ISL_3232901, EPI_ISL_3232902, EPI_ISL_3232903, EPI_ISL_3232904, EPI_ISL_3232905, EPI_ISL_3232906, EPI_ISL_3232907, EPI_ISL_3232908, EPI_ISL_3232909, EPI_ISL_3232910, EPI_ISL_3232911, EPI_ISL_3232912, EPI_ISL_3232913, EPI_ISL_3232914 | | | |
| see above | MMCRI | INSACOG-KA, NIMHANS | Chitra Pattabiraman, Anson Kunjumon George,Darshan Sreenivas, Ellango Ramasamy, Sony Sharma, Mahesh Kumar.C.S, Ananthapadmanabha Kotambail, Chetan G K, Gautham Arunachal Udupi, Anita S Desai, V Ravi |
| EPI_ISL_3232916, EPI_ISL_3232917, EPI_ISL_3232918, EPI_ISL_3232919, EPI_ISL_3232920, EPI_ISL_3232921, EPI_ISL_3232922, EPI_ISL_3232923 | NIMHANS | INSACOG-KA, NIMHANS | Chitra Pattabiraman, Anson Kunjumon George,Darshan Sreenivas, Ellango Ramasamy, Sony Sharma, Mahesh Kumar.C.S, Ananthapadmanabha Kotambail, Chetan G K, Gautham Arunachal Udupi, Anita S Desai, V Ravi |
| EPI_ISL_3232926, EPI_ISL_3232927, EPI_ISL_3232928 | COMMAND HOSPITAL | INSACOG-KA, NIMHANS | Chitra Pattabiraman, Anson Kunjumon George,Darshan Sreenivas, Ellango Ramasamy, Sony Sharma, Mahesh Kumar.C.S, Ananthapadmanabha Kotambail, Chetan G K, Gautham Arunachal Udupi, Anita S Desai, V Ravi |
| EPI_ISL_3232929 | RAILWAY HOSPITAL | INSACOG-KA, NIMHANS | Chitra Pattabiraman, Anson Kunjumon George,Darshan Sreenivas, Ellango Ramasamy, Sony Sharma, Mahesh Kumar.C.S, Ananthapadmanabha Kotambail, Chetan G K, Gautham Arunachal Udupi, Anita S Desai, V Ravi |
| EPI_ISL_3232930, EPI_ISL_3232931, EPI_ISL_3232932, EPI_ISL_3232933, EPI_ISL_3232934, EPI_ISL_3232935, EPI_ISL_3232936, EPI_ISL_3232937, EPI_ISL_3232938, EPI_ISL_3232939, EPI_ISL_3232940, EPI_ISL_3232941, EPI_ISL_3232942 | | | |
| see above | DSU BIDAR | INSACOG-KA, NIMHANS | Chitra Pattabiraman, Anson Kunjumon George,Darshan Sreenivas, Ellango Ramasamy, Sony Sharma, Mahesh Kumar.C.S, Ananthapadmanabha Kotambail, Chetan G K, Gautham Arunachal Udupi, Anita S Desai, V Ravi |
| EPI_ISL_3232943, EPI_ISL_3232944, EPI_ISL_3232945, EPI_ISL_3232946, EPI_ISL_3232947, EPI_ISL_3232948, EPI_ISL_3232949, EPI_ISL_3232950, EPI_ISL_3232951, EPI_ISL_3232952, EPI_ISL_3232953, EPI_ISL_3232954, EPI_ISL_3232955, EPI_ISL_3232956, EPI_ISL_3232957, EPI_ISL_3232958, EPI_ISL_3232959, EPI_ISL_3232960 | | | |
| see above | NIMHANS | INSACOG-KA, NIMHANS | Chitra Pattabiraman, Anson Kunjumon George,Darshan Sreenivas, Ellango Ramasamy, Sony Sharma, Mahesh Kumar.C.S, Ananthapadmanabha Kotambail, Chetan G K, Gautham Arunachal Udupi, Anita S Desai, V Ravi |
| EPI_ISL_3232962, EPI_ISL_3232963, EPI_ISL_3232964, EPI_ISL_3232965, EPI_ISL_3232966, EPI_ISL_3232967, EPI_ISL_3232970, EPI_ISL_3232971, EPI_ISL_3232972, EPI_ISL_3232973, EPI_ISL_3232974, EPI_ISL_3232975, EPI_ISL_3232976, EPI_ISL_3232977, EPI_ISL_3232978 | | | |
| see above | HIMS | INSACOG-KA, NIMHANS | Chitra Pattabiraman, Anson Kunjumon George,Darshan Sreenivas, Ellango Ramasamy, Sony Sharma, Mahesh Kumar.C.S, Ananthapadmanabha Kotambail, Chetan G K, Gautham Arunachal Udupi, Anita S Desai, V Ravi |
| EPI_ISL_3232979, EPI_ISL_3232980, EPI_ISL_3232981, EPI_ISL_3232982, EPI_ISL_3232983, EPI_ISL_3232984, EPI_ISL_3232985, EPI_ISL_3232986, EPI_ISL_3232987, EPI_ISL_3232988, EPI_ISL_3232989, EPI_ISL_3232990, EPI_ISL_3232991, EPI_ISL_3232992, EPI_ISL_3232993 | | | |
| see above | NIMHANS | INSACOG-KA, NIMHANS | Chitra Pattabiraman, Anson Kunjumon George,Darshan Sreenivas, Ellango Ramasamy, Sony Sharma, Mahesh Kumar.C.S, Ananthapadmanabha Kotambail, Chetan G K, Gautham Arunachal Udupi, Anita S Desai, V Ravi |
| EPI_ISL_3232994, EPI_ISL_3232995, EPI_ISL_3232996, EPI_ISL_3232997, EPI_ISL_3232998 | VRDL VIJAYAPURA | INSACOG-KA, NIMHANS | Chitra Pattabiraman, Anson Kunjumon George,Darshan Sreenivas, Ellango Ramasamy, Sony Sharma, Mahesh Kumar.C.S, Ananthapadmanabha Kotambail, Chetan G K, Gautham Arunachal Udupi, Anita S Desai, V Ravi |
| EPI_ISL_3232999 | NIMHANS | INSACOG-KA, NIMHANS | Chitra Pattabiraman, Anson Kunjumon George,Darshan Sreenivas, Ellango Ramasamy, Sony Sharma, Mahesh Kumar.C.S, Ananthapadmanabha Kotambail, Chetan G K, Gautham Arunachal Udupi, Anita S Desai, V Ravi |
| EPI_ISL_3233000 | RAILWAY HOSPITAL | INSACOG-KA, NIMHANS | Chitra Pattabiraman, Anson Kunjumon George,Darshan Sreenivas, Ellango Ramasamy, Sony Sharma, Mahesh Kumar.C.S, Ananthapadmanabha Kotambail, Chetan G K, Gautham Arunachal Udupi, Anita S Desai, V Ravi |

| | | | |
|---|---|---|---|
| EPI_ISL_3233001, EPI_ISL_3233002, EPI_ISL_3233003, EPI_ISL_3233004, EPI_ISL_3233005 | VRDL VIJAYAPURA | INSACOG-KA, NIMHANS | Chitra Pattabiraman, Anson Kunjumon George,Darshan Sreenivas, Ellango Ramasamy, Sony Sharma, Mahesh Kumar.C.S, Ananthapadmanabha Kotambail, Chetan G K, Gautham Arunachal Udupi, Anita S Desai, V Ravi |
| EPI_ISL_3233006, EPI_ISL_3233007, EPI_ISL_3233008, EPI_ISL_3233009, EPI_ISL_3233010, EPI_ISL_3233011, EPI_ISL_3233012 | DSU KALABURAGI | INSACOG-KA, NIMHANS | Chitra Pattabiraman, Anson Kunjumon George,Darshan Sreenivas, Ellango Ramasamy, Sony Sharma, Mahesh Kumar.C.S, Ananthapadmanabha Kotambail, Chetan G K, Gautham Arunachal Udupi, Anita S Desai, V Ravi |
| EPI_ISL_3233013, EPI_ISL_3233014, EPI_ISL_3233015, EPI_ISL_3233016, EPI_ISL_3233017, EPI_ISL_3233018, EPI_ISL_3233019, EPI_ISL_3233020, EPI_ISL_3233021, EPI_ISL_3233022, EPI_ISL_3233023, EPI_ISL_3233024 | | | |
| see above | NIMHANS | INSACOG-KA, NIMHANS | Chitra Pattabiraman, Anson Kunjumon George,Darshan Sreenivas, Ellango Ramasamy, Sony Sharma, Mahesh Kumar.C.S, Ananthapadmanabha Kotambail, Chetan G K, Gautham Arunachal Udupi, Anita S Desai, V Ravi |
| EPI_ISL_3233025 | COLOMBIA ASIA | INSACOG-KA, NIMHANS | Chitra Pattabiraman, Anson Kunjumon George,Darshan Sreenivas, Ellango Ramasamy, Sony Sharma, Mahesh Kumar.C.S, Ananthapadmanabha Kotambail, Chetan G K, Gautham Arunachal Udupi, Anita S Desai, V Ravi |
| EPI_ISL_3233026, EPI_ISL_3233027, EPI_ISL_3233028, EPI_ISL_3233029, EPI_ISL_3233030, EPI_ISL_3233031, EPI_ISL_3233032, EPI_ISL_3233033, EPI_ISL_3233034, EPI_ISL_3233035, EPI_ISL_3233036, EPI_ISL_3233037, EPI_ISL_3233038, EPI_ISL_3233039, EPI_ISL_3233040, EPI_ISL_3233041, EPI_ISL_3233042, EPI_ISL_3233043, EPI_ISL_3233044, EPI_ISL_3233045, EPI_ISL_3233046 | | | |
| see above | VICTORIA HOSPITAL | INSACOG-KA, NIMHANS | Chitra Pattabiraman, Anson Kunjumon George,Darshan Sreenivas, Ellango Ramasamy, Sony Sharma, Mahesh Kumar.C.S, Ananthapadmanabha Kotambail, Chetan G K, Gautham Arunachal Udupi, Anita S Desai, V Ravi |
| EPI_ISL_3233047, EPI_ISL_3233048, EPI_ISL_3233049, EPI_ISL_3233051, EPI_ISL_3233052, EPI_ISL_3233053, EPI_ISL_3233054, EPI_ISL_3233055, EPI_ISL_3233056, EPI_ISL_3233057, EPI_ISL_3233058, EPI_ISL_3233059 | | | |
| see above | OSPEDALE CIVILE TERAMO - CENTRO TRASFUSIONALE | Istituto Zooprofilattico Sperimentale dell’Abruzzo e Molise “G. Caporale” | Lorusso A, Marcacci M, Di Domenico M, Ancora M, Curini V, Di Lollo Valeria, Mangone I, Rinaldi A, Delli Compagni E, Scialabba S, Caporale M, Di Pasquale A, Cammà C, Puglia I, Calistri P, Savini G |
| EPI_ISL_3233060, EPI_ISL_3233061 | OSPEDALE SAN SALVATORE - MEDICINA DI LABORATORIO | Istituto Zooprofilattico Sperimentale dell’Abruzzo e Molise “G. Caporale” | Lorusso A, Marcacci M, Di Domenico M, Ancora M, Curini V, Di Lollo Valeria, Mangone I, Rinaldi A, Delli Compagni E, Scialabba S, Caporale M, Di Pasquale A, Cammà C, Puglia I, Calistri P, Savini G |
| EPI_ISL_3233063, EPI_ISL_3233065, EPI_ISL_3233066, EPI_ISL_3233067, EPI_ISL_3233068, EPI_ISL_3233069, EPI_ISL_3233070, EPI_ISL_3233071, EPI_ISL_3233072, EPI_ISL_3233073, EPI_ISL_3233074, EPI_ISL_3233075, EPI_ISL_3233076, EPI_ISL_3233077 | | | |
| see above | OSPEDALE CIVILE TERAMO - CENTRO TRASFUSIONALE | Istituto Zooprofilattico Sperimentale dell’Abruzzo e Molise “G. Caporale” | Lorusso A, Marcacci M, Di Domenico M, Ancora M, Curini V, Di Lollo Valeria, Mangone I, Rinaldi A, Delli Compagni E, Scialabba S, Caporale M, Di Pasquale A, Cammà C, Puglia I, Calistri P, Savini G |
| EPI_ISL_3233078, EPI_ISL_3233079, EPI_ISL_3233080, EPI_ISL_3233081, EPI_ISL_3233082 | OSPEDALE SAN SALVATORE - MEDICINA DI LABORATORIO | Istituto Zooprofilattico Sperimentale dell’Abruzzo e Molise “G. Caporale” | Lorusso A, Marcacci M, Di Domenico M, Ancora M, Curini V, Di Lollo Valeria, Mangone I, Rinaldi A, Delli Compagni E, Scialabba S, Caporale M, Di Pasquale A, Cammà C, Puglia I, Calistri P, Savini G |
| EPI_ISL_3233083, EPI_ISL_3233086, EPI_ISL_3233088 | PO AVEZZANO - U.O.C. Laboratorio Analisi | Istituto Zooprofilattico Sperimentale dell’Abruzzo e Molise “G. Caporale” | Lorusso A, Marcacci M, Di Domenico M, Ancora M, Curini V, Di Lollo Valeria, Mangone I, Rinaldi A, Delli Compagni E, Scialabba S, Caporale M, Di Pasquale A, Cammà C, Puglia I, Calistri P, Savini G |
| EPI_ISL_3233089, EPI_ISL_3233090, EPI_ISL_3233091, EPI_ISL_3233092, EPI_ISL_3233093, EPI_ISL_3233094, EPI_ISL_3233095, EPI_ISL_3233096, EPI_ISL_3233097, EPI_ISL_3233098 | OSPEDALE CIVILE TERAMO - CENTRO TRASFUSIONALE | Istituto Zooprofilattico Sperimentale dell’Abruzzo e Molise “G. Caporale” | Lorusso A, Marcacci M, Di Domenico M, Ancora M, Curini V, Di Lollo Valeria, Mangone I, Rinaldi A, Delli Compagni E, Scialabba S, Caporale M, Di Pasquale A, Cammà C, Puglia I, Calistri P, Savini G |
| EPI_ISL_3233102, EPI_ISL_3233103 | PO AVEZZANO - U.O.C. Laboratorio Analisi | Istituto Zooprofilattico Sperimentale dell’Abruzzo e Molise “G. Caporale” | Lorusso A, Marcacci M, Di Domenico M, Ancora M, Curini V, Di Lollo Valeria, Mangone I, Rinaldi A, Delli Compagni E, Scialabba S, Caporale M, Di Pasquale A, Cammà C, Puglia I, Calistri P, Savini G |
| EPI_ISL_3233104, EPI_ISL_3233105 | OSPEDALE SAN SALVATORE - MEDICINA DI LABORATORIO | Istituto Zooprofilattico Sperimentale dell’Abruzzo e Molise “G. Caporale” | Lorusso A, Marcacci M, Di Domenico M, Ancora M, Curini V, Di Lollo Valeria, Mangone I, Rinaldi A, Delli Compagni E, Scialabba S, Caporale M, Di Pasquale A, Cammà C, Puglia I, Calistri P, Savini G |
| EPI_ISL_3233109, EPI_ISL_3233110, EPI_ISL_3233111, EPI_ISL_3233112, EPI_ISL_3233113, EPI_ISL_3233114, EPI_ISL_3233115, EPI_ISL_3233117, EPI_ISL_3233118, EPI_ISL_3233119, EPI_ISL_3233120, EPI_ISL_3233121, EPI_ISL_3233122, EPI_ISL_3233123, EPI_ISL_3233124, EPI_ISL_3233125, EPI_ISL_3233126, EPI_ISL_3233127, EPI_ISL_3233128, EPI_ISL_3233129, EPI_ISL_3233130, EPI_ISL_3233131, EPI_ISL_3233132, EPI_ISL_3233133 | | | |
| see above | OSPEDALE CIVILE TERAMO - CENTRO TRASFUSIONALE | Istituto Zooprofilattico Sperimentale dell’Abruzzo e Molise “G. Caporale” | Lorusso A, Marcacci M, Di Domenico M, Ancora M, Curini V, Di Lollo Valeria, Mangone I, Rinaldi A, Delli Compagni E, Scialabba S, Caporale M, Di Pasquale A, Cammà C, Puglia I, Calistri P, Savini G |
| EPI_ISL_3233136, EPI_ISL_3233137, EPI_ISL_3233140 | PO AVEZZANO - U.O.C. Laboratorio Analisi | Istituto Zooprofilattico Sperimentale dell’Abruzzo e Molise “G. Caporale” | Lorusso A, Marcacci M, Di Domenico M, Ancora M, Curini V, Di Lollo Valeria, Mangone I, Rinaldi A, Delli Compagni E, Scialabba S, Caporale M, Di Pasquale A, Cammà C, Puglia I, Calistri P, Savini G |
| EPI_ISL_3233141, EPI_ISL_3233142, EPI_ISL_3233143 | OSPEDALE SAN SALVATORE - MEDICINA DI LABORATORIO | Istituto Zooprofilattico Sperimentale dell’Abruzzo e Molise “G. Caporale” | Lorusso A, Marcacci M, Di Domenico M, Ancora M, Curini V, Di Lollo Valeria, Mangone I, Rinaldi A, Delli Compagni E, Scialabba S, Caporale M, Di Pasquale A, Cammà C, Puglia I, Calistri P, Savini G |
| EPI_ISL_3233146 | OSPEDALE CIVILE TERAMO - CENTRO TRASFUSIONALE | Istituto Zooprofilattico Sperimentale dell’Abruzzo e Molise “G. Caporale” | Lorusso A, Marcacci M, Di Domenico M, Ancora M, Curini V, Di Lollo Valeria, Mangone I, Rinaldi A, Delli Compagni E, Scialabba S, Caporale M, Di Pasquale A, Cammà C, Puglia I, Calistri P, Savini G |
| EPI_ISL_3233147, EPI_ISL_3233148, EPI_ISL_3233150, EPI_ISL_3233151, EPI_ISL_3233152, EPI_ISL_3233154, EPI_ISL_3233155, EPI_ISL_3233156, EPI_ISL_3233157, EPI_ISL_3233158, EPI_ISL_3233159 | | | |
| see above | Heilig hart Lier | Imelda Ziekenhuis | Johan Frans, Hanne Valgaeren, Dagmar Obbels |
| EPI_ISL_3233160, EPI_ISL_3233161, EPI_ISL_3233162, EPI_ISL_3233164, EPI_ISL_3233165 | AZ Jan Portaels | Imelda Ziekenhuis | Johan Frans, Hanne Valgaeren, Dagmar Obbels |
| EPI_ISL_3233166 | Somedi | Imelda Ziekenhuis | Johan Frans, Hanne Valgaeren, Dagmar Obbels |
| EPI_ISL_3233167, EPI_ISL_3233168, EPI_ISL_3233169 | Imelda Ziekenhuis | Imelda Ziekenhuis | Johan Frans, Hanne Valgaeren, Dagmar Obbels |
| EPI_ISL_3233170, EPI_ISL_3233171, EPI_ISL_3233172, EPI_ISL_3233173 | Somedi | Imelda Ziekenhuis | Johan Frans, Hanne Valgaeren, Dagmar Obbels |
| EPI_ISL_3233174, EPI_ISL_3233175, EPI_ISL_3233176, EPI_ISL_3233177, EPI_ISL_3233178, EPI_ISL_3233179, EPI_ISL_3233180, EPI_ISL_3233181, EPI_ISL_3233182 | Imelda Ziekenhuis | Imelda Ziekenhuis | Johan Frans, Hanne Valgaeren, Dagmar Obbels |
| EPI_ISL_3233184, EPI_ISL_3233185, EPI_ISL_3233186, EPI_ISL_3233187, EPI_ISL_3233188, EPI_ISL_3233189, EPI_ISL_3233190, EPI_ISL_3233191, EPI_ISL_3233192, EPI_ISL_3233193, EPI_ISL_3233194, EPI_ISL_3233195, EPI_ISL_3233196, EPI_ISL_3233197, EPI_ISL_3233198, EPI_ISL_3233199, EPI_ISL_3233200, EPI_ISL_3233201, EPI_ISL_3233202, EPI_ISL_3233203, EPI_ISL_3233204, EPI_ISL_3233205, EPI_ISL_3233206, EPI_ISL_3233207, EPI_ISL_3233208, EPI_ISL_3233209, EPI_ISL_3233210, EPI_ISL_3233211, EPI_ISL_3233212, EPI_ISL_3233213, EPI_ISL_3233214, EPI_ISL_3233215, EPI_ISL_3233216, EPI_ISL_3233217, EPI_ISL_3233218, EPI_ISL_3233219, EPI_ISL_3233220, EPI_ISL_3233221, EPI_ISL_3233222, EPI_ISL_3233223, EPI_ISL_3233224, EPI_ISL_3233225, EPI_ISL_3233226, EPI_ISL_3233227, EPI_ISL_3233228, EPI_ISL_3233229, EPI_ISL_3233230, EPI_ISL_3233231 | | | |
| see above | Laboratory of Molecular Biology and Cancer Immunology, Faculty of Sciences, Lebanese University | Microbial Pathogenomics Lab - LAU | Sima tokajian, Georgi Merhi, Jad Koweyes, Bassam Badran, Fadi Abdel Sater, Rawan Makki, Tatiana Tohmeh, Nada Ghosn, Hamad Hassan |
| EPI_ISL_3233233, EPI_ISL_3233234, EPI_ISL_3233235, EPI_ISL_3233236, EPI_ISL_3233237, EPI_ISL_3233238, EPI_ISL_3233240, EPI_ISL_3233241, EPI_ISL_3233242, EPI_ISL_3233243, EPI_ISL_3233244, EPI_ISL_3233245, EPI_ISL_3233246, EPI_ISL_3233247, EPI_ISL_3233248, EPI_ISL_3233249, EPI_ISL_3233250, EPI_ISL_3233251, EPI_ISL_3233252 | | | |
| see above | Plateforme de testing Namuroise | Plateforme de testing Namuroise | Lesly Nyinkeu Kemamen; Nicolas Gilliard; Demars Aurore ; Maschietto Céline ; Nobis Chloé ; Otto Gaetan ; Denis Olivier ; Degosserie Jonathan ; Mullier François |
| EPI_ISL_3233253, EPI_ISL_3233254, EPI_ISL_3233255 | Diagnostyka. Laboratoria Medyczne. | 1. ViroGenetics - BSL3 Laboratory of Virology, Małopolska Centre of Biotechnology, Jagiellonian University; 2. genXone SA, Research & Development Laboratory | Mazur-Panasiuk,N., Grzegorz Nowicki, Wydmanski,W., Gromowski,T., Natalia Drweska-Matelska, Jakub Grabowski, Anna Brylak, Aleksandra Gidlewicz, Karol Szeszko, Maciej Sykulski, Łukasz Krych, Kowalski,M., Szulc,P., Sylwia Januszczak, Labaj,P.P., Michał Kaszuba, Pyrc,K. |
| EPI_ISL_3233256, EPI_ISL_3233257, EPI_ISL_3233259, EPI_ISL_3233260, EPI_ISL_3233261, EPI_ISL_3233262, EPI_ISL_3233263, EPI_ISL_3233264, EPI_ISL_3233265, EPI_ISL_3233266, EPI_ISL_3233267, EPI_ISL_3233268, EPI_ISL_3233269, EPI_ISL_3233270, EPI_ISL_3233271, EPI_ISL_3233272, EPI_ISL_3233273, EPI_ISL_3233274, EPI_ISL_3233275, EPI_ISL_3233276, EPI_ISL_3233277, EPI_ISL_3233278, | | | |

| | | | |
|---|---|---|---|
| EPI_ISL_3233279, EPI_ISL_3233281, EPI_ISL_3233282, EPI_ISL_3233283, EPI_ISL_3233284, EPI_ISL_3233285, EPI_ISL_3233286, EPI_ISL_3233287, EPI_ISL_3233288 | | | |
| see above | Dept. of Microbiology and Infection Control, Akershus University Hospital HF | Dept. of Microbiology and Infection Control, Akershus University Hospital HF | Hege Vangstein Aamot, Alexander Hesselberg Løvestad |
| EPI_ISL_3233328, EPI_ISL_3233329, EPI_ISL_3233330, EPI_ISL_3233331, EPI_ISL_3233332, EPI_ISL_3233333, EPI_ISL_3233334, EPI_ISL_3233335, EPI_ISL_3233336, EPI_ISL_3233337, EPI_ISL_3233338, EPI_ISL_3233339, EPI_ISL_3233340, EPI_ISL_3233341, EPI_ISL_3233342, EPI_ISL_3233343, EPI_ISL_3233344, EPI_ISL_3233345, EPI_ISL_3233346, EPI_ISL_3233347, EPI_ISL_3233348, EPI_ISL_3233350, EPI_ISL_3233351, EPI_ISL_3233352, EPI_ISL_3233353, EPI_ISL_3233354, EPI_ISL_3233355, EPI_ISL_3233356, EPI_ISL_3233357, EPI_ISL_3233358, EPI_ISL_3233359, EPI_ISL_3233360, EPI_ISL_3233361, EPI_ISL_3233362, EPI_ISL_3233363, EPI_ISL_3233364, EPI_ISL_3233365, EPI_ISL_3233366, EPI_ISL_3233368, EPI_ISL_3233369, EPI_ISL_3233370, EPI_ISL_3233371, EPI_ISL_3233372, EPI_ISL_3233373, EPI_ISL_3233374, EPI_ISL_3233375, EPI_ISL_3233376, EPI_ISL_3233377, EPI_ISL_3233378, EPI_ISL_3233379, EPI_ISL_3233380, EPI_ISL_3233381 | | | |
| see above | National Center of Infectious and Parasitic Diseases | National Center of Infectious and Parasitic Diseases | Alexiev et al |
| EPI_ISL_3233382, EPI_ISL_3233383, EPI_ISL_3233384, EPI_ISL_3233385, EPI_ISL_3233386, EPI_ISL_3233387, EPI_ISL_3233388, EPI_ISL_3233389 | AZ St.Jan Brugge-Oostende | AZ SINT-JAN BRUGGE | Marijke Reynders, Patrick Descheemaeker, Laurien Hoornaert, Thomas Van Landschoot, Jorn Hellemans |
| EPI_ISL_3233390, EPI_ISL_3233393, EPI_ISL_3233394, EPI_ISL_3233395, EPI_ISL_3233398, EPI_ISL_3233399, EPI_ISL_3233400, EPI_ISL_3233402, EPI_ISL_3233406, EPI_ISL_3233407, EPI_ISL_3233408, EPI_ISL_3233412, EPI_ISL_3233413, EPI_ISL_3233415, EPI_ISL_3233416, EPI_ISL_3233417 | | | |
| see above | Microbiology Department, University Hospital Donostia | Microbiology Department, University Hospital Donostia | Marimon JM, Montes M, Piñeiro L, Martin-Peñaranda T, Gomez M, Sorarrain A, Cilla G. |
| EPI_ISL_3233451, EPI_ISL_3233452, EPI_ISL_3233454, EPI_ISL_3233456, EPI_ISL_3233457, EPI_ISL_3233458, EPI_ISL_3233459, EPI_ISL_3233464, EPI_ISL_3233465, EPI_ISL_3233467, EPI_ISL_3233469, EPI_ISL_3233470, EPI_ISL_3233471, EPI_ISL_3233472 | | | |
| see above | Max von Pettenkofer Institute, Virology, National Reference Center for Retroviruses, LMU Munich | Laboratory for Functional Genome Analysis; Dept. Genomics; Gene Center of the LMU Munich | Max Muenchhoff; Stefan Krebs; Alexander Graf; Oliver Keppler; Helmut Blum |
| EPI_ISL_3233566, EPI_ISL_3233567, EPI_ISL_3233568, EPI_ISL_3233569, EPI_ISL_3233570, EPI_ISL_3233571, EPI_ISL_3233572, EPI_ISL_3233573, EPI_ISL_3233574, EPI_ISL_3233575, EPI_ISL_3233576, EPI_ISL_3233577, EPI_ISL_3233578, EPI_ISL_3233579, EPI_ISL_3233580, EPI_ISL_3233581, EPI_ISL_3233582, EPI_ISL_3233583, EPI_ISL_3233584, EPI_ISL_3233585 | | | |
| see above | National Platform bis UMONS/Jolimont | National Platform bis UMONS/Jolimont | François Dufrasne, Guillaume Bayon-Vicente, Florian Juszczak, Laetitia Gheysen, Eric Tarantino, Gautier Detry, Ruddy Wattiez |
| EPI_ISL_3233587, EPI_ISL_3233588, EPI_ISL_3233589, EPI_ISL_3233590 | ZOL | Jessa | Marijke Raymaekers et al. on behalf of the Jessa_cmdLab |
| EPI_ISL_3233591, EPI_ISL_3233592 | LKO | Jessa | Marijke Raymaekers et al. on behalf of the Jessa_cmdLab |
| EPI_ISL_3233593, EPI_ISL_3233594, EPI_ISL_3233595, EPI_ISL_3233596, EPI_ISL_3233597, EPI_ISL_3233598, EPI_ISL_3233599, EPI_ISL_3233600, EPI_ISL_3233601, EPI_ISL_3233602, EPI_ISL_3233603, EPI_ISL_3233604, EPI_ISL_3233605, EPI_ISL_3233606, EPI_ISL_3233607, EPI_ISL_3233608, EPI_ISL_3233609 | | | |
| see above | ZOL | Jessa | Marijke Raymaekers et al. on behalf of the Jessa_cmdLab |
| EPI_ISL_3233610, EPI_ISL_3233611, EPI_ISL_3233612, EPI_ISL_3233613, EPI_ISL_3233614, EPI_ISL_3233615, EPI_ISL_3233616, EPI_ISL_3233617, EPI_ISL_3233618, EPI_ISL_3233619, EPI_ISL_3233620 | | | |
| see above | Jessa | Jessa | Marijke Raymaekers et al. on behalf of the Jessa_cmdLab |
| EPI_ISL_3233698, EPI_ISL_3233701, EPI_ISL_3233702, EPI_ISL_3233704, EPI_ISL_3233707, EPI_ISL_3233717, EPI_ISL_3233718, EPI_ISL_3233721, EPI_ISL_3233723, EPI_ISL_3233724, EPI_ISL_3233725, EPI_ISL_3233726, EPI_ISL_3233728, EPI_ISL_3233730, EPI_ISL_3233733, EPI_ISL_3233734, EPI_ISL_3233738, EPI_ISL_3233743, EPI_ISL_3233744, EPI_ISL_3233745, EPI_ISL_3233748, EPI_ISL_3233750, EPI_ISL_3233751, EPI_ISL_3233752, EPI_ISL_3233753, EPI_ISL_3233754, EPI_ISL_3233755, EPI_ISL_3233756, EPI_ISL_3233757, EPI_ISL_3233758, EPI_ISL_3233759, EPI_ISL_3233760, EPI_ISL_3233761, EPI_ISL_3233762, EPI_ISL_3233763, EPI_ISL_3233765, EPI_ISL_3233770, EPI_ISL_3233771, EPI_ISL_3233773, EPI_ISL_3233774, EPI_ISL_3233775, EPI_ISL_3233776, EPI_ISL_3233777, EPI_ISL_3233778, EPI_ISL_3233779, EPI_ISL_3233780, EPI_ISL_3233781, EPI_ISL_3233782, EPI_ISL_3233783, EPI_ISL_3233784, EPI_ISL_3233785, EPI_ISL_3233787, EPI_ISL_3233788, EPI_ISL_3233789, EPI_ISL_3233790, EPI_ISL_3233791, EPI_ISL_3233792, EPI_ISL_3233793, EPI_ISL_3233794, EPI_ISL_3233795, EPI_ISL_3233796, EPI_ISL_3233798, EPI_ISL_3233799, EPI_ISL_3233801, EPI_ISL_3233802, EPI_ISL_3233803, EPI_ISL_3233804, EPI_ISL_3233805, EPI_ISL_3233806, EPI_ISL_3233807, EPI_ISL_3233808, EPI_ISL_3233809, EPI_ISL_3233810, EPI_ISL_3233811, EPI_ISL_3233812, EPI_ISL_3233815, EPI_ISL_3233816, EPI_ISL_3233818, EPI_ISL_3233819, EPI_ISL_3233820, EPI_ISL_3233821, EPI_ISL_3233822, EPI_ISL_3233823, EPI_ISL_3233824, EPI_ISL_3233825, EPI_ISL_3233827, EPI_ISL_3233830, EPI_ISL_3233833, EPI_ISL_3233834, EPI_ISL_3233835, EPI_ISL_3233836, EPI_ISL_3233838, EPI_ISL_3233839, EPI_ISL_3233840, EPI_ISL_3233843, EPI_ISL_3233844, EPI_ISL_3233845, EPI_ISL_3233846, EPI_ISL_3233847, EPI_ISL_3233848, EPI_ISL_3233849, EPI_ISL_3233850, EPI_ISL_3233851, EPI_ISL_3233853, EPI_ISL_3233854, EPI_ISL_3233855, EPI_ISL_3233856, EPI_ISL_3233858, EPI_ISL_3233860, EPI_ISL_3233861, EPI_ISL_3233862, EPI_ISL_3233863, EPI_ISL_3233864, EPI_ISL_3233865, EPI_ISL_3233866, EPI_ISL_3233867, EPI_ISL_3233868, EPI_ISL_3233869, EPI_ISL_3233870, EPI_ISL_3233871, EPI_ISL_3233872, EPI_ISL_3233873, EPI_ISL_3233874, EPI_ISL_3233875, EPI_ISL_3233876, EPI_ISL_3233877, EPI_ISL_3233879, EPI_ISL_3233880, EPI_ISL_3233881, EPI_ISL_3233882, EPI_ISL_3233883, EPI_ISL_3233884, EPI_ISL_3233885, EPI_ISL_3233886, EPI_ISL_3233887, EPI_ISL_3233888, EPI_ISL_3233889, EPI_ISL_3233890, EPI_ISL_3233891, EPI_ISL_3233892, EPI_ISL_3233893, EPI_ISL_3233894, EPI_ISL_3233895, EPI_ISL_3233896, EPI_ISL_3233897, EPI_ISL_3233898, EPI_ISL_3233899, EPI_ISL_3233900, EPI_ISL_3233901, EPI_ISL_3233902, EPI_ISL_3233904, EPI_ISL_3233905, EPI_ISL_3233906, EPI_ISL_3233907, EPI_ISL_3233909, EPI_ISL_3233910, EPI_ISL_3233911, EPI_ISL_3233913, EPI_ISL_3233914, EPI_ISL_3233915, EPI_ISL_3233916, EPI_ISL_3233918, EPI_ISL_3233919, EPI_ISL_3233920, EPI_ISL_3233921, EPI_ISL_3233922, EPI_ISL_3233925, EPI_ISL_3233926, EPI_ISL_3233927, EPI_ISL_3233929, EPI_ISL_3233930, EPI_ISL_3233932, EPI_ISL_3233933, EPI_ISL_3233936, EPI_ISL_3233937, EPI_ISL_3233938, EPI_ISL_3233939, EPI_ISL_3233940, EPI_ISL_3233941, EPI_ISL_3233942, EPI_ISL_3233943, EPI_ISL_3233944, EPI_ISL_3233945, EPI_ISL_3233946, EPI_ISL_3233947, EPI_ISL_3233948, EPI_ISL_3233949, EPI_ISL_3233950, EPI_ISL_3233951, EPI_ISL_3233952, EPI_ISL_3233953, EPI_ISL_3233954, EPI_ISL_3233955, EPI_ISL_3233956, EPI_ISL_3233957, EPI_ISL_3233958, EPI_ISL_3233959, EPI_ISL_3233960, EPI_ISL_3233961, EPI_ISL_3233962, EPI_ISL_3233963, EPI_ISL_3233964, EPI_ISL_3233965, EPI_ISL_3233966, EPI_ISL_3233967, EPI_ISL_3233968, EPI_ISL_3233969, EPI_ISL_3233972, EPI_ISL_3233973, EPI_ISL_3233975, EPI_ISL_3233976, EPI_ISL_3233977, EPI_ISL_3233978, EPI_ISL_3233979, EPI_ISL_3233980, EPI_ISL_3233981, EPI_ISL_3233982, EPI_ISL_3233984, EPI_ISL_3233985, EPI_ISL_3233986, EPI_ISL_3233987, EPI_ISL_3233988, EPI_ISL_3233990, EPI_ISL_3233991, EPI_ISL_3233992, EPI_ISL_3233993, EPI_ISL_3233995, EPI_ISL_3233997, EPI_ISL_3233998, EPI_ISL_3233999, EPI_ISL_3234000, EPI_ISL_3234001, EPI_ISL_3234002, EPI_ISL_3234003, EPI_ISL_3234004, EPI_ISL_3234005, EPI_ISL_3234007, EPI_ISL_3234009, EPI_ISL_3234010, EPI_ISL_3234011, EPI_ISL_3234012, EPI_ISL_3234013, EPI_ISL_3234014, EPI_ISL_3234015, EPI_ISL_3234016, EPI_ISL_3234017, EPI_ISL_3234018, EPI_ISL_3234019, EPI_ISL_3234020, EPI_ISL_3234021, EPI_ISL_3234022, EPI_ISL_3234023, EPI_ISL_3234024, EPI_ISL_3234025, EPI_ISL_3234026, EPI_ISL_3234027, EPI_ISL_3234028, EPI_ISL_3234029, EPI_ISL_3234030, EPI_ISL_3234031, EPI_ISL_3234032, EPI_ISL_3234033, EPI_ISL_3234034, EPI_ISL_3234035, EPI_ISL_3234036, EPI_ISL_3234037, EPI_ISL_3234038, EPI_ISL_3234039, EPI_ISL_3234040, EPI_ISL_3234041, EPI_ISL_3234042, EPI_ISL_3234043, EPI_ISL_3234046, EPI_ISL_3234049, EPI_ISL_3234052, EPI_ISL_3234053, EPI_ISL_3234054, EPI_ISL_3234055, EPI_ISL_3234056, EPI_ISL_3234057, EPI_ISL_3234058, EPI_ISL_3234060, EPI_ISL_3234061, EPI_ISL_3234062, EPI_ISL_3234063, EPI_ISL_3234064, EPI_ISL_3234066, EPI_ISL_3234070, EPI_ISL_3234071, EPI_ISL_3234072, EPI_ISL_3234073, EPI_ISL_3234074, EPI_ISL_3234075, EPI_ISL_3234076, EPI_ISL_3234077, EPI_ISL_3234078, EPI_ISL_3234079, EPI_ISL_3234080, EPI_ISL_3234081, EPI_ISL_3234082, EPI_ISL_3234083, EPI_ISL_3234084, EPI_ISL_3234085, EPI_ISL_3234086, EPI_ISL_3234087, EPI_ISL_3234088, EPI_ISL_3234089, EPI_ISL_3234090, EPI_ISL_3234091, EPI_ISL_3234092, EPI_ISL_3234093, EPI_ISL_3234094, EPI_ISL_3234095, EPI_ISL_3234096, EPI_ISL_3234097, EPI_ISL_3234098, EPI_ISL_3234099, EPI_ISL_3234100, EPI_ISL_3234101, EPI_ISL_3234102, EPI_ISL_3234103, EPI_ISL_3234104, EPI_ISL_3234105, EPI_ISL_3234106, EPI_ISL_3234107, EPI_ISL_3234109, EPI_ISL_3234110, EPI_ISL_3234111, EPI_ISL_3234113, EPI_ISL_3234114, EPI_ISL_3234115, EPI_ISL_3234116, EPI_ISL_3234117, EPI_ISL_3234118, EPI_ISL_3234119, EPI_ISL_3234120, EPI_ISL_3234121, EPI_ISL_3234122, EPI_ISL_3234123, EPI_ISL_3234124, EPI_ISL_3234126, EPI_ISL_3234127, EPI_ISL_3234128, EPI_ISL_3234129, EPI_ISL_3234130, EPI_ISL_3234131, EPI_ISL_3234132, EPI_ISL_3234133, EPI_ISL_3234134, EPI_ISL_3234135, EPI_ISL_3234138, EPI_ISL_3234139, EPI_ISL_3234140, EPI_ISL_3234141, EPI_ISL_3234142, EPI_ISL_3234143, EPI_ISL_3234144, EPI_ISL_3234147, EPI_ISL_3234148, EPI_ISL_3234149, EPI_ISL_3234150, EPI_ISL_3234151, EPI_ISL_3234152, EPI_ISL_3234153, EPI_ISL_3234154, EPI_ISL_3234155, EPI_ISL_3234156, EPI_ISL_3234157, EPI_ISL_3234158, EPI_ISL_3234159, EPI_ISL_3234160, EPI_ISL_3234164, EPI_ISL_3234166, EPI_ISL_3234167, EPI_ISL_3234168, EPI_ISL_3234169, EPI_ISL_3234170, EPI_ISL_3234171, EPI_ISL_3234172, EPI_ISL_3234173, EPI_ISL_3234174, EPI_ISL_3234176, EPI_ISL_3234177, EPI_ISL_3234178, EPI_ISL_3234180, EPI_ISL_3234181, EPI_ISL_3234182, EPI_ISL_3234183, EPI_ISL_3234184, EPI_ISL_3234185, EPI_ISL_3234186, EPI_ISL_3234187, EPI_ISL_3234188, EPI_ISL_3234189, EPI_ISL_3234190, EPI_ISL_3234191, EPI_ISL_3234192, EPI_ISL_3234193, EPI_ISL_3234194, EPI_ISL_3234195, EPI_ISL_3234196, EPI_ISL_3234197, EPI_ISL_3234198, EPI_ISL_3234199, EPI_ISL_3234200, EPI_ISL_3234202, EPI_ISL_3234203, EPI_ISL_3234204, EPI_ISL_3234205, EPI_ISL_3234206, EPI_ISL_3234207, EPI_ISL_3234208, EPI_ISL_3234209, EPI_ISL_3234210, EPI_ISL_3234211, EPI_ISL_3234212, EPI_ISL_3234213, EPI_ISL_3234214, EPI_ISL_3234215, EPI_ISL_3234216, EPI_ISL_3234217, EPI_ISL_3234218, EPI_ISL_3234219, EPI_ISL_3234220, EPI_ISL_3234221, EPI_ISL_3234222, EPI_ISL_3234223, EPI_ISL_3234224, EPI_ISL_3234225, EPI_ISL_3234226, EPI_ISL_3234227, EPI_ISL_3234228, EPI_ISL_3234229, EPI_ISL_3234230, EPI_ISL_3234231, EPI_ISL_3234232, EPI_ISL_3234233, EPI_ISL_3234234, EPI_ISL_3234235, EPI_ISL_3234236, EPI_ISL_3234237, EPI_ISL_3234238, EPI_ISL_3234239, EPI_ISL_3234240, EPI_ISL_3234241, EPI_ISL_3234242, EPI_ISL_3234243, EPI_ISL_3234247, EPI_ISL_3234248, EPI_ISL_3234249, EPI_ISL_3234250, EPI_ISL_3234251, EPI_ISL_3234252, EPI_ISL_3234253, EPI_ISL_3234254, EPI_ISL_3234255, EPI_ISL_3234256, EPI_ISL_3234257, EPI_ISL_3234258, EPI_ISL_3234259, EPI_ISL_3234260, EPI_ISL_3234261, EPI_ISL_3234262, EPI_ISL_3234263, EPI_ISL_3234264, EPI_ISL_3234265, EPI_ISL_3234266, EPI_ISL_3234267, EPI_ISL_3234268, EPI_ISL_3234269, EPI_ISL_3234270, EPI_ISL_3234272, EPI_ISL_3234273, EPI_ISL_3234274, EPI_ISL_3234275, EPI_ISL_3234277, EPI_ISL_3234278, EPI_ISL_3234279, EPI_ISL_3234280, EPI_ISL_3234281, EPI_ISL_3234282, EPI_ISL_3234283, EPI_ISL_3234284, EPI_ISL_3234285, EPI_ISL_3234286, EPI_ISL_3234287, EPI_ISL_3234289, EPI_ISL_3234290, EPI_ISL_3234291, EPI_ISL_3234292, EPI_ISL_3234293, EPI_ISL_3234294, EPI_ISL_3234295, EPI_ISL_3234296, EPI_ISL_3234297, EPI_ISL_3234298, EPI_ISL_3234299, EPI_ISL_3234300, EPI_ISL_3234301, EPI_ISL_3234302, EPI_ISL_3234303, EPI_ISL_3234304, EPI_ISL_3234305, EPI_ISL_3234307, EPI_ISL_3234308, EPI_ISL_3234309, EPI_ISL_3234310, EPI_ISL_3234311, EPI_ISL_3234314, EPI_ISL_3234315, EPI_ISL_3234316, EPI_ISL_3234317, EPI_ISL_3234318, EPI_ISL_3234319, EPI_ISL_3234320, EPI_ISL_3234321, EPI_ISL_3234323, EPI_ISL_3234325, EPI_ISL_3234326, EPI_ISL_3234327, EPI_ISL_3234328, EPI_ISL_3234329, EPI_ISL_3234330, EPI_ISL_3234331, EPI_ISL_3234332, EPI_ISL_3234333, EPI_ISL_3234334, EPI_ISL_3234335, EPI_ISL_3234336, EPI_ISL_3234337, EPI_ISL_3234338, EPI_ISL_3234339, EPI_ISL_3234340, EPI_ISL_3234341, EPI_ISL_3234342, EPI_ISL_3234343, EPI_ISL_3234344, EPI_ISL_3234345, EPI_ISL_3234346, EPI_ISL_3234347, EPI_ISL_3234348, EPI_ISL_3234349, EPI_ISL_3234350, EPI_ISL_3234351, EPI_ISL_3234352, EPI_ISL_3234353, EPI_ISL_3234354, EPI_ISL_3234355, EPI_ISL_3234356, EPI_ISL_3234357, EPI_ISL_3234358, EPI_ISL_3234359, EPI_ISL_3234360, EPI_ISL_3234361, EPI_ISL_3234362, EPI_ISL_3234363, EPI_ISL_3234364, EPI_ISL_3234365, EPI_ISL_3234366, EPI_ISL_3234367, EPI_ISL_3234368, EPI_ISL_3234369, EPI_ISL_3234370, EPI_ISL_3234371, EPI_ISL_3234372 | | | |
| see above | Public Health Ontario Laboratory | Public Health Ontario Laboratory | Vanessa G Allen, Philip Banh, Yao Chen, Richard de Borja, Alireza Eshaghi, Nahuel Fittipaldi, Christine Frantz, Jonathan B Gubbay, Jennifer L Guthrie, Lawrence Heisler, Esha Joshi, Michael Laszloffy, Aimin Li, Michael CY Li, Dean Maxwell, Sandeep Nagra, Samir N Patel, Jared Simpson, Karthikeyan Sivaraman, Ashleigh Sullivan, Yogi Sundaravadanam, Sarah Teatero, Andre Villegas, Matthew Watson, Sandra Zittermann |
| EPI_ISL_3234539 | CH AUCH | CHU Purpan - Laboratoire de Virologie - Institut Fédératif de Biologie | Latour J., Milhes M., Bulach T., Ranger N., Salin G., Nicot F., Tremeaux P., Donnadieu C., Izopet J. |
| EPI_ISL_3234540, EPI_ISL_3234541, EPI_ISL_3234542, EPI_ISL_3234543, EPI_ISL_3234544, EPI_ISL_3234545, EPI_ISL_3234546, EPI_ISL_3234547, EPI_ISL_3234548, EPI_ISL_3234549 | LABO BIO DOC | CHU Purpan - Laboratoire de Virologie - Institut Fédératif de Biologie | Latour J., Milhes M., Bulach T., Ranger N., Salin G., Nicot F., Tremeaux P., Donnadieu C., Izopet J. |
| EPI_ISL_3234550, EPI_ISL_3234551, EPI_ISL_3234553, EPI_ISL_3234555, EPI_ISL_3234557, EPI_ISL_3234558, EPI_ISL_3234559, EPI_ISL_3234562, EPI_ISL_3234563, EPI_ISL_3234564, EPI_ISL_3234565, EPI_ISL_3234566, EPI_ISL_3234567, EPI_ISL_3234568, EPI_ISL_3234569, EPI_ISL_3234570, EPI_ISL_3234571, EPI_ISL_3234572, EPI_ISL_3234573, EPI_ISL_3234574, EPI_ISL_3234575, EPI_ISL_3234576, EPI_ISL_3234577, EPI_ISL_3234578, EPI_ISL_3234579, EPI_ISL_3234580, EPI_ISL_3234581, EPI_ISL_3234583, EPI_ISL_3234584, EPI_ISL_3234585, EPI_ISL_3234586, EPI_ISL_3234587, EPI_ISL_3234588, EPI_ISL_3234589, EPI_ISL_3234591, EPI_ISL_3234592 | | | |
| see above | CHU Purpan - Laboratoire de Virologie - Institut Fédératif de Biologie | CHU Purpan - Laboratoire de Virologie - Institut Fédératif de Biologie | Latour J., Milhes M., Bulach T., Ranger N., Salin G., Nicot F., Tremeaux P., Donnadieu C., Izopet J. |
| EPI_ISL_3234593 | CH AUCH | CHU Purpan - Laboratoire de Virologie - Institut Fédératif de Biologie | Latour J., Milhes M., Bulach T., Ranger N., Salin G., Nicot F., Tremeaux P., Donnadieu C., Izopet J. |
| EPI_ISL_3234594, EPI_ISL_3234595, EPI_ISL_3234596, EPI_ISL_3234597, EPI_ISL_3234598 | CHU Purpan - Laboratoire de Virologie - Institut Fédératif de Biologie | CHU Purpan - Laboratoire de Virologie - Institut Fédératif de Biologie | Latour J., Milhes M., Bulach T., Ranger N., Salin G., Nicot F., Tremeaux P., Donnadieu C., Izopet J. |
| EPI_ISL_3234599, EPI_ISL_3234600, EPI_ISL_3234601, EPI_ISL_3234603, EPI_ISL_3234604, EPI_ISL_3234605, EPI_ISL_3234606, EPI_ISL_3234607, EPI_ISL_3234608 | CENTRE DE BIOLOGIE | CHU Purpan - Laboratoire de Virologie - Institut Fédératif de Biologie | Latour J., Milhes M., Bulach T., Ranger N., Salin G., Nicot F., Tremeaux P., Donnadieu C., Izopet J. |
| EPI_ISL_3234609 | CH ALBI | CHU Purpan - Laboratoire de Virologie - Institut Fédératif de Biologie | Latour J., Milhes M., Bulach T., Ranger N., Salin G., Nicot F., Tremeaux P., Donnadieu C., Izopet J. |
| EPI_ISL_3234611, EPI_ISL_3234612, EPI_ISL_3234613, EPI_ISL_3234614, EPI_ISL_3234615, EPI_ISL_3234616 | CENTRE DE BIOLOGIE | CHU Purpan - Laboratoire de Virologie - Institut Fédératif de Biologie | Latour J., Milhes M., Bulach T., Ranger N., Salin G., Nicot F., Tremeaux P., Donnadieu C., Izopet J. |
| EPI_ISL_3234617 | CH MONTAUBAN | CHU Purpan - Laboratoire de Virologie - Institut Fédératif de Biologie | Latour J., Milhes M., Bulach T., Ranger N., Salin G., Nicot F., Tremeaux P., Donnadieu C., Izopet J. |

| | | | |
|---|---|---|---|
| EPI_ISL_3234618 | HOPITAL INTERCOMUNAL CASTEL-MOISSAC | CHU Purpan - Laboratoire de Virologie - Institut Fédératif de Biologie | Latour J., Milhes M., Bulach T., Ranger N., Salin G., Nicot F., Tremeaux P., Donnadieu C., Izopet J. |
| EPI_ISL_3234619, EPI_ISL_3234620, EPI_ISL_3234621, EPI_ISL_3234622, EPI_ISL_3234623, EPI_ISL_3234624, EPI_ISL_3234625, EPI_ISL_3234626, EPI_ISL_3234627, EPI_ISL_3234628, EPI_ISL_3234629 | | | |
| see above | CENTRE DE BIOLOGIE | CHU Purpan - Laboratoire de Virologie - Institut Fédératif de Biologie | Latour J., Milhes M., Bulach T., Ranger N., Salin G., Nicot F., Tremeaux P., Donnadieu C., Izopet J. |
| EPI_ISL_3234630 | CH MONTAUBAN | CHU Purpan - Laboratoire de Virologie - Institut Fédératif de Biologie | Latour J., Milhes M., Bulach T., Ranger N., Salin G., Nicot F., Tremeaux P., Donnadieu C., Izopet J. |
| EPI_ISL_3234633, EPI_ISL_3234634, EPI_ISL_3234635, EPI_ISL_3234636, EPI_ISL_3234637, EPI_ISL_3234638, EPI_ISL_3234639, EPI_ISL_3234640, EPI_ISL_3234641, EPI_ISL_3234642, EPI_ISL_3234643, EPI_ISL_3234644, EPI_ISL_3234645, EPI_ISL_3234646, EPI_ISL_3234647, EPI_ISL_3234648, EPI_ISL_3234649, EPI_ISL_3234650, EPI_ISL_3234651, EPI_ISL_3234652, EPI_ISL_3234653, EPI_ISL_3234654, EPI_ISL_3234655 | | | |
| see above | LABO BIO DOC | CHU Purpan - Laboratoire de Virologie - Institut Fédératif de Biologie | Latour J., Milhes M., Bulach T., Ranger N., Salin G., Nicot F., Tremeaux P., Donnadieu C., Izopet J. |
| EPI_ISL_3234656, EPI_ISL_3234657, EPI_ISL_3234658 | CHU Purpan - Laboratoire de Virologie - Institut Fédératif de Biologie | CHU Purpan - Laboratoire de Virologie - Institut Fédératif de Biologie | Latour J., Milhes M., Bulach T., Ranger N., Salin G., Nicot F., Tremeaux P., Donnadieu C., Izopet J. |
| EPI_ISL_3234659 | GCS DE BIOLOGIE DU TERRITOIRE DE L'ARIEGE | CHU Purpan - Laboratoire de Virologie - Institut Fédératif de Biologie | Latour J., Milhes M., Bulach T., Ranger N., Salin G., Nicot F., Tremeaux P., Donnadieu C., Izopet J. |
| EPI_ISL_3234660, EPI_ISL_3234661, EPI_ISL_3234662 | CH ALBI | CHU Purpan - Laboratoire de Virologie - Institut Fédératif de Biologie | Latour J., Milhes M., Bulach T., Ranger N., Salin G., Nicot F., Tremeaux P., Donnadieu C., Izopet J. |
| EPI_ISL_3234663, EPI_ISL_3234664 | CH CARCASSONE | CHU Purpan - Laboratoire de Virologie - Institut Fédératif de Biologie | Latour J., Milhes M., Bulach T., Ranger N., Salin G., Nicot F., Tremeaux P., Donnadieu C., Izopet J. |
| EPI_ISL_3234666 | CHU Purpan - Laboratoire de Virologie - Institut Fédératif de Biologie | CHU Purpan - Laboratoire de Virologie - Institut Fédératif de Biologie | Latour J., Milhes M., Bulach T., Ranger N., Salin G., Nicot F., Tremeaux P., Donnadieu C., Izopet J. |
| EPI_ISL_3234667, EPI_ISL_3234668, EPI_ISL_3234669, EPI_ISL_3234673 | LABO BIO DOC | CHU Purpan - Laboratoire de Virologie - Institut Fédératif de Biologie | Latour J., Milhes M., Bulach T., Ranger N., Salin G., Nicot F., Tremeaux P., Donnadieu C., Izopet J. |
| EPI_ISL_3234674, EPI_ISL_3234675 | CHU Purpan - Laboratoire de Virologie - Institut Fédératif de Biologie | CHU Purpan - Laboratoire de Virologie - Institut Fédératif de Biologie | Latour J., Milhes M., Bulach T., Ranger N., Salin G., Nicot F., Tremeaux P., Donnadieu C., Izopet J. |
| EPI_ISL_3234676, EPI_ISL_3234677, EPI_ISL_3234678, EPI_ISL_3234679, EPI_ISL_3234680, EPI_ISL_3234681, EPI_ISL_3234682, EPI_ISL_3234685, EPI_ISL_3234686, EPI_ISL_3234687, EPI_ISL_3234688, EPI_ISL_3234689, EPI_ISL_3234690, EPI_ISL_3234691, EPI_ISL_3234692, EPI_ISL_3234693, EPI_ISL_3234694, EPI_ISL_3234695, EPI_ISL_3234696, EPI_ISL_3234697, EPI_ISL_3234698, EPI_ISL_3234699, EPI_ISL_3234700, EPI_ISL_3234702, EPI_ISL_3234703, EPI_ISL_3234705, EPI_ISL_3234706, EPI_ISL_3234707, EPI_ISL_3234708, EPI_ISL_3234709, EPI_ISL_3234710, EPI_ISL_3234711, EPI_ISL_3234712, EPI_ISL_3234713, EPI_ISL_3234714, EPI_ISL_3234715, EPI_ISL_3234716, EPI_ISL_3234717, EPI_ISL_3234719, EPI_ISL_3234721, EPI_ISL_3234722, EPI_ISL_3234723, EPI_ISL_3234724, EPI_ISL_3234725, EPI_ISL_3234726, EPI_ISL_3234727, EPI_ISL_3234728, EPI_ISL_3234729, EPI_ISL_3234730, EPI_ISL_3234731, EPI_ISL_3234732, EPI_ISL_3234733, EPI_ISL_3234734, EPI_ISL_3234735, EPI_ISL_3234736, EPI_ISL_3234737, EPI_ISL_3234738, EPI_ISL_3234739, EPI_ISL_3234741, EPI_ISL_3234742, EPI_ISL_3234743, EPI_ISL_3234744, EPI_ISL_3234745, EPI_ISL_3234746, EPI_ISL_3234747, EPI_ISL_3234748, EPI_ISL_3234749, EPI_ISL_3234750, EPI_ISL_3234751, EPI_ISL_3234752, EPI_ISL_3234753, EPI_ISL_3234754, EPI_ISL_3234755, EPI_ISL_3234756, EPI_ISL_3234757, EPI_ISL_3234758, EPI_ISL_3234759, EPI_ISL_3234761, EPI_ISL_3234762, EPI_ISL_3234764, EPI_ISL_3234765, EPI_ISL_3234766, EPI_ISL_3234767, EPI_ISL_3234768, EPI_ISL_3234769, EPI_ISL_3234770, EPI_ISL_3234772, EPI_ISL_3234773, EPI_ISL_3234774, EPI_ISL_3234775, EPI_ISL_3234776, EPI_ISL_3234777, EPI_ISL_3234778, EPI_ISL_3234779, EPI_ISL_3234780, EPI_ISL_3234781, EPI_ISL_3234782, EPI_ISL_3234784, EPI_ISL_3234785, EPI_ISL_3234787, EPI_ISL_3234788, EPI_ISL_3234789, EPI_ISL_3234790, EPI_ISL_3234791, EPI_ISL_3234794, EPI_ISL_3234795, EPI_ISL_3234796, EPI_ISL_3234797, EPI_ISL_3234798, EPI_ISL_3234799, EPI_ISL_3234800, EPI_ISL_3234801, EPI_ISL_3234802, EPI_ISL_3234803, EPI_ISL_3234804, EPI_ISL_3234805, EPI_ISL_3234806, EPI_ISL_3234807, EPI_ISL_3234808, EPI_ISL_3234809, EPI_ISL_3234811, EPI_ISL_3234812, EPI_ISL_3234813, EPI_ISL_3234815, EPI_ISL_3234816, EPI_ISL_3234817, EPI_ISL_3234819, EPI_ISL_3234820, EPI_ISL_3234821, EPI_ISL_3234823, EPI_ISL_3234824, EPI_ISL_3234826, EPI_ISL_3234827, EPI_ISL_3234828, EPI_ISL_3234829, EPI_ISL_3234830, EPI_ISL_3234831, EPI_ISL_3234832, EPI_ISL_3234833, EPI_ISL_3234835, EPI_ISL_3234836, EPI_ISL_3234837, EPI_ISL_3234838, EPI_ISL_3234839, EPI_ISL_3234841, EPI_ISL_3234842, EPI_ISL_3234843, EPI_ISL_3234844, EPI_ISL_3234845, EPI_ISL_3234846, EPI_ISL_3234847, EPI_ISL_3234848, EPI_ISL_3234849, EPI_ISL_3234850, EPI_ISL_3234851, EPI_ISL_3234852, EPI_ISL_3234853, EPI_ISL_3234854, EPI_ISL_3234855, EPI_ISL_3234856, EPI_ISL_3234857, EPI_ISL_3234858, EPI_ISL_3234859, EPI_ISL_3234860, EPI_ISL_3234861, EPI_ISL_3234862, EPI_ISL_3234863, EPI_ISL_3234864, EPI_ISL_3234865, EPI_ISL_3234866, EPI_ISL_3234867, EPI_ISL_3234868, EPI_ISL_3234869, EPI_ISL_3234870, EPI_ISL_3234871, EPI_ISL_3234872, EPI_ISL_3234873, EPI_ISL_3234874, EPI_ISL_3234875, EPI_ISL_3234876, EPI_ISL_3234877, EPI_ISL_3234878, EPI_ISL_3234879, EPI_ISL_3234880, EPI_ISL_3234881, EPI_ISL_3234882, EPI_ISL_3234883, EPI_ISL_3234884, EPI_ISL_3234885, EPI_ISL_3234886, EPI_ISL_3234887, EPI_ISL_3234888, EPI_ISL_3234889, EPI_ISL_3234890, EPI_ISL_3234892, EPI_ISL_3234893, EPI_ISL_3234894, EPI_ISL_3234896, EPI_ISL_3234897, EPI_ISL_3234898, EPI_ISL_3234899, EPI_ISL_3234900, EPI_ISL_3234901, EPI_ISL_3234902, EPI_ISL_3234903, EPI_ISL_3234904, EPI_ISL_3234905, EPI_ISL_3234907, EPI_ISL_3234908, EPI_ISL_3234909, EPI_ISL_3234910, EPI_ISL_3234911, EPI_ISL_3234912, EPI_ISL_3234913, EPI_ISL_3234914, EPI_ISL_3234915, EPI_ISL_3234916, EPI_ISL_3234917, EPI_ISL_3234918, EPI_ISL_3234919, EPI_ISL_3234920, EPI_ISL_3234921, EPI_ISL_3234922, EPI_ISL_3234923, EPI_ISL_3234924, EPI_ISL_3234925 | | | |
| see above | Wisconsin State Laboratory of Hygiene Communicable Disease Division | Wisconsin State Laboratory of Hygiene Communicable Disease Division | Abigail C. Shockey, Alicia J. Mooney, Erika M. Hanson, Tonya Danz, Richard Griesser, Sara Wagner, Kelsey R. Florek |
| EPI_ISL_3234929, EPI_ISL_3234930, EPI_ISL_3234931, EPI_ISL_3234933 | Microbiology Department, University Hospital Donostia | Microbiology Department, University Hospital Donostia | Marimon JM, Montes M, Piñeiro L, Martin-Peñaranda T, Gomez M, Sorarrain A, Cilla G. |
| EPI_ISL_3234934, EPI_ISL_3234935, EPI_ISL_3234936, EPI_ISL_3234937, EPI_ISL_3234938, EPI_ISL_3234939, EPI_ISL_3234940, EPI_ISL_3234941, EPI_ISL_3234942, EPI_ISL_3234943, EPI_ISL_3234944, EPI_ISL_3234945, EPI_ISL_3234946, EPI_ISL_3234947, EPI_ISL_3234948, EPI_ISL_3234949, EPI_ISL_3234950, EPI_ISL_3234951, EPI_ISL_3234952, EPI_ISL_3234953, EPI_ISL_3234955, EPI_ISL_3234956, EPI_ISL_3234957, EPI_ISL_3234958, EPI_ISL_3234959, EPI_ISL_3234960, EPI_ISL_3234961, EPI_ISL_3234962, EPI_ISL_3234963, EPI_ISL_3234964, EPI_ISL_3234965, EPI_ISL_3234966, EPI_ISL_3234967, EPI_ISL_3234968, EPI_ISL_3234969, EPI_ISL_3234970, EPI_ISL_3234971, EPI_ISL_3234972, EPI_ISL_3234973, EPI_ISL_3234974, EPI_ISL_3234975, EPI_ISL_3234976, EPI_ISL_3234977, EPI_ISL_3234978, EPI_ISL_3234979, EPI_ISL_3234980, EPI_ISL_3234981, EPI_ISL_3234982, EPI_ISL_3234983, EPI_ISL_3234984, EPI_ISL_3234985, EPI_ISL_3234986, EPI_ISL_3234987, EPI_ISL_3234988, EPI_ISL_3234989, EPI_ISL_3234990, EPI_ISL_3234991, EPI_ISL_3234992, EPI_ISL_3234993, EPI_ISL_3234994, EPI_ISL_3234995, EPI_ISL_3234996, EPI_ISL_3234997, EPI_ISL_3234998, EPI_ISL_3234999, EPI_ISL_3235000, EPI_ISL_3235001, EPI_ISL_3235002, EPI_ISL_3235003, EPI_ISL_3235004, EPI_ISL_3235005, EPI_ISL_3235006, EPI_ISL_3235007, EPI_ISL_3235009, EPI_ISL_3235010, EPI_ISL_3235011, EPI_ISL_3235012, EPI_ISL_3235013, EPI_ISL_3235014, EPI_ISL_3235015, EPI_ISL_3235016, EPI_ISL_3235017, EPI_ISL_3235018, EPI_ISL_3235019, EPI_ISL_3235020, EPI_ISL_3235021, EPI_ISL_3235022, EPI_ISL_3235023, EPI_ISL_3235024, EPI_ISL_3235025 | | | |
| see above | Nebraska Public Health Laboratory | NPHL COVID-19 Response Team | NPHL COVID-19 Response Team |
| EPI_ISL_3235026, EPI_ISL_3235027, EPI_ISL_3235028, EPI_ISL_3235029, EPI_ISL_3235030, EPI_ISL_3235031, EPI_ISL_3235032, EPI_ISL_3235033, EPI_ISL_3235034, EPI_ISL_3235035, EPI_ISL_3235036, EPI_ISL_3235037, EPI_ISL_3235038, EPI_ISL_3235040, EPI_ISL_3235041, EPI_ISL_3235042, EPI_ISL_3235043, EPI_ISL_3235044, EPI_ISL_3235045, EPI_ISL_3235046, EPI_ISL_3235049, EPI_ISL_3235050, EPI_ISL_3235051, EPI_ISL_3235052, EPI_ISL_3235053, EPI_ISL_3235054, EPI_ISL_3235055, EPI_ISL_3235057, EPI_ISL_3235058, EPI_ISL_3235061, EPI_ISL_3235062, EPI_ISL_3235063, EPI_ISL_3235064, EPI_ISL_3235065, EPI_ISL_3235066, EPI_ISL_3235067, EPI_ISL_3235068, EPI_ISL_3235069, EPI_ISL_3235070, EPI_ISL_3235071, EPI_ISL_3235072, EPI_ISL_3235073, EPI_ISL_3235074, EPI_ISL_3235075, EPI_ISL_3235076, EPI_ISL_3235078, EPI_ISL_3235079, EPI_ISL_3235080, EPI_ISL_3235081, EPI_ISL_3235082, EPI_ISL_3235083, EPI_ISL_3235084, EPI_ISL_3235085, EPI_ISL_3235086, EPI_ISL_3235087, EPI_ISL_3235088, EPI_ISL_3235089, EPI_ISL_3235090 | | | |
| see above | ASL Napoli 1 Centro | AMES Centro Polidiagnostico Strumentale S.r.l. | Giovanni Savarese, Raffaella Ruggiero, Eloisa Evangelista, Antonella Di Carlo, Luisa Circelli, Luigi D'Amore, Roberto Sirica,Maurizio D'Amora Antonio Fico |
| EPI_ISL_3235321, EPI_ISL_3235322, EPI_ISL_3235324, EPI_ISL_3235327, EPI_ISL_3235328, EPI_ISL_3235329, EPI_ISL_3235330, EPI_ISL_3235331, EPI_ISL_3235336, EPI_ISL_3235339, EPI_ISL_3235341, EPI_ISL_3235342, EPI_ISL_3235343, EPI_ISL_3235344, EPI_ISL_3235347, EPI_ISL_3235355, EPI_ISL_3235361 | | | |
| see above | Hackensack Medical Center | New York Genome Center | Michael Zody, Andre Corvelo, Dayna M. Oschwald, Amy Baldwin, Samantha Fennessey, Tom Maniatis, Liang Chen, Jose Mediavilla, Marcus Cunningham, Kaelea Composto, Kar Chow, David Perlin, Barry Kreiswirth |
| EPI_ISL_3235371, EPI_ISL_3235372, EPI_ISL_3235373, EPI_ISL_3235374, EPI_ISL_3235375, EPI_ISL_3235376, EPI_ISL_3235377, EPI_ISL_3235378, EPI_ISL_3235379, EPI_ISL_3235380, EPI_ISL_3235381, EPI_ISL_3235382, EPI_ISL_3235383, EPI_ISL_3235384, EPI_ISL_3235385, EPI_ISL_3235386, EPI_ISL_3235387, EPI_ISL_3235388, EPI_ISL_3235389, EPI_ISL_3235390, EPI_ISL_3235391, EPI_ISL_3235392, EPI_ISL_3235393, EPI_ISL_3235394, EPI_ISL_3235395, EPI_ISL_3235396, EPI_ISL_3235397, EPI_ISL_3235398, EPI_ISL_3235399, EPI_ISL_3235400, EPI_ISL_3235401, EPI_ISL_3235402, EPI_ISL_3235403, EPI_ISL_3235404, EPI_ISL_3235405, EPI_ISL_3235406, EPI_ISL_3235407, EPI_ISL_3235408, EPI_ISL_3235409, EPI_ISL_3235410, EPI_ISL_3235411, EPI_ISL_3235412, EPI_ISL_3235413, EPI_ISL_3235414, EPI_ISL_3235415, EPI_ISL_3235416, EPI_ISL_3235417, EPI_ISL_3235418, EPI_ISL_3235419, EPI_ISL_3235420, EPI_ISL_3235421, EPI_ISL_3235422, EPI_ISL_3235423, EPI_ISL_3235424, EPI_ISL_3235425, EPI_ISL_3235426, EPI_ISL_3235427, EPI_ISL_3235428, EPI_ISL_3235429, EPI_ISL_3235430, EPI_ISL_3235431, EPI_ISL_3235432, EPI_ISL_3235433, EPI_ISL_3235434, EPI_ISL_3235435, EPI_ISL_3235436, EPI_ISL_3235437, EPI_ISL_3235438, EPI_ISL_3235439, EPI_ISL_3235440, EPI_ISL_3235441, EPI_ISL_3235442, EPI_ISL_3235443, EPI_ISL_3235444, EPI_ISL_3235445, EPI_ISL_3235446, EPI_ISL_3235447, EPI_ISL_3235448, EPI_ISL_3235449, EPI_ISL_3235450, EPI_ISL_3235451, EPI_ISL_3235452, EPI_ISL_3235453, EPI_ISL_3235454, EPI_ISL_3235455, EPI_ISL_3235456, EPI_ISL_3235457, EPI_ISL_3235458, EPI_ISL_3235459, EPI_ISL_3235460, EPI_ISL_3235461, EPI_ISL_3235462, EPI_ISL_3235463, EPI_ISL_3235464, EPI_ISL_3235465, EPI_ISL_3235466, EPI_ISL_3235467, EPI_ISL_3235468, EPI_ISL_3235469, EPI_ISL_3235470, EPI_ISL_3235471, EPI_ISL_3235472, EPI_ISL_3235473, EPI_ISL_3235474, EPI_ISL_3235475, EPI_ISL_3235476, EPI_ISL_3235477, EPI_ISL_3235478, EPI_ISL_3235481, EPI_ISL_3235482, EPI_ISL_3235483, EPI_ISL_3235484, EPI_ISL_3235485, EPI_ISL_3235486, EPI_ISL_3235487, EPI_ISL_3235488, EPI_ISL_3235489, EPI_ISL_3235490, EPI_ISL_3235491, EPI_ISL_3235492, EPI_ISL_3235493, EPI_ISL_3235494, EPI_ISL_3235495, EPI_ISL_3235496, EPI_ISL_3235497, EPI_ISL_3235498, EPI_ISL_3235499, EPI_ISL_3235500, EPI_ISL_3235501, EPI_ISL_3235502, EPI_ISL_3235503, EPI_ISL_3235504, EPI_ISL_3235505, EPI_ISL_3235506, EPI_ISL_3235507, EPI_ISL_3235508, EPI_ISL_3235509, EPI_ISL_3235510, EPI_ISL_3235511, EPI_ISL_3235512, EPI_ISL_3235513, EPI_ISL_3235514, EPI_ISL_3235515, EPI_ISL_3235516, EPI_ISL_3235518, EPI_ISL_3235521, EPI_ISL_3235522, EPI_ISL_3235523, EPI_ISL_3235524, EPI_ISL_3235525, EPI_ISL_3235526, EPI_ISL_3235527, EPI_ISL_3235528, EPI_ISL_3235529 | | | |
| see above | Missouri State Public Health Laboratory | Missouri State Public Health Laboratory | Matthew Sinn, Joshua Barry, Ashley New |
| EPI_ISL_3235530, EPI_ISL_3235531, EPI_ISL_3235532, EPI_ISL_3235533, EPI_ISL_3235534, EPI_ISL_3235535, EPI_ISL_3235536, EPI_ISL_3235537, EPI_ISL_3235538, EPI_ISL_3235539, EPI_ISL_3235540, EPI_ISL_3235541, EPI_ISL_3235542, EPI_ISL_3235543, EPI_ISL_3235544, EPI_ISL_3235545, EPI_ISL_3235546, EPI_ISL_3235547, EPI_ISL_3235548, EPI_ISL_3235549, EPI_ISL_3235550, EPI_ISL_3235551, EPI_ISL_3235552, EPI_ISL_3235553, EPI_ISL_3235554, EPI_ISL_3235555, EPI_ISL_3235556, EPI_ISL_3235557, EPI_ISL_3235558, EPI_ISL_3235559, EPI_ISL_3235560 | | | |
| see above | Kansas Health and Environmental Lab | Kansas Health and Environmental Lab | Mike Grose, Jonathan Barnell, Ben Olsen, and Phil Adam |
| EPI_ISL_3235561, EPI_ISL_3235562, EPI_ISL_3235563, EPI_ISL_3235564, EPI_ISL_3235565, EPI_ISL_3235566, EPI_ISL_3235567, EPI_ISL_3235568, EPI_ISL_3235570, EPI_ISL_3235572, EPI_ISL_3235573, EPI_ISL_3235574, EPI_ISL_3235575, EPI_ISL_3235576, EPI_ISL_3235577, EPI_ISL_3235578, EPI_ISL_3235579, EPI_ISL_3235580, EPI_ISL_3235581, EPI_ISL_3235582, EPI_ISL_3235583, EPI_ISL_3235584, EPI_ISL_3235585, EPI_ISL_3235586, EPI_ISL_3235587, EPI_ISL_3235588, EPI_ISL_3235589, EPI_ISL_3235590, EPI_ISL_3235591, EPI_ISL_3235592, EPI_ISL_3235593, EPI_ISL_3235594, EPI_ISL_3235595, EPI_ISL_3235596, EPI_ISL_3235597, EPI_ISL_3235598, EPI_ISL_3235599, EPI_ISL_3235600, EPI_ISL_3235601, EPI_ISL_3235602, EPI_ISL_3235603, EPI_ISL_3235604, EPI_ISL_3235606, EPI_ISL_3235607, EPI_ISL_3235608, EPI_ISL_3235609, EPI_ISL_3235610, EPI_ISL_3235611, EPI_ISL_3235612, EPI_ISL_3235613, EPI_ISL_3235614, EPI_ISL_3235615, EPI_ISL_3235616, EPI_ISL_3235617, EPI_ISL_3235618, EPI_ISL_3235619, EPI_ISL_3235620, EPI_ISL_3235621, EPI_ISL_3235622, EPI_ISL_3235623, EPI_ISL_3235624, EPI_ISL_3235625, EPI_ISL_3235626, EPI_ISL_3235627, EPI_ISL_3235630, EPI_ISL_3235631, EPI_ISL_3235632, EPI_ISL_3235633, EPI_ISL_3235634, EPI_ISL_3235635, EPI_ISL_3235636, EPI_ISL_3235637, EPI_ISL_3235638, EPI_ISL_3235639, EPI_ISL_3235640, EPI_ISL_3235642, EPI_ISL_3235643, EPI_ISL_3235644 | | | |
| see above | MD PHL | Maryland Department of Health Laboratories Administration | Eric N. Keller, Jillian Loomis, Kwang Low, Terence L. Moore, Ami Patel, and Robert Myers |
| EPI_ISL_3235645 | Tripler Army Medical Center | Tripler Army Medical Center | Rachael Downham, Kendra Carter, Jonathan D'Ambrozio, Catherine Uyehara |
| EPI_ISL_3235646, EPI_ISL_3235647, EPI_ISL_3235648, EPI_ISL_3235649, EPI_ISL_3235650, EPI_ISL_3235651, EPI_ISL_3235652 | ALBANY MEDICAL CENTER | Wadsworth Center, New York State Department of Health | Kirsten St. George, Daryl M. Lamson, Alexis Russell, Matthew Shudt, Melissa A Leisner, Jonathan Plitnick, Catharine Prussing, Navjot Singh, John Kelly, Erasmus Schneider, Erica Lasek-Nesselquist |
| EPI_ISL_3235653 | BIO-REFERENCE LABORATORIES | Wadsworth Center, New York State Department of Health | Kirsten St. George, Daryl M. Lamson, Alexis Russell, Matthew Shudt, Melissa A Leisner, Jonathan Plitnick, Catharine Prussing, Navjot Singh, John Kelly, Erasmus Schneider, Erica Lasek-Nesselquist |
| EPI_ISL_3235654, EPI_ISL_3235655, EPI_ISL_3235656, EPI_ISL_3235657, EPI_ISL_3235659, EPI_ISL_3235660, EPI_ISL_3235661, EPI_ISL_3235662, EPI_ISL_3235663, EPI_ISL_3235664, EPI_ISL_3235665, EPI_ISL_3235666 | | | |
| see above | BOSTON HEART DIAGNOSTICS CORP | Wadsworth Center, New York State Department of Health | Kirsten St. George, Daryl M. Lamson, Alexis Russell, Matthew Shudt, Melissa A Leisner, Jonathan Plitnick, Catharine Prussing, Navjot Singh, John Kelly, Erasmus Schneider, Erica Lasek-Nesselquist |
| EPI_ISL_3235667, EPI_ISL_3235668, EPI_ISL_3235669, EPI_ISL_3235670, EPI_ISL_3235671, EPI_ISL_3235672, EPI_ISL_3235673, EPI_ISL_3235674, EPI_ISL_3235675, EPI_ISL_3235676, EPI_ISL_3235677, EPI_ISL_3235678, EPI_ISL_3235679, EPI_ISL_3235680, EPI_ISL_3235681, EPI_ISL_3235682, EPI_ISL_3235683, EPI_ISL_3235684, EPI_ISL_3235685, EPI_ISL_3235687, EPI_ISL_3235688, EPI_ISL_3235689, EPI_ISL_3235690, EPI_ISL_3235691, EPI_ISL_3235692, EPI_ISL_3235693, EPI_ISL_3235694, EPI_ISL_3235695, EPI_ISL_3235696, EPI_ISL_3235698, EPI_ISL_3235700, EPI_ISL_3235702, EPI_ISL_3235703, EPI_ISL_3235704, EPI_ISL_3235706, EPI_ISL_3235707 | | | |
| see above | NORTHWELL HEALTH LABORATORIES | Wadsworth Center, New York State Department of Health | Kirsten St. George, Daryl M. Lamson, Alexis Russell, Matthew Shudt, Melissa A Leisner, Jonathan Plitnick, Catharine Prussing, Navjot Singh, John Kelly, Erasmus Schneider, Erica Lasek-Nesselquist |
| EPI_ISL_3235708 | Quest Diagnostics | Wadsworth Center, New York State Department of Health | Kirsten St. George, Daryl M. Lamson, Alexis Russell, Matthew Shudt, Melissa A Leisner, Jonathan Plitnick, Catharine Prussing, Navjot Singh, John Kelly, Erasmus Schneider, Erica Lasek-Nesselquist |
| EPI_ISL_3235709, EPI_ISL_3235710 | Saratoga Hospital Laboratory | Wadsworth Center, New York State Department of Health | Kirsten St. George, Daryl M. Lamson, Alexis Russell, Matthew Shudt, Melissa A Leisner, Jonathan Plitnick, Catharine Prussing, Navjot Singh, John Kelly, Erasmus Schneider, Erica Lasek-Nesselquist |
| EPI_ISL_3235712, EPI_ISL_3235713, EPI_ISL_3235714, EPI_ISL_3235715, EPI_ISL_3235716, EPI_ISL_3235717, EPI_ISL_3235719, EPI_ISL_3235720, | STONY BROOK UNIVERSITY HOSPITAL | Wadsworth Center, New York State Department of Health | Kirsten St. George, Daryl M. Lamson, Alexis Russell, Matthew Shudt, Melissa A Leisner, Jonathan Plitnick, Catharine Prussing, Navjot Singh, John Kelly, Erasmus Schneider, Erica Lasek-Nesselquist |

| | | | |
|---|---|---|---|
| EPI_ISL_3235721 | | | |
| EPI_ISL_3235723 | SUNRISE MEDICAL LABORATORIES | Wadsworth Center, New York State Department of Health | Kirsten St. George, Daryl M. Lamson, Alexis Russell, Matthew Shudt, Melissa A Leisner, Jonathan Plitnick, Catharine Prussing, Navjot Singh, John Kelly, Erasmus Schneider, Erica Lasek-Nesselquist |
| EPI_ISL_3235724, EPI_ISL_3235725 | SUNY UPSTATE MEDICAL UNIVERSITY | Wadsworth Center, New York State Department of Health | Kirsten St. George, Daryl M. Lamson, Alexis Russell, Matthew Shudt, Melissa A Leisner, Jonathan Plitnick, Catharine Prussing, Navjot Singh, John Kelly, Erasmus Schneider, Erica Lasek-Nesselquist |
| EPI_ISL_3235727, EPI_ISL_3235728, EPI_ISL_3235729 | UNIVERSITY HOSPITAL- EASTERN LONG ISLAND HOSPITAL LABORATORY | Wadsworth Center, New York State Department of Health | Kirsten St. George, Daryl M. Lamson, Alexis Russell, Matthew Shudt, Melissa A Leisner, Jonathan Plitnick, Catharine Prussing, Navjot Singh, John Kelly, Erasmus Schneider, Erica Lasek-Nesselquist |
| EPI_ISL_3235730, EPI_ISL_3235731 | URMC LABS | Wadsworth Center, New York State Department of Health | Kirsten St. George, Daryl M. Lamson, Alexis Russell, Matthew Shudt, Melissa A Leisner, Jonathan Plitnick, Catharine Prussing, Navjot Singh, John Kelly, Erasmus Schneider, Erica Lasek-Nesselquist |
| EPI_ISL_3235732 | WellNow Urgent Care PC - Cicero | Wadsworth Center, New York State Department of Health | Kirsten St. George, Daryl M. Lamson, Alexis Russell, Matthew Shudt, Melissa A Leisner, Jonathan Plitnick, Catharine Prussing, Navjot Singh, John Kelly, Erasmus Schneider, Erica Lasek-Nesselquist |
| EPI_ISL_3236102, EPI_ISL_3236103, EPI_ISL_3236104, EPI_ISL_3236105, EPI_ISL_3236106, EPI_ISL_3236107, EPI_ISL_3236108, EPI_ISL_3236109, EPI_ISL_3236110, EPI_ISL_3236111, EPI_ISL_3236112, EPI_ISL_3236113, EPI_ISL_3236114, EPI_ISL_3236115, EPI_ISL_3236116, EPI_ISL_3236117, EPI_ISL_3236118, EPI_ISL_3236119 | | | |
| see above | Stanford Health Care | Stanford University School of Medicine, Clinical Virology Laboratory | Bernadette Troung, Malaya K. Sahoo, Becky Jiang, Nathan Hammond, James Zehnder, and Benjamin A. Pinsky |
| EPI_ISL_3236120, EPI_ISL_3236121, EPI_ISL_3236122, EPI_ISL_3236123, EPI_ISL_3236124, EPI_ISL_3236125, EPI_ISL_3236126, EPI_ISL_3236127, EPI_ISL_3236128, EPI_ISL_3236129, EPI_ISL_3236130, EPI_ISL_3236131, EPI_ISL_3236133, EPI_ISL_3236134, EPI_ISL_3236135, EPI_ISL_3236136, EPI_ISL_3236137, EPI_ISL_3236138, EPI_ISL_3236139 | | | |
| see above | Stanford Health Care | Stanford University School of Medicine, Clinical Virology Laboratory | Malaya K. Sahoo, Bernadette Troung, Becky Jiang, Nathan Hammond, James Zehnder, and Benjamin A. Pinsky |
| EPI_ISL_3236141, EPI_ISL_3236142, EPI_ISL_3236143, EPI_ISL_3236144, EPI_ISL_3236145, EPI_ISL_3236146, EPI_ISL_3236147, EPI_ISL_3236148, EPI_ISL_3236149, EPI_ISL_3236150, EPI_ISL_3236151, EPI_ISL_3236152, EPI_ISL_3236153, EPI_ISL_3236154, EPI_ISL_3236155, EPI_ISL_3236156, EPI_ISL_3236158, EPI_ISL_3236159, EPI_ISL_3236160, EPI_ISL_3236161 | | | |
| see above | Stanford Health Care | Stanford University School of Medicine, Clinical Virology Laboratory | Becky Jiang, Bernadette Troung, Malaya K. Sahoo, Nathan Hammond, James Zehnder, and Benjamin A. Pinsky |
| EPI_ISL_3236163, EPI_ISL_3236164, EPI_ISL_3236165, EPI_ISL_3236166, EPI_ISL_3236167, EPI_ISL_3236168, EPI_ISL_3236169, EPI_ISL_3236170, EPI_ISL_3236171, EPI_ISL_3236172, EPI_ISL_3236173, EPI_ISL_3236174, EPI_ISL_3236175, EPI_ISL_3236176, EPI_ISL_3236177, EPI_ISL_3236178, EPI_ISL_3236179, EPI_ISL_3236180, EPI_ISL_3236181, EPI_ISL_3236182, EPI_ISL_3236183, EPI_ISL_3236184, EPI_ISL_3236185 | | | |
| see above | Stanford Health Care | Stanford University School of Medicine, Clinical Virology Laboratory | Malaya K. Sahoo, Becky Jiang, Bernadette Troung, Nathan Hammond, James Zehnder, and Benjamin A. Pinsky |
| EPI_ISL_3236191, EPI_ISL_3236192, EPI_ISL_3236196, EPI_ISL_3236197, EPI_ISL_3236198, EPI_ISL_3236199 | Genome Information Research Center, Research Institute for Microbial Diseases, Osaka University | Genome Information Research Center, Research Institute for Microbial Diseases, Osaka University | Daisuke Motooka, Kentaro Tanaka, Hiroya Oki, and Shota Nakamura |
| EPI_ISL_3236287, EPI_ISL_3236288, EPI_ISL_3236289, EPI_ISL_3236290, EPI_ISL_3236291, EPI_ISL_3236292, EPI_ISL_3236293, EPI_ISL_3236294, EPI_ISL_3236295, EPI_ISL_3236296, EPI_ISL_3236297, EPI_ISL_3236298, EPI_ISL_3236299, EPI_ISL_3236300, EPI_ISL_3236301, EPI_ISL_3236302, EPI_ISL_3236303, EPI_ISL_3236304, EPI_ISL_3236305, EPI_ISL_3236306, EPI_ISL_3236307, EPI_ISL_3236308, EPI_ISL_3236309, EPI_ISL_3236310, EPI_ISL_3236311, EPI_ISL_3236312, EPI_ISL_3236313, EPI_ISL_3236314, EPI_ISL_3236315, EPI_ISL_3236316, EPI_ISL_3236317, EPI_ISL_3236318, EPI_ISL_3236319, EPI_ISL_3236320, EPI_ISL_3236321, EPI_ISL_3236322, EPI_ISL_3236323, EPI_ISL_3236324, EPI_ISL_3236325, EPI_ISL_3236326, EPI_ISL_3236327, EPI_ISL_3236328, EPI_ISL_3236329, EPI_ISL_3236330, EPI_ISL_3236331, EPI_ISL_3236332, EPI_ISL_3236334, EPI_ISL_3236335, EPI_ISL_3236336, EPI_ISL_3236338, EPI_ISL_3236339 | | | |
| see above | Hospital Universitari Vall d'Hebron - Vall d'Hebron Institut de Recerca | Hospital Universitari Vall d'Hebron - Vall d'Hebron Institut de Recerca | Cristina Andrés, Maria Piñana, Alejandra González-Sánchez, Damir Garcia-Cehic, Ariadna Rando, Juliana Esperalba, Maria Gema Codina, Carla Castillo, Karen García, Rodrigo Vásquez, Maria Carmen Martin, Tomàs Pumarola, Josep Quer, Andrés Antón |
| EPI_ISL_3236340, EPI_ISL_3236341, EPI_ISL_3236342, EPI_ISL_3236343, EPI_ISL_3236344, EPI_ISL_3236345, EPI_ISL_3236346, EPI_ISL_3236347, EPI_ISL_3236348, EPI_ISL_3236349, EPI_ISL_3236350, EPI_ISL_3236351, EPI_ISL_3236352, EPI_ISL_3236353, EPI_ISL_3236354, EPI_ISL_3236355, EPI_ISL_3236356, EPI_ISL_3236357, EPI_ISL_3236358 | | | |
| see above | Hospital Universitari Arnau de Vilanova | Hospital Universitari Vall d'Hebron - Vall d'Hebron Institut de Recerca | Cristina Andrés, Maria Piñana, Alejandra González-Sánchez, Damir Garcia-Cehic, Ariadna Rando, Juliana Esperalba, Maria Gema Codina, Carla Castillo, Karen García, Rodrigo Vásquez, Maria Carmen Martin, Tomàs Pumarola, Josep Quer, Andrés Antón |
| EPI_ISL_3236362, EPI_ISL_3236363, EPI_ISL_3236364, EPI_ISL_3236365, EPI_ISL_3236366, EPI_ISL_3236367, EPI_ISL_3236368, EPI_ISL_3236370, EPI_ISL_3236371, EPI_ISL_3236372, EPI_ISL_3236373, EPI_ISL_3236374, EPI_ISL_3236375, EPI_ISL_3236376, EPI_ISL_3236379, EPI_ISL_3236380, EPI_ISL_3236381, EPI_ISL_3236382, EPI_ISL_3236384, EPI_ISL_3236386, EPI_ISL_3236388, EPI_ISL_3236390, EPI_ISL_3236391, EPI_ISL_3236392, EPI_ISL_3236393, EPI_ISL_3236395, EPI_ISL_3236397, EPI_ISL_3236398, EPI_ISL_3236399, EPI_ISL_3236400, EPI_ISL_3236401, EPI_ISL_3236402, EPI_ISL_3236403, EPI_ISL_3236404, EPI_ISL_3236405, EPI_ISL_3236406, EPI_ISL_3236408, EPI_ISL_3236410, EPI_ISL_3236411, EPI_ISL_3236413, EPI_ISL_3236414, EPI_ISL_3236415, EPI_ISL_3236416, EPI_ISL_3236418, EPI_ISL_3236419, EPI_ISL_3236420, EPI_ISL_3236421, EPI_ISL_3236422, EPI_ISL_3236423, EPI_ISL_3236424, EPI_ISL_3236425, EPI_ISL_3236426, EPI_ISL_3236427, EPI_ISL_3236428, EPI_ISL_3236429, EPI_ISL_3236430, EPI_ISL_3236431, EPI_ISL_3236432, EPI_ISL_3236453, EPI_ISL_3236454, EPI_ISL_3236455, EPI_ISL_3236456, EPI_ISL_3236457, EPI_ISL_3236458, EPI_ISL_3236459, EPI_ISL_3236460, EPI_ISL_3236461, EPI_ISL_3236462, EPI_ISL_3236463, EPI_ISL_3236464, EPI_ISL_3236467, EPI_ISL_3236468, EPI_ISL_3236469, EPI_ISL_3236470, EPI_ISL_3236472, EPI_ISL_3236473, EPI_ISL_3236474, EPI_ISL_3236475, EPI_ISL_3236476, EPI_ISL_3236477, EPI_ISL_3236478, EPI_ISL_3236479, EPI_ISL_3236481, EPI_ISL_3236482, EPI_ISL_3236483, EPI_ISL_3236484, EPI_ISL_3236485, EPI_ISL_3236486, EPI_ISL_3236487, EPI_ISL_3236488, EPI_ISL_3236489, EPI_ISL_3236490, EPI_ISL_3236491, EPI_ISL_3236492, EPI_ISL_3236493, EPI_ISL_3236494 | | | |
| see above | Yale Clinical Virology Lab | Grubaugh Lab - Yale School of Public Health | Joseph Fauver, Mallery Breban, Isabel Ott, Tara Alpert, Mary Petrone, Anderson Brito, Chantal Vogels, Annie Watkins, Chaney Kalinich, Jessica Rothman, Kendall Billig, Tobias Koch, Marie L. Landry, Nathan Grubaugh |
| EPI_ISL_3236495, EPI_ISL_3236496, EPI_ISL_3236497, EPI_ISL_3236498, EPI_ISL_3236500, EPI_ISL_3236505, EPI_ISL_3236506, EPI_ISL_3236510, EPI_ISL_3236512, EPI_ISL_3236513, EPI_ISL_3236514, EPI_ISL_3236515, EPI_ISL_3236517, EPI_ISL_3236518, EPI_ISL_3236519, EPI_ISL_3236520 | | | |
| see above | VIDOH (Virgin Islands Department of Health) | Grubaugh Lab - Yale School of Public Health | Joseph Fauver, Mallery Breban, Isabel Ott, Tara Alpert, Mary Petrone, Anderson Brito, Chantal Vogels, Annie Watkins, Chaney Kalinich, Jessica Rothman, Kendall Billig, Tobias Koch, Jendai Richards, TaLesa Aderohunmu, Esther Ellis, Marlon Lawrence, Brett Ellis, Nathan Grubaugh |
| EPI_ISL_3236524, EPI_ISL_3236527, EPI_ISL_3236528, EPI_ISL_3236529, EPI_ISL_3236530, EPI_ISL_3236531, EPI_ISL_3236532, EPI_ISL_3236533, EPI_ISL_3236534, EPI_ISL_3236535, EPI_ISL_3236536, EPI_ISL_3236537, EPI_ISL_3236538, EPI_ISL_3236539, EPI_ISL_3236540, EPI_ISL_3236541, EPI_ISL_3236542, EPI_ISL_3236543, EPI_ISL_3236544, EPI_ISL_3236545, EPI_ISL_3236546, EPI_ISL_3236547, EPI_ISL_3236548, EPI_ISL_3236549, EPI_ISL_3236550, EPI_ISL_3236551, EPI_ISL_3236552, EPI_ISL_3236553, EPI_ISL_3236555, EPI_ISL_3236556, EPI_ISL_3236557, EPI_ISL_3236558, EPI_ISL_3236559, EPI_ISL_3236560, EPI_ISL_3236561, EPI_ISL_3236563, EPI_ISL_3236564, EPI_ISL_3236565, EPI_ISL_3236566, EPI_ISL_3236567, EPI_ISL_3236568, EPI_ISL_3236569, EPI_ISL_3236570, EPI_ISL_3236571, EPI_ISL_3236572 | | | |
| see above | UCC (Universidad Central del Caribe) | Grubaugh Lab - Yale School of Public Health | Joseph Fauver, Mallery Breban, Isabel Ott, Tara Alpert, Mary Petrone, Anderson Brito, Chantal Vogels, Annie Watkins, Chaney Kalinich, Jessica Rothman, Kendall Billig, Tobias Koch, Fabiola Cruz-Lopez, David Rivera Aponte, Fabiola Fontanet, Natalie Machargo, Andrea Firpo, Vianca Aponte, Ernesto Vazquez, Patricia Serrano, Tania Mitwalli, Karolane Gonzalez, Nathan Grubaugh |
| EPI_ISL_3236573, EPI_ISL_3236574, EPI_ISL_3236575, EPI_ISL_3236576, EPI_ISL_3236577, EPI_ISL_3236578, EPI_ISL_3236579, EPI_ISL_3236580, EPI_ISL_3236581, EPI_ISL_3236582, EPI_ISL_3236583, EPI_ISL_3236584 | | | |
| see above | Yale Pathology Lab | Grubaugh Lab - Yale School of Public Health | Joseph Fauver, Mallery Breban, Isabel Ott, Tara Alpert, Mary Petrone, Anderson Brito, Chantal Vogels, Annie Watkins, Chaney Kalinich, Jessica Rothman, Kendall Billig, Tobias Koch, Jianhui Wang, Chen Liu, Pei Hui, Nathan Grubaugh |
| EPI_ISL_3236586, EPI_ISL_3236587, EPI_ISL_3236588, EPI_ISL_3236589, EPI_ISL_3236590, EPI_ISL_3236591, EPI_ISL_3236593, EPI_ISL_3236594, EPI_ISL_3236595, EPI_ISL_3236596, EPI_ISL_3236597, EPI_ISL_3236598, EPI_ISL_3236600, EPI_ISL_3236601, EPI_ISL_3236602, EPI_ISL_3236603, EPI_ISL_3236604, EPI_ISL_3236605, EPI_ISL_3236606, EPI_ISL_3236607, EPI_ISL_3236608, EPI_ISL_3236609, EPI_ISL_3236610, EPI_ISL_3236611, EPI_ISL_3236612, EPI_ISL_3236613, EPI_ISL_3236615, EPI_ISL_3236616, EPI_ISL_3236617, EPI_ISL_3236618, EPI_ISL_3236619, EPI_ISL_3236620, EPI_ISL_3236621, EPI_ISL_3236622, EPI_ISL_3236624, EPI_ISL_3236625, EPI_ISL_3236626, EPI_ISL_3236627, EPI_ISL_3236628, EPI_ISL_3236629, EPI_ISL_3236630, EPI_ISL_3236631, EPI_ISL_3236632, EPI_ISL_3236633, EPI_ISL_3236634, EPI_ISL_3236636, EPI_ISL_3236637, EPI_ISL_3236638, EPI_ISL_3236639, EPI_ISL_3236640, EPI_ISL_3236641, EPI_ISL_3236642, EPI_ISL_3236643, EPI_ISL_3236644, EPI_ISL_3236645, EPI_ISL_3236646, EPI_ISL_3236647, EPI_ISL_3236648, EPI_ISL_3236649, EPI_ISL_3236651, EPI_ISL_3236652, EPI_ISL_3236653, EPI_ISL_3236654, EPI_ISL_3236655, EPI_ISL_3236656, EPI_ISL_3236657, EPI_ISL_3236658, EPI_ISL_3236659, EPI_ISL_3236660, EPI_ISL_3236661, EPI_ISL_3236662, EPI_ISL_3236663, EPI_ISL_3236664, EPI_ISL_3236665, EPI_ISL_3236666, EPI_ISL_3236667, EPI_ISL_3236668, EPI_ISL_3236669, EPI_ISL_3236670, EPI_ISL_3236671, EPI_ISL_3236672, EPI_ISL_3236673, EPI_ISL_3236674, EPI_ISL_3236675, EPI_ISL_3236676, EPI_ISL_3236677, EPI_ISL_3236678, EPI_ISL_3236679, EPI_ISL_3236680, EPI_ISL_3236681, EPI_ISL_3236682, EPI_ISL_3236683, EPI_ISL_3236684, EPI_ISL_3236685, EPI_ISL_3236686, EPI_ISL_3236687, EPI_ISL_3236688, EPI_ISL_3236689, EPI_ISL_3236690, EPI_ISL_3236691, EPI_ISL_3236692, EPI_ISL_3236693, EPI_ISL_3236695, EPI_ISL_3236696, EPI_ISL_3236697, EPI_ISL_3236698, EPI_ISL_3236699, EPI_ISL_3236700, EPI_ISL_3236701, EPI_ISL_3236702, EPI_ISL_3236703, EPI_ISL_3236704, EPI_ISL_3236706, EPI_ISL_3236707, EPI_ISL_3236708, EPI_ISL_3236709, EPI_ISL_3236710, EPI_ISL_3236711, EPI_ISL_3236712, EPI_ISL_3236713, EPI_ISL_3236714, EPI_ISL_3236715, EPI_ISL_3236716, EPI_ISL_3236717, EPI_ISL_3236718, EPI_ISL_3236719, EPI_ISL_3236720, EPI_ISL_3236721, EPI_ISL_3236722, EPI_ISL_3236723, EPI_ISL_3236724, EPI_ISL_3236725, EPI_ISL_3236726, EPI_ISL_3236727, EPI_ISL_3236728, EPI_ISL_3236729, EPI_ISL_3236730, EPI_ISL_3236731, EPI_ISL_3236732, EPI_ISL_3236733, EPI_ISL_3236734, EPI_ISL_3236736, EPI_ISL_3236737, EPI_ISL_3236739, EPI_ISL_3236740, EPI_ISL_3236741, EPI_ISL_3236743, EPI_ISL_3236744, EPI_ISL_3236745, EPI_ISL_3236746, EPI_ISL_3236747, EPI_ISL_3236748, EPI_ISL_3236749, EPI_ISL_3236750, EPI_ISL_3236751, EPI_ISL_3236752, EPI_ISL_3236753, EPI_ISL_3236755, EPI_ISL_3236756, EPI_ISL_3236757, EPI_ISL_3236758, EPI_ISL_3236759, EPI_ISL_3236760, EPI_ISL_3236761, EPI_ISL_3236762, EPI_ISL_3236763, EPI_ISL_3236764, EPI_ISL_3236765, EPI_ISL_3236766, EPI_ISL_3236767, EPI_ISL_3236768, EPI_ISL_3236769, EPI_ISL_3236770, EPI_ISL_3236771, EPI_ISL_3236772, EPI_ISL_3236773, EPI_ISL_3236774, EPI_ISL_3236775, EPI_ISL_3236776, EPI_ISL_3236777, EPI_ISL_3236779, EPI_ISL_3236780, EPI_ISL_3236781, EPI_ISL_3236782, EPI_ISL_3236783, EPI_ISL_3236784, EPI_ISL_3236785, EPI_ISL_3236787, EPI_ISL_3236788, EPI_ISL_3236789, EPI_ISL_3236790, EPI_ISL_3236791, EPI_ISL_3236792, EPI_ISL_3236793, EPI_ISL_3236794, EPI_ISL_3236795, EPI_ISL_3236796, EPI_ISL_3236797, EPI_ISL_3236798, EPI_ISL_3236799, EPI_ISL_3236800, EPI_ISL_3236801, EPI_ISL_3236802, EPI_ISL_3236803, EPI_ISL_3236804, EPI_ISL_3236805, EPI_ISL_3236806, EPI_ISL_3236807, EPI_ISL_3236808, EPI_ISL_3236810, EPI_ISL_3236811, EPI_ISL_3236812, EPI_ISL_3236813, EPI_ISL_3236814, EPI_ISL_3236815, EPI_ISL_3236816, EPI_ISL_3236817, EPI_ISL_3236818, EPI_ISL_3236819, EPI_ISL_3236820, EPI_ISL_3236821, EPI_ISL_3236822, EPI_ISL_3236824, EPI_ISL_3236825, EPI_ISL_3236826, EPI_ISL_3236827, EPI_ISL_3236829, EPI_ISL_3236830, EPI_ISL_3236831, EPI_ISL_3236832, EPI_ISL_3236833, EPI_ISL_3236834, EPI_ISL_3236835, EPI_ISL_3236836, EPI_ISL_3236837, EPI_ISL_3236838, EPI_ISL_3236840, EPI_ISL_3236841, EPI_ISL_3236842, EPI_ISL_3236843, EPI_ISL_3236844, EPI_ISL_3236845, EPI_ISL_3236846, EPI_ISL_3236847, EPI_ISL_3236848, EPI_ISL_3236849, EPI_ISL_3236850, EPI_ISL_3236851, EPI_ISL_3236852, EPI_ISL_3236853, EPI_ISL_3236854, EPI_ISL_3236855, EPI_ISL_3236856, EPI_ISL_3236857, EPI_ISL_3236858, EPI_ISL_3236859, EPI_ISL_3236860, EPI_ISL_3236861, EPI_ISL_3236862, EPI_ISL_3236863, EPI_ISL_3236864, EPI_ISL_3236865, EPI_ISL_3236866, EPI_ISL_3236867, EPI_ISL_3236868, EPI_ISL_3236869, EPI_ISL_3236870, EPI_ISL_3236871, EPI_ISL_3236873, EPI_ISL_3236874, EPI_ISL_3236875, EPI_ISL_3236876, EPI_ISL_3236877, EPI_ISL_3236878, EPI_ISL_3236880, EPI_ISL_3236881, EPI_ISL_3236882, EPI_ISL_3236883, EPI_ISL_3236884, EPI_ISL_3236885, EPI_ISL_3236886, EPI_ISL_3236887, EPI_ISL_3236888, EPI_ISL_3236889, EPI_ISL_3236890, EPI_ISL_3236891, EPI_ISL_3236892, EPI_ISL_3236893, EPI_ISL_3236894, EPI_ISL_3236895 | | | |
| see above | UW Virology Lab | UW Virology Lab | Pavitra Roychoudhury, Shah Mohamed Bakhash, Ricardo Perez, Hong Xie, Tien V. Nguyen, Maria Lukes, Sean Ellis, Nathan Breit, Robert J. Livingston, Meei-Li Huang, Keith R Jerome, Patrick Mathias, Alexander Greninger |
| EPI_ISL_3236931, EPI_ISL_3236932, EPI_ISL_3236933, EPI_ISL_3236934, EPI_ISL_3236935, EPI_ISL_3236936, EPI_ISL_3236937, EPI_ISL_3236938, EPI_ISL_3236939, EPI_ISL_3236940, EPI_ISL_3236941, EPI_ISL_3236942, EPI_ISL_3236943, EPI_ISL_3236944, EPI_ISL_3236945, EPI_ISL_3236946, EPI_ISL_3236947, EPI_ISL_3236948, EPI_ISL_3236949, EPI_ISL_3236950, EPI_ISL_3236952, EPI_ISL_3236954 | | | |
| see above | TAMBO MEMORIAL LABORATORY | National Institute for Communicable Diseases of the National Health Laboratory Service | Amoako DG, Everatt J, Scheepers C, Mohale T, Ntuli N, Mahlangu B, Mnguni A, Ismail A, Bhiman JN |
| EPI_ISL_3236956 | KOPANONG LABORATORY | National Institute for Communicable Diseases of the National Health Laboratory Service | Amoako DG, Everatt J, Scheepers C, Mohale T, Ntuli N, Mahlangu B, Mnguni A, Ismail A, Bhiman JN |
| EPI_ISL_3236957, EPI_ISL_3236958 | LANCET LABORATORY | National Institute for Communicable Diseases of the National Health Laboratory Service | Amoako DG, Everatt J, Scheepers C, Mohale T, Ntuli N, Mahlangu B, Mnguni A, Ismail A, Bhiman JN |
| EPI_ISL_3236960 | MANKWENG LABORATORY | National Institute for Communicable Diseases of the National Health Laboratory Service | Amoako DG, Everatt J, Scheepers C, Mohale T, Ntuli N, Mahlangu B, Mnguni A, Ismail A, Bhiman JN |
| EPI_ISL_3236961, EPI_ISL_3236963, EPI_ISL_3236964, EPI_ISL_3236965, EPI_ISL_3236966, EPI_ISL_3236967, EPI_ISL_3236968, EPI_ISL_3236969, EPI_ISL_3236970, EPI_ISL_3236971, EPI_ISL_3236973, EPI_ISL_3236974, EPI_ISL_3236975, EPI_ISL_3236977, EPI_ISL_3236979, EPI_ISL_3236980 | | | |
| see above | TSHEPONG LABORATORY | National Institute for Communicable Diseases of the National Health Laboratory Service | Amoako DG, Everatt J, Scheepers C, Mohale T, Ntuli N, Mahlangu B, Mnguni A, Ismail A, Bhiman JN |
| EPI_ISL_3236982, EPI_ISL_3236983, EPI_ISL_3236984, EPI_ISL_3236985 | RUSTENBURG LABORATORY | National Institute for Communicable Diseases of the National Health Laboratory Service | Amoako DG, Everatt J, Scheepers C, Mohale T, Ntuli N, Mahlangu B, Mnguni A, Ismail A, Bhiman JN |
| EPI_ISL_3236986, EPI_ISL_3236987, EPI_ISL_3236988 | Pathcare Vaal laboratory | National Institute for Communicable Diseases of the National Health Laboratory Service | Amoako DG, Everatt J, Scheepers C, Mohale T, Ntuli N, Mahlangu B, Mnguni A, Ismail A, Bhiman JN |
| EPI_ISL_3236989, EPI_ISL_3236990, EPI_ISL_3236991, EPI_ISL_3236992 | LANCET LABORATORY | National Institute for Communicable Diseases of the National Health Laboratory Service | Amoako DG, Everatt J, Scheepers C, Mohale T, Ntuli N, Mahlangu B, Mnguni A, Ismail A, Bhiman JN |
| EPI_ISL_3236993, EPI_ISL_3236995, EPI_ISL_3236997, EPI_ISL_3237001 | RUSTENBURG LABORATORY | National Institute for Communicable Diseases of the National Health Laboratory Service | Amoako DG, Everatt J, Scheepers C, Mohale T, Ntuli N, Mahlangu B, Mnguni A, Ismail A, Bhiman JN |
| EPI_ISL_3237004, EPI_ISL_3237005, EPI_ISL_3237006, EPI_ISL_3237007, EPI_ISL_3237008, EPI_ISL_3237009, EPI_ISL_3237010, EPI_ISL_3237011, EPI_ISL_3237012, EPI_ISL_3237013, EPI_ISL_3237014, EPI_ISL_3237015, EPI_ISL_3237016, EPI_ISL_3237017, EPI_ISL_3237018, EPI_ISL_3237019, EPI_ISL_3237020 | | | |

| | | | |
|---|---|---|---|
| see above | PORT ELIZABETH | National Institute for Communicable Diseases of the National Health Laboratory Service | Amoako DG, Everatt J, Scheepers C, Mohale T, Ntuli N, Mahlangu B, Mnguni A, Ismail A, Bhiman JN |
| EPI_ISL_3237021, EPI_ISL_3237022, EPI_ISL_3237023, EPI_ISL_3237024, EPI_ISL_3237025, EPI_ISL_3237026, EPI_ISL_3237027, EPI_ISL_3237028, EPI_ISL_3237029, EPI_ISL_3237030, EPI_ISL_3237031, EPI_ISL_3237032 | | | |
| see above | ROB FERREIRA LABORATORY | National Institute for Communicable Diseases of the National Health Laboratory Service | Amoako DG, Everatt J, Scheepers C, Mohale T, Ntuli N, Mahlangu B, Mnguni A, Ismail A, Bhiman JN |
| EPI_ISL_3237033, EPI_ISL_3237034 | LANCET LABORATORY | National Institute for Communicable Diseases of the National Health Laboratory Service | Amoako DG, Everatt J, Scheepers C, Mohale T, Ntuli N, Mahlangu B, Mnguni A, Ismail A, Bhiman JN |
| EPI_ISL_3237041, EPI_ISL_3237043 | University Pretoria | National Institute for Communicable Diseases of the National Health Laboratory Service | Amy Strydom, Adriano Mendes, Micheala Davids, Sim Mayaphi, Marietjie Venter, Amoako DG, Everatt J, Scheepers C, Mohale T, Ntuli N, Mahlangu B, Mnguni A, Ismail A, Bhiman JN |
| EPI_ISL_3237051, EPI_ISL_3237052, EPI_ISL_3237053 | TAMBO MEMORIAL LABORATORY | National Institute for Communicable Diseases of the National Health Laboratory Service | Amoako DG, Everatt J, Scheepers C, Mohale T, Ntuli N, Mahlangu B, Mnguni A, Ismail A, Bhiman JN |
| EPI_ISL_3237054 | BERTHA GXOWA LABORATORY | National Institute for Communicable Diseases of the National Health Laboratory Service | Amoako DG, Everatt J, Scheepers C, Mohale T, Ntuli N, Mahlangu B, Mnguni A, Ismail A, Bhiman JN |
| EPI_ISL_3237055, EPI_ISL_3237056, EPI_ISL_3237057, EPI_ISL_3237059, EPI_ISL_3237060, EPI_ISL_3237061, EPI_ISL_3237062, EPI_ISL_3237064, EPI_ISL_3237065, EPI_ISL_3237066, EPI_ISL_3237067, EPI_ISL_3237068, EPI_ISL_3237069, EPI_ISL_3237070, EPI_ISL_3237071, EPI_ISL_3237073, EPI_ISL_3237074, EPI_ISL_3237075, EPI_ISL_3237076, EPI_ISL_3237077, EPI_ISL_3237078, EPI_ISL_3237079, EPI_ISL_3237080, EPI_ISL_3237081, EPI_ISL_3237082, EPI_ISL_3237083, EPI_ISL_3237085, EPI_ISL_3237086, EPI_ISL_3237087, EPI_ISL_3237088, EPI_ISL_3237089, EPI_ISL_3237090, EPI_ISL_3237091, EPI_ISL_3237093, EPI_ISL_3237094, EPI_ISL_3237095, EPI_ISL_3237096, EPI_ISL_3237097, EPI_ISL_3237099, EPI_ISL_3237101, EPI_ISL_3237102, EPI_ISL_3237103 | | | |
| see above | TAMBO MEMORIAL LABORATORY | National Institute for Communicable Diseases of the National Health Laboratory Service | Amoako DG, Everatt J, Scheepers C, Mohale T, Ntuli N, Mahlangu B, Mnguni A, Ismail A, Bhiman JN |
| EPI_ISL_3237104, EPI_ISL_3237105, EPI_ISL_3237106 | KOPANONG LABORATORY | National Institute for Communicable Diseases of the National Health Laboratory Service | Amoako DG, Everatt J, Scheepers C, Mohale T, Ntuli N, Mahlangu B, Mnguni A, Ismail A, Bhiman JN |
| EPI_ISL_3237108, EPI_ISL_3237109, EPI_ISL_3237110, EPI_ISL_3237111, EPI_ISL_3237112 | LANCET LABORATORY | National Institute for Communicable Diseases of the National Health Laboratory Service | Amoako DG, Everatt J, Scheepers C, Mohale T, Ntuli N, Mahlangu B, Mnguni A, Ismail A, Bhiman JN |
| EPI_ISL_3237114 | MANKWENG LABORATORY | National Institute for Communicable Diseases of the National Health Laboratory Service | Amoako DG, Everatt J, Scheepers C, Mohale T, Ntuli N, Mahlangu B, Mnguni A, Ismail A, Bhiman JN |
| EPI_ISL_3237116, EPI_ISL_3237117, EPI_ISL_3237118, EPI_ISL_3237119, EPI_ISL_3237120, EPI_ISL_3237122, EPI_ISL_3237124, EPI_ISL_3237125, EPI_ISL_3237126, EPI_ISL_3237127, EPI_ISL_3237128, EPI_ISL_3237129, EPI_ISL_3237131, EPI_ISL_3237132, EPI_ISL_3237133, EPI_ISL_3237137, EPI_ISL_3237138, EPI_ISL_3237139, EPI_ISL_3237140, EPI_ISL_3237141, EPI_ISL_3237142, EPI_ISL_3237143, EPI_ISL_3237145, EPI_ISL_3237146, EPI_ISL_3237147, EPI_ISL_3237148, EPI_ISL_3237149 | | | |
| see above | TSHEPONG LABORATORY | National Institute for Communicable Diseases of the National Health Laboratory Service | Amoako DG, Everatt J, Scheepers C, Mohale T, Ntuli N, Mahlangu B, Mnguni A, Ismail A, Bhiman JN |
| EPI_ISL_3237150, EPI_ISL_3237151, EPI_ISL_3237152, EPI_ISL_3237153 | RUSTENBURG LABORATORY | National Institute for Communicable Diseases of the National Health Laboratory Service | Amoako DG, Everatt J, Scheepers C, Mohale T, Ntuli N, Mahlangu B, Mnguni A, Ismail A, Bhiman JN |
| EPI_ISL_3237154, EPI_ISL_3237155, EPI_ISL_3237156, EPI_ISL_3237158, EPI_ISL_3237159, EPI_ISL_3237160, EPI_ISL_3237161, EPI_ISL_3237162, EPI_ISL_3237163, EPI_ISL_3237164, EPI_ISL_3237166 | | | |
| see above | Pathcare Vaal laboratory | National Institute for Communicable Diseases of the National Health Laboratory Service | Amoako DG, Everatt J, Scheepers C, Mohale T, Ntuli N, Mahlangu B, Mnguni A, Ismail A, Bhiman JN |
| EPI_ISL_3237167, EPI_ISL_3237168, EPI_ISL_3237169, EPI_ISL_3237170, EPI_ISL_3237171, EPI_ISL_3237172 | LANCET LABORATORY | National Institute for Communicable Diseases of the National Health Laboratory Service | Amoako DG, Everatt J, Scheepers C, Mohale T, Ntuli N, Mahlangu B, Mnguni A, Ismail A, Bhiman JN |
| EPI_ISL_3237173, EPI_ISL_3237177, EPI_ISL_3237178, EPI_ISL_3237179, EPI_ISL_3237195 | RUSTENBURG LABORATORY | National Institute for Communicable Diseases of the National Health Laboratory Service | Amoako DG, Everatt J, Scheepers C, Mohale T, Ntuli N, Mahlangu B, Mnguni A, Ismail A, Bhiman JN |
| EPI_ISL_3237197, EPI_ISL_3237198, EPI_ISL_3237199, EPI_ISL_3237200, EPI_ISL_3237201, EPI_ISL_3237202, EPI_ISL_3237203, EPI_ISL_3237204, EPI_ISL_3237205, EPI_ISL_3237206, EPI_ISL_3237207, EPI_ISL_3237208, EPI_ISL_3237209, EPI_ISL_3237210, EPI_ISL_3237211, EPI_ISL_3237212, EPI_ISL_3237213, EPI_ISL_3237214, EPI_ISL_3237215, EPI_ISL_3237216, EPI_ISL_3237217, EPI_ISL_3237218, EPI_ISL_3237219, EPI_ISL_3237220, EPI_ISL_3237221 | | | |
| see above | PORT ELIZABETH | National Institute for Communicable Diseases of the National Health Laboratory Service | Amoako DG, Everatt J, Scheepers C, Mohale T, Ntuli N, Mahlangu B, Mnguni A, Ismail A, Bhiman JN |
| EPI_ISL_3237222, EPI_ISL_3237224, EPI_ISL_3237225, EPI_ISL_3237226, EPI_ISL_3237227, EPI_ISL_3237228, EPI_ISL_3237229, EPI_ISL_3237230, EPI_ISL_3237231, EPI_ISL_3237232, EPI_ISL_3237234 | | | |
| see above | ROB FERREIRA LABORATORY | National Institute for Communicable Diseases of the National Health Laboratory Service | Amoako DG, Everatt J, Scheepers C, Mohale T, Ntuli N, Mahlangu B, Mnguni A, Ismail A, Bhiman JN |
| EPI_ISL_3237235, EPI_ISL_3237236, EPI_ISL_3237238, EPI_ISL_3237239, EPI_ISL_3237240 | LANCET LABORATORY | National Institute for Communicable Diseases of the National Health Laboratory Service | Amoako DG, Everatt J, Scheepers C, Mohale T, Ntuli N, Mahlangu B, Mnguni A, Ismail A, Bhiman JN |
| EPI_ISL_3237243, EPI_ISL_3237245, EPI_ISL_3237250, EPI_ISL_3237251 | University Pretoria | National Institute for Communicable Diseases of the National Health Laboratory Service | Amy Strydom, Adriano Mendes, Micheala Davids, Sim Mayaphi, Marietjie Venter, Amoako DG, Everatt J, Scheepers C, Mohale T, Ntuli N, Mahlangu B, Mnguni A, Ismail A, Bhiman JN |
| EPI_ISL_3237254, EPI_ISL_3237257 | KOPANONG LABORATORY | National Institute for Communicable Diseases of the National Health Laboratory Service | Amoako DG, Everatt J, Scheepers C, Mohale T, Ntuli N, Mahlangu B, Mnguni A, Ismail A, Bhiman JN |
| EPI_ISL_3237266, EPI_ISL_3237267, EPI_ISL_3237268 | SYNLAB MVZ Weiden | Robert Koch Institute | unknown |
| EPI_ISL_3237269 | SYNLAB MVZ Trier | Robert Koch Institute | unknown |
| EPI_ISL_3237270, EPI_ISL_3237271 | SYNLAB MVZ Leverkusen | Robert Koch Institute | unknown |
| EPI_ISL_3237272 | SYNLAB MVZ Trier | Robert Koch Institute | unknown |
| EPI_ISL_3237273 | SYNLAB MVZ Leverkusen | Robert Koch Institute | unknown |
| EPI_ISL_3237274 | SYNLAB MVZ Weiden | Robert Koch Institute | unknown |
| EPI_ISL_3237275 | SYNLAB MVZ Leverkusen | Robert Koch Institute | unknown |
| EPI_ISL_3237276 | SYNLAB MVZ Weiden | Robert Koch Institute | unknown |
| EPI_ISL_3237277, EPI_ISL_3237278 | SYNLAB MVZ Leverkusen | Robert Koch Institute | unknown |
| EPI_ISL_3237279 | SYNLAB MVZ Trier | Robert Koch Institute | unknown |
| EPI_ISL_3237280 | SYNLAB MVZ Leinfelden-Echterdingen | Robert Koch Institute | unknown |
| EPI_ISL_3237281 | SYNLAB MVZ Weiden | Robert Koch Institute | unknown |
| EPI_ISL_3237282 | SYNLAB MVZ Trier | Robert Koch Institute | unknown |
| EPI_ISL_3237283 | SYNLAB MVZ Hamburg | Robert Koch Institute | unknown |
| EPI_ISL_3237284, EPI_ISL_3237285, EPI_ISL_3237286, EPI_ISL_3237287 | SYNLAB MVZ Weiden | Robert Koch Institute | unknown |
| EPI_ISL_3237288 | SYNLAB MVZ Leverkusen | Robert Koch Institute | unknown |

| | | | |
|---|---|---|---|
| EPI_ISL_3237289 | SYNLAB MVZ Leinfelden-Echterdingen | Robert Koch Institute | unknown |
| EPI_ISL_3237290 | Synlab MVZ Augsburg | Robert Koch Institute | unknown |
| EPI_ISL_3237291 | SYNLAB MVZ Weiden | Robert Koch Institute | unknown |
| EPI_ISL_3237292 | SYNLAB MVZ Leverkusen | Robert Koch Institute | unknown |
| EPI_ISL_3237293 | SYNLAB MVZ Leinfelden-Echterdingen | Robert Koch Institute | unknown |
| EPI_ISL_3237294 | SYNLAB MVZ Weiden | Robert Koch Institute | unknown |
| EPI_ISL_3237295 | SYNLAB Labor München Zentrum LMZ | Robert Koch Institute | unknown |
| EPI_ISL_3237296 | SYNLAB MVZ Leinfelden-Echterdingen | Robert Koch Institute | unknown |
| EPI_ISL_3237297 | SYNLAB MVZ Weiden | Robert Koch Institute | unknown |
| EPI_ISL_3237298 | SYNLAB MVZ Leinfelden-Echterdingen | Robert Koch Institute | unknown |
| EPI_ISL_3237299 | SYNLAB MVZ Leverkusen | Robert Koch Institute | unknown |
| EPI_ISL_3237300 | SYNLAB MVZ Trier | Robert Koch Institute | unknown |
| EPI_ISL_3237301 | SYNLAB MVZ Weiden | Robert Koch Institute | unknown |
| EPI_ISL_3237302 | SYNLAB MVZ Leinfelden-Echterdingen | Robert Koch Institute | unknown |
| EPI_ISL_3237303 | SYNLAB MVZ Dachau | Robert Koch Institute | unknown |
| EPI_ISL_3237304 | SYNLAB MVZ Weiden | Robert Koch Institute | unknown |
| EPI_ISL_3237305 | SYNLAB MVZ Dachau | Robert Koch Institute | unknown |
| EPI_ISL_3237306 | SYNLAB Jena Oncoscreen | Robert Koch Institute | unknown |
| EPI_ISL_3237307, EPI_ISL_3237308 | SYNLAB MVZ Weiden | Robert Koch Institute | unknown |
| EPI_ISL_3237309, EPI_ISL_3237310 | SYNLAB MVZ Leverkusen | Robert Koch Institute | unknown |
| EPI_ISL_3237311, EPI_ISL_3237312, EPI_ISL_3237313 | SYNLAB MVZ Weiden | Robert Koch Institute | unknown |
| EPI_ISL_3237314 | SYNLAB MVZ Leverkusen | Robert Koch Institute | unknown |
| EPI_ISL_3237315 | SYNLAB MVZ Weiden | Robert Koch Institute | unknown |
| EPI_ISL_3237316 | Synlab MVZ Augsburg | Robert Koch Institute | unknown |
| EPI_ISL_3237317 | SYNLAB MVZ Weiden | Robert Koch Institute | unknown |
| EPI_ISL_3237318 | SYNLAB MVZ Leinfelden-Echterdingen | Robert Koch Institute | unknown |
| EPI_ISL_3237319, EPI_ISL_3237320 | Synlab MVZ Augsburg | Robert Koch Institute | unknown |
| EPI_ISL_3237321, EPI_ISL_3237322 | SYNLAB MVZ Weiden | Robert Koch Institute | unknown |
| EPI_ISL_3237323, EPI_ISL_3237324 | Synlab MVZ Augsburg | Robert Koch Institute | unknown |
| EPI_ISL_3237325 | SYNLAB MVZ Hamburg | Robert Koch Institute | unknown |
| EPI_ISL_3237326 | SYNLAB MVZ Leverkusen | Robert Koch Institute | unknown |
| EPI_ISL_3237327, EPI_ISL_3237328 | SYNLAB MVZ Leinfelden-Echterdingen | Robert Koch Institute | unknown |
| EPI_ISL_3237329 | SYNLAB MVZ Weiden | Robert Koch Institute | unknown |
| EPI_ISL_3237330, EPI_ISL_3237331, EPI_ISL_3237332 | SYNLAB MVZ Leverkusen | Robert Koch Institute | unknown |
| EPI_ISL_3237333 | SYNLAB MVZ Weiden | Robert Koch Institute | unknown |
| EPI_ISL_3237334 | SYNLAB MVZ Leverkusen | Robert Koch Institute | unknown |
| EPI_ISL_3237335 | SYNLAB MVZ Leinfelden-Echterdingen | Robert Koch Institute | unknown |
| EPI_ISL_3237336, EPI_ISL_3237337 | SYNLAB MVZ Weiden | Robert Koch Institute | unknown |
| EPI_ISL_3237338, EPI_ISL_3237339, EPI_ISL_3237340 | SYNLAB MVZ Trier | Robert Koch Institute | unknown |
| EPI_ISL_3237341 | Synlab MVZ Augsburg | Robert Koch Institute | unknown |
| EPI_ISL_3237342 | SYNLAB MVZ Leinfelden-Echterdingen | Robert Koch Institute | unknown |
| EPI_ISL_3237343, EPI_ISL_3237344, EPI_ISL_3237345, EPI_ISL_3237346 | SYNLAB MVZ Weiden | Robert Koch Institute | unknown |
| EPI_ISL_3237347 | SYNLAB MVZ Trier | Robert Koch Institute | unknown |
| EPI_ISL_3237348 | SYNLAB MVZ Weiden | Robert Koch Institute | unknown |
| EPI_ISL_3237349 | SYNLAB MVZ Leverkusen | Robert Koch Institute | unknown |
| EPI_ISL_3237350 | Synlab MVZ Augsburg | Robert Koch Institute | unknown |
| EPI_ISL_3237351, EPI_ISL_3237352 | SYNLAB MVZ Weiden | Robert Koch Institute | unknown |
| EPI_ISL_3237353 | SYNLAB MVZ Trier | Robert Koch Institute | unknown |
| EPI_ISL_3237354 | SYNLAB MVZ Weiden | Robert Koch Institute | unknown |
| EPI_ISL_3237355, EPI_ISL_3237357, EPI_ISL_3237358 | SYNLAB MVZ Trier | Robert Koch Institute | unknown |
| EPI_ISL_3237359 | SYNLAB MVZ Weiden | Robert Koch Institute | unknown |
| EPI_ISL_3237360 | SYNLAB MVZ Leinfelden-Echterdingen | Robert Koch Institute | unknown |
| EPI_ISL_3237361 | SYNLAB MVZ Trier | Robert Koch Institute | unknown |
| EPI_ISL_3237362 | Synlab MVZ Augsburg | Robert Koch Institute | unknown |
| EPI_ISL_3237363 | SYNLAB MVZ Leverkusen | Robert Koch Institute | unknown |
| EPI_ISL_3237364 | SYNLAB MVZ Leinfelden-Echterdingen | Robert Koch Institute | unknown |
| EPI_ISL_3237365 | SYNLAB MVZ Weiden | Robert Koch Institute | unknown |
| EPI_ISL_3237366 | SYNLAB MVZ Leinfelden-Echterdingen | Robert Koch Institute | unknown |
| EPI_ISL_3237367 | SYNLAB MVZ Leverkusen | Robert Koch Institute | unknown |
| EPI_ISL_3237368 | SYNLAB MVZ Leinfelden-Echterdingen | Robert Koch Institute | unknown |
| EPI_ISL_3237369 | SYNLAB Jena Oncoscreen | Robert Koch Institute | unknown |
| EPI_ISL_3237370 | SYNLAB MVZ Leverkusen | Robert Koch Institute | unknown |
| EPI_ISL_3237371 | SYNLAB MVZ Leinfelden-Echterdingen | Robert Koch Institute | unknown |

| | | | |
|---|---|---|---|
| EPI_ISL_3237372 | SYNLAB MVZ Weiden | Robert Koch Institute | unknown |
| EPI_ISL_3237373, EPI_ISL_3237374 | SYNLAB MVZ Leverkusen | Robert Koch Institute | unknown |
| EPI_ISL_3237375 | SYNLAB MVZ Weiden | Robert Koch Institute | unknown |
| EPI_ISL_3237376 | SYNLAB MVZ Trier | Robert Koch Institute | unknown |
| EPI_ISL_3237377 | SYNLAB MVZ Leinfelden-Echterdingen | Robert Koch Institute | unknown |
| EPI_ISL_3237378 | Synlab MVZ Augsburg | Robert Koch Institute | unknown |
| EPI_ISL_3237379 | SYNLAB MVZ Leverkusen | Robert Koch Institute | unknown |
| EPI_ISL_3237380, EPI_ISL_3237381 | SYNLAB MVZ Weiden | Robert Koch Institute | unknown |
| EPI_ISL_3237382 | SYNLAB MVZ Leverkusen | Robert Koch Institute | unknown |
| EPI_ISL_3237383 | SYNLAB MVZ Weiden | Robert Koch Institute | unknown |
| EPI_ISL_3237384 | SYNLAB MVZ Leverkusen | Robert Koch Institute | unknown |
| EPI_ISL_3237385 | SYNLAB Labor München Zentrum LMZ | Robert Koch Institute | unknown |
| EPI_ISL_3237386 | SYNLAB MVZ Hamburg | Robert Koch Institute | unknown |
| EPI_ISL_3237387, EPI_ISL_3237388 | SYNLAB Labor München Zentrum LMZ | Robert Koch Institute | unknown |
| EPI_ISL_3237389 | SYNLAB MVZ Leverkusen | Robert Koch Institute | unknown |
| EPI_ISL_3237390 | SYNLAB MVZ Weiden | Robert Koch Institute | unknown |
| EPI_ISL_3237391, EPI_ISL_3237392 | SYNLAB MVZ Leinfelden-Echterdingen | Robert Koch Institute | unknown |
| EPI_ISL_3237393 | SYNLAB MVZ Weiden | Robert Koch Institute | unknown |
| EPI_ISL_3237394, EPI_ISL_3237395 | SYNLAB MVZ Leverkusen | Robert Koch Institute | unknown |
| EPI_ISL_3237396 | SYNLAB MVZ Leinfelden-Echterdingen | Robert Koch Institute | unknown |
| EPI_ISL_3237397 | SYNLAB Labor München Zentrum LMZ | Robert Koch Institute | unknown |
| EPI_ISL_3237398 | SYNLAB MVZ Weiden | Robert Koch Institute | unknown |
| EPI_ISL_3237399 | Synlab MVZ Augsburg | Robert Koch Institute | unknown |
| EPI_ISL_3237400 | SYNLAB MVZ Leinfelden-Echterdingen | Robert Koch Institute | unknown |
| EPI_ISL_3237401 | Synlab MVZ Augsburg | Robert Koch Institute | unknown |
| EPI_ISL_3237402 | SYNLAB MVZ Leverkusen | Robert Koch Institute | unknown |
| EPI_ISL_3237403 | SYNLAB MVZ Leinfelden-Echterdingen | Robert Koch Institute | unknown |
| EPI_ISL_3237404, EPI_ISL_3237405 | SYNLAB MVZ Weiden | Robert Koch Institute | unknown |
| EPI_ISL_3237406 | SYNLAB MVZ Trier | Robert Koch Institute | unknown |
| EPI_ISL_3237407, EPI_ISL_3237408 | SYNLAB MVZ Leverkusen | Robert Koch Institute | unknown |
| EPI_ISL_3237410 | SYNLAB MVZ Weiden | Robert Koch Institute | unknown |
| EPI_ISL_3237411, EPI_ISL_3237412 | SYNLAB MVZ Leverkusen | Robert Koch Institute | unknown |
| EPI_ISL_3237413 | SYNLAB MVZ Leinfelden-Echterdingen | Robert Koch Institute | unknown |
| EPI_ISL_3237414, EPI_ISL_3237416 | SYNLAB MVZ Leverkusen | Robert Koch Institute | unknown |
| EPI_ISL_3237417 | SYNLAB MVZ Weiden | Robert Koch Institute | unknown |
| EPI_ISL_3237418 | SYNLAB MVZ Leinfelden-Echterdingen | Robert Koch Institute | unknown |
| EPI_ISL_3237419 | SYNLAB MVZ Trier | Robert Koch Institute | unknown |
| EPI_ISL_3237420 | Synlab MVZ Augsburg | Robert Koch Institute | unknown |
| EPI_ISL_3237421 | SYNLAB MVZ Leinfelden-Echterdingen | Robert Koch Institute | unknown |
| EPI_ISL_3237422 | SYNLAB MVZ Leverkusen | Robert Koch Institute | unknown |
| EPI_ISL_3237423 | SYNLAB MVZ Weiden | Robert Koch Institute | unknown |
| EPI_ISL_3237424 | SYNLAB MVZ Trier | Robert Koch Institute | unknown |
| EPI_ISL_3237425 | SYNLAB MVZ Leinfelden-Echterdingen | Robert Koch Institute | unknown |
| EPI_ISL_3237426 | SYNLAB MVZ Dachau | Robert Koch Institute | unknown |
| EPI_ISL_3237427 | Synlab MVZ Augsburg | Robert Koch Institute | unknown |
| EPI_ISL_3237428 | SYNLAB MVZ Trier | Robert Koch Institute | unknown |
| EPI_ISL_3237429, EPI_ISL_3237430 | SYNLAB MVZ Weiden | Robert Koch Institute | unknown |
| EPI_ISL_3237432 | SYNLAB MVZ Leverkusen | Robert Koch Institute | unknown |
| EPI_ISL_3237433 | SYNLAB MVZ Weiden | Robert Koch Institute | unknown |
| EPI_ISL_3237434 | SYNLAB MVZ Leverkusen | Robert Koch Institute | unknown |
| EPI_ISL_3237435 | SYNLAB MVZ Trier | Robert Koch Institute | unknown |
| EPI_ISL_3237437 | SYNLAB MVZ Leinfelden-Echterdingen | Robert Koch Institute | unknown |
| EPI_ISL_3237438 | SYNLAB MVZ Leverkusen | Robert Koch Institute | unknown |
| EPI_ISL_3237440 | SYNLAB MVZ Trier | Robert Koch Institute | unknown |
| EPI_ISL_3237441 | SYNLAB Jena Oncoscreen | Robert Koch Institute | unknown |
| EPI_ISL_3237442 | SYNLAB MVZ Trier | Robert Koch Institute | unknown |
| EPI_ISL_3237443 | SYNLAB MVZ Leinfelden-Echterdingen | Robert Koch Institute | unknown |
| EPI_ISL_3237445, EPI_ISL_3237447 | SYNLAB MVZ Weiden | Robert Koch Institute | unknown |
| EPI_ISL_3237448, EPI_ISL_3237449 | SYNLAB MVZ Leverkusen | Robert Koch Institute | unknown |
| EPI_ISL_3237450, EPI_ISL_3237451 | SYNLAB MVZ Trier | Robert Koch Institute | unknown |
| EPI_ISL_3237452 | SYNLAB MVZ Weiden | Robert Koch Institute | unknown |
| EPI_ISL_3237453 | Synlab MVZ Augsburg | Robert Koch Institute | unknown |
| EPI_ISL_3237454 | SYNLAB MVZ Weiden | Robert Koch Institute | unknown |
| EPI_ISL_3237455 | SYNLAB MVZ Trier | Robert Koch Institute | unknown |
| EPI_ISL_3237456 | SYNLAB MVZ Weiden | Robert Koch Institute | unknown |
| EPI_ISL_3237457 | SYNLAB MVZ Trier | Robert Koch Institute | unknown |
| EPI_ISL_3237458 | SYNLAB MVZ Weiden | Robert Koch Institute | unknown |

| | | | |
|---|---|---|---|
| EPI_ISL_3237459 | SYNLAB MVZ Trier | Robert Koch Institute | unknown |
| EPI_ISL_3237460 | SYNLAB MVZ Leverkusen | Robert Koch Institute | unknown |
| EPI_ISL_3237461, EPI_ISL_3237462 | SYNLAB MVZ Weiden | Robert Koch Institute | unknown |
| EPI_ISL_3237463, EPI_ISL_3237464 | SYNLAB MVZ Leverkusen | Robert Koch Institute | unknown |
| EPI_ISL_3237465 | Synlab MVZ Augsburg | Robert Koch Institute | unknown |
| EPI_ISL_3237466 | SYNLAB MVZ Weiden | Robert Koch Institute | unknown |
| EPI_ISL_3237467 | SYNLAB MVZ Leinfelden-Echterdingen | Robert Koch Institute | unknown |
| EPI_ISL_3237468 | SYNLAB MVZ Weiden | Robert Koch Institute | unknown |
| EPI_ISL_3237469 | SYNLAB MVZ Leverkusen | Robert Koch Institute | unknown |
| EPI_ISL_3237470 | SYNLAB MVZ Trier | Robert Koch Institute | unknown |
| EPI_ISL_3237471 | Synlab MVZ Augsburg | Robert Koch Institute | unknown |
| EPI_ISL_3237473 | SYNLAB MVZ Weiden | Robert Koch Institute | unknown |
| EPI_ISL_3237474 | SYNLAB MVZ Leverkusen | Robert Koch Institute | unknown |
| EPI_ISL_3237475 | SYNLAB MVZ Weiden | Robert Koch Institute | unknown |
| EPI_ISL_3237476 | SYNLAB MVZ Leverkusen | Robert Koch Institute | unknown |
| EPI_ISL_3237477 | SYNLAB MVZ Trier | Robert Koch Institute | unknown |
| EPI_ISL_3237478 | SYNLAB MVZ Weiden | Robert Koch Institute | unknown |
| EPI_ISL_3237479 | SYNLAB MVZ Leinfelden-Echterdingen | Robert Koch Institute | unknown |
| EPI_ISL_3237480 | SYNLAB MVZ Berlin | Robert Koch Institute | unknown |
| EPI_ISL_3237481, EPI_ISL_3237482 | SYNLAB MVZ Leverkusen | Robert Koch Institute | unknown |
| EPI_ISL_3237483, EPI_ISL_3237484 | SYNLAB MVZ Leinfelden-Echterdingen | Robert Koch Institute | unknown |
| EPI_ISL_3237485 | SYNLAB MVZ Weiden | Robert Koch Institute | unknown |
| EPI_ISL_3237486 | SYNLAB MVZ Leverkusen | Robert Koch Institute | unknown |
| EPI_ISL_3237487 | SYNLAB MVZ Leinfelden-Echterdingen | Robert Koch Institute | unknown |
| EPI_ISL_3237488 | SYNLAB MVZ Weiden | Robert Koch Institute | unknown |
| EPI_ISL_3237489 | SYNLAB MVZ Trier | Robert Koch Institute | unknown |
| EPI_ISL_3237490 | SYNLAB MVZ Leinfelden-Echterdingen | Robert Koch Institute | unknown |
| EPI_ISL_3237491 | SYNLAB MVZ Weiden | Robert Koch Institute | unknown |
| EPI_ISL_3237492 | SYNLAB MVZ Trier | Robert Koch Institute | unknown |
| EPI_ISL_3237493 | SYNLAB MVZ Weiden | Robert Koch Institute | unknown |
| EPI_ISL_3237494 | Synlab MVZ Augsburg | Robert Koch Institute | unknown |
| EPI_ISL_3237495 | SYNLAB MVZ Weiden | Robert Koch Institute | unknown |
| EPI_ISL_3237496 | SYNLAB MVZ Leverkusen | Robert Koch Institute | unknown |
| EPI_ISL_3237497 | SYNLAB MVZ Berlin | Robert Koch Institute | unknown |
| EPI_ISL_3237498, EPI_ISL_3237499, EPI_ISL_3237500, EPI_ISL_3237501 | SYNLAB MVZ Weiden | Robert Koch Institute | unknown |
| EPI_ISL_3237502 | SYNLAB MVZ Leverkusen | Robert Koch Institute | unknown |
| EPI_ISL_3237503, EPI_ISL_3237504 | SYNLAB MVZ Weiden | Robert Koch Institute | unknown |
| EPI_ISL_3237505 | SYNLAB MVZ Trier | Robert Koch Institute | unknown |
| EPI_ISL_3237506 | SYNLAB MVZ Leinfelden-Echterdingen | Robert Koch Institute | unknown |
| EPI_ISL_3237507 | Landesamt für Verbraucherschutz des Landes Sachsen-Anhalt | Robert Koch Institute | unknown |
| EPI_ISL_3237508, EPI_ISL_3237509, EPI_ISL_3237510, EPI_ISL_3237511, EPI_ISL_3237512, EPI_ISL_3237513 | Robert Koch-Institut ZBS1 (Zentrum für biologische Gefahren und spezielle Pathogene hochpathogene Viren) | Robert Koch Institute | unknown |
| EPI_ISL_3237514, EPI_ISL_3237515 | Landesamt für Verbraucherschutz des Landes Sachsen-Anhalt | Robert Koch Institute | unknown |
| EPI_ISL_3237516, EPI_ISL_3237517, EPI_ISL_3237518, EPI_ISL_3237519, EPI_ISL_3237520, EPI_ISL_3237521, EPI_ISL_3237522, EPI_ISL_3237523 | Medizinisches Zentrallabor Altenburg GmbH & Co.KG | Robert Koch Institute | unknown |
| EPI_ISL_3237524 | IMD Labor Frankfurt | Robert Koch Institute | unknown |
| EPI_ISL_3237525 | Universitätsklinikum Bonn - Institut für Virologie | Robert Koch Institute | unknown |
| EPI_ISL_3237526, EPI_ISL_3237527 | IMD Labor Frankfurt | Robert Koch Institute | unknown |
| EPI_ISL_3237528 | Institut für Immunologie und Genetik Kaiserslautern: Medizinisches Labor Dr. med. B. Thiele | Robert Koch Institute | unknown |
| EPI_ISL_3237529 | IMD Labor Frankfurt | Robert Koch Institute | unknown |
| EPI_ISL_3237530 | Universitätsklinikum Bonn - Institut für Virologie | Robert Koch Institute | unknown |
| EPI_ISL_3237531 | Labormedizin Darmstadt | Robert Koch Institute | unknown |
| EPI_ISL_3237532 | IMD Labor Frankfurt | Robert Koch Institute | unknown |
| EPI_ISL_3237533 | Labormedizin Darmstadt | Robert Koch Institute | unknown |
| EPI_ISL_3237534, EPI_ISL_3237535 | IMD Labor Frankfurt | Robert Koch Institute | unknown |
| EPI_ISL_3237536 | Universitätsklinikum des Saarlandes - Institut für Virologie | Robert Koch Institute | unknown |
| EPI_ISL_3237537 | IMD Labor Frankfurt | Robert Koch Institute | unknown |
| EPI_ISL_3237539 | Universitätsklinikum Bonn - Institut für Virologie | Robert Koch Institute | unknown |
| EPI_ISL_3237540, EPI_ISL_3237541 | IMD Labor Frankfurt | Robert Koch Institute | unknown |
| EPI_ISL_3237542 | Universitätsklinikum Bonn - Institut für Virologie | Robert Koch Institute | unknown |
| EPI_ISL_3237543, EPI_ISL_3237545, | IMD Labor Frankfurt | Robert Koch Institute | unknown |

| | | | |
|---|---|---|---|
| EPI_ISL_3237546 | | | |
| EPI_ISL_3237547 | Institut für Immunologie und Genetik Kaiserslautern: Medizinisches Labor Dr. med. B. Thiele | Robert Koch Institute | unknown |
| EPI_ISL_3237548, EPI_ISL_3237549 | IMD Labor Frankfurt | Robert Koch Institute | unknown |
| EPI_ISL_3237550, EPI_ISL_3237551, EPI_ISL_3237552, EPI_ISL_3237553, EPI_ISL_3237554, EPI_ISL_3237555, EPI_ISL_3237556, EPI_ISL_3237557, EPI_ISL_3237558, EPI_ISL_3237559, EPI_ISL_3237560, EPI_ISL_3237561, EPI_ISL_3237562, EPI_ISL_3237563, EPI_ISL_3237564, EPI_ISL_3237565 | | | |
| see above | Klinikum Ernst von Bergmann gemeinnützige GmbH - stationärer Bereich | Robert Koch Institute | unknown |
| EPI_ISL_3237566, EPI_ISL_3237567, EPI_ISL_3237568, EPI_ISL_3237569, EPI_ISL_3237570 | MVZ Labor Dr. Limbach & Kollegen GbR | Robert Koch Institute | unknown |
| EPI_ISL_3237571 | Limbach - MVZ Labor Eveld & Kollegen Essen | Robert Koch Institute | unknown |
| EPI_ISL_3237572 | MVZ Labor Dr. Limbach & Kollegen GbR | Robert Koch Institute | unknown |
| EPI_ISL_3237573 | Limbach - MVZ Labor Eveld & Kollegen Essen | Robert Koch Institute | unknown |
| EPI_ISL_3237574, EPI_ISL_3237575, EPI_ISL_3237576, EPI_ISL_3237577, EPI_ISL_3237578, EPI_ISL_3237579 | MVZ Labor Dr. Limbach & Kollegen GbR | Robert Koch Institute | unknown |
| EPI_ISL_3237580 | Limbach - MVZ Labor Eveld & Kollegen Essen | Robert Koch Institute | unknown |
| EPI_ISL_3237581, EPI_ISL_3237582, EPI_ISL_3237583, EPI_ISL_3237584, EPI_ISL_3237585, EPI_ISL_3237586, EPI_ISL_3237587, EPI_ISL_3237588, EPI_ISL_3237589 | MVZ Labor Dr. Limbach & Kollegen GbR | Robert Koch Institute | unknown |
| EPI_ISL_3237591, EPI_ISL_3237592, EPI_ISL_3237593, EPI_ISL_3237594, EPI_ISL_3237596, EPI_ISL_3237597, EPI_ISL_3237598, EPI_ISL_3237600, EPI_ISL_3237601, EPI_ISL_3237602, EPI_ISL_3237604, EPI_ISL_3237605, EPI_ISL_3237606, EPI_ISL_3237607, EPI_ISL_3237608, EPI_ISL_3237609, EPI_ISL_3237610, EPI_ISL_3237612, EPI_ISL_3237614, EPI_ISL_3237615, EPI_ISL_3237616, EPI_ISL_3237617, EPI_ISL_3237620, EPI_ISL_3237622, EPI_ISL_3237623, EPI_ISL_3237625, EPI_ISL_3237626, EPI_ISL_3237627, EPI_ISL_3237630, EPI_ISL_3237633 | | | |
| see above | Diagnosticum - Labor Neukirchen | Robert Koch Institute | unknown |
| EPI_ISL_3237634, EPI_ISL_3237635, EPI_ISL_3237636, EPI_ISL_3237637, EPI_ISL_3237638, EPI_ISL_3237639, EPI_ISL_3237640, EPI_ISL_3237641, EPI_ISL_3237643, EPI_ISL_3237644, EPI_ISL_3237645, EPI_ISL_3237646, EPI_ISL_3237647, EPI_ISL_3237648, EPI_ISL_3237649, EPI_ISL_3237650, EPI_ISL_3237651 | | | |
| see above | MVZ für Laboratoriumsmedizin und Mikrobiologie Koblenz-Mittelrhein (Labor Koblenz) | Robert Koch Institute | unknown |
| EPI_ISL_3237652, EPI_ISL_3237655, EPI_ISL_3237656, EPI_ISL_3237657, EPI_ISL_3237658, EPI_ISL_3237659, EPI_ISL_3237660, EPI_ISL_3237661, EPI_ISL_3237662, EPI_ISL_3237664, EPI_ISL_3237665, EPI_ISL_3237666, EPI_ISL_3237667, EPI_ISL_3237668, EPI_ISL_3237669, EPI_ISL_3237670, EPI_ISL_3237672, EPI_ISL_3237675, EPI_ISL_3237676, EPI_ISL_3237678, EPI_ISL_3237679, EPI_ISL_3237681, EPI_ISL_3237682, EPI_ISL_3237683, EPI_ISL_3237684, EPI_ISL_3237685, EPI_ISL_3237686, EPI_ISL_3237687, EPI_ISL_3237688, EPI_ISL_3237689, EPI_ISL_3237690, EPI_ISL_3237692 | | | |
| see above | Labor ZOTZ\|KLIMAS; MVZ Düsseldorf-Centrum | Robert Koch Institute | unknown |
| EPI_ISL_3237695, EPI_ISL_3237696, EPI_ISL_3237697, EPI_ISL_3237698, EPI_ISL_3237699, EPI_ISL_3237700, EPI_ISL_3237701, EPI_ISL_3237702, EPI_ISL_3237703, EPI_ISL_3237705, EPI_ISL_3237706, EPI_ISL_3237707 | | | |
| see above | Helios Universitätsklinikum Wuppertal: Universität Witten/Herdecke: Institut für Medizinische Labordiagnostik | Robert Koch Institute | unknown |
| EPI_ISL_3237708, EPI_ISL_3237709, EPI_ISL_3237710, EPI_ISL_3237711, EPI_ISL_3237712, EPI_ISL_3237713, EPI_ISL_3237714, EPI_ISL_3237715, EPI_ISL_3237716, EPI_ISL_3237717, EPI_ISL_3237718, EPI_ISL_3237719, EPI_ISL_3237720, EPI_ISL_3237721, EPI_ISL_3237722, EPI_ISL_3237723, EPI_ISL_3237724, EPI_ISL_3237725, EPI_ISL_3237726, EPI_ISL_3237727, EPI_ISL_3237728, EPI_ISL_3237729, EPI_ISL_3237730, EPI_ISL_3237731, EPI_ISL_3237732, EPI_ISL_3237733, EPI_ISL_3237734, EPI_ISL_3237735, EPI_ISL_3237736, EPI_ISL_3237737, EPI_ISL_3237738, EPI_ISL_3237739, EPI_ISL_3237740, EPI_ISL_3237741, EPI_ISL_3237742, EPI_ISL_3237743, EPI_ISL_3237744, EPI_ISL_3237745, EPI_ISL_3237746, EPI_ISL_3237747 | | | |
| see above | MDI Limbach Berlin GmbH; MVZ Labor Berlin | Robert Koch Institute | unknown |
| EPI_ISL_3237748, EPI_ISL_3237749, EPI_ISL_3237750, EPI_ISL_3237751, EPI_ISL_3237752, EPI_ISL_3237753, EPI_ISL_3237754, EPI_ISL_3237755, EPI_ISL_3237756, EPI_ISL_3237757, EPI_ISL_3237758 | | | |
| see above | MVZ Dr. Eberhard & Partner Dortmund | Robert Koch Institute | unknown |
| EPI_ISL_3237759, EPI_ISL_3237760, EPI_ISL_3237761, EPI_ISL_3237762, EPI_ISL_3237763, EPI_ISL_3237764, EPI_ISL_3237765, EPI_ISL_3237766, EPI_ISL_3237767, EPI_ISL_3237768, EPI_ISL_3237769 | | | |
| see above | MDI Limbach Berlin GmbH; MVZ Labor Berlin | Robert Koch Institute | unknown |
| EPI_ISL_3237770 | Limbach - MVZ Gemeinschaftslabor Suhl | Robert Koch Institute | unknown |
| EPI_ISL_3237771 | MDI Limbach Berlin GmbH; MVZ Labor Berlin | Robert Koch Institute | unknown |
| EPI_ISL_3237772 | MVZ Labor Dr. Limbach & Kollegen GbR | Robert Koch Institute | unknown |
| EPI_ISL_3237773, EPI_ISL_3237774 | MDI Limbach Berlin GmbH; MVZ Labor Berlin | Robert Koch Institute | unknown |
| EPI_ISL_3237775, EPI_ISL_3237776 | MVZ Labor Dr. Limbach & Kollegen GbR | Robert Koch Institute | unknown |
| EPI_ISL_3237777, EPI_ISL_3237778, EPI_ISL_3237779 | MDI Limbach Berlin GmbH; MVZ Labor Berlin | Robert Koch Institute | unknown |
| EPI_ISL_3237780 | MVZ Labor Dr. Limbach & Kollegen GbR | Robert Koch Institute | unknown |
| EPI_ISL_3237781, EPI_ISL_3237782, EPI_ISL_3237783 | MDI Limbach Berlin GmbH; MVZ Labor Berlin | Robert Koch Institute | unknown |
| EPI_ISL_3237784 | MVZ Labor Dr. Limbach & Kollegen GbR | Robert Koch Institute | unknown |
| EPI_ISL_3237785, EPI_ISL_3237786, EPI_ISL_3237787, EPI_ISL_3237788, EPI_ISL_3237789 | MDI Limbach Berlin GmbH; MVZ Labor Berlin | Robert Koch Institute | unknown |
| EPI_ISL_3237790 | MVZ Labor Dr. Limbach & Kollegen GbR | Robert Koch Institute | unknown |
| EPI_ISL_3237791, EPI_ISL_3237792, EPI_ISL_3237793 | MDI Limbach Berlin GmbH; MVZ Labor Berlin | Robert Koch Institute | unknown |
| EPI_ISL_3237794, EPI_ISL_3237795, EPI_ISL_3237796, EPI_ISL_3237797, EPI_ISL_3237798, EPI_ISL_3237799, EPI_ISL_3237800, EPI_ISL_3237801, EPI_ISL_3237802, EPI_ISL_3237803, EPI_ISL_3237804, EPI_ISL_3237805, EPI_ISL_3237806, EPI_ISL_3237807, EPI_ISL_3237808, EPI_ISL_3237809, EPI_ISL_3237810, EPI_ISL_3237811, EPI_ISL_3237812 | | | |
| see above | Labor Mönchengladbach MVZ Dr. Stein + Kollegen GbR | Robert Koch Institute | unknown |
| EPI_ISL_3237813, EPI_ISL_3237814, EPI_ISL_3237815, EPI_ISL_3237816, EPI_ISL_3237817 | MVZ Labor Dr. Limbach & Kollegen GbR | Robert Koch Institute | unknown |
| EPI_ISL_3237818 | Labor Mönchengladbach MVZ Dr. Stein + Kollegen GbR | Robert Koch Institute | unknown |
| EPI_ISL_3237819, EPI_ISL_3237820 | MVZ Labor Dr. Limbach & Kollegen GbR | Robert Koch Institute | unknown |
| EPI_ISL_3237821 | MVZ für Laboratoriumsmedizin und Mikrobiologie Koblenz-Mittelrhein (Labor Koblenz) | Robert Koch Institute | unknown |
| EPI_ISL_3237822, EPI_ISL_3237823, EPI_ISL_3237824 | MVZ Labor Dr. Limbach & Kollegen GbR | Robert Koch Institute | unknown |

| | | | |
|---|---|---|---|
| EPI_ISL_3237825 | MVZ für Laboratoriumsmedizin und Mikrobiologie Koblenz-Mittelrhein (Labor Koblenz) | Robert Koch Institute | unknown |
| EPI_ISL_3237826 | MVZ Labor Dr. Limbach & Kollegen GbR | Robert Koch Institute | unknown |
| EPI_ISL_3237827 | MVZ für Laboratoriumsmedizin und Mikrobiologie Koblenz-Mittelrhein (Labor Koblenz) | Robert Koch Institute | unknown |
| EPI_ISL_3237828 | Labor Mönchengladbach MVZ Dr. Stein + Kollegen GbR | Robert Koch Institute | unknown |
| EPI_ISL_3237829 | MVZ für Laboratoriumsmedizin und Mikrobiologie Koblenz-Mittelrhein (Labor Koblenz) | Robert Koch Institute | unknown |
| EPI_ISL_3237830, EPI_ISL_3237831 | MVZ Labor Dr. Limbach & Kollegen GbR | Robert Koch Institute | unknown |
| EPI_ISL_3237832, EPI_ISL_3237833 | Labor Mönchengladbach MVZ Dr. Stein + Kollegen GbR | Robert Koch Institute | unknown |
| EPI_ISL_3237834 | MVZ für Laboratoriumsmedizin und Mikrobiologie Koblenz-Mittelrhein (Labor Koblenz) | Robert Koch Institute | unknown |
| EPI_ISL_3237835 | MDI Limbach Berlin GmbH; MVZ Labor Berlin | Robert Koch Institute | unknown |
| EPI_ISL_3237836 | MVZ für Laboratoriumsmedizin und Mikrobiologie Koblenz-Mittelrhein (Labor Koblenz) | Robert Koch Institute | unknown |
| EPI_ISL_3237837, EPI_ISL_3237838, EPI_ISL_3237839, EPI_ISL_3237840 | MDI Limbach Berlin GmbH; MVZ Labor Berlin | Robert Koch Institute | unknown |
| EPI_ISL_3237841, EPI_ISL_3237842 | MVZ für Laboratoriumsmedizin und Mikrobiologie Koblenz-Mittelrhein (Labor Koblenz) | Robert Koch Institute | unknown |
| EPI_ISL_3237843, EPI_ISL_3237844 | MDI Limbach Berlin GmbH; MVZ Labor Berlin | Robert Koch Institute | unknown |
| EPI_ISL_3237845 | MVZ Labor Dr. Reising-Ackermann und Kollegen | Robert Koch Institute | unknown |
| EPI_ISL_3237846 | MVZ für Laboratoriumsmedizin und Mikrobiologie Koblenz-Mittelrhein (Labor Koblenz) | Robert Koch Institute | unknown |
| EPI_ISL_3237847 | MDI Limbach Berlin GmbH; MVZ Labor Berlin | Robert Koch Institute | unknown |
| EPI_ISL_3237848 | MVZ für Laboratoriumsmedizin und Mikrobiologie Koblenz-Mittelrhein (Labor Koblenz) | Robert Koch Institute | unknown |
| EPI_ISL_3237849 | MDI Limbach Berlin GmbH; MVZ Labor Berlin | Robert Koch Institute | unknown |
| EPI_ISL_3237850 | MVZ für Laboratoriumsmedizin und Mikrobiologie Koblenz-Mittelrhein (Labor Koblenz) | Robert Koch Institute | unknown |
| EPI_ISL_3237851 | MDI Limbach Berlin GmbH; MVZ Labor Berlin | Robert Koch Institute | unknown |
| EPI_ISL_3237852, EPI_ISL_3237853, EPI_ISL_3237854, EPI_ISL_3237855, EPI_ISL_3237856, EPI_ISL_3237857, EPI_ISL_3237858, EPI_ISL_3237859, EPI_ISL_3237860 | Niedersächsisches Landesgesundheitsamt (NLGA) | Robert Koch Institute | unknown |
| EPI_ISL_3237861, EPI_ISL_3237862 | Labor für medizinische Mikrobiologie Bakteriologie Mykologie Virologie & Infektionsserologie - Labor Mölbis | Robert Koch Institute | unknown |
| EPI_ISL_3237863, EPI_ISL_3237864 | alphaomega Labor GbR (Medizinische Kooperationsgemeinschaft Dres. Ackermann/Tennert/Zaiß) | Robert Koch Institute | unknown |
| EPI_ISL_3237865, EPI_ISL_3237866, EPI_ISL_3237867, EPI_ISL_3237868, EPI_ISL_3237869, EPI_ISL_3237870, EPI_ISL_3237871, EPI_ISL_3237872, EPI_ISL_3237873, EPI_ISL_3237874, EPI_ISL_3237875, EPI_ISL_3237876, EPI_ISL_3237877, EPI_ISL_3237878, EPI_ISL_3237879, EPI_ISL_3237880, EPI_ISL_3237881, EPI_ISL_3237882, EPI_ISL_3237883, EPI_ISL_3237884, EPI_ISL_3237885, EPI_ISL_3237886, EPI_ISL_3237887, EPI_ISL_3237888, EPI_ISL_3237889, EPI_ISL_3237890, EPI_ISL_3237891, EPI_ISL_3237892, EPI_ISL_3237893, EPI_ISL_3237894, EPI_ISL_3237895, EPI_ISL_3237896, EPI_ISL_3237897, EPI_ISL_3237898, EPI_ISL_3237899, EPI_ISL_3237900, EPI_ISL_3237901, EPI_ISL_3237902, EPI_ISL_3237903, EPI_ISL_3237904, EPI_ISL_3237906, EPI_ISL_3237907, EPI_ISL_3237908, EPI_ISL_3237909, EPI_ISL_3237910, EPI_ISL_3237911, EPI_ISL_3237912, EPI_ISL_3237913, EPI_ISL_3237914, EPI_ISL_3237915, EPI_ISL_3237916, EPI_ISL_3237917, EPI_ISL_3237918, EPI_ISL_3237919 | | | |
| see above | Medizinische Laboratorien Düsseldorf | Robert Koch Institute | unknown |
| EPI_ISL_3237920, EPI_ISL_3237921, EPI_ISL_3237922, EPI_ISL_3237923, EPI_ISL_3237924, EPI_ISL_3237926, EPI_ISL_3237927, EPI_ISL_3237928, EPI_ISL_3237929, EPI_ISL_3237931 | Uniklinikum Regensburg; Institut für klinische Mikrobiologie und Hygiene | Robert Koch Institute | unknown |
| EPI_ISL_3237932, EPI_ISL_3237933, EPI_ISL_3237934, EPI_ISL_3237935, EPI_ISL_3237936, EPI_ISL_3237937, EPI_ISL_3237938, EPI_ISL_3237939, EPI_ISL_3237940, EPI_ISL_3237941, EPI_ISL_3237942, EPI_ISL_3237943, EPI_ISL_3237944, EPI_ISL_3237945, EPI_ISL_3237946 | | | |
| see above | Labor Dr. Heidrich & Kollegen MVZ GmbH Hamburg | Robert Koch Institute | unknown |
| EPI_ISL_3237947, EPI_ISL_3237948, EPI_ISL_3237949, EPI_ISL_3237950, EPI_ISL_3237951, EPI_ISL_3237952, EPI_ISL_3237953, EPI_ISL_3237954, EPI_ISL_3237955, EPI_ISL_3237956, EPI_ISL_3237957, EPI_ISL_3237958, EPI_ISL_3237959, EPI_ISL_3237960, EPI_ISL_3237961, EPI_ISL_3237962, EPI_ISL_3237964, EPI_ISL_3237965, EPI_ISL_3237966, EPI_ISL_3237967, EPI_ISL_3237968, EPI_ISL_3237969, EPI_ISL_3237970, EPI_ISL_3237971 | | | |
| see above | Landesgesundheitsamt Baden-Württemberg | Robert Koch Institute | unknown |
| EPI_ISL_3237972, EPI_ISL_3237973, EPI_ISL_3237974, EPI_ISL_3237975, EPI_ISL_3237977 | Klinikum Stuttgart | Robert Koch Institute | unknown |
| EPI_ISL_3237978 | LabKom - Labor Augsburg MVZ GmbH | Robert Koch Institute | unknown |
| EPI_ISL_3237979, EPI_ISL_3237980 | Bioscientia Labor Wermsdorf | Robert Koch Institute | unknown |
| EPI_ISL_3237981 | LabKom - Labor Augsburg MVZ GmbH | Robert Koch Institute | unknown |
| EPI_ISL_3237982 | Bioscientia Labor Wermsdorf | Robert Koch Institute | unknown |
| EPI_ISL_3237983 | LabKom - Labor Augsburg MVZ GmbH | Robert Koch Institute | unknown |
| EPI_ISL_3237984 | Labor 28 MVZ GmbH | Robert Koch Institute | unknown |
| EPI_ISL_3237985 | Sonic - Dr. Staber & Kollegen GmbH München | Robert Koch Institute | unknown |
| EPI_ISL_3237986 | Bioscientia Labor Wermsdorf | Robert Koch Institute | unknown |
| EPI_ISL_3237987 | LabKom - Labor Augsburg MVZ GmbH | Robert Koch Institute | unknown |
| EPI_ISL_3237988, EPI_ISL_3237989, EPI_ISL_3237990 | Bioscientia Labor Wermsdorf | Robert Koch Institute | unknown |
| EPI_ISL_3237991 | LabKom - Labor Augsburg MVZ GmbH | Robert Koch Institute | unknown |
| EPI_ISL_3237992 | Bioscientia Labor Wermsdorf | Robert Koch Institute | unknown |
| EPI_ISL_3237993 | LabKom - Labor Augsburg MVZ GmbH | Robert Koch Institute | unknown |
| EPI_ISL_3237994, EPI_ISL_3237995, EPI_ISL_3237996 | Bioscientia Labor Wermsdorf | Robert Koch Institute | unknown |

| | | | |
|---|---|---|---|
| EPI_ISL_3237997 | Labor 28 MVZ GmbH | Robert Koch Institute | unknown |
| EPI_ISL_3237998, EPI_ISL_3237999, EPI_ISL_3238000, EPI_ISL_3238001 | Bioscientia Labor Wermsdorf | Robert Koch Institute | unknown |
| EPI_ISL_3238002 | Bioscientia MVZ Labor Karlsruhe GmbH | Robert Koch Institute | unknown |
| EPI_ISL_3238003 | LabKom - Labor Augsburg MVZ GmbH | Robert Koch Institute | unknown |
| EPI_ISL_3238004 | Sonic - MVZ Medizinisches Labor Bremen GmbH | Robert Koch Institute | unknown |
| EPI_ISL_3238005 | Bioscientia Labor Wermsdorf | Robert Koch Institute | unknown |
| EPI_ISL_3238006, EPI_ISL_3238007 | LabKom - Labor Augsburg MVZ GmbH | Robert Koch Institute | unknown |
| EPI_ISL_3238008, EPI_ISL_3238009 | Bioscientia Labor Wermsdorf | Robert Koch Institute | unknown |
| EPI_ISL_3238010 | LabKom - Labor Augsburg MVZ GmbH | Robert Koch Institute | unknown |
| EPI_ISL_3238011, EPI_ISL_3238012, EPI_ISL_3238013, EPI_ISL_3238014 | Bioscientia Labor Wermsdorf | Robert Koch Institute | unknown |
| EPI_ISL_3238015 | LabKom - Labor Augsburg MVZ GmbH | Robert Koch Institute | unknown |
| EPI_ISL_3238016 | Sonic - Bioscientia - MVZ Labor Saar GmbH | Robert Koch Institute | unknown |
| EPI_ISL_3238017 | Bioscientia Labor Wermsdorf | Robert Koch Institute | unknown |
| EPI_ISL_3238018 | Sonic - Bioscientia - MVZ Labor Saar GmbH | Robert Koch Institute | unknown |
| EPI_ISL_3238019 | Bioscientia Labor Wermsdorf | Robert Koch Institute | unknown |
| EPI_ISL_3238021 | Universitätsklinikum Frankfurt - Institut für Medizinische Virologie | Robert Koch Institute | unknown |
| EPI_ISL_3238023, EPI_ISL_3238024, EPI_ISL_3238025, EPI_ISL_3238026 | Bioscientia Labor Wermsdorf | Robert Koch Institute | unknown |
| EPI_ISL_3238027 | Sonic - Labor Dr. von Froreich GmbH | Robert Koch Institute | unknown |
| EPI_ISL_3238028 | LabKom - Labor Augsburg MVZ GmbH | Robert Koch Institute | unknown |
| EPI_ISL_3238029 | Bioscientia Labor Wermsdorf | Robert Koch Institute | unknown |
| EPI_ISL_3238030 | LabKom - Labor an der Salzbrücke MVZ GmbH | Robert Koch Institute | unknown |
| EPI_ISL_3238031, EPI_ISL_3238032, EPI_ISL_3238033, EPI_ISL_3238034 | Bioscientia Labor Wermsdorf | Robert Koch Institute | unknown |
| EPI_ISL_3238035 | LabKom - Labor Augsburg MVZ GmbH | Robert Koch Institute | unknown |
| EPI_ISL_3238036 | Bioscientia Labor Wermsdorf | Robert Koch Institute | unknown |
| EPI_ISL_3238037, EPI_ISL_3238038 | LabKom - Labor Augsburg MVZ GmbH | Robert Koch Institute | unknown |
| EPI_ISL_3238039 | Sonic - Dr. Staber & Kollegen GmbH München | Robert Koch Institute | unknown |
| EPI_ISL_3238040 | Bioscientia Labor Wermsdorf | Robert Koch Institute | unknown |
| EPI_ISL_3238041 | Bioscientia MVZ Labor Karlsruhe GmbH | Robert Koch Institute | unknown |
| EPI_ISL_3238042 | Bioscientia Labor Wermsdorf | Robert Koch Institute | unknown |
| EPI_ISL_3238043 | Universitätsklinikum Frankfurt - Institut für Medizinische Virologie | Robert Koch Institute | unknown |
| EPI_ISL_3238044 | LabKom - Labor Augsburg MVZ GmbH | Robert Koch Institute | unknown |
| EPI_ISL_3238045, EPI_ISL_3238046, EPI_ISL_3238047, EPI_ISL_3238048, EPI_ISL_3238049 | Bioscientia Labor Wermsdorf | Robert Koch Institute | unknown |
| EPI_ISL_3238050 | Labor 28 MVZ GmbH | Robert Koch Institute | unknown |
| EPI_ISL_3238051 | Sonic - Bioscientia - MVZ Labor Saar GmbH | Robert Koch Institute | unknown |
| EPI_ISL_3238052 | Bioscientia Labor Wermsdorf | Robert Koch Institute | unknown |
| EPI_ISL_3238054 | Sonic - Bioscientia - MVZ Labor Saar GmbH | Robert Koch Institute | unknown |
| EPI_ISL_3238055, EPI_ISL_3238056, EPI_ISL_3238057, EPI_ISL_3238058 | Bioscientia Labor Wermsdorf | Robert Koch Institute | unknown |
| EPI_ISL_3238059 | Labor 28 MVZ GmbH | Robert Koch Institute | unknown |
| EPI_ISL_3238060 | Sonic - Labor Dr. von Froreich GmbH | Robert Koch Institute | unknown |
| EPI_ISL_3238061 | Bioscientia MVZ Labor Karlsruhe GmbH | Robert Koch Institute | unknown |
| EPI_ISL_3238062 | Bioscientia Labor Wermsdorf | Robert Koch Institute | unknown |
| EPI_ISL_3238063 | Sonic - Labor Dr. von Froreich GmbH | Robert Koch Institute | unknown |
| EPI_ISL_3238064 | Labor 28 MVZ GmbH | Robert Koch Institute | unknown |
| EPI_ISL_3238065 | LabKom - Labor Augsburg MVZ GmbH | Robert Koch Institute | unknown |
| EPI_ISL_3238066, EPI_ISL_3238067, EPI_ISL_3238068 | Bioscientia Labor Wermsdorf | Robert Koch Institute | unknown |
| EPI_ISL_3238069 | LabKom - Labor Augsburg MVZ GmbH | Robert Koch Institute | unknown |
| EPI_ISL_3238070 | Bioscientia Labor Wermsdorf | Robert Koch Institute | unknown |
| EPI_ISL_3238071 | Sonic - Bioscientia - MVZ Labor Saar GmbH | Robert Koch Institute | unknown |
| EPI_ISL_3238072, EPI_ISL_3238073 | Bioscientia Labor Wermsdorf | Robert Koch Institute | unknown |
| EPI_ISL_3238074 | LabKom - Labor Augsburg MVZ GmbH | Robert Koch Institute | unknown |
| EPI_ISL_3238075 | Sonic - MVZ Medizinisches Labor Bremen GmbH | Robert Koch Institute | unknown |
| EPI_ISL_3238076, EPI_ISL_3238077, EPI_ISL_3238079, EPI_ISL_3238080, EPI_ISL_3238081 | Bioscientia Labor Wermsdorf | Robert Koch Institute | unknown |
| EPI_ISL_3238082 | LabKom - Labor Augsburg MVZ GmbH | Robert Koch Institute | unknown |
| EPI_ISL_3238083, EPI_ISL_3238084 | Bioscientia Labor Wermsdorf | Robert Koch Institute | unknown |

| | | | |
|---|---|---|---|
| EPI_ISL_3238085 | Sonic - Bioscientia - MVZ Labor Saar GmbH | Robert Koch Institute | unknown |
| EPI_ISL_3238086 | Bioscientia Labor Wermsdorf | Robert Koch Institute | unknown |
| EPI_ISL_3238087 | Sonic - Dr. Staber & Kollegen GmbH München | Robert Koch Institute | unknown |
| EPI_ISL_3238088, EPI_ISL_3238089, EPI_ISL_3238090, EPI_ISL_3238091, EPI_ISL_3238092 | Bioscientia Labor Wermsdorf | Robert Koch Institute | unknown |
| EPI_ISL_3238093 | Labor 28 MVZ GmbH | Robert Koch Institute | unknown |
| EPI_ISL_3238094 | Bioscientia Labor Wermsdorf | Robert Koch Institute | unknown |
| EPI_ISL_3238095 | Labor 28 MVZ GmbH | Robert Koch Institute | unknown |
| EPI_ISL_3238096 | Sonic - Bioscientia - MVZ Labor Saar GmbH | Robert Koch Institute | unknown |
| EPI_ISL_3238097 | Bioscientia Labor Wermsdorf | Robert Koch Institute | unknown |
| EPI_ISL_3238099 | LabKom - Labor Augsburg MVZ GmbH | Robert Koch Institute | unknown |
| EPI_ISL_3238100 | Bioscientia Labor Wermsdorf | Robert Koch Institute | unknown |
| EPI_ISL_3238101 | LabKom - Labor Augsburg MVZ GmbH | Robert Koch Institute | unknown |
| EPI_ISL_3238102 | Bioscientia Labor Wermsdorf | Robert Koch Institute | unknown |
| EPI_ISL_3238103 | LabKom - Labor Augsburg MVZ GmbH | Robert Koch Institute | unknown |
| EPI_ISL_3238104, EPI_ISL_3238105, EPI_ISL_3238106 | Bioscientia Labor Wermsdorf | Robert Koch Institute | unknown |
| EPI_ISL_3238107 | LabKom - Labor Augsburg MVZ GmbH | Robert Koch Institute | unknown |
| EPI_ISL_3238108, EPI_ISL_3238109, EPI_ISL_3238110 | Bioscientia Labor Wermsdorf | Robert Koch Institute | unknown |
| EPI_ISL_3238111 | Bioscientia MVZ Labor Karlsruhe GmbH | Robert Koch Institute | unknown |
| EPI_ISL_3238112 | LabKom - Labor Augsburg MVZ GmbH | Robert Koch Institute | unknown |
| EPI_ISL_3238113, EPI_ISL_3238114 | Bioscientia Labor Wermsdorf | Robert Koch Institute | unknown |
| EPI_ISL_3238115 | Sonic - Bioscientia - MVZ Labor Saar GmbH | Robert Koch Institute | unknown |
| EPI_ISL_3238116, EPI_ISL_3238117, EPI_ISL_3238118 | Bioscientia Labor Wermsdorf | Robert Koch Institute | unknown |
| EPI_ISL_3238119 | Sonic - Dr. Staber & Kollegen GmbH München | Robert Koch Institute | unknown |
| EPI_ISL_3238120 | Sonic - Bioscientia - MVZ Labor Saar GmbH | Robert Koch Institute | unknown |
| EPI_ISL_3238121 | Sonic - Labor Dr. von Froreich GmbH | Robert Koch Institute | unknown |
| EPI_ISL_3238122, EPI_ISL_3238123, EPI_ISL_3238124, EPI_ISL_3238125, EPI_ISL_3238126, EPI_ISL_3238127 | Bioscientia Labor Wermsdorf | Robert Koch Institute | unknown |
| EPI_ISL_3238128 | Sonic - MVZ Medizinisches Labor Bremen GmbH | Robert Koch Institute | unknown |
| EPI_ISL_3238129 | LabKom - Labor Augsburg MVZ GmbH | Robert Koch Institute | unknown |
| EPI_ISL_3238130, EPI_ISL_3238131 | Bioscientia Labor Wermsdorf | Robert Koch Institute | unknown |
| EPI_ISL_3238132 | Robert-Bosch-Krankenhaus | Robert Koch Institute | unknown |
| EPI_ISL_3238133, EPI_ISL_3238134, EPI_ISL_3238135, EPI_ISL_3238136, EPI_ISL_3238138 | Bioscientia Labor Wermsdorf | Robert Koch Institute | unknown |
| EPI_ISL_3238139 | Sonic - MVZ Medizinisches Labor Bremen GmbH | Robert Koch Institute | unknown |
| EPI_ISL_3238140 | Sonic - Labor Dr. von Froreich GmbH | Robert Koch Institute | unknown |
| EPI_ISL_3238141 | Bioscientia Labor Wermsdorf | Robert Koch Institute | unknown |
| EPI_ISL_3238142, EPI_ISL_3238143 | LabKom - Labor Augsburg MVZ GmbH | Robert Koch Institute | unknown |
| EPI_ISL_3238144 | Labor 28 MVZ GmbH | Robert Koch Institute | unknown |
| EPI_ISL_3238145 | LabKom - Labor Augsburg MVZ GmbH | Robert Koch Institute | unknown |
| EPI_ISL_3238146 | Bioscientia Labor Wermsdorf | Robert Koch Institute | unknown |
| EPI_ISL_3238147 | Sonic - Dr. Staber & Kollegen GmbH München | Robert Koch Institute | unknown |
| EPI_ISL_3238148, EPI_ISL_3238149 | Bioscientia Labor Wermsdorf | Robert Koch Institute | unknown |
| EPI_ISL_3238150, EPI_ISL_3238151 | LabKom - Labor Augsburg MVZ GmbH | Robert Koch Institute | unknown |
| EPI_ISL_3238152, EPI_ISL_3238153 | Bioscientia Labor Wermsdorf | Robert Koch Institute | unknown |
| EPI_ISL_3238154 | Sonic - Labor Dr. von Froreich GmbH | Robert Koch Institute | unknown |
| EPI_ISL_3238155 | Labor 28 MVZ GmbH | Robert Koch Institute | unknown |
| EPI_ISL_3238156, EPI_ISL_3238157 | LabKom - Labor Augsburg MVZ GmbH | Robert Koch Institute | unknown |
| EPI_ISL_3238158, EPI_ISL_3238159 | Bioscientia Labor Wermsdorf | Robert Koch Institute | unknown |
| EPI_ISL_3238160, EPI_ISL_3238161 | LabKom - Labor Augsburg MVZ GmbH | Robert Koch Institute | unknown |
| EPI_ISL_3238162, EPI_ISL_3238163 | Bioscientia Labor Wermsdorf | Robert Koch Institute | unknown |
| EPI_ISL_3238164 | LabKom - Labor Augsburg MVZ GmbH | Robert Koch Institute | unknown |
| EPI_ISL_3238165 | Sonic - Labor Dr. von Froreich GmbH | Robert Koch Institute | unknown |
| EPI_ISL_3238166 | LabKom - Labor an der Salzbrücke MVZ GmbH | Robert Koch Institute | unknown |
| EPI_ISL_3238167, EPI_ISL_3238168 | Bioscientia Labor Wermsdorf | Robert Koch Institute | unknown |
| EPI_ISL_3238169 | Labor 28 MVZ GmbH | Robert Koch Institute | unknown |
| EPI_ISL_3238170 | Bioscientia Labor Wermsdorf | Robert Koch Institute | unknown |
| EPI_ISL_3238171 | Sonic - Bioscientia - MVZ Labor Saar GmbH | Robert Koch Institute | unknown |
| EPI_ISL_3238172 | LabKom - Labor Augsburg MVZ GmbH | Robert Koch Institute | unknown |

| | | | |
|---|---|---|---|
| EPI_ISL_3238173, EPI_ISL_3238174, EPI_ISL_3238175, EPI_ISL_3238176, EPI_ISL_3238177, EPI_ISL_3238178, EPI_ISL_3238179, EPI_ISL_3238180 | Bioscientia Labor Wermsdorf | Robert Koch Institute | unknown |
| EPI_ISL_3238181 | Sonic - Dr. Staber & Kollegen GmbH München | Robert Koch Institute | unknown |
| EPI_ISL_3238182, EPI_ISL_3238183, EPI_ISL_3238184 | LabKom - Labor Augsburg MVZ GmbH | Robert Koch Institute | unknown |
| EPI_ISL_3238185 | Labor 28 MVZ GmbH | Robert Koch Institute | unknown |
| EPI_ISL_3238186 | LabKom - Labor Augsburg MVZ GmbH | Robert Koch Institute | unknown |
| EPI_ISL_3238187, EPI_ISL_3238188 | Bioscientia Labor Wermsdorf | Robert Koch Institute | unknown |
| EPI_ISL_3238189 | LabKom - Labor an der Salzbrücke MVZ GmbH | Robert Koch Institute | unknown |
| EPI_ISL_3238190, EPI_ISL_3238191, EPI_ISL_3238192, EPI_ISL_3238193, EPI_ISL_3238194, EPI_ISL_3238195, EPI_ISL_3238196, EPI_ISL_3238197 | Bioscientia Labor Wermsdorf | Robert Koch Institute | unknown |
| EPI_ISL_3238198 | LabKom - Labor Augsburg MVZ GmbH | Robert Koch Institute | unknown |
| EPI_ISL_3238199, EPI_ISL_3238200, EPI_ISL_3238201 | Bioscientia Labor Wermsdorf | Robert Koch Institute | unknown |
| EPI_ISL_3238202 | Sonic - Bioscientia - MVZ Labor Saar GmbH | Robert Koch Institute | unknown |
| EPI_ISL_3238203 | Sonic - Dr. Staber & Kollegen GmbH München | Robert Koch Institute | unknown |
| EPI_ISL_3238204 | Sonic - Labor Dr. von Froreich GmbH | Robert Koch Institute | unknown |
| EPI_ISL_3238205, EPI_ISL_3238206 | Bioscientia Labor Wermsdorf | Robert Koch Institute | unknown |
| EPI_ISL_3238207 | Sonic - MVZ Medizinisches Labor Bremen GmbH | Robert Koch Institute | unknown |
| EPI_ISL_3238208 | Labor 28 MVZ GmbH | Robert Koch Institute | unknown |
| EPI_ISL_3238209, EPI_ISL_3238210, EPI_ISL_3238211, EPI_ISL_3238212 | Bioscientia Labor Wermsdorf | Robert Koch Institute | unknown |
| EPI_ISL_3238213 | Sonic - Bioscientia - MVZ Labor Saar GmbH | Robert Koch Institute | unknown |
| EPI_ISL_3238214 | Bioscientia Labor Wermsdorf | Robert Koch Institute | unknown |
| EPI_ISL_3238215, EPI_ISL_3238216, EPI_ISL_3238217, EPI_ISL_3238218, EPI_ISL_3238219, EPI_ISL_3238220, EPI_ISL_3238221, EPI_ISL_3238222, EPI_ISL_3238223, EPI_ISL_3238224, EPI_ISL_3238225, EPI_ISL_3238227, EPI_ISL_3238228, EPI_ISL_3238229, EPI_ISL_3238230, EPI_ISL_3238231, EPI_ISL_3238232, EPI_ISL_3238233, EPI_ISL_3238234, EPI_ISL_3238235, EPI_ISL_3238236, EPI_ISL_3238237, EPI_ISL_3238238, EPI_ISL_3238239, EPI_ISL_3238240, EPI_ISL_3238241, EPI_ISL_3238242, EPI_ISL_3238243, EPI_ISL_3238244, EPI_ISL_3238246, EPI_ISL_3238247, EPI_ISL_3238248, EPI_ISL_3238249, EPI_ISL_3238250, EPI_ISL_3238251, EPI_ISL_3238252, EPI_ISL_3238254, EPI_ISL_3238255, EPI_ISL_3238256, EPI_ISL_3238257, EPI_ISL_3238259, EPI_ISL_3238260, EPI_ISL_3238261, EPI_ISL_3238262, EPI_ISL_3238264, EPI_ISL_3238265, EPI_ISL_3238267, EPI_ISL_3238268, EPI_ISL_3238269, EPI_ISL_3238270, EPI_ISL_3238272, EPI_ISL_3238273, EPI_ISL_3238274, EPI_ISL_3238275, EPI_ISL_3238276, EPI_ISL_3238277, EPI_ISL_3238278, EPI_ISL_3238279, EPI_ISL_3238280, EPI_ISL_3238281, EPI_ISL_3238282, EPI_ISL_3238283, EPI_ISL_3238284, EPI_ISL_3238285, EPI_ISL_3238286, EPI_ISL_3238287, EPI_ISL_3238288, EPI_ISL_3238290, EPI_ISL_3238291, EPI_ISL_3238292, EPI_ISL_3238294, EPI_ISL_3238295, EPI_ISL_3238296, EPI_ISL_3238297, EPI_ISL_3238298, EPI_ISL_3238299, EPI_ISL_3238300, EPI_ISL_3238301, EPI_ISL_3238302, EPI_ISL_3238303, EPI_ISL_3238304, EPI_ISL_3238305, EPI_ISL_3238306, EPI_ISL_3238307, EPI_ISL_3238308, EPI_ISL_3238309, EPI_ISL_3238310, EPI_ISL_3238311, EPI_ISL_3238312, EPI_ISL_3238313, EPI_ISL_3238314, EPI_ISL_3238316, EPI_ISL_3238317, EPI_ISL_3238318, EPI_ISL_3238319, EPI_ISL_3238320, EPI_ISL_3238321, EPI_ISL_3238322, EPI_ISL_3238323, EPI_ISL_3238324, EPI_ISL_3238325, EPI_ISL_3238326, EPI_ISL_3238327, EPI_ISL_3238328, EPI_ISL_3238330, EPI_ISL_3238331, EPI_ISL_3238332, EPI_ISL_3238333, EPI_ISL_3238334, EPI_ISL_3238335, EPI_ISL_3238336, EPI_ISL_3238337, EPI_ISL_3238338, EPI_ISL_3238339, EPI_ISL_3238340, EPI_ISL_3238341, EPI_ISL_3238342, EPI_ISL_3238343, EPI_ISL_3238344, EPI_ISL_3238345, EPI_ISL_3238346, EPI_ISL_3238347, EPI_ISL_3238348 | | | |
| see above | CENTOGENE Frankfurt Laboratory: Niederlassung Industriepark Höchst | Robert Koch Institute | unknown |
| EPI_ISL_3238349, EPI_ISL_3238350, EPI_ISL_3238351 | Laborarztpraxis Osnabrück | Robert Koch Institute | unknown |
| EPI_ISL_3238352, EPI_ISL_3238353 | MVZ Dr. Eberhard & Partner Dortmund | Robert Koch Institute | unknown |
| EPI_ISL_3238354 | Laborarztpraxis Osnabrück | Robert Koch Institute | unknown |
| EPI_ISL_3238355 | MVZ Dr. Eberhard & Partner Dortmund | Robert Koch Institute | unknown |
| EPI_ISL_3238356 | labopart - Medizinische Laboratorien Dresden | Robert Koch Institute | unknown |
| EPI_ISL_3238357, EPI_ISL_3238358, EPI_ISL_3238359 | Laborarztpraxis Osnabrück | Robert Koch Institute | unknown |
| EPI_ISL_3238360, EPI_ISL_3238361, EPI_ISL_3238362 | MVZ Dr. Eberhard & Partner Dortmund | Robert Koch Institute | unknown |
| EPI_ISL_3238363 | Laborarztpraxis Osnabrück | Robert Koch Institute | unknown |
| EPI_ISL_3238364 | Medizinisches Labor Wahl Lüdenscheid | Robert Koch Institute | unknown |
| EPI_ISL_3238365 | Laborarztpraxis Osnabrück | Robert Koch Institute | unknown |
| EPI_ISL_3238366 | MVZ Dr. Eberhard & Partner Dortmund | Robert Koch Institute | unknown |
| EPI_ISL_3238367, EPI_ISL_3238368 | Laborarztpraxis Osnabrück | Robert Koch Institute | unknown |
| EPI_ISL_3238369 | MVZ Dr. Eberhard & Partner Dortmund | Robert Koch Institute | unknown |
| EPI_ISL_3238370 | labopart - Medizinische Laboratorien Dresden | Robert Koch Institute | unknown |
| EPI_ISL_3238371, EPI_ISL_3238372 | Laborarztpraxis Osnabrück | Robert Koch Institute | unknown |
| EPI_ISL_3238373 | labopart - Medizinische Laboratorien Dresden | Robert Koch Institute | unknown |
| EPI_ISL_3238374, EPI_ISL_3238375, EPI_ISL_3238376, EPI_ISL_3238377 | Laborarztpraxis Osnabrück | Robert Koch Institute | unknown |
| EPI_ISL_3238378 | MVZ Dr. Eberhard & Partner Dortmund | Robert Koch Institute | unknown |
| EPI_ISL_3238379, EPI_ISL_3238380, EPI_ISL_3238381 | Laborarztpraxis Osnabrück | Robert Koch Institute | unknown |
| EPI_ISL_3238382 | MVZ Dr. Eberhard & Partner Dortmund | Robert Koch Institute | unknown |
| EPI_ISL_3238383 | Laborarztpraxis Osnabrück | Robert Koch Institute | unknown |
| EPI_ISL_3238384 | MVZ Dr. Eberhard & Partner Dortmund | Robert Koch Institute | unknown |
| EPI_ISL_3238385, EPI_ISL_3238386, EPI_ISL_3238387, EPI_ISL_3238388, | Laborarztpraxis Osnabrück | Robert Koch Institute | unknown |

| | | | |
|---|---|---|---|
| EPI_ISL_3238389 | | | |
| EPI_ISL_3238390 | MVZ Dr. Eberhard & Partner Dortmund | Robert Koch Institute | unknown |
| EPI_ISL_3238391 | Laborarztpraxis Osnabrück | Robert Koch Institute | unknown |
| EPI_ISL_3238392, EPI_ISL_3238393, EPI_ISL_3238394, EPI_ISL_3238395, EPI_ISL_3238396, EPI_ISL_3238397 | Carl-Thiem-Klinikum Cottbus - Zentrum für Laboratoriumsmedizin Mikrobiologie und Krankenhaushygiene | Robert Koch Institute | unknown |
| EPI_ISL_3238398, EPI_ISL_3238399, EPI_ISL_3238400, EPI_ISL_3238401, EPI_ISL_3238402, EPI_ISL_3238403, EPI_ISL_3238404, EPI_ISL_3238405, EPI_ISL_3238406, EPI_ISL_3238407, EPI_ISL_3238408 | | | |
| see above | Labor Dr. Wisplinghoff - Köln | Robert Koch Institute | unknown |
| EPI_ISL_3238409, EPI_ISL_3238411, EPI_ISL_3238413, EPI_ISL_3238414, EPI_ISL_3238416 | Virologisches Institut des Universitätsklinikums Erlangen | Robert Koch Institute | unknown |
| EPI_ISL_3238417, EPI_ISL_3238418, EPI_ISL_3238419, EPI_ISL_3238420, EPI_ISL_3238421, EPI_ISL_3238422, EPI_ISL_3238423, EPI_ISL_3238424 | Labor Dr. Heidrich & Kollegen MVZ GmbH Hamburg | Robert Koch Institute | unknown |
| EPI_ISL_3238425, EPI_ISL_3238426, EPI_ISL_3238427, EPI_ISL_3238428, EPI_ISL_3238430, EPI_ISL_3238431, EPI_ISL_3238432, EPI_ISL_3238433, EPI_ISL_3238434, EPI_ISL_3238435, EPI_ISL_3238436, EPI_ISL_3238437, EPI_ISL_3238438, EPI_ISL_3238439, EPI_ISL_3238440, EPI_ISL_3238441, EPI_ISL_3238442, EPI_ISL_3238443, EPI_ISL_3238444, EPI_ISL_3238445, EPI_ISL_3238446, EPI_ISL_3238447, EPI_ISL_3238448, EPI_ISL_3238449, EPI_ISL_3238450, EPI_ISL_3238451, EPI_ISL_3238452, EPI_ISL_3238453, EPI_ISL_3238454 | | | |
| see above | MVZ Labor Krone GbR | Robert Koch Institute | unknown |
| EPI_ISL_3238455, EPI_ISL_3238456 | Thüringer Landesamtes für Verbraucherschutz | Robert Koch Institute | unknown |
| EPI_ISL_3238457, EPI_ISL_3238458 | Clemenshospital - Alexianer Clemenshospital Münster | Robert Koch Institute | unknown |
| EPI_ISL_3238461, EPI_ISL_3238462 | Bioscientia Labor Wermsdorf | Robert Koch Institute | unknown |
| EPI_ISL_3238463 | Bioscientia MVZ Labor Karlsruhe GmbH | Robert Koch Institute | unknown |
| EPI_ISL_3238464 | Bioscientia Labor Wermsdorf | Robert Koch Institute | unknown |
| EPI_ISL_3238465, EPI_ISL_3238466 | LabKom - Labor Augsburg MVZ GmbH | Robert Koch Institute | unknown |
| EPI_ISL_3238467, EPI_ISL_3238468 | Bioscientia Labor Wermsdorf | Robert Koch Institute | unknown |
| EPI_ISL_3238470 | LabKom - Labor Augsburg MVZ GmbH | Robert Koch Institute | unknown |
| EPI_ISL_3238471 | Bioscientia Labor Wermsdorf | Robert Koch Institute | unknown |
| EPI_ISL_3238472 | Sonic - Bioscientia - MVZ Labor Saar GmbH | Robert Koch Institute | unknown |
| EPI_ISL_3238473 | Bioscientia Labor Wermsdorf | Robert Koch Institute | unknown |
| EPI_ISL_3238474 | Sonic - Bioscientia - MVZ Labor Saar GmbH | Robert Koch Institute | unknown |
| EPI_ISL_3238475 | LabKom - Labor Augsburg MVZ GmbH | Robert Koch Institute | unknown |
| EPI_ISL_3238476, EPI_ISL_3238477 | Bioscientia MVZ Labor Karlsruhe GmbH | Robert Koch Institute | unknown |
| EPI_ISL_3238479, EPI_ISL_3238480 | LabKom - Labor Augsburg MVZ GmbH | Robert Koch Institute | unknown |
| EPI_ISL_3238481, EPI_ISL_3238482 | Bioscientia Labor Wermsdorf | Robert Koch Institute | unknown |
| EPI_ISL_3238483 | LabKom - Labor Augsburg MVZ GmbH | Robert Koch Institute | unknown |
| EPI_ISL_3238484 | Bioscientia Labor Wermsdorf | Robert Koch Institute | unknown |
| EPI_ISL_3238485, EPI_ISL_3238486 | LabKom - Labor Augsburg MVZ GmbH | Robert Koch Institute | unknown |
| EPI_ISL_3238487 | Bioscientia Labor Wermsdorf | Robert Koch Institute | unknown |
| EPI_ISL_3238488 | LabKom - Labor Augsburg MVZ GmbH | Robert Koch Institute | unknown |
| EPI_ISL_3238489 | Sonic - Bioscientia - MVZ Labor Saar GmbH | Robert Koch Institute | unknown |
| EPI_ISL_3238490 | LabKom - Labor Augsburg MVZ GmbH | Robert Koch Institute | unknown |
| EPI_ISL_3238491, EPI_ISL_3238492 | Bioscientia Labor Wermsdorf | Robert Koch Institute | unknown |
| EPI_ISL_3238493 | Sonic - Bioscientia - MVZ Labor Saar GmbH | Robert Koch Institute | unknown |
| EPI_ISL_3238494 | Sonic - Labor Staber Heilbronn | Robert Koch Institute | unknown |
| EPI_ISL_3238495 | Sonic - Bioscientia - MVZ Labor Saar GmbH | Robert Koch Institute | unknown |
| EPI_ISL_3238496 | Bioscientia Labor Wermsdorf | Robert Koch Institute | unknown |
| EPI_ISL_3238497 | LabKom - Labor Augsburg MVZ GmbH | Robert Koch Institute | unknown |
| EPI_ISL_3238498 | Bioscientia Labor Wermsdorf | Robert Koch Institute | unknown |
| EPI_ISL_3238499, EPI_ISL_3238500 | Sonic - Bioscientia - MVZ Labor Saar GmbH | Robert Koch Institute | unknown |
| EPI_ISL_3238501, EPI_ISL_3238502 | Bioscientia Labor Wermsdorf | Robert Koch Institute | unknown |
| EPI_ISL_3238503 | LabKom - Labor Augsburg MVZ GmbH | Robert Koch Institute | unknown |
| EPI_ISL_3238504 | Bioscientia Labor Wermsdorf | Robert Koch Institute | unknown |
| EPI_ISL_3238505, EPI_ISL_3238506, EPI_ISL_3238507 | Sonic - Bioscientia - MVZ Labor Saar GmbH | Robert Koch Institute | unknown |
| EPI_ISL_3238508 | LabKom - Labor Augsburg MVZ GmbH | Robert Koch Institute | unknown |
| EPI_ISL_3238509 | Bioscientia Labor Wermsdorf | Robert Koch Institute | unknown |
| EPI_ISL_3238510 | Bioscientia MVZ Labor Karlsruhe GmbH | Robert Koch Institute | unknown |
| EPI_ISL_3238511 | Bioscientia Labor Wermsdorf | Robert Koch Institute | unknown |
| EPI_ISL_3238512 | Bioscientia MVZ Labor Karlsruhe GmbH | Robert Koch Institute | unknown |
| EPI_ISL_3238513, EPI_ISL_3238514 | Bioscientia Labor Wermsdorf | Robert Koch Institute | unknown |
| EPI_ISL_3238515 | Sonic - Bioscientia - MVZ Labor Saar GmbH | Robert Koch Institute | unknown |
| EPI_ISL_3238516 | LabKom - Labor Augsburg MVZ GmbH | Robert Koch Institute | unknown |
| EPI_ISL_3238517 | Sonic - Bioscientia - MVZ Labor Saar GmbH | Robert Koch Institute | unknown |

| | | | |
|---|---|---|---|
| EPI_ISL_3238518, EPI_ISL_3238519 | Bioscientia Labor Wermsdorf | Robert Koch Institute | unknown |
| EPI_ISL_3238520 | LabKom - Labor Augsburg MVZ GmbH | Robert Koch Institute | unknown |
| EPI_ISL_3238521 | Sonic - Bioscientia - MVZ Labor Saar GmbH | Robert Koch Institute | unknown |
| EPI_ISL_3238522, EPI_ISL_3238523 | LabKom - Labor Augsburg MVZ GmbH | Robert Koch Institute | unknown |
| EPI_ISL_3238524 | Sonic - Bioscientia - MVZ Labor Saar GmbH | Robert Koch Institute | unknown |
| EPI_ISL_3238525 | Bioscientia MVZ Labor Karlsruhe GmbH | Robert Koch Institute | unknown |
| EPI_ISL_3238526 | Bioscientia Labor Wermsdorf | Robert Koch Institute | unknown |
| EPI_ISL_3238527 | LabKom - Labor Augsburg MVZ GmbH | Robert Koch Institute | unknown |
| EPI_ISL_3238528 | Bioscientia Labor Wermsdorf | Robert Koch Institute | unknown |
| EPI_ISL_3238529 | LabKom - Labor Augsburg MVZ GmbH | Robert Koch Institute | unknown |
| EPI_ISL_3238530, EPI_ISL_3238531 | Sonic - Bioscientia - MVZ Labor Saar GmbH | Robert Koch Institute | unknown |
| EPI_ISL_3238532, EPI_ISL_3238533, EPI_ISL_3238534 | Bioscientia Labor Wermsdorf | Robert Koch Institute | unknown |
| EPI_ISL_3238535, EPI_ISL_3238536, EPI_ISL_3238537 | LabKom - Labor Augsburg MVZ GmbH | Robert Koch Institute | unknown |
| EPI_ISL_3238538 | Sonic - Bioscientia - MVZ Labor Saar GmbH | Robert Koch Institute | unknown |
| EPI_ISL_3238541, EPI_ISL_3238543, EPI_ISL_3238544, EPI_ISL_3238545 | LabKom - Labor Augsburg MVZ GmbH | Robert Koch Institute | unknown |
| EPI_ISL_3238546, EPI_ISL_3238547, EPI_ISL_3238548 | Bioscientia Labor Wermsdorf | Robert Koch Institute | unknown |
| EPI_ISL_3238549 | Bioscientia MVZ Labor Karlsruhe GmbH | Robert Koch Institute | unknown |
| EPI_ISL_3238550 | Sonic - Bioscientia - MVZ Labor Saar GmbH | Robert Koch Institute | unknown |
| EPI_ISL_3238551 | Bioscientia Labor Wermsdorf | Robert Koch Institute | unknown |
| EPI_ISL_3238552 | Sonic - Bioscientia - MVZ Labor Saar GmbH | Robert Koch Institute | unknown |
| EPI_ISL_3238553 | Bioscientia Labor Wermsdorf | Robert Koch Institute | unknown |
| EPI_ISL_3238554 | Bioscientia MVZ Labor Karlsruhe GmbH | Robert Koch Institute | unknown |
| EPI_ISL_3238555 | LabKom - Labor Augsburg MVZ GmbH | Robert Koch Institute | unknown |
| EPI_ISL_3238556 | Sonic - Bioscientia - MVZ Labor Saar GmbH | Robert Koch Institute | unknown |
| EPI_ISL_3238557, EPI_ISL_3238558 | Bioscientia MVZ Labor Karlsruhe GmbH | Robert Koch Institute | unknown |
| EPI_ISL_3238559 | Sonic - Bioscientia - MVZ Labor Saar GmbH | Robert Koch Institute | unknown |
| EPI_ISL_3238560 | LabKom - Labor Augsburg MVZ GmbH | Robert Koch Institute | unknown |
| EPI_ISL_3238561 | Bioscientia MVZ Labor Karlsruhe GmbH | Robert Koch Institute | unknown |
| EPI_ISL_3238562 | Sonic - Bioscientia - MVZ Labor Saar GmbH | Robert Koch Institute | unknown |
| EPI_ISL_3238563 | Bioscientia Labor Wermsdorf | Robert Koch Institute | unknown |
| EPI_ISL_3238564 | LabKom - Labor Augsburg MVZ GmbH | Robert Koch Institute | unknown |
| EPI_ISL_3238565 | Bioscientia Labor Wermsdorf | Robert Koch Institute | unknown |
| EPI_ISL_3238566 | LabKom - Labor Augsburg MVZ GmbH | Robert Koch Institute | unknown |
| EPI_ISL_3238567 | Bioscientia Labor Wermsdorf | Robert Koch Institute | unknown |
| EPI_ISL_3238568 | LabKom - Labor Augsburg MVZ GmbH | Robert Koch Institute | unknown |
| EPI_ISL_3238569 | Bioscientia MVZ Labor Karlsruhe GmbH | Robert Koch Institute | unknown |
| EPI_ISL_3238570 | Sonic - Bioscientia - MVZ Labor Saar GmbH | Robert Koch Institute | unknown |
| EPI_ISL_3238571, EPI_ISL_3238572 | Bioscientia MVZ Labor Karlsruhe GmbH | Robert Koch Institute | unknown |
| EPI_ISL_3238573 | Bioscientia Labor Wermsdorf | Robert Koch Institute | unknown |
| EPI_ISL_3238574 | Sonic - Bioscientia - MVZ Labor Saar GmbH | Robert Koch Institute | unknown |
| EPI_ISL_3238575 | Bioscientia MVZ Labor Karlsruhe GmbH | Robert Koch Institute | unknown |
| EPI_ISL_3238576 | Bioscientia Labor Wermsdorf | Robert Koch Institute | unknown |
| EPI_ISL_3238577 | Bioscientia MVZ Labor Karlsruhe GmbH | Robert Koch Institute | unknown |
| EPI_ISL_3238578 | Sonic - Bioscientia - MVZ Labor Saar GmbH | Robert Koch Institute | unknown |
| EPI_ISL_3238579, EPI_ISL_3238580, EPI_ISL_3238581 | LabKom - Labor Augsburg MVZ GmbH | Robert Koch Institute | unknown |
| EPI_ISL_3238582 | Sonic - Dr. Staber & Kollegen GmbH München | Robert Koch Institute | unknown |
| EPI_ISL_3238583 | Bioscientia MVZ Labor Karlsruhe GmbH | Robert Koch Institute | unknown |
| EPI_ISL_3238584, EPI_ISL_3238585 | LabKom - Labor Augsburg MVZ GmbH | Robert Koch Institute | unknown |
| EPI_ISL_3238586, EPI_ISL_3238587 | Sonic - Bioscientia - MVZ Labor Saar GmbH | Robert Koch Institute | unknown |
| EPI_ISL_3238588 | Bioscientia Labor Wermsdorf | Robert Koch Institute | unknown |
| EPI_ISL_3238589 | LabKom - Labor Augsburg MVZ GmbH | Robert Koch Institute | unknown |
| EPI_ISL_3238590, EPI_ISL_3238591, EPI_ISL_3238592 | Bioscientia Labor Wermsdorf | Robert Koch Institute | unknown |
| EPI_ISL_3238593 | Sonic - Bioscientia - MVZ Labor Saar GmbH | Robert Koch Institute | unknown |
| EPI_ISL_3238594 | LabKom - Labor Augsburg MVZ GmbH | Robert Koch Institute | unknown |
| EPI_ISL_3238595 | Bioscientia Labor Wermsdorf | Robert Koch Institute | unknown |
| EPI_ISL_3238596 | LabKom - Labor Augsburg MVZ GmbH | Robert Koch Institute | unknown |
| EPI_ISL_3238597, EPI_ISL_3238598 | Bioscientia Labor Wermsdorf | Robert Koch Institute | unknown |
| EPI_ISL_3238599 | LabKom - Labor Augsburg MVZ GmbH | Robert Koch Institute | unknown |
| EPI_ISL_3238600, EPI_ISL_3238601, EPI_ISL_3238602, EPI_ISL_3238603 | Bioscientia Labor Wermsdorf | Robert Koch Institute | unknown |
| EPI_ISL_3238604 | Bioscientia MVZ Labor Karlsruhe GmbH | Robert Koch Institute | unknown |

| | | | |
|---|---|---|---|
| EPI_ISL_3238605 | Sonic - Bioscientia - MVZ Labor Saar GmbH | Robert Koch Institute | unknown |
| EPI_ISL_3238606 | LabKom - Labor Augsburg MVZ GmbH | Robert Koch Institute | unknown |
| EPI_ISL_3238607, EPI_ISL_3238608 | Bioscientia MVZ Labor Karlsruhe GmbH | Robert Koch Institute | unknown |
| EPI_ISL_3238609 | Sonic - Bioscientia - MVZ Labor Saar GmbH | Robert Koch Institute | unknown |
| EPI_ISL_3238610, EPI_ISL_3238611, EPI_ISL_3238612 | LabKom - Labor Augsburg MVZ GmbH | Robert Koch Institute | unknown |
| EPI_ISL_3238613 | Bioscientia MVZ Labor Karlsruhe GmbH | Robert Koch Institute | unknown |
| EPI_ISL_3238614, EPI_ISL_3238615 | Bioscientia Labor Wermsdorf | Robert Koch Institute | unknown |
| EPI_ISL_3238616 | Sonic - Bioscientia - MVZ Labor Saar GmbH | Robert Koch Institute | unknown |
| EPI_ISL_3238617 | Bioscientia MVZ Labor Karlsruhe GmbH | Robert Koch Institute | unknown |
| EPI_ISL_3238618 | Bioscientia Labor Wermsdorf | Robert Koch Institute | unknown |
| EPI_ISL_3238619 | LabKom - Labor Augsburg MVZ GmbH | Robert Koch Institute | unknown |
| EPI_ISL_3238620 | Sonic - Bioscientia - MVZ Labor Saar GmbH | Robert Koch Institute | unknown |
| EPI_ISL_3238621, EPI_ISL_3238622 | Bioscientia Labor Wermsdorf | Robert Koch Institute | unknown |
| EPI_ISL_3238623, EPI_ISL_3238624, EPI_ISL_3238625, EPI_ISL_3238626 | LabKom - Labor Augsburg MVZ GmbH | Robert Koch Institute | unknown |
| EPI_ISL_3238627 | Bioscientia Labor Wermsdorf | Robert Koch Institute | unknown |
| EPI_ISL_3238628 | LabKom - Labor Augsburg MVZ GmbH | Robert Koch Institute | unknown |
| EPI_ISL_3238629 | Sonic - Bioscientia - MVZ Labor Saar GmbH | Robert Koch Institute | unknown |
| EPI_ISL_3238630 | Bioscientia MVZ Labor Karlsruhe GmbH | Robert Koch Institute | unknown |
| EPI_ISL_3238631 | LabKom - Labor Augsburg MVZ GmbH | Robert Koch Institute | unknown |
| EPI_ISL_3238632 | Bioscientia Labor Wermsdorf | Robert Koch Institute | unknown |
| EPI_ISL_3238633 | Sonic - Dr. Staber & Kollegen GmbH München | Robert Koch Institute | unknown |
| EPI_ISL_3238634 | Bioscientia MVZ Labor Karlsruhe GmbH | Robert Koch Institute | unknown |
| EPI_ISL_3238635, EPI_ISL_3238636 | Sonic - Bioscientia - MVZ Labor Saar GmbH | Robert Koch Institute | unknown |
| EPI_ISL_3238638 | Bioscientia MVZ Labor Karlsruhe GmbH | Robert Koch Institute | unknown |
| EPI_ISL_3238639 | LabKom - Labor Augsburg MVZ GmbH | Robert Koch Institute | unknown |
| EPI_ISL_3238640 | Bioscientia Labor Wermsdorf | Robert Koch Institute | unknown |
| EPI_ISL_3238641, EPI_ISL_3238642 | Sonic - Bioscientia - MVZ Labor Saar GmbH | Robert Koch Institute | unknown |
| EPI_ISL_3238643, EPI_ISL_3238644, EPI_ISL_3238645 | Bioscientia Labor Wermsdorf | Robert Koch Institute | unknown |
| EPI_ISL_3238646 | Sonic - Bioscientia - MVZ Labor Saar GmbH | Robert Koch Institute | unknown |
| EPI_ISL_3238647 | Bioscientia Labor Wermsdorf | Robert Koch Institute | unknown |
| EPI_ISL_3238648 | LabKom - Labor Augsburg MVZ GmbH | Robert Koch Institute | unknown |
| EPI_ISL_3238649 | Bioscientia MVZ Labor Karlsruhe GmbH | Robert Koch Institute | unknown |
| EPI_ISL_3238650 | LabKom - Labor Augsburg MVZ GmbH | Robert Koch Institute | unknown |
| EPI_ISL_3238651, EPI_ISL_3238652, EPI_ISL_3238653 | Sonic - Bioscientia - MVZ Labor Saar GmbH | Robert Koch Institute | unknown |
| EPI_ISL_3238654 | Bioscientia Labor Wermsdorf | Robert Koch Institute | unknown |
| EPI_ISL_3238655 | Sonic - Bioscientia - MVZ Labor Saar GmbH | Robert Koch Institute | unknown |
| EPI_ISL_3238656 | LabKom - Labor Augsburg MVZ GmbH | Robert Koch Institute | unknown |
| EPI_ISL_3238657 | Bioscientia Labor Wermsdorf | Robert Koch Institute | unknown |
| EPI_ISL_3238658, EPI_ISL_3238659 | LabKom - Labor Augsburg MVZ GmbH | Robert Koch Institute | unknown |
| EPI_ISL_3238660 | Sonic - Bioscientia - MVZ Labor Saar GmbH | Robert Koch Institute | unknown |
| EPI_ISL_3238661, EPI_ISL_3238662, EPI_ISL_3238663 | LabKom - Labor Augsburg MVZ GmbH | Robert Koch Institute | unknown |
| EPI_ISL_3238664 | Sonic - Bioscientia - MVZ Labor Saar GmbH | Robert Koch Institute | unknown |
| EPI_ISL_3238665 | Bioscientia Labor Wermsdorf | Robert Koch Institute | unknown |
| EPI_ISL_3238666, EPI_ISL_3238667 | LabKom - Labor Augsburg MVZ GmbH | Robert Koch Institute | unknown |
| EPI_ISL_3238668, EPI_ISL_3238669 | Bioscientia Labor Wermsdorf | Robert Koch Institute | unknown |
| EPI_ISL_3238670 | Sonic - Bioscientia - MVZ Labor Saar GmbH | Robert Koch Institute | unknown |
| EPI_ISL_3238671, EPI_ISL_3238672 | LabKom - Labor Augsburg MVZ GmbH | Robert Koch Institute | unknown |
| EPI_ISL_3238673, EPI_ISL_3238674 | Bioscientia Labor Wermsdorf | Robert Koch Institute | unknown |
| EPI_ISL_3238675, EPI_ISL_3238676 | Bioscientia MVZ Labor Karlsruhe GmbH | Robert Koch Institute | unknown |
| EPI_ISL_3238677, EPI_ISL_3238678 | Bioscientia Labor Wermsdorf | Robert Koch Institute | unknown |
| EPI_ISL_3238679, EPI_ISL_3238680, EPI_ISL_3238681 | LabKom - Labor Augsburg MVZ GmbH | Robert Koch Institute | unknown |
| EPI_ISL_3238682 | Sonic - Bioscientia - MVZ Labor Saar GmbH | Robert Koch Institute | unknown |
| EPI_ISL_3238684 | Bioscientia MVZ Labor Karlsruhe GmbH | Robert Koch Institute | unknown |
| EPI_ISL_3238685 | Bioscientia Labor Wermsdorf | Robert Koch Institute | unknown |
| EPI_ISL_3238686 | LabKom - Labor Augsburg MVZ GmbH | Robert Koch Institute | unknown |
| EPI_ISL_3238687 | Bioscientia Labor Wermsdorf | Robert Koch Institute | unknown |
| EPI_ISL_3238688 | LabKom - Labor Augsburg MVZ GmbH | Robert Koch Institute | unknown |
| EPI_ISL_3238689 | Sonic - Bioscientia - MVZ Labor Saar GmbH | Robert Koch Institute | unknown |
| EPI_ISL_3238690 | Bioscientia Labor Wermsdorf | Robert Koch Institute | unknown |

| | | | |
|---|---|---|---|
| EPI_ISL_3238691 | Sonic - Bioscientia - MVZ Labor Saar GmbH | Robert Koch Institute | unknown |
| EPI_ISL_3238692, EPI_ISL_3238693 | Bioscientia Labor Wermsdorf | Robert Koch Institute | unknown |
| EPI_ISL_3238695 | Sonic - MVZ Medizinisches Labor Bremen GmbH | Robert Koch Institute | unknown |
| EPI_ISL_3238696, EPI_ISL_3238697 | Bioscientia Labor Wermsdorf | Robert Koch Institute | unknown |
| EPI_ISL_3238698 | LabKom - Labor Augsburg MVZ GmbH | Robert Koch Institute | unknown |
| EPI_ISL_3238699 | Bioscientia Labor Wermsdorf | Robert Koch Institute | unknown |
| EPI_ISL_3238700, EPI_ISL_3238701 | Labor 28 MVZ GmbH | Robert Koch Institute | unknown |
| EPI_ISL_3238702 | Bioscientia Labor Wermsdorf | Robert Koch Institute | unknown |
| EPI_ISL_3238703 | Bioscientia MVZ Labor Karlsruhe GmbH | Robert Koch Institute | unknown |
| EPI_ISL_3238704 | LabKom - Labor Augsburg MVZ GmbH | Robert Koch Institute | unknown |
| EPI_ISL_3238705, EPI_ISL_3238706 | Bioscientia Labor Wermsdorf | Robert Koch Institute | unknown |
| EPI_ISL_3238707 | LabKom - Labor Hannover MVZ GmbH | Robert Koch Institute | unknown |
| EPI_ISL_3238708 | LabKom - Labor Augsburg MVZ GmbH | Robert Koch Institute | unknown |
| EPI_ISL_3238709 | Labor 28 MVZ GmbH | Robert Koch Institute | unknown |
| EPI_ISL_3238711, EPI_ISL_3238712 | Bioscientia Labor Wermsdorf | Robert Koch Institute | unknown |
| EPI_ISL_3238713 | LabKom - Labor Augsburg MVZ GmbH | Robert Koch Institute | unknown |
| EPI_ISL_3238714 | Labor 28 MVZ GmbH | Robert Koch Institute | unknown |
| EPI_ISL_3238715 | Sonic - MVZ Medizinisches Labor Bremen GmbH | Robert Koch Institute | unknown |
| EPI_ISL_3238716, EPI_ISL_3238717 | Bioscientia MVZ Labor Karlsruhe GmbH | Robert Koch Institute | unknown |
| EPI_ISL_3238718 | Labor 28 MVZ GmbH | Robert Koch Institute | unknown |
| EPI_ISL_3238719 | LabKom - Labor Augsburg MVZ GmbH | Robert Koch Institute | unknown |
| EPI_ISL_3238720 | Sonic - MVZ Medizinisches Labor Bremen GmbH | Robert Koch Institute | unknown |
| EPI_ISL_3238721 | LabKom - Labor Augsburg MVZ GmbH | Robert Koch Institute | unknown |
| EPI_ISL_3238722 | Klinikum Fulda gAG | Robert Koch Institute | unknown |
| EPI_ISL_3238724 | Bioscientia Labor Wermsdorf | Robert Koch Institute | unknown |
| EPI_ISL_3238725, EPI_ISL_3238726 | Labor 28 MVZ GmbH | Robert Koch Institute | unknown |
| EPI_ISL_3238727 | LabKom - Labor Augsburg MVZ GmbH | Robert Koch Institute | unknown |
| EPI_ISL_3238729 | Labor 28 MVZ GmbH | Robert Koch Institute | unknown |
| EPI_ISL_3238732 | LabKom - Labor Augsburg MVZ GmbH | Robert Koch Institute | unknown |
| EPI_ISL_3238733 | Bioscientia MVZ Labor Karlsruhe GmbH | Robert Koch Institute | unknown |
| EPI_ISL_3238734 | Bioscientia Labor Wermsdorf | Robert Koch Institute | unknown |
| EPI_ISL_3238735 | Labor 28 MVZ GmbH | Robert Koch Institute | unknown |
| EPI_ISL_3238736 | LabKom - Labor Augsburg MVZ GmbH | Robert Koch Institute | unknown |
| EPI_ISL_3238737 | Labor 28 MVZ GmbH | Robert Koch Institute | unknown |
| EPI_ISL_3238738 | Bioscientia Labor Wermsdorf | Robert Koch Institute | unknown |
| EPI_ISL_3238739 | LabKom - Labor Augsburg MVZ GmbH | Robert Koch Institute | unknown |
| EPI_ISL_3238740, EPI_ISL_3238741 | Bioscientia Labor Wermsdorf | Robert Koch Institute | unknown |
| EPI_ISL_3238742, EPI_ISL_3238743 | LabKom - Labor Augsburg MVZ GmbH | Robert Koch Institute | unknown |
| EPI_ISL_3238744 | Labor 28 MVZ GmbH | Robert Koch Institute | unknown |
| EPI_ISL_3238745 | Bioscientia Labor Wermsdorf | Robert Koch Institute | unknown |
| EPI_ISL_3238746 | Labor 28 MVZ GmbH | Robert Koch Institute | unknown |
| EPI_ISL_3238747 | Sonic - MVZ Medizinisches Labor Bremen GmbH | Robert Koch Institute | unknown |
| EPI_ISL_3238748, EPI_ISL_3238749 | Labor 28 MVZ GmbH | Robert Koch Institute | unknown |
| EPI_ISL_3238750 | Bioscientia Labor Wermsdorf | Robert Koch Institute | unknown |
| EPI_ISL_3238751, EPI_ISL_3238752 | Labor 28 MVZ GmbH | Robert Koch Institute | unknown |
| EPI_ISL_3238753 | Bioscientia Labor Wermsdorf | Robert Koch Institute | unknown |
| EPI_ISL_3238754 | Labor 28 MVZ GmbH | Robert Koch Institute | unknown |
| EPI_ISL_3238755, EPI_ISL_3238756 | LabKom - Labor Augsburg MVZ GmbH | Robert Koch Institute | unknown |
| EPI_ISL_3238757 | Sonic - Dr. Staber & Kollegen GmbH München | Robert Koch Institute | unknown |
| EPI_ISL_3238758 | LabKom - Labor Augsburg MVZ GmbH | Robert Koch Institute | unknown |
| EPI_ISL_3238759 | Bioscientia Labor Wermsdorf | Robert Koch Institute | unknown |
| EPI_ISL_3238760 | LabKom - Labor Augsburg MVZ GmbH | Robert Koch Institute | unknown |
| EPI_ISL_3238761, EPI_ISL_3238762 | Bioscientia Labor Wermsdorf | Robert Koch Institute | unknown |
| EPI_ISL_3238763 | LabKom - Labor Augsburg MVZ GmbH | Robert Koch Institute | unknown |
| EPI_ISL_3238764 | Labor 28 MVZ GmbH | Robert Koch Institute | unknown |
| EPI_ISL_3238765 | Bioscientia Labor Wermsdorf | Robert Koch Institute | unknown |
| EPI_ISL_3238766 | Bioscientia MVZ Labor Karlsruhe GmbH | Robert Koch Institute | unknown |
| EPI_ISL_3238767 | LabKom - Labor Augsburg MVZ GmbH | Robert Koch Institute | unknown |
| EPI_ISL_3238768 | Sonic - MVZ Medizinisches Labor Bremen GmbH | Robert Koch Institute | unknown |
| EPI_ISL_3238769 | Labor 28 MVZ GmbH | Robert Koch Institute | unknown |
| EPI_ISL_3238770 | LabKom - Labor Augsburg MVZ GmbH | Robert Koch Institute | unknown |
| EPI_ISL_3238771, EPI_ISL_3238772 | Bioscientia Labor Wermsdorf | Robert Koch Institute | unknown |
| EPI_ISL_3238773 | Labor 28 MVZ GmbH | Robert Koch Institute | unknown |
| EPI_ISL_3238774, EPI_ISL_3238775 | Bioscientia Labor Wermsdorf | Robert Koch Institute | unknown |
| EPI_ISL_3238776 | LabKom - Labor Augsburg MVZ GmbH | Robert Koch Institute | unknown |
| EPI_ISL_3238777 | Labor 28 MVZ GmbH | Robert Koch Institute | unknown |

| | | | |
|---|---|---|---|
| EPI_ISL_3238778 | LabKom - Labor Augsburg MVZ GmbH | Robert Koch Institute | unknown |
| EPI_ISL_3238779 | Bioscientia MVZ Labor Karlsruhe GmbH | Robert Koch Institute | unknown |
| EPI_ISL_3238780, EPI_ISL_3238781 | LabKom - Labor Augsburg MVZ GmbH | Robert Koch Institute | unknown |
| EPI_ISL_3238782 | Labor 28 MVZ GmbH | Robert Koch Institute | unknown |
| EPI_ISL_3238783 | LabKom - Labor Hannover MVZ GmbH | Robert Koch Institute | unknown |
| EPI_ISL_3238784 | Sonic - MVZ Medizinisches Labor Bremen GmbH | Robert Koch Institute | unknown |
| EPI_ISL_3238785 | LabKom - Labor Augsburg MVZ GmbH | Robert Koch Institute | unknown |
| EPI_ISL_3238786 | Bioscientia Labor Wermsdorf | Robert Koch Institute | unknown |
| EPI_ISL_3238787 | LabKom - Labor Augsburg MVZ GmbH | Robert Koch Institute | unknown |
| EPI_ISL_3238788 | Sonic - MVZ Medizinisches Labor Bremen GmbH | Robert Koch Institute | unknown |
| EPI_ISL_3238789 | LabKom - Labor Augsburg MVZ GmbH | Robert Koch Institute | unknown |
| EPI_ISL_3238790, EPI_ISL_3238791 | Labor 28 MVZ GmbH | Robert Koch Institute | unknown |
| EPI_ISL_3238792, EPI_ISL_3238793, EPI_ISL_3238794, EPI_ISL_3238795, EPI_ISL_3238796 | LabKom - Labor Augsburg MVZ GmbH | Robert Koch Institute | unknown |
| EPI_ISL_3238797, EPI_ISL_3238799 | Labor 28 MVZ GmbH | Robert Koch Institute | unknown |
| EPI_ISL_3238800 | Bioscientia Labor Wermsdorf | Robert Koch Institute | unknown |
| EPI_ISL_3238801 | LabKom - Labor Augsburg MVZ GmbH | Robert Koch Institute | unknown |
| EPI_ISL_3238802 | Sonic - MVZ Medizinisches Labor Bremen GmbH | Robert Koch Institute | unknown |
| EPI_ISL_3238803 | Labor 28 MVZ GmbH | Robert Koch Institute | unknown |
| EPI_ISL_3238804, EPI_ISL_3238806, EPI_ISL_3238807 | Bioscientia Labor Wermsdorf | Robert Koch Institute | unknown |
| EPI_ISL_3238808 | Sonic - Labor Staber Bayreuth | Robert Koch Institute | unknown |
| EPI_ISL_3238809, EPI_ISL_3238810, EPI_ISL_3238811 | Bioscientia Labor Wermsdorf | Robert Koch Institute | unknown |
| EPI_ISL_3238813 | Sonic - MVZ Medizinisches Labor Bremen GmbH | Robert Koch Institute | unknown |
| EPI_ISL_3238814 | Sonic - Dr. Staber & Kollegen GmbH München | Robert Koch Institute | unknown |
| EPI_ISL_3238815 | LabKom - Labor Augsburg MVZ GmbH | Robert Koch Institute | unknown |
| EPI_ISL_3238816 | Labor 28 MVZ GmbH | Robert Koch Institute | unknown |
| EPI_ISL_3238818 | LabKom - Labor Augsburg MVZ GmbH | Robert Koch Institute | unknown |
| EPI_ISL_3238819 | Bioscientia MVZ Labor Karlsruhe GmbH | Robert Koch Institute | unknown |
| EPI_ISL_3238820, EPI_ISL_3238821 | Bioscientia Labor Wermsdorf | Robert Koch Institute | unknown |
| EPI_ISL_3238822 | Labor 28 MVZ GmbH | Robert Koch Institute | unknown |
| EPI_ISL_3238823 | Bioscientia Labor Wermsdorf | Robert Koch Institute | unknown |
| EPI_ISL_3238824 | LabKom - Labor an der Salzbrücke MVZ GmbH | Robert Koch Institute | unknown |
| EPI_ISL_3238825 | Sonic - MVZ Medizinisches Labor Bremen GmbH | Robert Koch Institute | unknown |
| EPI_ISL_3238826 | Bioscientia Labor Wermsdorf | Robert Koch Institute | unknown |
| EPI_ISL_3238827 | LabKom - Labor Augsburg MVZ GmbH | Robert Koch Institute | unknown |
| EPI_ISL_3238828 | Labor 28 MVZ GmbH | Robert Koch Institute | unknown |
| EPI_ISL_3238829 | LabKom - Labor Augsburg MVZ GmbH | Robert Koch Institute | unknown |
| EPI_ISL_3238830, EPI_ISL_3238831 | Labor 28 MVZ GmbH | Robert Koch Institute | unknown |
| EPI_ISL_3238832 | Bioscientia Labor Wermsdorf | Robert Koch Institute | unknown |
| EPI_ISL_3238833 | LabKom - Labor Augsburg MVZ GmbH | Robert Koch Institute | unknown |
| EPI_ISL_3238834 | Bioscientia Labor Wermsdorf | Robert Koch Institute | unknown |
| EPI_ISL_3238835, EPI_ISL_3238836, EPI_ISL_3238837 | LabKom - Labor Augsburg MVZ GmbH | Robert Koch Institute | unknown |
| EPI_ISL_3238838 | Labor 28 MVZ GmbH | Robert Koch Institute | unknown |
| EPI_ISL_3238839 | LabKom - Labor Augsburg MVZ GmbH | Robert Koch Institute | unknown |
| EPI_ISL_3238840 | Bioscientia Labor Wermsdorf | Robert Koch Institute | unknown |
| EPI_ISL_3238841, EPI_ISL_3238842 | LabKom - Labor Augsburg MVZ GmbH | Robert Koch Institute | unknown |
| EPI_ISL_3238843, EPI_ISL_3238844, EPI_ISL_3238845 | Bioscientia Labor Wermsdorf | Robert Koch Institute | unknown |
| EPI_ISL_3238846 | Labor 28 MVZ GmbH | Robert Koch Institute | unknown |
| EPI_ISL_3238847 | LabKom - Labor Augsburg MVZ GmbH | Robert Koch Institute | unknown |
| EPI_ISL_3238848 | Labor 28 MVZ GmbH | Robert Koch Institute | unknown |
| EPI_ISL_3238849 | Bioscientia Labor Wermsdorf | Robert Koch Institute | unknown |
| EPI_ISL_3238850, EPI_ISL_3238851 | Labor 28 MVZ GmbH | Robert Koch Institute | unknown |
| EPI_ISL_3238852 | LabKom - Labor Augsburg MVZ GmbH | Robert Koch Institute | unknown |
| EPI_ISL_3238853 | Labor 28 MVZ GmbH | Robert Koch Institute | unknown |
| EPI_ISL_3238854 | LabKom - Labor Augsburg MVZ GmbH | Robert Koch Institute | unknown |
| EPI_ISL_3238855 | Bioscientia MVZ Labor Karlsruhe GmbH | Robert Koch Institute | unknown |
| EPI_ISL_3238856 | Bioscientia Labor Wermsdorf | Robert Koch Institute | unknown |
| EPI_ISL_3238857 | LabKom - Labor Augsburg MVZ GmbH | Robert Koch Institute | unknown |
| EPI_ISL_3238858 | Bioscientia Labor Wermsdorf | Robert Koch Institute | unknown |
| EPI_ISL_3238859, EPI_ISL_3238860 | LabKom - Labor Augsburg MVZ GmbH | Robert Koch Institute | unknown |
| EPI_ISL_3238861 | Bioscientia Labor Wermsdorf | Robert Koch Institute | unknown |
| EPI_ISL_3238862, EPI_ISL_3238863 | LabKom - Labor Augsburg MVZ GmbH | Robert Koch Institute | unknown |
| EPI_ISL_3238864 | Bioscientia Labor Wermsdorf | Robert Koch Institute | unknown |

| | | | |
|---|---|---|---|
| EPI_ISL_3238865 | Labor 28 MVZ GmbH | Robert Koch Institute | unknown |
| EPI_ISL_3238866 | LabKom - Labor Hannover MVZ GmbH | Robert Koch Institute | unknown |
| EPI_ISL_3238867 | Bioscientia Labor Wermsdorf | Robert Koch Institute | unknown |
| EPI_ISL_3238868 | Labor 28 MVZ GmbH | Robert Koch Institute | unknown |
| EPI_ISL_3238869, EPI_ISL_3238870 | Sonic - MVZ Medizinisches Labor Bremen GmbH | Robert Koch Institute | unknown |
| EPI_ISL_3238871, EPI_ISL_3238872 | Bioscientia Labor Wermsdorf | Robert Koch Institute | unknown |
| EPI_ISL_3238873 | LabKom - Labor Augsburg MVZ GmbH | Robert Koch Institute | unknown |
| EPI_ISL_3238874, EPI_ISL_3238875 | Bioscientia Labor Wermsdorf | Robert Koch Institute | unknown |
| EPI_ISL_3238876 | Sonic - MVZ Medizinisches Labor Bremen GmbH | Robert Koch Institute | unknown |
| EPI_ISL_3238877 | LabKom - Labor Augsburg MVZ GmbH | Robert Koch Institute | unknown |
| EPI_ISL_3238878 | Bioscientia Labor Wermsdorf | Robert Koch Institute | unknown |
| EPI_ISL_3238879, EPI_ISL_3238880, EPI_ISL_3238881, EPI_ISL_3238882, EPI_ISL_3238883, EPI_ISL_3238884, EPI_ISL_3238885, EPI_ISL_3238887, EPI_ISL_3238888, EPI_ISL_3238889, EPI_ISL_3238890, EPI_ISL_3238891 | | | |
| see above | Eurofins MVZ Labor Gelsenkirchen | Robert Koch Institute | unknown |
| EPI_ISL_3238892 | SYNLAB MVZ Leverkusen | Robert Koch Institute | unknown |
| EPI_ISL_3238893 | SYNLAB Labor München Zentrum LMZ | Robert Koch Institute | unknown |
| EPI_ISL_3238894, EPI_ISL_3238895 | SYNLAB MVZ Leverkusen | Robert Koch Institute | unknown |
| EPI_ISL_3238896 | SYNLAB MVZ Berlin | Robert Koch Institute | unknown |
| EPI_ISL_3238897 | SYNLAB Labor München Zentrum LMZ | Robert Koch Institute | unknown |
| EPI_ISL_3238898 | SYNLAB MVZ Leverkusen | Robert Koch Institute | unknown |
| EPI_ISL_3238899 | SYNLAB Labor München Zentrum LMZ | Robert Koch Institute | unknown |
| EPI_ISL_3238900 | SYNLAB MVZ Leverkusen | Robert Koch Institute | unknown |
| EPI_ISL_3238901 | Synlab MVZ Augsburg | Robert Koch Institute | unknown |
| EPI_ISL_3238902, EPI_ISL_3238903, EPI_ISL_3238904 | SYNLAB Labor München Zentrum LMZ | Robert Koch Institute | unknown |
| EPI_ISL_3238905, EPI_ISL_3238906 | SYNLAB MVZ Leverkusen | Robert Koch Institute | unknown |
| EPI_ISL_3238907 | SYNLAB MVZ Trier | Robert Koch Institute | unknown |
| EPI_ISL_3238908, EPI_ISL_3238909 | SYNLAB MVZ Leverkusen | Robert Koch Institute | unknown |
| EPI_ISL_3238910 | SYNLAB Jena Oncoscreen | Robert Koch Institute | unknown |
| EPI_ISL_3238911, EPI_ISL_3238912, EPI_ISL_3238913 | SYNLAB MVZ Leverkusen | Robert Koch Institute | unknown |
| EPI_ISL_3238914 | SYNLAB MVZ Berlin | Robert Koch Institute | unknown |
| EPI_ISL_3238915 | SYNLAB MVZ Leverkusen | Robert Koch Institute | unknown |
| EPI_ISL_3238916 | Synlab MVZ Augsburg | Robert Koch Institute | unknown |
| EPI_ISL_3238917, EPI_ISL_3238918, EPI_ISL_3238919, EPI_ISL_3238921 | SYNLAB MVZ Leverkusen | Robert Koch Institute | unknown |
| EPI_ISL_3238922 | SYNLAB MVZ Leinfelden-Echterdingen | Robert Koch Institute | unknown |
| EPI_ISL_3238923 | SYNLAB MVZ Leverkusen | Robert Koch Institute | unknown |
| EPI_ISL_3238924 | SYNLAB MVZ Weiden | Robert Koch Institute | unknown |
| EPI_ISL_3238925, EPI_ISL_3238926 | SYNLAB MVZ Leverkusen | Robert Koch Institute | unknown |
| EPI_ISL_3238927 | Synlab MVZ Augsburg | Robert Koch Institute | unknown |
| EPI_ISL_3238928, EPI_ISL_3238929 | SYNLAB MVZ Leverkusen | Robert Koch Institute | unknown |
| EPI_ISL_3238930 | SYNLAB MVZ Weiden | Robert Koch Institute | unknown |
| EPI_ISL_3238931, EPI_ISL_3238932, EPI_ISL_3238933 | SYNLAB Labor München Zentrum LMZ | Robert Koch Institute | unknown |
| EPI_ISL_3238934 | SYNLAB MVZ Leinfelden-Echterdingen | Robert Koch Institute | unknown |
| EPI_ISL_3238935 | SYNLAB MVZ Berlin | Robert Koch Institute | unknown |
| EPI_ISL_3238936 | SYNLAB MVZ Leverkusen | Robert Koch Institute | unknown |
| EPI_ISL_3238937, EPI_ISL_3238938 | SYNLAB Labor München Zentrum LMZ | Robert Koch Institute | unknown |
| EPI_ISL_3238939, EPI_ISL_3238940 | SYNLAB MVZ Leverkusen | Robert Koch Institute | unknown |
| EPI_ISL_3238941 | SYNLAB MVZ Leinfelden-Echterdingen | Robert Koch Institute | unknown |
| EPI_ISL_3238942 | SYNLAB MVZ Weiden | Robert Koch Institute | unknown |
| EPI_ISL_3238943, EPI_ISL_3238944 | SYNLAB MVZ Leverkusen | Robert Koch Institute | unknown |
| EPI_ISL_3238945 | Synlab MVZ Augsburg | Robert Koch Institute | unknown |
| EPI_ISL_3238946 | SYNLAB MVZ Weiden | Robert Koch Institute | unknown |
| EPI_ISL_3238948 | SYNLAB MVZ Leverkusen | Robert Koch Institute | unknown |
| EPI_ISL_3238949 | SYNLAB MVZ Weiden | Robert Koch Institute | unknown |
| EPI_ISL_3238950 | SYNLAB MVZ Leverkusen | Robert Koch Institute | unknown |
| EPI_ISL_3238951 | SYNLAB Jena Oncoscreen | Robert Koch Institute | unknown |
| EPI_ISL_3238952 | SYNLAB MVZ Leinfelden-Echterdingen | Robert Koch Institute | unknown |
| EPI_ISL_3238953 | SYNLAB MVZ Trier | Robert Koch Institute | unknown |
| EPI_ISL_3238954, EPI_ISL_3238955 | SYNLAB Labor München Zentrum LMZ | Robert Koch Institute | unknown |
| EPI_ISL_3238956, EPI_ISL_3238957, EPI_ISL_3238958 | SYNLAB MVZ Leverkusen | Robert Koch Institute | unknown |
| EPI_ISL_3238959 | SYNLAB MVZ Berlin | Robert Koch Institute | unknown |
| EPI_ISL_3238960 | SYNLAB MVZ Leverkusen | Robert Koch Institute | unknown |

| | | | |
|---|---|---|---|
| EPI_ISL_3238962 | SYNLAB MVZ Leinfelden-Echterdingen | Robert Koch Institute | unknown |
| EPI_ISL_3238963 | SYNLAB MVZ Leverkusen | Robert Koch Institute | unknown |
| EPI_ISL_3238964 | SYNLAB MVZ Weiden | Robert Koch Institute | unknown |
| EPI_ISL_3238965,<br>EPI_ISL_3238966,<br>EPI_ISL_3238967,<br>EPI_ISL_3238968,<br>EPI_ISL_3238969,<br>EPI_ISL_3238970,<br>EPI_ISL_3238971 | SYNLAB MVZ Leverkusen | Robert Koch Institute | unknown |
| EPI_ISL_3238972 | SYNLAB MVZ Weiden | Robert Koch Institute | unknown |
| EPI_ISL_3238975 | SYNLAB MVZ Leverkusen | Robert Koch Institute | unknown |
| EPI_ISL_3238976 | SYNLAB MVZ Weiden | Robert Koch Institute | unknown |
| EPI_ISL_3238978 | SYNLAB MVZ Leverkusen | Robert Koch Institute | unknown |
| EPI_ISL_3238979 | SYNLAB MVZ Leinfelden-Echterdingen | Robert Koch Institute | unknown |
| EPI_ISL_3238980,<br>EPI_ISL_3238981 | SYNLAB MVZ Leverkusen | Robert Koch Institute | unknown |
| EPI_ISL_3238982 | Synlab MVZ Augsburg | Robert Koch Institute | unknown |
| EPI_ISL_3238983 | SYNLAB MVZ Leinfelden-Echterdingen | Robert Koch Institute | unknown |
| EPI_ISL_3238984 | Synlab MVZ Augsburg | Robert Koch Institute | unknown |
| EPI_ISL_3238985,<br>EPI_ISL_3238986 | SYNLAB MVZ Weiden | Robert Koch Institute | unknown |
| EPI_ISL_3238987 | SYNLAB MVZ Trier | Robert Koch Institute | unknown |
| EPI_ISL_3238988 | SYNLAB MVZ Leinfelden-Echterdingen | Robert Koch Institute | unknown |
| EPI_ISL_3238989 | SYNLAB MVZ Leverkusen | Robert Koch Institute | unknown |
| EPI_ISL_3238990 | SYNLAB MVZ Weiden | Robert Koch Institute | unknown |
| EPI_ISL_3238991 | SYNLAB MVZ Leinfelden-Echterdingen | Robert Koch Institute | unknown |
| EPI_ISL_3238992 | SYNLAB MVZ Leverkusen | Robert Koch Institute | unknown |
| EPI_ISL_3238993 | SYNLAB MVZ Weiden | Robert Koch Institute | unknown |
| EPI_ISL_3238994 | SYNLAB MVZ Leinfelden-Echterdingen | Robert Koch Institute | unknown |
| EPI_ISL_3238995 | SYNLAB MVZ Berlin | Robert Koch Institute | unknown |
| EPI_ISL_3238996,<br>EPI_ISL_3238997,<br>EPI_ISL_3238998,<br>EPI_ISL_3238999 | SYNLAB MVZ Leverkusen | Robert Koch Institute | unknown |
| EPI_ISL_3239001,<br>EPI_ISL_3239002 | SYNLAB MVZ Weiden | Robert Koch Institute | unknown |
| EPI_ISL_3239003 | SYNLAB MVZ Leverkusen | Robert Koch Institute | unknown |
| EPI_ISL_3239004,<br>EPI_ISL_3239005 | SYNLAB Labor München Zentrum LMZ | Robert Koch Institute | unknown |
| EPI_ISL_3239006 | SYNLAB MVZ Leverkusen | Robert Koch Institute | unknown |
| EPI_ISL_3239007 | SYNLAB Labor München Zentrum LMZ | Robert Koch Institute | unknown |
| EPI_ISL_3239008 | SYNLAB MVZ Weiden | Robert Koch Institute | unknown |
| EPI_ISL_3239009,<br>EPI_ISL_3239010 | SYNLAB MVZ Leinfelden-Echterdingen | Robert Koch Institute | unknown |
| EPI_ISL_3239011 | Synlab MVZ Augsburg | Robert Koch Institute | unknown |
| EPI_ISL_3239012 | SYNLAB Labor München Zentrum LMZ | Robert Koch Institute | unknown |
| EPI_ISL_3239013,<br>EPI_ISL_3239014 | SYNLAB MVZ Leverkusen | Robert Koch Institute | unknown |
| EPI_ISL_3239015 | SYNLAB Labor München Zentrum LMZ | Robert Koch Institute | unknown |
| EPI_ISL_3239017 | SYNLAB MVZ Trier | Robert Koch Institute | unknown |
| EPI_ISL_3239018,<br>EPI_ISL_3239019,<br>EPI_ISL_3239020 | SYNLAB MVZ Leverkusen | Robert Koch Institute | unknown |
| EPI_ISL_3239021 | SYNLAB MVZ Weiden | Robert Koch Institute | unknown |
| EPI_ISL_3239022,<br>EPI_ISL_3239023 | SYNLAB MVZ Leverkusen | Robert Koch Institute | unknown |
| EPI_ISL_3239024 | SYNLAB MVZ Trier | Robert Koch Institute | unknown |
| EPI_ISL_3239025 | SYNLAB MVZ Leverkusen | Robert Koch Institute | unknown |
| EPI_ISL_3239026 | SYNLAB MVZ Leinfelden-Echterdingen | Robert Koch Institute | unknown |
| EPI_ISL_3239027,<br>EPI_ISL_3239028 | SYNLAB MVZ Leverkusen | Robert Koch Institute | unknown |
| EPI_ISL_3239029 | SYNLAB MVZ Leinfelden-Echterdingen | Robert Koch Institute | unknown |
| EPI_ISL_3239030 | SYNLAB MVZ Dachau | Robert Koch Institute | unknown |
| EPI_ISL_3239031 | SYNLAB MVZ Weiden | Robert Koch Institute | unknown |
| EPI_ISL_3239033,<br>EPI_ISL_3239034 | SYNLAB MVZ Leverkusen | Robert Koch Institute | unknown |
| EPI_ISL_3239035 | SYNLAB MVZ Leinfelden-Echterdingen | Robert Koch Institute | unknown |
| EPI_ISL_3239036,<br>EPI_ISL_3239037,<br>EPI_ISL_3239038 | SYNLAB MVZ Weiden | Robert Koch Institute | unknown |
| EPI_ISL_3239039 | SYNLAB MVZ Leinfelden-Echterdingen | Robert Koch Institute | unknown |
| EPI_ISL_3239040 | SYNLAB MVZ Leverkusen | Robert Koch Institute | unknown |
| EPI_ISL_3239041 | SYNLAB MVZ Weiden | Robert Koch Institute | unknown |
| EPI_ISL_3239042 | SYNLAB MVZ Trier | Robert Koch Institute | unknown |
| EPI_ISL_3239043 | SYNLAB MVZ Leverkusen | Robert Koch Institute | unknown |
| EPI_ISL_3239044,<br>EPI_ISL_3239045,<br>EPI_ISL_3239046 | SYNLAB Labor München Zentrum LMZ | Robert Koch Institute | unknown |
| EPI_ISL_3239047 | SYNLAB MVZ Leverkusen | Robert Koch Institute | unknown |
| EPI_ISL_3239048 | SYNLAB MVZ Berlin | Robert Koch Institute | unknown |
| EPI_ISL_3239049 | SYNLAB MVZ Weiden | Robert Koch Institute | unknown |
| EPI_ISL_3239050 | SYNLAB Labor München Zentrum LMZ | Robert Koch Institute | unknown |
| EPI_ISL_3239051 | SYNLAB MVZ Trier | Robert Koch Institute | unknown |

| | | | |
|---|---|---|---|
| EPI_ISL_3239052 | Synlab MVZ Augsburg | Robert Koch Institute | unknown |
| EPI_ISL_3239053, EPI_ISL_3239054 | SYNLAB MVZ Leinfelden-Echterdingen | Robert Koch Institute | unknown |
| EPI_ISL_3239055, EPI_ISL_3239056 | SYNLAB MVZ Berlin | Robert Koch Institute | unknown |
| EPI_ISL_3239057 | SYNLAB MVZ Weiden | Robert Koch Institute | unknown |
| EPI_ISL_3239058, EPI_ISL_3239059 | SYNLAB MVZ Berlin | Robert Koch Institute | unknown |
| EPI_ISL_3239060 | SYNLAB MVZ Weiden | Robert Koch Institute | unknown |
| EPI_ISL_3239061 | SYNLAB MVZ Leinfelden-Echterdingen | Robert Koch Institute | unknown |
| EPI_ISL_3239062 | SYNLAB MVZ Leverkusen | Robert Koch Institute | unknown |
| EPI_ISL_3239063 | SYNLAB MVZ Weiden | Robert Koch Institute | unknown |
| EPI_ISL_3239064, EPI_ISL_3239066 | SYNLAB MVZ Leverkusen | Robert Koch Institute | unknown |
| EPI_ISL_3239067 | SYNLAB MVZ Weiden | Robert Koch Institute | unknown |
| EPI_ISL_3239068 | SYNLAB MVZ Leverkusen | Robert Koch Institute | unknown |
| EPI_ISL_3239069 | SYNLAB MVZ Trier | Robert Koch Institute | unknown |
| EPI_ISL_3239070 | SYNLAB MVZ Leverkusen | Robert Koch Institute | unknown |
| EPI_ISL_3239071 | SYNLAB MVZ Trier | Robert Koch Institute | unknown |
| EPI_ISL_3239073, EPI_ISL_3239074 | SYNLAB MVZ Weiden | Robert Koch Institute | unknown |
| EPI_ISL_3239076 | SYNLAB MVZ Trier | Robert Koch Institute | unknown |
| EPI_ISL_3239077, EPI_ISL_3239078 | SYNLAB MVZ Leinfelden-Echterdingen | Robert Koch Institute | unknown |
| EPI_ISL_3239079 | SYNLAB MVZ Trier | Robert Koch Institute | unknown |
| EPI_ISL_3239080, EPI_ISL_3239081 | SYNLAB MVZ Leverkusen | Robert Koch Institute | unknown |
| EPI_ISL_3239082 | SYNLAB MVZ Leinfelden-Echterdingen | Robert Koch Institute | unknown |
| EPI_ISL_3239083 | SYNLAB MVZ Leverkusen | Robert Koch Institute | unknown |
| EPI_ISL_3239084, EPI_ISL_3239085 | SYNLAB MVZ Trier | Robert Koch Institute | unknown |
| EPI_ISL_3239086, EPI_ISL_3239087 | SYNLAB MVZ Leverkusen | Robert Koch Institute | unknown |
| EPI_ISL_3239088, EPI_ISL_3239089 | SYNLAB MVZ Weiden | Robert Koch Institute | unknown |
| EPI_ISL_3239090, EPI_ISL_3239091 | SYNLAB MVZ Leverkusen | Robert Koch Institute | unknown |
| EPI_ISL_3239092, EPI_ISL_3239093 | SYNLAB MVZ Leinfelden-Echterdingen | Robert Koch Institute | unknown |
| EPI_ISL_3239094 | SYNLAB MVZ Weiden | Robert Koch Institute | unknown |
| EPI_ISL_3239095, EPI_ISL_3239096 | SYNLAB MVZ Leverkusen | Robert Koch Institute | unknown |
| EPI_ISL_3239097 | SYNLAB MVZ Leinfelden-Echterdingen | Robert Koch Institute | unknown |
| EPI_ISL_3239098 | SYNLAB MVZ Leverkusen | Robert Koch Institute | unknown |
| EPI_ISL_3239099 | SYNLAB Labor München Zentrum LMZ | Robert Koch Institute | unknown |
| EPI_ISL_3239100 | SYNLAB MVZ Leverkusen | Robert Koch Institute | unknown |
| EPI_ISL_3239101 | SYNLAB MVZ Berlin | Robert Koch Institute | unknown |
| EPI_ISL_3239102 | SYNLAB MVZ Weiden | Robert Koch Institute | unknown |
| EPI_ISL_3239103 | Synlab MVZ Augsburg | Robert Koch Institute | unknown |
| EPI_ISL_3239104, EPI_ISL_3239105, EPI_ISL_3239106 | SYNLAB MVZ Leverkusen | Robert Koch Institute | unknown |
| EPI_ISL_3239107, EPI_ISL_3239108 | SYNLAB MVZ Weiden | Robert Koch Institute | unknown |
| EPI_ISL_3239109, EPI_ISL_3239110 | SYNLAB MVZ Leinfelden-Echterdingen | Robert Koch Institute | unknown |
| EPI_ISL_3239111 | SYNLAB MVZ Leverkusen | Robert Koch Institute | unknown |
| EPI_ISL_3239112 | SYNLAB MVZ Berlin | Robert Koch Institute | unknown |
| EPI_ISL_3239113, EPI_ISL_3239114 | SYNLAB MVZ Weiden | Robert Koch Institute | unknown |
| EPI_ISL_3239116, EPI_ISL_3239117, EPI_ISL_3239118, EPI_ISL_3239119, EPI_ISL_3239120 | SYNLAB MVZ Leverkusen | Robert Koch Institute | unknown |
| EPI_ISL_3239121 | Synlab MVZ Augsburg | Robert Koch Institute | unknown |
| EPI_ISL_3239122, EPI_ISL_3239123 | SYNLAB MVZ Leverkusen | Robert Koch Institute | unknown |
| EPI_ISL_3239124 | Synlab MVZ Augsburg | Robert Koch Institute | unknown |
| EPI_ISL_3239125 | SYNLAB MVZ Weiden | Robert Koch Institute | unknown |
| EPI_ISL_3239126 | SYNLAB MVZ Leinfelden-Echterdingen | Robert Koch Institute | unknown |
| EPI_ISL_3239127 | SYNLAB MVZ Leverkusen | Robert Koch Institute | unknown |
| EPI_ISL_3239128 | SYNLAB MVZ Trier | Robert Koch Institute | unknown |
| EPI_ISL_3239129 | SYNLAB MVZ Leverkusen | Robert Koch Institute | unknown |
| EPI_ISL_3239130 | SYNLAB MVZ Weiden | Robert Koch Institute | unknown |
| EPI_ISL_3239131 | SYNLAB Jena Oncoscreen | Robert Koch Institute | unknown |
| EPI_ISL_3239132, EPI_ISL_3239133, EPI_ISL_3239134 | SYNLAB MVZ Leverkusen | Robert Koch Institute | unknown |
| EPI_ISL_3239135 | SYNLAB MVZ Weiden | Robert Koch Institute | unknown |
| EPI_ISL_3239136 | SYNLAB MVZ Leinfelden-Echterdingen | Robert Koch Institute | unknown |
| EPI_ISL_3239137 | SYNLAB MVZ Leverkusen | Robert Koch Institute | unknown |
| EPI_ISL_3239138 | SYNLAB MVZ Leinfelden-Echterdingen | Robert Koch Institute | unknown |
| EPI_ISL_3239139, | Universitätsmedizin Mannheim | Robert Koch Institute | unknown |

| | | | |
|---|---|---|---|
| EPI_ISL_3239140, EPI_ISL_3239141, EPI_ISL_3239142, EPI_ISL_3239143 | | | |
| EPI_ISL_3239144, EPI_ISL_3239145, EPI_ISL_3239147, EPI_ISL_3239148, EPI_ISL_3239149, EPI_ISL_3239150, EPI_ISL_3239151, EPI_ISL_3239152, EPI_ISL_3239153, EPI_ISL_3239154, EPI_ISL_3239155 | | | |
| see above | Universitätsklinikum Heidelberg | Robert Koch Institute | unknown |
| EPI_ISL_3239156, EPI_ISL_3239157, EPI_ISL_3239158, EPI_ISL_3239159 | Landesamt für Verbraucherschutz des Landes Sachsen-Anhalt | Robert Koch Institute | unknown |
| EPI_ISL_3239160 | IMD - Institut für Medizinische Diagnostik Berlin-Potsdam | Robert Koch Institute | unknown |
| EPI_ISL_3239161 | IMD - Labor Oderland | Robert Koch Institute | unknown |
| EPI_ISL_3239162 | IMD - Medizinisches Labor Rostock | Robert Koch Institute | unknown |
| EPI_ISL_3239163 | IMD - Labor Oderland | Robert Koch Institute | unknown |
| EPI_ISL_3239164 | IMD - MVZ Labor Greifswald | Robert Koch Institute | unknown |
| EPI_ISL_3239165 | IMD - MVZ Labor Martinsried | Robert Koch Institute | unknown |
| EPI_ISL_3239166, EPI_ISL_3239167 | IMD - Institut für Medizinische Diagnostik Berlin-Potsdam | Robert Koch Institute | unknown |
| EPI_ISL_3239168 | IMD - MVZ Labor Greifswald | Robert Koch Institute | unknown |
| EPI_ISL_3239169 | IMD - Labor Oderland | Robert Koch Institute | unknown |
| EPI_ISL_3239170, EPI_ISL_3239171 | IMD - MVZ Labor Greifswald | Robert Koch Institute | unknown |
| EPI_ISL_3239172, EPI_ISL_3239173 | IMD - MVZ Labor Martinsried | Robert Koch Institute | unknown |
| EPI_ISL_3239174, EPI_ISL_3239175, EPI_ISL_3239176, EPI_ISL_3239177, EPI_ISL_3239178, EPI_ISL_3239180, EPI_ISL_3239181, EPI_ISL_3239184, EPI_ISL_3239185, EPI_ISL_3239186, EPI_ISL_3239187, EPI_ISL_3239188, EPI_ISL_3239189, EPI_ISL_3239190, EPI_ISL_3239191, EPI_ISL_3239192, EPI_ISL_3239193, EPI_ISL_3239194, EPI_ISL_3239195, EPI_ISL_3239196 | | | |
| see above | Eurofins LifeCodexx GmbH | Robert Koch Institute | unknown |
| EPI_ISL_3239197, EPI_ISL_3239198, EPI_ISL_3239199, EPI_ISL_3239200, EPI_ISL_3239201, EPI_ISL_3239202, EPI_ISL_3239203, EPI_ISL_3239204, EPI_ISL_3239205, EPI_ISL_3239206, EPI_ISL_3239207, EPI_ISL_3239208, EPI_ISL_3239209 | | | |
| see above | Clemenshospital - Alexianer Clemenshospital Münster | Robert Koch Institute | unknown |
| EPI_ISL_3239210 | Landesuntersuchungsanstalt für das Gesundheits- und Veterinärwesen (LUA) Sachsen | Robert Koch Institute | unknown |
| EPI_ISL_3239211, EPI_ISL_3239212 | Uniklinikum Carl Gustav Carus an der TU Dresden; Institut für Virologie | Robert Koch Institute | unknown |
| EPI_ISL_3239213 | Landesuntersuchungsanstalt für das Gesundheits- und Veterinärwesen (LUA) Sachsen | Robert Koch Institute | unknown |
| EPI_ISL_3239214 | Uniklinikum Carl Gustav Carus an der TU Dresden; Institut für Virologie | Robert Koch Institute | unknown |
| EPI_ISL_3239215, EPI_ISL_3239217 | Medizinisches Labor Ostsachsen MVZ GbR Nebenbetriebsstätte Dresden | Robert Koch Institute | unknown |
| EPI_ISL_3239218 | Städtisches Klinikum Dresden Institut für Klinische Chemie und Laboratoriumsmedizin | Robert Koch Institute | unknown |
| EPI_ISL_3239219, EPI_ISL_3239220, EPI_ISL_3239221, EPI_ISL_3239222 | Medizinisches Labor Ostsachsen MVZ GbR Nebenbetriebsstätte Dresden | Robert Koch Institute | unknown |
| EPI_ISL_3239223 | Landesuntersuchungsanstalt für das Gesundheits- und Veterinärwesen (LUA) Sachsen | Robert Koch Institute | unknown |
| EPI_ISL_3239224, EPI_ISL_3239225, EPI_ISL_3239226, EPI_ISL_3239227, EPI_ISL_3239228, EPI_ISL_3239229, EPI_ISL_3239230, EPI_ISL_3239231, EPI_ISL_3239232 | Labor Blackholm MVZ | Robert Koch Institute | unknown |
| EPI_ISL_3239234, EPI_ISL_3239235, EPI_ISL_3239236 | MVZ Labor Dr. Quade & Kollegen GmbH | Robert Koch Institute | unknown |
| EPI_ISL_3239237 | Labor Blackholm MVZ | Robert Koch Institute | unknown |
| EPI_ISL_3239238 | MVZ Labor Dr. Quade & Kollegen GmbH | Robert Koch Institute | unknown |
| EPI_ISL_3239239, EPI_ISL_3239241, EPI_ISL_3239242 | Labor Blackholm MVZ | Robert Koch Institute | unknown |
| EPI_ISL_3239243, EPI_ISL_3239244 | MVZ Labor Dr. Quade & Kollegen GmbH | Robert Koch Institute | unknown |
| EPI_ISL_3239245, EPI_ISL_3239247, EPI_ISL_3239248, EPI_ISL_3239249 | Labor Blackholm MVZ | Robert Koch Institute | unknown |
| EPI_ISL_3239250, EPI_ISL_3239251, EPI_ISL_3239252, EPI_ISL_3239253 | MVZ Labor Dr. Quade & Kollegen GmbH | Robert Koch Institute | unknown |
| EPI_ISL_3239254, EPI_ISL_3239255, EPI_ISL_3239256 | Labor Blackholm MVZ | Robert Koch Institute | unknown |
| EPI_ISL_3239257 | MVZ Labor Dr. Quade & Kollegen GmbH | Robert Koch Institute | unknown |
| EPI_ISL_3239258, EPI_ISL_3239259, EPI_ISL_3239262, EPI_ISL_3239263, EPI_ISL_3239264, EPI_ISL_3239265, EPI_ISL_3239266, EPI_ISL_3239267, EPI_ISL_3239268, EPI_ISL_3239269, EPI_ISL_3239270, EPI_ISL_3239271, EPI_ISL_3239272, EPI_ISL_3239273, EPI_ISL_3239274, EPI_ISL_3239275, EPI_ISL_3239276, EPI_ISL_3239277, EPI_ISL_3239278, EPI_ISL_3239280, EPI_ISL_3239281, EPI_ISL_3239282, EPI_ISL_3239283, EPI_ISL_3239284, EPI_ISL_3239285 | | | |
| see above | Labor Becker & Kollegen (Standort München) | Robert Koch Institute | unknown |
| EPI_ISL_3239287 | SYNLAB MVZ Leverkusen | Robert Koch Institute | unknown |
| EPI_ISL_3239288 | SYNLAB MVZ Weiden | Robert Koch Institute | unknown |
| EPI_ISL_3239289 | SYNLAB MVZ Leverkusen | Robert Koch Institute | unknown |
| EPI_ISL_3239290, EPI_ISL_3239291 | SYNLAB MVZ Weiden | Robert Koch Institute | unknown |
| EPI_ISL_3239292 | SYNLAB MVZ Leverkusen | Robert Koch Institute | unknown |
| EPI_ISL_3239293, EPI_ISL_3239294 | SYNLAB MVZ Weiden | Robert Koch Institute | unknown |
| EPI_ISL_3239295, EPI_ISL_3239296 | SYNLAB MVZ Leverkusen | Robert Koch Institute | unknown |

| | | | |
|---|---|---|---|
| EPI_ISL_3239297 | SYNLAB MVZ Trier | Robert Koch Institute | unknown |
| EPI_ISL_3239298 | SYNLAB MVZ Leinfelden-Echterdingen | Robert Koch Institute | unknown |
| EPI_ISL_3239299 | SYNLAB Labor München Zentrum LMZ | Robert Koch Institute | unknown |
| EPI_ISL_3239300 | SYNLAB MVZ Weiden | Robert Koch Institute | unknown |
| EPI_ISL_3239301 | SYNLAB Labor München Zentrum LMZ | Robert Koch Institute | unknown |
| EPI_ISL_3239302 | SYNLAB MVZ Leinfelden-Echterdingen | Robert Koch Institute | unknown |
| EPI_ISL_3239303 | SYNLAB MVZ Leverkusen | Robert Koch Institute | unknown |
| EPI_ISL_3239304 | SYNLAB MVZ Weiden | Robert Koch Institute | unknown |
| EPI_ISL_3239306 | SYNLAB MVZ Trier | Robert Koch Institute | unknown |
| EPI_ISL_3239307, EPI_ISL_3239308 | SYNLAB MVZ Leverkusen | Robert Koch Institute | unknown |
| EPI_ISL_3239309 | SYNLAB MVZ Berlin | Robert Koch Institute | unknown |
| EPI_ISL_3239310 | SYNLAB MVZ Leverkusen | Robert Koch Institute | unknown |
| EPI_ISL_3239311 | SYNLAB MVZ Trier | Robert Koch Institute | unknown |
| EPI_ISL_3239312 | SYNLAB MVZ Leinfelden-Echterdingen | Robert Koch Institute | unknown |
| EPI_ISL_3239313 | SYNLAB MVZ Trier | Robert Koch Institute | unknown |
| EPI_ISL_3239314 | SYNLAB MVZ Leinfelden-Echterdingen | Robert Koch Institute | unknown |
| EPI_ISL_3239315 | SYNLAB MVZ Leverkusen | Robert Koch Institute | unknown |
| EPI_ISL_3239316 | SYNLAB Labor München Zentrum LMZ | Robert Koch Institute | unknown |
| EPI_ISL_3239317 | SYNLAB MVZ Weiden | Robert Koch Institute | unknown |
| EPI_ISL_3239318 | SYNLAB MVZ Trier | Robert Koch Institute | unknown |
| EPI_ISL_3239319, EPI_ISL_3239320 | SYNLAB Labor München Zentrum LMZ | Robert Koch Institute | unknown |
| EPI_ISL_3239322, EPI_ISL_3239323 | SYNLAB MVZ Leverkusen | Robert Koch Institute | unknown |
| EPI_ISL_3239324 | SYNLAB MVZ Weiden | Robert Koch Institute | unknown |
| EPI_ISL_3239325, EPI_ISL_3239326 | SYNLAB MVZ Leverkusen | Robert Koch Institute | unknown |
| EPI_ISL_3239327, EPI_ISL_3239328 | SYNLAB MVZ Weiden | Robert Koch Institute | unknown |
| EPI_ISL_3239329, EPI_ISL_3239330 | SYNLAB MVZ Leverkusen | Robert Koch Institute | unknown |
| EPI_ISL_3239331 | SYNLAB MVZ Weiden | Robert Koch Institute | unknown |
| EPI_ISL_3239332 | SYNLAB MVZ Leverkusen | Robert Koch Institute | unknown |
| EPI_ISL_3239333, EPI_ISL_3239334 | SYNLAB Labor München Zentrum LMZ | Robert Koch Institute | unknown |
| EPI_ISL_3239335, EPI_ISL_3239336, EPI_ISL_3239337 | SYNLAB MVZ Leverkusen | Robert Koch Institute | unknown |
| EPI_ISL_3239338 | SYNLAB MVZ Weiden | Robert Koch Institute | unknown |
| EPI_ISL_3239339 | SYNLAB MVZ Leverkusen | Robert Koch Institute | unknown |
| EPI_ISL_3239340, EPI_ISL_3239341 | SYNLAB MVZ Trier | Robert Koch Institute | unknown |
| EPI_ISL_3239342 | Synlab MVZ Augsburg | Robert Koch Institute | unknown |
| EPI_ISL_3239343 | SYNLAB MVZ Leverkusen | Robert Koch Institute | unknown |
| EPI_ISL_3239344, EPI_ISL_3239345 | SYNLAB MVZ Weiden | Robert Koch Institute | unknown |
| EPI_ISL_3239346 | SYNLAB MVZ Trier | Robert Koch Institute | unknown |
| EPI_ISL_3239347 | Synlab MVZ Augsburg | Robert Koch Institute | unknown |
| EPI_ISL_3239348, EPI_ISL_3239349 | SYNLAB MVZ Leverkusen | Robert Koch Institute | unknown |
| EPI_ISL_3239350 | SYNLAB MVZ Weiden | Robert Koch Institute | unknown |
| EPI_ISL_3239351 | SYNLAB MVZ Trier | Robert Koch Institute | unknown |
| EPI_ISL_3239353, EPI_ISL_3239354 | SYNLAB MVZ Leverkusen | Robert Koch Institute | unknown |
| EPI_ISL_3239355 | SYNLAB MVZ Weiden | Robert Koch Institute | unknown |
| EPI_ISL_3239356, EPI_ISL_3239357 | SYNLAB MVZ Leverkusen | Robert Koch Institute | unknown |
| EPI_ISL_3239358, EPI_ISL_3239359 | SYNLAB MVZ Weiden | Robert Koch Institute | unknown |
| EPI_ISL_3239360 | SYNLAB Labor München Zentrum LMZ | Robert Koch Institute | unknown |
| EPI_ISL_3239361 | SYNLAB MVZ Weiden | Robert Koch Institute | unknown |
| EPI_ISL_3239362, EPI_ISL_3239363, EPI_ISL_3239364 | SYNLAB MVZ Leverkusen | Robert Koch Institute | unknown |
| EPI_ISL_3239365 | SYNLAB MVZ Weiden | Robert Koch Institute | unknown |
| EPI_ISL_3239366 | SYNLAB MVZ Leverkusen | Robert Koch Institute | unknown |
| EPI_ISL_3239367, EPI_ISL_3239368 | SYNLAB MVZ Weiden | Robert Koch Institute | unknown |
| EPI_ISL_3239369 | SYNLAB MVZ Leverkusen | Robert Koch Institute | unknown |
| EPI_ISL_3239370 | SYNLAB MVZ Weiden | Robert Koch Institute | unknown |
| EPI_ISL_3239371, EPI_ISL_3239372, EPI_ISL_3239373, EPI_ISL_3239374 | SYNLAB MVZ Leverkusen | Robert Koch Institute | unknown |
| EPI_ISL_3239375 | SYNLAB MVZ Weiden | Robert Koch Institute | unknown |
| EPI_ISL_3239376 | SYNLAB MVZ Leverkusen | Robert Koch Institute | unknown |
| EPI_ISL_3239377 | Synlab MVZ Augsburg | Robert Koch Institute | unknown |
| EPI_ISL_3239378 | SYNLAB MVZ Leverkusen | Robert Koch Institute | unknown |
| EPI_ISL_3239379, EPI_ISL_3239380 | SYNLAB MVZ Trier | Robert Koch Institute | unknown |
| EPI_ISL_3239381, EPI_ISL_3239382 | SYNLAB MVZ Weiden | Robert Koch Institute | unknown |

| | | | |
|---|---|---|---|
| EPI_ISL_3239383 | SYNLAB MVZ Leverkusen | Robert Koch Institute | unknown |
| EPI_ISL_3239384 | SYNLAB Jena Oncoscreen | Robert Koch Institute | unknown |
| EPI_ISL_3239385, EPI_ISL_3239386 | SYNLAB MVZ Leverkusen | Robert Koch Institute | unknown |
| EPI_ISL_3239387 | SYNLAB MVZ Weiden | Robert Koch Institute | unknown |
| EPI_ISL_3239388, EPI_ISL_3239389 | SYNLAB MVZ Trier | Robert Koch Institute | unknown |
| EPI_ISL_3239391, EPI_ISL_3239392 | SYNLAB MVZ Weiden | Robert Koch Institute | unknown |
| EPI_ISL_3239393, EPI_ISL_3239394 | SYNLAB MVZ Leverkusen | Robert Koch Institute | unknown |
| EPI_ISL_3239395 | SYNLAB MVZ Weiden | Robert Koch Institute | unknown |
| EPI_ISL_3239396 | SYNLAB MVZ Trier | Robert Koch Institute | unknown |
| EPI_ISL_3239397 | SYNLAB MVZ Leverkusen | Robert Koch Institute | unknown |
| EPI_ISL_3239398, EPI_ISL_3239399, EPI_ISL_3239400 | SYNLAB MVZ Trier | Robert Koch Institute | unknown |
| EPI_ISL_3239401, EPI_ISL_3239402, EPI_ISL_3239403, EPI_ISL_3239404 | SYNLAB MVZ Weiden | Robert Koch Institute | unknown |
| EPI_ISL_3239405, EPI_ISL_3239406 | SYNLAB MVZ Leverkusen | Robert Koch Institute | unknown |
| EPI_ISL_3239407, EPI_ISL_3239408 | SYNLAB MVZ Trier | Robert Koch Institute | unknown |
| EPI_ISL_3239409 | SYNLAB MVZ Leverkusen | Robert Koch Institute | unknown |
| EPI_ISL_3239410 | SYNLAB MVZ Berlin | Robert Koch Institute | unknown |
| EPI_ISL_3239411 | SYNLAB Labor München Zentrum LMZ | Robert Koch Institute | unknown |
| EPI_ISL_3239412 | SYNLAB MVZ Leverkusen | Robert Koch Institute | unknown |
| EPI_ISL_3239413 | SYNLAB Jena Oncoscreen | Robert Koch Institute | unknown |
| EPI_ISL_3239414 | SYNLAB MVZ Leinfelden-Echterdingen | Robert Koch Institute | unknown |
| EPI_ISL_3239415, EPI_ISL_3239416, EPI_ISL_3239417 | SYNLAB MVZ Leverkusen | Robert Koch Institute | unknown |
| EPI_ISL_3239418 | SYNLAB MVZ Weiden | Robert Koch Institute | unknown |
| EPI_ISL_3239419 | Synlab MVZ Augsburg | Robert Koch Institute | unknown |
| EPI_ISL_3239420, EPI_ISL_3239421, EPI_ISL_3239422 | SYNLAB MVZ Leverkusen | Robert Koch Institute | unknown |
| EPI_ISL_3239423 | SYNLAB MVZ Hamburg | Robert Koch Institute | unknown |
| EPI_ISL_3239424 | SYNLAB MVZ Leverkusen | Robert Koch Institute | unknown |
| EPI_ISL_3239425 | Synlab MVZ Augsburg | Robert Koch Institute | unknown |
| EPI_ISL_3239426 | SYNLAB MVZ Berlin | Robert Koch Institute | unknown |
| EPI_ISL_3239427, EPI_ISL_3239428 | SYNLAB MVZ Leverkusen | Robert Koch Institute | unknown |
| EPI_ISL_3239429 | SYNLAB MVZ Weiden | Robert Koch Institute | unknown |
| EPI_ISL_3239430 | SYNLAB MVZ Trier | Robert Koch Institute | unknown |
| EPI_ISL_3239431 | SYNLAB MVZ Weiden | Robert Koch Institute | unknown |
| EPI_ISL_3239432, EPI_ISL_3239433, EPI_ISL_3239434, EPI_ISL_3239435 | SYNLAB MVZ Leverkusen | Robert Koch Institute | unknown |
| EPI_ISL_3239436 | SYNLAB MVZ Hamburg | Robert Koch Institute | unknown |
| EPI_ISL_3239437 | SYNLAB MVZ Leverkusen | Robert Koch Institute | unknown |
| EPI_ISL_3239438 | SYNLAB MVZ Trier | Robert Koch Institute | unknown |
| EPI_ISL_3239439, EPI_ISL_3239440 | SYNLAB MVZ Weiden | Robert Koch Institute | unknown |
| EPI_ISL_3239441 | SYNLAB MVZ Trier | Robert Koch Institute | unknown |
| EPI_ISL_3239442, EPI_ISL_3239443 | SYNLAB MVZ Leverkusen | Robert Koch Institute | unknown |
| EPI_ISL_3239444 | SYNLAB MVZ Trier | Robert Koch Institute | unknown |
| EPI_ISL_3239445, EPI_ISL_3239446, EPI_ISL_3239447 | SYNLAB MVZ Leverkusen | Robert Koch Institute | unknown |
| EPI_ISL_3239448 | SYNLAB MVZ Weiden | Robert Koch Institute | unknown |
| EPI_ISL_3239449 | Synlab MVZ Augsburg | Robert Koch Institute | unknown |
| EPI_ISL_3239450 | SYNLAB MVZ Trier | Robert Koch Institute | unknown |
| EPI_ISL_3239451 | SYNLAB MVZ Leverkusen | Robert Koch Institute | unknown |
| EPI_ISL_3239452 | SYNLAB MVZ Leinfelden-Echterdingen | Robert Koch Institute | unknown |
| EPI_ISL_3239453 | SYNLAB MVZ Leverkusen | Robert Koch Institute | unknown |
| EPI_ISL_3239454 | SYNLAB MVZ Weiden | Robert Koch Institute | unknown |
| EPI_ISL_3239455 | SYNLAB MVZ Trier | Robert Koch Institute | unknown |
| EPI_ISL_3239456, EPI_ISL_3239457, EPI_ISL_3239458 | SYNLAB MVZ Leverkusen | Robert Koch Institute | unknown |
| EPI_ISL_3239459 | SYNLAB MVZ Trier | Robert Koch Institute | unknown |
| EPI_ISL_3239460 | SYNLAB MVZ Leverkusen | Robert Koch Institute | unknown |
| EPI_ISL_3239461 | SYNLAB MVZ Weiden | Robert Koch Institute | unknown |
| EPI_ISL_3239462 | SYNLAB MVZ Leinfelden-Echterdingen | Robert Koch Institute | unknown |
| EPI_ISL_3239463 | SYNLAB MVZ Weiden | Robert Koch Institute | unknown |
| EPI_ISL_3239464, EPI_ISL_3239465, EPI_ISL_3239466, EPI_ISL_3239467, EPI_ISL_3239468, EPI_ISL_3239469 | SYNLAB MVZ Leverkusen | Robert Koch Institute | unknown |

| | | | |
|---|---|---|---|
| EPI_ISL_3239470 | Synlab MVZ Augsburg | Robert Koch Institute | unknown |
| EPI_ISL_3239471, EPI_ISL_3239472 | SYNLAB MVZ Leverkusen | Robert Koch Institute | unknown |
| EPI_ISL_3239473, EPI_ISL_3239474 | SYNLAB Labor München Zentrum LMZ | Robert Koch Institute | unknown |
| EPI_ISL_3239475 | SYNLAB MVZ Weiden | Robert Koch Institute | unknown |
| EPI_ISL_3239476 | SYNLAB MVZ Leverkusen | Robert Koch Institute | unknown |
| EPI_ISL_3239477, EPI_ISL_3239478 | SYNLAB MVZ Weiden | Robert Koch Institute | unknown |
| EPI_ISL_3239479, EPI_ISL_3239480 | SYNLAB MVZ Leverkusen | Robert Koch Institute | unknown |
| EPI_ISL_3239481 | SYNLAB MVZ Weiden | Robert Koch Institute | unknown |
| EPI_ISL_3239482, EPI_ISL_3239483 | SYNLAB MVZ Leverkusen | Robert Koch Institute | unknown |
| EPI_ISL_3239484 | SYNLAB MVZ Weiden | Robert Koch Institute | unknown |
| EPI_ISL_3239485, EPI_ISL_3239486, EPI_ISL_3239487 | SYNLAB MVZ Leverkusen | Robert Koch Institute | unknown |
| EPI_ISL_3239488, EPI_ISL_3239489 | Synlab MVZ Augsburg | Robert Koch Institute | unknown |
| EPI_ISL_3239490 | SYNLAB MVZ Leverkusen | Robert Koch Institute | unknown |
| EPI_ISL_3239492 | SYNLAB MVZ Weiden | Robert Koch Institute | unknown |
| EPI_ISL_3239493, EPI_ISL_3239494, EPI_ISL_3239495, EPI_ISL_3239496 | SYNLAB MVZ Leverkusen | Robert Koch Institute | unknown |
| EPI_ISL_3239497 | SYNLAB MVZ Dachau | Robert Koch Institute | unknown |
| EPI_ISL_3239498 | SYNLAB MVZ Weiden | Robert Koch Institute | unknown |
| EPI_ISL_3239499, EPI_ISL_3239500 | SYNLAB MVZ Leverkusen | Robert Koch Institute | unknown |
| EPI_ISL_3239501, EPI_ISL_3239502 | SYNLAB MVZ Weiden | Robert Koch Institute | unknown |
| EPI_ISL_3239503, EPI_ISL_3239504 | SYNLAB MVZ Leverkusen | Robert Koch Institute | unknown |
| EPI_ISL_3239505 | SYNLAB MVZ Weiden | Robert Koch Institute | unknown |
| EPI_ISL_3239506 | SYNLAB MVZ Leverkusen | Robert Koch Institute | unknown |
| EPI_ISL_3239507 | SYNLAB MVZ Dachau | Robert Koch Institute | unknown |
| EPI_ISL_3239508 | SYNLAB Jena Oncoscreen | Robert Koch Institute | unknown |
| EPI_ISL_3239509, EPI_ISL_3239510 | SYNLAB MVZ Weiden | Robert Koch Institute | unknown |
| EPI_ISL_3239511 | SYNLAB MVZ Leverkusen | Robert Koch Institute | unknown |
| EPI_ISL_3239512 | SYNLAB MVZ Weiden | Robert Koch Institute | unknown |
| EPI_ISL_3239513 | SYNLAB MVZ Leverkusen | Robert Koch Institute | unknown |
| EPI_ISL_3239514 | SYNLAB MVZ Trier | Robert Koch Institute | unknown |
| EPI_ISL_3239515 | SYNLAB MVZ Weiden | Robert Koch Institute | unknown |
| EPI_ISL_3239517 | SYNLAB MVZ Leverkusen | Robert Koch Institute | unknown |
| EPI_ISL_3239518 | SYNLAB MVZ Weiden | Robert Koch Institute | unknown |
| EPI_ISL_3239519 | Synlab MVZ Augsburg | Robert Koch Institute | unknown |
| EPI_ISL_3239520, EPI_ISL_3239522 | SYNLAB MVZ Leverkusen | Robert Koch Institute | unknown |
| EPI_ISL_3239523 | SYNLAB MVZ Weiden | Robert Koch Institute | unknown |
| EPI_ISL_3239524, EPI_ISL_3239525 | SYNLAB MVZ Leverkusen | Robert Koch Institute | unknown |
| EPI_ISL_3239526 | SYNLAB Labor München Zentrum LMZ | Robert Koch Institute | unknown |
| EPI_ISL_3239527 | SYNLAB MVZ Leverkusen | Robert Koch Institute | unknown |
| EPI_ISL_3239528 | SYNLAB MVZ Weiden | Robert Koch Institute | unknown |
| EPI_ISL_3239529 | SYNLAB MVZ Leverkusen | Robert Koch Institute | unknown |
| EPI_ISL_3239530 | SYNLAB MVZ Weiden | Robert Koch Institute | unknown |
| EPI_ISL_3239531 | SYNLAB Labor München Zentrum LMZ | Robert Koch Institute | unknown |
| EPI_ISL_3239532 | SYNLAB MVZ Leverkusen | Robert Koch Institute | unknown |
| EPI_ISL_3239533 | SYNLAB MVZ Trier | Robert Koch Institute | unknown |
| EPI_ISL_3239534 | SYNLAB MVZ Weiden | Robert Koch Institute | unknown |
| EPI_ISL_3239535, EPI_ISL_3239536 | SYNLAB MVZ Trier | Robert Koch Institute | unknown |
| EPI_ISL_3239537 | SYNLAB MVZ Berlin | Robert Koch Institute | unknown |
| EPI_ISL_3239538 | SYNLAB MVZ Leverkusen | Robert Koch Institute | unknown |
| EPI_ISL_3239539 | SYNLAB MVZ Weiden | Robert Koch Institute | unknown |
| EPI_ISL_3239540 | SYNLAB MVZ Leverkusen | Robert Koch Institute | unknown |
| EPI_ISL_3239541 | SYNLAB Labor München Zentrum LMZ | Robert Koch Institute | unknown |
| EPI_ISL_3239543 | SYNLAB MVZ Leverkusen | Robert Koch Institute | unknown |
| EPI_ISL_3239544, EPI_ISL_3239545 | SYNLAB Labor München Zentrum LMZ | Robert Koch Institute | unknown |
| EPI_ISL_3239546, EPI_ISL_3239547 | SYNLAB MVZ Leverkusen | Robert Koch Institute | unknown |
| EPI_ISL_3239548 | SYNLAB Labor München Zentrum LMZ | Robert Koch Institute | unknown |
| EPI_ISL_3239549 | SYNLAB MVZ Hamburg | Robert Koch Institute | unknown |
| EPI_ISL_3239550, EPI_ISL_3239551 | SYNLAB MVZ Leverkusen | Robert Koch Institute | unknown |
| EPI_ISL_3239552 | SYNLAB Labor München Zentrum LMZ | Robert Koch Institute | unknown |
| EPI_ISL_3239553 | SYNLAB MVZ Hamburg | Robert Koch Institute | unknown |
| EPI_ISL_3239554 | SYNLAB MVZ Leverkusen | Robert Koch Institute | unknown |
| EPI_ISL_3239555 | SYNLAB Labor München Zentrum LMZ | Robert Koch Institute | unknown |

| | | | |
|---|---|---|---|
| EPI_ISL_3239556 | SYNLAB MVZ Leverkusen | Robert Koch Institute | unknown |
| EPI_ISL_3239557 | SYNLAB Labor München Zentrum LMZ | Robert Koch Institute | unknown |
| EPI_ISL_3239558 | Dhulikhel Hospital, Kathmandu University Hospital | Molecular and Genomics Research Lab, Dhulikhel Hospital, Kathmandu University Hospital | Rajeev Shrestha, Nishan Katuwal, Navin Adhikari, Dipesh Tamrakar, Meghnath Dhimal, Pradip Gyanwali, Saroj Bhattarai, Surendra Kumar Madhup |
| EPI_ISL_3239559 | SYNLAB MVZ Weiden | Robert Koch Institute | unknown |
| EPI_ISL_3239560, EPI_ISL_3239561, EPI_ISL_3239562 | SYNLAB MVZ Leverkusen | Robert Koch Institute | unknown |
| EPI_ISL_3239563 | SYNLAB MVZ Weiden | Robert Koch Institute | unknown |
| EPI_ISL_3239564 | SYNLAB MVZ Trier | Robert Koch Institute | unknown |
| EPI_ISL_3239565, EPI_ISL_3239566, EPI_ISL_3239567 | SYNLAB MVZ Leverkusen | Robert Koch Institute | unknown |
| EPI_ISL_3239568 | SYNLAB Labor München Zentrum LMZ | Robert Koch Institute | unknown |
| EPI_ISL_3239569 | SYNLAB MVZ Weiden | Robert Koch Institute | unknown |
| EPI_ISL_3239570, EPI_ISL_3239571, EPI_ISL_3239572, EPI_ISL_3239573 | SYNLAB MVZ Leverkusen | Robert Koch Institute | unknown |
| EPI_ISL_3239574 | Synlab MVZ Augsburg | Robert Koch Institute | unknown |
| EPI_ISL_3239575, EPI_ISL_3239576 | SYNLAB MVZ Leverkusen | Robert Koch Institute | unknown |
| EPI_ISL_3239578, EPI_ISL_3239579 | SYNLAB MVZ Weiden | Robert Koch Institute | unknown |
| EPI_ISL_3239580, EPI_ISL_3239581 | SYNLAB MVZ Leverkusen | Robert Koch Institute | unknown |
| EPI_ISL_3239582 | SYNLAB MVZ Weiden | Robert Koch Institute | unknown |
| EPI_ISL_3239583, EPI_ISL_3239584 | SYNLAB MVZ Leverkusen | Robert Koch Institute | unknown |
| EPI_ISL_3239585 | SYNLAB MVZ Trier | Robert Koch Institute | unknown |
| EPI_ISL_3239586 | SYNLAB MVZ Weiden | Robert Koch Institute | unknown |
| EPI_ISL_3239587, EPI_ISL_3239588, EPI_ISL_3239589, EPI_ISL_3239590 | SYNLAB MVZ Leverkusen | Robert Koch Institute | unknown |
| EPI_ISL_3239591 | SYNLAB Jena Oncoscreen | Robert Koch Institute | unknown |
| EPI_ISL_3239592 | SYNLAB MVZ Leverkusen | Robert Koch Institute | unknown |
| EPI_ISL_3239593 | SYNLAB MVZ Trier | Robert Koch Institute | unknown |
| EPI_ISL_3239594, EPI_ISL_3239595 | SYNLAB MVZ Leverkusen | Robert Koch Institute | unknown |
| EPI_ISL_3239596 | SYNLAB MVZ Trier | Robert Koch Institute | unknown |
| EPI_ISL_3239597 | Synlab MVZ Augsburg | Robert Koch Institute | unknown |
| EPI_ISL_3239598 | SYNLAB MVZ Hamburg | Robert Koch Institute | unknown |
| EPI_ISL_3239599 | SYNLAB MVZ Dachau | Robert Koch Institute | unknown |
| EPI_ISL_3239600, EPI_ISL_3239601, EPI_ISL_3239602, EPI_ISL_3239603, EPI_ISL_3239604, EPI_ISL_3239605, EPI_ISL_3239606, EPI_ISL_3239607, EPI_ISL_3239608 | MDI Limbach Berlin GmbH; MVZ Labor Berlin | Robert Koch Institute | unknown |
| EPI_ISL_3239609 | Limbach - MVZ Gemeinschaftslabor Suhl | Robert Koch Institute | unknown |
| EPI_ISL_3239610 | MDI Limbach Berlin GmbH; MVZ Labor Berlin | Robert Koch Institute | unknown |
| EPI_ISL_3239611 | MVZ Medizinisches Labor Hannover GmbH | Robert Koch Institute | unknown |
| EPI_ISL_3239612, EPI_ISL_3239613, EPI_ISL_3239614, EPI_ISL_3239615, EPI_ISL_3239616, EPI_ISL_3239617, EPI_ISL_3239618, EPI_ISL_3239619, EPI_ISL_3239620, EPI_ISL_3239621, EPI_ISL_3239622, EPI_ISL_3239623, EPI_ISL_3239624, EPI_ISL_3239625 | | | |
| see above | MDI Limbach Berlin GmbH; MVZ Labor Berlin | Robert Koch Institute | unknown |
| EPI_ISL_3239626 | MVZ Labor Dr. Limbach & Kollegen GbR | Robert Koch Institute | unknown |
| EPI_ISL_3239627, EPI_ISL_3239628, EPI_ISL_3239629, EPI_ISL_3239630, EPI_ISL_3239631, EPI_ISL_3239632, EPI_ISL_3239633, EPI_ISL_3239634, EPI_ISL_3239635, EPI_ISL_3239636, EPI_ISL_3239637, EPI_ISL_3239638, EPI_ISL_3239639, EPI_ISL_3239640, EPI_ISL_3239641, EPI_ISL_3239642, EPI_ISL_3239643, EPI_ISL_3239644, EPI_ISL_3239645, EPI_ISL_3239646 | | | |
| see above | MDI Limbach Berlin GmbH; MVZ Labor Berlin | Robert Koch Institute | unknown |
| EPI_ISL_3239647 | SLK-Kliniken Heilbronn GmbH - Institut für Laboratoriumsmedizin | Robert Koch Institute | unknown |
| EPI_ISL_3239648 | MVZ Labor Dr. Reising-Ackermann und Kollegen | Robert Koch Institute | unknown |
| EPI_ISL_3239649, EPI_ISL_3239650, EPI_ISL_3239651, EPI_ISL_3239652, EPI_ISL_3239653, EPI_ISL_3239654 | MVZ Labor Dr. Limbach & Kollegen GbR | Robert Koch Institute | unknown |
| EPI_ISL_3239655, EPI_ISL_3239656, EPI_ISL_3239657, EPI_ISL_3239658, EPI_ISL_3239659 | MDI Limbach Berlin GmbH; MVZ Labor Berlin | Robert Koch Institute | unknown |
| EPI_ISL_3239660 | Labor Mönchengladbach MVZ Dr. Stein + Kollegen GbR | Robert Koch Institute | unknown |
| EPI_ISL_3239661, EPI_ISL_3239662 | MDI Limbach Berlin GmbH; MVZ Labor Berlin | Robert Koch Institute | unknown |
| EPI_ISL_3239663, EPI_ISL_3239664, EPI_ISL_3239666, EPI_ISL_3239667, EPI_ISL_3239668, EPI_ISL_3239669, EPI_ISL_3239670, EPI_ISL_3239671, EPI_ISL_3239672, EPI_ISL_3239673, EPI_ISL_3239674, EPI_ISL_3239675, EPI_ISL_3239676, EPI_ISL_3239677, EPI_ISL_3239678, EPI_ISL_3239679, EPI_ISL_3239680, EPI_ISL_3239681, EPI_ISL_3239682, EPI_ISL_3239683, EPI_ISL_3239684, EPI_ISL_3239685, EPI_ISL_3239686, EPI_ISL_3239687, EPI_ISL_3239688, EPI_ISL_3239689, EPI_ISL_3239690, EPI_ISL_3239691, EPI_ISL_3239692, EPI_ISL_3239693, EPI_ISL_3239694, EPI_ISL_3239696, EPI_ISL_3239697, EPI_ISL_3239699, EPI_ISL_3239700, EPI_ISL_3239701, EPI_ISL_3239702, EPI_ISL_3239703 | | | |
| see above | Diagnosticum - Labor Neukirchen | Robert Koch Institute | unknown |
| EPI_ISL_3239704, EPI_ISL_3239705, EPI_ISL_3239706, EPI_ISL_3239707, EPI_ISL_3239708, EPI_ISL_3239709, EPI_ISL_3239710, EPI_ISL_3239711, EPI_ISL_3239712, EPI_ISL_3239713, EPI_ISL_3239714, EPI_ISL_3239715, EPI_ISL_3239716, EPI_ISL_3239717, EPI_ISL_3239718, EPI_ISL_3239719, EPI_ISL_3239720, EPI_ISL_3239721, EPI_ISL_3239722, EPI_ISL_3239723, EPI_ISL_3239724 | | | |
| see above | Bayerisches Landesamt für Gesundheit und Lebensmittelsicherheit (LGL) | Robert Koch Institute | unknown |
| EPI_ISL_3239725 | MVZ Medizinisches Labor Hannover GmbH | Robert Koch Institute | unknown |
| EPI_ISL_3239726, EPI_ISL_3239727, EPI_ISL_3239728, EPI_ISL_3239729, EPI_ISL_3239730, | MVZ Labor Dr. Limbach & Kollegen GbR | Robert Koch Institute | unknown |

| | | | |
|---|---|---|---|
| EPI_ISL_3239731 | | | |
| EPI_ISL_3239732 | Limbach - MVZ Labor Eveld & Kollegen Essen | Robert Koch Institute | unknown |
| EPI_ISL_3239733 | MVZ Labor Dr. Limbach & Kollegen GbR | Robert Koch Institute | unknown |
| EPI_ISL_3239734 | Limbach - MVZ Labor Eveld & Kollegen Essen | Robert Koch Institute | unknown |
| EPI_ISL_3239735 | MVZ Labor Dr. Limbach & Kollegen GbR | Robert Koch Institute | unknown |
| EPI_ISL_3239736 | Limbach - MVZ Labor Eveld & Kollegen Essen | Robert Koch Institute | unknown |
| EPI_ISL_3239737, EPI_ISL_3239738 | MVZ Labor Dr. Limbach & Kollegen GbR | Robert Koch Institute | unknown |
| EPI_ISL_3239739 | Labor Prof. Dr. G. Enders MVZ GbR | Robert Koch Institute | unknown |
| EPI_ISL_3239740, EPI_ISL_3239741, EPI_ISL_3239742, EPI_ISL_3239743, EPI_ISL_3239744, EPI_ISL_3239745, EPI_ISL_3239746 | MVZ Labor Dr. Limbach & Kollegen GbR | Robert Koch Institute | unknown |
| EPI_ISL_3239747, EPI_ISL_3239748 | Labor Mönchengladbach MVZ Dr. Stein + Kollegen GbR | Robert Koch Institute | unknown |
| EPI_ISL_3239749, EPI_ISL_3239750 | MVZ Labor Dr. Limbach & Kollegen GbR | Robert Koch Institute | unknown |
| EPI_ISL_3239751 | Labor Mönchengladbach MVZ Dr. Stein + Kollegen GbR | Robert Koch Institute | unknown |
| EPI_ISL_3239752, EPI_ISL_3239753 | MVZ Labor Dr. Limbach & Kollegen GbR | Robert Koch Institute | unknown |
| EPI_ISL_3239754 | Labor Mönchengladbach MVZ Dr. Stein + Kollegen GbR | Robert Koch Institute | unknown |
| EPI_ISL_3239755 | MVZ Labor Dr. Limbach & Kollegen GbR | Robert Koch Institute | unknown |
| EPI_ISL_3239756, EPI_ISL_3239757, EPI_ISL_3239758, EPI_ISL_3239759 | Labor Mönchengladbach MVZ Dr. Stein + Kollegen GbR | Robert Koch Institute | unknown |
| EPI_ISL_3239760 | MVZ Labor Dr. Limbach & Kollegen GbR | Robert Koch Institute | unknown |
| EPI_ISL_3239761, EPI_ISL_3239762, EPI_ISL_3239763, EPI_ISL_3239764 | Labor Mönchengladbach MVZ Dr. Stein + Kollegen GbR | Robert Koch Institute | unknown |
| EPI_ISL_3239765 | MVZ Labor Dr. Limbach & Kollegen GbR | Robert Koch Institute | unknown |
| EPI_ISL_3239766, EPI_ISL_3239767 | Labor Mönchengladbach MVZ Dr. Stein + Kollegen GbR | Robert Koch Institute | unknown |
| EPI_ISL_3239768 | MVZ Labor Dr. Limbach & Kollegen GbR | Robert Koch Institute | unknown |
| EPI_ISL_3239769 | Labor Mönchengladbach MVZ Dr. Stein + Kollegen GbR | Robert Koch Institute | unknown |
| EPI_ISL_3239770, EPI_ISL_3239771, EPI_ISL_3239772, EPI_ISL_3239773, EPI_ISL_3239774, EPI_ISL_3239775, EPI_ISL_3239776, EPI_ISL_3239777, EPI_ISL_3239778, EPI_ISL_3239779, EPI_ISL_3239780, EPI_ISL_3239781, EPI_ISL_3239782, EPI_ISL_3239783, EPI_ISL_3239784, EPI_ISL_3239785, EPI_ISL_3239786, EPI_ISL_3239787, EPI_ISL_3239788, EPI_ISL_3239789, EPI_ISL_3239790, EPI_ISL_3239791, EPI_ISL_3239792 | | | |
| see above | MDI Limbach Berlin GmbH; MVZ Labor Berlin | Robert Koch Institute | unknown |
| EPI_ISL_3239793, EPI_ISL_3239794, EPI_ISL_3239795, EPI_ISL_3239796, EPI_ISL_3239797, EPI_ISL_3239798, EPI_ISL_3239799 | Laborarztpraxis Dres. med. Walther Weindel & Kollegen | Robert Koch Institute | unknown |
| EPI_ISL_3239800 | MVZ Labor Dr. Limbach & Kollegen GbR | Robert Koch Institute | unknown |
| EPI_ISL_3239801 | MDI Limbach Berlin GmbH; MVZ Labor Berlin | Robert Koch Institute | unknown |
| EPI_ISL_3239802 | MVZ Labor Dr. Limbach & Kollegen GbR | Robert Koch Institute | unknown |
| EPI_ISL_3239803, EPI_ISL_3239804, EPI_ISL_3239805, EPI_ISL_3239806 | MDI Limbach Berlin GmbH; MVZ Labor Berlin | Robert Koch Institute | unknown |
| EPI_ISL_3239807 | MVZ Labor Dr. Limbach & Kollegen GbR | Robert Koch Institute | unknown |
| EPI_ISL_3239808, EPI_ISL_3239809, EPI_ISL_3239810 | MDI Limbach Berlin GmbH; MVZ Labor Berlin | Robert Koch Institute | unknown |
| EPI_ISL_3239811 | MVZ Medizinisches Labor Hannover GmbH | Robert Koch Institute | unknown |
| EPI_ISL_3239812 | MDI Limbach Berlin GmbH; MVZ Labor Berlin | Robert Koch Institute | unknown |
| EPI_ISL_3239813 | Limbach - MVZ Gemeinschaftslabor Suhl | Robert Koch Institute | unknown |
| EPI_ISL_3239814 | MVZ Labor Dr. Limbach & Kollegen GbR | Robert Koch Institute | unknown |
| EPI_ISL_3239815 | MDI Limbach Berlin GmbH; MVZ Labor Berlin | Robert Koch Institute | unknown |
| EPI_ISL_3239816 | MVZ für Laboratoriumsmedizin und Mikrobiologie Koblenz-Mittelrhein (Labor Koblenz) | Robert Koch Institute | unknown |
| EPI_ISL_3239817, EPI_ISL_3239818, EPI_ISL_3239819, EPI_ISL_3239821, EPI_ISL_3239822, EPI_ISL_3239823 | MVZ Labor Dr. Limbach & Kollegen GbR | Robert Koch Institute | unknown |
| EPI_ISL_3239824 | Labor für medizinische Mikrobiologie Bakteriologie Mykologie Virologie & Infektionsserologie - Labor Mölbis | Robert Koch Institute | unknown |
| EPI_ISL_3239825, EPI_ISL_3239826, EPI_ISL_3239827, EPI_ISL_3239828, EPI_ISL_3239829, EPI_ISL_3239830, EPI_ISL_3239831, EPI_ISL_3239832, EPI_ISL_3239833, EPI_ISL_3239834 | alphaomega Labor GbR (Medizinische Kooperationsgemeinschaft Dres. Ackermann/Tennert/Zaiß) | Robert Koch Institute | unknown |
| EPI_ISL_3239835 | Labor für medizinische Mikrobiologie Bakteriologie Mykologie Virologie & Infektionsserologie - Labor Mölbis | Robert Koch Institute | unknown |
| EPI_ISL_3239836 | Institut für med. Mikrobiologie u. Virologie Klinikum Oldenburg | Robert Koch Institute | unknown |
| EPI_ISL_3239837, EPI_ISL_3239838, EPI_ISL_3239839 | Dhulikhel Hospital, Kathmandu University Hospital | Molecular and Genomics Research Lab, Dhulikhel Hospital, Kathmandu University Hospital | Rajeev Shrestha, Nishan Katuwal, Navin Adhikari, Dipesh Tamrakar, Meghnath Dhimal, Pradip Gyanwali, Saroj Bhattarai, Surendra Kumar Madhup |

| | | | |
|---|---|---|---|
| EPI_ISL_3239845 | A.S.L. CITTA' DI TORINO - OSPEDALE AMEDEO DI SAVOIA - SC MICROBIOLOGIA E VIROLOGIA - | Fondazione del Piemonte per l'Oncologia IRCCS | Antonino Sottile, Silvia Brossa, Paola Marino, Giorgio Giardina |
| EPI_ISL_3239846, EPI_ISL_3239847 | A.S.L. TO4 | Fondazione del Piemonte per l'Oncologia IRCCS | Antonino Sottile, Silvia Brossa, Paola Marino, Giorgio Giardina |
| EPI_ISL_3239848 | A.S.L. TO5 | Fondazione del Piemonte per l'Oncologia IRCCS | Antonino Sottile, Silvia Brossa, Paola Marino, Giorgio Giardina |
| EPI_ISL_3239849 | A.S.L. CITTA' DI TORINO - OSPEDALE AMEDEO DI SAVOIA - SC MICROBIOLOGIA E VIROLOGIA - | Fondazione del Piemonte per l'Oncologia IRCCS | Antonino Sottile, Silvia Brossa, Paola Marino, Giorgio Giardina |
| EPI_ISL_3239850, EPI_ISL_3239851, EPI_ISL_3239853, EPI_ISL_3239854, EPI_ISL_3239855, EPI_ISL_3239856, EPI_ISL_3239857, EPI_ISL_3239858, EPI_ISL_3239859, EPI_ISL_3239860, EPI_ISL_3239861, EPI_ISL_3239862, EPI_ISL_3239863, EPI_ISL_3239864, EPI_ISL_3239865, EPI_ISL_3239867, EPI_ISL_3239868, EPI_ISL_3239869, EPI_ISL_3239870, EPI_ISL_3239871, EPI_ISL_3239872, EPI_ISL_3239873, EPI_ISL_3239874, EPI_ISL_3239875, EPI_ISL_3239876, EPI_ISL_3239877, EPI_ISL_3239878, EPI_ISL_3239879, EPI_ISL_3239880, EPI_ISL_3239881, EPI_ISL_3239883, EPI_ISL_3239884, EPI_ISL_3239885, EPI_ISL_3239886, EPI_ISL_3239887, EPI_ISL_3239888, EPI_ISL_3239889, EPI_ISL_3239891, EPI_ISL_3239892, EPI_ISL_3239893, EPI_ISL_3239894 | | | |
| see above | A.S.L. NOVARA - ISTITUTO S.S. TRINITA' - BORGOMANERO - S.S. LABORATORIO ANALISI CHIMICO CLINICHE - | Fondazione del Piemonte per l'Oncologia IRCCS | Antonino Sottile, Silvia Brossa, Paola Marino, Giorgio Giardina |
| EPI_ISL_3239901, EPI_ISL_3239902, EPI_ISL_3239903, EPI_ISL_3239904, EPI_ISL_3239906 | Laboratorio de Virología HUCA | Laboratorio de Virología HUCA | Sandoval M, Castelló C, Gómez de Oña J, Boga JA, Rojo S, Alvarez-Arguelles ME, Abreu F, Costales I, Perez-Martínez Z, Martín-Rodríguez G, Coto E, Melón S |
| EPI_ISL_3239907 | OSPEDALE CIVILE TERAMO - CENTRO TRASFUSIONALE | Istituto Zooprofilattico Sperimentale dell'Abruzzo e Molise "G. Caporale" | Lorusso A, Marcacci M, Di Domenico M, Ancora M, Curini V, Di Lollo Valeria, Mangone I, Rinaldi A, Delli Compagni E, Scialabba S, Caporale M, Di Pasquale A, Cammà C, Puglia I, Calistri P, Savini G |
| EPI_ISL_3239908, EPI_ISL_3239909, EPI_ISL_3239913 | Laboratorio de Virología HUCA | Laboratorio de Virología HUCA | Sandoval M, Castelló C, Gómez de Oña J, Boga JA, Rojo S, Alvarez-Arguelles ME, Abreu F, Costales I, Perez-Martínez Z, Martín-Rodríguez G, Coto E, Melón S |
| EPI_ISL_3239916 | Department of Pathology & Laboratory Medicine | Centre for Research in Advanced Tropical Bioscience (Biotropic Centre) | Hajar Fauzan Ahmad |
| EPI_ISL_3239917 | National Platform bis UMONS/Jolimont | National Platform bis UMONS/Jolimont | François Dufrasne, Guillaume Bayon-Vicente, Florian Juszczak, Laetitia Gheysen, Eric Tarantino, Gautier Detry, Ruddy Wattiez |
| EPI_ISL_3239918 | National Platform bis UMONS/Jolimont | National Platform bis UMONS/Jolimont | Rue Institut Notre Dame de la Compassion, 1. La Louvière 7100. Belgium |
| EPI_ISL_3239919 | National Platform bis UMONS/Jolimont | National Platform bis UMONS/Jolimont | François Dufrasne, Guillaume Bayon-Vicente, Florian Juszczak, Laetitia Gheysen, Eric Tarantino, Gautier Detry, Ruddy Wattiez |
| EPI_ISL_3239920 | National Platform bis UMONS/Jolimont | National Platform bis UMONS/Jolimont | National Platform bis UMONS/Jolimont |
| EPI_ISL_3239921 | A.S.L. ALESSANDRIA | Fondazione del Piemonte per l'Oncologia IRCCS | Antonino Sottile, Silvia Brossa, Paola Marino, Giorgio Giardina |
| EPI_ISL_3239922, EPI_ISL_3239923 | A.S.L. NOVARA - ISTITUTO S.S. TRINITA' - BORGOMANERO - S.S. LABORATORIO ANALISI CHIMICO CLINICHE - | Fondazione del Piemonte per l'Oncologia IRCCS | Antonino Sottile, Silvia Brossa, Paola Marino, Giorgio Giardina |
| EPI_ISL_3239924 | A.S.L. BIELLA | Fondazione del Piemonte per l'Oncologia IRCCS | Antonino Sottile, Silvia Brossa, Paola Marino, Giorgio Giardina |
| EPI_ISL_3239925 | A.S.L. CUNEO 1 | Fondazione del Piemonte per l'Oncologia IRCCS | Antonino Sottile, Silvia Brossa, Paola Marino, Giorgio Giardina |
| EPI_ISL_3239926 | A.S.L. NOVARA - ISTITUTO S.S. TRINITA' - BORGOMANERO - S.S. LABORATORIO ANALISI CHIMICO CLINICHE - | Fondazione del Piemonte per l'Oncologia IRCCS | Antonino Sottile, Silvia Brossa, Paola Marino, Giorgio Giardina |
| EPI_ISL_3239928, EPI_ISL_3239929, EPI_ISL_3239930, EPI_ISL_3239931, EPI_ISL_3239932, EPI_ISL_3239933, EPI_ISL_3239934, EPI_ISL_3239935, EPI_ISL_3239936, EPI_ISL_3239937, EPI_ISL_3239938, EPI_ISL_3239939, EPI_ISL_3239940, EPI_ISL_3239941, EPI_ISL_3239943, EPI_ISL_3239945, EPI_ISL_3239946, EPI_ISL_3239947, EPI_ISL_3239948, EPI_ISL_3239949, EPI_ISL_3239950, EPI_ISL_3239951, EPI_ISL_3239953, EPI_ISL_3239954, EPI_ISL_3239955, EPI_ISL_3239956, EPI_ISL_3239957, EPI_ISL_3239958, EPI_ISL_3239959, EPI_ISL_3239960, EPI_ISL_3239961, EPI_ISL_3239962, EPI_ISL_3239963, EPI_ISL_3239964, EPI_ISL_3239965, EPI_ISL_3239966, EPI_ISL_3239967, EPI_ISL_3239968, EPI_ISL_3239969, EPI_ISL_3239970 | | | |
| see above | A.S.L. CUNEO 1 | Fondazione del Piemonte per l'Oncologia IRCCS | Antonino Sottile, Silvia Brossa, Paola Marino, Giorgio Giardina |
| EPI_ISL_3239971, EPI_ISL_3239972, EPI_ISL_3239973, EPI_ISL_3239974, EPI_ISL_3239975, EPI_ISL_3239976, EPI_ISL_3239977, EPI_ISL_3239978, EPI_ISL_3239979, EPI_ISL_3239980, EPI_ISL_3239981, EPI_ISL_3239982, EPI_ISL_3239983, EPI_ISL_3239984, EPI_ISL_3239985, EPI_ISL_3239986, EPI_ISL_3239987, EPI_ISL_3239988, EPI_ISL_3239989, EPI_ISL_3239990, EPI_ISL_3239991, EPI_ISL_3239992, EPI_ISL_3239993, EPI_ISL_3239994, EPI_ISL_3239995, EPI_ISL_3239996, EPI_ISL_3239997, EPI_ISL_3239998, EPI_ISL_3239999, EPI_ISL_3240000, EPI_ISL_3240001, EPI_ISL_3240002, EPI_ISL_3240003, EPI_ISL_3240004, EPI_ISL_3240005, EPI_ISL_3240006, EPI_ISL_3240007, EPI_ISL_3240008, EPI_ISL_3240009, EPI_ISL_3240010, EPI_ISL_3240011, EPI_ISL_3240012, EPI_ISL_3240013, EPI_ISL_3240014, EPI_ISL_3240015, EPI_ISL_3240016, EPI_ISL_3240017, EPI_ISL_3240018, EPI_ISL_3240019, EPI_ISL_3240020, EPI_ISL_3240021, EPI_ISL_3240022, EPI_ISL_3240023, EPI_ISL_3240024, EPI_ISL_3240025, EPI_ISL_3240026, EPI_ISL_3240027, EPI_ISL_3240028, EPI_ISL_3240029, EPI_ISL_3240030, EPI_ISL_3240031, EPI_ISL_3240032, EPI_ISL_3240033, EPI_ISL_3240034, EPI_ISL_3240035, EPI_ISL_3240036, EPI_ISL_3240037, EPI_ISL_3240038, EPI_ISL_3240039, EPI_ISL_3240040, EPI_ISL_3240041, EPI_ISL_3240042, EPI_ISL_3240043, EPI_ISL_3240044, EPI_ISL_3240045, EPI_ISL_3240046, EPI_ISL_3240047, EPI_ISL_3240048, EPI_ISL_3240049, EPI_ISL_3240050, EPI_ISL_3240051, EPI_ISL_3240052, EPI_ISL_3240053, EPI_ISL_3240054, EPI_ISL_3240055, EPI_ISL_3240056, EPI_ISL_3240057, EPI_ISL_3240058, EPI_ISL_3240059, EPI_ISL_3240060, EPI_ISL_3240061, EPI_ISL_3240062, EPI_ISL_3240063, EPI_ISL_3240064, EPI_ISL_3240065, EPI_ISL_3240066, EPI_ISL_3240067, EPI_ISL_3240068, EPI_ISL_3240069, EPI_ISL_3240070, EPI_ISL_3240071, EPI_ISL_3240072, EPI_ISL_3240073, EPI_ISL_3240074, EPI_ISL_3240075, EPI_ISL_3240076, EPI_ISL_3240077, EPI_ISL_3240078, EPI_ISL_3240079, EPI_ISL_3240080, EPI_ISL_3240081, EPI_ISL_3240082, EPI_ISL_3240083, EPI_ISL_3240084, EPI_ISL_3240085, EPI_ISL_3240086, EPI_ISL_3240087, EPI_ISL_3240088, EPI_ISL_3240089, EPI_ISL_3240090, EPI_ISL_3240091, EPI_ISL_3240092, EPI_ISL_3240093, EPI_ISL_3240094, EPI_ISL_3240095, EPI_ISL_3240096, EPI_ISL_3240097, EPI_ISL_3240098, EPI_ISL_3240099, EPI_ISL_3240100, EPI_ISL_3240101, EPI_ISL_3240102, EPI_ISL_3240103, EPI_ISL_3240104, EPI_ISL_3240105, EPI_ISL_3240106, EPI_ISL_3240107, EPI_ISL_3240108, EPI_ISL_3240109, EPI_ISL_3240110, EPI_ISL_3240111, EPI_ISL_3240112, EPI_ISL_3240113, EPI_ISL_3240114, EPI_ISL_3240115, EPI_ISL_3240116, EPI_ISL_3240117, EPI_ISL_3240118, EPI_ISL_3240119, EPI_ISL_3240120, EPI_ISL_3240121, EPI_ISL_3240122, EPI_ISL_3240123, EPI_ISL_3240124, EPI_ISL_3240125, EPI_ISL_3240126, EPI_ISL_3240127, EPI_ISL_3240128, EPI_ISL_3240129, EPI_ISL_3240130, EPI_ISL_3240131, EPI_ISL_3240132, EPI_ISL_3240133, EPI_ISL_3240134, EPI_ISL_3240135, EPI_ISL_3240136, EPI_ISL_3240137, EPI_ISL_3240138, EPI_ISL_3240139, EPI_ISL_3240140, EPI_ISL_3240141, EPI_ISL_3240142, EPI_ISL_3240143, EPI_ISL_3240144, EPI_ISL_3240145, EPI_ISL_3240146, EPI_ISL_3240147, EPI_ISL_3240148, EPI_ISL_3240149, EPI_ISL_3240150, EPI_ISL_3240151, EPI_ISL_3240152, EPI_ISL_3240153, EPI_ISL_3240154, EPI_ISL_3240155, EPI_ISL_3240156, EPI_ISL_3240157, EPI_ISL_3240158, EPI_ISL_3240159, EPI_ISL_3240160, EPI_ISL_3240161, EPI_ISL_3240162, EPI_ISL_3240163, EPI_ISL_3240164, EPI_ISL_3240165, EPI_ISL_3240166, EPI_ISL_3240167, EPI_ISL_3240168, EPI_ISL_3240169, EPI_ISL_3240170, EPI_ISL_3240171, EPI_ISL_3240172, EPI_ISL_3240173, EPI_ISL_3240174, EPI_ISL_3240175, EPI_ISL_3240176, EPI_ISL_3240177 | | | |
| see above | Department of Bacteria, Parasites and Fungi, Statens Serum Institut, Copenhagen, Denmark | Statens Serum Institut Bioinformatics and Microbial Genomics | Danish Covid-19 Genome Consortium |
| EPI_ISL_3240178 | Department of Clinical Microbiology, Odense University Hospital, Odense, Denmark | Statens Serum Institut Bioinformatics and Microbial Genomics | Danish Covid-19 Genome Consortium |
| EPI_ISL_3240179, EPI_ISL_3240180, EPI_ISL_3240181, EPI_ISL_3240182, EPI_ISL_3240183, EPI_ISL_3240184, EPI_ISL_3240185, EPI_ISL_3240186, EPI_ISL_3240187, EPI_ISL_3240188, EPI_ISL_3240189, EPI_ISL_3240190, EPI_ISL_3240191, EPI_ISL_3240192, EPI_ISL_3240193, EPI_ISL_3240194, EPI_ISL_3240195, EPI_ISL_3240196, EPI_ISL_3240197, EPI_ISL_3240198, EPI_ISL_3240199, EPI_ISL_3240200, EPI_ISL_3240201, EPI_ISL_3240202, EPI_ISL_3240203, EPI_ISL_3240204, EPI_ISL_3240205, EPI_ISL_3240206, EPI_ISL_3240207, EPI_ISL_3240208, EPI_ISL_3240209, EPI_ISL_3240210, EPI_ISL_3240211, EPI_ISL_3240212, EPI_ISL_3240213, EPI_ISL_3240214, EPI_ISL_3240215, EPI_ISL_3240216, EPI_ISL_3240217, EPI_ISL_3240218, EPI_ISL_3240219, EPI_ISL_3240220, EPI_ISL_3240222, EPI_ISL_3240223, EPI_ISL_3240224, EPI_ISL_3240225, EPI_ISL_3240226, EPI_ISL_3240227, EPI_ISL_3240228, EPI_ISL_3240229, EPI_ISL_3240230, EPI_ISL_3240231, EPI_ISL_3240232, EPI_ISL_3240233, EPI_ISL_3240234, EPI_ISL_3240235, EPI_ISL_3240236, EPI_ISL_3240237, EPI_ISL_3240238, EPI_ISL_3240239, EPI_ISL_3240240, EPI_ISL_3240241, EPI_ISL_3240242, EPI_ISL_3240243, EPI_ISL_3240244, EPI_ISL_3240245, EPI_ISL_3240246, EPI_ISL_3240247, EPI_ISL_3240248, EPI_ISL_3240249, EPI_ISL_3240250, EPI_ISL_3240251, EPI_ISL_3240252, EPI_ISL_3240253, EPI_ISL_3240254, EPI_ISL_3240255, EPI_ISL_3240256, EPI_ISL_3240257, EPI_ISL_3240258, EPI_ISL_3240259, EPI_ISL_3240260, EPI_ISL_3240261, EPI_ISL_3240262, EPI_ISL_3240263, EPI_ISL_3240264, EPI_ISL_3240265, EPI_ISL_3240266, EPI_ISL_3240267, EPI_ISL_3240268, EPI_ISL_3240269, EPI_ISL_3240270, EPI_ISL_3240271, EPI_ISL_3240272, EPI_ISL_3240273, EPI_ISL_3240274, EPI_ISL_3240275, EPI_ISL_3240276, EPI_ISL_3240277, EPI_ISL_3240278, EPI_ISL_3240279, EPI_ISL_3240280, EPI_ISL_3240281, EPI_ISL_3240282, EPI_ISL_3240283, EPI_ISL_3240284, EPI_ISL_3240285, EPI_ISL_3240286, EPI_ISL_3240287, EPI_ISL_3240288, EPI_ISL_3240289, EPI_ISL_3240290, EPI_ISL_3240291, EPI_ISL_3240293, EPI_ISL_3240294, EPI_ISL_3240295, EPI_ISL_3240296, EPI_ISL_3240297, EPI_ISL_3240298, EPI_ISL_3240299, EPI_ISL_3240300, EPI_ISL_3240301, EPI_ISL_3240302, EPI_ISL_3240303, EPI_ISL_3240304, EPI_ISL_3240305, EPI_ISL_3240306, EPI_ISL_3240307, EPI_ISL_3240308, EPI_ISL_3240309, EPI_ISL_3240310, EPI_ISL_3240311, EPI_ISL_3240312, EPI_ISL_3240313, EPI_ISL_3240314, EPI_ISL_3240315, EPI_ISL_3240316, EPI_ISL_3240317, EPI_ISL_3240318, EPI_ISL_3240319, EPI_ISL_3240320, EPI_ISL_3240321, EPI_ISL_3240322, EPI_ISL_3240323, EPI_ISL_3240324, EPI_ISL_3240325, EPI_ISL_3240326, EPI_ISL_3240327, EPI_ISL_3240328, EPI_ISL_3240329, EPI_ISL_3240330, EPI_ISL_3240331, EPI_ISL_3240332, EPI_ISL_3240333, EPI_ISL_3240334, EPI_ISL_3240335, EPI_ISL_3240336, EPI_ISL_3240337, EPI_ISL_3240338, EPI_ISL_3240339, EPI_ISL_3240340, EPI_ISL_3240341, EPI_ISL_3240342, EPI_ISL_3240343, EPI_ISL_3240344, EPI_ISL_3240345, EPI_ISL_3240346, EPI_ISL_3240347, EPI_ISL_3240348, EPI_ISL_3240349, EPI_ISL_3240350, EPI_ISL_3240351, EPI_ISL_3240352, EPI_ISL_3240353, EPI_ISL_3240354, EPI_ISL_3240355, EPI_ISL_3240356, EPI_ISL_3240357 | | | |
| see above | Department of Bacteria, Parasites and Fungi, Statens Serum Institut, Copenhagen, Denmark | Statens Serum Institut Bioinformatics and Microbial Genomics | Danish Covid-19 Genome Consortium |
| EPI_ISL_3240358 | Department of Clinical Microbiology, Odense University Hospital, Odense, Denmark | Statens Serum Institut Bioinformatics and Microbial Genomics | Danish Covid-19 Genome Consortium |
| EPI_ISL_3240359, EPI_ISL_3240360, EPI_ISL_3240361, EPI_ISL_3240362, EPI_ISL_3240363, EPI_ISL_3240364, EPI_ISL_3240365, EPI_ISL_3240366, EPI_ISL_3240367, EPI_ISL_3240368, EPI_ISL_3240369, EPI_ISL_3240370, EPI_ISL_3240371, EPI_ISL_3240372, EPI_ISL_3240373, EPI_ISL_3240374, EPI_ISL_3240375, EPI_ISL_3240376, EPI_ISL_3240377, EPI_ISL_3240378, EPI_ISL_3240379, EPI_ISL_3240380, EPI_ISL_3240381, EPI_ISL_3240382, EPI_ISL_3240383, EPI_ISL_3240384, EPI_ISL_3240385, EPI_ISL_3240386, EPI_ISL_3240387, EPI_ISL_3240388, EPI_ISL_3240389, EPI_ISL_3240390, EPI_ISL_3240391, EPI_ISL_3240392, EPI_ISL_3240393, EPI_ISL_3240394, EPI_ISL_3240395, EPI_ISL_3240396, EPI_ISL_3240397, EPI_ISL_3240398, EPI_ISL_3240399, EPI_ISL_3240400, EPI_ISL_3240401, EPI_ISL_3240402, EPI_ISL_3240403, EPI_ISL_3240404, EPI_ISL_3240405, EPI_ISL_3240406, EPI_ISL_3240407, EPI_ISL_3240408, EPI_ISL_3240409, EPI_ISL_3240410, EPI_ISL_3240411, EPI_ISL_3240412, EPI_ISL_3240413, EPI_ISL_3240414, EPI_ISL_3240415, EPI_ISL_3240416, EPI_ISL_3240417, EPI_ISL_3240418, EPI_ISL_3240419, EPI_ISL_3240420, EPI_ISL_3240421, EPI_ISL_3240422, EPI_ISL_3240423, EPI_ISL_3240424, EPI_ISL_3240425, EPI_ISL_3240426, EPI_ISL_3240427, EPI_ISL_3240428, EPI_ISL_3240429, EPI_ISL_3240430, EPI_ISL_3240431, EPI_ISL_3240432, EPI_ISL_3240433, EPI_ISL_3240434, EPI_ISL_3240435, EPI_ISL_3240436, EPI_ISL_3240437, EPI_ISL_3240438, EPI_ISL_3240439, EPI_ISL_3240440, EPI_ISL_3240441, EPI_ISL_3240442, EPI_ISL_3240443, EPI_ISL_3240444, EPI_ISL_3240445, EPI_ISL_3240446, EPI_ISL_3240447, EPI_ISL_3240448, EPI_ISL_3240449, EPI_ISL_3240450, EPI_ISL_3240451, EPI_ISL_3240452, EPI_ISL_3240453, EPI_ISL_3240454, EPI_ISL_3240455, EPI_ISL_3240456, EPI_ISL_3240457, EPI_ISL_3240458, EPI_ISL_3240459, EPI_ISL_3240460, EPI_ISL_3240461, EPI_ISL_3240462, EPI_ISL_3240463, EPI_ISL_3240464, EPI_ISL_3240465, EPI_ISL_3240466, EPI_ISL_3240467, EPI_ISL_3240468, EPI_ISL_3240469, EPI_ISL_3240470, EPI_ISL_3240471, EPI_ISL_3240472, EPI_ISL_3240473, EPI_ISL_3240474, EPI_ISL_3240475, EPI_ISL_3240476, EPI_ISL_3240477, EPI_ISL_3240478, EPI_ISL_3240479, EPI_ISL_3240480, EPI_ISL_3240481, EPI_ISL_3240482, EPI_ISL_3240483, EPI_ISL_3240484, EPI_ISL_3240485, EPI_ISL_3240486, EPI_ISL_3240487, EPI_ISL_3240488, EPI_ISL_3240489, EPI_ISL_3240490, EPI_ISL_3240491, EPI_ISL_3240492, EPI_ISL_3240493, EPI_ISL_3240494, EPI_ISL_3240495, EPI_ISL_3240496, EPI_ISL_3240497, EPI_ISL_3240498, EPI_ISL_3240499, EPI_ISL_3240500, EPI_ISL_3240501, EPI_ISL_3240502, EPI_ISL_3240503, EPI_ISL_3240504, EPI_ISL_3240505, EPI_ISL_3240506, EPI_ISL_3240507, EPI_ISL_3240508, EPI_ISL_3240509, EPI_ISL_3240510, EPI_ISL_3240511, EPI_ISL_3240512, EPI_ISL_3240513, EPI_ISL_3240514, EPI_ISL_3240515, EPI_ISL_3240516, EPI_ISL_3240517, EPI_ISL_3240518, EPI_ISL_3240519, EPI_ISL_3240520, EPI_ISL_3240521, EPI_ISL_3240522, EPI_ISL_3240523, EPI_ISL_3240524 | | | |
| see above | Department of Bacteria, Parasites and Fungi, Statens Serum Institut, Copenhagen, Denmark | Statens Serum Institut Bioinformatics and Microbial Genomics | Danish Covid-19 Genome Consortium |
| EPI_ISL_3240525 | Department of Clinical Microbiology, Odense University Hospital, Odense, Denmark | Statens Serum Institut Bioinformatics and Microbial Genomics | Danish Covid-19 Genome Consortium |
| EPI_ISL_3240526, EPI_ISL_3240527, EPI_ISL_3240528, EPI_ISL_3240529, EPI_ISL_3240530, EPI_ISL_3240531, EPI_ISL_3240532, EPI_ISL_3240533, EPI_ISL_3240534, EPI_ISL_3240535, EPI_ISL_3240536, EPI_ISL_3240537, EPI_ISL_3240538, EPI_ISL_3240539, EPI_ISL_3240540, EPI_ISL_3240541, EPI_ISL_3240542, EPI_ISL_3240543, EPI_ISL_3240544, EPI_ISL_3240545, EPI_ISL_3240546, EPI_ISL_3240547, EPI_ISL_3240548, EPI_ISL_3240549, EPI_ISL_3240550, EPI_ISL_3240551, EPI_ISL_3240552, EPI_ISL_3240553, EPI_ISL_3240554, EPI_ISL_3240555, EPI_ISL_3240556, EPI_ISL_3240557, EPI_ISL_3240558, EPI_ISL_3240559, EPI_ISL_3240560, EPI_ISL_3240562, EPI_ISL_3240563, EPI_ISL_3240564, EPI_ISL_3240565, EPI_ISL_3240566, EPI_ISL_3240567, EPI_ISL_3240568, EPI_ISL_3240569, EPI_ISL_3240570, EPI_ISL_3240571, EPI_ISL_3240572, EPI_ISL_3240573, EPI_ISL_3240574, EPI_ISL_3240575, EPI_ISL_3240576, EPI_ISL_3240577, EPI_ISL_3240578, EPI_ISL_3240579, EPI_ISL_3240580, EPI_ISL_3240581, EPI_ISL_3240582, EPI_ISL_3240583, EPI_ISL_3240584, EPI_ISL_3240585, EPI_ISL_3240586, EPI_ISL_3240587, EPI_ISL_3240588, EPI_ISL_3240589, EPI_ISL_3240591, EPI_ISL_3240592, EPI_ISL_3240593, EPI_ISL_3240594, EPI_ISL_3240595, EPI_ISL_3240596, EPI_ISL_3240597, EPI_ISL_3240598, EPI_ISL_3240599, EPI_ISL_3240600, EPI_ISL_3240601, EPI_ISL_3240602, EPI_ISL_3240603, EPI_ISL_3240604, EPI_ISL_3240605, EPI_ISL_3240606, EPI_ISL_3240607, EPI_ISL_3240608, EPI_ISL_3240609, EPI_ISL_3240610, EPI_ISL_3240611, EPI_ISL_3240612, EPI_ISL_3240613, EPI_ISL_3240615, EPI_ISL_3240616, EPI_ISL_3240617, EPI_ISL_3240618, EPI_ISL_3240619, EPI_ISL_3240620, EPI_ISL_3240621, EPI_ISL_3240622, EPI_ISL_3240623, EPI_ISL_3240624, EPI_ISL_3240625, EPI_ISL_3240626, EPI_ISL_3240627, EPI_ISL_3240628, EPI_ISL_3240629, EPI_ISL_3240630, EPI_ISL_3240631, EPI_ISL_3240632, EPI_ISL_3240633, EPI_ISL_3240634, EPI_ISL_3240635, EPI_ISL_3240636, EPI_ISL_3240637, EPI_ISL_3240638, EPI_ISL_3240639, EPI_ISL_3240640, EPI_ISL_3240641, EPI_ISL_3240642, EPI_ISL_3240643, EPI_ISL_3240644, EPI_ISL_3240645, EPI_ISL_3240646, EPI_ISL_3240647, EPI_ISL_3240648, EPI_ISL_3240649, EPI_ISL_3240650, EPI_ISL_3240651, EPI_ISL_3240652 | | | |
| see above | Department of Bacteria, Parasites and Fungi, Statens Serum Institut, Copenhagen, Denmark | Statens Serum Institut Bioinformatics and Microbial Genomics | Danish Covid-19 Genome Consortium |
| EPI_ISL_3240653 | Department of Clinical Microbiology, Odense University Hospital, Odense, Denmark | Statens Serum Institut Bioinformatics and Microbial Genomics | Danish Covid-19 Genome Consortium |
| EPI_ISL_3240654, EPI_ISL_3240655, EPI_ISL_3240656, EPI_ISL_3240657, EPI_ISL_3240658, EPI_ISL_3240659, EPI_ISL_3240660, EPI_ISL_3240661, EPI_ISL_3240662, EPI_ISL_3240663, EPI_ISL_3240664, EPI_ISL_3240665, EPI_ISL_3240666, EPI_ISL_3240667, EPI_ISL_3240668, EPI_ISL_3240669, EPI_ISL_3240670, EPI_ISL_3240671, EPI_ISL_3240672, EPI_ISL_3240673, EPI_ISL_3240674, EPI_ISL_3240675, EPI_ISL_3240676, EPI_ISL_3240677, EPI_ISL_3240678, EPI_ISL_3240679, EPI_ISL_3240680, EPI_ISL_3240681, EPI_ISL_3240682, EPI_ISL_3240683, EPI_ISL_3240684, EPI_ISL_3240685, EPI_ISL_3240686, EPI_ISL_3240687, EPI_ISL_3240689 | | | |
| see above | Department of Bacteria, Parasites and Fungi, Statens Serum Institut, Copenhagen, Denmark | Statens Serum Institut Bioinformatics and Microbial Genomics | Danish Covid-19 Genome Consortium |
| EPI_ISL_3240690 | Department of Clinical Microbiology, Odense University Hospital, Odense, Denmark | Statens Serum Institut Bioinformatics and Microbial Genomics | Danish Covid-19 Genome Consortium |
| EPI_ISL_3240691, EPI_ISL_3240692, EPI_ISL_3240693, EPI_ISL_3240694, EPI_ISL_3240695, EPI_ISL_3240696, EPI_ISL_3240697, EPI_ISL_3240698, EPI_ISL_3240699, EPI_ISL_3240700, EPI_ISL_3240701, EPI_ISL_3240702, EPI_ISL_3240703, EPI_ISL_3240704, EPI_ISL_3240705, EPI_ISL_3240706, EPI_ISL_3240707, EPI_ISL_3240708, EPI_ISL_3240709, EPI_ISL_3240710, EPI_ISL_3240711, EPI_ISL_3240712, EPI_ISL_3240713, EPI_ISL_3240714, EPI_ISL_3240715, EPI_ISL_3240716, EPI_ISL_3240717, EPI_ISL_3240718, EPI_ISL_3240719, EPI_ISL_3240720, EPI_ISL_3240721, EPI_ISL_3240722, EPI_ISL_3240723, EPI_ISL_3240724, EPI_ISL_3240725, EPI_ISL_3240726, EPI_ISL_3240727, EPI_ISL_3240728, EPI_ISL_3240729, EPI_ISL_3240730, EPI_ISL_3240731, EPI_ISL_3240732, EPI_ISL_3240734, EPI_ISL_3240735, | | | |

| | | | |
|---|---|---|---|
| EPI_ISL_3240736, EPI_ISL_3240737, EPI_ISL_3240738, EPI_ISL_3240739, EPI_ISL_3240740, EPI_ISL_3240741, EPI_ISL_3240742, EPI_ISL_3240743, EPI_ISL_3240744, EPI_ISL_3240745, EPI_ISL_3240746, EPI_ISL_3240747, EPI_ISL_3240748, EPI_ISL_3240749, EPI_ISL_3240750, EPI_ISL_3240751, EPI_ISL_3240752, EPI_ISL_3240753, EPI_ISL_3240754, EPI_ISL_3240755, EPI_ISL_3240756, EPI_ISL_3240757, EPI_ISL_3240758, EPI_ISL_3240759, EPI_ISL_3240760, EPI_ISL_3240761, EPI_ISL_3240762, EPI_ISL_3240763, EPI_ISL_3240764, EPI_ISL_3240765, EPI_ISL_3240766, EPI_ISL_3240767, EPI_ISL_3240768, EPI_ISL_3240769, EPI_ISL_3240770, EPI_ISL_3240771, EPI_ISL_3240772, EPI_ISL_3240773, EPI_ISL_3240774, EPI_ISL_3240775, EPI_ISL_3240776, EPI_ISL_3240777, EPI_ISL_3240778, EPI_ISL_3240779, EPI_ISL_3240780, EPI_ISL_3240781, EPI_ISL_3240782, EPI_ISL_3240783, EPI_ISL_3240784, EPI_ISL_3240785, EPI_ISL_3240786, EPI_ISL_3240787, EPI_ISL_3240788, EPI_ISL_3240789, EPI_ISL_3240790, EPI_ISL_3240791, EPI_ISL_3240792, EPI_ISL_3240793, EPI_ISL_3240794, EPI_ISL_3240795, EPI_ISL_3240796, EPI_ISL_3240797, EPI_ISL_3240798, EPI_ISL_3240799, EPI_ISL_3240800, EPI_ISL_3240801, EPI_ISL_3240802, EPI_ISL_3240803, EPI_ISL_3240804, EPI_ISL_3240805, EPI_ISL_3240806, EPI_ISL_3240807, EPI_ISL_3240808, EPI_ISL_3240809, EPI_ISL_3240810, EPI_ISL_3240811, EPI_ISL_3240812, EPI_ISL_3240813, EPI_ISL_3240814, EPI_ISL_3240815, EPI_ISL_3240816, EPI_ISL_3240817, EPI_ISL_3240818, EPI_ISL_3240819, EPI_ISL_3240820, EPI_ISL_3240821, EPI_ISL_3240822, EPI_ISL_3240823, EPI_ISL_3240824, EPI_ISL_3240825, EPI_ISL_3240826, EPI_ISL_3240827, EPI_ISL_3240828, EPI_ISL_3240829, EPI_ISL_3240830, EPI_ISL_3240831, EPI_ISL_3240832, EPI_ISL_3240833, EPI_ISL_3240834, EPI_ISL_3240835, EPI_ISL_3240836, EPI_ISL_3240837, EPI_ISL_3240838, EPI_ISL_3240839, EPI_ISL_3240840, EPI_ISL_3240841, EPI_ISL_3240842, EPI_ISL_3240843 | | | |
| see above | Department of Bacteria, Parasites and Fungi, Statens Serum Institut, Copenhagen, Denmark | Statens Serum Institut Bioinformatics and Microbial Genomics | Danish Covid-19 Genome Consortium |
| EPI_ISL_3240844 | Department of Clinical Microbiology, Odense University Hospital, Odense, Denmark | Statens Serum Institut Bioinformatics and Microbial Genomics | Danish Covid-19 Genome Consortium |
| EPI_ISL_3240845, EPI_ISL_3240846, EPI_ISL_3240847, EPI_ISL_3240848, EPI_ISL_3240849, EPI_ISL_3240850, EPI_ISL_3240851, EPI_ISL_3240852, EPI_ISL_3240853, EPI_ISL_3240854, EPI_ISL_3240855, EPI_ISL_3240856, EPI_ISL_3240857, EPI_ISL_3240858, EPI_ISL_3240859, EPI_ISL_3240860, EPI_ISL_3240861, EPI_ISL_3240862, EPI_ISL_3240863, EPI_ISL_3240864, EPI_ISL_3240865 | | | |
| see above | Department of Bacteria, Parasites and Fungi, Statens Serum Institut, Copenhagen, Denmark | Statens Serum Institut Bioinformatics and Microbial Genomics | Danish Covid-19 Genome Consortium |
| EPI_ISL_3240866 | Department of Clinical Microbiology, Odense University Hospital, Odense, Denmark | Statens Serum Institut Bioinformatics and Microbial Genomics | Danish Covid-19 Genome Consortium |
| EPI_ISL_3240867, EPI_ISL_3240868, EPI_ISL_3240869, EPI_ISL_3240870, EPI_ISL_3240871, EPI_ISL_3240872, EPI_ISL_3240873, EPI_ISL_3240874, EPI_ISL_3240875, EPI_ISL_3240876, EPI_ISL_3240877, EPI_ISL_3240878, EPI_ISL_3240879, EPI_ISL_3240880, EPI_ISL_3240881, EPI_ISL_3240882, EPI_ISL_3240883, EPI_ISL_3240884, EPI_ISL_3240885, EPI_ISL_3240886, EPI_ISL_3240887, EPI_ISL_3240888, EPI_ISL_3240889, EPI_ISL_3240890, EPI_ISL_3240891, EPI_ISL_3240892, EPI_ISL_3240893, EPI_ISL_3240894, EPI_ISL_3240895, EPI_ISL_3240896, EPI_ISL_3240897, EPI_ISL_3240898, EPI_ISL_3240899, EPI_ISL_3240900, EPI_ISL_3240901, EPI_ISL_3240902, EPI_ISL_3240903, EPI_ISL_3240904, EPI_ISL_3240905, EPI_ISL_3240906, EPI_ISL_3240907, EPI_ISL_3240908, EPI_ISL_3240909, EPI_ISL_3240910, EPI_ISL_3240911, EPI_ISL_3240912, EPI_ISL_3240913, EPI_ISL_3240914, EPI_ISL_3240915, EPI_ISL_3240916, EPI_ISL_3240917, EPI_ISL_3240918, EPI_ISL_3240919, EPI_ISL_3240920, EPI_ISL_3240921, EPI_ISL_3240922, EPI_ISL_3240923, EPI_ISL_3240924, EPI_ISL_3240925, EPI_ISL_3240926, EPI_ISL_3240927, EPI_ISL_3240928, EPI_ISL_3240929, EPI_ISL_3240930, EPI_ISL_3240931, EPI_ISL_3240932, EPI_ISL_3240933, EPI_ISL_3240934, EPI_ISL_3240935, EPI_ISL_3240936, EPI_ISL_3240937, EPI_ISL_3240938, EPI_ISL_3240939 | | | |
| see above | Department of Bacteria, Parasites and Fungi, Statens Serum Institut, Copenhagen, Denmark | Statens Serum Institut Bioinformatics and Microbial Genomics | Danish Covid-19 Genome Consortium |
| EPI_ISL_3240940 | Department of Clinical Microbiology, Odense University Hospital, Odense, Denmark | Statens Serum Institut Bioinformatics and Microbial Genomics | Danish Covid-19 Genome Consortium |
| EPI_ISL_3240941, EPI_ISL_3240942, EPI_ISL_3240943, EPI_ISL_3240944, EPI_ISL_3240945, EPI_ISL_3240946, EPI_ISL_3240947, EPI_ISL_3240948, EPI_ISL_3240949, EPI_ISL_3240950, EPI_ISL_3240951, EPI_ISL_3240952, EPI_ISL_3240953, EPI_ISL_3240954, EPI_ISL_3240955, EPI_ISL_3240956, EPI_ISL_3240957, EPI_ISL_3240958, EPI_ISL_3240959, EPI_ISL_3240960, EPI_ISL_3240961, EPI_ISL_3240962, EPI_ISL_3240963, EPI_ISL_3240964, EPI_ISL_3240965, EPI_ISL_3240966, EPI_ISL_3240967, EPI_ISL_3240968, EPI_ISL_3240969, EPI_ISL_3240970, EPI_ISL_3240971, EPI_ISL_3240972, EPI_ISL_3240973, EPI_ISL_3240974, EPI_ISL_3240975, EPI_ISL_3240976, EPI_ISL_3240977, EPI_ISL_3240978, EPI_ISL_3240979, EPI_ISL_3240980, EPI_ISL_3240981, EPI_ISL_3240982, EPI_ISL_3240983, EPI_ISL_3240984, EPI_ISL_3240985, EPI_ISL_3240986, EPI_ISL_3240987, EPI_ISL_3240988, EPI_ISL_3240989, EPI_ISL_3240990, EPI_ISL_3240991, EPI_ISL_3240992, EPI_ISL_3240993, EPI_ISL_3240994, EPI_ISL_3240995, EPI_ISL_3240996, EPI_ISL_3240997, EPI_ISL_3240998, EPI_ISL_3240999, EPI_ISL_3241000, EPI_ISL_3241001, EPI_ISL_3241002, EPI_ISL_3241003, EPI_ISL_3241004, EPI_ISL_3241005, EPI_ISL_3241006, EPI_ISL_3241007, EPI_ISL_3241008, EPI_ISL_3241009, EPI_ISL_3241010, EPI_ISL_3241011, EPI_ISL_3241012, EPI_ISL_3241013, EPI_ISL_3241014, EPI_ISL_3241015, EPI_ISL_3241016, EPI_ISL_3241017, EPI_ISL_3241018, EPI_ISL_3241019, EPI_ISL_3241020, EPI_ISL_3241021, EPI_ISL_3241022, EPI_ISL_3241023, EPI_ISL_3241024, EPI_ISL_3241025, EPI_ISL_3241026, EPI_ISL_3241027, EPI_ISL_3241028, EPI_ISL_3241029, EPI_ISL_3241030, EPI_ISL_3241031, EPI_ISL_3241032, EPI_ISL_3241033, EPI_ISL_3241034, EPI_ISL_3241035, EPI_ISL_3241036, EPI_ISL_3241037, EPI_ISL_3241038, EPI_ISL_3241039, EPI_ISL_3241040, EPI_ISL_3241041, EPI_ISL_3241042, EPI_ISL_3241043, EPI_ISL_3241044, EPI_ISL_3241045, EPI_ISL_3241046, EPI_ISL_3241047, EPI_ISL_3241048, EPI_ISL_3241049, EPI_ISL_3241050, EPI_ISL_3241051, EPI_ISL_3241052, EPI_ISL_3241053, EPI_ISL_3241054, EPI_ISL_3241055, EPI_ISL_3241056, EPI_ISL_3241057, EPI_ISL_3241058, EPI_ISL_3241059, EPI_ISL_3241060, EPI_ISL_3241061, EPI_ISL_3241062, EPI_ISL_3241063, EPI_ISL_3241064, EPI_ISL_3241065, EPI_ISL_3241066, EPI_ISL_3241067, EPI_ISL_3241068, EPI_ISL_3241069, EPI_ISL_3241070, EPI_ISL_3241071, EPI_ISL_3241072, EPI_ISL_3241073, EPI_ISL_3241074, EPI_ISL_3241075, EPI_ISL_3241076, EPI_ISL_3241077, EPI_ISL_3241078, EPI_ISL_3241079, EPI_ISL_3241080, EPI_ISL_3241081, EPI_ISL_3241082, EPI_ISL_3241083, EPI_ISL_3241084, EPI_ISL_3241085, EPI_ISL_3241086, EPI_ISL_3241087, EPI_ISL_3241088, EPI_ISL_3241089, EPI_ISL_3241090, EPI_ISL_3241091, EPI_ISL_3241092, EPI_ISL_3241093, EPI_ISL_3241094, EPI_ISL_3241095, EPI_ISL_3241096, EPI_ISL_3241097, EPI_ISL_3241098, EPI_ISL_3241099, EPI_ISL_3241100, EPI_ISL_3241101, EPI_ISL_3241102, EPI_ISL_3241103, EPI_ISL_3241104, EPI_ISL_3241105, EPI_ISL_3241106, EPI_ISL_3241107, EPI_ISL_3241108, EPI_ISL_3241109, EPI_ISL_3241110, EPI_ISL_3241111, EPI_ISL_3241112, EPI_ISL_3241113, EPI_ISL_3241114, EPI_ISL_3241115, EPI_ISL_3241116, EPI_ISL_3241117, EPI_ISL_3241118, EPI_ISL_3241119, EPI_ISL_3241120, EPI_ISL_3241121, EPI_ISL_3241122, EPI_ISL_3241123, EPI_ISL_3241124, EPI_ISL_3241125, EPI_ISL_3241126, EPI_ISL_3241127, EPI_ISL_3241128, EPI_ISL_3241129, EPI_ISL_3241130, EPI_ISL_3241131, EPI_ISL_3241132, EPI_ISL_3241133, EPI_ISL_3241134, EPI_ISL_3241135, EPI_ISL_3241136, EPI_ISL_3241137, EPI_ISL_3241138, EPI_ISL_3241139, EPI_ISL_3241140, EPI_ISL_3241141, EPI_ISL_3241142, EPI_ISL_3241143, EPI_ISL_3241144, EPI_ISL_3241145, EPI_ISL_3241146, EPI_ISL_3241148, EPI_ISL_3241149, EPI_ISL_3241150, EPI_ISL_3241151, EPI_ISL_3241152, EPI_ISL_3241153, EPI_ISL_3241154, EPI_ISL_3241155, EPI_ISL_3241156, EPI_ISL_3241157, EPI_ISL_3241158, EPI_ISL_3241159, EPI_ISL_3241160, EPI_ISL_3241161, EPI_ISL_3241162, EPI_ISL_3241163, EPI_ISL_3241164, EPI_ISL_3241165, EPI_ISL_3241166, EPI_ISL_3241167, EPI_ISL_3241168, EPI_ISL_3241169, EPI_ISL_3241170, EPI_ISL_3241171, EPI_ISL_3241172, EPI_ISL_3241173, EPI_ISL_3241174, EPI_ISL_3241175, EPI_ISL_3241176, EPI_ISL_3241177, EPI_ISL_3241178, EPI_ISL_3241179, EPI_ISL_3241180, EPI_ISL_3241181, EPI_ISL_3241182, EPI_ISL_3241183, EPI_ISL_3241184, EPI_ISL_3241185, EPI_ISL_3241186, EPI_ISL_3241187, EPI_ISL_3241188, EPI_ISL_3241189, EPI_ISL_3241190, EPI_ISL_3241191, EPI_ISL_3241192, EPI_ISL_3241193, EPI_ISL_3241194, EPI_ISL_3241195, EPI_ISL_3241196, EPI_ISL_3241197, EPI_ISL_3241198, EPI_ISL_3241199, EPI_ISL_3241201, EPI_ISL_3241202, EPI_ISL_3241203, EPI_ISL_3241204, EPI_ISL_3241205, EPI_ISL_3241206, EPI_ISL_3241207, EPI_ISL_3241208, EPI_ISL_3241209, EPI_ISL_3241210, EPI_ISL_3241211, EPI_ISL_3241212, EPI_ISL_3241213, EPI_ISL_3241214, EPI_ISL_3241215, EPI_ISL_3241216, EPI_ISL_3241217, EPI_ISL_3241218, EPI_ISL_3241219, EPI_ISL_3241220, EPI_ISL_3241221, EPI_ISL_3241222, EPI_ISL_3241223, EPI_ISL_3241224, EPI_ISL_3241225, EPI_ISL_3241226, EPI_ISL_3241227, EPI_ISL_3241228, EPI_ISL_3241229, EPI_ISL_3241230, EPI_ISL_3241231, EPI_ISL_3241232, EPI_ISL_3241233, EPI_ISL_3241234, EPI_ISL_3241235, EPI_ISL_3241236, EPI_ISL_3241237, EPI_ISL_3241238, EPI_ISL_3241239, EPI_ISL_3241240, EPI_ISL_3241241, EPI_ISL_3241242, EPI_ISL_3241243, EPI_ISL_3241244, EPI_ISL_3241245, EPI_ISL_3241246, EPI_ISL_3241247, EPI_ISL_3241248, EPI_ISL_3241249, EPI_ISL_3241250, EPI_ISL_3241251, EPI_ISL_3241252, EPI_ISL_3241253, EPI_ISL_3241254, EPI_ISL_3241255, EPI_ISL_3241256, EPI_ISL_3241257, EPI_ISL_3241258, EPI_ISL_3241259, EPI_ISL_3241260, EPI_ISL_3241261, EPI_ISL_3241262, EPI_ISL_3241263, EPI_ISL_3241264, EPI_ISL_3241265, EPI_ISL_3241266, EPI_ISL_3241267, EPI_ISL_3241268, EPI_ISL_3241269, EPI_ISL_3241270, EPI_ISL_3241271, EPI_ISL_3241272, EPI_ISL_3241273, EPI_ISL_3241274, EPI_ISL_3241275, EPI_ISL_3241276, EPI_ISL_3241277, EPI_ISL_3241278 | | | |
| see above | Department of Bacteria, Parasites and Fungi, Statens Serum Institut, Copenhagen, Denmark | Statens Serum Institut Bioinformatics and Microbial Genomics | Danish Covid-19 Genome Consortium |
| EPI_ISL_3241279 | Department of Clinical Microbiology, Odense University Hospital, Odense, Denmark | Statens Serum Institut Bioinformatics and Microbial Genomics | Danish Covid-19 Genome Consortium |
| EPI_ISL_3241280, EPI_ISL_3241281, EPI_ISL_3241282, EPI_ISL_3241283, EPI_ISL_3241284, EPI_ISL_3241285, EPI_ISL_3241286, EPI_ISL_3241287, EPI_ISL_3241288, EPI_ISL_3241289, EPI_ISL_3241290, EPI_ISL_3241291, EPI_ISL_3241292, EPI_ISL_3241293, EPI_ISL_3241294, EPI_ISL_3241295, EPI_ISL_3241296, EPI_ISL_3241297, EPI_ISL_3241298, EPI_ISL_3241299, EPI_ISL_3241300, EPI_ISL_3241301, EPI_ISL_3241302, EPI_ISL_3241303, EPI_ISL_3241304, EPI_ISL_3241305, EPI_ISL_3241306, EPI_ISL_3241307, EPI_ISL_3241308, EPI_ISL_3241309, EPI_ISL_3241310, EPI_ISL_3241311, EPI_ISL_3241312, EPI_ISL_3241313, EPI_ISL_3241314, EPI_ISL_3241315, EPI_ISL_3241316, EPI_ISL_3241317, EPI_ISL_3241318, EPI_ISL_3241319, EPI_ISL_3241320, EPI_ISL_3241321, EPI_ISL_3241322, EPI_ISL_3241323, EPI_ISL_3241324, EPI_ISL_3241325, EPI_ISL_3241326, EPI_ISL_3241327, EPI_ISL_3241328, EPI_ISL_3241329, EPI_ISL_3241330, EPI_ISL_3241331, EPI_ISL_3241332, EPI_ISL_3241333, EPI_ISL_3241334, EPI_ISL_3241335, EPI_ISL_3241336, EPI_ISL_3241337, EPI_ISL_3241338, EPI_ISL_3241339, EPI_ISL_3241340, EPI_ISL_3241341, EPI_ISL_3241342, EPI_ISL_3241343, EPI_ISL_3241344, EPI_ISL_3241345, EPI_ISL_3241346, EPI_ISL_3241347, EPI_ISL_3241348, EPI_ISL_3241349, EPI_ISL_3241350, EPI_ISL_3241351, EPI_ISL_3241352, EPI_ISL_3241353, EPI_ISL_3241354, EPI_ISL_3241355, EPI_ISL_3241356, EPI_ISL_3241357, EPI_ISL_3241358, EPI_ISL_3241359, EPI_ISL_3241360, EPI_ISL_3241361, EPI_ISL_3241362, EPI_ISL_3241363, EPI_ISL_3241364, EPI_ISL_3241365, EPI_ISL_3241366, EPI_ISL_3241367, EPI_ISL_3241368, EPI_ISL_3241369, EPI_ISL_3241370, EPI_ISL_3241371, EPI_ISL_3241372, EPI_ISL_3241373, EPI_ISL_3241374, EPI_ISL_3241375, EPI_ISL_3241376, EPI_ISL_3241377, EPI_ISL_3241378, EPI_ISL_3241379, EPI_ISL_3241380, EPI_ISL_3241381, EPI_ISL_3241382, EPI_ISL_3241383, EPI_ISL_3241384, EPI_ISL_3241385, EPI_ISL_3241386, EPI_ISL_3241387, EPI_ISL_3241388, EPI_ISL_3241389, EPI_ISL_3241390, EPI_ISL_3241391, EPI_ISL_3241392, EPI_ISL_3241393, EPI_ISL_3241394, EPI_ISL_3241395, EPI_ISL_3241396, EPI_ISL_3241397, EPI_ISL_3241398, EPI_ISL_3241399, EPI_ISL_3241400, EPI_ISL_3241401, EPI_ISL_3241402, EPI_ISL_3241403, EPI_ISL_3241404, EPI_ISL_3241405, EPI_ISL_3241406, EPI_ISL_3241407, EPI_ISL_3241408, EPI_ISL_3241409, EPI_ISL_3241410, EPI_ISL_3241411, EPI_ISL_3241412, EPI_ISL_3241413, EPI_ISL_3241414, EPI_ISL_3241415, EPI_ISL_3241416, EPI_ISL_3241417, EPI_ISL_3241418, EPI_ISL_3241419, EPI_ISL_3241421, EPI_ISL_3241422, EPI_ISL_3241423, EPI_ISL_3241424, EPI_ISL_3241425, EPI_ISL_3241426, EPI_ISL_3241427, EPI_ISL_3241428, EPI_ISL_3241429, EPI_ISL_3241430, EPI_ISL_3241431, EPI_ISL_3241432, EPI_ISL_3241433, EPI_ISL_3241434, EPI_ISL_3241435, EPI_ISL_3241436, EPI_ISL_3241437 | | | |
| see above | Department of Bacteria, Parasites and Fungi, Statens Serum Institut, Copenhagen, Denmark | Statens Serum Institut Bioinformatics and Microbial Genomics | Danish Covid-19 Genome Consortium |
| EPI_ISL_3241438 | Department of Clinical Microbiology, Odense University Hospital, Odense, Denmark | Statens Serum Institut Bioinformatics and Microbial Genomics | Danish Covid-19 Genome Consortium |
| EPI_ISL_3241439, EPI_ISL_3241440, EPI_ISL_3241441, EPI_ISL_3241442, EPI_ISL_3241443, EPI_ISL_3241444, EPI_ISL_3241445, EPI_ISL_3241446, EPI_ISL_3241447, EPI_ISL_3241448, EPI_ISL_3241449, EPI_ISL_3241450, EPI_ISL_3241451, EPI_ISL_3241452, EPI_ISL_3241453, EPI_ISL_3241454, EPI_ISL_3241455, EPI_ISL_3241456, EPI_ISL_3241457, EPI_ISL_3241458, EPI_ISL_3241459, EPI_ISL_3241460, EPI_ISL_3241461, EPI_ISL_3241462, EPI_ISL_3241463, EPI_ISL_3241464, EPI_ISL_3241465, EPI_ISL_3241466, EPI_ISL_3241467 | | | |
| see above | Department of Bacteria, Parasites and Fungi, Statens Serum Institut, Copenhagen, Denmark | Statens Serum Institut Bioinformatics and Microbial Genomics | Danish Covid-19 Genome Consortium |
| EPI_ISL_3241468 | Department of Clinical Microbiology, Odense University Hospital, Odense, Denmark | Statens Serum Institut Bioinformatics and Microbial Genomics | Danish Covid-19 Genome Consortium |
| EPI_ISL_3241469, EPI_ISL_3241470, EPI_ISL_3241471, EPI_ISL_3241472, EPI_ISL_3241473, EPI_ISL_3241474, EPI_ISL_3241475, EPI_ISL_3241476, EPI_ISL_3241477, EPI_ISL_3241478, EPI_ISL_3241479, EPI_ISL_3241480, EPI_ISL_3241481, EPI_ISL_3241482, EPI_ISL_3241483, EPI_ISL_3241484, EPI_ISL_3241485, EPI_ISL_3241486, EPI_ISL_3241487, EPI_ISL_3241488, EPI_ISL_3241489, EPI_ISL_3241490, EPI_ISL_3241491, EPI_ISL_3241492, EPI_ISL_3241493, EPI_ISL_3241494, EPI_ISL_3241495, EPI_ISL_3241496, EPI_ISL_3241497, EPI_ISL_3241498, EPI_ISL_3241499, EPI_ISL_3241500, EPI_ISL_3241501, EPI_ISL_3241502, EPI_ISL_3241503, EPI_ISL_3241504, EPI_ISL_3241505, EPI_ISL_3241506, EPI_ISL_3241507, EPI_ISL_3241508, EPI_ISL_3241509, EPI_ISL_3241510, EPI_ISL_3241511, EPI_ISL_3241512, EPI_ISL_3241513, EPI_ISL_3241514 | | | |
| see above | Department of Bacteria, Parasites and Fungi, Statens Serum Institut, Copenhagen, Denmark | Statens Serum Institut Bioinformatics and Microbial Genomics | Danish Covid-19 Genome Consortium |
| EPI_ISL_3241515 | Department of Clinical Microbiology, Odense University Hospital, Odense, Denmark | Statens Serum Institut Bioinformatics and Microbial Genomics | Danish Covid-19 Genome Consortium |
| EPI_ISL_3241516, EPI_ISL_3241517, EPI_ISL_3241518, EPI_ISL_3241519, EPI_ISL_3241520, EPI_ISL_3241521, EPI_ISL_3241522, EPI_ISL_3241523, EPI_ISL_3241524, EPI_ISL_3241525, EPI_ISL_3241526, EPI_ISL_3241527, EPI_ISL_3241528, EPI_ISL_3241529, EPI_ISL_3241530, EPI_ISL_3241531, EPI_ISL_3241532, EPI_ISL_3241533, EPI_ISL_3241534, EPI_ISL_3241535, EPI_ISL_3241536, EPI_ISL_3241537, EPI_ISL_3241538, EPI_ISL_3241539, EPI_ISL_3241540, EPI_ISL_3241541, EPI_ISL_3241542, EPI_ISL_3241543, EPI_ISL_3241544, EPI_ISL_3241545, EPI_ISL_3241546, EPI_ISL_3241547, EPI_ISL_3241548, EPI_ISL_3241549, EPI_ISL_3241550, EPI_ISL_3241551, EPI_ISL_3241552, EPI_ISL_3241553, EPI_ISL_3241554, EPI_ISL_3241555, EPI_ISL_3241556, EPI_ISL_3241557, EPI_ISL_3241558, EPI_ISL_3241559, EPI_ISL_3241560 | | | |
| see above | Department of Bacteria, Parasites and Fungi, Statens Serum Institut, Copenhagen, Denmark | Statens Serum Institut Bioinformatics and Microbial Genomics | Danish Covid-19 Genome Consortium |
| EPI_ISL_3241561 | Department of Pathology & Laboratory Medicine | Centre for Research in Advanced Tropical Bioscience (Biotropic Centre) | Ummu Afeera Zainulabid, Norhidayah Kamarudin, Mohd Nazli Kamarulzaman, How Soon Hin and Hajar Fauzan Ahmad |
| EPI_ISL_3241562, EPI_ISL_3241563, EPI_ISL_3241564, EPI_ISL_3241565, EPI_ISL_3241566, EPI_ISL_3241567, EPI_ISL_3241568, EPI_ISL_3241569, EPI_ISL_3241570, EPI_ISL_3241571, EPI_ISL_3241572, EPI_ISL_3241573, EPI_ISL_3241574, EPI_ISL_3241575, EPI_ISL_3241576, EPI_ISL_3241577, EPI_ISL_3241578, EPI_ISL_3241579 | | | |
| see above | Department of Bacteria, Parasites and Fungi, Statens Serum Institut, Copenhagen, Denmark | Statens Serum Institut Bioinformatics and Microbial Genomics | Danish Covid-19 Genome Consortium |
| EPI_ISL_3241580 | Department of Clinical Microbiology, Odense University Hospital, Odense, Denmark | Statens Serum Institut Bioinformatics and Microbial Genomics | Danish Covid-19 Genome Consortium |
| EPI_ISL_3241581, EPI_ISL_3241582, EPI_ISL_3241583, EPI_ISL_3241584, EPI_ISL_3241585, EPI_ISL_3241586, EPI_ISL_3241587, EPI_ISL_3241588, EPI_ISL_3241589, EPI_ISL_3241590, EPI_ISL_3241591, EPI_ISL_3241592, EPI_ISL_3241593, EPI_ISL_3241594, EPI_ISL_3241595, EPI_ISL_3241596, EPI_ISL_3241597, EPI_ISL_3241598, EPI_ISL_3241599, EPI_ISL_3241600, EPI_ISL_3241601, EPI_ISL_3241602, EPI_ISL_3241603, EPI_ISL_3241604, EPI_ISL_3241605, EPI_ISL_3241606 | | | |
| see above | Department of Bacteria, Parasites and Fungi, Statens Serum Institut, Copenhagen, Denmark | Statens Serum Institut Bioinformatics and Microbial Genomics | Danish Covid-19 Genome Consortium |
| EPI_ISL_3241611 | A.S.L. CUNEO 1 | Fondazione del Piemonte per l'Oncologia IRCCS | Antonino Sottile, Silvia Brossa, Paola Marino, Giorgio Giardina |
| EPI_ISL_3241612, EPI_ISL_3241613 | A.S.L. TO5 | Fondazione del Piemonte per l'Oncologia IRCCS | Antonino Sottile, Silvia Brossa, Paola Marino, Giorgio Giardina |
| EPI_ISL_3241614, EPI_ISL_3241615, EPI_ISL_3241616, EPI_ISL_3241617, EPI_ISL_3241618, EPI_ISL_3241619, EPI_ISL_3241620 | A.S.L. CITTA' DI TORINO-ASO MAURIZIANO | Fondazione del Piemonte per l'Oncologia IRCCS | Antonino Sottile, Silvia Brossa, Paola Marino, Giorgio Giardina |
| EPI_ISL_3241622, EPI_ISL_3241623, EPI_ISL_3241624, EPI_ISL_3241625, EPI_ISL_3241626, EPI_ISL_3241627, EPI_ISL_3241628, EPI_ISL_3241629, EPI_ISL_3241630, EPI_ISL_3241631, EPI_ISL_3241632, EPI_ISL_3241633, EPI_ISL_3241634, EPI_ISL_3241635, EPI_ISL_3241636, EPI_ISL_3241637, EPI_ISL_3241638, EPI_ISL_3241639, EPI_ISL_3241640, EPI_ISL_3241641, EPI_ISL_3241642, EPI_ISL_3241643, EPI_ISL_3241644, EPI_ISL_3241645, EPI_ISL_3241646, EPI_ISL_3241648, EPI_ISL_3241649, EPI_ISL_3241650, EPI_ISL_3241651, EPI_ISL_3241652, EPI_ISL_3241653, EPI_ISL_3241654 | | | |
| see above | A.S.L. NOVARA - ISTITUTO S.S. TRINITA' - BORGOMANERO - S.S. | Fondazione del Piemonte per l'Oncologia IRCCS | Antonino Sottile, Silvia Brossa, Paola Marino, Giorgio Giardina |

| | | | |
|---|---|---|---|
| | LABORATORIO ANALISI CHIMICO CLINICHE - | | |
| EPI_ISL_3241655 | A.S.L. TO4 | Fondazione del Piemonte per l'Oncologia IRCCS | Antonino Sottile, Silvia Brossa, Paola Marino, Giorgio Giardina |
| EPI_ISL_3241656 | A.S.L. NOVARA - ISTITUTO S.S. TRINITA' - BORGOMANERO - S.S. LABORATORIO ANALISI CHIMICO CLINICHE - | Fondazione del Piemonte per l'Oncologia IRCCS | Antonino Sottile, Silvia Brossa, Paola Marino, Giorgio Giardina |
| EPI_ISL_3241657, EPI_ISL_3241658 | A.S.L. TO4 | Fondazione del Piemonte per l'Oncologia IRCCS | Antonino Sottile, Silvia Brossa, Paola Marino, Giorgio Giardina |
| EPI_ISL_3241659 | A.S.L. NOVARA - ISTITUTO S.S. TRINITA' - BORGOMANERO - S.S. LABORATORIO ANALISI CHIMICO CLINICHE - | Fondazione del Piemonte per l'Oncologia IRCCS | Antonino Sottile, Silvia Brossa, Paola Marino, Giorgio Giardina |
| EPI_ISL_3241660, EPI_ISL_3241661, EPI_ISL_3241662, EPI_ISL_3241663, EPI_ISL_3241664 | A.S.L. TO4 | Fondazione del Piemonte per l'Oncologia IRCCS | Antonino Sottile, Silvia Brossa, Paola Marino, Giorgio Giardina |
| EPI_ISL_3241665, EPI_ISL_3241666 | A.S.L. VCO | Fondazione del Piemonte per l'Oncologia IRCCS | Antonino Sottile, Silvia Brossa, Paola Marino, Giorgio Giardina |
| EPI_ISL_3241667, EPI_ISL_3241668 | A.S.L. ASTI | Fondazione del Piemonte per l'Oncologia IRCCS | Antonino Sottile, Silvia Brossa, Paola Marino, Giorgio Giardina |
| EPI_ISL_3241669, EPI_ISL_3241670, EPI_ISL_3241671, EPI_ISL_3241672, EPI_ISL_3241675, EPI_ISL_3241676, EPI_ISL_3241677, EPI_ISL_3241678, EPI_ISL_3241679, EPI_ISL_3241680, EPI_ISL_3241681, EPI_ISL_3241682, EPI_ISL_3241683, EPI_ISL_3241684, EPI_ISL_3241685, EPI_ISL_3241686, EPI_ISL_3241687, EPI_ISL_3241688 | | | |
| see above | A.S.L. NOVARA - ISTITUTO S.S. TRINITA' - BORGOMANERO - S.S. LABORATORIO ANALISI CHIMICO CLINICHE - | Fondazione del Piemonte per l'Oncologia IRCCS | Antonino Sottile, Silvia Brossa, Paola Marino, Giorgio Giardina |
| EPI_ISL_3241689 | A.S.L. CUNEO 1 | Fondazione del Piemonte per l'Oncologia IRCCS | Antonino Sottile, Silvia Brossa, Paola Marino, Giorgio Giardina |
| EPI_ISL_3241690, EPI_ISL_3241691 | A.S.L. NOVARA - ISTITUTO S.S. TRINITA' - BORGOMANERO - S.S. LABORATORIO ANALISI CHIMICO CLINICHE - | Fondazione del Piemonte per l'Oncologia IRCCS | Antonino Sottile, Silvia Brossa, Paola Marino, Giorgio Giardina |
| EPI_ISL_3241692 | A.S.L. CUNEO 1 | Fondazione del Piemonte per l'Oncologia IRCCS | Antonino Sottile, Silvia Brossa, Paola Marino, Giorgio Giardina |
| EPI_ISL_3241693 | A.S.L. NOVARA - ISTITUTO S.S. TRINITA' - BORGOMANERO - S.S. LABORATORIO ANALISI CHIMICO CLINICHE - | Fondazione del Piemonte per l'Oncologia IRCCS | Antonino Sottile, Silvia Brossa, Paola Marino, Giorgio Giardina |
| EPI_ISL_3241694, EPI_ISL_3241695, EPI_ISL_3241696, EPI_ISL_3241697, EPI_ISL_3241698, EPI_ISL_3241700, EPI_ISL_3241701, EPI_ISL_3241702, EPI_ISL_3241704, EPI_ISL_3241705 | A.S.L. CUNEO 1 | Fondazione del Piemonte per l'Oncologia IRCCS | Antonino Sottile, Silvia Brossa, Paola Marino, Giorgio Giardina |
| EPI_ISL_3241706 | A.S.L. TO5 | Fondazione del Piemonte per l'Oncologia IRCCS | Antonino Sottile, Silvia Brossa, Paola Marino, Giorgio Giardina |
| EPI_ISL_3241707, EPI_ISL_3241708 | A.S.L. CUNEO 1 | Fondazione del Piemonte per l'Oncologia IRCCS | Antonino Sottile, Silvia Brossa, Paola Marino, Giorgio Giardina |
| EPI_ISL_3241709, EPI_ISL_3241710 | A.S.L. TO5 | Fondazione del Piemonte per l'Oncologia IRCCS | Antonino Sottile, Silvia Brossa, Paola Marino, Giorgio Giardina |
| EPI_ISL_3241713, EPI_ISL_3241714, EPI_ISL_3241715, EPI_ISL_3241716, EPI_ISL_3241717, EPI_ISL_3241718, EPI_ISL_3241719, EPI_ISL_3241720, EPI_ISL_3241721, EPI_ISL_3241722, EPI_ISL_3241723, EPI_ISL_3241724, EPI_ISL_3241725, EPI_ISL_3241726 | | | |
| see above | Clinical Molecular Microbiology Laboratory, UNC Hospitals | Jeremy Wang | Jeremy Wang, Shawn Hawken, Corbin Jones, Melissa Miller |
| EPI_ISL_3241728, EPI_ISL_3241729, EPI_ISL_3241734, EPI_ISL_3241735, EPI_ISL_3241736, EPI_ISL_3241737, EPI_ISL_3241738, EPI_ISL_3241739, EPI_ISL_3241740, EPI_ISL_3241741, EPI_ISL_3241742, EPI_ISL_3241743, EPI_ISL_3241744, EPI_ISL_3241745, EPI_ISL_3241746, EPI_ISL_3241747, EPI_ISL_3241748, EPI_ISL_3241749, EPI_ISL_3241750, EPI_ISL_3241751, EPI_ISL_3241752, EPI_ISL_3241753, EPI_ISL_3241754, EPI_ISL_3241755, EPI_ISL_3241756, EPI_ISL_3241757, EPI_ISL_3241758, EPI_ISL_3241759, EPI_ISL_3241760, EPI_ISL_3241761, EPI_ISL_3241762, EPI_ISL_3241763, EPI_ISL_3241766, EPI_ISL_3241767, EPI_ISL_3241768, EPI_ISL_3241769, EPI_ISL_3241770, EPI_ISL_3241771, EPI_ISL_3241772, EPI_ISL_3241775, EPI_ISL_3241776, EPI_ISL_3241777, EPI_ISL_3241778, EPI_ISL_3241779, EPI_ISL_3241780, EPI_ISL_3241781, EPI_ISL_3241782, EPI_ISL_3241783, EPI_ISL_3241784, EPI_ISL_3241785, EPI_ISL_3241786, EPI_ISL_3241787, EPI_ISL_3241788, EPI_ISL_3241789, EPI_ISL_3241790, EPI_ISL_3241791, EPI_ISL_3241792, EPI_ISL_3241793, EPI_ISL_3241794, EPI_ISL_3241795, EPI_ISL_3241796, EPI_ISL_3241797, EPI_ISL_3241798, EPI_ISL_3241799, EPI_ISL_3241800, EPI_ISL_3241801, EPI_ISL_3241802, EPI_ISL_3241803, EPI_ISL_3241804, EPI_ISL_3241805, EPI_ISL_3241806, EPI_ISL_3241807, EPI_ISL_3241808, EPI_ISL_3241809, EPI_ISL_3241810, EPI_ISL_3241814, EPI_ISL_3241815, EPI_ISL_3241816, EPI_ISL_3241817, EPI_ISL_3241818, EPI_ISL_3241819, EPI_ISL_3241820 | | | |
| see above | Microbiology Department. Complexo Hospitalario Universitario de Vigo | Microbiology Department. Complexo Hospitalario Universitario de Vigo | Alfaya N, Alonso I, Alvarez M, Cabrera JJ, Carballo R, Cores O, Cortizo S, del-Campo V, Martinez L, Mediero G, Perez S, Davina C, Potel C, Regueiro B, Rey S, Vasallo FJ |
| EPI_ISL_3241822 | COVID-19 Detection Lab, Chattogram Veterinary and Animal Sciences University | Genomic Research Lab, BCSIR | Md. Murshed Hasan Sarkar, Abu Sayeed Mohammad Mahmud, Mohammad Samir Uzzaman, Eshrar Osman, Md. Ahasan Habib, Shahina Akter, Tanjina Akhter Banu, Barna Goswami, Iffat Jahan, Mohammad Mohi Uddin, Md. Kamrul Islam, Goutam Buddha Das, Tridip Das, Tanvir Ahmad Nizami, Eaftekhar Ahmed Rana, Md. Sirazul Islam, Pronesh Dutta, Sharmin Chowdhury, Md. Paritosh Kumar Biswas, Md. Salim Khan |
| EPI_ISL_3241826 | Microbiology Department. Complexo Hospitalario Universitario de Vigo | Microbiology Department. Complexo Hospitalario Universitario de Vigo | Alfaya N, Alonso I, Alvarez M, Cabrera JJ, Carballo R, Cores O, Cortizo S, del-Campo V, Martinez L, Mediero G, Perez S, Davina C, Potel C, Regueiro B, Rey S, Vasallo FJ |
| EPI_ISL_3241828, EPI_ISL_3241829, EPI_ISL_3241831, EPI_ISL_3241832, EPI_ISL_3241833, EPI_ISL_3241834, EPI_ISL_3241835, EPI_ISL_3241836, EPI_ISL_3241837, EPI_ISL_3241838, EPI_ISL_3241839, EPI_ISL_3241840, EPI_ISL_3241841, EPI_ISL_3241842, EPI_ISL_3241843, EPI_ISL_3241844, EPI_ISL_3241845, EPI_ISL_3241846, EPI_ISL_3241847, EPI_ISL_3241848, EPI_ISL_3241849, EPI_ISL_3241850, EPI_ISL_3241851, EPI_ISL_3241852, EPI_ISL_3241853, EPI_ISL_3241854, EPI_ISL_3241855, EPI_ISL_3241856, EPI_ISL_3241857, EPI_ISL_3241858, EPI_ISL_3241859, EPI_ISL_3241860, EPI_ISL_3241861, EPI_ISL_3241862, EPI_ISL_3241863, EPI_ISL_3241864, EPI_ISL_3241865, EPI_ISL_3241866, EPI_ISL_3241867, EPI_ISL_3241868, EPI_ISL_3241869, EPI_ISL_3241870, EPI_ISL_3241871, EPI_ISL_3241872, EPI_ISL_3241873, EPI_ISL_3241874, EPI_ISL_3241875, EPI_ISL_3241876, EPI_ISL_3241878, EPI_ISL_3241879, EPI_ISL_3241880, EPI_ISL_3241881, EPI_ISL_3241882, EPI_ISL_3241883, EPI_ISL_3241884, EPI_ISL_3241885, EPI_ISL_3241886, EPI_ISL_3241887, EPI_ISL_3241888, EPI_ISL_3241889, EPI_ISL_3241890, EPI_ISL_3241891, EPI_ISL_3241892, EPI_ISL_3241893, EPI_ISL_3241894, EPI_ISL_3241895, EPI_ISL_3241896, EPI_ISL_3241897, EPI_ISL_3241898, EPI_ISL_3241899, EPI_ISL_3241900, EPI_ISL_3241901, EPI_ISL_3241902 | | | |
| see above | AZDelta | AZ Delta Medical Laboratories in Roeselare, Belgium | Geert Martens, Dieter De Smet, Frederik Van Hoecke, on behalf of AZ Delta COVID-19 Genomics core (member of Genomic surveillance of SARS-CoV-2 in Belgium network) |
| EPI_ISL_3241903, EPI_ISL_3241904, EPI_ISL_3241906, EPI_ISL_3241907, EPI_ISL_3241909, EPI_ISL_3241911, EPI_ISL_3241912, EPI_ISL_3241913, EPI_ISL_3241915, EPI_ISL_3241916 | MD PHL | MD PHL | Eric N. Keller, Jillian Loomis, Kwang Low, Terence L. Moore, Ami Patel, and Robert Myers |
| EPI_ISL_3241917 | FL Bureau of Public Health Laboratories-Miami | Centers for Disease Control and Prevention Division of Viral Diseases, Pathogen Discovery | Mili Sheth, Sarah Nobles, Jasmine Padilla, Alex Burgin, Meghan Bentz, Mark Burroughs, Peter Cook, Clinton R. Paden, Dhwani Batra, Krista Queen, Kristen Knipe, Dakota Howard, Yvette Unoarumhi, Matthew Schmerer, Ben L. Rambo-Martin, Kristine Lacek, Sam Shepard, Dave Wentworth, Vivien Dugan, Suxiang Tong, Justin Lee |
| EPI_ISL_3241918 | FL Bur. of Public Health Laboratories-Jacksonville | Centers for Disease Control and Prevention Division of Viral Diseases, Pathogen Discovery | Mili Sheth, Sarah Nobles, Jasmine Padilla, Alex Burgin, Meghan Bentz, Mark Burroughs, Peter Cook, Clinton R. Paden, Dhwani Batra, Krista Queen, Kristen Knipe, Dakota Howard, Yvette Unoarumhi, Matthew Schmerer, Ben L. Rambo-Martin, Kristine Lacek, Sam Shepard, Dave Wentworth, Vivien Dugan, Suxiang Tong, Justin Lee |
| EPI_ISL_3241919 | CT-Dr. Katherine A. Kelley State Public Health Lab | Centers for Disease Control and Prevention Division of Viral Diseases, Pathogen Discovery | Mili Sheth, Sarah Nobles, Jasmine Padilla, Alex Burgin, Meghan Bentz, Mark Burroughs, Peter Cook, Clinton R. Paden, Dhwani Batra, Krista Queen, Kristen Knipe, Dakota Howard, Yvette Unoarumhi, Matthew Schmerer, Ben L. Rambo-Martin, Kristine Lacek, Sam Shepard, Dave Wentworth, Vivien Dugan, Suxiang Tong, Justin Lee |
| EPI_ISL_3241920, EPI_ISL_3241921 | NE Public Health Laboratory | Centers for Disease Control and Prevention Division of Viral Diseases, Pathogen Discovery | Mili Sheth, Sarah Nobles, Jasmine Padilla, Alex Burgin, Meghan Bentz, Mark Burroughs, Peter Cook, Clinton R. Paden, Dhwani Batra, Krista Queen, Kristen Knipe, Dakota Howard, Yvette Unoarumhi, Matthew Schmerer, Ben L. Rambo-Martin, Kristine Lacek, Sam Shepard, Dave Wentworth, Vivien Dugan, Suxiang Tong, Justin Lee |
| EPI_ISL_3241922 | SD Public Health Laboratory | Centers for Disease Control and Prevention Division of Viral Diseases, Pathogen Discovery | Mili Sheth, Sarah Nobles, Jasmine Padilla, Alex Burgin, Meghan Bentz, Mark Burroughs, Peter Cook, Clinton R. Paden, Dhwani Batra, Krista Queen, Kristen Knipe, Dakota Howard, Yvette Unoarumhi, Matthew Schmerer, Ben L. Rambo-Martin, Kristine Lacek, Sam Shepard, Dave Wentworth, Vivien Dugan, Suxiang Tong, Justin Lee |
| EPI_ISL_3241923, EPI_ISL_3241924 | FL Bureau of Public Health Laboratories-Tampa | Centers for Disease Control and Prevention Division of Viral Diseases, Pathogen Discovery | Mili Sheth, Sarah Nobles, Jasmine Padilla, Alex Burgin, Meghan Bentz, Mark Burroughs, Peter Cook, Clinton R. Paden, Dhwani Batra, Krista Queen, Kristen Knipe, Dakota Howard, Yvette Unoarumhi, Matthew Schmerer, Ben L. Rambo-Martin, Kristine Lacek, Sam Shepard, Dave Wentworth, Vivien Dugan, Suxiang Tong, Justin Lee |
| EPI_ISL_3241925 | WY Public Health Laboratory | Centers for Disease Control and Prevention Division of Viral Diseases, Pathogen Discovery | Mili Sheth, Sarah Nobles, Jasmine Padilla, Alex Burgin, Meghan Bentz, Mark Burroughs, Peter Cook, Clinton R. Paden, Dhwani Batra, Krista Queen, Kristen Knipe, Dakota Howard, Yvette Unoarumhi, Matthew Schmerer, Ben L. Rambo-Martin, Kristine Lacek, Sam Shepard, Dave Wentworth, Vivien Dugan, Suxiang Tong, Justin Lee |
| EPI_ISL_3241926, EPI_ISL_3241927, EPI_ISL_3241928, EPI_ISL_3241929, EPI_ISL_3241930 | FL Bureau of Public Health Laboratories-Miami | Centers for Disease Control and Prevention Division of Viral Diseases, Pathogen Discovery | Mili Sheth, Sarah Nobles, Jasmine Padilla, Alex Burgin, Meghan Bentz, Mark Burroughs, Peter Cook, Clinton R. Paden, Dhwani Batra, Krista Queen, Kristen Knipe, Dakota Howard, Yvette Unoarumhi, Matthew Schmerer, Ben L. Rambo-Martin, Kristine Lacek, Sam Shepard, Dave Wentworth, Vivien Dugan, Suxiang Tong, Justin Lee |
| EPI_ISL_3241931, EPI_ISL_3241932, EPI_ISL_3241933, EPI_ISL_3241934, EPI_ISL_3241935, EPI_ISL_3241936, EPI_ISL_3241937, EPI_ISL_3241938, EPI_ISL_3241939, EPI_ISL_3241940, EPI_ISL_3241941, EPI_ISL_3241942, EPI_ISL_3241943, EPI_ISL_3241944, EPI_ISL_3241946, EPI_ISL_3241947, EPI_ISL_3241948 | | | |
| see above | FL Bur. of Public Health Laboratories-Jacksonville | Centers for Disease Control and Prevention Division of Viral Diseases, Pathogen Discovery | Mili Sheth, Sarah Nobles, Jasmine Padilla, Alex Burgin, Meghan Bentz, Mark Burroughs, Peter Cook, Clinton R. Paden, Dhwani Batra, Krista Queen, Kristen Knipe, Dakota Howard, Yvette Unoarumhi, Matthew Schmerer, Ben L. Rambo-Martin, Kristine Lacek, Sam Shepard, Dave Wentworth, Vivien Dugan, Suxiang Tong, Justin Lee |
| EPI_ISL_3241949, EPI_ISL_3241950, EPI_ISL_3241952, EPI_ISL_3241953 | LA Office of Public Health Laboratories | Centers for Disease Control and Prevention Division of Viral Diseases, Pathogen Discovery | Mili Sheth, Sarah Nobles, Jasmine Padilla, Alex Burgin, Meghan Bentz, Mark Burroughs, Peter Cook, Clinton R. Paden, Dhwani Batra, Krista Queen, Kristen Knipe, Dakota Howard, Yvette Unoarumhi, Matthew Schmerer, Ben L. Rambo-Martin, Kristine Lacek, Sam Shepard, Dave Wentworth, Vivien Dugan, Suxiang Tong, Justin Lee |

| | | | |
|---|---|---|---|
| EPI_ISL_3241967, EPI_ISL_3241968, EPI_ISL_3241969, EPI_ISL_3241970 | ME Health and Environmental Testing Laboratory | Centers for Disease Control and Prevention Division of Viral Diseases, Pathogen Discovery | Mili Sheth, Sarah Nobles, Jasmine Padilla, Alex Burgin, Meghan Bentz, Mark Burroughs, Peter Cook, Clinton R. Paden, Dhwani Batra, Krista Queen, Kristen Knipe, Dakota Howard, Yvette Unoarumhi, Matthew Schmerer, Ben L. Rambo-Martin, Kristine Lacek, Sam Shepard, Dave Wentworth, Vivien Dugan, Suxiang Tong, Justin Lee |
| EPI_ISL_3241971, EPI_ISL_3241973, EPI_ISL_3241974, EPI_ISL_3241975, EPI_ISL_3241976, EPI_ISL_3241978, EPI_ISL_3241979, EPI_ISL_3241980 | AR Dept. of Health-PHL, Molecular Diagnostics | Centers for Disease Control and Prevention Division of Viral Diseases, Pathogen Discovery | Mili Sheth, Sarah Nobles, Jasmine Padilla, Alex Burgin, Meghan Bentz, Mark Burroughs, Peter Cook, Clinton R. Paden, Dhwani Batra, Krista Queen, Kristen Knipe, Dakota Howard, Yvette Unoarumhi, Matthew Schmerer, Ben L. Rambo-Martin, Kristine Lacek, Sam Shepard, Dave Wentworth, Vivien Dugan, Suxiang Tong, Justin Lee |
| EPI_ISL_3241981, EPI_ISL_3241982, EPI_ISL_3241983, EPI_ISL_3241984, EPI_ISL_3241985, EPI_ISL_3241986 | NM Dept. Health, Scientific Laboratory Division | Centers for Disease Control and Prevention Division of Viral Diseases, Pathogen Discovery | Mili Sheth, Sarah Nobles, Jasmine Padilla, Alex Burgin, Meghan Bentz, Mark Burroughs, Peter Cook, Clinton R. Paden, Dhwani Batra, Krista Queen, Kristen Knipe, Dakota Howard, Yvette Unoarumhi, Matthew Schmerer, Ben L. Rambo-Martin, Kristine Lacek, Sam Shepard, Dave Wentworth, Vivien Dugan, Suxiang Tong, Justin Lee |
| EPI_ISL_3241987, EPI_ISL_3241989, EPI_ISL_3241990, EPI_ISL_3241994, EPI_ISL_3241998 | Guam Public Health Laboratory | Centers for Disease Control and Prevention Division of Viral Diseases, Pathogen Discovery | Mili Sheth, Sarah Nobles, Jasmine Padilla, Alex Burgin, Meghan Bentz, Mark Burroughs, Peter Cook, Clinton R. Paden, Dhwani Batra, Krista Queen, Kristen Knipe, Dakota Howard, Yvette Unoarumhi, Matthew Schmerer, Ben L. Rambo-Martin, Kristine Lacek, Sam Shepard, Dave Wentworth, Vivien Dugan, Suxiang Tong, Justin Lee |
| EPI_ISL_3242005, EPI_ISL_3242006, EPI_ISL_3242007, EPI_ISL_3242008, EPI_ISL_3242009, EPI_ISL_3242010, EPI_ISL_3242011, EPI_ISL_3242012, EPI_ISL_3242013, EPI_ISL_3242014, EPI_ISL_3242015, EPI_ISL_3242016 | | | |
| see above | IL Dept. of Public Health Springfield Laboratory | Centers for Disease Control and Prevention Division of Viral Diseases, Pathogen Discovery | Mili Sheth, Sarah Nobles, Jasmine Padilla, Alex Burgin, Meghan Bentz, Mark Burroughs, Peter Cook, Clinton R. Paden, Dhwani Batra, Krista Queen, Kristen Knipe, Dakota Howard, Yvette Unoarumhi, Matthew Schmerer, Ben L. Rambo-Martin, Kristine Lacek, Sam Shepard, Dave Wentworth, Vivien Dugan, Suxiang Tong, Justin Lee |
| EPI_ISL_3242018, EPI_ISL_3242019, EPI_ISL_3242020 | CT-Dr. Katherine A. Kelley State Public Health Lab | Centers for Disease Control and Prevention Division of Viral Diseases, Pathogen Discovery | Mili Sheth, Sarah Nobles, Jasmine Padilla, Alex Burgin, Meghan Bentz, Mark Burroughs, Peter Cook, Clinton R. Paden, Dhwani Batra, Krista Queen, Kristen Knipe, Dakota Howard, Yvette Unoarumhi, Matthew Schmerer, Ben L. Rambo-Martin, Kristine Lacek, Sam Shepard, Dave Wentworth, Vivien Dugan, Suxiang Tong, Justin Lee |
| EPI_ISL_3242021, EPI_ISL_3242022, EPI_ISL_3242023, EPI_ISL_3242024, EPI_ISL_3242026, EPI_ISL_3242027, EPI_ISL_3242028, EPI_ISL_3242029, EPI_ISL_3242030, EPI_ISL_3242032, EPI_ISL_3242033, EPI_ISL_3242034, EPI_ISL_3242035 | | | |
| see above | MI - Michigan Department of Health and Human Services - Bureau of Laboratories | Centers for Disease Control and Prevention Division of Viral Diseases, Pathogen Discovery | Mili Sheth, Sarah Nobles, Jasmine Padilla, Alex Burgin, Meghan Bentz, Mark Burroughs, Peter Cook, Clinton R. Paden, Dhwani Batra, Krista Queen, Kristen Knipe, Dakota Howard, Yvette Unoarumhi, Matthew Schmerer, Ben L. Rambo-Martin, Kristine Lacek, Sam Shepard, Dave Wentworth, Vivien Dugan, Suxiang Tong, Justin Lee |
| EPI_ISL_3242036, EPI_ISL_3242038, EPI_ISL_3242039, EPI_ISL_3242040 | TX-Houston Health Department Laboratory | Centers for Disease Control and Prevention Division of Viral Diseases, Pathogen Discovery | Mili Sheth, Sarah Nobles, Jasmine Padilla, Alex Burgin, Meghan Bentz, Mark Burroughs, Peter Cook, Clinton R. Paden, Dhwani Batra, Krista Queen, Kristen Knipe, Dakota Howard, Yvette Unoarumhi, Matthew Schmerer, Ben L. Rambo-Martin, Kristine Lacek, Sam Shepard, Dave Wentworth, Vivien Dugan, Suxiang Tong, Justin Lee |
| EPI_ISL_3242042, EPI_ISL_3242043, EPI_ISL_3242044, EPI_ISL_3242045, EPI_ISL_3242046, EPI_ISL_3242047 | NE Public Health Laboratory | Centers for Disease Control and Prevention Division of Viral Diseases, Pathogen Discovery | Mili Sheth, Sarah Nobles, Jasmine Padilla, Alex Burgin, Meghan Bentz, Mark Burroughs, Peter Cook, Clinton R. Paden, Dhwani Batra, Krista Queen, Kristen Knipe, Dakota Howard, Yvette Unoarumhi, Matthew Schmerer, Ben L. Rambo-Martin, Kristine Lacek, Sam Shepard, Dave Wentworth, Vivien Dugan, Suxiang Tong, Justin Lee |
| EPI_ISL_3242048, EPI_ISL_3242049, EPI_ISL_3242050, EPI_ISL_3242051, EPI_ISL_3242052, EPI_ISL_3242053, EPI_ISL_3242054, EPI_ISL_3242055, EPI_ISL_3242056, EPI_ISL_3242057, EPI_ISL_3242058, EPI_ISL_3242059, EPI_ISL_3242060, EPI_ISL_3242061 | | | |
| see above | SD Public Health Laboratory | Centers for Disease Control and Prevention Division of Viral Diseases, Pathogen Discovery | Mili Sheth, Sarah Nobles, Jasmine Padilla, Alex Burgin, Meghan Bentz, Mark Burroughs, Peter Cook, Clinton R. Paden, Dhwani Batra, Krista Queen, Kristen Knipe, Dakota Howard, Yvette Unoarumhi, Matthew Schmerer, Ben L. Rambo-Martin, Kristine Lacek, Sam Shepard, Dave Wentworth, Vivien Dugan, Suxiang Tong, Justin Lee |
| EPI_ISL_3242062 | MN PHL Division, Minnesota Department of Health | Centers for Disease Control and Prevention Division of Viral Diseases, Pathogen Discovery | Mili Sheth, Sarah Nobles, Jasmine Padilla, Alex Burgin, Meghan Bentz, Mark Burroughs, Peter Cook, Clinton R. Paden, Dhwani Batra, Krista Queen, Kristen Knipe, Dakota Howard, Yvette Unoarumhi, Matthew Schmerer, Ben L. Rambo-Martin, Kristine Lacek, Sam Shepard, Dave Wentworth, Vivien Dugan, Suxiang Tong, Justin Lee |
| EPI_ISL_3242063, EPI_ISL_3242065, EPI_ISL_3242070, EPI_ISL_3242071, EPI_ISL_3242072, EPI_ISL_3242073 | FL Bureau of Public Health Laboratories-Tampa | Centers for Disease Control and Prevention Division of Viral Diseases, Pathogen Discovery | Mili Sheth, Sarah Nobles, Jasmine Padilla, Alex Burgin, Meghan Bentz, Mark Burroughs, Peter Cook, Clinton R. Paden, Dhwani Batra, Krista Queen, Kristen Knipe, Dakota Howard, Yvette Unoarumhi, Matthew Schmerer, Ben L. Rambo-Martin, Kristine Lacek, Sam Shepard, Dave Wentworth, Vivien Dugan, Suxiang Tong, Justin Lee |
| EPI_ISL_3242075, EPI_ISL_3242076, EPI_ISL_3242079 | ND Dept. of Health Laboratory Services-Microbiology | Centers for Disease Control and Prevention Division of Viral Diseases, Pathogen Discovery | Mili Sheth, Sarah Nobles, Jasmine Padilla, Alex Burgin, Meghan Bentz, Mark Burroughs, Peter Cook, Clinton R. Paden, Dhwani Batra, Krista Queen, Kristen Knipe, Dakota Howard, Yvette Unoarumhi, Matthew Schmerer, Ben L. Rambo-Martin, Kristine Lacek, Sam Shepard, Dave Wentworth, Vivien Dugan, Suxiang Tong, Justin Lee |
| EPI_ISL_3242081, EPI_ISL_3242084, EPI_ISL_3242085, EPI_ISL_3242087, EPI_ISL_3242089 | AL Dept. of Public Health Bureau of Clinical Laboratories | Centers for Disease Control and Prevention Division of Viral Diseases, Pathogen Discovery | Mili Sheth, Sarah Nobles, Jasmine Padilla, Alex Burgin, Meghan Bentz, Mark Burroughs, Peter Cook, Clinton R. Paden, Dhwani Batra, Krista Queen, Kristen Knipe, Dakota Howard, Yvette Unoarumhi, Matthew Schmerer, Ben L. Rambo-Martin, Kristine Lacek, Sam Shepard, Dave Wentworth, Vivien Dugan, Suxiang Tong, Justin Lee |
| EPI_ISL_3242090, EPI_ISL_3242091, EPI_ISL_3242092, EPI_ISL_3242093, EPI_ISL_3242094, EPI_ISL_3242095, EPI_ISL_3242096, EPI_ISL_3242097 | WY Public Health Laboratory | Centers for Disease Control and Prevention Division of Viral Diseases, Pathogen Discovery | Mili Sheth, Sarah Nobles, Jasmine Padilla, Alex Burgin, Meghan Bentz, Mark Burroughs, Peter Cook, Clinton R. Paden, Dhwani Batra, Krista Queen, Kristen Knipe, Dakota Howard, Yvette Unoarumhi, Matthew Schmerer, Ben L. Rambo-Martin, Kristine Lacek, Sam Shepard, Dave Wentworth, Vivien Dugan, Suxiang Tong, Justin Lee |
| EPI_ISL_3242104, EPI_ISL_3242105, EPI_ISL_3242106, EPI_ISL_3242107, EPI_ISL_3242108 | RI State Health Laboratories | Centers for Disease Control and Prevention Division of Viral Diseases, Pathogen Discovery | Mili Sheth, Sarah Nobles, Jasmine Padilla, Alex Burgin, Meghan Bentz, Mark Burroughs, Peter Cook, Clinton R. Paden, Dhwani Batra, Krista Queen, Kristen Knipe, Dakota Howard, Yvette Unoarumhi, Matthew Schmerer, Ben L. Rambo-Martin, Kristine Lacek, Sam Shepard, Dave Wentworth, Vivien Dugan, Suxiang Tong, Justin Lee |
| EPI_ISL_3242109 | COVID-19 Detection Lab, Chattogram Veterinary and Animal Sciences University | Genomic Research Lab, BCSIR | Md. Murshed Hasan Sarkar, Abu Sayeed Mohammad Mahmud, Mohammad Samir Uzzaman, Eshrar Osman, Md. Ahasan Habib, Shahina Akter, Tanjina Akhter Banu, Barna Goswami, Iffat Jahan, Mohammad Mohi Uddin, Md. Kamrul Islam, Goutam Buddha Das, Tridip Das, Tanvir Ahmad Nizami, Eaftekhar Ahmed Rana, Md. Sirazul Islam, Pronesh Dutta, Sharmin Chowdhury, Md. Paritosh Kumar Biswas, Md. Salim Khan |
| EPI_ISL_3242110, EPI_ISL_3242111, EPI_ISL_3242112, EPI_ISL_3242113, EPI_ISL_3242114, EPI_ISL_3242115, EPI_ISL_3242116, EPI_ISL_3242117, EPI_ISL_3242118, EPI_ISL_3242119, EPI_ISL_3242120, EPI_ISL_3242121, EPI_ISL_3242122, EPI_ISL_3242123, EPI_ISL_3242124, EPI_ISL_3242125, EPI_ISL_3242126, EPI_ISL_3242127, EPI_ISL_3242128, EPI_ISL_3242129, EPI_ISL_3242130, EPI_ISL_3242131, EPI_ISL_3242132, EPI_ISL_3242133, EPI_ISL_3242134, EPI_ISL_3242135, EPI_ISL_3242136, EPI_ISL_3242137, EPI_ISL_3242138, EPI_ISL_3242139, EPI_ISL_3242140, EPI_ISL_3242141 | | | |
| see above | Kansas Health and Environmental Lab | Kansas Health and Environmental Lab | Mike Grose, Jonathan Barnell, Ben Olsen, and Phil Adam |
| EPI_ISL_3242142, EPI_ISL_3242143, EPI_ISL_3242144, EPI_ISL_3242145, EPI_ISL_3242146, EPI_ISL_3242147, EPI_ISL_3242148, EPI_ISL_3242149, EPI_ISL_3242150, EPI_ISL_3242151, EPI_ISL_3242152, EPI_ISL_3242153, EPI_ISL_3242155, EPI_ISL_3242156, EPI_ISL_3242157, EPI_ISL_3242158, EPI_ISL_3242159, EPI_ISL_3242160, EPI_ISL_3242161, EPI_ISL_3242162, EPI_ISL_3242163, EPI_ISL_3242165, EPI_ISL_3242166, EPI_ISL_3242167, EPI_ISL_3242169, EPI_ISL_3242170, EPI_ISL_3242171, EPI_ISL_3242172, EPI_ISL_3242173, EPI_ISL_3242174, EPI_ISL_3242175, EPI_ISL_3242176, EPI_ISL_3242177, EPI_ISL_3242178, EPI_ISL_3242179, EPI_ISL_3242180, EPI_ISL_3242181, EPI_ISL_3242182, EPI_ISL_3242183, EPI_ISL_3242184, EPI_ISL_3242185, EPI_ISL_3242186, EPI_ISL_3242187, EPI_ISL_3242188, EPI_ISL_3242189, EPI_ISL_3242190, EPI_ISL_3242191, EPI_ISL_3242192, EPI_ISL_3242193, EPI_ISL_3242194, EPI_ISL_3242195, EPI_ISL_3242196, EPI_ISL_3242197, EPI_ISL_3242198, EPI_ISL_3242199, EPI_ISL_3242200, EPI_ISL_3242201, EPI_ISL_3242202, EPI_ISL_3242203, EPI_ISL_3242204, EPI_ISL_3242205, EPI_ISL_3242206, EPI_ISL_3242207, EPI_ISL_3242208, EPI_ISL_3242209, EPI_ISL_3242210, EPI_ISL_3242211, EPI_ISL_3242213, EPI_ISL_3242214, EPI_ISL_3242215, EPI_ISL_3242216, EPI_ISL_3242217, EPI_ISL_3242218, EPI_ISL_3242219, EPI_ISL_3242220, EPI_ISL_3242221, EPI_ISL_3242222, EPI_ISL_3242223, EPI_ISL_3242224, EPI_ISL_3242225, EPI_ISL_3242226, EPI_ISL_3242227, EPI_ISL_3242228, EPI_ISL_3242229, EPI_ISL_3242230, EPI_ISL_3242231, EPI_ISL_3242232, EPI_ISL_3242233, EPI_ISL_3242234, EPI_ISL_3242236, EPI_ISL_3242237, EPI_ISL_3242238, EPI_ISL_3242239, EPI_ISL_3242240, EPI_ISL_3242241, EPI_ISL_3242242, EPI_ISL_3242243, EPI_ISL_3242244, EPI_ISL_3242245, EPI_ISL_3242246, EPI_ISL_3242247, EPI_ISL_3242248, EPI_ISL_3242249, EPI_ISL_3242250, EPI_ISL_3242251, EPI_ISL_3242252, EPI_ISL_3242253, EPI_ISL_3242254 | | | |
| see above | Ochsner Health | Bioinfoexperts | Amy Feehan, Judy Crabtree, Rebecca Rose, Ben Lain, Chris Huston, Samual Moot, David J. Nolan, Tessa LaFleur, Julia-Garcia-Diaz, Lucio Miele, Susanna L. Lamers |
| EPI_ISL_3242256, EPI_ISL_3242257, EPI_ISL_3242258, EPI_ISL_3242259, EPI_ISL_3242260, EPI_ISL_3242261, EPI_ISL_3242262, EPI_ISL_3242263, EPI_ISL_3242264, EPI_ISL_3242265, EPI_ISL_3242266, EPI_ISL_3242267, EPI_ISL_3242268, EPI_ISL_3242269, EPI_ISL_3242270, EPI_ISL_3242271, EPI_ISL_3242272, EPI_ISL_3242273, EPI_ISL_3242274 | | | |
| see above | Osmania Medical College | CDFD | Murali Bashyam, Asmita Gupta, Reelina Basu, Vinay Donipadi, Nagamani Kammili, Divya Vashisht, Ashwin Dalal |
| EPI_ISL_3242275, EPI_ISL_3242276, EPI_ISL_3242277, EPI_ISL_3242278, EPI_ISL_3242279, EPI_ISL_3242280, EPI_ISL_3242281, EPI_ISL_3242282, EPI_ISL_3242283, EPI_ISL_3242284, EPI_ISL_3242285, EPI_ISL_3242286, EPI_ISL_3242287, EPI_ISL_3242288, EPI_ISL_3242289, EPI_ISL_3242290, EPI_ISL_3242291, EPI_ISL_3242292, EPI_ISL_3242293, EPI_ISL_3242294, EPI_ISL_3242295, EPI_ISL_3242296 | | | |
| see above | Government Fever Hospital | CDFD | Murali Bashyam, Asmita Gupta, Reelina Basu, Vinay Donipadi, Nagamani Kammili, Divya Vashisht, Ashwin Dalal |
| EPI_ISL_3242297, EPI_ISL_3242298, EPI_ISL_3242299 | Goa Medical College | CDFD | Murali Bashyam, Asmita Gupta, Reelina Basu, Vinay Donipadi, Savio Rodrigues, Divya Vashisht, Ashwin Dalal |
| EPI_ISL_3242300, EPI_ISL_3242301, EPI_ISL_3242302, EPI_ISL_3242303, EPI_ISL_3242304, EPI_ISL_3242305, EPI_ISL_3242306, EPI_ISL_3242307, EPI_ISL_3242308, EPI_ISL_3242309, EPI_ISL_3242310, EPI_ISL_3242311, EPI_ISL_3242312, EPI_ISL_3242313 | | | |
| see above | Government Maternity Hospital | CDFD | Murali Bashyam, Asmita Gupta, Reelina Basu, Vinay Donipadi, Nagamani Kammili, Divya Vashisht, Ashwin Dalal |
| EPI_ISL_3242314, EPI_ISL_3242315, EPI_ISL_3242316, EPI_ISL_3242317, EPI_ISL_3242318, EPI_ISL_3242319, EPI_ISL_3242320, EPI_ISL_3242321, EPI_ISL_3242322, EPI_ISL_3242323, EPI_ISL_3242324, EPI_ISL_3242325, EPI_ISL_3242326, EPI_ISL_3242327, EPI_ISL_3242328, EPI_ISL_3242329, EPI_ISL_3242330, EPI_ISL_3242331, EPI_ISL_3242332, EPI_ISL_3242333, EPI_ISL_3242334, EPI_ISL_3242335, EPI_ISL_3242336, EPI_ISL_3242337, EPI_ISL_3242338, EPI_ISL_3242339, EPI_ISL_3242340, EPI_ISL_3242341, EPI_ISL_3242342, EPI_ISL_3242343, EPI_ISL_3242344, EPI_ISL_3242345, EPI_ISL_3242346, EPI_ISL_3242347, EPI_ISL_3242348, EPI_ISL_3242349, EPI_ISL_3242350, EPI_ISL_3242351, EPI_ISL_3242352, EPI_ISL_3242353, EPI_ISL_3242355, EPI_ISL_3242356, EPI_ISL_3242357, EPI_ISL_3242358, | | | |

| | | | |
|---|---|---|---|
| EPI_ISL_3242359, EPI_ISL_3242361, EPI_ISL_3242362, EPI_ISL_3242364, EPI_ISL_3242365, EPI_ISL_3242366, EPI_ISL_3242368, EPI_ISL_3242370, EPI_ISL_3242371, EPI_ISL_3242372, EPI_ISL_3242373, EPI_ISL_3242374, EPI_ISL_3242375, EPI_ISL_3242376, EPI_ISL_3242377, EPI_ISL_3242378, EPI_ISL_3242379, EPI_ISL_3242380, EPI_ISL_3242381, EPI_ISL_3242382, EPI_ISL_3242383, EPI_ISL_3242384, EPI_ISL_3242385, EPI_ISL_3242386, EPI_ISL_3242387, EPI_ISL_3242388, EPI_ISL_3242389, EPI_ISL_3242390 | | | |
| see above | Goa Medical College | CDFD | Murali Bashyam, Asmita Gupta, Reelina Basu, Vinay Donipadi, Savio Rodrigues, Divya Vashisht, Ashwin Dalal |
| EPI_ISL_3242391, EPI_ISL_3242392 | Government General Hospital | CDFD | Murali Bashyam, Asmita Gupta, Reelina Basu, Vinay Donipadi, Nagamani Kammili, Divya Vashisht, Ashwin Dalal |
| EPI_ISL_3242393, EPI_ISL_3242394, EPI_ISL_3242395, EPI_ISL_3242396, EPI_ISL_3242397, EPI_ISL_3242398, EPI_ISL_3242399, EPI_ISL_3242400 | Indian Institute of Science Education and Research | CDFD | Murali Bashyam, Asmita Gupta, Reelina Basu, Vinay Donipadi, Indranil Banerjee, Divya Vashisht, Ashwin Dalal |
| EPI_ISL_3242401, EPI_ISL_3242402, EPI_ISL_3242403, EPI_ISL_3242404, EPI_ISL_3242405, EPI_ISL_3242406, EPI_ISL_3242407, EPI_ISL_3242408, EPI_ISL_3242409, EPI_ISL_3242410, EPI_ISL_3242411, EPI_ISL_3242412, EPI_ISL_3242413, EPI_ISL_3242414, EPI_ISL_3242415 | | | |
| see above | Institute of Preventive Medicine | CDFD | Murali Bashyam, Asmita Gupta, Reelina Basu, Vinay Donipadi, Nagamani Kammili, Divya Vashisht, Ashwin Dalal |
| EPI_ISL_3242416 | COVID-19 Detection Lab, Chattogram Veterinary and Animal Sciences University | Genomic Research Lab, BCSIR | Md. Murshed Hasan Sarkar, Abu Sayeed Mohammad Mahmud, Mohammad Samir Uzzaman, Eshrar Osman, Md. Ahasan Habib, Shahina Akter, Tanjina Akhter Banu, Barna Goswami, Iffat Jahan, Mohammad Mohi Uddin, Md. Kamrul Islam, Goutam Buddha Das, Tridip Das, Tanvir Ahmad Nizami, Eaftekhar Ahmed Rana, Md. Sirazul Islam, Pronesh Dutta, Sharmin Chowdhury, Md. Paritosh Kumar Biswas, Md. Salim Khan |
| EPI_ISL_3242418, EPI_ISL_3242419, EPI_ISL_3242420, EPI_ISL_3242421 | Department of Health Technology and Informatics, The Hong Kong Polytechnic University | Department of Health Technology and Informatics, The Hong Kong Polytechnic University | Gilman Kit-Hang Siu, Lam-Kwong Lee, Kenneth Siu-Sing Leung, Jake Siu-Lun Leung, Wing-Hei Lo, Timothy Ting-Leung Ng, Chloe Toi-Mei Chan, Hoi-Ching Jim, Ivan Tak-Fai Wong, Kingsley King-Gee Tam, Hiu-Yin Lao, Denise Sze-Hang Wong, Alan Ka-Lun Wu, Miranda Chong-Yee Yau, Yvette Wai-Man Lai, Kitty Sau-Chun Fung, Sandy Ka-Yee Chau, Barry Kin-Chung Wong, Wing-Kin To, Kristine Luk, Alex Yat-Man Ho, Tak-Lun Que, Kam-Tong Yip, Wing Cheong Yam, David Ho-Keung Shum, Shea Ping Yip |
| EPI_ISL_3242444 | SC (UCO) Igiene e Sanità Pubblica, ASUGI, Trieste | ARGO Laboratorio Genomica ed Epigenomica | Licastro D, Dal Monego S, Degasperi M, Marcello A, Segat L, Piscianz E, Busetti M, Fontana F, Braida C, Barbone F, D'Agaro P |
| EPI_ISL_3242750, EPI_ISL_3242751, EPI_ISL_3242752, EPI_ISL_3242753, EPI_ISL_3242754, EPI_ISL_3242755, EPI_ISL_3242756, EPI_ISL_3242757, EPI_ISL_3242758, EPI_ISL_3242759, EPI_ISL_3242760, EPI_ISL_3242761, EPI_ISL_3242762, EPI_ISL_3242763, EPI_ISL_3242764, EPI_ISL_3242765, EPI_ISL_3242766, EPI_ISL_3242767, EPI_ISL_3242768, EPI_ISL_3242769, EPI_ISL_3242770, EPI_ISL_3242771, EPI_ISL_3242772, EPI_ISL_3242773, EPI_ISL_3242774, EPI_ISL_3242775, EPI_ISL_3242776, EPI_ISL_3242777, EPI_ISL_3242778, EPI_ISL_3242779, EPI_ISL_3242780, EPI_ISL_3242781, EPI_ISL_3242782, EPI_ISL_3242783 | | | |
| see above | Tampa General Hospital Esoteric Lab | Tampa General Hospital Esoteric Research & Development Lab | Jon Faughn, Marci O'Driscoll, Deanna Becker, Dominic Uy, Jorge Mecias-Frias, Katie Hurless, Vicki Healer, Amorce Lima, Suzane Silbert |
| EPI_ISL_3242784, EPI_ISL_3242785, EPI_ISL_3242786, EPI_ISL_3242787, EPI_ISL_3242788, EPI_ISL_3242789, EPI_ISL_3242791, EPI_ISL_3242792, EPI_ISL_3242793, EPI_ISL_3242794, EPI_ISL_3242795, EPI_ISL_3242797, EPI_ISL_3242798, EPI_ISL_3242799, EPI_ISL_3242800, EPI_ISL_3242801, EPI_ISL_3242802, EPI_ISL_3242803, EPI_ISL_3242804, EPI_ISL_3242805, EPI_ISL_3242806, EPI_ISL_3242807, EPI_ISL_3242808, EPI_ISL_3242814, EPI_ISL_3242815, EPI_ISL_3242816, EPI_ISL_3242817, EPI_ISL_3242821, EPI_ISL_3242822, EPI_ISL_3242823, EPI_ISL_3242824, EPI_ISL_3242825, EPI_ISL_3242828, EPI_ISL_3242829, EPI_ISL_3242831, EPI_ISL_3242832, EPI_ISL_3242833 | | | |
| see above | Wyoming Public Health Laboratory | Wyoming Public Health Laboratory | Jim Mildenberger, Taylor Fearing, Channing Weber, Ashley Norberg, Chayse Rowley, Marley Goetz, Brian Dominguez, Elliot Thomasson, Sam Britz, Cari Sloma, Robert Petit, and Rob Christensen |
| EPI_ISL_3242844, EPI_ISL_3242845, EPI_ISL_3242846, EPI_ISL_3242847, EPI_ISL_3242848, EPI_ISL_3242849, EPI_ISL_3242850, EPI_ISL_3242851, EPI_ISL_3242852, EPI_ISL_3242853, EPI_ISL_3242854, EPI_ISL_3242855, EPI_ISL_3242856, EPI_ISL_3242857, EPI_ISL_3242858, EPI_ISL_3242859, EPI_ISL_3242860, EPI_ISL_3242861, EPI_ISL_3242862, EPI_ISL_3242863, EPI_ISL_3242864, EPI_ISL_3242865, EPI_ISL_3242866, EPI_ISL_3242867, EPI_ISL_3242868, EPI_ISL_3242869, EPI_ISL_3242870, EPI_ISL_3242871, EPI_ISL_3242872, EPI_ISL_3242873, EPI_ISL_3242874, EPI_ISL_3242875, EPI_ISL_3242876, EPI_ISL_3242877, EPI_ISL_3242878, EPI_ISL_3242879, EPI_ISL_3242880, EPI_ISL_3242881, EPI_ISL_3242882, EPI_ISL_3242883, EPI_ISL_3242884, EPI_ISL_3242885, EPI_ISL_3242886, EPI_ISL_3242887, EPI_ISL_3242888, EPI_ISL_3242889, EPI_ISL_3242890, EPI_ISL_3242891, EPI_ISL_3242892, EPI_ISL_3242893, EPI_ISL_3242894, EPI_ISL_3242895, EPI_ISL_3242896, EPI_ISL_3242897, EPI_ISL_3242898, EPI_ISL_3242899, EPI_ISL_3242900, EPI_ISL_3242901, EPI_ISL_3242902, EPI_ISL_3242903, EPI_ISL_3242904, EPI_ISL_3242905, EPI_ISL_3242906, EPI_ISL_3242907, EPI_ISL_3242908, EPI_ISL_3242909, EPI_ISL_3242910, EPI_ISL_3242911, EPI_ISL_3242912, EPI_ISL_3242913, EPI_ISL_3242914, EPI_ISL_3242915, EPI_ISL_3242916 | | | |
| see above | AZDelta | AZ Delta Medical Laboratories in Roeselare, Belgium | Geert Martens, Dieter De Smet, Frederik Van Hoecke, on behalf of AZ Delta COVID-19 Genomics core (member of Genomic surveillance of SARS-CoV-2 in Belgium network) |
| EPI_ISL_3243000, EPI_ISL_3243001, EPI_ISL_3243002, EPI_ISL_3243003, EPI_ISL_3243004, EPI_ISL_3243006, EPI_ISL_3243007, EPI_ISL_3243008, EPI_ISL_3243009, EPI_ISL_3243010, EPI_ISL_3243011, EPI_ISL_3243012, EPI_ISL_3243013, EPI_ISL_3243015, EPI_ISL_3243016 | | | |
| see above | TXDSHS | TXDSHS | Rashmi Tuladhar, Bonnie Oh, Jenny Zhang, Maliha Rahman, Mayela Pedrueza, Anita Pokharel, Karen Bobier, Lorraine Rodriguez, Myong Koag, Chun Wang, Rachel Lee, Grace Kubin |
| EPI_ISL_3243023 | Applied DNA Sciences Inc | Applied DNA Sciences Inc | Huan Dong; Spencer William; Lucas Liu; Yuhua Sun |
| EPI_ISL_3243037, EPI_ISL_3243038, EPI_ISL_3243039, EPI_ISL_3243040, EPI_ISL_3243041, EPI_ISL_3243042, EPI_ISL_3243043, EPI_ISL_3243044, EPI_ISL_3243045, EPI_ISL_3243046, EPI_ISL_3243047, EPI_ISL_3243048, EPI_ISL_3243049, EPI_ISL_3243050, EPI_ISL_3243051, EPI_ISL_3243052, EPI_ISL_3243053, EPI_ISL_3243054, EPI_ISL_3243055, EPI_ISL_3243056, EPI_ISL_3243057, EPI_ISL_3243058, EPI_ISL_3243059, EPI_ISL_3243060, EPI_ISL_3243061, EPI_ISL_3243062, EPI_ISL_3243063, EPI_ISL_3243064, EPI_ISL_3243065, EPI_ISL_3243066, EPI_ISL_3243067, EPI_ISL_3243068, EPI_ISL_3243070, EPI_ISL_3243071, EPI_ISL_3243072, EPI_ISL_3243073 | | | |
| see above | Oregon State Public Health Laboratory | Oregon State Public Health Laboratory | Rafia Razzaque, Eugene Yeboah, Vanda Makris, Laura Tsaknaridis, John Fontana and Shane Sevey |
| EPI_ISL_3243114, EPI_ISL_3243115, EPI_ISL_3243118, EPI_ISL_3243120 | Hackensack Medical Center | New York Genome Center | Michael Zody, Andre Corvelo, Dayna M. Oschwald, Amy Baldwin, Samantha Fennessey, Tom Maniatis, Liang Chen, Jose Mediavilla, Marcus Cunningham, Kaelea Composto, Kar Chow, David Perlin, Barry Kreiswirth |
| EPI_ISL_3243196 | Osmania Medical College | CDFD | Murali Bashyam, Asmita Gupta, Reelina Basu, Vinay Donipadi, Nagamani Kammili, Divya Vashisht, Ashwin Dalal |
| EPI_ISL_3243201, EPI_ISL_3243203, EPI_ISL_3243204, EPI_ISL_3243205 | Indian Institute of Science Education and Research | CDFD | Murali Bashyam, Asmita Gupta, Reelina Basu, Vinay Donipadi, Indranil Banerjee, Divya Vashisht, Ashwin Dalal |
| EPI_ISL_3243206, EPI_ISL_3243207, EPI_ISL_3243208, EPI_ISL_3243209, EPI_ISL_3243210, EPI_ISL_3243211, EPI_ISL_3243212, EPI_ISL_3243214, EPI_ISL_3243215, EPI_ISL_3243216 | Gandhi Hospital | CDFD | Murali Bashyam, Asmita Gupta, Reelina Basu, Vinay Donipadi, Nagamani Kammili, Divya Vashisht, Ashwin Dalal |
| EPI_ISL_3243217, EPI_ISL_3243218 | Government Maternity Hospital | CDFD | Murali Bashyam, Asmita Gupta, Reelina Basu, Vinay Donipadi, Nagamani Kammili, Divya Vashisht, Ashwin Dalal |
| EPI_ISL_3243219 | Gandhi Hospital | CDFD | Murali Bashyam, Asmita Gupta, Reelina Basu, Vinay Donipadi, Nagamani Kammili, Divya Vashisht, Ashwin Dalal |
| EPI_ISL_3243220, EPI_ISL_3243221, EPI_ISL_3243222, EPI_ISL_3243223, EPI_ISL_3243224, EPI_ISL_3243225, EPI_ISL_3243226, EPI_ISL_3243227, EPI_ISL_3243228, EPI_ISL_3243229, EPI_ISL_3243230, EPI_ISL_3243231, EPI_ISL_3243232, EPI_ISL_3243233, EPI_ISL_3243234, EPI_ISL_3243235, EPI_ISL_3243236, EPI_ISL_3243237 | | | |
| see above | CDFD | CDFD | Murali Bashyam, Asmita Gupta, Reelina Basu, Vinay Donipadi, Divya Vashisht, Ashwin Dalal |
| EPI_ISL_3243238, EPI_ISL_3243239, EPI_ISL_3243240, EPI_ISL_3243241, EPI_ISL_3243242, EPI_ISL_3243243, EPI_ISL_3243244 | Government Fever Hospital | CDFD | Murali Bashyam, Asmita Gupta, Reelina Basu, Vinay Donipadi, Nagamani Kammili, Divya Vashisht, Ashwin Dalal |
| EPI_ISL_3243245, EPI_ISL_3243246, EPI_ISL_3243247, EPI_ISL_3243248, EPI_ISL_3243249, EPI_ISL_3243250, EPI_ISL_3243251, EPI_ISL_3243252 | Osmania Medical College | CDFD | Murali Bashyam, Asmita Gupta, Reelina Basu, Vinay Donipadi, Nagamani Kammili, Divya Vashisht, Ashwin Dalal |
| EPI_ISL_3243253, EPI_ISL_3243254, EPI_ISL_3243255, EPI_ISL_3243256, EPI_ISL_3243257, EPI_ISL_3243258, EPI_ISL_3243259, EPI_ISL_3243260, EPI_ISL_3243261, EPI_ISL_3243262, EPI_ISL_3243263, EPI_ISL_3243264, EPI_ISL_3243265, EPI_ISL_3243266, EPI_ISL_3243267, EPI_ISL_3243268, EPI_ISL_3243269, EPI_ISL_3243270, EPI_ISL_3243271, EPI_ISL_3243272, EPI_ISL_3243273, EPI_ISL_3243274, EPI_ISL_3243275, EPI_ISL_3243276, EPI_ISL_3243277, EPI_ISL_3243278, EPI_ISL_3243279, EPI_ISL_3243280, EPI_ISL_3243281, EPI_ISL_3243282, EPI_ISL_3243283, EPI_ISL_3243284, EPI_ISL_3243285, EPI_ISL_3243286, EPI_ISL_3243288, EPI_ISL_3243289 | | | |
| see above | Gandhi Hospital | CDFD | Murali Bashyam, Asmita Gupta, Reelina Basu, Vinay Donipadi, Nagamani Kammili, Divya Vashisht, Ashwin Dalal |
| EPI_ISL_3243290, EPI_ISL_3243291, EPI_ISL_3243292, EPI_ISL_3243293 | Government Fever Hospital | CDFD | Murali Bashyam, Asmita Gupta, Reelina Basu, Vinay Donipadi, Nagamani Kammili, Divya Vashisht, Ashwin Dalal |
| EPI_ISL_3243294 | Osmania Medical College | CDFD | Murali Bashyam, Asmita Gupta, Reelina Basu, Vinay Donipadi, Nagamani Kammili, Divya Vashisht, Ashwin Dalal |
| EPI_ISL_3243295 | Gandhi Hospital | CDFD | Murali Bashyam, Asmita Gupta, Reelina Basu, Vinay Donipadi, Nagamani Kammili, Divya Vashisht, Ashwin Dalal |
| EPI_ISL_3243296 | Government General Hospital | CDFD | Murali Bashyam, Asmita Gupta, Reelina Basu, Vinay Donipadi, Nagamani Kammili, Divya Vashisht, Ashwin Dalal |
| EPI_ISL_3243297, EPI_ISL_3243298, EPI_ISL_3243299, EPI_ISL_3243300, EPI_ISL_3243301, EPI_ISL_3243302, EPI_ISL_3243303, EPI_ISL_3243304, EPI_ISL_3243305, EPI_ISL_3243306, EPI_ISL_3243307 | | | |
| see above | Government Maternity Hospital | CDFD | Murali Bashyam, Asmita Gupta, Reelina Basu, Vinay Donipadi, Nagamani Kammili, Divya Vashisht, Ashwin Dalal |
| EPI_ISL_3243308, EPI_ISL_3243309, EPI_ISL_3243310, | Osmania Medical College | CDFD | Murali Bashyam, Asmita Gupta, Reelina Basu, Vinay Donipadi, Nagamani Kammili, Divya Vashisht, Ashwin Dalal |

| | | | |
|---|---|---|---|
| EPI_ISL_3243311 | | | |
| EPI_ISL_3243312, EPI_ISL_3243314, EPI_ISL_3243315, EPI_ISL_3243316 | Institute of Preventive Medicine | CDFD | Murali Bashyam, Asmita Gupta, Reelina Basu, Vinay Donipadi, Nagamani Kammili, Divya Vashisht, Ashwin Dalal |
| EPI_ISL_3243317 | Gandhi Hospital | CDFD | Murali Bashyam, Asmita Gupta, Reelina Basu, Vinay Donipadi, Nagamani Kammili, Divya Vashisht, Ashwin Dalal |
| EPI_ISL_3243318 | Osmania Medical College | CDFD | Murali Bashyam, Asmita Gupta, Reelina Basu, Vinay Donipadi, Nagamani Kammili, Divya Vashisht, Ashwin Dalal |
| EPI_ISL_3243319, EPI_ISL_3243320, EPI_ISL_3243321, EPI_ISL_3243322, EPI_ISL_3243323 | Institute of Preventive Medicine | CDFD | Murali Bashyam, Asmita Gupta, Reelina Basu, Vinay Donipadi, Nagamani Kammili, Divya Vashisht, Ashwin Dalal |
| EPI_ISL_3243324, EPI_ISL_3243325 | Gandhi Hospital | CDFD | Murali Bashyam, Asmita Gupta, Reelina Basu, Vinay Donipadi, Nagamani Kammili, Divya Vashisht, Ashwin Dalal |
| EPI_ISL_3243326, EPI_ISL_3243327 | Institute of Preventive Medicine | CDFD | Murali Bashyam, Asmita Gupta, Reelina Basu, Vinay Donipadi, Nagamani Kammili, Divya Vashisht, Ashwin Dalal |
| EPI_ISL_3243328, EPI_ISL_3243329, EPI_ISL_3243330 | Osmania Medical College | CDFD | Murali Bashyam, Asmita Gupta, Reelina Basu, Vinay Donipadi, Nagamani Kammili, Divya Vashisht, Ashwin Dalal |
| EPI_ISL_3243331, EPI_ISL_3243332, EPI_ISL_3243333 | Government General Hospital | CDFD | Murali Bashyam, Asmita Gupta, Reelina Basu, Vinay Donipadi, Nagamani Kammili, Divya Vashisht, Ashwin Dalal |
| EPI_ISL_3243334 | Osmania Medical College | CDFD | Murali Bashyam, Asmita Gupta, Reelina Basu, Vinay Donipadi, Nagamani Kammili, Divya Vashisht, Ashwin Dalal |
| EPI_ISL_3243335, EPI_ISL_3243336, EPI_ISL_3243337, EPI_ISL_3243338, EPI_ISL_3243339, EPI_ISL_3243340, EPI_ISL_3243341, EPI_ISL_3243343, EPI_ISL_3243344, EPI_ISL_3243345, EPI_ISL_3243346, EPI_ISL_3243347 | | | |
| see above | Government General Hospital | CDFD | Murali Bashyam, Asmita Gupta, Reelina Basu, Vinay Donipadi, Nagamani Kammili, Divya Vashisht, Ashwin Dalal |
| EPI_ISL_3243348, EPI_ISL_3243349, EPI_ISL_3243350, EPI_ISL_3243351, EPI_ISL_3243352, EPI_ISL_3243353 | Osmania Medical College | CDFD | Murali Bashyam, Asmita Gupta, Reelina Basu, Vinay Donipadi, Nagamani Kammili, Divya Vashisht, Ashwin Dalal |
| EPI_ISL_3243354, EPI_ISL_3243355, EPI_ISL_3243356, EPI_ISL_3243357 | Gandhi Hospital | CDFD | Murali Bashyam, Asmita Gupta, Reelina Basu, Vinay Donipadi, Nagamani Kammili, Divya Vashisht, Ashwin Dalal |
| EPI_ISL_3243358, EPI_ISL_3243359, EPI_ISL_3243360, EPI_ISL_3243361 | Government Fever Hospital | CDFD | Murali Bashyam, Asmita Gupta, Reelina Basu, Vinay Donipadi, Nagamani Kammili, Divya Vashisht, Ashwin Dalal |
| EPI_ISL_3243362, EPI_ISL_3243363, EPI_ISL_3243364 | Institute of Preventive Medicine | CDFD | Murali Bashyam, Asmita Gupta, Reelina Basu, Vinay Donipadi, Nagamani Kammili, Divya Vashisht, Ashwin Dalal |
| EPI_ISL_3243365, EPI_ISL_3243366, EPI_ISL_3243367, EPI_ISL_3243368, EPI_ISL_3243369, EPI_ISL_3243370, EPI_ISL_3243371, EPI_ISL_3243372, EPI_ISL_3243373, EPI_ISL_3243374 | Government General Hospital | CDFD | Murali Bashyam, Asmita Gupta, Reelina Basu, Vinay Donipadi, Nagamani Kammili, Divya Vashisht, Ashwin Dalal |
| EPI_ISL_3243375, EPI_ISL_3243376 | Osmania Medical College | CDFD | Murali Bashyam, Asmita Gupta, Reelina Basu, Vinay Donipadi, Nagamani Kammili, Divya Vashisht, Ashwin Dalal |
| EPI_ISL_3243377, EPI_ISL_3243378 | Government Maternity Hospital | CDFD | Murali Bashyam, Asmita Gupta, Reelina Basu, Vinay Donipadi, Nagamani Kammili, Divya Vashisht, Ashwin Dalal |
| EPI_ISL_3243379, EPI_ISL_3243380, EPI_ISL_3243381, EPI_ISL_3243382, EPI_ISL_3243383, EPI_ISL_3243384, EPI_ISL_3243385, EPI_ISL_3243386, EPI_ISL_3243387, EPI_ISL_3243389, EPI_ISL_3243390, EPI_ISL_3243391, EPI_ISL_3243392, EPI_ISL_3243393, EPI_ISL_3243394, EPI_ISL_3243395, EPI_ISL_3243396, EPI_ISL_3243397, EPI_ISL_3243398, EPI_ISL_3243400, EPI_ISL_3243401, EPI_ISL_3243402, EPI_ISL_3243403, EPI_ISL_3243404, EPI_ISL_3243405, EPI_ISL_3243406, EPI_ISL_3243407, EPI_ISL_3243408, EPI_ISL_3243409, EPI_ISL_3243410, EPI_ISL_3243411, EPI_ISL_3243412, EPI_ISL_3243413, EPI_ISL_3243414, EPI_ISL_3243415, EPI_ISL_3243416, EPI_ISL_3243417, EPI_ISL_3243418, EPI_ISL_3243420, EPI_ISL_3243421, EPI_ISL_3243422, EPI_ISL_3243423, EPI_ISL_3243424, EPI_ISL_3243425, EPI_ISL_3243426, EPI_ISL_3243427, EPI_ISL_3243428, EPI_ISL_3243429, EPI_ISL_3243430, EPI_ISL_3243431, EPI_ISL_3243432, EPI_ISL_3243433, EPI_ISL_3243435, EPI_ISL_3243436, EPI_ISL_3243437, EPI_ISL_3243438, EPI_ISL_3243439, EPI_ISL_3243441, EPI_ISL_3243442 | | | |
| see above | Orange County Public Health Laboratory | Orange County Public Health Laboratory | Angelica Torres, Julia Wolfe |
| EPI_ISL_3243768, EPI_ISL_3243799, EPI_ISL_3243801, EPI_ISL_3243825, EPI_ISL_3243854, EPI_ISL_3243862, EPI_ISL_3243884, EPI_ISL_3243933, EPI_ISL_3243995, EPI_ISL_3244121, EPI_ISL_3244124, EPI_ISL_3244131, EPI_ISL_3244132, EPI_ISL_3244138, EPI_ISL_3244207, EPI_ISL_3244214, EPI_ISL_3244228, EPI_ISL_3244229, EPI_ISL_3244243, EPI_ISL_3244256, EPI_ISL_3244263, EPI_ISL_3244304, EPI_ISL_3244305, EPI_ISL_3244350, EPI_ISL_3244361, EPI_ISL_3244377, EPI_ISL_3244385, EPI_ISL_3244413, EPI_ISL_3244422, EPI_ISL_3244428, EPI_ISL_3244459, EPI_ISL_3244486, EPI_ISL_3244491, EPI_ISL_3244513, EPI_ISL_3244520 | | | |
| see above | CDPH VBL | California Department of Public Health | Emily Smith on behalf of CDPH-COVIDNet and UCSD EXCITE lab |
| EPI_ISL_3244595, EPI_ISL_3244596, EPI_ISL_3244597, EPI_ISL_3244598, EPI_ISL_3244599 | Hospital Margarita Maza de Juárez | Microbial Genomics Laboratory | Consorcio Mexicano de Vigilancia Genómica (CoViGen-Mex). Authors (in alphabetical order): Julio Elias Alvarado-Yaah, Carlos F. Arias, Santiago Ávila-Ríos, Eduardo Becerril-Vargas, Víctor Hugo Borja-Aburto, Celia Boukadida, Cristóbal Cháidez-Quiróz, Juan Bautista Chale-Dzul, Ricardo Ciria Merce, Andreu Comas-García, Célida Duque Molina, Julissa Enciso-Ibarra, José Antonio Enciso-Moreno, Gloria Elena Espinosa-Ayala, Fernando Fontove-Herrera, Daniel Fregoso-Rueda, Victor Eduardo García-Arias, Alejandra García-Gasca, Bruno Gómez-Gil, Jean Pierre González, Irvin González-López, Concepción Grajales-Muñiz, Ricardo Grande, Rosa María Gutiérrez Rios, Jesús Hernández, Alejandra Hernández-Terán, Alfredo Herrera-Estrella, Carla Ivón Herrera-Najera, Pavel Isa, Verónica Mata-Haro, Daniel Lira Morales, Susana Lopez, Antonio Loza Román, Brenda Irasema Maldonado-Meza, Bernardo Martínez-Miguel, José Arturo Martínez-Orozco, Célida Martínez-Rodríguez, Margarita Matías-Florentino, Fidencio Mejía-Nepomuceno, María Guadalupe de Jesús Mireles-Rivera, Gloria María Molina-Salinas, Hector Montoya-Fuentes, , Mario Mújica-Sánchez, José Esteban Muñoz-Medina, José de Jesús Nuñez-Contreras, Alicia Ocaña-Mondragón, Luis Alberto Ochoa-Carrera, Hector Esteban Paz-Juárez, Marissa Perez-Garcia, Francisco Pulido, Helen Haydee Fernanda Ramírez-Plascencia, Jorge Salas-Hernández, Angel Gustavo Salas-Lais, Alejandro Sánchez-Flores, Clara Esperanza Santacruz-Tinoco, María Guadalupe Santiago-Mauricio, Selene Zárate, Nelly Sélem-Mojica, Blanca Taboada, Gloria Vazquez, Joel Armando Vázquez-Pérez. |
| EPI_ISL_3244601 | Imágenes Diagnósticas de Mazatlán | Microbial Genomics Laboratory | Consorcio Mexicano de Vigilancia Genómica (CoViGen-Mex). Authors (in alphabetical order): Julio Elias Alvarado-Yaah, Carlos F. Arias, Santiago Ávila-Ríos, Eduardo Becerril-Vargas, Víctor Hugo Borja-Aburto, Celia Boukadida, Cristóbal Cháidez-Quiróz, Juan Bautista Chale-Dzul, Ricardo Ciria Merce, Andreu Comas-García, Célida Duque Molina, Julissa Enciso-Ibarra, José Antonio Enciso-Moreno, Gloria Elena Espinosa-Ayala, Fernando Fontove-Herrera, Daniel Fregoso-Rueda, Victor Eduardo García-Arias, Alejandra García-Gasca, Bruno Gómez-Gil, Jean Pierre González, Irvin González-López, Concepción Grajales-Muñiz, Ricardo Grande, Rosa María Gutiérrez Rios, Jesús Hernández, Alejandra Hernández-Terán, Alfredo Herrera-Estrella, Carla Ivón Herrera-Najera, Pavel Isa, Verónica Mata-Haro, Daniel Lira Morales, Susana Lopez, Antonio Loza Román, Brenda Irasema Maldonado-Meza, Bernardo Martínez-Miguel, José Arturo Martínez-Orozco, Célida Martínez-Rodríguez, Margarita Matías-Florentino, Fidencio Mejía-Nepomuceno, María Guadalupe de Jesús Mireles-Rivera, Gloria María Molina-Salinas, Hector Montoya-Fuentes, , Mario Mújica-Sánchez, José Esteban Muñoz-Medina, José de Jesús Nuñez-Contreras, Alicia Ocaña-Mondragón, Luis Alberto Ochoa-Carrera, Hector Esteban Paz-Juárez, Marissa Perez-Garcia, Francisco Pulido, Helen Haydee Fernanda Ramírez-Plascencia, Jorge Salas-Hernández, Angel Gustavo Salas-Lais, Alejandro Sánchez-Flores, Clara Esperanza Santacruz-Tinoco, María Guadalupe Santiago-Mauricio, Selene Zárate, Nelly Sélem-Mojica, Blanca Taboada, Gloria Vazquez, Joel Armando Vázquez-Pérez. |
| EPI_ISL_3244602 | Hospital Sharp | Microbial Genomics Laboratory | Consorcio Mexicano de Vigilancia Genómica (CoViGen-Mex). Authors (in alphabetical order): Julio Elias Alvarado-Yaah, Carlos F. Arias, Santiago Ávila-Ríos, Eduardo Becerril-Vargas, Víctor Hugo Borja-Aburto, Celia Boukadida, Cristóbal Cháidez-Quiróz, Juan Bautista Chale-Dzul, Ricardo Ciria Merce, Andreu Comas-García, Célida Duque Molina, Julissa Enciso-Ibarra, José Antonio Enciso-Moreno, Gloria Elena Espinosa-Ayala, Fernando Fontove-Herrera, Daniel Fregoso-Rueda, Victor Eduardo García-Arias, Alejandra García-Gasca, Bruno Gómez-Gil, Jean Pierre González, Irvin González-López, Concepción Grajales-Muñiz, Ricardo Grande, Rosa María Gutiérrez Rios, Jesús Hernández, Alejandra Hernández-Terán, Alfredo Herrera-Estrella, Carla Ivón Herrera-Najera, Pavel Isa, Verónica Mata-Haro, Daniel Lira Morales, Susana Lopez, Antonio Loza Román, Brenda Irasema Maldonado-Meza, Bernardo Martínez-Miguel, José Arturo Martínez-Orozco, Célida Martínez-Rodríguez, Margarita Matías-Florentino, Fidencio Mejía-Nepomuceno, María Guadalupe de Jesús Mireles-Rivera, Gloria María Molina-Salinas, Hector Montoya-Fuentes, , Mario Mújica-Sánchez, José Esteban Muñoz-Medina, José de Jesús Nuñez-Contreras, Alicia Ocaña-Mondragón, Luis Alberto Ochoa-Carrera, Hector Esteban Paz-Juárez, Marissa Perez-Garcia, Francisco Pulido, Helen Haydee Fernanda Ramírez-Plascencia, Jorge Salas-Hernández, Angel Gustavo Salas-Lais, Alejandro Sánchez-Flores, Clara Esperanza Santacruz-Tinoco, María Guadalupe Santiago-Mauricio, Selene Zárate, Nelly Sélem-Mojica, Blanca Taboada, Gloria Vazquez, Joel Armando Vázquez-Pérez. |
| EPI_ISL_3244603, EPI_ISL_3244604, EPI_ISL_3244605, EPI_ISL_3244606, EPI_ISL_3244607, EPI_ISL_3244608, EPI_ISL_3244609, EPI_ISL_3244610, EPI_ISL_3244611, EPI_ISL_3244612, EPI_ISL_3244613, EPI_ISL_3244614, EPI_ISL_3244615, EPI_ISL_3244616, EPI_ISL_3244617, EPI_ISL_3244618, EPI_ISL_3244619, EPI_ISL_3244620, EPI_ISL_3244621, EPI_ISL_3244622, EPI_ISL_3244623, EPI_ISL_3244624, EPI_ISL_3244625, EPI_ISL_3244626, EPI_ISL_3244627, EPI_ISL_3244628, EPI_ISL_3244629, EPI_ISL_3244630, EPI_ISL_3244631, EPI_ISL_3244632, EPI_ISL_3244633, EPI_ISL_3244634, EPI_ISL_3244635, EPI_ISL_3244636, EPI_ISL_3244637, EPI_ISL_3244638, EPI_ISL_3244639, EPI_ISL_3244640, EPI_ISL_3244641, EPI_ISL_3244642, EPI_ISL_3244643, EPI_ISL_3244644, EPI_ISL_3244645, EPI_ISL_3244646, EPI_ISL_3244647, EPI_ISL_3244648, EPI_ISL_3244649, EPI_ISL_3244650, EPI_ISL_3244651, EPI_ISL_3244652, EPI_ISL_3244653, EPI_ISL_3244654, EPI_ISL_3244655, EPI_ISL_3244656, EPI_ISL_3244657, EPI_ISL_3244658, EPI_ISL_3244659, EPI_ISL_3244660, EPI_ISL_3244661, EPI_ISL_3244662, EPI_ISL_3244663, EPI_ISL_3244664, EPI_ISL_3244665, EPI_ISL_3244666, EPI_ISL_3244667, EPI_ISL_3244668, EPI_ISL_3244669, EPI_ISL_3244670, EPI_ISL_3244671, EPI_ISL_3244672, EPI_ISL_3244673, EPI_ISL_3244674, EPI_ISL_3244675, EPI_ISL_3244676, EPI_ISL_3244677, EPI_ISL_3244678, EPI_ISL_3244679, EPI_ISL_3244680, EPI_ISL_3244681, EPI_ISL_3244682, EPI_ISL_3244683, EPI_ISL_3244684, EPI_ISL_3244685, EPI_ISL_3244686, EPI_ISL_3244687, EPI_ISL_3244688, EPI_ISL_3244689, EPI_ISL_3244690, EPI_ISL_3244691, EPI_ISL_3244692, EPI_ISL_3244693, EPI_ISL_3244694, EPI_ISL_3244695, EPI_ISL_3244696, EPI_ISL_3244697, EPI_ISL_3244698, EPI_ISL_3244699, EPI_ISL_3244700, EPI_ISL_3244701, EPI_ISL_3244702, EPI_ISL_3244703, EPI_ISL_3244704, EPI_ISL_3244705, EPI_ISL_3244706, EPI_ISL_3244707, EPI_ISL_3244708, EPI_ISL_3244709, EPI_ISL_3244710, EPI_ISL_3244711, EPI_ISL_3244712, EPI_ISL_3244713, EPI_ISL_3244714, EPI_ISL_3244715, EPI_ISL_3244716, EPI_ISL_3244717, EPI_ISL_3244718, EPI_ISL_3244719, EPI_ISL_3244720, EPI_ISL_3244721, EPI_ISL_3244722, EPI_ISL_3244723, EPI_ISL_3244724, EPI_ISL_3244725, EPI_ISL_3244726, EPI_ISL_3244727, EPI_ISL_3244728, EPI_ISL_3244729, EPI_ISL_3244730, EPI_ISL_3244731, EPI_ISL_3244732, EPI_ISL_3244733, EPI_ISL_3244734, EPI_ISL_3244735, EPI_ISL_3244736, EPI_ISL_3244737, EPI_ISL_3244738, EPI_ISL_3244739, EPI_ISL_3244740, EPI_ISL_3244741, EPI_ISL_3244742, EPI_ISL_3244743, EPI_ISL_3244744, EPI_ISL_3244745, EPI_ISL_3244746, EPI_ISL_3244747, EPI_ISL_3244748, EPI_ISL_3244749, EPI_ISL_3244750, EPI_ISL_3244751, EPI_ISL_3244752, EPI_ISL_3244753, EPI_ISL_3244754, EPI_ISL_3244755, EPI_ISL_3244756, EPI_ISL_3244757, EPI_ISL_3244758, EPI_ISL_3244759, EPI_ISL_3244760, EPI_ISL_3244761, EPI_ISL_3244762, EPI_ISL_3244763, EPI_ISL_3244764, EPI_ISL_3244765, EPI_ISL_3244766, EPI_ISL_3244767, EPI_ISL_3244768, EPI_ISL_3244769, EPI_ISL_3244770, EPI_ISL_3244771, EPI_ISL_3244772, EPI_ISL_3244773, EPI_ISL_3244774, EPI_ISL_3244775, EPI_ISL_3244776, EPI_ISL_3244777, EPI_ISL_3244778, EPI_ISL_3244779 | | | |
| see above | ICMR-National Institute of Virology - INSACOG | NIV Influenza | Dr. Varsha Potdar and NIC Team |
| EPI_ISL_3244792, EPI_ISL_3244793, EPI_ISL_3244794, EPI_ISL_3244795, | UW Virology Lab | UW Virology Lab | Pavitra Roychoudhury, Shah Mohamed Bakhash, Ricardo Perez, Maria Lukes, Hong Xie, Tien V. Nguyen, Noah R. Baker, Sean Ellis, Meei-Li Huang, Keith R Jerome, Alexander Greninger |

| | | | |
|---|---|---|---|
| EPI_ISL_3244796, EPI_ISL_3244797 | | | |
| EPI_ISL_3244798, EPI_ISL_3244799, EPI_ISL_3244800, EPI_ISL_3244801, EPI_ISL_3244802, EPI_ISL_3244803, EPI_ISL_3244804, EPI_ISL_3244805, EPI_ISL_3244806, EPI_ISL_3244807, EPI_ISL_3244808, EPI_ISL_3244809, EPI_ISL_3244811, EPI_ISL_3244812, EPI_ISL_3244813, EPI_ISL_3244814, EPI_ISL_3244815, EPI_ISL_3244816, EPI_ISL_3244817, EPI_ISL_3244818, EPI_ISL_3244819, EPI_ISL_3244820, EPI_ISL_3244821, EPI_ISL_3244822, EPI_ISL_3244823, EPI_ISL_3244824, EPI_ISL_3244825, EPI_ISL_3244826, EPI_ISL_3244827, EPI_ISL_3244828, EPI_ISL_3244829, EPI_ISL_3244830, EPI_ISL_3244831, EPI_ISL_3244832, EPI_ISL_3244833, EPI_ISL_3244834, EPI_ISL_3244835, EPI_ISL_3244836, EPI_ISL_3244837, EPI_ISL_3244838, EPI_ISL_3244839, EPI_ISL_3244840, EPI_ISL_3244841, EPI_ISL_3244842, EPI_ISL_3244843, EPI_ISL_3244844, EPI_ISL_3244845, EPI_ISL_3244846, EPI_ISL_3244847, EPI_ISL_3244848, EPI_ISL_3244849, EPI_ISL_3244850, EPI_ISL_3244851, EPI_ISL_3244853, EPI_ISL_3244854, EPI_ISL_3244855, EPI_ISL_3244856, EPI_ISL_3244857, EPI_ISL_3244858, EPI_ISL_3244859, EPI_ISL_3244860, EPI_ISL_3244861, EPI_ISL_3244862, EPI_ISL_3244863, EPI_ISL_3244864, EPI_ISL_3244865, EPI_ISL_3244866, EPI_ISL_3244867, EPI_ISL_3244869, EPI_ISL_3244870, EPI_ISL_3244871, EPI_ISL_3244872, EPI_ISL_3244873, EPI_ISL_3244874, EPI_ISL_3244875, EPI_ISL_3244876, EPI_ISL_3244877, EPI_ISL_3244878, EPI_ISL_3244879, EPI_ISL_3244880, EPI_ISL_3244881, EPI_ISL_3244882, EPI_ISL_3244883, EPI_ISL_3244884, EPI_ISL_3244885, EPI_ISL_3244886, EPI_ISL_3244887 | | | |
| see above | UW Virology Lab | UW Virology Lab | Pavitra Roychoudhury, Shah Mohamed Bakhash, Ricardo Perez, Hong Xie, Tien V. Nguyen, Maria Lukes, Sean Ellis, Nathan Breit, Robert J. Livingston, Meei-Li Huang, Keith R Jerome, Patrick Mathias, Alexander Greninger |
| EPI_ISL_3244888, EPI_ISL_3244889, EPI_ISL_3244891, EPI_ISL_3244892, EPI_ISL_3244893, EPI_ISL_3244894, EPI_ISL_3244895, EPI_ISL_3244896, EPI_ISL_3244897, EPI_ISL_3244898, EPI_ISL_3244899, EPI_ISL_3244901, EPI_ISL_3244902, EPI_ISL_3244904, EPI_ISL_3244906, EPI_ISL_3244908, EPI_ISL_3244910, EPI_ISL_3244912, EPI_ISL_3244913, EPI_ISL_3244914, EPI_ISL_3244915, EPI_ISL_3244916, EPI_ISL_3244917, EPI_ISL_3244918, EPI_ISL_3244919, EPI_ISL_3244920, EPI_ISL_3244921, EPI_ISL_3244922, EPI_ISL_3244923, EPI_ISL_3244924, EPI_ISL_3244925, EPI_ISL_3244926, EPI_ISL_3244927, EPI_ISL_3244928, EPI_ISL_3244929, EPI_ISL_3244931, EPI_ISL_3244932, EPI_ISL_3244933, EPI_ISL_3244934, EPI_ISL_3244935, EPI_ISL_3244936, EPI_ISL_3244937, EPI_ISL_3244938, EPI_ISL_3244940, EPI_ISL_3244941, EPI_ISL_3244942, EPI_ISL_3244943, EPI_ISL_3244944, EPI_ISL_3244945, EPI_ISL_3244946, EPI_ISL_3244947, EPI_ISL_3244948, EPI_ISL_3244949, EPI_ISL_3244950, EPI_ISL_3244951, EPI_ISL_3244952, EPI_ISL_3244953, EPI_ISL_3244954, EPI_ISL_3244955, EPI_ISL_3244956, EPI_ISL_3244957, EPI_ISL_3244959, EPI_ISL_3244960, EPI_ISL_3244961, EPI_ISL_3244962, EPI_ISL_3244963, EPI_ISL_3244964, EPI_ISL_3244965, EPI_ISL_3244967, EPI_ISL_3244968, EPI_ISL_3244969, EPI_ISL_3244970, EPI_ISL_3244971, EPI_ISL_3244972, EPI_ISL_3244973, EPI_ISL_3244974, EPI_ISL_3244975, EPI_ISL_3244976, EPI_ISL_3244977, EPI_ISL_3244978, EPI_ISL_3244979, EPI_ISL_3244980, EPI_ISL_3244981, EPI_ISL_3244982, EPI_ISL_3244983, EPI_ISL_3244984, EPI_ISL_3244985, EPI_ISL_3244986, EPI_ISL_3244987, EPI_ISL_3244989, EPI_ISL_3244990, EPI_ISL_3244991, EPI_ISL_3244992, EPI_ISL_3244993, EPI_ISL_3244994, EPI_ISL_3244995, EPI_ISL_3244996, EPI_ISL_3244997, EPI_ISL_3244998, EPI_ISL_3244999, EPI_ISL_3245000, EPI_ISL_3245001, EPI_ISL_3245002, EPI_ISL_3245003, EPI_ISL_3245004, EPI_ISL_3245005, EPI_ISL_3245006, EPI_ISL_3245007, EPI_ISL_3245008, EPI_ISL_3245011, EPI_ISL_3245012, EPI_ISL_3245013, EPI_ISL_3245015, EPI_ISL_3245016, EPI_ISL_3245017, EPI_ISL_3245018, EPI_ISL_3245019, EPI_ISL_3245020, EPI_ISL_3245021, EPI_ISL_3245022, EPI_ISL_3245023, EPI_ISL_3245025, EPI_ISL_3245026, EPI_ISL_3245027, EPI_ISL_3245028, EPI_ISL_3245030, EPI_ISL_3245031, EPI_ISL_3245032, EPI_ISL_3245033, EPI_ISL_3245034, EPI_ISL_3245035, EPI_ISL_3245036, EPI_ISL_3245037, EPI_ISL_3245039, EPI_ISL_3245040, EPI_ISL_3245041, EPI_ISL_3245042, EPI_ISL_3245044, EPI_ISL_3245045, EPI_ISL_3245046, EPI_ISL_3245047, EPI_ISL_3245050, EPI_ISL_3245051, EPI_ISL_3245052, EPI_ISL_3245053, EPI_ISL_3245054, EPI_ISL_3245055, EPI_ISL_3245056, EPI_ISL_3245057, EPI_ISL_3245058, EPI_ISL_3245059, EPI_ISL_3245060, EPI_ISL_3245061, EPI_ISL_3245063, EPI_ISL_3245064, EPI_ISL_3245065, EPI_ISL_3245066, EPI_ISL_3245067, EPI_ISL_3245068, EPI_ISL_3245070, EPI_ISL_3245071, EPI_ISL_3245072, EPI_ISL_3245073, EPI_ISL_3245074, EPI_ISL_3245075, EPI_ISL_3245076, EPI_ISL_3245077, EPI_ISL_3245078, EPI_ISL_3245079, EPI_ISL_3245080, EPI_ISL_3245081, EPI_ISL_3245082, EPI_ISL_3245083, EPI_ISL_3245084, EPI_ISL_3245085, EPI_ISL_3245086, EPI_ISL_3245087, EPI_ISL_3245088, EPI_ISL_3245089, EPI_ISL_3245090, EPI_ISL_3245091, EPI_ISL_3245092, EPI_ISL_3245093, EPI_ISL_3245094, EPI_ISL_3245095, EPI_ISL_3245097, EPI_ISL_3245098, EPI_ISL_3245100 | | | |
| see above | UW Virology Lab | UW Virology Lab | Pavitra Roychoudhury, Shah Mohamed Bakhash, Ricardo Perez, Maria Lukes, Hong Xie, Tien V. Nguyen, Noah R. Baker, Sean Ellis, Meei-Li Huang, Keith R Jerome, Alexander Greninger |
| EPI_ISL_3245103, EPI_ISL_3245202, EPI_ISL_3245205, EPI_ISL_3245219, EPI_ISL_3245226, EPI_ISL_3245228, EPI_ISL_3245242, EPI_ISL_3245243, EPI_ISL_3245244, EPI_ISL_3245249, EPI_ISL_3245254, EPI_ISL_3245255, EPI_ISL_3245264, EPI_ISL_3245272, EPI_ISL_3245280, EPI_ISL_3245281, EPI_ISL_3245287, EPI_ISL_3245288, EPI_ISL_3245292, EPI_ISL_3245296, EPI_ISL_3245297, EPI_ISL_3245299, EPI_ISL_3245303, EPI_ISL_3245314, EPI_ISL_3245315, EPI_ISL_3245316, EPI_ISL_3245318, EPI_ISL_3245319, EPI_ISL_3245320, EPI_ISL_3245321, EPI_ISL_3245322, EPI_ISL_3245334, EPI_ISL_3245335, EPI_ISL_3245337, EPI_ISL_3245347, EPI_ISL_3245348, EPI_ISL_3245349 | | | |
| see above | Laboratorio Central Noel Nutels | Bioinformatics Laboratory / LNCC | Luiz G P de Almeida, Alessandra P Lamarca, Ronaldo da Silva F Jr, Liliane Cavalcante, Alexandra L Gerber, Ana Paula de C Guimaraes, Douglas Terra Machado, Erica Ramos dos Santos Nascimento, Fernanda Leitao dos Santos, Ricardo Jose Barbosa Salviano, Cintia Policarpo, Gleidson da Silva de Oliveira, Mario Sergio Ribeiro, Silvia Carvalho, Flavio Dias da Silva, Marcio Henrique de Oliveira Garcia, Leandro Magalhaes de Souza, Cristiane Gomes da Silva, Caio Luiz Pereira Ribeiro, Andrea Cony Cavalcanti, Claudia Maria Braga de Mello, Amilcar Tanuri, Ana Tereza R Vasconcelos |
| EPI_ISL_3245350, EPI_ISL_3245352, EPI_ISL_3245354, EPI_ISL_3245356, EPI_ISL_3245357, EPI_ISL_3245358, EPI_ISL_3245359, EPI_ISL_3245360, EPI_ISL_3245363, EPI_ISL_3245364, EPI_ISL_3245365, EPI_ISL_3245366, EPI_ISL_3245367, EPI_ISL_3245368, EPI_ISL_3245369, EPI_ISL_3245371, EPI_ISL_3245372, EPI_ISL_3245373, EPI_ISL_3245375, EPI_ISL_3245376, EPI_ISL_3245378, EPI_ISL_3245379, EPI_ISL_3245384, EPI_ISL_3245385, EPI_ISL_3245386, EPI_ISL_3245397, EPI_ISL_3245398, EPI_ISL_3245399, EPI_ISL_3245401, EPI_ISL_3245406, EPI_ISL_3245408, EPI_ISL_3245410, EPI_ISL_3245416, EPI_ISL_3245426, EPI_ISL_3245427, EPI_ISL_3245428, EPI_ISL_3245429, EPI_ISL_3245430, EPI_ISL_3245431, EPI_ISL_3245432, EPI_ISL_3245433, EPI_ISL_3245434, EPI_ISL_3245436, EPI_ISL_3245438, EPI_ISL_3245442, EPI_ISL_3245443, EPI_ISL_3245444, EPI_ISL_3245445, EPI_ISL_3245446, EPI_ISL_3245447, EPI_ISL_3245450, EPI_ISL_3245451, EPI_ISL_3245454, EPI_ISL_3245455, EPI_ISL_3245457, EPI_ISL_3245458, EPI_ISL_3245460, EPI_ISL_3245461, EPI_ISL_3245463 | | | |
| see above | Unidade de apoio ao diagnostico da COVID - UNADIG | Bioinformatics Laboratory / LNCC | Luiz G P de Almeida, Alessandra P Lamarca, Ronaldo da Silva F Jr, Liliane Cavalcante, Alexandra L Gerber, Ana Paula de C Guimaraes, Douglas Terra Machado, Erica Ramos dos Santos Nascimento, Fernanda Leitao dos Santos, Ricardo Jose Barbosa Salviano, Cintia Policarpo, Gleidson da Silva de Oliveira, Mario Sergio Ribeiro, Silvia Carvalho, Flavio Dias da Silva, Marcio Henrique de Oliveira Garcia, Leandro Magalhaes de Souza, Cristiane Gomes da Silva, Caio Luiz Pereira Ribeiro, Andrea Cony Cavalcanti, Claudia Maria Braga de Mello, Amilcar Tanuri, Ana Tereza R Vasconcelos |
| EPI_ISL_3245465, EPI_ISL_3245466, EPI_ISL_3245469, EPI_ISL_3245470, EPI_ISL_3245471, EPI_ISL_3245473, EPI_ISL_3245474, EPI_ISL_3245476, EPI_ISL_3245477, EPI_ISL_3245480, EPI_ISL_3245482, EPI_ISL_3245486, EPI_ISL_3245487, EPI_ISL_3245488, EPI_ISL_3245492, EPI_ISL_3245493, EPI_ISL_3245494, EPI_ISL_3245495, EPI_ISL_3245496, EPI_ISL_3245497, EPI_ISL_3245499, EPI_ISL_3245500, EPI_ISL_3245504, EPI_ISL_3245506, EPI_ISL_3245507, EPI_ISL_3245509, EPI_ISL_3245510, EPI_ISL_3245511, EPI_ISL_3245512, EPI_ISL_3245514, EPI_ISL_3245515, EPI_ISL_3245516, EPI_ISL_3245517, EPI_ISL_3245518, EPI_ISL_3245520, EPI_ISL_3245521, EPI_ISL_3245522, EPI_ISL_3245523, EPI_ISL_3245524, EPI_ISL_3245525, EPI_ISL_3245526, EPI_ISL_3245527, EPI_ISL_3245528, EPI_ISL_3245529, EPI_ISL_3245530, EPI_ISL_3245531, EPI_ISL_3245532, EPI_ISL_3245533, EPI_ISL_3245534, EPI_ISL_3245535, EPI_ISL_3245536, EPI_ISL_3245537, EPI_ISL_3245538, EPI_ISL_3245539, EPI_ISL_3245540, EPI_ISL_3245541, EPI_ISL_3245542, EPI_ISL_3245543, EPI_ISL_3245544, EPI_ISL_3245545, EPI_ISL_3245546, EPI_ISL_3245548, EPI_ISL_3245549, EPI_ISL_3245550, EPI_ISL_3245551, EPI_ISL_3245552, EPI_ISL_3245553, EPI_ISL_3245554, EPI_ISL_3245555, EPI_ISL_3245556, EPI_ISL_3245557, EPI_ISL_3245558, EPI_ISL_3245559, EPI_ISL_3245560, EPI_ISL_3245561, EPI_ISL_3245563, EPI_ISL_3245564, EPI_ISL_3245565, EPI_ISL_3245566, EPI_ISL_3245567, EPI_ISL_3245568, EPI_ISL_3245569, EPI_ISL_3245570, EPI_ISL_3245571, EPI_ISL_3245572, EPI_ISL_3245574, EPI_ISL_3245575, EPI_ISL_3245577, EPI_ISL_3245578, EPI_ISL_3245579, EPI_ISL_3245580, EPI_ISL_3245581, EPI_ISL_3245582, EPI_ISL_3245583, EPI_ISL_3245584, EPI_ISL_3245585, EPI_ISL_3245586, EPI_ISL_3245587, EPI_ISL_3245588, EPI_ISL_3245591, EPI_ISL_3245592, EPI_ISL_3245593, EPI_ISL_3245594, EPI_ISL_3245596, EPI_ISL_3245597, EPI_ISL_3245598, EPI_ISL_3245600, EPI_ISL_3245601, EPI_ISL_3245602, EPI_ISL_3245605, EPI_ISL_3245606, EPI_ISL_3245607, EPI_ISL_3245608, EPI_ISL_3245609, EPI_ISL_3245610, EPI_ISL_3245611, EPI_ISL_3245612, EPI_ISL_3245613, EPI_ISL_3245615, EPI_ISL_3245616, EPI_ISL_3245617, EPI_ISL_3245618, EPI_ISL_3245619, EPI_ISL_3245620, EPI_ISL_3245622, EPI_ISL_3245623, EPI_ISL_3245625, EPI_ISL_3245626, EPI_ISL_3245628, EPI_ISL_3245630, EPI_ISL_3245631, EPI_ISL_3245633, EPI_ISL_3245634, EPI_ISL_3245635, EPI_ISL_3245636, EPI_ISL_3245637, EPI_ISL_3245638, EPI_ISL_3245639, EPI_ISL_3245640, EPI_ISL_3245641, EPI_ISL_3245642, EPI_ISL_3245643, EPI_ISL_3245644, EPI_ISL_3245645, EPI_ISL_3245646, EPI_ISL_3245647, EPI_ISL_3245648, EPI_ISL_3245649, EPI_ISL_3245650, EPI_ISL_3245651, EPI_ISL_3245652, EPI_ISL_3245653, EPI_ISL_3245654, EPI_ISL_3245655, EPI_ISL_3245656, EPI_ISL_3245657, EPI_ISL_3245658, EPI_ISL_3245659, EPI_ISL_3245660, EPI_ISL_3245661, EPI_ISL_3245662, EPI_ISL_3245663, EPI_ISL_3245665, EPI_ISL_3245666, EPI_ISL_3245667, EPI_ISL_3245668, EPI_ISL_3245669 | | | |
| see above | NYU Langone Health | Departments of Pathology and Medicine, New York University School of Medicine | Adriana Heguy, Dacia Dimartino, Emily Guzman, Christian Marier, Peter Meyn, Sitharam Ramaswami, Gael Westby, Paul Zappile, Yutong Zhang, Paolo Cotzia, Guiqing Wang |
| EPI_ISL_3245670, EPI_ISL_3245671 | Hospital Universitari Vall d'Hebron - Vall d'Hebron Institut de Recerca | Hospital Universitari Vall d'Hebron - Vall d'Hebron Institut de Recerca | Cristina Andrés, Maria Piñana, Alejandra González-Sánchez, Damir Garcia-Cehic, Ariadna Rando, Juliana Esperalba, Maria Gema Codina, Carla Castillo, Karen García, Rodrigo Vásquez, Maria Carmen Martin, Tomàs Pumarola, Josep Quer, Andrés Antón |
| EPI_ISL_3245700, EPI_ISL_3245701, EPI_ISL_3245702, EPI_ISL_3245703, EPI_ISL_3245704, EPI_ISL_3245705, EPI_ISL_3245706, EPI_ISL_3245707, EPI_ISL_3245708, EPI_ISL_3245709, EPI_ISL_3245710, EPI_ISL_3245711, EPI_ISL_3245712, EPI_ISL_3245713, EPI_ISL_3245714, EPI_ISL_3245715, EPI_ISL_3245716, EPI_ISL_3245717, EPI_ISL_3245718, EPI_ISL_3245719, EPI_ISL_3245720 | | | |
| see above | North Dakota Department of Health, Public Health Laboratory | North Dakota Department of Health, Public Health Laboratory | Lisa Wingerter |
| EPI_ISL_3245742, EPI_ISL_3245743, EPI_ISL_3245744, EPI_ISL_3245745, EPI_ISL_3245746, EPI_ISL_3245748, EPI_ISL_3245749, EPI_ISL_3245750, EPI_ISL_3245751, EPI_ISL_3245752, EPI_ISL_3245754 | | | |
| see above | PathWest Laboratory Medicine WA | PathWest Laboratory Medicine WA Microbial Surveillance Unit | PathWest Laboratory Medicine WA Microbial Surveillance Unit |
| EPI_ISL_3245755, EPI_ISL_3245757, EPI_ISL_3245758, EPI_ISL_3245760, EPI_ISL_3245761, EPI_ISL_3245762, EPI_ISL_3245764, EPI_ISL_3245765, EPI_ISL_3245766, EPI_ISL_3245767, EPI_ISL_3245768, EPI_ISL_3245769, EPI_ISL_3245770 | | | |
| see above | Department of Microbiology, Yamanashi Institute for Public Health and Environment | Genome Analysis Center, Yamanashi Central Hospital | Yosuke Hirotsu |
| EPI_ISL_3245772, EPI_ISL_3245773, EPI_ISL_3245774, EPI_ISL_3245775, EPI_ISL_3245776, EPI_ISL_3245777, EPI_ISL_3245778, EPI_ISL_3245779, EPI_ISL_3245780, EPI_ISL_3245781, EPI_ISL_3245782, EPI_ISL_3245783, EPI_ISL_3245784, EPI_ISL_3245785 | | | |
| see above | Labo Analyses Med | National Reference Center for Viruses of Respiratory Infections, Institut Pasteur, Paris | Marion Barbet, Sylvie Behillil, Méline Bizard, Angela Brisebarre, Camille Capel, Vincent Enouf, Louise Lefrançois, Frédéric Lemoine, Christophe Malabat, Corinne Maufrais, Hub de Bioinformatique et Biostatistique, Etienne Simon-Lorière, Maud Vanpeene, Sylvie Van der Werf ,Vanessa Magin |
| EPI_ISL_3245786 | Hospital | National Reference Center for Viruses of Respiratory Infections, Institut Pasteur, Paris | Marion Barbet, Sylvie Behillil, Méline Bizard, Angela Brisebarre, Camille Capel, Vincent Enouf, Louise Lefrançois, Frédéric Lemoine, Christophe Malabat, Corinne Maufrais, Hub de Bioinformatique et Biostatistique, Etienne Simon-Lorière, Maud Vanpeene, Sylvie Van der Werf ,Anne-CéCile Hochart |
| EPI_ISL_3245787 | Research | National Reference Center for Viruses of Respiratory Infections, Institut Pasteur, Paris | Marion Barbet, Sylvie Behillil, Méline Bizard, Angela Brisebarre, Camille Capel, Vincent Enouf, Louise Lefrançois, Frédéric Lemoine, Christophe Malabat, Corinne Maufrais, Hub de Bioinformatique et Biostatistique, Etienne Simon-Lorière, Maud Vanpeene, Sylvie Van der Werf ,Marianne Leruez-Ville |
| EPI_ISL_3245788 | Hospital | National Reference Center for Viruses of Respiratory Infections, Institut Pasteur, Paris | Marion Barbet, Sylvie Behillil, Méline Bizard, Angela Brisebarre, Camille Capel, Vincent Enouf, Louise Lefrançois, Frédéric Lemoine, Christophe Malabat, Corinne Maufrais, Hub de Bioinformatique et Biostatistique, Etienne Simon-Lorière, Maud Vanpeene, Sylvie Van der Werf ,Guy Galeazzi |
| EPI_ISL_3245789, EPI_ISL_3245790, EPI_ISL_3245792, EPI_ISL_3245793 | Labo Analyses Med | National Reference Center for Viruses of Respiratory Infections, Institut Pasteur, Paris | Marion Barbet, Sylvie Behillil, Méline Bizard, Angela Brisebarre, Camille Capel, Vincent Enouf, Louise Lefrançois, Frédéric Lemoine, Christophe Malabat, Corinne Maufrais, Hub de Bioinformatique et Biostatistique, Etienne Simon-Lorière, Maud Vanpeene, Sylvie Van der Werf ,ValéRie Weber |
| EPI_ISL_3245810, EPI_ISL_3245812 | Outre mer | National Reference Center for Viruses of Respiratory Infections, Institut Pasteur, Paris | Marion Barbet, Sylvie Behillil, Méline Bizard, Angela Brisebarre, Camille Capel, Vincent Enouf, Louise Lefrançois, Frédéric Lemoine, Christophe Malabat, Corinne Maufrais, Hub de Bioinformatique et Biostatistique, Etienne Simon-Lorière, Maud Vanpeene, Sylvie Van der Werf ,Marie-HéLèNe Glaudon Louveau De La Guigneraye |
| EPI_ISL_3245820, EPI_ISL_3245821, EPI_ISL_3245822, EPI_ISL_3245823, EPI_ISL_3245824, EPI_ISL_3245825, EPI_ISL_3245826, EPI_ISL_3245827, EPI_ISL_3245828, EPI_ISL_3245829, EPI_ISL_3245830, EPI_ISL_3245831, EPI_ISL_3245832, EPI_ISL_3245833, EPI_ISL_3245834, EPI_ISL_3245835, EPI_ISL_3245836, EPI_ISL_3245837, EPI_ISL_3245838, EPI_ISL_3245839, EPI_ISL_3245840, EPI_ISL_3245841, EPI_ISL_3245842, EPI_ISL_3245843, EPI_ISL_3245844, EPI_ISL_3245845, EPI_ISL_3245846 | | | |
| see above | Hospital | National Reference Center for Viruses of Respiratory Infections, Institut Pasteur, Paris | Marion Barbet, Sylvie Behillil, Méline Bizard, Angela Brisebarre, Camille Capel, Vincent Enouf, Louise Lefrançois, Frédéric Lemoine, Christophe Malabat, Corinne Maufrais, Hub de Bioinformatique et Biostatistique, Etienne Simon-Lorière, Maud Vanpeene, Sylvie Van der Werf ,JéRome Guinard |
| EPI_ISL_3245847 | Hospital | National Reference Center for Viruses of Respiratory Infections, Institut Pasteur, Paris | Marion Barbet, Sylvie Behillil, Méline Bizard, Angela Brisebarre, Camille Capel, Vincent Enouf, Louise Lefrançois, Frédéric Lemoine, Christophe Malabat, Corinne Maufrais, Hub de Bioinformatique et Biostatistique, Etienne Simon-Lorière, Maud Vanpeene, Sylvie Van der Werf ,Catherine FranÇOis |
| EPI_ISL_3245848 | Hospital | National Reference Center for Viruses of Respiratory Infections, Institut Pasteur, Paris | Marion Barbet, Sylvie Behillil, Méline Bizard, Angela Brisebarre, Camille Capel, Vincent Enouf, Louise Lefrançois, Frédéric Lemoine, Christophe Malabat, Corinne Maufrais, Hub de Bioinformatique et Biostatistique, Etienne Simon-Lorière, Maud Vanpeene, Sylvie Van der Werf ,Pierre Alain Billy |
| EPI_ISL_3245849 | Hospital | National Reference Center for Viruses of Respiratory Infections, Institut Pasteur, Paris | Marion Barbet, Sylvie Behillil, Méline Bizard, Angela Brisebarre, Camille Capel, Vincent Enouf, Louise Lefrançois, Frédéric Lemoine, Christophe Malabat, Corinne Maufrais, Hub de Bioinformatique et Biostatistique, Etienne Simon-Lorière, Maud Vanpeene, Sylvie Van der Werf ,Pascale Martres |
| EPI_ISL_3245851 | Hospital | National Reference Center for Viruses of Respiratory Infections, Institut Pasteur, Paris | Marion Barbet, Sylvie Behillil, Méline Bizard, Angela Brisebarre, Camille Capel, Vincent Enouf, Louise Lefrançois, Frédéric Lemoine, Christophe Malabat, Corinne Maufrais, Hub de Bioinformatique et Biostatistique, Etienne Simon-Lorière, Maud Vanpeene, Sylvie Van der Werf ,ValéRie Serazin |
| EPI_ISL_3245852, EPI_ISL_3245853, EPI_ISL_3245854 | Hospital | National Reference Center for Viruses of Respiratory Infections, Institut Pasteur, Paris | Marion Barbet, Sylvie Behillil, Méline Bizard, Angela Brisebarre, Camille Capel, Vincent Enouf, Louise Lefrançois, Frédéric Lemoine, Christophe Malabat, Corinne Maufrais, Hub de Bioinformatique et Biostatistique, Etienne Simon-Lorière, Maud Vanpeene, Sylvie Van der Werf ,StéPhanie Marque Juillet |
| EPI_ISL_3245855, EPI_ISL_3245856, EPI_ISL_3245857, EPI_ISL_3245858, EPI_ISL_3245859 | Armies | National Reference Center for Viruses of Respiratory Infections, Institut Pasteur, Paris | Marion Barbet, Sylvie Behillil, Méline Bizard, Angela Brisebarre, Camille Capel, Vincent Enouf, Louise Lefrançois, Frédéric Lemoine, Christophe Malabat, Corinne Maufrais, Hub de Bioinformatique et Biostatistique, Etienne Simon-Lorière, Maud Vanpeene, Sylvie Van der Werf ,Marine Desroches |
| EPI_ISL_3245860, EPI_ISL_3245861, EPI_ISL_3245862, EPI_ISL_3245863, EPI_ISL_3245865, EPI_ISL_3245867 | Hospital | National Reference Center for Viruses of Respiratory Infections, Institut Pasteur, Paris | Marion Barbet, Sylvie Behillil, Méline Bizard, Angela Brisebarre, Camille Capel, Vincent Enouf, Louise Lefrançois, Frédéric Lemoine, Christophe Malabat, Corinne Maufrais, Hub de Bioinformatique et Biostatistique, Etienne Simon-Lorière, Maud Vanpeene, Sylvie Van der Werf ,JéRome Guinard |
| EPI_ISL_3245868 | Hospital | National Reference Center for Viruses of Respiratory Infections, Institut Pasteur, Paris | Marion Barbet, Sylvie Behillil, Méline Bizard, Angela Brisebarre, Camille Capel, Vincent Enouf, Louise Lefrançois, Frédéric Lemoine, Christophe Malabat, Corinne Maufrais, Hub de Bioinformatique et Biostatistique, Etienne Simon-Lorière, Maud Vanpeene, Sylvie Van der Werf ,Guy Galeazzi |
| EPI_ISL_3245869, EPI_ISL_3245870, EPI_ISL_3245873, EPI_ISL_3245874, EPI_ISL_3245875, | Labo Analyses Med | National Reference Center for Viruses of Respiratory Infections, Institut Pasteur, Paris | Marion Barbet, Sylvie Behillil, Méline Bizard, Angela Brisebarre, Camille Capel, Vincent Enouf, Louise Lefrançois, Frédéric Lemoine, Christophe Malabat, Corinne Maufrais, Hub de Bioinformatique et Biostatistique, Etienne Simon-Lorière, Maud Vanpeene, Sylvie Van der Werf ,Philippe Miara |

| | | | |
|---|---|---|---|
| EPI_ISL_3245876, EPI_ISL_3245877 | | | |
| EPI_ISL_3245896, EPI_ISL_3245897, EPI_ISL_3245898, EPI_ISL_3245899, EPI_ISL_3245900, EPI_ISL_3245903, EPI_ISL_3245904, EPI_ISL_3245905, EPI_ISL_3245906, EPI_ISL_3245907, EPI_ISL_3245909, EPI_ISL_3245910, EPI_ISL_3245911, EPI_ISL_3245912, EPI_ISL_3245913, EPI_ISL_3245914, EPI_ISL_3245915, EPI_ISL_3245916, EPI_ISL_3245917, EPI_ISL_3245918, EPI_ISL_3245919, EPI_ISL_3245920, EPI_ISL_3245921, EPI_ISL_3245922, EPI_ISL_3245923, EPI_ISL_3245924, EPI_ISL_3245925, EPI_ISL_3245927, EPI_ISL_3245928, EPI_ISL_3245929, EPI_ISL_3245930, EPI_ISL_3245931, EPI_ISL_3245932, EPI_ISL_3245933 | | | |
| see above | North Dakota Department of Health, Public Health Laboratory | North Dakota Department of Health, Public Health Laboratory | Lisa Wingerter |
| EPI_ISL_3245940 | Outre mer | National Reference Center for Viruses of Respiratory Infections, Institut Pasteur, Paris | Marion Barbet, Sylvie Behillil, Méline Bizard, Angela Brisebarre, Camille Capel, Vincent Enouf, Louise Lefrançois, Frédéric Lemoine, Christophe Malabat, Corinne Maufrais, Hub de bioinformatique et biostatistique, Etienne Simon-Lorière, Maud Vanpeene, Sylvie Van der Werf ,FrançOis Nestour |
| EPI_ISL_3245949, EPI_ISL_3245950, EPI_ISL_3245951, EPI_ISL_3245952, EPI_ISL_3245953 | Labo Analyses Med | National Reference Center for Viruses of Respiratory Infections, Institut Pasteur, Paris | Marion Barbet, Sylvie Behillil, Méline Bizard, Angela Brisebarre, Camille Capel, Vincent Enouf, Louise Lefrançois, Frédéric Lemoine, Christophe Malabat, Corinne Maufrais, Hub de bioinformatique et biostatistique, Etienne Simon-Lorière, Maud Vanpeene, Sylvie Van der Werf ,Gilles Abs |
| EPI_ISL_3245954, EPI_ISL_3245955, EPI_ISL_3245956, EPI_ISL_3245957, EPI_ISL_3245958, EPI_ISL_3245959 | Hospital | National Reference Center for Viruses of Respiratory Infections, Institut Pasteur, Paris | Marion Barbet, Sylvie Behillil, Méline Bizard, Angela Brisebarre, Camille Capel, Vincent Enouf, Louise Lefrançois, Frédéric Lemoine, Christophe Malabat, Corinne Maufrais, Hub de bioinformatique et biostatistique, Etienne Simon-Lorière, Maud Vanpeene, Sylvie Van der Werf ,Bruno Berdin |
| EPI_ISL_3245960 | Hospital | National Reference Center for Viruses of Respiratory Infections, Institut Pasteur, Paris | Marion Barbet, Sylvie Behillil, Méline Bizard, Angela Brisebarre, Camille Capel, Vincent Enouf, Louise Lefrançois, Frédéric Lemoine, Christophe Malabat, Corinne Maufrais, Hub de bioinformatique et biostatistique, Etienne Simon-Lorière, Maud Vanpeene, Sylvie Van der Werf ,Laurence Louvet |
| EPI_ISL_3245961, EPI_ISL_3245962 | Hospital | National Reference Center for Viruses of Respiratory Infections, Institut Pasteur, Paris | Marion Barbet, Sylvie Behillil, Méline Bizard, Angela Brisebarre, Camille Capel, Vincent Enouf, Louise Lefrançois, Frédéric Lemoine, Christophe Malabat, Corinne Maufrais, Hub de bioinformatique et biostatistique, Etienne Simon-Lorière, Maud Vanpeene, Sylvie Van der Werf ,ValéRie Serazin |
| EPI_ISL_3245963, EPI_ISL_3245964, EPI_ISL_3245965, EPI_ISL_3245966 | Hospital | National Reference Center for Viruses of Respiratory Infections, Institut Pasteur, Paris | Marion Barbet, Sylvie Behillil, Méline Bizard, Angela Brisebarre, Camille Capel, Vincent Enouf, Louise Lefrançois, Frédéric Lemoine, Christophe Malabat, Corinne Maufrais, Hub de bioinformatique et biostatistique, Etienne Simon-Lorière, Maud Vanpeene, Sylvie Van der Werf ,Pascale Martres |
| EPI_ISL_3245969, EPI_ISL_3245970 | Labo Analyses Med | National Reference Center for Viruses of Respiratory Infections, Institut Pasteur, Paris | Marion Barbet, Sylvie Behillil, Méline Bizard, Angela Brisebarre, Camille Capel, Vincent Enouf, Louise Lefrançois, Frédéric Lemoine, Christophe Malabat, Corinne Maufrais, Hub de bioinformatique et biostatistique, Etienne Simon-Lorière, Maud Vanpeene, Sylvie Van der Werf ,Philippe Miara |
| EPI_ISL_3245971, EPI_ISL_3245973, EPI_ISL_3245974, EPI_ISL_3245975, EPI_ISL_3245976, EPI_ISL_3245977 | Labo Analyses Med | National Reference Center for Viruses of Respiratory Infections, Institut Pasteur, Paris | Marion Barbet, Sylvie Behillil, Méline Bizard, Angela Brisebarre, Camille Capel, Vincent Enouf, Louise Lefrançois, Frédéric Lemoine, Christophe Malabat, Corinne Maufrais, Hub de bioinformatique et biostatistique, Etienne Simon-Lorière, Maud Vanpeene, Sylvie Van der Werf ,BéNéDicte Roquebert |
| EPI_ISL_3245978, EPI_ISL_3245979, EPI_ISL_3245980, EPI_ISL_3245981, EPI_ISL_3245982, EPI_ISL_3245983, EPI_ISL_3245984, EPI_ISL_3245985, EPI_ISL_3245986, EPI_ISL_3245987, EPI_ISL_3245988, EPI_ISL_3245989, EPI_ISL_3245990, EPI_ISL_3245991, EPI_ISL_3245992, EPI_ISL_3245993, EPI_ISL_3245994, EPI_ISL_3245995, EPI_ISL_3245996, EPI_ISL_3245997, EPI_ISL_3245998, EPI_ISL_3245999, EPI_ISL_3246000, EPI_ISL_3246001, EPI_ISL_3246002, EPI_ISL_3246003, EPI_ISL_3246004, EPI_ISL_3246005, EPI_ISL_3246006, EPI_ISL_3246007, EPI_ISL_3246008 | | | |
| see above | Labo Analyses Med | National Reference Center for Viruses of Respiratory Infections, Institut Pasteur, Paris | Marion Barbet, Sylvie Behillil, Méline Bizard, Angela Brisebarre, Camille Capel, Vincent Enouf, Louise Lefrançois, Frédéric Lemoine, Christophe Malabat, Corinne Maufrais, Hub de bioinformatique et biostatistique, Etienne Simon-Lorière, Maud Vanpeene, Sylvie Van der Werf ,GréGoire Potiron |
| EPI_ISL_3246010, EPI_ISL_3246020, EPI_ISL_3246024 | Arizona State University | Arizona State University | Ajeet Bains, LaRinda A. Holland, Matthew F. Smith, Peter T. Skidmore, Rabia Maqsood, Nicholas J. Mellor, Nathaniel Johnson, Joshua LaBaer, Vel Murugan, Efrem S. Lim |
| EPI_ISL_3246035, EPI_ISL_3246036, EPI_ISL_3246037, EPI_ISL_3246038, EPI_ISL_3246039, EPI_ISL_3246040, EPI_ISL_3246041, EPI_ISL_3246042, EPI_ISL_3246043, EPI_ISL_3246044, EPI_ISL_3246045, EPI_ISL_3246046, EPI_ISL_3246047 | | | |
| see above | Myanmar | National Institute of Health, Department of Medical Sciences, Ministry of Public Health, Thailand | Pilailuk Okada; Siripaporn Phuygun; Nuttida Thongpramul; Sittiporn Parnmen; Ratana Tacharoenmuang; Pakorn Piromtong; Natchaya Khiadsang; Thanutsapa Thanadachakul; Warawan Wongboot; sirikanda wimol; Sunthareeya Waicharoen; |
| EPI_ISL_3246049 | Labo Analyses Med | National Reference Center for Viruses of Respiratory Infections, Institut Pasteur, Paris | Marion Barbet, Sylvie Behillil, Méline Bizard, Angela Brisebarre, Camille Capel, Vincent Enouf, Louise Lefrançois, Frédéric Lemoine, Christophe Malabat, Corinne Maufrais, Hub de Bioinformatique et Biostatistique, Etienne Simon-Lorière, Maud Vanpeene, Sylvie Van der Werf ,Marie-HéLèNe Glaudon Louveau De La Guigneraye |
| EPI_ISL_3246050 | Hospital | National Reference Center for Viruses of Respiratory Infections, Institut Pasteur, Paris | Marion Barbet, Sylvie Behillil, Méline Bizard, Angela Brisebarre, Camille Capel, Vincent Enouf, Louise Lefrançois, Frédéric Lemoine, Christophe Malabat, Corinne Maufrais, Hub de Bioinformatique et Biostatistique, Etienne Simon-Lorière, Maud Vanpeene, Sylvie Van der Werf ,Marie-HéLèNe Glaudon Louveau De La Guigneraye |
| EPI_ISL_3246051 | Monterey County Public Health Laboratory | Monterey County Public Health Laboratory | Monterey County Public Health Laboratory |
| EPI_ISL_3246073, EPI_ISL_3246081, EPI_ISL_3246082 | Wyoming Public Health Laboratory | Wyoming Public Health Laboratory | Jim Mildenberger, Taylor Fearing, Channing Weber, Ashley Norberg, Chayse Rowley, Marley Goetz, Brian Dominguez, Elliot Thomasson, Sam Britz, Cari Sloma, Robert Petit, and Rob Christensen |
| EPI_ISL_3246085, EPI_ISL_3246086 | Hospital | National Reference Center for Viruses of Respiratory Infections, Institut Pasteur, Paris | Marion Barbet, Sylvie Behillil, Méline Bizard, Angela Brisebarre, Camille Capel, Vincent Enouf, Louise Lefrançois, Frédéric Lemoine, Christophe Malabat, Corinne Maufrais, Hub de Bioinformatique et Statistique, Etienne Simon-Lorière, Maud Vanpeene, Sylvie Van der Werf ,Guillaume Aubin (G) |
| EPI_ISL_3246088, EPI_ISL_3246090 | Monterey County Public Health Laboratory | Monterey County Public Health Laboratory | Monterey County Public Health Laboratory |
| EPI_ISL_3246091, EPI_ISL_3246092, EPI_ISL_3246093, EPI_ISL_3246094, EPI_ISL_3246095, EPI_ISL_3246096, EPI_ISL_3246097 | AFIP | Instituto Butantan | Dimas Tadeu Covas, Antonio Jorge Martins, Claudia Renata dos Santos Barros, David Schlesinger, Debora Botequio Moretti, Elaine Cristina Marqueze, Elaine Vieira Santos, Evandra Strazza Rodrigues, Heidge Fukumasu, Jayme Augusto de Souza-Neto, José Salvatore Leister Patané, Luiz Alcantara, Luiz Lehmann Coutinho, Maria Carolina Elias, Maurício Lacerda Nogueira, Rafael dos Santos Bezerra, Raul Machado Neto, Rejane Maria Tommasini Grotto, Ricardo Haddad, Sandra Coccuzzo Sampaio Vessoni, Simone Kashima, Svetoslav Nanev Slavov, Vincent Louis Viala |
| EPI_ISL_3246098, EPI_ISL_3246099, EPI_ISL_3246100, EPI_ISL_3246101, EPI_ISL_3246102 | BIOFAST | Instituto Butantan | Dimas Tadeu Covas, Antonio Jorge Martins, Claudia Renata dos Santos Barros, David Schlesinger, Debora Botequio Moretti, Elaine Cristina Marqueze, Elaine Vieira Santos, Evandra Strazza Rodrigues, Heidge Fukumasu, Jayme Augusto de Souza-Neto, José Salvatore Leister Patané, Luiz Alcantara, Luiz Lehmann Coutinho, Maria Carolina Elias, Maurício Lacerda Nogueira, Rafael dos Santos Bezerra, Raul Machado Neto, Rejane Maria Tommasini Grotto, Ricardo Haddad, Sandra Coccuzzo Sampaio Vessoni, Simone Kashima, Svetoslav Nanev Slavov, Vincent Louis Viala |
| EPI_ISL_3246103 | FACULDADE DE MEDICINA ABC | Instituto Butantan | Dimas Tadeu Covas, Antonio Jorge Martins, Claudia Renata dos Santos Barros, David Schlesinger, Debora Botequio Moretti, Elaine Cristina Marqueze, Elaine Vieira Santos, Evandra Strazza Rodrigues, Heidge Fukumasu, Jayme Augusto de Souza-Neto, José Salvatore Leister Patané, Luiz Alcantara, Luiz Lehmann Coutinho, Maria Carolina Elias, Maurício Lacerda Nogueira, Rafael dos Santos Bezerra, Raul Machado Neto, Rejane Maria Tommasini Grotto, Ricardo Haddad, Sandra Coccuzzo Sampaio Vessoni, Simone Kashima, Svetoslav Nanev Slavov, Vincent Louis Viala |
| EPI_ISL_3246104, EPI_ISL_3246105, EPI_ISL_3246106, EPI_ISL_3246107, EPI_ISL_3246108, EPI_ISL_3246109, EPI_ISL_3246110, EPI_ISL_3246111, EPI_ISL_3246112, EPI_ISL_3246113, EPI_ISL_3246114, EPI_ISL_3246115, EPI_ISL_3246116, EPI_ISL_3246117, EPI_ISL_3246118, EPI_ISL_3246119, EPI_ISL_3246120 | | | |
| see above | INSIDE DIAGNÓSTICOS | Instituto Butantan | Dimas Tadeu Covas, Antonio Jorge Martins, Claudia Renata dos Santos Barros, David Schlesinger, Debora Botequio Moretti, Elaine Cristina Marqueze, Elaine Vieira Santos, Evandra Strazza Rodrigues, Heidge Fukumasu, Jayme Augusto de Souza-Neto, José Salvatore Leister Patané, Luiz Alcantara, Luiz Lehmann Coutinho, Maria Carolina Elias, Maurício Lacerda Nogueira, Rafael dos Santos Bezerra, Raul Machado Neto, Rejane Maria Tommasini Grotto, Ricardo Haddad, Sandra Coccuzzo Sampaio Vessoni, Simone Kashima, Svetoslav Nanev Slavov, Vincent Louis Viala |
| EPI_ISL_3246121 | HOSPITAL GERAL JESUS TEIXEIRA DA COSTA GUAIANASES SAO PAULO | Instituto Butantan | Dimas Tadeu Covas, Antonio Jorge Martins, Claudia Renata dos Santos Barros, David Schlesinger, Debora Botequio Moretti, Elaine Cristina Marqueze, Elaine Vieira Santos, Evandra Strazza Rodrigues, Heidge Fukumasu, Jayme Augusto de Souza-Neto, José Salvatore Leister Patané, Luiz Alcantara, Luiz Lehmann Coutinho, Maria Carolina Elias, Maurício Lacerda Nogueira, Rafael dos Santos Bezerra, Raul Machado Neto, Rejane Maria Tommasini Grotto, Ricardo Haddad, Sandra Coccuzzo Sampaio Vessoni, Simone Kashima, Svetoslav Nanev Slavov, Vincent Louis Viala |
| EPI_ISL_3246122 | HOSPITAL DE CAMPANHA COVID 19 CAIEIRAS | Instituto Butantan | Dimas Tadeu Covas, Antonio Jorge Martins, Claudia Renata dos Santos Barros, David Schlesinger, Debora Botequio Moretti, Elaine Cristina Marqueze, Elaine Vieira Santos, Evandra Strazza Rodrigues, Heidge Fukumasu, Jayme Augusto de Souza-Neto, José Salvatore Leister Patané, Luiz Alcantara, Luiz Lehmann Coutinho, Maria Carolina Elias, Maurício Lacerda Nogueira, Rafael dos Santos Bezerra, Raul Machado Neto, Rejane Maria Tommasini Grotto, Ricardo Haddad, Sandra Coccuzzo Sampaio Vessoni, Simone Kashima, Svetoslav Nanev Slavov, Vincent Louis Viala |
| EPI_ISL_3246123, EPI_ISL_3246124 | UPA DR AKIRA TADA | Instituto Butantan | Dimas Tadeu Covas, Antonio Jorge Martins, Claudia Renata dos Santos Barros, David Schlesinger, Debora Botequio Moretti, Elaine Cristina Marqueze, Elaine Vieira Santos, Evandra Strazza Rodrigues, Heidge Fukumasu, Jayme Augusto de Souza-Neto, José Salvatore Leister Patané, Luiz Alcantara, Luiz Lehmann Coutinho, Maria Carolina Elias, Maurício Lacerda Nogueira, Rafael dos Santos Bezerra, Raul Machado Neto, Rejane Maria Tommasini Grotto, Ricardo Haddad, Sandra Coccuzzo Sampaio Vessoni, Simone Kashima, Svetoslav Nanev Slavov, Vincent Louis Viala |
| EPI_ISL_3246125, EPI_ISL_3246126, EPI_ISL_3246127, EPI_ISL_3246128, EPI_ISL_3246129, EPI_ISL_3246130, EPI_ISL_3246131 | VIGILANCIA SANITARIA E VIG EPIDEMIOLOGICA DE ITAPEVI | Instituto Butantan | Dimas Tadeu Covas, Antonio Jorge Martins, Claudia Renata dos Santos Barros, David Schlesinger, Debora Botequio Moretti, Elaine Cristina Marqueze, Elaine Vieira Santos, Evandra Strazza Rodrigues, Heidge Fukumasu, Jayme Augusto de Souza-Neto, José Salvatore Leister Patané, Luiz Alcantara, Luiz Lehmann Coutinho, Maria Carolina Elias, Maurício Lacerda Nogueira, Rafael dos Santos Bezerra, Raul Machado Neto, Rejane Maria Tommasini Grotto, Ricardo Haddad, Sandra Coccuzzo Sampaio Vessoni, Simone Kashima, Svetoslav Nanev Slavov, Vincent Louis Viala |
| EPI_ISL_3246132 | UPA DR FRANCO DA ROCHA | Instituto Butantan | Dimas Tadeu Covas, Antonio Jorge Martins, Claudia Renata dos Santos Barros, David Schlesinger, Debora Botequio Moretti, Elaine Cristina Marqueze, Elaine Vieira Santos, Evandra Strazza Rodrigues, Heidge Fukumasu, Jayme Augusto de Souza-Neto, José Salvatore Leister Patané, Luiz Alcantara, Luiz Lehmann Coutinho, Maria Carolina Elias, Maurício Lacerda Nogueira, Rafael dos Santos Bezerra, Raul Machado Neto, Rejane Maria Tommasini Grotto, Ricardo Haddad, Sandra Coccuzzo Sampaio Vessoni, Simone Kashima, Svetoslav Nanev Slavov, Vincent Louis Viala |
| EPI_ISL_3246133, EPI_ISL_3246134 | HOSPITAL MUNICIPAL DE SAO VICENTE | Instituto Butantan | Dimas Tadeu Covas, Antonio Jorge Martins, Claudia Renata dos Santos Barros, David Schlesinger, Debora Botequio Moretti, Elaine Cristina Marqueze, Elaine Vieira Santos, Evandra Strazza Rodrigues, Heidge Fukumasu, Jayme Augusto de Souza-Neto, José Salvatore Leister Patané, Luiz Alcantara, Luiz Lehmann Coutinho, Maria Carolina Elias, Maurício Lacerda Nogueira, Rafael dos Santos Bezerra, Raul Machado Neto, Rejane Maria Tommasini Grotto, Ricardo Haddad, Sandra |

| | | | |
|---|---|---|---|
| | | | Coccuzzo Sampaio Vessoni, Simone Kashima, Svetoslav Nanev Slavov, Vincent Louis Viala |
| EPI_ISL_3246135 | SECAO CENTRO DE DIAGNOSTICO SECEDI | Instituto Butantan | Dimas Tadeu Covas, Antonio Jorge Martins, Claudia Renata dos Santos Barros, David Schlesinger, Debora Botequio Moretti, Elaine Cristina Marqueze, Elaine Vieira Santos, Evandra Strazza Rodrigues, Heidge Fukumasu, Jayme Augusto de Souza-Neto, José Salvatore Leister Patané, Luiz Alcantara, Luiz Lehmann Coutinho, Maria Carolina Elias, Maurício Lacerda Nogueira, Rafael dos Santos Bezerra, Raul Machado Neto, Rejane Maria Tommasini Grotto, Ricardo Haddad, Sandra Coccuzzo Sampaio Vessoni, Simone Kashima, Svetoslav Nanev Slavov, Vincent Louis Viala |
| EPI_ISL_3246136, EPI_ISL_3246137 | CENTRO DE SAUDE DE IBIRAREMA | Instituto Butantan | Dimas Tadeu Covas, Antonio Jorge Martins, Claudia Renata dos Santos Barros, David Schlesinger, Debora Botequio Moretti, Elaine Cristina Marqueze, Elaine Vieira Santos, Evandra Strazza Rodrigues, Heidge Fukumasu, Jayme Augusto de Souza-Neto, José Salvatore Leister Patané, Luiz Alcantara, Luiz Lehmann Coutinho, Maria Carolina Elias, Maurício Lacerda Nogueira, Rafael dos Santos Bezerra, Raul Machado Neto, Rejane Maria Tommasini Grotto, Ricardo Haddad, Sandra Coccuzzo Sampaio Vessoni, Simone Kashima, Svetoslav Nanev Slavov, Vincent Louis Viala |
| EPI_ISL_3246138 | CENTRO DE ESPECIALIDDS MEDICAS IRMA LEOPOLDINA PIRASSUNUNGA | Instituto Butantan | Dimas Tadeu Covas, Antonio Jorge Martins, Claudia Renata dos Santos Barros, David Schlesinger, Debora Botequio Moretti, Elaine Cristina Marqueze, Elaine Vieira Santos, Evandra Strazza Rodrigues, Heidge Fukumasu, Jayme Augusto de Souza-Neto, José Salvatore Leister Patané, Luiz Alcantara, Luiz Lehmann Coutinho, Maria Carolina Elias, Maurício Lacerda Nogueira, Rafael dos Santos Bezerra, Raul Machado Neto, Rejane Maria Tommasini Grotto, Ricardo Haddad, Sandra Coccuzzo Sampaio Vessoni, Simone Kashima, Svetoslav Nanev Slavov, Vincent Louis Viala |
| EPI_ISL_3246139 | HOSPITAL DE CAMPANHA COVID 19 MUNICIPIO DE TAUBATE | Instituto Butantan | Dimas Tadeu Covas, Antonio Jorge Martins, Claudia Renata dos Santos Barros, David Schlesinger, Debora Botequio Moretti, Elaine Cristina Marqueze, Elaine Vieira Santos, Evandra Strazza Rodrigues, Heidge Fukumasu, Jayme Augusto de Souza-Neto, José Salvatore Leister Patané, Luiz Alcantara, Luiz Lehmann Coutinho, Maria Carolina Elias, Maurício Lacerda Nogueira, Rafael dos Santos Bezerra, Raul Machado Neto, Rejane Maria Tommasini Grotto, Ricardo Haddad, Sandra Coccuzzo Sampaio Vessoni, Simone Kashima, Svetoslav Nanev Slavov, Vincent Louis Viala |
| EPI_ISL_3246140 | UNIDADE DE PRONTO ATENDIMENTO CENTRAL DE CARAGUATATUBA | Instituto Butantan | Dimas Tadeu Covas, Antonio Jorge Martins, Claudia Renata dos Santos Barros, David Schlesinger, Debora Botequio Moretti, Elaine Cristina Marqueze, Elaine Vieira Santos, Evandra Strazza Rodrigues, Heidge Fukumasu, Jayme Augusto de Souza-Neto, José Salvatore Leister Patané, Luiz Alcantara, Luiz Lehmann Coutinho, Maria Carolina Elias, Maurício Lacerda Nogueira, Rafael dos Santos Bezerra, Raul Machado Neto, Rejane Maria Tommasini Grotto, Ricardo Haddad, Sandra Coccuzzo Sampaio Vessoni, Simone Kashima, Svetoslav Nanev Slavov, Vincent Louis Viala |
| EPI_ISL_3246141, EPI_ISL_3246142 | CENTRO DE APOIO EPIDEMIOLOGICO | Instituto Butantan | Dimas Tadeu Covas, Antonio Jorge Martins, Claudia Renata dos Santos Barros, David Schlesinger, Debora Botequio Moretti, Elaine Cristina Marqueze, Elaine Vieira Santos, Evandra Strazza Rodrigues, Heidge Fukumasu, Jayme Augusto de Souza-Neto, José Salvatore Leister Patané, Luiz Alcantara, Luiz Lehmann Coutinho, Maria Carolina Elias, Maurício Lacerda Nogueira, Rafael dos Santos Bezerra, Raul Machado Neto, Rejane Maria Tommasini Grotto, Ricardo Haddad, Sandra Coccuzzo Sampaio Vessoni, Simone Kashima, Svetoslav Nanev Slavov, Vincent Louis Viala |
| EPI_ISL_3246143 | UNIDADE BASICA DE SAUDE DE POTIM | Instituto Butantan | Dimas Tadeu Covas, Antonio Jorge Martins, Claudia Renata dos Santos Barros, David Schlesinger, Debora Botequio Moretti, Elaine Cristina Marqueze, Elaine Vieira Santos, Evandra Strazza Rodrigues, Heidge Fukumasu, Jayme Augusto de Souza-Neto, José Salvatore Leister Patané, Luiz Alcantara, Luiz Lehmann Coutinho, Maria Carolina Elias, Maurício Lacerda Nogueira, Rafael dos Santos Bezerra, Raul Machado Neto, Rejane Maria Tommasini Grotto, Ricardo Haddad, Sandra Coccuzzo Sampaio Vessoni, Simone Kashima, Svetoslav Nanev Slavov, Vincent Louis Viala |
| EPI_ISL_3246144 | HOSPITAL MUNICIPAL DR JOSE DE CARVALHO FLORENCE | Instituto Butantan | Dimas Tadeu Covas, Antonio Jorge Martins, Claudia Renata dos Santos Barros, David Schlesinger, Debora Botequio Moretti, Elaine Cristina Marqueze, Elaine Vieira Santos, Evandra Strazza Rodrigues, Heidge Fukumasu, Jayme Augusto de Souza-Neto, José Salvatore Leister Patané, Luiz Alcantara, Luiz Lehmann Coutinho, Maria Carolina Elias, Maurício Lacerda Nogueira, Rafael dos Santos Bezerra, Raul Machado Neto, Rejane Maria Tommasini Grotto, Ricardo Haddad, Sandra Coccuzzo Sampaio Vessoni, Simone Kashima, Svetoslav Nanev Slavov, Vincent Louis Viala |
| EPI_ISL_3246145, EPI_ISL_3246146 | SECRETARIA MUNICIPAL DA SAUDE DE GUARATINGUETA | Instituto Butantan | Dimas Tadeu Covas, Antonio Jorge Martins, Claudia Renata dos Santos Barros, David Schlesinger, Debora Botequio Moretti, Elaine Cristina Marqueze, Elaine Vieira Santos, Evandra Strazza Rodrigues, Heidge Fukumasu, Jayme Augusto de Souza-Neto, José Salvatore Leister Patané, Luiz Alcantara, Luiz Lehmann Coutinho, Maria Carolina Elias, Maurício Lacerda Nogueira, Rafael dos Santos Bezerra, Raul Machado Neto, Rejane Maria Tommasini Grotto, Ricardo Haddad, Sandra Coccuzzo Sampaio Vessoni, Simone Kashima, Svetoslav Nanev Slavov, Vincent Louis Viala |
| EPI_ISL_3246147 | HOSPITAL DE CAMPANHA COVID 19 MUNICIPIO DE TAUBATE | Instituto Butantan | Dimas Tadeu Covas, Antonio Jorge Martins, Claudia Renata dos Santos Barros, David Schlesinger, Debora Botequio Moretti, Elaine Cristina Marqueze, Elaine Vieira Santos, Evandra Strazza Rodrigues, Heidge Fukumasu, Jayme Augusto de Souza-Neto, José Salvatore Leister Patané, Luiz Alcantara, Luiz Lehmann Coutinho, Maria Carolina Elias, Maurício Lacerda Nogueira, Rafael dos Santos Bezerra, Raul Machado Neto, Rejane Maria Tommasini Grotto, Ricardo Haddad, Sandra Coccuzzo Sampaio Vessoni, Simone Kashima, Svetoslav Nanev Slavov, Vincent Louis Viala |
| EPI_ISL_3246148 | HOSPITAL MUNICIPAL DR JOSE DE CARVALHO FLORENCE | Instituto Butantan | Dimas Tadeu Covas, Antonio Jorge Martins, Claudia Renata dos Santos Barros, David Schlesinger, Debora Botequio Moretti, Elaine Cristina Marqueze, Elaine Vieira Santos, Evandra Strazza Rodrigues, Heidge Fukumasu, Jayme Augusto de Souza-Neto, José Salvatore Leister Patané, Luiz Alcantara, Luiz Lehmann Coutinho, Maria Carolina Elias, Maurício Lacerda Nogueira, Rafael dos Santos Bezerra, Raul Machado Neto, Rejane Maria Tommasini Grotto, Ricardo Haddad, Sandra Coccuzzo Sampaio Vessoni, Simone Kashima, Svetoslav Nanev Slavov, Vincent Louis Viala |
| EPI_ISL_3246149 | UNIDADE MISTA DE SAUDE ROSEIRA DR PAULO GUIMARAES CASTRO | Instituto Butantan | Dimas Tadeu Covas, Antonio Jorge Martins, Claudia Renata dos Santos Barros, David Schlesinger, Debora Botequio Moretti, Elaine Cristina Marqueze, Elaine Vieira Santos, Evandra Strazza Rodrigues, Heidge Fukumasu, Jayme Augusto de Souza-Neto, José Salvatore Leister Patané, Luiz Alcantara, Luiz Lehmann Coutinho, Maria Carolina Elias, Maurício Lacerda Nogueira, Rafael dos Santos Bezerra, Raul Machado Neto, Rejane Maria Tommasini Grotto, Ricardo Haddad, Sandra Coccuzzo Sampaio Vessoni, Simone Kashima, Svetoslav Nanev Slavov, Vincent Louis Viala |
| EPI_ISL_3246150 | SANTA CASA DE MISERICORDIA DE UBATUBA | Instituto Butantan | Dimas Tadeu Covas, Antonio Jorge Martins, Claudia Renata dos Santos Barros, David Schlesinger, Debora Botequio Moretti, Elaine Cristina Marqueze, Elaine Vieira Santos, Evandra Strazza Rodrigues, Heidge Fukumasu, Jayme Augusto de Souza-Neto, José Salvatore Leister Patané, Luiz Alcantara, Luiz Lehmann Coutinho, Maria Carolina Elias, Maurício Lacerda Nogueira, Rafael dos Santos Bezerra, Raul Machado Neto, Rejane Maria Tommasini Grotto, Ricardo Haddad, Sandra Coccuzzo Sampaio Vessoni, Simone Kashima, Svetoslav Nanev Slavov, Vincent Louis Viala |
| EPI_ISL_3246151 | UNIDADE MISTA DE SAUDE ROSEIRA DR PAULO GUIMARAES CASTRO | Instituto Butantan | Dimas Tadeu Covas, Antonio Jorge Martins, Claudia Renata dos Santos Barros, David Schlesinger, Debora Botequio Moretti, Elaine Cristina Marqueze, Elaine Vieira Santos, Evandra Strazza Rodrigues, Heidge Fukumasu, Jayme Augusto de Souza-Neto, José Salvatore Leister Patané, Luiz Alcantara, Luiz Lehmann Coutinho, Maria Carolina Elias, Maurício Lacerda Nogueira, Rafael dos Santos Bezerra, Raul Machado Neto, Rejane Maria Tommasini Grotto, Ricardo Haddad, Sandra Coccuzzo Sampaio Vessoni, Simone Kashima, Svetoslav Nanev Slavov, Vincent Louis Viala |
| EPI_ISL_3246156, EPI_ISL_3246166, EPI_ISL_3246169, EPI_ISL_3246175, EPI_ISL_3246182, EPI_ISL_3246190, EPI_ISL_3246192 | LATE - Laboratório de Técnicas Especiais - Hospital Israelita Albert Einstein | LATE - Laboratório de Técnicas Especiais - Hospital Israelita Albert Einstein | Deyvid Amgarten, Fernanda de Mello Malta, Raquel Riyuzo, Ana Paula Moreira Salles, Pedro Henrique Sebe Rodrigues, Alexandre Hideaki Takara, Anelisie da Silva Santos, Erick Gustavo Dorlass, Marcio Anunciacao Menezes, João Renato Rebello Pinho |
| EPI_ISL_3246198, EPI_ISL_3246199, EPI_ISL_3246200, EPI_ISL_3246201, EPI_ISL_3246202, EPI_ISL_3246203, EPI_ISL_3246204, EPI_ISL_3246205, EPI_ISL_3246206 | Plateforme de testing Namuroise | Plateforme de testing Namuroise | Lesly Nyinkeu Kemamen; Nicolas Gilliard; Demars Aurore ; Maschietto Céline ; Nobis Chloé ; Otto Gaetan ; Denis Olivier ; Degosserie Jonathan ; Mullier François |
| EPI_ISL_3246239 | Neuberg Supratech Refrence Laboratory, Surat | Gujarat Biotechnology Research Centre | Arpit Shukla,Bhadreshsinh Gohil,Nimesh Patel,Nitin Shukla,Sonal Sharma ,Zuber Saiyed,Ramesh Pandit,Janvi Raval,Zarna Patel,Nitin Savaliya,Dinesh Kumar,Twinkle Soni,Manthan Shah,Umang Mishra,Nitesh Shah,Chaitanya Joshi,Madhvi Joshi |
| EPI_ISL_3246240 | Neuberg Supratech Refrence Laboratory, Surat | Gujarat Biotechnology Research Centre | Bhadreshsinh Gohil,Nimesh Patel,Nitin Shukla,Sonal Sharma ,Zuber Saiyed,Ramesh Pandit,Janvi Raval,Zarna Patel,Nitin Savaliya,Dinesh Kumar,Twinkle Soni,Arpit Shukla,Manthan Shah,Umang Mishra,Nitesh Shah,Chaitanya Joshi,Madhvi Joshi |
| EPI_ISL_3246241 | Neuberg Supratech Refrence Laboratory, Surat | Gujarat Biotechnology Research Centre | Nimesh Patel,Nitin Shukla,Sonal Sharma ,Zuber Saiyed,Ramesh Pandit,Janvi Raval,Zarna Patel,Nitin Savaliya,Dinesh Kumar,Twinkle Soni,Arpit Shukla,Bhadreshsinh Gohil,Manthan Shah,Umang Mishra,Nitesh Shah,Chaitanya Joshi,Madhvi Joshi |
| EPI_ISL_3246242 | Neuberg Supratech Refrence Laboratory, Surat | Gujarat Biotechnology Research Centre | Nitin Shukla,Sonal Sharma ,Zuber Saiyed,Ramesh Pandit,Janvi Raval,Zarna Patel,Nitin Savaliya,Dinesh Kumar,Twinkle Soni,Arpit Shukla,Bhadreshsinh Gohil,Nimesh Patel,Manthan Shah,Umang Mishra,Nitesh Shah,Chaitanya Joshi,Madhvi Joshi |
| EPI_ISL_3246243 | Neuberg Supratech Refrence Laboratory, Surat | Gujarat Biotechnology Research Centre | Sonal Sharma ,Zuber Saiyed,Ramesh Pandit,Janvi Raval,Zarna Patel,Nitin Savaliya,Dinesh Kumar,Twinkle Soni,Arpit Shukla,Bhadreshsinh Gohil,Nimesh Patel,Nitin Shukla,Manthan Shah,Umang Mishra,Nitesh Shah,Chaitanya Joshi,Madhvi Joshi |
| EPI_ISL_3246244 | Neuberg Supratech Refrence Laboratory, Surat | Gujarat Biotechnology Research Centre | Zuber Saiyed,Ramesh Pandit,Janvi Raval,Zarna Patel,Nitin Savaliya,Dinesh Kumar,Twinkle Soni,Arpit Shukla,Bhadreshsinh Gohil,Nimesh Patel,Nitin Shukla,Sonal Sharma ,Manthan Shah,Umang Mishra,Nitesh Shah,Chaitanya Joshi,Madhvi Joshi |
| EPI_ISL_3246245 | Neuberg Supratech Refrence Laboratory, Surat | Gujarat Biotechnology Research Centre | Ramesh Pandit,Janvi Raval,Zarna Patel,Nitin Savaliya,Dinesh Kumar,Twinkle Soni,Arpit Shukla,Bhadreshsinh Gohil,Nimesh Patel,Nitin Shukla,Sonal Sharma ,Zuber Saiyed,Manthan Shah,Umang Mishra,Nitesh Shah,Chaitanya Joshi,Madhvi Joshi |
| EPI_ISL_3246246 | Neuberg Supratech Refrence Laboratory, Surat | Gujarat Biotechnology Research Centre | Janvi Raval,Zarna Patel,Nitin Savaliya,Dinesh Kumar,Twinkle Soni,Arpit Shukla,Bhadreshsinh Gohil,Nimesh Patel,Nitin Shukla,Sonal Sharma ,Zuber Saiyed,Ramesh Pandit,Manthan Shah,Umang Mishra,Nitesh Shah,Chaitanya Joshi,Madhvi Joshi |
| EPI_ISL_3246247 | Neuberg Supratech Refrence Laboratory, Surat | Gujarat Biotechnology Research Centre | Zarna Patel,Nitin Savaliya,Dinesh Kumar,Twinkle Soni,Arpit Shukla,Bhadreshsinh Gohil,Nimesh Patel,Nitin Shukla,Sonal Sharma ,Zuber Saiyed,Ramesh Pandit,Janvi Raval,Manthan Shah,Umang Mishra,Nitesh Shah,Chaitanya Joshi,Madhvi Joshi |
| EPI_ISL_3246248 | GMERS Medical College, Valsad | Gujarat Biotechnology Research Centre | Nitin Savaliya,Dinesh Kumar,Twinkle Soni,Arpit Shukla,Bhadreshsinh Gohil,Nimesh Patel,Nitin Shukla,Sonal Sharma ,Zuber Saiyed,Ramesh Pandit,Janvi Raval,Zarna Patel,Vicky Gandhi,Umang Mishra,Nitesh Shah,Chaitanya Joshi,Madhvi Joshi |
| EPI_ISL_3246249 | GMERS Medical College, Valsad | Gujarat Biotechnology Research Centre | Dinesh Kumar,Twinkle Soni,Arpit Shukla,Bhadreshsinh Gohil,Nimesh Patel,Nitin Shukla,Sonal Sharma ,Zuber Saiyed,Ramesh Pandit,Janvi Raval,Zarna Patel,Nitin Savaliya,Vicky Gandhi,Umang Mishra,Nitesh Shah,Chaitanya Joshi,Madhvi Joshi |
| EPI_ISL_3246250 | GMERS Medical College, Valsad | Gujarat Biotechnology Research Centre | Twinkle Soni,Arpit Shukla,Bhadreshsinh Gohil,Nimesh Patel,Nitin Shukla,Sonal Sharma ,Zuber Saiyed,Ramesh Pandit,Janvi Raval,Zarna Patel,Nitin Savaliya,Dinesh Kumar,Vicky Gandhi,Umang Mishra,Nitesh Shah,Chaitanya Joshi,Madhvi Joshi |
| EPI_ISL_3246251 | College of veterinary Science and A.H Dantiwada | Gujarat Biotechnology Research Centre | Arpit Shukla,Bhadreshsinh Gohil,Nimesh Patel,Nitin Shukla,Sonal Sharma ,Zuber Saiyed,Ramesh Pandit,Janvi Raval,Zarna Patel,Nitin Savaliya,Dinesh Kumar,Twinkle Soni,Hemlatta Sunsara,Umang Mishra,Nitesh Shah,Chaitanya Joshi,Madhvi Joshi |
| EPI_ISL_3246252 | Neuberg Supratech Refrence Laboratory, Surat | Gujarat Biotechnology Research Centre | Bhadreshsinh Gohil,Nimesh Patel,Nitin Shukla,Sonal Sharma ,Zuber Saiyed,Ramesh Pandit,Janvi Raval,Zarna Patel,Nitin Savaliya,Dinesh Kumar,Twinkle Soni,Arpit Shukla,Manthan Shah,Umang Mishra,Nitesh Shah,Chaitanya Joshi,Madhvi Joshi |
| EPI_ISL_3246253 | Neuberg Supratech Refrence Laboratory, Surat | Gujarat Biotechnology Research Centre | Nimesh Patel,Nitin Shukla,Sonal Sharma ,Zuber Saiyed,Ramesh Pandit,Janvi Raval,Zarna Patel,Nitin Savaliya,Dinesh Kumar,Twinkle Soni,Arpit Shukla,Bhadreshsinh Gohil,Manthan Shah,Umang Mishra,Nitesh Shah,Chaitanya Joshi,Madhvi Joshi |
| EPI_ISL_3246254 | Neuberg Supratech Refrence Laboratory, Surat | Gujarat Biotechnology Research Centre | Nitin Shukla,Sonal Sharma ,Zuber Saiyed,Ramesh Pandit,Janvi Raval,Zarna Patel,Nitin Savaliya,Dinesh Kumar,Twinkle Soni,Arpit Shukla,Bhadreshsinh Gohil,Nimesh Patel,Manthan Shah,Umang Mishra,Nitesh Shah,Chaitanya Joshi,Madhvi Joshi |
| EPI_ISL_3246255 | Neuberg Supratech Refrence Laboratory, Surat | Gujarat Biotechnology Research Centre | Sonal Sharma ,Zuber Saiyed,Ramesh Pandit,Janvi Raval,Zarna Patel,Nitin Savaliya,Dinesh Kumar,Twinkle Soni,Arpit Shukla,Bhadreshsinh Gohil,Nimesh Patel,Nitin Shukla,Manthan Shah,Umang Mishra,Nitesh Shah,Chaitanya Joshi,Madhvi Joshi |
| EPI_ISL_3246256 | Neuberg Supratech Refrence Laboratory, Surat | Gujarat Biotechnology Research Centre | Zuber Saiyed,Ramesh Pandit,Janvi Raval,Zarna Patel,Nitin Savaliya,Dinesh Kumar,Twinkle Soni,Arpit Shukla,Bhadreshsinh Gohil,Nimesh Patel,Nitin Shukla,Sonal Sharma ,Manthan Shah,Umang Mishra,Nitesh Shah,Chaitanya Joshi,Madhvi Joshi |
| EPI_ISL_3246257 | Supratech Micropath Laboratory Research Institute Pvt Ltd, Ahmedabad | Gujarat Biotechnology Research Centre | Ramesh Pandit,Janvi Raval,Zarna Patel,Nitin Savaliya,Dinesh Kumar,Twinkle Soni,Arpit Shukla,Bhadreshsinh Gohil,Nimesh Patel,Nitin Shukla,Sonal Sharma ,Zuber Saiyed,Mahendra Parikh,Umang Mishra,Nitesh Shah,Chaitanya Joshi,Madhvi Joshi |
| EPI_ISL_3246277, EPI_ISL_3246286, EPI_ISL_3246288, EPI_ISL_3246290, EPI_ISL_3246291, EPI_ISL_3246292, EPI_ISL_3246293, EPI_ISL_3246295, EPI_ISL_3246296, EPI_ISL_3246297, EPI_ISL_3246305, EPI_ISL_3246306, EPI_ISL_3246307, EPI_ISL_3246308, EPI_ISL_3246309, EPI_ISL_3246310, EPI_ISL_3246311, EPI_ISL_3246312, EPI_ISL_3246314, EPI_ISL_3246315, EPI_ISL_3246316, EPI_ISL_3246317, EPI_ISL_3246320, EPI_ISL_3246321, EPI_ISL_3246322, EPI_ISL_3246324, EPI_ISL_3246325, EPI_ISL_3246326, EPI_ISL_3246327, EPI_ISL_3246328 | | | |
| see above | Eurofins Diatherix | Hudsonalpha Genome Sequencing Center | Jane Grimwood, Melissa Williams, Lori H. Handley, Joshua Stough, Leslie Malone, Stefan Brzezinski, Ada Stewart, Teresa Jones, Jenell Webber, John Lovell, Jennifer Cart, and Jeremy Schmutz |
| EPI_ISL_3246329 | Fundacio Althaia-Manresa | Banc de Sang i Teixits | Francisco Vidal, Irene Corrales, Nina Borràs, Carlos Hobeich, Natália Comes, Lorena Ramírez, Noemí Gonzalez, Silvia Sauleda, Maria Glòria Soria |
| EPI_ISL_3246331 | CAP Sant Hipolit de Voltrega | Banc de Sang i Teixits | Francisco Vidal, Irene Corrales, Nina Borràs, Carlos Hobeich, Natália Comes, Lorena Ramírez, Noemí Gonzalez, Silvia Sauleda, Maria Glòria Soria |
| EPI_ISL_3246332 | CAP Vic Nord | Banc de Sang i Teixits | Francisco Vidal, Irene Corrales, Nina Borràs, Carlos Hobeich, Natália Comes, Lorena Ramírez, Noemí Gonzalez, Silvia Sauleda, Maria Glòria Soria |
| EPI_ISL_3246333 | CAP El Remei | Banc de Sang i Teixits | Francisco Vidal, Irene Corrales, Nina Borràs, Carlos Hobeich, Natália Comes, Lorena Ramírez, Noemí Gonzalez, Silvia Sauleda, Maria Glòria Soria |
| EPI_ISL_3246340 | CAP Torello | Banc de Sang i Teixits | Francisco Vidal, Irene Corrales, Nina Borràs, Carlos Hobeich, Natália Comes, Lorena Ramírez, Noemí Gonzalez, Silvia Sauleda, Maria Glòria Soria |
| EPI_ISL_3246344 | CAP Tona | Banc de Sang i Teixits | Francisco Vidal, Irene Corrales, Nina Borràs, Carlos Hobeich, Natália Comes, Lorena Ramírez, Noemí Gonzalez, Silvia Sauleda, Maria Glòria Soria |
| EPI_ISL_3246431, EPI_ISL_3246432, EPI_ISL_3246433, EPI_ISL_3246434, EPI_ISL_3246435, EPI_ISL_3246436, EPI_ISL_3246437, EPI_ISL_3246438, EPI_ISL_3246439, EPI_ISL_3246440, EPI_ISL_3246441, EPI_ISL_3246442, EPI_ISL_3246443, EPI_ISL_3246444, EPI_ISL_3246445, EPI_ISL_3246446, EPI_ISL_3246447, EPI_ISL_3246448, EPI_ISL_3246449, EPI_ISL_3246450, EPI_ISL_3246451, EPI_ISL_3246452, EPI_ISL_3246453, EPI_ISL_3246454, EPI_ISL_3246455, EPI_ISL_3246456, EPI_ISL_3246457, EPI_ISL_3246458, EPI_ISL_3246459, EPI_ISL_3246460, EPI_ISL_3246461, EPI_ISL_3246462, EPI_ISL_3246463, EPI_ISL_3246464, EPI_ISL_3246465, EPI_ISL_3246466, EPI_ISL_3246467, EPI_ISL_3246468, EPI_ISL_3246469, EPI_ISL_3246470, EPI_ISL_3246471, EPI_ISL_3246472, EPI_ISL_3246473, EPI_ISL_3246474, | | | |

| | | | |
|---|---|---|---|
| EPI_ISL_3246475, EPI_ISL_3246476, EPI_ISL_3246477, EPI_ISL_3246478, EPI_ISL_3246479, EPI_ISL_3246480, EPI_ISL_3246481, EPI_ISL_3246482, EPI_ISL_3246483, EPI_ISL_3246484, EPI_ISL_3246485, EPI_ISL_3246486, EPI_ISL_3246487, EPI_ISL_3246488, EPI_ISL_3246489, EPI_ISL_3246490, EPI_ISL_3246491, EPI_ISL_3246492, EPI_ISL_3246493, EPI_ISL_3246494, EPI_ISL_3246495, EPI_ISL_3246496, EPI_ISL_3246497, EPI_ISL_3246498, EPI_ISL_3246499, EPI_ISL_3246500, EPI_ISL_3246501, EPI_ISL_3246502, EPI_ISL_3246503, EPI_ISL_3246504, EPI_ISL_3246505, EPI_ISL_3246506, EPI_ISL_3246507, EPI_ISL_3246508, EPI_ISL_3246509, EPI_ISL_3246510, EPI_ISL_3246511, EPI_ISL_3246512, EPI_ISL_3246513, EPI_ISL_3246514, EPI_ISL_3246515, EPI_ISL_3246516, EPI_ISL_3246518, EPI_ISL_3246519, EPI_ISL_3246520, EPI_ISL_3246521 | | | |
| see above | Lifebrain Covid Labor GmbH | Dept. of Laboratory Medicine | Harald Esterbauer, Anna Gschaider, Petra Jurkowitsch, Fabian Konig, Sabina Plumer, Robert Strassl, Claudia Weber, Oswald Wagner |
| EPI_ISL_3246533, EPI_ISL_3246534, EPI_ISL_3246535, EPI_ISL_3246536, EPI_ISL_3246537, EPI_ISL_3246538, EPI_ISL_3246539, EPI_ISL_3246540, EPI_ISL_3246541, EPI_ISL_3246542, EPI_ISL_3246543, EPI_ISL_3246544, EPI_ISL_3246545, EPI_ISL_3246546, EPI_ISL_3246547, EPI_ISL_3246548, EPI_ISL_3246549, EPI_ISL_3246550, EPI_ISL_3246551, EPI_ISL_3246552, EPI_ISL_3246553, EPI_ISL_3246554, EPI_ISL_3246555, EPI_ISL_3246556, EPI_ISL_3246557, EPI_ISL_3246558, EPI_ISL_3246559, EPI_ISL_3246560, EPI_ISL_3246561, EPI_ISL_3246562, EPI_ISL_3246563, EPI_ISL_3246564, EPI_ISL_3246565, EPI_ISL_3246566, EPI_ISL_3246567, EPI_ISL_3246568, EPI_ISL_3246569, EPI_ISL_3246570, EPI_ISL_3246571, EPI_ISL_3246572, EPI_ISL_3246573, EPI_ISL_3246574, EPI_ISL_3246575, EPI_ISL_3246576, EPI_ISL_3246577, EPI_ISL_3246578, EPI_ISL_3246579, EPI_ISL_3246580, EPI_ISL_3246581, EPI_ISL_3246582, EPI_ISL_3246583, EPI_ISL_3246584, EPI_ISL_3246585, EPI_ISL_3246586, EPI_ISL_3246587, EPI_ISL_3246588, EPI_ISL_3246590 | | | |
| see above | Kansas Health and Environmental Lab | Kansas Health and Environmental Lab | Mike Grose, Jonathan Barnell, Amanda Bradley, and Phil Adam |
| EPI_ISL_3246604, EPI_ISL_3246605 | Marche en Famenne | Plateforme de testing Namuroise | Lesly Nyinkeu Kemamen; Nicolas Gilliard; Demars Aurore ; Maschietto Céline ; Nobis Chloé ; Otto Gaetan ; Denis Olivier ; Degosserie Jonathan ; Mullier François |
| EPI_ISL_3246606, EPI_ISL_3246607 | Plateforme de testing Namuroise | Plateforme de testing Namuroise | Lesly Nyinkeu Kemamen; Nicolas Gilliard; Demars Aurore ; Maschietto Céline ; Nobis Chloé ; Otto Gaetan ; Denis Olivier ; Degosserie Jonathan ; Mullier François |
| EPI_ISL_3246608 | Marche en Famenne | Plateforme de testing Namuroise | Lesly Nyinkeu Kemamen; Nicolas Gilliard; Demars Aurore ; Maschietto Céline ; Nobis Chloé ; Otto Gaetan ; Denis Olivier ; Degosserie Jonathan ; Mullier François |
| EPI_ISL_3246609, EPI_ISL_3246610 | Plateforme de testing Namuroise | Plateforme de testing Namuroise | Lesly Nyinkeu Kemamen; Nicolas Gilliard; Demars Aurore ; Maschietto Céline ; Nobis Chloé ; Otto Gaetan ; Denis Olivier ; Degosserie Jonathan ; Mullier François |
| EPI_ISL_3246611 | Marche en Famenne | Plateforme de testing Namuroise | Lesly Nyinkeu Kemamen; Nicolas Gilliard; Demars Aurore ; Maschietto Céline ; Nobis Chloé ; Otto Gaetan ; Denis Olivier ; Degosserie Jonathan ; Mullier François |
| EPI_ISL_3246612 | Plateforme de testing Namuroise | Plateforme de testing Namuroise | Lesly Nyinkeu Kemamen; Nicolas Gilliard; Demars Aurore ; Maschietto Céline ; Nobis Chloé ; Otto Gaetan ; Denis Olivier ; Degosserie Jonathan ; Mullier François |
| EPI_ISL_3246613 | Marche en Famenne | Plateforme de testing Namuroise | Lesly Nyinkeu Kemamen; Nicolas Gilliard; Demars Aurore ; Maschietto Céline ; Nobis Chloé ; Otto Gaetan ; Denis Olivier ; Degosserie Jonathan ; Mullier François |
| EPI_ISL_3246614, EPI_ISL_3246615, EPI_ISL_3246617, EPI_ISL_3246618 | Plateforme de testing Namuroise | Plateforme de testing Namuroise | Lesly Nyinkeu Kemamen; Nicolas Gilliard; Demars Aurore ; Maschietto Céline ; Nobis Chloé ; Otto Gaetan ; Denis Olivier ; Degosserie Jonathan ; Mullier François |
| EPI_ISL_3246619 | Marche en Famenne | Plateforme de testing Namuroise | Lesly Nyinkeu Kemamen; Nicolas Gilliard; Demars Aurore ; Maschietto Céline ; Nobis Chloé ; Otto Gaetan ; Denis Olivier ; Degosserie Jonathan ; Mullier François |
| EPI_ISL_3246620, EPI_ISL_3246621, EPI_ISL_3246622 | Plateforme de testing Namuroise | Plateforme de testing Namuroise | Lesly Nyinkeu Kemamen; Nicolas Gilliard; Demars Aurore ; Maschietto Céline ; Nobis Chloé ; Otto Gaetan ; Denis Olivier ; Degosserie Jonathan ; Mullier François |
| EPI_ISL_3246628, EPI_ISL_3246629, EPI_ISL_3246630, EPI_ISL_3246648, EPI_ISL_3246649, EPI_ISL_3246654, EPI_ISL_3246655, EPI_ISL_3246656, EPI_ISL_3246675 | Weill Cornell Medicine | New York Genome Center | Michael Zody, Andre Corvelo, Dayna M. Oschwald, Amy Baldwin, Samantha Fennessey, Tom Maniatis, Melissa Cushing, Olivier Elemento, Margaret Elizabeth Ross, Chris Mason, Priya Velu, Hanna Rennert, Arryn Craney, Lars F Westblade |
| EPI_ISL_3246684 | Jessa | Jessa | Severine Berden et al. on behalf of the Jessa_cmdLab |
| EPI_ISL_3246685, EPI_ISL_3246686, EPI_ISL_3246687, EPI_ISL_3246688, EPI_ISL_3246689, EPI_ISL_3246690, EPI_ISL_3246691, EPI_ISL_3246692, EPI_ISL_3246693, EPI_ISL_3246694, EPI_ISL_3246695, EPI_ISL_3246696, EPI_ISL_3246697, EPI_ISL_3246698, EPI_ISL_3246699, EPI_ISL_3246700, EPI_ISL_3246701, EPI_ISL_3246702, EPI_ISL_3246703 | | | |
| see above | ZOL | Jessa | Severine Berden et al. on behalf of the Jessa_cmdLab |
| EPI_ISL_3246704, EPI_ISL_3246705 | Jessa | Jessa | Severine Berden et al. on behalf of the Jessa_cmdLab |
| EPI_ISL_3246706, EPI_ISL_3246707 | LKO | Jessa | Severine Berden et al. on behalf of the Jessa_cmdLab |
| EPI_ISL_3246708, EPI_ISL_3246709, EPI_ISL_3246711, EPI_ISL_3246713, EPI_ISL_3246714, EPI_ISL_3246715, EPI_ISL_3246716, EPI_ISL_3246717, EPI_ISL_3246718, EPI_ISL_3246719, EPI_ISL_3246720, EPI_ISL_3246721, EPI_ISL_3246722, EPI_ISL_3246723, EPI_ISL_3246724, EPI_ISL_3246725, EPI_ISL_3246726, EPI_ISL_3246727 | | | |
| see above | Jessa | Jessa | Severine Berden et al. on behalf of the Jessa_cmdLab |
| EPI_ISL_3246728, EPI_ISL_3246729, EPI_ISL_3246730, EPI_ISL_3246731, EPI_ISL_3246732, EPI_ISL_3246733, EPI_ISL_3246734, EPI_ISL_3246735, EPI_ISL_3246736, EPI_ISL_3246737, EPI_ISL_3246738, EPI_ISL_3246739, EPI_ISL_3246740, EPI_ISL_3246741, EPI_ISL_3246742, EPI_ISL_3246743, EPI_ISL_3246744, EPI_ISL_3246745, EPI_ISL_3246746, EPI_ISL_3246747, EPI_ISL_3246749, EPI_ISL_3246750, EPI_ISL_3246751, EPI_ISL_3246752, EPI_ISL_3246753, EPI_ISL_3246754, EPI_ISL_3246755, EPI_ISL_3246756, EPI_ISL_3246757, EPI_ISL_3246758, EPI_ISL_3246759, EPI_ISL_3246760, EPI_ISL_3246762, EPI_ISL_3246763, EPI_ISL_3246764, EPI_ISL_3246765, EPI_ISL_3246766, EPI_ISL_3246767, EPI_ISL_3246768, EPI_ISL_3246769, EPI_ISL_3246770, EPI_ISL_3246771, EPI_ISL_3246772, EPI_ISL_3246773, EPI_ISL_3246774, EPI_ISL_3246775, EPI_ISL_3246776, EPI_ISL_3246777, EPI_ISL_3246778, EPI_ISL_3246779, EPI_ISL_3246780, EPI_ISL_3246781, EPI_ISL_3246782, EPI_ISL_3246783, EPI_ISL_3246784, EPI_ISL_3246785, EPI_ISL_3246786, EPI_ISL_3246787, EPI_ISL_3246788, EPI_ISL_3246789 | | | |
| see above | Hospital Universitari Vall d'Hebron - Vall d'Hebron Institut de Recerca | Hospital Universitari Vall d'Hebron - Vall d'Hebron Institut de Recerca | Cristina Andrés, Maria Piñana, Alejandra González-Sánchez, Damir Garcia-Cehic, Ariadna Rando, Juliana Esperalba, Maria Gema Codina, Carla Castillo, Karen García, Rodrigo Vásquez, Maria Carmen Martin, Tomàs Pumarola, Josep Quer, Andrés Antón |
| EPI_ISL_3246790, EPI_ISL_3246791, EPI_ISL_3246792, EPI_ISL_3246793, EPI_ISL_3246794, EPI_ISL_3246795, EPI_ISL_3246796, EPI_ISL_3246797, EPI_ISL_3246798, EPI_ISL_3246799, EPI_ISL_3246800, EPI_ISL_3246801, EPI_ISL_3246802, EPI_ISL_3246803, EPI_ISL_3246804, EPI_ISL_3246805, EPI_ISL_3246806, EPI_ISL_3246807, EPI_ISL_3246808, EPI_ISL_3246809, EPI_ISL_3246810, EPI_ISL_3246811, EPI_ISL_3246813, EPI_ISL_3246815, EPI_ISL_3246816, EPI_ISL_3246817, EPI_ISL_3246818, EPI_ISL_3246819 | | | |
| see above | IN State Department of Health Laboratory Services | IN State Department of Health Laboratory Services | Cassandra Campion, Jamie Yeadon, Brian Pope, Lixia Liu, Kyle Brownlee, Melissa Hindenlang, Mark Glazier |
| EPI_ISL_3246820, EPI_ISL_3246821, EPI_ISL_3246823, EPI_ISL_3246824, EPI_ISL_3246825, EPI_ISL_3246826, EPI_ISL_3246827, EPI_ISL_3246828, EPI_ISL_3246829, EPI_ISL_3246830, EPI_ISL_3246831, EPI_ISL_3246832, EPI_ISL_3246833, EPI_ISL_3246834, EPI_ISL_3246835, EPI_ISL_3246836, EPI_ISL_3246837, EPI_ISL_3246838, EPI_ISL_3246839, EPI_ISL_3246840, EPI_ISL_3246841, EPI_ISL_3246843, EPI_ISL_3246844, EPI_ISL_3246845, EPI_ISL_3246846, EPI_ISL_3246848, EPI_ISL_3246849 | | | |
| see above | Montana Public Health Laboratory | Montana Public Health Laboratory | Joy Ritter, Michelle Mozer, Carrie Biskupiak, Deborah Gibson, Michael Dills |
| EPI_ISL_3246860 | Arkansas Department of Health | Center for Global Health, University of New Mexico Health Sciences Center | Daryl Domman, Kurt Schwalm, Valerie Morley, Atul Kothari, Joshua L. Kennedy, Darrell Dinwiddie |
| EPI_ISL_3246862, EPI_ISL_3246872, EPI_ISL_3246873, EPI_ISL_3246874, EPI_ISL_3246880, EPI_ISL_3246891, EPI_ISL_3246894, EPI_ISL_3246895, EPI_ISL_3246896, EPI_ISL_3246897, EPI_ISL_3246898, EPI_ISL_3246899, EPI_ISL_3246903, EPI_ISL_3246904, EPI_ISL_3246910 | | | |
| see above | Baptist Health Medical Center | Center for Global Health, University of New Mexico Health Sciences Center | Daryl Domman, Kurt Schwalm, Valerie Morley, Amanda Novack, Dirk Haselow, Joshua L. Kennedy, Darrell Dinwiddie |
| EPI_ISL_3246917, EPI_ISL_3246918 | University of New Mexico Hospital | Center for Global Health, University of New Mexico Health Sciences Center | Daryl Domman, Justin Baca, Silas Bussman, Robert Taylor, Kurt Schwalm, Valerie Morley, Jon Femling, Darrell Dinwiddie |
| EPI_ISL_3246923, EPI_ISL_3246924, EPI_ISL_3246925, EPI_ISL_3246926, EPI_ISL_3246927, EPI_ISL_3246928, EPI_ISL_3246929, EPI_ISL_3246930, EPI_ISL_3246931, EPI_ISL_3246932, EPI_ISL_3246933, EPI_ISL_3246934, EPI_ISL_3246935, EPI_ISL_3246936, EPI_ISL_3246937, EPI_ISL_3246938, EPI_ISL_3246939, EPI_ISL_3246940, EPI_ISL_3246941, EPI_ISL_3246942, EPI_ISL_3246943, EPI_ISL_3246944, EPI_ISL_3246945, EPI_ISL_3246946, EPI_ISL_3246947, EPI_ISL_3246948, EPI_ISL_3246949, EPI_ISL_3246950, EPI_ISL_3246951, EPI_ISL_3246952, EPI_ISL_3246953, EPI_ISL_3246954, EPI_ISL_3246955, EPI_ISL_3246956, EPI_ISL_3246957, EPI_ISL_3246958, EPI_ISL_3246959, EPI_ISL_3246960, EPI_ISL_3246961, EPI_ISL_3246963, EPI_ISL_3246964, EPI_ISL_3246965, EPI_ISL_3246966, EPI_ISL_3246967, EPI_ISL_3246968, EPI_ISL_3246969, EPI_ISL_3246970, EPI_ISL_3246971, EPI_ISL_3246972, EPI_ISL_3246973, EPI_ISL_3246974, EPI_ISL_3246975, EPI_ISL_3246976, EPI_ISL_3246977, EPI_ISL_3246978, EPI_ISL_3246979, EPI_ISL_3246980 | | | |
| see above | North Dakota Department of Health, Public Health Laboratory | North Dakota Department of Health, Public Health Laboratory | Lisa Wingerter |
| EPI_ISL_3246981 | Tripler Army Medical Center Microbiology | Dept. of Clinical Investigation | Rachael Downham, Kendra Carter, Jonathan D'Ambrozio, Catherine Uyehara |
| EPI_ISL_3246982 | Tripler Army Medical Center Microbiology Department | Tripler Army Medical Center Microbiology Department | Rachael Downham, Kendra Carter, Jonathan D'Ambrozio, Catherine Uyehara |
| EPI_ISL_3246983, EPI_ISL_3246984, EPI_ISL_3246986, EPI_ISL_3246987, EPI_ISL_3246993, EPI_ISL_3247006, EPI_ISL_3247008 | Tripler Army Medical Center Microbiology Department | Department of Clinical Investigation | Rachael Downham, Kendra Carter, Jonathan D'Ambrozio, Catherine Uyehara |
| EPI_ISL_3247021 | Spital Zweisimmen STS AG | Institute for Infectious Diseases | Stefan Neuenschwander, Christian Baumann, Miguel A Terrazos Miani, Sonja Gempeler, Loïc Borcard, Cora Sägesser, Pascal Bittel, Peter Keller, Franziska Suter-Riniker, Stephen L Leib, Alban Ramette |
| EPI_ISL_3247022 | MCL Med. Laboratorien AG | Institute for Infectious Diseases | Stefan Neuenschwander, Christian Baumann, Miguel A Terrazos Miani, Sonja Gempeler, Loïc Borcard, Cora Sägesser, Pascal Bittel, Peter Keller, Franziska Suter-Riniker, Stephen L Leib, Alban Ramette |
| EPI_ISL_3247023, EPI_ISL_3247024, EPI_ISL_3247025, EPI_ISL_3247026, EPI_ISL_3247027 | Luzerner Kantonsspital | Institute for Infectious Diseases | Stefan Neuenschwander, Christian Baumann, Miguel A Terrazos Miani, Sonja Gempeler, Loïc Borcard, Cora Sägesser, Pascal Bittel, Peter Keller, Franziska Suter-Riniker, Stephen L Leib, Alban Ramette |
| EPI_ISL_3247028 | Spital Burgdorf | Institute for Infectious Diseases | Stefan Neuenschwander, Christian Baumann, Miguel A Terrazos Miani, Sonja Gempeler, Loïc Borcard, Cora Sägesser, Pascal Bittel, Peter Keller, Franziska Suter-Riniker, Stephen L Leib, Alban Ramette |
| EPI_ISL_3247029 | Bio Analytica AG | Institute for Infectious Diseases | Stefan Neuenschwander, Christian Baumann, Miguel A Terrazos Miani, Sonja Gempeler, Loïc Borcard, Cora Sägesser, Pascal Bittel, Peter Keller, Franziska Suter-Riniker, Stephen L Leib, Alban Ramette |
| EPI_ISL_3247030 | Luzerner Kantonsspital | Institute for Infectious Diseases | Stefan Neuenschwander, Christian Baumann, Miguel A Terrazos Miani, Sonja Gempeler, Loïc Borcard, Cora Sägesser, Pascal Bittel, Peter Keller, Franziska Suter-Riniker, Stephen L Leib, Alban Ramette |
| EPI_ISL_3247031 | Spital Riggisberg | Institute for Infectious Diseases | Stefan Neuenschwander, Christian Baumann, Miguel A Terrazos Miani, Sonja Gempeler, Loïc Borcard, Cora Sägesser, Pascal Bittel, Peter Keller, Franziska Suter-Riniker, Stephen L Leib, Alban Ramette |
| EPI_ISL_3247032, EPI_ISL_3247033 | Bakt. Institut Olten Bio AG | Institute for Infectious Diseases | Stefan Neuenschwander, Christian Baumann, Miguel A Terrazos Miani, Sonja Gempeler, Loïc Borcard, Cora Sägesser, Pascal Bittel, Peter Keller, Franziska Suter-Riniker, Stephen L Leib, Alban Ramette |
| EPI_ISL_3247034, EPI_ISL_3247035, | Inselspital Bern (Covid-Track) | Institute for Infectious Diseases | Stefan Neuenschwander, Christian Baumann, Miguel A Terrazos Miani, Sonja Gempeler, Loïc Borcard, Cora Sägesser, Pascal Bittel, Peter Keller, Franziska Suter-Riniker, Stephen L Leib, Alban Ramette |

| | | | |
|---|---|---|---|
| EPI_ISL_3247036, EPI_ISL_3247037 | | | |
| EPI_ISL_3247038 | Spital Riggisberg | Institute for Infectious Diseases | Stefan Neuenschwander, Christian Baumann, Miguel A Terrazos Miani, Sonja Gempeler, Loïc Borcard, Cora Sägesser, Pascal Bittel, Peter Keller, Franziska Suter-Riniker, Stephen L Leib, Alban Ramette |
| EPI_ISL_3247039, EPI_ISL_3247040, EPI_ISL_3247041, EPI_ISL_3247042, EPI_ISL_3247043, EPI_ISL_3247044, EPI_ISL_3247045, EPI_ISL_3247046, EPI_ISL_3247047, EPI_ISL_3247048, EPI_ISL_3247049, EPI_ISL_3247050, EPI_ISL_3247051, EPI_ISL_3247052, EPI_ISL_3247053, EPI_ISL_3247054, EPI_ISL_3247055, EPI_ISL_3247056, EPI_ISL_3247057, EPI_ISL_3247058 | | | |
| see above | Inselspital Bern (Covid-Track) | Institute for Infectious Diseases | Stefan Neuenschwander, Christian Baumann, Miguel A Terrazos Miani, Sonja Gempeler, Loïc Borcard, Cora Sägesser, Pascal Bittel, Peter Keller, Franziska Suter-Riniker, Stephen L Leib, Alban Ramette |
| EPI_ISL_3247059 | AZ Klina | AZ Klina | Carl Vael - Lynsey Berckmans |
| EPI_ISL_3247060 | Royal Hobart Hospital | Royal Hobart Hospital | Dr L. Cooley, Mr R Vanhaeften |
| EPI_ISL_3247062, EPI_ISL_3247068, EPI_ISL_3247069, EPI_ISL_3247071 | AZ Klina | AZ Klina | Carl Vael - Lynsey Berckmans |
| EPI_ISL_3247073 | AZ Klina | AZ Klina | Carl Vael, Lynsey Berckmans |
| EPI_ISL_3247074 | AZ Klina | AZ Klina | Carl Vael - Lynsey Berckmans |
| EPI_ISL_3247075, EPI_ISL_3247076, EPI_ISL_3247077, EPI_ISL_3247078, EPI_ISL_3247079, EPI_ISL_3247080, EPI_ISL_3247081, EPI_ISL_3247082, EPI_ISL_3247083, EPI_ISL_3247084, EPI_ISL_3247085, EPI_ISL_3247086, EPI_ISL_3247087, EPI_ISL_3247088, EPI_ISL_3247089, EPI_ISL_3247090, EPI_ISL_3247091, EPI_ISL_3247092, EPI_ISL_3247093, EPI_ISL_3247094, EPI_ISL_3247096, EPI_ISL_3247097, EPI_ISL_3247098, EPI_ISL_3247099, EPI_ISL_3247100, EPI_ISL_3247101, EPI_ISL_3247102, EPI_ISL_3247103, EPI_ISL_3247104, EPI_ISL_3247105, EPI_ISL_3247106, EPI_ISL_3247107, EPI_ISL_3247108, EPI_ISL_3247109, EPI_ISL_3247110, EPI_ISL_3247111, EPI_ISL_3247112, EPI_ISL_3247113, EPI_ISL_3247114, EPI_ISL_3247115, EPI_ISL_3247117, EPI_ISL_3247118, EPI_ISL_3247119, EPI_ISL_3247120, EPI_ISL_3247121, EPI_ISL_3247122, EPI_ISL_3247123, EPI_ISL_3247124, EPI_ISL_3247125, EPI_ISL_3247126, EPI_ISL_3247127, EPI_ISL_3247128, EPI_ISL_3247129, EPI_ISL_3247131, EPI_ISL_3247132, EPI_ISL_3247133, EPI_ISL_3247134, EPI_ISL_3247135, EPI_ISL_3247136, EPI_ISL_3247137, EPI_ISL_3247138, EPI_ISL_3247139, EPI_ISL_3247140, EPI_ISL_3247141, EPI_ISL_3247142, EPI_ISL_3247143, EPI_ISL_3247144, EPI_ISL_3247145, EPI_ISL_3247146, EPI_ISL_3247147, EPI_ISL_3247148, EPI_ISL_3247149, EPI_ISL_3247150, EPI_ISL_3247151, EPI_ISL_3247152, EPI_ISL_3247153 | | | |
| see above | Division of Medical Virology, National Health Laboratory Service (NHLS), Tygerberg Hospital / Stellenbosch University | Division of Medical Virology, National Health Laboratory Service (NHLS), Tygerberg Hospital / Stellenbosch University | Susan Engelbrecht, Tongai Maponga, Shannon Wilson, Tania Stander, Kamela Mahlakwane, Gert van Zyl, Wolfgang Preiser |
| EPI_ISL_3247154, EPI_ISL_3247155, EPI_ISL_3247156, EPI_ISL_3247157, EPI_ISL_3247158, EPI_ISL_3247159, EPI_ISL_3247160, EPI_ISL_3247161, EPI_ISL_3247162, EPI_ISL_3247167, EPI_ISL_3247168, EPI_ISL_3247169, EPI_ISL_3247170, EPI_ISL_3247171, EPI_ISL_3247172 | | | |
| see above | AZ Klina | AZ Klina | Carl Vael - Lynsey Berckmans |
| EPI_ISL_3247173 | Hospital del Oriente de Asturias | Laboratorio de Virología HUCA | Sandoval M, Castelló C, Gómez de Oña J, Boga JA, Rojo S, Alvarez-Arguelles ME, Abreu F, Costales I, Perez-Martínez Z, Martín-Rodríguez G, Coto E, Melón S |
| EPI_ISL_3247176 | Hospital Carmen y Severo Ochoa | Laboratorio de Virología HUCA | Sandoval M, Castelló C, Gómez de Oña J, Boga JA, Rojo S, Alvarez-Arguelles ME, Abreu F, Costales I, Perez-Martínez Z, Martín-Rodríguez G, Coto E, Melón S |
| EPI_ISL_3247178, EPI_ISL_3247179, EPI_ISL_3247180, EPI_ISL_3247181, EPI_ISL_3247182, EPI_ISL_3247183, EPI_ISL_3247184, EPI_ISL_3247185, EPI_ISL_3247186, EPI_ISL_3247187, EPI_ISL_3247188, EPI_ISL_3247189, EPI_ISL_3247190, EPI_ISL_3247191, EPI_ISL_3247192, EPI_ISL_3247193, EPI_ISL_3247194, EPI_ISL_3247195, EPI_ISL_3247196, EPI_ISL_3247197, EPI_ISL_3247198, EPI_ISL_3247199, EPI_ISL_3247200, EPI_ISL_3247201, EPI_ISL_3247202, EPI_ISL_3247203, EPI_ISL_3247204, EPI_ISL_3247205, EPI_ISL_3247206, EPI_ISL_3247207, EPI_ISL_3247208, EPI_ISL_3247209, EPI_ISL_3247210, EPI_ISL_3247211, EPI_ISL_3247212, EPI_ISL_3247213, EPI_ISL_3247214, EPI_ISL_3247215, EPI_ISL_3247216, EPI_ISL_3247217, EPI_ISL_3247218, EPI_ISL_3247219, EPI_ISL_3247220, EPI_ISL_3247221, EPI_ISL_3247222, EPI_ISL_3247223, EPI_ISL_3247224, EPI_ISL_3247225, EPI_ISL_3247226, EPI_ISL_3247227, EPI_ISL_3247228, EPI_ISL_3247229, EPI_ISL_3247230, EPI_ISL_3247231, EPI_ISL_3247232, EPI_ISL_3247233 | | | |
| see above | Queensland Health Forensic and Scientific Services | Queensland Health Forensic and Scientific Services | Q-PHIRE Genomics |
| EPI_ISL_3247235, EPI_ISL_3247236, EPI_ISL_3247237, EPI_ISL_3247238, EPI_ISL_3247239, EPI_ISL_3247241, EPI_ISL_3247242, EPI_ISL_3247243, EPI_ISL_3247245, EPI_ISL_3247246, EPI_ISL_3247247, EPI_ISL_3247248, EPI_ISL_3247249, EPI_ISL_3247250, EPI_ISL_3247251, EPI_ISL_3247252, EPI_ISL_3247253, EPI_ISL_3247254, EPI_ISL_3247255, EPI_ISL_3247256, EPI_ISL_3247257, EPI_ISL_3247258, EPI_ISL_3247259, EPI_ISL_3247260 | | | |
| see above | ZOTZ KLIMAS MVZ Düsseldorf-Centrum GbR ÜBAG für Labormedizin, Genetik, Zytologie, Pathologie | Center of Medical Microbiology, Virology, and Hospital Hygiene, University of Duesseldorf | Maximilian Damagnez;Alexander Dilthey;Patrick Finzer;Katrin Hoffmann;Torsten Houwaart;Lisanna Hülse;Malte Kohns Vasconcelos;Nadine Lübke;Jessica Nicolai;Klaus Pfeffer;Daniel Strelow;Jörg Timm;Andreas Walker;Tobias Wienemann;Rainer Zotz |
| EPI_ISL_3247261 | AZ Klina | AZ Klina | Carl Vael - Lynsey Berckmans |
| EPI_ISL_3247284 | Virology Unit, AOUP | Virology Unit, AOUP | Mauro Pistello, Annalisa Capria, Pietro Giorgio Spezia, Susanna Cappellini, Francesca Pasqualetti |
| EPI_ISL_3247285, EPI_ISL_3247286, EPI_ISL_3247287, EPI_ISL_3247288, EPI_ISL_3247289, EPI_ISL_3247290, EPI_ISL_3247292, EPI_ISL_3247293, EPI_ISL_3247294, EPI_ISL_3247295, EPI_ISL_3247296, EPI_ISL_3247297, EPI_ISL_3247298, EPI_ISL_3247299, EPI_ISL_3247300, EPI_ISL_3247301, EPI_ISL_3247302, EPI_ISL_3247303, EPI_ISL_3247304, EPI_ISL_3247306, EPI_ISL_3247307, EPI_ISL_3247308, EPI_ISL_3247309, EPI_ISL_3247310, EPI_ISL_3247311, EPI_ISL_3247312, EPI_ISL_3247313, EPI_ISL_3247314, EPI_ISL_3247315 | | | |
| see above | Clinical Virology | Clinical Bacteriology | Tim Roloff, Fanny Wegner, Helena MB Seth-Smith, Alfredo Mari, Karoline Leuzinger, Julia Bielicki, Manuel Battegay, Hans Hirsch, Adrian Egli |
| EPI_ISL_3247316, EPI_ISL_3247317, EPI_ISL_3247318, EPI_ISL_3247319, EPI_ISL_3247320, EPI_ISL_3247321, EPI_ISL_3247322, EPI_ISL_3247323 | Synlab Suisse SA | Clinical Bacteriology | Tim Roloff, Fanny Wegner, Helena MB Seth-Smith, Alfredo Mari, Karoline Leuzinger, Julia Bielicki, Manuel Battegay, Hans Hirsch, Adrian Egli |
| EPI_ISL_3247324, EPI_ISL_3247325, EPI_ISL_3247326, EPI_ISL_3247327, EPI_ISL_3247328, EPI_ISL_3247329, EPI_ISL_3247330, EPI_ISL_3247331 | Rothen Medizinische Laboratorien AG | Clinical Bacteriology | Tim Roloff, Fanny Wegner, Helena MB Seth-Smith, Alfredo Mari, Karoline Leuzinger, Julia Bielicki, Ingrid Steffen, Manuel Battegay, Hans Hirsch, Adrian Egli |
| EPI_ISL_3247332 | Hopital du Jura: Delemont | Clinical Bacteriology | Tim Roloff, Fanny Wegner, Helena MB Seth-Smith, Alfredo Mari, Karoline Leuzinger, Julia Bielicki, Manuel Battegay, Hans Hirsch, Adrian Egli |
| EPI_ISL_3247333, EPI_ISL_3247334, EPI_ISL_3247335, EPI_ISL_3247336, EPI_ISL_3247337, EPI_ISL_3247338, EPI_ISL_3247339, EPI_ISL_3247340, EPI_ISL_3247341, EPI_ISL_3247342, EPI_ISL_3247343, EPI_ISL_3247344, EPI_ISL_3247345, EPI_ISL_3247346, EPI_ISL_3247347, EPI_ISL_3247348, EPI_ISL_3247349 | | | |
| see above | Medics Labor AG | Clinical Bacteriology | Tim Roloff, Fanny Wegner, Helena MB Seth-Smith, Alfredo Mari, Karoline Leuzinger, Julia Bielicki, Manuel Battegay, Hans Hirsch, Adrian Egli |
| EPI_ISL_3247350 | Labor Team W Ag | Clinical Bacteriology | Tim Roloff, Fanny Wegner, Helena MB Seth-Smith, Alfredo Mari, Karoline Leuzinger, Julia Bielicki, Manuel Battegay, Hans Hirsch, Adrian Egli |
| EPI_ISL_3247351, EPI_ISL_3247352, EPI_ISL_3247353, EPI_ISL_3247354, EPI_ISL_3247355 | Synlab Suisse SA | Clinical Bacteriology | Tim Roloff, Fanny Wegner, Helena MB Seth-Smith, Alfredo Mari, Karoline Leuzinger, Julia Bielicki, Manuel Battegay, Hans Hirsch, Adrian Egli |
| EPI_ISL_3247356 | Clinical Virology | Clinical Bacteriology | Tim Roloff, Fanny Wegner, Helena MB Seth-Smith, Alfredo Mari, Karoline Leuzinger, Julia Bielicki, Manuel Battegay, Hans Hirsch, Adrian Egli |
| EPI_ISL_3247359 | Stadtspital Triemli | Institute of Medical Virology | Verena Kufner, Gabriela Ziltener, Maryam Zaheri, Stefan Schmutz, Annette Audigé, Maria Grünberg, Kevin Steiner, Jon Huder, Cyril Shah, Riccarda Capaul, Guido Bloemberg, Jürg Böni, Michael Huber, Alexandra Trkola |
| EPI_ISL_3247360 | UniversitätsSpital Zürich | Institute of Medical Virology | Verena Kufner, Gabriela Ziltener, Maryam Zaheri, Stefan Schmutz, Annette Audigé, Maria Grünberg, Kevin Steiner, Jon Huder, Cyril Shah, Riccarda Capaul, Guido Bloemberg, Jürg Böni, Michael Huber, Alexandra Trkola |
| EPI_ISL_3247361 | Stadtspital Triemli | Institute of Medical Virology | Verena Kufner, Gabriela Ziltener, Maryam Zaheri, Stefan Schmutz, Annette Audigé, Maria Grünberg, Kevin Steiner, Jon Huder, Cyril Shah, Riccarda Capaul, Guido Bloemberg, Jürg Böni, Michael Huber, Alexandra Trkola |
| EPI_ISL_3247362 | Medica | Institute of Medical Virology | Verena Kufner, Gabriela Ziltener, Maryam Zaheri, Stefan Schmutz, Annette Audigé, Maria Grünberg, Kevin Steiner, Jon Huder, Cyril Shah, Riccarda Capaul, Guido Bloemberg, Jürg Böni, Michael Huber, Alexandra Trkola |
| EPI_ISL_3247363 | Kantonsspital Baden AG | Institute of Medical Virology | Verena Kufner, Gabriela Ziltener, Maryam Zaheri, Stefan Schmutz, Annette Audigé, Maria Grünberg, Kevin Steiner, Jon Huder, Cyril Shah, Riccarda Capaul, Guido Bloemberg, Jürg Böni, Michael Huber, Alexandra Trkola |
| EPI_ISL_3247364 | Medica | Institute of Medical Virology | Verena Kufner, Gabriela Ziltener, Maryam Zaheri, Stefan Schmutz, Annette Audigé, Maria Grünberg, Kevin Steiner, Jon Huder, Cyril Shah, Riccarda Capaul, Guido Bloemberg, Jürg Böni, Michael Huber, Alexandra Trkola |
| EPI_ISL_3247365 | Analytica | Institute of Medical Virology | Verena Kufner, Gabriela Ziltener, Maryam Zaheri, Stefan Schmutz, Annette Audigé, Maria Grünberg, Kevin Steiner, Jon Huder, Cyril Shah, Riccarda Capaul, Guido Bloemberg, Jürg Böni, Michael Huber, Alexandra Trkola |
| EPI_ISL_3247366 | Zentrallabor Zürich | Institute of Medical Virology | Verena Kufner, Gabriela Ziltener, Maryam Zaheri, Stefan Schmutz, Annette Audigé, Maria Grünberg, Kevin Steiner, Jon Huder, Cyril Shah, Riccarda Capaul, Guido Bloemberg, Jürg Böni, Michael Huber, Alexandra Trkola |
| EPI_ISL_3247367 | Zentrum für Labormedizin | Institute of Medical Virology | Verena Kufner, Gabriela Ziltener, Maryam Zaheri, Stefan Schmutz, Annette Audigé, Maria Grünberg, Kevin Steiner, Jon Huder, Cyril Shah, Riccarda Capaul, Guido Bloemberg, Jürg Böni, Michael Huber, Alexandra Trkola |
| EPI_ISL_3247368 | Unilabs | Institute of Medical Virology | Verena Kufner, Gabriela Ziltener, Maryam Zaheri, Stefan Schmutz, Annette Audigé, Maria Grünberg, Kevin Steiner, Jon Huder, Cyril Shah, Riccarda Capaul, Guido Bloemberg, Jürg Böni, Michael Huber, Alexandra Trkola |
| EPI_ISL_3247369 | Kantonsspital Baden AG | Institute of Medical Virology | Verena Kufner, Gabriela Ziltener, Maryam Zaheri, Stefan Schmutz, Annette Audigé, Maria Grünberg, Kevin Steiner, Jon Huder, Cyril Shah, Riccarda Capaul, Guido Bloemberg, Jürg Böni, Michael Huber, Alexandra Trkola |
| EPI_ISL_3247370 | Medica | Institute of Medical Virology | Verena Kufner, Gabriela Ziltener, Maryam Zaheri, Stefan Schmutz, Annette Audigé, Maria Grünberg, Kevin Steiner, Jon Huder, Cyril Shah, Riccarda Capaul, Guido Bloemberg, Jürg Böni, Michael Huber, Alexandra Trkola |
| EPI_ISL_3247371 | Diamisanto GmbH | Institute of Medical Virology | Verena Kufner, Gabriela Ziltener, Maryam Zaheri, Stefan Schmutz, Annette Audigé, Maria Grünberg, Kevin Steiner, Jon Huder, Cyril Shah, Riccarda Capaul, Guido Bloemberg, Jürg Böni, Michael Huber, Alexandra Trkola |
| EPI_ISL_3247372 | Universität Zürich | Institute of Medical Virology | Verena Kufner, Gabriela Ziltener, Maryam Zaheri, Stefan Schmutz, Annette Audigé, Maria Grünberg, Kevin Steiner, Jon Huder, Cyril Shah, Riccarda Capaul, Guido Bloemberg, Jürg Böni, Michael Huber, Alexandra Trkola |
| EPI_ISL_3247373 | Diamisanto GmbH | Institute of Medical Virology | Verena Kufner, Gabriela Ziltener, Maryam Zaheri, Stefan Schmutz, Annette Audigé, Maria Grünberg, Kevin Steiner, Jon Huder, Cyril Shah, Riccarda Capaul, Guido Bloemberg, Jürg Böni, Michael Huber, Alexandra Trkola |
| EPI_ISL_3247374 | Zentrum für Labormedizin | Institute of Medical Virology | Verena Kufner, Gabriela Ziltener, Maryam Zaheri, Stefan Schmutz, Annette Audigé, Maria Grünberg, Kevin Steiner, Jon Huder, Cyril Shah, Riccarda Capaul, Guido Bloemberg, Jürg Böni, Michael Huber, Alexandra Trkola |
| EPI_ISL_3247375 | Kantonsspital Aarau | Institute of Medical Virology | Verena Kufner, Gabriela Ziltener, Maryam Zaheri, Stefan Schmutz, Annette Audigé, Maria Grünberg, Kevin Steiner, Jon Huder, Cyril Shah, Riccarda Capaul, Guido Bloemberg, Jürg Böni, Michael Huber, Alexandra Trkola |
| EPI_ISL_3247376, EPI_ISL_3247377, EPI_ISL_3247378, EPI_ISL_3247379, EPI_ISL_3247380, EPI_ISL_3247381 | UniversitätsSpital Zürich | Institute of Medical Virology | Verena Kufner, Gabriela Ziltener, Maryam Zaheri, Stefan Schmutz, Annette Audigé, Maria Grünberg, Kevin Steiner, Jon Huder, Cyril Shah, Riccarda Capaul, Guido Bloemberg, Jürg Böni, Michael Huber, Alexandra Trkola |
| EPI_ISL_3247383, EPI_ISL_3247384 | Universitäts-Kinderspital Zürich | Institute of Medical Virology | Verena Kufner, Gabriela Ziltener, Maryam Zaheri, Stefan Schmutz, Annette Audigé, Maria Grünberg, Kevin Steiner, Jon Huder, Cyril Shah, Riccarda Capaul, Guido Bloemberg, Jürg Böni, Michael Huber, Alexandra Trkola |
| EPI_ISL_3247385, | Zentrallabor Zürich | Institute of Medical Virology | Verena Kufner, Gabriela Ziltener, Maryam Zaheri, Stefan Schmutz, Annette Audigé, Maria Grünberg, Kevin Steiner, Jon Huder, Cyril Shah, Riccarda Capaul, Guido Bloemberg, Jürg Böni, Michael Huber, Alexandra Trkola |

| | | | |
|---|---|---|---|
| EPI_ISL_3247386, EPI_ISL_3247387, EPI_ISL_3247388, EPI_ISL_3247389 | | | |
| EPI_ISL_3247390, EPI_ISL_3247391, EPI_ISL_3247392, EPI_ISL_3247393, EPI_ISL_3247394 | Stadtspital Triemli | Institute of Medical Virology | Verena Kufner, Gabriela Ziltener, Maryam Zaheri, Stefan Schmutz, Annette Audigé, Maria Grünberg, Kevin Steiner, Jon Huder, Cyril Shah, Riccarda Capaul, Guido Bloemberg, Jürg Böni, Michael Huber, Alexandra Trkola |
| EPI_ISL_3247395, EPI_ISL_3247396, EPI_ISL_3247397, EPI_ISL_3247398 | Universität Zürich | Institute of Medical Virology | Verena Kufner, Gabriela Ziltener, Maryam Zaheri, Stefan Schmutz, Annette Audigé, Maria Grünberg, Kevin Steiner, Jon Huder, Cyril Shah, Riccarda Capaul, Guido Bloemberg, Jürg Böni, Michael Huber, Alexandra Trkola |
| EPI_ISL_3247401, EPI_ISL_3247402 | MD PHL | Maryland Department of Health Laboratories Administration | Eric N. Keller, Jillian Loomis, Kwang Low, Terence L. Moore, Ami Patel, and Robert Myers |
| EPI_ISL_3247488, EPI_ISL_3247489, EPI_ISL_3247491, EPI_ISL_3247492, EPI_ISL_3247493, EPI_ISL_3247494, EPI_ISL_3247495, EPI_ISL_3247496, EPI_ISL_3247497, EPI_ISL_3247498, EPI_ISL_3247499, EPI_ISL_3247500, EPI_ISL_3247501, EPI_ISL_3247502 | | | |
| see above | INNO Diagnostics Reference Laboratory | RCMI-Center for Research Resources, Ponce Research Institute | Vanessa Rivera-Amill, Andrea Arias-García, Raphael Sánchez-Torres |
| EPI_ISL_3247508, EPI_ISL_3247509, EPI_ISL_3247510, EPI_ISL_3247511, EPI_ISL_3247512, EPI_ISL_3247513, EPI_ISL_3247514, EPI_ISL_3247515, EPI_ISL_3247516, EPI_ISL_3247517, EPI_ISL_3247518, EPI_ISL_3247519, EPI_ISL_3247520, EPI_ISL_3247521, EPI_ISL_3247522, EPI_ISL_3247523, EPI_ISL_3247524, EPI_ISL_3247525, EPI_ISL_3247526 | | | |
| see above | ILSPPR | RCMI-Center for Research Resources, Ponce Research Institute | Vanessa Rivera-Amill, Andrea Arias-García, Raphael Sánchez-Torres |
| EPI_ISL_3247527, EPI_ISL_3247528, EPI_ISL_3247529 | Institute for Medical Research, Infectious Disease Research Centre, National Institutes of Health, Ministry of Health Malaysia | Institute for Medical Research, Infectious Disease Research Centre, National Institutes of Health, Ministry of Health Malaysia | Suppiah J, Kamel K, Mohd Zawawi Z, Azizan MA, Ramly N, Robert F, Anasir MI, Thayan R |
| EPI_ISL_3247530, EPI_ISL_3247531, EPI_ISL_3247532, EPI_ISL_3247533, EPI_ISL_3247534, EPI_ISL_3247535, EPI_ISL_3247536, EPI_ISL_3247537, EPI_ISL_3247539, EPI_ISL_3247540, EPI_ISL_3247541, EPI_ISL_3247542, EPI_ISL_3247545, EPI_ISL_3247546, EPI_ISL_3247547, EPI_ISL_3247548, EPI_ISL_3247549, EPI_ISL_3247550, EPI_ISL_3247551, EPI_ISL_3247554, EPI_ISL_3247555, EPI_ISL_3247556, EPI_ISL_3247557, EPI_ISL_3247558, EPI_ISL_3247559, EPI_ISL_3247563, EPI_ISL_3247565, EPI_ISL_3247566, EPI_ISL_3247567, EPI_ISL_3247569, EPI_ISL_3247570, EPI_ISL_3247571, EPI_ISL_3247572, EPI_ISL_3247573, EPI_ISL_3247574, EPI_ISL_3247575, EPI_ISL_3247576, EPI_ISL_3247577, EPI_ISL_3247578, EPI_ISL_3247579, EPI_ISL_3247580, EPI_ISL_3247581, EPI_ISL_3247582, EPI_ISL_3247583, EPI_ISL_3247584, EPI_ISL_3247585, EPI_ISL_3247586, EPI_ISL_3247587, EPI_ISL_3247588, EPI_ISL_3247589, EPI_ISL_3247590, EPI_ISL_3247591, EPI_ISL_3247592, EPI_ISL_3247593, EPI_ISL_3247594, EPI_ISL_3247595, EPI_ISL_3247596, EPI_ISL_3247597, EPI_ISL_3247598, EPI_ISL_3247599, EPI_ISL_3247600, EPI_ISL_3247601, EPI_ISL_3247602, EPI_ISL_3247603, EPI_ISL_3247604, EPI_ISL_3247605, EPI_ISL_3247608, EPI_ISL_3247609, EPI_ISL_3247610, EPI_ISL_3247611, EPI_ISL_3247612, EPI_ISL_3247613, EPI_ISL_3247614, EPI_ISL_3247615, EPI_ISL_3247616, EPI_ISL_3247617, EPI_ISL_3247618, EPI_ISL_3247619, EPI_ISL_3247620, EPI_ISL_3247621, EPI_ISL_3247622, EPI_ISL_3247623, EPI_ISL_3247624, EPI_ISL_3247625, EPI_ISL_3247626, EPI_ISL_3247627, EPI_ISL_3247628, EPI_ISL_3247629, EPI_ISL_3247630, EPI_ISL_3247631, EPI_ISL_3247632, EPI_ISL_3247633, EPI_ISL_3247634, EPI_ISL_3247635, EPI_ISL_3247636, EPI_ISL_3247637, EPI_ISL_3247640, EPI_ISL_3247641, EPI_ISL_3247642, EPI_ISL_3247643, EPI_ISL_3247644, EPI_ISL_3247645, EPI_ISL_3247646, EPI_ISL_3247647, EPI_ISL_3247648, EPI_ISL_3247649, EPI_ISL_3247650, EPI_ISL_3247651, EPI_ISL_3247652, EPI_ISL_3247653, EPI_ISL_3247654, EPI_ISL_3247655, EPI_ISL_3247656, EPI_ISL_3247657, EPI_ISL_3247658, EPI_ISL_3247659, EPI_ISL_3247660, EPI_ISL_3247661, EPI_ISL_3247662, EPI_ISL_3247663, EPI_ISL_3247664, EPI_ISL_3247665, EPI_ISL_3247666, EPI_ISL_3247667, EPI_ISL_3247668, EPI_ISL_3247670, EPI_ISL_3247671, EPI_ISL_3247672, EPI_ISL_3247673, EPI_ISL_3247674, EPI_ISL_3247675, EPI_ISL_3247676, EPI_ISL_3247677, EPI_ISL_3247678, EPI_ISL_3247679, EPI_ISL_3247680, EPI_ISL_3247681, EPI_ISL_3247682, EPI_ISL_3247683, EPI_ISL_3247684, EPI_ISL_3247685, EPI_ISL_3247686, EPI_ISL_3247687, EPI_ISL_3247688, EPI_ISL_3247689, EPI_ISL_3247690, EPI_ISL_3247691, EPI_ISL_3247692, EPI_ISL_3247693, EPI_ISL_3247694, EPI_ISL_3247695, EPI_ISL_3247696, EPI_ISL_3247697, EPI_ISL_3247698, EPI_ISL_3247699 | | | |
| see above | National Virus Reference Laboratory | National Virus Reference Laboratory | Zoe Yandle, Charlene Bennett, Gabriel Gonzalez, Michael Carr, Jonathan Dean, Cillian F De Gascun |
| EPI_ISL_3247700 | Institute for Medical Research, Infectious Disease Research Centre, National Institutes of Health, Ministry of Health Malaysia | Institute for Medical Research, Infectious Disease Research Centre, National Institutes of Health, Ministry of Health Malaysia | Suppiah J, Kamel K, Mohd Zawawi Z, Azizan MA, Ramly N, Robert F, Anasir MI, Thayan R |
| EPI_ISL_3247701, EPI_ISL_3247702, EPI_ISL_3247703, EPI_ISL_3247704, EPI_ISL_3247705, EPI_ISL_3247706, EPI_ISL_3247707, EPI_ISL_3247708, EPI_ISL_3247709, EPI_ISL_3247710, EPI_ISL_3247711, EPI_ISL_3247712, EPI_ISL_3247713, EPI_ISL_3247714, EPI_ISL_3247715, EPI_ISL_3247716, EPI_ISL_3247717, EPI_ISL_3247718, EPI_ISL_3247719, EPI_ISL_3247720, EPI_ISL_3247721, EPI_ISL_3247722, EPI_ISL_3247723, EPI_ISL_3247724, EPI_ISL_3247725, EPI_ISL_3247726, EPI_ISL_3247727, EPI_ISL_3247728, EPI_ISL_3247729, EPI_ISL_3247730, EPI_ISL_3247731, EPI_ISL_3247732, EPI_ISL_3247733, EPI_ISL_3247734, EPI_ISL_3247735, EPI_ISL_3247736, EPI_ISL_3247737, EPI_ISL_3247738, EPI_ISL_3247739, EPI_ISL_3247740, EPI_ISL_3247741, EPI_ISL_3247742, EPI_ISL_3247743, EPI_ISL_3247744, EPI_ISL_3247745, EPI_ISL_3247746, EPI_ISL_3247747, EPI_ISL_3247748, EPI_ISL_3247749, EPI_ISL_3247750, EPI_ISL_3247751, EPI_ISL_3247752, EPI_ISL_3247753, EPI_ISL_3247754, EPI_ISL_3247755, EPI_ISL_3247756, EPI_ISL_3247757, EPI_ISL_3247758, EPI_ISL_3247759, EPI_ISL_3247760, EPI_ISL_3247761, EPI_ISL_3247762, EPI_ISL_3247763, EPI_ISL_3247764, EPI_ISL_3247765, EPI_ISL_3247766, EPI_ISL_3247767 | | | |
| see above | National Virus Reference Laboratory | National Virus Reference Laboratory | Zoe Yandle, Charlene Bennett, Gabriel Gonzalez, Michael Carr, Jonathan Dean, Cillian F De Gascun |
| EPI_ISL_3247768, EPI_ISL_3247769 | Institute for Medical Research, Infectious Disease Research Centre, National Institutes of Health, Ministry of Health Malaysia | Institute for Medical Research, Infectious Disease Research Centre, National Institutes of Health, Ministry of Health Malaysia | Suppiah J, Kamel K, Mohd Zawawi Z, Azizan MA, Ramly N, Robert F, Anasir MI, Thayan R |
| EPI_ISL_3247773 | AZ Klina | AZ Klina | Carl Vael - Lynsey Berckmans |
| EPI_ISL_3247774, EPI_ISL_3247775, EPI_ISL_3247776 | Lighthouse Laboratory Plymouth | Wellcome Sanger Institute for the COVID-19 Genomics UK (COG-UK) Consortium | Lighthouse Laboratory Plymouth and Alex Alderton, Roberto Amato, Jeffrey Barrett, Sonia Goncalves, Ewan Harrison, David K. Jackson, Ian Johnston, Dominic Kwiatkowski, Cordelia Langford, John Sillitoe on behalf of the Wellcome Sanger Institute COVID-19 Surveillance Team |
| EPI_ISL_3247777 | Lighthouse Lab in Glasgow | Wellcome Sanger Institute for the COVID-19 Genomics UK (COG-UK) Consortium | Harper VanSteenhouse, Yumi Kasai, David Gray, Carol Clugston, Anna Dominiczak and Alex Alderton, Roberto Amato, Jeffrey Barrett, Sonia Goncalves, Ewan Harrison, David K. Jackson, Ian Johnston, Dominic Kwiatkowski, Cordelia Langford, John Sillitoe on behalf of the Wellcome Sanger Institute COVID-19 Surveillance Team |
| EPI_ISL_3247778, EPI_ISL_3247779, EPI_ISL_3247780, EPI_ISL_3247781, EPI_ISL_3247782, EPI_ISL_3247783, EPI_ISL_3247784, EPI_ISL_3247785 | Lighthouse Laboratory Plymouth | Wellcome Sanger Institute for the COVID-19 Genomics UK (COG-UK) Consortium | Lighthouse Laboratory Plymouth and Alex Alderton, Roberto Amato, Jeffrey Barrett, Sonia Goncalves, Ewan Harrison, David K. Jackson, Ian Johnston, Dominic Kwiatkowski, Cordelia Langford, John Sillitoe on behalf of the Wellcome Sanger Institute COVID-19 Surveillance Team |
| EPI_ISL_3247786 | Lighthouse Lab in Glasgow | Wellcome Sanger Institute for the COVID-19 Genomics UK (COG-UK) Consortium | Harper VanSteenhouse, Yumi Kasai, David Gray, Carol Clugston, Anna Dominiczak and Alex Alderton, Roberto Amato, Jeffrey Barrett, Sonia Goncalves, Ewan Harrison, David K. Jackson, Ian Johnston, Dominic Kwiatkowski, Cordelia Langford, John Sillitoe on behalf of the Wellcome Sanger Institute COVID-19 Surveillance Team |
| EPI_ISL_3247787 | Lighthouse Laboratory Plymouth | Wellcome Sanger Institute for the COVID-19 Genomics UK (COG-UK) Consortium | Lighthouse Laboratory Plymouth and Alex Alderton, Roberto Amato, Jeffrey Barrett, Sonia Goncalves, Ewan Harrison, David K. Jackson, Ian Johnston, Dominic Kwiatkowski, Cordelia Langford, John Sillitoe on behalf of the Wellcome Sanger Institute COVID-19 Surveillance Team |
| EPI_ISL_3247788 | Lighthouse Lab in Glasgow | Wellcome Sanger Institute for the COVID-19 Genomics UK (COG-UK) Consortium | Harper VanSteenhouse, Yumi Kasai, David Gray, Carol Clugston, Anna Dominiczak and Alex Alderton, Roberto Amato, Jeffrey Barrett, Sonia Goncalves, Ewan Harrison, David K. Jackson, Ian Johnston, Dominic Kwiatkowski, Cordelia Langford, John Sillitoe on behalf of the Wellcome Sanger Institute COVID-19 Surveillance Team |
| EPI_ISL_3247789, EPI_ISL_3247790, EPI_ISL_3247791 | Lighthouse Laboratory Plymouth | Wellcome Sanger Institute for the COVID-19 Genomics UK (COG-UK) Consortium | Lighthouse Laboratory Plymouth and Alex Alderton, Roberto Amato, Jeffrey Barrett, Sonia Goncalves, Ewan Harrison, David K. Jackson, Ian Johnston, Dominic Kwiatkowski, Cordelia Langford, John Sillitoe on behalf of the Wellcome Sanger Institute COVID-19 Surveillance Team |
| EPI_ISL_3247792 | Lighthouse Lab in Glasgow | Wellcome Sanger Institute for the COVID-19 Genomics UK (COG-UK) Consortium | Harper VanSteenhouse, Yumi Kasai, David Gray, Carol Clugston, Anna Dominiczak and Alex Alderton, Roberto Amato, Jeffrey Barrett, Sonia Goncalves, Ewan Harrison, David K. Jackson, Ian Johnston, Dominic Kwiatkowski, Cordelia Langford, John Sillitoe on behalf of the Wellcome Sanger Institute COVID-19 Surveillance Team |
| EPI_ISL_3247793, EPI_ISL_3247794, EPI_ISL_3247795, EPI_ISL_3247796, EPI_ISL_3247797 | Lighthouse Laboratory Plymouth | Wellcome Sanger Institute for the COVID-19 Genomics UK (COG-UK) Consortium | Lighthouse Laboratory Plymouth and Alex Alderton, Roberto Amato, Jeffrey Barrett, Sonia Goncalves, Ewan Harrison, David K. Jackson, Ian Johnston, Dominic Kwiatkowski, Cordelia Langford, John Sillitoe on behalf of the Wellcome Sanger Institute COVID-19 Surveillance Team |
| EPI_ISL_3247798, EPI_ISL_3247799 | Lighthouse Lab in Glasgow | Wellcome Sanger Institute for the COVID-19 Genomics UK (COG-UK) Consortium | Harper VanSteenhouse, Yumi Kasai, David Gray, Carol Clugston, Anna Dominiczak and Alex Alderton, Roberto Amato, Jeffrey Barrett, Sonia Goncalves, Ewan Harrison, David K. Jackson, Ian Johnston, Dominic Kwiatkowski, Cordelia Langford, John Sillitoe on behalf of the Wellcome Sanger Institute COVID-19 Surveillance Team |
| EPI_ISL_3247800, EPI_ISL_3247801, EPI_ISL_3247802, EPI_ISL_3247803, EPI_ISL_3247804, EPI_ISL_3247805, EPI_ISL_3247806, EPI_ISL_3247807, EPI_ISL_3247808, EPI_ISL_3247809, EPI_ISL_3247810 | | | |
| see above | Lighthouse Laboratory Plymouth | Wellcome Sanger Institute for the COVID-19 Genomics UK (COG-UK) Consortium | Lighthouse Laboratory Plymouth and Alex Alderton, Roberto Amato, Jeffrey Barrett, Sonia Goncalves, Ewan Harrison, David K. Jackson, Ian Johnston, Dominic Kwiatkowski, Cordelia Langford, John Sillitoe on behalf of the Wellcome Sanger Institute COVID-19 Surveillance Team |
| EPI_ISL_3247811 | Lighthouse Lab in Glasgow | Wellcome Sanger Institute for the COVID-19 Genomics UK (COG-UK) Consortium | Harper VanSteenhouse, Yumi Kasai, David Gray, Carol Clugston, Anna Dominiczak and Alex Alderton, Roberto Amato, Jeffrey Barrett, Sonia Goncalves, Ewan Harrison, David K. Jackson, Ian Johnston, Dominic Kwiatkowski, Cordelia Langford, John Sillitoe on behalf of the Wellcome Sanger Institute COVID-19 Surveillance Team |
| EPI_ISL_3247812, EPI_ISL_3247813, EPI_ISL_3247814, EPI_ISL_3247815, EPI_ISL_3247816 | Lighthouse Laboratory Plymouth | Wellcome Sanger Institute for the COVID-19 Genomics UK (COG-UK) Consortium | Lighthouse Laboratory Plymouth and Alex Alderton, Roberto Amato, Jeffrey Barrett, Sonia Goncalves, Ewan Harrison, David K. Jackson, Ian Johnston, Dominic Kwiatkowski, Cordelia Langford, John Sillitoe on behalf of the Wellcome Sanger Institute COVID-19 Surveillance Team |
| EPI_ISL_3247817 | Lighthouse Lab in Glasgow | Wellcome Sanger Institute for the COVID-19 Genomics UK (COG-UK) Consortium | Harper VanSteenhouse, Yumi Kasai, David Gray, Carol Clugston, Anna Dominiczak and Alex Alderton, Roberto Amato, Jeffrey Barrett, Sonia Goncalves, Ewan Harrison, David K. Jackson, Ian Johnston, Dominic Kwiatkowski, Cordelia Langford, John Sillitoe on behalf of the Wellcome Sanger Institute COVID-19 Surveillance Team |
| EPI_ISL_3247818, EPI_ISL_3247819, EPI_ISL_3247820, EPI_ISL_3247821 | Lighthouse Laboratory Plymouth | Wellcome Sanger Institute for the COVID-19 Genomics UK (COG-UK) Consortium | Lighthouse Laboratory Plymouth and Alex Alderton, Roberto Amato, Jeffrey Barrett, Sonia Goncalves, Ewan Harrison, David K. Jackson, Ian Johnston, Dominic Kwiatkowski, Cordelia Langford, John Sillitoe on behalf of the Wellcome Sanger Institute COVID-19 Surveillance Team |
| EPI_ISL_3247822 | Lighthouse Lab in Glasgow | Wellcome Sanger Institute for the COVID-19 Genomics UK (COG-UK) Consortium | Harper VanSteenhouse, Yumi Kasai, David Gray, Carol Clugston, Anna Dominiczak and Alex Alderton, Roberto Amato, Jeffrey Barrett, Sonia Goncalves, Ewan Harrison, David K. Jackson, Ian Johnston, Dominic Kwiatkowski, Cordelia Langford, John Sillitoe on behalf of the Wellcome Sanger Institute COVID-19 Surveillance Team |
| EPI_ISL_3247823, EPI_ISL_3247824 | Lighthouse Laboratory Plymouth | Wellcome Sanger Institute for the COVID-19 Genomics UK (COG-UK) Consortium | Lighthouse Laboratory Plymouth and Alex Alderton, Roberto Amato, Jeffrey Barrett, Sonia Goncalves, Ewan Harrison, David K. Jackson, Ian Johnston, Dominic Kwiatkowski, Cordelia Langford, John Sillitoe on behalf of the Wellcome Sanger Institute COVID-19 Surveillance Team |
| EPI_ISL_3247825 | Lighthouse Lab in Glasgow | Wellcome Sanger Institute for the COVID-19 Genomics UK (COG-UK) Consortium | Harper VanSteenhouse, Yumi Kasai, David Gray, Carol Clugston, Anna Dominiczak and Alex Alderton, Roberto Amato, Jeffrey Barrett, Sonia Goncalves, Ewan Harrison, David K. Jackson, Ian Johnston, Dominic Kwiatkowski, Cordelia Langford, John Sillitoe on behalf of the Wellcome Sanger Institute COVID-19 Surveillance Team |

| | | | |
|---|---|---|---|
| EPI_ISL_3247826, EPI_ISL_3247827, EPI_ISL_3247828, EPI_ISL_3247829, EPI_ISL_3247830, EPI_ISL_3247831, EPI_ISL_3247832, EPI_ISL_3247833, EPI_ISL_3247834, EPI_ISL_3247835, EPI_ISL_3247836, EPI_ISL_3247837, EPI_ISL_3247838, EPI_ISL_3247839, EPI_ISL_3247840, EPI_ISL_3247841, EPI_ISL_3247842, EPI_ISL_3247843, EPI_ISL_3247844 | | | |
| see above | Lighthouse Laboratory Plymouth | Wellcome Sanger Institute for the COVID-19 Genomics UK (COG-UK) Consortium | Lighthouse Laboratory Plymouth and Alex Alderton, Roberto Amato, Jeffrey Barrett, Sonia Goncalves, Ewan Harrison, David K. Jackson, Ian Johnston, Dominic Kwiatkowski, Cordelia Langford, John Sillitoe on behalf of the Wellcome Sanger Institute COVID-19 Surveillance Team |
| EPI_ISL_3247845, EPI_ISL_3247846 | Lighthouse Lab in Glasgow | Wellcome Sanger Institute for the COVID-19 Genomics UK (COG-UK) Consortium | Harper VanSteenhouse, Yumi Kasai, David Gray, Carol Clugston, Anna Dominiczak and Alex Alderton, Roberto Amato, Jeffrey Barrett, Sonia Goncalves, Ewan Harrison, David K. Jackson, Ian Johnston, Dominic Kwiatkowski, Cordelia Langford, John Sillitoe on behalf of the Wellcome Sanger Institute COVID-19 Surveillance Team |
| EPI_ISL_3247847, EPI_ISL_3247848, EPI_ISL_3247849, EPI_ISL_3247850, EPI_ISL_3247851, EPI_ISL_3247852, EPI_ISL_3247853, EPI_ISL_3247854 | Lighthouse Laboratory Plymouth | Wellcome Sanger Institute for the COVID-19 Genomics UK (COG-UK) Consortium | Lighthouse Laboratory Plymouth and Alex Alderton, Roberto Amato, Jeffrey Barrett, Sonia Goncalves, Ewan Harrison, David K. Jackson, Ian Johnston, Dominic Kwiatkowski, Cordelia Langford, John Sillitoe on behalf of the Wellcome Sanger Institute COVID-19 Surveillance Team |
| EPI_ISL_3247855 | Lighthouse Lab in Glasgow | Wellcome Sanger Institute for the COVID-19 Genomics UK (COG-UK) Consortium | Harper VanSteenhouse, Yumi Kasai, David Gray, Carol Clugston, Anna Dominiczak and Alex Alderton, Roberto Amato, Jeffrey Barrett, Sonia Goncalves, Ewan Harrison, David K. Jackson, Ian Johnston, Dominic Kwiatkowski, Cordelia Langford, John Sillitoe on behalf of the Wellcome Sanger Institute COVID-19 Surveillance Team |
| EPI_ISL_3247856, EPI_ISL_3247857, EPI_ISL_3247858 | Lighthouse Laboratory Plymouth | Wellcome Sanger Institute for the COVID-19 Genomics UK (COG-UK) Consortium | Lighthouse Laboratory Plymouth and Alex Alderton, Roberto Amato, Jeffrey Barrett, Sonia Goncalves, Ewan Harrison, David K. Jackson, Ian Johnston, Dominic Kwiatkowski, Cordelia Langford, John Sillitoe on behalf of the Wellcome Sanger Institute COVID-19 Surveillance Team |
| EPI_ISL_3247859 | Lighthouse Lab in Glasgow | Wellcome Sanger Institute for the COVID-19 Genomics UK (COG-UK) Consortium | Harper VanSteenhouse, Yumi Kasai, David Gray, Carol Clugston, Anna Dominiczak and Alex Alderton, Roberto Amato, Jeffrey Barrett, Sonia Goncalves, Ewan Harrison, David K. Jackson, Ian Johnston, Dominic Kwiatkowski, Cordelia Langford, John Sillitoe on behalf of the Wellcome Sanger Institute COVID-19 Surveillance Team |
| EPI_ISL_3247860, EPI_ISL_3247861, EPI_ISL_3247862, EPI_ISL_3247863, EPI_ISL_3247864, EPI_ISL_3247865, EPI_ISL_3247866, EPI_ISL_3247867, EPI_ISL_3247868, EPI_ISL_3247869, EPI_ISL_3247870, EPI_ISL_3247871 | | | |
| see above | Lighthouse Laboratory Plymouth | Wellcome Sanger Institute for the COVID-19 Genomics UK (COG-UK) Consortium | Lighthouse Laboratory Plymouth and Alex Alderton, Roberto Amato, Jeffrey Barrett, Sonia Goncalves, Ewan Harrison, David K. Jackson, Ian Johnston, Dominic Kwiatkowski, Cordelia Langford, John Sillitoe on behalf of the Wellcome Sanger Institute COVID-19 Surveillance Team |
| EPI_ISL_3247872, EPI_ISL_3247873 | Lighthouse Lab in Glasgow | Wellcome Sanger Institute for the COVID-19 Genomics UK (COG-UK) Consortium | Harper VanSteenhouse, Yumi Kasai, David Gray, Carol Clugston, Anna Dominiczak and Alex Alderton, Roberto Amato, Jeffrey Barrett, Sonia Goncalves, Ewan Harrison, David K. Jackson, Ian Johnston, Dominic Kwiatkowski, Cordelia Langford, John Sillitoe on behalf of the Wellcome Sanger Institute COVID-19 Surveillance Team |
| EPI_ISL_3247874, EPI_ISL_3247875, EPI_ISL_3247876, EPI_ISL_3247877, EPI_ISL_3247878, EPI_ISL_3247879, EPI_ISL_3247880, EPI_ISL_3247881 | Lighthouse Laboratory Plymouth | Wellcome Sanger Institute for the COVID-19 Genomics UK (COG-UK) Consortium | Lighthouse Laboratory Plymouth and Alex Alderton, Roberto Amato, Jeffrey Barrett, Sonia Goncalves, Ewan Harrison, David K. Jackson, Ian Johnston, Dominic Kwiatkowski, Cordelia Langford, John Sillitoe on behalf of the Wellcome Sanger Institute COVID-19 Surveillance Team |
| EPI_ISL_3247882 | Lighthouse Lab in Glasgow | Wellcome Sanger Institute for the COVID-19 Genomics UK (COG-UK) Consortium | Harper VanSteenhouse, Yumi Kasai, David Gray, Carol Clugston, Anna Dominiczak and Alex Alderton, Roberto Amato, Jeffrey Barrett, Sonia Goncalves, Ewan Harrison, David K. Jackson, Ian Johnston, Dominic Kwiatkowski, Cordelia Langford, John Sillitoe on behalf of the Wellcome Sanger Institute COVID-19 Surveillance Team |
| EPI_ISL_3247883, EPI_ISL_3247884, EPI_ISL_3247885, EPI_ISL_3247886, EPI_ISL_3247887 | Lighthouse Laboratory Plymouth | Wellcome Sanger Institute for the COVID-19 Genomics UK (COG-UK) Consortium | Lighthouse Laboratory Plymouth and Alex Alderton, Roberto Amato, Jeffrey Barrett, Sonia Goncalves, Ewan Harrison, David K. Jackson, Ian Johnston, Dominic Kwiatkowski, Cordelia Langford, John Sillitoe on behalf of the Wellcome Sanger Institute COVID-19 Surveillance Team |
| EPI_ISL_3247888 | Lighthouse Lab in Glasgow | Wellcome Sanger Institute for the COVID-19 Genomics UK (COG-UK) Consortium | Harper VanSteenhouse, Yumi Kasai, David Gray, Carol Clugston, Anna Dominiczak and Alex Alderton, Roberto Amato, Jeffrey Barrett, Sonia Goncalves, Ewan Harrison, David K. Jackson, Ian Johnston, Dominic Kwiatkowski, Cordelia Langford, John Sillitoe on behalf of the Wellcome Sanger Institute COVID-19 Surveillance Team |
| EPI_ISL_3247889, EPI_ISL_3247890, EPI_ISL_3247891, EPI_ISL_3247892, EPI_ISL_3247893, EPI_ISL_3247894, EPI_ISL_3247895, EPI_ISL_3247896 | Lighthouse Laboratory Plymouth | Wellcome Sanger Institute for the COVID-19 Genomics UK (COG-UK) Consortium | Lighthouse Laboratory Plymouth and Alex Alderton, Roberto Amato, Jeffrey Barrett, Sonia Goncalves, Ewan Harrison, David K. Jackson, Ian Johnston, Dominic Kwiatkowski, Cordelia Langford, John Sillitoe on behalf of the Wellcome Sanger Institute COVID-19 Surveillance Team |
| EPI_ISL_3247897, EPI_ISL_3247898 | Lighthouse Lab in Glasgow | Wellcome Sanger Institute for the COVID-19 Genomics UK (COG-UK) Consortium | Harper VanSteenhouse, Yumi Kasai, David Gray, Carol Clugston, Anna Dominiczak and Alex Alderton, Roberto Amato, Jeffrey Barrett, Sonia Goncalves, Ewan Harrison, David K. Jackson, Ian Johnston, Dominic Kwiatkowski, Cordelia Langford, John Sillitoe on behalf of the Wellcome Sanger Institute COVID-19 Surveillance Team |
| EPI_ISL_3247899, EPI_ISL_3247900, EPI_ISL_3247901, EPI_ISL_3247902 | Lighthouse Laboratory Plymouth | Wellcome Sanger Institute for the COVID-19 Genomics UK (COG-UK) Consortium | Lighthouse Laboratory Plymouth and Alex Alderton, Roberto Amato, Jeffrey Barrett, Sonia Goncalves, Ewan Harrison, David K. Jackson, Ian Johnston, Dominic Kwiatkowski, Cordelia Langford, John Sillitoe on behalf of the Wellcome Sanger Institute COVID-19 Surveillance Team |
| EPI_ISL_3247903 | Lighthouse Lab in Glasgow | Wellcome Sanger Institute for the COVID-19 Genomics UK (COG-UK) Consortium | Harper VanSteenhouse, Yumi Kasai, David Gray, Carol Clugston, Anna Dominiczak and Alex Alderton, Roberto Amato, Jeffrey Barrett, Sonia Goncalves, Ewan Harrison, David K. Jackson, Ian Johnston, Dominic Kwiatkowski, Cordelia Langford, John Sillitoe on behalf of the Wellcome Sanger Institute COVID-19 Surveillance Team |
| EPI_ISL_3247904, EPI_ISL_3247905, EPI_ISL_3247906 | Lighthouse Laboratory Plymouth | Wellcome Sanger Institute for the COVID-19 Genomics UK (COG-UK) Consortium | Lighthouse Laboratory Plymouth and Alex Alderton, Roberto Amato, Jeffrey Barrett, Sonia Goncalves, Ewan Harrison, David K. Jackson, Ian Johnston, Dominic Kwiatkowski, Cordelia Langford, John Sillitoe on behalf of the Wellcome Sanger Institute COVID-19 Surveillance Team |
| EPI_ISL_3247907 | Lighthouse Lab in Glasgow | Wellcome Sanger Institute for the COVID-19 Genomics UK (COG-UK) Consortium | Harper VanSteenhouse, Yumi Kasai, David Gray, Carol Clugston, Anna Dominiczak and Alex Alderton, Roberto Amato, Jeffrey Barrett, Sonia Goncalves, Ewan Harrison, David K. Jackson, Ian Johnston, Dominic Kwiatkowski, Cordelia Langford, John Sillitoe on behalf of the Wellcome Sanger Institute COVID-19 Surveillance Team |
| EPI_ISL_3247908, EPI_ISL_3247909 | Lighthouse Laboratory Plymouth | Wellcome Sanger Institute for the COVID-19 Genomics UK (COG-UK) Consortium | Lighthouse Laboratory Plymouth and Alex Alderton, Roberto Amato, Jeffrey Barrett, Sonia Goncalves, Ewan Harrison, David K. Jackson, Ian Johnston, Dominic Kwiatkowski, Cordelia Langford, John Sillitoe on behalf of the Wellcome Sanger Institute COVID-19 Surveillance Team |
| EPI_ISL_3247910 | Lighthouse Lab in Glasgow | Wellcome Sanger Institute for the COVID-19 Genomics UK (COG-UK) Consortium | Harper VanSteenhouse, Yumi Kasai, David Gray, Carol Clugston, Anna Dominiczak and Alex Alderton, Roberto Amato, Jeffrey Barrett, Sonia Goncalves, Ewan Harrison, David K. Jackson, Ian Johnston, Dominic Kwiatkowski, Cordelia Langford, John Sillitoe on behalf of the Wellcome Sanger Institute COVID-19 Surveillance Team |
| EPI_ISL_3247911, EPI_ISL_3247912, EPI_ISL_3247913, EPI_ISL_3247914 | Lighthouse Laboratory Plymouth | Wellcome Sanger Institute for the COVID-19 Genomics UK (COG-UK) Consortium | Lighthouse Laboratory Plymouth and Alex Alderton, Roberto Amato, Jeffrey Barrett, Sonia Goncalves, Ewan Harrison, David K. Jackson, Ian Johnston, Dominic Kwiatkowski, Cordelia Langford, John Sillitoe on behalf of the Wellcome Sanger Institute COVID-19 Surveillance Team |
| EPI_ISL_3247915 | Lighthouse Lab in Glasgow | Wellcome Sanger Institute for the COVID-19 Genomics UK (COG-UK) Consortium | Harper VanSteenhouse, Yumi Kasai, David Gray, Carol Clugston, Anna Dominiczak and Alex Alderton, Roberto Amato, Jeffrey Barrett, Sonia Goncalves, Ewan Harrison, David K. Jackson, Ian Johnston, Dominic Kwiatkowski, Cordelia Langford, John Sillitoe on behalf of the Wellcome Sanger Institute COVID-19 Surveillance Team |
| EPI_ISL_3247916, EPI_ISL_3247917, EPI_ISL_3247918, EPI_ISL_3247919, EPI_ISL_3247920, EPI_ISL_3247921, EPI_ISL_3247922, EPI_ISL_3247923, EPI_ISL_3247924, EPI_ISL_3247925, EPI_ISL_3247926, EPI_ISL_3247927, EPI_ISL_3247928, EPI_ISL_3247929, EPI_ISL_3247930, EPI_ISL_3247931, EPI_ISL_3247932, EPI_ISL_3247933, EPI_ISL_3247934, EPI_ISL_3247935, EPI_ISL_3247936, EPI_ISL_3247937, EPI_ISL_3247938, EPI_ISL_3247939, EPI_ISL_3247940, EPI_ISL_3247941, EPI_ISL_3247942 | | | |
| see above | Lighthouse Laboratory Plymouth | Wellcome Sanger Institute for the COVID-19 Genomics UK (COG-UK) Consortium | Lighthouse Laboratory Plymouth and Alex Alderton, Roberto Amato, Jeffrey Barrett, Sonia Goncalves, Ewan Harrison, David K. Jackson, Ian Johnston, Dominic Kwiatkowski, Cordelia Langford, John Sillitoe on behalf of the Wellcome Sanger Institute COVID-19 Surveillance Team |
| EPI_ISL_3247943 | Lighthouse Lab in Glasgow | Wellcome Sanger Institute for the COVID-19 Genomics UK (COG-UK) Consortium | Harper VanSteenhouse, Yumi Kasai, David Gray, Carol Clugston, Anna Dominiczak and Alex Alderton, Roberto Amato, Jeffrey Barrett, Sonia Goncalves, Ewan Harrison, David K. Jackson, Ian Johnston, Dominic Kwiatkowski, Cordelia Langford, John Sillitoe on behalf of the Wellcome Sanger Institute COVID-19 Surveillance Team |
| EPI_ISL_3247944, EPI_ISL_3247945, EPI_ISL_3247946, EPI_ISL_3247947, EPI_ISL_3247948 | Lighthouse Laboratory Plymouth | Wellcome Sanger Institute for the COVID-19 Genomics UK (COG-UK) Consortium | Lighthouse Laboratory Plymouth and Alex Alderton, Roberto Amato, Jeffrey Barrett, Sonia Goncalves, Ewan Harrison, David K. Jackson, Ian Johnston, Dominic Kwiatkowski, Cordelia Langford, John Sillitoe on behalf of the Wellcome Sanger Institute COVID-19 Surveillance Team |
| EPI_ISL_3247949, EPI_ISL_3247950 | Lighthouse Lab in Glasgow | Wellcome Sanger Institute for the COVID-19 Genomics UK (COG-UK) Consortium | Harper VanSteenhouse, Yumi Kasai, David Gray, Carol Clugston, Anna Dominiczak and Alex Alderton, Roberto Amato, Jeffrey Barrett, Sonia Goncalves, Ewan Harrison, David K. Jackson, Ian Johnston, Dominic Kwiatkowski, Cordelia Langford, John Sillitoe on behalf of the Wellcome Sanger Institute COVID-19 Surveillance Team |
| EPI_ISL_3247951, EPI_ISL_3247952, EPI_ISL_3247953 | Lighthouse Laboratory Plymouth | Wellcome Sanger Institute for the COVID-19 Genomics UK (COG-UK) Consortium | Lighthouse Laboratory Plymouth and Alex Alderton, Roberto Amato, Jeffrey Barrett, Sonia Goncalves, Ewan Harrison, David K. Jackson, Ian Johnston, Dominic Kwiatkowski, Cordelia Langford, John Sillitoe on behalf of the Wellcome Sanger Institute COVID-19 Surveillance Team |
| EPI_ISL_3247954, EPI_ISL_3247955 | Lighthouse Lab in Glasgow | Wellcome Sanger Institute for the COVID-19 Genomics UK (COG-UK) Consortium | Harper VanSteenhouse, Yumi Kasai, David Gray, Carol Clugston, Anna Dominiczak and Alex Alderton, Roberto Amato, Jeffrey Barrett, Sonia Goncalves, Ewan Harrison, David K. Jackson, Ian Johnston, Dominic Kwiatkowski, Cordelia Langford, John Sillitoe on behalf of the Wellcome Sanger Institute COVID-19 Surveillance Team |
| EPI_ISL_3247956, EPI_ISL_3247957, EPI_ISL_3247958, EPI_ISL_3247959 | Lighthouse Laboratory Plymouth | Wellcome Sanger Institute for the COVID-19 Genomics UK (COG-UK) Consortium | Lighthouse Laboratory Plymouth and Alex Alderton, Roberto Amato, Jeffrey Barrett, Sonia Goncalves, Ewan Harrison, David K. Jackson, Ian Johnston, Dominic Kwiatkowski, Cordelia Langford, John Sillitoe on behalf of the Wellcome Sanger Institute COVID-19 Surveillance Team |

| | | | |
|---|---|---|---|
| EPI_ISL_3247960, EPI_ISL_3247961 | Lighthouse Lab in Glasgow | Wellcome Sanger Institute for the COVID-19 Genomics UK (COG-UK) Consortium | Harper VanSteenhouse, Yumi Kasai, David Gray, Carol Clugston, Anna Dominiczak and Alex Alderton, Roberto Amato, Jeffrey Barrett, Sonia Goncalves, Ewan Harrison, David K. Jackson, Ian Johnston, Dominic Kwiatkowski, Cordelia Langford, John Sillitoe on behalf of the Wellcome Sanger Institute COVID-19 Surveillance Team |
| EPI_ISL_3247962, EPI_ISL_3247963, EPI_ISL_3247964, EPI_ISL_3247965 | Lighthouse Laboratory Plymouth | Wellcome Sanger Institute for the COVID-19 Genomics UK (COG-UK) Consortium | Lighthouse Laboratory Plymouth and Alex Alderton, Roberto Amato, Jeffrey Barrett, Sonia Goncalves, Ewan Harrison, David K. Jackson, Ian Johnston, Dominic Kwiatkowski, Cordelia Langford, John Sillitoe on behalf of the Wellcome Sanger Institute COVID-19 Surveillance Team |
| EPI_ISL_3247966 | Lighthouse Lab in Glasgow | Wellcome Sanger Institute for the COVID-19 Genomics UK (COG-UK) Consortium | Harper VanSteenhouse, Yumi Kasai, David Gray, Carol Clugston, Anna Dominiczak and Alex Alderton, Roberto Amato, Jeffrey Barrett, Sonia Goncalves, Ewan Harrison, David K. Jackson, Ian Johnston, Dominic Kwiatkowski, Cordelia Langford, John Sillitoe on behalf of the Wellcome Sanger Institute COVID-19 Surveillance Team |
| EPI_ISL_3247967, EPI_ISL_3247968, EPI_ISL_3247969 | Lighthouse Laboratory Plymouth | Wellcome Sanger Institute for the COVID-19 Genomics UK (COG-UK) Consortium | Lighthouse Laboratory Plymouth and Alex Alderton, Roberto Amato, Jeffrey Barrett, Sonia Goncalves, Ewan Harrison, David K. Jackson, Ian Johnston, Dominic Kwiatkowski, Cordelia Langford, John Sillitoe on behalf of the Wellcome Sanger Institute COVID-19 Surveillance Team |
| EPI_ISL_3247970, EPI_ISL_3247971 | Lighthouse Lab in Glasgow | Wellcome Sanger Institute for the COVID-19 Genomics UK (COG-UK) Consortium | Harper VanSteenhouse, Yumi Kasai, David Gray, Carol Clugston, Anna Dominiczak and Alex Alderton, Roberto Amato, Jeffrey Barrett, Sonia Goncalves, Ewan Harrison, David K. Jackson, Ian Johnston, Dominic Kwiatkowski, Cordelia Langford, John Sillitoe on behalf of the Wellcome Sanger Institute COVID-19 Surveillance Team |
| EPI_ISL_3247972, EPI_ISL_3247973, EPI_ISL_3247974 | Lighthouse Laboratory Plymouth | Wellcome Sanger Institute for the COVID-19 Genomics UK (COG-UK) Consortium | Lighthouse Laboratory Plymouth and Alex Alderton, Roberto Amato, Jeffrey Barrett, Sonia Goncalves, Ewan Harrison, David K. Jackson, Ian Johnston, Dominic Kwiatkowski, Cordelia Langford, John Sillitoe on behalf of the Wellcome Sanger Institute COVID-19 Surveillance Team |
| EPI_ISL_3247975 | Lighthouse Lab in Glasgow | Wellcome Sanger Institute for the COVID-19 Genomics UK (COG-UK) Consortium | Harper VanSteenhouse, Yumi Kasai, David Gray, Carol Clugston, Anna Dominiczak and Alex Alderton, Roberto Amato, Jeffrey Barrett, Sonia Goncalves, Ewan Harrison, David K. Jackson, Ian Johnston, Dominic Kwiatkowski, Cordelia Langford, John Sillitoe on behalf of the Wellcome Sanger Institute COVID-19 Surveillance Team |
| EPI_ISL_3247976, EPI_ISL_3247977, EPI_ISL_3247978, EPI_ISL_3247979, EPI_ISL_3247980, EPI_ISL_3247981, EPI_ISL_3247982, EPI_ISL_3247983, EPI_ISL_3247984 | Lighthouse Laboratory Plymouth | Wellcome Sanger Institute for the COVID-19 Genomics UK (COG-UK) Consortium | Lighthouse Laboratory Plymouth and Alex Alderton, Roberto Amato, Jeffrey Barrett, Sonia Goncalves, Ewan Harrison, David K. Jackson, Ian Johnston, Dominic Kwiatkowski, Cordelia Langford, John Sillitoe on behalf of the Wellcome Sanger Institute COVID-19 Surveillance Team |
| EPI_ISL_3247985 | Lighthouse Lab in Glasgow | Wellcome Sanger Institute for the COVID-19 Genomics UK (COG-UK) Consortium | Harper VanSteenhouse, Yumi Kasai, David Gray, Carol Clugston, Anna Dominiczak and Alex Alderton, Roberto Amato, Jeffrey Barrett, Sonia Goncalves, Ewan Harrison, David K. Jackson, Ian Johnston, Dominic Kwiatkowski, Cordelia Langford, John Sillitoe on behalf of the Wellcome Sanger Institute COVID-19 Surveillance Team |
| EPI_ISL_3247986, EPI_ISL_3247987 | Lighthouse Laboratory Plymouth | Wellcome Sanger Institute for the COVID-19 Genomics UK (COG-UK) Consortium | Lighthouse Laboratory Plymouth and Alex Alderton, Roberto Amato, Jeffrey Barrett, Sonia Goncalves, Ewan Harrison, David K. Jackson, Ian Johnston, Dominic Kwiatkowski, Cordelia Langford, John Sillitoe on behalf of the Wellcome Sanger Institute COVID-19 Surveillance Team |
| EPI_ISL_3247988 | Lighthouse Lab in Glasgow | Wellcome Sanger Institute for the COVID-19 Genomics UK (COG-UK) Consortium | Harper VanSteenhouse, Yumi Kasai, David Gray, Carol Clugston, Anna Dominiczak and Alex Alderton, Roberto Amato, Jeffrey Barrett, Sonia Goncalves, Ewan Harrison, David K. Jackson, Ian Johnston, Dominic Kwiatkowski, Cordelia Langford, John Sillitoe on behalf of the Wellcome Sanger Institute COVID-19 Surveillance Team |
| EPI_ISL_3247989 | Lighthouse Laboratory Plymouth | Wellcome Sanger Institute for the COVID-19 Genomics UK (COG-UK) Consortium | Lighthouse Laboratory Plymouth and Alex Alderton, Roberto Amato, Jeffrey Barrett, Sonia Goncalves, Ewan Harrison, David K. Jackson, Ian Johnston, Dominic Kwiatkowski, Cordelia Langford, John Sillitoe on behalf of the Wellcome Sanger Institute COVID-19 Surveillance Team |
| EPI_ISL_3247990 | Lighthouse Lab in Glasgow | Wellcome Sanger Institute for the COVID-19 Genomics UK (COG-UK) Consortium | Harper VanSteenhouse, Yumi Kasai, David Gray, Carol Clugston, Anna Dominiczak and Alex Alderton, Roberto Amato, Jeffrey Barrett, Sonia Goncalves, Ewan Harrison, David K. Jackson, Ian Johnston, Dominic Kwiatkowski, Cordelia Langford, John Sillitoe on behalf of the Wellcome Sanger Institute COVID-19 Surveillance Team |
| EPI_ISL_3247991 | Lighthouse Laboratory Plymouth | Wellcome Sanger Institute for the COVID-19 Genomics UK (COG-UK) Consortium | Lighthouse Laboratory Plymouth and Alex Alderton, Roberto Amato, Jeffrey Barrett, Sonia Goncalves, Ewan Harrison, David K. Jackson, Ian Johnston, Dominic Kwiatkowski, Cordelia Langford, John Sillitoe on behalf of the Wellcome Sanger Institute COVID-19 Surveillance Team |
| EPI_ISL_3247992 | Lighthouse Lab in Glasgow | Wellcome Sanger Institute for the COVID-19 Genomics UK (COG-UK) Consortium | Harper VanSteenhouse, Yumi Kasai, David Gray, Carol Clugston, Anna Dominiczak and Alex Alderton, Roberto Amato, Jeffrey Barrett, Sonia Goncalves, Ewan Harrison, David K. Jackson, Ian Johnston, Dominic Kwiatkowski, Cordelia Langford, John Sillitoe on behalf of the Wellcome Sanger Institute COVID-19 Surveillance Team |
| EPI_ISL_3247993, EPI_ISL_3247994, EPI_ISL_3247995, EPI_ISL_3247996, EPI_ISL_3247997 | Lighthouse Laboratory Plymouth | Wellcome Sanger Institute for the COVID-19 Genomics UK (COG-UK) Consortium | Lighthouse Laboratory Plymouth and Alex Alderton, Roberto Amato, Jeffrey Barrett, Sonia Goncalves, Ewan Harrison, David K. Jackson, Ian Johnston, Dominic Kwiatkowski, Cordelia Langford, John Sillitoe on behalf of the Wellcome Sanger Institute COVID-19 Surveillance Team |
| EPI_ISL_3247998 | Lighthouse Lab in Glasgow | Wellcome Sanger Institute for the COVID-19 Genomics UK (COG-UK) Consortium | Harper VanSteenhouse, Yumi Kasai, David Gray, Carol Clugston, Anna Dominiczak and Alex Alderton, Roberto Amato, Jeffrey Barrett, Sonia Goncalves, Ewan Harrison, David K. Jackson, Ian Johnston, Dominic Kwiatkowski, Cordelia Langford, John Sillitoe on behalf of the Wellcome Sanger Institute COVID-19 Surveillance Team |
| EPI_ISL_3247999 | Lighthouse Laboratory Plymouth | Wellcome Sanger Institute for the COVID-19 Genomics UK (COG-UK) Consortium | Lighthouse Laboratory Plymouth and Alex Alderton, Roberto Amato, Jeffrey Barrett, Sonia Goncalves, Ewan Harrison, David K. Jackson, Ian Johnston, Dominic Kwiatkowski, Cordelia Langford, John Sillitoe on behalf of the Wellcome Sanger Institute COVID-19 Surveillance Team |
| EPI_ISL_3248000 | Lighthouse Lab in Glasgow | Wellcome Sanger Institute for the COVID-19 Genomics UK (COG-UK) Consortium | Harper VanSteenhouse, Yumi Kasai, David Gray, Carol Clugston, Anna Dominiczak and Alex Alderton, Roberto Amato, Jeffrey Barrett, Sonia Goncalves, Ewan Harrison, David K. Jackson, Ian Johnston, Dominic Kwiatkowski, Cordelia Langford, John Sillitoe on behalf of the Wellcome Sanger Institute COVID-19 Surveillance Team |
| EPI_ISL_3248001, EPI_ISL_3248002, EPI_ISL_3248003, EPI_ISL_3248004 | Lighthouse Laboratory Plymouth | Wellcome Sanger Institute for the COVID-19 Genomics UK (COG-UK) Consortium | Lighthouse Laboratory Plymouth and Alex Alderton, Roberto Amato, Jeffrey Barrett, Sonia Goncalves, Ewan Harrison, David K. Jackson, Ian Johnston, Dominic Kwiatkowski, Cordelia Langford, John Sillitoe on behalf of the Wellcome Sanger Institute COVID-19 Surveillance Team |
| EPI_ISL_3248005 | Lighthouse Lab in Glasgow | Wellcome Sanger Institute for the COVID-19 Genomics UK (COG-UK) Consortium | Harper VanSteenhouse, Yumi Kasai, David Gray, Carol Clugston, Anna Dominiczak and Alex Alderton, Roberto Amato, Jeffrey Barrett, Sonia Goncalves, Ewan Harrison, David K. Jackson, Ian Johnston, Dominic Kwiatkowski, Cordelia Langford, John Sillitoe on behalf of the Wellcome Sanger Institute COVID-19 Surveillance Team |
| EPI_ISL_3248006, EPI_ISL_3248007, EPI_ISL_3248008, EPI_ISL_3248009, EPI_ISL_3248010, EPI_ISL_3248011, EPI_ISL_3248012, EPI_ISL_3248013, EPI_ISL_3248014, EPI_ISL_3248015, EPI_ISL_3248016, EPI_ISL_3248017 | | | |
| see above | Lighthouse Laboratory Plymouth | Wellcome Sanger Institute for the COVID-19 Genomics UK (COG-UK) Consortium | Lighthouse Laboratory Plymouth and Alex Alderton, Roberto Amato, Jeffrey Barrett, Sonia Goncalves, Ewan Harrison, David K. Jackson, Ian Johnston, Dominic Kwiatkowski, Cordelia Langford, John Sillitoe on behalf of the Wellcome Sanger Institute COVID-19 Surveillance Team |
| EPI_ISL_3248018 | Lighthouse Lab in Glasgow | Wellcome Sanger Institute for the COVID-19 Genomics UK (COG-UK) Consortium | Harper VanSteenhouse, Yumi Kasai, David Gray, Carol Clugston, Anna Dominiczak and Alex Alderton, Roberto Amato, Jeffrey Barrett, Sonia Goncalves, Ewan Harrison, David K. Jackson, Ian Johnston, Dominic Kwiatkowski, Cordelia Langford, John Sillitoe on behalf of the Wellcome Sanger Institute COVID-19 Surveillance Team |
| EPI_ISL_3248019, EPI_ISL_3248020, EPI_ISL_3248021, EPI_ISL_3248022, EPI_ISL_3248023 | Lighthouse Laboratory Plymouth | Wellcome Sanger Institute for the COVID-19 Genomics UK (COG-UK) Consortium | Lighthouse Laboratory Plymouth and Alex Alderton, Roberto Amato, Jeffrey Barrett, Sonia Goncalves, Ewan Harrison, David K. Jackson, Ian Johnston, Dominic Kwiatkowski, Cordelia Langford, John Sillitoe on behalf of the Wellcome Sanger Institute COVID-19 Surveillance Team |
| EPI_ISL_3248024 | Lighthouse Lab in Glasgow | Wellcome Sanger Institute for the COVID-19 Genomics UK (COG-UK) Consortium | Harper VanSteenhouse, Yumi Kasai, David Gray, Carol Clugston, Anna Dominiczak and Alex Alderton, Roberto Amato, Jeffrey Barrett, Sonia Goncalves, Ewan Harrison, David K. Jackson, Ian Johnston, Dominic Kwiatkowski, Cordelia Langford, John Sillitoe on behalf of the Wellcome Sanger Institute COVID-19 Surveillance Team |
| EPI_ISL_3248025, EPI_ISL_3248026, EPI_ISL_3248027, EPI_ISL_3248028, EPI_ISL_3248029 | Lighthouse Laboratory Plymouth | Wellcome Sanger Institute for the COVID-19 Genomics UK (COG-UK) Consortium | Lighthouse Laboratory Plymouth and Alex Alderton, Roberto Amato, Jeffrey Barrett, Sonia Goncalves, Ewan Harrison, David K. Jackson, Ian Johnston, Dominic Kwiatkowski, Cordelia Langford, John Sillitoe on behalf of the Wellcome Sanger Institute COVID-19 Surveillance Team |
| EPI_ISL_3248030 | Lighthouse Lab in Glasgow | Wellcome Sanger Institute for the COVID-19 Genomics UK (COG-UK) Consortium | Harper VanSteenhouse, Yumi Kasai, David Gray, Carol Clugston, Anna Dominiczak and Alex Alderton, Roberto Amato, Jeffrey Barrett, Sonia Goncalves, Ewan Harrison, David K. Jackson, Ian Johnston, Dominic Kwiatkowski, Cordelia Langford, John Sillitoe on behalf of the Wellcome Sanger Institute COVID-19 Surveillance Team |
| EPI_ISL_3248031, EPI_ISL_3248032, EPI_ISL_3248033, EPI_ISL_3248034, EPI_ISL_3248035 | Lighthouse Laboratory Plymouth | Wellcome Sanger Institute for the COVID-19 Genomics UK (COG-UK) Consortium | Lighthouse Laboratory Plymouth and Alex Alderton, Roberto Amato, Jeffrey Barrett, Sonia Goncalves, Ewan Harrison, David K. Jackson, Ian Johnston, Dominic Kwiatkowski, Cordelia Langford, John Sillitoe on behalf of the Wellcome Sanger Institute COVID-19 Surveillance Team |
| EPI_ISL_3248036 | Lighthouse Lab in Glasgow | Wellcome Sanger Institute for the COVID-19 Genomics UK (COG-UK) Consortium | Harper VanSteenhouse, Yumi Kasai, David Gray, Carol Clugston, Anna Dominiczak and Alex Alderton, Roberto Amato, Jeffrey Barrett, Sonia Goncalves, Ewan Harrison, David K. Jackson, Ian Johnston, Dominic Kwiatkowski, Cordelia Langford, John Sillitoe on behalf of the Wellcome Sanger Institute COVID-19 Surveillance Team |
| EPI_ISL_3248037 | Lighthouse Laboratory Plymouth | Wellcome Sanger Institute for the COVID-19 Genomics UK (COG-UK) Consortium | Lighthouse Laboratory Plymouth and Alex Alderton, Roberto Amato, Jeffrey Barrett, Sonia Goncalves, Ewan Harrison, David K. Jackson, Ian Johnston, Dominic Kwiatkowski, Cordelia Langford, John Sillitoe on behalf of the Wellcome Sanger Institute COVID-19 Surveillance Team |
| EPI_ISL_3248038 | Lighthouse Lab in Glasgow | Wellcome Sanger Institute for the COVID-19 Genomics UK (COG-UK) Consortium | Harper VanSteenhouse, Yumi Kasai, David Gray, Carol Clugston, Anna Dominiczak and Alex Alderton, Roberto Amato, Jeffrey Barrett, Sonia Goncalves, Ewan Harrison, David K. Jackson, Ian Johnston, Dominic Kwiatkowski, Cordelia Langford, John Sillitoe on behalf of the Wellcome Sanger Institute COVID-19 Surveillance Team |
| EPI_ISL_3248039, EPI_ISL_3248040, EPI_ISL_3248041 | Lighthouse Laboratory Plymouth | Wellcome Sanger Institute for the COVID-19 Genomics UK (COG-UK) Consortium | Lighthouse Laboratory Plymouth and Alex Alderton, Roberto Amato, Jeffrey Barrett, Sonia Goncalves, Ewan Harrison, David K. Jackson, Ian Johnston, Dominic Kwiatkowski, Cordelia Langford, John Sillitoe on behalf of the Wellcome Sanger Institute COVID-19 Surveillance Team |
| EPI_ISL_3248042 | Lighthouse Lab in Glasgow | Wellcome Sanger Institute for the COVID-19 Genomics UK (COG-UK) Consortium | Harper VanSteenhouse, Yumi Kasai, David Gray, Carol Clugston, Anna Dominiczak and Alex Alderton, Roberto Amato, Jeffrey Barrett, Sonia Goncalves, Ewan Harrison, David K. Jackson, Ian Johnston, Dominic Kwiatkowski, Cordelia Langford, John Sillitoe on behalf of the Wellcome Sanger Institute COVID-19 Surveillance Team |
| EPI_ISL_3248043, EPI_ISL_3248044 | Lighthouse Laboratory Plymouth | Wellcome Sanger Institute for the COVID-19 Genomics UK (COG-UK) Consortium | Lighthouse Laboratory Plymouth and Alex Alderton, Roberto Amato, Jeffrey Barrett, Sonia Goncalves, Ewan Harrison, David K. Jackson, Ian Johnston, Dominic Kwiatkowski, Cordelia Langford, John Sillitoe on behalf of the Wellcome Sanger Institute COVID-19 Surveillance Team |

| | | | |
|---|---|---|---|
| EPI_ISL_3248045 | Lighthouse Lab in Glasgow | Wellcome Sanger Institute for the COVID-19 Genomics UK (COG-UK) Consortium | Harper VanSteenhouse, Yumi Kasai, David Gray, Carol Clugston, Anna Dominiczak and Alex Alderton, Roberto Amato, Jeffrey Barrett, Sonia Goncalves, Ewan Harrison, David K. Jackson, Ian Johnston, Dominic Kwiatkowski, Cordelia Langford, John Sillitoe on behalf of the Wellcome Sanger Institute COVID-19 Surveillance Team |
| EPI_ISL_3248046, EPI_ISL_3248047 | Lighthouse Lab in Alderley Park | Wellcome Sanger Institute for the COVID-19 Genomics UK (COG-UK) Consortium | Jacquelyn Wynn, Mairead Hyland, The Lighthouse Lab in Alderley Park and Alex Alderton, Roberto Amato, Jeffrey Barrett, Sonia Goncalves, Ewan Harrison, David K. Jackson, Ian Johnston, Dominic Kwiatkowski, Cordelia Langford, John Sillitoe on behalf of the Wellcome Sanger Institute COVID-19 Surveillance Team |
| EPI_ISL_3248048 | Lighthouse Laboratory Plymouth | Wellcome Sanger Institute for the COVID-19 Genomics UK (COG-UK) Consortium | Lighthouse Laboratory Plymouth and Alex Alderton, Roberto Amato, Jeffrey Barrett, Sonia Goncalves, Ewan Harrison, David K. Jackson, Ian Johnston, Dominic Kwiatkowski, Cordelia Langford, John Sillitoe on behalf of the Wellcome Sanger Institute COVID-19 Surveillance Team |
| EPI_ISL_3248049 | Lighthouse Lab in Alderley Park | Wellcome Sanger Institute for the COVID-19 Genomics UK (COG-UK) Consortium | Jacquelyn Wynn, Mairead Hyland, The Lighthouse Lab in Alderley Park and Alex Alderton, Roberto Amato, Jeffrey Barrett, Sonia Goncalves, Ewan Harrison, David K. Jackson, Ian Johnston, Dominic Kwiatkowski, Cordelia Langford, John Sillitoe on behalf of the Wellcome Sanger Institute COVID-19 Surveillance Team |
| EPI_ISL_3248050, EPI_ISL_3248051, EPI_ISL_3248052, EPI_ISL_3248053, EPI_ISL_3248054 | Lighthouse Laboratory Plymouth | Wellcome Sanger Institute for the COVID-19 Genomics UK (COG-UK) Consortium | Lighthouse Laboratory Plymouth and Alex Alderton, Roberto Amato, Jeffrey Barrett, Sonia Goncalves, Ewan Harrison, David K. Jackson, Ian Johnston, Dominic Kwiatkowski, Cordelia Langford, John Sillitoe on behalf of the Wellcome Sanger Institute COVID-19 Surveillance Team |
| EPI_ISL_3248055, EPI_ISL_3248056 | Lighthouse Lab in Alderley Park | Wellcome Sanger Institute for the COVID-19 Genomics UK (COG-UK) Consortium | Jacquelyn Wynn, Mairead Hyland, The Lighthouse Lab in Alderley Park and Alex Alderton, Roberto Amato, Jeffrey Barrett, Sonia Goncalves, Ewan Harrison, David K. Jackson, Ian Johnston, Dominic Kwiatkowski, Cordelia Langford, John Sillitoe on behalf of the Wellcome Sanger Institute COVID-19 Surveillance Team |
| EPI_ISL_3248057, EPI_ISL_3248058, EPI_ISL_3248059, EPI_ISL_3248060 | Lighthouse Laboratory Plymouth | Wellcome Sanger Institute for the COVID-19 Genomics UK (COG-UK) Consortium | Lighthouse Laboratory Plymouth and Alex Alderton, Roberto Amato, Jeffrey Barrett, Sonia Goncalves, Ewan Harrison, David K. Jackson, Ian Johnston, Dominic Kwiatkowski, Cordelia Langford, John Sillitoe on behalf of the Wellcome Sanger Institute COVID-19 Surveillance Team |
| EPI_ISL_3248061 | Lighthouse Lab in Alderley Park | Wellcome Sanger Institute for the COVID-19 Genomics UK (COG-UK) Consortium | Jacquelyn Wynn, Mairead Hyland, The Lighthouse Lab in Alderley Park and Alex Alderton, Roberto Amato, Jeffrey Barrett, Sonia Goncalves, Ewan Harrison, David K. Jackson, Ian Johnston, Dominic Kwiatkowski, Cordelia Langford, John Sillitoe on behalf of the Wellcome Sanger Institute COVID-19 Surveillance Team |
| EPI_ISL_3248062 | Lighthouse Laboratory Plymouth | Wellcome Sanger Institute for the COVID-19 Genomics UK (COG-UK) Consortium | Lighthouse Laboratory Plymouth and Alex Alderton, Roberto Amato, Jeffrey Barrett, Sonia Goncalves, Ewan Harrison, David K. Jackson, Ian Johnston, Dominic Kwiatkowski, Cordelia Langford, John Sillitoe on behalf of the Wellcome Sanger Institute COVID-19 Surveillance Team |
| EPI_ISL_3248063 | Lighthouse Lab in Alderley Park | Wellcome Sanger Institute for the COVID-19 Genomics UK (COG-UK) Consortium | Jacquelyn Wynn, Mairead Hyland, The Lighthouse Lab in Alderley Park and Alex Alderton, Roberto Amato, Jeffrey Barrett, Sonia Goncalves, Ewan Harrison, David K. Jackson, Ian Johnston, Dominic Kwiatkowski, Cordelia Langford, John Sillitoe on behalf of the Wellcome Sanger Institute COVID-19 Surveillance Team |
| EPI_ISL_3248064, EPI_ISL_3248065, EPI_ISL_3248066, EPI_ISL_3248067, EPI_ISL_3248068, EPI_ISL_3248069 | Lighthouse Laboratory Plymouth | Wellcome Sanger Institute for the COVID-19 Genomics UK (COG-UK) Consortium | Lighthouse Laboratory Plymouth and Alex Alderton, Roberto Amato, Jeffrey Barrett, Sonia Goncalves, Ewan Harrison, David K. Jackson, Ian Johnston, Dominic Kwiatkowski, Cordelia Langford, John Sillitoe on behalf of the Wellcome Sanger Institute COVID-19 Surveillance Team |
| EPI_ISL_3248070, EPI_ISL_3248071, EPI_ISL_3248072, EPI_ISL_3248073, EPI_ISL_3248074, EPI_ISL_3248075, EPI_ISL_3248076, EPI_ISL_3248077, EPI_ISL_3248078, EPI_ISL_3248079 | Lighthouse Lab in Alderley Park | Wellcome Sanger Institute for the COVID-19 Genomics UK (COG-UK) Consortium | Jacquelyn Wynn, Mairead Hyland, The Lighthouse Lab in Alderley Park and Alex Alderton, Roberto Amato, Jeffrey Barrett, Sonia Goncalves, Ewan Harrison, David K. Jackson, Ian Johnston, Dominic Kwiatkowski, Cordelia Langford, John Sillitoe on behalf of the Wellcome Sanger Institute COVID-19 Surveillance Team |
| EPI_ISL_3248080 | Lighthouse Laboratory Plymouth | Wellcome Sanger Institute for the COVID-19 Genomics UK (COG-UK) Consortium | Lighthouse Laboratory Plymouth and Alex Alderton, Roberto Amato, Jeffrey Barrett, Sonia Goncalves, Ewan Harrison, David K. Jackson, Ian Johnston, Dominic Kwiatkowski, Cordelia Langford, John Sillitoe on behalf of the Wellcome Sanger Institute COVID-19 Surveillance Team |
| EPI_ISL_3248081 | Lighthouse Lab in Alderley Park | Wellcome Sanger Institute for the COVID-19 Genomics UK (COG-UK) Consortium | Jacquelyn Wynn, Mairead Hyland, The Lighthouse Lab in Alderley Park and Alex Alderton, Roberto Amato, Jeffrey Barrett, Sonia Goncalves, Ewan Harrison, David K. Jackson, Ian Johnston, Dominic Kwiatkowski, Cordelia Langford, John Sillitoe on behalf of the Wellcome Sanger Institute COVID-19 Surveillance Team |
| EPI_ISL_3248082, EPI_ISL_3248083, EPI_ISL_3248084 | Lighthouse Laboratory Plymouth | Wellcome Sanger Institute for the COVID-19 Genomics UK (COG-UK) Consortium | Lighthouse Laboratory Plymouth and Alex Alderton, Roberto Amato, Jeffrey Barrett, Sonia Goncalves, Ewan Harrison, David K. Jackson, Ian Johnston, Dominic Kwiatkowski, Cordelia Langford, John Sillitoe on behalf of the Wellcome Sanger Institute COVID-19 Surveillance Team |
| EPI_ISL_3248085 | Lighthouse Lab in Alderley Park | Wellcome Sanger Institute for the COVID-19 Genomics UK (COG-UK) Consortium | Jacquelyn Wynn, Mairead Hyland, The Lighthouse Lab in Alderley Park and Alex Alderton, Roberto Amato, Jeffrey Barrett, Sonia Goncalves, Ewan Harrison, David K. Jackson, Ian Johnston, Dominic Kwiatkowski, Cordelia Langford, John Sillitoe on behalf of the Wellcome Sanger Institute COVID-19 Surveillance Team |
| EPI_ISL_3248086 | Lighthouse Laboratory Plymouth | Wellcome Sanger Institute for the COVID-19 Genomics UK (COG-UK) Consortium | Lighthouse Laboratory Plymouth and Alex Alderton, Roberto Amato, Jeffrey Barrett, Sonia Goncalves, Ewan Harrison, David K. Jackson, Ian Johnston, Dominic Kwiatkowski, Cordelia Langford, John Sillitoe on behalf of the Wellcome Sanger Institute COVID-19 Surveillance Team |
| EPI_ISL_3248087 | Lighthouse Lab in Alderley Park | Wellcome Sanger Institute for the COVID-19 Genomics UK (COG-UK) Consortium | Jacquelyn Wynn, Mairead Hyland, The Lighthouse Lab in Alderley Park and Alex Alderton, Roberto Amato, Jeffrey Barrett, Sonia Goncalves, Ewan Harrison, David K. Jackson, Ian Johnston, Dominic Kwiatkowski, Cordelia Langford, John Sillitoe on behalf of the Wellcome Sanger Institute COVID-19 Surveillance Team |
| EPI_ISL_3248088, EPI_ISL_3248089, EPI_ISL_3248090, EPI_ISL_3248091 | Lighthouse Laboratory Plymouth | Wellcome Sanger Institute for the COVID-19 Genomics UK (COG-UK) Consortium | Lighthouse Laboratory Plymouth and Alex Alderton, Roberto Amato, Jeffrey Barrett, Sonia Goncalves, Ewan Harrison, David K. Jackson, Ian Johnston, Dominic Kwiatkowski, Cordelia Langford, John Sillitoe on behalf of the Wellcome Sanger Institute COVID-19 Surveillance Team |
| EPI_ISL_3248092, EPI_ISL_3248093, EPI_ISL_3248094, EPI_ISL_3248095 | Lighthouse Lab in Alderley Park | Wellcome Sanger Institute for the COVID-19 Genomics UK (COG-UK) Consortium | Jacquelyn Wynn, Mairead Hyland, The Lighthouse Lab in Alderley Park and Alex Alderton, Roberto Amato, Jeffrey Barrett, Sonia Goncalves, Ewan Harrison, David K. Jackson, Ian Johnston, Dominic Kwiatkowski, Cordelia Langford, John Sillitoe on behalf of the Wellcome Sanger Institute COVID-19 Surveillance Team |
| EPI_ISL_3248096, EPI_ISL_3248097, EPI_ISL_3248098, EPI_ISL_3248099, EPI_ISL_3248100, EPI_ISL_3248101 | Lighthouse Laboratory Plymouth | Wellcome Sanger Institute for the COVID-19 Genomics UK (COG-UK) Consortium | Lighthouse Laboratory Plymouth and Alex Alderton, Roberto Amato, Jeffrey Barrett, Sonia Goncalves, Ewan Harrison, David K. Jackson, Ian Johnston, Dominic Kwiatkowski, Cordelia Langford, John Sillitoe on behalf of the Wellcome Sanger Institute COVID-19 Surveillance Team |
| EPI_ISL_3248102 | Lighthouse Lab in Alderley Park | Wellcome Sanger Institute for the COVID-19 Genomics UK (COG-UK) Consortium | Jacquelyn Wynn, Mairead Hyland, The Lighthouse Lab in Alderley Park and Alex Alderton, Roberto Amato, Jeffrey Barrett, Sonia Goncalves, Ewan Harrison, David K. Jackson, Ian Johnston, Dominic Kwiatkowski, Cordelia Langford, John Sillitoe on behalf of the Wellcome Sanger Institute COVID-19 Surveillance Team |
| EPI_ISL_3248103 | Lighthouse Laboratory Plymouth | Wellcome Sanger Institute for the COVID-19 Genomics UK (COG-UK) Consortium | Lighthouse Laboratory Plymouth and Alex Alderton, Roberto Amato, Jeffrey Barrett, Sonia Goncalves, Ewan Harrison, David K. Jackson, Ian Johnston, Dominic Kwiatkowski, Cordelia Langford, John Sillitoe on behalf of the Wellcome Sanger Institute COVID-19 Surveillance Team |
| EPI_ISL_3248104, EPI_ISL_3248105, EPI_ISL_3248106 | Lighthouse Lab in Alderley Park | Wellcome Sanger Institute for the COVID-19 Genomics UK (COG-UK) Consortium | Jacquelyn Wynn, Mairead Hyland, The Lighthouse Lab in Alderley Park and Alex Alderton, Roberto Amato, Jeffrey Barrett, Sonia Goncalves, Ewan Harrison, David K. Jackson, Ian Johnston, Dominic Kwiatkowski, Cordelia Langford, John Sillitoe on behalf of the Wellcome Sanger Institute COVID-19 Surveillance Team |
| EPI_ISL_3248107, EPI_ISL_3248108, EPI_ISL_3248109 | Lighthouse Laboratory Plymouth | Wellcome Sanger Institute for the COVID-19 Genomics UK (COG-UK) Consortium | Lighthouse Laboratory Plymouth and Alex Alderton, Roberto Amato, Jeffrey Barrett, Sonia Goncalves, Ewan Harrison, David K. Jackson, Ian Johnston, Dominic Kwiatkowski, Cordelia Langford, John Sillitoe on behalf of the Wellcome Sanger Institute COVID-19 Surveillance Team |
| EPI_ISL_3248110 | Lighthouse Lab in Alderley Park | Wellcome Sanger Institute for the COVID-19 Genomics UK (COG-UK) Consortium | Jacquelyn Wynn, Mairead Hyland, The Lighthouse Lab in Alderley Park and Alex Alderton, Roberto Amato, Jeffrey Barrett, Sonia Goncalves, Ewan Harrison, David K. Jackson, Ian Johnston, Dominic Kwiatkowski, Cordelia Langford, John Sillitoe on behalf of the Wellcome Sanger Institute COVID-19 Surveillance Team |
| EPI_ISL_3248111 | Lighthouse Laboratory Plymouth | Wellcome Sanger Institute for the COVID-19 Genomics UK (COG-UK) Consortium | Lighthouse Laboratory Plymouth and Alex Alderton, Roberto Amato, Jeffrey Barrett, Sonia Goncalves, Ewan Harrison, David K. Jackson, Ian Johnston, Dominic Kwiatkowski, Cordelia Langford, John Sillitoe on behalf of the Wellcome Sanger Institute COVID-19 Surveillance Team |
| EPI_ISL_3248112 | Lighthouse Lab in Alderley Park | Wellcome Sanger Institute for the COVID-19 Genomics UK (COG-UK) Consortium | Jacquelyn Wynn, Mairead Hyland, The Lighthouse Lab in Alderley Park and Alex Alderton, Roberto Amato, Jeffrey Barrett, Sonia Goncalves, Ewan Harrison, David K. Jackson, Ian Johnston, Dominic Kwiatkowski, Cordelia Langford, John Sillitoe on behalf of the Wellcome Sanger Institute COVID-19 Surveillance Team |
| EPI_ISL_3248113, EPI_ISL_3248114 | Lighthouse Laboratory Plymouth | Wellcome Sanger Institute for the COVID-19 Genomics UK (COG-UK) Consortium | Lighthouse Laboratory Plymouth and Alex Alderton, Roberto Amato, Jeffrey Barrett, Sonia Goncalves, Ewan Harrison, David K. Jackson, Ian Johnston, Dominic Kwiatkowski, Cordelia Langford, John Sillitoe on behalf of the Wellcome Sanger Institute COVID-19 Surveillance Team |
| EPI_ISL_3248115, EPI_ISL_3248116 | Lighthouse Lab in Alderley Park | Wellcome Sanger Institute for the COVID-19 Genomics UK (COG-UK) Consortium | Jacquelyn Wynn, Mairead Hyland, The Lighthouse Lab in Alderley Park and Alex Alderton, Roberto Amato, Jeffrey Barrett, Sonia Goncalves, Ewan Harrison, David K. Jackson, Ian Johnston, Dominic Kwiatkowski, Cordelia Langford, John Sillitoe on behalf of the Wellcome Sanger Institute COVID-19 Surveillance Team |
| EPI_ISL_3248117 | Lighthouse Laboratory Plymouth | Wellcome Sanger Institute for the COVID-19 Genomics UK (COG-UK) Consortium | Lighthouse Laboratory Plymouth and Alex Alderton, Roberto Amato, Jeffrey Barrett, Sonia Goncalves, Ewan Harrison, David K. Jackson, Ian Johnston, Dominic Kwiatkowski, Cordelia Langford, John Sillitoe on behalf of the Wellcome Sanger Institute COVID-19 Surveillance Team |
| EPI_ISL_3248118 | Lighthouse Lab in Alderley Park | Wellcome Sanger Institute for the COVID-19 Genomics UK (COG-UK) Consortium | Jacquelyn Wynn, Mairead Hyland, The Lighthouse Lab in Alderley Park and Alex Alderton, Roberto Amato, Jeffrey Barrett, Sonia Goncalves, Ewan Harrison, David K. Jackson, Ian Johnston, Dominic Kwiatkowski, Cordelia Langford, John Sillitoe on behalf of the Wellcome Sanger Institute COVID-19 Surveillance Team |
| EPI_ISL_3248119 | Lighthouse Laboratory Plymouth | Wellcome Sanger Institute for the COVID-19 Genomics UK (COG-UK) Consortium | Lighthouse Laboratory Plymouth and Alex Alderton, Roberto Amato, Jeffrey Barrett, Sonia Goncalves, Ewan Harrison, David K. Jackson, Ian Johnston, Dominic Kwiatkowski, Cordelia Langford, John Sillitoe on behalf of the Wellcome Sanger Institute COVID-19 Surveillance Team |
| EPI_ISL_3248120, | Lighthouse Lab in Alderley Park | Wellcome Sanger Institute for the COVID-19 Genomics UK (COG-UK) | Jacquelyn Wynn, Mairead Hyland, The Lighthouse Lab in Alderley Park and Alex Alderton, Roberto Amato, Jeffrey Barrett, Sonia Goncalves, Ewan Harrison, David K. Jackson, Ian Johnston, Dominic Kwiatkowski, Cordelia Langford, John Sillitoe on |

| | | | |
|---|---|---|---|
| EPI_ISL_3248121, EPI_ISL_3248122 | | Consortium | behalf of the Wellcome Sanger Institute COVID-19 Surveillance Team |
| EPI_ISL_3248123, EPI_ISL_3248124 | Lighthouse Laboratory Plymouth | Wellcome Sanger Institute for the COVID-19 Genomics UK (COG-UK) Consortium | Lighthouse Laboratory Plymouth and Alex Alderton, Roberto Amato, Jeffrey Barrett, Sonia Goncalves, Ewan Harrison, David K. Jackson, Ian Johnston, Dominic Kwiatkowski, Cordelia Langford, John Sillitoe on behalf of the Wellcome Sanger Institute COVID-19 Surveillance Team |
| EPI_ISL_3248125 | Lighthouse Lab in Alderley Park | Wellcome Sanger Institute for the COVID-19 Genomics UK (COG-UK) Consortium | Jacquelyn Wynn, Mairead Hyland, The Lighthouse Lab in Alderley Park and Alex Alderton, Roberto Amato, Jeffrey Barrett, Sonia Goncalves, Ewan Harrison, David K. Jackson, Ian Johnston, Dominic Kwiatkowski, Cordelia Langford, John Sillitoe on behalf of the Wellcome Sanger Institute COVID-19 Surveillance Team |
| EPI_ISL_3248126, EPI_ISL_3248127 | Lighthouse Laboratory Plymouth | Wellcome Sanger Institute for the COVID-19 Genomics UK (COG-UK) Consortium | Lighthouse Laboratory Plymouth and Alex Alderton, Roberto Amato, Jeffrey Barrett, Sonia Goncalves, Ewan Harrison, David K. Jackson, Ian Johnston, Dominic Kwiatkowski, Cordelia Langford, John Sillitoe on behalf of the Wellcome Sanger Institute COVID-19 Surveillance Team |
| EPI_ISL_3248128 | Lighthouse Lab in Alderley Park | Wellcome Sanger Institute for the COVID-19 Genomics UK (COG-UK) Consortium | Jacquelyn Wynn, Mairead Hyland, The Lighthouse Lab in Alderley Park and Alex Alderton, Roberto Amato, Jeffrey Barrett, Sonia Goncalves, Ewan Harrison, David K. Jackson, Ian Johnston, Dominic Kwiatkowski, Cordelia Langford, John Sillitoe on behalf of the Wellcome Sanger Institute COVID-19 Surveillance Team |
| EPI_ISL_3248129 | Lighthouse Laboratory Plymouth | Wellcome Sanger Institute for the COVID-19 Genomics UK (COG-UK) Consortium | Lighthouse Laboratory Plymouth and Alex Alderton, Roberto Amato, Jeffrey Barrett, Sonia Goncalves, Ewan Harrison, David K. Jackson, Ian Johnston, Dominic Kwiatkowski, Cordelia Langford, John Sillitoe on behalf of the Wellcome Sanger Institute COVID-19 Surveillance Team |
| EPI_ISL_3248130 | Lighthouse Lab in Alderley Park | Wellcome Sanger Institute for the COVID-19 Genomics UK (COG-UK) Consortium | Jacquelyn Wynn, Mairead Hyland, The Lighthouse Lab in Alderley Park and Alex Alderton, Roberto Amato, Jeffrey Barrett, Sonia Goncalves, Ewan Harrison, David K. Jackson, Ian Johnston, Dominic Kwiatkowski, Cordelia Langford, John Sillitoe on behalf of the Wellcome Sanger Institute COVID-19 Surveillance Team |
| EPI_ISL_3248131, EPI_ISL_3248132 | Lighthouse Laboratory Plymouth | Wellcome Sanger Institute for the COVID-19 Genomics UK (COG-UK) Consortium | Lighthouse Laboratory Plymouth and Alex Alderton, Roberto Amato, Jeffrey Barrett, Sonia Goncalves, Ewan Harrison, David K. Jackson, Ian Johnston, Dominic Kwiatkowski, Cordelia Langford, John Sillitoe on behalf of the Wellcome Sanger Institute COVID-19 Surveillance Team |
| EPI_ISL_3248133 | Lighthouse Lab in Alderley Park | Wellcome Sanger Institute for the COVID-19 Genomics UK (COG-UK) Consortium | Jacquelyn Wynn, Mairead Hyland, The Lighthouse Lab in Alderley Park and Alex Alderton, Roberto Amato, Jeffrey Barrett, Sonia Goncalves, Ewan Harrison, David K. Jackson, Ian Johnston, Dominic Kwiatkowski, Cordelia Langford, John Sillitoe on behalf of the Wellcome Sanger Institute COVID-19 Surveillance Team |
| EPI_ISL_3248134 | Lighthouse Laboratory Plymouth | Wellcome Sanger Institute for the COVID-19 Genomics UK (COG-UK) Consortium | Lighthouse Laboratory Plymouth and Alex Alderton, Roberto Amato, Jeffrey Barrett, Sonia Goncalves, Ewan Harrison, David K. Jackson, Ian Johnston, Dominic Kwiatkowski, Cordelia Langford, John Sillitoe on behalf of the Wellcome Sanger Institute COVID-19 Surveillance Team |
| EPI_ISL_3248135, EPI_ISL_3248136, EPI_ISL_3248137 | Lighthouse Lab in Alderley Park | Wellcome Sanger Institute for the COVID-19 Genomics UK (COG-UK) Consortium | Jacquelyn Wynn, Mairead Hyland, The Lighthouse Lab in Alderley Park and Alex Alderton, Roberto Amato, Jeffrey Barrett, Sonia Goncalves, Ewan Harrison, David K. Jackson, Ian Johnston, Dominic Kwiatkowski, Cordelia Langford, John Sillitoe on behalf of the Wellcome Sanger Institute COVID-19 Surveillance Team |
| EPI_ISL_3248138 | Lighthouse Laboratory Plymouth | Wellcome Sanger Institute for the COVID-19 Genomics UK (COG-UK) Consortium | Lighthouse Laboratory Plymouth and Alex Alderton, Roberto Amato, Jeffrey Barrett, Sonia Goncalves, Ewan Harrison, David K. Jackson, Ian Johnston, Dominic Kwiatkowski, Cordelia Langford, John Sillitoe on behalf of the Wellcome Sanger Institute COVID-19 Surveillance Team |
| EPI_ISL_3248139, EPI_ISL_3248140, EPI_ISL_3248141, EPI_ISL_3248142, EPI_ISL_3248143, EPI_ISL_3248144, EPI_ISL_3248145, EPI_ISL_3248146, EPI_ISL_3248147 | Lighthouse Lab in Alderley Park | Wellcome Sanger Institute for the COVID-19 Genomics UK (COG-UK) Consortium | Jacquelyn Wynn, Mairead Hyland, The Lighthouse Lab in Alderley Park and Alex Alderton, Roberto Amato, Jeffrey Barrett, Sonia Goncalves, Ewan Harrison, David K. Jackson, Ian Johnston, Dominic Kwiatkowski, Cordelia Langford, John Sillitoe on behalf of the Wellcome Sanger Institute COVID-19 Surveillance Team |
| EPI_ISL_3248148, EPI_ISL_3248149 | Lighthouse Laboratory Plymouth | Wellcome Sanger Institute for the COVID-19 Genomics UK (COG-UK) Consortium | Lighthouse Laboratory Plymouth and Alex Alderton, Roberto Amato, Jeffrey Barrett, Sonia Goncalves, Ewan Harrison, David K. Jackson, Ian Johnston, Dominic Kwiatkowski, Cordelia Langford, John Sillitoe on behalf of the Wellcome Sanger Institute COVID-19 Surveillance Team |
| EPI_ISL_3248150 | Lighthouse Lab in Alderley Park | Wellcome Sanger Institute for the COVID-19 Genomics UK (COG-UK) Consortium | Jacquelyn Wynn, Mairead Hyland, The Lighthouse Lab in Alderley Park and Alex Alderton, Roberto Amato, Jeffrey Barrett, Sonia Goncalves, Ewan Harrison, David K. Jackson, Ian Johnston, Dominic Kwiatkowski, Cordelia Langford, John Sillitoe on behalf of the Wellcome Sanger Institute COVID-19 Surveillance Team |
| EPI_ISL_3248151 | Lighthouse Laboratory Plymouth | Wellcome Sanger Institute for the COVID-19 Genomics UK (COG-UK) Consortium | Lighthouse Laboratory Plymouth and Alex Alderton, Roberto Amato, Jeffrey Barrett, Sonia Goncalves, Ewan Harrison, David K. Jackson, Ian Johnston, Dominic Kwiatkowski, Cordelia Langford, John Sillitoe on behalf of the Wellcome Sanger Institute COVID-19 Surveillance Team |
| EPI_ISL_3248152, EPI_ISL_3248153, EPI_ISL_3248154 | Lighthouse Lab in Alderley Park | Wellcome Sanger Institute for the COVID-19 Genomics UK (COG-UK) Consortium | Jacquelyn Wynn, Mairead Hyland, The Lighthouse Lab in Alderley Park and Alex Alderton, Roberto Amato, Jeffrey Barrett, Sonia Goncalves, Ewan Harrison, David K. Jackson, Ian Johnston, Dominic Kwiatkowski, Cordelia Langford, John Sillitoe on behalf of the Wellcome Sanger Institute COVID-19 Surveillance Team |
| EPI_ISL_3248155, EPI_ISL_3248156, EPI_ISL_3248157, EPI_ISL_3248158 | Lighthouse Laboratory Plymouth | Wellcome Sanger Institute for the COVID-19 Genomics UK (COG-UK) Consortium | Lighthouse Laboratory Plymouth and Alex Alderton, Roberto Amato, Jeffrey Barrett, Sonia Goncalves, Ewan Harrison, David K. Jackson, Ian Johnston, Dominic Kwiatkowski, Cordelia Langford, John Sillitoe on behalf of the Wellcome Sanger Institute COVID-19 Surveillance Team |
| EPI_ISL_3248159, EPI_ISL_3248160 | Lighthouse Lab in Alderley Park | Wellcome Sanger Institute for the COVID-19 Genomics UK (COG-UK) Consortium | Jacquelyn Wynn, Mairead Hyland, The Lighthouse Lab in Alderley Park and Alex Alderton, Roberto Amato, Jeffrey Barrett, Sonia Goncalves, Ewan Harrison, David K. Jackson, Ian Johnston, Dominic Kwiatkowski, Cordelia Langford, John Sillitoe on behalf of the Wellcome Sanger Institute COVID-19 Surveillance Team |
| EPI_ISL_3248161 | Lighthouse Laboratory Plymouth | Wellcome Sanger Institute for the COVID-19 Genomics UK (COG-UK) Consortium | Lighthouse Laboratory Plymouth and Alex Alderton, Roberto Amato, Jeffrey Barrett, Sonia Goncalves, Ewan Harrison, David K. Jackson, Ian Johnston, Dominic Kwiatkowski, Cordelia Langford, John Sillitoe on behalf of the Wellcome Sanger Institute COVID-19 Surveillance Team |
| EPI_ISL_3248162 | Lighthouse Lab in Alderley Park | Wellcome Sanger Institute for the COVID-19 Genomics UK (COG-UK) Consortium | Jacquelyn Wynn, Mairead Hyland, The Lighthouse Lab in Alderley Park and Alex Alderton, Roberto Amato, Jeffrey Barrett, Sonia Goncalves, Ewan Harrison, David K. Jackson, Ian Johnston, Dominic Kwiatkowski, Cordelia Langford, John Sillitoe on behalf of the Wellcome Sanger Institute COVID-19 Surveillance Team |
| EPI_ISL_3248163, EPI_ISL_3248164, EPI_ISL_3248165 | Lighthouse Laboratory Plymouth | Wellcome Sanger Institute for the COVID-19 Genomics UK (COG-UK) Consortium | Lighthouse Laboratory Plymouth and Alex Alderton, Roberto Amato, Jeffrey Barrett, Sonia Goncalves, Ewan Harrison, David K. Jackson, Ian Johnston, Dominic Kwiatkowski, Cordelia Langford, John Sillitoe on behalf of the Wellcome Sanger Institute COVID-19 Surveillance Team |
| EPI_ISL_3248166 | Lighthouse Lab in Alderley Park | Wellcome Sanger Institute for the COVID-19 Genomics UK (COG-UK) Consortium | Jacquelyn Wynn, Mairead Hyland, The Lighthouse Lab in Alderley Park and Alex Alderton, Roberto Amato, Jeffrey Barrett, Sonia Goncalves, Ewan Harrison, David K. Jackson, Ian Johnston, Dominic Kwiatkowski, Cordelia Langford, John Sillitoe on behalf of the Wellcome Sanger Institute COVID-19 Surveillance Team |
| EPI_ISL_3248167 | Lighthouse Laboratory Plymouth | Wellcome Sanger Institute for the COVID-19 Genomics UK (COG-UK) Consortium | Lighthouse Laboratory Plymouth and Alex Alderton, Roberto Amato, Jeffrey Barrett, Sonia Goncalves, Ewan Harrison, David K. Jackson, Ian Johnston, Dominic Kwiatkowski, Cordelia Langford, John Sillitoe on behalf of the Wellcome Sanger Institute COVID-19 Surveillance Team |
| EPI_ISL_3248168, EPI_ISL_3248169 | Lighthouse Lab in Alderley Park | Wellcome Sanger Institute for the COVID-19 Genomics UK (COG-UK) Consortium | Jacquelyn Wynn, Mairead Hyland, The Lighthouse Lab in Alderley Park and Alex Alderton, Roberto Amato, Jeffrey Barrett, Sonia Goncalves, Ewan Harrison, David K. Jackson, Ian Johnston, Dominic Kwiatkowski, Cordelia Langford, John Sillitoe on behalf of the Wellcome Sanger Institute COVID-19 Surveillance Team |
| EPI_ISL_3248170, EPI_ISL_3248171 | Lighthouse Laboratory Plymouth | Wellcome Sanger Institute for the COVID-19 Genomics UK (COG-UK) Consortium | Lighthouse Laboratory Plymouth and Alex Alderton, Roberto Amato, Jeffrey Barrett, Sonia Goncalves, Ewan Harrison, David K. Jackson, Ian Johnston, Dominic Kwiatkowski, Cordelia Langford, John Sillitoe on behalf of the Wellcome Sanger Institute COVID-19 Surveillance Team |
| EPI_ISL_3248172, EPI_ISL_3248173 | Lighthouse Lab in Alderley Park | Wellcome Sanger Institute for the COVID-19 Genomics UK (COG-UK) Consortium | Jacquelyn Wynn, Mairead Hyland, The Lighthouse Lab in Alderley Park and Alex Alderton, Roberto Amato, Jeffrey Barrett, Sonia Goncalves, Ewan Harrison, David K. Jackson, Ian Johnston, Dominic Kwiatkowski, Cordelia Langford, John Sillitoe on behalf of the Wellcome Sanger Institute COVID-19 Surveillance Team |
| EPI_ISL_3248174 | Lighthouse Laboratory Plymouth | Wellcome Sanger Institute for the COVID-19 Genomics UK (COG-UK) Consortium | Lighthouse Laboratory Plymouth and Alex Alderton, Roberto Amato, Jeffrey Barrett, Sonia Goncalves, Ewan Harrison, David K. Jackson, Ian Johnston, Dominic Kwiatkowski, Cordelia Langford, John Sillitoe on behalf of the Wellcome Sanger Institute COVID-19 Surveillance Team |
| EPI_ISL_3248175 | Lighthouse Lab in Alderley Park | Wellcome Sanger Institute for the COVID-19 Genomics UK (COG-UK) Consortium | Jacquelyn Wynn, Mairead Hyland, The Lighthouse Lab in Alderley Park and Alex Alderton, Roberto Amato, Jeffrey Barrett, Sonia Goncalves, Ewan Harrison, David K. Jackson, Ian Johnston, Dominic Kwiatkowski, Cordelia Langford, John Sillitoe on behalf of the Wellcome Sanger Institute COVID-19 Surveillance Team |
| EPI_ISL_3248176, EPI_ISL_3248177, EPI_ISL_3248178, EPI_ISL_3248179, EPI_ISL_3248180, EPI_ISL_3248181 | Lighthouse Laboratory Plymouth | Wellcome Sanger Institute for the COVID-19 Genomics UK (COG-UK) Consortium | Lighthouse Laboratory Plymouth and Alex Alderton, Roberto Amato, Jeffrey Barrett, Sonia Goncalves, Ewan Harrison, David K. Jackson, Ian Johnston, Dominic Kwiatkowski, Cordelia Langford, John Sillitoe on behalf of the Wellcome Sanger Institute COVID-19 Surveillance Team |
| EPI_ISL_3248182, EPI_ISL_3248183, EPI_ISL_3248184 | Lighthouse Lab in Alderley Park | Wellcome Sanger Institute for the COVID-19 Genomics UK (COG-UK) Consortium | Jacquelyn Wynn, Mairead Hyland, The Lighthouse Lab in Alderley Park and Alex Alderton, Roberto Amato, Jeffrey Barrett, Sonia Goncalves, Ewan Harrison, David K. Jackson, Ian Johnston, Dominic Kwiatkowski, Cordelia Langford, John Sillitoe on behalf of the Wellcome Sanger Institute COVID-19 Surveillance Team |
| EPI_ISL_3248185 | Lighthouse Laboratory Plymouth | Wellcome Sanger Institute for the COVID-19 Genomics UK (COG-UK) Consortium | Lighthouse Laboratory Plymouth and Alex Alderton, Roberto Amato, Jeffrey Barrett, Sonia Goncalves, Ewan Harrison, David K. Jackson, Ian Johnston, Dominic Kwiatkowski, Cordelia Langford, John Sillitoe on behalf of the Wellcome Sanger Institute COVID-19 Surveillance Team |
| EPI_ISL_3248186, EPI_ISL_3248187, EPI_ISL_3248188 | Lighthouse Lab in Alderley Park | Wellcome Sanger Institute for the COVID-19 Genomics UK (COG-UK) Consortium | Jacquelyn Wynn, Mairead Hyland, The Lighthouse Lab in Alderley Park and Alex Alderton, Roberto Amato, Jeffrey Barrett, Sonia Goncalves, Ewan Harrison, David K. Jackson, Ian Johnston, Dominic Kwiatkowski, Cordelia Langford, John Sillitoe on behalf of the Wellcome Sanger Institute COVID-19 Surveillance Team |
| EPI_ISL_3248189 | Lighthouse Laboratory Plymouth | Wellcome Sanger Institute for the COVID-19 Genomics UK (COG-UK) Consortium | Lighthouse Laboratory Plymouth and Alex Alderton, Roberto Amato, Jeffrey Barrett, Sonia Goncalves, Ewan Harrison, David K. Jackson, Ian Johnston, Dominic Kwiatkowski, Cordelia Langford, John Sillitoe on behalf of the Wellcome Sanger Institute COVID-19 Surveillance Team |
| EPI_ISL_3248190 | Lighthouse Lab in Alderley Park | Wellcome Sanger Institute for the COVID-19 Genomics UK (COG-UK) Consortium | Jacquelyn Wynn, Mairead Hyland, The Lighthouse Lab in Alderley Park and Alex Alderton, Roberto Amato, Jeffrey Barrett, Sonia Goncalves, Ewan Harrison, David K. Jackson, Ian Johnston, Dominic Kwiatkowski, Cordelia Langford, John Sillitoe on behalf of the Wellcome Sanger Institute COVID-19 Surveillance Team |
| EPI_ISL_3248191 | Lighthouse Laboratory Plymouth | Wellcome Sanger Institute for the COVID-19 Genomics UK (COG-UK) Consortium | Lighthouse Laboratory Plymouth and Alex Alderton, Roberto Amato, Jeffrey Barrett, Sonia Goncalves, Ewan Harrison, David K. Jackson, Ian Johnston, Dominic Kwiatkowski, Cordelia Langford, John Sillitoe on behalf of the Wellcome Sanger Institute COVID-19 Surveillance Team |
| EPI_ISL_3248192 | Lighthouse Lab in Alderley Park | Wellcome Sanger Institute for the COVID-19 Genomics UK (COG-UK) Consortium | Jacquelyn Wynn, Mairead Hyland, The Lighthouse Lab in Alderley Park and Alex Alderton, Roberto Amato, Jeffrey Barrett, Sonia Goncalves, Ewan Harrison, David K. Jackson, Ian Johnston, Dominic Kwiatkowski, Cordelia Langford, John Sillitoe on behalf of the Wellcome Sanger Institute COVID-19 Surveillance Team |
| EPI_ISL_3248193 | Lighthouse Laboratory Plymouth | Wellcome Sanger Institute for the COVID-19 Genomics UK (COG-UK) Consortium | Lighthouse Laboratory Plymouth and Alex Alderton, Roberto Amato, Jeffrey Barrett, Sonia Goncalves, Ewan Harrison, David K. Jackson, Ian Johnston, Dominic Kwiatkowski, Cordelia Langford, John Sillitoe on behalf of the Wellcome Sanger Institute COVID-19 Surveillance Team |

| | | | |
|---|---|---|---|
| EPI_ISL_3248194, EPI_ISL_3248195 | Lighthouse Lab in Alderley Park | Wellcome Sanger Institute for the COVID-19 Genomics UK (COG-UK) Consortium | Jacquelyn Wynn, Mairead Hyland, The Lighthouse Lab in Alderley Park and Alex Alderton, Roberto Amato, Jeffrey Barrett, Sonia Goncalves, Ewan Harrison, David K. Jackson, Ian Johnston, Dominic Kwiatkowski, Cordelia Langford, John Sillitoe on behalf of the Wellcome Sanger Institute COVID-19 Surveillance Team |
| EPI_ISL_3248196 | Lighthouse Laboratory Plymouth | Wellcome Sanger Institute for the COVID-19 Genomics UK (COG-UK) Consortium | Lighthouse Laboratory Plymouth and Alex Alderton, Roberto Amato, Jeffrey Barrett, Sonia Goncalves, Ewan Harrison, David K. Jackson, Ian Johnston, Dominic Kwiatkowski, Cordelia Langford, John Sillitoe on behalf of the Wellcome Sanger Institute COVID-19 Surveillance Team |
| EPI_ISL_3248197, EPI_ISL_3248198 | Lighthouse Lab in Alderley Park | Wellcome Sanger Institute for the COVID-19 Genomics UK (COG-UK) Consortium | Jacquelyn Wynn, Mairead Hyland, The Lighthouse Lab in Alderley Park and Alex Alderton, Roberto Amato, Jeffrey Barrett, Sonia Goncalves, Ewan Harrison, David K. Jackson, Ian Johnston, Dominic Kwiatkowski, Cordelia Langford, John Sillitoe on behalf of the Wellcome Sanger Institute COVID-19 Surveillance Team |
| EPI_ISL_3248199 | Lighthouse Laboratory Plymouth | Wellcome Sanger Institute for the COVID-19 Genomics UK (COG-UK) Consortium | Lighthouse Laboratory Plymouth and Alex Alderton, Roberto Amato, Jeffrey Barrett, Sonia Goncalves, Ewan Harrison, David K. Jackson, Ian Johnston, Dominic Kwiatkowski, Cordelia Langford, John Sillitoe on behalf of the Wellcome Sanger Institute COVID-19 Surveillance Team |
| EPI_ISL_3248200, EPI_ISL_3248201 | Lighthouse Lab in Alderley Park | Wellcome Sanger Institute for the COVID-19 Genomics UK (COG-UK) Consortium | Jacquelyn Wynn, Mairead Hyland, The Lighthouse Lab in Alderley Park and Alex Alderton, Roberto Amato, Jeffrey Barrett, Sonia Goncalves, Ewan Harrison, David K. Jackson, Ian Johnston, Dominic Kwiatkowski, Cordelia Langford, John Sillitoe on behalf of the Wellcome Sanger Institute COVID-19 Surveillance Team |
| EPI_ISL_3248202, EPI_ISL_3248203, EPI_ISL_3248204 | Lighthouse Laboratory Plymouth | Wellcome Sanger Institute for the COVID-19 Genomics UK (COG-UK) Consortium | Lighthouse Laboratory Plymouth and Alex Alderton, Roberto Amato, Jeffrey Barrett, Sonia Goncalves, Ewan Harrison, David K. Jackson, Ian Johnston, Dominic Kwiatkowski, Cordelia Langford, John Sillitoe on behalf of the Wellcome Sanger Institute COVID-19 Surveillance Team |
| EPI_ISL_3248205, EPI_ISL_3248206 | Lighthouse Lab in Alderley Park | Wellcome Sanger Institute for the COVID-19 Genomics UK (COG-UK) Consortium | Jacquelyn Wynn, Mairead Hyland, The Lighthouse Lab in Alderley Park and Alex Alderton, Roberto Amato, Jeffrey Barrett, Sonia Goncalves, Ewan Harrison, David K. Jackson, Ian Johnston, Dominic Kwiatkowski, Cordelia Langford, John Sillitoe on behalf of the Wellcome Sanger Institute COVID-19 Surveillance Team |
| EPI_ISL_3248207, EPI_ISL_3248208, EPI_ISL_3248209 | Lighthouse Laboratory Plymouth | Wellcome Sanger Institute for the COVID-19 Genomics UK (COG-UK) Consortium | Lighthouse Laboratory Plymouth and Alex Alderton, Roberto Amato, Jeffrey Barrett, Sonia Goncalves, Ewan Harrison, David K. Jackson, Ian Johnston, Dominic Kwiatkowski, Cordelia Langford, John Sillitoe on behalf of the Wellcome Sanger Institute COVID-19 Surveillance Team |
| EPI_ISL_3248210 | Lighthouse Lab in Alderley Park | Wellcome Sanger Institute for the COVID-19 Genomics UK (COG-UK) Consortium | Jacquelyn Wynn, Mairead Hyland, The Lighthouse Lab in Alderley Park and Alex Alderton, Roberto Amato, Jeffrey Barrett, Sonia Goncalves, Ewan Harrison, David K. Jackson, Ian Johnston, Dominic Kwiatkowski, Cordelia Langford, John Sillitoe on behalf of the Wellcome Sanger Institute COVID-19 Surveillance Team |
| EPI_ISL_3248211 | Lighthouse Laboratory Plymouth | Wellcome Sanger Institute for the COVID-19 Genomics UK (COG-UK) Consortium | Lighthouse Laboratory Plymouth and Alex Alderton, Roberto Amato, Jeffrey Barrett, Sonia Goncalves, Ewan Harrison, David K. Jackson, Ian Johnston, Dominic Kwiatkowski, Cordelia Langford, John Sillitoe on behalf of the Wellcome Sanger Institute COVID-19 Surveillance Team |
| EPI_ISL_3248212 | Lighthouse Lab in Alderley Park | Wellcome Sanger Institute for the COVID-19 Genomics UK (COG-UK) Consortium | Jacquelyn Wynn, Mairead Hyland, The Lighthouse Lab in Alderley Park and Alex Alderton, Roberto Amato, Jeffrey Barrett, Sonia Goncalves, Ewan Harrison, David K. Jackson, Ian Johnston, Dominic Kwiatkowski, Cordelia Langford, John Sillitoe on behalf of the Wellcome Sanger Institute COVID-19 Surveillance Team |
| EPI_ISL_3248213, EPI_ISL_3248214, EPI_ISL_3248215, EPI_ISL_3248216 | Lighthouse Laboratory Plymouth | Wellcome Sanger Institute for the COVID-19 Genomics UK (COG-UK) Consortium | Lighthouse Laboratory Plymouth and Alex Alderton, Roberto Amato, Jeffrey Barrett, Sonia Goncalves, Ewan Harrison, David K. Jackson, Ian Johnston, Dominic Kwiatkowski, Cordelia Langford, John Sillitoe on behalf of the Wellcome Sanger Institute COVID-19 Surveillance Team |
| EPI_ISL_3248217, EPI_ISL_3248218 | Lighthouse Lab in Alderley Park | Wellcome Sanger Institute for the COVID-19 Genomics UK (COG-UK) Consortium | Jacquelyn Wynn, Mairead Hyland, The Lighthouse Lab in Alderley Park and Alex Alderton, Roberto Amato, Jeffrey Barrett, Sonia Goncalves, Ewan Harrison, David K. Jackson, Ian Johnston, Dominic Kwiatkowski, Cordelia Langford, John Sillitoe on behalf of the Wellcome Sanger Institute COVID-19 Surveillance Team |
| EPI_ISL_3248219 | Lighthouse Laboratory Plymouth | Wellcome Sanger Institute for the COVID-19 Genomics UK (COG-UK) Consortium | Lighthouse Laboratory Plymouth and Alex Alderton, Roberto Amato, Jeffrey Barrett, Sonia Goncalves, Ewan Harrison, David K. Jackson, Ian Johnston, Dominic Kwiatkowski, Cordelia Langford, John Sillitoe on behalf of the Wellcome Sanger Institute COVID-19 Surveillance Team |
| EPI_ISL_3248220 | Lighthouse Lab in Alderley Park | Wellcome Sanger Institute for the COVID-19 Genomics UK (COG-UK) Consortium | Jacquelyn Wynn, Mairead Hyland, The Lighthouse Lab in Alderley Park and Alex Alderton, Roberto Amato, Jeffrey Barrett, Sonia Goncalves, Ewan Harrison, David K. Jackson, Ian Johnston, Dominic Kwiatkowski, Cordelia Langford, John Sillitoe on behalf of the Wellcome Sanger Institute COVID-19 Surveillance Team |
| EPI_ISL_3248221, EPI_ISL_3248222, EPI_ISL_3248223, EPI_ISL_3248224 | Lighthouse Laboratory Plymouth | Wellcome Sanger Institute for the COVID-19 Genomics UK (COG-UK) Consortium | Lighthouse Laboratory Plymouth and Alex Alderton, Roberto Amato, Jeffrey Barrett, Sonia Goncalves, Ewan Harrison, David K. Jackson, Ian Johnston, Dominic Kwiatkowski, Cordelia Langford, John Sillitoe on behalf of the Wellcome Sanger Institute COVID-19 Surveillance Team |
| EPI_ISL_3248225, EPI_ISL_3248226, EPI_ISL_3248227 | Lighthouse Lab in Alderley Park | Wellcome Sanger Institute for the COVID-19 Genomics UK (COG-UK) Consortium | Jacquelyn Wynn, Mairead Hyland, The Lighthouse Lab in Alderley Park and Alex Alderton, Roberto Amato, Jeffrey Barrett, Sonia Goncalves, Ewan Harrison, David K. Jackson, Ian Johnston, Dominic Kwiatkowski, Cordelia Langford, John Sillitoe on behalf of the Wellcome Sanger Institute COVID-19 Surveillance Team |
| EPI_ISL_3248228, EPI_ISL_3248229, EPI_ISL_3248230 | Lighthouse Laboratory Plymouth | Wellcome Sanger Institute for the COVID-19 Genomics UK (COG-UK) Consortium | Lighthouse Laboratory Plymouth and Alex Alderton, Roberto Amato, Jeffrey Barrett, Sonia Goncalves, Ewan Harrison, David K. Jackson, Ian Johnston, Dominic Kwiatkowski, Cordelia Langford, John Sillitoe on behalf of the Wellcome Sanger Institute COVID-19 Surveillance Team |
| EPI_ISL_3248231, EPI_ISL_3248232, EPI_ISL_3248233, EPI_ISL_3248234 | Lighthouse Lab in Alderley Park | Wellcome Sanger Institute for the COVID-19 Genomics UK (COG-UK) Consortium | Jacquelyn Wynn, Mairead Hyland, The Lighthouse Lab in Alderley Park and Alex Alderton, Roberto Amato, Jeffrey Barrett, Sonia Goncalves, Ewan Harrison, David K. Jackson, Ian Johnston, Dominic Kwiatkowski, Cordelia Langford, John Sillitoe on behalf of the Wellcome Sanger Institute COVID-19 Surveillance Team |
| EPI_ISL_3248235 | Lighthouse Laboratory Plymouth | Wellcome Sanger Institute for the COVID-19 Genomics UK (COG-UK) Consortium | Lighthouse Laboratory Plymouth and Alex Alderton, Roberto Amato, Jeffrey Barrett, Sonia Goncalves, Ewan Harrison, David K. Jackson, Ian Johnston, Dominic Kwiatkowski, Cordelia Langford, John Sillitoe on behalf of the Wellcome Sanger Institute COVID-19 Surveillance Team |
| EPI_ISL_3248236, EPI_ISL_3248237, EPI_ISL_3248238 | Lighthouse Lab in Alderley Park | Wellcome Sanger Institute for the COVID-19 Genomics UK (COG-UK) Consortium | Jacquelyn Wynn, Mairead Hyland, The Lighthouse Lab in Alderley Park and Alex Alderton, Roberto Amato, Jeffrey Barrett, Sonia Goncalves, Ewan Harrison, David K. Jackson, Ian Johnston, Dominic Kwiatkowski, Cordelia Langford, John Sillitoe on behalf of the Wellcome Sanger Institute COVID-19 Surveillance Team |
| EPI_ISL_3248239, EPI_ISL_3248240 | Lighthouse Laboratory Plymouth | Wellcome Sanger Institute for the COVID-19 Genomics UK (COG-UK) Consortium | Lighthouse Laboratory Plymouth and Alex Alderton, Roberto Amato, Jeffrey Barrett, Sonia Goncalves, Ewan Harrison, David K. Jackson, Ian Johnston, Dominic Kwiatkowski, Cordelia Langford, John Sillitoe on behalf of the Wellcome Sanger Institute COVID-19 Surveillance Team |
| EPI_ISL_3248241, EPI_ISL_3248242, EPI_ISL_3248243, EPI_ISL_3248244, EPI_ISL_3248245 | Lighthouse Lab in Alderley Park | Wellcome Sanger Institute for the COVID-19 Genomics UK (COG-UK) Consortium | Jacquelyn Wynn, Mairead Hyland, The Lighthouse Lab in Alderley Park and Alex Alderton, Roberto Amato, Jeffrey Barrett, Sonia Goncalves, Ewan Harrison, David K. Jackson, Ian Johnston, Dominic Kwiatkowski, Cordelia Langford, John Sillitoe on behalf of the Wellcome Sanger Institute COVID-19 Surveillance Team |
| EPI_ISL_3248246 | Lighthouse Laboratory Plymouth | Wellcome Sanger Institute for the COVID-19 Genomics UK (COG-UK) Consortium | Lighthouse Laboratory Plymouth and Alex Alderton, Roberto Amato, Jeffrey Barrett, Sonia Goncalves, Ewan Harrison, David K. Jackson, Ian Johnston, Dominic Kwiatkowski, Cordelia Langford, John Sillitoe on behalf of the Wellcome Sanger Institute COVID-19 Surveillance Team |
| EPI_ISL_3248247 | Lighthouse Lab in Alderley Park | Wellcome Sanger Institute for the COVID-19 Genomics UK (COG-UK) Consortium | Jacquelyn Wynn, Mairead Hyland, The Lighthouse Lab in Alderley Park and Alex Alderton, Roberto Amato, Jeffrey Barrett, Sonia Goncalves, Ewan Harrison, David K. Jackson, Ian Johnston, Dominic Kwiatkowski, Cordelia Langford, John Sillitoe on behalf of the Wellcome Sanger Institute COVID-19 Surveillance Team |
| EPI_ISL_3248248 | Lighthouse Laboratory Plymouth | Wellcome Sanger Institute for the COVID-19 Genomics UK (COG-UK) Consortium | Lighthouse Laboratory Plymouth and Alex Alderton, Roberto Amato, Jeffrey Barrett, Sonia Goncalves, Ewan Harrison, David K. Jackson, Ian Johnston, Dominic Kwiatkowski, Cordelia Langford, John Sillitoe on behalf of the Wellcome Sanger Institute COVID-19 Surveillance Team |
| EPI_ISL_3248249 | Lighthouse Lab in Alderley Park | Wellcome Sanger Institute for the COVID-19 Genomics UK (COG-UK) Consortium | Jacquelyn Wynn, Mairead Hyland, The Lighthouse Lab in Alderley Park and Alex Alderton, Roberto Amato, Jeffrey Barrett, Sonia Goncalves, Ewan Harrison, David K. Jackson, Ian Johnston, Dominic Kwiatkowski, Cordelia Langford, John Sillitoe on behalf of the Wellcome Sanger Institute COVID-19 Surveillance Team |
| EPI_ISL_3248250, EPI_ISL_3248251 | Lighthouse Laboratory Plymouth | Wellcome Sanger Institute for the COVID-19 Genomics UK (COG-UK) Consortium | Lighthouse Laboratory Plymouth and Alex Alderton, Roberto Amato, Jeffrey Barrett, Sonia Goncalves, Ewan Harrison, David K. Jackson, Ian Johnston, Dominic Kwiatkowski, Cordelia Langford, John Sillitoe on behalf of the Wellcome Sanger Institute COVID-19 Surveillance Team |
| EPI_ISL_3248252, EPI_ISL_3248253, EPI_ISL_3248254, EPI_ISL_3248255, EPI_ISL_3248256, EPI_ISL_3248257 | Lighthouse Lab in Alderley Park | Wellcome Sanger Institute for the COVID-19 Genomics UK (COG-UK) Consortium | Jacquelyn Wynn, Mairead Hyland, The Lighthouse Lab in Alderley Park and Alex Alderton, Roberto Amato, Jeffrey Barrett, Sonia Goncalves, Ewan Harrison, David K. Jackson, Ian Johnston, Dominic Kwiatkowski, Cordelia Langford, John Sillitoe on behalf of the Wellcome Sanger Institute COVID-19 Surveillance Team |
| EPI_ISL_3248258, EPI_ISL_3248259, EPI_ISL_3248260 | Lighthouse Laboratory Plymouth | Wellcome Sanger Institute for the COVID-19 Genomics UK (COG-UK) Consortium | Lighthouse Laboratory Plymouth and Alex Alderton, Roberto Amato, Jeffrey Barrett, Sonia Goncalves, Ewan Harrison, David K. Jackson, Ian Johnston, Dominic Kwiatkowski, Cordelia Langford, John Sillitoe on behalf of the Wellcome Sanger Institute COVID-19 Surveillance Team |
| EPI_ISL_3248261 | Lighthouse Lab in Alderley Park | Wellcome Sanger Institute for the COVID-19 Genomics UK (COG-UK) Consortium | Jacquelyn Wynn, Mairead Hyland, The Lighthouse Lab in Alderley Park and Alex Alderton, Roberto Amato, Jeffrey Barrett, Sonia Goncalves, Ewan Harrison, David K. Jackson, Ian Johnston, Dominic Kwiatkowski, Cordelia Langford, John Sillitoe on behalf of the Wellcome Sanger Institute COVID-19 Surveillance Team |
| EPI_ISL_3248262 | Lighthouse Laboratory Plymouth | Wellcome Sanger Institute for the COVID-19 Genomics UK (COG-UK) Consortium | Lighthouse Laboratory Plymouth and Alex Alderton, Roberto Amato, Jeffrey Barrett, Sonia Goncalves, Ewan Harrison, David K. Jackson, Ian Johnston, Dominic Kwiatkowski, Cordelia Langford, John Sillitoe on behalf of the Wellcome Sanger Institute COVID-19 Surveillance Team |
| EPI_ISL_3248263, EPI_ISL_3248264, EPI_ISL_3248265, EPI_ISL_3248266 | Lighthouse Lab in Alderley Park | Wellcome Sanger Institute for the COVID-19 Genomics UK (COG-UK) Consortium | Jacquelyn Wynn, Mairead Hyland, The Lighthouse Lab in Alderley Park and Alex Alderton, Roberto Amato, Jeffrey Barrett, Sonia Goncalves, Ewan Harrison, David K. Jackson, Ian Johnston, Dominic Kwiatkowski, Cordelia Langford, John Sillitoe on behalf of the Wellcome Sanger Institute COVID-19 Surveillance Team |
| EPI_ISL_3248267 | Lighthouse Laboratory Plymouth | Wellcome Sanger Institute for the COVID-19 Genomics UK (COG-UK) Consortium | Lighthouse Laboratory Plymouth and Alex Alderton, Roberto Amato, Jeffrey Barrett, Sonia Goncalves, Ewan Harrison, David K. Jackson, Ian Johnston, Dominic Kwiatkowski, Cordelia Langford, John Sillitoe on behalf of the Wellcome Sanger Institute COVID-19 Surveillance Team |
| EPI_ISL_3248268, EPI_ISL_3248269 | Lighthouse Lab in Alderley Park | Wellcome Sanger Institute for the COVID-19 Genomics UK (COG-UK) Consortium | Jacquelyn Wynn, Mairead Hyland, The Lighthouse Lab in Alderley Park and Alex Alderton, Roberto Amato, Jeffrey Barrett, Sonia Goncalves, Ewan Harrison, David K. Jackson, Ian Johnston, Dominic Kwiatkowski, Cordelia Langford, John Sillitoe on behalf of the Wellcome Sanger Institute COVID-19 Surveillance Team |
| EPI_ISL_3248270, EPI_ISL_3248271 | Lighthouse Laboratory Plymouth | Wellcome Sanger Institute for the COVID-19 Genomics UK (COG-UK) Consortium | Lighthouse Laboratory Plymouth and Alex Alderton, Roberto Amato, Jeffrey Barrett, Sonia Goncalves, Ewan Harrison, David K. Jackson, Ian Johnston, Dominic Kwiatkowski, Cordelia Langford, John Sillitoe on behalf of the Wellcome Sanger Institute COVID-19 Surveillance Team |
| EPI_ISL_3248272 | Lighthouse Lab in Alderley Park | Wellcome Sanger Institute for the COVID-19 Genomics UK (COG-UK) | Jacquelyn Wynn, Mairead Hyland, The Lighthouse Lab in Alderley Park and Alex Alderton, Roberto Amato, Jeffrey Barrett, Sonia Goncalves, Ewan Harrison, David K. Jackson, Ian Johnston, Dominic Kwiatkowski, Cordelia Langford, John Sillitoe on |

| | | | |
|---|---|---|---|
| | | Consortium | behalf of the Wellcome Sanger Institute COVID-19 Surveillance Team |
| EPI_ISL_3248273 | Lighthouse Laboratory Plymouth | Wellcome Sanger Institute for the COVID-19 Genomics UK (COG-UK) Consortium | Lighthouse Laboratory Plymouth and Alex Alderton, Roberto Amato, Jeffrey Barrett, Sonia Goncalves, Ewan Harrison, David K. Jackson, Ian Johnston, Dominic Kwiatkowski, Cordelia Langford, John Sillitoe on behalf of the Wellcome Sanger Institute COVID-19 Surveillance Team |
| EPI_ISL_3248274, EPI_ISL_3248275, EPI_ISL_3248276, EPI_ISL_3248277 | Lighthouse Lab in Alderley Park | Wellcome Sanger Institute for the COVID-19 Genomics UK (COG-UK) Consortium | Jacquelyn Wynn, Mairead Hyland, The Lighthouse Lab in Alderley Park and Alex Alderton, Roberto Amato, Jeffrey Barrett, Sonia Goncalves, Ewan Harrison, David K. Jackson, Ian Johnston, Dominic Kwiatkowski, Cordelia Langford, John Sillitoe on behalf of the Wellcome Sanger Institute COVID-19 Surveillance Team |
| EPI_ISL_3248278 | Lighthouse Laboratory Plymouth | Wellcome Sanger Institute for the COVID-19 Genomics UK (COG-UK) Consortium | Lighthouse Laboratory Plymouth and Alex Alderton, Roberto Amato, Jeffrey Barrett, Sonia Goncalves, Ewan Harrison, David K. Jackson, Ian Johnston, Dominic Kwiatkowski, Cordelia Langford, John Sillitoe on behalf of the Wellcome Sanger Institute COVID-19 Surveillance Team |
| EPI_ISL_3248279 | Lighthouse Lab in Alderley Park | Wellcome Sanger Institute for the COVID-19 Genomics UK (COG-UK) Consortium | Jacquelyn Wynn, Mairead Hyland, The Lighthouse Lab in Alderley Park and Alex Alderton, Roberto Amato, Jeffrey Barrett, Sonia Goncalves, Ewan Harrison, David K. Jackson, Ian Johnston, Dominic Kwiatkowski, Cordelia Langford, John Sillitoe on behalf of the Wellcome Sanger Institute COVID-19 Surveillance Team |
| EPI_ISL_3248280, EPI_ISL_3248281 | Lighthouse Laboratory Plymouth | Wellcome Sanger Institute for the COVID-19 Genomics UK (COG-UK) Consortium | Lighthouse Laboratory Plymouth and Alex Alderton, Roberto Amato, Jeffrey Barrett, Sonia Goncalves, Ewan Harrison, David K. Jackson, Ian Johnston, Dominic Kwiatkowski, Cordelia Langford, John Sillitoe on behalf of the Wellcome Sanger Institute COVID-19 Surveillance Team |
| EPI_ISL_3248282, EPI_ISL_3248283, EPI_ISL_3248284, EPI_ISL_3248285, EPI_ISL_3248286, EPI_ISL_3248287, EPI_ISL_3248288, EPI_ISL_3248289, EPI_ISL_3248290, EPI_ISL_3248291 | Lighthouse Lab in Alderley Park | Wellcome Sanger Institute for the COVID-19 Genomics UK (COG-UK) Consortium | Jacquelyn Wynn, Mairead Hyland, The Lighthouse Lab in Alderley Park and Alex Alderton, Roberto Amato, Jeffrey Barrett, Sonia Goncalves, Ewan Harrison, David K. Jackson, Ian Johnston, Dominic Kwiatkowski, Cordelia Langford, John Sillitoe on behalf of the Wellcome Sanger Institute COVID-19 Surveillance Team |
| EPI_ISL_3248292, EPI_ISL_3248293 | Lighthouse Laboratory Plymouth | Wellcome Sanger Institute for the COVID-19 Genomics UK (COG-UK) Consortium | Lighthouse Laboratory Plymouth and Alex Alderton, Roberto Amato, Jeffrey Barrett, Sonia Goncalves, Ewan Harrison, David K. Jackson, Ian Johnston, Dominic Kwiatkowski, Cordelia Langford, John Sillitoe on behalf of the Wellcome Sanger Institute COVID-19 Surveillance Team |
| EPI_ISL_3248294 | Lighthouse Lab in Alderley Park | Wellcome Sanger Institute for the COVID-19 Genomics UK (COG-UK) Consortium | Jacquelyn Wynn, Mairead Hyland, The Lighthouse Lab in Alderley Park and Alex Alderton, Roberto Amato, Jeffrey Barrett, Sonia Goncalves, Ewan Harrison, David K. Jackson, Ian Johnston, Dominic Kwiatkowski, Cordelia Langford, John Sillitoe on behalf of the Wellcome Sanger Institute COVID-19 Surveillance Team |
| EPI_ISL_3248295, EPI_ISL_3248296, EPI_ISL_3248297, EPI_ISL_3248298, EPI_ISL_3248299 | Lighthouse Laboratory Plymouth | Wellcome Sanger Institute for the COVID-19 Genomics UK (COG-UK) Consortium | Lighthouse Laboratory Plymouth and Alex Alderton, Roberto Amato, Jeffrey Barrett, Sonia Goncalves, Ewan Harrison, David K. Jackson, Ian Johnston, Dominic Kwiatkowski, Cordelia Langford, John Sillitoe on behalf of the Wellcome Sanger Institute COVID-19 Surveillance Team |
| EPI_ISL_3248300 | Lighthouse Lab in Alderley Park | Wellcome Sanger Institute for the COVID-19 Genomics UK (COG-UK) Consortium | Jacquelyn Wynn, Mairead Hyland, The Lighthouse Lab in Alderley Park and Alex Alderton, Roberto Amato, Jeffrey Barrett, Sonia Goncalves, Ewan Harrison, David K. Jackson, Ian Johnston, Dominic Kwiatkowski, Cordelia Langford, John Sillitoe on behalf of the Wellcome Sanger Institute COVID-19 Surveillance Team |
| EPI_ISL_3248301 | Lighthouse Laboratory Plymouth | Wellcome Sanger Institute for the COVID-19 Genomics UK (COG-UK) Consortium | Lighthouse Laboratory Plymouth and Alex Alderton, Roberto Amato, Jeffrey Barrett, Sonia Goncalves, Ewan Harrison, David K. Jackson, Ian Johnston, Dominic Kwiatkowski, Cordelia Langford, John Sillitoe on behalf of the Wellcome Sanger Institute COVID-19 Surveillance Team |
| EPI_ISL_3248302 | Lighthouse Lab in Alderley Park | Wellcome Sanger Institute for the COVID-19 Genomics UK (COG-UK) Consortium | Jacquelyn Wynn, Mairead Hyland, The Lighthouse Lab in Alderley Park and Alex Alderton, Roberto Amato, Jeffrey Barrett, Sonia Goncalves, Ewan Harrison, David K. Jackson, Ian Johnston, Dominic Kwiatkowski, Cordelia Langford, John Sillitoe on behalf of the Wellcome Sanger Institute COVID-19 Surveillance Team |
| EPI_ISL_3248303 | AZ Klina | AZ Klina | Carl Vael - Lynsey Berckmans |
| EPI_ISL_3248304 | Lighthouse Lab in Alderley Park | Wellcome Sanger Institute for the COVID-19 Genomics UK (COG-UK) Consortium | Jacquelyn Wynn, Mairead Hyland, The Lighthouse Lab in Alderley Park and Alex Alderton, Roberto Amato, Jeffrey Barrett, Sonia Goncalves, Ewan Harrison, David K. Jackson, Ian Johnston, Dominic Kwiatkowski, Cordelia Langford, John Sillitoe on behalf of the Wellcome Sanger Institute COVID-19 Surveillance Team |
| EPI_ISL_3248305 | Lighthouse Laboratory Plymouth | Wellcome Sanger Institute for the COVID-19 Genomics UK (COG-UK) Consortium | Lighthouse Laboratory Plymouth and Alex Alderton, Roberto Amato, Jeffrey Barrett, Sonia Goncalves, Ewan Harrison, David K. Jackson, Ian Johnston, Dominic Kwiatkowski, Cordelia Langford, John Sillitoe on behalf of the Wellcome Sanger Institute COVID-19 Surveillance Team |
| EPI_ISL_3248306, EPI_ISL_3248307 | Lighthouse Lab in Alderley Park | Wellcome Sanger Institute for the COVID-19 Genomics UK (COG-UK) Consortium | Jacquelyn Wynn, Mairead Hyland, The Lighthouse Lab in Alderley Park and Alex Alderton, Roberto Amato, Jeffrey Barrett, Sonia Goncalves, Ewan Harrison, David K. Jackson, Ian Johnston, Dominic Kwiatkowski, Cordelia Langford, John Sillitoe on behalf of the Wellcome Sanger Institute COVID-19 Surveillance Team |
| EPI_ISL_3248308, EPI_ISL_3248309, EPI_ISL_3248310 | Lighthouse Laboratory Plymouth | Wellcome Sanger Institute for the COVID-19 Genomics UK (COG-UK) Consortium | Lighthouse Laboratory Plymouth and Alex Alderton, Roberto Amato, Jeffrey Barrett, Sonia Goncalves, Ewan Harrison, David K. Jackson, Ian Johnston, Dominic Kwiatkowski, Cordelia Langford, John Sillitoe on behalf of the Wellcome Sanger Institute COVID-19 Surveillance Team |
| EPI_ISL_3248311 | Lighthouse Lab in Alderley Park | Wellcome Sanger Institute for the COVID-19 Genomics UK (COG-UK) Consortium | Jacquelyn Wynn, Mairead Hyland, The Lighthouse Lab in Alderley Park and Alex Alderton, Roberto Amato, Jeffrey Barrett, Sonia Goncalves, Ewan Harrison, David K. Jackson, Ian Johnston, Dominic Kwiatkowski, Cordelia Langford, John Sillitoe on behalf of the Wellcome Sanger Institute COVID-19 Surveillance Team |
| EPI_ISL_3248312, EPI_ISL_3248313 | Lighthouse Laboratory Plymouth | Wellcome Sanger Institute for the COVID-19 Genomics UK (COG-UK) Consortium | Lighthouse Laboratory Plymouth and Alex Alderton, Roberto Amato, Jeffrey Barrett, Sonia Goncalves, Ewan Harrison, David K. Jackson, Ian Johnston, Dominic Kwiatkowski, Cordelia Langford, John Sillitoe on behalf of the Wellcome Sanger Institute COVID-19 Surveillance Team |
| EPI_ISL_3248314 | Lighthouse Lab in Alderley Park | Wellcome Sanger Institute for the COVID-19 Genomics UK (COG-UK) Consortium | Jacquelyn Wynn, Mairead Hyland, The Lighthouse Lab in Alderley Park and Alex Alderton, Roberto Amato, Jeffrey Barrett, Sonia Goncalves, Ewan Harrison, David K. Jackson, Ian Johnston, Dominic Kwiatkowski, Cordelia Langford, John Sillitoe on behalf of the Wellcome Sanger Institute COVID-19 Surveillance Team |
| EPI_ISL_3248315, EPI_ISL_3248316 | Lighthouse Laboratory Plymouth | Wellcome Sanger Institute for the COVID-19 Genomics UK (COG-UK) Consortium | Lighthouse Laboratory Plymouth and Alex Alderton, Roberto Amato, Jeffrey Barrett, Sonia Goncalves, Ewan Harrison, David K. Jackson, Ian Johnston, Dominic Kwiatkowski, Cordelia Langford, John Sillitoe on behalf of the Wellcome Sanger Institute COVID-19 Surveillance Team |
| EPI_ISL_3248317 | Lighthouse Lab in Alderley Park | Wellcome Sanger Institute for the COVID-19 Genomics UK (COG-UK) Consortium | Jacquelyn Wynn, Mairead Hyland, The Lighthouse Lab in Alderley Park and Alex Alderton, Roberto Amato, Jeffrey Barrett, Sonia Goncalves, Ewan Harrison, David K. Jackson, Ian Johnston, Dominic Kwiatkowski, Cordelia Langford, John Sillitoe on behalf of the Wellcome Sanger Institute COVID-19 Surveillance Team |
| EPI_ISL_3248318, EPI_ISL_3248319 | Lighthouse Laboratory Plymouth | Wellcome Sanger Institute for the COVID-19 Genomics UK (COG-UK) Consortium | Lighthouse Laboratory Plymouth and Alex Alderton, Roberto Amato, Jeffrey Barrett, Sonia Goncalves, Ewan Harrison, David K. Jackson, Ian Johnston, Dominic Kwiatkowski, Cordelia Langford, John Sillitoe on behalf of the Wellcome Sanger Institute COVID-19 Surveillance Team |
| EPI_ISL_3248320, EPI_ISL_3248321, EPI_ISL_3248322 | Lighthouse Lab in Alderley Park | Wellcome Sanger Institute for the COVID-19 Genomics UK (COG-UK) Consortium | Jacquelyn Wynn, Mairead Hyland, The Lighthouse Lab in Alderley Park and Alex Alderton, Roberto Amato, Jeffrey Barrett, Sonia Goncalves, Ewan Harrison, David K. Jackson, Ian Johnston, Dominic Kwiatkowski, Cordelia Langford, John Sillitoe on behalf of the Wellcome Sanger Institute COVID-19 Surveillance Team |
| EPI_ISL_3248323 | Lighthouse Laboratory Plymouth | Wellcome Sanger Institute for the COVID-19 Genomics UK (COG-UK) Consortium | Lighthouse Laboratory Plymouth and Alex Alderton, Roberto Amato, Jeffrey Barrett, Sonia Goncalves, Ewan Harrison, David K. Jackson, Ian Johnston, Dominic Kwiatkowski, Cordelia Langford, John Sillitoe on behalf of the Wellcome Sanger Institute COVID-19 Surveillance Team |
| EPI_ISL_3248324 | Lighthouse Lab in Alderley Park | Wellcome Sanger Institute for the COVID-19 Genomics UK (COG-UK) Consortium | Jacquelyn Wynn, Mairead Hyland, The Lighthouse Lab in Alderley Park and Alex Alderton, Roberto Amato, Jeffrey Barrett, Sonia Goncalves, Ewan Harrison, David K. Jackson, Ian Johnston, Dominic Kwiatkowski, Cordelia Langford, John Sillitoe on behalf of the Wellcome Sanger Institute COVID-19 Surveillance Team |
| EPI_ISL_3248325 | Lighthouse Laboratory Plymouth | Wellcome Sanger Institute for the COVID-19 Genomics UK (COG-UK) Consortium | Lighthouse Laboratory Plymouth and Alex Alderton, Roberto Amato, Jeffrey Barrett, Sonia Goncalves, Ewan Harrison, David K. Jackson, Ian Johnston, Dominic Kwiatkowski, Cordelia Langford, John Sillitoe on behalf of the Wellcome Sanger Institute COVID-19 Surveillance Team |
| EPI_ISL_3248326 | Lighthouse Lab in Alderley Park | Wellcome Sanger Institute for the COVID-19 Genomics UK (COG-UK) Consortium | Jacquelyn Wynn, Mairead Hyland, The Lighthouse Lab in Alderley Park and Alex Alderton, Roberto Amato, Jeffrey Barrett, Sonia Goncalves, Ewan Harrison, David K. Jackson, Ian Johnston, Dominic Kwiatkowski, Cordelia Langford, John Sillitoe on behalf of the Wellcome Sanger Institute COVID-19 Surveillance Team |
| EPI_ISL_3248327 | Lighthouse Laboratory Plymouth | Wellcome Sanger Institute for the COVID-19 Genomics UK (COG-UK) Consortium | Lighthouse Laboratory Plymouth and Alex Alderton, Roberto Amato, Jeffrey Barrett, Sonia Goncalves, Ewan Harrison, David K. Jackson, Ian Johnston, Dominic Kwiatkowski, Cordelia Langford, John Sillitoe on behalf of the Wellcome Sanger Institute COVID-19 Surveillance Team |
| EPI_ISL_3248328 | Lighthouse Lab in Alderley Park | Wellcome Sanger Institute for the COVID-19 Genomics UK (COG-UK) Consortium | Jacquelyn Wynn, Mairead Hyland, The Lighthouse Lab in Alderley Park and Alex Alderton, Roberto Amato, Jeffrey Barrett, Sonia Goncalves, Ewan Harrison, David K. Jackson, Ian Johnston, Dominic Kwiatkowski, Cordelia Langford, John Sillitoe on behalf of the Wellcome Sanger Institute COVID-19 Surveillance Team |
| EPI_ISL_3248329, EPI_ISL_3248330, EPI_ISL_3248331 | Lighthouse Laboratory Plymouth | Wellcome Sanger Institute for the COVID-19 Genomics UK (COG-UK) Consortium | Lighthouse Laboratory Plymouth and Alex Alderton, Roberto Amato, Jeffrey Barrett, Sonia Goncalves, Ewan Harrison, David K. Jackson, Ian Johnston, Dominic Kwiatkowski, Cordelia Langford, John Sillitoe on behalf of the Wellcome Sanger Institute COVID-19 Surveillance Team |
| EPI_ISL_3248332 | Lighthouse Lab in Alderley Park | Wellcome Sanger Institute for the COVID-19 Genomics UK (COG-UK) Consortium | Jacquelyn Wynn, Mairead Hyland, The Lighthouse Lab in Alderley Park and Alex Alderton, Roberto Amato, Jeffrey Barrett, Sonia Goncalves, Ewan Harrison, David K. Jackson, Ian Johnston, Dominic Kwiatkowski, Cordelia Langford, John Sillitoe on behalf of the Wellcome Sanger Institute COVID-19 Surveillance Team |
| EPI_ISL_3248333, EPI_ISL_3248334 | Lighthouse Laboratory Plymouth | Wellcome Sanger Institute for the COVID-19 Genomics UK (COG-UK) Consortium | Lighthouse Laboratory Plymouth and Alex Alderton, Roberto Amato, Jeffrey Barrett, Sonia Goncalves, Ewan Harrison, David K. Jackson, Ian Johnston, Dominic Kwiatkowski, Cordelia Langford, John Sillitoe on behalf of the Wellcome Sanger Institute COVID-19 Surveillance Team |
| EPI_ISL_3248335, EPI_ISL_3248336, EPI_ISL_3248337, EPI_ISL_3248338, EPI_ISL_3248339 | Lighthouse Lab in Alderley Park | Wellcome Sanger Institute for the COVID-19 Genomics UK (COG-UK) Consortium | Jacquelyn Wynn, Mairead Hyland, The Lighthouse Lab in Alderley Park and Alex Alderton, Roberto Amato, Jeffrey Barrett, Sonia Goncalves, Ewan Harrison, David K. Jackson, Ian Johnston, Dominic Kwiatkowski, Cordelia Langford, John Sillitoe on behalf of the Wellcome Sanger Institute COVID-19 Surveillance Team |
| EPI_ISL_3248340, EPI_ISL_3248341, EPI_ISL_3248342, EPI_ISL_3248343 | Lighthouse Laboratory Plymouth | Wellcome Sanger Institute for the COVID-19 Genomics UK (COG-UK) Consortium | Lighthouse Laboratory Plymouth and Alex Alderton, Roberto Amato, Jeffrey Barrett, Sonia Goncalves, Ewan Harrison, David K. Jackson, Ian Johnston, Dominic Kwiatkowski, Cordelia Langford, John Sillitoe on behalf of the Wellcome Sanger Institute COVID-19 Surveillance Team |
| EPI_ISL_3248344, EPI_ISL_3248345, EPI_ISL_3248346 | Lighthouse Lab in Alderley Park | Wellcome Sanger Institute for the COVID-19 Genomics UK (COG-UK) Consortium | Jacquelyn Wynn, Mairead Hyland, The Lighthouse Lab in Alderley Park and Alex Alderton, Roberto Amato, Jeffrey Barrett, Sonia Goncalves, Ewan Harrison, David K. Jackson, Ian Johnston, Dominic Kwiatkowski, Cordelia Langford, John Sillitoe on behalf of the Wellcome Sanger Institute COVID-19 Surveillance Team |

| | | | |
|---|---|---|---|
| EPI_ISL_3248347 | Lighthouse Laboratory Plymouth | Wellcome Sanger Institute for the COVID-19 Genomics UK (COG-UK) Consortium | Lighthouse Laboratory Plymouth and Alex Alderton, Roberto Amato, Jeffrey Barrett, Sonia Goncalves, Ewan Harrison, David K. Jackson, Ian Johnston, Dominic Kwiatkowski, Cordelia Langford, John Sillitoe on behalf of the Wellcome Sanger Institute COVID-19 Surveillance Team |
| EPI_ISL_3248348 | Lighthouse Lab in Alderley Park | Wellcome Sanger Institute for the COVID-19 Genomics UK (COG-UK) Consortium | Jacquelyn Wynn, Mairead Hyland, The Lighthouse Lab in Alderley Park and Alex Alderton, Roberto Amato, Jeffrey Barrett, Sonia Goncalves, Ewan Harrison, David K. Jackson, Ian Johnston, Dominic Kwiatkowski, Cordelia Langford, John Sillitoe on behalf of the Wellcome Sanger Institute COVID-19 Surveillance Team |
| EPI_ISL_3248349, EPI_ISL_3248350, EPI_ISL_3248351 | Lighthouse Laboratory Plymouth | Wellcome Sanger Institute for the COVID-19 Genomics UK (COG-UK) Consortium | Lighthouse Laboratory Plymouth and Alex Alderton, Roberto Amato, Jeffrey Barrett, Sonia Goncalves, Ewan Harrison, David K. Jackson, Ian Johnston, Dominic Kwiatkowski, Cordelia Langford, John Sillitoe on behalf of the Wellcome Sanger Institute COVID-19 Surveillance Team |
| EPI_ISL_3248352 | Lighthouse Lab in Alderley Park | Wellcome Sanger Institute for the COVID-19 Genomics UK (COG-UK) Consortium | Jacquelyn Wynn, Mairead Hyland, The Lighthouse Lab in Alderley Park and Alex Alderton, Roberto Amato, Jeffrey Barrett, Sonia Goncalves, Ewan Harrison, David K. Jackson, Ian Johnston, Dominic Kwiatkowski, Cordelia Langford, John Sillitoe on behalf of the Wellcome Sanger Institute COVID-19 Surveillance Team |
| EPI_ISL_3248353 | Lighthouse Laboratory Plymouth | Wellcome Sanger Institute for the COVID-19 Genomics UK (COG-UK) Consortium | Lighthouse Laboratory Plymouth and Alex Alderton, Roberto Amato, Jeffrey Barrett, Sonia Goncalves, Ewan Harrison, David K. Jackson, Ian Johnston, Dominic Kwiatkowski, Cordelia Langford, John Sillitoe on behalf of the Wellcome Sanger Institute COVID-19 Surveillance Team |
| EPI_ISL_3248354, EPI_ISL_3248355, EPI_ISL_3248356, EPI_ISL_3248357, EPI_ISL_3248359 | Lighthouse Lab in Alderley Park | Wellcome Sanger Institute for the COVID-19 Genomics UK (COG-UK) Consortium | Jacquelyn Wynn, Mairead Hyland, The Lighthouse Lab in Alderley Park and Alex Alderton, Roberto Amato, Jeffrey Barrett, Sonia Goncalves, Ewan Harrison, David K. Jackson, Ian Johnston, Dominic Kwiatkowski, Cordelia Langford, John Sillitoe on behalf of the Wellcome Sanger Institute COVID-19 Surveillance Team |
| EPI_ISL_3248360, EPI_ISL_3248361 | Lighthouse Laboratory Plymouth | Wellcome Sanger Institute for the COVID-19 Genomics UK (COG-UK) Consortium | Lighthouse Laboratory Plymouth and Alex Alderton, Roberto Amato, Jeffrey Barrett, Sonia Goncalves, Ewan Harrison, David K. Jackson, Ian Johnston, Dominic Kwiatkowski, Cordelia Langford, John Sillitoe on behalf of the Wellcome Sanger Institute COVID-19 Surveillance Team |
| EPI_ISL_3248362, EPI_ISL_3248363, EPI_ISL_3248364 | Lighthouse Lab in Alderley Park | Wellcome Sanger Institute for the COVID-19 Genomics UK (COG-UK) Consortium | Jacquelyn Wynn, Mairead Hyland, The Lighthouse Lab in Alderley Park and Alex Alderton, Roberto Amato, Jeffrey Barrett, Sonia Goncalves, Ewan Harrison, David K. Jackson, Ian Johnston, Dominic Kwiatkowski, Cordelia Langford, John Sillitoe on behalf of the Wellcome Sanger Institute COVID-19 Surveillance Team |
| EPI_ISL_3248365 | Lighthouse Laboratory Plymouth | Wellcome Sanger Institute for the COVID-19 Genomics UK (COG-UK) Consortium | Lighthouse Laboratory Plymouth and Alex Alderton, Roberto Amato, Jeffrey Barrett, Sonia Goncalves, Ewan Harrison, David K. Jackson, Ian Johnston, Dominic Kwiatkowski, Cordelia Langford, John Sillitoe on behalf of the Wellcome Sanger Institute COVID-19 Surveillance Team |
| EPI_ISL_3248366, EPI_ISL_3248367, EPI_ISL_3248368 | Lighthouse Lab in Alderley Park | Wellcome Sanger Institute for the COVID-19 Genomics UK (COG-UK) Consortium | Jacquelyn Wynn, Mairead Hyland, The Lighthouse Lab in Alderley Park and Alex Alderton, Roberto Amato, Jeffrey Barrett, Sonia Goncalves, Ewan Harrison, David K. Jackson, Ian Johnston, Dominic Kwiatkowski, Cordelia Langford, John Sillitoe on behalf of the Wellcome Sanger Institute COVID-19 Surveillance Team |
| EPI_ISL_3248369 | Lighthouse Laboratory Plymouth | Wellcome Sanger Institute for the COVID-19 Genomics UK (COG-UK) Consortium | Lighthouse Laboratory Plymouth and Alex Alderton, Roberto Amato, Jeffrey Barrett, Sonia Goncalves, Ewan Harrison, David K. Jackson, Ian Johnston, Dominic Kwiatkowski, Cordelia Langford, John Sillitoe on behalf of the Wellcome Sanger Institute COVID-19 Surveillance Team |
| EPI_ISL_3248370, EPI_ISL_3248371, EPI_ISL_3248372, EPI_ISL_3248373, EPI_ISL_3248374 | Lighthouse Lab in Glasgow | Wellcome Sanger Institute for the COVID-19 Genomics UK (COG-UK) Consortium | Harper VanSteenhouse, Yumi Kasai, David Gray, Carol Clugston, Anna Dominiczak and Alex Alderton, Roberto Amato, Jeffrey Barrett, Sonia Goncalves, Ewan Harrison, David K. Jackson, Ian Johnston, Dominic Kwiatkowski, Cordelia Langford, John Sillitoe on behalf of the Wellcome Sanger Institute COVID-19 Surveillance Team |
| EPI_ISL_3248375 | Lighthouse Lab in Alderley Park | Wellcome Sanger Institute for the COVID-19 Genomics UK (COG-UK) Consortium | Jacquelyn Wynn, Mairead Hyland, The Lighthouse Lab in Alderley Park and Alex Alderton, Roberto Amato, Jeffrey Barrett, Sonia Goncalves, Ewan Harrison, David K. Jackson, Ian Johnston, Dominic Kwiatkowski, Cordelia Langford, John Sillitoe on behalf of the Wellcome Sanger Institute COVID-19 Surveillance Team |
| EPI_ISL_3248376 | Lighthouse Lab in Glasgow | Wellcome Sanger Institute for the COVID-19 Genomics UK (COG-UK) Consortium | Harper VanSteenhouse, Yumi Kasai, David Gray, Carol Clugston, Anna Dominiczak and Alex Alderton, Roberto Amato, Jeffrey Barrett, Sonia Goncalves, Ewan Harrison, David K. Jackson, Ian Johnston, Dominic Kwiatkowski, Cordelia Langford, John Sillitoe on behalf of the Wellcome Sanger Institute COVID-19 Surveillance Team |
| EPI_ISL_3248377 | Lighthouse Lab in Alderley Park | Wellcome Sanger Institute for the COVID-19 Genomics UK (COG-UK) Consortium | Jacquelyn Wynn, Mairead Hyland, The Lighthouse Lab in Alderley Park and Alex Alderton, Roberto Amato, Jeffrey Barrett, Sonia Goncalves, Ewan Harrison, David K. Jackson, Ian Johnston, Dominic Kwiatkowski, Cordelia Langford, John Sillitoe on behalf of the Wellcome Sanger Institute COVID-19 Surveillance Team |
| EPI_ISL_3248378 | Lighthouse Lab in Glasgow | Wellcome Sanger Institute for the COVID-19 Genomics UK (COG-UK) Consortium | Harper VanSteenhouse, Yumi Kasai, David Gray, Carol Clugston, Anna Dominiczak and Alex Alderton, Roberto Amato, Jeffrey Barrett, Sonia Goncalves, Ewan Harrison, David K. Jackson, Ian Johnston, Dominic Kwiatkowski, Cordelia Langford, John Sillitoe on behalf of the Wellcome Sanger Institute COVID-19 Surveillance Team |
| EPI_ISL_3248379, EPI_ISL_3248380 | Lighthouse Lab in Alderley Park | Wellcome Sanger Institute for the COVID-19 Genomics UK (COG-UK) Consortium | Jacquelyn Wynn, Mairead Hyland, The Lighthouse Lab in Alderley Park and Alex Alderton, Roberto Amato, Jeffrey Barrett, Sonia Goncalves, Ewan Harrison, David K. Jackson, Ian Johnston, Dominic Kwiatkowski, Cordelia Langford, John Sillitoe on behalf of the Wellcome Sanger Institute COVID-19 Surveillance Team |
| EPI_ISL_3248381, EPI_ISL_3248382, EPI_ISL_3248383 | Lighthouse Lab in Glasgow | Wellcome Sanger Institute for the COVID-19 Genomics UK (COG-UK) Consortium | Harper VanSteenhouse, Yumi Kasai, David Gray, Carol Clugston, Anna Dominiczak and Alex Alderton, Roberto Amato, Jeffrey Barrett, Sonia Goncalves, Ewan Harrison, David K. Jackson, Ian Johnston, Dominic Kwiatkowski, Cordelia Langford, John Sillitoe on behalf of the Wellcome Sanger Institute COVID-19 Surveillance Team |
| EPI_ISL_3248384, EPI_ISL_3248385 | Lighthouse Lab in Alderley Park | Wellcome Sanger Institute for the COVID-19 Genomics UK (COG-UK) Consortium | Jacquelyn Wynn, Mairead Hyland, The Lighthouse Lab in Alderley Park and Alex Alderton, Roberto Amato, Jeffrey Barrett, Sonia Goncalves, Ewan Harrison, David K. Jackson, Ian Johnston, Dominic Kwiatkowski, Cordelia Langford, John Sillitoe on behalf of the Wellcome Sanger Institute COVID-19 Surveillance Team |
| EPI_ISL_3248386, EPI_ISL_3248387 | Lighthouse Lab in Glasgow | Wellcome Sanger Institute for the COVID-19 Genomics UK (COG-UK) Consortium | Harper VanSteenhouse, Yumi Kasai, David Gray, Carol Clugston, Anna Dominiczak and Alex Alderton, Roberto Amato, Jeffrey Barrett, Sonia Goncalves, Ewan Harrison, David K. Jackson, Ian Johnston, Dominic Kwiatkowski, Cordelia Langford, John Sillitoe on behalf of the Wellcome Sanger Institute COVID-19 Surveillance Team |
| EPI_ISL_3248388, EPI_ISL_3248389 | Lighthouse Lab in Alderley Park | Wellcome Sanger Institute for the COVID-19 Genomics UK (COG-UK) Consortium | Jacquelyn Wynn, Mairead Hyland, The Lighthouse Lab in Alderley Park and Alex Alderton, Roberto Amato, Jeffrey Barrett, Sonia Goncalves, Ewan Harrison, David K. Jackson, Ian Johnston, Dominic Kwiatkowski, Cordelia Langford, John Sillitoe on behalf of the Wellcome Sanger Institute COVID-19 Surveillance Team |
| EPI_ISL_3248390, EPI_ISL_3248391 | Lighthouse Lab in Glasgow | Wellcome Sanger Institute for the COVID-19 Genomics UK (COG-UK) Consortium | Harper VanSteenhouse, Yumi Kasai, David Gray, Carol Clugston, Anna Dominiczak and Alex Alderton, Roberto Amato, Jeffrey Barrett, Sonia Goncalves, Ewan Harrison, David K. Jackson, Ian Johnston, Dominic Kwiatkowski, Cordelia Langford, John Sillitoe on behalf of the Wellcome Sanger Institute COVID-19 Surveillance Team |
| EPI_ISL_3248392 | Lighthouse Lab in Alderley Park | Wellcome Sanger Institute for the COVID-19 Genomics UK (COG-UK) Consortium | Jacquelyn Wynn, Mairead Hyland, The Lighthouse Lab in Alderley Park and Alex Alderton, Roberto Amato, Jeffrey Barrett, Sonia Goncalves, Ewan Harrison, David K. Jackson, Ian Johnston, Dominic Kwiatkowski, Cordelia Langford, John Sillitoe on behalf of the Wellcome Sanger Institute COVID-19 Surveillance Team |
| EPI_ISL_3248393, EPI_ISL_3248394, EPI_ISL_3248395, EPI_ISL_3248396, EPI_ISL_3248397 | Lighthouse Lab in Glasgow | Wellcome Sanger Institute for the COVID-19 Genomics UK (COG-UK) Consortium | Harper VanSteenhouse, Yumi Kasai, David Gray, Carol Clugston, Anna Dominiczak and Alex Alderton, Roberto Amato, Jeffrey Barrett, Sonia Goncalves, Ewan Harrison, David K. Jackson, Ian Johnston, Dominic Kwiatkowski, Cordelia Langford, John Sillitoe on behalf of the Wellcome Sanger Institute COVID-19 Surveillance Team |
| EPI_ISL_3248398 | Lighthouse Lab in Alderley Park | Wellcome Sanger Institute for the COVID-19 Genomics UK (COG-UK) Consortium | Jacquelyn Wynn, Mairead Hyland, The Lighthouse Lab in Alderley Park and Alex Alderton, Roberto Amato, Jeffrey Barrett, Sonia Goncalves, Ewan Harrison, David K. Jackson, Ian Johnston, Dominic Kwiatkowski, Cordelia Langford, John Sillitoe on behalf of the Wellcome Sanger Institute COVID-19 Surveillance Team |
| EPI_ISL_3248399 | Lighthouse Lab in Glasgow | Wellcome Sanger Institute for the COVID-19 Genomics UK (COG-UK) Consortium | Harper VanSteenhouse, Yumi Kasai, David Gray, Carol Clugston, Anna Dominiczak and Alex Alderton, Roberto Amato, Jeffrey Barrett, Sonia Goncalves, Ewan Harrison, David K. Jackson, Ian Johnston, Dominic Kwiatkowski, Cordelia Langford, John Sillitoe on behalf of the Wellcome Sanger Institute COVID-19 Surveillance Team |
| EPI_ISL_3248400, EPI_ISL_3248401 | Lighthouse Lab in Alderley Park | Wellcome Sanger Institute for the COVID-19 Genomics UK (COG-UK) Consortium | Jacquelyn Wynn, Mairead Hyland, The Lighthouse Lab in Alderley Park and Alex Alderton, Roberto Amato, Jeffrey Barrett, Sonia Goncalves, Ewan Harrison, David K. Jackson, Ian Johnston, Dominic Kwiatkowski, Cordelia Langford, John Sillitoe on behalf of the Wellcome Sanger Institute COVID-19 Surveillance Team |
| EPI_ISL_3248402, EPI_ISL_3248403, EPI_ISL_3248404, EPI_ISL_3248405, EPI_ISL_3248406, EPI_ISL_3248407, EPI_ISL_3248408, EPI_ISL_3248409 | Lighthouse Lab in Glasgow | Wellcome Sanger Institute for the COVID-19 Genomics UK (COG-UK) Consortium | Harper VanSteenhouse, Yumi Kasai, David Gray, Carol Clugston, Anna Dominiczak and Alex Alderton, Roberto Amato, Jeffrey Barrett, Sonia Goncalves, Ewan Harrison, David K. Jackson, Ian Johnston, Dominic Kwiatkowski, Cordelia Langford, John Sillitoe on behalf of the Wellcome Sanger Institute COVID-19 Surveillance Team |
| EPI_ISL_3248410 | Lighthouse Lab in Alderley Park | Wellcome Sanger Institute for the COVID-19 Genomics UK (COG-UK) Consortium | Jacquelyn Wynn, Mairead Hyland, The Lighthouse Lab in Alderley Park and Alex Alderton, Roberto Amato, Jeffrey Barrett, Sonia Goncalves, Ewan Harrison, David K. Jackson, Ian Johnston, Dominic Kwiatkowski, Cordelia Langford, John Sillitoe on behalf of the Wellcome Sanger Institute COVID-19 Surveillance Team |
| EPI_ISL_3248411, EPI_ISL_3248412 | Lighthouse Lab in Glasgow | Wellcome Sanger Institute for the COVID-19 Genomics UK (COG-UK) Consortium | Harper VanSteenhouse, Yumi Kasai, David Gray, Carol Clugston, Anna Dominiczak and Alex Alderton, Roberto Amato, Jeffrey Barrett, Sonia Goncalves, Ewan Harrison, David K. Jackson, Ian Johnston, Dominic Kwiatkowski, Cordelia Langford, John Sillitoe on behalf of the Wellcome Sanger Institute COVID-19 Surveillance Team |
| EPI_ISL_3248413 | Lighthouse Lab in Alderley Park | Wellcome Sanger Institute for the COVID-19 Genomics UK (COG-UK) Consortium | Jacquelyn Wynn, Mairead Hyland, The Lighthouse Lab in Alderley Park and Alex Alderton, Roberto Amato, Jeffrey Barrett, Sonia Goncalves, Ewan Harrison, David K. Jackson, Ian Johnston, Dominic Kwiatkowski, Cordelia Langford, John Sillitoe on behalf of the Wellcome Sanger Institute COVID-19 Surveillance Team |
| EPI_ISL_3248414 | Lighthouse Lab in Glasgow | Wellcome Sanger Institute for the COVID-19 Genomics UK (COG-UK) Consortium | Harper VanSteenhouse, Yumi Kasai, David Gray, Carol Clugston, Anna Dominiczak and Alex Alderton, Roberto Amato, Jeffrey Barrett, Sonia Goncalves, Ewan Harrison, David K. Jackson, Ian Johnston, Dominic Kwiatkowski, Cordelia Langford, John Sillitoe on behalf of the Wellcome Sanger Institute COVID-19 Surveillance Team |
| EPI_ISL_3248415, EPI_ISL_3248416, EPI_ISL_3248417, EPI_ISL_3248418 | Lighthouse Lab in Alderley Park | Wellcome Sanger Institute for the COVID-19 Genomics UK (COG-UK) Consortium | Jacquelyn Wynn, Mairead Hyland, The Lighthouse Lab in Alderley Park and Alex Alderton, Roberto Amato, Jeffrey Barrett, Sonia Goncalves, Ewan Harrison, David K. Jackson, Ian Johnston, Dominic Kwiatkowski, Cordelia Langford, John Sillitoe on behalf of the Wellcome Sanger Institute COVID-19 Surveillance Team |
| EPI_ISL_3248419, EPI_ISL_3248420 | Lighthouse Lab in Glasgow | Wellcome Sanger Institute for the COVID-19 Genomics UK (COG-UK) Consortium | Harper VanSteenhouse, Yumi Kasai, David Gray, Carol Clugston, Anna Dominiczak and Alex Alderton, Roberto Amato, Jeffrey Barrett, Sonia Goncalves, Ewan Harrison, David K. Jackson, Ian Johnston, Dominic Kwiatkowski, Cordelia Langford, John Sillitoe on behalf of the Wellcome Sanger Institute COVID-19 Surveillance Team |
| EPI_ISL_3248421 | Lighthouse Lab in Alderley Park | Wellcome Sanger Institute for the COVID-19 Genomics UK (COG-UK) Consortium | Jacquelyn Wynn, Mairead Hyland, The Lighthouse Lab in Alderley Park and Alex Alderton, Roberto Amato, Jeffrey Barrett, Sonia Goncalves, Ewan Harrison, David K. Jackson, Ian Johnston, Dominic Kwiatkowski, Cordelia Langford, John Sillitoe on behalf of the Wellcome Sanger Institute COVID-19 Surveillance Team |
| EPI_ISL_3248422 | Lighthouse Lab in Glasgow | Wellcome Sanger Institute for the COVID-19 Genomics UK (COG-UK) Consortium | Harper VanSteenhouse, Yumi Kasai, David Gray, Carol Clugston, Anna Dominiczak and Alex Alderton, Roberto Amato, Jeffrey Barrett, Sonia Goncalves, Ewan Harrison, David K. Jackson, Ian Johnston, Dominic Kwiatkowski, Cordelia Langford, John Sillitoe on behalf of the Wellcome Sanger Institute COVID-19 Surveillance Team |
| EPI_ISL_3248423, | Lighthouse Lab in Alderley Park | Wellcome Sanger Institute for the COVID-19 Genomics UK (COG-UK) Consortium | Jacquelyn Wynn, Mairead Hyland, The Lighthouse Lab in Alderley Park and Alex Alderton, Roberto Amato, Jeffrey Barrett, Sonia Goncalves, Ewan Harrison, David K. Jackson, Ian Johnston, Dominic Kwiatkowski, Cordelia Langford, John Sillitoe on |

| | | | |
|---|---|---|---|
| EPI_ISL_3248424 | | Consortium | behalf of the Wellcome Sanger Institute COVID-19 Surveillance Team |
| EPI_ISL_3248425, EPI_ISL_3248426 | Lighthouse Lab in Glasgow | Wellcome Sanger Institute for the COVID-19 Genomics UK (COG-UK) Consortium | Harper VanSteenhouse, Yumi Kasai, David Gray, Carol Clugston, Anna Dominiczak and Alex Alderton, Roberto Amato, Jeffrey Barrett, Sonia Goncalves, Ewan Harrison, David K. Jackson, Ian Johnston, Dominic Kwiatkowski, Cordelia Langford, John Sillitoe on behalf of the Wellcome Sanger Institute COVID-19 Surveillance Team |
| EPI_ISL_3248427 | Lighthouse Lab in Alderley Park | Wellcome Sanger Institute for the COVID-19 Genomics UK (COG-UK) Consortium | Jacquelyn Wynn, Mairead Hyland, The Lighthouse Lab in Alderley Park and Alex Alderton, Roberto Amato, Jeffrey Barrett, Sonia Goncalves, Ewan Harrison, David K. Jackson, Ian Johnston, Dominic Kwiatkowski, Cordelia Langford, John Sillitoe on behalf of the Wellcome Sanger Institute COVID-19 Surveillance Team |
| EPI_ISL_3248428, EPI_ISL_3248429 | Lighthouse Lab in Glasgow | Wellcome Sanger Institute for the COVID-19 Genomics UK (COG-UK) Consortium | Harper VanSteenhouse, Yumi Kasai, David Gray, Carol Clugston, Anna Dominiczak and Alex Alderton, Roberto Amato, Jeffrey Barrett, Sonia Goncalves, Ewan Harrison, David K. Jackson, Ian Johnston, Dominic Kwiatkowski, Cordelia Langford, John Sillitoe on behalf of the Wellcome Sanger Institute COVID-19 Surveillance Team |
| EPI_ISL_3248430, EPI_ISL_3248431 | Lighthouse Lab in Alderley Park | Wellcome Sanger Institute for the COVID-19 Genomics UK (COG-UK) Consortium | Jacquelyn Wynn, Mairead Hyland, The Lighthouse Lab in Alderley Park and Alex Alderton, Roberto Amato, Jeffrey Barrett, Sonia Goncalves, Ewan Harrison, David K. Jackson, Ian Johnston, Dominic Kwiatkowski, Cordelia Langford, John Sillitoe on behalf of the Wellcome Sanger Institute COVID-19 Surveillance Team |
| EPI_ISL_3248432 | Lighthouse Lab in Glasgow | Wellcome Sanger Institute for the COVID-19 Genomics UK (COG-UK) Consortium | Harper VanSteenhouse, Yumi Kasai, David Gray, Carol Clugston, Anna Dominiczak and Alex Alderton, Roberto Amato, Jeffrey Barrett, Sonia Goncalves, Ewan Harrison, David K. Jackson, Ian Johnston, Dominic Kwiatkowski, Cordelia Langford, John Sillitoe on behalf of the Wellcome Sanger Institute COVID-19 Surveillance Team |
| EPI_ISL_3248433 | Lighthouse Lab in Alderley Park | Wellcome Sanger Institute for the COVID-19 Genomics UK (COG-UK) Consortium | Jacquelyn Wynn, Mairead Hyland, The Lighthouse Lab in Alderley Park and Alex Alderton, Roberto Amato, Jeffrey Barrett, Sonia Goncalves, Ewan Harrison, David K. Jackson, Ian Johnston, Dominic Kwiatkowski, Cordelia Langford, John Sillitoe on behalf of the Wellcome Sanger Institute COVID-19 Surveillance Team |
| EPI_ISL_3248434, EPI_ISL_3248435, EPI_ISL_3248436 | Lighthouse Lab in Glasgow | Wellcome Sanger Institute for the COVID-19 Genomics UK (COG-UK) Consortium | Harper VanSteenhouse, Yumi Kasai, David Gray, Carol Clugston, Anna Dominiczak and Alex Alderton, Roberto Amato, Jeffrey Barrett, Sonia Goncalves, Ewan Harrison, David K. Jackson, Ian Johnston, Dominic Kwiatkowski, Cordelia Langford, John Sillitoe on behalf of the Wellcome Sanger Institute COVID-19 Surveillance Team |
| EPI_ISL_3248437 | Lighthouse Lab in Alderley Park | Wellcome Sanger Institute for the COVID-19 Genomics UK (COG-UK) Consortium | Jacquelyn Wynn, Mairead Hyland, The Lighthouse Lab in Alderley Park and Alex Alderton, Roberto Amato, Jeffrey Barrett, Sonia Goncalves, Ewan Harrison, David K. Jackson, Ian Johnston, Dominic Kwiatkowski, Cordelia Langford, John Sillitoe on behalf of the Wellcome Sanger Institute COVID-19 Surveillance Team |
| EPI_ISL_3248438, EPI_ISL_3248439, EPI_ISL_3248440, EPI_ISL_3248441, EPI_ISL_3248442, EPI_ISL_3248443, EPI_ISL_3248444, EPI_ISL_3248445 | Lighthouse Lab in Glasgow | Wellcome Sanger Institute for the COVID-19 Genomics UK (COG-UK) Consortium | Harper VanSteenhouse, Yumi Kasai, David Gray, Carol Clugston, Anna Dominiczak and Alex Alderton, Roberto Amato, Jeffrey Barrett, Sonia Goncalves, Ewan Harrison, David K. Jackson, Ian Johnston, Dominic Kwiatkowski, Cordelia Langford, John Sillitoe on behalf of the Wellcome Sanger Institute COVID-19 Surveillance Team |
| EPI_ISL_3248446 | Lighthouse Lab in Alderley Park | Wellcome Sanger Institute for the COVID-19 Genomics UK (COG-UK) Consortium | Jacquelyn Wynn, Mairead Hyland, The Lighthouse Lab in Alderley Park and Alex Alderton, Roberto Amato, Jeffrey Barrett, Sonia Goncalves, Ewan Harrison, David K. Jackson, Ian Johnston, Dominic Kwiatkowski, Cordelia Langford, John Sillitoe on behalf of the Wellcome Sanger Institute COVID-19 Surveillance Team |
| EPI_ISL_3248447, EPI_ISL_3248448, EPI_ISL_3248449, EPI_ISL_3248450, EPI_ISL_3248451, EPI_ISL_3248452, EPI_ISL_3248453, EPI_ISL_3248454 | Lighthouse Lab in Glasgow | Wellcome Sanger Institute for the COVID-19 Genomics UK (COG-UK) Consortium | Harper VanSteenhouse, Yumi Kasai, David Gray, Carol Clugston, Anna Dominiczak and Alex Alderton, Roberto Amato, Jeffrey Barrett, Sonia Goncalves, Ewan Harrison, David K. Jackson, Ian Johnston, Dominic Kwiatkowski, Cordelia Langford, John Sillitoe on behalf of the Wellcome Sanger Institute COVID-19 Surveillance Team |
| EPI_ISL_3248455, EPI_ISL_3248456, EPI_ISL_3248457 | Lighthouse Lab in Alderley Park | Wellcome Sanger Institute for the COVID-19 Genomics UK (COG-UK) Consortium | Jacquelyn Wynn, Mairead Hyland, The Lighthouse Lab in Alderley Park and Alex Alderton, Roberto Amato, Jeffrey Barrett, Sonia Goncalves, Ewan Harrison, David K. Jackson, Ian Johnston, Dominic Kwiatkowski, Cordelia Langford, John Sillitoe on behalf of the Wellcome Sanger Institute COVID-19 Surveillance Team |
| EPI_ISL_3248458 | Lighthouse Lab in Glasgow | Wellcome Sanger Institute for the COVID-19 Genomics UK (COG-UK) Consortium | Harper VanSteenhouse, Yumi Kasai, David Gray, Carol Clugston, Anna Dominiczak and Alex Alderton, Roberto Amato, Jeffrey Barrett, Sonia Goncalves, Ewan Harrison, David K. Jackson, Ian Johnston, Dominic Kwiatkowski, Cordelia Langford, John Sillitoe on behalf of the Wellcome Sanger Institute COVID-19 Surveillance Team |
| EPI_ISL_3248459 | Lighthouse Lab in Alderley Park | Wellcome Sanger Institute for the COVID-19 Genomics UK (COG-UK) Consortium | Jacquelyn Wynn, Mairead Hyland, The Lighthouse Lab in Alderley Park and Alex Alderton, Roberto Amato, Jeffrey Barrett, Sonia Goncalves, Ewan Harrison, David K. Jackson, Ian Johnston, Dominic Kwiatkowski, Cordelia Langford, John Sillitoe on behalf of the Wellcome Sanger Institute COVID-19 Surveillance Team |
| EPI_ISL_3248460 | Lighthouse Lab in Glasgow | Wellcome Sanger Institute for the COVID-19 Genomics UK (COG-UK) Consortium | Harper VanSteenhouse, Yumi Kasai, David Gray, Carol Clugston, Anna Dominiczak and Alex Alderton, Roberto Amato, Jeffrey Barrett, Sonia Goncalves, Ewan Harrison, David K. Jackson, Ian Johnston, Dominic Kwiatkowski, Cordelia Langford, John Sillitoe on behalf of the Wellcome Sanger Institute COVID-19 Surveillance Team |
| EPI_ISL_3248461, EPI_ISL_3248462 | Lighthouse Lab in Alderley Park | Wellcome Sanger Institute for the COVID-19 Genomics UK (COG-UK) Consortium | Jacquelyn Wynn, Mairead Hyland, The Lighthouse Lab in Alderley Park and Alex Alderton, Roberto Amato, Jeffrey Barrett, Sonia Goncalves, Ewan Harrison, David K. Jackson, Ian Johnston, Dominic Kwiatkowski, Cordelia Langford, John Sillitoe on behalf of the Wellcome Sanger Institute COVID-19 Surveillance Team |
| EPI_ISL_3248463 | Lighthouse Lab in Glasgow | Wellcome Sanger Institute for the COVID-19 Genomics UK (COG-UK) Consortium | Harper VanSteenhouse, Yumi Kasai, David Gray, Carol Clugston, Anna Dominiczak and Alex Alderton, Roberto Amato, Jeffrey Barrett, Sonia Goncalves, Ewan Harrison, David K. Jackson, Ian Johnston, Dominic Kwiatkowski, Cordelia Langford, John Sillitoe on behalf of the Wellcome Sanger Institute COVID-19 Surveillance Team |
| EPI_ISL_3248464, EPI_ISL_3248465 | Lighthouse Lab in Alderley Park | Wellcome Sanger Institute for the COVID-19 Genomics UK (COG-UK) Consortium | Jacquelyn Wynn, Mairead Hyland, The Lighthouse Lab in Alderley Park and Alex Alderton, Roberto Amato, Jeffrey Barrett, Sonia Goncalves, Ewan Harrison, David K. Jackson, Ian Johnston, Dominic Kwiatkowski, Cordelia Langford, John Sillitoe on behalf of the Wellcome Sanger Institute COVID-19 Surveillance Team |
| EPI_ISL_3248466, EPI_ISL_3248467, EPI_ISL_3248468, EPI_ISL_3248469, EPI_ISL_3248470 | Lighthouse Lab in Glasgow | Wellcome Sanger Institute for the COVID-19 Genomics UK (COG-UK) Consortium | Harper VanSteenhouse, Yumi Kasai, David Gray, Carol Clugston, Anna Dominiczak and Alex Alderton, Roberto Amato, Jeffrey Barrett, Sonia Goncalves, Ewan Harrison, David K. Jackson, Ian Johnston, Dominic Kwiatkowski, Cordelia Langford, John Sillitoe on behalf of the Wellcome Sanger Institute COVID-19 Surveillance Team |
| EPI_ISL_3248471 | Lighthouse Lab in Alderley Park | Wellcome Sanger Institute for the COVID-19 Genomics UK (COG-UK) Consortium | Jacquelyn Wynn, Mairead Hyland, The Lighthouse Lab in Alderley Park and Alex Alderton, Roberto Amato, Jeffrey Barrett, Sonia Goncalves, Ewan Harrison, David K. Jackson, Ian Johnston, Dominic Kwiatkowski, Cordelia Langford, John Sillitoe on behalf of the Wellcome Sanger Institute COVID-19 Surveillance Team |
| EPI_ISL_3248472, EPI_ISL_3248473 | Lighthouse Lab in Glasgow | Wellcome Sanger Institute for the COVID-19 Genomics UK (COG-UK) Consortium | Harper VanSteenhouse, Yumi Kasai, David Gray, Carol Clugston, Anna Dominiczak and Alex Alderton, Roberto Amato, Jeffrey Barrett, Sonia Goncalves, Ewan Harrison, David K. Jackson, Ian Johnston, Dominic Kwiatkowski, Cordelia Langford, John Sillitoe on behalf of the Wellcome Sanger Institute COVID-19 Surveillance Team |
| EPI_ISL_3248474 | Lighthouse Lab in Alderley Park | Wellcome Sanger Institute for the COVID-19 Genomics UK (COG-UK) Consortium | Jacquelyn Wynn, Mairead Hyland, The Lighthouse Lab in Alderley Park and Alex Alderton, Roberto Amato, Jeffrey Barrett, Sonia Goncalves, Ewan Harrison, David K. Jackson, Ian Johnston, Dominic Kwiatkowski, Cordelia Langford, John Sillitoe on behalf of the Wellcome Sanger Institute COVID-19 Surveillance Team |
| EPI_ISL_3248475, EPI_ISL_3248476, EPI_ISL_3248477, EPI_ISL_3248478 | Lighthouse Lab in Glasgow | Wellcome Sanger Institute for the COVID-19 Genomics UK (COG-UK) Consortium | Harper VanSteenhouse, Yumi Kasai, David Gray, Carol Clugston, Anna Dominiczak and Alex Alderton, Roberto Amato, Jeffrey Barrett, Sonia Goncalves, Ewan Harrison, David K. Jackson, Ian Johnston, Dominic Kwiatkowski, Cordelia Langford, John Sillitoe on behalf of the Wellcome Sanger Institute COVID-19 Surveillance Team |
| EPI_ISL_3248479 | Lighthouse Lab in Alderley Park | Wellcome Sanger Institute for the COVID-19 Genomics UK (COG-UK) Consortium | Jacquelyn Wynn, Mairead Hyland, The Lighthouse Lab in Alderley Park and Alex Alderton, Roberto Amato, Jeffrey Barrett, Sonia Goncalves, Ewan Harrison, David K. Jackson, Ian Johnston, Dominic Kwiatkowski, Cordelia Langford, John Sillitoe on behalf of the Wellcome Sanger Institute COVID-19 Surveillance Team |
| EPI_ISL_3248480 | Lighthouse Lab in Glasgow | Wellcome Sanger Institute for the COVID-19 Genomics UK (COG-UK) Consortium | Harper VanSteenhouse, Yumi Kasai, David Gray, Carol Clugston, Anna Dominiczak and Alex Alderton, Roberto Amato, Jeffrey Barrett, Sonia Goncalves, Ewan Harrison, David K. Jackson, Ian Johnston, Dominic Kwiatkowski, Cordelia Langford, John Sillitoe on behalf of the Wellcome Sanger Institute COVID-19 Surveillance Team |
| EPI_ISL_3248481, EPI_ISL_3248482 | Lighthouse Lab in Alderley Park | Wellcome Sanger Institute for the COVID-19 Genomics UK (COG-UK) Consortium | Jacquelyn Wynn, Mairead Hyland, The Lighthouse Lab in Alderley Park and Alex Alderton, Roberto Amato, Jeffrey Barrett, Sonia Goncalves, Ewan Harrison, David K. Jackson, Ian Johnston, Dominic Kwiatkowski, Cordelia Langford, John Sillitoe on behalf of the Wellcome Sanger Institute COVID-19 Surveillance Team |
| EPI_ISL_3248483, EPI_ISL_3248484, EPI_ISL_3248485 | Lighthouse Lab in Glasgow | Wellcome Sanger Institute for the COVID-19 Genomics UK (COG-UK) Consortium | Harper VanSteenhouse, Yumi Kasai, David Gray, Carol Clugston, Anna Dominiczak and Alex Alderton, Roberto Amato, Jeffrey Barrett, Sonia Goncalves, Ewan Harrison, David K. Jackson, Ian Johnston, Dominic Kwiatkowski, Cordelia Langford, John Sillitoe on behalf of the Wellcome Sanger Institute COVID-19 Surveillance Team |
| EPI_ISL_3248486, EPI_ISL_3248487 | Lighthouse Lab in Alderley Park | Wellcome Sanger Institute for the COVID-19 Genomics UK (COG-UK) Consortium | Jacquelyn Wynn, Mairead Hyland, The Lighthouse Lab in Alderley Park and Alex Alderton, Roberto Amato, Jeffrey Barrett, Sonia Goncalves, Ewan Harrison, David K. Jackson, Ian Johnston, Dominic Kwiatkowski, Cordelia Langford, John Sillitoe on behalf of the Wellcome Sanger Institute COVID-19 Surveillance Team |
| EPI_ISL_3248488, EPI_ISL_3248489, EPI_ISL_3248490, EPI_ISL_3248491 | Lighthouse Lab in Glasgow | Wellcome Sanger Institute for the COVID-19 Genomics UK (COG-UK) Consortium | Harper VanSteenhouse, Yumi Kasai, David Gray, Carol Clugston, Anna Dominiczak and Alex Alderton, Roberto Amato, Jeffrey Barrett, Sonia Goncalves, Ewan Harrison, David K. Jackson, Ian Johnston, Dominic Kwiatkowski, Cordelia Langford, John Sillitoe on behalf of the Wellcome Sanger Institute COVID-19 Surveillance Team |
| EPI_ISL_3248492 | Lighthouse Lab in Alderley Park | Wellcome Sanger Institute for the COVID-19 Genomics UK (COG-UK) Consortium | Jacquelyn Wynn, Mairead Hyland, The Lighthouse Lab in Alderley Park and Alex Alderton, Roberto Amato, Jeffrey Barrett, Sonia Goncalves, Ewan Harrison, David K. Jackson, Ian Johnston, Dominic Kwiatkowski, Cordelia Langford, John Sillitoe on behalf of the Wellcome Sanger Institute COVID-19 Surveillance Team |
| EPI_ISL_3248493 | Lighthouse Lab in Glasgow | Wellcome Sanger Institute for the COVID-19 Genomics UK (COG-UK) Consortium | Harper VanSteenhouse, Yumi Kasai, David Gray, Carol Clugston, Anna Dominiczak and Alex Alderton, Roberto Amato, Jeffrey Barrett, Sonia Goncalves, Ewan Harrison, David K. Jackson, Ian Johnston, Dominic Kwiatkowski, Cordelia Langford, John Sillitoe on behalf of the Wellcome Sanger Institute COVID-19 Surveillance Team |
| EPI_ISL_3248494 | Lighthouse Lab in Alderley Park | Wellcome Sanger Institute for the COVID-19 Genomics UK (COG-UK) Consortium | Jacquelyn Wynn, Mairead Hyland, The Lighthouse Lab in Alderley Park and Alex Alderton, Roberto Amato, Jeffrey Barrett, Sonia Goncalves, Ewan Harrison, David K. Jackson, Ian Johnston, Dominic Kwiatkowski, Cordelia Langford, John Sillitoe on behalf of the Wellcome Sanger Institute COVID-19 Surveillance Team |
| EPI_ISL_3248495 | Lighthouse Lab in Glasgow | Wellcome Sanger Institute for the COVID-19 Genomics UK (COG-UK) Consortium | Harper VanSteenhouse, Yumi Kasai, David Gray, Carol Clugston, Anna Dominiczak and Alex Alderton, Roberto Amato, Jeffrey Barrett, Sonia Goncalves, Ewan Harrison, David K. Jackson, Ian Johnston, Dominic Kwiatkowski, Cordelia Langford, John Sillitoe on behalf of the Wellcome Sanger Institute COVID-19 Surveillance Team |
| EPI_ISL_3248496 | Lighthouse Lab in Alderley Park | Wellcome Sanger Institute for the COVID-19 Genomics UK (COG-UK) Consortium | Jacquelyn Wynn, Mairead Hyland, The Lighthouse Lab in Alderley Park and Alex Alderton, Roberto Amato, Jeffrey Barrett, Sonia Goncalves, Ewan Harrison, David K. Jackson, Ian Johnston, Dominic Kwiatkowski, Cordelia Langford, John Sillitoe on behalf of the Wellcome Sanger Institute COVID-19 Surveillance Team |
| EPI_ISL_3248497, EPI_ISL_3248498 | Lighthouse Lab in Glasgow | Wellcome Sanger Institute for the COVID-19 Genomics UK (COG-UK) Consortium | Harper VanSteenhouse, Yumi Kasai, David Gray, Carol Clugston, Anna Dominiczak and Alex Alderton, Roberto Amato, Jeffrey Barrett, Sonia Goncalves, Ewan Harrison, David K. Jackson, Ian Johnston, Dominic Kwiatkowski, Cordelia Langford, John Sillitoe on behalf of the Wellcome Sanger Institute COVID-19 Surveillance Team |
| EPI_ISL_3248499, | Lighthouse Lab in Alderley Park | Wellcome Sanger Institute for the COVID-19 Genomics UK (COG-UK) | Jacquelyn Wynn, Mairead Hyland, The Lighthouse Lab in Alderley Park and Alex Alderton, Roberto Amato, Jeffrey Barrett, Sonia Goncalves, Ewan Harrison, David K. Jackson, Ian Johnston, Dominic Kwiatkowski, Cordelia Langford, John Sillitoe on |

| | | | |
|---|---|---|---|
| EPI_ISL_3248500 | | Consortium | behalf of the Wellcome Sanger Institute COVID-19 Surveillance Team |
| EPI_ISL_3248501 | Lighthouse Lab in Glasgow | Wellcome Sanger Institute for the COVID-19 Genomics UK (COG-UK) Consortium | Harper VanSteenhouse, Yumi Kasai, David Gray, Carol Clugston, Anna Dominiczak and Alex Alderton, Roberto Amato, Jeffrey Barrett, Sonia Goncalves, Ewan Harrison, David K. Jackson, Ian Johnston, Dominic Kwiatkowski, Cordelia Langford, John Sillitoe on behalf of the Wellcome Sanger Institute COVID-19 Surveillance Team |
| EPI_ISL_3248502 | Lighthouse Lab in Alderley Park | Wellcome Sanger Institute for the COVID-19 Genomics UK (COG-UK) Consortium | Jacquelyn Wynn, Mairead Hyland, The Lighthouse Lab in Alderley Park and Alex Alderton, Roberto Amato, Jeffrey Barrett, Sonia Goncalves, Ewan Harrison, David K. Jackson, Ian Johnston, Dominic Kwiatkowski, Cordelia Langford, John Sillitoe on behalf of the Wellcome Sanger Institute COVID-19 Surveillance Team |
| EPI_ISL_3248503, EPI_ISL_3248504 | Lighthouse Lab in Glasgow | Wellcome Sanger Institute for the COVID-19 Genomics UK (COG-UK) Consortium | Harper VanSteenhouse, Yumi Kasai, David Gray, Carol Clugston, Anna Dominiczak and Alex Alderton, Roberto Amato, Jeffrey Barrett, Sonia Goncalves, Ewan Harrison, David K. Jackson, Ian Johnston, Dominic Kwiatkowski, Cordelia Langford, John Sillitoe on behalf of the Wellcome Sanger Institute COVID-19 Surveillance Team |
| EPI_ISL_3248505 | Lighthouse Lab in Alderley Park | Wellcome Sanger Institute for the COVID-19 Genomics UK (COG-UK) Consortium | Jacquelyn Wynn, Mairead Hyland, The Lighthouse Lab in Alderley Park and Alex Alderton, Roberto Amato, Jeffrey Barrett, Sonia Goncalves, Ewan Harrison, David K. Jackson, Ian Johnston, Dominic Kwiatkowski, Cordelia Langford, John Sillitoe on behalf of the Wellcome Sanger Institute COVID-19 Surveillance Team |
| EPI_ISL_3248506 | Lighthouse Lab in Glasgow | Wellcome Sanger Institute for the COVID-19 Genomics UK (COG-UK) Consortium | Harper VanSteenhouse, Yumi Kasai, David Gray, Carol Clugston, Anna Dominiczak and Alex Alderton, Roberto Amato, Jeffrey Barrett, Sonia Goncalves, Ewan Harrison, David K. Jackson, Ian Johnston, Dominic Kwiatkowski, Cordelia Langford, John Sillitoe on behalf of the Wellcome Sanger Institute COVID-19 Surveillance Team |
| EPI_ISL_3248507 | Lighthouse Lab in Alderley Park | Wellcome Sanger Institute for the COVID-19 Genomics UK (COG-UK) Consortium | Jacquelyn Wynn, Mairead Hyland, The Lighthouse Lab in Alderley Park and Alex Alderton, Roberto Amato, Jeffrey Barrett, Sonia Goncalves, Ewan Harrison, David K. Jackson, Ian Johnston, Dominic Kwiatkowski, Cordelia Langford, John Sillitoe on behalf of the Wellcome Sanger Institute COVID-19 Surveillance Team |
| EPI_ISL_3248508 | Lighthouse Lab in Glasgow | Wellcome Sanger Institute for the COVID-19 Genomics UK (COG-UK) Consortium | Harper VanSteenhouse, Yumi Kasai, David Gray, Carol Clugston, Anna Dominiczak and Alex Alderton, Roberto Amato, Jeffrey Barrett, Sonia Goncalves, Ewan Harrison, David K. Jackson, Ian Johnston, Dominic Kwiatkowski, Cordelia Langford, John Sillitoe on behalf of the Wellcome Sanger Institute COVID-19 Surveillance Team |
| EPI_ISL_3248509 | Lighthouse Lab in Alderley Park | Wellcome Sanger Institute for the COVID-19 Genomics UK (COG-UK) Consortium | Jacquelyn Wynn, Mairead Hyland, The Lighthouse Lab in Alderley Park and Alex Alderton, Roberto Amato, Jeffrey Barrett, Sonia Goncalves, Ewan Harrison, David K. Jackson, Ian Johnston, Dominic Kwiatkowski, Cordelia Langford, John Sillitoe on behalf of the Wellcome Sanger Institute COVID-19 Surveillance Team |
| EPI_ISL_3248510, EPI_ISL_3248511, EPI_ISL_3248512 | Lighthouse Lab in Glasgow | Wellcome Sanger Institute for the COVID-19 Genomics UK (COG-UK) Consortium | Harper VanSteenhouse, Yumi Kasai, David Gray, Carol Clugston, Anna Dominiczak and Alex Alderton, Roberto Amato, Jeffrey Barrett, Sonia Goncalves, Ewan Harrison, David K. Jackson, Ian Johnston, Dominic Kwiatkowski, Cordelia Langford, John Sillitoe on behalf of the Wellcome Sanger Institute COVID-19 Surveillance Team |
| EPI_ISL_3248513 | Lighthouse Lab in Alderley Park | Wellcome Sanger Institute for the COVID-19 Genomics UK (COG-UK) Consortium | Jacquelyn Wynn, Mairead Hyland, The Lighthouse Lab in Alderley Park and Alex Alderton, Roberto Amato, Jeffrey Barrett, Sonia Goncalves, Ewan Harrison, David K. Jackson, Ian Johnston, Dominic Kwiatkowski, Cordelia Langford, John Sillitoe on behalf of the Wellcome Sanger Institute COVID-19 Surveillance Team |
| EPI_ISL_3248514, EPI_ISL_3248515, EPI_ISL_3248516, EPI_ISL_3248517, EPI_ISL_3248518, EPI_ISL_3248519, EPI_ISL_3248520, EPI_ISL_3248521, EPI_ISL_3248522, EPI_ISL_3248523 | Lighthouse Lab in Glasgow | Wellcome Sanger Institute for the COVID-19 Genomics UK (COG-UK) Consortium | Harper VanSteenhouse, Yumi Kasai, David Gray, Carol Clugston, Anna Dominiczak and Alex Alderton, Roberto Amato, Jeffrey Barrett, Sonia Goncalves, Ewan Harrison, David K. Jackson, Ian Johnston, Dominic Kwiatkowski, Cordelia Langford, John Sillitoe on behalf of the Wellcome Sanger Institute COVID-19 Surveillance Team |
| EPI_ISL_3248524 | Lighthouse Lab in Alderley Park | Wellcome Sanger Institute for the COVID-19 Genomics UK (COG-UK) Consortium | Jacquelyn Wynn, Mairead Hyland, The Lighthouse Lab in Alderley Park and Alex Alderton, Roberto Amato, Jeffrey Barrett, Sonia Goncalves, Ewan Harrison, David K. Jackson, Ian Johnston, Dominic Kwiatkowski, Cordelia Langford, John Sillitoe on behalf of the Wellcome Sanger Institute COVID-19 Surveillance Team |
| EPI_ISL_3248525 | Lighthouse Lab in Glasgow | Wellcome Sanger Institute for the COVID-19 Genomics UK (COG-UK) Consortium | Harper VanSteenhouse, Yumi Kasai, David Gray, Carol Clugston, Anna Dominiczak and Alex Alderton, Roberto Amato, Jeffrey Barrett, Sonia Goncalves, Ewan Harrison, David K. Jackson, Ian Johnston, Dominic Kwiatkowski, Cordelia Langford, John Sillitoe on behalf of the Wellcome Sanger Institute COVID-19 Surveillance Team |
| EPI_ISL_3248526, EPI_ISL_3248527, EPI_ISL_3248528 | Lighthouse Lab in Alderley Park | Wellcome Sanger Institute for the COVID-19 Genomics UK (COG-UK) Consortium | Jacquelyn Wynn, Mairead Hyland, The Lighthouse Lab in Alderley Park and Alex Alderton, Roberto Amato, Jeffrey Barrett, Sonia Goncalves, Ewan Harrison, David K. Jackson, Ian Johnston, Dominic Kwiatkowski, Cordelia Langford, John Sillitoe on behalf of the Wellcome Sanger Institute COVID-19 Surveillance Team |
| EPI_ISL_3248529 | Lighthouse Lab in Glasgow | Wellcome Sanger Institute for the COVID-19 Genomics UK (COG-UK) Consortium | Harper VanSteenhouse, Yumi Kasai, David Gray, Carol Clugston, Anna Dominiczak and Alex Alderton, Roberto Amato, Jeffrey Barrett, Sonia Goncalves, Ewan Harrison, David K. Jackson, Ian Johnston, Dominic Kwiatkowski, Cordelia Langford, John Sillitoe on behalf of the Wellcome Sanger Institute COVID-19 Surveillance Team |
| EPI_ISL_3248530 | Lighthouse Lab in Alderley Park | Wellcome Sanger Institute for the COVID-19 Genomics UK (COG-UK) Consortium | Jacquelyn Wynn, Mairead Hyland, The Lighthouse Lab in Alderley Park and Alex Alderton, Roberto Amato, Jeffrey Barrett, Sonia Goncalves, Ewan Harrison, David K. Jackson, Ian Johnston, Dominic Kwiatkowski, Cordelia Langford, John Sillitoe on behalf of the Wellcome Sanger Institute COVID-19 Surveillance Team |
| EPI_ISL_3248531 | Lighthouse Lab in Glasgow | Wellcome Sanger Institute for the COVID-19 Genomics UK (COG-UK) Consortium | Harper VanSteenhouse, Yumi Kasai, David Gray, Carol Clugston, Anna Dominiczak and Alex Alderton, Roberto Amato, Jeffrey Barrett, Sonia Goncalves, Ewan Harrison, David K. Jackson, Ian Johnston, Dominic Kwiatkowski, Cordelia Langford, John Sillitoe on behalf of the Wellcome Sanger Institute COVID-19 Surveillance Team |
| EPI_ISL_3248532 | Lighthouse Lab in Alderley Park | Wellcome Sanger Institute for the COVID-19 Genomics UK (COG-UK) Consortium | Jacquelyn Wynn, Mairead Hyland, The Lighthouse Lab in Alderley Park and Alex Alderton, Roberto Amato, Jeffrey Barrett, Sonia Goncalves, Ewan Harrison, David K. Jackson, Ian Johnston, Dominic Kwiatkowski, Cordelia Langford, John Sillitoe on behalf of the Wellcome Sanger Institute COVID-19 Surveillance Team |
| EPI_ISL_3248533, EPI_ISL_3248534 | Lighthouse Lab in Glasgow | Wellcome Sanger Institute for the COVID-19 Genomics UK (COG-UK) Consortium | Harper VanSteenhouse, Yumi Kasai, David Gray, Carol Clugston, Anna Dominiczak and Alex Alderton, Roberto Amato, Jeffrey Barrett, Sonia Goncalves, Ewan Harrison, David K. Jackson, Ian Johnston, Dominic Kwiatkowski, Cordelia Langford, John Sillitoe on behalf of the Wellcome Sanger Institute COVID-19 Surveillance Team |
| EPI_ISL_3248535 | Lighthouse Lab in Alderley Park | Wellcome Sanger Institute for the COVID-19 Genomics UK (COG-UK) Consortium | Jacquelyn Wynn, Mairead Hyland, The Lighthouse Lab in Alderley Park and Alex Alderton, Roberto Amato, Jeffrey Barrett, Sonia Goncalves, Ewan Harrison, David K. Jackson, Ian Johnston, Dominic Kwiatkowski, Cordelia Langford, John Sillitoe on behalf of the Wellcome Sanger Institute COVID-19 Surveillance Team |
| EPI_ISL_3248536, EPI_ISL_3248537 | Lighthouse Lab in Glasgow | Wellcome Sanger Institute for the COVID-19 Genomics UK (COG-UK) Consortium | Harper VanSteenhouse, Yumi Kasai, David Gray, Carol Clugston, Anna Dominiczak and Alex Alderton, Roberto Amato, Jeffrey Barrett, Sonia Goncalves, Ewan Harrison, David K. Jackson, Ian Johnston, Dominic Kwiatkowski, Cordelia Langford, John Sillitoe on behalf of the Wellcome Sanger Institute COVID-19 Surveillance Team |
| EPI_ISL_3248538, EPI_ISL_3248539, EPI_ISL_3248540 | Lighthouse Lab in Alderley Park | Wellcome Sanger Institute for the COVID-19 Genomics UK (COG-UK) Consortium | Jacquelyn Wynn, Mairead Hyland, The Lighthouse Lab in Alderley Park and Alex Alderton, Roberto Amato, Jeffrey Barrett, Sonia Goncalves, Ewan Harrison, David K. Jackson, Ian Johnston, Dominic Kwiatkowski, Cordelia Langford, John Sillitoe on behalf of the Wellcome Sanger Institute COVID-19 Surveillance Team |
| EPI_ISL_3248541, EPI_ISL_3248542, EPI_ISL_3248543, EPI_ISL_3248544, EPI_ISL_3248545, EPI_ISL_3248546, EPI_ISL_3248547, EPI_ISL_3248548, EPI_ISL_3248549, EPI_ISL_3248550, EPI_ISL_3248551 | | | |
| see above | Lighthouse Lab in Glasgow | Wellcome Sanger Institute for the COVID-19 Genomics UK (COG-UK) Consortium | Harper VanSteenhouse, Yumi Kasai, David Gray, Carol Clugston, Anna Dominiczak and Alex Alderton, Roberto Amato, Jeffrey Barrett, Sonia Goncalves, Ewan Harrison, David K. Jackson, Ian Johnston, Dominic Kwiatkowski, Cordelia Langford, John Sillitoe on behalf of the Wellcome Sanger Institute COVID-19 Surveillance Team |
| EPI_ISL_3248552 | Lighthouse Lab in Alderley Park | Wellcome Sanger Institute for the COVID-19 Genomics UK (COG-UK) Consortium | Jacquelyn Wynn, Mairead Hyland, The Lighthouse Lab in Alderley Park and Alex Alderton, Roberto Amato, Jeffrey Barrett, Sonia Goncalves, Ewan Harrison, David K. Jackson, Ian Johnston, Dominic Kwiatkowski, Cordelia Langford, John Sillitoe on behalf of the Wellcome Sanger Institute COVID-19 Surveillance Team |
| EPI_ISL_3248553 | Lighthouse Lab in Glasgow | Wellcome Sanger Institute for the COVID-19 Genomics UK (COG-UK) Consortium | Harper VanSteenhouse, Yumi Kasai, David Gray, Carol Clugston, Anna Dominiczak and Alex Alderton, Roberto Amato, Jeffrey Barrett, Sonia Goncalves, Ewan Harrison, David K. Jackson, Ian Johnston, Dominic Kwiatkowski, Cordelia Langford, John Sillitoe on behalf of the Wellcome Sanger Institute COVID-19 Surveillance Team |
| EPI_ISL_3248554 | Lighthouse Lab in Alderley Park | Wellcome Sanger Institute for the COVID-19 Genomics UK (COG-UK) Consortium | Jacquelyn Wynn, Mairead Hyland, The Lighthouse Lab in Alderley Park and Alex Alderton, Roberto Amato, Jeffrey Barrett, Sonia Goncalves, Ewan Harrison, David K. Jackson, Ian Johnston, Dominic Kwiatkowski, Cordelia Langford, John Sillitoe on behalf of the Wellcome Sanger Institute COVID-19 Surveillance Team |
| EPI_ISL_3248555 | Lighthouse Lab in Glasgow | Wellcome Sanger Institute for the COVID-19 Genomics UK (COG-UK) Consortium | Harper VanSteenhouse, Yumi Kasai, David Gray, Carol Clugston, Anna Dominiczak and Alex Alderton, Roberto Amato, Jeffrey Barrett, Sonia Goncalves, Ewan Harrison, David K. Jackson, Ian Johnston, Dominic Kwiatkowski, Cordelia Langford, John Sillitoe on behalf of the Wellcome Sanger Institute COVID-19 Surveillance Team |
| EPI_ISL_3248556 | Lighthouse Lab in Alderley Park | Wellcome Sanger Institute for the COVID-19 Genomics UK (COG-UK) Consortium | Jacquelyn Wynn, Mairead Hyland, The Lighthouse Lab in Alderley Park and Alex Alderton, Roberto Amato, Jeffrey Barrett, Sonia Goncalves, Ewan Harrison, David K. Jackson, Ian Johnston, Dominic Kwiatkowski, Cordelia Langford, John Sillitoe on behalf of the Wellcome Sanger Institute COVID-19 Surveillance Team |
| EPI_ISL_3248557, EPI_ISL_3248558, EPI_ISL_3248559 | Lighthouse Lab in Glasgow | Wellcome Sanger Institute for the COVID-19 Genomics UK (COG-UK) Consortium | Harper VanSteenhouse, Yumi Kasai, David Gray, Carol Clugston, Anna Dominiczak and Alex Alderton, Roberto Amato, Jeffrey Barrett, Sonia Goncalves, Ewan Harrison, David K. Jackson, Ian Johnston, Dominic Kwiatkowski, Cordelia Langford, John Sillitoe on behalf of the Wellcome Sanger Institute COVID-19 Surveillance Team |
| EPI_ISL_3248560 | Lighthouse Lab in Alderley Park | Wellcome Sanger Institute for the COVID-19 Genomics UK (COG-UK) Consortium | Jacquelyn Wynn, Mairead Hyland, The Lighthouse Lab in Alderley Park and Alex Alderton, Roberto Amato, Jeffrey Barrett, Sonia Goncalves, Ewan Harrison, David K. Jackson, Ian Johnston, Dominic Kwiatkowski, Cordelia Langford, John Sillitoe on behalf of the Wellcome Sanger Institute COVID-19 Surveillance Team |
| EPI_ISL_3248561, EPI_ISL_3248562, EPI_ISL_3248563, EPI_ISL_3248564, EPI_ISL_3248565, EPI_ISL_3248566, EPI_ISL_3248567, EPI_ISL_3248568, EPI_ISL_3248569, EPI_ISL_3248570, EPI_ISL_3248571, EPI_ISL_3248572 | | | |
| see above | Lighthouse Lab in Glasgow | Wellcome Sanger Institute for the COVID-19 Genomics UK (COG-UK) Consortium | Harper VanSteenhouse, Yumi Kasai, David Gray, Carol Clugston, Anna Dominiczak and Alex Alderton, Roberto Amato, Jeffrey Barrett, Sonia Goncalves, Ewan Harrison, David K. Jackson, Ian Johnston, Dominic Kwiatkowski, Cordelia Langford, John Sillitoe on behalf of the Wellcome Sanger Institute COVID-19 Surveillance Team |
| EPI_ISL_3248573 | Lighthouse Lab in Alderley Park | Wellcome Sanger Institute for the COVID-19 Genomics UK (COG-UK) Consortium | Jacquelyn Wynn, Mairead Hyland, The Lighthouse Lab in Alderley Park and Alex Alderton, Roberto Amato, Jeffrey Barrett, Sonia Goncalves, Ewan Harrison, David K. Jackson, Ian Johnston, Dominic Kwiatkowski, Cordelia Langford, John Sillitoe on behalf of the Wellcome Sanger Institute COVID-19 Surveillance Team |
| EPI_ISL_3248574, EPI_ISL_3248575, EPI_ISL_3248576, EPI_ISL_3248577, EPI_ISL_3248578, EPI_ISL_3248579 | Lighthouse Lab in Glasgow | Wellcome Sanger Institute for the COVID-19 Genomics UK (COG-UK) Consortium | Harper VanSteenhouse, Yumi Kasai, David Gray, Carol Clugston, Anna Dominiczak and Alex Alderton, Roberto Amato, Jeffrey Barrett, Sonia Goncalves, Ewan Harrison, David K. Jackson, Ian Johnston, Dominic Kwiatkowski, Cordelia Langford, John Sillitoe on behalf of the Wellcome Sanger Institute COVID-19 Surveillance Team |
| EPI_ISL_3248580 | Lighthouse Lab in Alderley Park | Wellcome Sanger Institute for the COVID-19 Genomics UK (COG-UK) Consortium | Jacquelyn Wynn, Mairead Hyland, The Lighthouse Lab in Alderley Park and Alex Alderton, Roberto Amato, Jeffrey Barrett, Sonia Goncalves, Ewan Harrison, David K. Jackson, Ian Johnston, Dominic Kwiatkowski, Cordelia Langford, John Sillitoe on behalf of the Wellcome Sanger Institute COVID-19 Surveillance Team |
| EPI_ISL_3248581 | Lighthouse Lab in Glasgow | Wellcome Sanger Institute for the COVID-19 Genomics UK (COG-UK) Consortium | Harper VanSteenhouse, Yumi Kasai, David Gray, Carol Clugston, Anna Dominiczak and Alex Alderton, Roberto Amato, Jeffrey Barrett, Sonia Goncalves, Ewan Harrison, David K. Jackson, Ian Johnston, Dominic Kwiatkowski, Cordelia Langford, John Sillitoe on behalf of the Wellcome Sanger Institute COVID-19 Surveillance Team |
| EPI_ISL_3248582 | Lighthouse Lab in Alderley Park | Wellcome Sanger Institute for the COVID-19 Genomics UK (COG-UK) Consortium | Jacquelyn Wynn, Mairead Hyland, The Lighthouse Lab in Alderley Park and Alex Alderton, Roberto Amato, Jeffrey Barrett, Sonia Goncalves, Ewan Harrison, David K. Jackson, Ian Johnston, Dominic Kwiatkowski, Cordelia Langford, John Sillitoe on behalf of the Wellcome Sanger Institute COVID-19 Surveillance Team |
| EPI_ISL_3248583, EPI_ISL_3248584, | Lighthouse Lab in Glasgow | Wellcome Sanger Institute for the COVID-19 Genomics UK (COG-UK) Consortium | Harper VanSteenhouse, Yumi Kasai, David Gray, Carol Clugston, Anna Dominiczak and Alex Alderton, Roberto Amato, Jeffrey Barrett, Sonia Goncalves, Ewan Harrison, David K. Jackson, Ian Johnston, Dominic Kwiatkowski, Cordelia Langford, John Sillitoe on behalf of the Wellcome Sanger Institute COVID-19 Surveillance Team |

| | | | |
|---|---|---|---|
| EPI_ISL_3248585 | | | |
| EPI_ISL_3248586, EPI_ISL_3248587, EPI_ISL_3248588 | Lighthouse Lab in Alderley Park | Wellcome Sanger Institute for the COVID-19 Genomics UK (COG-UK) Consortium | Jacquelyn Wynn, Mairead Hyland, The Lighthouse Lab in Alderley Park and Alex Alderton, Roberto Amato, Jeffrey Barrett, Sonia Goncalves, Ewan Harrison, David K. Jackson, Ian Johnston, Dominic Kwiatkowski, Cordelia Langford, John Sillitoe on behalf of the Wellcome Sanger Institute COVID-19 Surveillance Team |
| EPI_ISL_3248589, EPI_ISL_3248590 | Lighthouse Lab in Glasgow | Wellcome Sanger Institute for the COVID-19 Genomics UK (COG-UK) Consortium | Harper VanSteenhouse, Yumi Kasai, David Gray, Carol Clugston, Anna Dominiczak and Alex Alderton, Roberto Amato, Jeffrey Barrett, Sonia Goncalves, Ewan Harrison, David K. Jackson, Ian Johnston, Dominic Kwiatkowski, Cordelia Langford, John Sillitoe on behalf of the Wellcome Sanger Institute COVID-19 Surveillance Team |
| EPI_ISL_3248591 | Lighthouse Lab in Alderley Park | Wellcome Sanger Institute for the COVID-19 Genomics UK (COG-UK) Consortium | Jacquelyn Wynn, Mairead Hyland, The Lighthouse Lab in Alderley Park and Alex Alderton, Roberto Amato, Jeffrey Barrett, Sonia Goncalves, Ewan Harrison, David K. Jackson, Ian Johnston, Dominic Kwiatkowski, Cordelia Langford, John Sillitoe on behalf of the Wellcome Sanger Institute COVID-19 Surveillance Team |
| EPI_ISL_3248592 | Lighthouse Lab in Glasgow | Wellcome Sanger Institute for the COVID-19 Genomics UK (COG-UK) Consortium | Harper VanSteenhouse, Yumi Kasai, David Gray, Carol Clugston, Anna Dominiczak and Alex Alderton, Roberto Amato, Jeffrey Barrett, Sonia Goncalves, Ewan Harrison, David K. Jackson, Ian Johnston, Dominic Kwiatkowski, Cordelia Langford, John Sillitoe on behalf of the Wellcome Sanger Institute COVID-19 Surveillance Team |
| EPI_ISL_3248593 | Lighthouse Lab in Alderley Park | Wellcome Sanger Institute for the COVID-19 Genomics UK (COG-UK) Consortium | Jacquelyn Wynn, Mairead Hyland, The Lighthouse Lab in Alderley Park and Alex Alderton, Roberto Amato, Jeffrey Barrett, Sonia Goncalves, Ewan Harrison, David K. Jackson, Ian Johnston, Dominic Kwiatkowski, Cordelia Langford, John Sillitoe on behalf of the Wellcome Sanger Institute COVID-19 Surveillance Team |
| EPI_ISL_3248594, EPI_ISL_3248595, EPI_ISL_3248596 | Lighthouse Lab in Glasgow | Wellcome Sanger Institute for the COVID-19 Genomics UK (COG-UK) Consortium | Harper VanSteenhouse, Yumi Kasai, David Gray, Carol Clugston, Anna Dominiczak and Alex Alderton, Roberto Amato, Jeffrey Barrett, Sonia Goncalves, Ewan Harrison, David K. Jackson, Ian Johnston, Dominic Kwiatkowski, Cordelia Langford, John Sillitoe on behalf of the Wellcome Sanger Institute COVID-19 Surveillance Team |
| EPI_ISL_3248597, EPI_ISL_3248598, EPI_ISL_3248599 | Lighthouse Lab in Alderley Park | Wellcome Sanger Institute for the COVID-19 Genomics UK (COG-UK) Consortium | Jacquelyn Wynn, Mairead Hyland, The Lighthouse Lab in Alderley Park and Alex Alderton, Roberto Amato, Jeffrey Barrett, Sonia Goncalves, Ewan Harrison, David K. Jackson, Ian Johnston, Dominic Kwiatkowski, Cordelia Langford, John Sillitoe on behalf of the Wellcome Sanger Institute COVID-19 Surveillance Team |
| EPI_ISL_3248600 | Lighthouse Lab in Glasgow | Wellcome Sanger Institute for the COVID-19 Genomics UK (COG-UK) Consortium | Harper VanSteenhouse, Yumi Kasai, David Gray, Carol Clugston, Anna Dominiczak and Alex Alderton, Roberto Amato, Jeffrey Barrett, Sonia Goncalves, Ewan Harrison, David K. Jackson, Ian Johnston, Dominic Kwiatkowski, Cordelia Langford, John Sillitoe on behalf of the Wellcome Sanger Institute COVID-19 Surveillance Team |
| EPI_ISL_3248601 | Lighthouse Lab in Alderley Park | Wellcome Sanger Institute for the COVID-19 Genomics UK (COG-UK) Consortium | Jacquelyn Wynn, Mairead Hyland, The Lighthouse Lab in Alderley Park and Alex Alderton, Roberto Amato, Jeffrey Barrett, Sonia Goncalves, Ewan Harrison, David K. Jackson, Ian Johnston, Dominic Kwiatkowski, Cordelia Langford, John Sillitoe on behalf of the Wellcome Sanger Institute COVID-19 Surveillance Team |
| EPI_ISL_3248602 | Lighthouse Lab in Glasgow | Wellcome Sanger Institute for the COVID-19 Genomics UK (COG-UK) Consortium | Harper VanSteenhouse, Yumi Kasai, David Gray, Carol Clugston, Anna Dominiczak and Alex Alderton, Roberto Amato, Jeffrey Barrett, Sonia Goncalves, Ewan Harrison, David K. Jackson, Ian Johnston, Dominic Kwiatkowski, Cordelia Langford, John Sillitoe on behalf of the Wellcome Sanger Institute COVID-19 Surveillance Team |
| EPI_ISL_3248603 | Lighthouse Lab in Alderley Park | Wellcome Sanger Institute for the COVID-19 Genomics UK (COG-UK) Consortium | Jacquelyn Wynn, Mairead Hyland, The Lighthouse Lab in Alderley Park and Alex Alderton, Roberto Amato, Jeffrey Barrett, Sonia Goncalves, Ewan Harrison, David K. Jackson, Ian Johnston, Dominic Kwiatkowski, Cordelia Langford, John Sillitoe on behalf of the Wellcome Sanger Institute COVID-19 Surveillance Team |
| EPI_ISL_3248604, EPI_ISL_3248605 | Lighthouse Lab in Glasgow | Wellcome Sanger Institute for the COVID-19 Genomics UK (COG-UK) Consortium | Harper VanSteenhouse, Yumi Kasai, David Gray, Carol Clugston, Anna Dominiczak and Alex Alderton, Roberto Amato, Jeffrey Barrett, Sonia Goncalves, Ewan Harrison, David K. Jackson, Ian Johnston, Dominic Kwiatkowski, Cordelia Langford, John Sillitoe on behalf of the Wellcome Sanger Institute COVID-19 Surveillance Team |
| EPI_ISL_3248606 | Lighthouse Lab in Alderley Park | Wellcome Sanger Institute for the COVID-19 Genomics UK (COG-UK) Consortium | Jacquelyn Wynn, Mairead Hyland, The Lighthouse Lab in Alderley Park and Alex Alderton, Roberto Amato, Jeffrey Barrett, Sonia Goncalves, Ewan Harrison, David K. Jackson, Ian Johnston, Dominic Kwiatkowski, Cordelia Langford, John Sillitoe on behalf of the Wellcome Sanger Institute COVID-19 Surveillance Team |
| EPI_ISL_3248607, EPI_ISL_3248608, EPI_ISL_3248609, EPI_ISL_3248610 | Lighthouse Lab in Glasgow | Wellcome Sanger Institute for the COVID-19 Genomics UK (COG-UK) Consortium | Harper VanSteenhouse, Yumi Kasai, David Gray, Carol Clugston, Anna Dominiczak and Alex Alderton, Roberto Amato, Jeffrey Barrett, Sonia Goncalves, Ewan Harrison, David K. Jackson, Ian Johnston, Dominic Kwiatkowski, Cordelia Langford, John Sillitoe on behalf of the Wellcome Sanger Institute COVID-19 Surveillance Team |
| EPI_ISL_3248611, EPI_ISL_3248612 | Lighthouse Lab in Alderley Park | Wellcome Sanger Institute for the COVID-19 Genomics UK (COG-UK) Consortium | Jacquelyn Wynn, Mairead Hyland, The Lighthouse Lab in Alderley Park and Alex Alderton, Roberto Amato, Jeffrey Barrett, Sonia Goncalves, Ewan Harrison, David K. Jackson, Ian Johnston, Dominic Kwiatkowski, Cordelia Langford, John Sillitoe on behalf of the Wellcome Sanger Institute COVID-19 Surveillance Team |
| EPI_ISL_3248613, EPI_ISL_3248614, EPI_ISL_3248615, EPI_ISL_3248616, EPI_ISL_3248617, EPI_ISL_3248618, EPI_ISL_3248619, EPI_ISL_3248620, EPI_ISL_3248621, EPI_ISL_3248622, EPI_ISL_3248623, EPI_ISL_3248624, EPI_ISL_3248625, EPI_ISL_3248626 | | | |
| see above | Lighthouse Lab in Glasgow | Wellcome Sanger Institute for the COVID-19 Genomics UK (COG-UK) Consortium | Harper VanSteenhouse, Yumi Kasai, David Gray, Carol Clugston, Anna Dominiczak and Alex Alderton, Roberto Amato, Jeffrey Barrett, Sonia Goncalves, Ewan Harrison, David K. Jackson, Ian Johnston, Dominic Kwiatkowski, Cordelia Langford, John Sillitoe on behalf of the Wellcome Sanger Institute COVID-19 Surveillance Team |
| EPI_ISL_3248627 | Lighthouse Lab in Alderley Park | Wellcome Sanger Institute for the COVID-19 Genomics UK (COG-UK) Consortium | Jacquelyn Wynn, Mairead Hyland, The Lighthouse Lab in Alderley Park and Alex Alderton, Roberto Amato, Jeffrey Barrett, Sonia Goncalves, Ewan Harrison, David K. Jackson, Ian Johnston, Dominic Kwiatkowski, Cordelia Langford, John Sillitoe on behalf of the Wellcome Sanger Institute COVID-19 Surveillance Team |
| EPI_ISL_3248628 | Lighthouse Lab in Glasgow | Wellcome Sanger Institute for the COVID-19 Genomics UK (COG-UK) Consortium | Harper VanSteenhouse, Yumi Kasai, David Gray, Carol Clugston, Anna Dominiczak and Alex Alderton, Roberto Amato, Jeffrey Barrett, Sonia Goncalves, Ewan Harrison, David K. Jackson, Ian Johnston, Dominic Kwiatkowski, Cordelia Langford, John Sillitoe on behalf of the Wellcome Sanger Institute COVID-19 Surveillance Team |
| EPI_ISL_3248629, EPI_ISL_3248630, EPI_ISL_3248631, EPI_ISL_3248632 | Lighthouse Lab in Alderley Park | Wellcome Sanger Institute for the COVID-19 Genomics UK (COG-UK) Consortium | Jacquelyn Wynn, Mairead Hyland, The Lighthouse Lab in Alderley Park and Alex Alderton, Roberto Amato, Jeffrey Barrett, Sonia Goncalves, Ewan Harrison, David K. Jackson, Ian Johnston, Dominic Kwiatkowski, Cordelia Langford, John Sillitoe on behalf of the Wellcome Sanger Institute COVID-19 Surveillance Team |
| EPI_ISL_3248633, EPI_ISL_3248634 | Lighthouse Lab in Glasgow | Wellcome Sanger Institute for the COVID-19 Genomics UK (COG-UK) Consortium | Harper VanSteenhouse, Yumi Kasai, David Gray, Carol Clugston, Anna Dominiczak and Alex Alderton, Roberto Amato, Jeffrey Barrett, Sonia Goncalves, Ewan Harrison, David K. Jackson, Ian Johnston, Dominic Kwiatkowski, Cordelia Langford, John Sillitoe on behalf of the Wellcome Sanger Institute COVID-19 Surveillance Team |
| EPI_ISL_3248635, EPI_ISL_3248636 | Lighthouse Lab in Alderley Park | Wellcome Sanger Institute for the COVID-19 Genomics UK (COG-UK) Consortium | Jacquelyn Wynn, Mairead Hyland, The Lighthouse Lab in Alderley Park and Alex Alderton, Roberto Amato, Jeffrey Barrett, Sonia Goncalves, Ewan Harrison, David K. Jackson, Ian Johnston, Dominic Kwiatkowski, Cordelia Langford, John Sillitoe on behalf of the Wellcome Sanger Institute COVID-19 Surveillance Team |
| EPI_ISL_3248637, EPI_ISL_3248638 | Berkshire and Surrey Pathology Services Lighthouse Laboratory | Wellcome Sanger Institute for the COVID-19 Genomics UK (COG-UK) Consortium | Berkshire and Surrey Pathology Services Lighthouse Laboratory and Alex Alderton, Roberto Amato, Jeffrey Barrett, Sonia Goncalves, Ewan Harrison, David K. Jackson, Ian Johnston, Dominic Kwiatkowski, Cordelia Langford, John Sillitoe on behalf of the Wellcome Sanger Institute COVID-19 Surveillance Team |
| EPI_ISL_3248639 | Lighthouse Lab in Alderley Park | Wellcome Sanger Institute for the COVID-19 Genomics UK (COG-UK) Consortium | Jacquelyn Wynn, Mairead Hyland, The Lighthouse Lab in Alderley Park and Alex Alderton, Roberto Amato, Jeffrey Barrett, Sonia Goncalves, Ewan Harrison, David K. Jackson, Ian Johnston, Dominic Kwiatkowski, Cordelia Langford, John Sillitoe on behalf of the Wellcome Sanger Institute COVID-19 Surveillance Team |
| EPI_ISL_3248640 | Lighthouse Lab in Glasgow | Wellcome Sanger Institute for the COVID-19 Genomics UK (COG-UK) Consortium | Harper VanSteenhouse, Yumi Kasai, David Gray, Carol Clugston, Anna Dominiczak and Alex Alderton, Roberto Amato, Jeffrey Barrett, Sonia Goncalves, Ewan Harrison, David K. Jackson, Ian Johnston, Dominic Kwiatkowski, Cordelia Langford, John Sillitoe on behalf of the Wellcome Sanger Institute COVID-19 Surveillance Team |
| EPI_ISL_3248641, EPI_ISL_3248642 | Berkshire and Surrey Pathology Services Lighthouse Laboratory | Wellcome Sanger Institute for the COVID-19 Genomics UK (COG-UK) Consortium | Berkshire and Surrey Pathology Services Lighthouse Laboratory and Alex Alderton, Roberto Amato, Jeffrey Barrett, Sonia Goncalves, Ewan Harrison, David K. Jackson, Ian Johnston, Dominic Kwiatkowski, Cordelia Langford, John Sillitoe on behalf of the Wellcome Sanger Institute COVID-19 Surveillance Team |
| EPI_ISL_3248643 | Lighthouse Lab in Milton Keynes | Wellcome Sanger Institute for the COVID-19 Genomics UK (COG-UK) Consortium | The Lighthouse Lab in Milton Keynes and Alex Alderton, Roberto Amato, Jeffrey Barrett, Sonia Goncalves, Ewan Harrison, David K. Jackson, Ian Johnston, Dominic Kwiatkowski, Cordelia Langford, John Sillitoe on behalf of the Wellcome Sanger Institute COVID-19 Surveillance Team |
| EPI_ISL_3248644 | Lighthouse Lab in Glasgow | Wellcome Sanger Institute for the COVID-19 Genomics UK (COG-UK) Consortium | Harper VanSteenhouse, Yumi Kasai, David Gray, Carol Clugston, Anna Dominiczak and Alex Alderton, Roberto Amato, Jeffrey Barrett, Sonia Goncalves, Ewan Harrison, David K. Jackson, Ian Johnston, Dominic Kwiatkowski, Cordelia Langford, John Sillitoe on behalf of the Wellcome Sanger Institute COVID-19 Surveillance Team |
| EPI_ISL_3248645 | Berkshire and Surrey Pathology Services Lighthouse Laboratory | Wellcome Sanger Institute for the COVID-19 Genomics UK (COG-UK) Consortium | Berkshire and Surrey Pathology Services Lighthouse Laboratory and Alex Alderton, Roberto Amato, Jeffrey Barrett, Sonia Goncalves, Ewan Harrison, David K. Jackson, Ian Johnston, Dominic Kwiatkowski, Cordelia Langford, John Sillitoe on behalf of the Wellcome Sanger Institute COVID-19 Surveillance Team |
| EPI_ISL_3248646 | Lighthouse Lab in Glasgow | Wellcome Sanger Institute for the COVID-19 Genomics UK (COG-UK) Consortium | Harper VanSteenhouse, Yumi Kasai, David Gray, Carol Clugston, Anna Dominiczak and Alex Alderton, Roberto Amato, Jeffrey Barrett, Sonia Goncalves, Ewan Harrison, David K. Jackson, Ian Johnston, Dominic Kwiatkowski, Cordelia Langford, John Sillitoe on behalf of the Wellcome Sanger Institute COVID-19 Surveillance Team |
| EPI_ISL_3248647 | Berkshire and Surrey Pathology Services Lighthouse Laboratory | Wellcome Sanger Institute for the COVID-19 Genomics UK (COG-UK) Consortium | Berkshire and Surrey Pathology Services Lighthouse Laboratory and Alex Alderton, Roberto Amato, Jeffrey Barrett, Sonia Goncalves, Ewan Harrison, David K. Jackson, Ian Johnston, Dominic Kwiatkowski, Cordelia Langford, John Sillitoe on behalf of the Wellcome Sanger Institute COVID-19 Surveillance Team |
| EPI_ISL_3248648 | Lighthouse Lab in Glasgow | Wellcome Sanger Institute for the COVID-19 Genomics UK (COG-UK) Consortium | Harper VanSteenhouse, Yumi Kasai, David Gray, Carol Clugston, Anna Dominiczak and Alex Alderton, Roberto Amato, Jeffrey Barrett, Sonia Goncalves, Ewan Harrison, David K. Jackson, Ian Johnston, Dominic Kwiatkowski, Cordelia Langford, John Sillitoe on behalf of the Wellcome Sanger Institute COVID-19 Surveillance Team |
| EPI_ISL_3248649, EPI_ISL_3248650 | Lighthouse Lab in Milton Keynes | Wellcome Sanger Institute for the COVID-19 Genomics UK (COG-UK) Consortium | The Lighthouse Lab in Milton Keynes and Alex Alderton, Roberto Amato, Jeffrey Barrett, Sonia Goncalves, Ewan Harrison, David K. Jackson, Ian Johnston, Dominic Kwiatkowski, Cordelia Langford, John Sillitoe on behalf of the Wellcome Sanger Institute COVID-19 Surveillance Team |
| EPI_ISL_3248651 | Lighthouse Lab in Alderley Park | Wellcome Sanger Institute for the COVID-19 Genomics UK (COG-UK) Consortium | Jacquelyn Wynn, Mairead Hyland, The Lighthouse Lab in Alderley Park and Alex Alderton, Roberto Amato, Jeffrey Barrett, Sonia Goncalves, Ewan Harrison, David K. Jackson, Ian Johnston, Dominic Kwiatkowski, Cordelia Langford, John Sillitoe on behalf of the Wellcome Sanger Institute COVID-19 Surveillance Team |
| EPI_ISL_3248652 | Lighthouse Lab in Milton Keynes | Wellcome Sanger Institute for the COVID-19 Genomics UK (COG-UK) Consortium | The Lighthouse Lab in Milton Keynes and Alex Alderton, Roberto Amato, Jeffrey Barrett, Sonia Goncalves, Ewan Harrison, David K. Jackson, Ian Johnston, Dominic Kwiatkowski, Cordelia Langford, John Sillitoe on behalf of the Wellcome Sanger Institute COVID-19 Surveillance Team |
| EPI_ISL_3248653 | Berkshire and Surrey Pathology Services Lighthouse Laboratory | Wellcome Sanger Institute for the COVID-19 Genomics UK (COG-UK) Consortium | Berkshire and Surrey Pathology Services Lighthouse Laboratory and Alex Alderton, Roberto Amato, Jeffrey Barrett, Sonia Goncalves, Ewan Harrison, David K. Jackson, Ian Johnston, Dominic Kwiatkowski, Cordelia Langford, John Sillitoe on behalf of the Wellcome Sanger Institute COVID-19 Surveillance Team |
| EPI_ISL_3248654 | Lighthouse Lab in Glasgow | Wellcome Sanger Institute for the COVID-19 Genomics UK (COG-UK) Consortium | Harper VanSteenhouse, Yumi Kasai, David Gray, Carol Clugston, Anna Dominiczak and Alex Alderton, Roberto Amato, Jeffrey Barrett, Sonia Goncalves, Ewan Harrison, David K. Jackson, Ian Johnston, Dominic Kwiatkowski, Cordelia Langford, John Sillitoe on behalf of the Wellcome Sanger Institute COVID-19 Surveillance Team |
| EPI_ISL_3248655 | Lighthouse Lab in Milton Keynes | Wellcome Sanger Institute for the COVID-19 Genomics UK (COG-UK) Consortium | The Lighthouse Lab in Milton Keynes and Alex Alderton, Roberto Amato, Jeffrey Barrett, Sonia Goncalves, Ewan Harrison, David K. Jackson, Ian Johnston, Dominic Kwiatkowski, Cordelia Langford, John Sillitoe on behalf of the Wellcome Sanger Institute COVID-19 Surveillance Team |
| EPI_ISL_3248656 | Lighthouse Lab in Alderley Park | Wellcome Sanger Institute for the COVID-19 Genomics UK (COG-UK) Consortium | Jacquelyn Wynn, Mairead Hyland, The Lighthouse Lab in Alderley Park and Alex Alderton, Roberto Amato, Jeffrey Barrett, Sonia Goncalves, Ewan Harrison, David K. Jackson, Ian Johnston, Dominic Kwiatkowski, Cordelia Langford, John Sillitoe on behalf of the Wellcome Sanger Institute COVID-19 Surveillance Team |
| EPI_ISL_3248657, EPI_ISL_3248658 | Lighthouse Lab in Milton Keynes | Wellcome Sanger Institute for the COVID-19 Genomics UK (COG-UK) Consortium | The Lighthouse Lab in Milton Keynes and Alex Alderton, Roberto Amato, Jeffrey Barrett, Sonia Goncalves, Ewan Harrison, David K. Jackson, Ian Johnston, Dominic Kwiatkowski, Cordelia Langford, John Sillitoe on behalf of the Wellcome Sanger Institute COVID-19 Surveillance Team |
| EPI_ISL_3248659, EPI_ISL_3248660 | Lighthouse Lab in Glasgow | Wellcome Sanger Institute for the COVID-19 Genomics UK (COG-UK) Consortium | Harper VanSteenhouse, Yumi Kasai, David Gray, Carol Clugston, Anna Dominiczak and Alex Alderton, Roberto Amato, Jeffrey Barrett, Sonia Goncalves, Ewan Harrison, David K. Jackson, Ian Johnston, Dominic Kwiatkowski, Cordelia Langford, John Sillitoe on behalf of the Wellcome Sanger Institute COVID-19 Surveillance Team |
| EPI_ISL_3248661, EPI_ISL_3248662 | Lighthouse Lab in Alderley Park | Wellcome Sanger Institute for the COVID-19 Genomics UK (COG-UK) Consortium | Jacquelyn Wynn, Mairead Hyland, The Lighthouse Lab in Alderley Park and Alex Alderton, Roberto Amato, Jeffrey Barrett, Sonia Goncalves, Ewan Harrison, David K. Jackson, Ian Johnston, Dominic Kwiatkowski, Cordelia Langford, John Sillitoe on behalf of the Wellcome Sanger Institute COVID-19 Surveillance Team |

| | | | |
|---|---|---|---|
| EPI_ISL_3248663 | Lighthouse Lab in Glasgow | Wellcome Sanger Institute for the COVID-19 Genomics UK (COG-UK) Consortium | Harper VanSteenhouse, Yumi Kasai, David Gray, Carol Clugston, Anna Dominiczak and Alex Alderton, Roberto Amato, Jeffrey Barrett, Sonia Goncalves, Ewan Harrison, David K. Jackson, Ian Johnston, Dominic Kwiatkowski, Cordelia Langford, John Sillitoe on behalf of the Wellcome Sanger Institute COVID-19 Surveillance Team |
| EPI_ISL_3248664 | Lighthouse Lab in Milton Keynes | Wellcome Sanger Institute for the COVID-19 Genomics UK (COG-UK) Consortium | The Lighthouse Lab in Milton Keynes and Alex Alderton, Roberto Amato, Jeffrey Barrett, Sonia Goncalves, Ewan Harrison, David K. Jackson, Ian Johnston, Dominic Kwiatkowski, Cordelia Langford, John Sillitoe on behalf of the Wellcome Sanger Institute COVID-19 Surveillance Team |
| EPI_ISL_3248665 | Lighthouse Lab in Glasgow | Wellcome Sanger Institute for the COVID-19 Genomics UK (COG-UK) Consortium | Harper VanSteenhouse, Yumi Kasai, David Gray, Carol Clugston, Anna Dominiczak and Alex Alderton, Roberto Amato, Jeffrey Barrett, Sonia Goncalves, Ewan Harrison, David K. Jackson, Ian Johnston, Dominic Kwiatkowski, Cordelia Langford, John Sillitoe on behalf of the Wellcome Sanger Institute COVID-19 Surveillance Team |
| EPI_ISL_3248666 | Lighthouse Lab in Alderley Park | Wellcome Sanger Institute for the COVID-19 Genomics UK (COG-UK) Consortium | Jacquelyn Wynn, Mairead Hyland, The Lighthouse Lab in Alderley Park and Alex Alderton, Roberto Amato, Jeffrey Barrett, Sonia Goncalves, Ewan Harrison, David K. Jackson, Ian Johnston, Dominic Kwiatkowski, Cordelia Langford, John Sillitoe on behalf of the Wellcome Sanger Institute COVID-19 Surveillance Team |
| EPI_ISL_3248667 | Lighthouse Lab in Milton Keynes | Wellcome Sanger Institute for the COVID-19 Genomics UK (COG-UK) Consortium | The Lighthouse Lab in Milton Keynes and Alex Alderton, Roberto Amato, Jeffrey Barrett, Sonia Goncalves, Ewan Harrison, David K. Jackson, Ian Johnston, Dominic Kwiatkowski, Cordelia Langford, John Sillitoe on behalf of the Wellcome Sanger Institute COVID-19 Surveillance Team |
| EPI_ISL_3248668, EPI_ISL_3248669 | Lighthouse Lab in Alderley Park | Wellcome Sanger Institute for the COVID-19 Genomics UK (COG-UK) Consortium | Jacquelyn Wynn, Mairead Hyland, The Lighthouse Lab in Alderley Park and Alex Alderton, Roberto Amato, Jeffrey Barrett, Sonia Goncalves, Ewan Harrison, David K. Jackson, Ian Johnston, Dominic Kwiatkowski, Cordelia Langford, John Sillitoe on behalf of the Wellcome Sanger Institute COVID-19 Surveillance Team |
| EPI_ISL_3248670, EPI_ISL_3248671, EPI_ISL_3248672 | Berkshire and Surrey Pathology Services Lighthouse Laboratory | Wellcome Sanger Institute for the COVID-19 Genomics UK (COG-UK) Consortium | Berkshire and Surrey Pathology Services Lighthouse Laboratory and Alex Alderton, Roberto Amato, Jeffrey Barrett, Sonia Goncalves, Ewan Harrison, David K. Jackson, Ian Johnston, Dominic Kwiatkowski, Cordelia Langford, John Sillitoe on behalf of the Wellcome Sanger Institute COVID-19 Surveillance Team |
| EPI_ISL_3248673 | Lighthouse Lab in Milton Keynes | Wellcome Sanger Institute for the COVID-19 Genomics UK (COG-UK) Consortium | The Lighthouse Lab in Milton Keynes and Alex Alderton, Roberto Amato, Jeffrey Barrett, Sonia Goncalves, Ewan Harrison, David K. Jackson, Ian Johnston, Dominic Kwiatkowski, Cordelia Langford, John Sillitoe on behalf of the Wellcome Sanger Institute COVID-19 Surveillance Team |
| EPI_ISL_3248674, EPI_ISL_3248675 | Lighthouse Lab in Alderley Park | Wellcome Sanger Institute for the COVID-19 Genomics UK (COG-UK) Consortium | Jacquelyn Wynn, Mairead Hyland, The Lighthouse Lab in Alderley Park and Alex Alderton, Roberto Amato, Jeffrey Barrett, Sonia Goncalves, Ewan Harrison, David K. Jackson, Ian Johnston, Dominic Kwiatkowski, Cordelia Langford, John Sillitoe on behalf of the Wellcome Sanger Institute COVID-19 Surveillance Team |
| EPI_ISL_3248676 | Berkshire and Surrey Pathology Services Lighthouse Laboratory | Wellcome Sanger Institute for the COVID-19 Genomics UK (COG-UK) Consortium | Berkshire and Surrey Pathology Services Lighthouse Laboratory and Alex Alderton, Roberto Amato, Jeffrey Barrett, Sonia Goncalves, Ewan Harrison, David K. Jackson, Ian Johnston, Dominic Kwiatkowski, Cordelia Langford, John Sillitoe on behalf of the Wellcome Sanger Institute COVID-19 Surveillance Team |
| EPI_ISL_3248677 | Lighthouse Lab in Glasgow | Wellcome Sanger Institute for the COVID-19 Genomics UK (COG-UK) Consortium | Harper VanSteenhouse, Yumi Kasai, David Gray, Carol Clugston, Anna Dominiczak and Alex Alderton, Roberto Amato, Jeffrey Barrett, Sonia Goncalves, Ewan Harrison, David K. Jackson, Ian Johnston, Dominic Kwiatkowski, Cordelia Langford, John Sillitoe on behalf of the Wellcome Sanger Institute COVID-19 Surveillance Team |
| EPI_ISL_3248678 | Lighthouse Lab in Alderley Park | Wellcome Sanger Institute for the COVID-19 Genomics UK (COG-UK) Consortium | Jacquelyn Wynn, Mairead Hyland, The Lighthouse Lab in Alderley Park and Alex Alderton, Roberto Amato, Jeffrey Barrett, Sonia Goncalves, Ewan Harrison, David K. Jackson, Ian Johnston, Dominic Kwiatkowski, Cordelia Langford, John Sillitoe on behalf of the Wellcome Sanger Institute COVID-19 Surveillance Team |
| EPI_ISL_3248679 | Berkshire and Surrey Pathology Services Lighthouse Laboratory | Wellcome Sanger Institute for the COVID-19 Genomics UK (COG-UK) Consortium | Berkshire and Surrey Pathology Services Lighthouse Laboratory and Alex Alderton, Roberto Amato, Jeffrey Barrett, Sonia Goncalves, Ewan Harrison, David K. Jackson, Ian Johnston, Dominic Kwiatkowski, Cordelia Langford, John Sillitoe on behalf of the Wellcome Sanger Institute COVID-19 Surveillance Team |
| EPI_ISL_3248680, EPI_ISL_3248681 | Lighthouse Lab in Glasgow | Wellcome Sanger Institute for the COVID-19 Genomics UK (COG-UK) Consortium | Harper VanSteenhouse, Yumi Kasai, David Gray, Carol Clugston, Anna Dominiczak and Alex Alderton, Roberto Amato, Jeffrey Barrett, Sonia Goncalves, Ewan Harrison, David K. Jackson, Ian Johnston, Dominic Kwiatkowski, Cordelia Langford, John Sillitoe on behalf of the Wellcome Sanger Institute COVID-19 Surveillance Team |
| EPI_ISL_3248682 | Berkshire and Surrey Pathology Services Lighthouse Laboratory | Wellcome Sanger Institute for the COVID-19 Genomics UK (COG-UK) Consortium | Berkshire and Surrey Pathology Services Lighthouse Laboratory and Alex Alderton, Roberto Amato, Jeffrey Barrett, Sonia Goncalves, Ewan Harrison, David K. Jackson, Ian Johnston, Dominic Kwiatkowski, Cordelia Langford, John Sillitoe on behalf of the Wellcome Sanger Institute COVID-19 Surveillance Team |
| EPI_ISL_3248683 | Lighthouse Lab in Milton Keynes | Wellcome Sanger Institute for the COVID-19 Genomics UK (COG-UK) Consortium | The Lighthouse Lab in Milton Keynes and Alex Alderton, Roberto Amato, Jeffrey Barrett, Sonia Goncalves, Ewan Harrison, David K. Jackson, Ian Johnston, Dominic Kwiatkowski, Cordelia Langford, John Sillitoe on behalf of the Wellcome Sanger Institute COVID-19 Surveillance Team |
| EPI_ISL_3248684 | Lighthouse Lab in Alderley Park | Wellcome Sanger Institute for the COVID-19 Genomics UK (COG-UK) Consortium | Jacquelyn Wynn, Mairead Hyland, The Lighthouse Lab in Alderley Park and Alex Alderton, Roberto Amato, Jeffrey Barrett, Sonia Goncalves, Ewan Harrison, David K. Jackson, Ian Johnston, Dominic Kwiatkowski, Cordelia Langford, John Sillitoe on behalf of the Wellcome Sanger Institute COVID-19 Surveillance Team |
| EPI_ISL_3248685 | Lighthouse Lab in Glasgow | Wellcome Sanger Institute for the COVID-19 Genomics UK (COG-UK) Consortium | Harper VanSteenhouse, Yumi Kasai, David Gray, Carol Clugston, Anna Dominiczak and Alex Alderton, Roberto Amato, Jeffrey Barrett, Sonia Goncalves, Ewan Harrison, David K. Jackson, Ian Johnston, Dominic Kwiatkowski, Cordelia Langford, John Sillitoe on behalf of the Wellcome Sanger Institute COVID-19 Surveillance Team |
| EPI_ISL_3248686 | Lighthouse Lab in Alderley Park | Wellcome Sanger Institute for the COVID-19 Genomics UK (COG-UK) Consortium | Jacquelyn Wynn, Mairead Hyland, The Lighthouse Lab in Alderley Park and Alex Alderton, Roberto Amato, Jeffrey Barrett, Sonia Goncalves, Ewan Harrison, David K. Jackson, Ian Johnston, Dominic Kwiatkowski, Cordelia Langford, John Sillitoe on behalf of the Wellcome Sanger Institute COVID-19 Surveillance Team |
| EPI_ISL_3248687 | Berkshire and Surrey Pathology Services Lighthouse Laboratory | Wellcome Sanger Institute for the COVID-19 Genomics UK (COG-UK) Consortium | Berkshire and Surrey Pathology Services Lighthouse Laboratory and Alex Alderton, Roberto Amato, Jeffrey Barrett, Sonia Goncalves, Ewan Harrison, David K. Jackson, Ian Johnston, Dominic Kwiatkowski, Cordelia Langford, John Sillitoe on behalf of the Wellcome Sanger Institute COVID-19 Surveillance Team |
| EPI_ISL_3248688, EPI_ISL_3248689 | Lighthouse Lab in Milton Keynes | Wellcome Sanger Institute for the COVID-19 Genomics UK (COG-UK) Consortium | The Lighthouse Lab in Milton Keynes and Alex Alderton, Roberto Amato, Jeffrey Barrett, Sonia Goncalves, Ewan Harrison, David K. Jackson, Ian Johnston, Dominic Kwiatkowski, Cordelia Langford, John Sillitoe on behalf of the Wellcome Sanger Institute COVID-19 Surveillance Team |
| EPI_ISL_3248690 | Berkshire and Surrey Pathology Services Lighthouse Laboratory | Wellcome Sanger Institute for the COVID-19 Genomics UK (COG-UK) Consortium | Berkshire and Surrey Pathology Services Lighthouse Laboratory and Alex Alderton, Roberto Amato, Jeffrey Barrett, Sonia Goncalves, Ewan Harrison, David K. Jackson, Ian Johnston, Dominic Kwiatkowski, Cordelia Langford, John Sillitoe on behalf of the Wellcome Sanger Institute COVID-19 Surveillance Team |
| EPI_ISL_3248691 | Lighthouse Lab in Glasgow | Wellcome Sanger Institute for the COVID-19 Genomics UK (COG-UK) Consortium | Harper VanSteenhouse, Yumi Kasai, David Gray, Carol Clugston, Anna Dominiczak and Alex Alderton, Roberto Amato, Jeffrey Barrett, Sonia Goncalves, Ewan Harrison, David K. Jackson, Ian Johnston, Dominic Kwiatkowski, Cordelia Langford, John Sillitoe on behalf of the Wellcome Sanger Institute COVID-19 Surveillance Team |
| EPI_ISL_3248692 | Berkshire and Surrey Pathology Services Lighthouse Laboratory | Wellcome Sanger Institute for the COVID-19 Genomics UK (COG-UK) Consortium | Berkshire and Surrey Pathology Services Lighthouse Laboratory and Alex Alderton, Roberto Amato, Jeffrey Barrett, Sonia Goncalves, Ewan Harrison, David K. Jackson, Ian Johnston, Dominic Kwiatkowski, Cordelia Langford, John Sillitoe on behalf of the Wellcome Sanger Institute COVID-19 Surveillance Team |
| EPI_ISL_3248693 | Lighthouse Lab in Alderley Park | Wellcome Sanger Institute for the COVID-19 Genomics UK (COG-UK) Consortium | Jacquelyn Wynn, Mairead Hyland, The Lighthouse Lab in Alderley Park and Alex Alderton, Roberto Amato, Jeffrey Barrett, Sonia Goncalves, Ewan Harrison, David K. Jackson, Ian Johnston, Dominic Kwiatkowski, Cordelia Langford, John Sillitoe on behalf of the Wellcome Sanger Institute COVID-19 Surveillance Team |
| EPI_ISL_3248694 | Berkshire and Surrey Pathology Services Lighthouse Laboratory | Wellcome Sanger Institute for the COVID-19 Genomics UK (COG-UK) Consortium | Berkshire and Surrey Pathology Services Lighthouse Laboratory and Alex Alderton, Roberto Amato, Jeffrey Barrett, Sonia Goncalves, Ewan Harrison, David K. Jackson, Ian Johnston, Dominic Kwiatkowski, Cordelia Langford, John Sillitoe on behalf of the Wellcome Sanger Institute COVID-19 Surveillance Team |
| EPI_ISL_3248695 | Lighthouse Lab in Milton Keynes | Wellcome Sanger Institute for the COVID-19 Genomics UK (COG-UK) Consortium | The Lighthouse Lab in Milton Keynes and Alex Alderton, Roberto Amato, Jeffrey Barrett, Sonia Goncalves, Ewan Harrison, David K. Jackson, Ian Johnston, Dominic Kwiatkowski, Cordelia Langford, John Sillitoe on behalf of the Wellcome Sanger Institute COVID-19 Surveillance Team |
| EPI_ISL_3248696 | Berkshire and Surrey Pathology Services Lighthouse Laboratory | Wellcome Sanger Institute for the COVID-19 Genomics UK (COG-UK) Consortium | Berkshire and Surrey Pathology Services Lighthouse Laboratory and Alex Alderton, Roberto Amato, Jeffrey Barrett, Sonia Goncalves, Ewan Harrison, David K. Jackson, Ian Johnston, Dominic Kwiatkowski, Cordelia Langford, John Sillitoe on behalf of the Wellcome Sanger Institute COVID-19 Surveillance Team |
| EPI_ISL_3248697 | Lighthouse Lab in Milton Keynes | Wellcome Sanger Institute for the COVID-19 Genomics UK (COG-UK) Consortium | The Lighthouse Lab in Milton Keynes and Alex Alderton, Roberto Amato, Jeffrey Barrett, Sonia Goncalves, Ewan Harrison, David K. Jackson, Ian Johnston, Dominic Kwiatkowski, Cordelia Langford, John Sillitoe on behalf of the Wellcome Sanger Institute COVID-19 Surveillance Team |
| EPI_ISL_3248698 | Lighthouse Lab in Alderley Park | Wellcome Sanger Institute for the COVID-19 Genomics UK (COG-UK) Consortium | Jacquelyn Wynn, Mairead Hyland, The Lighthouse Lab in Alderley Park and Alex Alderton, Roberto Amato, Jeffrey Barrett, Sonia Goncalves, Ewan Harrison, David K. Jackson, Ian Johnston, Dominic Kwiatkowski, Cordelia Langford, John Sillitoe on behalf of the Wellcome Sanger Institute COVID-19 Surveillance Team |
| EPI_ISL_3248699 | Berkshire and Surrey Pathology Services Lighthouse Laboratory | Wellcome Sanger Institute for the COVID-19 Genomics UK (COG-UK) Consortium | Berkshire and Surrey Pathology Services Lighthouse Laboratory and Alex Alderton, Roberto Amato, Jeffrey Barrett, Sonia Goncalves, Ewan Harrison, David K. Jackson, Ian Johnston, Dominic Kwiatkowski, Cordelia Langford, John Sillitoe on behalf of the Wellcome Sanger Institute COVID-19 Surveillance Team |
| EPI_ISL_3248700, EPI_ISL_3248701 | Lighthouse Lab in Milton Keynes | Wellcome Sanger Institute for the COVID-19 Genomics UK (COG-UK) Consortium | The Lighthouse Lab in Milton Keynes and Alex Alderton, Roberto Amato, Jeffrey Barrett, Sonia Goncalves, Ewan Harrison, David K. Jackson, Ian Johnston, Dominic Kwiatkowski, Cordelia Langford, John Sillitoe on behalf of the Wellcome Sanger Institute COVID-19 Surveillance Team |
| EPI_ISL_3248702 | Lighthouse Lab in Alderley Park | Wellcome Sanger Institute for the COVID-19 Genomics UK (COG-UK) Consortium | Jacquelyn Wynn, Mairead Hyland, The Lighthouse Lab in Alderley Park and Alex Alderton, Roberto Amato, Jeffrey Barrett, Sonia Goncalves, Ewan Harrison, David K. Jackson, Ian Johnston, Dominic Kwiatkowski, Cordelia Langford, John Sillitoe on behalf of the Wellcome Sanger Institute COVID-19 Surveillance Team |
| EPI_ISL_3248703 | Berkshire and Surrey Pathology Services Lighthouse Laboratory | Wellcome Sanger Institute for the COVID-19 Genomics UK (COG-UK) Consortium | Berkshire and Surrey Pathology Services Lighthouse Laboratory and Alex Alderton, Roberto Amato, Jeffrey Barrett, Sonia Goncalves, Ewan Harrison, David K. Jackson, Ian Johnston, Dominic Kwiatkowski, Cordelia Langford, John Sillitoe on behalf of the Wellcome Sanger Institute COVID-19 Surveillance Team |
| EPI_ISL_3248704 | Lighthouse Lab in Milton Keynes | Wellcome Sanger Institute for the COVID-19 Genomics UK (COG-UK) Consortium | The Lighthouse Lab in Milton Keynes and Alex Alderton, Roberto Amato, Jeffrey Barrett, Sonia Goncalves, Ewan Harrison, David K. Jackson, Ian Johnston, Dominic Kwiatkowski, Cordelia Langford, John Sillitoe on behalf of the Wellcome Sanger Institute COVID-19 Surveillance Team |
| EPI_ISL_3248705 | Berkshire and Surrey Pathology Services Lighthouse Laboratory | Wellcome Sanger Institute for the COVID-19 Genomics UK (COG-UK) Consortium | Berkshire and Surrey Pathology Services Lighthouse Laboratory and Alex Alderton, Roberto Amato, Jeffrey Barrett, Sonia Goncalves, Ewan Harrison, David K. Jackson, Ian Johnston, Dominic Kwiatkowski, Cordelia Langford, John Sillitoe on behalf of the Wellcome Sanger Institute COVID-19 Surveillance Team |
| EPI_ISL_3248706, EPI_ISL_3248707 | Lighthouse Lab in Alderley Park | Wellcome Sanger Institute for the COVID-19 Genomics UK (COG-UK) Consortium | Jacquelyn Wynn, Mairead Hyland, The Lighthouse Lab in Alderley Park and Alex Alderton, Roberto Amato, Jeffrey Barrett, Sonia Goncalves, Ewan Harrison, David K. Jackson, Ian Johnston, Dominic Kwiatkowski, Cordelia Langford, John Sillitoe on behalf of the Wellcome Sanger Institute COVID-19 Surveillance Team |
| EPI_ISL_3248708 | Lighthouse Lab in Glasgow | Wellcome Sanger Institute for the COVID-19 Genomics UK (COG-UK) Consortium | Harper VanSteenhouse, Yumi Kasai, David Gray, Carol Clugston, Anna Dominiczak and Alex Alderton, Roberto Amato, Jeffrey Barrett, Sonia Goncalves, Ewan Harrison, David K. Jackson, Ian Johnston, Dominic Kwiatkowski, Cordelia Langford, John Sillitoe on behalf of the Wellcome Sanger Institute COVID-19 Surveillance Team |
| EPI_ISL_3248709, EPI_ISL_3248710 | Berkshire and Surrey Pathology Services Lighthouse Laboratory | Wellcome Sanger Institute for the COVID-19 Genomics UK (COG-UK) Consortium | Berkshire and Surrey Pathology Services Lighthouse Laboratory and Alex Alderton, Roberto Amato, Jeffrey Barrett, Sonia Goncalves, Ewan Harrison, David K. Jackson, Ian Johnston, Dominic Kwiatkowski, Cordelia Langford, John Sillitoe on behalf of the Wellcome Sanger Institute COVID-19 Surveillance Team |
| EPI_ISL_3248711, EPI_ISL_3248712 | Lighthouse Lab in Milton Keynes | Wellcome Sanger Institute for the COVID-19 Genomics UK (COG-UK) Consortium | The Lighthouse Lab in Milton Keynes and Alex Alderton, Roberto Amato, Jeffrey Barrett, Sonia Goncalves, Ewan Harrison, David K. Jackson, Ian Johnston, Dominic Kwiatkowski, Cordelia Langford, John Sillitoe on behalf of the Wellcome Sanger Institute COVID-19 Surveillance Team |
| EPI_ISL_3248713 | Berkshire and Surrey Pathology Services Lighthouse Laboratory | Wellcome Sanger Institute for the COVID-19 Genomics UK (COG-UK) Consortium | Berkshire and Surrey Pathology Services Lighthouse Laboratory and Alex Alderton, Roberto Amato, Jeffrey Barrett, Sonia Goncalves, Ewan Harrison, David K. Jackson, Ian Johnston, Dominic Kwiatkowski, Cordelia Langford, John Sillitoe on behalf of the Wellcome Sanger Institute COVID-19 Surveillance Team |
| EPI_ISL_3248714 | Lighthouse Lab in Milton Keynes | Wellcome Sanger Institute for the COVID-19 Genomics UK (COG-UK) Consortium | The Lighthouse Lab in Milton Keynes and Alex Alderton, Roberto Amato, Jeffrey Barrett, Sonia Goncalves, Ewan Harrison, David K. Jackson, Ian Johnston, Dominic Kwiatkowski, Cordelia Langford, John Sillitoe on behalf of the Wellcome Sanger Institute COVID-19 Surveillance Team |
| EPI_ISL_3248715 | Berkshire and Surrey Pathology Services Lighthouse Laboratory | Wellcome Sanger Institute for the COVID-19 Genomics UK (COG-UK) Consortium | Berkshire and Surrey Pathology Services Lighthouse Laboratory and Alex Alderton, Roberto Amato, Jeffrey Barrett, Sonia Goncalves, Ewan Harrison, David K. Jackson, Ian Johnston, Dominic Kwiatkowski, Cordelia Langford, John Sillitoe on behalf of the Wellcome Sanger Institute COVID-19 Surveillance Team |
| EPI_ISL_3248716 | Lighthouse Lab in Milton Keynes | Wellcome Sanger Institute for the COVID-19 Genomics UK (COG-UK) Consortium | The Lighthouse Lab in Milton Keynes and Alex Alderton, Roberto Amato, Jeffrey Barrett, Sonia Goncalves, Ewan Harrison, David K. Jackson, Ian Johnston, Dominic Kwiatkowski, Cordelia Langford, John Sillitoe on behalf of the Wellcome Sanger Institute COVID-19 Surveillance Team |
| EPI_ISL_3248717 | Lighthouse Lab in Glasgow | Wellcome Sanger Institute for the COVID-19 Genomics UK (COG-UK) | Harper VanSteenhouse, Yumi Kasai, David Gray, Carol Clugston, Anna Dominiczak and Alex Alderton, Roberto Amato, Jeffrey Barrett, Sonia Goncalves, Ewan Harrison, David K. Jackson, Ian Johnston, Dominic Kwiatkowski, Cordelia Langford, |

| | | | |
|---|---|---|---|
| | | Consortium | John Sillitoe on behalf of the Wellcome Sanger Institute COVID-19 Surveillance Team |
| EPI_ISL_3248718 | Berkshire and Surrey Pathology Services Lighthouse Laboratory | Wellcome Sanger Institute for the COVID-19 Genomics UK (COG-UK) Consortium | Berkshire and Surrey Pathology Services Lighthouse Laboratory and Alex Alderton, Roberto Amato, Jeffrey Barrett, Sonia Goncalves, Ewan Harrison, David K. Jackson, Ian Johnston, Dominic Kwiatkowski, Cordelia Langford, John Sillitoe on behalf of the Wellcome Sanger Institute COVID-19 Surveillance Team |
| EPI_ISL_3248719 | Lighthouse Lab in Glasgow | Wellcome Sanger Institute for the COVID-19 Genomics UK (COG-UK) Consortium | Harper VanSteenhouse, Yumi Kasai, David Gray, Carol Clugston, Anna Dominiczak and Alex Alderton, Roberto Amato, Jeffrey Barrett, Sonia Goncalves, Ewan Harrison, David K. Jackson, Ian Johnston, Dominic Kwiatkowski, Cordelia Langford, John Sillitoe on behalf of the Wellcome Sanger Institute COVID-19 Surveillance Team |
| EPI_ISL_3248720 | Berkshire and Surrey Pathology Services Lighthouse Laboratory | Wellcome Sanger Institute for the COVID-19 Genomics UK (COG-UK) Consortium | Berkshire and Surrey Pathology Services Lighthouse Laboratory and Alex Alderton, Roberto Amato, Jeffrey Barrett, Sonia Goncalves, Ewan Harrison, David K. Jackson, Ian Johnston, Dominic Kwiatkowski, Cordelia Langford, John Sillitoe on behalf of the Wellcome Sanger Institute COVID-19 Surveillance Team |
| EPI_ISL_3248721, EPI_ISL_3248722 | Lighthouse Lab in Alderley Park | Wellcome Sanger Institute for the COVID-19 Genomics UK (COG-UK) Consortium | Jacquelyn Wynn, Mairead Hyland, The Lighthouse Lab in Alderley Park and Alex Alderton, Roberto Amato, Jeffrey Barrett, Sonia Goncalves, Ewan Harrison, David K. Jackson, Ian Johnston, Dominic Kwiatkowski, Cordelia Langford, John Sillitoe on behalf of the Wellcome Sanger Institute COVID-19 Surveillance Team |
| EPI_ISL_3248723, EPI_ISL_3248724 | Lighthouse Lab in Glasgow | Wellcome Sanger Institute for the COVID-19 Genomics UK (COG-UK) Consortium | Harper VanSteenhouse, Yumi Kasai, David Gray, Carol Clugston, Anna Dominiczak and Alex Alderton, Roberto Amato, Jeffrey Barrett, Sonia Goncalves, Ewan Harrison, David K. Jackson, Ian Johnston, Dominic Kwiatkowski, Cordelia Langford, John Sillitoe on behalf of the Wellcome Sanger Institute COVID-19 Surveillance Team |
| EPI_ISL_3248725 | Lighthouse Lab in Alderley Park | Wellcome Sanger Institute for the COVID-19 Genomics UK (COG-UK) Consortium | Jacquelyn Wynn, Mairead Hyland, The Lighthouse Lab in Alderley Park and Alex Alderton, Roberto Amato, Jeffrey Barrett, Sonia Goncalves, Ewan Harrison, David K. Jackson, Ian Johnston, Dominic Kwiatkowski, Cordelia Langford, John Sillitoe on behalf of the Wellcome Sanger Institute COVID-19 Surveillance Team |
| EPI_ISL_3248726 | Lighthouse Lab in Glasgow | Wellcome Sanger Institute for the COVID-19 Genomics UK (COG-UK) Consortium | Harper VanSteenhouse, Yumi Kasai, David Gray, Carol Clugston, Anna Dominiczak and Alex Alderton, Roberto Amato, Jeffrey Barrett, Sonia Goncalves, Ewan Harrison, David K. Jackson, Ian Johnston, Dominic Kwiatkowski, Cordelia Langford, John Sillitoe on behalf of the Wellcome Sanger Institute COVID-19 Surveillance Team |
| EPI_ISL_3248727 | Lighthouse Lab in Milton Keynes | Wellcome Sanger Institute for the COVID-19 Genomics UK (COG-UK) Consortium | The Lighthouse Lab in Milton Keynes and Alex Alderton, Roberto Amato, Jeffrey Barrett, Sonia Goncalves, Ewan Harrison, David K. Jackson, Ian Johnston, Dominic Kwiatkowski, Cordelia Langford, John Sillitoe on behalf of the Wellcome Sanger Institute COVID-19 Surveillance Team |
| EPI_ISL_3248728 | Lighthouse Lab in Glasgow | Wellcome Sanger Institute for the COVID-19 Genomics UK (COG-UK) Consortium | Harper VanSteenhouse, Yumi Kasai, David Gray, Carol Clugston, Anna Dominiczak and Alex Alderton, Roberto Amato, Jeffrey Barrett, Sonia Goncalves, Ewan Harrison, David K. Jackson, Ian Johnston, Dominic Kwiatkowski, Cordelia Langford, John Sillitoe on behalf of the Wellcome Sanger Institute COVID-19 Surveillance Team |
| EPI_ISL_3248729 | Lighthouse Lab in Milton Keynes | Wellcome Sanger Institute for the COVID-19 Genomics UK (COG-UK) Consortium | The Lighthouse Lab in Milton Keynes and Alex Alderton, Roberto Amato, Jeffrey Barrett, Sonia Goncalves, Ewan Harrison, David K. Jackson, Ian Johnston, Dominic Kwiatkowski, Cordelia Langford, John Sillitoe on behalf of the Wellcome Sanger Institute COVID-19 Surveillance Team |
| EPI_ISL_3248730 | Lighthouse Lab in Glasgow | Wellcome Sanger Institute for the COVID-19 Genomics UK (COG-UK) Consortium | Harper VanSteenhouse, Yumi Kasai, David Gray, Carol Clugston, Anna Dominiczak and Alex Alderton, Roberto Amato, Jeffrey Barrett, Sonia Goncalves, Ewan Harrison, David K. Jackson, Ian Johnston, Dominic Kwiatkowski, Cordelia Langford, John Sillitoe on behalf of the Wellcome Sanger Institute COVID-19 Surveillance Team |
| EPI_ISL_3248731, EPI_ISL_3248732, EPI_ISL_3248733, EPI_ISL_3248734 | Berkshire and Surrey Pathology Services Lighthouse Laboratory | Wellcome Sanger Institute for the COVID-19 Genomics UK (COG-UK) Consortium | Berkshire and Surrey Pathology Services Lighthouse Laboratory and Alex Alderton, Roberto Amato, Jeffrey Barrett, Sonia Goncalves, Ewan Harrison, David K. Jackson, Ian Johnston, Dominic Kwiatkowski, Cordelia Langford, John Sillitoe on behalf of the Wellcome Sanger Institute COVID-19 Surveillance Team |
| EPI_ISL_3248735 | Lighthouse Lab in Milton Keynes | Wellcome Sanger Institute for the COVID-19 Genomics UK (COG-UK) Consortium | The Lighthouse Lab in Milton Keynes and Alex Alderton, Roberto Amato, Jeffrey Barrett, Sonia Goncalves, Ewan Harrison, David K. Jackson, Ian Johnston, Dominic Kwiatkowski, Cordelia Langford, John Sillitoe on behalf of the Wellcome Sanger Institute COVID-19 Surveillance Team |
| EPI_ISL_3248736 | Lighthouse Lab in Glasgow | Wellcome Sanger Institute for the COVID-19 Genomics UK (COG-UK) Consortium | Harper VanSteenhouse, Yumi Kasai, David Gray, Carol Clugston, Anna Dominiczak and Alex Alderton, Roberto Amato, Jeffrey Barrett, Sonia Goncalves, Ewan Harrison, David K. Jackson, Ian Johnston, Dominic Kwiatkowski, Cordelia Langford, John Sillitoe on behalf of the Wellcome Sanger Institute COVID-19 Surveillance Team |
| EPI_ISL_3248737, EPI_ISL_3248738, EPI_ISL_3248739, EPI_ISL_3248740 | Lighthouse Lab in Alderley Park | Wellcome Sanger Institute for the COVID-19 Genomics UK (COG-UK) Consortium | Jacquelyn Wynn, Mairead Hyland, The Lighthouse Lab in Alderley Park and Alex Alderton, Roberto Amato, Jeffrey Barrett, Sonia Goncalves, Ewan Harrison, David K. Jackson, Ian Johnston, Dominic Kwiatkowski, Cordelia Langford, John Sillitoe on behalf of the Wellcome Sanger Institute COVID-19 Surveillance Team |
| EPI_ISL_3248741 | Berkshire and Surrey Pathology Services Lighthouse Laboratory | Wellcome Sanger Institute for the COVID-19 Genomics UK (COG-UK) Consortium | Berkshire and Surrey Pathology Services Lighthouse Laboratory and Alex Alderton, Roberto Amato, Jeffrey Barrett, Sonia Goncalves, Ewan Harrison, David K. Jackson, Ian Johnston, Dominic Kwiatkowski, Cordelia Langford, John Sillitoe on behalf of the Wellcome Sanger Institute COVID-19 Surveillance Team |
| EPI_ISL_3248742, EPI_ISL_3248743 | Lighthouse Lab in Milton Keynes | Wellcome Sanger Institute for the COVID-19 Genomics UK (COG-UK) Consortium | The Lighthouse Lab in Milton Keynes and Alex Alderton, Roberto Amato, Jeffrey Barrett, Sonia Goncalves, Ewan Harrison, David K. Jackson, Ian Johnston, Dominic Kwiatkowski, Cordelia Langford, John Sillitoe on behalf of the Wellcome Sanger Institute COVID-19 Surveillance Team |
| EPI_ISL_3248744 | Lighthouse Lab in Glasgow | Wellcome Sanger Institute for the COVID-19 Genomics UK (COG-UK) Consortium | Harper VanSteenhouse, Yumi Kasai, David Gray, Carol Clugston, Anna Dominiczak and Alex Alderton, Roberto Amato, Jeffrey Barrett, Sonia Goncalves, Ewan Harrison, David K. Jackson, Ian Johnston, Dominic Kwiatkowski, Cordelia Langford, John Sillitoe on behalf of the Wellcome Sanger Institute COVID-19 Surveillance Team |
| EPI_ISL_3248745 | Lighthouse Lab in Milton Keynes | Wellcome Sanger Institute for the COVID-19 Genomics UK (COG-UK) Consortium | The Lighthouse Lab in Milton Keynes and Alex Alderton, Roberto Amato, Jeffrey Barrett, Sonia Goncalves, Ewan Harrison, David K. Jackson, Ian Johnston, Dominic Kwiatkowski, Cordelia Langford, John Sillitoe on behalf of the Wellcome Sanger Institute COVID-19 Surveillance Team |
| EPI_ISL_3248746 | Lighthouse Lab in Glasgow | Wellcome Sanger Institute for the COVID-19 Genomics UK (COG-UK) Consortium | Harper VanSteenhouse, Yumi Kasai, David Gray, Carol Clugston, Anna Dominiczak and Alex Alderton, Roberto Amato, Jeffrey Barrett, Sonia Goncalves, Ewan Harrison, David K. Jackson, Ian Johnston, Dominic Kwiatkowski, Cordelia Langford, John Sillitoe on behalf of the Wellcome Sanger Institute COVID-19 Surveillance Team |
| EPI_ISL_3248747 | Berkshire and Surrey Pathology Services Lighthouse Laboratory | Wellcome Sanger Institute for the COVID-19 Genomics UK (COG-UK) Consortium | Berkshire and Surrey Pathology Services Lighthouse Laboratory and Alex Alderton, Roberto Amato, Jeffrey Barrett, Sonia Goncalves, Ewan Harrison, David K. Jackson, Ian Johnston, Dominic Kwiatkowski, Cordelia Langford, John Sillitoe on behalf of the Wellcome Sanger Institute COVID-19 Surveillance Team |
| EPI_ISL_3248748, EPI_ISL_3248749 | Lighthouse Lab in Milton Keynes | Wellcome Sanger Institute for the COVID-19 Genomics UK (COG-UK) Consortium | The Lighthouse Lab in Milton Keynes and Alex Alderton, Roberto Amato, Jeffrey Barrett, Sonia Goncalves, Ewan Harrison, David K. Jackson, Ian Johnston, Dominic Kwiatkowski, Cordelia Langford, John Sillitoe on behalf of the Wellcome Sanger Institute COVID-19 Surveillance Team |
| EPI_ISL_3248750 | Berkshire and Surrey Pathology Services Lighthouse Laboratory | Wellcome Sanger Institute for the COVID-19 Genomics UK (COG-UK) Consortium | Berkshire and Surrey Pathology Services Lighthouse Laboratory and Alex Alderton, Roberto Amato, Jeffrey Barrett, Sonia Goncalves, Ewan Harrison, David K. Jackson, Ian Johnston, Dominic Kwiatkowski, Cordelia Langford, John Sillitoe on behalf of the Wellcome Sanger Institute COVID-19 Surveillance Team |
| EPI_ISL_3248751, EPI_ISL_3248752 | Lighthouse Lab in Alderley Park | Wellcome Sanger Institute for the COVID-19 Genomics UK (COG-UK) Consortium | Jacquelyn Wynn, Mairead Hyland, The Lighthouse Lab in Alderley Park and Alex Alderton, Roberto Amato, Jeffrey Barrett, Sonia Goncalves, Ewan Harrison, David K. Jackson, Ian Johnston, Dominic Kwiatkowski, Cordelia Langford, John Sillitoe on behalf of the Wellcome Sanger Institute COVID-19 Surveillance Team |
| EPI_ISL_3248753 | Berkshire and Surrey Pathology Services Lighthouse Laboratory | Wellcome Sanger Institute for the COVID-19 Genomics UK (COG-UK) Consortium | Berkshire and Surrey Pathology Services Lighthouse Laboratory and Alex Alderton, Roberto Amato, Jeffrey Barrett, Sonia Goncalves, Ewan Harrison, David K. Jackson, Ian Johnston, Dominic Kwiatkowski, Cordelia Langford, John Sillitoe on behalf of the Wellcome Sanger Institute COVID-19 Surveillance Team |
| EPI_ISL_3248754 | Lighthouse Lab in Glasgow | Wellcome Sanger Institute for the COVID-19 Genomics UK (COG-UK) Consortium | Harper VanSteenhouse, Yumi Kasai, David Gray, Carol Clugston, Anna Dominiczak and Alex Alderton, Roberto Amato, Jeffrey Barrett, Sonia Goncalves, Ewan Harrison, David K. Jackson, Ian Johnston, Dominic Kwiatkowski, Cordelia Langford, John Sillitoe on behalf of the Wellcome Sanger Institute COVID-19 Surveillance Team |
| EPI_ISL_3248755 | Lighthouse Lab in Alderley Park | Wellcome Sanger Institute for the COVID-19 Genomics UK (COG-UK) Consortium | Jacquelyn Wynn, Mairead Hyland, The Lighthouse Lab in Alderley Park and Alex Alderton, Roberto Amato, Jeffrey Barrett, Sonia Goncalves, Ewan Harrison, David K. Jackson, Ian Johnston, Dominic Kwiatkowski, Cordelia Langford, John Sillitoe on behalf of the Wellcome Sanger Institute COVID-19 Surveillance Team |
| EPI_ISL_3248756, EPI_ISL_3248757 | Lighthouse Lab in Glasgow | Wellcome Sanger Institute for the COVID-19 Genomics UK (COG-UK) Consortium | Harper VanSteenhouse, Yumi Kasai, David Gray, Carol Clugston, Anna Dominiczak and Alex Alderton, Roberto Amato, Jeffrey Barrett, Sonia Goncalves, Ewan Harrison, David K. Jackson, Ian Johnston, Dominic Kwiatkowski, Cordelia Langford, John Sillitoe on behalf of the Wellcome Sanger Institute COVID-19 Surveillance Team |
| EPI_ISL_3248758, EPI_ISL_3248759, EPI_ISL_3248760 | Lighthouse Lab in Alderley Park | Wellcome Sanger Institute for the COVID-19 Genomics UK (COG-UK) Consortium | Jacquelyn Wynn, Mairead Hyland, The Lighthouse Lab in Alderley Park and Alex Alderton, Roberto Amato, Jeffrey Barrett, Sonia Goncalves, Ewan Harrison, David K. Jackson, Ian Johnston, Dominic Kwiatkowski, Cordelia Langford, John Sillitoe on behalf of the Wellcome Sanger Institute COVID-19 Surveillance Team |
| EPI_ISL_3248761 | Lighthouse Lab in Glasgow | Wellcome Sanger Institute for the COVID-19 Genomics UK (COG-UK) Consortium | Harper VanSteenhouse, Yumi Kasai, David Gray, Carol Clugston, Anna Dominiczak and Alex Alderton, Roberto Amato, Jeffrey Barrett, Sonia Goncalves, Ewan Harrison, David K. Jackson, Ian Johnston, Dominic Kwiatkowski, Cordelia Langford, John Sillitoe on behalf of the Wellcome Sanger Institute COVID-19 Surveillance Team |
| EPI_ISL_3248762 | Lighthouse Lab in Milton Keynes | Wellcome Sanger Institute for the COVID-19 Genomics UK (COG-UK) Consortium | The Lighthouse Lab in Milton Keynes and Alex Alderton, Roberto Amato, Jeffrey Barrett, Sonia Goncalves, Ewan Harrison, David K. Jackson, Ian Johnston, Dominic Kwiatkowski, Cordelia Langford, John Sillitoe on behalf of the Wellcome Sanger Institute COVID-19 Surveillance Team |
| EPI_ISL_3248763 | Lighthouse Lab in Glasgow | Wellcome Sanger Institute for the COVID-19 Genomics UK (COG-UK) Consortium | Harper VanSteenhouse, Yumi Kasai, David Gray, Carol Clugston, Anna Dominiczak and Alex Alderton, Roberto Amato, Jeffrey Barrett, Sonia Goncalves, Ewan Harrison, David K. Jackson, Ian Johnston, Dominic Kwiatkowski, Cordelia Langford, John Sillitoe on behalf of the Wellcome Sanger Institute COVID-19 Surveillance Team |
| EPI_ISL_3248764, EPI_ISL_3248765 | Lighthouse Lab in Milton Keynes | Wellcome Sanger Institute for the COVID-19 Genomics UK (COG-UK) Consortium | The Lighthouse Lab in Milton Keynes and Alex Alderton, Roberto Amato, Jeffrey Barrett, Sonia Goncalves, Ewan Harrison, David K. Jackson, Ian Johnston, Dominic Kwiatkowski, Cordelia Langford, John Sillitoe on behalf of the Wellcome Sanger Institute COVID-19 Surveillance Team |
| EPI_ISL_3248766 | Lighthouse Lab in Alderley Park | Wellcome Sanger Institute for the COVID-19 Genomics UK (COG-UK) Consortium | Jacquelyn Wynn, Mairead Hyland, The Lighthouse Lab in Alderley Park and Alex Alderton, Roberto Amato, Jeffrey Barrett, Sonia Goncalves, Ewan Harrison, David K. Jackson, Ian Johnston, Dominic Kwiatkowski, Cordelia Langford, John Sillitoe on behalf of the Wellcome Sanger Institute COVID-19 Surveillance Team |
| EPI_ISL_3248767 | Lighthouse Lab in Milton Keynes | Wellcome Sanger Institute for the COVID-19 Genomics UK (COG-UK) Consortium | The Lighthouse Lab in Milton Keynes and Alex Alderton, Roberto Amato, Jeffrey Barrett, Sonia Goncalves, Ewan Harrison, David K. Jackson, Ian Johnston, Dominic Kwiatkowski, Cordelia Langford, John Sillitoe on behalf of the Wellcome Sanger Institute COVID-19 Surveillance Team |
| EPI_ISL_3248768 | Lighthouse Lab in Glasgow | Wellcome Sanger Institute for the COVID-19 Genomics UK (COG-UK) Consortium | Harper VanSteenhouse, Yumi Kasai, David Gray, Carol Clugston, Anna Dominiczak and Alex Alderton, Roberto Amato, Jeffrey Barrett, Sonia Goncalves, Ewan Harrison, David K. Jackson, Ian Johnston, Dominic Kwiatkowski, Cordelia Langford, John Sillitoe on behalf of the Wellcome Sanger Institute COVID-19 Surveillance Team |
| EPI_ISL_3248769 | Berkshire and Surrey Pathology Services Lighthouse Laboratory | Wellcome Sanger Institute for the COVID-19 Genomics UK (COG-UK) Consortium | Berkshire and Surrey Pathology Services Lighthouse Laboratory and Alex Alderton, Roberto Amato, Jeffrey Barrett, Sonia Goncalves, Ewan Harrison, David K. Jackson, Ian Johnston, Dominic Kwiatkowski, Cordelia Langford, John Sillitoe on behalf of the Wellcome Sanger Institute COVID-19 Surveillance Team |
| EPI_ISL_3248770, EPI_ISL_3248771 | Lighthouse Lab in Glasgow | Wellcome Sanger Institute for the COVID-19 Genomics UK (COG-UK) Consortium | Harper VanSteenhouse, Yumi Kasai, David Gray, Carol Clugston, Anna Dominiczak and Alex Alderton, Roberto Amato, Jeffrey Barrett, Sonia Goncalves, Ewan Harrison, David K. Jackson, Ian Johnston, Dominic Kwiatkowski, Cordelia Langford, John Sillitoe on behalf of the Wellcome Sanger Institute COVID-19 Surveillance Team |
| EPI_ISL_3248772 | Berkshire and Surrey Pathology Services Lighthouse Laboratory | Wellcome Sanger Institute for the COVID-19 Genomics UK (COG-UK) Consortium | Berkshire and Surrey Pathology Services Lighthouse Laboratory and Alex Alderton, Roberto Amato, Jeffrey Barrett, Sonia Goncalves, Ewan Harrison, David K. Jackson, Ian Johnston, Dominic Kwiatkowski, Cordelia Langford, John Sillitoe on behalf of the Wellcome Sanger Institute COVID-19 Surveillance Team |
| EPI_ISL_3248773 | Lighthouse Lab in Glasgow | Wellcome Sanger Institute for the COVID-19 Genomics UK (COG-UK) Consortium | Harper VanSteenhouse, Yumi Kasai, David Gray, Carol Clugston, Anna Dominiczak and Alex Alderton, Roberto Amato, Jeffrey Barrett, Sonia Goncalves, Ewan Harrison, David K. Jackson, Ian Johnston, Dominic Kwiatkowski, Cordelia Langford, John Sillitoe on behalf of the Wellcome Sanger Institute COVID-19 Surveillance Team |
| EPI_ISL_3248774 | Lighthouse Lab in Milton Keynes | Wellcome Sanger Institute for the COVID-19 Genomics UK (COG-UK) Consortium | The Lighthouse Lab in Milton Keynes and Alex Alderton, Roberto Amato, Jeffrey Barrett, Sonia Goncalves, Ewan Harrison, David K. Jackson, Ian Johnston, Dominic Kwiatkowski, Cordelia Langford, John Sillitoe on behalf of the Wellcome Sanger Institute COVID-19 Surveillance Team |
| EPI_ISL_3248775 | Lighthouse Lab in Alderley Park | Wellcome Sanger Institute for the COVID-19 Genomics UK (COG-UK) Consortium | Jacquelyn Wynn, Mairead Hyland, The Lighthouse Lab in Alderley Park and Alex Alderton, Roberto Amato, Jeffrey Barrett, Sonia Goncalves, Ewan Harrison, David K. Jackson, Ian Johnston, Dominic Kwiatkowski, Cordelia Langford, John Sillitoe on behalf of the Wellcome Sanger Institute COVID-19 Surveillance Team |
| EPI_ISL_3248776 | Lighthouse Lab in Glasgow | Wellcome Sanger Institute for the COVID-19 Genomics UK (COG-UK) | Harper VanSteenhouse, Yumi Kasai, David Gray, Carol Clugston, Anna Dominiczak and Alex Alderton, Roberto Amato, Jeffrey Barrett, Sonia Goncalves, Ewan Harrison, David K. Jackson, Ian Johnston, Dominic Kwiatkowski, Cordelia Langford, |

| | | | |
|---|---|---|---|
| | | Consortium | John Sillitoe on behalf of the Wellcome Sanger Institute COVID-19 Surveillance Team |
| EPI_ISL_3248777 | Lighthouse Lab in Milton Keynes | Wellcome Sanger Institute for the COVID-19 Genomics UK (COG-UK) Consortium | The Lighthouse Lab in Milton Keynes and Alex Alderton, Roberto Amato, Jeffrey Barrett, Sonia Goncalves, Ewan Harrison, David K. Jackson, Ian Johnston, Dominic Kwiatkowski, Cordelia Langford, John Sillitoe on behalf of the Wellcome Sanger Institute COVID-19 Surveillance Team |
| EPI_ISL_3248778 | Berkshire and Surrey Pathology Services Lighthouse Laboratory | Wellcome Sanger Institute for the COVID-19 Genomics UK (COG-UK) Consortium | Berkshire and Surrey Pathology Services Lighthouse Laboratory and Alex Alderton, Roberto Amato, Jeffrey Barrett, Sonia Goncalves, Ewan Harrison, David K. Jackson, Ian Johnston, Dominic Kwiatkowski, Cordelia Langford, John Sillitoe on behalf of the Wellcome Sanger Institute COVID-19 Surveillance Team |
| EPI_ISL_3248779, EPI_ISL_3248780 | Lighthouse Lab in Glasgow | Wellcome Sanger Institute for the COVID-19 Genomics UK (COG-UK) Consortium | Harper VanSteenhouse, Yumi Kasai, David Gray, Carol Clugston, Anna Dominiczak and Alex Alderton, Roberto Amato, Jeffrey Barrett, Sonia Goncalves, Ewan Harrison, David K. Jackson, Ian Johnston, Dominic Kwiatkowski, Cordelia Langford, John Sillitoe on behalf of the Wellcome Sanger Institute COVID-19 Surveillance Team |
| EPI_ISL_3248781 | Berkshire and Surrey Pathology Services Lighthouse Laboratory | Wellcome Sanger Institute for the COVID-19 Genomics UK (COG-UK) Consortium | Berkshire and Surrey Pathology Services Lighthouse Laboratory and Alex Alderton, Roberto Amato, Jeffrey Barrett, Sonia Goncalves, Ewan Harrison, David K. Jackson, Ian Johnston, Dominic Kwiatkowski, Cordelia Langford, John Sillitoe on behalf of the Wellcome Sanger Institute COVID-19 Surveillance Team |
| EPI_ISL_3248782 | Lighthouse Lab in Alderley Park | Wellcome Sanger Institute for the COVID-19 Genomics UK (COG-UK) Consortium | Jacquelyn Wynn, Mairead Hyland, The Lighthouse Lab in Alderley Park and Alex Alderton, Roberto Amato, Jeffrey Barrett, Sonia Goncalves, Ewan Harrison, David K. Jackson, Ian Johnston, Dominic Kwiatkowski, Cordelia Langford, John Sillitoe on behalf of the Wellcome Sanger Institute COVID-19 Surveillance Team |
| EPI_ISL_3248783 | Berkshire and Surrey Pathology Services Lighthouse Laboratory | Wellcome Sanger Institute for the COVID-19 Genomics UK (COG-UK) Consortium | Berkshire and Surrey Pathology Services Lighthouse Laboratory and Alex Alderton, Roberto Amato, Jeffrey Barrett, Sonia Goncalves, Ewan Harrison, David K. Jackson, Ian Johnston, Dominic Kwiatkowski, Cordelia Langford, John Sillitoe on behalf of the Wellcome Sanger Institute COVID-19 Surveillance Team |
| EPI_ISL_3248784 | Lighthouse Lab in Alderley Park | Wellcome Sanger Institute for the COVID-19 Genomics UK (COG-UK) Consortium | Jacquelyn Wynn, Mairead Hyland, The Lighthouse Lab in Alderley Park and Alex Alderton, Roberto Amato, Jeffrey Barrett, Sonia Goncalves, Ewan Harrison, David K. Jackson, Ian Johnston, Dominic Kwiatkowski, Cordelia Langford, John Sillitoe on behalf of the Wellcome Sanger Institute COVID-19 Surveillance Team |
| EPI_ISL_3248785, EPI_ISL_3248786 | Berkshire and Surrey Pathology Services Lighthouse Laboratory | Wellcome Sanger Institute for the COVID-19 Genomics UK (COG-UK) Consortium | Berkshire and Surrey Pathology Services Lighthouse Laboratory and Alex Alderton, Roberto Amato, Jeffrey Barrett, Sonia Goncalves, Ewan Harrison, David K. Jackson, Ian Johnston, Dominic Kwiatkowski, Cordelia Langford, John Sillitoe on behalf of the Wellcome Sanger Institute COVID-19 Surveillance Team |
| EPI_ISL_3248787, EPI_ISL_3248788 | Lighthouse Lab in Milton Keynes | Wellcome Sanger Institute for the COVID-19 Genomics UK (COG-UK) Consortium | The Lighthouse Lab in Milton Keynes and Alex Alderton, Roberto Amato, Jeffrey Barrett, Sonia Goncalves, Ewan Harrison, David K. Jackson, Ian Johnston, Dominic Kwiatkowski, Cordelia Langford, John Sillitoe on behalf of the Wellcome Sanger Institute COVID-19 Surveillance Team |
| EPI_ISL_3248789 | Lighthouse Lab in Glasgow | Wellcome Sanger Institute for the COVID-19 Genomics UK (COG-UK) Consortium | Harper VanSteenhouse, Yumi Kasai, David Gray, Carol Clugston, Anna Dominiczak and Alex Alderton, Roberto Amato, Jeffrey Barrett, Sonia Goncalves, Ewan Harrison, David K. Jackson, Ian Johnston, Dominic Kwiatkowski, Cordelia Langford, John Sillitoe on behalf of the Wellcome Sanger Institute COVID-19 Surveillance Team |
| EPI_ISL_3248790 | Lighthouse Lab in Alderley Park | Wellcome Sanger Institute for the COVID-19 Genomics UK (COG-UK) Consortium | Jacquelyn Wynn, Mairead Hyland, The Lighthouse Lab in Alderley Park and Alex Alderton, Roberto Amato, Jeffrey Barrett, Sonia Goncalves, Ewan Harrison, David K. Jackson, Ian Johnston, Dominic Kwiatkowski, Cordelia Langford, John Sillitoe on behalf of the Wellcome Sanger Institute COVID-19 Surveillance Team |
| EPI_ISL_3248791 | Berkshire and Surrey Pathology Services Lighthouse Laboratory | Wellcome Sanger Institute for the COVID-19 Genomics UK (COG-UK) Consortium | Berkshire and Surrey Pathology Services Lighthouse Laboratory and Alex Alderton, Roberto Amato, Jeffrey Barrett, Sonia Goncalves, Ewan Harrison, David K. Jackson, Ian Johnston, Dominic Kwiatkowski, Cordelia Langford, John Sillitoe on behalf of the Wellcome Sanger Institute COVID-19 Surveillance Team |
| EPI_ISL_3248792 | Lighthouse Lab in Milton Keynes | Wellcome Sanger Institute for the COVID-19 Genomics UK (COG-UK) Consortium | The Lighthouse Lab in Milton Keynes and Alex Alderton, Roberto Amato, Jeffrey Barrett, Sonia Goncalves, Ewan Harrison, David K. Jackson, Ian Johnston, Dominic Kwiatkowski, Cordelia Langford, John Sillitoe on behalf of the Wellcome Sanger Institute COVID-19 Surveillance Team |
| EPI_ISL_3248793 | Lighthouse Lab in Alderley Park | Wellcome Sanger Institute for the COVID-19 Genomics UK (COG-UK) Consortium | Jacquelyn Wynn, Mairead Hyland, The Lighthouse Lab in Alderley Park and Alex Alderton, Roberto Amato, Jeffrey Barrett, Sonia Goncalves, Ewan Harrison, David K. Jackson, Ian Johnston, Dominic Kwiatkowski, Cordelia Langford, John Sillitoe on behalf of the Wellcome Sanger Institute COVID-19 Surveillance Team |
| EPI_ISL_3248794 | Lighthouse Lab in Glasgow | Wellcome Sanger Institute for the COVID-19 Genomics UK (COG-UK) Consortium | Harper VanSteenhouse, Yumi Kasai, David Gray, Carol Clugston, Anna Dominiczak and Alex Alderton, Roberto Amato, Jeffrey Barrett, Sonia Goncalves, Ewan Harrison, David K. Jackson, Ian Johnston, Dominic Kwiatkowski, Cordelia Langford, John Sillitoe on behalf of the Wellcome Sanger Institute COVID-19 Surveillance Team |
| EPI_ISL_3248795, EPI_ISL_3248796 | Berkshire and Surrey Pathology Services Lighthouse Laboratory | Wellcome Sanger Institute for the COVID-19 Genomics UK (COG-UK) Consortium | Berkshire and Surrey Pathology Services Lighthouse Laboratory and Alex Alderton, Roberto Amato, Jeffrey Barrett, Sonia Goncalves, Ewan Harrison, David K. Jackson, Ian Johnston, Dominic Kwiatkowski, Cordelia Langford, John Sillitoe on behalf of the Wellcome Sanger Institute COVID-19 Surveillance Team |
| EPI_ISL_3248797, EPI_ISL_3248798 | Lighthouse Lab in Alderley Park | Wellcome Sanger Institute for the COVID-19 Genomics UK (COG-UK) Consortium | Jacquelyn Wynn, Mairead Hyland, The Lighthouse Lab in Alderley Park and Alex Alderton, Roberto Amato, Jeffrey Barrett, Sonia Goncalves, Ewan Harrison, David K. Jackson, Ian Johnston, Dominic Kwiatkowski, Cordelia Langford, John Sillitoe on behalf of the Wellcome Sanger Institute COVID-19 Surveillance Team |
| EPI_ISL_3248799, EPI_ISL_3248800 | Lighthouse Lab in Glasgow | Wellcome Sanger Institute for the COVID-19 Genomics UK (COG-UK) Consortium | Harper VanSteenhouse, Yumi Kasai, David Gray, Carol Clugston, Anna Dominiczak and Alex Alderton, Roberto Amato, Jeffrey Barrett, Sonia Goncalves, Ewan Harrison, David K. Jackson, Ian Johnston, Dominic Kwiatkowski, Cordelia Langford, John Sillitoe on behalf of the Wellcome Sanger Institute COVID-19 Surveillance Team |
| EPI_ISL_3248801 | Berkshire and Surrey Pathology Services Lighthouse Laboratory | Wellcome Sanger Institute for the COVID-19 Genomics UK (COG-UK) Consortium | Berkshire and Surrey Pathology Services Lighthouse Laboratory and Alex Alderton, Roberto Amato, Jeffrey Barrett, Sonia Goncalves, Ewan Harrison, David K. Jackson, Ian Johnston, Dominic Kwiatkowski, Cordelia Langford, John Sillitoe on behalf of the Wellcome Sanger Institute COVID-19 Surveillance Team |
| EPI_ISL_3248802, EPI_ISL_3248803, EPI_ISL_3248804 | Lighthouse Lab in Milton Keynes | Wellcome Sanger Institute for the COVID-19 Genomics UK (COG-UK) Consortium | The Lighthouse Lab in Milton Keynes and Alex Alderton, Roberto Amato, Jeffrey Barrett, Sonia Goncalves, Ewan Harrison, David K. Jackson, Ian Johnston, Dominic Kwiatkowski, Cordelia Langford, John Sillitoe on behalf of the Wellcome Sanger Institute COVID-19 Surveillance Team |
| EPI_ISL_3248805 | Lighthouse Lab in Glasgow | Wellcome Sanger Institute for the COVID-19 Genomics UK (COG-UK) Consortium | Harper VanSteenhouse, Yumi Kasai, David Gray, Carol Clugston, Anna Dominiczak and Alex Alderton, Roberto Amato, Jeffrey Barrett, Sonia Goncalves, Ewan Harrison, David K. Jackson, Ian Johnston, Dominic Kwiatkowski, Cordelia Langford, John Sillitoe on behalf of the Wellcome Sanger Institute COVID-19 Surveillance Team |
| EPI_ISL_3248806, EPI_ISL_3248807 | Berkshire and Surrey Pathology Services Lighthouse Laboratory | Wellcome Sanger Institute for the COVID-19 Genomics UK (COG-UK) Consortium | Berkshire and Surrey Pathology Services Lighthouse Laboratory and Alex Alderton, Roberto Amato, Jeffrey Barrett, Sonia Goncalves, Ewan Harrison, David K. Jackson, Ian Johnston, Dominic Kwiatkowski, Cordelia Langford, John Sillitoe on behalf of the Wellcome Sanger Institute COVID-19 Surveillance Team |
| EPI_ISL_3248808 | Lighthouse Lab in Alderley Park | Wellcome Sanger Institute for the COVID-19 Genomics UK (COG-UK) Consortium | Jacquelyn Wynn, Mairead Hyland, The Lighthouse Lab in Alderley Park and Alex Alderton, Roberto Amato, Jeffrey Barrett, Sonia Goncalves, Ewan Harrison, David K. Jackson, Ian Johnston, Dominic Kwiatkowski, Cordelia Langford, John Sillitoe on behalf of the Wellcome Sanger Institute COVID-19 Surveillance Team |
| EPI_ISL_3248809 | Berkshire and Surrey Pathology Services Lighthouse Laboratory | Wellcome Sanger Institute for the COVID-19 Genomics UK (COG-UK) Consortium | Berkshire and Surrey Pathology Services Lighthouse Laboratory and Alex Alderton, Roberto Amato, Jeffrey Barrett, Sonia Goncalves, Ewan Harrison, David K. Jackson, Ian Johnston, Dominic Kwiatkowski, Cordelia Langford, John Sillitoe on behalf of the Wellcome Sanger Institute COVID-19 Surveillance Team |
| EPI_ISL_3248810 | Lighthouse Lab in Alderley Park | Wellcome Sanger Institute for the COVID-19 Genomics UK (COG-UK) Consortium | Jacquelyn Wynn, Mairead Hyland, The Lighthouse Lab in Alderley Park and Alex Alderton, Roberto Amato, Jeffrey Barrett, Sonia Goncalves, Ewan Harrison, David K. Jackson, Ian Johnston, Dominic Kwiatkowski, Cordelia Langford, John Sillitoe on behalf of the Wellcome Sanger Institute COVID-19 Surveillance Team |
| EPI_ISL_3248811 | Lighthouse Lab in Milton Keynes | Wellcome Sanger Institute for the COVID-19 Genomics UK (COG-UK) Consortium | The Lighthouse Lab in Milton Keynes and Alex Alderton, Roberto Amato, Jeffrey Barrett, Sonia Goncalves, Ewan Harrison, David K. Jackson, Ian Johnston, Dominic Kwiatkowski, Cordelia Langford, John Sillitoe on behalf of the Wellcome Sanger Institute COVID-19 Surveillance Team |
| EPI_ISL_3248812 | Lighthouse Lab in Glasgow | Wellcome Sanger Institute for the COVID-19 Genomics UK (COG-UK) Consortium | Harper VanSteenhouse, Yumi Kasai, David Gray, Carol Clugston, Anna Dominiczak and Alex Alderton, Roberto Amato, Jeffrey Barrett, Sonia Goncalves, Ewan Harrison, David K. Jackson, Ian Johnston, Dominic Kwiatkowski, Cordelia Langford, John Sillitoe on behalf of the Wellcome Sanger Institute COVID-19 Surveillance Team |
| EPI_ISL_3248813 | Berkshire and Surrey Pathology Services Lighthouse Laboratory | Wellcome Sanger Institute for the COVID-19 Genomics UK (COG-UK) Consortium | Berkshire and Surrey Pathology Services Lighthouse Laboratory and Alex Alderton, Roberto Amato, Jeffrey Barrett, Sonia Goncalves, Ewan Harrison, David K. Jackson, Ian Johnston, Dominic Kwiatkowski, Cordelia Langford, John Sillitoe on behalf of the Wellcome Sanger Institute COVID-19 Surveillance Team |
| EPI_ISL_3248814, EPI_ISL_3248815 | Lighthouse Lab in Alderley Park | Wellcome Sanger Institute for the COVID-19 Genomics UK (COG-UK) Consortium | Jacquelyn Wynn, Mairead Hyland, The Lighthouse Lab in Alderley Park and Alex Alderton, Roberto Amato, Jeffrey Barrett, Sonia Goncalves, Ewan Harrison, David K. Jackson, Ian Johnston, Dominic Kwiatkowski, Cordelia Langford, John Sillitoe on behalf of the Wellcome Sanger Institute COVID-19 Surveillance Team |
| EPI_ISL_3248816 | Berkshire and Surrey Pathology Services Lighthouse Laboratory | Wellcome Sanger Institute for the COVID-19 Genomics UK (COG-UK) Consortium | Berkshire and Surrey Pathology Services Lighthouse Laboratory and Alex Alderton, Roberto Amato, Jeffrey Barrett, Sonia Goncalves, Ewan Harrison, David K. Jackson, Ian Johnston, Dominic Kwiatkowski, Cordelia Langford, John Sillitoe on behalf of the Wellcome Sanger Institute COVID-19 Surveillance Team |
| EPI_ISL_3248817, EPI_ISL_3248818 | Lighthouse Lab in Milton Keynes | Wellcome Sanger Institute for the COVID-19 Genomics UK (COG-UK) Consortium | The Lighthouse Lab in Milton Keynes and Alex Alderton, Roberto Amato, Jeffrey Barrett, Sonia Goncalves, Ewan Harrison, David K. Jackson, Ian Johnston, Dominic Kwiatkowski, Cordelia Langford, John Sillitoe on behalf of the Wellcome Sanger Institute COVID-19 Surveillance Team |
| EPI_ISL_3248819 | Lighthouse Lab in Alderley Park | Wellcome Sanger Institute for the COVID-19 Genomics UK (COG-UK) Consortium | Jacquelyn Wynn, Mairead Hyland, The Lighthouse Lab in Alderley Park and Alex Alderton, Roberto Amato, Jeffrey Barrett, Sonia Goncalves, Ewan Harrison, David K. Jackson, Ian Johnston, Dominic Kwiatkowski, Cordelia Langford, John Sillitoe on behalf of the Wellcome Sanger Institute COVID-19 Surveillance Team |
| EPI_ISL_3248820 | Berkshire and Surrey Pathology Services Lighthouse Laboratory | Wellcome Sanger Institute for the COVID-19 Genomics UK (COG-UK) Consortium | Berkshire and Surrey Pathology Services Lighthouse Laboratory and Alex Alderton, Roberto Amato, Jeffrey Barrett, Sonia Goncalves, Ewan Harrison, David K. Jackson, Ian Johnston, Dominic Kwiatkowski, Cordelia Langford, John Sillitoe on behalf of the Wellcome Sanger Institute COVID-19 Surveillance Team |
| EPI_ISL_3248821 | Lighthouse Lab in Milton Keynes | Wellcome Sanger Institute for the COVID-19 Genomics UK (COG-UK) Consortium | The Lighthouse Lab in Milton Keynes and Alex Alderton, Roberto Amato, Jeffrey Barrett, Sonia Goncalves, Ewan Harrison, David K. Jackson, Ian Johnston, Dominic Kwiatkowski, Cordelia Langford, John Sillitoe on behalf of the Wellcome Sanger Institute COVID-19 Surveillance Team |
| EPI_ISL_3248822 | Lighthouse Lab in Glasgow | Wellcome Sanger Institute for the COVID-19 Genomics UK (COG-UK) Consortium | Harper VanSteenhouse, Yumi Kasai, David Gray, Carol Clugston, Anna Dominiczak and Alex Alderton, Roberto Amato, Jeffrey Barrett, Sonia Goncalves, Ewan Harrison, David K. Jackson, Ian Johnston, Dominic Kwiatkowski, Cordelia Langford, John Sillitoe on behalf of the Wellcome Sanger Institute COVID-19 Surveillance Team |
| EPI_ISL_3248823 | Berkshire and Surrey Pathology Services Lighthouse Laboratory | Wellcome Sanger Institute for the COVID-19 Genomics UK (COG-UK) Consortium | Berkshire and Surrey Pathology Services Lighthouse Laboratory and Alex Alderton, Roberto Amato, Jeffrey Barrett, Sonia Goncalves, Ewan Harrison, David K. Jackson, Ian Johnston, Dominic Kwiatkowski, Cordelia Langford, John Sillitoe on behalf of the Wellcome Sanger Institute COVID-19 Surveillance Team |
| EPI_ISL_3248824 | Lighthouse Lab in Alderley Park | Wellcome Sanger Institute for the COVID-19 Genomics UK (COG-UK) Consortium | Jacquelyn Wynn, Mairead Hyland, The Lighthouse Lab in Alderley Park and Alex Alderton, Roberto Amato, Jeffrey Barrett, Sonia Goncalves, Ewan Harrison, David K. Jackson, Ian Johnston, Dominic Kwiatkowski, Cordelia Langford, John Sillitoe on behalf of the Wellcome Sanger Institute COVID-19 Surveillance Team |
| EPI_ISL_3248825, EPI_ISL_3248826, EPI_ISL_3248827 | Lighthouse Lab in Milton Keynes | Wellcome Sanger Institute for the COVID-19 Genomics UK (COG-UK) Consortium | The Lighthouse Lab in Milton Keynes and Alex Alderton, Roberto Amato, Jeffrey Barrett, Sonia Goncalves, Ewan Harrison, David K. Jackson, Ian Johnston, Dominic Kwiatkowski, Cordelia Langford, John Sillitoe on behalf of the Wellcome Sanger Institute COVID-19 Surveillance Team |
| EPI_ISL_3248828 | Lighthouse Lab in Alderley Park | Wellcome Sanger Institute for the COVID-19 Genomics UK (COG-UK) Consortium | Jacquelyn Wynn, Mairead Hyland, The Lighthouse Lab in Alderley Park and Alex Alderton, Roberto Amato, Jeffrey Barrett, Sonia Goncalves, Ewan Harrison, David K. Jackson, Ian Johnston, Dominic Kwiatkowski, Cordelia Langford, John Sillitoe on behalf of the Wellcome Sanger Institute COVID-19 Surveillance Team |
| EPI_ISL_3248829, EPI_ISL_3248830 | Lighthouse Lab in Glasgow | Wellcome Sanger Institute for the COVID-19 Genomics UK (COG-UK) Consortium | Harper VanSteenhouse, Yumi Kasai, David Gray, Carol Clugston, Anna Dominiczak and Alex Alderton, Roberto Amato, Jeffrey Barrett, Sonia Goncalves, Ewan Harrison, David K. Jackson, Ian Johnston, Dominic Kwiatkowski, Cordelia Langford, John Sillitoe on behalf of the Wellcome Sanger Institute COVID-19 Surveillance Team |
| EPI_ISL_3248831 | Berkshire and Surrey Pathology Services Lighthouse Laboratory | Wellcome Sanger Institute for the COVID-19 Genomics UK (COG-UK) Consortium | Berkshire and Surrey Pathology Services Lighthouse Laboratory and Alex Alderton, Roberto Amato, Jeffrey Barrett, Sonia Goncalves, Ewan Harrison, David K. Jackson, Ian Johnston, Dominic Kwiatkowski, Cordelia Langford, John Sillitoe on behalf of the Wellcome Sanger Institute COVID-19 Surveillance Team |
| EPI_ISL_3248832 | Lighthouse Lab in Alderley Park | Wellcome Sanger Institute for the COVID-19 Genomics UK (COG-UK) Consortium | Jacquelyn Wynn, Mairead Hyland, The Lighthouse Lab in Alderley Park and Alex Alderton, Roberto Amato, Jeffrey Barrett, Sonia Goncalves, Ewan Harrison, David K. Jackson, Ian Johnston, Dominic Kwiatkowski, Cordelia Langford, John Sillitoe on behalf of the Wellcome Sanger Institute COVID-19 Surveillance Team |
| EPI_ISL_3248833, EPI_ISL_3248834 | Lighthouse Lab in Glasgow | Wellcome Sanger Institute for the COVID-19 Genomics UK (COG-UK) Consortium | Harper VanSteenhouse, Yumi Kasai, David Gray, Carol Clugston, Anna Dominiczak and Alex Alderton, Roberto Amato, Jeffrey Barrett, Sonia Goncalves, Ewan Harrison, David K. Jackson, Ian Johnston, Dominic Kwiatkowski, Cordelia Langford, John Sillitoe on behalf of the Wellcome Sanger Institute COVID-19 Surveillance Team |
| EPI_ISL_3248835 | Lighthouse Lab in Alderley Park | Wellcome Sanger Institute for the COVID-19 Genomics UK (COG-UK) | Jacquelyn Wynn, Mairead Hyland, The Lighthouse Lab in Alderley Park and Alex Alderton, Roberto Amato, Jeffrey Barrett, Sonia Goncalves, Ewan Harrison, David K. Jackson, Ian Johnston, Dominic Kwiatkowski, Cordelia Langford, John Sillitoe on |

| | | | |
|---|---|---|---|
| | | Consortium | behalf of the Wellcome Sanger Institute COVID-19 Surveillance Team |
| EPI_ISL_3248836 | Berkshire and Surrey Pathology Services Lighthouse Laboratory | Wellcome Sanger Institute for the COVID-19 Genomics UK (COG-UK) Consortium | Berkshire and Surrey Pathology Services Lighthouse Laboratory and Alex Alderton, Roberto Amato, Jeffrey Barrett, Sonia Goncalves, Ewan Harrison, David K. Jackson, Ian Johnston, Dominic Kwiatkowski, Cordelia Langford, John Sillitoe on behalf of the Wellcome Sanger Institute COVID-19 Surveillance Team |
| EPI_ISL_3248837 | Lighthouse Lab in Glasgow | Wellcome Sanger Institute for the COVID-19 Genomics UK (COG-UK) Consortium | Harper VanSteenhouse, Yumi Kasai, David Gray, Carol Clugston, Anna Dominiczak and Alex Alderton, Roberto Amato, Jeffrey Barrett, Sonia Goncalves, Ewan Harrison, David K. Jackson, Ian Johnston, Dominic Kwiatkowski, Cordelia Langford, John Sillitoe on behalf of the Wellcome Sanger Institute COVID-19 Surveillance Team |
| EPI_ISL_3248838 | Lighthouse Lab in Milton Keynes | Wellcome Sanger Institute for the COVID-19 Genomics UK (COG-UK) Consortium | The Lighthouse Lab in Milton Keynes and Alex Alderton, Roberto Amato, Jeffrey Barrett, Sonia Goncalves, Ewan Harrison, David K. Jackson, Ian Johnston, Dominic Kwiatkowski, Cordelia Langford, John Sillitoe on behalf of the Wellcome Sanger Institute COVID-19 Surveillance Team |
| EPI_ISL_3248839 | Lighthouse Lab in Alderley Park | Wellcome Sanger Institute for the COVID-19 Genomics UK (COG-UK) Consortium | Jacquelyn Wynn, Mairead Hyland, The Lighthouse Lab in Alderley Park and Alex Alderton, Roberto Amato, Jeffrey Barrett, Sonia Goncalves, Ewan Harrison, David K. Jackson, Ian Johnston, Dominic Kwiatkowski, Cordelia Langford, John Sillitoe on behalf of the Wellcome Sanger Institute COVID-19 Surveillance Team |
| EPI_ISL_3248840 | Lighthouse Lab in Milton Keynes | Wellcome Sanger Institute for the COVID-19 Genomics UK (COG-UK) Consortium | The Lighthouse Lab in Milton Keynes and Alex Alderton, Roberto Amato, Jeffrey Barrett, Sonia Goncalves, Ewan Harrison, David K. Jackson, Ian Johnston, Dominic Kwiatkowski, Cordelia Langford, John Sillitoe on behalf of the Wellcome Sanger Institute COVID-19 Surveillance Team |
| EPI_ISL_3248841 | Lighthouse Lab in Glasgow | Wellcome Sanger Institute for the COVID-19 Genomics UK (COG-UK) Consortium | Harper VanSteenhouse, Yumi Kasai, David Gray, Carol Clugston, Anna Dominiczak and Alex Alderton, Roberto Amato, Jeffrey Barrett, Sonia Goncalves, Ewan Harrison, David K. Jackson, Ian Johnston, Dominic Kwiatkowski, Cordelia Langford, John Sillitoe on behalf of the Wellcome Sanger Institute COVID-19 Surveillance Team |
| EPI_ISL_3248842 | Lighthouse Lab in Milton Keynes | Wellcome Sanger Institute for the COVID-19 Genomics UK (COG-UK) Consortium | The Lighthouse Lab in Milton Keynes and Alex Alderton, Roberto Amato, Jeffrey Barrett, Sonia Goncalves, Ewan Harrison, David K. Jackson, Ian Johnston, Dominic Kwiatkowski, Cordelia Langford, John Sillitoe on behalf of the Wellcome Sanger Institute COVID-19 Surveillance Team |
| EPI_ISL_3248843, EPI_ISL_3248844, EPI_ISL_3248845 | Lighthouse Lab in Alderley Park | Wellcome Sanger Institute for the COVID-19 Genomics UK (COG-UK) Consortium | Jacquelyn Wynn, Mairead Hyland, The Lighthouse Lab in Alderley Park and Alex Alderton, Roberto Amato, Jeffrey Barrett, Sonia Goncalves, Ewan Harrison, David K. Jackson, Ian Johnston, Dominic Kwiatkowski, Cordelia Langford, John Sillitoe on behalf of the Wellcome Sanger Institute COVID-19 Surveillance Team |
| EPI_ISL_3248846 | Berkshire and Surrey Pathology Services Lighthouse Laboratory | Wellcome Sanger Institute for the COVID-19 Genomics UK (COG-UK) Consortium | Berkshire and Surrey Pathology Services Lighthouse Laboratory and Alex Alderton, Roberto Amato, Jeffrey Barrett, Sonia Goncalves, Ewan Harrison, David K. Jackson, Ian Johnston, Dominic Kwiatkowski, Cordelia Langford, John Sillitoe on behalf of the Wellcome Sanger Institute COVID-19 Surveillance Team |
| EPI_ISL_3248847, EPI_ISL_3248848, EPI_ISL_3248849, EPI_ISL_3248850 | Lighthouse Lab in Glasgow | Wellcome Sanger Institute for the COVID-19 Genomics UK (COG-UK) Consortium | Harper VanSteenhouse, Yumi Kasai, David Gray, Carol Clugston, Anna Dominiczak and Alex Alderton, Roberto Amato, Jeffrey Barrett, Sonia Goncalves, Ewan Harrison, David K. Jackson, Ian Johnston, Dominic Kwiatkowski, Cordelia Langford, John Sillitoe on behalf of the Wellcome Sanger Institute COVID-19 Surveillance Team |
| EPI_ISL_3248851 | Lighthouse Lab in Alderley Park | Wellcome Sanger Institute for the COVID-19 Genomics UK (COG-UK) Consortium | Jacquelyn Wynn, Mairead Hyland, The Lighthouse Lab in Alderley Park and Alex Alderton, Roberto Amato, Jeffrey Barrett, Sonia Goncalves, Ewan Harrison, David K. Jackson, Ian Johnston, Dominic Kwiatkowski, Cordelia Langford, John Sillitoe on behalf of the Wellcome Sanger Institute COVID-19 Surveillance Team |
| EPI_ISL_3248852, EPI_ISL_3248853, EPI_ISL_3248854 | Lighthouse Lab in Milton Keynes | Wellcome Sanger Institute for the COVID-19 Genomics UK (COG-UK) Consortium | The Lighthouse Lab in Milton Keynes and Alex Alderton, Roberto Amato, Jeffrey Barrett, Sonia Goncalves, Ewan Harrison, David K. Jackson, Ian Johnston, Dominic Kwiatkowski, Cordelia Langford, John Sillitoe on behalf of the Wellcome Sanger Institute COVID-19 Surveillance Team |
| EPI_ISL_3248855, EPI_ISL_3248856 | Berkshire and Surrey Pathology Services Lighthouse Laboratory | Wellcome Sanger Institute for the COVID-19 Genomics UK (COG-UK) Consortium | Berkshire and Surrey Pathology Services Lighthouse Laboratory and Alex Alderton, Roberto Amato, Jeffrey Barrett, Sonia Goncalves, Ewan Harrison, David K. Jackson, Ian Johnston, Dominic Kwiatkowski, Cordelia Langford, John Sillitoe on behalf of the Wellcome Sanger Institute COVID-19 Surveillance Team |
| EPI_ISL_3248857 | Lighthouse Lab in Alderley Park | Wellcome Sanger Institute for the COVID-19 Genomics UK (COG-UK) Consortium | Jacquelyn Wynn, Mairead Hyland, The Lighthouse Lab in Alderley Park and Alex Alderton, Roberto Amato, Jeffrey Barrett, Sonia Goncalves, Ewan Harrison, David K. Jackson, Ian Johnston, Dominic Kwiatkowski, Cordelia Langford, John Sillitoe on behalf of the Wellcome Sanger Institute COVID-19 Surveillance Team |
| EPI_ISL_3248858, EPI_ISL_3248859, EPI_ISL_3248860 | Berkshire and Surrey Pathology Services Lighthouse Laboratory | Wellcome Sanger Institute for the COVID-19 Genomics UK (COG-UK) Consortium | Berkshire and Surrey Pathology Services Lighthouse Laboratory and Alex Alderton, Roberto Amato, Jeffrey Barrett, Sonia Goncalves, Ewan Harrison, David K. Jackson, Ian Johnston, Dominic Kwiatkowski, Cordelia Langford, John Sillitoe on behalf of the Wellcome Sanger Institute COVID-19 Surveillance Team |
| EPI_ISL_3248861 | Lighthouse Lab in Glasgow | Wellcome Sanger Institute for the COVID-19 Genomics UK (COG-UK) Consortium | Harper VanSteenhouse, Yumi Kasai, David Gray, Carol Clugston, Anna Dominiczak and Alex Alderton, Roberto Amato, Jeffrey Barrett, Sonia Goncalves, Ewan Harrison, David K. Jackson, Ian Johnston, Dominic Kwiatkowski, Cordelia Langford, John Sillitoe on behalf of the Wellcome Sanger Institute COVID-19 Surveillance Team |
| EPI_ISL_3248862 | Lighthouse Lab in Milton Keynes | Wellcome Sanger Institute for the COVID-19 Genomics UK (COG-UK) Consortium | The Lighthouse Lab in Milton Keynes and Alex Alderton, Roberto Amato, Jeffrey Barrett, Sonia Goncalves, Ewan Harrison, David K. Jackson, Ian Johnston, Dominic Kwiatkowski, Cordelia Langford, John Sillitoe on behalf of the Wellcome Sanger Institute COVID-19 Surveillance Team |
| EPI_ISL_3248863, EPI_ISL_3248864 | Berkshire and Surrey Pathology Services Lighthouse Laboratory | Wellcome Sanger Institute for the COVID-19 Genomics UK (COG-UK) Consortium | Berkshire and Surrey Pathology Services Lighthouse Laboratory and Alex Alderton, Roberto Amato, Jeffrey Barrett, Sonia Goncalves, Ewan Harrison, David K. Jackson, Ian Johnston, Dominic Kwiatkowski, Cordelia Langford, John Sillitoe on behalf of the Wellcome Sanger Institute COVID-19 Surveillance Team |
| EPI_ISL_3248865, EPI_ISL_3248866, EPI_ISL_3248867 | Lighthouse Lab in Alderley Park | Wellcome Sanger Institute for the COVID-19 Genomics UK (COG-UK) Consortium | Jacquelyn Wynn, Mairead Hyland, The Lighthouse Lab in Alderley Park and Alex Alderton, Roberto Amato, Jeffrey Barrett, Sonia Goncalves, Ewan Harrison, David K. Jackson, Ian Johnston, Dominic Kwiatkowski, Cordelia Langford, John Sillitoe on behalf of the Wellcome Sanger Institute COVID-19 Surveillance Team |
| EPI_ISL_3248868 | Lighthouse Lab in Milton Keynes | Wellcome Sanger Institute for the COVID-19 Genomics UK (COG-UK) Consortium | The Lighthouse Lab in Milton Keynes and Alex Alderton, Roberto Amato, Jeffrey Barrett, Sonia Goncalves, Ewan Harrison, David K. Jackson, Ian Johnston, Dominic Kwiatkowski, Cordelia Langford, John Sillitoe on behalf of the Wellcome Sanger Institute COVID-19 Surveillance Team |
| EPI_ISL_3248869 | Berkshire and Surrey Pathology Services Lighthouse Laboratory | Wellcome Sanger Institute for the COVID-19 Genomics UK (COG-UK) Consortium | Berkshire and Surrey Pathology Services Lighthouse Laboratory and Alex Alderton, Roberto Amato, Jeffrey Barrett, Sonia Goncalves, Ewan Harrison, David K. Jackson, Ian Johnston, Dominic Kwiatkowski, Cordelia Langford, John Sillitoe on behalf of the Wellcome Sanger Institute COVID-19 Surveillance Team |
| EPI_ISL_3248870, EPI_ISL_3248871 | Lighthouse Lab in Alderley Park | Wellcome Sanger Institute for the COVID-19 Genomics UK (COG-UK) Consortium | Jacquelyn Wynn, Mairead Hyland, The Lighthouse Lab in Alderley Park and Alex Alderton, Roberto Amato, Jeffrey Barrett, Sonia Goncalves, Ewan Harrison, David K. Jackson, Ian Johnston, Dominic Kwiatkowski, Cordelia Langford, John Sillitoe on behalf of the Wellcome Sanger Institute COVID-19 Surveillance Team |
| EPI_ISL_3248872 | Berkshire and Surrey Pathology Services Lighthouse Laboratory | Wellcome Sanger Institute for the COVID-19 Genomics UK (COG-UK) Consortium | Berkshire and Surrey Pathology Services Lighthouse Laboratory and Alex Alderton, Roberto Amato, Jeffrey Barrett, Sonia Goncalves, Ewan Harrison, David K. Jackson, Ian Johnston, Dominic Kwiatkowski, Cordelia Langford, John Sillitoe on behalf of the Wellcome Sanger Institute COVID-19 Surveillance Team |
| EPI_ISL_3248873, EPI_ISL_3248874 | Lighthouse Lab in Glasgow | Wellcome Sanger Institute for the COVID-19 Genomics UK (COG-UK) Consortium | Harper VanSteenhouse, Yumi Kasai, David Gray, Carol Clugston, Anna Dominiczak and Alex Alderton, Roberto Amato, Jeffrey Barrett, Sonia Goncalves, Ewan Harrison, David K. Jackson, Ian Johnston, Dominic Kwiatkowski, Cordelia Langford, John Sillitoe on behalf of the Wellcome Sanger Institute COVID-19 Surveillance Team |
| EPI_ISL_3248875 | Lighthouse Lab in Alderley Park | Wellcome Sanger Institute for the COVID-19 Genomics UK (COG-UK) Consortium | Jacquelyn Wynn, Mairead Hyland, The Lighthouse Lab in Alderley Park and Alex Alderton, Roberto Amato, Jeffrey Barrett, Sonia Goncalves, Ewan Harrison, David K. Jackson, Ian Johnston, Dominic Kwiatkowski, Cordelia Langford, John Sillitoe on behalf of the Wellcome Sanger Institute COVID-19 Surveillance Team |
| EPI_ISL_3248876, EPI_ISL_3248877, EPI_ISL_3248878 | Lighthouse Lab in Milton Keynes | Wellcome Sanger Institute for the COVID-19 Genomics UK (COG-UK) Consortium | The Lighthouse Lab in Milton Keynes and Alex Alderton, Roberto Amato, Jeffrey Barrett, Sonia Goncalves, Ewan Harrison, David K. Jackson, Ian Johnston, Dominic Kwiatkowski, Cordelia Langford, John Sillitoe on behalf of the Wellcome Sanger Institute COVID-19 Surveillance Team |
| EPI_ISL_3248879 | Lighthouse Lab in Glasgow | Wellcome Sanger Institute for the COVID-19 Genomics UK (COG-UK) Consortium | Harper VanSteenhouse, Yumi Kasai, David Gray, Carol Clugston, Anna Dominiczak and Alex Alderton, Roberto Amato, Jeffrey Barrett, Sonia Goncalves, Ewan Harrison, David K. Jackson, Ian Johnston, Dominic Kwiatkowski, Cordelia Langford, John Sillitoe on behalf of the Wellcome Sanger Institute COVID-19 Surveillance Team |
| EPI_ISL_3248880 | Berkshire and Surrey Pathology Services Lighthouse Laboratory | Wellcome Sanger Institute for the COVID-19 Genomics UK (COG-UK) Consortium | Berkshire and Surrey Pathology Services Lighthouse Laboratory and Alex Alderton, Roberto Amato, Jeffrey Barrett, Sonia Goncalves, Ewan Harrison, David K. Jackson, Ian Johnston, Dominic Kwiatkowski, Cordelia Langford, John Sillitoe on behalf of the Wellcome Sanger Institute COVID-19 Surveillance Team |
| EPI_ISL_3248881 | Lighthouse Lab in Alderley Park | Wellcome Sanger Institute for the COVID-19 Genomics UK (COG-UK) Consortium | Jacquelyn Wynn, Mairead Hyland, The Lighthouse Lab in Alderley Park and Alex Alderton, Roberto Amato, Jeffrey Barrett, Sonia Goncalves, Ewan Harrison, David K. Jackson, Ian Johnston, Dominic Kwiatkowski, Cordelia Langford, John Sillitoe on behalf of the Wellcome Sanger Institute COVID-19 Surveillance Team |
| EPI_ISL_3248882 | Lighthouse Lab in Milton Keynes | Wellcome Sanger Institute for the COVID-19 Genomics UK (COG-UK) Consortium | The Lighthouse Lab in Milton Keynes and Alex Alderton, Roberto Amato, Jeffrey Barrett, Sonia Goncalves, Ewan Harrison, David K. Jackson, Ian Johnston, Dominic Kwiatkowski, Cordelia Langford, John Sillitoe on behalf of the Wellcome Sanger Institute COVID-19 Surveillance Team |
| EPI_ISL_3248883 | Berkshire and Surrey Pathology Services Lighthouse Laboratory | Wellcome Sanger Institute for the COVID-19 Genomics UK (COG-UK) Consortium | Berkshire and Surrey Pathology Services Lighthouse Laboratory and Alex Alderton, Roberto Amato, Jeffrey Barrett, Sonia Goncalves, Ewan Harrison, David K. Jackson, Ian Johnston, Dominic Kwiatkowski, Cordelia Langford, John Sillitoe on behalf of the Wellcome Sanger Institute COVID-19 Surveillance Team |
| EPI_ISL_3248884 | Lighthouse Lab in Glasgow | Wellcome Sanger Institute for the COVID-19 Genomics UK (COG-UK) Consortium | Harper VanSteenhouse, Yumi Kasai, David Gray, Carol Clugston, Anna Dominiczak and Alex Alderton, Roberto Amato, Jeffrey Barrett, Sonia Goncalves, Ewan Harrison, David K. Jackson, Ian Johnston, Dominic Kwiatkowski, Cordelia Langford, John Sillitoe on behalf of the Wellcome Sanger Institute COVID-19 Surveillance Team |
| EPI_ISL_3248885 | Berkshire and Surrey Pathology Services Lighthouse Laboratory | Wellcome Sanger Institute for the COVID-19 Genomics UK (COG-UK) Consortium | Berkshire and Surrey Pathology Services Lighthouse Laboratory and Alex Alderton, Roberto Amato, Jeffrey Barrett, Sonia Goncalves, Ewan Harrison, David K. Jackson, Ian Johnston, Dominic Kwiatkowski, Cordelia Langford, John Sillitoe on behalf of the Wellcome Sanger Institute COVID-19 Surveillance Team |
| EPI_ISL_3248886 | Lighthouse Lab in Milton Keynes | Wellcome Sanger Institute for the COVID-19 Genomics UK (COG-UK) Consortium | The Lighthouse Lab in Milton Keynes and Alex Alderton, Roberto Amato, Jeffrey Barrett, Sonia Goncalves, Ewan Harrison, David K. Jackson, Ian Johnston, Dominic Kwiatkowski, Cordelia Langford, John Sillitoe on behalf of the Wellcome Sanger Institute COVID-19 Surveillance Team |
| EPI_ISL_3248887 | Lighthouse Lab in Glasgow | Wellcome Sanger Institute for the COVID-19 Genomics UK (COG-UK) Consortium | Harper VanSteenhouse, Yumi Kasai, David Gray, Carol Clugston, Anna Dominiczak and Alex Alderton, Roberto Amato, Jeffrey Barrett, Sonia Goncalves, Ewan Harrison, David K. Jackson, Ian Johnston, Dominic Kwiatkowski, Cordelia Langford, John Sillitoe on behalf of the Wellcome Sanger Institute COVID-19 Surveillance Team |
| EPI_ISL_3248888 | Lighthouse Lab in Alderley Park | Wellcome Sanger Institute for the COVID-19 Genomics UK (COG-UK) Consortium | Jacquelyn Wynn, Mairead Hyland, The Lighthouse Lab in Alderley Park and Alex Alderton, Roberto Amato, Jeffrey Barrett, Sonia Goncalves, Ewan Harrison, David K. Jackson, Ian Johnston, Dominic Kwiatkowski, Cordelia Langford, John Sillitoe on behalf of the Wellcome Sanger Institute COVID-19 Surveillance Team |
| EPI_ISL_3248889 | Berkshire and Surrey Pathology Services Lighthouse Laboratory | Wellcome Sanger Institute for the COVID-19 Genomics UK (COG-UK) Consortium | Berkshire and Surrey Pathology Services Lighthouse Laboratory and Alex Alderton, Roberto Amato, Jeffrey Barrett, Sonia Goncalves, Ewan Harrison, David K. Jackson, Ian Johnston, Dominic Kwiatkowski, Cordelia Langford, John Sillitoe on behalf of the Wellcome Sanger Institute COVID-19 Surveillance Team |
| EPI_ISL_3248890 | Lighthouse Lab in Glasgow | Wellcome Sanger Institute for the COVID-19 Genomics UK (COG-UK) Consortium | Harper VanSteenhouse, Yumi Kasai, David Gray, Carol Clugston, Anna Dominiczak and Alex Alderton, Roberto Amato, Jeffrey Barrett, Sonia Goncalves, Ewan Harrison, David K. Jackson, Ian Johnston, Dominic Kwiatkowski, Cordelia Langford, John Sillitoe on behalf of the Wellcome Sanger Institute COVID-19 Surveillance Team |
| EPI_ISL_3248891 | Lighthouse Lab in Milton Keynes | Wellcome Sanger Institute for the COVID-19 Genomics UK (COG-UK) Consortium | The Lighthouse Lab in Milton Keynes and Alex Alderton, Roberto Amato, Jeffrey Barrett, Sonia Goncalves, Ewan Harrison, David K. Jackson, Ian Johnston, Dominic Kwiatkowski, Cordelia Langford, John Sillitoe on behalf of the Wellcome Sanger Institute COVID-19 Surveillance Team |
| EPI_ISL_3248892 | Berkshire and Surrey Pathology Services Lighthouse Laboratory | Wellcome Sanger Institute for the COVID-19 Genomics UK (COG-UK) Consortium | Berkshire and Surrey Pathology Services Lighthouse Laboratory and Alex Alderton, Roberto Amato, Jeffrey Barrett, Sonia Goncalves, Ewan Harrison, David K. Jackson, Ian Johnston, Dominic Kwiatkowski, Cordelia Langford, John Sillitoe on behalf of the Wellcome Sanger Institute COVID-19 Surveillance Team |
| EPI_ISL_3248893 | Lighthouse Lab in Glasgow | Wellcome Sanger Institute for the COVID-19 Genomics UK (COG-UK) Consortium | Harper VanSteenhouse, Yumi Kasai, David Gray, Carol Clugston, Anna Dominiczak and Alex Alderton, Roberto Amato, Jeffrey Barrett, Sonia Goncalves, Ewan Harrison, David K. Jackson, Ian Johnston, Dominic Kwiatkowski, Cordelia Langford, John Sillitoe on behalf of the Wellcome Sanger Institute COVID-19 Surveillance Team |
| EPI_ISL_3248894 | Lighthouse Lab in Milton Keynes | Wellcome Sanger Institute for the COVID-19 Genomics UK (COG-UK) Consortium | The Lighthouse Lab in Milton Keynes and Alex Alderton, Roberto Amato, Jeffrey Barrett, Sonia Goncalves, Ewan Harrison, David K. Jackson, Ian Johnston, Dominic Kwiatkowski, Cordelia Langford, John Sillitoe on behalf of the Wellcome Sanger Institute COVID-19 Surveillance Team |

| | | | |
|---|---|---|---|
| EPI_ISL_3248895 | Berkshire and Surrey Pathology Services Lighthouse Laboratory | Wellcome Sanger Institute for the COVID-19 Genomics UK (COG-UK) Consortium | Berkshire and Surrey Pathology Services Lighthouse Laboratory and Alex Alderton, Roberto Amato, Jeffrey Barrett, Sonia Goncalves, Ewan Harrison, David K. Jackson, Ian Johnston, Dominic Kwiatkowski, Cordelia Langford, John Sillitoe on behalf of the Wellcome Sanger Institute COVID-19 Surveillance Team |
| EPI_ISL_3248896 | Lighthouse Lab in Alderley Park | Wellcome Sanger Institute for the COVID-19 Genomics UK (COG-UK) Consortium | Jacquelyn Wynn, Mairead Hyland, The Lighthouse Lab in Alderley Park and Alex Alderton, Roberto Amato, Jeffrey Barrett, Sonia Goncalves, Ewan Harrison, David K. Jackson, Ian Johnston, Dominic Kwiatkowski, Cordelia Langford, John Sillitoe on behalf of the Wellcome Sanger Institute COVID-19 Surveillance Team |
| EPI_ISL_3248897, EPI_ISL_3248898 | Lighthouse Lab in Glasgow | Wellcome Sanger Institute for the COVID-19 Genomics UK (COG-UK) Consortium | Harper VanSteenhouse, Yumi Kasai, David Gray, Carol Clugston, Anna Dominiczak and Alex Alderton, Roberto Amato, Jeffrey Barrett, Sonia Goncalves, Ewan Harrison, David K. Jackson, Ian Johnston, Dominic Kwiatkowski, Cordelia Langford, John Sillitoe on behalf of the Wellcome Sanger Institute COVID-19 Surveillance Team |
| EPI_ISL_3248899 | Berkshire and Surrey Pathology Services Lighthouse Laboratory | Wellcome Sanger Institute for the COVID-19 Genomics UK (COG-UK) Consortium | Berkshire and Surrey Pathology Services Lighthouse Laboratory and Alex Alderton, Roberto Amato, Jeffrey Barrett, Sonia Goncalves, Ewan Harrison, David K. Jackson, Ian Johnston, Dominic Kwiatkowski, Cordelia Langford, John Sillitoe on behalf of the Wellcome Sanger Institute COVID-19 Surveillance Team |
| EPI_ISL_3248900 | Lighthouse Lab in Alderley Park | Wellcome Sanger Institute for the COVID-19 Genomics UK (COG-UK) Consortium | Jacquelyn Wynn, Mairead Hyland, The Lighthouse Lab in Alderley Park and Alex Alderton, Roberto Amato, Jeffrey Barrett, Sonia Goncalves, Ewan Harrison, David K. Jackson, Ian Johnston, Dominic Kwiatkowski, Cordelia Langford, John Sillitoe on behalf of the Wellcome Sanger Institute COVID-19 Surveillance Team |
| EPI_ISL_3248901 | Lighthouse Lab in Milton Keynes | Wellcome Sanger Institute for the COVID-19 Genomics UK (COG-UK) Consortium | The Lighthouse Lab in Milton Keynes and Alex Alderton, Roberto Amato, Jeffrey Barrett, Sonia Goncalves, Ewan Harrison, David K. Jackson, Ian Johnston, Dominic Kwiatkowski, Cordelia Langford, John Sillitoe on behalf of the Wellcome Sanger Institute COVID-19 Surveillance Team |
| EPI_ISL_3248902 | Lighthouse Lab in Alderley Park | Wellcome Sanger Institute for the COVID-19 Genomics UK (COG-UK) Consortium | Jacquelyn Wynn, Mairead Hyland, The Lighthouse Lab in Alderley Park and Alex Alderton, Roberto Amato, Jeffrey Barrett, Sonia Goncalves, Ewan Harrison, David K. Jackson, Ian Johnston, Dominic Kwiatkowski, Cordelia Langford, John Sillitoe on behalf of the Wellcome Sanger Institute COVID-19 Surveillance Team |
| EPI_ISL_3248903 | Berkshire and Surrey Pathology Services Lighthouse Laboratory | Wellcome Sanger Institute for the COVID-19 Genomics UK (COG-UK) Consortium | Berkshire and Surrey Pathology Services Lighthouse Laboratory and Alex Alderton, Roberto Amato, Jeffrey Barrett, Sonia Goncalves, Ewan Harrison, David K. Jackson, Ian Johnston, Dominic Kwiatkowski, Cordelia Langford, John Sillitoe on behalf of the Wellcome Sanger Institute COVID-19 Surveillance Team |
| EPI_ISL_3248904, EPI_ISL_3248905, EPI_ISL_3248906 | Lighthouse Lab in Alderley Park | Wellcome Sanger Institute for the COVID-19 Genomics UK (COG-UK) Consortium | Jacquelyn Wynn, Mairead Hyland, The Lighthouse Lab in Alderley Park and Alex Alderton, Roberto Amato, Jeffrey Barrett, Sonia Goncalves, Ewan Harrison, David K. Jackson, Ian Johnston, Dominic Kwiatkowski, Cordelia Langford, John Sillitoe on behalf of the Wellcome Sanger Institute COVID-19 Surveillance Team |
| EPI_ISL_3248907 | Lighthouse Lab in Milton Keynes | Wellcome Sanger Institute for the COVID-19 Genomics UK (COG-UK) Consortium | The Lighthouse Lab in Milton Keynes and Alex Alderton, Roberto Amato, Jeffrey Barrett, Sonia Goncalves, Ewan Harrison, David K. Jackson, Ian Johnston, Dominic Kwiatkowski, Cordelia Langford, John Sillitoe on behalf of the Wellcome Sanger Institute COVID-19 Surveillance Team |
| EPI_ISL_3248908, EPI_ISL_3248909, EPI_ISL_3248910 | Lighthouse Lab in Alderley Park | Wellcome Sanger Institute for the COVID-19 Genomics UK (COG-UK) Consortium | Jacquelyn Wynn, Mairead Hyland, The Lighthouse Lab in Alderley Park and Alex Alderton, Roberto Amato, Jeffrey Barrett, Sonia Goncalves, Ewan Harrison, David K. Jackson, Ian Johnston, Dominic Kwiatkowski, Cordelia Langford, John Sillitoe on behalf of the Wellcome Sanger Institute COVID-19 Surveillance Team |
| EPI_ISL_3248911 | Lighthouse Lab in Glasgow | Wellcome Sanger Institute for the COVID-19 Genomics UK (COG-UK) Consortium | Harper VanSteenhouse, Yumi Kasai, David Gray, Carol Clugston, Anna Dominiczak and Alex Alderton, Roberto Amato, Jeffrey Barrett, Sonia Goncalves, Ewan Harrison, David K. Jackson, Ian Johnston, Dominic Kwiatkowski, Cordelia Langford, John Sillitoe on behalf of the Wellcome Sanger Institute COVID-19 Surveillance Team |
| EPI_ISL_3248912 | Lighthouse Lab in Milton Keynes | Wellcome Sanger Institute for the COVID-19 Genomics UK (COG-UK) Consortium | The Lighthouse Lab in Milton Keynes and Alex Alderton, Roberto Amato, Jeffrey Barrett, Sonia Goncalves, Ewan Harrison, David K. Jackson, Ian Johnston, Dominic Kwiatkowski, Cordelia Langford, John Sillitoe on behalf of the Wellcome Sanger Institute COVID-19 Surveillance Team |
| EPI_ISL_3248913 | Lighthouse Lab in Glasgow | Wellcome Sanger Institute for the COVID-19 Genomics UK (COG-UK) Consortium | Harper VanSteenhouse, Yumi Kasai, David Gray, Carol Clugston, Anna Dominiczak and Alex Alderton, Roberto Amato, Jeffrey Barrett, Sonia Goncalves, Ewan Harrison, David K. Jackson, Ian Johnston, Dominic Kwiatkowski, Cordelia Langford, John Sillitoe on behalf of the Wellcome Sanger Institute COVID-19 Surveillance Team |
| EPI_ISL_3248914 | Lighthouse Lab in Alderley Park | Wellcome Sanger Institute for the COVID-19 Genomics UK (COG-UK) Consortium | Jacquelyn Wynn, Mairead Hyland, The Lighthouse Lab in Alderley Park and Alex Alderton, Roberto Amato, Jeffrey Barrett, Sonia Goncalves, Ewan Harrison, David K. Jackson, Ian Johnston, Dominic Kwiatkowski, Cordelia Langford, John Sillitoe on behalf of the Wellcome Sanger Institute COVID-19 Surveillance Team |
| EPI_ISL_3248915 | Lighthouse Lab in Milton Keynes | Wellcome Sanger Institute for the COVID-19 Genomics UK (COG-UK) Consortium | The Lighthouse Lab in Milton Keynes and Alex Alderton, Roberto Amato, Jeffrey Barrett, Sonia Goncalves, Ewan Harrison, David K. Jackson, Ian Johnston, Dominic Kwiatkowski, Cordelia Langford, John Sillitoe on behalf of the Wellcome Sanger Institute COVID-19 Surveillance Team |
| EPI_ISL_3248916, EPI_ISL_3248917, EPI_ISL_3248918 | Lighthouse Lab in Alderley Park | Wellcome Sanger Institute for the COVID-19 Genomics UK (COG-UK) Consortium | Jacquelyn Wynn, Mairead Hyland, The Lighthouse Lab in Alderley Park and Alex Alderton, Roberto Amato, Jeffrey Barrett, Sonia Goncalves, Ewan Harrison, David K. Jackson, Ian Johnston, Dominic Kwiatkowski, Cordelia Langford, John Sillitoe on behalf of the Wellcome Sanger Institute COVID-19 Surveillance Team |
| EPI_ISL_3248919, EPI_ISL_3248920 | Lighthouse Lab in Milton Keynes | Wellcome Sanger Institute for the COVID-19 Genomics UK (COG-UK) Consortium | The Lighthouse Lab in Milton Keynes and Alex Alderton, Roberto Amato, Jeffrey Barrett, Sonia Goncalves, Ewan Harrison, David K. Jackson, Ian Johnston, Dominic Kwiatkowski, Cordelia Langford, John Sillitoe on behalf of the Wellcome Sanger Institute COVID-19 Surveillance Team |
| EPI_ISL_3248921, EPI_ISL_3248922 | Lighthouse Lab in Alderley Park | Wellcome Sanger Institute for the COVID-19 Genomics UK (COG-UK) Consortium | Jacquelyn Wynn, Mairead Hyland, The Lighthouse Lab in Alderley Park and Alex Alderton, Roberto Amato, Jeffrey Barrett, Sonia Goncalves, Ewan Harrison, David K. Jackson, Ian Johnston, Dominic Kwiatkowski, Cordelia Langford, John Sillitoe on behalf of the Wellcome Sanger Institute COVID-19 Surveillance Team |
| EPI_ISL_3248923, EPI_ISL_3248924 | Lighthouse Lab in Glasgow | Wellcome Sanger Institute for the COVID-19 Genomics UK (COG-UK) Consortium | Harper VanSteenhouse, Yumi Kasai, David Gray, Carol Clugston, Anna Dominiczak and Alex Alderton, Roberto Amato, Jeffrey Barrett, Sonia Goncalves, Ewan Harrison, David K. Jackson, Ian Johnston, Dominic Kwiatkowski, Cordelia Langford, John Sillitoe on behalf of the Wellcome Sanger Institute COVID-19 Surveillance Team |
| EPI_ISL_3248925 | Berkshire and Surrey Pathology Services Lighthouse Laboratory | Wellcome Sanger Institute for the COVID-19 Genomics UK (COG-UK) Consortium | Berkshire and Surrey Pathology Services Lighthouse Laboratory and Alex Alderton, Roberto Amato, Jeffrey Barrett, Sonia Goncalves, Ewan Harrison, David K. Jackson, Ian Johnston, Dominic Kwiatkowski, Cordelia Langford, John Sillitoe on behalf of the Wellcome Sanger Institute COVID-19 Surveillance Team |
| EPI_ISL_3248926 | Lighthouse Lab in Milton Keynes | Wellcome Sanger Institute for the COVID-19 Genomics UK (COG-UK) Consortium | The Lighthouse Lab in Milton Keynes and Alex Alderton, Roberto Amato, Jeffrey Barrett, Sonia Goncalves, Ewan Harrison, David K. Jackson, Ian Johnston, Dominic Kwiatkowski, Cordelia Langford, John Sillitoe on behalf of the Wellcome Sanger Institute COVID-19 Surveillance Team |
| EPI_ISL_3248927, EPI_ISL_3248928 | Lighthouse Lab in Alderley Park | Wellcome Sanger Institute for the COVID-19 Genomics UK (COG-UK) Consortium | Jacquelyn Wynn, Mairead Hyland, The Lighthouse Lab in Alderley Park and Alex Alderton, Roberto Amato, Jeffrey Barrett, Sonia Goncalves, Ewan Harrison, David K. Jackson, Ian Johnston, Dominic Kwiatkowski, Cordelia Langford, John Sillitoe on behalf of the Wellcome Sanger Institute COVID-19 Surveillance Team |
| EPI_ISL_3248929 | Lighthouse Lab in Glasgow | Wellcome Sanger Institute for the COVID-19 Genomics UK (COG-UK) Consortium | Harper VanSteenhouse, Yumi Kasai, David Gray, Carol Clugston, Anna Dominiczak and Alex Alderton, Roberto Amato, Jeffrey Barrett, Sonia Goncalves, Ewan Harrison, David K. Jackson, Ian Johnston, Dominic Kwiatkowski, Cordelia Langford, John Sillitoe on behalf of the Wellcome Sanger Institute COVID-19 Surveillance Team |
| EPI_ISL_3248930 | Lighthouse Lab in Milton Keynes | Wellcome Sanger Institute for the COVID-19 Genomics UK (COG-UK) Consortium | The Lighthouse Lab in Milton Keynes and Alex Alderton, Roberto Amato, Jeffrey Barrett, Sonia Goncalves, Ewan Harrison, David K. Jackson, Ian Johnston, Dominic Kwiatkowski, Cordelia Langford, John Sillitoe on behalf of the Wellcome Sanger Institute COVID-19 Surveillance Team |
| EPI_ISL_3248931 | Lighthouse Lab in Alderley Park | Wellcome Sanger Institute for the COVID-19 Genomics UK (COG-UK) Consortium | Jacquelyn Wynn, Mairead Hyland, The Lighthouse Lab in Alderley Park and Alex Alderton, Roberto Amato, Jeffrey Barrett, Sonia Goncalves, Ewan Harrison, David K. Jackson, Ian Johnston, Dominic Kwiatkowski, Cordelia Langford, John Sillitoe on behalf of the Wellcome Sanger Institute COVID-19 Surveillance Team |
| EPI_ISL_3248932 | Berkshire and Surrey Pathology Services Lighthouse Laboratory | Wellcome Sanger Institute for the COVID-19 Genomics UK (COG-UK) Consortium | Berkshire and Surrey Pathology Services Lighthouse Laboratory and Alex Alderton, Roberto Amato, Jeffrey Barrett, Sonia Goncalves, Ewan Harrison, David K. Jackson, Ian Johnston, Dominic Kwiatkowski, Cordelia Langford, John Sillitoe on behalf of the Wellcome Sanger Institute COVID-19 Surveillance Team |
| EPI_ISL_3248933, EPI_ISL_3248934 | Lighthouse Lab in Alderley Park | Wellcome Sanger Institute for the COVID-19 Genomics UK (COG-UK) Consortium | Jacquelyn Wynn, Mairead Hyland, The Lighthouse Lab in Alderley Park and Alex Alderton, Roberto Amato, Jeffrey Barrett, Sonia Goncalves, Ewan Harrison, David K. Jackson, Ian Johnston, Dominic Kwiatkowski, Cordelia Langford, John Sillitoe on behalf of the Wellcome Sanger Institute COVID-19 Surveillance Team |
| EPI_ISL_3248935 | Lighthouse Lab in Glasgow | Wellcome Sanger Institute for the COVID-19 Genomics UK (COG-UK) Consortium | Harper VanSteenhouse, Yumi Kasai, David Gray, Carol Clugston, Anna Dominiczak and Alex Alderton, Roberto Amato, Jeffrey Barrett, Sonia Goncalves, Ewan Harrison, David K. Jackson, Ian Johnston, Dominic Kwiatkowski, Cordelia Langford, John Sillitoe on behalf of the Wellcome Sanger Institute COVID-19 Surveillance Team |
| EPI_ISL_3248936 | Lighthouse Lab in Alderley Park | Wellcome Sanger Institute for the COVID-19 Genomics UK (COG-UK) Consortium | Jacquelyn Wynn, Mairead Hyland, The Lighthouse Lab in Alderley Park and Alex Alderton, Roberto Amato, Jeffrey Barrett, Sonia Goncalves, Ewan Harrison, David K. Jackson, Ian Johnston, Dominic Kwiatkowski, Cordelia Langford, John Sillitoe on behalf of the Wellcome Sanger Institute COVID-19 Surveillance Team |
| EPI_ISL_3248937 | Lighthouse Lab in Glasgow | Wellcome Sanger Institute for the COVID-19 Genomics UK (COG-UK) Consortium | Harper VanSteenhouse, Yumi Kasai, David Gray, Carol Clugston, Anna Dominiczak and Alex Alderton, Roberto Amato, Jeffrey Barrett, Sonia Goncalves, Ewan Harrison, David K. Jackson, Ian Johnston, Dominic Kwiatkowski, Cordelia Langford, John Sillitoe on behalf of the Wellcome Sanger Institute COVID-19 Surveillance Team |
| EPI_ISL_3248938, EPI_ISL_3248939 | Lighthouse Lab in Alderley Park | Wellcome Sanger Institute for the COVID-19 Genomics UK (COG-UK) Consortium | Jacquelyn Wynn, Mairead Hyland, The Lighthouse Lab in Alderley Park and Alex Alderton, Roberto Amato, Jeffrey Barrett, Sonia Goncalves, Ewan Harrison, David K. Jackson, Ian Johnston, Dominic Kwiatkowski, Cordelia Langford, John Sillitoe on behalf of the Wellcome Sanger Institute COVID-19 Surveillance Team |
| EPI_ISL_3248940 | Berkshire and Surrey Pathology Services Lighthouse Laboratory | Wellcome Sanger Institute for the COVID-19 Genomics UK (COG-UK) Consortium | Berkshire and Surrey Pathology Services Lighthouse Laboratory and Alex Alderton, Roberto Amato, Jeffrey Barrett, Sonia Goncalves, Ewan Harrison, David K. Jackson, Ian Johnston, Dominic Kwiatkowski, Cordelia Langford, John Sillitoe on behalf of the Wellcome Sanger Institute COVID-19 Surveillance Team |
| EPI_ISL_3248941 | Lighthouse Lab in Alderley Park | Wellcome Sanger Institute for the COVID-19 Genomics UK (COG-UK) Consortium | Jacquelyn Wynn, Mairead Hyland, The Lighthouse Lab in Alderley Park and Alex Alderton, Roberto Amato, Jeffrey Barrett, Sonia Goncalves, Ewan Harrison, David K. Jackson, Ian Johnston, Dominic Kwiatkowski, Cordelia Langford, John Sillitoe on behalf of the Wellcome Sanger Institute COVID-19 Surveillance Team |
| EPI_ISL_3248942, EPI_ISL_3248943, EPI_ISL_3248944 | Lighthouse Lab in Milton Keynes | Wellcome Sanger Institute for the COVID-19 Genomics UK (COG-UK) Consortium | The Lighthouse Lab in Milton Keynes and Alex Alderton, Roberto Amato, Jeffrey Barrett, Sonia Goncalves, Ewan Harrison, David K. Jackson, Ian Johnston, Dominic Kwiatkowski, Cordelia Langford, John Sillitoe on behalf of the Wellcome Sanger Institute COVID-19 Surveillance Team |
| EPI_ISL_3248945 | Lighthouse Lab in Alderley Park | Wellcome Sanger Institute for the COVID-19 Genomics UK (COG-UK) Consortium | Jacquelyn Wynn, Mairead Hyland, The Lighthouse Lab in Alderley Park and Alex Alderton, Roberto Amato, Jeffrey Barrett, Sonia Goncalves, Ewan Harrison, David K. Jackson, Ian Johnston, Dominic Kwiatkowski, Cordelia Langford, John Sillitoe on behalf of the Wellcome Sanger Institute COVID-19 Surveillance Team |
| EPI_ISL_3248946 | Lighthouse Lab in Glasgow | Wellcome Sanger Institute for the COVID-19 Genomics UK (COG-UK) Consortium | Harper VanSteenhouse, Yumi Kasai, David Gray, Carol Clugston, Anna Dominiczak and Alex Alderton, Roberto Amato, Jeffrey Barrett, Sonia Goncalves, Ewan Harrison, David K. Jackson, Ian Johnston, Dominic Kwiatkowski, Cordelia Langford, John Sillitoe on behalf of the Wellcome Sanger Institute COVID-19 Surveillance Team |
| EPI_ISL_3248947, EPI_ISL_3248948, EPI_ISL_3248949 | Lighthouse Lab in Milton Keynes | Wellcome Sanger Institute for the COVID-19 Genomics UK (COG-UK) Consortium | The Lighthouse Lab in Milton Keynes and Alex Alderton, Roberto Amato, Jeffrey Barrett, Sonia Goncalves, Ewan Harrison, David K. Jackson, Ian Johnston, Dominic Kwiatkowski, Cordelia Langford, John Sillitoe on behalf of the Wellcome Sanger Institute COVID-19 Surveillance Team |
| EPI_ISL_3248950, EPI_ISL_3248951 | Lighthouse Lab in Glasgow | Wellcome Sanger Institute for the COVID-19 Genomics UK (COG-UK) Consortium | Harper VanSteenhouse, Yumi Kasai, David Gray, Carol Clugston, Anna Dominiczak and Alex Alderton, Roberto Amato, Jeffrey Barrett, Sonia Goncalves, Ewan Harrison, David K. Jackson, Ian Johnston, Dominic Kwiatkowski, Cordelia Langford, John Sillitoe on behalf of the Wellcome Sanger Institute COVID-19 Surveillance Team |
| EPI_ISL_3248952 | Lighthouse Lab in Milton Keynes | Wellcome Sanger Institute for the COVID-19 Genomics UK (COG-UK) Consortium | The Lighthouse Lab in Milton Keynes and Alex Alderton, Roberto Amato, Jeffrey Barrett, Sonia Goncalves, Ewan Harrison, David K. Jackson, Ian Johnston, Dominic Kwiatkowski, Cordelia Langford, John Sillitoe on behalf of the Wellcome Sanger Institute COVID-19 Surveillance Team |
| EPI_ISL_3248953 | ASST MONZA | ASST MONZA | Sergio Maria Ivano Malandrin |
| EPI_ISL_3248954 | Lighthouse Lab in Milton Keynes | Wellcome Sanger Institute for the COVID-19 Genomics UK (COG-UK) Consortium | The Lighthouse Lab in Milton Keynes and Alex Alderton, Roberto Amato, Jeffrey Barrett, Sonia Goncalves, Ewan Harrison, David K. Jackson, Ian Johnston, Dominic Kwiatkowski, Cordelia Langford, John Sillitoe on behalf of the Wellcome Sanger Institute COVID-19 Surveillance Team |
| EPI_ISL_3248955 | Lighthouse Lab in Alderley Park | Wellcome Sanger Institute for the COVID-19 Genomics UK (COG-UK) Consortium | Jacquelyn Wynn, Mairead Hyland, The Lighthouse Lab in Alderley Park and Alex Alderton, Roberto Amato, Jeffrey Barrett, Sonia Goncalves, Ewan Harrison, David K. Jackson, Ian Johnston, Dominic Kwiatkowski, Cordelia Langford, John Sillitoe on behalf of the Wellcome Sanger Institute COVID-19 Surveillance Team |
| EPI_ISL_3248956 | ASST MONZA | ASST MONZA | Sergio Maria Ivano Malandrin |

| | | | |
|---|---|---|---|
| EPI_ISL_3248957, EPI_ISL_3248958 | Lighthouse Lab in Milton Keynes | Wellcome Sanger Institute for the COVID-19 Genomics UK (COG-UK) Consortium | The Lighthouse Lab in Milton Keynes and Alex Alderton, Roberto Amato, Jeffrey Barrett, Sonia Goncalves, Ewan Harrison, David K. Jackson, Ian Johnston, Dominic Kwiatkowski, Cordelia Langford, John Sillitoe on behalf of the Wellcome Sanger Institute COVID-19 Surveillance Team |
| EPI_ISL_3248959 | Lighthouse Lab in Alderley Park | Wellcome Sanger Institute for the COVID-19 Genomics UK (COG-UK) Consortium | Jacquelyn Wynn, Mairead Hyland, The Lighthouse Lab in Alderley Park and Alex Alderton, Roberto Amato, Jeffrey Barrett, Sonia Goncalves, Ewan Harrison, David K. Jackson, Ian Johnston, Dominic Kwiatkowski, Cordelia Langford, John Sillitoe on behalf of the Wellcome Sanger Institute COVID-19 Surveillance Team |
| EPI_ISL_3248960 | ASST MONZA | ASST MONZA | Sergio Maria Ivano Malandrin |
| EPI_ISL_3248961, EPI_ISL_3248962 | Lighthouse Lab in Milton Keynes | Wellcome Sanger Institute for the COVID-19 Genomics UK (COG-UK) Consortium | The Lighthouse Lab in Milton Keynes and Alex Alderton, Roberto Amato, Jeffrey Barrett, Sonia Goncalves, Ewan Harrison, David K. Jackson, Ian Johnston, Dominic Kwiatkowski, Cordelia Langford, John Sillitoe on behalf of the Wellcome Sanger Institute COVID-19 Surveillance Team |
| EPI_ISL_3248963 | Lighthouse Lab in Glasgow | Wellcome Sanger Institute for the COVID-19 Genomics UK (COG-UK) Consortium | Harper VanSteenhouse, Yumi Kasai, David Gray, Carol Clugston, Anna Dominiczak and Alex Alderton, Roberto Amato, Jeffrey Barrett, Sonia Goncalves, Ewan Harrison, David K. Jackson, Ian Johnston, Dominic Kwiatkowski, Cordelia Langford, John Sillitoe on behalf of the Wellcome Sanger Institute COVID-19 Surveillance Team |
| EPI_ISL_3248964 | ASST MONZA | ASST MONZA | Sergio Maria Ivano Malandrin |
| EPI_ISL_3248965 | Lighthouse Lab in Milton Keynes | Wellcome Sanger Institute for the COVID-19 Genomics UK (COG-UK) Consortium | The Lighthouse Lab in Milton Keynes and Alex Alderton, Roberto Amato, Jeffrey Barrett, Sonia Goncalves, Ewan Harrison, David K. Jackson, Ian Johnston, Dominic Kwiatkowski, Cordelia Langford, John Sillitoe on behalf of the Wellcome Sanger Institute COVID-19 Surveillance Team |
| EPI_ISL_3248966 | Lighthouse Lab in Glasgow | Wellcome Sanger Institute for the COVID-19 Genomics UK (COG-UK) Consortium | Harper VanSteenhouse, Yumi Kasai, David Gray, Carol Clugston, Anna Dominiczak and Alex Alderton, Roberto Amato, Jeffrey Barrett, Sonia Goncalves, Ewan Harrison, David K. Jackson, Ian Johnston, Dominic Kwiatkowski, Cordelia Langford, John Sillitoe on behalf of the Wellcome Sanger Institute COVID-19 Surveillance Team |
| EPI_ISL_3248967 | ASST MONZA | ASST MONZA | Sergio Maria Ivano Malandrin |
| EPI_ISL_3248968, EPI_ISL_3248969, EPI_ISL_3248970 | Lighthouse Lab in Milton Keynes | Wellcome Sanger Institute for the COVID-19 Genomics UK (COG-UK) Consortium | The Lighthouse Lab in Milton Keynes and Alex Alderton, Roberto Amato, Jeffrey Barrett, Sonia Goncalves, Ewan Harrison, David K. Jackson, Ian Johnston, Dominic Kwiatkowski, Cordelia Langford, John Sillitoe on behalf of the Wellcome Sanger Institute COVID-19 Surveillance Team |
| EPI_ISL_3248971 | ASST MONZA | ASST MONZA | Sergio Maria Ivano Malandrin |
| EPI_ISL_3248972 | Lighthouse Lab in Alderley Park | Wellcome Sanger Institute for the COVID-19 Genomics UK (COG-UK) Consortium | Jacquelyn Wynn, Mairead Hyland, The Lighthouse Lab in Alderley Park and Alex Alderton, Roberto Amato, Jeffrey Barrett, Sonia Goncalves, Ewan Harrison, David K. Jackson, Ian Johnston, Dominic Kwiatkowski, Cordelia Langford, John Sillitoe on behalf of the Wellcome Sanger Institute COVID-19 Surveillance Team |
| EPI_ISL_3248973, EPI_ISL_3248974 | Lighthouse Lab in Glasgow | Wellcome Sanger Institute for the COVID-19 Genomics UK (COG-UK) Consortium | Harper VanSteenhouse, Yumi Kasai, David Gray, Carol Clugston, Anna Dominiczak and Alex Alderton, Roberto Amato, Jeffrey Barrett, Sonia Goncalves, Ewan Harrison, David K. Jackson, Ian Johnston, Dominic Kwiatkowski, Cordelia Langford, John Sillitoe on behalf of the Wellcome Sanger Institute COVID-19 Surveillance Team |
| EPI_ISL_3248975 | ASST MONZA | ASST MONZA | Sergio Maria Ivano Malandrin |
| EPI_ISL_3248976, EPI_ISL_3248977 | Lighthouse Lab in Milton Keynes | Wellcome Sanger Institute for the COVID-19 Genomics UK (COG-UK) Consortium | The Lighthouse Lab in Milton Keynes and Alex Alderton, Roberto Amato, Jeffrey Barrett, Sonia Goncalves, Ewan Harrison, David K. Jackson, Ian Johnston, Dominic Kwiatkowski, Cordelia Langford, John Sillitoe on behalf of the Wellcome Sanger Institute COVID-19 Surveillance Team |
| EPI_ISL_3248978 | ASST MONZA | ASST MONZA | Sergio Maria Ivano Malandrin |
| EPI_ISL_3248979, EPI_ISL_3248980, EPI_ISL_3248981 | Lighthouse Lab in Milton Keynes | Wellcome Sanger Institute for the COVID-19 Genomics UK (COG-UK) Consortium | The Lighthouse Lab in Milton Keynes and Alex Alderton, Roberto Amato, Jeffrey Barrett, Sonia Goncalves, Ewan Harrison, David K. Jackson, Ian Johnston, Dominic Kwiatkowski, Cordelia Langford, John Sillitoe on behalf of the Wellcome Sanger Institute COVID-19 Surveillance Team |
| EPI_ISL_3248982 | ASST MONZA | ASST MONZA | Sergio Maria Ivano Malandrin |
| EPI_ISL_3248983 | Lighthouse Lab in Alderley Park | Wellcome Sanger Institute for the COVID-19 Genomics UK (COG-UK) Consortium | Jacquelyn Wynn, Mairead Hyland, The Lighthouse Lab in Alderley Park and Alex Alderton, Roberto Amato, Jeffrey Barrett, Sonia Goncalves, Ewan Harrison, David K. Jackson, Ian Johnston, Dominic Kwiatkowski, Cordelia Langford, John Sillitoe on behalf of the Wellcome Sanger Institute COVID-19 Surveillance Team |
| EPI_ISL_3248984 | Lighthouse Lab in Milton Keynes | Wellcome Sanger Institute for the COVID-19 Genomics UK (COG-UK) Consortium | The Lighthouse Lab in Milton Keynes and Alex Alderton, Roberto Amato, Jeffrey Barrett, Sonia Goncalves, Ewan Harrison, David K. Jackson, Ian Johnston, Dominic Kwiatkowski, Cordelia Langford, John Sillitoe on behalf of the Wellcome Sanger Institute COVID-19 Surveillance Team |
| EPI_ISL_3248985 | ASST MONZA | ASST MONZA | Sergio Maria Ivano Malandrin |
| EPI_ISL_3248986 | Lighthouse Lab in Milton Keynes | Wellcome Sanger Institute for the COVID-19 Genomics UK (COG-UK) Consortium | The Lighthouse Lab in Milton Keynes and Alex Alderton, Roberto Amato, Jeffrey Barrett, Sonia Goncalves, Ewan Harrison, David K. Jackson, Ian Johnston, Dominic Kwiatkowski, Cordelia Langford, John Sillitoe on behalf of the Wellcome Sanger Institute COVID-19 Surveillance Team |
| EPI_ISL_3248987 | Lighthouse Lab in Alderley Park | Wellcome Sanger Institute for the COVID-19 Genomics UK (COG-UK) Consortium | Jacquelyn Wynn, Mairead Hyland, The Lighthouse Lab in Alderley Park and Alex Alderton, Roberto Amato, Jeffrey Barrett, Sonia Goncalves, Ewan Harrison, David K. Jackson, Ian Johnston, Dominic Kwiatkowski, Cordelia Langford, John Sillitoe on behalf of the Wellcome Sanger Institute COVID-19 Surveillance Team |
| EPI_ISL_3248988 | Lighthouse Lab in Glasgow | Wellcome Sanger Institute for the COVID-19 Genomics UK (COG-UK) Consortium | Harper VanSteenhouse, Yumi Kasai, David Gray, Carol Clugston, Anna Dominiczak and Alex Alderton, Roberto Amato, Jeffrey Barrett, Sonia Goncalves, Ewan Harrison, David K. Jackson, Ian Johnston, Dominic Kwiatkowski, Cordelia Langford, John Sillitoe on behalf of the Wellcome Sanger Institute COVID-19 Surveillance Team |
| EPI_ISL_3248989 | ASST MONZA | ASST MONZA | Sergio Maria Ivano Malandrin |
| EPI_ISL_3248990, EPI_ISL_3248991 | Lighthouse Lab in Alderley Park | Wellcome Sanger Institute for the COVID-19 Genomics UK (COG-UK) Consortium | Jacquelyn Wynn, Mairead Hyland, The Lighthouse Lab in Alderley Park and Alex Alderton, Roberto Amato, Jeffrey Barrett, Sonia Goncalves, Ewan Harrison, David K. Jackson, Ian Johnston, Dominic Kwiatkowski, Cordelia Langford, John Sillitoe on behalf of the Wellcome Sanger Institute COVID-19 Surveillance Team |
| EPI_ISL_3248992 | ASST MONZA | ASST MONZA | Sergio Maria Ivano Malandrin |
| EPI_ISL_3248993 | Lighthouse Lab in Glasgow | Wellcome Sanger Institute for the COVID-19 Genomics UK (COG-UK) Consortium | Harper VanSteenhouse, Yumi Kasai, David Gray, Carol Clugston, Anna Dominiczak and Alex Alderton, Roberto Amato, Jeffrey Barrett, Sonia Goncalves, Ewan Harrison, David K. Jackson, Ian Johnston, Dominic Kwiatkowski, Cordelia Langford, John Sillitoe on behalf of the Wellcome Sanger Institute COVID-19 Surveillance Team |
| EPI_ISL_3248994 | Lighthouse Lab in Alderley Park | Wellcome Sanger Institute for the COVID-19 Genomics UK (COG-UK) Consortium | Jacquelyn Wynn, Mairead Hyland, The Lighthouse Lab in Alderley Park and Alex Alderton, Roberto Amato, Jeffrey Barrett, Sonia Goncalves, Ewan Harrison, David K. Jackson, Ian Johnston, Dominic Kwiatkowski, Cordelia Langford, John Sillitoe on behalf of the Wellcome Sanger Institute COVID-19 Surveillance Team |
| EPI_ISL_3248995 | Lighthouse Lab in Milton Keynes | Wellcome Sanger Institute for the COVID-19 Genomics UK (COG-UK) Consortium | The Lighthouse Lab in Milton Keynes and Alex Alderton, Roberto Amato, Jeffrey Barrett, Sonia Goncalves, Ewan Harrison, David K. Jackson, Ian Johnston, Dominic Kwiatkowski, Cordelia Langford, John Sillitoe on behalf of the Wellcome Sanger Institute COVID-19 Surveillance Team |
| EPI_ISL_3248996 | ASST MONZA | ASST MONZA | Sergio Maria Ivano Malandrin |
| EPI_ISL_3248997, EPI_ISL_3248998 | Lighthouse Lab in Milton Keynes | Wellcome Sanger Institute for the COVID-19 Genomics UK (COG-UK) Consortium | The Lighthouse Lab in Milton Keynes and Alex Alderton, Roberto Amato, Jeffrey Barrett, Sonia Goncalves, Ewan Harrison, David K. Jackson, Ian Johnston, Dominic Kwiatkowski, Cordelia Langford, John Sillitoe on behalf of the Wellcome Sanger Institute COVID-19 Surveillance Team |
| EPI_ISL_3248999 | ASST MONZA | ASST MONZA | Sergio Maria Ivano Malandrin |
| EPI_ISL_3249000 | Lighthouse Lab in Milton Keynes | Wellcome Sanger Institute for the COVID-19 Genomics UK (COG-UK) Consortium | The Lighthouse Lab in Milton Keynes and Alex Alderton, Roberto Amato, Jeffrey Barrett, Sonia Goncalves, Ewan Harrison, David K. Jackson, Ian Johnston, Dominic Kwiatkowski, Cordelia Langford, John Sillitoe on behalf of the Wellcome Sanger Institute COVID-19 Surveillance Team |
| EPI_ISL_3249001 | Lighthouse Lab in Glasgow | Wellcome Sanger Institute for the COVID-19 Genomics UK (COG-UK) Consortium | Harper VanSteenhouse, Yumi Kasai, David Gray, Carol Clugston, Anna Dominiczak and Alex Alderton, Roberto Amato, Jeffrey Barrett, Sonia Goncalves, Ewan Harrison, David K. Jackson, Ian Johnston, Dominic Kwiatkowski, Cordelia Langford, John Sillitoe on behalf of the Wellcome Sanger Institute COVID-19 Surveillance Team |
| EPI_ISL_3249002 | Lighthouse Lab in Milton Keynes | Wellcome Sanger Institute for the COVID-19 Genomics UK (COG-UK) Consortium | The Lighthouse Lab in Milton Keynes and Alex Alderton, Roberto Amato, Jeffrey Barrett, Sonia Goncalves, Ewan Harrison, David K. Jackson, Ian Johnston, Dominic Kwiatkowski, Cordelia Langford, John Sillitoe on behalf of the Wellcome Sanger Institute COVID-19 Surveillance Team |
| EPI_ISL_3249003 | ASST MONZA | ASST MONZA | Sergio Maria Ivano Malandrin |
| EPI_ISL_3249004 | Lighthouse Lab in Alderley Park | Wellcome Sanger Institute for the COVID-19 Genomics UK (COG-UK) Consortium | Jacquelyn Wynn, Mairead Hyland, The Lighthouse Lab in Alderley Park and Alex Alderton, Roberto Amato, Jeffrey Barrett, Sonia Goncalves, Ewan Harrison, David K. Jackson, Ian Johnston, Dominic Kwiatkowski, Cordelia Langford, John Sillitoe on behalf of the Wellcome Sanger Institute COVID-19 Surveillance Team |
| EPI_ISL_3249005 | Lighthouse Lab in Milton Keynes | Wellcome Sanger Institute for the COVID-19 Genomics UK (COG-UK) Consortium | The Lighthouse Lab in Milton Keynes and Alex Alderton, Roberto Amato, Jeffrey Barrett, Sonia Goncalves, Ewan Harrison, David K. Jackson, Ian Johnston, Dominic Kwiatkowski, Cordelia Langford, John Sillitoe on behalf of the Wellcome Sanger Institute COVID-19 Surveillance Team |
| EPI_ISL_3249006 | Lighthouse Lab in Glasgow | Wellcome Sanger Institute for the COVID-19 Genomics UK (COG-UK) Consortium | Harper VanSteenhouse, Yumi Kasai, David Gray, Carol Clugston, Anna Dominiczak and Alex Alderton, Roberto Amato, Jeffrey Barrett, Sonia Goncalves, Ewan Harrison, David K. Jackson, Ian Johnston, Dominic Kwiatkowski, Cordelia Langford, John Sillitoe on behalf of the Wellcome Sanger Institute COVID-19 Surveillance Team |
| EPI_ISL_3249007 | ASST MONZA | ASST MONZA | Sergio Maria Ivano Malandrin |
| EPI_ISL_3249008 | Lighthouse Lab in Alderley Park | Wellcome Sanger Institute for the COVID-19 Genomics UK (COG-UK) Consortium | Jacquelyn Wynn, Mairead Hyland, The Lighthouse Lab in Alderley Park and Alex Alderton, Roberto Amato, Jeffrey Barrett, Sonia Goncalves, Ewan Harrison, David K. Jackson, Ian Johnston, Dominic Kwiatkowski, Cordelia Langford, John Sillitoe on behalf of the Wellcome Sanger Institute COVID-19 Surveillance Team |
| EPI_ISL_3249009 | Lighthouse Lab in Milton Keynes | Wellcome Sanger Institute for the COVID-19 Genomics UK (COG-UK) Consortium | The Lighthouse Lab in Milton Keynes and Alex Alderton, Roberto Amato, Jeffrey Barrett, Sonia Goncalves, Ewan Harrison, David K. Jackson, Ian Johnston, Dominic Kwiatkowski, Cordelia Langford, John Sillitoe on behalf of the Wellcome Sanger Institute COVID-19 Surveillance Team |
| EPI_ISL_3249010 | Lighthouse Lab in Alderley Park | Wellcome Sanger Institute for the COVID-19 Genomics UK (COG-UK) Consortium | Jacquelyn Wynn, Mairead Hyland, The Lighthouse Lab in Alderley Park and Alex Alderton, Roberto Amato, Jeffrey Barrett, Sonia Goncalves, Ewan Harrison, David K. Jackson, Ian Johnston, Dominic Kwiatkowski, Cordelia Langford, John Sillitoe on behalf of the Wellcome Sanger Institute COVID-19 Surveillance Team |
| EPI_ISL_3249011 | ASST MONZA | ASST MONZA | Sergio Maria Ivano Malandrin |
| EPI_ISL_3249012 | Lighthouse Lab in Alderley Park | Wellcome Sanger Institute for the COVID-19 Genomics UK (COG-UK) Consortium | Jacquelyn Wynn, Mairead Hyland, The Lighthouse Lab in Alderley Park and Alex Alderton, Roberto Amato, Jeffrey Barrett, Sonia Goncalves, Ewan Harrison, David K. Jackson, Ian Johnston, Dominic Kwiatkowski, Cordelia Langford, John Sillitoe on behalf of the Wellcome Sanger Institute COVID-19 Surveillance Team |
| EPI_ISL_3249013 | Lighthouse Lab in Glasgow | Wellcome Sanger Institute for the COVID-19 Genomics UK (COG-UK) Consortium | Harper VanSteenhouse, Yumi Kasai, David Gray, Carol Clugston, Anna Dominiczak and Alex Alderton, Roberto Amato, Jeffrey Barrett, Sonia Goncalves, Ewan Harrison, David K. Jackson, Ian Johnston, Dominic Kwiatkowski, Cordelia Langford, John Sillitoe on behalf of the Wellcome Sanger Institute COVID-19 Surveillance Team |
| EPI_ISL_3249014 | ASST MONZA | ASST MONZA | Sergio Maria Ivano Malandrin |
| EPI_ISL_3249015 | Lighthouse Lab in Milton Keynes | Wellcome Sanger Institute for the COVID-19 Genomics UK (COG-UK) Consortium | The Lighthouse Lab in Milton Keynes and Alex Alderton, Roberto Amato, Jeffrey Barrett, Sonia Goncalves, Ewan Harrison, David K. Jackson, Ian Johnston, Dominic Kwiatkowski, Cordelia Langford, John Sillitoe on behalf of the Wellcome Sanger Institute COVID-19 Surveillance Team |
| EPI_ISL_3249016 | Lighthouse Lab in Glasgow | Wellcome Sanger Institute for the COVID-19 Genomics UK (COG-UK) Consortium | Harper VanSteenhouse, Yumi Kasai, David Gray, Carol Clugston, Anna Dominiczak and Alex Alderton, Roberto Amato, Jeffrey Barrett, Sonia Goncalves, Ewan Harrison, David K. Jackson, Ian Johnston, Dominic Kwiatkowski, Cordelia Langford, John Sillitoe on behalf of the Wellcome Sanger Institute COVID-19 Surveillance Team |
| EPI_ISL_3249017 | Lighthouse Lab in Milton Keynes | Wellcome Sanger Institute for the COVID-19 Genomics UK (COG-UK) | The Lighthouse Lab in Milton Keynes and Alex Alderton, Roberto Amato, Jeffrey Barrett, Sonia Goncalves, Ewan Harrison, David K. Jackson, Ian Johnston, Dominic Kwiatkowski, Cordelia Langford, John Sillitoe on behalf of the Wellcome Sanger |

| | | | |
|---|---|---|---|
| | | Consortium | Institute COVID-19 Surveillance Team |
| EPI_ISL_3249018 | ASST MONZA | ASST MONZA | Sergio Maria Ivano Malandrin |
| EPI_ISL_3249019, EPI_ISL_3249020 | Lighthouse Lab in Alderley Park | Wellcome Sanger Institute for the COVID-19 Genomics UK (COG-UK) Consortium | Jacquelyn Wynn, Mairead Hyland, The Lighthouse Lab in Alderley Park and Alex Alderton, Roberto Amato, Jeffrey Barrett, Sonia Goncalves, Ewan Harrison, David K. Jackson, Ian Johnston, Dominic Kwiatkowski, Cordelia Langford, John Sillitoe on behalf of the Wellcome Sanger Institute COVID-19 Surveillance Team |
| EPI_ISL_3249021 | ASST MONZA | ASST MONZA | Sergio Maria Ivano Malandrin |
| EPI_ISL_3249022, EPI_ISL_3249023, EPI_ISL_3249024 | Lighthouse Lab in Milton Keynes | Wellcome Sanger Institute for the COVID-19 Genomics UK (COG-UK) Consortium | The Lighthouse Lab in Milton Keynes and Alex Alderton, Roberto Amato, Jeffrey Barrett, Sonia Goncalves, Ewan Harrison, David K. Jackson, Ian Johnston, Dominic Kwiatkowski, Cordelia Langford, John Sillitoe on behalf of the Wellcome Sanger Institute COVID-19 Surveillance Team |
| EPI_ISL_3249025 | ASST MONZA | ASST MONZA | Sergio Maria Ivano Malandrin |
| EPI_ISL_3249026, EPI_ISL_3249027 | Lighthouse Lab in Milton Keynes | Wellcome Sanger Institute for the COVID-19 Genomics UK (COG-UK) Consortium | The Lighthouse Lab in Milton Keynes and Alex Alderton, Roberto Amato, Jeffrey Barrett, Sonia Goncalves, Ewan Harrison, David K. Jackson, Ian Johnston, Dominic Kwiatkowski, Cordelia Langford, John Sillitoe on behalf of the Wellcome Sanger Institute COVID-19 Surveillance Team |
| EPI_ISL_3249028 | ASST MONZA | ASST MONZA | Sergio Maria Ivano Malandrin |
| EPI_ISL_3249029 | Lighthouse Lab in Milton Keynes | Wellcome Sanger Institute for the COVID-19 Genomics UK (COG-UK) Consortium | The Lighthouse Lab in Milton Keynes and Alex Alderton, Roberto Amato, Jeffrey Barrett, Sonia Goncalves, Ewan Harrison, David K. Jackson, Ian Johnston, Dominic Kwiatkowski, Cordelia Langford, John Sillitoe on behalf of the Wellcome Sanger Institute COVID-19 Surveillance Team |
| EPI_ISL_3249030, EPI_ISL_3249032 | Lighthouse Lab in Alderley Park | Wellcome Sanger Institute for the COVID-19 Genomics UK (COG-UK) Consortium | Jacquelyn Wynn, Mairead Hyland, The Lighthouse Lab in Alderley Park and Alex Alderton, Roberto Amato, Jeffrey Barrett, Sonia Goncalves, Ewan Harrison, David K. Jackson, Ian Johnston, Dominic Kwiatkowski, Cordelia Langford, John Sillitoe on behalf of the Wellcome Sanger Institute COVID-19 Surveillance Team |
| EPI_ISL_3249033 | Lighthouse Lab in Milton Keynes | Wellcome Sanger Institute for the COVID-19 Genomics UK (COG-UK) Consortium | The Lighthouse Lab in Milton Keynes and Alex Alderton, Roberto Amato, Jeffrey Barrett, Sonia Goncalves, Ewan Harrison, David K. Jackson, Ian Johnston, Dominic Kwiatkowski, Cordelia Langford, John Sillitoe on behalf of the Wellcome Sanger Institute COVID-19 Surveillance Team |
| EPI_ISL_3249034 | Lighthouse Lab in Alderley Park | Wellcome Sanger Institute for the COVID-19 Genomics UK (COG-UK) Consortium | Jacquelyn Wynn, Mairead Hyland, The Lighthouse Lab in Alderley Park and Alex Alderton, Roberto Amato, Jeffrey Barrett, Sonia Goncalves, Ewan Harrison, David K. Jackson, Ian Johnston, Dominic Kwiatkowski, Cordelia Langford, John Sillitoe on behalf of the Wellcome Sanger Institute COVID-19 Surveillance Team |
| EPI_ISL_3249035 | ASST MONZA | ASST MONZA | Sergio Maria Ivano Malandrin |
| EPI_ISL_3249036, EPI_ISL_3249037 | Lighthouse Lab in Milton Keynes | Wellcome Sanger Institute for the COVID-19 Genomics UK (COG-UK) Consortium | The Lighthouse Lab in Milton Keynes and Alex Alderton, Roberto Amato, Jeffrey Barrett, Sonia Goncalves, Ewan Harrison, David K. Jackson, Ian Johnston, Dominic Kwiatkowski, Cordelia Langford, John Sillitoe on behalf of the Wellcome Sanger Institute COVID-19 Surveillance Team |
| EPI_ISL_3249038 | ASST MONZA | ASST MONZA | Sergio Maria Ivano Malandrin |
| EPI_ISL_3249039, EPI_ISL_3249040 | Lighthouse Lab in Milton Keynes | Wellcome Sanger Institute for the COVID-19 Genomics UK (COG-UK) Consortium | The Lighthouse Lab in Milton Keynes and Alex Alderton, Roberto Amato, Jeffrey Barrett, Sonia Goncalves, Ewan Harrison, David K. Jackson, Ian Johnston, Dominic Kwiatkowski, Cordelia Langford, John Sillitoe on behalf of the Wellcome Sanger Institute COVID-19 Surveillance Team |
| EPI_ISL_3249041 | ASST MONZA | ASST MONZA | Sergio Maria Ivano Malandrin |
| EPI_ISL_3249042 | Lighthouse Lab in Milton Keynes | Wellcome Sanger Institute for the COVID-19 Genomics UK (COG-UK) Consortium | The Lighthouse Lab in Milton Keynes and Alex Alderton, Roberto Amato, Jeffrey Barrett, Sonia Goncalves, Ewan Harrison, David K. Jackson, Ian Johnston, Dominic Kwiatkowski, Cordelia Langford, John Sillitoe on behalf of the Wellcome Sanger Institute COVID-19 Surveillance Team |
| EPI_ISL_3249043 | Lighthouse Lab in Alderley Park | Wellcome Sanger Institute for the COVID-19 Genomics UK (COG-UK) Consortium | Jacquelyn Wynn, Mairead Hyland, The Lighthouse Lab in Alderley Park and Alex Alderton, Roberto Amato, Jeffrey Barrett, Sonia Goncalves, Ewan Harrison, David K. Jackson, Ian Johnston, Dominic Kwiatkowski, Cordelia Langford, John Sillitoe on behalf of the Wellcome Sanger Institute COVID-19 Surveillance Team |
| EPI_ISL_3249044 | Lighthouse Lab in Milton Keynes | Wellcome Sanger Institute for the COVID-19 Genomics UK (COG-UK) Consortium | The Lighthouse Lab in Milton Keynes and Alex Alderton, Roberto Amato, Jeffrey Barrett, Sonia Goncalves, Ewan Harrison, David K. Jackson, Ian Johnston, Dominic Kwiatkowski, Cordelia Langford, John Sillitoe on behalf of the Wellcome Sanger Institute COVID-19 Surveillance Team |
| EPI_ISL_3249045 | ASST MONZA | ASST MONZA | Sergio Maria Ivano Malandrin |
| EPI_ISL_3249046 | Lighthouse Lab in Alderley Park | Wellcome Sanger Institute for the COVID-19 Genomics UK (COG-UK) Consortium | Jacquelyn Wynn, Mairead Hyland, The Lighthouse Lab in Alderley Park and Alex Alderton, Roberto Amato, Jeffrey Barrett, Sonia Goncalves, Ewan Harrison, David K. Jackson, Ian Johnston, Dominic Kwiatkowski, Cordelia Langford, John Sillitoe on behalf of the Wellcome Sanger Institute COVID-19 Surveillance Team |
| EPI_ISL_3249047 | Lighthouse Lab in Milton Keynes | Wellcome Sanger Institute for the COVID-19 Genomics UK (COG-UK) Consortium | The Lighthouse Lab in Milton Keynes and Alex Alderton, Roberto Amato, Jeffrey Barrett, Sonia Goncalves, Ewan Harrison, David K. Jackson, Ian Johnston, Dominic Kwiatkowski, Cordelia Langford, John Sillitoe on behalf of the Wellcome Sanger Institute COVID-19 Surveillance Team |
| EPI_ISL_3249048 | ASST MONZA | ASST MONZA | Sergio Maria Ivano Malandrin |
| EPI_ISL_3249049 | Lighthouse Lab in Glasgow | Wellcome Sanger Institute for the COVID-19 Genomics UK (COG-UK) Consortium | Harper VanSteenhouse, Yumi Kasai, David Gray, Carol Clugston, Anna Dominiczak and Alex Alderton, Roberto Amato, Jeffrey Barrett, Sonia Goncalves, Ewan Harrison, David K. Jackson, Ian Johnston, Dominic Kwiatkowski, Cordelia Langford, John Sillitoe on behalf of the Wellcome Sanger Institute COVID-19 Surveillance Team |
| EPI_ISL_3249050, EPI_ISL_3249051 | Lighthouse Lab in Milton Keynes | Wellcome Sanger Institute for the COVID-19 Genomics UK (COG-UK) Consortium | The Lighthouse Lab in Milton Keynes and Alex Alderton, Roberto Amato, Jeffrey Barrett, Sonia Goncalves, Ewan Harrison, David K. Jackson, Ian Johnston, Dominic Kwiatkowski, Cordelia Langford, John Sillitoe on behalf of the Wellcome Sanger Institute COVID-19 Surveillance Team |
| EPI_ISL_3249052 | ASST MONZA | ASST MONZA | Sergio Maria Ivano Malandrin |
| EPI_ISL_3249053, EPI_ISL_3249054 | Lighthouse Lab in Alderley Park | Wellcome Sanger Institute for the COVID-19 Genomics UK (COG-UK) Consortium | Jacquelyn Wynn, Mairead Hyland, The Lighthouse Lab in Alderley Park and Alex Alderton, Roberto Amato, Jeffrey Barrett, Sonia Goncalves, Ewan Harrison, David K. Jackson, Ian Johnston, Dominic Kwiatkowski, Cordelia Langford, John Sillitoe on behalf of the Wellcome Sanger Institute COVID-19 Surveillance Team |
| EPI_ISL_3249055 | Lighthouse Lab in Milton Keynes | Wellcome Sanger Institute for the COVID-19 Genomics UK (COG-UK) Consortium | The Lighthouse Lab in Milton Keynes and Alex Alderton, Roberto Amato, Jeffrey Barrett, Sonia Goncalves, Ewan Harrison, David K. Jackson, Ian Johnston, Dominic Kwiatkowski, Cordelia Langford, John Sillitoe on behalf of the Wellcome Sanger Institute COVID-19 Surveillance Team |
| EPI_ISL_3249056 | ASST MONZA | ASST MONZA | Sergio Maria Ivano Malandrin |
| EPI_ISL_3249057 | Lighthouse Lab in Milton Keynes | Wellcome Sanger Institute for the COVID-19 Genomics UK (COG-UK) Consortium | The Lighthouse Lab in Milton Keynes and Alex Alderton, Roberto Amato, Jeffrey Barrett, Sonia Goncalves, Ewan Harrison, David K. Jackson, Ian Johnston, Dominic Kwiatkowski, Cordelia Langford, John Sillitoe on behalf of the Wellcome Sanger Institute COVID-19 Surveillance Team |
| EPI_ISL_3249058 | Lighthouse Lab in Alderley Park | Wellcome Sanger Institute for the COVID-19 Genomics UK (COG-UK) Consortium | Jacquelyn Wynn, Mairead Hyland, The Lighthouse Lab in Alderley Park and Alex Alderton, Roberto Amato, Jeffrey Barrett, Sonia Goncalves, Ewan Harrison, David K. Jackson, Ian Johnston, Dominic Kwiatkowski, Cordelia Langford, John Sillitoe on behalf of the Wellcome Sanger Institute COVID-19 Surveillance Team |
| EPI_ISL_3249059 | ASST MONZA | ASST MONZA | Sergio Maria Ivano Malandrin |
| EPI_ISL_3249060 | Lighthouse Lab in Glasgow | Wellcome Sanger Institute for the COVID-19 Genomics UK (COG-UK) Consortium | Harper VanSteenhouse, Yumi Kasai, David Gray, Carol Clugston, Anna Dominiczak and Alex Alderton, Roberto Amato, Jeffrey Barrett, Sonia Goncalves, Ewan Harrison, David K. Jackson, Ian Johnston, Dominic Kwiatkowski, Cordelia Langford, John Sillitoe on behalf of the Wellcome Sanger Institute COVID-19 Surveillance Team |
| EPI_ISL_3249061 | Lighthouse Lab in Milton Keynes | Wellcome Sanger Institute for the COVID-19 Genomics UK (COG-UK) Consortium | The Lighthouse Lab in Milton Keynes and Alex Alderton, Roberto Amato, Jeffrey Barrett, Sonia Goncalves, Ewan Harrison, David K. Jackson, Ian Johnston, Dominic Kwiatkowski, Cordelia Langford, John Sillitoe on behalf of the Wellcome Sanger Institute COVID-19 Surveillance Team |
| EPI_ISL_3249062 | Lighthouse Lab in Alderley Park | Wellcome Sanger Institute for the COVID-19 Genomics UK (COG-UK) Consortium | Jacquelyn Wynn, Mairead Hyland, The Lighthouse Lab in Alderley Park and Alex Alderton, Roberto Amato, Jeffrey Barrett, Sonia Goncalves, Ewan Harrison, David K. Jackson, Ian Johnston, Dominic Kwiatkowski, Cordelia Langford, John Sillitoe on behalf of the Wellcome Sanger Institute COVID-19 Surveillance Team |
| EPI_ISL_3249063 | ASST MONZA | ASST MONZA | Sergio Maria Ivano Malandrin |
| EPI_ISL_3249064 | Lighthouse Lab in Glasgow | Wellcome Sanger Institute for the COVID-19 Genomics UK (COG-UK) Consortium | Harper VanSteenhouse, Yumi Kasai, David Gray, Carol Clugston, Anna Dominiczak and Alex Alderton, Roberto Amato, Jeffrey Barrett, Sonia Goncalves, Ewan Harrison, David K. Jackson, Ian Johnston, Dominic Kwiatkowski, Cordelia Langford, John Sillitoe on behalf of the Wellcome Sanger Institute COVID-19 Surveillance Team |
| EPI_ISL_3249065 | Lighthouse Lab in Milton Keynes | Wellcome Sanger Institute for the COVID-19 Genomics UK (COG-UK) Consortium | The Lighthouse Lab in Milton Keynes and Alex Alderton, Roberto Amato, Jeffrey Barrett, Sonia Goncalves, Ewan Harrison, David K. Jackson, Ian Johnston, Dominic Kwiatkowski, Cordelia Langford, John Sillitoe on behalf of the Wellcome Sanger Institute COVID-19 Surveillance Team |
| EPI_ISL_3249066 | ASST MONZA | ASST MONZA | Sergio Maria Ivano Malandrin |
| EPI_ISL_3249067, EPI_ISL_3249068 | Lighthouse Lab in Milton Keynes | Wellcome Sanger Institute for the COVID-19 Genomics UK (COG-UK) Consortium | The Lighthouse Lab in Milton Keynes and Alex Alderton, Roberto Amato, Jeffrey Barrett, Sonia Goncalves, Ewan Harrison, David K. Jackson, Ian Johnston, Dominic Kwiatkowski, Cordelia Langford, John Sillitoe on behalf of the Wellcome Sanger Institute COVID-19 Surveillance Team |
| EPI_ISL_3249069 | ASST MONZA | ASST MONZA | Sergio Maria Ivano Malandrin |
| EPI_ISL_3249070 | Lighthouse Lab in Alderley Park | Wellcome Sanger Institute for the COVID-19 Genomics UK (COG-UK) Consortium | Jacquelyn Wynn, Mairead Hyland, The Lighthouse Lab in Alderley Park and Alex Alderton, Roberto Amato, Jeffrey Barrett, Sonia Goncalves, Ewan Harrison, David K. Jackson, Ian Johnston, Dominic Kwiatkowski, Cordelia Langford, John Sillitoe on behalf of the Wellcome Sanger Institute COVID-19 Surveillance Team |
| EPI_ISL_3249071 | Lighthouse Lab in Glasgow | Wellcome Sanger Institute for the COVID-19 Genomics UK (COG-UK) Consortium | Harper VanSteenhouse, Yumi Kasai, David Gray, Carol Clugston, Anna Dominiczak and Alex Alderton, Roberto Amato, Jeffrey Barrett, Sonia Goncalves, Ewan Harrison, David K. Jackson, Ian Johnston, Dominic Kwiatkowski, Cordelia Langford, John Sillitoe on behalf of the Wellcome Sanger Institute COVID-19 Surveillance Team |
| EPI_ISL_3249072 | Lighthouse Lab in Alderley Park | Wellcome Sanger Institute for the COVID-19 Genomics UK (COG-UK) Consortium | Jacquelyn Wynn, Mairead Hyland, The Lighthouse Lab in Alderley Park and Alex Alderton, Roberto Amato, Jeffrey Barrett, Sonia Goncalves, Ewan Harrison, David K. Jackson, Ian Johnston, Dominic Kwiatkowski, Cordelia Langford, John Sillitoe on behalf of the Wellcome Sanger Institute COVID-19 Surveillance Team |
| EPI_ISL_3249073 | ASST MONZA | ASST MONZA | Sergio Maria Ivano Malandrin |
| EPI_ISL_3249074, EPI_ISL_3249075 | Lighthouse Lab in Milton Keynes | Wellcome Sanger Institute for the COVID-19 Genomics UK (COG-UK) Consortium | The Lighthouse Lab in Milton Keynes and Alex Alderton, Roberto Amato, Jeffrey Barrett, Sonia Goncalves, Ewan Harrison, David K. Jackson, Ian Johnston, Dominic Kwiatkowski, Cordelia Langford, John Sillitoe on behalf of the Wellcome Sanger Institute COVID-19 Surveillance Team |
| EPI_ISL_3249076 | ASST MONZA | ASST MONZA | Sergio Maria Ivano Malandrin |
| EPI_ISL_3249077 | Lighthouse Lab in Milton Keynes | Wellcome Sanger Institute for the COVID-19 Genomics UK (COG-UK) Consortium | The Lighthouse Lab in Milton Keynes and Alex Alderton, Roberto Amato, Jeffrey Barrett, Sonia Goncalves, Ewan Harrison, David K. Jackson, Ian Johnston, Dominic Kwiatkowski, Cordelia Langford, John Sillitoe on behalf of the Wellcome Sanger Institute COVID-19 Surveillance Team |
| EPI_ISL_3249078 | Lighthouse Lab in Alderley Park | Wellcome Sanger Institute for the COVID-19 Genomics UK (COG-UK) Consortium | Jacquelyn Wynn, Mairead Hyland, The Lighthouse Lab in Alderley Park and Alex Alderton, Roberto Amato, Jeffrey Barrett, Sonia Goncalves, Ewan Harrison, David K. Jackson, Ian Johnston, Dominic Kwiatkowski, Cordelia Langford, John Sillitoe on behalf of the Wellcome Sanger Institute COVID-19 Surveillance Team |
| EPI_ISL_3249079 | ASST MONZA | ASST MONZA | Sergio Maria Ivano Malandrin |
| EPI_ISL_3249080 | Lighthouse Lab in Milton Keynes | Wellcome Sanger Institute for the COVID-19 Genomics UK (COG-UK) Consortium | The Lighthouse Lab in Milton Keynes and Alex Alderton, Roberto Amato, Jeffrey Barrett, Sonia Goncalves, Ewan Harrison, David K. Jackson, Ian Johnston, Dominic Kwiatkowski, Cordelia Langford, John Sillitoe on behalf of the Wellcome Sanger Institute COVID-19 Surveillance Team |
| EPI_ISL_3249081 | Lighthouse Lab in Glasgow | Wellcome Sanger Institute for the COVID-19 Genomics UK (COG-UK) | Harper VanSteenhouse, Yumi Kasai, David Gray, Carol Clugston, Anna Dominiczak and Alex Alderton, Roberto Amato, Jeffrey Barrett, Sonia Goncalves, Ewan Harrison, David K. Jackson, Ian Johnston, Dominic Kwiatkowski, Cordelia Langford, |

| | | | |
|---|---|---|---|
| | | Consortium | John Sillitoe on behalf of the Wellcome Sanger Institute COVID-19 Surveillance Team |
| EPI_ISL_3249082 | Lighthouse Lab in Alderley Park | Wellcome Sanger Institute for the COVID-19 Genomics UK (COG-UK) Consortium | Jacquelyn Wynn, Mairead Hyland, The Lighthouse Lab in Alderley Park and Alex Alderton, Roberto Amato, Jeffrey Barrett, Sonia Goncalves, Ewan Harrison, David K. Jackson, Ian Johnston, Dominic Kwiatkowski, Cordelia Langford, John Sillitoe on behalf of the Wellcome Sanger Institute COVID-19 Surveillance Team |
| EPI_ISL_3249083 | ASST MONZA | ASST MONZA | Sergio Maria Ivano Malandrin |
| EPI_ISL_3249084 | Lighthouse Lab in Glasgow | Wellcome Sanger Institute for the COVID-19 Genomics UK (COG-UK) Consortium | Harper VanSteenhouse, Yumi Kasai, David Gray, Carol Clugston, Anna Dominiczak and Alex Alderton, Roberto Amato, Jeffrey Barrett, Sonia Goncalves, Ewan Harrison, David K. Jackson, Ian Johnston, Dominic Kwiatkowski, Cordelia Langford, John Sillitoe on behalf of the Wellcome Sanger Institute COVID-19 Surveillance Team |
| EPI_ISL_3249085 | Lighthouse Lab in Milton Keynes | Wellcome Sanger Institute for the COVID-19 Genomics UK (COG-UK) Consortium | The Lighthouse Lab in Milton Keynes and Alex Alderton, Roberto Amato, Jeffrey Barrett, Sonia Goncalves, Ewan Harrison, David K. Jackson, Ian Johnston, Dominic Kwiatkowski, Cordelia Langford, John Sillitoe on behalf of the Wellcome Sanger Institute COVID-19 Surveillance Team |
| EPI_ISL_3249086 | ASST MONZA | ASST MONZA | Sergio Maria Ivano Malandrin |
| EPI_ISL_3249087, EPI_ISL_3249088, EPI_ISL_3249089 | Lighthouse Lab in Alderley Park | Wellcome Sanger Institute for the COVID-19 Genomics UK (COG-UK) Consortium | Jacquelyn Wynn, Mairead Hyland, The Lighthouse Lab in Alderley Park and Alex Alderton, Roberto Amato, Jeffrey Barrett, Sonia Goncalves, Ewan Harrison, David K. Jackson, Ian Johnston, Dominic Kwiatkowski, Cordelia Langford, John Sillitoe on behalf of the Wellcome Sanger Institute COVID-19 Surveillance Team |
| EPI_ISL_3249090 | ASST MONZA | ASST MONZA | Sergio Maria Ivano Malandrin |
| EPI_ISL_3249091, EPI_ISL_3249092 | Lighthouse Lab in Alderley Park | Wellcome Sanger Institute for the COVID-19 Genomics UK (COG-UK) Consortium | Jacquelyn Wynn, Mairead Hyland, The Lighthouse Lab in Alderley Park and Alex Alderton, Roberto Amato, Jeffrey Barrett, Sonia Goncalves, Ewan Harrison, David K. Jackson, Ian Johnston, Dominic Kwiatkowski, Cordelia Langford, John Sillitoe on behalf of the Wellcome Sanger Institute COVID-19 Surveillance Team |
| EPI_ISL_3249093 | ASST MONZA | ASST MONZA | Sergio Maria Ivano Malandrin |
| EPI_ISL_3249094, EPI_ISL_3249095 | Lighthouse Lab in Glasgow | Wellcome Sanger Institute for the COVID-19 Genomics UK (COG-UK) Consortium | Harper VanSteenhouse, Yumi Kasai, David Gray, Carol Clugston, Anna Dominiczak and Alex Alderton, Roberto Amato, Jeffrey Barrett, Sonia Goncalves, Ewan Harrison, David K. Jackson, Ian Johnston, Dominic Kwiatkowski, Cordelia Langford, John Sillitoe on behalf of the Wellcome Sanger Institute COVID-19 Surveillance Team |
| EPI_ISL_3249096 | Lighthouse Lab in Milton Keynes | Wellcome Sanger Institute for the COVID-19 Genomics UK (COG-UK) Consortium | The Lighthouse Lab in Milton Keynes and Alex Alderton, Roberto Amato, Jeffrey Barrett, Sonia Goncalves, Ewan Harrison, David K. Jackson, Ian Johnston, Dominic Kwiatkowski, Cordelia Langford, John Sillitoe on behalf of the Wellcome Sanger Institute COVID-19 Surveillance Team |
| EPI_ISL_3249097 | ASST MONZA | ASST MONZA | Sergio Maria Ivano Malandrin |
| EPI_ISL_3249098, EPI_ISL_3249099 | Lighthouse Lab in Milton Keynes | Wellcome Sanger Institute for the COVID-19 Genomics UK (COG-UK) Consortium | The Lighthouse Lab in Milton Keynes and Alex Alderton, Roberto Amato, Jeffrey Barrett, Sonia Goncalves, Ewan Harrison, David K. Jackson, Ian Johnston, Dominic Kwiatkowski, Cordelia Langford, John Sillitoe on behalf of the Wellcome Sanger Institute COVID-19 Surveillance Team |
| EPI_ISL_3249101 | Lighthouse Lab in Glasgow | Wellcome Sanger Institute for the COVID-19 Genomics UK (COG-UK) Consortium | Harper VanSteenhouse, Yumi Kasai, David Gray, Carol Clugston, Anna Dominiczak and Alex Alderton, Roberto Amato, Jeffrey Barrett, Sonia Goncalves, Ewan Harrison, David K. Jackson, Ian Johnston, Dominic Kwiatkowski, Cordelia Langford, John Sillitoe on behalf of the Wellcome Sanger Institute COVID-19 Surveillance Team |
| EPI_ISL_3249102, EPI_ISL_3249103 | Lighthouse Lab in Milton Keynes | Wellcome Sanger Institute for the COVID-19 Genomics UK (COG-UK) Consortium | The Lighthouse Lab in Milton Keynes and Alex Alderton, Roberto Amato, Jeffrey Barrett, Sonia Goncalves, Ewan Harrison, David K. Jackson, Ian Johnston, Dominic Kwiatkowski, Cordelia Langford, John Sillitoe on behalf of the Wellcome Sanger Institute COVID-19 Surveillance Team |
| EPI_ISL_3249104 | ASST MONZA | ASST MONZA | Sergio Maria Ivano Malandrin |
| EPI_ISL_3249105 | Lighthouse Lab in Milton Keynes | Wellcome Sanger Institute for the COVID-19 Genomics UK (COG-UK) Consortium | The Lighthouse Lab in Milton Keynes and Alex Alderton, Roberto Amato, Jeffrey Barrett, Sonia Goncalves, Ewan Harrison, David K. Jackson, Ian Johnston, Dominic Kwiatkowski, Cordelia Langford, John Sillitoe on behalf of the Wellcome Sanger Institute COVID-19 Surveillance Team |
| EPI_ISL_3249106 | Lighthouse Lab in Glasgow | Wellcome Sanger Institute for the COVID-19 Genomics UK (COG-UK) Consortium | Harper VanSteenhouse, Yumi Kasai, David Gray, Carol Clugston, Anna Dominiczak and Alex Alderton, Roberto Amato, Jeffrey Barrett, Sonia Goncalves, Ewan Harrison, David K. Jackson, Ian Johnston, Dominic Kwiatkowski, Cordelia Langford, John Sillitoe on behalf of the Wellcome Sanger Institute COVID-19 Surveillance Team |
| EPI_ISL_3249107 | ASST MONZA | ASST MONZA | Sergio Maria Ivano Malandrin |
| EPI_ISL_3249108 | Lighthouse Lab in Alderley Park | Wellcome Sanger Institute for the COVID-19 Genomics UK (COG-UK) Consortium | Jacquelyn Wynn, Mairead Hyland, The Lighthouse Lab in Alderley Park and Alex Alderton, Roberto Amato, Jeffrey Barrett, Sonia Goncalves, Ewan Harrison, David K. Jackson, Ian Johnston, Dominic Kwiatkowski, Cordelia Langford, John Sillitoe on behalf of the Wellcome Sanger Institute COVID-19 Surveillance Team |
| EPI_ISL_3249109 | Lighthouse Lab in Milton Keynes | Wellcome Sanger Institute for the COVID-19 Genomics UK (COG-UK) Consortium | The Lighthouse Lab in Milton Keynes and Alex Alderton, Roberto Amato, Jeffrey Barrett, Sonia Goncalves, Ewan Harrison, David K. Jackson, Ian Johnston, Dominic Kwiatkowski, Cordelia Langford, John Sillitoe on behalf of the Wellcome Sanger Institute COVID-19 Surveillance Team |
| EPI_ISL_3249110 | ASST MONZA | ASST MONZA | Sergio Maria Ivano Malandrin |
| EPI_ISL_3249111 | Lighthouse Lab in Alderley Park | Wellcome Sanger Institute for the COVID-19 Genomics UK (COG-UK) Consortium | Jacquelyn Wynn, Mairead Hyland, The Lighthouse Lab in Alderley Park and Alex Alderton, Roberto Amato, Jeffrey Barrett, Sonia Goncalves, Ewan Harrison, David K. Jackson, Ian Johnston, Dominic Kwiatkowski, Cordelia Langford, John Sillitoe on behalf of the Wellcome Sanger Institute COVID-19 Surveillance Team |
| EPI_ISL_3249112 | Lighthouse Lab in Glasgow | Wellcome Sanger Institute for the COVID-19 Genomics UK (COG-UK) Consortium | Harper VanSteenhouse, Yumi Kasai, David Gray, Carol Clugston, Anna Dominiczak and Alex Alderton, Roberto Amato, Jeffrey Barrett, Sonia Goncalves, Ewan Harrison, David K. Jackson, Ian Johnston, Dominic Kwiatkowski, Cordelia Langford, John Sillitoe on behalf of the Wellcome Sanger Institute COVID-19 Surveillance Team |
| EPI_ISL_3249113 | Lighthouse Lab in Milton Keynes | Wellcome Sanger Institute for the COVID-19 Genomics UK (COG-UK) Consortium | The Lighthouse Lab in Milton Keynes and Alex Alderton, Roberto Amato, Jeffrey Barrett, Sonia Goncalves, Ewan Harrison, David K. Jackson, Ian Johnston, Dominic Kwiatkowski, Cordelia Langford, John Sillitoe on behalf of the Wellcome Sanger Institute COVID-19 Surveillance Team |
| EPI_ISL_3249114 | ASST MONZA | ASST MONZA | Sergio Maria Ivano Malandrin |
| EPI_ISL_3249115 | Lighthouse Lab in Alderley Park | Wellcome Sanger Institute for the COVID-19 Genomics UK (COG-UK) Consortium | Jacquelyn Wynn, Mairead Hyland, The Lighthouse Lab in Alderley Park and Alex Alderton, Roberto Amato, Jeffrey Barrett, Sonia Goncalves, Ewan Harrison, David K. Jackson, Ian Johnston, Dominic Kwiatkowski, Cordelia Langford, John Sillitoe on behalf of the Wellcome Sanger Institute COVID-19 Surveillance Team |
| EPI_ISL_3249116 | Lighthouse Lab in Milton Keynes | Wellcome Sanger Institute for the COVID-19 Genomics UK (COG-UK) Consortium | The Lighthouse Lab in Milton Keynes and Alex Alderton, Roberto Amato, Jeffrey Barrett, Sonia Goncalves, Ewan Harrison, David K. Jackson, Ian Johnston, Dominic Kwiatkowski, Cordelia Langford, John Sillitoe on behalf of the Wellcome Sanger Institute COVID-19 Surveillance Team |
| EPI_ISL_3249117 | Lighthouse Lab in Alderley Park | Wellcome Sanger Institute for the COVID-19 Genomics UK (COG-UK) Consortium | Jacquelyn Wynn, Mairead Hyland, The Lighthouse Lab in Alderley Park and Alex Alderton, Roberto Amato, Jeffrey Barrett, Sonia Goncalves, Ewan Harrison, David K. Jackson, Ian Johnston, Dominic Kwiatkowski, Cordelia Langford, John Sillitoe on behalf of the Wellcome Sanger Institute COVID-19 Surveillance Team |
| EPI_ISL_3249118 | ASST MONZA | ASST MONZA | Sergio Maria Ivano Malandrin |
| EPI_ISL_3249119 | Lighthouse Lab in Alderley Park | Wellcome Sanger Institute for the COVID-19 Genomics UK (COG-UK) Consortium | Jacquelyn Wynn, Mairead Hyland, The Lighthouse Lab in Alderley Park and Alex Alderton, Roberto Amato, Jeffrey Barrett, Sonia Goncalves, Ewan Harrison, David K. Jackson, Ian Johnston, Dominic Kwiatkowski, Cordelia Langford, John Sillitoe on behalf of the Wellcome Sanger Institute COVID-19 Surveillance Team |
| EPI_ISL_3249120 | Lighthouse Lab in Milton Keynes | Wellcome Sanger Institute for the COVID-19 Genomics UK (COG-UK) Consortium | The Lighthouse Lab in Milton Keynes and Alex Alderton, Roberto Amato, Jeffrey Barrett, Sonia Goncalves, Ewan Harrison, David K. Jackson, Ian Johnston, Dominic Kwiatkowski, Cordelia Langford, John Sillitoe on behalf of the Wellcome Sanger Institute COVID-19 Surveillance Team |
| EPI_ISL_3249121 | ASST MONZA | ASST MONZA | Sergio Maria Ivano Malandrin |
| EPI_ISL_3249122, EPI_ISL_3249123 | Lighthouse Lab in Milton Keynes | Wellcome Sanger Institute for the COVID-19 Genomics UK (COG-UK) Consortium | The Lighthouse Lab in Milton Keynes and Alex Alderton, Roberto Amato, Jeffrey Barrett, Sonia Goncalves, Ewan Harrison, David K. Jackson, Ian Johnston, Dominic Kwiatkowski, Cordelia Langford, John Sillitoe on behalf of the Wellcome Sanger Institute COVID-19 Surveillance Team |
| EPI_ISL_3249124 | Lighthouse Lab in Glasgow | Wellcome Sanger Institute for the COVID-19 Genomics UK (COG-UK) Consortium | Harper VanSteenhouse, Yumi Kasai, David Gray, Carol Clugston, Anna Dominiczak and Alex Alderton, Roberto Amato, Jeffrey Barrett, Sonia Goncalves, Ewan Harrison, David K. Jackson, Ian Johnston, Dominic Kwiatkowski, Cordelia Langford, John Sillitoe on behalf of the Wellcome Sanger Institute COVID-19 Surveillance Team |
| EPI_ISL_3249125 | ASST MONZA | ASST MONZA | Sergio Maria Ivano Malandrin |
| EPI_ISL_3249126 | Lighthouse Lab in Glasgow | Wellcome Sanger Institute for the COVID-19 Genomics UK (COG-UK) Consortium | Harper VanSteenhouse, Yumi Kasai, David Gray, Carol Clugston, Anna Dominiczak and Alex Alderton, Roberto Amato, Jeffrey Barrett, Sonia Goncalves, Ewan Harrison, David K. Jackson, Ian Johnston, Dominic Kwiatkowski, Cordelia Langford, John Sillitoe on behalf of the Wellcome Sanger Institute COVID-19 Surveillance Team |
| EPI_ISL_3249127 | Lighthouse Lab in Milton Keynes | Wellcome Sanger Institute for the COVID-19 Genomics UK (COG-UK) Consortium | The Lighthouse Lab in Milton Keynes and Alex Alderton, Roberto Amato, Jeffrey Barrett, Sonia Goncalves, Ewan Harrison, David K. Jackson, Ian Johnston, Dominic Kwiatkowski, Cordelia Langford, John Sillitoe on behalf of the Wellcome Sanger Institute COVID-19 Surveillance Team |
| EPI_ISL_3249128 | ASST MONZA | ASST MONZA | Sergio Maria Ivano Malandrin |
| EPI_ISL_3249129 | Lighthouse Lab in Glasgow | Wellcome Sanger Institute for the COVID-19 Genomics UK (COG-UK) Consortium | Harper VanSteenhouse, Yumi Kasai, David Gray, Carol Clugston, Anna Dominiczak and Alex Alderton, Roberto Amato, Jeffrey Barrett, Sonia Goncalves, Ewan Harrison, David K. Jackson, Ian Johnston, Dominic Kwiatkowski, Cordelia Langford, John Sillitoe on behalf of the Wellcome Sanger Institute COVID-19 Surveillance Team |
| EPI_ISL_3249130, EPI_ISL_3249131 | Lighthouse Lab in Milton Keynes | Wellcome Sanger Institute for the COVID-19 Genomics UK (COG-UK) Consortium | The Lighthouse Lab in Milton Keynes and Alex Alderton, Roberto Amato, Jeffrey Barrett, Sonia Goncalves, Ewan Harrison, David K. Jackson, Ian Johnston, Dominic Kwiatkowski, Cordelia Langford, John Sillitoe on behalf of the Wellcome Sanger Institute COVID-19 Surveillance Team |
| EPI_ISL_3249132 | ASST MONZA | ASST MONZA | Sergio Maria Ivano Malandrin |
| EPI_ISL_3249133 | Lighthouse Lab in Glasgow | Wellcome Sanger Institute for the COVID-19 Genomics UK (COG-UK) Consortium | Harper VanSteenhouse, Yumi Kasai, David Gray, Carol Clugston, Anna Dominiczak and Alex Alderton, Roberto Amato, Jeffrey Barrett, Sonia Goncalves, Ewan Harrison, David K. Jackson, Ian Johnston, Dominic Kwiatkowski, Cordelia Langford, John Sillitoe on behalf of the Wellcome Sanger Institute COVID-19 Surveillance Team |
| EPI_ISL_3249134 | Lighthouse Lab in Milton Keynes | Wellcome Sanger Institute for the COVID-19 Genomics UK (COG-UK) Consortium | The Lighthouse Lab in Milton Keynes and Alex Alderton, Roberto Amato, Jeffrey Barrett, Sonia Goncalves, Ewan Harrison, David K. Jackson, Ian Johnston, Dominic Kwiatkowski, Cordelia Langford, John Sillitoe on behalf of the Wellcome Sanger Institute COVID-19 Surveillance Team |
| EPI_ISL_3249135 | ASST MONZA | ASST MONZA | Sergio Maria Ivano Malandrin |
| EPI_ISL_3249136 | Lighthouse Lab in Alderley Park | Wellcome Sanger Institute for the COVID-19 Genomics UK (COG-UK) Consortium | Jacquelyn Wynn, Mairead Hyland, The Lighthouse Lab in Alderley Park and Alex Alderton, Roberto Amato, Jeffrey Barrett, Sonia Goncalves, Ewan Harrison, David K. Jackson, Ian Johnston, Dominic Kwiatkowski, Cordelia Langford, John Sillitoe on behalf of the Wellcome Sanger Institute COVID-19 Surveillance Team |
| EPI_ISL_3249137 | Lighthouse Lab in Milton Keynes | Wellcome Sanger Institute for the COVID-19 Genomics UK (COG-UK) Consortium | The Lighthouse Lab in Milton Keynes and Alex Alderton, Roberto Amato, Jeffrey Barrett, Sonia Goncalves, Ewan Harrison, David K. Jackson, Ian Johnston, Dominic Kwiatkowski, Cordelia Langford, John Sillitoe on behalf of the Wellcome Sanger Institute COVID-19 Surveillance Team |
| EPI_ISL_3249138 | ASST MONZA | ASST MONZA | Sergio Maria Ivano Malandrin |
| EPI_ISL_3249139 | Lighthouse Lab in Milton Keynes | Wellcome Sanger Institute for the COVID-19 Genomics UK (COG-UK) Consortium | The Lighthouse Lab in Milton Keynes and Alex Alderton, Roberto Amato, Jeffrey Barrett, Sonia Goncalves, Ewan Harrison, David K. Jackson, Ian Johnston, Dominic Kwiatkowski, Cordelia Langford, John Sillitoe on behalf of the Wellcome Sanger Institute COVID-19 Surveillance Team |
| EPI_ISL_3249140, EPI_ISL_3249141 | Lighthouse Lab in Glasgow | Wellcome Sanger Institute for the COVID-19 Genomics UK (COG-UK) Consortium | Harper VanSteenhouse, Yumi Kasai, David Gray, Carol Clugston, Anna Dominiczak and Alex Alderton, Roberto Amato, Jeffrey Barrett, Sonia Goncalves, Ewan Harrison, David K. Jackson, Ian Johnston, Dominic Kwiatkowski, Cordelia Langford, John Sillitoe on behalf of the Wellcome Sanger Institute COVID-19 Surveillance Team |
| EPI_ISL_3249142 | ASST MONZA | ASST MONZA | Sergio Maria Ivano Malandrin |

| | | | |
|---|---|---|---|
| EPI_ISL_3249143, EPI_ISL_3249144 | Lighthouse Lab in Alderley Park | Wellcome Sanger Institute for the COVID-19 Genomics UK (COG-UK) Consortium | Jacquelyn Wynn, Mairead Hyland, The Lighthouse Lab in Alderley Park and Alex Alderton, Roberto Amato, Jeffrey Barrett, Sonia Goncalves, Ewan Harrison, David K. Jackson, Ian Johnston, Dominic Kwiatkowski, Cordelia Langford, John Sillitoe on behalf of the Wellcome Sanger Institute COVID-19 Surveillance Team |
| EPI_ISL_3249145 | ASST MONZA | ASST MONZA | Sergio Maria Ivano Malandrin |
| EPI_ISL_3249146 | Lighthouse Lab in Glasgow | Wellcome Sanger Institute for the COVID-19 Genomics UK (COG-UK) Consortium | Harper VanSteenhouse, Yumi Kasai, David Gray, Carol Clugston, Anna Dominiczak and Alex Alderton, Roberto Amato, Jeffrey Barrett, Sonia Goncalves, Ewan Harrison, David K. Jackson, Ian Johnston, Dominic Kwiatkowski, Cordelia Langford, John Sillitoe on behalf of the Wellcome Sanger Institute COVID-19 Surveillance Team |
| EPI_ISL_3249147 | Lighthouse Lab in Alderley Park | Wellcome Sanger Institute for the COVID-19 Genomics UK (COG-UK) Consortium | Jacquelyn Wynn, Mairead Hyland, The Lighthouse Lab in Alderley Park and Alex Alderton, Roberto Amato, Jeffrey Barrett, Sonia Goncalves, Ewan Harrison, David K. Jackson, Ian Johnston, Dominic Kwiatkowski, Cordelia Langford, John Sillitoe on behalf of the Wellcome Sanger Institute COVID-19 Surveillance Team |
| EPI_ISL_3249148 | Lighthouse Lab in Milton Keynes | Wellcome Sanger Institute for the COVID-19 Genomics UK (COG-UK) Consortium | The Lighthouse Lab in Milton Keynes and Alex Alderton, Roberto Amato, Jeffrey Barrett, Sonia Goncalves, Ewan Harrison, David K. Jackson, Ian Johnston, Dominic Kwiatkowski, Cordelia Langford, John Sillitoe on behalf of the Wellcome Sanger Institute COVID-19 Surveillance Team |
| EPI_ISL_3249149 | ASST MONZA | ASST MONZA | Sergio Maria Ivano Malandrin |
| EPI_ISL_3249150 | Lighthouse Lab in Alderley Park | Wellcome Sanger Institute for the COVID-19 Genomics UK (COG-UK) Consortium | Jacquelyn Wynn, Mairead Hyland, The Lighthouse Lab in Alderley Park and Alex Alderton, Roberto Amato, Jeffrey Barrett, Sonia Goncalves, Ewan Harrison, David K. Jackson, Ian Johnston, Dominic Kwiatkowski, Cordelia Langford, John Sillitoe on behalf of the Wellcome Sanger Institute COVID-19 Surveillance Team |
| EPI_ISL_3249151 | Lighthouse Lab in Milton Keynes | Wellcome Sanger Institute for the COVID-19 Genomics UK (COG-UK) Consortium | The Lighthouse Lab in Milton Keynes and Alex Alderton, Roberto Amato, Jeffrey Barrett, Sonia Goncalves, Ewan Harrison, David K. Jackson, Ian Johnston, Dominic Kwiatkowski, Cordelia Langford, John Sillitoe on behalf of the Wellcome Sanger Institute COVID-19 Surveillance Team |
| EPI_ISL_3249152 | ASST MONZA | ASST MONZA | Sergio Maria Ivano Malandrin |
| EPI_ISL_3249153 | Lighthouse Lab in Milton Keynes | Wellcome Sanger Institute for the COVID-19 Genomics UK (COG-UK) Consortium | The Lighthouse Lab in Milton Keynes and Alex Alderton, Roberto Amato, Jeffrey Barrett, Sonia Goncalves, Ewan Harrison, David K. Jackson, Ian Johnston, Dominic Kwiatkowski, Cordelia Langford, John Sillitoe on behalf of the Wellcome Sanger Institute COVID-19 Surveillance Team |
| EPI_ISL_3249154 | Lighthouse Lab in Alderley Park | Wellcome Sanger Institute for the COVID-19 Genomics UK (COG-UK) Consortium | Jacquelyn Wynn, Mairead Hyland, The Lighthouse Lab in Alderley Park and Alex Alderton, Roberto Amato, Jeffrey Barrett, Sonia Goncalves, Ewan Harrison, David K. Jackson, Ian Johnston, Dominic Kwiatkowski, Cordelia Langford, John Sillitoe on behalf of the Wellcome Sanger Institute COVID-19 Surveillance Team |
| EPI_ISL_3249156 | Lighthouse Lab in Milton Keynes | Wellcome Sanger Institute for the COVID-19 Genomics UK (COG-UK) Consortium | The Lighthouse Lab in Milton Keynes and Alex Alderton, Roberto Amato, Jeffrey Barrett, Sonia Goncalves, Ewan Harrison, David K. Jackson, Ian Johnston, Dominic Kwiatkowski, Cordelia Langford, John Sillitoe on behalf of the Wellcome Sanger Institute COVID-19 Surveillance Team |
| EPI_ISL_3249157 | Lighthouse Lab in Alderley Park | Wellcome Sanger Institute for the COVID-19 Genomics UK (COG-UK) Consortium | Jacquelyn Wynn, Mairead Hyland, The Lighthouse Lab in Alderley Park and Alex Alderton, Roberto Amato, Jeffrey Barrett, Sonia Goncalves, Ewan Harrison, David K. Jackson, Ian Johnston, Dominic Kwiatkowski, Cordelia Langford, John Sillitoe on behalf of the Wellcome Sanger Institute COVID-19 Surveillance Team |
| EPI_ISL_3249158 | Lighthouse Lab in Milton Keynes | Wellcome Sanger Institute for the COVID-19 Genomics UK (COG-UK) Consortium | The Lighthouse Lab in Milton Keynes and Alex Alderton, Roberto Amato, Jeffrey Barrett, Sonia Goncalves, Ewan Harrison, David K. Jackson, Ian Johnston, Dominic Kwiatkowski, Cordelia Langford, John Sillitoe on behalf of the Wellcome Sanger Institute COVID-19 Surveillance Team |
| EPI_ISL_3249159, EPI_ISL_3249160 | Lighthouse Lab in Alderley Park | Wellcome Sanger Institute for the COVID-19 Genomics UK (COG-UK) Consortium | Jacquelyn Wynn, Mairead Hyland, The Lighthouse Lab in Alderley Park and Alex Alderton, Roberto Amato, Jeffrey Barrett, Sonia Goncalves, Ewan Harrison, David K. Jackson, Ian Johnston, Dominic Kwiatkowski, Cordelia Langford, John Sillitoe on behalf of the Wellcome Sanger Institute COVID-19 Surveillance Team |
| EPI_ISL_3249161 | Lighthouse Lab in Glasgow | Wellcome Sanger Institute for the COVID-19 Genomics UK (COG-UK) Consortium | Harper VanSteenhouse, Yumi Kasai, David Gray, Carol Clugston, Anna Dominiczak and Alex Alderton, Roberto Amato, Jeffrey Barrett, Sonia Goncalves, Ewan Harrison, David K. Jackson, Ian Johnston, Dominic Kwiatkowski, Cordelia Langford, John Sillitoe on behalf of the Wellcome Sanger Institute COVID-19 Surveillance Team |
| EPI_ISL_3249162, EPI_ISL_3249163, EPI_ISL_3249164, EPI_ISL_3249165, EPI_ISL_3249166, EPI_ISL_3249167, EPI_ISL_3249168, EPI_ISL_3249169 | Lighthouse Lab in Milton Keynes | Wellcome Sanger Institute for the COVID-19 Genomics UK (COG-UK) Consortium | The Lighthouse Lab in Milton Keynes and Alex Alderton, Roberto Amato, Jeffrey Barrett, Sonia Goncalves, Ewan Harrison, David K. Jackson, Ian Johnston, Dominic Kwiatkowski, Cordelia Langford, John Sillitoe on behalf of the Wellcome Sanger Institute COVID-19 Surveillance Team |
| EPI_ISL_3249170 | Lighthouse Lab in Glasgow | Wellcome Sanger Institute for the COVID-19 Genomics UK (COG-UK) Consortium | Harper VanSteenhouse, Yumi Kasai, David Gray, Carol Clugston, Anna Dominiczak and Alex Alderton, Roberto Amato, Jeffrey Barrett, Sonia Goncalves, Ewan Harrison, David K. Jackson, Ian Johnston, Dominic Kwiatkowski, Cordelia Langford, John Sillitoe on behalf of the Wellcome Sanger Institute COVID-19 Surveillance Team |
| EPI_ISL_3249171, EPI_ISL_3249172 | Lighthouse Lab in Milton Keynes | Wellcome Sanger Institute for the COVID-19 Genomics UK (COG-UK) Consortium | The Lighthouse Lab in Milton Keynes and Alex Alderton, Roberto Amato, Jeffrey Barrett, Sonia Goncalves, Ewan Harrison, David K. Jackson, Ian Johnston, Dominic Kwiatkowski, Cordelia Langford, John Sillitoe on behalf of the Wellcome Sanger Institute COVID-19 Surveillance Team |
| EPI_ISL_3249173 | Lighthouse Lab in Alderley Park | Wellcome Sanger Institute for the COVID-19 Genomics UK (COG-UK) Consortium | Jacquelyn Wynn, Mairead Hyland, The Lighthouse Lab in Alderley Park and Alex Alderton, Roberto Amato, Jeffrey Barrett, Sonia Goncalves, Ewan Harrison, David K. Jackson, Ian Johnston, Dominic Kwiatkowski, Cordelia Langford, John Sillitoe on behalf of the Wellcome Sanger Institute COVID-19 Surveillance Team |
| EPI_ISL_3249174 | Lighthouse Lab in Milton Keynes | Wellcome Sanger Institute for the COVID-19 Genomics UK (COG-UK) Consortium | The Lighthouse Lab in Milton Keynes and Alex Alderton, Roberto Amato, Jeffrey Barrett, Sonia Goncalves, Ewan Harrison, David K. Jackson, Ian Johnston, Dominic Kwiatkowski, Cordelia Langford, John Sillitoe on behalf of the Wellcome Sanger Institute COVID-19 Surveillance Team |
| EPI_ISL_3249175 | Lighthouse Lab in Glasgow | Wellcome Sanger Institute for the COVID-19 Genomics UK (COG-UK) Consortium | Harper VanSteenhouse, Yumi Kasai, David Gray, Carol Clugston, Anna Dominiczak and Alex Alderton, Roberto Amato, Jeffrey Barrett, Sonia Goncalves, Ewan Harrison, David K. Jackson, Ian Johnston, Dominic Kwiatkowski, Cordelia Langford, John Sillitoe on behalf of the Wellcome Sanger Institute COVID-19 Surveillance Team |
| EPI_ISL_3249176 | Lighthouse Lab in Alderley Park | Wellcome Sanger Institute for the COVID-19 Genomics UK (COG-UK) Consortium | Jacquelyn Wynn, Mairead Hyland, The Lighthouse Lab in Alderley Park and Alex Alderton, Roberto Amato, Jeffrey Barrett, Sonia Goncalves, Ewan Harrison, David K. Jackson, Ian Johnston, Dominic Kwiatkowski, Cordelia Langford, John Sillitoe on behalf of the Wellcome Sanger Institute COVID-19 Surveillance Team |
| EPI_ISL_3249177 | Lighthouse Lab in Milton Keynes | Wellcome Sanger Institute for the COVID-19 Genomics UK (COG-UK) Consortium | The Lighthouse Lab in Milton Keynes and Alex Alderton, Roberto Amato, Jeffrey Barrett, Sonia Goncalves, Ewan Harrison, David K. Jackson, Ian Johnston, Dominic Kwiatkowski, Cordelia Langford, John Sillitoe on behalf of the Wellcome Sanger Institute COVID-19 Surveillance Team |
| EPI_ISL_3249178 | Lighthouse Lab in Alderley Park | Wellcome Sanger Institute for the COVID-19 Genomics UK (COG-UK) Consortium | Jacquelyn Wynn, Mairead Hyland, The Lighthouse Lab in Alderley Park and Alex Alderton, Roberto Amato, Jeffrey Barrett, Sonia Goncalves, Ewan Harrison, David K. Jackson, Ian Johnston, Dominic Kwiatkowski, Cordelia Langford, John Sillitoe on behalf of the Wellcome Sanger Institute COVID-19 Surveillance Team |
| EPI_ISL_3249179, EPI_ISL_3249180 | Lighthouse Lab in Milton Keynes | Wellcome Sanger Institute for the COVID-19 Genomics UK (COG-UK) Consortium | The Lighthouse Lab in Milton Keynes and Alex Alderton, Roberto Amato, Jeffrey Barrett, Sonia Goncalves, Ewan Harrison, David K. Jackson, Ian Johnston, Dominic Kwiatkowski, Cordelia Langford, John Sillitoe on behalf of the Wellcome Sanger Institute COVID-19 Surveillance Team |
| EPI_ISL_3249181, EPI_ISL_3249182 | Lighthouse Lab in Alderley Park | Wellcome Sanger Institute for the COVID-19 Genomics UK (COG-UK) Consortium | Jacquelyn Wynn, Mairead Hyland, The Lighthouse Lab in Alderley Park and Alex Alderton, Roberto Amato, Jeffrey Barrett, Sonia Goncalves, Ewan Harrison, David K. Jackson, Ian Johnston, Dominic Kwiatkowski, Cordelia Langford, John Sillitoe on behalf of the Wellcome Sanger Institute COVID-19 Surveillance Team |
| EPI_ISL_3249183, EPI_ISL_3249184 | Lighthouse Lab in Milton Keynes | Wellcome Sanger Institute for the COVID-19 Genomics UK (COG-UK) Consortium | The Lighthouse Lab in Milton Keynes and Alex Alderton, Roberto Amato, Jeffrey Barrett, Sonia Goncalves, Ewan Harrison, David K. Jackson, Ian Johnston, Dominic Kwiatkowski, Cordelia Langford, John Sillitoe on behalf of the Wellcome Sanger Institute COVID-19 Surveillance Team |
| EPI_ISL_3249185 | Lighthouse Lab in Glasgow | Wellcome Sanger Institute for the COVID-19 Genomics UK (COG-UK) Consortium | Harper VanSteenhouse, Yumi Kasai, David Gray, Carol Clugston, Anna Dominiczak and Alex Alderton, Roberto Amato, Jeffrey Barrett, Sonia Goncalves, Ewan Harrison, David K. Jackson, Ian Johnston, Dominic Kwiatkowski, Cordelia Langford, John Sillitoe on behalf of the Wellcome Sanger Institute COVID-19 Surveillance Team |
| EPI_ISL_3249186, EPI_ISL_3249187 | Lighthouse Lab in Milton Keynes | Wellcome Sanger Institute for the COVID-19 Genomics UK (COG-UK) Consortium | The Lighthouse Lab in Milton Keynes and Alex Alderton, Roberto Amato, Jeffrey Barrett, Sonia Goncalves, Ewan Harrison, David K. Jackson, Ian Johnston, Dominic Kwiatkowski, Cordelia Langford, John Sillitoe on behalf of the Wellcome Sanger Institute COVID-19 Surveillance Team |
| EPI_ISL_3249188 | Lighthouse Lab in Glasgow | Wellcome Sanger Institute for the COVID-19 Genomics UK (COG-UK) Consortium | Harper VanSteenhouse, Yumi Kasai, David Gray, Carol Clugston, Anna Dominiczak and Alex Alderton, Roberto Amato, Jeffrey Barrett, Sonia Goncalves, Ewan Harrison, David K. Jackson, Ian Johnston, Dominic Kwiatkowski, Cordelia Langford, John Sillitoe on behalf of the Wellcome Sanger Institute COVID-19 Surveillance Team |
| EPI_ISL_3249189 | Lighthouse Lab in Milton Keynes | Wellcome Sanger Institute for the COVID-19 Genomics UK (COG-UK) Consortium | The Lighthouse Lab in Milton Keynes and Alex Alderton, Roberto Amato, Jeffrey Barrett, Sonia Goncalves, Ewan Harrison, David K. Jackson, Ian Johnston, Dominic Kwiatkowski, Cordelia Langford, John Sillitoe on behalf of the Wellcome Sanger Institute COVID-19 Surveillance Team |
| EPI_ISL_3249190, EPI_ISL_3249191 | Lighthouse Lab in Alderley Park | Wellcome Sanger Institute for the COVID-19 Genomics UK (COG-UK) Consortium | Jacquelyn Wynn, Mairead Hyland, The Lighthouse Lab in Alderley Park and Alex Alderton, Roberto Amato, Jeffrey Barrett, Sonia Goncalves, Ewan Harrison, David K. Jackson, Ian Johnston, Dominic Kwiatkowski, Cordelia Langford, John Sillitoe on behalf of the Wellcome Sanger Institute COVID-19 Surveillance Team |
| EPI_ISL_3249192, EPI_ISL_3249193 | Lighthouse Lab in Milton Keynes | Wellcome Sanger Institute for the COVID-19 Genomics UK (COG-UK) Consortium | The Lighthouse Lab in Milton Keynes and Alex Alderton, Roberto Amato, Jeffrey Barrett, Sonia Goncalves, Ewan Harrison, David K. Jackson, Ian Johnston, Dominic Kwiatkowski, Cordelia Langford, John Sillitoe on behalf of the Wellcome Sanger Institute COVID-19 Surveillance Team |
| EPI_ISL_3249194 | Lighthouse Lab in Glasgow | Wellcome Sanger Institute for the COVID-19 Genomics UK (COG-UK) Consortium | Harper VanSteenhouse, Yumi Kasai, David Gray, Carol Clugston, Anna Dominiczak and Alex Alderton, Roberto Amato, Jeffrey Barrett, Sonia Goncalves, Ewan Harrison, David K. Jackson, Ian Johnston, Dominic Kwiatkowski, Cordelia Langford, John Sillitoe on behalf of the Wellcome Sanger Institute COVID-19 Surveillance Team |
| EPI_ISL_3249195, EPI_ISL_3249196 | Lighthouse Lab in Milton Keynes | Wellcome Sanger Institute for the COVID-19 Genomics UK (COG-UK) Consortium | The Lighthouse Lab in Milton Keynes and Alex Alderton, Roberto Amato, Jeffrey Barrett, Sonia Goncalves, Ewan Harrison, David K. Jackson, Ian Johnston, Dominic Kwiatkowski, Cordelia Langford, John Sillitoe on behalf of the Wellcome Sanger Institute COVID-19 Surveillance Team |
| EPI_ISL_3249197 | Lighthouse Lab in Alderley Park | Wellcome Sanger Institute for the COVID-19 Genomics UK (COG-UK) Consortium | Jacquelyn Wynn, Mairead Hyland, The Lighthouse Lab in Alderley Park and Alex Alderton, Roberto Amato, Jeffrey Barrett, Sonia Goncalves, Ewan Harrison, David K. Jackson, Ian Johnston, Dominic Kwiatkowski, Cordelia Langford, John Sillitoe on behalf of the Wellcome Sanger Institute COVID-19 Surveillance Team |
| EPI_ISL_3249198, EPI_ISL_3249199, EPI_ISL_3249200 | Lighthouse Lab in Glasgow | Wellcome Sanger Institute for the COVID-19 Genomics UK (COG-UK) Consortium | Harper VanSteenhouse, Yumi Kasai, David Gray, Carol Clugston, Anna Dominiczak and Alex Alderton, Roberto Amato, Jeffrey Barrett, Sonia Goncalves, Ewan Harrison, David K. Jackson, Ian Johnston, Dominic Kwiatkowski, Cordelia Langford, John Sillitoe on behalf of the Wellcome Sanger Institute COVID-19 Surveillance Team |
| EPI_ISL_3249201 | Lighthouse Lab in Milton Keynes | Wellcome Sanger Institute for the COVID-19 Genomics UK (COG-UK) Consortium | The Lighthouse Lab in Milton Keynes and Alex Alderton, Roberto Amato, Jeffrey Barrett, Sonia Goncalves, Ewan Harrison, David K. Jackson, Ian Johnston, Dominic Kwiatkowski, Cordelia Langford, John Sillitoe on behalf of the Wellcome Sanger Institute COVID-19 Surveillance Team |
| EPI_ISL_3249202 | Lighthouse Lab in Alderley Park | Wellcome Sanger Institute for the COVID-19 Genomics UK (COG-UK) Consortium | Jacquelyn Wynn, Mairead Hyland, The Lighthouse Lab in Alderley Park and Alex Alderton, Roberto Amato, Jeffrey Barrett, Sonia Goncalves, Ewan Harrison, David K. Jackson, Ian Johnston, Dominic Kwiatkowski, Cordelia Langford, John Sillitoe on behalf of the Wellcome Sanger Institute COVID-19 Surveillance Team |
| EPI_ISL_3249203, EPI_ISL_3249204, EPI_ISL_3249205 | Lighthouse Lab in Milton Keynes | Wellcome Sanger Institute for the COVID-19 Genomics UK (COG-UK) Consortium | The Lighthouse Lab in Milton Keynes and Alex Alderton, Roberto Amato, Jeffrey Barrett, Sonia Goncalves, Ewan Harrison, David K. Jackson, Ian Johnston, Dominic Kwiatkowski, Cordelia Langford, John Sillitoe on behalf of the Wellcome Sanger Institute COVID-19 Surveillance Team |
| EPI_ISL_3249206 | Lighthouse Lab in Alderley Park | Wellcome Sanger Institute for the COVID-19 Genomics UK (COG-UK) Consortium | Jacquelyn Wynn, Mairead Hyland, The Lighthouse Lab in Alderley Park and Alex Alderton, Roberto Amato, Jeffrey Barrett, Sonia Goncalves, Ewan Harrison, David K. Jackson, Ian Johnston, Dominic Kwiatkowski, Cordelia Langford, John Sillitoe on behalf of the Wellcome Sanger Institute COVID-19 Surveillance Team |
| EPI_ISL_3249207, EPI_ISL_3249208, EPI_ISL_3249209, | Lighthouse Lab in Milton Keynes | Wellcome Sanger Institute for the COVID-19 Genomics UK (COG-UK) Consortium | The Lighthouse Lab in Milton Keynes and Alex Alderton, Roberto Amato, Jeffrey Barrett, Sonia Goncalves, Ewan Harrison, David K. Jackson, Ian Johnston, Dominic Kwiatkowski, Cordelia Langford, John Sillitoe on behalf of the Wellcome Sanger Institute COVID-19 Surveillance Team |

| | | | |
|---|---|---|---|
| EPI_ISL_3249210 | | | |
| EPI_ISL_3249211 | Lighthouse Lab in Alderley Park | Wellcome Sanger Institute for the COVID-19 Genomics UK (COG-UK) Consortium | Jacquelyn Wynn, Mairead Hyland, The Lighthouse Lab in Alderley Park and Alex Alderton, Roberto Amato, Jeffrey Barrett, Sonia Goncalves, Ewan Harrison, David K. Jackson, Ian Johnston, Dominic Kwiatkowski, Cordelia Langford, John Sillitoe on behalf of the Wellcome Sanger Institute COVID-19 Surveillance Team |
| EPI_ISL_3249212, EPI_ISL_3249213, EPI_ISL_3249214 | Lighthouse Lab in Milton Keynes | Wellcome Sanger Institute for the COVID-19 Genomics UK (COG-UK) Consortium | The Lighthouse Lab in Milton Keynes and Alex Alderton, Roberto Amato, Jeffrey Barrett, Sonia Goncalves, Ewan Harrison, David K. Jackson, Ian Johnston, Dominic Kwiatkowski, Cordelia Langford, John Sillitoe on behalf of the Wellcome Sanger Institute COVID-19 Surveillance Team |
| EPI_ISL_3249215, EPI_ISL_3249216 | Lighthouse Lab in Alderley Park | Wellcome Sanger Institute for the COVID-19 Genomics UK (COG-UK) Consortium | Jacquelyn Wynn, Mairead Hyland, The Lighthouse Lab in Alderley Park and Alex Alderton, Roberto Amato, Jeffrey Barrett, Sonia Goncalves, Ewan Harrison, David K. Jackson, Ian Johnston, Dominic Kwiatkowski, Cordelia Langford, John Sillitoe on behalf of the Wellcome Sanger Institute COVID-19 Surveillance Team |
| EPI_ISL_3249217, EPI_ISL_3249218, EPI_ISL_3249219, EPI_ISL_3249220 | Lighthouse Lab in Milton Keynes | Wellcome Sanger Institute for the COVID-19 Genomics UK (COG-UK) Consortium | The Lighthouse Lab in Milton Keynes and Alex Alderton, Roberto Amato, Jeffrey Barrett, Sonia Goncalves, Ewan Harrison, David K. Jackson, Ian Johnston, Dominic Kwiatkowski, Cordelia Langford, John Sillitoe on behalf of the Wellcome Sanger Institute COVID-19 Surveillance Team |
| EPI_ISL_3249221 | Lighthouse Lab in Glasgow | Wellcome Sanger Institute for the COVID-19 Genomics UK (COG-UK) Consortium | Harper VanSteenhouse, Yumi Kasai, David Gray, Carol Clugston, Anna Dominiczak and Alex Alderton, Roberto Amato, Jeffrey Barrett, Sonia Goncalves, Ewan Harrison, David K. Jackson, Ian Johnston, Dominic Kwiatkowski, Cordelia Langford, John Sillitoe on behalf of the Wellcome Sanger Institute COVID-19 Surveillance Team |
| EPI_ISL_3249222 | Lighthouse Lab in Alderley Park | Wellcome Sanger Institute for the COVID-19 Genomics UK (COG-UK) Consortium | Jacquelyn Wynn, Mairead Hyland, The Lighthouse Lab in Alderley Park and Alex Alderton, Roberto Amato, Jeffrey Barrett, Sonia Goncalves, Ewan Harrison, David K. Jackson, Ian Johnston, Dominic Kwiatkowski, Cordelia Langford, John Sillitoe on behalf of the Wellcome Sanger Institute COVID-19 Surveillance Team |
| EPI_ISL_3249223, EPI_ISL_3249224, EPI_ISL_3249225 | Lighthouse Lab in Milton Keynes | Wellcome Sanger Institute for the COVID-19 Genomics UK (COG-UK) Consortium | The Lighthouse Lab in Milton Keynes and Alex Alderton, Roberto Amato, Jeffrey Barrett, Sonia Goncalves, Ewan Harrison, David K. Jackson, Ian Johnston, Dominic Kwiatkowski, Cordelia Langford, John Sillitoe on behalf of the Wellcome Sanger Institute COVID-19 Surveillance Team |
| EPI_ISL_3249226 | Lighthouse Lab in Alderley Park | Wellcome Sanger Institute for the COVID-19 Genomics UK (COG-UK) Consortium | Jacquelyn Wynn, Mairead Hyland, The Lighthouse Lab in Alderley Park and Alex Alderton, Roberto Amato, Jeffrey Barrett, Sonia Goncalves, Ewan Harrison, David K. Jackson, Ian Johnston, Dominic Kwiatkowski, Cordelia Langford, John Sillitoe on behalf of the Wellcome Sanger Institute COVID-19 Surveillance Team |
| EPI_ISL_3249227 | Lighthouse Lab in Milton Keynes | Wellcome Sanger Institute for the COVID-19 Genomics UK (COG-UK) Consortium | The Lighthouse Lab in Milton Keynes and Alex Alderton, Roberto Amato, Jeffrey Barrett, Sonia Goncalves, Ewan Harrison, David K. Jackson, Ian Johnston, Dominic Kwiatkowski, Cordelia Langford, John Sillitoe on behalf of the Wellcome Sanger Institute COVID-19 Surveillance Team |
| EPI_ISL_3249228 | Lighthouse Lab in Glasgow | Wellcome Sanger Institute for the COVID-19 Genomics UK (COG-UK) Consortium | Harper VanSteenhouse, Yumi Kasai, David Gray, Carol Clugston, Anna Dominiczak and Alex Alderton, Roberto Amato, Jeffrey Barrett, Sonia Goncalves, Ewan Harrison, David K. Jackson, Ian Johnston, Dominic Kwiatkowski, Cordelia Langford, John Sillitoe on behalf of the Wellcome Sanger Institute COVID-19 Surveillance Team |
| EPI_ISL_3249229, EPI_ISL_3249230, EPI_ISL_3249231 | Lighthouse Lab in Milton Keynes | Wellcome Sanger Institute for the COVID-19 Genomics UK (COG-UK) Consortium | The Lighthouse Lab in Milton Keynes and Alex Alderton, Roberto Amato, Jeffrey Barrett, Sonia Goncalves, Ewan Harrison, David K. Jackson, Ian Johnston, Dominic Kwiatkowski, Cordelia Langford, John Sillitoe on behalf of the Wellcome Sanger Institute COVID-19 Surveillance Team |
| EPI_ISL_3249232 | Lighthouse Lab in Alderley Park | Wellcome Sanger Institute for the COVID-19 Genomics UK (COG-UK) Consortium | Jacquelyn Wynn, Mairead Hyland, The Lighthouse Lab in Alderley Park and Alex Alderton, Roberto Amato, Jeffrey Barrett, Sonia Goncalves, Ewan Harrison, David K. Jackson, Ian Johnston, Dominic Kwiatkowski, Cordelia Langford, John Sillitoe on behalf of the Wellcome Sanger Institute COVID-19 Surveillance Team |
| EPI_ISL_3249233 | Lighthouse Lab in Milton Keynes | Wellcome Sanger Institute for the COVID-19 Genomics UK (COG-UK) Consortium | The Lighthouse Lab in Milton Keynes and Alex Alderton, Roberto Amato, Jeffrey Barrett, Sonia Goncalves, Ewan Harrison, David K. Jackson, Ian Johnston, Dominic Kwiatkowski, Cordelia Langford, John Sillitoe on behalf of the Wellcome Sanger Institute COVID-19 Surveillance Team |
| EPI_ISL_3249234, EPI_ISL_3249235, EPI_ISL_3249236 | Lighthouse Lab in Alderley Park | Wellcome Sanger Institute for the COVID-19 Genomics UK (COG-UK) Consortium | Jacquelyn Wynn, Mairead Hyland, The Lighthouse Lab in Alderley Park and Alex Alderton, Roberto Amato, Jeffrey Barrett, Sonia Goncalves, Ewan Harrison, David K. Jackson, Ian Johnston, Dominic Kwiatkowski, Cordelia Langford, John Sillitoe on behalf of the Wellcome Sanger Institute COVID-19 Surveillance Team |
| EPI_ISL_3249237, EPI_ISL_3249238, EPI_ISL_3249239, EPI_ISL_3249240 | Lighthouse Lab in Milton Keynes | Wellcome Sanger Institute for the COVID-19 Genomics UK (COG-UK) Consortium | The Lighthouse Lab in Milton Keynes and Alex Alderton, Roberto Amato, Jeffrey Barrett, Sonia Goncalves, Ewan Harrison, David K. Jackson, Ian Johnston, Dominic Kwiatkowski, Cordelia Langford, John Sillitoe on behalf of the Wellcome Sanger Institute COVID-19 Surveillance Team |
| EPI_ISL_3249241 | Lighthouse Lab in Glasgow | Wellcome Sanger Institute for the COVID-19 Genomics UK (COG-UK) Consortium | Harper VanSteenhouse, Yumi Kasai, David Gray, Carol Clugston, Anna Dominiczak and Alex Alderton, Roberto Amato, Jeffrey Barrett, Sonia Goncalves, Ewan Harrison, David K. Jackson, Ian Johnston, Dominic Kwiatkowski, Cordelia Langford, John Sillitoe on behalf of the Wellcome Sanger Institute COVID-19 Surveillance Team |
| EPI_ISL_3249242 | Lighthouse Lab in Alderley Park | Wellcome Sanger Institute for the COVID-19 Genomics UK (COG-UK) Consortium | Jacquelyn Wynn, Mairead Hyland, The Lighthouse Lab in Alderley Park and Alex Alderton, Roberto Amato, Jeffrey Barrett, Sonia Goncalves, Ewan Harrison, David K. Jackson, Ian Johnston, Dominic Kwiatkowski, Cordelia Langford, John Sillitoe on behalf of the Wellcome Sanger Institute COVID-19 Surveillance Team |
| EPI_ISL_3249243, EPI_ISL_3249244 | Lighthouse Lab in Milton Keynes | Wellcome Sanger Institute for the COVID-19 Genomics UK (COG-UK) Consortium | The Lighthouse Lab in Milton Keynes and Alex Alderton, Roberto Amato, Jeffrey Barrett, Sonia Goncalves, Ewan Harrison, David K. Jackson, Ian Johnston, Dominic Kwiatkowski, Cordelia Langford, John Sillitoe on behalf of the Wellcome Sanger Institute COVID-19 Surveillance Team |
| EPI_ISL_3249245 | Lighthouse Lab in Glasgow | Wellcome Sanger Institute for the COVID-19 Genomics UK (COG-UK) Consortium | Harper VanSteenhouse, Yumi Kasai, David Gray, Carol Clugston, Anna Dominiczak and Alex Alderton, Roberto Amato, Jeffrey Barrett, Sonia Goncalves, Ewan Harrison, David K. Jackson, Ian Johnston, Dominic Kwiatkowski, Cordelia Langford, John Sillitoe on behalf of the Wellcome Sanger Institute COVID-19 Surveillance Team |
| EPI_ISL_3249246, EPI_ISL_3249247, EPI_ISL_3249248, EPI_ISL_3249249 | Lighthouse Lab in Milton Keynes | Wellcome Sanger Institute for the COVID-19 Genomics UK (COG-UK) Consortium | The Lighthouse Lab in Milton Keynes and Alex Alderton, Roberto Amato, Jeffrey Barrett, Sonia Goncalves, Ewan Harrison, David K. Jackson, Ian Johnston, Dominic Kwiatkowski, Cordelia Langford, John Sillitoe on behalf of the Wellcome Sanger Institute COVID-19 Surveillance Team |
| EPI_ISL_3249250 | Lighthouse Lab in Alderley Park | Wellcome Sanger Institute for the COVID-19 Genomics UK (COG-UK) Consortium | Jacquelyn Wynn, Mairead Hyland, The Lighthouse Lab in Alderley Park and Alex Alderton, Roberto Amato, Jeffrey Barrett, Sonia Goncalves, Ewan Harrison, David K. Jackson, Ian Johnston, Dominic Kwiatkowski, Cordelia Langford, John Sillitoe on behalf of the Wellcome Sanger Institute COVID-19 Surveillance Team |
| EPI_ISL_3249251 | Lighthouse Lab in Glasgow | Wellcome Sanger Institute for the COVID-19 Genomics UK (COG-UK) Consortium | Harper VanSteenhouse, Yumi Kasai, David Gray, Carol Clugston, Anna Dominiczak and Alex Alderton, Roberto Amato, Jeffrey Barrett, Sonia Goncalves, Ewan Harrison, David K. Jackson, Ian Johnston, Dominic Kwiatkowski, Cordelia Langford, John Sillitoe on behalf of the Wellcome Sanger Institute COVID-19 Surveillance Team |
| EPI_ISL_3249252, EPI_ISL_3249253 | Lighthouse Lab in Milton Keynes | Wellcome Sanger Institute for the COVID-19 Genomics UK (COG-UK) Consortium | The Lighthouse Lab in Milton Keynes and Alex Alderton, Roberto Amato, Jeffrey Barrett, Sonia Goncalves, Ewan Harrison, David K. Jackson, Ian Johnston, Dominic Kwiatkowski, Cordelia Langford, John Sillitoe on behalf of the Wellcome Sanger Institute COVID-19 Surveillance Team |
| EPI_ISL_3249254, EPI_ISL_3249255 | Lighthouse Lab in Alderley Park | Wellcome Sanger Institute for the COVID-19 Genomics UK (COG-UK) Consortium | Jacquelyn Wynn, Mairead Hyland, The Lighthouse Lab in Alderley Park and Alex Alderton, Roberto Amato, Jeffrey Barrett, Sonia Goncalves, Ewan Harrison, David K. Jackson, Ian Johnston, Dominic Kwiatkowski, Cordelia Langford, John Sillitoe on behalf of the Wellcome Sanger Institute COVID-19 Surveillance Team |
| EPI_ISL_3249256 | Lighthouse Lab in Glasgow | Wellcome Sanger Institute for the COVID-19 Genomics UK (COG-UK) Consortium | Harper VanSteenhouse, Yumi Kasai, David Gray, Carol Clugston, Anna Dominiczak and Alex Alderton, Roberto Amato, Jeffrey Barrett, Sonia Goncalves, Ewan Harrison, David K. Jackson, Ian Johnston, Dominic Kwiatkowski, Cordelia Langford, John Sillitoe on behalf of the Wellcome Sanger Institute COVID-19 Surveillance Team |
| EPI_ISL_3249257, EPI_ISL_3249258 | Lighthouse Lab in Milton Keynes | Wellcome Sanger Institute for the COVID-19 Genomics UK (COG-UK) Consortium | The Lighthouse Lab in Milton Keynes and Alex Alderton, Roberto Amato, Jeffrey Barrett, Sonia Goncalves, Ewan Harrison, David K. Jackson, Ian Johnston, Dominic Kwiatkowski, Cordelia Langford, John Sillitoe on behalf of the Wellcome Sanger Institute COVID-19 Surveillance Team |
| EPI_ISL_3249259, EPI_ISL_3249260 | Lighthouse Lab in Alderley Park | Wellcome Sanger Institute for the COVID-19 Genomics UK (COG-UK) Consortium | Jacquelyn Wynn, Mairead Hyland, The Lighthouse Lab in Alderley Park and Alex Alderton, Roberto Amato, Jeffrey Barrett, Sonia Goncalves, Ewan Harrison, David K. Jackson, Ian Johnston, Dominic Kwiatkowski, Cordelia Langford, John Sillitoe on behalf of the Wellcome Sanger Institute COVID-19 Surveillance Team |
| EPI_ISL_3249261 | Lighthouse Lab in Milton Keynes | Wellcome Sanger Institute for the COVID-19 Genomics UK (COG-UK) Consortium | The Lighthouse Lab in Milton Keynes and Alex Alderton, Roberto Amato, Jeffrey Barrett, Sonia Goncalves, Ewan Harrison, David K. Jackson, Ian Johnston, Dominic Kwiatkowski, Cordelia Langford, John Sillitoe on behalf of the Wellcome Sanger Institute COVID-19 Surveillance Team |
| EPI_ISL_3249262 | Lighthouse Lab in Glasgow | Wellcome Sanger Institute for the COVID-19 Genomics UK (COG-UK) Consortium | Harper VanSteenhouse, Yumi Kasai, David Gray, Carol Clugston, Anna Dominiczak and Alex Alderton, Roberto Amato, Jeffrey Barrett, Sonia Goncalves, Ewan Harrison, David K. Jackson, Ian Johnston, Dominic Kwiatkowski, Cordelia Langford, John Sillitoe on behalf of the Wellcome Sanger Institute COVID-19 Surveillance Team |
| EPI_ISL_3249263 | Lighthouse Lab in Alderley Park | Wellcome Sanger Institute for the COVID-19 Genomics UK (COG-UK) Consortium | Jacquelyn Wynn, Mairead Hyland, The Lighthouse Lab in Alderley Park and Alex Alderton, Roberto Amato, Jeffrey Barrett, Sonia Goncalves, Ewan Harrison, David K. Jackson, Ian Johnston, Dominic Kwiatkowski, Cordelia Langford, John Sillitoe on behalf of the Wellcome Sanger Institute COVID-19 Surveillance Team |
| EPI_ISL_3249264 | Lighthouse Lab in Milton Keynes | Wellcome Sanger Institute for the COVID-19 Genomics UK (COG-UK) Consortium | The Lighthouse Lab in Milton Keynes and Alex Alderton, Roberto Amato, Jeffrey Barrett, Sonia Goncalves, Ewan Harrison, David K. Jackson, Ian Johnston, Dominic Kwiatkowski, Cordelia Langford, John Sillitoe on behalf of the Wellcome Sanger Institute COVID-19 Surveillance Team |
| EPI_ISL_3249265, EPI_ISL_3249266 | Lighthouse Lab in Glasgow | Wellcome Sanger Institute for the COVID-19 Genomics UK (COG-UK) Consortium | Harper VanSteenhouse, Yumi Kasai, David Gray, Carol Clugston, Anna Dominiczak and Alex Alderton, Roberto Amato, Jeffrey Barrett, Sonia Goncalves, Ewan Harrison, David K. Jackson, Ian Johnston, Dominic Kwiatkowski, Cordelia Langford, John Sillitoe on behalf of the Wellcome Sanger Institute COVID-19 Surveillance Team |
| EPI_ISL_3249267 | Lighthouse Lab in Alderley Park | Wellcome Sanger Institute for the COVID-19 Genomics UK (COG-UK) Consortium | Jacquelyn Wynn, Mairead Hyland, The Lighthouse Lab in Alderley Park and Alex Alderton, Roberto Amato, Jeffrey Barrett, Sonia Goncalves, Ewan Harrison, David K. Jackson, Ian Johnston, Dominic Kwiatkowski, Cordelia Langford, John Sillitoe on behalf of the Wellcome Sanger Institute COVID-19 Surveillance Team |
| EPI_ISL_3249268, EPI_ISL_3249269 | Lighthouse Lab in Milton Keynes | Wellcome Sanger Institute for the COVID-19 Genomics UK (COG-UK) Consortium | The Lighthouse Lab in Milton Keynes and Alex Alderton, Roberto Amato, Jeffrey Barrett, Sonia Goncalves, Ewan Harrison, David K. Jackson, Ian Johnston, Dominic Kwiatkowski, Cordelia Langford, John Sillitoe on behalf of the Wellcome Sanger Institute COVID-19 Surveillance Team |
| EPI_ISL_3249270, EPI_ISL_3249271 | Lighthouse Lab in Glasgow | Wellcome Sanger Institute for the COVID-19 Genomics UK (COG-UK) Consortium | Harper VanSteenhouse, Yumi Kasai, David Gray, Carol Clugston, Anna Dominiczak and Alex Alderton, Roberto Amato, Jeffrey Barrett, Sonia Goncalves, Ewan Harrison, David K. Jackson, Ian Johnston, Dominic Kwiatkowski, Cordelia Langford, John Sillitoe on behalf of the Wellcome Sanger Institute COVID-19 Surveillance Team |
| EPI_ISL_3249272, EPI_ISL_3249273 | Lighthouse Lab in Alderley Park | Wellcome Sanger Institute for the COVID-19 Genomics UK (COG-UK) Consortium | Jacquelyn Wynn, Mairead Hyland, The Lighthouse Lab in Alderley Park and Alex Alderton, Roberto Amato, Jeffrey Barrett, Sonia Goncalves, Ewan Harrison, David K. Jackson, Ian Johnston, Dominic Kwiatkowski, Cordelia Langford, John Sillitoe on behalf of the Wellcome Sanger Institute COVID-19 Surveillance Team |
| EPI_ISL_3249274 | Lighthouse Lab in Milton Keynes | Wellcome Sanger Institute for the COVID-19 Genomics UK (COG-UK) Consortium | The Lighthouse Lab in Milton Keynes and Alex Alderton, Roberto Amato, Jeffrey Barrett, Sonia Goncalves, Ewan Harrison, David K. Jackson, Ian Johnston, Dominic Kwiatkowski, Cordelia Langford, John Sillitoe on behalf of the Wellcome Sanger Institute COVID-19 Surveillance Team |
| EPI_ISL_3249275, EPI_ISL_3249276 | Lighthouse Lab in Glasgow | Wellcome Sanger Institute for the COVID-19 Genomics UK (COG-UK) Consortium | Harper VanSteenhouse, Yumi Kasai, David Gray, Carol Clugston, Anna Dominiczak and Alex Alderton, Roberto Amato, Jeffrey Barrett, Sonia Goncalves, Ewan Harrison, David K. Jackson, Ian Johnston, Dominic Kwiatkowski, Cordelia Langford, John Sillitoe on behalf of the Wellcome Sanger Institute COVID-19 Surveillance Team |
| EPI_ISL_3249277 | Lighthouse Lab in Milton Keynes | Wellcome Sanger Institute for the COVID-19 Genomics UK (COG-UK) Consortium | The Lighthouse Lab in Milton Keynes and Alex Alderton, Roberto Amato, Jeffrey Barrett, Sonia Goncalves, Ewan Harrison, David K. Jackson, Ian Johnston, Dominic Kwiatkowski, Cordelia Langford, John Sillitoe on behalf of the Wellcome Sanger Institute COVID-19 Surveillance Team |
| EPI_ISL_3249278 | Lighthouse Lab in Alderley Park | Wellcome Sanger Institute for the COVID-19 Genomics UK (COG-UK) Consortium | Jacquelyn Wynn, Mairead Hyland, The Lighthouse Lab in Alderley Park and Alex Alderton, Roberto Amato, Jeffrey Barrett, Sonia Goncalves, Ewan Harrison, David K. Jackson, Ian Johnston, Dominic Kwiatkowski, Cordelia Langford, John Sillitoe on behalf of the Wellcome Sanger Institute COVID-19 Surveillance Team |
| EPI_ISL_3249279 | Lighthouse Lab in Milton Keynes | Wellcome Sanger Institute for the COVID-19 Genomics UK (COG-UK) | The Lighthouse Lab in Milton Keynes and Alex Alderton, Roberto Amato, Jeffrey Barrett, Sonia Goncalves, Ewan Harrison, David K. Jackson, Ian Johnston, Dominic Kwiatkowski, Cordelia Langford, John Sillitoe on behalf of the Wellcome Sanger |

| | | | |
|---|---|---|---|
| | | Consortium | Institute COVID-19 Surveillance Team |
| EPI_ISL_3249280 | Lighthouse Lab in Alderley Park | Wellcome Sanger Institute for the COVID-19 Genomics UK (COG-UK) Consortium | Jacquelyn Wynn, Mairead Hyland, The Lighthouse Lab in Alderley Park and Alex Alderton, Roberto Amato, Jeffrey Barrett, Sonia Goncalves, Ewan Harrison, David K. Jackson, Ian Johnston, Dominic Kwiatkowski, Cordelia Langford, John Sillitoe on behalf of the Wellcome Sanger Institute COVID-19 Surveillance Team |
| EPI_ISL_3249281 | Lighthouse Lab in Glasgow | Wellcome Sanger Institute for the COVID-19 Genomics UK (COG-UK) Consortium | Harper VanSteenhouse, Yumi Kasai, David Gray, Carol Clugston, Anna Dominiczak and Alex Alderton, Roberto Amato, Jeffrey Barrett, Sonia Goncalves, Ewan Harrison, David K. Jackson, Ian Johnston, Dominic Kwiatkowski, Cordelia Langford, John Sillitoe on behalf of the Wellcome Sanger Institute COVID-19 Surveillance Team |
| EPI_ISL_3249282 | Lighthouse Lab in Milton Keynes | Wellcome Sanger Institute for the COVID-19 Genomics UK (COG-UK) Consortium | The Lighthouse Lab in Milton Keynes and Alex Alderton, Roberto Amato, Jeffrey Barrett, Sonia Goncalves, Ewan Harrison, David K. Jackson, Ian Johnston, Dominic Kwiatkowski, Cordelia Langford, John Sillitoe on behalf of the Wellcome Sanger Institute COVID-19 Surveillance Team |
| EPI_ISL_3249283, EPI_ISL_3249284 | Lighthouse Lab in Alderley Park | Wellcome Sanger Institute for the COVID-19 Genomics UK (COG-UK) Consortium | Jacquelyn Wynn, Mairead Hyland, The Lighthouse Lab in Alderley Park and Alex Alderton, Roberto Amato, Jeffrey Barrett, Sonia Goncalves, Ewan Harrison, David K. Jackson, Ian Johnston, Dominic Kwiatkowski, Cordelia Langford, John Sillitoe on behalf of the Wellcome Sanger Institute COVID-19 Surveillance Team |
| EPI_ISL_3249285, EPI_ISL_3249286, EPI_ISL_3249287, EPI_ISL_3249288, EPI_ISL_3249289, EPI_ISL_3249290, EPI_ISL_3249291, EPI_ISL_3249292 | Lighthouse Lab in Milton Keynes | Wellcome Sanger Institute for the COVID-19 Genomics UK (COG-UK) Consortium | The Lighthouse Lab in Milton Keynes and Alex Alderton, Roberto Amato, Jeffrey Barrett, Sonia Goncalves, Ewan Harrison, David K. Jackson, Ian Johnston, Dominic Kwiatkowski, Cordelia Langford, John Sillitoe on behalf of the Wellcome Sanger Institute COVID-19 Surveillance Team |
| EPI_ISL_3249293, EPI_ISL_3249294 | Lighthouse Lab in Alderley Park | Wellcome Sanger Institute for the COVID-19 Genomics UK (COG-UK) Consortium | Jacquelyn Wynn, Mairead Hyland, The Lighthouse Lab in Alderley Park and Alex Alderton, Roberto Amato, Jeffrey Barrett, Sonia Goncalves, Ewan Harrison, David K. Jackson, Ian Johnston, Dominic Kwiatkowski, Cordelia Langford, John Sillitoe on behalf of the Wellcome Sanger Institute COVID-19 Surveillance Team |
| EPI_ISL_3249295 | Lighthouse Lab in Milton Keynes | Wellcome Sanger Institute for the COVID-19 Genomics UK (COG-UK) Consortium | The Lighthouse Lab in Milton Keynes and Alex Alderton, Roberto Amato, Jeffrey Barrett, Sonia Goncalves, Ewan Harrison, David K. Jackson, Ian Johnston, Dominic Kwiatkowski, Cordelia Langford, John Sillitoe on behalf of the Wellcome Sanger Institute COVID-19 Surveillance Team |
| EPI_ISL_3249296, EPI_ISL_3249297 | Lighthouse Lab in Alderley Park | Wellcome Sanger Institute for the COVID-19 Genomics UK (COG-UK) Consortium | Jacquelyn Wynn, Mairead Hyland, The Lighthouse Lab in Alderley Park and Alex Alderton, Roberto Amato, Jeffrey Barrett, Sonia Goncalves, Ewan Harrison, David K. Jackson, Ian Johnston, Dominic Kwiatkowski, Cordelia Langford, John Sillitoe on behalf of the Wellcome Sanger Institute COVID-19 Surveillance Team |
| EPI_ISL_3249298 | Lighthouse Lab in Milton Keynes | Wellcome Sanger Institute for the COVID-19 Genomics UK (COG-UK) Consortium | The Lighthouse Lab in Milton Keynes and Alex Alderton, Roberto Amato, Jeffrey Barrett, Sonia Goncalves, Ewan Harrison, David K. Jackson, Ian Johnston, Dominic Kwiatkowski, Cordelia Langford, John Sillitoe on behalf of the Wellcome Sanger Institute COVID-19 Surveillance Team |
| EPI_ISL_3249299 | Lighthouse Lab in Glasgow | Wellcome Sanger Institute for the COVID-19 Genomics UK (COG-UK) Consortium | Harper VanSteenhouse, Yumi Kasai, David Gray, Carol Clugston, Anna Dominiczak and Alex Alderton, Roberto Amato, Jeffrey Barrett, Sonia Goncalves, Ewan Harrison, David K. Jackson, Ian Johnston, Dominic Kwiatkowski, Cordelia Langford, John Sillitoe on behalf of the Wellcome Sanger Institute COVID-19 Surveillance Team |
| EPI_ISL_3249300, EPI_ISL_3249301, EPI_ISL_3249302 | Lighthouse Lab in Milton Keynes | Wellcome Sanger Institute for the COVID-19 Genomics UK (COG-UK) Consortium | The Lighthouse Lab in Milton Keynes and Alex Alderton, Roberto Amato, Jeffrey Barrett, Sonia Goncalves, Ewan Harrison, David K. Jackson, Ian Johnston, Dominic Kwiatkowski, Cordelia Langford, John Sillitoe on behalf of the Wellcome Sanger Institute COVID-19 Surveillance Team |
| EPI_ISL_3249303 | Lighthouse Lab in Glasgow | Wellcome Sanger Institute for the COVID-19 Genomics UK (COG-UK) Consortium | Harper VanSteenhouse, Yumi Kasai, David Gray, Carol Clugston, Anna Dominiczak and Alex Alderton, Roberto Amato, Jeffrey Barrett, Sonia Goncalves, Ewan Harrison, David K. Jackson, Ian Johnston, Dominic Kwiatkowski, Cordelia Langford, John Sillitoe on behalf of the Wellcome Sanger Institute COVID-19 Surveillance Team |
| EPI_ISL_3249304 | Lighthouse Lab in Milton Keynes | Wellcome Sanger Institute for the COVID-19 Genomics UK (COG-UK) Consortium | The Lighthouse Lab in Milton Keynes and Alex Alderton, Roberto Amato, Jeffrey Barrett, Sonia Goncalves, Ewan Harrison, David K. Jackson, Ian Johnston, Dominic Kwiatkowski, Cordelia Langford, John Sillitoe on behalf of the Wellcome Sanger Institute COVID-19 Surveillance Team |
| EPI_ISL_3249305 | Lighthouse Lab in Glasgow | Wellcome Sanger Institute for the COVID-19 Genomics UK (COG-UK) Consortium | Harper VanSteenhouse, Yumi Kasai, David Gray, Carol Clugston, Anna Dominiczak and Alex Alderton, Roberto Amato, Jeffrey Barrett, Sonia Goncalves, Ewan Harrison, David K. Jackson, Ian Johnston, Dominic Kwiatkowski, Cordelia Langford, John Sillitoe on behalf of the Wellcome Sanger Institute COVID-19 Surveillance Team |
| EPI_ISL_3249306, EPI_ISL_3249307 | Lighthouse Lab in Alderley Park | Wellcome Sanger Institute for the COVID-19 Genomics UK (COG-UK) Consortium | Jacquelyn Wynn, Mairead Hyland, The Lighthouse Lab in Alderley Park and Alex Alderton, Roberto Amato, Jeffrey Barrett, Sonia Goncalves, Ewan Harrison, David K. Jackson, Ian Johnston, Dominic Kwiatkowski, Cordelia Langford, John Sillitoe on behalf of the Wellcome Sanger Institute COVID-19 Surveillance Team |
| EPI_ISL_3249308, EPI_ISL_3249309 | Lighthouse Lab in Milton Keynes | Wellcome Sanger Institute for the COVID-19 Genomics UK (COG-UK) Consortium | The Lighthouse Lab in Milton Keynes and Alex Alderton, Roberto Amato, Jeffrey Barrett, Sonia Goncalves, Ewan Harrison, David K. Jackson, Ian Johnston, Dominic Kwiatkowski, Cordelia Langford, John Sillitoe on behalf of the Wellcome Sanger Institute COVID-19 Surveillance Team |
| EPI_ISL_3249310, EPI_ISL_3249311 | Lighthouse Lab in Alderley Park | Wellcome Sanger Institute for the COVID-19 Genomics UK (COG-UK) Consortium | Jacquelyn Wynn, Mairead Hyland, The Lighthouse Lab in Alderley Park and Alex Alderton, Roberto Amato, Jeffrey Barrett, Sonia Goncalves, Ewan Harrison, David K. Jackson, Ian Johnston, Dominic Kwiatkowski, Cordelia Langford, John Sillitoe on behalf of the Wellcome Sanger Institute COVID-19 Surveillance Team |
| EPI_ISL_3249312 | Lighthouse Lab in Glasgow | Wellcome Sanger Institute for the COVID-19 Genomics UK (COG-UK) Consortium | Harper VanSteenhouse, Yumi Kasai, David Gray, Carol Clugston, Anna Dominiczak and Alex Alderton, Roberto Amato, Jeffrey Barrett, Sonia Goncalves, Ewan Harrison, David K. Jackson, Ian Johnston, Dominic Kwiatkowski, Cordelia Langford, John Sillitoe on behalf of the Wellcome Sanger Institute COVID-19 Surveillance Team |
| EPI_ISL_3249313 | Lighthouse Lab in Milton Keynes | Wellcome Sanger Institute for the COVID-19 Genomics UK (COG-UK) Consortium | The Lighthouse Lab in Milton Keynes and Alex Alderton, Roberto Amato, Jeffrey Barrett, Sonia Goncalves, Ewan Harrison, David K. Jackson, Ian Johnston, Dominic Kwiatkowski, Cordelia Langford, John Sillitoe on behalf of the Wellcome Sanger Institute COVID-19 Surveillance Team |
| EPI_ISL_3249314 | Lighthouse Lab in Alderley Park | Wellcome Sanger Institute for the COVID-19 Genomics UK (COG-UK) Consortium | Jacquelyn Wynn, Mairead Hyland, The Lighthouse Lab in Alderley Park and Alex Alderton, Roberto Amato, Jeffrey Barrett, Sonia Goncalves, Ewan Harrison, David K. Jackson, Ian Johnston, Dominic Kwiatkowski, Cordelia Langford, John Sillitoe on behalf of the Wellcome Sanger Institute COVID-19 Surveillance Team |
| EPI_ISL_3249315 | Lighthouse Lab in Glasgow | Wellcome Sanger Institute for the COVID-19 Genomics UK (COG-UK) Consortium | Harper VanSteenhouse, Yumi Kasai, David Gray, Carol Clugston, Anna Dominiczak and Alex Alderton, Roberto Amato, Jeffrey Barrett, Sonia Goncalves, Ewan Harrison, David K. Jackson, Ian Johnston, Dominic Kwiatkowski, Cordelia Langford, John Sillitoe on behalf of the Wellcome Sanger Institute COVID-19 Surveillance Team |
| EPI_ISL_3249316, EPI_ISL_3249317, EPI_ISL_3249318, EPI_ISL_3249319, EPI_ISL_3249320 | Lighthouse Lab in Milton Keynes | Wellcome Sanger Institute for the COVID-19 Genomics UK (COG-UK) Consortium | The Lighthouse Lab in Milton Keynes and Alex Alderton, Roberto Amato, Jeffrey Barrett, Sonia Goncalves, Ewan Harrison, David K. Jackson, Ian Johnston, Dominic Kwiatkowski, Cordelia Langford, John Sillitoe on behalf of the Wellcome Sanger Institute COVID-19 Surveillance Team |
| EPI_ISL_3249321 | Lighthouse Lab in Glasgow | Wellcome Sanger Institute for the COVID-19 Genomics UK (COG-UK) Consortium | Harper VanSteenhouse, Yumi Kasai, David Gray, Carol Clugston, Anna Dominiczak and Alex Alderton, Roberto Amato, Jeffrey Barrett, Sonia Goncalves, Ewan Harrison, David K. Jackson, Ian Johnston, Dominic Kwiatkowski, Cordelia Langford, John Sillitoe on behalf of the Wellcome Sanger Institute COVID-19 Surveillance Team |
| EPI_ISL_3249322 | Lighthouse Lab in Alderley Park | Wellcome Sanger Institute for the COVID-19 Genomics UK (COG-UK) Consortium | Jacquelyn Wynn, Mairead Hyland, The Lighthouse Lab in Alderley Park and Alex Alderton, Roberto Amato, Jeffrey Barrett, Sonia Goncalves, Ewan Harrison, David K. Jackson, Ian Johnston, Dominic Kwiatkowski, Cordelia Langford, John Sillitoe on behalf of the Wellcome Sanger Institute COVID-19 Surveillance Team |
| EPI_ISL_3249323, EPI_ISL_3249324, EPI_ISL_3249325 | Lighthouse Lab in Milton Keynes | Wellcome Sanger Institute for the COVID-19 Genomics UK (COG-UK) Consortium | The Lighthouse Lab in Milton Keynes and Alex Alderton, Roberto Amato, Jeffrey Barrett, Sonia Goncalves, Ewan Harrison, David K. Jackson, Ian Johnston, Dominic Kwiatkowski, Cordelia Langford, John Sillitoe on behalf of the Wellcome Sanger Institute COVID-19 Surveillance Team |
| EPI_ISL_3249326 | Lighthouse Lab in Alderley Park | Wellcome Sanger Institute for the COVID-19 Genomics UK (COG-UK) Consortium | Jacquelyn Wynn, Mairead Hyland, The Lighthouse Lab in Alderley Park and Alex Alderton, Roberto Amato, Jeffrey Barrett, Sonia Goncalves, Ewan Harrison, David K. Jackson, Ian Johnston, Dominic Kwiatkowski, Cordelia Langford, John Sillitoe on behalf of the Wellcome Sanger Institute COVID-19 Surveillance Team |
| EPI_ISL_3249327 | Lighthouse Lab in Glasgow | Wellcome Sanger Institute for the COVID-19 Genomics UK (COG-UK) Consortium | Harper VanSteenhouse, Yumi Kasai, David Gray, Carol Clugston, Anna Dominiczak and Alex Alderton, Roberto Amato, Jeffrey Barrett, Sonia Goncalves, Ewan Harrison, David K. Jackson, Ian Johnston, Dominic Kwiatkowski, Cordelia Langford, John Sillitoe on behalf of the Wellcome Sanger Institute COVID-19 Surveillance Team |
| EPI_ISL_3249328, EPI_ISL_3249329 | Lighthouse Lab in Alderley Park | Wellcome Sanger Institute for the COVID-19 Genomics UK (COG-UK) Consortium | Jacquelyn Wynn, Mairead Hyland, The Lighthouse Lab in Alderley Park and Alex Alderton, Roberto Amato, Jeffrey Barrett, Sonia Goncalves, Ewan Harrison, David K. Jackson, Ian Johnston, Dominic Kwiatkowski, Cordelia Langford, John Sillitoe on behalf of the Wellcome Sanger Institute COVID-19 Surveillance Team |
| EPI_ISL_3249330 | Lighthouse Lab in Glasgow | Wellcome Sanger Institute for the COVID-19 Genomics UK (COG-UK) Consortium | Harper VanSteenhouse, Yumi Kasai, David Gray, Carol Clugston, Anna Dominiczak and Alex Alderton, Roberto Amato, Jeffrey Barrett, Sonia Goncalves, Ewan Harrison, David K. Jackson, Ian Johnston, Dominic Kwiatkowski, Cordelia Langford, John Sillitoe on behalf of the Wellcome Sanger Institute COVID-19 Surveillance Team |
| EPI_ISL_3249331 | Lighthouse Lab in Milton Keynes | Wellcome Sanger Institute for the COVID-19 Genomics UK (COG-UK) Consortium | The Lighthouse Lab in Milton Keynes and Alex Alderton, Roberto Amato, Jeffrey Barrett, Sonia Goncalves, Ewan Harrison, David K. Jackson, Ian Johnston, Dominic Kwiatkowski, Cordelia Langford, John Sillitoe on behalf of the Wellcome Sanger Institute COVID-19 Surveillance Team |
| EPI_ISL_3249332 | Berkshire and Surrey Pathology Services Lighthouse Laboratory | Wellcome Sanger Institute for the COVID-19 Genomics UK (COG-UK) Consortium | Berkshire and Surrey Pathology Services Lighthouse Laboratory and Alex Alderton, Roberto Amato, Jeffrey Barrett, Sonia Goncalves, Ewan Harrison, David K. Jackson, Ian Johnston, Dominic Kwiatkowski, Cordelia Langford, John Sillitoe on behalf of the Wellcome Sanger Institute COVID-19 Surveillance Team |
| EPI_ISL_3249333 | Lighthouse Lab in Alderley Park | Wellcome Sanger Institute for the COVID-19 Genomics UK (COG-UK) Consortium | Jacquelyn Wynn, Mairead Hyland, The Lighthouse Lab in Alderley Park and Alex Alderton, Roberto Amato, Jeffrey Barrett, Sonia Goncalves, Ewan Harrison, David K. Jackson, Ian Johnston, Dominic Kwiatkowski, Cordelia Langford, John Sillitoe on behalf of the Wellcome Sanger Institute COVID-19 Surveillance Team |
| EPI_ISL_3249334 | Berkshire and Surrey Pathology Services Lighthouse Laboratory | Wellcome Sanger Institute for the COVID-19 Genomics UK (COG-UK) Consortium | Berkshire and Surrey Pathology Services Lighthouse Laboratory and Alex Alderton, Roberto Amato, Jeffrey Barrett, Sonia Goncalves, Ewan Harrison, David K. Jackson, Ian Johnston, Dominic Kwiatkowski, Cordelia Langford, John Sillitoe on behalf of the Wellcome Sanger Institute COVID-19 Surveillance Team |
| EPI_ISL_3249335 | Lighthouse Lab in Alderley Park | Wellcome Sanger Institute for the COVID-19 Genomics UK (COG-UK) Consortium | Jacquelyn Wynn, Mairead Hyland, The Lighthouse Lab in Alderley Park and Alex Alderton, Roberto Amato, Jeffrey Barrett, Sonia Goncalves, Ewan Harrison, David K. Jackson, Ian Johnston, Dominic Kwiatkowski, Cordelia Langford, John Sillitoe on behalf of the Wellcome Sanger Institute COVID-19 Surveillance Team |
| EPI_ISL_3249336, EPI_ISL_3249337 | Lighthouse Lab in Milton Keynes | Wellcome Sanger Institute for the COVID-19 Genomics UK (COG-UK) Consortium | The Lighthouse Lab in Milton Keynes and Alex Alderton, Roberto Amato, Jeffrey Barrett, Sonia Goncalves, Ewan Harrison, David K. Jackson, Ian Johnston, Dominic Kwiatkowski, Cordelia Langford, John Sillitoe on behalf of the Wellcome Sanger Institute COVID-19 Surveillance Team |
| EPI_ISL_3249338, EPI_ISL_3249339, EPI_ISL_3249340 | Berkshire and Surrey Pathology Services Lighthouse Laboratory | Wellcome Sanger Institute for the COVID-19 Genomics UK (COG-UK) Consortium | Berkshire and Surrey Pathology Services Lighthouse Laboratory and Alex Alderton, Roberto Amato, Jeffrey Barrett, Sonia Goncalves, Ewan Harrison, David K. Jackson, Ian Johnston, Dominic Kwiatkowski, Cordelia Langford, John Sillitoe on behalf of the Wellcome Sanger Institute COVID-19 Surveillance Team |
| EPI_ISL_3249341, EPI_ISL_3249342 | Lighthouse Lab in Milton Keynes | Wellcome Sanger Institute for the COVID-19 Genomics UK (COG-UK) Consortium | The Lighthouse Lab in Milton Keynes and Alex Alderton, Roberto Amato, Jeffrey Barrett, Sonia Goncalves, Ewan Harrison, David K. Jackson, Ian Johnston, Dominic Kwiatkowski, Cordelia Langford, John Sillitoe on behalf of the Wellcome Sanger Institute COVID-19 Surveillance Team |
| EPI_ISL_3249343 | Lighthouse Lab in Alderley Park | Wellcome Sanger Institute for the COVID-19 Genomics UK (COG-UK) Consortium | Jacquelyn Wynn, Mairead Hyland, The Lighthouse Lab in Alderley Park and Alex Alderton, Roberto Amato, Jeffrey Barrett, Sonia Goncalves, Ewan Harrison, David K. Jackson, Ian Johnston, Dominic Kwiatkowski, Cordelia Langford, John Sillitoe on behalf of the Wellcome Sanger Institute COVID-19 Surveillance Team |
| EPI_ISL_3249344 | Berkshire and Surrey Pathology Services Lighthouse Laboratory | Wellcome Sanger Institute for the COVID-19 Genomics UK (COG-UK) Consortium | Berkshire and Surrey Pathology Services Lighthouse Laboratory and Alex Alderton, Roberto Amato, Jeffrey Barrett, Sonia Goncalves, Ewan Harrison, David K. Jackson, Ian Johnston, Dominic Kwiatkowski, Cordelia Langford, John Sillitoe on behalf of the Wellcome Sanger Institute COVID-19 Surveillance Team |
| EPI_ISL_3249345, | Lighthouse Lab in Milton Keynes | Wellcome Sanger Institute for the COVID-19 Genomics UK (COG-UK) | The Lighthouse Lab in Milton Keynes and Alex Alderton, Roberto Amato, Jeffrey Barrett, Sonia Goncalves, Ewan Harrison, David K. Jackson, Ian Johnston, Dominic Kwiatkowski, Cordelia Langford, John Sillitoe on behalf of the Wellcome Sanger |

| | | | |
|---|---|---|---|
| EPI_ISL_3249346 | | Consortium | Institute COVID-19 Surveillance Team |
| EPI_ISL_3249347 | Lighthouse Lab in Alderley Park | Wellcome Sanger Institute for the COVID-19 Genomics UK (COG-UK) Consortium | Jacquelyn Wynn, Mairead Hyland, The Lighthouse Lab in Alderley Park and Alex Alderton, Roberto Amato, Jeffrey Barrett, Sonia Goncalves, Ewan Harrison, David K. Jackson, Ian Johnston, Dominic Kwiatkowski, Cordelia Langford, John Sillitoe on behalf of the Wellcome Sanger Institute COVID-19 Surveillance Team |
| EPI_ISL_3249348, EPI_ISL_3249349, EPI_ISL_3249350 | Lighthouse Lab in Milton Keynes | Wellcome Sanger Institute for the COVID-19 Genomics UK (COG-UK) Consortium | The Lighthouse Lab in Milton Keynes and Alex Alderton, Roberto Amato, Jeffrey Barrett, Sonia Goncalves, Ewan Harrison, David K. Jackson, Ian Johnston, Dominic Kwiatkowski, Cordelia Langford, John Sillitoe on behalf of the Wellcome Sanger Institute COVID-19 Surveillance Team |
| EPI_ISL_3249351 | Berkshire and Surrey Pathology Services Lighthouse Laboratory | Wellcome Sanger Institute for the COVID-19 Genomics UK (COG-UK) Consortium | Berkshire and Surrey Pathology Services Lighthouse Laboratory and Alex Alderton, Roberto Amato, Jeffrey Barrett, Sonia Goncalves, Ewan Harrison, David K. Jackson, Ian Johnston, Dominic Kwiatkowski, Cordelia Langford, John Sillitoe on behalf of the Wellcome Sanger Institute COVID-19 Surveillance Team |
| EPI_ISL_3249352 | Lighthouse Lab in Alderley Park | Wellcome Sanger Institute for the COVID-19 Genomics UK (COG-UK) Consortium | Jacquelyn Wynn, Mairead Hyland, The Lighthouse Lab in Alderley Park and Alex Alderton, Roberto Amato, Jeffrey Barrett, Sonia Goncalves, Ewan Harrison, David K. Jackson, Ian Johnston, Dominic Kwiatkowski, Cordelia Langford, John Sillitoe on behalf of the Wellcome Sanger Institute COVID-19 Surveillance Team |
| EPI_ISL_3249353 | Lighthouse Lab in Milton Keynes | Wellcome Sanger Institute for the COVID-19 Genomics UK (COG-UK) Consortium | The Lighthouse Lab in Milton Keynes and Alex Alderton, Roberto Amato, Jeffrey Barrett, Sonia Goncalves, Ewan Harrison, David K. Jackson, Ian Johnston, Dominic Kwiatkowski, Cordelia Langford, John Sillitoe on behalf of the Wellcome Sanger Institute COVID-19 Surveillance Team |
| EPI_ISL_3249354 | Lighthouse Lab in Glasgow | Wellcome Sanger Institute for the COVID-19 Genomics UK (COG-UK) Consortium | Harper VanSteenhouse, Yumi Kasai, David Gray, Carol Clugston, Anna Dominiczak and Alex Alderton, Roberto Amato, Jeffrey Barrett, Sonia Goncalves, Ewan Harrison, David K. Jackson, Ian Johnston, Dominic Kwiatkowski, Cordelia Langford, John Sillitoe on behalf of the Wellcome Sanger Institute COVID-19 Surveillance Team |
| EPI_ISL_3249355 | Lighthouse Lab in Alderley Park | Wellcome Sanger Institute for the COVID-19 Genomics UK (COG-UK) Consortium | Jacquelyn Wynn, Mairead Hyland, The Lighthouse Lab in Alderley Park and Alex Alderton, Roberto Amato, Jeffrey Barrett, Sonia Goncalves, Ewan Harrison, David K. Jackson, Ian Johnston, Dominic Kwiatkowski, Cordelia Langford, John Sillitoe on behalf of the Wellcome Sanger Institute COVID-19 Surveillance Team |
| EPI_ISL_3249356 | Lighthouse Lab in Milton Keynes | Wellcome Sanger Institute for the COVID-19 Genomics UK (COG-UK) Consortium | The Lighthouse Lab in Milton Keynes and Alex Alderton, Roberto Amato, Jeffrey Barrett, Sonia Goncalves, Ewan Harrison, David K. Jackson, Ian Johnston, Dominic Kwiatkowski, Cordelia Langford, John Sillitoe on behalf of the Wellcome Sanger Institute COVID-19 Surveillance Team |
| EPI_ISL_3249357 | Lighthouse Lab in Alderley Park | Wellcome Sanger Institute for the COVID-19 Genomics UK (COG-UK) Consortium | Jacquelyn Wynn, Mairead Hyland, The Lighthouse Lab in Alderley Park and Alex Alderton, Roberto Amato, Jeffrey Barrett, Sonia Goncalves, Ewan Harrison, David K. Jackson, Ian Johnston, Dominic Kwiatkowski, Cordelia Langford, John Sillitoe on behalf of the Wellcome Sanger Institute COVID-19 Surveillance Team |
| EPI_ISL_3249358 | Lighthouse Lab in Milton Keynes | Wellcome Sanger Institute for the COVID-19 Genomics UK (COG-UK) Consortium | The Lighthouse Lab in Milton Keynes and Alex Alderton, Roberto Amato, Jeffrey Barrett, Sonia Goncalves, Ewan Harrison, David K. Jackson, Ian Johnston, Dominic Kwiatkowski, Cordelia Langford, John Sillitoe on behalf of the Wellcome Sanger Institute COVID-19 Surveillance Team |
| EPI_ISL_3249359 | Berkshire and Surrey Pathology Services Lighthouse Laboratory | Wellcome Sanger Institute for the COVID-19 Genomics UK (COG-UK) Consortium | Berkshire and Surrey Pathology Services Lighthouse Laboratory and Alex Alderton, Roberto Amato, Jeffrey Barrett, Sonia Goncalves, Ewan Harrison, David K. Jackson, Ian Johnston, Dominic Kwiatkowski, Cordelia Langford, John Sillitoe on behalf of the Wellcome Sanger Institute COVID-19 Surveillance Team |
| EPI_ISL_3249360 | Lighthouse Lab in Alderley Park | Wellcome Sanger Institute for the COVID-19 Genomics UK (COG-UK) Consortium | Jacquelyn Wynn, Mairead Hyland, The Lighthouse Lab in Alderley Park and Alex Alderton, Roberto Amato, Jeffrey Barrett, Sonia Goncalves, Ewan Harrison, David K. Jackson, Ian Johnston, Dominic Kwiatkowski, Cordelia Langford, John Sillitoe on behalf of the Wellcome Sanger Institute COVID-19 Surveillance Team |
| EPI_ISL_3249361 | Berkshire and Surrey Pathology Services Lighthouse Laboratory | Wellcome Sanger Institute for the COVID-19 Genomics UK (COG-UK) Consortium | Berkshire and Surrey Pathology Services Lighthouse Laboratory and Alex Alderton, Roberto Amato, Jeffrey Barrett, Sonia Goncalves, Ewan Harrison, David K. Jackson, Ian Johnston, Dominic Kwiatkowski, Cordelia Langford, John Sillitoe on behalf of the Wellcome Sanger Institute COVID-19 Surveillance Team |
| EPI_ISL_3249362 | Lighthouse Lab in Glasgow | Wellcome Sanger Institute for the COVID-19 Genomics UK (COG-UK) Consortium | Harper VanSteenhouse, Yumi Kasai, David Gray, Carol Clugston, Anna Dominiczak and Alex Alderton, Roberto Amato, Jeffrey Barrett, Sonia Goncalves, Ewan Harrison, David K. Jackson, Ian Johnston, Dominic Kwiatkowski, Cordelia Langford, John Sillitoe on behalf of the Wellcome Sanger Institute COVID-19 Surveillance Team |
| EPI_ISL_3249363 | Lighthouse Lab in Milton Keynes | Wellcome Sanger Institute for the COVID-19 Genomics UK (COG-UK) Consortium | The Lighthouse Lab in Milton Keynes and Alex Alderton, Roberto Amato, Jeffrey Barrett, Sonia Goncalves, Ewan Harrison, David K. Jackson, Ian Johnston, Dominic Kwiatkowski, Cordelia Langford, John Sillitoe on behalf of the Wellcome Sanger Institute COVID-19 Surveillance Team |
| EPI_ISL_3249364 | Lighthouse Lab in Glasgow | Wellcome Sanger Institute for the COVID-19 Genomics UK (COG-UK) Consortium | Harper VanSteenhouse, Yumi Kasai, David Gray, Carol Clugston, Anna Dominiczak and Alex Alderton, Roberto Amato, Jeffrey Barrett, Sonia Goncalves, Ewan Harrison, David K. Jackson, Ian Johnston, Dominic Kwiatkowski, Cordelia Langford, John Sillitoe on behalf of the Wellcome Sanger Institute COVID-19 Surveillance Team |
| EPI_ISL_3249365 | Berkshire and Surrey Pathology Services Lighthouse Laboratory | Wellcome Sanger Institute for the COVID-19 Genomics UK (COG-UK) Consortium | Berkshire and Surrey Pathology Services Lighthouse Laboratory and Alex Alderton, Roberto Amato, Jeffrey Barrett, Sonia Goncalves, Ewan Harrison, David K. Jackson, Ian Johnston, Dominic Kwiatkowski, Cordelia Langford, John Sillitoe on behalf of the Wellcome Sanger Institute COVID-19 Surveillance Team |
| EPI_ISL_3249366 | Lighthouse Lab in Milton Keynes | Wellcome Sanger Institute for the COVID-19 Genomics UK (COG-UK) Consortium | The Lighthouse Lab in Milton Keynes and Alex Alderton, Roberto Amato, Jeffrey Barrett, Sonia Goncalves, Ewan Harrison, David K. Jackson, Ian Johnston, Dominic Kwiatkowski, Cordelia Langford, John Sillitoe on behalf of the Wellcome Sanger Institute COVID-19 Surveillance Team |
| EPI_ISL_3249367 | Berkshire and Surrey Pathology Services Lighthouse Laboratory | Wellcome Sanger Institute for the COVID-19 Genomics UK (COG-UK) Consortium | Berkshire and Surrey Pathology Services Lighthouse Laboratory and Alex Alderton, Roberto Amato, Jeffrey Barrett, Sonia Goncalves, Ewan Harrison, David K. Jackson, Ian Johnston, Dominic Kwiatkowski, Cordelia Langford, John Sillitoe on behalf of the Wellcome Sanger Institute COVID-19 Surveillance Team |
| EPI_ISL_3249368 | Lighthouse Lab in Glasgow | Wellcome Sanger Institute for the COVID-19 Genomics UK (COG-UK) Consortium | Harper VanSteenhouse, Yumi Kasai, David Gray, Carol Clugston, Anna Dominiczak and Alex Alderton, Roberto Amato, Jeffrey Barrett, Sonia Goncalves, Ewan Harrison, David K. Jackson, Ian Johnston, Dominic Kwiatkowski, Cordelia Langford, John Sillitoe on behalf of the Wellcome Sanger Institute COVID-19 Surveillance Team |
| EPI_ISL_3249369 | Lighthouse Lab in Alderley Park | Wellcome Sanger Institute for the COVID-19 Genomics UK (COG-UK) Consortium | Jacquelyn Wynn, Mairead Hyland, The Lighthouse Lab in Alderley Park and Alex Alderton, Roberto Amato, Jeffrey Barrett, Sonia Goncalves, Ewan Harrison, David K. Jackson, Ian Johnston, Dominic Kwiatkowski, Cordelia Langford, John Sillitoe on behalf of the Wellcome Sanger Institute COVID-19 Surveillance Team |
| EPI_ISL_3249370 | Lighthouse Lab in Glasgow | Wellcome Sanger Institute for the COVID-19 Genomics UK (COG-UK) Consortium | Harper VanSteenhouse, Yumi Kasai, David Gray, Carol Clugston, Anna Dominiczak and Alex Alderton, Roberto Amato, Jeffrey Barrett, Sonia Goncalves, Ewan Harrison, David K. Jackson, Ian Johnston, Dominic Kwiatkowski, Cordelia Langford, John Sillitoe on behalf of the Wellcome Sanger Institute COVID-19 Surveillance Team |
| EPI_ISL_3249371, EPI_ISL_3249372, EPI_ISL_3249373 | Lighthouse Lab in Alderley Park | Wellcome Sanger Institute for the COVID-19 Genomics UK (COG-UK) Consortium | Jacquelyn Wynn, Mairead Hyland, The Lighthouse Lab in Alderley Park and Alex Alderton, Roberto Amato, Jeffrey Barrett, Sonia Goncalves, Ewan Harrison, David K. Jackson, Ian Johnston, Dominic Kwiatkowski, Cordelia Langford, John Sillitoe on behalf of the Wellcome Sanger Institute COVID-19 Surveillance Team |
| EPI_ISL_3249374 | Berkshire and Surrey Pathology Services Lighthouse Laboratory | Wellcome Sanger Institute for the COVID-19 Genomics UK (COG-UK) Consortium | Berkshire and Surrey Pathology Services Lighthouse Laboratory and Alex Alderton, Roberto Amato, Jeffrey Barrett, Sonia Goncalves, Ewan Harrison, David K. Jackson, Ian Johnston, Dominic Kwiatkowski, Cordelia Langford, John Sillitoe on behalf of the Wellcome Sanger Institute COVID-19 Surveillance Team |
| EPI_ISL_3249375, EPI_ISL_3249376, EPI_ISL_3249377, EPI_ISL_3249378, EPI_ISL_3249379 | Lighthouse Lab in Milton Keynes | Wellcome Sanger Institute for the COVID-19 Genomics UK (COG-UK) Consortium | The Lighthouse Lab in Milton Keynes and Alex Alderton, Roberto Amato, Jeffrey Barrett, Sonia Goncalves, Ewan Harrison, David K. Jackson, Ian Johnston, Dominic Kwiatkowski, Cordelia Langford, John Sillitoe on behalf of the Wellcome Sanger Institute COVID-19 Surveillance Team |
| EPI_ISL_3249380, EPI_ISL_3249381, EPI_ISL_3249382 | Lighthouse Lab in Alderley Park | Wellcome Sanger Institute for the COVID-19 Genomics UK (COG-UK) Consortium | Jacquelyn Wynn, Mairead Hyland, The Lighthouse Lab in Alderley Park and Alex Alderton, Roberto Amato, Jeffrey Barrett, Sonia Goncalves, Ewan Harrison, David K. Jackson, Ian Johnston, Dominic Kwiatkowski, Cordelia Langford, John Sillitoe on behalf of the Wellcome Sanger Institute COVID-19 Surveillance Team |
| EPI_ISL_3249383 | Berkshire and Surrey Pathology Services Lighthouse Laboratory | Wellcome Sanger Institute for the COVID-19 Genomics UK (COG-UK) Consortium | Berkshire and Surrey Pathology Services Lighthouse Laboratory and Alex Alderton, Roberto Amato, Jeffrey Barrett, Sonia Goncalves, Ewan Harrison, David K. Jackson, Ian Johnston, Dominic Kwiatkowski, Cordelia Langford, John Sillitoe on behalf of the Wellcome Sanger Institute COVID-19 Surveillance Team |
| EPI_ISL_3249384 | Lighthouse Lab in Glasgow | Wellcome Sanger Institute for the COVID-19 Genomics UK (COG-UK) Consortium | Harper VanSteenhouse, Yumi Kasai, David Gray, Carol Clugston, Anna Dominiczak and Alex Alderton, Roberto Amato, Jeffrey Barrett, Sonia Goncalves, Ewan Harrison, David K. Jackson, Ian Johnston, Dominic Kwiatkowski, Cordelia Langford, John Sillitoe on behalf of the Wellcome Sanger Institute COVID-19 Surveillance Team |
| EPI_ISL_3249385, EPI_ISL_3249386 | Lighthouse Lab in Milton Keynes | Wellcome Sanger Institute for the COVID-19 Genomics UK (COG-UK) Consortium | The Lighthouse Lab in Milton Keynes and Alex Alderton, Roberto Amato, Jeffrey Barrett, Sonia Goncalves, Ewan Harrison, David K. Jackson, Ian Johnston, Dominic Kwiatkowski, Cordelia Langford, John Sillitoe on behalf of the Wellcome Sanger Institute COVID-19 Surveillance Team |
| EPI_ISL_3249387 | Lighthouse Lab in Glasgow | Wellcome Sanger Institute for the COVID-19 Genomics UK (COG-UK) Consortium | Harper VanSteenhouse, Yumi Kasai, David Gray, Carol Clugston, Anna Dominiczak and Alex Alderton, Roberto Amato, Jeffrey Barrett, Sonia Goncalves, Ewan Harrison, David K. Jackson, Ian Johnston, Dominic Kwiatkowski, Cordelia Langford, John Sillitoe on behalf of the Wellcome Sanger Institute COVID-19 Surveillance Team |
| EPI_ISL_3249388 | Lighthouse Lab in Milton Keynes | Wellcome Sanger Institute for the COVID-19 Genomics UK (COG-UK) Consortium | The Lighthouse Lab in Milton Keynes and Alex Alderton, Roberto Amato, Jeffrey Barrett, Sonia Goncalves, Ewan Harrison, David K. Jackson, Ian Johnston, Dominic Kwiatkowski, Cordelia Langford, John Sillitoe on behalf of the Wellcome Sanger Institute COVID-19 Surveillance Team |
| EPI_ISL_3249389 | Berkshire and Surrey Pathology Services Lighthouse Laboratory | Wellcome Sanger Institute for the COVID-19 Genomics UK (COG-UK) Consortium | Berkshire and Surrey Pathology Services Lighthouse Laboratory and Alex Alderton, Roberto Amato, Jeffrey Barrett, Sonia Goncalves, Ewan Harrison, David K. Jackson, Ian Johnston, Dominic Kwiatkowski, Cordelia Langford, John Sillitoe on behalf of the Wellcome Sanger Institute COVID-19 Surveillance Team |
| EPI_ISL_3249390 | Lighthouse Lab in Alderley Park | Wellcome Sanger Institute for the COVID-19 Genomics UK (COG-UK) Consortium | Jacquelyn Wynn, Mairead Hyland, The Lighthouse Lab in Alderley Park and Alex Alderton, Roberto Amato, Jeffrey Barrett, Sonia Goncalves, Ewan Harrison, David K. Jackson, Ian Johnston, Dominic Kwiatkowski, Cordelia Langford, John Sillitoe on behalf of the Wellcome Sanger Institute COVID-19 Surveillance Team |
| EPI_ISL_3249391, EPI_ISL_3249392 | Berkshire and Surrey Pathology Services Lighthouse Laboratory | Wellcome Sanger Institute for the COVID-19 Genomics UK (COG-UK) Consortium | Berkshire and Surrey Pathology Services Lighthouse Laboratory and Alex Alderton, Roberto Amato, Jeffrey Barrett, Sonia Goncalves, Ewan Harrison, David K. Jackson, Ian Johnston, Dominic Kwiatkowski, Cordelia Langford, John Sillitoe on behalf of the Wellcome Sanger Institute COVID-19 Surveillance Team |
| EPI_ISL_3249393 | Lighthouse Lab in Alderley Park | Wellcome Sanger Institute for the COVID-19 Genomics UK (COG-UK) Consortium | Jacquelyn Wynn, Mairead Hyland, The Lighthouse Lab in Alderley Park and Alex Alderton, Roberto Amato, Jeffrey Barrett, Sonia Goncalves, Ewan Harrison, David K. Jackson, Ian Johnston, Dominic Kwiatkowski, Cordelia Langford, John Sillitoe on behalf of the Wellcome Sanger Institute COVID-19 Surveillance Team |
| EPI_ISL_3249394 | Lighthouse Lab in Milton Keynes | Wellcome Sanger Institute for the COVID-19 Genomics UK (COG-UK) Consortium | The Lighthouse Lab in Milton Keynes and Alex Alderton, Roberto Amato, Jeffrey Barrett, Sonia Goncalves, Ewan Harrison, David K. Jackson, Ian Johnston, Dominic Kwiatkowski, Cordelia Langford, John Sillitoe on behalf of the Wellcome Sanger Institute COVID-19 Surveillance Team |
| EPI_ISL_3249395 | Lighthouse Lab in Glasgow | Wellcome Sanger Institute for the COVID-19 Genomics UK (COG-UK) Consortium | Harper VanSteenhouse, Yumi Kasai, David Gray, Carol Clugston, Anna Dominiczak and Alex Alderton, Roberto Amato, Jeffrey Barrett, Sonia Goncalves, Ewan Harrison, David K. Jackson, Ian Johnston, Dominic Kwiatkowski, Cordelia Langford, John Sillitoe on behalf of the Wellcome Sanger Institute COVID-19 Surveillance Team |
| EPI_ISL_3249396 | Berkshire and Surrey Pathology Services Lighthouse Laboratory | Wellcome Sanger Institute for the COVID-19 Genomics UK (COG-UK) Consortium | Berkshire and Surrey Pathology Services Lighthouse Laboratory and Alex Alderton, Roberto Amato, Jeffrey Barrett, Sonia Goncalves, Ewan Harrison, David K. Jackson, Ian Johnston, Dominic Kwiatkowski, Cordelia Langford, John Sillitoe on behalf of the Wellcome Sanger Institute COVID-19 Surveillance Team |
| EPI_ISL_3249397 | Lighthouse Lab in Glasgow | Wellcome Sanger Institute for the COVID-19 Genomics UK (COG-UK) Consortium | Harper VanSteenhouse, Yumi Kasai, David Gray, Carol Clugston, Anna Dominiczak and Alex Alderton, Roberto Amato, Jeffrey Barrett, Sonia Goncalves, Ewan Harrison, David K. Jackson, Ian Johnston, Dominic Kwiatkowski, Cordelia Langford, John Sillitoe on behalf of the Wellcome Sanger Institute COVID-19 Surveillance Team |
| EPI_ISL_3249398 | Berkshire and Surrey Pathology Services Lighthouse Laboratory | Wellcome Sanger Institute for the COVID-19 Genomics UK (COG-UK) Consortium | Berkshire and Surrey Pathology Services Lighthouse Laboratory and Alex Alderton, Roberto Amato, Jeffrey Barrett, Sonia Goncalves, Ewan Harrison, David K. Jackson, Ian Johnston, Dominic Kwiatkowski, Cordelia Langford, John Sillitoe on behalf of the Wellcome Sanger Institute COVID-19 Surveillance Team |
| EPI_ISL_3249399 | Lighthouse Lab in Milton Keynes | Wellcome Sanger Institute for the COVID-19 Genomics UK (COG-UK) Consortium | The Lighthouse Lab in Milton Keynes and Alex Alderton, Roberto Amato, Jeffrey Barrett, Sonia Goncalves, Ewan Harrison, David K. Jackson, Ian Johnston, Dominic Kwiatkowski, Cordelia Langford, John Sillitoe on behalf of the Wellcome Sanger Institute COVID-19 Surveillance Team |
| EPI_ISL_3249400 | Lighthouse Lab in Glasgow | Wellcome Sanger Institute for the COVID-19 Genomics UK (COG-UK) Consortium | Harper VanSteenhouse, Yumi Kasai, David Gray, Carol Clugston, Anna Dominiczak and Alex Alderton, Roberto Amato, Jeffrey Barrett, Sonia Goncalves, Ewan Harrison, David K. Jackson, Ian Johnston, Dominic Kwiatkowski, Cordelia Langford, John Sillitoe on behalf of the Wellcome Sanger Institute COVID-19 Surveillance Team |

| | | | |
|---|---|---|---|
| EPI_ISL_3249401 | Berkshire and Surrey Pathology Services Lighthouse Laboratory | Wellcome Sanger Institute for the COVID-19 Genomics UK (COG-UK) Consortium | Berkshire and Surrey Pathology Services Lighthouse Laboratory and Alex Alderton, Roberto Amato, Jeffrey Barrett, Sonia Goncalves, Ewan Harrison, David K. Jackson, Ian Johnston, Dominic Kwiatkowski, Cordelia Langford, John Sillitoe on behalf of the Wellcome Sanger Institute COVID-19 Surveillance Team |
| EPI_ISL_3249402 | Lighthouse Lab in Alderley Park | Wellcome Sanger Institute for the COVID-19 Genomics UK (COG-UK) Consortium | Jacquelyn Wynn, Mairead Hyland, The Lighthouse Lab in Alderley Park and Alex Alderton, Roberto Amato, Jeffrey Barrett, Sonia Goncalves, Ewan Harrison, David K. Jackson, Ian Johnston, Dominic Kwiatkowski, Cordelia Langford, John Sillitoe on behalf of the Wellcome Sanger Institute COVID-19 Surveillance Team |
| EPI_ISL_3249403, EPI_ISL_3249404 | Lighthouse Lab in Glasgow | Wellcome Sanger Institute for the COVID-19 Genomics UK (COG-UK) Consortium | Harper VanSteenhouse, Yumi Kasai, David Gray, Carol Clugston, Anna Dominiczak and Alex Alderton, Roberto Amato, Jeffrey Barrett, Sonia Goncalves, Ewan Harrison, David K. Jackson, Ian Johnston, Dominic Kwiatkowski, Cordelia Langford, John Sillitoe on behalf of the Wellcome Sanger Institute COVID-19 Surveillance Team |
| EPI_ISL_3249405 | Berkshire and Surrey Pathology Services Lighthouse Laboratory | Wellcome Sanger Institute for the COVID-19 Genomics UK (COG-UK) Consortium | Berkshire and Surrey Pathology Services Lighthouse Laboratory and Alex Alderton, Roberto Amato, Jeffrey Barrett, Sonia Goncalves, Ewan Harrison, David K. Jackson, Ian Johnston, Dominic Kwiatkowski, Cordelia Langford, John Sillitoe on behalf of the Wellcome Sanger Institute COVID-19 Surveillance Team |
| EPI_ISL_3249406, EPI_ISL_3249407 | Lighthouse Lab in Alderley Park | Wellcome Sanger Institute for the COVID-19 Genomics UK (COG-UK) Consortium | Jacquelyn Wynn, Mairead Hyland, The Lighthouse Lab in Alderley Park and Alex Alderton, Roberto Amato, Jeffrey Barrett, Sonia Goncalves, Ewan Harrison, David K. Jackson, Ian Johnston, Dominic Kwiatkowski, Cordelia Langford, John Sillitoe on behalf of the Wellcome Sanger Institute COVID-19 Surveillance Team |
| EPI_ISL_3249408 | Lighthouse Lab in Glasgow | Wellcome Sanger Institute for the COVID-19 Genomics UK (COG-UK) Consortium | Harper VanSteenhouse, Yumi Kasai, David Gray, Carol Clugston, Anna Dominiczak and Alex Alderton, Roberto Amato, Jeffrey Barrett, Sonia Goncalves, Ewan Harrison, David K. Jackson, Ian Johnston, Dominic Kwiatkowski, Cordelia Langford, John Sillitoe on behalf of the Wellcome Sanger Institute COVID-19 Surveillance Team |
| EPI_ISL_3249409 | Lighthouse Lab in Alderley Park | Wellcome Sanger Institute for the COVID-19 Genomics UK (COG-UK) Consortium | Jacquelyn Wynn, Mairead Hyland, The Lighthouse Lab in Alderley Park and Alex Alderton, Roberto Amato, Jeffrey Barrett, Sonia Goncalves, Ewan Harrison, David K. Jackson, Ian Johnston, Dominic Kwiatkowski, Cordelia Langford, John Sillitoe on behalf of the Wellcome Sanger Institute COVID-19 Surveillance Team |
| EPI_ISL_3249410, EPI_ISL_3249411 | Lighthouse Lab in Glasgow | Wellcome Sanger Institute for the COVID-19 Genomics UK (COG-UK) Consortium | Harper VanSteenhouse, Yumi Kasai, David Gray, Carol Clugston, Anna Dominiczak and Alex Alderton, Roberto Amato, Jeffrey Barrett, Sonia Goncalves, Ewan Harrison, David K. Jackson, Ian Johnston, Dominic Kwiatkowski, Cordelia Langford, John Sillitoe on behalf of the Wellcome Sanger Institute COVID-19 Surveillance Team |
| EPI_ISL_3249412 | Lighthouse Lab in Alderley Park | Wellcome Sanger Institute for the COVID-19 Genomics UK (COG-UK) Consortium | Jacquelyn Wynn, Mairead Hyland, The Lighthouse Lab in Alderley Park and Alex Alderton, Roberto Amato, Jeffrey Barrett, Sonia Goncalves, Ewan Harrison, David K. Jackson, Ian Johnston, Dominic Kwiatkowski, Cordelia Langford, John Sillitoe on behalf of the Wellcome Sanger Institute COVID-19 Surveillance Team |
| EPI_ISL_3249413 | Lighthouse Lab in Milton Keynes | Wellcome Sanger Institute for the COVID-19 Genomics UK (COG-UK) Consortium | The Lighthouse Lab in Milton Keynes and Alex Alderton, Roberto Amato, Jeffrey Barrett, Sonia Goncalves, Ewan Harrison, David K. Jackson, Ian Johnston, Dominic Kwiatkowski, Cordelia Langford, John Sillitoe on behalf of the Wellcome Sanger Institute COVID-19 Surveillance Team |
| EPI_ISL_3249414 | Lighthouse Lab in Alderley Park | Wellcome Sanger Institute for the COVID-19 Genomics UK (COG-UK) Consortium | Jacquelyn Wynn, Mairead Hyland, The Lighthouse Lab in Alderley Park and Alex Alderton, Roberto Amato, Jeffrey Barrett, Sonia Goncalves, Ewan Harrison, David K. Jackson, Ian Johnston, Dominic Kwiatkowski, Cordelia Langford, John Sillitoe on behalf of the Wellcome Sanger Institute COVID-19 Surveillance Team |
| EPI_ISL_3249415 | Berkshire and Surrey Pathology Services Lighthouse Laboratory | Wellcome Sanger Institute for the COVID-19 Genomics UK (COG-UK) Consortium | Berkshire and Surrey Pathology Services Lighthouse Laboratory and Alex Alderton, Roberto Amato, Jeffrey Barrett, Sonia Goncalves, Ewan Harrison, David K. Jackson, Ian Johnston, Dominic Kwiatkowski, Cordelia Langford, John Sillitoe on behalf of the Wellcome Sanger Institute COVID-19 Surveillance Team |
| EPI_ISL_3249416 | Lighthouse Lab in Milton Keynes | Wellcome Sanger Institute for the COVID-19 Genomics UK (COG-UK) Consortium | The Lighthouse Lab in Milton Keynes and Alex Alderton, Roberto Amato, Jeffrey Barrett, Sonia Goncalves, Ewan Harrison, David K. Jackson, Ian Johnston, Dominic Kwiatkowski, Cordelia Langford, John Sillitoe on behalf of the Wellcome Sanger Institute COVID-19 Surveillance Team |
| EPI_ISL_3249417 | Lighthouse Lab in Alderley Park | Wellcome Sanger Institute for the COVID-19 Genomics UK (COG-UK) Consortium | Jacquelyn Wynn, Mairead Hyland, The Lighthouse Lab in Alderley Park and Alex Alderton, Roberto Amato, Jeffrey Barrett, Sonia Goncalves, Ewan Harrison, David K. Jackson, Ian Johnston, Dominic Kwiatkowski, Cordelia Langford, John Sillitoe on behalf of the Wellcome Sanger Institute COVID-19 Surveillance Team |
| EPI_ISL_3249418 | Berkshire and Surrey Pathology Services Lighthouse Laboratory | Wellcome Sanger Institute for the COVID-19 Genomics UK (COG-UK) Consortium | Berkshire and Surrey Pathology Services Lighthouse Laboratory and Alex Alderton, Roberto Amato, Jeffrey Barrett, Sonia Goncalves, Ewan Harrison, David K. Jackson, Ian Johnston, Dominic Kwiatkowski, Cordelia Langford, John Sillitoe on behalf of the Wellcome Sanger Institute COVID-19 Surveillance Team |
| EPI_ISL_3249419 | Lighthouse Lab in Milton Keynes | Wellcome Sanger Institute for the COVID-19 Genomics UK (COG-UK) Consortium | The Lighthouse Lab in Milton Keynes and Alex Alderton, Roberto Amato, Jeffrey Barrett, Sonia Goncalves, Ewan Harrison, David K. Jackson, Ian Johnston, Dominic Kwiatkowski, Cordelia Langford, John Sillitoe on behalf of the Wellcome Sanger Institute COVID-19 Surveillance Team |
| EPI_ISL_3249420 | Lighthouse Lab in Alderley Park | Wellcome Sanger Institute for the COVID-19 Genomics UK (COG-UK) Consortium | Jacquelyn Wynn, Mairead Hyland, The Lighthouse Lab in Alderley Park and Alex Alderton, Roberto Amato, Jeffrey Barrett, Sonia Goncalves, Ewan Harrison, David K. Jackson, Ian Johnston, Dominic Kwiatkowski, Cordelia Langford, John Sillitoe on behalf of the Wellcome Sanger Institute COVID-19 Surveillance Team |
| EPI_ISL_3249421 | Lighthouse Lab in Milton Keynes | Wellcome Sanger Institute for the COVID-19 Genomics UK (COG-UK) Consortium | The Lighthouse Lab in Milton Keynes and Alex Alderton, Roberto Amato, Jeffrey Barrett, Sonia Goncalves, Ewan Harrison, David K. Jackson, Ian Johnston, Dominic Kwiatkowski, Cordelia Langford, John Sillitoe on behalf of the Wellcome Sanger Institute COVID-19 Surveillance Team |
| EPI_ISL_3249422 | Berkshire and Surrey Pathology Services Lighthouse Laboratory | Wellcome Sanger Institute for the COVID-19 Genomics UK (COG-UK) Consortium | Berkshire and Surrey Pathology Services Lighthouse Laboratory and Alex Alderton, Roberto Amato, Jeffrey Barrett, Sonia Goncalves, Ewan Harrison, David K. Jackson, Ian Johnston, Dominic Kwiatkowski, Cordelia Langford, John Sillitoe on behalf of the Wellcome Sanger Institute COVID-19 Surveillance Team |
| EPI_ISL_3249423 | Lighthouse Lab in Milton Keynes | Wellcome Sanger Institute for the COVID-19 Genomics UK (COG-UK) Consortium | The Lighthouse Lab in Milton Keynes and Alex Alderton, Roberto Amato, Jeffrey Barrett, Sonia Goncalves, Ewan Harrison, David K. Jackson, Ian Johnston, Dominic Kwiatkowski, Cordelia Langford, John Sillitoe on behalf of the Wellcome Sanger Institute COVID-19 Surveillance Team |
| EPI_ISL_3249424 | Lighthouse Lab in Alderley Park | Wellcome Sanger Institute for the COVID-19 Genomics UK (COG-UK) Consortium | Jacquelyn Wynn, Mairead Hyland, The Lighthouse Lab in Alderley Park and Alex Alderton, Roberto Amato, Jeffrey Barrett, Sonia Goncalves, Ewan Harrison, David K. Jackson, Ian Johnston, Dominic Kwiatkowski, Cordelia Langford, John Sillitoe on behalf of the Wellcome Sanger Institute COVID-19 Surveillance Team |
| EPI_ISL_3249425 | Lighthouse Lab in Milton Keynes | Wellcome Sanger Institute for the COVID-19 Genomics UK (COG-UK) Consortium | The Lighthouse Lab in Milton Keynes and Alex Alderton, Roberto Amato, Jeffrey Barrett, Sonia Goncalves, Ewan Harrison, David K. Jackson, Ian Johnston, Dominic Kwiatkowski, Cordelia Langford, John Sillitoe on behalf of the Wellcome Sanger Institute COVID-19 Surveillance Team |
| EPI_ISL_3249426, EPI_ISL_3249427 | Berkshire and Surrey Pathology Services Lighthouse Laboratory | Wellcome Sanger Institute for the COVID-19 Genomics UK (COG-UK) Consortium | Berkshire and Surrey Pathology Services Lighthouse Laboratory and Alex Alderton, Roberto Amato, Jeffrey Barrett, Sonia Goncalves, Ewan Harrison, David K. Jackson, Ian Johnston, Dominic Kwiatkowski, Cordelia Langford, John Sillitoe on behalf of the Wellcome Sanger Institute COVID-19 Surveillance Team |
| EPI_ISL_3249428 | Lighthouse Lab in Milton Keynes | Wellcome Sanger Institute for the COVID-19 Genomics UK (COG-UK) Consortium | The Lighthouse Lab in Milton Keynes and Alex Alderton, Roberto Amato, Jeffrey Barrett, Sonia Goncalves, Ewan Harrison, David K. Jackson, Ian Johnston, Dominic Kwiatkowski, Cordelia Langford, John Sillitoe on behalf of the Wellcome Sanger Institute COVID-19 Surveillance Team |
| EPI_ISL_3249429 | Lighthouse Lab in Alderley Park | Wellcome Sanger Institute for the COVID-19 Genomics UK (COG-UK) Consortium | Jacquelyn Wynn, Mairead Hyland, The Lighthouse Lab in Alderley Park and Alex Alderton, Roberto Amato, Jeffrey Barrett, Sonia Goncalves, Ewan Harrison, David K. Jackson, Ian Johnston, Dominic Kwiatkowski, Cordelia Langford, John Sillitoe on behalf of the Wellcome Sanger Institute COVID-19 Surveillance Team |
| EPI_ISL_3249430 | Lighthouse Lab in Glasgow | Wellcome Sanger Institute for the COVID-19 Genomics UK (COG-UK) Consortium | Harper VanSteenhouse, Yumi Kasai, David Gray, Carol Clugston, Anna Dominiczak and Alex Alderton, Roberto Amato, Jeffrey Barrett, Sonia Goncalves, Ewan Harrison, David K. Jackson, Ian Johnston, Dominic Kwiatkowski, Cordelia Langford, John Sillitoe on behalf of the Wellcome Sanger Institute COVID-19 Surveillance Team |
| EPI_ISL_3249431 | Lighthouse Lab in Alderley Park | Wellcome Sanger Institute for the COVID-19 Genomics UK (COG-UK) Consortium | Jacquelyn Wynn, Mairead Hyland, The Lighthouse Lab in Alderley Park and Alex Alderton, Roberto Amato, Jeffrey Barrett, Sonia Goncalves, Ewan Harrison, David K. Jackson, Ian Johnston, Dominic Kwiatkowski, Cordelia Langford, John Sillitoe on behalf of the Wellcome Sanger Institute COVID-19 Surveillance Team |
| EPI_ISL_3249432, EPI_ISL_3249433, EPI_ISL_3249434 | Berkshire and Surrey Pathology Services Lighthouse Laboratory | Wellcome Sanger Institute for the COVID-19 Genomics UK (COG-UK) Consortium | Berkshire and Surrey Pathology Services Lighthouse Laboratory and Alex Alderton, Roberto Amato, Jeffrey Barrett, Sonia Goncalves, Ewan Harrison, David K. Jackson, Ian Johnston, Dominic Kwiatkowski, Cordelia Langford, John Sillitoe on behalf of the Wellcome Sanger Institute COVID-19 Surveillance Team |
| EPI_ISL_3249435, EPI_ISL_3249436, EPI_ISL_3249437 | Lighthouse Lab in Milton Keynes | Wellcome Sanger Institute for the COVID-19 Genomics UK (COG-UK) Consortium | The Lighthouse Lab in Milton Keynes and Alex Alderton, Roberto Amato, Jeffrey Barrett, Sonia Goncalves, Ewan Harrison, David K. Jackson, Ian Johnston, Dominic Kwiatkowski, Cordelia Langford, John Sillitoe on behalf of the Wellcome Sanger Institute COVID-19 Surveillance Team |
| EPI_ISL_3249438 | Berkshire and Surrey Pathology Services Lighthouse Laboratory | Wellcome Sanger Institute for the COVID-19 Genomics UK (COG-UK) Consortium | Berkshire and Surrey Pathology Services Lighthouse Laboratory and Alex Alderton, Roberto Amato, Jeffrey Barrett, Sonia Goncalves, Ewan Harrison, David K. Jackson, Ian Johnston, Dominic Kwiatkowski, Cordelia Langford, John Sillitoe on behalf of the Wellcome Sanger Institute COVID-19 Surveillance Team |
| EPI_ISL_3249439 | Lighthouse Lab in Alderley Park | Wellcome Sanger Institute for the COVID-19 Genomics UK (COG-UK) Consortium | Jacquelyn Wynn, Mairead Hyland, The Lighthouse Lab in Alderley Park and Alex Alderton, Roberto Amato, Jeffrey Barrett, Sonia Goncalves, Ewan Harrison, David K. Jackson, Ian Johnston, Dominic Kwiatkowski, Cordelia Langford, John Sillitoe on behalf of the Wellcome Sanger Institute COVID-19 Surveillance Team |
| EPI_ISL_3249440, EPI_ISL_3249441 | Lighthouse Lab in Milton Keynes | Wellcome Sanger Institute for the COVID-19 Genomics UK (COG-UK) Consortium | The Lighthouse Lab in Milton Keynes and Alex Alderton, Roberto Amato, Jeffrey Barrett, Sonia Goncalves, Ewan Harrison, David K. Jackson, Ian Johnston, Dominic Kwiatkowski, Cordelia Langford, John Sillitoe on behalf of the Wellcome Sanger Institute COVID-19 Surveillance Team |
| EPI_ISL_3249442 | Lighthouse Lab in Alderley Park | Wellcome Sanger Institute for the COVID-19 Genomics UK (COG-UK) Consortium | Jacquelyn Wynn, Mairead Hyland, The Lighthouse Lab in Alderley Park and Alex Alderton, Roberto Amato, Jeffrey Barrett, Sonia Goncalves, Ewan Harrison, David K. Jackson, Ian Johnston, Dominic Kwiatkowski, Cordelia Langford, John Sillitoe on behalf of the Wellcome Sanger Institute COVID-19 Surveillance Team |
| EPI_ISL_3249443 | Berkshire and Surrey Pathology Services Lighthouse Laboratory | Wellcome Sanger Institute for the COVID-19 Genomics UK (COG-UK) Consortium | Berkshire and Surrey Pathology Services Lighthouse Laboratory and Alex Alderton, Roberto Amato, Jeffrey Barrett, Sonia Goncalves, Ewan Harrison, David K. Jackson, Ian Johnston, Dominic Kwiatkowski, Cordelia Langford, John Sillitoe on behalf of the Wellcome Sanger Institute COVID-19 Surveillance Team |
| EPI_ISL_3249444 | Lighthouse Lab in Alderley Park | Wellcome Sanger Institute for the COVID-19 Genomics UK (COG-UK) Consortium | Jacquelyn Wynn, Mairead Hyland, The Lighthouse Lab in Alderley Park and Alex Alderton, Roberto Amato, Jeffrey Barrett, Sonia Goncalves, Ewan Harrison, David K. Jackson, Ian Johnston, Dominic Kwiatkowski, Cordelia Langford, John Sillitoe on behalf of the Wellcome Sanger Institute COVID-19 Surveillance Team |
| EPI_ISL_3249445 | Lighthouse Lab in Glasgow | Wellcome Sanger Institute for the COVID-19 Genomics UK (COG-UK) Consortium | Harper VanSteenhouse, Yumi Kasai, David Gray, Carol Clugston, Anna Dominiczak and Alex Alderton, Roberto Amato, Jeffrey Barrett, Sonia Goncalves, Ewan Harrison, David K. Jackson, Ian Johnston, Dominic Kwiatkowski, Cordelia Langford, John Sillitoe on behalf of the Wellcome Sanger Institute COVID-19 Surveillance Team |
| EPI_ISL_3249446 | Lighthouse Lab in Alderley Park | Wellcome Sanger Institute for the COVID-19 Genomics UK (COG-UK) Consortium | Jacquelyn Wynn, Mairead Hyland, The Lighthouse Lab in Alderley Park and Alex Alderton, Roberto Amato, Jeffrey Barrett, Sonia Goncalves, Ewan Harrison, David K. Jackson, Ian Johnston, Dominic Kwiatkowski, Cordelia Langford, John Sillitoe on behalf of the Wellcome Sanger Institute COVID-19 Surveillance Team |
| EPI_ISL_3249447, EPI_ISL_3249448 | Berkshire and Surrey Pathology Services Lighthouse Laboratory | Wellcome Sanger Institute for the COVID-19 Genomics UK (COG-UK) Consortium | Berkshire and Surrey Pathology Services Lighthouse Laboratory and Alex Alderton, Roberto Amato, Jeffrey Barrett, Sonia Goncalves, Ewan Harrison, David K. Jackson, Ian Johnston, Dominic Kwiatkowski, Cordelia Langford, John Sillitoe on behalf of the Wellcome Sanger Institute COVID-19 Surveillance Team |
| EPI_ISL_3249449 | Lighthouse Lab in Alderley Park | Wellcome Sanger Institute for the COVID-19 Genomics UK (COG-UK) Consortium | Jacquelyn Wynn, Mairead Hyland, The Lighthouse Lab in Alderley Park and Alex Alderton, Roberto Amato, Jeffrey Barrett, Sonia Goncalves, Ewan Harrison, David K. Jackson, Ian Johnston, Dominic Kwiatkowski, Cordelia Langford, John Sillitoe on behalf of the Wellcome Sanger Institute COVID-19 Surveillance Team |
| EPI_ISL_3249450 | Berkshire and Surrey Pathology Services Lighthouse Laboratory | Wellcome Sanger Institute for the COVID-19 Genomics UK (COG-UK) Consortium | Berkshire and Surrey Pathology Services Lighthouse Laboratory and Alex Alderton, Roberto Amato, Jeffrey Barrett, Sonia Goncalves, Ewan Harrison, David K. Jackson, Ian Johnston, Dominic Kwiatkowski, Cordelia Langford, John Sillitoe on behalf of the Wellcome Sanger Institute COVID-19 Surveillance Team |
| EPI_ISL_3249451 | Lighthouse Lab in Glasgow | Wellcome Sanger Institute for the COVID-19 Genomics UK (COG-UK) Consortium | Harper VanSteenhouse, Yumi Kasai, David Gray, Carol Clugston, Anna Dominiczak and Alex Alderton, Roberto Amato, Jeffrey Barrett, Sonia Goncalves, Ewan Harrison, David K. Jackson, Ian Johnston, Dominic Kwiatkowski, Cordelia Langford, John Sillitoe on behalf of the Wellcome Sanger Institute COVID-19 Surveillance Team |
| EPI_ISL_3249452 | Berkshire and Surrey Pathology Services Lighthouse Laboratory | Wellcome Sanger Institute for the COVID-19 Genomics UK (COG-UK) Consortium | Berkshire and Surrey Pathology Services Lighthouse Laboratory and Alex Alderton, Roberto Amato, Jeffrey Barrett, Sonia Goncalves, Ewan Harrison, David K. Jackson, Ian Johnston, Dominic Kwiatkowski, Cordelia Langford, John Sillitoe on behalf of the Wellcome Sanger Institute COVID-19 Surveillance Team |
| EPI_ISL_3249453 | Lighthouse Lab in Glasgow | Wellcome Sanger Institute for the COVID-19 Genomics UK (COG-UK) Consortium | Harper VanSteenhouse, Yumi Kasai, David Gray, Carol Clugston, Anna Dominiczak and Alex Alderton, Roberto Amato, Jeffrey Barrett, Sonia Goncalves, Ewan Harrison, David K. Jackson, Ian Johnston, Dominic Kwiatkowski, Cordelia Langford, John Sillitoe on behalf of the Wellcome Sanger Institute COVID-19 Surveillance Team |
| EPI_ISL_3249454 | Lighthouse Lab in Alderley Park | Wellcome Sanger Institute for the COVID-19 Genomics UK (COG-UK) Consortium | Jacquelyn Wynn, Mairead Hyland, The Lighthouse Lab in Alderley Park and Alex Alderton, Roberto Amato, Jeffrey Barrett, Sonia Goncalves, Ewan Harrison, David K. Jackson, Ian Johnston, Dominic Kwiatkowski, Cordelia Langford, John Sillitoe on behalf of the Wellcome Sanger Institute COVID-19 Surveillance Team |
| EPI_ISL_3249455 | Lighthouse Lab in Glasgow | Wellcome Sanger Institute for the COVID-19 Genomics UK (COG-UK) | Harper VanSteenhouse, Yumi Kasai, David Gray, Carol Clugston, Anna Dominiczak and Alex Alderton, Roberto Amato, Jeffrey Barrett, Sonia Goncalves, Ewan Harrison, David K. Jackson, Ian Johnston, Dominic Kwiatkowski, Cordelia Langford, |

| | | | |
|---|---|---|---|
| | | Consortium | John Sillitoe on behalf of the Wellcome Sanger Institute COVID-19 Surveillance Team |
| EPI_ISL_3249456 | Lighthouse Lab in Milton Keynes | Wellcome Sanger Institute for the COVID-19 Genomics UK (COG-UK) Consortium | The Lighthouse Lab in Milton Keynes and Alex Alderton, Roberto Amato, Jeffrey Barrett, Sonia Goncalves, Ewan Harrison, David K. Jackson, Ian Johnston, Dominic Kwiatkowski, Cordelia Langford, John Sillitoe on behalf of the Wellcome Sanger Institute COVID-19 Surveillance Team |
| EPI_ISL_3249457 | Berkshire and Surrey Pathology Services Lighthouse Laboratory | Wellcome Sanger Institute for the COVID-19 Genomics UK (COG-UK) Consortium | Berkshire and Surrey Pathology Services Lighthouse Laboratory and Alex Alderton, Roberto Amato, Jeffrey Barrett, Sonia Goncalves, Ewan Harrison, David K. Jackson, Ian Johnston, Dominic Kwiatkowski, Cordelia Langford, John Sillitoe on behalf of the Wellcome Sanger Institute COVID-19 Surveillance Team |
| EPI_ISL_3249458, EPI_ISL_3249459 | Lighthouse Lab in Milton Keynes | Wellcome Sanger Institute for the COVID-19 Genomics UK (COG-UK) Consortium | The Lighthouse Lab in Milton Keynes and Alex Alderton, Roberto Amato, Jeffrey Barrett, Sonia Goncalves, Ewan Harrison, David K. Jackson, Ian Johnston, Dominic Kwiatkowski, Cordelia Langford, John Sillitoe on behalf of the Wellcome Sanger Institute COVID-19 Surveillance Team |
| EPI_ISL_3249460, EPI_ISL_3249461 | Berkshire and Surrey Pathology Services Lighthouse Laboratory | Wellcome Sanger Institute for the COVID-19 Genomics UK (COG-UK) Consortium | Berkshire and Surrey Pathology Services Lighthouse Laboratory and Alex Alderton, Roberto Amato, Jeffrey Barrett, Sonia Goncalves, Ewan Harrison, David K. Jackson, Ian Johnston, Dominic Kwiatkowski, Cordelia Langford, John Sillitoe on behalf of the Wellcome Sanger Institute COVID-19 Surveillance Team |
| EPI_ISL_3249462 | Lighthouse Lab in Glasgow | Wellcome Sanger Institute for the COVID-19 Genomics UK (COG-UK) Consortium | Harper VanSteenhouse, Yumi Kasai, David Gray, Carol Clugston, Anna Dominiczak and Alex Alderton, Roberto Amato, Jeffrey Barrett, Sonia Goncalves, Ewan Harrison, David K. Jackson, Ian Johnston, Dominic Kwiatkowski, Cordelia Langford, John Sillitoe on behalf of the Wellcome Sanger Institute COVID-19 Surveillance Team |
| EPI_ISL_3249463, EPI_ISL_3249464, EPI_ISL_3249465 | Lighthouse Lab in Alderley Park | Wellcome Sanger Institute for the COVID-19 Genomics UK (COG-UK) Consortium | Jacquelyn Wynn, Mairead Hyland, The Lighthouse Lab in Alderley Park and Alex Alderton, Roberto Amato, Jeffrey Barrett, Sonia Goncalves, Ewan Harrison, David K. Jackson, Ian Johnston, Dominic Kwiatkowski, Cordelia Langford, John Sillitoe on behalf of the Wellcome Sanger Institute COVID-19 Surveillance Team |
| EPI_ISL_3249466 | Lighthouse Lab in Milton Keynes | Wellcome Sanger Institute for the COVID-19 Genomics UK (COG-UK) Consortium | The Lighthouse Lab in Milton Keynes and Alex Alderton, Roberto Amato, Jeffrey Barrett, Sonia Goncalves, Ewan Harrison, David K. Jackson, Ian Johnston, Dominic Kwiatkowski, Cordelia Langford, John Sillitoe on behalf of the Wellcome Sanger Institute COVID-19 Surveillance Team |
| EPI_ISL_3249467 | Berkshire and Surrey Pathology Services Lighthouse Laboratory | Wellcome Sanger Institute for the COVID-19 Genomics UK (COG-UK) Consortium | Berkshire and Surrey Pathology Services Lighthouse Laboratory and Alex Alderton, Roberto Amato, Jeffrey Barrett, Sonia Goncalves, Ewan Harrison, David K. Jackson, Ian Johnston, Dominic Kwiatkowski, Cordelia Langford, John Sillitoe on behalf of the Wellcome Sanger Institute COVID-19 Surveillance Team |
| EPI_ISL_3249468 | Lighthouse Lab in Milton Keynes | Wellcome Sanger Institute for the COVID-19 Genomics UK (COG-UK) Consortium | The Lighthouse Lab in Milton Keynes and Alex Alderton, Roberto Amato, Jeffrey Barrett, Sonia Goncalves, Ewan Harrison, David K. Jackson, Ian Johnston, Dominic Kwiatkowski, Cordelia Langford, John Sillitoe on behalf of the Wellcome Sanger Institute COVID-19 Surveillance Team |
| EPI_ISL_3249469, EPI_ISL_3249470 | Lighthouse Lab in Glasgow | Wellcome Sanger Institute for the COVID-19 Genomics UK (COG-UK) Consortium | Harper VanSteenhouse, Yumi Kasai, David Gray, Carol Clugston, Anna Dominiczak and Alex Alderton, Roberto Amato, Jeffrey Barrett, Sonia Goncalves, Ewan Harrison, David K. Jackson, Ian Johnston, Dominic Kwiatkowski, Cordelia Langford, John Sillitoe on behalf of the Wellcome Sanger Institute COVID-19 Surveillance Team |
| EPI_ISL_3249471 | Lighthouse Lab in Milton Keynes | Wellcome Sanger Institute for the COVID-19 Genomics UK (COG-UK) Consortium | The Lighthouse Lab in Milton Keynes and Alex Alderton, Roberto Amato, Jeffrey Barrett, Sonia Goncalves, Ewan Harrison, David K. Jackson, Ian Johnston, Dominic Kwiatkowski, Cordelia Langford, John Sillitoe on behalf of the Wellcome Sanger Institute COVID-19 Surveillance Team |
| EPI_ISL_3249472, EPI_ISL_3249473 | Lighthouse Lab in Alderley Park | Wellcome Sanger Institute for the COVID-19 Genomics UK (COG-UK) Consortium | Jacquelyn Wynn, Mairead Hyland, The Lighthouse Lab in Alderley Park and Alex Alderton, Roberto Amato, Jeffrey Barrett, Sonia Goncalves, Ewan Harrison, David K. Jackson, Ian Johnston, Dominic Kwiatkowski, Cordelia Langford, John Sillitoe on behalf of the Wellcome Sanger Institute COVID-19 Surveillance Team |
| EPI_ISL_3249474 | Berkshire and Surrey Pathology Services Lighthouse Laboratory | Wellcome Sanger Institute for the COVID-19 Genomics UK (COG-UK) Consortium | Berkshire and Surrey Pathology Services Lighthouse Laboratory and Alex Alderton, Roberto Amato, Jeffrey Barrett, Sonia Goncalves, Ewan Harrison, David K. Jackson, Ian Johnston, Dominic Kwiatkowski, Cordelia Langford, John Sillitoe on behalf of the Wellcome Sanger Institute COVID-19 Surveillance Team |
| EPI_ISL_3249475, EPI_ISL_3249476 | Lighthouse Lab in Milton Keynes | Wellcome Sanger Institute for the COVID-19 Genomics UK (COG-UK) Consortium | The Lighthouse Lab in Milton Keynes and Alex Alderton, Roberto Amato, Jeffrey Barrett, Sonia Goncalves, Ewan Harrison, David K. Jackson, Ian Johnston, Dominic Kwiatkowski, Cordelia Langford, John Sillitoe on behalf of the Wellcome Sanger Institute COVID-19 Surveillance Team |
| EPI_ISL_3249477 | Lighthouse Lab in Alderley Park | Wellcome Sanger Institute for the COVID-19 Genomics UK (COG-UK) Consortium | Jacquelyn Wynn, Mairead Hyland, The Lighthouse Lab in Alderley Park and Alex Alderton, Roberto Amato, Jeffrey Barrett, Sonia Goncalves, Ewan Harrison, David K. Jackson, Ian Johnston, Dominic Kwiatkowski, Cordelia Langford, John Sillitoe on behalf of the Wellcome Sanger Institute COVID-19 Surveillance Team |
| EPI_ISL_3249478 | Berkshire and Surrey Pathology Services Lighthouse Laboratory | Wellcome Sanger Institute for the COVID-19 Genomics UK (COG-UK) Consortium | Berkshire and Surrey Pathology Services Lighthouse Laboratory and Alex Alderton, Roberto Amato, Jeffrey Barrett, Sonia Goncalves, Ewan Harrison, David K. Jackson, Ian Johnston, Dominic Kwiatkowski, Cordelia Langford, John Sillitoe on behalf of the Wellcome Sanger Institute COVID-19 Surveillance Team |
| EPI_ISL_3249479 | Lighthouse Lab in Glasgow | Wellcome Sanger Institute for the COVID-19 Genomics UK (COG-UK) Consortium | Harper VanSteenhouse, Yumi Kasai, David Gray, Carol Clugston, Anna Dominiczak and Alex Alderton, Roberto Amato, Jeffrey Barrett, Sonia Goncalves, Ewan Harrison, David K. Jackson, Ian Johnston, Dominic Kwiatkowski, Cordelia Langford, John Sillitoe on behalf of the Wellcome Sanger Institute COVID-19 Surveillance Team |
| EPI_ISL_3249480, EPI_ISL_3249481 | Berkshire and Surrey Pathology Services Lighthouse Laboratory | Wellcome Sanger Institute for the COVID-19 Genomics UK (COG-UK) Consortium | Berkshire and Surrey Pathology Services Lighthouse Laboratory and Alex Alderton, Roberto Amato, Jeffrey Barrett, Sonia Goncalves, Ewan Harrison, David K. Jackson, Ian Johnston, Dominic Kwiatkowski, Cordelia Langford, John Sillitoe on behalf of the Wellcome Sanger Institute COVID-19 Surveillance Team |
| EPI_ISL_3249482 | Lighthouse Lab in Alderley Park | Wellcome Sanger Institute for the COVID-19 Genomics UK (COG-UK) Consortium | Jacquelyn Wynn, Mairead Hyland, The Lighthouse Lab in Alderley Park and Alex Alderton, Roberto Amato, Jeffrey Barrett, Sonia Goncalves, Ewan Harrison, David K. Jackson, Ian Johnston, Dominic Kwiatkowski, Cordelia Langford, John Sillitoe on behalf of the Wellcome Sanger Institute COVID-19 Surveillance Team |
| EPI_ISL_3249483 | Berkshire and Surrey Pathology Services Lighthouse Laboratory | Wellcome Sanger Institute for the COVID-19 Genomics UK (COG-UK) Consortium | Berkshire and Surrey Pathology Services Lighthouse Laboratory and Alex Alderton, Roberto Amato, Jeffrey Barrett, Sonia Goncalves, Ewan Harrison, David K. Jackson, Ian Johnston, Dominic Kwiatkowski, Cordelia Langford, John Sillitoe on behalf of the Wellcome Sanger Institute COVID-19 Surveillance Team |
| EPI_ISL_3249484 | Lighthouse Lab in Alderley Park | Wellcome Sanger Institute for the COVID-19 Genomics UK (COG-UK) Consortium | Jacquelyn Wynn, Mairead Hyland, The Lighthouse Lab in Alderley Park and Alex Alderton, Roberto Amato, Jeffrey Barrett, Sonia Goncalves, Ewan Harrison, David K. Jackson, Ian Johnston, Dominic Kwiatkowski, Cordelia Langford, John Sillitoe on behalf of the Wellcome Sanger Institute COVID-19 Surveillance Team |
| EPI_ISL_3249485, EPI_ISL_3249486, EPI_ISL_3249487 | Lighthouse Lab in Glasgow | Wellcome Sanger Institute for the COVID-19 Genomics UK (COG-UK) Consortium | Harper VanSteenhouse, Yumi Kasai, David Gray, Carol Clugston, Anna Dominiczak and Alex Alderton, Roberto Amato, Jeffrey Barrett, Sonia Goncalves, Ewan Harrison, David K. Jackson, Ian Johnston, Dominic Kwiatkowski, Cordelia Langford, John Sillitoe on behalf of the Wellcome Sanger Institute COVID-19 Surveillance Team |
| EPI_ISL_3249488 | Berkshire and Surrey Pathology Services Lighthouse Laboratory | Wellcome Sanger Institute for the COVID-19 Genomics UK (COG-UK) Consortium | Berkshire and Surrey Pathology Services Lighthouse Laboratory and Alex Alderton, Roberto Amato, Jeffrey Barrett, Sonia Goncalves, Ewan Harrison, David K. Jackson, Ian Johnston, Dominic Kwiatkowski, Cordelia Langford, John Sillitoe on behalf of the Wellcome Sanger Institute COVID-19 Surveillance Team |
| EPI_ISL_3249489 | Lighthouse Lab in Alderley Park | Wellcome Sanger Institute for the COVID-19 Genomics UK (COG-UK) Consortium | Jacquelyn Wynn, Mairead Hyland, The Lighthouse Lab in Alderley Park and Alex Alderton, Roberto Amato, Jeffrey Barrett, Sonia Goncalves, Ewan Harrison, David K. Jackson, Ian Johnston, Dominic Kwiatkowski, Cordelia Langford, John Sillitoe on behalf of the Wellcome Sanger Institute COVID-19 Surveillance Team |
| EPI_ISL_3249490 | Lighthouse Lab in Glasgow | Wellcome Sanger Institute for the COVID-19 Genomics UK (COG-UK) Consortium | Harper VanSteenhouse, Yumi Kasai, David Gray, Carol Clugston, Anna Dominiczak and Alex Alderton, Roberto Amato, Jeffrey Barrett, Sonia Goncalves, Ewan Harrison, David K. Jackson, Ian Johnston, Dominic Kwiatkowski, Cordelia Langford, John Sillitoe on behalf of the Wellcome Sanger Institute COVID-19 Surveillance Team |
| EPI_ISL_3249491 | Berkshire and Surrey Pathology Services Lighthouse Laboratory | Wellcome Sanger Institute for the COVID-19 Genomics UK (COG-UK) Consortium | Berkshire and Surrey Pathology Services Lighthouse Laboratory and Alex Alderton, Roberto Amato, Jeffrey Barrett, Sonia Goncalves, Ewan Harrison, David K. Jackson, Ian Johnston, Dominic Kwiatkowski, Cordelia Langford, John Sillitoe on behalf of the Wellcome Sanger Institute COVID-19 Surveillance Team |
| EPI_ISL_3249492 | Lighthouse Lab in Alderley Park | Wellcome Sanger Institute for the COVID-19 Genomics UK (COG-UK) Consortium | Jacquelyn Wynn, Mairead Hyland, The Lighthouse Lab in Alderley Park and Alex Alderton, Roberto Amato, Jeffrey Barrett, Sonia Goncalves, Ewan Harrison, David K. Jackson, Ian Johnston, Dominic Kwiatkowski, Cordelia Langford, John Sillitoe on behalf of the Wellcome Sanger Institute COVID-19 Surveillance Team |
| EPI_ISL_3249493 | Lighthouse Lab in Milton Keynes | Wellcome Sanger Institute for the COVID-19 Genomics UK (COG-UK) Consortium | The Lighthouse Lab in Milton Keynes and Alex Alderton, Roberto Amato, Jeffrey Barrett, Sonia Goncalves, Ewan Harrison, David K. Jackson, Ian Johnston, Dominic Kwiatkowski, Cordelia Langford, John Sillitoe on behalf of the Wellcome Sanger Institute COVID-19 Surveillance Team |
| EPI_ISL_3249494 | Lighthouse Lab in Alderley Park | Wellcome Sanger Institute for the COVID-19 Genomics UK (COG-UK) Consortium | Jacquelyn Wynn, Mairead Hyland, The Lighthouse Lab in Alderley Park and Alex Alderton, Roberto Amato, Jeffrey Barrett, Sonia Goncalves, Ewan Harrison, David K. Jackson, Ian Johnston, Dominic Kwiatkowski, Cordelia Langford, John Sillitoe on behalf of the Wellcome Sanger Institute COVID-19 Surveillance Team |
| EPI_ISL_3249495, EPI_ISL_3249496 | Berkshire and Surrey Pathology Services Lighthouse Laboratory | Wellcome Sanger Institute for the COVID-19 Genomics UK (COG-UK) Consortium | Berkshire and Surrey Pathology Services Lighthouse Laboratory and Alex Alderton, Roberto Amato, Jeffrey Barrett, Sonia Goncalves, Ewan Harrison, David K. Jackson, Ian Johnston, Dominic Kwiatkowski, Cordelia Langford, John Sillitoe on behalf of the Wellcome Sanger Institute COVID-19 Surveillance Team |
| EPI_ISL_3249497 | Lighthouse Lab in Glasgow | Wellcome Sanger Institute for the COVID-19 Genomics UK (COG-UK) Consortium | Harper VanSteenhouse, Yumi Kasai, David Gray, Carol Clugston, Anna Dominiczak and Alex Alderton, Roberto Amato, Jeffrey Barrett, Sonia Goncalves, Ewan Harrison, David K. Jackson, Ian Johnston, Dominic Kwiatkowski, Cordelia Langford, John Sillitoe on behalf of the Wellcome Sanger Institute COVID-19 Surveillance Team |
| EPI_ISL_3249498, EPI_ISL_3249499 | Berkshire and Surrey Pathology Services Lighthouse Laboratory | Wellcome Sanger Institute for the COVID-19 Genomics UK (COG-UK) Consortium | Berkshire and Surrey Pathology Services Lighthouse Laboratory and Alex Alderton, Roberto Amato, Jeffrey Barrett, Sonia Goncalves, Ewan Harrison, David K. Jackson, Ian Johnston, Dominic Kwiatkowski, Cordelia Langford, John Sillitoe on behalf of the Wellcome Sanger Institute COVID-19 Surveillance Team |
| EPI_ISL_3249500 | Lighthouse Lab in Glasgow | Wellcome Sanger Institute for the COVID-19 Genomics UK (COG-UK) Consortium | Harper VanSteenhouse, Yumi Kasai, David Gray, Carol Clugston, Anna Dominiczak and Alex Alderton, Roberto Amato, Jeffrey Barrett, Sonia Goncalves, Ewan Harrison, David K. Jackson, Ian Johnston, Dominic Kwiatkowski, Cordelia Langford, John Sillitoe on behalf of the Wellcome Sanger Institute COVID-19 Surveillance Team |
| EPI_ISL_3249501 | Lighthouse Lab in Alderley Park | Wellcome Sanger Institute for the COVID-19 Genomics UK (COG-UK) Consortium | Jacquelyn Wynn, Mairead Hyland, The Lighthouse Lab in Alderley Park and Alex Alderton, Roberto Amato, Jeffrey Barrett, Sonia Goncalves, Ewan Harrison, David K. Jackson, Ian Johnston, Dominic Kwiatkowski, Cordelia Langford, John Sillitoe on behalf of the Wellcome Sanger Institute COVID-19 Surveillance Team |
| EPI_ISL_3249502 | Lighthouse Lab in Milton Keynes | Wellcome Sanger Institute for the COVID-19 Genomics UK (COG-UK) Consortium | The Lighthouse Lab in Milton Keynes and Alex Alderton, Roberto Amato, Jeffrey Barrett, Sonia Goncalves, Ewan Harrison, David K. Jackson, Ian Johnston, Dominic Kwiatkowski, Cordelia Langford, John Sillitoe on behalf of the Wellcome Sanger Institute COVID-19 Surveillance Team |
| EPI_ISL_3249503 | Lighthouse Lab in Glasgow | Wellcome Sanger Institute for the COVID-19 Genomics UK (COG-UK) Consortium | Harper VanSteenhouse, Yumi Kasai, David Gray, Carol Clugston, Anna Dominiczak and Alex Alderton, Roberto Amato, Jeffrey Barrett, Sonia Goncalves, Ewan Harrison, David K. Jackson, Ian Johnston, Dominic Kwiatkowski, Cordelia Langford, John Sillitoe on behalf of the Wellcome Sanger Institute COVID-19 Surveillance Team |
| EPI_ISL_3249504 | Lighthouse Lab in Milton Keynes | Wellcome Sanger Institute for the COVID-19 Genomics UK (COG-UK) Consortium | The Lighthouse Lab in Milton Keynes and Alex Alderton, Roberto Amato, Jeffrey Barrett, Sonia Goncalves, Ewan Harrison, David K. Jackson, Ian Johnston, Dominic Kwiatkowski, Cordelia Langford, John Sillitoe on behalf of the Wellcome Sanger Institute COVID-19 Surveillance Team |
| EPI_ISL_3249505 | Berkshire and Surrey Pathology Services Lighthouse Laboratory | Wellcome Sanger Institute for the COVID-19 Genomics UK (COG-UK) Consortium | Berkshire and Surrey Pathology Services Lighthouse Laboratory and Alex Alderton, Roberto Amato, Jeffrey Barrett, Sonia Goncalves, Ewan Harrison, David K. Jackson, Ian Johnston, Dominic Kwiatkowski, Cordelia Langford, John Sillitoe on behalf of the Wellcome Sanger Institute COVID-19 Surveillance Team |
| EPI_ISL_3249506 | Lighthouse Lab in Milton Keynes | Wellcome Sanger Institute for the COVID-19 Genomics UK (COG-UK) Consortium | The Lighthouse Lab in Milton Keynes and Alex Alderton, Roberto Amato, Jeffrey Barrett, Sonia Goncalves, Ewan Harrison, David K. Jackson, Ian Johnston, Dominic Kwiatkowski, Cordelia Langford, John Sillitoe on behalf of the Wellcome Sanger Institute COVID-19 Surveillance Team |
| EPI_ISL_3249507 | Lighthouse Lab in Alderley Park | Wellcome Sanger Institute for the COVID-19 Genomics UK (COG-UK) Consortium | Jacquelyn Wynn, Mairead Hyland, The Lighthouse Lab in Alderley Park and Alex Alderton, Roberto Amato, Jeffrey Barrett, Sonia Goncalves, Ewan Harrison, David K. Jackson, Ian Johnston, Dominic Kwiatkowski, Cordelia Langford, John Sillitoe on behalf of the Wellcome Sanger Institute COVID-19 Surveillance Team |
| EPI_ISL_3249508 | Lighthouse Lab in Milton Keynes | Wellcome Sanger Institute for the COVID-19 Genomics UK (COG-UK) Consortium | The Lighthouse Lab in Milton Keynes and Alex Alderton, Roberto Amato, Jeffrey Barrett, Sonia Goncalves, Ewan Harrison, David K. Jackson, Ian Johnston, Dominic Kwiatkowski, Cordelia Langford, John Sillitoe on behalf of the Wellcome Sanger Institute COVID-19 Surveillance Team |
| EPI_ISL_3249509 | Lighthouse Lab in Alderley Park | Wellcome Sanger Institute for the COVID-19 Genomics UK (COG-UK) Consortium | Jacquelyn Wynn, Mairead Hyland, The Lighthouse Lab in Alderley Park and Alex Alderton, Roberto Amato, Jeffrey Barrett, Sonia Goncalves, Ewan Harrison, David K. Jackson, Ian Johnston, Dominic Kwiatkowski, Cordelia Langford, John Sillitoe on behalf of the Wellcome Sanger Institute COVID-19 Surveillance Team |
| EPI_ISL_3249510 | Berkshire and Surrey Pathology Services Lighthouse Laboratory | Wellcome Sanger Institute for the COVID-19 Genomics UK (COG-UK) Consortium | Berkshire and Surrey Pathology Services Lighthouse Laboratory and Alex Alderton, Roberto Amato, Jeffrey Barrett, Sonia Goncalves, Ewan Harrison, David K. Jackson, Ian Johnston, Dominic Kwiatkowski, Cordelia Langford, John Sillitoe on behalf of the Wellcome Sanger Institute COVID-19 Surveillance Team |
| EPI_ISL_3249511, EPI_ISL_3249512 | Lighthouse Lab in Milton Keynes | Wellcome Sanger Institute for the COVID-19 Genomics UK (COG-UK) Consortium | The Lighthouse Lab in Milton Keynes and Alex Alderton, Roberto Amato, Jeffrey Barrett, Sonia Goncalves, Ewan Harrison, David K. Jackson, Ian Johnston, Dominic Kwiatkowski, Cordelia Langford, John Sillitoe on behalf of the Wellcome Sanger Institute COVID-19 Surveillance Team |

| | | | |
|---|---|---|---|
| EPI_ISL_3249513 | Berkshire and Surrey Pathology Services Lighthouse Laboratory | Wellcome Sanger Institute for the COVID-19 Genomics UK (COG-UK) Consortium | Berkshire and Surrey Pathology Services Lighthouse Laboratory and Alex Alderton, Roberto Amato, Jeffrey Barrett, Sonia Goncalves, Ewan Harrison, David K. Jackson, Ian Johnston, Dominic Kwiatkowski, Cordelia Langford, John Sillitoe on behalf of the Wellcome Sanger Institute COVID-19 Surveillance Team |
| EPI_ISL_3249514, EPI_ISL_3249515, EPI_ISL_3249516 | Lighthouse Lab in Milton Keynes | Wellcome Sanger Institute for the COVID-19 Genomics UK (COG-UK) Consortium | The Lighthouse Lab in Milton Keynes and Alex Alderton, Roberto Amato, Jeffrey Barrett, Sonia Goncalves, Ewan Harrison, David K. Jackson, Ian Johnston, Dominic Kwiatkowski, Cordelia Langford, John Sillitoe on behalf of the Wellcome Sanger Institute COVID-19 Surveillance Team |
| EPI_ISL_3249517 | Lighthouse Lab in Alderley Park | Wellcome Sanger Institute for the COVID-19 Genomics UK (COG-UK) Consortium | Jacquelyn Wynn, Mairead Hyland, The Lighthouse Lab in Alderley Park and Alex Alderton, Roberto Amato, Jeffrey Barrett, Sonia Goncalves, Ewan Harrison, David K. Jackson, Ian Johnston, Dominic Kwiatkowski, Cordelia Langford, John Sillitoe on behalf of the Wellcome Sanger Institute COVID-19 Surveillance Team |
| EPI_ISL_3249518 | Lighthouse Lab in Glasgow | Wellcome Sanger Institute for the COVID-19 Genomics UK (COG-UK) Consortium | Harper VanSteenhouse, Yumi Kasai, David Gray, Carol Clugston, Anna Dominiczak and Alex Alderton, Roberto Amato, Jeffrey Barrett, Sonia Goncalves, Ewan Harrison, David K. Jackson, Ian Johnston, Dominic Kwiatkowski, Cordelia Langford, John Sillitoe on behalf of the Wellcome Sanger Institute COVID-19 Surveillance Team |
| EPI_ISL_3249519 | Lighthouse Lab in Alderley Park | Wellcome Sanger Institute for the COVID-19 Genomics UK (COG-UK) Consortium | Jacquelyn Wynn, Mairead Hyland, The Lighthouse Lab in Alderley Park and Alex Alderton, Roberto Amato, Jeffrey Barrett, Sonia Goncalves, Ewan Harrison, David K. Jackson, Ian Johnston, Dominic Kwiatkowski, Cordelia Langford, John Sillitoe on behalf of the Wellcome Sanger Institute COVID-19 Surveillance Team |
| EPI_ISL_3249520, EPI_ISL_3249521 | Lighthouse Lab in Glasgow | Wellcome Sanger Institute for the COVID-19 Genomics UK (COG-UK) Consortium | Harper VanSteenhouse, Yumi Kasai, David Gray, Carol Clugston, Anna Dominiczak and Alex Alderton, Roberto Amato, Jeffrey Barrett, Sonia Goncalves, Ewan Harrison, David K. Jackson, Ian Johnston, Dominic Kwiatkowski, Cordelia Langford, John Sillitoe on behalf of the Wellcome Sanger Institute COVID-19 Surveillance Team |
| EPI_ISL_3249522, EPI_ISL_3249523, EPI_ISL_3249524 | Lighthouse Lab in Alderley Park | Wellcome Sanger Institute for the COVID-19 Genomics UK (COG-UK) Consortium | Jacquelyn Wynn, Mairead Hyland, The Lighthouse Lab in Alderley Park and Alex Alderton, Roberto Amato, Jeffrey Barrett, Sonia Goncalves, Ewan Harrison, David K. Jackson, Ian Johnston, Dominic Kwiatkowski, Cordelia Langford, John Sillitoe on behalf of the Wellcome Sanger Institute COVID-19 Surveillance Team |
| EPI_ISL_3249525 | Lighthouse Lab in Milton Keynes | Wellcome Sanger Institute for the COVID-19 Genomics UK (COG-UK) Consortium | The Lighthouse Lab in Milton Keynes and Alex Alderton, Roberto Amato, Jeffrey Barrett, Sonia Goncalves, Ewan Harrison, David K. Jackson, Ian Johnston, Dominic Kwiatkowski, Cordelia Langford, John Sillitoe on behalf of the Wellcome Sanger Institute COVID-19 Surveillance Team |
| EPI_ISL_3249526 | Lighthouse Lab in Glasgow | Wellcome Sanger Institute for the COVID-19 Genomics UK (COG-UK) Consortium | Harper VanSteenhouse, Yumi Kasai, David Gray, Carol Clugston, Anna Dominiczak and Alex Alderton, Roberto Amato, Jeffrey Barrett, Sonia Goncalves, Ewan Harrison, David K. Jackson, Ian Johnston, Dominic Kwiatkowski, Cordelia Langford, John Sillitoe on behalf of the Wellcome Sanger Institute COVID-19 Surveillance Team |
| EPI_ISL_3249527, EPI_ISL_3249528 | Lighthouse Lab in Milton Keynes | Wellcome Sanger Institute for the COVID-19 Genomics UK (COG-UK) Consortium | The Lighthouse Lab in Milton Keynes and Alex Alderton, Roberto Amato, Jeffrey Barrett, Sonia Goncalves, Ewan Harrison, David K. Jackson, Ian Johnston, Dominic Kwiatkowski, Cordelia Langford, John Sillitoe on behalf of the Wellcome Sanger Institute COVID-19 Surveillance Team |
| EPI_ISL_3249529 | Berkshire and Surrey Pathology Services Lighthouse Laboratory | Wellcome Sanger Institute for the COVID-19 Genomics UK (COG-UK) Consortium | Berkshire and Surrey Pathology Services Lighthouse Laboratory and Alex Alderton, Roberto Amato, Jeffrey Barrett, Sonia Goncalves, Ewan Harrison, David K. Jackson, Ian Johnston, Dominic Kwiatkowski, Cordelia Langford, John Sillitoe on behalf of the Wellcome Sanger Institute COVID-19 Surveillance Team |
| EPI_ISL_3249530, EPI_ISL_3249531 | Lighthouse Lab in Alderley Park | Wellcome Sanger Institute for the COVID-19 Genomics UK (COG-UK) Consortium | Jacquelyn Wynn, Mairead Hyland, The Lighthouse Lab in Alderley Park and Alex Alderton, Roberto Amato, Jeffrey Barrett, Sonia Goncalves, Ewan Harrison, David K. Jackson, Ian Johnston, Dominic Kwiatkowski, Cordelia Langford, John Sillitoe on behalf of the Wellcome Sanger Institute COVID-19 Surveillance Team |
| EPI_ISL_3249532 | Berkshire and Surrey Pathology Services Lighthouse Laboratory | Wellcome Sanger Institute for the COVID-19 Genomics UK (COG-UK) Consortium | Berkshire and Surrey Pathology Services Lighthouse Laboratory and Alex Alderton, Roberto Amato, Jeffrey Barrett, Sonia Goncalves, Ewan Harrison, David K. Jackson, Ian Johnston, Dominic Kwiatkowski, Cordelia Langford, John Sillitoe on behalf of the Wellcome Sanger Institute COVID-19 Surveillance Team |
| EPI_ISL_3249533 | Lighthouse Lab in Glasgow | Wellcome Sanger Institute for the COVID-19 Genomics UK (COG-UK) Consortium | Harper VanSteenhouse, Yumi Kasai, David Gray, Carol Clugston, Anna Dominiczak and Alex Alderton, Roberto Amato, Jeffrey Barrett, Sonia Goncalves, Ewan Harrison, David K. Jackson, Ian Johnston, Dominic Kwiatkowski, Cordelia Langford, John Sillitoe on behalf of the Wellcome Sanger Institute COVID-19 Surveillance Team |
| EPI_ISL_3249534 | Lighthouse Lab in Alderley Park | Wellcome Sanger Institute for the COVID-19 Genomics UK (COG-UK) Consortium | Jacquelyn Wynn, Mairead Hyland, The Lighthouse Lab in Alderley Park and Alex Alderton, Roberto Amato, Jeffrey Barrett, Sonia Goncalves, Ewan Harrison, David K. Jackson, Ian Johnston, Dominic Kwiatkowski, Cordelia Langford, John Sillitoe on behalf of the Wellcome Sanger Institute COVID-19 Surveillance Team |
| EPI_ISL_3249535 | Lighthouse Lab in Milton Keynes | Wellcome Sanger Institute for the COVID-19 Genomics UK (COG-UK) Consortium | The Lighthouse Lab in Milton Keynes and Alex Alderton, Roberto Amato, Jeffrey Barrett, Sonia Goncalves, Ewan Harrison, David K. Jackson, Ian Johnston, Dominic Kwiatkowski, Cordelia Langford, John Sillitoe on behalf of the Wellcome Sanger Institute COVID-19 Surveillance Team |
| EPI_ISL_3249536, EPI_ISL_3249537 | Lighthouse Lab in Glasgow | Wellcome Sanger Institute for the COVID-19 Genomics UK (COG-UK) Consortium | Harper VanSteenhouse, Yumi Kasai, David Gray, Carol Clugston, Anna Dominiczak and Alex Alderton, Roberto Amato, Jeffrey Barrett, Sonia Goncalves, Ewan Harrison, David K. Jackson, Ian Johnston, Dominic Kwiatkowski, Cordelia Langford, John Sillitoe on behalf of the Wellcome Sanger Institute COVID-19 Surveillance Team |
| EPI_ISL_3249538, EPI_ISL_3249539 | Lighthouse Lab in Milton Keynes | Wellcome Sanger Institute for the COVID-19 Genomics UK (COG-UK) Consortium | The Lighthouse Lab in Milton Keynes and Alex Alderton, Roberto Amato, Jeffrey Barrett, Sonia Goncalves, Ewan Harrison, David K. Jackson, Ian Johnston, Dominic Kwiatkowski, Cordelia Langford, John Sillitoe on behalf of the Wellcome Sanger Institute COVID-19 Surveillance Team |
| EPI_ISL_3249540 | Berkshire and Surrey Pathology Services Lighthouse Laboratory | Wellcome Sanger Institute for the COVID-19 Genomics UK (COG-UK) Consortium | Berkshire and Surrey Pathology Services Lighthouse Laboratory and Alex Alderton, Roberto Amato, Jeffrey Barrett, Sonia Goncalves, Ewan Harrison, David K. Jackson, Ian Johnston, Dominic Kwiatkowski, Cordelia Langford, John Sillitoe on behalf of the Wellcome Sanger Institute COVID-19 Surveillance Team |
| EPI_ISL_3249541 | Lighthouse Lab in Alderley Park | Wellcome Sanger Institute for the COVID-19 Genomics UK (COG-UK) Consortium | Jacquelyn Wynn, Mairead Hyland, The Lighthouse Lab in Alderley Park and Alex Alderton, Roberto Amato, Jeffrey Barrett, Sonia Goncalves, Ewan Harrison, David K. Jackson, Ian Johnston, Dominic Kwiatkowski, Cordelia Langford, John Sillitoe on behalf of the Wellcome Sanger Institute COVID-19 Surveillance Team |
| EPI_ISL_3249542, EPI_ISL_3249543 | Lighthouse Lab in Milton Keynes | Wellcome Sanger Institute for the COVID-19 Genomics UK (COG-UK) Consortium | The Lighthouse Lab in Milton Keynes and Alex Alderton, Roberto Amato, Jeffrey Barrett, Sonia Goncalves, Ewan Harrison, David K. Jackson, Ian Johnston, Dominic Kwiatkowski, Cordelia Langford, John Sillitoe on behalf of the Wellcome Sanger Institute COVID-19 Surveillance Team |
| EPI_ISL_3249544 | Berkshire and Surrey Pathology Services Lighthouse Laboratory | Wellcome Sanger Institute for the COVID-19 Genomics UK (COG-UK) Consortium | Berkshire and Surrey Pathology Services Lighthouse Laboratory and Alex Alderton, Roberto Amato, Jeffrey Barrett, Sonia Goncalves, Ewan Harrison, David K. Jackson, Ian Johnston, Dominic Kwiatkowski, Cordelia Langford, John Sillitoe on behalf of the Wellcome Sanger Institute COVID-19 Surveillance Team |
| EPI_ISL_3249545, EPI_ISL_3249546, EPI_ISL_3249547, EPI_ISL_3249548, EPI_ISL_3249549 | Lighthouse Lab in Milton Keynes | Wellcome Sanger Institute for the COVID-19 Genomics UK (COG-UK) Consortium | The Lighthouse Lab in Milton Keynes and Alex Alderton, Roberto Amato, Jeffrey Barrett, Sonia Goncalves, Ewan Harrison, David K. Jackson, Ian Johnston, Dominic Kwiatkowski, Cordelia Langford, John Sillitoe on behalf of the Wellcome Sanger Institute COVID-19 Surveillance Team |
| EPI_ISL_3249550, EPI_ISL_3249551 | Lighthouse Lab in Alderley Park | Wellcome Sanger Institute for the COVID-19 Genomics UK (COG-UK) Consortium | Jacquelyn Wynn, Mairead Hyland, The Lighthouse Lab in Alderley Park and Alex Alderton, Roberto Amato, Jeffrey Barrett, Sonia Goncalves, Ewan Harrison, David K. Jackson, Ian Johnston, Dominic Kwiatkowski, Cordelia Langford, John Sillitoe on behalf of the Wellcome Sanger Institute COVID-19 Surveillance Team |
| EPI_ISL_3249552 | Berkshire and Surrey Pathology Services Lighthouse Laboratory | Wellcome Sanger Institute for the COVID-19 Genomics UK (COG-UK) Consortium | Berkshire and Surrey Pathology Services Lighthouse Laboratory and Alex Alderton, Roberto Amato, Jeffrey Barrett, Sonia Goncalves, Ewan Harrison, David K. Jackson, Ian Johnston, Dominic Kwiatkowski, Cordelia Langford, John Sillitoe on behalf of the Wellcome Sanger Institute COVID-19 Surveillance Team |
| EPI_ISL_3249553 | Lighthouse Lab in Milton Keynes | Wellcome Sanger Institute for the COVID-19 Genomics UK (COG-UK) Consortium | The Lighthouse Lab in Milton Keynes and Alex Alderton, Roberto Amato, Jeffrey Barrett, Sonia Goncalves, Ewan Harrison, David K. Jackson, Ian Johnston, Dominic Kwiatkowski, Cordelia Langford, John Sillitoe on behalf of the Wellcome Sanger Institute COVID-19 Surveillance Team |
| EPI_ISL_3249554 | Lighthouse Lab in Glasgow | Wellcome Sanger Institute for the COVID-19 Genomics UK (COG-UK) Consortium | Harper VanSteenhouse, Yumi Kasai, David Gray, Carol Clugston, Anna Dominiczak and Alex Alderton, Roberto Amato, Jeffrey Barrett, Sonia Goncalves, Ewan Harrison, David K. Jackson, Ian Johnston, Dominic Kwiatkowski, Cordelia Langford, John Sillitoe on behalf of the Wellcome Sanger Institute COVID-19 Surveillance Team |
| EPI_ISL_3249555 | Lighthouse Lab in Milton Keynes | Wellcome Sanger Institute for the COVID-19 Genomics UK (COG-UK) Consortium | The Lighthouse Lab in Milton Keynes and Alex Alderton, Roberto Amato, Jeffrey Barrett, Sonia Goncalves, Ewan Harrison, David K. Jackson, Ian Johnston, Dominic Kwiatkowski, Cordelia Langford, John Sillitoe on behalf of the Wellcome Sanger Institute COVID-19 Surveillance Team |
| EPI_ISL_3249556 | Lighthouse Lab in Alderley Park | Wellcome Sanger Institute for the COVID-19 Genomics UK (COG-UK) Consortium | Jacquelyn Wynn, Mairead Hyland, The Lighthouse Lab in Alderley Park and Alex Alderton, Roberto Amato, Jeffrey Barrett, Sonia Goncalves, Ewan Harrison, David K. Jackson, Ian Johnston, Dominic Kwiatkowski, Cordelia Langford, John Sillitoe on behalf of the Wellcome Sanger Institute COVID-19 Surveillance Team |
| EPI_ISL_3249557 | Lighthouse Lab in Milton Keynes | Wellcome Sanger Institute for the COVID-19 Genomics UK (COG-UK) Consortium | The Lighthouse Lab in Milton Keynes and Alex Alderton, Roberto Amato, Jeffrey Barrett, Sonia Goncalves, Ewan Harrison, David K. Jackson, Ian Johnston, Dominic Kwiatkowski, Cordelia Langford, John Sillitoe on behalf of the Wellcome Sanger Institute COVID-19 Surveillance Team |
| EPI_ISL_3249558 | Lighthouse Lab in Alderley Park | Wellcome Sanger Institute for the COVID-19 Genomics UK (COG-UK) Consortium | Jacquelyn Wynn, Mairead Hyland, The Lighthouse Lab in Alderley Park and Alex Alderton, Roberto Amato, Jeffrey Barrett, Sonia Goncalves, Ewan Harrison, David K. Jackson, Ian Johnston, Dominic Kwiatkowski, Cordelia Langford, John Sillitoe on behalf of the Wellcome Sanger Institute COVID-19 Surveillance Team |
| EPI_ISL_3249559 | Berkshire and Surrey Pathology Services Lighthouse Laboratory | Wellcome Sanger Institute for the COVID-19 Genomics UK (COG-UK) Consortium | Berkshire and Surrey Pathology Services Lighthouse Laboratory and Alex Alderton, Roberto Amato, Jeffrey Barrett, Sonia Goncalves, Ewan Harrison, David K. Jackson, Ian Johnston, Dominic Kwiatkowski, Cordelia Langford, John Sillitoe on behalf of the Wellcome Sanger Institute COVID-19 Surveillance Team |
| EPI_ISL_3249560 | Lighthouse Lab in Alderley Park | Wellcome Sanger Institute for the COVID-19 Genomics UK (COG-UK) Consortium | Jacquelyn Wynn, Mairead Hyland, The Lighthouse Lab in Alderley Park and Alex Alderton, Roberto Amato, Jeffrey Barrett, Sonia Goncalves, Ewan Harrison, David K. Jackson, Ian Johnston, Dominic Kwiatkowski, Cordelia Langford, John Sillitoe on behalf of the Wellcome Sanger Institute COVID-19 Surveillance Team |
| EPI_ISL_3249561 | Lighthouse Lab in Glasgow | Wellcome Sanger Institute for the COVID-19 Genomics UK (COG-UK) Consortium | Harper VanSteenhouse, Yumi Kasai, David Gray, Carol Clugston, Anna Dominiczak and Alex Alderton, Roberto Amato, Jeffrey Barrett, Sonia Goncalves, Ewan Harrison, David K. Jackson, Ian Johnston, Dominic Kwiatkowski, Cordelia Langford, John Sillitoe on behalf of the Wellcome Sanger Institute COVID-19 Surveillance Team |
| EPI_ISL_3249562, EPI_ISL_3249563 | Berkshire and Surrey Pathology Services Lighthouse Laboratory | Wellcome Sanger Institute for the COVID-19 Genomics UK (COG-UK) Consortium | Berkshire and Surrey Pathology Services Lighthouse Laboratory and Alex Alderton, Roberto Amato, Jeffrey Barrett, Sonia Goncalves, Ewan Harrison, David K. Jackson, Ian Johnston, Dominic Kwiatkowski, Cordelia Langford, John Sillitoe on behalf of the Wellcome Sanger Institute COVID-19 Surveillance Team |
| EPI_ISL_3249564 | Lighthouse Lab in Alderley Park | Wellcome Sanger Institute for the COVID-19 Genomics UK (COG-UK) Consortium | Jacquelyn Wynn, Mairead Hyland, The Lighthouse Lab in Alderley Park and Alex Alderton, Roberto Amato, Jeffrey Barrett, Sonia Goncalves, Ewan Harrison, David K. Jackson, Ian Johnston, Dominic Kwiatkowski, Cordelia Langford, John Sillitoe on behalf of the Wellcome Sanger Institute COVID-19 Surveillance Team |
| EPI_ISL_3249565 | Lighthouse Lab in Milton Keynes | Wellcome Sanger Institute for the COVID-19 Genomics UK (COG-UK) Consortium | The Lighthouse Lab in Milton Keynes and Alex Alderton, Roberto Amato, Jeffrey Barrett, Sonia Goncalves, Ewan Harrison, David K. Jackson, Ian Johnston, Dominic Kwiatkowski, Cordelia Langford, John Sillitoe on behalf of the Wellcome Sanger Institute COVID-19 Surveillance Team |
| EPI_ISL_3249566 | Lighthouse Lab in Glasgow | Wellcome Sanger Institute for the COVID-19 Genomics UK (COG-UK) Consortium | Harper VanSteenhouse, Yumi Kasai, David Gray, Carol Clugston, Anna Dominiczak and Alex Alderton, Roberto Amato, Jeffrey Barrett, Sonia Goncalves, Ewan Harrison, David K. Jackson, Ian Johnston, Dominic Kwiatkowski, Cordelia Langford, John Sillitoe on behalf of the Wellcome Sanger Institute COVID-19 Surveillance Team |
| EPI_ISL_3249567 | Lighthouse Lab in Milton Keynes | Wellcome Sanger Institute for the COVID-19 Genomics UK (COG-UK) Consortium | The Lighthouse Lab in Milton Keynes and Alex Alderton, Roberto Amato, Jeffrey Barrett, Sonia Goncalves, Ewan Harrison, David K. Jackson, Ian Johnston, Dominic Kwiatkowski, Cordelia Langford, John Sillitoe on behalf of the Wellcome Sanger Institute COVID-19 Surveillance Team |
| EPI_ISL_3249568 | Lighthouse Lab in Glasgow | Wellcome Sanger Institute for the COVID-19 Genomics UK (COG-UK) Consortium | Harper VanSteenhouse, Yumi Kasai, David Gray, Carol Clugston, Anna Dominiczak and Alex Alderton, Roberto Amato, Jeffrey Barrett, Sonia Goncalves, Ewan Harrison, David K. Jackson, Ian Johnston, Dominic Kwiatkowski, Cordelia Langford, John Sillitoe on behalf of the Wellcome Sanger Institute COVID-19 Surveillance Team |
| EPI_ISL_3249569, EPI_ISL_3249570 | Berkshire and Surrey Pathology Services Lighthouse Laboratory | Wellcome Sanger Institute for the COVID-19 Genomics UK (COG-UK) Consortium | Berkshire and Surrey Pathology Services Lighthouse Laboratory and Alex Alderton, Roberto Amato, Jeffrey Barrett, Sonia Goncalves, Ewan Harrison, David K. Jackson, Ian Johnston, Dominic Kwiatkowski, Cordelia Langford, John Sillitoe on behalf of the Wellcome Sanger Institute COVID-19 Surveillance Team |
| EPI_ISL_3249571 | Lighthouse Lab in Alderley Park | Wellcome Sanger Institute for the COVID-19 Genomics UK (COG-UK) Consortium | Jacquelyn Wynn, Mairead Hyland, The Lighthouse Lab in Alderley Park and Alex Alderton, Roberto Amato, Jeffrey Barrett, Sonia Goncalves, Ewan Harrison, David K. Jackson, Ian Johnston, Dominic Kwiatkowski, Cordelia Langford, John Sillitoe on behalf of the Wellcome Sanger Institute COVID-19 Surveillance Team |
| EPI_ISL_3249572 | Lighthouse Lab in Glasgow | Wellcome Sanger Institute for the COVID-19 Genomics UK (COG-UK) Consortium | Harper VanSteenhouse, Yumi Kasai, David Gray, Carol Clugston, Anna Dominiczak and Alex Alderton, Roberto Amato, Jeffrey Barrett, Sonia Goncalves, Ewan Harrison, David K. Jackson, Ian Johnston, Dominic Kwiatkowski, Cordelia Langford, John Sillitoe on behalf of the Wellcome Sanger Institute COVID-19 Surveillance Team |

| | | | |
|---|---|---|---|
| EPI_ISL_3249573 | Lighthouse Lab in Milton Keynes | Wellcome Sanger Institute for the COVID-19 Genomics UK (COG-UK) Consortium | The Lighthouse Lab in Milton Keynes and Alex Alderton, Roberto Amato, Jeffrey Barrett, Sonia Goncalves, Ewan Harrison, David K. Jackson, Ian Johnston, Dominic Kwiatkowski, Cordelia Langford, John Sillitoe on behalf of the Wellcome Sanger Institute COVID-19 Surveillance Team |
| EPI_ISL_3249574 | Lighthouse Lab in Alderley Park | Wellcome Sanger Institute for the COVID-19 Genomics UK (COG-UK) Consortium | Jacquelyn Wynn, Mairead Hyland, The Lighthouse Lab in Alderley Park and Alex Alderton, Roberto Amato, Jeffrey Barrett, Sonia Goncalves, Ewan Harrison, David K. Jackson, Ian Johnston, Dominic Kwiatkowski, Cordelia Langford, John Sillitoe on behalf of the Wellcome Sanger Institute COVID-19 Surveillance Team |
| EPI_ISL_3249575 | Lighthouse Lab in Glasgow | Wellcome Sanger Institute for the COVID-19 Genomics UK (COG-UK) Consortium | Harper VanSteenhouse, Yumi Kasai, David Gray, Carol Clugston, Anna Dominiczak and Alex Alderton, Roberto Amato, Jeffrey Barrett, Sonia Goncalves, Ewan Harrison, David K. Jackson, Ian Johnston, Dominic Kwiatkowski, Cordelia Langford, John Sillitoe on behalf of the Wellcome Sanger Institute COVID-19 Surveillance Team |
| EPI_ISL_3249576, EPI_ISL_3249577 | Berkshire and Surrey Pathology Services Lighthouse Laboratory | Wellcome Sanger Institute for the COVID-19 Genomics UK (COG-UK) Consortium | Berkshire and Surrey Pathology Services Lighthouse Laboratory and Alex Alderton, Roberto Amato, Jeffrey Barrett, Sonia Goncalves, Ewan Harrison, David K. Jackson, Ian Johnston, Dominic Kwiatkowski, Cordelia Langford, John Sillitoe on behalf of the Wellcome Sanger Institute COVID-19 Surveillance Team |
| EPI_ISL_3249578, EPI_ISL_3249579 | Lighthouse Lab in Alderley Park | Wellcome Sanger Institute for the COVID-19 Genomics UK (COG-UK) Consortium | Jacquelyn Wynn, Mairead Hyland, The Lighthouse Lab in Alderley Park and Alex Alderton, Roberto Amato, Jeffrey Barrett, Sonia Goncalves, Ewan Harrison, David K. Jackson, Ian Johnston, Dominic Kwiatkowski, Cordelia Langford, John Sillitoe on behalf of the Wellcome Sanger Institute COVID-19 Surveillance Team |
| EPI_ISL_3249580 | Berkshire and Surrey Pathology Services Lighthouse Laboratory | Wellcome Sanger Institute for the COVID-19 Genomics UK (COG-UK) Consortium | Berkshire and Surrey Pathology Services Lighthouse Laboratory and Alex Alderton, Roberto Amato, Jeffrey Barrett, Sonia Goncalves, Ewan Harrison, David K. Jackson, Ian Johnston, Dominic Kwiatkowski, Cordelia Langford, John Sillitoe on behalf of the Wellcome Sanger Institute COVID-19 Surveillance Team |
| EPI_ISL_3249581 | Lighthouse Lab in Alderley Park | Wellcome Sanger Institute for the COVID-19 Genomics UK (COG-UK) Consortium | Jacquelyn Wynn, Mairead Hyland, The Lighthouse Lab in Alderley Park and Alex Alderton, Roberto Amato, Jeffrey Barrett, Sonia Goncalves, Ewan Harrison, David K. Jackson, Ian Johnston, Dominic Kwiatkowski, Cordelia Langford, John Sillitoe on behalf of the Wellcome Sanger Institute COVID-19 Surveillance Team |
| EPI_ISL_3249582 | Berkshire and Surrey Pathology Services Lighthouse Laboratory | Wellcome Sanger Institute for the COVID-19 Genomics UK (COG-UK) Consortium | Berkshire and Surrey Pathology Services Lighthouse Laboratory and Alex Alderton, Roberto Amato, Jeffrey Barrett, Sonia Goncalves, Ewan Harrison, David K. Jackson, Ian Johnston, Dominic Kwiatkowski, Cordelia Langford, John Sillitoe on behalf of the Wellcome Sanger Institute COVID-19 Surveillance Team |
| EPI_ISL_3249583 | Lighthouse Lab in Milton Keynes | Wellcome Sanger Institute for the COVID-19 Genomics UK (COG-UK) Consortium | The Lighthouse Lab in Milton Keynes and Alex Alderton, Roberto Amato, Jeffrey Barrett, Sonia Goncalves, Ewan Harrison, David K. Jackson, Ian Johnston, Dominic Kwiatkowski, Cordelia Langford, John Sillitoe on behalf of the Wellcome Sanger Institute COVID-19 Surveillance Team |
| EPI_ISL_3249584 | Lighthouse Lab in Glasgow | Wellcome Sanger Institute for the COVID-19 Genomics UK (COG-UK) Consortium | Harper VanSteenhouse, Yumi Kasai, David Gray, Carol Clugston, Anna Dominiczak and Alex Alderton, Roberto Amato, Jeffrey Barrett, Sonia Goncalves, Ewan Harrison, David K. Jackson, Ian Johnston, Dominic Kwiatkowski, Cordelia Langford, John Sillitoe on behalf of the Wellcome Sanger Institute COVID-19 Surveillance Team |
| EPI_ISL_3249585 | Lighthouse Lab in Milton Keynes | Wellcome Sanger Institute for the COVID-19 Genomics UK (COG-UK) Consortium | The Lighthouse Lab in Milton Keynes and Alex Alderton, Roberto Amato, Jeffrey Barrett, Sonia Goncalves, Ewan Harrison, David K. Jackson, Ian Johnston, Dominic Kwiatkowski, Cordelia Langford, John Sillitoe on behalf of the Wellcome Sanger Institute COVID-19 Surveillance Team |
| EPI_ISL_3249586 | Berkshire and Surrey Pathology Services Lighthouse Laboratory | Wellcome Sanger Institute for the COVID-19 Genomics UK (COG-UK) Consortium | Berkshire and Surrey Pathology Services Lighthouse Laboratory and Alex Alderton, Roberto Amato, Jeffrey Barrett, Sonia Goncalves, Ewan Harrison, David K. Jackson, Ian Johnston, Dominic Kwiatkowski, Cordelia Langford, John Sillitoe on behalf of the Wellcome Sanger Institute COVID-19 Surveillance Team |
| EPI_ISL_3249587 | Lighthouse Lab in Milton Keynes | Wellcome Sanger Institute for the COVID-19 Genomics UK (COG-UK) Consortium | The Lighthouse Lab in Milton Keynes and Alex Alderton, Roberto Amato, Jeffrey Barrett, Sonia Goncalves, Ewan Harrison, David K. Jackson, Ian Johnston, Dominic Kwiatkowski, Cordelia Langford, John Sillitoe on behalf of the Wellcome Sanger Institute COVID-19 Surveillance Team |
| EPI_ISL_3249588 | Berkshire and Surrey Pathology Services Lighthouse Laboratory | Wellcome Sanger Institute for the COVID-19 Genomics UK (COG-UK) Consortium | Berkshire and Surrey Pathology Services Lighthouse Laboratory and Alex Alderton, Roberto Amato, Jeffrey Barrett, Sonia Goncalves, Ewan Harrison, David K. Jackson, Ian Johnston, Dominic Kwiatkowski, Cordelia Langford, John Sillitoe on behalf of the Wellcome Sanger Institute COVID-19 Surveillance Team |
| EPI_ISL_3249589, EPI_ISL_3249590 | Lighthouse Lab in Alderley Park | Wellcome Sanger Institute for the COVID-19 Genomics UK (COG-UK) Consortium | Jacquelyn Wynn, Mairead Hyland, The Lighthouse Lab in Alderley Park and Alex Alderton, Roberto Amato, Jeffrey Barrett, Sonia Goncalves, Ewan Harrison, David K. Jackson, Ian Johnston, Dominic Kwiatkowski, Cordelia Langford, John Sillitoe on behalf of the Wellcome Sanger Institute COVID-19 Surveillance Team |
| EPI_ISL_3249591 | Lighthouse Lab in Milton Keynes | Wellcome Sanger Institute for the COVID-19 Genomics UK (COG-UK) Consortium | The Lighthouse Lab in Milton Keynes and Alex Alderton, Roberto Amato, Jeffrey Barrett, Sonia Goncalves, Ewan Harrison, David K. Jackson, Ian Johnston, Dominic Kwiatkowski, Cordelia Langford, John Sillitoe on behalf of the Wellcome Sanger Institute COVID-19 Surveillance Team |
| EPI_ISL_3249592 | Lighthouse Lab in Alderley Park | Wellcome Sanger Institute for the COVID-19 Genomics UK (COG-UK) Consortium | Jacquelyn Wynn, Mairead Hyland, The Lighthouse Lab in Alderley Park and Alex Alderton, Roberto Amato, Jeffrey Barrett, Sonia Goncalves, Ewan Harrison, David K. Jackson, Ian Johnston, Dominic Kwiatkowski, Cordelia Langford, John Sillitoe on behalf of the Wellcome Sanger Institute COVID-19 Surveillance Team |
| EPI_ISL_3249593 | Lighthouse Lab in Milton Keynes | Wellcome Sanger Institute for the COVID-19 Genomics UK (COG-UK) Consortium | The Lighthouse Lab in Milton Keynes and Alex Alderton, Roberto Amato, Jeffrey Barrett, Sonia Goncalves, Ewan Harrison, David K. Jackson, Ian Johnston, Dominic Kwiatkowski, Cordelia Langford, John Sillitoe on behalf of the Wellcome Sanger Institute COVID-19 Surveillance Team |
| EPI_ISL_3249594 | Berkshire and Surrey Pathology Services Lighthouse Laboratory | Wellcome Sanger Institute for the COVID-19 Genomics UK (COG-UK) Consortium | Berkshire and Surrey Pathology Services Lighthouse Laboratory and Alex Alderton, Roberto Amato, Jeffrey Barrett, Sonia Goncalves, Ewan Harrison, David K. Jackson, Ian Johnston, Dominic Kwiatkowski, Cordelia Langford, John Sillitoe on behalf of the Wellcome Sanger Institute COVID-19 Surveillance Team |
| EPI_ISL_3249595, EPI_ISL_3249596, EPI_ISL_3249597 | Lighthouse Lab in Milton Keynes | Wellcome Sanger Institute for the COVID-19 Genomics UK (COG-UK) Consortium | The Lighthouse Lab in Milton Keynes and Alex Alderton, Roberto Amato, Jeffrey Barrett, Sonia Goncalves, Ewan Harrison, David K. Jackson, Ian Johnston, Dominic Kwiatkowski, Cordelia Langford, John Sillitoe on behalf of the Wellcome Sanger Institute COVID-19 Surveillance Team |
| EPI_ISL_3249598 | Lighthouse Lab in Alderley Park | Wellcome Sanger Institute for the COVID-19 Genomics UK (COG-UK) Consortium | Jacquelyn Wynn, Mairead Hyland, The Lighthouse Lab in Alderley Park and Alex Alderton, Roberto Amato, Jeffrey Barrett, Sonia Goncalves, Ewan Harrison, David K. Jackson, Ian Johnston, Dominic Kwiatkowski, Cordelia Langford, John Sillitoe on behalf of the Wellcome Sanger Institute COVID-19 Surveillance Team |
| EPI_ISL_3249599, EPI_ISL_3249600 | Lighthouse Lab in Milton Keynes | Wellcome Sanger Institute for the COVID-19 Genomics UK (COG-UK) Consortium | The Lighthouse Lab in Milton Keynes and Alex Alderton, Roberto Amato, Jeffrey Barrett, Sonia Goncalves, Ewan Harrison, David K. Jackson, Ian Johnston, Dominic Kwiatkowski, Cordelia Langford, John Sillitoe on behalf of the Wellcome Sanger Institute COVID-19 Surveillance Team |
| EPI_ISL_3249601 | Lighthouse Lab in Alderley Park | Wellcome Sanger Institute for the COVID-19 Genomics UK (COG-UK) Consortium | Jacquelyn Wynn, Mairead Hyland, The Lighthouse Lab in Alderley Park and Alex Alderton, Roberto Amato, Jeffrey Barrett, Sonia Goncalves, Ewan Harrison, David K. Jackson, Ian Johnston, Dominic Kwiatkowski, Cordelia Langford, John Sillitoe on behalf of the Wellcome Sanger Institute COVID-19 Surveillance Team |
| EPI_ISL_3249602 | Berkshire and Surrey Pathology Services Lighthouse Laboratory | Wellcome Sanger Institute for the COVID-19 Genomics UK (COG-UK) Consortium | Berkshire and Surrey Pathology Services Lighthouse Laboratory and Alex Alderton, Roberto Amato, Jeffrey Barrett, Sonia Goncalves, Ewan Harrison, David K. Jackson, Ian Johnston, Dominic Kwiatkowski, Cordelia Langford, John Sillitoe on behalf of the Wellcome Sanger Institute COVID-19 Surveillance Team |
| EPI_ISL_3249603 | Lighthouse Lab in Alderley Park | Wellcome Sanger Institute for the COVID-19 Genomics UK (COG-UK) Consortium | Jacquelyn Wynn, Mairead Hyland, The Lighthouse Lab in Alderley Park and Alex Alderton, Roberto Amato, Jeffrey Barrett, Sonia Goncalves, Ewan Harrison, David K. Jackson, Ian Johnston, Dominic Kwiatkowski, Cordelia Langford, John Sillitoe on behalf of the Wellcome Sanger Institute COVID-19 Surveillance Team |
| EPI_ISL_3249604 | Berkshire and Surrey Pathology Services Lighthouse Laboratory | Wellcome Sanger Institute for the COVID-19 Genomics UK (COG-UK) Consortium | Berkshire and Surrey Pathology Services Lighthouse Laboratory and Alex Alderton, Roberto Amato, Jeffrey Barrett, Sonia Goncalves, Ewan Harrison, David K. Jackson, Ian Johnston, Dominic Kwiatkowski, Cordelia Langford, John Sillitoe on behalf of the Wellcome Sanger Institute COVID-19 Surveillance Team |
| EPI_ISL_3249605, EPI_ISL_3249606 | Lighthouse Lab in Alderley Park | Wellcome Sanger Institute for the COVID-19 Genomics UK (COG-UK) Consortium | Jacquelyn Wynn, Mairead Hyland, The Lighthouse Lab in Alderley Park and Alex Alderton, Roberto Amato, Jeffrey Barrett, Sonia Goncalves, Ewan Harrison, David K. Jackson, Ian Johnston, Dominic Kwiatkowski, Cordelia Langford, John Sillitoe on behalf of the Wellcome Sanger Institute COVID-19 Surveillance Team |
| EPI_ISL_3249607 | Lighthouse Lab in Glasgow | Wellcome Sanger Institute for the COVID-19 Genomics UK (COG-UK) Consortium | Harper VanSteenhouse, Yumi Kasai, David Gray, Carol Clugston, Anna Dominiczak and Alex Alderton, Roberto Amato, Jeffrey Barrett, Sonia Goncalves, Ewan Harrison, David K. Jackson, Ian Johnston, Dominic Kwiatkowski, Cordelia Langford, John Sillitoe on behalf of the Wellcome Sanger Institute COVID-19 Surveillance Team |
| EPI_ISL_3249608 | Lighthouse Lab in Milton Keynes | Wellcome Sanger Institute for the COVID-19 Genomics UK (COG-UK) Consortium | The Lighthouse Lab in Milton Keynes and Alex Alderton, Roberto Amato, Jeffrey Barrett, Sonia Goncalves, Ewan Harrison, David K. Jackson, Ian Johnston, Dominic Kwiatkowski, Cordelia Langford, John Sillitoe on behalf of the Wellcome Sanger Institute COVID-19 Surveillance Team |
| EPI_ISL_3249609, EPI_ISL_3249610 | Berkshire and Surrey Pathology Services Lighthouse Laboratory | Wellcome Sanger Institute for the COVID-19 Genomics UK (COG-UK) Consortium | Berkshire and Surrey Pathology Services Lighthouse Laboratory and Alex Alderton, Roberto Amato, Jeffrey Barrett, Sonia Goncalves, Ewan Harrison, David K. Jackson, Ian Johnston, Dominic Kwiatkowski, Cordelia Langford, John Sillitoe on behalf of the Wellcome Sanger Institute COVID-19 Surveillance Team |
| EPI_ISL_3249611 | Lighthouse Lab in Glasgow | Wellcome Sanger Institute for the COVID-19 Genomics UK (COG-UK) Consortium | Harper VanSteenhouse, Yumi Kasai, David Gray, Carol Clugston, Anna Dominiczak and Alex Alderton, Roberto Amato, Jeffrey Barrett, Sonia Goncalves, Ewan Harrison, David K. Jackson, Ian Johnston, Dominic Kwiatkowski, Cordelia Langford, John Sillitoe on behalf of the Wellcome Sanger Institute COVID-19 Surveillance Team |
| EPI_ISL_3249612 | Lighthouse Lab in Milton Keynes | Wellcome Sanger Institute for the COVID-19 Genomics UK (COG-UK) Consortium | The Lighthouse Lab in Milton Keynes and Alex Alderton, Roberto Amato, Jeffrey Barrett, Sonia Goncalves, Ewan Harrison, David K. Jackson, Ian Johnston, Dominic Kwiatkowski, Cordelia Langford, John Sillitoe on behalf of the Wellcome Sanger Institute COVID-19 Surveillance Team |
| EPI_ISL_3249613, EPI_ISL_3249614, EPI_ISL_3249615 | Berkshire and Surrey Pathology Services Lighthouse Laboratory | Wellcome Sanger Institute for the COVID-19 Genomics UK (COG-UK) Consortium | Berkshire and Surrey Pathology Services Lighthouse Laboratory and Alex Alderton, Roberto Amato, Jeffrey Barrett, Sonia Goncalves, Ewan Harrison, David K. Jackson, Ian Johnston, Dominic Kwiatkowski, Cordelia Langford, John Sillitoe on behalf of the Wellcome Sanger Institute COVID-19 Surveillance Team |
| EPI_ISL_3249616 | Lighthouse Lab in Glasgow | Wellcome Sanger Institute for the COVID-19 Genomics UK (COG-UK) Consortium | Harper VanSteenhouse, Yumi Kasai, David Gray, Carol Clugston, Anna Dominiczak and Alex Alderton, Roberto Amato, Jeffrey Barrett, Sonia Goncalves, Ewan Harrison, David K. Jackson, Ian Johnston, Dominic Kwiatkowski, Cordelia Langford, John Sillitoe on behalf of the Wellcome Sanger Institute COVID-19 Surveillance Team |
| EPI_ISL_3249617 | Lighthouse Lab in Milton Keynes | Wellcome Sanger Institute for the COVID-19 Genomics UK (COG-UK) Consortium | The Lighthouse Lab in Milton Keynes and Alex Alderton, Roberto Amato, Jeffrey Barrett, Sonia Goncalves, Ewan Harrison, David K. Jackson, Ian Johnston, Dominic Kwiatkowski, Cordelia Langford, John Sillitoe on behalf of the Wellcome Sanger Institute COVID-19 Surveillance Team |
| EPI_ISL_3249618, EPI_ISL_3249619 | Lighthouse Lab in Glasgow | Wellcome Sanger Institute for the COVID-19 Genomics UK (COG-UK) Consortium | Harper VanSteenhouse, Yumi Kasai, David Gray, Carol Clugston, Anna Dominiczak and Alex Alderton, Roberto Amato, Jeffrey Barrett, Sonia Goncalves, Ewan Harrison, David K. Jackson, Ian Johnston, Dominic Kwiatkowski, Cordelia Langford, John Sillitoe on behalf of the Wellcome Sanger Institute COVID-19 Surveillance Team |
| EPI_ISL_3249620 | Lighthouse Lab in Milton Keynes | Wellcome Sanger Institute for the COVID-19 Genomics UK (COG-UK) Consortium | The Lighthouse Lab in Milton Keynes and Alex Alderton, Roberto Amato, Jeffrey Barrett, Sonia Goncalves, Ewan Harrison, David K. Jackson, Ian Johnston, Dominic Kwiatkowski, Cordelia Langford, John Sillitoe on behalf of the Wellcome Sanger Institute COVID-19 Surveillance Team |
| EPI_ISL_3249621, EPI_ISL_3249622 | Lighthouse Lab in Glasgow | Wellcome Sanger Institute for the COVID-19 Genomics UK (COG-UK) Consortium | Harper VanSteenhouse, Yumi Kasai, David Gray, Carol Clugston, Anna Dominiczak and Alex Alderton, Roberto Amato, Jeffrey Barrett, Sonia Goncalves, Ewan Harrison, David K. Jackson, Ian Johnston, Dominic Kwiatkowski, Cordelia Langford, John Sillitoe on behalf of the Wellcome Sanger Institute COVID-19 Surveillance Team |
| EPI_ISL_3249623, EPI_ISL_3249624 | Berkshire and Surrey Pathology Services Lighthouse Laboratory | Wellcome Sanger Institute for the COVID-19 Genomics UK (COG-UK) Consortium | Berkshire and Surrey Pathology Services Lighthouse Laboratory and Alex Alderton, Roberto Amato, Jeffrey Barrett, Sonia Goncalves, Ewan Harrison, David K. Jackson, Ian Johnston, Dominic Kwiatkowski, Cordelia Langford, John Sillitoe on behalf of the Wellcome Sanger Institute COVID-19 Surveillance Team |
| EPI_ISL_3249625 | Lighthouse Lab in Glasgow | Wellcome Sanger Institute for the COVID-19 Genomics UK (COG-UK) Consortium | Harper VanSteenhouse, Yumi Kasai, David Gray, Carol Clugston, Anna Dominiczak and Alex Alderton, Roberto Amato, Jeffrey Barrett, Sonia Goncalves, Ewan Harrison, David K. Jackson, Ian Johnston, Dominic Kwiatkowski, Cordelia Langford, John Sillitoe on behalf of the Wellcome Sanger Institute COVID-19 Surveillance Team |
| EPI_ISL_3249626 | Lighthouse Lab in Milton Keynes | Wellcome Sanger Institute for the COVID-19 Genomics UK (COG-UK) Consortium | The Lighthouse Lab in Milton Keynes and Alex Alderton, Roberto Amato, Jeffrey Barrett, Sonia Goncalves, Ewan Harrison, David K. Jackson, Ian Johnston, Dominic Kwiatkowski, Cordelia Langford, John Sillitoe on behalf of the Wellcome Sanger Institute COVID-19 Surveillance Team |
| EPI_ISL_3249627 | Berkshire and Surrey Pathology Services Lighthouse Laboratory | Wellcome Sanger Institute for the COVID-19 Genomics UK (COG-UK) Consortium | Berkshire and Surrey Pathology Services Lighthouse Laboratory and Alex Alderton, Roberto Amato, Jeffrey Barrett, Sonia Goncalves, Ewan Harrison, David K. Jackson, Ian Johnston, Dominic Kwiatkowski, Cordelia Langford, John Sillitoe on behalf of the Wellcome Sanger Institute COVID-19 Surveillance Team |
| EPI_ISL_3249628 | Lighthouse Lab in Milton Keynes | Wellcome Sanger Institute for the COVID-19 Genomics UK (COG-UK) Consortium | The Lighthouse Lab in Milton Keynes and Alex Alderton, Roberto Amato, Jeffrey Barrett, Sonia Goncalves, Ewan Harrison, David K. Jackson, Ian Johnston, Dominic Kwiatkowski, Cordelia Langford, John Sillitoe on behalf of the Wellcome Sanger Institute COVID-19 Surveillance Team |
| EPI_ISL_3249629 | Lighthouse Lab in Glasgow | Wellcome Sanger Institute for the COVID-19 Genomics UK (COG-UK) Consortium | Harper VanSteenhouse, Yumi Kasai, David Gray, Carol Clugston, Anna Dominiczak and Alex Alderton, Roberto Amato, Jeffrey Barrett, Sonia Goncalves, Ewan Harrison, David K. Jackson, Ian Johnston, Dominic Kwiatkowski, Cordelia Langford, John Sillitoe on behalf of the Wellcome Sanger Institute COVID-19 Surveillance Team |
| EPI_ISL_3249630 | Berkshire and Surrey Pathology Services Lighthouse Laboratory | Wellcome Sanger Institute for the COVID-19 Genomics UK (COG-UK) | Berkshire and Surrey Pathology Services Lighthouse Laboratory and Alex Alderton, Roberto Amato, Jeffrey Barrett, Sonia Goncalves, Ewan Harrison, David K. Jackson, Ian Johnston, Dominic Kwiatkowski, Cordelia Langford, John Sillitoe on behalf |

| | | | |
|---|---|---|---|
| | | Consortium | of the Wellcome Sanger Institute COVID-19 Surveillance Team |
| EPI_ISL_3249631, EPI_ISL_3249632 | Lighthouse Lab in Milton Keynes | Wellcome Sanger Institute for the COVID-19 Genomics UK (COG-UK) Consortium | The Lighthouse Lab in Milton Keynes and Alex Alderton, Roberto Amato, Jeffrey Barrett, Sonia Goncalves, Ewan Harrison, David K. Jackson, Ian Johnston, Dominic Kwiatkowski, Cordelia Langford, John Sillitoe on behalf of the Wellcome Sanger Institute COVID-19 Surveillance Team |
| EPI_ISL_3249633, EPI_ISL_3249634, EPI_ISL_3249635, EPI_ISL_3249636, EPI_ISL_3249637 | Berkshire and Surrey Pathology Services Lighthouse Laboratory | Wellcome Sanger Institute for the COVID-19 Genomics UK (COG-UK) Consortium | Berkshire and Surrey Pathology Services Lighthouse Laboratory and Alex Alderton, Roberto Amato, Jeffrey Barrett, Sonia Goncalves, Ewan Harrison, David K. Jackson, Ian Johnston, Dominic Kwiatkowski, Cordelia Langford, John Sillitoe on behalf of the Wellcome Sanger Institute COVID-19 Surveillance Team |
| EPI_ISL_3249638, EPI_ISL_3249639 | Lighthouse Lab in Milton Keynes | Wellcome Sanger Institute for the COVID-19 Genomics UK (COG-UK) Consortium | The Lighthouse Lab in Milton Keynes and Alex Alderton, Roberto Amato, Jeffrey Barrett, Sonia Goncalves, Ewan Harrison, David K. Jackson, Ian Johnston, Dominic Kwiatkowski, Cordelia Langford, John Sillitoe on behalf of the Wellcome Sanger Institute COVID-19 Surveillance Team |
| EPI_ISL_3249640, EPI_ISL_3249641, EPI_ISL_3249642, EPI_ISL_3249643 | Berkshire and Surrey Pathology Services Lighthouse Laboratory | Wellcome Sanger Institute for the COVID-19 Genomics UK (COG-UK) Consortium | Berkshire and Surrey Pathology Services Lighthouse Laboratory and Alex Alderton, Roberto Amato, Jeffrey Barrett, Sonia Goncalves, Ewan Harrison, David K. Jackson, Ian Johnston, Dominic Kwiatkowski, Cordelia Langford, John Sillitoe on behalf of the Wellcome Sanger Institute COVID-19 Surveillance Team |
| EPI_ISL_3249644 | Lighthouse Lab in Milton Keynes | Wellcome Sanger Institute for the COVID-19 Genomics UK (COG-UK) Consortium | The Lighthouse Lab in Milton Keynes and Alex Alderton, Roberto Amato, Jeffrey Barrett, Sonia Goncalves, Ewan Harrison, David K. Jackson, Ian Johnston, Dominic Kwiatkowski, Cordelia Langford, John Sillitoe on behalf of the Wellcome Sanger Institute COVID-19 Surveillance Team |
| EPI_ISL_3249645 | Lighthouse Lab in Glasgow | Wellcome Sanger Institute for the COVID-19 Genomics UK (COG-UK) Consortium | Harper VanSteenhouse, Yumi Kasai, David Gray, Carol Clugston, Anna Dominiczak and Alex Alderton, Roberto Amato, Jeffrey Barrett, Sonia Goncalves, Ewan Harrison, David K. Jackson, Ian Johnston, Dominic Kwiatkowski, Cordelia Langford, John Sillitoe on behalf of the Wellcome Sanger Institute COVID-19 Surveillance Team |
| EPI_ISL_3249646, EPI_ISL_3249647, EPI_ISL_3249648 | Berkshire and Surrey Pathology Services Lighthouse Laboratory | Wellcome Sanger Institute for the COVID-19 Genomics UK (COG-UK) Consortium | Berkshire and Surrey Pathology Services Lighthouse Laboratory and Alex Alderton, Roberto Amato, Jeffrey Barrett, Sonia Goncalves, Ewan Harrison, David K. Jackson, Ian Johnston, Dominic Kwiatkowski, Cordelia Langford, John Sillitoe on behalf of the Wellcome Sanger Institute COVID-19 Surveillance Team |
| EPI_ISL_3249649 | Lighthouse Lab in Glasgow | Wellcome Sanger Institute for the COVID-19 Genomics UK (COG-UK) Consortium | Harper VanSteenhouse, Yumi Kasai, David Gray, Carol Clugston, Anna Dominiczak and Alex Alderton, Roberto Amato, Jeffrey Barrett, Sonia Goncalves, Ewan Harrison, David K. Jackson, Ian Johnston, Dominic Kwiatkowski, Cordelia Langford, John Sillitoe on behalf of the Wellcome Sanger Institute COVID-19 Surveillance Team |
| EPI_ISL_3249650 | Lighthouse Lab in Milton Keynes | Wellcome Sanger Institute for the COVID-19 Genomics UK (COG-UK) Consortium | The Lighthouse Lab in Milton Keynes and Alex Alderton, Roberto Amato, Jeffrey Barrett, Sonia Goncalves, Ewan Harrison, David K. Jackson, Ian Johnston, Dominic Kwiatkowski, Cordelia Langford, John Sillitoe on behalf of the Wellcome Sanger Institute COVID-19 Surveillance Team |
| EPI_ISL_3249651, EPI_ISL_3249652, EPI_ISL_3249653 | Berkshire and Surrey Pathology Services Lighthouse Laboratory | Wellcome Sanger Institute for the COVID-19 Genomics UK (COG-UK) Consortium | Berkshire and Surrey Pathology Services Lighthouse Laboratory and Alex Alderton, Roberto Amato, Jeffrey Barrett, Sonia Goncalves, Ewan Harrison, David K. Jackson, Ian Johnston, Dominic Kwiatkowski, Cordelia Langford, John Sillitoe on behalf of the Wellcome Sanger Institute COVID-19 Surveillance Team |
| EPI_ISL_3249654, EPI_ISL_3249655 | Lighthouse Lab in Milton Keynes | Wellcome Sanger Institute for the COVID-19 Genomics UK (COG-UK) Consortium | The Lighthouse Lab in Milton Keynes and Alex Alderton, Roberto Amato, Jeffrey Barrett, Sonia Goncalves, Ewan Harrison, David K. Jackson, Ian Johnston, Dominic Kwiatkowski, Cordelia Langford, John Sillitoe on behalf of the Wellcome Sanger Institute COVID-19 Surveillance Team |
| EPI_ISL_3249656 | Berkshire and Surrey Pathology Services Lighthouse Laboratory | Wellcome Sanger Institute for the COVID-19 Genomics UK (COG-UK) Consortium | Berkshire and Surrey Pathology Services Lighthouse Laboratory and Alex Alderton, Roberto Amato, Jeffrey Barrett, Sonia Goncalves, Ewan Harrison, David K. Jackson, Ian Johnston, Dominic Kwiatkowski, Cordelia Langford, John Sillitoe on behalf of the Wellcome Sanger Institute COVID-19 Surveillance Team |
| EPI_ISL_3249657, EPI_ISL_3249658 | Lighthouse Lab in Milton Keynes | Wellcome Sanger Institute for the COVID-19 Genomics UK (COG-UK) Consortium | The Lighthouse Lab in Milton Keynes and Alex Alderton, Roberto Amato, Jeffrey Barrett, Sonia Goncalves, Ewan Harrison, David K. Jackson, Ian Johnston, Dominic Kwiatkowski, Cordelia Langford, John Sillitoe on behalf of the Wellcome Sanger Institute COVID-19 Surveillance Team |
| EPI_ISL_3249659 | Berkshire and Surrey Pathology Services Lighthouse Laboratory | Wellcome Sanger Institute for the COVID-19 Genomics UK (COG-UK) Consortium | Berkshire and Surrey Pathology Services Lighthouse Laboratory and Alex Alderton, Roberto Amato, Jeffrey Barrett, Sonia Goncalves, Ewan Harrison, David K. Jackson, Ian Johnston, Dominic Kwiatkowski, Cordelia Langford, John Sillitoe on behalf of the Wellcome Sanger Institute COVID-19 Surveillance Team |
| EPI_ISL_3249660 | Lighthouse Lab in Milton Keynes | Wellcome Sanger Institute for the COVID-19 Genomics UK (COG-UK) Consortium | The Lighthouse Lab in Milton Keynes and Alex Alderton, Roberto Amato, Jeffrey Barrett, Sonia Goncalves, Ewan Harrison, David K. Jackson, Ian Johnston, Dominic Kwiatkowski, Cordelia Langford, John Sillitoe on behalf of the Wellcome Sanger Institute COVID-19 Surveillance Team |
| EPI_ISL_3249661 | Berkshire and Surrey Pathology Services Lighthouse Laboratory | Wellcome Sanger Institute for the COVID-19 Genomics UK (COG-UK) Consortium | Berkshire and Surrey Pathology Services Lighthouse Laboratory and Alex Alderton, Roberto Amato, Jeffrey Barrett, Sonia Goncalves, Ewan Harrison, David K. Jackson, Ian Johnston, Dominic Kwiatkowski, Cordelia Langford, John Sillitoe on behalf of the Wellcome Sanger Institute COVID-19 Surveillance Team |
| EPI_ISL_3249662 | Lighthouse Lab in Glasgow | Wellcome Sanger Institute for the COVID-19 Genomics UK (COG-UK) Consortium | Harper VanSteenhouse, Yumi Kasai, David Gray, Carol Clugston, Anna Dominiczak and Alex Alderton, Roberto Amato, Jeffrey Barrett, Sonia Goncalves, Ewan Harrison, David K. Jackson, Ian Johnston, Dominic Kwiatkowski, Cordelia Langford, John Sillitoe on behalf of the Wellcome Sanger Institute COVID-19 Surveillance Team |
| EPI_ISL_3249663, EPI_ISL_3249664, EPI_ISL_3249665, EPI_ISL_3249666 | Berkshire and Surrey Pathology Services Lighthouse Laboratory | Wellcome Sanger Institute for the COVID-19 Genomics UK (COG-UK) Consortium | Berkshire and Surrey Pathology Services Lighthouse Laboratory and Alex Alderton, Roberto Amato, Jeffrey Barrett, Sonia Goncalves, Ewan Harrison, David K. Jackson, Ian Johnston, Dominic Kwiatkowski, Cordelia Langford, John Sillitoe on behalf of the Wellcome Sanger Institute COVID-19 Surveillance Team |
| EPI_ISL_3249667 | Lighthouse Lab in Glasgow | Wellcome Sanger Institute for the COVID-19 Genomics UK (COG-UK) Consortium | Harper VanSteenhouse, Yumi Kasai, David Gray, Carol Clugston, Anna Dominiczak and Alex Alderton, Roberto Amato, Jeffrey Barrett, Sonia Goncalves, Ewan Harrison, David K. Jackson, Ian Johnston, Dominic Kwiatkowski, Cordelia Langford, John Sillitoe on behalf of the Wellcome Sanger Institute COVID-19 Surveillance Team |
| EPI_ISL_3249668, EPI_ISL_3249669, EPI_ISL_3249670, EPI_ISL_3249671 | Berkshire and Surrey Pathology Services Lighthouse Laboratory | Wellcome Sanger Institute for the COVID-19 Genomics UK (COG-UK) Consortium | Berkshire and Surrey Pathology Services Lighthouse Laboratory and Alex Alderton, Roberto Amato, Jeffrey Barrett, Sonia Goncalves, Ewan Harrison, David K. Jackson, Ian Johnston, Dominic Kwiatkowski, Cordelia Langford, John Sillitoe on behalf of the Wellcome Sanger Institute COVID-19 Surveillance Team |
| EPI_ISL_3249672 | Lighthouse Lab in Milton Keynes | Wellcome Sanger Institute for the COVID-19 Genomics UK (COG-UK) Consortium | The Lighthouse Lab in Milton Keynes and Alex Alderton, Roberto Amato, Jeffrey Barrett, Sonia Goncalves, Ewan Harrison, David K. Jackson, Ian Johnston, Dominic Kwiatkowski, Cordelia Langford, John Sillitoe on behalf of the Wellcome Sanger Institute COVID-19 Surveillance Team |
| EPI_ISL_3249673, EPI_ISL_3249674, EPI_ISL_3249675 | Berkshire and Surrey Pathology Services Lighthouse Laboratory | Wellcome Sanger Institute for the COVID-19 Genomics UK (COG-UK) Consortium | Berkshire and Surrey Pathology Services Lighthouse Laboratory and Alex Alderton, Roberto Amato, Jeffrey Barrett, Sonia Goncalves, Ewan Harrison, David K. Jackson, Ian Johnston, Dominic Kwiatkowski, Cordelia Langford, John Sillitoe on behalf of the Wellcome Sanger Institute COVID-19 Surveillance Team |
| EPI_ISL_3249676 | Lighthouse Lab in Glasgow | Wellcome Sanger Institute for the COVID-19 Genomics UK (COG-UK) Consortium | Harper VanSteenhouse, Yumi Kasai, David Gray, Carol Clugston, Anna Dominiczak and Alex Alderton, Roberto Amato, Jeffrey Barrett, Sonia Goncalves, Ewan Harrison, David K. Jackson, Ian Johnston, Dominic Kwiatkowski, Cordelia Langford, John Sillitoe on behalf of the Wellcome Sanger Institute COVID-19 Surveillance Team |
| EPI_ISL_3249677 | Berkshire and Surrey Pathology Services Lighthouse Laboratory | Wellcome Sanger Institute for the COVID-19 Genomics UK (COG-UK) Consortium | Berkshire and Surrey Pathology Services Lighthouse Laboratory and Alex Alderton, Roberto Amato, Jeffrey Barrett, Sonia Goncalves, Ewan Harrison, David K. Jackson, Ian Johnston, Dominic Kwiatkowski, Cordelia Langford, John Sillitoe on behalf of the Wellcome Sanger Institute COVID-19 Surveillance Team |
| EPI_ISL_3249678 | Lighthouse Lab in Milton Keynes | Wellcome Sanger Institute for the COVID-19 Genomics UK (COG-UK) Consortium | The Lighthouse Lab in Milton Keynes and Alex Alderton, Roberto Amato, Jeffrey Barrett, Sonia Goncalves, Ewan Harrison, David K. Jackson, Ian Johnston, Dominic Kwiatkowski, Cordelia Langford, John Sillitoe on behalf of the Wellcome Sanger Institute COVID-19 Surveillance Team |
| EPI_ISL_3249679, EPI_ISL_3249680, EPI_ISL_3249681, EPI_ISL_3249682 | Berkshire and Surrey Pathology Services Lighthouse Laboratory | Wellcome Sanger Institute for the COVID-19 Genomics UK (COG-UK) Consortium | Berkshire and Surrey Pathology Services Lighthouse Laboratory and Alex Alderton, Roberto Amato, Jeffrey Barrett, Sonia Goncalves, Ewan Harrison, David K. Jackson, Ian Johnston, Dominic Kwiatkowski, Cordelia Langford, John Sillitoe on behalf of the Wellcome Sanger Institute COVID-19 Surveillance Team |
| EPI_ISL_3249683 | Lighthouse Lab in Milton Keynes | Wellcome Sanger Institute for the COVID-19 Genomics UK (COG-UK) Consortium | The Lighthouse Lab in Milton Keynes and Alex Alderton, Roberto Amato, Jeffrey Barrett, Sonia Goncalves, Ewan Harrison, David K. Jackson, Ian Johnston, Dominic Kwiatkowski, Cordelia Langford, John Sillitoe on behalf of the Wellcome Sanger Institute COVID-19 Surveillance Team |
| EPI_ISL_3249684, EPI_ISL_3249685, EPI_ISL_3249686, EPI_ISL_3249687, EPI_ISL_3249688, EPI_ISL_3249689 | Berkshire and Surrey Pathology Services Lighthouse Laboratory | Wellcome Sanger Institute for the COVID-19 Genomics UK (COG-UK) Consortium | Berkshire and Surrey Pathology Services Lighthouse Laboratory and Alex Alderton, Roberto Amato, Jeffrey Barrett, Sonia Goncalves, Ewan Harrison, David K. Jackson, Ian Johnston, Dominic Kwiatkowski, Cordelia Langford, John Sillitoe on behalf of the Wellcome Sanger Institute COVID-19 Surveillance Team |
| EPI_ISL_3249690 | Lighthouse Lab in Milton Keynes | Wellcome Sanger Institute for the COVID-19 Genomics UK (COG-UK) Consortium | The Lighthouse Lab in Milton Keynes and Alex Alderton, Roberto Amato, Jeffrey Barrett, Sonia Goncalves, Ewan Harrison, David K. Jackson, Ian Johnston, Dominic Kwiatkowski, Cordelia Langford, John Sillitoe on behalf of the Wellcome Sanger Institute COVID-19 Surveillance Team |
| EPI_ISL_3249691 | Berkshire and Surrey Pathology Services Lighthouse Laboratory | Wellcome Sanger Institute for the COVID-19 Genomics UK (COG-UK) Consortium | Berkshire and Surrey Pathology Services Lighthouse Laboratory and Alex Alderton, Roberto Amato, Jeffrey Barrett, Sonia Goncalves, Ewan Harrison, David K. Jackson, Ian Johnston, Dominic Kwiatkowski, Cordelia Langford, John Sillitoe on behalf of the Wellcome Sanger Institute COVID-19 Surveillance Team |
| EPI_ISL_3249692, EPI_ISL_3249693, EPI_ISL_3249694, EPI_ISL_3249695, EPI_ISL_3249696 | Lighthouse Lab in Milton Keynes | Wellcome Sanger Institute for the COVID-19 Genomics UK (COG-UK) Consortium | The Lighthouse Lab in Milton Keynes and Alex Alderton, Roberto Amato, Jeffrey Barrett, Sonia Goncalves, Ewan Harrison, David K. Jackson, Ian Johnston, Dominic Kwiatkowski, Cordelia Langford, John Sillitoe on behalf of the Wellcome Sanger Institute COVID-19 Surveillance Team |
| EPI_ISL_3249697 | Berkshire and Surrey Pathology Services Lighthouse Laboratory | Wellcome Sanger Institute for the COVID-19 Genomics UK (COG-UK) Consortium | Berkshire and Surrey Pathology Services Lighthouse Laboratory and Alex Alderton, Roberto Amato, Jeffrey Barrett, Sonia Goncalves, Ewan Harrison, David K. Jackson, Ian Johnston, Dominic Kwiatkowski, Cordelia Langford, John Sillitoe on behalf of the Wellcome Sanger Institute COVID-19 Surveillance Team |
| EPI_ISL_3249698 | Lighthouse Lab in Glasgow | Wellcome Sanger Institute for the COVID-19 Genomics UK (COG-UK) Consortium | Harper VanSteenhouse, Yumi Kasai, David Gray, Carol Clugston, Anna Dominiczak and Alex Alderton, Roberto Amato, Jeffrey Barrett, Sonia Goncalves, Ewan Harrison, David K. Jackson, Ian Johnston, Dominic Kwiatkowski, Cordelia Langford, John Sillitoe on behalf of the Wellcome Sanger Institute COVID-19 Surveillance Team |
| EPI_ISL_3249699 | Berkshire and Surrey Pathology Services Lighthouse Laboratory | Wellcome Sanger Institute for the COVID-19 Genomics UK (COG-UK) Consortium | Berkshire and Surrey Pathology Services Lighthouse Laboratory and Alex Alderton, Roberto Amato, Jeffrey Barrett, Sonia Goncalves, Ewan Harrison, David K. Jackson, Ian Johnston, Dominic Kwiatkowski, Cordelia Langford, John Sillitoe on behalf of the Wellcome Sanger Institute COVID-19 Surveillance Team |
| EPI_ISL_3249700 | Lighthouse Lab in Milton Keynes | Wellcome Sanger Institute for the COVID-19 Genomics UK (COG-UK) Consortium | The Lighthouse Lab in Milton Keynes and Alex Alderton, Roberto Amato, Jeffrey Barrett, Sonia Goncalves, Ewan Harrison, David K. Jackson, Ian Johnston, Dominic Kwiatkowski, Cordelia Langford, John Sillitoe on behalf of the Wellcome Sanger Institute COVID-19 Surveillance Team |
| EPI_ISL_3249701, | Berkshire and Surrey Pathology Services Lighthouse Laboratory | Wellcome Sanger Institute for the COVID-19 Genomics UK (COG-UK) | Berkshire and Surrey Pathology Services Lighthouse Laboratory and Alex Alderton, Roberto Amato, Jeffrey Barrett, Sonia Goncalves, Ewan Harrison, David K. Jackson, Ian Johnston, Dominic Kwiatkowski, Cordelia Langford, John Sillitoe on behalf |

| | | | |
|---|---|---|---|
| EPI_ISL_3249702 | | Consortium | of the Wellcome Sanger Institute COVID-19 Surveillance Team |
| EPI_ISL_3249703 | Lighthouse Lab in Glasgow | Wellcome Sanger Institute for the COVID-19 Genomics UK (COG-UK) Consortium | Harper VanSteenhouse, Yumi Kasai, David Gray, Carol Clugston, Anna Dominiczak and Alex Alderton, Roberto Amato, Jeffrey Barrett, Sonia Goncalves, Ewan Harrison, David K. Jackson, Ian Johnston, Dominic Kwiatkowski, Cordelia Langford, John Sillitoe on behalf of the Wellcome Sanger Institute COVID-19 Surveillance Team |
| EPI_ISL_3249704 | Berkshire and Surrey Pathology Services Lighthouse Laboratory | Wellcome Sanger Institute for the COVID-19 Genomics UK (COG-UK) Consortium | Berkshire and Surrey Pathology Services Lighthouse Laboratory and Alex Alderton, Roberto Amato, Jeffrey Barrett, Sonia Goncalves, Ewan Harrison, David K. Jackson, Ian Johnston, Dominic Kwiatkowski, Cordelia Langford, John Sillitoe on behalf of the Wellcome Sanger Institute COVID-19 Surveillance Team |
| EPI_ISL_3249705 | Lighthouse Lab in Milton Keynes | Wellcome Sanger Institute for the COVID-19 Genomics UK (COG-UK) Consortium | The Lighthouse Lab in Milton Keynes and Alex Alderton, Roberto Amato, Jeffrey Barrett, Sonia Goncalves, Ewan Harrison, David K. Jackson, Ian Johnston, Dominic Kwiatkowski, Cordelia Langford, John Sillitoe on behalf of the Wellcome Sanger Institute COVID-19 Surveillance Team |
| EPI_ISL_3249706 | Lighthouse Lab in Glasgow | Wellcome Sanger Institute for the COVID-19 Genomics UK (COG-UK) Consortium | Harper VanSteenhouse, Yumi Kasai, David Gray, Carol Clugston, Anna Dominiczak and Alex Alderton, Roberto Amato, Jeffrey Barrett, Sonia Goncalves, Ewan Harrison, David K. Jackson, Ian Johnston, Dominic Kwiatkowski, Cordelia Langford, John Sillitoe on behalf of the Wellcome Sanger Institute COVID-19 Surveillance Team |
| EPI_ISL_3249707 | Lighthouse Lab in Milton Keynes | Wellcome Sanger Institute for the COVID-19 Genomics UK (COG-UK) Consortium | The Lighthouse Lab in Milton Keynes and Alex Alderton, Roberto Amato, Jeffrey Barrett, Sonia Goncalves, Ewan Harrison, David K. Jackson, Ian Johnston, Dominic Kwiatkowski, Cordelia Langford, John Sillitoe on behalf of the Wellcome Sanger Institute COVID-19 Surveillance Team |
| EPI_ISL_3249708, EPI_ISL_3249709 | Berkshire and Surrey Pathology Services Lighthouse Laboratory | Wellcome Sanger Institute for the COVID-19 Genomics UK (COG-UK) Consortium | Berkshire and Surrey Pathology Services Lighthouse Laboratory and Alex Alderton, Roberto Amato, Jeffrey Barrett, Sonia Goncalves, Ewan Harrison, David K. Jackson, Ian Johnston, Dominic Kwiatkowski, Cordelia Langford, John Sillitoe on behalf of the Wellcome Sanger Institute COVID-19 Surveillance Team |
| EPI_ISL_3249710 | Lighthouse Lab in Milton Keynes | Wellcome Sanger Institute for the COVID-19 Genomics UK (COG-UK) Consortium | The Lighthouse Lab in Milton Keynes and Alex Alderton, Roberto Amato, Jeffrey Barrett, Sonia Goncalves, Ewan Harrison, David K. Jackson, Ian Johnston, Dominic Kwiatkowski, Cordelia Langford, John Sillitoe on behalf of the Wellcome Sanger Institute COVID-19 Surveillance Team |
| EPI_ISL_3249711, EPI_ISL_3249712 | Berkshire and Surrey Pathology Services Lighthouse Laboratory | Wellcome Sanger Institute for the COVID-19 Genomics UK (COG-UK) Consortium | Berkshire and Surrey Pathology Services Lighthouse Laboratory and Alex Alderton, Roberto Amato, Jeffrey Barrett, Sonia Goncalves, Ewan Harrison, David K. Jackson, Ian Johnston, Dominic Kwiatkowski, Cordelia Langford, John Sillitoe on behalf of the Wellcome Sanger Institute COVID-19 Surveillance Team |
| EPI_ISL_3249713, EPI_ISL_3249714 | Lighthouse Lab in Milton Keynes | Wellcome Sanger Institute for the COVID-19 Genomics UK (COG-UK) Consortium | The Lighthouse Lab in Milton Keynes and Alex Alderton, Roberto Amato, Jeffrey Barrett, Sonia Goncalves, Ewan Harrison, David K. Jackson, Ian Johnston, Dominic Kwiatkowski, Cordelia Langford, John Sillitoe on behalf of the Wellcome Sanger Institute COVID-19 Surveillance Team |
| EPI_ISL_3249715 | Lighthouse Lab in Glasgow | Wellcome Sanger Institute for the COVID-19 Genomics UK (COG-UK) Consortium | Harper VanSteenhouse, Yumi Kasai, David Gray, Carol Clugston, Anna Dominiczak and Alex Alderton, Roberto Amato, Jeffrey Barrett, Sonia Goncalves, Ewan Harrison, David K. Jackson, Ian Johnston, Dominic Kwiatkowski, Cordelia Langford, John Sillitoe on behalf of the Wellcome Sanger Institute COVID-19 Surveillance Team |
| EPI_ISL_3249716, EPI_ISL_3249717 | Berkshire and Surrey Pathology Services Lighthouse Laboratory | Wellcome Sanger Institute for the COVID-19 Genomics UK (COG-UK) Consortium | Berkshire and Surrey Pathology Services Lighthouse Laboratory and Alex Alderton, Roberto Amato, Jeffrey Barrett, Sonia Goncalves, Ewan Harrison, David K. Jackson, Ian Johnston, Dominic Kwiatkowski, Cordelia Langford, John Sillitoe on behalf of the Wellcome Sanger Institute COVID-19 Surveillance Team |
| EPI_ISL_3249718 | Lighthouse Lab in Milton Keynes | Wellcome Sanger Institute for the COVID-19 Genomics UK (COG-UK) Consortium | The Lighthouse Lab in Milton Keynes and Alex Alderton, Roberto Amato, Jeffrey Barrett, Sonia Goncalves, Ewan Harrison, David K. Jackson, Ian Johnston, Dominic Kwiatkowski, Cordelia Langford, John Sillitoe on behalf of the Wellcome Sanger Institute COVID-19 Surveillance Team |
| EPI_ISL_3249719 | Berkshire and Surrey Pathology Services Lighthouse Laboratory | Wellcome Sanger Institute for the COVID-19 Genomics UK (COG-UK) Consortium | Berkshire and Surrey Pathology Services Lighthouse Laboratory and Alex Alderton, Roberto Amato, Jeffrey Barrett, Sonia Goncalves, Ewan Harrison, David K. Jackson, Ian Johnston, Dominic Kwiatkowski, Cordelia Langford, John Sillitoe on behalf of the Wellcome Sanger Institute COVID-19 Surveillance Team |
| EPI_ISL_3249720, EPI_ISL_3249721 | Lighthouse Lab in Glasgow | Wellcome Sanger Institute for the COVID-19 Genomics UK (COG-UK) Consortium | Harper VanSteenhouse, Yumi Kasai, David Gray, Carol Clugston, Anna Dominiczak and Alex Alderton, Roberto Amato, Jeffrey Barrett, Sonia Goncalves, Ewan Harrison, David K. Jackson, Ian Johnston, Dominic Kwiatkowski, Cordelia Langford, John Sillitoe on behalf of the Wellcome Sanger Institute COVID-19 Surveillance Team |
| EPI_ISL_3249722 | Lighthouse Lab in Milton Keynes | Wellcome Sanger Institute for the COVID-19 Genomics UK (COG-UK) Consortium | The Lighthouse Lab in Milton Keynes and Alex Alderton, Roberto Amato, Jeffrey Barrett, Sonia Goncalves, Ewan Harrison, David K. Jackson, Ian Johnston, Dominic Kwiatkowski, Cordelia Langford, John Sillitoe on behalf of the Wellcome Sanger Institute COVID-19 Surveillance Team |
| EPI_ISL_3249723 | Lighthouse Lab in Glasgow | Wellcome Sanger Institute for the COVID-19 Genomics UK (COG-UK) Consortium | Harper VanSteenhouse, Yumi Kasai, David Gray, Carol Clugston, Anna Dominiczak and Alex Alderton, Roberto Amato, Jeffrey Barrett, Sonia Goncalves, Ewan Harrison, David K. Jackson, Ian Johnston, Dominic Kwiatkowski, Cordelia Langford, John Sillitoe on behalf of the Wellcome Sanger Institute COVID-19 Surveillance Team |
| EPI_ISL_3249724 | Lighthouse Lab in Milton Keynes | Wellcome Sanger Institute for the COVID-19 Genomics UK (COG-UK) Consortium | The Lighthouse Lab in Milton Keynes and Alex Alderton, Roberto Amato, Jeffrey Barrett, Sonia Goncalves, Ewan Harrison, David K. Jackson, Ian Johnston, Dominic Kwiatkowski, Cordelia Langford, John Sillitoe on behalf of the Wellcome Sanger Institute COVID-19 Surveillance Team |
| EPI_ISL_3249725 | Berkshire and Surrey Pathology Services Lighthouse Laboratory | Wellcome Sanger Institute for the COVID-19 Genomics UK (COG-UK) Consortium | Berkshire and Surrey Pathology Services Lighthouse Laboratory and Alex Alderton, Roberto Amato, Jeffrey Barrett, Sonia Goncalves, Ewan Harrison, David K. Jackson, Ian Johnston, Dominic Kwiatkowski, Cordelia Langford, John Sillitoe on behalf of the Wellcome Sanger Institute COVID-19 Surveillance Team |
| EPI_ISL_3249726 | Lighthouse Lab in Glasgow | Wellcome Sanger Institute for the COVID-19 Genomics UK (COG-UK) Consortium | Harper VanSteenhouse, Yumi Kasai, David Gray, Carol Clugston, Anna Dominiczak and Alex Alderton, Roberto Amato, Jeffrey Barrett, Sonia Goncalves, Ewan Harrison, David K. Jackson, Ian Johnston, Dominic Kwiatkowski, Cordelia Langford, John Sillitoe on behalf of the Wellcome Sanger Institute COVID-19 Surveillance Team |
| EPI_ISL_3249727 | Berkshire and Surrey Pathology Services Lighthouse Laboratory | Wellcome Sanger Institute for the COVID-19 Genomics UK (COG-UK) Consortium | Berkshire and Surrey Pathology Services Lighthouse Laboratory and Alex Alderton, Roberto Amato, Jeffrey Barrett, Sonia Goncalves, Ewan Harrison, David K. Jackson, Ian Johnston, Dominic Kwiatkowski, Cordelia Langford, John Sillitoe on behalf of the Wellcome Sanger Institute COVID-19 Surveillance Team |
| EPI_ISL_3249728, EPI_ISL_3249729, EPI_ISL_3249730, EPI_ISL_3249731 | Lighthouse Lab in Milton Keynes | Wellcome Sanger Institute for the COVID-19 Genomics UK (COG-UK) Consortium | The Lighthouse Lab in Milton Keynes and Alex Alderton, Roberto Amato, Jeffrey Barrett, Sonia Goncalves, Ewan Harrison, David K. Jackson, Ian Johnston, Dominic Kwiatkowski, Cordelia Langford, John Sillitoe on behalf of the Wellcome Sanger Institute COVID-19 Surveillance Team |
| EPI_ISL_3249732 | Berkshire and Surrey Pathology Services Lighthouse Laboratory | Wellcome Sanger Institute for the COVID-19 Genomics UK (COG-UK) Consortium | Berkshire and Surrey Pathology Services Lighthouse Laboratory and Alex Alderton, Roberto Amato, Jeffrey Barrett, Sonia Goncalves, Ewan Harrison, David K. Jackson, Ian Johnston, Dominic Kwiatkowski, Cordelia Langford, John Sillitoe on behalf of the Wellcome Sanger Institute COVID-19 Surveillance Team |
| EPI_ISL_3249733 | Lighthouse Lab in Glasgow | Wellcome Sanger Institute for the COVID-19 Genomics UK (COG-UK) Consortium | Harper VanSteenhouse, Yumi Kasai, David Gray, Carol Clugston, Anna Dominiczak and Alex Alderton, Roberto Amato, Jeffrey Barrett, Sonia Goncalves, Ewan Harrison, David K. Jackson, Ian Johnston, Dominic Kwiatkowski, Cordelia Langford, John Sillitoe on behalf of the Wellcome Sanger Institute COVID-19 Surveillance Team |
| EPI_ISL_3249734 | Lighthouse Lab in Milton Keynes | Wellcome Sanger Institute for the COVID-19 Genomics UK (COG-UK) Consortium | The Lighthouse Lab in Milton Keynes and Alex Alderton, Roberto Amato, Jeffrey Barrett, Sonia Goncalves, Ewan Harrison, David K. Jackson, Ian Johnston, Dominic Kwiatkowski, Cordelia Langford, John Sillitoe on behalf of the Wellcome Sanger Institute COVID-19 Surveillance Team |
| EPI_ISL_3249735, EPI_ISL_3249736, EPI_ISL_3249737 | Berkshire and Surrey Pathology Services Lighthouse Laboratory | Wellcome Sanger Institute for the COVID-19 Genomics UK (COG-UK) Consortium | Berkshire and Surrey Pathology Services Lighthouse Laboratory and Alex Alderton, Roberto Amato, Jeffrey Barrett, Sonia Goncalves, Ewan Harrison, David K. Jackson, Ian Johnston, Dominic Kwiatkowski, Cordelia Langford, John Sillitoe on behalf of the Wellcome Sanger Institute COVID-19 Surveillance Team |
| EPI_ISL_3249738 | Lighthouse Lab in Glasgow | Wellcome Sanger Institute for the COVID-19 Genomics UK (COG-UK) Consortium | Harper VanSteenhouse, Yumi Kasai, David Gray, Carol Clugston, Anna Dominiczak and Alex Alderton, Roberto Amato, Jeffrey Barrett, Sonia Goncalves, Ewan Harrison, David K. Jackson, Ian Johnston, Dominic Kwiatkowski, Cordelia Langford, John Sillitoe on behalf of the Wellcome Sanger Institute COVID-19 Surveillance Team |
| EPI_ISL_3249739 | Berkshire and Surrey Pathology Services Lighthouse Laboratory | Wellcome Sanger Institute for the COVID-19 Genomics UK (COG-UK) Consortium | Berkshire and Surrey Pathology Services Lighthouse Laboratory and Alex Alderton, Roberto Amato, Jeffrey Barrett, Sonia Goncalves, Ewan Harrison, David K. Jackson, Ian Johnston, Dominic Kwiatkowski, Cordelia Langford, John Sillitoe on behalf of the Wellcome Sanger Institute COVID-19 Surveillance Team |
| EPI_ISL_3249740, EPI_ISL_3249741 | Lighthouse Lab in Glasgow | Wellcome Sanger Institute for the COVID-19 Genomics UK (COG-UK) Consortium | Harper VanSteenhouse, Yumi Kasai, David Gray, Carol Clugston, Anna Dominiczak and Alex Alderton, Roberto Amato, Jeffrey Barrett, Sonia Goncalves, Ewan Harrison, David K. Jackson, Ian Johnston, Dominic Kwiatkowski, Cordelia Langford, John Sillitoe on behalf of the Wellcome Sanger Institute COVID-19 Surveillance Team |
| EPI_ISL_3249742 | Berkshire and Surrey Pathology Services Lighthouse Laboratory | Wellcome Sanger Institute for the COVID-19 Genomics UK (COG-UK) Consortium | Berkshire and Surrey Pathology Services Lighthouse Laboratory and Alex Alderton, Roberto Amato, Jeffrey Barrett, Sonia Goncalves, Ewan Harrison, David K. Jackson, Ian Johnston, Dominic Kwiatkowski, Cordelia Langford, John Sillitoe on behalf of the Wellcome Sanger Institute COVID-19 Surveillance Team |
| EPI_ISL_3249743 | Lighthouse Lab in Milton Keynes | Wellcome Sanger Institute for the COVID-19 Genomics UK (COG-UK) Consortium | The Lighthouse Lab in Milton Keynes and Alex Alderton, Roberto Amato, Jeffrey Barrett, Sonia Goncalves, Ewan Harrison, David K. Jackson, Ian Johnston, Dominic Kwiatkowski, Cordelia Langford, John Sillitoe on behalf of the Wellcome Sanger Institute COVID-19 Surveillance Team |
| EPI_ISL_3249744 | Berkshire and Surrey Pathology Services Lighthouse Laboratory | Wellcome Sanger Institute for the COVID-19 Genomics UK (COG-UK) Consortium | Berkshire and Surrey Pathology Services Lighthouse Laboratory and Alex Alderton, Roberto Amato, Jeffrey Barrett, Sonia Goncalves, Ewan Harrison, David K. Jackson, Ian Johnston, Dominic Kwiatkowski, Cordelia Langford, John Sillitoe on behalf of the Wellcome Sanger Institute COVID-19 Surveillance Team |
| EPI_ISL_3249745 | Lighthouse Lab in Milton Keynes | Wellcome Sanger Institute for the COVID-19 Genomics UK (COG-UK) Consortium | The Lighthouse Lab in Milton Keynes and Alex Alderton, Roberto Amato, Jeffrey Barrett, Sonia Goncalves, Ewan Harrison, David K. Jackson, Ian Johnston, Dominic Kwiatkowski, Cordelia Langford, John Sillitoe on behalf of the Wellcome Sanger Institute COVID-19 Surveillance Team |
| EPI_ISL_3249746 | Berkshire and Surrey Pathology Services Lighthouse Laboratory | Wellcome Sanger Institute for the COVID-19 Genomics UK (COG-UK) Consortium | Berkshire and Surrey Pathology Services Lighthouse Laboratory and Alex Alderton, Roberto Amato, Jeffrey Barrett, Sonia Goncalves, Ewan Harrison, David K. Jackson, Ian Johnston, Dominic Kwiatkowski, Cordelia Langford, John Sillitoe on behalf of the Wellcome Sanger Institute COVID-19 Surveillance Team |
| EPI_ISL_3249747 | Lighthouse Lab in Milton Keynes | Wellcome Sanger Institute for the COVID-19 Genomics UK (COG-UK) Consortium | The Lighthouse Lab in Milton Keynes and Alex Alderton, Roberto Amato, Jeffrey Barrett, Sonia Goncalves, Ewan Harrison, David K. Jackson, Ian Johnston, Dominic Kwiatkowski, Cordelia Langford, John Sillitoe on behalf of the Wellcome Sanger Institute COVID-19 Surveillance Team |
| EPI_ISL_3249748 | Lighthouse Lab in Glasgow | Wellcome Sanger Institute for the COVID-19 Genomics UK (COG-UK) Consortium | Harper VanSteenhouse, Yumi Kasai, David Gray, Carol Clugston, Anna Dominiczak and Alex Alderton, Roberto Amato, Jeffrey Barrett, Sonia Goncalves, Ewan Harrison, David K. Jackson, Ian Johnston, Dominic Kwiatkowski, Cordelia Langford, John Sillitoe on behalf of the Wellcome Sanger Institute COVID-19 Surveillance Team |
| EPI_ISL_3249749, EPI_ISL_3249750, EPI_ISL_3249751 | Berkshire and Surrey Pathology Services Lighthouse Laboratory | Wellcome Sanger Institute for the COVID-19 Genomics UK (COG-UK) Consortium | Berkshire and Surrey Pathology Services Lighthouse Laboratory and Alex Alderton, Roberto Amato, Jeffrey Barrett, Sonia Goncalves, Ewan Harrison, David K. Jackson, Ian Johnston, Dominic Kwiatkowski, Cordelia Langford, John Sillitoe on behalf of the Wellcome Sanger Institute COVID-19 Surveillance Team |
| EPI_ISL_3249752 | Lighthouse Lab in Glasgow | Wellcome Sanger Institute for the COVID-19 Genomics UK (COG-UK) Consortium | Harper VanSteenhouse, Yumi Kasai, David Gray, Carol Clugston, Anna Dominiczak and Alex Alderton, Roberto Amato, Jeffrey Barrett, Sonia Goncalves, Ewan Harrison, David K. Jackson, Ian Johnston, Dominic Kwiatkowski, Cordelia Langford, John Sillitoe on behalf of the Wellcome Sanger Institute COVID-19 Surveillance Team |
| EPI_ISL_3249753 | Lighthouse Lab in Milton Keynes | Wellcome Sanger Institute for the COVID-19 Genomics UK (COG-UK) Consortium | The Lighthouse Lab in Milton Keynes and Alex Alderton, Roberto Amato, Jeffrey Barrett, Sonia Goncalves, Ewan Harrison, David K. Jackson, Ian Johnston, Dominic Kwiatkowski, Cordelia Langford, John Sillitoe on behalf of the Wellcome Sanger Institute COVID-19 Surveillance Team |
| EPI_ISL_3249754 | Berkshire and Surrey Pathology Services Lighthouse Laboratory | Wellcome Sanger Institute for the COVID-19 Genomics UK (COG-UK) Consortium | Berkshire and Surrey Pathology Services Lighthouse Laboratory and Alex Alderton, Roberto Amato, Jeffrey Barrett, Sonia Goncalves, Ewan Harrison, David K. Jackson, Ian Johnston, Dominic Kwiatkowski, Cordelia Langford, John Sillitoe on behalf of the Wellcome Sanger Institute COVID-19 Surveillance Team |
| EPI_ISL_3249755 | Lighthouse Lab in Milton Keynes | Wellcome Sanger Institute for the COVID-19 Genomics UK (COG-UK) Consortium | The Lighthouse Lab in Milton Keynes and Alex Alderton, Roberto Amato, Jeffrey Barrett, Sonia Goncalves, Ewan Harrison, David K. Jackson, Ian Johnston, Dominic Kwiatkowski, Cordelia Langford, John Sillitoe on behalf of the Wellcome Sanger Institute COVID-19 Surveillance Team |
| EPI_ISL_3249756, EPI_ISL_3249757, EPI_ISL_3249758, EPI_ISL_3249759, EPI_ISL_3249760, EPI_ISL_3249761, EPI_ISL_3249762 | Berkshire and Surrey Pathology Services Lighthouse Laboratory | Wellcome Sanger Institute for the COVID-19 Genomics UK (COG-UK) Consortium | Berkshire and Surrey Pathology Services Lighthouse Laboratory and Alex Alderton, Roberto Amato, Jeffrey Barrett, Sonia Goncalves, Ewan Harrison, David K. Jackson, Ian Johnston, Dominic Kwiatkowski, Cordelia Langford, John Sillitoe on behalf of the Wellcome Sanger Institute COVID-19 Surveillance Team |

| | | | |
|---|---|---|---|
| EPI_ISL_3249763 | Lighthouse Lab in Glasgow | Wellcome Sanger Institute for the COVID-19 Genomics UK (COG-UK) Consortium | Harper VanSteenhouse, Yumi Kasai, David Gray, Carol Clugston, Anna Dominiczak and Alex Alderton, Roberto Amato, Jeffrey Barrett, Sonia Goncalves, Ewan Harrison, David K. Jackson, Ian Johnston, Dominic Kwiatkowski, Cordelia Langford, John Sillitoe on behalf of the Wellcome Sanger Institute COVID-19 Surveillance Team |
| EPI_ISL_3249764 | Berkshire and Surrey Pathology Services Lighthouse Laboratory | Wellcome Sanger Institute for the COVID-19 Genomics UK (COG-UK) Consortium | Berkshire and Surrey Pathology Services Lighthouse Laboratory and Alex Alderton, Roberto Amato, Jeffrey Barrett, Sonia Goncalves, Ewan Harrison, David K. Jackson, Ian Johnston, Dominic Kwiatkowski, Cordelia Langford, John Sillitoe on behalf of the Wellcome Sanger Institute COVID-19 Surveillance Team |
| EPI_ISL_3249765 | Lighthouse Lab in Glasgow | Wellcome Sanger Institute for the COVID-19 Genomics UK (COG-UK) Consortium | Harper VanSteenhouse, Yumi Kasai, David Gray, Carol Clugston, Anna Dominiczak and Alex Alderton, Roberto Amato, Jeffrey Barrett, Sonia Goncalves, Ewan Harrison, David K. Jackson, Ian Johnston, Dominic Kwiatkowski, Cordelia Langford, John Sillitoe on behalf of the Wellcome Sanger Institute COVID-19 Surveillance Team |
| EPI_ISL_3249766 | Berkshire and Surrey Pathology Services Lighthouse Laboratory | Wellcome Sanger Institute for the COVID-19 Genomics UK (COG-UK) Consortium | Berkshire and Surrey Pathology Services Lighthouse Laboratory and Alex Alderton, Roberto Amato, Jeffrey Barrett, Sonia Goncalves, Ewan Harrison, David K. Jackson, Ian Johnston, Dominic Kwiatkowski, Cordelia Langford, John Sillitoe on behalf of the Wellcome Sanger Institute COVID-19 Surveillance Team |
| EPI_ISL_3249767 | Lighthouse Lab in Glasgow | Wellcome Sanger Institute for the COVID-19 Genomics UK (COG-UK) Consortium | Harper VanSteenhouse, Yumi Kasai, David Gray, Carol Clugston, Anna Dominiczak and Alex Alderton, Roberto Amato, Jeffrey Barrett, Sonia Goncalves, Ewan Harrison, David K. Jackson, Ian Johnston, Dominic Kwiatkowski, Cordelia Langford, John Sillitoe on behalf of the Wellcome Sanger Institute COVID-19 Surveillance Team |
| EPI_ISL_3249768, EPI_ISL_3249769 | Lighthouse Lab in Milton Keynes | Wellcome Sanger Institute for the COVID-19 Genomics UK (COG-UK) Consortium | The Lighthouse Lab in Milton Keynes and Alex Alderton, Roberto Amato, Jeffrey Barrett, Sonia Goncalves, Ewan Harrison, David K. Jackson, Ian Johnston, Dominic Kwiatkowski, Cordelia Langford, John Sillitoe on behalf of the Wellcome Sanger Institute COVID-19 Surveillance Team |
| EPI_ISL_3249770, EPI_ISL_3249771, EPI_ISL_3249772 | Berkshire and Surrey Pathology Services Lighthouse Laboratory | Wellcome Sanger Institute for the COVID-19 Genomics UK (COG-UK) Consortium | Berkshire and Surrey Pathology Services Lighthouse Laboratory and Alex Alderton, Roberto Amato, Jeffrey Barrett, Sonia Goncalves, Ewan Harrison, David K. Jackson, Ian Johnston, Dominic Kwiatkowski, Cordelia Langford, John Sillitoe on behalf of the Wellcome Sanger Institute COVID-19 Surveillance Team |
| EPI_ISL_3249773 | Lighthouse Lab in Milton Keynes | Wellcome Sanger Institute for the COVID-19 Genomics UK (COG-UK) Consortium | The Lighthouse Lab in Milton Keynes and Alex Alderton, Roberto Amato, Jeffrey Barrett, Sonia Goncalves, Ewan Harrison, David K. Jackson, Ian Johnston, Dominic Kwiatkowski, Cordelia Langford, John Sillitoe on behalf of the Wellcome Sanger Institute COVID-19 Surveillance Team |
| EPI_ISL_3249774, EPI_ISL_3249775 | Berkshire and Surrey Pathology Services Lighthouse Laboratory | Wellcome Sanger Institute for the COVID-19 Genomics UK (COG-UK) Consortium | Berkshire and Surrey Pathology Services Lighthouse Laboratory and Alex Alderton, Roberto Amato, Jeffrey Barrett, Sonia Goncalves, Ewan Harrison, David K. Jackson, Ian Johnston, Dominic Kwiatkowski, Cordelia Langford, John Sillitoe on behalf of the Wellcome Sanger Institute COVID-19 Surveillance Team |
| EPI_ISL_3249776 | Lighthouse Lab in Milton Keynes | Wellcome Sanger Institute for the COVID-19 Genomics UK (COG-UK) Consortium | The Lighthouse Lab in Milton Keynes and Alex Alderton, Roberto Amato, Jeffrey Barrett, Sonia Goncalves, Ewan Harrison, David K. Jackson, Ian Johnston, Dominic Kwiatkowski, Cordelia Langford, John Sillitoe on behalf of the Wellcome Sanger Institute COVID-19 Surveillance Team |
| EPI_ISL_3249777, EPI_ISL_3249778, EPI_ISL_3249779 | Berkshire and Surrey Pathology Services Lighthouse Laboratory | Wellcome Sanger Institute for the COVID-19 Genomics UK (COG-UK) Consortium | Berkshire and Surrey Pathology Services Lighthouse Laboratory and Alex Alderton, Roberto Amato, Jeffrey Barrett, Sonia Goncalves, Ewan Harrison, David K. Jackson, Ian Johnston, Dominic Kwiatkowski, Cordelia Langford, John Sillitoe on behalf of the Wellcome Sanger Institute COVID-19 Surveillance Team |
| EPI_ISL_3249780, EPI_ISL_3249781, EPI_ISL_3249782 | Lighthouse Lab in Milton Keynes | Wellcome Sanger Institute for the COVID-19 Genomics UK (COG-UK) Consortium | The Lighthouse Lab in Milton Keynes and Alex Alderton, Roberto Amato, Jeffrey Barrett, Sonia Goncalves, Ewan Harrison, David K. Jackson, Ian Johnston, Dominic Kwiatkowski, Cordelia Langford, John Sillitoe on behalf of the Wellcome Sanger Institute COVID-19 Surveillance Team |
| EPI_ISL_3249783 | Lighthouse Lab in Glasgow | Wellcome Sanger Institute for the COVID-19 Genomics UK (COG-UK) Consortium | Harper VanSteenhouse, Yumi Kasai, David Gray, Carol Clugston, Anna Dominiczak and Alex Alderton, Roberto Amato, Jeffrey Barrett, Sonia Goncalves, Ewan Harrison, David K. Jackson, Ian Johnston, Dominic Kwiatkowski, Cordelia Langford, John Sillitoe on behalf of the Wellcome Sanger Institute COVID-19 Surveillance Team |
| EPI_ISL_3249784, EPI_ISL_3249785 | Berkshire and Surrey Pathology Services Lighthouse Laboratory | Wellcome Sanger Institute for the COVID-19 Genomics UK (COG-UK) Consortium | Berkshire and Surrey Pathology Services Lighthouse Laboratory and Alex Alderton, Roberto Amato, Jeffrey Barrett, Sonia Goncalves, Ewan Harrison, David K. Jackson, Ian Johnston, Dominic Kwiatkowski, Cordelia Langford, John Sillitoe on behalf of the Wellcome Sanger Institute COVID-19 Surveillance Team |
| EPI_ISL_3249786 | Lighthouse Lab in Glasgow | Wellcome Sanger Institute for the COVID-19 Genomics UK (COG-UK) Consortium | Harper VanSteenhouse, Yumi Kasai, David Gray, Carol Clugston, Anna Dominiczak and Alex Alderton, Roberto Amato, Jeffrey Barrett, Sonia Goncalves, Ewan Harrison, David K. Jackson, Ian Johnston, Dominic Kwiatkowski, Cordelia Langford, John Sillitoe on behalf of the Wellcome Sanger Institute COVID-19 Surveillance Team |
| EPI_ISL_3249787 | Berkshire and Surrey Pathology Services Lighthouse Laboratory | Wellcome Sanger Institute for the COVID-19 Genomics UK (COG-UK) Consortium | Berkshire and Surrey Pathology Services Lighthouse Laboratory and Alex Alderton, Roberto Amato, Jeffrey Barrett, Sonia Goncalves, Ewan Harrison, David K. Jackson, Ian Johnston, Dominic Kwiatkowski, Cordelia Langford, John Sillitoe on behalf of the Wellcome Sanger Institute COVID-19 Surveillance Team |
| EPI_ISL_3249788, EPI_ISL_3249789, EPI_ISL_3249790, EPI_ISL_3249791 | Lighthouse Lab in Milton Keynes | Wellcome Sanger Institute for the COVID-19 Genomics UK (COG-UK) Consortium | The Lighthouse Lab in Milton Keynes and Alex Alderton, Roberto Amato, Jeffrey Barrett, Sonia Goncalves, Ewan Harrison, David K. Jackson, Ian Johnston, Dominic Kwiatkowski, Cordelia Langford, John Sillitoe on behalf of the Wellcome Sanger Institute COVID-19 Surveillance Team |
| EPI_ISL_3249792 | Lighthouse Lab in Glasgow | Wellcome Sanger Institute for the COVID-19 Genomics UK (COG-UK) Consortium | Harper VanSteenhouse, Yumi Kasai, David Gray, Carol Clugston, Anna Dominiczak and Alex Alderton, Roberto Amato, Jeffrey Barrett, Sonia Goncalves, Ewan Harrison, David K. Jackson, Ian Johnston, Dominic Kwiatkowski, Cordelia Langford, John Sillitoe on behalf of the Wellcome Sanger Institute COVID-19 Surveillance Team |
| EPI_ISL_3249793 | Berkshire and Surrey Pathology Services Lighthouse Laboratory | Wellcome Sanger Institute for the COVID-19 Genomics UK (COG-UK) Consortium | Berkshire and Surrey Pathology Services Lighthouse Laboratory and Alex Alderton, Roberto Amato, Jeffrey Barrett, Sonia Goncalves, Ewan Harrison, David K. Jackson, Ian Johnston, Dominic Kwiatkowski, Cordelia Langford, John Sillitoe on behalf of the Wellcome Sanger Institute COVID-19 Surveillance Team |
| EPI_ISL_3249794 | Lighthouse Lab in Milton Keynes | Wellcome Sanger Institute for the COVID-19 Genomics UK (COG-UK) Consortium | The Lighthouse Lab in Milton Keynes and Alex Alderton, Roberto Amato, Jeffrey Barrett, Sonia Goncalves, Ewan Harrison, David K. Jackson, Ian Johnston, Dominic Kwiatkowski, Cordelia Langford, John Sillitoe on behalf of the Wellcome Sanger Institute COVID-19 Surveillance Team |
| EPI_ISL_3249795 | Lighthouse Lab in Glasgow | Wellcome Sanger Institute for the COVID-19 Genomics UK (COG-UK) Consortium | Harper VanSteenhouse, Yumi Kasai, David Gray, Carol Clugston, Anna Dominiczak and Alex Alderton, Roberto Amato, Jeffrey Barrett, Sonia Goncalves, Ewan Harrison, David K. Jackson, Ian Johnston, Dominic Kwiatkowski, Cordelia Langford, John Sillitoe on behalf of the Wellcome Sanger Institute COVID-19 Surveillance Team |
| EPI_ISL_3249796, EPI_ISL_3249797, EPI_ISL_3249798, EPI_ISL_3249799, EPI_ISL_3249800, EPI_ISL_3249801, EPI_ISL_3249802, EPI_ISL_3249803 | Berkshire and Surrey Pathology Services Lighthouse Laboratory | Wellcome Sanger Institute for the COVID-19 Genomics UK (COG-UK) Consortium | Berkshire and Surrey Pathology Services Lighthouse Laboratory and Alex Alderton, Roberto Amato, Jeffrey Barrett, Sonia Goncalves, Ewan Harrison, David K. Jackson, Ian Johnston, Dominic Kwiatkowski, Cordelia Langford, John Sillitoe on behalf of the Wellcome Sanger Institute COVID-19 Surveillance Team |
| EPI_ISL_3249804 | Lighthouse Lab in Milton Keynes | Wellcome Sanger Institute for the COVID-19 Genomics UK (COG-UK) Consortium | The Lighthouse Lab in Milton Keynes and Alex Alderton, Roberto Amato, Jeffrey Barrett, Sonia Goncalves, Ewan Harrison, David K. Jackson, Ian Johnston, Dominic Kwiatkowski, Cordelia Langford, John Sillitoe on behalf of the Wellcome Sanger Institute COVID-19 Surveillance Team |
| EPI_ISL_3249805 | Berkshire and Surrey Pathology Services Lighthouse Laboratory | Wellcome Sanger Institute for the COVID-19 Genomics UK (COG-UK) Consortium | Berkshire and Surrey Pathology Services Lighthouse Laboratory and Alex Alderton, Roberto Amato, Jeffrey Barrett, Sonia Goncalves, Ewan Harrison, David K. Jackson, Ian Johnston, Dominic Kwiatkowski, Cordelia Langford, John Sillitoe on behalf of the Wellcome Sanger Institute COVID-19 Surveillance Team |
| EPI_ISL_3249806 | Lighthouse Lab in Milton Keynes | Wellcome Sanger Institute for the COVID-19 Genomics UK (COG-UK) Consortium | The Lighthouse Lab in Milton Keynes and Alex Alderton, Roberto Amato, Jeffrey Barrett, Sonia Goncalves, Ewan Harrison, David K. Jackson, Ian Johnston, Dominic Kwiatkowski, Cordelia Langford, John Sillitoe on behalf of the Wellcome Sanger Institute COVID-19 Surveillance Team |
| EPI_ISL_3249807 | Berkshire and Surrey Pathology Services Lighthouse Laboratory | Wellcome Sanger Institute for the COVID-19 Genomics UK (COG-UK) Consortium | Berkshire and Surrey Pathology Services Lighthouse Laboratory and Alex Alderton, Roberto Amato, Jeffrey Barrett, Sonia Goncalves, Ewan Harrison, David K. Jackson, Ian Johnston, Dominic Kwiatkowski, Cordelia Langford, John Sillitoe on behalf of the Wellcome Sanger Institute COVID-19 Surveillance Team |
| EPI_ISL_3249808, EPI_ISL_3249809 | Lighthouse Lab in Milton Keynes | Wellcome Sanger Institute for the COVID-19 Genomics UK (COG-UK) Consortium | The Lighthouse Lab in Milton Keynes and Alex Alderton, Roberto Amato, Jeffrey Barrett, Sonia Goncalves, Ewan Harrison, David K. Jackson, Ian Johnston, Dominic Kwiatkowski, Cordelia Langford, John Sillitoe on behalf of the Wellcome Sanger Institute COVID-19 Surveillance Team |
| EPI_ISL_3249810, EPI_ISL_3249811, EPI_ISL_3249812, EPI_ISL_3249813, EPI_ISL_3249814, EPI_ISL_3249815 | Berkshire and Surrey Pathology Services Lighthouse Laboratory | Wellcome Sanger Institute for the COVID-19 Genomics UK (COG-UK) Consortium | Berkshire and Surrey Pathology Services Lighthouse Laboratory and Alex Alderton, Roberto Amato, Jeffrey Barrett, Sonia Goncalves, Ewan Harrison, David K. Jackson, Ian Johnston, Dominic Kwiatkowski, Cordelia Langford, John Sillitoe on behalf of the Wellcome Sanger Institute COVID-19 Surveillance Team |
| EPI_ISL_3249816, EPI_ISL_3249817 | Lighthouse Lab in Milton Keynes | Wellcome Sanger Institute for the COVID-19 Genomics UK (COG-UK) Consortium | The Lighthouse Lab in Milton Keynes and Alex Alderton, Roberto Amato, Jeffrey Barrett, Sonia Goncalves, Ewan Harrison, David K. Jackson, Ian Johnston, Dominic Kwiatkowski, Cordelia Langford, John Sillitoe on behalf of the Wellcome Sanger Institute COVID-19 Surveillance Team |
| EPI_ISL_3249818 | Lighthouse Lab in Glasgow | Wellcome Sanger Institute for the COVID-19 Genomics UK (COG-UK) Consortium | Harper VanSteenhouse, Yumi Kasai, David Gray, Carol Clugston, Anna Dominiczak and Alex Alderton, Roberto Amato, Jeffrey Barrett, Sonia Goncalves, Ewan Harrison, David K. Jackson, Ian Johnston, Dominic Kwiatkowski, Cordelia Langford, John Sillitoe on behalf of the Wellcome Sanger Institute COVID-19 Surveillance Team |
| EPI_ISL_3249819, EPI_ISL_3249820 | Lighthouse Lab in Milton Keynes | Wellcome Sanger Institute for the COVID-19 Genomics UK (COG-UK) Consortium | The Lighthouse Lab in Milton Keynes and Alex Alderton, Roberto Amato, Jeffrey Barrett, Sonia Goncalves, Ewan Harrison, David K. Jackson, Ian Johnston, Dominic Kwiatkowski, Cordelia Langford, John Sillitoe on behalf of the Wellcome Sanger Institute COVID-19 Surveillance Team |
| EPI_ISL_3249821, EPI_ISL_3249822, EPI_ISL_3249823 | Berkshire and Surrey Pathology Services Lighthouse Laboratory | Wellcome Sanger Institute for the COVID-19 Genomics UK (COG-UK) Consortium | Berkshire and Surrey Pathology Services Lighthouse Laboratory and Alex Alderton, Roberto Amato, Jeffrey Barrett, Sonia Goncalves, Ewan Harrison, David K. Jackson, Ian Johnston, Dominic Kwiatkowski, Cordelia Langford, John Sillitoe on behalf of the Wellcome Sanger Institute COVID-19 Surveillance Team |
| EPI_ISL_3249824 | Lighthouse Lab in Glasgow | Wellcome Sanger Institute for the COVID-19 Genomics UK (COG-UK) Consortium | Harper VanSteenhouse, Yumi Kasai, David Gray, Carol Clugston, Anna Dominiczak and Alex Alderton, Roberto Amato, Jeffrey Barrett, Sonia Goncalves, Ewan Harrison, David K. Jackson, Ian Johnston, Dominic Kwiatkowski, Cordelia Langford, John Sillitoe on behalf of the Wellcome Sanger Institute COVID-19 Surveillance Team |
| EPI_ISL_3249825 | Berkshire and Surrey Pathology Services Lighthouse Laboratory | Wellcome Sanger Institute for the COVID-19 Genomics UK (COG-UK) Consortium | Berkshire and Surrey Pathology Services Lighthouse Laboratory and Alex Alderton, Roberto Amato, Jeffrey Barrett, Sonia Goncalves, Ewan Harrison, David K. Jackson, Ian Johnston, Dominic Kwiatkowski, Cordelia Langford, John Sillitoe on behalf of the Wellcome Sanger Institute COVID-19 Surveillance Team |
| EPI_ISL_3249826 | Lighthouse Lab in Glasgow | Wellcome Sanger Institute for the COVID-19 Genomics UK (COG-UK) Consortium | Harper VanSteenhouse, Yumi Kasai, David Gray, Carol Clugston, Anna Dominiczak and Alex Alderton, Roberto Amato, Jeffrey Barrett, Sonia Goncalves, Ewan Harrison, David K. Jackson, Ian Johnston, Dominic Kwiatkowski, Cordelia Langford, John Sillitoe on behalf of the Wellcome Sanger Institute COVID-19 Surveillance Team |
| EPI_ISL_3249827, EPI_ISL_3249828, EPI_ISL_3249829 | Lighthouse Lab in Milton Keynes | Wellcome Sanger Institute for the COVID-19 Genomics UK (COG-UK) Consortium | The Lighthouse Lab in Milton Keynes and Alex Alderton, Roberto Amato, Jeffrey Barrett, Sonia Goncalves, Ewan Harrison, David K. Jackson, Ian Johnston, Dominic Kwiatkowski, Cordelia Langford, John Sillitoe on behalf of the Wellcome Sanger Institute COVID-19 Surveillance Team |
| EPI_ISL_3249830 | Berkshire and Surrey Pathology Services Lighthouse Laboratory | Wellcome Sanger Institute for the COVID-19 Genomics UK (COG-UK) Consortium | Berkshire and Surrey Pathology Services Lighthouse Laboratory and Alex Alderton, Roberto Amato, Jeffrey Barrett, Sonia Goncalves, Ewan Harrison, David K. Jackson, Ian Johnston, Dominic Kwiatkowski, Cordelia Langford, John Sillitoe on behalf of the Wellcome Sanger Institute COVID-19 Surveillance Team |
| EPI_ISL_3249831, EPI_ISL_3249832 | Lighthouse Lab in Milton Keynes | Wellcome Sanger Institute for the COVID-19 Genomics UK (COG-UK) Consortium | The Lighthouse Lab in Milton Keynes and Alex Alderton, Roberto Amato, Jeffrey Barrett, Sonia Goncalves, Ewan Harrison, David K. Jackson, Ian Johnston, Dominic Kwiatkowski, Cordelia Langford, John Sillitoe on behalf of the Wellcome Sanger Institute COVID-19 Surveillance Team |

| | | | |
|---|---|---|---|
| EPI_ISL_3249833, EPI_ISL_3249834 | Berkshire and Surrey Pathology Services Lighthouse Laboratory | Wellcome Sanger Institute for the COVID-19 Genomics UK (COG-UK) Consortium | Berkshire and Surrey Pathology Services Lighthouse Laboratory and Alex Alderton, Roberto Amato, Jeffrey Barrett, Sonia Goncalves, Ewan Harrison, David K. Jackson, Ian Johnston, Dominic Kwiatkowski, Cordelia Langford, John Sillitoe on behalf of the Wellcome Sanger Institute COVID-19 Surveillance Team |
| EPI_ISL_3249835, EPI_ISL_3249836, EPI_ISL_3249837 | Lighthouse Lab in Milton Keynes | Wellcome Sanger Institute for the COVID-19 Genomics UK (COG-UK) Consortium | The Lighthouse Lab in Milton Keynes and Alex Alderton, Roberto Amato, Jeffrey Barrett, Sonia Goncalves, Ewan Harrison, David K. Jackson, Ian Johnston, Dominic Kwiatkowski, Cordelia Langford, John Sillitoe on behalf of the Wellcome Sanger Institute COVID-19 Surveillance Team |
| EPI_ISL_3249838 | Lighthouse Lab in Glasgow | Wellcome Sanger Institute for the COVID-19 Genomics UK (COG-UK) Consortium | Harper VanSteenhouse, Yumi Kasai, David Gray, Carol Clugston, Anna Dominiczak and Alex Alderton, Roberto Amato, Jeffrey Barrett, Sonia Goncalves, Ewan Harrison, David K. Jackson, Ian Johnston, Dominic Kwiatkowski, Cordelia Langford, John Sillitoe on behalf of the Wellcome Sanger Institute COVID-19 Surveillance Team |
| EPI_ISL_3249839 | Lighthouse Lab in Milton Keynes | Wellcome Sanger Institute for the COVID-19 Genomics UK (COG-UK) Consortium | The Lighthouse Lab in Milton Keynes and Alex Alderton, Roberto Amato, Jeffrey Barrett, Sonia Goncalves, Ewan Harrison, David K. Jackson, Ian Johnston, Dominic Kwiatkowski, Cordelia Langford, John Sillitoe on behalf of the Wellcome Sanger Institute COVID-19 Surveillance Team |
| EPI_ISL_3249840, EPI_ISL_3249841, EPI_ISL_3249842, EPI_ISL_3249843, EPI_ISL_3249844, EPI_ISL_3249845, EPI_ISL_3249846, EPI_ISL_3249847 | Berkshire and Surrey Pathology Services Lighthouse Laboratory | Wellcome Sanger Institute for the COVID-19 Genomics UK (COG-UK) Consortium | Berkshire and Surrey Pathology Services Lighthouse Laboratory and Alex Alderton, Roberto Amato, Jeffrey Barrett, Sonia Goncalves, Ewan Harrison, David K. Jackson, Ian Johnston, Dominic Kwiatkowski, Cordelia Langford, John Sillitoe on behalf of the Wellcome Sanger Institute COVID-19 Surveillance Team |
| EPI_ISL_3249848 | Lighthouse Lab in Milton Keynes | Wellcome Sanger Institute for the COVID-19 Genomics UK (COG-UK) Consortium | The Lighthouse Lab in Milton Keynes and Alex Alderton, Roberto Amato, Jeffrey Barrett, Sonia Goncalves, Ewan Harrison, David K. Jackson, Ian Johnston, Dominic Kwiatkowski, Cordelia Langford, John Sillitoe on behalf of the Wellcome Sanger Institute COVID-19 Surveillance Team |
| EPI_ISL_3249849 | Berkshire and Surrey Pathology Services Lighthouse Laboratory | Wellcome Sanger Institute for the COVID-19 Genomics UK (COG-UK) Consortium | Berkshire and Surrey Pathology Services Lighthouse Laboratory and Alex Alderton, Roberto Amato, Jeffrey Barrett, Sonia Goncalves, Ewan Harrison, David K. Jackson, Ian Johnston, Dominic Kwiatkowski, Cordelia Langford, John Sillitoe on behalf of the Wellcome Sanger Institute COVID-19 Surveillance Team |
| EPI_ISL_3249850 | Lighthouse Lab in Milton Keynes | Wellcome Sanger Institute for the COVID-19 Genomics UK (COG-UK) Consortium | The Lighthouse Lab in Milton Keynes and Alex Alderton, Roberto Amato, Jeffrey Barrett, Sonia Goncalves, Ewan Harrison, David K. Jackson, Ian Johnston, Dominic Kwiatkowski, Cordelia Langford, John Sillitoe on behalf of the Wellcome Sanger Institute COVID-19 Surveillance Team |
| EPI_ISL_3249851 | Berkshire and Surrey Pathology Services Lighthouse Laboratory | Wellcome Sanger Institute for the COVID-19 Genomics UK (COG-UK) Consortium | Berkshire and Surrey Pathology Services Lighthouse Laboratory and Alex Alderton, Roberto Amato, Jeffrey Barrett, Sonia Goncalves, Ewan Harrison, David K. Jackson, Ian Johnston, Dominic Kwiatkowski, Cordelia Langford, John Sillitoe on behalf of the Wellcome Sanger Institute COVID-19 Surveillance Team |
| EPI_ISL_3249852 | Lighthouse Lab in Milton Keynes | Wellcome Sanger Institute for the COVID-19 Genomics UK (COG-UK) Consortium | The Lighthouse Lab in Milton Keynes and Alex Alderton, Roberto Amato, Jeffrey Barrett, Sonia Goncalves, Ewan Harrison, David K. Jackson, Ian Johnston, Dominic Kwiatkowski, Cordelia Langford, John Sillitoe on behalf of the Wellcome Sanger Institute COVID-19 Surveillance Team |
| EPI_ISL_3249853, EPI_ISL_3249854, EPI_ISL_3249855, EPI_ISL_3249856, EPI_ISL_3249857, EPI_ISL_3249858, EPI_ISL_3249859 | Berkshire and Surrey Pathology Services Lighthouse Laboratory | Wellcome Sanger Institute for the COVID-19 Genomics UK (COG-UK) Consortium | Berkshire and Surrey Pathology Services Lighthouse Laboratory and Alex Alderton, Roberto Amato, Jeffrey Barrett, Sonia Goncalves, Ewan Harrison, David K. Jackson, Ian Johnston, Dominic Kwiatkowski, Cordelia Langford, John Sillitoe on behalf of the Wellcome Sanger Institute COVID-19 Surveillance Team |
| EPI_ISL_3249860 | Lighthouse Lab in Milton Keynes | Wellcome Sanger Institute for the COVID-19 Genomics UK (COG-UK) Consortium | The Lighthouse Lab in Milton Keynes and Alex Alderton, Roberto Amato, Jeffrey Barrett, Sonia Goncalves, Ewan Harrison, David K. Jackson, Ian Johnston, Dominic Kwiatkowski, Cordelia Langford, John Sillitoe on behalf of the Wellcome Sanger Institute COVID-19 Surveillance Team |
| EPI_ISL_3249861, EPI_ISL_3249862 | Berkshire and Surrey Pathology Services Lighthouse Laboratory | Wellcome Sanger Institute for the COVID-19 Genomics UK (COG-UK) Consortium | Berkshire and Surrey Pathology Services Lighthouse Laboratory and Alex Alderton, Roberto Amato, Jeffrey Barrett, Sonia Goncalves, Ewan Harrison, David K. Jackson, Ian Johnston, Dominic Kwiatkowski, Cordelia Langford, John Sillitoe on behalf of the Wellcome Sanger Institute COVID-19 Surveillance Team |
| EPI_ISL_3249863 | Lighthouse Lab in Milton Keynes | Wellcome Sanger Institute for the COVID-19 Genomics UK (COG-UK) Consortium | The Lighthouse Lab in Milton Keynes and Alex Alderton, Roberto Amato, Jeffrey Barrett, Sonia Goncalves, Ewan Harrison, David K. Jackson, Ian Johnston, Dominic Kwiatkowski, Cordelia Langford, John Sillitoe on behalf of the Wellcome Sanger Institute COVID-19 Surveillance Team |
| EPI_ISL_3249864 | Lighthouse Lab in Glasgow | Wellcome Sanger Institute for the COVID-19 Genomics UK (COG-UK) Consortium | Harper VanSteenhouse, Yumi Kasai, David Gray, Carol Clugston, Anna Dominiczak and Alex Alderton, Roberto Amato, Jeffrey Barrett, Sonia Goncalves, Ewan Harrison, David K. Jackson, Ian Johnston, Dominic Kwiatkowski, Cordelia Langford, John Sillitoe on behalf of the Wellcome Sanger Institute COVID-19 Surveillance Team |
| EPI_ISL_3249865 | Berkshire and Surrey Pathology Services Lighthouse Laboratory | Wellcome Sanger Institute for the COVID-19 Genomics UK (COG-UK) Consortium | Berkshire and Surrey Pathology Services Lighthouse Laboratory and Alex Alderton, Roberto Amato, Jeffrey Barrett, Sonia Goncalves, Ewan Harrison, David K. Jackson, Ian Johnston, Dominic Kwiatkowski, Cordelia Langford, John Sillitoe on behalf of the Wellcome Sanger Institute COVID-19 Surveillance Team |
| EPI_ISL_3249866, EPI_ISL_3249867 | Lighthouse Lab in Milton Keynes | Wellcome Sanger Institute for the COVID-19 Genomics UK (COG-UK) Consortium | The Lighthouse Lab in Milton Keynes and Alex Alderton, Roberto Amato, Jeffrey Barrett, Sonia Goncalves, Ewan Harrison, David K. Jackson, Ian Johnston, Dominic Kwiatkowski, Cordelia Langford, John Sillitoe on behalf of the Wellcome Sanger Institute COVID-19 Surveillance Team |
| EPI_ISL_3249868 | Lighthouse Lab in Glasgow | Wellcome Sanger Institute for the COVID-19 Genomics UK (COG-UK) Consortium | Harper VanSteenhouse, Yumi Kasai, David Gray, Carol Clugston, Anna Dominiczak and Alex Alderton, Roberto Amato, Jeffrey Barrett, Sonia Goncalves, Ewan Harrison, David K. Jackson, Ian Johnston, Dominic Kwiatkowski, Cordelia Langford, John Sillitoe on behalf of the Wellcome Sanger Institute COVID-19 Surveillance Team |
| EPI_ISL_3249869 | Lighthouse Lab in Milton Keynes | Wellcome Sanger Institute for the COVID-19 Genomics UK (COG-UK) Consortium | The Lighthouse Lab in Milton Keynes and Alex Alderton, Roberto Amato, Jeffrey Barrett, Sonia Goncalves, Ewan Harrison, David K. Jackson, Ian Johnston, Dominic Kwiatkowski, Cordelia Langford, John Sillitoe on behalf of the Wellcome Sanger Institute COVID-19 Surveillance Team |
| EPI_ISL_3249870, EPI_ISL_3249871, EPI_ISL_3249872 | Berkshire and Surrey Pathology Services Lighthouse Laboratory | Wellcome Sanger Institute for the COVID-19 Genomics UK (COG-UK) Consortium | Berkshire and Surrey Pathology Services Lighthouse Laboratory and Alex Alderton, Roberto Amato, Jeffrey Barrett, Sonia Goncalves, Ewan Harrison, David K. Jackson, Ian Johnston, Dominic Kwiatkowski, Cordelia Langford, John Sillitoe on behalf of the Wellcome Sanger Institute COVID-19 Surveillance Team |
| EPI_ISL_3249873, EPI_ISL_3249874 | Lighthouse Lab in Milton Keynes | Wellcome Sanger Institute for the COVID-19 Genomics UK (COG-UK) Consortium | The Lighthouse Lab in Milton Keynes and Alex Alderton, Roberto Amato, Jeffrey Barrett, Sonia Goncalves, Ewan Harrison, David K. Jackson, Ian Johnston, Dominic Kwiatkowski, Cordelia Langford, John Sillitoe on behalf of the Wellcome Sanger Institute COVID-19 Surveillance Team |
| EPI_ISL_3249875, EPI_ISL_3249876 | Berkshire and Surrey Pathology Services Lighthouse Laboratory | Wellcome Sanger Institute for the COVID-19 Genomics UK (COG-UK) Consortium | Berkshire and Surrey Pathology Services Lighthouse Laboratory and Alex Alderton, Roberto Amato, Jeffrey Barrett, Sonia Goncalves, Ewan Harrison, David K. Jackson, Ian Johnston, Dominic Kwiatkowski, Cordelia Langford, John Sillitoe on behalf of the Wellcome Sanger Institute COVID-19 Surveillance Team |
| EPI_ISL_3249877 | Lighthouse Lab in Milton Keynes | Wellcome Sanger Institute for the COVID-19 Genomics UK (COG-UK) Consortium | The Lighthouse Lab in Milton Keynes and Alex Alderton, Roberto Amato, Jeffrey Barrett, Sonia Goncalves, Ewan Harrison, David K. Jackson, Ian Johnston, Dominic Kwiatkowski, Cordelia Langford, John Sillitoe on behalf of the Wellcome Sanger Institute COVID-19 Surveillance Team |
| EPI_ISL_3249878, EPI_ISL_3249879 | Lighthouse Lab in Glasgow | Wellcome Sanger Institute for the COVID-19 Genomics UK (COG-UK) Consortium | Harper VanSteenhouse, Yumi Kasai, David Gray, Carol Clugston, Anna Dominiczak and Alex Alderton, Roberto Amato, Jeffrey Barrett, Sonia Goncalves, Ewan Harrison, David K. Jackson, Ian Johnston, Dominic Kwiatkowski, Cordelia Langford, John Sillitoe on behalf of the Wellcome Sanger Institute COVID-19 Surveillance Team |
| EPI_ISL_3249880, EPI_ISL_3249881 | Berkshire and Surrey Pathology Services Lighthouse Laboratory | Wellcome Sanger Institute for the COVID-19 Genomics UK (COG-UK) Consortium | Berkshire and Surrey Pathology Services Lighthouse Laboratory and Alex Alderton, Roberto Amato, Jeffrey Barrett, Sonia Goncalves, Ewan Harrison, David K. Jackson, Ian Johnston, Dominic Kwiatkowski, Cordelia Langford, John Sillitoe on behalf of the Wellcome Sanger Institute COVID-19 Surveillance Team |
| EPI_ISL_3249882 | Lighthouse Lab in Milton Keynes | Wellcome Sanger Institute for the COVID-19 Genomics UK (COG-UK) Consortium | The Lighthouse Lab in Milton Keynes and Alex Alderton, Roberto Amato, Jeffrey Barrett, Sonia Goncalves, Ewan Harrison, David K. Jackson, Ian Johnston, Dominic Kwiatkowski, Cordelia Langford, John Sillitoe on behalf of the Wellcome Sanger Institute COVID-19 Surveillance Team |
| EPI_ISL_3249883, EPI_ISL_3249884, EPI_ISL_3249885 | Berkshire and Surrey Pathology Services Lighthouse Laboratory | Wellcome Sanger Institute for the COVID-19 Genomics UK (COG-UK) Consortium | Berkshire and Surrey Pathology Services Lighthouse Laboratory and Alex Alderton, Roberto Amato, Jeffrey Barrett, Sonia Goncalves, Ewan Harrison, David K. Jackson, Ian Johnston, Dominic Kwiatkowski, Cordelia Langford, John Sillitoe on behalf of the Wellcome Sanger Institute COVID-19 Surveillance Team |
| EPI_ISL_3249886, EPI_ISL_3249887 | Lighthouse Laboratory Plymouth | Wellcome Sanger Institute for the COVID-19 Genomics UK (COG-UK) Consortium | Lighthouse Laboratory Plymouth and Alex Alderton, Roberto Amato, Jeffrey Barrett, Sonia Goncalves, Ewan Harrison, David K. Jackson, Ian Johnston, Dominic Kwiatkowski, Cordelia Langford, John Sillitoe on behalf of the Wellcome Sanger Institute COVID-19 Surveillance Team |
| EPI_ISL_3249888 | Lighthouse Lab in Glasgow | Wellcome Sanger Institute for the COVID-19 Genomics UK (COG-UK) Consortium | Harper VanSteenhouse, Yumi Kasai, David Gray, Carol Clugston, Anna Dominiczak and Alex Alderton, Roberto Amato, Jeffrey Barrett, Sonia Goncalves, Ewan Harrison, David K. Jackson, Ian Johnston, Dominic Kwiatkowski, Cordelia Langford, John Sillitoe on behalf of the Wellcome Sanger Institute COVID-19 Surveillance Team |
| EPI_ISL_3249889, EPI_ISL_3249890, EPI_ISL_3249891, EPI_ISL_3249892, EPI_ISL_3249893, EPI_ISL_3249894, EPI_ISL_3249895, EPI_ISL_3249896, EPI_ISL_3249897 | Lighthouse Laboratory Plymouth | Wellcome Sanger Institute for the COVID-19 Genomics UK (COG-UK) Consortium | Lighthouse Laboratory Plymouth and Alex Alderton, Roberto Amato, Jeffrey Barrett, Sonia Goncalves, Ewan Harrison, David K. Jackson, Ian Johnston, Dominic Kwiatkowski, Cordelia Langford, John Sillitoe on behalf of the Wellcome Sanger Institute COVID-19 Surveillance Team |
| EPI_ISL_3249898 | AZ Klina | AZ Klina | Carl Vael - Lynsey Berckmans |
| EPI_ISL_3249899, EPI_ISL_3249900, EPI_ISL_3249901 | Lighthouse Laboratory Plymouth | Wellcome Sanger Institute for the COVID-19 Genomics UK (COG-UK) Consortium | Lighthouse Laboratory Plymouth and Alex Alderton, Roberto Amato, Jeffrey Barrett, Sonia Goncalves, Ewan Harrison, David K. Jackson, Ian Johnston, Dominic Kwiatkowski, Cordelia Langford, John Sillitoe on behalf of the Wellcome Sanger Institute COVID-19 Surveillance Team |
| EPI_ISL_3249902, EPI_ISL_3249903 | Lighthouse Lab in Glasgow | Wellcome Sanger Institute for the COVID-19 Genomics UK (COG-UK) Consortium | Harper VanSteenhouse, Yumi Kasai, David Gray, Carol Clugston, Anna Dominiczak and Alex Alderton, Roberto Amato, Jeffrey Barrett, Sonia Goncalves, Ewan Harrison, David K. Jackson, Ian Johnston, Dominic Kwiatkowski, Cordelia Langford, John Sillitoe on behalf of the Wellcome Sanger Institute COVID-19 Surveillance Team |
| EPI_ISL_3249904 | Lighthouse Laboratory Plymouth | Wellcome Sanger Institute for the COVID-19 Genomics UK (COG-UK) Consortium | Lighthouse Laboratory Plymouth and Alex Alderton, Roberto Amato, Jeffrey Barrett, Sonia Goncalves, Ewan Harrison, David K. Jackson, Ian Johnston, Dominic Kwiatkowski, Cordelia Langford, John Sillitoe on behalf of the Wellcome Sanger Institute COVID-19 Surveillance Team |
| EPI_ISL_3249905, EPI_ISL_3249906, EPI_ISL_3249907 | Lighthouse Lab in Glasgow | Wellcome Sanger Institute for the COVID-19 Genomics UK (COG-UK) Consortium | Harper VanSteenhouse, Yumi Kasai, David Gray, Carol Clugston, Anna Dominiczak and Alex Alderton, Roberto Amato, Jeffrey Barrett, Sonia Goncalves, Ewan Harrison, David K. Jackson, Ian Johnston, Dominic Kwiatkowski, Cordelia Langford, John Sillitoe on behalf of the Wellcome Sanger Institute COVID-19 Surveillance Team |
| EPI_ISL_3249908, EPI_ISL_3249909 | Lighthouse Laboratory Plymouth | Wellcome Sanger Institute for the COVID-19 Genomics UK (COG-UK) Consortium | Lighthouse Laboratory Plymouth and Alex Alderton, Roberto Amato, Jeffrey Barrett, Sonia Goncalves, Ewan Harrison, David K. Jackson, Ian Johnston, Dominic Kwiatkowski, Cordelia Langford, John Sillitoe on behalf of the Wellcome Sanger Institute COVID-19 Surveillance Team |

| | | | |
|---|---|---|---|
| EPI_ISL_3249910 | Lighthouse Lab in Glasgow | Wellcome Sanger Institute for the COVID-19 Genomics UK (COG-UK) Consortium | Harper VanSteenhouse, Yumi Kasai, David Gray, Carol Clugston, Anna Dominiczak and Alex Alderton, Roberto Amato, Jeffrey Barrett, Sonia Goncalves, Ewan Harrison, David K. Jackson, Ian Johnston, Dominic Kwiatkowski, Cordelia Langford, John Sillitoe on behalf of the Wellcome Sanger Institute COVID-19 Surveillance Team |
| EPI_ISL_3249911, EPI_ISL_3249912, EPI_ISL_3249913, EPI_ISL_3249914, EPI_ISL_3249915 | Lighthouse Laboratory Plymouth | Wellcome Sanger Institute for the COVID-19 Genomics UK (COG-UK) Consortium | Lighthouse Laboratory Plymouth and Alex Alderton, Roberto Amato, Jeffrey Barrett, Sonia Goncalves, Ewan Harrison, David K. Jackson, Ian Johnston, Dominic Kwiatkowski, Cordelia Langford, John Sillitoe on behalf of the Wellcome Sanger Institute COVID-19 Surveillance Team |
| EPI_ISL_3249916 | Lighthouse Lab in Glasgow | Wellcome Sanger Institute for the COVID-19 Genomics UK (COG-UK) Consortium | Harper VanSteenhouse, Yumi Kasai, David Gray, Carol Clugston, Anna Dominiczak and Alex Alderton, Roberto Amato, Jeffrey Barrett, Sonia Goncalves, Ewan Harrison, David K. Jackson, Ian Johnston, Dominic Kwiatkowski, Cordelia Langford, John Sillitoe on behalf of the Wellcome Sanger Institute COVID-19 Surveillance Team |
| EPI_ISL_3249917, EPI_ISL_3249918 | Lighthouse Laboratory Plymouth | Wellcome Sanger Institute for the COVID-19 Genomics UK (COG-UK) Consortium | Lighthouse Laboratory Plymouth and Alex Alderton, Roberto Amato, Jeffrey Barrett, Sonia Goncalves, Ewan Harrison, David K. Jackson, Ian Johnston, Dominic Kwiatkowski, Cordelia Langford, John Sillitoe on behalf of the Wellcome Sanger Institute COVID-19 Surveillance Team |
| EPI_ISL_3249919 | Lighthouse Lab in Glasgow | Wellcome Sanger Institute for the COVID-19 Genomics UK (COG-UK) Consortium | Harper VanSteenhouse, Yumi Kasai, David Gray, Carol Clugston, Anna Dominiczak and Alex Alderton, Roberto Amato, Jeffrey Barrett, Sonia Goncalves, Ewan Harrison, David K. Jackson, Ian Johnston, Dominic Kwiatkowski, Cordelia Langford, John Sillitoe on behalf of the Wellcome Sanger Institute COVID-19 Surveillance Team |
| EPI_ISL_3249920, EPI_ISL_3249921, EPI_ISL_3249922, EPI_ISL_3249923, EPI_ISL_3249924, EPI_ISL_3249925, EPI_ISL_3249926 | Lighthouse Laboratory Plymouth | Wellcome Sanger Institute for the COVID-19 Genomics UK (COG-UK) Consortium | Lighthouse Laboratory Plymouth and Alex Alderton, Roberto Amato, Jeffrey Barrett, Sonia Goncalves, Ewan Harrison, David K. Jackson, Ian Johnston, Dominic Kwiatkowski, Cordelia Langford, John Sillitoe on behalf of the Wellcome Sanger Institute COVID-19 Surveillance Team |
| EPI_ISL_3249927 | Lighthouse Lab in Glasgow | Wellcome Sanger Institute for the COVID-19 Genomics UK (COG-UK) Consortium | Harper VanSteenhouse, Yumi Kasai, David Gray, Carol Clugston, Anna Dominiczak and Alex Alderton, Roberto Amato, Jeffrey Barrett, Sonia Goncalves, Ewan Harrison, David K. Jackson, Ian Johnston, Dominic Kwiatkowski, Cordelia Langford, John Sillitoe on behalf of the Wellcome Sanger Institute COVID-19 Surveillance Team |
| EPI_ISL_3249928 | Lighthouse Laboratory Plymouth | Wellcome Sanger Institute for the COVID-19 Genomics UK (COG-UK) Consortium | Lighthouse Laboratory Plymouth and Alex Alderton, Roberto Amato, Jeffrey Barrett, Sonia Goncalves, Ewan Harrison, David K. Jackson, Ian Johnston, Dominic Kwiatkowski, Cordelia Langford, John Sillitoe on behalf of the Wellcome Sanger Institute COVID-19 Surveillance Team |
| EPI_ISL_3249929 | Lighthouse Lab in Glasgow | Wellcome Sanger Institute for the COVID-19 Genomics UK (COG-UK) Consortium | Harper VanSteenhouse, Yumi Kasai, David Gray, Carol Clugston, Anna Dominiczak and Alex Alderton, Roberto Amato, Jeffrey Barrett, Sonia Goncalves, Ewan Harrison, David K. Jackson, Ian Johnston, Dominic Kwiatkowski, Cordelia Langford, John Sillitoe on behalf of the Wellcome Sanger Institute COVID-19 Surveillance Team |
| EPI_ISL_3249930, EPI_ISL_3249931, EPI_ISL_3249932, EPI_ISL_3249933, EPI_ISL_3249934, EPI_ISL_3249935, EPI_ISL_3249936, EPI_ISL_3249937, EPI_ISL_3249938 | Lighthouse Laboratory Plymouth | Wellcome Sanger Institute for the COVID-19 Genomics UK (COG-UK) Consortium | Lighthouse Laboratory Plymouth and Alex Alderton, Roberto Amato, Jeffrey Barrett, Sonia Goncalves, Ewan Harrison, David K. Jackson, Ian Johnston, Dominic Kwiatkowski, Cordelia Langford, John Sillitoe on behalf of the Wellcome Sanger Institute COVID-19 Surveillance Team |
| EPI_ISL_3249939 | Lighthouse Lab in Glasgow | Wellcome Sanger Institute for the COVID-19 Genomics UK (COG-UK) Consortium | Harper VanSteenhouse, Yumi Kasai, David Gray, Carol Clugston, Anna Dominiczak and Alex Alderton, Roberto Amato, Jeffrey Barrett, Sonia Goncalves, Ewan Harrison, David K. Jackson, Ian Johnston, Dominic Kwiatkowski, Cordelia Langford, John Sillitoe on behalf of the Wellcome Sanger Institute COVID-19 Surveillance Team |
| EPI_ISL_3249940, EPI_ISL_3249941, EPI_ISL_3249942, EPI_ISL_3249943, EPI_ISL_3249944 | Lighthouse Laboratory Plymouth | Wellcome Sanger Institute for the COVID-19 Genomics UK (COG-UK) Consortium | Lighthouse Laboratory Plymouth and Alex Alderton, Roberto Amato, Jeffrey Barrett, Sonia Goncalves, Ewan Harrison, David K. Jackson, Ian Johnston, Dominic Kwiatkowski, Cordelia Langford, John Sillitoe on behalf of the Wellcome Sanger Institute COVID-19 Surveillance Team |
| EPI_ISL_3249945 | Lighthouse Lab in Glasgow | Wellcome Sanger Institute for the COVID-19 Genomics UK (COG-UK) Consortium | Harper VanSteenhouse, Yumi Kasai, David Gray, Carol Clugston, Anna Dominiczak and Alex Alderton, Roberto Amato, Jeffrey Barrett, Sonia Goncalves, Ewan Harrison, David K. Jackson, Ian Johnston, Dominic Kwiatkowski, Cordelia Langford, John Sillitoe on behalf of the Wellcome Sanger Institute COVID-19 Surveillance Team |
| EPI_ISL_3249946, EPI_ISL_3249947, EPI_ISL_3249948, EPI_ISL_3249949 | Lighthouse Laboratory Plymouth | Wellcome Sanger Institute for the COVID-19 Genomics UK (COG-UK) Consortium | Lighthouse Laboratory Plymouth and Alex Alderton, Roberto Amato, Jeffrey Barrett, Sonia Goncalves, Ewan Harrison, David K. Jackson, Ian Johnston, Dominic Kwiatkowski, Cordelia Langford, John Sillitoe on behalf of the Wellcome Sanger Institute COVID-19 Surveillance Team |
| EPI_ISL_3249950, EPI_ISL_3249951, EPI_ISL_3249952 | Lighthouse Lab in Glasgow | Wellcome Sanger Institute for the COVID-19 Genomics UK (COG-UK) Consortium | Harper VanSteenhouse, Yumi Kasai, David Gray, Carol Clugston, Anna Dominiczak and Alex Alderton, Roberto Amato, Jeffrey Barrett, Sonia Goncalves, Ewan Harrison, David K. Jackson, Ian Johnston, Dominic Kwiatkowski, Cordelia Langford, John Sillitoe on behalf of the Wellcome Sanger Institute COVID-19 Surveillance Team |
| EPI_ISL_3249953, EPI_ISL_3249954, EPI_ISL_3249955, EPI_ISL_3249956, EPI_ISL_3249957, EPI_ISL_3249958, EPI_ISL_3249959, EPI_ISL_3249960, EPI_ISL_3249961 | Lighthouse Laboratory Plymouth | Wellcome Sanger Institute for the COVID-19 Genomics UK (COG-UK) Consortium | Lighthouse Laboratory Plymouth and Alex Alderton, Roberto Amato, Jeffrey Barrett, Sonia Goncalves, Ewan Harrison, David K. Jackson, Ian Johnston, Dominic Kwiatkowski, Cordelia Langford, John Sillitoe on behalf of the Wellcome Sanger Institute COVID-19 Surveillance Team |
| EPI_ISL_3249962 | Lighthouse Lab in Glasgow | Wellcome Sanger Institute for the COVID-19 Genomics UK (COG-UK) Consortium | Harper VanSteenhouse, Yumi Kasai, David Gray, Carol Clugston, Anna Dominiczak and Alex Alderton, Roberto Amato, Jeffrey Barrett, Sonia Goncalves, Ewan Harrison, David K. Jackson, Ian Johnston, Dominic Kwiatkowski, Cordelia Langford, John Sillitoe on behalf of the Wellcome Sanger Institute COVID-19 Surveillance Team |
| EPI_ISL_3249963, EPI_ISL_3249964, EPI_ISL_3249965 | Lighthouse Laboratory Plymouth | Wellcome Sanger Institute for the COVID-19 Genomics UK (COG-UK) Consortium | Lighthouse Laboratory Plymouth and Alex Alderton, Roberto Amato, Jeffrey Barrett, Sonia Goncalves, Ewan Harrison, David K. Jackson, Ian Johnston, Dominic Kwiatkowski, Cordelia Langford, John Sillitoe on behalf of the Wellcome Sanger Institute COVID-19 Surveillance Team |
| EPI_ISL_3249966 | Lighthouse Lab in Glasgow | Wellcome Sanger Institute for the COVID-19 Genomics UK (COG-UK) Consortium | Harper VanSteenhouse, Yumi Kasai, David Gray, Carol Clugston, Anna Dominiczak and Alex Alderton, Roberto Amato, Jeffrey Barrett, Sonia Goncalves, Ewan Harrison, David K. Jackson, Ian Johnston, Dominic Kwiatkowski, Cordelia Langford, John Sillitoe on behalf of the Wellcome Sanger Institute COVID-19 Surveillance Team |
| EPI_ISL_3249967 | Lighthouse Laboratory Plymouth | Wellcome Sanger Institute for the COVID-19 Genomics UK (COG-UK) Consortium | Lighthouse Laboratory Plymouth and Alex Alderton, Roberto Amato, Jeffrey Barrett, Sonia Goncalves, Ewan Harrison, David K. Jackson, Ian Johnston, Dominic Kwiatkowski, Cordelia Langford, John Sillitoe on behalf of the Wellcome Sanger Institute COVID-19 Surveillance Team |
| EPI_ISL_3249968 | Lighthouse Lab in Glasgow | Wellcome Sanger Institute for the COVID-19 Genomics UK (COG-UK) Consortium | Harper VanSteenhouse, Yumi Kasai, David Gray, Carol Clugston, Anna Dominiczak and Alex Alderton, Roberto Amato, Jeffrey Barrett, Sonia Goncalves, Ewan Harrison, David K. Jackson, Ian Johnston, Dominic Kwiatkowski, Cordelia Langford, John Sillitoe on behalf of the Wellcome Sanger Institute COVID-19 Surveillance Team |
| EPI_ISL_3249969 | Lighthouse Laboratory Plymouth | Wellcome Sanger Institute for the COVID-19 Genomics UK (COG-UK) Consortium | Lighthouse Laboratory Plymouth and Alex Alderton, Roberto Amato, Jeffrey Barrett, Sonia Goncalves, Ewan Harrison, David K. Jackson, Ian Johnston, Dominic Kwiatkowski, Cordelia Langford, John Sillitoe on behalf of the Wellcome Sanger Institute COVID-19 Surveillance Team |
| EPI_ISL_3249970 | Lighthouse Lab in Glasgow | Wellcome Sanger Institute for the COVID-19 Genomics UK (COG-UK) Consortium | Harper VanSteenhouse, Yumi Kasai, David Gray, Carol Clugston, Anna Dominiczak and Alex Alderton, Roberto Amato, Jeffrey Barrett, Sonia Goncalves, Ewan Harrison, David K. Jackson, Ian Johnston, Dominic Kwiatkowski, Cordelia Langford, John Sillitoe on behalf of the Wellcome Sanger Institute COVID-19 Surveillance Team |
| EPI_ISL_3249971, EPI_ISL_3249972, EPI_ISL_3249973, EPI_ISL_3249974 | Lighthouse Laboratory Plymouth | Wellcome Sanger Institute for the COVID-19 Genomics UK (COG-UK) Consortium | Lighthouse Laboratory Plymouth and Alex Alderton, Roberto Amato, Jeffrey Barrett, Sonia Goncalves, Ewan Harrison, David K. Jackson, Ian Johnston, Dominic Kwiatkowski, Cordelia Langford, John Sillitoe on behalf of the Wellcome Sanger Institute COVID-19 Surveillance Team |
| EPI_ISL_3249975 | Lighthouse Lab in Glasgow | Wellcome Sanger Institute for the COVID-19 Genomics UK (COG-UK) Consortium | Harper VanSteenhouse, Yumi Kasai, David Gray, Carol Clugston, Anna Dominiczak and Alex Alderton, Roberto Amato, Jeffrey Barrett, Sonia Goncalves, Ewan Harrison, David K. Jackson, Ian Johnston, Dominic Kwiatkowski, Cordelia Langford, John Sillitoe on behalf of the Wellcome Sanger Institute COVID-19 Surveillance Team |
| EPI_ISL_3249976, EPI_ISL_3249977, EPI_ISL_3249978, EPI_ISL_3249979 | Lighthouse Laboratory Plymouth | Wellcome Sanger Institute for the COVID-19 Genomics UK (COG-UK) Consortium | Lighthouse Laboratory Plymouth and Alex Alderton, Roberto Amato, Jeffrey Barrett, Sonia Goncalves, Ewan Harrison, David K. Jackson, Ian Johnston, Dominic Kwiatkowski, Cordelia Langford, John Sillitoe on behalf of the Wellcome Sanger Institute COVID-19 Surveillance Team |
| EPI_ISL_3249980 | Lighthouse Lab in Glasgow | Wellcome Sanger Institute for the COVID-19 Genomics UK (COG-UK) Consortium | Harper VanSteenhouse, Yumi Kasai, David Gray, Carol Clugston, Anna Dominiczak and Alex Alderton, Roberto Amato, Jeffrey Barrett, Sonia Goncalves, Ewan Harrison, David K. Jackson, Ian Johnston, Dominic Kwiatkowski, Cordelia Langford, John Sillitoe on behalf of the Wellcome Sanger Institute COVID-19 Surveillance Team |
| EPI_ISL_3249981, EPI_ISL_3249982, EPI_ISL_3249983, EPI_ISL_3249984, EPI_ISL_3249985, EPI_ISL_3249986, EPI_ISL_3249987, EPI_ISL_3249988, EPI_ISL_3249989 | Lighthouse Laboratory Plymouth | Wellcome Sanger Institute for the COVID-19 Genomics UK (COG-UK) Consortium | Lighthouse Laboratory Plymouth and Alex Alderton, Roberto Amato, Jeffrey Barrett, Sonia Goncalves, Ewan Harrison, David K. Jackson, Ian Johnston, Dominic Kwiatkowski, Cordelia Langford, John Sillitoe on behalf of the Wellcome Sanger Institute COVID-19 Surveillance Team |
| EPI_ISL_3249990 | Lighthouse Lab in Glasgow | Wellcome Sanger Institute for the COVID-19 Genomics UK (COG-UK) | Harper VanSteenhouse, Yumi Kasai, David Gray, Carol Clugston, Anna Dominiczak and Alex Alderton, Roberto Amato, Jeffrey Barrett, Sonia Goncalves, Ewan Harrison, David K. Jackson, Ian Johnston, Dominic Kwiatkowski, Cordelia Langford, |

| | | | |
|---|---|---|---|
| | | Consortium | John Sillitoe on behalf of the Wellcome Sanger Institute COVID-19 Surveillance Team |
| EPI_ISL_3249991, EPI_ISL_3249992, EPI_ISL_3249993, EPI_ISL_3249994, EPI_ISL_3249995, EPI_ISL_3249996 | Lighthouse Laboratory Plymouth | Wellcome Sanger Institute for the COVID-19 Genomics UK (COG-UK) Consortium | Lighthouse Laboratory Plymouth and Alex Alderton, Roberto Amato, Jeffrey Barrett, Sonia Goncalves, Ewan Harrison, David K. Jackson, Ian Johnston, Dominic Kwiatkowski, Cordelia Langford, John Sillitoe on behalf of the Wellcome Sanger Institute COVID-19 Surveillance Team |
| EPI_ISL_3249997 | Lighthouse Lab in Glasgow | Wellcome Sanger Institute for the COVID-19 Genomics UK (COG-UK) Consortium | Harper VanSteenhouse, Yumi Kasai, David Gray, Carol Clugston, Anna Dominiczak and Alex Alderton, Roberto Amato, Jeffrey Barrett, Sonia Goncalves, Ewan Harrison, David K. Jackson, Ian Johnston, Dominic Kwiatkowski, Cordelia Langford, John Sillitoe on behalf of the Wellcome Sanger Institute COVID-19 Surveillance Team |
| EPI_ISL_3249998 | Lighthouse Laboratory Plymouth | Wellcome Sanger Institute for the COVID-19 Genomics UK (COG-UK) Consortium | Lighthouse Laboratory Plymouth and Alex Alderton, Roberto Amato, Jeffrey Barrett, Sonia Goncalves, Ewan Harrison, David K. Jackson, Ian Johnston, Dominic Kwiatkowski, Cordelia Langford, John Sillitoe on behalf of the Wellcome Sanger Institute COVID-19 Surveillance Team |
| EPI_ISL_3249999 | Lighthouse Lab in Glasgow | Wellcome Sanger Institute for the COVID-19 Genomics UK (COG-UK) Consortium | Harper VanSteenhouse, Yumi Kasai, David Gray, Carol Clugston, Anna Dominiczak and Alex Alderton, Roberto Amato, Jeffrey Barrett, Sonia Goncalves, Ewan Harrison, David K. Jackson, Ian Johnston, Dominic Kwiatkowski, Cordelia Langford, John Sillitoe on behalf of the Wellcome Sanger Institute COVID-19 Surveillance Team |
| EPI_ISL_3250000 | Lighthouse Laboratory Plymouth | Wellcome Sanger Institute for the COVID-19 Genomics UK (COG-UK) Consortium | Lighthouse Laboratory Plymouth and Alex Alderton, Roberto Amato, Jeffrey Barrett, Sonia Goncalves, Ewan Harrison, David K. Jackson, Ian Johnston, Dominic Kwiatkowski, Cordelia Langford, John Sillitoe on behalf of the Wellcome Sanger Institute COVID-19 Surveillance Team |
| EPI_ISL_3250001 | Lighthouse Lab in Glasgow | Wellcome Sanger Institute for the COVID-19 Genomics UK (COG-UK) Consortium | Harper VanSteenhouse, Yumi Kasai, David Gray, Carol Clugston, Anna Dominiczak and Alex Alderton, Roberto Amato, Jeffrey Barrett, Sonia Goncalves, Ewan Harrison, David K. Jackson, Ian Johnston, Dominic Kwiatkowski, Cordelia Langford, John Sillitoe on behalf of the Wellcome Sanger Institute COVID-19 Surveillance Team |
| EPI_ISL_3250002, EPI_ISL_3250003, EPI_ISL_3250004, EPI_ISL_3250005, EPI_ISL_3250006, EPI_ISL_3250007 | Lighthouse Laboratory Plymouth | Wellcome Sanger Institute for the COVID-19 Genomics UK (COG-UK) Consortium | Lighthouse Laboratory Plymouth and Alex Alderton, Roberto Amato, Jeffrey Barrett, Sonia Goncalves, Ewan Harrison, David K. Jackson, Ian Johnston, Dominic Kwiatkowski, Cordelia Langford, John Sillitoe on behalf of the Wellcome Sanger Institute COVID-19 Surveillance Team |
| EPI_ISL_3250008 | Lighthouse Lab in Glasgow | Wellcome Sanger Institute for the COVID-19 Genomics UK (COG-UK) Consortium | Harper VanSteenhouse, Yumi Kasai, David Gray, Carol Clugston, Anna Dominiczak and Alex Alderton, Roberto Amato, Jeffrey Barrett, Sonia Goncalves, Ewan Harrison, David K. Jackson, Ian Johnston, Dominic Kwiatkowski, Cordelia Langford, John Sillitoe on behalf of the Wellcome Sanger Institute COVID-19 Surveillance Team |
| EPI_ISL_3250009, EPI_ISL_3250010, EPI_ISL_3250011 | Lighthouse Laboratory Plymouth | Wellcome Sanger Institute for the COVID-19 Genomics UK (COG-UK) Consortium | Lighthouse Laboratory Plymouth and Alex Alderton, Roberto Amato, Jeffrey Barrett, Sonia Goncalves, Ewan Harrison, David K. Jackson, Ian Johnston, Dominic Kwiatkowski, Cordelia Langford, John Sillitoe on behalf of the Wellcome Sanger Institute COVID-19 Surveillance Team |
| EPI_ISL_3250012, EPI_ISL_3250013, EPI_ISL_3250014 | Lighthouse Lab in Glasgow | Wellcome Sanger Institute for the COVID-19 Genomics UK (COG-UK) Consortium | Harper VanSteenhouse, Yumi Kasai, David Gray, Carol Clugston, Anna Dominiczak and Alex Alderton, Roberto Amato, Jeffrey Barrett, Sonia Goncalves, Ewan Harrison, David K. Jackson, Ian Johnston, Dominic Kwiatkowski, Cordelia Langford, John Sillitoe on behalf of the Wellcome Sanger Institute COVID-19 Surveillance Team |
| EPI_ISL_3250015, EPI_ISL_3250016, EPI_ISL_3250017 | Lighthouse Laboratory Plymouth | Wellcome Sanger Institute for the COVID-19 Genomics UK (COG-UK) Consortium | Lighthouse Laboratory Plymouth and Alex Alderton, Roberto Amato, Jeffrey Barrett, Sonia Goncalves, Ewan Harrison, David K. Jackson, Ian Johnston, Dominic Kwiatkowski, Cordelia Langford, John Sillitoe on behalf of the Wellcome Sanger Institute COVID-19 Surveillance Team |
| EPI_ISL_3250018 | Lighthouse Lab in Glasgow | Wellcome Sanger Institute for the COVID-19 Genomics UK (COG-UK) Consortium | Harper VanSteenhouse, Yumi Kasai, David Gray, Carol Clugston, Anna Dominiczak and Alex Alderton, Roberto Amato, Jeffrey Barrett, Sonia Goncalves, Ewan Harrison, David K. Jackson, Ian Johnston, Dominic Kwiatkowski, Cordelia Langford, John Sillitoe on behalf of the Wellcome Sanger Institute COVID-19 Surveillance Team |
| EPI_ISL_3250019, EPI_ISL_3250020, EPI_ISL_3250021, EPI_ISL_3250022, EPI_ISL_3250023, EPI_ISL_3250024, EPI_ISL_3250025, EPI_ISL_3250026, EPI_ISL_3250027, EPI_ISL_3250028, EPI_ISL_3250029, EPI_ISL_3250030, EPI_ISL_3250031, EPI_ISL_3250032, EPI_ISL_3250033, EPI_ISL_3250034, EPI_ISL_3250035, EPI_ISL_3250036, EPI_ISL_3250037, EPI_ISL_3250038, EPI_ISL_3250039, EPI_ISL_3250040, EPI_ISL_3250041, EPI_ISL_3250042, EPI_ISL_3250043, EPI_ISL_3250044, EPI_ISL_3250045 | | | |
| see above | Lighthouse Laboratory Plymouth | Wellcome Sanger Institute for the COVID-19 Genomics UK (COG-UK) Consortium | Lighthouse Laboratory Plymouth and Alex Alderton, Roberto Amato, Jeffrey Barrett, Sonia Goncalves, Ewan Harrison, David K. Jackson, Ian Johnston, Dominic Kwiatkowski, Cordelia Langford, John Sillitoe on behalf of the Wellcome Sanger Institute COVID-19 Surveillance Team |
| EPI_ISL_3250046 | Lighthouse Lab in Glasgow | Wellcome Sanger Institute for the COVID-19 Genomics UK (COG-UK) Consortium | Harper VanSteenhouse, Yumi Kasai, David Gray, Carol Clugston, Anna Dominiczak and Alex Alderton, Roberto Amato, Jeffrey Barrett, Sonia Goncalves, Ewan Harrison, David K. Jackson, Ian Johnston, Dominic Kwiatkowski, Cordelia Langford, John Sillitoe on behalf of the Wellcome Sanger Institute COVID-19 Surveillance Team |
| EPI_ISL_3250047, EPI_ISL_3250048, EPI_ISL_3250049 | Lighthouse Laboratory Plymouth | Wellcome Sanger Institute for the COVID-19 Genomics UK (COG-UK) Consortium | Lighthouse Laboratory Plymouth and Alex Alderton, Roberto Amato, Jeffrey Barrett, Sonia Goncalves, Ewan Harrison, David K. Jackson, Ian Johnston, Dominic Kwiatkowski, Cordelia Langford, John Sillitoe on behalf of the Wellcome Sanger Institute COVID-19 Surveillance Team |
| EPI_ISL_3250050 | Lighthouse Lab in Glasgow | Wellcome Sanger Institute for the COVID-19 Genomics UK (COG-UK) Consortium | Harper VanSteenhouse, Yumi Kasai, David Gray, Carol Clugston, Anna Dominiczak and Alex Alderton, Roberto Amato, Jeffrey Barrett, Sonia Goncalves, Ewan Harrison, David K. Jackson, Ian Johnston, Dominic Kwiatkowski, Cordelia Langford, John Sillitoe on behalf of the Wellcome Sanger Institute COVID-19 Surveillance Team |
| EPI_ISL_3250051, EPI_ISL_3250052 | Lighthouse Laboratory Plymouth | Wellcome Sanger Institute for the COVID-19 Genomics UK (COG-UK) Consortium | Lighthouse Laboratory Plymouth and Alex Alderton, Roberto Amato, Jeffrey Barrett, Sonia Goncalves, Ewan Harrison, David K. Jackson, Ian Johnston, Dominic Kwiatkowski, Cordelia Langford, John Sillitoe on behalf of the Wellcome Sanger Institute COVID-19 Surveillance Team |
| EPI_ISL_3250053, EPI_ISL_3250054 | Lighthouse Lab in Glasgow | Wellcome Sanger Institute for the COVID-19 Genomics UK (COG-UK) Consortium | Harper VanSteenhouse, Yumi Kasai, David Gray, Carol Clugston, Anna Dominiczak and Alex Alderton, Roberto Amato, Jeffrey Barrett, Sonia Goncalves, Ewan Harrison, David K. Jackson, Ian Johnston, Dominic Kwiatkowski, Cordelia Langford, John Sillitoe on behalf of the Wellcome Sanger Institute COVID-19 Surveillance Team |
| EPI_ISL_3250055 | Lighthouse Laboratory Plymouth | Wellcome Sanger Institute for the COVID-19 Genomics UK (COG-UK) Consortium | Lighthouse Laboratory Plymouth and Alex Alderton, Roberto Amato, Jeffrey Barrett, Sonia Goncalves, Ewan Harrison, David K. Jackson, Ian Johnston, Dominic Kwiatkowski, Cordelia Langford, John Sillitoe on behalf of the Wellcome Sanger Institute COVID-19 Surveillance Team |
| EPI_ISL_3250056 | Lighthouse Lab in Glasgow | Wellcome Sanger Institute for the COVID-19 Genomics UK (COG-UK) Consortium | Harper VanSteenhouse, Yumi Kasai, David Gray, Carol Clugston, Anna Dominiczak and Alex Alderton, Roberto Amato, Jeffrey Barrett, Sonia Goncalves, Ewan Harrison, David K. Jackson, Ian Johnston, Dominic Kwiatkowski, Cordelia Langford, John Sillitoe on behalf of the Wellcome Sanger Institute COVID-19 Surveillance Team |
| EPI_ISL_3250057, EPI_ISL_3250058, EPI_ISL_3250059, EPI_ISL_3250060 | Lighthouse Laboratory Plymouth | Wellcome Sanger Institute for the COVID-19 Genomics UK (COG-UK) Consortium | Lighthouse Laboratory Plymouth and Alex Alderton, Roberto Amato, Jeffrey Barrett, Sonia Goncalves, Ewan Harrison, David K. Jackson, Ian Johnston, Dominic Kwiatkowski, Cordelia Langford, John Sillitoe on behalf of the Wellcome Sanger Institute COVID-19 Surveillance Team |
| EPI_ISL_3250061, EPI_ISL_3250062 | Lighthouse Lab in Glasgow | Wellcome Sanger Institute for the COVID-19 Genomics UK (COG-UK) Consortium | Harper VanSteenhouse, Yumi Kasai, David Gray, Carol Clugston, Anna Dominiczak and Alex Alderton, Roberto Amato, Jeffrey Barrett, Sonia Goncalves, Ewan Harrison, David K. Jackson, Ian Johnston, Dominic Kwiatkowski, Cordelia Langford, John Sillitoe on behalf of the Wellcome Sanger Institute COVID-19 Surveillance Team |
| EPI_ISL_3250063, EPI_ISL_3250064, EPI_ISL_3250065, EPI_ISL_3250066 | Lighthouse Laboratory Plymouth | Wellcome Sanger Institute for the COVID-19 Genomics UK (COG-UK) Consortium | Lighthouse Laboratory Plymouth and Alex Alderton, Roberto Amato, Jeffrey Barrett, Sonia Goncalves, Ewan Harrison, David K. Jackson, Ian Johnston, Dominic Kwiatkowski, Cordelia Langford, John Sillitoe on behalf of the Wellcome Sanger Institute COVID-19 Surveillance Team |
| EPI_ISL_3250067 | Lighthouse Lab in Glasgow | Wellcome Sanger Institute for the COVID-19 Genomics UK (COG-UK) Consortium | Harper VanSteenhouse, Yumi Kasai, David Gray, Carol Clugston, Anna Dominiczak and Alex Alderton, Roberto Amato, Jeffrey Barrett, Sonia Goncalves, Ewan Harrison, David K. Jackson, Ian Johnston, Dominic Kwiatkowski, Cordelia Langford, John Sillitoe on behalf of the Wellcome Sanger Institute COVID-19 Surveillance Team |
| EPI_ISL_3250068, EPI_ISL_3250069 | Lighthouse Laboratory Plymouth | Wellcome Sanger Institute for the COVID-19 Genomics UK (COG-UK) Consortium | Lighthouse Laboratory Plymouth and Alex Alderton, Roberto Amato, Jeffrey Barrett, Sonia Goncalves, Ewan Harrison, David K. Jackson, Ian Johnston, Dominic Kwiatkowski, Cordelia Langford, John Sillitoe on behalf of the Wellcome Sanger Institute COVID-19 Surveillance Team |
| EPI_ISL_3250070, EPI_ISL_3250071, EPI_ISL_3250072 | Lighthouse Lab in Glasgow | Wellcome Sanger Institute for the COVID-19 Genomics UK (COG-UK) Consortium | Harper VanSteenhouse, Yumi Kasai, David Gray, Carol Clugston, Anna Dominiczak and Alex Alderton, Roberto Amato, Jeffrey Barrett, Sonia Goncalves, Ewan Harrison, David K. Jackson, Ian Johnston, Dominic Kwiatkowski, Cordelia Langford, John Sillitoe on behalf of the Wellcome Sanger Institute COVID-19 Surveillance Team |
| EPI_ISL_3250073, EPI_ISL_3250074 | Lighthouse Laboratory Plymouth | Wellcome Sanger Institute for the COVID-19 Genomics UK (COG-UK) Consortium | Lighthouse Laboratory Plymouth and Alex Alderton, Roberto Amato, Jeffrey Barrett, Sonia Goncalves, Ewan Harrison, David K. Jackson, Ian Johnston, Dominic Kwiatkowski, Cordelia Langford, John Sillitoe on behalf of the Wellcome Sanger Institute COVID-19 Surveillance Team |
| EPI_ISL_3250075, EPI_ISL_3250076 | Lighthouse Lab in Glasgow | Wellcome Sanger Institute for the COVID-19 Genomics UK (COG-UK) Consortium | Harper VanSteenhouse, Yumi Kasai, David Gray, Carol Clugston, Anna Dominiczak and Alex Alderton, Roberto Amato, Jeffrey Barrett, Sonia Goncalves, Ewan Harrison, David K. Jackson, Ian Johnston, Dominic Kwiatkowski, Cordelia Langford, John Sillitoe on behalf of the Wellcome Sanger Institute COVID-19 Surveillance Team |
| EPI_ISL_3250077, EPI_ISL_3250078, EPI_ISL_3250079 | Lighthouse Laboratory Plymouth | Wellcome Sanger Institute for the COVID-19 Genomics UK (COG-UK) Consortium | Lighthouse Laboratory Plymouth and Alex Alderton, Roberto Amato, Jeffrey Barrett, Sonia Goncalves, Ewan Harrison, David K. Jackson, Ian Johnston, Dominic Kwiatkowski, Cordelia Langford, John Sillitoe on behalf of the Wellcome Sanger Institute COVID-19 Surveillance Team |
| EPI_ISL_3250080 | Lighthouse Lab in Glasgow | Wellcome Sanger Institute for the COVID-19 Genomics UK (COG-UK) Consortium | Harper VanSteenhouse, Yumi Kasai, David Gray, Carol Clugston, Anna Dominiczak and Alex Alderton, Roberto Amato, Jeffrey Barrett, Sonia Goncalves, Ewan Harrison, David K. Jackson, Ian Johnston, Dominic Kwiatkowski, Cordelia Langford, John Sillitoe on behalf of the Wellcome Sanger Institute COVID-19 Surveillance Team |
| EPI_ISL_3250081, EPI_ISL_3250082 | Lighthouse Laboratory Plymouth | Wellcome Sanger Institute for the COVID-19 Genomics UK (COG-UK) Consortium | Lighthouse Laboratory Plymouth and Alex Alderton, Roberto Amato, Jeffrey Barrett, Sonia Goncalves, Ewan Harrison, David K. Jackson, Ian Johnston, Dominic Kwiatkowski, Cordelia Langford, John Sillitoe on behalf of the Wellcome Sanger Institute COVID-19 Surveillance Team |
| EPI_ISL_3250083, EPI_ISL_3250084, EPI_ISL_3250085 | Lighthouse Lab in Glasgow | Wellcome Sanger Institute for the COVID-19 Genomics UK (COG-UK) Consortium | Harper VanSteenhouse, Yumi Kasai, David Gray, Carol Clugston, Anna Dominiczak and Alex Alderton, Roberto Amato, Jeffrey Barrett, Sonia Goncalves, Ewan Harrison, David K. Jackson, Ian Johnston, Dominic Kwiatkowski, Cordelia Langford, John Sillitoe on behalf of the Wellcome Sanger Institute COVID-19 Surveillance Team |
| EPI_ISL_3250086, EPI_ISL_3250087, EPI_ISL_3250088, EPI_ISL_3250089, EPI_ISL_3250090, EPI_ISL_3250091, EPI_ISL_3250092, EPI_ISL_3250093, EPI_ISL_3250094, | Lighthouse Laboratory Plymouth | Wellcome Sanger Institute for the COVID-19 Genomics UK (COG-UK) Consortium | Lighthouse Laboratory Plymouth and Alex Alderton, Roberto Amato, Jeffrey Barrett, Sonia Goncalves, Ewan Harrison, David K. Jackson, Ian Johnston, Dominic Kwiatkowski, Cordelia Langford, John Sillitoe on behalf of the Wellcome Sanger Institute COVID-19 Surveillance Team |

| | | | |
|---|---|---|---|
| EPI_ISL_3250095 | | | |
| EPI_ISL_3250096 | Lighthouse Lab in Glasgow | Wellcome Sanger Institute for the COVID-19 Genomics UK (COG-UK) Consortium | Harper VanSteenhouse, Yumi Kasai, David Gray, Carol Clugston, Anna Dominiczak and Alex Alderton, Roberto Amato, Jeffrey Barrett, Sonia Goncalves, Ewan Harrison, David K. Jackson, Ian Johnston, Dominic Kwiatkowski, Cordelia Langford, John Sillitoe on behalf of the Wellcome Sanger Institute COVID-19 Surveillance Team |
| EPI_ISL_3250097, EPI_ISL_3250098, EPI_ISL_3250099 | Lighthouse Laboratory Plymouth | Wellcome Sanger Institute for the COVID-19 Genomics UK (COG-UK) Consortium | Lighthouse Laboratory Plymouth and Alex Alderton, Roberto Amato, Jeffrey Barrett, Sonia Goncalves, Ewan Harrison, David K. Jackson, Ian Johnston, Dominic Kwiatkowski, Cordelia Langford, John Sillitoe on behalf of the Wellcome Sanger Institute COVID-19 Surveillance Team |
| EPI_ISL_3250100 | Lighthouse Lab in Glasgow | Wellcome Sanger Institute for the COVID-19 Genomics UK (COG-UK) Consortium | Harper VanSteenhouse, Yumi Kasai, David Gray, Carol Clugston, Anna Dominiczak and Alex Alderton, Roberto Amato, Jeffrey Barrett, Sonia Goncalves, Ewan Harrison, David K. Jackson, Ian Johnston, Dominic Kwiatkowski, Cordelia Langford, John Sillitoe on behalf of the Wellcome Sanger Institute COVID-19 Surveillance Team |
| EPI_ISL_3250101, EPI_ISL_3250102, EPI_ISL_3250103, EPI_ISL_3250104, EPI_ISL_3250105, EPI_ISL_3250106, EPI_ISL_3250107, EPI_ISL_3250108, EPI_ISL_3250109, EPI_ISL_3250110 | Lighthouse Laboratory Plymouth | Wellcome Sanger Institute for the COVID-19 Genomics UK (COG-UK) Consortium | Lighthouse Laboratory Plymouth and Alex Alderton, Roberto Amato, Jeffrey Barrett, Sonia Goncalves, Ewan Harrison, David K. Jackson, Ian Johnston, Dominic Kwiatkowski, Cordelia Langford, John Sillitoe on behalf of the Wellcome Sanger Institute COVID-19 Surveillance Team |
| EPI_ISL_3250111 | Lighthouse Lab in Glasgow | Wellcome Sanger Institute for the COVID-19 Genomics UK (COG-UK) Consortium | Harper VanSteenhouse, Yumi Kasai, David Gray, Carol Clugston, Anna Dominiczak and Alex Alderton, Roberto Amato, Jeffrey Barrett, Sonia Goncalves, Ewan Harrison, David K. Jackson, Ian Johnston, Dominic Kwiatkowski, Cordelia Langford, John Sillitoe on behalf of the Wellcome Sanger Institute COVID-19 Surveillance Team |
| EPI_ISL_3250112 | Lighthouse Laboratory Plymouth | Wellcome Sanger Institute for the COVID-19 Genomics UK (COG-UK) Consortium | Lighthouse Laboratory Plymouth and Alex Alderton, Roberto Amato, Jeffrey Barrett, Sonia Goncalves, Ewan Harrison, David K. Jackson, Ian Johnston, Dominic Kwiatkowski, Cordelia Langford, John Sillitoe on behalf of the Wellcome Sanger Institute COVID-19 Surveillance Team |
| EPI_ISL_3250113 | Lighthouse Lab in Glasgow | Wellcome Sanger Institute for the COVID-19 Genomics UK (COG-UK) Consortium | Harper VanSteenhouse, Yumi Kasai, David Gray, Carol Clugston, Anna Dominiczak and Alex Alderton, Roberto Amato, Jeffrey Barrett, Sonia Goncalves, Ewan Harrison, David K. Jackson, Ian Johnston, Dominic Kwiatkowski, Cordelia Langford, John Sillitoe on behalf of the Wellcome Sanger Institute COVID-19 Surveillance Team |
| EPI_ISL_3250114 | Lighthouse Laboratory Plymouth | Wellcome Sanger Institute for the COVID-19 Genomics UK (COG-UK) Consortium | Lighthouse Laboratory Plymouth and Alex Alderton, Roberto Amato, Jeffrey Barrett, Sonia Goncalves, Ewan Harrison, David K. Jackson, Ian Johnston, Dominic Kwiatkowski, Cordelia Langford, John Sillitoe on behalf of the Wellcome Sanger Institute COVID-19 Surveillance Team |
| EPI_ISL_3250115 | Lighthouse Lab in Glasgow | Wellcome Sanger Institute for the COVID-19 Genomics UK (COG-UK) Consortium | Harper VanSteenhouse, Yumi Kasai, David Gray, Carol Clugston, Anna Dominiczak and Alex Alderton, Roberto Amato, Jeffrey Barrett, Sonia Goncalves, Ewan Harrison, David K. Jackson, Ian Johnston, Dominic Kwiatkowski, Cordelia Langford, John Sillitoe on behalf of the Wellcome Sanger Institute COVID-19 Surveillance Team |
| EPI_ISL_3250116 | Lighthouse Laboratory Plymouth | Wellcome Sanger Institute for the COVID-19 Genomics UK (COG-UK) Consortium | Lighthouse Laboratory Plymouth and Alex Alderton, Roberto Amato, Jeffrey Barrett, Sonia Goncalves, Ewan Harrison, David K. Jackson, Ian Johnston, Dominic Kwiatkowski, Cordelia Langford, John Sillitoe on behalf of the Wellcome Sanger Institute COVID-19 Surveillance Team |
| EPI_ISL_3250117 | Lighthouse Lab in Glasgow | Wellcome Sanger Institute for the COVID-19 Genomics UK (COG-UK) Consortium | Harper VanSteenhouse, Yumi Kasai, David Gray, Carol Clugston, Anna Dominiczak and Alex Alderton, Roberto Amato, Jeffrey Barrett, Sonia Goncalves, Ewan Harrison, David K. Jackson, Ian Johnston, Dominic Kwiatkowski, Cordelia Langford, John Sillitoe on behalf of the Wellcome Sanger Institute COVID-19 Surveillance Team |
| EPI_ISL_3250118 | Lighthouse Laboratory Plymouth | Wellcome Sanger Institute for the COVID-19 Genomics UK (COG-UK) Consortium | Lighthouse Laboratory Plymouth and Alex Alderton, Roberto Amato, Jeffrey Barrett, Sonia Goncalves, Ewan Harrison, David K. Jackson, Ian Johnston, Dominic Kwiatkowski, Cordelia Langford, John Sillitoe on behalf of the Wellcome Sanger Institute COVID-19 Surveillance Team |
| EPI_ISL_3250119 | Lighthouse Lab in Glasgow | Wellcome Sanger Institute for the COVID-19 Genomics UK (COG-UK) Consortium | Harper VanSteenhouse, Yumi Kasai, David Gray, Carol Clugston, Anna Dominiczak and Alex Alderton, Roberto Amato, Jeffrey Barrett, Sonia Goncalves, Ewan Harrison, David K. Jackson, Ian Johnston, Dominic Kwiatkowski, Cordelia Langford, John Sillitoe on behalf of the Wellcome Sanger Institute COVID-19 Surveillance Team |
| EPI_ISL_3250120, EPI_ISL_3250121, EPI_ISL_3250122 | Lighthouse Laboratory Plymouth | Wellcome Sanger Institute for the COVID-19 Genomics UK (COG-UK) Consortium | Lighthouse Laboratory Plymouth and Alex Alderton, Roberto Amato, Jeffrey Barrett, Sonia Goncalves, Ewan Harrison, David K. Jackson, Ian Johnston, Dominic Kwiatkowski, Cordelia Langford, John Sillitoe on behalf of the Wellcome Sanger Institute COVID-19 Surveillance Team |
| EPI_ISL_3250123, EPI_ISL_3250124 | Lighthouse Lab in Glasgow | Wellcome Sanger Institute for the COVID-19 Genomics UK (COG-UK) Consortium | Harper VanSteenhouse, Yumi Kasai, David Gray, Carol Clugston, Anna Dominiczak and Alex Alderton, Roberto Amato, Jeffrey Barrett, Sonia Goncalves, Ewan Harrison, David K. Jackson, Ian Johnston, Dominic Kwiatkowski, Cordelia Langford, John Sillitoe on behalf of the Wellcome Sanger Institute COVID-19 Surveillance Team |
| EPI_ISL_3250125 | Lighthouse Laboratory Plymouth | Wellcome Sanger Institute for the COVID-19 Genomics UK (COG-UK) Consortium | Lighthouse Laboratory Plymouth and Alex Alderton, Roberto Amato, Jeffrey Barrett, Sonia Goncalves, Ewan Harrison, David K. Jackson, Ian Johnston, Dominic Kwiatkowski, Cordelia Langford, John Sillitoe on behalf of the Wellcome Sanger Institute COVID-19 Surveillance Team |
| EPI_ISL_3250126 | Lighthouse Lab in Glasgow | Wellcome Sanger Institute for the COVID-19 Genomics UK (COG-UK) Consortium | Harper VanSteenhouse, Yumi Kasai, David Gray, Carol Clugston, Anna Dominiczak and Alex Alderton, Roberto Amato, Jeffrey Barrett, Sonia Goncalves, Ewan Harrison, David K. Jackson, Ian Johnston, Dominic Kwiatkowski, Cordelia Langford, John Sillitoe on behalf of the Wellcome Sanger Institute COVID-19 Surveillance Team |
| EPI_ISL_3250127, EPI_ISL_3250128, EPI_ISL_3250129, EPI_ISL_3250130, EPI_ISL_3250131 | Lighthouse Laboratory Plymouth | Wellcome Sanger Institute for the COVID-19 Genomics UK (COG-UK) Consortium | Lighthouse Laboratory Plymouth and Alex Alderton, Roberto Amato, Jeffrey Barrett, Sonia Goncalves, Ewan Harrison, David K. Jackson, Ian Johnston, Dominic Kwiatkowski, Cordelia Langford, John Sillitoe on behalf of the Wellcome Sanger Institute COVID-19 Surveillance Team |
| EPI_ISL_3250132 | Lighthouse Lab in Glasgow | Wellcome Sanger Institute for the COVID-19 Genomics UK (COG-UK) Consortium | Harper VanSteenhouse, Yumi Kasai, David Gray, Carol Clugston, Anna Dominiczak and Alex Alderton, Roberto Amato, Jeffrey Barrett, Sonia Goncalves, Ewan Harrison, David K. Jackson, Ian Johnston, Dominic Kwiatkowski, Cordelia Langford, John Sillitoe on behalf of the Wellcome Sanger Institute COVID-19 Surveillance Team |
| EPI_ISL_3250133, EPI_ISL_3250134 | Lighthouse Laboratory Plymouth | Wellcome Sanger Institute for the COVID-19 Genomics UK (COG-UK) Consortium | Lighthouse Laboratory Plymouth and Alex Alderton, Roberto Amato, Jeffrey Barrett, Sonia Goncalves, Ewan Harrison, David K. Jackson, Ian Johnston, Dominic Kwiatkowski, Cordelia Langford, John Sillitoe on behalf of the Wellcome Sanger Institute COVID-19 Surveillance Team |
| EPI_ISL_3250135 | Lighthouse Lab in Glasgow | Wellcome Sanger Institute for the COVID-19 Genomics UK (COG-UK) Consortium | Harper VanSteenhouse, Yumi Kasai, David Gray, Carol Clugston, Anna Dominiczak and Alex Alderton, Roberto Amato, Jeffrey Barrett, Sonia Goncalves, Ewan Harrison, David K. Jackson, Ian Johnston, Dominic Kwiatkowski, Cordelia Langford, John Sillitoe on behalf of the Wellcome Sanger Institute COVID-19 Surveillance Team |
| EPI_ISL_3250136 | Lighthouse Laboratory Plymouth | Wellcome Sanger Institute for the COVID-19 Genomics UK (COG-UK) Consortium | Lighthouse Laboratory Plymouth and Alex Alderton, Roberto Amato, Jeffrey Barrett, Sonia Goncalves, Ewan Harrison, David K. Jackson, Ian Johnston, Dominic Kwiatkowski, Cordelia Langford, John Sillitoe on behalf of the Wellcome Sanger Institute COVID-19 Surveillance Team |
| EPI_ISL_3250137 | Lighthouse Lab in Glasgow | Wellcome Sanger Institute for the COVID-19 Genomics UK (COG-UK) Consortium | Harper VanSteenhouse, Yumi Kasai, David Gray, Carol Clugston, Anna Dominiczak and Alex Alderton, Roberto Amato, Jeffrey Barrett, Sonia Goncalves, Ewan Harrison, David K. Jackson, Ian Johnston, Dominic Kwiatkowski, Cordelia Langford, John Sillitoe on behalf of the Wellcome Sanger Institute COVID-19 Surveillance Team |
| EPI_ISL_3250138 | Lighthouse Laboratory Plymouth | Wellcome Sanger Institute for the COVID-19 Genomics UK (COG-UK) Consortium | Lighthouse Laboratory Plymouth and Alex Alderton, Roberto Amato, Jeffrey Barrett, Sonia Goncalves, Ewan Harrison, David K. Jackson, Ian Johnston, Dominic Kwiatkowski, Cordelia Langford, John Sillitoe on behalf of the Wellcome Sanger Institute COVID-19 Surveillance Team |
| EPI_ISL_3250139 | Lighthouse Lab in Glasgow | Wellcome Sanger Institute for the COVID-19 Genomics UK (COG-UK) Consortium | Harper VanSteenhouse, Yumi Kasai, David Gray, Carol Clugston, Anna Dominiczak and Alex Alderton, Roberto Amato, Jeffrey Barrett, Sonia Goncalves, Ewan Harrison, David K. Jackson, Ian Johnston, Dominic Kwiatkowski, Cordelia Langford, John Sillitoe on behalf of the Wellcome Sanger Institute COVID-19 Surveillance Team |
| EPI_ISL_3250140, EPI_ISL_3250141, EPI_ISL_3250142, EPI_ISL_3250143, EPI_ISL_3250144, EPI_ISL_3250145, EPI_ISL_3250146, EPI_ISL_3250147, EPI_ISL_3250148, EPI_ISL_3250149, EPI_ISL_3250150 | | | |
| see above | Lighthouse Laboratory Plymouth | Wellcome Sanger Institute for the COVID-19 Genomics UK (COG-UK) Consortium | Lighthouse Laboratory Plymouth and Alex Alderton, Roberto Amato, Jeffrey Barrett, Sonia Goncalves, Ewan Harrison, David K. Jackson, Ian Johnston, Dominic Kwiatkowski, Cordelia Langford, John Sillitoe on behalf of the Wellcome Sanger Institute COVID-19 Surveillance Team |
| EPI_ISL_3250151 | Lighthouse Lab in Glasgow | Wellcome Sanger Institute for the COVID-19 Genomics UK (COG-UK) Consortium | Harper VanSteenhouse, Yumi Kasai, David Gray, Carol Clugston, Anna Dominiczak and Alex Alderton, Roberto Amato, Jeffrey Barrett, Sonia Goncalves, Ewan Harrison, David K. Jackson, Ian Johnston, Dominic Kwiatkowski, Cordelia Langford, John Sillitoe on behalf of the Wellcome Sanger Institute COVID-19 Surveillance Team |
| EPI_ISL_3250152, EPI_ISL_3250153, EPI_ISL_3250154, EPI_ISL_3250155, EPI_ISL_3250156 | Lighthouse Laboratory Plymouth | Wellcome Sanger Institute for the COVID-19 Genomics UK (COG-UK) Consortium | Lighthouse Laboratory Plymouth and Alex Alderton, Roberto Amato, Jeffrey Barrett, Sonia Goncalves, Ewan Harrison, David K. Jackson, Ian Johnston, Dominic Kwiatkowski, Cordelia Langford, John Sillitoe on behalf of the Wellcome Sanger Institute COVID-19 Surveillance Team |
| EPI_ISL_3250157 | Lighthouse Lab in Glasgow | Wellcome Sanger Institute for the COVID-19 Genomics UK (COG-UK) Consortium | Harper VanSteenhouse, Yumi Kasai, David Gray, Carol Clugston, Anna Dominiczak and Alex Alderton, Roberto Amato, Jeffrey Barrett, Sonia Goncalves, Ewan Harrison, David K. Jackson, Ian Johnston, Dominic Kwiatkowski, Cordelia Langford, John Sillitoe on behalf of the Wellcome Sanger Institute COVID-19 Surveillance Team |
| EPI_ISL_3250158, EPI_ISL_3250159 | Lighthouse Laboratory Plymouth | Wellcome Sanger Institute for the COVID-19 Genomics UK (COG-UK) Consortium | Lighthouse Laboratory Plymouth and Alex Alderton, Roberto Amato, Jeffrey Barrett, Sonia Goncalves, Ewan Harrison, David K. Jackson, Ian Johnston, Dominic Kwiatkowski, Cordelia Langford, John Sillitoe on behalf of the Wellcome Sanger Institute COVID-19 Surveillance Team |
| EPI_ISL_3250160 | Lighthouse Lab in Glasgow | Wellcome Sanger Institute for the COVID-19 Genomics UK (COG-UK) Consortium | Harper VanSteenhouse, Yumi Kasai, David Gray, Carol Clugston, Anna Dominiczak and Alex Alderton, Roberto Amato, Jeffrey Barrett, Sonia Goncalves, Ewan Harrison, David K. Jackson, Ian Johnston, Dominic Kwiatkowski, Cordelia Langford, John Sillitoe on behalf of the Wellcome Sanger Institute COVID-19 Surveillance Team |
| EPI_ISL_3250161, EPI_ISL_3250162, EPI_ISL_3250163, EPI_ISL_3250164, EPI_ISL_3250165, EPI_ISL_3250166, EPI_ISL_3250167 | Lighthouse Laboratory Plymouth | Wellcome Sanger Institute for the COVID-19 Genomics UK (COG-UK) Consortium | Lighthouse Laboratory Plymouth and Alex Alderton, Roberto Amato, Jeffrey Barrett, Sonia Goncalves, Ewan Harrison, David K. Jackson, Ian Johnston, Dominic Kwiatkowski, Cordelia Langford, John Sillitoe on behalf of the Wellcome Sanger Institute COVID-19 Surveillance Team |
| EPI_ISL_3250168, EPI_ISL_3250169 | Lighthouse Lab in Glasgow | Wellcome Sanger Institute for the COVID-19 Genomics UK (COG-UK) Consortium | Harper VanSteenhouse, Yumi Kasai, David Gray, Carol Clugston, Anna Dominiczak and Alex Alderton, Roberto Amato, Jeffrey Barrett, Sonia Goncalves, Ewan Harrison, David K. Jackson, Ian Johnston, Dominic Kwiatkowski, Cordelia Langford, John Sillitoe on behalf of the Wellcome Sanger Institute COVID-19 Surveillance Team |
| EPI_ISL_3250170, EPI_ISL_3250171, EPI_ISL_3250172, EPI_ISL_3250173, EPI_ISL_3250174 | Lighthouse Laboratory Plymouth | Wellcome Sanger Institute for the COVID-19 Genomics UK (COG-UK) Consortium | Lighthouse Laboratory Plymouth and Alex Alderton, Roberto Amato, Jeffrey Barrett, Sonia Goncalves, Ewan Harrison, David K. Jackson, Ian Johnston, Dominic Kwiatkowski, Cordelia Langford, John Sillitoe on behalf of the Wellcome Sanger Institute COVID-19 Surveillance Team |

| | | | |
|---|---|---|---|
| EPI_ISL_3250175 | Lighthouse Lab in Glasgow | Wellcome Sanger Institute for the COVID-19 Genomics UK (COG-UK) Consortium | Harper VanSteenhouse, Yumi Kasai, David Gray, Carol Clugston, Anna Dominiczak and Alex Alderton, Roberto Amato, Jeffrey Barrett, Sonia Goncalves, Ewan Harrison, David K. Jackson, Ian Johnston, Dominic Kwiatkowski, Cordelia Langford, John Sillitoe on behalf of the Wellcome Sanger Institute COVID-19 Surveillance Team |
| EPI_ISL_3250176, EPI_ISL_3250177, EPI_ISL_3250178, EPI_ISL_3250179, EPI_ISL_3250180, EPI_ISL_3250181 | Lighthouse Laboratory Plymouth | Wellcome Sanger Institute for the COVID-19 Genomics UK (COG-UK) Consortium | Lighthouse Laboratory Plymouth and Alex Alderton, Roberto Amato, Jeffrey Barrett, Sonia Goncalves, Ewan Harrison, David K. Jackson, Ian Johnston, Dominic Kwiatkowski, Cordelia Langford, John Sillitoe on behalf of the Wellcome Sanger Institute COVID-19 Surveillance Team |
| EPI_ISL_3250182 | Lighthouse Lab in Glasgow | Wellcome Sanger Institute for the COVID-19 Genomics UK (COG-UK) Consortium | Harper VanSteenhouse, Yumi Kasai, David Gray, Carol Clugston, Anna Dominiczak and Alex Alderton, Roberto Amato, Jeffrey Barrett, Sonia Goncalves, Ewan Harrison, David K. Jackson, Ian Johnston, Dominic Kwiatkowski, Cordelia Langford, John Sillitoe on behalf of the Wellcome Sanger Institute COVID-19 Surveillance Team |
| EPI_ISL_3250183, EPI_ISL_3250184, EPI_ISL_3250185, EPI_ISL_3250186, EPI_ISL_3250187, EPI_ISL_3250188, EPI_ISL_3250189, EPI_ISL_3250190, EPI_ISL_3250191, EPI_ISL_3250192 | Lighthouse Laboratory Plymouth | Wellcome Sanger Institute for the COVID-19 Genomics UK (COG-UK) Consortium | Lighthouse Laboratory Plymouth and Alex Alderton, Roberto Amato, Jeffrey Barrett, Sonia Goncalves, Ewan Harrison, David K. Jackson, Ian Johnston, Dominic Kwiatkowski, Cordelia Langford, John Sillitoe on behalf of the Wellcome Sanger Institute COVID-19 Surveillance Team |
| EPI_ISL_3250193, EPI_ISL_3250194 | Lighthouse Lab in Glasgow | Wellcome Sanger Institute for the COVID-19 Genomics UK (COG-UK) Consortium | Harper VanSteenhouse, Yumi Kasai, David Gray, Carol Clugston, Anna Dominiczak and Alex Alderton, Roberto Amato, Jeffrey Barrett, Sonia Goncalves, Ewan Harrison, David K. Jackson, Ian Johnston, Dominic Kwiatkowski, Cordelia Langford, John Sillitoe on behalf of the Wellcome Sanger Institute COVID-19 Surveillance Team |
| EPI_ISL_3250195, EPI_ISL_3250196, EPI_ISL_3250197, EPI_ISL_3250198, EPI_ISL_3250199, EPI_ISL_3250200, EPI_ISL_3250201, EPI_ISL_3250202, EPI_ISL_3250203, EPI_ISL_3250204 | Lighthouse Laboratory Plymouth | Wellcome Sanger Institute for the COVID-19 Genomics UK (COG-UK) Consortium | Lighthouse Laboratory Plymouth and Alex Alderton, Roberto Amato, Jeffrey Barrett, Sonia Goncalves, Ewan Harrison, David K. Jackson, Ian Johnston, Dominic Kwiatkowski, Cordelia Langford, John Sillitoe on behalf of the Wellcome Sanger Institute COVID-19 Surveillance Team |
| EPI_ISL_3250205 | Lighthouse Lab in Glasgow | Wellcome Sanger Institute for the COVID-19 Genomics UK (COG-UK) Consortium | Harper VanSteenhouse, Yumi Kasai, David Gray, Carol Clugston, Anna Dominiczak and Alex Alderton, Roberto Amato, Jeffrey Barrett, Sonia Goncalves, Ewan Harrison, David K. Jackson, Ian Johnston, Dominic Kwiatkowski, Cordelia Langford, John Sillitoe on behalf of the Wellcome Sanger Institute COVID-19 Surveillance Team |
| EPI_ISL_3250206, EPI_ISL_3250207, EPI_ISL_3250208, EPI_ISL_3250209, EPI_ISL_3250210, EPI_ISL_3250211, EPI_ISL_3250212, EPI_ISL_3250213 | Lighthouse Laboratory Plymouth | Wellcome Sanger Institute for the COVID-19 Genomics UK (COG-UK) Consortium | Lighthouse Laboratory Plymouth and Alex Alderton, Roberto Amato, Jeffrey Barrett, Sonia Goncalves, Ewan Harrison, David K. Jackson, Ian Johnston, Dominic Kwiatkowski, Cordelia Langford, John Sillitoe on behalf of the Wellcome Sanger Institute COVID-19 Surveillance Team |
| EPI_ISL_3250214 | Lighthouse Lab in Glasgow | Wellcome Sanger Institute for the COVID-19 Genomics UK (COG-UK) Consortium | Harper VanSteenhouse, Yumi Kasai, David Gray, Carol Clugston, Anna Dominiczak and Alex Alderton, Roberto Amato, Jeffrey Barrett, Sonia Goncalves, Ewan Harrison, David K. Jackson, Ian Johnston, Dominic Kwiatkowski, Cordelia Langford, John Sillitoe on behalf of the Wellcome Sanger Institute COVID-19 Surveillance Team |
| EPI_ISL_3250215 | Lighthouse Laboratory Plymouth | Wellcome Sanger Institute for the COVID-19 Genomics UK (COG-UK) Consortium | Lighthouse Laboratory Plymouth and Alex Alderton, Roberto Amato, Jeffrey Barrett, Sonia Goncalves, Ewan Harrison, David K. Jackson, Ian Johnston, Dominic Kwiatkowski, Cordelia Langford, John Sillitoe on behalf of the Wellcome Sanger Institute COVID-19 Surveillance Team |
| EPI_ISL_3250216, EPI_ISL_3250217 | Lighthouse Lab in Glasgow | Wellcome Sanger Institute for the COVID-19 Genomics UK (COG-UK) Consortium | Harper VanSteenhouse, Yumi Kasai, David Gray, Carol Clugston, Anna Dominiczak and Alex Alderton, Roberto Amato, Jeffrey Barrett, Sonia Goncalves, Ewan Harrison, David K. Jackson, Ian Johnston, Dominic Kwiatkowski, Cordelia Langford, John Sillitoe on behalf of the Wellcome Sanger Institute COVID-19 Surveillance Team |
| EPI_ISL_3250218 | Lighthouse Laboratory Plymouth | Wellcome Sanger Institute for the COVID-19 Genomics UK (COG-UK) Consortium | Lighthouse Laboratory Plymouth and Alex Alderton, Roberto Amato, Jeffrey Barrett, Sonia Goncalves, Ewan Harrison, David K. Jackson, Ian Johnston, Dominic Kwiatkowski, Cordelia Langford, John Sillitoe on behalf of the Wellcome Sanger Institute COVID-19 Surveillance Team |
| EPI_ISL_3250219 | Lighthouse Lab in Glasgow | Wellcome Sanger Institute for the COVID-19 Genomics UK (COG-UK) Consortium | Harper VanSteenhouse, Yumi Kasai, David Gray, Carol Clugston, Anna Dominiczak and Alex Alderton, Roberto Amato, Jeffrey Barrett, Sonia Goncalves, Ewan Harrison, David K. Jackson, Ian Johnston, Dominic Kwiatkowski, Cordelia Langford, John Sillitoe on behalf of the Wellcome Sanger Institute COVID-19 Surveillance Team |
| EPI_ISL_3250220, EPI_ISL_3250221, EPI_ISL_3250222, EPI_ISL_3250223, EPI_ISL_3250224, EPI_ISL_3250225, EPI_ISL_3250226 | Lighthouse Laboratory Plymouth | Wellcome Sanger Institute for the COVID-19 Genomics UK (COG-UK) Consortium | Lighthouse Laboratory Plymouth and Alex Alderton, Roberto Amato, Jeffrey Barrett, Sonia Goncalves, Ewan Harrison, David K. Jackson, Ian Johnston, Dominic Kwiatkowski, Cordelia Langford, John Sillitoe on behalf of the Wellcome Sanger Institute COVID-19 Surveillance Team |
| EPI_ISL_3250227 | Lighthouse Lab in Glasgow | Wellcome Sanger Institute for the COVID-19 Genomics UK (COG-UK) Consortium | Harper VanSteenhouse, Yumi Kasai, David Gray, Carol Clugston, Anna Dominiczak and Alex Alderton, Roberto Amato, Jeffrey Barrett, Sonia Goncalves, Ewan Harrison, David K. Jackson, Ian Johnston, Dominic Kwiatkowski, Cordelia Langford, John Sillitoe on behalf of the Wellcome Sanger Institute COVID-19 Surveillance Team |
| EPI_ISL_3250228 | Lighthouse Laboratory Plymouth | Wellcome Sanger Institute for the COVID-19 Genomics UK (COG-UK) Consortium | Lighthouse Laboratory Plymouth and Alex Alderton, Roberto Amato, Jeffrey Barrett, Sonia Goncalves, Ewan Harrison, David K. Jackson, Ian Johnston, Dominic Kwiatkowski, Cordelia Langford, John Sillitoe on behalf of the Wellcome Sanger Institute COVID-19 Surveillance Team |
| EPI_ISL_3250229, EPI_ISL_3250230 | Lighthouse Lab in Glasgow | Wellcome Sanger Institute for the COVID-19 Genomics UK (COG-UK) Consortium | Harper VanSteenhouse, Yumi Kasai, David Gray, Carol Clugston, Anna Dominiczak and Alex Alderton, Roberto Amato, Jeffrey Barrett, Sonia Goncalves, Ewan Harrison, David K. Jackson, Ian Johnston, Dominic Kwiatkowski, Cordelia Langford, John Sillitoe on behalf of the Wellcome Sanger Institute COVID-19 Surveillance Team |
| EPI_ISL_3250231, EPI_ISL_3250232 | Lighthouse Laboratory Plymouth | Wellcome Sanger Institute for the COVID-19 Genomics UK (COG-UK) Consortium | Lighthouse Laboratory Plymouth and Alex Alderton, Roberto Amato, Jeffrey Barrett, Sonia Goncalves, Ewan Harrison, David K. Jackson, Ian Johnston, Dominic Kwiatkowski, Cordelia Langford, John Sillitoe on behalf of the Wellcome Sanger Institute COVID-19 Surveillance Team |
| EPI_ISL_3250233 | Lighthouse Lab in Glasgow | Wellcome Sanger Institute for the COVID-19 Genomics UK (COG-UK) Consortium | Harper VanSteenhouse, Yumi Kasai, David Gray, Carol Clugston, Anna Dominiczak and Alex Alderton, Roberto Amato, Jeffrey Barrett, Sonia Goncalves, Ewan Harrison, David K. Jackson, Ian Johnston, Dominic Kwiatkowski, Cordelia Langford, John Sillitoe on behalf of the Wellcome Sanger Institute COVID-19 Surveillance Team |
| EPI_ISL_3250234 | Lighthouse Laboratory Plymouth | Wellcome Sanger Institute for the COVID-19 Genomics UK (COG-UK) Consortium | Lighthouse Laboratory Plymouth and Alex Alderton, Roberto Amato, Jeffrey Barrett, Sonia Goncalves, Ewan Harrison, David K. Jackson, Ian Johnston, Dominic Kwiatkowski, Cordelia Langford, John Sillitoe on behalf of the Wellcome Sanger Institute COVID-19 Surveillance Team |
| EPI_ISL_3250235, EPI_ISL_3250236 | Lighthouse Lab in Glasgow | Wellcome Sanger Institute for the COVID-19 Genomics UK (COG-UK) Consortium | Harper VanSteenhouse, Yumi Kasai, David Gray, Carol Clugston, Anna Dominiczak and Alex Alderton, Roberto Amato, Jeffrey Barrett, Sonia Goncalves, Ewan Harrison, David K. Jackson, Ian Johnston, Dominic Kwiatkowski, Cordelia Langford, John Sillitoe on behalf of the Wellcome Sanger Institute COVID-19 Surveillance Team |
| EPI_ISL_3250237 | Lighthouse Laboratory Plymouth | Wellcome Sanger Institute for the COVID-19 Genomics UK (COG-UK) Consortium | Lighthouse Laboratory Plymouth and Alex Alderton, Roberto Amato, Jeffrey Barrett, Sonia Goncalves, Ewan Harrison, David K. Jackson, Ian Johnston, Dominic Kwiatkowski, Cordelia Langford, John Sillitoe on behalf of the Wellcome Sanger Institute COVID-19 Surveillance Team |
| EPI_ISL_3250238 | Lighthouse Lab in Alderley Park | Wellcome Sanger Institute for the COVID-19 Genomics UK (COG-UK) Consortium | Jacquelyn Wynn, Mairead Hyland, The Lighthouse Lab in Alderley Park and Alex Alderton, Roberto Amato, Jeffrey Barrett, Sonia Goncalves, Ewan Harrison, David K. Jackson, Ian Johnston, Dominic Kwiatkowski, Cordelia Langford, John Sillitoe on behalf of the Wellcome Sanger Institute COVID-19 Surveillance Team |
| EPI_ISL_3250239 | Lighthouse Lab in Glasgow | Wellcome Sanger Institute for the COVID-19 Genomics UK (COG-UK) Consortium | Harper VanSteenhouse, Yumi Kasai, David Gray, Carol Clugston, Anna Dominiczak and Alex Alderton, Roberto Amato, Jeffrey Barrett, Sonia Goncalves, Ewan Harrison, David K. Jackson, Ian Johnston, Dominic Kwiatkowski, Cordelia Langford, John Sillitoe on behalf of the Wellcome Sanger Institute COVID-19 Surveillance Team |
| EPI_ISL_3250240 | Lighthouse Laboratory Plymouth | Wellcome Sanger Institute for the COVID-19 Genomics UK (COG-UK) Consortium | Lighthouse Laboratory Plymouth and Alex Alderton, Roberto Amato, Jeffrey Barrett, Sonia Goncalves, Ewan Harrison, David K. Jackson, Ian Johnston, Dominic Kwiatkowski, Cordelia Langford, John Sillitoe on behalf of the Wellcome Sanger Institute COVID-19 Surveillance Team |
| EPI_ISL_3250241 | Lighthouse Lab in Alderley Park | Wellcome Sanger Institute for the COVID-19 Genomics UK (COG-UK) Consortium | Jacquelyn Wynn, Mairead Hyland, The Lighthouse Lab in Alderley Park and Alex Alderton, Roberto Amato, Jeffrey Barrett, Sonia Goncalves, Ewan Harrison, David K. Jackson, Ian Johnston, Dominic Kwiatkowski, Cordelia Langford, John Sillitoe on behalf of the Wellcome Sanger Institute COVID-19 Surveillance Team |
| EPI_ISL_3250242 | Lighthouse Lab in Glasgow | Wellcome Sanger Institute for the COVID-19 Genomics UK (COG-UK) Consortium | Harper VanSteenhouse, Yumi Kasai, David Gray, Carol Clugston, Anna Dominiczak and Alex Alderton, Roberto Amato, Jeffrey Barrett, Sonia Goncalves, Ewan Harrison, David K. Jackson, Ian Johnston, Dominic Kwiatkowski, Cordelia Langford, John Sillitoe on behalf of the Wellcome Sanger Institute COVID-19 Surveillance Team |
| EPI_ISL_3250243 | Lighthouse Lab in Alderley Park | Wellcome Sanger Institute for the COVID-19 Genomics UK (COG-UK) Consortium | Jacquelyn Wynn, Mairead Hyland, The Lighthouse Lab in Alderley Park and Alex Alderton, Roberto Amato, Jeffrey Barrett, Sonia Goncalves, Ewan Harrison, David K. Jackson, Ian Johnston, Dominic Kwiatkowski, Cordelia Langford, John Sillitoe on behalf of the Wellcome Sanger Institute COVID-19 Surveillance Team |
| EPI_ISL_3250244 | Lighthouse Lab in Glasgow | Wellcome Sanger Institute for the COVID-19 Genomics UK (COG-UK) Consortium | Harper VanSteenhouse, Yumi Kasai, David Gray, Carol Clugston, Anna Dominiczak and Alex Alderton, Roberto Amato, Jeffrey Barrett, Sonia Goncalves, Ewan Harrison, David K. Jackson, Ian Johnston, Dominic Kwiatkowski, Cordelia Langford, John Sillitoe on behalf of the Wellcome Sanger Institute COVID-19 Surveillance Team |
| EPI_ISL_3250245 | Lighthouse Laboratory Plymouth | Wellcome Sanger Institute for the COVID-19 Genomics UK (COG-UK) Consortium | Lighthouse Laboratory Plymouth and Alex Alderton, Roberto Amato, Jeffrey Barrett, Sonia Goncalves, Ewan Harrison, David K. Jackson, Ian Johnston, Dominic Kwiatkowski, Cordelia Langford, John Sillitoe on behalf of the Wellcome Sanger Institute COVID-19 Surveillance Team |
| EPI_ISL_3250246 | Lighthouse Lab in Alderley Park | Wellcome Sanger Institute for the COVID-19 Genomics UK (COG-UK) Consortium | Jacquelyn Wynn, Mairead Hyland, The Lighthouse Lab in Alderley Park and Alex Alderton, Roberto Amato, Jeffrey Barrett, Sonia Goncalves, Ewan Harrison, David K. Jackson, Ian Johnston, Dominic Kwiatkowski, Cordelia Langford, John Sillitoe on behalf of the Wellcome Sanger Institute COVID-19 Surveillance Team |
| EPI_ISL_3250247, EPI_ISL_3250248 | Lighthouse Lab in Glasgow | Wellcome Sanger Institute for the COVID-19 Genomics UK (COG-UK) Consortium | Harper VanSteenhouse, Yumi Kasai, David Gray, Carol Clugston, Anna Dominiczak and Alex Alderton, Roberto Amato, Jeffrey Barrett, Sonia Goncalves, Ewan Harrison, David K. Jackson, Ian Johnston, Dominic Kwiatkowski, Cordelia Langford, John Sillitoe on behalf of the Wellcome Sanger Institute COVID-19 Surveillance Team |

| | | | |
|---|---|---|---|
| EPI_ISL_3250249 | Lighthouse Laboratory Plymouth | Wellcome Sanger Institute for the COVID-19 Genomics UK (COG-UK) Consortium | Lighthouse Laboratory Plymouth and Alex Alderton, Roberto Amato, Jeffrey Barrett, Sonia Goncalves, Ewan Harrison, David K. Jackson, Ian Johnston, Dominic Kwiatkowski, Cordelia Langford, John Sillitoe on behalf of the Wellcome Sanger Institute COVID-19 Surveillance Team |
| EPI_ISL_3250250 | Lighthouse Lab in Alderley Park | Wellcome Sanger Institute for the COVID-19 Genomics UK (COG-UK) Consortium | Jacquelyn Wynn, Mairead Hyland, The Lighthouse Lab in Alderley Park and Alex Alderton, Roberto Amato, Jeffrey Barrett, Sonia Goncalves, Ewan Harrison, David K. Jackson, Ian Johnston, Dominic Kwiatkowski, Cordelia Langford, John Sillitoe on behalf of the Wellcome Sanger Institute COVID-19 Surveillance Team |
| EPI_ISL_3250251 | Lighthouse Lab in Glasgow | Wellcome Sanger Institute for the COVID-19 Genomics UK (COG-UK) Consortium | Harper VanSteenhouse, Yumi Kasai, David Gray, Carol Clugston, Anna Dominiczak and Alex Alderton, Roberto Amato, Jeffrey Barrett, Sonia Goncalves, Ewan Harrison, David K. Jackson, Ian Johnston, Dominic Kwiatkowski, Cordelia Langford, John Sillitoe on behalf of the Wellcome Sanger Institute COVID-19 Surveillance Team |
| EPI_ISL_3250252 | Lighthouse Laboratory Plymouth | Wellcome Sanger Institute for the COVID-19 Genomics UK (COG-UK) Consortium | Lighthouse Laboratory Plymouth and Alex Alderton, Roberto Amato, Jeffrey Barrett, Sonia Goncalves, Ewan Harrison, David K. Jackson, Ian Johnston, Dominic Kwiatkowski, Cordelia Langford, John Sillitoe on behalf of the Wellcome Sanger Institute COVID-19 Surveillance Team |
| EPI_ISL_3250253, EPI_ISL_3250254, EPI_ISL_3250255 | Lighthouse Lab in Glasgow | Wellcome Sanger Institute for the COVID-19 Genomics UK (COG-UK) Consortium | Harper VanSteenhouse, Yumi Kasai, David Gray, Carol Clugston, Anna Dominiczak and Alex Alderton, Roberto Amato, Jeffrey Barrett, Sonia Goncalves, Ewan Harrison, David K. Jackson, Ian Johnston, Dominic Kwiatkowski, Cordelia Langford, John Sillitoe on behalf of the Wellcome Sanger Institute COVID-19 Surveillance Team |
| EPI_ISL_3250256 | Lighthouse Laboratory Plymouth | Wellcome Sanger Institute for the COVID-19 Genomics UK (COG-UK) Consortium | Lighthouse Laboratory Plymouth and Alex Alderton, Roberto Amato, Jeffrey Barrett, Sonia Goncalves, Ewan Harrison, David K. Jackson, Ian Johnston, Dominic Kwiatkowski, Cordelia Langford, John Sillitoe on behalf of the Wellcome Sanger Institute COVID-19 Surveillance Team |
| EPI_ISL_3250257, EPI_ISL_3250258 | Lighthouse Lab in Alderley Park | Wellcome Sanger Institute for the COVID-19 Genomics UK (COG-UK) Consortium | Jacquelyn Wynn, Mairead Hyland, The Lighthouse Lab in Alderley Park and Alex Alderton, Roberto Amato, Jeffrey Barrett, Sonia Goncalves, Ewan Harrison, David K. Jackson, Ian Johnston, Dominic Kwiatkowski, Cordelia Langford, John Sillitoe on behalf of the Wellcome Sanger Institute COVID-19 Surveillance Team |
| EPI_ISL_3250259 | Lighthouse Laboratory Plymouth | Wellcome Sanger Institute for the COVID-19 Genomics UK (COG-UK) Consortium | Lighthouse Laboratory Plymouth and Alex Alderton, Roberto Amato, Jeffrey Barrett, Sonia Goncalves, Ewan Harrison, David K. Jackson, Ian Johnston, Dominic Kwiatkowski, Cordelia Langford, John Sillitoe on behalf of the Wellcome Sanger Institute COVID-19 Surveillance Team |
| EPI_ISL_3250260 | Lighthouse Lab in Alderley Park | Wellcome Sanger Institute for the COVID-19 Genomics UK (COG-UK) Consortium | Jacquelyn Wynn, Mairead Hyland, The Lighthouse Lab in Alderley Park and Alex Alderton, Roberto Amato, Jeffrey Barrett, Sonia Goncalves, Ewan Harrison, David K. Jackson, Ian Johnston, Dominic Kwiatkowski, Cordelia Langford, John Sillitoe on behalf of the Wellcome Sanger Institute COVID-19 Surveillance Team |
| EPI_ISL_3250261 | Lighthouse Laboratory Plymouth | Wellcome Sanger Institute for the COVID-19 Genomics UK (COG-UK) Consortium | Lighthouse Laboratory Plymouth and Alex Alderton, Roberto Amato, Jeffrey Barrett, Sonia Goncalves, Ewan Harrison, David K. Jackson, Ian Johnston, Dominic Kwiatkowski, Cordelia Langford, John Sillitoe on behalf of the Wellcome Sanger Institute COVID-19 Surveillance Team |
| EPI_ISL_3250262 | Lighthouse Lab in Alderley Park | Wellcome Sanger Institute for the COVID-19 Genomics UK (COG-UK) Consortium | Jacquelyn Wynn, Mairead Hyland, The Lighthouse Lab in Alderley Park and Alex Alderton, Roberto Amato, Jeffrey Barrett, Sonia Goncalves, Ewan Harrison, David K. Jackson, Ian Johnston, Dominic Kwiatkowski, Cordelia Langford, John Sillitoe on behalf of the Wellcome Sanger Institute COVID-19 Surveillance Team |
| EPI_ISL_3250263 | Lighthouse Laboratory Plymouth | Wellcome Sanger Institute for the COVID-19 Genomics UK (COG-UK) Consortium | Lighthouse Laboratory Plymouth and Alex Alderton, Roberto Amato, Jeffrey Barrett, Sonia Goncalves, Ewan Harrison, David K. Jackson, Ian Johnston, Dominic Kwiatkowski, Cordelia Langford, John Sillitoe on behalf of the Wellcome Sanger Institute COVID-19 Surveillance Team |
| EPI_ISL_3250264 | Lighthouse Lab in Glasgow | Wellcome Sanger Institute for the COVID-19 Genomics UK (COG-UK) Consortium | Harper VanSteenhouse, Yumi Kasai, David Gray, Carol Clugston, Anna Dominiczak and Alex Alderton, Roberto Amato, Jeffrey Barrett, Sonia Goncalves, Ewan Harrison, David K. Jackson, Ian Johnston, Dominic Kwiatkowski, Cordelia Langford, John Sillitoe on behalf of the Wellcome Sanger Institute COVID-19 Surveillance Team |
| EPI_ISL_3250265, EPI_ISL_3250266 | Lighthouse Lab in Alderley Park | Wellcome Sanger Institute for the COVID-19 Genomics UK (COG-UK) Consortium | Jacquelyn Wynn, Mairead Hyland, The Lighthouse Lab in Alderley Park and Alex Alderton, Roberto Amato, Jeffrey Barrett, Sonia Goncalves, Ewan Harrison, David K. Jackson, Ian Johnston, Dominic Kwiatkowski, Cordelia Langford, John Sillitoe on behalf of the Wellcome Sanger Institute COVID-19 Surveillance Team |
| EPI_ISL_3250267, EPI_ISL_3250268 | Lighthouse Lab in Glasgow | Wellcome Sanger Institute for the COVID-19 Genomics UK (COG-UK) Consortium | Harper VanSteenhouse, Yumi Kasai, David Gray, Carol Clugston, Anna Dominiczak and Alex Alderton, Roberto Amato, Jeffrey Barrett, Sonia Goncalves, Ewan Harrison, David K. Jackson, Ian Johnston, Dominic Kwiatkowski, Cordelia Langford, John Sillitoe on behalf of the Wellcome Sanger Institute COVID-19 Surveillance Team |
| EPI_ISL_3250269, EPI_ISL_3250270, EPI_ISL_3250271 | Lighthouse Lab in Alderley Park | Wellcome Sanger Institute for the COVID-19 Genomics UK (COG-UK) Consortium | Jacquelyn Wynn, Mairead Hyland, The Lighthouse Lab in Alderley Park and Alex Alderton, Roberto Amato, Jeffrey Barrett, Sonia Goncalves, Ewan Harrison, David K. Jackson, Ian Johnston, Dominic Kwiatkowski, Cordelia Langford, John Sillitoe on behalf of the Wellcome Sanger Institute COVID-19 Surveillance Team |
| EPI_ISL_3250272 | Lighthouse Lab in Glasgow | Wellcome Sanger Institute for the COVID-19 Genomics UK (COG-UK) Consortium | Harper VanSteenhouse, Yumi Kasai, David Gray, Carol Clugston, Anna Dominiczak and Alex Alderton, Roberto Amato, Jeffrey Barrett, Sonia Goncalves, Ewan Harrison, David K. Jackson, Ian Johnston, Dominic Kwiatkowski, Cordelia Langford, John Sillitoe on behalf of the Wellcome Sanger Institute COVID-19 Surveillance Team |
| EPI_ISL_3250273 | Lighthouse Laboratory Plymouth | Wellcome Sanger Institute for the COVID-19 Genomics UK (COG-UK) Consortium | Lighthouse Laboratory Plymouth and Alex Alderton, Roberto Amato, Jeffrey Barrett, Sonia Goncalves, Ewan Harrison, David K. Jackson, Ian Johnston, Dominic Kwiatkowski, Cordelia Langford, John Sillitoe on behalf of the Wellcome Sanger Institute COVID-19 Surveillance Team |
| EPI_ISL_3250274 | Lighthouse Lab in Glasgow | Wellcome Sanger Institute for the COVID-19 Genomics UK (COG-UK) Consortium | Harper VanSteenhouse, Yumi Kasai, David Gray, Carol Clugston, Anna Dominiczak and Alex Alderton, Roberto Amato, Jeffrey Barrett, Sonia Goncalves, Ewan Harrison, David K. Jackson, Ian Johnston, Dominic Kwiatkowski, Cordelia Langford, John Sillitoe on behalf of the Wellcome Sanger Institute COVID-19 Surveillance Team |
| EPI_ISL_3250275 | Lighthouse Laboratory Plymouth | Wellcome Sanger Institute for the COVID-19 Genomics UK (COG-UK) Consortium | Lighthouse Laboratory Plymouth and Alex Alderton, Roberto Amato, Jeffrey Barrett, Sonia Goncalves, Ewan Harrison, David K. Jackson, Ian Johnston, Dominic Kwiatkowski, Cordelia Langford, John Sillitoe on behalf of the Wellcome Sanger Institute COVID-19 Surveillance Team |
| EPI_ISL_3250276, EPI_ISL_3250277, EPI_ISL_3250278 | Lighthouse Lab in Glasgow | Wellcome Sanger Institute for the COVID-19 Genomics UK (COG-UK) Consortium | Harper VanSteenhouse, Yumi Kasai, David Gray, Carol Clugston, Anna Dominiczak and Alex Alderton, Roberto Amato, Jeffrey Barrett, Sonia Goncalves, Ewan Harrison, David K. Jackson, Ian Johnston, Dominic Kwiatkowski, Cordelia Langford, John Sillitoe on behalf of the Wellcome Sanger Institute COVID-19 Surveillance Team |
| EPI_ISL_3250279, EPI_ISL_3250280 | Lighthouse Lab in Alderley Park | Wellcome Sanger Institute for the COVID-19 Genomics UK (COG-UK) Consortium | Jacquelyn Wynn, Mairead Hyland, The Lighthouse Lab in Alderley Park and Alex Alderton, Roberto Amato, Jeffrey Barrett, Sonia Goncalves, Ewan Harrison, David K. Jackson, Ian Johnston, Dominic Kwiatkowski, Cordelia Langford, John Sillitoe on behalf of the Wellcome Sanger Institute COVID-19 Surveillance Team |
| EPI_ISL_3250281 | Lighthouse Lab in Glasgow | Wellcome Sanger Institute for the COVID-19 Genomics UK (COG-UK) Consortium | Harper VanSteenhouse, Yumi Kasai, David Gray, Carol Clugston, Anna Dominiczak and Alex Alderton, Roberto Amato, Jeffrey Barrett, Sonia Goncalves, Ewan Harrison, David K. Jackson, Ian Johnston, Dominic Kwiatkowski, Cordelia Langford, John Sillitoe on behalf of the Wellcome Sanger Institute COVID-19 Surveillance Team |
| EPI_ISL_3250282 | Lighthouse Laboratory Plymouth | Wellcome Sanger Institute for the COVID-19 Genomics UK (COG-UK) Consortium | Lighthouse Laboratory Plymouth and Alex Alderton, Roberto Amato, Jeffrey Barrett, Sonia Goncalves, Ewan Harrison, David K. Jackson, Ian Johnston, Dominic Kwiatkowski, Cordelia Langford, John Sillitoe on behalf of the Wellcome Sanger Institute COVID-19 Surveillance Team |
| EPI_ISL_3250283, EPI_ISL_3250284 | Lighthouse Lab in Alderley Park | Wellcome Sanger Institute for the COVID-19 Genomics UK (COG-UK) Consortium | Jacquelyn Wynn, Mairead Hyland, The Lighthouse Lab in Alderley Park and Alex Alderton, Roberto Amato, Jeffrey Barrett, Sonia Goncalves, Ewan Harrison, David K. Jackson, Ian Johnston, Dominic Kwiatkowski, Cordelia Langford, John Sillitoe on behalf of the Wellcome Sanger Institute COVID-19 Surveillance Team |
| EPI_ISL_3250285 | Lighthouse Laboratory Plymouth | Wellcome Sanger Institute for the COVID-19 Genomics UK (COG-UK) Consortium | Lighthouse Laboratory Plymouth and Alex Alderton, Roberto Amato, Jeffrey Barrett, Sonia Goncalves, Ewan Harrison, David K. Jackson, Ian Johnston, Dominic Kwiatkowski, Cordelia Langford, John Sillitoe on behalf of the Wellcome Sanger Institute COVID-19 Surveillance Team |
| EPI_ISL_3250286 | Lighthouse Lab in Glasgow | Wellcome Sanger Institute for the COVID-19 Genomics UK (COG-UK) Consortium | Harper VanSteenhouse, Yumi Kasai, David Gray, Carol Clugston, Anna Dominiczak and Alex Alderton, Roberto Amato, Jeffrey Barrett, Sonia Goncalves, Ewan Harrison, David K. Jackson, Ian Johnston, Dominic Kwiatkowski, Cordelia Langford, John Sillitoe on behalf of the Wellcome Sanger Institute COVID-19 Surveillance Team |
| EPI_ISL_3250287 | Lighthouse Lab in Alderley Park | Wellcome Sanger Institute for the COVID-19 Genomics UK (COG-UK) Consortium | Jacquelyn Wynn, Mairead Hyland, The Lighthouse Lab in Alderley Park and Alex Alderton, Roberto Amato, Jeffrey Barrett, Sonia Goncalves, Ewan Harrison, David K. Jackson, Ian Johnston, Dominic Kwiatkowski, Cordelia Langford, John Sillitoe on behalf of the Wellcome Sanger Institute COVID-19 Surveillance Team |
| EPI_ISL_3250288 | Lighthouse Lab in Glasgow | Wellcome Sanger Institute for the COVID-19 Genomics UK (COG-UK) Consortium | Harper VanSteenhouse, Yumi Kasai, David Gray, Carol Clugston, Anna Dominiczak and Alex Alderton, Roberto Amato, Jeffrey Barrett, Sonia Goncalves, Ewan Harrison, David K. Jackson, Ian Johnston, Dominic Kwiatkowski, Cordelia Langford, John Sillitoe on behalf of the Wellcome Sanger Institute COVID-19 Surveillance Team |
| EPI_ISL_3250289, EPI_ISL_3250290 | Lighthouse Laboratory Plymouth | Wellcome Sanger Institute for the COVID-19 Genomics UK (COG-UK) Consortium | Lighthouse Laboratory Plymouth and Alex Alderton, Roberto Amato, Jeffrey Barrett, Sonia Goncalves, Ewan Harrison, David K. Jackson, Ian Johnston, Dominic Kwiatkowski, Cordelia Langford, John Sillitoe on behalf of the Wellcome Sanger Institute COVID-19 Surveillance Team |
| EPI_ISL_3250291, EPI_ISL_3250292, EPI_ISL_3250293, EPI_ISL_3250294, EPI_ISL_3250295 | Lighthouse Lab in Glasgow | Wellcome Sanger Institute for the COVID-19 Genomics UK (COG-UK) Consortium | Harper VanSteenhouse, Yumi Kasai, David Gray, Carol Clugston, Anna Dominiczak and Alex Alderton, Roberto Amato, Jeffrey Barrett, Sonia Goncalves, Ewan Harrison, David K. Jackson, Ian Johnston, Dominic Kwiatkowski, Cordelia Langford, John Sillitoe on behalf of the Wellcome Sanger Institute COVID-19 Surveillance Team |
| EPI_ISL_3250296, EPI_ISL_3250297 | Lighthouse Laboratory Plymouth | Wellcome Sanger Institute for the COVID-19 Genomics UK (COG-UK) Consortium | Lighthouse Laboratory Plymouth and Alex Alderton, Roberto Amato, Jeffrey Barrett, Sonia Goncalves, Ewan Harrison, David K. Jackson, Ian Johnston, Dominic Kwiatkowski, Cordelia Langford, John Sillitoe on behalf of the Wellcome Sanger Institute COVID-19 Surveillance Team |
| EPI_ISL_3250298, EPI_ISL_3250299 | Lighthouse Lab in Glasgow | Wellcome Sanger Institute for the COVID-19 Genomics UK (COG-UK) Consortium | Harper VanSteenhouse, Yumi Kasai, David Gray, Carol Clugston, Anna Dominiczak and Alex Alderton, Roberto Amato, Jeffrey Barrett, Sonia Goncalves, Ewan Harrison, David K. Jackson, Ian Johnston, Dominic Kwiatkowski, Cordelia Langford, John Sillitoe on behalf of the Wellcome Sanger Institute COVID-19 Surveillance Team |
| EPI_ISL_3250300 | Lighthouse Laboratory Plymouth | Wellcome Sanger Institute for the COVID-19 Genomics UK (COG-UK) Consortium | Lighthouse Laboratory Plymouth and Alex Alderton, Roberto Amato, Jeffrey Barrett, Sonia Goncalves, Ewan Harrison, David K. Jackson, Ian Johnston, Dominic Kwiatkowski, Cordelia Langford, John Sillitoe on behalf of the Wellcome Sanger Institute COVID-19 Surveillance Team |
| EPI_ISL_3250301 | Lighthouse Lab in Glasgow | Wellcome Sanger Institute for the COVID-19 Genomics UK (COG-UK) Consortium | Harper VanSteenhouse, Yumi Kasai, David Gray, Carol Clugston, Anna Dominiczak and Alex Alderton, Roberto Amato, Jeffrey Barrett, Sonia Goncalves, Ewan Harrison, David K. Jackson, Ian Johnston, Dominic Kwiatkowski, Cordelia Langford, John Sillitoe on behalf of the Wellcome Sanger Institute COVID-19 Surveillance Team |
| EPI_ISL_3250302, EPI_ISL_3250303 | Lighthouse Laboratory Plymouth | Wellcome Sanger Institute for the COVID-19 Genomics UK (COG-UK) Consortium | Lighthouse Laboratory Plymouth and Alex Alderton, Roberto Amato, Jeffrey Barrett, Sonia Goncalves, Ewan Harrison, David K. Jackson, Ian Johnston, Dominic Kwiatkowski, Cordelia Langford, John Sillitoe on behalf of the Wellcome Sanger Institute COVID-19 Surveillance Team |
| EPI_ISL_3250304, EPI_ISL_3250305, EPI_ISL_3250306, EPI_ISL_3250307 | Lighthouse Lab in Alderley Park | Wellcome Sanger Institute for the COVID-19 Genomics UK (COG-UK) Consortium | Jacquelyn Wynn, Mairead Hyland, The Lighthouse Lab in Alderley Park and Alex Alderton, Roberto Amato, Jeffrey Barrett, Sonia Goncalves, Ewan Harrison, David K. Jackson, Ian Johnston, Dominic Kwiatkowski, Cordelia Langford, John Sillitoe on behalf of the Wellcome Sanger Institute COVID-19 Surveillance Team |
| EPI_ISL_3250308, EPI_ISL_3250309 | Lighthouse Laboratory Plymouth | Wellcome Sanger Institute for the COVID-19 Genomics UK (COG-UK) Consortium | Lighthouse Laboratory Plymouth and Alex Alderton, Roberto Amato, Jeffrey Barrett, Sonia Goncalves, Ewan Harrison, David K. Jackson, Ian Johnston, Dominic Kwiatkowski, Cordelia Langford, John Sillitoe on behalf of the Wellcome Sanger Institute COVID-19 Surveillance Team |
| EPI_ISL_3250310, EPI_ISL_3250311 | Lighthouse Lab in Alderley Park | Wellcome Sanger Institute for the COVID-19 Genomics UK (COG-UK) Consortium | Jacquelyn Wynn, Mairead Hyland, The Lighthouse Lab in Alderley Park and Alex Alderton, Roberto Amato, Jeffrey Barrett, Sonia Goncalves, Ewan Harrison, David K. Jackson, Ian Johnston, Dominic Kwiatkowski, Cordelia Langford, John Sillitoe on behalf of the Wellcome Sanger Institute COVID-19 Surveillance Team |
| EPI_ISL_3250312, EPI_ISL_3250313 | Lighthouse Laboratory Plymouth | Wellcome Sanger Institute for the COVID-19 Genomics UK (COG-UK) Consortium | Lighthouse Laboratory Plymouth and Alex Alderton, Roberto Amato, Jeffrey Barrett, Sonia Goncalves, Ewan Harrison, David K. Jackson, Ian Johnston, Dominic Kwiatkowski, Cordelia Langford, John Sillitoe on behalf of the Wellcome Sanger Institute COVID-19 Surveillance Team |
| EPI_ISL_3250314 | Lighthouse Lab in Alderley Park | Wellcome Sanger Institute for the COVID-19 Genomics UK (COG-UK) Consortium | Jacquelyn Wynn, Mairead Hyland, The Lighthouse Lab in Alderley Park and Alex Alderton, Roberto Amato, Jeffrey Barrett, Sonia Goncalves, Ewan Harrison, David K. Jackson, Ian Johnston, Dominic Kwiatkowski, Cordelia Langford, John Sillitoe on behalf of the Wellcome Sanger Institute COVID-19 Surveillance Team |
| EPI_ISL_3250315, EPI_ISL_3250316 | Lighthouse Laboratory Plymouth | Wellcome Sanger Institute for the COVID-19 Genomics UK (COG-UK) Consortium | Lighthouse Laboratory Plymouth and Alex Alderton, Roberto Amato, Jeffrey Barrett, Sonia Goncalves, Ewan Harrison, David K. Jackson, Ian Johnston, Dominic Kwiatkowski, Cordelia Langford, John Sillitoe on behalf of the Wellcome Sanger Institute COVID-19 Surveillance Team |

| | | | |
|---|---|---|---|
| EPI_ISL_3250317 | Lighthouse Lab in Alderley Park | Wellcome Sanger Institute for the COVID-19 Genomics UK (COG-UK) Consortium | Jacquelyn Wynn, Mairead Hyland, The Lighthouse Lab in Alderley Park and Alex Alderton, Roberto Amato, Jeffrey Barrett, Sonia Goncalves, Ewan Harrison, David K. Jackson, Ian Johnston, Dominic Kwiatkowski, Cordelia Langford, John Sillitoe on behalf of the Wellcome Sanger Institute COVID-19 Surveillance Team |
| EPI_ISL_3250318, EPI_ISL_3250319, EPI_ISL_3250320 | Lighthouse Lab in Glasgow | Wellcome Sanger Institute for the COVID-19 Genomics UK (COG-UK) Consortium | Harper VanSteenhouse, Yumi Kasai, David Gray, Carol Clugston, Anna Dominiczak and Alex Alderton, Roberto Amato, Jeffrey Barrett, Sonia Goncalves, Ewan Harrison, David K. Jackson, Ian Johnston, Dominic Kwiatkowski, Cordelia Langford, John Sillitoe on behalf of the Wellcome Sanger Institute COVID-19 Surveillance Team |
| EPI_ISL_3250321, EPI_ISL_3250322 | Lighthouse Lab in Alderley Park | Wellcome Sanger Institute for the COVID-19 Genomics UK (COG-UK) Consortium | Jacquelyn Wynn, Mairead Hyland, The Lighthouse Lab in Alderley Park and Alex Alderton, Roberto Amato, Jeffrey Barrett, Sonia Goncalves, Ewan Harrison, David K. Jackson, Ian Johnston, Dominic Kwiatkowski, Cordelia Langford, John Sillitoe on behalf of the Wellcome Sanger Institute COVID-19 Surveillance Team |
| EPI_ISL_3250323 | Lighthouse Laboratory Plymouth | Wellcome Sanger Institute for the COVID-19 Genomics UK (COG-UK) Consortium | Lighthouse Laboratory Plymouth and Alex Alderton, Roberto Amato, Jeffrey Barrett, Sonia Goncalves, Ewan Harrison, David K. Jackson, Ian Johnston, Dominic Kwiatkowski, Cordelia Langford, John Sillitoe on behalf of the Wellcome Sanger Institute COVID-19 Surveillance Team |
| EPI_ISL_3250324 | Lighthouse Lab in Alderley Park | Wellcome Sanger Institute for the COVID-19 Genomics UK (COG-UK) Consortium | Jacquelyn Wynn, Mairead Hyland, The Lighthouse Lab in Alderley Park and Alex Alderton, Roberto Amato, Jeffrey Barrett, Sonia Goncalves, Ewan Harrison, David K. Jackson, Ian Johnston, Dominic Kwiatkowski, Cordelia Langford, John Sillitoe on behalf of the Wellcome Sanger Institute COVID-19 Surveillance Team |
| EPI_ISL_3250325 | Lighthouse Lab in Glasgow | Wellcome Sanger Institute for the COVID-19 Genomics UK (COG-UK) Consortium | Harper VanSteenhouse, Yumi Kasai, David Gray, Carol Clugston, Anna Dominiczak and Alex Alderton, Roberto Amato, Jeffrey Barrett, Sonia Goncalves, Ewan Harrison, David K. Jackson, Ian Johnston, Dominic Kwiatkowski, Cordelia Langford, John Sillitoe on behalf of the Wellcome Sanger Institute COVID-19 Surveillance Team |
| EPI_ISL_3250326 | Lighthouse Laboratory Plymouth | Wellcome Sanger Institute for the COVID-19 Genomics UK (COG-UK) Consortium | Lighthouse Laboratory Plymouth and Alex Alderton, Roberto Amato, Jeffrey Barrett, Sonia Goncalves, Ewan Harrison, David K. Jackson, Ian Johnston, Dominic Kwiatkowski, Cordelia Langford, John Sillitoe on behalf of the Wellcome Sanger Institute COVID-19 Surveillance Team |
| EPI_ISL_3250327, EPI_ISL_3250328 | Lighthouse Lab in Glasgow | Wellcome Sanger Institute for the COVID-19 Genomics UK (COG-UK) Consortium | Harper VanSteenhouse, Yumi Kasai, David Gray, Carol Clugston, Anna Dominiczak and Alex Alderton, Roberto Amato, Jeffrey Barrett, Sonia Goncalves, Ewan Harrison, David K. Jackson, Ian Johnston, Dominic Kwiatkowski, Cordelia Langford, John Sillitoe on behalf of the Wellcome Sanger Institute COVID-19 Surveillance Team |
| EPI_ISL_3250329 | Lighthouse Lab in Alderley Park | Wellcome Sanger Institute for the COVID-19 Genomics UK (COG-UK) Consortium | Jacquelyn Wynn, Mairead Hyland, The Lighthouse Lab in Alderley Park and Alex Alderton, Roberto Amato, Jeffrey Barrett, Sonia Goncalves, Ewan Harrison, David K. Jackson, Ian Johnston, Dominic Kwiatkowski, Cordelia Langford, John Sillitoe on behalf of the Wellcome Sanger Institute COVID-19 Surveillance Team |
| EPI_ISL_3250330, EPI_ISL_3250331, EPI_ISL_3250332, EPI_ISL_3250333 | Lighthouse Lab in Glasgow | Wellcome Sanger Institute for the COVID-19 Genomics UK (COG-UK) Consortium | Harper VanSteenhouse, Yumi Kasai, David Gray, Carol Clugston, Anna Dominiczak and Alex Alderton, Roberto Amato, Jeffrey Barrett, Sonia Goncalves, Ewan Harrison, David K. Jackson, Ian Johnston, Dominic Kwiatkowski, Cordelia Langford, John Sillitoe on behalf of the Wellcome Sanger Institute COVID-19 Surveillance Team |
| EPI_ISL_3250334, EPI_ISL_3250335, EPI_ISL_3250336, EPI_ISL_3250337, EPI_ISL_3250338 | Lighthouse Laboratory Plymouth | Wellcome Sanger Institute for the COVID-19 Genomics UK (COG-UK) Consortium | Lighthouse Laboratory Plymouth and Alex Alderton, Roberto Amato, Jeffrey Barrett, Sonia Goncalves, Ewan Harrison, David K. Jackson, Ian Johnston, Dominic Kwiatkowski, Cordelia Langford, John Sillitoe on behalf of the Wellcome Sanger Institute COVID-19 Surveillance Team |
| EPI_ISL_3250339, EPI_ISL_3250340 | Lighthouse Lab in Glasgow | Wellcome Sanger Institute for the COVID-19 Genomics UK (COG-UK) Consortium | Harper VanSteenhouse, Yumi Kasai, David Gray, Carol Clugston, Anna Dominiczak and Alex Alderton, Roberto Amato, Jeffrey Barrett, Sonia Goncalves, Ewan Harrison, David K. Jackson, Ian Johnston, Dominic Kwiatkowski, Cordelia Langford, John Sillitoe on behalf of the Wellcome Sanger Institute COVID-19 Surveillance Team |
| EPI_ISL_3250341 | Lighthouse Lab in Alderley Park | Wellcome Sanger Institute for the COVID-19 Genomics UK (COG-UK) Consortium | Jacquelyn Wynn, Mairead Hyland, The Lighthouse Lab in Alderley Park and Alex Alderton, Roberto Amato, Jeffrey Barrett, Sonia Goncalves, Ewan Harrison, David K. Jackson, Ian Johnston, Dominic Kwiatkowski, Cordelia Langford, John Sillitoe on behalf of the Wellcome Sanger Institute COVID-19 Surveillance Team |
| EPI_ISL_3250342 | Lighthouse Lab in Glasgow | Wellcome Sanger Institute for the COVID-19 Genomics UK (COG-UK) Consortium | Harper VanSteenhouse, Yumi Kasai, David Gray, Carol Clugston, Anna Dominiczak and Alex Alderton, Roberto Amato, Jeffrey Barrett, Sonia Goncalves, Ewan Harrison, David K. Jackson, Ian Johnston, Dominic Kwiatkowski, Cordelia Langford, John Sillitoe on behalf of the Wellcome Sanger Institute COVID-19 Surveillance Team |
| EPI_ISL_3250343 | Lighthouse Lab in Alderley Park | Wellcome Sanger Institute for the COVID-19 Genomics UK (COG-UK) Consortium | Jacquelyn Wynn, Mairead Hyland, The Lighthouse Lab in Alderley Park and Alex Alderton, Roberto Amato, Jeffrey Barrett, Sonia Goncalves, Ewan Harrison, David K. Jackson, Ian Johnston, Dominic Kwiatkowski, Cordelia Langford, John Sillitoe on behalf of the Wellcome Sanger Institute COVID-19 Surveillance Team |
| EPI_ISL_3250344 | Lighthouse Lab in Glasgow | Wellcome Sanger Institute for the COVID-19 Genomics UK (COG-UK) Consortium | Harper VanSteenhouse, Yumi Kasai, David Gray, Carol Clugston, Anna Dominiczak and Alex Alderton, Roberto Amato, Jeffrey Barrett, Sonia Goncalves, Ewan Harrison, David K. Jackson, Ian Johnston, Dominic Kwiatkowski, Cordelia Langford, John Sillitoe on behalf of the Wellcome Sanger Institute COVID-19 Surveillance Team |
| EPI_ISL_3250345 | Lighthouse Lab in Alderley Park | Wellcome Sanger Institute for the COVID-19 Genomics UK (COG-UK) Consortium | Jacquelyn Wynn, Mairead Hyland, The Lighthouse Lab in Alderley Park and Alex Alderton, Roberto Amato, Jeffrey Barrett, Sonia Goncalves, Ewan Harrison, David K. Jackson, Ian Johnston, Dominic Kwiatkowski, Cordelia Langford, John Sillitoe on behalf of the Wellcome Sanger Institute COVID-19 Surveillance Team |
| EPI_ISL_3250346, EPI_ISL_3250347 | Lighthouse Lab in Glasgow | Wellcome Sanger Institute for the COVID-19 Genomics UK (COG-UK) Consortium | Harper VanSteenhouse, Yumi Kasai, David Gray, Carol Clugston, Anna Dominiczak and Alex Alderton, Roberto Amato, Jeffrey Barrett, Sonia Goncalves, Ewan Harrison, David K. Jackson, Ian Johnston, Dominic Kwiatkowski, Cordelia Langford, John Sillitoe on behalf of the Wellcome Sanger Institute COVID-19 Surveillance Team |
| EPI_ISL_3250348 | Lighthouse Laboratory Plymouth | Wellcome Sanger Institute for the COVID-19 Genomics UK (COG-UK) Consortium | Lighthouse Laboratory Plymouth and Alex Alderton, Roberto Amato, Jeffrey Barrett, Sonia Goncalves, Ewan Harrison, David K. Jackson, Ian Johnston, Dominic Kwiatkowski, Cordelia Langford, John Sillitoe on behalf of the Wellcome Sanger Institute COVID-19 Surveillance Team |
| EPI_ISL_3250349, EPI_ISL_3250350 | Lighthouse Lab in Alderley Park | Wellcome Sanger Institute for the COVID-19 Genomics UK (COG-UK) Consortium | Jacquelyn Wynn, Mairead Hyland, The Lighthouse Lab in Alderley Park and Alex Alderton, Roberto Amato, Jeffrey Barrett, Sonia Goncalves, Ewan Harrison, David K. Jackson, Ian Johnston, Dominic Kwiatkowski, Cordelia Langford, John Sillitoe on behalf of the Wellcome Sanger Institute COVID-19 Surveillance Team |
| EPI_ISL_3250351, EPI_ISL_3250352, EPI_ISL_3250353, EPI_ISL_3250354, EPI_ISL_3250355 | Lighthouse Laboratory Plymouth | Wellcome Sanger Institute for the COVID-19 Genomics UK (COG-UK) Consortium | Lighthouse Laboratory Plymouth and Alex Alderton, Roberto Amato, Jeffrey Barrett, Sonia Goncalves, Ewan Harrison, David K. Jackson, Ian Johnston, Dominic Kwiatkowski, Cordelia Langford, John Sillitoe on behalf of the Wellcome Sanger Institute COVID-19 Surveillance Team |
| EPI_ISL_3250356, EPI_ISL_3250357 | Lighthouse Lab in Glasgow | Wellcome Sanger Institute for the COVID-19 Genomics UK (COG-UK) Consortium | Harper VanSteenhouse, Yumi Kasai, David Gray, Carol Clugston, Anna Dominiczak and Alex Alderton, Roberto Amato, Jeffrey Barrett, Sonia Goncalves, Ewan Harrison, David K. Jackson, Ian Johnston, Dominic Kwiatkowski, Cordelia Langford, John Sillitoe on behalf of the Wellcome Sanger Institute COVID-19 Surveillance Team |
| EPI_ISL_3250358 | Lighthouse Laboratory Plymouth | Wellcome Sanger Institute for the COVID-19 Genomics UK (COG-UK) Consortium | Lighthouse Laboratory Plymouth and Alex Alderton, Roberto Amato, Jeffrey Barrett, Sonia Goncalves, Ewan Harrison, David K. Jackson, Ian Johnston, Dominic Kwiatkowski, Cordelia Langford, John Sillitoe on behalf of the Wellcome Sanger Institute COVID-19 Surveillance Team |
| EPI_ISL_3250359, EPI_ISL_3250360, EPI_ISL_3250361 | Lighthouse Lab in Alderley Park | Wellcome Sanger Institute for the COVID-19 Genomics UK (COG-UK) Consortium | Jacquelyn Wynn, Mairead Hyland, The Lighthouse Lab in Alderley Park and Alex Alderton, Roberto Amato, Jeffrey Barrett, Sonia Goncalves, Ewan Harrison, David K. Jackson, Ian Johnston, Dominic Kwiatkowski, Cordelia Langford, John Sillitoe on behalf of the Wellcome Sanger Institute COVID-19 Surveillance Team |
| EPI_ISL_3250362 | Lighthouse Laboratory Plymouth | Wellcome Sanger Institute for the COVID-19 Genomics UK (COG-UK) Consortium | Lighthouse Laboratory Plymouth and Alex Alderton, Roberto Amato, Jeffrey Barrett, Sonia Goncalves, Ewan Harrison, David K. Jackson, Ian Johnston, Dominic Kwiatkowski, Cordelia Langford, John Sillitoe on behalf of the Wellcome Sanger Institute COVID-19 Surveillance Team |
| EPI_ISL_3250363 | Lighthouse Lab in Glasgow | Wellcome Sanger Institute for the COVID-19 Genomics UK (COG-UK) Consortium | Harper VanSteenhouse, Yumi Kasai, David Gray, Carol Clugston, Anna Dominiczak and Alex Alderton, Roberto Amato, Jeffrey Barrett, Sonia Goncalves, Ewan Harrison, David K. Jackson, Ian Johnston, Dominic Kwiatkowski, Cordelia Langford, John Sillitoe on behalf of the Wellcome Sanger Institute COVID-19 Surveillance Team |
| EPI_ISL_3250364 | Lighthouse Lab in Alderley Park | Wellcome Sanger Institute for the COVID-19 Genomics UK (COG-UK) Consortium | Jacquelyn Wynn, Mairead Hyland, The Lighthouse Lab in Alderley Park and Alex Alderton, Roberto Amato, Jeffrey Barrett, Sonia Goncalves, Ewan Harrison, David K. Jackson, Ian Johnston, Dominic Kwiatkowski, Cordelia Langford, John Sillitoe on behalf of the Wellcome Sanger Institute COVID-19 Surveillance Team |
| EPI_ISL_3250365, EPI_ISL_3250366 | Lighthouse Lab in Glasgow | Wellcome Sanger Institute for the COVID-19 Genomics UK (COG-UK) Consortium | Harper VanSteenhouse, Yumi Kasai, David Gray, Carol Clugston, Anna Dominiczak and Alex Alderton, Roberto Amato, Jeffrey Barrett, Sonia Goncalves, Ewan Harrison, David K. Jackson, Ian Johnston, Dominic Kwiatkowski, Cordelia Langford, John Sillitoe on behalf of the Wellcome Sanger Institute COVID-19 Surveillance Team |
| EPI_ISL_3250367, EPI_ISL_3250368 | Lighthouse Lab in Alderley Park | Wellcome Sanger Institute for the COVID-19 Genomics UK (COG-UK) Consortium | Jacquelyn Wynn, Mairead Hyland, The Lighthouse Lab in Alderley Park and Alex Alderton, Roberto Amato, Jeffrey Barrett, Sonia Goncalves, Ewan Harrison, David K. Jackson, Ian Johnston, Dominic Kwiatkowski, Cordelia Langford, John Sillitoe on behalf of the Wellcome Sanger Institute COVID-19 Surveillance Team |
| EPI_ISL_3250369, EPI_ISL_3250370, EPI_ISL_3250371 | Lighthouse Lab in Glasgow | Wellcome Sanger Institute for the COVID-19 Genomics UK (COG-UK) Consortium | Harper VanSteenhouse, Yumi Kasai, David Gray, Carol Clugston, Anna Dominiczak and Alex Alderton, Roberto Amato, Jeffrey Barrett, Sonia Goncalves, Ewan Harrison, David K. Jackson, Ian Johnston, Dominic Kwiatkowski, Cordelia Langford, John Sillitoe on behalf of the Wellcome Sanger Institute COVID-19 Surveillance Team |
| EPI_ISL_3250372 | Lighthouse Laboratory Plymouth | Wellcome Sanger Institute for the COVID-19 Genomics UK (COG-UK) Consortium | Lighthouse Laboratory Plymouth and Alex Alderton, Roberto Amato, Jeffrey Barrett, Sonia Goncalves, Ewan Harrison, David K. Jackson, Ian Johnston, Dominic Kwiatkowski, Cordelia Langford, John Sillitoe on behalf of the Wellcome Sanger Institute COVID-19 Surveillance Team |
| EPI_ISL_3250373 | Lighthouse Lab in Alderley Park | Wellcome Sanger Institute for the COVID-19 Genomics UK (COG-UK) Consortium | Jacquelyn Wynn, Mairead Hyland, The Lighthouse Lab in Alderley Park and Alex Alderton, Roberto Amato, Jeffrey Barrett, Sonia Goncalves, Ewan Harrison, David K. Jackson, Ian Johnston, Dominic Kwiatkowski, Cordelia Langford, John Sillitoe on behalf of the Wellcome Sanger Institute COVID-19 Surveillance Team |
| EPI_ISL_3250374, EPI_ISL_3250375 | Lighthouse Lab in Glasgow | Wellcome Sanger Institute for the COVID-19 Genomics UK (COG-UK) Consortium | Harper VanSteenhouse, Yumi Kasai, David Gray, Carol Clugston, Anna Dominiczak and Alex Alderton, Roberto Amato, Jeffrey Barrett, Sonia Goncalves, Ewan Harrison, David K. Jackson, Ian Johnston, Dominic Kwiatkowski, Cordelia Langford, John Sillitoe on behalf of the Wellcome Sanger Institute COVID-19 Surveillance Team |
| EPI_ISL_3250376 | Lighthouse Laboratory Plymouth | Wellcome Sanger Institute for the COVID-19 Genomics UK (COG-UK) Consortium | Lighthouse Laboratory Plymouth and Alex Alderton, Roberto Amato, Jeffrey Barrett, Sonia Goncalves, Ewan Harrison, David K. Jackson, Ian Johnston, Dominic Kwiatkowski, Cordelia Langford, John Sillitoe on behalf of the Wellcome Sanger Institute COVID-19 Surveillance Team |
| EPI_ISL_3250377, EPI_ISL_3250378, EPI_ISL_3250379 | Lighthouse Lab in Glasgow | Wellcome Sanger Institute for the COVID-19 Genomics UK (COG-UK) Consortium | Harper VanSteenhouse, Yumi Kasai, David Gray, Carol Clugston, Anna Dominiczak and Alex Alderton, Roberto Amato, Jeffrey Barrett, Sonia Goncalves, Ewan Harrison, David K. Jackson, Ian Johnston, Dominic Kwiatkowski, Cordelia Langford, John Sillitoe on behalf of the Wellcome Sanger Institute COVID-19 Surveillance Team |
| EPI_ISL_3250380 | Lighthouse Lab in Alderley Park | Wellcome Sanger Institute for the COVID-19 Genomics UK (COG-UK) Consortium | Jacquelyn Wynn, Mairead Hyland, The Lighthouse Lab in Alderley Park and Alex Alderton, Roberto Amato, Jeffrey Barrett, Sonia Goncalves, Ewan Harrison, David K. Jackson, Ian Johnston, Dominic Kwiatkowski, Cordelia Langford, John Sillitoe on behalf of the Wellcome Sanger Institute COVID-19 Surveillance Team |
| EPI_ISL_3250381, EPI_ISL_3250382, EPI_ISL_3250383 | Lighthouse Lab in Glasgow | Wellcome Sanger Institute for the COVID-19 Genomics UK (COG-UK) Consortium | Harper VanSteenhouse, Yumi Kasai, David Gray, Carol Clugston, Anna Dominiczak and Alex Alderton, Roberto Amato, Jeffrey Barrett, Sonia Goncalves, Ewan Harrison, David K. Jackson, Ian Johnston, Dominic Kwiatkowski, Cordelia Langford, John Sillitoe on behalf of the Wellcome Sanger Institute COVID-19 Surveillance Team |
| EPI_ISL_3250384, EPI_ISL_3250385 | Lighthouse Laboratory Plymouth | Wellcome Sanger Institute for the COVID-19 Genomics UK (COG-UK) Consortium | Lighthouse Laboratory Plymouth and Alex Alderton, Roberto Amato, Jeffrey Barrett, Sonia Goncalves, Ewan Harrison, David K. Jackson, Ian Johnston, Dominic Kwiatkowski, Cordelia Langford, John Sillitoe on behalf of the Wellcome Sanger Institute COVID-19 Surveillance Team |
| EPI_ISL_3250386, EPI_ISL_3250387, EPI_ISL_3250388, EPI_ISL_3250389, | Lighthouse Lab in Glasgow | Wellcome Sanger Institute for the COVID-19 Genomics UK (COG-UK) Consortium | Harper VanSteenhouse, Yumi Kasai, David Gray, Carol Clugston, Anna Dominiczak and Alex Alderton, Roberto Amato, Jeffrey Barrett, Sonia Goncalves, Ewan Harrison, David K. Jackson, Ian Johnston, Dominic Kwiatkowski, Cordelia Langford, John Sillitoe on behalf of the Wellcome Sanger Institute COVID-19 Surveillance Team |

| | | | |
|---|---|---|---|
| EPI_ISL_3250390, EPI_ISL_3250391 | | | |
| EPI_ISL_3250392 | AZ Klina | AZ Klina | Carl Vael - Lynsey Berckmans |
| EPI_ISL_3250393 | Lighthouse Lab in Glasgow | Wellcome Sanger Institute for the COVID-19 Genomics UK (COG-UK) Consortium | Harper VanSteenhouse, Yumi Kasai, David Gray, Carol Clugston, Anna Dominiczak and Alex Alderton, Roberto Amato, Jeffrey Barrett, Sonia Goncalves, Ewan Harrison, David K. Jackson, Ian Johnston, Dominic Kwiatkowski, Cordelia Langford, John Sillitoe on behalf of the Wellcome Sanger Institute COVID-19 Surveillance Team |
| EPI_ISL_3250394 | Lighthouse Lab in Alderley Park | Wellcome Sanger Institute for the COVID-19 Genomics UK (COG-UK) Consortium | Jacquelyn Wynn, Mairead Hyland, The Lighthouse Lab in Alderley Park and Alex Alderton, Roberto Amato, Jeffrey Barrett, Sonia Goncalves, Ewan Harrison, David K. Jackson, Ian Johnston, Dominic Kwiatkowski, Cordelia Langford, John Sillitoe on behalf of the Wellcome Sanger Institute COVID-19 Surveillance Team |
| EPI_ISL_3250395 | Lighthouse Lab in Glasgow | Wellcome Sanger Institute for the COVID-19 Genomics UK (COG-UK) Consortium | Harper VanSteenhouse, Yumi Kasai, David Gray, Carol Clugston, Anna Dominiczak and Alex Alderton, Roberto Amato, Jeffrey Barrett, Sonia Goncalves, Ewan Harrison, David K. Jackson, Ian Johnston, Dominic Kwiatkowski, Cordelia Langford, John Sillitoe on behalf of the Wellcome Sanger Institute COVID-19 Surveillance Team |
| EPI_ISL_3250396 | Lighthouse Lab in Alderley Park | Wellcome Sanger Institute for the COVID-19 Genomics UK (COG-UK) Consortium | Jacquelyn Wynn, Mairead Hyland, The Lighthouse Lab in Alderley Park and Alex Alderton, Roberto Amato, Jeffrey Barrett, Sonia Goncalves, Ewan Harrison, David K. Jackson, Ian Johnston, Dominic Kwiatkowski, Cordelia Langford, John Sillitoe on behalf of the Wellcome Sanger Institute COVID-19 Surveillance Team |
| EPI_ISL_3250397, EPI_ISL_3250398 | Lighthouse Laboratory Plymouth | Wellcome Sanger Institute for the COVID-19 Genomics UK (COG-UK) Consortium | Lighthouse Laboratory Plymouth and Alex Alderton, Roberto Amato, Jeffrey Barrett, Sonia Goncalves, Ewan Harrison, David K. Jackson, Ian Johnston, Dominic Kwiatkowski, Cordelia Langford, John Sillitoe on behalf of the Wellcome Sanger Institute COVID-19 Surveillance Team |
| EPI_ISL_3250399 | Lighthouse Lab in Alderley Park | Wellcome Sanger Institute for the COVID-19 Genomics UK (COG-UK) Consortium | Jacquelyn Wynn, Mairead Hyland, The Lighthouse Lab in Alderley Park and Alex Alderton, Roberto Amato, Jeffrey Barrett, Sonia Goncalves, Ewan Harrison, David K. Jackson, Ian Johnston, Dominic Kwiatkowski, Cordelia Langford, John Sillitoe on behalf of the Wellcome Sanger Institute COVID-19 Surveillance Team |
| EPI_ISL_3250400, EPI_ISL_3250401, EPI_ISL_3250402, EPI_ISL_3250403 | Lighthouse Laboratory Plymouth | Wellcome Sanger Institute for the COVID-19 Genomics UK (COG-UK) Consortium | Lighthouse Laboratory Plymouth and Alex Alderton, Roberto Amato, Jeffrey Barrett, Sonia Goncalves, Ewan Harrison, David K. Jackson, Ian Johnston, Dominic Kwiatkowski, Cordelia Langford, John Sillitoe on behalf of the Wellcome Sanger Institute COVID-19 Surveillance Team |
| EPI_ISL_3250404 | Lighthouse Lab in Glasgow | Wellcome Sanger Institute for the COVID-19 Genomics UK (COG-UK) Consortium | Harper VanSteenhouse, Yumi Kasai, David Gray, Carol Clugston, Anna Dominiczak and Alex Alderton, Roberto Amato, Jeffrey Barrett, Sonia Goncalves, Ewan Harrison, David K. Jackson, Ian Johnston, Dominic Kwiatkowski, Cordelia Langford, John Sillitoe on behalf of the Wellcome Sanger Institute COVID-19 Surveillance Team |
| EPI_ISL_3250405 | Lighthouse Laboratory Plymouth | Wellcome Sanger Institute for the COVID-19 Genomics UK (COG-UK) Consortium | Lighthouse Laboratory Plymouth and Alex Alderton, Roberto Amato, Jeffrey Barrett, Sonia Goncalves, Ewan Harrison, David K. Jackson, Ian Johnston, Dominic Kwiatkowski, Cordelia Langford, John Sillitoe on behalf of the Wellcome Sanger Institute COVID-19 Surveillance Team |
| EPI_ISL_3250406 | Lighthouse Lab in Glasgow | Wellcome Sanger Institute for the COVID-19 Genomics UK (COG-UK) Consortium | Harper VanSteenhouse, Yumi Kasai, David Gray, Carol Clugston, Anna Dominiczak and Alex Alderton, Roberto Amato, Jeffrey Barrett, Sonia Goncalves, Ewan Harrison, David K. Jackson, Ian Johnston, Dominic Kwiatkowski, Cordelia Langford, John Sillitoe on behalf of the Wellcome Sanger Institute COVID-19 Surveillance Team |
| EPI_ISL_3250407, EPI_ISL_3250408 | Lighthouse Lab in Alderley Park | Wellcome Sanger Institute for the COVID-19 Genomics UK (COG-UK) Consortium | Jacquelyn Wynn, Mairead Hyland, The Lighthouse Lab in Alderley Park and Alex Alderton, Roberto Amato, Jeffrey Barrett, Sonia Goncalves, Ewan Harrison, David K. Jackson, Ian Johnston, Dominic Kwiatkowski, Cordelia Langford, John Sillitoe on behalf of the Wellcome Sanger Institute COVID-19 Surveillance Team |
| EPI_ISL_3250409, EPI_ISL_3250410 | Lighthouse Lab in Glasgow | Wellcome Sanger Institute for the COVID-19 Genomics UK (COG-UK) Consortium | Harper VanSteenhouse, Yumi Kasai, David Gray, Carol Clugston, Anna Dominiczak and Alex Alderton, Roberto Amato, Jeffrey Barrett, Sonia Goncalves, Ewan Harrison, David K. Jackson, Ian Johnston, Dominic Kwiatkowski, Cordelia Langford, John Sillitoe on behalf of the Wellcome Sanger Institute COVID-19 Surveillance Team |
| EPI_ISL_3250411 | Lighthouse Laboratory Plymouth | Wellcome Sanger Institute for the COVID-19 Genomics UK (COG-UK) Consortium | Lighthouse Laboratory Plymouth and Alex Alderton, Roberto Amato, Jeffrey Barrett, Sonia Goncalves, Ewan Harrison, David K. Jackson, Ian Johnston, Dominic Kwiatkowski, Cordelia Langford, John Sillitoe on behalf of the Wellcome Sanger Institute COVID-19 Surveillance Team |
| EPI_ISL_3250412 | Lighthouse Lab in Alderley Park | Wellcome Sanger Institute for the COVID-19 Genomics UK (COG-UK) Consortium | Jacquelyn Wynn, Mairead Hyland, The Lighthouse Lab in Alderley Park and Alex Alderton, Roberto Amato, Jeffrey Barrett, Sonia Goncalves, Ewan Harrison, David K. Jackson, Ian Johnston, Dominic Kwiatkowski, Cordelia Langford, John Sillitoe on behalf of the Wellcome Sanger Institute COVID-19 Surveillance Team |
| EPI_ISL_3250413, EPI_ISL_3250414 | Lighthouse Lab in Glasgow | Wellcome Sanger Institute for the COVID-19 Genomics UK (COG-UK) Consortium | Harper VanSteenhouse, Yumi Kasai, David Gray, Carol Clugston, Anna Dominiczak and Alex Alderton, Roberto Amato, Jeffrey Barrett, Sonia Goncalves, Ewan Harrison, David K. Jackson, Ian Johnston, Dominic Kwiatkowski, Cordelia Langford, John Sillitoe on behalf of the Wellcome Sanger Institute COVID-19 Surveillance Team |
| EPI_ISL_3250415, EPI_ISL_3250416, EPI_ISL_3250417 | Lighthouse Lab in Alderley Park | Wellcome Sanger Institute for the COVID-19 Genomics UK (COG-UK) Consortium | Jacquelyn Wynn, Mairead Hyland, The Lighthouse Lab in Alderley Park and Alex Alderton, Roberto Amato, Jeffrey Barrett, Sonia Goncalves, Ewan Harrison, David K. Jackson, Ian Johnston, Dominic Kwiatkowski, Cordelia Langford, John Sillitoe on behalf of the Wellcome Sanger Institute COVID-19 Surveillance Team |
| EPI_ISL_3250418, EPI_ISL_3250419 | Lighthouse Lab in Glasgow | Wellcome Sanger Institute for the COVID-19 Genomics UK (COG-UK) Consortium | Harper VanSteenhouse, Yumi Kasai, David Gray, Carol Clugston, Anna Dominiczak and Alex Alderton, Roberto Amato, Jeffrey Barrett, Sonia Goncalves, Ewan Harrison, David K. Jackson, Ian Johnston, Dominic Kwiatkowski, Cordelia Langford, John Sillitoe on behalf of the Wellcome Sanger Institute COVID-19 Surveillance Team |
| EPI_ISL_3250420, EPI_ISL_3250421 | Lighthouse Laboratory Plymouth | Wellcome Sanger Institute for the COVID-19 Genomics UK (COG-UK) Consortium | Lighthouse Laboratory Plymouth and Alex Alderton, Roberto Amato, Jeffrey Barrett, Sonia Goncalves, Ewan Harrison, David K. Jackson, Ian Johnston, Dominic Kwiatkowski, Cordelia Langford, John Sillitoe on behalf of the Wellcome Sanger Institute COVID-19 Surveillance Team |
| EPI_ISL_3250422, EPI_ISL_3250423 | Lighthouse Lab in Alderley Park | Wellcome Sanger Institute for the COVID-19 Genomics UK (COG-UK) Consortium | Jacquelyn Wynn, Mairead Hyland, The Lighthouse Lab in Alderley Park and Alex Alderton, Roberto Amato, Jeffrey Barrett, Sonia Goncalves, Ewan Harrison, David K. Jackson, Ian Johnston, Dominic Kwiatkowski, Cordelia Langford, John Sillitoe on behalf of the Wellcome Sanger Institute COVID-19 Surveillance Team |
| EPI_ISL_3250424 | Lighthouse Laboratory Plymouth | Wellcome Sanger Institute for the COVID-19 Genomics UK (COG-UK) Consortium | Lighthouse Laboratory Plymouth and Alex Alderton, Roberto Amato, Jeffrey Barrett, Sonia Goncalves, Ewan Harrison, David K. Jackson, Ian Johnston, Dominic Kwiatkowski, Cordelia Langford, John Sillitoe on behalf of the Wellcome Sanger Institute COVID-19 Surveillance Team |
| EPI_ISL_3250425 | Lighthouse Lab in Glasgow | Wellcome Sanger Institute for the COVID-19 Genomics UK (COG-UK) Consortium | Harper VanSteenhouse, Yumi Kasai, David Gray, Carol Clugston, Anna Dominiczak and Alex Alderton, Roberto Amato, Jeffrey Barrett, Sonia Goncalves, Ewan Harrison, David K. Jackson, Ian Johnston, Dominic Kwiatkowski, Cordelia Langford, John Sillitoe on behalf of the Wellcome Sanger Institute COVID-19 Surveillance Team |
| EPI_ISL_3250426, EPI_ISL_3250427 | Lighthouse Laboratory Plymouth | Wellcome Sanger Institute for the COVID-19 Genomics UK (COG-UK) Consortium | Lighthouse Laboratory Plymouth and Alex Alderton, Roberto Amato, Jeffrey Barrett, Sonia Goncalves, Ewan Harrison, David K. Jackson, Ian Johnston, Dominic Kwiatkowski, Cordelia Langford, John Sillitoe on behalf of the Wellcome Sanger Institute COVID-19 Surveillance Team |
| EPI_ISL_3250428 | Lighthouse Lab in Glasgow | Wellcome Sanger Institute for the COVID-19 Genomics UK (COG-UK) Consortium | Harper VanSteenhouse, Yumi Kasai, David Gray, Carol Clugston, Anna Dominiczak and Alex Alderton, Roberto Amato, Jeffrey Barrett, Sonia Goncalves, Ewan Harrison, David K. Jackson, Ian Johnston, Dominic Kwiatkowski, Cordelia Langford, John Sillitoe on behalf of the Wellcome Sanger Institute COVID-19 Surveillance Team |
| EPI_ISL_3250429 | Lighthouse Laboratory Plymouth | Wellcome Sanger Institute for the COVID-19 Genomics UK (COG-UK) Consortium | Lighthouse Laboratory Plymouth and Alex Alderton, Roberto Amato, Jeffrey Barrett, Sonia Goncalves, Ewan Harrison, David K. Jackson, Ian Johnston, Dominic Kwiatkowski, Cordelia Langford, John Sillitoe on behalf of the Wellcome Sanger Institute COVID-19 Surveillance Team |
| EPI_ISL_3250430, EPI_ISL_3250431, EPI_ISL_3250432 | Lighthouse Lab in Glasgow | Wellcome Sanger Institute for the COVID-19 Genomics UK (COG-UK) Consortium | Harper VanSteenhouse, Yumi Kasai, David Gray, Carol Clugston, Anna Dominiczak and Alex Alderton, Roberto Amato, Jeffrey Barrett, Sonia Goncalves, Ewan Harrison, David K. Jackson, Ian Johnston, Dominic Kwiatkowski, Cordelia Langford, John Sillitoe on behalf of the Wellcome Sanger Institute COVID-19 Surveillance Team |
| EPI_ISL_3250433 | Lighthouse Lab in Alderley Park | Wellcome Sanger Institute for the COVID-19 Genomics UK (COG-UK) Consortium | Jacquelyn Wynn, Mairead Hyland, The Lighthouse Lab in Alderley Park and Alex Alderton, Roberto Amato, Jeffrey Barrett, Sonia Goncalves, Ewan Harrison, David K. Jackson, Ian Johnston, Dominic Kwiatkowski, Cordelia Langford, John Sillitoe on behalf of the Wellcome Sanger Institute COVID-19 Surveillance Team |
| EPI_ISL_3250434 | Lighthouse Lab in Glasgow | Wellcome Sanger Institute for the COVID-19 Genomics UK (COG-UK) Consortium | Harper VanSteenhouse, Yumi Kasai, David Gray, Carol Clugston, Anna Dominiczak and Alex Alderton, Roberto Amato, Jeffrey Barrett, Sonia Goncalves, Ewan Harrison, David K. Jackson, Ian Johnston, Dominic Kwiatkowski, Cordelia Langford, John Sillitoe on behalf of the Wellcome Sanger Institute COVID-19 Surveillance Team |
| EPI_ISL_3250435, EPI_ISL_3250436 | Lighthouse Laboratory Plymouth | Wellcome Sanger Institute for the COVID-19 Genomics UK (COG-UK) Consortium | Lighthouse Laboratory Plymouth and Alex Alderton, Roberto Amato, Jeffrey Barrett, Sonia Goncalves, Ewan Harrison, David K. Jackson, Ian Johnston, Dominic Kwiatkowski, Cordelia Langford, John Sillitoe on behalf of the Wellcome Sanger Institute COVID-19 Surveillance Team |
| EPI_ISL_3250437, EPI_ISL_3250438, EPI_ISL_3250439, EPI_ISL_3250440, EPI_ISL_3250441 | Lighthouse Lab in Glasgow | Wellcome Sanger Institute for the COVID-19 Genomics UK (COG-UK) Consortium | Harper VanSteenhouse, Yumi Kasai, David Gray, Carol Clugston, Anna Dominiczak and Alex Alderton, Roberto Amato, Jeffrey Barrett, Sonia Goncalves, Ewan Harrison, David K. Jackson, Ian Johnston, Dominic Kwiatkowski, Cordelia Langford, John Sillitoe on behalf of the Wellcome Sanger Institute COVID-19 Surveillance Team |
| EPI_ISL_3250442 | Lighthouse Lab in Alderley Park | Wellcome Sanger Institute for the COVID-19 Genomics UK (COG-UK) Consortium | Jacquelyn Wynn, Mairead Hyland, The Lighthouse Lab in Alderley Park and Alex Alderton, Roberto Amato, Jeffrey Barrett, Sonia Goncalves, Ewan Harrison, David K. Jackson, Ian Johnston, Dominic Kwiatkowski, Cordelia Langford, John Sillitoe on behalf of the Wellcome Sanger Institute COVID-19 Surveillance Team |
| EPI_ISL_3250443 | Lighthouse Lab in Glasgow | Wellcome Sanger Institute for the COVID-19 Genomics UK (COG-UK) Consortium | Harper VanSteenhouse, Yumi Kasai, David Gray, Carol Clugston, Anna Dominiczak and Alex Alderton, Roberto Amato, Jeffrey Barrett, Sonia Goncalves, Ewan Harrison, David K. Jackson, Ian Johnston, Dominic Kwiatkowski, Cordelia Langford, John Sillitoe on behalf of the Wellcome Sanger Institute COVID-19 Surveillance Team |
| EPI_ISL_3250444 | Lighthouse Laboratory Plymouth | Wellcome Sanger Institute for the COVID-19 Genomics UK (COG-UK) Consortium | Lighthouse Laboratory Plymouth and Alex Alderton, Roberto Amato, Jeffrey Barrett, Sonia Goncalves, Ewan Harrison, David K. Jackson, Ian Johnston, Dominic Kwiatkowski, Cordelia Langford, John Sillitoe on behalf of the Wellcome Sanger Institute COVID-19 Surveillance Team |
| EPI_ISL_3250445 | Lighthouse Lab in Alderley Park | Wellcome Sanger Institute for the COVID-19 Genomics UK (COG-UK) Consortium | Jacquelyn Wynn, Mairead Hyland, The Lighthouse Lab in Alderley Park and Alex Alderton, Roberto Amato, Jeffrey Barrett, Sonia Goncalves, Ewan Harrison, David K. Jackson, Ian Johnston, Dominic Kwiatkowski, Cordelia Langford, John Sillitoe on behalf of the Wellcome Sanger Institute COVID-19 Surveillance Team |
| EPI_ISL_3250446, EPI_ISL_3250447 | Lighthouse Lab in Glasgow | Wellcome Sanger Institute for the COVID-19 Genomics UK (COG-UK) Consortium | Harper VanSteenhouse, Yumi Kasai, David Gray, Carol Clugston, Anna Dominiczak and Alex Alderton, Roberto Amato, Jeffrey Barrett, Sonia Goncalves, Ewan Harrison, David K. Jackson, Ian Johnston, Dominic Kwiatkowski, Cordelia Langford, John Sillitoe on behalf of the Wellcome Sanger Institute COVID-19 Surveillance Team |
| EPI_ISL_3250448 | Lighthouse Lab in Alderley Park | Wellcome Sanger Institute for the COVID-19 Genomics UK (COG-UK) Consortium | Jacquelyn Wynn, Mairead Hyland, The Lighthouse Lab in Alderley Park and Alex Alderton, Roberto Amato, Jeffrey Barrett, Sonia Goncalves, Ewan Harrison, David K. Jackson, Ian Johnston, Dominic Kwiatkowski, Cordelia Langford, John Sillitoe on behalf of the Wellcome Sanger Institute COVID-19 Surveillance Team |
| EPI_ISL_3250449 | Lighthouse Lab in Glasgow | Wellcome Sanger Institute for the COVID-19 Genomics UK (COG-UK) Consortium | Harper VanSteenhouse, Yumi Kasai, David Gray, Carol Clugston, Anna Dominiczak and Alex Alderton, Roberto Amato, Jeffrey Barrett, Sonia Goncalves, Ewan Harrison, David K. Jackson, Ian Johnston, Dominic Kwiatkowski, Cordelia Langford, John Sillitoe on behalf of the Wellcome Sanger Institute COVID-19 Surveillance Team |
| EPI_ISL_3250450 | Lighthouse Lab in Alderley Park | Wellcome Sanger Institute for the COVID-19 Genomics UK (COG-UK) Consortium | Jacquelyn Wynn, Mairead Hyland, The Lighthouse Lab in Alderley Park and Alex Alderton, Roberto Amato, Jeffrey Barrett, Sonia Goncalves, Ewan Harrison, David K. Jackson, Ian Johnston, Dominic Kwiatkowski, Cordelia Langford, John Sillitoe on behalf of the Wellcome Sanger Institute COVID-19 Surveillance Team |
| EPI_ISL_3250451, EPI_ISL_3250452 | Lighthouse Lab in Glasgow | Wellcome Sanger Institute for the COVID-19 Genomics UK (COG-UK) Consortium | Harper VanSteenhouse, Yumi Kasai, David Gray, Carol Clugston, Anna Dominiczak and Alex Alderton, Roberto Amato, Jeffrey Barrett, Sonia Goncalves, Ewan Harrison, David K. Jackson, Ian Johnston, Dominic Kwiatkowski, Cordelia Langford, John Sillitoe on behalf of the Wellcome Sanger Institute COVID-19 Surveillance Team |
| EPI_ISL_3250453, EPI_ISL_3250454 | Lighthouse Laboratory Plymouth | Wellcome Sanger Institute for the COVID-19 Genomics UK (COG-UK) Consortium | Lighthouse Laboratory Plymouth and Alex Alderton, Roberto Amato, Jeffrey Barrett, Sonia Goncalves, Ewan Harrison, David K. Jackson, Ian Johnston, Dominic Kwiatkowski, Cordelia Langford, John Sillitoe on behalf of the Wellcome Sanger Institute COVID-19 Surveillance Team |
| EPI_ISL_3250455 | Lighthouse Lab in Glasgow | Wellcome Sanger Institute for the COVID-19 Genomics UK (COG-UK) Consortium | Harper VanSteenhouse, Yumi Kasai, David Gray, Carol Clugston, Anna Dominiczak and Alex Alderton, Roberto Amato, Jeffrey Barrett, Sonia Goncalves, Ewan Harrison, David K. Jackson, Ian Johnston, Dominic Kwiatkowski, Cordelia Langford, John Sillitoe on behalf of the Wellcome Sanger Institute COVID-19 Surveillance Team |
| EPI_ISL_3250456 | Lighthouse Lab in Alderley Park | Wellcome Sanger Institute for the COVID-19 Genomics UK (COG-UK) | Jacquelyn Wynn, Mairead Hyland, The Lighthouse Lab in Alderley Park and Alex Alderton, Roberto Amato, Jeffrey Barrett, Sonia Goncalves, Ewan Harrison, David K. Jackson, Ian Johnston, Dominic Kwiatkowski, Cordelia Langford, John Sillitoe on |

| | | | |
|---|---|---|---|
| | | Consortium | behalf of the Wellcome Sanger Institute COVID-19 Surveillance Team |
| EPI_ISL_3250457, EPI_ISL_3250458, EPI_ISL_3250459 | Lighthouse Laboratory Plymouth | Wellcome Sanger Institute for the COVID-19 Genomics UK (COG-UK) Consortium | Lighthouse Laboratory Plymouth and Alex Alderton, Roberto Amato, Jeffrey Barrett, Sonia Goncalves, Ewan Harrison, David K. Jackson, Ian Johnston, Dominic Kwiatkowski, Cordelia Langford, John Sillitoe on behalf of the Wellcome Sanger Institute COVID-19 Surveillance Team |
| EPI_ISL_3250460, EPI_ISL_3250461, EPI_ISL_3250462, EPI_ISL_3250463, EPI_ISL_3250464 | Lighthouse Lab in Glasgow | Wellcome Sanger Institute for the COVID-19 Genomics UK (COG-UK) Consortium | Harper VanSteenhouse, Yumi Kasai, David Gray, Carol Clugston, Anna Dominiczak and Alex Alderton, Roberto Amato, Jeffrey Barrett, Sonia Goncalves, Ewan Harrison, David K. Jackson, Ian Johnston, Dominic Kwiatkowski, Cordelia Langford, John Sillitoe on behalf of the Wellcome Sanger Institute COVID-19 Surveillance Team |
| EPI_ISL_3250465 | Lighthouse Lab in Alderley Park | Wellcome Sanger Institute for the COVID-19 Genomics UK (COG-UK) Consortium | Jacquelyn Wynn, Mairead Hyland, The Lighthouse Lab in Alderley Park and Alex Alderton, Roberto Amato, Jeffrey Barrett, Sonia Goncalves, Ewan Harrison, David K. Jackson, Ian Johnston, Dominic Kwiatkowski, Cordelia Langford, John Sillitoe on behalf of the Wellcome Sanger Institute COVID-19 Surveillance Team |
| EPI_ISL_3250466, EPI_ISL_3250467 | Lighthouse Laboratory Plymouth | Wellcome Sanger Institute for the COVID-19 Genomics UK (COG-UK) Consortium | Lighthouse Laboratory Plymouth and Alex Alderton, Roberto Amato, Jeffrey Barrett, Sonia Goncalves, Ewan Harrison, David K. Jackson, Ian Johnston, Dominic Kwiatkowski, Cordelia Langford, John Sillitoe on behalf of the Wellcome Sanger Institute COVID-19 Surveillance Team |
| EPI_ISL_3250468, EPI_ISL_3250469 | Lighthouse Lab in Alderley Park | Wellcome Sanger Institute for the COVID-19 Genomics UK (COG-UK) Consortium | Jacquelyn Wynn, Mairead Hyland, The Lighthouse Lab in Alderley Park and Alex Alderton, Roberto Amato, Jeffrey Barrett, Sonia Goncalves, Ewan Harrison, David K. Jackson, Ian Johnston, Dominic Kwiatkowski, Cordelia Langford, John Sillitoe on behalf of the Wellcome Sanger Institute COVID-19 Surveillance Team |
| EPI_ISL_3250470 | Lighthouse Laboratory Plymouth | Wellcome Sanger Institute for the COVID-19 Genomics UK (COG-UK) Consortium | Lighthouse Laboratory Plymouth and Alex Alderton, Roberto Amato, Jeffrey Barrett, Sonia Goncalves, Ewan Harrison, David K. Jackson, Ian Johnston, Dominic Kwiatkowski, Cordelia Langford, John Sillitoe on behalf of the Wellcome Sanger Institute COVID-19 Surveillance Team |
| EPI_ISL_3250471 | Lighthouse Lab in Glasgow | Wellcome Sanger Institute for the COVID-19 Genomics UK (COG-UK) Consortium | Harper VanSteenhouse, Yumi Kasai, David Gray, Carol Clugston, Anna Dominiczak and Alex Alderton, Roberto Amato, Jeffrey Barrett, Sonia Goncalves, Ewan Harrison, David K. Jackson, Ian Johnston, Dominic Kwiatkowski, Cordelia Langford, John Sillitoe on behalf of the Wellcome Sanger Institute COVID-19 Surveillance Team |
| EPI_ISL_3250472 | Lighthouse Lab in Alderley Park | Wellcome Sanger Institute for the COVID-19 Genomics UK (COG-UK) Consortium | Jacquelyn Wynn, Mairead Hyland, The Lighthouse Lab in Alderley Park and Alex Alderton, Roberto Amato, Jeffrey Barrett, Sonia Goncalves, Ewan Harrison, David K. Jackson, Ian Johnston, Dominic Kwiatkowski, Cordelia Langford, John Sillitoe on behalf of the Wellcome Sanger Institute COVID-19 Surveillance Team |
| EPI_ISL_3250473, EPI_ISL_3250474, EPI_ISL_3250475 | Lighthouse Lab in Glasgow | Wellcome Sanger Institute for the COVID-19 Genomics UK (COG-UK) Consortium | Harper VanSteenhouse, Yumi Kasai, David Gray, Carol Clugston, Anna Dominiczak and Alex Alderton, Roberto Amato, Jeffrey Barrett, Sonia Goncalves, Ewan Harrison, David K. Jackson, Ian Johnston, Dominic Kwiatkowski, Cordelia Langford, John Sillitoe on behalf of the Wellcome Sanger Institute COVID-19 Surveillance Team |
| EPI_ISL_3250476 | Lighthouse Laboratory Plymouth | Wellcome Sanger Institute for the COVID-19 Genomics UK (COG-UK) Consortium | Lighthouse Laboratory Plymouth and Alex Alderton, Roberto Amato, Jeffrey Barrett, Sonia Goncalves, Ewan Harrison, David K. Jackson, Ian Johnston, Dominic Kwiatkowski, Cordelia Langford, John Sillitoe on behalf of the Wellcome Sanger Institute COVID-19 Surveillance Team |
| EPI_ISL_3250477, EPI_ISL_3250478 | Lighthouse Lab in Glasgow | Wellcome Sanger Institute for the COVID-19 Genomics UK (COG-UK) Consortium | Harper VanSteenhouse, Yumi Kasai, David Gray, Carol Clugston, Anna Dominiczak and Alex Alderton, Roberto Amato, Jeffrey Barrett, Sonia Goncalves, Ewan Harrison, David K. Jackson, Ian Johnston, Dominic Kwiatkowski, Cordelia Langford, John Sillitoe on behalf of the Wellcome Sanger Institute COVID-19 Surveillance Team |
| EPI_ISL_3250479, EPI_ISL_3250480 | Lighthouse Laboratory Plymouth | Wellcome Sanger Institute for the COVID-19 Genomics UK (COG-UK) Consortium | Lighthouse Laboratory Plymouth and Alex Alderton, Roberto Amato, Jeffrey Barrett, Sonia Goncalves, Ewan Harrison, David K. Jackson, Ian Johnston, Dominic Kwiatkowski, Cordelia Langford, John Sillitoe on behalf of the Wellcome Sanger Institute COVID-19 Surveillance Team |
| EPI_ISL_3250481 | Lighthouse Lab in Glasgow | Wellcome Sanger Institute for the COVID-19 Genomics UK (COG-UK) Consortium | Harper VanSteenhouse, Yumi Kasai, David Gray, Carol Clugston, Anna Dominiczak and Alex Alderton, Roberto Amato, Jeffrey Barrett, Sonia Goncalves, Ewan Harrison, David K. Jackson, Ian Johnston, Dominic Kwiatkowski, Cordelia Langford, John Sillitoe on behalf of the Wellcome Sanger Institute COVID-19 Surveillance Team |
| EPI_ISL_3250482 | Lighthouse Laboratory Plymouth | Wellcome Sanger Institute for the COVID-19 Genomics UK (COG-UK) Consortium | Lighthouse Laboratory Plymouth and Alex Alderton, Roberto Amato, Jeffrey Barrett, Sonia Goncalves, Ewan Harrison, David K. Jackson, Ian Johnston, Dominic Kwiatkowski, Cordelia Langford, John Sillitoe on behalf of the Wellcome Sanger Institute COVID-19 Surveillance Team |
| EPI_ISL_3250483 | Lighthouse Lab in Glasgow | Wellcome Sanger Institute for the COVID-19 Genomics UK (COG-UK) Consortium | Harper VanSteenhouse, Yumi Kasai, David Gray, Carol Clugston, Anna Dominiczak and Alex Alderton, Roberto Amato, Jeffrey Barrett, Sonia Goncalves, Ewan Harrison, David K. Jackson, Ian Johnston, Dominic Kwiatkowski, Cordelia Langford, John Sillitoe on behalf of the Wellcome Sanger Institute COVID-19 Surveillance Team |
| EPI_ISL_3250484 | Lighthouse Laboratory Plymouth | Wellcome Sanger Institute for the COVID-19 Genomics UK (COG-UK) Consortium | Lighthouse Laboratory Plymouth and Alex Alderton, Roberto Amato, Jeffrey Barrett, Sonia Goncalves, Ewan Harrison, David K. Jackson, Ian Johnston, Dominic Kwiatkowski, Cordelia Langford, John Sillitoe on behalf of the Wellcome Sanger Institute COVID-19 Surveillance Team |
| EPI_ISL_3250485 | Lighthouse Lab in Alderley Park | Wellcome Sanger Institute for the COVID-19 Genomics UK (COG-UK) Consortium | Jacquelyn Wynn, Mairead Hyland, The Lighthouse Lab in Alderley Park and Alex Alderton, Roberto Amato, Jeffrey Barrett, Sonia Goncalves, Ewan Harrison, David K. Jackson, Ian Johnston, Dominic Kwiatkowski, Cordelia Langford, John Sillitoe on behalf of the Wellcome Sanger Institute COVID-19 Surveillance Team |
| EPI_ISL_3250486, EPI_ISL_3250487 | Lighthouse Laboratory Plymouth | Wellcome Sanger Institute for the COVID-19 Genomics UK (COG-UK) Consortium | Lighthouse Laboratory Plymouth and Alex Alderton, Roberto Amato, Jeffrey Barrett, Sonia Goncalves, Ewan Harrison, David K. Jackson, Ian Johnston, Dominic Kwiatkowski, Cordelia Langford, John Sillitoe on behalf of the Wellcome Sanger Institute COVID-19 Surveillance Team |
| EPI_ISL_3250488 | Lighthouse Lab in Glasgow | Wellcome Sanger Institute for the COVID-19 Genomics UK (COG-UK) Consortium | Harper VanSteenhouse, Yumi Kasai, David Gray, Carol Clugston, Anna Dominiczak and Alex Alderton, Roberto Amato, Jeffrey Barrett, Sonia Goncalves, Ewan Harrison, David K. Jackson, Ian Johnston, Dominic Kwiatkowski, Cordelia Langford, John Sillitoe on behalf of the Wellcome Sanger Institute COVID-19 Surveillance Team |
| EPI_ISL_3250489, EPI_ISL_3250490 | Lighthouse Lab in Alderley Park | Wellcome Sanger Institute for the COVID-19 Genomics UK (COG-UK) Consortium | Jacquelyn Wynn, Mairead Hyland, The Lighthouse Lab in Alderley Park and Alex Alderton, Roberto Amato, Jeffrey Barrett, Sonia Goncalves, Ewan Harrison, David K. Jackson, Ian Johnston, Dominic Kwiatkowski, Cordelia Langford, John Sillitoe on behalf of the Wellcome Sanger Institute COVID-19 Surveillance Team |
| EPI_ISL_3250491 | Lighthouse Laboratory Plymouth | Wellcome Sanger Institute for the COVID-19 Genomics UK (COG-UK) Consortium | Lighthouse Laboratory Plymouth and Alex Alderton, Roberto Amato, Jeffrey Barrett, Sonia Goncalves, Ewan Harrison, David K. Jackson, Ian Johnston, Dominic Kwiatkowski, Cordelia Langford, John Sillitoe on behalf of the Wellcome Sanger Institute COVID-19 Surveillance Team |
| EPI_ISL_3250492, EPI_ISL_3250493, EPI_ISL_3250494 | Lighthouse Lab in Glasgow | Wellcome Sanger Institute for the COVID-19 Genomics UK (COG-UK) Consortium | Harper VanSteenhouse, Yumi Kasai, David Gray, Carol Clugston, Anna Dominiczak and Alex Alderton, Roberto Amato, Jeffrey Barrett, Sonia Goncalves, Ewan Harrison, David K. Jackson, Ian Johnston, Dominic Kwiatkowski, Cordelia Langford, John Sillitoe on behalf of the Wellcome Sanger Institute COVID-19 Surveillance Team |
| EPI_ISL_3250495, EPI_ISL_3250496 | Lighthouse Laboratory Plymouth | Wellcome Sanger Institute for the COVID-19 Genomics UK (COG-UK) Consortium | Lighthouse Laboratory Plymouth and Alex Alderton, Roberto Amato, Jeffrey Barrett, Sonia Goncalves, Ewan Harrison, David K. Jackson, Ian Johnston, Dominic Kwiatkowski, Cordelia Langford, John Sillitoe on behalf of the Wellcome Sanger Institute COVID-19 Surveillance Team |
| EPI_ISL_3250497 | Lighthouse Lab in Alderley Park | Wellcome Sanger Institute for the COVID-19 Genomics UK (COG-UK) Consortium | Jacquelyn Wynn, Mairead Hyland, The Lighthouse Lab in Alderley Park and Alex Alderton, Roberto Amato, Jeffrey Barrett, Sonia Goncalves, Ewan Harrison, David K. Jackson, Ian Johnston, Dominic Kwiatkowski, Cordelia Langford, John Sillitoe on behalf of the Wellcome Sanger Institute COVID-19 Surveillance Team |
| EPI_ISL_3250498, EPI_ISL_3250499 | Lighthouse Lab in Glasgow | Wellcome Sanger Institute for the COVID-19 Genomics UK (COG-UK) Consortium | Harper VanSteenhouse, Yumi Kasai, David Gray, Carol Clugston, Anna Dominiczak and Alex Alderton, Roberto Amato, Jeffrey Barrett, Sonia Goncalves, Ewan Harrison, David K. Jackson, Ian Johnston, Dominic Kwiatkowski, Cordelia Langford, John Sillitoe on behalf of the Wellcome Sanger Institute COVID-19 Surveillance Team |
| EPI_ISL_3250500 | Lighthouse Lab in Alderley Park | Wellcome Sanger Institute for the COVID-19 Genomics UK (COG-UK) Consortium | Jacquelyn Wynn, Mairead Hyland, The Lighthouse Lab in Alderley Park and Alex Alderton, Roberto Amato, Jeffrey Barrett, Sonia Goncalves, Ewan Harrison, David K. Jackson, Ian Johnston, Dominic Kwiatkowski, Cordelia Langford, John Sillitoe on behalf of the Wellcome Sanger Institute COVID-19 Surveillance Team |
| EPI_ISL_3250501 | Lighthouse Lab in Glasgow | Wellcome Sanger Institute for the COVID-19 Genomics UK (COG-UK) Consortium | Harper VanSteenhouse, Yumi Kasai, David Gray, Carol Clugston, Anna Dominiczak and Alex Alderton, Roberto Amato, Jeffrey Barrett, Sonia Goncalves, Ewan Harrison, David K. Jackson, Ian Johnston, Dominic Kwiatkowski, Cordelia Langford, John Sillitoe on behalf of the Wellcome Sanger Institute COVID-19 Surveillance Team |
| EPI_ISL_3250502 | Lighthouse Lab in Alderley Park | Wellcome Sanger Institute for the COVID-19 Genomics UK (COG-UK) Consortium | Jacquelyn Wynn, Mairead Hyland, The Lighthouse Lab in Alderley Park and Alex Alderton, Roberto Amato, Jeffrey Barrett, Sonia Goncalves, Ewan Harrison, David K. Jackson, Ian Johnston, Dominic Kwiatkowski, Cordelia Langford, John Sillitoe on behalf of the Wellcome Sanger Institute COVID-19 Surveillance Team |
| EPI_ISL_3250503, EPI_ISL_3250504 | Lighthouse Lab in Glasgow | Wellcome Sanger Institute for the COVID-19 Genomics UK (COG-UK) Consortium | Harper VanSteenhouse, Yumi Kasai, David Gray, Carol Clugston, Anna Dominiczak and Alex Alderton, Roberto Amato, Jeffrey Barrett, Sonia Goncalves, Ewan Harrison, David K. Jackson, Ian Johnston, Dominic Kwiatkowski, Cordelia Langford, John Sillitoe on behalf of the Wellcome Sanger Institute COVID-19 Surveillance Team |
| EPI_ISL_3250505 | Lighthouse Laboratory Plymouth | Wellcome Sanger Institute for the COVID-19 Genomics UK (COG-UK) Consortium | Lighthouse Laboratory Plymouth and Alex Alderton, Roberto Amato, Jeffrey Barrett, Sonia Goncalves, Ewan Harrison, David K. Jackson, Ian Johnston, Dominic Kwiatkowski, Cordelia Langford, John Sillitoe on behalf of the Wellcome Sanger Institute COVID-19 Surveillance Team |
| EPI_ISL_3250506 | Lighthouse Lab in Alderley Park | Wellcome Sanger Institute for the COVID-19 Genomics UK (COG-UK) Consortium | Jacquelyn Wynn, Mairead Hyland, The Lighthouse Lab in Alderley Park and Alex Alderton, Roberto Amato, Jeffrey Barrett, Sonia Goncalves, Ewan Harrison, David K. Jackson, Ian Johnston, Dominic Kwiatkowski, Cordelia Langford, John Sillitoe on behalf of the Wellcome Sanger Institute COVID-19 Surveillance Team |
| EPI_ISL_3250507 | Lighthouse Lab in Glasgow | Wellcome Sanger Institute for the COVID-19 Genomics UK (COG-UK) Consortium | Harper VanSteenhouse, Yumi Kasai, David Gray, Carol Clugston, Anna Dominiczak and Alex Alderton, Roberto Amato, Jeffrey Barrett, Sonia Goncalves, Ewan Harrison, David K. Jackson, Ian Johnston, Dominic Kwiatkowski, Cordelia Langford, John Sillitoe on behalf of the Wellcome Sanger Institute COVID-19 Surveillance Team |
| EPI_ISL_3250508 | Lighthouse Lab in Alderley Park | Wellcome Sanger Institute for the COVID-19 Genomics UK (COG-UK) Consortium | Jacquelyn Wynn, Mairead Hyland, The Lighthouse Lab in Alderley Park and Alex Alderton, Roberto Amato, Jeffrey Barrett, Sonia Goncalves, Ewan Harrison, David K. Jackson, Ian Johnston, Dominic Kwiatkowski, Cordelia Langford, John Sillitoe on behalf of the Wellcome Sanger Institute COVID-19 Surveillance Team |
| EPI_ISL_3250509 | Lighthouse Laboratory Plymouth | Wellcome Sanger Institute for the COVID-19 Genomics UK (COG-UK) Consortium | Lighthouse Laboratory Plymouth and Alex Alderton, Roberto Amato, Jeffrey Barrett, Sonia Goncalves, Ewan Harrison, David K. Jackson, Ian Johnston, Dominic Kwiatkowski, Cordelia Langford, John Sillitoe on behalf of the Wellcome Sanger Institute COVID-19 Surveillance Team |
| EPI_ISL_3250510 | Lighthouse Lab in Alderley Park | Wellcome Sanger Institute for the COVID-19 Genomics UK (COG-UK) Consortium | Jacquelyn Wynn, Mairead Hyland, The Lighthouse Lab in Alderley Park and Alex Alderton, Roberto Amato, Jeffrey Barrett, Sonia Goncalves, Ewan Harrison, David K. Jackson, Ian Johnston, Dominic Kwiatkowski, Cordelia Langford, John Sillitoe on behalf of the Wellcome Sanger Institute COVID-19 Surveillance Team |
| EPI_ISL_3250511, EPI_ISL_3250512 | Lighthouse Lab in Glasgow | Wellcome Sanger Institute for the COVID-19 Genomics UK (COG-UK) Consortium | Harper VanSteenhouse, Yumi Kasai, David Gray, Carol Clugston, Anna Dominiczak and Alex Alderton, Roberto Amato, Jeffrey Barrett, Sonia Goncalves, Ewan Harrison, David K. Jackson, Ian Johnston, Dominic Kwiatkowski, Cordelia Langford, John Sillitoe on behalf of the Wellcome Sanger Institute COVID-19 Surveillance Team |
| EPI_ISL_3250513, EPI_ISL_3250514, EPI_ISL_3250515, EPI_ISL_3250516 | Lighthouse Lab in Alderley Park | Wellcome Sanger Institute for the COVID-19 Genomics UK (COG-UK) Consortium | Jacquelyn Wynn, Mairead Hyland, The Lighthouse Lab in Alderley Park and Alex Alderton, Roberto Amato, Jeffrey Barrett, Sonia Goncalves, Ewan Harrison, David K. Jackson, Ian Johnston, Dominic Kwiatkowski, Cordelia Langford, John Sillitoe on behalf of the Wellcome Sanger Institute COVID-19 Surveillance Team |
| EPI_ISL_3250517, EPI_ISL_3250518 | Lighthouse Lab in Glasgow | Wellcome Sanger Institute for the COVID-19 Genomics UK (COG-UK) Consortium | Harper VanSteenhouse, Yumi Kasai, David Gray, Carol Clugston, Anna Dominiczak and Alex Alderton, Roberto Amato, Jeffrey Barrett, Sonia Goncalves, Ewan Harrison, David K. Jackson, Ian Johnston, Dominic Kwiatkowski, Cordelia Langford, John Sillitoe on behalf of the Wellcome Sanger Institute COVID-19 Surveillance Team |
| EPI_ISL_3250519 | Lighthouse Laboratory Plymouth | Wellcome Sanger Institute for the COVID-19 Genomics UK (COG-UK) Consortium | Lighthouse Laboratory Plymouth and Alex Alderton, Roberto Amato, Jeffrey Barrett, Sonia Goncalves, Ewan Harrison, David K. Jackson, Ian Johnston, Dominic Kwiatkowski, Cordelia Langford, John Sillitoe on behalf of the Wellcome Sanger Institute COVID-19 Surveillance Team |
| EPI_ISL_3250520 | Lighthouse Lab in Alderley Park | Wellcome Sanger Institute for the COVID-19 Genomics UK (COG-UK) Consortium | Jacquelyn Wynn, Mairead Hyland, The Lighthouse Lab in Alderley Park and Alex Alderton, Roberto Amato, Jeffrey Barrett, Sonia Goncalves, Ewan Harrison, David K. Jackson, Ian Johnston, Dominic Kwiatkowski, Cordelia Langford, John Sillitoe on behalf of the Wellcome Sanger Institute COVID-19 Surveillance Team |
| EPI_ISL_3250521 | Lighthouse Lab in Glasgow | Wellcome Sanger Institute for the COVID-19 Genomics UK (COG-UK) Consortium | Harper VanSteenhouse, Yumi Kasai, David Gray, Carol Clugston, Anna Dominiczak and Alex Alderton, Roberto Amato, Jeffrey Barrett, Sonia Goncalves, Ewan Harrison, David K. Jackson, Ian Johnston, Dominic Kwiatkowski, Cordelia Langford, John Sillitoe on behalf of the Wellcome Sanger Institute COVID-19 Surveillance Team |
| EPI_ISL_3250522, | Lighthouse Lab in Alderley Park | Wellcome Sanger Institute for the COVID-19 Genomics UK (COG-UK) | Jacquelyn Wynn, Mairead Hyland, The Lighthouse Lab in Alderley Park and Alex Alderton, Roberto Amato, Jeffrey Barrett, Sonia Goncalves, Ewan Harrison, David K. Jackson, Ian Johnston, Dominic Kwiatkowski, Cordelia Langford, John Sillitoe on |

| | | | |
|---|---|---|---|
| EPI_ISL_3250523 | | Consortium | behalf of the Wellcome Sanger Institute COVID-19 Surveillance Team |
| EPI_ISL_3250524, EPI_ISL_3250525 | Lighthouse Lab in Glasgow | Wellcome Sanger Institute for the COVID-19 Genomics UK (COG-UK) Consortium | Harper VanSteenhouse, Yumi Kasai, David Gray, Carol Clugston, Anna Dominiczak and Alex Alderton, Roberto Amato, Jeffrey Barrett, Sonia Goncalves, Ewan Harrison, David K. Jackson, Ian Johnston, Dominic Kwiatkowski, Cordelia Langford, John Sillitoe on behalf of the Wellcome Sanger Institute COVID-19 Surveillance Team |
| EPI_ISL_3250526 | Lighthouse Lab in Alderley Park | Wellcome Sanger Institute for the COVID-19 Genomics UK (COG-UK) Consortium | Jacquelyn Wynn, Mairead Hyland, The Lighthouse Lab in Alderley Park and Alex Alderton, Roberto Amato, Jeffrey Barrett, Sonia Goncalves, Ewan Harrison, David K. Jackson, Ian Johnston, Dominic Kwiatkowski, Cordelia Langford, John Sillitoe on behalf of the Wellcome Sanger Institute COVID-19 Surveillance Team |
| EPI_ISL_3250527, EPI_ISL_3250528 | Lighthouse Laboratory Plymouth | Wellcome Sanger Institute for the COVID-19 Genomics UK (COG-UK) Consortium | Lighthouse Laboratory Plymouth and Alex Alderton, Roberto Amato, Jeffrey Barrett, Sonia Goncalves, Ewan Harrison, David K. Jackson, Ian Johnston, Dominic Kwiatkowski, Cordelia Langford, John Sillitoe on behalf of the Wellcome Sanger Institute COVID-19 Surveillance Team |
| EPI_ISL_3250529 | Lighthouse Lab in Alderley Park | Wellcome Sanger Institute for the COVID-19 Genomics UK (COG-UK) Consortium | Jacquelyn Wynn, Mairead Hyland, The Lighthouse Lab in Alderley Park and Alex Alderton, Roberto Amato, Jeffrey Barrett, Sonia Goncalves, Ewan Harrison, David K. Jackson, Ian Johnston, Dominic Kwiatkowski, Cordelia Langford, John Sillitoe on behalf of the Wellcome Sanger Institute COVID-19 Surveillance Team |
| EPI_ISL_3250530, EPI_ISL_3250531 | Lighthouse Laboratory Plymouth | Wellcome Sanger Institute for the COVID-19 Genomics UK (COG-UK) Consortium | Lighthouse Laboratory Plymouth and Alex Alderton, Roberto Amato, Jeffrey Barrett, Sonia Goncalves, Ewan Harrison, David K. Jackson, Ian Johnston, Dominic Kwiatkowski, Cordelia Langford, John Sillitoe on behalf of the Wellcome Sanger Institute COVID-19 Surveillance Team |
| EPI_ISL_3250532, EPI_ISL_3250533, EPI_ISL_3250534 | Lighthouse Lab in Glasgow | Wellcome Sanger Institute for the COVID-19 Genomics UK (COG-UK) Consortium | Harper VanSteenhouse, Yumi Kasai, David Gray, Carol Clugston, Anna Dominiczak and Alex Alderton, Roberto Amato, Jeffrey Barrett, Sonia Goncalves, Ewan Harrison, David K. Jackson, Ian Johnston, Dominic Kwiatkowski, Cordelia Langford, John Sillitoe on behalf of the Wellcome Sanger Institute COVID-19 Surveillance Team |
| EPI_ISL_3250535 | Berkshire and Surrey Pathology Services Lighthouse Laboratory | Wellcome Sanger Institute for the COVID-19 Genomics UK (COG-UK) Consortium | Berkshire and Surrey Pathology Services Lighthouse Laboratory and Alex Alderton, Roberto Amato, Jeffrey Barrett, Sonia Goncalves, Ewan Harrison, David K. Jackson, Ian Johnston, Dominic Kwiatkowski, Cordelia Langford, John Sillitoe on behalf of the Wellcome Sanger Institute COVID-19 Surveillance Team |
| EPI_ISL_3250536 | Lighthouse Laboratory Plymouth | Wellcome Sanger Institute for the COVID-19 Genomics UK (COG-UK) Consortium | Lighthouse Laboratory Plymouth and Alex Alderton, Roberto Amato, Jeffrey Barrett, Sonia Goncalves, Ewan Harrison, David K. Jackson, Ian Johnston, Dominic Kwiatkowski, Cordelia Langford, John Sillitoe on behalf of the Wellcome Sanger Institute COVID-19 Surveillance Team |
| EPI_ISL_3250537, EPI_ISL_3250538 | Berkshire and Surrey Pathology Services Lighthouse Laboratory | Wellcome Sanger Institute for the COVID-19 Genomics UK (COG-UK) Consortium | Berkshire and Surrey Pathology Services Lighthouse Laboratory and Alex Alderton, Roberto Amato, Jeffrey Barrett, Sonia Goncalves, Ewan Harrison, David K. Jackson, Ian Johnston, Dominic Kwiatkowski, Cordelia Langford, John Sillitoe on behalf of the Wellcome Sanger Institute COVID-19 Surveillance Team |
| EPI_ISL_3250539 | Lighthouse Lab in Milton Keynes | Wellcome Sanger Institute for the COVID-19 Genomics UK (COG-UK) Consortium | The Lighthouse Lab in Milton Keynes and Alex Alderton, Roberto Amato, Jeffrey Barrett, Sonia Goncalves, Ewan Harrison, David K. Jackson, Ian Johnston, Dominic Kwiatkowski, Cordelia Langford, John Sillitoe on behalf of the Wellcome Sanger Institute COVID-19 Surveillance Team |
| EPI_ISL_3250540, EPI_ISL_3250541 | Berkshire and Surrey Pathology Services Lighthouse Laboratory | Wellcome Sanger Institute for the COVID-19 Genomics UK (COG-UK) Consortium | Berkshire and Surrey Pathology Services Lighthouse Laboratory and Alex Alderton, Roberto Amato, Jeffrey Barrett, Sonia Goncalves, Ewan Harrison, David K. Jackson, Ian Johnston, Dominic Kwiatkowski, Cordelia Langford, John Sillitoe on behalf of the Wellcome Sanger Institute COVID-19 Surveillance Team |
| EPI_ISL_3250542 | Lighthouse Lab in Milton Keynes | Wellcome Sanger Institute for the COVID-19 Genomics UK (COG-UK) Consortium | The Lighthouse Lab in Milton Keynes and Alex Alderton, Roberto Amato, Jeffrey Barrett, Sonia Goncalves, Ewan Harrison, David K. Jackson, Ian Johnston, Dominic Kwiatkowski, Cordelia Langford, John Sillitoe on behalf of the Wellcome Sanger Institute COVID-19 Surveillance Team |
| EPI_ISL_3250543, EPI_ISL_3250544, EPI_ISL_3250545, EPI_ISL_3250546 | Berkshire and Surrey Pathology Services Lighthouse Laboratory | Wellcome Sanger Institute for the COVID-19 Genomics UK (COG-UK) Consortium | Berkshire and Surrey Pathology Services Lighthouse Laboratory and Alex Alderton, Roberto Amato, Jeffrey Barrett, Sonia Goncalves, Ewan Harrison, David K. Jackson, Ian Johnston, Dominic Kwiatkowski, Cordelia Langford, John Sillitoe on behalf of the Wellcome Sanger Institute COVID-19 Surveillance Team |
| EPI_ISL_3250547, EPI_ISL_3250548, EPI_ISL_3250549, EPI_ISL_3250550, EPI_ISL_3250551, EPI_ISL_3250552, EPI_ISL_3250553, EPI_ISL_3250554, EPI_ISL_3250555, EPI_ISL_3250556, EPI_ISL_3250557, EPI_ISL_3250558, EPI_ISL_3250559, EPI_ISL_3250560, EPI_ISL_3250561, EPI_ISL_3250562 | | | |
| see above | Lighthouse Lab in Glasgow | Wellcome Sanger Institute for the COVID-19 Genomics UK (COG-UK) Consortium | Harper VanSteenhouse, Yumi Kasai, David Gray, Carol Clugston, Anna Dominiczak and Alex Alderton, Roberto Amato, Jeffrey Barrett, Sonia Goncalves, Ewan Harrison, David K. Jackson, Ian Johnston, Dominic Kwiatkowski, Cordelia Langford, John Sillitoe on behalf of the Wellcome Sanger Institute COVID-19 Surveillance Team |
| EPI_ISL_3250563 | Lighthouse Lab in Alderley Park | Wellcome Sanger Institute for the COVID-19 Genomics UK (COG-UK) Consortium | Jacquelyn Wynn, Mairead Hyland, The Lighthouse Lab in Alderley Park and Alex Alderton, Roberto Amato, Jeffrey Barrett, Sonia Goncalves, Ewan Harrison, David K. Jackson, Ian Johnston, Dominic Kwiatkowski, Cordelia Langford, John Sillitoe on behalf of the Wellcome Sanger Institute COVID-19 Surveillance Team |
| EPI_ISL_3250564, EPI_ISL_3250565, EPI_ISL_3250566, EPI_ISL_3250567, EPI_ISL_3250568, EPI_ISL_3250569, EPI_ISL_3250570, EPI_ISL_3250571, EPI_ISL_3250572, EPI_ISL_3250573, EPI_ISL_3250574, EPI_ISL_3250575, EPI_ISL_3250576, EPI_ISL_3250577, EPI_ISL_3250578, EPI_ISL_3250579, EPI_ISL_3250580, EPI_ISL_3250581, EPI_ISL_3250582, EPI_ISL_3250583, EPI_ISL_3250584, EPI_ISL_3250585, EPI_ISL_3250586, EPI_ISL_3250587, EPI_ISL_3250588 | | | |
| see above | Lighthouse Lab in Glasgow | Wellcome Sanger Institute for the COVID-19 Genomics UK (COG-UK) Consortium | Harper VanSteenhouse, Yumi Kasai, David Gray, Carol Clugston, Anna Dominiczak and Alex Alderton, Roberto Amato, Jeffrey Barrett, Sonia Goncalves, Ewan Harrison, David K. Jackson, Ian Johnston, Dominic Kwiatkowski, Cordelia Langford, John Sillitoe on behalf of the Wellcome Sanger Institute COVID-19 Surveillance Team |
| EPI_ISL_3250589, EPI_ISL_3250590, EPI_ISL_3250591, EPI_ISL_3250592, EPI_ISL_3250593, EPI_ISL_3250594, EPI_ISL_3250595, EPI_ISL_3250596, EPI_ISL_3250597, EPI_ISL_3250598, EPI_ISL_3250599, EPI_ISL_3250600, EPI_ISL_3250601, EPI_ISL_3250602, EPI_ISL_3250603, EPI_ISL_3250604, EPI_ISL_3250605, EPI_ISL_3250606, EPI_ISL_3250607, EPI_ISL_3250608, EPI_ISL_3250609, EPI_ISL_3250610, EPI_ISL_3250611, EPI_ISL_3250612, EPI_ISL_3250613, EPI_ISL_3250614, EPI_ISL_3250615, EPI_ISL_3250616, EPI_ISL_3250617, EPI_ISL_3250618, EPI_ISL_3250619, EPI_ISL_3250620, EPI_ISL_3250621, EPI_ISL_3250622, EPI_ISL_3250623, EPI_ISL_3250624, EPI_ISL_3250625, EPI_ISL_3250626, EPI_ISL_3250627, EPI_ISL_3250628, EPI_ISL_3250629, EPI_ISL_3250630, EPI_ISL_3250631, EPI_ISL_3250632, EPI_ISL_3250633, EPI_ISL_3250634, EPI_ISL_3250635, EPI_ISL_3250636, EPI_ISL_3250637, EPI_ISL_3250638, EPI_ISL_3250639, EPI_ISL_3250640, EPI_ISL_3250641, EPI_ISL_3250642, EPI_ISL_3250643, EPI_ISL_3250644, EPI_ISL_3250645, EPI_ISL_3250646, EPI_ISL_3250647, EPI_ISL_3250648, EPI_ISL_3250649, EPI_ISL_3250650, EPI_ISL_3250651, EPI_ISL_3250652, EPI_ISL_3250653, EPI_ISL_3250654, EPI_ISL_3250655, EPI_ISL_3250656, EPI_ISL_3250657, EPI_ISL_3250658, EPI_ISL_3250659, EPI_ISL_3250660, EPI_ISL_3250661, EPI_ISL_3250662, EPI_ISL_3250663, EPI_ISL_3250664, EPI_ISL_3250665, EPI_ISL_3250666, EPI_ISL_3250667, EPI_ISL_3250668, EPI_ISL_3250669, EPI_ISL_3250670, EPI_ISL_3250671, EPI_ISL_3250672, EPI_ISL_3250673, EPI_ISL_3250674 | | | |
| see above | Lighthouse Lab in Milton Keynes | Wellcome Sanger Institute for the COVID-19 Genomics UK (COG-UK) Consortium | The Lighthouse Lab in Milton Keynes and Alex Alderton, Roberto Amato, Jeffrey Barrett, Sonia Goncalves, Ewan Harrison, David K. Jackson, Ian Johnston, Dominic Kwiatkowski, Cordelia Langford, John Sillitoe on behalf of the Wellcome Sanger Institute COVID-19 Surveillance Team |
| EPI_ISL_3250675, EPI_ISL_3250676, EPI_ISL_3250677, EPI_ISL_3250678, EPI_ISL_3250679, EPI_ISL_3250680, EPI_ISL_3250681, EPI_ISL_3250682, EPI_ISL_3250683, EPI_ISL_3250684, EPI_ISL_3250685 | | | |
| see above | Lighthouse Lab in Alderley Park | Wellcome Sanger Institute for the COVID-19 Genomics UK (COG-UK) Consortium | Jacquelyn Wynn, Mairead Hyland, The Lighthouse Lab in Alderley Park and Alex Alderton, Roberto Amato, Jeffrey Barrett, Sonia Goncalves, Ewan Harrison, David K. Jackson, Ian Johnston, Dominic Kwiatkowski, Cordelia Langford, John Sillitoe on behalf of the Wellcome Sanger Institute COVID-19 Surveillance Team |
| EPI_ISL_3250686, EPI_ISL_3250688 | Berkshire and Surrey Pathology Services Lighthouse Laboratory | Wellcome Sanger Institute for the COVID-19 Genomics UK (COG-UK) Consortium | Berkshire and Surrey Pathology Services Lighthouse Laboratory and Alex Alderton, Roberto Amato, Jeffrey Barrett, Sonia Goncalves, Ewan Harrison, David K. Jackson, Ian Johnston, Dominic Kwiatkowski, Cordelia Langford, John Sillitoe on behalf of the Wellcome Sanger Institute COVID-19 Surveillance Team |
| EPI_ISL_3250689 | Lighthouse Lab in Alderley Park | Wellcome Sanger Institute for the COVID-19 Genomics UK (COG-UK) Consortium | Jacquelyn Wynn, Mairead Hyland, The Lighthouse Lab in Alderley Park and Alex Alderton, Roberto Amato, Jeffrey Barrett, Sonia Goncalves, Ewan Harrison, David K. Jackson, Ian Johnston, Dominic Kwiatkowski, Cordelia Langford, John Sillitoe on behalf of the Wellcome Sanger Institute COVID-19 Surveillance Team |
| EPI_ISL_3250690 | Berkshire and Surrey Pathology Services Lighthouse Laboratory | Wellcome Sanger Institute for the COVID-19 Genomics UK (COG-UK) Consortium | Berkshire and Surrey Pathology Services Lighthouse Laboratory and Alex Alderton, Roberto Amato, Jeffrey Barrett, Sonia Goncalves, Ewan Harrison, David K. Jackson, Ian Johnston, Dominic Kwiatkowski, Cordelia Langford, John Sillitoe on behalf of the Wellcome Sanger Institute COVID-19 Surveillance Team |
| EPI_ISL_3250691, EPI_ISL_3250692 | Lighthouse Lab in Alderley Park | Wellcome Sanger Institute for the COVID-19 Genomics UK (COG-UK) Consortium | Jacquelyn Wynn, Mairead Hyland, The Lighthouse Lab in Alderley Park and Alex Alderton, Roberto Amato, Jeffrey Barrett, Sonia Goncalves, Ewan Harrison, David K. Jackson, Ian Johnston, Dominic Kwiatkowski, Cordelia Langford, John Sillitoe on behalf of the Wellcome Sanger Institute COVID-19 Surveillance Team |
| EPI_ISL_3250693, EPI_ISL_3250694 | Berkshire and Surrey Pathology Services Lighthouse Laboratory | Wellcome Sanger Institute for the COVID-19 Genomics UK (COG-UK) Consortium | Berkshire and Surrey Pathology Services Lighthouse Laboratory and Alex Alderton, Roberto Amato, Jeffrey Barrett, Sonia Goncalves, Ewan Harrison, David K. Jackson, Ian Johnston, Dominic Kwiatkowski, Cordelia Langford, John Sillitoe on behalf of the Wellcome Sanger Institute COVID-19 Surveillance Team |
| EPI_ISL_3250695, EPI_ISL_3250696 | Lighthouse Lab in Alderley Park | Wellcome Sanger Institute for the COVID-19 Genomics UK (COG-UK) Consortium | Jacquelyn Wynn, Mairead Hyland, The Lighthouse Lab in Alderley Park and Alex Alderton, Roberto Amato, Jeffrey Barrett, Sonia Goncalves, Ewan Harrison, David K. Jackson, Ian Johnston, Dominic Kwiatkowski, Cordelia Langford, John Sillitoe on behalf of the Wellcome Sanger Institute COVID-19 Surveillance Team |
| EPI_ISL_3250697, EPI_ISL_3250698, EPI_ISL_3250699, EPI_ISL_3250700, EPI_ISL_3250701, EPI_ISL_3250702, EPI_ISL_3250703, EPI_ISL_3250704, EPI_ISL_3250705, EPI_ISL_3250706, EPI_ISL_3250707, EPI_ISL_3250708, EPI_ISL_3250709, EPI_ISL_3250710, EPI_ISL_3250711, EPI_ISL_3250712, EPI_ISL_3250713, EPI_ISL_3250714, EPI_ISL_3250715, EPI_ISL_3250716, EPI_ISL_3250717, EPI_ISL_3250718, EPI_ISL_3250719, EPI_ISL_3250720, EPI_ISL_3250721, EPI_ISL_3250722, EPI_ISL_3250723, EPI_ISL_3250724, EPI_ISL_3250725, EPI_ISL_3250726, EPI_ISL_3250727, EPI_ISL_3250728, EPI_ISL_3250729, EPI_ISL_3250730, EPI_ISL_3250731, EPI_ISL_3250732, EPI_ISL_3250733, EPI_ISL_3250734, EPI_ISL_3250735, EPI_ISL_3250736, EPI_ISL_3250737, EPI_ISL_3250738, EPI_ISL_3250739, EPI_ISL_3250740, EPI_ISL_3250741, EPI_ISL_3250742, EPI_ISL_3250743, EPI_ISL_3250744, EPI_ISL_3250745, EPI_ISL_3250746, EPI_ISL_3250747, EPI_ISL_3250748, EPI_ISL_3250749, EPI_ISL_3250750, EPI_ISL_3250751, EPI_ISL_3250752, EPI_ISL_3250753, EPI_ISL_3250754, EPI_ISL_3250755, EPI_ISL_3250756, EPI_ISL_3250757, EPI_ISL_3250758, EPI_ISL_3250759, EPI_ISL_3250760, EPI_ISL_3250761, EPI_ISL_3250762, EPI_ISL_3250763, EPI_ISL_3250764, EPI_ISL_3250765, EPI_ISL_3250766, EPI_ISL_3250767, EPI_ISL_3250768, EPI_ISL_3250769, EPI_ISL_3250770, EPI_ISL_3250771, EPI_ISL_3250772, EPI_ISL_3250773, EPI_ISL_3250774 | | | |
| see above | Lighthouse Lab in Milton Keynes | Wellcome Sanger Institute for the COVID-19 Genomics UK (COG-UK) Consortium | The Lighthouse Lab in Milton Keynes and Alex Alderton, Roberto Amato, Jeffrey Barrett, Sonia Goncalves, Ewan Harrison, David K. Jackson, Ian Johnston, Dominic Kwiatkowski, Cordelia Langford, John Sillitoe on behalf of the Wellcome Sanger Institute COVID-19 Surveillance Team |
| EPI_ISL_3250775 | Lighthouse Lab in Glasgow | Wellcome Sanger Institute for the COVID-19 Genomics UK (COG-UK) Consortium | Harper VanSteenhouse, Yumi Kasai, David Gray, Carol Clugston, Anna Dominiczak and Alex Alderton, Roberto Amato, Jeffrey Barrett, Sonia Goncalves, Ewan Harrison, David K. Jackson, Ian Johnston, Dominic Kwiatkowski, Cordelia Langford, John Sillitoe on behalf of the Wellcome Sanger Institute COVID-19 Surveillance Team |
| EPI_ISL_3250776, EPI_ISL_3250777, EPI_ISL_3250778, EPI_ISL_3250779, EPI_ISL_3250780, EPI_ISL_3250781, EPI_ISL_3250782, EPI_ISL_3250783, EPI_ISL_3250784, EPI_ISL_3250785, EPI_ISL_3250786, EPI_ISL_3250787, EPI_ISL_3250788, EPI_ISL_3250789, EPI_ISL_3250790, EPI_ISL_3250791, EPI_ISL_3250792, EPI_ISL_3250793, EPI_ISL_3250794, EPI_ISL_3250795, EPI_ISL_3250796, EPI_ISL_3250797, EPI_ISL_3250798, EPI_ISL_3250799, EPI_ISL_3250800, EPI_ISL_3250801, EPI_ISL_3250802, EPI_ISL_3250803, EPI_ISL_3250804, EPI_ISL_3250805, EPI_ISL_3250806, EPI_ISL_3250807, EPI_ISL_3250808, EPI_ISL_3250809, EPI_ISL_3250810, EPI_ISL_3250811, EPI_ISL_3250812, EPI_ISL_3250813, EPI_ISL_3250814, EPI_ISL_3250815, EPI_ISL_3250816, EPI_ISL_3250817, EPI_ISL_3250818, EPI_ISL_3250819, EPI_ISL_3250820, EPI_ISL_3250821, EPI_ISL_3250822, EPI_ISL_3250823, EPI_ISL_3250824, EPI_ISL_3250825, EPI_ISL_3250826, EPI_ISL_3250827, EPI_ISL_3250828, EPI_ISL_3250829, EPI_ISL_3250830, EPI_ISL_3250831, EPI_ISL_3250832, EPI_ISL_3250833, EPI_ISL_3250834, EPI_ISL_3250835, EPI_ISL_3250836, EPI_ISL_3250837, EPI_ISL_3250838, EPI_ISL_3250839, EPI_ISL_3250840, EPI_ISL_3250841, EPI_ISL_3250842, EPI_ISL_3250843, EPI_ISL_3250844, EPI_ISL_3250845, EPI_ISL_3250846 | | | |
| see above | Lighthouse Lab in Milton Keynes | Wellcome Sanger Institute for the COVID-19 Genomics UK (COG-UK) Consortium | The Lighthouse Lab in Milton Keynes and Alex Alderton, Roberto Amato, Jeffrey Barrett, Sonia Goncalves, Ewan Harrison, David K. Jackson, Ian Johnston, Dominic Kwiatkowski, Cordelia Langford, John Sillitoe on behalf of the Wellcome Sanger Institute COVID-19 Surveillance Team |
| EPI_ISL_3250847 | AZ Klina | AZ Klina | Carl Vael - Lynsey Berckmans |
| EPI_ISL_3250848, EPI_ISL_3250849, EPI_ISL_3250850, EPI_ISL_3250851, EPI_ISL_3250852, EPI_ISL_3250853, EPI_ISL_3250854, EPI_ISL_3250855, EPI_ISL_3250856, EPI_ISL_3250857, EPI_ISL_3250858, EPI_ISL_3250859, EPI_ISL_3250860, EPI_ISL_3250861, EPI_ISL_3250862, EPI_ISL_3250863, EPI_ISL_3250864, EPI_ISL_3250865, EPI_ISL_3250866, EPI_ISL_3250867, EPI_ISL_3250868, EPI_ISL_3250869, EPI_ISL_3250870, EPI_ISL_3250871, EPI_ISL_3250872, EPI_ISL_3250873, EPI_ISL_3250874, EPI_ISL_3250875, EPI_ISL_3250876, EPI_ISL_3250877, EPI_ISL_3250878, EPI_ISL_3250879, EPI_ISL_3250880, EPI_ISL_3250881, EPI_ISL_3250882, EPI_ISL_3250883, EPI_ISL_3250884, EPI_ISL_3250885, EPI_ISL_3250886, EPI_ISL_3250887, EPI_ISL_3250888, EPI_ISL_3250889, EPI_ISL_3250890, EPI_ISL_3250891, EPI_ISL_3250892, EPI_ISL_3250893, EPI_ISL_3250894, EPI_ISL_3250895, EPI_ISL_3250896, EPI_ISL_3250897, EPI_ISL_3250898, EPI_ISL_3250899, EPI_ISL_3250900, EPI_ISL_3250901, EPI_ISL_3250902, EPI_ISL_3250903, EPI_ISL_3250904, EPI_ISL_3250905, EPI_ISL_3250906, EPI_ISL_3250907, EPI_ISL_3250908, EPI_ISL_3250909, EPI_ISL_3250910, EPI_ISL_3250911, EPI_ISL_3250912, EPI_ISL_3250913, EPI_ISL_3250914, EPI_ISL_3250915, EPI_ISL_3250916, EPI_ISL_3250917, EPI_ISL_3250918, EPI_ISL_3250919, EPI_ISL_3250920, EPI_ISL_3250921, EPI_ISL_3250922, EPI_ISL_3250923, EPI_ISL_3250924, EPI_ISL_3250925, EPI_ISL_3250926, EPI_ISL_3250927, EPI_ISL_3250928, EPI_ISL_3250929, EPI_ISL_3250930, EPI_ISL_3250931, EPI_ISL_3250932, EPI_ISL_3250933, EPI_ISL_3250934, EPI_ISL_3250935, EPI_ISL_3250936, EPI_ISL_3250937, EPI_ISL_3250938, EPI_ISL_3250939, EPI_ISL_3250940, EPI_ISL_3250941, EPI_ISL_3250942, EPI_ISL_3250943, EPI_ISL_3250944, EPI_ISL_3250945, EPI_ISL_3250946, EPI_ISL_3250947, EPI_ISL_3250948, EPI_ISL_3250949, EPI_ISL_3250950, EPI_ISL_3250951, EPI_ISL_3250952, EPI_ISL_3250953, EPI_ISL_3250954, EPI_ISL_3250955, EPI_ISL_3250956, EPI_ISL_3250957, EPI_ISL_3250958, EPI_ISL_3250959, EPI_ISL_3250960, EPI_ISL_3250961, EPI_ISL_3250962, EPI_ISL_3250963, EPI_ISL_3250964, EPI_ISL_3250965, EPI_ISL_3250966, EPI_ISL_3250967, EPI_ISL_3250968, EPI_ISL_3250969, EPI_ISL_3250970, EPI_ISL_3250971, EPI_ISL_3250972, EPI_ISL_3250973, EPI_ISL_3250974, EPI_ISL_3250975, EPI_ISL_3250976, EPI_ISL_3250977, EPI_ISL_3250978, EPI_ISL_3250979, EPI_ISL_3250980, EPI_ISL_3250981, EPI_ISL_3250982, EPI_ISL_3250983, EPI_ISL_3250984, EPI_ISL_3250985, EPI_ISL_3250986, EPI_ISL_3250987, EPI_ISL_3250988, EPI_ISL_3250989, EPI_ISL_3250990, EPI_ISL_3250991, EPI_ISL_3250992, EPI_ISL_3250993, EPI_ISL_3250994, EPI_ISL_3250995, EPI_ISL_3250996, EPI_ISL_3250997, EPI_ISL_3250998, EPI_ISL_3250999, EPI_ISL_3251000, EPI_ISL_3251001, EPI_ISL_3251002, EPI_ISL_3251003, EPI_ISL_3251004, EPI_ISL_3251005, EPI_ISL_3251006, EPI_ISL_3251007, EPI_ISL_3251008, EPI_ISL_3251009, EPI_ISL_3251010, EPI_ISL_3251011, EPI_ISL_3251012, EPI_ISL_3251013, EPI_ISL_3251014, EPI_ISL_3251015, EPI_ISL_3251016, EPI_ISL_3251017, EPI_ISL_3251018, EPI_ISL_3251019, EPI_ISL_3251020, EPI_ISL_3251021, EPI_ISL_3251022, EPI_ISL_3251023, EPI_ISL_3251024, EPI_ISL_3251025, EPI_ISL_3251026, EPI_ISL_3251027, EPI_ISL_3251028, EPI_ISL_3251029, EPI_ISL_3251030, EPI_ISL_3251031, EPI_ISL_3251032, EPI_ISL_3251033, EPI_ISL_3251034, EPI_ISL_3251035, EPI_ISL_3251036, EPI_ISL_3251037, EPI_ISL_3251038, EPI_ISL_3251039, EPI_ISL_3251040, EPI_ISL_3251041, EPI_ISL_3251042, EPI_ISL_3251043, EPI_ISL_3251044, EPI_ISL_3251045, EPI_ISL_3251046, EPI_ISL_3251047, EPI_ISL_3251048, EPI_ISL_3251049, EPI_ISL_3251050, EPI_ISL_3251051, EPI_ISL_3251052, EPI_ISL_3251053, EPI_ISL_3251054, EPI_ISL_3251055, EPI_ISL_3251056, EPI_ISL_3251057, EPI_ISL_3251058, EPI_ISL_3251059, EPI_ISL_3251060, EPI_ISL_3251061, EPI_ISL_3251062, EPI_ISL_3251063, EPI_ISL_3251064, EPI_ISL_3251065, EPI_ISL_3251066, EPI_ISL_3251067, EPI_ISL_3251068, EPI_ISL_3251069, EPI_ISL_3251070, EPI_ISL_3251071, EPI_ISL_3251072, EPI_ISL_3251073, EPI_ISL_3251074, EPI_ISL_3251075, EPI_ISL_3251076, EPI_ISL_3251077, EPI_ISL_3251078, EPI_ISL_3251079, EPI_ISL_3251080, EPI_ISL_3251081, EPI_ISL_3251082, EPI_ISL_3251083, EPI_ISL_3251084, EPI_ISL_3251085, EPI_ISL_3251086, EPI_ISL_3251087, EPI_ISL_3251088, EPI_ISL_3251089, EPI_ISL_3251090, EPI_ISL_3251091, EPI_ISL_3251092, EPI_ISL_3251093, EPI_ISL_3251094, EPI_ISL_3251095, EPI_ISL_3251096, EPI_ISL_3251097, EPI_ISL_3251098, EPI_ISL_3251099, EPI_ISL_3251100, EPI_ISL_3251101, EPI_ISL_3251102, EPI_ISL_3251103, EPI_ISL_3251104, EPI_ISL_3251105, EPI_ISL_3251106, EPI_ISL_3251107, EPI_ISL_3251108, EPI_ISL_3251109, EPI_ISL_3251110, EPI_ISL_3251111, EPI_ISL_3251112, EPI_ISL_3251113, EPI_ISL_3251114, EPI_ISL_3251115, EPI_ISL_3251116, EPI_ISL_3251117, EPI_ISL_3251118, EPI_ISL_3251119, EPI_ISL_3251120, EPI_ISL_3251121, EPI_ISL_3251122, EPI_ISL_3251123, EPI_ISL_3251124, EPI_ISL_3251125, EPI_ISL_3251126, EPI_ISL_3251127, EPI_ISL_3251128, EPI_ISL_3251129, EPI_ISL_3251130, EPI_ISL_3251131, EPI_ISL_3251132, EPI_ISL_3251133, EPI_ISL_3251134, EPI_ISL_3251135, EPI_ISL_3251136, EPI_ISL_3251137, EPI_ISL_3251138, EPI_ISL_3251139, EPI_ISL_3251140, EPI_ISL_3251141, EPI_ISL_3251142, EPI_ISL_3251143, EPI_ISL_3251144, EPI_ISL_3251145, EPI_ISL_3251146, EPI_ISL_3251147, EPI_ISL_3251148, EPI_ISL_3251149, EPI_ISL_3251150, EPI_ISL_3251151, EPI_ISL_3251152, EPI_ISL_3251153, EPI_ISL_3251154, EPI_ISL_3251155, EPI_ISL_3251156, EPI_ISL_3251157, EPI_ISL_3251158, EPI_ISL_3251159, EPI_ISL_3251160, EPI_ISL_3251161, EPI_ISL_3251162, EPI_ISL_3251163, EPI_ISL_3251164, EPI_ISL_3251165, EPI_ISL_3251166, EPI_ISL_3251167, EPI_ISL_3251168, EPI_ISL_3251169, EPI_ISL_3251170, EPI_ISL_3251171, EPI_ISL_3251172, EPI_ISL_3251173, EPI_ISL_3251174, EPI_ISL_3251175, EPI_ISL_3251176, EPI_ISL_3251177, EPI_ISL_3251178, EPI_ISL_3251179, EPI_ISL_3251180, EPI_ISL_3251181, EPI_ISL_3251182, EPI_ISL_3251183, EPI_ISL_3251184, EPI_ISL_3251185, EPI_ISL_3251187, EPI_ISL_3251188, EPI_ISL_3251189, EPI_ISL_3251190, EPI_ISL_3251191, EPI_ISL_3251192, EPI_ISL_3251193, EPI_ISL_3251195, EPI_ISL_3251196, EPI_ISL_3251197, EPI_ISL_3251198, EPI_ISL_3251199, EPI_ISL_3251200, EPI_ISL_3251201, EPI_ISL_3251202, EPI_ISL_3251203, EPI_ISL_3251204, EPI_ISL_3251205, EPI_ISL_3251206, EPI_ISL_3251207, EPI_ISL_3251208, EPI_ISL_3251209, EPI_ISL_3251210, EPI_ISL_3251211, EPI_ISL_3251212, EPI_ISL_3251213, EPI_ISL_3251214, EPI_ISL_3251215, EPI_ISL_3251216, EPI_ISL_3251217, EPI_ISL_3251218, EPI_ISL_3251219, EPI_ISL_3251220, EPI_ISL_3251221, EPI_ISL_3251222, EPI_ISL_3251223, EPI_ISL_3251224, EPI_ISL_3251225, EPI_ISL_3251226, EPI_ISL_3251227, EPI_ISL_3251228, EPI_ISL_3251229, EPI_ISL_3251230, EPI_ISL_3251231, EPI_ISL_3251232, EPI_ISL_3251233, EPI_ISL_3251234, EPI_ISL_3251235, EPI_ISL_3251236, EPI_ISL_3251237, EPI_ISL_3251238, EPI_ISL_3251239, EPI_ISL_3251240 | | | |
| see above | Lighthouse Lab in Milton Keynes | Wellcome Sanger Institute for the COVID-19 Genomics UK (COG-UK) | The Lighthouse Lab in Milton Keynes and Alex Alderton, Roberto Amato, Jeffrey Barrett, Sonia Goncalves, Ewan Harrison, David K. Jackson, Ian Johnston, Dominic Kwiatkowski, Cordelia Langford, John Sillitoe on behalf of the Wellcome Sanger |

| | | | |
|---|---|---|---|
| | | Consortium | Institute COVID-19 Surveillance Team |
| EPI_ISL_3251242, EPI_ISL_3251243, EPI_ISL_3251244, EPI_ISL_3251245, EPI_ISL_3251246 | AZ Klina | AZ Klina | Carl Vael - Lynsey Berckmans |
| EPI_ISL_3251247, EPI_ISL_3251248, EPI_ISL_3251249, EPI_ISL_3251250, EPI_ISL_3251251, EPI_ISL_3251252, EPI_ISL_3251253, EPI_ISL_3251254, EPI_ISL_3251255, EPI_ISL_3251256, EPI_ISL_3251257, EPI_ISL_3251258, EPI_ISL_3251259, EPI_ISL_3251260, EPI_ISL_3251261, EPI_ISL_3251262, EPI_ISL_3251263, EPI_ISL_3251264, EPI_ISL_3251265, EPI_ISL_3251266, EPI_ISL_3251267, EPI_ISL_3251268, EPI_ISL_3251269, EPI_ISL_3251270, EPI_ISL_3251271, EPI_ISL_3251272, EPI_ISL_3251273, EPI_ISL_3251274, EPI_ISL_3251275, EPI_ISL_3251276, EPI_ISL_3251277, EPI_ISL_3251278, EPI_ISL_3251279, EPI_ISL_3251280, EPI_ISL_3251281, EPI_ISL_3251282, EPI_ISL_3251283, EPI_ISL_3251284, EPI_ISL_3251285, EPI_ISL_3251286, EPI_ISL_3251287, EPI_ISL_3251288, EPI_ISL_3251289, EPI_ISL_3251290, EPI_ISL_3251291, EPI_ISL_3251292, EPI_ISL_3251293, EPI_ISL_3251294, EPI_ISL_3251295, EPI_ISL_3251296, EPI_ISL_3251297, EPI_ISL_3251298, EPI_ISL_3251299, EPI_ISL_3251300, EPI_ISL_3251301, EPI_ISL_3251302, EPI_ISL_3251303, EPI_ISL_3251304, EPI_ISL_3251305, EPI_ISL_3251306, EPI_ISL_3251307 | | | |
| see above | Department of Bacteria, Parasites and Fungi, Statens Serum Institut, Copenhagen, Denmark | Statens Serum Institut Bioinformatics and Microbial Genomics | Danish Covid-19 Genome Consortium |
| EPI_ISL_3251308 | Molekylær Medicinsk Afdeling, Aarhus University Hospital, Aarhus, Denmark | Statens Serum Institut Bioinformatics and Microbial Genomics | Danish Covid-19 Genome Consortium |
| EPI_ISL_3251309, EPI_ISL_3251310, EPI_ISL_3251311, EPI_ISL_3251312, EPI_ISL_3251313, EPI_ISL_3251314 | Department of Bacteria, Parasites and Fungi, Statens Serum Institut, Copenhagen, Denmark | Statens Serum Institut Bioinformatics and Microbial Genomics | Danish Covid-19 Genome Consortium |
| EPI_ISL_3251315 | AZ Klina | AZ Klina | Carl Vael - Lynsey Berckmans |
| EPI_ISL_3251316, EPI_ISL_3251317, EPI_ISL_3251318, EPI_ISL_3251319, EPI_ISL_3251320, EPI_ISL_3251321, EPI_ISL_3251322, EPI_ISL_3251323, EPI_ISL_3251324, EPI_ISL_3251325, EPI_ISL_3251326, EPI_ISL_3251327, EPI_ISL_3251328, EPI_ISL_3251329, EPI_ISL_3251330, EPI_ISL_3251331, EPI_ISL_3251332, EPI_ISL_3251333, EPI_ISL_3251334, EPI_ISL_3251335, EPI_ISL_3251336, EPI_ISL_3251337, EPI_ISL_3251338, EPI_ISL_3251339, EPI_ISL_3251340, EPI_ISL_3251341 | | | |
| see above | Department of Bacteria, Parasites and Fungi, Statens Serum Institut, Copenhagen, Denmark | Statens Serum Institut Bioinformatics and Microbial Genomics | Danish Covid-19 Genome Consortium |
| EPI_ISL_3251342 | Department of Clinical Microbiology and Center for Genomic Medicine, Rigshospitalet, Copenhagen, Denmark | Statens Serum Institut Bioinformatics and Microbial Genomics | Danish Covid-19 Genome Consortium |
| EPI_ISL_3251343, EPI_ISL_3251344, EPI_ISL_3251345, EPI_ISL_3251346, EPI_ISL_3251347, EPI_ISL_3251348, EPI_ISL_3251349, EPI_ISL_3251350, EPI_ISL_3251351, EPI_ISL_3251352, EPI_ISL_3251353, EPI_ISL_3251354, EPI_ISL_3251355, EPI_ISL_3251356, EPI_ISL_3251357, EPI_ISL_3251358, EPI_ISL_3251359, EPI_ISL_3251360, EPI_ISL_3251361, EPI_ISL_3251362, EPI_ISL_3251363, EPI_ISL_3251364, EPI_ISL_3251365, EPI_ISL_3251366, EPI_ISL_3251367, EPI_ISL_3251368, EPI_ISL_3251369, EPI_ISL_3251370, EPI_ISL_3251371, EPI_ISL_3251372, EPI_ISL_3251373, EPI_ISL_3251375, EPI_ISL_3251376, EPI_ISL_3251377, EPI_ISL_3251378, EPI_ISL_3251379, EPI_ISL_3251380, EPI_ISL_3251381, EPI_ISL_3251382, EPI_ISL_3251383, EPI_ISL_3251384, EPI_ISL_3251385, EPI_ISL_3251386, EPI_ISL_3251387, EPI_ISL_3251388, EPI_ISL_3251389, EPI_ISL_3251390, EPI_ISL_3251391, EPI_ISL_3251392, EPI_ISL_3251393, EPI_ISL_3251394, EPI_ISL_3251395, EPI_ISL_3251396, EPI_ISL_3251397, EPI_ISL_3251398, EPI_ISL_3251399, EPI_ISL_3251400, EPI_ISL_3251401, EPI_ISL_3251402, EPI_ISL_3251403, EPI_ISL_3251404, EPI_ISL_3251405, EPI_ISL_3251406, EPI_ISL_3251407, EPI_ISL_3251408, EPI_ISL_3251409, EPI_ISL_3251410, EPI_ISL_3251411, EPI_ISL_3251412, EPI_ISL_3251413, EPI_ISL_3251414, EPI_ISL_3251415, EPI_ISL_3251416 | | | |
| see above | Department of Bacteria, Parasites and Fungi, Statens Serum Institut, Copenhagen, Denmark | Statens Serum Institut Bioinformatics and Microbial Genomics | Danish Covid-19 Genome Consortium |
| EPI_ISL_3251417 | Molekylær Medicinsk Afdeling, Aarhus University Hospital, Aarhus, Denmark | Statens Serum Institut Bioinformatics and Microbial Genomics | Danish Covid-19 Genome Consortium |
| EPI_ISL_3251418, EPI_ISL_3251419, EPI_ISL_3251421, EPI_ISL_3251422, EPI_ISL_3251423, EPI_ISL_3251424, EPI_ISL_3251425, EPI_ISL_3251426, EPI_ISL_3251427, EPI_ISL_3251428, EPI_ISL_3251429 | | | |
| see above | Department of Bacteria, Parasites and Fungi, Statens Serum Institut, Copenhagen, Denmark | Statens Serum Institut Bioinformatics and Microbial Genomics | Danish Covid-19 Genome Consortium |
| EPI_ISL_3251430 | Molekylær Medicinsk Afdeling, Aarhus University Hospital, Aarhus, Denmark | Statens Serum Institut Bioinformatics and Microbial Genomics | Danish Covid-19 Genome Consortium |
| EPI_ISL_3251431, EPI_ISL_3251432, EPI_ISL_3251433, EPI_ISL_3251434, EPI_ISL_3251435, EPI_ISL_3251436, EPI_ISL_3251437, EPI_ISL_3251438 | Department of Bacteria, Parasites and Fungi, Statens Serum Institut, Copenhagen, Denmark | Statens Serum Institut Bioinformatics and Microbial Genomics | Danish Covid-19 Genome Consortium |
| EPI_ISL_3251439 | Laboratorium Analiz Lekarskich ALAB Rzeszow | Wojewodzka Stacja Sanitarno-Epidemiologiczna w Rzeszowie, Laboratorium Diagnostyki Medycznej | Marzena Baranowska, Katarzyna Wilk, Karolina Ostrowska, Anna Nowakowska |
| EPI_ISL_3251440, EPI_ISL_3251441 | Department of Bacteria, Parasites and Fungi, Statens Serum Institut, Copenhagen, Denmark | Statens Serum Institut Bioinformatics and Microbial Genomics | Danish Covid-19 Genome Consortium |
| EPI_ISL_3251442 | Centrum Medyczne w Lancucie Dzial Diagnostyki Laboratoryjnej | Wojewodzka Stacja Sanitarno-Epidemiologiczna w Rzeszowie, Laboratorium Diagnostyki Medycznej | Marzena Baranowska, Katarzyna Wilk, Karolina Ostrowska, Anna Nowakowska |
| EPI_ISL_3251443, EPI_ISL_3251445, EPI_ISL_3251447 | Department of Bacteria, Parasites and Fungi, Statens Serum Institut, Copenhagen, Denmark | Statens Serum Institut Bioinformatics and Microbial Genomics | Danish Covid-19 Genome Consortium |
| EPI_ISL_3251448 | Laboratoria Medyczne OptiMed | Wojewodzka Stacja Sanitarno-Epidemiologiczna w Rzeszowie, Laboratorium Diagnostyki Medycznej | Marzena Baranowska, Katarzyna Wilk, Karolina Ostrowska, Anna Nowakowska |
| EPI_ISL_3251449 | Department of Bacteria, Parasites and Fungi, Statens Serum Institut, Copenhagen, Denmark | Statens Serum Institut Bioinformatics and Microbial Genomics | Danish Covid-19 Genome Consortium |
| EPI_ISL_3251450 | Laboratorium Medyczne GenLab | Wojewodzka Stacja Sanitarno-Epidemiologiczna w Rzeszowie, Laboratorium Diagnostyki Medycznej | Marzena Baranowska, Katarzyna Wilk, Karolina Ostrowska, Anna Nowakowska |
| EPI_ISL_3251451 | Department of Bacteria, Parasites and Fungi, Statens Serum Institut, Copenhagen, Denmark | Statens Serum Institut Bioinformatics and Microbial Genomics | Danish Covid-19 Genome Consortium |
| EPI_ISL_3251452 | Szpital Wojewodzki w Lomzy | Wojewodzka Stacja Sanitarno-Epidemiologiczna w Rzeszowie, Laboratorium Diagnostyki Medycznej | Marzena Baranowska, Katarzyna Wilk, Karolina Ostrowska, Anna Nowakowska |
| EPI_ISL_3251453 | Department of Bacteria, Parasites and Fungi, Statens Serum Institut, Copenhagen, Denmark | Statens Serum Institut Bioinformatics and Microbial Genomics | Danish Covid-19 Genome Consortium |
| EPI_ISL_3251454 | Uniwersytecki Szpital Kliniczny w Bialymstoku Zaklad Laboratoryjnej Diagnostyki Klinicznej | Wojewodzka Stacja Sanitarno-Epidemiologiczna w Rzeszowie, Laboratorium Diagnostyki Medycznej | Marzena Baranowska, Katarzyna Wilk, Karolina Ostrowska, Anna Nowakowska |
| EPI_ISL_3251455, EPI_ISL_3251456 | Department of Bacteria, Parasites and Fungi, Statens Serum Institut, Copenhagen, Denmark | Statens Serum Institut Bioinformatics and Microbial Genomics | Danish Covid-19 Genome Consortium |
| EPI_ISL_3251457 | Regionalne Centrum Krwiodawstwa i Krwiolecznictwa z Bialegostoku | Wojewodzka Stacja Sanitarno-Epidemiologiczna w Rzeszowie, Laboratorium Diagnostyki Medycznej | Marzena Baranowska, Katarzyna Wilk, Karolina Ostrowska, Anna Nowakowska |
| EPI_ISL_3251458 | Department of Bacteria, Parasites and Fungi, Statens Serum Institut, Copenhagen, Denmark | Statens Serum Institut Bioinformatics and Microbial Genomics | Danish Covid-19 Genome Consortium |
| EPI_ISL_3251459 | WSSE w Warszawie | Wojewodzka Stacja Sanitarno-Epidemiologiczna w Rzeszowie, Laboratorium Diagnostyki Medycznej | Marzena Baranowska, Katarzyna Wilk, Karolina Ostrowska, Anna Nowakowska |
| EPI_ISL_3251460, EPI_ISL_3251462 | Department of Bacteria, Parasites and Fungi, Statens Serum Institut, Copenhagen, Denmark | Statens Serum Institut Bioinformatics and Microbial Genomics | Danish Covid-19 Genome Consortium |
| EPI_ISL_3251463 | Cellis Laboratorium | Wojewodzka Stacja Sanitarno-Epidemiologiczna w Rzeszowie, Laboratorium Diagnostyki Medycznej | Marzena Baranowska, Katarzyna Wilk, Karolina Ostrowska, Anna Nowakowska |
| EPI_ISL_3251464 | Department of Bacteria, Parasites and Fungi, Statens Serum Institut, Copenhagen, Denmark | Statens Serum Institut Bioinformatics and Microbial Genomics | Danish Covid-19 Genome Consortium |
| EPI_ISL_3251465 | NZOZ MediGen Daignostyka Molekularna | Wojewodzka Stacja Sanitarno-Epidemiologiczna w Rzeszowie, Laboratorium Diagnostyki Medycznej | Marzena Baranowska, Katarzyna Wilk, Karolina Ostrowska, Anna Nowakowska |
| EPI_ISL_3251466 | Department of Bacteria, Parasites and Fungi, Statens Serum Institut, Copenhagen, Denmark | Statens Serum Institut Bioinformatics and Microbial Genomics | Danish Covid-19 Genome Consortium |
| EPI_ISL_3251467 | Medyczne Laboratorium Diagnostyka | Wojewodzka Stacja Sanitarno-Epidemiologiczna w Rzeszowie, Laboratorium Diagnostyki Medycznej | Marzena Baranowska, Katarzyna Wilk, Karolina Ostrowska, Anna Nowakowska |
| EPI_ISL_3251468, EPI_ISL_3251469 | Department of Bacteria, Parasites and Fungi, Statens Serum Institut, Copenhagen, Denmark | Statens Serum Institut Bioinformatics and Microbial Genomics | Danish Covid-19 Genome Consortium |
| EPI_ISL_3251470 | Medyczne Laboratorium Diagnostyka | Wojewodzka Stacja Sanitarno-Epidemiologiczna w Rzeszowie, | Marzena Baranowska, Katarzyna Wilk, Karolina Ostrowska, Anna Nowakowska |

| | | | |
|---|---|---|---|
| | | Laboratorium Diagnostyki Medycznej | |
| EPI_ISL_3251471 | Department of Bacteria, Parasites and Fungi, Statens Serum Institut, Copenhagen, Denmark | Statens Serum Institut Bioinformatics and Microbial Genomics | Danish Covid-19 Genome Consortium |
| EPI_ISL_3251472 | Medyczne Laboratorium Diagnostyka | Wojewodzka Stacja Sanitarno-Epidemiologiczna w Rzeszowie, Laboratorium Diagnostyki Medycznej | Marzena Baranowska, Katarzyna Wilk, Karolina Ostrowska, Anna Nowakowska |
| EPI_ISL_3251473 | Department of Bacteria, Parasites and Fungi, Statens Serum Institut, Copenhagen, Denmark | Statens Serum Institut Bioinformatics and Microbial Genomics | Danish Covid-19 Genome Consortium |
| EPI_ISL_3251474 | Laboratorium Mikrobiologiczne RSS w Grudziadzu | Wojewodzka Stacja Sanitarno-Epidemiologiczna w Rzeszowie, Laboratorium Diagnostyki Medycznej | Marzena Baranowska, Katarzyna Wilk, Karolina Ostrowska, Anna Nowakowska |
| EPI_ISL_3251475 | Department of Bacteria, Parasites and Fungi, Statens Serum Institut, Copenhagen, Denmark | Statens Serum Institut Bioinformatics and Microbial Genomics | Danish Covid-19 Genome Consortium |
| EPI_ISL_3251476 | Kujawsko-Pomorskie Centrum Pulmonologii Zaklad Diagnostyki Mikrobiologicznej | Wojewodzka Stacja Sanitarno-Epidemiologiczna w Rzeszowie, Laboratorium Diagnostyki Medycznej | Marzena Baranowska, Katarzyna Wilk, Karolina Ostrowska, Anna Nowakowska |
| EPI_ISL_3251477 | Department of Bacteria, Parasites and Fungi, Statens Serum Institut, Copenhagen, Denmark | Statens Serum Institut Bioinformatics and Microbial Genomics | Danish Covid-19 Genome Consortium |
| EPI_ISL_3251478 | Kujawsko-Pomorskie Centrum Pulmonologii Zaklad Diagnostyki Mikrobiologicznej | Wojewodzka Stacja Sanitarno-Epidemiologiczna w Rzeszowie, Laboratorium Diagnostyki Medycznej | Marzena Baranowska, Katarzyna Wilk, Karolina Ostrowska, Anna Nowakowska |
| EPI_ISL_3251479 | Department of Bacteria, Parasites and Fungi, Statens Serum Institut, Copenhagen, Denmark | Statens Serum Institut Bioinformatics and Microbial Genomics | Danish Covid-19 Genome Consortium |
| EPI_ISL_3251480 | Laboratorium Mikrobiologiczne RSS w Grudziadzu | Wojewodzka Stacja Sanitarno-Epidemiologiczna w Rzeszowie, Laboratorium Diagnostyki Medycznej | Marzena Baranowska, Katarzyna Wilk, Karolina Ostrowska, Anna Nowakowska |
| EPI_ISL_3251481 | Department of Bacteria, Parasites and Fungi, Statens Serum Institut, Copenhagen, Denmark | Statens Serum Institut Bioinformatics and Microbial Genomics | Danish Covid-19 Genome Consortium |
| EPI_ISL_3251482 | Laboratorium Mikrobiologiczne RSS w Grudziadzu | Wojewodzka Stacja Sanitarno-Epidemiologiczna w Rzeszowie, Laboratorium Diagnostyki Medycznej | Marzena Baranowska, Katarzyna Wilk, Karolina Ostrowska, Anna Nowakowska |
| EPI_ISL_3251483, EPI_ISL_3251484 | Department of Bacteria, Parasites and Fungi, Statens Serum Institut, Copenhagen, Denmark | Statens Serum Institut Bioinformatics and Microbial Genomics | Danish Covid-19 Genome Consortium |
| EPI_ISL_3251485 | Kujawsko-Pomorskie Centrum Pulmonologii Zaklad Diagnostyki Mikrobiologicznej | Wojewodzka Stacja Sanitarno-Epidemiologiczna w Rzeszowie, Laboratorium Diagnostyki Medycznej | Marzena Baranowska, Katarzyna Wilk, Karolina Ostrowska, Anna Nowakowska |
| EPI_ISL_3251486 | Department of Bacteria, Parasites and Fungi, Statens Serum Institut, Copenhagen, Denmark | Statens Serum Institut Bioinformatics and Microbial Genomics | Danish Covid-19 Genome Consortium |
| EPI_ISL_3251487 | IMMQuest | Wojewodzka Stacja Sanitarno-Epidemiologiczna w Rzeszowie, Laboratorium Diagnostyki Medycznej | Marzena Baranowska, Katarzyna Wilk, Karolina Ostrowska, Anna Nowakowska |
| EPI_ISL_3251488 | Department of Bacteria, Parasites and Fungi, Statens Serum Institut, Copenhagen, Denmark | Statens Serum Institut Bioinformatics and Microbial Genomics | Danish Covid-19 Genome Consortium |
| EPI_ISL_3251489 | Diagnostyka | Wojewodzka Stacja Sanitarno-Epidemiologiczna w Rzeszowie, Laboratorium Diagnostyki Medycznej | Marzena Baranowska, Katarzyna Wilk, Karolina Ostrowska, Anna Nowakowska |
| EPI_ISL_3251490 | Department of Bacteria, Parasites and Fungi, Statens Serum Institut, Copenhagen, Denmark | Statens Serum Institut Bioinformatics and Microbial Genomics | Danish Covid-19 Genome Consortium |
| EPI_ISL_3251491 | Diagnostyka | Wojewodzka Stacja Sanitarno-Epidemiologiczna w Rzeszowie, Laboratorium Diagnostyki Medycznej | Marzena Baranowska, Katarzyna Wilk, Karolina Ostrowska, Anna Nowakowska |
| EPI_ISL_3251492 | Department of Bacteria, Parasites and Fungi, Statens Serum Institut, Copenhagen, Denmark | Statens Serum Institut Bioinformatics and Microbial Genomics | Danish Covid-19 Genome Consortium |
| EPI_ISL_3251493 | Laboratorium Biologii Molekularnej USK | Wojewodzka Stacja Sanitarno-Epidemiologiczna w Rzeszowie, Laboratorium Diagnostyki Medycznej | Marzena Baranowska, Katarzyna Wilk, Karolina Ostrowska, Anna Nowakowska |
| EPI_ISL_3251494 | Department of Bacteria, Parasites and Fungi, Statens Serum Institut, Copenhagen, Denmark | Statens Serum Institut Bioinformatics and Microbial Genomics | Danish Covid-19 Genome Consortium |
| EPI_ISL_3251495 | Laboratorium Biologii Molekularnej USK | Wojewodzka Stacja Sanitarno-Epidemiologiczna w Rzeszowie, Laboratorium Diagnostyki Medycznej | Marzena Baranowska, Katarzyna Wilk, Karolina Ostrowska, Anna Nowakowska |
| EPI_ISL_3251496, EPI_ISL_3251497 | Department of Bacteria, Parasites and Fungi, Statens Serum Institut, Copenhagen, Denmark | Statens Serum Institut Bioinformatics and Microbial Genomics | Danish Covid-19 Genome Consortium |
| EPI_ISL_3251498 | Laboratorium Biologii Molekularnej USK | Wojewodzka Stacja Sanitarno-Epidemiologiczna w Rzeszowie, Laboratorium Diagnostyki Medycznej | Marzena Baranowska, Katarzyna Wilk, Karolina Ostrowska, Anna Nowakowska |
| EPI_ISL_3251499 | Department of Bacteria, Parasites and Fungi, Statens Serum Institut, Copenhagen, Denmark | Statens Serum Institut Bioinformatics and Microbial Genomics | Danish Covid-19 Genome Consortium |
| EPI_ISL_3251500 | Laboratorium Biologii Molekularnej USK | Wojewodzka Stacja Sanitarno-Epidemiologiczna w Rzeszowie, Laboratorium Diagnostyki Medycznej | Marzena Baranowska, Katarzyna Wilk, Karolina Ostrowska, Anna Nowakowska |
| EPI_ISL_3251501 | Department of Bacteria, Parasites and Fungi, Statens Serum Institut, Copenhagen, Denmark | Statens Serum Institut Bioinformatics and Microbial Genomics | Danish Covid-19 Genome Consortium |
| EPI_ISL_3251502 | Centrum Medyczne MEDYK Zaklad Diagnostyki Medycznej | Wojewodzka Stacja Sanitarno-Epidemiologiczna w Rzeszowie, Laboratorium Diagnostyki Medycznej | Marzena Baranowska, Katarzyna Wilk, Karolina Ostrowska, Anna Nowakowska |
| EPI_ISL_3251503 | Laboratorium Diagnostyczne Centrum Medyczne LuxMed | Wojewodzka Stacja Sanitarno-Epidemiologiczna w Rzeszowie, Laboratorium Diagnostyki Medycznej | Marzena Baranowska, Katarzyna Wilk, Karolina Ostrowska, Anna Nowakowska |
| EPI_ISL_3251504 | Department of Bacteria, Parasites and Fungi, Statens Serum Institut, Copenhagen, Denmark | Statens Serum Institut Bioinformatics and Microbial Genomics | Danish Covid-19 Genome Consortium |
| EPI_ISL_3251505 | Zaklad Diagnostyki Laboratoryjnej Koagulologii i Mikrobiologii | Wojewodzka Stacja Sanitarno-Epidemiologiczna w Rzeszowie, Laboratorium Diagnostyki Medycznej | Marzena Baranowska, Katarzyna Wilk, Karolina Ostrowska, Anna Nowakowska |
| EPI_ISL_3251506 | Department of Bacteria, Parasites and Fungi, Statens Serum Institut, Copenhagen, Denmark | Statens Serum Institut Bioinformatics and Microbial Genomics | Danish Covid-19 Genome Consortium |
| EPI_ISL_3251507 | Zaklad Diagnostyki Laboratoryjnej Koagulologii i Mikrobiologii | Wojewodzka Stacja Sanitarno-Epidemiologiczna w Rzeszowie, Laboratorium Diagnostyki Medycznej | Marzena Baranowska, Katarzyna Wilk, Karolina Ostrowska, Anna Nowakowska |
| EPI_ISL_3251508 | Department of Bacteria, Parasites and Fungi, Statens Serum Institut, Copenhagen, Denmark | Statens Serum Institut Bioinformatics and Microbial Genomics | Danish Covid-19 Genome Consortium |
| EPI_ISL_3251509 | MakLab Laboratorium Diagnostyczne | Wojewodzka Stacja Sanitarno-Epidemiologiczna w Rzeszowie, Laboratorium Diagnostyki Medycznej | Marzena Baranowska, Katarzyna Wilk, Karolina Ostrowska, Anna Nowakowska |
| EPI_ISL_3251510 | Department of Bacteria, Parasites and Fungi, Statens Serum Institut, Copenhagen, Denmark | Statens Serum Institut Bioinformatics and Microbial Genomics | Danish Covid-19 Genome Consortium |
| EPI_ISL_3251511 | MakLab Laboratorium Diagnostyczne | Wojewodzka Stacja Sanitarno-Epidemiologiczna w Rzeszowie, Laboratorium Diagnostyki Medycznej | Marzena Baranowska, Katarzyna Wilk, Karolina Ostrowska, Anna Nowakowska |
| EPI_ISL_3251512, EPI_ISL_3251513 | Department of Bacteria, Parasites and Fungi, Statens Serum Institut, Copenhagen, Denmark | Statens Serum Institut Bioinformatics and Microbial Genomics | Danish Covid-19 Genome Consortium |
| EPI_ISL_3251514 | Wojewodzki Szpital Specjalistyczny Laboratorium Mikrobiologiczne | Wojewodzka Stacja Sanitarno-Epidemiologiczna w Rzeszowie, Laboratorium Diagnostyki Medycznej | Marzena Baranowska, Katarzyna Wilk, Karolina Ostrowska, Anna Nowakowska |
| EPI_ISL_3251515 | Department of Bacteria, Parasites and Fungi, Statens Serum Institut, Copenhagen, Denmark | Statens Serum Institut Bioinformatics and Microbial Genomics | Danish Covid-19 Genome Consortium |
| EPI_ISL_3251516 | Laboratorium Genetyczne Alab Plus | Wojewodzka Stacja Sanitarno-Epidemiologiczna w Rzeszowie, Laboratorium Diagnostyki Medycznej | Marzena Baranowska, Katarzyna Wilk, Karolina Ostrowska, Anna Nowakowska |
| EPI_ISL_3251517 | Department of Bacteria, Parasites and Fungi, Statens Serum Institut, Copenhagen, Denmark | Statens Serum Institut Bioinformatics and Microbial Genomics | Danish Covid-19 Genome Consortium |
| EPI_ISL_3251518 | Laboratorium Diagnostyczne Centrum Medyczne LuxMed | Wojewodzka Stacja Sanitarno-Epidemiologiczna w Rzeszowie, | Marzena Baranowska, Katarzyna Wilk, Karolina Ostrowska, Anna Nowakowska |

| | | | |
|---|---|---|---|
| | | Laboratorium Diagnostyki Medycznej | |
| EPI_ISL_3251519, EPI_ISL_3251521 | Department of Bacteria, Parasites and Fungi, Statens Serum Institut, Copenhagen, Denmark | Statens Serum Institut Bioinformatics and Microbial Genomics | Danish Covid-19 Genome Consortium |
| EPI_ISL_3251522 | WSSE Lublin Laboratorium | Wojewodzka Stacja Sanitarno-Epidemiologiczna w Rzeszowie, Laboratorium Diagnostyki Medycznej | Marzena Baranowska, Katarzyna Wilk, Karolina Ostrowska, Anna Nowakowska |
| EPI_ISL_3251523 | Department of Bacteria, Parasites and Fungi, Statens Serum Institut, Copenhagen, Denmark | Statens Serum Institut Bioinformatics and Microbial Genomics | Danish Covid-19 Genome Consortium |
| EPI_ISL_3251524 | Centrum Medyczne MEDYK Zaklad Diagnostyki Medycznej | Wojewodzka Stacja Sanitarno-Epidemiologiczna w Rzeszowie, Laboratorium Diagnostyki Medycznej | Marzena Baranowska, Katarzyna Wilk, Karolina Ostrowska, Anna Nowakowska |
| EPI_ISL_3251525 | Department of Bacteria, Parasites and Fungi, Statens Serum Institut, Copenhagen, Denmark | Statens Serum Institut Bioinformatics and Microbial Genomics | Danish Covid-19 Genome Consortium |
| EPI_ISL_3251526 | WSSE Lublin Laboratorium | Wojewodzka Stacja Sanitarno-Epidemiologiczna w Rzeszowie, Laboratorium Diagnostyki Medycznej | Marzena Baranowska, Katarzyna Wilk, Karolina Ostrowska, Anna Nowakowska |
| EPI_ISL_3251527 | Department of Bacteria, Parasites and Fungi, Statens Serum Institut, Copenhagen, Denmark | Statens Serum Institut Bioinformatics and Microbial Genomics | Danish Covid-19 Genome Consortium |
| EPI_ISL_3251528 | SP WSZ w Szczecinie Zaklad Diagnostyki Laboratoryjnej | Wojewodzka Stacja Sanitarno-Epidemiologiczna w Rzeszowie, Laboratorium Diagnostyki Medycznej | Marzena Baranowska, Katarzyna Wilk, Karolina Ostrowska, Anna Nowakowska |
| EPI_ISL_3251529 | Department of Bacteria, Parasites and Fungi, Statens Serum Institut, Copenhagen, Denmark | Statens Serum Institut Bioinformatics and Microbial Genomics | Danish Covid-19 Genome Consortium |
| EPI_ISL_3251530 | GenXOne | Wojewodzka Stacja Sanitarno-Epidemiologiczna w Rzeszowie, Laboratorium Diagnostyki Medycznej | Marzena Baranowska, Katarzyna Wilk, Karolina Ostrowska, Anna Nowakowska |
| EPI_ISL_3251531, EPI_ISL_3251532 | Department of Bacteria, Parasites and Fungi, Statens Serum Institut, Copenhagen, Denmark | Statens Serum Institut Bioinformatics and Microbial Genomics | Danish Covid-19 Genome Consortium |
| EPI_ISL_3251533 | GenXOne | Wojewodzka Stacja Sanitarno-Epidemiologiczna w Rzeszowie, Laboratorium Diagnostyki Medycznej | Marzena Baranowska, Katarzyna Wilk, Karolina Ostrowska, Anna Nowakowska |
| EPI_ISL_3251534 | Department of Bacteria, Parasites and Fungi, Statens Serum Institut, Copenhagen, Denmark | Statens Serum Institut Bioinformatics and Microbial Genomics | Danish Covid-19 Genome Consortium |
| EPI_ISL_3251535 | GenXOne | Wojewodzka Stacja Sanitarno-Epidemiologiczna w Rzeszowie, Laboratorium Diagnostyki Medycznej | Marzena Baranowska, Katarzyna Wilk, Karolina Ostrowska, Anna Nowakowska |
| EPI_ISL_3251536 | Department of Bacteria, Parasites and Fungi, Statens Serum Institut, Copenhagen, Denmark | Statens Serum Institut Bioinformatics and Microbial Genomics | Danish Covid-19 Genome Consortium |
| EPI_ISL_3251537 | GenXOne | Wojewodzka Stacja Sanitarno-Epidemiologiczna w Rzeszowie, Laboratorium Diagnostyki Medycznej | Marzena Baranowska, Katarzyna Wilk, Karolina Ostrowska, Anna Nowakowska |
| EPI_ISL_3251538 | Department of Bacteria, Parasites and Fungi, Statens Serum Institut, Copenhagen, Denmark | Statens Serum Institut Bioinformatics and Microbial Genomics | Danish Covid-19 Genome Consortium |
| EPI_ISL_3251539 | GenXOne | Wojewodzka Stacja Sanitarno-Epidemiologiczna w Rzeszowie, Laboratorium Diagnostyki Medycznej | Marzena Baranowska, Katarzyna Wilk, Karolina Ostrowska, Anna Nowakowska |
| EPI_ISL_3251540 | Department of Bacteria, Parasites and Fungi, Statens Serum Institut, Copenhagen, Denmark | Statens Serum Institut Bioinformatics and Microbial Genomics | Danish Covid-19 Genome Consortium |
| EPI_ISL_3251541 | GenXOne | Wojewodzka Stacja Sanitarno-Epidemiologiczna w Rzeszowie, Laboratorium Diagnostyki Medycznej | Marzena Baranowska, Katarzyna Wilk, Karolina Ostrowska, Anna Nowakowska |
| EPI_ISL_3251542 | Department of Bacteria, Parasites and Fungi, Statens Serum Institut, Copenhagen, Denmark | Statens Serum Institut Bioinformatics and Microbial Genomics | Danish Covid-19 Genome Consortium |
| EPI_ISL_3251543 | Centrum Badan DNA | Wojewodzka Stacja Sanitarno-Epidemiologiczna w Rzeszowie, Laboratorium Diagnostyki Medycznej | Marzena Baranowska, Katarzyna Wilk, Karolina Ostrowska, Anna Nowakowska |
| EPI_ISL_3251544 | Department of Bacteria, Parasites and Fungi, Statens Serum Institut, Copenhagen, Denmark | Statens Serum Institut Bioinformatics and Microbial Genomics | Danish Covid-19 Genome Consortium |
| EPI_ISL_3251545 | Centrum Badan DNA | Wojewodzka Stacja Sanitarno-Epidemiologiczna w Rzeszowie, Laboratorium Diagnostyki Medycznej | Marzena Baranowska, Katarzyna Wilk, Karolina Ostrowska, Anna Nowakowska |
| EPI_ISL_3251546 | Department of Bacteria, Parasites and Fungi, Statens Serum Institut, Copenhagen, Denmark | Statens Serum Institut Bioinformatics and Microbial Genomics | Danish Covid-19 Genome Consortium |
| EPI_ISL_3251547 | Centrum Medyczne MEDYK Zaklad Diagnostyki Medycznej | Wojewodzka Stacja Sanitarno-Epidemiologiczna w Rzeszowie, Laboratorium Diagnostyki Medycznej | Marzena Baranowska, Katarzyna Wilk, Karolina Ostrowska, Anna Nowakowska |
| EPI_ISL_3251548, EPI_ISL_3251549 | Department of Bacteria, Parasites and Fungi, Statens Serum Institut, Copenhagen, Denmark | Statens Serum Institut Bioinformatics and Microbial Genomics | Danish Covid-19 Genome Consortium |
| EPI_ISL_3251550 | Laboratorium Diagnostyczno-analityczne Osrodek Profilaktyki i Epidemiologii Nowotworow im. A. Pienkowskiej | Wojewodzka Stacja Sanitarno-Epidemiologiczna w Rzeszowie, Laboratorium Diagnostyki Medycznej | Marzena Baranowska, Katarzyna Wilk, Karolina Ostrowska, Anna Nowakowska |
| EPI_ISL_3251551 | WSZ im. dr R. Ostrzyckiego w Koninie Zaklad Mikrobiologii Klinicznej | Wojewodzka Stacja Sanitarno-Epidemiologiczna w Rzeszowie, Laboratorium Diagnostyki Medycznej | Marzena Baranowska, Katarzyna Wilk, Karolina Ostrowska, Anna Nowakowska |
| EPI_ISL_3251552, EPI_ISL_3251553 | Department of Bacteria, Parasites and Fungi, Statens Serum Institut, Copenhagen, Denmark | Statens Serum Institut Bioinformatics and Microbial Genomics | Danish Covid-19 Genome Consortium |
| EPI_ISL_3251554 | WSZ im. dr R. Ostrzyckiego w Koninie Zaklad Mikrobiologii Klinicznej | Wojewodzka Stacja Sanitarno-Epidemiologiczna w Rzeszowie, Laboratorium Diagnostyki Medycznej | Marzena Baranowska, Katarzyna Wilk, Karolina Ostrowska, Anna Nowakowska |
| EPI_ISL_3251555 | Department of Bacteria, Parasites and Fungi, Statens Serum Institut, Copenhagen, Denmark | Statens Serum Institut Bioinformatics and Microbial Genomics | Danish Covid-19 Genome Consortium |
| EPI_ISL_3251556 | SP-ZOZ w Krotoszynie Pracownia Biologii Molekularnej Laboratorium Centralne | Wojewodzka Stacja Sanitarno-Epidemiologiczna w Rzeszowie, Laboratorium Diagnostyki Medycznej | Marzena Baranowska, Katarzyna Wilk, Karolina Ostrowska, Anna Nowakowska |
| EPI_ISL_3251557 | Laboratoria Medyczne Bruss grupa ALAB Medyczne Laboratorium Diagnostyczne Pracownia Genetyki | Wojewodzka Stacja Sanitarno-Epidemiologiczna w Rzeszowie, Laboratorium Diagnostyki Medycznej | Marzena Baranowska, Katarzyna Wilk, Karolina Ostrowska, Anna Nowakowska |
| EPI_ISL_3251558, EPI_ISL_3251559 | Department of Bacteria, Parasites and Fungi, Statens Serum Institut, Copenhagen, Denmark | Statens Serum Institut Bioinformatics and Microbial Genomics | Danish Covid-19 Genome Consortium |
| EPI_ISL_3251560 | Medyczne Laboratoria Diagnostyczne INVICTA | Wojewodzka Stacja Sanitarno-Epidemiologiczna w Rzeszowie, Laboratorium Diagnostyki Medycznej | Marzena Baranowska, Katarzyna Wilk, Karolina Ostrowska, Anna Nowakowska |
| EPI_ISL_3251561 | Department of Bacteria, Parasites and Fungi, Statens Serum Institut, Copenhagen, Denmark | Statens Serum Institut Bioinformatics and Microbial Genomics | Danish Covid-19 Genome Consortium |
| EPI_ISL_3251562 | Medyczne Laboratoria Diagnostyczne INVICTA | Wojewodzka Stacja Sanitarno-Epidemiologiczna w Rzeszowie, Laboratorium Diagnostyki Medycznej | Marzena Baranowska, Katarzyna Wilk, Karolina Ostrowska, Anna Nowakowska |
| EPI_ISL_3251563 | Department of Bacteria, Parasites and Fungi, Statens Serum Institut, Copenhagen, Denmark | Statens Serum Institut Bioinformatics and Microbial Genomics | Danish Covid-19 Genome Consortium |
| EPI_ISL_3251564 | Medyczne Laboratoria Diagnostyczne INVICTA | Wojewodzka Stacja Sanitarno-Epidemiologiczna w Rzeszowie, Laboratorium Diagnostyki Medycznej | Marzena Baranowska, Katarzyna Wilk, Karolina Ostrowska, Anna Nowakowska |
| EPI_ISL_3251565 | Department of Bacteria, Parasites and Fungi, Statens Serum Institut, Copenhagen, Denmark | Statens Serum Institut Bioinformatics and Microbial Genomics | Danish Covid-19 Genome Consortium |
| EPI_ISL_3251566 | Medyczne Laboratoria Diagnostyczne INVICTA | Wojewodzka Stacja Sanitarno-Epidemiologiczna w Rzeszowie, Laboratorium Diagnostyki Medycznej | Marzena Baranowska, Katarzyna Wilk, Karolina Ostrowska, Anna Nowakowska |
| EPI_ISL_3251567, EPI_ISL_3251569, EPI_ISL_3251570 | Department of Bacteria, Parasites and Fungi, Statens Serum Institut, Copenhagen, Denmark | Statens Serum Institut Bioinformatics and Microbial Genomics | Danish Covid-19 Genome Consortium |
| EPI_ISL_3251571 | Laboratorium Analiz Lekarskich ALAB Tarnobrzeg | Wojewodzka Stacja Sanitarno-Epidemiologiczna w Rzeszowie, Laboratorium Diagnostyki Medycznej | Marzena Baranowska, Katarzyna Wilk, Karolina Ostrowska, Anna Nowakowska |

| | | | |
|---|---|---|---|
| EPI_ISL_3251572, EPI_ISL_3251574 | Department of Bacteria, Parasites and Fungi, Statens Serum Institut, Copenhagen, Denmark | Statens Serum Institut Bioinformatics and Microbial Genomics | Danish Covid-19 Genome Consortium |
| EPI_ISL_3251575 | WSSE Gdansk Laboratorium | Wojewodzka Stacja Sanitarno-Epidemiologiczna w Rzeszowie, Laboratorium Diagnostyki Medycznej | Marzena Baranowska, Katarzyna Wilk, Karolina Ostrowska, Anna Nowakowska |
| EPI_ISL_3251576 | Department of Bacteria, Parasites and Fungi, Statens Serum Institut, Copenhagen, Denmark | Statens Serum Institut Bioinformatics and Microbial Genomics | Danish Covid-19 Genome Consortium |
| EPI_ISL_3251577 | Uniwersyteckie Centrum Kliniczne | Wojewodzka Stacja Sanitarno-Epidemiologiczna w Rzeszowie, Laboratorium Diagnostyki Medycznej | Marzena Baranowska, Katarzyna Wilk, Karolina Ostrowska, Anna Nowakowska |
| EPI_ISL_3251578 | Department of Bacteria, Parasites and Fungi, Statens Serum Institut, Copenhagen, Denmark | Statens Serum Institut Bioinformatics and Microbial Genomics | Danish Covid-19 Genome Consortium |
| EPI_ISL_3251579 | WSSE Gdansk Laboratorium | Wojewodzka Stacja Sanitarno-Epidemiologiczna w Rzeszowie, Laboratorium Diagnostyki Medycznej | Marzena Baranowska, Katarzyna Wilk, Karolina Ostrowska, Anna Nowakowska |
| EPI_ISL_3251580 | Department of Bacteria, Parasites and Fungi, Statens Serum Institut, Copenhagen, Denmark | Statens Serum Institut Bioinformatics and Microbial Genomics | Danish Covid-19 Genome Consortium |
| EPI_ISL_3251581 | Laboratoria Medyczne Bruss grupa ALAB Medyczne Laboratorium Diagnostyczne Pracownia Genetyki | Wojewodzka Stacja Sanitarno-Epidemiologiczna w Rzeszowie, Laboratorium Diagnostyki Medycznej | Marzena Baranowska, Katarzyna Wilk, Karolina Ostrowska, Anna Nowakowska |
| EPI_ISL_3251582, EPI_ISL_3251583 | Department of Bacteria, Parasites and Fungi, Statens Serum Institut, Copenhagen, Denmark | Statens Serum Institut Bioinformatics and Microbial Genomics | Danish Covid-19 Genome Consortium |
| EPI_ISL_3251584 | WSSE Gdansk Laboratorium | Wojewodzka Stacja Sanitarno-Epidemiologiczna w Rzeszowie, Laboratorium Diagnostyki Medycznej | Marzena Baranowska, Katarzyna Wilk, Karolina Ostrowska, Anna Nowakowska |
| EPI_ISL_3251585 | Laboratoria Medyczne Bruss grupa ALAB Medyczne Laboratorium Diagnostyczne Pracownia Genetyki | Wojewodzka Stacja Sanitarno-Epidemiologiczna w Rzeszowie, Laboratorium Diagnostyki Medycznej | Marzena Baranowska, Katarzyna Wilk, Karolina Ostrowska, Anna Nowakowska |
| EPI_ISL_3251586, EPI_ISL_3251587, EPI_ISL_3251589 | Department of Bacteria, Parasites and Fungi, Statens Serum Institut, Copenhagen, Denmark | Statens Serum Institut Bioinformatics and Microbial Genomics | Danish Covid-19 Genome Consortium |
| EPI_ISL_3251590 | Centrum Medycyny Klinicznej dr n. med. Karol Majewski | Wojewodzka Stacja Sanitarno-Epidemiologiczna w Rzeszowie, Laboratorium Diagnostyki Medycznej | Marzena Baranowska, Katarzyna Wilk, Karolina Ostrowska, Anna Nowakowska |
| EPI_ISL_3251591 | Department of Bacteria, Parasites and Fungi, Statens Serum Institut, Copenhagen, Denmark | Statens Serum Institut Bioinformatics and Microbial Genomics | Danish Covid-19 Genome Consortium |
| EPI_ISL_3251592 | Centrum Medycyny Klinicznej dr n. med. Karol Majewski | Wojewodzka Stacja Sanitarno-Epidemiologiczna w Rzeszowie, Laboratorium Diagnostyki Medycznej | Marzena Baranowska, Katarzyna Wilk, Karolina Ostrowska, Anna Nowakowska |
| EPI_ISL_3251593 | Department of Bacteria, Parasites and Fungi, Statens Serum Institut, Copenhagen, Denmark | Statens Serum Institut Bioinformatics and Microbial Genomics | Danish Covid-19 Genome Consortium |
| EPI_ISL_3251594 | Laboratorium Analiz Lekarskich ALAB Tarnobrzeg | Wojewodzka Stacja Sanitarno-Epidemiologiczna w Rzeszowie, Laboratorium Diagnostyki Medycznej | Marzena Baranowska, Katarzyna Wilk, Karolina Ostrowska, Anna Nowakowska |
| EPI_ISL_3251595 | Department of Bacteria, Parasites and Fungi, Statens Serum Institut, Copenhagen, Denmark | Statens Serum Institut Bioinformatics and Microbial Genomics | Danish Covid-19 Genome Consortium |
| EPI_ISL_3251596 | Centrum Medycyny Klinicznej dr n. med. Karol Majewski | Wojewodzka Stacja Sanitarno-Epidemiologiczna w Rzeszowie, Laboratorium Diagnostyki Medycznej | Marzena Baranowska, Katarzyna Wilk, Karolina Ostrowska, Anna Nowakowska |
| EPI_ISL_3251597 | Department of Clinical Microbiology and Center for Genomic Medicine, Rigshospitalet, Copenhagen, Denmark | Statens Serum Institut Bioinformatics and Microbial Genomics | Danish Covid-19 Genome Consortium |
| EPI_ISL_3251598 | Department of Bacteria, Parasites and Fungi, Statens Serum Institut, Copenhagen, Denmark | Statens Serum Institut Bioinformatics and Microbial Genomics | Danish Covid-19 Genome Consortium |
| EPI_ISL_3251599 | Centrum Medycyny Klinicznej dr n. med. Karol Majewski | Wojewodzka Stacja Sanitarno-Epidemiologiczna w Rzeszowie, Laboratorium Diagnostyki Medycznej | Marzena Baranowska, Katarzyna Wilk, Karolina Ostrowska, Anna Nowakowska |
| EPI_ISL_3251600 | Department of Bacteria, Parasites and Fungi, Statens Serum Institut, Copenhagen, Denmark | Statens Serum Institut Bioinformatics and Microbial Genomics | Danish Covid-19 Genome Consortium |
| EPI_ISL_3251601 | Wojewodzki Szpital Specjalistyczny im. Marii Sklodowskiej-Curie w Zgierzu Zaklad Diagnostyki Laboratoryjnej | Wojewodzka Stacja Sanitarno-Epidemiologiczna w Rzeszowie, Laboratorium Diagnostyki Medycznej | Marzena Baranowska, Katarzyna Wilk, Karolina Ostrowska, Anna Nowakowska |
| EPI_ISL_3251602, EPI_ISL_3251604, EPI_ISL_3251606 | Department of Bacteria, Parasites and Fungi, Statens Serum Institut, Copenhagen, Denmark | Statens Serum Institut Bioinformatics and Microbial Genomics | Danish Covid-19 Genome Consortium |
| EPI_ISL_3251607 | Szpital Uniwersytecki w Krakowie | Wojewodzka Stacja Sanitarno-Epidemiologiczna w Rzeszowie, Laboratorium Diagnostyki Medycznej | Marzena Baranowska, Katarzyna Wilk, Karolina Ostrowska, Anna Nowakowska |
| EPI_ISL_3251608, EPI_ISL_3251610 | Department of Bacteria, Parasites and Fungi, Statens Serum Institut, Copenhagen, Denmark | Statens Serum Institut Bioinformatics and Microbial Genomics | Danish Covid-19 Genome Consortium |
| EPI_ISL_3251611 | Szpital Uniwersytecki w Krakowie | Wojewodzka Stacja Sanitarno-Epidemiologiczna w Rzeszowie, Laboratorium Diagnostyki Medycznej | Marzena Baranowska, Katarzyna Wilk, Karolina Ostrowska, Anna Nowakowska |
| EPI_ISL_3251612 | Department of Bacteria, Parasites and Fungi, Statens Serum Institut, Copenhagen, Denmark | Statens Serum Institut Bioinformatics and Microbial Genomics | Danish Covid-19 Genome Consortium |
| EPI_ISL_3251613 | WSSE w Krakowie | Wojewodzka Stacja Sanitarno-Epidemiologiczna w Rzeszowie, Laboratorium Diagnostyki Medycznej | Marzena Baranowska, Katarzyna Wilk, Karolina Ostrowska, Anna Nowakowska |
| EPI_ISL_3251614, EPI_ISL_3251615 | Department of Bacteria, Parasites and Fungi, Statens Serum Institut, Copenhagen, Denmark | Statens Serum Institut Bioinformatics and Microbial Genomics | Danish Covid-19 Genome Consortium |
| EPI_ISL_3251616 | WSSE w Krakowie | Wojewodzka Stacja Sanitarno-Epidemiologiczna w Rzeszowie, Laboratorium Diagnostyki Medycznej | Marzena Baranowska, Katarzyna Wilk, Karolina Ostrowska, Anna Nowakowska |
| EPI_ISL_3251617 | Department of Bacteria, Parasites and Fungi, Statens Serum Institut, Copenhagen, Denmark | Statens Serum Institut Bioinformatics and Microbial Genomics | Danish Covid-19 Genome Consortium |
| EPI_ISL_3251618 | Laboratorium Analiz Lekarskich ALAB Rzeszow | Wojewodzka Stacja Sanitarno-Epidemiologiczna w Rzeszowie, Laboratorium Diagnostyki Medycznej | Marzena Baranowska, Katarzyna Wilk, Karolina Ostrowska, Anna Nowakowska |
| EPI_ISL_3251619 | Department of Bacteria, Parasites and Fungi, Statens Serum Institut, Copenhagen, Denmark | Statens Serum Institut Bioinformatics and Microbial Genomics | Danish Covid-19 Genome Consortium |
| EPI_ISL_3251620 | Szpital Uniwersytecki w Krakowie | Wojewodzka Stacja Sanitarno-Epidemiologiczna w Rzeszowie, Laboratorium Diagnostyki Medycznej | Marzena Baranowska, Katarzyna Wilk, Karolina Ostrowska, Anna Nowakowska |
| EPI_ISL_3251621 | Department of Bacteria, Parasites and Fungi, Statens Serum Institut, Copenhagen, Denmark | Statens Serum Institut Bioinformatics and Microbial Genomics | Danish Covid-19 Genome Consortium |
| EPI_ISL_3251622 | Szpital Uniwersytecki w Krakowie | Wojewodzka Stacja Sanitarno-Epidemiologiczna w Rzeszowie, Laboratorium Diagnostyki Medycznej | Marzena Baranowska, Katarzyna Wilk, Karolina Ostrowska, Anna Nowakowska |
| EPI_ISL_3251623 | Department of Bacteria, Parasites and Fungi, Statens Serum Institut, Copenhagen, Denmark | Statens Serum Institut Bioinformatics and Microbial Genomics | Danish Covid-19 Genome Consortium |
| EPI_ISL_3251624 | Szpital Uniwersytecki w Krakowie | Wojewodzka Stacja Sanitarno-Epidemiologiczna w Rzeszowie, Laboratorium Diagnostyki Medycznej | Marzena Baranowska, Katarzyna Wilk, Karolina Ostrowska, Anna Nowakowska |
| EPI_ISL_3251625 | Department of Bacteria, Parasites and Fungi, Statens Serum Institut, Copenhagen, Denmark | Statens Serum Institut Bioinformatics and Microbial Genomics | Danish Covid-19 Genome Consortium |
| EPI_ISL_3251626 | WSSE w Krakowie | Wojewodzka Stacja Sanitarno-Epidemiologiczna w Rzeszowie, Laboratorium Diagnostyki Medycznej | Marzena Baranowska, Katarzyna Wilk, Karolina Ostrowska, Anna Nowakowska |
| EPI_ISL_3251627 | Department of Bacteria, Parasites and Fungi, Statens Serum Institut, Copenhagen, Denmark | Statens Serum Institut Bioinformatics and Microbial Genomics | Danish Covid-19 Genome Consortium |
| EPI_ISL_3251628 | Szpital Uniwersytecki w Krakowie | Wojewodzka Stacja Sanitarno-Epidemiologiczna w Rzeszowie, Laboratorium Diagnostyki Medycznej | Marzena Baranowska, Katarzyna Wilk, Karolina Ostrowska, Anna Nowakowska |

| | | | |
|---|---|---|---|
| EPI_ISL_3251629 | Department of Bacteria, Parasites and Fungi, Statens Serum Institut, Copenhagen, Denmark | Statens Serum Institut Bioinformatics and Microbial Genomics | Danish Covid-19 Genome Consortium |
| EPI_ISL_3251630 | Szpital Uniwersytecki w Krakowie | Wojewodzka Stacja Sanitarno-Epidemiologiczna w Rzeszowie, Laboratorium Diagnostyki Medycznej | Marzena Baranowska, Katarzyna Wilk, Karolina Ostrowska, Anna Nowakowska |
| EPI_ISL_3251631 | Department of Bacteria, Parasites and Fungi, Statens Serum Institut, Copenhagen, Denmark | Statens Serum Institut Bioinformatics and Microbial Genomics | Danish Covid-19 Genome Consortium |
| EPI_ISL_3251632 | Szpital Specjalistyczny im. Edmunda Biernackiego w Mielcu Laboratorium Mikrobiologiczne z Pracownia Cytologiczna | Wojewodzka Stacja Sanitarno-Epidemiologiczna w Rzeszowie, Laboratorium Diagnostyki Medycznej | Marzena Baranowska, Katarzyna Wilk, Karolina Ostrowska, Anna Nowakowska |
| EPI_ISL_3251633, EPI_ISL_3251634, EPI_ISL_3251635, EPI_ISL_3251636, EPI_ISL_3251637, EPI_ISL_3251638, EPI_ISL_3251639, EPI_ISL_3251640, EPI_ISL_3251641, EPI_ISL_3251642, EPI_ISL_3251643, EPI_ISL_3251644, EPI_ISL_3251645, EPI_ISL_3251646, EPI_ISL_3251647 | | | |
| see above | Department of Bacteria, Parasites and Fungi, Statens Serum Institut, Copenhagen, Denmark | Statens Serum Institut Bioinformatics and Microbial Genomics | Danish Covid-19 Genome Consortium |
| EPI_ISL_3251648 | Department of Clinical Microbiology, Odense University Hospital, Odense, Denmark | Statens Serum Institut Bioinformatics and Microbial Genomics | Danish Covid-19 Genome Consortium |
| EPI_ISL_3251649, EPI_ISL_3251650, EPI_ISL_3251651, EPI_ISL_3251652, EPI_ISL_3251653, EPI_ISL_3251654, EPI_ISL_3251655, EPI_ISL_3251656, EPI_ISL_3251657, EPI_ISL_3251658, EPI_ISL_3251659, EPI_ISL_3251660, EPI_ISL_3251661, EPI_ISL_3251662, EPI_ISL_3251663, EPI_ISL_3251664, EPI_ISL_3251665, EPI_ISL_3251666, EPI_ISL_3251667, EPI_ISL_3251668, EPI_ISL_3251669, EPI_ISL_3251670, EPI_ISL_3251671, EPI_ISL_3251672, EPI_ISL_3251673, EPI_ISL_3251674, EPI_ISL_3251675, EPI_ISL_3251676, EPI_ISL_3251677, EPI_ISL_3251678, EPI_ISL_3251679, EPI_ISL_3251680, EPI_ISL_3251681, EPI_ISL_3251682, EPI_ISL_3251683, EPI_ISL_3251684, EPI_ISL_3251685, EPI_ISL_3251686, EPI_ISL_3251687, EPI_ISL_3251688, EPI_ISL_3251689, EPI_ISL_3251690, EPI_ISL_3251691, EPI_ISL_3251692, EPI_ISL_3251693, EPI_ISL_3251694, EPI_ISL_3251695, EPI_ISL_3251696, EPI_ISL_3251697, EPI_ISL_3251698, EPI_ISL_3251699, EPI_ISL_3251700, EPI_ISL_3251701, EPI_ISL_3251702, EPI_ISL_3251703, EPI_ISL_3251704, EPI_ISL_3251705, EPI_ISL_3251706, EPI_ISL_3251707, EPI_ISL_3251708, EPI_ISL_3251709, EPI_ISL_3251710, EPI_ISL_3251711, EPI_ISL_3251712, EPI_ISL_3251713, EPI_ISL_3251714, EPI_ISL_3251715, EPI_ISL_3251716, EPI_ISL_3251717, EPI_ISL_3251718, EPI_ISL_3251719, EPI_ISL_3251720, EPI_ISL_3251721, EPI_ISL_3251722, EPI_ISL_3251723, EPI_ISL_3251724, EPI_ISL_3251725, EPI_ISL_3251726, EPI_ISL_3251727, EPI_ISL_3251728, EPI_ISL_3251729, EPI_ISL_3251730, EPI_ISL_3251731, EPI_ISL_3251732, EPI_ISL_3251733, EPI_ISL_3251734, EPI_ISL_3251735, EPI_ISL_3251736, EPI_ISL_3251737, EPI_ISL_3251738, EPI_ISL_3251739, EPI_ISL_3251740, EPI_ISL_3251741 | | | |
| see above | Department of Bacteria, Parasites and Fungi, Statens Serum Institut, Copenhagen, Denmark | Statens Serum Institut Bioinformatics and Microbial Genomics | Danish Covid-19 Genome Consortium |
| EPI_ISL_3251742 | Molekylær Medicinsk Afdeling, Aarhus University Hospital, Aarhus, Denmark | Statens Serum Institut Bioinformatics and Microbial Genomics | Danish Covid-19 Genome Consortium |
| EPI_ISL_3251743, EPI_ISL_3251744, EPI_ISL_3251745, EPI_ISL_3251746, EPI_ISL_3251747, EPI_ISL_3251748, EPI_ISL_3251749, EPI_ISL_3251750, EPI_ISL_3251751, EPI_ISL_3251752, EPI_ISL_3251753, EPI_ISL_3251754, EPI_ISL_3251755, EPI_ISL_3251756, EPI_ISL_3251757, EPI_ISL_3251758, EPI_ISL_3251759, EPI_ISL_3251760, EPI_ISL_3251761, EPI_ISL_3251762, EPI_ISL_3251763, EPI_ISL_3251764, EPI_ISL_3251765, EPI_ISL_3251766, EPI_ISL_3251767, EPI_ISL_3251768, EPI_ISL_3251769, EPI_ISL_3251770, EPI_ISL_3251771, EPI_ISL_3251772, EPI_ISL_3251773, EPI_ISL_3251774, EPI_ISL_3251775, EPI_ISL_3251776, EPI_ISL_3251777, EPI_ISL_3251778, EPI_ISL_3251779 | | | |
| see above | Department of Bacteria, Parasites and Fungi, Statens Serum Institut, Copenhagen, Denmark | Statens Serum Institut Bioinformatics and Microbial Genomics | Danish Covid-19 Genome Consortium |
| EPI_ISL_3251780 | Molekylær Medicinsk Afdeling, Aarhus University Hospital, Aarhus, Denmark | Statens Serum Institut Bioinformatics and Microbial Genomics | Danish Covid-19 Genome Consortium |
| EPI_ISL_3251781, EPI_ISL_3251782, EPI_ISL_3251783, EPI_ISL_3251784, EPI_ISL_3251785, EPI_ISL_3251786, EPI_ISL_3251787, EPI_ISL_3251788, EPI_ISL_3251789, EPI_ISL_3251790, EPI_ISL_3251791, EPI_ISL_3251792, EPI_ISL_3251793, EPI_ISL_3251794, EPI_ISL_3251795, EPI_ISL_3251796, EPI_ISL_3251797, EPI_ISL_3251798, EPI_ISL_3251799, EPI_ISL_3251800 | | | |
| see above | Department of Bacteria, Parasites and Fungi, Statens Serum Institut, Copenhagen, Denmark | Statens Serum Institut Bioinformatics and Microbial Genomics | Danish Covid-19 Genome Consortium |
| EPI_ISL_3251801 | Department of Clinical Microbiology and Center for Genomic Medicine, Rigshospitalet, Copenhagen, Denmark | Statens Serum Institut Bioinformatics and Microbial Genomics | Danish Covid-19 Genome Consortium |
| EPI_ISL_3251802, EPI_ISL_3251803, EPI_ISL_3251804, EPI_ISL_3251805, EPI_ISL_3251806, EPI_ISL_3251807, EPI_ISL_3251808, EPI_ISL_3251809, EPI_ISL_3251810, EPI_ISL_3251811, EPI_ISL_3251812, EPI_ISL_3251813, EPI_ISL_3251814, EPI_ISL_3251815, EPI_ISL_3251816, EPI_ISL_3251817, EPI_ISL_3251818, EPI_ISL_3251819, EPI_ISL_3251820, EPI_ISL_3251821, EPI_ISL_3251822, EPI_ISL_3251823, EPI_ISL_3251824, EPI_ISL_3251825, EPI_ISL_3251826, EPI_ISL_3251827, EPI_ISL_3251828, EPI_ISL_3251829, EPI_ISL_3251830, EPI_ISL_3251831, EPI_ISL_3251832, EPI_ISL_3251833, EPI_ISL_3251834, EPI_ISL_3251835, EPI_ISL_3251836 | | | |
| see above | Department of Bacteria, Parasites and Fungi, Statens Serum Institut, Copenhagen, Denmark | Statens Serum Institut Bioinformatics and Microbial Genomics | Danish Covid-19 Genome Consortium |
| EPI_ISL_3251837 | Molekylær Medicinsk Afdeling, Aarhus University Hospital, Aarhus, Denmark | Statens Serum Institut Bioinformatics and Microbial Genomics | Danish Covid-19 Genome Consortium |
| EPI_ISL_3251838, EPI_ISL_3251839, EPI_ISL_3251840, EPI_ISL_3251842, EPI_ISL_3251843, EPI_ISL_3251844, EPI_ISL_3251845, EPI_ISL_3251846, EPI_ISL_3251847, EPI_ISL_3251848, EPI_ISL_3251849, EPI_ISL_3251850, EPI_ISL_3251851, EPI_ISL_3251852, EPI_ISL_3251853, EPI_ISL_3251854, EPI_ISL_3251855, EPI_ISL_3251856, EPI_ISL_3251857, EPI_ISL_3251858, EPI_ISL_3251859, EPI_ISL_3251860, EPI_ISL_3251861, EPI_ISL_3251862, EPI_ISL_3251863, EPI_ISL_3251864, EPI_ISL_3251865, EPI_ISL_3251866, EPI_ISL_3251867, EPI_ISL_3251868, EPI_ISL_3251869, EPI_ISL_3251870, EPI_ISL_3251871, EPI_ISL_3251872, EPI_ISL_3251873, EPI_ISL_3251874, EPI_ISL_3251875, EPI_ISL_3251876, EPI_ISL_3251877, EPI_ISL_3251878, EPI_ISL_3251879, EPI_ISL_3251880, EPI_ISL_3251881, EPI_ISL_3251882, EPI_ISL_3251883, EPI_ISL_3251884, EPI_ISL_3251885, EPI_ISL_3251886, EPI_ISL_3251887, EPI_ISL_3251888, EPI_ISL_3251889, EPI_ISL_3251890, EPI_ISL_3251891, EPI_ISL_3251892, EPI_ISL_3251893, EPI_ISL_3251894, EPI_ISL_3251895, EPI_ISL_3251896, EPI_ISL_3251897, EPI_ISL_3251898, EPI_ISL_3251899, EPI_ISL_3251900, EPI_ISL_3251901, EPI_ISL_3251902, EPI_ISL_3251903, EPI_ISL_3251904, EPI_ISL_3251905, EPI_ISL_3251906, EPI_ISL_3251907, EPI_ISL_3251908, EPI_ISL_3251909, EPI_ISL_3251910, EPI_ISL_3251911, EPI_ISL_3251912, EPI_ISL_3251913, EPI_ISL_3251914, EPI_ISL_3251915, EPI_ISL_3251916, EPI_ISL_3251917, EPI_ISL_3251918, EPI_ISL_3251919, EPI_ISL_3251920, EPI_ISL_3251921, EPI_ISL_3251922, EPI_ISL_3251923, EPI_ISL_3251924, EPI_ISL_3251925, EPI_ISL_3251926, EPI_ISL_3251927, EPI_ISL_3251928, EPI_ISL_3251930, EPI_ISL_3251931, EPI_ISL_3251932, EPI_ISL_3251933, EPI_ISL_3251934, EPI_ISL_3251935, EPI_ISL_3251936, EPI_ISL_3251937, EPI_ISL_3251938, EPI_ISL_3251939, EPI_ISL_3251940, EPI_ISL_3251941, EPI_ISL_3251942, EPI_ISL_3251943, EPI_ISL_3251944, EPI_ISL_3251945, EPI_ISL_3251946, EPI_ISL_3251947, EPI_ISL_3251948, EPI_ISL_3251949, EPI_ISL_3251950, EPI_ISL_3251951, EPI_ISL_3251952, EPI_ISL_3251953, EPI_ISL_3251954 | | | |
| see above | Department of Bacteria, Parasites and Fungi, Statens Serum Institut, Copenhagen, Denmark | Statens Serum Institut Bioinformatics and Microbial Genomics | Danish Covid-19 Genome Consortium |
| EPI_ISL_3251955 | Department of Clinical Microbiology, Odense University Hospital, Odense, Denmark | Statens Serum Institut Bioinformatics and Microbial Genomics | Danish Covid-19 Genome Consortium |
| EPI_ISL_3251956, EPI_ISL_3251957, EPI_ISL_3251958, EPI_ISL_3251959, EPI_ISL_3251960, EPI_ISL_3251961, EPI_ISL_3251962, EPI_ISL_3251963, EPI_ISL_3251964, EPI_ISL_3251965, EPI_ISL_3251966, EPI_ISL_3251967, EPI_ISL_3251968, EPI_ISL_3251969, EPI_ISL_3251970, EPI_ISL_3251971, EPI_ISL_3251972, EPI_ISL_3251973, EPI_ISL_3251974, EPI_ISL_3251975, EPI_ISL_3251976, EPI_ISL_3251977, EPI_ISL_3251978, EPI_ISL_3251979, EPI_ISL_3251980, EPI_ISL_3251981, EPI_ISL_3251982, EPI_ISL_3251983, EPI_ISL_3251984, EPI_ISL_3251985, EPI_ISL_3251986, EPI_ISL_3251987, EPI_ISL_3251988, EPI_ISL_3251989, EPI_ISL_3251990, EPI_ISL_3251991, EPI_ISL_3251992, EPI_ISL_3251994, EPI_ISL_3251995, EPI_ISL_3251996, EPI_ISL_3251997, EPI_ISL_3251998 | | | |
| see above | Department of Bacteria, Parasites and Fungi, Statens Serum Institut, Copenhagen, Denmark | Statens Serum Institut Bioinformatics and Microbial Genomics | Danish Covid-19 Genome Consortium |
| EPI_ISL_3251999 | Molekylær Medicinsk Afdeling, Aarhus University Hospital, Aarhus, Denmark | Statens Serum Institut Bioinformatics and Microbial Genomics | Danish Covid-19 Genome Consortium |
| EPI_ISL_3252000, EPI_ISL_3252001, EPI_ISL_3252002, EPI_ISL_3252003, EPI_ISL_3252004, EPI_ISL_3252005, EPI_ISL_3252006, EPI_ISL_3252007, EPI_ISL_3252008, EPI_ISL_3252009, EPI_ISL_3252010, EPI_ISL_3252011, EPI_ISL_3252012, EPI_ISL_3252013, EPI_ISL_3252014, EPI_ISL_3252015, EPI_ISL_3252016, EPI_ISL_3252017, EPI_ISL_3252018, EPI_ISL_3252019, EPI_ISL_3252020, EPI_ISL_3252021, EPI_ISL_3252022, EPI_ISL_3252023, EPI_ISL_3252024, EPI_ISL_3252025, EPI_ISL_3252026 | | | |
| see above | Department of Bacteria, Parasites and Fungi, Statens Serum Institut, Copenhagen, Denmark | Statens Serum Institut Bioinformatics and Microbial Genomics | Danish Covid-19 Genome Consortium |
| EPI_ISL_3252027 | Molekylær Medicinsk Afdeling, Aarhus University Hospital, Aarhus, Denmark | Statens Serum Institut Bioinformatics and Microbial Genomics | Danish Covid-19 Genome Consortium |
| EPI_ISL_3252028, EPI_ISL_3252029, EPI_ISL_3252030, EPI_ISL_3252031, EPI_ISL_3252032, EPI_ISL_3252033, EPI_ISL_3252034, EPI_ISL_3252035, EPI_ISL_3252036, EPI_ISL_3252037, EPI_ISL_3252038, EPI_ISL_3252039, EPI_ISL_3252040, EPI_ISL_3252041, EPI_ISL_3252042, EPI_ISL_3252043, EPI_ISL_3252044, EPI_ISL_3252045, EPI_ISL_3252046, EPI_ISL_3252047, EPI_ISL_3252048, EPI_ISL_3252049, EPI_ISL_3252050, EPI_ISL_3252051 | | | |
| see above | Department of Bacteria, Parasites and Fungi, Statens Serum Institut, Copenhagen, Denmark | Statens Serum Institut Bioinformatics and Microbial Genomics | Danish Covid-19 Genome Consortium |
| EPI_ISL_3252052 | Molekylær Medicinsk Afdeling, Aarhus University Hospital, Aarhus, Denmark | Statens Serum Institut Bioinformatics and Microbial Genomics | Danish Covid-19 Genome Consortium |
| EPI_ISL_3252053 | Department of Clinical Microbiology, Odense University Hospital, Odense, Denmark | Statens Serum Institut Bioinformatics and Microbial Genomics | Danish Covid-19 Genome Consortium |
| EPI_ISL_3252054, EPI_ISL_3252055, EPI_ISL_3252056, EPI_ISL_3252058, EPI_ISL_3252059, EPI_ISL_3252060, EPI_ISL_3252061, EPI_ISL_3252062, EPI_ISL_3252063, EPI_ISL_3252064, EPI_ISL_3252065, EPI_ISL_3252066, EPI_ISL_3252067, EPI_ISL_3252068, EPI_ISL_3252069, EPI_ISL_3252070, EPI_ISL_3252071, EPI_ISL_3252072, EPI_ISL_3252073, EPI_ISL_3252074, EPI_ISL_3252075, EPI_ISL_3252076, EPI_ISL_3252077, EPI_ISL_3252078, EPI_ISL_3252079, EPI_ISL_3252080, EPI_ISL_3252081, EPI_ISL_3252082, EPI_ISL_3252083, EPI_ISL_3252084, EPI_ISL_3252085, EPI_ISL_3252086, EPI_ISL_3252087, EPI_ISL_3252088, EPI_ISL_3252089, EPI_ISL_3252090, EPI_ISL_3252091, EPI_ISL_3252092, EPI_ISL_3252093, EPI_ISL_3252094, EPI_ISL_3252095, EPI_ISL_3252096, EPI_ISL_3252097, EPI_ISL_3252098, EPI_ISL_3252099, EPI_ISL_3252100, EPI_ISL_3252101, EPI_ISL_3252102, EPI_ISL_3252103, EPI_ISL_3252104, EPI_ISL_3252105, EPI_ISL_3252106, EPI_ISL_3252107, EPI_ISL_3252108, EPI_ISL_3252109, EPI_ISL_3252110, EPI_ISL_3252111, EPI_ISL_3252112, EPI_ISL_3252113, EPI_ISL_3252114, EPI_ISL_3252115, EPI_ISL_3252116, EPI_ISL_3252117, EPI_ISL_3252118, EPI_ISL_3252119, EPI_ISL_3252120, EPI_ISL_3252121, EPI_ISL_3252122, EPI_ISL_3252123, EPI_ISL_3252124, EPI_ISL_3252125, EPI_ISL_3252126, EPI_ISL_3252127, EPI_ISL_3252128, EPI_ISL_3252129, EPI_ISL_3252130, EPI_ISL_3252131, EPI_ISL_3252132, EPI_ISL_3252133, EPI_ISL_3252134, EPI_ISL_3252135, EPI_ISL_3252136, EPI_ISL_3252137, EPI_ISL_3252138, EPI_ISL_3252139, EPI_ISL_3252140, EPI_ISL_3252141, EPI_ISL_3252142, EPI_ISL_3252143, EPI_ISL_3252144, EPI_ISL_3252145, EPI_ISL_3252146, EPI_ISL_3252147, EPI_ISL_3252148, EPI_ISL_3252149, EPI_ISL_3252150, EPI_ISL_3252151, EPI_ISL_3252152, EPI_ISL_3252153, EPI_ISL_3252154, EPI_ISL_3252155, EPI_ISL_3252156, EPI_ISL_3252157, EPI_ISL_3252158, EPI_ISL_3252159, EPI_ISL_3252160, EPI_ISL_3252161, EPI_ISL_3252162, EPI_ISL_3252163, EPI_ISL_3252164, EPI_ISL_3252165, EPI_ISL_3252166, EPI_ISL_3252167, EPI_ISL_3252168, EPI_ISL_3252169, EPI_ISL_3252170, EPI_ISL_3252171, EPI_ISL_3252172, EPI_ISL_3252173, EPI_ISL_3252174, EPI_ISL_3252175, EPI_ISL_3252176, EPI_ISL_3252177, EPI_ISL_3252178, EPI_ISL_3252179, EPI_ISL_3252180, EPI_ISL_3252181, EPI_ISL_3252182, EPI_ISL_3252183, EPI_ISL_3252184, EPI_ISL_3252185, EPI_ISL_3252186, EPI_ISL_3252187, EPI_ISL_3252188, EPI_ISL_3252189, EPI_ISL_3252190, EPI_ISL_3252191, EPI_ISL_3252192, EPI_ISL_3252193, EPI_ISL_3252194, EPI_ISL_3252195, EPI_ISL_3252196, EPI_ISL_3252197, EPI_ISL_3252198, EPI_ISL_3252199, EPI_ISL_3252200, EPI_ISL_3252201, EPI_ISL_3252202, EPI_ISL_3252203 | | | |
| see above | Department of Bacteria, Parasites and Fungi, Statens Serum Institut, Copenhagen, Denmark | Statens Serum Institut Bioinformatics and Microbial Genomics | Danish Covid-19 Genome Consortium |
| EPI_ISL_3252204 | Molekylær Medicinsk Afdeling, Aarhus University Hospital, Aarhus, Denmark | Statens Serum Institut Bioinformatics and Microbial Genomics | Danish Covid-19 Genome Consortium |
| EPI_ISL_3252205, EPI_ISL_3252206, EPI_ISL_3252207, EPI_ISL_3252208, EPI_ISL_3252209, EPI_ISL_3252210, EPI_ISL_3252211, EPI_ISL_3252212, EPI_ISL_3252213, EPI_ISL_3252214, EPI_ISL_3252215, EPI_ISL_3252217, EPI_ISL_3252218, EPI_ISL_3252219, EPI_ISL_3252220, EPI_ISL_3252221, EPI_ISL_3252222, EPI_ISL_3252223, EPI_ISL_3252224, EPI_ISL_3252225, EPI_ISL_3252226, EPI_ISL_3252227, EPI_ISL_3252228, EPI_ISL_3252229, EPI_ISL_3252230, EPI_ISL_3252231, EPI_ISL_3252232, EPI_ISL_3252233, EPI_ISL_3252234, EPI_ISL_3252235, EPI_ISL_3252236, EPI_ISL_3252237, EPI_ISL_3252238, EPI_ISL_3252239, EPI_ISL_3252240, EPI_ISL_3252241, EPI_ISL_3252242, EPI_ISL_3252243, EPI_ISL_3252244, EPI_ISL_3252245, EPI_ISL_3252246, EPI_ISL_3252247, EPI_ISL_3252248, EPI_ISL_3252249, EPI_ISL_3252250, EPI_ISL_3252251, EPI_ISL_3252252, EPI_ISL_3252253, EPI_ISL_3252254, EPI_ISL_3252255, EPI_ISL_3252256, EPI_ISL_3252257, EPI_ISL_3252258, EPI_ISL_3252259, EPI_ISL_3252260, EPI_ISL_3252261, EPI_ISL_3252262, EPI_ISL_3252263, EPI_ISL_3252264, EPI_ISL_3252265, EPI_ISL_3252266, EPI_ISL_3252267, EPI_ISL_3252268, EPI_ISL_3252269, EPI_ISL_3252270, EPI_ISL_3252271, EPI_ISL_3252272, EPI_ISL_3252274, EPI_ISL_3252275, EPI_ISL_3252276, EPI_ISL_3252277, EPI_ISL_3252278, EPI_ISL_3252279, EPI_ISL_3252280, EPI_ISL_3252281, EPI_ISL_3252282, EPI_ISL_3252283, EPI_ISL_3252284, EPI_ISL_3252285, EPI_ISL_3252286, EPI_ISL_3252287, EPI_ISL_3252288, EPI_ISL_3252289, EPI_ISL_3252290, EPI_ISL_3252291, EPI_ISL_3252292, EPI_ISL_3252293, EPI_ISL_3252294, EPI_ISL_3252295, EPI_ISL_3252296, EPI_ISL_3252297, EPI_ISL_3252298, EPI_ISL_3252299, EPI_ISL_3252300, EPI_ISL_3252301, EPI_ISL_3252302, EPI_ISL_3252303, EPI_ISL_3252304, EPI_ISL_3252305, EPI_ISL_3252306 | | | |
| see above | Department of Bacteria, Parasites and Fungi, Statens Serum Institut, Copenhagen, Denmark | Statens Serum Institut Bioinformatics and Microbial Genomics | Danish Covid-19 Genome Consortium |
| EPI_ISL_3252307 | Department of Clinical Microbiology and Center for Genomic Medicine, Rigshospitalet, Copenhagen, Denmark | Statens Serum Institut Bioinformatics and Microbial Genomics | Danish Covid-19 Genome Consortium |
| EPI_ISL_3252308, EPI_ISL_3252309, EPI_ISL_3252310, EPI_ISL_3252311, EPI_ISL_3252312, EPI_ISL_3252313, EPI_ISL_3252314, EPI_ISL_3252315, EPI_ISL_3252316, EPI_ISL_3252317, EPI_ISL_3252318, EPI_ISL_3252319, EPI_ISL_3252320, EPI_ISL_3252321, EPI_ISL_3252322, EPI_ISL_3252323, EPI_ISL_3252324, EPI_ISL_3252325, EPI_ISL_3252326, EPI_ISL_3252328, EPI_ISL_3252329, EPI_ISL_3252330, EPI_ISL_3252331, EPI_ISL_3252332, EPI_ISL_3252333, EPI_ISL_3252334, EPI_ISL_3252335, EPI_ISL_3252336, EPI_ISL_3252337, EPI_ISL_3252338, EPI_ISL_3252339, EPI_ISL_3252340, EPI_ISL_3252341, EPI_ISL_3252342, EPI_ISL_3252343, EPI_ISL_3252344, EPI_ISL_3252345, EPI_ISL_3252346, EPI_ISL_3252347, EPI_ISL_3252348, EPI_ISL_3252349, EPI_ISL_3252350, EPI_ISL_3252351, EPI_ISL_3252352, EPI_ISL_3252353 | | | |

| | | | |
|---|---|---|---|
| see above | Department of Bacteria, Parasites and Fungi, Statens Serum Institut, Copenhagen, Denmark | Statens Serum Institut Bioinformatics and Microbial Genomics | Danish Covid-19 Genome Consortium |
| EPI_ISL_3252354 | Molekylær Medicinsk Afdeling, Aarhus University Hospital, Aarhus, Denmark | Statens Serum Institut Bioinformatics and Microbial Genomics | Danish Covid-19 Genome Consortium |
| EPI_ISL_3252355, EPI_ISL_3252356, EPI_ISL_3252357, EPI_ISL_3252358, EPI_ISL_3252359, EPI_ISL_3252360, EPI_ISL_3252361, EPI_ISL_3252362, EPI_ISL_3252363, EPI_ISL_3252364, EPI_ISL_3252365, EPI_ISL_3252366, EPI_ISL_3252367, EPI_ISL_3252368, EPI_ISL_3252369, EPI_ISL_3252370, EPI_ISL_3252371, EPI_ISL_3252372, EPI_ISL_3252373, EPI_ISL_3252374, EPI_ISL_3252375, EPI_ISL_3252376, EPI_ISL_3252377, EPI_ISL_3252378, EPI_ISL_3252379, EPI_ISL_3252380, EPI_ISL_3252381, EPI_ISL_3252382, EPI_ISL_3252383, EPI_ISL_3252384, EPI_ISL_3252385, EPI_ISL_3252386, EPI_ISL_3252387, EPI_ISL_3252388, EPI_ISL_3252389, EPI_ISL_3252390, EPI_ISL_3252391, EPI_ISL_3252392, EPI_ISL_3252393, EPI_ISL_3252394, EPI_ISL_3252395, EPI_ISL_3252396, EPI_ISL_3252397, EPI_ISL_3252398, EPI_ISL_3252399, EPI_ISL_3252400, EPI_ISL_3252401, EPI_ISL_3252402, EPI_ISL_3252403, EPI_ISL_3252404, EPI_ISL_3252405, EPI_ISL_3252406, EPI_ISL_3252407, EPI_ISL_3252408, EPI_ISL_3252409, EPI_ISL_3252410, EPI_ISL_3252411, EPI_ISL_3252412, EPI_ISL_3252413, EPI_ISL_3252414, EPI_ISL_3252416, EPI_ISL_3252417, EPI_ISL_3252418, EPI_ISL_3252419, EPI_ISL_3252420, EPI_ISL_3252421, EPI_ISL_3252422, EPI_ISL_3252423, EPI_ISL_3252424, EPI_ISL_3252425, EPI_ISL_3252426, EPI_ISL_3252427, EPI_ISL_3252428, EPI_ISL_3252429, EPI_ISL_3252430, EPI_ISL_3252431, EPI_ISL_3252432, EPI_ISL_3252433, EPI_ISL_3252434, EPI_ISL_3252435, EPI_ISL_3252436, EPI_ISL_3252437, EPI_ISL_3252438, EPI_ISL_3252439, EPI_ISL_3252440, EPI_ISL_3252441, EPI_ISL_3252442, EPI_ISL_3252443, EPI_ISL_3252444, EPI_ISL_3252445, EPI_ISL_3252446, EPI_ISL_3252447, EPI_ISL_3252448, EPI_ISL_3252449, EPI_ISL_3252450, EPI_ISL_3252451, EPI_ISL_3252452, EPI_ISL_3252453, EPI_ISL_3252454, EPI_ISL_3252455, EPI_ISL_3252456, EPI_ISL_3252457, EPI_ISL_3252458, EPI_ISL_3252459, EPI_ISL_3252460, EPI_ISL_3252461, EPI_ISL_3252462, EPI_ISL_3252463, EPI_ISL_3252464, EPI_ISL_3252465, EPI_ISL_3252466, EPI_ISL_3252467, EPI_ISL_3252468, EPI_ISL_3252469, EPI_ISL_3252470, EPI_ISL_3252471, EPI_ISL_3252472, EPI_ISL_3252473, EPI_ISL_3252474, EPI_ISL_3252475, EPI_ISL_3252476, EPI_ISL_3252477, EPI_ISL_3252478, EPI_ISL_3252479, EPI_ISL_3252480, EPI_ISL_3252481, EPI_ISL_3252482, EPI_ISL_3252483, EPI_ISL_3252484, EPI_ISL_3252485, EPI_ISL_3252486, EPI_ISL_3252487, EPI_ISL_3252488, EPI_ISL_3252489, EPI_ISL_3252490, EPI_ISL_3252491, EPI_ISL_3252492, EPI_ISL_3252493, EPI_ISL_3252494, EPI_ISL_3252495, EPI_ISL_3252496, EPI_ISL_3252497, EPI_ISL_3252498, EPI_ISL_3252499, EPI_ISL_3252500, EPI_ISL_3252501, EPI_ISL_3252502, EPI_ISL_3252503, EPI_ISL_3252504, EPI_ISL_3252505, EPI_ISL_3252506, EPI_ISL_3252507, EPI_ISL_3252508, EPI_ISL_3252509, EPI_ISL_3252510, EPI_ISL_3252511, EPI_ISL_3252512, EPI_ISL_3252513 | | | |
| see above | Department of Bacteria, Parasites and Fungi, Statens Serum Institut, Copenhagen, Denmark | Statens Serum Institut Bioinformatics and Microbial Genomics | Danish Covid-19 Genome Consortium |
| EPI_ISL_3252514 | Molekylær Medicinsk Afdeling, Aarhus University Hospital, Aarhus, Denmark | Statens Serum Institut Bioinformatics and Microbial Genomics | Danish Covid-19 Genome Consortium |
| EPI_ISL_3252515, EPI_ISL_3252516, EPI_ISL_3252517, EPI_ISL_3252518, EPI_ISL_3252519, EPI_ISL_3252520, EPI_ISL_3252521, EPI_ISL_3252522, EPI_ISL_3252523, EPI_ISL_3252524, EPI_ISL_3252525, EPI_ISL_3252526, EPI_ISL_3252527, EPI_ISL_3252528, EPI_ISL_3252529, EPI_ISL_3252530, EPI_ISL_3252531, EPI_ISL_3252532, EPI_ISL_3252533, EPI_ISL_3252534, EPI_ISL_3252535, EPI_ISL_3252537, EPI_ISL_3252538, EPI_ISL_3252539, EPI_ISL_3252540, EPI_ISL_3252542, EPI_ISL_3252543, EPI_ISL_3252544, EPI_ISL_3252545, EPI_ISL_3252546, EPI_ISL_3252547, EPI_ISL_3252548, EPI_ISL_3252549, EPI_ISL_3252550, EPI_ISL_3252551, EPI_ISL_3252552, EPI_ISL_3252553, EPI_ISL_3252554, EPI_ISL_3252555, EPI_ISL_3252556, EPI_ISL_3252557, EPI_ISL_3252558, EPI_ISL_3252559, EPI_ISL_3252560, EPI_ISL_3252561, EPI_ISL_3252562, EPI_ISL_3252563, EPI_ISL_3252564, EPI_ISL_3252565, EPI_ISL_3252566, EPI_ISL_3252567, EPI_ISL_3252568, EPI_ISL_3252569, EPI_ISL_3252570, EPI_ISL_3252571, EPI_ISL_3252572, EPI_ISL_3252573, EPI_ISL_3252574, EPI_ISL_3252575, EPI_ISL_3252577, EPI_ISL_3252578, EPI_ISL_3252579, EPI_ISL_3252580, EPI_ISL_3252581, EPI_ISL_3252582, EPI_ISL_3252583, EPI_ISL_3252584, EPI_ISL_3252585, EPI_ISL_3252586, EPI_ISL_3252587, EPI_ISL_3252588, EPI_ISL_3252589, EPI_ISL_3252590, EPI_ISL_3252591, EPI_ISL_3252592, EPI_ISL_3252593, EPI_ISL_3252594, EPI_ISL_3252595, EPI_ISL_3252596, EPI_ISL_3252597, EPI_ISL_3252598, EPI_ISL_3252599, EPI_ISL_3252600, EPI_ISL_3252601, EPI_ISL_3252602, EPI_ISL_3252603, EPI_ISL_3252604, EPI_ISL_3252605, EPI_ISL_3252606, EPI_ISL_3252607, EPI_ISL_3252608, EPI_ISL_3252609, EPI_ISL_3252610, EPI_ISL_3252611, EPI_ISL_3252612, EPI_ISL_3252613, EPI_ISL_3252614, EPI_ISL_3252615, EPI_ISL_3252616, EPI_ISL_3252617, EPI_ISL_3252618, EPI_ISL_3252619, EPI_ISL_3252620, EPI_ISL_3252621, EPI_ISL_3252622, EPI_ISL_3252623, EPI_ISL_3252624, EPI_ISL_3252625, EPI_ISL_3252626, EPI_ISL_3252627, EPI_ISL_3252628, EPI_ISL_3252629, EPI_ISL_3252630, EPI_ISL_3252631, EPI_ISL_3252632, EPI_ISL_3252633, EPI_ISL_3252634, EPI_ISL_3252635, EPI_ISL_3252636, EPI_ISL_3252637, EPI_ISL_3252638, EPI_ISL_3252639, EPI_ISL_3252640, EPI_ISL_3252641, EPI_ISL_3252642, EPI_ISL_3252643, EPI_ISL_3252644, EPI_ISL_3252645, EPI_ISL_3252646, EPI_ISL_3252647, EPI_ISL_3252648, EPI_ISL_3252649, EPI_ISL_3252650, EPI_ISL_3252651, EPI_ISL_3252652, EPI_ISL_3252653, EPI_ISL_3252654, EPI_ISL_3252655, EPI_ISL_3252656, EPI_ISL_3252657, EPI_ISL_3252658, EPI_ISL_3252659, EPI_ISL_3252660, EPI_ISL_3252661, EPI_ISL_3252662, EPI_ISL_3252663, EPI_ISL_3252664, EPI_ISL_3252665, EPI_ISL_3252666, EPI_ISL_3252667, EPI_ISL_3252668, EPI_ISL_3252669, EPI_ISL_3252670, EPI_ISL_3252671, EPI_ISL_3252672, EPI_ISL_3252673, EPI_ISL_3252674, EPI_ISL_3252675, EPI_ISL_3252676, EPI_ISL_3252677, EPI_ISL_3252678, EPI_ISL_3252679, EPI_ISL_3252680, EPI_ISL_3252681, EPI_ISL_3252682, EPI_ISL_3252683, EPI_ISL_3252684, EPI_ISL_3252685, EPI_ISL_3252686, EPI_ISL_3252687, EPI_ISL_3252688, EPI_ISL_3252689, EPI_ISL_3252690, EPI_ISL_3252691, EPI_ISL_3252692, EPI_ISL_3252693, EPI_ISL_3252694, EPI_ISL_3252695, EPI_ISL_3252696, EPI_ISL_3252697, EPI_ISL_3252698, EPI_ISL_3252699, EPI_ISL_3252700, EPI_ISL_3252701, EPI_ISL_3252702, EPI_ISL_3252703, EPI_ISL_3252704, EPI_ISL_3252705, EPI_ISL_3252706, EPI_ISL_3252707, EPI_ISL_3252708, EPI_ISL_3252709, EPI_ISL_3252710, EPI_ISL_3252711, EPI_ISL_3252712, EPI_ISL_3252713, EPI_ISL_3252714, EPI_ISL_3252715, EPI_ISL_3252716, EPI_ISL_3252717, EPI_ISL_3252718, EPI_ISL_3252719, EPI_ISL_3252720, EPI_ISL_3252721, EPI_ISL_3252722, EPI_ISL_3252723, EPI_ISL_3252724, EPI_ISL_3252725, EPI_ISL_3252726, EPI_ISL_3252727, EPI_ISL_3252728, EPI_ISL_3252729, EPI_ISL_3252730, EPI_ISL_3252731, EPI_ISL_3252732, EPI_ISL_3252733, EPI_ISL_3252734, EPI_ISL_3252735, EPI_ISL_3252736, EPI_ISL_3252737, EPI_ISL_3252738, EPI_ISL_3252739, EPI_ISL_3252740, EPI_ISL_3252741, EPI_ISL_3252742, EPI_ISL_3252743, EPI_ISL_3252744, EPI_ISL_3252745, EPI_ISL_3252746, EPI_ISL_3252747, EPI_ISL_3252748, EPI_ISL_3252749, EPI_ISL_3252750, EPI_ISL_3252751, EPI_ISL_3252752 | | | |
| see above | Department of Bacteria, Parasites and Fungi, Statens Serum Institut, Copenhagen, Denmark | Statens Serum Institut Bioinformatics and Microbial Genomics | Danish Covid-19 Genome Consortium |
| EPI_ISL_3252753 | Molekylær Medicinsk Afdeling, Aarhus University Hospital, Aarhus, Denmark | Statens Serum Institut Bioinformatics and Microbial Genomics | Danish Covid-19 Genome Consortium |
| EPI_ISL_3252754 | Department of Bacteria, Parasites and Fungi, Statens Serum Institut, Copenhagen, Denmark | Statens Serum Institut Bioinformatics and Microbial Genomics | Danish Covid-19 Genome Consortium |
| EPI_ISL_3252755 | Molekylær Medicinsk Afdeling, Aarhus University Hospital, Aarhus, Denmark | Statens Serum Institut Bioinformatics and Microbial Genomics | Danish Covid-19 Genome Consortium |
| EPI_ISL_3252756, EPI_ISL_3252757, EPI_ISL_3252758, EPI_ISL_3252760, EPI_ISL_3252761, EPI_ISL_3252762, EPI_ISL_3252763, EPI_ISL_3252764, EPI_ISL_3252765, EPI_ISL_3252766, EPI_ISL_3252767, EPI_ISL_3252768, EPI_ISL_3252769, EPI_ISL_3252770, EPI_ISL_3252771, EPI_ISL_3252772, EPI_ISL_3252773, EPI_ISL_3252774, EPI_ISL_3252775, EPI_ISL_3252776, EPI_ISL_3252777, EPI_ISL_3252778, EPI_ISL_3252779, EPI_ISL_3252780, EPI_ISL_3252781, EPI_ISL_3252782, EPI_ISL_3252783, EPI_ISL_3252784, EPI_ISL_3252785, EPI_ISL_3252786, EPI_ISL_3252787, EPI_ISL_3252788, EPI_ISL_3252789, EPI_ISL_3252790, EPI_ISL_3252791, EPI_ISL_3252792, EPI_ISL_3252793, EPI_ISL_3252794, EPI_ISL_3252795, EPI_ISL_3252796, EPI_ISL_3252797, EPI_ISL_3252798, EPI_ISL_3252799, EPI_ISL_3252800, EPI_ISL_3252801, EPI_ISL_3252802, EPI_ISL_3252803, EPI_ISL_3252804, EPI_ISL_3252805, EPI_ISL_3252806, EPI_ISL_3252807, EPI_ISL_3252808, EPI_ISL_3252809, EPI_ISL_3252810, EPI_ISL_3252811, EPI_ISL_3252812, EPI_ISL_3252813, EPI_ISL_3252814, EPI_ISL_3252815, EPI_ISL_3252816 | | | |
| see above | Department of Bacteria, Parasites and Fungi, Statens Serum Institut, Copenhagen, Denmark | Statens Serum Institut Bioinformatics and Microbial Genomics | Danish Covid-19 Genome Consortium |
| EPI_ISL_3252817 | Molekylær Medicinsk Afdeling, Aarhus University Hospital, Aarhus, Denmark | Statens Serum Institut Bioinformatics and Microbial Genomics | Danish Covid-19 Genome Consortium |
| EPI_ISL_3252818, EPI_ISL_3252819, EPI_ISL_3252820, EPI_ISL_3252821, EPI_ISL_3252822, EPI_ISL_3252823, EPI_ISL_3252825, EPI_ISL_3252826, EPI_ISL_3252827, EPI_ISL_3252828, EPI_ISL_3252829, EPI_ISL_3252830, EPI_ISL_3252831, EPI_ISL_3252832, EPI_ISL_3252833, EPI_ISL_3252834, EPI_ISL_3252835, EPI_ISL_3252836, EPI_ISL_3252837, EPI_ISL_3252838, EPI_ISL_3252839, EPI_ISL_3252840, EPI_ISL_3252841, EPI_ISL_3252842, EPI_ISL_3252843, EPI_ISL_3252844, EPI_ISL_3252845, EPI_ISL_3252846, EPI_ISL_3252847, EPI_ISL_3252848, EPI_ISL_3252849, EPI_ISL_3252850, EPI_ISL_3252851, EPI_ISL_3252852, EPI_ISL_3252853, EPI_ISL_3252854, EPI_ISL_3252855, EPI_ISL_3252856, EPI_ISL_3252857, EPI_ISL_3252858, EPI_ISL_3252859, EPI_ISL_3252860, EPI_ISL_3252861, EPI_ISL_3252862, EPI_ISL_3252863, EPI_ISL_3252864, EPI_ISL_3252865, EPI_ISL_3252866, EPI_ISL_3252867, EPI_ISL_3252868, EPI_ISL_3252869, EPI_ISL_3252870, EPI_ISL_3252871, EPI_ISL_3252872, EPI_ISL_3252873, EPI_ISL_3252874, EPI_ISL_3252875, EPI_ISL_3252876, EPI_ISL_3252877, EPI_ISL_3252878, EPI_ISL_3252879, EPI_ISL_3252880, EPI_ISL_3252881, EPI_ISL_3252882 | | | |
| see above | Department of Bacteria, Parasites and Fungi, Statens Serum Institut, Copenhagen, Denmark | Statens Serum Institut Bioinformatics and Microbial Genomics | Danish Covid-19 Genome Consortium |
| EPI_ISL_3252883 | Molekylær Medicinsk Afdeling, Aarhus University Hospital, Aarhus, Denmark | Statens Serum Institut Bioinformatics and Microbial Genomics | Danish Covid-19 Genome Consortium |
| EPI_ISL_3252884, EPI_ISL_3252885, EPI_ISL_3252886, EPI_ISL_3252887, EPI_ISL_3252888, EPI_ISL_3252889, EPI_ISL_3252890, EPI_ISL_3252891, EPI_ISL_3252892, EPI_ISL_3252893, EPI_ISL_3252894, EPI_ISL_3252895, EPI_ISL_3252896, EPI_ISL_3252897, EPI_ISL_3252898, EPI_ISL_3252899, EPI_ISL_3252900, EPI_ISL_3252901, EPI_ISL_3252902, EPI_ISL_3252903, EPI_ISL_3252904, EPI_ISL_3252905, EPI_ISL_3252906, EPI_ISL_3252907, EPI_ISL_3252908, EPI_ISL_3252909, EPI_ISL_3252910, EPI_ISL_3252911, EPI_ISL_3252912, EPI_ISL_3252913, EPI_ISL_3252914, EPI_ISL_3252915, EPI_ISL_3252916, EPI_ISL_3252917, EPI_ISL_3252918, EPI_ISL_3252919, EPI_ISL_3252920, EPI_ISL_3252921, EPI_ISL_3252922, EPI_ISL_3252923, EPI_ISL_3252924, EPI_ISL_3252925, EPI_ISL_3252926, EPI_ISL_3252927, EPI_ISL_3252928, EPI_ISL_3252929, EPI_ISL_3252930, EPI_ISL_3252931, EPI_ISL_3252932, EPI_ISL_3252933, EPI_ISL_3252934, EPI_ISL_3252935, EPI_ISL_3252936, EPI_ISL_3252937, EPI_ISL_3252938, EPI_ISL_3252939, EPI_ISL_3252940, EPI_ISL_3252941, EPI_ISL_3252942, EPI_ISL_3252943, EPI_ISL_3252944, EPI_ISL_3252945, EPI_ISL_3252946, EPI_ISL_3252947, EPI_ISL_3252948, EPI_ISL_3252949, EPI_ISL_3252950, EPI_ISL_3252951, EPI_ISL_3252952, EPI_ISL_3252953, EPI_ISL_3252954, EPI_ISL_3252955, EPI_ISL_3252956, EPI_ISL_3252957, EPI_ISL_3252958, EPI_ISL_3252959, EPI_ISL_3252960, EPI_ISL_3252961, EPI_ISL_3252962, EPI_ISL_3252963, EPI_ISL_3252964, EPI_ISL_3252965, EPI_ISL_3252966, EPI_ISL_3252967, EPI_ISL_3252968, EPI_ISL_3252969, EPI_ISL_3252970, EPI_ISL_3252971 | | | |
| see above | Department of Bacteria, Parasites and Fungi, Statens Serum Institut, Copenhagen, Denmark | Statens Serum Institut Bioinformatics and Microbial Genomics | Danish Covid-19 Genome Consortium |
| EPI_ISL_3252982, EPI_ISL_3252983, EPI_ISL_3252984, EPI_ISL_3252985, EPI_ISL_3252986, EPI_ISL_3252987, EPI_ISL_3252988, EPI_ISL_3252989, EPI_ISL_3252990, EPI_ISL_3252991, EPI_ISL_3252992, EPI_ISL_3252993, EPI_ISL_3252994, EPI_ISL_3252995, EPI_ISL_3252996, EPI_ISL_3252997, EPI_ISL_3252998, EPI_ISL_3252999, EPI_ISL_3253000, EPI_ISL_3253001, EPI_ISL_3253002, EPI_ISL_3253003, EPI_ISL_3253004, EPI_ISL_3253005, EPI_ISL_3253006, EPI_ISL_3253007, EPI_ISL_3253008, EPI_ISL_3253009, EPI_ISL_3253010, EPI_ISL_3253011, EPI_ISL_3253012, EPI_ISL_3253013, EPI_ISL_3253014, EPI_ISL_3253015, EPI_ISL_3253016, EPI_ISL_3253017, EPI_ISL_3253018, EPI_ISL_3253019, EPI_ISL_3253020, EPI_ISL_3253021, EPI_ISL_3253022, EPI_ISL_3253023, EPI_ISL_3253024, EPI_ISL_3253025, EPI_ISL_3253026, EPI_ISL_3253027, EPI_ISL_3253028, EPI_ISL_3253029, EPI_ISL_3253030, EPI_ISL_3253031, EPI_ISL_3253032, EPI_ISL_3253033, EPI_ISL_3253034, EPI_ISL_3253035, EPI_ISL_3253036, EPI_ISL_3253037, EPI_ISL_3253038, EPI_ISL_3253039, EPI_ISL_3253040, EPI_ISL_3253041, EPI_ISL_3253042, EPI_ISL_3253043, EPI_ISL_3253044, EPI_ISL_3253045, EPI_ISL_3253046, EPI_ISL_3253047, EPI_ISL_3253048, EPI_ISL_3253049, EPI_ISL_3253050, EPI_ISL_3253051, EPI_ISL_3253052, EPI_ISL_3253053, EPI_ISL_3253054, EPI_ISL_3253055, EPI_ISL_3253056, EPI_ISL_3253057, EPI_ISL_3253058, EPI_ISL_3253059, EPI_ISL_3253060, EPI_ISL_3253061, EPI_ISL_3253062, EPI_ISL_3253063, EPI_ISL_3253064, EPI_ISL_3253065, EPI_ISL_3253066, EPI_ISL_3253067, EPI_ISL_3253068, EPI_ISL_3253069, EPI_ISL_3253070, EPI_ISL_3253071, EPI_ISL_3253072, EPI_ISL_3253073, EPI_ISL_3253074, EPI_ISL_3253075, EPI_ISL_3253076, EPI_ISL_3253077, EPI_ISL_3253078, EPI_ISL_3253079, EPI_ISL_3253080, EPI_ISL_3253081, EPI_ISL_3253082, EPI_ISL_3253083, EPI_ISL_3253084, EPI_ISL_3253085, EPI_ISL_3253086, EPI_ISL_3253087, EPI_ISL_3253088, EPI_ISL_3253089, EPI_ISL_3253090, EPI_ISL_3253091, EPI_ISL_3253092, EPI_ISL_3253093, EPI_ISL_3253094, EPI_ISL_3253095, EPI_ISL_3253096, EPI_ISL_3253097, EPI_ISL_3253098, EPI_ISL_3253099, EPI_ISL_3253100, EPI_ISL_3253101, EPI_ISL_3253102, EPI_ISL_3253103, EPI_ISL_3253104, EPI_ISL_3253105, EPI_ISL_3253106, EPI_ISL_3253107, EPI_ISL_3253108, EPI_ISL_3253109, EPI_ISL_3253110, EPI_ISL_3253111, EPI_ISL_3253112, EPI_ISL_3253113, EPI_ISL_3253114, EPI_ISL_3253115, EPI_ISL_3253116, EPI_ISL_3253117, EPI_ISL_3253118, EPI_ISL_3253119, EPI_ISL_3253120, EPI_ISL_3253121, EPI_ISL_3253122, EPI_ISL_3253123, EPI_ISL_3253124, EPI_ISL_3253125, EPI_ISL_3253126, EPI_ISL_3253127, EPI_ISL_3253128, EPI_ISL_3253129, EPI_ISL_3253130, EPI_ISL_3253131, EPI_ISL_3253132, EPI_ISL_3253133, EPI_ISL_3253134, EPI_ISL_3253135, EPI_ISL_3253136, EPI_ISL_3253137, EPI_ISL_3253138, EPI_ISL_3253139, EPI_ISL_3253140, EPI_ISL_3253141, EPI_ISL_3253142, EPI_ISL_3253143, EPI_ISL_3253144, EPI_ISL_3253145, EPI_ISL_3253146, EPI_ISL_3253147, EPI_ISL_3253148, EPI_ISL_3253149, EPI_ISL_3253150, EPI_ISL_3253151, EPI_ISL_3253152, EPI_ISL_3253153, EPI_ISL_3253154, EPI_ISL_3253155, EPI_ISL_3253156, EPI_ISL_3253157, EPI_ISL_3253158, EPI_ISL_3253159, EPI_ISL_3253160, EPI_ISL_3253161, EPI_ISL_3253162, EPI_ISL_3253163, EPI_ISL_3253164, EPI_ISL_3253165, EPI_ISL_3253166, EPI_ISL_3253167, EPI_ISL_3253168, EPI_ISL_3253169, EPI_ISL_3253170, EPI_ISL_3253171, EPI_ISL_3253172, EPI_ISL_3253173, EPI_ISL_3253174, EPI_ISL_3253175, EPI_ISL_3253176, EPI_ISL_3253177, EPI_ISL_3253178, EPI_ISL_3253179, EPI_ISL_3253180, EPI_ISL_3253181, EPI_ISL_3253182, EPI_ISL_3253183, EPI_ISL_3253184, EPI_ISL_3253185, EPI_ISL_3253186, EPI_ISL_3253187, EPI_ISL_3253188, EPI_ISL_3253189, EPI_ISL_3253190, EPI_ISL_3253191, EPI_ISL_3253192, EPI_ISL_3253193, EPI_ISL_3253194, EPI_ISL_3253195, EPI_ISL_3253196, EPI_ISL_3253197, EPI_ISL_3253198, EPI_ISL_3253199, EPI_ISL_3253200, EPI_ISL_3253201, EPI_ISL_3253202, EPI_ISL_3253203, EPI_ISL_3253204, EPI_ISL_3253205, EPI_ISL_3253206, EPI_ISL_3253207, EPI_ISL_3253208, EPI_ISL_3253209, EPI_ISL_3253210, EPI_ISL_3253211, EPI_ISL_3253212, EPI_ISL_3253213, EPI_ISL_3253214, EPI_ISL_3253215, EPI_ISL_3253216, EPI_ISL_3253217, EPI_ISL_3253218, EPI_ISL_3253219, EPI_ISL_3253220, EPI_ISL_3253221, EPI_ISL_3253222, EPI_ISL_3253223, EPI_ISL_3253224, EPI_ISL_3253225, EPI_ISL_3253226, EPI_ISL_3253232, EPI_ISL_3253233, EPI_ISL_3253234, EPI_ISL_3253235, EPI_ISL_3253236, EPI_ISL_3253237, EPI_ISL_3253238, EPI_ISL_3253239, EPI_ISL_3253240, EPI_ISL_3253241, EPI_ISL_3253242, EPI_ISL_3253243, EPI_ISL_3253244, EPI_ISL_3253245, EPI_ISL_3253246, EPI_ISL_3253248, EPI_ISL_3253250, EPI_ISL_3253251, EPI_ISL_3253253 | | | |
| see above | National Virus Reference Laboratory | National Virus Reference Laboratory | Zoe Yandle, Charlene Bennett, Gabriel Gonzalez, Michael Carr, Jonathan Dean, Cillian F De Gascun |
| EPI_ISL_3253298, EPI_ISL_3253299, EPI_ISL_3253300, EPI_ISL_3253301, EPI_ISL_3253302, EPI_ISL_3253303, EPI_ISL_3253304, EPI_ISL_3253305, EPI_ISL_3253306, EPI_ISL_3253308, EPI_ISL_3253309, EPI_ISL_3253310, EPI_ISL_3253312, EPI_ISL_3253313, EPI_ISL_3253314, EPI_ISL_3253315, EPI_ISL_3253316, EPI_ISL_3253317, EPI_ISL_3253318, EPI_ISL_3253319, EPI_ISL_3253320, EPI_ISL_3253321 | | | |
| see above | Wyoming Public Health Laboratory | Wyoming Public Health Laboratory | Jim Mildenberger, Taylor Fearing, Channing Weber, Ashley Norberg, Chayse Rowley, Marley Goetz, Brian Dominguez, Elliot Thomasson, Sam Britz, Cari Sloma, Robert Petit, and Rob Christensen |
| EPI_ISL_3253324 | A.S.L. CUNEO 1 | Fondazione del Piemonte per l'Oncologia IRCCS | Antonino Sottile, Silvia Brossa, Paola Marino, Giorgio Giardina |
| EPI_ISL_3253325 | A.S.L. CITTA' DI TORINO - OSPEDALE AMEDEO DI SAVOIA - SC MICROBIOLOGIA E VIROLOGIA - | Fondazione del Piemonte per l'Oncologia IRCCS | Antonino Sottile, Silvia Brossa, Paola Marino, Giorgio Giardina |
| EPI_ISL_3253326 | A.S.L. CITTA' DI TORINO-OSPEDALE SAN LUIGI- ORBASSANO | Fondazione del Piemonte per l'Oncologia IRCCS | Antonino Sottile, Silvia Brossa, Paola Marino, Giorgio Giardina |
| EPI_ISL_3253327, EPI_ISL_3253331, EPI_ISL_3253332, EPI_ISL_3253333, EPI_ISL_3253334, EPI_ISL_3253335, EPI_ISL_3253336, EPI_ISL_3253337, EPI_ISL_3253338, EPI_ISL_3253339, EPI_ISL_3253340, EPI_ISL_3253341, EPI_ISL_3253342, EPI_ISL_3253343, EPI_ISL_3253344, EPI_ISL_3253345, EPI_ISL_3253346, EPI_ISL_3253348, EPI_ISL_3253349, EPI_ISL_3253350, EPI_ISL_3253351, EPI_ISL_3253352, EPI_ISL_3253353, EPI_ISL_3253354, EPI_ISL_3253355, EPI_ISL_3253356, EPI_ISL_3253357, EPI_ISL_3253358, EPI_ISL_3253359, EPI_ISL_3253360 | | | |
| see above | A.S.L. CUNEO 1 | Fondazione del Piemonte per l'Oncologia IRCCS | Antonino Sottile, Silvia Brossa, Paola Marino, Giorgio Giardina |
| EPI_ISL_3253361, EPI_ISL_3253362, EPI_ISL_3253363, EPI_ISL_3253364 | A.S.L. VERCELLI | Fondazione del Piemonte per l'Oncologia IRCCS | Antonino Sottile, Silvia Brossa, Paola Marino, Giorgio Giardina |
| EPI_ISL_3253365 | A.S.L. CUNEO 1 | Fondazione del Piemonte per l'Oncologia IRCCS | Antonino Sottile, Silvia Brossa, Paola Marino, Giorgio Giardina |
| EPI_ISL_3253366, EPI_ISL_3253367, EPI_ISL_3253368, EPI_ISL_3253369, EPI_ISL_3253370, EPI_ISL_3253371, EPI_ISL_3253372, EPI_ISL_3253373, EPI_ISL_3253374 | A.S.L. VERCELLI | Fondazione del Piemonte per l'Oncologia IRCCS | Antonino Sottile, Silvia Brossa, Paola Marino, Giorgio Giardina |
| EPI_ISL_3253375, EPI_ISL_3253376, EPI_ISL_3253377, EPI_ISL_3253378, EPI_ISL_3253379, EPI_ISL_3253380, EPI_ISL_3253381, EPI_ISL_3253382, EPI_ISL_3253383, EPI_ISL_3253384, EPI_ISL_3253385, EPI_ISL_3253386, EPI_ISL_3253387, EPI_ISL_3253388, EPI_ISL_3253389, EPI_ISL_3253390, EPI_ISL_3253391, EPI_ISL_3253392, EPI_ISL_3253393, EPI_ISL_3253395, EPI_ISL_3253396, EPI_ISL_3253397, EPI_ISL_3253398, EPI_ISL_3253399, EPI_ISL_3253400, EPI_ISL_3253401, EPI_ISL_3253402, EPI_ISL_3253403, EPI_ISL_3253404, EPI_ISL_3253405, EPI_ISL_3253406, EPI_ISL_3253407, EPI_ISL_3253408, EPI_ISL_3253410, EPI_ISL_3253411, EPI_ISL_3253412, EPI_ISL_3253413, EPI_ISL_3253414, EPI_ISL_3253415, EPI_ISL_3253416, EPI_ISL_3253417, EPI_ISL_3253418, EPI_ISL_3253419, EPI_ISL_3253420, EPI_ISL_3253421, EPI_ISL_3253422, EPI_ISL_3253423 | | | |
| see above | A.S.L. NOVARA | Fondazione del Piemonte per l'Oncologia IRCCS | Antonino Sottile, Silvia Brossa, Paola Marino, Giorgio Giardina |
| EPI_ISL_3253424, | AZ Klina | AZ Klina | Carl Vael - Lynsey Berckmans |

| | | | |
|---|---|---|---|
| EPI_ISL_3253425 | | | |
| EPI_ISL_3253426 | Laboratoire de Recherche et d'Analyses Médicales de la Gendarmerie Royale | Laboratoire de Recherche et d'Analyses Médicales de la Gendarmerie Royale | Sanaâ LEMRISS, Amal SOUIRI, Hajar LEMRISS, Saâd EL KABBAJ |
| EPI_ISL_3253427, EPI_ISL_3253428 | AZ Klina | AZ Klina | Carl Vael - Lynsey Berckmans |
| EPI_ISL_3253429 | NCSLPH | NCSLPH | Chase K, Miller MC, Greene S, Glover W |
| EPI_ISL_3253430 | AZ Klina | AZ Klina | Carl Vael - Lynsey Berckmans |
| EPI_ISL_3253431 | NCSLPH | NCSLPH | Chase K, Miller MC, Greene S, Glover W |
| EPI_ISL_3253432 | AZ Klina | AZ Klina | Carl Vael - Lynsey Berckmans |
| EPI_ISL_3253433 | NCSLPH | NCSLPH | Chase K, Miller MC, Greene S, Glover W |
| EPI_ISL_3253434, EPI_ISL_3253551, EPI_ISL_3253795 | AZ Klina | AZ Klina | Carl Vael - Lynsey Berckmans |
| EPI_ISL_3253871 | NCSLPH | NCSLPH | Chase K, Miller MC, Greene S, Glover W |
| EPI_ISL_3254030, EPI_ISL_3254213 | AZ Klina | AZ Klina | Carl Vael - Lynsey Berckmans |
| EPI_ISL_3254214, EPI_ISL_3254215, EPI_ISL_3254216 | NCSLPH | NCSLPH | Chase K, Miller MC, Greene S, Glover W |
| EPI_ISL_3254217 | DOHMH Riverside | New York City Public Health Laboratory | Jade Wang, et al. |
| EPI_ISL_3254218 | DOHMH Jamaica | New York City Public Health Laboratory | Jade Wang, et al. |
| EPI_ISL_3254221, EPI_ISL_3254222, EPI_ISL_3254223 | DOHMH Crown Heights | New York City Public Health Laboratory | Jade Wang, et al. |
| EPI_ISL_3254224 | DOHMH Riverside | New York City Public Health Laboratory | Jade Wang, et al. |
| EPI_ISL_3254225 | DOHMH Chelsea | New York City Public Health Laboratory | Jade Wang, et al. |
| EPI_ISL_3254226 | DOHMH Fort Greene | New York City Public Health Laboratory | Jade Wang, et al. |
| EPI_ISL_3254227 | DOHMH PHL | New York City Public Health Laboratory | Jade Wang, et al. |
| EPI_ISL_3254228 | DOHMH Corona | New York City Public Health Laboratory | Jade Wang, et al. |
| EPI_ISL_3254229 | DOHMH Chelsea | New York City Public Health Laboratory | Jade Wang, et al. |
| EPI_ISL_3254230 | DOHMH Crown Heights | New York City Public Health Laboratory | Jade Wang, et al. |
| EPI_ISL_3254231, EPI_ISL_3254232 | DOHMH Fort Greene | New York City Public Health Laboratory | Jade Wang, et al. |
| EPI_ISL_3254233 | DOHMH Jamaica | New York City Public Health Laboratory | Jade Wang, et al. |
| EPI_ISL_3254234, EPI_ISL_3254235 | DOHMH PHL | New York City Public Health Laboratory | Jade Wang, et al. |
| EPI_ISL_3254237, EPI_ISL_3254238, EPI_ISL_3254239, EPI_ISL_3254240 | Quest Diagnostics | New York City Public Health Laboratory | Jade Wang, et al. |
| EPI_ISL_3254241, EPI_ISL_3254242 | DOHMH Crown Heights | New York City Public Health Laboratory | Jade Wang, et al. |
| EPI_ISL_3254243 | DOHMH PHL | New York City Public Health Laboratory | Jade Wang, et al. |
| EPI_ISL_3254245 | DOHMH Riverside | New York City Public Health Laboratory | Jade Wang, et al. |
| EPI_ISL_3254246 | DOHMH Chelsea | New York City Public Health Laboratory | Jade Wang, et al. |
| EPI_ISL_3254247 | DOHMH PHL | New York City Public Health Laboratory | Jade Wang, et al. |
| EPI_ISL_3254248 | DOHMH Chelsea | New York City Public Health Laboratory | Jade Wang, et al. |
| EPI_ISL_3254249 | DOHMH Fort Greene | New York City Public Health Laboratory | Jade Wang, et al. |
| EPI_ISL_3254250 | DOHMH Corona | New York City Public Health Laboratory | Jade Wang, et al. |
| EPI_ISL_3254251 | DOHMH Crown Heights | New York City Public Health Laboratory | Jade Wang, et al. |
| EPI_ISL_3254252 | DOHMH Morrisania | New York City Public Health Laboratory | Jade Wang, et al. |
| EPI_ISL_3254253 | DOHMH PHL | New York City Public Health Laboratory | Jade Wang, et al. |
| EPI_ISL_3254254 | DOHMH Jamaica | New York City Public Health Laboratory | Jade Wang, et al. |
| EPI_ISL_3254255 | DOHMH Crown Heights | New York City Public Health Laboratory | Jade Wang, et al. |
| EPI_ISL_3254256, EPI_ISL_3254257, EPI_ISL_3254258, EPI_ISL_3254259 | DOHMH PHL | New York City Public Health Laboratory | Jade Wang, et al. |
| EPI_ISL_3254260, EPI_ISL_3254261 | DOHMH Riverside | New York City Public Health Laboratory | Jade Wang, et al. |
| EPI_ISL_3254262 | DOHMH Chelsea | New York City Public Health Laboratory | Jade Wang, et al. |
| EPI_ISL_3254263, EPI_ISL_3254264 | DOHMH Fort Greene | New York City Public Health Laboratory | Jade Wang, et al. |
| EPI_ISL_3254265, EPI_ISL_3254266, EPI_ISL_3254267, EPI_ISL_3254268 | DOHMH Jamaica | New York City Public Health Laboratory | Jade Wang, et al. |
| EPI_ISL_3254269, EPI_ISL_3254270, EPI_ISL_3254271, EPI_ISL_3254272, EPI_ISL_3254273, EPI_ISL_3254274, EPI_ISL_3254275 | DOHMH PHL | New York City Public Health Laboratory | Jade Wang, et al. |
| EPI_ISL_3254276, EPI_ISL_3254277 | Centers Lab NJ LLC | New York City Public Health Laboratory | Jade Wang, et al. |
| EPI_ISL_3254279, EPI_ISL_3254280, EPI_ISL_3254281, EPI_ISL_3254282, EPI_ISL_3254283, EPI_ISL_3254284 | Quest Diagnostics | New York City Public Health Laboratory | Jade Wang, et al. |
| EPI_ISL_3254286, EPI_ISL_3254287, EPI_ISL_3254288 | Lincoln Diagnostics | New York City Public Health Laboratory | Jade Wang, et al. |
| EPI_ISL_3254289 | NCSLPH | NCSLPH | Chase K, Miller MC, Greene S, Glover W |
| EPI_ISL_3254290 | DOHMH Crown Heights | New York City Public Health Laboratory | Jade Wang, et al. |
| EPI_ISL_3254291, | DOHMH Jamaica | New York City Public Health Laboratory | Jade Wang, et al. |

| | | | |
|---|---|---|---|
| EPI_ISL_3254292, EPI_ISL_3254293, EPI_ISL_3254294, EPI_ISL_3254295, EPI_ISL_3254296, EPI_ISL_3254297 | | | |
| EPI_ISL_3254298, EPI_ISL_3254299 | DOHMH Riverside | New York City Public Health Laboratory | Jade Wang, et al. |
| EPI_ISL_3254300 | DOHMH Corona | New York City Public Health Laboratory | Jade Wang, et al. |
| EPI_ISL_3254301, EPI_ISL_3254302 | DOHMH Chelsea | New York City Public Health Laboratory | Jade Wang, et al. |
| EPI_ISL_3254303, EPI_ISL_3254304, EPI_ISL_3254305 | DOHMH Morrisania | New York City Public Health Laboratory | Jade Wang, et al. |
| EPI_ISL_3254306 | DOHMH Chelsea | New York City Public Health Laboratory | Jade Wang, et al. |
| EPI_ISL_3254314, EPI_ISL_3254315, EPI_ISL_3254367, EPI_ISL_3254527, EPI_ISL_3254685, EPI_ISL_3254686, EPI_ISL_3254687 | NCSLPH | NCSLPH | Chase K, Miller MC, Greene S, Glover W |
| EPI_ISL_3254689, EPI_ISL_3254694, EPI_ISL_3254695, EPI_ISL_3254696, EPI_ISL_3254697, EPI_ISL_3254700, EPI_ISL_3254701, EPI_ISL_3254705, EPI_ISL_3254709, EPI_ISL_3254710, EPI_ISL_3254711, EPI_ISL_3254712, EPI_ISL_3254715, EPI_ISL_3254716, EPI_ISL_3254731, EPI_ISL_3254732, EPI_ISL_3254733, EPI_ISL_3254734, EPI_ISL_3254735, EPI_ISL_3254736, EPI_ISL_3254737, EPI_ISL_3254738, EPI_ISL_3254739, EPI_ISL_3254740, EPI_ISL_3254741, EPI_ISL_3254742, EPI_ISL_3254743, EPI_ISL_3254744, EPI_ISL_3254745, EPI_ISL_3254746, EPI_ISL_3254747, EPI_ISL_3254748, EPI_ISL_3254749 | | | |
| see above | Illinois Department of Public Health | Gagnon Lab, Southern Illinois University | Keith Gagnon |
| EPI_ISL_3254751 | Simple Laboratories | Gagnon Lab, Southern Illinois University | Keith Gagnon |
| EPI_ISL_3254752, EPI_ISL_3254753, EPI_ISL_3254754, EPI_ISL_3254755, EPI_ISL_3254756, EPI_ISL_3254782, EPI_ISL_3254789 | Illinois Department of Public Health | Gagnon Lab, Southern Illinois University | Keith Gagnon |
| EPI_ISL_3254790 | NCSLPH | NCSLPH | Chase K, Miller MC, Greene S, Glover W |
| EPI_ISL_3254793, EPI_ISL_3254795, EPI_ISL_3254798, EPI_ISL_3254800, EPI_ISL_3254801, EPI_ISL_3254802, EPI_ISL_3254807, EPI_ISL_3254808, EPI_ISL_3254810, EPI_ISL_3254812, EPI_ISL_3254813, EPI_ISL_3254816 | | | |
| see above | Illinois Department of Public Health | Gagnon Lab, Southern Illinois University | Keith Gagnon |
| EPI_ISL_3254821 | Simple Laboratories | Gagnon Lab, Southern Illinois University | Keith Gagnon |
| EPI_ISL_3254823, EPI_ISL_3254824, EPI_ISL_3254825, EPI_ISL_3254826, EPI_ISL_3254827, EPI_ISL_3254828 | Illinois Department of Public Health | Gagnon Lab, Southern Illinois University | Keith Gagnon |
| EPI_ISL_3254829 | Simple Laboratories | Gagnon Lab, Southern Illinois University | Keith Gagnon |
| EPI_ISL_3254961, EPI_ISL_3254962 | NCSLPH | NCSLPH | Chase K, Miller MC, Greene S, Glover W |
| EPI_ISL_3254963, EPI_ISL_3254964, EPI_ISL_3254965, EPI_ISL_3254966, EPI_ISL_3254967, EPI_ISL_3254968, EPI_ISL_3254969, EPI_ISL_3254970, EPI_ISL_3254971, EPI_ISL_3254972, EPI_ISL_3254973, EPI_ISL_3254974, EPI_ISL_3254975, EPI_ISL_3254976, EPI_ISL_3254977, EPI_ISL_3254978, EPI_ISL_3254979, EPI_ISL_3254980, EPI_ISL_3254981, EPI_ISL_3254982, EPI_ISL_3254983, EPI_ISL_3254984, EPI_ISL_3254985, EPI_ISL_3254987, EPI_ISL_3254989, EPI_ISL_3254990, EPI_ISL_3254991, EPI_ISL_3254992, EPI_ISL_3254993, EPI_ISL_3254994, EPI_ISL_3254995, EPI_ISL_3254996, EPI_ISL_3254997, EPI_ISL_3254998, EPI_ISL_3254999, EPI_ISL_3255000 | | | |
| see above | AZDelta | AZ Delta Medical Laboratories in Roeselare, Belgium | Geert Martens, Dieter De Smet, Frederik Van Hoecke, on behalf of AZ Delta COVID-19 Genomics core (member of Genomic surveillance of SARS-CoV-2 in Belgium network) |
| EPI_ISL_3255001, EPI_ISL_3255002, EPI_ISL_3255003, EPI_ISL_3255004, EPI_ISL_3255005 | NCSLPH | NCSLPH | Chase K, Miller MC, Greene S, Glover W |
| EPI_ISL_3255006 | A.S.L. TO3 | Fondazione del Piemonte per l'Oncologia IRCCS | Antonino Sottile, Silvia Brossa, Paola Marino, Giorgio Giardina |
| EPI_ISL_3255007 | A.S.L. NOVARA | Fondazione del Piemonte per l'Oncologia IRCCS | Antonino Sottile, Silvia Brossa, Paola Marino, Giorgio Giardina |
| EPI_ISL_3255008 | A.S.L. TO5 | Fondazione del Piemonte per l'Oncologia IRCCS | Antonino Sottile, Silvia Brossa, Paola Marino, Giorgio Giardina |
| EPI_ISL_3255009 | A.S.L. TO4 | Fondazione del Piemonte per l'Oncologia IRCCS | Antonino Sottile, Silvia Brossa, Paola Marino, Giorgio Giardina |
| EPI_ISL_3255010 | A.S.L. TO5 | Fondazione del Piemonte per l'Oncologia IRCCS | Antonino Sottile, Silvia Brossa, Paola Marino, Giorgio Giardina |
| EPI_ISL_3255011 | A.S.L. CITTA' DI TORINO - OSPEDALE AMEDEO DI SAVOIA - SC MICROBIOLOGIA E VIROLOGIA - | Fondazione del Piemonte per l'Oncologia IRCCS | Antonino Sottile, Silvia Brossa, Paola Marino, Giorgio Giardina |
| EPI_ISL_3255012 | A.S.L. TO5 | Fondazione del Piemonte per l'Oncologia IRCCS | Antonino Sottile, Silvia Brossa, Paola Marino, Giorgio Giardina |
| EPI_ISL_3255013 | A.S.L. CUNEO 1 | Fondazione del Piemonte per l'Oncologia IRCCS | Antonino Sottile, Silvia Brossa, Paola Marino, Giorgio Giardina |
| EPI_ISL_3255014, EPI_ISL_3255015 | A.S.L. TO4 | Fondazione del Piemonte per l'Oncologia IRCCS | Antonino Sottile, Silvia Brossa, Paola Marino, Giorgio Giardina |
| EPI_ISL_3255016, EPI_ISL_3255017, EPI_ISL_3255018, EPI_ISL_3255019, EPI_ISL_3255020, EPI_ISL_3255021, EPI_ISL_3255022, EPI_ISL_3255023 | A.S.L. CUNEO 1 | Fondazione del Piemonte per l'Oncologia IRCCS | Antonino Sottile, Silvia Brossa, Paola Marino, Giorgio Giardina |
| EPI_ISL_3255024 | A.S.L. TO3 | Fondazione del Piemonte per l'Oncologia IRCCS | Antonino Sottile, Silvia Brossa, Paola Marino, Giorgio Giardina |
| EPI_ISL_3255025, EPI_ISL_3255026, EPI_ISL_3255027 | A.S.L. CUNEO 1 | Fondazione del Piemonte per l'Oncologia IRCCS | Antonino Sottile, Silvia Brossa, Paola Marino, Giorgio Giardina |
| EPI_ISL_3255028, EPI_ISL_3255029, EPI_ISL_3255030, EPI_ISL_3255031, EPI_ISL_3255032, EPI_ISL_3255033, EPI_ISL_3255034, EPI_ISL_3255035 | A.S.L. TO3 | Fondazione del Piemonte per l'Oncologia IRCCS | Antonino Sottile, Silvia Brossa, Paola Marino, Giorgio Giardina |
| EPI_ISL_3255036, EPI_ISL_3255037 | A.S.L. CUNEO 1 | Fondazione del Piemonte per l'Oncologia IRCCS | Antonino Sottile, Silvia Brossa, Paola Marino, Giorgio Giardina |
| EPI_ISL_3255038 | A.S.L. TO3 | Fondazione del Piemonte per l'Oncologia IRCCS | Antonino Sottile, Silvia Brossa, Paola Marino, Giorgio Giardina |
| EPI_ISL_3255039, EPI_ISL_3255040, EPI_ISL_3255041, EPI_ISL_3255042 | A.S.L. CUNEO 1 | Fondazione del Piemonte per l'Oncologia IRCCS | Antonino Sottile, Silvia Brossa, Paola Marino, Giorgio Giardina |
| EPI_ISL_3255043, | NCSLPH | NCSLPH | Chase K, Miller MC, Greene S, Glover W |

| | | | |
|---|---|---|---|
| EPI_ISL_3255044, EPI_ISL_3255045 | | | |
| EPI_ISL_3255047 | Hospital General Universitario de Ciudad Real | Hospital General Universitario de Ciudad Real | José Martínez-Alarcón, Soledad Illescas Fernández-Bermejo, Marta Torres-Narbona, Cristina Colmenarejo, Lidia García-Agudo |
| EPI_ISL_3255048 | Hospital General de Valdepeñas | Hospital General Universitario de Ciudad Real | José Martínez-Alarcón, Soledad Illescas Fernández-Bermejo, Marta Torres-Narbona, Cristina Colmenarejo, Lidia García-Agudo |
| EPI_ISL_3255049, EPI_ISL_3255050 | Hospital General Universitario de Ciudad Real | Hospital General Universitario de Ciudad Real | José Martínez-Alarcón, Soledad Illescas Fernández-Bermejo, Marta Torres-Narbona, Cristina Colmenarejo, Lidia García-Agudo |
| EPI_ISL_3255051 | Hospital Virgen De Altagracia | Hospital General Universitario de Ciudad Real | José Martínez-Alarcón, Soledad Illescas Fernández-Bermejo, Marta Torres-Narbona, Cristina Colmenarejo, Lidia García-Agudo |
| EPI_ISL_3255052 | Hospital General de Valdepeñas | Hospital General Universitario de Ciudad Real | José Martínez-Alarcón, Soledad Illescas Fernández-Bermejo, Marta Torres-Narbona, Cristina Colmenarejo, Lidia García-Agudo |
| EPI_ISL_3255053 | Hospital General Universitario de Ciudad Real | Hospital General Universitario de Ciudad Real | José Martínez-Alarcón, Soledad Illescas Fernández-Bermejo, Marta Torres-Narbona, Cristina Colmenarejo, Lidia García-Agudo |
| EPI_ISL_3255054 | Hospital General de Valdepeñas | Hospital General Universitario de Ciudad Real | José Martínez-Alarcón, Soledad Illescas Fernández-Bermejo, Marta Torres-Narbona, Cristina Colmenarejo, Lidia García-Agudo |
| EPI_ISL_3255288, EPI_ISL_3255289, EPI_ISL_3255290, EPI_ISL_3255291, EPI_ISL_3255292, EPI_ISL_3255293, EPI_ISL_3255294, EPI_ISL_3255295, EPI_ISL_3255296, EPI_ISL_3255297, EPI_ISL_3255298, EPI_ISL_3255299, EPI_ISL_3255300, EPI_ISL_3255301, EPI_ISL_3255302, EPI_ISL_3255303, EPI_ISL_3255304, EPI_ISL_3255305, EPI_ISL_3255306, EPI_ISL_3255307, EPI_ISL_3255308, EPI_ISL_3255309, EPI_ISL_3255310, EPI_ISL_3255311, EPI_ISL_3255312, EPI_ISL_3255313, EPI_ISL_3255314, EPI_ISL_3255315, EPI_ISL_3255316, EPI_ISL_3255317, EPI_ISL_3255318, EPI_ISL_3255319, EPI_ISL_3255320, EPI_ISL_3255321, EPI_ISL_3255322, EPI_ISL_3255323, EPI_ISL_3255324, EPI_ISL_3255325, EPI_ISL_3255326, EPI_ISL_3255327, EPI_ISL_3255328, EPI_ISL_3255329, EPI_ISL_3255330, EPI_ISL_3255331, EPI_ISL_3255332, EPI_ISL_3255333, EPI_ISL_3255334, EPI_ISL_3255335, EPI_ISL_3255336, EPI_ISL_3255337, EPI_ISL_3255338, EPI_ISL_3255339, EPI_ISL_3255340, EPI_ISL_3255341, EPI_ISL_3255342, EPI_ISL_3255343, EPI_ISL_3255344, EPI_ISL_3255345, EPI_ISL_3255346, EPI_ISL_3255347, EPI_ISL_3255348, EPI_ISL_3255349, EPI_ISL_3255350, EPI_ISL_3255351, EPI_ISL_3255352, EPI_ISL_3255353, EPI_ISL_3255354, EPI_ISL_3255355, EPI_ISL_3255356, EPI_ISL_3255357, EPI_ISL_3255358, EPI_ISL_3255359, EPI_ISL_3255360, EPI_ISL_3255361, EPI_ISL_3255362, EPI_ISL_3255363, EPI_ISL_3255364, EPI_ISL_3255365, EPI_ISL_3255366, EPI_ISL_3255367, EPI_ISL_3255368, EPI_ISL_3255369, EPI_ISL_3255370, EPI_ISL_3255371, EPI_ISL_3255372, EPI_ISL_3255373, EPI_ISL_3255374, EPI_ISL_3255375, EPI_ISL_3255376, EPI_ISL_3255377, EPI_ISL_3255378, EPI_ISL_3255379, EPI_ISL_3255380, EPI_ISL_3255381, EPI_ISL_3255382, EPI_ISL_3255383, EPI_ISL_3255384, EPI_ISL_3255385, EPI_ISL_3255386, EPI_ISL_3255387, EPI_ISL_3255388, EPI_ISL_3255389, EPI_ISL_3255390, EPI_ISL_3255391, EPI_ISL_3255392, EPI_ISL_3255393, EPI_ISL_3255394, EPI_ISL_3255395, EPI_ISL_3255396, EPI_ISL_3255397, EPI_ISL_3255398, EPI_ISL_3255399, EPI_ISL_3255400, EPI_ISL_3255401, EPI_ISL_3255402, EPI_ISL_3255403, EPI_ISL_3255404, EPI_ISL_3255405, EPI_ISL_3255406, EPI_ISL_3255407, EPI_ISL_3255408, EPI_ISL_3255409, EPI_ISL_3255410, EPI_ISL_3255411, EPI_ISL_3255412, EPI_ISL_3255414, EPI_ISL_3255415, EPI_ISL_3255416, EPI_ISL_3255417, EPI_ISL_3255418, EPI_ISL_3255419, EPI_ISL_3255420, EPI_ISL_3255421, EPI_ISL_3255422, EPI_ISL_3255423, EPI_ISL_3255424, EPI_ISL_3255425 | | | |
| see above | Azienda Sanitaria Locale di Piacenza - Presidio Ospedaliero - Laboratorio di Microbiologia | Istituto Zooprofilattico Sperimentale della Lombardia e dell'Emilia Romagna (IZSLER), Risk Analysis and Genomic Epidemiology Unit | Lo Cascio Giuliana, Schiavo Roberta, Peroni Irene, Marina Morganti, Ilaria Menozzi, Erika Scaltriti, Stefano Pongolini |
| EPI_ISL_3255548, EPI_ISL_3255556, EPI_ISL_3255559, EPI_ISL_3255563, EPI_ISL_3255566, EPI_ISL_3255567, EPI_ISL_3255575, EPI_ISL_3255580, EPI_ISL_3255582, EPI_ISL_3255584, EPI_ISL_3255585, EPI_ISL_3255586, EPI_ISL_3255587, EPI_ISL_3255588, EPI_ISL_3255589, EPI_ISL_3255590, EPI_ISL_3255591, EPI_ISL_3255593, EPI_ISL_3255595, EPI_ISL_3255597, EPI_ISL_3255598, EPI_ISL_3255599, EPI_ISL_3255601, EPI_ISL_3255602, EPI_ISL_3255603, EPI_ISL_3255604, EPI_ISL_3255605, EPI_ISL_3255606, EPI_ISL_3255609, EPI_ISL_3255613, EPI_ISL_3255616, EPI_ISL_3255617, EPI_ISL_3255621, EPI_ISL_3255622, EPI_ISL_3255627, EPI_ISL_3255643, EPI_ISL_3255648, EPI_ISL_3255649, EPI_ISL_3255652, EPI_ISL_3255653, EPI_ISL_3255655, EPI_ISL_3255656, EPI_ISL_3255657, EPI_ISL_3255658, EPI_ISL_3255659, EPI_ISL_3255660, EPI_ISL_3255662, EPI_ISL_3255667, EPI_ISL_3255671, EPI_ISL_3255675, EPI_ISL_3255677, EPI_ISL_3255686, EPI_ISL_3255690, EPI_ISL_3255696, EPI_ISL_3255699, EPI_ISL_3255700, EPI_ISL_3255704, EPI_ISL_3255715, EPI_ISL_3255721, EPI_ISL_3255723, EPI_ISL_3255725, EPI_ISL_3255726, EPI_ISL_3255729, EPI_ISL_3255731, EPI_ISL_3255732, EPI_ISL_3255733, EPI_ISL_3255735, EPI_ISL_3255737, EPI_ISL_3255738, EPI_ISL_3255739, EPI_ISL_3255740, EPI_ISL_3255741, EPI_ISL_3255742, EPI_ISL_3255744, EPI_ISL_3255745, EPI_ISL_3255746, EPI_ISL_3255761, EPI_ISL_3255763, EPI_ISL_3255766, EPI_ISL_3255767, EPI_ISL_3255769, EPI_ISL_3255770, EPI_ISL_3255775, EPI_ISL_3255776, EPI_ISL_3255777, EPI_ISL_3255778, EPI_ISL_3255779, EPI_ISL_3255784, EPI_ISL_3255785, EPI_ISL_3255786, EPI_ISL_3255787, EPI_ISL_3255789, EPI_ISL_3255802, EPI_ISL_3255804, EPI_ISL_3255805, EPI_ISL_3255806, EPI_ISL_3255807, EPI_ISL_3255808, EPI_ISL_3255810, EPI_ISL_3255812, EPI_ISL_3255813, EPI_ISL_3255814, EPI_ISL_3255815, EPI_ISL_3255816, EPI_ISL_3255818, EPI_ISL_3255819, EPI_ISL_3255820, EPI_ISL_3255823, EPI_ISL_3255824, EPI_ISL_3255826, EPI_ISL_3255829, EPI_ISL_3255830, EPI_ISL_3255835, EPI_ISL_3255843, EPI_ISL_3255844, EPI_ISL_3255849, EPI_ISL_3255852, EPI_ISL_3255855, EPI_ISL_3255858, EPI_ISL_3255866, EPI_ISL_3255875, EPI_ISL_3255877, EPI_ISL_3255880, EPI_ISL_3255881, EPI_ISL_3255884, EPI_ISL_3255885, EPI_ISL_3255887, EPI_ISL_3255888, EPI_ISL_3255890, EPI_ISL_3255891, EPI_ISL_3255893, EPI_ISL_3255895, EPI_ISL_3255897, EPI_ISL_3255898, EPI_ISL_3255907, EPI_ISL_3255908, EPI_ISL_3255909, EPI_ISL_3255911, EPI_ISL_3255913, EPI_ISL_3255915, EPI_ISL_3255918, EPI_ISL_3255920, EPI_ISL_3255921, EPI_ISL_3255936, EPI_ISL_3255938, EPI_ISL_3255940, EPI_ISL_3255942, EPI_ISL_3255949, EPI_ISL_3255955, EPI_ISL_3255959, EPI_ISL_3255960, EPI_ISL_3255961, EPI_ISL_3255966, EPI_ISL_3255972 | | | |
| see above | SK-Roy Romanow Provincial Laboratory | National Microbiology Laboratory (NML) | Anna Majer, Shari Tyson, Grace Seo, Philip Mabon, Elsie Grudeski, Rhiannon Huzarewich, Russell Mandes, Anneliese Landgraff, Jennifer Tanner, Natalie Knox, Morag Graham, Gary Van Domselaar, Ryan McDonald, Keith MacKenzie, Meredith Faires, Kara Loos, Rachel DePaulo, Alanna Senecal, Amanda Lang, Jessica Minion, Nathalie Bastien, Yan Li, Timothy Booth, Darian Hole, Madison Chapel, Kirsten Biggar, CanCOGeN's metadata curation team, Public Health Agency of Canada CanCOGeN team |
| EPI_ISL_3255984, EPI_ISL_3255985, EPI_ISL_3255987, EPI_ISL_3255988, EPI_ISL_3255989, EPI_ISL_3255990, EPI_ISL_3255991, EPI_ISL_3255992, EPI_ISL_3255993, EPI_ISL_3255994, EPI_ISL_3255995, EPI_ISL_3255996, EPI_ISL_3255997, EPI_ISL_3255998, EPI_ISL_3255999, EPI_ISL_3256000, EPI_ISL_3256002, EPI_ISL_3256003, EPI_ISL_3256004, EPI_ISL_3256005, EPI_ISL_3256009, EPI_ISL_3256010, EPI_ISL_3256011, EPI_ISL_3256012, EPI_ISL_3256013, EPI_ISL_3256014, EPI_ISL_3256015, EPI_ISL_3256016, EPI_ISL_3256017, EPI_ISL_3256018, EPI_ISL_3256019, EPI_ISL_3256020, EPI_ISL_3256021, EPI_ISL_3256022, EPI_ISL_3256023, EPI_ISL_3256024, EPI_ISL_3256025 | | | |
| see above | Molecular Diagnostics Laboratory | National Reference Laboratory, Nigeria Centre for Disease Control | Dr Ndodo Nnaemeka, Olusola Anuoluwapo Akanbi, Chimaobi Chukwu, Dr Omoare Adesuyi, Celestina Obiekea, Anthony Ahumibe, Catherine Okoi, Nwando Mba, Dr Chikwe Ihekweazu |
| EPI_ISL_3256026, EPI_ISL_3256027, EPI_ISL_3256028, EPI_ISL_3256029, EPI_ISL_3256030, EPI_ISL_3256031, EPI_ISL_3256032, EPI_ISL_3256033, EPI_ISL_3256034, EPI_ISL_3256035, EPI_ISL_3256036, EPI_ISL_3256037, EPI_ISL_3256038, EPI_ISL_3256039, EPI_ISL_3256040, EPI_ISL_3256041, EPI_ISL_3256042, EPI_ISL_3256043, EPI_ISL_3256044, EPI_ISL_3256045, EPI_ISL_3256046, EPI_ISL_3256047, EPI_ISL_3256048, EPI_ISL_3256049, EPI_ISL_3256050, EPI_ISL_3256051, EPI_ISL_3256052, EPI_ISL_3256053, EPI_ISL_3256054, EPI_ISL_3256055, EPI_ISL_3256056, EPI_ISL_3256057, EPI_ISL_3256058, EPI_ISL_3256059, EPI_ISL_3256060, EPI_ISL_3256061, EPI_ISL_3256062, EPI_ISL_3256063, EPI_ISL_3256064, EPI_ISL_3256065, EPI_ISL_3256066, EPI_ISL_3256067, EPI_ISL_3256068, EPI_ISL_3256069, EPI_ISL_3256070, EPI_ISL_3256071, EPI_ISL_3256072, EPI_ISL_3256073, EPI_ISL_3256074, EPI_ISL_3256075, EPI_ISL_3256076, EPI_ISL_3256077, EPI_ISL_3256078, EPI_ISL_3256079, EPI_ISL_3256080, EPI_ISL_3256081, EPI_ISL_3256082, EPI_ISL_3256083, EPI_ISL_3256084, EPI_ISL_3256086, EPI_ISL_3256087, EPI_ISL_3256088, EPI_ISL_3256089, EPI_ISL_3256090, EPI_ISL_3256091, EPI_ISL_3256092, EPI_ISL_3256093, EPI_ISL_3256094, EPI_ISL_3256095, EPI_ISL_3256096, EPI_ISL_3256097, EPI_ISL_3256098, EPI_ISL_3256099, EPI_ISL_3256100, EPI_ISL_3256101, EPI_ISL_3256102, EPI_ISL_3256103, EPI_ISL_3256104, EPI_ISL_3256105, EPI_ISL_3256106, EPI_ISL_3256107, EPI_ISL_3256108, EPI_ISL_3256109, EPI_ISL_3256110, EPI_ISL_3256111, EPI_ISL_3256112, EPI_ISL_3256113, EPI_ISL_3256114, EPI_ISL_3256115, EPI_ISL_3256116, EPI_ISL_3256117, EPI_ISL_3256118, EPI_ISL_3256119, EPI_ISL_3256120, EPI_ISL_3256121, EPI_ISL_3256122, EPI_ISL_3256123, EPI_ISL_3256124, EPI_ISL_3256125, EPI_ISL_3256126, EPI_ISL_3256127, EPI_ISL_3256128, EPI_ISL_3256129, EPI_ISL_3256130, EPI_ISL_3256131, EPI_ISL_3256132, EPI_ISL_3256133, EPI_ISL_3256134, EPI_ISL_3256135, EPI_ISL_3256136, EPI_ISL_3256137, EPI_ISL_3256138, EPI_ISL_3256139, EPI_ISL_3256140, EPI_ISL_3256141, EPI_ISL_3256142, EPI_ISL_3256143, EPI_ISL_3256144, EPI_ISL_3256145, EPI_ISL_3256146, EPI_ISL_3256147, EPI_ISL_3256148, EPI_ISL_3256149, EPI_ISL_3256150, EPI_ISL_3256151, EPI_ISL_3256152, EPI_ISL_3256153, EPI_ISL_3256154, EPI_ISL_3256155, EPI_ISL_3256156, EPI_ISL_3256157, EPI_ISL_3256158, EPI_ISL_3256159, EPI_ISL_3256160, EPI_ISL_3256161, EPI_ISL_3256162, EPI_ISL_3256163, EPI_ISL_3256164, EPI_ISL_3256165, EPI_ISL_3256166, EPI_ISL_3256167, EPI_ISL_3256168, EPI_ISL_3256169, EPI_ISL_3256170, EPI_ISL_3256171, EPI_ISL_3256172, EPI_ISL_3256173, EPI_ISL_3256174, EPI_ISL_3256175, EPI_ISL_3256176, EPI_ISL_3256177, EPI_ISL_3256178, EPI_ISL_3256179, EPI_ISL_3256180, EPI_ISL_3256181, EPI_ISL_3256182, EPI_ISL_3256183, EPI_ISL_3256184, EPI_ISL_3256185, EPI_ISL_3256186, EPI_ISL_3256187, EPI_ISL_3256188, EPI_ISL_3256189, EPI_ISL_3256190, EPI_ISL_3256191, EPI_ISL_3256192, EPI_ISL_3256193, EPI_ISL_3256194, EPI_ISL_3256195, EPI_ISL_3256196, EPI_ISL_3256197, EPI_ISL_3256198, EPI_ISL_3256199, EPI_ISL_3256200, EPI_ISL_3256201, EPI_ISL_3256202, EPI_ISL_3256203, EPI_ISL_3256204, EPI_ISL_3256205, EPI_ISL_3256206, EPI_ISL_3256207, EPI_ISL_3256208, EPI_ISL_3256209, EPI_ISL_3256210, EPI_ISL_3256211, EPI_ISL_3256212, EPI_ISL_3256213, EPI_ISL_3256214, EPI_ISL_3256215, EPI_ISL_3256216, EPI_ISL_3256218, EPI_ISL_3256219, EPI_ISL_3256220, EPI_ISL_3256221, EPI_ISL_3256222, EPI_ISL_3256224, EPI_ISL_3256225, EPI_ISL_3256226, EPI_ISL_3256227, EPI_ISL_3256228, EPI_ISL_3256229, EPI_ISL_3256230, EPI_ISL_3256231, EPI_ISL_3256232, EPI_ISL_3256233, EPI_ISL_3256234, EPI_ISL_3256235, EPI_ISL_3256236, EPI_ISL_3256237, EPI_ISL_3256238, EPI_ISL_3256239, EPI_ISL_3256240, EPI_ISL_3256241, EPI_ISL_3256242, EPI_ISL_3256243, EPI_ISL_3256244, EPI_ISL_3256245, EPI_ISL_3256246, EPI_ISL_3256247, EPI_ISL_3256248, EPI_ISL_3256249, EPI_ISL_3256251, EPI_ISL_3256252, EPI_ISL_3256253, EPI_ISL_3256254, EPI_ISL_3256255, EPI_ISL_3256256, EPI_ISL_3256257, EPI_ISL_3256258, EPI_ISL_3256259, EPI_ISL_3256260, EPI_ISL_3256261, EPI_ISL_3256262, EPI_ISL_3256263, EPI_ISL_3256264, EPI_ISL_3256265, EPI_ISL_3256266, EPI_ISL_3256268, EPI_ISL_3256269, EPI_ISL_3256270, EPI_ISL_3256271, EPI_ISL_3256272, EPI_ISL_3256273, EPI_ISL_3256274, EPI_ISL_3256275, EPI_ISL_3256276, EPI_ISL_3256277, EPI_ISL_3256278, EPI_ISL_3256279, EPI_ISL_3256280, EPI_ISL_3256281, EPI_ISL_3256284, EPI_ISL_3256285, EPI_ISL_3256294, EPI_ISL_3256295, EPI_ISL_3256296, EPI_ISL_3256298, EPI_ISL_3256299, EPI_ISL_3256300, EPI_ISL_3256301, EPI_ISL_3256302, EPI_ISL_3256303, EPI_ISL_3256304, EPI_ISL_3256305, EPI_ISL_3256306, EPI_ISL_3256307, EPI_ISL_3256308, EPI_ISL_3256309, EPI_ISL_3256310, EPI_ISL_3256311, EPI_ISL_3256312, EPI_ISL_3256313, EPI_ISL_3256314, EPI_ISL_3256315, EPI_ISL_3256316, EPI_ISL_3256317, EPI_ISL_3256318, EPI_ISL_3256319, EPI_ISL_3256320, EPI_ISL_3256321, EPI_ISL_3256322, EPI_ISL_3256323, EPI_ISL_3256324, EPI_ISL_3256325, EPI_ISL_3256326, EPI_ISL_3256327, EPI_ISL_3256328, EPI_ISL_3256329, EPI_ISL_3256330, EPI_ISL_3256331, EPI_ISL_3256332, EPI_ISL_3256333, EPI_ISL_3256334, EPI_ISL_3256335, EPI_ISL_3256336, EPI_ISL_3256337, EPI_ISL_3256338, EPI_ISL_3256339, EPI_ISL_3256340, EPI_ISL_3256341, EPI_ISL_3256342, EPI_ISL_3256343, EPI_ISL_3256344, EPI_ISL_3256345, EPI_ISL_3256346, EPI_ISL_3256347, EPI_ISL_3256348, EPI_ISL_3256349, EPI_ISL_3256350, EPI_ISL_3256351, EPI_ISL_3256352, EPI_ISL_3256353, EPI_ISL_3256354, EPI_ISL_3256355, EPI_ISL_3256356, EPI_ISL_3256357, EPI_ISL_3256358, EPI_ISL_3256359, EPI_ISL_3256360, EPI_ISL_3256361, EPI_ISL_3256362, EPI_ISL_3256363, EPI_ISL_3256364, EPI_ISL_3256365, EPI_ISL_3256366, EPI_ISL_3256367, EPI_ISL_3256368, EPI_ISL_3256369, EPI_ISL_3256370, EPI_ISL_3256371, EPI_ISL_3256372, EPI_ISL_3256373, EPI_ISL_3256374, EPI_ISL_3256375, EPI_ISL_3256376, EPI_ISL_3256377, EPI_ISL_3256378, EPI_ISL_3256379, EPI_ISL_3256380, EPI_ISL_3256381, EPI_ISL_3256382, EPI_ISL_3256383, EPI_ISL_3256384, EPI_ISL_3256385, EPI_ISL_3256386, EPI_ISL_3256387, EPI_ISL_3256388, EPI_ISL_3256389, EPI_ISL_3256390, EPI_ISL_3256391, EPI_ISL_3256392, EPI_ISL_3256393, EPI_ISL_3256394, EPI_ISL_3256395, EPI_ISL_3256396, EPI_ISL_3256397, EPI_ISL_3256398, EPI_ISL_3256399, EPI_ISL_3256400, EPI_ISL_3256401, EPI_ISL_3256402, EPI_ISL_3256403, EPI_ISL_3256404, EPI_ISL_3256405, EPI_ISL_3256406, EPI_ISL_3256407, EPI_ISL_3256408, EPI_ISL_3256409, EPI_ISL_3256410, EPI_ISL_3256411, EPI_ISL_3256412, EPI_ISL_3256413, EPI_ISL_3256414, EPI_ISL_3256415, EPI_ISL_3256416, EPI_ISL_3256417 | | | |
| see above | Arkansas Public Health Laboratory, Arkansas Department of Health | University of Minnesota Genomics Center | Daryl M. Gohl, John Garbe, Jaquelyn Kuriger-Laber, Corbin Dirkx |
| EPI_ISL_3256418, EPI_ISL_3256419, EPI_ISL_3256420, EPI_ISL_3256421, EPI_ISL_3256422, EPI_ISL_3256423, EPI_ISL_3256424, EPI_ISL_3256425, EPI_ISL_3256426, EPI_ISL_3256427, EPI_ISL_3256428, EPI_ISL_3256429, EPI_ISL_3256430, EPI_ISL_3256431, EPI_ISL_3256432, EPI_ISL_3256433, EPI_ISL_3256434, EPI_ISL_3256435, EPI_ISL_3256436, EPI_ISL_3256437, EPI_ISL_3256438, EPI_ISL_3256439, EPI_ISL_3256440, EPI_ISL_3256441, EPI_ISL_3256442, EPI_ISL_3256443, EPI_ISL_3256444, EPI_ISL_3256445, EPI_ISL_3256446 | | | |
| see above | Montana Public Health Laboratory | Montana Public Health Laboratory | Joy Ritter, Michelle Mozer, Carrie Biskupiak, Deborah Gibson, Michael Dills |
| EPI_ISL_3256536, EPI_ISL_3256553, EPI_ISL_3256560, EPI_ISL_3256563, EPI_ISL_3256575, EPI_ISL_3256580, EPI_ISL_3256875, EPI_ISL_3256934 | Ministry of Health Turkey | Ministry of Health Turkey | Fatma Bayrakdar, Yasemin Coşgun, Süleyman Yalcin, Gülay Korukluoğlu |
| EPI_ISL_3257364, EPI_ISL_3257365, EPI_ISL_3257366, EPI_ISL_3257367, EPI_ISL_3257368, EPI_ISL_3257369, EPI_ISL_3257370, EPI_ISL_3257371, EPI_ISL_3257372, EPI_ISL_3257374, EPI_ISL_3257375, EPI_ISL_3257376, EPI_ISL_3257377, EPI_ISL_3257378, EPI_ISL_3257379, EPI_ISL_3257380, EPI_ISL_3257381, EPI_ISL_3257382, EPI_ISL_3257383, EPI_ISL_3257384, EPI_ISL_3257385, EPI_ISL_3257386, EPI_ISL_3257387, EPI_ISL_3257388, EPI_ISL_3257390, EPI_ISL_3257391, EPI_ISL_3257392, EPI_ISL_3257393, EPI_ISL_3257394, EPI_ISL_3257395, EPI_ISL_3257396, EPI_ISL_3257397, EPI_ISL_3257398, EPI_ISL_3257399, EPI_ISL_3257400, EPI_ISL_3257401, EPI_ISL_3257402, EPI_ISL_3257403, EPI_ISL_3257404, EPI_ISL_3257405, EPI_ISL_3257406, EPI_ISL_3257407, EPI_ISL_3257408, EPI_ISL_3257410, EPI_ISL_3257411, EPI_ISL_3257412, EPI_ISL_3257413, EPI_ISL_3257414, EPI_ISL_3257415, EPI_ISL_3257416, EPI_ISL_3257417, EPI_ISL_3257418, EPI_ISL_3257419, EPI_ISL_3257420, EPI_ISL_3257421, EPI_ISL_3257422, EPI_ISL_3257423, EPI_ISL_3257424, EPI_ISL_3257425, EPI_ISL_3257511, EPI_ISL_3257512, EPI_ISL_3257524, EPI_ISL_3257525, EPI_ISL_3257527, EPI_ISL_3257528, EPI_ISL_3257529, EPI_ISL_3257530, EPI_ISL_3257531, EPI_ISL_3257533, EPI_ISL_3257534, EPI_ISL_3257535, EPI_ISL_3257536, EPI_ISL_3257537, EPI_ISL_3257539, EPI_ISL_3257540, EPI_ISL_3257541, EPI_ISL_3257542, EPI_ISL_3257543, EPI_ISL_3257544, EPI_ISL_3257549, EPI_ISL_3257550, EPI_ISL_3257551, EPI_ISL_3257552, EPI_ISL_3257553, EPI_ISL_3257554, EPI_ISL_3257555, EPI_ISL_3257556, EPI_ISL_3257557, EPI_ISL_3257558, EPI_ISL_3257559, EPI_ISL_3257560, EPI_ISL_3257561, EPI_ISL_3257562, EPI_ISL_3257563, EPI_ISL_3257571, EPI_ISL_3257572, EPI_ISL_3257573, EPI_ISL_3257574, EPI_ISL_3257575, EPI_ISL_3257576, EPI_ISL_3257577, EPI_ISL_3257578, EPI_ISL_3257579, EPI_ISL_3257580, EPI_ISL_3257581, EPI_ISL_3257582, EPI_ISL_3257586, EPI_ISL_3257587, EPI_ISL_3257588, EPI_ISL_3257589, EPI_ISL_3257590, EPI_ISL_3257591, EPI_ISL_3257592, EPI_ISL_3257593, EPI_ISL_3257594, EPI_ISL_3257595, EPI_ISL_3257596, EPI_ISL_3257597, EPI_ISL_3257598, EPI_ISL_3257599, EPI_ISL_3257600, EPI_ISL_3257601, EPI_ISL_3257602, EPI_ISL_3257603, EPI_ISL_3257604, EPI_ISL_3257605, EPI_ISL_3257606, EPI_ISL_3257607, EPI_ISL_3257608, EPI_ISL_3257609, EPI_ISL_3257610, EPI_ISL_3257611, EPI_ISL_3257612, EPI_ISL_3257613, EPI_ISL_3257614, EPI_ISL_3257615, EPI_ISL_3257616, EPI_ISL_3257617, EPI_ISL_3257637, EPI_ISL_3257638, EPI_ISL_3257639, EPI_ISL_3257640, EPI_ISL_3257641, EPI_ISL_3257642, EPI_ISL_3257643, EPI_ISL_3257644, EPI_ISL_3257645, EPI_ISL_3257646, EPI_ISL_3257647, EPI_ISL_3257648, EPI_ISL_3257649, EPI_ISL_3257650, EPI_ISL_3257651, EPI_ISL_3257652, EPI_ISL_3257653, EPI_ISL_3257654, EPI_ISL_3257655, EPI_ISL_3257656, EPI_ISL_3257657, EPI_ISL_3257658, EPI_ISL_3257659, EPI_ISL_3257660, EPI_ISL_3257661, EPI_ISL_3257662, EPI_ISL_3257663, EPI_ISL_3257664, EPI_ISL_3257665, EPI_ISL_3257666, EPI_ISL_3257667, EPI_ISL_3257668, EPI_ISL_3257669, EPI_ISL_3257670, EPI_ISL_3257671, EPI_ISL_3257672, EPI_ISL_3257673, EPI_ISL_3257674, EPI_ISL_3257675, EPI_ISL_3257676, EPI_ISL_3257677, EPI_ISL_3257678, EPI_ISL_3257679, EPI_ISL_3257680, EPI_ISL_3257681, EPI_ISL_3257682, EPI_ISL_3257683, EPI_ISL_3257684, EPI_ISL_3257685, EPI_ISL_3257686, EPI_ISL_3257687, EPI_ISL_3257688, EPI_ISL_3257689, EPI_ISL_3257690, EPI_ISL_3257691, EPI_ISL_3257692, EPI_ISL_3257693, EPI_ISL_3257694, EPI_ISL_3257695, EPI_ISL_3257696, EPI_ISL_3257697, EPI_ISL_3257698, EPI_ISL_3257699, EPI_ISL_3257700, EPI_ISL_3257701, EPI_ISL_3257702, EPI_ISL_3257703, EPI_ISL_3257704, EPI_ISL_3257705, EPI_ISL_3257706, EPI_ISL_3257707, EPI_ISL_3257708, EPI_ISL_3257709, EPI_ISL_3257710, EPI_ISL_3257711, EPI_ISL_3257712, EPI_ISL_3257713, EPI_ISL_3257714, EPI_ISL_3257715, EPI_ISL_3257716, EPI_ISL_3257717, EPI_ISL_3257718, EPI_ISL_3257719, EPI_ISL_3257720, EPI_ISL_3257721, EPI_ISL_3257722, EPI_ISL_3257723, EPI_ISL_3257724, EPI_ISL_3257725, EPI_ISL_3257726, EPI_ISL_3257727, EPI_ISL_3257728, EPI_ISL_3257729, EPI_ISL_3257730, EPI_ISL_3257731, EPI_ISL_3257732, EPI_ISL_3257733, EPI_ISL_3257734, EPI_ISL_3257735, EPI_ISL_3257736, EPI_ISL_3257737, EPI_ISL_3257738, EPI_ISL_3257739, EPI_ISL_3257740, EPI_ISL_3257741, EPI_ISL_3257742, EPI_ISL_3257743, EPI_ISL_3257746, EPI_ISL_3257747, EPI_ISL_3257748, EPI_ISL_3257749, EPI_ISL_3257750, EPI_ISL_3257751, EPI_ISL_3257752, EPI_ISL_3257753, EPI_ISL_3257754, EPI_ISL_3257755, EPI_ISL_3257756, EPI_ISL_3257762, EPI_ISL_3257763, EPI_ISL_3257764, EPI_ISL_3257765, EPI_ISL_3257766, EPI_ISL_3257767, EPI_ISL_3257768, EPI_ISL_3257769, EPI_ISL_3257770, EPI_ISL_3257771, EPI_ISL_3257772, EPI_ISL_3257773, EPI_ISL_3257774, EPI_ISL_3257775, EPI_ISL_3257776, EPI_ISL_3257777, EPI_ISL_3257778, EPI_ISL_3257779, EPI_ISL_3257780, EPI_ISL_3257781, EPI_ISL_3257782, EPI_ISL_3257783, EPI_ISL_3257784, EPI_ISL_3257785, EPI_ISL_3257786, EPI_ISL_3257787, EPI_ISL_3257788, EPI_ISL_3257789, EPI_ISL_3257790, EPI_ISL_3257791, EPI_ISL_3257792, EPI_ISL_3257793, EPI_ISL_3257794, EPI_ISL_3257795, EPI_ISL_3257796, EPI_ISL_3257798, EPI_ISL_3257799, EPI_ISL_3257800, EPI_ISL_3257801, EPI_ISL_3257802, EPI_ISL_3257803, EPI_ISL_3257804, EPI_ISL_3257805, EPI_ISL_3257806, EPI_ISL_3257807, EPI_ISL_3257808, EPI_ISL_3257809, EPI_ISL_3257810, EPI_ISL_3257811, EPI_ISL_3257812, EPI_ISL_3257813, EPI_ISL_3257814, EPI_ISL_3257815, EPI_ISL_3257816, EPI_ISL_3257817, EPI_ISL_3257818, EPI_ISL_3257819, EPI_ISL_3257820, EPI_ISL_3257821, EPI_ISL_3257822, EPI_ISL_3257823, EPI_ISL_3257824, EPI_ISL_3257825, EPI_ISL_3257826, EPI_ISL_3257827, EPI_ISL_3257828, EPI_ISL_3257829, EPI_ISL_3257830, EPI_ISL_3257831, EPI_ISL_3257832, EPI_ISL_3257833, EPI_ISL_3257834, EPI_ISL_3257835, EPI_ISL_3257836, EPI_ISL_3257837, EPI_ISL_3257838, EPI_ISL_3257839, EPI_ISL_3257840, EPI_ISL_3257841, EPI_ISL_3257842, EPI_ISL_3257843, EPI_ISL_3257844, EPI_ISL_3257845, EPI_ISL_3257846, EPI_ISL_3257847, EPI_ISL_3257848, EPI_ISL_3257849, EPI_ISL_3257850, EPI_ISL_3257851, EPI_ISL_3257852, EPI_ISL_3257853, EPI_ISL_3257854, EPI_ISL_3257855, EPI_ISL_3257856, EPI_ISL_3257857, EPI_ISL_3257858, EPI_ISL_3257859, EPI_ISL_3257860, EPI_ISL_3257861, EPI_ISL_3257862, EPI_ISL_3257863, EPI_ISL_3257864, EPI_ISL_3257865, EPI_ISL_3257866, EPI_ISL_3257867, EPI_ISL_3257868, EPI_ISL_3257869, EPI_ISL_3257870, EPI_ISL_3257871, EPI_ISL_3257872, EPI_ISL_3257873, EPI_ISL_3257874, EPI_ISL_3257875, EPI_ISL_3257876, EPI_ISL_3257877, EPI_ISL_3257878, EPI_ISL_3257879, EPI_ISL_3257880, EPI_ISL_3257881, EPI_ISL_3257882, EPI_ISL_3257883, EPI_ISL_3257884, EPI_ISL_3257885, EPI_ISL_3257886, EPI_ISL_3257887, EPI_ISL_3257888, EPI_ISL_3257889, EPI_ISL_3257890, EPI_ISL_3257891, EPI_ISL_3257892, EPI_ISL_3257893, EPI_ISL_3257894, EPI_ISL_3257895, EPI_ISL_3257896, EPI_ISL_3257897, EPI_ISL_3257898, EPI_ISL_3257899, EPI_ISL_3257900, EPI_ISL_3257901, EPI_ISL_3257902, EPI_ISL_3257903, EPI_ISL_3257904, EPI_ISL_3257905, EPI_ISL_3257906, EPI_ISL_3257907, EPI_ISL_3257908, EPI_ISL_3257909, EPI_ISL_3257910, EPI_ISL_3257911, EPI_ISL_3257912, EPI_ISL_3257913, EPI_ISL_3257914, EPI_ISL_3257915, EPI_ISL_3257916, EPI_ISL_3257917, EPI_ISL_3257918, EPI_ISL_3257919, EPI_ISL_3257920, EPI_ISL_3257921, EPI_ISL_3257922, EPI_ISL_3257923, EPI_ISL_3257924, EPI_ISL_3257925, EPI_ISL_3257926, EPI_ISL_3257927, EPI_ISL_3257928, EPI_ISL_3257929, EPI_ISL_3257930, EPI_ISL_3257931, EPI_ISL_3257932, EPI_ISL_3257933, EPI_ISL_3257934, EPI_ISL_3257935, EPI_ISL_3257936, EPI_ISL_3257937, EPI_ISL_3257938, EPI_ISL_3257939, EPI_ISL_3257940, EPI_ISL_3257941, EPI_ISL_3257942, EPI_ISL_3257943, EPI_ISL_3257944, EPI_ISL_3257945, EPI_ISL_3257946, EPI_ISL_3257947, EPI_ISL_3257948, EPI_ISL_3257949, EPI_ISL_3257950, EPI_ISL_3257951, EPI_ISL_3257952, EPI_ISL_3257953, EPI_ISL_3257954, EPI_ISL_3257955, EPI_ISL_3257956, EPI_ISL_3257957, EPI_ISL_3257958, EPI_ISL_3257959, EPI_ISL_3257960, EPI_ISL_3257961, EPI_ISL_3257962, EPI_ISL_3257963, EPI_ISL_3257964, EPI_ISL_3257965, EPI_ISL_3257966, EPI_ISL_3257967, EPI_ISL_3257968, EPI_ISL_3257969, EPI_ISL_3257970, EPI_ISL_3257971, EPI_ISL_3257972, EPI_ISL_3257973, EPI_ISL_3257974, EPI_ISL_3257975, EPI_ISL_3257976, EPI_ISL_3257977, EPI_ISL_3257978, EPI_ISL_3257979, EPI_ISL_3257980, EPI_ISL_3257981, EPI_ISL_3257982, EPI_ISL_3257983, EPI_ISL_3257984, EPI_ISL_3257985, EPI_ISL_3257986, EPI_ISL_3257987, EPI_ISL_3257988, EPI_ISL_3257989, EPI_ISL_3257990, EPI_ISL_3257991, EPI_ISL_3257992, EPI_ISL_3257993, EPI_ISL_3257994, EPI_ISL_3257995, EPI_ISL_3257996, EPI_ISL_3257997, EPI_ISL_3257998, EPI_ISL_3257999, EPI_ISL_3258000, EPI_ISL_3258001, EPI_ISL_3258002, EPI_ISL_3258003, EPI_ISL_3258004, EPI_ISL_3258005, EPI_ISL_3258006, EPI_ISL_3258007, EPI_ISL_3258008, EPI_ISL_3258009, EPI_ISL_3258010, EPI_ISL_3258011, EPI_ISL_3258012, EPI_ISL_3258013, EPI_ISL_3258014, EPI_ISL_3258015, EPI_ISL_3258016, EPI_ISL_3258017, EPI_ISL_3258018, EPI_ISL_3258019, EPI_ISL_3258020, EPI_ISL_3258021, EPI_ISL_3258022, EPI_ISL_3258023, EPI_ISL_3258024, EPI_ISL_3258025, EPI_ISL_3258026, EPI_ISL_3258027, EPI_ISL_3258028, EPI_ISL_3258029, EPI_ISL_3258030, EPI_ISL_3258031, EPI_ISL_3258032, EPI_ISL_3258033, EPI_ISL_3258034, EPI_ISL_3258035, EPI_ISL_3258036, EPI_ISL_3258037, EPI_ISL_3258038, EPI_ISL_3258039, EPI_ISL_3258040, EPI_ISL_3258041, EPI_ISL_3258042, EPI_ISL_3258043, EPI_ISL_3258044, EPI_ISL_3258045, EPI_ISL_3258046, EPI_ISL_3258047, EPI_ISL_3258048, EPI_ISL_3258049, EPI_ISL_3258050, EPI_ISL_3258051, EPI_ISL_3258052, EPI_ISL_3258053, EPI_ISL_3258054, EPI_ISL_3258055, EPI_ISL_3258056, EPI_ISL_3258057, EPI_ISL_3258058, EPI_ISL_3258059, EPI_ISL_3258060, EPI_ISL_3258285, EPI_ISL_3258286, EPI_ISL_3258287, EPI_ISL_3258288, EPI_ISL_3258289, EPI_ISL_3258290, EPI_ISL_3258291, EPI_ISL_3258292, EPI_ISL_3258293, EPI_ISL_3258294, EPI_ISL_3258295, EPI_ISL_3258296, EPI_ISL_3258297, EPI_ISL_3258298, EPI_ISL_3258299, EPI_ISL_3258300, EPI_ISL_3258301, EPI_ISL_3258302, EPI_ISL_3258303, EPI_ISL_3258304, EPI_ISL_3258305, EPI_ISL_3258306, EPI_ISL_3258307, EPI_ISL_3258308, EPI_ISL_3258309, EPI_ISL_3258310, EPI_ISL_3258311, EPI_ISL_3258312, EPI_ISL_3258313, EPI_ISL_3258314, EPI_ISL_3258315, EPI_ISL_3258316, EPI_ISL_3258317, EPI_ISL_3258318, EPI_ISL_3258319, EPI_ISL_3258320, EPI_ISL_3258321, EPI_ISL_3258322, EPI_ISL_3258323, EPI_ISL_3258324, EPI_ISL_3258325, EPI_ISL_3258326, EPI_ISL_3258327, EPI_ISL_3258328, EPI_ISL_3258329, EPI_ISL_3258330, EPI_ISL_3258331, EPI_ISL_3258332, EPI_ISL_3258333, EPI_ISL_3258334, EPI_ISL_3258335, EPI_ISL_3258336, EPI_ISL_3258337, EPI_ISL_3258338, EPI_ISL_3258339, EPI_ISL_3258340, EPI_ISL_3258341, EPI_ISL_3258342, EPI_ISL_3258343, EPI_ISL_3258344, EPI_ISL_3258345, EPI_ISL_3258346, EPI_ISL_3258347, EPI_ISL_3258348, EPI_ISL_3258349, EPI_ISL_3258350, EPI_ISL_3258351, EPI_ISL_3258352, EPI_ISL_3258353, EPI_ISL_3258354, EPI_ISL_3258355, EPI_ISL_3258356, EPI_ISL_3258357, EPI_ISL_3258358, EPI_ISL_3258359, EPI_ISL_3258360, EPI_ISL_3258361, EPI_ISL_3258362, EPI_ISL_3258363, EPI_ISL_3258364, EPI_ISL_3258365, EPI_ISL_3258366, EPI_ISL_3258367, EPI_ISL_3258368, EPI_ISL_3258369, EPI_ISL_3258370, EPI_ISL_3258371, EPI_ISL_3258372, EPI_ISL_3258373, EPI_ISL_3258374, EPI_ISL_3258375, EPI_ISL_3258376, EPI_ISL_3258377, EPI_ISL_3258378, EPI_ISL_3258379, EPI_ISL_3258380, EPI_ISL_3258381, EPI_ISL_3258382, EPI_ISL_3258383, EPI_ISL_3258384, EPI_ISL_3258385, EPI_ISL_3258386, EPI_ISL_3258387, EPI_ISL_3258388, EPI_ISL_3258389, EPI_ISL_3258390, EPI_ISL_3258391, EPI_ISL_3258392, EPI_ISL_3258393, EPI_ISL_3258394, EPI_ISL_3258395, EPI_ISL_3258396, EPI_ISL_3258397, EPI_ISL_3258398, EPI_ISL_3258399, EPI_ISL_3258400, EPI_ISL_3258401, EPI_ISL_3258402, EPI_ISL_3258403, EPI_ISL_3258404, EPI_ISL_3258405, EPI_ISL_3258406, EPI_ISL_3258407, EPI_ISL_3258408, EPI_ISL_3258409, EPI_ISL_3258410, EPI_ISL_3258411, EPI_ISL_3258412, EPI_ISL_3258413, EPI_ISL_3258414, EPI_ISL_3258415, EPI_ISL_3258416, EPI_ISL_3258417, EPI_ISL_3258418, EPI_ISL_3258419, EPI_ISL_3258420, EPI_ISL_3258421, EPI_ISL_3258422, EPI_ISL_3258423, EPI_ISL_3258424, EPI_ISL_3258425, EPI_ISL_3258426, EPI_ISL_3258427, EPI_ISL_3258428, EPI_ISL_3258429, EPI_ISL_3258430, EPI_ISL_3258431, EPI_ISL_3258432, EPI_ISL_3258433, EPI_ISL_3258434, EPI_ISL_3258435, EPI_ISL_3258436, EPI_ISL_3258437, EPI_ISL_3258438, EPI_ISL_3258439, EPI_ISL_3258440, EPI_ISL_3258441, EPI_ISL_3258442, EPI_ISL_3258443, EPI_ISL_3258444, EPI_ISL_3258445, EPI_ISL_3258446, EPI_ISL_3258447, EPI_ISL_3258448, EPI_ISL_3258449, EPI_ISL_3258450, EPI_ISL_3258451, EPI_ISL_3258452, EPI_ISL_3258453, EPI_ISL_3258454, EPI_ISL_3258455, EPI_ISL_3258456, EPI_ISL_3258457, EPI_ISL_3258458, EPI_ISL_3258459, EPI_ISL_3258460, EPI_ISL_3258461, EPI_ISL_3258462, EPI_ISL_3258463, EPI_ISL_3258464, EPI_ISL_3258465, EPI_ISL_3258466, EPI_ISL_3258467, EPI_ISL_3258468, EPI_ISL_3258469, EPI_ISL_3258470, EPI_ISL_3258471, EPI_ISL_3258472, EPI_ISL_3258473, EPI_ISL_3258474, EPI_ISL_3258475, EPI_ISL_3258476, EPI_ISL_3258477, EPI_ISL_3258478, EPI_ISL_3258479, EPI_ISL_3258480, EPI_ISL_3258481, EPI_ISL_3258482, EPI_ISL_3258483, EPI_ISL_3258484, EPI_ISL_3258485, EPI_ISL_3258486, EPI_ISL_3258487, EPI_ISL_3258488, EPI_ISL_3258489, EPI_ISL_3258490, EPI_ISL_3258491, EPI_ISL_3258492, EPI_ISL_3258493, EPI_ISL_3258494, EPI_ISL_3258495, EPI_ISL_3258496, EPI_ISL_3258497, EPI_ISL_3258498, EPI_ISL_3258499, EPI_ISL_3258500, EPI_ISL_3258501, EPI_ISL_3258502, EPI_ISL_3258503, EPI_ISL_3258504, EPI_ISL_3258505, EPI_ISL_3258506, EPI_ISL_3258507, EPI_ISL_3258508, EPI_ISL_3258509, EPI_ISL_3258510, EPI_ISL_3258511, EPI_ISL_3258512, EPI_ISL_3258513, EPI_ISL_3258514, EPI_ISL_3258515, EPI_ISL_3258516, EPI_ISL_3258517, EPI_ISL_3258518, EPI_ISL_3258519, EPI_ISL_3258520, EPI_ISL_3258521, EPI_ISL_3258522, EPI_ISL_3258523, EPI_ISL_3258524, EPI_ISL_3258525, EPI_ISL_3258526, EPI_ISL_3258527, EPI_ISL_3258528, EPI_ISL_3258529, EPI_ISL_3258530, EPI_ISL_3258531, EPI_ISL_3258532, EPI_ISL_3258533, EPI_ISL_3258534, EPI_ISL_3258535, EPI_ISL_3258536, EPI_ISL_3258537, EPI_ISL_3258538, EPI_ISL_3258539, EPI_ISL_3258540, EPI_ISL_3258541, EPI_ISL_3258542, EPI_ISL_3258543, EPI_ISL_3258544, EPI_ISL_3258545, EPI_ISL_3258546, EPI_ISL_3258547, EPI_ISL_3258548, EPI_ISL_3258549, EPI_ISL_3258550, EPI_ISL_3258551, EPI_ISL_3258552, EPI_ISL_3258553, EPI_ISL_3258554, EPI_ISL_3258555, EPI_ISL_3258556, EPI_ISL_3258557, EPI_ISL_3258558, EPI_ISL_3258559, EPI_ISL_3258560, EPI_ISL_3258561, EPI_ISL_3258562, EPI_ISL_3258563, EPI_ISL_3258564, EPI_ISL_3258565, EPI_ISL_3258566, | | | |

| | | | |
|---|---|---|---|
| EPI_ISL_3258567, EPI_ISL_3258568, EPI_ISL_3258569, EPI_ISL_3258570, EPI_ISL_3258571, EPI_ISL_3258572, EPI_ISL_3258573, EPI_ISL_3258574, EPI_ISL_3258575, EPI_ISL_3258576, EPI_ISL_3258577, EPI_ISL_3258578, EPI_ISL_3258579, EPI_ISL_3258580, EPI_ISL_3258581, EPI_ISL_3258582, EPI_ISL_3258583, EPI_ISL_3258584, EPI_ISL_3258585, EPI_ISL_3258586, EPI_ISL_3258587, EPI_ISL_3258588, EPI_ISL_3258589, EPI_ISL_3258590, EPI_ISL_3258591, EPI_ISL_3258592, EPI_ISL_3258593, EPI_ISL_3258594, EPI_ISL_3258595, EPI_ISL_3258596, EPI_ISL_3258597, EPI_ISL_3258598, EPI_ISL_3258599, EPI_ISL_3258600, EPI_ISL_3258601, EPI_ISL_3258602, EPI_ISL_3258603, EPI_ISL_3258604, EPI_ISL_3258605, EPI_ISL_3258606, EPI_ISL_3258607, EPI_ISL_3258608, EPI_ISL_3258609, EPI_ISL_3258610, EPI_ISL_3258611, EPI_ISL_3258612, EPI_ISL_3258613, EPI_ISL_3258614, EPI_ISL_3258615, EPI_ISL_3258616, EPI_ISL_3258617, EPI_ISL_3258618, EPI_ISL_3258619, EPI_ISL_3258620, EPI_ISL_3258621, EPI_ISL_3258622, EPI_ISL_3258623, EPI_ISL_3258624, EPI_ISL_3258625, EPI_ISL_3258626, EPI_ISL_3258627, EPI_ISL_3258628, EPI_ISL_3258629, EPI_ISL_3258630, EPI_ISL_3258631, EPI_ISL_3258632, EPI_ISL_3258633, EPI_ISL_3258634, EPI_ISL_3258635, EPI_ISL_3258636, EPI_ISL_3258637, EPI_ISL_3258638, EPI_ISL_3258639, EPI_ISL_3258640, EPI_ISL_3258641, EPI_ISL_3258642, EPI_ISL_3258643, EPI_ISL_3258644, EPI_ISL_3258645, EPI_ISL_3258646, EPI_ISL_3258647, EPI_ISL_3258648, EPI_ISL_3258649, EPI_ISL_3258650, EPI_ISL_3258651, EPI_ISL_3258652, EPI_ISL_3258653, EPI_ISL_3258654, EPI_ISL_3258655, EPI_ISL_3258656, EPI_ISL_3258657, EPI_ISL_3258658, EPI_ISL_3258659, EPI_ISL_3258660, EPI_ISL_3258661, EPI_ISL_3258662, EPI_ISL_3258663, EPI_ISL_3258664, EPI_ISL_3258665, EPI_ISL_3258666, EPI_ISL_3258667, EPI_ISL_3258668, EPI_ISL_3258669, EPI_ISL_3258670, EPI_ISL_3258671, EPI_ISL_3258672, EPI_ISL_3258673, EPI_ISL_3258674, EPI_ISL_3258675, EPI_ISL_3258676, EPI_ISL_3258677, EPI_ISL_3258678, EPI_ISL_3258679, EPI_ISL_3258680, EPI_ISL_3258681, EPI_ISL_3258682, EPI_ISL_3258683, EPI_ISL_3258684, EPI_ISL_3258685, EPI_ISL_3258686, EPI_ISL_3258687, EPI_ISL_3258688, EPI_ISL_3258689, EPI_ISL_3258690, EPI_ISL_3258691, EPI_ISL_3258692, EPI_ISL_3258693, EPI_ISL_3258694, EPI_ISL_3258695, EPI_ISL_3258696, EPI_ISL_3258697, EPI_ISL_3258698, EPI_ISL_3258699, EPI_ISL_3258700, EPI_ISL_3258701, EPI_ISL_3258702, EPI_ISL_3258703, EPI_ISL_3258704, EPI_ISL_3258705, EPI_ISL_3258706, EPI_ISL_3258707, EPI_ISL_3258708, EPI_ISL_3258709, EPI_ISL_3258710, EPI_ISL_3258711, EPI_ISL_3258712, EPI_ISL_3258713, EPI_ISL_3258714, EPI_ISL_3258715, EPI_ISL_3258716, EPI_ISL_3258717, EPI_ISL_3258718, EPI_ISL_3258719, EPI_ISL_3258720, EPI_ISL_3258721, EPI_ISL_3258722, EPI_ISL_3258723, EPI_ISL_3258724, EPI_ISL_3258725, EPI_ISL_3258726, EPI_ISL_3258727, EPI_ISL_3258728, EPI_ISL_3258729, EPI_ISL_3258730, EPI_ISL_3258731, EPI_ISL_3258732, EPI_ISL_3258733, EPI_ISL_3258734, EPI_ISL_3258735, EPI_ISL_3258736, EPI_ISL_3258737, EPI_ISL_3258738, EPI_ISL_3258739, EPI_ISL_3258740, EPI_ISL_3258741, EPI_ISL_3258742, EPI_ISL_3258743, EPI_ISL_3258744, EPI_ISL_3258745, EPI_ISL_3258746, EPI_ISL_3258747, EPI_ISL_3258748, EPI_ISL_3258749, EPI_ISL_3258750, EPI_ISL_3258751, EPI_ISL_3258752, EPI_ISL_3258753, EPI_ISL_3258754, EPI_ISL_3258755, EPI_ISL_3258756, EPI_ISL_3258757, EPI_ISL_3258758, EPI_ISL_3258759, EPI_ISL_3258760, EPI_ISL_3258761, EPI_ISL_3258762, EPI_ISL_3258763, EPI_ISL_3258764, EPI_ISL_3258765, EPI_ISL_3258766, EPI_ISL_3258767, EPI_ISL_3258768, EPI_ISL_3258769, EPI_ISL_3258770, EPI_ISL_3258771, EPI_ISL_3258772, EPI_ISL_3258773, EPI_ISL_3258774, EPI_ISL_3258775, EPI_ISL_3258776, EPI_ISL_3258777, EPI_ISL_3258778, EPI_ISL_3258779, EPI_ISL_3258780, EPI_ISL_3258781, EPI_ISL_3258782, EPI_ISL_3258783, EPI_ISL_3258784, EPI_ISL_3258785, EPI_ISL_3258786, EPI_ISL_3258787, EPI_ISL_3258788, EPI_ISL_3258789, EPI_ISL_3258790, EPI_ISL_3258791, EPI_ISL_3258792, EPI_ISL_3258793, EPI_ISL_3258794, EPI_ISL_3258795, EPI_ISL_3258796, EPI_ISL_3258797, EPI_ISL_3258798, EPI_ISL_3258799, EPI_ISL_3258800, EPI_ISL_3258801, EPI_ISL_3258802, EPI_ISL_3258803, EPI_ISL_3258804, EPI_ISL_3258805, EPI_ISL_3258806, EPI_ISL_3258807, EPI_ISL_3258808, EPI_ISL_3258809, EPI_ISL_3258810, EPI_ISL_3258811, EPI_ISL_3258812, EPI_ISL_3258813, EPI_ISL_3258814, EPI_ISL_3258815, EPI_ISL_3258816, EPI_ISL_3258817, EPI_ISL_3258818, EPI_ISL_3258819, EPI_ISL_3258820, EPI_ISL_3258821, EPI_ISL_3258822, EPI_ISL_3258823, EPI_ISL_3258824, EPI_ISL_3258825, EPI_ISL_3258826, EPI_ISL_3258827, EPI_ISL_3258828, EPI_ISL_3258829, EPI_ISL_3258830, EPI_ISL_3258831, EPI_ISL_3258832, EPI_ISL_3258833, EPI_ISL_3258834, EPI_ISL_3258835, EPI_ISL_3258836, EPI_ISL_3258837, EPI_ISL_3258838, EPI_ISL_3258839, EPI_ISL_3258840, EPI_ISL_3258841, EPI_ISL_3258842, EPI_ISL_3258843, EPI_ISL_3258844, EPI_ISL_3258845, EPI_ISL_3258846, EPI_ISL_3258847, EPI_ISL_3258848, EPI_ISL_3258849, EPI_ISL_3258850, EPI_ISL_3258851, EPI_ISL_3258852, EPI_ISL_3258853, EPI_ISL_3258854, EPI_ISL_3258855, EPI_ISL_3258856, EPI_ISL_3258857, EPI_ISL_3258858, EPI_ISL_3258859, EPI_ISL_3258860, EPI_ISL_3258861, EPI_ISL_3258862, EPI_ISL_3258863, EPI_ISL_3258864, EPI_ISL_3258865, EPI_ISL_3258866, EPI_ISL_3258867, EPI_ISL_3258868, EPI_ISL_3258869, EPI_ISL_3258870, EPI_ISL_3258871, EPI_ISL_3258872, EPI_ISL_3258873, EPI_ISL_3258874, EPI_ISL_3258875, EPI_ISL_3258876, EPI_ISL_3258877, EPI_ISL_3258878, EPI_ISL_3258879, EPI_ISL_3258880, EPI_ISL_3258881, EPI_ISL_3258882, EPI_ISL_3258883, EPI_ISL_3258884, EPI_ISL_3258885, EPI_ISL_3258886, EPI_ISL_3258887, EPI_ISL_3258888, EPI_ISL_3258889, EPI_ISL_3258890, EPI_ISL_3258891, EPI_ISL_3258892, EPI_ISL_3258893, EPI_ISL_3258894, EPI_ISL_3258895, EPI_ISL_3258896, EPI_ISL_3258897, EPI_ISL_3258898, EPI_ISL_3258899, EPI_ISL_3258900, EPI_ISL_3258901, EPI_ISL_3258902, EPI_ISL_3258903, EPI_ISL_3258904, EPI_ISL_3258905, EPI_ISL_3258906, EPI_ISL_3258907, EPI_ISL_3258908, EPI_ISL_3258909, EPI_ISL_3258910, EPI_ISL_3258911, EPI_ISL_3258912, EPI_ISL_3258913, EPI_ISL_3258914, EPI_ISL_3258915, EPI_ISL_3258916, EPI_ISL_3258917, EPI_ISL_3258918, EPI_ISL_3258919, EPI_ISL_3258920, EPI_ISL_3258921, EPI_ISL_3258922, EPI_ISL_3258923, EPI_ISL_3258924, EPI_ISL_3258925, EPI_ISL_3258926, EPI_ISL_3258927, EPI_ISL_3258928, EPI_ISL_3258929, EPI_ISL_3258930, EPI_ISL_3258931, EPI_ISL_3258932, EPI_ISL_3258933, EPI_ISL_3258934, EPI_ISL_3258935, EPI_ISL_3258936, EPI_ISL_3258937, EPI_ISL_3258938, EPI_ISL_3258939, EPI_ISL_3258940, EPI_ISL_3258941, EPI_ISL_3258942, EPI_ISL_3258943, EPI_ISL_3258944, EPI_ISL_3258945, EPI_ISL_3258946, EPI_ISL_3258947, EPI_ISL_3258948, EPI_ISL_3258949, EPI_ISL_3258950, EPI_ISL_3258951, EPI_ISL_3258952, EPI_ISL_3258953, EPI_ISL_3258954, EPI_ISL_3258955, EPI_ISL_3258956, EPI_ISL_3258957, EPI_ISL_3258958, EPI_ISL_3258959, EPI_ISL_3258960, EPI_ISL_3258961, EPI_ISL_3258962, EPI_ISL_3258963, EPI_ISL_3258964, EPI_ISL_3258965, EPI_ISL_3258966, EPI_ISL_3258967, EPI_ISL_3258968, EPI_ISL_3258969, EPI_ISL_3258970, EPI_ISL_3258971, EPI_ISL_3258972, EPI_ISL_3258973, EPI_ISL_3258974, EPI_ISL_3258975, EPI_ISL_3258976, EPI_ISL_3258977, EPI_ISL_3258978, EPI_ISL_3258979, EPI_ISL_3258980, EPI_ISL_3258981, EPI_ISL_3258982, EPI_ISL_3258983, EPI_ISL_3258984, EPI_ISL_3258985, EPI_ISL_3258986, EPI_ISL_3258987, EPI_ISL_3258988, EPI_ISL_3258989, EPI_ISL_3258990, EPI_ISL_3258991, EPI_ISL_3258992, EPI_ISL_3258993, EPI_ISL_3258994, EPI_ISL_3258995, EPI_ISL_3258996, EPI_ISL_3258997, EPI_ISL_3258998, EPI_ISL_3258999, EPI_ISL_3259000, EPI_ISL_3259001, EPI_ISL_3259002, EPI_ISL_3259003, EPI_ISL_3259004, EPI_ISL_3259005, EPI_ISL_3259006, EPI_ISL_3259007, EPI_ISL_3259008, EPI_ISL_3259009, EPI_ISL_3259010, EPI_ISL_3259011, EPI_ISL_3259012, EPI_ISL_3259013, EPI_ISL_3259014, EPI_ISL_3259015, EPI_ISL_3259016, EPI_ISL_3259017, EPI_ISL_3259018, EPI_ISL_3259019, EPI_ISL_3259020, EPI_ISL_3259021, EPI_ISL_3259022, EPI_ISL_3259023, EPI_ISL_3259024, EPI_ISL_3259025, EPI_ISL_3259026, EPI_ISL_3259027, EPI_ISL_3259028, EPI_ISL_3259029, EPI_ISL_3259030, EPI_ISL_3259031, EPI_ISL_3259032, EPI_ISL_3259033, EPI_ISL_3259034, EPI_ISL_3259035, EPI_ISL_3259036, EPI_ISL_3259037, EPI_ISL_3259038, EPI_ISL_3259039, EPI_ISL_3259040, EPI_ISL_3259041, EPI_ISL_3259042, EPI_ISL_3259043, EPI_ISL_3259044, EPI_ISL_3259045, EPI_ISL_3259046, EPI_ISL_3259047, EPI_ISL_3259048, EPI_ISL_3259049, EPI_ISL_3259050, EPI_ISL_3259051, EPI_ISL_3259052, EPI_ISL_3259053, EPI_ISL_3259054, EPI_ISL_3259055, EPI_ISL_3259056, EPI_ISL_3259057, EPI_ISL_3259058, EPI_ISL_3259059, EPI_ISL_3259060, EPI_ISL_3259061, EPI_ISL_3259062, EPI_ISL_3259063, EPI_ISL_3259064, EPI_ISL_3259065, EPI_ISL_3259066, EPI_ISL_3259067, EPI_ISL_3259068, EPI_ISL_3259069, EPI_ISL_3259070, EPI_ISL_3259071, EPI_ISL_3259072, EPI_ISL_3259073, EPI_ISL_3259074, EPI_ISL_3259075, EPI_ISL_3259076, EPI_ISL_3259077, EPI_ISL_3259078, EPI_ISL_3259079, EPI_ISL_3259080, EPI_ISL_3259081, EPI_ISL_3259082, EPI_ISL_3259083, EPI_ISL_3259084, EPI_ISL_3259085, EPI_ISL_3259086, EPI_ISL_3259087, EPI_ISL_3259088, EPI_ISL_3259089, EPI_ISL_3259090, EPI_ISL_3259091, EPI_ISL_3259092, EPI_ISL_3259093, EPI_ISL_3259094, EPI_ISL_3259095, EPI_ISL_3259096, EPI_ISL_3259097, EPI_ISL_3259098, EPI_ISL_3259099, EPI_ISL_3259100, EPI_ISL_3259101, EPI_ISL_3259102, EPI_ISL_3259103, EPI_ISL_3259104, EPI_ISL_3259105, EPI_ISL_3259106, EPI_ISL_3259107, EPI_ISL_3259108, EPI_ISL_3259109, EPI_ISL_3259110, EPI_ISL_3259111, EPI_ISL_3259112, EPI_ISL_3259113, EPI_ISL_3259114, EPI_ISL_3259115, EPI_ISL_3259116, EPI_ISL_3259117, EPI_ISL_3259118, EPI_ISL_3259119, EPI_ISL_3259120, EPI_ISL_3259121, EPI_ISL_3259122, EPI_ISL_3259123, EPI_ISL_3259124, EPI_ISL_3259125, EPI_ISL_3259126, EPI_ISL_3259127, EPI_ISL_3259128, EPI_ISL_3259129, EPI_ISL_3259130, EPI_ISL_3259131, EPI_ISL_3259132, EPI_ISL_3259133, EPI_ISL_3259134, EPI_ISL_3259135, EPI_ISL_3259136, EPI_ISL_3259137, EPI_ISL_3259138, EPI_ISL_3259139, EPI_ISL_3259140, EPI_ISL_3259141, EPI_ISL_3259142, EPI_ISL_3259143, EPI_ISL_3259144, EPI_ISL_3259145, EPI_ISL_3259146, EPI_ISL_3259147, EPI_ISL_3259148, EPI_ISL_3259149, EPI_ISL_3259150, EPI_ISL_3259151, EPI_ISL_3259152, EPI_ISL_3259153, EPI_ISL_3259154, EPI_ISL_3259155, EPI_ISL_3259156, EPI_ISL_3259157, EPI_ISL_3259158, EPI_ISL_3259159, EPI_ISL_3259160, EPI_ISL_3259161, EPI_ISL_3259162, EPI_ISL_3259163, EPI_ISL_3259164, EPI_ISL_3259165, EPI_ISL_3259166, EPI_ISL_3259167, EPI_ISL_3259168, EPI_ISL_3259169, EPI_ISL_3259170, EPI_ISL_3259171, EPI_ISL_3259172, EPI_ISL_3259173, EPI_ISL_3259174, EPI_ISL_3259175, EPI_ISL_3259176, EPI_ISL_3259177, EPI_ISL_3259178, EPI_ISL_3259179, EPI_ISL_3259180, EPI_ISL_3259181, EPI_ISL_3259182, EPI_ISL_3259183, EPI_ISL_3259184, EPI_ISL_3259185, EPI_ISL_3259186, EPI_ISL_3259187, EPI_ISL_3259188, EPI_ISL_3259189, EPI_ISL_3259190, EPI_ISL_3259191, EPI_ISL_3259192, EPI_ISL_3259193, EPI_ISL_3259194, EPI_ISL_3259195, EPI_ISL_3259196, EPI_ISL_3259197, EPI_ISL_3259198, EPI_ISL_3259199, EPI_ISL_3259200, EPI_ISL_3259201, EPI_ISL_3259202, EPI_ISL_3259203, EPI_ISL_3259204, EPI_ISL_3259205, EPI_ISL_3259206, EPI_ISL_3259207, EPI_ISL_3259208, EPI_ISL_3259209, EPI_ISL_3259210, EPI_ISL_3259211, EPI_ISL_3259212, EPI_ISL_3259213, EPI_ISL_3259214, EPI_ISL_3259215, EPI_ISL_3259216, EPI_ISL_3259217, EPI_ISL_3259218, EPI_ISL_3259219, EPI_ISL_3259220, EPI_ISL_3259221, EPI_ISL_3259222, EPI_ISL_3259223, EPI_ISL_3259224, EPI_ISL_3259225, EPI_ISL_3259226, EPI_ISL_3259227, EPI_ISL_3259228, EPI_ISL_3259229, EPI_ISL_3259230, EPI_ISL_3259231, EPI_ISL_3259232, EPI_ISL_3259233, EPI_ISL_3259234, EPI_ISL_3259235, EPI_ISL_3259236, EPI_ISL_3259237, EPI_ISL_3259238, EPI_ISL_3259239, EPI_ISL_3259240, EPI_ISL_3259241, EPI_ISL_3259242, EPI_ISL_3259243, EPI_ISL_3259244, EPI_ISL_3259245, EPI_ISL_3259246, EPI_ISL_3259247, EPI_ISL_3259248, EPI_ISL_3259249, EPI_ISL_3259250, EPI_ISL_3259251, EPI_ISL_3259252, EPI_ISL_3259253, EPI_ISL_3259254, EPI_ISL_3259255, EPI_ISL_3259256, EPI_ISL_3259257, EPI_ISL_3259258, EPI_ISL_3259259, EPI_ISL_3259260, EPI_ISL_3259261, EPI_ISL_3259262, EPI_ISL_3259263, EPI_ISL_3259264, EPI_ISL_3259265, EPI_ISL_3259266, EPI_ISL_3259267, EPI_ISL_3259268, EPI_ISL_3259269, EPI_ISL_3259270, EPI_ISL_3259271, EPI_ISL_3259272, EPI_ISL_3259273, EPI_ISL_3259274, EPI_ISL_3259275, EPI_ISL_3259276, EPI_ISL_3259277, EPI_ISL_3259278, EPI_ISL_3259279, EPI_ISL_3259280, EPI_ISL_3259281, EPI_ISL_3259282, EPI_ISL_3259283, EPI_ISL_3259284, EPI_ISL_3259285, EPI_ISL_3259286, EPI_ISL_3259287, EPI_ISL_3259288, EPI_ISL_3259289, EPI_ISL_3259290, EPI_ISL_3259291, EPI_ISL_3259292, EPI_ISL_3259293, EPI_ISL_3259294, EPI_ISL_3259295, EPI_ISL_3259296, EPI_ISL_3259297, EPI_ISL_3259298, EPI_ISL_3259299, EPI_ISL_3259300, EPI_ISL_3259301, EPI_ISL_3259302, EPI_ISL_3259303, EPI_ISL_3259304, EPI_ISL_3259305, EPI_ISL_3259306, EPI_ISL_3259307, EPI_ISL_3259308, EPI_ISL_3259309, EPI_ISL_3259310, EPI_ISL_3259311, EPI_ISL_3259312, EPI_ISL_3259313, EPI_ISL_3259314, EPI_ISL_3259315, EPI_ISL_3259316, EPI_ISL_3259317, EPI_ISL_3259318, EPI_ISL_3259319, EPI_ISL_3259320, EPI_ISL_3259321, EPI_ISL_3259322, EPI_ISL_3259323, EPI_ISL_3259324, EPI_ISL_3259325, EPI_ISL_3259326, EPI_ISL_3259327, EPI_ISL_3259328, EPI_ISL_3259329, EPI_ISL_3259330, EPI_ISL_3259331, EPI_ISL_3259332, EPI_ISL_3259333, EPI_ISL_3259334, EPI_ISL_3259335, EPI_ISL_3259336, EPI_ISL_3259337, EPI_ISL_3259338, EPI_ISL_3259339, EPI_ISL_3259340, EPI_ISL_3259341, EPI_ISL_3259342, EPI_ISL_3259343, EPI_ISL_3259344, EPI_ISL_3259345, EPI_ISL_3259346, EPI_ISL_3259347, EPI_ISL_3259348, EPI_ISL_3259349 | | | |
| see above | Dutch COVID-19 response team | National Institute for Public Health and the Environment (RIVM) | Adam Meijer, Harry Vennema, Dirk Eggink, Jeroen Cremer, Sharon van den Brink, Bas van der Veer, AnneMarie van den Brandt, Lisa Wijsman, Kim Freriks, Ryanne Jaarsma, Euníce Then, Lynn Aarts, Sanne Bos, Melissa van Tuil, Florian Zwagemaker, Dennis Schmitz, Annelies Kroneman, Karim Hajji, Chantal Reusken, on behalf of the national COVID-19 response team |
| EPI_ISL_3259387, EPI_ISL_3259388, EPI_ISL_3259389, EPI_ISL_3259390, EPI_ISL_3259391, EPI_ISL_3259392, EPI_ISL_3259393, EPI_ISL_3259394, EPI_ISL_3259395, EPI_ISL_3259396, EPI_ISL_3259397, EPI_ISL_3259398, EPI_ISL_3259399, EPI_ISL_3259400, EPI_ISL_3259401, EPI_ISL_3259402, EPI_ISL_3259403, EPI_ISL_3259404, EPI_ISL_3259405, EPI_ISL_3259406, EPI_ISL_3259407, EPI_ISL_3259408, EPI_ISL_3259409, EPI_ISL_3259410, EPI_ISL_3259411, EPI_ISL_3259412, EPI_ISL_3259413, EPI_ISL_3259415, EPI_ISL_3259416, EPI_ISL_3259417, EPI_ISL_3259418, EPI_ISL_3259419, EPI_ISL_3259420, EPI_ISL_3259421, EPI_ISL_3259422, EPI_ISL_3259423, EPI_ISL_3259424, EPI_ISL_3259425, EPI_ISL_3259426, EPI_ISL_3259427, EPI_ISL_3259428, EPI_ISL_3259429, EPI_ISL_3259430, EPI_ISL_3259431, EPI_ISL_3259432, EPI_ISL_3259433, EPI_ISL_3259434, EPI_ISL_3259435, EPI_ISL_3259436, EPI_ISL_3259437, EPI_ISL_3259438, EPI_ISL_3259439, EPI_ISL_3259440, EPI_ISL_3259441, EPI_ISL_3259442, EPI_ISL_3259443, EPI_ISL_3259444, EPI_ISL_3259445, EPI_ISL_3259446, EPI_ISL_3259447, EPI_ISL_3259448, EPI_ISL_3259449, EPI_ISL_3259450, EPI_ISL_3259451, EPI_ISL_3259452, EPI_ISL_3259453, EPI_ISL_3259454, EPI_ISL_3259455, EPI_ISL_3259456, EPI_ISL_3259457, EPI_ISL_3259458, EPI_ISL_3259459, EPI_ISL_3259460, EPI_ISL_3259461, EPI_ISL_3259462, EPI_ISL_3259463, EPI_ISL_3259464, EPI_ISL_3259465, EPI_ISL_3259466, EPI_ISL_3259467, EPI_ISL_3259468, EPI_ISL_3259469, EPI_ISL_3259470, EPI_ISL_3259471, EPI_ISL_3259472, EPI_ISL_3259473, EPI_ISL_3259474, EPI_ISL_3259475, EPI_ISL_3259476, EPI_ISL_3259477, EPI_ISL_3259478, EPI_ISL_3259479, EPI_ISL_3259480, EPI_ISL_3259481, EPI_ISL_3259482, EPI_ISL_3259483, EPI_ISL_3259484, EPI_ISL_3259485, EPI_ISL_3259486, EPI_ISL_3259487, EPI_ISL_3259488, EPI_ISL_3259489, EPI_ISL_3259490, EPI_ISL_3259491, EPI_ISL_3259492, EPI_ISL_3259493, EPI_ISL_3259494, EPI_ISL_3259495, EPI_ISL_3259496, EPI_ISL_3259497, EPI_ISL_3259498, EPI_ISL_3259499, EPI_ISL_3259500, EPI_ISL_3259501, EPI_ISL_3259502, EPI_ISL_3259503, EPI_ISL_3259504, EPI_ISL_3259505, EPI_ISL_3259506, EPI_ISL_3259507, EPI_ISL_3259508, EPI_ISL_3259509, EPI_ISL_3259510, EPI_ISL_3259511, EPI_ISL_3259512, EPI_ISL_3259513, EPI_ISL_3259514, EPI_ISL_3259515, EPI_ISL_3259516, EPI_ISL_3259517, EPI_ISL_3259518, EPI_ISL_3259519, EPI_ISL_3259520, EPI_ISL_3259521, EPI_ISL_3259522, EPI_ISL_3259523, EPI_ISL_3259524, EPI_ISL_3259525, EPI_ISL_3259526, EPI_ISL_3259527, EPI_ISL_3259528, EPI_ISL_3259529, EPI_ISL_3259530 | | | |
| see above | Ochsner Health | Bioinfoexperts | Amy Feehan, Judy Crabtree, Rebecca Rose, Ben Lain, Chris Huston, Samual Moot, David J. Nolan, Tessa LaFleur, Julia-Garcia-Diaz, Lucio Miele, Susanna L. Lamers |
| EPI_ISL_3259533, EPI_ISL_3259534 | Ohio Department of Health Laboratory | Ohio Department of Health Laboratory | Holmes, Jennifer; Eric Brandt, Keoni Omura, Glen McGillivary, Caitlin McDonnell, Jade Mowery, Stephanie Mccracken, Tyler Payne, Kirtana Ramadugu, Erica Leasure, Brent Lee, Kelsey Florek, Heather Blankenship, Allison Black, Quanta Brown, and Tammy Bannerman |
| EPI_ISL_3259535 | National Institute of Infectious Diseases-Prof. Dr. Matei Bals Molecular Diagnostics Laboratory | National Institute of Infectious Diseases-Prof. Dr. Matei Bals Molecular Diagnostics Laboratory | Corina Casangiu, Leontina Banica, Marius Surleac, Ovidiu Vlaicu, Petre Milu, Robert Hohan, Simona Paraschiv, Dan Otelea |
| EPI_ISL_3259536 | Department of Public Health Bucharest | National Institute of Infectious Diseases-Prof. Dr. Matei Bals Molecular Diagnostics Laboratory | Corina Casangiu, Leontina Banica, Marius Surleac, Ovidiu Vlaicu, Petre Milu, Robert Hohan, Simona Paraschiv, Dan Otelea |
| EPI_ISL_3259537, EPI_ISL_3259538 | National Institute of Infectious Diseases-Prof. Dr. Matei Bals Molecular Diagnostics Laboratory | National Institute of Infectious Diseases-Prof. Dr. Matei Bals Molecular Diagnostics Laboratory | Corina Casangiu, Leontina Banica, Marius Surleac, Ovidiu Vlaicu, Petre Milu, Robert Hohan, Simona Paraschiv, Dan Otelea |
| EPI_ISL_3259539, EPI_ISL_3259540, EPI_ISL_3259541, EPI_ISL_3259542 | Department of Public Health Bucharest | National Institute of Infectious Diseases-Prof. Dr. Matei Bals Molecular Diagnostics Laboratory | Corina Casangiu, Leontina Banica, Marius Surleac, Ovidiu Vlaicu, Petre Milu, Robert Hohan, Simona Paraschiv, Dan Otelea |
| EPI_ISL_3259543 | Department of Public Health Constanta | National Institute of Infectious Diseases-Prof. Dr. Matei Bals Molecular Diagnostics Laboratory | Corina Casangiu, Leontina Banica, Marius Surleac, Ovidiu Vlaicu, Petre Milu, Robert Hohan, Simona Paraschiv, Dan Otelea |
| EPI_ISL_3259544 | Department of Public Health Vrancea | National Institute of Infectious Diseases-Prof. Dr. Matei Bals Molecular Diagnostics Laboratory | Corina Casangiu, Leontina Banica, Marius Surleac, Ovidiu Vlaicu, Petre Milu, Robert Hohan, Simona Paraschiv, Dan Otelea |
| EPI_ISL_3259545, EPI_ISL_3259546 | Department of Public Health Constanta | National Institute of Infectious Diseases-Prof. Dr. Matei Bals Molecular Diagnostics Laboratory | Corina Casangiu, Leontina Banica, Marius Surleac, Ovidiu Vlaicu, Petre Milu, Robert Hohan, Simona Paraschiv, Dan Otelea |
| EPI_ISL_3259554, EPI_ISL_3259555 | Department of Public Health Brasov | National Institute of Infectious Diseases-Prof. Dr. Matei Bals Molecular Diagnostics Laboratory | Corina Casangiu, Leontina Banica, Marius Surleac, Ovidiu Vlaicu, Petre Milu, Robert Hohan, Simona Paraschiv, Dan Otelea |
| EPI_ISL_3259556 | Department of Public Health Sibiu | National Institute of Infectious Diseases-Prof. Dr. Matei Bals Molecular Diagnostics Laboratory | Corina Casangiu, Leontina Banica, Marius Surleac, Ovidiu Vlaicu, Petre Milu, Robert Hohan, Simona Paraschiv, Dan Otelea |
| EPI_ISL_3259559 | Department of Public Health Vaslui | National Institute of Infectious Diseases-Prof. Dr. Matei Bals Molecular Diagnostics Laboratory | Corina Casangiu, Leontina Banica, Marius Surleac, Ovidiu Vlaicu, Petre Milu, Robert Hohan, Simona Paraschiv, Dan Otelea |
| EPI_ISL_3259562 | Department of Public Health Neamt | National Institute of Infectious Diseases-Prof. Dr. Matei Bals Molecular Diagnostics Laboratory | Corina Casangiu, Leontina Banica, Marius Surleac, Ovidiu Vlaicu, Petre Milu, Robert Hohan, Simona Paraschiv, Dan Otelea |
| EPI_ISL_3259563, EPI_ISL_3259564, EPI_ISL_3259565, EPI_ISL_3259566, EPI_ISL_3259567, EPI_ISL_3259568, EPI_ISL_3259569, EPI_ISL_3259570 | National Institute of Infectious Diseases-Prof. Dr. Matei Bals Molecular Diagnostics Laboratory | National Institute of Infectious Diseases-Prof. Dr. Matei Bals Molecular Diagnostics Laboratory | Corina Casangiu, Leontina Banica, Marius Surleac, Ovidiu Vlaicu, Petre Milu, Robert Hohan, Simona Paraschiv, Dan Otelea |
| EPI_ISL_3259571 | Emergency County Hospital Timisoara | National Institute of Infectious Diseases-Prof. Dr. Matei Bals Molecular Diagnostics Laboratory | Corina Casangiu, Leontina Banica, Marius Surleac, Ovidiu Vlaicu, Petre Milu, Robert Hohan, Simona Paraschiv, Dan Otelea |
| EPI_ISL_3259572 | Department of Public Health Galati | National Institute of Infectious Diseases-Prof. Dr. Matei Bals Molecular Diagnostics Laboratory | Corina Casangiu, Leontina Banica, Marius Surleac, Ovidiu Vlaicu, Petre Milu, Robert Hohan, Simona Paraschiv, Dan Otelea |
| EPI_ISL_3259573 | National Institute of Infectious Diseases-Prof. Dr. Matei Bals Molecular Diagnostics Laboratory | National Institute of Infectious Diseases-Prof. Dr. Matei Bals Molecular Diagnostics Laboratory | Corina Casangiu, Leontina Banica, Marius Surleac, Ovidiu Vlaicu, Petre Milu, Robert Hohan, Simona Paraschiv, Dan Otelea |
| EPI_ISL_3259574, EPI_ISL_3259575, EPI_ISL_3259576, EPI_ISL_3259577, EPI_ISL_3259578, EPI_ISL_3259579, EPI_ISL_3259580, EPI_ISL_3259581, EPI_ISL_3259582, EPI_ISL_3259583, EPI_ISL_3259584, EPI_ISL_3259585, EPI_ISL_3259586, EPI_ISL_3259587, EPI_ISL_3259588, EPI_ISL_3259589, EPI_ISL_3259590, EPI_ISL_3259591 | | | |
| see above | Department of Public Health Iasi | National Institute of Infectious Diseases-Prof. Dr. Matei Bals Molecular Diagnostics Laboratory | Corina Casangiu, Leontina Banica, Marius Surleac, Ovidiu Vlaicu, Petre Milu, Robert Hohan, Simona Paraschiv, Dan Otelea |
| EPI_ISL_3259592, EPI_ISL_3259593, EPI_ISL_3259594, EPI_ISL_3259595 | Department of Public Health Constanta | National Institute of Infectious Diseases-Prof. Dr. Matei Bals Molecular Diagnostics Laboratory | Corina Casangiu, Leontina Banica, Marius Surleac, Ovidiu Vlaicu, Petre Milu, Robert Hohan, Simona Paraschiv, Dan Otelea |
| EPI_ISL_3259596, EPI_ISL_3259597, EPI_ISL_3259598, EPI_ISL_3259599, EPI_ISL_3259600, EPI_ISL_3259601, | National Institute of Infectious Diseases-Prof. Dr. Matei Bals Molecular Diagnostics Laboratory | National Institute of Infectious Diseases-Prof. Dr. Matei Bals Molecular Diagnostics Laboratory | Corina Casangiu, Leontina Banica, Marius Surleac, Ovidiu Vlaicu, Petre Milu, Robert Hohan, Simona Paraschiv, Dan Otelea |

| | | | |
|---|---|---|---|
| EPI_ISL_3259602, EPI_ISL_3259603, EPI_ISL_3259604 | | | |
| EPI_ISL_3259605 | Emergency County Hospital Timisoara | National Institute of Infectious Diseases-Prof. Dr. Matei Bals Molecular Diagnostics Laboratory | Corina Casangiu, Leontina Banica, Marius Surleac, Ovidiu Vlaicu, Petre Milu, Robert Hohan, Simona Paraschiv, Dan Otelea |
| EPI_ISL_3259606 | National Institute of Infectious Diseases-Prof. Dr. Matei Bals Molecular Diagnostics Laboratory | National Institute of Infectious Diseases-Prof. Dr. Matei Bals Molecular Diagnostics Laboratory | Corina Casangiu, Leontina Banica, Marius Surleac, Ovidiu Vlaicu, Petre Milu, Robert Hohan, Simona Paraschiv, Dan Otelea |
| EPI_ISL_3259607, EPI_ISL_3259608, EPI_ISL_3259609, EPI_ISL_3259610, EPI_ISL_3259611, EPI_ISL_3259612, EPI_ISL_3259613, EPI_ISL_3259614, EPI_ISL_3259615, EPI_ISL_3259616, EPI_ISL_3259617, EPI_ISL_3259618, EPI_ISL_3259619, EPI_ISL_3259620, EPI_ISL_3259622, EPI_ISL_3259623 | | | |
| see above | Department of Public Health Iasi | National Institute of Infectious Diseases-Prof. Dr. Matei Bals Molecular Diagnostics Laboratory | Corina Casangiu, Leontina Banica, Marius Surleac, Ovidiu Vlaicu, Petre Milu, Robert Hohan, Simona Paraschiv, Dan Otelea |
| EPI_ISL_3259624, EPI_ISL_3259625, EPI_ISL_3259626, EPI_ISL_3259627, EPI_ISL_3259628, EPI_ISL_3259629, EPI_ISL_3259630, EPI_ISL_3259631, EPI_ISL_3259632 | Victor Babes Hospital | National Institute of Infectious Diseases-Prof. Dr. Matei Bals Molecular Diagnostics Laboratory | Corina Casangiu, Leontina Banica, Marius Surleac, Ovidiu Vlaicu, Petre Milu, Robert Hohan, Simona Paraschiv, Dan Otelea |
| EPI_ISL_3259633, EPI_ISL_3259634, EPI_ISL_3259635, EPI_ISL_3259636 | Department of Public Health Ilfov | National Institute of Infectious Diseases-Prof. Dr. Matei Bals Molecular Diagnostics Laboratory | Corina Casangiu, Leontina Banica, Marius Surleac, Ovidiu Vlaicu, Petre Milu, Robert Hohan, Simona Paraschiv, Dan Otelea |
| EPI_ISL_3259637 | National Institute of Infectious Diseases-Prof. Dr. Matei Bals Molecular Diagnostics Laboratory | National Institute of Infectious Diseases-Prof. Dr. Matei Bals Molecular Diagnostics Laboratory | Corina Casangiu, Leontina Banica, Marius Surleac, Ovidiu Vlaicu, Petre Milu, Robert Hohan, Simona Paraschiv, Dan Otelea |
| EPI_ISL_3259638, EPI_ISL_3259639 | Department of Public Health Ilfov | National Institute of Infectious Diseases-Prof. Dr. Matei Bals Molecular Diagnostics Laboratory | Corina Casangiu, Leontina Banica, Marius Surleac, Ovidiu Vlaicu, Petre Milu, Robert Hohan, Simona Paraschiv, Dan Otelea |
| EPI_ISL_3259640, EPI_ISL_3259641 | National Institute of Infectious Diseases-Prof. Dr. Matei Bals Molecular Diagnostics Laboratory | National Institute of Infectious Diseases-Prof. Dr. Matei Bals Molecular Diagnostics Laboratory | Corina Casangiu, Leontina Banica, Marius Surleac, Ovidiu Vlaicu, Petre Milu, Robert Hohan, Simona Paraschiv, Dan Otelea |
| EPI_ISL_3259642 | Department of Public Health Ilfov | National Institute of Infectious Diseases-Prof. Dr. Matei Bals Molecular Diagnostics Laboratory | Corina Casangiu, Leontina Banica, Marius Surleac, Ovidiu Vlaicu, Petre Milu, Robert Hohan, Simona Paraschiv, Dan Otelea |
| EPI_ISL_3259643 | National Institute of Infectious Diseases-Prof. Dr. Matei Bals Molecular Diagnostics Laboratory | National Institute of Infectious Diseases-Prof. Dr. Matei Bals Molecular Diagnostics Laboratory | Corina Casangiu, Leontina Banica, Marius Surleac, Ovidiu Vlaicu, Petre Milu, Robert Hohan, Simona Paraschiv, Dan Otelea |
| EPI_ISL_3259644 | Ohio Department of Health Laboratory | Ohio Department of Health Laboratory | Holmes, Jennifer; Eric Brandt, Keoni Omura, Glen McGillivary, Caitlin McDonnell, Jade Mowery, Stephanie Mccracken, Tyler Payne, Kirtana Ramadugu, Erica Leasure, Brent Lee, Kelsey Florek, Heather Blankenship, Allison Black, Quanta Brown, and Tammy Bannerman |
| EPI_ISL_3259645, EPI_ISL_3259646, EPI_ISL_3259647, EPI_ISL_3259648, EPI_ISL_3259649, EPI_ISL_3259650, EPI_ISL_3259651, EPI_ISL_3259652, EPI_ISL_3259653, EPI_ISL_3259654, EPI_ISL_3259655, EPI_ISL_3259656, EPI_ISL_3259657, EPI_ISL_3259658, EPI_ISL_3259659, EPI_ISL_3259660, EPI_ISL_3259661, EPI_ISL_3259662, EPI_ISL_3259663, EPI_ISL_3259664, EPI_ISL_3259665, EPI_ISL_3259666, EPI_ISL_3259667, EPI_ISL_3259668, EPI_ISL_3259670, EPI_ISL_3259671, EPI_ISL_3259672, EPI_ISL_3259673, EPI_ISL_3259674, EPI_ISL_3259675 | | | |
| see above | Montana Public Health Laboratory | Montana Public Health Laboratory | Joy Ritter, Michelle Mozer, Carrie Biskupiak, Deborah Gibson, Michael Dills |
| EPI_ISL_3259676 | Ohio Department of Health Laboratory | Ohio Department of Health Laboratory | Holmes, Jennifer; Eric Brandt, Keoni Omura, Glen McGillivary, Caitlin McDonnell, Jade Mowery, Stephanie Mccracken, Tyler Payne, Kirtana Ramadugu, Erica Leasure, Brent Lee, Kelsey Florek, Heather Blankenship, Allison Black, Quanta Brown, and Tammy Bannerman |
| EPI_ISL_3259855, EPI_ISL_3259856, EPI_ISL_3259857 | EHNV | Laboratory of genomics and metagenomics | Trestan Pillonel, Damien Jacot, Sébastien Aeby, Gilbert Greub, Claire Bertelli |
| EPI_ISL_3259858, EPI_ISL_3259859, EPI_ISL_3259860 | CHUV | Laboratory of genomics and metagenomics | Trestan Pillonel, Damien Jacot, Sébastien Aeby, Gilbert Greub, Claire Bertelli |
| EPI_ISL_3259861 | SYNLAB | Laboratory of genomics and metagenomics | Trestan Pillonel, Damien Jacot, Sébastien Aeby, Gilbert Greub, Claire Bertelli |
| EPI_ISL_3259862, EPI_ISL_3259863, EPI_ISL_3259864 | CHUV | Laboratory of genomics and metagenomics | Trestan Pillonel, Damien Jacot, Sébastien Aeby, Gilbert Greub, Claire Bertelli |
| EPI_ISL_3259865 | EHNV | Laboratory of genomics and metagenomics | Trestan Pillonel, Damien Jacot, Sébastien Aeby, Gilbert Greub, Claire Bertelli |
| EPI_ISL_3259866 | CHUV | Laboratory of genomics and metagenomics | Trestan Pillonel, Damien Jacot, Sébastien Aeby, Gilbert Greub, Claire Bertelli |
| EPI_ISL_3259867 | EHNV | Laboratory of genomics and metagenomics | Trestan Pillonel, Damien Jacot, Sébastien Aeby, Gilbert Greub, Claire Bertelli |
| EPI_ISL_3259868 | CHUV | Laboratory of genomics and metagenomics | Trestan Pillonel, Damien Jacot, Sébastien Aeby, Gilbert Greub, Claire Bertelli |
| EPI_ISL_3259869, EPI_ISL_3259870, EPI_ISL_3259871, EPI_ISL_3259872, EPI_ISL_3259873, EPI_ISL_3259874, EPI_ISL_3259875, EPI_ISL_3259876, EPI_ISL_3259877, EPI_ISL_3259878, EPI_ISL_3259879, EPI_ISL_3259880, EPI_ISL_3259881, EPI_ISL_3259882, EPI_ISL_3259883, EPI_ISL_3259884, EPI_ISL_3259885, EPI_ISL_3259886, EPI_ISL_3259887, EPI_ISL_3259888, EPI_ISL_3259889, EPI_ISL_3259890, EPI_ISL_3259891, EPI_ISL_3259892, EPI_ISL_3259893, EPI_ISL_3259894, EPI_ISL_3259895, EPI_ISL_3259896, EPI_ISL_3259898, EPI_ISL_3259899, EPI_ISL_3259900, EPI_ISL_3259901, EPI_ISL_3259902, EPI_ISL_3259903, EPI_ISL_3259904, EPI_ISL_3259905, EPI_ISL_3259906, EPI_ISL_3259907, EPI_ISL_3259908, EPI_ISL_3259909, EPI_ISL_3259910, EPI_ISL_3259911, EPI_ISL_3259912, EPI_ISL_3259913, EPI_ISL_3259914, EPI_ISL_3259915, EPI_ISL_3259916, EPI_ISL_3259917, EPI_ISL_3259918, EPI_ISL_3259919, EPI_ISL_3259920, EPI_ISL_3259921, EPI_ISL_3259922, EPI_ISL_3259923, EPI_ISL_3259924, EPI_ISL_3259925, EPI_ISL_3259926, EPI_ISL_3259928, EPI_ISL_3259929, EPI_ISL_3259930 | | | |
| see above | EHNV | Laboratory of genomics and metagenomics | Trestan Pillonel, Damien Jacot, Sébastien Aeby, Gilbert Greub, Claire Bertelli |
| EPI_ISL_3259931, EPI_ISL_3259935 | CHUV | Laboratory of genomics and metagenomics | Trestan Pillonel, Damien Jacot, Sébastien Aeby, Gilbert Greub, Claire Bertelli |
| EPI_ISL_3259936 | VIDYMED LAUSANNE | Laboratory of genomics and metagenomics | Trestan Pillonel, Damien Jacot, Sébastien Aeby, Gilbert Greub, Claire Bertelli |
| EPI_ISL_3259938 | CHUV | Laboratory of genomics and metagenomics | Trestan Pillonel, Damien Jacot, Sébastien Aeby, Gilbert Greub, Claire Bertelli |
| EPI_ISL_3259939 | VIDYMED LAUSANNE | Laboratory of genomics and metagenomics | Trestan Pillonel, Damien Jacot, Sébastien Aeby, Gilbert Greub, Claire Bertelli |
| EPI_ISL_3259940, EPI_ISL_3259941 | CHUV | Laboratory of genomics and metagenomics | Trestan Pillonel, Damien Jacot, Sébastien Aeby, Gilbert Greub, Claire Bertelli |
| EPI_ISL_3259942 | VIDYMED LAUSANNE | Laboratory of genomics and metagenomics | Trestan Pillonel, Damien Jacot, Sébastien Aeby, Gilbert Greub, Claire Bertelli |
| EPI_ISL_3259943 | CHUV | Laboratory of genomics and metagenomics | Trestan Pillonel, Damien Jacot, Sébastien Aeby, Gilbert Greub, Claire Bertelli |
| EPI_ISL_3259944, EPI_ISL_3259945 | VIDYMED LAUSANNE | Laboratory of genomics and metagenomics | Trestan Pillonel, Damien Jacot, Sébastien Aeby, Gilbert Greub, Claire Bertelli |
| EPI_ISL_3259946 | CHUV | Laboratory of genomics and metagenomics | Trestan Pillonel, Damien Jacot, Sébastien Aeby, Gilbert Greub, Claire Bertelli |
| EPI_ISL_3259947 | Kantonsspital Baden AG | Institute of Medical Virology | Daniel Ehrsam, Isabel Stürmer, Catharine Aquino, Joel Wirz, Weihong Qi, Hubert Rehrauer, Verena Kufner, Gabriela Ziltener, Maryam Zaheri, Stefan Schmutz, Annette Audigé, Maria Grünberg, Kevin Steiner, Jon Huder, Cyril Shah, Riccarda Capaul, Guido Bloemberg, Jürg Böni, Michael Huber, Alexandra Trkola |
| EPI_ISL_3259948, EPI_ISL_3259949 | Universität Zürich | Institute of Medical Virology | Daniel Ehrsam, Isabel Stürmer, Catharine Aquino, Joel Wirz, Weihong Qi, Hubert Rehrauer, Verena Kufner, Gabriela Ziltener, Maryam Zaheri, Stefan Schmutz, Annette Audigé, Maria Grünberg, Kevin Steiner, Jon Huder, Cyril Shah, Riccarda Capaul, Guido Bloemberg, Jürg Böni, Michael Huber, Alexandra Trkola |
| EPI_ISL_3259951, EPI_ISL_3259952 | Stadtspital Triemli | Institute of Medical Virology | Daniel Ehrsam, Isabel Stürmer, Catharine Aquino, Joel Wirz, Weihong Qi, Hubert Rehrauer, Verena Kufner, Gabriela Ziltener, Maryam Zaheri, Stefan Schmutz, Annette Audigé, Maria Grünberg, Kevin Steiner, Jon Huder, Cyril Shah, Riccarda Capaul, Guido Bloemberg, Jürg Böni, Michael Huber, Alexandra Trkola |
| EPI_ISL_3259953, EPI_ISL_3259954, EPI_ISL_3259955, EPI_ISL_3259956, EPI_ISL_3259957, EPI_ISL_3259959, EPI_ISL_3259960, EPI_ISL_3259961, EPI_ISL_3259962, EPI_ISL_3259963, EPI_ISL_3259964, EPI_ISL_3259965, EPI_ISL_3259966, EPI_ISL_3259967, EPI_ISL_3259968 | | | |
| see above | Universität Zürich | Institute of Medical Virology | Daniel Ehrsam, Isabel Stürmer, Catharine Aquino, Joel Wirz, Weihong Qi, Hubert Rehrauer, Verena Kufner, Gabriela Ziltener, Maryam Zaheri, Stefan Schmutz, Annette Audigé, Maria Grünberg, Kevin Steiner, Jon Huder, Cyril Shah, Riccarda Capaul, Guido Bloemberg, Jürg Böni, Michael Huber, Alexandra Trkola |
| EPI_ISL_3259969, EPI_ISL_3259970 | Stadtspital Triemli | Institute of Medical Virology | Daniel Ehrsam, Isabel Stürmer, Catharine Aquino, Joel Wirz, Weihong Qi, Hubert Rehrauer, Verena Kufner, Gabriela Ziltener, Maryam Zaheri, Stefan Schmutz, Annette Audigé, Maria Grünberg, Kevin Steiner, Jon Huder, Cyril Shah, Riccarda Capaul, Guido Bloemberg, Jürg Böni, Michael Huber, Alexandra Trkola |
| EPI_ISL_3259971, EPI_ISL_3259972 | Universität Zürich | Institute of Medical Virology | Daniel Ehrsam, Isabel Stürmer, Catharine Aquino, Joel Wirz, Weihong Qi, Hubert Rehrauer, Verena Kufner, Gabriela Ziltener, Maryam Zaheri, Stefan Schmutz, Annette Audigé, Maria Grünberg, Kevin Steiner, Jon Huder, Cyril Shah, Riccarda Capaul, Guido Bloemberg, Jürg Böni, Michael Huber, Alexandra Trkola |
| EPI_ISL_3259973, EPI_ISL_3259974, EPI_ISL_3259975, EPI_ISL_3259976, EPI_ISL_3259977, EPI_ISL_3259978 | UniversitätsSpital Zürich | Institute of Medical Virology | Daniel Ehrsam, Isabel Stürmer, Catharine Aquino, Joel Wirz, Weihong Qi, Hubert Rehrauer, Verena Kufner, Gabriela Ziltener, Maryam Zaheri, Stefan Schmutz, Annette Audigé, Maria Grünberg, Kevin Steiner, Jon Huder, Cyril Shah, Riccarda Capaul, Guido Bloemberg, Jürg Böni, Michael Huber, Alexandra Trkola |
| EPI_ISL_3259979, EPI_ISL_3259980, EPI_ISL_3259981, EPI_ISL_3259982, EPI_ISL_3259983, EPI_ISL_3259984, EPI_ISL_3259985, EPI_ISL_3259986, EPI_ISL_3259987, EPI_ISL_3259988, EPI_ISL_3259989, EPI_ISL_3259990, EPI_ISL_3259991, EPI_ISL_3259992, EPI_ISL_3259993, EPI_ISL_3259994 | | | |
| see above | Stadtspital Triemli | Institute of Medical Virology | Daniel Ehrsam, Isabel Stürmer, Catharine Aquino, Joel Wirz, Weihong Qi, Hubert Rehrauer, Verena Kufner, Gabriela Ziltener, Maryam Zaheri, Stefan Schmutz, Annette Audigé, Maria Grünberg, Kevin Steiner, Jon Huder, Cyril Shah, Riccarda |

| | | | |
|---|---|---|---|
| | | | Capaul, Guido Bloemberg, Jürg Böni, Michael Huber, Alexandra Trkola |
| EPI_ISL_3259995, EPI_ISL_3259996, EPI_ISL_3259997, EPI_ISL_3259998, EPI_ISL_3259999, EPI_ISL_3260000, EPI_ISL_3260001, EPI_ISL_3260002 | UniversitätsSpital Zürich | Institute of Medical Virology | Daniel Ehrsam, Isabel Stürmer, Catharine Aquino, Joel Wirz, Weihong Qi, Hubert Rehrauer, Verena Kufner, Gabriela Ziltener, Maryam Zaheri, Stefan Schmutz, Annette Audigé, Maria Grünberg, Kevin Steiner, Jon Huder, Cyril Shah, Riccarda Capaul, Guido Bloemberg, Jürg Böni, Michael Huber, Alexandra Trkola |
| EPI_ISL_3260003, EPI_ISL_3260004 | Universitäts-Kinderspital Zürich | Institute of Medical Virology | Daniel Ehrsam, Isabel Stürmer, Catharine Aquino, Joel Wirz, Weihong Qi, Hubert Rehrauer, Verena Kufner, Gabriela Ziltener, Maryam Zaheri, Stefan Schmutz, Annette Audigé, Maria Grünberg, Kevin Steiner, Jon Huder, Cyril Shah, Riccarda Capaul, Guido Bloemberg, Jürg Böni, Michael Huber, Alexandra Trkola |
| EPI_ISL_3260005, EPI_ISL_3260006, EPI_ISL_3260007 | UniversitätsSpital Zürich | Institute of Medical Virology | Daniel Ehrsam, Isabel Stürmer, Catharine Aquino, Joel Wirz, Weihong Qi, Hubert Rehrauer, Verena Kufner, Gabriela Ziltener, Maryam Zaheri, Stefan Schmutz, Annette Audigé, Maria Grünberg, Kevin Steiner, Jon Huder, Cyril Shah, Riccarda Capaul, Guido Bloemberg, Jürg Böni, Michael Huber, Alexandra Trkola |
| EPI_ISL_3260008, EPI_ISL_3260009, EPI_ISL_3260010, EPI_ISL_3260011, EPI_ISL_3260012, EPI_ISL_3260013, EPI_ISL_3260014, EPI_ISL_3260015, EPI_ISL_3260016, EPI_ISL_3260017, EPI_ISL_3260018, EPI_ISL_3260019, EPI_ISL_3260020, EPI_ISL_3260021, EPI_ISL_3260022, EPI_ISL_3260023, EPI_ISL_3260024, EPI_ISL_3260025, EPI_ISL_3260026 | | | |
| see above | Stadtspital Triemli | Institute of Medical Virology | Daniel Ehrsam, Isabel Stürmer, Catharine Aquino, Joel Wirz, Weihong Qi, Hubert Rehrauer, Verena Kufner, Gabriela Ziltener, Maryam Zaheri, Stefan Schmutz, Annette Audigé, Maria Grünberg, Kevin Steiner, Jon Huder, Cyril Shah, Riccarda Capaul, Guido Bloemberg, Jürg Böni, Michael Huber, Alexandra Trkola |
| EPI_ISL_3260027 | Kinderspital Zürich | Institute of Medical Virology | Daniel Ehrsam, Isabel Stürmer, Catharine Aquino, Joel Wirz, Weihong Qi, Hubert Rehrauer, Verena Kufner, Gabriela Ziltener, Maryam Zaheri, Stefan Schmutz, Annette Audigé, Maria Grünberg, Kevin Steiner, Jon Huder, Cyril Shah, Riccarda Capaul, Guido Bloemberg, Jürg Böni, Michael Huber, Alexandra Trkola |
| EPI_ISL_3260028, EPI_ISL_3260029, EPI_ISL_3260030, EPI_ISL_3260031, EPI_ISL_3260032, EPI_ISL_3260034, EPI_ISL_3260035, EPI_ISL_3260036, EPI_ISL_3260037, EPI_ISL_3260038, EPI_ISL_3260039, EPI_ISL_3260040, EPI_ISL_3260041, EPI_ISL_3260042, EPI_ISL_3260043, EPI_ISL_3260044, EPI_ISL_3260045, EPI_ISL_3260046, EPI_ISL_3260047, EPI_ISL_3260048, EPI_ISL_3260049, EPI_ISL_3260050, EPI_ISL_3260051, EPI_ISL_3260052, EPI_ISL_3260053, EPI_ISL_3260054, EPI_ISL_3260056, EPI_ISL_3260057, EPI_ISL_3260058, EPI_ISL_3260059, EPI_ISL_3260060, EPI_ISL_3260061, EPI_ISL_3260062, EPI_ISL_3260063, EPI_ISL_3260065, EPI_ISL_3260066, EPI_ISL_3260067, EPI_ISL_3260068, EPI_ISL_3260069, EPI_ISL_3260070, EPI_ISL_3260071, EPI_ISL_3260072, EPI_ISL_3260073, EPI_ISL_3260074, EPI_ISL_3260075, EPI_ISL_3260076, EPI_ISL_3260077, EPI_ISL_3260078, EPI_ISL_3260079, EPI_ISL_3260080, EPI_ISL_3260081, EPI_ISL_3260082, EPI_ISL_3260083, EPI_ISL_3260084, EPI_ISL_3260085, EPI_ISL_3260086, EPI_ISL_3260087, EPI_ISL_3260088, EPI_ISL_3260089, EPI_ISL_3260090 | | | |
| see above | Michigan Department of Health and Human Services, Bureau of Laboratories | Michigan Department of Health and Human Services, Bureau of Laboratories | Blankenship HM, Riner D, Soehnlen MK |
| EPI_ISL_3260113 | Minnesota Department of Health, Public Health Laboratory | Minnesota Department of Health, Public Health Laboratory | Alexandra Lorentz, Jacob Garfin, Matt Plumb, and Xiong Wang |
| EPI_ISL_3260114, EPI_ISL_3260116, EPI_ISL_3260117, EPI_ISL_3260118, EPI_ISL_3260119, EPI_ISL_3260120, EPI_ISL_3260121, EPI_ISL_3260122, EPI_ISL_3260123, EPI_ISL_3260124, EPI_ISL_3260125, EPI_ISL_3260126, EPI_ISL_3260127, EPI_ISL_3260128, EPI_ISL_3260129, EPI_ISL_3260130, EPI_ISL_3260131, EPI_ISL_3260132, EPI_ISL_3260133, EPI_ISL_3260134, EPI_ISL_3260135, EPI_ISL_3260136, EPI_ISL_3260137, EPI_ISL_3260138, EPI_ISL_3260139, EPI_ISL_3260140, EPI_ISL_3260141, EPI_ISL_3260142, EPI_ISL_3260143, EPI_ISL_3260144, EPI_ISL_3260145, EPI_ISL_3260146, EPI_ISL_3260147, EPI_ISL_3260148, EPI_ISL_3260149, EPI_ISL_3260150, EPI_ISL_3260151, EPI_ISL_3260152, EPI_ISL_3260153, EPI_ISL_3260154, EPI_ISL_3260155, EPI_ISL_3260156, EPI_ISL_3260157, EPI_ISL_3260158, EPI_ISL_3260159, EPI_ISL_3260160, EPI_ISL_3260161, EPI_ISL_3260162, EPI_ISL_3260163, EPI_ISL_3260164, EPI_ISL_3260165, EPI_ISL_3260166, EPI_ISL_3260167, EPI_ISL_3260168, EPI_ISL_3260169, EPI_ISL_3260170, EPI_ISL_3260171, EPI_ISL_3260172, EPI_ISL_3260173, EPI_ISL_3260174 | | | |
| see above | Centracare Laboratory Services | Minnesota Department of Health, Public Health Laboratory | Alexandra Lorentz, Jacob Garfin, Matt Plumb, and Xiong Wang |
| EPI_ISL_3260175, EPI_ISL_3260176, EPI_ISL_3260177, EPI_ISL_3260178, EPI_ISL_3260179, EPI_ISL_3260180, EPI_ISL_3260181, EPI_ISL_3260182, EPI_ISL_3260183, EPI_ISL_3260184, EPI_ISL_3260185, EPI_ISL_3260187, EPI_ISL_3260188, EPI_ISL_3260189, EPI_ISL_3260190, EPI_ISL_3260191, EPI_ISL_3260192, EPI_ISL_3260193, EPI_ISL_3260194, EPI_ISL_3260195, EPI_ISL_3260196, EPI_ISL_3260197, EPI_ISL_3260198, EPI_ISL_3260199, EPI_ISL_3260200 | | | |
| see above | Minnesota Department of Health, Public Health Laboratory | Minnesota Department of Health, Public Health Laboratory | Alexandra Lorentz, Jacob Garfin, Matt Plumb, and Xiong Wang |
| EPI_ISL_3260201, EPI_ISL_3260202, EPI_ISL_3260203, EPI_ISL_3260204, EPI_ISL_3260205, EPI_ISL_3260206, EPI_ISL_3260207, EPI_ISL_3260208, EPI_ISL_3260209, EPI_ISL_3260210, EPI_ISL_3260211, EPI_ISL_3260212, EPI_ISL_3260213, EPI_ISL_3260214 | | | |
| see above | Centracare Laboratory Services | Minnesota Department of Health, Public Health Laboratory | Alexandra Lorentz, Jacob Garfin, Matt Plumb, and Xiong Wang |
| EPI_ISL_3260215, EPI_ISL_3260216, EPI_ISL_3260217, EPI_ISL_3260218, EPI_ISL_3260219, EPI_ISL_3260220, EPI_ISL_3260221, EPI_ISL_3260222, EPI_ISL_3260224, EPI_ISL_3260225, EPI_ISL_3260226, EPI_ISL_3260227, EPI_ISL_3260228, EPI_ISL_3260229, EPI_ISL_3260230, EPI_ISL_3260232, EPI_ISL_3260233, EPI_ISL_3260234, EPI_ISL_3260235, EPI_ISL_3260236, EPI_ISL_3260237, EPI_ISL_3260238, EPI_ISL_3260240, EPI_ISL_3260241, EPI_ISL_3260242, EPI_ISL_3260243, EPI_ISL_3260244, EPI_ISL_3260246, EPI_ISL_3260248, EPI_ISL_3260251, EPI_ISL_3260252, EPI_ISL_3260253, EPI_ISL_3260254, EPI_ISL_3260255, EPI_ISL_3260256, EPI_ISL_3260257, EPI_ISL_3260258, EPI_ISL_3260259, EPI_ISL_3260260, EPI_ISL_3260261, EPI_ISL_3260262, EPI_ISL_3260263, EPI_ISL_3260264, EPI_ISL_3260265, EPI_ISL_3260266, EPI_ISL_3260267, EPI_ISL_3260268, EPI_ISL_3260269, EPI_ISL_3260270, EPI_ISL_3260271, EPI_ISL_3260272, EPI_ISL_3260273, EPI_ISL_3260274, EPI_ISL_3260275, EPI_ISL_3260276, EPI_ISL_3260277 | | | |
| see above | Minnesota Department of Health, Public Health Laboratory | Minnesota Department of Health, Public Health Laboratory | Alexandra Lorentz, Jacob Garfin, Matt Plumb, and Xiong Wang |
| EPI_ISL_3260278, EPI_ISL_3260279, EPI_ISL_3260280, EPI_ISL_3260281, EPI_ISL_3260282, EPI_ISL_3260283, EPI_ISL_3260284, EPI_ISL_3260285, EPI_ISL_3260286, EPI_ISL_3260287, EPI_ISL_3260288, EPI_ISL_3260289, EPI_ISL_3260290, EPI_ISL_3260291, EPI_ISL_3260292 | | | |
| see above | Centracare Laboratory Services | Minnesota Department of Health, Public Health Laboratory | Alexandra Lorentz, Jacob Garfin, Matt Plumb, and Xiong Wang |
| EPI_ISL_3260294, EPI_ISL_3260297, EPI_ISL_3260298, EPI_ISL_3260299, EPI_ISL_3260300, EPI_ISL_3260301, EPI_ISL_3260302, EPI_ISL_3260303, EPI_ISL_3260304, EPI_ISL_3260305, EPI_ISL_3260306, EPI_ISL_3260307, EPI_ISL_3260308, EPI_ISL_3260309, EPI_ISL_3260310, EPI_ISL_3260311, EPI_ISL_3260312, EPI_ISL_3260313, EPI_ISL_3260314, EPI_ISL_3260315, EPI_ISL_3260316, EPI_ISL_3260317, EPI_ISL_3260318, EPI_ISL_3260319, EPI_ISL_3260320, EPI_ISL_3260321 | | | |
| see above | IN State Department of Health Laboratory Services | IN State Department of Health Laboratory Services | Cassandra Campion, Jamie Yeadon, Brian Pope, Lixia Liu, Kyle Brownlee, Melissa Hindenlang, Mark Glazier |
| EPI_ISL_3260322, EPI_ISL_3260323, EPI_ISL_3260324, EPI_ISL_3260325, EPI_ISL_3260326, EPI_ISL_3260327, EPI_ISL_3260328, EPI_ISL_3260329, EPI_ISL_3260330, EPI_ISL_3260331, EPI_ISL_3260332, EPI_ISL_3260333, EPI_ISL_3260334, EPI_ISL_3260335, EPI_ISL_3260336, EPI_ISL_3260337, EPI_ISL_3260338, EPI_ISL_3260339, EPI_ISL_3260340, EPI_ISL_3260341, EPI_ISL_3260342, EPI_ISL_3260343, EPI_ISL_3260344, EPI_ISL_3260345, EPI_ISL_3260346, EPI_ISL_3260347, EPI_ISL_3260348, EPI_ISL_3260349, EPI_ISL_3260350, EPI_ISL_3260351, EPI_ISL_3260352, EPI_ISL_3260353, EPI_ISL_3260354, EPI_ISL_3260355 | | | |
| see above | SYNLAB | GIGA Medical Genomics | Keith Durkin, Maria Artesi, Bouchra Boujemla, Nathalie Renotte, Nadine Cambisano, Sébastien Bontems, Cécile Meex, Claire Gourzonès, Olivier Ek, Laurent Gillet, Marie-Pierre Hayette, Vincent Bours |
| EPI_ISL_3260356, EPI_ISL_3260357, EPI_ISL_3260358, EPI_ISL_3260359, EPI_ISL_3260360, EPI_ISL_3260361, EPI_ISL_3260362, EPI_ISL_3260363 | Department of Clinical Microbiology | GIGA Medical Genomics | Keith Durkin, Maria Artesi, Bouchra Boujemla, Nathalie Renotte, Nadine Cambisano, Sébastien Bontems, Cécile Meex, Claire Gourzonès, Olivier Ek, Laurent Gillet, Marie-Pierre Hayette, Vincent Bours |
| EPI_ISL_3260364, EPI_ISL_3260365, EPI_ISL_3260366, EPI_ISL_3260367, EPI_ISL_3260368, EPI_ISL_3260369, EPI_ISL_3260370, EPI_ISL_3260371, EPI_ISL_3260372, EPI_ISL_3260373, EPI_ISL_3260374, EPI_ISL_3260375, EPI_ISL_3260376, EPI_ISL_3260377, EPI_ISL_3260378, EPI_ISL_3260379, EPI_ISL_3260380, EPI_ISL_3260381, EPI_ISL_3260382, EPI_ISL_3260383, EPI_ISL_3260384, EPI_ISL_3260385, EPI_ISL_3260386 | | | |
| see above | University of Liège COVID-19 testing center | GIGA Medical Genomics | Keith Durkin, Maria Artesi, Bouchra Boujemla, Nathalie Renotte, Nadine Cambisano, Sébastien Bontems, Cécile Meex, Claire Gourzonès, Olivier Ek, Laurent Gillet, Marie-Pierre Hayette, Vincent Bours |
| EPI_ISL_3260387, EPI_ISL_3260388 | C.H.C MontLégia | GIGA Medical Genomics | Keith Durkin, Maria Artesi, Bouchra Boujemla, Nathalie Renotte, Nadine Cambisano, Sébastien Bontems, Cécile Meex, Claire Gourzonès, Olivier Ek, Laurent Gillet, Marie-Pierre Hayette, Vincent Bours |
| EPI_ISL_3260389 | University of Liège COVID-19 testing center | GIGA Medical Genomics | Keith Durkin, Maria Artesi, Bouchra Boujemla, Nathalie Renotte, Nadine Cambisano, Sébastien Bontems, Cécile Meex, Claire Gourzonès, Olivier Ek, Laurent Gillet, Marie-Pierre Hayette, Vincent Bours |
| EPI_ISL_3260390, EPI_ISL_3260391 | Department of Clinical Microbiology | GIGA Medical Genomics | Keith Durkin, Maria Artesi, Bouchra Boujemla, Nathalie Renotte, Nadine Cambisano, Sébastien Bontems, Cécile Meex, Claire Gourzonès, Olivier Ek, Laurent Gillet, Marie-Pierre Hayette, Vincent Bours |
| EPI_ISL_3260392 | SYNLAB | GIGA Medical Genomics | Keith Durkin, Maria Artesi, Bouchra Boujemla, Nathalie Renotte, Nadine Cambisano, Sébastien Bontems, Cécile Meex, Claire Gourzonès, Olivier Ek, Laurent Gillet, Marie-Pierre Hayette, Vincent Bours |
| EPI_ISL_3260393 | Department of Clinical Microbiology | GIGA Medical Genomics | Keith Durkin, Maria Artesi, Bouchra Boujemla, Nathalie Renotte, Nadine Cambisano, Sébastien Bontems, Cécile Meex, Claire Gourzonès, Olivier Ek, Laurent Gillet, Marie-Pierre Hayette, Vincent Bours |
| EPI_ISL_3260394 | C.H.C MontLégia | GIGA Medical Genomics | Keith Durkin, Maria Artesi, Bouchra Boujemla, Nathalie Renotte, Nadine Cambisano, Sébastien Bontems, Cécile Meex, Claire Gourzonès, Olivier Ek, Laurent Gillet, Marie-Pierre Hayette, Vincent Bours |
| EPI_ISL_3260395 | Department of Clinical Microbiology | GIGA Medical Genomics | Keith Durkin, Maria Artesi, Bouchra Boujemla, Nathalie Renotte, Nadine Cambisano, Sébastien Bontems, Cécile Meex, Claire Gourzonès, Olivier Ek, Laurent Gillet, Marie-Pierre Hayette, Vincent Bours |
| EPI_ISL_3260396, EPI_ISL_3260397 | University of Liège COVID-19 testing center | GIGA Medical Genomics | Keith Durkin, Maria Artesi, Bouchra Boujemla, Nathalie Renotte, Nadine Cambisano, Sébastien Bontems, Cécile Meex, Claire Gourzonès, Olivier Ek, Laurent Gillet, Marie-Pierre Hayette, Vincent Bours |
| EPI_ISL_3260398, EPI_ISL_3260399, EPI_ISL_3260400 | SYNLAB | GIGA Medical Genomics | Keith Durkin, Maria Artesi, Bouchra Boujemla, Nathalie Renotte, Nadine Cambisano, Sébastien Bontems, Cécile Meex, Claire Gourzonès, Olivier Ek, Laurent Gillet, Marie-Pierre Hayette, Vincent Bours |
| EPI_ISL_3260401, EPI_ISL_3260402 | Department of Clinical Microbiology | GIGA Medical Genomics | Keith Durkin, Maria Artesi, Bouchra Boujemla, Nathalie Renotte, Nadine Cambisano, Sébastien Bontems, Cécile Meex, Claire Gourzonès, Olivier Ek, Laurent Gillet, Marie-Pierre Hayette, Vincent Bours |
| EPI_ISL_3260403 | University of Liège COVID-19 testing center | GIGA Medical Genomics | Keith Durkin, Maria Artesi, Bouchra Boujemla, Nathalie Renotte, Nadine Cambisano, Sébastien Bontems, Cécile Meex, Claire Gourzonès, Olivier Ek, Laurent Gillet, Marie-Pierre Hayette, Vincent Bours |
| EPI_ISL_3260404, EPI_ISL_3260405 | SYNLAB | GIGA Medical Genomics | Keith Durkin, Maria Artesi, Bouchra Boujemla, Nathalie Renotte, Nadine Cambisano, Sébastien Bontems, Cécile Meex, Claire Gourzonès, Olivier Ek, Laurent Gillet, Marie-Pierre Hayette, Vincent Bours |
| EPI_ISL_3260406, EPI_ISL_3260407 | University of Liège COVID-19 testing center | GIGA Medical Genomics | Keith Durkin, Maria Artesi, Bouchra Boujemla, Nathalie Renotte, Nadine Cambisano, Sébastien Bontems, Cécile Meex, Claire Gourzonès, Olivier Ek, Laurent Gillet, Marie-Pierre Hayette, Vincent Bours |
| EPI_ISL_3260408 | SYNLAB | GIGA Medical Genomics | Keith Durkin, Maria Artesi, Bouchra Boujemla, Nathalie Renotte, Nadine Cambisano, Sébastien Bontems, Cécile Meex, Claire Gourzonès, Olivier Ek, Laurent Gillet, Marie-Pierre Hayette, Vincent Bours |
| EPI_ISL_3260409, EPI_ISL_3260410, EPI_ISL_3260411, EPI_ISL_3260412, EPI_ISL_3260413, EPI_ISL_3260414, EPI_ISL_3260415, EPI_ISL_3260416, EPI_ISL_3260417, EPI_ISL_3260418, EPI_ISL_3260419, EPI_ISL_3260420, EPI_ISL_3260421, EPI_ISL_3260422, EPI_ISL_3260423, EPI_ISL_3260425, EPI_ISL_3260426, EPI_ISL_3260427, EPI_ISL_3260428, EPI_ISL_3260429, EPI_ISL_3260430, EPI_ISL_3260431, EPI_ISL_3260432, EPI_ISL_3260433, EPI_ISL_3260434, EPI_ISL_3260435, EPI_ISL_3260436, EPI_ISL_3260437, EPI_ISL_3260438, EPI_ISL_3260439, EPI_ISL_3260440, EPI_ISL_3260441, EPI_ISL_3260442, EPI_ISL_3260443, EPI_ISL_3260444, EPI_ISL_3260445, EPI_ISL_3260446, EPI_ISL_3260447, EPI_ISL_3260448, EPI_ISL_3260449, EPI_ISL_3260450, EPI_ISL_3260451, EPI_ISL_3260452, EPI_ISL_3260453, EPI_ISL_3260454, EPI_ISL_3260455, EPI_ISL_3260457, EPI_ISL_3260458, EPI_ISL_3260459, EPI_ISL_3260460, EPI_ISL_3260461, EPI_ISL_3260463, EPI_ISL_3260464, EPI_ISL_3260465, EPI_ISL_3260466, EPI_ISL_3260467, EPI_ISL_3260468, EPI_ISL_3260469, EPI_ISL_3260470, EPI_ISL_3260472, EPI_ISL_3260473, EPI_ISL_3260475, EPI_ISL_3260476, EPI_ISL_3260478, EPI_ISL_3260479, EPI_ISL_3260480, EPI_ISL_3260481, EPI_ISL_3260482, EPI_ISL_3260483, EPI_ISL_3260484, EPI_ISL_3260485, EPI_ISL_3260486, EPI_ISL_3260487, EPI_ISL_3260488, EPI_ISL_3260489, EPI_ISL_3260490, EPI_ISL_3260491, EPI_ISL_3260492, EPI_ISL_3260493, EPI_ISL_3260494, EPI_ISL_3260495, EPI_ISL_3260496, EPI_ISL_3260497, EPI_ISL_3260498, EPI_ISL_3260499, EPI_ISL_3260500, EPI_ISL_3260501, EPI_ISL_3260502, EPI_ISL_3260503, EPI_ISL_3260504, EPI_ISL_3260505, EPI_ISL_3260507, EPI_ISL_3260508, EPI_ISL_3260509, EPI_ISL_3260510, EPI_ISL_3260511, EPI_ISL_3260512, EPI_ISL_3260514, EPI_ISL_3260515, EPI_ISL_3260516, EPI_ISL_3260517, EPI_ISL_3260518, EPI_ISL_3260519, EPI_ISL_3260520, EPI_ISL_3260521, EPI_ISL_3260522, EPI_ISL_3260523, EPI_ISL_3260524, EPI_ISL_3260525, EPI_ISL_3260526, EPI_ISL_3260527, EPI_ISL_3260528, EPI_ISL_3260529, EPI_ISL_3260530, EPI_ISL_3260531, EPI_ISL_3260532, EPI_ISL_3260533, EPI_ISL_3260534, EPI_ISL_3260535, EPI_ISL_3260536, EPI_ISL_3260537, EPI_ISL_3260538, EPI_ISL_3260539, EPI_ISL_3260540, EPI_ISL_3260541, EPI_ISL_3260542, EPI_ISL_3260543, EPI_ISL_3260544, EPI_ISL_3260545, EPI_ISL_3260546, EPI_ISL_3260547, EPI_ISL_3260548, EPI_ISL_3260549, EPI_ISL_3260550, EPI_ISL_3260551, EPI_ISL_3260552, EPI_ISL_3260553, EPI_ISL_3260554, EPI_ISL_3260555, EPI_ISL_3260556, EPI_ISL_3260557, EPI_ISL_3260558, EPI_ISL_3260560, EPI_ISL_3260561, EPI_ISL_3260562, EPI_ISL_3260563, EPI_ISL_3260564, EPI_ISL_3260565, EPI_ISL_3260566, EPI_ISL_3260567, EPI_ISL_3260568, EPI_ISL_3260569 | | | |
| see above | Arizona State University | Arizona State University | Ajeet Bains, LaRinda A. Holland, Matthew F. Smith, Peter T. Skidmore, Rabia Maqsood, Nicholas J. Mellor, Nathaniel Johnson, Joshua LaBaer, Vel Murugan, Efrem S. Lim |
| EPI_ISL_3260572, EPI_ISL_3260576 | Ohio Department of Health Laboratory | Ohio Department of Health Laboratory | Holmes, Jennifer; Eric Brandt, Keoni Omura, Glen McGillivary, Caitlin McDonnell, Jade Mowery, Stephanie Mccracken, Tyler Payne, Kirtana Ramadugu, Erica Leasure, Brent Lee, Kelsey Florek, Heather Blankenship, Allison Black, Quanta Brown, and Tammy Bannerman |
| EPI_ISL_3260577, EPI_ISL_3260578, EPI_ISL_3260579, EPI_ISL_3260580, EPI_ISL_3260581, EPI_ISL_3260582, EPI_ISL_3260583, EPI_ISL_3260584, EPI_ISL_3260585, EPI_ISL_3260586, EPI_ISL_3260587, EPI_ISL_3260588, EPI_ISL_3260589, EPI_ISL_3260590, EPI_ISL_3260591, EPI_ISL_3260592, EPI_ISL_3260593, EPI_ISL_3260594, EPI_ISL_3260595, EPI_ISL_3260596, EPI_ISL_3260597, EPI_ISL_3260598, EPI_ISL_3260599, EPI_ISL_3260600, EPI_ISL_3260601, EPI_ISL_3260602, EPI_ISL_3260603, EPI_ISL_3260604, EPI_ISL_3260605, EPI_ISL_3260606 | | | |

| | | | |
|---|---|---|---|
| see above | Nebraska Public Health Laboratory | NPHL COVID-19 Response Team | NPHL COVID-19 Response Team |
| EPI_ISL_3260607, EPI_ISL_3260608 | KEMRI-Wellcome Trust Research Programme,Kilifi | KEMRI-Wellcome Trust Research Programme,Kilifi | Githinji G.,Ochola I.,Agoti C.,Mohamed K.S.,de Laurent Z.,Mburu M.W.,Morobe J.,Lambisia A., Makori T., Edidah O., Ndwiga L. |
| EPI_ISL_3260610, EPI_ISL_3260611, EPI_ISL_3260612, EPI_ISL_3260613, EPI_ISL_3260614, EPI_ISL_3260615, EPI_ISL_3260616, EPI_ISL_3260617, EPI_ISL_3260618, EPI_ISL_3260619, EPI_ISL_3260620, EPI_ISL_3260621, EPI_ISL_3260622, EPI_ISL_3260623, EPI_ISL_3260624, EPI_ISL_3260625, EPI_ISL_3260626, EPI_ISL_3260627, EPI_ISL_3260628, EPI_ISL_3260629, EPI_ISL_3260630, EPI_ISL_3260631, EPI_ISL_3260632, EPI_ISL_3260633, EPI_ISL_3260634, EPI_ISL_3260635, EPI_ISL_3260636, EPI_ISL_3260637, EPI_ISL_3260638, EPI_ISL_3260639, EPI_ISL_3260640, EPI_ISL_3260641 | | | |
| see above | Nebraska Public Health Laboratory | NPHL COVID-19 Response Team | NPHL COVID-19 Response Team |
| EPI_ISL_3260655, EPI_ISL_3260656, EPI_ISL_3260657, EPI_ISL_3260658, EPI_ISL_3260659, EPI_ISL_3260660, EPI_ISL_3260661, EPI_ISL_3260662, EPI_ISL_3260663, EPI_ISL_3260664, EPI_ISL_3260665, EPI_ISL_3260667, EPI_ISL_3260668, EPI_ISL_3260669, EPI_ISL_3260670, EPI_ISL_3260671, EPI_ISL_3260672, EPI_ISL_3260674, EPI_ISL_3260676, EPI_ISL_3260677, EPI_ISL_3260678, EPI_ISL_3260679, EPI_ISL_3260680, EPI_ISL_3260681, EPI_ISL_3260682, EPI_ISL_3260683, EPI_ISL_3260684, EPI_ISL_3260685, EPI_ISL_3260686, EPI_ISL_3260687, EPI_ISL_3260688, EPI_ISL_3260689 | | | |
| see above | Dept. of Microbiology and Infection Control, Akershus University Hospital HF | Dept. of Microbiology and Infection Control, Akershus University Hospital HF | Hege Vangstein Aamot, Alexander Hesselberg Løvestad |
| EPI_ISL_3260694 | Institute of Microbiology, Universidad San Francisco de Quito | Institute of Microbiology, Universidad San Francisco de Quito | Sully Márquez, Juan José Guadalupe, Belén Prado-Vivar, Erika Muñoz, Monica Becerra-Wong, Fernanda Zurita, Bernardo Gutiérrez, Maria Nataly Velastegui Peralta, Verónica Barragán, Patricio Rojas-Silva, Gabriel Trueba, Michelle Grunauer, Paúl Cárdenas |
| EPI_ISL_3260696 | Institute of Microbiology, Universidad San Francisco de Quito | Institute of Microbiology, Universidad San Francisco de Quito | Fernanda Zurita, Sully Márquez, Juan José Guadalupe, Belén Prado-Vivar, Monica Becerra-Wong, Erika Muñoz, Bernardo Gutiérrez, Andrea Pinos, Verónica Barragán, Patricio Rojas-Silva, Gabriel Trueba, Michelle Grunauer, Paúl Cárdenas |
| EPI_ISL_3260699 | Institute of Microbiology, Universidad San Francisco de Quito | Institute of Microbiology, Universidad San Francisco de Quito | Monica Becerra-Wong, Erika Muñoz, Sully Márquez, Juan José Guadalupe, Belén Prado-Vivar, Fernanda Zurita, Bernardo Gutiérrez, Andrea Pinos, Verónica Barragán, Patricio Rojas-Silva, Gabriel Trueba, Michelle Grunauer, Paúl Cárdenas |
| EPI_ISL_3260712, EPI_ISL_3260714, EPI_ISL_3260715, EPI_ISL_3260716, EPI_ISL_3260717, EPI_ISL_3260718, EPI_ISL_3260719, EPI_ISL_3260720, EPI_ISL_3260723, EPI_ISL_3260724, EPI_ISL_3260725, EPI_ISL_3260727 | | | |
| see above | Department of Microbiology, Yamanashi Institute for Public Health and Environment | Genome Analysis Center, Yamanashi Central Hospital | Yosuke Hirotsu |
| EPI_ISL_3260754, EPI_ISL_3260755, EPI_ISL_3260756, EPI_ISL_3260757, EPI_ISL_3260758, EPI_ISL_3260759, EPI_ISL_3260760, EPI_ISL_3260761, EPI_ISL_3260762, EPI_ISL_3260763, EPI_ISL_3260764, EPI_ISL_3260765, EPI_ISL_3260766, EPI_ISL_3260767, EPI_ISL_3260768, EPI_ISL_3260769, EPI_ISL_3260770, EPI_ISL_3260771, EPI_ISL_3260772, EPI_ISL_3260773, EPI_ISL_3260774, EPI_ISL_3260775, EPI_ISL_3260776, EPI_ISL_3260777, EPI_ISL_3260778, EPI_ISL_3260779, EPI_ISL_3260780, EPI_ISL_3260781, EPI_ISL_3260782, EPI_ISL_3260783, EPI_ISL_3260784, EPI_ISL_3260785, EPI_ISL_3260786, EPI_ISL_3260787, EPI_ISL_3260788, EPI_ISL_3260789, EPI_ISL_3260790, EPI_ISL_3260791, EPI_ISL_3260792, EPI_ISL_3260793, EPI_ISL_3260794, EPI_ISL_3260795, EPI_ISL_3260796, EPI_ISL_3260797, EPI_ISL_3260798 | | | |
| see above | Servicio Microbiología Hospital La Paz | Servicio Microbiología Hospital La Paz | Fernando Lázaro, Rubén Cáceres, Iván Bloise, Jesús Mingorance Cruz, Elie Dahdouh |
| EPI_ISL_3260890 | HOSPITAL UNIVERSITARIO VIRGEN DE LA ARRIXACA | Instituto de Salud Carlos III | Iglesias-Caballero, M. Sandonís,V. Vázquez-Morón, S. Camarero, S. Pozo, F. Casas, I. Jiménez, P. Zaballos, A. Monzón, S. Varona, S. Cuesta, I.MORENO PARRADO, LAURA |
| EPI_ISL_3260898, EPI_ISL_3260899, EPI_ISL_3260900, EPI_ISL_3260901, EPI_ISL_3260902, EPI_ISL_3260903, EPI_ISL_3260904, EPI_ISL_3260905, EPI_ISL_3260906, EPI_ISL_3260907, EPI_ISL_3260908, EPI_ISL_3260909, EPI_ISL_3260910, EPI_ISL_3260911, EPI_ISL_3260912, EPI_ISL_3260913, EPI_ISL_3260914, EPI_ISL_3260915, EPI_ISL_3260916, EPI_ISL_3260917, EPI_ISL_3260918, EPI_ISL_3260919, EPI_ISL_3260920, EPI_ISL_3260921, EPI_ISL_3260923, EPI_ISL_3260924, EPI_ISL_3260925, EPI_ISL_3260926, EPI_ISL_3260927, EPI_ISL_3260928, EPI_ISL_3260929, EPI_ISL_3260930, EPI_ISL_3260931, EPI_ISL_3260932, EPI_ISL_3260933, EPI_ISL_3260934, EPI_ISL_3260935, EPI_ISL_3260936, EPI_ISL_3260937, EPI_ISL_3260938, EPI_ISL_3260939, EPI_ISL_3260940, EPI_ISL_3260941, EPI_ISL_3260942, EPI_ISL_3260943, EPI_ISL_3260944, EPI_ISL_3260945, EPI_ISL_3260946, EPI_ISL_3260947, EPI_ISL_3260948, EPI_ISL_3260949, EPI_ISL_3260950, EPI_ISL_3260951, EPI_ISL_3260952, EPI_ISL_3260953, EPI_ISL_3260954, EPI_ISL_3260955, EPI_ISL_3260956, EPI_ISL_3260957, EPI_ISL_3260958, EPI_ISL_3260959, EPI_ISL_3260960, EPI_ISL_3260961, EPI_ISL_3260962, EPI_ISL_3260963, EPI_ISL_3260964, EPI_ISL_3260965, EPI_ISL_3260966, EPI_ISL_3260967, EPI_ISL_3260968, EPI_ISL_3260969, EPI_ISL_3260970, EPI_ISL_3260971, EPI_ISL_3260972 | | | |
| see above | Dipartimento di Medicina di Laboratorio, Azienda sanitaria universitaria Friuli Centrale (ASU FC) | Dipartimento di Medicina di Laboratorio, Azienda sanitaria universitaria Friuli Centrale (ASU FC) | Corrado Pipan, Catia Mio, Chiara Dal Secco, Stefania Marzinotto, Francesco Curcio |
| EPI_ISL_3260978, EPI_ISL_3260980, EPI_ISL_3260981, EPI_ISL_3260982, EPI_ISL_3260983, EPI_ISL_3260984, EPI_ISL_3260989, EPI_ISL_3260990, EPI_ISL_3260991, EPI_ISL_3260992, EPI_ISL_3260993, EPI_ISL_3260995, EPI_ISL_3260996, EPI_ISL_3260997, EPI_ISL_3260998, EPI_ISL_3261001, EPI_ISL_3261002, EPI_ISL_3261003, EPI_ISL_3261004, EPI_ISL_3261007, EPI_ISL_3261008, EPI_ISL_3261009, EPI_ISL_3261012, EPI_ISL_3261014, EPI_ISL_3261015, EPI_ISL_3261017, EPI_ISL_3261018, EPI_ISL_3261019 | | | |
| see above | Servizo de Microbioloxía. Complexo Hospitalario de Santiago de Compostela | Servizo de Microbioloxía. Complexo Hospitalario de Santiago de Compostela | Xosé Costa, Antonio Aguilera, Gema Barbeito, Amparo Coira, Carlos García_Riestra, Manuela Hernández, José Llovo, Teresa Lopez_Valiño, Laura Millán, Daniel Navarro, Rocío trastoy, Mercedes Treviño, María Luisa Pérez_del_Molino |
| EPI_ISL_3261020, EPI_ISL_3261021, EPI_ISL_3261022, EPI_ISL_3261023, EPI_ISL_3261024, EPI_ISL_3261025 | CHU Pontchaillou | CHU Pontchaillou | GROLHIER Claire, DENOUAL Florent, ETCHEVERRY Amandine, SASSI Mohamed, JAGLINE Steven, FEBREAU Christine, PRONIER Charlotte, GALIBERT Marie Dominique, DE TAYRAC Marie, THIBAULT Vincent |
| EPI_ISL_3261027, EPI_ISL_3261028 | AZ Jan Portaels | Imelda Ziekenhuis | Johan Frans, Dagmar Obbels, Hanne Valgaeren |
| EPI_ISL_3261029, EPI_ISL_3261030, EPI_ISL_3261031, EPI_ISL_3261032, EPI_ISL_3261033, EPI_ISL_3261034, EPI_ISL_3261035, EPI_ISL_3261036 | Imelda Ziekenhuis | Imelda Ziekenhuis | Johan Frans, Dagmar Obbels, Hanne Valgaeren |
| EPI_ISL_3261040 | SC (UCO) Igiene e Sanità Pubblica, ASUGI, Trieste | ARGO Laboratorio Genomica ed Epigenomica | Licastro D, Dal Monego S, Degasperi M, Marcello A, Maggiore A, Busetti M, Segat L, Piscianz E, Braida C, Barbone F, D'Agaro P |
| EPI_ISL_3261116, EPI_ISL_3261117, EPI_ISL_3261119 | URMC LABS | URMC LABS | Ghinwa Dumyati, Dwight Hardy, Andrew Cameron, Stepan Voljehn, Kelly DeLary |
| EPI_ISL_3261130, EPI_ISL_3261132, EPI_ISL_3261133, EPI_ISL_3261134, EPI_ISL_3261135, EPI_ISL_3261136, EPI_ISL_3261137, EPI_ISL_3261138, EPI_ISL_3261141, EPI_ISL_3261143, EPI_ISL_3261145, EPI_ISL_3261146, EPI_ISL_3261147, EPI_ISL_3261148, EPI_ISL_3261149, EPI_ISL_3261150, EPI_ISL_3261151, EPI_ISL_3261152, EPI_ISL_3261153, EPI_ISL_3261154, EPI_ISL_3261155, EPI_ISL_3261156, EPI_ISL_3261157, EPI_ISL_3261158, EPI_ISL_3261159, EPI_ISL_3261160, EPI_ISL_3261161, EPI_ISL_3261162, EPI_ISL_3261163, EPI_ISL_3261164, EPI_ISL_3261165, EPI_ISL_3261166, EPI_ISL_3261167, EPI_ISL_3261168, EPI_ISL_3261169, EPI_ISL_3261170, EPI_ISL_3261171, EPI_ISL_3261172, EPI_ISL_3261173, EPI_ISL_3261174, EPI_ISL_3261175, EPI_ISL_3261176, EPI_ISL_3261177, EPI_ISL_3261178, EPI_ISL_3261179, EPI_ISL_3261180, EPI_ISL_3261182, EPI_ISL_3261183, EPI_ISL_3261184, EPI_ISL_3261185 | | | |
| see above | Arcispedale Santa Maria Nuova, Autoimmunità, Allergologia e Biotecnologie Innovative | Istituto Zooprofilattico Sperimentale della Lombardia e dell'Emilia Romagna (IZSLER), Risk Analysis and Genomic Epidemiology Unit | Alessandro Zerbini, Lucia Belloni, Stefania Croci, Marina Morganti, Ilaria Menozzi, Erika Scaltriti, Stefano Pongolini |
| EPI_ISL_3261186 | Tampa General Hospital Esoteric Lab | Tampa General Hospital Esoteric Research & Development Lab | Jon Faughn, Deanna Becker, Dominic Uy, Jorge Mecias-Frias, Katie Hurless, Vicki Healer, Amorce Lima, Suzane Silbert |
| EPI_ISL_3261187 | Tampa General Hospital Esoteric Lab | Tampa General Hospital Esoteric Research & Development Lab | Jon Faughn, Marci O'Driscoll, Deanna Becker, Dominic Uy, Jorge Mecias-Frias, Katie Hurless, Vicki Healer, Amorce Lima, Suzane Silbert |
| EPI_ISL_3261188, EPI_ISL_3261189, EPI_ISL_3261190, EPI_ISL_3261191, EPI_ISL_3261192, EPI_ISL_3261193, EPI_ISL_3261194 | Hospital Universitari de Bellvitge | Microbiology Department | Sara Marti, Aida Gonzalez-Diaz, Yolanda Hernandez, Laura Calatayud, Jordi Niubó, Carmen Ardanuy, Jordi Camara, M Angeles Domínguez |
| EPI_ISL_3261195, EPI_ISL_3261196, EPI_ISL_3261197, EPI_ISL_3261198, EPI_ISL_3261199, EPI_ISL_3261200, EPI_ISL_3261201, EPI_ISL_3261202 | CLILAB | Microbiology Department | Sara Marti, Aida Gonzalez-Diaz, Yolanda Hernandez, Laura Calatayud, Jordi Niubó, Carmen Ardanuy, Jordi Camara, M Angeles Domínguez |
| EPI_ISL_3261203, EPI_ISL_3261204, EPI_ISL_3261205, EPI_ISL_3261206 | Hospital Universitari de Bellvitge | Microbiology Department | Sara Marti, Aida Gonzalez-Diaz, Yolanda Hernandez, Laura Calatayud, Jordi Niubó, Carmen Ardanuy, Jordi Camara, M Angeles Domínguez |
| EPI_ISL_3261207, EPI_ISL_3261208, EPI_ISL_3261210 | CLILAB | Microbiology Department | Sara Marti, Aida Gonzalez-Diaz, Yolanda Hernandez, Laura Calatayud, Jordi Niubó, Carmen Ardanuy, Jordi Camara, M Angeles Domínguez |
| EPI_ISL_3261211, EPI_ISL_3261212, EPI_ISL_3261213, EPI_ISL_3261214, EPI_ISL_3261215, EPI_ISL_3261216, EPI_ISL_3261217, EPI_ISL_3261218, EPI_ISL_3261219, EPI_ISL_3261221, EPI_ISL_3261222, EPI_ISL_3261223, EPI_ISL_3261224, EPI_ISL_3261225, EPI_ISL_3261226, EPI_ISL_3261227, EPI_ISL_3261228, EPI_ISL_3261229, EPI_ISL_3261230, EPI_ISL_3261231, EPI_ISL_3261232, EPI_ISL_3261233, EPI_ISL_3261234, EPI_ISL_3261235, EPI_ISL_3261236, EPI_ISL_3261237, EPI_ISL_3261238, EPI_ISL_3261239, EPI_ISL_3261240, EPI_ISL_3261241, EPI_ISL_3261242, EPI_ISL_3261243, EPI_ISL_3261244, EPI_ISL_3261245, EPI_ISL_3261246, EPI_ISL_3261247, EPI_ISL_3261248, EPI_ISL_3261249, EPI_ISL_3261250, EPI_ISL_3261251, EPI_ISL_3261252, EPI_ISL_3261253, EPI_ISL_3261254, EPI_ISL_3261255, EPI_ISL_3261256, EPI_ISL_3261257, EPI_ISL_3261258, EPI_ISL_3261259, EPI_ISL_3261260, EPI_ISL_3261261, EPI_ISL_3261262, EPI_ISL_3261263, EPI_ISL_3261264 | | | |
| see above | Hospital Universitari de Bellvitge | Microbiology Department | Sara Marti, Aida Gonzalez-Diaz, Yolanda Hernandez, Laura Calatayud, Jordi Niubó, Carmen Ardanuy, Jordi Camara, M Angeles Domínguez |
| EPI_ISL_3261265, EPI_ISL_3261266, EPI_ISL_3261268, EPI_ISL_3261269, EPI_ISL_3261270, EPI_ISL_3261271, EPI_ISL_3261272, EPI_ISL_3261273, EPI_ISL_3261274, EPI_ISL_3261275, EPI_ISL_3261277, EPI_ISL_3261278, EPI_ISL_3261279, EPI_ISL_3261280, EPI_ISL_3261281, EPI_ISL_3261282, EPI_ISL_3261283, EPI_ISL_3261284, EPI_ISL_3261285, EPI_ISL_3261286, EPI_ISL_3261287, EPI_ISL_3261289, EPI_ISL_3261290, EPI_ISL_3261291, EPI_ISL_3261292, EPI_ISL_3261293, EPI_ISL_3261294, EPI_ISL_3261295, EPI_ISL_3261296, EPI_ISL_3261297, EPI_ISL_3261298, EPI_ISL_3261299, EPI_ISL_3261300, EPI_ISL_3261301, EPI_ISL_3261302, EPI_ISL_3261303, EPI_ISL_3261304, EPI_ISL_3261305, EPI_ISL_3261306, EPI_ISL_3261307, EPI_ISL_3261308, EPI_ISL_3261309, EPI_ISL_3261310, EPI_ISL_3261311, EPI_ISL_3261312, EPI_ISL_3261313, EPI_ISL_3261314, EPI_ISL_3261315, EPI_ISL_3261316, EPI_ISL_3261317, EPI_ISL_3261318, EPI_ISL_3261319, EPI_ISL_3261320, EPI_ISL_3261321, EPI_ISL_3261322, EPI_ISL_3261323, EPI_ISL_3261326, EPI_ISL_3261328, EPI_ISL_3261329, EPI_ISL_3261332, EPI_ISL_3261333, EPI_ISL_3261334, EPI_ISL_3261335, EPI_ISL_3261336, EPI_ISL_3261337, EPI_ISL_3261338, EPI_ISL_3261339, EPI_ISL_3261340, EPI_ISL_3261341, EPI_ISL_3261342, EPI_ISL_3261343, EPI_ISL_3261344, EPI_ISL_3261346, EPI_ISL_3261347, EPI_ISL_3261348, EPI_ISL_3261349, EPI_ISL_3261350, EPI_ISL_3261351, EPI_ISL_3261352, EPI_ISL_3261353, EPI_ISL_3261354, EPI_ISL_3261355, EPI_ISL_3261356, EPI_ISL_3261357, EPI_ISL_3261358, EPI_ISL_3261359, EPI_ISL_3261360, EPI_ISL_3261361, EPI_ISL_3261362, EPI_ISL_3261363, EPI_ISL_3261364, EPI_ISL_3261365, EPI_ISL_3261366, EPI_ISL_3261367, EPI_ISL_3261368, EPI_ISL_3261369, EPI_ISL_3261370, EPI_ISL_3261371, EPI_ISL_3261372, EPI_ISL_3261374, EPI_ISL_3261375, EPI_ISL_3261376, EPI_ISL_3261377, EPI_ISL_3261378, EPI_ISL_3261379, EPI_ISL_3261380, EPI_ISL_3261381, EPI_ISL_3261382, EPI_ISL_3261383, EPI_ISL_3261384, EPI_ISL_3261385, EPI_ISL_3261386, EPI_ISL_3261387, EPI_ISL_3261388, EPI_ISL_3261389, EPI_ISL_3261390, EPI_ISL_3261391, EPI_ISL_3261392, EPI_ISL_3261393, EPI_ISL_3261394, EPI_ISL_3261395, EPI_ISL_3261396, EPI_ISL_3261397, EPI_ISL_3261398, EPI_ISL_3261399, EPI_ISL_3261402, EPI_ISL_3261403, EPI_ISL_3261404, EPI_ISL_3261406, EPI_ISL_3261407, EPI_ISL_3261408, EPI_ISL_3261409, EPI_ISL_3261410, EPI_ISL_3261411, EPI_ISL_3261412, EPI_ISL_3261413, EPI_ISL_3261414, EPI_ISL_3261415, EPI_ISL_3261416, EPI_ISL_3261417, EPI_ISL_3261418, EPI_ISL_3261419, EPI_ISL_3261421, EPI_ISL_3261422, EPI_ISL_3261423, EPI_ISL_3261424, EPI_ISL_3261425, EPI_ISL_3261426, EPI_ISL_3261428, EPI_ISL_3261429, EPI_ISL_3261430, EPI_ISL_3261431, EPI_ISL_3261433, EPI_ISL_3261434, EPI_ISL_3261435, EPI_ISL_3261436, EPI_ISL_3261437, EPI_ISL_3261438, EPI_ISL_3261439, EPI_ISL_3261441, EPI_ISL_3261442, EPI_ISL_3261443, EPI_ISL_3261445, EPI_ISL_3261446, EPI_ISL_3261447, EPI_ISL_3261448, EPI_ISL_3261449, EPI_ISL_3261450, EPI_ISL_3261451, EPI_ISL_3261453, EPI_ISL_3261455, EPI_ISL_3261457, EPI_ISL_3261458, EPI_ISL_3261459, EPI_ISL_3261460, EPI_ISL_3261461, EPI_ISL_3261462, EPI_ISL_3261463, EPI_ISL_3261464, EPI_ISL_3261465, EPI_ISL_3261466, EPI_ISL_3261467, EPI_ISL_3261468, EPI_ISL_3261469, EPI_ISL_3261471, EPI_ISL_3261472, EPI_ISL_3261473, EPI_ISL_3261474, EPI_ISL_3261475, EPI_ISL_3261476, EPI_ISL_3261477, EPI_ISL_3261478, EPI_ISL_3261480, EPI_ISL_3261481, EPI_ISL_3261482, EPI_ISL_3261483, EPI_ISL_3261484, EPI_ISL_3261486, EPI_ISL_3261487, EPI_ISL_3261488, EPI_ISL_3261489, EPI_ISL_3261490, EPI_ISL_3261491, EPI_ISL_3261492, EPI_ISL_3261493, EPI_ISL_3261494, EPI_ISL_3261495, EPI_ISL_3261496, EPI_ISL_3261497, EPI_ISL_3261498, EPI_ISL_3261499, EPI_ISL_3261500, EPI_ISL_3261501, EPI_ISL_3261502, EPI_ISL_3261503, EPI_ISL_3261504, EPI_ISL_3261505, EPI_ISL_3261506, EPI_ISL_3261507, EPI_ISL_3261508, | | | |

| | | | |
|---|---|---|---|
| EPI_ISL_3261509 | | | |
| see above | Sharp HealthCare Laboratory | Andersen lab at Scripps Research | SEARCH Alliance San Diego with Aaron Harding, Jacquelyn Berumen, Cathy Woerle, Liam McGinnis, Art Mendoza, Omid Bakhtar |
| EPI_ISL_3261511 | Clinique Saint-Pierre Ottignies | UCLouvain/IREC/MBLG | Jean Ruelle, Bertrand Bearzatto, Benoit Kabamba Mukadi |
| EPI_ISL_3261512 | Sharp HealthCare Laboratory | Andersen lab at Scripps Research | SEARCH Alliance San Diego with Aaron Harding, Jacquelyn Berumen, Cathy Woerle, Liam McGinnis, Art Mendoza, Omid Bakhtar |
| EPI_ISL_3261513 | Clinique Saint-Pierre Ottignies | UCLouvain/IREC/MBLG | Jean Ruelle, Bertrand Bearzatto, Benoit Kabamba Mukadi |
| EPI_ISL_3261514 | Sharp HealthCare Laboratory | Andersen lab at Scripps Research | SEARCH Alliance San Diego with Aaron Harding, Jacquelyn Berumen, Cathy Woerle, Liam McGinnis, Art Mendoza, Omid Bakhtar |
| EPI_ISL_3261515, EPI_ISL_3261516 | Clinique Saint-Pierre Ottignies | UCLouvain/IREC/MBLG | Jean Ruelle, Bertrand Bearzatto, Benoit Kabamba Mukadi |
| EPI_ISL_3261517 | Sharp HealthCare Laboratory | Andersen lab at Scripps Research | SEARCH Alliance San Diego with Aaron Harding, Jacquelyn Berumen, Cathy Woerle, Liam McGinnis, Art Mendoza, Omid Bakhtar |
| EPI_ISL_3261518, EPI_ISL_3261519 | Clinique Saint-Pierre Ottignies | UCLouvain/IREC/MBLG | Jean Ruelle, Bertrand Bearzatto, Benoit Kabamba Mukadi |
| EPI_ISL_3261520 | Sharp HealthCare Laboratory | Andersen lab at Scripps Research | SEARCH Alliance San Diego with Aaron Harding, Jacquelyn Berumen, Cathy Woerle, Liam McGinnis, Art Mendoza, Omid Bakhtar |
| EPI_ISL_3261521, EPI_ISL_3261522 | Clinique Saint-Pierre Ottignies | UCLouvain/IREC/MBLG | Jean Ruelle, Bertrand Bearzatto, Benoit Kabamba Mukadi |
| EPI_ISL_3261523 | Sharp HealthCare Laboratory | Andersen lab at Scripps Research | SEARCH Alliance San Diego with Aaron Harding, Jacquelyn Berumen, Cathy Woerle, Liam McGinnis, Art Mendoza, Omid Bakhtar |
| EPI_ISL_3261524, EPI_ISL_3261525 | Clinique Saint-Pierre Ottignies | UCLouvain/IREC/MBLG | Jean Ruelle, Bertrand Bearzatto, Benoit Kabamba Mukadi |
| EPI_ISL_3261526 | Sharp HealthCare Laboratory | Andersen lab at Scripps Research | SEARCH Alliance San Diego with Aaron Harding, Jacquelyn Berumen, Cathy Woerle, Liam McGinnis, Art Mendoza, Omid Bakhtar |
| EPI_ISL_3261527, EPI_ISL_3261528 | Clinique Saint-Pierre Ottignies | UCLouvain/IREC/MBLG | Jean Ruelle, Bertrand Bearzatto, Benoit Kabamba Mukadi |
| EPI_ISL_3261529 | Sharp HealthCare Laboratory | Andersen lab at Scripps Research | SEARCH Alliance San Diego with Aaron Harding, Jacquelyn Berumen, Cathy Woerle, Liam McGinnis, Art Mendoza, Omid Bakhtar |
| EPI_ISL_3261530 | Clinique Saint-Pierre Ottignies | UCLouvain/IREC/MBLG | Jean Ruelle, Bertrand Bearzatto, Benoit Kabamba Mukadi |
| EPI_ISL_3261531 | Cliniques universitaires Saint-Luc | UCLouvain/IREC/MBLG | Jean Ruelle, Bertrand Bearzatto, Benoit Kabamba Mukadi |
| EPI_ISL_3261532 | Sharp HealthCare Laboratory | Andersen lab at Scripps Research | SEARCH Alliance San Diego with Aaron Harding, Jacquelyn Berumen, Cathy Woerle, Liam McGinnis, Art Mendoza, Omid Bakhtar |
| EPI_ISL_3261533 | Cliniques universitaires Saint-Luc | UCLouvain/IREC/MBLG | Jean Ruelle, Bertrand Bearzatto, Benoit Kabamba Mukadi |
| EPI_ISL_3261534 | Sharp HealthCare Laboratory | Andersen lab at Scripps Research | SEARCH Alliance San Diego with Aaron Harding, Jacquelyn Berumen, Cathy Woerle, Liam McGinnis, Art Mendoza, Omid Bakhtar |
| EPI_ISL_3261535, EPI_ISL_3261536, EPI_ISL_3261537 | Cliniques universitaires Saint-Luc | UCLouvain/IREC/MBLG | Jean Ruelle, Bertrand Bearzatto, Benoit Kabamba Mukadi |
| EPI_ISL_3261538 | Sharp HealthCare Laboratory | Andersen lab at Scripps Research | SEARCH Alliance San Diego with Aaron Harding, Jacquelyn Berumen, Cathy Woerle, Liam McGinnis, Art Mendoza, Omid Bakhtar |
| EPI_ISL_3261539 | Cliniques universitaires Saint-Luc | UCLouvain/IREC/MBLG | Jean Ruelle, Bertrand Bearzatto, Benoit Kabamba Mukadi |
| EPI_ISL_3261540 | Sharp HealthCare Laboratory | Andersen lab at Scripps Research | SEARCH Alliance San Diego with Aaron Harding, Jacquelyn Berumen, Cathy Woerle, Liam McGinnis, Art Mendoza, Omid Bakhtar |
| EPI_ISL_3261541, EPI_ISL_3261542 | Cliniques universitaires Saint-Luc | UCLouvain/IREC/MBLG | Jean Ruelle, Bertrand Bearzatto, Benoit Kabamba Mukadi |
| EPI_ISL_3261543 | Sharp HealthCare Laboratory | Andersen lab at Scripps Research | SEARCH Alliance San Diego with Aaron Harding, Jacquelyn Berumen, Cathy Woerle, Liam McGinnis, Art Mendoza, Omid Bakhtar |
| EPI_ISL_3261544, EPI_ISL_3261545 | Cliniques universitaires Saint-Luc | UCLouvain/IREC/MBLG | Jean Ruelle, Bertrand Bearzatto, Benoit Kabamba Mukadi |
| EPI_ISL_3261546 | Sharp HealthCare Laboratory | Andersen lab at Scripps Research | SEARCH Alliance San Diego with Aaron Harding, Jacquelyn Berumen, Cathy Woerle, Liam McGinnis, Art Mendoza, Omid Bakhtar |
| EPI_ISL_3261547, EPI_ISL_3261548 | Cliniques universitaires Saint-Luc | UCLouvain/IREC/MBLG | Jean Ruelle, Bertrand Bearzatto, Benoit Kabamba Mukadi |
| EPI_ISL_3261549 | Sharp HealthCare Laboratory | Andersen lab at Scripps Research | SEARCH Alliance San Diego with Aaron Harding, Jacquelyn Berumen, Cathy Woerle, Liam McGinnis, Art Mendoza, Omid Bakhtar |
| EPI_ISL_3261550, EPI_ISL_3261551 | Cliniques universitaires Saint-Luc | UCLouvain/IREC/MBLG | Jean Ruelle, Bertrand Bearzatto, Benoit Kabamba Mukadi |
| EPI_ISL_3261552 | Sharp HealthCare Laboratory | Andersen lab at Scripps Research | SEARCH Alliance San Diego with Aaron Harding, Jacquelyn Berumen, Cathy Woerle, Liam McGinnis, Art Mendoza, Omid Bakhtar |
| EPI_ISL_3261553, EPI_ISL_3261554 | Cliniques universitaires Saint-Luc | UCLouvain/IREC/MBLG | Jean Ruelle, Bertrand Bearzatto, Benoit Kabamba Mukadi |
| EPI_ISL_3261555 | Sharp HealthCare Laboratory | Andersen lab at Scripps Research | SEARCH Alliance San Diego with Aaron Harding, Jacquelyn Berumen, Cathy Woerle, Liam McGinnis, Art Mendoza, Omid Bakhtar |
| EPI_ISL_3261556, EPI_ISL_3261557 | Cliniques universitaires Saint-Luc | UCLouvain/IREC/MBLG | Jean Ruelle, Bertrand Bearzatto, Benoit Kabamba Mukadi |
| EPI_ISL_3261558 | Sharp HealthCare Laboratory | Andersen lab at Scripps Research | SEARCH Alliance San Diego with Aaron Harding, Jacquelyn Berumen, Cathy Woerle, Liam McGinnis, Art Mendoza, Omid Bakhtar |
| EPI_ISL_3261559, EPI_ISL_3261560 | Cliniques universitaires Saint-Luc | UCLouvain/IREC/MBLG | Jean Ruelle, Bertrand Bearzatto, Benoit Kabamba Mukadi |
| EPI_ISL_3261561 | Sharp HealthCare Laboratory | Andersen lab at Scripps Research | SEARCH Alliance San Diego with Aaron Harding, Jacquelyn Berumen, Cathy Woerle, Liam McGinnis, Art Mendoza, Omid Bakhtar |
| EPI_ISL_3261562, EPI_ISL_3261563 | Cliniques universitaires Saint-Luc | UCLouvain/IREC/MBLG | Jean Ruelle, Bertrand Bearzatto, Benoit Kabamba Mukadi |
| EPI_ISL_3261564 | Sharp HealthCare Laboratory | Andersen lab at Scripps Research | SEARCH Alliance San Diego with Aaron Harding, Jacquelyn Berumen, Cathy Woerle, Liam McGinnis, Art Mendoza, Omid Bakhtar |
| EPI_ISL_3261565, EPI_ISL_3261566 | Cliniques universitaires Saint-Luc | UCLouvain/IREC/MBLG | Jean Ruelle, Bertrand Bearzatto, Benoit Kabamba Mukadi |
| EPI_ISL_3261567 | Sharp HealthCare Laboratory | Andersen lab at Scripps Research | SEARCH Alliance San Diego with Aaron Harding, Jacquelyn Berumen, Cathy Woerle, Liam McGinnis, Art Mendoza, Omid Bakhtar |
| EPI_ISL_3261568, EPI_ISL_3261569 | Cliniques universitaires Saint-Luc | UCLouvain/IREC/MBLG | Jean Ruelle, Bertrand Bearzatto, Benoit Kabamba Mukadi |
| EPI_ISL_3261570 | Sharp HealthCare Laboratory | Andersen lab at Scripps Research | SEARCH Alliance San Diego with Aaron Harding, Jacquelyn Berumen, Cathy Woerle, Liam McGinnis, Art Mendoza, Omid Bakhtar |
| EPI_ISL_3261571, EPI_ISL_3261572 | Cliniques universitaires Saint-Luc | UCLouvain/IREC/MBLG | Jean Ruelle, Bertrand Bearzatto, Benoit Kabamba Mukadi |
| EPI_ISL_3261573 | Sharp HealthCare Laboratory | Andersen lab at Scripps Research | SEARCH Alliance San Diego with Aaron Harding, Jacquelyn Berumen, Cathy Woerle, Liam McGinnis, Art Mendoza, Omid Bakhtar |
| EPI_ISL_3261574 | Cliniques universitaires Saint-Luc | UCLouvain/IREC/MBLG | Jean Ruelle, Bertrand Bearzatto, Benoit Kabamba Mukadi |
| EPI_ISL_3261575 | Sharp HealthCare Laboratory | Andersen lab at Scripps Research | SEARCH Alliance San Diego with Aaron Harding, Jacquelyn Berumen, Cathy Woerle, Liam McGinnis, Art Mendoza, Omid Bakhtar |
| EPI_ISL_3261576 | Cliniques universitaires Saint-Luc | UCLouvain/IREC/MBLG | Jean Ruelle, Bertrand Bearzatto, Benoit Kabamba Mukadi |
| EPI_ISL_3261577 | Sharp HealthCare Laboratory | Andersen lab at Scripps Research | SEARCH Alliance San Diego with Aaron Harding, Jacquelyn Berumen, Cathy Woerle, Liam McGinnis, Art Mendoza, Omid Bakhtar |
| EPI_ISL_3261578, EPI_ISL_3261579 | Cliniques universitaires Saint-Luc | UCLouvain/IREC/MBLG | Jean Ruelle, Bertrand Bearzatto, Benoit Kabamba Mukadi |
| EPI_ISL_3261580 | Sharp HealthCare Laboratory | Andersen lab at Scripps Research | SEARCH Alliance San Diego with Aaron Harding, Jacquelyn Berumen, Cathy Woerle, Liam McGinnis, Art Mendoza, Omid Bakhtar |
| EPI_ISL_3261581, EPI_ISL_3261582 | Cliniques universitaires Saint-Luc | UCLouvain/IREC/MBLG | Jean Ruelle, Bertrand Bearzatto, Benoit Kabamba Mukadi |
| EPI_ISL_3261583 | Sharp HealthCare Laboratory | Andersen lab at Scripps Research | SEARCH Alliance San Diego with Aaron Harding, Jacquelyn Berumen, Cathy Woerle, Liam McGinnis, Art Mendoza, Omid Bakhtar |
| EPI_ISL_3261584, EPI_ISL_3261585 | Cliniques universitaires Saint-Luc | UCLouvain/IREC/MBLG | Jean Ruelle, Bertrand Bearzatto, Benoit Kabamba Mukadi |
| EPI_ISL_3261586 | Sharp HealthCare Laboratory | Andersen lab at Scripps Research | SEARCH Alliance San Diego with Aaron Harding, Jacquelyn Berumen, Cathy Woerle, Liam McGinnis, Art Mendoza, Omid Bakhtar |
| EPI_ISL_3261587, EPI_ISL_3261589, EPI_ISL_3261590 | Cliniques universitaires Saint-Luc | UCLouvain/IREC/MBLG | Jean Ruelle, Bertrand Bearzatto, Benoit Kabamba Mukadi |
| EPI_ISL_3261591 | Sharp HealthCare Laboratory | Andersen lab at Scripps Research | SEARCH Alliance San Diego with Aaron Harding, Jacquelyn Berumen, Cathy Woerle, Liam McGinnis, Art Mendoza, Omid Bakhtar |
| EPI_ISL_3261592, EPI_ISL_3261593, | Cliniques universitaires Saint-Luc | UCLouvain/IREC/MBLG | Jean Ruelle, Bertrand Bearzatto, Benoit Kabamba Mukadi |

| | | | |
|---|---|---|---|
| EPI_ISL_3261594 | | | |
| EPI_ISL_3261595 | Sharp HealthCare Laboratory | Andersen lab at Scripps Research | SEARCH Alliance San Diego with Aaron Harding, Jacquelyn Berumen, Cathy Woerle, Liam McGinnis, Art Mendoza, Omid Bakhtar |
| EPI_ISL_3261596 | Cliniques universitaires Saint-Luc | UCLouvain/IREC/MBLG | Jean Ruelle, Bertrand Bearzatto, Benoit Kabamba Mukadi |
| EPI_ISL_3261597 | Sharp HealthCare Laboratory | Andersen lab at Scripps Research | SEARCH Alliance San Diego with Aaron Harding, Jacquelyn Berumen, Cathy Woerle, Liam McGinnis, Art Mendoza, Omid Bakhtar |
| EPI_ISL_3261598, EPI_ISL_3261599 | Cliniques universitaires Saint-Luc | UCLouvain/IREC/MBLG | Jean Ruelle, Bertrand Bearzatto, Benoit Kabamba Mukadi |
| EPI_ISL_3261600 | Sharp HealthCare Laboratory | Andersen lab at Scripps Research | SEARCH Alliance San Diego with Aaron Harding, Jacquelyn Berumen, Cathy Woerle, Liam McGinnis, Art Mendoza, Omid Bakhtar |
| EPI_ISL_3261601, EPI_ISL_3261602 | Cliniques universitaires Saint-Luc | UCLouvain/IREC/MBLG | Jean Ruelle, Bertrand Bearzatto, Benoit Kabamba Mukadi |
| EPI_ISL_3261603 | Scripps Medical Laboratory | Andersen lab at Scripps Research | SEARCH Alliance San Diego with Michael Quigley, Ellen Stefanski, Ian Mchardy |
| EPI_ISL_3261604, EPI_ISL_3261605 | Cliniques universitaires Saint-Luc | UCLouvain/IREC/MBLG | Jean Ruelle, Bertrand Bearzatto, Benoit Kabamba Mukadi |
| EPI_ISL_3261606 | Scripps Medical Laboratory | Andersen lab at Scripps Research | SEARCH Alliance San Diego with Michael Quigley, Ellen Stefanski, Ian Mchardy |
| EPI_ISL_3261607 | Cliniques universitaires Saint-Luc | UCLouvain/IREC/MBLG | Jean Ruelle, Bertrand Bearzatto, Benoit Kabamba Mukadi |
| EPI_ISL_3261608 | Scripps Medical Laboratory | Andersen lab at Scripps Research | SEARCH Alliance San Diego with Michael Quigley, Ellen Stefanski, Ian Mchardy |
| EPI_ISL_3261609, EPI_ISL_3261610 | Cliniques universitaires Saint-Luc | UCLouvain/IREC/MBLG | Jean Ruelle, Bertrand Bearzatto, Benoit Kabamba Mukadi |
| EPI_ISL_3261611 | Scripps Medical Laboratory | Andersen lab at Scripps Research | SEARCH Alliance San Diego with Michael Quigley, Ellen Stefanski, Ian Mchardy |
| EPI_ISL_3261612, EPI_ISL_3261613 | Cliniques universitaires Saint-Luc | UCLouvain/IREC/MBLG | Jean Ruelle, Bertrand Bearzatto, Benoit Kabamba Mukadi |
| EPI_ISL_3261614 | Scripps Medical Laboratory | Andersen lab at Scripps Research | SEARCH Alliance San Diego with Michael Quigley, Ellen Stefanski, Ian Mchardy |
| EPI_ISL_3261615, EPI_ISL_3261616 | Cliniques universitaires Saint-Luc | UCLouvain/IREC/MBLG | Jean Ruelle, Bertrand Bearzatto, Benoit Kabamba Mukadi |
| EPI_ISL_3261617 | Scripps Medical Laboratory | Andersen lab at Scripps Research | SEARCH Alliance San Diego with Michael Quigley, Ellen Stefanski, Ian Mchardy |
| EPI_ISL_3261618, EPI_ISL_3261619 | Cliniques universitaires Saint-Luc | UCLouvain/IREC/MBLG | Jean Ruelle, Bertrand Bearzatto, Benoit Kabamba Mukadi |
| EPI_ISL_3261620 | Scripps Medical Laboratory | Andersen lab at Scripps Research | SEARCH Alliance San Diego with Michael Quigley, Ellen Stefanski, Ian Mchardy |
| EPI_ISL_3261621 | Cliniques universitaires Saint-Luc | UCLouvain/IREC/MBLG | Jean Ruelle, Bertrand Bearzatto, Benoit Kabamba Mukadi |
| EPI_ISL_3261622 | Scripps Medical Laboratory | Andersen lab at Scripps Research | SEARCH Alliance San Diego with Michael Quigley, Ellen Stefanski, Ian Mchardy |
| EPI_ISL_3261623, EPI_ISL_3261624 | Cliniques universitaires Saint-Luc | UCLouvain/IREC/MBLG | Jean Ruelle, Bertrand Bearzatto, Benoit Kabamba Mukadi |
| EPI_ISL_3261625 | Scripps Medical Laboratory | Andersen lab at Scripps Research | SEARCH Alliance San Diego with Michael Quigley, Ellen Stefanski, Ian Mchardy |
| EPI_ISL_3261626, EPI_ISL_3261627 | Cliniques universitaires Saint-Luc | UCLouvain/IREC/MBLG | Jean Ruelle, Bertrand Bearzatto, Benoit Kabamba Mukadi |
| EPI_ISL_3261628 | Scripps Medical Laboratory | Andersen lab at Scripps Research | SEARCH Alliance San Diego with Michael Quigley, Ellen Stefanski, Ian Mchardy |
| EPI_ISL_3261629 | Cliniques universitaires Saint-Luc | UCLouvain/IREC/MBLG | Jean Ruelle, Bertrand Bearzatto, Benoit Kabamba Mukadi |
| EPI_ISL_3261630 | Scripps Medical Laboratory | Andersen lab at Scripps Research | SEARCH Alliance San Diego with Michael Quigley, Ellen Stefanski, Ian Mchardy |
| EPI_ISL_3261631, EPI_ISL_3261632 | Cliniques universitaires Saint-Luc | UCLouvain/IREC/MBLG | Jean Ruelle, Bertrand Bearzatto, Benoit Kabamba Mukadi |
| EPI_ISL_3261633 | Scripps Medical Laboratory | Andersen lab at Scripps Research | SEARCH Alliance San Diego with Michael Quigley, Ellen Stefanski, Ian Mchardy |
| EPI_ISL_3261634, EPI_ISL_3261635 | Cliniques universitaires Saint-Luc | UCLouvain/IREC/MBLG | Jean Ruelle, Bertrand Bearzatto, Benoit Kabamba Mukadi |
| EPI_ISL_3261636 | Scripps Medical Laboratory | Andersen lab at Scripps Research | SEARCH Alliance San Diego with Michael Quigley, Ellen Stefanski, Ian Mchardy |
| EPI_ISL_3261637 | Cliniques universitaires Saint-Luc | UCLouvain/IREC/MBLG | Jean Ruelle, Bertrand Bearzatto, Benoit Kabamba Mukadi |
| EPI_ISL_3261638 | Scripps Medical Laboratory | Andersen lab at Scripps Research | SEARCH Alliance San Diego with Michael Quigley, Ellen Stefanski, Ian Mchardy |
| EPI_ISL_3261639, EPI_ISL_3261640 | Cliniques universitaires Saint-Luc | UCLouvain/IREC/MBLG | Jean Ruelle, Bertrand Bearzatto, Benoit Kabamba Mukadi |
| EPI_ISL_3261641 | Scripps Medical Laboratory | Andersen lab at Scripps Research | SEARCH Alliance San Diego with Michael Quigley, Ellen Stefanski, Ian Mchardy |
| EPI_ISL_3261642, EPI_ISL_3261643 | Cliniques universitaires Saint-Luc | UCLouvain/IREC/MBLG | Jean Ruelle, Bertrand Bearzatto, Benoit Kabamba Mukadi |
| EPI_ISL_3261644 | Scripps Medical Laboratory | Andersen lab at Scripps Research | SEARCH Alliance San Diego with Michael Quigley, Ellen Stefanski, Ian Mchardy |
| EPI_ISL_3261645, EPI_ISL_3261646 | Cliniques universitaires Saint-Luc | UCLouvain/IREC/MBLG | Jean Ruelle, Bertrand Bearzatto, Benoit Kabamba Mukadi |
| EPI_ISL_3261647, EPI_ISL_3261648, EPI_ISL_3261649, EPI_ISL_3261650, EPI_ISL_3261651, EPI_ISL_3261652, EPI_ISL_3261653, EPI_ISL_3261654, EPI_ISL_3261655, EPI_ISL_3261656, EPI_ISL_3261658, EPI_ISL_3261659, EPI_ISL_3261660, EPI_ISL_3261661, EPI_ISL_3261662, EPI_ISL_3261663, EPI_ISL_3261664, EPI_ISL_3261665, EPI_ISL_3261666, EPI_ISL_3261668, EPI_ISL_3261669, EPI_ISL_3261670, EPI_ISL_3261671, EPI_ISL_3261672, EPI_ISL_3261673, EPI_ISL_3261674, EPI_ISL_3261675, EPI_ISL_3261676, EPI_ISL_3261677, EPI_ISL_3261678, EPI_ISL_3261679, EPI_ISL_3261680, EPI_ISL_3261681, EPI_ISL_3261682, EPI_ISL_3261683, EPI_ISL_3261684, EPI_ISL_3261685, EPI_ISL_3261686, EPI_ISL_3261687, EPI_ISL_3261688, EPI_ISL_3261689, EPI_ISL_3261690, EPI_ISL_3261693, EPI_ISL_3261695, EPI_ISL_3261696, EPI_ISL_3261697, EPI_ISL_3261698, EPI_ISL_3261699, EPI_ISL_3261700, EPI_ISL_3261701, EPI_ISL_3261703, EPI_ISL_3261704, EPI_ISL_3261705, EPI_ISL_3261706, EPI_ISL_3261707, EPI_ISL_3261708, EPI_ISL_3261710, EPI_ISL_3261711, EPI_ISL_3261712, EPI_ISL_3261713, EPI_ISL_3261714, EPI_ISL_3261716, EPI_ISL_3261718, EPI_ISL_3261719, EPI_ISL_3261720, EPI_ISL_3261722, EPI_ISL_3261727, EPI_ISL_3261728, EPI_ISL_3261729, EPI_ISL_3261730, EPI_ISL_3261731, EPI_ISL_3261732, EPI_ISL_3261733, EPI_ISL_3261734, EPI_ISL_3261735, EPI_ISL_3261736, EPI_ISL_3261737, EPI_ISL_3261738, EPI_ISL_3261739, EPI_ISL_3261741, EPI_ISL_3261744, EPI_ISL_3261745 | | | |
| see above | Scripps Medical Laboratory | Andersen lab at Scripps Research | SEARCH Alliance San Diego with Michael Quigley, Ellen Stefanski, Ian Mchardy |
| EPI_ISL_3261854 | Microbiology laboratory, National Center for Global Health and Medicine Hospital | Pathogen Genomics Center, National Institute of Infectious Diseases | Tsuyoshi Sekizuka, Kentaro Itokawa, Rina Tanaka, Masanori Hashino, Makoto Kuroda, Harutaka Katano, Yuichiro Hirata, Shun Iida, Shinji Saito, Tadaki Suzuki, Norio Ohmagari |
| EPI_ISL_3261864, EPI_ISL_3261865, EPI_ISL_3261866, EPI_ISL_3261867, EPI_ISL_3261868, EPI_ISL_3261869, EPI_ISL_3261870, EPI_ISL_3261871, EPI_ISL_3261872, EPI_ISL_3261873, EPI_ISL_3261874, EPI_ISL_3261875, EPI_ISL_3261877, EPI_ISL_3261878, EPI_ISL_3261879, EPI_ISL_3261880, EPI_ISL_3261881, EPI_ISL_3261882, EPI_ISL_3261883, EPI_ISL_3261884, EPI_ISL_3261885, EPI_ISL_3261886, EPI_ISL_3261887, EPI_ISL_3261888, EPI_ISL_3261889, EPI_ISL_3261890, EPI_ISL_3261891, EPI_ISL_3261892, EPI_ISL_3261893, EPI_ISL_3261894, EPI_ISL_3261895, EPI_ISL_3261896, EPI_ISL_3261897, EPI_ISL_3261898, EPI_ISL_3261899, EPI_ISL_3261900, EPI_ISL_3261901, EPI_ISL_3261902, EPI_ISL_3261903, EPI_ISL_3261904, EPI_ISL_3261905 | | | |
| see above | NHLS_VIRO | KRISP, KZN Research Innovation and Sequencing Platform | Giandhari Jennifer, Pillay Sureshnee, Yajna Ramphal, Naidoo Yeshnee, Tshabuila Derek,Tegally Houriiyah, San James, Wilkinson Eduan, de Oliveira Tulio |
| EPI_ISL_3261906, EPI_ISL_3261907, EPI_ISL_3261908, EPI_ISL_3261909, EPI_ISL_3261910, EPI_ISL_3261911, EPI_ISL_3261912, EPI_ISL_3261913 | NHLS Port Elizabeth Laboratory | KRISP, KZN Research Innovation and Sequencing Platform | Giandhari Jennifer, Pillay Sureshnee, Yajna Ramphal, Naidoo Yeshnee, Tshabuila Derek,Tegally Houriiyah, San James, Wilkinson Eduan, de Oliveira Tulio |
| EPI_ISL_3261914, EPI_ISL_3261915, EPI_ISL_3261916, EPI_ISL_3261917, EPI_ISL_3261919, EPI_ISL_3261920, EPI_ISL_3261921, EPI_ISL_3261922, EPI_ISL_3261923 | NHLS_JHB (Ship 4) | KRISP, KZN Research Innovation and Sequencing Platform | Giandhari Jennifer, Pillay Sureshnee, Yajna Ramphal, Naidoo Yeshnee, Tshabuila Derek,Tegally Houriiyah, San James, Wilkinson Eduan, de Oliveira Tulio |
| EPI_ISL_3261924, EPI_ISL_3261925, EPI_ISL_3261926, EPI_ISL_3261927, EPI_ISL_3261928, EPI_ISL_3261929, EPI_ISL_3261930, EPI_ISL_3261931, EPI_ISL_3261932, EPI_ISL_3261933, EPI_ISL_3261934, EPI_ISL_3261935, EPI_ISL_3261936, EPI_ISL_3261938, EPI_ISL_3261939, EPI_ISL_3261940, EPI_ISL_3261941, EPI_ISL_3261942, EPI_ISL_3261943, EPI_ISL_3261944, EPI_ISL_3261945, EPI_ISL_3261946, EPI_ISL_3261947, EPI_ISL_3261948, EPI_ISL_3261949, EPI_ISL_3261950, EPI_ISL_3261951, EPI_ISL_3261952, EPI_ISL_3261953, EPI_ISL_3261954, EPI_ISL_3261955, EPI_ISL_3261956, EPI_ISL_3261958, EPI_ISL_3261959, EPI_ISL_3261960, EPI_ISL_3261961, EPI_ISL_3261962, EPI_ISL_3261963, EPI_ISL_3261964, EPI_ISL_3261965, EPI_ISL_3261966, EPI_ISL_3261967, EPI_ISL_3261968, EPI_ISL_3261969, EPI_ISL_3261971, EPI_ISL_3261972, EPI_ISL_3261973, EPI_ISL_3261974, EPI_ISL_3261975, EPI_ISL_3261976, EPI_ISL_3261977, EPI_ISL_3261978, EPI_ISL_3261979, EPI_ISL_3261980, EPI_ISL_3261981, EPI_ISL_3261982, EPI_ISL_3261983 | | | |
| see above | NHLS_VIRO | KRISP, KZN Research Innovation and Sequencing Platform | Giandhari Jennifer, Pillay Sureshnee, Yajna Ramphal, Naidoo Yeshnee, Tshabuila Derek,Tegally Houriiyah, San James, Wilkinson Eduan, de Oliveira Tulio |
| EPI_ISL_3261984, EPI_ISL_3261985, EPI_ISL_3261986, EPI_ISL_3261987, EPI_ISL_3261988, EPI_ISL_3261989, EPI_ISL_3261990, EPI_ISL_3261991, EPI_ISL_3261992, EPI_ISL_3261993, EPI_ISL_3261994, EPI_ISL_3261995, EPI_ISL_3261996, EPI_ISL_3261997, EPI_ISL_3261998, EPI_ISL_3261999, EPI_ISL_3262000, EPI_ISL_3262001, EPI_ISL_3262002, EPI_ISL_3262003, EPI_ISL_3262004, EPI_ISL_3262005, EPI_ISL_3262006, EPI_ISL_3262007, EPI_ISL_3262008, EPI_ISL_3262009, EPI_ISL_3262010, EPI_ISL_3262011, EPI_ISL_3262012, EPI_ISL_3262013, EPI_ISL_3262014, EPI_ISL_3262015, EPI_ISL_3262016, EPI_ISL_3262017, EPI_ISL_3262018, EPI_ISL_3262019, EPI_ISL_3262020, EPI_ISL_3262021, EPI_ISL_3262022, EPI_ISL_3262023, EPI_ISL_3262024, EPI_ISL_3262025, EPI_ISL_3262026, EPI_ISL_3262027, EPI_ISL_3262028, EPI_ISL_3262029, EPI_ISL_3262030, EPI_ISL_3262031, EPI_ISL_3262032, EPI_ISL_3262033, EPI_ISL_3262034, EPI_ISL_3262035, EPI_ISL_3262036, EPI_ISL_3262037, EPI_ISL_3262038, EPI_ISL_3262039, EPI_ISL_3262040, EPI_ISL_3262041, EPI_ISL_3262042, EPI_ISL_3262043, EPI_ISL_3262044, EPI_ISL_3262045, EPI_ISL_3262046, EPI_ISL_3262047, EPI_ISL_3262048, EPI_ISL_3262049, EPI_ISL_3262050, EPI_ISL_3262051, EPI_ISL_3262052, EPI_ISL_3262053, EPI_ISL_3262054, EPI_ISL_3262055, EPI_ISL_3262056, EPI_ISL_3262057, EPI_ISL_3262058, EPI_ISL_3262059, EPI_ISL_3262060, EPI_ISL_3262061, EPI_ISL_3262062, EPI_ISL_3262063, EPI_ISL_3262064, EPI_ISL_3262065, EPI_ISL_3262066, EPI_ISL_3262067, EPI_ISL_3262068, EPI_ISL_3262069, EPI_ISL_3262070, EPI_ISL_3262071, | | | |

| | | | |
|---|---|---|---|
| EPI_ISL_3262072, EPI_ISL_3262073, EPI_ISL_3262074, EPI_ISL_3262075, EPI_ISL_3262076, EPI_ISL_3262077, EPI_ISL_3262078, EPI_ISL_3262079, EPI_ISL_3262080, EPI_ISL_3262081, EPI_ISL_3262082, EPI_ISL_3262083, EPI_ISL_3262084, EPI_ISL_3262085, EPI_ISL_3262086, EPI_ISL_3262087, EPI_ISL_3262088, EPI_ISL_3262089, EPI_ISL_3262090, EPI_ISL_3262091, EPI_ISL_3262092, EPI_ISL_3262093, EPI_ISL_3262094, EPI_ISL_3262095, EPI_ISL_3262096, EPI_ISL_3262097, EPI_ISL_3262098, EPI_ISL_3262099, EPI_ISL_3262100, EPI_ISL_3262101, EPI_ISL_3262102, EPI_ISL_3262103, EPI_ISL_3262104, EPI_ISL_3262105, EPI_ISL_3262106, EPI_ISL_3262107, EPI_ISL_3262108, EPI_ISL_3262109, EPI_ISL_3262110, EPI_ISL_3262111, EPI_ISL_3262112, EPI_ISL_3262113, EPI_ISL_3262114, EPI_ISL_3262115, EPI_ISL_3262116, EPI_ISL_3262117 | | | |
| see above | National Public Health Laboratory, National Centre for Infectious Diseases | National Public Health Laboratory, National Centre for Infectious Diseases | Tze Minn Mak, Katherine Ching, Zhenyang Zhou, Royce Ang, Lin Cui, Raymond Tzer Pin Lin |
| EPI_ISL_3262119, EPI_ISL_3262120, EPI_ISL_3262121, EPI_ISL_3262123, EPI_ISL_3262126, EPI_ISL_3262128, EPI_ISL_3262129, EPI_ISL_3262133, EPI_ISL_3262136, EPI_ISL_3262138 | SI «Public Health Center of MHU» | The Institute of Molecular Biology and Genetics of NASU | M.Tukalo et al. |
| EPI_ISL_3262140, EPI_ISL_3262141, EPI_ISL_3262142, EPI_ISL_3262143, EPI_ISL_3262144, EPI_ISL_3262145, EPI_ISL_3262146, EPI_ISL_3262147, EPI_ISL_3262148, EPI_ISL_3262149, EPI_ISL_3262150, EPI_ISL_3262151, EPI_ISL_3262152, EPI_ISL_3262153, EPI_ISL_3262154, EPI_ISL_3262155, EPI_ISL_3262156, EPI_ISL_3262157, EPI_ISL_3262158, EPI_ISL_3262159, EPI_ISL_3262160, EPI_ISL_3262161, EPI_ISL_3262162, EPI_ISL_3262163, EPI_ISL_3262164, EPI_ISL_3262165, EPI_ISL_3262166, EPI_ISL_3262168, EPI_ISL_3262169, EPI_ISL_3262170, EPI_ISL_3262171, EPI_ISL_3262172, EPI_ISL_3262173, EPI_ISL_3262174, EPI_ISL_3262175, EPI_ISL_3262176, EPI_ISL_3262177, EPI_ISL_3262178, EPI_ISL_3262179, EPI_ISL_3262180, EPI_ISL_3262181, EPI_ISL_3262182, EPI_ISL_3262183, EPI_ISL_3262184, EPI_ISL_3262185, EPI_ISL_3262186, EPI_ISL_3262187, EPI_ISL_3262188, EPI_ISL_3262189, EPI_ISL_3262190, EPI_ISL_3262191, EPI_ISL_3262192, EPI_ISL_3262193, EPI_ISL_3262194, EPI_ISL_3262195, EPI_ISL_3262196, EPI_ISL_3262197, EPI_ISL_3262198, EPI_ISL_3262199, EPI_ISL_3262200, EPI_ISL_3262201, EPI_ISL_3262202, EPI_ISL_3262203, EPI_ISL_3262204, EPI_ISL_3262205, EPI_ISL_3262206, EPI_ISL_3262207, EPI_ISL_3262208, EPI_ISL_3262209 | | | |
| see above | UMC Groningen, Clinical Virology, Department of Medical Microbiology and Infection Prevention | UMC Groningen, Clinical Virology, Department of Medical Microbiology and Infection Prevention | Hubert Niesters, Alexander Friedrich, Erley Lizarazo-Forero, Monika Fliss, Lilli Gard, Sigrid Rosema, Coretta Van Leer-Buter, Xuewei Zhou, Marjolein Knoester |
| EPI_ISL_3262211, EPI_ISL_3262212, EPI_ISL_3262213, EPI_ISL_3262214, EPI_ISL_3262215, EPI_ISL_3262216, EPI_ISL_3262217, EPI_ISL_3262218, EPI_ISL_3262219, EPI_ISL_3262220, EPI_ISL_3262221, EPI_ISL_3262222, EPI_ISL_3262223, EPI_ISL_3262224, EPI_ISL_3262225, EPI_ISL_3262226, EPI_ISL_3262227, EPI_ISL_3262228, EPI_ISL_3262229, EPI_ISL_3262230, EPI_ISL_3262231, EPI_ISL_3262232, EPI_ISL_3262233, EPI_ISL_3262234, EPI_ISL_3262235, EPI_ISL_3262236, EPI_ISL_3262237, EPI_ISL_3262238, EPI_ISL_3262239, EPI_ISL_3262240, EPI_ISL_3262241, EPI_ISL_3262242, EPI_ISL_3262243, EPI_ISL_3262244, EPI_ISL_3262245, EPI_ISL_3262246, EPI_ISL_3262247, EPI_ISL_3262248, EPI_ISL_3262249, EPI_ISL_3262250, EPI_ISL_3262251, EPI_ISL_3262252, EPI_ISL_3262253, EPI_ISL_3262254, EPI_ISL_3262255, EPI_ISL_3262256, EPI_ISL_3262257, EPI_ISL_3262258, EPI_ISL_3262259, EPI_ISL_3262260, EPI_ISL_3262261, EPI_ISL_3262262, EPI_ISL_3262263, EPI_ISL_3262264, EPI_ISL_3262265, EPI_ISL_3262266, EPI_ISL_3262267, EPI_ISL_3262268, EPI_ISL_3262269, EPI_ISL_3262270, EPI_ISL_3262271, EPI_ISL_3262272, EPI_ISL_3262273, EPI_ISL_3262274, EPI_ISL_3262275, EPI_ISL_3262276, EPI_ISL_3262277, EPI_ISL_3262278, EPI_ISL_3262279, EPI_ISL_3262280, EPI_ISL_3262281, EPI_ISL_3262282, EPI_ISL_3262283, EPI_ISL_3262284, EPI_ISL_3262285, EPI_ISL_3262286, EPI_ISL_3262287, EPI_ISL_3262288, EPI_ISL_3262289, EPI_ISL_3262290, EPI_ISL_3262291, EPI_ISL_3262292, EPI_ISL_3262293, EPI_ISL_3262294, EPI_ISL_3262295, EPI_ISL_3262296, EPI_ISL_3262297, EPI_ISL_3262298, EPI_ISL_3262299, EPI_ISL_3262300, EPI_ISL_3262301, EPI_ISL_3262302, EPI_ISL_3262303, EPI_ISL_3262304, EPI_ISL_3262305, EPI_ISL_3262306, EPI_ISL_3262307, EPI_ISL_3262308, EPI_ISL_3262309, EPI_ISL_3262310, EPI_ISL_3262311, EPI_ISL_3262312, EPI_ISL_3262313, EPI_ISL_3262314, EPI_ISL_3262315, EPI_ISL_3262316, EPI_ISL_3262317, EPI_ISL_3262318, EPI_ISL_3262319, EPI_ISL_3262320, EPI_ISL_3262321, EPI_ISL_3262322, EPI_ISL_3262323, EPI_ISL_3262324, EPI_ISL_3262325, EPI_ISL_3262326, EPI_ISL_3262327, EPI_ISL_3262328, EPI_ISL_3262329, EPI_ISL_3262330, EPI_ISL_3262331, EPI_ISL_3262332, EPI_ISL_3262333, EPI_ISL_3262334, EPI_ISL_3262335, EPI_ISL_3262336, EPI_ISL_3262337, EPI_ISL_3262338, EPI_ISL_3262339, EPI_ISL_3262340, EPI_ISL_3262341, EPI_ISL_3262342, EPI_ISL_3262343, EPI_ISL_3262344, EPI_ISL_3262345, EPI_ISL_3262346, EPI_ISL_3262347, EPI_ISL_3262348, EPI_ISL_3262349, EPI_ISL_3262350, EPI_ISL_3262351, EPI_ISL_3262352, EPI_ISL_3262353, EPI_ISL_3262354, EPI_ISL_3262355, EPI_ISL_3262356, EPI_ISL_3262357, EPI_ISL_3262358, EPI_ISL_3262359, EPI_ISL_3262360, EPI_ISL_3262361, EPI_ISL_3262362, EPI_ISL_3262363, EPI_ISL_3262364, EPI_ISL_3262365, EPI_ISL_3262366, EPI_ISL_3262367, EPI_ISL_3262368, EPI_ISL_3262369, EPI_ISL_3262370, EPI_ISL_3262371, EPI_ISL_3262372, EPI_ISL_3262373, EPI_ISL_3262374, EPI_ISL_3262375, EPI_ISL_3262376, EPI_ISL_3262377, EPI_ISL_3262378, EPI_ISL_3262379, EPI_ISL_3262380, EPI_ISL_3262381, EPI_ISL_3262382, EPI_ISL_3262383, EPI_ISL_3262384, EPI_ISL_3262385, EPI_ISL_3262386, EPI_ISL_3262387, EPI_ISL_3262388, EPI_ISL_3262389, EPI_ISL_3262390, EPI_ISL_3262391, EPI_ISL_3262392, EPI_ISL_3262393, EPI_ISL_3262394, EPI_ISL_3262395, EPI_ISL_3262396, EPI_ISL_3262397, EPI_ISL_3262398, EPI_ISL_3262399, EPI_ISL_3262400, EPI_ISL_3262401, EPI_ISL_3262402, EPI_ISL_3262403, EPI_ISL_3262404, EPI_ISL_3262405, EPI_ISL_3262406, EPI_ISL_3262407, EPI_ISL_3262408, EPI_ISL_3262409, EPI_ISL_3262410, EPI_ISL_3262411, EPI_ISL_3262412, EPI_ISL_3262413, EPI_ISL_3262414, EPI_ISL_3262415, EPI_ISL_3262416, EPI_ISL_3262417, EPI_ISL_3262418, EPI_ISL_3262419, EPI_ISL_3262420, EPI_ISL_3262421 | | | |
| see above | University Hospitals of Geneva, Laboratory of Virology | HUG, Laboratory of Virology and the Health2030 Genome Center | Samuel Cordey, Aline Mamin, Francisco Perez, Ana Rita Goncalves, Laurent Kaiser, Lorenzo Cerutti, Cedric Howald, Henri Pegeot, Melyssa Elies, Deborah Penet, Keith Harshman, Ioannis Xenarios |
| EPI_ISL_3262633, EPI_ISL_3262634, EPI_ISL_3262635, EPI_ISL_3262636, EPI_ISL_3262637, EPI_ISL_3262638, EPI_ISL_3262639, EPI_ISL_3262640, EPI_ISL_3262641, EPI_ISL_3262642, EPI_ISL_3262643, EPI_ISL_3262644, EPI_ISL_3262645, EPI_ISL_3262646, EPI_ISL_3262647, EPI_ISL_3262648, EPI_ISL_3262649, EPI_ISL_3262650, EPI_ISL_3262651, EPI_ISL_3262652, EPI_ISL_3262653, EPI_ISL_3262654, EPI_ISL_3262655, EPI_ISL_3262657, EPI_ISL_3262658, EPI_ISL_3262659, EPI_ISL_3262660, EPI_ISL_3262661, EPI_ISL_3262662 | | | |
| see above | Kansas Health and Environmental Lab | Kansas Health and Environmental Lab | Mike Grose, Jonathan Barnell, Amanda Bradley, and Phil Adam |
| EPI_ISL_3262723, EPI_ISL_3262724, EPI_ISL_3262725, EPI_ISL_3262726, EPI_ISL_3262727, EPI_ISL_3262728, EPI_ISL_3262729, EPI_ISL_3262730, EPI_ISL_3262731, EPI_ISL_3262732, EPI_ISL_3262734, EPI_ISL_3262735, EPI_ISL_3262736, EPI_ISL_3262737, EPI_ISL_3262738, EPI_ISL_3262739, EPI_ISL_3262740, EPI_ISL_3262741, EPI_ISL_3262742, EPI_ISL_3262743, EPI_ISL_3262744, EPI_ISL_3262745, EPI_ISL_3262746, EPI_ISL_3262747, EPI_ISL_3262748, EPI_ISL_3262750, EPI_ISL_3262751, EPI_ISL_3262752, EPI_ISL_3262753, EPI_ISL_3262754, EPI_ISL_3262755, EPI_ISL_3262756, EPI_ISL_3262757, EPI_ISL_3262758, EPI_ISL_3262759, EPI_ISL_3262761, EPI_ISL_3262762, EPI_ISL_3262763, EPI_ISL_3262764, EPI_ISL_3262765, EPI_ISL_3262766, EPI_ISL_3262767, EPI_ISL_3262768, EPI_ISL_3262769, EPI_ISL_3262771, EPI_ISL_3262772, EPI_ISL_3262773, EPI_ISL_3262774, EPI_ISL_3262775, EPI_ISL_3262776, EPI_ISL_3262777, EPI_ISL_3262778, EPI_ISL_3262779, EPI_ISL_3262780, EPI_ISL_3262781, EPI_ISL_3262782, EPI_ISL_3262786, EPI_ISL_3262787 | | | |
| see above | Arizona State Public Health Laboratory | Arizona State Public Health Laboratory | Trung Huynh, Jessica Escobar, Katherine Fullerton, Nobuko Fukushima, Matthew Contursi, Stacy White, Linda Getsinger, Victor Waddell |
| EPI_ISL_3262788, EPI_ISL_3262789, EPI_ISL_3262790, EPI_ISL_3262791, EPI_ISL_3262792, EPI_ISL_3262793, EPI_ISL_3262794, EPI_ISL_3262795, EPI_ISL_3262796, EPI_ISL_3262797, EPI_ISL_3262798, EPI_ISL_3262799, EPI_ISL_3262800, EPI_ISL_3262801, EPI_ISL_3262802, EPI_ISL_3262804, EPI_ISL_3262805, EPI_ISL_3262806, EPI_ISL_3262807, EPI_ISL_3262808, EPI_ISL_3262810, EPI_ISL_3262811, EPI_ISL_3262812, EPI_ISL_3262813, EPI_ISL_3262814, EPI_ISL_3262815, EPI_ISL_3262816, EPI_ISL_3262818, EPI_ISL_3262819, EPI_ISL_3262820, EPI_ISL_3262821, EPI_ISL_3262822, EPI_ISL_3262823, EPI_ISL_3262824, EPI_ISL_3262825, EPI_ISL_3262826, EPI_ISL_3262827, EPI_ISL_3262828, EPI_ISL_3262829, EPI_ISL_3262830, EPI_ISL_3262831, EPI_ISL_3262832, EPI_ISL_3262833, EPI_ISL_3262834, EPI_ISL_3262835, EPI_ISL_3262836, EPI_ISL_3262837, EPI_ISL_3262838, EPI_ISL_3262839, EPI_ISL_3262840, EPI_ISL_3262841, EPI_ISL_3262842, EPI_ISL_3262843, EPI_ISL_3262844, EPI_ISL_3262845, EPI_ISL_3262846, EPI_ISL_3262847, EPI_ISL_3262848, EPI_ISL_3262849, EPI_ISL_3262850, EPI_ISL_3262851, EPI_ISL_3262852, EPI_ISL_3262853, EPI_ISL_3262854, EPI_ISL_3262855, EPI_ISL_3262856, EPI_ISL_3262857, EPI_ISL_3262858 | | | |
| see above | Oregon State Public Health Laboratory | Oregon State Public Health Laboratory | Rafia Razzaque, Eugene Yeboah, Vanda Makris, Laura Tsaknaridis, John Fontana and Shane Sevey |
| EPI_ISL_3262888, EPI_ISL_3262889, EPI_ISL_3262891, EPI_ISL_3262892, EPI_ISL_3262893, EPI_ISL_3262894, EPI_ISL_3262895, EPI_ISL_3262896, EPI_ISL_3262897, EPI_ISL_3262898, EPI_ISL_3262899, EPI_ISL_3262900, EPI_ISL_3262901, EPI_ISL_3262902, EPI_ISL_3262903, EPI_ISL_3262904, EPI_ISL_3262905, EPI_ISL_3262906, EPI_ISL_3262907, EPI_ISL_3262908, EPI_ISL_3262909, EPI_ISL_3262910, EPI_ISL_3262911, EPI_ISL_3262912, EPI_ISL_3262913, EPI_ISL_3262914, EPI_ISL_3262915, EPI_ISL_3262916 | | | |
| see above | State Hygienic Laboratory at the University of Iowa | State Hygienic Laboratory at the University of Iowa | Valerie Reeb, Erik Twait, Anna Yakos, Mariah Knutson, Kris Eveland, Kristen Zanon, Jeff, Benfer, Davis Rieckenberg, Wes Hottel, Alankar Kampoowale |
| EPI_ISL_3262920, EPI_ISL_3262921, EPI_ISL_3262922, EPI_ISL_3262923, EPI_ISL_3262924, EPI_ISL_3262925, EPI_ISL_3262926, EPI_ISL_3262927, EPI_ISL_3262928, EPI_ISL_3262929, EPI_ISL_3262930, EPI_ISL_3262931, EPI_ISL_3262932, EPI_ISL_3262933, EPI_ISL_3262934, EPI_ISL_3262935, EPI_ISL_3262936, EPI_ISL_3262937, EPI_ISL_3262938, EPI_ISL_3262939, EPI_ISL_3262940, EPI_ISL_3262941, EPI_ISL_3262942, EPI_ISL_3262943, EPI_ISL_3262944, EPI_ISL_3262945, EPI_ISL_3262946, EPI_ISL_3262947, EPI_ISL_3262948, EPI_ISL_3262949, EPI_ISL_3262950, EPI_ISL_3262951, EPI_ISL_3262952, EPI_ISL_3262953, EPI_ISL_3262954, EPI_ISL_3262955, EPI_ISL_3262956, EPI_ISL_3262957, EPI_ISL_3262958, EPI_ISL_3262959, EPI_ISL_3262960, EPI_ISL_3262961, EPI_ISL_3262962, EPI_ISL_3262963, EPI_ISL_3262964, EPI_ISL_3262965, EPI_ISL_3262966, EPI_ISL_3262967, EPI_ISL_3262968, EPI_ISL_3262969, EPI_ISL_3262970, EPI_ISL_3262971, EPI_ISL_3262972, EPI_ISL_3262973, EPI_ISL_3262974, EPI_ISL_3262975, EPI_ISL_3262976, EPI_ISL_3262977, EPI_ISL_3262978, EPI_ISL_3262979, EPI_ISL_3262980, EPI_ISL_3262981, EPI_ISL_3262982, EPI_ISL_3262983, EPI_ISL_3262984, EPI_ISL_3262985, EPI_ISL_3262986, EPI_ISL_3262987, EPI_ISL_3262988, EPI_ISL_3262989, EPI_ISL_3262990, EPI_ISL_3262991, EPI_ISL_3262992, EPI_ISL_3262993, EPI_ISL_3262994, EPI_ISL_3262995, EPI_ISL_3262996, EPI_ISL_3262997, EPI_ISL_3262998, EPI_ISL_3262999, EPI_ISL_3263001, EPI_ISL_3263002, EPI_ISL_3263003, EPI_ISL_3263004, EPI_ISL_3263005, EPI_ISL_3263006, EPI_ISL_3263007, EPI_ISL_3263008, EPI_ISL_3263009, EPI_ISL_3263010, EPI_ISL_3263011, EPI_ISL_3263012, EPI_ISL_3263013, EPI_ISL_3263014, EPI_ISL_3263015, EPI_ISL_3263016, EPI_ISL_3263017, EPI_ISL_3263018, EPI_ISL_3263019, EPI_ISL_3263020, EPI_ISL_3263021, EPI_ISL_3263022, EPI_ISL_3263023, EPI_ISL_3263024, EPI_ISL_3263025, EPI_ISL_3263026, EPI_ISL_3263027, EPI_ISL_3263029, EPI_ISL_3263030, EPI_ISL_3263031, EPI_ISL_3263032, EPI_ISL_3263033, EPI_ISL_3263034, EPI_ISL_3263035, EPI_ISL_3263036, EPI_ISL_3263038, EPI_ISL_3263039, EPI_ISL_3263040, EPI_ISL_3263041, EPI_ISL_3263042, EPI_ISL_3263043, EPI_ISL_3263045, EPI_ISL_3263046, EPI_ISL_3263050, EPI_ISL_3263053, EPI_ISL_3263054, EPI_ISL_3263055, EPI_ISL_3263056, EPI_ISL_3263057, EPI_ISL_3263058, EPI_ISL_3263059, EPI_ISL_3263060, EPI_ISL_3263061, EPI_ISL_3263062, EPI_ISL_3263063, EPI_ISL_3263065, EPI_ISL_3263066, EPI_ISL_3263067, EPI_ISL_3263068, EPI_ISL_3263069, EPI_ISL_3263070, EPI_ISL_3263071, EPI_ISL_3263072, EPI_ISL_3263073, EPI_ISL_3263074, EPI_ISL_3263075, EPI_ISL_3263076, EPI_ISL_3263077, EPI_ISL_3263078, EPI_ISL_3263079, EPI_ISL_3263080, EPI_ISL_3263081, EPI_ISL_3263082, EPI_ISL_3263083, EPI_ISL_3263084, EPI_ISL_3263085, EPI_ISL_3263086, EPI_ISL_3263087, EPI_ISL_3263088, EPI_ISL_3263089, EPI_ISL_3263090, EPI_ISL_3263092, EPI_ISL_3263093, EPI_ISL_3263094, EPI_ISL_3263095, EPI_ISL_3263096, EPI_ISL_3263098, EPI_ISL_3263099, EPI_ISL_3263100, EPI_ISL_3263102, EPI_ISL_3263103, EPI_ISL_3263104, EPI_ISL_3263105, EPI_ISL_3263106, EPI_ISL_3263107, EPI_ISL_3263108, EPI_ISL_3263109, EPI_ISL_3263110, EPI_ISL_3263111, EPI_ISL_3263112, EPI_ISL_3263114, EPI_ISL_3263115, EPI_ISL_3263116, EPI_ISL_3263117, EPI_ISL_3263118, EPI_ISL_3263119, EPI_ISL_3263123, EPI_ISL_3263124, EPI_ISL_3263125, EPI_ISL_3263126, EPI_ISL_3263127, EPI_ISL_3263128, EPI_ISL_3263132, EPI_ISL_3263136, EPI_ISL_3263139, EPI_ISL_3263140, EPI_ISL_3263141, EPI_ISL_3263142, EPI_ISL_3263145, EPI_ISL_3263148, EPI_ISL_3263149, EPI_ISL_3263150, EPI_ISL_3263152, EPI_ISL_3263153, EPI_ISL_3263157, EPI_ISL_3263158, EPI_ISL_3263159, EPI_ISL_3263162, EPI_ISL_3263165, EPI_ISL_3263166, EPI_ISL_3263168, EPI_ISL_3263170, EPI_ISL_3263172, EPI_ISL_3263173, EPI_ISL_3263174, EPI_ISL_3263177, EPI_ISL_3263178, EPI_ISL_3263179, EPI_ISL_3263180, EPI_ISL_3263181, EPI_ISL_3263183, EPI_ISL_3263184, EPI_ISL_3263185, EPI_ISL_3263186, EPI_ISL_3263187, EPI_ISL_3263188, EPI_ISL_3263189, EPI_ISL_3263190, EPI_ISL_3263193, EPI_ISL_3263194, EPI_ISL_3263195, EPI_ISL_3263198, EPI_ISL_3263199, EPI_ISL_3263201, EPI_ISL_3263202, EPI_ISL_3263206, EPI_ISL_3263208, EPI_ISL_3263211, EPI_ISL_3263214, EPI_ISL_3263215, EPI_ISL_3263216, EPI_ISL_3263217, EPI_ISL_3263219, EPI_ISL_3263221, EPI_ISL_3263222, EPI_ISL_3263225, EPI_ISL_3263228, EPI_ISL_3263231, EPI_ISL_3263234, EPI_ISL_3263237, EPI_ISL_3263239, EPI_ISL_3263242, EPI_ISL_3263243, EPI_ISL_3263244, EPI_ISL_3263245, EPI_ISL_3263246, EPI_ISL_3263247, EPI_ISL_3263249, EPI_ISL_3263250, EPI_ISL_3263254, EPI_ISL_3263255, EPI_ISL_3263256, EPI_ISL_3263257, EPI_ISL_3263260, EPI_ISL_3263262, EPI_ISL_3263263, EPI_ISL_3263265, EPI_ISL_3263266, EPI_ISL_3263267, EPI_ISL_3263269, EPI_ISL_3263270, EPI_ISL_3263271, EPI_ISL_3263272, EPI_ISL_3263273, EPI_ISL_3263274, EPI_ISL_3263275, EPI_ISL_3263276, EPI_ISL_3263279, EPI_ISL_3263280, EPI_ISL_3263281, EPI_ISL_3263283, EPI_ISL_3263285, EPI_ISL_3263286, EPI_ISL_3263287, EPI_ISL_3263288, EPI_ISL_3263289, EPI_ISL_3263291, EPI_ISL_3263292, EPI_ISL_3263293, EPI_ISL_3263294, EPI_ISL_3263296, EPI_ISL_3263297, EPI_ISL_3263298, EPI_ISL_3263299, EPI_ISL_3263300, EPI_ISL_3263301, EPI_ISL_3263302, EPI_ISL_3263303, EPI_ISL_3263304, EPI_ISL_3263305, EPI_ISL_3263306, EPI_ISL_3263307, EPI_ISL_3263308, EPI_ISL_3263309, EPI_ISL_3263310, EPI_ISL_3263311, EPI_ISL_3263312, EPI_ISL_3263313, EPI_ISL_3263314, EPI_ISL_3263315, EPI_ISL_3263316, EPI_ISL_3263317, EPI_ISL_3263319, EPI_ISL_3263320, EPI_ISL_3263322, EPI_ISL_3263323, EPI_ISL_3263324, EPI_ISL_3263325, EPI_ISL_3263326, EPI_ISL_3263327, EPI_ISL_3263328, EPI_ISL_3263329, EPI_ISL_3263330, EPI_ISL_3263331, EPI_ISL_3263332, EPI_ISL_3263333, EPI_ISL_3263335, EPI_ISL_3263336, EPI_ISL_3263337, EPI_ISL_3263338, EPI_ISL_3263340, EPI_ISL_3263341, EPI_ISL_3263342, EPI_ISL_3263343, EPI_ISL_3263345, EPI_ISL_3263346, EPI_ISL_3263347, EPI_ISL_3263348, EPI_ISL_3263349, EPI_ISL_3263350, EPI_ISL_3263351, EPI_ISL_3263352, EPI_ISL_3263353, EPI_ISL_3263354, EPI_ISL_3263355, EPI_ISL_3263356, EPI_ISL_3263358, EPI_ISL_3263359, EPI_ISL_3263361, EPI_ISL_3263362, EPI_ISL_3263363, EPI_ISL_3263364, EPI_ISL_3263365, EPI_ISL_3263367, EPI_ISL_3263368, EPI_ISL_3263369, EPI_ISL_3263370, EPI_ISL_3263371, EPI_ISL_3263372, EPI_ISL_3263373, EPI_ISL_3263375, EPI_ISL_3263376, EPI_ISL_3263378, EPI_ISL_3263379, EPI_ISL_3263380, EPI_ISL_3263382, EPI_ISL_3263383, EPI_ISL_3263384, EPI_ISL_3263385, EPI_ISL_3263386, EPI_ISL_3263387, EPI_ISL_3263389, EPI_ISL_3263390, EPI_ISL_3263391, EPI_ISL_3263393, EPI_ISL_3263395, EPI_ISL_3263396, EPI_ISL_3263398, EPI_ISL_3263399, EPI_ISL_3263400, EPI_ISL_3263401, EPI_ISL_3263402, EPI_ISL_3263403, EPI_ISL_3263404, EPI_ISL_3263405, EPI_ISL_3263406, EPI_ISL_3263407, EPI_ISL_3263409, EPI_ISL_3263410, EPI_ISL_3263411, EPI_ISL_3263412, EPI_ISL_3263413, EPI_ISL_3263414, EPI_ISL_3263417, EPI_ISL_3263419, EPI_ISL_3263420, EPI_ISL_3263421, EPI_ISL_3263422, EPI_ISL_3263423, EPI_ISL_3263424, EPI_ISL_3263425, EPI_ISL_3263426, EPI_ISL_3263428, EPI_ISL_3263429, EPI_ISL_3263431, EPI_ISL_3263432, EPI_ISL_3263434, EPI_ISL_3263435, EPI_ISL_3263436, EPI_ISL_3263437, EPI_ISL_3263438, EPI_ISL_3263440, EPI_ISL_3263441, EPI_ISL_3263443, EPI_ISL_3263444, EPI_ISL_3263445, EPI_ISL_3263446, EPI_ISL_3263447, EPI_ISL_3263448, EPI_ISL_3263449, EPI_ISL_3263451, EPI_ISL_3263454, EPI_ISL_3263455, EPI_ISL_3263456, EPI_ISL_3263458, EPI_ISL_3263460, EPI_ISL_3263461, EPI_ISL_3263462, EPI_ISL_3263463, EPI_ISL_3263464, EPI_ISL_3263466, EPI_ISL_3263467, EPI_ISL_3263469, EPI_ISL_3263470, EPI_ISL_3263472, EPI_ISL_3263473, EPI_ISL_3263474, EPI_ISL_3263475, EPI_ISL_3263476, EPI_ISL_3263479, EPI_ISL_3263480, EPI_ISL_3263481, EPI_ISL_3263482, EPI_ISL_3263483, EPI_ISL_3263484, EPI_ISL_3263485, EPI_ISL_3263486, EPI_ISL_3263488, EPI_ISL_3263489, EPI_ISL_3263490, EPI_ISL_3263492, EPI_ISL_3263493, EPI_ISL_3263494, EPI_ISL_3263495, EPI_ISL_3263496, EPI_ISL_3263497, EPI_ISL_3263498, EPI_ISL_3263500, EPI_ISL_3263501, EPI_ISL_3263502, EPI_ISL_3263503, EPI_ISL_3263504, EPI_ISL_3263505, EPI_ISL_3263506, EPI_ISL_3263508, EPI_ISL_3263509, EPI_ISL_3263511, EPI_ISL_3263512, EPI_ISL_3263513, EPI_ISL_3263514, EPI_ISL_3263515, EPI_ISL_3263516, EPI_ISL_3263517, EPI_ISL_3263518, EPI_ISL_3263519, EPI_ISL_3263521, EPI_ISL_3263523, EPI_ISL_3263524, EPI_ISL_3263525, EPI_ISL_3263527, EPI_ISL_3263528, EPI_ISL_3263529, EPI_ISL_3263530, EPI_ISL_3263531, EPI_ISL_3263532, EPI_ISL_3263533, EPI_ISL_3263534, EPI_ISL_3263535, EPI_ISL_3263537, EPI_ISL_3263538, EPI_ISL_3263539, EPI_ISL_3263541, EPI_ISL_3263542, EPI_ISL_3263543, EPI_ISL_3263544, EPI_ISL_3263546, EPI_ISL_3263551, EPI_ISL_3263553, EPI_ISL_3263555, EPI_ISL_3263557, EPI_ISL_3263558, EPI_ISL_3263559, EPI_ISL_3263560, EPI_ISL_3263561, EPI_ISL_3263562, EPI_ISL_3263563, EPI_ISL_3263564, EPI_ISL_3263565, EPI_ISL_3263566, EPI_ISL_3263567, EPI_ISL_3263568, EPI_ISL_3263569, EPI_ISL_3263571, EPI_ISL_3263572, EPI_ISL_3263573, EPI_ISL_3263574, EPI_ISL_3263575, EPI_ISL_3263576, EPI_ISL_3263577, EPI_ISL_3263578, EPI_ISL_3263579, EPI_ISL_3263580, EPI_ISL_3263581, EPI_ISL_3263582, EPI_ISL_3263583, EPI_ISL_3263584, EPI_ISL_3263585, EPI_ISL_3263587, EPI_ISL_3263588, EPI_ISL_3263589, EPI_ISL_3263590, EPI_ISL_3263591, EPI_ISL_3263592, EPI_ISL_3263593, EPI_ISL_3263594, EPI_ISL_3263595, EPI_ISL_3263596, EPI_ISL_3263597, EPI_ISL_3263598, EPI_ISL_3263599, EPI_ISL_3263600, EPI_ISL_3263602, EPI_ISL_3263603, EPI_ISL_3263604, EPI_ISL_3263605, EPI_ISL_3263609, EPI_ISL_3263611, EPI_ISL_3263612, EPI_ISL_3263613, EPI_ISL_3263614, EPI_ISL_3263615, EPI_ISL_3263616, EPI_ISL_3263617, EPI_ISL_3263618, EPI_ISL_3263619, EPI_ISL_3263622, EPI_ISL_3263624, EPI_ISL_3263625, EPI_ISL_3263626, EPI_ISL_3263629, EPI_ISL_3263630, EPI_ISL_3263631, EPI_ISL_3263632, EPI_ISL_3263633, EPI_ISL_3263635, EPI_ISL_3263636, EPI_ISL_3263638, EPI_ISL_3263639, EPI_ISL_3263640, EPI_ISL_3263642, EPI_ISL_3263644, EPI_ISL_3263645, EPI_ISL_3263646, EPI_ISL_3263647, EPI_ISL_3263648, EPI_ISL_3263649, EPI_ISL_3263650, EPI_ISL_3263651, EPI_ISL_3263652, EPI_ISL_3263653, EPI_ISL_3263654, EPI_ISL_3263655, EPI_ISL_3263656, EPI_ISL_3263658, EPI_ISL_3263662, EPI_ISL_3263663, EPI_ISL_3263664, EPI_ISL_3263665, EPI_ISL_3263666, EPI_ISL_3263667, EPI_ISL_3263668, EPI_ISL_3263669, EPI_ISL_3263670, EPI_ISL_3263671, EPI_ISL_3263672, EPI_ISL_3263674, EPI_ISL_3263675, EPI_ISL_3263677, EPI_ISL_3263679, EPI_ISL_3263680, EPI_ISL_3263681, EPI_ISL_3263685, EPI_ISL_3263686, EPI_ISL_3263688, EPI_ISL_3263689, EPI_ISL_3263690, EPI_ISL_3263691, EPI_ISL_3263692, EPI_ISL_3263693, EPI_ISL_3263696, EPI_ISL_3263697, EPI_ISL_3263699, EPI_ISL_3263700, EPI_ISL_3263701, EPI_ISL_3263702, EPI_ISL_3263703, EPI_ISL_3263704, EPI_ISL_3263705, EPI_ISL_3263706, EPI_ISL_3263707, EPI_ISL_3263708, EPI_ISL_3263712, EPI_ISL_3263713, EPI_ISL_3263714, EPI_ISL_3263716, EPI_ISL_3263717, EPI_ISL_3263718, EPI_ISL_3263719, EPI_ISL_3263720, EPI_ISL_3263721, EPI_ISL_3263722, EPI_ISL_3263724, EPI_ISL_3263725, EPI_ISL_3263726, EPI_ISL_3263727, EPI_ISL_3263728, EPI_ISL_3263729, EPI_ISL_3263730, EPI_ISL_3263731, EPI_ISL_3263733, EPI_ISL_3263734, EPI_ISL_3263735, EPI_ISL_3263736, EPI_ISL_3263737, EPI_ISL_3263738, EPI_ISL_3263739, EPI_ISL_3263740, EPI_ISL_3263741, EPI_ISL_3263742, EPI_ISL_3263743, EPI_ISL_3263744, EPI_ISL_3263745, EPI_ISL_3263746, EPI_ISL_3263747, EPI_ISL_3263748, EPI_ISL_3263749, EPI_ISL_3263750, EPI_ISL_3263751, EPI_ISL_3263752, EPI_ISL_3263753, EPI_ISL_3263754, EPI_ISL_3263755, EPI_ISL_3263757, EPI_ISL_3263758, EPI_ISL_3263759, EPI_ISL_3263760, EPI_ISL_3263761, EPI_ISL_3263762, EPI_ISL_3263763, EPI_ISL_3263764, EPI_ISL_3263765, EPI_ISL_3263766, EPI_ISL_3263767, EPI_ISL_3263768, EPI_ISL_3263769, EPI_ISL_3263770, EPI_ISL_3263771, EPI_ISL_3263772, EPI_ISL_3263773, EPI_ISL_3263774, EPI_ISL_3263775, EPI_ISL_3263776, EPI_ISL_3263777, EPI_ISL_3263778, EPI_ISL_3263779, EPI_ISL_3263780, EPI_ISL_3263781, EPI_ISL_3263782, EPI_ISL_3263783, EPI_ISL_3263784, EPI_ISL_3263786, EPI_ISL_3263788, EPI_ISL_3263789, EPI_ISL_3263790, EPI_ISL_3263791, EPI_ISL_3263793, EPI_ISL_3263795, EPI_ISL_3263796, EPI_ISL_3263797, EPI_ISL_3263798, EPI_ISL_3263799, EPI_ISL_3263800, EPI_ISL_3263802, EPI_ISL_3263803, EPI_ISL_3263804, EPI_ISL_3263806, EPI_ISL_3263807, EPI_ISL_3263808, EPI_ISL_3263809, EPI_ISL_3263810, EPI_ISL_3263812, EPI_ISL_3263813, EPI_ISL_3263815, EPI_ISL_3263816, EPI_ISL_3263818, EPI_ISL_3263819, EPI_ISL_3263820, EPI_ISL_3263821, EPI_ISL_3263822, EPI_ISL_3263823, EPI_ISL_3263824, EPI_ISL_3263825, EPI_ISL_3263826, EPI_ISL_3263827, EPI_ISL_3263828, EPI_ISL_3263829, EPI_ISL_3263830, EPI_ISL_3263831, EPI_ISL_3263832, EPI_ISL_3263833, EPI_ISL_3263834, EPI_ISL_3263835, EPI_ISL_3263836, EPI_ISL_3263837, EPI_ISL_3263838, EPI_ISL_3263839, EPI_ISL_3263840, EPI_ISL_3263841, EPI_ISL_3263842, EPI_ISL_3263843, EPI_ISL_3263845, EPI_ISL_3263846, EPI_ISL_3263847, EPI_ISL_3263848, EPI_ISL_3263849, EPI_ISL_3263850, EPI_ISL_3263852, EPI_ISL_3263853, EPI_ISL_3263854, EPI_ISL_3263855, EPI_ISL_3263856, EPI_ISL_3263857, EPI_ISL_3263858, EPI_ISL_3263859, EPI_ISL_3263860, EPI_ISL_3263862, EPI_ISL_3263863, EPI_ISL_3263864, EPI_ISL_3263866, EPI_ISL_3263867, EPI_ISL_3263868, EPI_ISL_3263869, EPI_ISL_3263870, EPI_ISL_3263871, EPI_ISL_3263872, EPI_ISL_3263873, EPI_ISL_3263874, EPI_ISL_3263875, EPI_ISL_3263876, EPI_ISL_3263878, EPI_ISL_3263879, EPI_ISL_3263880, EPI_ISL_3263881, EPI_ISL_3263882, EPI_ISL_3263883, EPI_ISL_3263884, EPI_ISL_3263885, EPI_ISL_3263886, EPI_ISL_3263887, EPI_ISL_3263888, EPI_ISL_3263889, EPI_ISL_3263890, EPI_ISL_3263891, EPI_ISL_3263893, EPI_ISL_3263894, EPI_ISL_3263895, EPI_ISL_3263897, EPI_ISL_3263898, EPI_ISL_3263899, EPI_ISL_3263900, EPI_ISL_3263901, EPI_ISL_3263902, EPI_ISL_3263903, EPI_ISL_3263904, EPI_ISL_3263905, EPI_ISL_3263906, EPI_ISL_3263907, EPI_ISL_3263908, EPI_ISL_3263909, EPI_ISL_3263911, EPI_ISL_3263912, EPI_ISL_3263914, EPI_ISL_3263916, EPI_ISL_3263917, EPI_ISL_3263919, EPI_ISL_3263920, EPI_ISL_3263922, EPI_ISL_3263923, EPI_ISL_3263924, EPI_ISL_3263925, EPI_ISL_3263926, EPI_ISL_3263929, EPI_ISL_3263930, EPI_ISL_3263931, EPI_ISL_3263932, EPI_ISL_3263933, EPI_ISL_3263937 | | | |
| see above | Utah Public Health Laboratory | Utah Public Health Laboratory | Erin L. Young, Kelly F. Oakeson, Olinto Linares-Perdomo |
| EPI_ISL_3264957, EPI_ISL_3264958, EPI_ISL_3264959, EPI_ISL_3264961, EPI_ISL_3264964, EPI_ISL_3264965, EPI_ISL_3264966, EPI_ISL_3264967, EPI_ISL_3264968, EPI_ISL_3264969, EPI_ISL_3264970, EPI_ISL_3264971, EPI_ISL_3264972, EPI_ISL_3264974, EPI_ISL_3264975, EPI_ISL_3264977, EPI_ISL_3264978, EPI_ISL_3264979, EPI_ISL_3264980, EPI_ISL_3264981, EPI_ISL_3264982, EPI_ISL_3264983, EPI_ISL_3264984, EPI_ISL_3264985, EPI_ISL_3264986, EPI_ISL_3264987, EPI_ISL_3264988, EPI_ISL_3264989, EPI_ISL_3264990, EPI_ISL_3264991, EPI_ISL_3264992, EPI_ISL_3264993, EPI_ISL_3264994, EPI_ISL_3264995, EPI_ISL_3264996, EPI_ISL_3264997, EPI_ISL_3264998, EPI_ISL_3264999, EPI_ISL_3265000, EPI_ISL_3265002, EPI_ISL_3265003, EPI_ISL_3265004, EPI_ISL_3265005, EPI_ISL_3265006, EPI_ISL_3265007, EPI_ISL_3265008, EPI_ISL_3265009, EPI_ISL_3265010, EPI_ISL_3265011, EPI_ISL_3265012, EPI_ISL_3265013, EPI_ISL_3265014, EPI_ISL_3265015, EPI_ISL_3265016, EPI_ISL_3265017, EPI_ISL_3265018, EPI_ISL_3265019, EPI_ISL_3265020, EPI_ISL_3265021, EPI_ISL_3265022, EPI_ISL_3265023, EPI_ISL_3265024, EPI_ISL_3265025, EPI_ISL_3265026, EPI_ISL_3265027, EPI_ISL_3265029, EPI_ISL_3265030, EPI_ISL_3265031, EPI_ISL_3265032, EPI_ISL_3265033, EPI_ISL_3265034, EPI_ISL_3265035, EPI_ISL_3265036, EPI_ISL_3265037, EPI_ISL_3265038, EPI_ISL_3265039, EPI_ISL_3265040, EPI_ISL_3265041, EPI_ISL_3265042 | | | |
| see above | JIPMER Puducherry | inStem NCBS – INSACOG | Uma Ramakrishnan Dasaradhi Palakodeti Aswin SaiNarain |
| EPI_ISL_3265043, EPI_ISL_3265044, EPI_ISL_3265045, EPI_ISL_3265046, EPI_ISL_3265047, EPI_ISL_3265048, EPI_ISL_3265049 | Dr. Madhusudhan, Wenlock District Hospital | inStem NCBS – INSACOG | Uma Ramakrishnan Dasaradhi Palakodeti Aswin SaiNarain |
| EPI_ISL_3265050, EPI_ISL_3265051, EPI_ISL_3265052, EPI_ISL_3265053, EPI_ISL_3265054, EPI_ISL_3265055, EPI_ISL_3265056, EPI_ISL_3265057, EPI_ISL_3265058, EPI_ISL_3265059, EPI_ISL_3265060, EPI_ISL_3265061, EPI_ISL_3265062, EPI_ISL_3265063, EPI_ISL_3265064, EPI_ISL_3265065, EPI_ISL_3265066, EPI_ISL_3265067, EPI_ISL_3265068, EPI_ISL_3265069, EPI_ISL_3265070, EPI_ISL_3265071, EPI_ISL_3265072, EPI_ISL_3265073, EPI_ISL_3265074, EPI_ISL_3265075, EPI_ISL_3265076, EPI_ISL_3265078, EPI_ISL_3265079, EPI_ISL_3265080, EPI_ISL_3265081, EPI_ISL_3265082, EPI_ISL_3265083, EPI_ISL_3265084, EPI_ISL_3265085, EPI_ISL_3265087, EPI_ISL_3265088, EPI_ISL_3265089, EPI_ISL_3265090, EPI_ISL_3265091, EPI_ISL_3265092, EPI_ISL_3265093, EPI_ISL_3265094, EPI_ISL_3265095, EPI_ISL_3265096, EPI_ISL_3265097, EPI_ISL_3265098, EPI_ISL_3265099, EPI_ISL_3265100, EPI_ISL_3265101, EPI_ISL_3265102, EPI_ISL_3265103, EPI_ISL_3265104, EPI_ISL_3265105, EPI_ISL_3265106, EPI_ISL_3265107, EPI_ISL_3265108, EPI_ISL_3265109, EPI_ISL_3265110, EPI_ISL_3265111, EPI_ISL_3265112, EPI_ISL_3265113, EPI_ISL_3265114, EPI_ISL_3265116, EPI_ISL_3265117, EPI_ISL_3265118, | | | |

| | | | |
|---|---|---|---|
| EPI_ISL_3265119, EPI_ISL_3265120, EPI_ISL_3265121, EPI_ISL_3265122, EPI_ISL_3265123, EPI_ISL_3265124, EPI_ISL_3265125, EPI_ISL_3265126, EPI_ISL_3265127, EPI_ISL_3265128, EPI_ISL_3265129, EPI_ISL_3265130, EPI_ISL_3265132, EPI_ISL_3265133, EPI_ISL_3265134, EPI_ISL_3265135, EPI_ISL_3265136, EPI_ISL_3265137, EPI_ISL_3265138, EPI_ISL_3265139, EPI_ISL_3265140, EPI_ISL_3265141, EPI_ISL_3265142, EPI_ISL_3265143, EPI_ISL_3265144, EPI_ISL_3265145, EPI_ISL_3265146, EPI_ISL_3265147, EPI_ISL_3265148, EPI_ISL_3265149, EPI_ISL_3265150, EPI_ISL_3265151, EPI_ISL_3265152, EPI_ISL_3265153, EPI_ISL_3265154, EPI_ISL_3265155, EPI_ISL_3265156, EPI_ISL_3265157, EPI_ISL_3265158, EPI_ISL_3265159, EPI_ISL_3265160, EPI_ISL_3265161, EPI_ISL_3265162, EPI_ISL_3265163, EPI_ISL_3265164, EPI_ISL_3265165, EPI_ISL_3265166, EPI_ISL_3265167, EPI_ISL_3265168, EPI_ISL_3265169, EPI_ISL_3265170, EPI_ISL_3265172, EPI_ISL_3265173, EPI_ISL_3265175, EPI_ISL_3265176, EPI_ISL_3265177, EPI_ISL_3265178, EPI_ISL_3265179, EPI_ISL_3265180, EPI_ISL_3265181, EPI_ISL_3265182, EPI_ISL_3265183, EPI_ISL_3265184, EPI_ISL_3265185, EPI_ISL_3265186, EPI_ISL_3265187, EPI_ISL_3265188, EPI_ISL_3265189, EPI_ISL_3265190, EPI_ISL_3265191, EPI_ISL_3265192, EPI_ISL_3265193, EPI_ISL_3265194, EPI_ISL_3265195, EPI_ISL_3265196, EPI_ISL_3265197, EPI_ISL_3265198, EPI_ISL_3265199, EPI_ISL_3265200, EPI_ISL_3265201, EPI_ISL_3265202, EPI_ISL_3265203, EPI_ISL_3265204, EPI_ISL_3265205, EPI_ISL_3265206, EPI_ISL_3265207, EPI_ISL_3265208, EPI_ISL_3265209, EPI_ISL_3265210, EPI_ISL_3265211, EPI_ISL_3265212, EPI_ISL_3265213, EPI_ISL_3265215, EPI_ISL_3265216, EPI_ISL_3265217, EPI_ISL_3265218, EPI_ISL_3265219, EPI_ISL_3265220, EPI_ISL_3265221, EPI_ISL_3265222, EPI_ISL_3265223, EPI_ISL_3265224, EPI_ISL_3265225, EPI_ISL_3265226, EPI_ISL_3265227, EPI_ISL_3265228, EPI_ISL_3265229, EPI_ISL_3265230, EPI_ISL_3265231, EPI_ISL_3265232, EPI_ISL_3265233, EPI_ISL_3265234, EPI_ISL_3265235, EPI_ISL_3265236, EPI_ISL_3265237, EPI_ISL_3265238, EPI_ISL_3265239, EPI_ISL_3265240, EPI_ISL_3265241, EPI_ISL_3265242, EPI_ISL_3265243, EPI_ISL_3265244, EPI_ISL_3265245, EPI_ISL_3265246, EPI_ISL_3265247, EPI_ISL_3265248, EPI_ISL_3265249, EPI_ISL_3265250, EPI_ISL_3265251, EPI_ISL_3265252, EPI_ISL_3265253, EPI_ISL_3265254, EPI_ISL_3265255, EPI_ISL_3265256, EPI_ISL_3265257, EPI_ISL_3265258, EPI_ISL_3265259, EPI_ISL_3265260, EPI_ISL_3265261, EPI_ISL_3265262, EPI_ISL_3265263, EPI_ISL_3265264, EPI_ISL_3265265, EPI_ISL_3265266, EPI_ISL_3265267, EPI_ISL_3265268, EPI_ISL_3265269, EPI_ISL_3265270, EPI_ISL_3265271, EPI_ISL_3265272, EPI_ISL_3265273, EPI_ISL_3265274, EPI_ISL_3265275, EPI_ISL_3265276, EPI_ISL_3265277, EPI_ISL_3265278, EPI_ISL_3265279, EPI_ISL_3265280, EPI_ISL_3265281, EPI_ISL_3265282, EPI_ISL_3265283, EPI_ISL_3265284, EPI_ISL_3265285, EPI_ISL_3265286, EPI_ISL_3265287, EPI_ISL_3265288, EPI_ISL_3265290, EPI_ISL_3265291, EPI_ISL_3265292, EPI_ISL_3265293, EPI_ISL_3265294, EPI_ISL_3265295, EPI_ISL_3265296, EPI_ISL_3265297, EPI_ISL_3265298, EPI_ISL_3265299, EPI_ISL_3265300, EPI_ISL_3265301, EPI_ISL_3265302, EPI_ISL_3265303, EPI_ISL_3265304, EPI_ISL_3265305, EPI_ISL_3265306, EPI_ISL_3265307, EPI_ISL_3265308, EPI_ISL_3265309, EPI_ISL_3265310, EPI_ISL_3265311, EPI_ISL_3265312, EPI_ISL_3265313, EPI_ISL_3265314, EPI_ISL_3265315, EPI_ISL_3265316, EPI_ISL_3265317, EPI_ISL_3265318, EPI_ISL_3265319, EPI_ISL_3265320, EPI_ISL_3265321, EPI_ISL_3265322, EPI_ISL_3265324, EPI_ISL_3265325, EPI_ISL_3265326, EPI_ISL_3265327, EPI_ISL_3265328, EPI_ISL_3265329 | | | |
| see above | JIPMER Puducherry | inStem NCBS – INSACOG | Uma Ramakrishnan Dasaradhi Palakodeti Aswin SaiNarain |
| EPI_ISL_3265330, EPI_ISL_3265331, EPI_ISL_3265332, EPI_ISL_3265333, EPI_ISL_3265334, EPI_ISL_3265335, EPI_ISL_3265336, EPI_ISL_3265337, EPI_ISL_3265338, EPI_ISL_3265339 | LESP Hidalgo | Instituto de Diagnostico y Referencia Epidemiologicos (INDRE) | Claudia Wong-Arambula, Abril Rodriguez-Maldonado, Vanessa Rivero-Arredondo, Ariadna Medina-Benitez, Joaquin Quiroz-Mercado, Sergio Rangel-Guerrero, Maribel Gonzalez-Villa, Natividad Cruz-Ortiz, Tatiana Nunez-Garcia, Gisela Barrera-Badillo, Lucia Hernandez-Rivas, Irma Lopez-Martinez, Ernesto Ramirez-Gonzalez. |
| EPI_ISL_3265340 | LESP Ciudad de Mexico | Instituto de Diagnostico y Referencia Epidemiologicos (INDRE) | Claudia Wong-Arambula, Abril Rodriguez-Maldonado, Vanessa Rivero-Arredondo, Ariadna Medina-Benitez, Joaquin Quiroz-Mercado, Sergio Rangel-Guerrero, Maribel Gonzalez-Villa, Natividad Cruz-Ortiz, Tatiana Nunez-Garcia, Gisela Barrera-Badillo, Lucia Hernandez-Rivas, Irma Lopez-Martinez, Ernesto Ramirez-Gonzalez. |
| EPI_ISL_3265341, EPI_ISL_3265342, EPI_ISL_3265343, EPI_ISL_3265344, EPI_ISL_3265345, EPI_ISL_3265346, EPI_ISL_3265347 | LESP Hidalgo | Instituto de Diagnostico y Referencia Epidemiologicos (INDRE) | Claudia Wong-Arambula, Abril Rodriguez-Maldonado, Vanessa Rivero-Arredondo, Ariadna Medina-Benitez, Joaquin Quiroz-Mercado, Sergio Rangel-Guerrero, Maribel Gonzalez-Villa, Natividad Cruz-Ortiz, Tatiana Nunez-Garcia, Gisela Barrera-Badillo, Lucia Hernandez-Rivas, Irma Lopez-Martinez, Ernesto Ramirez-Gonzalez. |
| EPI_ISL_3265348 | LESP Estado de Mexico | Instituto de Diagnostico y Referencia Epidemiologicos (INDRE) | Claudia Wong-Arambula, Abril Rodriguez-Maldonado, Vanessa Rivero-Arredondo, Ariadna Medina-Benitez, Joaquin Quiroz-Mercado, Sergio Rangel-Guerrero, Maribel Gonzalez-Villa, Natividad Cruz-Ortiz, Tatiana Nunez-Garcia, Gisela Barrera-Badillo, Lucia Hernandez-Rivas, Irma Lopez-Martinez, Ernesto Ramirez-Gonzalez. |
| EPI_ISL_3265349 | LESP Queretaro | Instituto de Diagnostico y Referencia Epidemiologicos (INDRE) | Claudia Wong-Arambula, Abril Rodriguez-Maldonado, Vanessa Rivero-Arredondo, Ariadna Medina-Benitez, Joaquin Quiroz-Mercado, Sergio Rangel-Guerrero, Maribel Gonzalez-Villa, Natividad Cruz-Ortiz, Tatiana Nunez-Garcia, Gisela Barrera-Badillo, Lucia Hernandez-Rivas, Irma Lopez-Martinez, Ernesto Ramirez-Gonzalez. |
| EPI_ISL_3265350 | LESP Guanajuato | Instituto de Diagnostico y Referencia Epidemiologicos (INDRE) | Claudia Wong-Arambula, Abril Rodriguez-Maldonado, Vanessa Rivero-Arredondo, Ariadna Medina-Benitez, Joaquin Quiroz-Mercado, Sergio Rangel-Guerrero, Maribel Gonzalez-Villa, Natividad Cruz-Ortiz, Tatiana Nunez-Garcia, Gisela Barrera-Badillo, Lucia Hernandez-Rivas, Irma Lopez-Martinez, Ernesto Ramirez-Gonzalez. |
| EPI_ISL_3265351, EPI_ISL_3265352 | LESP Estado de Mexico | Instituto de Diagnostico y Referencia Epidemiologicos (INDRE) | Claudia Wong-Arambula, Abril Rodriguez-Maldonado, Vanessa Rivero-Arredondo, Ariadna Medina-Benitez, Joaquin Quiroz-Mercado, Sergio Rangel-Guerrero, Maribel Gonzalez-Villa, Natividad Cruz-Ortiz, Tatiana Nunez-Garcia, Gisela Barrera-Badillo, Lucia Hernandez-Rivas, Irma Lopez-Martinez, Ernesto Ramirez-Gonzalez. |
| EPI_ISL_3265353, EPI_ISL_3265354 | LESP Queretaro | Instituto de Diagnostico y Referencia Epidemiologicos (INDRE) | Claudia Wong-Arambula, Abril Rodriguez-Maldonado, Vanessa Rivero-Arredondo, Ariadna Medina-Benitez, Joaquin Quiroz-Mercado, Sergio Rangel-Guerrero, Maribel Gonzalez-Villa, Natividad Cruz-Ortiz, Tatiana Nunez-Garcia, Gisela Barrera-Badillo, Lucia Hernandez-Rivas, Irma Lopez-Martinez, Ernesto Ramirez-Gonzalez. |
| EPI_ISL_3265355, EPI_ISL_3265356, EPI_ISL_3265357, EPI_ISL_3265358, EPI_ISL_3265359, EPI_ISL_3265360, EPI_ISL_3265361, EPI_ISL_3265362, EPI_ISL_3265363 | LESP Estado de Mexico | Instituto de Diagnostico y Referencia Epidemiologicos (INDRE) | Claudia Wong-Arambula, Abril Rodriguez-Maldonado, Vanessa Rivero-Arredondo, Ariadna Medina-Benitez, Joaquin Quiroz-Mercado, Sergio Rangel-Guerrero, Maribel Gonzalez-Villa, Natividad Cruz-Ortiz, Tatiana Nunez-Garcia, Gisela Barrera-Badillo, Lucia Hernandez-Rivas, Irma Lopez-Martinez, Ernesto Ramirez-Gonzalez. |
| EPI_ISL_3265364 | LESP Guerrero | Instituto de Diagnostico y Referencia Epidemiologicos (INDRE) | Claudia Wong-Arambula, Abril Rodriguez-Maldonado, Vanessa Rivero-Arredondo, Ariadna Medina-Benitez, Joaquin Quiroz-Mercado, Sergio Rangel-Guerrero, Maribel Gonzalez-Villa, Natividad Cruz-Ortiz, Tatiana Nunez-Garcia, Gisela Barrera-Badillo, Lucia Hernandez-Rivas, Irma Lopez-Martinez, Ernesto Ramirez-Gonzalez. |
| EPI_ISL_3265365 | LESP Estado de Mexico | Instituto de Diagnostico y Referencia Epidemiologicos (INDRE) | Claudia Wong-Arambula, Abril Rodriguez-Maldonado, Vanessa Rivero-Arredondo, Ariadna Medina-Benitez, Joaquin Quiroz-Mercado, Sergio Rangel-Guerrero, Maribel Gonzalez-Villa, Natividad Cruz-Ortiz, Tatiana Nunez-Garcia, Gisela Barrera-Badillo, Lucia Hernandez-Rivas, Irma Lopez-Martinez, Ernesto Ramirez-Gonzalez. |
| EPI_ISL_3265366, EPI_ISL_3265367 | LESP Ciudad de Mexico | Instituto de Diagnostico y Referencia Epidemiologicos (INDRE) | Claudia Wong-Arambula, Abril Rodriguez-Maldonado, Vanessa Rivero-Arredondo, Ariadna Medina-Benitez, Joaquin Quiroz-Mercado, Sergio Rangel-Guerrero, Maribel Gonzalez-Villa, Natividad Cruz-Ortiz, Tatiana Nunez-Garcia, Gisela Barrera-Badillo, Lucia Hernandez-Rivas, Irma Lopez-Martinez, Ernesto Ramirez-Gonzalez. |
| EPI_ISL_3265368, EPI_ISL_3265369, EPI_ISL_3265370, EPI_ISL_3265371, EPI_ISL_3265372, EPI_ISL_3265373 | LESP Estado de Mexico | Instituto de Diagnostico y Referencia Epidemiologicos (INDRE) | Claudia Wong-Arambula, Abril Rodriguez-Maldonado, Vanessa Rivero-Arredondo, Ariadna Medina-Benitez, Joaquin Quiroz-Mercado, Sergio Rangel-Guerrero, Maribel Gonzalez-Villa, Natividad Cruz-Ortiz, Tatiana Nunez-Garcia, Gisela Barrera-Badillo, Lucia Hernandez-Rivas, Irma Lopez-Martinez, Ernesto Ramirez-Gonzalez. |
| EPI_ISL_3265374 | LESP Jalisco | Instituto de Diagnostico y Referencia Epidemiologicos (INDRE) | Claudia Wong-Arambula, Abril Rodriguez-Maldonado, Vanessa Rivero-Arredondo, Ariadna Medina-Benitez, Joaquin Quiroz-Mercado, Sergio Rangel-Guerrero, Maribel Gonzalez-Villa, Natividad Cruz-Ortiz, Tatiana Nunez-Garcia, Gisela Barrera-Badillo, Lucia Hernandez-Rivas, Irma Lopez-Martinez, Ernesto Ramirez-Gonzalez. |
| EPI_ISL_3265375, EPI_ISL_3265376, EPI_ISL_3265377, EPI_ISL_3265378, EPI_ISL_3265379, EPI_ISL_3265380, EPI_ISL_3265381, EPI_ISL_3265382, EPI_ISL_3265383, EPI_ISL_3265384 | LESP Estado de Mexico | Instituto de Diagnostico y Referencia Epidemiologicos (INDRE) | Claudia Wong-Arambula, Abril Rodriguez-Maldonado, Vanessa Rivero-Arredondo, Ariadna Medina-Benitez, Joaquin Quiroz-Mercado, Sergio Rangel-Guerrero, Maribel Gonzalez-Villa, Natividad Cruz-Ortiz, Tatiana Nunez-Garcia, Gisela Barrera-Badillo, Lucia Hernandez-Rivas, Irma Lopez-Martinez, Ernesto Ramirez-Gonzalez. |
| EPI_ISL_3265385 | LESP Hidalgo | Instituto de Diagnostico y Referencia Epidemiologicos (INDRE) | Claudia Wong-Arambula, Abril Rodriguez-Maldonado, Vanessa Rivero-Arredondo, Ariadna Medina-Benitez, Joaquin Quiroz-Mercado, Sergio Rangel-Guerrero, Maribel Gonzalez-Villa, Natividad Cruz-Ortiz, Tatiana Nunez-Garcia, Gisela Barrera-Badillo, Lucia Hernandez-Rivas, Irma Lopez-Martinez, Ernesto Ramirez-Gonzalez. |
| EPI_ISL_3265386, EPI_ISL_3265387, EPI_ISL_3265388, EPI_ISL_3265389, EPI_ISL_3265390, EPI_ISL_3265391, EPI_ISL_3265392, EPI_ISL_3265393, EPI_ISL_3265394, EPI_ISL_3265395, EPI_ISL_3265396, EPI_ISL_3265397 | | | |
| see above | LESP Estado de Mexico | Instituto de Diagnostico y Referencia Epidemiologicos (INDRE) | Claudia Wong-Arambula, Abril Rodriguez-Maldonado, Vanessa Rivero-Arredondo, Ariadna Medina-Benitez, Joaquin Quiroz-Mercado, Sergio Rangel-Guerrero, Maribel Gonzalez-Villa, Natividad Cruz-Ortiz, Tatiana Nunez-Garcia, Gisela Barrera-Badillo, Lucia Hernandez-Rivas, Irma Lopez-Martinez, Ernesto Ramirez-Gonzalez. |
| EPI_ISL_3265398 | LESP Colima | Instituto de Diagnostico y Referencia Epidemiologicos (INDRE) | Claudia Wong-Arambula, Abril Rodriguez-Maldonado, Vanessa Rivero-Arredondo, Ariadna Medina-Benitez, Joaquin Quiroz-Mercado, Sergio Rangel-Guerrero, Maribel Gonzalez-Villa, Natividad Cruz-Ortiz, Tatiana Nunez-Garcia, Gisela Barrera-Badillo, Lucia Hernandez-Rivas, Irma Lopez-Martinez, Ernesto Ramirez-Gonzalez. |
| EPI_ISL_3265399 | LESP Ciudad de Mexico | Instituto de Diagnostico y Referencia Epidemiologicos (INDRE) | Claudia Wong-Arambula, Abril Rodriguez-Maldonado, Vanessa Rivero-Arredondo, Ariadna Medina-Benitez, Joaquin Quiroz-Mercado, Sergio Rangel-Guerrero, Maribel Gonzalez-Villa, Natividad Cruz-Ortiz, Tatiana Nunez-Garcia, Gisela Barrera-Badillo, Lucia Hernandez-Rivas, Irma Lopez-Martinez, Ernesto Ramirez-Gonzalez. |
| EPI_ISL_3265400, EPI_ISL_3265401, EPI_ISL_3265402, EPI_ISL_3265403, EPI_ISL_3265404, EPI_ISL_3265405, EPI_ISL_3265406, EPI_ISL_3265407, EPI_ISL_3265408, EPI_ISL_3265409, EPI_ISL_3265410, EPI_ISL_3265411 | | | |
| see above | LESP Aguascalientes | Instituto de Diagnostico y Referencia Epidemiologicos (INDRE) | Claudia Wong-Arambula, Abril Rodriguez-Maldonado, Vanessa Rivero-Arredondo, Ariadna Medina-Benitez, Joaquin Quiroz-Mercado, Sergio Rangel-Guerrero, Maribel Gonzalez-Villa, Natividad Cruz-Ortiz, Tatiana Nunez-Garcia, Gisela Barrera-Badillo, Lucia Hernandez-Rivas, Irma Lopez-Martinez, Ernesto Ramirez-Gonzalez. |
| EPI_ISL_3265412 | LESP Campeche | Instituto de Diagnostico y Referencia Epidemiologicos (INDRE) | Claudia Wong-Arambula, Abril Rodriguez-Maldonado, Vanessa Rivero-Arredondo, Ariadna Medina-Benitez, Joaquin Quiroz-Mercado, Sergio Rangel-Guerrero, Maribel Gonzalez-Villa, Natividad Cruz-Ortiz, Tatiana Nunez-Garcia, Gisela Barrera-Badillo, Lucia Hernandez-Rivas, Irma Lopez-Martinez, Ernesto Ramirez-Gonzalez. |
| EPI_ISL_3265413, EPI_ISL_3265414, EPI_ISL_3265415, EPI_ISL_3265416, EPI_ISL_3265417, EPI_ISL_3265418, EPI_ISL_3265419 | LESP Sinaloa | Instituto de Diagnostico y Referencia Epidemiologicos (INDRE) | Claudia Wong-Arambula, Abril Rodriguez-Maldonado, Vanessa Rivero-Arredondo, Ariadna Medina-Benitez, Joaquin Quiroz-Mercado, Sergio Rangel-Guerrero, Maribel Gonzalez-Villa, Natividad Cruz-Ortiz, Tatiana Nunez-Garcia, Gisela Barrera-Badillo, Lucia Hernandez-Rivas, Irma Lopez-Martinez, Ernesto Ramirez-Gonzalez. |
| EPI_ISL_3265420, EPI_ISL_3265421, EPI_ISL_3265422, EPI_ISL_3265423, EPI_ISL_3265424, EPI_ISL_3265425, EPI_ISL_3265426, EPI_ISL_3265427, EPI_ISL_3265428, EPI_ISL_3265429, EPI_ISL_3265430, EPI_ISL_3265431, EPI_ISL_3265432, EPI_ISL_3265433, EPI_ISL_3265434, EPI_ISL_3265435, EPI_ISL_3265436, EPI_ISL_3265437, EPI_ISL_3265438, EPI_ISL_3265439, EPI_ISL_3265440, EPI_ISL_3265441, EPI_ISL_3265442, EPI_ISL_3265443, EPI_ISL_3265444, EPI_ISL_3265445, EPI_ISL_3265446, EPI_ISL_3265447, EPI_ISL_3265448, EPI_ISL_3265449, EPI_ISL_3265450 | | | |
| see above | LESP Baja California Sur | Instituto de Diagnostico y Referencia Epidemiologicos (INDRE) | Claudia Wong-Arambula, Abril Rodriguez-Maldonado, Vanessa Rivero-Arredondo, Ariadna Medina-Benitez, Joaquin Quiroz-Mercado, Sergio Rangel-Guerrero, Maribel Gonzalez-Villa, Natividad Cruz-Ortiz, Tatiana Nunez-Garcia, Gisela Barrera-Badillo, Lucia Hernandez-Rivas, Irma Lopez-Martinez, Ernesto Ramirez-Gonzalez. |

| | | | |
|---|---|---|---|
| EPI_ISL_3265451, EPI_ISL_3265452, EPI_ISL_3265453, EPI_ISL_3265454 | LESP Sonora | Instituto de Diagnostico y Referencia Epidemiologicos (INDRE) | Claudia Wong-Arambula, Abril Rodriguez-Maldonado, Vanessa Rivero-Arredondo, Ariadna Medina-Benitez, Joaquin Quiroz-Mercado, Sergio Rangel-Guerrero, Maribel Gonzalez-Villa, Natividad Cruz-Ortiz, Tatiana Nunez-Garcia, Gisela Barrera-Badillo, Lucia Hernandez-Rivas, Irma Lopez-Martinez, Ernesto Ramirez-Gonzalez. |
| EPI_ISL_3265455 | LESP Colima | Instituto de Diagnostico y Referencia Epidemiologicos (INDRE) | Claudia Wong-Arambula, Abril Rodriguez-Maldonado, Vanessa Rivero-Arredondo, Ariadna Medina-Benitez, Joaquin Quiroz-Mercado, Sergio Rangel-Guerrero, Maribel Gonzalez-Villa, Natividad Cruz-Ortiz, Tatiana Nunez-Garcia, Gisela Barrera-Badillo, Lucia Hernandez-Rivas, Irma Lopez-Martinez, Ernesto Ramirez-Gonzalez. |
| EPI_ISL_3265456 | LESP Michoacan | Instituto de Diagnostico y Referencia Epidemiologicos (INDRE) | Claudia Wong-Arambula, Abril Rodriguez-Maldonado, Vanessa Rivero-Arredondo, Ariadna Medina-Benitez, Joaquin Quiroz-Mercado, Sergio Rangel-Guerrero, Maribel Gonzalez-Villa, Natividad Cruz-Ortiz, Tatiana Nunez-Garcia, Gisela Barrera-Badillo, Lucia Hernandez-Rivas, Irma Lopez-Martinez, Ernesto Ramirez-Gonzalez. |
| EPI_ISL_3265457 | LESP Colima | Instituto de Diagnostico y Referencia Epidemiologicos (INDRE) | Claudia Wong-Arambula, Abril Rodriguez-Maldonado, Vanessa Rivero-Arredondo, Ariadna Medina-Benitez, Joaquin Quiroz-Mercado, Sergio Rangel-Guerrero, Maribel Gonzalez-Villa, Natividad Cruz-Ortiz, Tatiana Nunez-Garcia, Gisela Barrera-Badillo, Lucia Hernandez-Rivas, Irma Lopez-Martinez, Ernesto Ramirez-Gonzalez. |
| EPI_ISL_3265458, EPI_ISL_3265459, EPI_ISL_3265460, EPI_ISL_3265461, EPI_ISL_3265462, EPI_ISL_3265463 | LESP Ciudad de Mexico | Instituto de Diagnostico y Referencia Epidemiologicos (INDRE) | Claudia Wong-Arambula, Abril Rodriguez-Maldonado, Vanessa Rivero-Arredondo, Ariadna Medina-Benitez, Joaquin Quiroz-Mercado, Sergio Rangel-Guerrero, Maribel Gonzalez-Villa, Natividad Cruz-Ortiz, Tatiana Nunez-Garcia, Gisela Barrera-Badillo, Lucia Hernandez-Rivas, Irma Lopez-Martinez, Ernesto Ramirez-Gonzalez. |
| EPI_ISL_3265464, EPI_ISL_3265465, EPI_ISL_3265466 | LESP Hidalgo | Instituto de Diagnostico y Referencia Epidemiologicos (INDRE) | Claudia Wong-Arambula, Abril Rodriguez-Maldonado, Vanessa Rivero-Arredondo, Ariadna Medina-Benitez, Joaquin Quiroz-Mercado, Sergio Rangel-Guerrero, Maribel Gonzalez-Villa, Natividad Cruz-Ortiz, Tatiana Nunez-Garcia, Gisela Barrera-Badillo, Lucia Hernandez-Rivas, Irma Lopez-Martinez, Ernesto Ramirez-Gonzalez. |
| EPI_ISL_3265467, EPI_ISL_3265468, EPI_ISL_3265469 | LESP Estado de Mexico | Instituto de Diagnostico y Referencia Epidemiologicos (INDRE) | Claudia Wong-Arambula, Abril Rodriguez-Maldonado, Vanessa Rivero-Arredondo, Ariadna Medina-Benitez, Joaquin Quiroz-Mercado, Sergio Rangel-Guerrero, Maribel Gonzalez-Villa, Natividad Cruz-Ortiz, Tatiana Nunez-Garcia, Gisela Barrera-Badillo, Lucia Hernandez-Rivas, Irma Lopez-Martinez, Ernesto Ramirez-Gonzalez. |
| EPI_ISL_3265470 | LESP San Luis Potosi | Instituto de Diagnostico y Referencia Epidemiologicos (INDRE) | Claudia Wong-Arambula, Abril Rodriguez-Maldonado, Vanessa Rivero-Arredondo, Ariadna Medina-Benitez, Joaquin Quiroz-Mercado, Sergio Rangel-Guerrero, Maribel Gonzalez-Villa, Natividad Cruz-Ortiz, Tatiana Nunez-Garcia, Gisela Barrera-Badillo, Lucia Hernandez-Rivas, Irma Lopez-Martinez, Ernesto Ramirez-Gonzalez. |
| EPI_ISL_3265471 | LESP Morelos | Instituto de Diagnostico y Referencia Epidemiologicos (INDRE) | Claudia Wong-Arambula, Abril Rodriguez-Maldonado, Vanessa Rivero-Arredondo, Ariadna Medina-Benitez, Joaquin Quiroz-Mercado, Sergio Rangel-Guerrero, Maribel Gonzalez-Villa, Natividad Cruz-Ortiz, Tatiana Nunez-Garcia, Gisela Barrera-Badillo, Lucia Hernandez-Rivas, Irma Lopez-Martinez, Ernesto Ramirez-Gonzalez. |
| EPI_ISL_3265472, EPI_ISL_3265473, EPI_ISL_3265474 | LESP Guanajuato | Instituto de Diagnostico y Referencia Epidemiologicos (INDRE) | Claudia Wong-Arambula, Abril Rodriguez-Maldonado, Vanessa Rivero-Arredondo, Ariadna Medina-Benitez, Joaquin Quiroz-Mercado, Sergio Rangel-Guerrero, Maribel Gonzalez-Villa, Natividad Cruz-Ortiz, Tatiana Nunez-Garcia, Gisela Barrera-Badillo, Lucia Hernandez-Rivas, Irma Lopez-Martinez, Ernesto Ramirez-Gonzalez. |
| EPI_ISL_3265475, EPI_ISL_3265476, EPI_ISL_3265477, EPI_ISL_3265478, EPI_ISL_3265479 | LESP Sinaloa | Instituto de Diagnostico y Referencia Epidemiologicos (INDRE) | Claudia Wong-Arambula, Abril Rodriguez-Maldonado, Vanessa Rivero-Arredondo, Ariadna Medina-Benitez, Joaquin Quiroz-Mercado, Sergio Rangel-Guerrero, Maribel Gonzalez-Villa, Natividad Cruz-Ortiz, Tatiana Nunez-Garcia, Gisela Barrera-Badillo, Lucia Hernandez-Rivas, Irma Lopez-Martinez, Ernesto Ramirez-Gonzalez. |
| EPI_ISL_3265480, EPI_ISL_3265481, EPI_ISL_3265482, EPI_ISL_3265483, EPI_ISL_3265484, EPI_ISL_3265485 | LESP Aguascalientes | Instituto de Diagnostico y Referencia Epidemiologicos (INDRE) | Claudia Wong-Arambula, Abril Rodriguez-Maldonado, Vanessa Rivero-Arredondo, Ariadna Medina-Benitez, Joaquin Quiroz-Mercado, Sergio Rangel-Guerrero, Maribel Gonzalez-Villa, Natividad Cruz-Ortiz, Tatiana Nunez-Garcia, Gisela Barrera-Badillo, Lucia Hernandez-Rivas, Irma Lopez-Martinez, Ernesto Ramirez-Gonzalez. |
| EPI_ISL_3265486, EPI_ISL_3265487, EPI_ISL_3265488, EPI_ISL_3265489, EPI_ISL_3265490, EPI_ISL_3265491, EPI_ISL_3265492, EPI_ISL_3265493, EPI_ISL_3265494, EPI_ISL_3265495, EPI_ISL_3265496, EPI_ISL_3265497, EPI_ISL_3265498, EPI_ISL_3265499, EPI_ISL_3265500, EPI_ISL_3265501, EPI_ISL_3265502, EPI_ISL_3265503, EPI_ISL_3265504, EPI_ISL_3265505, EPI_ISL_3265506, EPI_ISL_3265507, EPI_ISL_3265508 | | | |
| see above | LESP Veracruz | Instituto de Diagnostico y Referencia Epidemiologicos (INDRE) | Claudia Wong-Arambula, Abril Rodriguez-Maldonado, Vanessa Rivero-Arredondo, Ariadna Medina-Benitez, Joaquin Quiroz-Mercado, Sergio Rangel-Guerrero, Maribel Gonzalez-Villa, Natividad Cruz-Ortiz, Tatiana Nunez-Garcia, Gisela Barrera-Badillo, Lucia Hernandez-Rivas, Irma Lopez-Martinez, Ernesto Ramirez-Gonzalez. |
| EPI_ISL_3265509, EPI_ISL_3265510, EPI_ISL_3265511, EPI_ISL_3265512 | LESP Durango | Instituto de Diagnostico y Referencia Epidemiologicos (INDRE) | Claudia Wong-Arambula, Abril Rodriguez-Maldonado, Vanessa Rivero-Arredondo, Ariadna Medina-Benitez, Joaquin Quiroz-Mercado, Sergio Rangel-Guerrero, Maribel Gonzalez-Villa, Natividad Cruz-Ortiz, Tatiana Nunez-Garcia, Gisela Barrera-Badillo, Lucia Hernandez-Rivas, Irma Lopez-Martinez, Ernesto Ramirez-Gonzalez. |
| EPI_ISL_3265513 | LESP Nayarit | Instituto de Diagnostico y Referencia Epidemiologicos (INDRE) | Claudia Wong-Arambula, Abril Rodriguez-Maldonado, Vanessa Rivero-Arredondo, Ariadna Medina-Benitez, Joaquin Quiroz-Mercado, Sergio Rangel-Guerrero, Maribel Gonzalez-Villa, Natividad Cruz-Ortiz, Tatiana Nunez-Garcia, Gisela Barrera-Badillo, Lucia Hernandez-Rivas, Irma Lopez-Martinez, Ernesto Ramirez-Gonzalez. |
| EPI_ISL_3265514, EPI_ISL_3265515 | LESP Tamaulipas | Instituto de Diagnostico y Referencia Epidemiologicos (INDRE) | Claudia Wong-Arambula, Abril Rodriguez-Maldonado, Vanessa Rivero-Arredondo, Ariadna Medina-Benitez, Joaquin Quiroz-Mercado, Sergio Rangel-Guerrero, Maribel Gonzalez-Villa, Natividad Cruz-Ortiz, Tatiana Nunez-Garcia, Gisela Barrera-Badillo, Lucia Hernandez-Rivas, Irma Lopez-Martinez, Ernesto Ramirez-Gonzalez. |
| EPI_ISL_3265516, EPI_ISL_3265517, EPI_ISL_3265518, EPI_ISL_3265519 | LESP Michoacan | Instituto de Diagnostico y Referencia Epidemiologicos (INDRE) | Claudia Wong-Arambula, Abril Rodriguez-Maldonado, Vanessa Rivero-Arredondo, Ariadna Medina-Benitez, Joaquin Quiroz-Mercado, Sergio Rangel-Guerrero, Maribel Gonzalez-Villa, Natividad Cruz-Ortiz, Tatiana Nunez-Garcia, Gisela Barrera-Badillo, Lucia Hernandez-Rivas, Irma Lopez-Martinez, Ernesto Ramirez-Gonzalez. |
| EPI_ISL_3265520, EPI_ISL_3265521, EPI_ISL_3265522, EPI_ISL_3265523, EPI_ISL_3265524, EPI_ISL_3265525, EPI_ISL_3265526, EPI_ISL_3265527, EPI_ISL_3265528, EPI_ISL_3265529, EPI_ISL_3265530 | | | |
| see above | LESP Baja California Sur | Instituto de Diagnostico y Referencia Epidemiologicos (INDRE) | Claudia Wong-Arambula, Abril Rodriguez-Maldonado, Vanessa Rivero-Arredondo, Ariadna Medina-Benitez, Joaquin Quiroz-Mercado, Sergio Rangel-Guerrero, Maribel Gonzalez-Villa, Natividad Cruz-Ortiz, Tatiana Nunez-Garcia, Gisela Barrera-Badillo, Lucia Hernandez-Rivas, Irma Lopez-Martinez, Ernesto Ramirez-Gonzalez. |
| EPI_ISL_3265531 | LESP Estado de Mexico | Instituto de Diagnostico y Referencia Epidemiologicos (INDRE) | Claudia Wong-Arambula, Abril Rodriguez-Maldonado, Vanessa Rivero-Arredondo, Ariadna Medina-Benitez, Joaquin Quiroz-Mercado, Sergio Rangel-Guerrero, Maribel Gonzalez-Villa, Natividad Cruz-Ortiz, Tatiana Nunez-Garcia, Gisela Barrera-Badillo, Lucia Hernandez-Rivas, Irma Lopez-Martinez, Ernesto Ramirez-Gonzalez. |
| EPI_ISL_3265532, EPI_ISL_3265533 | LESP Jalisco | Instituto de Diagnostico y Referencia Epidemiologicos (INDRE) | Claudia Wong-Arambula, Abril Rodriguez-Maldonado, Vanessa Rivero-Arredondo, Ariadna Medina-Benitez, Joaquin Quiroz-Mercado, Sergio Rangel-Guerrero, Maribel Gonzalez-Villa, Natividad Cruz-Ortiz, Tatiana Nunez-Garcia, Gisela Barrera-Badillo, Lucia Hernandez-Rivas, Irma Lopez-Martinez, Ernesto Ramirez-Gonzalez. |
| EPI_ISL_3265534, EPI_ISL_3265535, EPI_ISL_3265536, EPI_ISL_3265537, EPI_ISL_3265538 | LESP Colima | Instituto de Diagnostico y Referencia Epidemiologicos (INDRE) | Claudia Wong-Arambula, Abril Rodriguez-Maldonado, Vanessa Rivero-Arredondo, Ariadna Medina-Benitez, Joaquin Quiroz-Mercado, Sergio Rangel-Guerrero, Maribel Gonzalez-Villa, Natividad Cruz-Ortiz, Tatiana Nunez-Garcia, Gisela Barrera-Badillo, Lucia Hernandez-Rivas, Irma Lopez-Martinez, Ernesto Ramirez-Gonzalez. |
| EPI_ISL_3265539, EPI_ISL_3265540, EPI_ISL_3265541, EPI_ISL_3265542 | LESP Sinaloa | Instituto de Diagnostico y Referencia Epidemiologicos (INDRE) | Claudia Wong-Arambula, Abril Rodriguez-Maldonado, Vanessa Rivero-Arredondo, Ariadna Medina-Benitez, Joaquin Quiroz-Mercado, Sergio Rangel-Guerrero, Maribel Gonzalez-Villa, Natividad Cruz-Ortiz, Tatiana Nunez-Garcia, Gisela Barrera-Badillo, Lucia Hernandez-Rivas, Irma Lopez-Martinez, Ernesto Ramirez-Gonzalez. |
| EPI_ISL_3265543 | LESP Estado de Mexico | Instituto de Diagnostico y Referencia Epidemiologicos (INDRE) | Claudia Wong-Arambula, Abril Rodriguez-Maldonado, Vanessa Rivero-Arredondo, Ariadna Medina-Benitez, Joaquin Quiroz-Mercado, Sergio Rangel-Guerrero, Maribel Gonzalez-Villa, Natividad Cruz-Ortiz, Tatiana Nunez-Garcia, Gisela Barrera-Badillo, Lucia Hernandez-Rivas, Irma Lopez-Martinez, Ernesto Ramirez-Gonzalez. |
| EPI_ISL_3265544, EPI_ISL_3265545, EPI_ISL_3265546 | LESP Quintana Roo | Instituto de Diagnostico y Referencia Epidemiologicos (INDRE) | Claudia Wong-Arambula, Abril Rodriguez-Maldonado, Vanessa Rivero-Arredondo, Ariadna Medina-Benitez, Joaquin Quiroz-Mercado, Sergio Rangel-Guerrero, Maribel Gonzalez-Villa, Natividad Cruz-Ortiz, Tatiana Nunez-Garcia, Gisela Barrera-Badillo, Lucia Hernandez-Rivas, Irma Lopez-Martinez, Ernesto Ramirez-Gonzalez. |
| EPI_ISL_3265547 | LESP Campeche | Instituto de Diagnostico y Referencia Epidemiologicos (INDRE) | Claudia Wong-Arambula, Abril Rodriguez-Maldonado, Vanessa Rivero-Arredondo, Ariadna Medina-Benitez, Joaquin Quiroz-Mercado, Sergio Rangel-Guerrero, Maribel Gonzalez-Villa, Natividad Cruz-Ortiz, Tatiana Nunez-Garcia, Gisela Barrera-Badillo, Lucia Hernandez-Rivas, Irma Lopez-Martinez, Ernesto Ramirez-Gonzalez. |
| EPI_ISL_3265548 | LESP Ciudad de Mexico | Instituto de Diagnostico y Referencia Epidemiologicos (INDRE) | Claudia Wong-Arambula, Abril Rodriguez-Maldonado, Vanessa Rivero-Arredondo, Ariadna Medina-Benitez, Joaquin Quiroz-Mercado, Sergio Rangel-Guerrero, Maribel Gonzalez-Villa, Natividad Cruz-Ortiz, Tatiana Nunez-Garcia, Gisela Barrera-Badillo, Lucia Hernandez-Rivas, Irma Lopez-Martinez, Ernesto Ramirez-Gonzalez. |
| EPI_ISL_3265549 | LESP Campeche | Instituto de Diagnostico y Referencia Epidemiologicos (INDRE) | Claudia Wong-Arambula, Abril Rodriguez-Maldonado, Vanessa Rivero-Arredondo, Ariadna Medina-Benitez, Joaquin Quiroz-Mercado, Sergio Rangel-Guerrero, Maribel Gonzalez-Villa, Natividad Cruz-Ortiz, Tatiana Nunez-Garcia, Gisela Barrera-Badillo, Lucia Hernandez-Rivas, Irma Lopez-Martinez, Ernesto Ramirez-Gonzalez. |
| EPI_ISL_3265550 | LESP Sonora | Instituto de Diagnostico y Referencia Epidemiologicos (INDRE) | Claudia Wong-Arambula, Abril Rodriguez-Maldonado, Vanessa Rivero-Arredondo, Ariadna Medina-Benitez, Joaquin Quiroz-Mercado, Sergio Rangel-Guerrero, Maribel Gonzalez-Villa, Natividad Cruz-Ortiz, Tatiana Nunez-Garcia, Gisela Barrera-Badillo, Lucia Hernandez-Rivas, Irma Lopez-Martinez, Ernesto Ramirez-Gonzalez. |
| EPI_ISL_3265551 | LESP Coahuila | Instituto de Diagnostico y Referencia Epidemiologicos (INDRE) | Claudia Wong-Arambula, Abril Rodriguez-Maldonado, Vanessa Rivero-Arredondo, Ariadna Medina-Benitez, Joaquin Quiroz-Mercado, Sergio Rangel-Guerrero, Maribel Gonzalez-Villa, Natividad Cruz-Ortiz, Tatiana Nunez-Garcia, Gisela Barrera-Badillo, Lucia Hernandez-Rivas, Irma Lopez-Martinez, Ernesto Ramirez-Gonzalez. |
| EPI_ISL_3265552 | LESP Ciudad de Mexico | Instituto de Diagnostico y Referencia Epidemiologicos (INDRE) | Claudia Wong-Arambula, Abril Rodriguez-Maldonado, Vanessa Rivero-Arredondo, Ariadna Medina-Benitez, Joaquin Quiroz-Mercado, Sergio Rangel-Guerrero, Maribel Gonzalez-Villa, Natividad Cruz-Ortiz, Tatiana Nunez-Garcia, Gisela Barrera-Badillo, Lucia Hernandez-Rivas, Irma Lopez-Martinez, Ernesto Ramirez-Gonzalez. |
| EPI_ISL_3265553 | LESP Campeche | Instituto de Diagnostico y Referencia Epidemiologicos (INDRE) | Claudia Wong-Arambula, Abril Rodriguez-Maldonado, Vanessa Rivero-Arredondo, Ariadna Medina-Benitez, Joaquin Quiroz-Mercado, Sergio Rangel-Guerrero, Maribel Gonzalez-Villa, Natividad Cruz-Ortiz, Tatiana Nunez-Garcia, Gisela Barrera-Badillo, Lucia Hernandez-Rivas, Irma Lopez-Martinez, Ernesto Ramirez-Gonzalez. |
| EPI_ISL_3265554 | LESP Baja California | Instituto de Diagnostico y Referencia Epidemiologicos (INDRE) | Claudia Wong-Arambula, Abril Rodriguez-Maldonado, Vanessa Rivero-Arredondo, Ariadna Medina-Benitez, Joaquin Quiroz-Mercado, Sergio Rangel-Guerrero, Maribel Gonzalez-Villa, Natividad Cruz-Ortiz, Tatiana Nunez-Garcia, Gisela Barrera-Badillo, Lucia Hernandez-Rivas, Irma Lopez-Martinez, Ernesto Ramirez-Gonzalez. |
| EPI_ISL_3265555 | LESP Sonora | Instituto de Diagnostico y Referencia Epidemiologicos (INDRE) | Claudia Wong-Arambula, Abril Rodriguez-Maldonado, Vanessa Rivero-Arredondo, Ariadna Medina-Benitez, Joaquin Quiroz-Mercado, Sergio Rangel-Guerrero, Maribel Gonzalez-Villa, Natividad Cruz-Ortiz, Tatiana Nunez-Garcia, Gisela Barrera-Badillo, Lucia Hernandez-Rivas, Irma Lopez-Martinez, Ernesto Ramirez-Gonzalez. |

| | | | |
|---|---|---|---|
| EPI_ISL_3265556 | LESP Campeche | Instituto de Diagnostico y Referencia Epidemiologicos (INDRE) | Claudia Wong-Arambula, Abril Rodriguez-Maldonado, Vanessa Rivero-Arredondo, Ariadna Medina-Benitez, Joaquin Quiroz-Mercado, Sergio Rangel-Guerrero, Maribel Gonzalez-Villa, Natividad Cruz-Ortiz, Tatiana Nunez-Garcia, Gisela Barrera-Badillo, Lucia Hernandez-Rivas, Irma Lopez-Martinez, Ernesto Ramirez-Gonzalez. |
| EPI_ISL_3265557 | LESP Baja California | Instituto de Diagnostico y Referencia Epidemiologicos (INDRE) | Claudia Wong-Arambula, Abril Rodriguez-Maldonado, Vanessa Rivero-Arredondo, Ariadna Medina-Benitez, Joaquin Quiroz-Mercado, Sergio Rangel-Guerrero, Maribel Gonzalez-Villa, Natividad Cruz-Ortiz, Tatiana Nunez-Garcia, Gisela Barrera-Badillo, Lucia Hernandez-Rivas, Irma Lopez-Martinez, Ernesto Ramirez-Gonzalez. |
| EPI_ISL_3265558 | LESP Sonora | Instituto de Diagnostico y Referencia Epidemiologicos (INDRE) | Claudia Wong-Arambula, Abril Rodriguez-Maldonado, Vanessa Rivero-Arredondo, Ariadna Medina-Benitez, Joaquin Quiroz-Mercado, Sergio Rangel-Guerrero, Maribel Gonzalez-Villa, Natividad Cruz-Ortiz, Tatiana Nunez-Garcia, Gisela Barrera-Badillo, Lucia Hernandez-Rivas, Irma Lopez-Martinez, Ernesto Ramirez-Gonzalez. |
| EPI_ISL_3265559 | LESP Coahuila | Instituto de Diagnostico y Referencia Epidemiologicos (INDRE) | Claudia Wong-Arambula, Abril Rodriguez-Maldonado, Vanessa Rivero-Arredondo, Ariadna Medina-Benitez, Joaquin Quiroz-Mercado, Sergio Rangel-Guerrero, Maribel Gonzalez-Villa, Natividad Cruz-Ortiz, Tatiana Nunez-Garcia, Gisela Barrera-Badillo, Lucia Hernandez-Rivas, Irma Lopez-Martinez, Ernesto Ramirez-Gonzalez. |
| EPI_ISL_3265560 | LESP Sonora | Instituto de Diagnostico y Referencia Epidemiologicos (INDRE) | Claudia Wong-Arambula, Abril Rodriguez-Maldonado, Vanessa Rivero-Arredondo, Ariadna Medina-Benitez, Joaquin Quiroz-Mercado, Sergio Rangel-Guerrero, Maribel Gonzalez-Villa, Natividad Cruz-Ortiz, Tatiana Nunez-Garcia, Gisela Barrera-Badillo, Lucia Hernandez-Rivas, Irma Lopez-Martinez, Ernesto Ramirez-Gonzalez. |
| EPI_ISL_3265561, EPI_ISL_3265562, EPI_ISL_3265563 | LESP Campeche | Instituto de Diagnostico y Referencia Epidemiologicos (INDRE) | Claudia Wong-Arambula, Abril Rodriguez-Maldonado, Vanessa Rivero-Arredondo, Ariadna Medina-Benitez, Joaquin Quiroz-Mercado, Sergio Rangel-Guerrero, Maribel Gonzalez-Villa, Natividad Cruz-Ortiz, Tatiana Nunez-Garcia, Gisela Barrera-Badillo, Lucia Hernandez-Rivas, Irma Lopez-Martinez, Ernesto Ramirez-Gonzalez. |
| EPI_ISL_3265564 | LESP Baja California | Instituto de Diagnostico y Referencia Epidemiologicos (INDRE) | Claudia Wong-Arambula, Abril Rodriguez-Maldonado, Vanessa Rivero-Arredondo, Ariadna Medina-Benitez, Joaquin Quiroz-Mercado, Sergio Rangel-Guerrero, Maribel Gonzalez-Villa, Natividad Cruz-Ortiz, Tatiana Nunez-Garcia, Gisela Barrera-Badillo, Lucia Hernandez-Rivas, Irma Lopez-Martinez, Ernesto Ramirez-Gonzalez. |
| EPI_ISL_3265565 | LESP Estado de Mexico | Instituto de Diagnostico y Referencia Epidemiologicos (INDRE) | Claudia Wong-Arambula, Abril Rodriguez-Maldonado, Vanessa Rivero-Arredondo, Ariadna Medina-Benitez, Joaquin Quiroz-Mercado, Sergio Rangel-Guerrero, Maribel Gonzalez-Villa, Natividad Cruz-Ortiz, Tatiana Nunez-Garcia, Gisela Barrera-Badillo, Lucia Hernandez-Rivas, Irma Lopez-Martinez, Ernesto Ramirez-Gonzalez. |
| EPI_ISL_3265678, EPI_ISL_3265679, EPI_ISL_3265680, EPI_ISL_3265684, EPI_ISL_3265685, EPI_ISL_3265686, EPI_ISL_3265687, EPI_ISL_3265688, EPI_ISL_3265689, EPI_ISL_3265690, EPI_ISL_3265691, EPI_ISL_3265692, EPI_ISL_3265693, EPI_ISL_3265694, EPI_ISL_3265695, EPI_ISL_3265696, EPI_ISL_3265697, EPI_ISL_3265698, EPI_ISL_3265699, EPI_ISL_3265700, EPI_ISL_3265701, EPI_ISL_3265702, EPI_ISL_3265704, EPI_ISL_3265705, EPI_ISL_3265706, EPI_ISL_3265714, EPI_ISL_3265725, EPI_ISL_3265726, EPI_ISL_3265727, EPI_ISL_3265728, EPI_ISL_3265729, EPI_ISL_3265730, EPI_ISL_3265731, EPI_ISL_3265732, EPI_ISL_3265733, EPI_ISL_3265734, EPI_ISL_3265735, EPI_ISL_3265736, EPI_ISL_3265737, EPI_ISL_3265738, EPI_ISL_3265739, EPI_ISL_3265740, EPI_ISL_3265741, EPI_ISL_3265742, EPI_ISL_3265743, EPI_ISL_3265744, EPI_ISL_3265745, EPI_ISL_3265746, EPI_ISL_3265747, EPI_ISL_3265748, EPI_ISL_3265749, EPI_ISL_3265750, EPI_ISL_3265751, EPI_ISL_3265752, EPI_ISL_3265753, EPI_ISL_3265754, EPI_ISL_3265755, EPI_ISL_3265756, EPI_ISL_3265757, EPI_ISL_3265758, EPI_ISL_3265759, EPI_ISL_3265760, EPI_ISL_3265761, EPI_ISL_3265762, EPI_ISL_3265763, EPI_ISL_3265764, EPI_ISL_3265765, EPI_ISL_3265766, EPI_ISL_3265767, EPI_ISL_3265769, EPI_ISL_3265770, EPI_ISL_3265771, EPI_ISL_3265772, EPI_ISL_3265774, EPI_ISL_3265775, EPI_ISL_3265776, EPI_ISL_3265777, EPI_ISL_3265778, EPI_ISL_3265779, EPI_ISL_3265781, EPI_ISL_3265782, EPI_ISL_3265784, EPI_ISL_3265786, EPI_ISL_3265787, EPI_ISL_3265788, EPI_ISL_3265789, EPI_ISL_3265790, EPI_ISL_3265791, EPI_ISL_3265792, EPI_ISL_3265794, EPI_ISL_3265795, EPI_ISL_3265796, EPI_ISL_3265797, EPI_ISL_3265798, EPI_ISL_3265800, EPI_ISL_3265801, EPI_ISL_3265802, EPI_ISL_3265803, EPI_ISL_3265804, EPI_ISL_3265806, EPI_ISL_3265807, EPI_ISL_3265808, EPI_ISL_3265809, EPI_ISL_3265811, EPI_ISL_3265812, EPI_ISL_3265813, EPI_ISL_3265814, EPI_ISL_3265815, EPI_ISL_3265816, EPI_ISL_3265817, EPI_ISL_3265818, EPI_ISL_3265819, EPI_ISL_3265820, EPI_ISL_3265821, EPI_ISL_3265822, EPI_ISL_3265823, EPI_ISL_3265824, EPI_ISL_3265826, EPI_ISL_3265827, EPI_ISL_3265828, EPI_ISL_3265830, EPI_ISL_3265831, EPI_ISL_3265832, EPI_ISL_3265833, EPI_ISL_3265834, EPI_ISL_3265836, EPI_ISL_3265837, EPI_ISL_3265838, EPI_ISL_3265840, EPI_ISL_3265841, EPI_ISL_3265842, EPI_ISL_3265844, EPI_ISL_3265845, EPI_ISL_3265846, EPI_ISL_3265847, EPI_ISL_3265848, EPI_ISL_3265849, EPI_ISL_3265850, EPI_ISL_3265852, EPI_ISL_3265853, EPI_ISL_3265854, EPI_ISL_3265855, EPI_ISL_3265856, EPI_ISL_3265857, EPI_ISL_3265858, EPI_ISL_3265859, EPI_ISL_3265860, EPI_ISL_3265862, EPI_ISL_3265863, EPI_ISL_3265864, EPI_ISL_3265865, EPI_ISL_3265866, EPI_ISL_3265867, EPI_ISL_3265868, EPI_ISL_3265869, EPI_ISL_3265870, EPI_ISL_3265871, EPI_ISL_3265872, EPI_ISL_3265873, EPI_ISL_3265874, EPI_ISL_3265875, EPI_ISL_3265876, EPI_ISL_3265877, EPI_ISL_3265878, EPI_ISL_3265879, EPI_ISL_3265880, EPI_ISL_3265881, EPI_ISL_3265882, EPI_ISL_3265883, EPI_ISL_3265884, EPI_ISL_3265885, EPI_ISL_3265886, EPI_ISL_3265887, EPI_ISL_3265889, EPI_ISL_3265890, EPI_ISL_3265891, EPI_ISL_3265892, EPI_ISL_3265894, EPI_ISL_3265896, EPI_ISL_3265898, EPI_ISL_3265900, EPI_ISL_3265902, EPI_ISL_3265903, EPI_ISL_3265904, EPI_ISL_3265905, EPI_ISL_3265906, EPI_ISL_3265907, EPI_ISL_3265908, EPI_ISL_3265909, EPI_ISL_3265910, EPI_ISL_3265911, EPI_ISL_3265912, EPI_ISL_3265916, EPI_ISL_3265917, EPI_ISL_3265918, EPI_ISL_3265920, EPI_ISL_3265921, EPI_ISL_3265922, EPI_ISL_3265923, EPI_ISL_3265924, EPI_ISL_3265925, EPI_ISL_3265926, EPI_ISL_3265927, EPI_ISL_3265928, EPI_ISL_3265929, EPI_ISL_3265930, EPI_ISL_3265931, EPI_ISL_3265933, EPI_ISL_3265934, EPI_ISL_3265935, EPI_ISL_3265936, EPI_ISL_3265937, EPI_ISL_3265938, EPI_ISL_3265939, EPI_ISL_3265940, EPI_ISL_3265941, EPI_ISL_3265942, EPI_ISL_3265943, EPI_ISL_3265944, EPI_ISL_3265945, EPI_ISL_3265946, EPI_ISL_3265947, EPI_ISL_3265948, EPI_ISL_3265949, EPI_ISL_3265950, EPI_ISL_3265951, EPI_ISL_3265952, EPI_ISL_3265953, EPI_ISL_3265954, EPI_ISL_3265955, EPI_ISL_3265969, EPI_ISL_3265981, EPI_ISL_3265982, EPI_ISL_3265983, EPI_ISL_3265984, EPI_ISL_3265985, EPI_ISL_3265986, EPI_ISL_3265988, EPI_ISL_3265989, EPI_ISL_3265990, EPI_ISL_3265991, EPI_ISL_3265994, EPI_ISL_3265995, EPI_ISL_3265996, EPI_ISL_3265997, EPI_ISL_3265999, EPI_ISL_3266000, EPI_ISL_3266001, EPI_ISL_3266002, EPI_ISL_3266004, EPI_ISL_3266005, EPI_ISL_3266006, EPI_ISL_3266014, EPI_ISL_3266015, EPI_ISL_3266016, EPI_ISL_3266017, EPI_ISL_3266018, EPI_ISL_3266023, EPI_ISL_3266025, EPI_ISL_3266028, EPI_ISL_3266029, EPI_ISL_3266030, EPI_ISL_3266031 | | | |
| see above | Servicio de Microbiología Clínica (Complejo Hospitalario de Navarra, Pamplona) | Centro de Secuenciación NASERTIC | Carmen Ezpeleta Baquedano, Ana Navascués, Ana Miqueleiz |
| EPI_ISL_3266032 | Muar Clinic | Institute for Medical Research, Infectious Disease Research Centre, National Institutes of Health, Ministry of Health Malaysia | Suppiah J, Kamel K, Mohd Zawawi Z, Azizan MA, Ramly N, Robert F, Anasir MI, Thayan R |
| EPI_ISL_3266033, EPI_ISL_3266034, EPI_ISL_3266035, EPI_ISL_3266036, EPI_ISL_3266037, EPI_ISL_3266038, EPI_ISL_3266040, EPI_ISL_3266041, EPI_ISL_3266042 | State Hygienic Laboratory at the University of Iowa | State Hygienic Laboratory at the University of Iowa | Valerie Reeb, Erik Twait, Anna Yakos, Mariah Knutson, Kris Eveland, Kristen Zanon, Jeff, Benfer, Davis Rieckenberg, Wes Hottel, Alankar Kampoowale |
| EPI_ISL_3266043 | University of Iowa Hospitals and Clinics | State Hygienic Laboratory at the University of Iowa | Valerie Reeb, Erik Twait, Anna Yakos, Mariah Knutson, Kris Eveland, Kristen Zanon, Jeff, Benfer, Davis Rieckenberg, Wes Hottel, Alankar Kampoowale |
| EPI_ISL_3266044, EPI_ISL_3266045, EPI_ISL_3266046, EPI_ISL_3266047, EPI_ISL_3266048, EPI_ISL_3266049, EPI_ISL_3266050, EPI_ISL_3266051, EPI_ISL_3266052, EPI_ISL_3266053, EPI_ISL_3266055, EPI_ISL_3266056, EPI_ISL_3266057, EPI_ISL_3266058, EPI_ISL_3266059, EPI_ISL_3266060, EPI_ISL_3266061 | | | |
| see above | State Hygienic Laboratory at the University of Iowa | State Hygienic Laboratory at the University of Iowa | Valerie Reeb, Erik Twait, Anna Yakos, Mariah Knutson, Kris Eveland, Kristen Zanon, Jeff, Benfer, Davis Rieckenberg, Wes Hottel, Alankar Kampoowale |
| EPI_ISL_3266062 | Asia Muar Clinic | Institute for Medical Research, Infectious Disease Research Centre, National Institutes of Health, Ministry of Health Malaysia | Suppiah J, Kamel K, Mohd Zawawi Z, Azizan MA, Ramly N, Robert F, Anasir MI, Thayan R |
| EPI_ISL_3266063, EPI_ISL_3266064, EPI_ISL_3266065 | Melaka Hospital | Institute for Medical Research, Infectious Disease Research Centre, National Institutes of Health, Ministry of Health Malaysia | Suppiah J, Kamel K, Mohd Zawawi Z, Azizan MA, Ramly N, Robert F, Anasir MI, Thayan R |
| EPI_ISL_3266066, EPI_ISL_3266068 | Multidisciplinary Research Unit, Institute of Medical Sciences, Banaras Hindu University, Varanasi - 221005 | Multidisciplinary Research Unit, Institute of Medical Sciences, Banaras Hindu University, Varanasi - 221005 | Chetan Sahni, Royana Singh, Priyoneel Basu, Nitish Kumar Singh, Saumya Singh, Pranav Singh, Deepa Devadas, Gunjan Rai, Sarita Chowdhary, Saurabh Singh, Sangeeta Rai, Umesh Chaudhary, Ashish Ashish, Abhay Kumar Yadav, Manpreet Kaur, Ajay Kumar Yadav, Shivangi Mishra, Meghna Wadi, Shani Vishwakarma, Shivam Tiwari, Meghna Wadi, Nargis Khanam, Janhavi Yadav, Ragini Mishra, Surendra Pratap Mishra, Tribhuwan Mohan Mohapatra |
| EPI_ISL_3266069, EPI_ISL_3266070 | Multidisciplinary Research Unit, Institute of Medical Sciences, Banaras Hindu University, Varanasi - 221005 | Multidisciplinary Research Unit, Institute of Medical Sciences, Banaras Hindu University, Varanasi - 221005 | Priyoneel Basu, Royana Singh, Chetan Sahni, Nitish Kumar Singh, Ashish Ashish, Abhay Kumar Yadav, Manpreet Kaur, Saumya Singh, Deepa Devadas, Gunjan Rai, Sarita Chowdhary, Saurabh Singh, Sangeeta Rai, Umesh Chaudhary, Ajay Kumar Yadav, Shivangi Mishra, Meghna Wadi, Shani Vishwakarma, Shivam Tiwari, Meghna Wadi, Nargis Khanam, Janhavi Yadav, Ragini Mishra, Surendra Pratap Mishra, Tribhuwan Mohan Mohapatra |
| EPI_ISL_3266126, EPI_ISL_3266127, EPI_ISL_3266128, EPI_ISL_3266129, EPI_ISL_3266130, EPI_ISL_3266131, EPI_ISL_3266132, EPI_ISL_3266133, EPI_ISL_3266134, EPI_ISL_3266135, EPI_ISL_3266136, EPI_ISL_3266137, EPI_ISL_3266138, EPI_ISL_3266139, EPI_ISL_3266140, EPI_ISL_3266141, EPI_ISL_3266142, EPI_ISL_3266143, EPI_ISL_3266144, EPI_ISL_3266145, EPI_ISL_3266146, EPI_ISL_3266147, EPI_ISL_3266148, EPI_ISL_3266149, EPI_ISL_3266150, EPI_ISL_3266151, EPI_ISL_3266152, EPI_ISL_3266153, EPI_ISL_3266154, EPI_ISL_3266155, EPI_ISL_3266156, EPI_ISL_3266157, EPI_ISL_3266158, EPI_ISL_3266159, EPI_ISL_3266160, EPI_ISL_3266161, EPI_ISL_3266162, EPI_ISL_3266163, EPI_ISL_3266164, EPI_ISL_3266165, EPI_ISL_3266166, EPI_ISL_3266167, EPI_ISL_3266168, EPI_ISL_3266169, EPI_ISL_3266170, EPI_ISL_3266171, EPI_ISL_3266172, EPI_ISL_3266173, EPI_ISL_3266174, EPI_ISL_3266175, EPI_ISL_3266176, EPI_ISL_3266177, EPI_ISL_3266178, EPI_ISL_3266179, EPI_ISL_3266180, EPI_ISL_3266181, EPI_ISL_3266182, EPI_ISL_3266183, EPI_ISL_3266184, EPI_ISL_3266185, EPI_ISL_3266186, EPI_ISL_3266187, EPI_ISL_3266188, EPI_ISL_3266189, EPI_ISL_3266190, EPI_ISL_3266191, EPI_ISL_3266192, EPI_ISL_3266193, EPI_ISL_3266194, EPI_ISL_3266195, EPI_ISL_3266196, EPI_ISL_3266197, EPI_ISL_3266198, EPI_ISL_3266199, EPI_ISL_3266200, EPI_ISL_3266201, EPI_ISL_3266202, EPI_ISL_3266203, EPI_ISL_3266204, EPI_ISL_3266205, EPI_ISL_3266206, EPI_ISL_3266207, EPI_ISL_3266208, EPI_ISL_3266209 | | | |
| see above | National Public Health Laboratory, National Centre for Infectious Diseases | National Public Health Laboratory, National Centre for Infectious Diseases | Tze Minn Mak, Katherine Ching, Zhenyang Zhou, Royce Ang, Lin Cui, Raymond Tzer Pin Lin |
| EPI_ISL_3266235 | AZDelta | AZ Delta Medical Laboratories in Roeselare, Belgium | Geert Martens, Dieter De Smet, Frederik Van Hoecke, on behalf of AZ Delta COVID-19 Genomics core (member of Genomic surveillance of SARS-CoV-2 in Belgium network) |
| EPI_ISL_3266243, EPI_ISL_3266244, EPI_ISL_3266245, EPI_ISL_3266246, EPI_ISL_3266247, EPI_ISL_3266248, EPI_ISL_3266249, EPI_ISL_3266250, EPI_ISL_3266251, EPI_ISL_3266253, EPI_ISL_3266254, EPI_ISL_3266255, EPI_ISL_3266256, EPI_ISL_3266257, EPI_ISL_3266258 | | | |
| see above | Medical Laboratories Duesseldorf | Center of Medical Microbiology, Virology, and Hospital Hygiene, University of Duesseldorf | Maximilian Damagnez;Alexander Dilthey;Angelika Helmer;Torsten Houwaart;Lisanna Hülse;Christian Lange;Malte Kohns Vasconcelos;Nadine Lübke;Jessica Nicolai;Klaus Pfeffer;Daniel Strelow;Jörg Timm;Andreas Walker;Tobias Wienemann |
| EPI_ISL_3266266, EPI_ISL_3266295, EPI_ISL_3266296, EPI_ISL_3266300, EPI_ISL_3266306, EPI_ISL_3266307, EPI_ISL_3266309, EPI_ISL_3266313, EPI_ISL_3266333, EPI_ISL_3266334, EPI_ISL_3266335, EPI_ISL_3266336, EPI_ISL_3266337, EPI_ISL_3266338, EPI_ISL_3266339, EPI_ISL_3266346, EPI_ISL_3266348 | | | |
| see above | U.O. Microbiologia Laboratorio Unico Centro Servizi - AUSL della Romagna | U.O. Microbiologia, Laboratorio Unico Centro Servizi - AUSL della Romagna | Giorgio Dirani |
| EPI_ISL_3266358, EPI_ISL_3266359, EPI_ISL_3266361, EPI_ISL_3266364, EPI_ISL_3266365, EPI_ISL_3266379, EPI_ISL_3266389, EPI_ISL_3266390, EPI_ISL_3266392, EPI_ISL_3266393, EPI_ISL_3266395, EPI_ISL_3266398, EPI_ISL_3266402, EPI_ISL_3266403, EPI_ISL_3266404, EPI_ISL_3266405, EPI_ISL_3266406, EPI_ISL_3266407, EPI_ISL_3266408, EPI_ISL_3266409, EPI_ISL_3266410, EPI_ISL_3266411, EPI_ISL_3266414, EPI_ISL_3266415, EPI_ISL_3266416, EPI_ISL_3266417, EPI_ISL_3266418, EPI_ISL_3266419 | | | |
| see above | Università di Parma, Laboratorio di Igiene e Sanità Pubblica | Istituto Zooprofilattico Sperimentale della Lombardia e dell'Emilia Romagna (IZSLER), Risk Analysis and Genomic Epidemiology Unit | Maria Eugenia Colucci, Licia Veronesi, Paola Affanni, Marina Morganti, Ilaria Menozzi, Erika Scaltriti, Stefano Pongolini |
| EPI_ISL_3266442, EPI_ISL_3266446, EPI_ISL_3266448, EPI_ISL_3266449, EPI_ISL_3266450, EPI_ISL_3266451 | U.O. Microbiologia Laboratorio Unico Centro Servizi - AUSL della Romagna | U.O. Microbiologia, Laboratorio Unico Centro Servizi - AUSL della Romagna | Giorgio Dirani |
| EPI_ISL_3266454 | Innlandet Hospital Trust, Division Lillehammer, Department for Medical Microbiology | Norwegian Institute of Public Health, Department of Virology | Kathrine Stene-Johansen, Kamilla Heddeland Instefjord, Hilde Elshaug, Garcia Llorente Ignacio, Jon Bråte, Engebretsen Serina Beate,Pedersen Benedikte Nevjen, Line Victoria Moen, Debech Nadia, Atiya R Ali,Marie Paulsen Madsen, Rasmus Riis Kopperud, Hilde Vollan, Karoline Bragstad, Olav Hungnes |
| EPI_ISL_3266455 | Vestfold Hospital, Toensberg Department of Microbiology | Norwegian Institute of Public Health, Department of Virology | Kathrine Stene-Johansen, Kamilla Heddeland Instefjord, Hilde Elshaug, Garcia Llorente Ignacio, Jon Bråte, Engebretsen Serina Beate,Pedersen Benedikte Nevjen, Line Victoria Moen, Debech Nadia, Atiya R Ali,Marie Paulsen Madsen, Rasmus Riis Kopperud, Hilde Vollan, Karoline Bragstad, Olav Hungnes |
| EPI_ISL_3266456 | Hospital of Southern Norway - Kristiansand, Department of Medical Microbiology | Norwegian Institute of Public Health, Department of Virology | Kathrine Stene-Johansen, Kamilla Heddeland Instefjord, Hilde Elshaug, Garcia Llorente Ignacio, Jon Bråte, Engebretsen Serina Beate,Pedersen Benedikte Nevjen, Line Victoria Moen, Debech Nadia, Atiya R Ali,Marie Paulsen Madsen, Rasmus Riis Kopperud, Hilde Vollan, Karoline Bragstad, Olav Hungnes |
| EPI_ISL_3266457, EPI_ISL_3266460 | Nordland Hospital - Bodo, Laboratory Department, Molecular Biology Unit | Norwegian Institute of Public Health, Department of Virology | Kathrine Stene-Johansen, Kamilla Heddeland Instefjord, Hilde Elshaug, Garcia Llorente Ignacio, Jon Bråte, Engebretsen Serina Beate,Pedersen Benedikte Nevjen, Line Victoria Moen, Debech Nadia, Atiya R Ali,Marie Paulsen Madsen, Rasmus Riis Kopperud, Hilde Vollan, Karoline Bragstad, Olav Hungnes |
| EPI_ISL_3266462 | Innlandet Hospital Trust, Division Lillehammer, Department for Medical Microbiology | Norwegian Institute of Public Health, Department of Virology | Kathrine Stene-Johansen, Kamilla Heddeland Instefjord, Hilde Elshaug, Garcia Llorente Ignacio, Jon Bråte, Engebretsen Serina Beate,Pedersen Benedikte Nevjen, Line Victoria Moen, Debech Nadia, Atiya R Ali,Marie Paulsen Madsen, Rasmus Riis Kopperud, Hilde Vollan, Karoline Bragstad, Olav Hungnes |
| EPI_ISL_3266465 | Hospital of Southern Norway - Kristiansand, Department of Medical Microbiology | Norwegian Institute of Public Health, Department of Virology | Kathrine Stene-Johansen, Kamilla Heddeland Instefjord, Hilde Elshaug, Garcia Llorente Ignacio, Jon Bråte, Engebretsen Serina Beate,Pedersen Benedikte Nevjen, Line Victoria Moen, Debech Nadia, Atiya R Ali,Marie Paulsen Madsen, Rasmus Riis Kopperud, Hilde Vollan, Karoline Bragstad, Olav Hungnes |
| EPI_ISL_3266468 | Department of Medical Microbiology - section Molde, Molde Hospital | Norwegian Institute of Public Health, Department of Virology | Kathrine Stene-Johansen, Kamilla Heddeland Instefjord, Hilde Elshaug, Garcia Llorente Ignacio, Jon Bråte, Engebretsen Serina Beate,Pedersen Benedikte Nevjen, Line Victoria Moen, Debech Nadia, Atiya R Ali,Marie Paulsen Madsen, Rasmus Riis Kopperud, Hilde Vollan, Karoline Bragstad, Olav Hungnes |
| EPI_ISL_3266469, EPI_ISL_3266470 | Nordland Hospital - Bodo, Laboratory Department, Molecular Biology Unit | Norwegian Institute of Public Health, Department of Virology | Kathrine Stene-Johansen, Kamilla Heddeland Instefjord, Hilde Elshaug, Garcia Llorente Ignacio, Jon Bråte, Engebretsen Serina Beate,Pedersen Benedikte Nevjen, Line Victoria Moen, Debech Nadia, Atiya R Ali,Marie Paulsen Madsen, Rasmus Riis Kopperud, Hilde Vollan, Karoline Bragstad, Olav Hungnes |

| | | | |
|---|---|---|---|
| EPI_ISL_3266471 | Department of Medical Microbiology - section Molde, Molde Hospital | Norwegian Institute of Public Health, Department of Virology | Kathrine Stene-Johansen, Kamilla Heddeland Instefjord, Hilde Elshaug, Garcia Llorente Ignacio, Jon Bråte, Engebretsen Serina Beate,Pedersen Benedikte Nevjen, Line Victoria Moen, Debech Nadia, Atiya R Ali,Marie Paulsen Madsen, Rasmus Riis Kopperud, Hilde Vollan, Karoline Bragstad, Olav Hungnes |
| EPI_ISL_3266472 | Nordland Hospital - Bodo, Laboratory Department, Molecular Biology Unit | Norwegian Institute of Public Health, Department of Virology | Kathrine Stene-Johansen, Kamilla Heddeland Instefjord, Hilde Elshaug, Garcia Llorente Ignacio, Jon Bråte, Engebretsen Serina Beate,Pedersen Benedikte Nevjen, Line Victoria Moen, Debech Nadia, Atiya R Ali,Marie Paulsen Madsen, Rasmus Riis Kopperud, Hilde Vollan, Karoline Bragstad, Olav Hungnes |
| EPI_ISL_3266473 | Department of Medical Microbiology, St. Olavs hospital | Norwegian Institute of Public Health, Department of Virology | Kathrine Stene-Johansen, Kamilla Heddeland Instefjord, Hilde Elshaug, Garcia Llorente Ignacio, Jon Bråte, Engebretsen Serina Beate,Pedersen Benedikte Nevjen, Line Victoria Moen, Debech Nadia, Atiya R Ali,Marie Paulsen Madsen, Rasmus Riis Kopperud, Hilde Vollan, Karoline Bragstad, Olav Hungnes |
| EPI_ISL_3266474 | Department of Medical Microbiology - section Molde, Molde Hospital | Norwegian Institute of Public Health, Department of Virology | Kathrine Stene-Johansen, Kamilla Heddeland Instefjord, Hilde Elshaug, Garcia Llorente Ignacio, Jon Bråte, Engebretsen Serina Beate,Pedersen Benedikte Nevjen, Line Victoria Moen, Debech Nadia, Atiya R Ali,Marie Paulsen Madsen, Rasmus Riis Kopperud, Hilde Vollan, Karoline Bragstad, Olav Hungnes |
| EPI_ISL_3266475, EPI_ISL_3266476 | Hospital of Southern Norway - Kristiansand, Department of Medical Microbiology | Norwegian Institute of Public Health, Department of Virology | Kathrine Stene-Johansen, Kamilla Heddeland Instefjord, Hilde Elshaug, Garcia Llorente Ignacio, Jon Bråte, Engebretsen Serina Beate,Pedersen Benedikte Nevjen, Line Victoria Moen, Debech Nadia, Atiya R Ali,Marie Paulsen Madsen, Rasmus Riis Kopperud, Hilde Vollan, Karoline Bragstad, Olav Hungnes |
| EPI_ISL_3266478 | Furst Medical Laboratory | Norwegian Institute of Public Health, Department of Virology | Kathrine Stene-Johansen, Kamilla Heddeland Instefjord, Hilde Elshaug, Garcia Llorente Ignacio, Jon Bråte, Engebretsen Serina Beate,Pedersen Benedikte Nevjen, Line Victoria Moen, Debech Nadia, Atiya R Ali,Marie Paulsen Madsen, Rasmus Riis Kopperud, Hilde Vollan, Karoline Bragstad, Olav Hungnes |
| EPI_ISL_3266479 | Haukeland University Hospital, Dept. of Microbiology | Norwegian Institute of Public Health, Department of Virology | Kathrine Stene-Johansen, Kamilla Heddeland Instefjord, Hilde Elshaug, Garcia Llorente Ignacio, Jon Bråte, Engebretsen Serina Beate,Pedersen Benedikte Nevjen, Line Victoria Moen, Debech Nadia, Atiya R Ali,Marie Paulsen Madsen, Rasmus Riis Kopperud, Hilde Vollan, Karoline Bragstad, Olav Hungnes |
| EPI_ISL_3266481 | Furst Medical Laboratory | Norwegian Institute of Public Health, Department of Virology | Kathrine Stene-Johansen, Kamilla Heddeland Instefjord, Hilde Elshaug, Garcia Llorente Ignacio, Jon Bråte, Engebretsen Serina Beate,Pedersen Benedikte Nevjen, Line Victoria Moen, Debech Nadia, Atiya R Ali,Marie Paulsen Madsen, Rasmus Riis Kopperud, Hilde Vollan, Karoline Bragstad, Olav Hungnes |
| EPI_ISL_3266482 | Ostfold Hospital Trust - Kalnes, Centre for Laboratory Medicine, Section for gene technology and infection serology | Norwegian Institute of Public Health, Department of Virology | Kathrine Stene-Johansen, Kamilla Heddeland Instefjord, Hilde Elshaug, Garcia Llorente Ignacio, Jon Bråte, Engebretsen Serina Beate,Pedersen Benedikte Nevjen, Line Victoria Moen, Debech Nadia, Atiya R Ali,Marie Paulsen Madsen, Rasmus Riis Kopperud, Hilde Vollan, Karoline Bragstad, Olav Hungnes |
| EPI_ISL_3266483 | Haukeland University Hospital, Dept. of Microbiology | Norwegian Institute of Public Health, Department of Virology | Kathrine Stene-Johansen, Kamilla Heddeland Instefjord, Hilde Elshaug, Garcia Llorente Ignacio, Jon Bråte, Engebretsen Serina Beate,Pedersen Benedikte Nevjen, Line Victoria Moen, Debech Nadia, Atiya R Ali,Marie Paulsen Madsen, Rasmus Riis Kopperud, Hilde Vollan, Karoline Bragstad, Olav Hungnes |
| EPI_ISL_3266485 | Medical Microbiology Unit, Department for Laboratory Medicine, Drammen Hospital, Vestre Viken Health Trust, | Norwegian Institute of Public Health, Department of Virology | Kathrine Stene-Johansen, Kamilla Heddeland Instefjord, Hilde Elshaug, Garcia Llorente Ignacio, Jon Bråte, Engebretsen Serina Beate,Pedersen Benedikte Nevjen, Line Victoria Moen, Debech Nadia, Atiya R Ali,Marie Paulsen Madsen, Rasmus Riis Kopperud, Hilde Vollan, Karoline Bragstad, Olav Hungnes |
| EPI_ISL_3266487, EPI_ISL_3266488 | Haukeland University Hospital, Dept. of Microbiology | Norwegian Institute of Public Health, Department of Virology | Kathrine Stene-Johansen, Kamilla Heddeland Instefjord, Hilde Elshaug, Garcia Llorente Ignacio, Jon Bråte, Engebretsen Serina Beate,Pedersen Benedikte Nevjen, Line Victoria Moen, Debech Nadia, Atiya R Ali,Marie Paulsen Madsen, Rasmus Riis Kopperud, Hilde Vollan, Karoline Bragstad, Olav Hungnes |
| EPI_ISL_3266490 | Medical Microbiology Unit, Department for Laboratory Medicine, Drammen Hospital, Vestre Viken Health Trust, | Norwegian Institute of Public Health, Department of Virology | Kathrine Stene-Johansen, Kamilla Heddeland Instefjord, Hilde Elshaug, Garcia Llorente Ignacio, Jon Bråte, Engebretsen Serina Beate,Pedersen Benedikte Nevjen, Line Victoria Moen, Debech Nadia, Atiya R Ali,Marie Paulsen Madsen, Rasmus Riis Kopperud, Hilde Vollan, Karoline Bragstad, Olav Hungnes |
| EPI_ISL_3266491 | Unilabs Laboratory Medicine | Norwegian Institute of Public Health, Department of Virology | Kathrine Stene-Johansen, Kamilla Heddeland Instefjord, Hilde Elshaug, Garcia Llorente Ignacio, Jon Bråte, Engebretsen Serina Beate,Pedersen Benedikte Nevjen, Line Victoria Moen, Debech Nadia, Atiya R Ali,Marie Paulsen Madsen, Rasmus Riis Kopperud, Hilde Vollan, Karoline Bragstad, Olav Hungnes |
| EPI_ISL_3266492, EPI_ISL_3266493, EPI_ISL_3266494, EPI_ISL_3266495, EPI_ISL_3266496 | Haukeland University Hospital, Dept. of Microbiology | Norwegian Institute of Public Health, Department of Virology | Kathrine Stene-Johansen, Kamilla Heddeland Instefjord, Hilde Elshaug, Garcia Llorente Ignacio, Jon Bråte, Engebretsen Serina Beate,Pedersen Benedikte Nevjen, Line Victoria Moen, Debech Nadia, Atiya R Ali,Marie Paulsen Madsen, Rasmus Riis Kopperud, Hilde Vollan, Karoline Bragstad, Olav Hungnes |
| EPI_ISL_3266497 | Ostfold Hospital Trust - Kalnes, Centre for Laboratory Medicine, Section for gene technology and infection serology | Norwegian Institute of Public Health, Department of Virology | Kathrine Stene-Johansen, Kamilla Heddeland Instefjord, Hilde Elshaug, Garcia Llorente Ignacio, Jon Bråte, Engebretsen Serina Beate,Pedersen Benedikte Nevjen, Line Victoria Moen, Debech Nadia, Atiya R Ali,Marie Paulsen Madsen, Rasmus Riis Kopperud, Hilde Vollan, Karoline Bragstad, Olav Hungnes |
| EPI_ISL_3266499 | Medical Microbiology Unit, Department for Laboratory Medicine, Drammen Hospital, Vestre Viken Health Trust, | Norwegian Institute of Public Health, Department of Virology | Kathrine Stene-Johansen, Kamilla Heddeland Instefjord, Hilde Elshaug, Garcia Llorente Ignacio, Jon Bråte, Engebretsen Serina Beate,Pedersen Benedikte Nevjen, Line Victoria Moen, Debech Nadia, Atiya R Ali,Marie Paulsen Madsen, Rasmus Riis Kopperud, Hilde Vollan, Karoline Bragstad, Olav Hungnes |
| EPI_ISL_3266500, EPI_ISL_3266501 | Haukeland University Hospital, Dept. of Microbiology | Norwegian Institute of Public Health, Department of Virology | Kathrine Stene-Johansen, Kamilla Heddeland Instefjord, Hilde Elshaug, Garcia Llorente Ignacio, Jon Bråte, Engebretsen Serina Beate,Pedersen Benedikte Nevjen, Line Victoria Moen, Debech Nadia, Atiya R Ali,Marie Paulsen Madsen, Rasmus Riis Kopperud, Hilde Vollan, Karoline Bragstad, Olav Hungnes |
| EPI_ISL_3266502 | Unilabs Laboratory Medicine | Norwegian Institute of Public Health, Department of Virology | Kathrine Stene-Johansen, Kamilla Heddeland Instefjord, Hilde Elshaug, Garcia Llorente Ignacio, Jon Bråte, Engebretsen Serina Beate,Pedersen Benedikte Nevjen, Line Victoria Moen, Debech Nadia, Atiya R Ali,Marie Paulsen Madsen, Rasmus Riis Kopperud, Hilde Vollan, Karoline Bragstad, Olav Hungnes |
| EPI_ISL_3266503 | Ostfold Hospital Trust - Kalnes, Centre for Laboratory Medicine, Section for gene technology and infection serology | Norwegian Institute of Public Health, Department of Virology | Kathrine Stene-Johansen, Kamilla Heddeland Instefjord, Hilde Elshaug, Garcia Llorente Ignacio, Jon Bråte, Engebretsen Serina Beate,Pedersen Benedikte Nevjen, Line Victoria Moen, Debech Nadia, Atiya R Ali,Marie Paulsen Madsen, Rasmus Riis Kopperud, Hilde Vollan, Karoline Bragstad, Olav Hungnes |
| EPI_ISL_3266504 | Furst Medical Laboratory | Norwegian Institute of Public Health, Department of Virology | Kathrine Stene-Johansen, Kamilla Heddeland Instefjord, Hilde Elshaug, Garcia Llorente Ignacio, Jon Bråte, Engebretsen Serina Beate,Pedersen Benedikte Nevjen, Line Victoria Moen, Debech Nadia, Atiya R Ali,Marie Paulsen Madsen, Rasmus Riis Kopperud, Hilde Vollan, Karoline Bragstad, Olav Hungnes |
| EPI_ISL_3266505 | Haukeland University Hospital, Dept. of Microbiology | Norwegian Institute of Public Health, Department of Virology | Kathrine Stene-Johansen, Kamilla Heddeland Instefjord, Hilde Elshaug, Garcia Llorente Ignacio, Jon Bråte, Engebretsen Serina Beate,Pedersen Benedikte Nevjen, Line Victoria Moen, Debech Nadia, Atiya R Ali,Marie Paulsen Madsen, Rasmus Riis Kopperud, Hilde Vollan, Karoline Bragstad, Olav Hungnes |
| EPI_ISL_3266506 | Medical Microbiology Unit, Department for Laboratory Medicine, Drammen Hospital, Vestre Viken Health Trust, | Norwegian Institute of Public Health, Department of Virology | Kathrine Stene-Johansen, Kamilla Heddeland Instefjord, Hilde Elshaug, Garcia Llorente Ignacio, Jon Bråte, Engebretsen Serina Beate,Pedersen Benedikte Nevjen, Line Victoria Moen, Debech Nadia, Atiya R Ali,Marie Paulsen Madsen, Rasmus Riis Kopperud, Hilde Vollan, Karoline Bragstad, Olav Hungnes |
| EPI_ISL_3266507 | Haukeland University Hospital, Dept. of Microbiology | Norwegian Institute of Public Health, Department of Virology | Kathrine Stene-Johansen, Kamilla Heddeland Instefjord, Hilde Elshaug, Garcia Llorente Ignacio, Jon Bråte, Engebretsen Serina Beate,Pedersen Benedikte Nevjen, Line Victoria Moen, Debech Nadia, Atiya R Ali,Marie Paulsen Madsen, Rasmus Riis Kopperud, Hilde Vollan, Karoline Bragstad, Olav Hungnes |
| EPI_ISL_3266508 | Medical Microbiology Unit, Department for Laboratory Medicine, Drammen Hospital, Vestre Viken Health Trust, | Norwegian Institute of Public Health, Department of Virology | Kathrine Stene-Johansen, Kamilla Heddeland Instefjord, Hilde Elshaug, Garcia Llorente Ignacio, Jon Bråte, Engebretsen Serina Beate,Pedersen Benedikte Nevjen, Line Victoria Moen, Debech Nadia, Atiya R Ali,Marie Paulsen Madsen, Rasmus Riis Kopperud, Hilde Vollan, Karoline Bragstad, Olav Hungnes |
| EPI_ISL_3266510, EPI_ISL_3266511 | Oslo University Hospital, Department of Medical Microbiology | Norwegian Institute of Public Health, Department of Virology | Kathrine Stene-Johansen, Kamilla Heddeland Instefjord, Hilde Elshaug, Garcia Llorente Ignacio, Jon Bråte, Engebretsen Serina Beate,Pedersen Benedikte Nevjen, Line Victoria Moen, Debech Nadia, Atiya R Ali,Marie Paulsen Madsen, Rasmus Riis Kopperud, Hilde Vollan, Karoline Bragstad, Olav Hungnes |
| EPI_ISL_3266512 | Furst Medical Laboratory | Norwegian Institute of Public Health, Department of Virology | Kathrine Stene-Johansen, Kamilla Heddeland Instefjord, Hilde Elshaug, Garcia Llorente Ignacio, Jon Bråte, Engebretsen Serina Beate,Pedersen Benedikte Nevjen, Line Victoria Moen, Debech Nadia, Atiya R Ali,Marie Paulsen Madsen, Rasmus Riis Kopperud, Hilde Vollan, Karoline Bragstad, Olav Hungnes |
| EPI_ISL_3266513, EPI_ISL_3266514, EPI_ISL_3266515 | Nordland Hospital - Bodo, Laboratory Department, Molecular Biology Unit | Norwegian Institute of Public Health, Department of Virology | Kathrine Stene-Johansen, Kamilla Heddeland Instefjord, Hilde Elshaug, Garcia Llorente Ignacio, Jon Bråte, Engebretsen Serina Beate,Pedersen Benedikte Nevjen, Line Victoria Moen, Debech Nadia, Atiya R Ali,Marie Paulsen Madsen, Rasmus Riis Kopperud, Hilde Vollan, Karoline Bragstad, Olav Hungnes |
| EPI_ISL_3266516 | Oslo University Hospital, Department of Medical Microbiology | Norwegian Institute of Public Health, Department of Virology | Kathrine Stene-Johansen, Kamilla Heddeland Instefjord, Hilde Elshaug, Garcia Llorente Ignacio, Jon Bråte, Engebretsen Serina Beate,Pedersen Benedikte Nevjen, Line Victoria Moen, Debech Nadia, Atiya R Ali,Marie Paulsen Madsen, Rasmus Riis Kopperud, Hilde Vollan, Karoline Bragstad, Olav Hungnes |
| EPI_ISL_3266517 | Medical Microbiology Unit, Department for Laboratory Medicine, Drammen Hospital, Vestre Viken Health Trust, | Norwegian Institute of Public Health, Department of Virology | Kathrine Stene-Johansen, Kamilla Heddeland Instefjord, Hilde Elshaug, Garcia Llorente Ignacio, Jon Bråte, Engebretsen Serina Beate,Pedersen Benedikte Nevjen, Line Victoria Moen, Debech Nadia, Atiya R Ali,Marie Paulsen Madsen, Rasmus Riis Kopperud, Hilde Vollan, Karoline Bragstad, Olav Hungnes |
| EPI_ISL_3266520 | Furst Medical Laboratory | Norwegian Institute of Public Health, Department of Virology | Kathrine Stene-Johansen, Kamilla Heddeland Instefjord, Hilde Elshaug, Garcia Llorente Ignacio, Jon Bråte, Engebretsen Serina Beate,Pedersen Benedikte Nevjen, Line Victoria Moen, Debech Nadia, Atiya R Ali,Marie Paulsen Madsen, Rasmus Riis Kopperud, Hilde Vollan, Karoline Bragstad, Olav Hungnes |
| EPI_ISL_3266521, EPI_ISL_3266522 | University Hospital of Northern Norway, Department for Microbiology and Infectious Disease Control | Norwegian Institute of Public Health, Department of Virology | Kathrine Stene-Johansen, Kamilla Heddeland Instefjord, Hilde Elshaug, Garcia Llorente Ignacio, Jon Bråte, Engebretsen Serina Beate,Pedersen Benedikte Nevjen, Line Victoria Moen, Debech Nadia, Atiya R Ali,Marie Paulsen Madsen, Rasmus Riis Kopperud, Hilde Vollan, Karoline Bragstad, Olav Hungnes |
| EPI_ISL_3266524 | Innlandet Hospital Trust, Division Lillehammer, Department for Medical Microbiology | Norwegian Institute of Public Health, Department of Virology | Kathrine Stene-Johansen, Kamilla Heddeland Instefjord, Hilde Elshaug, Garcia Llorente Ignacio, Jon Bråte, Engebretsen Serina Beate,Pedersen Benedikte Nevjen, Line Victoria Moen, Debech Nadia, Atiya R Ali,Marie Paulsen Madsen, Rasmus Riis Kopperud, Hilde Vollan, Karoline Bragstad, Olav Hungnes |
| EPI_ISL_3266525 | Hospital of Southern Norway - Kristiansand, Department of Medical Microbiology | Norwegian Institute of Public Health, Department of Virology | Kathrine Stene-Johansen, Kamilla Heddeland Instefjord, Hilde Elshaug, Garcia Llorente Ignacio, Jon Bråte, Engebretsen Serina Beate,Pedersen Benedikte Nevjen, Line Victoria Moen, Debech Nadia, Atiya R Ali,Marie Paulsen Madsen, Rasmus Riis Kopperud, Hilde Vollan, Karoline Bragstad, Olav Hungnes |
| EPI_ISL_3266528 | Nordland Hospital - Bodo, Laboratory Department, Molecular Biology Unit | Norwegian Institute of Public Health, Department of Virology | Kathrine Stene-Johansen, Kamilla Heddeland Instefjord, Hilde Elshaug, Garcia Llorente Ignacio, Jon Bråte, Engebretsen Serina Beate,Pedersen Benedikte Nevjen, Line Victoria Moen, Debech Nadia, Atiya R Ali,Marie Paulsen Madsen, Rasmus Riis Kopperud, Hilde Vollan, Karoline Bragstad, Olav Hungnes |
| EPI_ISL_3266530 | Department of Medical Microbiology, St. Olavs hospital | Norwegian Institute of Public Health, Department of Virology | Kathrine Stene-Johansen, Kamilla Heddeland Instefjord, Hilde Elshaug, Garcia Llorente Ignacio, Jon Bråte, Engebretsen Serina Beate,Pedersen Benedikte Nevjen, Line Victoria Moen, Debech Nadia, Atiya R Ali,Marie Paulsen Madsen, Rasmus Riis Kopperud, Hilde Vollan, Karoline Bragstad, Olav Hungnes |
| EPI_ISL_3266531 | Nordland Hospital - Bodo, Laboratory Department, Molecular Biology Unit | Norwegian Institute of Public Health, Department of Virology | Kathrine Stene-Johansen, Kamilla Heddeland Instefjord, Hilde Elshaug, Garcia Llorente Ignacio, Jon Bråte, Engebretsen Serina Beate,Pedersen Benedikte Nevjen, Line Victoria Moen, Debech Nadia, Atiya R Ali,Marie Paulsen Madsen, Rasmus Riis Kopperud, Hilde Vollan, Karoline Bragstad, Olav Hungnes |
| EPI_ISL_3266532 | Department of Medical Microbiology - section Molde, Molde Hospital | Norwegian Institute of Public Health, Department of Virology | Kathrine Stene-Johansen, Kamilla Heddeland Instefjord, Hilde Elshaug, Garcia Llorente Ignacio, Jon Bråte, Engebretsen Serina Beate,Pedersen Benedikte Nevjen, Line Victoria Moen, Debech Nadia, Atiya R Ali,Marie Paulsen Madsen, Rasmus Riis Kopperud, Hilde Vollan, Karoline Bragstad, Olav Hungnes |
| EPI_ISL_3266533 | Hospital of Southern Norway - Kristiansand, Department of Medical Microbiology | Norwegian Institute of Public Health, Department of Virology | Kathrine Stene-Johansen, Kamilla Heddeland Instefjord, Hilde Elshaug, Garcia Llorente Ignacio, Jon Bråte, Engebretsen Serina Beate,Pedersen Benedikte Nevjen, Line Victoria Moen, Debech Nadia, Atiya R Ali,Marie Paulsen Madsen, Rasmus Riis Kopperud, Hilde Vollan, Karoline Bragstad, Olav Hungnes |
| EPI_ISL_3266535, EPI_ISL_3266536, EPI_ISL_3266537, EPI_ISL_3266542, EPI_ISL_3266544, EPI_ISL_3266545, EPI_ISL_3266550, EPI_ISL_3266551, EPI_ISL_3266552, EPI_ISL_3266553, EPI_ISL_3266554, EPI_ISL_3266556, EPI_ISL_3266562, EPI_ISL_3266563, EPI_ISL_3266566 | | | |
| see above | U.O. Microbiologia Laboratorio Unico Centro Servizi - AUSL della Romagna | U.O. Microbiologia, Laboratorio Unico Centro Servizi - AUSL della Romagna | Giorgio Dirani |
| EPI_ISL_3266574, EPI_ISL_3266575, EPI_ISL_3266576, EPI_ISL_3266577, EPI_ISL_3266578, EPI_ISL_3266579, EPI_ISL_3266580, | Platform BIS UZA/UAntwerpen | Labo Klinische Biologie, UZA | Marie Le Mercier, Jasmine Coppens, Kathleen Holemans, Ines Verbesselt, Basil Britto Xavier, Christine Lammens, Veerle Matheeussen, Herman Goossens |

| | | | |
|---|---|---|---|
| EPI_ISL_3266581, EPI_ISL_3266582 | | | |
| EPI_ISL_3266583 | LABO LKO - LMC | Labo Klinische Biologie, UZA | Marie Le Mercier, Jasmine Coppens, Kathleen Holemans, Ines Verbesselt, Basil Britto Xavier, Christine Lammens, Veerle Matheeussen, Herman Goossens |
| EPI_ISL_3266584, EPI_ISL_3266585 | Algemeen Klinisch Labo | Labo Klinische Biologie, UZA | Marie Le Mercier, Jasmine Coppens, Kathleen Holemans, Ines Verbesselt, Basil Britto Xavier, Christine Lammens, Veerle Matheeussen, Herman Goossens |
| EPI_ISL_3266587, EPI_ISL_3266588, EPI_ISL_3266589, EPI_ISL_3266590 | Centrum voor Medische Analyse | Labo Klinische Biologie, UZA | Marie Le Mercier, Jasmine Coppens, Kathleen Holemans, Ines Verbesselt, Basil Britto Xavier, Christine Lammens, Veerle Matheeussen, Herman Goossens |
| EPI_ISL_3266591, EPI_ISL_3266592, EPI_ISL_3266593, EPI_ISL_3266594, EPI_ISL_3266595, EPI_ISL_3266596, EPI_ISL_3266597 | Platform BIS UZA/UAntwerpen | Labo Klinische Biologie, UZA | Marie Le Mercier, Jasmine Coppens, Kathleen Holemans, Ines Verbesselt, Basil Britto Xavier, Christine Lammens, Veerle Matheeussen, Herman Goossens |
| EPI_ISL_3266598, EPI_ISL_3266599, EPI_ISL_3266600, EPI_ISL_3266601 | Centrum voor Medische Analyse | Labo Klinische Biologie, UZA | Marie Le Mercier, Jasmine Coppens, Kathleen Holemans, Ines Verbesselt, Basil Britto Xavier, Christine Lammens, Veerle Matheeussen, Herman Goossens |
| EPI_ISL_3266602, EPI_ISL_3266603, EPI_ISL_3266604, EPI_ISL_3266605, EPI_ISL_3266606, EPI_ISL_3266607, EPI_ISL_3266608, EPI_ISL_3266609, EPI_ISL_3266610, EPI_ISL_3266611, EPI_ISL_3266612, EPI_ISL_3266613, EPI_ISL_3266614, EPI_ISL_3266615, EPI_ISL_3266616, EPI_ISL_3266617, EPI_ISL_3266618, EPI_ISL_3266619, EPI_ISL_3266620, EPI_ISL_3266621, EPI_ISL_3266622, EPI_ISL_3266623, EPI_ISL_3266624, EPI_ISL_3266625, EPI_ISL_3266626, EPI_ISL_3266627, EPI_ISL_3266628, EPI_ISL_3266629, EPI_ISL_3266630, EPI_ISL_3266631, EPI_ISL_3266632, EPI_ISL_3266633, EPI_ISL_3266634, EPI_ISL_3266635, EPI_ISL_3266636, EPI_ISL_3266637, EPI_ISL_3266638 | | | |
| see above | Platform BIS UZA/UAntwerpen | Labo Klinische Biologie, UZA | Marie Le Mercier, Jasmine Coppens, Kathleen Holemans, Ines Verbesselt, Basil Britto Xavier, Christine Lammens, Veerle Matheeussen, Herman Goossens |
| EPI_ISL_3266639, EPI_ISL_3266640 | Labo Klinische Biologie, UZA | Labo Klinische Biologie, UZA | Marie Le Mercier, Jasmine Coppens, Kathleen Holemans, Ines Verbesselt, Basil Britto Xavier, Christine Lammens, Veerle Matheeussen, Herman Goossens |
| EPI_ISL_3266641, EPI_ISL_3266642, EPI_ISL_3266643, EPI_ISL_3266644, EPI_ISL_3266645, EPI_ISL_3266646, EPI_ISL_3266647, EPI_ISL_3266648, EPI_ISL_3266649, EPI_ISL_3266650, EPI_ISL_3266651, EPI_ISL_3266652, EPI_ISL_3266653, EPI_ISL_3266654, EPI_ISL_3266655, EPI_ISL_3266656, EPI_ISL_3266657, EPI_ISL_3266658, EPI_ISL_3266659, EPI_ISL_3266660 | | | |
| see above | Algemeen Medisch Labo | Labo Klinische Biologie, UZA | Marie Le Mercier, Jasmine Coppens, Kathleen Holemans, Ines Verbesselt, Basil Britto Xavier, Christine Lammens, Veerle Matheeussen, Herman Goossens |
| EPI_ISL_3266661, EPI_ISL_3266662, EPI_ISL_3266663, EPI_ISL_3266664, EPI_ISL_3266665, EPI_ISL_3266666, EPI_ISL_3266668, EPI_ISL_3266669, EPI_ISL_3266671, EPI_ISL_3266672, EPI_ISL_3266674, EPI_ISL_3266676, EPI_ISL_3266681, EPI_ISL_3266683, EPI_ISL_3266684, EPI_ISL_3266686, EPI_ISL_3266688, EPI_ISL_3266689, EPI_ISL_3266690, EPI_ISL_3266692, EPI_ISL_3266693, EPI_ISL_3266694, EPI_ISL_3266695, EPI_ISL_3266697, EPI_ISL_3266698, EPI_ISL_3266700, EPI_ISL_3266701, EPI_ISL_3266702, EPI_ISL_3266706, EPI_ISL_3266707, EPI_ISL_3266709, EPI_ISL_3266710, EPI_ISL_3266711, EPI_ISL_3266712, EPI_ISL_3266713, EPI_ISL_3266714, EPI_ISL_3266715, EPI_ISL_3266716, EPI_ISL_3266717, EPI_ISL_3266718, EPI_ISL_3266719, EPI_ISL_3266722, EPI_ISL_3266723, EPI_ISL_3266726, EPI_ISL_3266727, EPI_ISL_3266728, EPI_ISL_3266729, EPI_ISL_3266730, EPI_ISL_3266731, EPI_ISL_3266732, EPI_ISL_3266733, EPI_ISL_3266734, EPI_ISL_3266735, EPI_ISL_3266739, EPI_ISL_3266742, EPI_ISL_3266744, EPI_ISL_3266747, EPI_ISL_3266748, EPI_ISL_3266750, EPI_ISL_3266751, EPI_ISL_3266752, EPI_ISL_3266753, EPI_ISL_3266755, EPI_ISL_3266757, EPI_ISL_3266758, EPI_ISL_3266760, EPI_ISL_3266764, EPI_ISL_3266768, EPI_ISL_3266769, EPI_ISL_3266770, EPI_ISL_3266772, EPI_ISL_3266773, EPI_ISL_3266774, EPI_ISL_3266775, EPI_ISL_3266776, EPI_ISL_3266778, EPI_ISL_3266779, EPI_ISL_3266780, EPI_ISL_3266781, EPI_ISL_3266783, EPI_ISL_3266786, EPI_ISL_3266787, EPI_ISL_3266788, EPI_ISL_3266789, EPI_ISL_3266790, EPI_ISL_3266791, EPI_ISL_3266792, EPI_ISL_3266793, EPI_ISL_3266794, EPI_ISL_3266795, EPI_ISL_3266796, EPI_ISL_3266797, EPI_ISL_3266798, EPI_ISL_3266799, EPI_ISL_3266800, EPI_ISL_3266801, EPI_ISL_3266802, EPI_ISL_3266803, EPI_ISL_3266804, EPI_ISL_3266805, EPI_ISL_3266806, EPI_ISL_3266807, EPI_ISL_3266808, EPI_ISL_3266809, EPI_ISL_3266810, EPI_ISL_3266811, EPI_ISL_3266812, EPI_ISL_3266813, EPI_ISL_3266814, EPI_ISL_3266815, EPI_ISL_3266816, EPI_ISL_3266817, EPI_ISL_3266818, EPI_ISL_3266819, EPI_ISL_3266820, EPI_ISL_3266821, EPI_ISL_3266822, EPI_ISL_3266823, EPI_ISL_3266824, EPI_ISL_3266825, EPI_ISL_3266826, EPI_ISL_3266828, EPI_ISL_3266830 | | | |
| see above | U.O. Microbiologia Laboratorio Unico Centro Servizi - AUSL della Romagna | U.O. Microbiologia, Laboratorio Unico Centro Servizi - AUSL della Romagna | Giorgio Dirani |
| EPI_ISL_3266831, EPI_ISL_3266832, EPI_ISL_3266833, EPI_ISL_3266834, EPI_ISL_3266851, EPI_ISL_3266852, EPI_ISL_3266853, EPI_ISL_3266854, EPI_ISL_3266855, EPI_ISL_3266856, EPI_ISL_3266857, EPI_ISL_3266860, EPI_ISL_3266861, EPI_ISL_3266863 | | | |
| see above | Dutch COVID-19 response team | National Institute for Public Health and the Environment (RIVM) | Adam Meijer, Harry Vennema, Dirk Eggink, Jeroen Cremer, Sharon van den Brink, Bas van der Veer, AnneMarie van den Brandt, Lisa Wijsman, Kim Freriks, Ryanne Jaarsma, Euníce Then, Lynn Aarts, Sanne Bos, Melissa van Tuil, Florian Zwagemaker, Dennis Schmitz, Annelies Kroneman, Karim Hajji, Chantal Reusken, on behalf of the national COVID-19 response team |
| EPI_ISL_3266864, EPI_ISL_3266865, EPI_ISL_3266866, EPI_ISL_3266867, EPI_ISL_3266868, EPI_ISL_3266870, EPI_ISL_3266871, EPI_ISL_3266872, EPI_ISL_3266873, EPI_ISL_3266874, EPI_ISL_3266876, EPI_ISL_3266877, EPI_ISL_3266878, EPI_ISL_3266879, EPI_ISL_3266880, EPI_ISL_3266881, EPI_ISL_3266882, EPI_ISL_3266883, EPI_ISL_3266884, EPI_ISL_3266885, EPI_ISL_3266886, EPI_ISL_3266888, EPI_ISL_3266889, EPI_ISL_3266890, EPI_ISL_3266891, EPI_ISL_3266892, EPI_ISL_3266893, EPI_ISL_3266894, EPI_ISL_3266895, EPI_ISL_3266896, EPI_ISL_3266897, EPI_ISL_3266898, EPI_ISL_3266899, EPI_ISL_3266900 | | | |
| see above | U.O. Microbiologia Laboratorio Unico Centro Servizi - AUSL della Romagna | U.O. Microbiologia, Laboratorio Unico Centro Servizi - AUSL della Romagna | Giorgio Dirani |
| EPI_ISL_3266903, EPI_ISL_3266904, EPI_ISL_3266905, EPI_ISL_3266906, EPI_ISL_3266907, EPI_ISL_3266908, EPI_ISL_3266909, EPI_ISL_3266910, EPI_ISL_3266911, EPI_ISL_3266912, EPI_ISL_3266913, EPI_ISL_3266914, EPI_ISL_3266915, EPI_ISL_3266922, EPI_ISL_3266923, EPI_ISL_3266924, EPI_ISL_3266925, EPI_ISL_3266926, EPI_ISL_3266927, EPI_ISL_3266928, EPI_ISL_3266929, EPI_ISL_3266930, EPI_ISL_3266931, EPI_ISL_3266932, EPI_ISL_3266933 | | | |
| see above | Microbiology Department. Complexo Hospitalario Universitario de Vigo | Microbiology Department. Complexo Hospitalario Universitario de Vigo | Alfaya N, Alonso I, Alvarez M, Cabrera JJ, Carballo R, Cores O, Cortizo S, del-Campo V, Martinez L, Mediero G, Perez S, Davina C, Potel C, Regueiro B, Rey S, Vasallo FJ |
| EPI_ISL_3266934, EPI_ISL_3266935, EPI_ISL_3266936, EPI_ISL_3266937, EPI_ISL_3266938, EPI_ISL_3266939, EPI_ISL_3266940, EPI_ISL_3266941, EPI_ISL_3266942, EPI_ISL_3266943, EPI_ISL_3266944, EPI_ISL_3266945, EPI_ISL_3266946, EPI_ISL_3266947, EPI_ISL_3266949, EPI_ISL_3266952, EPI_ISL_3266953, EPI_ISL_3266954, EPI_ISL_3266955, EPI_ISL_3266956, EPI_ISL_3266957, EPI_ISL_3266958, EPI_ISL_3266959, EPI_ISL_3266960, EPI_ISL_3266961, EPI_ISL_3266962, EPI_ISL_3266963, EPI_ISL_3266964, EPI_ISL_3266965, EPI_ISL_3266966, EPI_ISL_3266967, EPI_ISL_3266968, EPI_ISL_3266969, EPI_ISL_3266970, EPI_ISL_3266971, EPI_ISL_3266972 | | | |
| see above | U.O. Microbiologia Laboratorio Unico Centro Servizi - AUSL della Romagna | U.O. Microbiologia, Laboratorio Unico Centro Servizi - AUSL della Romagna | Giorgio Dirani |
| EPI_ISL_3266973, EPI_ISL_3266976, EPI_ISL_3266977, EPI_ISL_3266978, EPI_ISL_3266979, EPI_ISL_3266980, EPI_ISL_3266981, EPI_ISL_3266982, EPI_ISL_3266983, EPI_ISL_3266984, EPI_ISL_3266985, EPI_ISL_3266986, EPI_ISL_3266987, EPI_ISL_3266988, EPI_ISL_3266989, EPI_ISL_3266990, EPI_ISL_3266991, EPI_ISL_3266992 | | | |
| see above | Wyoming Public Health Laboratory | Wyoming Public Health Laboratory | Jim Mildenberger, Taylor Fearing, Channing Weber, Ashley Norberg, Chayse Rowley, Marley Goetz, Brian Dominguez, Elliot Thomasson, Sam Britz, Cari Sloma, Robert Petit, and Rob Christensen |
| EPI_ISL_3267053, EPI_ISL_3267054, EPI_ISL_3267055, EPI_ISL_3267056, EPI_ISL_3267057, EPI_ISL_3267058, EPI_ISL_3267059, EPI_ISL_3267060, EPI_ISL_3267061, EPI_ISL_3267062, EPI_ISL_3267063, EPI_ISL_3267064, EPI_ISL_3267065 | | | |
| see above | LABO BIO DOC | CHU Purpan - Laboratoire de Virologie - Institut Fédératif de Biologie | Latour J., Jeanne N., Ranger N., Carcenac R., Harter A., Boyer P., Ferrer V., Tremeaux R., Izopet J. |
| EPI_ISL_3267066, EPI_ISL_3267067, EPI_ISL_3267068 | HOPITAL NOSTRA SENYORA MERITXELL | CHU Purpan - Laboratoire de Virologie - Institut Fédératif de Biologie | Latour J., Jeanne N., Ranger N., Carcenac R., Harter A., Boyer P., Ferrer V., Tremeaux R., Izopet J. |
| EPI_ISL_3267069, EPI_ISL_3267070, EPI_ISL_3267071, EPI_ISL_3267072, EPI_ISL_3267073, EPI_ISL_3267074, EPI_ISL_3267075, EPI_ISL_3267076 | CH PERPIGNAN | CHU Purpan - Laboratoire de Virologie - Institut Fédératif de Biologie | Latour J., Jeanne N., Ranger N., Carcenac R., Harter A., Boyer P., Ferrer V., Tremeaux R., Izopet J. |
| EPI_ISL_3267077, EPI_ISL_3267078 | CHU Purpan - Laboratoire de Virologie - Institut Fédératif de Biologie | CHU Purpan - Laboratoire de Virologie - Institut Fédératif de Biologie | Latour J., Jeanne N., Ranger N., Carcenac R., Harter A., Boyer P., Ferrer V., Tremeaux R., Izopet J. |
| EPI_ISL_3267079 | CH AUCH | CHU Purpan - Laboratoire de Virologie - Institut Fédératif de Biologie | Latour J., Jeanne N., Ranger N., Carcenac R., Harter A., Boyer P., Ferrer V., Tremeaux R., Izopet J. |
| EPI_ISL_3267080 | CEDIBIO-UNILABS OUSTALOUS | CHU Purpan - Laboratoire de Virologie - Institut Fédératif de Biologie | Latour J., Jeanne N., Ranger N., Carcenac R., Harter A., Boyer P., Ferrer V., Tremeaux R., Izopet J. |
| EPI_ISL_3267081, EPI_ISL_3267082 | CH AUCH | CHU Purpan - Laboratoire de Virologie - Institut Fédératif de Biologie | Latour J., Jeanne N., Ranger N., Carcenac R., Harter A., Boyer P., Ferrer V., Tremeaux R., Izopet J. |
| EPI_ISL_3267083 | CEDIBIO-UNILABS ST ORENS ARCADES | CHU Purpan - Laboratoire de Virologie - Institut Fédératif de Biologie | Latour J., Jeanne N., Ranger N., Carcenac R., Harter A., Boyer P., Ferrer V., Tremeaux R., Izopet J. |
| EPI_ISL_3267084, EPI_ISL_3267085 | CEDIBIO-UNILABS CASTANET DELHERM | CHU Purpan - Laboratoire de Virologie - Institut Fédératif de Biologie | Latour J., Jeanne N., Ranger N., Carcenac R., Harter A., Boyer P., Ferrer V., Tremeaux R., Izopet J. |
| EPI_ISL_3267086 | CHU Purpan - Laboratoire de Virologie - Institut Fédératif de Biologie | CHU Purpan - Laboratoire de Virologie - Institut Fédératif de Biologie | Latour J., Jeanne N., Ranger N., Carcenac R., Harter A., Boyer P., Ferrer V., Tremeaux R., Izopet J. |
| EPI_ISL_3267087 | CEDIBIO-UNILABS CASTANET DELHERM | CHU Purpan - Laboratoire de Virologie - Institut Fédératif de Biologie | Latour J., Jeanne N., Ranger N., Carcenac R., Harter A., Boyer P., Ferrer V., Tremeaux R., Izopet J. |
| EPI_ISL_3267088, EPI_ISL_3267089, EPI_ISL_3267090, EPI_ISL_3267091 | CEDIBIO UNILABS | CHU Purpan - Laboratoire de Virologie - Institut Fédératif de Biologie | Latour J., Jeanne N., Ranger N., Carcenac R., Harter A., Boyer P., Ferrer V., Tremeaux R., Izopet J. |
| EPI_ISL_3267092 | GCS DE BIOLOGIE DU TERRITOIRE DE L'ARIEGE | CHU Purpan - Laboratoire de Virologie - Institut Fédératif de Biologie | Latour J., Jeanne N., Ranger N., Carcenac R., Harter A., Boyer P., Ferrer V., Tremeaux R., Izopet J. |
| EPI_ISL_3267093, EPI_ISL_3267094, EPI_ISL_3267095 | CH ALBI | CHU Purpan - Laboratoire de Virologie - Institut Fédératif de Biologie | Latour J., Jeanne N., Ranger N., Carcenac R., Harter A., Boyer P., Ferrer V., Tremeaux R., Izopet J. |
| EPI_ISL_3267096 | CH MONTAUBAN | CHU Purpan - Laboratoire de Virologie - Institut Fédératif de Biologie | Latour J., Jeanne N., Ranger N., Carcenac R., Harter A., Boyer P., Ferrer V., Tremeaux R., Izopet J. |
| EPI_ISL_3267097 | CEDIBIO UNILABS | CHU Purpan - Laboratoire de Virologie - Institut Fédératif de Biologie | Latour J., Jeanne N., Ranger N., Carcenac R., Harter A., Boyer P., Ferrer V., Tremeaux R., Izopet J. |
| EPI_ISL_3267105 | LABO BIO DOC | CHU Purpan - Laboratoire de Virologie - Institut Fédératif de Biologie | Latour J., Jeanne N., Ranger N., Carcenac R., Harter A., Boyer P., Ferrer V., Tremeaux R., Izopet J. |
| EPI_ISL_3267106, EPI_ISL_3267107 | CH PERPIGNAN | CHU Purpan - Laboratoire de Virologie - Institut Fédératif de Biologie | Latour J., Jeanne N., Ranger N., Carcenac R., Harter A., Boyer P., Ferrer V., Tremeaux R., Izopet J. |
| EPI_ISL_3267108 | LABO BIO DOC | CHU Purpan - Laboratoire de Virologie - Institut Fédératif de Biologie | Latour J., Jeanne N., Ranger N., Carcenac R., Harter A., Boyer P., Ferrer V., Tremeaux R., Izopet J. |

| | | | |
|---|---|---|---|
| EPI_ISL_3267109 | CH PERPIGNAN | CHU Purpan - Laboratoire de Virologie - Institut Fédératif de Biologie | Latour J., Jeanne N., Ranger N., Carcenac R., Harter A., Boyer P., Ferrer V., Tremeaux R., Izopet J. |
| EPI_ISL_3267110 | CH MONTAUBAN | CHU Purpan - Laboratoire de Virologie - Institut Fédératif de Biologie | Latour J., Jeanne N., Ranger N., Carcenac R., Harter A., Boyer P., Ferrer V., Tremeaux R., Izopet J. |
| EPI_ISL_3267111 | LABO BIO DOC | CHU Purpan - Laboratoire de Virologie - Institut Fédératif de Biologie | Latour J., Jeanne N., Ranger N., Carcenac R., Harter A., Boyer P., Ferrer V., Tremeaux R., Izopet J. |
| EPI_ISL_3267112 | CEDIBIO UNILABS | CHU Purpan - Laboratoire de Virologie - Institut Fédératif de Biologie | Latour J., Jeanne N., Ranger N., Carcenac R., Harter A., Boyer P., Ferrer V., Tremeaux R., Izopet J. |
| EPI_ISL_3267113, EPI_ISL_3267114 | CHU Purpan - Laboratoire de Virologie - Institut Fédératif de Biologie | CHU Purpan - Laboratoire de Virologie - Institut Fédératif de Biologie | Latour J., Jeanne N., Ranger N., Carcenac R., Harter A., Boyer P., Ferrer V., Tremeaux R., Izopet J. |
| EPI_ISL_3267115 | CH ALBI | CHU Purpan - Laboratoire de Virologie - Institut Fédératif de Biologie | Latour J., Jeanne N., Ranger N., Carcenac R., Harter A., Boyer P., Ferrer V., Tremeaux R., Izopet J. |
| EPI_ISL_3267116, EPI_ISL_3267117 | CH MONTAUBAN | CHU Purpan - Laboratoire de Virologie - Institut Fédératif de Biologie | Latour J., Jeanne N., Ranger N., Carcenac R., Harter A., Boyer P., Ferrer V., Tremeaux R., Izopet J. |
| EPI_ISL_3267118 | CHU Purpan - Laboratoire de Virologie - Institut Fédératif de Biologie | CHU Purpan - Laboratoire de Virologie - Institut Fédératif de Biologie | Latour J., Jeanne N., Ranger N., Carcenac R., Harter A., Boyer P., Ferrer V., Tremeaux R., Izopet J. |
| EPI_ISL_3267119 | LABO BIO DOC | CHU Purpan - Laboratoire de Virologie - Institut Fédératif de Biologie | Latour J., Jeanne N., Ranger N., Carcenac R., Harter A., Boyer P., Ferrer V., Tremeaux R., Izopet J. |
| EPI_ISL_3267127 | CH MONTAUBAN | CHU Purpan - Laboratoire de Virologie - Institut Fédératif de Biologie | Latour J., Jeanne N., Ranger N., Carcenac R., Harter A., Boyer P., Ferrer V., Tremeaux R., Izopet J. |
| EPI_ISL_3267128 | CEDIBIO-UNILABS OUSTALOUS | CHU Purpan - Laboratoire de Virologie - Institut Fédératif de Biologie | Latour J., Jeanne N., Ranger N., Carcenac R., Harter A., Boyer P., Ferrer V., Tremeaux R., Izopet J. |
| EPI_ISL_3267129 | CHU Purpan - Laboratoire de Virologie - Institut Fédératif de Biologie | CHU Purpan - Laboratoire de Virologie - Institut Fédératif de Biologie | Latour J., Jeanne N., Ranger N., Carcenac R., Harter A., Boyer P., Ferrer V., Tremeaux R., Izopet J. |
| EPI_ISL_3267130 | CEDIBIO-UNILABS CASTANET DELHERM | CHU Purpan - Laboratoire de Virologie - Institut Fédératif de Biologie | Latour J., Jeanne N., Ranger N., Carcenac R., Harter A., Boyer P., Ferrer V., Tremeaux R., Izopet J. |
| EPI_ISL_3267131 | CH CAHORS | CHU Purpan - Laboratoire de Virologie - Institut Fédératif de Biologie | Latour J., Jeanne N., Ranger N., Carcenac R., Harter A., Boyer P., Ferrer V., Tremeaux R., Izopet J. |
| EPI_ISL_3267132 | CH PERPIGNAN | CHU Purpan - Laboratoire de Virologie - Institut Fédératif de Biologie | Latour J., Jeanne N., Ranger N., Carcenac R., Harter A., Boyer P., Ferrer V., Tremeaux R., Izopet J. |
| EPI_ISL_3267133 | CH CARCASSONE | CHU Purpan - Laboratoire de Virologie - Institut Fédératif de Biologie | Latour J., Jeanne N., Ranger N., Carcenac R., Harter A., Boyer P., Ferrer V., Tremeaux R., Izopet J. |
| EPI_ISL_3267134, EPI_ISL_3267135, EPI_ISL_3267136 | LABO BIO DOC | CHU Purpan - Laboratoire de Virologie - Institut Fédératif de Biologie | Latour J., Jeanne N., Ranger N., Carcenac R., Harter A., Boyer P., Ferrer V., Tremeaux R., Izopet J. |
| EPI_ISL_3267137 | CENTRE HOSPITALIER DE BIGORRE | CHU Purpan - Laboratoire de Virologie - Institut Fédératif de Biologie | Latour J., Jeanne N., Ranger N., Carcenac R., Harter A., Boyer P., Ferrer V., Tremeaux R., Izopet J. |
| EPI_ISL_3267138, EPI_ISL_3267139 | LABO BIO DOC | CHU Purpan - Laboratoire de Virologie - Institut Fédératif de Biologie | Latour J., Jeanne N., Ranger N., Carcenac R., Harter A., Boyer P., Ferrer V., Tremeaux R., Izopet J. |
| EPI_ISL_3267140 | HOPITAL NOSTRA SENYORA MERITXELL | CHU Purpan - Laboratoire de Virologie - Institut Fédératif de Biologie | Latour J., Jeanne N., Ranger N., Carcenac R., Harter A., Boyer P., Ferrer V., Tremeaux R., Izopet J. |
| EPI_ISL_3267141, EPI_ISL_3267142, EPI_ISL_3267143 | CH PERPIGNAN | CHU Purpan - Laboratoire de Virologie - Institut Fédératif de Biologie | Latour J., Jeanne N., Ranger N., Carcenac R., Harter A., Boyer P., Ferrer V., Tremeaux R., Izopet J. |
| EPI_ISL_3267144 | CEDIBIO UNILABS | CHU Purpan - Laboratoire de Virologie - Institut Fédératif de Biologie | Latour J., Jeanne N., Ranger N., Carcenac R., Harter A., Boyer P., Ferrer V., Tremeaux R., Izopet J. |
| EPI_ISL_3267145 | CH MONTAUBAN | CHU Purpan - Laboratoire de Virologie - Institut Fédératif de Biologie | Latour J., Jeanne N., Ranger N., Carcenac R., Harter A., Boyer P., Ferrer V., Tremeaux R., Izopet J. |
| EPI_ISL_3267146, EPI_ISL_3267147, EPI_ISL_3267148, EPI_ISL_3267149, EPI_ISL_3267150, EPI_ISL_3267151, EPI_ISL_3267152, EPI_ISL_3267153, EPI_ISL_3267155, EPI_ISL_3267156, EPI_ISL_3267160, EPI_ISL_3267161, EPI_ISL_3267162, EPI_ISL_3267163, EPI_ISL_3267164, EPI_ISL_3267165, EPI_ISL_3267166, EPI_ISL_3267167, EPI_ISL_3267168, EPI_ISL_3267169, EPI_ISL_3267170, EPI_ISL_3267171, EPI_ISL_3267172, EPI_ISL_3267173, EPI_ISL_3267174, EPI_ISL_3267175, EPI_ISL_3267176, EPI_ISL_3267177, EPI_ISL_3267178, EPI_ISL_3267179, EPI_ISL_3267181, EPI_ISL_3267182, EPI_ISL_3267183, EPI_ISL_3267184, EPI_ISL_3267185 | | | |
| see above | U.O. Microbiologia Laboratorio Unico Centro Servizi - AUSL della Romagna | U.O. Microbiologia, Laboratorio Unico Centro Servizi - AUSL della Romagna | Giorgio Dirani |
| EPI_ISL_3267187 | RSUD dr. Soegiri Lamongan | Institute of Tropical Disease, Universitas Airlangga | Rima R Prasetya, Krisnoadi Rahardjo, Aldise M Nastri, Jezzy R Dewantari, M Chaidir Annas, Gatot Soegiarto, Laksmi Wulandari, Resti Yudhawati, Yasuko Mori, Soetjipto, Kazufumi Shimizu, Maria I Lusida |
| EPI_ISL_3267188, EPI_ISL_3267189, EPI_ISL_3267190, EPI_ISL_3267191, EPI_ISL_3267192, EPI_ISL_3267193, EPI_ISL_3267194, EPI_ISL_3267196, EPI_ISL_3267197, EPI_ISL_3267198, EPI_ISL_3267199, EPI_ISL_3267200, EPI_ISL_3267201, EPI_ISL_3267202, EPI_ISL_3267205, EPI_ISL_3267206, EPI_ISL_3267207, EPI_ISL_3267208, EPI_ISL_3267209, EPI_ISL_3267210, EPI_ISL_3267211, EPI_ISL_3267212, EPI_ISL_3267213, EPI_ISL_3267215, EPI_ISL_3267218, EPI_ISL_3267219, EPI_ISL_3267221, EPI_ISL_3267222, EPI_ISL_3267223, EPI_ISL_3267225, EPI_ISL_3267226, EPI_ISL_3267228, EPI_ISL_3267229, EPI_ISL_3267230, EPI_ISL_3267235, EPI_ISL_3267236, EPI_ISL_3267239, EPI_ISL_3267242, EPI_ISL_3267243, EPI_ISL_3267245, EPI_ISL_3267247, EPI_ISL_3267248, EPI_ISL_3267249, EPI_ISL_3267250, EPI_ISL_3267252, EPI_ISL_3267253, EPI_ISL_3267254, EPI_ISL_3267255, EPI_ISL_3267256, EPI_ISL_3267257, EPI_ISL_3267258, EPI_ISL_3267259, EPI_ISL_3267260, EPI_ISL_3267261, EPI_ISL_3267262, EPI_ISL_3267263, EPI_ISL_3267264, EPI_ISL_3267265, EPI_ISL_3267266, EPI_ISL_3267267, EPI_ISL_3267268, EPI_ISL_3267269, EPI_ISL_3267270, EPI_ISL_3267271, EPI_ISL_3267272, EPI_ISL_3267273, EPI_ISL_3267274, EPI_ISL_3267275, EPI_ISL_3267276, EPI_ISL_3267277, EPI_ISL_3267278, EPI_ISL_3267279, EPI_ISL_3267280, EPI_ISL_3267281, EPI_ISL_3267282, EPI_ISL_3267284, EPI_ISL_3267285, EPI_ISL_3267288, EPI_ISL_3267289, EPI_ISL_3267290, EPI_ISL_3267291, EPI_ISL_3267292, EPI_ISL_3267293, EPI_ISL_3267294, EPI_ISL_3267295, EPI_ISL_3267296, EPI_ISL_3267297, EPI_ISL_3267298, EPI_ISL_3267299, EPI_ISL_3267300, EPI_ISL_3267301, EPI_ISL_3267302, EPI_ISL_3267303, EPI_ISL_3267304, EPI_ISL_3267305, EPI_ISL_3267306, EPI_ISL_3267307, EPI_ISL_3267308, EPI_ISL_3267309, EPI_ISL_3267310, EPI_ISL_3267312, EPI_ISL_3267313, EPI_ISL_3267314, EPI_ISL_3267315, EPI_ISL_3267316, EPI_ISL_3267317, EPI_ISL_3267318, EPI_ISL_3267319, EPI_ISL_3267320, EPI_ISL_3267321, EPI_ISL_3267322, EPI_ISL_3267323, EPI_ISL_3267324, EPI_ISL_3267325, EPI_ISL_3267326, EPI_ISL_3267327, EPI_ISL_3267328, EPI_ISL_3267329, EPI_ISL_3267330, EPI_ISL_3267331, EPI_ISL_3267332, EPI_ISL_3267333, EPI_ISL_3267334, EPI_ISL_3267335, EPI_ISL_3267336, EPI_ISL_3267337, EPI_ISL_3267338, EPI_ISL_3267339, EPI_ISL_3267340, EPI_ISL_3267341, EPI_ISL_3267342, EPI_ISL_3267343, EPI_ISL_3267344, EPI_ISL_3267345, EPI_ISL_3267346, EPI_ISL_3267347, EPI_ISL_3267348, EPI_ISL_3267349, EPI_ISL_3267350, EPI_ISL_3267351, EPI_ISL_3267352, EPI_ISL_3267353, EPI_ISL_3267354, EPI_ISL_3267355, EPI_ISL_3267356, EPI_ISL_3267357, EPI_ISL_3267358, EPI_ISL_3267359, EPI_ISL_3267360, EPI_ISL_3267361, EPI_ISL_3267362, EPI_ISL_3267363, EPI_ISL_3267364, EPI_ISL_3267365, EPI_ISL_3267366, EPI_ISL_3267367, EPI_ISL_3267368, EPI_ISL_3267369, EPI_ISL_3267370, EPI_ISL_3267371, EPI_ISL_3267372, EPI_ISL_3267375, EPI_ISL_3267379, EPI_ISL_3267380, EPI_ISL_3267381, EPI_ISL_3267382, EPI_ISL_3267383, EPI_ISL_3267384, EPI_ISL_3267385, EPI_ISL_3267386, EPI_ISL_3267387, EPI_ISL_3267388, EPI_ISL_3267389, EPI_ISL_3267391, EPI_ISL_3267392, EPI_ISL_3267393, EPI_ISL_3267394, EPI_ISL_3267395, EPI_ISL_3267396, EPI_ISL_3267397, EPI_ISL_3267398, EPI_ISL_3267399, EPI_ISL_3267401, EPI_ISL_3267402, EPI_ISL_3267403, EPI_ISL_3267404, EPI_ISL_3267405, EPI_ISL_3267407, EPI_ISL_3267408, EPI_ISL_3267409 | | | |
| see above | INSACOG Surveillance | INSACOG at CSIR Institute of Genomics and Integrative Biology | INSACOG |
| EPI_ISL_3267411, EPI_ISL_3267412, EPI_ISL_3267413, EPI_ISL_3267414, EPI_ISL_3267415, EPI_ISL_3267416, EPI_ISL_3267417, EPI_ISL_3267418, EPI_ISL_3267419, EPI_ISL_3267420, EPI_ISL_3267421, EPI_ISL_3267422, EPI_ISL_3267423, EPI_ISL_3267424, EPI_ISL_3267425, EPI_ISL_3267426, EPI_ISL_3267427, EPI_ISL_3267428, EPI_ISL_3267429, EPI_ISL_3267430, EPI_ISL_3267431, EPI_ISL_3267432, EPI_ISL_3267433, EPI_ISL_3267434, EPI_ISL_3267435, EPI_ISL_3267436, EPI_ISL_3267437, EPI_ISL_3267438, EPI_ISL_3267439, EPI_ISL_3267440, EPI_ISL_3267441, EPI_ISL_3267442, EPI_ISL_3267443, EPI_ISL_3267444, EPI_ISL_3267445, EPI_ISL_3267446, EPI_ISL_3267447, EPI_ISL_3267448, EPI_ISL_3267449, EPI_ISL_3267450, EPI_ISL_3267451, EPI_ISL_3267452, EPI_ISL_3267453, EPI_ISL_3267454, EPI_ISL_3267455, EPI_ISL_3267456, EPI_ISL_3267457, EPI_ISL_3267458, EPI_ISL_3267459, EPI_ISL_3267460, EPI_ISL_3267461, EPI_ISL_3267462, EPI_ISL_3267463, EPI_ISL_3267464, EPI_ISL_3267465, EPI_ISL_3267466, EPI_ISL_3267467, EPI_ISL_3267468, EPI_ISL_3267469, EPI_ISL_3267470, EPI_ISL_3267471, EPI_ISL_3267472, EPI_ISL_3267473, EPI_ISL_3267474, EPI_ISL_3267475, EPI_ISL_3267476, EPI_ISL_3267477, EPI_ISL_3267478, EPI_ISL_3267479, EPI_ISL_3267480, EPI_ISL_3267481, EPI_ISL_3267482, EPI_ISL_3267483, EPI_ISL_3267484, EPI_ISL_3267485, EPI_ISL_3267486, EPI_ISL_3267487, EPI_ISL_3267488, EPI_ISL_3267489, EPI_ISL_3267490, EPI_ISL_3267491, EPI_ISL_3267492, EPI_ISL_3267493, EPI_ISL_3267494, EPI_ISL_3267495, EPI_ISL_3267496, EPI_ISL_3267497, EPI_ISL_3267498, EPI_ISL_3267499, EPI_ISL_3267500, EPI_ISL_3267501, EPI_ISL_3267502, EPI_ISL_3267503, EPI_ISL_3267504, EPI_ISL_3267505, EPI_ISL_3267506, EPI_ISL_3267507, EPI_ISL_3267508, EPI_ISL_3267509, EPI_ISL_3267510, EPI_ISL_3267511, EPI_ISL_3267512, EPI_ISL_3267513, EPI_ISL_3267514, EPI_ISL_3267515, EPI_ISL_3267516, EPI_ISL_3267517, EPI_ISL_3267518, EPI_ISL_3267519, EPI_ISL_3267520, EPI_ISL_3267521, EPI_ISL_3267522, EPI_ISL_3267523, EPI_ISL_3267524, EPI_ISL_3267525, EPI_ISL_3267526, EPI_ISL_3267527, EPI_ISL_3267528, EPI_ISL_3267529, EPI_ISL_3267530, EPI_ISL_3267531, EPI_ISL_3267532, EPI_ISL_3267533, EPI_ISL_3267534, EPI_ISL_3267535, EPI_ISL_3267536, EPI_ISL_3267537, EPI_ISL_3267538, EPI_ISL_3267539, EPI_ISL_3267540, EPI_ISL_3267541, EPI_ISL_3267542, EPI_ISL_3267543, EPI_ISL_3267544, EPI_ISL_3267545, EPI_ISL_3267546, EPI_ISL_3267547, EPI_ISL_3267548, EPI_ISL_3267549, EPI_ISL_3267550, EPI_ISL_3267551, EPI_ISL_3267552, EPI_ISL_3267553, EPI_ISL_3267554, EPI_ISL_3267555, EPI_ISL_3267556, EPI_ISL_3267557, EPI_ISL_3267558, EPI_ISL_3267563 | | | |
| see above | Pathology and Laboratory Medicine Institute, Cleveland Clinic, Ohio, USA | Pathology and Laboratory Medicine Institute, Cleveland Clinic, Ohio, USA | Jessica Spildener, Concetta Peck, Jennifer Starbuck, Joy Nakitandwe, Kristen McDonnell, David Plunkett, Zheng Jin Tu, Jay Brock, Yu-Wei Cheng, Gary Procop, Daniel Rhoads, Daniel H. Farkas, David Bosler |
| EPI_ISL_3267722, EPI_ISL_3267723, EPI_ISL_3267724, EPI_ISL_3267725, EPI_ISL_3267726, EPI_ISL_3267727, EPI_ISL_3267728, EPI_ISL_3267729, EPI_ISL_3267730, EPI_ISL_3267732, EPI_ISL_3267733, EPI_ISL_3267734, EPI_ISL_3267735, EPI_ISL_3267736, EPI_ISL_3267737, EPI_ISL_3267738, EPI_ISL_3267739, EPI_ISL_3267740, EPI_ISL_3267741, EPI_ISL_3267742, EPI_ISL_3267743, EPI_ISL_3267744, EPI_ISL_3267745, EPI_ISL_3267746, EPI_ISL_3267747, EPI_ISL_3267748, EPI_ISL_3267749, EPI_ISL_3267750, EPI_ISL_3267752, EPI_ISL_3267753, EPI_ISL_3267754, EPI_ISL_3267755, EPI_ISL_3267756 | | | |
| see above | Sisonke - BARC SA | CERI, Centre for Epidemic Response and Innovation, Stellenbosch Univeristy & KRISP, KZN Research Innovation and Sequencing Platform | Giandhari Jennifer, Pillay Sureshnee, Yajna Ramphal, Naidoo Yeshnee, Tshabuila Derek,Tegally Houriiyah, San James, Wilkinson Eduan, de Oliveira Tulio |
| EPI_ISL_3267758, EPI_ISL_3267759 | CHUV | Laboratory of genomics and metagenomics | Trestan Pillonel, Damien Jacot, Sébastien Aeby, Gilbert Greub, Claire Bertelli |
| EPI_ISL_3267760 | VIDYMED LAUSANNE | Laboratory of genomics and metagenomics | Trestan Pillonel, Damien Jacot, Sébastien Aeby, Gilbert Greub, Claire Bertelli |
| EPI_ISL_3267761 | CHUV | Laboratory of genomics and metagenomics | Trestan Pillonel, Damien Jacot, Sébastien Aeby, Gilbert Greub, Claire Bertelli |
| EPI_ISL_3267762 | VIDYMED LAUSANNE | Laboratory of genomics and metagenomics | Trestan Pillonel, Damien Jacot, Sébastien Aeby, Gilbert Greub, Claire Bertelli |
| EPI_ISL_3267763 | EHNV | Laboratory of genomics and metagenomics | Trestan Pillonel, Damien Jacot, Sébastien Aeby, Gilbert Greub, Claire Bertelli |
| EPI_ISL_3267764 | CHUV | Laboratory of genomics and metagenomics | Trestan Pillonel, Damien Jacot, Sébastien Aeby, Gilbert Greub, Claire Bertelli |
| EPI_ISL_3267765 | VIDYMED LAUSANNE | Laboratory of genomics and metagenomics | Trestan Pillonel, Damien Jacot, Sébastien Aeby, Gilbert Greub, Claire Bertelli |
| EPI_ISL_3267766 | CHUV | Laboratory of genomics and metagenomics | Trestan Pillonel, Damien Jacot, Sébastien Aeby, Gilbert Greub, Claire Bertelli |
| EPI_ISL_3267767 | EHNV | Laboratory of genomics and metagenomics | Trestan Pillonel, Damien Jacot, Sébastien Aeby, Gilbert Greub, Claire Bertelli |
| EPI_ISL_3267768 | VIDYMED LAUSANNE | Laboratory of genomics and metagenomics | Trestan Pillonel, Damien Jacot, Sébastien Aeby, Gilbert Greub, Claire Bertelli |
| EPI_ISL_3267769 | EHNV | Laboratory of genomics and metagenomics | Trestan Pillonel, Damien Jacot, Sébastien Aeby, Gilbert Greub, Claire Bertelli |
| EPI_ISL_3267770 | VIDYMED LAUSANNE | Laboratory of genomics and metagenomics | Trestan Pillonel, Damien Jacot, Sébastien Aeby, Gilbert Greub, Claire Bertelli |
| EPI_ISL_3267771 | EHNV | Laboratory of genomics and metagenomics | Trestan Pillonel, Damien Jacot, Sébastien Aeby, Gilbert Greub, Claire Bertelli |
| EPI_ISL_3267772, EPI_ISL_3267773 | VIDYMED LAUSANNE | Laboratory of genomics and metagenomics | Trestan Pillonel, Damien Jacot, Sébastien Aeby, Gilbert Greub, Claire Bertelli |
| EPI_ISL_3267774, EPI_ISL_3267775 | EHNV | Laboratory of genomics and metagenomics | Trestan Pillonel, Damien Jacot, Sébastien Aeby, Gilbert Greub, Claire Bertelli |
| EPI_ISL_3267776, EPI_ISL_3267777, EPI_ISL_3267778 | CHUV | Laboratory of genomics and metagenomics | Trestan Pillonel, Damien Jacot, Sébastien Aeby, Gilbert Greub, Claire Bertelli |
| EPI_ISL_3267779 | VIDYMED LAUSANNE | Laboratory of genomics and metagenomics | Trestan Pillonel, Damien Jacot, Sébastien Aeby, Gilbert Greub, Claire Bertelli |
| EPI_ISL_3267780 | CHUV | Laboratory of genomics and metagenomics | Trestan Pillonel, Damien Jacot, Sébastien Aeby, Gilbert Greub, Claire Bertelli |
| EPI_ISL_3267781, EPI_ISL_3267782, EPI_ISL_3267783 | EHNV | Laboratory of genomics and metagenomics | Trestan Pillonel, Damien Jacot, Sébastien Aeby, Gilbert Greub, Claire Bertelli |

| | | | |
|---|---|---|---|
| EPI_ISL_3267784 | VIDYMED LAUSANNE | Laboratory of genomics and metagenomics | Trestan Pillonel, Damien Jacot, Sébastien Aeby, Gilbert Greub, Claire Bertelli |
| EPI_ISL_3267785 | EHNV | Laboratory of genomics and metagenomics | Trestan Pillonel, Damien Jacot, Sébastien Aeby, Gilbert Greub, Claire Bertelli |
| EPI_ISL_3267787, EPI_ISL_3267788 | CHUV | Laboratory of genomics and metagenomics | Trestan Pillonel, Damien Jacot, Sébastien Aeby, Gilbert Greub, Claire Bertelli |
| EPI_ISL_3267789 | EHNV | Laboratory of genomics and metagenomics | Trestan Pillonel, Damien Jacot, Sébastien Aeby, Gilbert Greub, Claire Bertelli |
| EPI_ISL_3267790 | CHUV | Laboratory of genomics and metagenomics | Trestan Pillonel, Damien Jacot, Sébastien Aeby, Gilbert Greub, Claire Bertelli |
| EPI_ISL_3267791, EPI_ISL_3267792 | EHNV | Laboratory of genomics and metagenomics | Trestan Pillonel, Damien Jacot, Sébastien Aeby, Gilbert Greub, Claire Bertelli |
| EPI_ISL_3267793, EPI_ISL_3267794 | CHUV | Laboratory of genomics and metagenomics | Trestan Pillonel, Damien Jacot, Sébastien Aeby, Gilbert Greub, Claire Bertelli |
| EPI_ISL_3267795 | EHC MORGES | Laboratory of genomics and metagenomics | Trestan Pillonel, Damien Jacot, Sébastien Aeby, Gilbert Greub, Claire Bertelli |
| EPI_ISL_3267796 | EHNV | Laboratory of genomics and metagenomics | Trestan Pillonel, Damien Jacot, Sébastien Aeby, Gilbert Greub, Claire Bertelli |
| EPI_ISL_3267797 | VIDYMED LAUSANNE | Laboratory of genomics and metagenomics | Trestan Pillonel, Damien Jacot, Sébastien Aeby, Gilbert Greub, Claire Bertelli |
| EPI_ISL_3267798 | SYNLAB | Laboratory of genomics and metagenomics | Trestan Pillonel, Damien Jacot, Sébastien Aeby, Gilbert Greub, Claire Bertelli |
| EPI_ISL_3267799 | CHUV | Laboratory of genomics and metagenomics | Trestan Pillonel, Damien Jacot, Sébastien Aeby, Gilbert Greub, Claire Bertelli |
| EPI_ISL_3267800 | VIDYMED LAUSANNE | Laboratory of genomics and metagenomics | Trestan Pillonel, Damien Jacot, Sébastien Aeby, Gilbert Greub, Claire Bertelli |
| EPI_ISL_3267801 | CHUV | Laboratory of genomics and metagenomics | Trestan Pillonel, Damien Jacot, Sébastien Aeby, Gilbert Greub, Claire Bertelli |
| EPI_ISL_3267803 | VIDYMED LAUSANNE | Laboratory of genomics and metagenomics | Trestan Pillonel, Damien Jacot, Sébastien Aeby, Gilbert Greub, Claire Bertelli |
| EPI_ISL_3267804, EPI_ISL_3267805, EPI_ISL_3267806 | EHNV | Laboratory of genomics and metagenomics | Trestan Pillonel, Damien Jacot, Sébastien Aeby, Gilbert Greub, Claire Bertelli |
| EPI_ISL_3267807 | VIDYMED LAUSANNE | Laboratory of genomics and metagenomics | Trestan Pillonel, Damien Jacot, Sébastien Aeby, Gilbert Greub, Claire Bertelli |
| EPI_ISL_3267808, EPI_ISL_3267809, EPI_ISL_3267810 | CHUV | Laboratory of genomics and metagenomics | Trestan Pillonel, Damien Jacot, Sébastien Aeby, Gilbert Greub, Claire Bertelli |
| EPI_ISL_3267811 | VIDYMED LAUSANNE | Laboratory of genomics and metagenomics | Trestan Pillonel, Damien Jacot, Sébastien Aeby, Gilbert Greub, Claire Bertelli |
| EPI_ISL_3267813 | EHNV | Laboratory of genomics and metagenomics | Trestan Pillonel, Damien Jacot, Sébastien Aeby, Gilbert Greub, Claire Bertelli |
| EPI_ISL_3267815 | CHUV | Laboratory of genomics and metagenomics | Trestan Pillonel, Damien Jacot, Sébastien Aeby, Gilbert Greub, Claire Bertelli |
| EPI_ISL_3267816, EPI_ISL_3267817 | VIDYMED LAUSANNE | Laboratory of genomics and metagenomics | Trestan Pillonel, Damien Jacot, Sébastien Aeby, Gilbert Greub, Claire Bertelli |
| EPI_ISL_3267819, EPI_ISL_3267820 | EHNV | Laboratory of genomics and metagenomics | Trestan Pillonel, Damien Jacot, Sébastien Aeby, Gilbert Greub, Claire Bertelli |
| EPI_ISL_3267821 | VIDYMED LAUSANNE | Laboratory of genomics and metagenomics | Trestan Pillonel, Damien Jacot, Sébastien Aeby, Gilbert Greub, Claire Bertelli |
| EPI_ISL_3267822 | CHUV | Laboratory of genomics and metagenomics | Trestan Pillonel, Damien Jacot, Sébastien Aeby, Gilbert Greub, Claire Bertelli |
| EPI_ISL_3267823 | VIDYMED LAUSANNE | Laboratory of genomics and metagenomics | Trestan Pillonel, Damien Jacot, Sébastien Aeby, Gilbert Greub, Claire Bertelli |
| EPI_ISL_3267824 | CHUV | Laboratory of genomics and metagenomics | Trestan Pillonel, Damien Jacot, Sébastien Aeby, Gilbert Greub, Claire Bertelli |
| EPI_ISL_3267826 | VIDYMED LAUSANNE | Laboratory of genomics and metagenomics | Trestan Pillonel, Damien Jacot, Sébastien Aeby, Gilbert Greub, Claire Bertelli |
| EPI_ISL_3267827, EPI_ISL_3267828, EPI_ISL_3267829, EPI_ISL_3267830, EPI_ISL_3267831, EPI_ISL_3267832, EPI_ISL_3267833, EPI_ISL_3267834, EPI_ISL_3267835, EPI_ISL_3267836, EPI_ISL_3267837, EPI_ISL_3267838, EPI_ISL_3267839, EPI_ISL_3267840, EPI_ISL_3267841, EPI_ISL_3267842, EPI_ISL_3267843, EPI_ISL_3267844, EPI_ISL_3267845, EPI_ISL_3267846, EPI_ISL_3267847, EPI_ISL_3267848, EPI_ISL_3267849, EPI_ISL_3267850, EPI_ISL_3267851, EPI_ISL_3267852, EPI_ISL_3267853, EPI_ISL_3267854, EPI_ISL_3267855, EPI_ISL_3267856, EPI_ISL_3267857, EPI_ISL_3267858 | | | |
| see above | Institute for Infectious Diseases | Institute for Infectious Diseases | Stefan Neuenschwander, Christian Baumann, Miguel A Terrazos Miani, Cora Sägesser, Pascal Bittel, Peter Keller, Franziska Suter-Riniker, Stephen L Leib, Alban Ramette |
| EPI_ISL_3267859, EPI_ISL_3267860 | Instituto Adolfo Lutz Central | Instituto Adolfo Lutz, Interdiciplinary Procedures Center, Strategic Laboratory | Claudio Tavares Sacchi, Claudia Regina Gonçalves, Erica Valessa Ramos Gomes, Karoline Rodrigues Campos, Caio Vinicius Dias Lopes, Leonardo Tadeu de Araujo, Marlon Benedito Nascimento Santos |
| EPI_ISL_3267862, EPI_ISL_3267865, EPI_ISL_3267867, EPI_ISL_3267868, EPI_ISL_3267869, EPI_ISL_3267870, EPI_ISL_3267871, EPI_ISL_3267872 | INTERLAB | Omics Sciences Laboratory | Derly Andrade Molina, Gabriel Morey León, Darlyn Amaya, Rubén Armas González, Katheryn Sacheri, Daniela Bravo Prieto, Elena Romero Cevallos, Esteban Fernández, Karen Muñoz, Juan Carlos Fernández Cadena |
| EPI_ISL_3267873, EPI_ISL_3267874, EPI_ISL_3267875, EPI_ISL_3267877, EPI_ISL_3267878, EPI_ISL_3267879, EPI_ISL_3267881, EPI_ISL_3267882, EPI_ISL_3267888, EPI_ISL_3267889, EPI_ISL_3267890, EPI_ISL_3267891, EPI_ISL_3267892, EPI_ISL_3267893, EPI_ISL_3267894, EPI_ISL_3267895, EPI_ISL_3267896, EPI_ISL_3267897, EPI_ISL_3267898, EPI_ISL_3267899, EPI_ISL_3267900, EPI_ISL_3267901, EPI_ISL_3267902, EPI_ISL_3267903, EPI_ISL_3267904, EPI_ISL_3267905 | | | |
| see above | TXDSHS | TXDSHS | Rashmi Tuladhar, Bonnie Oh, Jenny Zhang, Maliha Rahman, Mayela Pedrueza, Anita Pokharel, Karen Bobier, Lorraine Rodriguez, Myong Koag, Chun Wang, Rachel Lee, Grace Kubin |
| EPI_ISL_3268026, EPI_ISL_3268028, EPI_ISL_3268029, EPI_ISL_3268030, EPI_ISL_3268031, EPI_ISL_3268032, EPI_ISL_3268033, EPI_ISL_3268034, EPI_ISL_3268035, EPI_ISL_3268036, EPI_ISL_3268037, EPI_ISL_3268038, EPI_ISL_3268039, EPI_ISL_3268040, EPI_ISL_3268041, EPI_ISL_3268042, EPI_ISL_3268043, EPI_ISL_3268045, EPI_ISL_3268048, EPI_ISL_3268050, EPI_ISL_3268053, EPI_ISL_3268054, EPI_ISL_3268055, EPI_ISL_3268056, EPI_ISL_3268057, EPI_ISL_3268058, EPI_ISL_3268059, EPI_ISL_3268060, EPI_ISL_3268063, EPI_ISL_3268064, EPI_ISL_3268065, EPI_ISL_3268066, EPI_ISL_3268067, EPI_ISL_3268068, EPI_ISL_3268069, EPI_ISL_3268070, EPI_ISL_3268071, EPI_ISL_3268072, EPI_ISL_3268073, EPI_ISL_3268075, EPI_ISL_3268076, EPI_ISL_3268077, EPI_ISL_3268078, EPI_ISL_3268079, EPI_ISL_3268080, EPI_ISL_3268084, EPI_ISL_3268085, EPI_ISL_3268087, EPI_ISL_3268088, EPI_ISL_3268089, EPI_ISL_3268090, EPI_ISL_3268091, EPI_ISL_3268092, EPI_ISL_3268093, EPI_ISL_3268094, EPI_ISL_3268095, EPI_ISL_3268096, EPI_ISL_3268097, EPI_ISL_3268098, EPI_ISL_3268099, EPI_ISL_3268128, EPI_ISL_3268129, EPI_ISL_3268130, EPI_ISL_3268131, EPI_ISL_3268132, EPI_ISL_3268133, EPI_ISL_3268134, EPI_ISL_3268135, EPI_ISL_3268136, EPI_ISL_3268138, EPI_ISL_3268140, EPI_ISL_3268141, EPI_ISL_3268142, EPI_ISL_3268143, EPI_ISL_3268144, EPI_ISL_3268146, EPI_ISL_3268148, EPI_ISL_3268149, EPI_ISL_3268150, EPI_ISL_3268151, EPI_ISL_3268152, EPI_ISL_3268153, EPI_ISL_3268155, EPI_ISL_3268156, EPI_ISL_3268157, EPI_ISL_3268158, EPI_ISL_3268159, EPI_ISL_3268160, EPI_ISL_3268162, EPI_ISL_3268163, EPI_ISL_3268164, EPI_ISL_3268165, EPI_ISL_3268166, EPI_ISL_3268167, EPI_ISL_3268168, EPI_ISL_3268169, EPI_ISL_3268170, EPI_ISL_3268171, EPI_ISL_3268172, EPI_ISL_3268173, EPI_ISL_3268174, EPI_ISL_3268175, EPI_ISL_3268176, EPI_ISL_3268177, EPI_ISL_3268178, EPI_ISL_3268179, EPI_ISL_3268180, EPI_ISL_3268181, EPI_ISL_3268182, EPI_ISL_3268183, EPI_ISL_3268184, EPI_ISL_3268185, EPI_ISL_3268186, EPI_ISL_3268187, EPI_ISL_3268188, EPI_ISL_3268191, EPI_ISL_3268192, EPI_ISL_3268193, EPI_ISL_3268194, EPI_ISL_3268195, EPI_ISL_3268196 | | | |
| see above | WACCBIP, University of Ghana, Accra, Ghana | WACCBIP, University of Ghana, Volta Road, Legon-Accra, Ghana | Collins M. Morang'a, Joyce M. Ngoi, Dominic S. Y. Amuzu, Vincent Appiah, Evelyn B. Quansah, Samirah Saiid, Bright K. Yemi, Israel Osei-Wusu, Deborah N. A. Mettle, Philip M. Soglo, Violette V. M'cormack, Paul Owusu-Oduro, Samuel Armoo, Oliver Commey, Sylvester Dassah, Victor Asoala, Richard Odame Phillips, Michael Owusu, Frederick M. Tei-Maya, Joe K. Mutungi, Nicaise T. Ndam, William K. Ampofo, Yaw Bediako, Peter K. Quashie, Lucas N. Amenga-Etego, and Gordon A. Awandare, |
| EPI_ISL_3268197, EPI_ISL_3268198, EPI_ISL_3268199, EPI_ISL_3268200, EPI_ISL_3268201, EPI_ISL_3268202, EPI_ISL_3268203, EPI_ISL_3268204, EPI_ISL_3268205, EPI_ISL_3268206, EPI_ISL_3268207, EPI_ISL_3268208, EPI_ISL_3268209, EPI_ISL_3268210, EPI_ISL_3268211, EPI_ISL_3268212, EPI_ISL_3268213, EPI_ISL_3268214, EPI_ISL_3268215, EPI_ISL_3268216, EPI_ISL_3268217, EPI_ISL_3268218, EPI_ISL_3268219, EPI_ISL_3268220, EPI_ISL_3268221, EPI_ISL_3268222, EPI_ISL_3268223, EPI_ISL_3268224, EPI_ISL_3268225, EPI_ISL_3268227 | | | |
| see above | Montana Public Health Laboratory | Montana Public Health Laboratory | Joy Ritter, Michelle Mozer, Carrie Biskupiak, Deborah Gibson, Michael Dills |
| EPI_ISL_3268228, EPI_ISL_3268229, EPI_ISL_3268230, EPI_ISL_3268231, EPI_ISL_3268232, EPI_ISL_3268233, EPI_ISL_3268234, EPI_ISL_3268235, EPI_ISL_3268236, EPI_ISL_3268237, EPI_ISL_3268238, EPI_ISL_3268239, EPI_ISL_3268240, EPI_ISL_3268241, EPI_ISL_3268242, EPI_ISL_3268243, EPI_ISL_3268244, EPI_ISL_3268245, EPI_ISL_3268246, EPI_ISL_3268247, EPI_ISL_3268248, EPI_ISL_3268249, EPI_ISL_3268250, EPI_ISL_3268251, EPI_ISL_3268252, EPI_ISL_3268253, EPI_ISL_3268254, EPI_ISL_3268255, EPI_ISL_3268256 | | | |
| see above | Kansas Health and Environmental Lab | Kansas Health and Environmental Lab | Mike Grose, Jonathan Barnell, Amanda Bradley, and Phil Adam |
| EPI_ISL_3268257, EPI_ISL_3268258, EPI_ISL_3268259, EPI_ISL_3268260, EPI_ISL_3268261, EPI_ISL_3268262, EPI_ISL_3268263, EPI_ISL_3268264, EPI_ISL_3268265, EPI_ISL_3268266, EPI_ISL_3268267, EPI_ISL_3268268, EPI_ISL_3268269, EPI_ISL_3268270, EPI_ISL_3268271, EPI_ISL_3268272, EPI_ISL_3268273, EPI_ISL_3268274, EPI_ISL_3268275, EPI_ISL_3268276, EPI_ISL_3268277, EPI_ISL_3268278, EPI_ISL_3268279, EPI_ISL_3268280, EPI_ISL_3268281, EPI_ISL_3268282, EPI_ISL_3268283, EPI_ISL_3268284, EPI_ISL_3268288, EPI_ISL_3268289, EPI_ISL_3268290, EPI_ISL_3268292, EPI_ISL_3268293, EPI_ISL_3268294, EPI_ISL_3268295, EPI_ISL_3268296, EPI_ISL_3268297, EPI_ISL_3268298, EPI_ISL_3268299, EPI_ISL_3268300, EPI_ISL_3268301, EPI_ISL_3268302, EPI_ISL_3268303, EPI_ISL_3268306, EPI_ISL_3268307, EPI_ISL_3268308, EPI_ISL_3268309, EPI_ISL_3268310, EPI_ISL_3268311, EPI_ISL_3268312, EPI_ISL_3268313, EPI_ISL_3268314, EPI_ISL_3268315, EPI_ISL_3268316, EPI_ISL_3268317, EPI_ISL_3268318, EPI_ISL_3268320, EPI_ISL_3268321, EPI_ISL_3268322 | | | |
| see above | North Dakota Department of Health, Public Health Laboratory | North Dakota Department of Health, Public Health Laboratory | Lisa Wingerter |
| EPI_ISL_3268323, EPI_ISL_3268324, EPI_ISL_3268325, EPI_ISL_3268326, EPI_ISL_3268327, EPI_ISL_3268328, EPI_ISL_3268329, EPI_ISL_3268330, EPI_ISL_3268331, EPI_ISL_3268332, EPI_ISL_3268333, EPI_ISL_3268334, EPI_ISL_3268335, EPI_ISL_3268336, EPI_ISL_3268337, EPI_ISL_3268338, EPI_ISL_3268339, EPI_ISL_3268340, EPI_ISL_3268341, EPI_ISL_3268342, EPI_ISL_3268343, EPI_ISL_3268344, EPI_ISL_3268345, EPI_ISL_3268346, EPI_ISL_3268347, EPI_ISL_3268348, EPI_ISL_3268349, EPI_ISL_3268350, EPI_ISL_3268351, EPI_ISL_3268352, EPI_ISL_3268353, EPI_ISL_3268354, EPI_ISL_3268355, EPI_ISL_3268356, EPI_ISL_3268357, EPI_ISL_3268358, EPI_ISL_3268359, EPI_ISL_3268360, EPI_ISL_3268362, EPI_ISL_3268363, EPI_ISL_3268364, EPI_ISL_3268365, EPI_ISL_3268366, EPI_ISL_3268367 | | | |
| see above | Tampa General Hospital Esoteric Lab | Tampa General Hospital Esoteric Research & Development Lab | Jon Faughn, Deanna Becker, Dominic Uy, Jorge Mecias-Frias, Katie Hurless, Vicki Healer, Amorce Lima, Suzane Silbert |
| EPI_ISL_3268371, EPI_ISL_3268389 | Instituto Nacional de Enfermedades Respiratorias (INER) | Centro de Investigación en Enfermedades Infecciosas (CIENI), Instituto Nacional de Enfermedades Respiratorias (INER) | Consorcio Mexicano de Vigilancia Genómica (CoViGen-Mex). Authors (in alphabetical order): Julio Elias Alvarado-Yaah, Carlos F. Arias, Santiago Ávila-Ríos, Eduardo Becerril-Vargas, Víctor Hugo Borja-Aburto, Celia Boukadida, Cristóbal Cháidez-Quiróz, Juan Bautista Chale-Dzul, Ricardo Ciria Merce, Andreu Comas-García, Célida Duque Molina, Julissa Enciso-Ibarra, José Antonio Enciso-Moreno, Gloria Elena Espinosa-Ayala, Fernando Fontove-Herrera, Daniel Fregoso-Rueda, Victor Eduardo Garcia-Arias, Alejandra Garcia-Gasca, Bruno Gomez-Gil, Jean Pierre González, Irvin González-López, Concepción Grajales-Muñiz, Ricardo Grande, Rosa María Gutierrez Rios, Alejandra Hernández-Terán, Alfredo Herrera-Estrella, Carla Ivón Herrera-Najera, Pavel Isa, Daniel Lira Morales, Susana Lopez, Antonio Loza Román, Brenda Irasema Maldonado-Meza, Bernardo Martínez-Miguel, José Arturo Martínez-Orozco, Célida Martínez- Rodríguez, Margarita Matías-Florentino, Fidencio Mejía-Nepomuceno, María Guadalupe de Jesús Mireles-Rivera, Gloria María Molina-Salinas, Hector Montoya-Fuentes, Mario Mújica-Sánchez, José Esteban Muñoz-Medina, José de Jesús Nuñez-Contreras, Alicia Ocaña-Mondragón, Luis Alberto Ochoa-Carrera, Hector Esteban Paz-Juárez, Marissa Pérez-García, Francisco Pulido, Helen Haydee Fernanda Ramírez-Plascencia, Jorge Salas-Hernández, Angel Gustavo Salas-Lais, Alejandro Sanchez-Flores, Clara Esperanza Santacruz-Tinoco, María Guadalupe Santiago-Mauricio, Selene Zárate, Nelly Sélem-Mojica, Blanca Taboada, Kathia Elizabeth Tapia-Diaz, Gloria Vazquez, Joel Armando Vázquez-Pérez. |
| EPI_ISL_3268390 | Instituto Nacional de Cancerología | Centro de Investigación en Enfermedades Infecciosas (CIENI), Instituto Nacional de Enfermedades Respiratorias (INER) | Consorcio Mexicano de Vigilancia Genómica (CoViGen-Mex). Authors (in alphabetical order): Julio Elias Alvarado-Yaah, Carlos F. Arias, Santiago Ávila-Ríos, Eduardo Becerril-Vargas, Víctor Hugo Borja-Aburto, Celia Boukadida, Cristóbal Cháidez-Quiróz, Juan Bautista Chale-Dzul, Ricardo Ciria Merce, Andreu Comas-García, Célida Duque Molina, Julissa Enciso-Ibarra, José Antonio Enciso-Moreno, Gloria Elena Espinosa-Ayala, Fernando Fontove-Herrera, Daniel Fregoso-Rueda, Victor Eduardo Garcia-Arias, Alejandra Garcia-Gasca, Bruno Gomez-Gil, Jean Pierre González, Irvin González-López, Concepción Grajales-Muñiz, Ricardo Grande, Rosa María Gutierrez Rios, Alejandra Hernández-Terán, Alfredo Herrera-Estrella, Carla Ivón Herrera-Najera, Pavel Isa, Daniel Lira Morales, Susana Lopez, Antonio Loza Román, Brenda Irasema Maldonado-Meza, Bernardo Martínez-Miguel, José Arturo Martínez-Orozco, Célida Martínez- Rodríguez, Margarita Matías-Florentino, Fidencio Mejía-Nepomuceno, María Guadalupe de Jesús Mireles-Rivera, Gloria María Molina-Salinas, Hector Montoya-Fuentes, Mario Mújica-Sánchez, José Esteban Muñoz-Medina, José de Jesús Nuñez-Contreras, Alicia Ocaña-Mondragón, Luis Alberto Ochoa-Carrera, Hector Esteban Paz-Juárez, Marissa Pérez-García, Francisco Pulido, Helen Haydee Fernanda Ramírez-Plascencia, Jorge Salas-Hernández, Angel Gustavo Salas-Lais, Alejandro Sanchez-Flores, Clara Esperanza Santacruz-Tinoco, María Guadalupe Santiago-Mauricio, Selene Zárate, Nelly Sélem-Mojica, Blanca Taboada, Kathia Elizabeth Tapia-Diaz, Gloria Vazquez, Joel Armando Vázquez-Pérez. |
| EPI_ISL_3268391, EPI_ISL_3268392, EPI_ISL_3268393, EPI_ISL_3268394, EPI_ISL_3268395, EPI_ISL_3268396, EPI_ISL_3268397, EPI_ISL_3268398, EPI_ISL_3268399, EPI_ISL_3268400, EPI_ISL_3268401, EPI_ISL_3268402, EPI_ISL_3268403, EPI_ISL_3268404, EPI_ISL_3268405, EPI_ISL_3268406, EPI_ISL_3268407, EPI_ISL_3268408, EPI_ISL_3268409, EPI_ISL_3268410, EPI_ISL_3268411, EPI_ISL_3268412, | | | |

EPI_ISL_3268413, EPI_ISL_3268414, EPI_ISL_3268415, EPI_ISL_3268416

| | | | |
|---|---|---|---|
| see above | Instituto Nacional de Enfermedades Respiratorias (INER) | Centro de Investigación en Enfermedades Infecciosas (CIENI), Instituto Nacional de Enfermedades Respiratorias (INER) | Consorcio Mexicano de Vigilancia Genómica (CoViGen-Mex). Authors (in alphabetical order): Julio Elias Alvarado-Yaah, Carlos F. Arias, Santiago Ávila-Ríos, Eduardo Becerril-Vargas, Víctor Hugo Borja-Aburto, Celia Boukadida, Cristóbal Cháidez-Quiróz, Juan Bautista Chale-Dzul, Ricardo Ciria Merce, Andreu Comas-García, Célida Duque Molina, Julissa Enciso-Ibarra, José Antonio Enciso-Moreno, Gloria Elena Espinosa-Ayala, Fernando Fontove-Herrera, Daniel Fregoso-Rueda, Victor Eduardo Garcia-Arias, Alejandra Garcia-Gasca, Bruno Gomez-Gil, Jean Pierre González, Irvin González-López, Concepción Grajales-Muñiz, Ricardo Grande, Rosa María Gutierrez Rios, Alejandra Hernández-Terán, Alfredo Herrera-Estrella, Carla Ivón Herrera-Najera, Pavel Isa, Daniel Lira Morales, Susana Lopez, Antonio Loza Román, Brenda Irasema Maldonado-Meza, Bernardo Martínez-Miguel, José Arturo Martínez-Orozco, Célida Martínez- Rodríguez, Margarita Matías-Florentino, Fidencio Mejía-Nepomuceno, María Guadalupe de Jesús Mireles-Rivera, Gloria María Molina-Salinas, Hector Montoya-Fuentes, Mario Mújica-Sánchez, José Esteban Muñoz-Medina, José de Jesús Nuñez-Contreras, Alicia Ocaña-Mondragón, Luis Alberto Ochoa-Carrera, Hector Esteban Paz-Juárez, Marissa Pérez-García, Francisco Pulido, Helen Haydee Fernanda Ramírez-Plascencia, Jorge Salas-Hernández, Angel Gustavo Salas-Lais, Alejandro Sanchez-Flores, Clara Esperanza Santacruz-Tinoco, María Guadalupe Santiago-Mauricio, Selene Zárate, Nelly Sélem-Mojica, Blanca Taboada, Kathia Elizabeth Tapia-Diaz, Gloria Vazquez, Joel Armando Vázquez-Pérez. |
| EPI_ISL_3268417 | Instituto Nacional de Cancerología | Centro de Investigación en Enfermedades Infecciosas (CIENI), Instituto Nacional de Enfermedades Respiratorias (INER) | Consorcio Mexicano de Vigilancia Genómica (CoViGen-Mex). Authors (in alphabetical order): Julio Elias Alvarado-Yaah, Carlos F. Arias, Santiago Ávila-Ríos, Eduardo Becerril-Vargas, Víctor Hugo Borja-Aburto, Celia Boukadida, Cristóbal Cháidez-Quiróz, Juan Bautista Chale-Dzul, Ricardo Ciria Merce, Andreu Comas-García, Célida Duque Molina, Julissa Enciso-Ibarra, José Antonio Enciso-Moreno, Gloria Elena Espinosa-Ayala, Fernando Fontove-Herrera, Daniel Fregoso-Rueda, Victor Eduardo Garcia-Arias, Alejandra Garcia-Gasca, Bruno Gomez-Gil, Jean Pierre González, Irvin González-López, Concepción Grajales-Muñiz, Ricardo Grande, Rosa María Gutierrez Rios, Alejandra Hernández-Terán, Alfredo Herrera-Estrella, Carla Ivón Herrera-Najera, Pavel Isa, Daniel Lira Morales, Susana Lopez, Antonio Loza Román, Brenda Irasema Maldonado-Meza, Bernardo Martínez-Miguel, José Arturo Martínez-Orozco, Célida Martínez- Rodríguez, Margarita Matías-Florentino, Fidencio Mejía-Nepomuceno, María Guadalupe de Jesús Mireles-Rivera, Gloria María Molina-Salinas, Hector Montoya-Fuentes, Mario Mújica-Sánchez, José Esteban Muñoz-Medina, José de Jesús Nuñez-Contreras, Alicia Ocaña-Mondragón, Luis Alberto Ochoa-Carrera, Hector Esteban Paz-Juárez, Marissa Pérez-García, Francisco Pulido, Helen Haydee Fernanda Ramírez-Plascencia, Jorge Salas-Hernández, Angel Gustavo Salas-Lais, Alejandro Sanchez-Flores, Clara Esperanza Santacruz-Tinoco, María Guadalupe Santiago-Mauricio, Selene Zárate, Nelly Sélem-Mojica, Blanca Taboada, Kathia Elizabeth Tapia-Diaz, Gloria Vazquez, Joel Armando Vázquez-Pérez. |
| EPI_ISL_3268418, EPI_ISL_3268419, EPI_ISL_3268420, EPI_ISL_3268421 | Instituto Nacional de Enfermedades Respiratorias (INER) | Centro de Investigación en Enfermedades Infecciosas (CIENI), Instituto Nacional de Enfermedades Respiratorias (INER) | Consorcio Mexicano de Vigilancia Genómica (CoViGen-Mex). Authors (in alphabetical order): Julio Elias Alvarado-Yaah, Carlos F. Arias, Santiago Ávila-Ríos, Eduardo Becerril-Vargas, Víctor Hugo Borja-Aburto, Celia Boukadida, Cristóbal Cháidez-Quiróz, Juan Bautista Chale-Dzul, Ricardo Ciria Merce, Andreu Comas-García, Célida Duque Molina, Julissa Enciso-Ibarra, José Antonio Enciso-Moreno, Gloria Elena Espinosa-Ayala, Fernando Fontove-Herrera, Daniel Fregoso-Rueda, Victor Eduardo Garcia-Arias, Alejandra Garcia-Gasca, Bruno Gomez-Gil, Jean Pierre González, Irvin González-López, Concepción Grajales-Muñiz, Ricardo Grande, Rosa María Gutierrez Rios, Alejandra Hernández-Terán, Alfredo Herrera-Estrella, Carla Ivón Herrera-Najera, Pavel Isa, Daniel Lira Morales, Susana Lopez, Antonio Loza Román, Brenda Irasema Maldonado-Meza, Bernardo Martínez-Miguel, José Arturo Martínez-Orozco, Célida Martínez- Rodríguez, Margarita Matías-Florentino, Fidencio Mejía-Nepomuceno, María Guadalupe de Jesús Mireles-Rivera, Gloria María Molina-Salinas, Hector Montoya-Fuentes, Mario Mújica-Sánchez, José Esteban Muñoz-Medina, José de Jesús Nuñez-Contreras, Alicia Ocaña-Mondragón, Luis Alberto Ochoa-Carrera, Hector Esteban Paz-Juárez, Marissa Pérez-García, Francisco Pulido, Helen Haydee Fernanda Ramírez-Plascencia, Jorge Salas-Hernández, Angel Gustavo Salas-Lais, Alejandro Sanchez-Flores, Clara Esperanza Santacruz-Tinoco, María Guadalupe Santiago-Mauricio, Selene Zárate, Nelly Sélem-Mojica, Blanca Taboada, Kathia Elizabeth Tapia-Diaz, Gloria Vazquez, Joel Armando Vázquez-Pérez. |
| EPI_ISL_3268422 | Instituto Nacional de Cancerología | Centro de Investigación en Enfermedades Infecciosas (CIENI), Instituto Nacional de Enfermedades Respiratorias (INER) | Consorcio Mexicano de Vigilancia Genómica (CoViGen-Mex). Authors (in alphabetical order): Julio Elias Alvarado-Yaah, Carlos F. Arias, Santiago Ávila-Ríos, Eduardo Becerril-Vargas, Víctor Hugo Borja-Aburto, Celia Boukadida, Cristóbal Cháidez-Quiróz, Juan Bautista Chale-Dzul, Ricardo Ciria Merce, Andreu Comas-García, Célida Duque Molina, Julissa Enciso-Ibarra, José Antonio Enciso-Moreno, Gloria Elena Espinosa-Ayala, Fernando Fontove-Herrera, Daniel Fregoso-Rueda, Victor Eduardo Garcia-Arias, Alejandra Garcia-Gasca, Bruno Gomez-Gil, Jean Pierre González, Irvin González-López, Concepción Grajales-Muñiz, Ricardo Grande, Rosa María Gutierrez Rios, Alejandra Hernández-Terán, Alfredo Herrera-Estrella, Carla Ivón Herrera-Najera, Pavel Isa, Daniel Lira Morales, Susana Lopez, Antonio Loza Román, Brenda Irasema Maldonado-Meza, Bernardo Martínez-Miguel, José Arturo Martínez-Orozco, Célida Martínez- Rodríguez, Margarita Matías-Florentino, Fidencio Mejía-Nepomuceno, María Guadalupe de Jesús Mireles-Rivera, Gloria María Molina-Salinas, Hector Montoya-Fuentes, Mario Mújica-Sánchez, José Esteban Muñoz-Medina, José de Jesús Nuñez-Contreras, Alicia Ocaña-Mondragón, Luis Alberto Ochoa-Carrera, Hector Esteban Paz-Juárez, Marissa Pérez-García, Francisco Pulido, Helen Haydee Fernanda Ramírez-Plascencia, Jorge Salas-Hernández, Angel Gustavo Salas-Lais, Alejandro Sanchez-Flores, Clara Esperanza Santacruz-Tinoco, María Guadalupe Santiago-Mauricio, Selene Zárate, Nelly Sélem-Mojica, Blanca Taboada, Kathia Elizabeth Tapia-Diaz, Gloria Vazquez, Joel Armando Vázquez-Pérez. |
| EPI_ISL_3268423, EPI_ISL_3268424, EPI_ISL_3268425 | Instituto Nacional de Enfermedades Respiratorias (INER) | Centro de Investigación en Enfermedades Infecciosas (CIENI), Instituto Nacional de Enfermedades Respiratorias (INER) | Consorcio Mexicano de Vigilancia Genómica (CoViGen-Mex). Authors (in alphabetical order): Julio Elias Alvarado-Yaah, Carlos F. Arias, Santiago Ávila-Ríos, Eduardo Becerril-Vargas, Víctor Hugo Borja-Aburto, Celia Boukadida, Cristóbal Cháidez-Quiróz, Juan Bautista Chale-Dzul, Ricardo Ciria Merce, Andreu Comas-García, Célida Duque Molina, Julissa Enciso-Ibarra, José Antonio Enciso-Moreno, Gloria Elena Espinosa-Ayala, Fernando Fontove-Herrera, Daniel Fregoso-Rueda, Victor Eduardo Garcia-Arias, Alejandra Garcia-Gasca, Bruno Gomez-Gil, Jean Pierre González, Irvin González-López, Concepción Grajales-Muñiz, Ricardo Grande, Rosa María Gutierrez Rios, Alejandra Hernández-Terán, Alfredo Herrera-Estrella, Carla Ivón Herrera-Najera, Pavel Isa, Daniel Lira Morales, Susana Lopez, Antonio Loza Román, Brenda Irasema Maldonado-Meza, Bernardo Martínez-Miguel, José Arturo Martínez-Orozco, Célida Martínez- Rodríguez, Margarita Matías-Florentino, Fidencio Mejía-Nepomuceno, María Guadalupe de Jesús Mireles-Rivera, Gloria María Molina-Salinas, Hector Montoya-Fuentes, Mario Mújica-Sánchez, José Esteban Muñoz-Medina, José de Jesús Nuñez-Contreras, Alicia Ocaña-Mondragón, Luis Alberto Ochoa-Carrera, Hector Esteban Paz-Juárez, Marissa Pérez-García, Francisco Pulido, Helen Haydee Fernanda Ramírez-Plascencia, Jorge Salas-Hernández, Angel Gustavo Salas-Lais, Alejandro Sanchez-Flores, Clara Esperanza Santacruz-Tinoco, María Guadalupe Santiago-Mauricio, Selene Zárate, Nelly Sélem-Mojica, Blanca Taboada, Kathia Elizabeth Tapia-Diaz, Gloria Vazquez, Joel Armando Vázquez-Pérez. |
| EPI_ISL_3268426 | Instituto Nacional de Cancerología | Centro de Investigación en Enfermedades Infecciosas (CIENI), Instituto Nacional de Enfermedades Respiratorias (INER) | Consorcio Mexicano de Vigilancia Genómica (CoViGen-Mex). Authors (in alphabetical order): Julio Elias Alvarado-Yaah, Carlos F. Arias, Santiago Ávila-Ríos, Eduardo Becerril-Vargas, Víctor Hugo Borja-Aburto, Celia Boukadida, Cristóbal Cháidez-Quiróz, Juan Bautista Chale-Dzul, Ricardo Ciria Merce, Andreu Comas-García, Célida Duque Molina, Julissa Enciso-Ibarra, José Antonio Enciso-Moreno, Gloria Elena Espinosa-Ayala, Fernando Fontove-Herrera, Daniel Fregoso-Rueda, Victor Eduardo Garcia-Arias, Alejandra Garcia-Gasca, Bruno Gomez-Gil, Jean Pierre González, Irvin González-López, Concepción Grajales-Muñiz, Ricardo Grande, Rosa María Gutierrez Rios, Alejandra Hernández-Terán, Alfredo Herrera-Estrella, Carla Ivón Herrera-Najera, Pavel Isa, Daniel Lira Morales, Susana Lopez, Antonio Loza Román, Brenda Irasema Maldonado-Meza, Bernardo Martínez-Miguel, José Arturo Martínez-Orozco, Célida Martínez- Rodríguez, Margarita Matías-Florentino, Fidencio Mejía-Nepomuceno, María Guadalupe de Jesús Mireles-Rivera, Gloria María Molina-Salinas, Hector Montoya-Fuentes, Mario Mújica-Sánchez, José Esteban Muñoz-Medina, José de Jesús Nuñez-Contreras, Alicia Ocaña-Mondragón, Luis Alberto Ochoa-Carrera, Hector Esteban Paz-Juárez, Marissa Pérez-García, Francisco Pulido, Helen Haydee Fernanda Ramírez-Plascencia, Jorge Salas-Hernández, Angel Gustavo Salas-Lais, Alejandro Sanchez-Flores, Clara Esperanza Santacruz-Tinoco, María Guadalupe Santiago-Mauricio, Selene Zárate, Nelly Sélem-Mojica, Blanca Taboada, Kathia Elizabeth Tapia-Diaz, Gloria Vazquez, Joel Armando Vázquez-Pérez. |
| EPI_ISL_3268427 | Instituto Nacional de Enfermedades Respiratorias (INER) | Centro de Investigación en Enfermedades Infecciosas (CIENI), Instituto Nacional de Enfermedades Respiratorias (INER) | Consorcio Mexicano de Vigilancia Genómica (CoViGen-Mex). Authors (in alphabetical order): Julio Elias Alvarado-Yaah, Carlos F. Arias, Santiago Ávila-Ríos, Eduardo Becerril-Vargas, Víctor Hugo Borja-Aburto, Celia Boukadida, Cristóbal Cháidez-Quiróz, Juan Bautista Chale-Dzul, Ricardo Ciria Merce, Andreu Comas-García, Célida Duque Molina, Julissa Enciso-Ibarra, José Antonio Enciso-Moreno, Gloria Elena Espinosa-Ayala, Fernando Fontove-Herrera, Daniel Fregoso-Rueda, Victor Eduardo Garcia-Arias, Alejandra Garcia-Gasca, Bruno Gomez-Gil, Jean Pierre González, Irvin González-López, Concepción Grajales-Muñiz, Ricardo Grande, Rosa María Gutierrez Rios, Alejandra Hernández-Terán, Alfredo Herrera-Estrella, Carla Ivón Herrera-Najera, Pavel Isa, Daniel Lira Morales, Susana Lopez, Antonio Loza Román, Brenda Irasema Maldonado-Meza, Bernardo Martínez-Miguel, José Arturo Martínez-Orozco, Célida Martínez- Rodríguez, Margarita Matías-Florentino, Fidencio Mejía-Nepomuceno, María Guadalupe de Jesús Mireles-Rivera, Gloria María Molina-Salinas, Hector Montoya-Fuentes, Mario Mújica-Sánchez, José Esteban Muñoz-Medina, José de Jesús Nuñez-Contreras, Alicia Ocaña-Mondragón, Luis Alberto Ochoa-Carrera, Hector Esteban Paz-Juárez, Marissa Pérez-García, Francisco Pulido, Helen Haydee Fernanda Ramírez-Plascencia, Jorge Salas-Hernández, Angel Gustavo Salas-Lais, Alejandro Sanchez-Flores, Clara Esperanza Santacruz-Tinoco, María Guadalupe Santiago-Mauricio, Selene Zárate, Nelly Sélem-Mojica, Blanca Taboada, Kathia Elizabeth Tapia-Diaz, Gloria Vazquez, Joel Armando Vázquez-Pérez. |
| EPI_ISL_3268428 | Instituto Nacional de Cancerología | Centro de Investigación en Enfermedades Infecciosas (CIENI), Instituto Nacional de Enfermedades Respiratorias (INER) | Consorcio Mexicano de Vigilancia Genómica (CoViGen-Mex). Authors (in alphabetical order): Julio Elias Alvarado-Yaah, Carlos F. Arias, Santiago Ávila-Ríos, Eduardo Becerril-Vargas, Víctor Hugo Borja-Aburto, Celia Boukadida, Cristóbal Cháidez-Quiróz, Juan Bautista Chale-Dzul, Ricardo Ciria Merce, Andreu Comas-García, Célida Duque Molina, Julissa Enciso-Ibarra, José Antonio Enciso-Moreno, Gloria Elena Espinosa-Ayala, Fernando Fontove-Herrera, Daniel Fregoso-Rueda, Victor Eduardo Garcia-Arias, Alejandra Garcia-Gasca, Bruno Gomez-Gil, Jean Pierre González, Irvin González-López, Concepción Grajales-Muñiz, Ricardo Grande, Rosa María Gutierrez Rios, Alejandra Hernández-Terán, Alfredo Herrera-Estrella, Carla Ivón Herrera-Najera, Pavel Isa, Daniel Lira Morales, Susana Lopez, Antonio Loza Román, Brenda Irasema Maldonado-Meza, Bernardo Martínez-Miguel, José Arturo Martínez-Orozco, Célida Martínez- Rodríguez, Margarita Matías-Florentino, Fidencio Mejía-Nepomuceno, María Guadalupe de Jesús Mireles-Rivera, Gloria María Molina-Salinas, Hector Montoya-Fuentes, Mario Mújica-Sánchez, José Esteban Muñoz-Medina, José de Jesús Nuñez-Contreras, Alicia Ocaña-Mondragón, Luis Alberto Ochoa-Carrera, Hector Esteban Paz-Juárez, Marissa Pérez-García, Francisco Pulido, Helen Haydee Fernanda Ramírez-Plascencia, Jorge Salas-Hernández, Angel Gustavo Salas-Lais, Alejandro Sanchez-Flores, Clara Esperanza Santacruz-Tinoco, María Guadalupe Santiago-Mauricio, Selene Zárate, Nelly Sélem-Mojica, Blanca Taboada, Kathia Elizabeth Tapia-Diaz, Gloria Vazquez, Joel Armando Vázquez-Pérez. |

EPI_ISL_3268429, EPI_ISL_3268430, EPI_ISL_3268431, EPI_ISL_3268432, EPI_ISL_3268433, EPI_ISL_3268434, EPI_ISL_3268435, EPI_ISL_3268436, EPI_ISL_3268437, EPI_ISL_3268438, EPI_ISL_3268439, EPI_ISL_3268440, EPI_ISL_3268441, EPI_ISL_3268442, EPI_ISL_3268443, EPI_ISL_3268444, EPI_ISL_3268445, EPI_ISL_3268446, EPI_ISL_3268447, EPI_ISL_3268448, EPI_ISL_3268449, EPI_ISL_3268450, EPI_ISL_3268451, EPI_ISL_3268452

| | | | |
|---|---|---|---|
| see above | Instituto Nacional de Enfermedades Respiratorias (INER) | Centro de Investigación en Enfermedades Infecciosas (CIENI), Instituto Nacional de Enfermedades Respiratorias (INER) | Consorcio Mexicano de Vigilancia Genómica (CoViGen-Mex). Authors (in alphabetical order): Julio Elias Alvarado-Yaah, Carlos F. Arias, Santiago Ávila-Ríos, Eduardo Becerril-Vargas, Víctor Hugo Borja-Aburto, Celia Boukadida, Cristóbal Cháidez-Quiróz, Juan Bautista Chale-Dzul, Ricardo Ciria Merce, Andreu Comas-García, Célida Duque Molina, Julissa Enciso-Ibarra, José Antonio Enciso-Moreno, Gloria Elena Espinosa-Ayala, Fernando Fontove-Herrera, Daniel Fregoso-Rueda, Victor Eduardo Garcia-Arias, Alejandra Garcia-Gasca, Bruno Gomez-Gil, Jean Pierre González, Irvin González-López, Concepción Grajales-Muñiz, Ricardo Grande, Rosa María Gutierrez Rios, Alejandra Hernández-Terán, Alfredo Herrera-Estrella, Carla Ivón Herrera-Najera, Pavel Isa, Daniel Lira Morales, Susana Lopez, Antonio Loza Román, Brenda Irasema Maldonado-Meza, Bernardo Martínez-Miguel, José Arturo Martínez-Orozco, Célida Martínez- Rodríguez, Margarita Matías-Florentino, Fidencio Mejía-Nepomuceno, María Guadalupe de Jesús Mireles-Rivera, Gloria María Molina-Salinas, Hector Montoya-Fuentes, Mario Mújica-Sánchez, José Esteban Muñoz-Medina, José de Jesús Nuñez-Contreras, Alicia Ocaña-Mondragón, Luis Alberto Ochoa-Carrera, Hector Esteban Paz-Juárez, Marissa Pérez-García, Francisco Pulido, Helen Haydee Fernanda Ramírez-Plascencia, Jorge Salas-Hernández, Angel Gustavo Salas-Lais, Alejandro Sanchez-Flores, Clara Esperanza Santacruz-Tinoco, María Guadalupe Santiago-Mauricio, Selene Zárate, Nelly Sélem-Mojica, Blanca Taboada, Kathia Elizabeth Tapia-Diaz, Gloria Vazquez, Joel Armando Vázquez-Pérez. |
| EPI_ISL_3268453, EPI_ISL_3268454 | Instituto Nacional de Cancerología | Centro de Investigación en Enfermedades Infecciosas (CIENI), Instituto Nacional de Enfermedades Respiratorias (INER) | Consorcio Mexicano de Vigilancia Genómica (CoViGen-Mex). Authors (in alphabetical order): Julio Elias Alvarado-Yaah, Carlos F. Arias, Santiago Ávila-Ríos, Eduardo Becerril-Vargas, Víctor Hugo Borja-Aburto, Celia Boukadida, Cristóbal Cháidez-Quiróz, Juan Bautista Chale-Dzul, Ricardo Ciria Merce, Andreu Comas-García, Célida Duque Molina, Julissa Enciso-Ibarra, José Antonio Enciso-Moreno, Gloria Elena Espinosa-Ayala, Fernando Fontove-Herrera, Daniel Fregoso-Rueda, Victor Eduardo Garcia-Arias, Alejandra Garcia-Gasca, Bruno Gomez-Gil, Jean Pierre González, Irvin González-López, Concepción Grajales-Muñiz, Ricardo Grande, Rosa María Gutierrez Rios, Alejandra Hernández-Terán, Alfredo Herrera-Estrella, Carla Ivón Herrera-Najera, Pavel Isa, Daniel Lira Morales, Susana Lopez, Antonio Loza Román, Brenda Irasema Maldonado-Meza, Bernardo Martínez-Miguel, José Arturo Martínez-Orozco, Célida Martínez- Rodríguez, Margarita Matías-Florentino, Fidencio Mejía-Nepomuceno, María Guadalupe de Jesús Mireles-Rivera, Gloria María Molina-Salinas, Hector Montoya-Fuentes, Mario Mújica-Sánchez, José Esteban Muñoz-Medina, José de Jesús Nuñez-Contreras, Alicia Ocaña-Mondragón, Luis Alberto Ochoa-Carrera, Hector Esteban Paz-Juárez, Marissa Pérez-García, Francisco Pulido, Helen Haydee Fernanda Ramírez-Plascencia, Jorge Salas-Hernández, Angel Gustavo Salas-Lais, Alejandro Sanchez-Flores, Clara Esperanza Santacruz-Tinoco, María Guadalupe Santiago-Mauricio, Selene Zárate, Nelly Sélem-Mojica, Blanca Taboada, Kathia Elizabeth Tapia-Diaz, Gloria Vazquez, Joel Armando Vázquez-Pérez. |
| EPI_ISL_3268455, EPI_ISL_3268456 | Instituto Nacional de Enfermedades Respiratorias (INER) | Centro de Investigación en Enfermedades Infecciosas (CIENI), Instituto Nacional de Enfermedades Respiratorias (INER) | Consorcio Mexicano de Vigilancia Genómica (CoViGen-Mex). Authors (in alphabetical order): Julio Elias Alvarado-Yaah, Carlos F. Arias, Santiago Ávila-Ríos, Eduardo Becerril-Vargas, Víctor Hugo Borja-Aburto, Celia Boukadida, Cristóbal Cháidez-Quiróz, Juan Bautista Chale-Dzul, Ricardo Ciria Merce, Andreu Comas-García, Célida Duque Molina, Julissa Enciso-Ibarra, José Antonio Enciso-Moreno, Gloria Elena Espinosa-Ayala, Fernando Fontove-Herrera, Daniel Fregoso-Rueda, Victor Eduardo Garcia-Arias, Alejandra Garcia-Gasca, Bruno Gomez-Gil, Jean Pierre González, Irvin González-López, Concepción Grajales-Muñiz, Ricardo Grande, Rosa María Gutierrez Rios, Alejandra Hernández-Terán, Alfredo Herrera-Estrella, Carla Ivón Herrera-Najera, Pavel Isa, Daniel Lira Morales, Susana Lopez, Antonio Loza Román, Brenda Irasema Maldonado-Meza, Bernardo Martínez-Miguel, José Arturo Martínez-Orozco, Célida Martínez- Rodríguez, Margarita Matías-Florentino, Fidencio Mejía-Nepomuceno, María Guadalupe de Jesús Mireles-Rivera, Gloria María Molina-Salinas, Hector Montoya-Fuentes, Mario Mújica-Sánchez, José Esteban Muñoz-Medina, José de Jesús Nuñez-Contreras, Alicia Ocaña-Mondragón, Luis Alberto Ochoa-Carrera, Hector Esteban Paz-Juárez, Marissa Pérez-García, Francisco Pulido, Helen Haydee Fernanda Ramírez-Plascencia, Jorge Salas-Hernández, Angel Gustavo Salas-Lais, Alejandro Sanchez-Flores, Clara Esperanza Santacruz-Tinoco, María Guadalupe Santiago-Mauricio, Selene Zárate, Nelly Sélem-Mojica, Blanca Taboada, Kathia Elizabeth Tapia-Diaz, Gloria Vazquez, Joel Armando Vázquez-Pérez. |

EPI_ISL_3268458, EPI_ISL_3268459, EPI_ISL_3268460, EPI_ISL_3268461, EPI_ISL_3268462, EPI_ISL_3268463, EPI_ISL_3268464, EPI_ISL_3268465, EPI_ISL_3268466, EPI_ISL_3268467, EPI_ISL_3268469, EPI_ISL_3268470, EPI_ISL_3268471, EPI_ISL_3268472, EPI_ISL_3268473, EPI_ISL_3268474, EPI_ISL_3268475, EPI_ISL_3268476, EPI_ISL_3268477, EPI_ISL_3268478, EPI_ISL_3268480, EPI_ISL_3268481, EPI_ISL_3268482, EPI_ISL_3268483, EPI_ISL_3268484, EPI_ISL_3268485, EPI_ISL_3268486, EPI_ISL_3268487, EPI_ISL_3268488, EPI_ISL_3268489, EPI_ISL_3268490, EPI_ISL_3268491, EPI_ISL_3268492, EPI_ISL_3268493, EPI_ISL_3268494, EPI_ISL_3268495, EPI_ISL_3268496, EPI_ISL_3268497, EPI_ISL_3268498, EPI_ISL_3268499, EPI_ISL_3268500, EPI_ISL_3268501, EPI_ISL_3268502, EPI_ISL_3268503, EPI_ISL_3268504, EPI_ISL_3268505, EPI_ISL_3268507, EPI_ISL_3268508, EPI_ISL_3268509, EPI_ISL_3268510, EPI_ISL_3268511, EPI_ISL_3268512, EPI_ISL_3268513, EPI_ISL_3268514, EPI_ISL_3268515, EPI_ISL_3268516, EPI_ISL_3268517, EPI_ISL_3268518, EPI_ISL_3268519, EPI_ISL_3268520, EPI_ISL_3268521, EPI_ISL_3268522, EPI_ISL_3268523, EPI_ISL_3268524, EPI_ISL_3268525, EPI_ISL_3268527, EPI_ISL_3268528, EPI_ISL_3268529, EPI_ISL_3268531, EPI_ISL_3268533, EPI_ISL_3268534, EPI_ISL_3268535, EPI_ISL_3268536, EPI_ISL_3268538, EPI_ISL_3268539, EPI_ISL_3268540, EPI_ISL_3268541, EPI_ISL_3268542, EPI_ISL_3268543, EPI_ISL_3268544, EPI_ISL_3268545, EPI_ISL_3268546, EPI_ISL_3268547, EPI_ISL_3268548, EPI_ISL_3268549, EPI_ISL_3268550, EPI_ISL_3268551, EPI_ISL_3268553, EPI_ISL_3268554, EPI_ISL_3268555, EPI_ISL_3268556, EPI_ISL_3268557, EPI_ISL_3268558, EPI_ISL_3268560, EPI_ISL_3268561, EPI_ISL_3268562, EPI_ISL_3268563, EPI_ISL_3268564, EPI_ISL_3268565, EPI_ISL_3268566, EPI_ISL_3268568, EPI_ISL_3268569, EPI_ISL_3268570, EPI_ISL_3268572, EPI_ISL_3268573, EPI_ISL_3268574, EPI_ISL_3268575, EPI_ISL_3268576, EPI_ISL_3268578, EPI_ISL_3268579, EPI_ISL_3268580, EPI_ISL_3268581, EPI_ISL_3268582, EPI_ISL_3268583, EPI_ISL_3268585, EPI_ISL_3268586, EPI_ISL_3268587, EPI_ISL_3268588, EPI_ISL_3268589, EPI_ISL_3268590, EPI_ISL_3268591, EPI_ISL_3268592, EPI_ISL_3268593, EPI_ISL_3268594, EPI_ISL_3268595, EPI_ISL_3268596, EPI_ISL_3268597, EPI_ISL_3268598, EPI_ISL_3268599, EPI_ISL_3268600, EPI_ISL_3268602, EPI_ISL_3268603, EPI_ISL_3268604, EPI_ISL_3268605, EPI_ISL_3268606, EPI_ISL_3268607, EPI_ISL_3268608, EPI_ISL_3268610, EPI_ISL_3268611, EPI_ISL_3268612, EPI_ISL_3268613, EPI_ISL_3268616, EPI_ISL_3268617, EPI_ISL_3268618, EPI_ISL_3268619, EPI_ISL_3268620, EPI_ISL_3268621, EPI_ISL_3268622, EPI_ISL_3268623, EPI_ISL_3268624, EPI_ISL_3268625, EPI_ISL_3268626, EPI_ISL_3268627, EPI_ISL_3268628, EPI_ISL_3268629, EPI_ISL_3268630, EPI_ISL_3268631, EPI_ISL_3268633, EPI_ISL_3268634, EPI_ISL_3268635, EPI_ISL_3268636, EPI_ISL_3268637, EPI_ISL_3268638, EPI_ISL_3268639, EPI_ISL_3268640, EPI_ISL_3268641, EPI_ISL_3268642, EPI_ISL_3268643, EPI_ISL_3268644, EPI_ISL_3268645, EPI_ISL_3268646, EPI_ISL_3268647, EPI_ISL_3268648, EPI_ISL_3268649, EPI_ISL_3268650, EPI_ISL_3268651, EPI_ISL_3268652, EPI_ISL_3268653, EPI_ISL_3268654, EPI_ISL_3268655, EPI_ISL_3268656, EPI_ISL_3268657, EPI_ISL_3268658, EPI_ISL_3268659, EPI_ISL_3268660, EPI_ISL_3268661, EPI_ISL_3268662, EPI_ISL_3268663, EPI_ISL_3268664, EPI_ISL_3268665, EPI_ISL_3268666, EPI_ISL_3268667, EPI_ISL_3268668, EPI_ISL_3268669, EPI_ISL_3268670, EPI_ISL_3268671, EPI_ISL_3268672, EPI_ISL_3268673, EPI_ISL_3268674, EPI_ISL_3268675, EPI_ISL_3268676, EPI_ISL_3268677, EPI_ISL_3268678, EPI_ISL_3268679, EPI_ISL_3268680, EPI_ISL_3268681, EPI_ISL_3268682, EPI_ISL_3268683, EPI_ISL_3268684, EPI_ISL_3268685, EPI_ISL_3268686, EPI_ISL_3268687, EPI_ISL_3268688, EPI_ISL_3268689, EPI_ISL_3268691, EPI_ISL_3268692, EPI_ISL_3268693, EPI_ISL_3268694, EPI_ISL_3268695, EPI_ISL_3268696, EPI_ISL_3268697, EPI_ISL_3268698, EPI_ISL_3268699, EPI_ISL_3268700, EPI_ISL_3268702, EPI_ISL_3268703, EPI_ISL_3268704, EPI_ISL_3268705, EPI_ISL_3268707, EPI_ISL_3268709, EPI_ISL_3268710, EPI_ISL_3268711, EPI_ISL_3268712, EPI_ISL_3268713, EPI_ISL_3268714, EPI_ISL_3268715, EPI_ISL_3268716, EPI_ISL_3268717, EPI_ISL_3268718, EPI_ISL_3268719, EPI_ISL_3268720, EPI_ISL_3268721, EPI_ISL_3268722, EPI_ISL_3268723, EPI_ISL_3268724, EPI_ISL_3268725, EPI_ISL_3268726, EPI_ISL_3268727, EPI_ISL_3268728, EPI_ISL_3268729, EPI_ISL_3268730, EPI_ISL_3268731, EPI_ISL_3268732, EPI_ISL_3268733, EPI_ISL_3268734, EPI_ISL_3268735, EPI_ISL_3268736, EPI_ISL_3268737, EPI_ISL_3268738, EPI_ISL_3268739, EPI_ISL_3268741, EPI_ISL_3268742, EPI_ISL_3268743, EPI_ISL_3268744, EPI_ISL_3268745, EPI_ISL_3268747, EPI_ISL_3268748, EPI_ISL_3268749, EPI_ISL_3268750, EPI_ISL_3268751, EPI_ISL_3268752, EPI_ISL_3268754, EPI_ISL_3268755, EPI_ISL_3268756, EPI_ISL_3268757, EPI_ISL_3268758, EPI_ISL_3268759, EPI_ISL_3268760, EPI_ISL_3268761, EPI_ISL_3268763, EPI_ISL_3268764, EPI_ISL_3268765, EPI_ISL_3268766, EPI_ISL_3268767, EPI_ISL_3268768, EPI_ISL_3268769, EPI_ISL_3268770, EPI_ISL_3268771, EPI_ISL_3268772, EPI_ISL_3268773, EPI_ISL_3268774, EPI_ISL_3268775, EPI_ISL_3268776, EPI_ISL_3268777, EPI_ISL_3268778, EPI_ISL_3268779, EPI_ISL_3268781, EPI_ISL_3268782, EPI_ISL_3268783, EPI_ISL_3268784, EPI_ISL_3268785, EPI_ISL_3268787, EPI_ISL_3268788, EPI_ISL_3268789, EPI_ISL_3268790, EPI_ISL_3268791, EPI_ISL_3268792, EPI_ISL_3268793, EPI_ISL_3268794, EPI_ISL_3268795, EPI_ISL_3268797, EPI_ISL_3268798, EPI_ISL_3268799, EPI_ISL_3268800, EPI_ISL_3268801, EPI_ISL_3268802, EPI_ISL_3268803, EPI_ISL_3268804, EPI_ISL_3268806, EPI_ISL_3268807, EPI_ISL_3268808, EPI_ISL_3268809, EPI_ISL_3268810, EPI_ISL_3268811, EPI_ISL_3268812, EPI_ISL_3268813, EPI_ISL_3268814, EPI_ISL_3268815, EPI_ISL_3268816, EPI_ISL_3268817, EPI_ISL_3268818, EPI_ISL_3268819, EPI_ISL_3268820, EPI_ISL_3268821, EPI_ISL_3268822, EPI_ISL_3268823, EPI_ISL_3268824, EPI_ISL_3268825, EPI_ISL_3268826, EPI_ISL_3268828, EPI_ISL_3268829, EPI_ISL_3268830, EPI_ISL_3268831, EPI_ISL_3268832, EPI_ISL_3268834, EPI_ISL_3268836, EPI_ISL_3268837, EPI_ISL_3268838, EPI_ISL_3268839, EPI_ISL_3268840, EPI_ISL_3268841, EPI_ISL_3268842, EPI_ISL_3268843, EPI_ISL_3268844, EPI_ISL_3268846, EPI_ISL_3268848, EPI_ISL_3268849, EPI_ISL_3268850, EPI_ISL_3268851, EPI_ISL_3268852, EPI_ISL_3268853, EPI_ISL_3268854, EPI_ISL_3268855, EPI_ISL_3268856, EPI_ISL_3268857, EPI_ISL_3268858, EPI_ISL_3268859, EPI_ISL_3268860, EPI_ISL_3268863, EPI_ISL_3268865, EPI_ISL_3268867, EPI_ISL_3268869, EPI_ISL_3268870, EPI_ISL_3268871, EPI_ISL_3268872, EPI_ISL_3268873, EPI_ISL_3268874, EPI_ISL_3268875, EPI_ISL_3268876, EPI_ISL_3268877, EPI_ISL_3268878, EPI_ISL_3268879, EPI_ISL_3268880, EPI_ISL_3268881, EPI_ISL_3268882, EPI_ISL_3268883, EPI_ISL_3268884, EPI_ISL_3268885, EPI_ISL_3268886, EPI_ISL_3268887, EPI_ISL_3268888, EPI_ISL_3268889, EPI_ISL_3268890, EPI_ISL_3268891, EPI_ISL_3268892, EPI_ISL_3268893, EPI_ISL_3268894, EPI_ISL_3268895, EPI_ISL_3268896, EPI_ISL_3268897, EPI_ISL_3268898, EPI_ISL_3268899, EPI_ISL_3268900, EPI_ISL_3268901, EPI_ISL_3268902, EPI_ISL_3268904, EPI_ISL_3268905, EPI_ISL_3268906, EPI_ISL_3268908, EPI_ISL_3268909, EPI_ISL_3268910, EPI_ISL_3268911, EPI_ISL_3268912, EPI_ISL_3268914, EPI_ISL_3268915, EPI_ISL_3268917, EPI_ISL_3268918, EPI_ISL_3268919, EPI_ISL_3268920, EPI_ISL_3268921, EPI_ISL_3268923, EPI_ISL_3268924, EPI_ISL_3268925, EPI_ISL_3268926, EPI_ISL_3268927, EPI_ISL_3268928, EPI_ISL_3268929, EPI_ISL_3268930, EPI_ISL_3268931, EPI_ISL_3268932, EPI_ISL_3268935, EPI_ISL_3268938, EPI_ISL_3268939, EPI_ISL_3268940, EPI_ISL_3268941, EPI_ISL_3268942, EPI_ISL_3268943, EPI_ISL_3268944, EPI_ISL_3268946, EPI_ISL_3268947, EPI_ISL_3268948, EPI_ISL_3268949, EPI_ISL_3268950, EPI_ISL_3268953, EPI_ISL_3268954, EPI_ISL_3268955, EPI_ISL_3268956, EPI_ISL_3268957, EPI_ISL_3268958

| | | | |
|---|---|---|---|
| see above | Utah Public Health Laboratory | Utah Public Health Laboratory | Olinto Linares-Perdomo, Erin L. Young, Kelly F. Oakeson |

EPI_ISL_3269460, EPI_ISL_3269461, EPI_ISL_3269462, EPI_ISL_3269463, EPI_ISL_3269464, EPI_ISL_3269465, EPI_ISL_3269466, EPI_ISL_3269467, EPI_ISL_3269468, EPI_ISL_3269469, EPI_ISL_3269470, EPI_ISL_3269471, EPI_ISL_3269472, EPI_ISL_3269473, EPI_ISL_3269474, EPI_ISL_3269475, EPI_ISL_3269476, EPI_ISL_3269477, EPI_ISL_3269478, EPI_ISL_3269479, EPI_ISL_3269480, EPI_ISL_3269481,

| | | | |
|---|---|---|---|
| EPI_ISL_3269482, EPI_ISL_3269483, EPI_ISL_3269484, EPI_ISL_3269485, EPI_ISL_3269486, EPI_ISL_3269487, EPI_ISL_3269488, EPI_ISL_3269490, EPI_ISL_3269491, EPI_ISL_3269492, EPI_ISL_3269494, EPI_ISL_3269495, EPI_ISL_3269496, EPI_ISL_3269497, EPI_ISL_3269498, EPI_ISL_3269499, EPI_ISL_3269500, EPI_ISL_3269501, EPI_ISL_3269502, EPI_ISL_3269503, EPI_ISL_3269504, EPI_ISL_3269505, EPI_ISL_3269506, EPI_ISL_3269507, EPI_ISL_3269508, EPI_ISL_3269509, EPI_ISL_3269510, EPI_ISL_3269511, EPI_ISL_3269512, EPI_ISL_3269513, EPI_ISL_3269514, EPI_ISL_3269515, EPI_ISL_3269516, EPI_ISL_3269517, EPI_ISL_3269518, EPI_ISL_3269519, EPI_ISL_3269520, EPI_ISL_3269521, EPI_ISL_3269522, EPI_ISL_3269523, EPI_ISL_3269524, EPI_ISL_3269525, EPI_ISL_3269526, EPI_ISL_3269527, EPI_ISL_3269528, EPI_ISL_3269529, EPI_ISL_3269530, EPI_ISL_3269531, EPI_ISL_3269533, EPI_ISL_3269534, EPI_ISL_3269535, EPI_ISL_3269536, EPI_ISL_3269537, EPI_ISL_3269538, EPI_ISL_3269539, EPI_ISL_3269540, EPI_ISL_3269541, EPI_ISL_3269542, EPI_ISL_3269543, EPI_ISL_3269544, EPI_ISL_3269545, EPI_ISL_3269546, EPI_ISL_3269547, EPI_ISL_3269548, EPI_ISL_3269549, EPI_ISL_3269550, EPI_ISL_3269551, EPI_ISL_3269552, EPI_ISL_3269553, EPI_ISL_3269554, EPI_ISL_3269555, EPI_ISL_3269556, EPI_ISL_3269557, EPI_ISL_3269558, EPI_ISL_3269559, EPI_ISL_3269560, EPI_ISL_3269561, EPI_ISL_3269562, EPI_ISL_3269563, EPI_ISL_3269564, EPI_ISL_3269565, EPI_ISL_3269566, EPI_ISL_3269567, EPI_ISL_3269568, EPI_ISL_3269569, EPI_ISL_3269570, EPI_ISL_3269571, EPI_ISL_3269572, EPI_ISL_3269573, EPI_ISL_3269574, EPI_ISL_3269575, EPI_ISL_3269576, EPI_ISL_3269577, EPI_ISL_3269578, EPI_ISL_3269579, EPI_ISL_3269580, EPI_ISL_3269581, EPI_ISL_3269582, EPI_ISL_3269583, EPI_ISL_3269584, EPI_ISL_3269585, EPI_ISL_3269586, EPI_ISL_3269587, EPI_ISL_3269588, EPI_ISL_3269589, EPI_ISL_3269590, EPI_ISL_3269591, EPI_ISL_3269592, EPI_ISL_3269593, EPI_ISL_3269594, EPI_ISL_3269595, EPI_ISL_3269596, EPI_ISL_3269597, EPI_ISL_3269598, EPI_ISL_3269599, EPI_ISL_3269600, EPI_ISL_3269601 | | | |
| see above | University of Mississippi Medical Center, Department of Pathology | University of Mississippi Medical Center, Molecular and Genomics Core Facility | Ashley C. Johnson, Wenjie Wu, Ithiel J. Frame, Krishna K. Ayyalasomayajula, Michael R. Garrett, D. Ashley Robinson |
| EPI_ISL_3269606 | CHC Andrée Rosemon | Institut Pasteur de la Guyane | Anne Lavergne, Dominique Rousset, Antoine Enfissi, Arielle Salmier |
| EPI_ISL_3269609 | CNR Institut Pasteur de la Guyane | Institut Pasteur de la Guyane | Anne Lavergne, Dominique Rousset, Antoine Enfissi, Arielle Salmier |
| EPI_ISL_3269614 | CH de l'Ouest Guyanais Frank Joly | Institut Pasteur de la Guyane | Anne Lavergne, Dominique Rousset, Antoine Enfissi, Arielle Salmier |
| EPI_ISL_3269623, EPI_ISL_3269624, EPI_ISL_3269625, EPI_ISL_3269626, EPI_ISL_3269627, EPI_ISL_3269628, EPI_ISL_3269629, EPI_ISL_3269630, EPI_ISL_3269632, EPI_ISL_3269633, EPI_ISL_3269634, EPI_ISL_3269635, EPI_ISL_3269636, EPI_ISL_3269637, EPI_ISL_3269638, EPI_ISL_3269639, EPI_ISL_3269640, EPI_ISL_3269641, EPI_ISL_3269642, EPI_ISL_3269643, EPI_ISL_3269644, EPI_ISL_3269645, EPI_ISL_3269646, EPI_ISL_3269647, EPI_ISL_3269648, EPI_ISL_3269649, EPI_ISL_3269650, EPI_ISL_3269651, EPI_ISL_3269652, EPI_ISL_3269653, EPI_ISL_3269654, EPI_ISL_3269655, EPI_ISL_3269656, EPI_ISL_3269657, EPI_ISL_3269658, EPI_ISL_3269659, EPI_ISL_3269660, EPI_ISL_3269661, EPI_ISL_3269662, EPI_ISL_3269730, EPI_ISL_3269731, EPI_ISL_3269733, EPI_ISL_3269770, EPI_ISL_3269868, EPI_ISL_3269869, EPI_ISL_3269870, EPI_ISL_3269873, EPI_ISL_3269874, EPI_ISL_3269875, EPI_ISL_3269877, EPI_ISL_3269878, EPI_ISL_3269879, EPI_ISL_3269880, EPI_ISL_3269881, EPI_ISL_3269882, EPI_ISL_3269883, EPI_ISL_3269884, EPI_ISL_3269885, EPI_ISL_3269886, EPI_ISL_3269887, EPI_ISL_3269888, EPI_ISL_3269889, EPI_ISL_3269890, EPI_ISL_3269891, EPI_ISL_3269892, EPI_ISL_3269893, EPI_ISL_3269894, EPI_ISL_3269895, EPI_ISL_3269896, EPI_ISL_3269898, EPI_ISL_3269899, EPI_ISL_3269900, EPI_ISL_3269901, EPI_ISL_3269902, EPI_ISL_3269903, EPI_ISL_3269905, EPI_ISL_3269906, EPI_ISL_3269907, EPI_ISL_3269908, EPI_ISL_3269909, EPI_ISL_3269910, EPI_ISL_3269911, EPI_ISL_3269912, EPI_ISL_3269913, EPI_ISL_3269914, EPI_ISL_3269916, EPI_ISL_3269917, EPI_ISL_3269918, EPI_ISL_3269919, EPI_ISL_3269921, EPI_ISL_3269923, EPI_ISL_3269924, EPI_ISL_3269925, EPI_ISL_3269926, EPI_ISL_3269927, EPI_ISL_3269928, EPI_ISL_3269929, EPI_ISL_3269930, EPI_ISL_3269931, EPI_ISL_3269932, EPI_ISL_3269933, EPI_ISL_3269934, EPI_ISL_3269935, EPI_ISL_3269936, EPI_ISL_3269937, EPI_ISL_3269938, EPI_ISL_3269939, EPI_ISL_3269940, EPI_ISL_3269941, EPI_ISL_3269942, EPI_ISL_3269943, EPI_ISL_3269944, EPI_ISL_3269945, EPI_ISL_3269946, EPI_ISL_3269947, EPI_ISL_3269948, EPI_ISL_3269949, EPI_ISL_3269951, EPI_ISL_3269952, EPI_ISL_3269953, EPI_ISL_3269956, EPI_ISL_3269957, EPI_ISL_3269960, EPI_ISL_3269961, EPI_ISL_3269962, EPI_ISL_3269963, EPI_ISL_3269964, EPI_ISL_3269965, EPI_ISL_3269966, EPI_ISL_3269967, EPI_ISL_3269969, EPI_ISL_3269970, EPI_ISL_3269971, EPI_ISL_3269972, EPI_ISL_3269974, EPI_ISL_3269975, EPI_ISL_3269976, EPI_ISL_3269977, EPI_ISL_3269978, EPI_ISL_3269979, EPI_ISL_3269980, EPI_ISL_3269981, EPI_ISL_3269982, EPI_ISL_3269983, EPI_ISL_3269984, EPI_ISL_3269985, EPI_ISL_3269986, EPI_ISL_3269987, EPI_ISL_3269989, EPI_ISL_3269990, EPI_ISL_3269991, EPI_ISL_3269992, EPI_ISL_3269994, EPI_ISL_3269995, EPI_ISL_3269996, EPI_ISL_3269997, EPI_ISL_3269998, EPI_ISL_3269999, EPI_ISL_3270000, EPI_ISL_3270001, EPI_ISL_3270002, EPI_ISL_3270003, EPI_ISL_3270005, EPI_ISL_3270006, EPI_ISL_3270007, EPI_ISL_3270008, EPI_ISL_3270009, EPI_ISL_3270010, EPI_ISL_3270011, EPI_ISL_3270012, EPI_ISL_3270013, EPI_ISL_3270014, EPI_ISL_3270015, EPI_ISL_3270016, EPI_ISL_3270017, EPI_ISL_3270018, EPI_ISL_3270019, EPI_ISL_3270020, EPI_ISL_3270021, EPI_ISL_3270022, EPI_ISL_3270023, EPI_ISL_3270024, EPI_ISL_3270025, EPI_ISL_3270026, EPI_ISL_3270027, EPI_ISL_3270028, EPI_ISL_3270029, EPI_ISL_3270030, EPI_ISL_3270031, EPI_ISL_3270032, EPI_ISL_3270033, EPI_ISL_3270034, EPI_ISL_3270035, EPI_ISL_3270036, EPI_ISL_3270037, EPI_ISL_3270038, EPI_ISL_3270039, EPI_ISL_3270040, EPI_ISL_3270041, EPI_ISL_3270042, EPI_ISL_3270043, EPI_ISL_3270044, EPI_ISL_3270045, EPI_ISL_3270046, EPI_ISL_3270047, EPI_ISL_3270048, EPI_ISL_3270049, EPI_ISL_3270050, EPI_ISL_3270051, EPI_ISL_3270052, EPI_ISL_3270053, EPI_ISL_3270054, EPI_ISL_3270055, EPI_ISL_3270056, EPI_ISL_3270057, EPI_ISL_3270058, EPI_ISL_3270059, EPI_ISL_3270060, EPI_ISL_3270061, EPI_ISL_3270062, EPI_ISL_3270063, EPI_ISL_3270064, EPI_ISL_3270065, EPI_ISL_3270066, EPI_ISL_3270067, EPI_ISL_3270068, EPI_ISL_3270069, EPI_ISL_3270070, EPI_ISL_3270071, EPI_ISL_3270072, EPI_ISL_3270073, EPI_ISL_3270074, EPI_ISL_3270075, EPI_ISL_3270076, EPI_ISL_3270077, EPI_ISL_3270078, EPI_ISL_3270079, EPI_ISL_3270080, EPI_ISL_3270081, EPI_ISL_3270082, EPI_ISL_3270083, EPI_ISL_3270084 | | | |
| see above | Utah Public Health Laboratory | Utah Public Health Laboratory | Olinto Linares-Perdomo, Erin L. Young, Kelly F. Oakeson |
| EPI_ISL_3270091, EPI_ISL_3270095, EPI_ISL_3270098, EPI_ISL_3270099, EPI_ISL_3270100, EPI_ISL_3270103, EPI_ISL_3270107, EPI_ISL_3270109, EPI_ISL_3270112, EPI_ISL_3270114, EPI_ISL_3270117, EPI_ISL_3270118, EPI_ISL_3270119, EPI_ISL_3270120, EPI_ISL_3270121, EPI_ISL_3270122, EPI_ISL_3270124, EPI_ISL_3270130, EPI_ISL_3270131, EPI_ISL_3270132, EPI_ISL_3270133, EPI_ISL_3270134, EPI_ISL_3270141, EPI_ISL_3270143, EPI_ISL_3270146, EPI_ISL_3270148, EPI_ISL_3270150, EPI_ISL_3270152, EPI_ISL_3270153, EPI_ISL_3270154, EPI_ISL_3270156, EPI_ISL_3270158, EPI_ISL_3270170, EPI_ISL_3270174, EPI_ISL_3270183, EPI_ISL_3270188, EPI_ISL_3270194, EPI_ISL_3270199, EPI_ISL_3270200, EPI_ISL_3270203, EPI_ISL_3270205, EPI_ISL_3270209, EPI_ISL_3270215, EPI_ISL_3270217, EPI_ISL_3270219, EPI_ISL_3270226, EPI_ISL_3270228, EPI_ISL_3270233, EPI_ISL_3270235, EPI_ISL_3270242, EPI_ISL_3270243, EPI_ISL_3270269, EPI_ISL_3270274, EPI_ISL_3270279, EPI_ISL_3270281, EPI_ISL_3270283, EPI_ISL_3270289, EPI_ISL_3270292, EPI_ISL_3270293, EPI_ISL_3270298, EPI_ISL_3270302, EPI_ISL_3270307, EPI_ISL_3270309, EPI_ISL_3270310, EPI_ISL_3270320, EPI_ISL_3270322, EPI_ISL_3270325, EPI_ISL_3270335, EPI_ISL_3270338, EPI_ISL_3270339, EPI_ISL_3270348, EPI_ISL_3270360, EPI_ISL_3270363, EPI_ISL_3270364, EPI_ISL_3270369, EPI_ISL_3270373, EPI_ISL_3270375, EPI_ISL_3270378, EPI_ISL_3270381, EPI_ISL_3270390, EPI_ISL_3270391, EPI_ISL_3270399, EPI_ISL_3270405, EPI_ISL_3270410, EPI_ISL_3270415, EPI_ISL_3270416, EPI_ISL_3270425, EPI_ISL_3270426, EPI_ISL_3270427, EPI_ISL_3270428 | | | |
| see above | Curative Labs | Curative Labs | Mikhail Hanewich-Hollatz, J.Cesar Ignacio-Espinoza, Elias L. Salfati, Nabjot Sandhu, George Way, Zhiyi Xie, Eugenia Khorosheva, Sophia Quasem, Janet Chen, Vladimir Slepnev |
| EPI_ISL_3270429, EPI_ISL_3270430, EPI_ISL_3270431, EPI_ISL_3270432, EPI_ISL_3270433, EPI_ISL_3270434, EPI_ISL_3270435, EPI_ISL_3270436, EPI_ISL_3270437, EPI_ISL_3270438, EPI_ISL_3270439, EPI_ISL_3270440, EPI_ISL_3270441, EPI_ISL_3270442, EPI_ISL_3270443, EPI_ISL_3270444, EPI_ISL_3270445, EPI_ISL_3270446, EPI_ISL_3270447, EPI_ISL_3270448, EPI_ISL_3270449, EPI_ISL_3270450, EPI_ISL_3270451, EPI_ISL_3270452, EPI_ISL_3270453, EPI_ISL_3270454, EPI_ISL_3270455, EPI_ISL_3270456, EPI_ISL_3270457, EPI_ISL_3270458, EPI_ISL_3270459, EPI_ISL_3270460, EPI_ISL_3270461, EPI_ISL_3270462, EPI_ISL_3270464, EPI_ISL_3270465, EPI_ISL_3270466, EPI_ISL_3270467, EPI_ISL_3270468, EPI_ISL_3270476, EPI_ISL_3270477, EPI_ISL_3270478, EPI_ISL_3270479, EPI_ISL_3270480, EPI_ISL_3270481, EPI_ISL_3270482, EPI_ISL_3270483, EPI_ISL_3270484, EPI_ISL_3270485, EPI_ISL_3270486, EPI_ISL_3270487, EPI_ISL_3270488, EPI_ISL_3270489, EPI_ISL_3270490, EPI_ISL_3270491, EPI_ISL_3270492, EPI_ISL_3270493, EPI_ISL_3270494, EPI_ISL_3270495, EPI_ISL_3270496, EPI_ISL_3270497, EPI_ISL_3270498, EPI_ISL_3270499, EPI_ISL_3270500, EPI_ISL_3270501, EPI_ISL_3270502, EPI_ISL_3270503, EPI_ISL_3270504, EPI_ISL_3270505, EPI_ISL_3270506, EPI_ISL_3270507, EPI_ISL_3270508, EPI_ISL_3270509, EPI_ISL_3270510 | | | |
| see above | Alaska State Virology Laboratory | Alaska State Virology Laboratory | Stephanie DeRonde, Elva House, Jacob Zidek, Lisa Smith, Ph.D., Jack Chen, Ph.D. |
| EPI_ISL_3270511, EPI_ISL_3270512 | Wyoming Public Health Laboratory | Wyoming Public Health Laboratory | Jim Mildenberger, Taylor Fearing, Channing Weber, Ashley Norberg, Chayse Rowley, Marley Goetz, Brian Dominguez, Elliot Thomasson, Sam Britz, Cari Sloma, Robert Petit, and Rob Christensen |
| EPI_ISL_3270544, EPI_ISL_3270545, EPI_ISL_3270546, EPI_ISL_3270547, EPI_ISL_3270548, EPI_ISL_3270549, EPI_ISL_3270550, EPI_ISL_3270551, EPI_ISL_3270552, EPI_ISL_3270553, EPI_ISL_3270554, EPI_ISL_3270555, EPI_ISL_3270556, EPI_ISL_3270557, EPI_ISL_3270558, EPI_ISL_3270559, EPI_ISL_3270560, EPI_ISL_3270561, EPI_ISL_3270562, EPI_ISL_3270563, EPI_ISL_3270565, EPI_ISL_3270566, EPI_ISL_3270567, EPI_ISL_3270568, EPI_ISL_3270569, EPI_ISL_3270570, EPI_ISL_3270571, EPI_ISL_3270574, EPI_ISL_3270575, EPI_ISL_3270576, EPI_ISL_3270577, EPI_ISL_3270578, EPI_ISL_3270580, EPI_ISL_3270581, EPI_ISL_3270582, EPI_ISL_3270583, EPI_ISL_3270584, EPI_ISL_3270585, EPI_ISL_3270586, EPI_ISL_3270587, EPI_ISL_3270588, EPI_ISL_3270589, EPI_ISL_3270590, EPI_ISL_3270591, EPI_ISL_3270592, EPI_ISL_3270593, EPI_ISL_3270594, EPI_ISL_3270595, EPI_ISL_3270596, EPI_ISL_3270597, EPI_ISL_3270598, EPI_ISL_3270599, EPI_ISL_3270600, EPI_ISL_3270601, EPI_ISL_3270602, EPI_ISL_3270603, EPI_ISL_3270605, EPI_ISL_3270606, EPI_ISL_3270607, EPI_ISL_3270608, EPI_ISL_3270609, EPI_ISL_3270610, EPI_ISL_3270611, EPI_ISL_3270612, EPI_ISL_3270613, EPI_ISL_3270615, EPI_ISL_3270616, EPI_ISL_3270617 | | | |
| see above | U.O. Microbiologia Laboratorio Unico Centro Servizi - AUSL della Romagna | U.O. Microbiologia, Laboratorio Unico Centro Servizi - AUSL della Romagna | Giorgio Dirani |
| EPI_ISL_3270707, EPI_ISL_3270708, EPI_ISL_3270709, EPI_ISL_3270713, EPI_ISL_3270714, EPI_ISL_3270715, EPI_ISL_3270717, EPI_ISL_3270718, EPI_ISL_3270719, EPI_ISL_3270720, EPI_ISL_3270721, EPI_ISL_3270723, EPI_ISL_3270724, EPI_ISL_3270725, EPI_ISL_3270727, EPI_ISL_3270728, EPI_ISL_3270730, EPI_ISL_3270733, EPI_ISL_3270738, EPI_ISL_3270741, EPI_ISL_3270742, EPI_ISL_3270743, EPI_ISL_3270744, EPI_ISL_3270747, EPI_ISL_3270748, EPI_ISL_3270749, EPI_ISL_3270751, EPI_ISL_3270755, EPI_ISL_3270763, EPI_ISL_3270764 | | | |
| see above | Università Federico II - Dipartimento di scienze mediche traslazionali - Napoli | Telethon Institute of Genetics and Medicine (TIGEM) | Antonio Grimaldi Patrizia Annunziata Francesco Panariello Teresa Giuliano Michele Cennamo Valentina Bouche Chiara Colantuono Lucio Di Filippo Mariano Fiorenza Anna Manfredi Marcello Salvi Giuseppe Portella Andrea Ballabio Davide Cacchiarelli |
| EPI_ISL_3272351, EPI_ISL_3272352, EPI_ISL_3272353, EPI_ISL_3272354, EPI_ISL_3272355, EPI_ISL_3272356, EPI_ISL_3272359, EPI_ISL_3272360, EPI_ISL_3272363, EPI_ISL_3272364, EPI_ISL_3272366, EPI_ISL_3272368, EPI_ISL_3272369, EPI_ISL_3272370, EPI_ISL_3272372, EPI_ISL_3272373, EPI_ISL_3272375, EPI_ISL_3272376, EPI_ISL_3272377, EPI_ISL_3272378, EPI_ISL_3272379, EPI_ISL_3272380, EPI_ISL_3272381, EPI_ISL_3272382, EPI_ISL_3272383, EPI_ISL_3272384, EPI_ISL_3272385, EPI_ISL_3272386, EPI_ISL_3272387, EPI_ISL_3272388, EPI_ISL_3272389, EPI_ISL_3272390, EPI_ISL_3272391, EPI_ISL_3272392, EPI_ISL_3272393, EPI_ISL_3272394, EPI_ISL_3272395, EPI_ISL_3272396, EPI_ISL_3272397, EPI_ISL_3272398, EPI_ISL_3272399, EPI_ISL_3272400, EPI_ISL_3272401, EPI_ISL_3272402, EPI_ISL_3272403, EPI_ISL_3272404, EPI_ISL_3272407, EPI_ISL_3272408, EPI_ISL_3272409, EPI_ISL_3272410, EPI_ISL_3272411, EPI_ISL_3272413, EPI_ISL_3272414, EPI_ISL_3272415, EPI_ISL_3272416, EPI_ISL_3272417, EPI_ISL_3272418, EPI_ISL_3272420, EPI_ISL_3272421, EPI_ISL_3272422, EPI_ISL_3272423, EPI_ISL_3272424, EPI_ISL_3272425, EPI_ISL_3272427, EPI_ISL_3272428, EPI_ISL_3272429, EPI_ISL_3272430, EPI_ISL_3272431, EPI_ISL_3272432, EPI_ISL_3272433, EPI_ISL_3272434, EPI_ISL_3272435, EPI_ISL_3272436, EPI_ISL_3272437, EPI_ISL_3272438, EPI_ISL_3272439, EPI_ISL_3272440, EPI_ISL_3272441, EPI_ISL_3272442, EPI_ISL_3272443, EPI_ISL_3272444, EPI_ISL_3272445, EPI_ISL_3272446, EPI_ISL_3272447, EPI_ISL_3272448, EPI_ISL_3272449, EPI_ISL_3272452, EPI_ISL_3272454, EPI_ISL_3272457, EPI_ISL_3272460, EPI_ISL_3272463, EPI_ISL_3272466, EPI_ISL_3272469, EPI_ISL_3272471, EPI_ISL_3272474, EPI_ISL_3272477, EPI_ISL_3272480, EPI_ISL_3272483, EPI_ISL_3272485, EPI_ISL_3272489, EPI_ISL_3272491, EPI_ISL_3272494, EPI_ISL_3272500, EPI_ISL_3272503, EPI_ISL_3272506, EPI_ISL_3272509, EPI_ISL_3272512, EPI_ISL_3272515, EPI_ISL_3272518, EPI_ISL_3272521, EPI_ISL_3272523, EPI_ISL_3272527, EPI_ISL_3272530, EPI_ISL_3272972, EPI_ISL_3273142, EPI_ISL_3273147, EPI_ISL_3273175, EPI_ISL_3273176, EPI_ISL_3273180, EPI_ISL_3273187, EPI_ISL_3273193, EPI_ISL_3273206, EPI_ISL_3273222 | | | |
| see above | Idaho Bureau of Laboratories | Boise VA Medical Center, PALMS | Matthew Burns, Aimee Ceniseros, Christian Loera, Dr. Christopher Ball, Ying Pei, Cheri Lamb McFarlane, James Razor, Emily Bartlett |
| EPI_ISL_3273233, EPI_ISL_3273236, EPI_ISL_3273237, EPI_ISL_3273239, EPI_ISL_3273240, EPI_ISL_3273241, EPI_ISL_3273243, EPI_ISL_3273244, EPI_ISL_3273245, EPI_ISL_3273246, EPI_ISL_3273249, EPI_ISL_3273250, EPI_ISL_3273251, EPI_ISL_3273253, EPI_ISL_3273256, EPI_ISL_3273257, EPI_ISL_3273258, EPI_ISL_3273259, EPI_ISL_3273261, EPI_ISL_3273263, EPI_ISL_3273264, EPI_ISL_3273266, EPI_ISL_3273270, EPI_ISL_3273274, EPI_ISL_3273275, EPI_ISL_3273276, EPI_ISL_3273277, EPI_ISL_3273279, EPI_ISL_3273280, EPI_ISL_3273282, EPI_ISL_3273283, EPI_ISL_3273285, EPI_ISL_3273288, EPI_ISL_3273289, EPI_ISL_3273290, EPI_ISL_3273291, EPI_ISL_3273294, EPI_ISL_3273295, EPI_ISL_3273297, EPI_ISL_3273299, EPI_ISL_3273300, EPI_ISL_3273301, EPI_ISL_3273304, EPI_ISL_3273305, EPI_ISL_3273310, EPI_ISL_3273311, EPI_ISL_3273312, EPI_ISL_3273313, EPI_ISL_3273314, EPI_ISL_3273316, EPI_ISL_3273317, EPI_ISL_3273318, EPI_ISL_3273354, EPI_ISL_3273361, EPI_ISL_3273366, EPI_ISL_3273368, EPI_ISL_3273370, EPI_ISL_3273372, EPI_ISL_3273373, EPI_ISL_3273376, EPI_ISL_3273378, EPI_ISL_3273379, EPI_ISL_3273381, EPI_ISL_3273382, EPI_ISL_3273383, EPI_ISL_3273384, EPI_ISL_3273386, EPI_ISL_3273387, EPI_ISL_3273389, EPI_ISL_3273390, EPI_ISL_3273391, EPI_ISL_3273392, EPI_ISL_3273393, EPI_ISL_3273394, EPI_ISL_3273395, EPI_ISL_3273399, EPI_ISL_3273400, EPI_ISL_3273401, EPI_ISL_3273402, EPI_ISL_3273404, EPI_ISL_3273405, EPI_ISL_3273406, EPI_ISL_3273407, EPI_ISL_3273408, EPI_ISL_3273409, EPI_ISL_3273410, EPI_ISL_3273411, EPI_ISL_3273412, EPI_ISL_3273413, EPI_ISL_3273414, EPI_ISL_3273415, EPI_ISL_3273416, EPI_ISL_3273417, EPI_ISL_3273418, EPI_ISL_3273419, EPI_ISL_3273421, EPI_ISL_3273422, EPI_ISL_3273423, EPI_ISL_3273424, EPI_ISL_3273425, EPI_ISL_3273426, EPI_ISL_3273427, EPI_ISL_3273428, EPI_ISL_3273430, EPI_ISL_3273431, EPI_ISL_3273433, EPI_ISL_3273435, EPI_ISL_3273436, EPI_ISL_3273438, EPI_ISL_3273439, EPI_ISL_3273440, EPI_ISL_3273441, EPI_ISL_3273445, EPI_ISL_3273446, EPI_ISL_3273447, EPI_ISL_3273448, EPI_ISL_3273449, EPI_ISL_3273450, EPI_ISL_3273451, EPI_ISL_3273452, EPI_ISL_3273453, EPI_ISL_3273455, EPI_ISL_3273456, EPI_ISL_3273457, EPI_ISL_3273458, EPI_ISL_3273461, EPI_ISL_3273462, EPI_ISL_3273463, EPI_ISL_3273464, EPI_ISL_3273465, EPI_ISL_3273466, EPI_ISL_3273468, EPI_ISL_3273470, EPI_ISL_3273472, EPI_ISL_3273474, EPI_ISL_3273475, EPI_ISL_3273476, EPI_ISL_3273477, EPI_ISL_3273478, EPI_ISL_3273479, EPI_ISL_3273480, EPI_ISL_3273482, EPI_ISL_3273485, EPI_ISL_3273486, EPI_ISL_3273488, EPI_ISL_3273491, EPI_ISL_3273493, EPI_ISL_3273494, EPI_ISL_3273496, EPI_ISL_3273499, EPI_ISL_3273502, EPI_ISL_3273504, EPI_ISL_3273507, EPI_ISL_3273509, EPI_ISL_3273510, EPI_ISL_3273512, EPI_ISL_3273514, EPI_ISL_3273517, EPI_ISL_3273519, EPI_ISL_3273523, EPI_ISL_3273524, EPI_ISL_3273526, EPI_ISL_3273529, EPI_ISL_3273530, EPI_ISL_3273532, EPI_ISL_3273535, EPI_ISL_3273537, EPI_ISL_3273538, EPI_ISL_3273542, EPI_ISL_3273543, EPI_ISL_3273544, EPI_ISL_3273545, EPI_ISL_3273546, EPI_ISL_3273547, EPI_ISL_3273549, EPI_ISL_3273552, EPI_ISL_3273553, EPI_ISL_3273555, EPI_ISL_3273556, EPI_ISL_3273558, EPI_ISL_3273559, EPI_ISL_3273562, EPI_ISL_3273565, EPI_ISL_3273568, EPI_ISL_3273569, EPI_ISL_3273570, EPI_ISL_3273572, EPI_ISL_3273574, EPI_ISL_3273575, EPI_ISL_3273576, EPI_ISL_3273578, EPI_ISL_3273579, EPI_ISL_3273581, EPI_ISL_3273583, EPI_ISL_3273584, EPI_ISL_3273586, EPI_ISL_3273587, EPI_ISL_3273589, EPI_ISL_3273590, EPI_ISL_3273592, EPI_ISL_3273593, EPI_ISL_3273596, EPI_ISL_3273598, EPI_ISL_3273600, EPI_ISL_3273603, EPI_ISL_3273604, EPI_ISL_3273606, EPI_ISL_3273607, EPI_ISL_3273608, EPI_ISL_3273610, EPI_ISL_3273611, EPI_ISL_3273612, EPI_ISL_3273614, EPI_ISL_3273616, EPI_ISL_3273618, EPI_ISL_3273619, EPI_ISL_3273621, EPI_ISL_3273622, EPI_ISL_3273624, EPI_ISL_3273626 | | | |
| see above | Curative Labs | Curative Labs | Mikhail Hanewich-Hollatz, J.Cesar Ignacio-Espinoza, Elias L. Salfati, Nabjot Sandhu, George Way, Zhiyi Xie, Eugenia Khorosheva, Sophia Quasem, Janet Chen, Vladimir Slepnev |
| EPI_ISL_3274088, EPI_ISL_3274089, EPI_ISL_3274090, EPI_ISL_3274091, EPI_ISL_3274092 | INSPI-CRN DE INFLUENZA Y OTROS VIRUS RESPIRATORIOS | NIC-INSPI | Alfredo Bruno, Maritza Olmedo, Michelle Páez, Jimmy Garcés, Johanna Laines, Lizbeth Patiño, Manuel Gonzalez, Domenica de Mora. |
| EPI_ISL_3274093 | INSPI-CRN DE INFLUENZA Y OTROS VIRUS RESPIRATORIOS | NIC-INSPI | Alfredo Bruno, Maritza Olmedo, Michelle Páez, Jimmy Garcés, Johanna Laines, Marcela Mejía, Milton Medina, Lizbeth Patiño, Manuel Gonzalez, Domenica de Mora. |
| EPI_ISL_3274094, EPI_ISL_3274095, EPI_ISL_3274096, EPI_ISL_3274097 | INSPI-CRN DE INFLUENZA Y OTROS VIRUS RESPIRATORIOS | NIC-INSPI | Alfredo Bruno, Maritza Olmedo, Michelle Páez, Jimmy Garcés, Johanna Laines, Lizbeth Patiño, Manuel Gonzalez, Domenica de Mora. |
| EPI_ISL_3274098 | INSPI-CRN DE INFLUENZA Y OTROS VIRUS RESPIRATORIOS | NIC-INSPI | Alfredo Bruno, Maritza Olmedo, Michelle Páez, Jimmy Garcés, Johanna Laines, Marcela Mejía, Milton Medina, Lizbeth Patiño, Manuel Gonzalez, Domenica de Mora. |
| EPI_ISL_3274099 | INSPI-CRN DE INFLUENZA Y OTROS VIRUS RESPIRATORIOS | NIC-INSPI | Alfredo Bruno, Maritza Olmedo, Michelle Páez, Jimmy Garcés, Johanna Laines, Lizbeth Patiño, Manuel Gonzalez, Domenica de Mora. |
| EPI_ISL_3274100, EPI_ISL_3274101, EPI_ISL_3274102 | INSPI-CRN DE INFLUENZA Y OTROS VIRUS RESPIRATORIOS | NIC-INSPI | Alfredo Bruno, Maritza Olmedo, Michelle Páez, Jimmy Garcés, Johanna Laines, Mayra Wilca, Maria Angelica Becerra, Daniel Ramos, Lizbeth Patiño, Manuel Gonzalez, Domenica de Mora. |
| EPI_ISL_3274103, EPI_ISL_3274104, EPI_ISL_3274105, EPI_ISL_3274106 | INSPI-CRN DE INFLUENZA Y OTROS VIRUS RESPIRATORIOS | NIC-INSPI | Alfredo Bruno, Maritza Olmedo, Michelle Páez, Jimmy Garcés, Johanna Laines, Lizbeth Patiño, Manuel Gonzalez, Domenica de Mora. |
| EPI_ISL_3274107, EPI_ISL_3274108 | INSPI-CRN DE INFLUENZA Y OTROS VIRUS RESPIRATORIOS | NIC-INSPI | Alfredo Bruno, Maritza Olmedo, Michelle Páez, Jimmy Garcés, Johanna Laines, Mayra Wilca, Maria Angelica Becerra, Daniel Ramos, Lizbeth Patiño, Manuel Gonzalez, Domenica de Mora. |
| EPI_ISL_3274109, EPI_ISL_3274110 | INSPI-CRN DE INFLUENZA Y OTROS VIRUS RESPIRATORIOS | NIC-INSPI | Alfredo Bruno, Maritza Olmedo, Michelle Páez, Jimmy Garcés, Johanna Laines, Lizbeth Patiño, Manuel Gonzalez, Domenica de Mora. |
| EPI_ISL_3274111 | INSPI-CRN DE INFLUENZA Y OTROS VIRUS RESPIRATORIOS | NIC-INSPI | Alfredo Bruno, Maritza Olmedo, Michelle Páez, Jimmy Garcés, Johanna Laines, Mayra Wilca, Maria Angelica Becerra, Daniel Ramos, Lizbeth Patiño, Manuel Gonzalez, Domenica de Mora. |
| EPI_ISL_3274112, EPI_ISL_3274113 | INSPI-CRN DE INFLUENZA Y OTROS VIRUS RESPIRATORIOS | NIC-INSPI | Alfredo Bruno, Maritza Olmedo, Michelle Páez, Jimmy Garcés, Johanna Laines, Lizbeth Patiño, Manuel Gonzalez, Domenica de Mora. |
| EPI_ISL_3274114, EPI_ISL_3274115 | INSPI-CRN DE INFLUENZA Y OTROS VIRUS RESPIRATORIOS | NIC-INSPI | Alfredo Bruno, Maritza Olmedo, Michelle Páez, Jimmy Garcés, Johanna Laines, Mayra Wilca, Maria Angelica Becerra, Daniel Ramos, Lizbeth Patiño, Manuel Gonzalez, Domenica de Mora. |
| EPI_ISL_3274116 | INSPI-CRN DE INFLUENZA Y OTROS VIRUS RESPIRATORIOS | NIC-INSPI | Alfredo Bruno, Maritza Olmedo, Michelle Páez, Jimmy Garcés, Johanna Laines, Lizbeth Patiño, Manuel Gonzalez, Domenica de Mora. |
| EPI_ISL_3274117 | INSPI-CRN DE INFLUENZA Y OTROS VIRUS RESPIRATORIOS | NIC-INSPI | Alfredo Bruno, Maritza Olmedo, Michelle Páez, Jimmy Garcés, Johanna Laines, Mayra Wilca, Maria Angelica Becerra, Daniel Ramos, Lizbeth Patiño, Manuel Gonzalez, Domenica de Mora. |
| EPI_ISL_3274166, | Arkansas Children's Hospital | Center for Global Health, University of New Mexico Health Sciences | Daryl Domman, Kurt Schwalm, Valerie Morley, Amanda Novack, Dirk Haselow, Joshua L. Kennedy, Darrell Dinwiddie |

| | | | |
|---|---|---|---|
| EPI_ISL_3274167, EPI_ISL_3274168, EPI_ISL_3274169, EPI_ISL_3274172, EPI_ISL_3274174, EPI_ISL_3274175, EPI_ISL_3274176 | | Center | |
| EPI_ISL_3274182, EPI_ISL_3274183, EPI_ISL_3274184, EPI_ISL_3274185, EPI_ISL_3274186, EPI_ISL_3274187, EPI_ISL_3274188, EPI_ISL_3274189, EPI_ISL_3274190, EPI_ISL_3274191, EPI_ISL_3274192, EPI_ISL_3274194, EPI_ISL_3274195, EPI_ISL_3274196, EPI_ISL_3274197, EPI_ISL_3274198, EPI_ISL_3274199, EPI_ISL_3274201, EPI_ISL_3274202, EPI_ISL_3274204, EPI_ISL_3274205, EPI_ISL_3274206, EPI_ISL_3274207, EPI_ISL_3274208, EPI_ISL_3274209, EPI_ISL_3274210, EPI_ISL_3274211, EPI_ISL_3274212, EPI_ISL_3274213, EPI_ISL_3274214, EPI_ISL_3274215, EPI_ISL_3274216, EPI_ISL_3274218, EPI_ISL_3274219, EPI_ISL_3274220, EPI_ISL_3274221, EPI_ISL_3274222, EPI_ISL_3274223, EPI_ISL_3274224, EPI_ISL_3274225, EPI_ISL_3274226, EPI_ISL_3274227, EPI_ISL_3274230, EPI_ISL_3274232, EPI_ISL_3274234, EPI_ISL_3274235, EPI_ISL_3274236, EPI_ISL_3274237, EPI_ISL_3274238, EPI_ISL_3274239, EPI_ISL_3274240, EPI_ISL_3274241, EPI_ISL_3274242, EPI_ISL_3274243, EPI_ISL_3274244, EPI_ISL_3274245 | | | |
| see above | Baptist Health Medical Center | Center for Global Health, University of New Mexico Health Sciences Center | Daryl Domman, Kurt Schwalm, Valerie Morley, Amanda Novack, Dirk Haselow, Joshua L. Kennedy, Darrell Dinwiddie |
| EPI_ISL_3274313 | Virology Laboratory, Scientific Department, Army Medical Center | Virology Laboratory, Scientific Department, Army Medical Center | Silvia Fillo, Riccardo De Sanctis, Antonella Fortunato, Anella Monte, Rossella Brandi, Giulia Campoli, Marzia Cavalli, Lucia Nicosia, Anna Anselmo, Vanessa Vera Fain, Francesco Giordani, Giandomenico Cerreto, Filippo Molinari, Giancarlo Petralito, Florigio Lista. |
| EPI_ISL_3274314 | Virology Laboratory, Scientific Department, Army Medical Center | Virology Laboratory, Scientific Department, Army Medical Center | Silvia Fillo, Riccardo De Sanctis, Antonella Fortunato, Anella Monte, Rossella Brandi, Giulia Campoli, Marzia Cavalli, Lucia Nicosia, Anna Anselmo, Vanessa Vera Fain, Francesco Giordani, Giandomenico Cerreto, Filippo Molinari, Giancarlo Petralito, Filippo Moramarco, Florigio Lista |
| EPI_ISL_3274315 | Virology Laboratory, Scientific Department, Army Medical Center | Virology Laboratory, Scientific Department, Army Medical Center | Silvia Fillo, Riccardo De Sanctis, Antonella Fortunato, Anella Monte, Rossella Brandi, Giulia Campoli, Marzia Cavalli, Lucia Nicosia, Anna Anselmo, Vanessa Vera Fain, Francesco Giordani, Giandomenico Cerreto, Filippo Molinari, Giancarlo Petralito, Florigio Lista, Maria Anna Spinelli, Margherita De Santis |
| EPI_ISL_3274316, EPI_ISL_3274317 | Virology Laboratory, Scientific Department, Army Medical Center | Virology Laboratory, Scientific Department, Army Medical Center | Silvia Fillo, Riccardo De Sanctis, Antonella Fortunato, Anella Monte, Rossella Brandi, Giulia Campoli, Marzia Cavalli, Lucia Nicosia, Anna Anselmo, Vanessa Vera Fain, Francesco Giordani, Giandomenico Cerreto, Filippo Molinari, Giancarlo Petralito, Florigio Lista. |
| EPI_ISL_3274348 | HOSPITAL DR. FERNANDO ESCALANTE PRADILLA [PEREZ ZELEDON/SAN JOSE] | Inciensa, Instituto Costarricense de Investigación y Enseñanza en Nutrición y Salud | Francisco Duarte, Hebleen Porras, Claudio Soto-Garita, Estela Cordero, Adriana Godínez, Melany Calderón, José Luis Vargas, Mariela Gutiérrez, Joselyn Prado, Caterina Guzmán, Nazareth Ruíz & Fabricio Aguilar |
| EPI_ISL_3274349 | AREA DE SALUD MATA REDONDA-HOSPITAL - CLINICA DR. MORENO CAÑAS [SAN JOSE/SAN JOSE] | Inciensa, Instituto Costarricense de Investigación y Enseñanza en Nutrición y Salud | Francisco Duarte, Hebleen Porras, Claudio Soto-Garita, Estela Cordero, Adriana Godínez, Melany Calderón, José Luis Vargas, Mariela Gutiérrez, Joselyn Prado, Caterina Guzmán, Nazareth Ruíz & Ricardo González |
| EPI_ISL_3274402, EPI_ISL_3274407, EPI_ISL_3274411, EPI_ISL_3274420, EPI_ISL_3274423 | INSPI-CRN DE INFLUENZA Y OTROS VIRUS RESPIRATORIOS | NIC-INSPI | Alfredo Bruno , Maritza Olmedo, Michelle Páez, Jimmy Garcés, Johanna Laines, Lizbeth Patiño, Manuel Gonzalez, Domenica de Mora. |
| EPI_ISL_3274433 | INSPI-CRN DE INFLUENZA Y OTROS VIRUS RESPIRATORIOS | NIC-INSPI | Alfredo Bruno, Maritza Olmedo, Michelle Páez, Jimmy Garcés, Johanna Laines, Maria Angelica Becerra, Daniel Ramos, Lizbeth Patiño, Manuel Gonzalez, Domenica de Mora. |
| EPI_ISL_3274437, EPI_ISL_3274445 | INSPI-CRN DE INFLUENZA Y OTROS VIRUS RESPIRATORIOS | NIC-INSPI | Alfredo Bruno , Maritza Olmedo, Michelle Páez, Jimmy Garcés, Johanna Laines, Lizbeth Patiño, Manuel Gonzalez, Domenica de Mora. |
| EPI_ISL_3274454 | INSPI-CRN DE INFLUENZA Y OTROS VIRUS RESPIRATORIOS | NIC-INSPI | Alfredo Bruno, Maritza Olmedo, Michelle Páez, Jimmy Garcés, Johanna Laines, Marcela Mejía, Lizbeth Patiño, Manuel Gonzalez, Domenica de Mora. |
| EPI_ISL_3274455, EPI_ISL_3274459, EPI_ISL_3274463, EPI_ISL_3274466, EPI_ISL_3274473, EPI_ISL_3274477, EPI_ISL_3274479, EPI_ISL_3274482, EPI_ISL_3274488, EPI_ISL_3274491, EPI_ISL_3274510, EPI_ISL_3274515, EPI_ISL_3274518, EPI_ISL_3274521, EPI_ISL_3274526, EPI_ISL_3274536, EPI_ISL_3274541, EPI_ISL_3274547, EPI_ISL_3274551, EPI_ISL_3274555, EPI_ISL_3274561 | | | |
| see above | INSPI-CRN DE INFLUENZA Y OTROS VIRUS RESPIRATORIOS | NIC-INSPI | Alfredo Bruno , Maritza Olmedo, Michelle Páez, Jimmy Garcés, Johanna Laines, Lizbeth Patiño, Manuel Gonzalez, Domenica de Mora. |
| EPI_ISL_3274566 | INSPI-CRN DE INFLUENZA Y OTROS VIRUS RESPIRATORIOS | NIC-INSPI | Alfredo Bruno, Maritza Olmedo, Michelle Páez, Jimmy Garcés, Johanna Laines, Marcela Mejía, Lizbeth Patiño, Manuel Gonzalez, Domenica de Mora. |
| EPI_ISL_3274571, EPI_ISL_3274582, EPI_ISL_3274586, EPI_ISL_3274587, EPI_ISL_3274595, EPI_ISL_3274607, EPI_ISL_3274610, EPI_ISL_3274618 | INSPI-CRN DE INFLUENZA Y OTROS VIRUS RESPIRATORIOS | NIC-INSPI | Alfredo Bruno , Maritza Olmedo, Michelle Páez, Jimmy Garcés, Johanna Laines, Lizbeth Patiño, Manuel Gonzalez, Domenica de Mora. |
| EPI_ISL_3274643, EPI_ISL_3274666, EPI_ISL_3274670, EPI_ISL_3274672 | TriCore Reference Laboratories | Center for Global Health, University of New Mexico Health Sciences Center | Daryl Domman, Kurt Schwalm, Valerie Morley, Cecilia Thompson, Kendra Pesko, Karissa Culbreath, Darrell Dinwiddie |
| EPI_ISL_3274747, EPI_ISL_3274750, EPI_ISL_3274764 | HLAGYN - Laboratorio de Imunologia de Transplantes de Goias | HLAGYN - Laboratorio de Imunologia de Transplantes de Goias | Fernando Antonio Vinhal dos Santos, Erika Lopes Rocha Batista, Alessandro Leonardo Alvares Magalhaes, Frederico Rodrigues Vinhal, Sabrina Sara Moreira Duarte, Lucas Carlos Gomes Pereira, Kamila Oliveira Reis De Freitas. |
| EPI_ISL_3274790 | HLAGYN - Laboratorio de Imunologia de Transplantes de Goias | Laboratorio INAC | Fernando Antonio Vinhal dos Santos, Erika Lopes Rocha Batista, Alessandro Leonardo Alvares Magalhaes, Frederico Rodrigues Vinhal, Sabrina Sara Moreira Duarte, Lucas Carlos Gomes Pereira, Kamila Oliveira Reis De Freitas. |
| EPI_ISL_3274791, EPI_ISL_3274792, EPI_ISL_3274793, EPI_ISL_3274794, EPI_ISL_3274795, EPI_ISL_3274796, EPI_ISL_3274797, EPI_ISL_3274798, EPI_ISL_3274799, EPI_ISL_3274800, EPI_ISL_3274801, EPI_ISL_3274802, EPI_ISL_3274803, EPI_ISL_3274805, EPI_ISL_3274806, EPI_ISL_3274807, EPI_ISL_3274808, EPI_ISL_3274809, EPI_ISL_3274810, EPI_ISL_3274811, EPI_ISL_3274812, EPI_ISL_3274813, EPI_ISL_3274814, EPI_ISL_3274815, EPI_ISL_3274816, EPI_ISL_3274817, EPI_ISL_3274818, EPI_ISL_3274819, EPI_ISL_3274820, EPI_ISL_3274821, EPI_ISL_3274822, EPI_ISL_3274823, EPI_ISL_3274824, EPI_ISL_3274825, EPI_ISL_3274826, EPI_ISL_3274827, EPI_ISL_3274828, EPI_ISL_3274829 | | | |
| see above | Fondazione IRCCS Ca' Granda Ospedale Maggiore Policlinico | Fondazione IRCCS Ca' Granda Ospedale Maggiore Policlinico | Ferruccio Ceriotti, Sara Uceda Renteria |
| EPI_ISL_3274831, EPI_ISL_3274832, EPI_ISL_3274833, EPI_ISL_3274834, EPI_ISL_3274835, EPI_ISL_3274836, EPI_ISL_3274837, EPI_ISL_3274838, EPI_ISL_3274839, EPI_ISL_3274840, EPI_ISL_3274841, EPI_ISL_3274842, EPI_ISL_3274843, EPI_ISL_3274844, EPI_ISL_3274845, EPI_ISL_3274846, EPI_ISL_3274847, EPI_ISL_3274848, EPI_ISL_3274849, EPI_ISL_3274851, EPI_ISL_3274852, EPI_ISL_3274853, EPI_ISL_3274854, EPI_ISL_3274855, EPI_ISL_3274856, EPI_ISL_3274857, EPI_ISL_3274858, EPI_ISL_3274859, EPI_ISL_3274860, EPI_ISL_3274861, EPI_ISL_3274862, EPI_ISL_3274863, EPI_ISL_3274864, EPI_ISL_3274865, EPI_ISL_3274866, EPI_ISL_3274867, EPI_ISL_3274868, EPI_ISL_3274869, EPI_ISL_3274870, EPI_ISL_3274871, EPI_ISL_3274872, EPI_ISL_3274873, EPI_ISL_3274874, EPI_ISL_3274875, EPI_ISL_3274876, EPI_ISL_3274877, EPI_ISL_3274878, EPI_ISL_3274879, EPI_ISL_3274880, EPI_ISL_3274881, EPI_ISL_3274882, EPI_ISL_3274883, EPI_ISL_3274884, EPI_ISL_3274886, EPI_ISL_3274887, EPI_ISL_3274888, EPI_ISL_3274889, EPI_ISL_3274890, EPI_ISL_3274891, EPI_ISL_3274892, EPI_ISL_3274893, EPI_ISL_3274894, EPI_ISL_3274895, EPI_ISL_3274896, EPI_ISL_3274897, EPI_ISL_3274898, EPI_ISL_3274899, EPI_ISL_3274900, EPI_ISL_3274901, EPI_ISL_3274902, EPI_ISL_3274903, EPI_ISL_3274904, EPI_ISL_3274905, EPI_ISL_3274906, EPI_ISL_3274907, EPI_ISL_3274908, EPI_ISL_3274909, EPI_ISL_3274910, EPI_ISL_3274911, EPI_ISL_3274912, EPI_ISL_3274913, EPI_ISL_3274914, EPI_ISL_3274915, EPI_ISL_3274916, EPI_ISL_3274917, EPI_ISL_3274918, EPI_ISL_3274919, EPI_ISL_3274920, EPI_ISL_3274921, EPI_ISL_3274922, EPI_ISL_3274923, EPI_ISL_3274924, EPI_ISL_3274925, EPI_ISL_3274926, EPI_ISL_3274927, EPI_ISL_3274928, EPI_ISL_3274929, EPI_ISL_3274930, EPI_ISL_3274931, EPI_ISL_3274932, EPI_ISL_3274933, EPI_ISL_3274934, EPI_ISL_3274935, EPI_ISL_3274936, EPI_ISL_3274937, EPI_ISL_3274938, EPI_ISL_3274939, EPI_ISL_3274940, EPI_ISL_3274941, EPI_ISL_3274942, EPI_ISL_3274943, EPI_ISL_3274944, EPI_ISL_3274945, EPI_ISL_3274946, EPI_ISL_3274947, EPI_ISL_3274948, EPI_ISL_3274949, EPI_ISL_3274950, EPI_ISL_3274951, EPI_ISL_3274952, EPI_ISL_3274953, EPI_ISL_3274954, EPI_ISL_3274955, EPI_ISL_3274956, EPI_ISL_3274957, EPI_ISL_3274958, EPI_ISL_3274959, EPI_ISL_3274960, EPI_ISL_3274961, EPI_ISL_3274962, EPI_ISL_3274964, EPI_ISL_3274965, EPI_ISL_3274966, EPI_ISL_3274967, EPI_ISL_3274968, EPI_ISL_3274969, EPI_ISL_3274970, EPI_ISL_3274971, EPI_ISL_3274972, EPI_ISL_3274973, EPI_ISL_3274974, EPI_ISL_3274975, EPI_ISL_3274976, EPI_ISL_3274977, EPI_ISL_3274978, EPI_ISL_3274979, EPI_ISL_3274980, EPI_ISL_3274981, EPI_ISL_3274982, EPI_ISL_3274983, EPI_ISL_3274984, EPI_ISL_3274985, EPI_ISL_3274986, EPI_ISL_3274987, EPI_ISL_3274988, EPI_ISL_3274989, EPI_ISL_3274990, EPI_ISL_3274991, EPI_ISL_3274992, EPI_ISL_3274993, EPI_ISL_3274994, EPI_ISL_3274995, EPI_ISL_3274996, EPI_ISL_3274997, EPI_ISL_3274998, EPI_ISL_3274999, EPI_ISL_3275000, EPI_ISL_3275001, EPI_ISL_3275002, EPI_ISL_3275003, EPI_ISL_3275004, EPI_ISL_3275005, EPI_ISL_3275006, EPI_ISL_3275007, EPI_ISL_3275008, EPI_ISL_3275009, EPI_ISL_3275010, EPI_ISL_3275011, EPI_ISL_3275012, EPI_ISL_3275013, EPI_ISL_3275014, EPI_ISL_3275015, EPI_ISL_3275016, EPI_ISL_3275017, EPI_ISL_3275018, EPI_ISL_3275019, EPI_ISL_3275020, EPI_ISL_3275021, EPI_ISL_3275022, EPI_ISL_3275023, EPI_ISL_3275024, EPI_ISL_3275025, EPI_ISL_3275026, EPI_ISL_3275027, EPI_ISL_3275028, EPI_ISL_3275029, EPI_ISL_3275030, EPI_ISL_3275031, EPI_ISL_3275032, EPI_ISL_3275033, EPI_ISL_3275034, EPI_ISL_3275035, EPI_ISL_3275036, EPI_ISL_3275037 | | | |
| see above | Clinical Microbiology, Infection Prevention and Control | Section for Molecular Diagnostics | Björn Hallström, Jonas Björkman |
| EPI_ISL_3275038 | BTKLPP Kelas 1 Manado | National Institute of Health Research and Development | Vivi Setiawaty, Hana Apsari Pawestri, Subangkit, Kartika Dewi Puspa, Arie Ardiansyah Nugraha, Hartanti Dian Ikawati, Krisna Nur Andriana Pangesti, Triyani Soekarso, Nelly Puspandari |
| EPI_ISL_3275039, EPI_ISL_3275040, EPI_ISL_3275043, EPI_ISL_3275046, EPI_ISL_3275047, EPI_ISL_3275050, EPI_ISL_3275051 | San Diego County Public Health Laboratory | San Diego County Public Health Laboratory | Ashleigh Murphy Schafer, Brett Austin |
| EPI_ISL_3275106, EPI_ISL_3275107, EPI_ISL_3275108, EPI_ISL_3275109, EPI_ISL_3275111, EPI_ISL_3275112, EPI_ISL_3275113, EPI_ISL_3275114, EPI_ISL_3275115, EPI_ISL_3275120, EPI_ISL_3275121, EPI_ISL_3275122, EPI_ISL_3275123, EPI_ISL_3275124, EPI_ISL_3275126, EPI_ISL_3275127, EPI_ISL_3275128, EPI_ISL_3275129, EPI_ISL_3275131, EPI_ISL_3275133, EPI_ISL_3275135, EPI_ISL_3275137, EPI_ISL_3275138 | | | |
| see above | Idaho Bureau of Laboratories | Center for Global Health, University of New Mexico Health Sciences Center | Daryl Domman, Kurt Schwalm, Valerie Morley, Matthew Burns, Robert Voermans, Christopher Ball, Darrell Dinwiddie |
| EPI_ISL_3275144, EPI_ISL_3275145 | Arizona State University | Arizona State University | Ajeet Bains, LaRinda A. Holland, Matthew F. Smith, Peter T. Skidmore, Rabia Maqsood, Nicholas J. Mellor, Nathaniel Johnson, Joshua LaBaer, Vel Murugan, Efrem S. Lim |
| EPI_ISL_3275166 | AA. Rasdhoo | Indira Gandhi Memorial Hospital | Dr. Milza Abdul Muhsin; Ms. Aminath Nazfa ; Ms. Aishath Shuhudha; Ms. Fathimath Zimna ;Mr. Ibrahim Nishan Ahmed |
| EPI_ISL_3275167 | K. Himmafushi | Indira Gandhi Memorial Hospital | Dr. Milza Abdul Muhsin; Ms. Aminath Nazfa ; Ms. Aishath Shuhudha; Ms. Fathimath Zimna ;Mr. Ibrahim Nishan Ahmed |
| EPI_ISL_3275169 | AA. Mathiveri | Indira Gandhi Memorial Hospital | Dr. Milza Abdul Muhsin; Ms. Aminath Nazfa ; Ms. Aishath Shuhudha; Ms. Fathimath Zimna ;Mr. Ibrahim Nishan Ahmed |
| EPI_ISL_3275170, EPI_ISL_3275172 | K. Maafushi | Indira Gandhi Memorial Hospital | Dr. Milza Abdul Muhsin; Ms. Aminath Nazfa ; Ms. Aishath Shuhudha; Ms. Fathimath Zimna ;Mr. Ibrahim Nishan Ahmed |
| EPI_ISL_3275173 | F. Feeali | Indira Gandhi Memorial Hospital | Dr. Milza Abdul Muhsin; Ms. Aminath Nazfa ; Ms. Aishath Shuhudha; Ms. Fathimath Zimna ;Mr. Ibrahim Nishan Ahmed |
| EPI_ISL_3275174 | Indira Gandhi Memorial Hospital | Indira Gandhi Memorial Hospital | Dr. Milza Abdul Muhsin; Ms. Aminath Nazfa ; Ms. Aishath Shuhudha; Ms. Fathimath Zimna ;Mr. Ibrahim Nishan Ahmed |
| EPI_ISL_3275352, EPI_ISL_3275353, EPI_ISL_3275354, EPI_ISL_3275355, EPI_ISL_3275356, EPI_ISL_3275357, EPI_ISL_3275358, EPI_ISL_3275359, EPI_ISL_3275360, EPI_ISL_3275366, EPI_ISL_3275367, EPI_ISL_3275368, EPI_ISL_3275370, EPI_ISL_3275372, EPI_ISL_3275373, EPI_ISL_3275375 | | | |
| see above | NHLS_VIRO Eastern Cape | KRISP, KZN Research Innovation and Sequencing Platform | Laguda-Akingba O, Giandhari Jennifer, Pillay Sureshnee, Yajna Ramphal, Naidoo Yeshnee, Tshabuila Derek,Tegally Houriiyah, San James, Wilkinson Eduan, de Oliveira Tulio |
| EPI_ISL_3275394, EPI_ISL_3275397, EPI_ISL_3275398, EPI_ISL_3275399, EPI_ISL_3275400, EPI_ISL_3275401, EPI_ISL_3275402, EPI_ISL_3275403, EPI_ISL_3275404, EPI_ISL_3275405, EPI_ISL_3275407, EPI_ISL_3275408, EPI_ISL_3275409, EPI_ISL_3275410, EPI_ISL_3275411, EPI_ISL_3275412, EPI_ISL_3275413, EPI_ISL_3275414, EPI_ISL_3275415, EPI_ISL_3275416, EPI_ISL_3275417, EPI_ISL_3275418, EPI_ISL_3275420, EPI_ISL_3275450, EPI_ISL_3275451, EPI_ISL_3275455, EPI_ISL_3275456, EPI_ISL_3275457, EPI_ISL_3275458, EPI_ISL_3275459, EPI_ISL_3275460, EPI_ISL_3275461, EPI_ISL_3275462, EPI_ISL_3275463, EPI_ISL_3275464, EPI_ISL_3275465, EPI_ISL_3275466, EPI_ISL_3275467, EPI_ISL_3275468, EPI_ISL_3275469, EPI_ISL_3275470, EPI_ISL_3275471, EPI_ISL_3275472, EPI_ISL_3275473, EPI_ISL_3275474, EPI_ISL_3275475, EPI_ISL_3275476, EPI_ISL_3275477, EPI_ISL_3275478, EPI_ISL_3275479, EPI_ISL_3275481, EPI_ISL_3275482, EPI_ISL_3275483, EPI_ISL_3275484, EPI_ISL_3275485, EPI_ISL_3275486 | | | |
| see above | Allergy, Immunology and Cell Biology Unit (AICBU) | Allergy, Immunology and Cell Biology Unit (AICBU) | Chandima Jeewandara, Deshni Jayathilaka, Dumni Gunasinghe, Tibutius Thanesh, Dinuka Ariyarathne, Ayesha Wijesinghe, Diyanath Ranasinghe, Gathsaurie Neelika Malavige |

| | | | |
|---|---|---|---|
| EPI_ISL_3275487, EPI_ISL_3275488, EPI_ISL_3275489, EPI_ISL_3275490, EPI_ISL_3275491, EPI_ISL_3275492, EPI_ISL_3275493, EPI_ISL_3275494, EPI_ISL_3275495, EPI_ISL_3275496, EPI_ISL_3275497, EPI_ISL_3275498, EPI_ISL_3275499, EPI_ISL_3275500, EPI_ISL_3275501, EPI_ISL_3275502, EPI_ISL_3275503, EPI_ISL_3275504, EPI_ISL_3275505, EPI_ISL_3275506, EPI_ISL_3275507, EPI_ISL_3275508, EPI_ISL_3275509, EPI_ISL_3275510, EPI_ISL_3275511, EPI_ISL_3275512, EPI_ISL_3275513, EPI_ISL_3275514, EPI_ISL_3275515, EPI_ISL_3275516, EPI_ISL_3275517, EPI_ISL_3275518, EPI_ISL_3275519, EPI_ISL_3275520, EPI_ISL_3275521, EPI_ISL_3275522, EPI_ISL_3275523, EPI_ISL_3275524, EPI_ISL_3275525, EPI_ISL_3275526, EPI_ISL_3275527, EPI_ISL_3275528, EPI_ISL_3275529, EPI_ISL_3275530, EPI_ISL_3275531, EPI_ISL_3275532, EPI_ISL_3275533, EPI_ISL_3275534, EPI_ISL_3275535, EPI_ISL_3275536, EPI_ISL_3275537, EPI_ISL_3275538, EPI_ISL_3275539, EPI_ISL_3275540, EPI_ISL_3275541, EPI_ISL_3275542, EPI_ISL_3275543, EPI_ISL_3275544, EPI_ISL_3275545, EPI_ISL_3275546, EPI_ISL_3275547, EPI_ISL_3275548, EPI_ISL_3275549, EPI_ISL_3275550, EPI_ISL_3275551, EPI_ISL_3275552, EPI_ISL_3275553, EPI_ISL_3275554, EPI_ISL_3275555, EPI_ISL_3275556, EPI_ISL_3275557, EPI_ISL_3275558, EPI_ISL_3275559, EPI_ISL_3275560, EPI_ISL_3275561, EPI_ISL_3275562, EPI_ISL_3275563, EPI_ISL_3275564, EPI_ISL_3275565, EPI_ISL_3275566, EPI_ISL_3275567, EPI_ISL_3275568, EPI_ISL_3275569, EPI_ISL_3275570, EPI_ISL_3275571, EPI_ISL_3275572, EPI_ISL_3275573, EPI_ISL_3275574, EPI_ISL_3275575, EPI_ISL_3275576, EPI_ISL_3275577, EPI_ISL_3275578, EPI_ISL_3275579, EPI_ISL_3275580, EPI_ISL_3275581, EPI_ISL_3275582, EPI_ISL_3275583, EPI_ISL_3275584, EPI_ISL_3275585, EPI_ISL_3275586, EPI_ISL_3275587, EPI_ISL_3275588, EPI_ISL_3275589, EPI_ISL_3275590, EPI_ISL_3275591, EPI_ISL_3275592, EPI_ISL_3275593, EPI_ISL_3275594, EPI_ISL_3275595, EPI_ISL_3275596, EPI_ISL_3275597, EPI_ISL_3275598, EPI_ISL_3275599, EPI_ISL_3275600, EPI_ISL_3275601, EPI_ISL_3275602, EPI_ISL_3275603, EPI_ISL_3275604, EPI_ISL_3275605, EPI_ISL_3275606, EPI_ISL_3275607, EPI_ISL_3275608, EPI_ISL_3275609, EPI_ISL_3275610, EPI_ISL_3275611, EPI_ISL_3275612, EPI_ISL_3275613, EPI_ISL_3275614, EPI_ISL_3275615, EPI_ISL_3275616, EPI_ISL_3275617, EPI_ISL_3275618, EPI_ISL_3275619 | | | |
| see above | Dianalabs SA | Genesupport | Tanguy ARAUD, Geraldine Jost, Katia Jaton, Nadia Liassine |
| EPI_ISL_3275620, EPI_ISL_3275621 | Polyanalytic SA | Genesupport | Tanguy ARAUD, Geraldine Jost, Katia Jaton, Nadia Liassine |
| EPI_ISL_3275622, EPI_ISL_3275623, EPI_ISL_3275624, EPI_ISL_3275625, EPI_ISL_3275626 | Dianalabs-Valais SA | Genesupport | Tanguy ARAUD, Geraldine Jost, Katia Jaton, Nadia Liassine |
| EPI_ISL_3275627, EPI_ISL_3275628, EPI_ISL_3275629, EPI_ISL_3275630, EPI_ISL_3275631, EPI_ISL_3275632, EPI_ISL_3275633, EPI_ISL_3275634, EPI_ISL_3275635, EPI_ISL_3275636, EPI_ISL_3275637, EPI_ISL_3275638, EPI_ISL_3275639, EPI_ISL_3275640, EPI_ISL_3275641, EPI_ISL_3275642, EPI_ISL_3275643, EPI_ISL_3275644, EPI_ISL_3275645, EPI_ISL_3275646, EPI_ISL_3275647, EPI_ISL_3275648 | | | |
| see above | Dianalabs SA | Genesupport | Tanguy ARAUD, Geraldine Jost, Katia Jaton, Nadia Liassine |
| EPI_ISL_3275649, EPI_ISL_3275650, EPI_ISL_3275651, EPI_ISL_3275652, EPI_ISL_3275653 | Proxilab SA | Genesupport | Tanguy ARAUD, Geraldine Jost, Katia Jaton, Nadia Liassine |
| EPI_ISL_3275654, EPI_ISL_3275655, EPI_ISL_3275656, EPI_ISL_3275657, EPI_ISL_3275658, EPI_ISL_3275659, EPI_ISL_3275660, EPI_ISL_3275661, EPI_ISL_3275662, EPI_ISL_3275663, EPI_ISL_3275664, EPI_ISL_3275665, EPI_ISL_3275666, EPI_ISL_3275667, EPI_ISL_3275668, EPI_ISL_3275669, EPI_ISL_3275670, EPI_ISL_3275671, EPI_ISL_3275672, EPI_ISL_3275673, EPI_ISL_3275675, EPI_ISL_3275676, EPI_ISL_3275677 | | | |
| see above | Plateforme de testing Namuroise | Plateforme de testing Namuroise | Lesly Nyinkeu Kemamen; Nicolas Gilliard; Demars Aurore ; Maschietto Céline ; Nobis Chloé ; Otto Gaetan ; Denis Olivier ; Degosserie Jonathan ; Mullier François |
| EPI_ISL_3275695 | Dr. Madhusudhan, MBBD, MD, ICMR VRDL, Wenlock District Hospital | inStem NCBS – INSACOG | Uma Ramakrishnan Dasaradhi Palakodeti Aswin SaiNarain |
| EPI_ISL_3275696, EPI_ISL_3275697, EPI_ISL_3275698 | Dr S Raju, Director of Public Health and Preventive Medicine | inStem NCBS – INSACOG | Uma Ramakrishnan Dasaradhi Palakodeti Aswin SaiNarain |
| EPI_ISL_3275699 | VRDL, Wenlock District Hospital | inStem NCBS – INSACOG | Uma Ramakrishnan Dasaradhi Palakodeti Aswin SaiNarain |
| EPI_ISL_3275700, EPI_ISL_3275701, EPI_ISL_3275702, EPI_ISL_3275703 | Dr S Raju, Director of Public Health and Preventive Medicine | inStem NCBS – INSACOG | Uma Ramakrishnan Dasaradhi Palakodeti Aswin SaiNarain |
| EPI_ISL_3275704 | Dr. Madhusudhan, MBBD, MD, ICMR VRDL, Wenlock District Hospital | inStem NCBS – INSACOG | Uma Ramakrishnan Dasaradhi Palakodeti Aswin SaiNarain |
| EPI_ISL_3275705 | Dr S Raju, Director of Public Health and Preventive Medicine | inStem NCBS – INSACOG | Uma Ramakrishnan Dasaradhi Palakodeti Aswin SaiNarain |
| EPI_ISL_3275706 | Dr. Madhusudhan, MBBD, MD, ICMR VRDL, Wenlock District Hospital | inStem NCBS – INSACOG | Uma Ramakrishnan Dasaradhi Palakodeti Aswin SaiNarain |
| EPI_ISL_3275707, EPI_ISL_3275708, EPI_ISL_3275709, EPI_ISL_3275710 | Dr S Raju, Director of Public Health and Preventive Medicine | inStem NCBS – INSACOG | Uma Ramakrishnan Dasaradhi Palakodeti Aswin SaiNarain |
| EPI_ISL_3275711 | VRDL, Wenlock District Hospital | inStem NCBS – INSACOG | Uma Ramakrishnan Dasaradhi Palakodeti Aswin SaiNarain |
| EPI_ISL_3275712 | Dr S Raju, Director of Public Health and Preventive Medicine | inStem NCBS – INSACOG | Uma Ramakrishnan Dasaradhi Palakodeti Aswin SaiNarain |
| EPI_ISL_3275713 | VRDL, Wenlock District Hospital | inStem NCBS – INSACOG | Uma Ramakrishnan Dasaradhi Palakodeti Aswin SaiNarain |
| EPI_ISL_3275714 | Dr. Madhusudhan, MBBD, MD, ICMR VRDL, Wenlock District Hospital | inStem NCBS – INSACOG | Uma Ramakrishnan Dasaradhi Palakodeti Aswin SaiNarain |
| EPI_ISL_3275715 | Dr S Raju, Director of Public Health and Preventive Medicine | inStem NCBS – INSACOG | Uma Ramakrishnan Dasaradhi Palakodeti Aswin SaiNarain |
| EPI_ISL_3275716 | Dr. Madhusudhan, MBBD, MD, ICMR VRDL, Wenlock District Hospital | inStem NCBS – INSACOG | Uma Ramakrishnan Dasaradhi Palakodeti Aswin SaiNarain |
| EPI_ISL_3275717, EPI_ISL_3275719 | Dr S Raju, Director of Public Health and Preventive Medicine | inStem NCBS – INSACOG | Uma Ramakrishnan Dasaradhi Palakodeti Aswin SaiNarain |
| EPI_ISL_3275720 | Dr. Madhusudhan, MBBD, MD, ICMR VRDL, Wenlock District Hospital | inStem NCBS – INSACOG | Uma Ramakrishnan Dasaradhi Palakodeti Aswin SaiNarain |
| EPI_ISL_3275721 | Dr S Raju, Director of Public Health and Preventive Medicine | inStem NCBS – INSACOG | Uma Ramakrishnan Dasaradhi Palakodeti Aswin SaiNarain |
| EPI_ISL_3275722 | Dr. Madhusudhan, MBBD, MD, ICMR VRDL, Wenlock District Hospital | inStem NCBS – INSACOG | Uma Ramakrishnan Dasaradhi Palakodeti Aswin SaiNarain |
| EPI_ISL_3275723, EPI_ISL_3275724, EPI_ISL_3275725 | Dr S Raju, Director of Public Health and Preventive Medicine | inStem NCBS – INSACOG | Uma Ramakrishnan Dasaradhi Palakodeti Aswin SaiNarain |
| EPI_ISL_3275726 | Dr. Madhusudhan, MBBD, MD, ICMR VRDL, Wenlock District Hospital | inStem NCBS – INSACOG | Uma Ramakrishnan Dasaradhi Palakodeti Aswin SaiNarain |
| EPI_ISL_3275727, EPI_ISL_3275730 | Dr S Raju, Director of Public Health and Preventive Medicine | inStem NCBS – INSACOG | Uma Ramakrishnan Dasaradhi Palakodeti Aswin SaiNarain |
| EPI_ISL_3275731 | Department of Microbiology, Belagavi Institute of Medical Sciences | inStem NCBS – INSACOG | Uma Ramakrishnan Dasaradhi Palakodeti Aswin SaiNarain |
| EPI_ISL_3275732, EPI_ISL_3275733 | Dr S Raju, Director of Public Health and Preventive Medicine | inStem NCBS – INSACOG | Uma Ramakrishnan Dasaradhi Palakodeti Aswin SaiNarain |
| EPI_ISL_3275734 | JIPMER Puducherry | inStem NCBS – INSACOG | Uma Ramakrishnan Dasaradhi Palakodeti Aswin SaiNarain |
| EPI_ISL_3275735 | Department of Microbiology, Belagavi Institute of Medical Sciences | inStem NCBS – INSACOG | Uma Ramakrishnan Dasaradhi Palakodeti Aswin SaiNarain |
| EPI_ISL_3275736, EPI_ISL_3275737 | Dr S Raju, Director of Public Health and Preventive Medicine | inStem NCBS – INSACOG | Uma Ramakrishnan Dasaradhi Palakodeti Aswin SaiNarain |
| EPI_ISL_3275738 | JIPMER Puducherry | inStem NCBS – INSACOG | Uma Ramakrishnan Dasaradhi Palakodeti Aswin SaiNarain |
| EPI_ISL_3275739, EPI_ISL_3275740, EPI_ISL_3275741, EPI_ISL_3275742 | Dr S Raju, Director of Public Health and Preventive Medicine | inStem NCBS – INSACOG | Uma Ramakrishnan Dasaradhi Palakodeti Aswin SaiNarain |
| EPI_ISL_3275743 | JIPMER Puducherry | inStem NCBS – INSACOG | Uma Ramakrishnan Dasaradhi Palakodeti Aswin SaiNarain |
| EPI_ISL_3275744, EPI_ISL_3275745, EPI_ISL_3275746, EPI_ISL_3275747 | Dr S Raju, Director of Public Health and Preventive Medicine | inStem NCBS – INSACOG | Uma Ramakrishnan Dasaradhi Palakodeti Aswin SaiNarain |
| EPI_ISL_3275748 | JIPMER Puducherry | inStem NCBS – INSACOG | Uma Ramakrishnan Dasaradhi Palakodeti Aswin SaiNarain |
| EPI_ISL_3275749, EPI_ISL_3275750 | Dr S Raju, Director of Public Health and Preventive Medicine | inStem NCBS – INSACOG | Uma Ramakrishnan Dasaradhi Palakodeti Aswin SaiNarain |
| EPI_ISL_3275751 | JIPMER Puducherry | inStem NCBS – INSACOG | Uma Ramakrishnan Dasaradhi Palakodeti Aswin SaiNarain |
| EPI_ISL_3275752, EPI_ISL_3275753 | Dr S Raju, Director of Public Health and Preventive Medicine | inStem NCBS – INSACOG | Uma Ramakrishnan Dasaradhi Palakodeti Aswin SaiNarain |

| | | | |
|---|---|---|---|
| EPI_ISL_3275754 | JIPMER Puducherry | inStem NCBS – INSACOG | Uma Ramakrishnan Dasaradhi Palakodeti Aswin SaiNarain |
| EPI_ISL_3275755 | Dr S Raju, Director of Public Health and Preventive Medicine | inStem NCBS – INSACOG | Uma Ramakrishnan Dasaradhi Palakodeti Aswin SaiNarain |
| EPI_ISL_3275756 | Dr. Madhusudhan, MBBD, MD, ICMR VRDL, Wenlock District Hospital | inStem NCBS – INSACOG | Uma Ramakrishnan Dasaradhi Palakodeti Aswin SaiNarain |
| EPI_ISL_3275758, EPI_ISL_3275759, EPI_ISL_3275760 | Dr S Raju, Director of Public Health and Preventive Medicine | inStem NCBS – INSACOG | Uma Ramakrishnan Dasaradhi Palakodeti Aswin SaiNarain |
| EPI_ISL_3275761 | Dr. Madhusudhan, MBBD, MD, ICMR VRDL, Wenlock District Hospital | inStem NCBS – INSACOG | Uma Ramakrishnan Dasaradhi Palakodeti Aswin SaiNarain |
| EPI_ISL_3275762, EPI_ISL_3275763, EPI_ISL_3275764 | Dr S Raju, Director of Public Health and Preventive Medicine | inStem NCBS – INSACOG | Uma Ramakrishnan Dasaradhi Palakodeti Aswin SaiNarain |
| EPI_ISL_3275765, EPI_ISL_3275766 | JIPMER Puducherry | inStem NCBS – INSACOG | Uma Ramakrishnan Dasaradhi Palakodeti Aswin SaiNarain |
| EPI_ISL_3275767 | Dr S Raju, Director of Public Health and Preventive Medicine | inStem NCBS – INSACOG | Uma Ramakrishnan Dasaradhi Palakodeti Aswin SaiNarain |
| EPI_ISL_3275768 | Dr. Madhusudhan, MBBD, MD, ICMR VRDL, Wenlock District Hospital | inStem NCBS – INSACOG | Uma Ramakrishnan Dasaradhi Palakodeti Aswin SaiNarain |
| EPI_ISL_3275770 | JIPMER Puducherry | inStem NCBS – INSACOG | Uma Ramakrishnan Dasaradhi Palakodeti Aswin SaiNarain |
| EPI_ISL_3275771, EPI_ISL_3275772 | Dr S Raju, Director of Public Health and Preventive Medicine | inStem NCBS – INSACOG | Uma Ramakrishnan Dasaradhi Palakodeti Aswin SaiNarain |
| EPI_ISL_3275773, EPI_ISL_3275774 | Dr. Madhusudhan, MBBD, MD, ICMR VRDL, Wenlock District Hospital | inStem NCBS – INSACOG | Uma Ramakrishnan Dasaradhi Palakodeti Aswin SaiNarain |
| EPI_ISL_3275775 | Dr S Raju, Director of Public Health and Preventive Medicine | inStem NCBS – INSACOG | Uma Ramakrishnan Dasaradhi Palakodeti Aswin SaiNarain |
| EPI_ISL_3275776 | JIPMER Puducherry | inStem NCBS – INSACOG | Uma Ramakrishnan Dasaradhi Palakodeti Aswin SaiNarain |
| EPI_ISL_3275777 | Dr. Madhusudhan, MBBD, MD, ICMR VRDL, Wenlock District Hospital | inStem NCBS – INSACOG | Uma Ramakrishnan Dasaradhi Palakodeti Aswin SaiNarain |
| EPI_ISL_3275778 | Dr S Raju, Director of Public Health and Preventive Medicine | inStem NCBS – INSACOG | Uma Ramakrishnan Dasaradhi Palakodeti Aswin SaiNarain |
| EPI_ISL_3275779 | Dr. Madhusudhan, MBBD, MD, ICMR VRDL, Wenlock District Hospital | inStem NCBS – INSACOG | Uma Ramakrishnan Dasaradhi Palakodeti Aswin SaiNarain |
| EPI_ISL_3275780 | Dr S Raju, Director of Public Health and Preventive Medicine | inStem NCBS – INSACOG | Uma Ramakrishnan Dasaradhi Palakodeti Aswin SaiNarain |
| EPI_ISL_3275781 | Dr. Madhusudhan, MBBD, MD, ICMR VRDL, Wenlock District Hospital | inStem NCBS – INSACOG | Uma Ramakrishnan Dasaradhi Palakodeti Aswin SaiNarain |
| EPI_ISL_3275782, EPI_ISL_3275783 | Dr S Raju, Director of Public Health and Preventive Medicine | inStem NCBS – INSACOG | Uma Ramakrishnan Dasaradhi Palakodeti Aswin SaiNarain |
| EPI_ISL_3275784, EPI_ISL_3275785 | Dr. Madhusudhan, MBBD, MD, ICMR VRDL, Wenlock District Hospital | inStem NCBS – INSACOG | Uma Ramakrishnan Dasaradhi Palakodeti Aswin SaiNarain |
| EPI_ISL_3275786 | Dr S Raju, Director of Public Health and Preventive Medicine | inStem NCBS – INSACOG | Uma Ramakrishnan Dasaradhi Palakodeti Aswin SaiNarain |
| EPI_ISL_3275787, EPI_ISL_3275788 | JIPMER Puducherry | inStem NCBS – INSACOG | Uma Ramakrishnan Dasaradhi Palakodeti Aswin SaiNarain |
| EPI_ISL_3275789 | Dr. Madhusudhan, MBBD, MD, ICMR VRDL, Wenlock District Hospital | inStem NCBS – INSACOG | Uma Ramakrishnan Dasaradhi Palakodeti Aswin SaiNarain |
| EPI_ISL_3275791 | JIPMER Puducherry | inStem NCBS – INSACOG | Uma Ramakrishnan Dasaradhi Palakodeti Aswin SaiNarain |
| EPI_ISL_3275792 | Dr. Madhusudhan, MBBD, MD, ICMR VRDL, Wenlock District Hospital | inStem NCBS – INSACOG | Uma Ramakrishnan Dasaradhi Palakodeti Aswin SaiNarain |
| EPI_ISL_3275793, EPI_ISL_3275794 | Dr S Raju, Director of Public Health and Preventive Medicine | inStem NCBS – INSACOG | Uma Ramakrishnan Dasaradhi Palakodeti Aswin SaiNarain |
| EPI_ISL_3275795 | JIPMER Puducherry | inStem NCBS – INSACOG | Uma Ramakrishnan Dasaradhi Palakodeti Aswin SaiNarain |
| EPI_ISL_3275797 | Dr S Raju, Director of Public Health and Preventive Medicine | inStem NCBS – INSACOG | Uma Ramakrishnan Dasaradhi Palakodeti Aswin SaiNarain |
| EPI_ISL_3275798, EPI_ISL_3275799 | Dr. Madhusudhan, MBBD, MD, ICMR VRDL, Wenlock District Hospital | inStem NCBS – INSACOG | Uma Ramakrishnan Dasaradhi Palakodeti Aswin SaiNarain |
| EPI_ISL_3275800, EPI_ISL_3275801 | Dr S Raju, Director of Public Health and Preventive Medicine | inStem NCBS – INSACOG | Uma Ramakrishnan Dasaradhi Palakodeti Aswin SaiNarain |
| EPI_ISL_3275802 | JIPMER Puducherry | inStem NCBS – INSACOG | Uma Ramakrishnan Dasaradhi Palakodeti Aswin SaiNarain |
| EPI_ISL_3275803, EPI_ISL_3275805 | Dr S Raju, Director of Public Health and Preventive Medicine | inStem NCBS – INSACOG | Uma Ramakrishnan Dasaradhi Palakodeti Aswin SaiNarain |
| EPI_ISL_3275806, EPI_ISL_3275807, EPI_ISL_3275808 | Dr. Madhusudhan, MBBD, MD, ICMR VRDL, Wenlock District Hospital | inStem NCBS – INSACOG | Uma Ramakrishnan Dasaradhi Palakodeti Aswin SaiNarain |
| EPI_ISL_3275809 | JIPMER Puducherry | inStem NCBS – INSACOG | Uma Ramakrishnan Dasaradhi Palakodeti Aswin SaiNarain |
| EPI_ISL_3275810, EPI_ISL_3275811 | Dr S Raju, Director of Public Health and Preventive Medicine | inStem NCBS – INSACOG | Uma Ramakrishnan Dasaradhi Palakodeti Aswin SaiNarain |
| EPI_ISL_3275812 | JIPMER Puducherry | inStem NCBS – INSACOG | Uma Ramakrishnan Dasaradhi Palakodeti Aswin SaiNarain |
| EPI_ISL_3275813, EPI_ISL_3275814 | Dr. Madhusudhan, MBBD, MD, ICMR VRDL, Wenlock District Hospital | inStem NCBS – INSACOG | Uma Ramakrishnan Dasaradhi Palakodeti Aswin SaiNarain |
| EPI_ISL_3275815 | VRDL, Wenlock District Hospital | inStem NCBS – INSACOG | Uma Ramakrishnan Dasaradhi Palakodeti Aswin SaiNarain |
| EPI_ISL_3275816 | JIPMER Puducherry | inStem NCBS – INSACOG | Uma Ramakrishnan Dasaradhi Palakodeti Aswin SaiNarain |
| EPI_ISL_3275817 | Dr. Madhusudhan, MBBD, MD, ICMR VRDL, Wenlock District Hospital | inStem NCBS – INSACOG | Uma Ramakrishnan Dasaradhi Palakodeti Aswin SaiNarain |
| EPI_ISL_3275818 | Dr S Raju, Director of Public Health and Preventive Medicine | inStem NCBS – INSACOG | Uma Ramakrishnan Dasaradhi Palakodeti Aswin SaiNarain |
| EPI_ISL_3275820 | JIPMER Puducherry | inStem NCBS – INSACOG | Uma Ramakrishnan Dasaradhi Palakodeti Aswin SaiNarain |
| EPI_ISL_3275821 | Dr. Madhusudhan, MBBD, MD, ICMR VRDL, Wenlock District Hospital | inStem NCBS – INSACOG | Uma Ramakrishnan Dasaradhi Palakodeti Aswin SaiNarain |
| EPI_ISL_3275822, EPI_ISL_3275823, EPI_ISL_3275824, EPI_ISL_3275825 | Dr S Raju, Director of Public Health and Preventive Medicine | inStem NCBS – INSACOG | Uma Ramakrishnan Dasaradhi Palakodeti Aswin SaiNarain |
| EPI_ISL_3275826 | JIPMER Puducherry | inStem NCBS – INSACOG | Uma Ramakrishnan Dasaradhi Palakodeti Aswin SaiNarain |
| EPI_ISL_3275827 | Dr S Raju, Director of Public Health and Preventive Medicine | inStem NCBS – INSACOG | Uma Ramakrishnan Dasaradhi Palakodeti Aswin SaiNarain |
| EPI_ISL_3275828 | JIPMER Puducherry | inStem NCBS – INSACOG | Uma Ramakrishnan Dasaradhi Palakodeti Aswin SaiNarain |
| EPI_ISL_3275829 | Dr S Raju, Director of Public Health and Preventive Medicine | inStem NCBS – INSACOG | Uma Ramakrishnan Dasaradhi Palakodeti Aswin SaiNarain |
| EPI_ISL_3275830 | Dr. Madhusudhan, MBBD, MD, ICMR VRDL, Wenlock District Hospital | inStem NCBS – INSACOG | Uma Ramakrishnan Dasaradhi Palakodeti Aswin SaiNarain |
| EPI_ISL_3275831 | Dr S Raju, Director of Public Health and Preventive Medicine | inStem NCBS – INSACOG | Uma Ramakrishnan Dasaradhi Palakodeti Aswin SaiNarain |
| EPI_ISL_3275832, EPI_ISL_3275833 | JIPMER Puducherry | inStem NCBS – INSACOG | Uma Ramakrishnan Dasaradhi Palakodeti Aswin SaiNarain |
| EPI_ISL_3275834, EPI_ISL_3275835, | Dr S Raju, Director of Public Health and Preventive Medicine | inStem NCBS – INSACOG | Uma Ramakrishnan Dasaradhi Palakodeti Aswin SaiNarain |

| | | | |
|---|---|---|---|
| EPI_ISL_3275836 | | | |
| EPI_ISL_3275837 | JIPMER Puducherry | inStem NCBS – INSACOG | Uma Ramakrishnan Dasaradhi Palakodeti Aswin SaiNarain |
| EPI_ISL_3275838, EPI_ISL_3275839, EPI_ISL_3275840 | Dr S Raju, Director of Public Health and Preventive Medicine | inStem NCBS – INSACOG | Uma Ramakrishnan Dasaradhi Palakodeti Aswin SaiNarain |
| EPI_ISL_3275841 | Department of Microbiology, Belagavi Institute of Medical Sciences | inStem NCBS – INSACOG | Uma Ramakrishnan Dasaradhi Palakodeti Aswin SaiNarain |
| EPI_ISL_3275842, EPI_ISL_3275843 | JIPMER Puducherry | inStem NCBS – INSACOG | Uma Ramakrishnan Dasaradhi Palakodeti Aswin SaiNarain |
| EPI_ISL_3275844 | Dr. Madhusudhan, MBBD, MD, ICMR VRDL, Wenlock District Hospital | inStem NCBS – INSACOG | Uma Ramakrishnan Dasaradhi Palakodeti Aswin SaiNarain |
| EPI_ISL_3275845, EPI_ISL_3275846, EPI_ISL_3275847, EPI_ISL_3275848 | Dr S Raju, Director of Public Health and Preventive Medicine | inStem NCBS – INSACOG | Uma Ramakrishnan Dasaradhi Palakodeti Aswin SaiNarain |
| EPI_ISL_3275849 | JIPMER Puducherry | inStem NCBS – INSACOG | Uma Ramakrishnan Dasaradhi Palakodeti Aswin SaiNarain |
| EPI_ISL_3275850 | Dr S Raju, Director of Public Health and Preventive Medicine | inStem NCBS – INSACOG | Uma Ramakrishnan Dasaradhi Palakodeti Aswin SaiNarain |
| EPI_ISL_3275851 | Dr. Madhusudhan, MBBD, MD, ICMR VRDL, Wenlock District Hospital | inStem NCBS – INSACOG | Uma Ramakrishnan Dasaradhi Palakodeti Aswin SaiNarain |
| EPI_ISL_3275852 | Dr S Raju, Director of Public Health and Preventive Medicine | inStem NCBS – INSACOG | Uma Ramakrishnan Dasaradhi Palakodeti Aswin SaiNarain |
| EPI_ISL_3275853 | Dr. Madhusudhan, MBBD, MD, ICMR VRDL, Wenlock District Hospital | inStem NCBS – INSACOG | Uma Ramakrishnan Dasaradhi Palakodeti Aswin SaiNarain |
| EPI_ISL_3275854, EPI_ISL_3275855, EPI_ISL_3275856, EPI_ISL_3275857, EPI_ISL_3275858 | Dr S Raju, Director of Public Health and Preventive Medicine | inStem NCBS – INSACOG | Uma Ramakrishnan Dasaradhi Palakodeti Aswin SaiNarain |
| EPI_ISL_3275859 | JIPMER Puducherry | inStem NCBS – INSACOG | Uma Ramakrishnan Dasaradhi Palakodeti Aswin SaiNarain |
| EPI_ISL_3275860, EPI_ISL_3275861 | Dr S Raju, Director of Public Health and Preventive Medicine | inStem NCBS – INSACOG | Uma Ramakrishnan Dasaradhi Palakodeti Aswin SaiNarain |
| EPI_ISL_3275862 | Dr. Madhusudhan, MBBD, MD, ICMR VRDL, Wenlock District Hospital | inStem NCBS – INSACOG | Uma Ramakrishnan Dasaradhi Palakodeti Aswin SaiNarain |
| EPI_ISL_3275863, EPI_ISL_3275864, EPI_ISL_3275865 | Dr S Raju, Director of Public Health and Preventive Medicine | inStem NCBS – INSACOG | Uma Ramakrishnan Dasaradhi Palakodeti Aswin SaiNarain |
| EPI_ISL_3275866, EPI_ISL_3275867 | JIPMER Puducherry | inStem NCBS – INSACOG | Uma Ramakrishnan Dasaradhi Palakodeti Aswin SaiNarain |
| EPI_ISL_3275868, EPI_ISL_3275869 | Dr S Raju, Director of Public Health and Preventive Medicine | inStem NCBS – INSACOG | Uma Ramakrishnan Dasaradhi Palakodeti Aswin SaiNarain |
| EPI_ISL_3275870 | Dr. Madhusudhan, MBBD, MD, ICMR VRDL, Wenlock District Hospital | inStem NCBS – INSACOG | Uma Ramakrishnan Dasaradhi Palakodeti Aswin SaiNarain |
| EPI_ISL_3275871 | Department of Microbiology, Belagavi Institute of Medical Sciences | inStem NCBS – INSACOG | Uma Ramakrishnan Dasaradhi Palakodeti Aswin SaiNarain |
| EPI_ISL_3275872, EPI_ISL_3275873, EPI_ISL_3275875, EPI_ISL_3275876 | Dr S Raju, Director of Public Health and Preventive Medicine | inStem NCBS – INSACOG | Uma Ramakrishnan Dasaradhi Palakodeti Aswin SaiNarain |
| EPI_ISL_3275877 | Dr. Madhusudhan, MBBD, MD, ICMR VRDL, Wenlock District Hospital | inStem NCBS – INSACOG | Uma Ramakrishnan Dasaradhi Palakodeti Aswin SaiNarain |
| EPI_ISL_3275878, EPI_ISL_3275881, EPI_ISL_3275887 | Dr S Raju, Director of Public Health and Preventive Medicine | inStem NCBS – INSACOG | Uma Ramakrishnan Dasaradhi Palakodeti Aswin SaiNarain |
| EPI_ISL_3275888 | Dr. Madhusudhan, MBBD, MD, ICMR VRDL, Wenlock District Hospital | inStem NCBS – INSACOG | Uma Ramakrishnan Dasaradhi Palakodeti Aswin SaiNarain |
| EPI_ISL_3275889, EPI_ISL_3275890 | Dr S Raju, Director of Public Health and Preventive Medicine | inStem NCBS – INSACOG | Uma Ramakrishnan Dasaradhi Palakodeti Aswin SaiNarain |
| EPI_ISL_3275891 | VRDL, Wenlock District Hospital | inStem NCBS – INSACOG | Uma Ramakrishnan Dasaradhi Palakodeti Aswin SaiNarain |
| EPI_ISL_3275892 | Department of Microbiology, Belagavi Institute of Medical Sciences | inStem NCBS – INSACOG | Uma Ramakrishnan Dasaradhi Palakodeti Aswin SaiNarain |
| EPI_ISL_3275893 | Dr. Madhusudhan, MBBD, MD, ICMR VRDL, Wenlock District Hospital | inStem NCBS – INSACOG | Uma Ramakrishnan Dasaradhi Palakodeti Aswin SaiNarain |
| EPI_ISL_3275895, EPI_ISL_3275896, EPI_ISL_3275897 | Dr S Raju, Director of Public Health and Preventive Medicine | inStem NCBS – INSACOG | Uma Ramakrishnan Dasaradhi Palakodeti Aswin SaiNarain |
| EPI_ISL_3275898 | VRDL, Wenlock District Hospital | inStem NCBS – INSACOG | Uma Ramakrishnan Dasaradhi Palakodeti Aswin SaiNarain |
| EPI_ISL_3275899 | Dr. Madhusudhan, MBBD, MD, ICMR VRDL, Wenlock District Hospital | inStem NCBS – INSACOG | Uma Ramakrishnan Dasaradhi Palakodeti Aswin SaiNarain |
| EPI_ISL_3275900 | Dr S Raju, Director of Public Health and Preventive Medicine | inStem NCBS – INSACOG | Uma Ramakrishnan Dasaradhi Palakodeti Aswin SaiNarain |
| EPI_ISL_3275901 | Dr. Madhusudhan, MBBD, MD, ICMR VRDL, Wenlock District Hospital | inStem NCBS – INSACOG | Uma Ramakrishnan Dasaradhi Palakodeti Aswin SaiNarain |
| EPI_ISL_3275902 | Dr S Raju, Director of Public Health and Preventive Medicine | inStem NCBS – INSACOG | Uma Ramakrishnan Dasaradhi Palakodeti Aswin SaiNarain |
| EPI_ISL_3275903 | VRDL, Wenlock District Hospital | inStem NCBS – INSACOG | Uma Ramakrishnan Dasaradhi Palakodeti Aswin SaiNarain |
| EPI_ISL_3275904, EPI_ISL_3275905 | Dr S Raju, Director of Public Health and Preventive Medicine | inStem NCBS – INSACOG | Uma Ramakrishnan Dasaradhi Palakodeti Aswin SaiNarain |
| EPI_ISL_3275906 | JIPMER Puducherry | inStem NCBS – INSACOG | Uma Ramakrishnan Dasaradhi Palakodeti Aswin SaiNarain |
| EPI_ISL_3275907, EPI_ISL_3275908, EPI_ISL_3275909, EPI_ISL_3275910, EPI_ISL_3275911 | Dr S Raju, Director of Public Health and Preventive Medicine | inStem NCBS – INSACOG | Uma Ramakrishnan Dasaradhi Palakodeti Aswin SaiNarain |
| EPI_ISL_3275912 | JIPMER Puducherry | inStem NCBS – INSACOG | Uma Ramakrishnan Dasaradhi Palakodeti Aswin SaiNarain |
| EPI_ISL_3275913, EPI_ISL_3275914, EPI_ISL_3275915, EPI_ISL_3275916 | Dr S Raju, Director of Public Health and Preventive Medicine | inStem NCBS – INSACOG | Uma Ramakrishnan Dasaradhi Palakodeti Aswin SaiNarain |
| EPI_ISL_3275917 | Dr. Madhusudhan, MBBD, MD, ICMR VRDL, Wenlock District Hospital | inStem NCBS – INSACOG | Uma Ramakrishnan Dasaradhi Palakodeti Aswin SaiNarain |
| EPI_ISL_3275918, EPI_ISL_3275951 | Dr S Raju, Director of Public Health and Preventive Medicine | inStem NCBS – INSACOG | Uma Ramakrishnan Dasaradhi Palakodeti Aswin SaiNarain |
| EPI_ISL_3275952 | VRDL, Wenlock District Hospital | inStem NCBS – INSACOG | Uma Ramakrishnan Dasaradhi Palakodeti Aswin SaiNarain |

| | | | |
|---|---|---|---|
| EPI_ISL_3275953, EPI_ISL_3275954 | Dr S Raju, Director of Public Health and Preventive Medicine | inStem NCBS – INSACOG | Uma Ramakrishnan Dasaradhi Palakodeti Aswin SaiNarain |
| EPI_ISL_3275955 | JIPMER Puducherry | inStem NCBS – INSACOG | Uma Ramakrishnan Dasaradhi Palakodeti Aswin SaiNarain |
| EPI_ISL_3275956, EPI_ISL_3275957, EPI_ISL_3275958, EPI_ISL_3275959, EPI_ISL_3275960, EPI_ISL_3275961 | Dr S Raju, Director of Public Health and Preventive Medicine | inStem NCBS – INSACOG | Uma Ramakrishnan Dasaradhi Palakodeti Aswin SaiNarain |
| EPI_ISL_3275962 | Department of Microbiology, Belagavi Institute of Medical Sciences | inStem NCBS – INSACOG | Uma Ramakrishnan Dasaradhi Palakodeti Aswin SaiNarain |
| EPI_ISL_3275963, EPI_ISL_3275964, EPI_ISL_3275965, EPI_ISL_3275966, EPI_ISL_3275967, EPI_ISL_3275968, EPI_ISL_3275970, EPI_ISL_3275971 | Dr S Raju, Director of Public Health and Preventive Medicine | inStem NCBS – INSACOG | Uma Ramakrishnan Dasaradhi Palakodeti Aswin SaiNarain |
| EPI_ISL_3275972 | JIPMER Puducherry | inStem NCBS – INSACOG | Uma Ramakrishnan Dasaradhi Palakodeti Aswin SaiNarain |
| EPI_ISL_3275973 | Dr S Raju, Director of Public Health and Preventive Medicine | inStem NCBS – INSACOG | Uma Ramakrishnan Dasaradhi Palakodeti Aswin SaiNarain |
| EPI_ISL_3275974 | VRDL, Wenlock District Hospital | inStem NCBS – INSACOG | Uma Ramakrishnan Dasaradhi Palakodeti Aswin SaiNarain |
| EPI_ISL_3275975 | Dr. Madhusudhan, MBBD, MD, ICMR VRDL, Wenlock District Hospital | inStem NCBS – INSACOG | Uma Ramakrishnan Dasaradhi Palakodeti Aswin SaiNarain |
| EPI_ISL_3275976 | Dr S Raju, Director of Public Health and Preventive Medicine | inStem NCBS – INSACOG | Uma Ramakrishnan Dasaradhi Palakodeti Aswin SaiNarain |
| EPI_ISL_3275977 | JIPMER Puducherry | inStem NCBS – INSACOG | Uma Ramakrishnan Dasaradhi Palakodeti Aswin SaiNarain |
| EPI_ISL_3275978 | Dr S Raju, Director of Public Health and Preventive Medicine | inStem NCBS – INSACOG | Uma Ramakrishnan Dasaradhi Palakodeti Aswin SaiNarain |
| EPI_ISL_3275979 | JIPMER Puducherry | inStem NCBS – INSACOG | Uma Ramakrishnan Dasaradhi Palakodeti Aswin SaiNarain |
| EPI_ISL_3275980, EPI_ISL_3275981, EPI_ISL_3275982 | Dr S Raju, Director of Public Health and Preventive Medicine | inStem NCBS – INSACOG | Uma Ramakrishnan Dasaradhi Palakodeti Aswin SaiNarain |
| EPI_ISL_3275983 | Dr. Madhusudhan, MBBD, MD, ICMR VRDL, Wenlock District Hospital | inStem NCBS – INSACOG | Uma Ramakrishnan Dasaradhi Palakodeti Aswin SaiNarain |
| EPI_ISL_3275984, EPI_ISL_3275985, EPI_ISL_3275986, EPI_ISL_3275987, EPI_ISL_3275988, EPI_ISL_3275989, EPI_ISL_3275990 | Dr S Raju, Director of Public Health and Preventive Medicine | inStem NCBS – INSACOG | Uma Ramakrishnan Dasaradhi Palakodeti Aswin SaiNarain |
| EPI_ISL_3275991 | JIPMER Puducherry | inStem NCBS – INSACOG | Uma Ramakrishnan Dasaradhi Palakodeti Aswin SaiNarain |
| EPI_ISL_3275992 | Dr S Raju, Director of Public Health and Preventive Medicine | inStem NCBS – INSACOG | Uma Ramakrishnan Dasaradhi Palakodeti Aswin SaiNarain |
| EPI_ISL_3275993, EPI_ISL_3275994, EPI_ISL_3275995, EPI_ISL_3275996, EPI_ISL_3275997, EPI_ISL_3275998 | Dr S raju, Director of public heath and preventive medicine | inStem NCBS – INSACOG | Uma Ramakrishnan Dasaradhi Palakodeti Aswin SaiNarain |
| EPI_ISL_3275999 | Kodigehalli PHC | inStem NCBS – INSACOG | Uma Ramakrishnan Dasaradhi Palakodeti Aswin SaiNarain |
| EPI_ISL_3276000, EPI_ISL_3276001, EPI_ISL_3276002, EPI_ISL_3276003, EPI_ISL_3276004, EPI_ISL_3276005 | Dr S raju, Director of public heath and preventive medicine | inStem NCBS – INSACOG | Uma Ramakrishnan Dasaradhi Palakodeti Aswin SaiNarain |
| EPI_ISL_3276006, EPI_ISL_3276007 | Kodigehalli PHC | inStem NCBS – INSACOG | Uma Ramakrishnan Dasaradhi Palakodeti Aswin SaiNarain |
| EPI_ISL_3276008, EPI_ISL_3276009, EPI_ISL_3276010, EPI_ISL_3276011, EPI_ISL_3276012, EPI_ISL_3276013, EPI_ISL_3276014, EPI_ISL_3276015 | Dr S raju, Director of public heath and preventive medicine | inStem NCBS – INSACOG | Uma Ramakrishnan Dasaradhi Palakodeti Aswin SaiNarain |
| EPI_ISL_3276016 | Kodigehalli PHC | inStem NCBS – INSACOG | Uma Ramakrishnan Dasaradhi Palakodeti Aswin SaiNarain |
| EPI_ISL_3276017, EPI_ISL_3276018, EPI_ISL_3276019, EPI_ISL_3276020, EPI_ISL_3276021, EPI_ISL_3276022, EPI_ISL_3276023, EPI_ISL_3276024, EPI_ISL_3276025, EPI_ISL_3276026 | Dr S raju, Director of public heath and preventive medicine | inStem NCBS – INSACOG | Uma Ramakrishnan Dasaradhi Palakodeti Aswin SaiNarain |
| EPI_ISL_3276027 | Kodigehalli PHC | inStem NCBS – INSACOG | Uma Ramakrishnan Dasaradhi Palakodeti Aswin SaiNarain |
| EPI_ISL_3276028, EPI_ISL_3276029, EPI_ISL_3276030, EPI_ISL_3276031, EPI_ISL_3276032 | Dr S raju, Director of public heath and preventive medicine | inStem NCBS – INSACOG | Uma Ramakrishnan Dasaradhi Palakodeti Aswin SaiNarain |
| EPI_ISL_3276033, EPI_ISL_3276034 | Kodigehalli PHC | inStem NCBS – INSACOG | Uma Ramakrishnan Dasaradhi Palakodeti Aswin SaiNarain |
| EPI_ISL_3276035, EPI_ISL_3276036, EPI_ISL_3276037, EPI_ISL_3276038, EPI_ISL_3276039, EPI_ISL_3276040, EPI_ISL_3276041 | Dr S raju, Director of public heath and preventive medicine | inStem NCBS – INSACOG | Uma Ramakrishnan Dasaradhi Palakodeti Aswin SaiNarain |
| EPI_ISL_3276042 | Kodigehalli PHC | inStem NCBS – INSACOG | Uma Ramakrishnan Dasaradhi Palakodeti Aswin SaiNarain |
| EPI_ISL_3276043 | Dr S raju, Director of public heath and preventive medicine | inStem NCBS – INSACOG | Uma Ramakrishnan Dasaradhi Palakodeti Aswin SaiNarain |
| EPI_ISL_3276044 | Sahakara Nagar PHC | inStem NCBS – INSACOG | Uma Ramakrishnan Dasaradhi Palakodeti Aswin SaiNarain |

EPI_ISL_3276045, EPI_ISL_3276046, EPI_ISL_3276047, EPI_ISL_3276048, EPI_ISL_3276049, EPI_ISL_3276050, EPI_ISL_3276051, EPI_ISL_3276052, EPI_ISL_3276053, EPI_ISL_3276054, EPI_ISL_3276055, EPI_ISL_3276056, EPI_ISL_3276057, EPI_ISL_3276058, EPI_ISL_3276059, EPI_ISL_3276060, EPI_ISL_3276061, EPI_ISL_3276062, EPI_ISL_3276063

| | | | |
|---|---|---|---|
| see above | Dr S raju, Director of public heath and preventive medicine | inStem NCBS – INSACOG | Uma Ramakrishnan Dasaradhi Palakodeti Aswin SaiNarain |
| EPI_ISL_3276064, EPI_ISL_3276065 | Kodigehalli PHC | inStem NCBS – INSACOG | Uma Ramakrishnan Dasaradhi Palakodeti Aswin SaiNarain |
| EPI_ISL_3276066, EPI_ISL_3276067, EPI_ISL_3276068, EPI_ISL_3276069, EPI_ISL_3276070, EPI_ISL_3276071, EPI_ISL_3276072 | Dr S raju, Director of public heath and preventive medicine | inStem NCBS – INSACOG | Uma Ramakrishnan Dasaradhi Palakodeti Aswin SaiNarain |
| EPI_ISL_3276073 | Kodigehalli PHC | inStem NCBS – INSACOG | Uma Ramakrishnan Dasaradhi Palakodeti Aswin SaiNarain |
| EPI_ISL_3276074, EPI_ISL_3276075, EPI_ISL_3276076, EPI_ISL_3276077, EPI_ISL_3276078, EPI_ISL_3276079, EPI_ISL_3276080, EPI_ISL_3276081, EPI_ISL_3276082, EPI_ISL_3276083, EPI_ISL_3276084, EPI_ISL_3276085, EPI_ISL_3276086, EPI_ISL_3276087, EPI_ISL_3276088, EPI_ISL_3276089, EPI_ISL_3276090, EPI_ISL_3276091 | | | |
| see above | Dr S raju, Director of public heath and preventive medicine | inStem NCBS – INSACOG | Uma Ramakrishnan Dasaradhi Palakodeti Aswin SaiNarain |
| EPI_ISL_3276092 | Kodigehalli PHC | inStem NCBS – INSACOG | Uma Ramakrishnan Dasaradhi Palakodeti Aswin SaiNarain |
| EPI_ISL_3276093, EPI_ISL_3276094, EPI_ISL_3276095, EPI_ISL_3276096, EPI_ISL_3276097, EPI_ISL_3276098, EPI_ISL_3276099, EPI_ISL_3276100 | Dr S raju, Director of public heath and preventive medicine | inStem NCBS – INSACOG | Uma Ramakrishnan Dasaradhi Palakodeti Aswin SaiNarain |
| EPI_ISL_3276101 | Sahakara Nagar PHC | inStem NCBS – INSACOG | Uma Ramakrishnan Dasaradhi Palakodeti Aswin SaiNarain |
| EPI_ISL_3276102 | Dr S raju, Director of public heath and preventive medicine | inStem NCBS – INSACOG | Uma Ramakrishnan Dasaradhi Palakodeti Aswin SaiNarain |
| EPI_ISL_3276103 | Kodigehalli PHC | inStem NCBS – INSACOG | Uma Ramakrishnan Dasaradhi Palakodeti Aswin SaiNarain |
| EPI_ISL_3276104, EPI_ISL_3276105, EPI_ISL_3276106, EPI_ISL_3276107, EPI_ISL_3276108, EPI_ISL_3276109, EPI_ISL_3276110, EPI_ISL_3276111 | Dr S raju, Director of public heath and preventive medicine | inStem NCBS – INSACOG | Uma Ramakrishnan Dasaradhi Palakodeti Aswin SaiNarain |
| EPI_ISL_3276112 | Kodigehalli PHC | inStem NCBS – INSACOG | Uma Ramakrishnan Dasaradhi Palakodeti Aswin SaiNarain |
| EPI_ISL_3276113, EPI_ISL_3276114, EPI_ISL_3276115, EPI_ISL_3276116, EPI_ISL_3276117, EPI_ISL_3276118, EPI_ISL_3276119, EPI_ISL_3276120 | Dr S raju, Director of public heath and preventive medicine | inStem NCBS – INSACOG | Uma Ramakrishnan Dasaradhi Palakodeti Aswin SaiNarain |
| EPI_ISL_3276121, EPI_ISL_3276122, EPI_ISL_3276123 | Kodigehalli PHC | inStem NCBS – INSACOG | Uma Ramakrishnan Dasaradhi Palakodeti Aswin SaiNarain |
| EPI_ISL_3276124, EPI_ISL_3276125, EPI_ISL_3276126 | Dr S raju, Director of public heath and preventive medicine | inStem NCBS – INSACOG | Uma Ramakrishnan Dasaradhi Palakodeti Aswin SaiNarain |
| EPI_ISL_3276127 | Kodigehalli PHC | inStem NCBS – INSACOG | Uma Ramakrishnan Dasaradhi Palakodeti Aswin SaiNarain |
| EPI_ISL_3276128, EPI_ISL_3276130, EPI_ISL_3276131, EPI_ISL_3276132, EPI_ISL_3276133, EPI_ISL_3276134, EPI_ISL_3276135, EPI_ISL_3276136, EPI_ISL_3276137, EPI_ISL_3276138, EPI_ISL_3276139, EPI_ISL_3276140 | | | |
| see above | Dr S raju, Director of public heath and preventive medicine | inStem NCBS – INSACOG | Uma Ramakrishnan Dasaradhi Palakodeti Aswin SaiNarain |
| EPI_ISL_3276141 | Kodigehalli PHC | inStem NCBS – INSACOG | Uma Ramakrishnan Dasaradhi Palakodeti Aswin SaiNarain |
| EPI_ISL_3276142 | Dr S raju, Director of public heath and preventive medicine | inStem NCBS – INSACOG | Uma Ramakrishnan Dasaradhi Palakodeti Aswin SaiNarain |
| EPI_ISL_3276143 | Kodigehalli PHC | inStem NCBS – INSACOG | Uma Ramakrishnan Dasaradhi Palakodeti Aswin SaiNarain |
| EPI_ISL_3276144, EPI_ISL_3276145, EPI_ISL_3276146 | Dr S raju, Director of public heath and preventive medicine | inStem NCBS – INSACOG | Uma Ramakrishnan Dasaradhi Palakodeti Aswin SaiNarain |
| EPI_ISL_3276147 | Sahakara Nagar PHC | inStem NCBS – INSACOG | Uma Ramakrishnan Dasaradhi Palakodeti Aswin SaiNarain |
| EPI_ISL_3276148, EPI_ISL_3276149, EPI_ISL_3276150, EPI_ISL_3276151 | Dr S raju, Director of public heath and preventive medicine | inStem NCBS – INSACOG | Uma Ramakrishnan Dasaradhi Palakodeti Aswin SaiNarain |
| EPI_ISL_3276152 | Sahakara Nagar PHC | inStem NCBS – INSACOG | Uma Ramakrishnan Dasaradhi Palakodeti Aswin SaiNarain |
| EPI_ISL_3276153, EPI_ISL_3276154 | Dr S raju, Director of public heath and preventive medicine | inStem NCBS – INSACOG | Uma Ramakrishnan Dasaradhi Palakodeti Aswin SaiNarain |
| EPI_ISL_3276155 | Kodigehalli PHC | inStem NCBS – INSACOG | Uma Ramakrishnan Dasaradhi Palakodeti Aswin SaiNarain |
| EPI_ISL_3276156, EPI_ISL_3276173, EPI_ISL_3276174, EPI_ISL_3276175, EPI_ISL_3276176 | Dr S raju, Director of public heath and preventive medicine | inStem NCBS – INSACOG | Uma Ramakrishnan Dasaradhi Palakodeti Aswin SaiNarain |
| EPI_ISL_3276177, EPI_ISL_3276178, EPI_ISL_3276179 | Kodigehalli PHC | inStem NCBS – INSACOG | Uma Ramakrishnan Dasaradhi Palakodeti Aswin SaiNarain |
| EPI_ISL_3276180 | Dr S raju, Director of public heath and preventive medicine | inStem NCBS – INSACOG | Uma Ramakrishnan Dasaradhi Palakodeti Aswin SaiNarain |
| EPI_ISL_3276181 | Kodigehalli PHC | inStem NCBS – INSACOG | Uma Ramakrishnan Dasaradhi Palakodeti Aswin SaiNarain |
| EPI_ISL_3276182, EPI_ISL_3276183, EPI_ISL_3276184 | Dr S raju, Director of public heath and preventive medicine | inStem NCBS – INSACOG | Uma Ramakrishnan Dasaradhi Palakodeti Aswin SaiNarain |
| EPI_ISL_3276185 | Kodigehalli PHC | inStem NCBS – INSACOG | Uma Ramakrishnan Dasaradhi Palakodeti Aswin SaiNarain |
| EPI_ISL_3276186 | Dr S raju, Director of public heath and preventive medicine | inStem NCBS – INSACOG | Uma Ramakrishnan Dasaradhi Palakodeti Aswin SaiNarain |
| EPI_ISL_3276187 | Kodigehalli PHC | inStem NCBS – INSACOG | Uma Ramakrishnan Dasaradhi Palakodeti Aswin SaiNarain |
| EPI_ISL_3276188, EPI_ISL_3276189, EPI_ISL_3276190, EPI_ISL_3276191, EPI_ISL_3276192, EPI_ISL_3276193, EPI_ISL_3276194, EPI_ISL_3276195, EPI_ISL_3276196, EPI_ISL_3276197, EPI_ISL_3276198, EPI_ISL_3276199, EPI_ISL_3276200, EPI_ISL_3276201 | | | |
| see above | Dr S raju, Director of public heath and preventive medicine | inStem NCBS – INSACOG | Uma Ramakrishnan Dasaradhi Palakodeti Aswin SaiNarain |
| EPI_ISL_3276202, EPI_ISL_3276203 | Kodigehalli PHC | inStem NCBS – INSACOG | Uma Ramakrishnan Dasaradhi Palakodeti Aswin SaiNarain |
| EPI_ISL_3276204, EPI_ISL_3276205, EPI_ISL_3276206, EPI_ISL_3276207, EPI_ISL_3276208, EPI_ISL_3276209 | Dr S raju, Director of public heath and preventive medicine | inStem NCBS – INSACOG | Uma Ramakrishnan Dasaradhi Palakodeti Aswin SaiNarain |

| | | | |
|---|---|---|---|
| EPI_ISL_3276210 | Kodigehalli PHC | inStem NCBS – INSACOG | Uma Ramakrishnan Dasaradhi Palakodeti Aswin SaiNarain |
| EPI_ISL_3276211, EPI_ISL_3276212 | Dr S raju, Director of public heath and preventive medicine | inStem NCBS – INSACOG | Uma Ramakrishnan Dasaradhi Palakodeti Aswin SaiNarain |
| EPI_ISL_3276213 | Kodigehalli PHC | inStem NCBS – INSACOG | Uma Ramakrishnan Dasaradhi Palakodeti Aswin SaiNarain |
| EPI_ISL_3276214, EPI_ISL_3276215, EPI_ISL_3276216 | Dr S raju, Director of public heath and preventive medicine | inStem NCBS – INSACOG | Uma Ramakrishnan Dasaradhi Palakodeti Aswin SaiNarain |
| EPI_ISL_3276217 | Kodigehalli PHC | inStem NCBS – INSACOG | Uma Ramakrishnan Dasaradhi Palakodeti Aswin SaiNarain |
| EPI_ISL_3276218, EPI_ISL_3276219, EPI_ISL_3276220 | Dr S raju, Director of public heath and preventive medicine | inStem NCBS – INSACOG | Uma Ramakrishnan Dasaradhi Palakodeti Aswin SaiNarain |
| EPI_ISL_3276221 | Kodigehalli PHC | inStem NCBS – INSACOG | Uma Ramakrishnan Dasaradhi Palakodeti Aswin SaiNarain |
| EPI_ISL_3276222, EPI_ISL_3276223, EPI_ISL_3276224, EPI_ISL_3276225, EPI_ISL_3276226, EPI_ISL_3276227, EPI_ISL_3276228, EPI_ISL_3276229, EPI_ISL_3276230, EPI_ISL_3276231, EPI_ISL_3276232, EPI_ISL_3276233, EPI_ISL_3276234, EPI_ISL_3276235, EPI_ISL_3276236 | | | |
| see above | Dr S raju, Director of public heath and preventive medicine | inStem NCBS – INSACOG | Uma Ramakrishnan Dasaradhi Palakodeti Aswin SaiNarain |
| EPI_ISL_3276237 | Kodigehalli PHC | inStem NCBS – INSACOG | Uma Ramakrishnan Dasaradhi Palakodeti Aswin SaiNarain |
| EPI_ISL_3276238 | Sahakara Nagar PHC | inStem NCBS – INSACOG | Uma Ramakrishnan Dasaradhi Palakodeti Aswin SaiNarain |
| EPI_ISL_3276239 | Kodigehalli PHC | inStem NCBS – INSACOG | Uma Ramakrishnan Dasaradhi Palakodeti Aswin SaiNarain |
| EPI_ISL_3276240, EPI_ISL_3276241, EPI_ISL_3276242, EPI_ISL_3276243, EPI_ISL_3276244, EPI_ISL_3276245, EPI_ISL_3276246, EPI_ISL_3276247, EPI_ISL_3276248, EPI_ISL_3276249, EPI_ISL_3276250, EPI_ISL_3276251, EPI_ISL_3276252, EPI_ISL_3276253, EPI_ISL_3276254, EPI_ISL_3276255, EPI_ISL_3276256, EPI_ISL_3276257, EPI_ISL_3276258, EPI_ISL_3276259, EPI_ISL_3276260, EPI_ISL_3276261, EPI_ISL_3276262, EPI_ISL_3276263, EPI_ISL_3276264, EPI_ISL_3276265, EPI_ISL_3276266, EPI_ISL_3276267, EPI_ISL_3276268 | | | |
| see above | Dr S raju, Director of public heath and preventive medicine | inStem NCBS – INSACOG | Uma Ramakrishnan Dasaradhi Palakodeti Aswin SaiNarain |
| EPI_ISL_3276269, EPI_ISL_3276270, EPI_ISL_3276271, EPI_ISL_3276272, EPI_ISL_3276273, EPI_ISL_3276274, EPI_ISL_3276275, EPI_ISL_3276276, EPI_ISL_3276277, EPI_ISL_3276278 | Kodigehalli PHC | inStem NCBS – INSACOG | Uma Ramakrishnan Dasaradhi Palakodeti Aswin SaiNarain |
| EPI_ISL_3276279, EPI_ISL_3276280, EPI_ISL_3276281, EPI_ISL_3276282, EPI_ISL_3276283, EPI_ISL_3276284, EPI_ISL_3276285, EPI_ISL_3276286, EPI_ISL_3276287, EPI_ISL_3276288, EPI_ISL_3276289, EPI_ISL_3276290, EPI_ISL_3276291, EPI_ISL_3276292, EPI_ISL_3276293 | | | |
| see above | Sahakara Nagar PHC | inStem NCBS – INSACOG | Uma Ramakrishnan Dasaradhi Palakodeti Aswin SaiNarain |
| EPI_ISL_3276294, EPI_ISL_3276295, EPI_ISL_3276296, EPI_ISL_3276297, EPI_ISL_3276298, EPI_ISL_3276299, EPI_ISL_3276300, EPI_ISL_3276301, EPI_ISL_3276302, EPI_ISL_3276303, EPI_ISL_3276304, EPI_ISL_3276305, EPI_ISL_3276306, EPI_ISL_3276307, EPI_ISL_3276308, EPI_ISL_3276309, EPI_ISL_3276310, EPI_ISL_3276311, EPI_ISL_3276312, EPI_ISL_3276313 | | | |
| see above | Kodigehalli PHC | inStem NCBS – INSACOG | Uma Ramakrishnan Dasaradhi Palakodeti Aswin SaiNarain |
| EPI_ISL_3276314, EPI_ISL_3276315 | Sahakara Nagar PHC | inStem NCBS – INSACOG | Uma Ramakrishnan Dasaradhi Palakodeti Aswin SaiNarain |
| EPI_ISL_3276316, EPI_ISL_3276317, EPI_ISL_3276318, EPI_ISL_3276319, EPI_ISL_3276320, EPI_ISL_3276321, EPI_ISL_3276322, EPI_ISL_3276323, EPI_ISL_3276324, EPI_ISL_3276325 | Kodigehalli PHC | inStem NCBS – INSACOG | Uma Ramakrishnan Dasaradhi Palakodeti Aswin SaiNarain |
| EPI_ISL_3276347, EPI_ISL_3276348, EPI_ISL_3276349, EPI_ISL_3276350, EPI_ISL_3276351, EPI_ISL_3276352, EPI_ISL_3276353, EPI_ISL_3276354, EPI_ISL_3276355, EPI_ISL_3276356, EPI_ISL_3276357, EPI_ISL_3276358, EPI_ISL_3276359, EPI_ISL_3276360, EPI_ISL_3276362, EPI_ISL_3276363, EPI_ISL_3276364, EPI_ISL_3276365, EPI_ISL_3276366, EPI_ISL_3276367, EPI_ISL_3276368, EPI_ISL_3276369, EPI_ISL_3276370, EPI_ISL_3276371, EPI_ISL_3276372, EPI_ISL_3276373, EPI_ISL_3276374, EPI_ISL_3276375, EPI_ISL_3276376, EPI_ISL_3276377, EPI_ISL_3276378, EPI_ISL_3276379, EPI_ISL_3276380, EPI_ISL_3276381, EPI_ISL_3276383, EPI_ISL_3276384, EPI_ISL_3276385, EPI_ISL_3276386, EPI_ISL_3276387, EPI_ISL_3276388, EPI_ISL_3276389, EPI_ISL_3276390, EPI_ISL_3276391, EPI_ISL_3276392, EPI_ISL_3276393, EPI_ISL_3276394, EPI_ISL_3276395, EPI_ISL_3276396, EPI_ISL_3276397, EPI_ISL_3276399, EPI_ISL_3276400, EPI_ISL_3276402, EPI_ISL_3276403, EPI_ISL_3276404, EPI_ISL_3276405, EPI_ISL_3276406, EPI_ISL_3276407, EPI_ISL_3276408, EPI_ISL_3276409, EPI_ISL_3276410, EPI_ISL_3276411, EPI_ISL_3276412, EPI_ISL_3276413, EPI_ISL_3276415, EPI_ISL_3276416, EPI_ISL_3276417, EPI_ISL_3276418, EPI_ISL_3276419, EPI_ISL_3276420, EPI_ISL_3276421, EPI_ISL_3276422, EPI_ISL_3276423, EPI_ISL_3276424, EPI_ISL_3276425, EPI_ISL_3276426, EPI_ISL_3276427, EPI_ISL_3276428, EPI_ISL_3276429, EPI_ISL_3276433, EPI_ISL_3276434, EPI_ISL_3276435, EPI_ISL_3276436, EPI_ISL_3276437, EPI_ISL_3276438, EPI_ISL_3276439, EPI_ISL_3276441, EPI_ISL_3276442, EPI_ISL_3276443, EPI_ISL_3276444, EPI_ISL_3276445, EPI_ISL_3276446, EPI_ISL_3276447, EPI_ISL_3276448, EPI_ISL_3276449, EPI_ISL_3276450, EPI_ISL_3276451, EPI_ISL_3276452, EPI_ISL_3276453, EPI_ISL_3276454, EPI_ISL_3276455, EPI_ISL_3276456, EPI_ISL_3276457, EPI_ISL_3276458, EPI_ISL_3276459, EPI_ISL_3276460, EPI_ISL_3276461, EPI_ISL_3276462, EPI_ISL_3276463, EPI_ISL_3276464, EPI_ISL_3276465, EPI_ISL_3276466, EPI_ISL_3276467, EPI_ISL_3276468, EPI_ISL_3276469, EPI_ISL_3276470, EPI_ISL_3276471, EPI_ISL_3276472, EPI_ISL_3276473, EPI_ISL_3276474, EPI_ISL_3276487 | | | |
| see above | Lab. Microbiologia e Virologia Cotugno A.O. dei Colli | TIGEM | Antonio Grimaldi Patrizia Annunziata Francesco Panariello Biancamaria Pierri Claudia Tiberio Teresa Giuliano Valentina Bouche Chiara Colantuono Maria Concetta Cuomo Denise Di Concilio Lucio Di Filippo Anna Manfredi Marcello Salvi Antonio Limone Luigi Atripaldi Pellegrino Cerino Andrea Ballabio Davide Cacchiarelli |
| EPI_ISL_3276654, EPI_ISL_3276657, EPI_ISL_3276658, EPI_ISL_3276659, EPI_ISL_3276660 | Saint Francis Medical Center | Los Angeles County Public Health Laboratories | P. Hemarajata et al. |
| EPI_ISL_3276662, EPI_ISL_3276663, EPI_ISL_3276664, EPI_ISL_3276665, EPI_ISL_3276666, EPI_ISL_3276667, EPI_ISL_3276668, EPI_ISL_3276669, EPI_ISL_3276670 | USC Clinical Lab | Los Angeles County Public Health Laboratories | P. Hemarajata et al. |
| EPI_ISL_3276672, EPI_ISL_3276673 | Torrance Memorial Medical Center | Los Angeles County Public Health Laboratories | P. Hemarajata et al. |
| EPI_ISL_3276674 | Quest Diagnostics West Hills Medical Center | Los Angeles County Public Health Laboratories | P. Hemarajata et al. |
| EPI_ISL_3276675, EPI_ISL_3276676, EPI_ISL_3276677, EPI_ISL_3276678 | Sun Clinical Laboratory | Los Angeles County Public Health Laboratories | P. Hemarajata et al. |
| EPI_ISL_3276681, EPI_ISL_3276683 | Pomona Valley Hospital Medical Center | Los Angeles County Public Health Laboratories | P. Hemarajata et al. |
| EPI_ISL_3276684 | TOPA Diagnostics | Los Angeles County Public Health Laboratories | P. Hemarajata et al. |
| EPI_ISL_3276687 | California Hospital Medical Center | Los Angeles County Public Health Laboratories | P. Hemarajata et al. |
| EPI_ISL_3276688 | Huntington Memorial Hospital | Los Angeles County Public Health Laboratories | P. Hemarajata et al. |
| EPI_ISL_3276690, EPI_ISL_3276691, EPI_ISL_3276694, EPI_ISL_3276695, EPI_ISL_3276696, EPI_ISL_3276698, EPI_ISL_3276699, EPI_ISL_3276700, EPI_ISL_3276701 | Torrance Memorial Medical Center | Los Angeles County Public Health Laboratories | P. Hemarajata et al. |
| EPI_ISL_3276703 | Pomona Valley Hospital Medical Center | Los Angeles County Public Health Laboratories | P. Hemarajata et al. |
| EPI_ISL_3276704, EPI_ISL_3276705 | California Hospital Medical Center | Los Angeles County Public Health Laboratories | P. Hemarajata et al. |
| EPI_ISL_3276706, EPI_ISL_3276707 | Los Angeles County Public Health Laboratories | Los Angeles County Public Health Laboratories | P. Hemarajata et al. |

| | | | |
|---|---|---|---|
| EPI_ISL_3276708, EPI_ISL_3276709, EPI_ISL_3276711, EPI_ISL_3276712, EPI_ISL_3276713, EPI_ISL_3276714, EPI_ISL_3276717 | Huntington Memorial Hospital | Los Angeles County Public Health Laboratories | P. Hemarajata et al. |
| EPI_ISL_3276718, EPI_ISL_3276719, EPI_ISL_3276720, EPI_ISL_3276721, EPI_ISL_3276722, EPI_ISL_3276723 | Los Angeles County Public Health Laboratories | Los Angeles County Public Health Laboratories | P. Hemarajata et al. |
| EPI_ISL_3276724 | Torrance Memorial Medical Center | Los Angeles County Public Health Laboratories | P. Hemarajata et al. |
| EPI_ISL_3276725, EPI_ISL_3276726, EPI_ISL_3276727 | Los Angeles County Public Health Laboratories | Los Angeles County Public Health Laboratories | P. Hemarajata et al. |
| EPI_ISL_3276728 | PIH Health Whittier Hospital | Los Angeles County Public Health Laboratories | P. Hemarajata et al. |
| EPI_ISL_3276729 | Los Angeles County Public Health Laboratories | Los Angeles County Public Health Laboratories | P. Hemarajata et al. |
| EPI_ISL_3276730 | Quest Diagnostics West Hills Medical Center | Los Angeles County Public Health Laboratories | P. Hemarajata et al. |
| EPI_ISL_3276731, EPI_ISL_3276732 | Los Angeles County Public Health Laboratories | Los Angeles County Public Health Laboratories | P. Hemarajata et al. |
| EPI_ISL_3276733, EPI_ISL_3276735, EPI_ISL_3276736, EPI_ISL_3276738, EPI_ISL_3276739 | Pomona Valley Hospital Medical Center | Los Angeles County Public Health Laboratories | P. Hemarajata et al. |
| EPI_ISL_3276740, EPI_ISL_3276741, EPI_ISL_3276742, EPI_ISL_3276743, EPI_ISL_3276744 | Huntington Memorial Hospital | Los Angeles County Public Health Laboratories | P. Hemarajata et al. |
| EPI_ISL_3276746, EPI_ISL_3276748, EPI_ISL_3276750, EPI_ISL_3276751, EPI_ISL_3276752, EPI_ISL_3276753, EPI_ISL_3276754, EPI_ISL_3276755, EPI_ISL_3276756, EPI_ISL_3276757, EPI_ISL_3276758, EPI_ISL_3276759, EPI_ISL_3276760 | | | |
| see above | Torrance Memorial Medical Center | Los Angeles County Public Health Laboratories | P. Hemarajata et al. |
| EPI_ISL_3276761, EPI_ISL_3276762, EPI_ISL_3276764, EPI_ISL_3276765, EPI_ISL_3276766, EPI_ISL_3276767, EPI_ISL_3276768 | Centinela Hospital Medical Center | Los Angeles County Public Health Laboratories | P. Hemarajata et al. |
| EPI_ISL_3276770, EPI_ISL_3276771, EPI_ISL_3276772, EPI_ISL_3276773, EPI_ISL_3276774, EPI_ISL_3276775, EPI_ISL_3276776, EPI_ISL_3276777, EPI_ISL_3276778 | USC Clinical Lab | Los Angeles County Public Health Laboratories | P. Hemarajata et al. |
| EPI_ISL_3276779 | Martin Luther King, Jr. Outpatient Center | Los Angeles County Public Health Laboratories | P. Hemarajata et al. |
| EPI_ISL_3276812, EPI_ISL_3276813 | Providence Saint John's Health Center | Los Angeles County Public Health Laboratories | P. Hemarajata et al. |
| EPI_ISL_3276814, EPI_ISL_3276816, EPI_ISL_3276817, EPI_ISL_3276818, EPI_ISL_3276819, EPI_ISL_3276820, EPI_ISL_3276821, EPI_ISL_3276822, EPI_ISL_3276823, EPI_ISL_3276824, EPI_ISL_3276825, EPI_ISL_3276827 | | | |
| see above | Martin Luther King Jr. Community Hospital | Los Angeles County Public Health Laboratories | P. Hemarajata et al. |
| EPI_ISL_3276828, EPI_ISL_3276829, EPI_ISL_3276830, EPI_ISL_3276832, EPI_ISL_3276833 | Palmdale Regional Medical Center | Los Angeles County Public Health Laboratories | P. Hemarajata et al. |
| EPI_ISL_3276835, EPI_ISL_3276837, EPI_ISL_3276838 | Martin Luther King, Jr. Outpatient Center | Los Angeles County Public Health Laboratories | P. Hemarajata et al. |
| EPI_ISL_3276839 | MLK COMMUNITY HOSPITAL | Los Angeles County Public Health Laboratories | P. Hemarajata et al. |
| EPI_ISL_3276840 | Martin Luther King, Jr. Outpatient Center | Los Angeles County Public Health Laboratories | P. Hemarajata et al. |
| EPI_ISL_3276875 | Clinical Virology Laboratory, Institute of Liver and Biliary Sciences | ILBS on behalf of 'Proyecto Argentino Interinstitucional de genomica de SARS-CoV-2' (PAIS Consortium) | Ekta Gupta, Varun Suroliya, Diptanu Paul, Amit Pandey, Reshu Agarwal, Chhagan Bihari Sharma, Shiv Kumar Sarin |
| EPI_ISL_3276964, EPI_ISL_3276972, EPI_ISL_3276984, EPI_ISL_3276988, EPI_ISL_3276995, EPI_ISL_3276998, EPI_ISL_3277001, EPI_ISL_3277035, EPI_ISL_3277036, EPI_ISL_3277037, EPI_ISL_3277039, EPI_ISL_3277041, EPI_ISL_3277042, EPI_ISL_3277043, EPI_ISL_3277045, EPI_ISL_3277046, EPI_ISL_3277047, EPI_ISL_3277048, EPI_ISL_3277049, EPI_ISL_3277050, EPI_ISL_3277051, EPI_ISL_3277053, EPI_ISL_3277054, EPI_ISL_3277055, EPI_ISL_3277056, EPI_ISL_3277057, EPI_ISL_3277058, EPI_ISL_3277059, EPI_ISL_3277060, EPI_ISL_3277063, EPI_ISL_3277064, EPI_ISL_3277065, EPI_ISL_3277066, EPI_ISL_3277067, EPI_ISL_3277068, EPI_ISL_3277070, EPI_ISL_3277072, EPI_ISL_3277073, EPI_ISL_3277074, EPI_ISL_3277075, EPI_ISL_3277077, EPI_ISL_3277078, EPI_ISL_3277079, EPI_ISL_3277080, EPI_ISL_3277081, EPI_ISL_3277082, EPI_ISL_3277083, EPI_ISL_3277084, EPI_ISL_3277085, EPI_ISL_3277087, EPI_ISL_3277088, EPI_ISL_3277089, EPI_ISL_3277090, EPI_ISL_3277091, EPI_ISL_3277093, EPI_ISL_3277094, EPI_ISL_3277095, EPI_ISL_3277097, EPI_ISL_3277098, EPI_ISL_3277099, EPI_ISL_3277100, EPI_ISL_3277101, EPI_ISL_3277102, EPI_ISL_3277104, EPI_ISL_3277105, EPI_ISL_3277106, EPI_ISL_3277107, EPI_ISL_3277108, EPI_ISL_3277109, EPI_ISL_3277110, EPI_ISL_3277111, EPI_ISL_3277112, EPI_ISL_3277113, EPI_ISL_3277114, EPI_ISL_3277115, EPI_ISL_3277116, EPI_ISL_3277117, EPI_ISL_3277118, EPI_ISL_3277119, EPI_ISL_3277120, EPI_ISL_3277121, EPI_ISL_3277122, EPI_ISL_3277123, EPI_ISL_3277124, EPI_ISL_3277125, EPI_ISL_3277126, EPI_ISL_3277127, EPI_ISL_3277128, EPI_ISL_3277129, EPI_ISL_3277131, EPI_ISL_3277132, EPI_ISL_3277133, EPI_ISL_3277134, EPI_ISL_3277135, EPI_ISL_3277136, EPI_ISL_3277137, EPI_ISL_3277138, EPI_ISL_3277139 | | | |
| see above | Clinical Virology Laboratory, Institute of Liver and Biliary Sciences | ILBS | Ekta Gupta, Varun Suroliya, Diptanu Paul, Amit Pandey, Reshu Agarwal, Chhagan Bihari Sharma, Shiv Kumar Sarin |
| EPI_ISL_3277396 | Institute of Virology, Biomedical Research Center of the Slovak Academy of Sciences, Bratislava | Faculty of Mathematics, Physics and Informatics, Comenius University, Bratislava | Viktoria Cabanova, Kristina Borsova, Brona Brejova, Veronika Vanova, Sabina Fumacova Havlikova, Juraj Kopacek, Martina Lickova, Lubomira Lukacikova, Martina Nebohacova, Monika Slavikova, Tomas Vinar, Jozef Nosek, Boris Klempa |
| EPI_ISL_3277397, EPI_ISL_3277398 | Institute of Virology, Biomedical Research Center of the Slovak Academy of Sciences, Bratislava | Faculty of Mathematics, Physics and Informatics, Comenius University, Bratislava | Kristina Borsova, Viktoria Cabanova, Brona Brejova, Veronika Vanova, Sabina Fumacova Havlikova, Juraj Kopacek, Martina Lickova, Lubomira Lukacikova, Martina Nebohacova, Monika Slavikova, Tomas Vinar, Boris Klempa, Jozef Nosek |
| EPI_ISL_3277399 | Institute of Virology, Biomedical Research Center of the Slovak Academy of Sciences, Bratislava | Faculty of Mathematics, Physics and Informatics, Comenius University, Bratislava | Viktoria Cabanova, Kristina Borsova, Brona Brejova, Veronika Vanova, Sabina Fumacova Havlikova, Juraj Kopacek, Martina Lickova, Lubomira Lukacikova, Martina Nebohacova, Monika Slavikova, Tomas Vinar, Jozef Nosek, Boris Klempa |
| EPI_ISL_3277400 | Institute of Virology, Biomedical Research Center of the Slovak Academy of Sciences, Bratislava | Faculty of Mathematics, Physics and Informatics, Comenius University, Bratislava | Brona Brejova, Viktoria Cabanova, Kristina Borsova, Veronika Vanova, Sabina Fumacova Havlikova, Juraj Kopacek, Martina Lickova, Lubomira Lukacikova, Martina Nebohacova, Monika Slavikova, Tomas Vinar, Jozef Nosek, Boris Klempa |
| EPI_ISL_3277401 | Institute of Virology, Biomedical Research Center of the Slovak Academy of Sciences, Bratislava | Faculty of Mathematics, Physics and Informatics, Comenius University, Bratislava | Kristina Borsova, Viktoria Cabanova, Brona Brejova, Veronika Vanova, Sabina Fumacova Havlikova, Juraj Kopacek, Martina Lickova, Lubomira Lukacikova, Martina Nebohacova, Monika Slavikova, Tomas Vinar, Boris Klempa, Jozef Nosek |
| EPI_ISL_3277402, EPI_ISL_3277403, EPI_ISL_3277404, EPI_ISL_3277405 | Institute of Virology, Biomedical Research Center of the Slovak Academy of Sciences, Bratislava | Faculty of Mathematics, Physics and Informatics, Comenius University, Bratislava | Viktoria Cabanova, Kristina Borsova, Brona Brejova, Veronika Vanova, Sabina Fumacova Havlikova, Juraj Kopacek, Martina Lickova, Lubomira Lukacikova, Martina Nebohacova, Monika Slavikova, Tomas Vinar, Jozef Nosek, Boris Klempa |
| EPI_ISL_3277406 | Institute of Virology, Biomedical Research Center of the Slovak Academy of Sciences, Bratislava | Faculty of Mathematics, Physics and Informatics, Comenius University, Bratislava | Kristina Borsova, Viktoria Cabanova, Brona Brejova, Veronika Vanova, Sabina Fumacova Havlikova, Juraj Kopacek, Martina Lickova, Lubomira Lukacikova, Martina Nebohacova, Monika Slavikova, Tomas Vinar, Boris Klempa, Jozef Nosek |
| EPI_ISL_3277407 | Institute of Virology, Biomedical Research Center of the Slovak Academy of Sciences, Bratislava | Faculty of Mathematics, Physics and Informatics, Comenius University, Bratislava | Brona Brejova, Viktoria Cabanova, Kristina Borsova, Veronika Vanova, Sabina Fumacova Havlikova, Juraj Kopacek, Martina Lickova, Lubomira Lukacikova, Martina Nebohacova, Monika Slavikova, Tomas Vinar, Jozef Nosek, Boris Klempa |
| EPI_ISL_3277409, EPI_ISL_3277410 | Institute of Virology, Biomedical Research Center of the Slovak Academy of Sciences, Bratislava | Faculty of Mathematics, Physics and Informatics, Comenius University, Bratislava | Viktoria Cabanova, Kristina Borsova, Brona Brejova, Veronika Vanova, Sabina Fumacova Havlikova, Juraj Kopacek, Martina Lickova, Lubomira Lukacikova, Martina Nebohacova, Monika Slavikova, Tomas Vinar, Jozef Nosek, Boris Klempa |
| EPI_ISL_3277411 | Institute of Virology, Biomedical Research Center of the Slovak | Faculty of Mathematics, Physics and Informatics, Comenius | Brona Brejova, Viktoria Cabanova, Kristina Borsova, Veronika Vanova, Sabina Fumacova Havlikova, Juraj Kopacek, Martina Lickova, Lubomira Lukacikova, Martina Nebohacova, Monika Slavikova, Tomas Vinar, Jozef Nosek, Boris Klempa |

| | | | |
|---|---|---|---|
| | Academy of Sciences, Bratislava | University, Bratislava | |
| EPI_ISL_3277412 | Institute of Virology, Biomedical Research Center of the Slovak Academy of Sciences, Bratislava | Faculty of Mathematics, Physics and Informatics, Comenius University, Bratislava | Kristina Borsova, Viktoria Cabanova, Brona Brejova, Veronika Vanova, Sabina Fumacova Havlikova, Juraj Kopacek, Martina Lickova, Lubomira Lukacikova, Martina Nebohacova, Monika Slavikova, Tomas Vinar, Boris Klempa, Jozef Nosek |
| EPI_ISL_3277413 | Institute of Virology, Biomedical Research Center of the Slovak Academy of Sciences, Bratislava | Faculty of Mathematics, Physics and Informatics, Comenius University, Bratislava | Brona Brejova, Viktoria Cabanova, Kristina Borsova, Veronika Vanova, Sabina Fumacova Havlikova, Juraj Kopacek, Martina Lickova, Lubomira Lukacikova, Martina Nebohacova, Monika Slavikova, Tomas Vinar, Jozef Nosek, Boris Klempa |
| EPI_ISL_3277414 | Institute of Virology, Biomedical Research Center of the Slovak Academy of Sciences, Bratislava | Faculty of Mathematics, Physics and Informatics, Comenius University, Bratislava | Viktoria Cabanova, Kristina Borsova, Brona Brejova, Veronika Vanova, Sabina Fumacova Havlikova, Juraj Kopacek, Martina Lickova, Lubomira Lukacikova, Martina Nebohacova, Monika Slavikova, Tomas Vinar, Jozef Nosek, Boris Klempa |
| EPI_ISL_3277415 | Institute of Virology, Biomedical Research Center of the Slovak Academy of Sciences, Bratislava | Faculty of Mathematics, Physics and Informatics, Comenius University, Bratislava | Kristina Borsova, Viktoria Cabanova, Brona Brejova, Veronika Vanova, Sabina Fumacova Havlikova, Juraj Kopacek, Martina Lickova, Lubomira Lukacikova, Martina Nebohacova, Monika Slavikova, Tomas Vinar, Boris Klempa, Jozef Nosek |
| EPI_ISL_3277416 | Institute of Virology, Biomedical Research Center of the Slovak Academy of Sciences, Bratislava | Faculty of Mathematics, Physics and Informatics, Comenius University, Bratislava | Viktoria Cabanova, Kristina Borsova, Brona Brejova, Veronika Vanova, Sabina Fumacova Havlikova, Juraj Kopacek, Martina Lickova, Lubomira Lukacikova, Martina Nebohacova, Monika Slavikova, Tomas Vinar, Jozef Nosek, Boris Klempa |
| EPI_ISL_3277417 | Institute of Virology, Biomedical Research Center of the Slovak Academy of Sciences, Bratislava | Faculty of Mathematics, Physics and Informatics, Comenius University, Bratislava | Kristina Borsova, Viktoria Cabanova, Brona Brejova, Veronika Vanova, Sabina Fumacova Havlikova, Juraj Kopacek, Martina Lickova, Lubomira Lukacikova, Martina Nebohacova, Monika Slavikova, Tomas Vinar, Boris Klempa, Jozef Nosek |
| EPI_ISL_3277418 | Institute of Virology, Biomedical Research Center of the Slovak Academy of Sciences, Bratislava | Faculty of Mathematics, Physics and Informatics, Comenius University, Bratislava | Viktoria Cabanova, Kristina Borsova, Brona Brejova, Veronika Vanova, Sabina Fumacova Havlikova, Juraj Kopacek, Martina Lickova, Lubomira Lukacikova, Martina Nebohacova, Monika Slavikova, Tomas Vinar, Jozef Nosek, Boris Klempa |
| EPI_ISL_3277419 | Institute of Virology, Biomedical Research Center of the Slovak Academy of Sciences, Bratislava | Faculty of Mathematics, Physics and Informatics, Comenius University, Bratislava | Brona Brejova, Viktoria Cabanova, Kristina Borsova, Veronika Vanova, Sabina Fumacova Havlikova, Juraj Kopacek, Martina Lickova, Lubomira Lukacikova, Martina Nebohacova, Monika Slavikova, Tomas Vinar, Jozef Nosek, Boris Klempa |
| EPI_ISL_3277420, EPI_ISL_3277421, EPI_ISL_3277422 | Institute of Virology, Biomedical Research Center of the Slovak Academy of Sciences, Bratislava | Faculty of Mathematics, Physics and Informatics, Comenius University, Bratislava | Viktoria Cabanova, Kristina Borsova, Brona Brejova, Veronika Vanova, Sabina Fumacova Havlikova, Juraj Kopacek, Martina Lickova, Lubomira Lukacikova, Martina Nebohacova, Monika Slavikova, Tomas Vinar, Jozef Nosek, Boris Klempa |
| EPI_ISL_3277423, EPI_ISL_3277424 | Institute of Virology, Biomedical Research Center of the Slovak Academy of Sciences, Bratislava | Faculty of Mathematics, Physics and Informatics, Comenius University, Bratislava | Kristina Borsova, Viktoria Cabanova, Brona Brejova, Veronika Vanova, Sabina Fumacova Havlikova, Juraj Kopacek, Martina Lickova, Lubomira Lukacikova, Martina Nebohacova, Monika Slavikova, Tomas Vinar, Boris Klempa, Jozef Nosek |
| EPI_ISL_3277425, EPI_ISL_3277426 | Institute of Virology, Biomedical Research Center of the Slovak Academy of Sciences, Bratislava | Faculty of Mathematics, Physics and Informatics, Comenius University, Bratislava | Brona Brejova, Viktoria Cabanova, Kristina Borsova, Veronika Vanova, Sabina Fumacova Havlikova, Juraj Kopacek, Martina Lickova, Lubomira Lukacikova, Martina Nebohacova, Monika Slavikova, Tomas Vinar, Jozef Nosek, Boris Klempa |
| EPI_ISL_3277427, EPI_ISL_3277428, EPI_ISL_3277429 | Institute of Virology, Biomedical Research Center of the Slovak Academy of Sciences, Bratislava | Faculty of Mathematics, Physics and Informatics, Comenius University, Bratislava | Viktoria Cabanova, Kristina Borsova, Brona Brejova, Veronika Vanova, Sabina Fumacova Havlikova, Juraj Kopacek, Martina Lickova, Lubomira Lukacikova, Martina Nebohacova, Monika Slavikova, Tomas Vinar, Jozef Nosek, Boris Klempa |
| EPI_ISL_3277430 | Institute of Virology, Biomedical Research Center of the Slovak Academy of Sciences, Bratislava | Faculty of Mathematics, Physics and Informatics, Comenius University, Bratislava | Kristina Borsova, Viktoria Cabanova, Brona Brejova, Veronika Vanova, Sabina Fumacova Havlikova, Juraj Kopacek, Martina Lickova, Lubomira Lukacikova, Martina Nebohacova, Monika Slavikova, Tomas Vinar, Boris Klempa, Jozef Nosek |
| EPI_ISL_3277431, EPI_ISL_3277432 | Institute of Virology, Biomedical Research Center of the Slovak Academy of Sciences, Bratislava | Faculty of Mathematics, Physics and Informatics, Comenius University, Bratislava | Brona Brejova, Viktoria Cabanova, Kristina Borsova, Veronika Vanova, Sabina Fumacova Havlikova, Juraj Kopacek, Martina Lickova, Lubomira Lukacikova, Martina Nebohacova, Monika Slavikova, Tomas Vinar, Jozef Nosek, Boris Klempa |
| EPI_ISL_3277433 | Institute of Virology, Biomedical Research Center of the Slovak Academy of Sciences, Bratislava | Faculty of Mathematics, Physics and Informatics, Comenius University, Bratislava | Kristina Borsova, Viktoria Cabanova, Brona Brejova, Veronika Vanova, Sabina Fumacova Havlikova, Juraj Kopacek, Martina Lickova, Lubomira Lukacikova, Martina Nebohacova, Monika Slavikova, Tomas Vinar, Boris Klempa, Jozef Nosek |
| EPI_ISL_3277434, EPI_ISL_3277435, EPI_ISL_3277436, EPI_ISL_3277437, EPI_ISL_3277438, EPI_ISL_3277439, EPI_ISL_3277440 | Institute of Virology, Biomedical Research Center of the Slovak Academy of Sciences, Bratislava | Faculty of Mathematics, Physics and Informatics, Comenius University, Bratislava | Viktoria Cabanova, Kristina Borsova, Brona Brejova, Veronika Vanova, Sabina Fumacova Havlikova, Juraj Kopacek, Martina Lickova, Lubomira Lukacikova, Martina Nebohacova, Monika Slavikova, Tomas Vinar, Jozef Nosek, Boris Klempa |
| EPI_ISL_3277441 | Institute of Virology, Biomedical Research Center of the Slovak Academy of Sciences, Bratislava | Faculty of Mathematics, Physics and Informatics, Comenius University, Bratislava | Kristina Borsova, Viktoria Cabanova, Brona Brejova, Veronika Vanova, Sabina Fumacova Havlikova, Juraj Kopacek, Martina Lickova, Lubomira Lukacikova, Martina Nebohacova, Monika Slavikova, Tomas Vinar, Boris Klempa, Jozef Nosek |
| EPI_ISL_3277442 | Institute of Virology, Biomedical Research Center of the Slovak Academy of Sciences, Bratislava | Faculty of Mathematics, Physics and Informatics, Comenius University, Bratislava | Brona Brejova, Viktoria Cabanova, Kristina Borsova, Veronika Vanova, Sabina Fumacova Havlikova, Juraj Kopacek, Martina Lickova, Lubomira Lukacikova, Martina Nebohacova, Monika Slavikova, Tomas Vinar, Jozef Nosek, Boris Klempa |
| EPI_ISL_3277443 | Institute of Virology, Biomedical Research Center of the Slovak Academy of Sciences, Bratislava | Faculty of Mathematics, Physics and Informatics, Comenius University, Bratislava | Kristina Borsova, Viktoria Cabanova, Brona Brejova, Veronika Vanova, Sabina Fumacova Havlikova, Juraj Kopacek, Martina Lickova, Lubomira Lukacikova, Martina Nebohacova, Monika Slavikova, Tomas Vinar, Boris Klempa, Jozef Nosek |
| EPI_ISL_3277444 | Institute of Virology, Biomedical Research Center of the Slovak Academy of Sciences, Bratislava | Faculty of Mathematics, Physics and Informatics, Comenius University, Bratislava | Brona Brejova, Viktoria Cabanova, Kristina Borsova, Veronika Vanova, Sabina Fumacova Havlikova, Juraj Kopacek, Martina Lickova, Lubomira Lukacikova, Martina Nebohacova, Monika Slavikova, Tomas Vinar, Jozef Nosek, Boris Klempa |
| EPI_ISL_3277445 | Institute of Virology, Biomedical Research Center of the Slovak Academy of Sciences, Bratislava | Faculty of Mathematics, Physics and Informatics, Comenius University, Bratislava | Viktoria Cabanova, Kristina Borsova, Brona Brejova, Veronika Vanova, Sabina Fumacova Havlikova, Juraj Kopacek, Martina Lickova, Lubomira Lukacikova, Martina Nebohacova, Monika Slavikova, Tomas Vinar, Jozef Nosek, Boris Klempa |
| EPI_ISL_3277447, EPI_ISL_3277448, EPI_ISL_3277449 | Institute of Virology, Biomedical Research Center of the Slovak Academy of Sciences, Bratislava | Faculty of Mathematics, Physics and Informatics, Comenius University, Bratislava | Brona Brejova, Viktoria Cabanova, Kristina Borsova, Veronika Vanova, Sabina Fumacova Havlikova, Juraj Kopacek, Martina Lickova, Lubomira Lukacikova, Martina Nebohacova, Monika Slavikova, Tomas Vinar, Jozef Nosek, Boris Klempa |
| EPI_ISL_3277450 | Institute of Virology, Biomedical Research Center of the Slovak Academy of Sciences, Bratislava | Faculty of Mathematics, Physics and Informatics, Comenius University, Bratislava | Viktoria Cabanova, Kristina Borsova, Brona Brejova, Veronika Vanova, Sabina Fumacova Havlikova, Juraj Kopacek, Martina Lickova, Lubomira Lukacikova, Martina Nebohacova, Monika Slavikova, Tomas Vinar, Jozef Nosek, Boris Klempa |
| EPI_ISL_3277451, EPI_ISL_3277452 | Institute of Virology, Biomedical Research Center of the Slovak Academy of Sciences, Bratislava | Faculty of Mathematics, Physics and Informatics, Comenius University, Bratislava | Kristina Borsova, Viktoria Cabanova, Brona Brejova, Veronika Vanova, Sabina Fumacova Havlikova, Juraj Kopacek, Martina Lickova, Lubomira Lukacikova, Martina Nebohacova, Monika Slavikova, Tomas Vinar, Boris Klempa, Jozef Nosek |
| EPI_ISL_3277453, EPI_ISL_3277454 | Institute of Virology, Biomedical Research Center of the Slovak Academy of Sciences, Bratislava | Faculty of Mathematics, Physics and Informatics, Comenius University, Bratislava | Viktoria Cabanova, Kristina Borsova, Brona Brejova, Veronika Vanova, Sabina Fumacova Havlikova, Juraj Kopacek, Martina Lickova, Lubomira Lukacikova, Martina Nebohacova, Monika Slavikova, Tomas Vinar, Jozef Nosek, Boris Klempa |
| EPI_ISL_3277455 | Institute of Virology, Biomedical Research Center of the Slovak Academy of Sciences, Bratislava | Faculty of Mathematics, Physics and Informatics, Comenius University, Bratislava | Kristina Borsova, Viktoria Cabanova, Brona Brejova, Veronika Vanova, Sabina Fumacova Havlikova, Juraj Kopacek, Martina Lickova, Lubomira Lukacikova, Martina Nebohacova, Monika Slavikova, Tomas Vinar, Boris Klempa, Jozef Nosek |
| EPI_ISL_3277456 | Institute of Virology, Biomedical Research Center of the Slovak Academy of Sciences, Bratislava | Faculty of Mathematics, Physics and Informatics, Comenius University, Bratislava | Viktoria Cabanova, Kristina Borsova, Brona Brejova, Veronika Vanova, Sabina Fumacova Havlikova, Juraj Kopacek, Martina Lickova, Lubomira Lukacikova, Martina Nebohacova, Monika Slavikova, Tomas Vinar, Jozef Nosek, Boris Klempa |
| EPI_ISL_3277457 | Institute of Virology, Biomedical Research Center of the Slovak Academy of Sciences, Bratislava | Faculty of Mathematics, Physics and Informatics, Comenius University, Bratislava | Brona Brejova, Viktoria Cabanova, Kristina Borsova, Veronika Vanova, Sabina Fumacova Havlikova, Juraj Kopacek, Martina Lickova, Lubomira Lukacikova, Martina Nebohacova, Monika Slavikova, Tomas Vinar, Jozef Nosek, Boris Klempa |
| EPI_ISL_3277458 | Institute of Virology, Biomedical Research Center of the Slovak Academy of Sciences, Bratislava | Faculty of Mathematics, Physics and Informatics, Comenius University, Bratislava | Kristina Borsova, Viktoria Cabanova, Brona Brejova, Veronika Vanova, Sabina Fumacova Havlikova, Juraj Kopacek, Martina Lickova, Lubomira Lukacikova, Martina Nebohacova, Monika Slavikova, Tomas Vinar, Boris Klempa, Jozef Nosek |
| EPI_ISL_3277459 | Institute of Virology, Biomedical Research Center of the Slovak Academy of Sciences, Bratislava | Faculty of Mathematics, Physics and Informatics, Comenius University, Bratislava | Brona Brejova, Viktoria Cabanova, Kristina Borsova, Veronika Vanova, Sabina Fumacova Havlikova, Juraj Kopacek, Martina Lickova, Lubomira Lukacikova, Martina Nebohacova, Monika Slavikova, Tomas Vinar, Jozef Nosek, Boris Klempa |
| EPI_ISL_3277460 | Institute of Virology, Biomedical Research Center of the Slovak Academy of Sciences, Bratislava | Faculty of Mathematics, Physics and Informatics, Comenius University, Bratislava | Viktoria Cabanova, Kristina Borsova, Brona Brejova, Veronika Vanova, Sabina Fumacova Havlikova, Juraj Kopacek, Martina Lickova, Lubomira Lukacikova, Martina Nebohacova, Monika Slavikova, Tomas Vinar, Jozef Nosek, Boris Klempa |
| EPI_ISL_3277461 | Institute of Virology, Biomedical Research Center of the Slovak Academy of Sciences, Bratislava | Faculty of Mathematics, Physics and Informatics, Comenius University, Bratislava | Kristina Borsova, Viktoria Cabanova, Brona Brejova, Veronika Vanova, Sabina Fumacova Havlikova, Juraj Kopacek, Martina Lickova, Lubomira Lukacikova, Martina Nebohacova, Monika Slavikova, Tomas Vinar, Boris Klempa, Jozef Nosek |
| EPI_ISL_3277462 | Institute of Virology, Biomedical Research Center of the Slovak Academy of Sciences, Bratislava | Faculty of Mathematics, Physics and Informatics, Comenius University, Bratislava | Brona Brejova, Viktoria Cabanova, Kristina Borsova, Veronika Vanova, Sabina Fumacova Havlikova, Juraj Kopacek, Martina Lickova, Lubomira Lukacikova, Martina Nebohacova, Monika Slavikova, Tomas Vinar, Jozef Nosek, Boris Klempa |
| EPI_ISL_3277463 | Institute of Virology, Biomedical Research Center of the Slovak Academy of Sciences, Bratislava | Faculty of Mathematics, Physics and Informatics, Comenius University, Bratislava | Viktoria Cabanova, Kristina Borsova, Brona Brejova, Veronika Vanova, Sabina Fumacova Havlikova, Juraj Kopacek, Martina Lickova, Lubomira Lukacikova, Martina Nebohacova, Monika Slavikova, Tomas Vinar, Jozef Nosek, Boris Klempa |
| EPI_ISL_3277464, EPI_ISL_3277465 | Institute of Virology, Biomedical Research Center of the Slovak Academy of Sciences, Bratislava | Faculty of Mathematics, Physics and Informatics, Comenius University, Bratislava | Kristina Borsova, Viktoria Cabanova, Brona Brejova, Veronika Vanova, Sabina Fumacova Havlikova, Juraj Kopacek, Martina Lickova, Lubomira Lukacikova, Martina Nebohacova, Monika Slavikova, Tomas Vinar, Boris Klempa, Jozef Nosek |
| EPI_ISL_3277466 | Institute of Virology, Biomedical Research Center of the Slovak Academy of Sciences, Bratislava | Faculty of Mathematics, Physics and Informatics, Comenius University, Bratislava | Viktoria Cabanova, Kristina Borsova, Brona Brejova, Veronika Vanova, Sabina Fumacova Havlikova, Juraj Kopacek, Martina Lickova, Lubomira Lukacikova, Martina Nebohacova, Monika Slavikova, Tomas Vinar, Jozef Nosek, Boris Klempa |
| EPI_ISL_3277467, EPI_ISL_3277468 | Institute of Virology, Biomedical Research Center of the Slovak Academy of Sciences, Bratislava | Faculty of Mathematics, Physics and Informatics, Comenius University, Bratislava | Kristina Borsova, Viktoria Cabanova, Brona Brejova, Veronika Vanova, Sabina Fumacova Havlikova, Juraj Kopacek, Martina Lickova, Lubomira Lukacikova, Martina Nebohacova, Monika Slavikova, Tomas Vinar, Boris Klempa, Jozef Nosek |
| EPI_ISL_3277469, EPI_ISL_3277470 | Institute of Virology, Biomedical Research Center of the Slovak Academy of Sciences, Bratislava | Faculty of Mathematics, Physics and Informatics, Comenius University, Bratislava | Brona Brejova, Viktoria Cabanova, Kristina Borsova, Veronika Vanova, Sabina Fumacova Havlikova, Juraj Kopacek, Martina Lickova, Lubomira Lukacikova, Martina Nebohacova, Monika Slavikova, Tomas Vinar, Jozef Nosek, Boris Klempa |
| EPI_ISL_3277471 | Institute of Virology, Biomedical Research Center of the Slovak Academy of Sciences, Bratislava | Faculty of Mathematics, Physics and Informatics, Comenius University, Bratislava | Kristina Borsova, Viktoria Cabanova, Brona Brejova, Veronika Vanova, Sabina Fumacova Havlikova, Juraj Kopacek, Martina Lickova, Lubomira Lukacikova, Martina Nebohacova, Monika Slavikova, Tomas Vinar, Boris Klempa, Jozef Nosek |
| EPI_ISL_3277472, EPI_ISL_3277473, EPI_ISL_3277474 | Institute of Virology, Biomedical Research Center of the Slovak Academy of Sciences, Bratislava | Faculty of Mathematics, Physics and Informatics, Comenius University, Bratislava | Brona Brejova, Viktoria Cabanova, Kristina Borsova, Veronika Vanova, Sabina Fumacova Havlikova, Juraj Kopacek, Martina Lickova, Lubomira Lukacikova, Martina Nebohacova, Monika Slavikova, Tomas Vinar, Jozef Nosek, Boris Klempa |

| | | | |
|---|---|---|---|
| EPI_ISL_3277475 | Institute of Virology, Biomedical Research Center of the Slovak Academy of Sciences, Bratislava | Faculty of Mathematics, Physics and Informatics, Comenius University, Bratislava | Kristina Borsova, Viktoria Cabanova, Brona Brejova, Veronika Vanova, Sabina Fumacova Havlikova, Juraj Kopacek, Martina Lickova, Lubomira Lukacikova, Martina Nebohacova, Monika Slavikova, Tomas Vinar, Boris Klempa, Jozef Nosek |
| EPI_ISL_3277476 | Institute of Virology, Biomedical Research Center of the Slovak Academy of Sciences, Bratislava | Faculty of Mathematics, Physics and Informatics, Comenius University, Bratislava | Viktoria Cabanova, Kristina Borsova, Brona Brejova, Veronika Vanova, Sabina Fumacova Havlikova, Juraj Kopacek, Martina Lickova, Lubomira Lukacikova, Martina Nebohacova, Monika Slavikova, Tomas Vinar, Jozef Nosek, Boris Klempa |
| EPI_ISL_3277477 | Institute of Virology, Biomedical Research Center of the Slovak Academy of Sciences, Bratislava | Faculty of Mathematics, Physics and Informatics, Comenius University, Bratislava | Brona Brejova, Viktoria Cabanova, Kristina Borsova, Veronika Vanova, Sabina Fumacova Havlikova, Juraj Kopacek, Martina Lickova, Lubomira Lukacikova, Martina Nebohacova, Monika Slavikova, Tomas Vinar, Jozef Nosek, Boris Klempa |
| EPI_ISL_3277478 | Institute of Virology, Biomedical Research Center of the Slovak Academy of Sciences, Bratislava | Faculty of Mathematics, Physics and Informatics, Comenius University, Bratislava | Viktoria Cabanova, Kristina Borsova, Brona Brejova, Veronika Vanova, Sabina Fumacova Havlikova, Juraj Kopacek, Martina Lickova, Lubomira Lukacikova, Martina Nebohacova, Monika Slavikova, Tomas Vinar, Jozef Nosek, Boris Klempa |
| EPI_ISL_3277479 | Institute of Virology, Biomedical Research Center of the Slovak Academy of Sciences, Bratislava | Faculty of Mathematics, Physics and Informatics, Comenius University, Bratislava | Kristina Borsova, Viktoria Cabanova, Brona Brejova, Veronika Vanova, Sabina Fumacova Havlikova, Juraj Kopacek, Martina Lickova, Lubomira Lukacikova, Martina Nebohacova, Monika Slavikova, Tomas Vinar, Boris Klempa, Jozef Nosek |
| EPI_ISL_3277480, EPI_ISL_3277481 | Institute of Virology, Biomedical Research Center of the Slovak Academy of Sciences, Bratislava | Faculty of Mathematics, Physics and Informatics, Comenius University, Bratislava | Brona Brejova, Viktoria Cabanova, Kristina Borsova, Veronika Vanova, Sabina Fumacova Havlikova, Juraj Kopacek, Martina Lickova, Lubomira Lukacikova, Martina Nebohacova, Monika Slavikova, Tomas Vinar, Jozef Nosek, Boris Klempa |
| EPI_ISL_3277482 | Institute of Virology, Biomedical Research Center of the Slovak Academy of Sciences, Bratislava | Faculty of Mathematics, Physics and Informatics, Comenius University, Bratislava | Kristina Borsova, Viktoria Cabanova, Brona Brejova, Veronika Vanova, Sabina Fumacova Havlikova, Juraj Kopacek, Martina Lickova, Lubomira Lukacikova, Martina Nebohacova, Monika Slavikova, Tomas Vinar, Boris Klempa, Jozef Nosek |
| EPI_ISL_3277484 | CHC Andrée Rosemon | Institut Pasteur de la Guyane | Anne Lavergne, Dominique Rousset, Antoine Enfissi, Arielle Salmier |
| EPI_ISL_3277485, EPI_ISL_3277490, EPI_ISL_3277491, EPI_ISL_3277495 | CNR Institut Pasteur de la Guyane | Institut Pasteur de la Guyane | Anne Lavergne, Dominique Rousset, Antoine Enfissi, Arielle Salmier |
| EPI_ISL_3277497 | CHC Andrée Rosemon | Institut Pasteur de la Guyane | Anne Lavergne, Dominique Rousset, Antoine Enfissi, Arielle Salmier |
| EPI_ISL_3277502, EPI_ISL_3277503 | Laboratoire Carage | Institut Pasteur de la Guyane | Anne Lavergne, Dominique Rousset, Antoine Enfissi, Arielle Salmier |
| EPI_ISL_3277521 | BTKL PP Kelas I Batam | Indonesian Institute of Sciences (LIPI)/National Agency of Research and Innovation (BRIN) and National Institute of Health Research and Development (NIHRD) | Syam B. Iryanto, Masrukhin, Yuliawati, Listiana Oktavia, Indriawati, Subangkit, Hana Apsari Pawestri, Vivi Setiawaty, Anik B. Dharmayanthi, Anggia Prasetyoputri, Sugiyono Saputra, Ratih A. Ningrum |
| EPI_ISL_3277544, EPI_ISL_3277545, EPI_ISL_3277546, EPI_ISL_3277547, EPI_ISL_3277548, EPI_ISL_3277549, EPI_ISL_3277550, EPI_ISL_3277551, EPI_ISL_3277552, EPI_ISL_3277553, EPI_ISL_3277554, EPI_ISL_3277555, EPI_ISL_3277556, EPI_ISL_3277557 | | | |
| see above | Centro Medico ABC | Instituto Nacional de Medicina Genomica | Cedro-Tanda A, Escobar-Arrazola MA, Ramirez-Vega O, Roxana Trejo-González, Cesar Lara, Jacomine Reyes-Carrasco, Daniel Aguirre M, Rangel-DeLeon D, Munguia-Garza P, Hidalgo-Miranda A, Mendoza-Vargas A, Reyes-Grajeda JP, Herrera-Montalvo LA. |
| EPI_ISL_3277558 | Instituto Nacional de Medicina Genomica | Instituto Nacional de Medicina Genomica | Cedro-Tanda A, Escobar-Arrazola MA, Ramirez-Vega O, Rangel-DeLeon D, Munguia-Garza P, Hidalgo-Miranda A, Mendoza-Vargas A, Reyes-Grajeda JP, Herrera-Montalvo LA. |
| EPI_ISL_3277559 | Instituto Nacional de Cancerología | Instituto Nacional de Medicina Genomica | Cedro-Tanda A, Escobar-Arrazola MA, Ramirez-Vega O, Rangel-DeLeon D, Munguia-Garza P, Hidalgo-Miranda A, Mendoza-Vargas A, Reyes-Grajeda JP, Herrera-Montalvo LA. |
| EPI_ISL_3277560, EPI_ISL_3277561, EPI_ISL_3277562, EPI_ISL_3277563, EPI_ISL_3277564, EPI_ISL_3277565, EPI_ISL_3277566, EPI_ISL_3277567, EPI_ISL_3277568 | INCMNSZ | Instituto Nacional de Medicina Genomica | Cedro-Tanda A, Escobar-Arrazola MA, Ramirez-Vega O, Rangel-DeLeon D, Munguia-Garza P, Hidalgo-Miranda A, Mendoza-Vargas A, Reyes-Grajeda JP, Herrera-Montalvo LA. |
| EPI_ISL_3277569, EPI_ISL_3277570, EPI_ISL_3277571, EPI_ISL_3277572, EPI_ISL_3277573, EPI_ISL_3277574, EPI_ISL_3277575, EPI_ISL_3277576, EPI_ISL_3277578, EPI_ISL_3277579, EPI_ISL_3277580, EPI_ISL_3277581, EPI_ISL_3277582, EPI_ISL_3277583, EPI_ISL_3277584, EPI_ISL_3277585, EPI_ISL_3277586, EPI_ISL_3277587, EPI_ISL_3277588, EPI_ISL_3277589, EPI_ISL_3277590 | | | |
| see above | Instituto Nacional de Medicina Genomica | Instituto Nacional de Medicina Genomica | Cedro-Tanda A, Escobar-Arrazola MA, Ramirez-Vega O, Rangel-DeLeon D, Munguia-Garza P, Hidalgo-Miranda A, Mendoza-Vargas A, Reyes-Grajeda JP, Herrera-Montalvo LA. |
| EPI_ISL_3277591, EPI_ISL_3277592, EPI_ISL_3277593, EPI_ISL_3277594, EPI_ISL_3277595, EPI_ISL_3277596, EPI_ISL_3277597, EPI_ISL_3277598 | INCMNSZ | Instituto Nacional de Medicina Genomica | Cedro-Tanda A, Escobar-Arrazola MA, Ramirez-Vega O, Rangel-DeLeon D, Munguia-Garza P, Hidalgo-Miranda A, Mendoza-Vargas A, Reyes-Grajeda JP, Herrera-Montalvo LA. |
| EPI_ISL_3277599, EPI_ISL_3277600 | Instituto Nacional de Medicina Genomica | Instituto Nacional de Medicina Genomica | Cedro-Tanda A, Escobar-Arrazola MA, Ramirez-Vega O, Rangel-DeLeon D, Munguia-Garza P, Hidalgo-Miranda A, Mendoza-Vargas A, Reyes-Grajeda JP, Herrera-Montalvo LA. |
| EPI_ISL_3277601, EPI_ISL_3277602, EPI_ISL_3277603, EPI_ISL_3277604, EPI_ISL_3277605, EPI_ISL_3277607, EPI_ISL_3277615, EPI_ISL_3277617, EPI_ISL_3277619, EPI_ISL_3277620, EPI_ISL_3277621, EPI_ISL_3277622, EPI_ISL_3277623, EPI_ISL_3277626, EPI_ISL_3277627, EPI_ISL_3277628, EPI_ISL_3277629, EPI_ISL_3277630, EPI_ISL_3277631, EPI_ISL_3277632, EPI_ISL_3277634, EPI_ISL_3277635, EPI_ISL_3277636, EPI_ISL_3277637, EPI_ISL_3277639, EPI_ISL_3277640, EPI_ISL_3277641, EPI_ISL_3277643, EPI_ISL_3277644, EPI_ISL_3277645, EPI_ISL_3277646, EPI_ISL_3277647, EPI_ISL_3277648, EPI_ISL_3277649, EPI_ISL_3277651, EPI_ISL_3277652, EPI_ISL_3277653, EPI_ISL_3277655, EPI_ISL_3277657, EPI_ISL_3277659, EPI_ISL_3277662, EPI_ISL_3277663, EPI_ISL_3277664, EPI_ISL_3277665, EPI_ISL_3277666, EPI_ISL_3277667, EPI_ISL_3277668, EPI_ISL_3277669, EPI_ISL_3277670, EPI_ISL_3277671, EPI_ISL_3277672, EPI_ISL_3277673, EPI_ISL_3277674, EPI_ISL_3277675, EPI_ISL_3277676, EPI_ISL_3277677, EPI_ISL_3277680, EPI_ISL_3277681, EPI_ISL_3277682, EPI_ISL_3277683, EPI_ISL_3277684, EPI_ISL_3277685, EPI_ISL_3277686, EPI_ISL_3277687, EPI_ISL_3277688, EPI_ISL_3277691, EPI_ISL_3277692, EPI_ISL_3277693, EPI_ISL_3277694, EPI_ISL_3277696, EPI_ISL_3277697, EPI_ISL_3277698, EPI_ISL_3277699, EPI_ISL_3277703, EPI_ISL_3277704, EPI_ISL_3277705, EPI_ISL_3277706, EPI_ISL_3277707, EPI_ISL_3277709, EPI_ISL_3277710, EPI_ISL_3277711, EPI_ISL_3277712, EPI_ISL_3277713, EPI_ISL_3277714, EPI_ISL_3277715, EPI_ISL_3277716, EPI_ISL_3277717, EPI_ISL_3277718, EPI_ISL_3277719, EPI_ISL_3277720, EPI_ISL_3277722, EPI_ISL_3277723, EPI_ISL_3277724, EPI_ISL_3277725, EPI_ISL_3277726, EPI_ISL_3277727, EPI_ISL_3277728, EPI_ISL_3277729, EPI_ISL_3277730, EPI_ISL_3277731, EPI_ISL_3277732, EPI_ISL_3277733, EPI_ISL_3277734, EPI_ISL_3277735, EPI_ISL_3277736, EPI_ISL_3277737, EPI_ISL_3277738, EPI_ISL_3277739, EPI_ISL_3277740, EPI_ISL_3277741, EPI_ISL_3277742, EPI_ISL_3277743, EPI_ISL_3277744, EPI_ISL_3277745, EPI_ISL_3277746, EPI_ISL_3277747, EPI_ISL_3277748, EPI_ISL_3277749, EPI_ISL_3277750, EPI_ISL_3277751, EPI_ISL_3277752, EPI_ISL_3277753, EPI_ISL_3277754, EPI_ISL_3277755, EPI_ISL_3277756, EPI_ISL_3277758, EPI_ISL_3277759, EPI_ISL_3277760, EPI_ISL_3277761, EPI_ISL_3277762, EPI_ISL_3277766, EPI_ISL_3277767, EPI_ISL_3277768, EPI_ISL_3277769, EPI_ISL_3277771, EPI_ISL_3277773, EPI_ISL_3277774, EPI_ISL_3277775, EPI_ISL_3277776, EPI_ISL_3277777, EPI_ISL_3277778, EPI_ISL_3277779, EPI_ISL_3277780, EPI_ISL_3277782, EPI_ISL_3277783, EPI_ISL_3277784, EPI_ISL_3277785, EPI_ISL_3277786, EPI_ISL_3277787, EPI_ISL_3277788, EPI_ISL_3277789, EPI_ISL_3277790, EPI_ISL_3277791, EPI_ISL_3277792, EPI_ISL_3277793, EPI_ISL_3277794, EPI_ISL_3277795, EPI_ISL_3277796, EPI_ISL_3277797 | | | |
| see above | Salud Digna | Instituto Nacional de Medicina Genomica | Cedro-Tanda A, Escobar-Arrazola MA, Ramirez-Vega O, Abraham Campos-Romero, Moreno-Camacho José Luis, Rodriguez-Gallegos Jorge, Luna-Ruiz Marco, Rangel-DeLeon D, Munguia-Garza P, Hidalgo-Miranda A, Mendoza-Vargas A, Reyes-Grajeda JP, Herrera-Montalvo LA. |
| EPI_ISL_3277798, EPI_ISL_3277799, EPI_ISL_3277800, EPI_ISL_3277801, EPI_ISL_3277802, EPI_ISL_3277803, EPI_ISL_3277804, EPI_ISL_3277805, EPI_ISL_3277806, EPI_ISL_3277807, EPI_ISL_3277808, EPI_ISL_3277809, EPI_ISL_3277810, EPI_ISL_3277811, EPI_ISL_3277812, EPI_ISL_3277813, EPI_ISL_3277814, EPI_ISL_3277815, EPI_ISL_3277816, EPI_ISL_3277817, EPI_ISL_3277818, EPI_ISL_3277819, EPI_ISL_3277820, EPI_ISL_3277821, EPI_ISL_3277822 | | | |
| see above | Instituto Nacional de Medicina Genomica | Instituto Nacional de Medicina Genomica | Cedro-Tanda A, Escobar-Arrazola MA, Ramirez-Vega O, Rangel-DeLeon D, Munguia-Garza P, Hidalgo-Miranda A, Mendoza-Vargas A, Reyes-Grajeda JP, Herrera-Montalvo LA. |
| EPI_ISL_3277823, EPI_ISL_3277824, EPI_ISL_3277825, EPI_ISL_3277826, EPI_ISL_3277827, EPI_ISL_3277828, EPI_ISL_3277829 | INCMNSZ | Instituto Nacional de Medicina Genomica | Cedro-Tanda A, Escobar-Arrazola MA, Ramirez-Vega O, Rangel-DeLeon D, Munguia-Garza P, Hidalgo-Miranda A, Mendoza-Vargas A, Reyes-Grajeda JP, Herrera-Montalvo LA. |
| EPI_ISL_3277830, EPI_ISL_3277831, EPI_ISL_3277832, EPI_ISL_3277833, EPI_ISL_3277834, EPI_ISL_3277835, EPI_ISL_3277836, EPI_ISL_3277837, EPI_ISL_3277838, EPI_ISL_3277839, EPI_ISL_3277840, EPI_ISL_3277841 | | | |
| see above | Instituto Nacional de Medicina Genomica | Instituto Nacional de Medicina Genomica | Cedro-Tanda A, Escobar-Arrazola MA, Ramirez-Vega O, Rangel-DeLeon D, Munguia-Garza P, Hidalgo-Miranda A, Mendoza-Vargas A, Reyes-Grajeda JP, Herrera-Montalvo LA. |
| EPI_ISL_3277842, EPI_ISL_3277843, EPI_ISL_3277844, EPI_ISL_3277845, EPI_ISL_3277846 | INCMNSZ | Instituto Nacional de Medicina Genomica | Cedro-Tanda A, Escobar-Arrazola MA, Ramirez-Vega O, Rangel-DeLeon D, Munguia-Garza P, Hidalgo-Miranda A, Mendoza-Vargas A, Reyes-Grajeda JP, Herrera-Montalvo LA. |
| EPI_ISL_3277847, EPI_ISL_3277848 | Instituto Nacional de Cancerología | Instituto Nacional de Medicina Genomica | Cedro-Tanda A, Escobar-Arrazola MA, Ramirez-Vega O, Rangel-DeLeon D, Munguia-Garza P, Hidalgo-Miranda A, Mendoza-Vargas A, Reyes-Grajeda JP, Herrera-Montalvo LA. |
| EPI_ISL_3277849, EPI_ISL_3277850, EPI_ISL_3277851, EPI_ISL_3277852, EPI_ISL_3277853, EPI_ISL_3277854, EPI_ISL_3277855, EPI_ISL_3277856, EPI_ISL_3277857, EPI_ISL_3277858, EPI_ISL_3277859, EPI_ISL_3277860, EPI_ISL_3277861, EPI_ISL_3277862, EPI_ISL_3277863, EPI_ISL_3277864, EPI_ISL_3277865, EPI_ISL_3277866, EPI_ISL_3277867, EPI_ISL_3277868, EPI_ISL_3277869, EPI_ISL_3277871, EPI_ISL_3277872, EPI_ISL_3277873, EPI_ISL_3277874, EPI_ISL_3277875, EPI_ISL_3277876, EPI_ISL_3277877, EPI_ISL_3277878, EPI_ISL_3277879, EPI_ISL_3277880, EPI_ISL_3277881, EPI_ISL_3277882, EPI_ISL_3277883 | | | |
| see above | Salud Digna | Instituto Nacional de Medicina Genomica | Cedro-Tanda A, Escobar-Arrazola MA, Ramirez-Vega O, Abraham Campos-Romero, Moreno-Camacho José Luis, Rodriguez-Gallegos Jorge, Luna-Ruiz Marco, Rangel-DeLeon D, Munguia-Garza P, Hidalgo-Miranda A, Mendoza-Vargas A, Reyes-Grajeda JP, Herrera-Montalvo LA. |
| EPI_ISL_3278291 | Sultanah Nur Zahirah Hospital | Institute for Medical Research, Infectious Disease Research Centre, National Institutes of Health, Ministry of Health Malaysia | Suppiah J, Kamel K, Mohd Zawawi Z, Azizan MA, Ramly N, Robert F, Anasir MI, Thayan R |
| EPI_ISL_3278292 | Shah Alam Hospital | Institute for Medical Research, Infectious Disease Research Centre, National Institutes of Health, Ministry of Health Malaysia | Suppiah J, Kamel K, Mohd Zawawi Z, Azizan MA, Ramly N, Robert F, Anasir MI, Thayan R |
| EPI_ISL_3278296 | RSUD Dr. W.Z. Johanes kupang | National Institute of Health Research and Development | Vivi Setiawaty, Hana Apsari Pawestri, Subangkit, Kartika Dewi Puspa, Arie Ardiansyah Nugraha, Hartanti Dian Ikawati, Krisna Pangesti, Triyani Soekarso, Nelly Puspandari |
| EPI_ISL_3278301, EPI_ISL_3278302 | Sultanah Bahiyah Hospital | Institute for Medical Research, Infectious Disease Research Centre, National Institutes of Health, Ministry of Health Malaysia | Suppiah J, Kamel K, Mohd Zawawi Z, Azizan MA, Ramly N, Robert F, Anasir MI, Thayan R |
| EPI_ISL_3278304, EPI_ISL_3278305, EPI_ISL_3278306, EPI_ISL_3278308, EPI_ISL_3278310, EPI_ISL_3278311, EPI_ISL_3278312, EPI_ISL_3278313, EPI_ISL_3278314, EPI_ISL_3278315, EPI_ISL_3278316, EPI_ISL_3278317, EPI_ISL_3278318, EPI_ISL_3278319, EPI_ISL_3278320, EPI_ISL_3278321, EPI_ISL_3278322, EPI_ISL_3278324, EPI_ISL_3278325, EPI_ISL_3278326, EPI_ISL_3278327, EPI_ISL_3278328, EPI_ISL_3278329, EPI_ISL_3278330, EPI_ISL_3278331, EPI_ISL_3278334, EPI_ISL_3278335, EPI_ISL_3278336, EPI_ISL_3278337, EPI_ISL_3278338, EPI_ISL_3278339, EPI_ISL_3278340, EPI_ISL_3278341, EPI_ISL_3278342, EPI_ISL_3278343, EPI_ISL_3278344, EPI_ISL_3278345, EPI_ISL_3278346, EPI_ISL_3278350, EPI_ISL_3278351, EPI_ISL_3278352, EPI_ISL_3278353, EPI_ISL_3278354, EPI_ISL_3278355, EPI_ISL_3278356, EPI_ISL_3278357, EPI_ISL_3278358, EPI_ISL_3278359, EPI_ISL_3278361, EPI_ISL_3278362, EPI_ISL_3278363, EPI_ISL_3278364, EPI_ISL_3278365, EPI_ISL_3278366, EPI_ISL_3278367, EPI_ISL_3278369, EPI_ISL_3278370, EPI_ISL_3278372, EPI_ISL_3278373, EPI_ISL_3278374, EPI_ISL_3278375, EPI_ISL_3278376, EPI_ISL_3278377, EPI_ISL_3278378, EPI_ISL_3278379, EPI_ISL_3278380, | | | |

EPI_ISL_3278381, EPI_ISL_3278382, EPI_ISL_3278383, EPI_ISL_3278384, EPI_ISL_3278385, EPI_ISL_3278386, EPI_ISL_3278387, EPI_ISL_3278388, EPI_ISL_3278389, EPI_ISL_3278390, EPI_ISL_3278391, EPI_ISL_3278392, EPI_ISL_3278393, EPI_ISL_3278394, EPI_ISL_3278395, EPI_ISL_3278396, EPI_ISL_3278397, EPI_ISL_3278398, EPI_ISL_3278399, EPI_ISL_3278400, EPI_ISL_3278401, EPI_ISL_3278402, EPI_ISL_3278403, EPI_ISL_3278404, EPI_ISL_3278405, EPI_ISL_3278406, EPI_ISL_3278407, EPI_ISL_3278408, EPI_ISL_3278409, EPI_ISL_3278410, EPI_ISL_3278412, EPI_ISL_3278414, EPI_ISL_3278415, EPI_ISL_3278416, EPI_ISL_3278417, EPI_ISL_3278418, EPI_ISL_3278419, EPI_ISL_3278420, EPI_ISL_3278421, EPI_ISL_3278423, EPI_ISL_3278424, EPI_ISL_3278425, EPI_ISL_3278426, EPI_ISL_3278427, EPI_ISL_3278428, EPI_ISL_3278429, EPI_ISL_3278430, EPI_ISL_3278432, EPI_ISL_3278433, EPI_ISL_3278434, EPI_ISL_3278435, EPI_ISL_3278436, EPI_ISL_3278437, EPI_ISL_3278438, EPI_ISL_3278439, EPI_ISL_3278440, EPI_ISL_3278441, EPI_ISL_3278442, EPI_ISL_3278444, EPI_ISL_3278445, EPI_ISL_3278446, EPI_ISL_3278447, EPI_ISL_3278448, EPI_ISL_3278449, EPI_ISL_3278451, EPI_ISL_3278452, EPI_ISL_3278453, EPI_ISL_3278454, EPI_ISL_3278455, EPI_ISL_3278456, EPI_ISL_3278458, EPI_ISL_3278459, EPI_ISL_3278461, EPI_ISL_3278462, EPI_ISL_3278463, EPI_ISL_3278464, EPI_ISL_3278465, EPI_ISL_3278466, EPI_ISL_3278467, EPI_ISL_3278468, EPI_ISL_3278469, EPI_ISL_3278470, EPI_ISL_3278471, EPI_ISL_3278472, EPI_ISL_3278473, EPI_ISL_3278474, EPI_ISL_3278475, EPI_ISL_3278476, EPI_ISL_3278477, EPI_ISL_3278478, EPI_ISL_3278479, EPI_ISL_3278480, EPI_ISL_3278481, EPI_ISL_3278482, EPI_ISL_3278483, EPI_ISL_3278484, EPI_ISL_3278485, EPI_ISL_3278486, EPI_ISL_3278487, EPI_ISL_3278488, EPI_ISL_3278489, EPI_ISL_3278490, EPI_ISL_3278491, EPI_ISL_3278492, EPI_ISL_3278493, EPI_ISL_3278494, EPI_ISL_3278495, EPI_ISL_3278496, EPI_ISL_3278497, EPI_ISL_3278498, EPI_ISL_3278499, EPI_ISL_3278500, EPI_ISL_3278501, EPI_ISL_3278502, EPI_ISL_3278503, EPI_ISL_3278504, EPI_ISL_3278505, EPI_ISL_3278506, EPI_ISL_3278507, EPI_ISL_3278509, EPI_ISL_3278510, EPI_ISL_3278511, EPI_ISL_3278512, EPI_ISL_3278513, EPI_ISL_3278514, EPI_ISL_3278515, EPI_ISL_3278516, EPI_ISL_3278517, EPI_ISL_3278518, EPI_ISL_3278519, EPI_ISL_3278520, EPI_ISL_3278521, EPI_ISL_3278522, EPI_ISL_3278523, EPI_ISL_3278524, EPI_ISL_3278525, EPI_ISL_3278526, EPI_ISL_3278528, EPI_ISL_3278529, EPI_ISL_3278530, EPI_ISL_3278531, EPI_ISL_3278532, EPI_ISL_3278533, EPI_ISL_3278534, EPI_ISL_3278535, EPI_ISL_3278536, EPI_ISL_3278537, EPI_ISL_3278538, EPI_ISL_3278539, EPI_ISL_3278540, EPI_ISL_3278541, EPI_ISL_3278542, EPI_ISL_3278543, EPI_ISL_3278544, EPI_ISL_3278545, EPI_ISL_3278546, EPI_ISL_3278547, EPI_ISL_3278548, EPI_ISL_3278549, EPI_ISL_3278550, EPI_ISL_3278551, EPI_ISL_3278554, EPI_ISL_3278555, EPI_ISL_3278556, EPI_ISL_3278557, EPI_ISL_3278558, EPI_ISL_3278559, EPI_ISL_3278560, EPI_ISL_3278562, EPI_ISL_3278563, EPI_ISL_3278565, EPI_ISL_3278566, EPI_ISL_3278567, EPI_ISL_3278568, EPI_ISL_3278569, EPI_ISL_3278570, EPI_ISL_3278571, EPI_ISL_3278572, EPI_ISL_3278573, EPI_ISL_3278574, EPI_ISL_3278575, EPI_ISL_3278576, EPI_ISL_3278577, EPI_ISL_3278578, EPI_ISL_3278579, EPI_ISL_3278580, EPI_ISL_3278581, EPI_ISL_3278582, EPI_ISL_3278583, EPI_ISL_3278584, EPI_ISL_3278585, EPI_ISL_3278586, EPI_ISL_3278587, EPI_ISL_3278588, EPI_ISL_3278590, EPI_ISL_3278591, EPI_ISL_3278593, EPI_ISL_3278594, EPI_ISL_3278595, EPI_ISL_3278597, EPI_ISL_3278598, EPI_ISL_3278599, EPI_ISL_3278600, EPI_ISL_3278601, EPI_ISL_3278603, EPI_ISL_3278604, EPI_ISL_3278606, EPI_ISL_3278607, EPI_ISL_3278608, EPI_ISL_3278609, EPI_ISL_3278610, EPI_ISL_3278611, EPI_ISL_3278612, EPI_ISL_3278614, EPI_ISL_3278615, EPI_ISL_3278616, EPI_ISL_3278617, EPI_ISL_3278619, EPI_ISL_3278620, EPI_ISL_3278623, EPI_ISL_3278625, EPI_ISL_3278626, EPI_ISL_3278628, EPI_ISL_3278629, EPI_ISL_3278630, EPI_ISL_3278631, EPI_ISL_3278632, EPI_ISL_3278633, EPI_ISL_3278634, EPI_ISL_3278635, EPI_ISL_3278636, EPI_ISL_3278637, EPI_ISL_3278638, EPI_ISL_3278639, EPI_ISL_3278640, EPI_ISL_3278641, EPI_ISL_3278642, EPI_ISL_3278643, EPI_ISL_3278644, EPI_ISL_3278645, EPI_ISL_3278646, EPI_ISL_3278647, EPI_ISL_3278648, EPI_ISL_3278649, EPI_ISL_3278652, EPI_ISL_3278653, EPI_ISL_3278654, EPI_ISL_3278655, EPI_ISL_3278656, EPI_ISL_3278658, EPI_ISL_3278659, EPI_ISL_3278660, EPI_ISL_3278661, EPI_ISL_3278662, EPI_ISL_3278663, EPI_ISL_3278664, EPI_ISL_3278665, EPI_ISL_3278666, EPI_ISL_3278667, EPI_ISL_3278668, EPI_ISL_3278669, EPI_ISL_3278670, EPI_ISL_3278671, EPI_ISL_3278672, EPI_ISL_3278673, EPI_ISL_3278674, EPI_ISL_3278675, EPI_ISL_3278676, EPI_ISL_3278677, EPI_ISL_3278678, EPI_ISL_3278679, EPI_ISL_3278680, EPI_ISL_3278681, EPI_ISL_3278682, EPI_ISL_3278683, EPI_ISL_3278684, EPI_ISL_3278685, EPI_ISL_3278686, EPI_ISL_3278687, EPI_ISL_3278688, EPI_ISL_3278689, EPI_ISL_3278690, EPI_ISL_3278691, EPI_ISL_3278692, EPI_ISL_3278693, EPI_ISL_3278694, EPI_ISL_3278695, EPI_ISL_3278696, EPI_ISL_3278697, EPI_ISL_3278698, EPI_ISL_3278699, EPI_ISL_3278700, EPI_ISL_3278701, EPI_ISL_3278702, EPI_ISL_3278703, EPI_ISL_3278704, EPI_ISL_3278705, EPI_ISL_3278706, EPI_ISL_3278707, EPI_ISL_3278708, EPI_ISL_3278709, EPI_ISL_3278710, EPI_ISL_3278711, EPI_ISL_3278712, EPI_ISL_3278713, EPI_ISL_3278714, EPI_ISL_3278715, EPI_ISL_3278716, EPI_ISL_3278718, EPI_ISL_3278721, EPI_ISL_3278723, EPI_ISL_3278724, EPI_ISL_3278726, EPI_ISL_3278727, EPI_ISL_3278728, EPI_ISL_3278729, EPI_ISL_3278730, EPI_ISL_3278731, EPI_ISL_3278732, EPI_ISL_3278733, EPI_ISL_3278734, EPI_ISL_3278735, EPI_ISL_3278736, EPI_ISL_3278737, EPI_ISL_3278739, EPI_ISL_3278740, EPI_ISL_3278741, EPI_ISL_3278743, EPI_ISL_3278744, EPI_ISL_3278745, EPI_ISL_3278746, EPI_ISL_3278747, EPI_ISL_3278748, EPI_ISL_3278749, EPI_ISL_3278750, EPI_ISL_3278752, EPI_ISL_3278753, EPI_ISL_3278754, EPI_ISL_3278755, EPI_ISL_3278756, EPI_ISL_3278757, EPI_ISL_3278758, EPI_ISL_3278759, EPI_ISL_3278760, EPI_ISL_3278761, EPI_ISL_3278762, EPI_ISL_3278763, EPI_ISL_3278764, EPI_ISL_3278765, EPI_ISL_3278766, EPI_ISL_3278767, EPI_ISL_3278768, EPI_ISL_3278769, EPI_ISL_3278770, EPI_ISL_3278771, EPI_ISL_3278772, EPI_ISL_3278773, EPI_ISL_3278774, EPI_ISL_3278775, EPI_ISL_3278776, EPI_ISL_3278778, EPI_ISL_3278779, EPI_ISL_3278780, EPI_ISL_3278781, EPI_ISL_3278782, EPI_ISL_3278783, EPI_ISL_3278784, EPI_ISL_3278785, EPI_ISL_3278786, EPI_ISL_3278787, EPI_ISL_3278788, EPI_ISL_3278789, EPI_ISL_3278790, EPI_ISL_3278791, EPI_ISL_3278793, EPI_ISL_3278794, EPI_ISL_3278795, EPI_ISL_3278796, EPI_ISL_3278797, EPI_ISL_3278798, EPI_ISL_3278799, EPI_ISL_3278800, EPI_ISL_3278801, EPI_ISL_3278802, EPI_ISL_3278803, EPI_ISL_3278804, EPI_ISL_3278805, EPI_ISL_3278806, EPI_ISL_3278807, EPI_ISL_3278808, EPI_ISL_3278809, EPI_ISL_3278810, EPI_ISL_3278811, EPI_ISL_3278812, EPI_ISL_3278813, EPI_ISL_3278814, EPI_ISL_3278815, EPI_ISL_3278816, EPI_ISL_3278817, EPI_ISL_3278818, EPI_ISL_3278819, EPI_ISL_3278820, EPI_ISL_3278821, EPI_ISL_3278822, EPI_ISL_3278823, EPI_ISL_3278824, EPI_ISL_3278825, EPI_ISL_3278826, EPI_ISL_3278827, EPI_ISL_3278828, EPI_ISL_3278829, EPI_ISL_3278830, EPI_ISL_3278831, EPI_ISL_3278832, EPI_ISL_3278833, EPI_ISL_3278834, EPI_ISL_3278835, EPI_ISL_3278836, EPI_ISL_3278837, EPI_ISL_3278838, EPI_ISL_3278839, EPI_ISL_3278841, EPI_ISL_3278842, EPI_ISL_3278844, EPI_ISL_3278845, EPI_ISL_3278846, EPI_ISL_3278847, EPI_ISL_3278849, EPI_ISL_3278850, EPI_ISL_3278851, EPI_ISL_3278852, EPI_ISL_3278853, EPI_ISL_3278854, EPI_ISL_3278855, EPI_ISL_3278856, EPI_ISL_3278857, EPI_ISL_3278858, EPI_ISL_3278859, EPI_ISL_3278860, EPI_ISL_3278861, EPI_ISL_3278862, EPI_ISL_3278863, EPI_ISL_3278864, EPI_ISL_3278865, EPI_ISL_3278866, EPI_ISL_3278867, EPI_ISL_3278868, EPI_ISL_3278869, EPI_ISL_3278870, EPI_ISL_3278871, EPI_ISL_3278872, EPI_ISL_3278873, EPI_ISL_3278874, EPI_ISL_3278877, EPI_ISL_3278878, EPI_ISL_3278880, EPI_ISL_3278881, EPI_ISL_3278882, EPI_ISL_3278883, EPI_ISL_3278884, EPI_ISL_3278885, EPI_ISL_3278886, EPI_ISL_3278887, EPI_ISL_3278888, EPI_ISL_3278889, EPI_ISL_3278890, EPI_ISL_3278891, EPI_ISL_3278892, EPI_ISL_3278893, EPI_ISL_3278894, EPI_ISL_3278895, EPI_ISL_3278896, EPI_ISL_3278897, EPI_ISL_3278899, EPI_ISL_3278900, EPI_ISL_3278901, EPI_ISL_3278902, EPI_ISL_3278904, EPI_ISL_3278905, EPI_ISL_3278906, EPI_ISL_3278907, EPI_ISL_3278908, EPI_ISL_3278910, EPI_ISL_3278912, EPI_ISL_3278913, EPI_ISL_3278914, EPI_ISL_3278915, EPI_ISL_3278916, EPI_ISL_3278917, EPI_ISL_3278918, EPI_ISL_3278919, EPI_ISL_3278920, EPI_ISL_3278921, EPI_ISL_3278922, EPI_ISL_3278923, EPI_ISL_3278925, EPI_ISL_3278927, EPI_ISL_3278928, EPI_ISL_3278929, EPI_ISL_3278930, EPI_ISL_3278931, EPI_ISL_3278932, EPI_ISL_3278933, EPI_ISL_3278934, EPI_ISL_3278936, EPI_ISL_3278937, EPI_ISL_3278938, EPI_ISL_3278939, EPI_ISL_3278940, EPI_ISL_3278942, EPI_ISL_3278943, EPI_ISL_3278944, EPI_ISL_3278945, EPI_ISL_3278946, EPI_ISL_3278947, EPI_ISL_3278948, EPI_ISL_3278949, EPI_ISL_3278950, EPI_ISL_3278951, EPI_ISL_3278952, EPI_ISL_3278953, EPI_ISL_3278954, EPI_ISL_3278955, EPI_ISL_3278956, EPI_ISL_3278957, EPI_ISL_3278958, EPI_ISL_3278959, EPI_ISL_3278960, EPI_ISL_3278961, EPI_ISL_3278962, EPI_ISL_3278963, EPI_ISL_3278964, EPI_ISL_3278965, EPI_ISL_3278967, EPI_ISL_3278968, EPI_ISL_3278969, EPI_ISL_3278970, EPI_ISL_3278971, EPI_ISL_3278972, EPI_ISL_3278973, EPI_ISL_3278974, EPI_ISL_3278977, EPI_ISL_3278978, EPI_ISL_3278980, EPI_ISL_3278981, EPI_ISL_3278982, EPI_ISL_3278983, EPI_ISL_3278984, EPI_ISL_3278985, EPI_ISL_3278986, EPI_ISL_3278987, EPI_ISL_3278988, EPI_ISL_3278989, EPI_ISL_3278990, EPI_ISL_3278991, EPI_ISL_3278992, EPI_ISL_3278993, EPI_ISL_3278994, EPI_ISL_3278995, EPI_ISL_3278996, EPI_ISL_3278997, EPI_ISL_3278998, EPI_ISL_3278999, EPI_ISL_3279000, EPI_ISL_3279001, EPI_ISL_3279002, EPI_ISL_3279003, EPI_ISL_3279004, EPI_ISL_3279005, EPI_ISL_3279006, EPI_ISL_3279007, EPI_ISL_3279008, EPI_ISL_3279009, EPI_ISL_3279012, EPI_ISL_3279013, EPI_ISL_3279015, EPI_ISL_3279016, EPI_ISL_3279017, EPI_ISL_3279019, EPI_ISL_3279020, EPI_ISL_3279021, EPI_ISL_3279022, EPI_ISL_3279023, EPI_ISL_3279024, EPI_ISL_3279025, EPI_ISL_3279026, EPI_ISL_3279027, EPI_ISL_3279028, EPI_ISL_3279029, EPI_ISL_3279030, EPI_ISL_3279031, EPI_ISL_3279032, EPI_ISL_3279033, EPI_ISL_3279034, EPI_ISL_3279035, EPI_ISL_3279036, EPI_ISL_3279037, EPI_ISL_3279038, EPI_ISL_3279039, EPI_ISL_3279040, EPI_ISL_3279041, EPI_ISL_3279042, EPI_ISL_3279043, EPI_ISL_3279044, EPI_ISL_3279045, EPI_ISL_3279046, EPI_ISL_3279047, EPI_ISL_3279048, EPI_ISL_3279049, EPI_ISL_3279050, EPI_ISL_3279051, EPI_ISL_3279053, EPI_ISL_3279054, EPI_ISL_3279055, EPI_ISL_3279057, EPI_ISL_3279058, EPI_ISL_3279060, EPI_ISL_3279061, EPI_ISL_3279062, EPI_ISL_3279063, EPI_ISL_3279064, EPI_ISL_3279065, EPI_ISL_3279066, EPI_ISL_3279067, EPI_ISL_3279068, EPI_ISL_3279069, EPI_ISL_3279070, EPI_ISL_3279071, EPI_ISL_3279072, EPI_ISL_3279073, EPI_ISL_3279074, EPI_ISL_3279075, EPI_ISL_3279076, EPI_ISL_3279077, EPI_ISL_3279078, EPI_ISL_3279079, EPI_ISL_3279080, EPI_ISL_3279081, EPI_ISL_3279082, EPI_ISL_3279083, EPI_ISL_3279084, EPI_ISL_3279085, EPI_ISL_3279086, EPI_ISL_3279087, EPI_ISL_3279088, EPI_ISL_3279089, EPI_ISL_3279090, EPI_ISL_3279091, EPI_ISL_3279092, EPI_ISL_3279093, EPI_ISL_3279094, EPI_ISL_3279095, EPI_ISL_3279096, EPI_ISL_3279098, EPI_ISL_3279099, EPI_ISL_3279101, EPI_ISL_3279102, EPI_ISL_3279103, EPI_ISL_3279104, EPI_ISL_3279105, EPI_ISL_3279106, EPI_ISL_3279108, EPI_ISL_3279109, EPI_ISL_3279110, EPI_ISL_3279111, EPI_ISL_3279112, EPI_ISL_3279113, EPI_ISL_3279114, EPI_ISL_3279115, EPI_ISL_3279116, EPI_ISL_3279117, EPI_ISL_3279118, EPI_ISL_3279119, EPI_ISL_3279120, EPI_ISL_3279121, EPI_ISL_3279122, EPI_ISL_3279123, EPI_ISL_3279124, EPI_ISL_3279125, EPI_ISL_3279126, EPI_ISL_3279127, EPI_ISL_3279128, EPI_ISL_3279129, EPI_ISL_3279130, EPI_ISL_3279131, EPI_ISL_3279132, EPI_ISL_3279133, EPI_ISL_3279134, EPI_ISL_3279135, EPI_ISL_3279136, EPI_ISL_3279137, EPI_ISL_3279138, EPI_ISL_3279139, EPI_ISL_3279141, EPI_ISL_3279142, EPI_ISL_3279145, EPI_ISL_3279146, EPI_ISL_3279148, EPI_ISL_3279149, EPI_ISL_3279150, EPI_ISL_3279151, EPI_ISL_3279152, EPI_ISL_3279153, EPI_ISL_3279155, EPI_ISL_3279156, EPI_ISL_3279157, EPI_ISL_3279158, EPI_ISL_3279159, EPI_ISL_3279160, EPI_ISL_3279161, EPI_ISL_3279162, EPI_ISL_3279163, EPI_ISL_3279164, EPI_ISL_3279166, EPI_ISL_3279167, EPI_ISL_3279168, EPI_ISL_3279169, EPI_ISL_3279170, EPI_ISL_3279172, EPI_ISL_3279173, EPI_ISL_3279174, EPI_ISL_3279175, EPI_ISL_3279177, EPI_ISL_3279178, EPI_ISL_3279179, EPI_ISL_3279180, EPI_ISL_3279181, EPI_ISL_3279182, EPI_ISL_3279183, EPI_ISL_3279184, EPI_ISL_3279185, EPI_ISL_3279186, EPI_ISL_3279187, EPI_ISL_3279188, EPI_ISL_3279189, EPI_ISL_3279190, EPI_ISL_3279191, EPI_ISL_3279192, EPI_ISL_3279193, EPI_ISL_3279194, EPI_ISL_3279195, EPI_ISL_3279196, EPI_ISL_3279197, EPI_ISL_3279198, EPI_ISL_3279200, EPI_ISL_3279201, EPI_ISL_3279202, EPI_ISL_3279203, EPI_ISL_3279204, EPI_ISL_3279205, EPI_ISL_3279206, EPI_ISL_3279207, EPI_ISL_3279208, EPI_ISL_3279210, EPI_ISL_3279212, EPI_ISL_3279216, EPI_ISL_3279217, EPI_ISL_3279218, EPI_ISL_3279220, EPI_ISL_3279221, EPI_ISL_3279224, EPI_ISL_3279225, EPI_ISL_3279226, EPI_ISL_3279227, EPI_ISL_3279228, EPI_ISL_3279229, EPI_ISL_3279230, EPI_ISL_3279232, EPI_ISL_3279233, EPI_ISL_3279234, EPI_ISL_3279235, EPI_ISL_3279236, EPI_ISL_3279237, EPI_ISL_3279238, EPI_ISL_3279239, EPI_ISL_3279241, EPI_ISL_3279242, EPI_ISL_3279243, EPI_ISL_3279244, EPI_ISL_3279245, EPI_ISL_3279246, EPI_ISL_3279248, EPI_ISL_3279249, EPI_ISL_3279250, EPI_ISL_3279251, EPI_ISL_3279252, EPI_ISL_3279253, EPI_ISL_3279254, EPI_ISL_3279255, EPI_ISL_3279256, EPI_ISL_3279257, EPI_ISL_3279258, EPI_ISL_3279259, EPI_ISL_3279261, EPI_ISL_3279263, EPI_ISL_3279265, EPI_ISL_3279267, EPI_ISL_3279268, EPI_ISL_3279270, EPI_ISL_3279271, EPI_ISL_3279272, EPI_ISL_3279273, EPI_ISL_3279274, EPI_ISL_3279275, EPI_ISL_3279276, EPI_ISL_3279278, EPI_ISL_3279279, EPI_ISL_3279280, EPI_ISL_3279281, EPI_ISL_3279282, EPI_ISL_3279283, EPI_ISL_3279284, EPI_ISL_3279285, EPI_ISL_3279286, EPI_ISL_3279287, EPI_ISL_3279288, EPI_ISL_3279289, EPI_ISL_3279290, EPI_ISL_3279291, EPI_ISL_3279292, EPI_ISL_3279293, EPI_ISL_3279294, EPI_ISL_3279295, EPI_ISL_3279296, EPI_ISL_3279297, EPI_ISL_3279298, EPI_ISL_3279300, EPI_ISL_3279301, EPI_ISL_3279302, EPI_ISL_3279303, EPI_ISL_3279305, EPI_ISL_3279306, EPI_ISL_3279307, EPI_ISL_3279308, EPI_ISL_3279309, EPI_ISL_3279310, EPI_ISL_3279311, EPI_ISL_3279312, EPI_ISL_3279313, EPI_ISL_3279315, EPI_ISL_3279316, EPI_ISL_3279317, EPI_ISL_3279319, EPI_ISL_3279320, EPI_ISL_3279322, EPI_ISL_3279323, EPI_ISL_3279324, EPI_ISL_3279325, EPI_ISL_3279326, EPI_ISL_3279329, EPI_ISL_3279330, EPI_ISL_3279331, EPI_ISL_3279334, EPI_ISL_3279335, EPI_ISL_3279336, EPI_ISL_3279337, EPI_ISL_3279339, EPI_ISL_3279340, EPI_ISL_3279341, EPI_ISL_3279342, EPI_ISL_3279344, EPI_ISL_3279347, EPI_ISL_3279348, EPI_ISL_3279350, EPI_ISL_3279356, EPI_ISL_3279357, EPI_ISL_3279361, EPI_ISL_3279362, EPI_ISL_3279363, EPI_ISL_3279364, EPI_ISL_3279365, EPI_ISL_3279367, EPI_ISL_3279368, EPI_ISL_3279370, EPI_ISL_3279371, EPI_ISL_3279372, EPI_ISL_3279373, EPI_ISL_3279374, EPI_ISL_3279375, EPI_ISL_3279376, EPI_ISL_3279377, EPI_ISL_3279378, EPI_ISL_3279379, EPI_ISL_3279380, EPI_ISL_3279381, EPI_ISL_3279383, EPI_ISL_3279385, EPI_ISL_3279386, EPI_ISL_3279388, EPI_ISL_3279389, EPI_ISL_3279390, EPI_ISL_3279391, EPI_ISL_3279393, EPI_ISL_3279394, EPI_ISL_3279395, EPI_ISL_3279396, EPI_ISL_3279397, EPI_ISL_3279398, EPI_ISL_3279399, EPI_ISL_3279401, EPI_ISL_3279402, EPI_ISL_3279403, EPI_ISL_3279404, EPI_ISL_3279405, EPI_ISL_3279406, EPI_ISL_3279407, EPI_ISL_3279408

| | | | |
|---|---|---|---|
| see above | Israel Central Virology laboratory | Israel National Consortium for SARS-CoV-2 sequencing | Neta Zuckerman, Efrat Dahan Bucris, Michal Mandelboim, Dana Bar-Ilan, Miranda Geva, Omer Asraf, Oran Erster, Efrat Glick-Saar, Omri Nayshool, Gideon Rechavi, Danit Sofer, Ella Mendelson, Orna Mor |
| EPI_ISL_3279413 | RSUD dr. Soedarsono | Institute of Tropical Disease, Universitas Airlangga | Kazufumi Shimizu, Krisnoadi Rahardjo, Aldise M Nastri, Jezzy R Dewantari, Rima R Prasetya, M Burhan, Gatot Soegiarto, Laksmi Wulandari, Resti Yudhawati, Yasuko Mori, Soetjipto, Maria I Lusida |
| EPI_ISL_3279414 | RS Universitas Brawijaya | Institute of Tropical Disease, Universitas Airlangga | Krisnoadi Rahardjo, Aldise M Nastri, Jezzy R Dewantari, Rima R Prasetya, Sri Andarini, Gatot Soegiarto, Laksmi Wulandari, Resti Yudhawati, Yasuko Mori, Soetjipto, Kazufumi Shimizu, Maria I Lusida |
| EPI_ISL_3279415, EPI_ISL_3279416, EPI_ISL_3279417 | Department of Bacteria, Parasites and Fungi, Statens Serum Institut, Copenhagen, Denmark | Statens Serum Institut Bioinformatics and Microbial Genomics | Danish Covid-19 Genome Consortium |
| EPI_ISL_3279418 | Department of Clinical Microbiology, Odense University Hospital, Odense, Denmark | Statens Serum Institut Bioinformatics and Microbial Genomics | Danish Covid-19 Genome Consortium |

EPI_ISL_3279419, EPI_ISL_3279420, EPI_ISL_3279421, EPI_ISL_3279422, EPI_ISL_3279423, EPI_ISL_3279424, EPI_ISL_3279425, EPI_ISL_3279426, EPI_ISL_3279427, EPI_ISL_3279428, EPI_ISL_3279429, EPI_ISL_3279430, EPI_ISL_3279431, EPI_ISL_3279432, EPI_ISL_3279433, EPI_ISL_3279434, EPI_ISL_3279435, EPI_ISL_3279436, EPI_ISL_3279437, EPI_ISL_3279438, EPI_ISL_3279439, EPI_ISL_3279440, EPI_ISL_3279441, EPI_ISL_3279442, EPI_ISL_3279443, EPI_ISL_3279444, EPI_ISL_3279445, EPI_ISL_3279446, EPI_ISL_3279447, EPI_ISL_3279448, EPI_ISL_3279449, EPI_ISL_3279450, EPI_ISL_3279451, EPI_ISL_3279452, EPI_ISL_3279453, EPI_ISL_3279454, EPI_ISL_3279455, EPI_ISL_3279456, EPI_ISL_3279457, EPI_ISL_3279458, EPI_ISL_3279459, EPI_ISL_3279460, EPI_ISL_3279461, EPI_ISL_3279462, EPI_ISL_3279463, EPI_ISL_3279464, EPI_ISL_3279465, EPI_ISL_3279466, EPI_ISL_3279467, EPI_ISL_3279468, EPI_ISL_3279469, EPI_ISL_3279470, EPI_ISL_3279471, EPI_ISL_3279472, EPI_ISL_3279473, EPI_ISL_3279474, EPI_ISL_3279475, EPI_ISL_3279476, EPI_ISL_3279477, EPI_ISL_3279478, EPI_ISL_3279479, EPI_ISL_3279480, EPI_ISL_3279481, EPI_ISL_3279482, EPI_ISL_3279483, EPI_ISL_3279484, EPI_ISL_3279485, EPI_ISL_3279486, EPI_ISL_3279487, EPI_ISL_3279488, EPI_ISL_3279489

| | | | |
|---|---|---|---|
| see above | Department of Bacteria, Parasites and Fungi, Statens Serum Institut, Copenhagen, Denmark | Statens Serum Institut Bioinformatics and Microbial Genomics | Danish Covid-19 Genome Consortium |
| EPI_ISL_3279490 | Department of Clinical Microbiology, Odense University Hospital, Odense, Denmark | Statens Serum Institut Bioinformatics and Microbial Genomics | Danish Covid-19 Genome Consortium |

EPI_ISL_3279491, EPI_ISL_3279492, EPI_ISL_3279493, EPI_ISL_3279494, EPI_ISL_3279495, EPI_ISL_3279496, EPI_ISL_3279497, EPI_ISL_3279499, EPI_ISL_3279500, EPI_ISL_3279501, EPI_ISL_3279502, EPI_ISL_3279503, EPI_ISL_3279504, EPI_ISL_3279505, EPI_ISL_3279506, EPI_ISL_3279507, EPI_ISL_3279508, EPI_ISL_3279509, EPI_ISL_3279510, EPI_ISL_3279511, EPI_ISL_3279512, EPI_ISL_3279513, EPI_ISL_3279514, EPI_ISL_3279515, EPI_ISL_3279516, EPI_ISL_3279517, EPI_ISL_3279518, EPI_ISL_3279519, EPI_ISL_3279520, EPI_ISL_3279521, EPI_ISL_3279522, EPI_ISL_3279523, EPI_ISL_3279524, EPI_ISL_3279525, EPI_ISL_3279526, EPI_ISL_3279527, EPI_ISL_3279528, EPI_ISL_3279529, EPI_ISL_3279530, EPI_ISL_3279531, EPI_ISL_3279532, EPI_ISL_3279534, EPI_ISL_3279535, EPI_ISL_3279536, EPI_ISL_3279537, EPI_ISL_3279538, EPI_ISL_3279539, EPI_ISL_3279540, EPI_ISL_3279541, EPI_ISL_3279542, EPI_ISL_3279544, EPI_ISL_3279545, EPI_ISL_3279546, EPI_ISL_3279547, EPI_ISL_3279548, EPI_ISL_3279549, EPI_ISL_3279550, EPI_ISL_3279551, EPI_ISL_3279552, EPI_ISL_3279553, EPI_ISL_3279554, EPI_ISL_3279555, EPI_ISL_3279556, EPI_ISL_3279557, EPI_ISL_3279558, EPI_ISL_3279559, EPI_ISL_3279560, EPI_ISL_3279561, EPI_ISL_3279562, EPI_ISL_3279563, EPI_ISL_3279564, EPI_ISL_3279565, EPI_ISL_3279566

| | | | |
|---|---|---|---|
| see above | Department of Bacteria, Parasites and Fungi, Statens Serum Institut, Copenhagen, Denmark | Statens Serum Institut Bioinformatics and Microbial Genomics | Danish Covid-19 Genome Consortium |
| EPI_ISL_3279567 | Department of Clinical Microbiology, Odense University Hospital, Odense, Denmark | Statens Serum Institut Bioinformatics and Microbial Genomics | Danish Covid-19 Genome Consortium |

EPI_ISL_3279568, EPI_ISL_3279569, EPI_ISL_3279570, EPI_ISL_3279571, EPI_ISL_3279572, EPI_ISL_3279573, EPI_ISL_3279575, EPI_ISL_3279576, EPI_ISL_3279577, EPI_ISL_3279578, EPI_ISL_3279579, EPI_ISL_3279580, EPI_ISL_3279581, EPI_ISL_3279582, EPI_ISL_3279583, EPI_ISL_3279584, EPI_ISL_3279587, EPI_ISL_3279588, EPI_ISL_3279589, EPI_ISL_3279590

| | | | |
|---|---|---|---|
| see above | Department of Bacteria, Parasites and Fungi, Statens Serum Institut, Copenhagen, Denmark | Statens Serum Institut Bioinformatics and Microbial Genomics | Danish Covid-19 Genome Consortium |
| EPI_ISL_3279591 | Department of Clinical Microbiology, Odense University Hospital, Odense, Denmark | Statens Serum Institut Bioinformatics and Microbial Genomics | Danish Covid-19 Genome Consortium |

EPI_ISL_3279592, EPI_ISL_3279593, EPI_ISL_3279594, EPI_ISL_3279595, EPI_ISL_3279596, EPI_ISL_3279597, EPI_ISL_3279598, EPI_ISL_3279599, EPI_ISL_3279600, EPI_ISL_3279601, EPI_ISL_3279602, EPI_ISL_3279603, EPI_ISL_3279604, EPI_ISL_3279605, EPI_ISL_3279606, EPI_ISL_3279607, EPI_ISL_3279608, EPI_ISL_3279609, EPI_ISL_3279610, EPI_ISL_3279611, EPI_ISL_3279612, EPI_ISL_3279613, EPI_ISL_3279614, EPI_ISL_3279615, EPI_ISL_3279616, EPI_ISL_3279617, EPI_ISL_3279618, EPI_ISL_3279619, EPI_ISL_3279620, EPI_ISL_3279621, EPI_ISL_3279622, EPI_ISL_3279623, EPI_ISL_3279624, EPI_ISL_3279625, EPI_ISL_3279626, EPI_ISL_3279627, EPI_ISL_3279628, EPI_ISL_3279629, EPI_ISL_3279630, EPI_ISL_3279631, EPI_ISL_3279632, EPI_ISL_3279633, EPI_ISL_3279634, EPI_ISL_3279635, EPI_ISL_3279636, EPI_ISL_3279637, EPI_ISL_3279638, EPI_ISL_3279639, EPI_ISL_3279640, EPI_ISL_3279641, EPI_ISL_3279642, EPI_ISL_3279643, EPI_ISL_3279644, EPI_ISL_3279645, EPI_ISL_3279646, EPI_ISL_3279647, EPI_ISL_3279648, EPI_ISL_3279649, EPI_ISL_3279651, EPI_ISL_3279652, EPI_ISL_3279653, EPI_ISL_3279654, EPI_ISL_3279655, EPI_ISL_3279656, EPI_ISL_3279657, EPI_ISL_3279658, EPI_ISL_3279659, EPI_ISL_3279660, EPI_ISL_3279661, EPI_ISL_3279662, EPI_ISL_3279663, EPI_ISL_3279664, EPI_ISL_3279665, EPI_ISL_3279666, EPI_ISL_3279667, EPI_ISL_3279668, EPI_ISL_3279669, EPI_ISL_3279670, EPI_ISL_3279671, EPI_ISL_3279672, EPI_ISL_3279673, EPI_ISL_3279674, EPI_ISL_3279675, EPI_ISL_3279676, EPI_ISL_3279678, EPI_ISL_3279679, EPI_ISL_3279680, EPI_ISL_3279681, EPI_ISL_3279682, EPI_ISL_3279683, EPI_ISL_3279684, EPI_ISL_3279685, EPI_ISL_3279686, EPI_ISL_3279687, EPI_ISL_3279688, EPI_ISL_3279689, EPI_ISL_3279690, EPI_ISL_3279691, EPI_ISL_3279692, EPI_ISL_3279693, EPI_ISL_3279694, EPI_ISL_3279695, EPI_ISL_3279696, EPI_ISL_3279697, EPI_ISL_3279698, EPI_ISL_3279699, EPI_ISL_3279700, EPI_ISL_3279701

| | | | |
|---|---|---|---|
| see above | Department of Bacteria, Parasites and Fungi, Statens Serum Institut, Copenhagen, Denmark | Statens Serum Institut Bioinformatics and Microbial Genomics | Danish Covid-19 Genome Consortium |
| EPI_ISL_3279702 | Department of Clinical Microbiology, Odense University Hospital, Odense, Denmark | Statens Serum Institut Bioinformatics and Microbial Genomics | Danish Covid-19 Genome Consortium |

EPI_ISL_3279703, EPI_ISL_3279704, EPI_ISL_3279705, EPI_ISL_3279706, EPI_ISL_3279707, EPI_ISL_3279708, EPI_ISL_3279709, EPI_ISL_3279710, EPI_ISL_3279711, EPI_ISL_3279712, EPI_ISL_3279713, EPI_ISL_3279714, EPI_ISL_3279715, EPI_ISL_3279716, EPI_ISL_3279717, EPI_ISL_3279718, EPI_ISL_3279719, EPI_ISL_3279720, EPI_ISL_3279721, EPI_ISL_3279722, EPI_ISL_3279723, EPI_ISL_3279724, EPI_ISL_3279725, EPI_ISL_3279726, EPI_ISL_3279727, EPI_ISL_3279728, EPI_ISL_3279729, EPI_ISL_3279730, EPI_ISL_3279731, EPI_ISL_3279732, EPI_ISL_3279733, EPI_ISL_3279734, EPI_ISL_3279735, EPI_ISL_3279736, EPI_ISL_3279737, EPI_ISL_3279738, EPI_ISL_3279739, EPI_ISL_3279740, EPI_ISL_3279741, EPI_ISL_3279742, EPI_ISL_3279743, EPI_ISL_3279744, EPI_ISL_3279746, EPI_ISL_3279747, EPI_ISL_3279748, EPI_ISL_3279749, EPI_ISL_3279750, EPI_ISL_3279751, EPI_ISL_3279752, EPI_ISL_3279753, EPI_ISL_3279754, EPI_ISL_3279755, EPI_ISL_3279756, EPI_ISL_3279757, EPI_ISL_3279758, EPI_ISL_3279759, EPI_ISL_3279760, EPI_ISL_3279761, EPI_ISL_3279762, EPI_ISL_3279763, EPI_ISL_3279764, EPI_ISL_3279765, EPI_ISL_3279766, EPI_ISL_3279767, EPI_ISL_3279768, EPI_ISL_3279770, EPI_ISL_3279771, EPI_ISL_3279772, EPI_ISL_3279773, EPI_ISL_3279774, EPI_ISL_3279775, EPI_ISL_3279776, EPI_ISL_3279777, EPI_ISL_3279778, EPI_ISL_3279779, EPI_ISL_3279780, EPI_ISL_3279781, EPI_ISL_3279782, EPI_ISL_3279783, EPI_ISL_3279784, EPI_ISL_3279785, EPI_ISL_3279786, EPI_ISL_3279787, EPI_ISL_3279788, EPI_ISL_3279789, EPI_ISL_3279790, EPI_ISL_3279791, EPI_ISL_3279792, EPI_ISL_3279793, EPI_ISL_3279794, EPI_ISL_3279795, EPI_ISL_3279796, EPI_ISL_3279797, EPI_ISL_3279798

| | | | |
|---|---|---|---|
| see above | Department of Bacteria, Parasites and Fungi, Statens Serum Institut, Copenhagen, Denmark | Statens Serum Institut Bioinformatics and Microbial Genomics | Danish Covid-19 Genome Consortium |
| EPI_ISL_3279799 | Department of Clinical Microbiology, Odense University Hospital, Odense, Denmark | Statens Serum Institut Bioinformatics and Microbial Genomics | Danish Covid-19 Genome Consortium |

EPI_ISL_3279800, EPI_ISL_3279801, EPI_ISL_3279802, EPI_ISL_3279803, EPI_ISL_3279804, EPI_ISL_3279805, EPI_ISL_3279806, EPI_ISL_3279807, EPI_ISL_3279808, EPI_ISL_3279809, EPI_ISL_3279810, EPI_ISL_3279811, EPI_ISL_3279812, EPI_ISL_3279813, EPI_ISL_3279814, EPI_ISL_3279815, EPI_ISL_3279816, EPI_ISL_3279817, EPI_ISL_3279818, EPI_ISL_3279819, EPI_ISL_3279820, EPI_ISL_3279821, EPI_ISL_3279822, EPI_ISL_3279823, EPI_ISL_3279824, EPI_ISL_3279825, EPI_ISL_3279826, EPI_ISL_3279827, EPI_ISL_3279828, EPI_ISL_3279829, EPI_ISL_3279830, EPI_ISL_3279831, EPI_ISL_3279832, EPI_ISL_3279833, EPI_ISL_3279834, EPI_ISL_3279835, EPI_ISL_3279836, EPI_ISL_3279837, EPI_ISL_3279838, EPI_ISL_3279839, EPI_ISL_3279840, EPI_ISL_3279841, EPI_ISL_3279842, EPI_ISL_3279843, EPI_ISL_3279844, EPI_ISL_3279845, EPI_ISL_3279846, EPI_ISL_3279847, EPI_ISL_3279848, EPI_ISL_3279849, EPI_ISL_3279850, EPI_ISL_3279851, EPI_ISL_3279852, EPI_ISL_3279853, EPI_ISL_3279854, EPI_ISL_3279855, EPI_ISL_3279856, EPI_ISL_3279857, EPI_ISL_3279858, EPI_ISL_3279859, EPI_ISL_3279860, EPI_ISL_3279861, EPI_ISL_3279862, EPI_ISL_3279863, EPI_ISL_3279864, EPI_ISL_3279865, EPI_ISL_3279866, EPI_ISL_3279867, EPI_ISL_3279868, EPI_ISL_3279869, EPI_ISL_3279870, EPI_ISL_3279871, EPI_ISL_3279872, EPI_ISL_3279873, EPI_ISL_3279874, EPI_ISL_3279875, EPI_ISL_3279876, EPI_ISL_3279877, EPI_ISL_3279878, EPI_ISL_3279879, EPI_ISL_3279880, EPI_ISL_3279881, EPI_ISL_3279882, EPI_ISL_3279883, EPI_ISL_3279884, EPI_ISL_3279885, EPI_ISL_3279886, EPI_ISL_3279887, EPI_ISL_3279889, EPI_ISL_3279890, EPI_ISL_3279891, EPI_ISL_3279892, EPI_ISL_3279893, EPI_ISL_3279894, EPI_ISL_3279895, EPI_ISL_3279896, EPI_ISL_3279897, EPI_ISL_3279898, EPI_ISL_3279899, EPI_ISL_3279900, EPI_ISL_3279901, EPI_ISL_3279902, EPI_ISL_3279903, EPI_ISL_3279904, EPI_ISL_3279905, EPI_ISL_3279906, EPI_ISL_3279907, EPI_ISL_3279908, EPI_ISL_3279909, EPI_ISL_3279910, EPI_ISL_3279911, EPI_ISL_3279912, EPI_ISL_3279913, EPI_ISL_3279914, EPI_ISL_3279915, EPI_ISL_3279916, EPI_ISL_3279917, EPI_ISL_3279918, EPI_ISL_3279919, EPI_ISL_3279920, EPI_ISL_3279921, EPI_ISL_3279922, EPI_ISL_3279923, EPI_ISL_3279924, EPI_ISL_3279925, EPI_ISL_3279926, EPI_ISL_3279927, EPI_ISL_3279928, EPI_ISL_3279929, EPI_ISL_3279930, EPI_ISL_3279931, EPI_ISL_3279932, EPI_ISL_3279933, EPI_ISL_3279934, EPI_ISL_3279935, EPI_ISL_3279936, EPI_ISL_3279937, EPI_ISL_3279938, EPI_ISL_3279939, EPI_ISL_3279940, EPI_ISL_3279941, EPI_ISL_3279942, EPI_ISL_3279943, EPI_ISL_3279944, EPI_ISL_3279945, EPI_ISL_3279946, EPI_ISL_3279947, EPI_ISL_3279948, EPI_ISL_3279949, EPI_ISL_3279950, EPI_ISL_3279951, EPI_ISL_3279952, EPI_ISL_3279953, EPI_ISL_3279954, EPI_ISL_3279955, EPI_ISL_3279956, EPI_ISL_3279957, EPI_ISL_3279958, EPI_ISL_3279959, EPI_ISL_3279960, EPI_ISL_3279961, EPI_ISL_3279962, EPI_ISL_3279963, EPI_ISL_3279964, EPI_ISL_3279965, EPI_ISL_3279966, EPI_ISL_3279967, EPI_ISL_3279968, EPI_ISL_3279969, EPI_ISL_3279970, EPI_ISL_3279971, EPI_ISL_3279972, EPI_ISL_3279973, EPI_ISL_3279974, EPI_ISL_3279975, EPI_ISL_3279976, EPI_ISL_3279977, EPI_ISL_3279978, EPI_ISL_3279979, EPI_ISL_3279980, EPI_ISL_3279981, EPI_ISL_3279982, EPI_ISL_3279983, EPI_ISL_3279984, EPI_ISL_3279985, EPI_ISL_3279986, EPI_ISL_3279987, EPI_ISL_3279988, EPI_ISL_3279989, EPI_ISL_3279990, EPI_ISL_3279991, EPI_ISL_3279992, EPI_ISL_3279993, EPI_ISL_3279994, EPI_ISL_3279995, EPI_ISL_3279996, EPI_ISL_3279997, EPI_ISL_3279998, EPI_ISL_3279999, EPI_ISL_3280000, EPI_ISL_3280001, EPI_ISL_3280002, EPI_ISL_3280003, EPI_ISL_3280004, EPI_ISL_3280005, EPI_ISL_3280006, EPI_ISL_3280007, EPI_ISL_3280008, EPI_ISL_3280009, EPI_ISL_3280010, EPI_ISL_3280011, EPI_ISL_3280012, EPI_ISL_3280013, EPI_ISL_3280014, EPI_ISL_3280015, EPI_ISL_3280016, EPI_ISL_3280017, EPI_ISL_3280018, EPI_ISL_3280019, EPI_ISL_3280020, EPI_ISL_3280021, EPI_ISL_3280022, EPI_ISL_3280023, EPI_ISL_3280024, EPI_ISL_3280025, EPI_ISL_3280026, EPI_ISL_3280027, EPI_ISL_3280028, EPI_ISL_3280029, EPI_ISL_3280030, EPI_ISL_3280031, EPI_ISL_3280032, EPI_ISL_3280033, EPI_ISL_3280034, EPI_ISL_3280035, EPI_ISL_3280036, EPI_ISL_3280037, EPI_ISL_3280038, EPI_ISL_3280039, EPI_ISL_3280040, EPI_ISL_3280041, EPI_ISL_3280042, EPI_ISL_3280043

| | | | |
|---|---|---|---|
| see above | Department of Bacteria, Parasites and Fungi, Statens Serum Institut, Copenhagen, Denmark | Statens Serum Institut Bioinformatics and Microbial Genomics | Danish Covid-19 Genome Consortium |
| EPI_ISL_3280044 | Department of Clinical Microbiology, Odense University Hospital, Odense, Denmark | Statens Serum Institut Bioinformatics and Microbial Genomics | Danish Covid-19 Genome Consortium |
| EPI_ISL_3280045, EPI_ISL_3280046, EPI_ISL_3280047, EPI_ISL_3280048, EPI_ISL_3280049 | Department of Bacteria, Parasites and Fungi, Statens Serum Institut, Copenhagen, Denmark | Statens Serum Institut Bioinformatics and Microbial Genomics | Danish Covid-19 Genome Consortium |
| EPI_ISL_3280050 | Department of Clinical Microbiology, Odense University Hospital, Odense, Denmark | Statens Serum Institut Bioinformatics and Microbial Genomics | Danish Covid-19 Genome Consortium |
| EPI_ISL_3280051, EPI_ISL_3280053, EPI_ISL_3280054, EPI_ISL_3280055, EPI_ISL_3280056, EPI_ISL_3280057, EPI_ISL_3280058, EPI_ISL_3280059, EPI_ISL_3280060, EPI_ISL_3280061, EPI_ISL_3280062, EPI_ISL_3280063, EPI_ISL_3280064, EPI_ISL_3280065, EPI_ISL_3280066, EPI_ISL_3280067, EPI_ISL_3280068, EPI_ISL_3280069, EPI_ISL_3280070, EPI_ISL_3280071, EPI_ISL_3280072, EPI_ISL_3280073, EPI_ISL_3280074, EPI_ISL_3280075, EPI_ISL_3280076, EPI_ISL_3280077, EPI_ISL_3280078, EPI_ISL_3280079, EPI_ISL_3280080, EPI_ISL_3280081, EPI_ISL_3280082, EPI_ISL_3280083, EPI_ISL_3280084, EPI_ISL_3280085, EPI_ISL_3280086, EPI_ISL_3280087, EPI_ISL_3280088, EPI_ISL_3280089, EPI_ISL_3280090 | | | |
| see above | Department of Bacteria, Parasites and Fungi, Statens Serum Institut, Copenhagen, Denmark | Statens Serum Institut Bioinformatics and Microbial Genomics | Danish Covid-19 Genome Consortium |
| EPI_ISL_3280091 | Department of Clinical Microbiology, Odense University Hospital, Odense, Denmark | Statens Serum Institut Bioinformatics and Microbial Genomics | Danish Covid-19 Genome Consortium |
| EPI_ISL_3280092, EPI_ISL_3280093, EPI_ISL_3280094, EPI_ISL_3280095, EPI_ISL_3280096, EPI_ISL_3280097, EPI_ISL_3280098, EPI_ISL_3280099, EPI_ISL_3280100, EPI_ISL_3280101, EPI_ISL_3280102, EPI_ISL_3280103, EPI_ISL_3280104, EPI_ISL_3280105, EPI_ISL_3280106, EPI_ISL_3280107, EPI_ISL_3280108, EPI_ISL_3280109, EPI_ISL_3280110, EPI_ISL_3280111, EPI_ISL_3280112, EPI_ISL_3280113, EPI_ISL_3280114, EPI_ISL_3280115, EPI_ISL_3280116, EPI_ISL_3280117, EPI_ISL_3280118, EPI_ISL_3280119, EPI_ISL_3280120, EPI_ISL_3280121, EPI_ISL_3280122, EPI_ISL_3280123, EPI_ISL_3280124, EPI_ISL_3280125, EPI_ISL_3280127, EPI_ISL_3280128, EPI_ISL_3280129, EPI_ISL_3280130, EPI_ISL_3280131, EPI_ISL_3280132, EPI_ISL_3280133, EPI_ISL_3280134, EPI_ISL_3280135, EPI_ISL_3280136, EPI_ISL_3280137, EPI_ISL_3280138, EPI_ISL_3280139, EPI_ISL_3280140, EPI_ISL_3280141, EPI_ISL_3280142, EPI_ISL_3280143, EPI_ISL_3280144, EPI_ISL_3280145, EPI_ISL_3280146, EPI_ISL_3280147, EPI_ISL_3280148, EPI_ISL_3280149, EPI_ISL_3280150, EPI_ISL_3280151, EPI_ISL_3280152, EPI_ISL_3280153, EPI_ISL_3280154, EPI_ISL_3280155, EPI_ISL_3280156, EPI_ISL_3280157, EPI_ISL_3280158 | | | |
| see above | Department of Bacteria, Parasites and Fungi, Statens Serum Institut, Copenhagen, Denmark | Statens Serum Institut Bioinformatics and Microbial Genomics | Danish Covid-19 Genome Consortium |
| EPI_ISL_3280159 | Department of Clinical Microbiology, Odense University Hospital, Odense, Denmark | Statens Serum Institut Bioinformatics and Microbial Genomics | Danish Covid-19 Genome Consortium |
| EPI_ISL_3280160, EPI_ISL_3280161, EPI_ISL_3280162, EPI_ISL_3280163 | Department of Bacteria, Parasites and Fungi, Statens Serum Institut, Copenhagen, Denmark | Statens Serum Institut Bioinformatics and Microbial Genomics | Danish Covid-19 Genome Consortium |
| EPI_ISL_3280164 | Department of Clinical Microbiology, Odense University Hospital, Odense, Denmark | Statens Serum Institut Bioinformatics and Microbial Genomics | Danish Covid-19 Genome Consortium |
| EPI_ISL_3280165, EPI_ISL_3280166, EPI_ISL_3280167, EPI_ISL_3280168, EPI_ISL_3280169, EPI_ISL_3280170, EPI_ISL_3280171, EPI_ISL_3280172 | Department of Bacteria, Parasites and Fungi, Statens Serum Institut, Copenhagen, Denmark | Statens Serum Institut Bioinformatics and Microbial Genomics | Danish Covid-19 Genome Consortium |
| EPI_ISL_3280173 | Department of Clinical Microbiology, Odense University Hospital, Odense, Denmark | Statens Serum Institut Bioinformatics and Microbial Genomics | Danish Covid-19 Genome Consortium |
| EPI_ISL_3280174, EPI_ISL_3280175, EPI_ISL_3280176, EPI_ISL_3280177, EPI_ISL_3280178, EPI_ISL_3280179, EPI_ISL_3280180, EPI_ISL_3280181, EPI_ISL_3280182, EPI_ISL_3280183, EPI_ISL_3280184, EPI_ISL_3280185, EPI_ISL_3280186, EPI_ISL_3280187, EPI_ISL_3280188, EPI_ISL_3280189, EPI_ISL_3280190, EPI_ISL_3280191, EPI_ISL_3280192, EPI_ISL_3280193, EPI_ISL_3280194, EPI_ISL_3280195, EPI_ISL_3280196, EPI_ISL_3280197, EPI_ISL_3280198, EPI_ISL_3280199, EPI_ISL_3280200, EPI_ISL_3280201, EPI_ISL_3280202, EPI_ISL_3280203, EPI_ISL_3280204, EPI_ISL_3280205, EPI_ISL_3280206, EPI_ISL_3280207, EPI_ISL_3280208, EPI_ISL_3280209, EPI_ISL_3280210, EPI_ISL_3280211, EPI_ISL_3280212, EPI_ISL_3280213, EPI_ISL_3280214, EPI_ISL_3280215, EPI_ISL_3280216, EPI_ISL_3280217, EPI_ISL_3280218, EPI_ISL_3280219, EPI_ISL_3280220, EPI_ISL_3280221, EPI_ISL_3280222, EPI_ISL_3280223, EPI_ISL_3280224, EPI_ISL_3280225, EPI_ISL_3280226, EPI_ISL_3280227, EPI_ISL_3280228, EPI_ISL_3280229, EPI_ISL_3280230, EPI_ISL_3280231, EPI_ISL_3280232, EPI_ISL_3280233, EPI_ISL_3280234, EPI_ISL_3280235, EPI_ISL_3280236, EPI_ISL_3280237, EPI_ISL_3280238, EPI_ISL_3280239, EPI_ISL_3280240, EPI_ISL_3280241, EPI_ISL_3280242, EPI_ISL_3280244, EPI_ISL_3280245, EPI_ISL_3280246, EPI_ISL_3280247, EPI_ISL_3280248, EPI_ISL_3280249, EPI_ISL_3280250, EPI_ISL_3280251, EPI_ISL_3280252, EPI_ISL_3280253, EPI_ISL_3280254, EPI_ISL_3280255, EPI_ISL_3280256, EPI_ISL_3280257, EPI_ISL_3280258, EPI_ISL_3280259, EPI_ISL_3280260, EPI_ISL_3280261, EPI_ISL_3280262, EPI_ISL_3280263, EPI_ISL_3280264, EPI_ISL_3280265, EPI_ISL_3280266, EPI_ISL_3280267, EPI_ISL_3280268, EPI_ISL_3280269, EPI_ISL_3280270, EPI_ISL_3280271, EPI_ISL_3280272, EPI_ISL_3280273, EPI_ISL_3280274, EPI_ISL_3280275, EPI_ISL_3280276, EPI_ISL_3280277, EPI_ISL_3280278, EPI_ISL_3280279, EPI_ISL_3280280, EPI_ISL_3280281, EPI_ISL_3280282, EPI_ISL_3280283, EPI_ISL_3280284, EPI_ISL_3280285, EPI_ISL_3280286, EPI_ISL_3280287, EPI_ISL_3280288, EPI_ISL_3280289, EPI_ISL_3280290, EPI_ISL_3280291, EPI_ISL_3280292, EPI_ISL_3280293, EPI_ISL_3280294, EPI_ISL_3280295, EPI_ISL_3280296, EPI_ISL_3280297, EPI_ISL_3280298, EPI_ISL_3280299, EPI_ISL_3280300, EPI_ISL_3280301, EPI_ISL_3280302, EPI_ISL_3280303, EPI_ISL_3280304, EPI_ISL_3280305, EPI_ISL_3280306, EPI_ISL_3280307, EPI_ISL_3280308, EPI_ISL_3280309, EPI_ISL_3280310 | | | |
| see above | Department of Bacteria, Parasites and Fungi, Statens Serum Institut, Copenhagen, Denmark | Statens Serum Institut Bioinformatics and Microbial Genomics | Danish Covid-19 Genome Consortium |
| EPI_ISL_3280311 | Department of Clinical Microbiology, Odense University Hospital, Odense, Denmark | Statens Serum Institut Bioinformatics and Microbial Genomics | Danish Covid-19 Genome Consortium |
| EPI_ISL_3280312, EPI_ISL_3280313, EPI_ISL_3280314, EPI_ISL_3280315, EPI_ISL_3280316, EPI_ISL_3280317, EPI_ISL_3280319, EPI_ISL_3280320, EPI_ISL_3280321, EPI_ISL_3280322, EPI_ISL_3280323, EPI_ISL_3280324, EPI_ISL_3280325, EPI_ISL_3280326, EPI_ISL_3280327, EPI_ISL_3280328, EPI_ISL_3280329, EPI_ISL_3280330, EPI_ISL_3280331, EPI_ISL_3280332, EPI_ISL_3280333, EPI_ISL_3280334, EPI_ISL_3280335, EPI_ISL_3280336, EPI_ISL_3280337, EPI_ISL_3280338, EPI_ISL_3280339, EPI_ISL_3280340, EPI_ISL_3280341, EPI_ISL_3280342, EPI_ISL_3280343, EPI_ISL_3280344, EPI_ISL_3280345, EPI_ISL_3280346, EPI_ISL_3280347, EPI_ISL_3280348, EPI_ISL_3280349, EPI_ISL_3280350, EPI_ISL_3280351, EPI_ISL_3280352, EPI_ISL_3280353, EPI_ISL_3280354, EPI_ISL_3280355, EPI_ISL_3280356, EPI_ISL_3280357, EPI_ISL_3280358, EPI_ISL_3280359, EPI_ISL_3280360, EPI_ISL_3280361, EPI_ISL_3280362, EPI_ISL_3280363, EPI_ISL_3280364, EPI_ISL_3280365, EPI_ISL_3280366, EPI_ISL_3280367, EPI_ISL_3280368, EPI_ISL_3280369, EPI_ISL_3280370, EPI_ISL_3280371, EPI_ISL_3280372, EPI_ISL_3280373, EPI_ISL_3280374, EPI_ISL_3280375, EPI_ISL_3280376 | | | |
| see above | Department of Bacteria, Parasites and Fungi, Statens Serum Institut, Copenhagen, Denmark | Statens Serum Institut Bioinformatics and Microbial Genomics | Danish Covid-19 Genome Consortium |
| EPI_ISL_3280377 | Department of Clinical Microbiology, Odense University Hospital, Odense, Denmark | Statens Serum Institut Bioinformatics and Microbial Genomics | Danish Covid-19 Genome Consortium |
| EPI_ISL_3280378, EPI_ISL_3280379, EPI_ISL_3280380, EPI_ISL_3280381, EPI_ISL_3280382, EPI_ISL_3280383, EPI_ISL_3280384, EPI_ISL_3280385, EPI_ISL_3280386, EPI_ISL_3280387, EPI_ISL_3280388, EPI_ISL_3280389, EPI_ISL_3280390, EPI_ISL_3280391, EPI_ISL_3280392, EPI_ISL_3280393, EPI_ISL_3280394, EPI_ISL_3280395, EPI_ISL_3280396, EPI_ISL_3280397, EPI_ISL_3280398, EPI_ISL_3280399, EPI_ISL_3280400, EPI_ISL_3280401, EPI_ISL_3280402, EPI_ISL_3280403, EPI_ISL_3280404, EPI_ISL_3280405, EPI_ISL_3280406, EPI_ISL_3280407, EPI_ISL_3280409, EPI_ISL_3280410, EPI_ISL_3280411, EPI_ISL_3280412, EPI_ISL_3280413, EPI_ISL_3280414, EPI_ISL_3280415, EPI_ISL_3280416, EPI_ISL_3280417, EPI_ISL_3280418, EPI_ISL_3280419, EPI_ISL_3280420, EPI_ISL_3280421, EPI_ISL_3280422, EPI_ISL_3280424, EPI_ISL_3280425, EPI_ISL_3280426, EPI_ISL_3280427, EPI_ISL_3280428, EPI_ISL_3280429, EPI_ISL_3280430, EPI_ISL_3280431, EPI_ISL_3280432, EPI_ISL_3280433, EPI_ISL_3280434, EPI_ISL_3280435, EPI_ISL_3280436, EPI_ISL_3280437, EPI_ISL_3280438, EPI_ISL_3280439, EPI_ISL_3280440, EPI_ISL_3280441, EPI_ISL_3280442, EPI_ISL_3280443, EPI_ISL_3280444, EPI_ISL_3280445, EPI_ISL_3280446, EPI_ISL_3280447, EPI_ISL_3280448, EPI_ISL_3280449, EPI_ISL_3280450, EPI_ISL_3280451, EPI_ISL_3280452, EPI_ISL_3280453, EPI_ISL_3280454, EPI_ISL_3280455, EPI_ISL_3280456, EPI_ISL_3280457, EPI_ISL_3280458, EPI_ISL_3280459, EPI_ISL_3280460, EPI_ISL_3280461, EPI_ISL_3280462, EPI_ISL_3280463, EPI_ISL_3280464, EPI_ISL_3280465, EPI_ISL_3280466, EPI_ISL_3280467, EPI_ISL_3280468, EPI_ISL_3280469, EPI_ISL_3280470, EPI_ISL_3280471, EPI_ISL_3280472, EPI_ISL_3280473, EPI_ISL_3280474, EPI_ISL_3280475, EPI_ISL_3280476, EPI_ISL_3280477, EPI_ISL_3280478, EPI_ISL_3280479, EPI_ISL_3280480, EPI_ISL_3280481, EPI_ISL_3280482, EPI_ISL_3280483, EPI_ISL_3280484, EPI_ISL_3280485, EPI_ISL_3280486, EPI_ISL_3280487 | | | |
| see above | Department of Bacteria, Parasites and Fungi, Statens Serum Institut, Copenhagen, Denmark | Statens Serum Institut Bioinformatics and Microbial Genomics | Danish Covid-19 Genome Consortium |
| EPI_ISL_3280488 | Department of Clinical Microbiology, Odense University Hospital, Odense, Denmark | Statens Serum Institut Bioinformatics and Microbial Genomics | Danish Covid-19 Genome Consortium |
| EPI_ISL_3280489, EPI_ISL_3280490, EPI_ISL_3280491, EPI_ISL_3280492, EPI_ISL_3280493, EPI_ISL_3280494, EPI_ISL_3280495, EPI_ISL_3280496, EPI_ISL_3280497, EPI_ISL_3280498, EPI_ISL_3280499, EPI_ISL_3280500, EPI_ISL_3280501, EPI_ISL_3280502, EPI_ISL_3280503, EPI_ISL_3280504, EPI_ISL_3280505, EPI_ISL_3280506, EPI_ISL_3280507, EPI_ISL_3280508, EPI_ISL_3280509, EPI_ISL_3280510, EPI_ISL_3280512, EPI_ISL_3280513, EPI_ISL_3280514, EPI_ISL_3280515, EPI_ISL_3280516, EPI_ISL_3280517, EPI_ISL_3280518, EPI_ISL_3280519, EPI_ISL_3280520, EPI_ISL_3280521, EPI_ISL_3280522, EPI_ISL_3280523, EPI_ISL_3280524, EPI_ISL_3280525, EPI_ISL_3280526, EPI_ISL_3280527, EPI_ISL_3280528 | | | |
| see above | Department of Bacteria, Parasites and Fungi, Statens Serum Institut, Copenhagen, Denmark | Statens Serum Institut Bioinformatics and Microbial Genomics | Danish Covid-19 Genome Consortium |
| EPI_ISL_3280529 | Department of Clinical Microbiology, Odense University Hospital, Odense, Denmark | Statens Serum Institut Bioinformatics and Microbial Genomics | Danish Covid-19 Genome Consortium |
| EPI_ISL_3280530, EPI_ISL_3280531, EPI_ISL_3280532, EPI_ISL_3280533, EPI_ISL_3280534, EPI_ISL_3280535, EPI_ISL_3280536, EPI_ISL_3280537, EPI_ISL_3280538, EPI_ISL_3280539, EPI_ISL_3280540, EPI_ISL_3280541, EPI_ISL_3280542, EPI_ISL_3280543, EPI_ISL_3280544, EPI_ISL_3280545, EPI_ISL_3280546, EPI_ISL_3280547, EPI_ISL_3280548, EPI_ISL_3280549, EPI_ISL_3280550, EPI_ISL_3280551, EPI_ISL_3280552, EPI_ISL_3280553, EPI_ISL_3280554, EPI_ISL_3280555, EPI_ISL_3280556, EPI_ISL_3280557, EPI_ISL_3280558, EPI_ISL_3280559 | | | |
| see above | Department of Bacteria, Parasites and Fungi, Statens Serum Institut, Copenhagen, Denmark | Statens Serum Institut Bioinformatics and Microbial Genomics | Danish Covid-19 Genome Consortium |
| EPI_ISL_3280560 | Department of Clinical Microbiology, Odense University Hospital, Odense, Denmark | Statens Serum Institut Bioinformatics and Microbial Genomics | Danish Covid-19 Genome Consortium |
| EPI_ISL_3280561, EPI_ISL_3280562, EPI_ISL_3280563, EPI_ISL_3280564, EPI_ISL_3280565, EPI_ISL_3280566, EPI_ISL_3280567, EPI_ISL_3280568, EPI_ISL_3280569, EPI_ISL_3280570, EPI_ISL_3280571, EPI_ISL_3280572, EPI_ISL_3280573, EPI_ISL_3280574, EPI_ISL_3280575, EPI_ISL_3280576, EPI_ISL_3280577, EPI_ISL_3280578, EPI_ISL_3280579, EPI_ISL_3280580, EPI_ISL_3280581, EPI_ISL_3280582, EPI_ISL_3280583, EPI_ISL_3280584, EPI_ISL_3280585, EPI_ISL_3280586, EPI_ISL_3280587, EPI_ISL_3280588, EPI_ISL_3280589, EPI_ISL_3280590, EPI_ISL_3280591, EPI_ISL_3280592, EPI_ISL_3280593, EPI_ISL_3280594, EPI_ISL_3280595, EPI_ISL_3280596, EPI_ISL_3280597, EPI_ISL_3280598, EPI_ISL_3280599, EPI_ISL_3280600, EPI_ISL_3280601, EPI_ISL_3280602, EPI_ISL_3280603, EPI_ISL_3280604, EPI_ISL_3280605, EPI_ISL_3280606, EPI_ISL_3280607, EPI_ISL_3280608, EPI_ISL_3280609, EPI_ISL_3280610, EPI_ISL_3280611, EPI_ISL_3280612, EPI_ISL_3280613, EPI_ISL_3280614, EPI_ISL_3280615, EPI_ISL_3280616, EPI_ISL_3280617, EPI_ISL_3280618, EPI_ISL_3280619, EPI_ISL_3280620, EPI_ISL_3280621, EPI_ISL_3280622, EPI_ISL_3280623, EPI_ISL_3280624, EPI_ISL_3280625, EPI_ISL_3280626, EPI_ISL_3280627, EPI_ISL_3280628, EPI_ISL_3280629, EPI_ISL_3280630, EPI_ISL_3280631, EPI_ISL_3280632, EPI_ISL_3280633, EPI_ISL_3280634, EPI_ISL_3280635, EPI_ISL_3280636, EPI_ISL_3280637, EPI_ISL_3280638, EPI_ISL_3280639, EPI_ISL_3280640, EPI_ISL_3280641, EPI_ISL_3280642, EPI_ISL_3280643, EPI_ISL_3280644, EPI_ISL_3280645, EPI_ISL_3280646, EPI_ISL_3280647, EPI_ISL_3280648, EPI_ISL_3280649, EPI_ISL_3280650, EPI_ISL_3280651, EPI_ISL_3280652, EPI_ISL_3280653, EPI_ISL_3280654, EPI_ISL_3280655, EPI_ISL_3280656, EPI_ISL_3280657, EPI_ISL_3280658, EPI_ISL_3280659, EPI_ISL_3280660, EPI_ISL_3280661, EPI_ISL_3280662, EPI_ISL_3280663, EPI_ISL_3280664, EPI_ISL_3280665, EPI_ISL_3280667, EPI_ISL_3280668, EPI_ISL_3280669, EPI_ISL_3280670, EPI_ISL_3280671, EPI_ISL_3280672, EPI_ISL_3280673, EPI_ISL_3280674, EPI_ISL_3280675, EPI_ISL_3280676, EPI_ISL_3280677, EPI_ISL_3280678, EPI_ISL_3280679, EPI_ISL_3280680, EPI_ISL_3280681, EPI_ISL_3280682, EPI_ISL_3280683, EPI_ISL_3280684, EPI_ISL_3280687, EPI_ISL_3280688, EPI_ISL_3280689, EPI_ISL_3280690, EPI_ISL_3280691, EPI_ISL_3280692, EPI_ISL_3280693, EPI_ISL_3280694, EPI_ISL_3280695, EPI_ISL_3280696, EPI_ISL_3280697, EPI_ISL_3280698, EPI_ISL_3280699, EPI_ISL_3280700, EPI_ISL_3280701, EPI_ISL_3280702, EPI_ISL_3280703, EPI_ISL_3280704, EPI_ISL_3280705, EPI_ISL_3280706 | | | |
| see above | Department of Bacteria, Parasites and Fungi, Statens Serum Institut, Copenhagen, Denmark | Statens Serum Institut Bioinformatics and Microbial Genomics | Danish Covid-19 Genome Consortium |
| EPI_ISL_3280707 | Department of Clinical Microbiology, Odense University Hospital, Odense, Denmark | Statens Serum Institut Bioinformatics and Microbial Genomics | Danish Covid-19 Genome Consortium |
| EPI_ISL_3280708, EPI_ISL_3280709, EPI_ISL_3280710, EPI_ISL_3280711, EPI_ISL_3280712, EPI_ISL_3280713, EPI_ISL_3280714, EPI_ISL_3280715, EPI_ISL_3280716, EPI_ISL_3280717, EPI_ISL_3280718, EPI_ISL_3280719, EPI_ISL_3280720, EPI_ISL_3280721, EPI_ISL_3280722, EPI_ISL_3280723, EPI_ISL_3280724, EPI_ISL_3280725, EPI_ISL_3280726, EPI_ISL_3280727, EPI_ISL_3280728, EPI_ISL_3280729, EPI_ISL_3280730, EPI_ISL_3280731, EPI_ISL_3280732, EPI_ISL_3280733, EPI_ISL_3280734, EPI_ISL_3280735, EPI_ISL_3280736, EPI_ISL_3280737, EPI_ISL_3280738, EPI_ISL_3280739, EPI_ISL_3280740, EPI_ISL_3280741, EPI_ISL_3280742, EPI_ISL_3280743, EPI_ISL_3280744, EPI_ISL_3280745, EPI_ISL_3280746, EPI_ISL_3280747, EPI_ISL_3280748, EPI_ISL_3280749, EPI_ISL_3280750, EPI_ISL_3280751, EPI_ISL_3280752, EPI_ISL_3280753, EPI_ISL_3280754, EPI_ISL_3280755, EPI_ISL_3280756, EPI_ISL_3280757, EPI_ISL_3280758, EPI_ISL_3280759, EPI_ISL_3280760, EPI_ISL_3280761, EPI_ISL_3280762, EPI_ISL_3280763, EPI_ISL_3280764, EPI_ISL_3280765, EPI_ISL_3280766, EPI_ISL_3280767, EPI_ISL_3280768, EPI_ISL_3280769, EPI_ISL_3280770, EPI_ISL_3280771, EPI_ISL_3280772, EPI_ISL_3280773, EPI_ISL_3280774, EPI_ISL_3280775, EPI_ISL_3280776, EPI_ISL_3280777, EPI_ISL_3280778, EPI_ISL_3280779, EPI_ISL_3280780, EPI_ISL_3280781, EPI_ISL_3280782, EPI_ISL_3280783, EPI_ISL_3280784, EPI_ISL_3280785, EPI_ISL_3280786, EPI_ISL_3280787, EPI_ISL_3280788, EPI_ISL_3280789, EPI_ISL_3280790, EPI_ISL_3280791, EPI_ISL_3280792, EPI_ISL_3280793, EPI_ISL_3280794, EPI_ISL_3280795 | | | |
| see above | Department of Bacteria, Parasites and Fungi, Statens Serum | Statens Serum Institut Bioinformatics and Microbial Genomics | Danish Covid-19 Genome Consortium |

| | | | |
|---|---|---|---|
| | Institut, Copenhagen, Denmark | | |
| EPI_ISL_3280796 | Department of Clinical Microbiology, Odense University Hospital, Odense, Denmark | Statens Serum Institut Bioinformatics and Microbial Genomics | Danish Covid-19 Genome Consortium |
| EPI_ISL_3280797, EPI_ISL_3280798, EPI_ISL_3280799, EPI_ISL_3280800, EPI_ISL_3280801, EPI_ISL_3280802, EPI_ISL_3280803, EPI_ISL_3280804, EPI_ISL_3280805, EPI_ISL_3280806, EPI_ISL_3280807, EPI_ISL_3280808, EPI_ISL_3280809, EPI_ISL_3280810, EPI_ISL_3280811 | | | |
| see above | Department of Bacteria, Parasites and Fungi, Statens Serum Institut, Copenhagen, Denmark | Statens Serum Institut Bioinformatics and Microbial Genomics | Danish Covid-19 Genome Consortium |
| EPI_ISL_3280812, EPI_ISL_3280813, EPI_ISL_3280814, EPI_ISL_3280815, EPI_ISL_3280816, EPI_ISL_3280817, EPI_ISL_3280818, EPI_ISL_3280819, EPI_ISL_3280820, EPI_ISL_3280821, EPI_ISL_3280822, EPI_ISL_3280823, EPI_ISL_3280824, EPI_ISL_3280825, EPI_ISL_3280826, EPI_ISL_3280827, EPI_ISL_3280828, EPI_ISL_3280829, EPI_ISL_3280830, EPI_ISL_3280831, EPI_ISL_3280832, EPI_ISL_3280833, EPI_ISL_3280834, EPI_ISL_3280835, EPI_ISL_3280836, EPI_ISL_3280837, EPI_ISL_3280838, EPI_ISL_3280839 | | | |
| see above | New South Wales Health Pathology Royal Prince Alfred Hospital | Microbiology RPAH | Foster, C.; Au, J.; Ruiz Silva, M.; Deveson, I.; Bull, R.; Van Hal, S.; Rawlinson, W. |
| EPI_ISL_3280840, EPI_ISL_3280841, EPI_ISL_3280842, EPI_ISL_3280844, EPI_ISL_3280845, EPI_ISL_3280846, EPI_ISL_3280848, EPI_ISL_3280849, EPI_ISL_3280850, EPI_ISL_3280851, EPI_ISL_3280852, EPI_ISL_3280853, EPI_ISL_3280854, EPI_ISL_3280855, EPI_ISL_3280856, EPI_ISL_3280857, EPI_ISL_3280858, EPI_ISL_3280859, EPI_ISL_3280861, EPI_ISL_3280862, EPI_ISL_3280863, EPI_ISL_3280864, EPI_ISL_3280865, EPI_ISL_3280866, EPI_ISL_3280867, EPI_ISL_3280869, EPI_ISL_3280870, EPI_ISL_3280871, EPI_ISL_3280872 | | | |
| see above | Laboratorio di Riferimento Regionale della Sicilia Occidentale per l'Emergenza COVID-19 | Laboratorio di Riferimento Regionale della Sicilia Occidentale per l'Emergenza COVID-19 | Fabio Tramuto, Carmelo Massimo Maida, Daniela Di Naro, Giulia Randazzo, Walter Mazzucco, Giorgio Graziano, Vincenzo Restivo, Claudio Costantino, Francesco Vitale |
| EPI_ISL_3280874 | RM1/IRCCS IDI | National Institute for Infectious Diseases (INMI) L. Spallanzani I.R.C.C.S | E Giombini, B Bartolini, M Rueca, G Bonfiglio, O Butera, CEM Gruber, F Santini, F Messina, A Di Caro, MR Capobianchi |
| EPI_ISL_3280875 | RM2/A.U. Tor Vergata | National Institute for Infectious Diseases (INMI) L. Spallanzani I.R.C.C.S | E Giombini, B Bartolini, M Rueca, G Bonfiglio, O Butera, CEM Gruber, F Santini, F Messina, A Di Caro, MR Capobianchi |
| EPI_ISL_3280876, EPI_ISL_3280877, EPI_ISL_3280878 | RM1/IRCCS IDI | National Institute for Infectious Diseases (INMI) L. Spallanzani I.R.C.C.S | E Giombini, B Bartolini, M Rueca, G Bonfiglio, O Butera, CEM Gruber, F Santini, F Messina, A Di Caro, MR Capobianchi |
| EPI_ISL_3280879 | RM3/Osp. G.Grassi | National Institute for Infectious Diseases (INMI) L. Spallanzani I.R.C.C.S | E Giombini, B Bartolini, M Rueca, G Bonfiglio, O Butera, CEM Gruber, F Santini, F Messina, A Di Caro, MR Capobianchi |
| EPI_ISL_3280880 | RM1/IRCCS IDI | National Institute for Infectious Diseases (INMI) L. Spallanzani I.R.C.C.S | E Giombini, B Bartolini, M Rueca, G Bonfiglio, O Butera, CEM Gruber, F Santini, F Messina, A Di Caro, MR Capobianchi |
| EPI_ISL_3280881 | RM2/A.U. Tor Vergata | National Institute for Infectious Diseases (INMI) L. Spallanzani I.R.C.C.S | E Giombini, B Bartolini, M Rueca, G Bonfiglio, O Butera, CEM Gruber, F Santini, F Messina, A Di Caro, MR Capobianchi |
| EPI_ISL_3280882, EPI_ISL_3280883, EPI_ISL_3280884 | RM1/IRCCS IDI | National Institute for Infectious Diseases (INMI) L. Spallanzani I.R.C.C.S | E Giombini, B Bartolini, M Rueca, G Bonfiglio, O Butera, CEM Gruber, F Santini, F Messina, A Di Caro, MR Capobianchi |
| EPI_ISL_3280885 | RM2/A.U. Tor Vergata | National Institute for Infectious Diseases (INMI) L. Spallanzani I.R.C.C.S | E Giombini, B Bartolini, M Rueca, G Bonfiglio, O Butera, CEM Gruber, F Santini, F Messina, A Di Caro, MR Capobianchi |
| EPI_ISL_3280886 | RM2/Osp. S.Eugenio | National Institute for Infectious Diseases (INMI) L. Spallanzani I.R.C.C.S | E Giombini, B Bartolini, M Rueca, G Bonfiglio, O Butera, CEM Gruber, F Santini, F Messina, A Di Caro, MR Capobianchi |
| EPI_ISL_3280887 | RM2/A.U. Tor Vergata | National Institute for Infectious Diseases (INMI) L. Spallanzani I.R.C.C.S | E Giombini, B Bartolini, M Rueca, G Bonfiglio, O Butera, CEM Gruber, F Santini, F Messina, A Di Caro, MR Capobianchi |
| EPI_ISL_3280888 | RM1/IRCCS IDI | National Institute for Infectious Diseases (INMI) L. Spallanzani I.R.C.C.S | E Giombini, B Bartolini, M Rueca, G Bonfiglio, O Butera, CEM Gruber, F Santini, F Messina, A Di Caro, MR Capobianchi |
| EPI_ISL_3280889, EPI_ISL_3280890 | RM2/A.U. Tor Vergata | National Institute for Infectious Diseases (INMI) L. Spallanzani I.R.C.C.S | E Giombini, B Bartolini, M Rueca, G Bonfiglio, O Butera, CEM Gruber, F Santini, F Messina, A Di Caro, MR Capobianchi |
| EPI_ISL_3280892, EPI_ISL_3280893 | RM1/IRCCS IDI | National Institute for Infectious Diseases (INMI) L. Spallanzani I.R.C.C.S | E Giombini, B Bartolini, M Rueca, G Bonfiglio, O Butera, CEM Gruber, F Santini, F Messina, A Di Caro, MR Capobianchi |
| EPI_ISL_3280894, EPI_ISL_3280895 | LT/Osp. S.Maria Goretti | National Institute for Infectious Diseases (INMI) L. Spallanzani I.R.C.C.S | E Giombini, B Bartolini, M Rueca, G Bonfiglio, O Butera, CEM Gruber, F Santini, F Messina, A Di Caro, MR Capobianchi |
| EPI_ISL_3280896 | RM2/A.U. Tor Vergata | National Institute for Infectious Diseases (INMI) L. Spallanzani I.R.C.C.S | E Giombini, B Bartolini, M Rueca, G Bonfiglio, O Butera, CEM Gruber, F Santini, F Messina, A Di Caro, MR Capobianchi |
| EPI_ISL_3280897 | RM1/IRCCS IDI | National Institute for Infectious Diseases (INMI) L. Spallanzani I.R.C.C.S | E Giombini, B Bartolini, M Rueca, G Bonfiglio, O Butera, CEM Gruber, F Santini, F Messina, A Di Caro, MR Capobianchi |
| EPI_ISL_3280898, EPI_ISL_3280901, EPI_ISL_3280902, EPI_ISL_3280903, EPI_ISL_3280904, EPI_ISL_3280905, EPI_ISL_3280906, EPI_ISL_3280907, EPI_ISL_3280908, EPI_ISL_3280909 | RM2/A.U. Tor Vergata | National Institute for Infectious Diseases (INMI) L. Spallanzani I.R.C.C.S | E Giombini, B Bartolini, M Rueca, G Bonfiglio, O Butera, CEM Gruber, F Santini, F Messina, A Di Caro, MR Capobianchi |
| EPI_ISL_3280911, EPI_ISL_3280912 | LT/Osp. S.Maria Goretti | National Institute for Infectious Diseases (INMI) L. Spallanzani I.R.C.C.S | E Giombini, B Bartolini, M Rueca, G Bonfiglio, O Butera, CEM Gruber, F Santini, F Messina, A Di Caro, MR Capobianchi |
| EPI_ISL_3280913, EPI_ISL_3280914 | RM2/Osp. S.Eugenio | National Institute for Infectious Diseases (INMI) L. Spallanzani I.R.C.C.S | E Giombini, B Bartolini, M Rueca, G Bonfiglio, O Butera, CEM Gruber, F Santini, F Messina, A Di Caro, MR Capobianchi |
| EPI_ISL_3280915 | VT/Osp. Di Belcolle | National Institute for Infectious Diseases (INMI) L. Spallanzani I.R.C.C.S | E Giombini, B Bartolini, M Rueca, G Bonfiglio, O Butera, CEM Gruber, F Santini, F Messina, A Di Caro, MR Capobianchi |
| EPI_ISL_3280916 | RM2/Osp. S.Eugenio | National Institute for Infectious Diseases (INMI) L. Spallanzani I.R.C.C.S | E Giombini, B Bartolini, M Rueca, G Bonfiglio, O Butera, CEM Gruber, F Santini, F Messina, A Di Caro, MR Capobianchi |
| EPI_ISL_3280917 | VT/Osp. Di Belcolle | National Institute for Infectious Diseases (INMI) L. Spallanzani I.R.C.C.S | E Giombini, B Bartolini, M Rueca, G Bonfiglio, O Butera, CEM Gruber, F Santini, F Messina, A Di Caro, MR Capobianchi |
| EPI_ISL_3280918 | RM2/Osp. S.Eugenio | National Institute for Infectious Diseases (INMI) L. Spallanzani I.R.C.C.S | E Giombini, B Bartolini, M Rueca, G Bonfiglio, O Butera, CEM Gruber, F Santini, F Messina, A Di Caro, MR Capobianchi |
| EPI_ISL_3280919 | VT/Osp. Di Belcolle | National Institute for Infectious Diseases (INMI) L. Spallanzani I.R.C.C.S | E Giombini, B Bartolini, M Rueca, G Bonfiglio, O Butera, CEM Gruber, F Santini, F Messina, A Di Caro, MR Capobianchi |
| EPI_ISL_3280920 | RM2/Osp. S.Eugenio | National Institute for Infectious Diseases (INMI) L. Spallanzani I.R.C.C.S | E Giombini, B Bartolini, M Rueca, G Bonfiglio, O Butera, CEM Gruber, F Santini, F Messina, A Di Caro, MR Capobianchi |
| EPI_ISL_3280921, EPI_ISL_3280922, EPI_ISL_3280923, EPI_ISL_3280924 | VT/Osp. Di Belcolle | National Institute for Infectious Diseases (INMI) L. Spallanzani I.R.C.C.S | E Giombini, B Bartolini, M Rueca, G Bonfiglio, O Butera, CEM Gruber, F Santini, F Messina, A Di Caro, MR Capobianchi |
| EPI_ISL_3280927, EPI_ISL_3280928, EPI_ISL_3280929, EPI_ISL_3280930, EPI_ISL_3280931, EPI_ISL_3280932, EPI_ISL_3280933, EPI_ISL_3280934, EPI_ISL_3280935, EPI_ISL_3280936, EPI_ISL_3280937, EPI_ISL_3280938, EPI_ISL_3280939, EPI_ISL_3280940, EPI_ISL_3280941, EPI_ISL_3280942, EPI_ISL_3280943, EPI_ISL_3280944, EPI_ISL_3280945, EPI_ISL_3280946, EPI_ISL_3280947, EPI_ISL_3280948, EPI_ISL_3280949, EPI_ISL_3280950, EPI_ISL_3280951, EPI_ISL_3280952, EPI_ISL_3280953, EPI_ISL_3280954, EPI_ISL_3280955, EPI_ISL_3280956, EPI_ISL_3280957, EPI_ISL_3280958, EPI_ISL_3280959, EPI_ISL_3280960, EPI_ISL_3280961, EPI_ISL_3280962, EPI_ISL_3280963, EPI_ISL_3280964, EPI_ISL_3280965, EPI_ISL_3280966, EPI_ISL_3280967, EPI_ISL_3280968, EPI_ISL_3280969, EPI_ISL_3280970, EPI_ISL_3280971, EPI_ISL_3280972, EPI_ISL_3280973, EPI_ISL_3280974, EPI_ISL_3280975, EPI_ISL_3280976, EPI_ISL_3280977, EPI_ISL_3280978, EPI_ISL_3280979, EPI_ISL_3280980, EPI_ISL_3280981, EPI_ISL_3280982, EPI_ISL_3280983, EPI_ISL_3280984, EPI_ISL_3280985, EPI_ISL_3280986, EPI_ISL_3280987, EPI_ISL_3280988, EPI_ISL_3280989, EPI_ISL_3280990, EPI_ISL_3280991, EPI_ISL_3280992, EPI_ISL_3280993, EPI_ISL_3280994, EPI_ISL_3280995, EPI_ISL_3280996, EPI_ISL_3280997, EPI_ISL_3280998, EPI_ISL_3280999, EPI_ISL_3281000, EPI_ISL_3281001, EPI_ISL_3281002, EPI_ISL_3281003, EPI_ISL_3281004, EPI_ISL_3281005, EPI_ISL_3281006, EPI_ISL_3281007, EPI_ISL_3281008, EPI_ISL_3281009, EPI_ISL_3281010, EPI_ISL_3281011, EPI_ISL_3281012, EPI_ISL_3281013 | | | |
| see above | New South Wales Health Pathology Royal Prince Alfred Hospital | Microbiology RPAH | Foster, C.; Au, J.; Ruiz Silva, M.; Deveson, I.; Bull, R.; Van Hal, S.; Rawlinson, W. |
| EPI_ISL_3281014, EPI_ISL_3281015, EPI_ISL_3281016 | National Public Health Laboratory | Institute for Medical Research, Infectious Disease Research Centre, National Institutes of Health, Ministry of Health Malaysia | Suppiah J, Kamel K, Mohd Zawawi Z, Azizan MA, Ramly N, Robert F, Anasir MI, Thayan R |
| EPI_ISL_3281017, EPI_ISL_3281018 | St Vincent's Pathology (SydPath) | NSW Health Pathology - Institute of Clinical Pathology and Medical Research; Westmead Hospital; University of Sydney | Arnott A., Draper J., Gall M., Martinez E., Rockett R., Sintchenko V., on behalf of ICPMR |
| EPI_ISL_3281019 | Histopath | NSW Health Pathology - Institute of Clinical Pathology and Medical Research; Westmead Hospital; University of Sydney | Arnott A., Draper J., Gall M., Martinez E., Rockett R., Sintchenko V., on behalf of ICPMR |
| EPI_ISL_3281020, EPI_ISL_3281021 | Laverty Pathology | NSW Health Pathology - Institute of Clinical Pathology and Medical Research; Westmead Hospital; University of Sydney | Arnott A., Draper J., Gall M., Martinez E., Rockett R., Sintchenko V., on behalf of ICPMR |
| EPI_ISL_3281022 | Histopath | NSW Health Pathology - Institute of Clinical Pathology and Medical | Arnott A., Draper J., Gall M., Martinez E., Rockett R., Sintchenko V., on behalf of ICPMR |

| | | | |
|---|---|---|---|
| | | Research; Westmead Hospital; University of Sydney | |
| EPI_ISL_3281023 | Laverty Pathology | NSW Health Pathology - Institute of Clinical Pathology and Medical Research; Westmead Hospital; University of Sydney | Arnott A., Draper J., Gall M., Martinez E., Rockett R., Sintchenko V., on behalf of ICPMR |
| EPI_ISL_3281024, EPI_ISL_3281025 | Histopath | NSW Health Pathology - Institute of Clinical Pathology and Medical Research; Westmead Hospital; University of Sydney | Arnott A., Draper J., Gall M., Martinez E., Rockett R., Sintchenko V., on behalf of ICPMR |
| EPI_ISL_3281026 | Laverty Pathology | NSW Health Pathology - Institute of Clinical Pathology and Medical Research; Westmead Hospital; University of Sydney | Arnott A., Draper J., Gall M., Martinez E., Rockett R., Sintchenko V., on behalf of ICPMR |
| EPI_ISL_3281027 | Sydney South West Pathology Service (SSWPS) - Liverpool Hospital - NSW Health Pathology | NSW Health Pathology - Institute of Clinical Pathology and Medical Research; Westmead Hospital; University of Sydney | Arnott A., Draper J., Gall M., Martinez E., Rockett R., Sintchenko V., on behalf of ICPMR |
| EPI_ISL_3281028 | Laverty Pathology | NSW Health Pathology - Institute of Clinical Pathology and Medical Research; Westmead Hospital; University of Sydney | Arnott A., Draper J., Gall M., Martinez E., Rockett R., Sintchenko V., on behalf of ICPMR |
| EPI_ISL_3281029, EPI_ISL_3281030 | St Vincent's Pathology (SydPath) | NSW Health Pathology - Institute of Clinical Pathology and Medical Research; Westmead Hospital; University of Sydney | Arnott A., Draper J., Gall M., Martinez E., Rockett R., Sintchenko V., on behalf of ICPMR |
| EPI_ISL_3281031 | Histopath | NSW Health Pathology - Institute of Clinical Pathology and Medical Research; Westmead Hospital; University of Sydney | Arnott A., Draper J., Gall M., Martinez E., Rockett R., Sintchenko V., on behalf of ICPMR |
| EPI_ISL_3281032 | Sydney South West Pathology Service (SSWPS) - Liverpool Hospital - NSW Health Pathology | NSW Health Pathology - Institute of Clinical Pathology and Medical Research; Westmead Hospital; University of Sydney | Arnott A., Draper J., Gall M., Martinez E., Rockett R., Sintchenko V., on behalf of ICPMR |
| EPI_ISL_3281033 | Douglass Hanly Moir Pathology | NSW Health Pathology - Institute of Clinical Pathology and Medical Research; Westmead Hospital; University of Sydney | Arnott A., Draper J., Gall M., Martinez E., Rockett R., Sintchenko V., on behalf of ICPMR |
| EPI_ISL_3281034, EPI_ISL_3281035 | Laverty Pathology | NSW Health Pathology - Institute of Clinical Pathology and Medical Research; Westmead Hospital; University of Sydney | Arnott A., Draper J., Gall M., Martinez E., Rockett R., Sintchenko V., on behalf of ICPMR |
| EPI_ISL_3281036 | Pathology North - Gosford Hospital - NSW Health Pathology | NSW Health Pathology - Institute of Clinical Pathology and Medical Research; Westmead Hospital; University of Sydney | Arnott A., Draper J., Gall M., Martinez E., Rockett R., Sintchenko V., on behalf of ICPMR |
| EPI_ISL_3281037, EPI_ISL_3281038 | Sydney South West Pathology Service (SSWPS) - Concord Repatriation General Hospital - NSW Health Pathology | NSW Health Pathology - Institute of Clinical Pathology and Medical Research; Westmead Hospital; University of Sydney | Arnott A., Draper J., Gall M., Martinez E., Rockett R., Sintchenko V., on behalf of ICPMR |
| EPI_ISL_3281039 | Austech Medical Laboratories | NSW Health Pathology - Institute of Clinical Pathology and Medical Research; Westmead Hospital; University of Sydney | Arnott A., Draper J., Gall M., Martinez E., Rockett R., Sintchenko V., on behalf of ICPMR |
| EPI_ISL_3281040, EPI_ISL_3281041, EPI_ISL_3281042 | Laverty Pathology | NSW Health Pathology - Institute of Clinical Pathology and Medical Research; Westmead Hospital; University of Sydney | Arnott A., Draper J., Gall M., Martinez E., Rockett R., Sintchenko V., on behalf of ICPMR |
| EPI_ISL_3281043 | Pathology North - Royal North Shore Hospital - NSW Health Pathology | NSW Health Pathology - Institute of Clinical Pathology and Medical Research; Westmead Hospital; University of Sydney | Arnott A., Draper J., Gall M., Martinez E., Rockett R., Sintchenko V., on behalf of ICPMR |
| EPI_ISL_3281044, EPI_ISL_3281045, EPI_ISL_3281046 | Histopath | NSW Health Pathology - Institute of Clinical Pathology and Medical Research; Westmead Hospital; University of Sydney | Arnott A., Draper J., Gall M., Martinez E., Rockett R., Sintchenko V., on behalf of ICPMR |
| EPI_ISL_3281047, EPI_ISL_3281048, EPI_ISL_3281049 | Sydney South West Pathology Service (SSWPS) - Liverpool Hospital - NSW Health Pathology | NSW Health Pathology - Institute of Clinical Pathology and Medical Research; Westmead Hospital; University of Sydney | Arnott A., Draper J., Gall M., Martinez E., Rockett R., Sintchenko V., on behalf of ICPMR |
| EPI_ISL_3281050 | St Vincent's Pathology (SydPath) | NSW Health Pathology - Institute of Clinical Pathology and Medical Research; Westmead Hospital; University of Sydney | Arnott A., Draper J., Gall M., Martinez E., Rockett R., Sintchenko V., on behalf of ICPMR |
| EPI_ISL_3281051 | Australian Clinical Labs (formerly Healthscope Pathology) | NSW Health Pathology - Institute of Clinical Pathology and Medical Research; Westmead Hospital; University of Sydney | Arnott A., Draper J., Gall M., Martinez E., Rockett R., Sintchenko V., on behalf of ICPMR |
| EPI_ISL_3281052 | Medlab Pathology | NSW Health Pathology - Institute of Clinical Pathology and Medical Research; Westmead Hospital; University of Sydney | Arnott A., Draper J., Gall M., Martinez E., Rockett R., Sintchenko V., on behalf of ICPMR |
| EPI_ISL_3281053 | Pathology West - NSW Health Pathology | NSW Health Pathology - Institute of Clinical Pathology and Medical Research; Westmead Hospital; University of Sydney | Arnott A., Draper J., Gall M., Martinez E., Rockett R., Sintchenko V., on behalf of ICPMR |
| EPI_ISL_3281054 | Histopath | NSW Health Pathology - Institute of Clinical Pathology and Medical Research; Westmead Hospital; University of Sydney | Arnott A., Draper J., Gall M., Martinez E., Rockett R., Sintchenko V., on behalf of ICPMR |
| EPI_ISL_3281055 | 4Cyte Pathology | NSW Health Pathology - Institute of Clinical Pathology and Medical Research; Westmead Hospital; University of Sydney | Arnott A., Draper J., Gall M., Martinez E., Rockett R., Sintchenko V., on behalf of ICPMR |
| EPI_ISL_3281056 | Sydney South West Pathology Service (SSWPS) - Liverpool Hospital - NSW Health Pathology | NSW Health Pathology - Institute of Clinical Pathology and Medical Research; Westmead Hospital; University of Sydney | Arnott A., Draper J., Gall M., Martinez E., Rockett R., Sintchenko V., on behalf of ICPMR |
| EPI_ISL_3281057, EPI_ISL_3281058 | Laverty Pathology | NSW Health Pathology - Institute of Clinical Pathology and Medical Research; Westmead Hospital; University of Sydney | Arnott A., Draper J., Gall M., Martinez E., Rockett R., Sintchenko V., on behalf of ICPMR |
| EPI_ISL_3281059 | Pathology North - Royal North Shore Hospital - NSW Health Pathology | NSW Health Pathology - Institute of Clinical Pathology and Medical Research; Westmead Hospital; University of Sydney | Arnott A., Draper J., Gall M., Martinez E., Rockett R., Sintchenko V., on behalf of ICPMR |
| EPI_ISL_3281060, EPI_ISL_3281061, EPI_ISL_3281062, EPI_ISL_3281063, EPI_ISL_3281064, EPI_ISL_3281065, EPI_ISL_3281066, EPI_ISL_3281067, EPI_ISL_3281068, EPI_ISL_3281069, EPI_ISL_3281070, EPI_ISL_3281071, EPI_ISL_3281072, EPI_ISL_3281073, EPI_ISL_3281074, EPI_ISL_3281075, EPI_ISL_3281076, EPI_ISL_3281077, EPI_ISL_3281078, EPI_ISL_3281079, EPI_ISL_3281080, EPI_ISL_3281081, EPI_ISL_3281082, EPI_ISL_3281083, EPI_ISL_3281084, EPI_ISL_3281085, EPI_ISL_3281086, EPI_ISL_3281087, EPI_ISL_3281088, EPI_ISL_3281089, EPI_ISL_3281090, EPI_ISL_3281091, EPI_ISL_3281092, EPI_ISL_3281093, EPI_ISL_3281094, EPI_ISL_3281095, EPI_ISL_3281096, EPI_ISL_3281097, EPI_ISL_3281098, EPI_ISL_3281099, EPI_ISL_3281100, EPI_ISL_3281101, EPI_ISL_3281102, EPI_ISL_3281103, EPI_ISL_3281104, EPI_ISL_3281105, EPI_ISL_3281106, EPI_ISL_3281107, EPI_ISL_3281108, EPI_ISL_3281109, EPI_ISL_3281110, EPI_ISL_3281111, EPI_ISL_3281112, EPI_ISL_3281113, EPI_ISL_3281114, EPI_ISL_3281115, EPI_ISL_3281116, EPI_ISL_3281117, EPI_ISL_3281118, EPI_ISL_3281119, EPI_ISL_3281120, EPI_ISL_3281121, EPI_ISL_3281122, EPI_ISL_3281123, EPI_ISL_3281124, EPI_ISL_3281125, EPI_ISL_3281126, EPI_ISL_3281127, EPI_ISL_3281128, EPI_ISL_3281129, EPI_ISL_3281130, EPI_ISL_3281131, EPI_ISL_3281132, EPI_ISL_3281133, EPI_ISL_3281134, EPI_ISL_3281135, EPI_ISL_3281136 | | | |
| see above | Histopath | NSW Health Pathology - Institute of Clinical Pathology and Medical Research; Westmead Hospital; University of Sydney | Arnott A., Draper J., Gall M., Martinez E., Rockett R., Sintchenko V., on behalf of ICPMR |
| EPI_ISL_3281137, EPI_ISL_3281138 | Sydney South West Pathology Service (SSWPS) - Concord Repatriation General Hospital - NSW Health Pathology | NSW Health Pathology - Institute of Clinical Pathology and Medical Research; Westmead Hospital; University of Sydney | Arnott A., Draper J., Gall M., Martinez E., Rockett R., Sintchenko V., on behalf of ICPMR |
| EPI_ISL_3281139, EPI_ISL_3281140, EPI_ISL_3281141, EPI_ISL_3281142, EPI_ISL_3281143, EPI_ISL_3281144, EPI_ISL_3281145, EPI_ISL_3281146, EPI_ISL_3281147, EPI_ISL_3281148, EPI_ISL_3281149, EPI_ISL_3281150, EPI_ISL_3281151, EPI_ISL_3281152, EPI_ISL_3281153, EPI_ISL_3281154, EPI_ISL_3281155, EPI_ISL_3281156, EPI_ISL_3281157, EPI_ISL_3281158, EPI_ISL_3281159, EPI_ISL_3281160, EPI_ISL_3281161 | | | |
| see above | Medlab Pathology | NSW Health Pathology - Institute of Clinical Pathology and Medical Research; Westmead Hospital; University of Sydney | Arnott A., Draper J., Gall M., Martinez E., Rockett R., Sintchenko V., on behalf of ICPMR |
| EPI_ISL_3281162, EPI_ISL_3281163, EPI_ISL_3281164, EPI_ISL_3281165, EPI_ISL_3281166, EPI_ISL_3281167, EPI_ISL_3281168, EPI_ISL_3281169, EPI_ISL_3281170, EPI_ISL_3281171, EPI_ISL_3281172, EPI_ISL_3281173 | | | |
| see above | 4Cyte Pathology | NSW Health Pathology - Institute of Clinical Pathology and Medical Research; Westmead Hospital; University of Sydney | Arnott A., Draper J., Gall M., Martinez E., Rockett R., Sintchenko V., on behalf of ICPMR |
| EPI_ISL_3281174, EPI_ISL_3281175, EPI_ISL_3281176, EPI_ISL_3281177, EPI_ISL_3281178, EPI_ISL_3281179, EPI_ISL_3281180, EPI_ISL_3281181, EPI_ISL_3281182, EPI_ISL_3281183 | Douglass Hanly Moir Pathology | NSW Health Pathology - Institute of Clinical Pathology and Medical Research; Westmead Hospital; University of Sydney | Arnott A., Draper J., Gall M., Martinez E., Rockett R., Sintchenko V., on behalf of ICPMR |
| EPI_ISL_3281184, EPI_ISL_3281185, EPI_ISL_3281186, EPI_ISL_3281187, EPI_ISL_3281188, EPI_ISL_3281189, EPI_ISL_3281190, EPI_ISL_3281191, EPI_ISL_3281192, EPI_ISL_3281193, EPI_ISL_3281194, EPI_ISL_3281195, EPI_ISL_3281196, EPI_ISL_3281197, EPI_ISL_3281198, EPI_ISL_3281199, EPI_ISL_3281200, EPI_ISL_3281201, EPI_ISL_3281202, EPI_ISL_3281203, EPI_ISL_3281204, EPI_ISL_3281205, EPI_ISL_3281206, EPI_ISL_3281207, EPI_ISL_3281208 | | | |
| see above | Laverty Pathology | NSW Health Pathology - Institute of Clinical Pathology and Medical Research; Westmead Hospital; University of Sydney | Arnott A., Draper J., Gall M., Martinez E., Rockett R., Sintchenko V., on behalf of ICPMR |
| EPI_ISL_3281209, EPI_ISL_3281210, EPI_ISL_3281211, EPI_ISL_3281212, EPI_ISL_3281213, EPI_ISL_3281214, EPI_ISL_3281215, EPI_ISL_3281216, EPI_ISL_3281217, EPI_ISL_3281218, EPI_ISL_3281219, EPI_ISL_3281220, EPI_ISL_3281221, EPI_ISL_3281222, EPI_ISL_3281223, EPI_ISL_3281224, EPI_ISL_3281225 | | | |
| see above | 4Cyte Pathology | NSW Health Pathology - Institute of Clinical Pathology and Medical Research; Westmead Hospital; University of Sydney | Arnott A., Draper J., Gall M., Martinez E., Rockett R., Sintchenko V., on behalf of ICPMR |
| EPI_ISL_3281226, EPI_ISL_3281227, EPI_ISL_3281228, EPI_ISL_3281229, | Australian Clinical Labs (formerly Healthscope Pathology) | NSW Health Pathology - Institute of Clinical Pathology and Medical Research; Westmead Hospital; University of Sydney | Arnott A., Draper J., Gall M., Martinez E., Rockett R., Sintchenko V., on behalf of ICPMR |

| | | | |
|---|---|---|---|
| EPI_ISL_3281230, EPI_ISL_3281231 | | | |
| EPI_ISL_3281232 | Pathology West - NSW Health Pathology | NSW Health Pathology - Institute of Clinical Pathology and Medical Research; Westmead Hospital; University of Sydney | Arnott A., Draper J., Gall M., Martinez E., Rockett R., Sintchenko V., on behalf of ICPMR |
| EPI_ISL_3281233, EPI_ISL_3281234, EPI_ISL_3281235 | Sydney South West Pathology Service (SSWPS) - Concord Repatriation General Hospital - NSW Health Pathology | NSW Health Pathology - Institute of Clinical Pathology and Medical Research; Westmead Hospital; University of Sydney | Arnott A., Draper J., Gall M., Martinez E., Rockett R., Sintchenko V., on behalf of ICPMR |
| EPI_ISL_3281236, EPI_ISL_3281237, EPI_ISL_3281238 | Medlab Pathology | NSW Health Pathology - Institute of Clinical Pathology and Medical Research; Westmead Hospital; University of Sydney | Arnott A., Draper J., Gall M., Martinez E., Rockett R., Sintchenko V., on behalf of ICPMR |
| EPI_ISL_3281239 | ACT Pathology | NSW Health Pathology - Institute of Clinical Pathology and Medical Research; Westmead Hospital; University of Sydney | Arnott A., Draper J., Gall M., Martinez E., Rockett R., Sintchenko V., on behalf of ICPMR |
| EPI_ISL_3281240 | Medlab Pathology | NSW Health Pathology - Institute of Clinical Pathology and Medical Research; Westmead Hospital; University of Sydney | Arnott A., Draper J., Gall M., Martinez E., Rockett R., Sintchenko V., on behalf of ICPMR |
| EPI_ISL_3281241 | Pathology North - Royal North Shore Hospital - NSW Health Pathology | NSW Health Pathology - Institute of Clinical Pathology and Medical Research; Westmead Hospital; University of Sydney | Arnott A., Draper J., Gall M., Martinez E., Rockett R., Sintchenko V., on behalf of ICPMR |
| EPI_ISL_3281242, EPI_ISL_3281243, EPI_ISL_3281244, EPI_ISL_3281245, EPI_ISL_3281246, EPI_ISL_3281247, EPI_ISL_3281248, EPI_ISL_3281249, EPI_ISL_3281250, EPI_ISL_3281251, EPI_ISL_3281252, EPI_ISL_3281253, EPI_ISL_3281254, EPI_ISL_3281255, EPI_ISL_3281256, EPI_ISL_3281257, EPI_ISL_3281258, EPI_ISL_3281259, EPI_ISL_3281260, EPI_ISL_3281261, EPI_ISL_3281262, EPI_ISL_3281263, EPI_ISL_3281264, EPI_ISL_3281265, EPI_ISL_3281266, EPI_ISL_3281267, EPI_ISL_3281268, EPI_ISL_3281269, EPI_ISL_3281270, EPI_ISL_3281271, EPI_ISL_3281272, EPI_ISL_3281273, EPI_ISL_3281274, EPI_ISL_3281275, EPI_ISL_3281276, EPI_ISL_3281277, EPI_ISL_3281278, EPI_ISL_3281279, EPI_ISL_3281280, EPI_ISL_3281281, EPI_ISL_3281282, EPI_ISL_3281283, EPI_ISL_3281284 | | | |
| see above | Histopath | NSW Health Pathology - Institute of Clinical Pathology and Medical Research; Westmead Hospital; University of Sydney | Arnott A., Draper J., Gall M., Martinez E., Rockett R., Sintchenko V., on behalf of ICPMR |
| EPI_ISL_3281285, EPI_ISL_3281286, EPI_ISL_3281287, EPI_ISL_3281288, EPI_ISL_3281289, EPI_ISL_3281290, EPI_ISL_3281291, EPI_ISL_3281292, EPI_ISL_3281293, EPI_ISL_3281294, EPI_ISL_3281295, EPI_ISL_3281296, EPI_ISL_3281297, EPI_ISL_3281298, EPI_ISL_3281299, EPI_ISL_3281300, EPI_ISL_3281301, EPI_ISL_3281302 | | | |
| see above | 4Cyte Pathology | NSW Health Pathology - Institute of Clinical Pathology and Medical Research; Westmead Hospital; University of Sydney | Arnott A., Draper J., Gall M., Martinez E., Rockett R., Sintchenko V., on behalf of ICPMR |
| EPI_ISL_3281303, EPI_ISL_3281304, EPI_ISL_3281305 | Australian Clinical Labs (formerly Healthscope Pathology) | NSW Health Pathology - Institute of Clinical Pathology and Medical Research; Westmead Hospital; University of Sydney | Arnott A., Draper J., Gall M., Martinez E., Rockett R., Sintchenko V., on behalf of ICPMR |
| EPI_ISL_3281306, EPI_ISL_3281307, EPI_ISL_3281308, EPI_ISL_3281309, EPI_ISL_3281310, EPI_ISL_3281311, EPI_ISL_3281312, EPI_ISL_3281313, EPI_ISL_3281314, EPI_ISL_3281315, EPI_ISL_3281316, EPI_ISL_3281317, EPI_ISL_3281318, EPI_ISL_3281319, EPI_ISL_3281320, EPI_ISL_3281321, EPI_ISL_3281322 | | | |
| see above | The Children's Hospital at Westmead | NSW Health Pathology - Institute of Clinical Pathology and Medical Research; Westmead Hospital; University of Sydney | Arnott A., Draper J., Gall M., Martinez E., Rockett R., Sintchenko V., on behalf of ICPMR |
| EPI_ISL_3281323, EPI_ISL_3281324, EPI_ISL_3281325, EPI_ISL_3281326, EPI_ISL_3281327, EPI_ISL_3281328, EPI_ISL_3281329, EPI_ISL_3281330, EPI_ISL_3281331, EPI_ISL_3281332, EPI_ISL_3281333, EPI_ISL_3281334, EPI_ISL_3281335 | | | |
| see above | Medlab Pathology | NSW Health Pathology - Institute of Clinical Pathology and Medical Research; Westmead Hospital; University of Sydney | Arnott A., Draper J., Gall M., Martinez E., Rockett R., Sintchenko V., on behalf of ICPMR |
| EPI_ISL_3281336, EPI_ISL_3281337 | Pathology North - Royal North Shore Hospital - NSW Health Pathology | NSW Health Pathology - Institute of Clinical Pathology and Medical Research; Westmead Hospital; University of Sydney | Arnott A., Draper J., Gall M., Martinez E., Rockett R., Sintchenko V., on behalf of ICPMR |
| EPI_ISL_3281338, EPI_ISL_3281339, EPI_ISL_3281340 | Sydney South West Pathology Service (SSWPS) - Concord Repatriation General Hospital - NSW Health Pathology | NSW Health Pathology - Institute of Clinical Pathology and Medical Research; Westmead Hospital; University of Sydney | Arnott A., Draper J., Gall M., Martinez E., Rockett R., Sintchenko V., on behalf of ICPMR |
| EPI_ISL_3281341, EPI_ISL_3281342, EPI_ISL_3281343, EPI_ISL_3281344, EPI_ISL_3281345, EPI_ISL_3281346, EPI_ISL_3281347, EPI_ISL_3281348, EPI_ISL_3281349, EPI_ISL_3281350, EPI_ISL_3281351, EPI_ISL_3281352, EPI_ISL_3281353, EPI_ISL_3281354, EPI_ISL_3281355, EPI_ISL_3281356, EPI_ISL_3281357, EPI_ISL_3281358, EPI_ISL_3281359, EPI_ISL_3281360, EPI_ISL_3281361, EPI_ISL_3281362, EPI_ISL_3281363, EPI_ISL_3281364, EPI_ISL_3281365, EPI_ISL_3281366 | | | |
| see above | Histopath | NSW Health Pathology - Institute of Clinical Pathology and Medical Research; Westmead Hospital; University of Sydney | Arnott A., Draper J., Gall M., Martinez E., Rockett R., Sintchenko V., on behalf of ICPMR |
| EPI_ISL_3281367 | JSS Medical Lab SDN BHD | Institute for Medical Research, Infectious Disease Research Centre, National Institutes of Health, Ministry of Health Malaysia | Suppiah J, Kamel K, Mohd Zawawi Z, Azizan MA, Ramly N, Robert F, Anasir MI, Thayan R |
| EPI_ISL_3281368, EPI_ISL_3281369, EPI_ISL_3281370, EPI_ISL_3281371, EPI_ISL_3281372, EPI_ISL_3281373, EPI_ISL_3281374, EPI_ISL_3281375, EPI_ISL_3281376, EPI_ISL_3281377, EPI_ISL_3281378, EPI_ISL_3281379, EPI_ISL_3281380, EPI_ISL_3281381, EPI_ISL_3281382, EPI_ISL_3281383, EPI_ISL_3281384, EPI_ISL_3281385, EPI_ISL_3281386, EPI_ISL_3281387, EPI_ISL_3281388, EPI_ISL_3281389, EPI_ISL_3281390, EPI_ISL_3281391, EPI_ISL_3281392, EPI_ISL_3281393, EPI_ISL_3281394, EPI_ISL_3281395 | | | |
| see above | Histopath | NSW Health Pathology - Institute of Clinical Pathology and Medical Research; Westmead Hospital; University of Sydney | Arnott A., Draper J., Gall M., Martinez E., Rockett R., Sintchenko V., on behalf of ICPMR |
| EPI_ISL_3281396, EPI_ISL_3281397, EPI_ISL_3281398, EPI_ISL_3281399, EPI_ISL_3281400, EPI_ISL_3281401 | Pathology West - NSW Health Pathology | NSW Health Pathology - Institute of Clinical Pathology and Medical Research; Westmead Hospital; University of Sydney | Arnott A., Draper J., Gall M., Martinez E., Rockett R., Sintchenko V., on behalf of ICPMR |
| EPI_ISL_3281402, EPI_ISL_3281403, EPI_ISL_3281404, EPI_ISL_3281405, EPI_ISL_3281406, EPI_ISL_3281407, EPI_ISL_3281408 | Laverty Pathology | NSW Health Pathology - Institute of Clinical Pathology and Medical Research; Westmead Hospital; University of Sydney | Arnott A., Draper J., Gall M., Martinez E., Rockett R., Sintchenko V., on behalf of ICPMR |
| EPI_ISL_3281409, EPI_ISL_3281410, EPI_ISL_3281411 | Medlab Pathology | NSW Health Pathology - Institute of Clinical Pathology and Medical Research; Westmead Hospital; University of Sydney | Arnott A., Draper J., Gall M., Martinez E., Rockett R., Sintchenko V., on behalf of ICPMR |
| EPI_ISL_3281412 | Pathology West - NSW Health Pathology | NSW Health Pathology - Institute of Clinical Pathology and Medical Research; Westmead Hospital; University of Sydney | Arnott A., Draper J., Gall M., Martinez E., Rockett R., Sintchenko V., on behalf of ICPMR |
| EPI_ISL_3281413 | National Cancer Institute | Institute for Medical Research, Infectious Disease Research Centre, National Institutes of Health, Ministry of Health Malaysia | Suppiah J, Kamel K, Mohd Zawawi Z, Azizan MA, Ramly N, Robert F, Anasir MI, Thayan R |
| EPI_ISL_3281415, EPI_ISL_3281416, EPI_ISL_3281417 | Microbiology Department, Laboratori Clínic Metropolitana Nord. Hospital Universitari Germans Trias i Pujol | Can Ruti SARS-CoV-2 Sequencing Hub (HUGTiP/IrsiCaixa/IGTP) | Marc Noguera-Julian, Pilar Armengol, Ignacio Blanco, Antoni E Bordoy, Francesc Catala-Moll, Pere-Joan Cardona, Maria Casadellà, Cristina Casañ, Gemma Clara, Bonaventura Clotet, Montserrat Giménez, Mercedes Guerrero, Anna Not, David Panisello, Roger Paredes, Mariona Parera, Alexia París, Verónica Saludes, Alba Sánchez, Laia Soler, Lauro Sumoy, and Elisa Martró on behalf of the Can Ruti SARS-CoV-2 Sequencing Hub |
| EPI_ISL_3281418 | Microbiologia CATLAB | Can Ruti SARS-CoV-2 Sequencing Hub (HUGTiP/IrsiCaixa/IGTP) | Marc Noguera-Julian, Pilar Armengol, Ignacio Blanco, Antoni E Bordoy, Francesc Catala-Moll, Pere-Joan Cardona, Maria Casadellà, Cristina Casañ, Gemma Clara, Bonaventura Clotet, Montserrat Giménez, Mercedes Guerrero, Anna Not, David Panisello, Roger Paredes, Mariona Parera, Alexia París, Verónica Saludes, Alba Sánchez, Laia Soler, Lauro Sumoy, and Elisa Martró on behalf of the Can Ruti SARS-CoV-2 Sequencing Hub |
| EPI_ISL_3281419, EPI_ISL_3281420, EPI_ISL_3281421, EPI_ISL_3281422, EPI_ISL_3281423, EPI_ISL_3281424, EPI_ISL_3281425, EPI_ISL_3281427, EPI_ISL_3281428, EPI_ISL_3281430, EPI_ISL_3281431, EPI_ISL_3281432, EPI_ISL_3281433, EPI_ISL_3281434, EPI_ISL_3281435, EPI_ISL_3281436, EPI_ISL_3281437, EPI_ISL_3281438, EPI_ISL_3281439, EPI_ISL_3281440, EPI_ISL_3281441, EPI_ISL_3281442, EPI_ISL_3281443, EPI_ISL_3281444, EPI_ISL_3281445, EPI_ISL_3281446, EPI_ISL_3281447, EPI_ISL_3281448, EPI_ISL_3281449, EPI_ISL_3281450, EPI_ISL_3281451, EPI_ISL_3281452, EPI_ISL_3281453, EPI_ISL_3281454, EPI_ISL_3281455, EPI_ISL_3281456, EPI_ISL_3281457, EPI_ISL_3281458, EPI_ISL_3281459, EPI_ISL_3281460, EPI_ISL_3281461, EPI_ISL_3281462, EPI_ISL_3281463, EPI_ISL_3281464, EPI_ISL_3281465, EPI_ISL_3281466, EPI_ISL_3281467, EPI_ISL_3281468, EPI_ISL_3281469, EPI_ISL_3281470, EPI_ISL_3281471, EPI_ISL_3281472, EPI_ISL_3281473, EPI_ISL_3281474, EPI_ISL_3281475, EPI_ISL_3281476 | | | |
| see above | Microbiology Department, Laboratori Clínic Metropolitana Nord. Hospital Universitari Germans Trias i Pujol | Can Ruti SARS-CoV-2 Sequencing Hub (HUGTiP/IrsiCaixa/IGTP) | Marc Noguera-Julian, Pilar Armengol, Ignacio Blanco, Antoni E Bordoy, Francesc Catala-Moll, Pere-Joan Cardona, Maria Casadellà, Cristina Casañ, Gemma Clara, Bonaventura Clotet, Montserrat Giménez, Mercedes Guerrero, Anna Not, David Panisello, Roger Paredes, Mariona Parera, Alexia París, Verónica Saludes, Alba Sánchez, Laia Soler, Lauro Sumoy, and Elisa Martró on behalf of the Can Ruti SARS-CoV-2 Sequencing Hub |
| EPI_ISL_3281477 | Microbiologia CATLAB | Can Ruti SARS-CoV-2 Sequencing Hub (HUGTiP/IrsiCaixa/IGTP) | Marc Noguera-Julian, Pilar Armengol, Ignacio Blanco, Antoni E Bordoy, Francesc Catala-Moll, Pere-Joan Cardona, Maria Casadellà, Cristina Casañ, Gemma Clara, Bonaventura Clotet, Montserrat Giménez, Mercedes Guerrero, Anna Not, David Panisello, Roger Paredes, Mariona Parera, Alexia París, Verónica Saludes, Alba Sánchez, Laia Soler, Lauro Sumoy, and Elisa Martró on behalf of the Can Ruti SARS-CoV-2 Sequencing Hub |
| EPI_ISL_3281478, EPI_ISL_3281479, EPI_ISL_3281480, EPI_ISL_3281481, EPI_ISL_3281482, EPI_ISL_3281483, EPI_ISL_3281484, EPI_ISL_3281485, EPI_ISL_3281486, EPI_ISL_3281487, EPI_ISL_3281490, EPI_ISL_3281491, EPI_ISL_3281492, EPI_ISL_3281493, EPI_ISL_3281494, EPI_ISL_3281495, EPI_ISL_3281496, EPI_ISL_3281497 | | | |
| see above | Microbiology Department, Laboratori Clínic Metropolitana Nord. Hospital Universitari Germans Trias i Pujol | Can Ruti SARS-CoV-2 Sequencing Hub (HUGTiP/IrsiCaixa/IGTP) | Marc Noguera-Julian, Pilar Armengol, Ignacio Blanco, Antoni E Bordoy, Francesc Catala-Moll, Pere-Joan Cardona, Maria Casadellà, Cristina Casañ, Gemma Clara, Bonaventura Clotet, Montserrat Giménez, Mercedes Guerrero, Anna Not, David Panisello, Roger Paredes, Mariona Parera, Alexia París, Verónica Saludes, Alba Sánchez, Laia Soler, Lauro Sumoy, and Elisa Martró on behalf of the Can Ruti SARS-CoV-2 Sequencing Hub |
| EPI_ISL_3281499 | Microbiologia CATLAB | Can Ruti SARS-CoV-2 Sequencing Hub (HUGTiP/IrsiCaixa/IGTP) | Marc Noguera-Julian, Pilar Armengol, Ignacio Blanco, Antoni E Bordoy, Francesc Catala-Moll, Pere-Joan Cardona, Maria Casadellà, Cristina Casañ, Gemma Clara, Bonaventura Clotet, Montserrat Giménez, Mercedes Guerrero, Anna Not, David Panisello, Roger Paredes, Mariona Parera, Alexia París, Verónica Saludes, Alba Sánchez, Laia Soler, Lauro Sumoy, and Elisa Martró on behalf of the Can Ruti SARS-CoV-2 Sequencing Hub |
| EPI_ISL_3281500, EPI_ISL_3281501, EPI_ISL_3281502, EPI_ISL_3281503, EPI_ISL_3281504, EPI_ISL_3281505, EPI_ISL_3281506, EPI_ISL_3281507, EPI_ISL_3281508 | Microbiology Department, Laboratori Clínic Metropolitana Nord. Hospital Universitari Germans Trias i Pujol | Can Ruti SARS-CoV-2 Sequencing Hub (HUGTiP/IrsiCaixa/IGTP) | Marc Noguera-Julian, Pilar Armengol, Ignacio Blanco, Antoni E Bordoy, Francesc Catala-Moll, Pere-Joan Cardona, Maria Casadellà, Cristina Casañ, Gemma Clara, Bonaventura Clotet, Montserrat Giménez, Mercedes Guerrero, Anna Not, David Panisello, Roger Paredes, Mariona Parera, Alexia París, Verónica Saludes, Alba Sánchez, Laia Soler, Lauro Sumoy, and Elisa Martró on behalf of the Can Ruti SARS-CoV-2 Sequencing Hub |
| EPI_ISL_3281509, EPI_ISL_3281510, EPI_ISL_3281511, EPI_ISL_3281512, EPI_ISL_3281513, EPI_ISL_3281514, EPI_ISL_3281515, EPI_ISL_3281516, EPI_ISL_3281517, EPI_ISL_3281518, EPI_ISL_3281519 | | | |

| | | | |
|---|---|---|---|
| see above | Althaia. Xarxa Assistencial Universitària de Manresa | IrsiCaixa | Marc Noguera-Julian, Mariona Parera, Maria Casadellà, Pilar Armengol, Francesc Catala-Moll, Roger Paredes, Bonaventura Clotet Gloria Trujillo, Rafel Perez Vidal, Jaume Trapé, Carolina Gonzalez-Fernandez, Miquel Micó, Antonia Flor, Roger Paredes, Eulalia Grau, Bonaventura Clotet |
| EPI_ISL_3281522, EPI_ISL_3281523, EPI_ISL_3281524, EPI_ISL_3281525, EPI_ISL_3281526, EPI_ISL_3281527, EPI_ISL_3281528, EPI_ISL_3281529, EPI_ISL_3281532, EPI_ISL_3281533, EPI_ISL_3281537, EPI_ISL_3281538, EPI_ISL_3281539, EPI_ISL_3281540, EPI_ISL_3281541, EPI_ISL_3281542, EPI_ISL_3281543, EPI_ISL_3281545, EPI_ISL_3281546, EPI_ISL_3281547, EPI_ISL_3281548, EPI_ISL_3281549, EPI_ISL_3281550, EPI_ISL_3281551, EPI_ISL_3281553, EPI_ISL_3281554, EPI_ISL_3281555, EPI_ISL_3281556, EPI_ISL_3281557, EPI_ISL_3281558, EPI_ISL_3281559, EPI_ISL_3281560, EPI_ISL_3281561, EPI_ISL_3281562, EPI_ISL_3281563, EPI_ISL_3281564, EPI_ISL_3281565, EPI_ISL_3281566, EPI_ISL_3281567, EPI_ISL_3281568, EPI_ISL_3281569, EPI_ISL_3281570, EPI_ISL_3281571, EPI_ISL_3281572, EPI_ISL_3281573, EPI_ISL_3281574, EPI_ISL_3281575, EPI_ISL_3281576, EPI_ISL_3281578, EPI_ISL_3281579, EPI_ISL_3281580, EPI_ISL_3281581 | | | |
| see above | IVY3 Central Lab, Vanderbilt University Medical Center | Lauring Lab, University of Michigan, Department of Microbiology and Immunology | Gilbert |
| EPI_ISL_3281585, EPI_ISL_3281586, EPI_ISL_3281587, EPI_ISL_3281588, EPI_ISL_3281589, EPI_ISL_3281590, EPI_ISL_3281591, EPI_ISL_3281592, EPI_ISL_3281593, EPI_ISL_3281594, EPI_ISL_3281595, EPI_ISL_3281596, EPI_ISL_3281597, EPI_ISL_3281598 | | | |
| see above | National Institute for Communicable Diseases of the National Health Laboratory Service | National Institute for Communicable Diseases of the National Health Laboratory Service | Amoako DG, Everatt J, Scheepers C, Mohale T, Ntuli N, Mahlangu B, Mnguni A, Ismail A, Bhiman JN |
| EPI_ISL_3281607 | Lighthouse Lab in Glasgow | Wellcome Sanger Institute for the COVID-19 Genomics UK (COG-UK) Consortium | Harper VanSteenhouse, Yumi Kasai, David Gray, Carol Clugston, Anna Dominiczak and Alex Alderton, Roberto Amato, Jeffrey Barrett, Sonia Goncalves, Ewan Harrison, David K. Jackson, Ian Johnston, Dominic Kwiatkowski, Cordelia Langford, John Sillitoe on behalf of the Wellcome Sanger Institute COVID-19 Surveillance Team |
| EPI_ISL_3281608 | Lighthouse Lab in Milton Keynes | Wellcome Sanger Institute for the COVID-19 Genomics UK (COG-UK) Consortium | The Lighthouse Lab in Milton Keynes and Alex Alderton, Roberto Amato, Jeffrey Barrett, Sonia Goncalves, Ewan Harrison, David K. Jackson, Ian Johnston, Dominic Kwiatkowski, Cordelia Langford, John Sillitoe on behalf of the Wellcome Sanger Institute COVID-19 Surveillance Team |
| EPI_ISL_3281609, EPI_ISL_3281610 | Lighthouse Lab in Glasgow | Wellcome Sanger Institute for the COVID-19 Genomics UK (COG-UK) Consortium | Harper VanSteenhouse, Yumi Kasai, David Gray, Carol Clugston, Anna Dominiczak and Alex Alderton, Roberto Amato, Jeffrey Barrett, Sonia Goncalves, Ewan Harrison, David K. Jackson, Ian Johnston, Dominic Kwiatkowski, Cordelia Langford, John Sillitoe on behalf of the Wellcome Sanger Institute COVID-19 Surveillance Team |
| EPI_ISL_3281611 | Lighthouse Lab in Alderley Park | Wellcome Sanger Institute for the COVID-19 Genomics UK (COG-UK) Consortium | Jacquelyn Wynn, Mairead Hyland, The Lighthouse Lab in Alderley Park and Alex Alderton, Roberto Amato, Jeffrey Barrett, Sonia Goncalves, Ewan Harrison, David K. Jackson, Ian Johnston, Dominic Kwiatkowski, Cordelia Langford, John Sillitoe on behalf of the Wellcome Sanger Institute COVID-19 Surveillance Team |
| EPI_ISL_3281612 | Lighthouse Lab in Milton Keynes | Wellcome Sanger Institute for the COVID-19 Genomics UK (COG-UK) Consortium | The Lighthouse Lab in Milton Keynes and Alex Alderton, Roberto Amato, Jeffrey Barrett, Sonia Goncalves, Ewan Harrison, David K. Jackson, Ian Johnston, Dominic Kwiatkowski, Cordelia Langford, John Sillitoe on behalf of the Wellcome Sanger Institute COVID-19 Surveillance Team |
| EPI_ISL_3281613 | Lighthouse Lab in Glasgow | Wellcome Sanger Institute for the COVID-19 Genomics UK (COG-UK) Consortium | Harper VanSteenhouse, Yumi Kasai, David Gray, Carol Clugston, Anna Dominiczak and Alex Alderton, Roberto Amato, Jeffrey Barrett, Sonia Goncalves, Ewan Harrison, David K. Jackson, Ian Johnston, Dominic Kwiatkowski, Cordelia Langford, John Sillitoe on behalf of the Wellcome Sanger Institute COVID-19 Surveillance Team |
| EPI_ISL_3281614 | Lighthouse Lab in Alderley Park | Wellcome Sanger Institute for the COVID-19 Genomics UK (COG-UK) Consortium | Jacquelyn Wynn, Mairead Hyland, The Lighthouse Lab in Alderley Park and Alex Alderton, Roberto Amato, Jeffrey Barrett, Sonia Goncalves, Ewan Harrison, David K. Jackson, Ian Johnston, Dominic Kwiatkowski, Cordelia Langford, John Sillitoe on behalf of the Wellcome Sanger Institute COVID-19 Surveillance Team |
| EPI_ISL_3281615 | Lighthouse Lab in Glasgow | Wellcome Sanger Institute for the COVID-19 Genomics UK (COG-UK) Consortium | Harper VanSteenhouse, Yumi Kasai, David Gray, Carol Clugston, Anna Dominiczak and Alex Alderton, Roberto Amato, Jeffrey Barrett, Sonia Goncalves, Ewan Harrison, David K. Jackson, Ian Johnston, Dominic Kwiatkowski, Cordelia Langford, John Sillitoe on behalf of the Wellcome Sanger Institute COVID-19 Surveillance Team |
| EPI_ISL_3281616 | Lighthouse Lab in Milton Keynes | Wellcome Sanger Institute for the COVID-19 Genomics UK (COG-UK) Consortium | The Lighthouse Lab in Milton Keynes and Alex Alderton, Roberto Amato, Jeffrey Barrett, Sonia Goncalves, Ewan Harrison, David K. Jackson, Ian Johnston, Dominic Kwiatkowski, Cordelia Langford, John Sillitoe on behalf of the Wellcome Sanger Institute COVID-19 Surveillance Team |
| EPI_ISL_3281617, EPI_ISL_3281618 | Lighthouse Lab in Alderley Park | Wellcome Sanger Institute for the COVID-19 Genomics UK (COG-UK) Consortium | Jacquelyn Wynn, Mairead Hyland, The Lighthouse Lab in Alderley Park and Alex Alderton, Roberto Amato, Jeffrey Barrett, Sonia Goncalves, Ewan Harrison, David K. Jackson, Ian Johnston, Dominic Kwiatkowski, Cordelia Langford, John Sillitoe on behalf of the Wellcome Sanger Institute COVID-19 Surveillance Team |
| EPI_ISL_3281619 | Lighthouse Lab in Glasgow | Wellcome Sanger Institute for the COVID-19 Genomics UK (COG-UK) Consortium | Harper VanSteenhouse, Yumi Kasai, David Gray, Carol Clugston, Anna Dominiczak and Alex Alderton, Roberto Amato, Jeffrey Barrett, Sonia Goncalves, Ewan Harrison, David K. Jackson, Ian Johnston, Dominic Kwiatkowski, Cordelia Langford, John Sillitoe on behalf of the Wellcome Sanger Institute COVID-19 Surveillance Team |
| EPI_ISL_3281620, EPI_ISL_3281621, EPI_ISL_3281622, EPI_ISL_3281623 | Lighthouse Lab in Milton Keynes | Wellcome Sanger Institute for the COVID-19 Genomics UK (COG-UK) Consortium | The Lighthouse Lab in Milton Keynes and Alex Alderton, Roberto Amato, Jeffrey Barrett, Sonia Goncalves, Ewan Harrison, David K. Jackson, Ian Johnston, Dominic Kwiatkowski, Cordelia Langford, John Sillitoe on behalf of the Wellcome Sanger Institute COVID-19 Surveillance Team |
| EPI_ISL_3281624, EPI_ISL_3281625 | Lighthouse Lab in Glasgow | Wellcome Sanger Institute for the COVID-19 Genomics UK (COG-UK) Consortium | Harper VanSteenhouse, Yumi Kasai, David Gray, Carol Clugston, Anna Dominiczak and Alex Alderton, Roberto Amato, Jeffrey Barrett, Sonia Goncalves, Ewan Harrison, David K. Jackson, Ian Johnston, Dominic Kwiatkowski, Cordelia Langford, John Sillitoe on behalf of the Wellcome Sanger Institute COVID-19 Surveillance Team |
| EPI_ISL_3281626, EPI_ISL_3281627 | Lighthouse Lab in Alderley Park | Wellcome Sanger Institute for the COVID-19 Genomics UK (COG-UK) Consortium | Jacquelyn Wynn, Mairead Hyland, The Lighthouse Lab in Alderley Park and Alex Alderton, Roberto Amato, Jeffrey Barrett, Sonia Goncalves, Ewan Harrison, David K. Jackson, Ian Johnston, Dominic Kwiatkowski, Cordelia Langford, John Sillitoe on behalf of the Wellcome Sanger Institute COVID-19 Surveillance Team |
| EPI_ISL_3281628 | Lighthouse Lab in Milton Keynes | Wellcome Sanger Institute for the COVID-19 Genomics UK (COG-UK) Consortium | The Lighthouse Lab in Milton Keynes and Alex Alderton, Roberto Amato, Jeffrey Barrett, Sonia Goncalves, Ewan Harrison, David K. Jackson, Ian Johnston, Dominic Kwiatkowski, Cordelia Langford, John Sillitoe on behalf of the Wellcome Sanger Institute COVID-19 Surveillance Team |
| EPI_ISL_3281629 | Lighthouse Lab in Glasgow | Wellcome Sanger Institute for the COVID-19 Genomics UK (COG-UK) Consortium | Harper VanSteenhouse, Yumi Kasai, David Gray, Carol Clugston, Anna Dominiczak and Alex Alderton, Roberto Amato, Jeffrey Barrett, Sonia Goncalves, Ewan Harrison, David K. Jackson, Ian Johnston, Dominic Kwiatkowski, Cordelia Langford, John Sillitoe on behalf of the Wellcome Sanger Institute COVID-19 Surveillance Team |
| EPI_ISL_3281630 | Lighthouse Lab in Alderley Park | Wellcome Sanger Institute for the COVID-19 Genomics UK (COG-UK) Consortium | Jacquelyn Wynn, Mairead Hyland, The Lighthouse Lab in Alderley Park and Alex Alderton, Roberto Amato, Jeffrey Barrett, Sonia Goncalves, Ewan Harrison, David K. Jackson, Ian Johnston, Dominic Kwiatkowski, Cordelia Langford, John Sillitoe on behalf of the Wellcome Sanger Institute COVID-19 Surveillance Team |
| EPI_ISL_3281631 | Lighthouse Lab in Milton Keynes | Wellcome Sanger Institute for the COVID-19 Genomics UK (COG-UK) Consortium | The Lighthouse Lab in Milton Keynes and Alex Alderton, Roberto Amato, Jeffrey Barrett, Sonia Goncalves, Ewan Harrison, David K. Jackson, Ian Johnston, Dominic Kwiatkowski, Cordelia Langford, John Sillitoe on behalf of the Wellcome Sanger Institute COVID-19 Surveillance Team |
| EPI_ISL_3281632, EPI_ISL_3281633 | Lighthouse Lab in Alderley Park | Wellcome Sanger Institute for the COVID-19 Genomics UK (COG-UK) Consortium | Jacquelyn Wynn, Mairead Hyland, The Lighthouse Lab in Alderley Park and Alex Alderton, Roberto Amato, Jeffrey Barrett, Sonia Goncalves, Ewan Harrison, David K. Jackson, Ian Johnston, Dominic Kwiatkowski, Cordelia Langford, John Sillitoe on behalf of the Wellcome Sanger Institute COVID-19 Surveillance Team |
| EPI_ISL_3281634 | Lighthouse Lab in Milton Keynes | Wellcome Sanger Institute for the COVID-19 Genomics UK (COG-UK) Consortium | The Lighthouse Lab in Milton Keynes and Alex Alderton, Roberto Amato, Jeffrey Barrett, Sonia Goncalves, Ewan Harrison, David K. Jackson, Ian Johnston, Dominic Kwiatkowski, Cordelia Langford, John Sillitoe on behalf of the Wellcome Sanger Institute COVID-19 Surveillance Team |
| EPI_ISL_3281635 | Lighthouse Lab in Glasgow | Wellcome Sanger Institute for the COVID-19 Genomics UK (COG-UK) Consortium | Harper VanSteenhouse, Yumi Kasai, David Gray, Carol Clugston, Anna Dominiczak and Alex Alderton, Roberto Amato, Jeffrey Barrett, Sonia Goncalves, Ewan Harrison, David K. Jackson, Ian Johnston, Dominic Kwiatkowski, Cordelia Langford, John Sillitoe on behalf of the Wellcome Sanger Institute COVID-19 Surveillance Team |
| EPI_ISL_3281636, EPI_ISL_3281637, EPI_ISL_3281638 | Lighthouse Lab in Alderley Park | Wellcome Sanger Institute for the COVID-19 Genomics UK (COG-UK) Consortium | Jacquelyn Wynn, Mairead Hyland, The Lighthouse Lab in Alderley Park and Alex Alderton, Roberto Amato, Jeffrey Barrett, Sonia Goncalves, Ewan Harrison, David K. Jackson, Ian Johnston, Dominic Kwiatkowski, Cordelia Langford, John Sillitoe on behalf of the Wellcome Sanger Institute COVID-19 Surveillance Team |
| EPI_ISL_3281639 | Lighthouse Lab in Glasgow | Wellcome Sanger Institute for the COVID-19 Genomics UK (COG-UK) Consortium | Harper VanSteenhouse, Yumi Kasai, David Gray, Carol Clugston, Anna Dominiczak and Alex Alderton, Roberto Amato, Jeffrey Barrett, Sonia Goncalves, Ewan Harrison, David K. Jackson, Ian Johnston, Dominic Kwiatkowski, Cordelia Langford, John Sillitoe on behalf of the Wellcome Sanger Institute COVID-19 Surveillance Team |
| EPI_ISL_3281640, EPI_ISL_3281641 | Lighthouse Lab in Milton Keynes | Wellcome Sanger Institute for the COVID-19 Genomics UK (COG-UK) Consortium | The Lighthouse Lab in Milton Keynes and Alex Alderton, Roberto Amato, Jeffrey Barrett, Sonia Goncalves, Ewan Harrison, David K. Jackson, Ian Johnston, Dominic Kwiatkowski, Cordelia Langford, John Sillitoe on behalf of the Wellcome Sanger Institute COVID-19 Surveillance Team |
| EPI_ISL_3281642 | Lighthouse Lab in Glasgow | Wellcome Sanger Institute for the COVID-19 Genomics UK (COG-UK) Consortium | Harper VanSteenhouse, Yumi Kasai, David Gray, Carol Clugston, Anna Dominiczak and Alex Alderton, Roberto Amato, Jeffrey Barrett, Sonia Goncalves, Ewan Harrison, David K. Jackson, Ian Johnston, Dominic Kwiatkowski, Cordelia Langford, John Sillitoe on behalf of the Wellcome Sanger Institute COVID-19 Surveillance Team |
| EPI_ISL_3281643, EPI_ISL_3281644, EPI_ISL_3281645, EPI_ISL_3281646 | Lighthouse Lab in Milton Keynes | Wellcome Sanger Institute for the COVID-19 Genomics UK (COG-UK) Consortium | The Lighthouse Lab in Milton Keynes and Alex Alderton, Roberto Amato, Jeffrey Barrett, Sonia Goncalves, Ewan Harrison, David K. Jackson, Ian Johnston, Dominic Kwiatkowski, Cordelia Langford, John Sillitoe on behalf of the Wellcome Sanger Institute COVID-19 Surveillance Team |
| EPI_ISL_3281647, EPI_ISL_3281648 | Lighthouse Lab in Glasgow | Wellcome Sanger Institute for the COVID-19 Genomics UK (COG-UK) Consortium | Harper VanSteenhouse, Yumi Kasai, David Gray, Carol Clugston, Anna Dominiczak and Alex Alderton, Roberto Amato, Jeffrey Barrett, Sonia Goncalves, Ewan Harrison, David K. Jackson, Ian Johnston, Dominic Kwiatkowski, Cordelia Langford, John Sillitoe on behalf of the Wellcome Sanger Institute COVID-19 Surveillance Team |
| EPI_ISL_3281649 | Lighthouse Lab in Milton Keynes | Wellcome Sanger Institute for the COVID-19 Genomics UK (COG-UK) Consortium | The Lighthouse Lab in Milton Keynes and Alex Alderton, Roberto Amato, Jeffrey Barrett, Sonia Goncalves, Ewan Harrison, David K. Jackson, Ian Johnston, Dominic Kwiatkowski, Cordelia Langford, John Sillitoe on behalf of the Wellcome Sanger Institute COVID-19 Surveillance Team |
| EPI_ISL_3281650 | Lighthouse Lab in Alderley Park | Wellcome Sanger Institute for the COVID-19 Genomics UK (COG-UK) Consortium | Jacquelyn Wynn, Mairead Hyland, The Lighthouse Lab in Alderley Park and Alex Alderton, Roberto Amato, Jeffrey Barrett, Sonia Goncalves, Ewan Harrison, David K. Jackson, Ian Johnston, Dominic Kwiatkowski, Cordelia Langford, John Sillitoe on behalf of the Wellcome Sanger Institute COVID-19 Surveillance Team |
| EPI_ISL_3281651, EPI_ISL_3281652 | Lighthouse Lab in Glasgow | Wellcome Sanger Institute for the COVID-19 Genomics UK (COG-UK) Consortium | Harper VanSteenhouse, Yumi Kasai, David Gray, Carol Clugston, Anna Dominiczak and Alex Alderton, Roberto Amato, Jeffrey Barrett, Sonia Goncalves, Ewan Harrison, David K. Jackson, Ian Johnston, Dominic Kwiatkowski, Cordelia Langford, John Sillitoe on behalf of the Wellcome Sanger Institute COVID-19 Surveillance Team |
| EPI_ISL_3281653 | Lighthouse Lab in Milton Keynes | Wellcome Sanger Institute for the COVID-19 Genomics UK (COG-UK) Consortium | The Lighthouse Lab in Milton Keynes and Alex Alderton, Roberto Amato, Jeffrey Barrett, Sonia Goncalves, Ewan Harrison, David K. Jackson, Ian Johnston, Dominic Kwiatkowski, Cordelia Langford, John Sillitoe on behalf of the Wellcome Sanger Institute COVID-19 Surveillance Team |
| EPI_ISL_3281654, EPI_ISL_3281655 | Lighthouse Lab in Glasgow | Wellcome Sanger Institute for the COVID-19 Genomics UK (COG-UK) Consortium | Harper VanSteenhouse, Yumi Kasai, David Gray, Carol Clugston, Anna Dominiczak and Alex Alderton, Roberto Amato, Jeffrey Barrett, Sonia Goncalves, Ewan Harrison, David K. Jackson, Ian Johnston, Dominic Kwiatkowski, Cordelia Langford, John Sillitoe on behalf of the Wellcome Sanger Institute COVID-19 Surveillance Team |
| EPI_ISL_3281656, EPI_ISL_3281657 | Lighthouse Lab in Alderley Park | Wellcome Sanger Institute for the COVID-19 Genomics UK (COG-UK) Consortium | Jacquelyn Wynn, Mairead Hyland, The Lighthouse Lab in Alderley Park and Alex Alderton, Roberto Amato, Jeffrey Barrett, Sonia Goncalves, Ewan Harrison, David K. Jackson, Ian Johnston, Dominic Kwiatkowski, Cordelia Langford, John Sillitoe on behalf of the Wellcome Sanger Institute COVID-19 Surveillance Team |
| EPI_ISL_3281658, EPI_ISL_3281659 | Lighthouse Lab in Glasgow | Wellcome Sanger Institute for the COVID-19 Genomics UK (COG-UK) Consortium | Harper VanSteenhouse, Yumi Kasai, David Gray, Carol Clugston, Anna Dominiczak and Alex Alderton, Roberto Amato, Jeffrey Barrett, Sonia Goncalves, Ewan Harrison, David K. Jackson, Ian Johnston, Dominic Kwiatkowski, Cordelia Langford, John Sillitoe on behalf of the Wellcome Sanger Institute COVID-19 Surveillance Team |
| EPI_ISL_3281660 | Lighthouse Lab in Alderley Park | Wellcome Sanger Institute for the COVID-19 Genomics UK (COG-UK) Consortium | Jacquelyn Wynn, Mairead Hyland, The Lighthouse Lab in Alderley Park and Alex Alderton, Roberto Amato, Jeffrey Barrett, Sonia Goncalves, Ewan Harrison, David K. Jackson, Ian Johnston, Dominic Kwiatkowski, Cordelia Langford, John Sillitoe on behalf of the Wellcome Sanger Institute COVID-19 Surveillance Team |
| EPI_ISL_3281661, EPI_ISL_3281662 | Lighthouse Lab in Milton Keynes | Wellcome Sanger Institute for the COVID-19 Genomics UK (COG-UK) Consortium | The Lighthouse Lab in Milton Keynes and Alex Alderton, Roberto Amato, Jeffrey Barrett, Sonia Goncalves, Ewan Harrison, David K. Jackson, Ian Johnston, Dominic Kwiatkowski, Cordelia Langford, John Sillitoe on behalf of the Wellcome Sanger Institute COVID-19 Surveillance Team |
| EPI_ISL_3281663, EPI_ISL_3281664, EPI_ISL_3281665 | Lighthouse Lab in Alderley Park | Wellcome Sanger Institute for the COVID-19 Genomics UK (COG-UK) Consortium | Jacquelyn Wynn, Mairead Hyland, The Lighthouse Lab in Alderley Park and Alex Alderton, Roberto Amato, Jeffrey Barrett, Sonia Goncalves, Ewan Harrison, David K. Jackson, Ian Johnston, Dominic Kwiatkowski, Cordelia Langford, John Sillitoe on behalf of the Wellcome Sanger Institute COVID-19 Surveillance Team |
| EPI_ISL_3281666 | Lighthouse Lab in Milton Keynes | Wellcome Sanger Institute for the COVID-19 Genomics UK (COG-UK) | The Lighthouse Lab in Milton Keynes and Alex Alderton, Roberto Amato, Jeffrey Barrett, Sonia Goncalves, Ewan Harrison, David K. Jackson, Ian Johnston, Dominic Kwiatkowski, Cordelia Langford, John Sillitoe on behalf of the Wellcome Sanger |

| | | | |
|---|---|---|---|
| | | Consortium | Institute COVID-19 Surveillance Team |
| EPI_ISL_3281667 | Lighthouse Lab in Alderley Park | Wellcome Sanger Institute for the COVID-19 Genomics UK (COG-UK) Consortium | Jacquelyn Wynn, Mairead Hyland, The Lighthouse Lab in Alderley Park and Alex Alderton, Roberto Amato, Jeffrey Barrett, Sonia Goncalves, Ewan Harrison, David K. Jackson, Ian Johnston, Dominic Kwiatkowski, Cordelia Langford, John Sillitoe on behalf of the Wellcome Sanger Institute COVID-19 Surveillance Team |
| EPI_ISL_3281668 | Lighthouse Lab in Glasgow | Wellcome Sanger Institute for the COVID-19 Genomics UK (COG-UK) Consortium | Harper VanSteenhouse, Yumi Kasai, David Gray, Carol Clugston, Anna Dominiczak and Alex Alderton, Roberto Amato, Jeffrey Barrett, Sonia Goncalves, Ewan Harrison, David K. Jackson, Ian Johnston, Dominic Kwiatkowski, Cordelia Langford, John Sillitoe on behalf of the Wellcome Sanger Institute COVID-19 Surveillance Team |
| EPI_ISL_3281669 | Lighthouse Lab in Alderley Park | Wellcome Sanger Institute for the COVID-19 Genomics UK (COG-UK) Consortium | Jacquelyn Wynn, Mairead Hyland, The Lighthouse Lab in Alderley Park and Alex Alderton, Roberto Amato, Jeffrey Barrett, Sonia Goncalves, Ewan Harrison, David K. Jackson, Ian Johnston, Dominic Kwiatkowski, Cordelia Langford, John Sillitoe on behalf of the Wellcome Sanger Institute COVID-19 Surveillance Team |
| EPI_ISL_3281670 | Lighthouse Lab in Milton Keynes | Wellcome Sanger Institute for the COVID-19 Genomics UK (COG-UK) Consortium | The Lighthouse Lab in Milton Keynes and Alex Alderton, Roberto Amato, Jeffrey Barrett, Sonia Goncalves, Ewan Harrison, David K. Jackson, Ian Johnston, Dominic Kwiatkowski, Cordelia Langford, John Sillitoe on behalf of the Wellcome Sanger Institute COVID-19 Surveillance Team |
| EPI_ISL_3281671 | Lighthouse Lab in Glasgow | Wellcome Sanger Institute for the COVID-19 Genomics UK (COG-UK) Consortium | Harper VanSteenhouse, Yumi Kasai, David Gray, Carol Clugston, Anna Dominiczak and Alex Alderton, Roberto Amato, Jeffrey Barrett, Sonia Goncalves, Ewan Harrison, David K. Jackson, Ian Johnston, Dominic Kwiatkowski, Cordelia Langford, John Sillitoe on behalf of the Wellcome Sanger Institute COVID-19 Surveillance Team |
| EPI_ISL_3281672, EPI_ISL_3281673 | Lighthouse Lab in Alderley Park | Wellcome Sanger Institute for the COVID-19 Genomics UK (COG-UK) Consortium | Jacquelyn Wynn, Mairead Hyland, The Lighthouse Lab in Alderley Park and Alex Alderton, Roberto Amato, Jeffrey Barrett, Sonia Goncalves, Ewan Harrison, David K. Jackson, Ian Johnston, Dominic Kwiatkowski, Cordelia Langford, John Sillitoe on behalf of the Wellcome Sanger Institute COVID-19 Surveillance Team |
| EPI_ISL_3281674 | Lighthouse Lab in Milton Keynes | Wellcome Sanger Institute for the COVID-19 Genomics UK (COG-UK) Consortium | The Lighthouse Lab in Milton Keynes and Alex Alderton, Roberto Amato, Jeffrey Barrett, Sonia Goncalves, Ewan Harrison, David K. Jackson, Ian Johnston, Dominic Kwiatkowski, Cordelia Langford, John Sillitoe on behalf of the Wellcome Sanger Institute COVID-19 Surveillance Team |
| EPI_ISL_3281675 | Lighthouse Lab in Glasgow | Wellcome Sanger Institute for the COVID-19 Genomics UK (COG-UK) Consortium | Harper VanSteenhouse, Yumi Kasai, David Gray, Carol Clugston, Anna Dominiczak and Alex Alderton, Roberto Amato, Jeffrey Barrett, Sonia Goncalves, Ewan Harrison, David K. Jackson, Ian Johnston, Dominic Kwiatkowski, Cordelia Langford, John Sillitoe on behalf of the Wellcome Sanger Institute COVID-19 Surveillance Team |
| EPI_ISL_3281676, EPI_ISL_3281677 | Lighthouse Lab in Alderley Park | Wellcome Sanger Institute for the COVID-19 Genomics UK (COG-UK) Consortium | Jacquelyn Wynn, Mairead Hyland, The Lighthouse Lab in Alderley Park and Alex Alderton, Roberto Amato, Jeffrey Barrett, Sonia Goncalves, Ewan Harrison, David K. Jackson, Ian Johnston, Dominic Kwiatkowski, Cordelia Langford, John Sillitoe on behalf of the Wellcome Sanger Institute COVID-19 Surveillance Team |
| EPI_ISL_3281678 | Lighthouse Lab in Glasgow | Wellcome Sanger Institute for the COVID-19 Genomics UK (COG-UK) Consortium | Harper VanSteenhouse, Yumi Kasai, David Gray, Carol Clugston, Anna Dominiczak and Alex Alderton, Roberto Amato, Jeffrey Barrett, Sonia Goncalves, Ewan Harrison, David K. Jackson, Ian Johnston, Dominic Kwiatkowski, Cordelia Langford, John Sillitoe on behalf of the Wellcome Sanger Institute COVID-19 Surveillance Team |
| EPI_ISL_3281679, EPI_ISL_3281680 | Lighthouse Lab in Alderley Park | Wellcome Sanger Institute for the COVID-19 Genomics UK (COG-UK) Consortium | Jacquelyn Wynn, Mairead Hyland, The Lighthouse Lab in Alderley Park and Alex Alderton, Roberto Amato, Jeffrey Barrett, Sonia Goncalves, Ewan Harrison, David K. Jackson, Ian Johnston, Dominic Kwiatkowski, Cordelia Langford, John Sillitoe on behalf of the Wellcome Sanger Institute COVID-19 Surveillance Team |
| EPI_ISL_3281681 | Lighthouse Lab in Milton Keynes | Wellcome Sanger Institute for the COVID-19 Genomics UK (COG-UK) Consortium | The Lighthouse Lab in Milton Keynes and Alex Alderton, Roberto Amato, Jeffrey Barrett, Sonia Goncalves, Ewan Harrison, David K. Jackson, Ian Johnston, Dominic Kwiatkowski, Cordelia Langford, John Sillitoe on behalf of the Wellcome Sanger Institute COVID-19 Surveillance Team |
| EPI_ISL_3281682, EPI_ISL_3281683, EPI_ISL_3281684 | Lighthouse Lab in Glasgow | Wellcome Sanger Institute for the COVID-19 Genomics UK (COG-UK) Consortium | Harper VanSteenhouse, Yumi Kasai, David Gray, Carol Clugston, Anna Dominiczak and Alex Alderton, Roberto Amato, Jeffrey Barrett, Sonia Goncalves, Ewan Harrison, David K. Jackson, Ian Johnston, Dominic Kwiatkowski, Cordelia Langford, John Sillitoe on behalf of the Wellcome Sanger Institute COVID-19 Surveillance Team |
| EPI_ISL_3281685 | Lighthouse Lab in Alderley Park | Wellcome Sanger Institute for the COVID-19 Genomics UK (COG-UK) Consortium | Jacquelyn Wynn, Mairead Hyland, The Lighthouse Lab in Alderley Park and Alex Alderton, Roberto Amato, Jeffrey Barrett, Sonia Goncalves, Ewan Harrison, David K. Jackson, Ian Johnston, Dominic Kwiatkowski, Cordelia Langford, John Sillitoe on behalf of the Wellcome Sanger Institute COVID-19 Surveillance Team |
| EPI_ISL_3281686 | Lighthouse Lab in Milton Keynes | Wellcome Sanger Institute for the COVID-19 Genomics UK (COG-UK) Consortium | The Lighthouse Lab in Milton Keynes and Alex Alderton, Roberto Amato, Jeffrey Barrett, Sonia Goncalves, Ewan Harrison, David K. Jackson, Ian Johnston, Dominic Kwiatkowski, Cordelia Langford, John Sillitoe on behalf of the Wellcome Sanger Institute COVID-19 Surveillance Team |
| EPI_ISL_3281687 | Lighthouse Lab in Alderley Park | Wellcome Sanger Institute for the COVID-19 Genomics UK (COG-UK) Consortium | Jacquelyn Wynn, Mairead Hyland, The Lighthouse Lab in Alderley Park and Alex Alderton, Roberto Amato, Jeffrey Barrett, Sonia Goncalves, Ewan Harrison, David K. Jackson, Ian Johnston, Dominic Kwiatkowski, Cordelia Langford, John Sillitoe on behalf of the Wellcome Sanger Institute COVID-19 Surveillance Team |
| EPI_ISL_3281688 | Lighthouse Lab in Milton Keynes | Wellcome Sanger Institute for the COVID-19 Genomics UK (COG-UK) Consortium | The Lighthouse Lab in Milton Keynes and Alex Alderton, Roberto Amato, Jeffrey Barrett, Sonia Goncalves, Ewan Harrison, David K. Jackson, Ian Johnston, Dominic Kwiatkowski, Cordelia Langford, John Sillitoe on behalf of the Wellcome Sanger Institute COVID-19 Surveillance Team |
| EPI_ISL_3281689 | Lighthouse Lab in Alderley Park | Wellcome Sanger Institute for the COVID-19 Genomics UK (COG-UK) Consortium | Jacquelyn Wynn, Mairead Hyland, The Lighthouse Lab in Alderley Park and Alex Alderton, Roberto Amato, Jeffrey Barrett, Sonia Goncalves, Ewan Harrison, David K. Jackson, Ian Johnston, Dominic Kwiatkowski, Cordelia Langford, John Sillitoe on behalf of the Wellcome Sanger Institute COVID-19 Surveillance Team |
| EPI_ISL_3281690 | Lighthouse Lab in Glasgow | Wellcome Sanger Institute for the COVID-19 Genomics UK (COG-UK) Consortium | Harper VanSteenhouse, Yumi Kasai, David Gray, Carol Clugston, Anna Dominiczak and Alex Alderton, Roberto Amato, Jeffrey Barrett, Sonia Goncalves, Ewan Harrison, David K. Jackson, Ian Johnston, Dominic Kwiatkowski, Cordelia Langford, John Sillitoe on behalf of the Wellcome Sanger Institute COVID-19 Surveillance Team |
| EPI_ISL_3281691, EPI_ISL_3281692 | Lighthouse Lab in Milton Keynes | Wellcome Sanger Institute for the COVID-19 Genomics UK (COG-UK) Consortium | The Lighthouse Lab in Milton Keynes and Alex Alderton, Roberto Amato, Jeffrey Barrett, Sonia Goncalves, Ewan Harrison, David K. Jackson, Ian Johnston, Dominic Kwiatkowski, Cordelia Langford, John Sillitoe on behalf of the Wellcome Sanger Institute COVID-19 Surveillance Team |
| EPI_ISL_3281693 | Lighthouse Lab in Glasgow | Wellcome Sanger Institute for the COVID-19 Genomics UK (COG-UK) Consortium | Harper VanSteenhouse, Yumi Kasai, David Gray, Carol Clugston, Anna Dominiczak and Alex Alderton, Roberto Amato, Jeffrey Barrett, Sonia Goncalves, Ewan Harrison, David K. Jackson, Ian Johnston, Dominic Kwiatkowski, Cordelia Langford, John Sillitoe on behalf of the Wellcome Sanger Institute COVID-19 Surveillance Team |
| EPI_ISL_3281694, EPI_ISL_3281695, EPI_ISL_3281696 | Lighthouse Lab in Alderley Park | Wellcome Sanger Institute for the COVID-19 Genomics UK (COG-UK) Consortium | Jacquelyn Wynn, Mairead Hyland, The Lighthouse Lab in Alderley Park and Alex Alderton, Roberto Amato, Jeffrey Barrett, Sonia Goncalves, Ewan Harrison, David K. Jackson, Ian Johnston, Dominic Kwiatkowski, Cordelia Langford, John Sillitoe on behalf of the Wellcome Sanger Institute COVID-19 Surveillance Team |
| EPI_ISL_3281697, EPI_ISL_3281698 | Lighthouse Lab in Milton Keynes | Wellcome Sanger Institute for the COVID-19 Genomics UK (COG-UK) Consortium | The Lighthouse Lab in Milton Keynes and Alex Alderton, Roberto Amato, Jeffrey Barrett, Sonia Goncalves, Ewan Harrison, David K. Jackson, Ian Johnston, Dominic Kwiatkowski, Cordelia Langford, John Sillitoe on behalf of the Wellcome Sanger Institute COVID-19 Surveillance Team |
| EPI_ISL_3281699 | Lighthouse Lab in Glasgow | Wellcome Sanger Institute for the COVID-19 Genomics UK (COG-UK) Consortium | Harper VanSteenhouse, Yumi Kasai, David Gray, Carol Clugston, Anna Dominiczak and Alex Alderton, Roberto Amato, Jeffrey Barrett, Sonia Goncalves, Ewan Harrison, David K. Jackson, Ian Johnston, Dominic Kwiatkowski, Cordelia Langford, John Sillitoe on behalf of the Wellcome Sanger Institute COVID-19 Surveillance Team |
| EPI_ISL_3281700 | Lighthouse Lab in Alderley Park | Wellcome Sanger Institute for the COVID-19 Genomics UK (COG-UK) Consortium | Jacquelyn Wynn, Mairead Hyland, The Lighthouse Lab in Alderley Park and Alex Alderton, Roberto Amato, Jeffrey Barrett, Sonia Goncalves, Ewan Harrison, David K. Jackson, Ian Johnston, Dominic Kwiatkowski, Cordelia Langford, John Sillitoe on behalf of the Wellcome Sanger Institute COVID-19 Surveillance Team |
| EPI_ISL_3281701, EPI_ISL_3281702 | Lighthouse Lab in Milton Keynes | Wellcome Sanger Institute for the COVID-19 Genomics UK (COG-UK) Consortium | The Lighthouse Lab in Milton Keynes and Alex Alderton, Roberto Amato, Jeffrey Barrett, Sonia Goncalves, Ewan Harrison, David K. Jackson, Ian Johnston, Dominic Kwiatkowski, Cordelia Langford, John Sillitoe on behalf of the Wellcome Sanger Institute COVID-19 Surveillance Team |
| EPI_ISL_3281703 | Lighthouse Lab in Glasgow | Wellcome Sanger Institute for the COVID-19 Genomics UK (COG-UK) Consortium | Harper VanSteenhouse, Yumi Kasai, David Gray, Carol Clugston, Anna Dominiczak and Alex Alderton, Roberto Amato, Jeffrey Barrett, Sonia Goncalves, Ewan Harrison, David K. Jackson, Ian Johnston, Dominic Kwiatkowski, Cordelia Langford, John Sillitoe on behalf of the Wellcome Sanger Institute COVID-19 Surveillance Team |
| EPI_ISL_3281704 | Lighthouse Lab in Alderley Park | Wellcome Sanger Institute for the COVID-19 Genomics UK (COG-UK) Consortium | Jacquelyn Wynn, Mairead Hyland, The Lighthouse Lab in Alderley Park and Alex Alderton, Roberto Amato, Jeffrey Barrett, Sonia Goncalves, Ewan Harrison, David K. Jackson, Ian Johnston, Dominic Kwiatkowski, Cordelia Langford, John Sillitoe on behalf of the Wellcome Sanger Institute COVID-19 Surveillance Team |
| EPI_ISL_3281705 | Lighthouse Lab in Milton Keynes | Wellcome Sanger Institute for the COVID-19 Genomics UK (COG-UK) Consortium | The Lighthouse Lab in Milton Keynes and Alex Alderton, Roberto Amato, Jeffrey Barrett, Sonia Goncalves, Ewan Harrison, David K. Jackson, Ian Johnston, Dominic Kwiatkowski, Cordelia Langford, John Sillitoe on behalf of the Wellcome Sanger Institute COVID-19 Surveillance Team |
| EPI_ISL_3281706 | Lighthouse Lab in Glasgow | Wellcome Sanger Institute for the COVID-19 Genomics UK (COG-UK) Consortium | Harper VanSteenhouse, Yumi Kasai, David Gray, Carol Clugston, Anna Dominiczak and Alex Alderton, Roberto Amato, Jeffrey Barrett, Sonia Goncalves, Ewan Harrison, David K. Jackson, Ian Johnston, Dominic Kwiatkowski, Cordelia Langford, John Sillitoe on behalf of the Wellcome Sanger Institute COVID-19 Surveillance Team |
| EPI_ISL_3281707 | Lighthouse Lab in Milton Keynes | Wellcome Sanger Institute for the COVID-19 Genomics UK (COG-UK) Consortium | The Lighthouse Lab in Milton Keynes and Alex Alderton, Roberto Amato, Jeffrey Barrett, Sonia Goncalves, Ewan Harrison, David K. Jackson, Ian Johnston, Dominic Kwiatkowski, Cordelia Langford, John Sillitoe on behalf of the Wellcome Sanger Institute COVID-19 Surveillance Team |
| EPI_ISL_3281708 | Lighthouse Lab in Alderley Park | Wellcome Sanger Institute for the COVID-19 Genomics UK (COG-UK) Consortium | Jacquelyn Wynn, Mairead Hyland, The Lighthouse Lab in Alderley Park and Alex Alderton, Roberto Amato, Jeffrey Barrett, Sonia Goncalves, Ewan Harrison, David K. Jackson, Ian Johnston, Dominic Kwiatkowski, Cordelia Langford, John Sillitoe on behalf of the Wellcome Sanger Institute COVID-19 Surveillance Team |
| EPI_ISL_3281709 | Lighthouse Lab in Glasgow | Wellcome Sanger Institute for the COVID-19 Genomics UK (COG-UK) Consortium | Harper VanSteenhouse, Yumi Kasai, David Gray, Carol Clugston, Anna Dominiczak and Alex Alderton, Roberto Amato, Jeffrey Barrett, Sonia Goncalves, Ewan Harrison, David K. Jackson, Ian Johnston, Dominic Kwiatkowski, Cordelia Langford, John Sillitoe on behalf of the Wellcome Sanger Institute COVID-19 Surveillance Team |
| EPI_ISL_3281710 | Lighthouse Lab in Alderley Park | Wellcome Sanger Institute for the COVID-19 Genomics UK (COG-UK) Consortium | Jacquelyn Wynn, Mairead Hyland, The Lighthouse Lab in Alderley Park and Alex Alderton, Roberto Amato, Jeffrey Barrett, Sonia Goncalves, Ewan Harrison, David K. Jackson, Ian Johnston, Dominic Kwiatkowski, Cordelia Langford, John Sillitoe on behalf of the Wellcome Sanger Institute COVID-19 Surveillance Team |
| EPI_ISL_3281711, EPI_ISL_3281712 | Lighthouse Lab in Glasgow | Wellcome Sanger Institute for the COVID-19 Genomics UK (COG-UK) Consortium | Harper VanSteenhouse, Yumi Kasai, David Gray, Carol Clugston, Anna Dominiczak and Alex Alderton, Roberto Amato, Jeffrey Barrett, Sonia Goncalves, Ewan Harrison, David K. Jackson, Ian Johnston, Dominic Kwiatkowski, Cordelia Langford, John Sillitoe on behalf of the Wellcome Sanger Institute COVID-19 Surveillance Team |
| EPI_ISL_3281713 | Lighthouse Lab in Alderley Park | Wellcome Sanger Institute for the COVID-19 Genomics UK (COG-UK) Consortium | Jacquelyn Wynn, Mairead Hyland, The Lighthouse Lab in Alderley Park and Alex Alderton, Roberto Amato, Jeffrey Barrett, Sonia Goncalves, Ewan Harrison, David K. Jackson, Ian Johnston, Dominic Kwiatkowski, Cordelia Langford, John Sillitoe on behalf of the Wellcome Sanger Institute COVID-19 Surveillance Team |
| EPI_ISL_3281714 | Lighthouse Lab in Milton Keynes | Wellcome Sanger Institute for the COVID-19 Genomics UK (COG-UK) Consortium | The Lighthouse Lab in Milton Keynes and Alex Alderton, Roberto Amato, Jeffrey Barrett, Sonia Goncalves, Ewan Harrison, David K. Jackson, Ian Johnston, Dominic Kwiatkowski, Cordelia Langford, John Sillitoe on behalf of the Wellcome Sanger Institute COVID-19 Surveillance Team |
| EPI_ISL_3281715, EPI_ISL_3281716, EPI_ISL_3281717 | Lighthouse Lab in Alderley Park | Wellcome Sanger Institute for the COVID-19 Genomics UK (COG-UK) Consortium | Jacquelyn Wynn, Mairead Hyland, The Lighthouse Lab in Alderley Park and Alex Alderton, Roberto Amato, Jeffrey Barrett, Sonia Goncalves, Ewan Harrison, David K. Jackson, Ian Johnston, Dominic Kwiatkowski, Cordelia Langford, John Sillitoe on behalf of the Wellcome Sanger Institute COVID-19 Surveillance Team |
| EPI_ISL_3281718 | Lighthouse Lab in Milton Keynes | Wellcome Sanger Institute for the COVID-19 Genomics UK (COG-UK) Consortium | The Lighthouse Lab in Milton Keynes and Alex Alderton, Roberto Amato, Jeffrey Barrett, Sonia Goncalves, Ewan Harrison, David K. Jackson, Ian Johnston, Dominic Kwiatkowski, Cordelia Langford, John Sillitoe on behalf of the Wellcome Sanger Institute COVID-19 Surveillance Team |
| EPI_ISL_3281719 | Lighthouse Lab in Glasgow | Wellcome Sanger Institute for the COVID-19 Genomics UK (COG-UK) Consortium | Harper VanSteenhouse, Yumi Kasai, David Gray, Carol Clugston, Anna Dominiczak and Alex Alderton, Roberto Amato, Jeffrey Barrett, Sonia Goncalves, Ewan Harrison, David K. Jackson, Ian Johnston, Dominic Kwiatkowski, Cordelia Langford, John Sillitoe on behalf of the Wellcome Sanger Institute COVID-19 Surveillance Team |
| EPI_ISL_3281720 | Lighthouse Lab in Alderley Park | Wellcome Sanger Institute for the COVID-19 Genomics UK (COG-UK) Consortium | Jacquelyn Wynn, Mairead Hyland, The Lighthouse Lab in Alderley Park and Alex Alderton, Roberto Amato, Jeffrey Barrett, Sonia Goncalves, Ewan Harrison, David K. Jackson, Ian Johnston, Dominic Kwiatkowski, Cordelia Langford, John Sillitoe on behalf of the Wellcome Sanger Institute COVID-19 Surveillance Team |
| EPI_ISL_3281721, EPI_ISL_3281722 | Lighthouse Lab in Glasgow | Wellcome Sanger Institute for the COVID-19 Genomics UK (COG-UK) Consortium | Harper VanSteenhouse, Yumi Kasai, David Gray, Carol Clugston, Anna Dominiczak and Alex Alderton, Roberto Amato, Jeffrey Barrett, Sonia Goncalves, Ewan Harrison, David K. Jackson, Ian Johnston, Dominic Kwiatkowski, Cordelia Langford, John Sillitoe on behalf of the Wellcome Sanger Institute COVID-19 Surveillance Team |
| EPI_ISL_3281723 | Lighthouse Lab in Milton Keynes | Wellcome Sanger Institute for the COVID-19 Genomics UK (COG-UK) Consortium | The Lighthouse Lab in Milton Keynes and Alex Alderton, Roberto Amato, Jeffrey Barrett, Sonia Goncalves, Ewan Harrison, David K. Jackson, Ian Johnston, Dominic Kwiatkowski, Cordelia Langford, John Sillitoe on behalf of the Wellcome Sanger Institute COVID-19 Surveillance Team |
| EPI_ISL_3281724, | Lighthouse Lab in Glasgow | Wellcome Sanger Institute for the COVID-19 Genomics UK (COG-UK) | Harper VanSteenhouse, Yumi Kasai, David Gray, Carol Clugston, Anna Dominiczak and Alex Alderton, Roberto Amato, Jeffrey Barrett, Sonia Goncalves, Ewan Harrison, David K. Jackson, Ian Johnston, Dominic Kwiatkowski, Cordelia Langford, |

| | | | |
|---|---|---|---|
| EPI_ISL_3281725, EPI_ISL_3281726, EPI_ISL_3281727 | | Consortium | John Sillitoe on behalf of the Wellcome Sanger Institute COVID-19 Surveillance Team |
| EPI_ISL_3281728 | Lighthouse Lab in Alderley Park | Wellcome Sanger Institute for the COVID-19 Genomics UK (COG-UK) Consortium | Jacquelyn Wynn, Mairead Hyland, The Lighthouse Lab in Alderley Park and Alex Alderton, Roberto Amato, Jeffrey Barrett, Sonia Goncalves, Ewan Harrison, David K. Jackson, Ian Johnston, Dominic Kwiatkowski, Cordelia Langford, John Sillitoe on behalf of the Wellcome Sanger Institute COVID-19 Surveillance Team |
| EPI_ISL_3281729 | Lighthouse Lab in Glasgow | Wellcome Sanger Institute for the COVID-19 Genomics UK (COG-UK) Consortium | Harper VanSteenhouse, Yumi Kasai, David Gray, Carol Clugston, Anna Dominiczak and Alex Alderton, Roberto Amato, Jeffrey Barrett, Sonia Goncalves, Ewan Harrison, David K. Jackson, Ian Johnston, Dominic Kwiatkowski, Cordelia Langford, John Sillitoe on behalf of the Wellcome Sanger Institute COVID-19 Surveillance Team |
| EPI_ISL_3281730 | Lighthouse Lab in Milton Keynes | Wellcome Sanger Institute for the COVID-19 Genomics UK (COG-UK) Consortium | The Lighthouse Lab in Milton Keynes and Alex Alderton, Roberto Amato, Jeffrey Barrett, Sonia Goncalves, Ewan Harrison, David K. Jackson, Ian Johnston, Dominic Kwiatkowski, Cordelia Langford, John Sillitoe on behalf of the Wellcome Sanger Institute COVID-19 Surveillance Team |
| EPI_ISL_3281731 | Lighthouse Lab in Glasgow | Wellcome Sanger Institute for the COVID-19 Genomics UK (COG-UK) Consortium | Harper VanSteenhouse, Yumi Kasai, David Gray, Carol Clugston, Anna Dominiczak and Alex Alderton, Roberto Amato, Jeffrey Barrett, Sonia Goncalves, Ewan Harrison, David K. Jackson, Ian Johnston, Dominic Kwiatkowski, Cordelia Langford, John Sillitoe on behalf of the Wellcome Sanger Institute COVID-19 Surveillance Team |
| EPI_ISL_3281732, EPI_ISL_3281733 | Lighthouse Lab in Alderley Park | Wellcome Sanger Institute for the COVID-19 Genomics UK (COG-UK) Consortium | Jacquelyn Wynn, Mairead Hyland, The Lighthouse Lab in Alderley Park and Alex Alderton, Roberto Amato, Jeffrey Barrett, Sonia Goncalves, Ewan Harrison, David K. Jackson, Ian Johnston, Dominic Kwiatkowski, Cordelia Langford, John Sillitoe on behalf of the Wellcome Sanger Institute COVID-19 Surveillance Team |
| EPI_ISL_3281734 | Lighthouse Lab in Glasgow | Wellcome Sanger Institute for the COVID-19 Genomics UK (COG-UK) Consortium | Harper VanSteenhouse, Yumi Kasai, David Gray, Carol Clugston, Anna Dominiczak and Alex Alderton, Roberto Amato, Jeffrey Barrett, Sonia Goncalves, Ewan Harrison, David K. Jackson, Ian Johnston, Dominic Kwiatkowski, Cordelia Langford, John Sillitoe on behalf of the Wellcome Sanger Institute COVID-19 Surveillance Team |
| EPI_ISL_3281735 | Lighthouse Lab in Alderley Park | Wellcome Sanger Institute for the COVID-19 Genomics UK (COG-UK) Consortium | Jacquelyn Wynn, Mairead Hyland, The Lighthouse Lab in Alderley Park and Alex Alderton, Roberto Amato, Jeffrey Barrett, Sonia Goncalves, Ewan Harrison, David K. Jackson, Ian Johnston, Dominic Kwiatkowski, Cordelia Langford, John Sillitoe on behalf of the Wellcome Sanger Institute COVID-19 Surveillance Team |
| EPI_ISL_3281736 | Lighthouse Lab in Milton Keynes | Wellcome Sanger Institute for the COVID-19 Genomics UK (COG-UK) Consortium | The Lighthouse Lab in Milton Keynes and Alex Alderton, Roberto Amato, Jeffrey Barrett, Sonia Goncalves, Ewan Harrison, David K. Jackson, Ian Johnston, Dominic Kwiatkowski, Cordelia Langford, John Sillitoe on behalf of the Wellcome Sanger Institute COVID-19 Surveillance Team |
| EPI_ISL_3281737 | Lighthouse Lab in Glasgow | Wellcome Sanger Institute for the COVID-19 Genomics UK (COG-UK) Consortium | Harper VanSteenhouse, Yumi Kasai, David Gray, Carol Clugston, Anna Dominiczak and Alex Alderton, Roberto Amato, Jeffrey Barrett, Sonia Goncalves, Ewan Harrison, David K. Jackson, Ian Johnston, Dominic Kwiatkowski, Cordelia Langford, John Sillitoe on behalf of the Wellcome Sanger Institute COVID-19 Surveillance Team |
| EPI_ISL_3281738 | Lighthouse Lab in Alderley Park | Wellcome Sanger Institute for the COVID-19 Genomics UK (COG-UK) Consortium | Jacquelyn Wynn, Mairead Hyland, The Lighthouse Lab in Alderley Park and Alex Alderton, Roberto Amato, Jeffrey Barrett, Sonia Goncalves, Ewan Harrison, David K. Jackson, Ian Johnston, Dominic Kwiatkowski, Cordelia Langford, John Sillitoe on behalf of the Wellcome Sanger Institute COVID-19 Surveillance Team |
| EPI_ISL_3281739, EPI_ISL_3281740 | Lighthouse Lab in Glasgow | Wellcome Sanger Institute for the COVID-19 Genomics UK (COG-UK) Consortium | Harper VanSteenhouse, Yumi Kasai, David Gray, Carol Clugston, Anna Dominiczak and Alex Alderton, Roberto Amato, Jeffrey Barrett, Sonia Goncalves, Ewan Harrison, David K. Jackson, Ian Johnston, Dominic Kwiatkowski, Cordelia Langford, John Sillitoe on behalf of the Wellcome Sanger Institute COVID-19 Surveillance Team |
| EPI_ISL_3281741 | Lighthouse Lab in Milton Keynes | Wellcome Sanger Institute for the COVID-19 Genomics UK (COG-UK) Consortium | The Lighthouse Lab in Milton Keynes and Alex Alderton, Roberto Amato, Jeffrey Barrett, Sonia Goncalves, Ewan Harrison, David K. Jackson, Ian Johnston, Dominic Kwiatkowski, Cordelia Langford, John Sillitoe on behalf of the Wellcome Sanger Institute COVID-19 Surveillance Team |
| EPI_ISL_3281742 | Lighthouse Lab in Glasgow | Wellcome Sanger Institute for the COVID-19 Genomics UK (COG-UK) Consortium | Harper VanSteenhouse, Yumi Kasai, David Gray, Carol Clugston, Anna Dominiczak and Alex Alderton, Roberto Amato, Jeffrey Barrett, Sonia Goncalves, Ewan Harrison, David K. Jackson, Ian Johnston, Dominic Kwiatkowski, Cordelia Langford, John Sillitoe on behalf of the Wellcome Sanger Institute COVID-19 Surveillance Team |
| EPI_ISL_3281743 | Lighthouse Lab in Alderley Park | Wellcome Sanger Institute for the COVID-19 Genomics UK (COG-UK) Consortium | Jacquelyn Wynn, Mairead Hyland, The Lighthouse Lab in Alderley Park and Alex Alderton, Roberto Amato, Jeffrey Barrett, Sonia Goncalves, Ewan Harrison, David K. Jackson, Ian Johnston, Dominic Kwiatkowski, Cordelia Langford, John Sillitoe on behalf of the Wellcome Sanger Institute COVID-19 Surveillance Team |
| EPI_ISL_3281744 | Lighthouse Lab in Glasgow | Wellcome Sanger Institute for the COVID-19 Genomics UK (COG-UK) Consortium | Harper VanSteenhouse, Yumi Kasai, David Gray, Carol Clugston, Anna Dominiczak and Alex Alderton, Roberto Amato, Jeffrey Barrett, Sonia Goncalves, Ewan Harrison, David K. Jackson, Ian Johnston, Dominic Kwiatkowski, Cordelia Langford, John Sillitoe on behalf of the Wellcome Sanger Institute COVID-19 Surveillance Team |
| EPI_ISL_3281745 | Lighthouse Lab in Milton Keynes | Wellcome Sanger Institute for the COVID-19 Genomics UK (COG-UK) Consortium | The Lighthouse Lab in Milton Keynes and Alex Alderton, Roberto Amato, Jeffrey Barrett, Sonia Goncalves, Ewan Harrison, David K. Jackson, Ian Johnston, Dominic Kwiatkowski, Cordelia Langford, John Sillitoe on behalf of the Wellcome Sanger Institute COVID-19 Surveillance Team |
| EPI_ISL_3281746 | Lighthouse Lab in Alderley Park | Wellcome Sanger Institute for the COVID-19 Genomics UK (COG-UK) Consortium | Jacquelyn Wynn, Mairead Hyland, The Lighthouse Lab in Alderley Park and Alex Alderton, Roberto Amato, Jeffrey Barrett, Sonia Goncalves, Ewan Harrison, David K. Jackson, Ian Johnston, Dominic Kwiatkowski, Cordelia Langford, John Sillitoe on behalf of the Wellcome Sanger Institute COVID-19 Surveillance Team |
| EPI_ISL_3281747, EPI_ISL_3281748 | Lighthouse Lab in Milton Keynes | Wellcome Sanger Institute for the COVID-19 Genomics UK (COG-UK) Consortium | The Lighthouse Lab in Milton Keynes and Alex Alderton, Roberto Amato, Jeffrey Barrett, Sonia Goncalves, Ewan Harrison, David K. Jackson, Ian Johnston, Dominic Kwiatkowski, Cordelia Langford, John Sillitoe on behalf of the Wellcome Sanger Institute COVID-19 Surveillance Team |
| EPI_ISL_3281749, EPI_ISL_3281750 | Lighthouse Lab in Glasgow | Wellcome Sanger Institute for the COVID-19 Genomics UK (COG-UK) Consortium | Harper VanSteenhouse, Yumi Kasai, David Gray, Carol Clugston, Anna Dominiczak and Alex Alderton, Roberto Amato, Jeffrey Barrett, Sonia Goncalves, Ewan Harrison, David K. Jackson, Ian Johnston, Dominic Kwiatkowski, Cordelia Langford, John Sillitoe on behalf of the Wellcome Sanger Institute COVID-19 Surveillance Team |
| EPI_ISL_3281751 | Lighthouse Lab in Alderley Park | Wellcome Sanger Institute for the COVID-19 Genomics UK (COG-UK) Consortium | Jacquelyn Wynn, Mairead Hyland, The Lighthouse Lab in Alderley Park and Alex Alderton, Roberto Amato, Jeffrey Barrett, Sonia Goncalves, Ewan Harrison, David K. Jackson, Ian Johnston, Dominic Kwiatkowski, Cordelia Langford, John Sillitoe on behalf of the Wellcome Sanger Institute COVID-19 Surveillance Team |
| EPI_ISL_3281752 | Lighthouse Lab in Glasgow | Wellcome Sanger Institute for the COVID-19 Genomics UK (COG-UK) Consortium | Harper VanSteenhouse, Yumi Kasai, David Gray, Carol Clugston, Anna Dominiczak and Alex Alderton, Roberto Amato, Jeffrey Barrett, Sonia Goncalves, Ewan Harrison, David K. Jackson, Ian Johnston, Dominic Kwiatkowski, Cordelia Langford, John Sillitoe on behalf of the Wellcome Sanger Institute COVID-19 Surveillance Team |
| EPI_ISL_3281753, EPI_ISL_3281754 | Lighthouse Lab in Milton Keynes | Wellcome Sanger Institute for the COVID-19 Genomics UK (COG-UK) Consortium | The Lighthouse Lab in Milton Keynes and Alex Alderton, Roberto Amato, Jeffrey Barrett, Sonia Goncalves, Ewan Harrison, David K. Jackson, Ian Johnston, Dominic Kwiatkowski, Cordelia Langford, John Sillitoe on behalf of the Wellcome Sanger Institute COVID-19 Surveillance Team |
| EPI_ISL_3281755, EPI_ISL_3281756 | Lighthouse Lab in Alderley Park | Wellcome Sanger Institute for the COVID-19 Genomics UK (COG-UK) Consortium | Jacquelyn Wynn, Mairead Hyland, The Lighthouse Lab in Alderley Park and Alex Alderton, Roberto Amato, Jeffrey Barrett, Sonia Goncalves, Ewan Harrison, David K. Jackson, Ian Johnston, Dominic Kwiatkowski, Cordelia Langford, John Sillitoe on behalf of the Wellcome Sanger Institute COVID-19 Surveillance Team |
| EPI_ISL_3281757 | Lighthouse Lab in Glasgow | Wellcome Sanger Institute for the COVID-19 Genomics UK (COG-UK) Consortium | Harper VanSteenhouse, Yumi Kasai, David Gray, Carol Clugston, Anna Dominiczak and Alex Alderton, Roberto Amato, Jeffrey Barrett, Sonia Goncalves, Ewan Harrison, David K. Jackson, Ian Johnston, Dominic Kwiatkowski, Cordelia Langford, John Sillitoe on behalf of the Wellcome Sanger Institute COVID-19 Surveillance Team |
| EPI_ISL_3281758, EPI_ISL_3281759 | Lighthouse Lab in Milton Keynes | Wellcome Sanger Institute for the COVID-19 Genomics UK (COG-UK) Consortium | The Lighthouse Lab in Milton Keynes and Alex Alderton, Roberto Amato, Jeffrey Barrett, Sonia Goncalves, Ewan Harrison, David K. Jackson, Ian Johnston, Dominic Kwiatkowski, Cordelia Langford, John Sillitoe on behalf of the Wellcome Sanger Institute COVID-19 Surveillance Team |
| EPI_ISL_3281760 | Lighthouse Lab in Glasgow | Wellcome Sanger Institute for the COVID-19 Genomics UK (COG-UK) Consortium | Harper VanSteenhouse, Yumi Kasai, David Gray, Carol Clugston, Anna Dominiczak and Alex Alderton, Roberto Amato, Jeffrey Barrett, Sonia Goncalves, Ewan Harrison, David K. Jackson, Ian Johnston, Dominic Kwiatkowski, Cordelia Langford, John Sillitoe on behalf of the Wellcome Sanger Institute COVID-19 Surveillance Team |
| EPI_ISL_3281761 | Lighthouse Lab in Milton Keynes | Wellcome Sanger Institute for the COVID-19 Genomics UK (COG-UK) Consortium | The Lighthouse Lab in Milton Keynes and Alex Alderton, Roberto Amato, Jeffrey Barrett, Sonia Goncalves, Ewan Harrison, David K. Jackson, Ian Johnston, Dominic Kwiatkowski, Cordelia Langford, John Sillitoe on behalf of the Wellcome Sanger Institute COVID-19 Surveillance Team |
| EPI_ISL_3281762, EPI_ISL_3281763, EPI_ISL_3281764 | Lighthouse Lab in Alderley Park | Wellcome Sanger Institute for the COVID-19 Genomics UK (COG-UK) Consortium | Jacquelyn Wynn, Mairead Hyland, The Lighthouse Lab in Alderley Park and Alex Alderton, Roberto Amato, Jeffrey Barrett, Sonia Goncalves, Ewan Harrison, David K. Jackson, Ian Johnston, Dominic Kwiatkowski, Cordelia Langford, John Sillitoe on behalf of the Wellcome Sanger Institute COVID-19 Surveillance Team |
| EPI_ISL_3281765, EPI_ISL_3281766 | Lighthouse Lab in Glasgow | Wellcome Sanger Institute for the COVID-19 Genomics UK (COG-UK) Consortium | Harper VanSteenhouse, Yumi Kasai, David Gray, Carol Clugston, Anna Dominiczak and Alex Alderton, Roberto Amato, Jeffrey Barrett, Sonia Goncalves, Ewan Harrison, David K. Jackson, Ian Johnston, Dominic Kwiatkowski, Cordelia Langford, John Sillitoe on behalf of the Wellcome Sanger Institute COVID-19 Surveillance Team |
| EPI_ISL_3281767 | Lighthouse Lab in Milton Keynes | Wellcome Sanger Institute for the COVID-19 Genomics UK (COG-UK) Consortium | The Lighthouse Lab in Milton Keynes and Alex Alderton, Roberto Amato, Jeffrey Barrett, Sonia Goncalves, Ewan Harrison, David K. Jackson, Ian Johnston, Dominic Kwiatkowski, Cordelia Langford, John Sillitoe on behalf of the Wellcome Sanger Institute COVID-19 Surveillance Team |
| EPI_ISL_3281768 | Lighthouse Lab in Alderley Park | Wellcome Sanger Institute for the COVID-19 Genomics UK (COG-UK) Consortium | Jacquelyn Wynn, Mairead Hyland, The Lighthouse Lab in Alderley Park and Alex Alderton, Roberto Amato, Jeffrey Barrett, Sonia Goncalves, Ewan Harrison, David K. Jackson, Ian Johnston, Dominic Kwiatkowski, Cordelia Langford, John Sillitoe on behalf of the Wellcome Sanger Institute COVID-19 Surveillance Team |
| EPI_ISL_3281769 | Lighthouse Lab in Glasgow | Wellcome Sanger Institute for the COVID-19 Genomics UK (COG-UK) Consortium | Harper VanSteenhouse, Yumi Kasai, David Gray, Carol Clugston, Anna Dominiczak and Alex Alderton, Roberto Amato, Jeffrey Barrett, Sonia Goncalves, Ewan Harrison, David K. Jackson, Ian Johnston, Dominic Kwiatkowski, Cordelia Langford, John Sillitoe on behalf of the Wellcome Sanger Institute COVID-19 Surveillance Team |
| EPI_ISL_3281770 | Lighthouse Lab in Milton Keynes | Wellcome Sanger Institute for the COVID-19 Genomics UK (COG-UK) Consortium | The Lighthouse Lab in Milton Keynes and Alex Alderton, Roberto Amato, Jeffrey Barrett, Sonia Goncalves, Ewan Harrison, David K. Jackson, Ian Johnston, Dominic Kwiatkowski, Cordelia Langford, John Sillitoe on behalf of the Wellcome Sanger Institute COVID-19 Surveillance Team |
| EPI_ISL_3281771 | Lighthouse Lab in Alderley Park | Wellcome Sanger Institute for the COVID-19 Genomics UK (COG-UK) Consortium | Jacquelyn Wynn, Mairead Hyland, The Lighthouse Lab in Alderley Park and Alex Alderton, Roberto Amato, Jeffrey Barrett, Sonia Goncalves, Ewan Harrison, David K. Jackson, Ian Johnston, Dominic Kwiatkowski, Cordelia Langford, John Sillitoe on behalf of the Wellcome Sanger Institute COVID-19 Surveillance Team |
| EPI_ISL_3281772 | Lighthouse Lab in Glasgow | Wellcome Sanger Institute for the COVID-19 Genomics UK (COG-UK) Consortium | Harper VanSteenhouse, Yumi Kasai, David Gray, Carol Clugston, Anna Dominiczak and Alex Alderton, Roberto Amato, Jeffrey Barrett, Sonia Goncalves, Ewan Harrison, David K. Jackson, Ian Johnston, Dominic Kwiatkowski, Cordelia Langford, John Sillitoe on behalf of the Wellcome Sanger Institute COVID-19 Surveillance Team |
| EPI_ISL_3281773 | Lighthouse Lab in Alderley Park | Wellcome Sanger Institute for the COVID-19 Genomics UK (COG-UK) Consortium | Jacquelyn Wynn, Mairead Hyland, The Lighthouse Lab in Alderley Park and Alex Alderton, Roberto Amato, Jeffrey Barrett, Sonia Goncalves, Ewan Harrison, David K. Jackson, Ian Johnston, Dominic Kwiatkowski, Cordelia Langford, John Sillitoe on behalf of the Wellcome Sanger Institute COVID-19 Surveillance Team |
| EPI_ISL_3281774, EPI_ISL_3281775, EPI_ISL_3281776 | Lighthouse Lab in Milton Keynes | Wellcome Sanger Institute for the COVID-19 Genomics UK (COG-UK) Consortium | The Lighthouse Lab in Milton Keynes and Alex Alderton, Roberto Amato, Jeffrey Barrett, Sonia Goncalves, Ewan Harrison, David K. Jackson, Ian Johnston, Dominic Kwiatkowski, Cordelia Langford, John Sillitoe on behalf of the Wellcome Sanger Institute COVID-19 Surveillance Team |
| EPI_ISL_3281777 | Lighthouse Lab in Glasgow | Wellcome Sanger Institute for the COVID-19 Genomics UK (COG-UK) Consortium | Harper VanSteenhouse, Yumi Kasai, David Gray, Carol Clugston, Anna Dominiczak and Alex Alderton, Roberto Amato, Jeffrey Barrett, Sonia Goncalves, Ewan Harrison, David K. Jackson, Ian Johnston, Dominic Kwiatkowski, Cordelia Langford, John Sillitoe on behalf of the Wellcome Sanger Institute COVID-19 Surveillance Team |
| EPI_ISL_3281778 | Lighthouse Lab in Milton Keynes | Wellcome Sanger Institute for the COVID-19 Genomics UK (COG-UK) Consortium | The Lighthouse Lab in Milton Keynes and Alex Alderton, Roberto Amato, Jeffrey Barrett, Sonia Goncalves, Ewan Harrison, David K. Jackson, Ian Johnston, Dominic Kwiatkowski, Cordelia Langford, John Sillitoe on behalf of the Wellcome Sanger Institute COVID-19 Surveillance Team |
| EPI_ISL_3281779 | Lighthouse Lab in Glasgow | Wellcome Sanger Institute for the COVID-19 Genomics UK (COG-UK) Consortium | Harper VanSteenhouse, Yumi Kasai, David Gray, Carol Clugston, Anna Dominiczak and Alex Alderton, Roberto Amato, Jeffrey Barrett, Sonia Goncalves, Ewan Harrison, David K. Jackson, Ian Johnston, Dominic Kwiatkowski, Cordelia Langford, John Sillitoe on behalf of the Wellcome Sanger Institute COVID-19 Surveillance Team |
| EPI_ISL_3281780, EPI_ISL_3281781 | Lighthouse Lab in Milton Keynes | Wellcome Sanger Institute for the COVID-19 Genomics UK (COG-UK) Consortium | The Lighthouse Lab in Milton Keynes and Alex Alderton, Roberto Amato, Jeffrey Barrett, Sonia Goncalves, Ewan Harrison, David K. Jackson, Ian Johnston, Dominic Kwiatkowski, Cordelia Langford, John Sillitoe on behalf of the Wellcome Sanger Institute COVID-19 Surveillance Team |
| EPI_ISL_3281782 | Lighthouse Lab in Glasgow | Wellcome Sanger Institute for the COVID-19 Genomics UK (COG-UK) Consortium | Harper VanSteenhouse, Yumi Kasai, David Gray, Carol Clugston, Anna Dominiczak and Alex Alderton, Roberto Amato, Jeffrey Barrett, Sonia Goncalves, Ewan Harrison, David K. Jackson, Ian Johnston, Dominic Kwiatkowski, Cordelia Langford, John Sillitoe on behalf of the Wellcome Sanger Institute COVID-19 Surveillance Team |
| EPI_ISL_3281783 | Lighthouse Lab in Alderley Park | Wellcome Sanger Institute for the COVID-19 Genomics UK (COG-UK) | Jacquelyn Wynn, Mairead Hyland, The Lighthouse Lab in Alderley Park and Alex Alderton, Roberto Amato, Jeffrey Barrett, Sonia Goncalves, Ewan Harrison, David K. Jackson, Ian Johnston, Dominic Kwiatkowski, Cordelia Langford, John Sillitoe on |

| | | | |
|---|---|---|---|
| | | Consortium | behalf of the Wellcome Sanger Institute COVID-19 Surveillance Team |
| EPI_ISL_3281784 | Lighthouse Lab in Glasgow | Wellcome Sanger Institute for the COVID-19 Genomics UK (COG-UK) Consortium | Harper VanSteenhouse, Yumi Kasai, David Gray, Carol Clugston, Anna Dominiczak and Alex Alderton, Roberto Amato, Jeffrey Barrett, Sonia Goncalves, Ewan Harrison, David K. Jackson, Ian Johnston, Dominic Kwiatkowski, Cordelia Langford, John Sillitoe on behalf of the Wellcome Sanger Institute COVID-19 Surveillance Team |
| EPI_ISL_3281785, EPI_ISL_3281786 | Lighthouse Lab in Alderley Park | Wellcome Sanger Institute for the COVID-19 Genomics UK (COG-UK) Consortium | Jacquelyn Wynn, Mairead Hyland, The Lighthouse Lab in Alderley Park and Alex Alderton, Roberto Amato, Jeffrey Barrett, Sonia Goncalves, Ewan Harrison, David K. Jackson, Ian Johnston, Dominic Kwiatkowski, Cordelia Langford, John Sillitoe on behalf of the Wellcome Sanger Institute COVID-19 Surveillance Team |
| EPI_ISL_3281787 | Lighthouse Lab in Glasgow | Wellcome Sanger Institute for the COVID-19 Genomics UK (COG-UK) Consortium | Harper VanSteenhouse, Yumi Kasai, David Gray, Carol Clugston, Anna Dominiczak and Alex Alderton, Roberto Amato, Jeffrey Barrett, Sonia Goncalves, Ewan Harrison, David K. Jackson, Ian Johnston, Dominic Kwiatkowski, Cordelia Langford, John Sillitoe on behalf of the Wellcome Sanger Institute COVID-19 Surveillance Team |
| EPI_ISL_3281788 | Lighthouse Lab in Alderley Park | Wellcome Sanger Institute for the COVID-19 Genomics UK (COG-UK) Consortium | Jacquelyn Wynn, Mairead Hyland, The Lighthouse Lab in Alderley Park and Alex Alderton, Roberto Amato, Jeffrey Barrett, Sonia Goncalves, Ewan Harrison, David K. Jackson, Ian Johnston, Dominic Kwiatkowski, Cordelia Langford, John Sillitoe on behalf of the Wellcome Sanger Institute COVID-19 Surveillance Team |
| EPI_ISL_3281789 | Lighthouse Lab in Glasgow | Wellcome Sanger Institute for the COVID-19 Genomics UK (COG-UK) Consortium | Harper VanSteenhouse, Yumi Kasai, David Gray, Carol Clugston, Anna Dominiczak and Alex Alderton, Roberto Amato, Jeffrey Barrett, Sonia Goncalves, Ewan Harrison, David K. Jackson, Ian Johnston, Dominic Kwiatkowski, Cordelia Langford, John Sillitoe on behalf of the Wellcome Sanger Institute COVID-19 Surveillance Team |
| EPI_ISL_3281790, EPI_ISL_3281791 | Lighthouse Lab in Alderley Park | Wellcome Sanger Institute for the COVID-19 Genomics UK (COG-UK) Consortium | Jacquelyn Wynn, Mairead Hyland, The Lighthouse Lab in Alderley Park and Alex Alderton, Roberto Amato, Jeffrey Barrett, Sonia Goncalves, Ewan Harrison, David K. Jackson, Ian Johnston, Dominic Kwiatkowski, Cordelia Langford, John Sillitoe on behalf of the Wellcome Sanger Institute COVID-19 Surveillance Team |
| EPI_ISL_3281792, EPI_ISL_3281793, EPI_ISL_3281794, EPI_ISL_3281795 | Lighthouse Lab in Milton Keynes | Wellcome Sanger Institute for the COVID-19 Genomics UK (COG-UK) Consortium | The Lighthouse Lab in Milton Keynes and Alex Alderton, Roberto Amato, Jeffrey Barrett, Sonia Goncalves, Ewan Harrison, David K. Jackson, Ian Johnston, Dominic Kwiatkowski, Cordelia Langford, John Sillitoe on behalf of the Wellcome Sanger Institute COVID-19 Surveillance Team |
| EPI_ISL_3281796 | Lighthouse Lab in Glasgow | Wellcome Sanger Institute for the COVID-19 Genomics UK (COG-UK) Consortium | Harper VanSteenhouse, Yumi Kasai, David Gray, Carol Clugston, Anna Dominiczak and Alex Alderton, Roberto Amato, Jeffrey Barrett, Sonia Goncalves, Ewan Harrison, David K. Jackson, Ian Johnston, Dominic Kwiatkowski, Cordelia Langford, John Sillitoe on behalf of the Wellcome Sanger Institute COVID-19 Surveillance Team |
| EPI_ISL_3281797 | Lighthouse Lab in Milton Keynes | Wellcome Sanger Institute for the COVID-19 Genomics UK (COG-UK) Consortium | The Lighthouse Lab in Milton Keynes and Alex Alderton, Roberto Amato, Jeffrey Barrett, Sonia Goncalves, Ewan Harrison, David K. Jackson, Ian Johnston, Dominic Kwiatkowski, Cordelia Langford, John Sillitoe on behalf of the Wellcome Sanger Institute COVID-19 Surveillance Team |
| EPI_ISL_3281798, EPI_ISL_3281799 | Lighthouse Lab in Glasgow | Wellcome Sanger Institute for the COVID-19 Genomics UK (COG-UK) Consortium | Harper VanSteenhouse, Yumi Kasai, David Gray, Carol Clugston, Anna Dominiczak and Alex Alderton, Roberto Amato, Jeffrey Barrett, Sonia Goncalves, Ewan Harrison, David K. Jackson, Ian Johnston, Dominic Kwiatkowski, Cordelia Langford, John Sillitoe on behalf of the Wellcome Sanger Institute COVID-19 Surveillance Team |
| EPI_ISL_3281800, EPI_ISL_3281801 | Lighthouse Lab in Milton Keynes | Wellcome Sanger Institute for the COVID-19 Genomics UK (COG-UK) Consortium | The Lighthouse Lab in Milton Keynes and Alex Alderton, Roberto Amato, Jeffrey Barrett, Sonia Goncalves, Ewan Harrison, David K. Jackson, Ian Johnston, Dominic Kwiatkowski, Cordelia Langford, John Sillitoe on behalf of the Wellcome Sanger Institute COVID-19 Surveillance Team |
| EPI_ISL_3281802, EPI_ISL_3281803 | Lighthouse Lab in Alderley Park | Wellcome Sanger Institute for the COVID-19 Genomics UK (COG-UK) Consortium | Jacquelyn Wynn, Mairead Hyland, The Lighthouse Lab in Alderley Park and Alex Alderton, Roberto Amato, Jeffrey Barrett, Sonia Goncalves, Ewan Harrison, David K. Jackson, Ian Johnston, Dominic Kwiatkowski, Cordelia Langford, John Sillitoe on behalf of the Wellcome Sanger Institute COVID-19 Surveillance Team |
| EPI_ISL_3281804, EPI_ISL_3281805 | Lighthouse Lab in Milton Keynes | Wellcome Sanger Institute for the COVID-19 Genomics UK (COG-UK) Consortium | The Lighthouse Lab in Milton Keynes and Alex Alderton, Roberto Amato, Jeffrey Barrett, Sonia Goncalves, Ewan Harrison, David K. Jackson, Ian Johnston, Dominic Kwiatkowski, Cordelia Langford, John Sillitoe on behalf of the Wellcome Sanger Institute COVID-19 Surveillance Team |
| EPI_ISL_3281806 | Lighthouse Lab in Glasgow | Wellcome Sanger Institute for the COVID-19 Genomics UK (COG-UK) Consortium | Harper VanSteenhouse, Yumi Kasai, David Gray, Carol Clugston, Anna Dominiczak and Alex Alderton, Roberto Amato, Jeffrey Barrett, Sonia Goncalves, Ewan Harrison, David K. Jackson, Ian Johnston, Dominic Kwiatkowski, Cordelia Langford, John Sillitoe on behalf of the Wellcome Sanger Institute COVID-19 Surveillance Team |
| EPI_ISL_3281807, EPI_ISL_3281808 | Lighthouse Lab in Alderley Park | Wellcome Sanger Institute for the COVID-19 Genomics UK (COG-UK) Consortium | Jacquelyn Wynn, Mairead Hyland, The Lighthouse Lab in Alderley Park and Alex Alderton, Roberto Amato, Jeffrey Barrett, Sonia Goncalves, Ewan Harrison, David K. Jackson, Ian Johnston, Dominic Kwiatkowski, Cordelia Langford, John Sillitoe on behalf of the Wellcome Sanger Institute COVID-19 Surveillance Team |
| EPI_ISL_3281809 | Lighthouse Lab in Milton Keynes | Wellcome Sanger Institute for the COVID-19 Genomics UK (COG-UK) Consortium | The Lighthouse Lab in Milton Keynes and Alex Alderton, Roberto Amato, Jeffrey Barrett, Sonia Goncalves, Ewan Harrison, David K. Jackson, Ian Johnston, Dominic Kwiatkowski, Cordelia Langford, John Sillitoe on behalf of the Wellcome Sanger Institute COVID-19 Surveillance Team |
| EPI_ISL_3281810 | Lighthouse Lab in Glasgow | Wellcome Sanger Institute for the COVID-19 Genomics UK (COG-UK) Consortium | Harper VanSteenhouse, Yumi Kasai, David Gray, Carol Clugston, Anna Dominiczak and Alex Alderton, Roberto Amato, Jeffrey Barrett, Sonia Goncalves, Ewan Harrison, David K. Jackson, Ian Johnston, Dominic Kwiatkowski, Cordelia Langford, John Sillitoe on behalf of the Wellcome Sanger Institute COVID-19 Surveillance Team |
| EPI_ISL_3281811, EPI_ISL_3281812 | Lighthouse Lab in Milton Keynes | Wellcome Sanger Institute for the COVID-19 Genomics UK (COG-UK) Consortium | The Lighthouse Lab in Milton Keynes and Alex Alderton, Roberto Amato, Jeffrey Barrett, Sonia Goncalves, Ewan Harrison, David K. Jackson, Ian Johnston, Dominic Kwiatkowski, Cordelia Langford, John Sillitoe on behalf of the Wellcome Sanger Institute COVID-19 Surveillance Team |
| EPI_ISL_3281813 | Lighthouse Lab in Glasgow | Wellcome Sanger Institute for the COVID-19 Genomics UK (COG-UK) Consortium | Harper VanSteenhouse, Yumi Kasai, David Gray, Carol Clugston, Anna Dominiczak and Alex Alderton, Roberto Amato, Jeffrey Barrett, Sonia Goncalves, Ewan Harrison, David K. Jackson, Ian Johnston, Dominic Kwiatkowski, Cordelia Langford, John Sillitoe on behalf of the Wellcome Sanger Institute COVID-19 Surveillance Team |
| EPI_ISL_3281814 | Lighthouse Lab in Alderley Park | Wellcome Sanger Institute for the COVID-19 Genomics UK (COG-UK) Consortium | Jacquelyn Wynn, Mairead Hyland, The Lighthouse Lab in Alderley Park and Alex Alderton, Roberto Amato, Jeffrey Barrett, Sonia Goncalves, Ewan Harrison, David K. Jackson, Ian Johnston, Dominic Kwiatkowski, Cordelia Langford, John Sillitoe on behalf of the Wellcome Sanger Institute COVID-19 Surveillance Team |
| EPI_ISL_3281815 | Lighthouse Lab in Milton Keynes | Wellcome Sanger Institute for the COVID-19 Genomics UK (COG-UK) Consortium | The Lighthouse Lab in Milton Keynes and Alex Alderton, Roberto Amato, Jeffrey Barrett, Sonia Goncalves, Ewan Harrison, David K. Jackson, Ian Johnston, Dominic Kwiatkowski, Cordelia Langford, John Sillitoe on behalf of the Wellcome Sanger Institute COVID-19 Surveillance Team |
| EPI_ISL_3281816 | Lighthouse Lab in Glasgow | Wellcome Sanger Institute for the COVID-19 Genomics UK (COG-UK) Consortium | Harper VanSteenhouse, Yumi Kasai, David Gray, Carol Clugston, Anna Dominiczak and Alex Alderton, Roberto Amato, Jeffrey Barrett, Sonia Goncalves, Ewan Harrison, David K. Jackson, Ian Johnston, Dominic Kwiatkowski, Cordelia Langford, John Sillitoe on behalf of the Wellcome Sanger Institute COVID-19 Surveillance Team |
| EPI_ISL_3281817 | Lighthouse Lab in Alderley Park | Wellcome Sanger Institute for the COVID-19 Genomics UK (COG-UK) Consortium | Jacquelyn Wynn, Mairead Hyland, The Lighthouse Lab in Alderley Park and Alex Alderton, Roberto Amato, Jeffrey Barrett, Sonia Goncalves, Ewan Harrison, David K. Jackson, Ian Johnston, Dominic Kwiatkowski, Cordelia Langford, John Sillitoe on behalf of the Wellcome Sanger Institute COVID-19 Surveillance Team |
| EPI_ISL_3281818 | Lighthouse Lab in Milton Keynes | Wellcome Sanger Institute for the COVID-19 Genomics UK (COG-UK) Consortium | The Lighthouse Lab in Milton Keynes and Alex Alderton, Roberto Amato, Jeffrey Barrett, Sonia Goncalves, Ewan Harrison, David K. Jackson, Ian Johnston, Dominic Kwiatkowski, Cordelia Langford, John Sillitoe on behalf of the Wellcome Sanger Institute COVID-19 Surveillance Team |
| EPI_ISL_3281819, EPI_ISL_3281820 | Lighthouse Lab in Alderley Park | Wellcome Sanger Institute for the COVID-19 Genomics UK (COG-UK) Consortium | Jacquelyn Wynn, Mairead Hyland, The Lighthouse Lab in Alderley Park and Alex Alderton, Roberto Amato, Jeffrey Barrett, Sonia Goncalves, Ewan Harrison, David K. Jackson, Ian Johnston, Dominic Kwiatkowski, Cordelia Langford, John Sillitoe on behalf of the Wellcome Sanger Institute COVID-19 Surveillance Team |
| EPI_ISL_3281821 | Lighthouse Lab in Glasgow | Wellcome Sanger Institute for the COVID-19 Genomics UK (COG-UK) Consortium | Harper VanSteenhouse, Yumi Kasai, David Gray, Carol Clugston, Anna Dominiczak and Alex Alderton, Roberto Amato, Jeffrey Barrett, Sonia Goncalves, Ewan Harrison, David K. Jackson, Ian Johnston, Dominic Kwiatkowski, Cordelia Langford, John Sillitoe on behalf of the Wellcome Sanger Institute COVID-19 Surveillance Team |
| EPI_ISL_3281822, EPI_ISL_3281823 | Lighthouse Lab in Alderley Park | Wellcome Sanger Institute for the COVID-19 Genomics UK (COG-UK) Consortium | Jacquelyn Wynn, Mairead Hyland, The Lighthouse Lab in Alderley Park and Alex Alderton, Roberto Amato, Jeffrey Barrett, Sonia Goncalves, Ewan Harrison, David K. Jackson, Ian Johnston, Dominic Kwiatkowski, Cordelia Langford, John Sillitoe on behalf of the Wellcome Sanger Institute COVID-19 Surveillance Team |
| EPI_ISL_3281824, EPI_ISL_3281825 | Lighthouse Lab in Glasgow | Wellcome Sanger Institute for the COVID-19 Genomics UK (COG-UK) Consortium | Harper VanSteenhouse, Yumi Kasai, David Gray, Carol Clugston, Anna Dominiczak and Alex Alderton, Roberto Amato, Jeffrey Barrett, Sonia Goncalves, Ewan Harrison, David K. Jackson, Ian Johnston, Dominic Kwiatkowski, Cordelia Langford, John Sillitoe on behalf of the Wellcome Sanger Institute COVID-19 Surveillance Team |
| EPI_ISL_3281826 | Lighthouse Lab in Milton Keynes | Wellcome Sanger Institute for the COVID-19 Genomics UK (COG-UK) Consortium | The Lighthouse Lab in Milton Keynes and Alex Alderton, Roberto Amato, Jeffrey Barrett, Sonia Goncalves, Ewan Harrison, David K. Jackson, Ian Johnston, Dominic Kwiatkowski, Cordelia Langford, John Sillitoe on behalf of the Wellcome Sanger Institute COVID-19 Surveillance Team |
| EPI_ISL_3281827 | Lighthouse Lab in Alderley Park | Wellcome Sanger Institute for the COVID-19 Genomics UK (COG-UK) Consortium | Jacquelyn Wynn, Mairead Hyland, The Lighthouse Lab in Alderley Park and Alex Alderton, Roberto Amato, Jeffrey Barrett, Sonia Goncalves, Ewan Harrison, David K. Jackson, Ian Johnston, Dominic Kwiatkowski, Cordelia Langford, John Sillitoe on behalf of the Wellcome Sanger Institute COVID-19 Surveillance Team |
| EPI_ISL_3281828, EPI_ISL_3281829 | Lighthouse Lab in Glasgow | Wellcome Sanger Institute for the COVID-19 Genomics UK (COG-UK) Consortium | Harper VanSteenhouse, Yumi Kasai, David Gray, Carol Clugston, Anna Dominiczak and Alex Alderton, Roberto Amato, Jeffrey Barrett, Sonia Goncalves, Ewan Harrison, David K. Jackson, Ian Johnston, Dominic Kwiatkowski, Cordelia Langford, John Sillitoe on behalf of the Wellcome Sanger Institute COVID-19 Surveillance Team |
| EPI_ISL_3281830, EPI_ISL_3281831 | Lighthouse Lab in Alderley Park | Wellcome Sanger Institute for the COVID-19 Genomics UK (COG-UK) Consortium | Jacquelyn Wynn, Mairead Hyland, The Lighthouse Lab in Alderley Park and Alex Alderton, Roberto Amato, Jeffrey Barrett, Sonia Goncalves, Ewan Harrison, David K. Jackson, Ian Johnston, Dominic Kwiatkowski, Cordelia Langford, John Sillitoe on behalf of the Wellcome Sanger Institute COVID-19 Surveillance Team |
| EPI_ISL_3281832 | Lighthouse Lab in Glasgow | Wellcome Sanger Institute for the COVID-19 Genomics UK (COG-UK) Consortium | Harper VanSteenhouse, Yumi Kasai, David Gray, Carol Clugston, Anna Dominiczak and Alex Alderton, Roberto Amato, Jeffrey Barrett, Sonia Goncalves, Ewan Harrison, David K. Jackson, Ian Johnston, Dominic Kwiatkowski, Cordelia Langford, John Sillitoe on behalf of the Wellcome Sanger Institute COVID-19 Surveillance Team |
| EPI_ISL_3281833 | Lighthouse Lab in Alderley Park | Wellcome Sanger Institute for the COVID-19 Genomics UK (COG-UK) Consortium | Jacquelyn Wynn, Mairead Hyland, The Lighthouse Lab in Alderley Park and Alex Alderton, Roberto Amato, Jeffrey Barrett, Sonia Goncalves, Ewan Harrison, David K. Jackson, Ian Johnston, Dominic Kwiatkowski, Cordelia Langford, John Sillitoe on behalf of the Wellcome Sanger Institute COVID-19 Surveillance Team |
| EPI_ISL_3281834, EPI_ISL_3281835, EPI_ISL_3281836, EPI_ISL_3281837 | Lighthouse Lab in Glasgow | Wellcome Sanger Institute for the COVID-19 Genomics UK (COG-UK) Consortium | Harper VanSteenhouse, Yumi Kasai, David Gray, Carol Clugston, Anna Dominiczak and Alex Alderton, Roberto Amato, Jeffrey Barrett, Sonia Goncalves, Ewan Harrison, David K. Jackson, Ian Johnston, Dominic Kwiatkowski, Cordelia Langford, John Sillitoe on behalf of the Wellcome Sanger Institute COVID-19 Surveillance Team |
| EPI_ISL_3281838 | Lighthouse Lab in Milton Keynes | Wellcome Sanger Institute for the COVID-19 Genomics UK (COG-UK) Consortium | The Lighthouse Lab in Milton Keynes and Alex Alderton, Roberto Amato, Jeffrey Barrett, Sonia Goncalves, Ewan Harrison, David K. Jackson, Ian Johnston, Dominic Kwiatkowski, Cordelia Langford, John Sillitoe on behalf of the Wellcome Sanger Institute COVID-19 Surveillance Team |
| EPI_ISL_3281839 | Lighthouse Lab in Alderley Park | Wellcome Sanger Institute for the COVID-19 Genomics UK (COG-UK) Consortium | Jacquelyn Wynn, Mairead Hyland, The Lighthouse Lab in Alderley Park and Alex Alderton, Roberto Amato, Jeffrey Barrett, Sonia Goncalves, Ewan Harrison, David K. Jackson, Ian Johnston, Dominic Kwiatkowski, Cordelia Langford, John Sillitoe on behalf of the Wellcome Sanger Institute COVID-19 Surveillance Team |
| EPI_ISL_3281840, EPI_ISL_3281841 | Lighthouse Lab in Glasgow | Wellcome Sanger Institute for the COVID-19 Genomics UK (COG-UK) Consortium | Harper VanSteenhouse, Yumi Kasai, David Gray, Carol Clugston, Anna Dominiczak and Alex Alderton, Roberto Amato, Jeffrey Barrett, Sonia Goncalves, Ewan Harrison, David K. Jackson, Ian Johnston, Dominic Kwiatkowski, Cordelia Langford, John Sillitoe on behalf of the Wellcome Sanger Institute COVID-19 Surveillance Team |
| EPI_ISL_3281842 | Lighthouse Lab in Alderley Park | Wellcome Sanger Institute for the COVID-19 Genomics UK (COG-UK) Consortium | Jacquelyn Wynn, Mairead Hyland, The Lighthouse Lab in Alderley Park and Alex Alderton, Roberto Amato, Jeffrey Barrett, Sonia Goncalves, Ewan Harrison, David K. Jackson, Ian Johnston, Dominic Kwiatkowski, Cordelia Langford, John Sillitoe on behalf of the Wellcome Sanger Institute COVID-19 Surveillance Team |
| EPI_ISL_3281843, EPI_ISL_3281844 | Lighthouse Lab in Milton Keynes | Wellcome Sanger Institute for the COVID-19 Genomics UK (COG-UK) Consortium | The Lighthouse Lab in Milton Keynes and Alex Alderton, Roberto Amato, Jeffrey Barrett, Sonia Goncalves, Ewan Harrison, David K. Jackson, Ian Johnston, Dominic Kwiatkowski, Cordelia Langford, John Sillitoe on behalf of the Wellcome Sanger Institute COVID-19 Surveillance Team |
| EPI_ISL_3281845 | Lighthouse Lab in Glasgow | Wellcome Sanger Institute for the COVID-19 Genomics UK (COG-UK) Consortium | Harper VanSteenhouse, Yumi Kasai, David Gray, Carol Clugston, Anna Dominiczak and Alex Alderton, Roberto Amato, Jeffrey Barrett, Sonia Goncalves, Ewan Harrison, David K. Jackson, Ian Johnston, Dominic Kwiatkowski, Cordelia Langford, John Sillitoe on behalf of the Wellcome Sanger Institute COVID-19 Surveillance Team |
| EPI_ISL_3281846, EPI_ISL_3281847, EPI_ISL_3281848 | Lighthouse Lab in Milton Keynes | Wellcome Sanger Institute for the COVID-19 Genomics UK (COG-UK) Consortium | The Lighthouse Lab in Milton Keynes and Alex Alderton, Roberto Amato, Jeffrey Barrett, Sonia Goncalves, Ewan Harrison, David K. Jackson, Ian Johnston, Dominic Kwiatkowski, Cordelia Langford, John Sillitoe on behalf of the Wellcome Sanger Institute COVID-19 Surveillance Team |
| EPI_ISL_3281849, | Lighthouse Laboratory Plymouth | Wellcome Sanger Institute for the COVID-19 Genomics UK (COG-UK) | Lighthouse Laboratory Plymouth and Alex Alderton, Roberto Amato, Jeffrey Barrett, Sonia Goncalves, Ewan Harrison, David K. Jackson, Ian Johnston, Dominic Kwiatkowski, Cordelia Langford, John Sillitoe on behalf of the Wellcome Sanger |

| | | | |
|---|---|---|---|
| EPI_ISL_3281850, EPI_ISL_3281851, EPI_ISL_3281852 | | Consortium | Institute COVID-19 Surveillance Team |
| EPI_ISL_3281853 | Lighthouse Lab in Glasgow | Wellcome Sanger Institute for the COVID-19 Genomics UK (COG-UK) Consortium | Harper VanSteenhouse, Yumi Kasai, David Gray, Carol Clugston, Anna Dominiczak and Alex Alderton, Roberto Amato, Jeffrey Barrett, Sonia Goncalves, Ewan Harrison, David K. Jackson, Ian Johnston, Dominic Kwiatkowski, Cordelia Langford, John Sillitoe on behalf of the Wellcome Sanger Institute COVID-19 Surveillance Team |
| EPI_ISL_3281854 | Berkshire and Surrey Pathology Services Lighthouse Laboratory | Wellcome Sanger Institute for the COVID-19 Genomics UK (COG-UK) Consortium | Berkshire and Surrey Pathology Services Lighthouse Laboratory and Alex Alderton, Roberto Amato, Jeffrey Barrett, Sonia Goncalves, Ewan Harrison, David K. Jackson, Ian Johnston, Dominic Kwiatkowski, Cordelia Langford, John Sillitoe on behalf of the Wellcome Sanger Institute COVID-19 Surveillance Team |
| EPI_ISL_3281855 | Lighthouse Laboratory Plymouth | Wellcome Sanger Institute for the COVID-19 Genomics UK (COG-UK) Consortium | Lighthouse Laboratory Plymouth and Alex Alderton, Roberto Amato, Jeffrey Barrett, Sonia Goncalves, Ewan Harrison, David K. Jackson, Ian Johnston, Dominic Kwiatkowski, Cordelia Langford, John Sillitoe on behalf of the Wellcome Sanger Institute COVID-19 Surveillance Team |
| EPI_ISL_3281856 | Lighthouse Lab in Glasgow | Wellcome Sanger Institute for the COVID-19 Genomics UK (COG-UK) Consortium | Harper VanSteenhouse, Yumi Kasai, David Gray, Carol Clugston, Anna Dominiczak and Alex Alderton, Roberto Amato, Jeffrey Barrett, Sonia Goncalves, Ewan Harrison, David K. Jackson, Ian Johnston, Dominic Kwiatkowski, Cordelia Langford, John Sillitoe on behalf of the Wellcome Sanger Institute COVID-19 Surveillance Team |
| EPI_ISL_3281857, EPI_ISL_3281858, EPI_ISL_3281859, EPI_ISL_3281860 | Lighthouse Laboratory Plymouth | Wellcome Sanger Institute for the COVID-19 Genomics UK (COG-UK) Consortium | Lighthouse Laboratory Plymouth and Alex Alderton, Roberto Amato, Jeffrey Barrett, Sonia Goncalves, Ewan Harrison, David K. Jackson, Ian Johnston, Dominic Kwiatkowski, Cordelia Langford, John Sillitoe on behalf of the Wellcome Sanger Institute COVID-19 Surveillance Team |
| EPI_ISL_3281861 | Berkshire and Surrey Pathology Services Lighthouse Laboratory | Wellcome Sanger Institute for the COVID-19 Genomics UK (COG-UK) Consortium | Berkshire and Surrey Pathology Services Lighthouse Laboratory and Alex Alderton, Roberto Amato, Jeffrey Barrett, Sonia Goncalves, Ewan Harrison, David K. Jackson, Ian Johnston, Dominic Kwiatkowski, Cordelia Langford, John Sillitoe on behalf of the Wellcome Sanger Institute COVID-19 Surveillance Team |
| EPI_ISL_3281862, EPI_ISL_3281863, EPI_ISL_3281864 | Lighthouse Laboratory Plymouth | Wellcome Sanger Institute for the COVID-19 Genomics UK (COG-UK) Consortium | Lighthouse Laboratory Plymouth and Alex Alderton, Roberto Amato, Jeffrey Barrett, Sonia Goncalves, Ewan Harrison, David K. Jackson, Ian Johnston, Dominic Kwiatkowski, Cordelia Langford, John Sillitoe on behalf of the Wellcome Sanger Institute COVID-19 Surveillance Team |
| EPI_ISL_3281865 | Lighthouse Lab in Glasgow | Wellcome Sanger Institute for the COVID-19 Genomics UK (COG-UK) Consortium | Harper VanSteenhouse, Yumi Kasai, David Gray, Carol Clugston, Anna Dominiczak and Alex Alderton, Roberto Amato, Jeffrey Barrett, Sonia Goncalves, Ewan Harrison, David K. Jackson, Ian Johnston, Dominic Kwiatkowski, Cordelia Langford, John Sillitoe on behalf of the Wellcome Sanger Institute COVID-19 Surveillance Team |
| EPI_ISL_3281866, EPI_ISL_3281867, EPI_ISL_3281868, EPI_ISL_3281869, EPI_ISL_3281870 | Lighthouse Laboratory Plymouth | Wellcome Sanger Institute for the COVID-19 Genomics UK (COG-UK) Consortium | Lighthouse Laboratory Plymouth and Alex Alderton, Roberto Amato, Jeffrey Barrett, Sonia Goncalves, Ewan Harrison, David K. Jackson, Ian Johnston, Dominic Kwiatkowski, Cordelia Langford, John Sillitoe on behalf of the Wellcome Sanger Institute COVID-19 Surveillance Team |
| EPI_ISL_3281871 | Berkshire and Surrey Pathology Services Lighthouse Laboratory | Wellcome Sanger Institute for the COVID-19 Genomics UK (COG-UK) Consortium | Berkshire and Surrey Pathology Services Lighthouse Laboratory and Alex Alderton, Roberto Amato, Jeffrey Barrett, Sonia Goncalves, Ewan Harrison, David K. Jackson, Ian Johnston, Dominic Kwiatkowski, Cordelia Langford, John Sillitoe on behalf of the Wellcome Sanger Institute COVID-19 Surveillance Team |
| EPI_ISL_3281872 | Lighthouse Lab in Glasgow | Wellcome Sanger Institute for the COVID-19 Genomics UK (COG-UK) Consortium | Harper VanSteenhouse, Yumi Kasai, David Gray, Carol Clugston, Anna Dominiczak and Alex Alderton, Roberto Amato, Jeffrey Barrett, Sonia Goncalves, Ewan Harrison, David K. Jackson, Ian Johnston, Dominic Kwiatkowski, Cordelia Langford, John Sillitoe on behalf of the Wellcome Sanger Institute COVID-19 Surveillance Team |
| EPI_ISL_3281873 | Lighthouse Laboratory Plymouth | Wellcome Sanger Institute for the COVID-19 Genomics UK (COG-UK) Consortium | Lighthouse Laboratory Plymouth and Alex Alderton, Roberto Amato, Jeffrey Barrett, Sonia Goncalves, Ewan Harrison, David K. Jackson, Ian Johnston, Dominic Kwiatkowski, Cordelia Langford, John Sillitoe on behalf of the Wellcome Sanger Institute COVID-19 Surveillance Team |
| EPI_ISL_3281874 | Berkshire and Surrey Pathology Services Lighthouse Laboratory | Wellcome Sanger Institute for the COVID-19 Genomics UK (COG-UK) Consortium | Berkshire and Surrey Pathology Services Lighthouse Laboratory and Alex Alderton, Roberto Amato, Jeffrey Barrett, Sonia Goncalves, Ewan Harrison, David K. Jackson, Ian Johnston, Dominic Kwiatkowski, Cordelia Langford, John Sillitoe on behalf of the Wellcome Sanger Institute COVID-19 Surveillance Team |
| EPI_ISL_3281875 | Lighthouse Lab in Glasgow | Wellcome Sanger Institute for the COVID-19 Genomics UK (COG-UK) Consortium | Harper VanSteenhouse, Yumi Kasai, David Gray, Carol Clugston, Anna Dominiczak and Alex Alderton, Roberto Amato, Jeffrey Barrett, Sonia Goncalves, Ewan Harrison, David K. Jackson, Ian Johnston, Dominic Kwiatkowski, Cordelia Langford, John Sillitoe on behalf of the Wellcome Sanger Institute COVID-19 Surveillance Team |
| EPI_ISL_3281876, EPI_ISL_3281877 | Lighthouse Laboratory Plymouth | Wellcome Sanger Institute for the COVID-19 Genomics UK (COG-UK) Consortium | Lighthouse Laboratory Plymouth and Alex Alderton, Roberto Amato, Jeffrey Barrett, Sonia Goncalves, Ewan Harrison, David K. Jackson, Ian Johnston, Dominic Kwiatkowski, Cordelia Langford, John Sillitoe on behalf of the Wellcome Sanger Institute COVID-19 Surveillance Team |
| EPI_ISL_3281878, EPI_ISL_3281879 | Berkshire and Surrey Pathology Services Lighthouse Laboratory | Wellcome Sanger Institute for the COVID-19 Genomics UK (COG-UK) Consortium | Berkshire and Surrey Pathology Services Lighthouse Laboratory and Alex Alderton, Roberto Amato, Jeffrey Barrett, Sonia Goncalves, Ewan Harrison, David K. Jackson, Ian Johnston, Dominic Kwiatkowski, Cordelia Langford, John Sillitoe on behalf of the Wellcome Sanger Institute COVID-19 Surveillance Team |
| EPI_ISL_3281880 | Lighthouse Laboratory Plymouth | Wellcome Sanger Institute for the COVID-19 Genomics UK (COG-UK) Consortium | Lighthouse Laboratory Plymouth and Alex Alderton, Roberto Amato, Jeffrey Barrett, Sonia Goncalves, Ewan Harrison, David K. Jackson, Ian Johnston, Dominic Kwiatkowski, Cordelia Langford, John Sillitoe on behalf of the Wellcome Sanger Institute COVID-19 Surveillance Team |
| EPI_ISL_3281881 | Lighthouse Lab in Glasgow | Wellcome Sanger Institute for the COVID-19 Genomics UK (COG-UK) Consortium | Harper VanSteenhouse, Yumi Kasai, David Gray, Carol Clugston, Anna Dominiczak and Alex Alderton, Roberto Amato, Jeffrey Barrett, Sonia Goncalves, Ewan Harrison, David K. Jackson, Ian Johnston, Dominic Kwiatkowski, Cordelia Langford, John Sillitoe on behalf of the Wellcome Sanger Institute COVID-19 Surveillance Team |
| EPI_ISL_3281882, EPI_ISL_3281883, EPI_ISL_3281884 | Lighthouse Laboratory Plymouth | Wellcome Sanger Institute for the COVID-19 Genomics UK (COG-UK) Consortium | Lighthouse Laboratory Plymouth and Alex Alderton, Roberto Amato, Jeffrey Barrett, Sonia Goncalves, Ewan Harrison, David K. Jackson, Ian Johnston, Dominic Kwiatkowski, Cordelia Langford, John Sillitoe on behalf of the Wellcome Sanger Institute COVID-19 Surveillance Team |
| EPI_ISL_3281885 | Berkshire and Surrey Pathology Services Lighthouse Laboratory | Wellcome Sanger Institute for the COVID-19 Genomics UK (COG-UK) Consortium | Berkshire and Surrey Pathology Services Lighthouse Laboratory and Alex Alderton, Roberto Amato, Jeffrey Barrett, Sonia Goncalves, Ewan Harrison, David K. Jackson, Ian Johnston, Dominic Kwiatkowski, Cordelia Langford, John Sillitoe on behalf of the Wellcome Sanger Institute COVID-19 Surveillance Team |
| EPI_ISL_3281886, EPI_ISL_3281887, EPI_ISL_3281888 | Lighthouse Laboratory Plymouth | Wellcome Sanger Institute for the COVID-19 Genomics UK (COG-UK) Consortium | Lighthouse Laboratory Plymouth and Alex Alderton, Roberto Amato, Jeffrey Barrett, Sonia Goncalves, Ewan Harrison, David K. Jackson, Ian Johnston, Dominic Kwiatkowski, Cordelia Langford, John Sillitoe on behalf of the Wellcome Sanger Institute COVID-19 Surveillance Team |
| EPI_ISL_3281889 | Berkshire and Surrey Pathology Services Lighthouse Laboratory | Wellcome Sanger Institute for the COVID-19 Genomics UK (COG-UK) Consortium | Berkshire and Surrey Pathology Services Lighthouse Laboratory and Alex Alderton, Roberto Amato, Jeffrey Barrett, Sonia Goncalves, Ewan Harrison, David K. Jackson, Ian Johnston, Dominic Kwiatkowski, Cordelia Langford, John Sillitoe on behalf of the Wellcome Sanger Institute COVID-19 Surveillance Team |
| EPI_ISL_3281890 | Lighthouse Laboratory Plymouth | Wellcome Sanger Institute for the COVID-19 Genomics UK (COG-UK) Consortium | Lighthouse Laboratory Plymouth and Alex Alderton, Roberto Amato, Jeffrey Barrett, Sonia Goncalves, Ewan Harrison, David K. Jackson, Ian Johnston, Dominic Kwiatkowski, Cordelia Langford, John Sillitoe on behalf of the Wellcome Sanger Institute COVID-19 Surveillance Team |
| EPI_ISL_3281891 | Lighthouse Lab in Glasgow | Wellcome Sanger Institute for the COVID-19 Genomics UK (COG-UK) Consortium | Harper VanSteenhouse, Yumi Kasai, David Gray, Carol Clugston, Anna Dominiczak and Alex Alderton, Roberto Amato, Jeffrey Barrett, Sonia Goncalves, Ewan Harrison, David K. Jackson, Ian Johnston, Dominic Kwiatkowski, Cordelia Langford, John Sillitoe on behalf of the Wellcome Sanger Institute COVID-19 Surveillance Team |
| EPI_ISL_3281892 | Berkshire and Surrey Pathology Services Lighthouse Laboratory | Wellcome Sanger Institute for the COVID-19 Genomics UK (COG-UK) Consortium | Berkshire and Surrey Pathology Services Lighthouse Laboratory and Alex Alderton, Roberto Amato, Jeffrey Barrett, Sonia Goncalves, Ewan Harrison, David K. Jackson, Ian Johnston, Dominic Kwiatkowski, Cordelia Langford, John Sillitoe on behalf of the Wellcome Sanger Institute COVID-19 Surveillance Team |
| EPI_ISL_3281893 | Lighthouse Laboratory Plymouth | Wellcome Sanger Institute for the COVID-19 Genomics UK (COG-UK) Consortium | Lighthouse Laboratory Plymouth and Alex Alderton, Roberto Amato, Jeffrey Barrett, Sonia Goncalves, Ewan Harrison, David K. Jackson, Ian Johnston, Dominic Kwiatkowski, Cordelia Langford, John Sillitoe on behalf of the Wellcome Sanger Institute COVID-19 Surveillance Team |
| EPI_ISL_3281894 | Berkshire and Surrey Pathology Services Lighthouse Laboratory | Wellcome Sanger Institute for the COVID-19 Genomics UK (COG-UK) Consortium | Berkshire and Surrey Pathology Services Lighthouse Laboratory and Alex Alderton, Roberto Amato, Jeffrey Barrett, Sonia Goncalves, Ewan Harrison, David K. Jackson, Ian Johnston, Dominic Kwiatkowski, Cordelia Langford, John Sillitoe on behalf of the Wellcome Sanger Institute COVID-19 Surveillance Team |
| EPI_ISL_3281895 | Lighthouse Laboratory Plymouth | Wellcome Sanger Institute for the COVID-19 Genomics UK (COG-UK) Consortium | Lighthouse Laboratory Plymouth and Alex Alderton, Roberto Amato, Jeffrey Barrett, Sonia Goncalves, Ewan Harrison, David K. Jackson, Ian Johnston, Dominic Kwiatkowski, Cordelia Langford, John Sillitoe on behalf of the Wellcome Sanger Institute COVID-19 Surveillance Team |
| EPI_ISL_3281896 | Berkshire and Surrey Pathology Services Lighthouse Laboratory | Wellcome Sanger Institute for the COVID-19 Genomics UK (COG-UK) Consortium | Berkshire and Surrey Pathology Services Lighthouse Laboratory and Alex Alderton, Roberto Amato, Jeffrey Barrett, Sonia Goncalves, Ewan Harrison, David K. Jackson, Ian Johnston, Dominic Kwiatkowski, Cordelia Langford, John Sillitoe on behalf of the Wellcome Sanger Institute COVID-19 Surveillance Team |
| EPI_ISL_3281897 | Lighthouse Laboratory Plymouth | Wellcome Sanger Institute for the COVID-19 Genomics UK (COG-UK) Consortium | Lighthouse Laboratory Plymouth and Alex Alderton, Roberto Amato, Jeffrey Barrett, Sonia Goncalves, Ewan Harrison, David K. Jackson, Ian Johnston, Dominic Kwiatkowski, Cordelia Langford, John Sillitoe on behalf of the Wellcome Sanger Institute COVID-19 Surveillance Team |
| EPI_ISL_3281898 | Lighthouse Lab in Glasgow | Wellcome Sanger Institute for the COVID-19 Genomics UK (COG-UK) Consortium | Harper VanSteenhouse, Yumi Kasai, David Gray, Carol Clugston, Anna Dominiczak and Alex Alderton, Roberto Amato, Jeffrey Barrett, Sonia Goncalves, Ewan Harrison, David K. Jackson, Ian Johnston, Dominic Kwiatkowski, Cordelia Langford, John Sillitoe on behalf of the Wellcome Sanger Institute COVID-19 Surveillance Team |
| EPI_ISL_3281899, EPI_ISL_3281900 | Lighthouse Laboratory Plymouth | Wellcome Sanger Institute for the COVID-19 Genomics UK (COG-UK) Consortium | Lighthouse Laboratory Plymouth and Alex Alderton, Roberto Amato, Jeffrey Barrett, Sonia Goncalves, Ewan Harrison, David K. Jackson, Ian Johnston, Dominic Kwiatkowski, Cordelia Langford, John Sillitoe on behalf of the Wellcome Sanger Institute COVID-19 Surveillance Team |
| EPI_ISL_3281901 | Berkshire and Surrey Pathology Services Lighthouse Laboratory | Wellcome Sanger Institute for the COVID-19 Genomics UK (COG-UK) Consortium | Berkshire and Surrey Pathology Services Lighthouse Laboratory and Alex Alderton, Roberto Amato, Jeffrey Barrett, Sonia Goncalves, Ewan Harrison, David K. Jackson, Ian Johnston, Dominic Kwiatkowski, Cordelia Langford, John Sillitoe on behalf of the Wellcome Sanger Institute COVID-19 Surveillance Team |
| EPI_ISL_3281902, EPI_ISL_3281903, EPI_ISL_3281904, EPI_ISL_3281905 | Lighthouse Laboratory Plymouth | Wellcome Sanger Institute for the COVID-19 Genomics UK (COG-UK) Consortium | Lighthouse Laboratory Plymouth and Alex Alderton, Roberto Amato, Jeffrey Barrett, Sonia Goncalves, Ewan Harrison, David K. Jackson, Ian Johnston, Dominic Kwiatkowski, Cordelia Langford, John Sillitoe on behalf of the Wellcome Sanger Institute COVID-19 Surveillance Team |
| EPI_ISL_3281906, EPI_ISL_3281907 | Berkshire and Surrey Pathology Services Lighthouse Laboratory | Wellcome Sanger Institute for the COVID-19 Genomics UK (COG-UK) Consortium | Berkshire and Surrey Pathology Services Lighthouse Laboratory and Alex Alderton, Roberto Amato, Jeffrey Barrett, Sonia Goncalves, Ewan Harrison, David K. Jackson, Ian Johnston, Dominic Kwiatkowski, Cordelia Langford, John Sillitoe on behalf of the Wellcome Sanger Institute COVID-19 Surveillance Team |
| EPI_ISL_3281908 | Lighthouse Laboratory Plymouth | Wellcome Sanger Institute for the COVID-19 Genomics UK (COG-UK) Consortium | Lighthouse Laboratory Plymouth and Alex Alderton, Roberto Amato, Jeffrey Barrett, Sonia Goncalves, Ewan Harrison, David K. Jackson, Ian Johnston, Dominic Kwiatkowski, Cordelia Langford, John Sillitoe on behalf of the Wellcome Sanger Institute COVID-19 Surveillance Team |
| EPI_ISL_3281909, EPI_ISL_3281910 | Lighthouse Lab in Glasgow | Wellcome Sanger Institute for the COVID-19 Genomics UK (COG-UK) Consortium | Harper VanSteenhouse, Yumi Kasai, David Gray, Carol Clugston, Anna Dominiczak and Alex Alderton, Roberto Amato, Jeffrey Barrett, Sonia Goncalves, Ewan Harrison, David K. Jackson, Ian Johnston, Dominic Kwiatkowski, Cordelia Langford, John Sillitoe on behalf of the Wellcome Sanger Institute COVID-19 Surveillance Team |
| EPI_ISL_3281911 | Lighthouse Laboratory Plymouth | Wellcome Sanger Institute for the COVID-19 Genomics UK (COG-UK) Consortium | Lighthouse Laboratory Plymouth and Alex Alderton, Roberto Amato, Jeffrey Barrett, Sonia Goncalves, Ewan Harrison, David K. Jackson, Ian Johnston, Dominic Kwiatkowski, Cordelia Langford, John Sillitoe on behalf of the Wellcome Sanger Institute COVID-19 Surveillance Team |
| EPI_ISL_3281912, EPI_ISL_3281913 | Berkshire and Surrey Pathology Services Lighthouse Laboratory | Wellcome Sanger Institute for the COVID-19 Genomics UK (COG-UK) Consortium | Berkshire and Surrey Pathology Services Lighthouse Laboratory and Alex Alderton, Roberto Amato, Jeffrey Barrett, Sonia Goncalves, Ewan Harrison, David K. Jackson, Ian Johnston, Dominic Kwiatkowski, Cordelia Langford, John Sillitoe on behalf of the Wellcome Sanger Institute COVID-19 Surveillance Team |
| EPI_ISL_3281914, | Lighthouse Laboratory Plymouth | Wellcome Sanger Institute for the COVID-19 Genomics UK (COG-UK) | Lighthouse Laboratory Plymouth and Alex Alderton, Roberto Amato, Jeffrey Barrett, Sonia Goncalves, Ewan Harrison, David K. Jackson, Ian Johnston, Dominic Kwiatkowski, Cordelia Langford, John Sillitoe on behalf of the Wellcome Sanger |

| | | | |
|---|---|---|---|
| EPI_ISL_3281915, EPI_ISL_3281916, EPI_ISL_3281917 | | Consortium | Institute COVID-19 Surveillance Team |
| EPI_ISL_3281918 | Lighthouse Lab in Glasgow | Wellcome Sanger Institute for the COVID-19 Genomics UK (COG-UK) Consortium | Harper VanSteenhouse, Yumi Kasai, David Gray, Carol Clugston, Anna Dominiczak and Alex Alderton, Roberto Amato, Jeffrey Barrett, Sonia Goncalves, Ewan Harrison, David K. Jackson, Ian Johnston, Dominic Kwiatkowski, Cordelia Langford, John Sillitoe on behalf of the Wellcome Sanger Institute COVID-19 Surveillance Team |
| EPI_ISL_3281919 | Berkshire and Surrey Pathology Services Lighthouse Laboratory | Wellcome Sanger Institute for the COVID-19 Genomics UK (COG-UK) Consortium | Berkshire and Surrey Pathology Services Lighthouse Laboratory and Alex Alderton, Roberto Amato, Jeffrey Barrett, Sonia Goncalves, Ewan Harrison, David K. Jackson, Ian Johnston, Dominic Kwiatkowski, Cordelia Langford, John Sillitoe on behalf of the Wellcome Sanger Institute COVID-19 Surveillance Team |
| EPI_ISL_3281920 | Lighthouse Laboratory Plymouth | Wellcome Sanger Institute for the COVID-19 Genomics UK (COG-UK) Consortium | Lighthouse Laboratory Plymouth and Alex Alderton, Roberto Amato, Jeffrey Barrett, Sonia Goncalves, Ewan Harrison, David K. Jackson, Ian Johnston, Dominic Kwiatkowski, Cordelia Langford, John Sillitoe on behalf of the Wellcome Sanger Institute COVID-19 Surveillance Team |
| EPI_ISL_3281921 | Lighthouse Lab in Glasgow | Wellcome Sanger Institute for the COVID-19 Genomics UK (COG-UK) Consortium | Harper VanSteenhouse, Yumi Kasai, David Gray, Carol Clugston, Anna Dominiczak and Alex Alderton, Roberto Amato, Jeffrey Barrett, Sonia Goncalves, Ewan Harrison, David K. Jackson, Ian Johnston, Dominic Kwiatkowski, Cordelia Langford, John Sillitoe on behalf of the Wellcome Sanger Institute COVID-19 Surveillance Team |
| EPI_ISL_3281922 | Berkshire and Surrey Pathology Services Lighthouse Laboratory | Wellcome Sanger Institute for the COVID-19 Genomics UK (COG-UK) Consortium | Berkshire and Surrey Pathology Services Lighthouse Laboratory and Alex Alderton, Roberto Amato, Jeffrey Barrett, Sonia Goncalves, Ewan Harrison, David K. Jackson, Ian Johnston, Dominic Kwiatkowski, Cordelia Langford, John Sillitoe on behalf of the Wellcome Sanger Institute COVID-19 Surveillance Team |
| EPI_ISL_3281923, EPI_ISL_3281924, EPI_ISL_3281925, EPI_ISL_3281926, EPI_ISL_3281927, EPI_ISL_3281928 | Lighthouse Laboratory Plymouth | Wellcome Sanger Institute for the COVID-19 Genomics UK (COG-UK) Consortium | Lighthouse Laboratory Plymouth and Alex Alderton, Roberto Amato, Jeffrey Barrett, Sonia Goncalves, Ewan Harrison, David K. Jackson, Ian Johnston, Dominic Kwiatkowski, Cordelia Langford, John Sillitoe on behalf of the Wellcome Sanger Institute COVID-19 Surveillance Team |
| EPI_ISL_3281929, EPI_ISL_3281930 | Berkshire and Surrey Pathology Services Lighthouse Laboratory | Wellcome Sanger Institute for the COVID-19 Genomics UK (COG-UK) Consortium | Berkshire and Surrey Pathology Services Lighthouse Laboratory and Alex Alderton, Roberto Amato, Jeffrey Barrett, Sonia Goncalves, Ewan Harrison, David K. Jackson, Ian Johnston, Dominic Kwiatkowski, Cordelia Langford, John Sillitoe on behalf of the Wellcome Sanger Institute COVID-19 Surveillance Team |
| EPI_ISL_3281931 | Lighthouse Laboratory Plymouth | Wellcome Sanger Institute for the COVID-19 Genomics UK (COG-UK) Consortium | Lighthouse Laboratory Plymouth and Alex Alderton, Roberto Amato, Jeffrey Barrett, Sonia Goncalves, Ewan Harrison, David K. Jackson, Ian Johnston, Dominic Kwiatkowski, Cordelia Langford, John Sillitoe on behalf of the Wellcome Sanger Institute COVID-19 Surveillance Team |
| EPI_ISL_3281932 | Berkshire and Surrey Pathology Services Lighthouse Laboratory | Wellcome Sanger Institute for the COVID-19 Genomics UK (COG-UK) Consortium | Berkshire and Surrey Pathology Services Lighthouse Laboratory and Alex Alderton, Roberto Amato, Jeffrey Barrett, Sonia Goncalves, Ewan Harrison, David K. Jackson, Ian Johnston, Dominic Kwiatkowski, Cordelia Langford, John Sillitoe on behalf of the Wellcome Sanger Institute COVID-19 Surveillance Team |
| EPI_ISL_3281933 | Lighthouse Laboratory Plymouth | Wellcome Sanger Institute for the COVID-19 Genomics UK (COG-UK) Consortium | Lighthouse Laboratory Plymouth and Alex Alderton, Roberto Amato, Jeffrey Barrett, Sonia Goncalves, Ewan Harrison, David K. Jackson, Ian Johnston, Dominic Kwiatkowski, Cordelia Langford, John Sillitoe on behalf of the Wellcome Sanger Institute COVID-19 Surveillance Team |
| EPI_ISL_3281934 | Berkshire and Surrey Pathology Services Lighthouse Laboratory | Wellcome Sanger Institute for the COVID-19 Genomics UK (COG-UK) Consortium | Berkshire and Surrey Pathology Services Lighthouse Laboratory and Alex Alderton, Roberto Amato, Jeffrey Barrett, Sonia Goncalves, Ewan Harrison, David K. Jackson, Ian Johnston, Dominic Kwiatkowski, Cordelia Langford, John Sillitoe on behalf of the Wellcome Sanger Institute COVID-19 Surveillance Team |
| EPI_ISL_3281935, EPI_ISL_3281936, EPI_ISL_3281937, EPI_ISL_3281938 | Lighthouse Laboratory Plymouth | Wellcome Sanger Institute for the COVID-19 Genomics UK (COG-UK) Consortium | Lighthouse Laboratory Plymouth and Alex Alderton, Roberto Amato, Jeffrey Barrett, Sonia Goncalves, Ewan Harrison, David K. Jackson, Ian Johnston, Dominic Kwiatkowski, Cordelia Langford, John Sillitoe on behalf of the Wellcome Sanger Institute COVID-19 Surveillance Team |
| EPI_ISL_3281939 | Lighthouse Lab in Glasgow | Wellcome Sanger Institute for the COVID-19 Genomics UK (COG-UK) Consortium | Harper VanSteenhouse, Yumi Kasai, David Gray, Carol Clugston, Anna Dominiczak and Alex Alderton, Roberto Amato, Jeffrey Barrett, Sonia Goncalves, Ewan Harrison, David K. Jackson, Ian Johnston, Dominic Kwiatkowski, Cordelia Langford, John Sillitoe on behalf of the Wellcome Sanger Institute COVID-19 Surveillance Team |
| EPI_ISL_3281940 | Lighthouse Laboratory Plymouth | Wellcome Sanger Institute for the COVID-19 Genomics UK (COG-UK) Consortium | Lighthouse Laboratory Plymouth and Alex Alderton, Roberto Amato, Jeffrey Barrett, Sonia Goncalves, Ewan Harrison, David K. Jackson, Ian Johnston, Dominic Kwiatkowski, Cordelia Langford, John Sillitoe on behalf of the Wellcome Sanger Institute COVID-19 Surveillance Team |
| EPI_ISL_3281941 | Lighthouse Lab in Glasgow | Wellcome Sanger Institute for the COVID-19 Genomics UK (COG-UK) Consortium | Harper VanSteenhouse, Yumi Kasai, David Gray, Carol Clugston, Anna Dominiczak and Alex Alderton, Roberto Amato, Jeffrey Barrett, Sonia Goncalves, Ewan Harrison, David K. Jackson, Ian Johnston, Dominic Kwiatkowski, Cordelia Langford, John Sillitoe on behalf of the Wellcome Sanger Institute COVID-19 Surveillance Team |
| EPI_ISL_3281942 | Berkshire and Surrey Pathology Services Lighthouse Laboratory | Wellcome Sanger Institute for the COVID-19 Genomics UK (COG-UK) Consortium | Berkshire and Surrey Pathology Services Lighthouse Laboratory and Alex Alderton, Roberto Amato, Jeffrey Barrett, Sonia Goncalves, Ewan Harrison, David K. Jackson, Ian Johnston, Dominic Kwiatkowski, Cordelia Langford, John Sillitoe on behalf of the Wellcome Sanger Institute COVID-19 Surveillance Team |
| EPI_ISL_3281943, EPI_ISL_3281944, EPI_ISL_3281945, EPI_ISL_3281946 | Lighthouse Laboratory Plymouth | Wellcome Sanger Institute for the COVID-19 Genomics UK (COG-UK) Consortium | Lighthouse Laboratory Plymouth and Alex Alderton, Roberto Amato, Jeffrey Barrett, Sonia Goncalves, Ewan Harrison, David K. Jackson, Ian Johnston, Dominic Kwiatkowski, Cordelia Langford, John Sillitoe on behalf of the Wellcome Sanger Institute COVID-19 Surveillance Team |
| EPI_ISL_3281947 | Lighthouse Lab in Glasgow | Wellcome Sanger Institute for the COVID-19 Genomics UK (COG-UK) Consortium | Harper VanSteenhouse, Yumi Kasai, David Gray, Carol Clugston, Anna Dominiczak and Alex Alderton, Roberto Amato, Jeffrey Barrett, Sonia Goncalves, Ewan Harrison, David K. Jackson, Ian Johnston, Dominic Kwiatkowski, Cordelia Langford, John Sillitoe on behalf of the Wellcome Sanger Institute COVID-19 Surveillance Team |
| EPI_ISL_3281948, EPI_ISL_3281949, EPI_ISL_3281950, EPI_ISL_3281951 | Lighthouse Laboratory Plymouth | Wellcome Sanger Institute for the COVID-19 Genomics UK (COG-UK) Consortium | Lighthouse Laboratory Plymouth and Alex Alderton, Roberto Amato, Jeffrey Barrett, Sonia Goncalves, Ewan Harrison, David K. Jackson, Ian Johnston, Dominic Kwiatkowski, Cordelia Langford, John Sillitoe on behalf of the Wellcome Sanger Institute COVID-19 Surveillance Team |
| EPI_ISL_3281952 | Berkshire and Surrey Pathology Services Lighthouse Laboratory | Wellcome Sanger Institute for the COVID-19 Genomics UK (COG-UK) Consortium | Berkshire and Surrey Pathology Services Lighthouse Laboratory and Alex Alderton, Roberto Amato, Jeffrey Barrett, Sonia Goncalves, Ewan Harrison, David K. Jackson, Ian Johnston, Dominic Kwiatkowski, Cordelia Langford, John Sillitoe on behalf of the Wellcome Sanger Institute COVID-19 Surveillance Team |
| EPI_ISL_3281953, EPI_ISL_3281954, EPI_ISL_3281955, EPI_ISL_3281956, EPI_ISL_3281957 | Lighthouse Laboratory Plymouth | Wellcome Sanger Institute for the COVID-19 Genomics UK (COG-UK) Consortium | Lighthouse Laboratory Plymouth and Alex Alderton, Roberto Amato, Jeffrey Barrett, Sonia Goncalves, Ewan Harrison, David K. Jackson, Ian Johnston, Dominic Kwiatkowski, Cordelia Langford, John Sillitoe on behalf of the Wellcome Sanger Institute COVID-19 Surveillance Team |
| EPI_ISL_3281958 | Berkshire and Surrey Pathology Services Lighthouse Laboratory | Wellcome Sanger Institute for the COVID-19 Genomics UK (COG-UK) Consortium | Berkshire and Surrey Pathology Services Lighthouse Laboratory and Alex Alderton, Roberto Amato, Jeffrey Barrett, Sonia Goncalves, Ewan Harrison, David K. Jackson, Ian Johnston, Dominic Kwiatkowski, Cordelia Langford, John Sillitoe on behalf of the Wellcome Sanger Institute COVID-19 Surveillance Team |
| EPI_ISL_3281959, EPI_ISL_3281960 | Lighthouse Laboratory Plymouth | Wellcome Sanger Institute for the COVID-19 Genomics UK (COG-UK) Consortium | Lighthouse Laboratory Plymouth and Alex Alderton, Roberto Amato, Jeffrey Barrett, Sonia Goncalves, Ewan Harrison, David K. Jackson, Ian Johnston, Dominic Kwiatkowski, Cordelia Langford, John Sillitoe on behalf of the Wellcome Sanger Institute COVID-19 Surveillance Team |
| EPI_ISL_3281961 | Berkshire and Surrey Pathology Services Lighthouse Laboratory | Wellcome Sanger Institute for the COVID-19 Genomics UK (COG-UK) Consortium | Berkshire and Surrey Pathology Services Lighthouse Laboratory and Alex Alderton, Roberto Amato, Jeffrey Barrett, Sonia Goncalves, Ewan Harrison, David K. Jackson, Ian Johnston, Dominic Kwiatkowski, Cordelia Langford, John Sillitoe on behalf of the Wellcome Sanger Institute COVID-19 Surveillance Team |
| EPI_ISL_3281962, EPI_ISL_3281963, EPI_ISL_3281964, EPI_ISL_3281965 | Lighthouse Laboratory Plymouth | Wellcome Sanger Institute for the COVID-19 Genomics UK (COG-UK) Consortium | Lighthouse Laboratory Plymouth and Alex Alderton, Roberto Amato, Jeffrey Barrett, Sonia Goncalves, Ewan Harrison, David K. Jackson, Ian Johnston, Dominic Kwiatkowski, Cordelia Langford, John Sillitoe on behalf of the Wellcome Sanger Institute COVID-19 Surveillance Team |
| EPI_ISL_3281966, EPI_ISL_3281967 | Berkshire and Surrey Pathology Services Lighthouse Laboratory | Wellcome Sanger Institute for the COVID-19 Genomics UK (COG-UK) Consortium | Berkshire and Surrey Pathology Services Lighthouse Laboratory and Alex Alderton, Roberto Amato, Jeffrey Barrett, Sonia Goncalves, Ewan Harrison, David K. Jackson, Ian Johnston, Dominic Kwiatkowski, Cordelia Langford, John Sillitoe on behalf of the Wellcome Sanger Institute COVID-19 Surveillance Team |
| EPI_ISL_3281968 | Lighthouse Laboratory Plymouth | Wellcome Sanger Institute for the COVID-19 Genomics UK (COG-UK) Consortium | Lighthouse Laboratory Plymouth and Alex Alderton, Roberto Amato, Jeffrey Barrett, Sonia Goncalves, Ewan Harrison, David K. Jackson, Ian Johnston, Dominic Kwiatkowski, Cordelia Langford, John Sillitoe on behalf of the Wellcome Sanger Institute COVID-19 Surveillance Team |
| EPI_ISL_3281969 | Berkshire and Surrey Pathology Services Lighthouse Laboratory | Wellcome Sanger Institute for the COVID-19 Genomics UK (COG-UK) Consortium | Berkshire and Surrey Pathology Services Lighthouse Laboratory and Alex Alderton, Roberto Amato, Jeffrey Barrett, Sonia Goncalves, Ewan Harrison, David K. Jackson, Ian Johnston, Dominic Kwiatkowski, Cordelia Langford, John Sillitoe on behalf of the Wellcome Sanger Institute COVID-19 Surveillance Team |
| EPI_ISL_3281970, EPI_ISL_3281971, EPI_ISL_3281972, EPI_ISL_3281973, EPI_ISL_3281974, EPI_ISL_3281975, EPI_ISL_3281976 | Lighthouse Laboratory Plymouth | Wellcome Sanger Institute for the COVID-19 Genomics UK (COG-UK) Consortium | Lighthouse Laboratory Plymouth and Alex Alderton, Roberto Amato, Jeffrey Barrett, Sonia Goncalves, Ewan Harrison, David K. Jackson, Ian Johnston, Dominic Kwiatkowski, Cordelia Langford, John Sillitoe on behalf of the Wellcome Sanger Institute COVID-19 Surveillance Team |
| EPI_ISL_3281977 | Berkshire and Surrey Pathology Services Lighthouse Laboratory | Wellcome Sanger Institute for the COVID-19 Genomics UK (COG-UK) Consortium | Berkshire and Surrey Pathology Services Lighthouse Laboratory and Alex Alderton, Roberto Amato, Jeffrey Barrett, Sonia Goncalves, Ewan Harrison, David K. Jackson, Ian Johnston, Dominic Kwiatkowski, Cordelia Langford, John Sillitoe on behalf of the Wellcome Sanger Institute COVID-19 Surveillance Team |
| EPI_ISL_3281978 | Lighthouse Laboratory Plymouth | Wellcome Sanger Institute for the COVID-19 Genomics UK (COG-UK) Consortium | Lighthouse Laboratory Plymouth and Alex Alderton, Roberto Amato, Jeffrey Barrett, Sonia Goncalves, Ewan Harrison, David K. Jackson, Ian Johnston, Dominic Kwiatkowski, Cordelia Langford, John Sillitoe on behalf of the Wellcome Sanger Institute COVID-19 Surveillance Team |
| EPI_ISL_3281979, EPI_ISL_3281980, EPI_ISL_3281981 | Berkshire and Surrey Pathology Services Lighthouse Laboratory | Wellcome Sanger Institute for the COVID-19 Genomics UK (COG-UK) Consortium | Berkshire and Surrey Pathology Services Lighthouse Laboratory and Alex Alderton, Roberto Amato, Jeffrey Barrett, Sonia Goncalves, Ewan Harrison, David K. Jackson, Ian Johnston, Dominic Kwiatkowski, Cordelia Langford, John Sillitoe on behalf of the Wellcome Sanger Institute COVID-19 Surveillance Team |
| EPI_ISL_3281982 | Lighthouse Laboratory Plymouth | Wellcome Sanger Institute for the COVID-19 Genomics UK (COG-UK) Consortium | Lighthouse Laboratory Plymouth and Alex Alderton, Roberto Amato, Jeffrey Barrett, Sonia Goncalves, Ewan Harrison, David K. Jackson, Ian Johnston, Dominic Kwiatkowski, Cordelia Langford, John Sillitoe on behalf of the Wellcome Sanger Institute COVID-19 Surveillance Team |
| EPI_ISL_3281983 | Berkshire and Surrey Pathology Services Lighthouse Laboratory | Wellcome Sanger Institute for the COVID-19 Genomics UK (COG-UK) Consortium | Berkshire and Surrey Pathology Services Lighthouse Laboratory and Alex Alderton, Roberto Amato, Jeffrey Barrett, Sonia Goncalves, Ewan Harrison, David K. Jackson, Ian Johnston, Dominic Kwiatkowski, Cordelia Langford, John Sillitoe on behalf of the Wellcome Sanger Institute COVID-19 Surveillance Team |
| EPI_ISL_3281984, EPI_ISL_3281985, | Lighthouse Laboratory Plymouth | Wellcome Sanger Institute for the COVID-19 Genomics UK (COG-UK) Consortium | Lighthouse Laboratory Plymouth and Alex Alderton, Roberto Amato, Jeffrey Barrett, Sonia Goncalves, Ewan Harrison, David K. Jackson, Ian Johnston, Dominic Kwiatkowski, Cordelia Langford, John Sillitoe on behalf of the Wellcome Sanger Institute COVID-19 Surveillance Team |

| | | | |
|---|---|---|---|
| EPI_ISL_3281986 | | | |
| EPI_ISL_3281987 | Lighthouse Lab in Glasgow | Wellcome Sanger Institute for the COVID-19 Genomics UK (COG-UK) Consortium | Harper VanSteenhouse, Yumi Kasai, David Gray, Carol Clugston, Anna Dominiczak and Alex Alderton, Roberto Amato, Jeffrey Barrett, Sonia Goncalves, Ewan Harrison, David K. Jackson, Ian Johnston, Dominic Kwiatkowski, Cordelia Langford, John Sillitoe on behalf of the Wellcome Sanger Institute COVID-19 Surveillance Team |
| EPI_ISL_3281988, EPI_ISL_3281989 | Berkshire and Surrey Pathology Services Lighthouse Laboratory | Wellcome Sanger Institute for the COVID-19 Genomics UK (COG-UK) Consortium | Berkshire and Surrey Pathology Services Lighthouse Laboratory and Alex Alderton, Roberto Amato, Jeffrey Barrett, Sonia Goncalves, Ewan Harrison, David K. Jackson, Ian Johnston, Dominic Kwiatkowski, Cordelia Langford, John Sillitoe on behalf of the Wellcome Sanger Institute COVID-19 Surveillance Team |
| EPI_ISL_3281990 | Lighthouse Laboratory Plymouth | Wellcome Sanger Institute for the COVID-19 Genomics UK (COG-UK) Consortium | Lighthouse Laboratory Plymouth and Alex Alderton, Roberto Amato, Jeffrey Barrett, Sonia Goncalves, Ewan Harrison, David K. Jackson, Ian Johnston, Dominic Kwiatkowski, Cordelia Langford, John Sillitoe on behalf of the Wellcome Sanger Institute COVID-19 Surveillance Team |
| EPI_ISL_3281991 | Lighthouse Lab in Glasgow | Wellcome Sanger Institute for the COVID-19 Genomics UK (COG-UK) Consortium | Harper VanSteenhouse, Yumi Kasai, David Gray, Carol Clugston, Anna Dominiczak and Alex Alderton, Roberto Amato, Jeffrey Barrett, Sonia Goncalves, Ewan Harrison, David K. Jackson, Ian Johnston, Dominic Kwiatkowski, Cordelia Langford, John Sillitoe on behalf of the Wellcome Sanger Institute COVID-19 Surveillance Team |
| EPI_ISL_3281992 | Berkshire and Surrey Pathology Services Lighthouse Laboratory | Wellcome Sanger Institute for the COVID-19 Genomics UK (COG-UK) Consortium | Berkshire and Surrey Pathology Services Lighthouse Laboratory and Alex Alderton, Roberto Amato, Jeffrey Barrett, Sonia Goncalves, Ewan Harrison, David K. Jackson, Ian Johnston, Dominic Kwiatkowski, Cordelia Langford, John Sillitoe on behalf of the Wellcome Sanger Institute COVID-19 Surveillance Team |
| EPI_ISL_3281993, EPI_ISL_3281994 | Lighthouse Lab in Glasgow | Wellcome Sanger Institute for the COVID-19 Genomics UK (COG-UK) Consortium | Harper VanSteenhouse, Yumi Kasai, David Gray, Carol Clugston, Anna Dominiczak and Alex Alderton, Roberto Amato, Jeffrey Barrett, Sonia Goncalves, Ewan Harrison, David K. Jackson, Ian Johnston, Dominic Kwiatkowski, Cordelia Langford, John Sillitoe on behalf of the Wellcome Sanger Institute COVID-19 Surveillance Team |
| EPI_ISL_3281995 | Berkshire and Surrey Pathology Services Lighthouse Laboratory | Wellcome Sanger Institute for the COVID-19 Genomics UK (COG-UK) Consortium | Berkshire and Surrey Pathology Services Lighthouse Laboratory and Alex Alderton, Roberto Amato, Jeffrey Barrett, Sonia Goncalves, Ewan Harrison, David K. Jackson, Ian Johnston, Dominic Kwiatkowski, Cordelia Langford, John Sillitoe on behalf of the Wellcome Sanger Institute COVID-19 Surveillance Team |
| EPI_ISL_3281996, EPI_ISL_3281997 | Lighthouse Laboratory Plymouth | Wellcome Sanger Institute for the COVID-19 Genomics UK (COG-UK) Consortium | Lighthouse Laboratory Plymouth and Alex Alderton, Roberto Amato, Jeffrey Barrett, Sonia Goncalves, Ewan Harrison, David K. Jackson, Ian Johnston, Dominic Kwiatkowski, Cordelia Langford, John Sillitoe on behalf of the Wellcome Sanger Institute COVID-19 Surveillance Team |
| EPI_ISL_3281998, EPI_ISL_3281999, EPI_ISL_3282000, EPI_ISL_3282001 | Berkshire and Surrey Pathology Services Lighthouse Laboratory | Wellcome Sanger Institute for the COVID-19 Genomics UK (COG-UK) Consortium | Berkshire and Surrey Pathology Services Lighthouse Laboratory and Alex Alderton, Roberto Amato, Jeffrey Barrett, Sonia Goncalves, Ewan Harrison, David K. Jackson, Ian Johnston, Dominic Kwiatkowski, Cordelia Langford, John Sillitoe on behalf of the Wellcome Sanger Institute COVID-19 Surveillance Team |
| EPI_ISL_3282002, EPI_ISL_3282003 | Lighthouse Laboratory Plymouth | Wellcome Sanger Institute for the COVID-19 Genomics UK (COG-UK) Consortium | Lighthouse Laboratory Plymouth and Alex Alderton, Roberto Amato, Jeffrey Barrett, Sonia Goncalves, Ewan Harrison, David K. Jackson, Ian Johnston, Dominic Kwiatkowski, Cordelia Langford, John Sillitoe on behalf of the Wellcome Sanger Institute COVID-19 Surveillance Team |
| EPI_ISL_3282004 | Berkshire and Surrey Pathology Services Lighthouse Laboratory | Wellcome Sanger Institute for the COVID-19 Genomics UK (COG-UK) Consortium | Berkshire and Surrey Pathology Services Lighthouse Laboratory and Alex Alderton, Roberto Amato, Jeffrey Barrett, Sonia Goncalves, Ewan Harrison, David K. Jackson, Ian Johnston, Dominic Kwiatkowski, Cordelia Langford, John Sillitoe on behalf of the Wellcome Sanger Institute COVID-19 Surveillance Team |
| EPI_ISL_3282005 | Lighthouse Laboratory Plymouth | Wellcome Sanger Institute for the COVID-19 Genomics UK (COG-UK) Consortium | Lighthouse Laboratory Plymouth and Alex Alderton, Roberto Amato, Jeffrey Barrett, Sonia Goncalves, Ewan Harrison, David K. Jackson, Ian Johnston, Dominic Kwiatkowski, Cordelia Langford, John Sillitoe on behalf of the Wellcome Sanger Institute COVID-19 Surveillance Team |
| EPI_ISL_3282006 | Berkshire and Surrey Pathology Services Lighthouse Laboratory | Wellcome Sanger Institute for the COVID-19 Genomics UK (COG-UK) Consortium | Berkshire and Surrey Pathology Services Lighthouse Laboratory and Alex Alderton, Roberto Amato, Jeffrey Barrett, Sonia Goncalves, Ewan Harrison, David K. Jackson, Ian Johnston, Dominic Kwiatkowski, Cordelia Langford, John Sillitoe on behalf of the Wellcome Sanger Institute COVID-19 Surveillance Team |
| EPI_ISL_3282007 | Lighthouse Laboratory Plymouth | Wellcome Sanger Institute for the COVID-19 Genomics UK (COG-UK) Consortium | Lighthouse Laboratory Plymouth and Alex Alderton, Roberto Amato, Jeffrey Barrett, Sonia Goncalves, Ewan Harrison, David K. Jackson, Ian Johnston, Dominic Kwiatkowski, Cordelia Langford, John Sillitoe on behalf of the Wellcome Sanger Institute COVID-19 Surveillance Team |
| EPI_ISL_3282008 | Lighthouse Lab in Glasgow | Wellcome Sanger Institute for the COVID-19 Genomics UK (COG-UK) Consortium | Harper VanSteenhouse, Yumi Kasai, David Gray, Carol Clugston, Anna Dominiczak and Alex Alderton, Roberto Amato, Jeffrey Barrett, Sonia Goncalves, Ewan Harrison, David K. Jackson, Ian Johnston, Dominic Kwiatkowski, Cordelia Langford, John Sillitoe on behalf of the Wellcome Sanger Institute COVID-19 Surveillance Team |
| EPI_ISL_3282009, EPI_ISL_3282010, EPI_ISL_3282011 | Lighthouse Laboratory Plymouth | Wellcome Sanger Institute for the COVID-19 Genomics UK (COG-UK) Consortium | Lighthouse Laboratory Plymouth and Alex Alderton, Roberto Amato, Jeffrey Barrett, Sonia Goncalves, Ewan Harrison, David K. Jackson, Ian Johnston, Dominic Kwiatkowski, Cordelia Langford, John Sillitoe on behalf of the Wellcome Sanger Institute COVID-19 Surveillance Team |
| EPI_ISL_3282012 | Berkshire and Surrey Pathology Services Lighthouse Laboratory | Wellcome Sanger Institute for the COVID-19 Genomics UK (COG-UK) Consortium | Berkshire and Surrey Pathology Services Lighthouse Laboratory and Alex Alderton, Roberto Amato, Jeffrey Barrett, Sonia Goncalves, Ewan Harrison, David K. Jackson, Ian Johnston, Dominic Kwiatkowski, Cordelia Langford, John Sillitoe on behalf of the Wellcome Sanger Institute COVID-19 Surveillance Team |
| EPI_ISL_3282013, EPI_ISL_3282014 | Lighthouse Laboratory Plymouth | Wellcome Sanger Institute for the COVID-19 Genomics UK (COG-UK) Consortium | Lighthouse Laboratory Plymouth and Alex Alderton, Roberto Amato, Jeffrey Barrett, Sonia Goncalves, Ewan Harrison, David K. Jackson, Ian Johnston, Dominic Kwiatkowski, Cordelia Langford, John Sillitoe on behalf of the Wellcome Sanger Institute COVID-19 Surveillance Team |
| EPI_ISL_3282015 | Berkshire and Surrey Pathology Services Lighthouse Laboratory | Wellcome Sanger Institute for the COVID-19 Genomics UK (COG-UK) Consortium | Berkshire and Surrey Pathology Services Lighthouse Laboratory and Alex Alderton, Roberto Amato, Jeffrey Barrett, Sonia Goncalves, Ewan Harrison, David K. Jackson, Ian Johnston, Dominic Kwiatkowski, Cordelia Langford, John Sillitoe on behalf of the Wellcome Sanger Institute COVID-19 Surveillance Team |
| EPI_ISL_3282016, EPI_ISL_3282017, EPI_ISL_3282018 | Lighthouse Laboratory Plymouth | Wellcome Sanger Institute for the COVID-19 Genomics UK (COG-UK) Consortium | Lighthouse Laboratory Plymouth and Alex Alderton, Roberto Amato, Jeffrey Barrett, Sonia Goncalves, Ewan Harrison, David K. Jackson, Ian Johnston, Dominic Kwiatkowski, Cordelia Langford, John Sillitoe on behalf of the Wellcome Sanger Institute COVID-19 Surveillance Team |
| EPI_ISL_3282019, EPI_ISL_3282020, EPI_ISL_3282021 | Lighthouse Lab in Glasgow | Wellcome Sanger Institute for the COVID-19 Genomics UK (COG-UK) Consortium | Harper VanSteenhouse, Yumi Kasai, David Gray, Carol Clugston, Anna Dominiczak and Alex Alderton, Roberto Amato, Jeffrey Barrett, Sonia Goncalves, Ewan Harrison, David K. Jackson, Ian Johnston, Dominic Kwiatkowski, Cordelia Langford, John Sillitoe on behalf of the Wellcome Sanger Institute COVID-19 Surveillance Team |
| EPI_ISL_3282022 | Lighthouse Laboratory Plymouth | Wellcome Sanger Institute for the COVID-19 Genomics UK (COG-UK) Consortium | Lighthouse Laboratory Plymouth and Alex Alderton, Roberto Amato, Jeffrey Barrett, Sonia Goncalves, Ewan Harrison, David K. Jackson, Ian Johnston, Dominic Kwiatkowski, Cordelia Langford, John Sillitoe on behalf of the Wellcome Sanger Institute COVID-19 Surveillance Team |
| EPI_ISL_3282023 | Berkshire and Surrey Pathology Services Lighthouse Laboratory | Wellcome Sanger Institute for the COVID-19 Genomics UK (COG-UK) Consortium | Berkshire and Surrey Pathology Services Lighthouse Laboratory and Alex Alderton, Roberto Amato, Jeffrey Barrett, Sonia Goncalves, Ewan Harrison, David K. Jackson, Ian Johnston, Dominic Kwiatkowski, Cordelia Langford, John Sillitoe on behalf of the Wellcome Sanger Institute COVID-19 Surveillance Team |
| EPI_ISL_3282024, EPI_ISL_3282025 | Lighthouse Laboratory Plymouth | Wellcome Sanger Institute for the COVID-19 Genomics UK (COG-UK) Consortium | Lighthouse Laboratory Plymouth and Alex Alderton, Roberto Amato, Jeffrey Barrett, Sonia Goncalves, Ewan Harrison, David K. Jackson, Ian Johnston, Dominic Kwiatkowski, Cordelia Langford, John Sillitoe on behalf of the Wellcome Sanger Institute COVID-19 Surveillance Team |
| EPI_ISL_3282026 | Berkshire and Surrey Pathology Services Lighthouse Laboratory | Wellcome Sanger Institute for the COVID-19 Genomics UK (COG-UK) Consortium | Berkshire and Surrey Pathology Services Lighthouse Laboratory and Alex Alderton, Roberto Amato, Jeffrey Barrett, Sonia Goncalves, Ewan Harrison, David K. Jackson, Ian Johnston, Dominic Kwiatkowski, Cordelia Langford, John Sillitoe on behalf of the Wellcome Sanger Institute COVID-19 Surveillance Team |
| EPI_ISL_3282027, EPI_ISL_3282028 | Lighthouse Laboratory Plymouth | Wellcome Sanger Institute for the COVID-19 Genomics UK (COG-UK) Consortium | Lighthouse Laboratory Plymouth and Alex Alderton, Roberto Amato, Jeffrey Barrett, Sonia Goncalves, Ewan Harrison, David K. Jackson, Ian Johnston, Dominic Kwiatkowski, Cordelia Langford, John Sillitoe on behalf of the Wellcome Sanger Institute COVID-19 Surveillance Team |
| EPI_ISL_3282029 | Berkshire and Surrey Pathology Services Lighthouse Laboratory | Wellcome Sanger Institute for the COVID-19 Genomics UK (COG-UK) Consortium | Berkshire and Surrey Pathology Services Lighthouse Laboratory and Alex Alderton, Roberto Amato, Jeffrey Barrett, Sonia Goncalves, Ewan Harrison, David K. Jackson, Ian Johnston, Dominic Kwiatkowski, Cordelia Langford, John Sillitoe on behalf of the Wellcome Sanger Institute COVID-19 Surveillance Team |
| EPI_ISL_3282030 | Lighthouse Laboratory Plymouth | Wellcome Sanger Institute for the COVID-19 Genomics UK (COG-UK) Consortium | Lighthouse Laboratory Plymouth and Alex Alderton, Roberto Amato, Jeffrey Barrett, Sonia Goncalves, Ewan Harrison, David K. Jackson, Ian Johnston, Dominic Kwiatkowski, Cordelia Langford, John Sillitoe on behalf of the Wellcome Sanger Institute COVID-19 Surveillance Team |
| EPI_ISL_3282031, EPI_ISL_3282032 | Lighthouse Lab in Glasgow | Wellcome Sanger Institute for the COVID-19 Genomics UK (COG-UK) Consortium | Harper VanSteenhouse, Yumi Kasai, David Gray, Carol Clugston, Anna Dominiczak and Alex Alderton, Roberto Amato, Jeffrey Barrett, Sonia Goncalves, Ewan Harrison, David K. Jackson, Ian Johnston, Dominic Kwiatkowski, Cordelia Langford, John Sillitoe on behalf of the Wellcome Sanger Institute COVID-19 Surveillance Team |
| EPI_ISL_3282033, EPI_ISL_3282034, EPI_ISL_3282035 | Lighthouse Laboratory Plymouth | Wellcome Sanger Institute for the COVID-19 Genomics UK (COG-UK) Consortium | Lighthouse Laboratory Plymouth and Alex Alderton, Roberto Amato, Jeffrey Barrett, Sonia Goncalves, Ewan Harrison, David K. Jackson, Ian Johnston, Dominic Kwiatkowski, Cordelia Langford, John Sillitoe on behalf of the Wellcome Sanger Institute COVID-19 Surveillance Team |
| EPI_ISL_3282036 | Lighthouse Lab in Glasgow | Wellcome Sanger Institute for the COVID-19 Genomics UK (COG-UK) Consortium | Harper VanSteenhouse, Yumi Kasai, David Gray, Carol Clugston, Anna Dominiczak and Alex Alderton, Roberto Amato, Jeffrey Barrett, Sonia Goncalves, Ewan Harrison, David K. Jackson, Ian Johnston, Dominic Kwiatkowski, Cordelia Langford, John Sillitoe on behalf of the Wellcome Sanger Institute COVID-19 Surveillance Team |
| EPI_ISL_3282037 | Lighthouse Laboratory Plymouth | Wellcome Sanger Institute for the COVID-19 Genomics UK (COG-UK) Consortium | Lighthouse Laboratory Plymouth and Alex Alderton, Roberto Amato, Jeffrey Barrett, Sonia Goncalves, Ewan Harrison, David K. Jackson, Ian Johnston, Dominic Kwiatkowski, Cordelia Langford, John Sillitoe on behalf of the Wellcome Sanger Institute COVID-19 Surveillance Team |
| EPI_ISL_3282038 | Lighthouse Lab in Glasgow | Wellcome Sanger Institute for the COVID-19 Genomics UK (COG-UK) Consortium | Harper VanSteenhouse, Yumi Kasai, David Gray, Carol Clugston, Anna Dominiczak and Alex Alderton, Roberto Amato, Jeffrey Barrett, Sonia Goncalves, Ewan Harrison, David K. Jackson, Ian Johnston, Dominic Kwiatkowski, Cordelia Langford, John Sillitoe on behalf of the Wellcome Sanger Institute COVID-19 Surveillance Team |
| EPI_ISL_3282039 | Lighthouse Laboratory Plymouth | Wellcome Sanger Institute for the COVID-19 Genomics UK (COG-UK) Consortium | Lighthouse Laboratory Plymouth and Alex Alderton, Roberto Amato, Jeffrey Barrett, Sonia Goncalves, Ewan Harrison, David K. Jackson, Ian Johnston, Dominic Kwiatkowski, Cordelia Langford, John Sillitoe on behalf of the Wellcome Sanger Institute COVID-19 Surveillance Team |
| EPI_ISL_3282040 | Berkshire and Surrey Pathology Services Lighthouse Laboratory | Wellcome Sanger Institute for the COVID-19 Genomics UK (COG-UK) Consortium | Berkshire and Surrey Pathology Services Lighthouse Laboratory and Alex Alderton, Roberto Amato, Jeffrey Barrett, Sonia Goncalves, Ewan Harrison, David K. Jackson, Ian Johnston, Dominic Kwiatkowski, Cordelia Langford, John Sillitoe on behalf of the Wellcome Sanger Institute COVID-19 Surveillance Team |
| EPI_ISL_3282041, EPI_ISL_3282042 | Lighthouse Laboratory Plymouth | Wellcome Sanger Institute for the COVID-19 Genomics UK (COG-UK) Consortium | Lighthouse Laboratory Plymouth and Alex Alderton, Roberto Amato, Jeffrey Barrett, Sonia Goncalves, Ewan Harrison, David K. Jackson, Ian Johnston, Dominic Kwiatkowski, Cordelia Langford, John Sillitoe on behalf of the Wellcome Sanger Institute COVID-19 Surveillance Team |
| EPI_ISL_3282043, EPI_ISL_3282044, EPI_ISL_3282045 | Berkshire and Surrey Pathology Services Lighthouse Laboratory | Wellcome Sanger Institute for the COVID-19 Genomics UK (COG-UK) Consortium | Berkshire and Surrey Pathology Services Lighthouse Laboratory and Alex Alderton, Roberto Amato, Jeffrey Barrett, Sonia Goncalves, Ewan Harrison, David K. Jackson, Ian Johnston, Dominic Kwiatkowski, Cordelia Langford, John Sillitoe on behalf of the Wellcome Sanger Institute COVID-19 Surveillance Team |
| EPI_ISL_3282046, EPI_ISL_3282047, EPI_ISL_3282048 | Lighthouse Laboratory Plymouth | Wellcome Sanger Institute for the COVID-19 Genomics UK (COG-UK) Consortium | Lighthouse Laboratory Plymouth and Alex Alderton, Roberto Amato, Jeffrey Barrett, Sonia Goncalves, Ewan Harrison, David K. Jackson, Ian Johnston, Dominic Kwiatkowski, Cordelia Langford, John Sillitoe on behalf of the Wellcome Sanger Institute COVID-19 Surveillance Team |
| EPI_ISL_3282049 | Lighthouse Lab in Glasgow | Wellcome Sanger Institute for the COVID-19 Genomics UK (COG-UK) Consortium | Harper VanSteenhouse, Yumi Kasai, David Gray, Carol Clugston, Anna Dominiczak and Alex Alderton, Roberto Amato, Jeffrey Barrett, Sonia Goncalves, Ewan Harrison, David K. Jackson, Ian Johnston, Dominic Kwiatkowski, Cordelia Langford, John Sillitoe on behalf of the Wellcome Sanger Institute COVID-19 Surveillance Team |
| EPI_ISL_3282050 | Berkshire and Surrey Pathology Services Lighthouse Laboratory | Wellcome Sanger Institute for the COVID-19 Genomics UK (COG-UK) Consortium | Berkshire and Surrey Pathology Services Lighthouse Laboratory and Alex Alderton, Roberto Amato, Jeffrey Barrett, Sonia Goncalves, Ewan Harrison, David K. Jackson, Ian Johnston, Dominic Kwiatkowski, Cordelia Langford, John Sillitoe on behalf of the Wellcome Sanger Institute COVID-19 Surveillance Team |
| EPI_ISL_3282051 | Lighthouse Laboratory Plymouth | Wellcome Sanger Institute for the COVID-19 Genomics UK (COG-UK) Consortium | Lighthouse Laboratory Plymouth and Alex Alderton, Roberto Amato, Jeffrey Barrett, Sonia Goncalves, Ewan Harrison, David K. Jackson, Ian Johnston, Dominic Kwiatkowski, Cordelia Langford, John Sillitoe on behalf of the Wellcome Sanger Institute COVID-19 Surveillance Team |
| EPI_ISL_3282052 | Lighthouse Lab in Glasgow | Wellcome Sanger Institute for the COVID-19 Genomics UK (COG-UK) | Harper VanSteenhouse, Yumi Kasai, David Gray, Carol Clugston, Anna Dominiczak and Alex Alderton, Roberto Amato, Jeffrey Barrett, Sonia Goncalves, Ewan Harrison, David K. Jackson, Ian Johnston, Dominic Kwiatkowski, Cordelia Langford, |

| | | | |
|---|---|---|---|
| | | Consortium | John Sillitoe on behalf of the Wellcome Sanger Institute COVID-19 Surveillance Team |
| EPI_ISL_3282053 | Lighthouse Laboratory Plymouth | Wellcome Sanger Institute for the COVID-19 Genomics UK (COG-UK) Consortium | Lighthouse Laboratory Plymouth and Alex Alderton, Roberto Amato, Jeffrey Barrett, Sonia Goncalves, Ewan Harrison, David K. Jackson, Ian Johnston, Dominic Kwiatkowski, Cordelia Langford, John Sillitoe on behalf of the Wellcome Sanger Institute COVID-19 Surveillance Team |
| EPI_ISL_3282054 | Berkshire and Surrey Pathology Services Lighthouse Laboratory | Wellcome Sanger Institute for the COVID-19 Genomics UK (COG-UK) Consortium | Berkshire and Surrey Pathology Services Lighthouse Laboratory and Alex Alderton, Roberto Amato, Jeffrey Barrett, Sonia Goncalves, Ewan Harrison, David K. Jackson, Ian Johnston, Dominic Kwiatkowski, Cordelia Langford, John Sillitoe on behalf of the Wellcome Sanger Institute COVID-19 Surveillance Team |
| EPI_ISL_3282055, EPI_ISL_3282056 | Lighthouse Lab in Glasgow | Wellcome Sanger Institute for the COVID-19 Genomics UK (COG-UK) Consortium | Harper VanSteenhouse, Yumi Kasai, David Gray, Carol Clugston, Anna Dominiczak and Alex Alderton, Roberto Amato, Jeffrey Barrett, Sonia Goncalves, Ewan Harrison, David K. Jackson, Ian Johnston, Dominic Kwiatkowski, Cordelia Langford, John Sillitoe on behalf of the Wellcome Sanger Institute COVID-19 Surveillance Team |
| EPI_ISL_3282057, EPI_ISL_3282058, EPI_ISL_3282059 | Lighthouse Laboratory Plymouth | Wellcome Sanger Institute for the COVID-19 Genomics UK (COG-UK) Consortium | Lighthouse Laboratory Plymouth and Alex Alderton, Roberto Amato, Jeffrey Barrett, Sonia Goncalves, Ewan Harrison, David K. Jackson, Ian Johnston, Dominic Kwiatkowski, Cordelia Langford, John Sillitoe on behalf of the Wellcome Sanger Institute COVID-19 Surveillance Team |
| EPI_ISL_3282060 | Lighthouse Lab in Glasgow | Wellcome Sanger Institute for the COVID-19 Genomics UK (COG-UK) Consortium | Harper VanSteenhouse, Yumi Kasai, David Gray, Carol Clugston, Anna Dominiczak and Alex Alderton, Roberto Amato, Jeffrey Barrett, Sonia Goncalves, Ewan Harrison, David K. Jackson, Ian Johnston, Dominic Kwiatkowski, Cordelia Langford, John Sillitoe on behalf of the Wellcome Sanger Institute COVID-19 Surveillance Team |
| EPI_ISL_3282061, EPI_ISL_3282062, EPI_ISL_3282063, EPI_ISL_3282064, EPI_ISL_3282065, EPI_ISL_3282066, EPI_ISL_3282067, EPI_ISL_3282068 | Lighthouse Laboratory Plymouth | Wellcome Sanger Institute for the COVID-19 Genomics UK (COG-UK) Consortium | Lighthouse Laboratory Plymouth and Alex Alderton, Roberto Amato, Jeffrey Barrett, Sonia Goncalves, Ewan Harrison, David K. Jackson, Ian Johnston, Dominic Kwiatkowski, Cordelia Langford, John Sillitoe on behalf of the Wellcome Sanger Institute COVID-19 Surveillance Team |
| EPI_ISL_3282069 | Lighthouse Lab in Glasgow | Wellcome Sanger Institute for the COVID-19 Genomics UK (COG-UK) Consortium | Harper VanSteenhouse, Yumi Kasai, David Gray, Carol Clugston, Anna Dominiczak and Alex Alderton, Roberto Amato, Jeffrey Barrett, Sonia Goncalves, Ewan Harrison, David K. Jackson, Ian Johnston, Dominic Kwiatkowski, Cordelia Langford, John Sillitoe on behalf of the Wellcome Sanger Institute COVID-19 Surveillance Team |
| EPI_ISL_3282070, EPI_ISL_3282071, EPI_ISL_3282072 | Lighthouse Laboratory Plymouth | Wellcome Sanger Institute for the COVID-19 Genomics UK (COG-UK) Consortium | Lighthouse Laboratory Plymouth and Alex Alderton, Roberto Amato, Jeffrey Barrett, Sonia Goncalves, Ewan Harrison, David K. Jackson, Ian Johnston, Dominic Kwiatkowski, Cordelia Langford, John Sillitoe on behalf of the Wellcome Sanger Institute COVID-19 Surveillance Team |
| EPI_ISL_3282073 | Lighthouse Lab in Glasgow | Wellcome Sanger Institute for the COVID-19 Genomics UK (COG-UK) Consortium | Harper VanSteenhouse, Yumi Kasai, David Gray, Carol Clugston, Anna Dominiczak and Alex Alderton, Roberto Amato, Jeffrey Barrett, Sonia Goncalves, Ewan Harrison, David K. Jackson, Ian Johnston, Dominic Kwiatkowski, Cordelia Langford, John Sillitoe on behalf of the Wellcome Sanger Institute COVID-19 Surveillance Team |
| EPI_ISL_3282074 | Berkshire and Surrey Pathology Services Lighthouse Laboratory | Wellcome Sanger Institute for the COVID-19 Genomics UK (COG-UK) Consortium | Berkshire and Surrey Pathology Services Lighthouse Laboratory and Alex Alderton, Roberto Amato, Jeffrey Barrett, Sonia Goncalves, Ewan Harrison, David K. Jackson, Ian Johnston, Dominic Kwiatkowski, Cordelia Langford, John Sillitoe on behalf of the Wellcome Sanger Institute COVID-19 Surveillance Team |
| EPI_ISL_3282075 | Lighthouse Laboratory Plymouth | Wellcome Sanger Institute for the COVID-19 Genomics UK (COG-UK) Consortium | Lighthouse Laboratory Plymouth and Alex Alderton, Roberto Amato, Jeffrey Barrett, Sonia Goncalves, Ewan Harrison, David K. Jackson, Ian Johnston, Dominic Kwiatkowski, Cordelia Langford, John Sillitoe on behalf of the Wellcome Sanger Institute COVID-19 Surveillance Team |
| EPI_ISL_3282076 | Berkshire and Surrey Pathology Services Lighthouse Laboratory | Wellcome Sanger Institute for the COVID-19 Genomics UK (COG-UK) Consortium | Berkshire and Surrey Pathology Services Lighthouse Laboratory and Alex Alderton, Roberto Amato, Jeffrey Barrett, Sonia Goncalves, Ewan Harrison, David K. Jackson, Ian Johnston, Dominic Kwiatkowski, Cordelia Langford, John Sillitoe on behalf of the Wellcome Sanger Institute COVID-19 Surveillance Team |
| EPI_ISL_3282077 | Lighthouse Laboratory Plymouth | Wellcome Sanger Institute for the COVID-19 Genomics UK (COG-UK) Consortium | Lighthouse Laboratory Plymouth and Alex Alderton, Roberto Amato, Jeffrey Barrett, Sonia Goncalves, Ewan Harrison, David K. Jackson, Ian Johnston, Dominic Kwiatkowski, Cordelia Langford, John Sillitoe on behalf of the Wellcome Sanger Institute COVID-19 Surveillance Team |
| EPI_ISL_3282078 | Berkshire and Surrey Pathology Services Lighthouse Laboratory | Wellcome Sanger Institute for the COVID-19 Genomics UK (COG-UK) Consortium | Berkshire and Surrey Pathology Services Lighthouse Laboratory and Alex Alderton, Roberto Amato, Jeffrey Barrett, Sonia Goncalves, Ewan Harrison, David K. Jackson, Ian Johnston, Dominic Kwiatkowski, Cordelia Langford, John Sillitoe on behalf of the Wellcome Sanger Institute COVID-19 Surveillance Team |
| EPI_ISL_3282079 | Lighthouse Lab in Glasgow | Wellcome Sanger Institute for the COVID-19 Genomics UK (COG-UK) Consortium | Harper VanSteenhouse, Yumi Kasai, David Gray, Carol Clugston, Anna Dominiczak and Alex Alderton, Roberto Amato, Jeffrey Barrett, Sonia Goncalves, Ewan Harrison, David K. Jackson, Ian Johnston, Dominic Kwiatkowski, Cordelia Langford, John Sillitoe on behalf of the Wellcome Sanger Institute COVID-19 Surveillance Team |
| EPI_ISL_3282080 | Berkshire and Surrey Pathology Services Lighthouse Laboratory | Wellcome Sanger Institute for the COVID-19 Genomics UK (COG-UK) Consortium | Berkshire and Surrey Pathology Services Lighthouse Laboratory and Alex Alderton, Roberto Amato, Jeffrey Barrett, Sonia Goncalves, Ewan Harrison, David K. Jackson, Ian Johnston, Dominic Kwiatkowski, Cordelia Langford, John Sillitoe on behalf of the Wellcome Sanger Institute COVID-19 Surveillance Team |
| EPI_ISL_3282081, EPI_ISL_3282082, EPI_ISL_3282083 | Lighthouse Laboratory Plymouth | Wellcome Sanger Institute for the COVID-19 Genomics UK (COG-UK) Consortium | Lighthouse Laboratory Plymouth and Alex Alderton, Roberto Amato, Jeffrey Barrett, Sonia Goncalves, Ewan Harrison, David K. Jackson, Ian Johnston, Dominic Kwiatkowski, Cordelia Langford, John Sillitoe on behalf of the Wellcome Sanger Institute COVID-19 Surveillance Team |
| EPI_ISL_3282084 | Berkshire and Surrey Pathology Services Lighthouse Laboratory | Wellcome Sanger Institute for the COVID-19 Genomics UK (COG-UK) Consortium | Berkshire and Surrey Pathology Services Lighthouse Laboratory and Alex Alderton, Roberto Amato, Jeffrey Barrett, Sonia Goncalves, Ewan Harrison, David K. Jackson, Ian Johnston, Dominic Kwiatkowski, Cordelia Langford, John Sillitoe on behalf of the Wellcome Sanger Institute COVID-19 Surveillance Team |
| EPI_ISL_3282085 | Lighthouse Laboratory Plymouth | Wellcome Sanger Institute for the COVID-19 Genomics UK (COG-UK) Consortium | Lighthouse Laboratory Plymouth and Alex Alderton, Roberto Amato, Jeffrey Barrett, Sonia Goncalves, Ewan Harrison, David K. Jackson, Ian Johnston, Dominic Kwiatkowski, Cordelia Langford, John Sillitoe on behalf of the Wellcome Sanger Institute COVID-19 Surveillance Team |
| EPI_ISL_3282086 | Lighthouse Lab in Glasgow | Wellcome Sanger Institute for the COVID-19 Genomics UK (COG-UK) Consortium | Harper VanSteenhouse, Yumi Kasai, David Gray, Carol Clugston, Anna Dominiczak and Alex Alderton, Roberto Amato, Jeffrey Barrett, Sonia Goncalves, Ewan Harrison, David K. Jackson, Ian Johnston, Dominic Kwiatkowski, Cordelia Langford, John Sillitoe on behalf of the Wellcome Sanger Institute COVID-19 Surveillance Team |
| EPI_ISL_3282087, EPI_ISL_3282088, EPI_ISL_3282089 | Lighthouse Laboratory Plymouth | Wellcome Sanger Institute for the COVID-19 Genomics UK (COG-UK) Consortium | Lighthouse Laboratory Plymouth and Alex Alderton, Roberto Amato, Jeffrey Barrett, Sonia Goncalves, Ewan Harrison, David K. Jackson, Ian Johnston, Dominic Kwiatkowski, Cordelia Langford, John Sillitoe on behalf of the Wellcome Sanger Institute COVID-19 Surveillance Team |
| EPI_ISL_3282090 | Lighthouse Lab in Glasgow | Wellcome Sanger Institute for the COVID-19 Genomics UK (COG-UK) Consortium | Harper VanSteenhouse, Yumi Kasai, David Gray, Carol Clugston, Anna Dominiczak and Alex Alderton, Roberto Amato, Jeffrey Barrett, Sonia Goncalves, Ewan Harrison, David K. Jackson, Ian Johnston, Dominic Kwiatkowski, Cordelia Langford, John Sillitoe on behalf of the Wellcome Sanger Institute COVID-19 Surveillance Team |
| EPI_ISL_3282091, EPI_ISL_3282092 | Lighthouse Laboratory Plymouth | Wellcome Sanger Institute for the COVID-19 Genomics UK (COG-UK) Consortium | Lighthouse Laboratory Plymouth and Alex Alderton, Roberto Amato, Jeffrey Barrett, Sonia Goncalves, Ewan Harrison, David K. Jackson, Ian Johnston, Dominic Kwiatkowski, Cordelia Langford, John Sillitoe on behalf of the Wellcome Sanger Institute COVID-19 Surveillance Team |
| EPI_ISL_3282093 | Lighthouse Lab in Glasgow | Wellcome Sanger Institute for the COVID-19 Genomics UK (COG-UK) Consortium | Harper VanSteenhouse, Yumi Kasai, David Gray, Carol Clugston, Anna Dominiczak and Alex Alderton, Roberto Amato, Jeffrey Barrett, Sonia Goncalves, Ewan Harrison, David K. Jackson, Ian Johnston, Dominic Kwiatkowski, Cordelia Langford, John Sillitoe on behalf of the Wellcome Sanger Institute COVID-19 Surveillance Team |
| EPI_ISL_3282094 | Lighthouse Laboratory Plymouth | Wellcome Sanger Institute for the COVID-19 Genomics UK (COG-UK) Consortium | Lighthouse Laboratory Plymouth and Alex Alderton, Roberto Amato, Jeffrey Barrett, Sonia Goncalves, Ewan Harrison, David K. Jackson, Ian Johnston, Dominic Kwiatkowski, Cordelia Langford, John Sillitoe on behalf of the Wellcome Sanger Institute COVID-19 Surveillance Team |
| EPI_ISL_3282095 | Berkshire and Surrey Pathology Services Lighthouse Laboratory | Wellcome Sanger Institute for the COVID-19 Genomics UK (COG-UK) Consortium | Berkshire and Surrey Pathology Services Lighthouse Laboratory and Alex Alderton, Roberto Amato, Jeffrey Barrett, Sonia Goncalves, Ewan Harrison, David K. Jackson, Ian Johnston, Dominic Kwiatkowski, Cordelia Langford, John Sillitoe on behalf of the Wellcome Sanger Institute COVID-19 Surveillance Team |
| EPI_ISL_3282096 | Lighthouse Laboratory Plymouth | Wellcome Sanger Institute for the COVID-19 Genomics UK (COG-UK) Consortium | Lighthouse Laboratory Plymouth and Alex Alderton, Roberto Amato, Jeffrey Barrett, Sonia Goncalves, Ewan Harrison, David K. Jackson, Ian Johnston, Dominic Kwiatkowski, Cordelia Langford, John Sillitoe on behalf of the Wellcome Sanger Institute COVID-19 Surveillance Team |
| EPI_ISL_3282097 | Berkshire and Surrey Pathology Services Lighthouse Laboratory | Wellcome Sanger Institute for the COVID-19 Genomics UK (COG-UK) Consortium | Berkshire and Surrey Pathology Services Lighthouse Laboratory and Alex Alderton, Roberto Amato, Jeffrey Barrett, Sonia Goncalves, Ewan Harrison, David K. Jackson, Ian Johnston, Dominic Kwiatkowski, Cordelia Langford, John Sillitoe on behalf of the Wellcome Sanger Institute COVID-19 Surveillance Team |
| EPI_ISL_3282098, EPI_ISL_3282099 | Lighthouse Laboratory Plymouth | Wellcome Sanger Institute for the COVID-19 Genomics UK (COG-UK) Consortium | Lighthouse Laboratory Plymouth and Alex Alderton, Roberto Amato, Jeffrey Barrett, Sonia Goncalves, Ewan Harrison, David K. Jackson, Ian Johnston, Dominic Kwiatkowski, Cordelia Langford, John Sillitoe on behalf of the Wellcome Sanger Institute COVID-19 Surveillance Team |
| EPI_ISL_3282100 | Lighthouse Lab in Glasgow | Wellcome Sanger Institute for the COVID-19 Genomics UK (COG-UK) Consortium | Harper VanSteenhouse, Yumi Kasai, David Gray, Carol Clugston, Anna Dominiczak and Alex Alderton, Roberto Amato, Jeffrey Barrett, Sonia Goncalves, Ewan Harrison, David K. Jackson, Ian Johnston, Dominic Kwiatkowski, Cordelia Langford, John Sillitoe on behalf of the Wellcome Sanger Institute COVID-19 Surveillance Team |
| EPI_ISL_3282101 | Lighthouse Laboratory Plymouth | Wellcome Sanger Institute for the COVID-19 Genomics UK (COG-UK) Consortium | Lighthouse Laboratory Plymouth and Alex Alderton, Roberto Amato, Jeffrey Barrett, Sonia Goncalves, Ewan Harrison, David K. Jackson, Ian Johnston, Dominic Kwiatkowski, Cordelia Langford, John Sillitoe on behalf of the Wellcome Sanger Institute COVID-19 Surveillance Team |
| EPI_ISL_3282102 | Lighthouse Lab in Glasgow | Wellcome Sanger Institute for the COVID-19 Genomics UK (COG-UK) Consortium | Harper VanSteenhouse, Yumi Kasai, David Gray, Carol Clugston, Anna Dominiczak and Alex Alderton, Roberto Amato, Jeffrey Barrett, Sonia Goncalves, Ewan Harrison, David K. Jackson, Ian Johnston, Dominic Kwiatkowski, Cordelia Langford, John Sillitoe on behalf of the Wellcome Sanger Institute COVID-19 Surveillance Team |
| EPI_ISL_3282103 | Lighthouse Laboratory Plymouth | Wellcome Sanger Institute for the COVID-19 Genomics UK (COG-UK) Consortium | Lighthouse Laboratory Plymouth and Alex Alderton, Roberto Amato, Jeffrey Barrett, Sonia Goncalves, Ewan Harrison, David K. Jackson, Ian Johnston, Dominic Kwiatkowski, Cordelia Langford, John Sillitoe on behalf of the Wellcome Sanger Institute COVID-19 Surveillance Team |
| EPI_ISL_3282104, EPI_ISL_3282105 | Berkshire and Surrey Pathology Services Lighthouse Laboratory | Wellcome Sanger Institute for the COVID-19 Genomics UK (COG-UK) Consortium | Berkshire and Surrey Pathology Services Lighthouse Laboratory and Alex Alderton, Roberto Amato, Jeffrey Barrett, Sonia Goncalves, Ewan Harrison, David K. Jackson, Ian Johnston, Dominic Kwiatkowski, Cordelia Langford, John Sillitoe on behalf of the Wellcome Sanger Institute COVID-19 Surveillance Team |
| EPI_ISL_3282106, EPI_ISL_3282107, EPI_ISL_3282108 | Lighthouse Laboratory Plymouth | Wellcome Sanger Institute for the COVID-19 Genomics UK (COG-UK) Consortium | Lighthouse Laboratory Plymouth and Alex Alderton, Roberto Amato, Jeffrey Barrett, Sonia Goncalves, Ewan Harrison, David K. Jackson, Ian Johnston, Dominic Kwiatkowski, Cordelia Langford, John Sillitoe on behalf of the Wellcome Sanger Institute COVID-19 Surveillance Team |
| EPI_ISL_3282109 | Berkshire and Surrey Pathology Services Lighthouse Laboratory | Wellcome Sanger Institute for the COVID-19 Genomics UK (COG-UK) Consortium | Berkshire and Surrey Pathology Services Lighthouse Laboratory and Alex Alderton, Roberto Amato, Jeffrey Barrett, Sonia Goncalves, Ewan Harrison, David K. Jackson, Ian Johnston, Dominic Kwiatkowski, Cordelia Langford, John Sillitoe on behalf of the Wellcome Sanger Institute COVID-19 Surveillance Team |
| EPI_ISL_3282110, EPI_ISL_3282111 | Lighthouse Laboratory Plymouth | Wellcome Sanger Institute for the COVID-19 Genomics UK (COG-UK) Consortium | Lighthouse Laboratory Plymouth and Alex Alderton, Roberto Amato, Jeffrey Barrett, Sonia Goncalves, Ewan Harrison, David K. Jackson, Ian Johnston, Dominic Kwiatkowski, Cordelia Langford, John Sillitoe on behalf of the Wellcome Sanger Institute COVID-19 Surveillance Team |
| EPI_ISL_3282112 | Berkshire and Surrey Pathology Services Lighthouse Laboratory | Wellcome Sanger Institute for the COVID-19 Genomics UK (COG-UK) Consortium | Berkshire and Surrey Pathology Services Lighthouse Laboratory and Alex Alderton, Roberto Amato, Jeffrey Barrett, Sonia Goncalves, Ewan Harrison, David K. Jackson, Ian Johnston, Dominic Kwiatkowski, Cordelia Langford, John Sillitoe on behalf of the Wellcome Sanger Institute COVID-19 Surveillance Team |
| EPI_ISL_3282113 | Lighthouse Laboratory Plymouth | Wellcome Sanger Institute for the COVID-19 Genomics UK (COG-UK) Consortium | Lighthouse Laboratory Plymouth and Alex Alderton, Roberto Amato, Jeffrey Barrett, Sonia Goncalves, Ewan Harrison, David K. Jackson, Ian Johnston, Dominic Kwiatkowski, Cordelia Langford, John Sillitoe on behalf of the Wellcome Sanger Institute COVID-19 Surveillance Team |
| EPI_ISL_3282114 | Berkshire and Surrey Pathology Services Lighthouse Laboratory | Wellcome Sanger Institute for the COVID-19 Genomics UK (COG-UK) Consortium | Berkshire and Surrey Pathology Services Lighthouse Laboratory and Alex Alderton, Roberto Amato, Jeffrey Barrett, Sonia Goncalves, Ewan Harrison, David K. Jackson, Ian Johnston, Dominic Kwiatkowski, Cordelia Langford, John Sillitoe on behalf of the Wellcome Sanger Institute COVID-19 Surveillance Team |

| | | | |
|---|---|---|---|
| EPI_ISL_3282115 | Lighthouse Laboratory Plymouth | Wellcome Sanger Institute for the COVID-19 Genomics UK (COG-UK) Consortium | Lighthouse Laboratory Plymouth and Alex Alderton, Roberto Amato, Jeffrey Barrett, Sonia Goncalves, Ewan Harrison, David K. Jackson, Ian Johnston, Dominic Kwiatkowski, Cordelia Langford, John Sillitoe on behalf of the Wellcome Sanger Institute COVID-19 Surveillance Team |
| EPI_ISL_3282116, EPI_ISL_3282117 | Berkshire and Surrey Pathology Services Lighthouse Laboratory | Wellcome Sanger Institute for the COVID-19 Genomics UK (COG-UK) Consortium | Berkshire and Surrey Pathology Services Lighthouse Laboratory and Alex Alderton, Roberto Amato, Jeffrey Barrett, Sonia Goncalves, Ewan Harrison, David K. Jackson, Ian Johnston, Dominic Kwiatkowski, Cordelia Langford, John Sillitoe on behalf of the Wellcome Sanger Institute COVID-19 Surveillance Team |
| EPI_ISL_3282118 | Lighthouse Lab in Glasgow | Wellcome Sanger Institute for the COVID-19 Genomics UK (COG-UK) Consortium | Harper VanSteenhouse, Yumi Kasai, David Gray, Carol Clugston, Anna Dominiczak and Alex Alderton, Roberto Amato, Jeffrey Barrett, Sonia Goncalves, Ewan Harrison, David K. Jackson, Ian Johnston, Dominic Kwiatkowski, Cordelia Langford, John Sillitoe on behalf of the Wellcome Sanger Institute COVID-19 Surveillance Team |
| EPI_ISL_3282119 | Lighthouse Laboratory Plymouth | Wellcome Sanger Institute for the COVID-19 Genomics UK (COG-UK) Consortium | Lighthouse Laboratory Plymouth and Alex Alderton, Roberto Amato, Jeffrey Barrett, Sonia Goncalves, Ewan Harrison, David K. Jackson, Ian Johnston, Dominic Kwiatkowski, Cordelia Langford, John Sillitoe on behalf of the Wellcome Sanger Institute COVID-19 Surveillance Team |
| EPI_ISL_3282120 | Berkshire and Surrey Pathology Services Lighthouse Laboratory | Wellcome Sanger Institute for the COVID-19 Genomics UK (COG-UK) Consortium | Berkshire and Surrey Pathology Services Lighthouse Laboratory and Alex Alderton, Roberto Amato, Jeffrey Barrett, Sonia Goncalves, Ewan Harrison, David K. Jackson, Ian Johnston, Dominic Kwiatkowski, Cordelia Langford, John Sillitoe on behalf of the Wellcome Sanger Institute COVID-19 Surveillance Team |
| EPI_ISL_3282121 | Lighthouse Lab in Glasgow | Wellcome Sanger Institute for the COVID-19 Genomics UK (COG-UK) Consortium | Harper VanSteenhouse, Yumi Kasai, David Gray, Carol Clugston, Anna Dominiczak and Alex Alderton, Roberto Amato, Jeffrey Barrett, Sonia Goncalves, Ewan Harrison, David K. Jackson, Ian Johnston, Dominic Kwiatkowski, Cordelia Langford, John Sillitoe on behalf of the Wellcome Sanger Institute COVID-19 Surveillance Team |
| EPI_ISL_3282122, EPI_ISL_3282123, EPI_ISL_3282124 | Lighthouse Laboratory Plymouth | Wellcome Sanger Institute for the COVID-19 Genomics UK (COG-UK) Consortium | Lighthouse Laboratory Plymouth and Alex Alderton, Roberto Amato, Jeffrey Barrett, Sonia Goncalves, Ewan Harrison, David K. Jackson, Ian Johnston, Dominic Kwiatkowski, Cordelia Langford, John Sillitoe on behalf of the Wellcome Sanger Institute COVID-19 Surveillance Team |
| EPI_ISL_3282125 | Berkshire and Surrey Pathology Services Lighthouse Laboratory | Wellcome Sanger Institute for the COVID-19 Genomics UK (COG-UK) Consortium | Berkshire and Surrey Pathology Services Lighthouse Laboratory and Alex Alderton, Roberto Amato, Jeffrey Barrett, Sonia Goncalves, Ewan Harrison, David K. Jackson, Ian Johnston, Dominic Kwiatkowski, Cordelia Langford, John Sillitoe on behalf of the Wellcome Sanger Institute COVID-19 Surveillance Team |
| EPI_ISL_3282126 | Lighthouse Lab in Milton Keynes | Wellcome Sanger Institute for the COVID-19 Genomics UK (COG-UK) Consortium | The Lighthouse Lab in Milton Keynes and Alex Alderton, Roberto Amato, Jeffrey Barrett, Sonia Goncalves, Ewan Harrison, David K. Jackson, Ian Johnston, Dominic Kwiatkowski, Cordelia Langford, John Sillitoe on behalf of the Wellcome Sanger Institute COVID-19 Surveillance Team |
| EPI_ISL_3282127 | Berkshire and Surrey Pathology Services Lighthouse Laboratory | Wellcome Sanger Institute for the COVID-19 Genomics UK (COG-UK) Consortium | Berkshire and Surrey Pathology Services Lighthouse Laboratory and Alex Alderton, Roberto Amato, Jeffrey Barrett, Sonia Goncalves, Ewan Harrison, David K. Jackson, Ian Johnston, Dominic Kwiatkowski, Cordelia Langford, John Sillitoe on behalf of the Wellcome Sanger Institute COVID-19 Surveillance Team |
| EPI_ISL_3282128, EPI_ISL_3282129 | Lighthouse Lab in Alderley Park | Wellcome Sanger Institute for the COVID-19 Genomics UK (COG-UK) Consortium | Jacquelyn Wynn, Mairead Hyland, The Lighthouse Lab in Alderley Park and Alex Alderton, Roberto Amato, Jeffrey Barrett, Sonia Goncalves, Ewan Harrison, David K. Jackson, Ian Johnston, Dominic Kwiatkowski, Cordelia Langford, John Sillitoe on behalf of the Wellcome Sanger Institute COVID-19 Surveillance Team |
| EPI_ISL_3282130, EPI_ISL_3282131 | Lighthouse Lab in Milton Keynes | Wellcome Sanger Institute for the COVID-19 Genomics UK (COG-UK) Consortium | The Lighthouse Lab in Milton Keynes and Alex Alderton, Roberto Amato, Jeffrey Barrett, Sonia Goncalves, Ewan Harrison, David K. Jackson, Ian Johnston, Dominic Kwiatkowski, Cordelia Langford, John Sillitoe on behalf of the Wellcome Sanger Institute COVID-19 Surveillance Team |
| EPI_ISL_3282132 | Berkshire and Surrey Pathology Services Lighthouse Laboratory | Wellcome Sanger Institute for the COVID-19 Genomics UK (COG-UK) Consortium | Berkshire and Surrey Pathology Services Lighthouse Laboratory and Alex Alderton, Roberto Amato, Jeffrey Barrett, Sonia Goncalves, Ewan Harrison, David K. Jackson, Ian Johnston, Dominic Kwiatkowski, Cordelia Langford, John Sillitoe on behalf of the Wellcome Sanger Institute COVID-19 Surveillance Team |
| EPI_ISL_3282133, EPI_ISL_3282134 | Lighthouse Lab in Alderley Park | Wellcome Sanger Institute for the COVID-19 Genomics UK (COG-UK) Consortium | Jacquelyn Wynn, Mairead Hyland, The Lighthouse Lab in Alderley Park and Alex Alderton, Roberto Amato, Jeffrey Barrett, Sonia Goncalves, Ewan Harrison, David K. Jackson, Ian Johnston, Dominic Kwiatkowski, Cordelia Langford, John Sillitoe on behalf of the Wellcome Sanger Institute COVID-19 Surveillance Team |
| EPI_ISL_3282135 | Berkshire and Surrey Pathology Services Lighthouse Laboratory | Wellcome Sanger Institute for the COVID-19 Genomics UK (COG-UK) Consortium | Berkshire and Surrey Pathology Services Lighthouse Laboratory and Alex Alderton, Roberto Amato, Jeffrey Barrett, Sonia Goncalves, Ewan Harrison, David K. Jackson, Ian Johnston, Dominic Kwiatkowski, Cordelia Langford, John Sillitoe on behalf of the Wellcome Sanger Institute COVID-19 Surveillance Team |
| EPI_ISL_3282136 | Lighthouse Lab in Alderley Park | Wellcome Sanger Institute for the COVID-19 Genomics UK (COG-UK) Consortium | Jacquelyn Wynn, Mairead Hyland, The Lighthouse Lab in Alderley Park and Alex Alderton, Roberto Amato, Jeffrey Barrett, Sonia Goncalves, Ewan Harrison, David K. Jackson, Ian Johnston, Dominic Kwiatkowski, Cordelia Langford, John Sillitoe on behalf of the Wellcome Sanger Institute COVID-19 Surveillance Team |
| EPI_ISL_3282137 | Berkshire and Surrey Pathology Services Lighthouse Laboratory | Wellcome Sanger Institute for the COVID-19 Genomics UK (COG-UK) Consortium | Berkshire and Surrey Pathology Services Lighthouse Laboratory and Alex Alderton, Roberto Amato, Jeffrey Barrett, Sonia Goncalves, Ewan Harrison, David K. Jackson, Ian Johnston, Dominic Kwiatkowski, Cordelia Langford, John Sillitoe on behalf of the Wellcome Sanger Institute COVID-19 Surveillance Team |
| EPI_ISL_3282138 | Lighthouse Lab in Alderley Park | Wellcome Sanger Institute for the COVID-19 Genomics UK (COG-UK) Consortium | Jacquelyn Wynn, Mairead Hyland, The Lighthouse Lab in Alderley Park and Alex Alderton, Roberto Amato, Jeffrey Barrett, Sonia Goncalves, Ewan Harrison, David K. Jackson, Ian Johnston, Dominic Kwiatkowski, Cordelia Langford, John Sillitoe on behalf of the Wellcome Sanger Institute COVID-19 Surveillance Team |
| EPI_ISL_3282139 | Lighthouse Lab in Milton Keynes | Wellcome Sanger Institute for the COVID-19 Genomics UK (COG-UK) Consortium | The Lighthouse Lab in Milton Keynes and Alex Alderton, Roberto Amato, Jeffrey Barrett, Sonia Goncalves, Ewan Harrison, David K. Jackson, Ian Johnston, Dominic Kwiatkowski, Cordelia Langford, John Sillitoe on behalf of the Wellcome Sanger Institute COVID-19 Surveillance Team |
| EPI_ISL_3282140 | Berkshire and Surrey Pathology Services Lighthouse Laboratory | Wellcome Sanger Institute for the COVID-19 Genomics UK (COG-UK) Consortium | Berkshire and Surrey Pathology Services Lighthouse Laboratory and Alex Alderton, Roberto Amato, Jeffrey Barrett, Sonia Goncalves, Ewan Harrison, David K. Jackson, Ian Johnston, Dominic Kwiatkowski, Cordelia Langford, John Sillitoe on behalf of the Wellcome Sanger Institute COVID-19 Surveillance Team |
| EPI_ISL_3282141 | Lighthouse Lab in Alderley Park | Wellcome Sanger Institute for the COVID-19 Genomics UK (COG-UK) Consortium | Jacquelyn Wynn, Mairead Hyland, The Lighthouse Lab in Alderley Park and Alex Alderton, Roberto Amato, Jeffrey Barrett, Sonia Goncalves, Ewan Harrison, David K. Jackson, Ian Johnston, Dominic Kwiatkowski, Cordelia Langford, John Sillitoe on behalf of the Wellcome Sanger Institute COVID-19 Surveillance Team |
| EPI_ISL_3282142, EPI_ISL_3282143 | Lighthouse Lab in Milton Keynes | Wellcome Sanger Institute for the COVID-19 Genomics UK (COG-UK) Consortium | The Lighthouse Lab in Milton Keynes and Alex Alderton, Roberto Amato, Jeffrey Barrett, Sonia Goncalves, Ewan Harrison, David K. Jackson, Ian Johnston, Dominic Kwiatkowski, Cordelia Langford, John Sillitoe on behalf of the Wellcome Sanger Institute COVID-19 Surveillance Team |
| EPI_ISL_3282144 | Berkshire and Surrey Pathology Services Lighthouse Laboratory | Wellcome Sanger Institute for the COVID-19 Genomics UK (COG-UK) Consortium | Berkshire and Surrey Pathology Services Lighthouse Laboratory and Alex Alderton, Roberto Amato, Jeffrey Barrett, Sonia Goncalves, Ewan Harrison, David K. Jackson, Ian Johnston, Dominic Kwiatkowski, Cordelia Langford, John Sillitoe on behalf of the Wellcome Sanger Institute COVID-19 Surveillance Team |
| EPI_ISL_3282145 | Lighthouse Lab in Alderley Park | Wellcome Sanger Institute for the COVID-19 Genomics UK (COG-UK) Consortium | Jacquelyn Wynn, Mairead Hyland, The Lighthouse Lab in Alderley Park and Alex Alderton, Roberto Amato, Jeffrey Barrett, Sonia Goncalves, Ewan Harrison, David K. Jackson, Ian Johnston, Dominic Kwiatkowski, Cordelia Langford, John Sillitoe on behalf of the Wellcome Sanger Institute COVID-19 Surveillance Team |
| EPI_ISL_3282146 | Lighthouse Lab in Milton Keynes | Wellcome Sanger Institute for the COVID-19 Genomics UK (COG-UK) Consortium | The Lighthouse Lab in Milton Keynes and Alex Alderton, Roberto Amato, Jeffrey Barrett, Sonia Goncalves, Ewan Harrison, David K. Jackson, Ian Johnston, Dominic Kwiatkowski, Cordelia Langford, John Sillitoe on behalf of the Wellcome Sanger Institute COVID-19 Surveillance Team |
| EPI_ISL_3282147 | Lighthouse Lab in Alderley Park | Wellcome Sanger Institute for the COVID-19 Genomics UK (COG-UK) Consortium | Jacquelyn Wynn, Mairead Hyland, The Lighthouse Lab in Alderley Park and Alex Alderton, Roberto Amato, Jeffrey Barrett, Sonia Goncalves, Ewan Harrison, David K. Jackson, Ian Johnston, Dominic Kwiatkowski, Cordelia Langford, John Sillitoe on behalf of the Wellcome Sanger Institute COVID-19 Surveillance Team |
| EPI_ISL_3282148 | Lighthouse Lab in Milton Keynes | Wellcome Sanger Institute for the COVID-19 Genomics UK (COG-UK) Consortium | The Lighthouse Lab in Milton Keynes and Alex Alderton, Roberto Amato, Jeffrey Barrett, Sonia Goncalves, Ewan Harrison, David K. Jackson, Ian Johnston, Dominic Kwiatkowski, Cordelia Langford, John Sillitoe on behalf of the Wellcome Sanger Institute COVID-19 Surveillance Team |
| EPI_ISL_3282149, EPI_ISL_3282150, EPI_ISL_3282151, EPI_ISL_3282152, EPI_ISL_3282153 | Lighthouse Lab in Alderley Park | Wellcome Sanger Institute for the COVID-19 Genomics UK (COG-UK) Consortium | Jacquelyn Wynn, Mairead Hyland, The Lighthouse Lab in Alderley Park and Alex Alderton, Roberto Amato, Jeffrey Barrett, Sonia Goncalves, Ewan Harrison, David K. Jackson, Ian Johnston, Dominic Kwiatkowski, Cordelia Langford, John Sillitoe on behalf of the Wellcome Sanger Institute COVID-19 Surveillance Team |
| EPI_ISL_3282154, EPI_ISL_3282155 | Lighthouse Lab in Milton Keynes | Wellcome Sanger Institute for the COVID-19 Genomics UK (COG-UK) Consortium | The Lighthouse Lab in Milton Keynes and Alex Alderton, Roberto Amato, Jeffrey Barrett, Sonia Goncalves, Ewan Harrison, David K. Jackson, Ian Johnston, Dominic Kwiatkowski, Cordelia Langford, John Sillitoe on behalf of the Wellcome Sanger Institute COVID-19 Surveillance Team |
| EPI_ISL_3282156 | Lighthouse Lab in Alderley Park | Wellcome Sanger Institute for the COVID-19 Genomics UK (COG-UK) Consortium | Jacquelyn Wynn, Mairead Hyland, The Lighthouse Lab in Alderley Park and Alex Alderton, Roberto Amato, Jeffrey Barrett, Sonia Goncalves, Ewan Harrison, David K. Jackson, Ian Johnston, Dominic Kwiatkowski, Cordelia Langford, John Sillitoe on behalf of the Wellcome Sanger Institute COVID-19 Surveillance Team |
| EPI_ISL_3282157, EPI_ISL_3282158 | Lighthouse Lab in Milton Keynes | Wellcome Sanger Institute for the COVID-19 Genomics UK (COG-UK) Consortium | The Lighthouse Lab in Milton Keynes and Alex Alderton, Roberto Amato, Jeffrey Barrett, Sonia Goncalves, Ewan Harrison, David K. Jackson, Ian Johnston, Dominic Kwiatkowski, Cordelia Langford, John Sillitoe on behalf of the Wellcome Sanger Institute COVID-19 Surveillance Team |
| EPI_ISL_3282159 | Lighthouse Lab in Alderley Park | Wellcome Sanger Institute for the COVID-19 Genomics UK (COG-UK) Consortium | Jacquelyn Wynn, Mairead Hyland, The Lighthouse Lab in Alderley Park and Alex Alderton, Roberto Amato, Jeffrey Barrett, Sonia Goncalves, Ewan Harrison, David K. Jackson, Ian Johnston, Dominic Kwiatkowski, Cordelia Langford, John Sillitoe on behalf of the Wellcome Sanger Institute COVID-19 Surveillance Team |
| EPI_ISL_3282160 | Lighthouse Lab in Milton Keynes | Wellcome Sanger Institute for the COVID-19 Genomics UK (COG-UK) Consortium | The Lighthouse Lab in Milton Keynes and Alex Alderton, Roberto Amato, Jeffrey Barrett, Sonia Goncalves, Ewan Harrison, David K. Jackson, Ian Johnston, Dominic Kwiatkowski, Cordelia Langford, John Sillitoe on behalf of the Wellcome Sanger Institute COVID-19 Surveillance Team |
| EPI_ISL_3282161, EPI_ISL_3282162 | Lighthouse Lab in Alderley Park | Wellcome Sanger Institute for the COVID-19 Genomics UK (COG-UK) Consortium | Jacquelyn Wynn, Mairead Hyland, The Lighthouse Lab in Alderley Park and Alex Alderton, Roberto Amato, Jeffrey Barrett, Sonia Goncalves, Ewan Harrison, David K. Jackson, Ian Johnston, Dominic Kwiatkowski, Cordelia Langford, John Sillitoe on behalf of the Wellcome Sanger Institute COVID-19 Surveillance Team |
| EPI_ISL_3282163, EPI_ISL_3282164 | Lighthouse Lab in Milton Keynes | Wellcome Sanger Institute for the COVID-19 Genomics UK (COG-UK) Consortium | The Lighthouse Lab in Milton Keynes and Alex Alderton, Roberto Amato, Jeffrey Barrett, Sonia Goncalves, Ewan Harrison, David K. Jackson, Ian Johnston, Dominic Kwiatkowski, Cordelia Langford, John Sillitoe on behalf of the Wellcome Sanger Institute COVID-19 Surveillance Team |
| EPI_ISL_3282165 | Lighthouse Lab in Alderley Park | Wellcome Sanger Institute for the COVID-19 Genomics UK (COG-UK) Consortium | Jacquelyn Wynn, Mairead Hyland, The Lighthouse Lab in Alderley Park and Alex Alderton, Roberto Amato, Jeffrey Barrett, Sonia Goncalves, Ewan Harrison, David K. Jackson, Ian Johnston, Dominic Kwiatkowski, Cordelia Langford, John Sillitoe on behalf of the Wellcome Sanger Institute COVID-19 Surveillance Team |
| EPI_ISL_3282166 | Lighthouse Lab in Milton Keynes | Wellcome Sanger Institute for the COVID-19 Genomics UK (COG-UK) Consortium | The Lighthouse Lab in Milton Keynes and Alex Alderton, Roberto Amato, Jeffrey Barrett, Sonia Goncalves, Ewan Harrison, David K. Jackson, Ian Johnston, Dominic Kwiatkowski, Cordelia Langford, John Sillitoe on behalf of the Wellcome Sanger Institute COVID-19 Surveillance Team |
| EPI_ISL_3282167, EPI_ISL_3282168 | Lighthouse Lab in Alderley Park | Wellcome Sanger Institute for the COVID-19 Genomics UK (COG-UK) Consortium | Jacquelyn Wynn, Mairead Hyland, The Lighthouse Lab in Alderley Park and Alex Alderton, Roberto Amato, Jeffrey Barrett, Sonia Goncalves, Ewan Harrison, David K. Jackson, Ian Johnston, Dominic Kwiatkowski, Cordelia Langford, John Sillitoe on behalf of the Wellcome Sanger Institute COVID-19 Surveillance Team |
| EPI_ISL_3282169 | Lighthouse Lab in Milton Keynes | Wellcome Sanger Institute for the COVID-19 Genomics UK (COG-UK) Consortium | The Lighthouse Lab in Milton Keynes and Alex Alderton, Roberto Amato, Jeffrey Barrett, Sonia Goncalves, Ewan Harrison, David K. Jackson, Ian Johnston, Dominic Kwiatkowski, Cordelia Langford, John Sillitoe on behalf of the Wellcome Sanger Institute COVID-19 Surveillance Team |
| EPI_ISL_3282170, EPI_ISL_3282171, EPI_ISL_3282172, EPI_ISL_3282173 | Lighthouse Lab in Alderley Park | Wellcome Sanger Institute for the COVID-19 Genomics UK (COG-UK) Consortium | Jacquelyn Wynn, Mairead Hyland, The Lighthouse Lab in Alderley Park and Alex Alderton, Roberto Amato, Jeffrey Barrett, Sonia Goncalves, Ewan Harrison, David K. Jackson, Ian Johnston, Dominic Kwiatkowski, Cordelia Langford, John Sillitoe on behalf of the Wellcome Sanger Institute COVID-19 Surveillance Team |
| EPI_ISL_3282174 | Lighthouse Lab in Milton Keynes | Wellcome Sanger Institute for the COVID-19 Genomics UK (COG-UK) Consortium | The Lighthouse Lab in Milton Keynes and Alex Alderton, Roberto Amato, Jeffrey Barrett, Sonia Goncalves, Ewan Harrison, David K. Jackson, Ian Johnston, Dominic Kwiatkowski, Cordelia Langford, John Sillitoe on behalf of the Wellcome Sanger Institute COVID-19 Surveillance Team |
| EPI_ISL_3282175 | Berkshire and Surrey Pathology Services Lighthouse Laboratory | Wellcome Sanger Institute for the COVID-19 Genomics UK (COG-UK) Consortium | Berkshire and Surrey Pathology Services Lighthouse Laboratory and Alex Alderton, Roberto Amato, Jeffrey Barrett, Sonia Goncalves, Ewan Harrison, David K. Jackson, Ian Johnston, Dominic Kwiatkowski, Cordelia Langford, John Sillitoe on behalf of the Wellcome Sanger Institute COVID-19 Surveillance Team |
| EPI_ISL_3282176, EPI_ISL_3282177 | Lighthouse Lab in Milton Keynes | Wellcome Sanger Institute for the COVID-19 Genomics UK (COG-UK) Consortium | The Lighthouse Lab in Milton Keynes and Alex Alderton, Roberto Amato, Jeffrey Barrett, Sonia Goncalves, Ewan Harrison, David K. Jackson, Ian Johnston, Dominic Kwiatkowski, Cordelia Langford, John Sillitoe on behalf of the Wellcome Sanger Institute COVID-19 Surveillance Team |

| | | | |
|---|---|---|---|
| EPI_ISL_3282178, EPI_ISL_3282179 | Lighthouse Lab in Alderley Park | Wellcome Sanger Institute for the COVID-19 Genomics UK (COG-UK) Consortium | Jacquelyn Wynn, Mairead Hyland, The Lighthouse Lab in Alderley Park and Alex Alderton, Roberto Amato, Jeffrey Barrett, Sonia Goncalves, Ewan Harrison, David K. Jackson, Ian Johnston, Dominic Kwiatkowski, Cordelia Langford, John Sillitoe on behalf of the Wellcome Sanger Institute COVID-19 Surveillance Team |
| EPI_ISL_3282180 | Berkshire and Surrey Pathology Services Lighthouse Laboratory | Wellcome Sanger Institute for the COVID-19 Genomics UK (COG-UK) Consortium | Berkshire and Surrey Pathology Services Lighthouse Laboratory and Alex Alderton, Roberto Amato, Jeffrey Barrett, Sonia Goncalves, Ewan Harrison, David K. Jackson, Ian Johnston, Dominic Kwiatkowski, Cordelia Langford, John Sillitoe on behalf of the Wellcome Sanger Institute COVID-19 Surveillance Team |
| EPI_ISL_3282181 | Lighthouse Lab in Alderley Park | Wellcome Sanger Institute for the COVID-19 Genomics UK (COG-UK) Consortium | Jacquelyn Wynn, Mairead Hyland, The Lighthouse Lab in Alderley Park and Alex Alderton, Roberto Amato, Jeffrey Barrett, Sonia Goncalves, Ewan Harrison, David K. Jackson, Ian Johnston, Dominic Kwiatkowski, Cordelia Langford, John Sillitoe on behalf of the Wellcome Sanger Institute COVID-19 Surveillance Team |
| EPI_ISL_3282182 | Lighthouse Lab in Milton Keynes | Wellcome Sanger Institute for the COVID-19 Genomics UK (COG-UK) Consortium | The Lighthouse Lab in Milton Keynes and Alex Alderton, Roberto Amato, Jeffrey Barrett, Sonia Goncalves, Ewan Harrison, David K. Jackson, Ian Johnston, Dominic Kwiatkowski, Cordelia Langford, John Sillitoe on behalf of the Wellcome Sanger Institute COVID-19 Surveillance Team |
| EPI_ISL_3282183, EPI_ISL_3282184, EPI_ISL_3282185, EPI_ISL_3282186, EPI_ISL_3282187, EPI_ISL_3282188 | Lighthouse Lab in Alderley Park | Wellcome Sanger Institute for the COVID-19 Genomics UK (COG-UK) Consortium | Jacquelyn Wynn, Mairead Hyland, The Lighthouse Lab in Alderley Park and Alex Alderton, Roberto Amato, Jeffrey Barrett, Sonia Goncalves, Ewan Harrison, David K. Jackson, Ian Johnston, Dominic Kwiatkowski, Cordelia Langford, John Sillitoe on behalf of the Wellcome Sanger Institute COVID-19 Surveillance Team |
| EPI_ISL_3282189, EPI_ISL_3282190, EPI_ISL_3282191 | Lighthouse Lab in Milton Keynes | Wellcome Sanger Institute for the COVID-19 Genomics UK (COG-UK) Consortium | The Lighthouse Lab in Milton Keynes and Alex Alderton, Roberto Amato, Jeffrey Barrett, Sonia Goncalves, Ewan Harrison, David K. Jackson, Ian Johnston, Dominic Kwiatkowski, Cordelia Langford, John Sillitoe on behalf of the Wellcome Sanger Institute COVID-19 Surveillance Team |
| EPI_ISL_3282192, EPI_ISL_3282193, EPI_ISL_3282194 | Lighthouse Lab in Alderley Park | Wellcome Sanger Institute for the COVID-19 Genomics UK (COG-UK) Consortium | Jacquelyn Wynn, Mairead Hyland, The Lighthouse Lab in Alderley Park and Alex Alderton, Roberto Amato, Jeffrey Barrett, Sonia Goncalves, Ewan Harrison, David K. Jackson, Ian Johnston, Dominic Kwiatkowski, Cordelia Langford, John Sillitoe on behalf of the Wellcome Sanger Institute COVID-19 Surveillance Team |
| EPI_ISL_3282195 | Lighthouse Lab in Milton Keynes | Wellcome Sanger Institute for the COVID-19 Genomics UK (COG-UK) Consortium | The Lighthouse Lab in Milton Keynes and Alex Alderton, Roberto Amato, Jeffrey Barrett, Sonia Goncalves, Ewan Harrison, David K. Jackson, Ian Johnston, Dominic Kwiatkowski, Cordelia Langford, John Sillitoe on behalf of the Wellcome Sanger Institute COVID-19 Surveillance Team |
| EPI_ISL_3282196, EPI_ISL_3282197 | Lighthouse Lab in Alderley Park | Wellcome Sanger Institute for the COVID-19 Genomics UK (COG-UK) Consortium | Jacquelyn Wynn, Mairead Hyland, The Lighthouse Lab in Alderley Park and Alex Alderton, Roberto Amato, Jeffrey Barrett, Sonia Goncalves, Ewan Harrison, David K. Jackson, Ian Johnston, Dominic Kwiatkowski, Cordelia Langford, John Sillitoe on behalf of the Wellcome Sanger Institute COVID-19 Surveillance Team |
| EPI_ISL_3282198 | Berkshire and Surrey Pathology Services Lighthouse Laboratory | Wellcome Sanger Institute for the COVID-19 Genomics UK (COG-UK) Consortium | Berkshire and Surrey Pathology Services Lighthouse Laboratory and Alex Alderton, Roberto Amato, Jeffrey Barrett, Sonia Goncalves, Ewan Harrison, David K. Jackson, Ian Johnston, Dominic Kwiatkowski, Cordelia Langford, John Sillitoe on behalf of the Wellcome Sanger Institute COVID-19 Surveillance Team |
| EPI_ISL_3282199 | Lighthouse Lab in Alderley Park | Wellcome Sanger Institute for the COVID-19 Genomics UK (COG-UK) Consortium | Jacquelyn Wynn, Mairead Hyland, The Lighthouse Lab in Alderley Park and Alex Alderton, Roberto Amato, Jeffrey Barrett, Sonia Goncalves, Ewan Harrison, David K. Jackson, Ian Johnston, Dominic Kwiatkowski, Cordelia Langford, John Sillitoe on behalf of the Wellcome Sanger Institute COVID-19 Surveillance Team |
| EPI_ISL_3282200, EPI_ISL_3282201 | Berkshire and Surrey Pathology Services Lighthouse Laboratory | Wellcome Sanger Institute for the COVID-19 Genomics UK (COG-UK) Consortium | Berkshire and Surrey Pathology Services Lighthouse Laboratory and Alex Alderton, Roberto Amato, Jeffrey Barrett, Sonia Goncalves, Ewan Harrison, David K. Jackson, Ian Johnston, Dominic Kwiatkowski, Cordelia Langford, John Sillitoe on behalf of the Wellcome Sanger Institute COVID-19 Surveillance Team |
| EPI_ISL_3282202, EPI_ISL_3282203, EPI_ISL_3282204 | Lighthouse Lab in Alderley Park | Wellcome Sanger Institute for the COVID-19 Genomics UK (COG-UK) Consortium | Jacquelyn Wynn, Mairead Hyland, The Lighthouse Lab in Alderley Park and Alex Alderton, Roberto Amato, Jeffrey Barrett, Sonia Goncalves, Ewan Harrison, David K. Jackson, Ian Johnston, Dominic Kwiatkowski, Cordelia Langford, John Sillitoe on behalf of the Wellcome Sanger Institute COVID-19 Surveillance Team |
| EPI_ISL_3282205, EPI_ISL_3282206 | Lighthouse Lab in Milton Keynes | Wellcome Sanger Institute for the COVID-19 Genomics UK (COG-UK) Consortium | The Lighthouse Lab in Milton Keynes and Alex Alderton, Roberto Amato, Jeffrey Barrett, Sonia Goncalves, Ewan Harrison, David K. Jackson, Ian Johnston, Dominic Kwiatkowski, Cordelia Langford, John Sillitoe on behalf of the Wellcome Sanger Institute COVID-19 Surveillance Team |
| EPI_ISL_3282207 | Lighthouse Lab in Alderley Park | Wellcome Sanger Institute for the COVID-19 Genomics UK (COG-UK) Consortium | Jacquelyn Wynn, Mairead Hyland, The Lighthouse Lab in Alderley Park and Alex Alderton, Roberto Amato, Jeffrey Barrett, Sonia Goncalves, Ewan Harrison, David K. Jackson, Ian Johnston, Dominic Kwiatkowski, Cordelia Langford, John Sillitoe on behalf of the Wellcome Sanger Institute COVID-19 Surveillance Team |
| EPI_ISL_3282208, EPI_ISL_3282209, EPI_ISL_3282210 | Lighthouse Lab in Milton Keynes | Wellcome Sanger Institute for the COVID-19 Genomics UK (COG-UK) Consortium | The Lighthouse Lab in Milton Keynes and Alex Alderton, Roberto Amato, Jeffrey Barrett, Sonia Goncalves, Ewan Harrison, David K. Jackson, Ian Johnston, Dominic Kwiatkowski, Cordelia Langford, John Sillitoe on behalf of the Wellcome Sanger Institute COVID-19 Surveillance Team |
| EPI_ISL_3282211 | Lighthouse Lab in Alderley Park | Wellcome Sanger Institute for the COVID-19 Genomics UK (COG-UK) Consortium | Jacquelyn Wynn, Mairead Hyland, The Lighthouse Lab in Alderley Park and Alex Alderton, Roberto Amato, Jeffrey Barrett, Sonia Goncalves, Ewan Harrison, David K. Jackson, Ian Johnston, Dominic Kwiatkowski, Cordelia Langford, John Sillitoe on behalf of the Wellcome Sanger Institute COVID-19 Surveillance Team |
| EPI_ISL_3282212 | Berkshire and Surrey Pathology Services Lighthouse Laboratory | Wellcome Sanger Institute for the COVID-19 Genomics UK (COG-UK) Consortium | Berkshire and Surrey Pathology Services Lighthouse Laboratory and Alex Alderton, Roberto Amato, Jeffrey Barrett, Sonia Goncalves, Ewan Harrison, David K. Jackson, Ian Johnston, Dominic Kwiatkowski, Cordelia Langford, John Sillitoe on behalf of the Wellcome Sanger Institute COVID-19 Surveillance Team |
| EPI_ISL_3282213 | Lighthouse Lab in Milton Keynes | Wellcome Sanger Institute for the COVID-19 Genomics UK (COG-UK) Consortium | The Lighthouse Lab in Milton Keynes and Alex Alderton, Roberto Amato, Jeffrey Barrett, Sonia Goncalves, Ewan Harrison, David K. Jackson, Ian Johnston, Dominic Kwiatkowski, Cordelia Langford, John Sillitoe on behalf of the Wellcome Sanger Institute COVID-19 Surveillance Team |
| EPI_ISL_3282215 | Berkshire and Surrey Pathology Services Lighthouse Laboratory | Wellcome Sanger Institute for the COVID-19 Genomics UK (COG-UK) Consortium | Berkshire and Surrey Pathology Services Lighthouse Laboratory and Alex Alderton, Roberto Amato, Jeffrey Barrett, Sonia Goncalves, Ewan Harrison, David K. Jackson, Ian Johnston, Dominic Kwiatkowski, Cordelia Langford, John Sillitoe on behalf of the Wellcome Sanger Institute COVID-19 Surveillance Team |
| EPI_ISL_3282216, EPI_ISL_3282217, EPI_ISL_3282218, EPI_ISL_3282219, EPI_ISL_3282220 | Lighthouse Lab in Alderley Park | Wellcome Sanger Institute for the COVID-19 Genomics UK (COG-UK) Consortium | Jacquelyn Wynn, Mairead Hyland, The Lighthouse Lab in Alderley Park and Alex Alderton, Roberto Amato, Jeffrey Barrett, Sonia Goncalves, Ewan Harrison, David K. Jackson, Ian Johnston, Dominic Kwiatkowski, Cordelia Langford, John Sillitoe on behalf of the Wellcome Sanger Institute COVID-19 Surveillance Team |
| EPI_ISL_3282221 | Lighthouse Lab in Milton Keynes | Wellcome Sanger Institute for the COVID-19 Genomics UK (COG-UK) Consortium | The Lighthouse Lab in Milton Keynes and Alex Alderton, Roberto Amato, Jeffrey Barrett, Sonia Goncalves, Ewan Harrison, David K. Jackson, Ian Johnston, Dominic Kwiatkowski, Cordelia Langford, John Sillitoe on behalf of the Wellcome Sanger Institute COVID-19 Surveillance Team |
| EPI_ISL_3282222, EPI_ISL_3282223, EPI_ISL_3282224, EPI_ISL_3282225 | Lighthouse Lab in Alderley Park | Wellcome Sanger Institute for the COVID-19 Genomics UK (COG-UK) Consortium | Jacquelyn Wynn, Mairead Hyland, The Lighthouse Lab in Alderley Park and Alex Alderton, Roberto Amato, Jeffrey Barrett, Sonia Goncalves, Ewan Harrison, David K. Jackson, Ian Johnston, Dominic Kwiatkowski, Cordelia Langford, John Sillitoe on behalf of the Wellcome Sanger Institute COVID-19 Surveillance Team |
| EPI_ISL_3282226 | Lighthouse Lab in Milton Keynes | Wellcome Sanger Institute for the COVID-19 Genomics UK (COG-UK) Consortium | The Lighthouse Lab in Milton Keynes and Alex Alderton, Roberto Amato, Jeffrey Barrett, Sonia Goncalves, Ewan Harrison, David K. Jackson, Ian Johnston, Dominic Kwiatkowski, Cordelia Langford, John Sillitoe on behalf of the Wellcome Sanger Institute COVID-19 Surveillance Team |
| EPI_ISL_3282227, EPI_ISL_3282228 | Lighthouse Lab in Alderley Park | Wellcome Sanger Institute for the COVID-19 Genomics UK (COG-UK) Consortium | Jacquelyn Wynn, Mairead Hyland, The Lighthouse Lab in Alderley Park and Alex Alderton, Roberto Amato, Jeffrey Barrett, Sonia Goncalves, Ewan Harrison, David K. Jackson, Ian Johnston, Dominic Kwiatkowski, Cordelia Langford, John Sillitoe on behalf of the Wellcome Sanger Institute COVID-19 Surveillance Team |
| EPI_ISL_3282229, EPI_ISL_3282230 | Lighthouse Lab in Milton Keynes | Wellcome Sanger Institute for the COVID-19 Genomics UK (COG-UK) Consortium | The Lighthouse Lab in Milton Keynes and Alex Alderton, Roberto Amato, Jeffrey Barrett, Sonia Goncalves, Ewan Harrison, David K. Jackson, Ian Johnston, Dominic Kwiatkowski, Cordelia Langford, John Sillitoe on behalf of the Wellcome Sanger Institute COVID-19 Surveillance Team |
| EPI_ISL_3282231 | Lighthouse Lab in Alderley Park | Wellcome Sanger Institute for the COVID-19 Genomics UK (COG-UK) Consortium | Jacquelyn Wynn, Mairead Hyland, The Lighthouse Lab in Alderley Park and Alex Alderton, Roberto Amato, Jeffrey Barrett, Sonia Goncalves, Ewan Harrison, David K. Jackson, Ian Johnston, Dominic Kwiatkowski, Cordelia Langford, John Sillitoe on behalf of the Wellcome Sanger Institute COVID-19 Surveillance Team |
| EPI_ISL_3282232 | Lighthouse Lab in Milton Keynes | Wellcome Sanger Institute for the COVID-19 Genomics UK (COG-UK) Consortium | The Lighthouse Lab in Milton Keynes and Alex Alderton, Roberto Amato, Jeffrey Barrett, Sonia Goncalves, Ewan Harrison, David K. Jackson, Ian Johnston, Dominic Kwiatkowski, Cordelia Langford, John Sillitoe on behalf of the Wellcome Sanger Institute COVID-19 Surveillance Team |
| EPI_ISL_3282233, EPI_ISL_3282234, EPI_ISL_3282235 | Lighthouse Lab in Alderley Park | Wellcome Sanger Institute for the COVID-19 Genomics UK (COG-UK) Consortium | Jacquelyn Wynn, Mairead Hyland, The Lighthouse Lab in Alderley Park and Alex Alderton, Roberto Amato, Jeffrey Barrett, Sonia Goncalves, Ewan Harrison, David K. Jackson, Ian Johnston, Dominic Kwiatkowski, Cordelia Langford, John Sillitoe on behalf of the Wellcome Sanger Institute COVID-19 Surveillance Team |
| EPI_ISL_3282236 | Lighthouse Lab in Milton Keynes | Wellcome Sanger Institute for the COVID-19 Genomics UK (COG-UK) Consortium | The Lighthouse Lab in Milton Keynes and Alex Alderton, Roberto Amato, Jeffrey Barrett, Sonia Goncalves, Ewan Harrison, David K. Jackson, Ian Johnston, Dominic Kwiatkowski, Cordelia Langford, John Sillitoe on behalf of the Wellcome Sanger Institute COVID-19 Surveillance Team |
| EPI_ISL_3282237 | Berkshire and Surrey Pathology Services Lighthouse Laboratory | Wellcome Sanger Institute for the COVID-19 Genomics UK (COG-UK) Consortium | Berkshire and Surrey Pathology Services Lighthouse Laboratory and Alex Alderton, Roberto Amato, Jeffrey Barrett, Sonia Goncalves, Ewan Harrison, David K. Jackson, Ian Johnston, Dominic Kwiatkowski, Cordelia Langford, John Sillitoe on behalf of the Wellcome Sanger Institute COVID-19 Surveillance Team |
| EPI_ISL_3282238, EPI_ISL_3282239, EPI_ISL_3282240, EPI_ISL_3282241 | Lighthouse Lab in Milton Keynes | Wellcome Sanger Institute for the COVID-19 Genomics UK (COG-UK) Consortium | The Lighthouse Lab in Milton Keynes and Alex Alderton, Roberto Amato, Jeffrey Barrett, Sonia Goncalves, Ewan Harrison, David K. Jackson, Ian Johnston, Dominic Kwiatkowski, Cordelia Langford, John Sillitoe on behalf of the Wellcome Sanger Institute COVID-19 Surveillance Team |
| EPI_ISL_3282242, EPI_ISL_3282243 | Lighthouse Lab in Alderley Park | Wellcome Sanger Institute for the COVID-19 Genomics UK (COG-UK) Consortium | Jacquelyn Wynn, Mairead Hyland, The Lighthouse Lab in Alderley Park and Alex Alderton, Roberto Amato, Jeffrey Barrett, Sonia Goncalves, Ewan Harrison, David K. Jackson, Ian Johnston, Dominic Kwiatkowski, Cordelia Langford, John Sillitoe on behalf of the Wellcome Sanger Institute COVID-19 Surveillance Team |
| EPI_ISL_3282244, EPI_ISL_3282245 | Lighthouse Lab in Milton Keynes | Wellcome Sanger Institute for the COVID-19 Genomics UK (COG-UK) Consortium | The Lighthouse Lab in Milton Keynes and Alex Alderton, Roberto Amato, Jeffrey Barrett, Sonia Goncalves, Ewan Harrison, David K. Jackson, Ian Johnston, Dominic Kwiatkowski, Cordelia Langford, John Sillitoe on behalf of the Wellcome Sanger Institute COVID-19 Surveillance Team |
| EPI_ISL_3282246, EPI_ISL_3282247 | Lighthouse Lab in Alderley Park | Wellcome Sanger Institute for the COVID-19 Genomics UK (COG-UK) Consortium | Jacquelyn Wynn, Mairead Hyland, The Lighthouse Lab in Alderley Park and Alex Alderton, Roberto Amato, Jeffrey Barrett, Sonia Goncalves, Ewan Harrison, David K. Jackson, Ian Johnston, Dominic Kwiatkowski, Cordelia Langford, John Sillitoe on behalf of the Wellcome Sanger Institute COVID-19 Surveillance Team |
| EPI_ISL_3282248 | Berkshire and Surrey Pathology Services Lighthouse Laboratory | Wellcome Sanger Institute for the COVID-19 Genomics UK (COG-UK) Consortium | Berkshire and Surrey Pathology Services Lighthouse Laboratory and Alex Alderton, Roberto Amato, Jeffrey Barrett, Sonia Goncalves, Ewan Harrison, David K. Jackson, Ian Johnston, Dominic Kwiatkowski, Cordelia Langford, John Sillitoe on behalf of the Wellcome Sanger Institute COVID-19 Surveillance Team |
| EPI_ISL_3282249 | Lighthouse Lab in Alderley Park | Wellcome Sanger Institute for the COVID-19 Genomics UK (COG-UK) Consortium | Jacquelyn Wynn, Mairead Hyland, The Lighthouse Lab in Alderley Park and Alex Alderton, Roberto Amato, Jeffrey Barrett, Sonia Goncalves, Ewan Harrison, David K. Jackson, Ian Johnston, Dominic Kwiatkowski, Cordelia Langford, John Sillitoe on behalf of the Wellcome Sanger Institute COVID-19 Surveillance Team |
| EPI_ISL_3282250 | Berkshire and Surrey Pathology Services Lighthouse Laboratory | Wellcome Sanger Institute for the COVID-19 Genomics UK (COG-UK) Consortium | Berkshire and Surrey Pathology Services Lighthouse Laboratory and Alex Alderton, Roberto Amato, Jeffrey Barrett, Sonia Goncalves, Ewan Harrison, David K. Jackson, Ian Johnston, Dominic Kwiatkowski, Cordelia Langford, John Sillitoe on behalf of the Wellcome Sanger Institute COVID-19 Surveillance Team |
| EPI_ISL_3282251 | Lighthouse Lab in Alderley Park | Wellcome Sanger Institute for the COVID-19 Genomics UK (COG-UK) | Jacquelyn Wynn, Mairead Hyland, The Lighthouse Lab in Alderley Park and Alex Alderton, Roberto Amato, Jeffrey Barrett, Sonia Goncalves, Ewan Harrison, David K. Jackson, Ian Johnston, Dominic Kwiatkowski, Cordelia Langford, John Sillitoe on |

| | | | |
|---|---|---|---|
| | | Consortium | behalf of the Wellcome Sanger Institute COVID-19 Surveillance Team |
| EPI_ISL_3282252 | Lighthouse Lab in Milton Keynes | Wellcome Sanger Institute for the COVID-19 Genomics UK (COG-UK) Consortium | The Lighthouse Lab in Milton Keynes and Alex Alderton, Roberto Amato, Jeffrey Barrett, Sonia Goncalves, Ewan Harrison, David K. Jackson, Ian Johnston, Dominic Kwiatkowski, Cordelia Langford, John Sillitoe on behalf of the Wellcome Sanger Institute COVID-19 Surveillance Team |
| EPI_ISL_3282253, EPI_ISL_3282254 | Lighthouse Lab in Alderley Park | Wellcome Sanger Institute for the COVID-19 Genomics UK (COG-UK) Consortium | Jacquelyn Wynn, Mairead Hyland, The Lighthouse Lab in Alderley Park and Alex Alderton, Roberto Amato, Jeffrey Barrett, Sonia Goncalves, Ewan Harrison, David K. Jackson, Ian Johnston, Dominic Kwiatkowski, Cordelia Langford, John Sillitoe on behalf of the Wellcome Sanger Institute COVID-19 Surveillance Team |
| EPI_ISL_3282255 | Berkshire and Surrey Pathology Services Lighthouse Laboratory | Wellcome Sanger Institute for the COVID-19 Genomics UK (COG-UK) Consortium | Berkshire and Surrey Pathology Services Lighthouse Laboratory and Alex Alderton, Roberto Amato, Jeffrey Barrett, Sonia Goncalves, Ewan Harrison, David K. Jackson, Ian Johnston, Dominic Kwiatkowski, Cordelia Langford, John Sillitoe on behalf of the Wellcome Sanger Institute COVID-19 Surveillance Team |
| EPI_ISL_3282256 | Lighthouse Lab in Alderley Park | Wellcome Sanger Institute for the COVID-19 Genomics UK (COG-UK) Consortium | Jacquelyn Wynn, Mairead Hyland, The Lighthouse Lab in Alderley Park and Alex Alderton, Roberto Amato, Jeffrey Barrett, Sonia Goncalves, Ewan Harrison, David K. Jackson, Ian Johnston, Dominic Kwiatkowski, Cordelia Langford, John Sillitoe on behalf of the Wellcome Sanger Institute COVID-19 Surveillance Team |
| EPI_ISL_3282257 | Lighthouse Lab in Milton Keynes | Wellcome Sanger Institute for the COVID-19 Genomics UK (COG-UK) Consortium | The Lighthouse Lab in Milton Keynes and Alex Alderton, Roberto Amato, Jeffrey Barrett, Sonia Goncalves, Ewan Harrison, David K. Jackson, Ian Johnston, Dominic Kwiatkowski, Cordelia Langford, John Sillitoe on behalf of the Wellcome Sanger Institute COVID-19 Surveillance Team |
| EPI_ISL_3282258 | Lighthouse Lab in Alderley Park | Wellcome Sanger Institute for the COVID-19 Genomics UK (COG-UK) Consortium | Jacquelyn Wynn, Mairead Hyland, The Lighthouse Lab in Alderley Park and Alex Alderton, Roberto Amato, Jeffrey Barrett, Sonia Goncalves, Ewan Harrison, David K. Jackson, Ian Johnston, Dominic Kwiatkowski, Cordelia Langford, John Sillitoe on behalf of the Wellcome Sanger Institute COVID-19 Surveillance Team |
| EPI_ISL_3282259 | Berkshire and Surrey Pathology Services Lighthouse Laboratory | Wellcome Sanger Institute for the COVID-19 Genomics UK (COG-UK) Consortium | Berkshire and Surrey Pathology Services Lighthouse Laboratory and Alex Alderton, Roberto Amato, Jeffrey Barrett, Sonia Goncalves, Ewan Harrison, David K. Jackson, Ian Johnston, Dominic Kwiatkowski, Cordelia Langford, John Sillitoe on behalf of the Wellcome Sanger Institute COVID-19 Surveillance Team |
| EPI_ISL_3282260 | Lighthouse Lab in Alderley Park | Wellcome Sanger Institute for the COVID-19 Genomics UK (COG-UK) Consortium | Jacquelyn Wynn, Mairead Hyland, The Lighthouse Lab in Alderley Park and Alex Alderton, Roberto Amato, Jeffrey Barrett, Sonia Goncalves, Ewan Harrison, David K. Jackson, Ian Johnston, Dominic Kwiatkowski, Cordelia Langford, John Sillitoe on behalf of the Wellcome Sanger Institute COVID-19 Surveillance Team |
| EPI_ISL_3282261 | Lighthouse Lab in Milton Keynes | Wellcome Sanger Institute for the COVID-19 Genomics UK (COG-UK) Consortium | The Lighthouse Lab in Milton Keynes and Alex Alderton, Roberto Amato, Jeffrey Barrett, Sonia Goncalves, Ewan Harrison, David K. Jackson, Ian Johnston, Dominic Kwiatkowski, Cordelia Langford, John Sillitoe on behalf of the Wellcome Sanger Institute COVID-19 Surveillance Team |
| EPI_ISL_3282262, EPI_ISL_3282263, EPI_ISL_3282264, EPI_ISL_3282265 | Lighthouse Lab in Alderley Park | Wellcome Sanger Institute for the COVID-19 Genomics UK (COG-UK) Consortium | Jacquelyn Wynn, Mairead Hyland, The Lighthouse Lab in Alderley Park and Alex Alderton, Roberto Amato, Jeffrey Barrett, Sonia Goncalves, Ewan Harrison, David K. Jackson, Ian Johnston, Dominic Kwiatkowski, Cordelia Langford, John Sillitoe on behalf of the Wellcome Sanger Institute COVID-19 Surveillance Team |
| EPI_ISL_3282266 | Berkshire and Surrey Pathology Services Lighthouse Laboratory | Wellcome Sanger Institute for the COVID-19 Genomics UK (COG-UK) Consortium | Berkshire and Surrey Pathology Services Lighthouse Laboratory and Alex Alderton, Roberto Amato, Jeffrey Barrett, Sonia Goncalves, Ewan Harrison, David K. Jackson, Ian Johnston, Dominic Kwiatkowski, Cordelia Langford, John Sillitoe on behalf of the Wellcome Sanger Institute COVID-19 Surveillance Team |
| EPI_ISL_3282267 | Lighthouse Lab in Alderley Park | Wellcome Sanger Institute for the COVID-19 Genomics UK (COG-UK) Consortium | Jacquelyn Wynn, Mairead Hyland, The Lighthouse Lab in Alderley Park and Alex Alderton, Roberto Amato, Jeffrey Barrett, Sonia Goncalves, Ewan Harrison, David K. Jackson, Ian Johnston, Dominic Kwiatkowski, Cordelia Langford, John Sillitoe on behalf of the Wellcome Sanger Institute COVID-19 Surveillance Team |
| EPI_ISL_3282268 | Lighthouse Lab in Milton Keynes | Wellcome Sanger Institute for the COVID-19 Genomics UK (COG-UK) Consortium | The Lighthouse Lab in Milton Keynes and Alex Alderton, Roberto Amato, Jeffrey Barrett, Sonia Goncalves, Ewan Harrison, David K. Jackson, Ian Johnston, Dominic Kwiatkowski, Cordelia Langford, John Sillitoe on behalf of the Wellcome Sanger Institute COVID-19 Surveillance Team |
| EPI_ISL_3282269, EPI_ISL_3282270, EPI_ISL_3282272, EPI_ISL_3282273, EPI_ISL_3282274 | Lighthouse Lab in Alderley Park | Wellcome Sanger Institute for the COVID-19 Genomics UK (COG-UK) Consortium | Jacquelyn Wynn, Mairead Hyland, The Lighthouse Lab in Alderley Park and Alex Alderton, Roberto Amato, Jeffrey Barrett, Sonia Goncalves, Ewan Harrison, David K. Jackson, Ian Johnston, Dominic Kwiatkowski, Cordelia Langford, John Sillitoe on behalf of the Wellcome Sanger Institute COVID-19 Surveillance Team |
| EPI_ISL_3282275, EPI_ISL_3282276, EPI_ISL_3282277 | Lighthouse Lab in Milton Keynes | Wellcome Sanger Institute for the COVID-19 Genomics UK (COG-UK) Consortium | The Lighthouse Lab in Milton Keynes and Alex Alderton, Roberto Amato, Jeffrey Barrett, Sonia Goncalves, Ewan Harrison, David K. Jackson, Ian Johnston, Dominic Kwiatkowski, Cordelia Langford, John Sillitoe on behalf of the Wellcome Sanger Institute COVID-19 Surveillance Team |
| EPI_ISL_3282278 | Lighthouse Lab in Alderley Park | Wellcome Sanger Institute for the COVID-19 Genomics UK (COG-UK) Consortium | Jacquelyn Wynn, Mairead Hyland, The Lighthouse Lab in Alderley Park and Alex Alderton, Roberto Amato, Jeffrey Barrett, Sonia Goncalves, Ewan Harrison, David K. Jackson, Ian Johnston, Dominic Kwiatkowski, Cordelia Langford, John Sillitoe on behalf of the Wellcome Sanger Institute COVID-19 Surveillance Team |
| EPI_ISL_3282279 | Berkshire and Surrey Pathology Services Lighthouse Laboratory | Wellcome Sanger Institute for the COVID-19 Genomics UK (COG-UK) Consortium | Berkshire and Surrey Pathology Services Lighthouse Laboratory and Alex Alderton, Roberto Amato, Jeffrey Barrett, Sonia Goncalves, Ewan Harrison, David K. Jackson, Ian Johnston, Dominic Kwiatkowski, Cordelia Langford, John Sillitoe on behalf of the Wellcome Sanger Institute COVID-19 Surveillance Team |
| EPI_ISL_3282280, EPI_ISL_3282281, EPI_ISL_3282282 | Lighthouse Lab in Alderley Park | Wellcome Sanger Institute for the COVID-19 Genomics UK (COG-UK) Consortium | Jacquelyn Wynn, Mairead Hyland, The Lighthouse Lab in Alderley Park and Alex Alderton, Roberto Amato, Jeffrey Barrett, Sonia Goncalves, Ewan Harrison, David K. Jackson, Ian Johnston, Dominic Kwiatkowski, Cordelia Langford, John Sillitoe on behalf of the Wellcome Sanger Institute COVID-19 Surveillance Team |
| EPI_ISL_3282283 | Lighthouse Lab in Milton Keynes | Wellcome Sanger Institute for the COVID-19 Genomics UK (COG-UK) Consortium | The Lighthouse Lab in Milton Keynes and Alex Alderton, Roberto Amato, Jeffrey Barrett, Sonia Goncalves, Ewan Harrison, David K. Jackson, Ian Johnston, Dominic Kwiatkowski, Cordelia Langford, John Sillitoe on behalf of the Wellcome Sanger Institute COVID-19 Surveillance Team |
| EPI_ISL_3282284 | Lighthouse Lab in Alderley Park | Wellcome Sanger Institute for the COVID-19 Genomics UK (COG-UK) Consortium | Jacquelyn Wynn, Mairead Hyland, The Lighthouse Lab in Alderley Park and Alex Alderton, Roberto Amato, Jeffrey Barrett, Sonia Goncalves, Ewan Harrison, David K. Jackson, Ian Johnston, Dominic Kwiatkowski, Cordelia Langford, John Sillitoe on behalf of the Wellcome Sanger Institute COVID-19 Surveillance Team |
| EPI_ISL_3282285 | Lighthouse Lab in Milton Keynes | Wellcome Sanger Institute for the COVID-19 Genomics UK (COG-UK) Consortium | The Lighthouse Lab in Milton Keynes and Alex Alderton, Roberto Amato, Jeffrey Barrett, Sonia Goncalves, Ewan Harrison, David K. Jackson, Ian Johnston, Dominic Kwiatkowski, Cordelia Langford, John Sillitoe on behalf of the Wellcome Sanger Institute COVID-19 Surveillance Team |
| EPI_ISL_3282286 | Berkshire and Surrey Pathology Services Lighthouse Laboratory | Wellcome Sanger Institute for the COVID-19 Genomics UK (COG-UK) Consortium | Berkshire and Surrey Pathology Services Lighthouse Laboratory and Alex Alderton, Roberto Amato, Jeffrey Barrett, Sonia Goncalves, Ewan Harrison, David K. Jackson, Ian Johnston, Dominic Kwiatkowski, Cordelia Langford, John Sillitoe on behalf of the Wellcome Sanger Institute COVID-19 Surveillance Team |
| EPI_ISL_3282287 | Lighthouse Lab in Alderley Park | Wellcome Sanger Institute for the COVID-19 Genomics UK (COG-UK) Consortium | Jacquelyn Wynn, Mairead Hyland, The Lighthouse Lab in Alderley Park and Alex Alderton, Roberto Amato, Jeffrey Barrett, Sonia Goncalves, Ewan Harrison, David K. Jackson, Ian Johnston, Dominic Kwiatkowski, Cordelia Langford, John Sillitoe on behalf of the Wellcome Sanger Institute COVID-19 Surveillance Team |
| EPI_ISL_3282288, EPI_ISL_3282289 | Lighthouse Lab in Milton Keynes | Wellcome Sanger Institute for the COVID-19 Genomics UK (COG-UK) Consortium | The Lighthouse Lab in Milton Keynes and Alex Alderton, Roberto Amato, Jeffrey Barrett, Sonia Goncalves, Ewan Harrison, David K. Jackson, Ian Johnston, Dominic Kwiatkowski, Cordelia Langford, John Sillitoe on behalf of the Wellcome Sanger Institute COVID-19 Surveillance Team |
| EPI_ISL_3282290 | Lighthouse Lab in Alderley Park | Wellcome Sanger Institute for the COVID-19 Genomics UK (COG-UK) Consortium | Jacquelyn Wynn, Mairead Hyland, The Lighthouse Lab in Alderley Park and Alex Alderton, Roberto Amato, Jeffrey Barrett, Sonia Goncalves, Ewan Harrison, David K. Jackson, Ian Johnston, Dominic Kwiatkowski, Cordelia Langford, John Sillitoe on behalf of the Wellcome Sanger Institute COVID-19 Surveillance Team |
| EPI_ISL_3282291 | Berkshire and Surrey Pathology Services Lighthouse Laboratory | Wellcome Sanger Institute for the COVID-19 Genomics UK (COG-UK) Consortium | Berkshire and Surrey Pathology Services Lighthouse Laboratory and Alex Alderton, Roberto Amato, Jeffrey Barrett, Sonia Goncalves, Ewan Harrison, David K. Jackson, Ian Johnston, Dominic Kwiatkowski, Cordelia Langford, John Sillitoe on behalf of the Wellcome Sanger Institute COVID-19 Surveillance Team |
| EPI_ISL_3282292 | Lighthouse Lab in Alderley Park | Wellcome Sanger Institute for the COVID-19 Genomics UK (COG-UK) Consortium | Jacquelyn Wynn, Mairead Hyland, The Lighthouse Lab in Alderley Park and Alex Alderton, Roberto Amato, Jeffrey Barrett, Sonia Goncalves, Ewan Harrison, David K. Jackson, Ian Johnston, Dominic Kwiatkowski, Cordelia Langford, John Sillitoe on behalf of the Wellcome Sanger Institute COVID-19 Surveillance Team |
| EPI_ISL_3282293 | Lighthouse Lab in Milton Keynes | Wellcome Sanger Institute for the COVID-19 Genomics UK (COG-UK) Consortium | The Lighthouse Lab in Milton Keynes and Alex Alderton, Roberto Amato, Jeffrey Barrett, Sonia Goncalves, Ewan Harrison, David K. Jackson, Ian Johnston, Dominic Kwiatkowski, Cordelia Langford, John Sillitoe on behalf of the Wellcome Sanger Institute COVID-19 Surveillance Team |
| EPI_ISL_3282294, EPI_ISL_3282295 | Lighthouse Lab in Alderley Park | Wellcome Sanger Institute for the COVID-19 Genomics UK (COG-UK) Consortium | Jacquelyn Wynn, Mairead Hyland, The Lighthouse Lab in Alderley Park and Alex Alderton, Roberto Amato, Jeffrey Barrett, Sonia Goncalves, Ewan Harrison, David K. Jackson, Ian Johnston, Dominic Kwiatkowski, Cordelia Langford, John Sillitoe on behalf of the Wellcome Sanger Institute COVID-19 Surveillance Team |
| EPI_ISL_3282296 | Berkshire and Surrey Pathology Services Lighthouse Laboratory | Wellcome Sanger Institute for the COVID-19 Genomics UK (COG-UK) Consortium | Berkshire and Surrey Pathology Services Lighthouse Laboratory and Alex Alderton, Roberto Amato, Jeffrey Barrett, Sonia Goncalves, Ewan Harrison, David K. Jackson, Ian Johnston, Dominic Kwiatkowski, Cordelia Langford, John Sillitoe on behalf of the Wellcome Sanger Institute COVID-19 Surveillance Team |
| EPI_ISL_3282297 | Lighthouse Lab in Milton Keynes | Wellcome Sanger Institute for the COVID-19 Genomics UK (COG-UK) Consortium | The Lighthouse Lab in Milton Keynes and Alex Alderton, Roberto Amato, Jeffrey Barrett, Sonia Goncalves, Ewan Harrison, David K. Jackson, Ian Johnston, Dominic Kwiatkowski, Cordelia Langford, John Sillitoe on behalf of the Wellcome Sanger Institute COVID-19 Surveillance Team |
| EPI_ISL_3282298, EPI_ISL_3282299, EPI_ISL_3282300, EPI_ISL_3282301, EPI_ISL_3282302 | Lighthouse Lab in Alderley Park | Wellcome Sanger Institute for the COVID-19 Genomics UK (COG-UK) Consortium | Jacquelyn Wynn, Mairead Hyland, The Lighthouse Lab in Alderley Park and Alex Alderton, Roberto Amato, Jeffrey Barrett, Sonia Goncalves, Ewan Harrison, David K. Jackson, Ian Johnston, Dominic Kwiatkowski, Cordelia Langford, John Sillitoe on behalf of the Wellcome Sanger Institute COVID-19 Surveillance Team |
| EPI_ISL_3282303 | Lighthouse Lab in Milton Keynes | Wellcome Sanger Institute for the COVID-19 Genomics UK (COG-UK) Consortium | The Lighthouse Lab in Milton Keynes and Alex Alderton, Roberto Amato, Jeffrey Barrett, Sonia Goncalves, Ewan Harrison, David K. Jackson, Ian Johnston, Dominic Kwiatkowski, Cordelia Langford, John Sillitoe on behalf of the Wellcome Sanger Institute COVID-19 Surveillance Team |
| EPI_ISL_3282304, EPI_ISL_3282305, EPI_ISL_3282306, EPI_ISL_3282307, EPI_ISL_3282308, EPI_ISL_3282309 | Lighthouse Lab in Alderley Park | Wellcome Sanger Institute for the COVID-19 Genomics UK (COG-UK) Consortium | Jacquelyn Wynn, Mairead Hyland, The Lighthouse Lab in Alderley Park and Alex Alderton, Roberto Amato, Jeffrey Barrett, Sonia Goncalves, Ewan Harrison, David K. Jackson, Ian Johnston, Dominic Kwiatkowski, Cordelia Langford, John Sillitoe on behalf of the Wellcome Sanger Institute COVID-19 Surveillance Team |
| EPI_ISL_3282310 | Lighthouse Lab in Milton Keynes | Wellcome Sanger Institute for the COVID-19 Genomics UK (COG-UK) Consortium | The Lighthouse Lab in Milton Keynes and Alex Alderton, Roberto Amato, Jeffrey Barrett, Sonia Goncalves, Ewan Harrison, David K. Jackson, Ian Johnston, Dominic Kwiatkowski, Cordelia Langford, John Sillitoe on behalf of the Wellcome Sanger Institute COVID-19 Surveillance Team |
| EPI_ISL_3282311, EPI_ISL_3282312 | Lighthouse Lab in Alderley Park | Wellcome Sanger Institute for the COVID-19 Genomics UK (COG-UK) Consortium | Jacquelyn Wynn, Mairead Hyland, The Lighthouse Lab in Alderley Park and Alex Alderton, Roberto Amato, Jeffrey Barrett, Sonia Goncalves, Ewan Harrison, David K. Jackson, Ian Johnston, Dominic Kwiatkowski, Cordelia Langford, John Sillitoe on behalf of the Wellcome Sanger Institute COVID-19 Surveillance Team |
| EPI_ISL_3282313, EPI_ISL_3282314 | Berkshire and Surrey Pathology Services Lighthouse Laboratory | Wellcome Sanger Institute for the COVID-19 Genomics UK (COG-UK) Consortium | Berkshire and Surrey Pathology Services Lighthouse Laboratory and Alex Alderton, Roberto Amato, Jeffrey Barrett, Sonia Goncalves, Ewan Harrison, David K. Jackson, Ian Johnston, Dominic Kwiatkowski, Cordelia Langford, John Sillitoe on behalf of the Wellcome Sanger Institute COVID-19 Surveillance Team |
| EPI_ISL_3282315 | Lighthouse Lab in Alderley Park | Wellcome Sanger Institute for the COVID-19 Genomics UK (COG-UK) Consortium | Jacquelyn Wynn, Mairead Hyland, The Lighthouse Lab in Alderley Park and Alex Alderton, Roberto Amato, Jeffrey Barrett, Sonia Goncalves, Ewan Harrison, David K. Jackson, Ian Johnston, Dominic Kwiatkowski, Cordelia Langford, John Sillitoe on behalf of the Wellcome Sanger Institute COVID-19 Surveillance Team |
| EPI_ISL_3282316 | Lighthouse Lab in Milton Keynes | Wellcome Sanger Institute for the COVID-19 Genomics UK (COG-UK) Consortium | The Lighthouse Lab in Milton Keynes and Alex Alderton, Roberto Amato, Jeffrey Barrett, Sonia Goncalves, Ewan Harrison, David K. Jackson, Ian Johnston, Dominic Kwiatkowski, Cordelia Langford, John Sillitoe on behalf of the Wellcome Sanger Institute COVID-19 Surveillance Team |

| | | | |
|---|---|---|---|
| EPI_ISL_3282317 | Lighthouse Lab in Alderley Park | Wellcome Sanger Institute for the COVID-19 Genomics UK (COG-UK) Consortium | Jacquelyn Wynn, Mairead Hyland, The Lighthouse Lab in Alderley Park and Alex Alderton, Roberto Amato, Jeffrey Barrett, Sonia Goncalves, Ewan Harrison, David K. Jackson, Ian Johnston, Dominic Kwiatkowski, Cordelia Langford, John Sillitoe on behalf of the Wellcome Sanger Institute COVID-19 Surveillance Team |
| EPI_ISL_3282318 | Berkshire and Surrey Pathology Services Lighthouse Laboratory | Wellcome Sanger Institute for the COVID-19 Genomics UK (COG-UK) Consortium | Berkshire and Surrey Pathology Services Lighthouse Laboratory and Alex Alderton, Roberto Amato, Jeffrey Barrett, Sonia Goncalves, Ewan Harrison, David K. Jackson, Ian Johnston, Dominic Kwiatkowski, Cordelia Langford, John Sillitoe on behalf of the Wellcome Sanger Institute COVID-19 Surveillance Team |
| EPI_ISL_3282319 | Lighthouse Lab in Alderley Park | Wellcome Sanger Institute for the COVID-19 Genomics UK (COG-UK) Consortium | Jacquelyn Wynn, Mairead Hyland, The Lighthouse Lab in Alderley Park and Alex Alderton, Roberto Amato, Jeffrey Barrett, Sonia Goncalves, Ewan Harrison, David K. Jackson, Ian Johnston, Dominic Kwiatkowski, Cordelia Langford, John Sillitoe on behalf of the Wellcome Sanger Institute COVID-19 Surveillance Team |
| EPI_ISL_3282320 | Berkshire and Surrey Pathology Services Lighthouse Laboratory | Wellcome Sanger Institute for the COVID-19 Genomics UK (COG-UK) Consortium | Berkshire and Surrey Pathology Services Lighthouse Laboratory and Alex Alderton, Roberto Amato, Jeffrey Barrett, Sonia Goncalves, Ewan Harrison, David K. Jackson, Ian Johnston, Dominic Kwiatkowski, Cordelia Langford, John Sillitoe on behalf of the Wellcome Sanger Institute COVID-19 Surveillance Team |
| EPI_ISL_3282321, EPI_ISL_3282322 | Lighthouse Lab in Alderley Park | Wellcome Sanger Institute for the COVID-19 Genomics UK (COG-UK) Consortium | Jacquelyn Wynn, Mairead Hyland, The Lighthouse Lab in Alderley Park and Alex Alderton, Roberto Amato, Jeffrey Barrett, Sonia Goncalves, Ewan Harrison, David K. Jackson, Ian Johnston, Dominic Kwiatkowski, Cordelia Langford, John Sillitoe on behalf of the Wellcome Sanger Institute COVID-19 Surveillance Team |
| EPI_ISL_3282323 | Berkshire and Surrey Pathology Services Lighthouse Laboratory | Wellcome Sanger Institute for the COVID-19 Genomics UK (COG-UK) Consortium | Berkshire and Surrey Pathology Services Lighthouse Laboratory and Alex Alderton, Roberto Amato, Jeffrey Barrett, Sonia Goncalves, Ewan Harrison, David K. Jackson, Ian Johnston, Dominic Kwiatkowski, Cordelia Langford, John Sillitoe on behalf of the Wellcome Sanger Institute COVID-19 Surveillance Team |
| EPI_ISL_3282324 | Lighthouse Lab in Alderley Park | Wellcome Sanger Institute for the COVID-19 Genomics UK (COG-UK) Consortium | Jacquelyn Wynn, Mairead Hyland, The Lighthouse Lab in Alderley Park and Alex Alderton, Roberto Amato, Jeffrey Barrett, Sonia Goncalves, Ewan Harrison, David K. Jackson, Ian Johnston, Dominic Kwiatkowski, Cordelia Langford, John Sillitoe on behalf of the Wellcome Sanger Institute COVID-19 Surveillance Team |
| EPI_ISL_3282325 | Berkshire and Surrey Pathology Services Lighthouse Laboratory | Wellcome Sanger Institute for the COVID-19 Genomics UK (COG-UK) Consortium | Berkshire and Surrey Pathology Services Lighthouse Laboratory and Alex Alderton, Roberto Amato, Jeffrey Barrett, Sonia Goncalves, Ewan Harrison, David K. Jackson, Ian Johnston, Dominic Kwiatkowski, Cordelia Langford, John Sillitoe on behalf of the Wellcome Sanger Institute COVID-19 Surveillance Team |
| EPI_ISL_3282326, EPI_ISL_3282327, EPI_ISL_3282328 | Lighthouse Lab in Alderley Park | Wellcome Sanger Institute for the COVID-19 Genomics UK (COG-UK) Consortium | Jacquelyn Wynn, Mairead Hyland, The Lighthouse Lab in Alderley Park and Alex Alderton, Roberto Amato, Jeffrey Barrett, Sonia Goncalves, Ewan Harrison, David K. Jackson, Ian Johnston, Dominic Kwiatkowski, Cordelia Langford, John Sillitoe on behalf of the Wellcome Sanger Institute COVID-19 Surveillance Team |
| EPI_ISL_3282329, EPI_ISL_3282330 | Lighthouse Lab in Milton Keynes | Wellcome Sanger Institute for the COVID-19 Genomics UK (COG-UK) Consortium | The Lighthouse Lab in Milton Keynes and Alex Alderton, Roberto Amato, Jeffrey Barrett, Sonia Goncalves, Ewan Harrison, David K. Jackson, Ian Johnston, Dominic Kwiatkowski, Cordelia Langford, John Sillitoe on behalf of the Wellcome Sanger Institute COVID-19 Surveillance Team |
| EPI_ISL_3282331 | Lighthouse Lab in Alderley Park | Wellcome Sanger Institute for the COVID-19 Genomics UK (COG-UK) Consortium | Jacquelyn Wynn, Mairead Hyland, The Lighthouse Lab in Alderley Park and Alex Alderton, Roberto Amato, Jeffrey Barrett, Sonia Goncalves, Ewan Harrison, David K. Jackson, Ian Johnston, Dominic Kwiatkowski, Cordelia Langford, John Sillitoe on behalf of the Wellcome Sanger Institute COVID-19 Surveillance Team |
| EPI_ISL_3282332, EPI_ISL_3282333, EPI_ISL_3282334 | Berkshire and Surrey Pathology Services Lighthouse Laboratory | Wellcome Sanger Institute for the COVID-19 Genomics UK (COG-UK) Consortium | Berkshire and Surrey Pathology Services Lighthouse Laboratory and Alex Alderton, Roberto Amato, Jeffrey Barrett, Sonia Goncalves, Ewan Harrison, David K. Jackson, Ian Johnston, Dominic Kwiatkowski, Cordelia Langford, John Sillitoe on behalf of the Wellcome Sanger Institute COVID-19 Surveillance Team |
| EPI_ISL_3282335 | Lighthouse Lab in Alderley Park | Wellcome Sanger Institute for the COVID-19 Genomics UK (COG-UK) Consortium | Jacquelyn Wynn, Mairead Hyland, The Lighthouse Lab in Alderley Park and Alex Alderton, Roberto Amato, Jeffrey Barrett, Sonia Goncalves, Ewan Harrison, David K. Jackson, Ian Johnston, Dominic Kwiatkowski, Cordelia Langford, John Sillitoe on behalf of the Wellcome Sanger Institute COVID-19 Surveillance Team |
| EPI_ISL_3282336 | Lighthouse Lab in Milton Keynes | Wellcome Sanger Institute for the COVID-19 Genomics UK (COG-UK) Consortium | The Lighthouse Lab in Milton Keynes and Alex Alderton, Roberto Amato, Jeffrey Barrett, Sonia Goncalves, Ewan Harrison, David K. Jackson, Ian Johnston, Dominic Kwiatkowski, Cordelia Langford, John Sillitoe on behalf of the Wellcome Sanger Institute COVID-19 Surveillance Team |
| EPI_ISL_3282337 | Berkshire and Surrey Pathology Services Lighthouse Laboratory | Wellcome Sanger Institute for the COVID-19 Genomics UK (COG-UK) Consortium | Berkshire and Surrey Pathology Services Lighthouse Laboratory and Alex Alderton, Roberto Amato, Jeffrey Barrett, Sonia Goncalves, Ewan Harrison, David K. Jackson, Ian Johnston, Dominic Kwiatkowski, Cordelia Langford, John Sillitoe on behalf of the Wellcome Sanger Institute COVID-19 Surveillance Team |
| EPI_ISL_3282338, EPI_ISL_3282339 | Lighthouse Lab in Alderley Park | Wellcome Sanger Institute for the COVID-19 Genomics UK (COG-UK) Consortium | Jacquelyn Wynn, Mairead Hyland, The Lighthouse Lab in Alderley Park and Alex Alderton, Roberto Amato, Jeffrey Barrett, Sonia Goncalves, Ewan Harrison, David K. Jackson, Ian Johnston, Dominic Kwiatkowski, Cordelia Langford, John Sillitoe on behalf of the Wellcome Sanger Institute COVID-19 Surveillance Team |
| EPI_ISL_3282340 | Berkshire and Surrey Pathology Services Lighthouse Laboratory | Wellcome Sanger Institute for the COVID-19 Genomics UK (COG-UK) Consortium | Berkshire and Surrey Pathology Services Lighthouse Laboratory and Alex Alderton, Roberto Amato, Jeffrey Barrett, Sonia Goncalves, Ewan Harrison, David K. Jackson, Ian Johnston, Dominic Kwiatkowski, Cordelia Langford, John Sillitoe on behalf of the Wellcome Sanger Institute COVID-19 Surveillance Team |
| EPI_ISL_3282341 | Lighthouse Lab in Alderley Park | Wellcome Sanger Institute for the COVID-19 Genomics UK (COG-UK) Consortium | Jacquelyn Wynn, Mairead Hyland, The Lighthouse Lab in Alderley Park and Alex Alderton, Roberto Amato, Jeffrey Barrett, Sonia Goncalves, Ewan Harrison, David K. Jackson, Ian Johnston, Dominic Kwiatkowski, Cordelia Langford, John Sillitoe on behalf of the Wellcome Sanger Institute COVID-19 Surveillance Team |
| EPI_ISL_3282342 | Berkshire and Surrey Pathology Services Lighthouse Laboratory | Wellcome Sanger Institute for the COVID-19 Genomics UK (COG-UK) Consortium | Berkshire and Surrey Pathology Services Lighthouse Laboratory and Alex Alderton, Roberto Amato, Jeffrey Barrett, Sonia Goncalves, Ewan Harrison, David K. Jackson, Ian Johnston, Dominic Kwiatkowski, Cordelia Langford, John Sillitoe on behalf of the Wellcome Sanger Institute COVID-19 Surveillance Team |
| EPI_ISL_3282343 | Lighthouse Lab in Milton Keynes | Wellcome Sanger Institute for the COVID-19 Genomics UK (COG-UK) Consortium | The Lighthouse Lab in Milton Keynes and Alex Alderton, Roberto Amato, Jeffrey Barrett, Sonia Goncalves, Ewan Harrison, David K. Jackson, Ian Johnston, Dominic Kwiatkowski, Cordelia Langford, John Sillitoe on behalf of the Wellcome Sanger Institute COVID-19 Surveillance Team |
| EPI_ISL_3282344, EPI_ISL_3282345 | Lighthouse Lab in Alderley Park | Wellcome Sanger Institute for the COVID-19 Genomics UK (COG-UK) Consortium | Jacquelyn Wynn, Mairead Hyland, The Lighthouse Lab in Alderley Park and Alex Alderton, Roberto Amato, Jeffrey Barrett, Sonia Goncalves, Ewan Harrison, David K. Jackson, Ian Johnston, Dominic Kwiatkowski, Cordelia Langford, John Sillitoe on behalf of the Wellcome Sanger Institute COVID-19 Surveillance Team |
| EPI_ISL_3282346 | Lighthouse Lab in Milton Keynes | Wellcome Sanger Institute for the COVID-19 Genomics UK (COG-UK) Consortium | The Lighthouse Lab in Milton Keynes and Alex Alderton, Roberto Amato, Jeffrey Barrett, Sonia Goncalves, Ewan Harrison, David K. Jackson, Ian Johnston, Dominic Kwiatkowski, Cordelia Langford, John Sillitoe on behalf of the Wellcome Sanger Institute COVID-19 Surveillance Team |
| EPI_ISL_3282347 | Lighthouse Lab in Alderley Park | Wellcome Sanger Institute for the COVID-19 Genomics UK (COG-UK) Consortium | Jacquelyn Wynn, Mairead Hyland, The Lighthouse Lab in Alderley Park and Alex Alderton, Roberto Amato, Jeffrey Barrett, Sonia Goncalves, Ewan Harrison, David K. Jackson, Ian Johnston, Dominic Kwiatkowski, Cordelia Langford, John Sillitoe on behalf of the Wellcome Sanger Institute COVID-19 Surveillance Team |
| EPI_ISL_3282348 | Lighthouse Lab in Milton Keynes | Wellcome Sanger Institute for the COVID-19 Genomics UK (COG-UK) Consortium | The Lighthouse Lab in Milton Keynes and Alex Alderton, Roberto Amato, Jeffrey Barrett, Sonia Goncalves, Ewan Harrison, David K. Jackson, Ian Johnston, Dominic Kwiatkowski, Cordelia Langford, John Sillitoe on behalf of the Wellcome Sanger Institute COVID-19 Surveillance Team |
| EPI_ISL_3282349 | Berkshire and Surrey Pathology Services Lighthouse Laboratory | Wellcome Sanger Institute for the COVID-19 Genomics UK (COG-UK) Consortium | Berkshire and Surrey Pathology Services Lighthouse Laboratory and Alex Alderton, Roberto Amato, Jeffrey Barrett, Sonia Goncalves, Ewan Harrison, David K. Jackson, Ian Johnston, Dominic Kwiatkowski, Cordelia Langford, John Sillitoe on behalf of the Wellcome Sanger Institute COVID-19 Surveillance Team |
| EPI_ISL_3282350, EPI_ISL_3282351 | Lighthouse Lab in Alderley Park | Wellcome Sanger Institute for the COVID-19 Genomics UK (COG-UK) Consortium | Jacquelyn Wynn, Mairead Hyland, The Lighthouse Lab in Alderley Park and Alex Alderton, Roberto Amato, Jeffrey Barrett, Sonia Goncalves, Ewan Harrison, David K. Jackson, Ian Johnston, Dominic Kwiatkowski, Cordelia Langford, John Sillitoe on behalf of the Wellcome Sanger Institute COVID-19 Surveillance Team |
| EPI_ISL_3282352 | Lighthouse Lab in Milton Keynes | Wellcome Sanger Institute for the COVID-19 Genomics UK (COG-UK) Consortium | The Lighthouse Lab in Milton Keynes and Alex Alderton, Roberto Amato, Jeffrey Barrett, Sonia Goncalves, Ewan Harrison, David K. Jackson, Ian Johnston, Dominic Kwiatkowski, Cordelia Langford, John Sillitoe on behalf of the Wellcome Sanger Institute COVID-19 Surveillance Team |
| EPI_ISL_3282353, EPI_ISL_3282354, EPI_ISL_3282355, EPI_ISL_3282356, EPI_ISL_3282357 | Lighthouse Lab in Alderley Park | Wellcome Sanger Institute for the COVID-19 Genomics UK (COG-UK) Consortium | Jacquelyn Wynn, Mairead Hyland, The Lighthouse Lab in Alderley Park and Alex Alderton, Roberto Amato, Jeffrey Barrett, Sonia Goncalves, Ewan Harrison, David K. Jackson, Ian Johnston, Dominic Kwiatkowski, Cordelia Langford, John Sillitoe on behalf of the Wellcome Sanger Institute COVID-19 Surveillance Team |
| EPI_ISL_3282358 | Lighthouse Lab in Milton Keynes | Wellcome Sanger Institute for the COVID-19 Genomics UK (COG-UK) Consortium | The Lighthouse Lab in Milton Keynes and Alex Alderton, Roberto Amato, Jeffrey Barrett, Sonia Goncalves, Ewan Harrison, David K. Jackson, Ian Johnston, Dominic Kwiatkowski, Cordelia Langford, John Sillitoe on behalf of the Wellcome Sanger Institute COVID-19 Surveillance Team |
| EPI_ISL_3282359, EPI_ISL_3282360 | Berkshire and Surrey Pathology Services Lighthouse Laboratory | Wellcome Sanger Institute for the COVID-19 Genomics UK (COG-UK) Consortium | Berkshire and Surrey Pathology Services Lighthouse Laboratory and Alex Alderton, Roberto Amato, Jeffrey Barrett, Sonia Goncalves, Ewan Harrison, David K. Jackson, Ian Johnston, Dominic Kwiatkowski, Cordelia Langford, John Sillitoe on behalf of the Wellcome Sanger Institute COVID-19 Surveillance Team |
| EPI_ISL_3282361, EPI_ISL_3282362 | Lighthouse Lab in Milton Keynes | Wellcome Sanger Institute for the COVID-19 Genomics UK (COG-UK) Consortium | The Lighthouse Lab in Milton Keynes and Alex Alderton, Roberto Amato, Jeffrey Barrett, Sonia Goncalves, Ewan Harrison, David K. Jackson, Ian Johnston, Dominic Kwiatkowski, Cordelia Langford, John Sillitoe on behalf of the Wellcome Sanger Institute COVID-19 Surveillance Team |
| EPI_ISL_3282363, EPI_ISL_3282364 | Lighthouse Lab in Alderley Park | Wellcome Sanger Institute for the COVID-19 Genomics UK (COG-UK) Consortium | Jacquelyn Wynn, Mairead Hyland, The Lighthouse Lab in Alderley Park and Alex Alderton, Roberto Amato, Jeffrey Barrett, Sonia Goncalves, Ewan Harrison, David K. Jackson, Ian Johnston, Dominic Kwiatkowski, Cordelia Langford, John Sillitoe on behalf of the Wellcome Sanger Institute COVID-19 Surveillance Team |
| EPI_ISL_3282365 | Lighthouse Lab in Milton Keynes | Wellcome Sanger Institute for the COVID-19 Genomics UK (COG-UK) Consortium | The Lighthouse Lab in Milton Keynes and Alex Alderton, Roberto Amato, Jeffrey Barrett, Sonia Goncalves, Ewan Harrison, David K. Jackson, Ian Johnston, Dominic Kwiatkowski, Cordelia Langford, John Sillitoe on behalf of the Wellcome Sanger Institute COVID-19 Surveillance Team |
| EPI_ISL_3282366, EPI_ISL_3282367, EPI_ISL_3282368, EPI_ISL_3282369, EPI_ISL_3282370, EPI_ISL_3282371, EPI_ISL_3282372, EPI_ISL_3282373, EPI_ISL_3282374, EPI_ISL_3282375 | Lighthouse Lab in Alderley Park | Wellcome Sanger Institute for the COVID-19 Genomics UK (COG-UK) Consortium | Jacquelyn Wynn, Mairead Hyland, The Lighthouse Lab in Alderley Park and Alex Alderton, Roberto Amato, Jeffrey Barrett, Sonia Goncalves, Ewan Harrison, David K. Jackson, Ian Johnston, Dominic Kwiatkowski, Cordelia Langford, John Sillitoe on behalf of the Wellcome Sanger Institute COVID-19 Surveillance Team |
| EPI_ISL_3282376 | Lighthouse Lab in Milton Keynes | Wellcome Sanger Institute for the COVID-19 Genomics UK (COG-UK) Consortium | The Lighthouse Lab in Milton Keynes and Alex Alderton, Roberto Amato, Jeffrey Barrett, Sonia Goncalves, Ewan Harrison, David K. Jackson, Ian Johnston, Dominic Kwiatkowski, Cordelia Langford, John Sillitoe on behalf of the Wellcome Sanger Institute COVID-19 Surveillance Team |
| EPI_ISL_3282377, EPI_ISL_3282378 | Lighthouse Lab in Alderley Park | Wellcome Sanger Institute for the COVID-19 Genomics UK (COG-UK) Consortium | Jacquelyn Wynn, Mairead Hyland, The Lighthouse Lab in Alderley Park and Alex Alderton, Roberto Amato, Jeffrey Barrett, Sonia Goncalves, Ewan Harrison, David K. Jackson, Ian Johnston, Dominic Kwiatkowski, Cordelia Langford, John Sillitoe on behalf of the Wellcome Sanger Institute COVID-19 Surveillance Team |
| EPI_ISL_3282379 | Lighthouse Lab in Milton Keynes | Wellcome Sanger Institute for the COVID-19 Genomics UK (COG-UK) Consortium | The Lighthouse Lab in Milton Keynes and Alex Alderton, Roberto Amato, Jeffrey Barrett, Sonia Goncalves, Ewan Harrison, David K. Jackson, Ian Johnston, Dominic Kwiatkowski, Cordelia Langford, John Sillitoe on behalf of the Wellcome Sanger Institute COVID-19 Surveillance Team |
| EPI_ISL_3282380, EPI_ISL_3282381, EPI_ISL_3282382 | Lighthouse Lab in Alderley Park | Wellcome Sanger Institute for the COVID-19 Genomics UK (COG-UK) Consortium | Jacquelyn Wynn, Mairead Hyland, The Lighthouse Lab in Alderley Park and Alex Alderton, Roberto Amato, Jeffrey Barrett, Sonia Goncalves, Ewan Harrison, David K. Jackson, Ian Johnston, Dominic Kwiatkowski, Cordelia Langford, John Sillitoe on behalf of the Wellcome Sanger Institute COVID-19 Surveillance Team |
| EPI_ISL_3282383 | Lighthouse Lab in Milton Keynes | Wellcome Sanger Institute for the COVID-19 Genomics UK (COG-UK) Consortium | The Lighthouse Lab in Milton Keynes and Alex Alderton, Roberto Amato, Jeffrey Barrett, Sonia Goncalves, Ewan Harrison, David K. Jackson, Ian Johnston, Dominic Kwiatkowski, Cordelia Langford, John Sillitoe on behalf of the Wellcome Sanger Institute COVID-19 Surveillance Team |
| EPI_ISL_3282384 | Lighthouse Lab in Alderley Park | Wellcome Sanger Institute for the COVID-19 Genomics UK (COG-UK) | Jacquelyn Wynn, Mairead Hyland, The Lighthouse Lab in Alderley Park and Alex Alderton, Roberto Amato, Jeffrey Barrett, Sonia Goncalves, Ewan Harrison, David K. Jackson, Ian Johnston, Dominic Kwiatkowski, Cordelia Langford, John Sillitoe on |

| | | | |
|---|---|---|---|
| | | Consortium | behalf of the Wellcome Sanger Institute COVID-19 Surveillance Team |
| EPI_ISL_3282385, EPI_ISL_3282386 | Lighthouse Lab in Milton Keynes | Wellcome Sanger Institute for the COVID-19 Genomics UK (COG-UK) Consortium | The Lighthouse Lab in Milton Keynes and Alex Alderton, Roberto Amato, Jeffrey Barrett, Sonia Goncalves, Ewan Harrison, David K. Jackson, Ian Johnston, Dominic Kwiatkowski, Cordelia Langford, John Sillitoe on behalf of the Wellcome Sanger Institute COVID-19 Surveillance Team |
| EPI_ISL_3282387, EPI_ISL_3282388, EPI_ISL_3282389 | Lighthouse Lab in Alderley Park | Wellcome Sanger Institute for the COVID-19 Genomics UK (COG-UK) Consortium | Jacquelyn Wynn, Mairead Hyland, The Lighthouse Lab in Alderley Park and Alex Alderton, Roberto Amato, Jeffrey Barrett, Sonia Goncalves, Ewan Harrison, David K. Jackson, Ian Johnston, Dominic Kwiatkowski, Cordelia Langford, John Sillitoe on behalf of the Wellcome Sanger Institute COVID-19 Surveillance Team |
| EPI_ISL_3282390 | Berkshire and Surrey Pathology Services Lighthouse Laboratory | Wellcome Sanger Institute for the COVID-19 Genomics UK (COG-UK) Consortium | Berkshire and Surrey Pathology Services Lighthouse Laboratory and Alex Alderton, Roberto Amato, Jeffrey Barrett, Sonia Goncalves, Ewan Harrison, David K. Jackson, Ian Johnston, Dominic Kwiatkowski, Cordelia Langford, John Sillitoe on behalf of the Wellcome Sanger Institute COVID-19 Surveillance Team |
| EPI_ISL_3282391, EPI_ISL_3282392, EPI_ISL_3282393, EPI_ISL_3282394, EPI_ISL_3282395 | Lighthouse Lab in Alderley Park | Wellcome Sanger Institute for the COVID-19 Genomics UK (COG-UK) Consortium | Jacquelyn Wynn, Mairead Hyland, The Lighthouse Lab in Alderley Park and Alex Alderton, Roberto Amato, Jeffrey Barrett, Sonia Goncalves, Ewan Harrison, David K. Jackson, Ian Johnston, Dominic Kwiatkowski, Cordelia Langford, John Sillitoe on behalf of the Wellcome Sanger Institute COVID-19 Surveillance Team |
| EPI_ISL_3282396, EPI_ISL_3282397, EPI_ISL_3282398 | Berkshire and Surrey Pathology Services Lighthouse Laboratory | Wellcome Sanger Institute for the COVID-19 Genomics UK (COG-UK) Consortium | Berkshire and Surrey Pathology Services Lighthouse Laboratory and Alex Alderton, Roberto Amato, Jeffrey Barrett, Sonia Goncalves, Ewan Harrison, David K. Jackson, Ian Johnston, Dominic Kwiatkowski, Cordelia Langford, John Sillitoe on behalf of the Wellcome Sanger Institute COVID-19 Surveillance Team |
| EPI_ISL_3282399 | Lighthouse Lab in Alderley Park | Wellcome Sanger Institute for the COVID-19 Genomics UK (COG-UK) Consortium | Jacquelyn Wynn, Mairead Hyland, The Lighthouse Lab in Alderley Park and Alex Alderton, Roberto Amato, Jeffrey Barrett, Sonia Goncalves, Ewan Harrison, David K. Jackson, Ian Johnston, Dominic Kwiatkowski, Cordelia Langford, John Sillitoe on behalf of the Wellcome Sanger Institute COVID-19 Surveillance Team |
| EPI_ISL_3282400, EPI_ISL_3282401 | Lighthouse Lab in Milton Keynes | Wellcome Sanger Institute for the COVID-19 Genomics UK (COG-UK) Consortium | The Lighthouse Lab in Milton Keynes and Alex Alderton, Roberto Amato, Jeffrey Barrett, Sonia Goncalves, Ewan Harrison, David K. Jackson, Ian Johnston, Dominic Kwiatkowski, Cordelia Langford, John Sillitoe on behalf of the Wellcome Sanger Institute COVID-19 Surveillance Team |
| EPI_ISL_3282402, EPI_ISL_3282403, EPI_ISL_3282404 | Lighthouse Lab in Alderley Park | Wellcome Sanger Institute for the COVID-19 Genomics UK (COG-UK) Consortium | Jacquelyn Wynn, Mairead Hyland, The Lighthouse Lab in Alderley Park and Alex Alderton, Roberto Amato, Jeffrey Barrett, Sonia Goncalves, Ewan Harrison, David K. Jackson, Ian Johnston, Dominic Kwiatkowski, Cordelia Langford, John Sillitoe on behalf of the Wellcome Sanger Institute COVID-19 Surveillance Team |
| EPI_ISL_3282405 | Berkshire and Surrey Pathology Services Lighthouse Laboratory | Wellcome Sanger Institute for the COVID-19 Genomics UK (COG-UK) Consortium | Berkshire and Surrey Pathology Services Lighthouse Laboratory and Alex Alderton, Roberto Amato, Jeffrey Barrett, Sonia Goncalves, Ewan Harrison, David K. Jackson, Ian Johnston, Dominic Kwiatkowski, Cordelia Langford, John Sillitoe on behalf of the Wellcome Sanger Institute COVID-19 Surveillance Team |
| EPI_ISL_3282406, EPI_ISL_3282407 | Lighthouse Lab in Alderley Park | Wellcome Sanger Institute for the COVID-19 Genomics UK (COG-UK) Consortium | Jacquelyn Wynn, Mairead Hyland, The Lighthouse Lab in Alderley Park and Alex Alderton, Roberto Amato, Jeffrey Barrett, Sonia Goncalves, Ewan Harrison, David K. Jackson, Ian Johnston, Dominic Kwiatkowski, Cordelia Langford, John Sillitoe on behalf of the Wellcome Sanger Institute COVID-19 Surveillance Team |
| EPI_ISL_3282408 | Lighthouse Lab in Milton Keynes | Wellcome Sanger Institute for the COVID-19 Genomics UK (COG-UK) Consortium | The Lighthouse Lab in Milton Keynes and Alex Alderton, Roberto Amato, Jeffrey Barrett, Sonia Goncalves, Ewan Harrison, David K. Jackson, Ian Johnston, Dominic Kwiatkowski, Cordelia Langford, John Sillitoe on behalf of the Wellcome Sanger Institute COVID-19 Surveillance Team |
| EPI_ISL_3282409 | Lighthouse Lab in Alderley Park | Wellcome Sanger Institute for the COVID-19 Genomics UK (COG-UK) Consortium | Jacquelyn Wynn, Mairead Hyland, The Lighthouse Lab in Alderley Park and Alex Alderton, Roberto Amato, Jeffrey Barrett, Sonia Goncalves, Ewan Harrison, David K. Jackson, Ian Johnston, Dominic Kwiatkowski, Cordelia Langford, John Sillitoe on behalf of the Wellcome Sanger Institute COVID-19 Surveillance Team |
| EPI_ISL_3282410 | Berkshire and Surrey Pathology Services Lighthouse Laboratory | Wellcome Sanger Institute for the COVID-19 Genomics UK (COG-UK) Consortium | Berkshire and Surrey Pathology Services Lighthouse Laboratory and Alex Alderton, Roberto Amato, Jeffrey Barrett, Sonia Goncalves, Ewan Harrison, David K. Jackson, Ian Johnston, Dominic Kwiatkowski, Cordelia Langford, John Sillitoe on behalf of the Wellcome Sanger Institute COVID-19 Surveillance Team |
| EPI_ISL_3282411 | Lighthouse Lab in Milton Keynes | Wellcome Sanger Institute for the COVID-19 Genomics UK (COG-UK) Consortium | The Lighthouse Lab in Milton Keynes and Alex Alderton, Roberto Amato, Jeffrey Barrett, Sonia Goncalves, Ewan Harrison, David K. Jackson, Ian Johnston, Dominic Kwiatkowski, Cordelia Langford, John Sillitoe on behalf of the Wellcome Sanger Institute COVID-19 Surveillance Team |
| EPI_ISL_3282412 | Lighthouse Lab in Alderley Park | Wellcome Sanger Institute for the COVID-19 Genomics UK (COG-UK) Consortium | Jacquelyn Wynn, Mairead Hyland, The Lighthouse Lab in Alderley Park and Alex Alderton, Roberto Amato, Jeffrey Barrett, Sonia Goncalves, Ewan Harrison, David K. Jackson, Ian Johnston, Dominic Kwiatkowski, Cordelia Langford, John Sillitoe on behalf of the Wellcome Sanger Institute COVID-19 Surveillance Team |
| EPI_ISL_3282413 | Berkshire and Surrey Pathology Services Lighthouse Laboratory | Wellcome Sanger Institute for the COVID-19 Genomics UK (COG-UK) Consortium | Berkshire and Surrey Pathology Services Lighthouse Laboratory and Alex Alderton, Roberto Amato, Jeffrey Barrett, Sonia Goncalves, Ewan Harrison, David K. Jackson, Ian Johnston, Dominic Kwiatkowski, Cordelia Langford, John Sillitoe on behalf of the Wellcome Sanger Institute COVID-19 Surveillance Team |
| EPI_ISL_3282414 | Lighthouse Lab in Alderley Park | Wellcome Sanger Institute for the COVID-19 Genomics UK (COG-UK) Consortium | Jacquelyn Wynn, Mairead Hyland, The Lighthouse Lab in Alderley Park and Alex Alderton, Roberto Amato, Jeffrey Barrett, Sonia Goncalves, Ewan Harrison, David K. Jackson, Ian Johnston, Dominic Kwiatkowski, Cordelia Langford, John Sillitoe on behalf of the Wellcome Sanger Institute COVID-19 Surveillance Team |
| EPI_ISL_3282415 | Lighthouse Lab in Milton Keynes | Wellcome Sanger Institute for the COVID-19 Genomics UK (COG-UK) Consortium | The Lighthouse Lab in Milton Keynes and Alex Alderton, Roberto Amato, Jeffrey Barrett, Sonia Goncalves, Ewan Harrison, David K. Jackson, Ian Johnston, Dominic Kwiatkowski, Cordelia Langford, John Sillitoe on behalf of the Wellcome Sanger Institute COVID-19 Surveillance Team |
| EPI_ISL_3282416 | Lighthouse Lab in Alderley Park | Wellcome Sanger Institute for the COVID-19 Genomics UK (COG-UK) Consortium | Jacquelyn Wynn, Mairead Hyland, The Lighthouse Lab in Alderley Park and Alex Alderton, Roberto Amato, Jeffrey Barrett, Sonia Goncalves, Ewan Harrison, David K. Jackson, Ian Johnston, Dominic Kwiatkowski, Cordelia Langford, John Sillitoe on behalf of the Wellcome Sanger Institute COVID-19 Surveillance Team |
| EPI_ISL_3282417 | Lighthouse Lab in Milton Keynes | Wellcome Sanger Institute for the COVID-19 Genomics UK (COG-UK) Consortium | The Lighthouse Lab in Milton Keynes and Alex Alderton, Roberto Amato, Jeffrey Barrett, Sonia Goncalves, Ewan Harrison, David K. Jackson, Ian Johnston, Dominic Kwiatkowski, Cordelia Langford, John Sillitoe on behalf of the Wellcome Sanger Institute COVID-19 Surveillance Team |
| EPI_ISL_3282418, EPI_ISL_3282419 | Lighthouse Lab in Alderley Park | Wellcome Sanger Institute for the COVID-19 Genomics UK (COG-UK) Consortium | Jacquelyn Wynn, Mairead Hyland, The Lighthouse Lab in Alderley Park and Alex Alderton, Roberto Amato, Jeffrey Barrett, Sonia Goncalves, Ewan Harrison, David K. Jackson, Ian Johnston, Dominic Kwiatkowski, Cordelia Langford, John Sillitoe on behalf of the Wellcome Sanger Institute COVID-19 Surveillance Team |
| EPI_ISL_3282420 | Berkshire and Surrey Pathology Services Lighthouse Laboratory | Wellcome Sanger Institute for the COVID-19 Genomics UK (COG-UK) Consortium | Berkshire and Surrey Pathology Services Lighthouse Laboratory and Alex Alderton, Roberto Amato, Jeffrey Barrett, Sonia Goncalves, Ewan Harrison, David K. Jackson, Ian Johnston, Dominic Kwiatkowski, Cordelia Langford, John Sillitoe on behalf of the Wellcome Sanger Institute COVID-19 Surveillance Team |
| EPI_ISL_3282421, EPI_ISL_3282422 | Lighthouse Lab in Milton Keynes | Wellcome Sanger Institute for the COVID-19 Genomics UK (COG-UK) Consortium | The Lighthouse Lab in Milton Keynes and Alex Alderton, Roberto Amato, Jeffrey Barrett, Sonia Goncalves, Ewan Harrison, David K. Jackson, Ian Johnston, Dominic Kwiatkowski, Cordelia Langford, John Sillitoe on behalf of the Wellcome Sanger Institute COVID-19 Surveillance Team |
| EPI_ISL_3282423 | Lighthouse Lab in Alderley Park | Wellcome Sanger Institute for the COVID-19 Genomics UK (COG-UK) Consortium | Jacquelyn Wynn, Mairead Hyland, The Lighthouse Lab in Alderley Park and Alex Alderton, Roberto Amato, Jeffrey Barrett, Sonia Goncalves, Ewan Harrison, David K. Jackson, Ian Johnston, Dominic Kwiatkowski, Cordelia Langford, John Sillitoe on behalf of the Wellcome Sanger Institute COVID-19 Surveillance Team |
| EPI_ISL_3282424, EPI_ISL_3282425 | Berkshire and Surrey Pathology Services Lighthouse Laboratory | Wellcome Sanger Institute for the COVID-19 Genomics UK (COG-UK) Consortium | Berkshire and Surrey Pathology Services Lighthouse Laboratory and Alex Alderton, Roberto Amato, Jeffrey Barrett, Sonia Goncalves, Ewan Harrison, David K. Jackson, Ian Johnston, Dominic Kwiatkowski, Cordelia Langford, John Sillitoe on behalf of the Wellcome Sanger Institute COVID-19 Surveillance Team |
| EPI_ISL_3282426 | Lighthouse Lab in Milton Keynes | Wellcome Sanger Institute for the COVID-19 Genomics UK (COG-UK) Consortium | The Lighthouse Lab in Milton Keynes and Alex Alderton, Roberto Amato, Jeffrey Barrett, Sonia Goncalves, Ewan Harrison, David K. Jackson, Ian Johnston, Dominic Kwiatkowski, Cordelia Langford, John Sillitoe on behalf of the Wellcome Sanger Institute COVID-19 Surveillance Team |
| EPI_ISL_3282427, EPI_ISL_3282428, EPI_ISL_3282429 | Lighthouse Lab in Alderley Park | Wellcome Sanger Institute for the COVID-19 Genomics UK (COG-UK) Consortium | Jacquelyn Wynn, Mairead Hyland, The Lighthouse Lab in Alderley Park and Alex Alderton, Roberto Amato, Jeffrey Barrett, Sonia Goncalves, Ewan Harrison, David K. Jackson, Ian Johnston, Dominic Kwiatkowski, Cordelia Langford, John Sillitoe on behalf of the Wellcome Sanger Institute COVID-19 Surveillance Team |
| EPI_ISL_3282430 | Lighthouse Lab in Milton Keynes | Wellcome Sanger Institute for the COVID-19 Genomics UK (COG-UK) Consortium | The Lighthouse Lab in Milton Keynes and Alex Alderton, Roberto Amato, Jeffrey Barrett, Sonia Goncalves, Ewan Harrison, David K. Jackson, Ian Johnston, Dominic Kwiatkowski, Cordelia Langford, John Sillitoe on behalf of the Wellcome Sanger Institute COVID-19 Surveillance Team |
| EPI_ISL_3282431 | Lighthouse Lab in Alderley Park | Wellcome Sanger Institute for the COVID-19 Genomics UK (COG-UK) Consortium | Jacquelyn Wynn, Mairead Hyland, The Lighthouse Lab in Alderley Park and Alex Alderton, Roberto Amato, Jeffrey Barrett, Sonia Goncalves, Ewan Harrison, David K. Jackson, Ian Johnston, Dominic Kwiatkowski, Cordelia Langford, John Sillitoe on behalf of the Wellcome Sanger Institute COVID-19 Surveillance Team |
| EPI_ISL_3282432 | Lighthouse Lab in Milton Keynes | Wellcome Sanger Institute for the COVID-19 Genomics UK (COG-UK) Consortium | The Lighthouse Lab in Milton Keynes and Alex Alderton, Roberto Amato, Jeffrey Barrett, Sonia Goncalves, Ewan Harrison, David K. Jackson, Ian Johnston, Dominic Kwiatkowski, Cordelia Langford, John Sillitoe on behalf of the Wellcome Sanger Institute COVID-19 Surveillance Team |
| EPI_ISL_3282433 | Lighthouse Lab in Alderley Park | Wellcome Sanger Institute for the COVID-19 Genomics UK (COG-UK) Consortium | Jacquelyn Wynn, Mairead Hyland, The Lighthouse Lab in Alderley Park and Alex Alderton, Roberto Amato, Jeffrey Barrett, Sonia Goncalves, Ewan Harrison, David K. Jackson, Ian Johnston, Dominic Kwiatkowski, Cordelia Langford, John Sillitoe on behalf of the Wellcome Sanger Institute COVID-19 Surveillance Team |
| EPI_ISL_3282434 | Lighthouse Lab in Milton Keynes | Wellcome Sanger Institute for the COVID-19 Genomics UK (COG-UK) Consortium | The Lighthouse Lab in Milton Keynes and Alex Alderton, Roberto Amato, Jeffrey Barrett, Sonia Goncalves, Ewan Harrison, David K. Jackson, Ian Johnston, Dominic Kwiatkowski, Cordelia Langford, John Sillitoe on behalf of the Wellcome Sanger Institute COVID-19 Surveillance Team |
| EPI_ISL_3282435 | Lighthouse Lab in Glasgow | Wellcome Sanger Institute for the COVID-19 Genomics UK (COG-UK) Consortium | Harper VanSteenhouse, Yumi Kasai, David Gray, Carol Clugston, Anna Dominiczak and Alex Alderton, Roberto Amato, Jeffrey Barrett, Sonia Goncalves, Ewan Harrison, David K. Jackson, Ian Johnston, Dominic Kwiatkowski, Cordelia Langford, John Sillitoe on behalf of the Wellcome Sanger Institute COVID-19 Surveillance Team |
| EPI_ISL_3282436 | Lighthouse Lab in Milton Keynes | Wellcome Sanger Institute for the COVID-19 Genomics UK (COG-UK) Consortium | The Lighthouse Lab in Milton Keynes and Alex Alderton, Roberto Amato, Jeffrey Barrett, Sonia Goncalves, Ewan Harrison, David K. Jackson, Ian Johnston, Dominic Kwiatkowski, Cordelia Langford, John Sillitoe on behalf of the Wellcome Sanger Institute COVID-19 Surveillance Team |
| EPI_ISL_3282437 | Lighthouse Lab in Alderley Park | Wellcome Sanger Institute for the COVID-19 Genomics UK (COG-UK) Consortium | Jacquelyn Wynn, Mairead Hyland, The Lighthouse Lab in Alderley Park and Alex Alderton, Roberto Amato, Jeffrey Barrett, Sonia Goncalves, Ewan Harrison, David K. Jackson, Ian Johnston, Dominic Kwiatkowski, Cordelia Langford, John Sillitoe on behalf of the Wellcome Sanger Institute COVID-19 Surveillance Team |
| EPI_ISL_3282438 | Lighthouse Lab in Milton Keynes | Wellcome Sanger Institute for the COVID-19 Genomics UK (COG-UK) Consortium | The Lighthouse Lab in Milton Keynes and Alex Alderton, Roberto Amato, Jeffrey Barrett, Sonia Goncalves, Ewan Harrison, David K. Jackson, Ian Johnston, Dominic Kwiatkowski, Cordelia Langford, John Sillitoe on behalf of the Wellcome Sanger Institute COVID-19 Surveillance Team |
| EPI_ISL_3282439, EPI_ISL_3282440 | Lighthouse Lab in Glasgow | Wellcome Sanger Institute for the COVID-19 Genomics UK (COG-UK) Consortium | Harper VanSteenhouse, Yumi Kasai, David Gray, Carol Clugston, Anna Dominiczak and Alex Alderton, Roberto Amato, Jeffrey Barrett, Sonia Goncalves, Ewan Harrison, David K. Jackson, Ian Johnston, Dominic Kwiatkowski, Cordelia Langford, John Sillitoe on behalf of the Wellcome Sanger Institute COVID-19 Surveillance Team |
| EPI_ISL_3282441 | Lighthouse Lab in Alderley Park | Wellcome Sanger Institute for the COVID-19 Genomics UK (COG-UK) Consortium | Jacquelyn Wynn, Mairead Hyland, The Lighthouse Lab in Alderley Park and Alex Alderton, Roberto Amato, Jeffrey Barrett, Sonia Goncalves, Ewan Harrison, David K. Jackson, Ian Johnston, Dominic Kwiatkowski, Cordelia Langford, John Sillitoe on behalf of the Wellcome Sanger Institute COVID-19 Surveillance Team |
| EPI_ISL_3282442 | Lighthouse Lab in Glasgow | Wellcome Sanger Institute for the COVID-19 Genomics UK (COG-UK) Consortium | Harper VanSteenhouse, Yumi Kasai, David Gray, Carol Clugston, Anna Dominiczak and Alex Alderton, Roberto Amato, Jeffrey Barrett, Sonia Goncalves, Ewan Harrison, David K. Jackson, Ian Johnston, Dominic Kwiatkowski, Cordelia Langford, John Sillitoe on behalf of the Wellcome Sanger Institute COVID-19 Surveillance Team |
| EPI_ISL_3282443, EPI_ISL_3282444 | Lighthouse Lab in Alderley Park | Wellcome Sanger Institute for the COVID-19 Genomics UK (COG-UK) Consortium | Jacquelyn Wynn, Mairead Hyland, The Lighthouse Lab in Alderley Park and Alex Alderton, Roberto Amato, Jeffrey Barrett, Sonia Goncalves, Ewan Harrison, David K. Jackson, Ian Johnston, Dominic Kwiatkowski, Cordelia Langford, John Sillitoe on behalf of the Wellcome Sanger Institute COVID-19 Surveillance Team |
| EPI_ISL_3282445, EPI_ISL_3282446, EPI_ISL_3282447 | Lighthouse Lab in Glasgow | Wellcome Sanger Institute for the COVID-19 Genomics UK (COG-UK) Consortium | Harper VanSteenhouse, Yumi Kasai, David Gray, Carol Clugston, Anna Dominiczak and Alex Alderton, Roberto Amato, Jeffrey Barrett, Sonia Goncalves, Ewan Harrison, David K. Jackson, Ian Johnston, Dominic Kwiatkowski, Cordelia Langford, John Sillitoe on behalf of the Wellcome Sanger Institute COVID-19 Surveillance Team |

| | | | |
|---|---|---|---|
| EPI_ISL_3282448 | Lighthouse Lab in Alderley Park | Wellcome Sanger Institute for the COVID-19 Genomics UK (COG-UK) Consortium | Jacquelyn Wynn, Mairead Hyland, The Lighthouse Lab in Alderley Park and Alex Alderton, Roberto Amato, Jeffrey Barrett, Sonia Goncalves, Ewan Harrison, David K. Jackson, Ian Johnston, Dominic Kwiatkowski, Cordelia Langford, John Sillitoe on behalf of the Wellcome Sanger Institute COVID-19 Surveillance Team |
| EPI_ISL_3282449, EPI_ISL_3282450, EPI_ISL_3282451, EPI_ISL_3282452, EPI_ISL_3282453 | Lighthouse Lab in Glasgow | Wellcome Sanger Institute for the COVID-19 Genomics UK (COG-UK) Consortium | Harper VanSteenhouse, Yumi Kasai, David Gray, Carol Clugston, Anna Dominiczak and Alex Alderton, Roberto Amato, Jeffrey Barrett, Sonia Goncalves, Ewan Harrison, David K. Jackson, Ian Johnston, Dominic Kwiatkowski, Cordelia Langford, John Sillitoe on behalf of the Wellcome Sanger Institute COVID-19 Surveillance Team |
| EPI_ISL_3282454 | Lighthouse Lab in Alderley Park | Wellcome Sanger Institute for the COVID-19 Genomics UK (COG-UK) Consortium | Jacquelyn Wynn, Mairead Hyland, The Lighthouse Lab in Alderley Park and Alex Alderton, Roberto Amato, Jeffrey Barrett, Sonia Goncalves, Ewan Harrison, David K. Jackson, Ian Johnston, Dominic Kwiatkowski, Cordelia Langford, John Sillitoe on behalf of the Wellcome Sanger Institute COVID-19 Surveillance Team |
| EPI_ISL_3282455, EPI_ISL_3282456 | Lighthouse Lab in Glasgow | Wellcome Sanger Institute for the COVID-19 Genomics UK (COG-UK) Consortium | Harper VanSteenhouse, Yumi Kasai, David Gray, Carol Clugston, Anna Dominiczak and Alex Alderton, Roberto Amato, Jeffrey Barrett, Sonia Goncalves, Ewan Harrison, David K. Jackson, Ian Johnston, Dominic Kwiatkowski, Cordelia Langford, John Sillitoe on behalf of the Wellcome Sanger Institute COVID-19 Surveillance Team |
| EPI_ISL_3282457, EPI_ISL_3282458 | Lighthouse Lab in Milton Keynes | Wellcome Sanger Institute for the COVID-19 Genomics UK (COG-UK) Consortium | The Lighthouse Lab in Milton Keynes and Alex Alderton, Roberto Amato, Jeffrey Barrett, Sonia Goncalves, Ewan Harrison, David K. Jackson, Ian Johnston, Dominic Kwiatkowski, Cordelia Langford, John Sillitoe on behalf of the Wellcome Sanger Institute COVID-19 Surveillance Team |
| EPI_ISL_3282459, EPI_ISL_3282460, EPI_ISL_3282461 | Lighthouse Lab in Glasgow | Wellcome Sanger Institute for the COVID-19 Genomics UK (COG-UK) Consortium | Harper VanSteenhouse, Yumi Kasai, David Gray, Carol Clugston, Anna Dominiczak and Alex Alderton, Roberto Amato, Jeffrey Barrett, Sonia Goncalves, Ewan Harrison, David K. Jackson, Ian Johnston, Dominic Kwiatkowski, Cordelia Langford, John Sillitoe on behalf of the Wellcome Sanger Institute COVID-19 Surveillance Team |
| EPI_ISL_3282462 | Lighthouse Lab in Milton Keynes | Wellcome Sanger Institute for the COVID-19 Genomics UK (COG-UK) Consortium | The Lighthouse Lab in Milton Keynes and Alex Alderton, Roberto Amato, Jeffrey Barrett, Sonia Goncalves, Ewan Harrison, David K. Jackson, Ian Johnston, Dominic Kwiatkowski, Cordelia Langford, John Sillitoe on behalf of the Wellcome Sanger Institute COVID-19 Surveillance Team |
| EPI_ISL_3282463 | Lighthouse Lab in Alderley Park | Wellcome Sanger Institute for the COVID-19 Genomics UK (COG-UK) Consortium | Jacquelyn Wynn, Mairead Hyland, The Lighthouse Lab in Alderley Park and Alex Alderton, Roberto Amato, Jeffrey Barrett, Sonia Goncalves, Ewan Harrison, David K. Jackson, Ian Johnston, Dominic Kwiatkowski, Cordelia Langford, John Sillitoe on behalf of the Wellcome Sanger Institute COVID-19 Surveillance Team |
| EPI_ISL_3282464, EPI_ISL_3282465 | Lighthouse Lab in Glasgow | Wellcome Sanger Institute for the COVID-19 Genomics UK (COG-UK) Consortium | Harper VanSteenhouse, Yumi Kasai, David Gray, Carol Clugston, Anna Dominiczak and Alex Alderton, Roberto Amato, Jeffrey Barrett, Sonia Goncalves, Ewan Harrison, David K. Jackson, Ian Johnston, Dominic Kwiatkowski, Cordelia Langford, John Sillitoe on behalf of the Wellcome Sanger Institute COVID-19 Surveillance Team |
| EPI_ISL_3282466, EPI_ISL_3282467, EPI_ISL_3282468 | Lighthouse Lab in Alderley Park | Wellcome Sanger Institute for the COVID-19 Genomics UK (COG-UK) Consortium | Jacquelyn Wynn, Mairead Hyland, The Lighthouse Lab in Alderley Park and Alex Alderton, Roberto Amato, Jeffrey Barrett, Sonia Goncalves, Ewan Harrison, David K. Jackson, Ian Johnston, Dominic Kwiatkowski, Cordelia Langford, John Sillitoe on behalf of the Wellcome Sanger Institute COVID-19 Surveillance Team |
| EPI_ISL_3282469 | Lighthouse Lab in Glasgow | Wellcome Sanger Institute for the COVID-19 Genomics UK (COG-UK) Consortium | Harper VanSteenhouse, Yumi Kasai, David Gray, Carol Clugston, Anna Dominiczak and Alex Alderton, Roberto Amato, Jeffrey Barrett, Sonia Goncalves, Ewan Harrison, David K. Jackson, Ian Johnston, Dominic Kwiatkowski, Cordelia Langford, John Sillitoe on behalf of the Wellcome Sanger Institute COVID-19 Surveillance Team |
| EPI_ISL_3282470 | Lighthouse Lab in Milton Keynes | Wellcome Sanger Institute for the COVID-19 Genomics UK (COG-UK) Consortium | The Lighthouse Lab in Milton Keynes and Alex Alderton, Roberto Amato, Jeffrey Barrett, Sonia Goncalves, Ewan Harrison, David K. Jackson, Ian Johnston, Dominic Kwiatkowski, Cordelia Langford, John Sillitoe on behalf of the Wellcome Sanger Institute COVID-19 Surveillance Team |
| EPI_ISL_3282471 | Lighthouse Lab in Glasgow | Wellcome Sanger Institute for the COVID-19 Genomics UK (COG-UK) Consortium | Harper VanSteenhouse, Yumi Kasai, David Gray, Carol Clugston, Anna Dominiczak and Alex Alderton, Roberto Amato, Jeffrey Barrett, Sonia Goncalves, Ewan Harrison, David K. Jackson, Ian Johnston, Dominic Kwiatkowski, Cordelia Langford, John Sillitoe on behalf of the Wellcome Sanger Institute COVID-19 Surveillance Team |
| EPI_ISL_3282472 | Lighthouse Lab in Alderley Park | Wellcome Sanger Institute for the COVID-19 Genomics UK (COG-UK) Consortium | Jacquelyn Wynn, Mairead Hyland, The Lighthouse Lab in Alderley Park and Alex Alderton, Roberto Amato, Jeffrey Barrett, Sonia Goncalves, Ewan Harrison, David K. Jackson, Ian Johnston, Dominic Kwiatkowski, Cordelia Langford, John Sillitoe on behalf of the Wellcome Sanger Institute COVID-19 Surveillance Team |
| EPI_ISL_3282473, EPI_ISL_3282474, EPI_ISL_3282475 | Lighthouse Lab in Milton Keynes | Wellcome Sanger Institute for the COVID-19 Genomics UK (COG-UK) Consortium | The Lighthouse Lab in Milton Keynes and Alex Alderton, Roberto Amato, Jeffrey Barrett, Sonia Goncalves, Ewan Harrison, David K. Jackson, Ian Johnston, Dominic Kwiatkowski, Cordelia Langford, John Sillitoe on behalf of the Wellcome Sanger Institute COVID-19 Surveillance Team |
| EPI_ISL_3282476 | Lighthouse Lab in Glasgow | Wellcome Sanger Institute for the COVID-19 Genomics UK (COG-UK) Consortium | Harper VanSteenhouse, Yumi Kasai, David Gray, Carol Clugston, Anna Dominiczak and Alex Alderton, Roberto Amato, Jeffrey Barrett, Sonia Goncalves, Ewan Harrison, David K. Jackson, Ian Johnston, Dominic Kwiatkowski, Cordelia Langford, John Sillitoe on behalf of the Wellcome Sanger Institute COVID-19 Surveillance Team |
| EPI_ISL_3282477 | Lighthouse Lab in Alderley Park | Wellcome Sanger Institute for the COVID-19 Genomics UK (COG-UK) Consortium | Jacquelyn Wynn, Mairead Hyland, The Lighthouse Lab in Alderley Park and Alex Alderton, Roberto Amato, Jeffrey Barrett, Sonia Goncalves, Ewan Harrison, David K. Jackson, Ian Johnston, Dominic Kwiatkowski, Cordelia Langford, John Sillitoe on behalf of the Wellcome Sanger Institute COVID-19 Surveillance Team |
| EPI_ISL_3282478 | Lighthouse Lab in Glasgow | Wellcome Sanger Institute for the COVID-19 Genomics UK (COG-UK) Consortium | Harper VanSteenhouse, Yumi Kasai, David Gray, Carol Clugston, Anna Dominiczak and Alex Alderton, Roberto Amato, Jeffrey Barrett, Sonia Goncalves, Ewan Harrison, David K. Jackson, Ian Johnston, Dominic Kwiatkowski, Cordelia Langford, John Sillitoe on behalf of the Wellcome Sanger Institute COVID-19 Surveillance Team |
| EPI_ISL_3282479 | Lighthouse Lab in Milton Keynes | Wellcome Sanger Institute for the COVID-19 Genomics UK (COG-UK) Consortium | The Lighthouse Lab in Milton Keynes and Alex Alderton, Roberto Amato, Jeffrey Barrett, Sonia Goncalves, Ewan Harrison, David K. Jackson, Ian Johnston, Dominic Kwiatkowski, Cordelia Langford, John Sillitoe on behalf of the Wellcome Sanger Institute COVID-19 Surveillance Team |
| EPI_ISL_3282480 | Lighthouse Lab in Alderley Park | Wellcome Sanger Institute for the COVID-19 Genomics UK (COG-UK) Consortium | Jacquelyn Wynn, Mairead Hyland, The Lighthouse Lab in Alderley Park and Alex Alderton, Roberto Amato, Jeffrey Barrett, Sonia Goncalves, Ewan Harrison, David K. Jackson, Ian Johnston, Dominic Kwiatkowski, Cordelia Langford, John Sillitoe on behalf of the Wellcome Sanger Institute COVID-19 Surveillance Team |
| EPI_ISL_3282481, EPI_ISL_3282482 | Lighthouse Lab in Glasgow | Wellcome Sanger Institute for the COVID-19 Genomics UK (COG-UK) Consortium | Harper VanSteenhouse, Yumi Kasai, David Gray, Carol Clugston, Anna Dominiczak and Alex Alderton, Roberto Amato, Jeffrey Barrett, Sonia Goncalves, Ewan Harrison, David K. Jackson, Ian Johnston, Dominic Kwiatkowski, Cordelia Langford, John Sillitoe on behalf of the Wellcome Sanger Institute COVID-19 Surveillance Team |
| EPI_ISL_3282483, EPI_ISL_3282484, EPI_ISL_3282485, EPI_ISL_3282486 | Lighthouse Lab in Alderley Park | Wellcome Sanger Institute for the COVID-19 Genomics UK (COG-UK) Consortium | Jacquelyn Wynn, Mairead Hyland, The Lighthouse Lab in Alderley Park and Alex Alderton, Roberto Amato, Jeffrey Barrett, Sonia Goncalves, Ewan Harrison, David K. Jackson, Ian Johnston, Dominic Kwiatkowski, Cordelia Langford, John Sillitoe on behalf of the Wellcome Sanger Institute COVID-19 Surveillance Team |
| EPI_ISL_3282487 | Lighthouse Lab in Glasgow | Wellcome Sanger Institute for the COVID-19 Genomics UK (COG-UK) Consortium | Harper VanSteenhouse, Yumi Kasai, David Gray, Carol Clugston, Anna Dominiczak and Alex Alderton, Roberto Amato, Jeffrey Barrett, Sonia Goncalves, Ewan Harrison, David K. Jackson, Ian Johnston, Dominic Kwiatkowski, Cordelia Langford, John Sillitoe on behalf of the Wellcome Sanger Institute COVID-19 Surveillance Team |
| EPI_ISL_3282488 | Lighthouse Lab in Milton Keynes | Wellcome Sanger Institute for the COVID-19 Genomics UK (COG-UK) Consortium | The Lighthouse Lab in Milton Keynes and Alex Alderton, Roberto Amato, Jeffrey Barrett, Sonia Goncalves, Ewan Harrison, David K. Jackson, Ian Johnston, Dominic Kwiatkowski, Cordelia Langford, John Sillitoe on behalf of the Wellcome Sanger Institute COVID-19 Surveillance Team |
| EPI_ISL_3282489 | Lighthouse Lab in Glasgow | Wellcome Sanger Institute for the COVID-19 Genomics UK (COG-UK) Consortium | Harper VanSteenhouse, Yumi Kasai, David Gray, Carol Clugston, Anna Dominiczak and Alex Alderton, Roberto Amato, Jeffrey Barrett, Sonia Goncalves, Ewan Harrison, David K. Jackson, Ian Johnston, Dominic Kwiatkowski, Cordelia Langford, John Sillitoe on behalf of the Wellcome Sanger Institute COVID-19 Surveillance Team |
| EPI_ISL_3282490, EPI_ISL_3282491, EPI_ISL_3282492 | Lighthouse Lab in Alderley Park | Wellcome Sanger Institute for the COVID-19 Genomics UK (COG-UK) Consortium | Jacquelyn Wynn, Mairead Hyland, The Lighthouse Lab in Alderley Park and Alex Alderton, Roberto Amato, Jeffrey Barrett, Sonia Goncalves, Ewan Harrison, David K. Jackson, Ian Johnston, Dominic Kwiatkowski, Cordelia Langford, John Sillitoe on behalf of the Wellcome Sanger Institute COVID-19 Surveillance Team |
| EPI_ISL_3282493, EPI_ISL_3282494 | Lighthouse Lab in Glasgow | Wellcome Sanger Institute for the COVID-19 Genomics UK (COG-UK) Consortium | Harper VanSteenhouse, Yumi Kasai, David Gray, Carol Clugston, Anna Dominiczak and Alex Alderton, Roberto Amato, Jeffrey Barrett, Sonia Goncalves, Ewan Harrison, David K. Jackson, Ian Johnston, Dominic Kwiatkowski, Cordelia Langford, John Sillitoe on behalf of the Wellcome Sanger Institute COVID-19 Surveillance Team |
| EPI_ISL_3282495 | Lighthouse Lab in Milton Keynes | Wellcome Sanger Institute for the COVID-19 Genomics UK (COG-UK) Consortium | The Lighthouse Lab in Milton Keynes and Alex Alderton, Roberto Amato, Jeffrey Barrett, Sonia Goncalves, Ewan Harrison, David K. Jackson, Ian Johnston, Dominic Kwiatkowski, Cordelia Langford, John Sillitoe on behalf of the Wellcome Sanger Institute COVID-19 Surveillance Team |
| EPI_ISL_3282496, EPI_ISL_3282497, EPI_ISL_3282498, EPI_ISL_3282499, EPI_ISL_3282500 | Lighthouse Lab in Glasgow | Wellcome Sanger Institute for the COVID-19 Genomics UK (COG-UK) Consortium | Harper VanSteenhouse, Yumi Kasai, David Gray, Carol Clugston, Anna Dominiczak and Alex Alderton, Roberto Amato, Jeffrey Barrett, Sonia Goncalves, Ewan Harrison, David K. Jackson, Ian Johnston, Dominic Kwiatkowski, Cordelia Langford, John Sillitoe on behalf of the Wellcome Sanger Institute COVID-19 Surveillance Team |
| EPI_ISL_3282501 | Lighthouse Lab in Milton Keynes | Wellcome Sanger Institute for the COVID-19 Genomics UK (COG-UK) Consortium | The Lighthouse Lab in Milton Keynes and Alex Alderton, Roberto Amato, Jeffrey Barrett, Sonia Goncalves, Ewan Harrison, David K. Jackson, Ian Johnston, Dominic Kwiatkowski, Cordelia Langford, John Sillitoe on behalf of the Wellcome Sanger Institute COVID-19 Surveillance Team |
| EPI_ISL_3282502 | Lighthouse Lab in Glasgow | Wellcome Sanger Institute for the COVID-19 Genomics UK (COG-UK) Consortium | Harper VanSteenhouse, Yumi Kasai, David Gray, Carol Clugston, Anna Dominiczak and Alex Alderton, Roberto Amato, Jeffrey Barrett, Sonia Goncalves, Ewan Harrison, David K. Jackson, Ian Johnston, Dominic Kwiatkowski, Cordelia Langford, John Sillitoe on behalf of the Wellcome Sanger Institute COVID-19 Surveillance Team |
| EPI_ISL_3282503, EPI_ISL_3282504 | Lighthouse Lab in Milton Keynes | Wellcome Sanger Institute for the COVID-19 Genomics UK (COG-UK) Consortium | The Lighthouse Lab in Milton Keynes and Alex Alderton, Roberto Amato, Jeffrey Barrett, Sonia Goncalves, Ewan Harrison, David K. Jackson, Ian Johnston, Dominic Kwiatkowski, Cordelia Langford, John Sillitoe on behalf of the Wellcome Sanger Institute COVID-19 Surveillance Team |
| EPI_ISL_3282505 | Lighthouse Lab in Glasgow | Wellcome Sanger Institute for the COVID-19 Genomics UK (COG-UK) Consortium | Harper VanSteenhouse, Yumi Kasai, David Gray, Carol Clugston, Anna Dominiczak and Alex Alderton, Roberto Amato, Jeffrey Barrett, Sonia Goncalves, Ewan Harrison, David K. Jackson, Ian Johnston, Dominic Kwiatkowski, Cordelia Langford, John Sillitoe on behalf of the Wellcome Sanger Institute COVID-19 Surveillance Team |
| EPI_ISL_3282506 | Lighthouse Lab in Alderley Park | Wellcome Sanger Institute for the COVID-19 Genomics UK (COG-UK) Consortium | Jacquelyn Wynn, Mairead Hyland, The Lighthouse Lab in Alderley Park and Alex Alderton, Roberto Amato, Jeffrey Barrett, Sonia Goncalves, Ewan Harrison, David K. Jackson, Ian Johnston, Dominic Kwiatkowski, Cordelia Langford, John Sillitoe on behalf of the Wellcome Sanger Institute COVID-19 Surveillance Team |
| EPI_ISL_3282507 | Lighthouse Lab in Glasgow | Wellcome Sanger Institute for the COVID-19 Genomics UK (COG-UK) Consortium | Harper VanSteenhouse, Yumi Kasai, David Gray, Carol Clugston, Anna Dominiczak and Alex Alderton, Roberto Amato, Jeffrey Barrett, Sonia Goncalves, Ewan Harrison, David K. Jackson, Ian Johnston, Dominic Kwiatkowski, Cordelia Langford, John Sillitoe on behalf of the Wellcome Sanger Institute COVID-19 Surveillance Team |
| EPI_ISL_3282508 | Lighthouse Lab in Alderley Park | Wellcome Sanger Institute for the COVID-19 Genomics UK (COG-UK) Consortium | Jacquelyn Wynn, Mairead Hyland, The Lighthouse Lab in Alderley Park and Alex Alderton, Roberto Amato, Jeffrey Barrett, Sonia Goncalves, Ewan Harrison, David K. Jackson, Ian Johnston, Dominic Kwiatkowski, Cordelia Langford, John Sillitoe on behalf of the Wellcome Sanger Institute COVID-19 Surveillance Team |
| EPI_ISL_3282509 | Lighthouse Lab in Glasgow | Wellcome Sanger Institute for the COVID-19 Genomics UK (COG-UK) Consortium | Harper VanSteenhouse, Yumi Kasai, David Gray, Carol Clugston, Anna Dominiczak and Alex Alderton, Roberto Amato, Jeffrey Barrett, Sonia Goncalves, Ewan Harrison, David K. Jackson, Ian Johnston, Dominic Kwiatkowski, Cordelia Langford, John Sillitoe on behalf of the Wellcome Sanger Institute COVID-19 Surveillance Team |
| EPI_ISL_3282510 | Lighthouse Lab in Milton Keynes | Wellcome Sanger Institute for the COVID-19 Genomics UK (COG-UK) Consortium | The Lighthouse Lab in Milton Keynes and Alex Alderton, Roberto Amato, Jeffrey Barrett, Sonia Goncalves, Ewan Harrison, David K. Jackson, Ian Johnston, Dominic Kwiatkowski, Cordelia Langford, John Sillitoe on behalf of the Wellcome Sanger Institute COVID-19 Surveillance Team |
| EPI_ISL_3282511, EPI_ISL_3282512, EPI_ISL_3282513 | Lighthouse Lab in Glasgow | Wellcome Sanger Institute for the COVID-19 Genomics UK (COG-UK) Consortium | Harper VanSteenhouse, Yumi Kasai, David Gray, Carol Clugston, Anna Dominiczak and Alex Alderton, Roberto Amato, Jeffrey Barrett, Sonia Goncalves, Ewan Harrison, David K. Jackson, Ian Johnston, Dominic Kwiatkowski, Cordelia Langford, John Sillitoe on behalf of the Wellcome Sanger Institute COVID-19 Surveillance Team |
| EPI_ISL_3282514 | Lighthouse Lab in Milton Keynes | Wellcome Sanger Institute for the COVID-19 Genomics UK (COG-UK) Consortium | The Lighthouse Lab in Milton Keynes and Alex Alderton, Roberto Amato, Jeffrey Barrett, Sonia Goncalves, Ewan Harrison, David K. Jackson, Ian Johnston, Dominic Kwiatkowski, Cordelia Langford, John Sillitoe on behalf of the Wellcome Sanger Institute COVID-19 Surveillance Team |

| | | | |
|---|---|---|---|
| EPI_ISL_3282515 | Lighthouse Lab in Glasgow | Wellcome Sanger Institute for the COVID-19 Genomics UK (COG-UK) Consortium | Harper VanSteenhouse, Yumi Kasai, David Gray, Carol Clugston, Anna Dominiczak and Alex Alderton, Roberto Amato, Jeffrey Barrett, Sonia Goncalves, Ewan Harrison, David K. Jackson, Ian Johnston, Dominic Kwiatkowski, Cordelia Langford, John Sillitoe on behalf of the Wellcome Sanger Institute COVID-19 Surveillance Team |
| EPI_ISL_3282516 | Lighthouse Lab in Milton Keynes | Wellcome Sanger Institute for the COVID-19 Genomics UK (COG-UK) Consortium | The Lighthouse Lab in Milton Keynes and Alex Alderton, Roberto Amato, Jeffrey Barrett, Sonia Goncalves, Ewan Harrison, David K. Jackson, Ian Johnston, Dominic Kwiatkowski, Cordelia Langford, John Sillitoe on behalf of the Wellcome Sanger Institute COVID-19 Surveillance Team |
| EPI_ISL_3282517 | Lighthouse Lab in Alderley Park | Wellcome Sanger Institute for the COVID-19 Genomics UK (COG-UK) Consortium | Jacquelyn Wynn, Mairead Hyland, The Lighthouse Lab in Alderley Park and Alex Alderton, Roberto Amato, Jeffrey Barrett, Sonia Goncalves, Ewan Harrison, David K. Jackson, Ian Johnston, Dominic Kwiatkowski, Cordelia Langford, John Sillitoe on behalf of the Wellcome Sanger Institute COVID-19 Surveillance Team |
| EPI_ISL_3282518 | Lighthouse Lab in Milton Keynes | Wellcome Sanger Institute for the COVID-19 Genomics UK (COG-UK) Consortium | The Lighthouse Lab in Milton Keynes and Alex Alderton, Roberto Amato, Jeffrey Barrett, Sonia Goncalves, Ewan Harrison, David K. Jackson, Ian Johnston, Dominic Kwiatkowski, Cordelia Langford, John Sillitoe on behalf of the Wellcome Sanger Institute COVID-19 Surveillance Team |
| EPI_ISL_3282519 | Lighthouse Lab in Glasgow | Wellcome Sanger Institute for the COVID-19 Genomics UK (COG-UK) Consortium | Harper VanSteenhouse, Yumi Kasai, David Gray, Carol Clugston, Anna Dominiczak and Alex Alderton, Roberto Amato, Jeffrey Barrett, Sonia Goncalves, Ewan Harrison, David K. Jackson, Ian Johnston, Dominic Kwiatkowski, Cordelia Langford, John Sillitoe on behalf of the Wellcome Sanger Institute COVID-19 Surveillance Team |
| EPI_ISL_3282520 | Lighthouse Lab in Milton Keynes | Wellcome Sanger Institute for the COVID-19 Genomics UK (COG-UK) Consortium | The Lighthouse Lab in Milton Keynes and Alex Alderton, Roberto Amato, Jeffrey Barrett, Sonia Goncalves, Ewan Harrison, David K. Jackson, Ian Johnston, Dominic Kwiatkowski, Cordelia Langford, John Sillitoe on behalf of the Wellcome Sanger Institute COVID-19 Surveillance Team |
| EPI_ISL_3282521 | Lighthouse Lab in Glasgow | Wellcome Sanger Institute for the COVID-19 Genomics UK (COG-UK) Consortium | Harper VanSteenhouse, Yumi Kasai, David Gray, Carol Clugston, Anna Dominiczak and Alex Alderton, Roberto Amato, Jeffrey Barrett, Sonia Goncalves, Ewan Harrison, David K. Jackson, Ian Johnston, Dominic Kwiatkowski, Cordelia Langford, John Sillitoe on behalf of the Wellcome Sanger Institute COVID-19 Surveillance Team |
| EPI_ISL_3282522, EPI_ISL_3282523 | Lighthouse Lab in Alderley Park | Wellcome Sanger Institute for the COVID-19 Genomics UK (COG-UK) Consortium | Jacquelyn Wynn, Mairead Hyland, The Lighthouse Lab in Alderley Park and Alex Alderton, Roberto Amato, Jeffrey Barrett, Sonia Goncalves, Ewan Harrison, David K. Jackson, Ian Johnston, Dominic Kwiatkowski, Cordelia Langford, John Sillitoe on behalf of the Wellcome Sanger Institute COVID-19 Surveillance Team |
| EPI_ISL_3282524 | Lighthouse Lab in Milton Keynes | Wellcome Sanger Institute for the COVID-19 Genomics UK (COG-UK) Consortium | The Lighthouse Lab in Milton Keynes and Alex Alderton, Roberto Amato, Jeffrey Barrett, Sonia Goncalves, Ewan Harrison, David K. Jackson, Ian Johnston, Dominic Kwiatkowski, Cordelia Langford, John Sillitoe on behalf of the Wellcome Sanger Institute COVID-19 Surveillance Team |
| EPI_ISL_3282525, EPI_ISL_3282526, EPI_ISL_3282527, EPI_ISL_3282528 | Lighthouse Lab in Glasgow | Wellcome Sanger Institute for the COVID-19 Genomics UK (COG-UK) Consortium | Harper VanSteenhouse, Yumi Kasai, David Gray, Carol Clugston, Anna Dominiczak and Alex Alderton, Roberto Amato, Jeffrey Barrett, Sonia Goncalves, Ewan Harrison, David K. Jackson, Ian Johnston, Dominic Kwiatkowski, Cordelia Langford, John Sillitoe on behalf of the Wellcome Sanger Institute COVID-19 Surveillance Team |
| EPI_ISL_3282529 | Lighthouse Lab in Alderley Park | Wellcome Sanger Institute for the COVID-19 Genomics UK (COG-UK) Consortium | Jacquelyn Wynn, Mairead Hyland, The Lighthouse Lab in Alderley Park and Alex Alderton, Roberto Amato, Jeffrey Barrett, Sonia Goncalves, Ewan Harrison, David K. Jackson, Ian Johnston, Dominic Kwiatkowski, Cordelia Langford, John Sillitoe on behalf of the Wellcome Sanger Institute COVID-19 Surveillance Team |
| EPI_ISL_3282530 | Lighthouse Lab in Glasgow | Wellcome Sanger Institute for the COVID-19 Genomics UK (COG-UK) Consortium | Harper VanSteenhouse, Yumi Kasai, David Gray, Carol Clugston, Anna Dominiczak and Alex Alderton, Roberto Amato, Jeffrey Barrett, Sonia Goncalves, Ewan Harrison, David K. Jackson, Ian Johnston, Dominic Kwiatkowski, Cordelia Langford, John Sillitoe on behalf of the Wellcome Sanger Institute COVID-19 Surveillance Team |
| EPI_ISL_3282531, EPI_ISL_3282532 | Lighthouse Lab in Milton Keynes | Wellcome Sanger Institute for the COVID-19 Genomics UK (COG-UK) Consortium | The Lighthouse Lab in Milton Keynes and Alex Alderton, Roberto Amato, Jeffrey Barrett, Sonia Goncalves, Ewan Harrison, David K. Jackson, Ian Johnston, Dominic Kwiatkowski, Cordelia Langford, John Sillitoe on behalf of the Wellcome Sanger Institute COVID-19 Surveillance Team |
| EPI_ISL_3282533, EPI_ISL_3282534 | Lighthouse Lab in Glasgow | Wellcome Sanger Institute for the COVID-19 Genomics UK (COG-UK) Consortium | Harper VanSteenhouse, Yumi Kasai, David Gray, Carol Clugston, Anna Dominiczak and Alex Alderton, Roberto Amato, Jeffrey Barrett, Sonia Goncalves, Ewan Harrison, David K. Jackson, Ian Johnston, Dominic Kwiatkowski, Cordelia Langford, John Sillitoe on behalf of the Wellcome Sanger Institute COVID-19 Surveillance Team |
| EPI_ISL_3282535 | Lighthouse Lab in Alderley Park | Wellcome Sanger Institute for the COVID-19 Genomics UK (COG-UK) Consortium | Jacquelyn Wynn, Mairead Hyland, The Lighthouse Lab in Alderley Park and Alex Alderton, Roberto Amato, Jeffrey Barrett, Sonia Goncalves, Ewan Harrison, David K. Jackson, Ian Johnston, Dominic Kwiatkowski, Cordelia Langford, John Sillitoe on behalf of the Wellcome Sanger Institute COVID-19 Surveillance Team |
| EPI_ISL_3282536 | Lighthouse Lab in Glasgow | Wellcome Sanger Institute for the COVID-19 Genomics UK (COG-UK) Consortium | Harper VanSteenhouse, Yumi Kasai, David Gray, Carol Clugston, Anna Dominiczak and Alex Alderton, Roberto Amato, Jeffrey Barrett, Sonia Goncalves, Ewan Harrison, David K. Jackson, Ian Johnston, Dominic Kwiatkowski, Cordelia Langford, John Sillitoe on behalf of the Wellcome Sanger Institute COVID-19 Surveillance Team |
| EPI_ISL_3282537 | Lighthouse Lab in Alderley Park | Wellcome Sanger Institute for the COVID-19 Genomics UK (COG-UK) Consortium | Jacquelyn Wynn, Mairead Hyland, The Lighthouse Lab in Alderley Park and Alex Alderton, Roberto Amato, Jeffrey Barrett, Sonia Goncalves, Ewan Harrison, David K. Jackson, Ian Johnston, Dominic Kwiatkowski, Cordelia Langford, John Sillitoe on behalf of the Wellcome Sanger Institute COVID-19 Surveillance Team |
| EPI_ISL_3282538 | Lighthouse Lab in Glasgow | Wellcome Sanger Institute for the COVID-19 Genomics UK (COG-UK) Consortium | Harper VanSteenhouse, Yumi Kasai, David Gray, Carol Clugston, Anna Dominiczak and Alex Alderton, Roberto Amato, Jeffrey Barrett, Sonia Goncalves, Ewan Harrison, David K. Jackson, Ian Johnston, Dominic Kwiatkowski, Cordelia Langford, John Sillitoe on behalf of the Wellcome Sanger Institute COVID-19 Surveillance Team |
| EPI_ISL_3282539, EPI_ISL_3282540 | Lighthouse Lab in Milton Keynes | Wellcome Sanger Institute for the COVID-19 Genomics UK (COG-UK) Consortium | The Lighthouse Lab in Milton Keynes and Alex Alderton, Roberto Amato, Jeffrey Barrett, Sonia Goncalves, Ewan Harrison, David K. Jackson, Ian Johnston, Dominic Kwiatkowski, Cordelia Langford, John Sillitoe on behalf of the Wellcome Sanger Institute COVID-19 Surveillance Team |
| EPI_ISL_3282541 | Lighthouse Lab in Alderley Park | Wellcome Sanger Institute for the COVID-19 Genomics UK (COG-UK) Consortium | Jacquelyn Wynn, Mairead Hyland, The Lighthouse Lab in Alderley Park and Alex Alderton, Roberto Amato, Jeffrey Barrett, Sonia Goncalves, Ewan Harrison, David K. Jackson, Ian Johnston, Dominic Kwiatkowski, Cordelia Langford, John Sillitoe on behalf of the Wellcome Sanger Institute COVID-19 Surveillance Team |
| EPI_ISL_3282542 | Lighthouse Lab in Milton Keynes | Wellcome Sanger Institute for the COVID-19 Genomics UK (COG-UK) Consortium | The Lighthouse Lab in Milton Keynes and Alex Alderton, Roberto Amato, Jeffrey Barrett, Sonia Goncalves, Ewan Harrison, David K. Jackson, Ian Johnston, Dominic Kwiatkowski, Cordelia Langford, John Sillitoe on behalf of the Wellcome Sanger Institute COVID-19 Surveillance Team |
| EPI_ISL_3282543, EPI_ISL_3282544, EPI_ISL_3282545 | Lighthouse Lab in Glasgow | Wellcome Sanger Institute for the COVID-19 Genomics UK (COG-UK) Consortium | Harper VanSteenhouse, Yumi Kasai, David Gray, Carol Clugston, Anna Dominiczak and Alex Alderton, Roberto Amato, Jeffrey Barrett, Sonia Goncalves, Ewan Harrison, David K. Jackson, Ian Johnston, Dominic Kwiatkowski, Cordelia Langford, John Sillitoe on behalf of the Wellcome Sanger Institute COVID-19 Surveillance Team |
| EPI_ISL_3282546 | Lighthouse Lab in Alderley Park | Wellcome Sanger Institute for the COVID-19 Genomics UK (COG-UK) Consortium | Jacquelyn Wynn, Mairead Hyland, The Lighthouse Lab in Alderley Park and Alex Alderton, Roberto Amato, Jeffrey Barrett, Sonia Goncalves, Ewan Harrison, David K. Jackson, Ian Johnston, Dominic Kwiatkowski, Cordelia Langford, John Sillitoe on behalf of the Wellcome Sanger Institute COVID-19 Surveillance Team |
| EPI_ISL_3282547 | Lighthouse Lab in Milton Keynes | Wellcome Sanger Institute for the COVID-19 Genomics UK (COG-UK) Consortium | The Lighthouse Lab in Milton Keynes and Alex Alderton, Roberto Amato, Jeffrey Barrett, Sonia Goncalves, Ewan Harrison, David K. Jackson, Ian Johnston, Dominic Kwiatkowski, Cordelia Langford, John Sillitoe on behalf of the Wellcome Sanger Institute COVID-19 Surveillance Team |
| EPI_ISL_3282548 | Lighthouse Lab in Glasgow | Wellcome Sanger Institute for the COVID-19 Genomics UK (COG-UK) Consortium | Harper VanSteenhouse, Yumi Kasai, David Gray, Carol Clugston, Anna Dominiczak and Alex Alderton, Roberto Amato, Jeffrey Barrett, Sonia Goncalves, Ewan Harrison, David K. Jackson, Ian Johnston, Dominic Kwiatkowski, Cordelia Langford, John Sillitoe on behalf of the Wellcome Sanger Institute COVID-19 Surveillance Team |
| EPI_ISL_3282549, EPI_ISL_3282550 | Lighthouse Lab in Alderley Park | Wellcome Sanger Institute for the COVID-19 Genomics UK (COG-UK) Consortium | Jacquelyn Wynn, Mairead Hyland, The Lighthouse Lab in Alderley Park and Alex Alderton, Roberto Amato, Jeffrey Barrett, Sonia Goncalves, Ewan Harrison, David K. Jackson, Ian Johnston, Dominic Kwiatkowski, Cordelia Langford, John Sillitoe on behalf of the Wellcome Sanger Institute COVID-19 Surveillance Team |
| EPI_ISL_3282551 | Lighthouse Lab in Glasgow | Wellcome Sanger Institute for the COVID-19 Genomics UK (COG-UK) Consortium | Harper VanSteenhouse, Yumi Kasai, David Gray, Carol Clugston, Anna Dominiczak and Alex Alderton, Roberto Amato, Jeffrey Barrett, Sonia Goncalves, Ewan Harrison, David K. Jackson, Ian Johnston, Dominic Kwiatkowski, Cordelia Langford, John Sillitoe on behalf of the Wellcome Sanger Institute COVID-19 Surveillance Team |
| EPI_ISL_3282552, EPI_ISL_3282553 | Lighthouse Lab in Milton Keynes | Wellcome Sanger Institute for the COVID-19 Genomics UK (COG-UK) Consortium | The Lighthouse Lab in Milton Keynes and Alex Alderton, Roberto Amato, Jeffrey Barrett, Sonia Goncalves, Ewan Harrison, David K. Jackson, Ian Johnston, Dominic Kwiatkowski, Cordelia Langford, John Sillitoe on behalf of the Wellcome Sanger Institute COVID-19 Surveillance Team |
| EPI_ISL_3282554 | Lighthouse Lab in Alderley Park | Wellcome Sanger Institute for the COVID-19 Genomics UK (COG-UK) Consortium | Jacquelyn Wynn, Mairead Hyland, The Lighthouse Lab in Alderley Park and Alex Alderton, Roberto Amato, Jeffrey Barrett, Sonia Goncalves, Ewan Harrison, David K. Jackson, Ian Johnston, Dominic Kwiatkowski, Cordelia Langford, John Sillitoe on behalf of the Wellcome Sanger Institute COVID-19 Surveillance Team |
| EPI_ISL_3282555, EPI_ISL_3282556, EPI_ISL_3282557, EPI_ISL_3282558 | Lighthouse Lab in Glasgow | Wellcome Sanger Institute for the COVID-19 Genomics UK (COG-UK) Consortium | Harper VanSteenhouse, Yumi Kasai, David Gray, Carol Clugston, Anna Dominiczak and Alex Alderton, Roberto Amato, Jeffrey Barrett, Sonia Goncalves, Ewan Harrison, David K. Jackson, Ian Johnston, Dominic Kwiatkowski, Cordelia Langford, John Sillitoe on behalf of the Wellcome Sanger Institute COVID-19 Surveillance Team |
| EPI_ISL_3282559 | Lighthouse Lab in Milton Keynes | Wellcome Sanger Institute for the COVID-19 Genomics UK (COG-UK) Consortium | The Lighthouse Lab in Milton Keynes and Alex Alderton, Roberto Amato, Jeffrey Barrett, Sonia Goncalves, Ewan Harrison, David K. Jackson, Ian Johnston, Dominic Kwiatkowski, Cordelia Langford, John Sillitoe on behalf of the Wellcome Sanger Institute COVID-19 Surveillance Team |
| EPI_ISL_3282560 | Lighthouse Lab in Glasgow | Wellcome Sanger Institute for the COVID-19 Genomics UK (COG-UK) Consortium | Harper VanSteenhouse, Yumi Kasai, David Gray, Carol Clugston, Anna Dominiczak and Alex Alderton, Roberto Amato, Jeffrey Barrett, Sonia Goncalves, Ewan Harrison, David K. Jackson, Ian Johnston, Dominic Kwiatkowski, Cordelia Langford, John Sillitoe on behalf of the Wellcome Sanger Institute COVID-19 Surveillance Team |
| EPI_ISL_3282561 | Lighthouse Lab in Milton Keynes | Wellcome Sanger Institute for the COVID-19 Genomics UK (COG-UK) Consortium | The Lighthouse Lab in Milton Keynes and Alex Alderton, Roberto Amato, Jeffrey Barrett, Sonia Goncalves, Ewan Harrison, David K. Jackson, Ian Johnston, Dominic Kwiatkowski, Cordelia Langford, John Sillitoe on behalf of the Wellcome Sanger Institute COVID-19 Surveillance Team |
| EPI_ISL_3282562, EPI_ISL_3282563 | Lighthouse Lab in Glasgow | Wellcome Sanger Institute for the COVID-19 Genomics UK (COG-UK) Consortium | Harper VanSteenhouse, Yumi Kasai, David Gray, Carol Clugston, Anna Dominiczak and Alex Alderton, Roberto Amato, Jeffrey Barrett, Sonia Goncalves, Ewan Harrison, David K. Jackson, Ian Johnston, Dominic Kwiatkowski, Cordelia Langford, John Sillitoe on behalf of the Wellcome Sanger Institute COVID-19 Surveillance Team |
| EPI_ISL_3282564, EPI_ISL_3282565, EPI_ISL_3282566 | Lighthouse Lab in Alderley Park | Wellcome Sanger Institute for the COVID-19 Genomics UK (COG-UK) Consortium | Jacquelyn Wynn, Mairead Hyland, The Lighthouse Lab in Alderley Park and Alex Alderton, Roberto Amato, Jeffrey Barrett, Sonia Goncalves, Ewan Harrison, David K. Jackson, Ian Johnston, Dominic Kwiatkowski, Cordelia Langford, John Sillitoe on behalf of the Wellcome Sanger Institute COVID-19 Surveillance Team |
| EPI_ISL_3282567 | Lighthouse Lab in Milton Keynes | Wellcome Sanger Institute for the COVID-19 Genomics UK (COG-UK) Consortium | The Lighthouse Lab in Milton Keynes and Alex Alderton, Roberto Amato, Jeffrey Barrett, Sonia Goncalves, Ewan Harrison, David K. Jackson, Ian Johnston, Dominic Kwiatkowski, Cordelia Langford, John Sillitoe on behalf of the Wellcome Sanger Institute COVID-19 Surveillance Team |
| EPI_ISL_3282568 | Lighthouse Lab in Alderley Park | Wellcome Sanger Institute for the COVID-19 Genomics UK (COG-UK) Consortium | Jacquelyn Wynn, Mairead Hyland, The Lighthouse Lab in Alderley Park and Alex Alderton, Roberto Amato, Jeffrey Barrett, Sonia Goncalves, Ewan Harrison, David K. Jackson, Ian Johnston, Dominic Kwiatkowski, Cordelia Langford, John Sillitoe on behalf of the Wellcome Sanger Institute COVID-19 Surveillance Team |
| EPI_ISL_3282569 | Lighthouse Lab in Milton Keynes | Wellcome Sanger Institute for the COVID-19 Genomics UK (COG-UK) Consortium | The Lighthouse Lab in Milton Keynes and Alex Alderton, Roberto Amato, Jeffrey Barrett, Sonia Goncalves, Ewan Harrison, David K. Jackson, Ian Johnston, Dominic Kwiatkowski, Cordelia Langford, John Sillitoe on behalf of the Wellcome Sanger Institute COVID-19 Surveillance Team |
| EPI_ISL_3282570, EPI_ISL_3282571, EPI_ISL_3282572 | Lighthouse Lab in Glasgow | Wellcome Sanger Institute for the COVID-19 Genomics UK (COG-UK) Consortium | Harper VanSteenhouse, Yumi Kasai, David Gray, Carol Clugston, Anna Dominiczak and Alex Alderton, Roberto Amato, Jeffrey Barrett, Sonia Goncalves, Ewan Harrison, David K. Jackson, Ian Johnston, Dominic Kwiatkowski, Cordelia Langford, John Sillitoe on behalf of the Wellcome Sanger Institute COVID-19 Surveillance Team |
| EPI_ISL_3282573, EPI_ISL_3282574 | Lighthouse Lab in Milton Keynes | Wellcome Sanger Institute for the COVID-19 Genomics UK (COG-UK) Consortium | The Lighthouse Lab in Milton Keynes and Alex Alderton, Roberto Amato, Jeffrey Barrett, Sonia Goncalves, Ewan Harrison, David K. Jackson, Ian Johnston, Dominic Kwiatkowski, Cordelia Langford, John Sillitoe on behalf of the Wellcome Sanger Institute COVID-19 Surveillance Team |
| EPI_ISL_3282575 | Lighthouse Lab in Alderley Park | Wellcome Sanger Institute for the COVID-19 Genomics UK (COG-UK) Consortium | Jacquelyn Wynn, Mairead Hyland, The Lighthouse Lab in Alderley Park and Alex Alderton, Roberto Amato, Jeffrey Barrett, Sonia Goncalves, Ewan Harrison, David K. Jackson, Ian Johnston, Dominic Kwiatkowski, Cordelia Langford, John Sillitoe on behalf of the Wellcome Sanger Institute COVID-19 Surveillance Team |
| EPI_ISL_3282576, EPI_ISL_3282577, EPI_ISL_3282578 | Lighthouse Lab in Milton Keynes | Wellcome Sanger Institute for the COVID-19 Genomics UK (COG-UK) Consortium | The Lighthouse Lab in Milton Keynes and Alex Alderton, Roberto Amato, Jeffrey Barrett, Sonia Goncalves, Ewan Harrison, David K. Jackson, Ian Johnston, Dominic Kwiatkowski, Cordelia Langford, John Sillitoe on behalf of the Wellcome Sanger Institute COVID-19 Surveillance Team |

| | | | |
|---|---|---|---|
| EPI_ISL_3282579 | Lighthouse Lab in Alderley Park | Wellcome Sanger Institute for the COVID-19 Genomics UK (COG-UK) Consortium | Jacquelyn Wynn, Mairead Hyland, The Lighthouse Lab in Alderley Park and Alex Alderton, Roberto Amato, Jeffrey Barrett, Sonia Goncalves, Ewan Harrison, David K. Jackson, Ian Johnston, Dominic Kwiatkowski, Cordelia Langford, John Sillitoe on behalf of the Wellcome Sanger Institute COVID-19 Surveillance Team |
| EPI_ISL_3282580 | Lighthouse Lab in Glasgow | Wellcome Sanger Institute for the COVID-19 Genomics UK (COG-UK) Consortium | Harper VanSteenhouse, Yumi Kasai, David Gray, Carol Clugston, Anna Dominiczak and Alex Alderton, Roberto Amato, Jeffrey Barrett, Sonia Goncalves, Ewan Harrison, David K. Jackson, Ian Johnston, Dominic Kwiatkowski, Cordelia Langford, John Sillitoe on behalf of the Wellcome Sanger Institute COVID-19 Surveillance Team |
| EPI_ISL_3282581 | Lighthouse Lab in Alderley Park | Wellcome Sanger Institute for the COVID-19 Genomics UK (COG-UK) Consortium | Jacquelyn Wynn, Mairead Hyland, The Lighthouse Lab in Alderley Park and Alex Alderton, Roberto Amato, Jeffrey Barrett, Sonia Goncalves, Ewan Harrison, David K. Jackson, Ian Johnston, Dominic Kwiatkowski, Cordelia Langford, John Sillitoe on behalf of the Wellcome Sanger Institute COVID-19 Surveillance Team |
| EPI_ISL_3282582 | Lighthouse Lab in Glasgow | Wellcome Sanger Institute for the COVID-19 Genomics UK (COG-UK) Consortium | Harper VanSteenhouse, Yumi Kasai, David Gray, Carol Clugston, Anna Dominiczak and Alex Alderton, Roberto Amato, Jeffrey Barrett, Sonia Goncalves, Ewan Harrison, David K. Jackson, Ian Johnston, Dominic Kwiatkowski, Cordelia Langford, John Sillitoe on behalf of the Wellcome Sanger Institute COVID-19 Surveillance Team |
| EPI_ISL_3282583 | Lighthouse Lab in Milton Keynes | Wellcome Sanger Institute for the COVID-19 Genomics UK (COG-UK) Consortium | The Lighthouse Lab in Milton Keynes and Alex Alderton, Roberto Amato, Jeffrey Barrett, Sonia Goncalves, Ewan Harrison, David K. Jackson, Ian Johnston, Dominic Kwiatkowski, Cordelia Langford, John Sillitoe on behalf of the Wellcome Sanger Institute COVID-19 Surveillance Team |
| EPI_ISL_3282584, EPI_ISL_3282585, EPI_ISL_3282586 | Lighthouse Lab in Glasgow | Wellcome Sanger Institute for the COVID-19 Genomics UK (COG-UK) Consortium | Harper VanSteenhouse, Yumi Kasai, David Gray, Carol Clugston, Anna Dominiczak and Alex Alderton, Roberto Amato, Jeffrey Barrett, Sonia Goncalves, Ewan Harrison, David K. Jackson, Ian Johnston, Dominic Kwiatkowski, Cordelia Langford, John Sillitoe on behalf of the Wellcome Sanger Institute COVID-19 Surveillance Team |
| EPI_ISL_3282587 | Lighthouse Lab in Alderley Park | Wellcome Sanger Institute for the COVID-19 Genomics UK (COG-UK) Consortium | Jacquelyn Wynn, Mairead Hyland, The Lighthouse Lab in Alderley Park and Alex Alderton, Roberto Amato, Jeffrey Barrett, Sonia Goncalves, Ewan Harrison, David K. Jackson, Ian Johnston, Dominic Kwiatkowski, Cordelia Langford, John Sillitoe on behalf of the Wellcome Sanger Institute COVID-19 Surveillance Team |
| EPI_ISL_3282588, EPI_ISL_3282589 | Lighthouse Lab in Glasgow | Wellcome Sanger Institute for the COVID-19 Genomics UK (COG-UK) Consortium | Harper VanSteenhouse, Yumi Kasai, David Gray, Carol Clugston, Anna Dominiczak and Alex Alderton, Roberto Amato, Jeffrey Barrett, Sonia Goncalves, Ewan Harrison, David K. Jackson, Ian Johnston, Dominic Kwiatkowski, Cordelia Langford, John Sillitoe on behalf of the Wellcome Sanger Institute COVID-19 Surveillance Team |
| EPI_ISL_3282590, EPI_ISL_3282591 | Lighthouse Lab in Milton Keynes | Wellcome Sanger Institute for the COVID-19 Genomics UK (COG-UK) Consortium | The Lighthouse Lab in Milton Keynes and Alex Alderton, Roberto Amato, Jeffrey Barrett, Sonia Goncalves, Ewan Harrison, David K. Jackson, Ian Johnston, Dominic Kwiatkowski, Cordelia Langford, John Sillitoe on behalf of the Wellcome Sanger Institute COVID-19 Surveillance Team |
| EPI_ISL_3282592 | Lighthouse Lab in Alderley Park | Wellcome Sanger Institute for the COVID-19 Genomics UK (COG-UK) Consortium | Jacquelyn Wynn, Mairead Hyland, The Lighthouse Lab in Alderley Park and Alex Alderton, Roberto Amato, Jeffrey Barrett, Sonia Goncalves, Ewan Harrison, David K. Jackson, Ian Johnston, Dominic Kwiatkowski, Cordelia Langford, John Sillitoe on behalf of the Wellcome Sanger Institute COVID-19 Surveillance Team |
| EPI_ISL_3282593 | Lighthouse Lab in Glasgow | Wellcome Sanger Institute for the COVID-19 Genomics UK (COG-UK) Consortium | Harper VanSteenhouse, Yumi Kasai, David Gray, Carol Clugston, Anna Dominiczak and Alex Alderton, Roberto Amato, Jeffrey Barrett, Sonia Goncalves, Ewan Harrison, David K. Jackson, Ian Johnston, Dominic Kwiatkowski, Cordelia Langford, John Sillitoe on behalf of the Wellcome Sanger Institute COVID-19 Surveillance Team |
| EPI_ISL_3282594, EPI_ISL_3282595 | Lighthouse Lab in Alderley Park | Wellcome Sanger Institute for the COVID-19 Genomics UK (COG-UK) Consortium | Jacquelyn Wynn, Mairead Hyland, The Lighthouse Lab in Alderley Park and Alex Alderton, Roberto Amato, Jeffrey Barrett, Sonia Goncalves, Ewan Harrison, David K. Jackson, Ian Johnston, Dominic Kwiatkowski, Cordelia Langford, John Sillitoe on behalf of the Wellcome Sanger Institute COVID-19 Surveillance Team |
| EPI_ISL_3282596, EPI_ISL_3282597 | Lighthouse Lab in Milton Keynes | Wellcome Sanger Institute for the COVID-19 Genomics UK (COG-UK) Consortium | The Lighthouse Lab in Milton Keynes and Alex Alderton, Roberto Amato, Jeffrey Barrett, Sonia Goncalves, Ewan Harrison, David K. Jackson, Ian Johnston, Dominic Kwiatkowski, Cordelia Langford, John Sillitoe on behalf of the Wellcome Sanger Institute COVID-19 Surveillance Team |
| EPI_ISL_3282598 | Lighthouse Lab in Glasgow | Wellcome Sanger Institute for the COVID-19 Genomics UK (COG-UK) Consortium | Harper VanSteenhouse, Yumi Kasai, David Gray, Carol Clugston, Anna Dominiczak and Alex Alderton, Roberto Amato, Jeffrey Barrett, Sonia Goncalves, Ewan Harrison, David K. Jackson, Ian Johnston, Dominic Kwiatkowski, Cordelia Langford, John Sillitoe on behalf of the Wellcome Sanger Institute COVID-19 Surveillance Team |
| EPI_ISL_3282599 | Lighthouse Lab in Milton Keynes | Wellcome Sanger Institute for the COVID-19 Genomics UK (COG-UK) Consortium | The Lighthouse Lab in Milton Keynes and Alex Alderton, Roberto Amato, Jeffrey Barrett, Sonia Goncalves, Ewan Harrison, David K. Jackson, Ian Johnston, Dominic Kwiatkowski, Cordelia Langford, John Sillitoe on behalf of the Wellcome Sanger Institute COVID-19 Surveillance Team |
| EPI_ISL_3282600 | Lighthouse Lab in Glasgow | Wellcome Sanger Institute for the COVID-19 Genomics UK (COG-UK) Consortium | Harper VanSteenhouse, Yumi Kasai, David Gray, Carol Clugston, Anna Dominiczak and Alex Alderton, Roberto Amato, Jeffrey Barrett, Sonia Goncalves, Ewan Harrison, David K. Jackson, Ian Johnston, Dominic Kwiatkowski, Cordelia Langford, John Sillitoe on behalf of the Wellcome Sanger Institute COVID-19 Surveillance Team |
| EPI_ISL_3282601 | Lighthouse Lab in Alderley Park | Wellcome Sanger Institute for the COVID-19 Genomics UK (COG-UK) Consortium | Jacquelyn Wynn, Mairead Hyland, The Lighthouse Lab in Alderley Park and Alex Alderton, Roberto Amato, Jeffrey Barrett, Sonia Goncalves, Ewan Harrison, David K. Jackson, Ian Johnston, Dominic Kwiatkowski, Cordelia Langford, John Sillitoe on behalf of the Wellcome Sanger Institute COVID-19 Surveillance Team |
| EPI_ISL_3282602, EPI_ISL_3282603, EPI_ISL_3282604, EPI_ISL_3282605 | Lighthouse Lab in Glasgow | Wellcome Sanger Institute for the COVID-19 Genomics UK (COG-UK) Consortium | Harper VanSteenhouse, Yumi Kasai, David Gray, Carol Clugston, Anna Dominiczak and Alex Alderton, Roberto Amato, Jeffrey Barrett, Sonia Goncalves, Ewan Harrison, David K. Jackson, Ian Johnston, Dominic Kwiatkowski, Cordelia Langford, John Sillitoe on behalf of the Wellcome Sanger Institute COVID-19 Surveillance Team |
| EPI_ISL_3282606 | Lighthouse Lab in Milton Keynes | Wellcome Sanger Institute for the COVID-19 Genomics UK (COG-UK) Consortium | The Lighthouse Lab in Milton Keynes and Alex Alderton, Roberto Amato, Jeffrey Barrett, Sonia Goncalves, Ewan Harrison, David K. Jackson, Ian Johnston, Dominic Kwiatkowski, Cordelia Langford, John Sillitoe on behalf of the Wellcome Sanger Institute COVID-19 Surveillance Team |
| EPI_ISL_3282607 | Lighthouse Lab in Alderley Park | Wellcome Sanger Institute for the COVID-19 Genomics UK (COG-UK) Consortium | Jacquelyn Wynn, Mairead Hyland, The Lighthouse Lab in Alderley Park and Alex Alderton, Roberto Amato, Jeffrey Barrett, Sonia Goncalves, Ewan Harrison, David K. Jackson, Ian Johnston, Dominic Kwiatkowski, Cordelia Langford, John Sillitoe on behalf of the Wellcome Sanger Institute COVID-19 Surveillance Team |
| EPI_ISL_3282608, EPI_ISL_3282609, EPI_ISL_3282610, EPI_ISL_3282611 | Lighthouse Lab in Milton Keynes | Wellcome Sanger Institute for the COVID-19 Genomics UK (COG-UK) Consortium | The Lighthouse Lab in Milton Keynes and Alex Alderton, Roberto Amato, Jeffrey Barrett, Sonia Goncalves, Ewan Harrison, David K. Jackson, Ian Johnston, Dominic Kwiatkowski, Cordelia Langford, John Sillitoe on behalf of the Wellcome Sanger Institute COVID-19 Surveillance Team |
| EPI_ISL_3282612 | Lighthouse Lab in Glasgow | Wellcome Sanger Institute for the COVID-19 Genomics UK (COG-UK) Consortium | Harper VanSteenhouse, Yumi Kasai, David Gray, Carol Clugston, Anna Dominiczak and Alex Alderton, Roberto Amato, Jeffrey Barrett, Sonia Goncalves, Ewan Harrison, David K. Jackson, Ian Johnston, Dominic Kwiatkowski, Cordelia Langford, John Sillitoe on behalf of the Wellcome Sanger Institute COVID-19 Surveillance Team |
| EPI_ISL_3282613, EPI_ISL_3282614 | Lighthouse Lab in Alderley Park | Wellcome Sanger Institute for the COVID-19 Genomics UK (COG-UK) Consortium | Jacquelyn Wynn, Mairead Hyland, The Lighthouse Lab in Alderley Park and Alex Alderton, Roberto Amato, Jeffrey Barrett, Sonia Goncalves, Ewan Harrison, David K. Jackson, Ian Johnston, Dominic Kwiatkowski, Cordelia Langford, John Sillitoe on behalf of the Wellcome Sanger Institute COVID-19 Surveillance Team |
| EPI_ISL_3282615 | Lighthouse Lab in Glasgow | Wellcome Sanger Institute for the COVID-19 Genomics UK (COG-UK) Consortium | Harper VanSteenhouse, Yumi Kasai, David Gray, Carol Clugston, Anna Dominiczak and Alex Alderton, Roberto Amato, Jeffrey Barrett, Sonia Goncalves, Ewan Harrison, David K. Jackson, Ian Johnston, Dominic Kwiatkowski, Cordelia Langford, John Sillitoe on behalf of the Wellcome Sanger Institute COVID-19 Surveillance Team |
| EPI_ISL_3282616 | Lighthouse Lab in Milton Keynes | Wellcome Sanger Institute for the COVID-19 Genomics UK (COG-UK) Consortium | The Lighthouse Lab in Milton Keynes and Alex Alderton, Roberto Amato, Jeffrey Barrett, Sonia Goncalves, Ewan Harrison, David K. Jackson, Ian Johnston, Dominic Kwiatkowski, Cordelia Langford, John Sillitoe on behalf of the Wellcome Sanger Institute COVID-19 Surveillance Team |
| EPI_ISL_3282617 | Lighthouse Lab in Glasgow | Wellcome Sanger Institute for the COVID-19 Genomics UK (COG-UK) Consortium | Harper VanSteenhouse, Yumi Kasai, David Gray, Carol Clugston, Anna Dominiczak and Alex Alderton, Roberto Amato, Jeffrey Barrett, Sonia Goncalves, Ewan Harrison, David K. Jackson, Ian Johnston, Dominic Kwiatkowski, Cordelia Langford, John Sillitoe on behalf of the Wellcome Sanger Institute COVID-19 Surveillance Team |
| EPI_ISL_3282618 | Lighthouse Lab in Alderley Park | Wellcome Sanger Institute for the COVID-19 Genomics UK (COG-UK) Consortium | Jacquelyn Wynn, Mairead Hyland, The Lighthouse Lab in Alderley Park and Alex Alderton, Roberto Amato, Jeffrey Barrett, Sonia Goncalves, Ewan Harrison, David K. Jackson, Ian Johnston, Dominic Kwiatkowski, Cordelia Langford, John Sillitoe on behalf of the Wellcome Sanger Institute COVID-19 Surveillance Team |
| EPI_ISL_3282619 | Lighthouse Lab in Glasgow | Wellcome Sanger Institute for the COVID-19 Genomics UK (COG-UK) Consortium | Harper VanSteenhouse, Yumi Kasai, David Gray, Carol Clugston, Anna Dominiczak and Alex Alderton, Roberto Amato, Jeffrey Barrett, Sonia Goncalves, Ewan Harrison, David K. Jackson, Ian Johnston, Dominic Kwiatkowski, Cordelia Langford, John Sillitoe on behalf of the Wellcome Sanger Institute COVID-19 Surveillance Team |
| EPI_ISL_3282620, EPI_ISL_3282621 | Lighthouse Lab in Alderley Park | Wellcome Sanger Institute for the COVID-19 Genomics UK (COG-UK) Consortium | Jacquelyn Wynn, Mairead Hyland, The Lighthouse Lab in Alderley Park and Alex Alderton, Roberto Amato, Jeffrey Barrett, Sonia Goncalves, Ewan Harrison, David K. Jackson, Ian Johnston, Dominic Kwiatkowski, Cordelia Langford, John Sillitoe on behalf of the Wellcome Sanger Institute COVID-19 Surveillance Team |
| EPI_ISL_3282622 | Lighthouse Lab in Milton Keynes | Wellcome Sanger Institute for the COVID-19 Genomics UK (COG-UK) Consortium | The Lighthouse Lab in Milton Keynes and Alex Alderton, Roberto Amato, Jeffrey Barrett, Sonia Goncalves, Ewan Harrison, David K. Jackson, Ian Johnston, Dominic Kwiatkowski, Cordelia Langford, John Sillitoe on behalf of the Wellcome Sanger Institute COVID-19 Surveillance Team |
| EPI_ISL_3282623, EPI_ISL_3282624 | Lighthouse Lab in Glasgow | Wellcome Sanger Institute for the COVID-19 Genomics UK (COG-UK) Consortium | Harper VanSteenhouse, Yumi Kasai, David Gray, Carol Clugston, Anna Dominiczak and Alex Alderton, Roberto Amato, Jeffrey Barrett, Sonia Goncalves, Ewan Harrison, David K. Jackson, Ian Johnston, Dominic Kwiatkowski, Cordelia Langford, John Sillitoe on behalf of the Wellcome Sanger Institute COVID-19 Surveillance Team |
| EPI_ISL_3282625, EPI_ISL_3282626 | Lighthouse Lab in Alderley Park | Wellcome Sanger Institute for the COVID-19 Genomics UK (COG-UK) Consortium | Jacquelyn Wynn, Mairead Hyland, The Lighthouse Lab in Alderley Park and Alex Alderton, Roberto Amato, Jeffrey Barrett, Sonia Goncalves, Ewan Harrison, David K. Jackson, Ian Johnston, Dominic Kwiatkowski, Cordelia Langford, John Sillitoe on behalf of the Wellcome Sanger Institute COVID-19 Surveillance Team |
| EPI_ISL_3282627, EPI_ISL_3282628 | Lighthouse Lab in Glasgow | Wellcome Sanger Institute for the COVID-19 Genomics UK (COG-UK) Consortium | Harper VanSteenhouse, Yumi Kasai, David Gray, Carol Clugston, Anna Dominiczak and Alex Alderton, Roberto Amato, Jeffrey Barrett, Sonia Goncalves, Ewan Harrison, David K. Jackson, Ian Johnston, Dominic Kwiatkowski, Cordelia Langford, John Sillitoe on behalf of the Wellcome Sanger Institute COVID-19 Surveillance Team |
| EPI_ISL_3282629 | Lighthouse Lab in Milton Keynes | Wellcome Sanger Institute for the COVID-19 Genomics UK (COG-UK) Consortium | The Lighthouse Lab in Milton Keynes and Alex Alderton, Roberto Amato, Jeffrey Barrett, Sonia Goncalves, Ewan Harrison, David K. Jackson, Ian Johnston, Dominic Kwiatkowski, Cordelia Langford, John Sillitoe on behalf of the Wellcome Sanger Institute COVID-19 Surveillance Team |
| EPI_ISL_3282630 | Lighthouse Lab in Glasgow | Wellcome Sanger Institute for the COVID-19 Genomics UK (COG-UK) Consortium | Harper VanSteenhouse, Yumi Kasai, David Gray, Carol Clugston, Anna Dominiczak and Alex Alderton, Roberto Amato, Jeffrey Barrett, Sonia Goncalves, Ewan Harrison, David K. Jackson, Ian Johnston, Dominic Kwiatkowski, Cordelia Langford, John Sillitoe on behalf of the Wellcome Sanger Institute COVID-19 Surveillance Team |
| EPI_ISL_3282631 | Lighthouse Lab in Milton Keynes | Wellcome Sanger Institute for the COVID-19 Genomics UK (COG-UK) Consortium | The Lighthouse Lab in Milton Keynes and Alex Alderton, Roberto Amato, Jeffrey Barrett, Sonia Goncalves, Ewan Harrison, David K. Jackson, Ian Johnston, Dominic Kwiatkowski, Cordelia Langford, John Sillitoe on behalf of the Wellcome Sanger Institute COVID-19 Surveillance Team |
| EPI_ISL_3282632 | Lighthouse Lab in Glasgow | Wellcome Sanger Institute for the COVID-19 Genomics UK (COG-UK) Consortium | Harper VanSteenhouse, Yumi Kasai, David Gray, Carol Clugston, Anna Dominiczak and Alex Alderton, Roberto Amato, Jeffrey Barrett, Sonia Goncalves, Ewan Harrison, David K. Jackson, Ian Johnston, Dominic Kwiatkowski, Cordelia Langford, John Sillitoe on behalf of the Wellcome Sanger Institute COVID-19 Surveillance Team |
| EPI_ISL_3282633 | Lighthouse Lab in Alderley Park | Wellcome Sanger Institute for the COVID-19 Genomics UK (COG-UK) Consortium | Jacquelyn Wynn, Mairead Hyland, The Lighthouse Lab in Alderley Park and Alex Alderton, Roberto Amato, Jeffrey Barrett, Sonia Goncalves, Ewan Harrison, David K. Jackson, Ian Johnston, Dominic Kwiatkowski, Cordelia Langford, John Sillitoe on behalf of the Wellcome Sanger Institute COVID-19 Surveillance Team |
| EPI_ISL_3282634 | Lighthouse Lab in Milton Keynes | Wellcome Sanger Institute for the COVID-19 Genomics UK (COG-UK) Consortium | The Lighthouse Lab in Milton Keynes and Alex Alderton, Roberto Amato, Jeffrey Barrett, Sonia Goncalves, Ewan Harrison, David K. Jackson, Ian Johnston, Dominic Kwiatkowski, Cordelia Langford, John Sillitoe on behalf of the Wellcome Sanger Institute COVID-19 Surveillance Team |
| EPI_ISL_3282635 | Lighthouse Lab in Glasgow | Wellcome Sanger Institute for the COVID-19 Genomics UK (COG-UK) Consortium | Harper VanSteenhouse, Yumi Kasai, David Gray, Carol Clugston, Anna Dominiczak and Alex Alderton, Roberto Amato, Jeffrey Barrett, Sonia Goncalves, Ewan Harrison, David K. Jackson, Ian Johnston, Dominic Kwiatkowski, Cordelia Langford, John Sillitoe on behalf of the Wellcome Sanger Institute COVID-19 Surveillance Team |
| EPI_ISL_3282636 | Lighthouse Lab in Milton Keynes | Wellcome Sanger Institute for the COVID-19 Genomics UK (COG-UK) Consortium | The Lighthouse Lab in Milton Keynes and Alex Alderton, Roberto Amato, Jeffrey Barrett, Sonia Goncalves, Ewan Harrison, David K. Jackson, Ian Johnston, Dominic Kwiatkowski, Cordelia Langford, John Sillitoe on behalf of the Wellcome Sanger Institute COVID-19 Surveillance Team |
| EPI_ISL_3282637, EPI_ISL_3282638, EPI_ISL_3282639 | Lighthouse Lab in Alderley Park | Wellcome Sanger Institute for the COVID-19 Genomics UK (COG-UK) Consortium | Jacquelyn Wynn, Mairead Hyland, The Lighthouse Lab in Alderley Park and Alex Alderton, Roberto Amato, Jeffrey Barrett, Sonia Goncalves, Ewan Harrison, David K. Jackson, Ian Johnston, Dominic Kwiatkowski, Cordelia Langford, John Sillitoe on behalf of the Wellcome Sanger Institute COVID-19 Surveillance Team |
| EPI_ISL_3282640, EPI_ISL_3282641 | Lighthouse Lab in Glasgow | Wellcome Sanger Institute for the COVID-19 Genomics UK (COG-UK) Consortium | Harper VanSteenhouse, Yumi Kasai, David Gray, Carol Clugston, Anna Dominiczak and Alex Alderton, Roberto Amato, Jeffrey Barrett, Sonia Goncalves, Ewan Harrison, David K. Jackson, Ian Johnston, Dominic Kwiatkowski, Cordelia Langford, John Sillitoe on behalf of the Wellcome Sanger Institute COVID-19 Surveillance Team |

| | | | |
|---|---|---|---|
| EPI_ISL_3282642 | Lighthouse Lab in Milton Keynes | Wellcome Sanger Institute for the COVID-19 Genomics UK (COG-UK) Consortium | The Lighthouse Lab in Milton Keynes and Alex Alderton, Roberto Amato, Jeffrey Barrett, Sonia Goncalves, Ewan Harrison, David K. Jackson, Ian Johnston, Dominic Kwiatkowski, Cordelia Langford, John Sillitoe on behalf of the Wellcome Sanger Institute COVID-19 Surveillance Team |
| EPI_ISL_3282643, EPI_ISL_3282644, EPI_ISL_3282645 | Lighthouse Lab in Glasgow | Wellcome Sanger Institute for the COVID-19 Genomics UK (COG-UK) Consortium | Harper VanSteenhouse, Yumi Kasai, David Gray, Carol Clugston, Anna Dominiczak and Alex Alderton, Roberto Amato, Jeffrey Barrett, Sonia Goncalves, Ewan Harrison, David K. Jackson, Ian Johnston, Dominic Kwiatkowski, Cordelia Langford, John Sillitoe on behalf of the Wellcome Sanger Institute COVID-19 Surveillance Team |
| EPI_ISL_3282646 | Lighthouse Lab in Alderley Park | Wellcome Sanger Institute for the COVID-19 Genomics UK (COG-UK) Consortium | Jacquelyn Wynn, Mairead Hyland, The Lighthouse Lab in Alderley Park and Alex Alderton, Roberto Amato, Jeffrey Barrett, Sonia Goncalves, Ewan Harrison, David K. Jackson, Ian Johnston, Dominic Kwiatkowski, Cordelia Langford, John Sillitoe on behalf of the Wellcome Sanger Institute COVID-19 Surveillance Team |
| EPI_ISL_3282647, EPI_ISL_3282648, EPI_ISL_3282649 | Lighthouse Lab in Milton Keynes | Wellcome Sanger Institute for the COVID-19 Genomics UK (COG-UK) Consortium | The Lighthouse Lab in Milton Keynes and Alex Alderton, Roberto Amato, Jeffrey Barrett, Sonia Goncalves, Ewan Harrison, David K. Jackson, Ian Johnston, Dominic Kwiatkowski, Cordelia Langford, John Sillitoe on behalf of the Wellcome Sanger Institute COVID-19 Surveillance Team |
| EPI_ISL_3282650 | Lighthouse Lab in Glasgow | Wellcome Sanger Institute for the COVID-19 Genomics UK (COG-UK) Consortium | Harper VanSteenhouse, Yumi Kasai, David Gray, Carol Clugston, Anna Dominiczak and Alex Alderton, Roberto Amato, Jeffrey Barrett, Sonia Goncalves, Ewan Harrison, David K. Jackson, Ian Johnston, Dominic Kwiatkowski, Cordelia Langford, John Sillitoe on behalf of the Wellcome Sanger Institute COVID-19 Surveillance Team |
| EPI_ISL_3282651 | Lighthouse Lab in Milton Keynes | Wellcome Sanger Institute for the COVID-19 Genomics UK (COG-UK) Consortium | The Lighthouse Lab in Milton Keynes and Alex Alderton, Roberto Amato, Jeffrey Barrett, Sonia Goncalves, Ewan Harrison, David K. Jackson, Ian Johnston, Dominic Kwiatkowski, Cordelia Langford, John Sillitoe on behalf of the Wellcome Sanger Institute COVID-19 Surveillance Team |
| EPI_ISL_3282652, EPI_ISL_3282653 | Lighthouse Lab in Glasgow | Wellcome Sanger Institute for the COVID-19 Genomics UK (COG-UK) Consortium | Harper VanSteenhouse, Yumi Kasai, David Gray, Carol Clugston, Anna Dominiczak and Alex Alderton, Roberto Amato, Jeffrey Barrett, Sonia Goncalves, Ewan Harrison, David K. Jackson, Ian Johnston, Dominic Kwiatkowski, Cordelia Langford, John Sillitoe on behalf of the Wellcome Sanger Institute COVID-19 Surveillance Team |
| EPI_ISL_3282654 | Lighthouse Lab in Milton Keynes | Wellcome Sanger Institute for the COVID-19 Genomics UK (COG-UK) Consortium | The Lighthouse Lab in Milton Keynes and Alex Alderton, Roberto Amato, Jeffrey Barrett, Sonia Goncalves, Ewan Harrison, David K. Jackson, Ian Johnston, Dominic Kwiatkowski, Cordelia Langford, John Sillitoe on behalf of the Wellcome Sanger Institute COVID-19 Surveillance Team |
| EPI_ISL_3282655, EPI_ISL_3282656 | Lighthouse Lab in Alderley Park | Wellcome Sanger Institute for the COVID-19 Genomics UK (COG-UK) Consortium | Jacquelyn Wynn, Mairead Hyland, The Lighthouse Lab in Alderley Park and Alex Alderton, Roberto Amato, Jeffrey Barrett, Sonia Goncalves, Ewan Harrison, David K. Jackson, Ian Johnston, Dominic Kwiatkowski, Cordelia Langford, John Sillitoe on behalf of the Wellcome Sanger Institute COVID-19 Surveillance Team |
| EPI_ISL_3282657, EPI_ISL_3282658 | Lighthouse Lab in Milton Keynes | Wellcome Sanger Institute for the COVID-19 Genomics UK (COG-UK) Consortium | The Lighthouse Lab in Milton Keynes and Alex Alderton, Roberto Amato, Jeffrey Barrett, Sonia Goncalves, Ewan Harrison, David K. Jackson, Ian Johnston, Dominic Kwiatkowski, Cordelia Langford, John Sillitoe on behalf of the Wellcome Sanger Institute COVID-19 Surveillance Team |
| EPI_ISL_3282659 | Lighthouse Lab in Glasgow | Wellcome Sanger Institute for the COVID-19 Genomics UK (COG-UK) Consortium | Harper VanSteenhouse, Yumi Kasai, David Gray, Carol Clugston, Anna Dominiczak and Alex Alderton, Roberto Amato, Jeffrey Barrett, Sonia Goncalves, Ewan Harrison, David K. Jackson, Ian Johnston, Dominic Kwiatkowski, Cordelia Langford, John Sillitoe on behalf of the Wellcome Sanger Institute COVID-19 Surveillance Team |
| EPI_ISL_3282660 | Lighthouse Lab in Alderley Park | Wellcome Sanger Institute for the COVID-19 Genomics UK (COG-UK) Consortium | Jacquelyn Wynn, Mairead Hyland, The Lighthouse Lab in Alderley Park and Alex Alderton, Roberto Amato, Jeffrey Barrett, Sonia Goncalves, Ewan Harrison, David K. Jackson, Ian Johnston, Dominic Kwiatkowski, Cordelia Langford, John Sillitoe on behalf of the Wellcome Sanger Institute COVID-19 Surveillance Team |
| EPI_ISL_3282661, EPI_ISL_3282662, EPI_ISL_3282663 | Lighthouse Lab in Glasgow | Wellcome Sanger Institute for the COVID-19 Genomics UK (COG-UK) Consortium | Harper VanSteenhouse, Yumi Kasai, David Gray, Carol Clugston, Anna Dominiczak and Alex Alderton, Roberto Amato, Jeffrey Barrett, Sonia Goncalves, Ewan Harrison, David K. Jackson, Ian Johnston, Dominic Kwiatkowski, Cordelia Langford, John Sillitoe on behalf of the Wellcome Sanger Institute COVID-19 Surveillance Team |
| EPI_ISL_3282664 | Lighthouse Lab in Milton Keynes | Wellcome Sanger Institute for the COVID-19 Genomics UK (COG-UK) Consortium | The Lighthouse Lab in Milton Keynes and Alex Alderton, Roberto Amato, Jeffrey Barrett, Sonia Goncalves, Ewan Harrison, David K. Jackson, Ian Johnston, Dominic Kwiatkowski, Cordelia Langford, John Sillitoe on behalf of the Wellcome Sanger Institute COVID-19 Surveillance Team |
| EPI_ISL_3282665 | Lighthouse Lab in Alderley Park | Wellcome Sanger Institute for the COVID-19 Genomics UK (COG-UK) Consortium | Jacquelyn Wynn, Mairead Hyland, The Lighthouse Lab in Alderley Park and Alex Alderton, Roberto Amato, Jeffrey Barrett, Sonia Goncalves, Ewan Harrison, David K. Jackson, Ian Johnston, Dominic Kwiatkowski, Cordelia Langford, John Sillitoe on behalf of the Wellcome Sanger Institute COVID-19 Surveillance Team |
| EPI_ISL_3282666, EPI_ISL_3282667 | Lighthouse Lab in Glasgow | Wellcome Sanger Institute for the COVID-19 Genomics UK (COG-UK) Consortium | Harper VanSteenhouse, Yumi Kasai, David Gray, Carol Clugston, Anna Dominiczak and Alex Alderton, Roberto Amato, Jeffrey Barrett, Sonia Goncalves, Ewan Harrison, David K. Jackson, Ian Johnston, Dominic Kwiatkowski, Cordelia Langford, John Sillitoe on behalf of the Wellcome Sanger Institute COVID-19 Surveillance Team |
| EPI_ISL_3282668 | Lighthouse Lab in Alderley Park | Wellcome Sanger Institute for the COVID-19 Genomics UK (COG-UK) Consortium | Jacquelyn Wynn, Mairead Hyland, The Lighthouse Lab in Alderley Park and Alex Alderton, Roberto Amato, Jeffrey Barrett, Sonia Goncalves, Ewan Harrison, David K. Jackson, Ian Johnston, Dominic Kwiatkowski, Cordelia Langford, John Sillitoe on behalf of the Wellcome Sanger Institute COVID-19 Surveillance Team |
| EPI_ISL_3282669 | Lighthouse Lab in Glasgow | Wellcome Sanger Institute for the COVID-19 Genomics UK (COG-UK) Consortium | Harper VanSteenhouse, Yumi Kasai, David Gray, Carol Clugston, Anna Dominiczak and Alex Alderton, Roberto Amato, Jeffrey Barrett, Sonia Goncalves, Ewan Harrison, David K. Jackson, Ian Johnston, Dominic Kwiatkowski, Cordelia Langford, John Sillitoe on behalf of the Wellcome Sanger Institute COVID-19 Surveillance Team |
| EPI_ISL_3282670 | Lighthouse Lab in Alderley Park | Wellcome Sanger Institute for the COVID-19 Genomics UK (COG-UK) Consortium | Jacquelyn Wynn, Mairead Hyland, The Lighthouse Lab in Alderley Park and Alex Alderton, Roberto Amato, Jeffrey Barrett, Sonia Goncalves, Ewan Harrison, David K. Jackson, Ian Johnston, Dominic Kwiatkowski, Cordelia Langford, John Sillitoe on behalf of the Wellcome Sanger Institute COVID-19 Surveillance Team |
| EPI_ISL_3282671 | Lighthouse Lab in Milton Keynes | Wellcome Sanger Institute for the COVID-19 Genomics UK (COG-UK) Consortium | The Lighthouse Lab in Milton Keynes and Alex Alderton, Roberto Amato, Jeffrey Barrett, Sonia Goncalves, Ewan Harrison, David K. Jackson, Ian Johnston, Dominic Kwiatkowski, Cordelia Langford, John Sillitoe on behalf of the Wellcome Sanger Institute COVID-19 Surveillance Team |
| EPI_ISL_3282672 | Lighthouse Lab in Glasgow | Wellcome Sanger Institute for the COVID-19 Genomics UK (COG-UK) Consortium | Harper VanSteenhouse, Yumi Kasai, David Gray, Carol Clugston, Anna Dominiczak and Alex Alderton, Roberto Amato, Jeffrey Barrett, Sonia Goncalves, Ewan Harrison, David K. Jackson, Ian Johnston, Dominic Kwiatkowski, Cordelia Langford, John Sillitoe on behalf of the Wellcome Sanger Institute COVID-19 Surveillance Team |
| EPI_ISL_3282673, EPI_ISL_3282674, EPI_ISL_3282675 | Lighthouse Lab in Alderley Park | Wellcome Sanger Institute for the COVID-19 Genomics UK (COG-UK) Consortium | Jacquelyn Wynn, Mairead Hyland, The Lighthouse Lab in Alderley Park and Alex Alderton, Roberto Amato, Jeffrey Barrett, Sonia Goncalves, Ewan Harrison, David K. Jackson, Ian Johnston, Dominic Kwiatkowski, Cordelia Langford, John Sillitoe on behalf of the Wellcome Sanger Institute COVID-19 Surveillance Team |
| EPI_ISL_3282676 | Lighthouse Lab in Glasgow | Wellcome Sanger Institute for the COVID-19 Genomics UK (COG-UK) Consortium | Harper VanSteenhouse, Yumi Kasai, David Gray, Carol Clugston, Anna Dominiczak and Alex Alderton, Roberto Amato, Jeffrey Barrett, Sonia Goncalves, Ewan Harrison, David K. Jackson, Ian Johnston, Dominic Kwiatkowski, Cordelia Langford, John Sillitoe on behalf of the Wellcome Sanger Institute COVID-19 Surveillance Team |
| EPI_ISL_3282677, EPI_ISL_3282678 | Lighthouse Lab in Alderley Park | Wellcome Sanger Institute for the COVID-19 Genomics UK (COG-UK) Consortium | Jacquelyn Wynn, Mairead Hyland, The Lighthouse Lab in Alderley Park and Alex Alderton, Roberto Amato, Jeffrey Barrett, Sonia Goncalves, Ewan Harrison, David K. Jackson, Ian Johnston, Dominic Kwiatkowski, Cordelia Langford, John Sillitoe on behalf of the Wellcome Sanger Institute COVID-19 Surveillance Team |
| EPI_ISL_3282679, EPI_ISL_3282680 | Lighthouse Lab in Glasgow | Wellcome Sanger Institute for the COVID-19 Genomics UK (COG-UK) Consortium | Harper VanSteenhouse, Yumi Kasai, David Gray, Carol Clugston, Anna Dominiczak and Alex Alderton, Roberto Amato, Jeffrey Barrett, Sonia Goncalves, Ewan Harrison, David K. Jackson, Ian Johnston, Dominic Kwiatkowski, Cordelia Langford, John Sillitoe on behalf of the Wellcome Sanger Institute COVID-19 Surveillance Team |
| EPI_ISL_3282681 | Lighthouse Lab in Alderley Park | Wellcome Sanger Institute for the COVID-19 Genomics UK (COG-UK) Consortium | Jacquelyn Wynn, Mairead Hyland, The Lighthouse Lab in Alderley Park and Alex Alderton, Roberto Amato, Jeffrey Barrett, Sonia Goncalves, Ewan Harrison, David K. Jackson, Ian Johnston, Dominic Kwiatkowski, Cordelia Langford, John Sillitoe on behalf of the Wellcome Sanger Institute COVID-19 Surveillance Team |
| EPI_ISL_3282682 | Lighthouse Lab in Milton Keynes | Wellcome Sanger Institute for the COVID-19 Genomics UK (COG-UK) Consortium | The Lighthouse Lab in Milton Keynes and Alex Alderton, Roberto Amato, Jeffrey Barrett, Sonia Goncalves, Ewan Harrison, David K. Jackson, Ian Johnston, Dominic Kwiatkowski, Cordelia Langford, John Sillitoe on behalf of the Wellcome Sanger Institute COVID-19 Surveillance Team |
| EPI_ISL_3282683 | Lighthouse Lab in Alderley Park | Wellcome Sanger Institute for the COVID-19 Genomics UK (COG-UK) Consortium | Jacquelyn Wynn, Mairead Hyland, The Lighthouse Lab in Alderley Park and Alex Alderton, Roberto Amato, Jeffrey Barrett, Sonia Goncalves, Ewan Harrison, David K. Jackson, Ian Johnston, Dominic Kwiatkowski, Cordelia Langford, John Sillitoe on behalf of the Wellcome Sanger Institute COVID-19 Surveillance Team |
| EPI_ISL_3282684 | Lighthouse Lab in Glasgow | Wellcome Sanger Institute for the COVID-19 Genomics UK (COG-UK) Consortium | Harper VanSteenhouse, Yumi Kasai, David Gray, Carol Clugston, Anna Dominiczak and Alex Alderton, Roberto Amato, Jeffrey Barrett, Sonia Goncalves, Ewan Harrison, David K. Jackson, Ian Johnston, Dominic Kwiatkowski, Cordelia Langford, John Sillitoe on behalf of the Wellcome Sanger Institute COVID-19 Surveillance Team |
| EPI_ISL_3282685 | Lighthouse Lab in Alderley Park | Wellcome Sanger Institute for the COVID-19 Genomics UK (COG-UK) Consortium | Jacquelyn Wynn, Mairead Hyland, The Lighthouse Lab in Alderley Park and Alex Alderton, Roberto Amato, Jeffrey Barrett, Sonia Goncalves, Ewan Harrison, David K. Jackson, Ian Johnston, Dominic Kwiatkowski, Cordelia Langford, John Sillitoe on behalf of the Wellcome Sanger Institute COVID-19 Surveillance Team |
| EPI_ISL_3282686, EPI_ISL_3282687 | Lighthouse Lab in Glasgow | Wellcome Sanger Institute for the COVID-19 Genomics UK (COG-UK) Consortium | Harper VanSteenhouse, Yumi Kasai, David Gray, Carol Clugston, Anna Dominiczak and Alex Alderton, Roberto Amato, Jeffrey Barrett, Sonia Goncalves, Ewan Harrison, David K. Jackson, Ian Johnston, Dominic Kwiatkowski, Cordelia Langford, John Sillitoe on behalf of the Wellcome Sanger Institute COVID-19 Surveillance Team |
| EPI_ISL_3282688 | Lighthouse Lab in Alderley Park | Wellcome Sanger Institute for the COVID-19 Genomics UK (COG-UK) Consortium | Jacquelyn Wynn, Mairead Hyland, The Lighthouse Lab in Alderley Park and Alex Alderton, Roberto Amato, Jeffrey Barrett, Sonia Goncalves, Ewan Harrison, David K. Jackson, Ian Johnston, Dominic Kwiatkowski, Cordelia Langford, John Sillitoe on behalf of the Wellcome Sanger Institute COVID-19 Surveillance Team |
| EPI_ISL_3282689, EPI_ISL_3282690 | Lighthouse Lab in Milton Keynes | Wellcome Sanger Institute for the COVID-19 Genomics UK (COG-UK) Consortium | The Lighthouse Lab in Milton Keynes and Alex Alderton, Roberto Amato, Jeffrey Barrett, Sonia Goncalves, Ewan Harrison, David K. Jackson, Ian Johnston, Dominic Kwiatkowski, Cordelia Langford, John Sillitoe on behalf of the Wellcome Sanger Institute COVID-19 Surveillance Team |
| EPI_ISL_3282691 | Lighthouse Lab in Alderley Park | Wellcome Sanger Institute for the COVID-19 Genomics UK (COG-UK) Consortium | Jacquelyn Wynn, Mairead Hyland, The Lighthouse Lab in Alderley Park and Alex Alderton, Roberto Amato, Jeffrey Barrett, Sonia Goncalves, Ewan Harrison, David K. Jackson, Ian Johnston, Dominic Kwiatkowski, Cordelia Langford, John Sillitoe on behalf of the Wellcome Sanger Institute COVID-19 Surveillance Team |
| EPI_ISL_3282692, EPI_ISL_3282693, EPI_ISL_3282694, EPI_ISL_3282695, EPI_ISL_3282696, EPI_ISL_3282697 | Lighthouse Lab in Milton Keynes | Wellcome Sanger Institute for the COVID-19 Genomics UK (COG-UK) Consortium | The Lighthouse Lab in Milton Keynes and Alex Alderton, Roberto Amato, Jeffrey Barrett, Sonia Goncalves, Ewan Harrison, David K. Jackson, Ian Johnston, Dominic Kwiatkowski, Cordelia Langford, John Sillitoe on behalf of the Wellcome Sanger Institute COVID-19 Surveillance Team |
| EPI_ISL_3282698, EPI_ISL_3282699 | Lighthouse Lab in Glasgow | Wellcome Sanger Institute for the COVID-19 Genomics UK (COG-UK) Consortium | Harper VanSteenhouse, Yumi Kasai, David Gray, Carol Clugston, Anna Dominiczak and Alex Alderton, Roberto Amato, Jeffrey Barrett, Sonia Goncalves, Ewan Harrison, David K. Jackson, Ian Johnston, Dominic Kwiatkowski, Cordelia Langford, John Sillitoe on behalf of the Wellcome Sanger Institute COVID-19 Surveillance Team |
| EPI_ISL_3282700 | Lighthouse Lab in Alderley Park | Wellcome Sanger Institute for the COVID-19 Genomics UK (COG-UK) Consortium | Jacquelyn Wynn, Mairead Hyland, The Lighthouse Lab in Alderley Park and Alex Alderton, Roberto Amato, Jeffrey Barrett, Sonia Goncalves, Ewan Harrison, David K. Jackson, Ian Johnston, Dominic Kwiatkowski, Cordelia Langford, John Sillitoe on behalf of the Wellcome Sanger Institute COVID-19 Surveillance Team |
| EPI_ISL_3282701, EPI_ISL_3282702 | Lighthouse Lab in Glasgow | Wellcome Sanger Institute for the COVID-19 Genomics UK (COG-UK) Consortium | Harper VanSteenhouse, Yumi Kasai, David Gray, Carol Clugston, Anna Dominiczak and Alex Alderton, Roberto Amato, Jeffrey Barrett, Sonia Goncalves, Ewan Harrison, David K. Jackson, Ian Johnston, Dominic Kwiatkowski, Cordelia Langford, John Sillitoe on behalf of the Wellcome Sanger Institute COVID-19 Surveillance Team |
| EPI_ISL_3282703 | Lighthouse Lab in Milton Keynes | Wellcome Sanger Institute for the COVID-19 Genomics UK (COG-UK) Consortium | The Lighthouse Lab in Milton Keynes and Alex Alderton, Roberto Amato, Jeffrey Barrett, Sonia Goncalves, Ewan Harrison, David K. Jackson, Ian Johnston, Dominic Kwiatkowski, Cordelia Langford, John Sillitoe on behalf of the Wellcome Sanger Institute COVID-19 Surveillance Team |
| EPI_ISL_3282704, EPI_ISL_3282705, EPI_ISL_3282706 | Lighthouse Lab in Glasgow | Wellcome Sanger Institute for the COVID-19 Genomics UK (COG-UK) Consortium | Harper VanSteenhouse, Yumi Kasai, David Gray, Carol Clugston, Anna Dominiczak and Alex Alderton, Roberto Amato, Jeffrey Barrett, Sonia Goncalves, Ewan Harrison, David K. Jackson, Ian Johnston, Dominic Kwiatkowski, Cordelia Langford, John Sillitoe on behalf of the Wellcome Sanger Institute COVID-19 Surveillance Team |
| EPI_ISL_3282707, EPI_ISL_3282708 | Lighthouse Lab in Alderley Park | Wellcome Sanger Institute for the COVID-19 Genomics UK (COG-UK) Consortium | Jacquelyn Wynn, Mairead Hyland, The Lighthouse Lab in Alderley Park and Alex Alderton, Roberto Amato, Jeffrey Barrett, Sonia Goncalves, Ewan Harrison, David K. Jackson, Ian Johnston, Dominic Kwiatkowski, Cordelia Langford, John Sillitoe on behalf of the Wellcome Sanger Institute COVID-19 Surveillance Team |

| | | | |
|---|---|---|---|
| EPI_ISL_3282709 | Lighthouse Lab in Glasgow | Wellcome Sanger Institute for the COVID-19 Genomics UK (COG-UK) Consortium | Harper VanSteenhouse, Yumi Kasai, David Gray, Carol Clugston, Anna Dominiczak and Alex Alderton, Roberto Amato, Jeffrey Barrett, Sonia Goncalves, Ewan Harrison, David K. Jackson, Ian Johnston, Dominic Kwiatkowski, Cordelia Langford, John Sillitoe on behalf of the Wellcome Sanger Institute COVID-19 Surveillance Team |
| EPI_ISL_3282710, EPI_ISL_3282711 | Lighthouse Laboratory Plymouth | Wellcome Sanger Institute for the COVID-19 Genomics UK (COG-UK) Consortium | Lighthouse Laboratory Plymouth and Alex Alderton, Roberto Amato, Jeffrey Barrett, Sonia Goncalves, Ewan Harrison, David K. Jackson, Ian Johnston, Dominic Kwiatkowski, Cordelia Langford, John Sillitoe on behalf of the Wellcome Sanger Institute COVID-19 Surveillance Team |
| EPI_ISL_3282712, EPI_ISL_3282713, EPI_ISL_3282714 | Lighthouse Lab in Alderley Park | Wellcome Sanger Institute for the COVID-19 Genomics UK (COG-UK) Consortium | Jacquelyn Wynn, Mairead Hyland, The Lighthouse Lab in Alderley Park and Alex Alderton, Roberto Amato, Jeffrey Barrett, Sonia Goncalves, Ewan Harrison, David K. Jackson, Ian Johnston, Dominic Kwiatkowski, Cordelia Langford, John Sillitoe on behalf of the Wellcome Sanger Institute COVID-19 Surveillance Team |
| EPI_ISL_3282715 | Lighthouse Laboratory Plymouth | Wellcome Sanger Institute for the COVID-19 Genomics UK (COG-UK) Consortium | Lighthouse Laboratory Plymouth and Alex Alderton, Roberto Amato, Jeffrey Barrett, Sonia Goncalves, Ewan Harrison, David K. Jackson, Ian Johnston, Dominic Kwiatkowski, Cordelia Langford, John Sillitoe on behalf of the Wellcome Sanger Institute COVID-19 Surveillance Team |
| EPI_ISL_3282716 | Lighthouse Lab in Alderley Park | Wellcome Sanger Institute for the COVID-19 Genomics UK (COG-UK) Consortium | Jacquelyn Wynn, Mairead Hyland, The Lighthouse Lab in Alderley Park and Alex Alderton, Roberto Amato, Jeffrey Barrett, Sonia Goncalves, Ewan Harrison, David K. Jackson, Ian Johnston, Dominic Kwiatkowski, Cordelia Langford, John Sillitoe on behalf of the Wellcome Sanger Institute COVID-19 Surveillance Team |
| EPI_ISL_3282717 | Lighthouse Laboratory Plymouth | Wellcome Sanger Institute for the COVID-19 Genomics UK (COG-UK) Consortium | Lighthouse Laboratory Plymouth and Alex Alderton, Roberto Amato, Jeffrey Barrett, Sonia Goncalves, Ewan Harrison, David K. Jackson, Ian Johnston, Dominic Kwiatkowski, Cordelia Langford, John Sillitoe on behalf of the Wellcome Sanger Institute COVID-19 Surveillance Team |
| EPI_ISL_3282718 | Lighthouse Lab in Alderley Park | Wellcome Sanger Institute for the COVID-19 Genomics UK (COG-UK) Consortium | Jacquelyn Wynn, Mairead Hyland, The Lighthouse Lab in Alderley Park and Alex Alderton, Roberto Amato, Jeffrey Barrett, Sonia Goncalves, Ewan Harrison, David K. Jackson, Ian Johnston, Dominic Kwiatkowski, Cordelia Langford, John Sillitoe on behalf of the Wellcome Sanger Institute COVID-19 Surveillance Team |
| EPI_ISL_3282719 | Lighthouse Lab in Glasgow | Wellcome Sanger Institute for the COVID-19 Genomics UK (COG-UK) Consortium | Harper VanSteenhouse, Yumi Kasai, David Gray, Carol Clugston, Anna Dominiczak and Alex Alderton, Roberto Amato, Jeffrey Barrett, Sonia Goncalves, Ewan Harrison, David K. Jackson, Ian Johnston, Dominic Kwiatkowski, Cordelia Langford, John Sillitoe on behalf of the Wellcome Sanger Institute COVID-19 Surveillance Team |
| EPI_ISL_3282720 | Lighthouse Laboratory Plymouth | Wellcome Sanger Institute for the COVID-19 Genomics UK (COG-UK) Consortium | Lighthouse Laboratory Plymouth and Alex Alderton, Roberto Amato, Jeffrey Barrett, Sonia Goncalves, Ewan Harrison, David K. Jackson, Ian Johnston, Dominic Kwiatkowski, Cordelia Langford, John Sillitoe on behalf of the Wellcome Sanger Institute COVID-19 Surveillance Team |
| EPI_ISL_3282721 | Lighthouse Lab in Alderley Park | Wellcome Sanger Institute for the COVID-19 Genomics UK (COG-UK) Consortium | Jacquelyn Wynn, Mairead Hyland, The Lighthouse Lab in Alderley Park and Alex Alderton, Roberto Amato, Jeffrey Barrett, Sonia Goncalves, Ewan Harrison, David K. Jackson, Ian Johnston, Dominic Kwiatkowski, Cordelia Langford, John Sillitoe on behalf of the Wellcome Sanger Institute COVID-19 Surveillance Team |
| EPI_ISL_3282722 | Lighthouse Laboratory Plymouth | Wellcome Sanger Institute for the COVID-19 Genomics UK (COG-UK) Consortium | Lighthouse Laboratory Plymouth and Alex Alderton, Roberto Amato, Jeffrey Barrett, Sonia Goncalves, Ewan Harrison, David K. Jackson, Ian Johnston, Dominic Kwiatkowski, Cordelia Langford, John Sillitoe on behalf of the Wellcome Sanger Institute COVID-19 Surveillance Team |
| EPI_ISL_3282723 | Lighthouse Lab in Glasgow | Wellcome Sanger Institute for the COVID-19 Genomics UK (COG-UK) Consortium | Harper VanSteenhouse, Yumi Kasai, David Gray, Carol Clugston, Anna Dominiczak and Alex Alderton, Roberto Amato, Jeffrey Barrett, Sonia Goncalves, Ewan Harrison, David K. Jackson, Ian Johnston, Dominic Kwiatkowski, Cordelia Langford, John Sillitoe on behalf of the Wellcome Sanger Institute COVID-19 Surveillance Team |
| EPI_ISL_3282724, EPI_ISL_3282725, EPI_ISL_3282726 | Lighthouse Lab in Alderley Park | Wellcome Sanger Institute for the COVID-19 Genomics UK (COG-UK) Consortium | Jacquelyn Wynn, Mairead Hyland, The Lighthouse Lab in Alderley Park and Alex Alderton, Roberto Amato, Jeffrey Barrett, Sonia Goncalves, Ewan Harrison, David K. Jackson, Ian Johnston, Dominic Kwiatkowski, Cordelia Langford, John Sillitoe on behalf of the Wellcome Sanger Institute COVID-19 Surveillance Team |
| EPI_ISL_3282727, EPI_ISL_3282728 | Lighthouse Lab in Glasgow | Wellcome Sanger Institute for the COVID-19 Genomics UK (COG-UK) Consortium | Harper VanSteenhouse, Yumi Kasai, David Gray, Carol Clugston, Anna Dominiczak and Alex Alderton, Roberto Amato, Jeffrey Barrett, Sonia Goncalves, Ewan Harrison, David K. Jackson, Ian Johnston, Dominic Kwiatkowski, Cordelia Langford, John Sillitoe on behalf of the Wellcome Sanger Institute COVID-19 Surveillance Team |
| EPI_ISL_3282729, EPI_ISL_3282730 | Lighthouse Lab in Alderley Park | Wellcome Sanger Institute for the COVID-19 Genomics UK (COG-UK) Consortium | Jacquelyn Wynn, Mairead Hyland, The Lighthouse Lab in Alderley Park and Alex Alderton, Roberto Amato, Jeffrey Barrett, Sonia Goncalves, Ewan Harrison, David K. Jackson, Ian Johnston, Dominic Kwiatkowski, Cordelia Langford, John Sillitoe on behalf of the Wellcome Sanger Institute COVID-19 Surveillance Team |
| EPI_ISL_3282731 | Lighthouse Laboratory Plymouth | Wellcome Sanger Institute for the COVID-19 Genomics UK (COG-UK) Consortium | Lighthouse Laboratory Plymouth and Alex Alderton, Roberto Amato, Jeffrey Barrett, Sonia Goncalves, Ewan Harrison, David K. Jackson, Ian Johnston, Dominic Kwiatkowski, Cordelia Langford, John Sillitoe on behalf of the Wellcome Sanger Institute COVID-19 Surveillance Team |
| EPI_ISL_3282732, EPI_ISL_3282733 | Lighthouse Lab in Alderley Park | Wellcome Sanger Institute for the COVID-19 Genomics UK (COG-UK) Consortium | Jacquelyn Wynn, Mairead Hyland, The Lighthouse Lab in Alderley Park and Alex Alderton, Roberto Amato, Jeffrey Barrett, Sonia Goncalves, Ewan Harrison, David K. Jackson, Ian Johnston, Dominic Kwiatkowski, Cordelia Langford, John Sillitoe on behalf of the Wellcome Sanger Institute COVID-19 Surveillance Team |
| EPI_ISL_3282734 | Lighthouse Laboratory Plymouth | Wellcome Sanger Institute for the COVID-19 Genomics UK (COG-UK) Consortium | Lighthouse Laboratory Plymouth and Alex Alderton, Roberto Amato, Jeffrey Barrett, Sonia Goncalves, Ewan Harrison, David K. Jackson, Ian Johnston, Dominic Kwiatkowski, Cordelia Langford, John Sillitoe on behalf of the Wellcome Sanger Institute COVID-19 Surveillance Team |
| EPI_ISL_3282735, EPI_ISL_3282736 | Lighthouse Lab in Alderley Park | Wellcome Sanger Institute for the COVID-19 Genomics UK (COG-UK) Consortium | Jacquelyn Wynn, Mairead Hyland, The Lighthouse Lab in Alderley Park and Alex Alderton, Roberto Amato, Jeffrey Barrett, Sonia Goncalves, Ewan Harrison, David K. Jackson, Ian Johnston, Dominic Kwiatkowski, Cordelia Langford, John Sillitoe on behalf of the Wellcome Sanger Institute COVID-19 Surveillance Team |
| EPI_ISL_3282737 | Lighthouse Laboratory Plymouth | Wellcome Sanger Institute for the COVID-19 Genomics UK (COG-UK) Consortium | Lighthouse Laboratory Plymouth and Alex Alderton, Roberto Amato, Jeffrey Barrett, Sonia Goncalves, Ewan Harrison, David K. Jackson, Ian Johnston, Dominic Kwiatkowski, Cordelia Langford, John Sillitoe on behalf of the Wellcome Sanger Institute COVID-19 Surveillance Team |
| EPI_ISL_3282738, EPI_ISL_3282739 | Lighthouse Lab in Alderley Park | Wellcome Sanger Institute for the COVID-19 Genomics UK (COG-UK) Consortium | Jacquelyn Wynn, Mairead Hyland, The Lighthouse Lab in Alderley Park and Alex Alderton, Roberto Amato, Jeffrey Barrett, Sonia Goncalves, Ewan Harrison, David K. Jackson, Ian Johnston, Dominic Kwiatkowski, Cordelia Langford, John Sillitoe on behalf of the Wellcome Sanger Institute COVID-19 Surveillance Team |
| EPI_ISL_3282740 | Lighthouse Lab in Glasgow | Wellcome Sanger Institute for the COVID-19 Genomics UK (COG-UK) Consortium | Harper VanSteenhouse, Yumi Kasai, David Gray, Carol Clugston, Anna Dominiczak and Alex Alderton, Roberto Amato, Jeffrey Barrett, Sonia Goncalves, Ewan Harrison, David K. Jackson, Ian Johnston, Dominic Kwiatkowski, Cordelia Langford, John Sillitoe on behalf of the Wellcome Sanger Institute COVID-19 Surveillance Team |
| EPI_ISL_3282741 | Lighthouse Laboratory Plymouth | Wellcome Sanger Institute for the COVID-19 Genomics UK (COG-UK) Consortium | Lighthouse Laboratory Plymouth and Alex Alderton, Roberto Amato, Jeffrey Barrett, Sonia Goncalves, Ewan Harrison, David K. Jackson, Ian Johnston, Dominic Kwiatkowski, Cordelia Langford, John Sillitoe on behalf of the Wellcome Sanger Institute COVID-19 Surveillance Team |
| EPI_ISL_3282742, EPI_ISL_3282743, EPI_ISL_3282744, EPI_ISL_3282745, EPI_ISL_3282746, EPI_ISL_3282747, EPI_ISL_3282748 | Lighthouse Lab in Alderley Park | Wellcome Sanger Institute for the COVID-19 Genomics UK (COG-UK) Consortium | Jacquelyn Wynn, Mairead Hyland, The Lighthouse Lab in Alderley Park and Alex Alderton, Roberto Amato, Jeffrey Barrett, Sonia Goncalves, Ewan Harrison, David K. Jackson, Ian Johnston, Dominic Kwiatkowski, Cordelia Langford, John Sillitoe on behalf of the Wellcome Sanger Institute COVID-19 Surveillance Team |
| EPI_ISL_3282749, EPI_ISL_3282750 | Lighthouse Lab in Glasgow | Wellcome Sanger Institute for the COVID-19 Genomics UK (COG-UK) Consortium | Harper VanSteenhouse, Yumi Kasai, David Gray, Carol Clugston, Anna Dominiczak and Alex Alderton, Roberto Amato, Jeffrey Barrett, Sonia Goncalves, Ewan Harrison, David K. Jackson, Ian Johnston, Dominic Kwiatkowski, Cordelia Langford, John Sillitoe on behalf of the Wellcome Sanger Institute COVID-19 Surveillance Team |
| EPI_ISL_3282751, EPI_ISL_3282752 | Lighthouse Laboratory Plymouth | Wellcome Sanger Institute for the COVID-19 Genomics UK (COG-UK) Consortium | Lighthouse Laboratory Plymouth and Alex Alderton, Roberto Amato, Jeffrey Barrett, Sonia Goncalves, Ewan Harrison, David K. Jackson, Ian Johnston, Dominic Kwiatkowski, Cordelia Langford, John Sillitoe on behalf of the Wellcome Sanger Institute COVID-19 Surveillance Team |
| EPI_ISL_3282753, EPI_ISL_3282754 | Lighthouse Lab in Alderley Park | Wellcome Sanger Institute for the COVID-19 Genomics UK (COG-UK) Consortium | Jacquelyn Wynn, Mairead Hyland, The Lighthouse Lab in Alderley Park and Alex Alderton, Roberto Amato, Jeffrey Barrett, Sonia Goncalves, Ewan Harrison, David K. Jackson, Ian Johnston, Dominic Kwiatkowski, Cordelia Langford, John Sillitoe on behalf of the Wellcome Sanger Institute COVID-19 Surveillance Team |
| EPI_ISL_3282755 | Lighthouse Lab in Glasgow | Wellcome Sanger Institute for the COVID-19 Genomics UK (COG-UK) Consortium | Harper VanSteenhouse, Yumi Kasai, David Gray, Carol Clugston, Anna Dominiczak and Alex Alderton, Roberto Amato, Jeffrey Barrett, Sonia Goncalves, Ewan Harrison, David K. Jackson, Ian Johnston, Dominic Kwiatkowski, Cordelia Langford, John Sillitoe on behalf of the Wellcome Sanger Institute COVID-19 Surveillance Team |
| EPI_ISL_3282756, EPI_ISL_3282757 | Lighthouse Laboratory Plymouth | Wellcome Sanger Institute for the COVID-19 Genomics UK (COG-UK) Consortium | Lighthouse Laboratory Plymouth and Alex Alderton, Roberto Amato, Jeffrey Barrett, Sonia Goncalves, Ewan Harrison, David K. Jackson, Ian Johnston, Dominic Kwiatkowski, Cordelia Langford, John Sillitoe on behalf of the Wellcome Sanger Institute COVID-19 Surveillance Team |
| EPI_ISL_3282758 | Lighthouse Lab in Alderley Park | Wellcome Sanger Institute for the COVID-19 Genomics UK (COG-UK) Consortium | Jacquelyn Wynn, Mairead Hyland, The Lighthouse Lab in Alderley Park and Alex Alderton, Roberto Amato, Jeffrey Barrett, Sonia Goncalves, Ewan Harrison, David K. Jackson, Ian Johnston, Dominic Kwiatkowski, Cordelia Langford, John Sillitoe on behalf of the Wellcome Sanger Institute COVID-19 Surveillance Team |
| EPI_ISL_3282759 | Lighthouse Lab in Glasgow | Wellcome Sanger Institute for the COVID-19 Genomics UK (COG-UK) Consortium | Harper VanSteenhouse, Yumi Kasai, David Gray, Carol Clugston, Anna Dominiczak and Alex Alderton, Roberto Amato, Jeffrey Barrett, Sonia Goncalves, Ewan Harrison, David K. Jackson, Ian Johnston, Dominic Kwiatkowski, Cordelia Langford, John Sillitoe on behalf of the Wellcome Sanger Institute COVID-19 Surveillance Team |
| EPI_ISL_3282760, EPI_ISL_3282761 | Lighthouse Laboratory Plymouth | Wellcome Sanger Institute for the COVID-19 Genomics UK (COG-UK) Consortium | Lighthouse Laboratory Plymouth and Alex Alderton, Roberto Amato, Jeffrey Barrett, Sonia Goncalves, Ewan Harrison, David K. Jackson, Ian Johnston, Dominic Kwiatkowski, Cordelia Langford, John Sillitoe on behalf of the Wellcome Sanger Institute COVID-19 Surveillance Team |
| EPI_ISL_3282762 | Lighthouse Lab in Glasgow | Wellcome Sanger Institute for the COVID-19 Genomics UK (COG-UK) Consortium | Harper VanSteenhouse, Yumi Kasai, David Gray, Carol Clugston, Anna Dominiczak and Alex Alderton, Roberto Amato, Jeffrey Barrett, Sonia Goncalves, Ewan Harrison, David K. Jackson, Ian Johnston, Dominic Kwiatkowski, Cordelia Langford, John Sillitoe on behalf of the Wellcome Sanger Institute COVID-19 Surveillance Team |
| EPI_ISL_3282763 | Lighthouse Laboratory Plymouth | Wellcome Sanger Institute for the COVID-19 Genomics UK (COG-UK) Consortium | Lighthouse Laboratory Plymouth and Alex Alderton, Roberto Amato, Jeffrey Barrett, Sonia Goncalves, Ewan Harrison, David K. Jackson, Ian Johnston, Dominic Kwiatkowski, Cordelia Langford, John Sillitoe on behalf of the Wellcome Sanger Institute COVID-19 Surveillance Team |
| EPI_ISL_3282764 | Lighthouse Lab in Alderley Park | Wellcome Sanger Institute for the COVID-19 Genomics UK (COG-UK) Consortium | Jacquelyn Wynn, Mairead Hyland, The Lighthouse Lab in Alderley Park and Alex Alderton, Roberto Amato, Jeffrey Barrett, Sonia Goncalves, Ewan Harrison, David K. Jackson, Ian Johnston, Dominic Kwiatkowski, Cordelia Langford, John Sillitoe on behalf of the Wellcome Sanger Institute COVID-19 Surveillance Team |
| EPI_ISL_3282765 | Lighthouse Lab in Glasgow | Wellcome Sanger Institute for the COVID-19 Genomics UK (COG-UK) Consortium | Harper VanSteenhouse, Yumi Kasai, David Gray, Carol Clugston, Anna Dominiczak and Alex Alderton, Roberto Amato, Jeffrey Barrett, Sonia Goncalves, Ewan Harrison, David K. Jackson, Ian Johnston, Dominic Kwiatkowski, Cordelia Langford, John Sillitoe on behalf of the Wellcome Sanger Institute COVID-19 Surveillance Team |
| EPI_ISL_3282766 | Lighthouse Lab in Alderley Park | Wellcome Sanger Institute for the COVID-19 Genomics UK (COG-UK) Consortium | Jacquelyn Wynn, Mairead Hyland, The Lighthouse Lab in Alderley Park and Alex Alderton, Roberto Amato, Jeffrey Barrett, Sonia Goncalves, Ewan Harrison, David K. Jackson, Ian Johnston, Dominic Kwiatkowski, Cordelia Langford, John Sillitoe on behalf of the Wellcome Sanger Institute COVID-19 Surveillance Team |
| EPI_ISL_3282767 | Lighthouse Laboratory Plymouth | Wellcome Sanger Institute for the COVID-19 Genomics UK (COG-UK) Consortium | Lighthouse Laboratory Plymouth and Alex Alderton, Roberto Amato, Jeffrey Barrett, Sonia Goncalves, Ewan Harrison, David K. Jackson, Ian Johnston, Dominic Kwiatkowski, Cordelia Langford, John Sillitoe on behalf of the Wellcome Sanger Institute COVID-19 Surveillance Team |
| EPI_ISL_3282768 | Lighthouse Lab in Alderley Park | Wellcome Sanger Institute for the COVID-19 Genomics UK (COG-UK) Consortium | Jacquelyn Wynn, Mairead Hyland, The Lighthouse Lab in Alderley Park and Alex Alderton, Roberto Amato, Jeffrey Barrett, Sonia Goncalves, Ewan Harrison, David K. Jackson, Ian Johnston, Dominic Kwiatkowski, Cordelia Langford, John Sillitoe on behalf of the Wellcome Sanger Institute COVID-19 Surveillance Team |
| EPI_ISL_3282769 | Lighthouse Laboratory Plymouth | Wellcome Sanger Institute for the COVID-19 Genomics UK (COG-UK) Consortium | Lighthouse Laboratory Plymouth and Alex Alderton, Roberto Amato, Jeffrey Barrett, Sonia Goncalves, Ewan Harrison, David K. Jackson, Ian Johnston, Dominic Kwiatkowski, Cordelia Langford, John Sillitoe on behalf of the Wellcome Sanger Institute COVID-19 Surveillance Team |
| EPI_ISL_3282770 | Lighthouse Lab in Glasgow | Wellcome Sanger Institute for the COVID-19 Genomics UK (COG-UK) Consortium | Harper VanSteenhouse, Yumi Kasai, David Gray, Carol Clugston, Anna Dominiczak and Alex Alderton, Roberto Amato, Jeffrey Barrett, Sonia Goncalves, Ewan Harrison, David K. Jackson, Ian Johnston, Dominic Kwiatkowski, Cordelia Langford, John Sillitoe on behalf of the Wellcome Sanger Institute COVID-19 Surveillance Team |
| EPI_ISL_3282771 | Lighthouse Laboratory Plymouth | Wellcome Sanger Institute for the COVID-19 Genomics UK (COG-UK) Consortium | Lighthouse Laboratory Plymouth and Alex Alderton, Roberto Amato, Jeffrey Barrett, Sonia Goncalves, Ewan Harrison, David K. Jackson, Ian Johnston, Dominic Kwiatkowski, Cordelia Langford, John Sillitoe on behalf of the Wellcome Sanger Institute COVID-19 Surveillance Team |
| EPI_ISL_3282772, | Lighthouse Lab in Alderley Park | Wellcome Sanger Institute for the COVID-19 Genomics UK (COG-UK) | Jacquelyn Wynn, Mairead Hyland, The Lighthouse Lab in Alderley Park and Alex Alderton, Roberto Amato, Jeffrey Barrett, Sonia Goncalves, Ewan Harrison, David K. Jackson, Ian Johnston, Dominic Kwiatkowski, Cordelia Langford, John Sillitoe on |

| | | | |
|---|---|---|---|
| EPI_ISL_3282773, EPI_ISL_3282774 | | Consortium | behalf of the Wellcome Sanger Institute COVID-19 Surveillance Team |
| EPI_ISL_3282775 | Lighthouse Lab in Glasgow | Wellcome Sanger Institute for the COVID-19 Genomics UK (COG-UK) Consortium | Harper VanSteenhouse, Yumi Kasai, David Gray, Carol Clugston, Anna Dominiczak and Alex Alderton, Roberto Amato, Jeffrey Barrett, Sonia Goncalves, Ewan Harrison, David K. Jackson, Ian Johnston, Dominic Kwiatkowski, Cordelia Langford, John Sillitoe on behalf of the Wellcome Sanger Institute COVID-19 Surveillance Team |
| EPI_ISL_3282776 | Lighthouse Laboratory Plymouth | Wellcome Sanger Institute for the COVID-19 Genomics UK (COG-UK) Consortium | Lighthouse Laboratory Plymouth and Alex Alderton, Roberto Amato, Jeffrey Barrett, Sonia Goncalves, Ewan Harrison, David K. Jackson, Ian Johnston, Dominic Kwiatkowski, Cordelia Langford, John Sillitoe on behalf of the Wellcome Sanger Institute COVID-19 Surveillance Team |
| EPI_ISL_3282777 | Lighthouse Lab in Glasgow | Wellcome Sanger Institute for the COVID-19 Genomics UK (COG-UK) Consortium | Harper VanSteenhouse, Yumi Kasai, David Gray, Carol Clugston, Anna Dominiczak and Alex Alderton, Roberto Amato, Jeffrey Barrett, Sonia Goncalves, Ewan Harrison, David K. Jackson, Ian Johnston, Dominic Kwiatkowski, Cordelia Langford, John Sillitoe on behalf of the Wellcome Sanger Institute COVID-19 Surveillance Team |
| EPI_ISL_3282778, EPI_ISL_3282779, EPI_ISL_3282780, EPI_ISL_3282781 | Lighthouse Laboratory Plymouth | Wellcome Sanger Institute for the COVID-19 Genomics UK (COG-UK) Consortium | Lighthouse Laboratory Plymouth and Alex Alderton, Roberto Amato, Jeffrey Barrett, Sonia Goncalves, Ewan Harrison, David K. Jackson, Ian Johnston, Dominic Kwiatkowski, Cordelia Langford, John Sillitoe on behalf of the Wellcome Sanger Institute COVID-19 Surveillance Team |
| EPI_ISL_3282782 | Lighthouse Lab in Glasgow | Wellcome Sanger Institute for the COVID-19 Genomics UK (COG-UK) Consortium | Harper VanSteenhouse, Yumi Kasai, David Gray, Carol Clugston, Anna Dominiczak and Alex Alderton, Roberto Amato, Jeffrey Barrett, Sonia Goncalves, Ewan Harrison, David K. Jackson, Ian Johnston, Dominic Kwiatkowski, Cordelia Langford, John Sillitoe on behalf of the Wellcome Sanger Institute COVID-19 Surveillance Team |
| EPI_ISL_3282783, EPI_ISL_3282784 | Lighthouse Lab in Alderley Park | Wellcome Sanger Institute for the COVID-19 Genomics UK (COG-UK) Consortium | Jacquelyn Wynn, Mairead Hyland, The Lighthouse Lab in Alderley Park and Alex Alderton, Roberto Amato, Jeffrey Barrett, Sonia Goncalves, Ewan Harrison, David K. Jackson, Ian Johnston, Dominic Kwiatkowski, Cordelia Langford, John Sillitoe on behalf of the Wellcome Sanger Institute COVID-19 Surveillance Team |
| EPI_ISL_3282785 | Lighthouse Laboratory Plymouth | Wellcome Sanger Institute for the COVID-19 Genomics UK (COG-UK) Consortium | Lighthouse Laboratory Plymouth and Alex Alderton, Roberto Amato, Jeffrey Barrett, Sonia Goncalves, Ewan Harrison, David K. Jackson, Ian Johnston, Dominic Kwiatkowski, Cordelia Langford, John Sillitoe on behalf of the Wellcome Sanger Institute COVID-19 Surveillance Team |
| EPI_ISL_3282786, EPI_ISL_3282787 | Lighthouse Lab in Glasgow | Wellcome Sanger Institute for the COVID-19 Genomics UK (COG-UK) Consortium | Harper VanSteenhouse, Yumi Kasai, David Gray, Carol Clugston, Anna Dominiczak and Alex Alderton, Roberto Amato, Jeffrey Barrett, Sonia Goncalves, Ewan Harrison, David K. Jackson, Ian Johnston, Dominic Kwiatkowski, Cordelia Langford, John Sillitoe on behalf of the Wellcome Sanger Institute COVID-19 Surveillance Team |
| EPI_ISL_3282788, EPI_ISL_3282789 | Lighthouse Lab in Alderley Park | Wellcome Sanger Institute for the COVID-19 Genomics UK (COG-UK) Consortium | Jacquelyn Wynn, Mairead Hyland, The Lighthouse Lab in Alderley Park and Alex Alderton, Roberto Amato, Jeffrey Barrett, Sonia Goncalves, Ewan Harrison, David K. Jackson, Ian Johnston, Dominic Kwiatkowski, Cordelia Langford, John Sillitoe on behalf of the Wellcome Sanger Institute COVID-19 Surveillance Team |
| EPI_ISL_3282790 | Lighthouse Lab in Glasgow | Wellcome Sanger Institute for the COVID-19 Genomics UK (COG-UK) Consortium | Harper VanSteenhouse, Yumi Kasai, David Gray, Carol Clugston, Anna Dominiczak and Alex Alderton, Roberto Amato, Jeffrey Barrett, Sonia Goncalves, Ewan Harrison, David K. Jackson, Ian Johnston, Dominic Kwiatkowski, Cordelia Langford, John Sillitoe on behalf of the Wellcome Sanger Institute COVID-19 Surveillance Team |
| EPI_ISL_3282791, EPI_ISL_3282792, EPI_ISL_3282793 | Lighthouse Lab in Alderley Park | Wellcome Sanger Institute for the COVID-19 Genomics UK (COG-UK) Consortium | Jacquelyn Wynn, Mairead Hyland, The Lighthouse Lab in Alderley Park and Alex Alderton, Roberto Amato, Jeffrey Barrett, Sonia Goncalves, Ewan Harrison, David K. Jackson, Ian Johnston, Dominic Kwiatkowski, Cordelia Langford, John Sillitoe on behalf of the Wellcome Sanger Institute COVID-19 Surveillance Team |
| EPI_ISL_3282794 | Lighthouse Laboratory Plymouth | Wellcome Sanger Institute for the COVID-19 Genomics UK (COG-UK) Consortium | Lighthouse Laboratory Plymouth and Alex Alderton, Roberto Amato, Jeffrey Barrett, Sonia Goncalves, Ewan Harrison, David K. Jackson, Ian Johnston, Dominic Kwiatkowski, Cordelia Langford, John Sillitoe on behalf of the Wellcome Sanger Institute COVID-19 Surveillance Team |
| EPI_ISL_3282795, EPI_ISL_3282796 | Lighthouse Lab in Alderley Park | Wellcome Sanger Institute for the COVID-19 Genomics UK (COG-UK) Consortium | Jacquelyn Wynn, Mairead Hyland, The Lighthouse Lab in Alderley Park and Alex Alderton, Roberto Amato, Jeffrey Barrett, Sonia Goncalves, Ewan Harrison, David K. Jackson, Ian Johnston, Dominic Kwiatkowski, Cordelia Langford, John Sillitoe on behalf of the Wellcome Sanger Institute COVID-19 Surveillance Team |
| EPI_ISL_3282797 | Lighthouse Laboratory Plymouth | Wellcome Sanger Institute for the COVID-19 Genomics UK (COG-UK) Consortium | Lighthouse Laboratory Plymouth and Alex Alderton, Roberto Amato, Jeffrey Barrett, Sonia Goncalves, Ewan Harrison, David K. Jackson, Ian Johnston, Dominic Kwiatkowski, Cordelia Langford, John Sillitoe on behalf of the Wellcome Sanger Institute COVID-19 Surveillance Team |
| EPI_ISL_3282798 | Lighthouse Lab in Glasgow | Wellcome Sanger Institute for the COVID-19 Genomics UK (COG-UK) Consortium | Harper VanSteenhouse, Yumi Kasai, David Gray, Carol Clugston, Anna Dominiczak and Alex Alderton, Roberto Amato, Jeffrey Barrett, Sonia Goncalves, Ewan Harrison, David K. Jackson, Ian Johnston, Dominic Kwiatkowski, Cordelia Langford, John Sillitoe on behalf of the Wellcome Sanger Institute COVID-19 Surveillance Team |
| EPI_ISL_3282799, EPI_ISL_3282800, EPI_ISL_3282801, EPI_ISL_3282802, EPI_ISL_3282803 | Lighthouse Lab in Alderley Park | Wellcome Sanger Institute for the COVID-19 Genomics UK (COG-UK) Consortium | Jacquelyn Wynn, Mairead Hyland, The Lighthouse Lab in Alderley Park and Alex Alderton, Roberto Amato, Jeffrey Barrett, Sonia Goncalves, Ewan Harrison, David K. Jackson, Ian Johnston, Dominic Kwiatkowski, Cordelia Langford, John Sillitoe on behalf of the Wellcome Sanger Institute COVID-19 Surveillance Team |
| EPI_ISL_3282804, EPI_ISL_3282805, EPI_ISL_3282806 | Lighthouse Lab in Glasgow | Wellcome Sanger Institute for the COVID-19 Genomics UK (COG-UK) Consortium | Harper VanSteenhouse, Yumi Kasai, David Gray, Carol Clugston, Anna Dominiczak and Alex Alderton, Roberto Amato, Jeffrey Barrett, Sonia Goncalves, Ewan Harrison, David K. Jackson, Ian Johnston, Dominic Kwiatkowski, Cordelia Langford, John Sillitoe on behalf of the Wellcome Sanger Institute COVID-19 Surveillance Team |
| EPI_ISL_3282807 | Lighthouse Laboratory Plymouth | Wellcome Sanger Institute for the COVID-19 Genomics UK (COG-UK) Consortium | Lighthouse Laboratory Plymouth and Alex Alderton, Roberto Amato, Jeffrey Barrett, Sonia Goncalves, Ewan Harrison, David K. Jackson, Ian Johnston, Dominic Kwiatkowski, Cordelia Langford, John Sillitoe on behalf of the Wellcome Sanger Institute COVID-19 Surveillance Team |
| EPI_ISL_3282808 | Lighthouse Lab in Glasgow | Wellcome Sanger Institute for the COVID-19 Genomics UK (COG-UK) Consortium | Harper VanSteenhouse, Yumi Kasai, David Gray, Carol Clugston, Anna Dominiczak and Alex Alderton, Roberto Amato, Jeffrey Barrett, Sonia Goncalves, Ewan Harrison, David K. Jackson, Ian Johnston, Dominic Kwiatkowski, Cordelia Langford, John Sillitoe on behalf of the Wellcome Sanger Institute COVID-19 Surveillance Team |
| EPI_ISL_3282809 | Lighthouse Lab in Alderley Park | Wellcome Sanger Institute for the COVID-19 Genomics UK (COG-UK) Consortium | Jacquelyn Wynn, Mairead Hyland, The Lighthouse Lab in Alderley Park and Alex Alderton, Roberto Amato, Jeffrey Barrett, Sonia Goncalves, Ewan Harrison, David K. Jackson, Ian Johnston, Dominic Kwiatkowski, Cordelia Langford, John Sillitoe on behalf of the Wellcome Sanger Institute COVID-19 Surveillance Team |
| EPI_ISL_3282810 | Lighthouse Lab in Glasgow | Wellcome Sanger Institute for the COVID-19 Genomics UK (COG-UK) Consortium | Harper VanSteenhouse, Yumi Kasai, David Gray, Carol Clugston, Anna Dominiczak and Alex Alderton, Roberto Amato, Jeffrey Barrett, Sonia Goncalves, Ewan Harrison, David K. Jackson, Ian Johnston, Dominic Kwiatkowski, Cordelia Langford, John Sillitoe on behalf of the Wellcome Sanger Institute COVID-19 Surveillance Team |
| EPI_ISL_3282811, EPI_ISL_3282812, EPI_ISL_3282813 | Lighthouse Lab in Alderley Park | Wellcome Sanger Institute for the COVID-19 Genomics UK (COG-UK) Consortium | Jacquelyn Wynn, Mairead Hyland, The Lighthouse Lab in Alderley Park and Alex Alderton, Roberto Amato, Jeffrey Barrett, Sonia Goncalves, Ewan Harrison, David K. Jackson, Ian Johnston, Dominic Kwiatkowski, Cordelia Langford, John Sillitoe on behalf of the Wellcome Sanger Institute COVID-19 Surveillance Team |
| EPI_ISL_3282814 | Lighthouse Laboratory Plymouth | Wellcome Sanger Institute for the COVID-19 Genomics UK (COG-UK) Consortium | Lighthouse Laboratory Plymouth and Alex Alderton, Roberto Amato, Jeffrey Barrett, Sonia Goncalves, Ewan Harrison, David K. Jackson, Ian Johnston, Dominic Kwiatkowski, Cordelia Langford, John Sillitoe on behalf of the Wellcome Sanger Institute COVID-19 Surveillance Team |
| EPI_ISL_3282815, EPI_ISL_3282816, EPI_ISL_3282817, EPI_ISL_3282818 | Lighthouse Lab in Alderley Park | Wellcome Sanger Institute for the COVID-19 Genomics UK (COG-UK) Consortium | Jacquelyn Wynn, Mairead Hyland, The Lighthouse Lab in Alderley Park and Alex Alderton, Roberto Amato, Jeffrey Barrett, Sonia Goncalves, Ewan Harrison, David K. Jackson, Ian Johnston, Dominic Kwiatkowski, Cordelia Langford, John Sillitoe on behalf of the Wellcome Sanger Institute COVID-19 Surveillance Team |
| EPI_ISL_3282819 | Lighthouse Laboratory Plymouth | Wellcome Sanger Institute for the COVID-19 Genomics UK (COG-UK) Consortium | Lighthouse Laboratory Plymouth and Alex Alderton, Roberto Amato, Jeffrey Barrett, Sonia Goncalves, Ewan Harrison, David K. Jackson, Ian Johnston, Dominic Kwiatkowski, Cordelia Langford, John Sillitoe on behalf of the Wellcome Sanger Institute COVID-19 Surveillance Team |
| EPI_ISL_3282820 | Lighthouse Lab in Glasgow | Wellcome Sanger Institute for the COVID-19 Genomics UK (COG-UK) Consortium | Harper VanSteenhouse, Yumi Kasai, David Gray, Carol Clugston, Anna Dominiczak and Alex Alderton, Roberto Amato, Jeffrey Barrett, Sonia Goncalves, Ewan Harrison, David K. Jackson, Ian Johnston, Dominic Kwiatkowski, Cordelia Langford, John Sillitoe on behalf of the Wellcome Sanger Institute COVID-19 Surveillance Team |
| EPI_ISL_3282821, EPI_ISL_3282822, EPI_ISL_3282823 | Lighthouse Lab in Alderley Park | Wellcome Sanger Institute for the COVID-19 Genomics UK (COG-UK) Consortium | Jacquelyn Wynn, Mairead Hyland, The Lighthouse Lab in Alderley Park and Alex Alderton, Roberto Amato, Jeffrey Barrett, Sonia Goncalves, Ewan Harrison, David K. Jackson, Ian Johnston, Dominic Kwiatkowski, Cordelia Langford, John Sillitoe on behalf of the Wellcome Sanger Institute COVID-19 Surveillance Team |
| EPI_ISL_3282824 | Lighthouse Laboratory Plymouth | Wellcome Sanger Institute for the COVID-19 Genomics UK (COG-UK) Consortium | Lighthouse Laboratory Plymouth and Alex Alderton, Roberto Amato, Jeffrey Barrett, Sonia Goncalves, Ewan Harrison, David K. Jackson, Ian Johnston, Dominic Kwiatkowski, Cordelia Langford, John Sillitoe on behalf of the Wellcome Sanger Institute COVID-19 Surveillance Team |
| EPI_ISL_3282825, EPI_ISL_3282826 | Lighthouse Lab in Alderley Park | Wellcome Sanger Institute for the COVID-19 Genomics UK (COG-UK) Consortium | Jacquelyn Wynn, Mairead Hyland, The Lighthouse Lab in Alderley Park and Alex Alderton, Roberto Amato, Jeffrey Barrett, Sonia Goncalves, Ewan Harrison, David K. Jackson, Ian Johnston, Dominic Kwiatkowski, Cordelia Langford, John Sillitoe on behalf of the Wellcome Sanger Institute COVID-19 Surveillance Team |
| EPI_ISL_3282827 | Lighthouse Laboratory Plymouth | Wellcome Sanger Institute for the COVID-19 Genomics UK (COG-UK) Consortium | Lighthouse Laboratory Plymouth and Alex Alderton, Roberto Amato, Jeffrey Barrett, Sonia Goncalves, Ewan Harrison, David K. Jackson, Ian Johnston, Dominic Kwiatkowski, Cordelia Langford, John Sillitoe on behalf of the Wellcome Sanger Institute COVID-19 Surveillance Team |
| EPI_ISL_3282828 | Lighthouse Lab in Alderley Park | Wellcome Sanger Institute for the COVID-19 Genomics UK (COG-UK) Consortium | Jacquelyn Wynn, Mairead Hyland, The Lighthouse Lab in Alderley Park and Alex Alderton, Roberto Amato, Jeffrey Barrett, Sonia Goncalves, Ewan Harrison, David K. Jackson, Ian Johnston, Dominic Kwiatkowski, Cordelia Langford, John Sillitoe on behalf of the Wellcome Sanger Institute COVID-19 Surveillance Team |
| EPI_ISL_3282829, EPI_ISL_3282830 | Lighthouse Laboratory Plymouth | Wellcome Sanger Institute for the COVID-19 Genomics UK (COG-UK) Consortium | Lighthouse Laboratory Plymouth and Alex Alderton, Roberto Amato, Jeffrey Barrett, Sonia Goncalves, Ewan Harrison, David K. Jackson, Ian Johnston, Dominic Kwiatkowski, Cordelia Langford, John Sillitoe on behalf of the Wellcome Sanger Institute COVID-19 Surveillance Team |
| EPI_ISL_3282831 | Lighthouse Lab in Alderley Park | Wellcome Sanger Institute for the COVID-19 Genomics UK (COG-UK) Consortium | Jacquelyn Wynn, Mairead Hyland, The Lighthouse Lab in Alderley Park and Alex Alderton, Roberto Amato, Jeffrey Barrett, Sonia Goncalves, Ewan Harrison, David K. Jackson, Ian Johnston, Dominic Kwiatkowski, Cordelia Langford, John Sillitoe on behalf of the Wellcome Sanger Institute COVID-19 Surveillance Team |
| EPI_ISL_3282832 | Lighthouse Lab in Glasgow | Wellcome Sanger Institute for the COVID-19 Genomics UK (COG-UK) Consortium | Harper VanSteenhouse, Yumi Kasai, David Gray, Carol Clugston, Anna Dominiczak and Alex Alderton, Roberto Amato, Jeffrey Barrett, Sonia Goncalves, Ewan Harrison, David K. Jackson, Ian Johnston, Dominic Kwiatkowski, Cordelia Langford, John Sillitoe on behalf of the Wellcome Sanger Institute COVID-19 Surveillance Team |
| EPI_ISL_3282833 | Lighthouse Lab in Alderley Park | Wellcome Sanger Institute for the COVID-19 Genomics UK (COG-UK) Consortium | Jacquelyn Wynn, Mairead Hyland, The Lighthouse Lab in Alderley Park and Alex Alderton, Roberto Amato, Jeffrey Barrett, Sonia Goncalves, Ewan Harrison, David K. Jackson, Ian Johnston, Dominic Kwiatkowski, Cordelia Langford, John Sillitoe on behalf of the Wellcome Sanger Institute COVID-19 Surveillance Team |
| EPI_ISL_3282834 | Lighthouse Lab in Glasgow | Wellcome Sanger Institute for the COVID-19 Genomics UK (COG-UK) Consortium | Harper VanSteenhouse, Yumi Kasai, David Gray, Carol Clugston, Anna Dominiczak and Alex Alderton, Roberto Amato, Jeffrey Barrett, Sonia Goncalves, Ewan Harrison, David K. Jackson, Ian Johnston, Dominic Kwiatkowski, Cordelia Langford, John Sillitoe on behalf of the Wellcome Sanger Institute COVID-19 Surveillance Team |
| EPI_ISL_3282835, EPI_ISL_3282836 | Lighthouse Lab in Alderley Park | Wellcome Sanger Institute for the COVID-19 Genomics UK (COG-UK) Consortium | Jacquelyn Wynn, Mairead Hyland, The Lighthouse Lab in Alderley Park and Alex Alderton, Roberto Amato, Jeffrey Barrett, Sonia Goncalves, Ewan Harrison, David K. Jackson, Ian Johnston, Dominic Kwiatkowski, Cordelia Langford, John Sillitoe on behalf of the Wellcome Sanger Institute COVID-19 Surveillance Team |
| EPI_ISL_3282837 | Lighthouse Laboratory Plymouth | Wellcome Sanger Institute for the COVID-19 Genomics UK (COG-UK) Consortium | Lighthouse Laboratory Plymouth and Alex Alderton, Roberto Amato, Jeffrey Barrett, Sonia Goncalves, Ewan Harrison, David K. Jackson, Ian Johnston, Dominic Kwiatkowski, Cordelia Langford, John Sillitoe on behalf of the Wellcome Sanger Institute COVID-19 Surveillance Team |
| EPI_ISL_3282838, EPI_ISL_3282839 | Lighthouse Lab in Alderley Park | Wellcome Sanger Institute for the COVID-19 Genomics UK (COG-UK) Consortium | Jacquelyn Wynn, Mairead Hyland, The Lighthouse Lab in Alderley Park and Alex Alderton, Roberto Amato, Jeffrey Barrett, Sonia Goncalves, Ewan Harrison, David K. Jackson, Ian Johnston, Dominic Kwiatkowski, Cordelia Langford, John Sillitoe on behalf of the Wellcome Sanger Institute COVID-19 Surveillance Team |
| EPI_ISL_3282840, EPI_ISL_3282841 | Lighthouse Lab in Glasgow | Wellcome Sanger Institute for the COVID-19 Genomics UK (COG-UK) Consortium | Harper VanSteenhouse, Yumi Kasai, David Gray, Carol Clugston, Anna Dominiczak and Alex Alderton, Roberto Amato, Jeffrey Barrett, Sonia Goncalves, Ewan Harrison, David K. Jackson, Ian Johnston, Dominic Kwiatkowski, Cordelia Langford, John Sillitoe on behalf of the Wellcome Sanger Institute COVID-19 Surveillance Team |

| | | | |
|---|---|---|---|
| EPI_ISL_3282842 | Lighthouse Lab in Alderley Park | Wellcome Sanger Institute for the COVID-19 Genomics UK (COG-UK) Consortium | Jacquelyn Wynn, Mairead Hyland, The Lighthouse Lab in Alderley Park and Alex Alderton, Roberto Amato, Jeffrey Barrett, Sonia Goncalves, Ewan Harrison, David K. Jackson, Ian Johnston, Dominic Kwiatkowski, Cordelia Langford, John Sillitoe on behalf of the Wellcome Sanger Institute COVID-19 Surveillance Team |
| EPI_ISL_3282843, EPI_ISL_3282844, EPI_ISL_3282845, EPI_ISL_3282846 | Lighthouse Laboratory Plymouth | Wellcome Sanger Institute for the COVID-19 Genomics UK (COG-UK) Consortium | Lighthouse Laboratory Plymouth and Alex Alderton, Roberto Amato, Jeffrey Barrett, Sonia Goncalves, Ewan Harrison, David K. Jackson, Ian Johnston, Dominic Kwiatkowski, Cordelia Langford, John Sillitoe on behalf of the Wellcome Sanger Institute COVID-19 Surveillance Team |
| EPI_ISL_3282847, EPI_ISL_3282848 | Lighthouse Lab in Glasgow | Wellcome Sanger Institute for the COVID-19 Genomics UK (COG-UK) Consortium | Harper VanSteenhouse, Yumi Kasai, David Gray, Carol Clugston, Anna Dominiczak and Alex Alderton, Roberto Amato, Jeffrey Barrett, Sonia Goncalves, Ewan Harrison, David K. Jackson, Ian Johnston, Dominic Kwiatkowski, Cordelia Langford, John Sillitoe on behalf of the Wellcome Sanger Institute COVID-19 Surveillance Team |
| EPI_ISL_3282849 | Lighthouse Lab in Alderley Park | Wellcome Sanger Institute for the COVID-19 Genomics UK (COG-UK) Consortium | Jacquelyn Wynn, Mairead Hyland, The Lighthouse Lab in Alderley Park and Alex Alderton, Roberto Amato, Jeffrey Barrett, Sonia Goncalves, Ewan Harrison, David K. Jackson, Ian Johnston, Dominic Kwiatkowski, Cordelia Langford, John Sillitoe on behalf of the Wellcome Sanger Institute COVID-19 Surveillance Team |
| EPI_ISL_3282850 | Lighthouse Lab in Glasgow | Wellcome Sanger Institute for the COVID-19 Genomics UK (COG-UK) Consortium | Harper VanSteenhouse, Yumi Kasai, David Gray, Carol Clugston, Anna Dominiczak and Alex Alderton, Roberto Amato, Jeffrey Barrett, Sonia Goncalves, Ewan Harrison, David K. Jackson, Ian Johnston, Dominic Kwiatkowski, Cordelia Langford, John Sillitoe on behalf of the Wellcome Sanger Institute COVID-19 Surveillance Team |
| EPI_ISL_3282851, EPI_ISL_3282852, EPI_ISL_3282853 | Lighthouse Laboratory Plymouth | Wellcome Sanger Institute for the COVID-19 Genomics UK (COG-UK) Consortium | Lighthouse Laboratory Plymouth and Alex Alderton, Roberto Amato, Jeffrey Barrett, Sonia Goncalves, Ewan Harrison, David K. Jackson, Ian Johnston, Dominic Kwiatkowski, Cordelia Langford, John Sillitoe on behalf of the Wellcome Sanger Institute COVID-19 Surveillance Team |
| EPI_ISL_3282854 | Lighthouse Lab in Alderley Park | Wellcome Sanger Institute for the COVID-19 Genomics UK (COG-UK) Consortium | Jacquelyn Wynn, Mairead Hyland, The Lighthouse Lab in Alderley Park and Alex Alderton, Roberto Amato, Jeffrey Barrett, Sonia Goncalves, Ewan Harrison, David K. Jackson, Ian Johnston, Dominic Kwiatkowski, Cordelia Langford, John Sillitoe on behalf of the Wellcome Sanger Institute COVID-19 Surveillance Team |
| EPI_ISL_3282855 | Lighthouse Lab in Glasgow | Wellcome Sanger Institute for the COVID-19 Genomics UK (COG-UK) Consortium | Harper VanSteenhouse, Yumi Kasai, David Gray, Carol Clugston, Anna Dominiczak and Alex Alderton, Roberto Amato, Jeffrey Barrett, Sonia Goncalves, Ewan Harrison, David K. Jackson, Ian Johnston, Dominic Kwiatkowski, Cordelia Langford, John Sillitoe on behalf of the Wellcome Sanger Institute COVID-19 Surveillance Team |
| EPI_ISL_3282856, EPI_ISL_3282857 | Lighthouse Laboratory Plymouth | Wellcome Sanger Institute for the COVID-19 Genomics UK (COG-UK) Consortium | Lighthouse Laboratory Plymouth and Alex Alderton, Roberto Amato, Jeffrey Barrett, Sonia Goncalves, Ewan Harrison, David K. Jackson, Ian Johnston, Dominic Kwiatkowski, Cordelia Langford, John Sillitoe on behalf of the Wellcome Sanger Institute COVID-19 Surveillance Team |
| EPI_ISL_3282858 | Lighthouse Lab in Alderley Park | Wellcome Sanger Institute for the COVID-19 Genomics UK (COG-UK) Consortium | Jacquelyn Wynn, Mairead Hyland, The Lighthouse Lab in Alderley Park and Alex Alderton, Roberto Amato, Jeffrey Barrett, Sonia Goncalves, Ewan Harrison, David K. Jackson, Ian Johnston, Dominic Kwiatkowski, Cordelia Langford, John Sillitoe on behalf of the Wellcome Sanger Institute COVID-19 Surveillance Team |
| EPI_ISL_3282859, EPI_ISL_3282860 | Lighthouse Laboratory Plymouth | Wellcome Sanger Institute for the COVID-19 Genomics UK (COG-UK) Consortium | Lighthouse Laboratory Plymouth and Alex Alderton, Roberto Amato, Jeffrey Barrett, Sonia Goncalves, Ewan Harrison, David K. Jackson, Ian Johnston, Dominic Kwiatkowski, Cordelia Langford, John Sillitoe on behalf of the Wellcome Sanger Institute COVID-19 Surveillance Team |
| EPI_ISL_3282861 | Lighthouse Lab in Alderley Park | Wellcome Sanger Institute for the COVID-19 Genomics UK (COG-UK) Consortium | Jacquelyn Wynn, Mairead Hyland, The Lighthouse Lab in Alderley Park and Alex Alderton, Roberto Amato, Jeffrey Barrett, Sonia Goncalves, Ewan Harrison, David K. Jackson, Ian Johnston, Dominic Kwiatkowski, Cordelia Langford, John Sillitoe on behalf of the Wellcome Sanger Institute COVID-19 Surveillance Team |
| EPI_ISL_3282862 | Lighthouse Lab in Glasgow | Wellcome Sanger Institute for the COVID-19 Genomics UK (COG-UK) Consortium | Harper VanSteenhouse, Yumi Kasai, David Gray, Carol Clugston, Anna Dominiczak and Alex Alderton, Roberto Amato, Jeffrey Barrett, Sonia Goncalves, Ewan Harrison, David K. Jackson, Ian Johnston, Dominic Kwiatkowski, Cordelia Langford, John Sillitoe on behalf of the Wellcome Sanger Institute COVID-19 Surveillance Team |
| EPI_ISL_3282863 | Lighthouse Lab in Alderley Park | Wellcome Sanger Institute for the COVID-19 Genomics UK (COG-UK) Consortium | Jacquelyn Wynn, Mairead Hyland, The Lighthouse Lab in Alderley Park and Alex Alderton, Roberto Amato, Jeffrey Barrett, Sonia Goncalves, Ewan Harrison, David K. Jackson, Ian Johnston, Dominic Kwiatkowski, Cordelia Langford, John Sillitoe on behalf of the Wellcome Sanger Institute COVID-19 Surveillance Team |
| EPI_ISL_3282864 | Lighthouse Lab in Glasgow | Wellcome Sanger Institute for the COVID-19 Genomics UK (COG-UK) Consortium | Harper VanSteenhouse, Yumi Kasai, David Gray, Carol Clugston, Anna Dominiczak and Alex Alderton, Roberto Amato, Jeffrey Barrett, Sonia Goncalves, Ewan Harrison, David K. Jackson, Ian Johnston, Dominic Kwiatkowski, Cordelia Langford, John Sillitoe on behalf of the Wellcome Sanger Institute COVID-19 Surveillance Team |
| EPI_ISL_3282865 | Lighthouse Lab in Alderley Park | Wellcome Sanger Institute for the COVID-19 Genomics UK (COG-UK) Consortium | Jacquelyn Wynn, Mairead Hyland, The Lighthouse Lab in Alderley Park and Alex Alderton, Roberto Amato, Jeffrey Barrett, Sonia Goncalves, Ewan Harrison, David K. Jackson, Ian Johnston, Dominic Kwiatkowski, Cordelia Langford, John Sillitoe on behalf of the Wellcome Sanger Institute COVID-19 Surveillance Team |
| EPI_ISL_3282866 | Lighthouse Lab in Glasgow | Wellcome Sanger Institute for the COVID-19 Genomics UK (COG-UK) Consortium | Harper VanSteenhouse, Yumi Kasai, David Gray, Carol Clugston, Anna Dominiczak and Alex Alderton, Roberto Amato, Jeffrey Barrett, Sonia Goncalves, Ewan Harrison, David K. Jackson, Ian Johnston, Dominic Kwiatkowski, Cordelia Langford, John Sillitoe on behalf of the Wellcome Sanger Institute COVID-19 Surveillance Team |
| EPI_ISL_3282867, EPI_ISL_3282868, EPI_ISL_3282869, EPI_ISL_3282870, EPI_ISL_3282871 | Lighthouse Lab in Alderley Park | Wellcome Sanger Institute for the COVID-19 Genomics UK (COG-UK) Consortium | Jacquelyn Wynn, Mairead Hyland, The Lighthouse Lab in Alderley Park and Alex Alderton, Roberto Amato, Jeffrey Barrett, Sonia Goncalves, Ewan Harrison, David K. Jackson, Ian Johnston, Dominic Kwiatkowski, Cordelia Langford, John Sillitoe on behalf of the Wellcome Sanger Institute COVID-19 Surveillance Team |
| EPI_ISL_3282872, EPI_ISL_3282873 | Lighthouse Lab in Glasgow | Wellcome Sanger Institute for the COVID-19 Genomics UK (COG-UK) Consortium | Harper VanSteenhouse, Yumi Kasai, David Gray, Carol Clugston, Anna Dominiczak and Alex Alderton, Roberto Amato, Jeffrey Barrett, Sonia Goncalves, Ewan Harrison, David K. Jackson, Ian Johnston, Dominic Kwiatkowski, Cordelia Langford, John Sillitoe on behalf of the Wellcome Sanger Institute COVID-19 Surveillance Team |
| EPI_ISL_3282874 | Lighthouse Lab in Alderley Park | Wellcome Sanger Institute for the COVID-19 Genomics UK (COG-UK) Consortium | Jacquelyn Wynn, Mairead Hyland, The Lighthouse Lab in Alderley Park and Alex Alderton, Roberto Amato, Jeffrey Barrett, Sonia Goncalves, Ewan Harrison, David K. Jackson, Ian Johnston, Dominic Kwiatkowski, Cordelia Langford, John Sillitoe on behalf of the Wellcome Sanger Institute COVID-19 Surveillance Team |
| EPI_ISL_3282875 | Lighthouse Lab in Glasgow | Wellcome Sanger Institute for the COVID-19 Genomics UK (COG-UK) Consortium | Harper VanSteenhouse, Yumi Kasai, David Gray, Carol Clugston, Anna Dominiczak and Alex Alderton, Roberto Amato, Jeffrey Barrett, Sonia Goncalves, Ewan Harrison, David K. Jackson, Ian Johnston, Dominic Kwiatkowski, Cordelia Langford, John Sillitoe on behalf of the Wellcome Sanger Institute COVID-19 Surveillance Team |
| EPI_ISL_3282876 | Lighthouse Lab in Alderley Park | Wellcome Sanger Institute for the COVID-19 Genomics UK (COG-UK) Consortium | Jacquelyn Wynn, Mairead Hyland, The Lighthouse Lab in Alderley Park and Alex Alderton, Roberto Amato, Jeffrey Barrett, Sonia Goncalves, Ewan Harrison, David K. Jackson, Ian Johnston, Dominic Kwiatkowski, Cordelia Langford, John Sillitoe on behalf of the Wellcome Sanger Institute COVID-19 Surveillance Team |
| EPI_ISL_3282877, EPI_ISL_3282878 | Lighthouse Laboratory Plymouth | Wellcome Sanger Institute for the COVID-19 Genomics UK (COG-UK) Consortium | Lighthouse Laboratory Plymouth and Alex Alderton, Roberto Amato, Jeffrey Barrett, Sonia Goncalves, Ewan Harrison, David K. Jackson, Ian Johnston, Dominic Kwiatkowski, Cordelia Langford, John Sillitoe on behalf of the Wellcome Sanger Institute COVID-19 Surveillance Team |
| EPI_ISL_3282879 | Lighthouse Lab in Alderley Park | Wellcome Sanger Institute for the COVID-19 Genomics UK (COG-UK) Consortium | Jacquelyn Wynn, Mairead Hyland, The Lighthouse Lab in Alderley Park and Alex Alderton, Roberto Amato, Jeffrey Barrett, Sonia Goncalves, Ewan Harrison, David K. Jackson, Ian Johnston, Dominic Kwiatkowski, Cordelia Langford, John Sillitoe on behalf of the Wellcome Sanger Institute COVID-19 Surveillance Team |
| EPI_ISL_3282880 | Lighthouse Laboratory Plymouth | Wellcome Sanger Institute for the COVID-19 Genomics UK (COG-UK) Consortium | Lighthouse Laboratory Plymouth and Alex Alderton, Roberto Amato, Jeffrey Barrett, Sonia Goncalves, Ewan Harrison, David K. Jackson, Ian Johnston, Dominic Kwiatkowski, Cordelia Langford, John Sillitoe on behalf of the Wellcome Sanger Institute COVID-19 Surveillance Team |
| EPI_ISL_3282881 | Lighthouse Lab in Alderley Park | Wellcome Sanger Institute for the COVID-19 Genomics UK (COG-UK) Consortium | Jacquelyn Wynn, Mairead Hyland, The Lighthouse Lab in Alderley Park and Alex Alderton, Roberto Amato, Jeffrey Barrett, Sonia Goncalves, Ewan Harrison, David K. Jackson, Ian Johnston, Dominic Kwiatkowski, Cordelia Langford, John Sillitoe on behalf of the Wellcome Sanger Institute COVID-19 Surveillance Team |
| EPI_ISL_3282882, EPI_ISL_3282883 | Lighthouse Lab in Glasgow | Wellcome Sanger Institute for the COVID-19 Genomics UK (COG-UK) Consortium | Harper VanSteenhouse, Yumi Kasai, David Gray, Carol Clugston, Anna Dominiczak and Alex Alderton, Roberto Amato, Jeffrey Barrett, Sonia Goncalves, Ewan Harrison, David K. Jackson, Ian Johnston, Dominic Kwiatkowski, Cordelia Langford, John Sillitoe on behalf of the Wellcome Sanger Institute COVID-19 Surveillance Team |
| EPI_ISL_3282884 | Lighthouse Lab in Alderley Park | Wellcome Sanger Institute for the COVID-19 Genomics UK (COG-UK) Consortium | Jacquelyn Wynn, Mairead Hyland, The Lighthouse Lab in Alderley Park and Alex Alderton, Roberto Amato, Jeffrey Barrett, Sonia Goncalves, Ewan Harrison, David K. Jackson, Ian Johnston, Dominic Kwiatkowski, Cordelia Langford, John Sillitoe on behalf of the Wellcome Sanger Institute COVID-19 Surveillance Team |
| EPI_ISL_3282885 | Lighthouse Laboratory Plymouth | Wellcome Sanger Institute for the COVID-19 Genomics UK (COG-UK) Consortium | Lighthouse Laboratory Plymouth and Alex Alderton, Roberto Amato, Jeffrey Barrett, Sonia Goncalves, Ewan Harrison, David K. Jackson, Ian Johnston, Dominic Kwiatkowski, Cordelia Langford, John Sillitoe on behalf of the Wellcome Sanger Institute COVID-19 Surveillance Team |
| EPI_ISL_3282886 | Lighthouse Lab in Alderley Park | Wellcome Sanger Institute for the COVID-19 Genomics UK (COG-UK) Consortium | Jacquelyn Wynn, Mairead Hyland, The Lighthouse Lab in Alderley Park and Alex Alderton, Roberto Amato, Jeffrey Barrett, Sonia Goncalves, Ewan Harrison, David K. Jackson, Ian Johnston, Dominic Kwiatkowski, Cordelia Langford, John Sillitoe on behalf of the Wellcome Sanger Institute COVID-19 Surveillance Team |
| EPI_ISL_3282887 | Lighthouse Lab in Glasgow | Wellcome Sanger Institute for the COVID-19 Genomics UK (COG-UK) Consortium | Harper VanSteenhouse, Yumi Kasai, David Gray, Carol Clugston, Anna Dominiczak and Alex Alderton, Roberto Amato, Jeffrey Barrett, Sonia Goncalves, Ewan Harrison, David K. Jackson, Ian Johnston, Dominic Kwiatkowski, Cordelia Langford, John Sillitoe on behalf of the Wellcome Sanger Institute COVID-19 Surveillance Team |
| EPI_ISL_3282888, EPI_ISL_3282889, EPI_ISL_3282890 | Lighthouse Lab in Alderley Park | Wellcome Sanger Institute for the COVID-19 Genomics UK (COG-UK) Consortium | Jacquelyn Wynn, Mairead Hyland, The Lighthouse Lab in Alderley Park and Alex Alderton, Roberto Amato, Jeffrey Barrett, Sonia Goncalves, Ewan Harrison, David K. Jackson, Ian Johnston, Dominic Kwiatkowski, Cordelia Langford, John Sillitoe on behalf of the Wellcome Sanger Institute COVID-19 Surveillance Team |
| EPI_ISL_3282891 | Lighthouse Laboratory Plymouth | Wellcome Sanger Institute for the COVID-19 Genomics UK (COG-UK) Consortium | Lighthouse Laboratory Plymouth and Alex Alderton, Roberto Amato, Jeffrey Barrett, Sonia Goncalves, Ewan Harrison, David K. Jackson, Ian Johnston, Dominic Kwiatkowski, Cordelia Langford, John Sillitoe on behalf of the Wellcome Sanger Institute COVID-19 Surveillance Team |
| EPI_ISL_3282892 | Lighthouse Lab in Glasgow | Wellcome Sanger Institute for the COVID-19 Genomics UK (COG-UK) Consortium | Harper VanSteenhouse, Yumi Kasai, David Gray, Carol Clugston, Anna Dominiczak and Alex Alderton, Roberto Amato, Jeffrey Barrett, Sonia Goncalves, Ewan Harrison, David K. Jackson, Ian Johnston, Dominic Kwiatkowski, Cordelia Langford, John Sillitoe on behalf of the Wellcome Sanger Institute COVID-19 Surveillance Team |
| EPI_ISL_3282893, EPI_ISL_3282894, EPI_ISL_3282895 | Lighthouse Lab in Alderley Park | Wellcome Sanger Institute for the COVID-19 Genomics UK (COG-UK) Consortium | Jacquelyn Wynn, Mairead Hyland, The Lighthouse Lab in Alderley Park and Alex Alderton, Roberto Amato, Jeffrey Barrett, Sonia Goncalves, Ewan Harrison, David K. Jackson, Ian Johnston, Dominic Kwiatkowski, Cordelia Langford, John Sillitoe on behalf of the Wellcome Sanger Institute COVID-19 Surveillance Team |
| EPI_ISL_3282896 | Lighthouse Lab in Glasgow | Wellcome Sanger Institute for the COVID-19 Genomics UK (COG-UK) Consortium | Harper VanSteenhouse, Yumi Kasai, David Gray, Carol Clugston, Anna Dominiczak and Alex Alderton, Roberto Amato, Jeffrey Barrett, Sonia Goncalves, Ewan Harrison, David K. Jackson, Ian Johnston, Dominic Kwiatkowski, Cordelia Langford, John Sillitoe on behalf of the Wellcome Sanger Institute COVID-19 Surveillance Team |
| EPI_ISL_3282897 | Lighthouse Lab in Alderley Park | Wellcome Sanger Institute for the COVID-19 Genomics UK (COG-UK) Consortium | Jacquelyn Wynn, Mairead Hyland, The Lighthouse Lab in Alderley Park and Alex Alderton, Roberto Amato, Jeffrey Barrett, Sonia Goncalves, Ewan Harrison, David K. Jackson, Ian Johnston, Dominic Kwiatkowski, Cordelia Langford, John Sillitoe on behalf of the Wellcome Sanger Institute COVID-19 Surveillance Team |
| EPI_ISL_3282898 | Lighthouse Lab in Glasgow | Wellcome Sanger Institute for the COVID-19 Genomics UK (COG-UK) Consortium | Harper VanSteenhouse, Yumi Kasai, David Gray, Carol Clugston, Anna Dominiczak and Alex Alderton, Roberto Amato, Jeffrey Barrett, Sonia Goncalves, Ewan Harrison, David K. Jackson, Ian Johnston, Dominic Kwiatkowski, Cordelia Langford, John Sillitoe on behalf of the Wellcome Sanger Institute COVID-19 Surveillance Team |
| EPI_ISL_3282899 | Lighthouse Lab in Alderley Park | Wellcome Sanger Institute for the COVID-19 Genomics UK (COG-UK) Consortium | Jacquelyn Wynn, Mairead Hyland, The Lighthouse Lab in Alderley Park and Alex Alderton, Roberto Amato, Jeffrey Barrett, Sonia Goncalves, Ewan Harrison, David K. Jackson, Ian Johnston, Dominic Kwiatkowski, Cordelia Langford, John Sillitoe on behalf of the Wellcome Sanger Institute COVID-19 Surveillance Team |
| EPI_ISL_3282900, EPI_ISL_3282901 | Lighthouse Lab in Glasgow | Wellcome Sanger Institute for the COVID-19 Genomics UK (COG-UK) Consortium | Harper VanSteenhouse, Yumi Kasai, David Gray, Carol Clugston, Anna Dominiczak and Alex Alderton, Roberto Amato, Jeffrey Barrett, Sonia Goncalves, Ewan Harrison, David K. Jackson, Ian Johnston, Dominic Kwiatkowski, Cordelia Langford, John Sillitoe on behalf of the Wellcome Sanger Institute COVID-19 Surveillance Team |
| EPI_ISL_3282902 | Lighthouse Laboratory Plymouth | Wellcome Sanger Institute for the COVID-19 Genomics UK (COG-UK) Consortium | Lighthouse Laboratory Plymouth and Alex Alderton, Roberto Amato, Jeffrey Barrett, Sonia Goncalves, Ewan Harrison, David K. Jackson, Ian Johnston, Dominic Kwiatkowski, Cordelia Langford, John Sillitoe on behalf of the Wellcome Sanger Institute COVID-19 Surveillance Team |
| EPI_ISL_3282903 | Lighthouse Lab in Glasgow | Wellcome Sanger Institute for the COVID-19 Genomics UK (COG-UK) | Harper VanSteenhouse, Yumi Kasai, David Gray, Carol Clugston, Anna Dominiczak and Alex Alderton, Roberto Amato, Jeffrey Barrett, Sonia Goncalves, Ewan Harrison, David K. Jackson, Ian Johnston, Dominic Kwiatkowski, Cordelia Langford, |

| | | | |
|---|---|---|---|
| | | Consortium | John Sillitoe on behalf of the Wellcome Sanger Institute COVID-19 Surveillance Team |
| EPI_ISL_3282904 | Lighthouse Lab in Alderley Park | Wellcome Sanger Institute for the COVID-19 Genomics UK (COG-UK) Consortium | Jacquelyn Wynn, Mairead Hyland, The Lighthouse Lab in Alderley Park and Alex Alderton, Roberto Amato, Jeffrey Barrett, Sonia Goncalves, Ewan Harrison, David K. Jackson, Ian Johnston, Dominic Kwiatkowski, Cordelia Langford, John Sillitoe on behalf of the Wellcome Sanger Institute COVID-19 Surveillance Team |
| EPI_ISL_3282905 | Lighthouse Laboratory Plymouth | Wellcome Sanger Institute for the COVID-19 Genomics UK (COG-UK) Consortium | Lighthouse Laboratory Plymouth and Alex Alderton, Roberto Amato, Jeffrey Barrett, Sonia Goncalves, Ewan Harrison, David K. Jackson, Ian Johnston, Dominic Kwiatkowski, Cordelia Langford, John Sillitoe on behalf of the Wellcome Sanger Institute COVID-19 Surveillance Team |
| EPI_ISL_3282906 | Lighthouse Lab in Alderley Park | Wellcome Sanger Institute for the COVID-19 Genomics UK (COG-UK) Consortium | Jacquelyn Wynn, Mairead Hyland, The Lighthouse Lab in Alderley Park and Alex Alderton, Roberto Amato, Jeffrey Barrett, Sonia Goncalves, Ewan Harrison, David K. Jackson, Ian Johnston, Dominic Kwiatkowski, Cordelia Langford, John Sillitoe on behalf of the Wellcome Sanger Institute COVID-19 Surveillance Team |
| EPI_ISL_3282907 | Lighthouse Lab in Glasgow | Wellcome Sanger Institute for the COVID-19 Genomics UK (COG-UK) Consortium | Harper VanSteenhouse, Yumi Kasai, David Gray, Carol Clugston, Anna Dominiczak and Alex Alderton, Roberto Amato, Jeffrey Barrett, Sonia Goncalves, Ewan Harrison, David K. Jackson, Ian Johnston, Dominic Kwiatkowski, Cordelia Langford, John Sillitoe on behalf of the Wellcome Sanger Institute COVID-19 Surveillance Team |
| EPI_ISL_3282908, EPI_ISL_3282909 | Lighthouse Lab in Alderley Park | Wellcome Sanger Institute for the COVID-19 Genomics UK (COG-UK) Consortium | Jacquelyn Wynn, Mairead Hyland, The Lighthouse Lab in Alderley Park and Alex Alderton, Roberto Amato, Jeffrey Barrett, Sonia Goncalves, Ewan Harrison, David K. Jackson, Ian Johnston, Dominic Kwiatkowski, Cordelia Langford, John Sillitoe on behalf of the Wellcome Sanger Institute COVID-19 Surveillance Team |
| EPI_ISL_3282910 | Lighthouse Lab in Glasgow | Wellcome Sanger Institute for the COVID-19 Genomics UK (COG-UK) Consortium | Harper VanSteenhouse, Yumi Kasai, David Gray, Carol Clugston, Anna Dominiczak and Alex Alderton, Roberto Amato, Jeffrey Barrett, Sonia Goncalves, Ewan Harrison, David K. Jackson, Ian Johnston, Dominic Kwiatkowski, Cordelia Langford, John Sillitoe on behalf of the Wellcome Sanger Institute COVID-19 Surveillance Team |
| EPI_ISL_3282911 | Lighthouse Lab in Alderley Park | Wellcome Sanger Institute for the COVID-19 Genomics UK (COG-UK) Consortium | Jacquelyn Wynn, Mairead Hyland, The Lighthouse Lab in Alderley Park and Alex Alderton, Roberto Amato, Jeffrey Barrett, Sonia Goncalves, Ewan Harrison, David K. Jackson, Ian Johnston, Dominic Kwiatkowski, Cordelia Langford, John Sillitoe on behalf of the Wellcome Sanger Institute COVID-19 Surveillance Team |
| EPI_ISL_3282912 | Lighthouse Lab in Glasgow | Wellcome Sanger Institute for the COVID-19 Genomics UK (COG-UK) Consortium | Harper VanSteenhouse, Yumi Kasai, David Gray, Carol Clugston, Anna Dominiczak and Alex Alderton, Roberto Amato, Jeffrey Barrett, Sonia Goncalves, Ewan Harrison, David K. Jackson, Ian Johnston, Dominic Kwiatkowski, Cordelia Langford, John Sillitoe on behalf of the Wellcome Sanger Institute COVID-19 Surveillance Team |
| EPI_ISL_3282913, EPI_ISL_3282914, EPI_ISL_3282915 | Lighthouse Lab in Alderley Park | Wellcome Sanger Institute for the COVID-19 Genomics UK (COG-UK) Consortium | Jacquelyn Wynn, Mairead Hyland, The Lighthouse Lab in Alderley Park and Alex Alderton, Roberto Amato, Jeffrey Barrett, Sonia Goncalves, Ewan Harrison, David K. Jackson, Ian Johnston, Dominic Kwiatkowski, Cordelia Langford, John Sillitoe on behalf of the Wellcome Sanger Institute COVID-19 Surveillance Team |
| EPI_ISL_3282916 | Lighthouse Laboratory Plymouth | Wellcome Sanger Institute for the COVID-19 Genomics UK (COG-UK) Consortium | Lighthouse Laboratory Plymouth and Alex Alderton, Roberto Amato, Jeffrey Barrett, Sonia Goncalves, Ewan Harrison, David K. Jackson, Ian Johnston, Dominic Kwiatkowski, Cordelia Langford, John Sillitoe on behalf of the Wellcome Sanger Institute COVID-19 Surveillance Team |
| EPI_ISL_3282917, EPI_ISL_3282918 | Lighthouse Lab in Alderley Park | Wellcome Sanger Institute for the COVID-19 Genomics UK (COG-UK) Consortium | Jacquelyn Wynn, Mairead Hyland, The Lighthouse Lab in Alderley Park and Alex Alderton, Roberto Amato, Jeffrey Barrett, Sonia Goncalves, Ewan Harrison, David K. Jackson, Ian Johnston, Dominic Kwiatkowski, Cordelia Langford, John Sillitoe on behalf of the Wellcome Sanger Institute COVID-19 Surveillance Team |
| EPI_ISL_3282919 | Lighthouse Lab in Glasgow | Wellcome Sanger Institute for the COVID-19 Genomics UK (COG-UK) Consortium | Harper VanSteenhouse, Yumi Kasai, David Gray, Carol Clugston, Anna Dominiczak and Alex Alderton, Roberto Amato, Jeffrey Barrett, Sonia Goncalves, Ewan Harrison, David K. Jackson, Ian Johnston, Dominic Kwiatkowski, Cordelia Langford, John Sillitoe on behalf of the Wellcome Sanger Institute COVID-19 Surveillance Team |
| EPI_ISL_3282920 | Lighthouse Laboratory Plymouth | Wellcome Sanger Institute for the COVID-19 Genomics UK (COG-UK) Consortium | Lighthouse Laboratory Plymouth and Alex Alderton, Roberto Amato, Jeffrey Barrett, Sonia Goncalves, Ewan Harrison, David K. Jackson, Ian Johnston, Dominic Kwiatkowski, Cordelia Langford, John Sillitoe on behalf of the Wellcome Sanger Institute COVID-19 Surveillance Team |
| EPI_ISL_3282921 | Lighthouse Lab in Glasgow | Wellcome Sanger Institute for the COVID-19 Genomics UK (COG-UK) Consortium | Harper VanSteenhouse, Yumi Kasai, David Gray, Carol Clugston, Anna Dominiczak and Alex Alderton, Roberto Amato, Jeffrey Barrett, Sonia Goncalves, Ewan Harrison, David K. Jackson, Ian Johnston, Dominic Kwiatkowski, Cordelia Langford, John Sillitoe on behalf of the Wellcome Sanger Institute COVID-19 Surveillance Team |
| EPI_ISL_3282922 | Lighthouse Lab in Alderley Park | Wellcome Sanger Institute for the COVID-19 Genomics UK (COG-UK) Consortium | Jacquelyn Wynn, Mairead Hyland, The Lighthouse Lab in Alderley Park and Alex Alderton, Roberto Amato, Jeffrey Barrett, Sonia Goncalves, Ewan Harrison, David K. Jackson, Ian Johnston, Dominic Kwiatkowski, Cordelia Langford, John Sillitoe on behalf of the Wellcome Sanger Institute COVID-19 Surveillance Team |
| EPI_ISL_3282923 | Lighthouse Laboratory Plymouth | Wellcome Sanger Institute for the COVID-19 Genomics UK (COG-UK) Consortium | Lighthouse Laboratory Plymouth and Alex Alderton, Roberto Amato, Jeffrey Barrett, Sonia Goncalves, Ewan Harrison, David K. Jackson, Ian Johnston, Dominic Kwiatkowski, Cordelia Langford, John Sillitoe on behalf of the Wellcome Sanger Institute COVID-19 Surveillance Team |
| EPI_ISL_3282924, EPI_ISL_3282925, EPI_ISL_3282926 | Lighthouse Lab in Alderley Park | Wellcome Sanger Institute for the COVID-19 Genomics UK (COG-UK) Consortium | Jacquelyn Wynn, Mairead Hyland, The Lighthouse Lab in Alderley Park and Alex Alderton, Roberto Amato, Jeffrey Barrett, Sonia Goncalves, Ewan Harrison, David K. Jackson, Ian Johnston, Dominic Kwiatkowski, Cordelia Langford, John Sillitoe on behalf of the Wellcome Sanger Institute COVID-19 Surveillance Team |
| EPI_ISL_3282927 | Lighthouse Lab in Glasgow | Wellcome Sanger Institute for the COVID-19 Genomics UK (COG-UK) Consortium | Harper VanSteenhouse, Yumi Kasai, David Gray, Carol Clugston, Anna Dominiczak and Alex Alderton, Roberto Amato, Jeffrey Barrett, Sonia Goncalves, Ewan Harrison, David K. Jackson, Ian Johnston, Dominic Kwiatkowski, Cordelia Langford, John Sillitoe on behalf of the Wellcome Sanger Institute COVID-19 Surveillance Team |
| EPI_ISL_3282928, EPI_ISL_3282929, EPI_ISL_3282930, EPI_ISL_3282931 | Lighthouse Lab in Alderley Park | Wellcome Sanger Institute for the COVID-19 Genomics UK (COG-UK) Consortium | Jacquelyn Wynn, Mairead Hyland, The Lighthouse Lab in Alderley Park and Alex Alderton, Roberto Amato, Jeffrey Barrett, Sonia Goncalves, Ewan Harrison, David K. Jackson, Ian Johnston, Dominic Kwiatkowski, Cordelia Langford, John Sillitoe on behalf of the Wellcome Sanger Institute COVID-19 Surveillance Team |
| EPI_ISL_3282932, EPI_ISL_3282933 | Lighthouse Laboratory Plymouth | Wellcome Sanger Institute for the COVID-19 Genomics UK (COG-UK) Consortium | Lighthouse Laboratory Plymouth and Alex Alderton, Roberto Amato, Jeffrey Barrett, Sonia Goncalves, Ewan Harrison, David K. Jackson, Ian Johnston, Dominic Kwiatkowski, Cordelia Langford, John Sillitoe on behalf of the Wellcome Sanger Institute COVID-19 Surveillance Team |
| EPI_ISL_3282934 | Lighthouse Lab in Glasgow | Wellcome Sanger Institute for the COVID-19 Genomics UK (COG-UK) Consortium | Harper VanSteenhouse, Yumi Kasai, David Gray, Carol Clugston, Anna Dominiczak and Alex Alderton, Roberto Amato, Jeffrey Barrett, Sonia Goncalves, Ewan Harrison, David K. Jackson, Ian Johnston, Dominic Kwiatkowski, Cordelia Langford, John Sillitoe on behalf of the Wellcome Sanger Institute COVID-19 Surveillance Team |
| EPI_ISL_3282935, EPI_ISL_3282936, EPI_ISL_3282937 | Lighthouse Laboratory Plymouth | Wellcome Sanger Institute for the COVID-19 Genomics UK (COG-UK) Consortium | Lighthouse Laboratory Plymouth and Alex Alderton, Roberto Amato, Jeffrey Barrett, Sonia Goncalves, Ewan Harrison, David K. Jackson, Ian Johnston, Dominic Kwiatkowski, Cordelia Langford, John Sillitoe on behalf of the Wellcome Sanger Institute COVID-19 Surveillance Team |
| EPI_ISL_3282938, EPI_ISL_3282939, EPI_ISL_3282940, EPI_ISL_3282941 | Lighthouse Lab in Alderley Park | Wellcome Sanger Institute for the COVID-19 Genomics UK (COG-UK) Consortium | Jacquelyn Wynn, Mairead Hyland, The Lighthouse Lab in Alderley Park and Alex Alderton, Roberto Amato, Jeffrey Barrett, Sonia Goncalves, Ewan Harrison, David K. Jackson, Ian Johnston, Dominic Kwiatkowski, Cordelia Langford, John Sillitoe on behalf of the Wellcome Sanger Institute COVID-19 Surveillance Team |
| EPI_ISL_3282942 | Lighthouse Lab in Glasgow | Wellcome Sanger Institute for the COVID-19 Genomics UK (COG-UK) Consortium | Harper VanSteenhouse, Yumi Kasai, David Gray, Carol Clugston, Anna Dominiczak and Alex Alderton, Roberto Amato, Jeffrey Barrett, Sonia Goncalves, Ewan Harrison, David K. Jackson, Ian Johnston, Dominic Kwiatkowski, Cordelia Langford, John Sillitoe on behalf of the Wellcome Sanger Institute COVID-19 Surveillance Team |
| EPI_ISL_3282943, EPI_ISL_3282944 | Lighthouse Laboratory Plymouth | Wellcome Sanger Institute for the COVID-19 Genomics UK (COG-UK) Consortium | Lighthouse Laboratory Plymouth and Alex Alderton, Roberto Amato, Jeffrey Barrett, Sonia Goncalves, Ewan Harrison, David K. Jackson, Ian Johnston, Dominic Kwiatkowski, Cordelia Langford, John Sillitoe on behalf of the Wellcome Sanger Institute COVID-19 Surveillance Team |
| EPI_ISL_3282945, EPI_ISL_3282946 | Lighthouse Lab in Alderley Park | Wellcome Sanger Institute for the COVID-19 Genomics UK (COG-UK) Consortium | Jacquelyn Wynn, Mairead Hyland, The Lighthouse Lab in Alderley Park and Alex Alderton, Roberto Amato, Jeffrey Barrett, Sonia Goncalves, Ewan Harrison, David K. Jackson, Ian Johnston, Dominic Kwiatkowski, Cordelia Langford, John Sillitoe on behalf of the Wellcome Sanger Institute COVID-19 Surveillance Team |
| EPI_ISL_3282947, EPI_ISL_3282948 | Lighthouse Lab in Glasgow | Wellcome Sanger Institute for the COVID-19 Genomics UK (COG-UK) Consortium | Harper VanSteenhouse, Yumi Kasai, David Gray, Carol Clugston, Anna Dominiczak and Alex Alderton, Roberto Amato, Jeffrey Barrett, Sonia Goncalves, Ewan Harrison, David K. Jackson, Ian Johnston, Dominic Kwiatkowski, Cordelia Langford, John Sillitoe on behalf of the Wellcome Sanger Institute COVID-19 Surveillance Team |
| EPI_ISL_3282949 | Lighthouse Lab in Alderley Park | Wellcome Sanger Institute for the COVID-19 Genomics UK (COG-UK) Consortium | Jacquelyn Wynn, Mairead Hyland, The Lighthouse Lab in Alderley Park and Alex Alderton, Roberto Amato, Jeffrey Barrett, Sonia Goncalves, Ewan Harrison, David K. Jackson, Ian Johnston, Dominic Kwiatkowski, Cordelia Langford, John Sillitoe on behalf of the Wellcome Sanger Institute COVID-19 Surveillance Team |
| EPI_ISL_3282950, EPI_ISL_3282951 | Lighthouse Lab in Glasgow | Wellcome Sanger Institute for the COVID-19 Genomics UK (COG-UK) Consortium | Harper VanSteenhouse, Yumi Kasai, David Gray, Carol Clugston, Anna Dominiczak and Alex Alderton, Roberto Amato, Jeffrey Barrett, Sonia Goncalves, Ewan Harrison, David K. Jackson, Ian Johnston, Dominic Kwiatkowski, Cordelia Langford, John Sillitoe on behalf of the Wellcome Sanger Institute COVID-19 Surveillance Team |
| EPI_ISL_3282952 | Lighthouse Laboratory Plymouth | Wellcome Sanger Institute for the COVID-19 Genomics UK (COG-UK) Consortium | Lighthouse Laboratory Plymouth and Alex Alderton, Roberto Amato, Jeffrey Barrett, Sonia Goncalves, Ewan Harrison, David K. Jackson, Ian Johnston, Dominic Kwiatkowski, Cordelia Langford, John Sillitoe on behalf of the Wellcome Sanger Institute COVID-19 Surveillance Team |
| EPI_ISL_3282953 | Lighthouse Lab in Glasgow | Wellcome Sanger Institute for the COVID-19 Genomics UK (COG-UK) Consortium | Harper VanSteenhouse, Yumi Kasai, David Gray, Carol Clugston, Anna Dominiczak and Alex Alderton, Roberto Amato, Jeffrey Barrett, Sonia Goncalves, Ewan Harrison, David K. Jackson, Ian Johnston, Dominic Kwiatkowski, Cordelia Langford, John Sillitoe on behalf of the Wellcome Sanger Institute COVID-19 Surveillance Team |
| EPI_ISL_3282954, EPI_ISL_3282955 | Lighthouse Lab in Alderley Park | Wellcome Sanger Institute for the COVID-19 Genomics UK (COG-UK) Consortium | Jacquelyn Wynn, Mairead Hyland, The Lighthouse Lab in Alderley Park and Alex Alderton, Roberto Amato, Jeffrey Barrett, Sonia Goncalves, Ewan Harrison, David K. Jackson, Ian Johnston, Dominic Kwiatkowski, Cordelia Langford, John Sillitoe on behalf of the Wellcome Sanger Institute COVID-19 Surveillance Team |
| EPI_ISL_3282956 | Lighthouse Laboratory Plymouth | Wellcome Sanger Institute for the COVID-19 Genomics UK (COG-UK) Consortium | Lighthouse Laboratory Plymouth and Alex Alderton, Roberto Amato, Jeffrey Barrett, Sonia Goncalves, Ewan Harrison, David K. Jackson, Ian Johnston, Dominic Kwiatkowski, Cordelia Langford, John Sillitoe on behalf of the Wellcome Sanger Institute COVID-19 Surveillance Team |
| EPI_ISL_3282957, EPI_ISL_3282958, EPI_ISL_3282959, EPI_ISL_3282960 | Lighthouse Lab in Alderley Park | Wellcome Sanger Institute for the COVID-19 Genomics UK (COG-UK) Consortium | Jacquelyn Wynn, Mairead Hyland, The Lighthouse Lab in Alderley Park and Alex Alderton, Roberto Amato, Jeffrey Barrett, Sonia Goncalves, Ewan Harrison, David K. Jackson, Ian Johnston, Dominic Kwiatkowski, Cordelia Langford, John Sillitoe on behalf of the Wellcome Sanger Institute COVID-19 Surveillance Team |
| EPI_ISL_3282961 | Lighthouse Lab in Glasgow | Wellcome Sanger Institute for the COVID-19 Genomics UK (COG-UK) Consortium | Harper VanSteenhouse, Yumi Kasai, David Gray, Carol Clugston, Anna Dominiczak and Alex Alderton, Roberto Amato, Jeffrey Barrett, Sonia Goncalves, Ewan Harrison, David K. Jackson, Ian Johnston, Dominic Kwiatkowski, Cordelia Langford, John Sillitoe on behalf of the Wellcome Sanger Institute COVID-19 Surveillance Team |
| EPI_ISL_3282962, EPI_ISL_3282963 | Lighthouse Lab in Alderley Park | Wellcome Sanger Institute for the COVID-19 Genomics UK (COG-UK) Consortium | Jacquelyn Wynn, Mairead Hyland, The Lighthouse Lab in Alderley Park and Alex Alderton, Roberto Amato, Jeffrey Barrett, Sonia Goncalves, Ewan Harrison, David K. Jackson, Ian Johnston, Dominic Kwiatkowski, Cordelia Langford, John Sillitoe on behalf of the Wellcome Sanger Institute COVID-19 Surveillance Team |
| EPI_ISL_3282964 | Lighthouse Laboratory Plymouth | Wellcome Sanger Institute for the COVID-19 Genomics UK (COG-UK) Consortium | Lighthouse Laboratory Plymouth and Alex Alderton, Roberto Amato, Jeffrey Barrett, Sonia Goncalves, Ewan Harrison, David K. Jackson, Ian Johnston, Dominic Kwiatkowski, Cordelia Langford, John Sillitoe on behalf of the Wellcome Sanger Institute COVID-19 Surveillance Team |
| EPI_ISL_3282965 | Lighthouse Lab in Glasgow | Wellcome Sanger Institute for the COVID-19 Genomics UK (COG-UK) Consortium | Harper VanSteenhouse, Yumi Kasai, David Gray, Carol Clugston, Anna Dominiczak and Alex Alderton, Roberto Amato, Jeffrey Barrett, Sonia Goncalves, Ewan Harrison, David K. Jackson, Ian Johnston, Dominic Kwiatkowski, Cordelia Langford, John Sillitoe on behalf of the Wellcome Sanger Institute COVID-19 Surveillance Team |
| EPI_ISL_3282966, EPI_ISL_3282967, EPI_ISL_3282968, EPI_ISL_3282969, EPI_ISL_3282970, EPI_ISL_3282971, | Lighthouse Lab in Alderley Park | Wellcome Sanger Institute for the COVID-19 Genomics UK (COG-UK) Consortium | Jacquelyn Wynn, Mairead Hyland, The Lighthouse Lab in Alderley Park and Alex Alderton, Roberto Amato, Jeffrey Barrett, Sonia Goncalves, Ewan Harrison, David K. Jackson, Ian Johnston, Dominic Kwiatkowski, Cordelia Langford, John Sillitoe on behalf of the Wellcome Sanger Institute COVID-19 Surveillance Team |

| | | | |
|---|---|---|---|
| EPI_ISL_3282972 | | | |
| EPI_ISL_3282973 | Lighthouse Laboratory Plymouth | Wellcome Sanger Institute for the COVID-19 Genomics UK (COG-UK) Consortium | Lighthouse Laboratory Plymouth and Alex Alderton, Roberto Amato, Jeffrey Barrett, Sonia Goncalves, Ewan Harrison, David K. Jackson, Ian Johnston, Dominic Kwiatkowski, Cordelia Langford, John Sillitoe on behalf of the Wellcome Sanger Institute COVID-19 Surveillance Team |
| EPI_ISL_3282974 | Lighthouse Lab in Glasgow | Wellcome Sanger Institute for the COVID-19 Genomics UK (COG-UK) Consortium | Harper VanSteenhouse, Yumi Kasai, David Gray, Carol Clugston, Anna Dominiczak and Alex Alderton, Roberto Amato, Jeffrey Barrett, Sonia Goncalves, Ewan Harrison, David K. Jackson, Ian Johnston, Dominic Kwiatkowski, Cordelia Langford, John Sillitoe on behalf of the Wellcome Sanger Institute COVID-19 Surveillance Team |
| EPI_ISL_3282975 | Lighthouse Laboratory Plymouth | Wellcome Sanger Institute for the COVID-19 Genomics UK (COG-UK) Consortium | Lighthouse Laboratory Plymouth and Alex Alderton, Roberto Amato, Jeffrey Barrett, Sonia Goncalves, Ewan Harrison, David K. Jackson, Ian Johnston, Dominic Kwiatkowski, Cordelia Langford, John Sillitoe on behalf of the Wellcome Sanger Institute COVID-19 Surveillance Team |
| EPI_ISL_3282976 | Lighthouse Lab in Glasgow | Wellcome Sanger Institute for the COVID-19 Genomics UK (COG-UK) Consortium | Harper VanSteenhouse, Yumi Kasai, David Gray, Carol Clugston, Anna Dominiczak and Alex Alderton, Roberto Amato, Jeffrey Barrett, Sonia Goncalves, Ewan Harrison, David K. Jackson, Ian Johnston, Dominic Kwiatkowski, Cordelia Langford, John Sillitoe on behalf of the Wellcome Sanger Institute COVID-19 Surveillance Team |
| EPI_ISL_3282977 | Lighthouse Lab in Alderley Park | Wellcome Sanger Institute for the COVID-19 Genomics UK (COG-UK) Consortium | Jacquelyn Wynn, Mairead Hyland, The Lighthouse Lab in Alderley Park and Alex Alderton, Roberto Amato, Jeffrey Barrett, Sonia Goncalves, Ewan Harrison, David K. Jackson, Ian Johnston, Dominic Kwiatkowski, Cordelia Langford, John Sillitoe on behalf of the Wellcome Sanger Institute COVID-19 Surveillance Team |
| EPI_ISL_3282978 | Lighthouse Laboratory Plymouth | Wellcome Sanger Institute for the COVID-19 Genomics UK (COG-UK) Consortium | Lighthouse Laboratory Plymouth and Alex Alderton, Roberto Amato, Jeffrey Barrett, Sonia Goncalves, Ewan Harrison, David K. Jackson, Ian Johnston, Dominic Kwiatkowski, Cordelia Langford, John Sillitoe on behalf of the Wellcome Sanger Institute COVID-19 Surveillance Team |
| EPI_ISL_3282979 | Lighthouse Lab in Glasgow | Wellcome Sanger Institute for the COVID-19 Genomics UK (COG-UK) Consortium | Harper VanSteenhouse, Yumi Kasai, David Gray, Carol Clugston, Anna Dominiczak and Alex Alderton, Roberto Amato, Jeffrey Barrett, Sonia Goncalves, Ewan Harrison, David K. Jackson, Ian Johnston, Dominic Kwiatkowski, Cordelia Langford, John Sillitoe on behalf of the Wellcome Sanger Institute COVID-19 Surveillance Team |
| EPI_ISL_3282980, EPI_ISL_3282981, EPI_ISL_3282982 | Lighthouse Lab in Alderley Park | Wellcome Sanger Institute for the COVID-19 Genomics UK (COG-UK) Consortium | Jacquelyn Wynn, Mairead Hyland, The Lighthouse Lab in Alderley Park and Alex Alderton, Roberto Amato, Jeffrey Barrett, Sonia Goncalves, Ewan Harrison, David K. Jackson, Ian Johnston, Dominic Kwiatkowski, Cordelia Langford, John Sillitoe on behalf of the Wellcome Sanger Institute COVID-19 Surveillance Team |
| EPI_ISL_3282983 | Lighthouse Laboratory Plymouth | Wellcome Sanger Institute for the COVID-19 Genomics UK (COG-UK) Consortium | Lighthouse Laboratory Plymouth and Alex Alderton, Roberto Amato, Jeffrey Barrett, Sonia Goncalves, Ewan Harrison, David K. Jackson, Ian Johnston, Dominic Kwiatkowski, Cordelia Langford, John Sillitoe on behalf of the Wellcome Sanger Institute COVID-19 Surveillance Team |
| EPI_ISL_3282984, EPI_ISL_3282985, EPI_ISL_3282986, EPI_ISL_3282987, EPI_ISL_3282988 | Lighthouse Lab in Alderley Park | Wellcome Sanger Institute for the COVID-19 Genomics UK (COG-UK) Consortium | Jacquelyn Wynn, Mairead Hyland, The Lighthouse Lab in Alderley Park and Alex Alderton, Roberto Amato, Jeffrey Barrett, Sonia Goncalves, Ewan Harrison, David K. Jackson, Ian Johnston, Dominic Kwiatkowski, Cordelia Langford, John Sillitoe on behalf of the Wellcome Sanger Institute COVID-19 Surveillance Team |
| EPI_ISL_3282989, EPI_ISL_3282990 | Lighthouse Laboratory Plymouth | Wellcome Sanger Institute for the COVID-19 Genomics UK (COG-UK) Consortium | Lighthouse Laboratory Plymouth and Alex Alderton, Roberto Amato, Jeffrey Barrett, Sonia Goncalves, Ewan Harrison, David K. Jackson, Ian Johnston, Dominic Kwiatkowski, Cordelia Langford, John Sillitoe on behalf of the Wellcome Sanger Institute COVID-19 Surveillance Team |
| EPI_ISL_3282991 | Lighthouse Lab in Alderley Park | Wellcome Sanger Institute for the COVID-19 Genomics UK (COG-UK) Consortium | Jacquelyn Wynn, Mairead Hyland, The Lighthouse Lab in Alderley Park and Alex Alderton, Roberto Amato, Jeffrey Barrett, Sonia Goncalves, Ewan Harrison, David K. Jackson, Ian Johnston, Dominic Kwiatkowski, Cordelia Langford, John Sillitoe on behalf of the Wellcome Sanger Institute COVID-19 Surveillance Team |
| EPI_ISL_3282992 | Lighthouse Lab in Glasgow | Wellcome Sanger Institute for the COVID-19 Genomics UK (COG-UK) Consortium | Harper VanSteenhouse, Yumi Kasai, David Gray, Carol Clugston, Anna Dominiczak and Alex Alderton, Roberto Amato, Jeffrey Barrett, Sonia Goncalves, Ewan Harrison, David K. Jackson, Ian Johnston, Dominic Kwiatkowski, Cordelia Langford, John Sillitoe on behalf of the Wellcome Sanger Institute COVID-19 Surveillance Team |
| EPI_ISL_3282993, EPI_ISL_3282994, EPI_ISL_3282995, EPI_ISL_3282996 | Lighthouse Lab in Alderley Park | Wellcome Sanger Institute for the COVID-19 Genomics UK (COG-UK) Consortium | Jacquelyn Wynn, Mairead Hyland, The Lighthouse Lab in Alderley Park and Alex Alderton, Roberto Amato, Jeffrey Barrett, Sonia Goncalves, Ewan Harrison, David K. Jackson, Ian Johnston, Dominic Kwiatkowski, Cordelia Langford, John Sillitoe on behalf of the Wellcome Sanger Institute COVID-19 Surveillance Team |
| EPI_ISL_3282997, EPI_ISL_3282998 | Lighthouse Laboratory Plymouth | Wellcome Sanger Institute for the COVID-19 Genomics UK (COG-UK) Consortium | Lighthouse Laboratory Plymouth and Alex Alderton, Roberto Amato, Jeffrey Barrett, Sonia Goncalves, Ewan Harrison, David K. Jackson, Ian Johnston, Dominic Kwiatkowski, Cordelia Langford, John Sillitoe on behalf of the Wellcome Sanger Institute COVID-19 Surveillance Team |
| EPI_ISL_3282999 | Lighthouse Lab in Alderley Park | Wellcome Sanger Institute for the COVID-19 Genomics UK (COG-UK) Consortium | Jacquelyn Wynn, Mairead Hyland, The Lighthouse Lab in Alderley Park and Alex Alderton, Roberto Amato, Jeffrey Barrett, Sonia Goncalves, Ewan Harrison, David K. Jackson, Ian Johnston, Dominic Kwiatkowski, Cordelia Langford, John Sillitoe on behalf of the Wellcome Sanger Institute COVID-19 Surveillance Team |
| EPI_ISL_3283000 | Lighthouse Lab in Glasgow | Wellcome Sanger Institute for the COVID-19 Genomics UK (COG-UK) Consortium | Harper VanSteenhouse, Yumi Kasai, David Gray, Carol Clugston, Anna Dominiczak and Alex Alderton, Roberto Amato, Jeffrey Barrett, Sonia Goncalves, Ewan Harrison, David K. Jackson, Ian Johnston, Dominic Kwiatkowski, Cordelia Langford, John Sillitoe on behalf of the Wellcome Sanger Institute COVID-19 Surveillance Team |
| EPI_ISL_3283001 | Lighthouse Laboratory Plymouth | Wellcome Sanger Institute for the COVID-19 Genomics UK (COG-UK) Consortium | Lighthouse Laboratory Plymouth and Alex Alderton, Roberto Amato, Jeffrey Barrett, Sonia Goncalves, Ewan Harrison, David K. Jackson, Ian Johnston, Dominic Kwiatkowski, Cordelia Langford, John Sillitoe on behalf of the Wellcome Sanger Institute COVID-19 Surveillance Team |
| EPI_ISL_3283002, EPI_ISL_3283003 | Lighthouse Lab in Alderley Park | Wellcome Sanger Institute for the COVID-19 Genomics UK (COG-UK) Consortium | Jacquelyn Wynn, Mairead Hyland, The Lighthouse Lab in Alderley Park and Alex Alderton, Roberto Amato, Jeffrey Barrett, Sonia Goncalves, Ewan Harrison, David K. Jackson, Ian Johnston, Dominic Kwiatkowski, Cordelia Langford, John Sillitoe on behalf of the Wellcome Sanger Institute COVID-19 Surveillance Team |
| EPI_ISL_3283004 | Lighthouse Laboratory Plymouth | Wellcome Sanger Institute for the COVID-19 Genomics UK (COG-UK) Consortium | Lighthouse Laboratory Plymouth and Alex Alderton, Roberto Amato, Jeffrey Barrett, Sonia Goncalves, Ewan Harrison, David K. Jackson, Ian Johnston, Dominic Kwiatkowski, Cordelia Langford, John Sillitoe on behalf of the Wellcome Sanger Institute COVID-19 Surveillance Team |
| EPI_ISL_3283005 | Lighthouse Lab in Alderley Park | Wellcome Sanger Institute for the COVID-19 Genomics UK (COG-UK) Consortium | Jacquelyn Wynn, Mairead Hyland, The Lighthouse Lab in Alderley Park and Alex Alderton, Roberto Amato, Jeffrey Barrett, Sonia Goncalves, Ewan Harrison, David K. Jackson, Ian Johnston, Dominic Kwiatkowski, Cordelia Langford, John Sillitoe on behalf of the Wellcome Sanger Institute COVID-19 Surveillance Team |
| EPI_ISL_3283006, EPI_ISL_3283007 | Lighthouse Lab in Glasgow | Wellcome Sanger Institute for the COVID-19 Genomics UK (COG-UK) Consortium | Harper VanSteenhouse, Yumi Kasai, David Gray, Carol Clugston, Anna Dominiczak and Alex Alderton, Roberto Amato, Jeffrey Barrett, Sonia Goncalves, Ewan Harrison, David K. Jackson, Ian Johnston, Dominic Kwiatkowski, Cordelia Langford, John Sillitoe on behalf of the Wellcome Sanger Institute COVID-19 Surveillance Team |
| EPI_ISL_3283008, EPI_ISL_3283009, EPI_ISL_3283010, EPI_ISL_3283011 | Lighthouse Lab in Alderley Park | Wellcome Sanger Institute for the COVID-19 Genomics UK (COG-UK) Consortium | Jacquelyn Wynn, Mairead Hyland, The Lighthouse Lab in Alderley Park and Alex Alderton, Roberto Amato, Jeffrey Barrett, Sonia Goncalves, Ewan Harrison, David K. Jackson, Ian Johnston, Dominic Kwiatkowski, Cordelia Langford, John Sillitoe on behalf of the Wellcome Sanger Institute COVID-19 Surveillance Team |
| EPI_ISL_3283012, EPI_ISL_3283013 | Lighthouse Laboratory Plymouth | Wellcome Sanger Institute for the COVID-19 Genomics UK (COG-UK) Consortium | Lighthouse Laboratory Plymouth and Alex Alderton, Roberto Amato, Jeffrey Barrett, Sonia Goncalves, Ewan Harrison, David K. Jackson, Ian Johnston, Dominic Kwiatkowski, Cordelia Langford, John Sillitoe on behalf of the Wellcome Sanger Institute COVID-19 Surveillance Team |
| EPI_ISL_3283014 | Lighthouse Lab in Glasgow | Wellcome Sanger Institute for the COVID-19 Genomics UK (COG-UK) Consortium | Harper VanSteenhouse, Yumi Kasai, David Gray, Carol Clugston, Anna Dominiczak and Alex Alderton, Roberto Amato, Jeffrey Barrett, Sonia Goncalves, Ewan Harrison, David K. Jackson, Ian Johnston, Dominic Kwiatkowski, Cordelia Langford, John Sillitoe on behalf of the Wellcome Sanger Institute COVID-19 Surveillance Team |
| EPI_ISL_3283015 | Lighthouse Laboratory Plymouth | Wellcome Sanger Institute for the COVID-19 Genomics UK (COG-UK) Consortium | Lighthouse Laboratory Plymouth and Alex Alderton, Roberto Amato, Jeffrey Barrett, Sonia Goncalves, Ewan Harrison, David K. Jackson, Ian Johnston, Dominic Kwiatkowski, Cordelia Langford, John Sillitoe on behalf of the Wellcome Sanger Institute COVID-19 Surveillance Team |
| EPI_ISL_3283016 | Lighthouse Lab in Glasgow | Wellcome Sanger Institute for the COVID-19 Genomics UK (COG-UK) Consortium | Harper VanSteenhouse, Yumi Kasai, David Gray, Carol Clugston, Anna Dominiczak and Alex Alderton, Roberto Amato, Jeffrey Barrett, Sonia Goncalves, Ewan Harrison, David K. Jackson, Ian Johnston, Dominic Kwiatkowski, Cordelia Langford, John Sillitoe on behalf of the Wellcome Sanger Institute COVID-19 Surveillance Team |
| EPI_ISL_3283017 | Lighthouse Lab in Alderley Park | Wellcome Sanger Institute for the COVID-19 Genomics UK (COG-UK) Consortium | Jacquelyn Wynn, Mairead Hyland, The Lighthouse Lab in Alderley Park and Alex Alderton, Roberto Amato, Jeffrey Barrett, Sonia Goncalves, Ewan Harrison, David K. Jackson, Ian Johnston, Dominic Kwiatkowski, Cordelia Langford, John Sillitoe on behalf of the Wellcome Sanger Institute COVID-19 Surveillance Team |
| EPI_ISL_3283018, EPI_ISL_3283019 | Lighthouse Lab in Glasgow | Wellcome Sanger Institute for the COVID-19 Genomics UK (COG-UK) Consortium | Harper VanSteenhouse, Yumi Kasai, David Gray, Carol Clugston, Anna Dominiczak and Alex Alderton, Roberto Amato, Jeffrey Barrett, Sonia Goncalves, Ewan Harrison, David K. Jackson, Ian Johnston, Dominic Kwiatkowski, Cordelia Langford, John Sillitoe on behalf of the Wellcome Sanger Institute COVID-19 Surveillance Team |
| EPI_ISL_3283020 | Lighthouse Lab in Alderley Park | Wellcome Sanger Institute for the COVID-19 Genomics UK (COG-UK) Consortium | Jacquelyn Wynn, Mairead Hyland, The Lighthouse Lab in Alderley Park and Alex Alderton, Roberto Amato, Jeffrey Barrett, Sonia Goncalves, Ewan Harrison, David K. Jackson, Ian Johnston, Dominic Kwiatkowski, Cordelia Langford, John Sillitoe on behalf of the Wellcome Sanger Institute COVID-19 Surveillance Team |
| EPI_ISL_3283021 | Lighthouse Laboratory Plymouth | Wellcome Sanger Institute for the COVID-19 Genomics UK (COG-UK) Consortium | Lighthouse Laboratory Plymouth and Alex Alderton, Roberto Amato, Jeffrey Barrett, Sonia Goncalves, Ewan Harrison, David K. Jackson, Ian Johnston, Dominic Kwiatkowski, Cordelia Langford, John Sillitoe on behalf of the Wellcome Sanger Institute COVID-19 Surveillance Team |
| EPI_ISL_3283022 | Lighthouse Lab in Alderley Park | Wellcome Sanger Institute for the COVID-19 Genomics UK (COG-UK) Consortium | Jacquelyn Wynn, Mairead Hyland, The Lighthouse Lab in Alderley Park and Alex Alderton, Roberto Amato, Jeffrey Barrett, Sonia Goncalves, Ewan Harrison, David K. Jackson, Ian Johnston, Dominic Kwiatkowski, Cordelia Langford, John Sillitoe on behalf of the Wellcome Sanger Institute COVID-19 Surveillance Team |
| EPI_ISL_3283023 | Lighthouse Lab in Glasgow | Wellcome Sanger Institute for the COVID-19 Genomics UK (COG-UK) Consortium | Harper VanSteenhouse, Yumi Kasai, David Gray, Carol Clugston, Anna Dominiczak and Alex Alderton, Roberto Amato, Jeffrey Barrett, Sonia Goncalves, Ewan Harrison, David K. Jackson, Ian Johnston, Dominic Kwiatkowski, Cordelia Langford, John Sillitoe on behalf of the Wellcome Sanger Institute COVID-19 Surveillance Team |
| EPI_ISL_3283024 | Lighthouse Laboratory Plymouth | Wellcome Sanger Institute for the COVID-19 Genomics UK (COG-UK) Consortium | Lighthouse Laboratory Plymouth and Alex Alderton, Roberto Amato, Jeffrey Barrett, Sonia Goncalves, Ewan Harrison, David K. Jackson, Ian Johnston, Dominic Kwiatkowski, Cordelia Langford, John Sillitoe on behalf of the Wellcome Sanger Institute COVID-19 Surveillance Team |
| EPI_ISL_3283025, EPI_ISL_3283026 | Lighthouse Lab in Glasgow | Wellcome Sanger Institute for the COVID-19 Genomics UK (COG-UK) Consortium | Harper VanSteenhouse, Yumi Kasai, David Gray, Carol Clugston, Anna Dominiczak and Alex Alderton, Roberto Amato, Jeffrey Barrett, Sonia Goncalves, Ewan Harrison, David K. Jackson, Ian Johnston, Dominic Kwiatkowski, Cordelia Langford, John Sillitoe on behalf of the Wellcome Sanger Institute COVID-19 Surveillance Team |
| EPI_ISL_3283027 | Lighthouse Lab in Alderley Park | Wellcome Sanger Institute for the COVID-19 Genomics UK (COG-UK) Consortium | Jacquelyn Wynn, Mairead Hyland, The Lighthouse Lab in Alderley Park and Alex Alderton, Roberto Amato, Jeffrey Barrett, Sonia Goncalves, Ewan Harrison, David K. Jackson, Ian Johnston, Dominic Kwiatkowski, Cordelia Langford, John Sillitoe on behalf of the Wellcome Sanger Institute COVID-19 Surveillance Team |
| EPI_ISL_3283028 | Lighthouse Lab in Glasgow | Wellcome Sanger Institute for the COVID-19 Genomics UK (COG-UK) Consortium | Harper VanSteenhouse, Yumi Kasai, David Gray, Carol Clugston, Anna Dominiczak and Alex Alderton, Roberto Amato, Jeffrey Barrett, Sonia Goncalves, Ewan Harrison, David K. Jackson, Ian Johnston, Dominic Kwiatkowski, Cordelia Langford, John Sillitoe on behalf of the Wellcome Sanger Institute COVID-19 Surveillance Team |
| EPI_ISL_3283029, EPI_ISL_3283030, EPI_ISL_3283031, EPI_ISL_3283032 | Lighthouse Lab in Alderley Park | Wellcome Sanger Institute for the COVID-19 Genomics UK (COG-UK) Consortium | Jacquelyn Wynn, Mairead Hyland, The Lighthouse Lab in Alderley Park and Alex Alderton, Roberto Amato, Jeffrey Barrett, Sonia Goncalves, Ewan Harrison, David K. Jackson, Ian Johnston, Dominic Kwiatkowski, Cordelia Langford, John Sillitoe on behalf of the Wellcome Sanger Institute COVID-19 Surveillance Team |
| EPI_ISL_3283033, EPI_ISL_3283034 | Lighthouse Lab in Glasgow | Wellcome Sanger Institute for the COVID-19 Genomics UK (COG-UK) Consortium | Harper VanSteenhouse, Yumi Kasai, David Gray, Carol Clugston, Anna Dominiczak and Alex Alderton, Roberto Amato, Jeffrey Barrett, Sonia Goncalves, Ewan Harrison, David K. Jackson, Ian Johnston, Dominic Kwiatkowski, Cordelia Langford, John Sillitoe on behalf of the Wellcome Sanger Institute COVID-19 Surveillance Team |
| EPI_ISL_3283035, EPI_ISL_3283036, | Lighthouse Lab in Alderley Park | Wellcome Sanger Institute for the COVID-19 Genomics UK (COG-UK) Consortium | Jacquelyn Wynn, Mairead Hyland, The Lighthouse Lab in Alderley Park and Alex Alderton, Roberto Amato, Jeffrey Barrett, Sonia Goncalves, Ewan Harrison, David K. Jackson, Ian Johnston, Dominic Kwiatkowski, Cordelia Langford, John Sillitoe on behalf of the Wellcome Sanger Institute COVID-19 Surveillance Team |

| | | | |
|---|---|---|---|
| EPI_ISL_3283037 | | | |
| EPI_ISL_3283038 | Lighthouse Lab in Glasgow | Wellcome Sanger Institute for the COVID-19 Genomics UK (COG-UK) Consortium | Harper VanSteenhouse, Yumi Kasai, David Gray, Carol Clugston, Anna Dominiczak and Alex Alderton, Roberto Amato, Jeffrey Barrett, Sonia Goncalves, Ewan Harrison, David K. Jackson, Ian Johnston, Dominic Kwiatkowski, Cordelia Langford, John Sillitoe on behalf of the Wellcome Sanger Institute COVID-19 Surveillance Team |
| EPI_ISL_3283039, EPI_ISL_3283040 | Lighthouse Lab in Alderley Park | Wellcome Sanger Institute for the COVID-19 Genomics UK (COG-UK) Consortium | Jacquelyn Wynn, Mairead Hyland, The Lighthouse Lab in Alderley Park and Alex Alderton, Roberto Amato, Jeffrey Barrett, Sonia Goncalves, Ewan Harrison, David K. Jackson, Ian Johnston, Dominic Kwiatkowski, Cordelia Langford, John Sillitoe on behalf of the Wellcome Sanger Institute COVID-19 Surveillance Team |
| EPI_ISL_3283041, EPI_ISL_3283042 | Lighthouse Laboratory Plymouth | Wellcome Sanger Institute for the COVID-19 Genomics UK (COG-UK) Consortium | Lighthouse Laboratory Plymouth and Alex Alderton, Roberto Amato, Jeffrey Barrett, Sonia Goncalves, Ewan Harrison, David K. Jackson, Ian Johnston, Dominic Kwiatkowski, Cordelia Langford, John Sillitoe on behalf of the Wellcome Sanger Institute COVID-19 Surveillance Team |
| EPI_ISL_3283043 | Lighthouse Lab in Alderley Park | Wellcome Sanger Institute for the COVID-19 Genomics UK (COG-UK) Consortium | Jacquelyn Wynn, Mairead Hyland, The Lighthouse Lab in Alderley Park and Alex Alderton, Roberto Amato, Jeffrey Barrett, Sonia Goncalves, Ewan Harrison, David K. Jackson, Ian Johnston, Dominic Kwiatkowski, Cordelia Langford, John Sillitoe on behalf of the Wellcome Sanger Institute COVID-19 Surveillance Team |
| EPI_ISL_3283044, EPI_ISL_3283045 | Lighthouse Laboratory Plymouth | Wellcome Sanger Institute for the COVID-19 Genomics UK (COG-UK) Consortium | Lighthouse Laboratory Plymouth and Alex Alderton, Roberto Amato, Jeffrey Barrett, Sonia Goncalves, Ewan Harrison, David K. Jackson, Ian Johnston, Dominic Kwiatkowski, Cordelia Langford, John Sillitoe on behalf of the Wellcome Sanger Institute COVID-19 Surveillance Team |
| EPI_ISL_3283046 | Lighthouse Lab in Glasgow | Wellcome Sanger Institute for the COVID-19 Genomics UK (COG-UK) Consortium | Harper VanSteenhouse, Yumi Kasai, David Gray, Carol Clugston, Anna Dominiczak and Alex Alderton, Roberto Amato, Jeffrey Barrett, Sonia Goncalves, Ewan Harrison, David K. Jackson, Ian Johnston, Dominic Kwiatkowski, Cordelia Langford, John Sillitoe on behalf of the Wellcome Sanger Institute COVID-19 Surveillance Team |
| EPI_ISL_3283047, EPI_ISL_3283048, EPI_ISL_3283049 | Lighthouse Lab in Alderley Park | Wellcome Sanger Institute for the COVID-19 Genomics UK (COG-UK) Consortium | Jacquelyn Wynn, Mairead Hyland, The Lighthouse Lab in Alderley Park and Alex Alderton, Roberto Amato, Jeffrey Barrett, Sonia Goncalves, Ewan Harrison, David K. Jackson, Ian Johnston, Dominic Kwiatkowski, Cordelia Langford, John Sillitoe on behalf of the Wellcome Sanger Institute COVID-19 Surveillance Team |
| EPI_ISL_3283050, EPI_ISL_3283051, EPI_ISL_3283052, EPI_ISL_3283053, EPI_ISL_3283054, EPI_ISL_3283055 | Lighthouse Lab in Glasgow | Wellcome Sanger Institute for the COVID-19 Genomics UK (COG-UK) Consortium | Harper VanSteenhouse, Yumi Kasai, David Gray, Carol Clugston, Anna Dominiczak and Alex Alderton, Roberto Amato, Jeffrey Barrett, Sonia Goncalves, Ewan Harrison, David K. Jackson, Ian Johnston, Dominic Kwiatkowski, Cordelia Langford, John Sillitoe on behalf of the Wellcome Sanger Institute COVID-19 Surveillance Team |
| EPI_ISL_3283056 | Lighthouse Laboratory Plymouth | Wellcome Sanger Institute for the COVID-19 Genomics UK (COG-UK) Consortium | Lighthouse Laboratory Plymouth and Alex Alderton, Roberto Amato, Jeffrey Barrett, Sonia Goncalves, Ewan Harrison, David K. Jackson, Ian Johnston, Dominic Kwiatkowski, Cordelia Langford, John Sillitoe on behalf of the Wellcome Sanger Institute COVID-19 Surveillance Team |
| EPI_ISL_3283057 | Lighthouse Lab in Glasgow | Wellcome Sanger Institute for the COVID-19 Genomics UK (COG-UK) Consortium | Harper VanSteenhouse, Yumi Kasai, David Gray, Carol Clugston, Anna Dominiczak and Alex Alderton, Roberto Amato, Jeffrey Barrett, Sonia Goncalves, Ewan Harrison, David K. Jackson, Ian Johnston, Dominic Kwiatkowski, Cordelia Langford, John Sillitoe on behalf of the Wellcome Sanger Institute COVID-19 Surveillance Team |
| EPI_ISL_3283058, EPI_ISL_3283059 | Lighthouse Lab in Alderley Park | Wellcome Sanger Institute for the COVID-19 Genomics UK (COG-UK) Consortium | Jacquelyn Wynn, Mairead Hyland, The Lighthouse Lab in Alderley Park and Alex Alderton, Roberto Amato, Jeffrey Barrett, Sonia Goncalves, Ewan Harrison, David K. Jackson, Ian Johnston, Dominic Kwiatkowski, Cordelia Langford, John Sillitoe on behalf of the Wellcome Sanger Institute COVID-19 Surveillance Team |
| EPI_ISL_3283060 | Lighthouse Lab in Glasgow | Wellcome Sanger Institute for the COVID-19 Genomics UK (COG-UK) Consortium | Harper VanSteenhouse, Yumi Kasai, David Gray, Carol Clugston, Anna Dominiczak and Alex Alderton, Roberto Amato, Jeffrey Barrett, Sonia Goncalves, Ewan Harrison, David K. Jackson, Ian Johnston, Dominic Kwiatkowski, Cordelia Langford, John Sillitoe on behalf of the Wellcome Sanger Institute COVID-19 Surveillance Team |
| EPI_ISL_3283061 | Lighthouse Laboratory Plymouth | Wellcome Sanger Institute for the COVID-19 Genomics UK (COG-UK) Consortium | Lighthouse Laboratory Plymouth and Alex Alderton, Roberto Amato, Jeffrey Barrett, Sonia Goncalves, Ewan Harrison, David K. Jackson, Ian Johnston, Dominic Kwiatkowski, Cordelia Langford, John Sillitoe on behalf of the Wellcome Sanger Institute COVID-19 Surveillance Team |
| EPI_ISL_3283062 | Lighthouse Lab in Glasgow | Wellcome Sanger Institute for the COVID-19 Genomics UK (COG-UK) Consortium | Harper VanSteenhouse, Yumi Kasai, David Gray, Carol Clugston, Anna Dominiczak and Alex Alderton, Roberto Amato, Jeffrey Barrett, Sonia Goncalves, Ewan Harrison, David K. Jackson, Ian Johnston, Dominic Kwiatkowski, Cordelia Langford, John Sillitoe on behalf of the Wellcome Sanger Institute COVID-19 Surveillance Team |
| EPI_ISL_3283063 | Lighthouse Lab in Alderley Park | Wellcome Sanger Institute for the COVID-19 Genomics UK (COG-UK) Consortium | Jacquelyn Wynn, Mairead Hyland, The Lighthouse Lab in Alderley Park and Alex Alderton, Roberto Amato, Jeffrey Barrett, Sonia Goncalves, Ewan Harrison, David K. Jackson, Ian Johnston, Dominic Kwiatkowski, Cordelia Langford, John Sillitoe on behalf of the Wellcome Sanger Institute COVID-19 Surveillance Team |
| EPI_ISL_3283064 | Lighthouse Laboratory Plymouth | Wellcome Sanger Institute for the COVID-19 Genomics UK (COG-UK) Consortium | Lighthouse Laboratory Plymouth and Alex Alderton, Roberto Amato, Jeffrey Barrett, Sonia Goncalves, Ewan Harrison, David K. Jackson, Ian Johnston, Dominic Kwiatkowski, Cordelia Langford, John Sillitoe on behalf of the Wellcome Sanger Institute COVID-19 Surveillance Team |
| EPI_ISL_3283065 | Lighthouse Lab in Alderley Park | Wellcome Sanger Institute for the COVID-19 Genomics UK (COG-UK) Consortium | Jacquelyn Wynn, Mairead Hyland, The Lighthouse Lab in Alderley Park and Alex Alderton, Roberto Amato, Jeffrey Barrett, Sonia Goncalves, Ewan Harrison, David K. Jackson, Ian Johnston, Dominic Kwiatkowski, Cordelia Langford, John Sillitoe on behalf of the Wellcome Sanger Institute COVID-19 Surveillance Team |
| EPI_ISL_3283066 | Lighthouse Lab in Glasgow | Wellcome Sanger Institute for the COVID-19 Genomics UK (COG-UK) Consortium | Harper VanSteenhouse, Yumi Kasai, David Gray, Carol Clugston, Anna Dominiczak and Alex Alderton, Roberto Amato, Jeffrey Barrett, Sonia Goncalves, Ewan Harrison, David K. Jackson, Ian Johnston, Dominic Kwiatkowski, Cordelia Langford, John Sillitoe on behalf of the Wellcome Sanger Institute COVID-19 Surveillance Team |
| EPI_ISL_3283067, EPI_ISL_3283068 | Lighthouse Lab in Alderley Park | Wellcome Sanger Institute for the COVID-19 Genomics UK (COG-UK) Consortium | Jacquelyn Wynn, Mairead Hyland, The Lighthouse Lab in Alderley Park and Alex Alderton, Roberto Amato, Jeffrey Barrett, Sonia Goncalves, Ewan Harrison, David K. Jackson, Ian Johnston, Dominic Kwiatkowski, Cordelia Langford, John Sillitoe on behalf of the Wellcome Sanger Institute COVID-19 Surveillance Team |
| EPI_ISL_3283069, EPI_ISL_3283070 | Lighthouse Lab in Glasgow | Wellcome Sanger Institute for the COVID-19 Genomics UK (COG-UK) Consortium | Harper VanSteenhouse, Yumi Kasai, David Gray, Carol Clugston, Anna Dominiczak and Alex Alderton, Roberto Amato, Jeffrey Barrett, Sonia Goncalves, Ewan Harrison, David K. Jackson, Ian Johnston, Dominic Kwiatkowski, Cordelia Langford, John Sillitoe on behalf of the Wellcome Sanger Institute COVID-19 Surveillance Team |
| EPI_ISL_3283071 | Lighthouse Laboratory Plymouth | Wellcome Sanger Institute for the COVID-19 Genomics UK (COG-UK) Consortium | Lighthouse Laboratory Plymouth and Alex Alderton, Roberto Amato, Jeffrey Barrett, Sonia Goncalves, Ewan Harrison, David K. Jackson, Ian Johnston, Dominic Kwiatkowski, Cordelia Langford, John Sillitoe on behalf of the Wellcome Sanger Institute COVID-19 Surveillance Team |
| EPI_ISL_3283072 | Lighthouse Lab in Glasgow | Wellcome Sanger Institute for the COVID-19 Genomics UK (COG-UK) Consortium | Harper VanSteenhouse, Yumi Kasai, David Gray, Carol Clugston, Anna Dominiczak and Alex Alderton, Roberto Amato, Jeffrey Barrett, Sonia Goncalves, Ewan Harrison, David K. Jackson, Ian Johnston, Dominic Kwiatkowski, Cordelia Langford, John Sillitoe on behalf of the Wellcome Sanger Institute COVID-19 Surveillance Team |
| EPI_ISL_3283073, EPI_ISL_3283074, EPI_ISL_3283075 | Lighthouse Laboratory Plymouth | Wellcome Sanger Institute for the COVID-19 Genomics UK (COG-UK) Consortium | Lighthouse Laboratory Plymouth and Alex Alderton, Roberto Amato, Jeffrey Barrett, Sonia Goncalves, Ewan Harrison, David K. Jackson, Ian Johnston, Dominic Kwiatkowski, Cordelia Langford, John Sillitoe on behalf of the Wellcome Sanger Institute COVID-19 Surveillance Team |
| EPI_ISL_3283076, EPI_ISL_3283077, EPI_ISL_3283078 | Lighthouse Lab in Glasgow | Wellcome Sanger Institute for the COVID-19 Genomics UK (COG-UK) Consortium | Harper VanSteenhouse, Yumi Kasai, David Gray, Carol Clugston, Anna Dominiczak and Alex Alderton, Roberto Amato, Jeffrey Barrett, Sonia Goncalves, Ewan Harrison, David K. Jackson, Ian Johnston, Dominic Kwiatkowski, Cordelia Langford, John Sillitoe on behalf of the Wellcome Sanger Institute COVID-19 Surveillance Team |
| EPI_ISL_3283079 | Lighthouse Laboratory Plymouth | Wellcome Sanger Institute for the COVID-19 Genomics UK (COG-UK) Consortium | Lighthouse Laboratory Plymouth and Alex Alderton, Roberto Amato, Jeffrey Barrett, Sonia Goncalves, Ewan Harrison, David K. Jackson, Ian Johnston, Dominic Kwiatkowski, Cordelia Langford, John Sillitoe on behalf of the Wellcome Sanger Institute COVID-19 Surveillance Team |
| EPI_ISL_3283080, EPI_ISL_3283081 | Lighthouse Lab in Glasgow | Wellcome Sanger Institute for the COVID-19 Genomics UK (COG-UK) Consortium | Harper VanSteenhouse, Yumi Kasai, David Gray, Carol Clugston, Anna Dominiczak and Alex Alderton, Roberto Amato, Jeffrey Barrett, Sonia Goncalves, Ewan Harrison, David K. Jackson, Ian Johnston, Dominic Kwiatkowski, Cordelia Langford, John Sillitoe on behalf of the Wellcome Sanger Institute COVID-19 Surveillance Team |
| EPI_ISL_3283082, EPI_ISL_3283083 | Lighthouse Lab in Alderley Park | Wellcome Sanger Institute for the COVID-19 Genomics UK (COG-UK) Consortium | Jacquelyn Wynn, Mairead Hyland, The Lighthouse Lab in Alderley Park and Alex Alderton, Roberto Amato, Jeffrey Barrett, Sonia Goncalves, Ewan Harrison, David K. Jackson, Ian Johnston, Dominic Kwiatkowski, Cordelia Langford, John Sillitoe on behalf of the Wellcome Sanger Institute COVID-19 Surveillance Team |
| EPI_ISL_3283084, EPI_ISL_3283085, EPI_ISL_3283086 | Lighthouse Lab in Glasgow | Wellcome Sanger Institute for the COVID-19 Genomics UK (COG-UK) Consortium | Harper VanSteenhouse, Yumi Kasai, David Gray, Carol Clugston, Anna Dominiczak and Alex Alderton, Roberto Amato, Jeffrey Barrett, Sonia Goncalves, Ewan Harrison, David K. Jackson, Ian Johnston, Dominic Kwiatkowski, Cordelia Langford, John Sillitoe on behalf of the Wellcome Sanger Institute COVID-19 Surveillance Team |
| EPI_ISL_3283087 | Lighthouse Lab in Alderley Park | Wellcome Sanger Institute for the COVID-19 Genomics UK (COG-UK) Consortium | Jacquelyn Wynn, Mairead Hyland, The Lighthouse Lab in Alderley Park and Alex Alderton, Roberto Amato, Jeffrey Barrett, Sonia Goncalves, Ewan Harrison, David K. Jackson, Ian Johnston, Dominic Kwiatkowski, Cordelia Langford, John Sillitoe on behalf of the Wellcome Sanger Institute COVID-19 Surveillance Team |
| EPI_ISL_3283088 | Lighthouse Lab in Glasgow | Wellcome Sanger Institute for the COVID-19 Genomics UK (COG-UK) Consortium | Harper VanSteenhouse, Yumi Kasai, David Gray, Carol Clugston, Anna Dominiczak and Alex Alderton, Roberto Amato, Jeffrey Barrett, Sonia Goncalves, Ewan Harrison, David K. Jackson, Ian Johnston, Dominic Kwiatkowski, Cordelia Langford, John Sillitoe on behalf of the Wellcome Sanger Institute COVID-19 Surveillance Team |
| EPI_ISL_3283089, EPI_ISL_3283090 | Lighthouse Laboratory Plymouth | Wellcome Sanger Institute for the COVID-19 Genomics UK (COG-UK) Consortium | Lighthouse Laboratory Plymouth and Alex Alderton, Roberto Amato, Jeffrey Barrett, Sonia Goncalves, Ewan Harrison, David K. Jackson, Ian Johnston, Dominic Kwiatkowski, Cordelia Langford, John Sillitoe on behalf of the Wellcome Sanger Institute COVID-19 Surveillance Team |
| EPI_ISL_3283091, EPI_ISL_3283092, EPI_ISL_3283093 | Lighthouse Lab in Glasgow | Wellcome Sanger Institute for the COVID-19 Genomics UK (COG-UK) Consortium | Harper VanSteenhouse, Yumi Kasai, David Gray, Carol Clugston, Anna Dominiczak and Alex Alderton, Roberto Amato, Jeffrey Barrett, Sonia Goncalves, Ewan Harrison, David K. Jackson, Ian Johnston, Dominic Kwiatkowski, Cordelia Langford, John Sillitoe on behalf of the Wellcome Sanger Institute COVID-19 Surveillance Team |
| EPI_ISL_3283094, EPI_ISL_3283095 | Lighthouse Lab in Alderley Park | Wellcome Sanger Institute for the COVID-19 Genomics UK (COG-UK) Consortium | Jacquelyn Wynn, Mairead Hyland, The Lighthouse Lab in Alderley Park and Alex Alderton, Roberto Amato, Jeffrey Barrett, Sonia Goncalves, Ewan Harrison, David K. Jackson, Ian Johnston, Dominic Kwiatkowski, Cordelia Langford, John Sillitoe on behalf of the Wellcome Sanger Institute COVID-19 Surveillance Team |
| EPI_ISL_3283096 | Lighthouse Laboratory Plymouth | Wellcome Sanger Institute for the COVID-19 Genomics UK (COG-UK) Consortium | Lighthouse Laboratory Plymouth and Alex Alderton, Roberto Amato, Jeffrey Barrett, Sonia Goncalves, Ewan Harrison, David K. Jackson, Ian Johnston, Dominic Kwiatkowski, Cordelia Langford, John Sillitoe on behalf of the Wellcome Sanger Institute COVID-19 Surveillance Team |
| EPI_ISL_3283097, EPI_ISL_3283098 | Lighthouse Lab in Glasgow | Wellcome Sanger Institute for the COVID-19 Genomics UK (COG-UK) Consortium | Harper VanSteenhouse, Yumi Kasai, David Gray, Carol Clugston, Anna Dominiczak and Alex Alderton, Roberto Amato, Jeffrey Barrett, Sonia Goncalves, Ewan Harrison, David K. Jackson, Ian Johnston, Dominic Kwiatkowski, Cordelia Langford, John Sillitoe on behalf of the Wellcome Sanger Institute COVID-19 Surveillance Team |
| EPI_ISL_3283099 | Lighthouse Laboratory Plymouth | Wellcome Sanger Institute for the COVID-19 Genomics UK (COG-UK) Consortium | Lighthouse Laboratory Plymouth and Alex Alderton, Roberto Amato, Jeffrey Barrett, Sonia Goncalves, Ewan Harrison, David K. Jackson, Ian Johnston, Dominic Kwiatkowski, Cordelia Langford, John Sillitoe on behalf of the Wellcome Sanger Institute COVID-19 Surveillance Team |
| EPI_ISL_3283100, EPI_ISL_3283101 | Lighthouse Lab in Alderley Park | Wellcome Sanger Institute for the COVID-19 Genomics UK (COG-UK) Consortium | Jacquelyn Wynn, Mairead Hyland, The Lighthouse Lab in Alderley Park and Alex Alderton, Roberto Amato, Jeffrey Barrett, Sonia Goncalves, Ewan Harrison, David K. Jackson, Ian Johnston, Dominic Kwiatkowski, Cordelia Langford, John Sillitoe on behalf of the Wellcome Sanger Institute COVID-19 Surveillance Team |
| EPI_ISL_3283102 | Lighthouse Laboratory Plymouth | Wellcome Sanger Institute for the COVID-19 Genomics UK (COG-UK) Consortium | Lighthouse Laboratory Plymouth and Alex Alderton, Roberto Amato, Jeffrey Barrett, Sonia Goncalves, Ewan Harrison, David K. Jackson, Ian Johnston, Dominic Kwiatkowski, Cordelia Langford, John Sillitoe on behalf of the Wellcome Sanger Institute COVID-19 Surveillance Team |
| EPI_ISL_3283103 | Lighthouse Lab in Alderley Park | Wellcome Sanger Institute for the COVID-19 Genomics UK (COG-UK) Consortium | Jacquelyn Wynn, Mairead Hyland, The Lighthouse Lab in Alderley Park and Alex Alderton, Roberto Amato, Jeffrey Barrett, Sonia Goncalves, Ewan Harrison, David K. Jackson, Ian Johnston, Dominic Kwiatkowski, Cordelia Langford, John Sillitoe on behalf of the Wellcome Sanger Institute COVID-19 Surveillance Team |
| EPI_ISL_3283104 | Lighthouse Laboratory Plymouth | Wellcome Sanger Institute for the COVID-19 Genomics UK (COG-UK) Consortium | Lighthouse Laboratory Plymouth and Alex Alderton, Roberto Amato, Jeffrey Barrett, Sonia Goncalves, Ewan Harrison, David K. Jackson, Ian Johnston, Dominic Kwiatkowski, Cordelia Langford, John Sillitoe on behalf of the Wellcome Sanger Institute COVID-19 Surveillance Team |
| EPI_ISL_3283105 | Lighthouse Lab in Alderley Park | Wellcome Sanger Institute for the COVID-19 Genomics UK (COG-UK) Consortium | Jacquelyn Wynn, Mairead Hyland, The Lighthouse Lab in Alderley Park and Alex Alderton, Roberto Amato, Jeffrey Barrett, Sonia Goncalves, Ewan Harrison, David K. Jackson, Ian Johnston, Dominic Kwiatkowski, Cordelia Langford, John Sillitoe on behalf of the Wellcome Sanger Institute COVID-19 Surveillance Team |

| | | | |
|---|---|---|---|
| EPI_ISL_3283106 | Lighthouse Lab in Glasgow | Wellcome Sanger Institute for the COVID-19 Genomics UK (COG-UK) Consortium | Harper VanSteenhouse, Yumi Kasai, David Gray, Carol Clugston, Anna Dominiczak and Alex Alderton, Roberto Amato, Jeffrey Barrett, Sonia Goncalves, Ewan Harrison, David K. Jackson, Ian Johnston, Dominic Kwiatkowski, Cordelia Langford, John Sillitoe on behalf of the Wellcome Sanger Institute COVID-19 Surveillance Team |
| EPI_ISL_3283107 | Lighthouse Laboratory Plymouth | Wellcome Sanger Institute for the COVID-19 Genomics UK (COG-UK) Consortium | Lighthouse Laboratory Plymouth and Alex Alderton, Roberto Amato, Jeffrey Barrett, Sonia Goncalves, Ewan Harrison, David K. Jackson, Ian Johnston, Dominic Kwiatkowski, Cordelia Langford, John Sillitoe on behalf of the Wellcome Sanger Institute COVID-19 Surveillance Team |
| EPI_ISL_3283108 | Lighthouse Lab in Glasgow | Wellcome Sanger Institute for the COVID-19 Genomics UK (COG-UK) Consortium | Harper VanSteenhouse, Yumi Kasai, David Gray, Carol Clugston, Anna Dominiczak and Alex Alderton, Roberto Amato, Jeffrey Barrett, Sonia Goncalves, Ewan Harrison, David K. Jackson, Ian Johnston, Dominic Kwiatkowski, Cordelia Langford, John Sillitoe on behalf of the Wellcome Sanger Institute COVID-19 Surveillance Team |
| EPI_ISL_3283109 | Lighthouse Laboratory Plymouth | Wellcome Sanger Institute for the COVID-19 Genomics UK (COG-UK) Consortium | Lighthouse Laboratory Plymouth and Alex Alderton, Roberto Amato, Jeffrey Barrett, Sonia Goncalves, Ewan Harrison, David K. Jackson, Ian Johnston, Dominic Kwiatkowski, Cordelia Langford, John Sillitoe on behalf of the Wellcome Sanger Institute COVID-19 Surveillance Team |
| EPI_ISL_3283110 | Lighthouse Lab in Glasgow | Wellcome Sanger Institute for the COVID-19 Genomics UK (COG-UK) Consortium | Harper VanSteenhouse, Yumi Kasai, David Gray, Carol Clugston, Anna Dominiczak and Alex Alderton, Roberto Amato, Jeffrey Barrett, Sonia Goncalves, Ewan Harrison, David K. Jackson, Ian Johnston, Dominic Kwiatkowski, Cordelia Langford, John Sillitoe on behalf of the Wellcome Sanger Institute COVID-19 Surveillance Team |
| EPI_ISL_3283111 | Lighthouse Laboratory Plymouth | Wellcome Sanger Institute for the COVID-19 Genomics UK (COG-UK) Consortium | Lighthouse Laboratory Plymouth and Alex Alderton, Roberto Amato, Jeffrey Barrett, Sonia Goncalves, Ewan Harrison, David K. Jackson, Ian Johnston, Dominic Kwiatkowski, Cordelia Langford, John Sillitoe on behalf of the Wellcome Sanger Institute COVID-19 Surveillance Team |
| EPI_ISL_3283112, EPI_ISL_3283113 | Lighthouse Lab in Alderley Park | Wellcome Sanger Institute for the COVID-19 Genomics UK (COG-UK) Consortium | Jacquelyn Wynn, Mairead Hyland, The Lighthouse Lab in Alderley Park and Alex Alderton, Roberto Amato, Jeffrey Barrett, Sonia Goncalves, Ewan Harrison, David K. Jackson, Ian Johnston, Dominic Kwiatkowski, Cordelia Langford, John Sillitoe on behalf of the Wellcome Sanger Institute COVID-19 Surveillance Team |
| EPI_ISL_3283114 | Lighthouse Laboratory Plymouth | Wellcome Sanger Institute for the COVID-19 Genomics UK (COG-UK) Consortium | Lighthouse Laboratory Plymouth and Alex Alderton, Roberto Amato, Jeffrey Barrett, Sonia Goncalves, Ewan Harrison, David K. Jackson, Ian Johnston, Dominic Kwiatkowski, Cordelia Langford, John Sillitoe on behalf of the Wellcome Sanger Institute COVID-19 Surveillance Team |
| EPI_ISL_3283115, EPI_ISL_3283116 | Lighthouse Lab in Alderley Park | Wellcome Sanger Institute for the COVID-19 Genomics UK (COG-UK) Consortium | Jacquelyn Wynn, Mairead Hyland, The Lighthouse Lab in Alderley Park and Alex Alderton, Roberto Amato, Jeffrey Barrett, Sonia Goncalves, Ewan Harrison, David K. Jackson, Ian Johnston, Dominic Kwiatkowski, Cordelia Langford, John Sillitoe on behalf of the Wellcome Sanger Institute COVID-19 Surveillance Team |
| EPI_ISL_3283117 | Lighthouse Laboratory Plymouth | Wellcome Sanger Institute for the COVID-19 Genomics UK (COG-UK) Consortium | Lighthouse Laboratory Plymouth and Alex Alderton, Roberto Amato, Jeffrey Barrett, Sonia Goncalves, Ewan Harrison, David K. Jackson, Ian Johnston, Dominic Kwiatkowski, Cordelia Langford, John Sillitoe on behalf of the Wellcome Sanger Institute COVID-19 Surveillance Team |
| EPI_ISL_3283118, EPI_ISL_3283119 | Lighthouse Lab in Glasgow | Wellcome Sanger Institute for the COVID-19 Genomics UK (COG-UK) Consortium | Harper VanSteenhouse, Yumi Kasai, David Gray, Carol Clugston, Anna Dominiczak and Alex Alderton, Roberto Amato, Jeffrey Barrett, Sonia Goncalves, Ewan Harrison, David K. Jackson, Ian Johnston, Dominic Kwiatkowski, Cordelia Langford, John Sillitoe on behalf of the Wellcome Sanger Institute COVID-19 Surveillance Team |
| EPI_ISL_3283120, EPI_ISL_3283121 | Lighthouse Lab in Alderley Park | Wellcome Sanger Institute for the COVID-19 Genomics UK (COG-UK) Consortium | Jacquelyn Wynn, Mairead Hyland, The Lighthouse Lab in Alderley Park and Alex Alderton, Roberto Amato, Jeffrey Barrett, Sonia Goncalves, Ewan Harrison, David K. Jackson, Ian Johnston, Dominic Kwiatkowski, Cordelia Langford, John Sillitoe on behalf of the Wellcome Sanger Institute COVID-19 Surveillance Team |
| EPI_ISL_3283122, EPI_ISL_3283123 | Lighthouse Laboratory Plymouth | Wellcome Sanger Institute for the COVID-19 Genomics UK (COG-UK) Consortium | Lighthouse Laboratory Plymouth and Alex Alderton, Roberto Amato, Jeffrey Barrett, Sonia Goncalves, Ewan Harrison, David K. Jackson, Ian Johnston, Dominic Kwiatkowski, Cordelia Langford, John Sillitoe on behalf of the Wellcome Sanger Institute COVID-19 Surveillance Team |
| EPI_ISL_3283124, EPI_ISL_3283125, EPI_ISL_3283126 | Lighthouse Lab in Alderley Park | Wellcome Sanger Institute for the COVID-19 Genomics UK (COG-UK) Consortium | Jacquelyn Wynn, Mairead Hyland, The Lighthouse Lab in Alderley Park and Alex Alderton, Roberto Amato, Jeffrey Barrett, Sonia Goncalves, Ewan Harrison, David K. Jackson, Ian Johnston, Dominic Kwiatkowski, Cordelia Langford, John Sillitoe on behalf of the Wellcome Sanger Institute COVID-19 Surveillance Team |
| EPI_ISL_3283127 | Lighthouse Lab in Glasgow | Wellcome Sanger Institute for the COVID-19 Genomics UK (COG-UK) Consortium | Harper VanSteenhouse, Yumi Kasai, David Gray, Carol Clugston, Anna Dominiczak and Alex Alderton, Roberto Amato, Jeffrey Barrett, Sonia Goncalves, Ewan Harrison, David K. Jackson, Ian Johnston, Dominic Kwiatkowski, Cordelia Langford, John Sillitoe on behalf of the Wellcome Sanger Institute COVID-19 Surveillance Team |
| EPI_ISL_3283128 | Lighthouse Laboratory Plymouth | Wellcome Sanger Institute for the COVID-19 Genomics UK (COG-UK) Consortium | Lighthouse Laboratory Plymouth and Alex Alderton, Roberto Amato, Jeffrey Barrett, Sonia Goncalves, Ewan Harrison, David K. Jackson, Ian Johnston, Dominic Kwiatkowski, Cordelia Langford, John Sillitoe on behalf of the Wellcome Sanger Institute COVID-19 Surveillance Team |
| EPI_ISL_3283129 | Lighthouse Lab in Glasgow | Wellcome Sanger Institute for the COVID-19 Genomics UK (COG-UK) Consortium | Harper VanSteenhouse, Yumi Kasai, David Gray, Carol Clugston, Anna Dominiczak and Alex Alderton, Roberto Amato, Jeffrey Barrett, Sonia Goncalves, Ewan Harrison, David K. Jackson, Ian Johnston, Dominic Kwiatkowski, Cordelia Langford, John Sillitoe on behalf of the Wellcome Sanger Institute COVID-19 Surveillance Team |
| EPI_ISL_3283130 | Lighthouse Laboratory Plymouth | Wellcome Sanger Institute for the COVID-19 Genomics UK (COG-UK) Consortium | Lighthouse Laboratory Plymouth and Alex Alderton, Roberto Amato, Jeffrey Barrett, Sonia Goncalves, Ewan Harrison, David K. Jackson, Ian Johnston, Dominic Kwiatkowski, Cordelia Langford, John Sillitoe on behalf of the Wellcome Sanger Institute COVID-19 Surveillance Team |
| EPI_ISL_3283131 | Lighthouse Lab in Alderley Park | Wellcome Sanger Institute for the COVID-19 Genomics UK (COG-UK) Consortium | Jacquelyn Wynn, Mairead Hyland, The Lighthouse Lab in Alderley Park and Alex Alderton, Roberto Amato, Jeffrey Barrett, Sonia Goncalves, Ewan Harrison, David K. Jackson, Ian Johnston, Dominic Kwiatkowski, Cordelia Langford, John Sillitoe on behalf of the Wellcome Sanger Institute COVID-19 Surveillance Team |
| EPI_ISL_3283132, EPI_ISL_3283133 | Lighthouse Lab in Glasgow | Wellcome Sanger Institute for the COVID-19 Genomics UK (COG-UK) Consortium | Harper VanSteenhouse, Yumi Kasai, David Gray, Carol Clugston, Anna Dominiczak and Alex Alderton, Roberto Amato, Jeffrey Barrett, Sonia Goncalves, Ewan Harrison, David K. Jackson, Ian Johnston, Dominic Kwiatkowski, Cordelia Langford, John Sillitoe on behalf of the Wellcome Sanger Institute COVID-19 Surveillance Team |
| EPI_ISL_3283134 | Lighthouse Lab in Alderley Park | Wellcome Sanger Institute for the COVID-19 Genomics UK (COG-UK) Consortium | Jacquelyn Wynn, Mairead Hyland, The Lighthouse Lab in Alderley Park and Alex Alderton, Roberto Amato, Jeffrey Barrett, Sonia Goncalves, Ewan Harrison, David K. Jackson, Ian Johnston, Dominic Kwiatkowski, Cordelia Langford, John Sillitoe on behalf of the Wellcome Sanger Institute COVID-19 Surveillance Team |
| EPI_ISL_3283135 | Lighthouse Lab in Glasgow | Wellcome Sanger Institute for the COVID-19 Genomics UK (COG-UK) Consortium | Harper VanSteenhouse, Yumi Kasai, David Gray, Carol Clugston, Anna Dominiczak and Alex Alderton, Roberto Amato, Jeffrey Barrett, Sonia Goncalves, Ewan Harrison, David K. Jackson, Ian Johnston, Dominic Kwiatkowski, Cordelia Langford, John Sillitoe on behalf of the Wellcome Sanger Institute COVID-19 Surveillance Team |
| EPI_ISL_3283136 | Lighthouse Lab in Alderley Park | Wellcome Sanger Institute for the COVID-19 Genomics UK (COG-UK) Consortium | Jacquelyn Wynn, Mairead Hyland, The Lighthouse Lab in Alderley Park and Alex Alderton, Roberto Amato, Jeffrey Barrett, Sonia Goncalves, Ewan Harrison, David K. Jackson, Ian Johnston, Dominic Kwiatkowski, Cordelia Langford, John Sillitoe on behalf of the Wellcome Sanger Institute COVID-19 Surveillance Team |
| EPI_ISL_3283137 | Lighthouse Lab in Glasgow | Wellcome Sanger Institute for the COVID-19 Genomics UK (COG-UK) Consortium | Harper VanSteenhouse, Yumi Kasai, David Gray, Carol Clugston, Anna Dominiczak and Alex Alderton, Roberto Amato, Jeffrey Barrett, Sonia Goncalves, Ewan Harrison, David K. Jackson, Ian Johnston, Dominic Kwiatkowski, Cordelia Langford, John Sillitoe on behalf of the Wellcome Sanger Institute COVID-19 Surveillance Team |
| EPI_ISL_3283138 | Lighthouse Laboratory Plymouth | Wellcome Sanger Institute for the COVID-19 Genomics UK (COG-UK) Consortium | Lighthouse Laboratory Plymouth and Alex Alderton, Roberto Amato, Jeffrey Barrett, Sonia Goncalves, Ewan Harrison, David K. Jackson, Ian Johnston, Dominic Kwiatkowski, Cordelia Langford, John Sillitoe on behalf of the Wellcome Sanger Institute COVID-19 Surveillance Team |
| EPI_ISL_3283139, EPI_ISL_3283140, EPI_ISL_3283141 | Lighthouse Lab in Glasgow | Wellcome Sanger Institute for the COVID-19 Genomics UK (COG-UK) Consortium | Harper VanSteenhouse, Yumi Kasai, David Gray, Carol Clugston, Anna Dominiczak and Alex Alderton, Roberto Amato, Jeffrey Barrett, Sonia Goncalves, Ewan Harrison, David K. Jackson, Ian Johnston, Dominic Kwiatkowski, Cordelia Langford, John Sillitoe on behalf of the Wellcome Sanger Institute COVID-19 Surveillance Team |
| EPI_ISL_3283142 | Lighthouse Laboratory Plymouth | Wellcome Sanger Institute for the COVID-19 Genomics UK (COG-UK) Consortium | Lighthouse Laboratory Plymouth and Alex Alderton, Roberto Amato, Jeffrey Barrett, Sonia Goncalves, Ewan Harrison, David K. Jackson, Ian Johnston, Dominic Kwiatkowski, Cordelia Langford, John Sillitoe on behalf of the Wellcome Sanger Institute COVID-19 Surveillance Team |
| EPI_ISL_3283143, EPI_ISL_3283144, EPI_ISL_3283145 | Lighthouse Lab in Glasgow | Wellcome Sanger Institute for the COVID-19 Genomics UK (COG-UK) Consortium | Harper VanSteenhouse, Yumi Kasai, David Gray, Carol Clugston, Anna Dominiczak and Alex Alderton, Roberto Amato, Jeffrey Barrett, Sonia Goncalves, Ewan Harrison, David K. Jackson, Ian Johnston, Dominic Kwiatkowski, Cordelia Langford, John Sillitoe on behalf of the Wellcome Sanger Institute COVID-19 Surveillance Team |
| EPI_ISL_3283146 | Lighthouse Lab in Alderley Park | Wellcome Sanger Institute for the COVID-19 Genomics UK (COG-UK) Consortium | Jacquelyn Wynn, Mairead Hyland, The Lighthouse Lab in Alderley Park and Alex Alderton, Roberto Amato, Jeffrey Barrett, Sonia Goncalves, Ewan Harrison, David K. Jackson, Ian Johnston, Dominic Kwiatkowski, Cordelia Langford, John Sillitoe on behalf of the Wellcome Sanger Institute COVID-19 Surveillance Team |
| EPI_ISL_3283147 | Lighthouse Laboratory Plymouth | Wellcome Sanger Institute for the COVID-19 Genomics UK (COG-UK) Consortium | Lighthouse Laboratory Plymouth and Alex Alderton, Roberto Amato, Jeffrey Barrett, Sonia Goncalves, Ewan Harrison, David K. Jackson, Ian Johnston, Dominic Kwiatkowski, Cordelia Langford, John Sillitoe on behalf of the Wellcome Sanger Institute COVID-19 Surveillance Team |
| EPI_ISL_3283148 | Lighthouse Lab in Alderley Park | Wellcome Sanger Institute for the COVID-19 Genomics UK (COG-UK) Consortium | Jacquelyn Wynn, Mairead Hyland, The Lighthouse Lab in Alderley Park and Alex Alderton, Roberto Amato, Jeffrey Barrett, Sonia Goncalves, Ewan Harrison, David K. Jackson, Ian Johnston, Dominic Kwiatkowski, Cordelia Langford, John Sillitoe on behalf of the Wellcome Sanger Institute COVID-19 Surveillance Team |
| EPI_ISL_3283149, EPI_ISL_3283150, EPI_ISL_3283151 | Lighthouse Lab in Glasgow | Wellcome Sanger Institute for the COVID-19 Genomics UK (COG-UK) Consortium | Harper VanSteenhouse, Yumi Kasai, David Gray, Carol Clugston, Anna Dominiczak and Alex Alderton, Roberto Amato, Jeffrey Barrett, Sonia Goncalves, Ewan Harrison, David K. Jackson, Ian Johnston, Dominic Kwiatkowski, Cordelia Langford, John Sillitoe on behalf of the Wellcome Sanger Institute COVID-19 Surveillance Team |
| EPI_ISL_3283152, EPI_ISL_3283153 | Lighthouse Lab in Alderley Park | Wellcome Sanger Institute for the COVID-19 Genomics UK (COG-UK) Consortium | Jacquelyn Wynn, Mairead Hyland, The Lighthouse Lab in Alderley Park and Alex Alderton, Roberto Amato, Jeffrey Barrett, Sonia Goncalves, Ewan Harrison, David K. Jackson, Ian Johnston, Dominic Kwiatkowski, Cordelia Langford, John Sillitoe on behalf of the Wellcome Sanger Institute COVID-19 Surveillance Team |
| EPI_ISL_3283154, EPI_ISL_3283155 | Lighthouse Lab in Glasgow | Wellcome Sanger Institute for the COVID-19 Genomics UK (COG-UK) Consortium | Harper VanSteenhouse, Yumi Kasai, David Gray, Carol Clugston, Anna Dominiczak and Alex Alderton, Roberto Amato, Jeffrey Barrett, Sonia Goncalves, Ewan Harrison, David K. Jackson, Ian Johnston, Dominic Kwiatkowski, Cordelia Langford, John Sillitoe on behalf of the Wellcome Sanger Institute COVID-19 Surveillance Team |
| EPI_ISL_3283156, EPI_ISL_3283157, EPI_ISL_3283158 | Lighthouse Lab in Alderley Park | Wellcome Sanger Institute for the COVID-19 Genomics UK (COG-UK) Consortium | Jacquelyn Wynn, Mairead Hyland, The Lighthouse Lab in Alderley Park and Alex Alderton, Roberto Amato, Jeffrey Barrett, Sonia Goncalves, Ewan Harrison, David K. Jackson, Ian Johnston, Dominic Kwiatkowski, Cordelia Langford, John Sillitoe on behalf of the Wellcome Sanger Institute COVID-19 Surveillance Team |
| EPI_ISL_3283159 | Lighthouse Laboratory Plymouth | Wellcome Sanger Institute for the COVID-19 Genomics UK (COG-UK) Consortium | Lighthouse Laboratory Plymouth and Alex Alderton, Roberto Amato, Jeffrey Barrett, Sonia Goncalves, Ewan Harrison, David K. Jackson, Ian Johnston, Dominic Kwiatkowski, Cordelia Langford, John Sillitoe on behalf of the Wellcome Sanger Institute COVID-19 Surveillance Team |
| EPI_ISL_3283160, EPI_ISL_3283161 | Lighthouse Lab in Glasgow | Wellcome Sanger Institute for the COVID-19 Genomics UK (COG-UK) Consortium | Harper VanSteenhouse, Yumi Kasai, David Gray, Carol Clugston, Anna Dominiczak and Alex Alderton, Roberto Amato, Jeffrey Barrett, Sonia Goncalves, Ewan Harrison, David K. Jackson, Ian Johnston, Dominic Kwiatkowski, Cordelia Langford, John Sillitoe on behalf of the Wellcome Sanger Institute COVID-19 Surveillance Team |
| EPI_ISL_3283162 | Lighthouse Laboratory Plymouth | Wellcome Sanger Institute for the COVID-19 Genomics UK (COG-UK) Consortium | Lighthouse Laboratory Plymouth and Alex Alderton, Roberto Amato, Jeffrey Barrett, Sonia Goncalves, Ewan Harrison, David K. Jackson, Ian Johnston, Dominic Kwiatkowski, Cordelia Langford, John Sillitoe on behalf of the Wellcome Sanger Institute COVID-19 Surveillance Team |
| EPI_ISL_3283163 | Lighthouse Lab in Glasgow | Wellcome Sanger Institute for the COVID-19 Genomics UK (COG-UK) Consortium | Harper VanSteenhouse, Yumi Kasai, David Gray, Carol Clugston, Anna Dominiczak and Alex Alderton, Roberto Amato, Jeffrey Barrett, Sonia Goncalves, Ewan Harrison, David K. Jackson, Ian Johnston, Dominic Kwiatkowski, Cordelia Langford, John Sillitoe on behalf of the Wellcome Sanger Institute COVID-19 Surveillance Team |
| EPI_ISL_3283164 | Lighthouse Laboratory Plymouth | Wellcome Sanger Institute for the COVID-19 Genomics UK (COG-UK) Consortium | Lighthouse Laboratory Plymouth and Alex Alderton, Roberto Amato, Jeffrey Barrett, Sonia Goncalves, Ewan Harrison, David K. Jackson, Ian Johnston, Dominic Kwiatkowski, Cordelia Langford, John Sillitoe on behalf of the Wellcome Sanger Institute COVID-19 Surveillance Team |
| EPI_ISL_3283165 | Lighthouse Lab in Glasgow | Wellcome Sanger Institute for the COVID-19 Genomics UK (COG-UK) Consortium | Harper VanSteenhouse, Yumi Kasai, David Gray, Carol Clugston, Anna Dominiczak and Alex Alderton, Roberto Amato, Jeffrey Barrett, Sonia Goncalves, Ewan Harrison, David K. Jackson, Ian Johnston, Dominic Kwiatkowski, Cordelia Langford, John Sillitoe on behalf of the Wellcome Sanger Institute COVID-19 Surveillance Team |
| EPI_ISL_3283166 | Lighthouse Laboratory Plymouth | Wellcome Sanger Institute for the COVID-19 Genomics UK (COG-UK) Consortium | Lighthouse Laboratory Plymouth and Alex Alderton, Roberto Amato, Jeffrey Barrett, Sonia Goncalves, Ewan Harrison, David K. Jackson, Ian Johnston, Dominic Kwiatkowski, Cordelia Langford, John Sillitoe on behalf of the Wellcome Sanger Institute COVID-19 Surveillance Team |
| EPI_ISL_3283167 | Lighthouse Lab in Glasgow | Wellcome Sanger Institute for the COVID-19 Genomics UK (COG-UK) Consortium | Harper VanSteenhouse, Yumi Kasai, David Gray, Carol Clugston, Anna Dominiczak and Alex Alderton, Roberto Amato, Jeffrey Barrett, Sonia Goncalves, Ewan Harrison, David K. Jackson, Ian Johnston, Dominic Kwiatkowski, Cordelia Langford, John Sillitoe on behalf of the Wellcome Sanger Institute COVID-19 Surveillance Team |

| | | | |
|---|---|---|---|
| EPI_ISL_3283168 | Lighthouse Lab in Alderley Park | Wellcome Sanger Institute for the COVID-19 Genomics UK (COG-UK) Consortium | Jacquelyn Wynn, Mairead Hyland, The Lighthouse Lab in Alderley Park and Alex Alderton, Roberto Amato, Jeffrey Barrett, Sonia Goncalves, Ewan Harrison, David K. Jackson, Ian Johnston, Dominic Kwiatkowski, Cordelia Langford, John Sillitoe on behalf of the Wellcome Sanger Institute COVID-19 Surveillance Team |
| EPI_ISL_3283169 | Lighthouse Lab in Glasgow | Wellcome Sanger Institute for the COVID-19 Genomics UK (COG-UK) Consortium | Harper VanSteenhouse, Yumi Kasai, David Gray, Carol Clugston, Anna Dominiczak and Alex Alderton, Roberto Amato, Jeffrey Barrett, Sonia Goncalves, Ewan Harrison, David K. Jackson, Ian Johnston, Dominic Kwiatkowski, Cordelia Langford, John Sillitoe on behalf of the Wellcome Sanger Institute COVID-19 Surveillance Team |
| EPI_ISL_3283170, EPI_ISL_3283171 | Lighthouse Laboratory Plymouth | Wellcome Sanger Institute for the COVID-19 Genomics UK (COG-UK) Consortium | Lighthouse Laboratory Plymouth and Alex Alderton, Roberto Amato, Jeffrey Barrett, Sonia Goncalves, Ewan Harrison, David K. Jackson, Ian Johnston, Dominic Kwiatkowski, Cordelia Langford, John Sillitoe on behalf of the Wellcome Sanger Institute COVID-19 Surveillance Team |
| EPI_ISL_3283172, EPI_ISL_3283173 | Lighthouse Lab in Glasgow | Wellcome Sanger Institute for the COVID-19 Genomics UK (COG-UK) Consortium | Harper VanSteenhouse, Yumi Kasai, David Gray, Carol Clugston, Anna Dominiczak and Alex Alderton, Roberto Amato, Jeffrey Barrett, Sonia Goncalves, Ewan Harrison, David K. Jackson, Ian Johnston, Dominic Kwiatkowski, Cordelia Langford, John Sillitoe on behalf of the Wellcome Sanger Institute COVID-19 Surveillance Team |
| EPI_ISL_3283174, EPI_ISL_3283175 | Lighthouse Lab in Alderley Park | Wellcome Sanger Institute for the COVID-19 Genomics UK (COG-UK) Consortium | Jacquelyn Wynn, Mairead Hyland, The Lighthouse Lab in Alderley Park and Alex Alderton, Roberto Amato, Jeffrey Barrett, Sonia Goncalves, Ewan Harrison, David K. Jackson, Ian Johnston, Dominic Kwiatkowski, Cordelia Langford, John Sillitoe on behalf of the Wellcome Sanger Institute COVID-19 Surveillance Team |
| EPI_ISL_3283176 | Lighthouse Lab in Glasgow | Wellcome Sanger Institute for the COVID-19 Genomics UK (COG-UK) Consortium | Harper VanSteenhouse, Yumi Kasai, David Gray, Carol Clugston, Anna Dominiczak and Alex Alderton, Roberto Amato, Jeffrey Barrett, Sonia Goncalves, Ewan Harrison, David K. Jackson, Ian Johnston, Dominic Kwiatkowski, Cordelia Langford, John Sillitoe on behalf of the Wellcome Sanger Institute COVID-19 Surveillance Team |
| EPI_ISL_3283177, EPI_ISL_3283178 | Lighthouse Laboratory Plymouth | Wellcome Sanger Institute for the COVID-19 Genomics UK (COG-UK) Consortium | Lighthouse Laboratory Plymouth and Alex Alderton, Roberto Amato, Jeffrey Barrett, Sonia Goncalves, Ewan Harrison, David K. Jackson, Ian Johnston, Dominic Kwiatkowski, Cordelia Langford, John Sillitoe on behalf of the Wellcome Sanger Institute COVID-19 Surveillance Team |
| EPI_ISL_3283179 | Lighthouse Lab in Glasgow | Wellcome Sanger Institute for the COVID-19 Genomics UK (COG-UK) Consortium | Harper VanSteenhouse, Yumi Kasai, David Gray, Carol Clugston, Anna Dominiczak and Alex Alderton, Roberto Amato, Jeffrey Barrett, Sonia Goncalves, Ewan Harrison, David K. Jackson, Ian Johnston, Dominic Kwiatkowski, Cordelia Langford, John Sillitoe on behalf of the Wellcome Sanger Institute COVID-19 Surveillance Team |
| EPI_ISL_3283180 | Lighthouse Lab in Alderley Park | Wellcome Sanger Institute for the COVID-19 Genomics UK (COG-UK) Consortium | Jacquelyn Wynn, Mairead Hyland, The Lighthouse Lab in Alderley Park and Alex Alderton, Roberto Amato, Jeffrey Barrett, Sonia Goncalves, Ewan Harrison, David K. Jackson, Ian Johnston, Dominic Kwiatkowski, Cordelia Langford, John Sillitoe on behalf of the Wellcome Sanger Institute COVID-19 Surveillance Team |
| EPI_ISL_3283181 | Lighthouse Lab in Glasgow | Wellcome Sanger Institute for the COVID-19 Genomics UK (COG-UK) Consortium | Harper VanSteenhouse, Yumi Kasai, David Gray, Carol Clugston, Anna Dominiczak and Alex Alderton, Roberto Amato, Jeffrey Barrett, Sonia Goncalves, Ewan Harrison, David K. Jackson, Ian Johnston, Dominic Kwiatkowski, Cordelia Langford, John Sillitoe on behalf of the Wellcome Sanger Institute COVID-19 Surveillance Team |
| EPI_ISL_3283182 | Lighthouse Laboratory Plymouth | Wellcome Sanger Institute for the COVID-19 Genomics UK (COG-UK) Consortium | Lighthouse Laboratory Plymouth and Alex Alderton, Roberto Amato, Jeffrey Barrett, Sonia Goncalves, Ewan Harrison, David K. Jackson, Ian Johnston, Dominic Kwiatkowski, Cordelia Langford, John Sillitoe on behalf of the Wellcome Sanger Institute COVID-19 Surveillance Team |
| EPI_ISL_3283183, EPI_ISL_3283184, EPI_ISL_3283185 | Lighthouse Lab in Glasgow | Wellcome Sanger Institute for the COVID-19 Genomics UK (COG-UK) Consortium | Harper VanSteenhouse, Yumi Kasai, David Gray, Carol Clugston, Anna Dominiczak and Alex Alderton, Roberto Amato, Jeffrey Barrett, Sonia Goncalves, Ewan Harrison, David K. Jackson, Ian Johnston, Dominic Kwiatkowski, Cordelia Langford, John Sillitoe on behalf of the Wellcome Sanger Institute COVID-19 Surveillance Team |
| EPI_ISL_3283186, EPI_ISL_3283187 | Lighthouse Laboratory Plymouth | Wellcome Sanger Institute for the COVID-19 Genomics UK (COG-UK) Consortium | Lighthouse Laboratory Plymouth and Alex Alderton, Roberto Amato, Jeffrey Barrett, Sonia Goncalves, Ewan Harrison, David K. Jackson, Ian Johnston, Dominic Kwiatkowski, Cordelia Langford, John Sillitoe on behalf of the Wellcome Sanger Institute COVID-19 Surveillance Team |
| EPI_ISL_3283188 | Lighthouse Lab in Alderley Park | Wellcome Sanger Institute for the COVID-19 Genomics UK (COG-UK) Consortium | Jacquelyn Wynn, Mairead Hyland, The Lighthouse Lab in Alderley Park and Alex Alderton, Roberto Amato, Jeffrey Barrett, Sonia Goncalves, Ewan Harrison, David K. Jackson, Ian Johnston, Dominic Kwiatkowski, Cordelia Langford, John Sillitoe on behalf of the Wellcome Sanger Institute COVID-19 Surveillance Team |
| EPI_ISL_3283189 | Lighthouse Lab in Glasgow | Wellcome Sanger Institute for the COVID-19 Genomics UK (COG-UK) Consortium | Harper VanSteenhouse, Yumi Kasai, David Gray, Carol Clugston, Anna Dominiczak and Alex Alderton, Roberto Amato, Jeffrey Barrett, Sonia Goncalves, Ewan Harrison, David K. Jackson, Ian Johnston, Dominic Kwiatkowski, Cordelia Langford, John Sillitoe on behalf of the Wellcome Sanger Institute COVID-19 Surveillance Team |
| EPI_ISL_3283190 | Lighthouse Lab in Alderley Park | Wellcome Sanger Institute for the COVID-19 Genomics UK (COG-UK) Consortium | Jacquelyn Wynn, Mairead Hyland, The Lighthouse Lab in Alderley Park and Alex Alderton, Roberto Amato, Jeffrey Barrett, Sonia Goncalves, Ewan Harrison, David K. Jackson, Ian Johnston, Dominic Kwiatkowski, Cordelia Langford, John Sillitoe on behalf of the Wellcome Sanger Institute COVID-19 Surveillance Team |
| EPI_ISL_3283191 | Lighthouse Laboratory Plymouth | Wellcome Sanger Institute for the COVID-19 Genomics UK (COG-UK) Consortium | Lighthouse Laboratory Plymouth and Alex Alderton, Roberto Amato, Jeffrey Barrett, Sonia Goncalves, Ewan Harrison, David K. Jackson, Ian Johnston, Dominic Kwiatkowski, Cordelia Langford, John Sillitoe on behalf of the Wellcome Sanger Institute COVID-19 Surveillance Team |
| EPI_ISL_3283192, EPI_ISL_3283193, EPI_ISL_3283194, EPI_ISL_3283195 | Lighthouse Lab in Glasgow | Wellcome Sanger Institute for the COVID-19 Genomics UK (COG-UK) Consortium | Harper VanSteenhouse, Yumi Kasai, David Gray, Carol Clugston, Anna Dominiczak and Alex Alderton, Roberto Amato, Jeffrey Barrett, Sonia Goncalves, Ewan Harrison, David K. Jackson, Ian Johnston, Dominic Kwiatkowski, Cordelia Langford, John Sillitoe on behalf of the Wellcome Sanger Institute COVID-19 Surveillance Team |
| EPI_ISL_3283196 | Lighthouse Lab in Alderley Park | Wellcome Sanger Institute for the COVID-19 Genomics UK (COG-UK) Consortium | Jacquelyn Wynn, Mairead Hyland, The Lighthouse Lab in Alderley Park and Alex Alderton, Roberto Amato, Jeffrey Barrett, Sonia Goncalves, Ewan Harrison, David K. Jackson, Ian Johnston, Dominic Kwiatkowski, Cordelia Langford, John Sillitoe on behalf of the Wellcome Sanger Institute COVID-19 Surveillance Team |
| EPI_ISL_3283197 | Lighthouse Lab in Glasgow | Wellcome Sanger Institute for the COVID-19 Genomics UK (COG-UK) Consortium | Harper VanSteenhouse, Yumi Kasai, David Gray, Carol Clugston, Anna Dominiczak and Alex Alderton, Roberto Amato, Jeffrey Barrett, Sonia Goncalves, Ewan Harrison, David K. Jackson, Ian Johnston, Dominic Kwiatkowski, Cordelia Langford, John Sillitoe on behalf of the Wellcome Sanger Institute COVID-19 Surveillance Team |
| EPI_ISL_3283198, EPI_ISL_3283199, EPI_ISL_3283200 | Lighthouse Laboratory Plymouth | Wellcome Sanger Institute for the COVID-19 Genomics UK (COG-UK) Consortium | Lighthouse Laboratory Plymouth and Alex Alderton, Roberto Amato, Jeffrey Barrett, Sonia Goncalves, Ewan Harrison, David K. Jackson, Ian Johnston, Dominic Kwiatkowski, Cordelia Langford, John Sillitoe on behalf of the Wellcome Sanger Institute COVID-19 Surveillance Team |
| EPI_ISL_3283201, EPI_ISL_3283202 | Lighthouse Lab in Glasgow | Wellcome Sanger Institute for the COVID-19 Genomics UK (COG-UK) Consortium | Harper VanSteenhouse, Yumi Kasai, David Gray, Carol Clugston, Anna Dominiczak and Alex Alderton, Roberto Amato, Jeffrey Barrett, Sonia Goncalves, Ewan Harrison, David K. Jackson, Ian Johnston, Dominic Kwiatkowski, Cordelia Langford, John Sillitoe on behalf of the Wellcome Sanger Institute COVID-19 Surveillance Team |
| EPI_ISL_3283203 | Lighthouse Laboratory Plymouth | Wellcome Sanger Institute for the COVID-19 Genomics UK (COG-UK) Consortium | Lighthouse Laboratory Plymouth and Alex Alderton, Roberto Amato, Jeffrey Barrett, Sonia Goncalves, Ewan Harrison, David K. Jackson, Ian Johnston, Dominic Kwiatkowski, Cordelia Langford, John Sillitoe on behalf of the Wellcome Sanger Institute COVID-19 Surveillance Team |
| EPI_ISL_3283204, EPI_ISL_3283205 | Lighthouse Lab in Alderley Park | Wellcome Sanger Institute for the COVID-19 Genomics UK (COG-UK) Consortium | Jacquelyn Wynn, Mairead Hyland, The Lighthouse Lab in Alderley Park and Alex Alderton, Roberto Amato, Jeffrey Barrett, Sonia Goncalves, Ewan Harrison, David K. Jackson, Ian Johnston, Dominic Kwiatkowski, Cordelia Langford, John Sillitoe on behalf of the Wellcome Sanger Institute COVID-19 Surveillance Team |
| EPI_ISL_3283206, EPI_ISL_3283207, EPI_ISL_3283208, EPI_ISL_3283209 | Lighthouse Lab in Glasgow | Wellcome Sanger Institute for the COVID-19 Genomics UK (COG-UK) Consortium | Harper VanSteenhouse, Yumi Kasai, David Gray, Carol Clugston, Anna Dominiczak and Alex Alderton, Roberto Amato, Jeffrey Barrett, Sonia Goncalves, Ewan Harrison, David K. Jackson, Ian Johnston, Dominic Kwiatkowski, Cordelia Langford, John Sillitoe on behalf of the Wellcome Sanger Institute COVID-19 Surveillance Team |
| EPI_ISL_3283210 | Lighthouse Laboratory Plymouth | Wellcome Sanger Institute for the COVID-19 Genomics UK (COG-UK) Consortium | Lighthouse Laboratory Plymouth and Alex Alderton, Roberto Amato, Jeffrey Barrett, Sonia Goncalves, Ewan Harrison, David K. Jackson, Ian Johnston, Dominic Kwiatkowski, Cordelia Langford, John Sillitoe on behalf of the Wellcome Sanger Institute COVID-19 Surveillance Team |
| EPI_ISL_3283211 | Lighthouse Lab in Glasgow | Wellcome Sanger Institute for the COVID-19 Genomics UK (COG-UK) Consortium | Harper VanSteenhouse, Yumi Kasai, David Gray, Carol Clugston, Anna Dominiczak and Alex Alderton, Roberto Amato, Jeffrey Barrett, Sonia Goncalves, Ewan Harrison, David K. Jackson, Ian Johnston, Dominic Kwiatkowski, Cordelia Langford, John Sillitoe on behalf of the Wellcome Sanger Institute COVID-19 Surveillance Team |
| EPI_ISL_3283212, EPI_ISL_3283213, EPI_ISL_3283214 | Lighthouse Lab in Alderley Park | Wellcome Sanger Institute for the COVID-19 Genomics UK (COG-UK) Consortium | Jacquelyn Wynn, Mairead Hyland, The Lighthouse Lab in Alderley Park and Alex Alderton, Roberto Amato, Jeffrey Barrett, Sonia Goncalves, Ewan Harrison, David K. Jackson, Ian Johnston, Dominic Kwiatkowski, Cordelia Langford, John Sillitoe on behalf of the Wellcome Sanger Institute COVID-19 Surveillance Team |
| EPI_ISL_3283215, EPI_ISL_3283216 | Lighthouse Laboratory Plymouth | Wellcome Sanger Institute for the COVID-19 Genomics UK (COG-UK) Consortium | Lighthouse Laboratory Plymouth and Alex Alderton, Roberto Amato, Jeffrey Barrett, Sonia Goncalves, Ewan Harrison, David K. Jackson, Ian Johnston, Dominic Kwiatkowski, Cordelia Langford, John Sillitoe on behalf of the Wellcome Sanger Institute COVID-19 Surveillance Team |
| EPI_ISL_3283217 | Lighthouse Lab in Alderley Park | Wellcome Sanger Institute for the COVID-19 Genomics UK (COG-UK) Consortium | Jacquelyn Wynn, Mairead Hyland, The Lighthouse Lab in Alderley Park and Alex Alderton, Roberto Amato, Jeffrey Barrett, Sonia Goncalves, Ewan Harrison, David K. Jackson, Ian Johnston, Dominic Kwiatkowski, Cordelia Langford, John Sillitoe on behalf of the Wellcome Sanger Institute COVID-19 Surveillance Team |
| EPI_ISL_3283218, EPI_ISL_3283219 | Lighthouse Laboratory Plymouth | Wellcome Sanger Institute for the COVID-19 Genomics UK (COG-UK) Consortium | Lighthouse Laboratory Plymouth and Alex Alderton, Roberto Amato, Jeffrey Barrett, Sonia Goncalves, Ewan Harrison, David K. Jackson, Ian Johnston, Dominic Kwiatkowski, Cordelia Langford, John Sillitoe on behalf of the Wellcome Sanger Institute COVID-19 Surveillance Team |
| EPI_ISL_3283220 | Lighthouse Lab in Alderley Park | Wellcome Sanger Institute for the COVID-19 Genomics UK (COG-UK) Consortium | Jacquelyn Wynn, Mairead Hyland, The Lighthouse Lab in Alderley Park and Alex Alderton, Roberto Amato, Jeffrey Barrett, Sonia Goncalves, Ewan Harrison, David K. Jackson, Ian Johnston, Dominic Kwiatkowski, Cordelia Langford, John Sillitoe on behalf of the Wellcome Sanger Institute COVID-19 Surveillance Team |
| EPI_ISL_3283221 | Lighthouse Lab in Glasgow | Wellcome Sanger Institute for the COVID-19 Genomics UK (COG-UK) Consortium | Harper VanSteenhouse, Yumi Kasai, David Gray, Carol Clugston, Anna Dominiczak and Alex Alderton, Roberto Amato, Jeffrey Barrett, Sonia Goncalves, Ewan Harrison, David K. Jackson, Ian Johnston, Dominic Kwiatkowski, Cordelia Langford, John Sillitoe on behalf of the Wellcome Sanger Institute COVID-19 Surveillance Team |
| EPI_ISL_3283222, EPI_ISL_3283223 | Lighthouse Laboratory Plymouth | Wellcome Sanger Institute for the COVID-19 Genomics UK (COG-UK) Consortium | Lighthouse Laboratory Plymouth and Alex Alderton, Roberto Amato, Jeffrey Barrett, Sonia Goncalves, Ewan Harrison, David K. Jackson, Ian Johnston, Dominic Kwiatkowski, Cordelia Langford, John Sillitoe on behalf of the Wellcome Sanger Institute COVID-19 Surveillance Team |
| EPI_ISL_3283224 | Lighthouse Lab in Glasgow | Wellcome Sanger Institute for the COVID-19 Genomics UK (COG-UK) Consortium | Harper VanSteenhouse, Yumi Kasai, David Gray, Carol Clugston, Anna Dominiczak and Alex Alderton, Roberto Amato, Jeffrey Barrett, Sonia Goncalves, Ewan Harrison, David K. Jackson, Ian Johnston, Dominic Kwiatkowski, Cordelia Langford, John Sillitoe on behalf of the Wellcome Sanger Institute COVID-19 Surveillance Team |
| EPI_ISL_3283225, EPI_ISL_3283226, EPI_ISL_3283227 | Lighthouse Laboratory Plymouth | Wellcome Sanger Institute for the COVID-19 Genomics UK (COG-UK) Consortium | Lighthouse Laboratory Plymouth and Alex Alderton, Roberto Amato, Jeffrey Barrett, Sonia Goncalves, Ewan Harrison, David K. Jackson, Ian Johnston, Dominic Kwiatkowski, Cordelia Langford, John Sillitoe on behalf of the Wellcome Sanger Institute COVID-19 Surveillance Team |
| EPI_ISL_3283228 | Lighthouse Lab in Alderley Park | Wellcome Sanger Institute for the COVID-19 Genomics UK (COG-UK) Consortium | Jacquelyn Wynn, Mairead Hyland, The Lighthouse Lab in Alderley Park and Alex Alderton, Roberto Amato, Jeffrey Barrett, Sonia Goncalves, Ewan Harrison, David K. Jackson, Ian Johnston, Dominic Kwiatkowski, Cordelia Langford, John Sillitoe on behalf of the Wellcome Sanger Institute COVID-19 Surveillance Team |
| EPI_ISL_3283229 | Lighthouse Lab in Glasgow | Wellcome Sanger Institute for the COVID-19 Genomics UK (COG-UK) Consortium | Harper VanSteenhouse, Yumi Kasai, David Gray, Carol Clugston, Anna Dominiczak and Alex Alderton, Roberto Amato, Jeffrey Barrett, Sonia Goncalves, Ewan Harrison, David K. Jackson, Ian Johnston, Dominic Kwiatkowski, Cordelia Langford, John Sillitoe on behalf of the Wellcome Sanger Institute COVID-19 Surveillance Team |
| EPI_ISL_3283230 | Lighthouse Lab in Alderley Park | Wellcome Sanger Institute for the COVID-19 Genomics UK (COG-UK) Consortium | Jacquelyn Wynn, Mairead Hyland, The Lighthouse Lab in Alderley Park and Alex Alderton, Roberto Amato, Jeffrey Barrett, Sonia Goncalves, Ewan Harrison, David K. Jackson, Ian Johnston, Dominic Kwiatkowski, Cordelia Langford, John Sillitoe on behalf of the Wellcome Sanger Institute COVID-19 Surveillance Team |
| EPI_ISL_3283231 | Lighthouse Lab in Glasgow | Wellcome Sanger Institute for the COVID-19 Genomics UK (COG-UK) Consortium | Harper VanSteenhouse, Yumi Kasai, David Gray, Carol Clugston, Anna Dominiczak and Alex Alderton, Roberto Amato, Jeffrey Barrett, Sonia Goncalves, Ewan Harrison, David K. Jackson, Ian Johnston, Dominic Kwiatkowski, Cordelia Langford, John Sillitoe on behalf of the Wellcome Sanger Institute COVID-19 Surveillance Team |
| EPI_ISL_3283232 | Lighthouse Laboratory Plymouth | Wellcome Sanger Institute for the COVID-19 Genomics UK (COG-UK) Consortium | Lighthouse Laboratory Plymouth and Alex Alderton, Roberto Amato, Jeffrey Barrett, Sonia Goncalves, Ewan Harrison, David K. Jackson, Ian Johnston, Dominic Kwiatkowski, Cordelia Langford, John Sillitoe on behalf of the Wellcome Sanger Institute COVID-19 Surveillance Team |
| EPI_ISL_3283233 | Lighthouse Lab in Glasgow | Wellcome Sanger Institute for the COVID-19 Genomics UK (COG-UK) Consortium | Harper VanSteenhouse, Yumi Kasai, David Gray, Carol Clugston, Anna Dominiczak and Alex Alderton, Roberto Amato, Jeffrey Barrett, Sonia Goncalves, Ewan Harrison, David K. Jackson, Ian Johnston, Dominic Kwiatkowski, Cordelia Langford, John Sillitoe on behalf of the Wellcome Sanger Institute COVID-19 Surveillance Team |

| | | | |
|---|---|---|---|
| EPI_ISL_3283234, EPI_ISL_3283235 | Lighthouse Lab in Alderley Park | Wellcome Sanger Institute for the COVID-19 Genomics UK (COG-UK) Consortium | Jacquelyn Wynn, Mairead Hyland, The Lighthouse Lab in Alderley Park and Alex Alderton, Roberto Amato, Jeffrey Barrett, Sonia Goncalves, Ewan Harrison, David K. Jackson, Ian Johnston, Dominic Kwiatkowski, Cordelia Langford, John Sillitoe on behalf of the Wellcome Sanger Institute COVID-19 Surveillance Team |
| EPI_ISL_3283236, EPI_ISL_3283237 | Lighthouse Lab in Glasgow | Wellcome Sanger Institute for the COVID-19 Genomics UK (COG-UK) Consortium | Harper VanSteenhouse, Yumi Kasai, David Gray, Carol Clugston, Anna Dominiczak and Alex Alderton, Roberto Amato, Jeffrey Barrett, Sonia Goncalves, Ewan Harrison, David K. Jackson, Ian Johnston, Dominic Kwiatkowski, Cordelia Langford, John Sillitoe on behalf of the Wellcome Sanger Institute COVID-19 Surveillance Team |
| EPI_ISL_3283238 | Lighthouse Laboratory Plymouth | Wellcome Sanger Institute for the COVID-19 Genomics UK (COG-UK) Consortium | Lighthouse Laboratory Plymouth and Alex Alderton, Roberto Amato, Jeffrey Barrett, Sonia Goncalves, Ewan Harrison, David K. Jackson, Ian Johnston, Dominic Kwiatkowski, Cordelia Langford, John Sillitoe on behalf of the Wellcome Sanger Institute COVID-19 Surveillance Team |
| EPI_ISL_3283239, EPI_ISL_3283240, EPI_ISL_3283241 | Lighthouse Lab in Glasgow | Wellcome Sanger Institute for the COVID-19 Genomics UK (COG-UK) Consortium | Harper VanSteenhouse, Yumi Kasai, David Gray, Carol Clugston, Anna Dominiczak and Alex Alderton, Roberto Amato, Jeffrey Barrett, Sonia Goncalves, Ewan Harrison, David K. Jackson, Ian Johnston, Dominic Kwiatkowski, Cordelia Langford, John Sillitoe on behalf of the Wellcome Sanger Institute COVID-19 Surveillance Team |
| EPI_ISL_3283242 | Lighthouse Laboratory Plymouth | Wellcome Sanger Institute for the COVID-19 Genomics UK (COG-UK) Consortium | Lighthouse Laboratory Plymouth and Alex Alderton, Roberto Amato, Jeffrey Barrett, Sonia Goncalves, Ewan Harrison, David K. Jackson, Ian Johnston, Dominic Kwiatkowski, Cordelia Langford, John Sillitoe on behalf of the Wellcome Sanger Institute COVID-19 Surveillance Team |
| EPI_ISL_3283243 | Lighthouse Lab in Alderley Park | Wellcome Sanger Institute for the COVID-19 Genomics UK (COG-UK) Consortium | Jacquelyn Wynn, Mairead Hyland, The Lighthouse Lab in Alderley Park and Alex Alderton, Roberto Amato, Jeffrey Barrett, Sonia Goncalves, Ewan Harrison, David K. Jackson, Ian Johnston, Dominic Kwiatkowski, Cordelia Langford, John Sillitoe on behalf of the Wellcome Sanger Institute COVID-19 Surveillance Team |
| EPI_ISL_3283244, EPI_ISL_3283245 | Lighthouse Lab in Glasgow | Wellcome Sanger Institute for the COVID-19 Genomics UK (COG-UK) Consortium | Harper VanSteenhouse, Yumi Kasai, David Gray, Carol Clugston, Anna Dominiczak and Alex Alderton, Roberto Amato, Jeffrey Barrett, Sonia Goncalves, Ewan Harrison, David K. Jackson, Ian Johnston, Dominic Kwiatkowski, Cordelia Langford, John Sillitoe on behalf of the Wellcome Sanger Institute COVID-19 Surveillance Team |
| EPI_ISL_3283246 | Lighthouse Laboratory Plymouth | Wellcome Sanger Institute for the COVID-19 Genomics UK (COG-UK) Consortium | Lighthouse Laboratory Plymouth and Alex Alderton, Roberto Amato, Jeffrey Barrett, Sonia Goncalves, Ewan Harrison, David K. Jackson, Ian Johnston, Dominic Kwiatkowski, Cordelia Langford, John Sillitoe on behalf of the Wellcome Sanger Institute COVID-19 Surveillance Team |
| EPI_ISL_3283247 | Lighthouse Lab in Alderley Park | Wellcome Sanger Institute for the COVID-19 Genomics UK (COG-UK) Consortium | Jacquelyn Wynn, Mairead Hyland, The Lighthouse Lab in Alderley Park and Alex Alderton, Roberto Amato, Jeffrey Barrett, Sonia Goncalves, Ewan Harrison, David K. Jackson, Ian Johnston, Dominic Kwiatkowski, Cordelia Langford, John Sillitoe on behalf of the Wellcome Sanger Institute COVID-19 Surveillance Team |
| EPI_ISL_3283248 | Lighthouse Laboratory Plymouth | Wellcome Sanger Institute for the COVID-19 Genomics UK (COG-UK) Consortium | Lighthouse Laboratory Plymouth and Alex Alderton, Roberto Amato, Jeffrey Barrett, Sonia Goncalves, Ewan Harrison, David K. Jackson, Ian Johnston, Dominic Kwiatkowski, Cordelia Langford, John Sillitoe on behalf of the Wellcome Sanger Institute COVID-19 Surveillance Team |
| EPI_ISL_3283249 | Lighthouse Lab in Alderley Park | Wellcome Sanger Institute for the COVID-19 Genomics UK (COG-UK) Consortium | Jacquelyn Wynn, Mairead Hyland, The Lighthouse Lab in Alderley Park and Alex Alderton, Roberto Amato, Jeffrey Barrett, Sonia Goncalves, Ewan Harrison, David K. Jackson, Ian Johnston, Dominic Kwiatkowski, Cordelia Langford, John Sillitoe on behalf of the Wellcome Sanger Institute COVID-19 Surveillance Team |
| EPI_ISL_3283250 | Lighthouse Lab in Glasgow | Wellcome Sanger Institute for the COVID-19 Genomics UK (COG-UK) Consortium | Harper VanSteenhouse, Yumi Kasai, David Gray, Carol Clugston, Anna Dominiczak and Alex Alderton, Roberto Amato, Jeffrey Barrett, Sonia Goncalves, Ewan Harrison, David K. Jackson, Ian Johnston, Dominic Kwiatkowski, Cordelia Langford, John Sillitoe on behalf of the Wellcome Sanger Institute COVID-19 Surveillance Team |
| EPI_ISL_3283251 | Lighthouse Laboratory Plymouth | Wellcome Sanger Institute for the COVID-19 Genomics UK (COG-UK) Consortium | Lighthouse Laboratory Plymouth and Alex Alderton, Roberto Amato, Jeffrey Barrett, Sonia Goncalves, Ewan Harrison, David K. Jackson, Ian Johnston, Dominic Kwiatkowski, Cordelia Langford, John Sillitoe on behalf of the Wellcome Sanger Institute COVID-19 Surveillance Team |
| EPI_ISL_3283252 | Lighthouse Lab in Alderley Park | Wellcome Sanger Institute for the COVID-19 Genomics UK (COG-UK) Consortium | Jacquelyn Wynn, Mairead Hyland, The Lighthouse Lab in Alderley Park and Alex Alderton, Roberto Amato, Jeffrey Barrett, Sonia Goncalves, Ewan Harrison, David K. Jackson, Ian Johnston, Dominic Kwiatkowski, Cordelia Langford, John Sillitoe on behalf of the Wellcome Sanger Institute COVID-19 Surveillance Team |
| EPI_ISL_3283253 | Lighthouse Lab in Glasgow | Wellcome Sanger Institute for the COVID-19 Genomics UK (COG-UK) Consortium | Harper VanSteenhouse, Yumi Kasai, David Gray, Carol Clugston, Anna Dominiczak and Alex Alderton, Roberto Amato, Jeffrey Barrett, Sonia Goncalves, Ewan Harrison, David K. Jackson, Ian Johnston, Dominic Kwiatkowski, Cordelia Langford, John Sillitoe on behalf of the Wellcome Sanger Institute COVID-19 Surveillance Team |
| EPI_ISL_3283254, EPI_ISL_3283255 | Lighthouse Laboratory Plymouth | Wellcome Sanger Institute for the COVID-19 Genomics UK (COG-UK) Consortium | Lighthouse Laboratory Plymouth and Alex Alderton, Roberto Amato, Jeffrey Barrett, Sonia Goncalves, Ewan Harrison, David K. Jackson, Ian Johnston, Dominic Kwiatkowski, Cordelia Langford, John Sillitoe on behalf of the Wellcome Sanger Institute COVID-19 Surveillance Team |
| EPI_ISL_3283256 | Lighthouse Lab in Alderley Park | Wellcome Sanger Institute for the COVID-19 Genomics UK (COG-UK) Consortium | Jacquelyn Wynn, Mairead Hyland, The Lighthouse Lab in Alderley Park and Alex Alderton, Roberto Amato, Jeffrey Barrett, Sonia Goncalves, Ewan Harrison, David K. Jackson, Ian Johnston, Dominic Kwiatkowski, Cordelia Langford, John Sillitoe on behalf of the Wellcome Sanger Institute COVID-19 Surveillance Team |
| EPI_ISL_3283257 | Lighthouse Laboratory Plymouth | Wellcome Sanger Institute for the COVID-19 Genomics UK (COG-UK) Consortium | Lighthouse Laboratory Plymouth and Alex Alderton, Roberto Amato, Jeffrey Barrett, Sonia Goncalves, Ewan Harrison, David K. Jackson, Ian Johnston, Dominic Kwiatkowski, Cordelia Langford, John Sillitoe on behalf of the Wellcome Sanger Institute COVID-19 Surveillance Team |
| EPI_ISL_3283258 | Lighthouse Lab in Glasgow | Wellcome Sanger Institute for the COVID-19 Genomics UK (COG-UK) Consortium | Harper VanSteenhouse, Yumi Kasai, David Gray, Carol Clugston, Anna Dominiczak and Alex Alderton, Roberto Amato, Jeffrey Barrett, Sonia Goncalves, Ewan Harrison, David K. Jackson, Ian Johnston, Dominic Kwiatkowski, Cordelia Langford, John Sillitoe on behalf of the Wellcome Sanger Institute COVID-19 Surveillance Team |
| EPI_ISL_3283259 | Lighthouse Laboratory Plymouth | Wellcome Sanger Institute for the COVID-19 Genomics UK (COG-UK) Consortium | Lighthouse Laboratory Plymouth and Alex Alderton, Roberto Amato, Jeffrey Barrett, Sonia Goncalves, Ewan Harrison, David K. Jackson, Ian Johnston, Dominic Kwiatkowski, Cordelia Langford, John Sillitoe on behalf of the Wellcome Sanger Institute COVID-19 Surveillance Team |
| EPI_ISL_3283260 | Lighthouse Lab in Glasgow | Wellcome Sanger Institute for the COVID-19 Genomics UK (COG-UK) Consortium | Harper VanSteenhouse, Yumi Kasai, David Gray, Carol Clugston, Anna Dominiczak and Alex Alderton, Roberto Amato, Jeffrey Barrett, Sonia Goncalves, Ewan Harrison, David K. Jackson, Ian Johnston, Dominic Kwiatkowski, Cordelia Langford, John Sillitoe on behalf of the Wellcome Sanger Institute COVID-19 Surveillance Team |
| EPI_ISL_3283261 | Lighthouse Lab in Alderley Park | Wellcome Sanger Institute for the COVID-19 Genomics UK (COG-UK) Consortium | Jacquelyn Wynn, Mairead Hyland, The Lighthouse Lab in Alderley Park and Alex Alderton, Roberto Amato, Jeffrey Barrett, Sonia Goncalves, Ewan Harrison, David K. Jackson, Ian Johnston, Dominic Kwiatkowski, Cordelia Langford, John Sillitoe on behalf of the Wellcome Sanger Institute COVID-19 Surveillance Team |
| EPI_ISL_3283262 | Lighthouse Laboratory Plymouth | Wellcome Sanger Institute for the COVID-19 Genomics UK (COG-UK) Consortium | Lighthouse Laboratory Plymouth and Alex Alderton, Roberto Amato, Jeffrey Barrett, Sonia Goncalves, Ewan Harrison, David K. Jackson, Ian Johnston, Dominic Kwiatkowski, Cordelia Langford, John Sillitoe on behalf of the Wellcome Sanger Institute COVID-19 Surveillance Team |
| EPI_ISL_3283263 | Lighthouse Lab in Glasgow | Wellcome Sanger Institute for the COVID-19 Genomics UK (COG-UK) Consortium | Harper VanSteenhouse, Yumi Kasai, David Gray, Carol Clugston, Anna Dominiczak and Alex Alderton, Roberto Amato, Jeffrey Barrett, Sonia Goncalves, Ewan Harrison, David K. Jackson, Ian Johnston, Dominic Kwiatkowski, Cordelia Langford, John Sillitoe on behalf of the Wellcome Sanger Institute COVID-19 Surveillance Team |
| EPI_ISL_3283264 | Lighthouse Laboratory Plymouth | Wellcome Sanger Institute for the COVID-19 Genomics UK (COG-UK) Consortium | Lighthouse Laboratory Plymouth and Alex Alderton, Roberto Amato, Jeffrey Barrett, Sonia Goncalves, Ewan Harrison, David K. Jackson, Ian Johnston, Dominic Kwiatkowski, Cordelia Langford, John Sillitoe on behalf of the Wellcome Sanger Institute COVID-19 Surveillance Team |
| EPI_ISL_3283265, EPI_ISL_3283266, EPI_ISL_3283267, EPI_ISL_3283268 | Lighthouse Lab in Glasgow | Wellcome Sanger Institute for the COVID-19 Genomics UK (COG-UK) Consortium | Harper VanSteenhouse, Yumi Kasai, David Gray, Carol Clugston, Anna Dominiczak and Alex Alderton, Roberto Amato, Jeffrey Barrett, Sonia Goncalves, Ewan Harrison, David K. Jackson, Ian Johnston, Dominic Kwiatkowski, Cordelia Langford, John Sillitoe on behalf of the Wellcome Sanger Institute COVID-19 Surveillance Team |
| EPI_ISL_3283269 | Lighthouse Laboratory Plymouth | Wellcome Sanger Institute for the COVID-19 Genomics UK (COG-UK) Consortium | Lighthouse Laboratory Plymouth and Alex Alderton, Roberto Amato, Jeffrey Barrett, Sonia Goncalves, Ewan Harrison, David K. Jackson, Ian Johnston, Dominic Kwiatkowski, Cordelia Langford, John Sillitoe on behalf of the Wellcome Sanger Institute COVID-19 Surveillance Team |
| EPI_ISL_3283270 | Lighthouse Lab in Glasgow | Wellcome Sanger Institute for the COVID-19 Genomics UK (COG-UK) Consortium | Harper VanSteenhouse, Yumi Kasai, David Gray, Carol Clugston, Anna Dominiczak and Alex Alderton, Roberto Amato, Jeffrey Barrett, Sonia Goncalves, Ewan Harrison, David K. Jackson, Ian Johnston, Dominic Kwiatkowski, Cordelia Langford, John Sillitoe on behalf of the Wellcome Sanger Institute COVID-19 Surveillance Team |
| EPI_ISL_3283271, EPI_ISL_3283272 | Lighthouse Laboratory Plymouth | Wellcome Sanger Institute for the COVID-19 Genomics UK (COG-UK) Consortium | Lighthouse Laboratory Plymouth and Alex Alderton, Roberto Amato, Jeffrey Barrett, Sonia Goncalves, Ewan Harrison, David K. Jackson, Ian Johnston, Dominic Kwiatkowski, Cordelia Langford, John Sillitoe on behalf of the Wellcome Sanger Institute COVID-19 Surveillance Team |
| EPI_ISL_3283273 | Lighthouse Lab in Glasgow | Wellcome Sanger Institute for the COVID-19 Genomics UK (COG-UK) Consortium | Harper VanSteenhouse, Yumi Kasai, David Gray, Carol Clugston, Anna Dominiczak and Alex Alderton, Roberto Amato, Jeffrey Barrett, Sonia Goncalves, Ewan Harrison, David K. Jackson, Ian Johnston, Dominic Kwiatkowski, Cordelia Langford, John Sillitoe on behalf of the Wellcome Sanger Institute COVID-19 Surveillance Team |
| EPI_ISL_3283274 | Lighthouse Laboratory Plymouth | Wellcome Sanger Institute for the COVID-19 Genomics UK (COG-UK) Consortium | Lighthouse Laboratory Plymouth and Alex Alderton, Roberto Amato, Jeffrey Barrett, Sonia Goncalves, Ewan Harrison, David K. Jackson, Ian Johnston, Dominic Kwiatkowski, Cordelia Langford, John Sillitoe on behalf of the Wellcome Sanger Institute COVID-19 Surveillance Team |
| EPI_ISL_3283275, EPI_ISL_3283276 | Lighthouse Lab in Alderley Park | Wellcome Sanger Institute for the COVID-19 Genomics UK (COG-UK) Consortium | Jacquelyn Wynn, Mairead Hyland, The Lighthouse Lab in Alderley Park and Alex Alderton, Roberto Amato, Jeffrey Barrett, Sonia Goncalves, Ewan Harrison, David K. Jackson, Ian Johnston, Dominic Kwiatkowski, Cordelia Langford, John Sillitoe on behalf of the Wellcome Sanger Institute COVID-19 Surveillance Team |
| EPI_ISL_3283277 | Lighthouse Lab in Glasgow | Wellcome Sanger Institute for the COVID-19 Genomics UK (COG-UK) Consortium | Harper VanSteenhouse, Yumi Kasai, David Gray, Carol Clugston, Anna Dominiczak and Alex Alderton, Roberto Amato, Jeffrey Barrett, Sonia Goncalves, Ewan Harrison, David K. Jackson, Ian Johnston, Dominic Kwiatkowski, Cordelia Langford, John Sillitoe on behalf of the Wellcome Sanger Institute COVID-19 Surveillance Team |
| EPI_ISL_3283278, EPI_ISL_3283279, EPI_ISL_3283280 | Lighthouse Laboratory Plymouth | Wellcome Sanger Institute for the COVID-19 Genomics UK (COG-UK) Consortium | Lighthouse Laboratory Plymouth and Alex Alderton, Roberto Amato, Jeffrey Barrett, Sonia Goncalves, Ewan Harrison, David K. Jackson, Ian Johnston, Dominic Kwiatkowski, Cordelia Langford, John Sillitoe on behalf of the Wellcome Sanger Institute COVID-19 Surveillance Team |
| EPI_ISL_3283281, EPI_ISL_3283282 | Lighthouse Lab in Alderley Park | Wellcome Sanger Institute for the COVID-19 Genomics UK (COG-UK) Consortium | Jacquelyn Wynn, Mairead Hyland, The Lighthouse Lab in Alderley Park and Alex Alderton, Roberto Amato, Jeffrey Barrett, Sonia Goncalves, Ewan Harrison, David K. Jackson, Ian Johnston, Dominic Kwiatkowski, Cordelia Langford, John Sillitoe on behalf of the Wellcome Sanger Institute COVID-19 Surveillance Team |
| EPI_ISL_3283283 | Lighthouse Lab in Glasgow | Wellcome Sanger Institute for the COVID-19 Genomics UK (COG-UK) Consortium | Harper VanSteenhouse, Yumi Kasai, David Gray, Carol Clugston, Anna Dominiczak and Alex Alderton, Roberto Amato, Jeffrey Barrett, Sonia Goncalves, Ewan Harrison, David K. Jackson, Ian Johnston, Dominic Kwiatkowski, Cordelia Langford, John Sillitoe on behalf of the Wellcome Sanger Institute COVID-19 Surveillance Team |
| EPI_ISL_3283284 | Lighthouse Laboratory Plymouth | Wellcome Sanger Institute for the COVID-19 Genomics UK (COG-UK) Consortium | Lighthouse Laboratory Plymouth and Alex Alderton, Roberto Amato, Jeffrey Barrett, Sonia Goncalves, Ewan Harrison, David K. Jackson, Ian Johnston, Dominic Kwiatkowski, Cordelia Langford, John Sillitoe on behalf of the Wellcome Sanger Institute COVID-19 Surveillance Team |
| EPI_ISL_3283285 | Lighthouse Lab in Alderley Park | Wellcome Sanger Institute for the COVID-19 Genomics UK (COG-UK) Consortium | Jacquelyn Wynn, Mairead Hyland, The Lighthouse Lab in Alderley Park and Alex Alderton, Roberto Amato, Jeffrey Barrett, Sonia Goncalves, Ewan Harrison, David K. Jackson, Ian Johnston, Dominic Kwiatkowski, Cordelia Langford, John Sillitoe on behalf of the Wellcome Sanger Institute COVID-19 Surveillance Team |
| EPI_ISL_3283286, EPI_ISL_3283287 | Lighthouse Laboratory Plymouth | Wellcome Sanger Institute for the COVID-19 Genomics UK (COG-UK) Consortium | Lighthouse Laboratory Plymouth and Alex Alderton, Roberto Amato, Jeffrey Barrett, Sonia Goncalves, Ewan Harrison, David K. Jackson, Ian Johnston, Dominic Kwiatkowski, Cordelia Langford, John Sillitoe on behalf of the Wellcome Sanger Institute COVID-19 Surveillance Team |
| EPI_ISL_3283288, EPI_ISL_3283289 | Lighthouse Lab in Alderley Park | Wellcome Sanger Institute for the COVID-19 Genomics UK (COG-UK) Consortium | Jacquelyn Wynn, Mairead Hyland, The Lighthouse Lab in Alderley Park and Alex Alderton, Roberto Amato, Jeffrey Barrett, Sonia Goncalves, Ewan Harrison, David K. Jackson, Ian Johnston, Dominic Kwiatkowski, Cordelia Langford, John Sillitoe on behalf of the Wellcome Sanger Institute COVID-19 Surveillance Team |
| EPI_ISL_3283290, EPI_ISL_3283291, EPI_ISL_3283292, EPI_ISL_3283293 | Lighthouse Lab in Glasgow | Wellcome Sanger Institute for the COVID-19 Genomics UK (COG-UK) Consortium | Harper VanSteenhouse, Yumi Kasai, David Gray, Carol Clugston, Anna Dominiczak and Alex Alderton, Roberto Amato, Jeffrey Barrett, Sonia Goncalves, Ewan Harrison, David K. Jackson, Ian Johnston, Dominic Kwiatkowski, Cordelia Langford, John Sillitoe on behalf of the Wellcome Sanger Institute COVID-19 Surveillance Team |
| EPI_ISL_3283294 | Lighthouse Laboratory Plymouth | Wellcome Sanger Institute for the COVID-19 Genomics UK (COG-UK) Consortium | Lighthouse Laboratory Plymouth and Alex Alderton, Roberto Amato, Jeffrey Barrett, Sonia Goncalves, Ewan Harrison, David K. Jackson, Ian Johnston, Dominic Kwiatkowski, Cordelia Langford, John Sillitoe on behalf of the Wellcome Sanger Institute COVID-19 Surveillance Team |
| EPI_ISL_3283295 | Lighthouse Lab in Alderley Park | Wellcome Sanger Institute for the COVID-19 Genomics UK (COG-UK) | Jacquelyn Wynn, Mairead Hyland, The Lighthouse Lab in Alderley Park and Alex Alderton, Roberto Amato, Jeffrey Barrett, Sonia Goncalves, Ewan Harrison, David K. Jackson, Ian Johnston, Dominic Kwiatkowski, Cordelia Langford, John Sillitoe on |

| | | | |
|---|---|---|---|
| | | Consortium | behalf of the Wellcome Sanger Institute COVID-19 Surveillance Team |
| EPI_ISL_3283296 | Lighthouse Laboratory Plymouth | Wellcome Sanger Institute for the COVID-19 Genomics UK (COG-UK) Consortium | Lighthouse Laboratory Plymouth and Alex Alderton, Roberto Amato, Jeffrey Barrett, Sonia Goncalves, Ewan Harrison, David K. Jackson, Ian Johnston, Dominic Kwiatkowski, Cordelia Langford, John Sillitoe on behalf of the Wellcome Sanger Institute COVID-19 Surveillance Team |
| EPI_ISL_3283297 | Lighthouse Lab in Alderley Park | Wellcome Sanger Institute for the COVID-19 Genomics UK (COG-UK) Consortium | Jacquelyn Wynn, Mairead Hyland, The Lighthouse Lab in Alderley Park and Alex Alderton, Roberto Amato, Jeffrey Barrett, Sonia Goncalves, Ewan Harrison, David K. Jackson, Ian Johnston, Dominic Kwiatkowski, Cordelia Langford, John Sillitoe on behalf of the Wellcome Sanger Institute COVID-19 Surveillance Team |
| EPI_ISL_3283298 | Lighthouse Lab in Glasgow | Wellcome Sanger Institute for the COVID-19 Genomics UK (COG-UK) Consortium | Harper VanSteenhouse, Yumi Kasai, David Gray, Carol Clugston, Anna Dominiczak and Alex Alderton, Roberto Amato, Jeffrey Barrett, Sonia Goncalves, Ewan Harrison, David K. Jackson, Ian Johnston, Dominic Kwiatkowski, Cordelia Langford, John Sillitoe on behalf of the Wellcome Sanger Institute COVID-19 Surveillance Team |
| EPI_ISL_3283299 | Lighthouse Laboratory Plymouth | Wellcome Sanger Institute for the COVID-19 Genomics UK (COG-UK) Consortium | Lighthouse Laboratory Plymouth and Alex Alderton, Roberto Amato, Jeffrey Barrett, Sonia Goncalves, Ewan Harrison, David K. Jackson, Ian Johnston, Dominic Kwiatkowski, Cordelia Langford, John Sillitoe on behalf of the Wellcome Sanger Institute COVID-19 Surveillance Team |
| EPI_ISL_3283300, EPI_ISL_3283301, EPI_ISL_3283302, EPI_ISL_3283303, EPI_ISL_3283304 | Lighthouse Lab in Glasgow | Wellcome Sanger Institute for the COVID-19 Genomics UK (COG-UK) Consortium | Harper VanSteenhouse, Yumi Kasai, David Gray, Carol Clugston, Anna Dominiczak and Alex Alderton, Roberto Amato, Jeffrey Barrett, Sonia Goncalves, Ewan Harrison, David K. Jackson, Ian Johnston, Dominic Kwiatkowski, Cordelia Langford, John Sillitoe on behalf of the Wellcome Sanger Institute COVID-19 Surveillance Team |
| EPI_ISL_3283305, EPI_ISL_3283306 | Lighthouse Lab in Alderley Park | Wellcome Sanger Institute for the COVID-19 Genomics UK (COG-UK) Consortium | Jacquelyn Wynn, Mairead Hyland, The Lighthouse Lab in Alderley Park and Alex Alderton, Roberto Amato, Jeffrey Barrett, Sonia Goncalves, Ewan Harrison, David K. Jackson, Ian Johnston, Dominic Kwiatkowski, Cordelia Langford, John Sillitoe on behalf of the Wellcome Sanger Institute COVID-19 Surveillance Team |
| EPI_ISL_3283307 | Lighthouse Lab in Glasgow | Wellcome Sanger Institute for the COVID-19 Genomics UK (COG-UK) Consortium | Harper VanSteenhouse, Yumi Kasai, David Gray, Carol Clugston, Anna Dominiczak and Alex Alderton, Roberto Amato, Jeffrey Barrett, Sonia Goncalves, Ewan Harrison, David K. Jackson, Ian Johnston, Dominic Kwiatkowski, Cordelia Langford, John Sillitoe on behalf of the Wellcome Sanger Institute COVID-19 Surveillance Team |
| EPI_ISL_3283308 | Lighthouse Lab in Alderley Park | Wellcome Sanger Institute for the COVID-19 Genomics UK (COG-UK) Consortium | Jacquelyn Wynn, Mairead Hyland, The Lighthouse Lab in Alderley Park and Alex Alderton, Roberto Amato, Jeffrey Barrett, Sonia Goncalves, Ewan Harrison, David K. Jackson, Ian Johnston, Dominic Kwiatkowski, Cordelia Langford, John Sillitoe on behalf of the Wellcome Sanger Institute COVID-19 Surveillance Team |
| EPI_ISL_3283309 | Lighthouse Lab in Glasgow | Wellcome Sanger Institute for the COVID-19 Genomics UK (COG-UK) Consortium | Harper VanSteenhouse, Yumi Kasai, David Gray, Carol Clugston, Anna Dominiczak and Alex Alderton, Roberto Amato, Jeffrey Barrett, Sonia Goncalves, Ewan Harrison, David K. Jackson, Ian Johnston, Dominic Kwiatkowski, Cordelia Langford, John Sillitoe on behalf of the Wellcome Sanger Institute COVID-19 Surveillance Team |
| EPI_ISL_3283310 | Lighthouse Laboratory Plymouth | Wellcome Sanger Institute for the COVID-19 Genomics UK (COG-UK) Consortium | Lighthouse Laboratory Plymouth and Alex Alderton, Roberto Amato, Jeffrey Barrett, Sonia Goncalves, Ewan Harrison, David K. Jackson, Ian Johnston, Dominic Kwiatkowski, Cordelia Langford, John Sillitoe on behalf of the Wellcome Sanger Institute COVID-19 Surveillance Team |
| EPI_ISL_3283311 | Lighthouse Lab in Glasgow | Wellcome Sanger Institute for the COVID-19 Genomics UK (COG-UK) Consortium | Harper VanSteenhouse, Yumi Kasai, David Gray, Carol Clugston, Anna Dominiczak and Alex Alderton, Roberto Amato, Jeffrey Barrett, Sonia Goncalves, Ewan Harrison, David K. Jackson, Ian Johnston, Dominic Kwiatkowski, Cordelia Langford, John Sillitoe on behalf of the Wellcome Sanger Institute COVID-19 Surveillance Team |
| EPI_ISL_3283312 | Lighthouse Laboratory Plymouth | Wellcome Sanger Institute for the COVID-19 Genomics UK (COG-UK) Consortium | Lighthouse Laboratory Plymouth and Alex Alderton, Roberto Amato, Jeffrey Barrett, Sonia Goncalves, Ewan Harrison, David K. Jackson, Ian Johnston, Dominic Kwiatkowski, Cordelia Langford, John Sillitoe on behalf of the Wellcome Sanger Institute COVID-19 Surveillance Team |
| EPI_ISL_3283313 | Lighthouse Lab in Glasgow | Wellcome Sanger Institute for the COVID-19 Genomics UK (COG-UK) Consortium | Harper VanSteenhouse, Yumi Kasai, David Gray, Carol Clugston, Anna Dominiczak and Alex Alderton, Roberto Amato, Jeffrey Barrett, Sonia Goncalves, Ewan Harrison, David K. Jackson, Ian Johnston, Dominic Kwiatkowski, Cordelia Langford, John Sillitoe on behalf of the Wellcome Sanger Institute COVID-19 Surveillance Team |
| EPI_ISL_3283314, EPI_ISL_3283315 | Lighthouse Lab in Alderley Park | Wellcome Sanger Institute for the COVID-19 Genomics UK (COG-UK) Consortium | Jacquelyn Wynn, Mairead Hyland, The Lighthouse Lab in Alderley Park and Alex Alderton, Roberto Amato, Jeffrey Barrett, Sonia Goncalves, Ewan Harrison, David K. Jackson, Ian Johnston, Dominic Kwiatkowski, Cordelia Langford, John Sillitoe on behalf of the Wellcome Sanger Institute COVID-19 Surveillance Team |
| EPI_ISL_3283316 | Lighthouse Lab in Milton Keynes | Wellcome Sanger Institute for the COVID-19 Genomics UK (COG-UK) Consortium | The Lighthouse Lab in Milton Keynes and Alex Alderton, Roberto Amato, Jeffrey Barrett, Sonia Goncalves, Ewan Harrison, David K. Jackson, Ian Johnston, Dominic Kwiatkowski, Cordelia Langford, John Sillitoe on behalf of the Wellcome Sanger Institute COVID-19 Surveillance Team |
| EPI_ISL_3283317 | Lighthouse Lab in Alderley Park | Wellcome Sanger Institute for the COVID-19 Genomics UK (COG-UK) Consortium | Jacquelyn Wynn, Mairead Hyland, The Lighthouse Lab in Alderley Park and Alex Alderton, Roberto Amato, Jeffrey Barrett, Sonia Goncalves, Ewan Harrison, David K. Jackson, Ian Johnston, Dominic Kwiatkowski, Cordelia Langford, John Sillitoe on behalf of the Wellcome Sanger Institute COVID-19 Surveillance Team |
| EPI_ISL_3283318, EPI_ISL_3283319, EPI_ISL_3283320, EPI_ISL_3283321 | Lighthouse Lab in Milton Keynes | Wellcome Sanger Institute for the COVID-19 Genomics UK (COG-UK) Consortium | The Lighthouse Lab in Milton Keynes and Alex Alderton, Roberto Amato, Jeffrey Barrett, Sonia Goncalves, Ewan Harrison, David K. Jackson, Ian Johnston, Dominic Kwiatkowski, Cordelia Langford, John Sillitoe on behalf of the Wellcome Sanger Institute COVID-19 Surveillance Team |
| EPI_ISL_3283322, EPI_ISL_3283323 | Lighthouse Lab in Alderley Park | Wellcome Sanger Institute for the COVID-19 Genomics UK (COG-UK) Consortium | Jacquelyn Wynn, Mairead Hyland, The Lighthouse Lab in Alderley Park and Alex Alderton, Roberto Amato, Jeffrey Barrett, Sonia Goncalves, Ewan Harrison, David K. Jackson, Ian Johnston, Dominic Kwiatkowski, Cordelia Langford, John Sillitoe on behalf of the Wellcome Sanger Institute COVID-19 Surveillance Team |
| EPI_ISL_3283324, EPI_ISL_3283325 | Lighthouse Lab in Milton Keynes | Wellcome Sanger Institute for the COVID-19 Genomics UK (COG-UK) Consortium | The Lighthouse Lab in Milton Keynes and Alex Alderton, Roberto Amato, Jeffrey Barrett, Sonia Goncalves, Ewan Harrison, David K. Jackson, Ian Johnston, Dominic Kwiatkowski, Cordelia Langford, John Sillitoe on behalf of the Wellcome Sanger Institute COVID-19 Surveillance Team |
| EPI_ISL_3283326, EPI_ISL_3283327, EPI_ISL_3283328 | Lighthouse Lab in Alderley Park | Wellcome Sanger Institute for the COVID-19 Genomics UK (COG-UK) Consortium | Jacquelyn Wynn, Mairead Hyland, The Lighthouse Lab in Alderley Park and Alex Alderton, Roberto Amato, Jeffrey Barrett, Sonia Goncalves, Ewan Harrison, David K. Jackson, Ian Johnston, Dominic Kwiatkowski, Cordelia Langford, John Sillitoe on behalf of the Wellcome Sanger Institute COVID-19 Surveillance Team |
| EPI_ISL_3283329, EPI_ISL_3283330 | Lighthouse Lab in Milton Keynes | Wellcome Sanger Institute for the COVID-19 Genomics UK (COG-UK) Consortium | The Lighthouse Lab in Milton Keynes and Alex Alderton, Roberto Amato, Jeffrey Barrett, Sonia Goncalves, Ewan Harrison, David K. Jackson, Ian Johnston, Dominic Kwiatkowski, Cordelia Langford, John Sillitoe on behalf of the Wellcome Sanger Institute COVID-19 Surveillance Team |
| EPI_ISL_3283331 | Lighthouse Lab in Alderley Park | Wellcome Sanger Institute for the COVID-19 Genomics UK (COG-UK) Consortium | Jacquelyn Wynn, Mairead Hyland, The Lighthouse Lab in Alderley Park and Alex Alderton, Roberto Amato, Jeffrey Barrett, Sonia Goncalves, Ewan Harrison, David K. Jackson, Ian Johnston, Dominic Kwiatkowski, Cordelia Langford, John Sillitoe on behalf of the Wellcome Sanger Institute COVID-19 Surveillance Team |
| EPI_ISL_3283332, EPI_ISL_3283333, EPI_ISL_3283334, EPI_ISL_3283335, EPI_ISL_3283336, EPI_ISL_3283337, EPI_ISL_3283338 | Lighthouse Lab in Milton Keynes | Wellcome Sanger Institute for the COVID-19 Genomics UK (COG-UK) Consortium | The Lighthouse Lab in Milton Keynes and Alex Alderton, Roberto Amato, Jeffrey Barrett, Sonia Goncalves, Ewan Harrison, David K. Jackson, Ian Johnston, Dominic Kwiatkowski, Cordelia Langford, John Sillitoe on behalf of the Wellcome Sanger Institute COVID-19 Surveillance Team |
| EPI_ISL_3283339, EPI_ISL_3283340 | Lighthouse Lab in Alderley Park | Wellcome Sanger Institute for the COVID-19 Genomics UK (COG-UK) Consortium | Jacquelyn Wynn, Mairead Hyland, The Lighthouse Lab in Alderley Park and Alex Alderton, Roberto Amato, Jeffrey Barrett, Sonia Goncalves, Ewan Harrison, David K. Jackson, Ian Johnston, Dominic Kwiatkowski, Cordelia Langford, John Sillitoe on behalf of the Wellcome Sanger Institute COVID-19 Surveillance Team |
| EPI_ISL_3283341 | Lighthouse Lab in Milton Keynes | Wellcome Sanger Institute for the COVID-19 Genomics UK (COG-UK) Consortium | The Lighthouse Lab in Milton Keynes and Alex Alderton, Roberto Amato, Jeffrey Barrett, Sonia Goncalves, Ewan Harrison, David K. Jackson, Ian Johnston, Dominic Kwiatkowski, Cordelia Langford, John Sillitoe on behalf of the Wellcome Sanger Institute COVID-19 Surveillance Team |
| EPI_ISL_3283342 | Lighthouse Lab in Alderley Park | Wellcome Sanger Institute for the COVID-19 Genomics UK (COG-UK) Consortium | Jacquelyn Wynn, Mairead Hyland, The Lighthouse Lab in Alderley Park and Alex Alderton, Roberto Amato, Jeffrey Barrett, Sonia Goncalves, Ewan Harrison, David K. Jackson, Ian Johnston, Dominic Kwiatkowski, Cordelia Langford, John Sillitoe on behalf of the Wellcome Sanger Institute COVID-19 Surveillance Team |
| EPI_ISL_3283343 | Lighthouse Lab in Milton Keynes | Wellcome Sanger Institute for the COVID-19 Genomics UK (COG-UK) Consortium | The Lighthouse Lab in Milton Keynes and Alex Alderton, Roberto Amato, Jeffrey Barrett, Sonia Goncalves, Ewan Harrison, David K. Jackson, Ian Johnston, Dominic Kwiatkowski, Cordelia Langford, John Sillitoe on behalf of the Wellcome Sanger Institute COVID-19 Surveillance Team |
| EPI_ISL_3283344, EPI_ISL_3283345, EPI_ISL_3283346 | Lighthouse Lab in Alderley Park | Wellcome Sanger Institute for the COVID-19 Genomics UK (COG-UK) Consortium | Jacquelyn Wynn, Mairead Hyland, The Lighthouse Lab in Alderley Park and Alex Alderton, Roberto Amato, Jeffrey Barrett, Sonia Goncalves, Ewan Harrison, David K. Jackson, Ian Johnston, Dominic Kwiatkowski, Cordelia Langford, John Sillitoe on behalf of the Wellcome Sanger Institute COVID-19 Surveillance Team |
| EPI_ISL_3283347, EPI_ISL_3283348 | Lighthouse Lab in Milton Keynes | Wellcome Sanger Institute for the COVID-19 Genomics UK (COG-UK) Consortium | The Lighthouse Lab in Milton Keynes and Alex Alderton, Roberto Amato, Jeffrey Barrett, Sonia Goncalves, Ewan Harrison, David K. Jackson, Ian Johnston, Dominic Kwiatkowski, Cordelia Langford, John Sillitoe on behalf of the Wellcome Sanger Institute COVID-19 Surveillance Team |
| EPI_ISL_3283349 | Lighthouse Lab in Alderley Park | Wellcome Sanger Institute for the COVID-19 Genomics UK (COG-UK) Consortium | Jacquelyn Wynn, Mairead Hyland, The Lighthouse Lab in Alderley Park and Alex Alderton, Roberto Amato, Jeffrey Barrett, Sonia Goncalves, Ewan Harrison, David K. Jackson, Ian Johnston, Dominic Kwiatkowski, Cordelia Langford, John Sillitoe on behalf of the Wellcome Sanger Institute COVID-19 Surveillance Team |
| EPI_ISL_3283350, EPI_ISL_3283351, EPI_ISL_3283352 | Lighthouse Lab in Milton Keynes | Wellcome Sanger Institute for the COVID-19 Genomics UK (COG-UK) Consortium | The Lighthouse Lab in Milton Keynes and Alex Alderton, Roberto Amato, Jeffrey Barrett, Sonia Goncalves, Ewan Harrison, David K. Jackson, Ian Johnston, Dominic Kwiatkowski, Cordelia Langford, John Sillitoe on behalf of the Wellcome Sanger Institute COVID-19 Surveillance Team |
| EPI_ISL_3283353 | Lighthouse Lab in Alderley Park | Wellcome Sanger Institute for the COVID-19 Genomics UK (COG-UK) Consortium | Jacquelyn Wynn, Mairead Hyland, The Lighthouse Lab in Alderley Park and Alex Alderton, Roberto Amato, Jeffrey Barrett, Sonia Goncalves, Ewan Harrison, David K. Jackson, Ian Johnston, Dominic Kwiatkowski, Cordelia Langford, John Sillitoe on behalf of the Wellcome Sanger Institute COVID-19 Surveillance Team |
| EPI_ISL_3283354, EPI_ISL_3283355, EPI_ISL_3283356, EPI_ISL_3283357 | Lighthouse Lab in Milton Keynes | Wellcome Sanger Institute for the COVID-19 Genomics UK (COG-UK) Consortium | The Lighthouse Lab in Milton Keynes and Alex Alderton, Roberto Amato, Jeffrey Barrett, Sonia Goncalves, Ewan Harrison, David K. Jackson, Ian Johnston, Dominic Kwiatkowski, Cordelia Langford, John Sillitoe on behalf of the Wellcome Sanger Institute COVID-19 Surveillance Team |
| EPI_ISL_3283358 | Lighthouse Lab in Alderley Park | Wellcome Sanger Institute for the COVID-19 Genomics UK (COG-UK) Consortium | Jacquelyn Wynn, Mairead Hyland, The Lighthouse Lab in Alderley Park and Alex Alderton, Roberto Amato, Jeffrey Barrett, Sonia Goncalves, Ewan Harrison, David K. Jackson, Ian Johnston, Dominic Kwiatkowski, Cordelia Langford, John Sillitoe on behalf of the Wellcome Sanger Institute COVID-19 Surveillance Team |
| EPI_ISL_3283359 | Lighthouse Lab in Milton Keynes | Wellcome Sanger Institute for the COVID-19 Genomics UK (COG-UK) Consortium | The Lighthouse Lab in Milton Keynes and Alex Alderton, Roberto Amato, Jeffrey Barrett, Sonia Goncalves, Ewan Harrison, David K. Jackson, Ian Johnston, Dominic Kwiatkowski, Cordelia Langford, John Sillitoe on behalf of the Wellcome Sanger Institute COVID-19 Surveillance Team |
| EPI_ISL_3283360, EPI_ISL_3283361, EPI_ISL_3283362, EPI_ISL_3283363, EPI_ISL_3283364 | Lighthouse Lab in Alderley Park | Wellcome Sanger Institute for the COVID-19 Genomics UK (COG-UK) Consortium | Jacquelyn Wynn, Mairead Hyland, The Lighthouse Lab in Alderley Park and Alex Alderton, Roberto Amato, Jeffrey Barrett, Sonia Goncalves, Ewan Harrison, David K. Jackson, Ian Johnston, Dominic Kwiatkowski, Cordelia Langford, John Sillitoe on behalf of the Wellcome Sanger Institute COVID-19 Surveillance Team |
| EPI_ISL_3283365, EPI_ISL_3283366, EPI_ISL_3283367 | Lighthouse Lab in Milton Keynes | Wellcome Sanger Institute for the COVID-19 Genomics UK (COG-UK) Consortium | The Lighthouse Lab in Milton Keynes and Alex Alderton, Roberto Amato, Jeffrey Barrett, Sonia Goncalves, Ewan Harrison, David K. Jackson, Ian Johnston, Dominic Kwiatkowski, Cordelia Langford, John Sillitoe on behalf of the Wellcome Sanger Institute COVID-19 Surveillance Team |

| | | | |
|---|---|---|---|
| EPI_ISL_3283368, EPI_ISL_3283369 | Lighthouse Lab in Alderley Park | Wellcome Sanger Institute for the COVID-19 Genomics UK (COG-UK) Consortium | Jacquelyn Wynn, Mairead Hyland, The Lighthouse Lab in Alderley Park and Alex Alderton, Roberto Amato, Jeffrey Barrett, Sonia Goncalves, Ewan Harrison, David K. Jackson, Ian Johnston, Dominic Kwiatkowski, Cordelia Langford, John Sillitoe on behalf of the Wellcome Sanger Institute COVID-19 Surveillance Team |
| EPI_ISL_3283370 | Lighthouse Lab in Milton Keynes | Wellcome Sanger Institute for the COVID-19 Genomics UK (COG-UK) Consortium | The Lighthouse Lab in Milton Keynes and Alex Alderton, Roberto Amato, Jeffrey Barrett, Sonia Goncalves, Ewan Harrison, David K. Jackson, Ian Johnston, Dominic Kwiatkowski, Cordelia Langford, John Sillitoe on behalf of the Wellcome Sanger Institute COVID-19 Surveillance Team |
| EPI_ISL_3283371 | Lighthouse Lab in Alderley Park | Wellcome Sanger Institute for the COVID-19 Genomics UK (COG-UK) Consortium | Jacquelyn Wynn, Mairead Hyland, The Lighthouse Lab in Alderley Park and Alex Alderton, Roberto Amato, Jeffrey Barrett, Sonia Goncalves, Ewan Harrison, David K. Jackson, Ian Johnston, Dominic Kwiatkowski, Cordelia Langford, John Sillitoe on behalf of the Wellcome Sanger Institute COVID-19 Surveillance Team |
| EPI_ISL_3283372, EPI_ISL_3283373 | Lighthouse Lab in Milton Keynes | Wellcome Sanger Institute for the COVID-19 Genomics UK (COG-UK) Consortium | The Lighthouse Lab in Milton Keynes and Alex Alderton, Roberto Amato, Jeffrey Barrett, Sonia Goncalves, Ewan Harrison, David K. Jackson, Ian Johnston, Dominic Kwiatkowski, Cordelia Langford, John Sillitoe on behalf of the Wellcome Sanger Institute COVID-19 Surveillance Team |
| EPI_ISL_3283374, EPI_ISL_3283375, EPI_ISL_3283376 | Lighthouse Lab in Alderley Park | Wellcome Sanger Institute for the COVID-19 Genomics UK (COG-UK) Consortium | Jacquelyn Wynn, Mairead Hyland, The Lighthouse Lab in Alderley Park and Alex Alderton, Roberto Amato, Jeffrey Barrett, Sonia Goncalves, Ewan Harrison, David K. Jackson, Ian Johnston, Dominic Kwiatkowski, Cordelia Langford, John Sillitoe on behalf of the Wellcome Sanger Institute COVID-19 Surveillance Team |
| EPI_ISL_3283377, EPI_ISL_3283378, EPI_ISL_3283379 | Lighthouse Lab in Milton Keynes | Wellcome Sanger Institute for the COVID-19 Genomics UK (COG-UK) Consortium | The Lighthouse Lab in Milton Keynes and Alex Alderton, Roberto Amato, Jeffrey Barrett, Sonia Goncalves, Ewan Harrison, David K. Jackson, Ian Johnston, Dominic Kwiatkowski, Cordelia Langford, John Sillitoe on behalf of the Wellcome Sanger Institute COVID-19 Surveillance Team |
| EPI_ISL_3283380, EPI_ISL_3283381 | Lighthouse Lab in Alderley Park | Wellcome Sanger Institute for the COVID-19 Genomics UK (COG-UK) Consortium | Jacquelyn Wynn, Mairead Hyland, The Lighthouse Lab in Alderley Park and Alex Alderton, Roberto Amato, Jeffrey Barrett, Sonia Goncalves, Ewan Harrison, David K. Jackson, Ian Johnston, Dominic Kwiatkowski, Cordelia Langford, John Sillitoe on behalf of the Wellcome Sanger Institute COVID-19 Surveillance Team |
| EPI_ISL_3283382, EPI_ISL_3283383 | Lighthouse Lab in Milton Keynes | Wellcome Sanger Institute for the COVID-19 Genomics UK (COG-UK) Consortium | The Lighthouse Lab in Milton Keynes and Alex Alderton, Roberto Amato, Jeffrey Barrett, Sonia Goncalves, Ewan Harrison, David K. Jackson, Ian Johnston, Dominic Kwiatkowski, Cordelia Langford, John Sillitoe on behalf of the Wellcome Sanger Institute COVID-19 Surveillance Team |
| EPI_ISL_3283384 | Lighthouse Lab in Alderley Park | Wellcome Sanger Institute for the COVID-19 Genomics UK (COG-UK) Consortium | Jacquelyn Wynn, Mairead Hyland, The Lighthouse Lab in Alderley Park and Alex Alderton, Roberto Amato, Jeffrey Barrett, Sonia Goncalves, Ewan Harrison, David K. Jackson, Ian Johnston, Dominic Kwiatkowski, Cordelia Langford, John Sillitoe on behalf of the Wellcome Sanger Institute COVID-19 Surveillance Team |
| EPI_ISL_3283385, EPI_ISL_3283386, EPI_ISL_3283387 | Lighthouse Lab in Milton Keynes | Wellcome Sanger Institute for the COVID-19 Genomics UK (COG-UK) Consortium | The Lighthouse Lab in Milton Keynes and Alex Alderton, Roberto Amato, Jeffrey Barrett, Sonia Goncalves, Ewan Harrison, David K. Jackson, Ian Johnston, Dominic Kwiatkowski, Cordelia Langford, John Sillitoe on behalf of the Wellcome Sanger Institute COVID-19 Surveillance Team |
| EPI_ISL_3283388 | Lighthouse Lab in Alderley Park | Wellcome Sanger Institute for the COVID-19 Genomics UK (COG-UK) Consortium | Jacquelyn Wynn, Mairead Hyland, The Lighthouse Lab in Alderley Park and Alex Alderton, Roberto Amato, Jeffrey Barrett, Sonia Goncalves, Ewan Harrison, David K. Jackson, Ian Johnston, Dominic Kwiatkowski, Cordelia Langford, John Sillitoe on behalf of the Wellcome Sanger Institute COVID-19 Surveillance Team |
| EPI_ISL_3283389, EPI_ISL_3283390, EPI_ISL_3283391 | Lighthouse Lab in Milton Keynes | Wellcome Sanger Institute for the COVID-19 Genomics UK (COG-UK) Consortium | The Lighthouse Lab in Milton Keynes and Alex Alderton, Roberto Amato, Jeffrey Barrett, Sonia Goncalves, Ewan Harrison, David K. Jackson, Ian Johnston, Dominic Kwiatkowski, Cordelia Langford, John Sillitoe on behalf of the Wellcome Sanger Institute COVID-19 Surveillance Team |
| EPI_ISL_3283392, EPI_ISL_3283393, EPI_ISL_3283394, EPI_ISL_3283395 | Lighthouse Lab in Alderley Park | Wellcome Sanger Institute for the COVID-19 Genomics UK (COG-UK) Consortium | Jacquelyn Wynn, Mairead Hyland, The Lighthouse Lab in Alderley Park and Alex Alderton, Roberto Amato, Jeffrey Barrett, Sonia Goncalves, Ewan Harrison, David K. Jackson, Ian Johnston, Dominic Kwiatkowski, Cordelia Langford, John Sillitoe on behalf of the Wellcome Sanger Institute COVID-19 Surveillance Team |
| EPI_ISL_3283396, EPI_ISL_3283397 | Lighthouse Lab in Milton Keynes | Wellcome Sanger Institute for the COVID-19 Genomics UK (COG-UK) Consortium | The Lighthouse Lab in Milton Keynes and Alex Alderton, Roberto Amato, Jeffrey Barrett, Sonia Goncalves, Ewan Harrison, David K. Jackson, Ian Johnston, Dominic Kwiatkowski, Cordelia Langford, John Sillitoe on behalf of the Wellcome Sanger Institute COVID-19 Surveillance Team |
| EPI_ISL_3283398, EPI_ISL_3283399 | Lighthouse Lab in Alderley Park | Wellcome Sanger Institute for the COVID-19 Genomics UK (COG-UK) Consortium | Jacquelyn Wynn, Mairead Hyland, The Lighthouse Lab in Alderley Park and Alex Alderton, Roberto Amato, Jeffrey Barrett, Sonia Goncalves, Ewan Harrison, David K. Jackson, Ian Johnston, Dominic Kwiatkowski, Cordelia Langford, John Sillitoe on behalf of the Wellcome Sanger Institute COVID-19 Surveillance Team |
| EPI_ISL_3283400, EPI_ISL_3283401 | Lighthouse Lab in Milton Keynes | Wellcome Sanger Institute for the COVID-19 Genomics UK (COG-UK) Consortium | The Lighthouse Lab in Milton Keynes and Alex Alderton, Roberto Amato, Jeffrey Barrett, Sonia Goncalves, Ewan Harrison, David K. Jackson, Ian Johnston, Dominic Kwiatkowski, Cordelia Langford, John Sillitoe on behalf of the Wellcome Sanger Institute COVID-19 Surveillance Team |
| EPI_ISL_3283402, EPI_ISL_3283403 | Lighthouse Lab in Alderley Park | Wellcome Sanger Institute for the COVID-19 Genomics UK (COG-UK) Consortium | Jacquelyn Wynn, Mairead Hyland, The Lighthouse Lab in Alderley Park and Alex Alderton, Roberto Amato, Jeffrey Barrett, Sonia Goncalves, Ewan Harrison, David K. Jackson, Ian Johnston, Dominic Kwiatkowski, Cordelia Langford, John Sillitoe on behalf of the Wellcome Sanger Institute COVID-19 Surveillance Team |
| EPI_ISL_3283404 | Lighthouse Lab in Milton Keynes | Wellcome Sanger Institute for the COVID-19 Genomics UK (COG-UK) Consortium | The Lighthouse Lab in Milton Keynes and Alex Alderton, Roberto Amato, Jeffrey Barrett, Sonia Goncalves, Ewan Harrison, David K. Jackson, Ian Johnston, Dominic Kwiatkowski, Cordelia Langford, John Sillitoe on behalf of the Wellcome Sanger Institute COVID-19 Surveillance Team |
| EPI_ISL_3283405, EPI_ISL_3283406 | Lighthouse Lab in Alderley Park | Wellcome Sanger Institute for the COVID-19 Genomics UK (COG-UK) Consortium | Jacquelyn Wynn, Mairead Hyland, The Lighthouse Lab in Alderley Park and Alex Alderton, Roberto Amato, Jeffrey Barrett, Sonia Goncalves, Ewan Harrison, David K. Jackson, Ian Johnston, Dominic Kwiatkowski, Cordelia Langford, John Sillitoe on behalf of the Wellcome Sanger Institute COVID-19 Surveillance Team |
| EPI_ISL_3283407, EPI_ISL_3283408, EPI_ISL_3283409 | Lighthouse Lab in Milton Keynes | Wellcome Sanger Institute for the COVID-19 Genomics UK (COG-UK) Consortium | The Lighthouse Lab in Milton Keynes and Alex Alderton, Roberto Amato, Jeffrey Barrett, Sonia Goncalves, Ewan Harrison, David K. Jackson, Ian Johnston, Dominic Kwiatkowski, Cordelia Langford, John Sillitoe on behalf of the Wellcome Sanger Institute COVID-19 Surveillance Team |
| EPI_ISL_3283410, EPI_ISL_3283411 | Lighthouse Lab in Alderley Park | Wellcome Sanger Institute for the COVID-19 Genomics UK (COG-UK) Consortium | Jacquelyn Wynn, Mairead Hyland, The Lighthouse Lab in Alderley Park and Alex Alderton, Roberto Amato, Jeffrey Barrett, Sonia Goncalves, Ewan Harrison, David K. Jackson, Ian Johnston, Dominic Kwiatkowski, Cordelia Langford, John Sillitoe on behalf of the Wellcome Sanger Institute COVID-19 Surveillance Team |
| EPI_ISL_3283412, EPI_ISL_3283413 | Lighthouse Lab in Milton Keynes | Wellcome Sanger Institute for the COVID-19 Genomics UK (COG-UK) Consortium | The Lighthouse Lab in Milton Keynes and Alex Alderton, Roberto Amato, Jeffrey Barrett, Sonia Goncalves, Ewan Harrison, David K. Jackson, Ian Johnston, Dominic Kwiatkowski, Cordelia Langford, John Sillitoe on behalf of the Wellcome Sanger Institute COVID-19 Surveillance Team |
| EPI_ISL_3283414, EPI_ISL_3283415 | Lighthouse Lab in Alderley Park | Wellcome Sanger Institute for the COVID-19 Genomics UK (COG-UK) Consortium | Jacquelyn Wynn, Mairead Hyland, The Lighthouse Lab in Alderley Park and Alex Alderton, Roberto Amato, Jeffrey Barrett, Sonia Goncalves, Ewan Harrison, David K. Jackson, Ian Johnston, Dominic Kwiatkowski, Cordelia Langford, John Sillitoe on behalf of the Wellcome Sanger Institute COVID-19 Surveillance Team |
| EPI_ISL_3283416 | Lighthouse Lab in Milton Keynes | Wellcome Sanger Institute for the COVID-19 Genomics UK (COG-UK) Consortium | The Lighthouse Lab in Milton Keynes and Alex Alderton, Roberto Amato, Jeffrey Barrett, Sonia Goncalves, Ewan Harrison, David K. Jackson, Ian Johnston, Dominic Kwiatkowski, Cordelia Langford, John Sillitoe on behalf of the Wellcome Sanger Institute COVID-19 Surveillance Team |
| EPI_ISL_3283417, EPI_ISL_3283418, EPI_ISL_3283419, EPI_ISL_3283420 | Lighthouse Lab in Alderley Park | Wellcome Sanger Institute for the COVID-19 Genomics UK (COG-UK) Consortium | Jacquelyn Wynn, Mairead Hyland, The Lighthouse Lab in Alderley Park and Alex Alderton, Roberto Amato, Jeffrey Barrett, Sonia Goncalves, Ewan Harrison, David K. Jackson, Ian Johnston, Dominic Kwiatkowski, Cordelia Langford, John Sillitoe on behalf of the Wellcome Sanger Institute COVID-19 Surveillance Team |
| EPI_ISL_3283421, EPI_ISL_3283422, EPI_ISL_3283423 | Lighthouse Lab in Milton Keynes | Wellcome Sanger Institute for the COVID-19 Genomics UK (COG-UK) Consortium | The Lighthouse Lab in Milton Keynes and Alex Alderton, Roberto Amato, Jeffrey Barrett, Sonia Goncalves, Ewan Harrison, David K. Jackson, Ian Johnston, Dominic Kwiatkowski, Cordelia Langford, John Sillitoe on behalf of the Wellcome Sanger Institute COVID-19 Surveillance Team |
| EPI_ISL_3283424 | Lighthouse Lab in Alderley Park | Wellcome Sanger Institute for the COVID-19 Genomics UK (COG-UK) Consortium | Jacquelyn Wynn, Mairead Hyland, The Lighthouse Lab in Alderley Park and Alex Alderton, Roberto Amato, Jeffrey Barrett, Sonia Goncalves, Ewan Harrison, David K. Jackson, Ian Johnston, Dominic Kwiatkowski, Cordelia Langford, John Sillitoe on behalf of the Wellcome Sanger Institute COVID-19 Surveillance Team |
| EPI_ISL_3283425, EPI_ISL_3283426 | Lighthouse Lab in Milton Keynes | Wellcome Sanger Institute for the COVID-19 Genomics UK (COG-UK) Consortium | The Lighthouse Lab in Milton Keynes and Alex Alderton, Roberto Amato, Jeffrey Barrett, Sonia Goncalves, Ewan Harrison, David K. Jackson, Ian Johnston, Dominic Kwiatkowski, Cordelia Langford, John Sillitoe on behalf of the Wellcome Sanger Institute COVID-19 Surveillance Team |
| EPI_ISL_3283427 | Lighthouse Lab in Alderley Park | Wellcome Sanger Institute for the COVID-19 Genomics UK (COG-UK) Consortium | Jacquelyn Wynn, Mairead Hyland, The Lighthouse Lab in Alderley Park and Alex Alderton, Roberto Amato, Jeffrey Barrett, Sonia Goncalves, Ewan Harrison, David K. Jackson, Ian Johnston, Dominic Kwiatkowski, Cordelia Langford, John Sillitoe on behalf of the Wellcome Sanger Institute COVID-19 Surveillance Team |
| EPI_ISL_3283428 | Lighthouse Lab in Milton Keynes | Wellcome Sanger Institute for the COVID-19 Genomics UK (COG-UK) Consortium | The Lighthouse Lab in Milton Keynes and Alex Alderton, Roberto Amato, Jeffrey Barrett, Sonia Goncalves, Ewan Harrison, David K. Jackson, Ian Johnston, Dominic Kwiatkowski, Cordelia Langford, John Sillitoe on behalf of the Wellcome Sanger Institute COVID-19 Surveillance Team |
| EPI_ISL_3283429, EPI_ISL_3283430, EPI_ISL_3283431 | Lighthouse Lab in Alderley Park | Wellcome Sanger Institute for the COVID-19 Genomics UK (COG-UK) Consortium | Jacquelyn Wynn, Mairead Hyland, The Lighthouse Lab in Alderley Park and Alex Alderton, Roberto Amato, Jeffrey Barrett, Sonia Goncalves, Ewan Harrison, David K. Jackson, Ian Johnston, Dominic Kwiatkowski, Cordelia Langford, John Sillitoe on behalf of the Wellcome Sanger Institute COVID-19 Surveillance Team |
| EPI_ISL_3283432, EPI_ISL_3283433, EPI_ISL_3283434, EPI_ISL_3283435 | Lighthouse Lab in Milton Keynes | Wellcome Sanger Institute for the COVID-19 Genomics UK (COG-UK) Consortium | The Lighthouse Lab in Milton Keynes and Alex Alderton, Roberto Amato, Jeffrey Barrett, Sonia Goncalves, Ewan Harrison, David K. Jackson, Ian Johnston, Dominic Kwiatkowski, Cordelia Langford, John Sillitoe on behalf of the Wellcome Sanger Institute COVID-19 Surveillance Team |
| EPI_ISL_3283436 | Lighthouse Lab in Alderley Park | Wellcome Sanger Institute for the COVID-19 Genomics UK (COG-UK) Consortium | Jacquelyn Wynn, Mairead Hyland, The Lighthouse Lab in Alderley Park and Alex Alderton, Roberto Amato, Jeffrey Barrett, Sonia Goncalves, Ewan Harrison, David K. Jackson, Ian Johnston, Dominic Kwiatkowski, Cordelia Langford, John Sillitoe on behalf of the Wellcome Sanger Institute COVID-19 Surveillance Team |
| EPI_ISL_3283437 | Lighthouse Lab in Milton Keynes | Wellcome Sanger Institute for the COVID-19 Genomics UK (COG-UK) Consortium | The Lighthouse Lab in Milton Keynes and Alex Alderton, Roberto Amato, Jeffrey Barrett, Sonia Goncalves, Ewan Harrison, David K. Jackson, Ian Johnston, Dominic Kwiatkowski, Cordelia Langford, John Sillitoe on behalf of the Wellcome Sanger Institute COVID-19 Surveillance Team |
| EPI_ISL_3283438, EPI_ISL_3283439, EPI_ISL_3283440 | Lighthouse Lab in Alderley Park | Wellcome Sanger Institute for the COVID-19 Genomics UK (COG-UK) Consortium | Jacquelyn Wynn, Mairead Hyland, The Lighthouse Lab in Alderley Park and Alex Alderton, Roberto Amato, Jeffrey Barrett, Sonia Goncalves, Ewan Harrison, David K. Jackson, Ian Johnston, Dominic Kwiatkowski, Cordelia Langford, John Sillitoe on behalf of the Wellcome Sanger Institute COVID-19 Surveillance Team |
| EPI_ISL_3283441 | Lighthouse Lab in Milton Keynes | Wellcome Sanger Institute for the COVID-19 Genomics UK (COG-UK) Consortium | The Lighthouse Lab in Milton Keynes and Alex Alderton, Roberto Amato, Jeffrey Barrett, Sonia Goncalves, Ewan Harrison, David K. Jackson, Ian Johnston, Dominic Kwiatkowski, Cordelia Langford, John Sillitoe on behalf of the Wellcome Sanger Institute COVID-19 Surveillance Team |
| EPI_ISL_3283442, EPI_ISL_3283443, EPI_ISL_3283444, EPI_ISL_3283445, EPI_ISL_3283446, EPI_ISL_3283447, EPI_ISL_3283448, | Lighthouse Lab in Alderley Park | Wellcome Sanger Institute for the COVID-19 Genomics UK (COG-UK) Consortium | Jacquelyn Wynn, Mairead Hyland, The Lighthouse Lab in Alderley Park and Alex Alderton, Roberto Amato, Jeffrey Barrett, Sonia Goncalves, Ewan Harrison, David K. Jackson, Ian Johnston, Dominic Kwiatkowski, Cordelia Langford, John Sillitoe on behalf of the Wellcome Sanger Institute COVID-19 Surveillance Team |

| | | | |
|---|---|---|---|
| EPI_ISL_3283449 | | | |
| EPI_ISL_3283450 | Lighthouse Lab in Milton Keynes | Wellcome Sanger Institute for the COVID-19 Genomics UK (COG-UK) Consortium | The Lighthouse Lab in Milton Keynes and Alex Alderton, Roberto Amato, Jeffrey Barrett, Sonia Goncalves, Ewan Harrison, David K. Jackson, Ian Johnston, Dominic Kwiatkowski, Cordelia Langford, John Sillitoe on behalf of the Wellcome Sanger Institute COVID-19 Surveillance Team |
| EPI_ISL_3283451 | Lighthouse Lab in Alderley Park | Wellcome Sanger Institute for the COVID-19 Genomics UK (COG-UK) Consortium | Jacquelyn Wynn, Mairead Hyland, The Lighthouse Lab in Alderley Park and Alex Alderton, Roberto Amato, Jeffrey Barrett, Sonia Goncalves, Ewan Harrison, David K. Jackson, Ian Johnston, Dominic Kwiatkowski, Cordelia Langford, John Sillitoe on behalf of the Wellcome Sanger Institute COVID-19 Surveillance Team |
| EPI_ISL_3283452 | Lighthouse Lab in Milton Keynes | Wellcome Sanger Institute for the COVID-19 Genomics UK (COG-UK) Consortium | The Lighthouse Lab in Milton Keynes and Alex Alderton, Roberto Amato, Jeffrey Barrett, Sonia Goncalves, Ewan Harrison, David K. Jackson, Ian Johnston, Dominic Kwiatkowski, Cordelia Langford, John Sillitoe on behalf of the Wellcome Sanger Institute COVID-19 Surveillance Team |
| EPI_ISL_3283453 | Lighthouse Lab in Alderley Park | Wellcome Sanger Institute for the COVID-19 Genomics UK (COG-UK) Consortium | Jacquelyn Wynn, Mairead Hyland, The Lighthouse Lab in Alderley Park and Alex Alderton, Roberto Amato, Jeffrey Barrett, Sonia Goncalves, Ewan Harrison, David K. Jackson, Ian Johnston, Dominic Kwiatkowski, Cordelia Langford, John Sillitoe on behalf of the Wellcome Sanger Institute COVID-19 Surveillance Team |
| EPI_ISL_3283454, EPI_ISL_3283455 | Lighthouse Lab in Milton Keynes | Wellcome Sanger Institute for the COVID-19 Genomics UK (COG-UK) Consortium | The Lighthouse Lab in Milton Keynes and Alex Alderton, Roberto Amato, Jeffrey Barrett, Sonia Goncalves, Ewan Harrison, David K. Jackson, Ian Johnston, Dominic Kwiatkowski, Cordelia Langford, John Sillitoe on behalf of the Wellcome Sanger Institute COVID-19 Surveillance Team |
| EPI_ISL_3283456 | Lighthouse Lab in Alderley Park | Wellcome Sanger Institute for the COVID-19 Genomics UK (COG-UK) Consortium | Jacquelyn Wynn, Mairead Hyland, The Lighthouse Lab in Alderley Park and Alex Alderton, Roberto Amato, Jeffrey Barrett, Sonia Goncalves, Ewan Harrison, David K. Jackson, Ian Johnston, Dominic Kwiatkowski, Cordelia Langford, John Sillitoe on behalf of the Wellcome Sanger Institute COVID-19 Surveillance Team |
| EPI_ISL_3283457 | Lighthouse Lab in Milton Keynes | Wellcome Sanger Institute for the COVID-19 Genomics UK (COG-UK) Consortium | The Lighthouse Lab in Milton Keynes and Alex Alderton, Roberto Amato, Jeffrey Barrett, Sonia Goncalves, Ewan Harrison, David K. Jackson, Ian Johnston, Dominic Kwiatkowski, Cordelia Langford, John Sillitoe on behalf of the Wellcome Sanger Institute COVID-19 Surveillance Team |
| EPI_ISL_3283458, EPI_ISL_3283459 | Lighthouse Lab in Alderley Park | Wellcome Sanger Institute for the COVID-19 Genomics UK (COG-UK) Consortium | Jacquelyn Wynn, Mairead Hyland, The Lighthouse Lab in Alderley Park and Alex Alderton, Roberto Amato, Jeffrey Barrett, Sonia Goncalves, Ewan Harrison, David K. Jackson, Ian Johnston, Dominic Kwiatkowski, Cordelia Langford, John Sillitoe on behalf of the Wellcome Sanger Institute COVID-19 Surveillance Team |
| EPI_ISL_3283460, EPI_ISL_3283461 | Lighthouse Lab in Milton Keynes | Wellcome Sanger Institute for the COVID-19 Genomics UK (COG-UK) Consortium | The Lighthouse Lab in Milton Keynes and Alex Alderton, Roberto Amato, Jeffrey Barrett, Sonia Goncalves, Ewan Harrison, David K. Jackson, Ian Johnston, Dominic Kwiatkowski, Cordelia Langford, John Sillitoe on behalf of the Wellcome Sanger Institute COVID-19 Surveillance Team |
| EPI_ISL_3283462, EPI_ISL_3283463, EPI_ISL_3283464 | Lighthouse Lab in Alderley Park | Wellcome Sanger Institute for the COVID-19 Genomics UK (COG-UK) Consortium | Jacquelyn Wynn, Mairead Hyland, The Lighthouse Lab in Alderley Park and Alex Alderton, Roberto Amato, Jeffrey Barrett, Sonia Goncalves, Ewan Harrison, David K. Jackson, Ian Johnston, Dominic Kwiatkowski, Cordelia Langford, John Sillitoe on behalf of the Wellcome Sanger Institute COVID-19 Surveillance Team |
| EPI_ISL_3283465 | Lighthouse Lab in Milton Keynes | Wellcome Sanger Institute for the COVID-19 Genomics UK (COG-UK) Consortium | The Lighthouse Lab in Milton Keynes and Alex Alderton, Roberto Amato, Jeffrey Barrett, Sonia Goncalves, Ewan Harrison, David K. Jackson, Ian Johnston, Dominic Kwiatkowski, Cordelia Langford, John Sillitoe on behalf of the Wellcome Sanger Institute COVID-19 Surveillance Team |
| EPI_ISL_3283466 | Lighthouse Lab in Alderley Park | Wellcome Sanger Institute for the COVID-19 Genomics UK (COG-UK) Consortium | Jacquelyn Wynn, Mairead Hyland, The Lighthouse Lab in Alderley Park and Alex Alderton, Roberto Amato, Jeffrey Barrett, Sonia Goncalves, Ewan Harrison, David K. Jackson, Ian Johnston, Dominic Kwiatkowski, Cordelia Langford, John Sillitoe on behalf of the Wellcome Sanger Institute COVID-19 Surveillance Team |
| EPI_ISL_3283467 | Lighthouse Lab in Milton Keynes | Wellcome Sanger Institute for the COVID-19 Genomics UK (COG-UK) Consortium | The Lighthouse Lab in Milton Keynes and Alex Alderton, Roberto Amato, Jeffrey Barrett, Sonia Goncalves, Ewan Harrison, David K. Jackson, Ian Johnston, Dominic Kwiatkowski, Cordelia Langford, John Sillitoe on behalf of the Wellcome Sanger Institute COVID-19 Surveillance Team |
| EPI_ISL_3283468, EPI_ISL_3283469 | Lighthouse Lab in Alderley Park | Wellcome Sanger Institute for the COVID-19 Genomics UK (COG-UK) Consortium | Jacquelyn Wynn, Mairead Hyland, The Lighthouse Lab in Alderley Park and Alex Alderton, Roberto Amato, Jeffrey Barrett, Sonia Goncalves, Ewan Harrison, David K. Jackson, Ian Johnston, Dominic Kwiatkowski, Cordelia Langford, John Sillitoe on behalf of the Wellcome Sanger Institute COVID-19 Surveillance Team |
| EPI_ISL_3283470, EPI_ISL_3283471, EPI_ISL_3283472 | Lighthouse Lab in Milton Keynes | Wellcome Sanger Institute for the COVID-19 Genomics UK (COG-UK) Consortium | The Lighthouse Lab in Milton Keynes and Alex Alderton, Roberto Amato, Jeffrey Barrett, Sonia Goncalves, Ewan Harrison, David K. Jackson, Ian Johnston, Dominic Kwiatkowski, Cordelia Langford, John Sillitoe on behalf of the Wellcome Sanger Institute COVID-19 Surveillance Team |
| EPI_ISL_3283473 | Lighthouse Lab in Alderley Park | Wellcome Sanger Institute for the COVID-19 Genomics UK (COG-UK) Consortium | Jacquelyn Wynn, Mairead Hyland, The Lighthouse Lab in Alderley Park and Alex Alderton, Roberto Amato, Jeffrey Barrett, Sonia Goncalves, Ewan Harrison, David K. Jackson, Ian Johnston, Dominic Kwiatkowski, Cordelia Langford, John Sillitoe on behalf of the Wellcome Sanger Institute COVID-19 Surveillance Team |
| EPI_ISL_3283474 | Lighthouse Lab in Milton Keynes | Wellcome Sanger Institute for the COVID-19 Genomics UK (COG-UK) Consortium | The Lighthouse Lab in Milton Keynes and Alex Alderton, Roberto Amato, Jeffrey Barrett, Sonia Goncalves, Ewan Harrison, David K. Jackson, Ian Johnston, Dominic Kwiatkowski, Cordelia Langford, John Sillitoe on behalf of the Wellcome Sanger Institute COVID-19 Surveillance Team |
| EPI_ISL_3283475, EPI_ISL_3283476, EPI_ISL_3283477 | Lighthouse Lab in Alderley Park | Wellcome Sanger Institute for the COVID-19 Genomics UK (COG-UK) Consortium | Jacquelyn Wynn, Mairead Hyland, The Lighthouse Lab in Alderley Park and Alex Alderton, Roberto Amato, Jeffrey Barrett, Sonia Goncalves, Ewan Harrison, David K. Jackson, Ian Johnston, Dominic Kwiatkowski, Cordelia Langford, John Sillitoe on behalf of the Wellcome Sanger Institute COVID-19 Surveillance Team |
| EPI_ISL_3283478 | Lighthouse Lab in Milton Keynes | Wellcome Sanger Institute for the COVID-19 Genomics UK (COG-UK) Consortium | The Lighthouse Lab in Milton Keynes and Alex Alderton, Roberto Amato, Jeffrey Barrett, Sonia Goncalves, Ewan Harrison, David K. Jackson, Ian Johnston, Dominic Kwiatkowski, Cordelia Langford, John Sillitoe on behalf of the Wellcome Sanger Institute COVID-19 Surveillance Team |
| EPI_ISL_3283479, EPI_ISL_3283480, EPI_ISL_3283481, EPI_ISL_3283482, EPI_ISL_3283483, EPI_ISL_3283484, EPI_ISL_3283485, EPI_ISL_3283486, EPI_ISL_3283487 | Lighthouse Lab in Alderley Park | Wellcome Sanger Institute for the COVID-19 Genomics UK (COG-UK) Consortium | Jacquelyn Wynn, Mairead Hyland, The Lighthouse Lab in Alderley Park and Alex Alderton, Roberto Amato, Jeffrey Barrett, Sonia Goncalves, Ewan Harrison, David K. Jackson, Ian Johnston, Dominic Kwiatkowski, Cordelia Langford, John Sillitoe on behalf of the Wellcome Sanger Institute COVID-19 Surveillance Team |
| EPI_ISL_3283488, EPI_ISL_3283489, EPI_ISL_3283490, EPI_ISL_3283491, EPI_ISL_3283492 | Lighthouse Lab in Milton Keynes | Wellcome Sanger Institute for the COVID-19 Genomics UK (COG-UK) Consortium | The Lighthouse Lab in Milton Keynes and Alex Alderton, Roberto Amato, Jeffrey Barrett, Sonia Goncalves, Ewan Harrison, David K. Jackson, Ian Johnston, Dominic Kwiatkowski, Cordelia Langford, John Sillitoe on behalf of the Wellcome Sanger Institute COVID-19 Surveillance Team |
| EPI_ISL_3283493, EPI_ISL_3283494 | Lighthouse Lab in Alderley Park | Wellcome Sanger Institute for the COVID-19 Genomics UK (COG-UK) Consortium | Jacquelyn Wynn, Mairead Hyland, The Lighthouse Lab in Alderley Park and Alex Alderton, Roberto Amato, Jeffrey Barrett, Sonia Goncalves, Ewan Harrison, David K. Jackson, Ian Johnston, Dominic Kwiatkowski, Cordelia Langford, John Sillitoe on behalf of the Wellcome Sanger Institute COVID-19 Surveillance Team |
| EPI_ISL_3283495 | Lighthouse Lab in Milton Keynes | Wellcome Sanger Institute for the COVID-19 Genomics UK (COG-UK) Consortium | The Lighthouse Lab in Milton Keynes and Alex Alderton, Roberto Amato, Jeffrey Barrett, Sonia Goncalves, Ewan Harrison, David K. Jackson, Ian Johnston, Dominic Kwiatkowski, Cordelia Langford, John Sillitoe on behalf of the Wellcome Sanger Institute COVID-19 Surveillance Team |
| EPI_ISL_3283496, EPI_ISL_3283497 | Lighthouse Lab in Alderley Park | Wellcome Sanger Institute for the COVID-19 Genomics UK (COG-UK) Consortium | Jacquelyn Wynn, Mairead Hyland, The Lighthouse Lab in Alderley Park and Alex Alderton, Roberto Amato, Jeffrey Barrett, Sonia Goncalves, Ewan Harrison, David K. Jackson, Ian Johnston, Dominic Kwiatkowski, Cordelia Langford, John Sillitoe on behalf of the Wellcome Sanger Institute COVID-19 Surveillance Team |
| EPI_ISL_3283498 | Lighthouse Lab in Milton Keynes | Wellcome Sanger Institute for the COVID-19 Genomics UK (COG-UK) Consortium | The Lighthouse Lab in Milton Keynes and Alex Alderton, Roberto Amato, Jeffrey Barrett, Sonia Goncalves, Ewan Harrison, David K. Jackson, Ian Johnston, Dominic Kwiatkowski, Cordelia Langford, John Sillitoe on behalf of the Wellcome Sanger Institute COVID-19 Surveillance Team |
| EPI_ISL_3283499, EPI_ISL_3283500, EPI_ISL_3283501, EPI_ISL_3283502 | Lighthouse Lab in Alderley Park | Wellcome Sanger Institute for the COVID-19 Genomics UK (COG-UK) Consortium | Jacquelyn Wynn, Mairead Hyland, The Lighthouse Lab in Alderley Park and Alex Alderton, Roberto Amato, Jeffrey Barrett, Sonia Goncalves, Ewan Harrison, David K. Jackson, Ian Johnston, Dominic Kwiatkowski, Cordelia Langford, John Sillitoe on behalf of the Wellcome Sanger Institute COVID-19 Surveillance Team |
| EPI_ISL_3283503, EPI_ISL_3283504 | Lighthouse Lab in Milton Keynes | Wellcome Sanger Institute for the COVID-19 Genomics UK (COG-UK) Consortium | The Lighthouse Lab in Milton Keynes and Alex Alderton, Roberto Amato, Jeffrey Barrett, Sonia Goncalves, Ewan Harrison, David K. Jackson, Ian Johnston, Dominic Kwiatkowski, Cordelia Langford, John Sillitoe on behalf of the Wellcome Sanger Institute COVID-19 Surveillance Team |
| EPI_ISL_3283505 | Lighthouse Lab in Alderley Park | Wellcome Sanger Institute for the COVID-19 Genomics UK (COG-UK) Consortium | Jacquelyn Wynn, Mairead Hyland, The Lighthouse Lab in Alderley Park and Alex Alderton, Roberto Amato, Jeffrey Barrett, Sonia Goncalves, Ewan Harrison, David K. Jackson, Ian Johnston, Dominic Kwiatkowski, Cordelia Langford, John Sillitoe on behalf of the Wellcome Sanger Institute COVID-19 Surveillance Team |
| EPI_ISL_3283506, EPI_ISL_3283507 | Lighthouse Lab in Milton Keynes | Wellcome Sanger Institute for the COVID-19 Genomics UK (COG-UK) Consortium | The Lighthouse Lab in Milton Keynes and Alex Alderton, Roberto Amato, Jeffrey Barrett, Sonia Goncalves, Ewan Harrison, David K. Jackson, Ian Johnston, Dominic Kwiatkowski, Cordelia Langford, John Sillitoe on behalf of the Wellcome Sanger Institute COVID-19 Surveillance Team |
| EPI_ISL_3283508 | Lighthouse Lab in Alderley Park | Wellcome Sanger Institute for the COVID-19 Genomics UK (COG-UK) Consortium | Jacquelyn Wynn, Mairead Hyland, The Lighthouse Lab in Alderley Park and Alex Alderton, Roberto Amato, Jeffrey Barrett, Sonia Goncalves, Ewan Harrison, David K. Jackson, Ian Johnston, Dominic Kwiatkowski, Cordelia Langford, John Sillitoe on behalf of the Wellcome Sanger Institute COVID-19 Surveillance Team |
| EPI_ISL_3283509 | Lighthouse Lab in Milton Keynes | Wellcome Sanger Institute for the COVID-19 Genomics UK (COG-UK) Consortium | The Lighthouse Lab in Milton Keynes and Alex Alderton, Roberto Amato, Jeffrey Barrett, Sonia Goncalves, Ewan Harrison, David K. Jackson, Ian Johnston, Dominic Kwiatkowski, Cordelia Langford, John Sillitoe on behalf of the Wellcome Sanger Institute COVID-19 Surveillance Team |
| EPI_ISL_3283510 | Lighthouse Lab in Alderley Park | Wellcome Sanger Institute for the COVID-19 Genomics UK (COG-UK) Consortium | Jacquelyn Wynn, Mairead Hyland, The Lighthouse Lab in Alderley Park and Alex Alderton, Roberto Amato, Jeffrey Barrett, Sonia Goncalves, Ewan Harrison, David K. Jackson, Ian Johnston, Dominic Kwiatkowski, Cordelia Langford, John Sillitoe on behalf of the Wellcome Sanger Institute COVID-19 Surveillance Team |
| EPI_ISL_3283511 | Lighthouse Lab in Milton Keynes | Wellcome Sanger Institute for the COVID-19 Genomics UK (COG-UK) Consortium | The Lighthouse Lab in Milton Keynes and Alex Alderton, Roberto Amato, Jeffrey Barrett, Sonia Goncalves, Ewan Harrison, David K. Jackson, Ian Johnston, Dominic Kwiatkowski, Cordelia Langford, John Sillitoe on behalf of the Wellcome Sanger Institute COVID-19 Surveillance Team |
| EPI_ISL_3283512, EPI_ISL_3283513 | Lighthouse Lab in Alderley Park | Wellcome Sanger Institute for the COVID-19 Genomics UK (COG-UK) Consortium | Jacquelyn Wynn, Mairead Hyland, The Lighthouse Lab in Alderley Park and Alex Alderton, Roberto Amato, Jeffrey Barrett, Sonia Goncalves, Ewan Harrison, David K. Jackson, Ian Johnston, Dominic Kwiatkowski, Cordelia Langford, John Sillitoe on behalf of the Wellcome Sanger Institute COVID-19 Surveillance Team |
| EPI_ISL_3283514, EPI_ISL_3283515 | Lighthouse Lab in Milton Keynes | Wellcome Sanger Institute for the COVID-19 Genomics UK (COG-UK) Consortium | The Lighthouse Lab in Milton Keynes and Alex Alderton, Roberto Amato, Jeffrey Barrett, Sonia Goncalves, Ewan Harrison, David K. Jackson, Ian Johnston, Dominic Kwiatkowski, Cordelia Langford, John Sillitoe on behalf of the Wellcome Sanger Institute COVID-19 Surveillance Team |
| EPI_ISL_3283516 | Lighthouse Lab in Alderley Park | Wellcome Sanger Institute for the COVID-19 Genomics UK (COG-UK) Consortium | Jacquelyn Wynn, Mairead Hyland, The Lighthouse Lab in Alderley Park and Alex Alderton, Roberto Amato, Jeffrey Barrett, Sonia Goncalves, Ewan Harrison, David K. Jackson, Ian Johnston, Dominic Kwiatkowski, Cordelia Langford, John Sillitoe on behalf of the Wellcome Sanger Institute COVID-19 Surveillance Team |
| EPI_ISL_3283517 | Lighthouse Lab in Milton Keynes | Wellcome Sanger Institute for the COVID-19 Genomics UK (COG-UK) Consortium | The Lighthouse Lab in Milton Keynes and Alex Alderton, Roberto Amato, Jeffrey Barrett, Sonia Goncalves, Ewan Harrison, David K. Jackson, Ian Johnston, Dominic Kwiatkowski, Cordelia Langford, John Sillitoe on behalf of the Wellcome Sanger Institute COVID-19 Surveillance Team |
| EPI_ISL_3283518 | Lighthouse Lab in Alderley Park | Wellcome Sanger Institute for the COVID-19 Genomics UK (COG-UK) Consortium | Jacquelyn Wynn, Mairead Hyland, The Lighthouse Lab in Alderley Park and Alex Alderton, Roberto Amato, Jeffrey Barrett, Sonia Goncalves, Ewan Harrison, David K. Jackson, Ian Johnston, Dominic Kwiatkowski, Cordelia Langford, John Sillitoe on behalf of the Wellcome Sanger Institute COVID-19 Surveillance Team |
| EPI_ISL_3283519, | Lighthouse Lab in Milton Keynes | Wellcome Sanger Institute for the COVID-19 Genomics UK (COG-UK) | The Lighthouse Lab in Milton Keynes and Alex Alderton, Roberto Amato, Jeffrey Barrett, Sonia Goncalves, Ewan Harrison, David K. Jackson, Ian Johnston, Dominic Kwiatkowski, Cordelia Langford, John Sillitoe on behalf of the Wellcome Sanger |

| | | | |
|---|---|---|---|
| EPI_ISL_3283520 | | Consortium | Institute COVID-19 Surveillance Team |
| EPI_ISL_3283521 | Lighthouse Lab in Alderley Park | Wellcome Sanger Institute for the COVID-19 Genomics UK (COG-UK) Consortium | Jacquelyn Wynn, Mairead Hyland, The Lighthouse Lab in Alderley Park and Alex Alderton, Roberto Amato, Jeffrey Barrett, Sonia Goncalves, Ewan Harrison, David K. Jackson, Ian Johnston, Dominic Kwiatkowski, Cordelia Langford, John Sillitoe on behalf of the Wellcome Sanger Institute COVID-19 Surveillance Team |
| EPI_ISL_3283522, EPI_ISL_3283523, EPI_ISL_3283524 | Lighthouse Lab in Milton Keynes | Wellcome Sanger Institute for the COVID-19 Genomics UK (COG-UK) Consortium | The Lighthouse Lab in Milton Keynes and Alex Alderton, Roberto Amato, Jeffrey Barrett, Sonia Goncalves, Ewan Harrison, David K. Jackson, Ian Johnston, Dominic Kwiatkowski, Cordelia Langford, John Sillitoe on behalf of the Wellcome Sanger Institute COVID-19 Surveillance Team |
| EPI_ISL_3283525 | Lighthouse Lab in Alderley Park | Wellcome Sanger Institute for the COVID-19 Genomics UK (COG-UK) Consortium | Jacquelyn Wynn, Mairead Hyland, The Lighthouse Lab in Alderley Park and Alex Alderton, Roberto Amato, Jeffrey Barrett, Sonia Goncalves, Ewan Harrison, David K. Jackson, Ian Johnston, Dominic Kwiatkowski, Cordelia Langford, John Sillitoe on behalf of the Wellcome Sanger Institute COVID-19 Surveillance Team |
| EPI_ISL_3283526 | Lighthouse Lab in Milton Keynes | Wellcome Sanger Institute for the COVID-19 Genomics UK (COG-UK) Consortium | The Lighthouse Lab in Milton Keynes and Alex Alderton, Roberto Amato, Jeffrey Barrett, Sonia Goncalves, Ewan Harrison, David K. Jackson, Ian Johnston, Dominic Kwiatkowski, Cordelia Langford, John Sillitoe on behalf of the Wellcome Sanger Institute COVID-19 Surveillance Team |
| EPI_ISL_3283527, EPI_ISL_3283528, EPI_ISL_3283529 | Lighthouse Lab in Alderley Park | Wellcome Sanger Institute for the COVID-19 Genomics UK (COG-UK) Consortium | Jacquelyn Wynn, Mairead Hyland, The Lighthouse Lab in Alderley Park and Alex Alderton, Roberto Amato, Jeffrey Barrett, Sonia Goncalves, Ewan Harrison, David K. Jackson, Ian Johnston, Dominic Kwiatkowski, Cordelia Langford, John Sillitoe on behalf of the Wellcome Sanger Institute COVID-19 Surveillance Team |
| EPI_ISL_3283530, EPI_ISL_3283531 | Lighthouse Lab in Milton Keynes | Wellcome Sanger Institute for the COVID-19 Genomics UK (COG-UK) Consortium | The Lighthouse Lab in Milton Keynes and Alex Alderton, Roberto Amato, Jeffrey Barrett, Sonia Goncalves, Ewan Harrison, David K. Jackson, Ian Johnston, Dominic Kwiatkowski, Cordelia Langford, John Sillitoe on behalf of the Wellcome Sanger Institute COVID-19 Surveillance Team |
| EPI_ISL_3283532 | Lighthouse Lab in Alderley Park | Wellcome Sanger Institute for the COVID-19 Genomics UK (COG-UK) Consortium | Jacquelyn Wynn, Mairead Hyland, The Lighthouse Lab in Alderley Park and Alex Alderton, Roberto Amato, Jeffrey Barrett, Sonia Goncalves, Ewan Harrison, David K. Jackson, Ian Johnston, Dominic Kwiatkowski, Cordelia Langford, John Sillitoe on behalf of the Wellcome Sanger Institute COVID-19 Surveillance Team |
| EPI_ISL_3283533, EPI_ISL_3283534, EPI_ISL_3283535, EPI_ISL_3283536, EPI_ISL_3283537, EPI_ISL_3283538, EPI_ISL_3283539, EPI_ISL_3283540, EPI_ISL_3283541 | Lighthouse Lab in Milton Keynes | Wellcome Sanger Institute for the COVID-19 Genomics UK (COG-UK) Consortium | The Lighthouse Lab in Milton Keynes and Alex Alderton, Roberto Amato, Jeffrey Barrett, Sonia Goncalves, Ewan Harrison, David K. Jackson, Ian Johnston, Dominic Kwiatkowski, Cordelia Langford, John Sillitoe on behalf of the Wellcome Sanger Institute COVID-19 Surveillance Team |
| EPI_ISL_3283542 | Lighthouse Lab in Alderley Park | Wellcome Sanger Institute for the COVID-19 Genomics UK (COG-UK) Consortium | Jacquelyn Wynn, Mairead Hyland, The Lighthouse Lab in Alderley Park and Alex Alderton, Roberto Amato, Jeffrey Barrett, Sonia Goncalves, Ewan Harrison, David K. Jackson, Ian Johnston, Dominic Kwiatkowski, Cordelia Langford, John Sillitoe on behalf of the Wellcome Sanger Institute COVID-19 Surveillance Team |
| EPI_ISL_3283543, EPI_ISL_3283544 | Lighthouse Lab in Milton Keynes | Wellcome Sanger Institute for the COVID-19 Genomics UK (COG-UK) Consortium | The Lighthouse Lab in Milton Keynes and Alex Alderton, Roberto Amato, Jeffrey Barrett, Sonia Goncalves, Ewan Harrison, David K. Jackson, Ian Johnston, Dominic Kwiatkowski, Cordelia Langford, John Sillitoe on behalf of the Wellcome Sanger Institute COVID-19 Surveillance Team |
| EPI_ISL_3283545, EPI_ISL_3283546 | Lighthouse Lab in Alderley Park | Wellcome Sanger Institute for the COVID-19 Genomics UK (COG-UK) Consortium | Jacquelyn Wynn, Mairead Hyland, The Lighthouse Lab in Alderley Park and Alex Alderton, Roberto Amato, Jeffrey Barrett, Sonia Goncalves, Ewan Harrison, David K. Jackson, Ian Johnston, Dominic Kwiatkowski, Cordelia Langford, John Sillitoe on behalf of the Wellcome Sanger Institute COVID-19 Surveillance Team |
| EPI_ISL_3283547 | Lighthouse Lab in Milton Keynes | Wellcome Sanger Institute for the COVID-19 Genomics UK (COG-UK) Consortium | The Lighthouse Lab in Milton Keynes and Alex Alderton, Roberto Amato, Jeffrey Barrett, Sonia Goncalves, Ewan Harrison, David K. Jackson, Ian Johnston, Dominic Kwiatkowski, Cordelia Langford, John Sillitoe on behalf of the Wellcome Sanger Institute COVID-19 Surveillance Team |
| EPI_ISL_3283548 | Lighthouse Lab in Alderley Park | Wellcome Sanger Institute for the COVID-19 Genomics UK (COG-UK) Consortium | Jacquelyn Wynn, Mairead Hyland, The Lighthouse Lab in Alderley Park and Alex Alderton, Roberto Amato, Jeffrey Barrett, Sonia Goncalves, Ewan Harrison, David K. Jackson, Ian Johnston, Dominic Kwiatkowski, Cordelia Langford, John Sillitoe on behalf of the Wellcome Sanger Institute COVID-19 Surveillance Team |
| EPI_ISL_3283549 | Lighthouse Lab in Milton Keynes | Wellcome Sanger Institute for the COVID-19 Genomics UK (COG-UK) Consortium | The Lighthouse Lab in Milton Keynes and Alex Alderton, Roberto Amato, Jeffrey Barrett, Sonia Goncalves, Ewan Harrison, David K. Jackson, Ian Johnston, Dominic Kwiatkowski, Cordelia Langford, John Sillitoe on behalf of the Wellcome Sanger Institute COVID-19 Surveillance Team |
| EPI_ISL_3283550, EPI_ISL_3283551, EPI_ISL_3283552 | Lighthouse Lab in Alderley Park | Wellcome Sanger Institute for the COVID-19 Genomics UK (COG-UK) Consortium | Jacquelyn Wynn, Mairead Hyland, The Lighthouse Lab in Alderley Park and Alex Alderton, Roberto Amato, Jeffrey Barrett, Sonia Goncalves, Ewan Harrison, David K. Jackson, Ian Johnston, Dominic Kwiatkowski, Cordelia Langford, John Sillitoe on behalf of the Wellcome Sanger Institute COVID-19 Surveillance Team |
| EPI_ISL_3283553, EPI_ISL_3283554, EPI_ISL_3283555 | Lighthouse Lab in Milton Keynes | Wellcome Sanger Institute for the COVID-19 Genomics UK (COG-UK) Consortium | The Lighthouse Lab in Milton Keynes and Alex Alderton, Roberto Amato, Jeffrey Barrett, Sonia Goncalves, Ewan Harrison, David K. Jackson, Ian Johnston, Dominic Kwiatkowski, Cordelia Langford, John Sillitoe on behalf of the Wellcome Sanger Institute COVID-19 Surveillance Team |
| EPI_ISL_3283556 | Lighthouse Lab in Alderley Park | Wellcome Sanger Institute for the COVID-19 Genomics UK (COG-UK) Consortium | Jacquelyn Wynn, Mairead Hyland, The Lighthouse Lab in Alderley Park and Alex Alderton, Roberto Amato, Jeffrey Barrett, Sonia Goncalves, Ewan Harrison, David K. Jackson, Ian Johnston, Dominic Kwiatkowski, Cordelia Langford, John Sillitoe on behalf of the Wellcome Sanger Institute COVID-19 Surveillance Team |
| EPI_ISL_3283557 | Lighthouse Lab in Milton Keynes | Wellcome Sanger Institute for the COVID-19 Genomics UK (COG-UK) Consortium | The Lighthouse Lab in Milton Keynes and Alex Alderton, Roberto Amato, Jeffrey Barrett, Sonia Goncalves, Ewan Harrison, David K. Jackson, Ian Johnston, Dominic Kwiatkowski, Cordelia Langford, John Sillitoe on behalf of the Wellcome Sanger Institute COVID-19 Surveillance Team |
| EPI_ISL_3283558 | Lighthouse Lab in Alderley Park | Wellcome Sanger Institute for the COVID-19 Genomics UK (COG-UK) Consortium | Jacquelyn Wynn, Mairead Hyland, The Lighthouse Lab in Alderley Park and Alex Alderton, Roberto Amato, Jeffrey Barrett, Sonia Goncalves, Ewan Harrison, David K. Jackson, Ian Johnston, Dominic Kwiatkowski, Cordelia Langford, John Sillitoe on behalf of the Wellcome Sanger Institute COVID-19 Surveillance Team |
| EPI_ISL_3283559 | Lighthouse Lab in Milton Keynes | Wellcome Sanger Institute for the COVID-19 Genomics UK (COG-UK) Consortium | The Lighthouse Lab in Milton Keynes and Alex Alderton, Roberto Amato, Jeffrey Barrett, Sonia Goncalves, Ewan Harrison, David K. Jackson, Ian Johnston, Dominic Kwiatkowski, Cordelia Langford, John Sillitoe on behalf of the Wellcome Sanger Institute COVID-19 Surveillance Team |
| EPI_ISL_3283560, EPI_ISL_3283561 | Lighthouse Lab in Alderley Park | Wellcome Sanger Institute for the COVID-19 Genomics UK (COG-UK) Consortium | Jacquelyn Wynn, Mairead Hyland, The Lighthouse Lab in Alderley Park and Alex Alderton, Roberto Amato, Jeffrey Barrett, Sonia Goncalves, Ewan Harrison, David K. Jackson, Ian Johnston, Dominic Kwiatkowski, Cordelia Langford, John Sillitoe on behalf of the Wellcome Sanger Institute COVID-19 Surveillance Team |
| EPI_ISL_3283562, EPI_ISL_3283563 | Lighthouse Lab in Milton Keynes | Wellcome Sanger Institute for the COVID-19 Genomics UK (COG-UK) Consortium | The Lighthouse Lab in Milton Keynes and Alex Alderton, Roberto Amato, Jeffrey Barrett, Sonia Goncalves, Ewan Harrison, David K. Jackson, Ian Johnston, Dominic Kwiatkowski, Cordelia Langford, John Sillitoe on behalf of the Wellcome Sanger Institute COVID-19 Surveillance Team |
| EPI_ISL_3283564 | Lighthouse Lab in Alderley Park | Wellcome Sanger Institute for the COVID-19 Genomics UK (COG-UK) Consortium | Jacquelyn Wynn, Mairead Hyland, The Lighthouse Lab in Alderley Park and Alex Alderton, Roberto Amato, Jeffrey Barrett, Sonia Goncalves, Ewan Harrison, David K. Jackson, Ian Johnston, Dominic Kwiatkowski, Cordelia Langford, John Sillitoe on behalf of the Wellcome Sanger Institute COVID-19 Surveillance Team |
| EPI_ISL_3283565 | Lighthouse Lab in Milton Keynes | Wellcome Sanger Institute for the COVID-19 Genomics UK (COG-UK) Consortium | The Lighthouse Lab in Milton Keynes and Alex Alderton, Roberto Amato, Jeffrey Barrett, Sonia Goncalves, Ewan Harrison, David K. Jackson, Ian Johnston, Dominic Kwiatkowski, Cordelia Langford, John Sillitoe on behalf of the Wellcome Sanger Institute COVID-19 Surveillance Team |
| EPI_ISL_3283566, EPI_ISL_3283567, EPI_ISL_3283568, EPI_ISL_3283569, EPI_ISL_3283570, EPI_ISL_3283571, EPI_ISL_3283572, EPI_ISL_3283573 | Lighthouse Lab in Alderley Park | Wellcome Sanger Institute for the COVID-19 Genomics UK (COG-UK) Consortium | Jacquelyn Wynn, Mairead Hyland, The Lighthouse Lab in Alderley Park and Alex Alderton, Roberto Amato, Jeffrey Barrett, Sonia Goncalves, Ewan Harrison, David K. Jackson, Ian Johnston, Dominic Kwiatkowski, Cordelia Langford, John Sillitoe on behalf of the Wellcome Sanger Institute COVID-19 Surveillance Team |
| EPI_ISL_3283574 | Lighthouse Lab in Milton Keynes | Wellcome Sanger Institute for the COVID-19 Genomics UK (COG-UK) Consortium | The Lighthouse Lab in Milton Keynes and Alex Alderton, Roberto Amato, Jeffrey Barrett, Sonia Goncalves, Ewan Harrison, David K. Jackson, Ian Johnston, Dominic Kwiatkowski, Cordelia Langford, John Sillitoe on behalf of the Wellcome Sanger Institute COVID-19 Surveillance Team |
| EPI_ISL_3283575 | Lighthouse Lab in Alderley Park | Wellcome Sanger Institute for the COVID-19 Genomics UK (COG-UK) Consortium | Jacquelyn Wynn, Mairead Hyland, The Lighthouse Lab in Alderley Park and Alex Alderton, Roberto Amato, Jeffrey Barrett, Sonia Goncalves, Ewan Harrison, David K. Jackson, Ian Johnston, Dominic Kwiatkowski, Cordelia Langford, John Sillitoe on behalf of the Wellcome Sanger Institute COVID-19 Surveillance Team |
| EPI_ISL_3283576 | Lighthouse Lab in Milton Keynes | Wellcome Sanger Institute for the COVID-19 Genomics UK (COG-UK) Consortium | The Lighthouse Lab in Milton Keynes and Alex Alderton, Roberto Amato, Jeffrey Barrett, Sonia Goncalves, Ewan Harrison, David K. Jackson, Ian Johnston, Dominic Kwiatkowski, Cordelia Langford, John Sillitoe on behalf of the Wellcome Sanger Institute COVID-19 Surveillance Team |
| EPI_ISL_3283577, EPI_ISL_3283578 | Lighthouse Lab in Alderley Park | Wellcome Sanger Institute for the COVID-19 Genomics UK (COG-UK) Consortium | Jacquelyn Wynn, Mairead Hyland, The Lighthouse Lab in Alderley Park and Alex Alderton, Roberto Amato, Jeffrey Barrett, Sonia Goncalves, Ewan Harrison, David K. Jackson, Ian Johnston, Dominic Kwiatkowski, Cordelia Langford, John Sillitoe on behalf of the Wellcome Sanger Institute COVID-19 Surveillance Team |
| EPI_ISL_3283579 | Lighthouse Lab in Milton Keynes | Wellcome Sanger Institute for the COVID-19 Genomics UK (COG-UK) Consortium | The Lighthouse Lab in Milton Keynes and Alex Alderton, Roberto Amato, Jeffrey Barrett, Sonia Goncalves, Ewan Harrison, David K. Jackson, Ian Johnston, Dominic Kwiatkowski, Cordelia Langford, John Sillitoe on behalf of the Wellcome Sanger Institute COVID-19 Surveillance Team |
| EPI_ISL_3283580, EPI_ISL_3283581, EPI_ISL_3283582 | Lighthouse Lab in Alderley Park | Wellcome Sanger Institute for the COVID-19 Genomics UK (COG-UK) Consortium | Jacquelyn Wynn, Mairead Hyland, The Lighthouse Lab in Alderley Park and Alex Alderton, Roberto Amato, Jeffrey Barrett, Sonia Goncalves, Ewan Harrison, David K. Jackson, Ian Johnston, Dominic Kwiatkowski, Cordelia Langford, John Sillitoe on behalf of the Wellcome Sanger Institute COVID-19 Surveillance Team |
| EPI_ISL_3283583, EPI_ISL_3283584, EPI_ISL_3283585, EPI_ISL_3283586, EPI_ISL_3283587 | Lighthouse Lab in Milton Keynes | Wellcome Sanger Institute for the COVID-19 Genomics UK (COG-UK) Consortium | The Lighthouse Lab in Milton Keynes and Alex Alderton, Roberto Amato, Jeffrey Barrett, Sonia Goncalves, Ewan Harrison, David K. Jackson, Ian Johnston, Dominic Kwiatkowski, Cordelia Langford, John Sillitoe on behalf of the Wellcome Sanger Institute COVID-19 Surveillance Team |
| EPI_ISL_3283588, EPI_ISL_3283589, EPI_ISL_3283590, EPI_ISL_3283591, EPI_ISL_3283592 | Lighthouse Lab in Alderley Park | Wellcome Sanger Institute for the COVID-19 Genomics UK (COG-UK) Consortium | Jacquelyn Wynn, Mairead Hyland, The Lighthouse Lab in Alderley Park and Alex Alderton, Roberto Amato, Jeffrey Barrett, Sonia Goncalves, Ewan Harrison, David K. Jackson, Ian Johnston, Dominic Kwiatkowski, Cordelia Langford, John Sillitoe on behalf of the Wellcome Sanger Institute COVID-19 Surveillance Team |
| EPI_ISL_3283593 | Lighthouse Lab in Milton Keynes | Wellcome Sanger Institute for the COVID-19 Genomics UK (COG-UK) Consortium | The Lighthouse Lab in Milton Keynes and Alex Alderton, Roberto Amato, Jeffrey Barrett, Sonia Goncalves, Ewan Harrison, David K. Jackson, Ian Johnston, Dominic Kwiatkowski, Cordelia Langford, John Sillitoe on behalf of the Wellcome Sanger Institute COVID-19 Surveillance Team |
| EPI_ISL_3283594, EPI_ISL_3283595, | Lighthouse Lab in Alderley Park | Wellcome Sanger Institute for the COVID-19 Genomics UK (COG-UK) Consortium | Jacquelyn Wynn, Mairead Hyland, The Lighthouse Lab in Alderley Park and Alex Alderton, Roberto Amato, Jeffrey Barrett, Sonia Goncalves, Ewan Harrison, David K. Jackson, Ian Johnston, Dominic Kwiatkowski, Cordelia Langford, John Sillitoe on behalf of the Wellcome Sanger Institute COVID-19 Surveillance Team |

| | | | |
|---|---|---|---|
| EPI_ISL_3283596 | | | |
| EPI_ISL_3283597, EPI_ISL_3283598, EPI_ISL_3283599, EPI_ISL_3283600 | Lighthouse Lab in Milton Keynes | Wellcome Sanger Institute for the COVID-19 Genomics UK (COG-UK) Consortium | The Lighthouse Lab in Milton Keynes and Alex Alderton, Roberto Amato, Jeffrey Barrett, Sonia Goncalves, Ewan Harrison, David K. Jackson, Ian Johnston, Dominic Kwiatkowski, Cordelia Langford, John Sillitoe on behalf of the Wellcome Sanger Institute COVID-19 Surveillance Team |
| EPI_ISL_3283601, EPI_ISL_3283602 | Lighthouse Lab in Alderley Park | Wellcome Sanger Institute for the COVID-19 Genomics UK (COG-UK) Consortium | Jacquelyn Wynn, Mairead Hyland, The Lighthouse Lab in Alderley Park and Alex Alderton, Roberto Amato, Jeffrey Barrett, Sonia Goncalves, Ewan Harrison, David K. Jackson, Ian Johnston, Dominic Kwiatkowski, Cordelia Langford, John Sillitoe on behalf of the Wellcome Sanger Institute COVID-19 Surveillance Team |
| EPI_ISL_3283603, EPI_ISL_3283604 | Lighthouse Lab in Milton Keynes | Wellcome Sanger Institute for the COVID-19 Genomics UK (COG-UK) Consortium | The Lighthouse Lab in Milton Keynes and Alex Alderton, Roberto Amato, Jeffrey Barrett, Sonia Goncalves, Ewan Harrison, David K. Jackson, Ian Johnston, Dominic Kwiatkowski, Cordelia Langford, John Sillitoe on behalf of the Wellcome Sanger Institute COVID-19 Surveillance Team |
| EPI_ISL_3283605, EPI_ISL_3283606, EPI_ISL_3283607, EPI_ISL_3283608 | Lighthouse Lab in Alderley Park | Wellcome Sanger Institute for the COVID-19 Genomics UK (COG-UK) Consortium | Jacquelyn Wynn, Mairead Hyland, The Lighthouse Lab in Alderley Park and Alex Alderton, Roberto Amato, Jeffrey Barrett, Sonia Goncalves, Ewan Harrison, David K. Jackson, Ian Johnston, Dominic Kwiatkowski, Cordelia Langford, John Sillitoe on behalf of the Wellcome Sanger Institute COVID-19 Surveillance Team |
| EPI_ISL_3283609, EPI_ISL_3283610, EPI_ISL_3283611 | Lighthouse Lab in Milton Keynes | Wellcome Sanger Institute for the COVID-19 Genomics UK (COG-UK) Consortium | The Lighthouse Lab in Milton Keynes and Alex Alderton, Roberto Amato, Jeffrey Barrett, Sonia Goncalves, Ewan Harrison, David K. Jackson, Ian Johnston, Dominic Kwiatkowski, Cordelia Langford, John Sillitoe on behalf of the Wellcome Sanger Institute COVID-19 Surveillance Team |
| EPI_ISL_3283612, EPI_ISL_3283613 | Lighthouse Lab in Alderley Park | Wellcome Sanger Institute for the COVID-19 Genomics UK (COG-UK) Consortium | Jacquelyn Wynn, Mairead Hyland, The Lighthouse Lab in Alderley Park and Alex Alderton, Roberto Amato, Jeffrey Barrett, Sonia Goncalves, Ewan Harrison, David K. Jackson, Ian Johnston, Dominic Kwiatkowski, Cordelia Langford, John Sillitoe on behalf of the Wellcome Sanger Institute COVID-19 Surveillance Team |
| EPI_ISL_3283614, EPI_ISL_3283615, EPI_ISL_3283616, EPI_ISL_3283617 | Lighthouse Lab in Milton Keynes | Wellcome Sanger Institute for the COVID-19 Genomics UK (COG-UK) Consortium | The Lighthouse Lab in Milton Keynes and Alex Alderton, Roberto Amato, Jeffrey Barrett, Sonia Goncalves, Ewan Harrison, David K. Jackson, Ian Johnston, Dominic Kwiatkowski, Cordelia Langford, John Sillitoe on behalf of the Wellcome Sanger Institute COVID-19 Surveillance Team |
| EPI_ISL_3283618 | Lighthouse Lab in Alderley Park | Wellcome Sanger Institute for the COVID-19 Genomics UK (COG-UK) Consortium | Jacquelyn Wynn, Mairead Hyland, The Lighthouse Lab in Alderley Park and Alex Alderton, Roberto Amato, Jeffrey Barrett, Sonia Goncalves, Ewan Harrison, David K. Jackson, Ian Johnston, Dominic Kwiatkowski, Cordelia Langford, John Sillitoe on behalf of the Wellcome Sanger Institute COVID-19 Surveillance Team |
| EPI_ISL_3283619 | Lighthouse Lab in Milton Keynes | Wellcome Sanger Institute for the COVID-19 Genomics UK (COG-UK) Consortium | The Lighthouse Lab in Milton Keynes and Alex Alderton, Roberto Amato, Jeffrey Barrett, Sonia Goncalves, Ewan Harrison, David K. Jackson, Ian Johnston, Dominic Kwiatkowski, Cordelia Langford, John Sillitoe on behalf of the Wellcome Sanger Institute COVID-19 Surveillance Team |
| EPI_ISL_3283620, EPI_ISL_3283621, EPI_ISL_3283622 | Lighthouse Lab in Alderley Park | Wellcome Sanger Institute for the COVID-19 Genomics UK (COG-UK) Consortium | Jacquelyn Wynn, Mairead Hyland, The Lighthouse Lab in Alderley Park and Alex Alderton, Roberto Amato, Jeffrey Barrett, Sonia Goncalves, Ewan Harrison, David K. Jackson, Ian Johnston, Dominic Kwiatkowski, Cordelia Langford, John Sillitoe on behalf of the Wellcome Sanger Institute COVID-19 Surveillance Team |
| EPI_ISL_3283623 | Lighthouse Lab in Milton Keynes | Wellcome Sanger Institute for the COVID-19 Genomics UK (COG-UK) Consortium | The Lighthouse Lab in Milton Keynes and Alex Alderton, Roberto Amato, Jeffrey Barrett, Sonia Goncalves, Ewan Harrison, David K. Jackson, Ian Johnston, Dominic Kwiatkowski, Cordelia Langford, John Sillitoe on behalf of the Wellcome Sanger Institute COVID-19 Surveillance Team |
| EPI_ISL_3283624, EPI_ISL_3283625 | Lighthouse Lab in Alderley Park | Wellcome Sanger Institute for the COVID-19 Genomics UK (COG-UK) Consortium | Jacquelyn Wynn, Mairead Hyland, The Lighthouse Lab in Alderley Park and Alex Alderton, Roberto Amato, Jeffrey Barrett, Sonia Goncalves, Ewan Harrison, David K. Jackson, Ian Johnston, Dominic Kwiatkowski, Cordelia Langford, John Sillitoe on behalf of the Wellcome Sanger Institute COVID-19 Surveillance Team |
| EPI_ISL_3283626, EPI_ISL_3283627 | Lighthouse Lab in Milton Keynes | Wellcome Sanger Institute for the COVID-19 Genomics UK (COG-UK) Consortium | The Lighthouse Lab in Milton Keynes and Alex Alderton, Roberto Amato, Jeffrey Barrett, Sonia Goncalves, Ewan Harrison, David K. Jackson, Ian Johnston, Dominic Kwiatkowski, Cordelia Langford, John Sillitoe on behalf of the Wellcome Sanger Institute COVID-19 Surveillance Team |
| EPI_ISL_3283628, EPI_ISL_3283629 | Lighthouse Lab in Alderley Park | Wellcome Sanger Institute for the COVID-19 Genomics UK (COG-UK) Consortium | Jacquelyn Wynn, Mairead Hyland, The Lighthouse Lab in Alderley Park and Alex Alderton, Roberto Amato, Jeffrey Barrett, Sonia Goncalves, Ewan Harrison, David K. Jackson, Ian Johnston, Dominic Kwiatkowski, Cordelia Langford, John Sillitoe on behalf of the Wellcome Sanger Institute COVID-19 Surveillance Team |
| EPI_ISL_3283630 | Lighthouse Lab in Milton Keynes | Wellcome Sanger Institute for the COVID-19 Genomics UK (COG-UK) Consortium | The Lighthouse Lab in Milton Keynes and Alex Alderton, Roberto Amato, Jeffrey Barrett, Sonia Goncalves, Ewan Harrison, David K. Jackson, Ian Johnston, Dominic Kwiatkowski, Cordelia Langford, John Sillitoe on behalf of the Wellcome Sanger Institute COVID-19 Surveillance Team |
| EPI_ISL_3283631 | Lighthouse Lab in Glasgow | Wellcome Sanger Institute for the COVID-19 Genomics UK (COG-UK) Consortium | Harper VanSteenhouse, Yumi Kasai, David Gray, Carol Clugston, Anna Dominiczak and Alex Alderton, Roberto Amato, Jeffrey Barrett, Sonia Goncalves, Ewan Harrison, David K. Jackson, Ian Johnston, Dominic Kwiatkowski, Cordelia Langford, John Sillitoe on behalf of the Wellcome Sanger Institute COVID-19 Surveillance Team |
| EPI_ISL_3283632, EPI_ISL_3283633, EPI_ISL_3283634, EPI_ISL_3283635 | Lighthouse Lab in Milton Keynes | Wellcome Sanger Institute for the COVID-19 Genomics UK (COG-UK) Consortium | The Lighthouse Lab in Milton Keynes and Alex Alderton, Roberto Amato, Jeffrey Barrett, Sonia Goncalves, Ewan Harrison, David K. Jackson, Ian Johnston, Dominic Kwiatkowski, Cordelia Langford, John Sillitoe on behalf of the Wellcome Sanger Institute COVID-19 Surveillance Team |
| EPI_ISL_3283636 | Lighthouse Lab in Glasgow | Wellcome Sanger Institute for the COVID-19 Genomics UK (COG-UK) Consortium | Harper VanSteenhouse, Yumi Kasai, David Gray, Carol Clugston, Anna Dominiczak and Alex Alderton, Roberto Amato, Jeffrey Barrett, Sonia Goncalves, Ewan Harrison, David K. Jackson, Ian Johnston, Dominic Kwiatkowski, Cordelia Langford, John Sillitoe on behalf of the Wellcome Sanger Institute COVID-19 Surveillance Team |
| EPI_ISL_3283637 | Lighthouse Lab in Milton Keynes | Wellcome Sanger Institute for the COVID-19 Genomics UK (COG-UK) Consortium | The Lighthouse Lab in Milton Keynes and Alex Alderton, Roberto Amato, Jeffrey Barrett, Sonia Goncalves, Ewan Harrison, David K. Jackson, Ian Johnston, Dominic Kwiatkowski, Cordelia Langford, John Sillitoe on behalf of the Wellcome Sanger Institute COVID-19 Surveillance Team |
| EPI_ISL_3283638 | Lighthouse Lab in Glasgow | Wellcome Sanger Institute for the COVID-19 Genomics UK (COG-UK) Consortium | Harper VanSteenhouse, Yumi Kasai, David Gray, Carol Clugston, Anna Dominiczak and Alex Alderton, Roberto Amato, Jeffrey Barrett, Sonia Goncalves, Ewan Harrison, David K. Jackson, Ian Johnston, Dominic Kwiatkowski, Cordelia Langford, John Sillitoe on behalf of the Wellcome Sanger Institute COVID-19 Surveillance Team |
| EPI_ISL_3283639, EPI_ISL_3283640 | Lighthouse Lab in Milton Keynes | Wellcome Sanger Institute for the COVID-19 Genomics UK (COG-UK) Consortium | The Lighthouse Lab in Milton Keynes and Alex Alderton, Roberto Amato, Jeffrey Barrett, Sonia Goncalves, Ewan Harrison, David K. Jackson, Ian Johnston, Dominic Kwiatkowski, Cordelia Langford, John Sillitoe on behalf of the Wellcome Sanger Institute COVID-19 Surveillance Team |
| EPI_ISL_3283641 | Lighthouse Lab in Glasgow | Wellcome Sanger Institute for the COVID-19 Genomics UK (COG-UK) Consortium | Harper VanSteenhouse, Yumi Kasai, David Gray, Carol Clugston, Anna Dominiczak and Alex Alderton, Roberto Amato, Jeffrey Barrett, Sonia Goncalves, Ewan Harrison, David K. Jackson, Ian Johnston, Dominic Kwiatkowski, Cordelia Langford, John Sillitoe on behalf of the Wellcome Sanger Institute COVID-19 Surveillance Team |
| EPI_ISL_3283642, EPI_ISL_3283643, EPI_ISL_3283644 | Lighthouse Lab in Milton Keynes | Wellcome Sanger Institute for the COVID-19 Genomics UK (COG-UK) Consortium | The Lighthouse Lab in Milton Keynes and Alex Alderton, Roberto Amato, Jeffrey Barrett, Sonia Goncalves, Ewan Harrison, David K. Jackson, Ian Johnston, Dominic Kwiatkowski, Cordelia Langford, John Sillitoe on behalf of the Wellcome Sanger Institute COVID-19 Surveillance Team |
| EPI_ISL_3283645, EPI_ISL_3283646 | Lighthouse Lab in Glasgow | Wellcome Sanger Institute for the COVID-19 Genomics UK (COG-UK) Consortium | Harper VanSteenhouse, Yumi Kasai, David Gray, Carol Clugston, Anna Dominiczak and Alex Alderton, Roberto Amato, Jeffrey Barrett, Sonia Goncalves, Ewan Harrison, David K. Jackson, Ian Johnston, Dominic Kwiatkowski, Cordelia Langford, John Sillitoe on behalf of the Wellcome Sanger Institute COVID-19 Surveillance Team |
| EPI_ISL_3283647 | Lighthouse Lab in Milton Keynes | Wellcome Sanger Institute for the COVID-19 Genomics UK (COG-UK) Consortium | The Lighthouse Lab in Milton Keynes and Alex Alderton, Roberto Amato, Jeffrey Barrett, Sonia Goncalves, Ewan Harrison, David K. Jackson, Ian Johnston, Dominic Kwiatkowski, Cordelia Langford, John Sillitoe on behalf of the Wellcome Sanger Institute COVID-19 Surveillance Team |
| EPI_ISL_3283648, EPI_ISL_3283649 | Lighthouse Lab in Glasgow | Wellcome Sanger Institute for the COVID-19 Genomics UK (COG-UK) Consortium | Harper VanSteenhouse, Yumi Kasai, David Gray, Carol Clugston, Anna Dominiczak and Alex Alderton, Roberto Amato, Jeffrey Barrett, Sonia Goncalves, Ewan Harrison, David K. Jackson, Ian Johnston, Dominic Kwiatkowski, Cordelia Langford, John Sillitoe on behalf of the Wellcome Sanger Institute COVID-19 Surveillance Team |
| EPI_ISL_3283650 | Lighthouse Lab in Milton Keynes | Wellcome Sanger Institute for the COVID-19 Genomics UK (COG-UK) Consortium | The Lighthouse Lab in Milton Keynes and Alex Alderton, Roberto Amato, Jeffrey Barrett, Sonia Goncalves, Ewan Harrison, David K. Jackson, Ian Johnston, Dominic Kwiatkowski, Cordelia Langford, John Sillitoe on behalf of the Wellcome Sanger Institute COVID-19 Surveillance Team |
| EPI_ISL_3283651 | Lighthouse Lab in Glasgow | Wellcome Sanger Institute for the COVID-19 Genomics UK (COG-UK) Consortium | Harper VanSteenhouse, Yumi Kasai, David Gray, Carol Clugston, Anna Dominiczak and Alex Alderton, Roberto Amato, Jeffrey Barrett, Sonia Goncalves, Ewan Harrison, David K. Jackson, Ian Johnston, Dominic Kwiatkowski, Cordelia Langford, John Sillitoe on behalf of the Wellcome Sanger Institute COVID-19 Surveillance Team |
| EPI_ISL_3283652, EPI_ISL_3283653, EPI_ISL_3283654, EPI_ISL_3283655, EPI_ISL_3283656 | Lighthouse Lab in Milton Keynes | Wellcome Sanger Institute for the COVID-19 Genomics UK (COG-UK) Consortium | The Lighthouse Lab in Milton Keynes and Alex Alderton, Roberto Amato, Jeffrey Barrett, Sonia Goncalves, Ewan Harrison, David K. Jackson, Ian Johnston, Dominic Kwiatkowski, Cordelia Langford, John Sillitoe on behalf of the Wellcome Sanger Institute COVID-19 Surveillance Team |
| EPI_ISL_3283657 | Lighthouse Lab in Glasgow | Wellcome Sanger Institute for the COVID-19 Genomics UK (COG-UK) Consortium | Harper VanSteenhouse, Yumi Kasai, David Gray, Carol Clugston, Anna Dominiczak and Alex Alderton, Roberto Amato, Jeffrey Barrett, Sonia Goncalves, Ewan Harrison, David K. Jackson, Ian Johnston, Dominic Kwiatkowski, Cordelia Langford, John Sillitoe on behalf of the Wellcome Sanger Institute COVID-19 Surveillance Team |
| EPI_ISL_3283658, EPI_ISL_3283659, EPI_ISL_3283660, EPI_ISL_3283661 | Lighthouse Lab in Milton Keynes | Wellcome Sanger Institute for the COVID-19 Genomics UK (COG-UK) Consortium | The Lighthouse Lab in Milton Keynes and Alex Alderton, Roberto Amato, Jeffrey Barrett, Sonia Goncalves, Ewan Harrison, David K. Jackson, Ian Johnston, Dominic Kwiatkowski, Cordelia Langford, John Sillitoe on behalf of the Wellcome Sanger Institute COVID-19 Surveillance Team |
| EPI_ISL_3283662 | Lighthouse Lab in Glasgow | Wellcome Sanger Institute for the COVID-19 Genomics UK (COG-UK) Consortium | Harper VanSteenhouse, Yumi Kasai, David Gray, Carol Clugston, Anna Dominiczak and Alex Alderton, Roberto Amato, Jeffrey Barrett, Sonia Goncalves, Ewan Harrison, David K. Jackson, Ian Johnston, Dominic Kwiatkowski, Cordelia Langford, John Sillitoe on behalf of the Wellcome Sanger Institute COVID-19 Surveillance Team |
| EPI_ISL_3283663, EPI_ISL_3283664 | Lighthouse Lab in Milton Keynes | Wellcome Sanger Institute for the COVID-19 Genomics UK (COG-UK) Consortium | The Lighthouse Lab in Milton Keynes and Alex Alderton, Roberto Amato, Jeffrey Barrett, Sonia Goncalves, Ewan Harrison, David K. Jackson, Ian Johnston, Dominic Kwiatkowski, Cordelia Langford, John Sillitoe on behalf of the Wellcome Sanger Institute COVID-19 Surveillance Team |
| EPI_ISL_3283665, EPI_ISL_3283666 | Lighthouse Lab in Glasgow | Wellcome Sanger Institute for the COVID-19 Genomics UK (COG-UK) Consortium | Harper VanSteenhouse, Yumi Kasai, David Gray, Carol Clugston, Anna Dominiczak and Alex Alderton, Roberto Amato, Jeffrey Barrett, Sonia Goncalves, Ewan Harrison, David K. Jackson, Ian Johnston, Dominic Kwiatkowski, Cordelia Langford, John Sillitoe on behalf of the Wellcome Sanger Institute COVID-19 Surveillance Team |
| EPI_ISL_3283667, EPI_ISL_3283668 | Lighthouse Lab in Milton Keynes | Wellcome Sanger Institute for the COVID-19 Genomics UK (COG-UK) Consortium | The Lighthouse Lab in Milton Keynes and Alex Alderton, Roberto Amato, Jeffrey Barrett, Sonia Goncalves, Ewan Harrison, David K. Jackson, Ian Johnston, Dominic Kwiatkowski, Cordelia Langford, John Sillitoe on behalf of the Wellcome Sanger Institute COVID-19 Surveillance Team |
| EPI_ISL_3283669, EPI_ISL_3283670 | Lighthouse Lab in Glasgow | Wellcome Sanger Institute for the COVID-19 Genomics UK (COG-UK) Consortium | Harper VanSteenhouse, Yumi Kasai, David Gray, Carol Clugston, Anna Dominiczak and Alex Alderton, Roberto Amato, Jeffrey Barrett, Sonia Goncalves, Ewan Harrison, David K. Jackson, Ian Johnston, Dominic Kwiatkowski, Cordelia Langford, John Sillitoe on behalf of the Wellcome Sanger Institute COVID-19 Surveillance Team |
| EPI_ISL_3283671, EPI_ISL_3283672, EPI_ISL_3283673 | Lighthouse Lab in Milton Keynes | Wellcome Sanger Institute for the COVID-19 Genomics UK (COG-UK) Consortium | The Lighthouse Lab in Milton Keynes and Alex Alderton, Roberto Amato, Jeffrey Barrett, Sonia Goncalves, Ewan Harrison, David K. Jackson, Ian Johnston, Dominic Kwiatkowski, Cordelia Langford, John Sillitoe on behalf of the Wellcome Sanger Institute COVID-19 Surveillance Team |
| EPI_ISL_3283674 | Lighthouse Lab in Glasgow | Wellcome Sanger Institute for the COVID-19 Genomics UK (COG-UK) | Harper VanSteenhouse, Yumi Kasai, David Gray, Carol Clugston, Anna Dominiczak and Alex Alderton, Roberto Amato, Jeffrey Barrett, Sonia Goncalves, Ewan Harrison, David K. Jackson, Ian Johnston, Dominic Kwiatkowski, Cordelia Langford, |

| | | | |
|---|---|---|---|
| | | Consortium | John Sillitoe on behalf of the Wellcome Sanger Institute COVID-19 Surveillance Team |
| EPI_ISL_3283675, EPI_ISL_3283676 | Lighthouse Lab in Milton Keynes | Wellcome Sanger Institute for the COVID-19 Genomics UK (COG-UK) Consortium | The Lighthouse Lab in Milton Keynes and Alex Alderton, Roberto Amato, Jeffrey Barrett, Sonia Goncalves, Ewan Harrison, David K. Jackson, Ian Johnston, Dominic Kwiatkowski, Cordelia Langford, John Sillitoe on behalf of the Wellcome Sanger Institute COVID-19 Surveillance Team |
| EPI_ISL_3283677 | Lighthouse Lab in Glasgow | Wellcome Sanger Institute for the COVID-19 Genomics UK (COG-UK) Consortium | Harper VanSteenhouse, Yumi Kasai, David Gray, Carol Clugston, Anna Dominiczak and Alex Alderton, Roberto Amato, Jeffrey Barrett, Sonia Goncalves, Ewan Harrison, David K. Jackson, Ian Johnston, Dominic Kwiatkowski, Cordelia Langford, John Sillitoe on behalf of the Wellcome Sanger Institute COVID-19 Surveillance Team |
| EPI_ISL_3283678, EPI_ISL_3283679 | Lighthouse Lab in Milton Keynes | Wellcome Sanger Institute for the COVID-19 Genomics UK (COG-UK) Consortium | The Lighthouse Lab in Milton Keynes and Alex Alderton, Roberto Amato, Jeffrey Barrett, Sonia Goncalves, Ewan Harrison, David K. Jackson, Ian Johnston, Dominic Kwiatkowski, Cordelia Langford, John Sillitoe on behalf of the Wellcome Sanger Institute COVID-19 Surveillance Team |
| EPI_ISL_3283680 | Lighthouse Lab in Alderley Park | Wellcome Sanger Institute for the COVID-19 Genomics UK (COG-UK) Consortium | Jacquelyn Wynn, Mairead Hyland, The Lighthouse Lab in Alderley Park and Alex Alderton, Roberto Amato, Jeffrey Barrett, Sonia Goncalves, Ewan Harrison, David K. Jackson, Ian Johnston, Dominic Kwiatkowski, Cordelia Langford, John Sillitoe on behalf of the Wellcome Sanger Institute COVID-19 Surveillance Team |
| EPI_ISL_3283681, EPI_ISL_3283682, EPI_ISL_3283683 | Lighthouse Lab in Milton Keynes | Wellcome Sanger Institute for the COVID-19 Genomics UK (COG-UK) Consortium | The Lighthouse Lab in Milton Keynes and Alex Alderton, Roberto Amato, Jeffrey Barrett, Sonia Goncalves, Ewan Harrison, David K. Jackson, Ian Johnston, Dominic Kwiatkowski, Cordelia Langford, John Sillitoe on behalf of the Wellcome Sanger Institute COVID-19 Surveillance Team |
| EPI_ISL_3283684, EPI_ISL_3283685 | Lighthouse Lab in Glasgow | Wellcome Sanger Institute for the COVID-19 Genomics UK (COG-UK) Consortium | Harper VanSteenhouse, Yumi Kasai, David Gray, Carol Clugston, Anna Dominiczak and Alex Alderton, Roberto Amato, Jeffrey Barrett, Sonia Goncalves, Ewan Harrison, David K. Jackson, Ian Johnston, Dominic Kwiatkowski, Cordelia Langford, John Sillitoe on behalf of the Wellcome Sanger Institute COVID-19 Surveillance Team |
| EPI_ISL_3283686 | Lighthouse Lab in Milton Keynes | Wellcome Sanger Institute for the COVID-19 Genomics UK (COG-UK) Consortium | The Lighthouse Lab in Milton Keynes and Alex Alderton, Roberto Amato, Jeffrey Barrett, Sonia Goncalves, Ewan Harrison, David K. Jackson, Ian Johnston, Dominic Kwiatkowski, Cordelia Langford, John Sillitoe on behalf of the Wellcome Sanger Institute COVID-19 Surveillance Team |
| EPI_ISL_3283687 | Lighthouse Lab in Glasgow | Wellcome Sanger Institute for the COVID-19 Genomics UK (COG-UK) Consortium | Harper VanSteenhouse, Yumi Kasai, David Gray, Carol Clugston, Anna Dominiczak and Alex Alderton, Roberto Amato, Jeffrey Barrett, Sonia Goncalves, Ewan Harrison, David K. Jackson, Ian Johnston, Dominic Kwiatkowski, Cordelia Langford, John Sillitoe on behalf of the Wellcome Sanger Institute COVID-19 Surveillance Team |
| EPI_ISL_3283688 | Lighthouse Lab in Milton Keynes | Wellcome Sanger Institute for the COVID-19 Genomics UK (COG-UK) Consortium | The Lighthouse Lab in Milton Keynes and Alex Alderton, Roberto Amato, Jeffrey Barrett, Sonia Goncalves, Ewan Harrison, David K. Jackson, Ian Johnston, Dominic Kwiatkowski, Cordelia Langford, John Sillitoe on behalf of the Wellcome Sanger Institute COVID-19 Surveillance Team |
| EPI_ISL_3283689 | Lighthouse Lab in Glasgow | Wellcome Sanger Institute for the COVID-19 Genomics UK (COG-UK) Consortium | Harper VanSteenhouse, Yumi Kasai, David Gray, Carol Clugston, Anna Dominiczak and Alex Alderton, Roberto Amato, Jeffrey Barrett, Sonia Goncalves, Ewan Harrison, David K. Jackson, Ian Johnston, Dominic Kwiatkowski, Cordelia Langford, John Sillitoe on behalf of the Wellcome Sanger Institute COVID-19 Surveillance Team |
| EPI_ISL_3283690 | Lighthouse Lab in Milton Keynes | Wellcome Sanger Institute for the COVID-19 Genomics UK (COG-UK) Consortium | The Lighthouse Lab in Milton Keynes and Alex Alderton, Roberto Amato, Jeffrey Barrett, Sonia Goncalves, Ewan Harrison, David K. Jackson, Ian Johnston, Dominic Kwiatkowski, Cordelia Langford, John Sillitoe on behalf of the Wellcome Sanger Institute COVID-19 Surveillance Team |
| EPI_ISL_3283691, EPI_ISL_3283692, EPI_ISL_3283693, EPI_ISL_3283694 | Lighthouse Lab in Glasgow | Wellcome Sanger Institute for the COVID-19 Genomics UK (COG-UK) Consortium | Harper VanSteenhouse, Yumi Kasai, David Gray, Carol Clugston, Anna Dominiczak and Alex Alderton, Roberto Amato, Jeffrey Barrett, Sonia Goncalves, Ewan Harrison, David K. Jackson, Ian Johnston, Dominic Kwiatkowski, Cordelia Langford, John Sillitoe on behalf of the Wellcome Sanger Institute COVID-19 Surveillance Team |
| EPI_ISL_3283695 | Lighthouse Lab in Milton Keynes | Wellcome Sanger Institute for the COVID-19 Genomics UK (COG-UK) Consortium | The Lighthouse Lab in Milton Keynes and Alex Alderton, Roberto Amato, Jeffrey Barrett, Sonia Goncalves, Ewan Harrison, David K. Jackson, Ian Johnston, Dominic Kwiatkowski, Cordelia Langford, John Sillitoe on behalf of the Wellcome Sanger Institute COVID-19 Surveillance Team |
| EPI_ISL_3283696, EPI_ISL_3283697, EPI_ISL_3283698 | Lighthouse Lab in Glasgow | Wellcome Sanger Institute for the COVID-19 Genomics UK (COG-UK) Consortium | Harper VanSteenhouse, Yumi Kasai, David Gray, Carol Clugston, Anna Dominiczak and Alex Alderton, Roberto Amato, Jeffrey Barrett, Sonia Goncalves, Ewan Harrison, David K. Jackson, Ian Johnston, Dominic Kwiatkowski, Cordelia Langford, John Sillitoe on behalf of the Wellcome Sanger Institute COVID-19 Surveillance Team |
| EPI_ISL_3283699, EPI_ISL_3283700, EPI_ISL_3283701 | Lighthouse Lab in Milton Keynes | Wellcome Sanger Institute for the COVID-19 Genomics UK (COG-UK) Consortium | The Lighthouse Lab in Milton Keynes and Alex Alderton, Roberto Amato, Jeffrey Barrett, Sonia Goncalves, Ewan Harrison, David K. Jackson, Ian Johnston, Dominic Kwiatkowski, Cordelia Langford, John Sillitoe on behalf of the Wellcome Sanger Institute COVID-19 Surveillance Team |
| EPI_ISL_3283702 | Lighthouse Lab in Glasgow | Wellcome Sanger Institute for the COVID-19 Genomics UK (COG-UK) Consortium | Harper VanSteenhouse, Yumi Kasai, David Gray, Carol Clugston, Anna Dominiczak and Alex Alderton, Roberto Amato, Jeffrey Barrett, Sonia Goncalves, Ewan Harrison, David K. Jackson, Ian Johnston, Dominic Kwiatkowski, Cordelia Langford, John Sillitoe on behalf of the Wellcome Sanger Institute COVID-19 Surveillance Team |
| EPI_ISL_3283703, EPI_ISL_3283704, EPI_ISL_3283705, EPI_ISL_3283706 | Lighthouse Lab in Milton Keynes | Wellcome Sanger Institute for the COVID-19 Genomics UK (COG-UK) Consortium | The Lighthouse Lab in Milton Keynes and Alex Alderton, Roberto Amato, Jeffrey Barrett, Sonia Goncalves, Ewan Harrison, David K. Jackson, Ian Johnston, Dominic Kwiatkowski, Cordelia Langford, John Sillitoe on behalf of the Wellcome Sanger Institute COVID-19 Surveillance Team |
| EPI_ISL_3283707 | Lighthouse Lab in Glasgow | Wellcome Sanger Institute for the COVID-19 Genomics UK (COG-UK) Consortium | Harper VanSteenhouse, Yumi Kasai, David Gray, Carol Clugston, Anna Dominiczak and Alex Alderton, Roberto Amato, Jeffrey Barrett, Sonia Goncalves, Ewan Harrison, David K. Jackson, Ian Johnston, Dominic Kwiatkowski, Cordelia Langford, John Sillitoe on behalf of the Wellcome Sanger Institute COVID-19 Surveillance Team |
| EPI_ISL_3283708 | Lighthouse Lab in Milton Keynes | Wellcome Sanger Institute for the COVID-19 Genomics UK (COG-UK) Consortium | The Lighthouse Lab in Milton Keynes and Alex Alderton, Roberto Amato, Jeffrey Barrett, Sonia Goncalves, Ewan Harrison, David K. Jackson, Ian Johnston, Dominic Kwiatkowski, Cordelia Langford, John Sillitoe on behalf of the Wellcome Sanger Institute COVID-19 Surveillance Team |
| EPI_ISL_3283709 | Lighthouse Lab in Glasgow | Wellcome Sanger Institute for the COVID-19 Genomics UK (COG-UK) Consortium | Harper VanSteenhouse, Yumi Kasai, David Gray, Carol Clugston, Anna Dominiczak and Alex Alderton, Roberto Amato, Jeffrey Barrett, Sonia Goncalves, Ewan Harrison, David K. Jackson, Ian Johnston, Dominic Kwiatkowski, Cordelia Langford, John Sillitoe on behalf of the Wellcome Sanger Institute COVID-19 Surveillance Team |
| EPI_ISL_3283710, EPI_ISL_3283711 | Lighthouse Lab in Milton Keynes | Wellcome Sanger Institute for the COVID-19 Genomics UK (COG-UK) Consortium | The Lighthouse Lab in Milton Keynes and Alex Alderton, Roberto Amato, Jeffrey Barrett, Sonia Goncalves, Ewan Harrison, David K. Jackson, Ian Johnston, Dominic Kwiatkowski, Cordelia Langford, John Sillitoe on behalf of the Wellcome Sanger Institute COVID-19 Surveillance Team |
| EPI_ISL_3283712 | Lighthouse Lab in Glasgow | Wellcome Sanger Institute for the COVID-19 Genomics UK (COG-UK) Consortium | Harper VanSteenhouse, Yumi Kasai, David Gray, Carol Clugston, Anna Dominiczak and Alex Alderton, Roberto Amato, Jeffrey Barrett, Sonia Goncalves, Ewan Harrison, David K. Jackson, Ian Johnston, Dominic Kwiatkowski, Cordelia Langford, John Sillitoe on behalf of the Wellcome Sanger Institute COVID-19 Surveillance Team |
| EPI_ISL_3283713, EPI_ISL_3283714 | Lighthouse Lab in Milton Keynes | Wellcome Sanger Institute for the COVID-19 Genomics UK (COG-UK) Consortium | The Lighthouse Lab in Milton Keynes and Alex Alderton, Roberto Amato, Jeffrey Barrett, Sonia Goncalves, Ewan Harrison, David K. Jackson, Ian Johnston, Dominic Kwiatkowski, Cordelia Langford, John Sillitoe on behalf of the Wellcome Sanger Institute COVID-19 Surveillance Team |
| EPI_ISL_3283715 | Lighthouse Lab in Glasgow | Wellcome Sanger Institute for the COVID-19 Genomics UK (COG-UK) Consortium | Harper VanSteenhouse, Yumi Kasai, David Gray, Carol Clugston, Anna Dominiczak and Alex Alderton, Roberto Amato, Jeffrey Barrett, Sonia Goncalves, Ewan Harrison, David K. Jackson, Ian Johnston, Dominic Kwiatkowski, Cordelia Langford, John Sillitoe on behalf of the Wellcome Sanger Institute COVID-19 Surveillance Team |
| EPI_ISL_3283716, EPI_ISL_3283717, EPI_ISL_3283718, EPI_ISL_3283719, EPI_ISL_3283720 | Lighthouse Lab in Milton Keynes | Wellcome Sanger Institute for the COVID-19 Genomics UK (COG-UK) Consortium | The Lighthouse Lab in Milton Keynes and Alex Alderton, Roberto Amato, Jeffrey Barrett, Sonia Goncalves, Ewan Harrison, David K. Jackson, Ian Johnston, Dominic Kwiatkowski, Cordelia Langford, John Sillitoe on behalf of the Wellcome Sanger Institute COVID-19 Surveillance Team |
| EPI_ISL_3283721 | Lighthouse Lab in Alderley Park | Wellcome Sanger Institute for the COVID-19 Genomics UK (COG-UK) Consortium | Jacquelyn Wynn, Mairead Hyland, The Lighthouse Lab in Alderley Park and Alex Alderton, Roberto Amato, Jeffrey Barrett, Sonia Goncalves, Ewan Harrison, David K. Jackson, Ian Johnston, Dominic Kwiatkowski, Cordelia Langford, John Sillitoe on behalf of the Wellcome Sanger Institute COVID-19 Surveillance Team |
| EPI_ISL_3283722 | Lighthouse Lab in Glasgow | Wellcome Sanger Institute for the COVID-19 Genomics UK (COG-UK) Consortium | Harper VanSteenhouse, Yumi Kasai, David Gray, Carol Clugston, Anna Dominiczak and Alex Alderton, Roberto Amato, Jeffrey Barrett, Sonia Goncalves, Ewan Harrison, David K. Jackson, Ian Johnston, Dominic Kwiatkowski, Cordelia Langford, John Sillitoe on behalf of the Wellcome Sanger Institute COVID-19 Surveillance Team |
| EPI_ISL_3283723 | Lighthouse Lab in Milton Keynes | Wellcome Sanger Institute for the COVID-19 Genomics UK (COG-UK) Consortium | The Lighthouse Lab in Milton Keynes and Alex Alderton, Roberto Amato, Jeffrey Barrett, Sonia Goncalves, Ewan Harrison, David K. Jackson, Ian Johnston, Dominic Kwiatkowski, Cordelia Langford, John Sillitoe on behalf of the Wellcome Sanger Institute COVID-19 Surveillance Team |
| EPI_ISL_3283724, EPI_ISL_3283725 | Lighthouse Lab in Glasgow | Wellcome Sanger Institute for the COVID-19 Genomics UK (COG-UK) Consortium | Harper VanSteenhouse, Yumi Kasai, David Gray, Carol Clugston, Anna Dominiczak and Alex Alderton, Roberto Amato, Jeffrey Barrett, Sonia Goncalves, Ewan Harrison, David K. Jackson, Ian Johnston, Dominic Kwiatkowski, Cordelia Langford, John Sillitoe on behalf of the Wellcome Sanger Institute COVID-19 Surveillance Team |
| EPI_ISL_3283726, EPI_ISL_3283727, EPI_ISL_3283728, EPI_ISL_3283729, EPI_ISL_3283730, EPI_ISL_3283731 | Lighthouse Lab in Milton Keynes | Wellcome Sanger Institute for the COVID-19 Genomics UK (COG-UK) Consortium | The Lighthouse Lab in Milton Keynes and Alex Alderton, Roberto Amato, Jeffrey Barrett, Sonia Goncalves, Ewan Harrison, David K. Jackson, Ian Johnston, Dominic Kwiatkowski, Cordelia Langford, John Sillitoe on behalf of the Wellcome Sanger Institute COVID-19 Surveillance Team |
| EPI_ISL_3283732 | Lighthouse Lab in Glasgow | Wellcome Sanger Institute for the COVID-19 Genomics UK (COG-UK) Consortium | Harper VanSteenhouse, Yumi Kasai, David Gray, Carol Clugston, Anna Dominiczak and Alex Alderton, Roberto Amato, Jeffrey Barrett, Sonia Goncalves, Ewan Harrison, David K. Jackson, Ian Johnston, Dominic Kwiatkowski, Cordelia Langford, John Sillitoe on behalf of the Wellcome Sanger Institute COVID-19 Surveillance Team |
| EPI_ISL_3283733, EPI_ISL_3283734, EPI_ISL_3283735, EPI_ISL_3283736, EPI_ISL_3283737 | Lighthouse Lab in Milton Keynes | Wellcome Sanger Institute for the COVID-19 Genomics UK (COG-UK) Consortium | The Lighthouse Lab in Milton Keynes and Alex Alderton, Roberto Amato, Jeffrey Barrett, Sonia Goncalves, Ewan Harrison, David K. Jackson, Ian Johnston, Dominic Kwiatkowski, Cordelia Langford, John Sillitoe on behalf of the Wellcome Sanger Institute COVID-19 Surveillance Team |
| EPI_ISL_3283738 | Lighthouse Lab in Glasgow | Wellcome Sanger Institute for the COVID-19 Genomics UK (COG-UK) Consortium | Harper VanSteenhouse, Yumi Kasai, David Gray, Carol Clugston, Anna Dominiczak and Alex Alderton, Roberto Amato, Jeffrey Barrett, Sonia Goncalves, Ewan Harrison, David K. Jackson, Ian Johnston, Dominic Kwiatkowski, Cordelia Langford, John Sillitoe on behalf of the Wellcome Sanger Institute COVID-19 Surveillance Team |
| EPI_ISL_3283739, EPI_ISL_3283740, EPI_ISL_3283741 | Lighthouse Lab in Milton Keynes | Wellcome Sanger Institute for the COVID-19 Genomics UK (COG-UK) Consortium | The Lighthouse Lab in Milton Keynes and Alex Alderton, Roberto Amato, Jeffrey Barrett, Sonia Goncalves, Ewan Harrison, David K. Jackson, Ian Johnston, Dominic Kwiatkowski, Cordelia Langford, John Sillitoe on behalf of the Wellcome Sanger Institute COVID-19 Surveillance Team |
| EPI_ISL_3283742, EPI_ISL_3283743, EPI_ISL_3283744, EPI_ISL_3283745 | Lighthouse Lab in Glasgow | Wellcome Sanger Institute for the COVID-19 Genomics UK (COG-UK) Consortium | Harper VanSteenhouse, Yumi Kasai, David Gray, Carol Clugston, Anna Dominiczak and Alex Alderton, Roberto Amato, Jeffrey Barrett, Sonia Goncalves, Ewan Harrison, David K. Jackson, Ian Johnston, Dominic Kwiatkowski, Cordelia Langford, John Sillitoe on behalf of the Wellcome Sanger Institute COVID-19 Surveillance Team |
| EPI_ISL_3283746, EPI_ISL_3283747 | Lighthouse Lab in Milton Keynes | Wellcome Sanger Institute for the COVID-19 Genomics UK (COG-UK) Consortium | The Lighthouse Lab in Milton Keynes and Alex Alderton, Roberto Amato, Jeffrey Barrett, Sonia Goncalves, Ewan Harrison, David K. Jackson, Ian Johnston, Dominic Kwiatkowski, Cordelia Langford, John Sillitoe on behalf of the Wellcome Sanger Institute COVID-19 Surveillance Team |
| EPI_ISL_3283748, | Lighthouse Laboratory Plymouth | Wellcome Sanger Institute for the COVID-19 Genomics UK (COG-UK) | Lighthouse Laboratory Plymouth and Alex Alderton, Roberto Amato, Jeffrey Barrett, Sonia Goncalves, Ewan Harrison, David K. Jackson, Ian Johnston, Dominic Kwiatkowski, Cordelia Langford, John Sillitoe on behalf of the Wellcome Sanger |

| | | | |
|---|---|---|---|
| EPI_ISL_3283749, EPI_ISL_3283750, EPI_ISL_3283751, EPI_ISL_3283752 | | Consortium | Institute COVID-19 Surveillance Team |
| EPI_ISL_3283753 | Berkshire and Surrey Pathology Services Lighthouse Laboratory | Wellcome Sanger Institute for the COVID-19 Genomics UK (COG-UK) Consortium | Berkshire and Surrey Pathology Services Lighthouse Laboratory and Alex Alderton, Roberto Amato, Jeffrey Barrett, Sonia Goncalves, Ewan Harrison, David K. Jackson, Ian Johnston, Dominic Kwiatkowski, Cordelia Langford, John Sillitoe on behalf of the Wellcome Sanger Institute COVID-19 Surveillance Team |
| EPI_ISL_3283754 | Lighthouse Laboratory Plymouth | Wellcome Sanger Institute for the COVID-19 Genomics UK (COG-UK) Consortium | Lighthouse Laboratory Plymouth and Alex Alderton, Roberto Amato, Jeffrey Barrett, Sonia Goncalves, Ewan Harrison, David K. Jackson, Ian Johnston, Dominic Kwiatkowski, Cordelia Langford, John Sillitoe on behalf of the Wellcome Sanger Institute COVID-19 Surveillance Team |
| EPI_ISL_3283755 | Berkshire and Surrey Pathology Services Lighthouse Laboratory | Wellcome Sanger Institute for the COVID-19 Genomics UK (COG-UK) Consortium | Berkshire and Surrey Pathology Services Lighthouse Laboratory and Alex Alderton, Roberto Amato, Jeffrey Barrett, Sonia Goncalves, Ewan Harrison, David K. Jackson, Ian Johnston, Dominic Kwiatkowski, Cordelia Langford, John Sillitoe on behalf of the Wellcome Sanger Institute COVID-19 Surveillance Team |
| EPI_ISL_3283756 | Lighthouse Laboratory Plymouth | Wellcome Sanger Institute for the COVID-19 Genomics UK (COG-UK) Consortium | Lighthouse Laboratory Plymouth and Alex Alderton, Roberto Amato, Jeffrey Barrett, Sonia Goncalves, Ewan Harrison, David K. Jackson, Ian Johnston, Dominic Kwiatkowski, Cordelia Langford, John Sillitoe on behalf of the Wellcome Sanger Institute COVID-19 Surveillance Team |
| EPI_ISL_3283757 | Berkshire and Surrey Pathology Services Lighthouse Laboratory | Wellcome Sanger Institute for the COVID-19 Genomics UK (COG-UK) Consortium | Berkshire and Surrey Pathology Services Lighthouse Laboratory and Alex Alderton, Roberto Amato, Jeffrey Barrett, Sonia Goncalves, Ewan Harrison, David K. Jackson, Ian Johnston, Dominic Kwiatkowski, Cordelia Langford, John Sillitoe on behalf of the Wellcome Sanger Institute COVID-19 Surveillance Team |
| EPI_ISL_3283758, EPI_ISL_3283759, EPI_ISL_3283760, EPI_ISL_3283761 | Lighthouse Laboratory Plymouth | Wellcome Sanger Institute for the COVID-19 Genomics UK (COG-UK) Consortium | Lighthouse Laboratory Plymouth and Alex Alderton, Roberto Amato, Jeffrey Barrett, Sonia Goncalves, Ewan Harrison, David K. Jackson, Ian Johnston, Dominic Kwiatkowski, Cordelia Langford, John Sillitoe on behalf of the Wellcome Sanger Institute COVID-19 Surveillance Team |
| EPI_ISL_3283762 | Berkshire and Surrey Pathology Services Lighthouse Laboratory | Wellcome Sanger Institute for the COVID-19 Genomics UK (COG-UK) Consortium | Berkshire and Surrey Pathology Services Lighthouse Laboratory and Alex Alderton, Roberto Amato, Jeffrey Barrett, Sonia Goncalves, Ewan Harrison, David K. Jackson, Ian Johnston, Dominic Kwiatkowski, Cordelia Langford, John Sillitoe on behalf of the Wellcome Sanger Institute COVID-19 Surveillance Team |
| EPI_ISL_3283763, EPI_ISL_3283764, EPI_ISL_3283765, EPI_ISL_3283766 | Lighthouse Laboratory Plymouth | Wellcome Sanger Institute for the COVID-19 Genomics UK (COG-UK) Consortium | Lighthouse Laboratory Plymouth and Alex Alderton, Roberto Amato, Jeffrey Barrett, Sonia Goncalves, Ewan Harrison, David K. Jackson, Ian Johnston, Dominic Kwiatkowski, Cordelia Langford, John Sillitoe on behalf of the Wellcome Sanger Institute COVID-19 Surveillance Team |
| EPI_ISL_3283767 | Berkshire and Surrey Pathology Services Lighthouse Laboratory | Wellcome Sanger Institute for the COVID-19 Genomics UK (COG-UK) Consortium | Berkshire and Surrey Pathology Services Lighthouse Laboratory and Alex Alderton, Roberto Amato, Jeffrey Barrett, Sonia Goncalves, Ewan Harrison, David K. Jackson, Ian Johnston, Dominic Kwiatkowski, Cordelia Langford, John Sillitoe on behalf of the Wellcome Sanger Institute COVID-19 Surveillance Team |
| EPI_ISL_3283768 | Lighthouse Laboratory Plymouth | Wellcome Sanger Institute for the COVID-19 Genomics UK (COG-UK) Consortium | Lighthouse Laboratory Plymouth and Alex Alderton, Roberto Amato, Jeffrey Barrett, Sonia Goncalves, Ewan Harrison, David K. Jackson, Ian Johnston, Dominic Kwiatkowski, Cordelia Langford, John Sillitoe on behalf of the Wellcome Sanger Institute COVID-19 Surveillance Team |
| EPI_ISL_3283769 | Berkshire and Surrey Pathology Services Lighthouse Laboratory | Wellcome Sanger Institute for the COVID-19 Genomics UK (COG-UK) Consortium | Berkshire and Surrey Pathology Services Lighthouse Laboratory and Alex Alderton, Roberto Amato, Jeffrey Barrett, Sonia Goncalves, Ewan Harrison, David K. Jackson, Ian Johnston, Dominic Kwiatkowski, Cordelia Langford, John Sillitoe on behalf of the Wellcome Sanger Institute COVID-19 Surveillance Team |
| EPI_ISL_3283770, EPI_ISL_3283771 | Lighthouse Laboratory Plymouth | Wellcome Sanger Institute for the COVID-19 Genomics UK (COG-UK) Consortium | Lighthouse Laboratory Plymouth and Alex Alderton, Roberto Amato, Jeffrey Barrett, Sonia Goncalves, Ewan Harrison, David K. Jackson, Ian Johnston, Dominic Kwiatkowski, Cordelia Langford, John Sillitoe on behalf of the Wellcome Sanger Institute COVID-19 Surveillance Team |
| EPI_ISL_3283772 | Berkshire and Surrey Pathology Services Lighthouse Laboratory | Wellcome Sanger Institute for the COVID-19 Genomics UK (COG-UK) Consortium | Berkshire and Surrey Pathology Services Lighthouse Laboratory and Alex Alderton, Roberto Amato, Jeffrey Barrett, Sonia Goncalves, Ewan Harrison, David K. Jackson, Ian Johnston, Dominic Kwiatkowski, Cordelia Langford, John Sillitoe on behalf of the Wellcome Sanger Institute COVID-19 Surveillance Team |
| EPI_ISL_3283773, EPI_ISL_3283774, EPI_ISL_3283775, EPI_ISL_3283776, EPI_ISL_3283777, EPI_ISL_3283778, EPI_ISL_3283779, EPI_ISL_3283780, EPI_ISL_3283781, EPI_ISL_3283782, EPI_ISL_3283783 | | | |
| see above | Lighthouse Laboratory Plymouth | Wellcome Sanger Institute for the COVID-19 Genomics UK (COG-UK) Consortium | Lighthouse Laboratory Plymouth and Alex Alderton, Roberto Amato, Jeffrey Barrett, Sonia Goncalves, Ewan Harrison, David K. Jackson, Ian Johnston, Dominic Kwiatkowski, Cordelia Langford, John Sillitoe on behalf of the Wellcome Sanger Institute COVID-19 Surveillance Team |
| EPI_ISL_3283784 | Berkshire and Surrey Pathology Services Lighthouse Laboratory | Wellcome Sanger Institute for the COVID-19 Genomics UK (COG-UK) Consortium | Berkshire and Surrey Pathology Services Lighthouse Laboratory and Alex Alderton, Roberto Amato, Jeffrey Barrett, Sonia Goncalves, Ewan Harrison, David K. Jackson, Ian Johnston, Dominic Kwiatkowski, Cordelia Langford, John Sillitoe on behalf of the Wellcome Sanger Institute COVID-19 Surveillance Team |
| EPI_ISL_3283785, EPI_ISL_3283786, EPI_ISL_3283787, EPI_ISL_3283788 | Lighthouse Laboratory Plymouth | Wellcome Sanger Institute for the COVID-19 Genomics UK (COG-UK) Consortium | Lighthouse Laboratory Plymouth and Alex Alderton, Roberto Amato, Jeffrey Barrett, Sonia Goncalves, Ewan Harrison, David K. Jackson, Ian Johnston, Dominic Kwiatkowski, Cordelia Langford, John Sillitoe on behalf of the Wellcome Sanger Institute COVID-19 Surveillance Team |
| EPI_ISL_3283789, EPI_ISL_3283790 | Berkshire and Surrey Pathology Services Lighthouse Laboratory | Wellcome Sanger Institute for the COVID-19 Genomics UK (COG-UK) Consortium | Berkshire and Surrey Pathology Services Lighthouse Laboratory and Alex Alderton, Roberto Amato, Jeffrey Barrett, Sonia Goncalves, Ewan Harrison, David K. Jackson, Ian Johnston, Dominic Kwiatkowski, Cordelia Langford, John Sillitoe on behalf of the Wellcome Sanger Institute COVID-19 Surveillance Team |
| EPI_ISL_3283791, EPI_ISL_3283792, EPI_ISL_3283793, EPI_ISL_3283794, EPI_ISL_3283795, EPI_ISL_3283796 | Lighthouse Laboratory Plymouth | Wellcome Sanger Institute for the COVID-19 Genomics UK (COG-UK) Consortium | Lighthouse Laboratory Plymouth and Alex Alderton, Roberto Amato, Jeffrey Barrett, Sonia Goncalves, Ewan Harrison, David K. Jackson, Ian Johnston, Dominic Kwiatkowski, Cordelia Langford, John Sillitoe on behalf of the Wellcome Sanger Institute COVID-19 Surveillance Team |
| EPI_ISL_3283797 | Berkshire and Surrey Pathology Services Lighthouse Laboratory | Wellcome Sanger Institute for the COVID-19 Genomics UK (COG-UK) Consortium | Berkshire and Surrey Pathology Services Lighthouse Laboratory and Alex Alderton, Roberto Amato, Jeffrey Barrett, Sonia Goncalves, Ewan Harrison, David K. Jackson, Ian Johnston, Dominic Kwiatkowski, Cordelia Langford, John Sillitoe on behalf of the Wellcome Sanger Institute COVID-19 Surveillance Team |
| EPI_ISL_3283798, EPI_ISL_3283799 | Lighthouse Laboratory Plymouth | Wellcome Sanger Institute for the COVID-19 Genomics UK (COG-UK) Consortium | Lighthouse Laboratory Plymouth and Alex Alderton, Roberto Amato, Jeffrey Barrett, Sonia Goncalves, Ewan Harrison, David K. Jackson, Ian Johnston, Dominic Kwiatkowski, Cordelia Langford, John Sillitoe on behalf of the Wellcome Sanger Institute COVID-19 Surveillance Team |
| EPI_ISL_3283800 | Berkshire and Surrey Pathology Services Lighthouse Laboratory | Wellcome Sanger Institute for the COVID-19 Genomics UK (COG-UK) Consortium | Berkshire and Surrey Pathology Services Lighthouse Laboratory and Alex Alderton, Roberto Amato, Jeffrey Barrett, Sonia Goncalves, Ewan Harrison, David K. Jackson, Ian Johnston, Dominic Kwiatkowski, Cordelia Langford, John Sillitoe on behalf of the Wellcome Sanger Institute COVID-19 Surveillance Team |
| EPI_ISL_3283801, EPI_ISL_3283802, EPI_ISL_3283803, EPI_ISL_3283804, EPI_ISL_3283805, EPI_ISL_3283806, EPI_ISL_3283807, EPI_ISL_3283808 | Lighthouse Laboratory Plymouth | Wellcome Sanger Institute for the COVID-19 Genomics UK (COG-UK) Consortium | Lighthouse Laboratory Plymouth and Alex Alderton, Roberto Amato, Jeffrey Barrett, Sonia Goncalves, Ewan Harrison, David K. Jackson, Ian Johnston, Dominic Kwiatkowski, Cordelia Langford, John Sillitoe on behalf of the Wellcome Sanger Institute COVID-19 Surveillance Team |
| EPI_ISL_3283809 | Berkshire and Surrey Pathology Services Lighthouse Laboratory | Wellcome Sanger Institute for the COVID-19 Genomics UK (COG-UK) Consortium | Berkshire and Surrey Pathology Services Lighthouse Laboratory and Alex Alderton, Roberto Amato, Jeffrey Barrett, Sonia Goncalves, Ewan Harrison, David K. Jackson, Ian Johnston, Dominic Kwiatkowski, Cordelia Langford, John Sillitoe on behalf of the Wellcome Sanger Institute COVID-19 Surveillance Team |
| EPI_ISL_3283810 | Lighthouse Laboratory Plymouth | Wellcome Sanger Institute for the COVID-19 Genomics UK (COG-UK) Consortium | Lighthouse Laboratory Plymouth and Alex Alderton, Roberto Amato, Jeffrey Barrett, Sonia Goncalves, Ewan Harrison, David K. Jackson, Ian Johnston, Dominic Kwiatkowski, Cordelia Langford, John Sillitoe on behalf of the Wellcome Sanger Institute COVID-19 Surveillance Team |
| EPI_ISL_3283811, EPI_ISL_3283812, EPI_ISL_3283813 | Berkshire and Surrey Pathology Services Lighthouse Laboratory | Wellcome Sanger Institute for the COVID-19 Genomics UK (COG-UK) Consortium | Berkshire and Surrey Pathology Services Lighthouse Laboratory and Alex Alderton, Roberto Amato, Jeffrey Barrett, Sonia Goncalves, Ewan Harrison, David K. Jackson, Ian Johnston, Dominic Kwiatkowski, Cordelia Langford, John Sillitoe on behalf of the Wellcome Sanger Institute COVID-19 Surveillance Team |
| EPI_ISL_3283814, EPI_ISL_3283815, EPI_ISL_3283816, EPI_ISL_3283817, EPI_ISL_3283818, EPI_ISL_3283819 | Lighthouse Laboratory Plymouth | Wellcome Sanger Institute for the COVID-19 Genomics UK (COG-UK) Consortium | Lighthouse Laboratory Plymouth and Alex Alderton, Roberto Amato, Jeffrey Barrett, Sonia Goncalves, Ewan Harrison, David K. Jackson, Ian Johnston, Dominic Kwiatkowski, Cordelia Langford, John Sillitoe on behalf of the Wellcome Sanger Institute COVID-19 Surveillance Team |
| EPI_ISL_3283820 | Berkshire and Surrey Pathology Services Lighthouse Laboratory | Wellcome Sanger Institute for the COVID-19 Genomics UK (COG-UK) Consortium | Berkshire and Surrey Pathology Services Lighthouse Laboratory and Alex Alderton, Roberto Amato, Jeffrey Barrett, Sonia Goncalves, Ewan Harrison, David K. Jackson, Ian Johnston, Dominic Kwiatkowski, Cordelia Langford, John Sillitoe on behalf of the Wellcome Sanger Institute COVID-19 Surveillance Team |
| EPI_ISL_3283821, EPI_ISL_3283822, EPI_ISL_3283823 | Lighthouse Laboratory Plymouth | Wellcome Sanger Institute for the COVID-19 Genomics UK (COG-UK) Consortium | Lighthouse Laboratory Plymouth and Alex Alderton, Roberto Amato, Jeffrey Barrett, Sonia Goncalves, Ewan Harrison, David K. Jackson, Ian Johnston, Dominic Kwiatkowski, Cordelia Langford, John Sillitoe on behalf of the Wellcome Sanger Institute COVID-19 Surveillance Team |
| EPI_ISL_3283824 | Berkshire and Surrey Pathology Services Lighthouse Laboratory | Wellcome Sanger Institute for the COVID-19 Genomics UK (COG-UK) Consortium | Berkshire and Surrey Pathology Services Lighthouse Laboratory and Alex Alderton, Roberto Amato, Jeffrey Barrett, Sonia Goncalves, Ewan Harrison, David K. Jackson, Ian Johnston, Dominic Kwiatkowski, Cordelia Langford, John Sillitoe on behalf of the Wellcome Sanger Institute COVID-19 Surveillance Team |
| EPI_ISL_3283825 | Lighthouse Laboratory Plymouth | Wellcome Sanger Institute for the COVID-19 Genomics UK (COG-UK) Consortium | Lighthouse Laboratory Plymouth and Alex Alderton, Roberto Amato, Jeffrey Barrett, Sonia Goncalves, Ewan Harrison, David K. Jackson, Ian Johnston, Dominic Kwiatkowski, Cordelia Langford, John Sillitoe on behalf of the Wellcome Sanger Institute COVID-19 Surveillance Team |
| EPI_ISL_3283826 | Berkshire and Surrey Pathology Services Lighthouse Laboratory | Wellcome Sanger Institute for the COVID-19 Genomics UK (COG-UK) Consortium | Berkshire and Surrey Pathology Services Lighthouse Laboratory and Alex Alderton, Roberto Amato, Jeffrey Barrett, Sonia Goncalves, Ewan Harrison, David K. Jackson, Ian Johnston, Dominic Kwiatkowski, Cordelia Langford, John Sillitoe on behalf of the Wellcome Sanger Institute COVID-19 Surveillance Team |
| EPI_ISL_3283827 | Lighthouse Laboratory Plymouth | Wellcome Sanger Institute for the COVID-19 Genomics UK (COG-UK) Consortium | Lighthouse Laboratory Plymouth and Alex Alderton, Roberto Amato, Jeffrey Barrett, Sonia Goncalves, Ewan Harrison, David K. Jackson, Ian Johnston, Dominic Kwiatkowski, Cordelia Langford, John Sillitoe on behalf of the Wellcome Sanger Institute COVID-19 Surveillance Team |
| EPI_ISL_3283828 | Berkshire and Surrey Pathology Services Lighthouse Laboratory | Wellcome Sanger Institute for the COVID-19 Genomics UK (COG-UK) Consortium | Berkshire and Surrey Pathology Services Lighthouse Laboratory and Alex Alderton, Roberto Amato, Jeffrey Barrett, Sonia Goncalves, Ewan Harrison, David K. Jackson, Ian Johnston, Dominic Kwiatkowski, Cordelia Langford, John Sillitoe on behalf of the Wellcome Sanger Institute COVID-19 Surveillance Team |

| | | | |
|---|---|---|---|
| EPI_ISL_3283829, EPI_ISL_3283830, EPI_ISL_3283831, EPI_ISL_3283832, EPI_ISL_3283833, EPI_ISL_3283834 | Lighthouse Laboratory Plymouth | Wellcome Sanger Institute for the COVID-19 Genomics UK (COG-UK) Consortium | Lighthouse Laboratory Plymouth and Alex Alderton, Roberto Amato, Jeffrey Barrett, Sonia Goncalves, Ewan Harrison, David K. Jackson, Ian Johnston, Dominic Kwiatkowski, Cordelia Langford, John Sillitoe on behalf of the Wellcome Sanger Institute COVID-19 Surveillance Team |
| EPI_ISL_3283835 | Berkshire and Surrey Pathology Services Lighthouse Laboratory | Wellcome Sanger Institute for the COVID-19 Genomics UK (COG-UK) Consortium | Berkshire and Surrey Pathology Services Lighthouse Laboratory and Alex Alderton, Roberto Amato, Jeffrey Barrett, Sonia Goncalves, Ewan Harrison, David K. Jackson, Ian Johnston, Dominic Kwiatkowski, Cordelia Langford, John Sillitoe on behalf of the Wellcome Sanger Institute COVID-19 Surveillance Team |
| EPI_ISL_3283836, EPI_ISL_3283837, EPI_ISL_3283838, EPI_ISL_3283839, EPI_ISL_3283840, EPI_ISL_3283841, EPI_ISL_3283842, EPI_ISL_3283843, EPI_ISL_3283844, EPI_ISL_3283845, EPI_ISL_3283846, EPI_ISL_3283847 | | | |
| see above | Lighthouse Laboratory Plymouth | Wellcome Sanger Institute for the COVID-19 Genomics UK (COG-UK) Consortium | Lighthouse Laboratory Plymouth and Alex Alderton, Roberto Amato, Jeffrey Barrett, Sonia Goncalves, Ewan Harrison, David K. Jackson, Ian Johnston, Dominic Kwiatkowski, Cordelia Langford, John Sillitoe on behalf of the Wellcome Sanger Institute COVID-19 Surveillance Team |
| EPI_ISL_3283848, EPI_ISL_3283849, EPI_ISL_3283850 | Berkshire and Surrey Pathology Services Lighthouse Laboratory | Wellcome Sanger Institute for the COVID-19 Genomics UK (COG-UK) Consortium | Berkshire and Surrey Pathology Services Lighthouse Laboratory and Alex Alderton, Roberto Amato, Jeffrey Barrett, Sonia Goncalves, Ewan Harrison, David K. Jackson, Ian Johnston, Dominic Kwiatkowski, Cordelia Langford, John Sillitoe on behalf of the Wellcome Sanger Institute COVID-19 Surveillance Team |
| EPI_ISL_3283851, EPI_ISL_3283852 | Lighthouse Laboratory Plymouth | Wellcome Sanger Institute for the COVID-19 Genomics UK (COG-UK) Consortium | Lighthouse Laboratory Plymouth and Alex Alderton, Roberto Amato, Jeffrey Barrett, Sonia Goncalves, Ewan Harrison, David K. Jackson, Ian Johnston, Dominic Kwiatkowski, Cordelia Langford, John Sillitoe on behalf of the Wellcome Sanger Institute COVID-19 Surveillance Team |
| EPI_ISL_3283853 | Berkshire and Surrey Pathology Services Lighthouse Laboratory | Wellcome Sanger Institute for the COVID-19 Genomics UK (COG-UK) Consortium | Berkshire and Surrey Pathology Services Lighthouse Laboratory and Alex Alderton, Roberto Amato, Jeffrey Barrett, Sonia Goncalves, Ewan Harrison, David K. Jackson, Ian Johnston, Dominic Kwiatkowski, Cordelia Langford, John Sillitoe on behalf of the Wellcome Sanger Institute COVID-19 Surveillance Team |
| EPI_ISL_3283854, EPI_ISL_3283855 | Lighthouse Laboratory Plymouth | Wellcome Sanger Institute for the COVID-19 Genomics UK (COG-UK) Consortium | Lighthouse Laboratory Plymouth and Alex Alderton, Roberto Amato, Jeffrey Barrett, Sonia Goncalves, Ewan Harrison, David K. Jackson, Ian Johnston, Dominic Kwiatkowski, Cordelia Langford, John Sillitoe on behalf of the Wellcome Sanger Institute COVID-19 Surveillance Team |
| EPI_ISL_3283856, EPI_ISL_3283857 | Berkshire and Surrey Pathology Services Lighthouse Laboratory | Wellcome Sanger Institute for the COVID-19 Genomics UK (COG-UK) Consortium | Berkshire and Surrey Pathology Services Lighthouse Laboratory and Alex Alderton, Roberto Amato, Jeffrey Barrett, Sonia Goncalves, Ewan Harrison, David K. Jackson, Ian Johnston, Dominic Kwiatkowski, Cordelia Langford, John Sillitoe on behalf of the Wellcome Sanger Institute COVID-19 Surveillance Team |
| EPI_ISL_3283858 | Lighthouse Laboratory Plymouth | Wellcome Sanger Institute for the COVID-19 Genomics UK (COG-UK) Consortium | Lighthouse Laboratory Plymouth and Alex Alderton, Roberto Amato, Jeffrey Barrett, Sonia Goncalves, Ewan Harrison, David K. Jackson, Ian Johnston, Dominic Kwiatkowski, Cordelia Langford, John Sillitoe on behalf of the Wellcome Sanger Institute COVID-19 Surveillance Team |
| EPI_ISL_3283859 | Berkshire and Surrey Pathology Services Lighthouse Laboratory | Wellcome Sanger Institute for the COVID-19 Genomics UK (COG-UK) Consortium | Berkshire and Surrey Pathology Services Lighthouse Laboratory and Alex Alderton, Roberto Amato, Jeffrey Barrett, Sonia Goncalves, Ewan Harrison, David K. Jackson, Ian Johnston, Dominic Kwiatkowski, Cordelia Langford, John Sillitoe on behalf of the Wellcome Sanger Institute COVID-19 Surveillance Team |
| EPI_ISL_3283860, EPI_ISL_3283861 | Lighthouse Laboratory Plymouth | Wellcome Sanger Institute for the COVID-19 Genomics UK (COG-UK) Consortium | Lighthouse Laboratory Plymouth and Alex Alderton, Roberto Amato, Jeffrey Barrett, Sonia Goncalves, Ewan Harrison, David K. Jackson, Ian Johnston, Dominic Kwiatkowski, Cordelia Langford, John Sillitoe on behalf of the Wellcome Sanger Institute COVID-19 Surveillance Team |
| EPI_ISL_3283862 | Berkshire and Surrey Pathology Services Lighthouse Laboratory | Wellcome Sanger Institute for the COVID-19 Genomics UK (COG-UK) Consortium | Berkshire and Surrey Pathology Services Lighthouse Laboratory and Alex Alderton, Roberto Amato, Jeffrey Barrett, Sonia Goncalves, Ewan Harrison, David K. Jackson, Ian Johnston, Dominic Kwiatkowski, Cordelia Langford, John Sillitoe on behalf of the Wellcome Sanger Institute COVID-19 Surveillance Team |
| EPI_ISL_3283863, EPI_ISL_3283864, EPI_ISL_3283865, EPI_ISL_3283866, EPI_ISL_3283867 | Lighthouse Laboratory Plymouth | Wellcome Sanger Institute for the COVID-19 Genomics UK (COG-UK) Consortium | Lighthouse Laboratory Plymouth and Alex Alderton, Roberto Amato, Jeffrey Barrett, Sonia Goncalves, Ewan Harrison, David K. Jackson, Ian Johnston, Dominic Kwiatkowski, Cordelia Langford, John Sillitoe on behalf of the Wellcome Sanger Institute COVID-19 Surveillance Team |
| EPI_ISL_3283868 | Berkshire and Surrey Pathology Services Lighthouse Laboratory | Wellcome Sanger Institute for the COVID-19 Genomics UK (COG-UK) Consortium | Berkshire and Surrey Pathology Services Lighthouse Laboratory and Alex Alderton, Roberto Amato, Jeffrey Barrett, Sonia Goncalves, Ewan Harrison, David K. Jackson, Ian Johnston, Dominic Kwiatkowski, Cordelia Langford, John Sillitoe on behalf of the Wellcome Sanger Institute COVID-19 Surveillance Team |
| EPI_ISL_3283869, EPI_ISL_3283870 | Lighthouse Laboratory Plymouth | Wellcome Sanger Institute for the COVID-19 Genomics UK (COG-UK) Consortium | Lighthouse Laboratory Plymouth and Alex Alderton, Roberto Amato, Jeffrey Barrett, Sonia Goncalves, Ewan Harrison, David K. Jackson, Ian Johnston, Dominic Kwiatkowski, Cordelia Langford, John Sillitoe on behalf of the Wellcome Sanger Institute COVID-19 Surveillance Team |
| EPI_ISL_3283871 | Berkshire and Surrey Pathology Services Lighthouse Laboratory | Wellcome Sanger Institute for the COVID-19 Genomics UK (COG-UK) Consortium | Berkshire and Surrey Pathology Services Lighthouse Laboratory and Alex Alderton, Roberto Amato, Jeffrey Barrett, Sonia Goncalves, Ewan Harrison, David K. Jackson, Ian Johnston, Dominic Kwiatkowski, Cordelia Langford, John Sillitoe on behalf of the Wellcome Sanger Institute COVID-19 Surveillance Team |
| EPI_ISL_3283872 | Lighthouse Laboratory Plymouth | Wellcome Sanger Institute for the COVID-19 Genomics UK (COG-UK) Consortium | Lighthouse Laboratory Plymouth and Alex Alderton, Roberto Amato, Jeffrey Barrett, Sonia Goncalves, Ewan Harrison, David K. Jackson, Ian Johnston, Dominic Kwiatkowski, Cordelia Langford, John Sillitoe on behalf of the Wellcome Sanger Institute COVID-19 Surveillance Team |
| EPI_ISL_3283873 | Berkshire and Surrey Pathology Services Lighthouse Laboratory | Wellcome Sanger Institute for the COVID-19 Genomics UK (COG-UK) Consortium | Berkshire and Surrey Pathology Services Lighthouse Laboratory and Alex Alderton, Roberto Amato, Jeffrey Barrett, Sonia Goncalves, Ewan Harrison, David K. Jackson, Ian Johnston, Dominic Kwiatkowski, Cordelia Langford, John Sillitoe on behalf of the Wellcome Sanger Institute COVID-19 Surveillance Team |
| EPI_ISL_3283874, EPI_ISL_3283875, EPI_ISL_3283876 | Lighthouse Laboratory Plymouth | Wellcome Sanger Institute for the COVID-19 Genomics UK (COG-UK) Consortium | Lighthouse Laboratory Plymouth and Alex Alderton, Roberto Amato, Jeffrey Barrett, Sonia Goncalves, Ewan Harrison, David K. Jackson, Ian Johnston, Dominic Kwiatkowski, Cordelia Langford, John Sillitoe on behalf of the Wellcome Sanger Institute COVID-19 Surveillance Team |
| EPI_ISL_3283877 | Berkshire and Surrey Pathology Services Lighthouse Laboratory | Wellcome Sanger Institute for the COVID-19 Genomics UK (COG-UK) Consortium | Berkshire and Surrey Pathology Services Lighthouse Laboratory and Alex Alderton, Roberto Amato, Jeffrey Barrett, Sonia Goncalves, Ewan Harrison, David K. Jackson, Ian Johnston, Dominic Kwiatkowski, Cordelia Langford, John Sillitoe on behalf of the Wellcome Sanger Institute COVID-19 Surveillance Team |
| EPI_ISL_3283878, EPI_ISL_3283879, EPI_ISL_3283880, EPI_ISL_3283881, EPI_ISL_3283882, EPI_ISL_3283883, EPI_ISL_3283884 | Lighthouse Laboratory Plymouth | Wellcome Sanger Institute for the COVID-19 Genomics UK (COG-UK) Consortium | Lighthouse Laboratory Plymouth and Alex Alderton, Roberto Amato, Jeffrey Barrett, Sonia Goncalves, Ewan Harrison, David K. Jackson, Ian Johnston, Dominic Kwiatkowski, Cordelia Langford, John Sillitoe on behalf of the Wellcome Sanger Institute COVID-19 Surveillance Team |
| EPI_ISL_3283885 | Berkshire and Surrey Pathology Services Lighthouse Laboratory | Wellcome Sanger Institute for the COVID-19 Genomics UK (COG-UK) Consortium | Berkshire and Surrey Pathology Services Lighthouse Laboratory and Alex Alderton, Roberto Amato, Jeffrey Barrett, Sonia Goncalves, Ewan Harrison, David K. Jackson, Ian Johnston, Dominic Kwiatkowski, Cordelia Langford, John Sillitoe on behalf of the Wellcome Sanger Institute COVID-19 Surveillance Team |
| EPI_ISL_3283886, EPI_ISL_3283887, EPI_ISL_3283888 | Lighthouse Laboratory Plymouth | Wellcome Sanger Institute for the COVID-19 Genomics UK (COG-UK) Consortium | Lighthouse Laboratory Plymouth and Alex Alderton, Roberto Amato, Jeffrey Barrett, Sonia Goncalves, Ewan Harrison, David K. Jackson, Ian Johnston, Dominic Kwiatkowski, Cordelia Langford, John Sillitoe on behalf of the Wellcome Sanger Institute COVID-19 Surveillance Team |
| EPI_ISL_3283889 | Berkshire and Surrey Pathology Services Lighthouse Laboratory | Wellcome Sanger Institute for the COVID-19 Genomics UK (COG-UK) Consortium | Berkshire and Surrey Pathology Services Lighthouse Laboratory and Alex Alderton, Roberto Amato, Jeffrey Barrett, Sonia Goncalves, Ewan Harrison, David K. Jackson, Ian Johnston, Dominic Kwiatkowski, Cordelia Langford, John Sillitoe on behalf of the Wellcome Sanger Institute COVID-19 Surveillance Team |
| EPI_ISL_3283890 | Lighthouse Laboratory Plymouth | Wellcome Sanger Institute for the COVID-19 Genomics UK (COG-UK) Consortium | Lighthouse Laboratory Plymouth and Alex Alderton, Roberto Amato, Jeffrey Barrett, Sonia Goncalves, Ewan Harrison, David K. Jackson, Ian Johnston, Dominic Kwiatkowski, Cordelia Langford, John Sillitoe on behalf of the Wellcome Sanger Institute COVID-19 Surveillance Team |
| EPI_ISL_3283891 | Berkshire and Surrey Pathology Services Lighthouse Laboratory | Wellcome Sanger Institute for the COVID-19 Genomics UK (COG-UK) Consortium | Berkshire and Surrey Pathology Services Lighthouse Laboratory and Alex Alderton, Roberto Amato, Jeffrey Barrett, Sonia Goncalves, Ewan Harrison, David K. Jackson, Ian Johnston, Dominic Kwiatkowski, Cordelia Langford, John Sillitoe on behalf of the Wellcome Sanger Institute COVID-19 Surveillance Team |
| EPI_ISL_3283892, EPI_ISL_3283893, EPI_ISL_3283894, EPI_ISL_3283895, EPI_ISL_3283896 | Lighthouse Laboratory Plymouth | Wellcome Sanger Institute for the COVID-19 Genomics UK (COG-UK) Consortium | Lighthouse Laboratory Plymouth and Alex Alderton, Roberto Amato, Jeffrey Barrett, Sonia Goncalves, Ewan Harrison, David K. Jackson, Ian Johnston, Dominic Kwiatkowski, Cordelia Langford, John Sillitoe on behalf of the Wellcome Sanger Institute COVID-19 Surveillance Team |
| EPI_ISL_3283897 | Berkshire and Surrey Pathology Services Lighthouse Laboratory | Wellcome Sanger Institute for the COVID-19 Genomics UK (COG-UK) Consortium | Berkshire and Surrey Pathology Services Lighthouse Laboratory and Alex Alderton, Roberto Amato, Jeffrey Barrett, Sonia Goncalves, Ewan Harrison, David K. Jackson, Ian Johnston, Dominic Kwiatkowski, Cordelia Langford, John Sillitoe on behalf of the Wellcome Sanger Institute COVID-19 Surveillance Team |
| EPI_ISL_3283898 | Lighthouse Laboratory Plymouth | Wellcome Sanger Institute for the COVID-19 Genomics UK (COG-UK) Consortium | Lighthouse Laboratory Plymouth and Alex Alderton, Roberto Amato, Jeffrey Barrett, Sonia Goncalves, Ewan Harrison, David K. Jackson, Ian Johnston, Dominic Kwiatkowski, Cordelia Langford, John Sillitoe on behalf of the Wellcome Sanger Institute COVID-19 Surveillance Team |
| EPI_ISL_3283899 | Berkshire and Surrey Pathology Services Lighthouse Laboratory | Wellcome Sanger Institute for the COVID-19 Genomics UK (COG-UK) Consortium | Berkshire and Surrey Pathology Services Lighthouse Laboratory and Alex Alderton, Roberto Amato, Jeffrey Barrett, Sonia Goncalves, Ewan Harrison, David K. Jackson, Ian Johnston, Dominic Kwiatkowski, Cordelia Langford, John Sillitoe on behalf of the Wellcome Sanger Institute COVID-19 Surveillance Team |
| EPI_ISL_3283900, EPI_ISL_3283901, EPI_ISL_3283902, EPI_ISL_3283903 | Lighthouse Laboratory Plymouth | Wellcome Sanger Institute for the COVID-19 Genomics UK (COG-UK) Consortium | Lighthouse Laboratory Plymouth and Alex Alderton, Roberto Amato, Jeffrey Barrett, Sonia Goncalves, Ewan Harrison, David K. Jackson, Ian Johnston, Dominic Kwiatkowski, Cordelia Langford, John Sillitoe on behalf of the Wellcome Sanger Institute COVID-19 Surveillance Team |
| EPI_ISL_3283904 | Berkshire and Surrey Pathology Services Lighthouse Laboratory | Wellcome Sanger Institute for the COVID-19 Genomics UK (COG-UK) Consortium | Berkshire and Surrey Pathology Services Lighthouse Laboratory and Alex Alderton, Roberto Amato, Jeffrey Barrett, Sonia Goncalves, Ewan Harrison, David K. Jackson, Ian Johnston, Dominic Kwiatkowski, Cordelia Langford, John Sillitoe on behalf of the Wellcome Sanger Institute COVID-19 Surveillance Team |
| EPI_ISL_3283905, EPI_ISL_3283906, EPI_ISL_3283907, EPI_ISL_3283908, EPI_ISL_3283909, EPI_ISL_3283910, EPI_ISL_3283911, EPI_ISL_3283912 | Lighthouse Laboratory Plymouth | Wellcome Sanger Institute for the COVID-19 Genomics UK (COG-UK) Consortium | Lighthouse Laboratory Plymouth and Alex Alderton, Roberto Amato, Jeffrey Barrett, Sonia Goncalves, Ewan Harrison, David K. Jackson, Ian Johnston, Dominic Kwiatkowski, Cordelia Langford, John Sillitoe on behalf of the Wellcome Sanger Institute COVID-19 Surveillance Team |
| EPI_ISL_3283913 | Berkshire and Surrey Pathology Services Lighthouse Laboratory | Wellcome Sanger Institute for the COVID-19 Genomics UK (COG-UK) Consortium | Berkshire and Surrey Pathology Services Lighthouse Laboratory and Alex Alderton, Roberto Amato, Jeffrey Barrett, Sonia Goncalves, Ewan Harrison, David K. Jackson, Ian Johnston, Dominic Kwiatkowski, Cordelia Langford, John Sillitoe on behalf of the Wellcome Sanger Institute COVID-19 Surveillance Team |

| | | | |
|---|---|---|---|
| EPI_ISL_3283914, EPI_ISL_3283915, EPI_ISL_3283916, EPI_ISL_3283917, EPI_ISL_3283918, EPI_ISL_3283919, EPI_ISL_3283920, EPI_ISL_3283921, EPI_ISL_3283922 | Lighthouse Laboratory Plymouth | Wellcome Sanger Institute for the COVID-19 Genomics UK (COG-UK) Consortium | Lighthouse Laboratory Plymouth and Alex Alderton, Roberto Amato, Jeffrey Barrett, Sonia Goncalves, Ewan Harrison, David K. Jackson, Ian Johnston, Dominic Kwiatkowski, Cordelia Langford, John Sillitoe on behalf of the Wellcome Sanger Institute COVID-19 Surveillance Team |
| EPI_ISL_3283923 | Berkshire and Surrey Pathology Services Lighthouse Laboratory | Wellcome Sanger Institute for the COVID-19 Genomics UK (COG-UK) Consortium | Berkshire and Surrey Pathology Services Lighthouse Laboratory and Alex Alderton, Roberto Amato, Jeffrey Barrett, Sonia Goncalves, Ewan Harrison, David K. Jackson, Ian Johnston, Dominic Kwiatkowski, Cordelia Langford, John Sillitoe on behalf of the Wellcome Sanger Institute COVID-19 Surveillance Team |
| EPI_ISL_3283924 | Lighthouse Laboratory Plymouth | Wellcome Sanger Institute for the COVID-19 Genomics UK (COG-UK) Consortium | Lighthouse Laboratory Plymouth and Alex Alderton, Roberto Amato, Jeffrey Barrett, Sonia Goncalves, Ewan Harrison, David K. Jackson, Ian Johnston, Dominic Kwiatkowski, Cordelia Langford, John Sillitoe on behalf of the Wellcome Sanger Institute COVID-19 Surveillance Team |
| EPI_ISL_3283925 | Berkshire and Surrey Pathology Services Lighthouse Laboratory | Wellcome Sanger Institute for the COVID-19 Genomics UK (COG-UK) Consortium | Berkshire and Surrey Pathology Services Lighthouse Laboratory and Alex Alderton, Roberto Amato, Jeffrey Barrett, Sonia Goncalves, Ewan Harrison, David K. Jackson, Ian Johnston, Dominic Kwiatkowski, Cordelia Langford, John Sillitoe on behalf of the Wellcome Sanger Institute COVID-19 Surveillance Team |
| EPI_ISL_3283926, EPI_ISL_3283927, EPI_ISL_3283928, EPI_ISL_3283929, EPI_ISL_3283930, EPI_ISL_3283931 | Lighthouse Laboratory Plymouth | Wellcome Sanger Institute for the COVID-19 Genomics UK (COG-UK) Consortium | Lighthouse Laboratory Plymouth and Alex Alderton, Roberto Amato, Jeffrey Barrett, Sonia Goncalves, Ewan Harrison, David K. Jackson, Ian Johnston, Dominic Kwiatkowski, Cordelia Langford, John Sillitoe on behalf of the Wellcome Sanger Institute COVID-19 Surveillance Team |
| EPI_ISL_3283932, EPI_ISL_3283933 | Berkshire and Surrey Pathology Services Lighthouse Laboratory | Wellcome Sanger Institute for the COVID-19 Genomics UK (COG-UK) Consortium | Berkshire and Surrey Pathology Services Lighthouse Laboratory and Alex Alderton, Roberto Amato, Jeffrey Barrett, Sonia Goncalves, Ewan Harrison, David K. Jackson, Ian Johnston, Dominic Kwiatkowski, Cordelia Langford, John Sillitoe on behalf of the Wellcome Sanger Institute COVID-19 Surveillance Team |
| EPI_ISL_3283934 | Lighthouse Laboratory Plymouth | Wellcome Sanger Institute for the COVID-19 Genomics UK (COG-UK) Consortium | Lighthouse Laboratory Plymouth and Alex Alderton, Roberto Amato, Jeffrey Barrett, Sonia Goncalves, Ewan Harrison, David K. Jackson, Ian Johnston, Dominic Kwiatkowski, Cordelia Langford, John Sillitoe on behalf of the Wellcome Sanger Institute COVID-19 Surveillance Team |
| EPI_ISL_3283935 | Berkshire and Surrey Pathology Services Lighthouse Laboratory | Wellcome Sanger Institute for the COVID-19 Genomics UK (COG-UK) Consortium | Berkshire and Surrey Pathology Services Lighthouse Laboratory and Alex Alderton, Roberto Amato, Jeffrey Barrett, Sonia Goncalves, Ewan Harrison, David K. Jackson, Ian Johnston, Dominic Kwiatkowski, Cordelia Langford, John Sillitoe on behalf of the Wellcome Sanger Institute COVID-19 Surveillance Team |
| EPI_ISL_3283936, EPI_ISL_3283937, EPI_ISL_3283938, EPI_ISL_3283939, EPI_ISL_3283940, EPI_ISL_3283941, EPI_ISL_3283942 | Lighthouse Laboratory Plymouth | Wellcome Sanger Institute for the COVID-19 Genomics UK (COG-UK) Consortium | Lighthouse Laboratory Plymouth and Alex Alderton, Roberto Amato, Jeffrey Barrett, Sonia Goncalves, Ewan Harrison, David K. Jackson, Ian Johnston, Dominic Kwiatkowski, Cordelia Langford, John Sillitoe on behalf of the Wellcome Sanger Institute COVID-19 Surveillance Team |
| EPI_ISL_3283943 | Berkshire and Surrey Pathology Services Lighthouse Laboratory | Wellcome Sanger Institute for the COVID-19 Genomics UK (COG-UK) Consortium | Berkshire and Surrey Pathology Services Lighthouse Laboratory and Alex Alderton, Roberto Amato, Jeffrey Barrett, Sonia Goncalves, Ewan Harrison, David K. Jackson, Ian Johnston, Dominic Kwiatkowski, Cordelia Langford, John Sillitoe on behalf of the Wellcome Sanger Institute COVID-19 Surveillance Team |
| EPI_ISL_3283944 | Lighthouse Laboratory Plymouth | Wellcome Sanger Institute for the COVID-19 Genomics UK (COG-UK) Consortium | Lighthouse Laboratory Plymouth and Alex Alderton, Roberto Amato, Jeffrey Barrett, Sonia Goncalves, Ewan Harrison, David K. Jackson, Ian Johnston, Dominic Kwiatkowski, Cordelia Langford, John Sillitoe on behalf of the Wellcome Sanger Institute COVID-19 Surveillance Team |
| EPI_ISL_3283945, EPI_ISL_3283946 | Berkshire and Surrey Pathology Services Lighthouse Laboratory | Wellcome Sanger Institute for the COVID-19 Genomics UK (COG-UK) Consortium | Berkshire and Surrey Pathology Services Lighthouse Laboratory and Alex Alderton, Roberto Amato, Jeffrey Barrett, Sonia Goncalves, Ewan Harrison, David K. Jackson, Ian Johnston, Dominic Kwiatkowski, Cordelia Langford, John Sillitoe on behalf of the Wellcome Sanger Institute COVID-19 Surveillance Team |
| EPI_ISL_3283947, EPI_ISL_3283948, EPI_ISL_3283949, EPI_ISL_3283950, EPI_ISL_3283951, EPI_ISL_3283952, EPI_ISL_3283953 | Lighthouse Laboratory Plymouth | Wellcome Sanger Institute for the COVID-19 Genomics UK (COG-UK) Consortium | Lighthouse Laboratory Plymouth and Alex Alderton, Roberto Amato, Jeffrey Barrett, Sonia Goncalves, Ewan Harrison, David K. Jackson, Ian Johnston, Dominic Kwiatkowski, Cordelia Langford, John Sillitoe on behalf of the Wellcome Sanger Institute COVID-19 Surveillance Team |
| EPI_ISL_3283954 | Berkshire and Surrey Pathology Services Lighthouse Laboratory | Wellcome Sanger Institute for the COVID-19 Genomics UK (COG-UK) Consortium | Berkshire and Surrey Pathology Services Lighthouse Laboratory and Alex Alderton, Roberto Amato, Jeffrey Barrett, Sonia Goncalves, Ewan Harrison, David K. Jackson, Ian Johnston, Dominic Kwiatkowski, Cordelia Langford, John Sillitoe on behalf of the Wellcome Sanger Institute COVID-19 Surveillance Team |
| EPI_ISL_3283955 | Lighthouse Laboratory Plymouth | Wellcome Sanger Institute for the COVID-19 Genomics UK (COG-UK) Consortium | Lighthouse Laboratory Plymouth and Alex Alderton, Roberto Amato, Jeffrey Barrett, Sonia Goncalves, Ewan Harrison, David K. Jackson, Ian Johnston, Dominic Kwiatkowski, Cordelia Langford, John Sillitoe on behalf of the Wellcome Sanger Institute COVID-19 Surveillance Team |
| EPI_ISL_3283956, EPI_ISL_3283957 | Berkshire and Surrey Pathology Services Lighthouse Laboratory | Wellcome Sanger Institute for the COVID-19 Genomics UK (COG-UK) Consortium | Berkshire and Surrey Pathology Services Lighthouse Laboratory and Alex Alderton, Roberto Amato, Jeffrey Barrett, Sonia Goncalves, Ewan Harrison, David K. Jackson, Ian Johnston, Dominic Kwiatkowski, Cordelia Langford, John Sillitoe on behalf of the Wellcome Sanger Institute COVID-19 Surveillance Team |
| EPI_ISL_3283958 | Lighthouse Laboratory Plymouth | Wellcome Sanger Institute for the COVID-19 Genomics UK (COG-UK) Consortium | Lighthouse Laboratory Plymouth and Alex Alderton, Roberto Amato, Jeffrey Barrett, Sonia Goncalves, Ewan Harrison, David K. Jackson, Ian Johnston, Dominic Kwiatkowski, Cordelia Langford, John Sillitoe on behalf of the Wellcome Sanger Institute COVID-19 Surveillance Team |
| EPI_ISL_3283959 | Berkshire and Surrey Pathology Services Lighthouse Laboratory | Wellcome Sanger Institute for the COVID-19 Genomics UK (COG-UK) Consortium | Berkshire and Surrey Pathology Services Lighthouse Laboratory and Alex Alderton, Roberto Amato, Jeffrey Barrett, Sonia Goncalves, Ewan Harrison, David K. Jackson, Ian Johnston, Dominic Kwiatkowski, Cordelia Langford, John Sillitoe on behalf of the Wellcome Sanger Institute COVID-19 Surveillance Team |
| EPI_ISL_3283960, EPI_ISL_3283961, EPI_ISL_3283962, EPI_ISL_3283963, EPI_ISL_3283964, EPI_ISL_3283965, EPI_ISL_3283966, EPI_ISL_3283967, EPI_ISL_3283968, EPI_ISL_3283969 | Lighthouse Laboratory Plymouth | Wellcome Sanger Institute for the COVID-19 Genomics UK (COG-UK) Consortium | Lighthouse Laboratory Plymouth and Alex Alderton, Roberto Amato, Jeffrey Barrett, Sonia Goncalves, Ewan Harrison, David K. Jackson, Ian Johnston, Dominic Kwiatkowski, Cordelia Langford, John Sillitoe on behalf of the Wellcome Sanger Institute COVID-19 Surveillance Team |
| EPI_ISL_3283970 | Berkshire and Surrey Pathology Services Lighthouse Laboratory | Wellcome Sanger Institute for the COVID-19 Genomics UK (COG-UK) Consortium | Berkshire and Surrey Pathology Services Lighthouse Laboratory and Alex Alderton, Roberto Amato, Jeffrey Barrett, Sonia Goncalves, Ewan Harrison, David K. Jackson, Ian Johnston, Dominic Kwiatkowski, Cordelia Langford, John Sillitoe on behalf of the Wellcome Sanger Institute COVID-19 Surveillance Team |
| EPI_ISL_3283971, EPI_ISL_3283972, EPI_ISL_3283973, EPI_ISL_3283974, EPI_ISL_3283975, EPI_ISL_3283976, EPI_ISL_3283977 | Lighthouse Laboratory Plymouth | Wellcome Sanger Institute for the COVID-19 Genomics UK (COG-UK) Consortium | Lighthouse Laboratory Plymouth and Alex Alderton, Roberto Amato, Jeffrey Barrett, Sonia Goncalves, Ewan Harrison, David K. Jackson, Ian Johnston, Dominic Kwiatkowski, Cordelia Langford, John Sillitoe on behalf of the Wellcome Sanger Institute COVID-19 Surveillance Team |
| EPI_ISL_3283978, EPI_ISL_3283979 | Berkshire and Surrey Pathology Services Lighthouse Laboratory | Wellcome Sanger Institute for the COVID-19 Genomics UK (COG-UK) Consortium | Berkshire and Surrey Pathology Services Lighthouse Laboratory and Alex Alderton, Roberto Amato, Jeffrey Barrett, Sonia Goncalves, Ewan Harrison, David K. Jackson, Ian Johnston, Dominic Kwiatkowski, Cordelia Langford, John Sillitoe on behalf of the Wellcome Sanger Institute COVID-19 Surveillance Team |
| EPI_ISL_3283980 | Lighthouse Laboratory Plymouth | Wellcome Sanger Institute for the COVID-19 Genomics UK (COG-UK) Consortium | Lighthouse Laboratory Plymouth and Alex Alderton, Roberto Amato, Jeffrey Barrett, Sonia Goncalves, Ewan Harrison, David K. Jackson, Ian Johnston, Dominic Kwiatkowski, Cordelia Langford, John Sillitoe on behalf of the Wellcome Sanger Institute COVID-19 Surveillance Team |
| EPI_ISL_3283981 | Berkshire and Surrey Pathology Services Lighthouse Laboratory | Wellcome Sanger Institute for the COVID-19 Genomics UK (COG-UK) Consortium | Berkshire and Surrey Pathology Services Lighthouse Laboratory and Alex Alderton, Roberto Amato, Jeffrey Barrett, Sonia Goncalves, Ewan Harrison, David K. Jackson, Ian Johnston, Dominic Kwiatkowski, Cordelia Langford, John Sillitoe on behalf of the Wellcome Sanger Institute COVID-19 Surveillance Team |
| EPI_ISL_3283982, EPI_ISL_3283983, EPI_ISL_3283984, EPI_ISL_3283985 | Lighthouse Laboratory Plymouth | Wellcome Sanger Institute for the COVID-19 Genomics UK (COG-UK) Consortium | Lighthouse Laboratory Plymouth and Alex Alderton, Roberto Amato, Jeffrey Barrett, Sonia Goncalves, Ewan Harrison, David K. Jackson, Ian Johnston, Dominic Kwiatkowski, Cordelia Langford, John Sillitoe on behalf of the Wellcome Sanger Institute COVID-19 Surveillance Team |
| EPI_ISL_3283986 | Berkshire and Surrey Pathology Services Lighthouse Laboratory | Wellcome Sanger Institute for the COVID-19 Genomics UK (COG-UK) Consortium | Berkshire and Surrey Pathology Services Lighthouse Laboratory and Alex Alderton, Roberto Amato, Jeffrey Barrett, Sonia Goncalves, Ewan Harrison, David K. Jackson, Ian Johnston, Dominic Kwiatkowski, Cordelia Langford, John Sillitoe on behalf of the Wellcome Sanger Institute COVID-19 Surveillance Team |
| EPI_ISL_3283987, EPI_ISL_3283988, EPI_ISL_3283989 | Lighthouse Laboratory Plymouth | Wellcome Sanger Institute for the COVID-19 Genomics UK (COG-UK) Consortium | Lighthouse Laboratory Plymouth and Alex Alderton, Roberto Amato, Jeffrey Barrett, Sonia Goncalves, Ewan Harrison, David K. Jackson, Ian Johnston, Dominic Kwiatkowski, Cordelia Langford, John Sillitoe on behalf of the Wellcome Sanger Institute COVID-19 Surveillance Team |
| EPI_ISL_3283990 | Berkshire and Surrey Pathology Services Lighthouse Laboratory | Wellcome Sanger Institute for the COVID-19 Genomics UK (COG-UK) Consortium | Berkshire and Surrey Pathology Services Lighthouse Laboratory and Alex Alderton, Roberto Amato, Jeffrey Barrett, Sonia Goncalves, Ewan Harrison, David K. Jackson, Ian Johnston, Dominic Kwiatkowski, Cordelia Langford, John Sillitoe on behalf of the Wellcome Sanger Institute COVID-19 Surveillance Team |
| EPI_ISL_3283991 | Lighthouse Laboratory Plymouth | Wellcome Sanger Institute for the COVID-19 Genomics UK (COG-UK) Consortium | Lighthouse Laboratory Plymouth and Alex Alderton, Roberto Amato, Jeffrey Barrett, Sonia Goncalves, Ewan Harrison, David K. Jackson, Ian Johnston, Dominic Kwiatkowski, Cordelia Langford, John Sillitoe on behalf of the Wellcome Sanger Institute COVID-19 Surveillance Team |
| EPI_ISL_3283992 | Berkshire and Surrey Pathology Services Lighthouse Laboratory | Wellcome Sanger Institute for the COVID-19 Genomics UK (COG-UK) Consortium | Berkshire and Surrey Pathology Services Lighthouse Laboratory and Alex Alderton, Roberto Amato, Jeffrey Barrett, Sonia Goncalves, Ewan Harrison, David K. Jackson, Ian Johnston, Dominic Kwiatkowski, Cordelia Langford, John Sillitoe on behalf of the Wellcome Sanger Institute COVID-19 Surveillance Team |

| | | | |
|---|---|---|---|
| EPI_ISL_3283993, EPI_ISL_3283994, EPI_ISL_3283995, EPI_ISL_3283996, EPI_ISL_3283997, EPI_ISL_3283998, EPI_ISL_3283999 | Lighthouse Laboratory Plymouth | Wellcome Sanger Institute for the COVID-19 Genomics UK (COG-UK) Consortium | Lighthouse Laboratory Plymouth and Alex Alderton, Roberto Amato, Jeffrey Barrett, Sonia Goncalves, Ewan Harrison, David K. Jackson, Ian Johnston, Dominic Kwiatkowski, Cordelia Langford, John Sillitoe on behalf of the Wellcome Sanger Institute COVID-19 Surveillance Team |
| EPI_ISL_3284000 | Berkshire and Surrey Pathology Services Lighthouse Laboratory | Wellcome Sanger Institute for the COVID-19 Genomics UK (COG-UK) Consortium | Berkshire and Surrey Pathology Services Lighthouse Laboratory and Alex Alderton, Roberto Amato, Jeffrey Barrett, Sonia Goncalves, Ewan Harrison, David K. Jackson, Ian Johnston, Dominic Kwiatkowski, Cordelia Langford, John Sillitoe on behalf of the Wellcome Sanger Institute COVID-19 Surveillance Team |
| EPI_ISL_3284001, EPI_ISL_3284002, EPI_ISL_3284003, EPI_ISL_3284004 | Lighthouse Laboratory Plymouth | Wellcome Sanger Institute for the COVID-19 Genomics UK (COG-UK) Consortium | Lighthouse Laboratory Plymouth and Alex Alderton, Roberto Amato, Jeffrey Barrett, Sonia Goncalves, Ewan Harrison, David K. Jackson, Ian Johnston, Dominic Kwiatkowski, Cordelia Langford, John Sillitoe on behalf of the Wellcome Sanger Institute COVID-19 Surveillance Team |
| EPI_ISL_3284005 | Berkshire and Surrey Pathology Services Lighthouse Laboratory | Wellcome Sanger Institute for the COVID-19 Genomics UK (COG-UK) Consortium | Berkshire and Surrey Pathology Services Lighthouse Laboratory and Alex Alderton, Roberto Amato, Jeffrey Barrett, Sonia Goncalves, Ewan Harrison, David K. Jackson, Ian Johnston, Dominic Kwiatkowski, Cordelia Langford, John Sillitoe on behalf of the Wellcome Sanger Institute COVID-19 Surveillance Team |
| EPI_ISL_3284006, EPI_ISL_3284007, EPI_ISL_3284008, EPI_ISL_3284009, EPI_ISL_3284010, EPI_ISL_3284011, EPI_ISL_3284012, EPI_ISL_3284013 | Lighthouse Laboratory Plymouth | Wellcome Sanger Institute for the COVID-19 Genomics UK (COG-UK) Consortium | Lighthouse Laboratory Plymouth and Alex Alderton, Roberto Amato, Jeffrey Barrett, Sonia Goncalves, Ewan Harrison, David K. Jackson, Ian Johnston, Dominic Kwiatkowski, Cordelia Langford, John Sillitoe on behalf of the Wellcome Sanger Institute COVID-19 Surveillance Team |
| EPI_ISL_3284014 | Berkshire and Surrey Pathology Services Lighthouse Laboratory | Wellcome Sanger Institute for the COVID-19 Genomics UK (COG-UK) Consortium | Berkshire and Surrey Pathology Services Lighthouse Laboratory and Alex Alderton, Roberto Amato, Jeffrey Barrett, Sonia Goncalves, Ewan Harrison, David K. Jackson, Ian Johnston, Dominic Kwiatkowski, Cordelia Langford, John Sillitoe on behalf of the Wellcome Sanger Institute COVID-19 Surveillance Team |
| EPI_ISL_3284015, EPI_ISL_3284016, EPI_ISL_3284017 | Lighthouse Laboratory Plymouth | Wellcome Sanger Institute for the COVID-19 Genomics UK (COG-UK) Consortium | Lighthouse Laboratory Plymouth and Alex Alderton, Roberto Amato, Jeffrey Barrett, Sonia Goncalves, Ewan Harrison, David K. Jackson, Ian Johnston, Dominic Kwiatkowski, Cordelia Langford, John Sillitoe on behalf of the Wellcome Sanger Institute COVID-19 Surveillance Team |
| EPI_ISL_3284018 | Berkshire and Surrey Pathology Services Lighthouse Laboratory | Wellcome Sanger Institute for the COVID-19 Genomics UK (COG-UK) Consortium | Berkshire and Surrey Pathology Services Lighthouse Laboratory and Alex Alderton, Roberto Amato, Jeffrey Barrett, Sonia Goncalves, Ewan Harrison, David K. Jackson, Ian Johnston, Dominic Kwiatkowski, Cordelia Langford, John Sillitoe on behalf of the Wellcome Sanger Institute COVID-19 Surveillance Team |
| EPI_ISL_3284019, EPI_ISL_3284020, EPI_ISL_3284021, EPI_ISL_3284022, EPI_ISL_3284023 | Lighthouse Laboratory Plymouth | Wellcome Sanger Institute for the COVID-19 Genomics UK (COG-UK) Consortium | Lighthouse Laboratory Plymouth and Alex Alderton, Roberto Amato, Jeffrey Barrett, Sonia Goncalves, Ewan Harrison, David K. Jackson, Ian Johnston, Dominic Kwiatkowski, Cordelia Langford, John Sillitoe on behalf of the Wellcome Sanger Institute COVID-19 Surveillance Team |
| EPI_ISL_3284024 | Berkshire and Surrey Pathology Services Lighthouse Laboratory | Wellcome Sanger Institute for the COVID-19 Genomics UK (COG-UK) Consortium | Berkshire and Surrey Pathology Services Lighthouse Laboratory and Alex Alderton, Roberto Amato, Jeffrey Barrett, Sonia Goncalves, Ewan Harrison, David K. Jackson, Ian Johnston, Dominic Kwiatkowski, Cordelia Langford, John Sillitoe on behalf of the Wellcome Sanger Institute COVID-19 Surveillance Team |
| EPI_ISL_3284025, EPI_ISL_3284026, EPI_ISL_3284027 | Lighthouse Laboratory Plymouth | Wellcome Sanger Institute for the COVID-19 Genomics UK (COG-UK) Consortium | Lighthouse Laboratory Plymouth and Alex Alderton, Roberto Amato, Jeffrey Barrett, Sonia Goncalves, Ewan Harrison, David K. Jackson, Ian Johnston, Dominic Kwiatkowski, Cordelia Langford, John Sillitoe on behalf of the Wellcome Sanger Institute COVID-19 Surveillance Team |
| EPI_ISL_3284028 | Berkshire and Surrey Pathology Services Lighthouse Laboratory | Wellcome Sanger Institute for the COVID-19 Genomics UK (COG-UK) Consortium | Berkshire and Surrey Pathology Services Lighthouse Laboratory and Alex Alderton, Roberto Amato, Jeffrey Barrett, Sonia Goncalves, Ewan Harrison, David K. Jackson, Ian Johnston, Dominic Kwiatkowski, Cordelia Langford, John Sillitoe on behalf of the Wellcome Sanger Institute COVID-19 Surveillance Team |
| EPI_ISL_3284029, EPI_ISL_3284030 | Lighthouse Laboratory Plymouth | Wellcome Sanger Institute for the COVID-19 Genomics UK (COG-UK) Consortium | Lighthouse Laboratory Plymouth and Alex Alderton, Roberto Amato, Jeffrey Barrett, Sonia Goncalves, Ewan Harrison, David K. Jackson, Ian Johnston, Dominic Kwiatkowski, Cordelia Langford, John Sillitoe on behalf of the Wellcome Sanger Institute COVID-19 Surveillance Team |
| EPI_ISL_3284031 | Berkshire and Surrey Pathology Services Lighthouse Laboratory | Wellcome Sanger Institute for the COVID-19 Genomics UK (COG-UK) Consortium | Berkshire and Surrey Pathology Services Lighthouse Laboratory and Alex Alderton, Roberto Amato, Jeffrey Barrett, Sonia Goncalves, Ewan Harrison, David K. Jackson, Ian Johnston, Dominic Kwiatkowski, Cordelia Langford, John Sillitoe on behalf of the Wellcome Sanger Institute COVID-19 Surveillance Team |
| EPI_ISL_3284032, EPI_ISL_3284033, EPI_ISL_3284034, EPI_ISL_3284035, EPI_ISL_3284036, EPI_ISL_3284037, EPI_ISL_3284038, EPI_ISL_3284039 | Lighthouse Laboratory Plymouth | Wellcome Sanger Institute for the COVID-19 Genomics UK (COG-UK) Consortium | Lighthouse Laboratory Plymouth and Alex Alderton, Roberto Amato, Jeffrey Barrett, Sonia Goncalves, Ewan Harrison, David K. Jackson, Ian Johnston, Dominic Kwiatkowski, Cordelia Langford, John Sillitoe on behalf of the Wellcome Sanger Institute COVID-19 Surveillance Team |
| EPI_ISL_3284040 | Berkshire and Surrey Pathology Services Lighthouse Laboratory | Wellcome Sanger Institute for the COVID-19 Genomics UK (COG-UK) Consortium | Berkshire and Surrey Pathology Services Lighthouse Laboratory and Alex Alderton, Roberto Amato, Jeffrey Barrett, Sonia Goncalves, Ewan Harrison, David K. Jackson, Ian Johnston, Dominic Kwiatkowski, Cordelia Langford, John Sillitoe on behalf of the Wellcome Sanger Institute COVID-19 Surveillance Team |
| EPI_ISL_3284041, EPI_ISL_3284042, EPI_ISL_3284043, EPI_ISL_3284044 | Lighthouse Laboratory Plymouth | Wellcome Sanger Institute for the COVID-19 Genomics UK (COG-UK) Consortium | Lighthouse Laboratory Plymouth and Alex Alderton, Roberto Amato, Jeffrey Barrett, Sonia Goncalves, Ewan Harrison, David K. Jackson, Ian Johnston, Dominic Kwiatkowski, Cordelia Langford, John Sillitoe on behalf of the Wellcome Sanger Institute COVID-19 Surveillance Team |
| EPI_ISL_3284045 | Berkshire and Surrey Pathology Services Lighthouse Laboratory | Wellcome Sanger Institute for the COVID-19 Genomics UK (COG-UK) Consortium | Berkshire and Surrey Pathology Services Lighthouse Laboratory and Alex Alderton, Roberto Amato, Jeffrey Barrett, Sonia Goncalves, Ewan Harrison, David K. Jackson, Ian Johnston, Dominic Kwiatkowski, Cordelia Langford, John Sillitoe on behalf of the Wellcome Sanger Institute COVID-19 Surveillance Team |
| EPI_ISL_3284046, EPI_ISL_3284047, EPI_ISL_3284048, EPI_ISL_3284049, EPI_ISL_3284050, EPI_ISL_3284051, EPI_ISL_3284052 | Lighthouse Laboratory Plymouth | Wellcome Sanger Institute for the COVID-19 Genomics UK (COG-UK) Consortium | Lighthouse Laboratory Plymouth and Alex Alderton, Roberto Amato, Jeffrey Barrett, Sonia Goncalves, Ewan Harrison, David K. Jackson, Ian Johnston, Dominic Kwiatkowski, Cordelia Langford, John Sillitoe on behalf of the Wellcome Sanger Institute COVID-19 Surveillance Team |
| EPI_ISL_3284053 | Berkshire and Surrey Pathology Services Lighthouse Laboratory | Wellcome Sanger Institute for the COVID-19 Genomics UK (COG-UK) Consortium | Berkshire and Surrey Pathology Services Lighthouse Laboratory and Alex Alderton, Roberto Amato, Jeffrey Barrett, Sonia Goncalves, Ewan Harrison, David K. Jackson, Ian Johnston, Dominic Kwiatkowski, Cordelia Langford, John Sillitoe on behalf of the Wellcome Sanger Institute COVID-19 Surveillance Team |
| EPI_ISL_3284054, EPI_ISL_3284055, EPI_ISL_3284056, EPI_ISL_3284057, EPI_ISL_3284058 | Lighthouse Laboratory Plymouth | Wellcome Sanger Institute for the COVID-19 Genomics UK (COG-UK) Consortium | Lighthouse Laboratory Plymouth and Alex Alderton, Roberto Amato, Jeffrey Barrett, Sonia Goncalves, Ewan Harrison, David K. Jackson, Ian Johnston, Dominic Kwiatkowski, Cordelia Langford, John Sillitoe on behalf of the Wellcome Sanger Institute COVID-19 Surveillance Team |
| EPI_ISL_3284059 | Berkshire and Surrey Pathology Services Lighthouse Laboratory | Wellcome Sanger Institute for the COVID-19 Genomics UK (COG-UK) Consortium | Berkshire and Surrey Pathology Services Lighthouse Laboratory and Alex Alderton, Roberto Amato, Jeffrey Barrett, Sonia Goncalves, Ewan Harrison, David K. Jackson, Ian Johnston, Dominic Kwiatkowski, Cordelia Langford, John Sillitoe on behalf of the Wellcome Sanger Institute COVID-19 Surveillance Team |
| EPI_ISL_3284060, EPI_ISL_3284061 | Lighthouse Laboratory Plymouth | Wellcome Sanger Institute for the COVID-19 Genomics UK (COG-UK) Consortium | Lighthouse Laboratory Plymouth and Alex Alderton, Roberto Amato, Jeffrey Barrett, Sonia Goncalves, Ewan Harrison, David K. Jackson, Ian Johnston, Dominic Kwiatkowski, Cordelia Langford, John Sillitoe on behalf of the Wellcome Sanger Institute COVID-19 Surveillance Team |
| EPI_ISL_3284062 | Berkshire and Surrey Pathology Services Lighthouse Laboratory | Wellcome Sanger Institute for the COVID-19 Genomics UK (COG-UK) Consortium | Berkshire and Surrey Pathology Services Lighthouse Laboratory and Alex Alderton, Roberto Amato, Jeffrey Barrett, Sonia Goncalves, Ewan Harrison, David K. Jackson, Ian Johnston, Dominic Kwiatkowski, Cordelia Langford, John Sillitoe on behalf of the Wellcome Sanger Institute COVID-19 Surveillance Team |
| EPI_ISL_3284063, EPI_ISL_3284064, EPI_ISL_3284065, EPI_ISL_3284066, EPI_ISL_3284067, EPI_ISL_3284068, EPI_ISL_3284069, EPI_ISL_3284070, EPI_ISL_3284071, EPI_ISL_3284072, EPI_ISL_3284073 | | | |
| see above | Lighthouse Laboratory Plymouth | Wellcome Sanger Institute for the COVID-19 Genomics UK (COG-UK) Consortium | Lighthouse Laboratory Plymouth and Alex Alderton, Roberto Amato, Jeffrey Barrett, Sonia Goncalves, Ewan Harrison, David K. Jackson, Ian Johnston, Dominic Kwiatkowski, Cordelia Langford, John Sillitoe on behalf of the Wellcome Sanger Institute COVID-19 Surveillance Team |
| EPI_ISL_3284074, EPI_ISL_3284075, EPI_ISL_3284076, EPI_ISL_3284077 | Berkshire and Surrey Pathology Services Lighthouse Laboratory | Wellcome Sanger Institute for the COVID-19 Genomics UK (COG-UK) Consortium | Berkshire and Surrey Pathology Services Lighthouse Laboratory and Alex Alderton, Roberto Amato, Jeffrey Barrett, Sonia Goncalves, Ewan Harrison, David K. Jackson, Ian Johnston, Dominic Kwiatkowski, Cordelia Langford, John Sillitoe on behalf of the Wellcome Sanger Institute COVID-19 Surveillance Team |
| EPI_ISL_3284078 | Lighthouse Laboratory Plymouth | Wellcome Sanger Institute for the COVID-19 Genomics UK (COG-UK) Consortium | Lighthouse Laboratory Plymouth and Alex Alderton, Roberto Amato, Jeffrey Barrett, Sonia Goncalves, Ewan Harrison, David K. Jackson, Ian Johnston, Dominic Kwiatkowski, Cordelia Langford, John Sillitoe on behalf of the Wellcome Sanger Institute COVID-19 Surveillance Team |
| EPI_ISL_3284079, EPI_ISL_3284080, EPI_ISL_3284081, EPI_ISL_3284082, EPI_ISL_3284083, EPI_ISL_3284084, EPI_ISL_3284085, EPI_ISL_3284086, EPI_ISL_3284087, EPI_ISL_3284088, EPI_ISL_3284089, EPI_ISL_3284090, EPI_ISL_3284091, EPI_ISL_3284092, EPI_ISL_3284093 | | | |
| see above | Berkshire and Surrey Pathology Services Lighthouse Laboratory | Wellcome Sanger Institute for the COVID-19 Genomics UK (COG-UK) Consortium | Berkshire and Surrey Pathology Services Lighthouse Laboratory and Alex Alderton, Roberto Amato, Jeffrey Barrett, Sonia Goncalves, Ewan Harrison, David K. Jackson, Ian Johnston, Dominic Kwiatkowski, Cordelia Langford, John Sillitoe on behalf of the Wellcome Sanger Institute COVID-19 Surveillance Team |
| EPI_ISL_3284094, EPI_ISL_3284095 | Lighthouse Laboratory Plymouth | Wellcome Sanger Institute for the COVID-19 Genomics UK (COG-UK) Consortium | Lighthouse Laboratory Plymouth and Alex Alderton, Roberto Amato, Jeffrey Barrett, Sonia Goncalves, Ewan Harrison, David K. Jackson, Ian Johnston, Dominic Kwiatkowski, Cordelia Langford, John Sillitoe on behalf of the Wellcome Sanger Institute COVID-19 Surveillance Team |
| EPI_ISL_3284096, | Berkshire and Surrey Pathology Services Lighthouse Laboratory | Wellcome Sanger Institute for the COVID-19 Genomics UK (COG-UK) | Berkshire and Surrey Pathology Services Lighthouse Laboratory and Alex Alderton, Roberto Amato, Jeffrey Barrett, Sonia Goncalves, Ewan Harrison, David K. Jackson, Ian Johnston, Dominic Kwiatkowski, Cordelia Langford, John Sillitoe on behalf |

| | | | |
|---|---|---|---|
| EPI_ISL_3284097, EPI_ISL_3284098, EPI_ISL_3284099, EPI_ISL_3284100, EPI_ISL_3284101, EPI_ISL_3284102, EPI_ISL_3284103 | | Consortium | of the Wellcome Sanger Institute COVID-19 Surveillance Team |
| EPI_ISL_3284104 | Lighthouse Laboratory Plymouth | Wellcome Sanger Institute for the COVID-19 Genomics UK (COG-UK) Consortium | Lighthouse Laboratory Plymouth and Alex Alderton, Roberto Amato, Jeffrey Barrett, Sonia Goncalves, Ewan Harrison, David K. Jackson, Ian Johnston, Dominic Kwiatkowski, Cordelia Langford, John Sillitoe on behalf of the Wellcome Sanger Institute COVID-19 Surveillance Team |
| EPI_ISL_3284105, EPI_ISL_3284106, EPI_ISL_3284107 | Berkshire and Surrey Pathology Services Lighthouse Laboratory | Wellcome Sanger Institute for the COVID-19 Genomics UK (COG-UK) Consortium | Berkshire and Surrey Pathology Services Lighthouse Laboratory and Alex Alderton, Roberto Amato, Jeffrey Barrett, Sonia Goncalves, Ewan Harrison, David K. Jackson, Ian Johnston, Dominic Kwiatkowski, Cordelia Langford, John Sillitoe on behalf of the Wellcome Sanger Institute COVID-19 Surveillance Team |
| EPI_ISL_3284108 | Lighthouse Laboratory Plymouth | Wellcome Sanger Institute for the COVID-19 Genomics UK (COG-UK) Consortium | Lighthouse Laboratory Plymouth and Alex Alderton, Roberto Amato, Jeffrey Barrett, Sonia Goncalves, Ewan Harrison, David K. Jackson, Ian Johnston, Dominic Kwiatkowski, Cordelia Langford, John Sillitoe on behalf of the Wellcome Sanger Institute COVID-19 Surveillance Team |
| EPI_ISL_3284109, EPI_ISL_3284110 | Berkshire and Surrey Pathology Services Lighthouse Laboratory | Wellcome Sanger Institute for the COVID-19 Genomics UK (COG-UK) Consortium | Berkshire and Surrey Pathology Services Lighthouse Laboratory and Alex Alderton, Roberto Amato, Jeffrey Barrett, Sonia Goncalves, Ewan Harrison, David K. Jackson, Ian Johnston, Dominic Kwiatkowski, Cordelia Langford, John Sillitoe on behalf of the Wellcome Sanger Institute COVID-19 Surveillance Team |
| EPI_ISL_3284111 | Lighthouse Laboratory Plymouth | Wellcome Sanger Institute for the COVID-19 Genomics UK (COG-UK) Consortium | Lighthouse Laboratory Plymouth and Alex Alderton, Roberto Amato, Jeffrey Barrett, Sonia Goncalves, Ewan Harrison, David K. Jackson, Ian Johnston, Dominic Kwiatkowski, Cordelia Langford, John Sillitoe on behalf of the Wellcome Sanger Institute COVID-19 Surveillance Team |
| EPI_ISL_3284112, EPI_ISL_3284113, EPI_ISL_3284114, EPI_ISL_3284115, EPI_ISL_3284116, EPI_ISL_3284117, EPI_ISL_3284118 | Berkshire and Surrey Pathology Services Lighthouse Laboratory | Wellcome Sanger Institute for the COVID-19 Genomics UK (COG-UK) Consortium | Berkshire and Surrey Pathology Services Lighthouse Laboratory and Alex Alderton, Roberto Amato, Jeffrey Barrett, Sonia Goncalves, Ewan Harrison, David K. Jackson, Ian Johnston, Dominic Kwiatkowski, Cordelia Langford, John Sillitoe on behalf of the Wellcome Sanger Institute COVID-19 Surveillance Team |
| EPI_ISL_3284119, EPI_ISL_3284120 | Lighthouse Laboratory Plymouth | Wellcome Sanger Institute for the COVID-19 Genomics UK (COG-UK) Consortium | Lighthouse Laboratory Plymouth and Alex Alderton, Roberto Amato, Jeffrey Barrett, Sonia Goncalves, Ewan Harrison, David K. Jackson, Ian Johnston, Dominic Kwiatkowski, Cordelia Langford, John Sillitoe on behalf of the Wellcome Sanger Institute COVID-19 Surveillance Team |
| EPI_ISL_3284121, EPI_ISL_3284122 | Berkshire and Surrey Pathology Services Lighthouse Laboratory | Wellcome Sanger Institute for the COVID-19 Genomics UK (COG-UK) Consortium | Berkshire and Surrey Pathology Services Lighthouse Laboratory and Alex Alderton, Roberto Amato, Jeffrey Barrett, Sonia Goncalves, Ewan Harrison, David K. Jackson, Ian Johnston, Dominic Kwiatkowski, Cordelia Langford, John Sillitoe on behalf of the Wellcome Sanger Institute COVID-19 Surveillance Team |
| EPI_ISL_3284123 | Lighthouse Laboratory Plymouth | Wellcome Sanger Institute for the COVID-19 Genomics UK (COG-UK) Consortium | Lighthouse Laboratory Plymouth and Alex Alderton, Roberto Amato, Jeffrey Barrett, Sonia Goncalves, Ewan Harrison, David K. Jackson, Ian Johnston, Dominic Kwiatkowski, Cordelia Langford, John Sillitoe on behalf of the Wellcome Sanger Institute COVID-19 Surveillance Team |
| EPI_ISL_3284124, EPI_ISL_3284125, EPI_ISL_3284126 | Berkshire and Surrey Pathology Services Lighthouse Laboratory | Wellcome Sanger Institute for the COVID-19 Genomics UK (COG-UK) Consortium | Berkshire and Surrey Pathology Services Lighthouse Laboratory and Alex Alderton, Roberto Amato, Jeffrey Barrett, Sonia Goncalves, Ewan Harrison, David K. Jackson, Ian Johnston, Dominic Kwiatkowski, Cordelia Langford, John Sillitoe on behalf of the Wellcome Sanger Institute COVID-19 Surveillance Team |
| EPI_ISL_3284127, EPI_ISL_3284128 | Lighthouse Laboratory Plymouth | Wellcome Sanger Institute for the COVID-19 Genomics UK (COG-UK) Consortium | Lighthouse Laboratory Plymouth and Alex Alderton, Roberto Amato, Jeffrey Barrett, Sonia Goncalves, Ewan Harrison, David K. Jackson, Ian Johnston, Dominic Kwiatkowski, Cordelia Langford, John Sillitoe on behalf of the Wellcome Sanger Institute COVID-19 Surveillance Team |
| EPI_ISL_3284129, EPI_ISL_3284130, EPI_ISL_3284131, EPI_ISL_3284132 | Berkshire and Surrey Pathology Services Lighthouse Laboratory | Wellcome Sanger Institute for the COVID-19 Genomics UK (COG-UK) Consortium | Berkshire and Surrey Pathology Services Lighthouse Laboratory and Alex Alderton, Roberto Amato, Jeffrey Barrett, Sonia Goncalves, Ewan Harrison, David K. Jackson, Ian Johnston, Dominic Kwiatkowski, Cordelia Langford, John Sillitoe on behalf of the Wellcome Sanger Institute COVID-19 Surveillance Team |
| EPI_ISL_3284133 | Lighthouse Laboratory Plymouth | Wellcome Sanger Institute for the COVID-19 Genomics UK (COG-UK) Consortium | Lighthouse Laboratory Plymouth and Alex Alderton, Roberto Amato, Jeffrey Barrett, Sonia Goncalves, Ewan Harrison, David K. Jackson, Ian Johnston, Dominic Kwiatkowski, Cordelia Langford, John Sillitoe on behalf of the Wellcome Sanger Institute COVID-19 Surveillance Team |
| EPI_ISL_3284134, EPI_ISL_3284135, EPI_ISL_3284136, EPI_ISL_3284137 | Berkshire and Surrey Pathology Services Lighthouse Laboratory | Wellcome Sanger Institute for the COVID-19 Genomics UK (COG-UK) Consortium | Berkshire and Surrey Pathology Services Lighthouse Laboratory and Alex Alderton, Roberto Amato, Jeffrey Barrett, Sonia Goncalves, Ewan Harrison, David K. Jackson, Ian Johnston, Dominic Kwiatkowski, Cordelia Langford, John Sillitoe on behalf of the Wellcome Sanger Institute COVID-19 Surveillance Team |
| EPI_ISL_3284138 | Lighthouse Laboratory Plymouth | Wellcome Sanger Institute for the COVID-19 Genomics UK (COG-UK) Consortium | Lighthouse Laboratory Plymouth and Alex Alderton, Roberto Amato, Jeffrey Barrett, Sonia Goncalves, Ewan Harrison, David K. Jackson, Ian Johnston, Dominic Kwiatkowski, Cordelia Langford, John Sillitoe on behalf of the Wellcome Sanger Institute COVID-19 Surveillance Team |
| EPI_ISL_3284139, EPI_ISL_3284140, EPI_ISL_3284141, EPI_ISL_3284142, EPI_ISL_3284143, EPI_ISL_3284144, EPI_ISL_3284145, EPI_ISL_3284146, EPI_ISL_3284147, EPI_ISL_3284148 | Berkshire and Surrey Pathology Services Lighthouse Laboratory | Wellcome Sanger Institute for the COVID-19 Genomics UK (COG-UK) Consortium | Berkshire and Surrey Pathology Services Lighthouse Laboratory and Alex Alderton, Roberto Amato, Jeffrey Barrett, Sonia Goncalves, Ewan Harrison, David K. Jackson, Ian Johnston, Dominic Kwiatkowski, Cordelia Langford, John Sillitoe on behalf of the Wellcome Sanger Institute COVID-19 Surveillance Team |
| EPI_ISL_3284149 | Lighthouse Laboratory Plymouth | Wellcome Sanger Institute for the COVID-19 Genomics UK (COG-UK) Consortium | Lighthouse Laboratory Plymouth and Alex Alderton, Roberto Amato, Jeffrey Barrett, Sonia Goncalves, Ewan Harrison, David K. Jackson, Ian Johnston, Dominic Kwiatkowski, Cordelia Langford, John Sillitoe on behalf of the Wellcome Sanger Institute COVID-19 Surveillance Team |
| EPI_ISL_3284150 | Berkshire and Surrey Pathology Services Lighthouse Laboratory | Wellcome Sanger Institute for the COVID-19 Genomics UK (COG-UK) Consortium | Berkshire and Surrey Pathology Services Lighthouse Laboratory and Alex Alderton, Roberto Amato, Jeffrey Barrett, Sonia Goncalves, Ewan Harrison, David K. Jackson, Ian Johnston, Dominic Kwiatkowski, Cordelia Langford, John Sillitoe on behalf of the Wellcome Sanger Institute COVID-19 Surveillance Team |
| EPI_ISL_3284151 | Lighthouse Laboratory Plymouth | Wellcome Sanger Institute for the COVID-19 Genomics UK (COG-UK) Consortium | Lighthouse Laboratory Plymouth and Alex Alderton, Roberto Amato, Jeffrey Barrett, Sonia Goncalves, Ewan Harrison, David K. Jackson, Ian Johnston, Dominic Kwiatkowski, Cordelia Langford, John Sillitoe on behalf of the Wellcome Sanger Institute COVID-19 Surveillance Team |
| EPI_ISL_3284152, EPI_ISL_3284153, EPI_ISL_3284154, EPI_ISL_3284155 | Berkshire and Surrey Pathology Services Lighthouse Laboratory | Wellcome Sanger Institute for the COVID-19 Genomics UK (COG-UK) Consortium | Berkshire and Surrey Pathology Services Lighthouse Laboratory and Alex Alderton, Roberto Amato, Jeffrey Barrett, Sonia Goncalves, Ewan Harrison, David K. Jackson, Ian Johnston, Dominic Kwiatkowski, Cordelia Langford, John Sillitoe on behalf of the Wellcome Sanger Institute COVID-19 Surveillance Team |
| EPI_ISL_3284156 | Lighthouse Laboratory Plymouth | Wellcome Sanger Institute for the COVID-19 Genomics UK (COG-UK) Consortium | Lighthouse Laboratory Plymouth and Alex Alderton, Roberto Amato, Jeffrey Barrett, Sonia Goncalves, Ewan Harrison, David K. Jackson, Ian Johnston, Dominic Kwiatkowski, Cordelia Langford, John Sillitoe on behalf of the Wellcome Sanger Institute COVID-19 Surveillance Team |
| EPI_ISL_3284157 | Berkshire and Surrey Pathology Services Lighthouse Laboratory | Wellcome Sanger Institute for the COVID-19 Genomics UK (COG-UK) Consortium | Berkshire and Surrey Pathology Services Lighthouse Laboratory and Alex Alderton, Roberto Amato, Jeffrey Barrett, Sonia Goncalves, Ewan Harrison, David K. Jackson, Ian Johnston, Dominic Kwiatkowski, Cordelia Langford, John Sillitoe on behalf of the Wellcome Sanger Institute COVID-19 Surveillance Team |
| EPI_ISL_3284158, EPI_ISL_3284159 | Lighthouse Laboratory Plymouth | Wellcome Sanger Institute for the COVID-19 Genomics UK (COG-UK) Consortium | Lighthouse Laboratory Plymouth and Alex Alderton, Roberto Amato, Jeffrey Barrett, Sonia Goncalves, Ewan Harrison, David K. Jackson, Ian Johnston, Dominic Kwiatkowski, Cordelia Langford, John Sillitoe on behalf of the Wellcome Sanger Institute COVID-19 Surveillance Team |
| EPI_ISL_3284160 | Berkshire and Surrey Pathology Services Lighthouse Laboratory | Wellcome Sanger Institute for the COVID-19 Genomics UK (COG-UK) Consortium | Berkshire and Surrey Pathology Services Lighthouse Laboratory and Alex Alderton, Roberto Amato, Jeffrey Barrett, Sonia Goncalves, Ewan Harrison, David K. Jackson, Ian Johnston, Dominic Kwiatkowski, Cordelia Langford, John Sillitoe on behalf of the Wellcome Sanger Institute COVID-19 Surveillance Team |
| EPI_ISL_3284161 | Lighthouse Laboratory Plymouth | Wellcome Sanger Institute for the COVID-19 Genomics UK (COG-UK) Consortium | Lighthouse Laboratory Plymouth and Alex Alderton, Roberto Amato, Jeffrey Barrett, Sonia Goncalves, Ewan Harrison, David K. Jackson, Ian Johnston, Dominic Kwiatkowski, Cordelia Langford, John Sillitoe on behalf of the Wellcome Sanger Institute COVID-19 Surveillance Team |
| EPI_ISL_3284162, EPI_ISL_3284163, EPI_ISL_3284164, EPI_ISL_3284165, EPI_ISL_3284166, EPI_ISL_3284167, EPI_ISL_3284168, EPI_ISL_3284169 | Berkshire and Surrey Pathology Services Lighthouse Laboratory | Wellcome Sanger Institute for the COVID-19 Genomics UK (COG-UK) Consortium | Berkshire and Surrey Pathology Services Lighthouse Laboratory and Alex Alderton, Roberto Amato, Jeffrey Barrett, Sonia Goncalves, Ewan Harrison, David K. Jackson, Ian Johnston, Dominic Kwiatkowski, Cordelia Langford, John Sillitoe on behalf of the Wellcome Sanger Institute COVID-19 Surveillance Team |
| EPI_ISL_3284170 | Lighthouse Laboratory Plymouth | Wellcome Sanger Institute for the COVID-19 Genomics UK (COG-UK) Consortium | Lighthouse Laboratory Plymouth and Alex Alderton, Roberto Amato, Jeffrey Barrett, Sonia Goncalves, Ewan Harrison, David K. Jackson, Ian Johnston, Dominic Kwiatkowski, Cordelia Langford, John Sillitoe on behalf of the Wellcome Sanger Institute COVID-19 Surveillance Team |
| EPI_ISL_3284171, EPI_ISL_3284172, EPI_ISL_3284173, EPI_ISL_3284174, EPI_ISL_3284175 | Berkshire and Surrey Pathology Services Lighthouse Laboratory | Wellcome Sanger Institute for the COVID-19 Genomics UK (COG-UK) Consortium | Berkshire and Surrey Pathology Services Lighthouse Laboratory and Alex Alderton, Roberto Amato, Jeffrey Barrett, Sonia Goncalves, Ewan Harrison, David K. Jackson, Ian Johnston, Dominic Kwiatkowski, Cordelia Langford, John Sillitoe on behalf of the Wellcome Sanger Institute COVID-19 Surveillance Team |
| EPI_ISL_3284176 | Lighthouse Laboratory Plymouth | Wellcome Sanger Institute for the COVID-19 Genomics UK (COG-UK) Consortium | Lighthouse Laboratory Plymouth and Alex Alderton, Roberto Amato, Jeffrey Barrett, Sonia Goncalves, Ewan Harrison, David K. Jackson, Ian Johnston, Dominic Kwiatkowski, Cordelia Langford, John Sillitoe on behalf of the Wellcome Sanger Institute COVID-19 Surveillance Team |
| EPI_ISL_3284177, EPI_ISL_3284178, | Berkshire and Surrey Pathology Services Lighthouse Laboratory | Wellcome Sanger Institute for the COVID-19 Genomics UK (COG-UK) Consortium | Berkshire and Surrey Pathology Services Lighthouse Laboratory and Alex Alderton, Roberto Amato, Jeffrey Barrett, Sonia Goncalves, Ewan Harrison, David K. Jackson, Ian Johnston, Dominic Kwiatkowski, Cordelia Langford, John Sillitoe on behalf of the Wellcome Sanger Institute COVID-19 Surveillance Team |

| | | | |
|---|---|---|---|
| EPI_ISL_3284179 | | | |
| EPI_ISL_3284180 | Lighthouse Laboratory Plymouth | Wellcome Sanger Institute for the COVID-19 Genomics UK (COG-UK) Consortium | Lighthouse Laboratory Plymouth and Alex Alderton, Roberto Amato, Jeffrey Barrett, Sonia Goncalves, Ewan Harrison, David K. Jackson, Ian Johnston, Dominic Kwiatkowski, Cordelia Langford, John Sillitoe on behalf of the Wellcome Sanger Institute COVID-19 Surveillance Team |
| EPI_ISL_3284181, EPI_ISL_3284182, EPI_ISL_3284183 | Berkshire and Surrey Pathology Services Lighthouse Laboratory | Wellcome Sanger Institute for the COVID-19 Genomics UK (COG-UK) Consortium | Berkshire and Surrey Pathology Services Lighthouse Laboratory and Alex Alderton, Roberto Amato, Jeffrey Barrett, Sonia Goncalves, Ewan Harrison, David K. Jackson, Ian Johnston, Dominic Kwiatkowski, Cordelia Langford, John Sillitoe on behalf of the Wellcome Sanger Institute COVID-19 Surveillance Team |
| EPI_ISL_3284184 | Lighthouse Laboratory Plymouth | Wellcome Sanger Institute for the COVID-19 Genomics UK (COG-UK) Consortium | Lighthouse Laboratory Plymouth and Alex Alderton, Roberto Amato, Jeffrey Barrett, Sonia Goncalves, Ewan Harrison, David K. Jackson, Ian Johnston, Dominic Kwiatkowski, Cordelia Langford, John Sillitoe on behalf of the Wellcome Sanger Institute COVID-19 Surveillance Team |
| EPI_ISL_3284185, EPI_ISL_3284186, EPI_ISL_3284187, EPI_ISL_3284188 | Berkshire and Surrey Pathology Services Lighthouse Laboratory | Wellcome Sanger Institute for the COVID-19 Genomics UK (COG-UK) Consortium | Berkshire and Surrey Pathology Services Lighthouse Laboratory and Alex Alderton, Roberto Amato, Jeffrey Barrett, Sonia Goncalves, Ewan Harrison, David K. Jackson, Ian Johnston, Dominic Kwiatkowski, Cordelia Langford, John Sillitoe on behalf of the Wellcome Sanger Institute COVID-19 Surveillance Team |
| EPI_ISL_3284189 | Lighthouse Laboratory Plymouth | Wellcome Sanger Institute for the COVID-19 Genomics UK (COG-UK) Consortium | Lighthouse Laboratory Plymouth and Alex Alderton, Roberto Amato, Jeffrey Barrett, Sonia Goncalves, Ewan Harrison, David K. Jackson, Ian Johnston, Dominic Kwiatkowski, Cordelia Langford, John Sillitoe on behalf of the Wellcome Sanger Institute COVID-19 Surveillance Team |
| EPI_ISL_3284190, EPI_ISL_3284191, EPI_ISL_3284192, EPI_ISL_3284193, EPI_ISL_3284194, EPI_ISL_3284195, EPI_ISL_3284196 | Berkshire and Surrey Pathology Services Lighthouse Laboratory | Wellcome Sanger Institute for the COVID-19 Genomics UK (COG-UK) Consortium | Berkshire and Surrey Pathology Services Lighthouse Laboratory and Alex Alderton, Roberto Amato, Jeffrey Barrett, Sonia Goncalves, Ewan Harrison, David K. Jackson, Ian Johnston, Dominic Kwiatkowski, Cordelia Langford, John Sillitoe on behalf of the Wellcome Sanger Institute COVID-19 Surveillance Team |
| EPI_ISL_3284197 | Lighthouse Laboratory Plymouth | Wellcome Sanger Institute for the COVID-19 Genomics UK (COG-UK) Consortium | Lighthouse Laboratory Plymouth and Alex Alderton, Roberto Amato, Jeffrey Barrett, Sonia Goncalves, Ewan Harrison, David K. Jackson, Ian Johnston, Dominic Kwiatkowski, Cordelia Langford, John Sillitoe on behalf of the Wellcome Sanger Institute COVID-19 Surveillance Team |
| EPI_ISL_3284198 | Berkshire and Surrey Pathology Services Lighthouse Laboratory | Wellcome Sanger Institute for the COVID-19 Genomics UK (COG-UK) Consortium | Berkshire and Surrey Pathology Services Lighthouse Laboratory and Alex Alderton, Roberto Amato, Jeffrey Barrett, Sonia Goncalves, Ewan Harrison, David K. Jackson, Ian Johnston, Dominic Kwiatkowski, Cordelia Langford, John Sillitoe on behalf of the Wellcome Sanger Institute COVID-19 Surveillance Team |
| EPI_ISL_3284199 | Lighthouse Laboratory Plymouth | Wellcome Sanger Institute for the COVID-19 Genomics UK (COG-UK) Consortium | Lighthouse Laboratory Plymouth and Alex Alderton, Roberto Amato, Jeffrey Barrett, Sonia Goncalves, Ewan Harrison, David K. Jackson, Ian Johnston, Dominic Kwiatkowski, Cordelia Langford, John Sillitoe on behalf of the Wellcome Sanger Institute COVID-19 Surveillance Team |
| EPI_ISL_3284200 | Berkshire and Surrey Pathology Services Lighthouse Laboratory | Wellcome Sanger Institute for the COVID-19 Genomics UK (COG-UK) Consortium | Berkshire and Surrey Pathology Services Lighthouse Laboratory and Alex Alderton, Roberto Amato, Jeffrey Barrett, Sonia Goncalves, Ewan Harrison, David K. Jackson, Ian Johnston, Dominic Kwiatkowski, Cordelia Langford, John Sillitoe on behalf of the Wellcome Sanger Institute COVID-19 Surveillance Team |
| EPI_ISL_3284201 | Lighthouse Laboratory Plymouth | Wellcome Sanger Institute for the COVID-19 Genomics UK (COG-UK) Consortium | Lighthouse Laboratory Plymouth and Alex Alderton, Roberto Amato, Jeffrey Barrett, Sonia Goncalves, Ewan Harrison, David K. Jackson, Ian Johnston, Dominic Kwiatkowski, Cordelia Langford, John Sillitoe on behalf of the Wellcome Sanger Institute COVID-19 Surveillance Team |
| EPI_ISL_3284202, EPI_ISL_3284203 | Berkshire and Surrey Pathology Services Lighthouse Laboratory | Wellcome Sanger Institute for the COVID-19 Genomics UK (COG-UK) Consortium | Berkshire and Surrey Pathology Services Lighthouse Laboratory and Alex Alderton, Roberto Amato, Jeffrey Barrett, Sonia Goncalves, Ewan Harrison, David K. Jackson, Ian Johnston, Dominic Kwiatkowski, Cordelia Langford, John Sillitoe on behalf of the Wellcome Sanger Institute COVID-19 Surveillance Team |
| EPI_ISL_3284204, EPI_ISL_3284205, EPI_ISL_3284206 | Lighthouse Laboratory Plymouth | Wellcome Sanger Institute for the COVID-19 Genomics UK (COG-UK) Consortium | Lighthouse Laboratory Plymouth and Alex Alderton, Roberto Amato, Jeffrey Barrett, Sonia Goncalves, Ewan Harrison, David K. Jackson, Ian Johnston, Dominic Kwiatkowski, Cordelia Langford, John Sillitoe on behalf of the Wellcome Sanger Institute COVID-19 Surveillance Team |
| EPI_ISL_3284207, EPI_ISL_3284208 | Berkshire and Surrey Pathology Services Lighthouse Laboratory | Wellcome Sanger Institute for the COVID-19 Genomics UK (COG-UK) Consortium | Berkshire and Surrey Pathology Services Lighthouse Laboratory and Alex Alderton, Roberto Amato, Jeffrey Barrett, Sonia Goncalves, Ewan Harrison, David K. Jackson, Ian Johnston, Dominic Kwiatkowski, Cordelia Langford, John Sillitoe on behalf of the Wellcome Sanger Institute COVID-19 Surveillance Team |
| EPI_ISL_3284209, EPI_ISL_3284210, EPI_ISL_3284211, EPI_ISL_3284212, EPI_ISL_3284213 | Lighthouse Laboratory Plymouth | Wellcome Sanger Institute for the COVID-19 Genomics UK (COG-UK) Consortium | Lighthouse Laboratory Plymouth and Alex Alderton, Roberto Amato, Jeffrey Barrett, Sonia Goncalves, Ewan Harrison, David K. Jackson, Ian Johnston, Dominic Kwiatkowski, Cordelia Langford, John Sillitoe on behalf of the Wellcome Sanger Institute COVID-19 Surveillance Team |
| EPI_ISL_3284214, EPI_ISL_3284215, EPI_ISL_3284216 | Berkshire and Surrey Pathology Services Lighthouse Laboratory | Wellcome Sanger Institute for the COVID-19 Genomics UK (COG-UK) Consortium | Berkshire and Surrey Pathology Services Lighthouse Laboratory and Alex Alderton, Roberto Amato, Jeffrey Barrett, Sonia Goncalves, Ewan Harrison, David K. Jackson, Ian Johnston, Dominic Kwiatkowski, Cordelia Langford, John Sillitoe on behalf of the Wellcome Sanger Institute COVID-19 Surveillance Team |
| EPI_ISL_3284217 | Lighthouse Laboratory Plymouth | Wellcome Sanger Institute for the COVID-19 Genomics UK (COG-UK) Consortium | Lighthouse Laboratory Plymouth and Alex Alderton, Roberto Amato, Jeffrey Barrett, Sonia Goncalves, Ewan Harrison, David K. Jackson, Ian Johnston, Dominic Kwiatkowski, Cordelia Langford, John Sillitoe on behalf of the Wellcome Sanger Institute COVID-19 Surveillance Team |
| EPI_ISL_3284218, EPI_ISL_3284219, EPI_ISL_3284220, EPI_ISL_3284221, EPI_ISL_3284222 | Berkshire and Surrey Pathology Services Lighthouse Laboratory | Wellcome Sanger Institute for the COVID-19 Genomics UK (COG-UK) Consortium | Berkshire and Surrey Pathology Services Lighthouse Laboratory and Alex Alderton, Roberto Amato, Jeffrey Barrett, Sonia Goncalves, Ewan Harrison, David K. Jackson, Ian Johnston, Dominic Kwiatkowski, Cordelia Langford, John Sillitoe on behalf of the Wellcome Sanger Institute COVID-19 Surveillance Team |
| EPI_ISL_3284223, EPI_ISL_3284224, EPI_ISL_3284225 | Lighthouse Laboratory Plymouth | Wellcome Sanger Institute for the COVID-19 Genomics UK (COG-UK) Consortium | Lighthouse Laboratory Plymouth and Alex Alderton, Roberto Amato, Jeffrey Barrett, Sonia Goncalves, Ewan Harrison, David K. Jackson, Ian Johnston, Dominic Kwiatkowski, Cordelia Langford, John Sillitoe on behalf of the Wellcome Sanger Institute COVID-19 Surveillance Team |
| EPI_ISL_3284226, EPI_ISL_3284227 | Berkshire and Surrey Pathology Services Lighthouse Laboratory | Wellcome Sanger Institute for the COVID-19 Genomics UK (COG-UK) Consortium | Berkshire and Surrey Pathology Services Lighthouse Laboratory and Alex Alderton, Roberto Amato, Jeffrey Barrett, Sonia Goncalves, Ewan Harrison, David K. Jackson, Ian Johnston, Dominic Kwiatkowski, Cordelia Langford, John Sillitoe on behalf of the Wellcome Sanger Institute COVID-19 Surveillance Team |
| EPI_ISL_3284228 | Lighthouse Laboratory Plymouth | Wellcome Sanger Institute for the COVID-19 Genomics UK (COG-UK) Consortium | Lighthouse Laboratory Plymouth and Alex Alderton, Roberto Amato, Jeffrey Barrett, Sonia Goncalves, Ewan Harrison, David K. Jackson, Ian Johnston, Dominic Kwiatkowski, Cordelia Langford, John Sillitoe on behalf of the Wellcome Sanger Institute COVID-19 Surveillance Team |
| EPI_ISL_3284229, EPI_ISL_3284230 | Berkshire and Surrey Pathology Services Lighthouse Laboratory | Wellcome Sanger Institute for the COVID-19 Genomics UK (COG-UK) Consortium | Berkshire and Surrey Pathology Services Lighthouse Laboratory and Alex Alderton, Roberto Amato, Jeffrey Barrett, Sonia Goncalves, Ewan Harrison, David K. Jackson, Ian Johnston, Dominic Kwiatkowski, Cordelia Langford, John Sillitoe on behalf of the Wellcome Sanger Institute COVID-19 Surveillance Team |
| EPI_ISL_3284231 | Lighthouse Laboratory Plymouth | Wellcome Sanger Institute for the COVID-19 Genomics UK (COG-UK) Consortium | Lighthouse Laboratory Plymouth and Alex Alderton, Roberto Amato, Jeffrey Barrett, Sonia Goncalves, Ewan Harrison, David K. Jackson, Ian Johnston, Dominic Kwiatkowski, Cordelia Langford, John Sillitoe on behalf of the Wellcome Sanger Institute COVID-19 Surveillance Team |
| EPI_ISL_3284232, EPI_ISL_3284233 | Berkshire and Surrey Pathology Services Lighthouse Laboratory | Wellcome Sanger Institute for the COVID-19 Genomics UK (COG-UK) Consortium | Berkshire and Surrey Pathology Services Lighthouse Laboratory and Alex Alderton, Roberto Amato, Jeffrey Barrett, Sonia Goncalves, Ewan Harrison, David K. Jackson, Ian Johnston, Dominic Kwiatkowski, Cordelia Langford, John Sillitoe on behalf of the Wellcome Sanger Institute COVID-19 Surveillance Team |
| EPI_ISL_3284234 | Lighthouse Laboratory Plymouth | Wellcome Sanger Institute for the COVID-19 Genomics UK (COG-UK) Consortium | Lighthouse Laboratory Plymouth and Alex Alderton, Roberto Amato, Jeffrey Barrett, Sonia Goncalves, Ewan Harrison, David K. Jackson, Ian Johnston, Dominic Kwiatkowski, Cordelia Langford, John Sillitoe on behalf of the Wellcome Sanger Institute COVID-19 Surveillance Team |
| EPI_ISL_3284235 | Berkshire and Surrey Pathology Services Lighthouse Laboratory | Wellcome Sanger Institute for the COVID-19 Genomics UK (COG-UK) Consortium | Berkshire and Surrey Pathology Services Lighthouse Laboratory and Alex Alderton, Roberto Amato, Jeffrey Barrett, Sonia Goncalves, Ewan Harrison, David K. Jackson, Ian Johnston, Dominic Kwiatkowski, Cordelia Langford, John Sillitoe on behalf of the Wellcome Sanger Institute COVID-19 Surveillance Team |
| EPI_ISL_3284236 | Lighthouse Laboratory Plymouth | Wellcome Sanger Institute for the COVID-19 Genomics UK (COG-UK) Consortium | Lighthouse Laboratory Plymouth and Alex Alderton, Roberto Amato, Jeffrey Barrett, Sonia Goncalves, Ewan Harrison, David K. Jackson, Ian Johnston, Dominic Kwiatkowski, Cordelia Langford, John Sillitoe on behalf of the Wellcome Sanger Institute COVID-19 Surveillance Team |
| EPI_ISL_3284237, EPI_ISL_3284238, EPI_ISL_3284239, EPI_ISL_3284240 | Berkshire and Surrey Pathology Services Lighthouse Laboratory | Wellcome Sanger Institute for the COVID-19 Genomics UK (COG-UK) Consortium | Berkshire and Surrey Pathology Services Lighthouse Laboratory and Alex Alderton, Roberto Amato, Jeffrey Barrett, Sonia Goncalves, Ewan Harrison, David K. Jackson, Ian Johnston, Dominic Kwiatkowski, Cordelia Langford, John Sillitoe on behalf of the Wellcome Sanger Institute COVID-19 Surveillance Team |
| EPI_ISL_3284241 | Lighthouse Laboratory Plymouth | Wellcome Sanger Institute for the COVID-19 Genomics UK (COG-UK) Consortium | Lighthouse Laboratory Plymouth and Alex Alderton, Roberto Amato, Jeffrey Barrett, Sonia Goncalves, Ewan Harrison, David K. Jackson, Ian Johnston, Dominic Kwiatkowski, Cordelia Langford, John Sillitoe on behalf of the Wellcome Sanger Institute COVID-19 Surveillance Team |
| EPI_ISL_3284242, EPI_ISL_3284243, EPI_ISL_3284244, EPI_ISL_3284245, EPI_ISL_3284246, EPI_ISL_3284247, EPI_ISL_3284248, EPI_ISL_3284249, EPI_ISL_3284250, EPI_ISL_3284251, EPI_ISL_3284252, EPI_ISL_3284253, EPI_ISL_3284254, EPI_ISL_3284255 | | | |
| see above | Berkshire and Surrey Pathology Services Lighthouse Laboratory | Wellcome Sanger Institute for the COVID-19 Genomics UK (COG-UK) Consortium | Berkshire and Surrey Pathology Services Lighthouse Laboratory and Alex Alderton, Roberto Amato, Jeffrey Barrett, Sonia Goncalves, Ewan Harrison, David K. Jackson, Ian Johnston, Dominic Kwiatkowski, Cordelia Langford, John Sillitoe on behalf of the Wellcome Sanger Institute COVID-19 Surveillance Team |
| EPI_ISL_3284256 | Lighthouse Laboratory Plymouth | Wellcome Sanger Institute for the COVID-19 Genomics UK (COG-UK) Consortium | Lighthouse Laboratory Plymouth and Alex Alderton, Roberto Amato, Jeffrey Barrett, Sonia Goncalves, Ewan Harrison, David K. Jackson, Ian Johnston, Dominic Kwiatkowski, Cordelia Langford, John Sillitoe on behalf of the Wellcome Sanger Institute COVID-19 Surveillance Team |
| EPI_ISL_3284257 | Berkshire and Surrey Pathology Services Lighthouse Laboratory | Wellcome Sanger Institute for the COVID-19 Genomics UK (COG-UK) Consortium | Berkshire and Surrey Pathology Services Lighthouse Laboratory and Alex Alderton, Roberto Amato, Jeffrey Barrett, Sonia Goncalves, Ewan Harrison, David K. Jackson, Ian Johnston, Dominic Kwiatkowski, Cordelia Langford, John Sillitoe on behalf of the Wellcome Sanger Institute COVID-19 Surveillance Team |
| EPI_ISL_3284258, EPI_ISL_3284259, EPI_ISL_3284260 | Lighthouse Laboratory Plymouth | Wellcome Sanger Institute for the COVID-19 Genomics UK (COG-UK) Consortium | Lighthouse Laboratory Plymouth and Alex Alderton, Roberto Amato, Jeffrey Barrett, Sonia Goncalves, Ewan Harrison, David K. Jackson, Ian Johnston, Dominic Kwiatkowski, Cordelia Langford, John Sillitoe on behalf of the Wellcome Sanger Institute COVID-19 Surveillance Team |
| EPI_ISL_3284261 | Berkshire and Surrey Pathology Services Lighthouse Laboratory | Wellcome Sanger Institute for the COVID-19 Genomics UK (COG-UK) Consortium | Berkshire and Surrey Pathology Services Lighthouse Laboratory and Alex Alderton, Roberto Amato, Jeffrey Barrett, Sonia Goncalves, Ewan Harrison, David K. Jackson, Ian Johnston, Dominic Kwiatkowski, Cordelia Langford, John Sillitoe on behalf of the Wellcome Sanger Institute COVID-19 Surveillance Team |
| EPI_ISL_3284262 | Lighthouse Laboratory Plymouth | Wellcome Sanger Institute for the COVID-19 Genomics UK (COG-UK) Consortium | Lighthouse Laboratory Plymouth and Alex Alderton, Roberto Amato, Jeffrey Barrett, Sonia Goncalves, Ewan Harrison, David K. Jackson, Ian Johnston, Dominic Kwiatkowski, Cordelia Langford, John Sillitoe on behalf of the Wellcome Sanger Institute COVID-19 Surveillance Team |
| EPI_ISL_3284263 | Berkshire and Surrey Pathology Services Lighthouse Laboratory | Wellcome Sanger Institute for the COVID-19 Genomics UK (COG-UK) Consortium | Berkshire and Surrey Pathology Services Lighthouse Laboratory and Alex Alderton, Roberto Amato, Jeffrey Barrett, Sonia Goncalves, Ewan Harrison, David K. Jackson, Ian Johnston, Dominic Kwiatkowski, Cordelia Langford, John Sillitoe on behalf of the Wellcome Sanger Institute COVID-19 Surveillance Team |

| | | | |
|---|---|---|---|
| EPI_ISL_3284264 | Lighthouse Laboratory Plymouth | Wellcome Sanger Institute for the COVID-19 Genomics UK (COG-UK) Consortium | Lighthouse Laboratory Plymouth and Alex Alderton, Roberto Amato, Jeffrey Barrett, Sonia Goncalves, Ewan Harrison, David K. Jackson, Ian Johnston, Dominic Kwiatkowski, Cordelia Langford, John Sillitoe on behalf of the Wellcome Sanger Institute COVID-19 Surveillance Team |
| EPI_ISL_3284265, EPI_ISL_3284266, EPI_ISL_3284267, EPI_ISL_3284268 | Berkshire and Surrey Pathology Services Lighthouse Laboratory | Wellcome Sanger Institute for the COVID-19 Genomics UK (COG-UK) Consortium | Berkshire and Surrey Pathology Services Lighthouse Laboratory and Alex Alderton, Roberto Amato, Jeffrey Barrett, Sonia Goncalves, Ewan Harrison, David K. Jackson, Ian Johnston, Dominic Kwiatkowski, Cordelia Langford, John Sillitoe on behalf of the Wellcome Sanger Institute COVID-19 Surveillance Team |
| EPI_ISL_3284269 | Lighthouse Laboratory Plymouth | Wellcome Sanger Institute for the COVID-19 Genomics UK (COG-UK) Consortium | Lighthouse Laboratory Plymouth and Alex Alderton, Roberto Amato, Jeffrey Barrett, Sonia Goncalves, Ewan Harrison, David K. Jackson, Ian Johnston, Dominic Kwiatkowski, Cordelia Langford, John Sillitoe on behalf of the Wellcome Sanger Institute COVID-19 Surveillance Team |
| EPI_ISL_3284270 | Berkshire and Surrey Pathology Services Lighthouse Laboratory | Wellcome Sanger Institute for the COVID-19 Genomics UK (COG-UK) Consortium | Berkshire and Surrey Pathology Services Lighthouse Laboratory and Alex Alderton, Roberto Amato, Jeffrey Barrett, Sonia Goncalves, Ewan Harrison, David K. Jackson, Ian Johnston, Dominic Kwiatkowski, Cordelia Langford, John Sillitoe on behalf of the Wellcome Sanger Institute COVID-19 Surveillance Team |
| EPI_ISL_3284271 | Lighthouse Laboratory Plymouth | Wellcome Sanger Institute for the COVID-19 Genomics UK (COG-UK) Consortium | Lighthouse Laboratory Plymouth and Alex Alderton, Roberto Amato, Jeffrey Barrett, Sonia Goncalves, Ewan Harrison, David K. Jackson, Ian Johnston, Dominic Kwiatkowski, Cordelia Langford, John Sillitoe on behalf of the Wellcome Sanger Institute COVID-19 Surveillance Team |
| EPI_ISL_3284272 | Berkshire and Surrey Pathology Services Lighthouse Laboratory | Wellcome Sanger Institute for the COVID-19 Genomics UK (COG-UK) Consortium | Berkshire and Surrey Pathology Services Lighthouse Laboratory and Alex Alderton, Roberto Amato, Jeffrey Barrett, Sonia Goncalves, Ewan Harrison, David K. Jackson, Ian Johnston, Dominic Kwiatkowski, Cordelia Langford, John Sillitoe on behalf of the Wellcome Sanger Institute COVID-19 Surveillance Team |
| EPI_ISL_3284273 | Lighthouse Laboratory Plymouth | Wellcome Sanger Institute for the COVID-19 Genomics UK (COG-UK) Consortium | Lighthouse Laboratory Plymouth and Alex Alderton, Roberto Amato, Jeffrey Barrett, Sonia Goncalves, Ewan Harrison, David K. Jackson, Ian Johnston, Dominic Kwiatkowski, Cordelia Langford, John Sillitoe on behalf of the Wellcome Sanger Institute COVID-19 Surveillance Team |
| EPI_ISL_3284274 | Berkshire and Surrey Pathology Services Lighthouse Laboratory | Wellcome Sanger Institute for the COVID-19 Genomics UK (COG-UK) Consortium | Berkshire and Surrey Pathology Services Lighthouse Laboratory and Alex Alderton, Roberto Amato, Jeffrey Barrett, Sonia Goncalves, Ewan Harrison, David K. Jackson, Ian Johnston, Dominic Kwiatkowski, Cordelia Langford, John Sillitoe on behalf of the Wellcome Sanger Institute COVID-19 Surveillance Team |
| EPI_ISL_3284275, EPI_ISL_3284276, EPI_ISL_3284277 | Lighthouse Laboratory Plymouth | Wellcome Sanger Institute for the COVID-19 Genomics UK (COG-UK) Consortium | Lighthouse Laboratory Plymouth and Alex Alderton, Roberto Amato, Jeffrey Barrett, Sonia Goncalves, Ewan Harrison, David K. Jackson, Ian Johnston, Dominic Kwiatkowski, Cordelia Langford, John Sillitoe on behalf of the Wellcome Sanger Institute COVID-19 Surveillance Team |
| EPI_ISL_3284278, EPI_ISL_3284279 | Berkshire and Surrey Pathology Services Lighthouse Laboratory | Wellcome Sanger Institute for the COVID-19 Genomics UK (COG-UK) Consortium | Berkshire and Surrey Pathology Services Lighthouse Laboratory and Alex Alderton, Roberto Amato, Jeffrey Barrett, Sonia Goncalves, Ewan Harrison, David K. Jackson, Ian Johnston, Dominic Kwiatkowski, Cordelia Langford, John Sillitoe on behalf of the Wellcome Sanger Institute COVID-19 Surveillance Team |
| EPI_ISL_3284280, EPI_ISL_3284281 | Lighthouse Laboratory Plymouth | Wellcome Sanger Institute for the COVID-19 Genomics UK (COG-UK) Consortium | Lighthouse Laboratory Plymouth and Alex Alderton, Roberto Amato, Jeffrey Barrett, Sonia Goncalves, Ewan Harrison, David K. Jackson, Ian Johnston, Dominic Kwiatkowski, Cordelia Langford, John Sillitoe on behalf of the Wellcome Sanger Institute COVID-19 Surveillance Team |
| EPI_ISL_3284282, EPI_ISL_3284283, EPI_ISL_3284284, EPI_ISL_3284285 | Berkshire and Surrey Pathology Services Lighthouse Laboratory | Wellcome Sanger Institute for the COVID-19 Genomics UK (COG-UK) Consortium | Berkshire and Surrey Pathology Services Lighthouse Laboratory and Alex Alderton, Roberto Amato, Jeffrey Barrett, Sonia Goncalves, Ewan Harrison, David K. Jackson, Ian Johnston, Dominic Kwiatkowski, Cordelia Langford, John Sillitoe on behalf of the Wellcome Sanger Institute COVID-19 Surveillance Team |
| EPI_ISL_3284286 | Lighthouse Laboratory Plymouth | Wellcome Sanger Institute for the COVID-19 Genomics UK (COG-UK) Consortium | Lighthouse Laboratory Plymouth and Alex Alderton, Roberto Amato, Jeffrey Barrett, Sonia Goncalves, Ewan Harrison, David K. Jackson, Ian Johnston, Dominic Kwiatkowski, Cordelia Langford, John Sillitoe on behalf of the Wellcome Sanger Institute COVID-19 Surveillance Team |
| EPI_ISL_3284287 | Berkshire and Surrey Pathology Services Lighthouse Laboratory | Wellcome Sanger Institute for the COVID-19 Genomics UK (COG-UK) Consortium | Berkshire and Surrey Pathology Services Lighthouse Laboratory and Alex Alderton, Roberto Amato, Jeffrey Barrett, Sonia Goncalves, Ewan Harrison, David K. Jackson, Ian Johnston, Dominic Kwiatkowski, Cordelia Langford, John Sillitoe on behalf of the Wellcome Sanger Institute COVID-19 Surveillance Team |
| EPI_ISL_3284288 | Lighthouse Laboratory Plymouth | Wellcome Sanger Institute for the COVID-19 Genomics UK (COG-UK) Consortium | Lighthouse Laboratory Plymouth and Alex Alderton, Roberto Amato, Jeffrey Barrett, Sonia Goncalves, Ewan Harrison, David K. Jackson, Ian Johnston, Dominic Kwiatkowski, Cordelia Langford, John Sillitoe on behalf of the Wellcome Sanger Institute COVID-19 Surveillance Team |
| EPI_ISL_3284289, EPI_ISL_3284290, EPI_ISL_3284291, EPI_ISL_3284292, EPI_ISL_3284293 | Berkshire and Surrey Pathology Services Lighthouse Laboratory | Wellcome Sanger Institute for the COVID-19 Genomics UK (COG-UK) Consortium | Berkshire and Surrey Pathology Services Lighthouse Laboratory and Alex Alderton, Roberto Amato, Jeffrey Barrett, Sonia Goncalves, Ewan Harrison, David K. Jackson, Ian Johnston, Dominic Kwiatkowski, Cordelia Langford, John Sillitoe on behalf of the Wellcome Sanger Institute COVID-19 Surveillance Team |
| EPI_ISL_3284294, EPI_ISL_3284295, EPI_ISL_3284296, EPI_ISL_3284297, EPI_ISL_3284298 | Lighthouse Laboratory Plymouth | Wellcome Sanger Institute for the COVID-19 Genomics UK (COG-UK) Consortium | Lighthouse Laboratory Plymouth and Alex Alderton, Roberto Amato, Jeffrey Barrett, Sonia Goncalves, Ewan Harrison, David K. Jackson, Ian Johnston, Dominic Kwiatkowski, Cordelia Langford, John Sillitoe on behalf of the Wellcome Sanger Institute COVID-19 Surveillance Team |
| EPI_ISL_3284299, EPI_ISL_3284300 | Berkshire and Surrey Pathology Services Lighthouse Laboratory | Wellcome Sanger Institute for the COVID-19 Genomics UK (COG-UK) Consortium | Berkshire and Surrey Pathology Services Lighthouse Laboratory and Alex Alderton, Roberto Amato, Jeffrey Barrett, Sonia Goncalves, Ewan Harrison, David K. Jackson, Ian Johnston, Dominic Kwiatkowski, Cordelia Langford, John Sillitoe on behalf of the Wellcome Sanger Institute COVID-19 Surveillance Team |
| EPI_ISL_3284301, EPI_ISL_3284302, EPI_ISL_3284303 | Lighthouse Laboratory Plymouth | Wellcome Sanger Institute for the COVID-19 Genomics UK (COG-UK) Consortium | Lighthouse Laboratory Plymouth and Alex Alderton, Roberto Amato, Jeffrey Barrett, Sonia Goncalves, Ewan Harrison, David K. Jackson, Ian Johnston, Dominic Kwiatkowski, Cordelia Langford, John Sillitoe on behalf of the Wellcome Sanger Institute COVID-19 Surveillance Team |
| EPI_ISL_3284304 | Berkshire and Surrey Pathology Services Lighthouse Laboratory | Wellcome Sanger Institute for the COVID-19 Genomics UK (COG-UK) Consortium | Berkshire and Surrey Pathology Services Lighthouse Laboratory and Alex Alderton, Roberto Amato, Jeffrey Barrett, Sonia Goncalves, Ewan Harrison, David K. Jackson, Ian Johnston, Dominic Kwiatkowski, Cordelia Langford, John Sillitoe on behalf of the Wellcome Sanger Institute COVID-19 Surveillance Team |
| EPI_ISL_3284305 | Lighthouse Laboratory Plymouth | Wellcome Sanger Institute for the COVID-19 Genomics UK (COG-UK) Consortium | Lighthouse Laboratory Plymouth and Alex Alderton, Roberto Amato, Jeffrey Barrett, Sonia Goncalves, Ewan Harrison, David K. Jackson, Ian Johnston, Dominic Kwiatkowski, Cordelia Langford, John Sillitoe on behalf of the Wellcome Sanger Institute COVID-19 Surveillance Team |
| EPI_ISL_3284306, EPI_ISL_3284307, EPI_ISL_3284308, EPI_ISL_3284309 | Berkshire and Surrey Pathology Services Lighthouse Laboratory | Wellcome Sanger Institute for the COVID-19 Genomics UK (COG-UK) Consortium | Berkshire and Surrey Pathology Services Lighthouse Laboratory and Alex Alderton, Roberto Amato, Jeffrey Barrett, Sonia Goncalves, Ewan Harrison, David K. Jackson, Ian Johnston, Dominic Kwiatkowski, Cordelia Langford, John Sillitoe on behalf of the Wellcome Sanger Institute COVID-19 Surveillance Team |
| EPI_ISL_3284310 | Lighthouse Laboratory Plymouth | Wellcome Sanger Institute for the COVID-19 Genomics UK (COG-UK) Consortium | Lighthouse Laboratory Plymouth and Alex Alderton, Roberto Amato, Jeffrey Barrett, Sonia Goncalves, Ewan Harrison, David K. Jackson, Ian Johnston, Dominic Kwiatkowski, Cordelia Langford, John Sillitoe on behalf of the Wellcome Sanger Institute COVID-19 Surveillance Team |
| EPI_ISL_3284311, EPI_ISL_3284312, EPI_ISL_3284313 | Berkshire and Surrey Pathology Services Lighthouse Laboratory | Wellcome Sanger Institute for the COVID-19 Genomics UK (COG-UK) Consortium | Berkshire and Surrey Pathology Services Lighthouse Laboratory and Alex Alderton, Roberto Amato, Jeffrey Barrett, Sonia Goncalves, Ewan Harrison, David K. Jackson, Ian Johnston, Dominic Kwiatkowski, Cordelia Langford, John Sillitoe on behalf of the Wellcome Sanger Institute COVID-19 Surveillance Team |
| EPI_ISL_3284314 | Lighthouse Laboratory Plymouth | Wellcome Sanger Institute for the COVID-19 Genomics UK (COG-UK) Consortium | Lighthouse Laboratory Plymouth and Alex Alderton, Roberto Amato, Jeffrey Barrett, Sonia Goncalves, Ewan Harrison, David K. Jackson, Ian Johnston, Dominic Kwiatkowski, Cordelia Langford, John Sillitoe on behalf of the Wellcome Sanger Institute COVID-19 Surveillance Team |
| EPI_ISL_3284315, EPI_ISL_3284316, EPI_ISL_3284317, EPI_ISL_3284318 | Berkshire and Surrey Pathology Services Lighthouse Laboratory | Wellcome Sanger Institute for the COVID-19 Genomics UK (COG-UK) Consortium | Berkshire and Surrey Pathology Services Lighthouse Laboratory and Alex Alderton, Roberto Amato, Jeffrey Barrett, Sonia Goncalves, Ewan Harrison, David K. Jackson, Ian Johnston, Dominic Kwiatkowski, Cordelia Langford, John Sillitoe on behalf of the Wellcome Sanger Institute COVID-19 Surveillance Team |
| EPI_ISL_3284319 | Lighthouse Laboratory Plymouth | Wellcome Sanger Institute for the COVID-19 Genomics UK (COG-UK) Consortium | Lighthouse Laboratory Plymouth and Alex Alderton, Roberto Amato, Jeffrey Barrett, Sonia Goncalves, Ewan Harrison, David K. Jackson, Ian Johnston, Dominic Kwiatkowski, Cordelia Langford, John Sillitoe on behalf of the Wellcome Sanger Institute COVID-19 Surveillance Team |
| EPI_ISL_3284320 | Berkshire and Surrey Pathology Services Lighthouse Laboratory | Wellcome Sanger Institute for the COVID-19 Genomics UK (COG-UK) Consortium | Berkshire and Surrey Pathology Services Lighthouse Laboratory and Alex Alderton, Roberto Amato, Jeffrey Barrett, Sonia Goncalves, Ewan Harrison, David K. Jackson, Ian Johnston, Dominic Kwiatkowski, Cordelia Langford, John Sillitoe on behalf of the Wellcome Sanger Institute COVID-19 Surveillance Team |
| EPI_ISL_3284321 | Lighthouse Laboratory Plymouth | Wellcome Sanger Institute for the COVID-19 Genomics UK (COG-UK) Consortium | Lighthouse Laboratory Plymouth and Alex Alderton, Roberto Amato, Jeffrey Barrett, Sonia Goncalves, Ewan Harrison, David K. Jackson, Ian Johnston, Dominic Kwiatkowski, Cordelia Langford, John Sillitoe on behalf of the Wellcome Sanger Institute COVID-19 Surveillance Team |
| EPI_ISL_3284322, EPI_ISL_3284323 | Berkshire and Surrey Pathology Services Lighthouse Laboratory | Wellcome Sanger Institute for the COVID-19 Genomics UK (COG-UK) Consortium | Berkshire and Surrey Pathology Services Lighthouse Laboratory and Alex Alderton, Roberto Amato, Jeffrey Barrett, Sonia Goncalves, Ewan Harrison, David K. Jackson, Ian Johnston, Dominic Kwiatkowski, Cordelia Langford, John Sillitoe on behalf of the Wellcome Sanger Institute COVID-19 Surveillance Team |
| EPI_ISL_3284324, EPI_ISL_3284325, EPI_ISL_3284326 | Lighthouse Laboratory Plymouth | Wellcome Sanger Institute for the COVID-19 Genomics UK (COG-UK) Consortium | Lighthouse Laboratory Plymouth and Alex Alderton, Roberto Amato, Jeffrey Barrett, Sonia Goncalves, Ewan Harrison, David K. Jackson, Ian Johnston, Dominic Kwiatkowski, Cordelia Langford, John Sillitoe on behalf of the Wellcome Sanger Institute COVID-19 Surveillance Team |
| EPI_ISL_3284327 | Berkshire and Surrey Pathology Services Lighthouse Laboratory | Wellcome Sanger Institute for the COVID-19 Genomics UK (COG-UK) Consortium | Berkshire and Surrey Pathology Services Lighthouse Laboratory and Alex Alderton, Roberto Amato, Jeffrey Barrett, Sonia Goncalves, Ewan Harrison, David K. Jackson, Ian Johnston, Dominic Kwiatkowski, Cordelia Langford, John Sillitoe on behalf of the Wellcome Sanger Institute COVID-19 Surveillance Team |
| EPI_ISL_3284328 | Lighthouse Laboratory Plymouth | Wellcome Sanger Institute for the COVID-19 Genomics UK (COG-UK) Consortium | Lighthouse Laboratory Plymouth and Alex Alderton, Roberto Amato, Jeffrey Barrett, Sonia Goncalves, Ewan Harrison, David K. Jackson, Ian Johnston, Dominic Kwiatkowski, Cordelia Langford, John Sillitoe on behalf of the Wellcome Sanger Institute COVID-19 Surveillance Team |
| EPI_ISL_3284329, EPI_ISL_3284330 | Berkshire and Surrey Pathology Services Lighthouse Laboratory | Wellcome Sanger Institute for the COVID-19 Genomics UK (COG-UK) Consortium | Berkshire and Surrey Pathology Services Lighthouse Laboratory and Alex Alderton, Roberto Amato, Jeffrey Barrett, Sonia Goncalves, Ewan Harrison, David K. Jackson, Ian Johnston, Dominic Kwiatkowski, Cordelia Langford, John Sillitoe on behalf of the Wellcome Sanger Institute COVID-19 Surveillance Team |
| EPI_ISL_3284331 | Lighthouse Laboratory Plymouth | Wellcome Sanger Institute for the COVID-19 Genomics UK (COG-UK) Consortium | Lighthouse Laboratory Plymouth and Alex Alderton, Roberto Amato, Jeffrey Barrett, Sonia Goncalves, Ewan Harrison, David K. Jackson, Ian Johnston, Dominic Kwiatkowski, Cordelia Langford, John Sillitoe on behalf of the Wellcome Sanger Institute COVID-19 Surveillance Team |
| EPI_ISL_3284332, EPI_ISL_3284333, EPI_ISL_3284334, EPI_ISL_3284335 | Berkshire and Surrey Pathology Services Lighthouse Laboratory | Wellcome Sanger Institute for the COVID-19 Genomics UK (COG-UK) Consortium | Berkshire and Surrey Pathology Services Lighthouse Laboratory and Alex Alderton, Roberto Amato, Jeffrey Barrett, Sonia Goncalves, Ewan Harrison, David K. Jackson, Ian Johnston, Dominic Kwiatkowski, Cordelia Langford, John Sillitoe on behalf of the Wellcome Sanger Institute COVID-19 Surveillance Team |
| EPI_ISL_3284336 | Lighthouse Laboratory Plymouth | Wellcome Sanger Institute for the COVID-19 Genomics UK (COG-UK) | Lighthouse Laboratory Plymouth and Alex Alderton, Roberto Amato, Jeffrey Barrett, Sonia Goncalves, Ewan Harrison, David K. Jackson, Ian Johnston, Dominic Kwiatkowski, Cordelia Langford, John Sillitoe on behalf of the Wellcome Sanger |

| | | | |
|---|---|---|---|
| | | Consortium | Institute COVID-19 Surveillance Team |
| EPI_ISL_3284337 | Berkshire and Surrey Pathology Services Lighthouse Laboratory | Wellcome Sanger Institute for the COVID-19 Genomics UK (COG-UK) Consortium | Berkshire and Surrey Pathology Services Lighthouse Laboratory and Alex Alderton, Roberto Amato, Jeffrey Barrett, Sonia Goncalves, Ewan Harrison, David K. Jackson, Ian Johnston, Dominic Kwiatkowski, Cordelia Langford, John Sillitoe on behalf of the Wellcome Sanger Institute COVID-19 Surveillance Team |
| EPI_ISL_3284338 | Lighthouse Laboratory Plymouth | Wellcome Sanger Institute for the COVID-19 Genomics UK (COG-UK) Consortium | Lighthouse Laboratory Plymouth and Alex Alderton, Roberto Amato, Jeffrey Barrett, Sonia Goncalves, Ewan Harrison, David K. Jackson, Ian Johnston, Dominic Kwiatkowski, Cordelia Langford, John Sillitoe on behalf of the Wellcome Sanger Institute COVID-19 Surveillance Team |
| EPI_ISL_3284339, EPI_ISL_3284340, EPI_ISL_3284341, EPI_ISL_3284342, EPI_ISL_3284343 | Berkshire and Surrey Pathology Services Lighthouse Laboratory | Wellcome Sanger Institute for the COVID-19 Genomics UK (COG-UK) Consortium | Berkshire and Surrey Pathology Services Lighthouse Laboratory and Alex Alderton, Roberto Amato, Jeffrey Barrett, Sonia Goncalves, Ewan Harrison, David K. Jackson, Ian Johnston, Dominic Kwiatkowski, Cordelia Langford, John Sillitoe on behalf of the Wellcome Sanger Institute COVID-19 Surveillance Team |
| EPI_ISL_3284344, EPI_ISL_3284345, EPI_ISL_3284346 | Lighthouse Laboratory Plymouth | Wellcome Sanger Institute for the COVID-19 Genomics UK (COG-UK) Consortium | Lighthouse Laboratory Plymouth and Alex Alderton, Roberto Amato, Jeffrey Barrett, Sonia Goncalves, Ewan Harrison, David K. Jackson, Ian Johnston, Dominic Kwiatkowski, Cordelia Langford, John Sillitoe on behalf of the Wellcome Sanger Institute COVID-19 Surveillance Team |
| EPI_ISL_3284347, EPI_ISL_3284348, EPI_ISL_3284349, EPI_ISL_3284350, EPI_ISL_3284351, EPI_ISL_3284352 | Berkshire and Surrey Pathology Services Lighthouse Laboratory | Wellcome Sanger Institute for the COVID-19 Genomics UK (COG-UK) Consortium | Berkshire and Surrey Pathology Services Lighthouse Laboratory and Alex Alderton, Roberto Amato, Jeffrey Barrett, Sonia Goncalves, Ewan Harrison, David K. Jackson, Ian Johnston, Dominic Kwiatkowski, Cordelia Langford, John Sillitoe on behalf of the Wellcome Sanger Institute COVID-19 Surveillance Team |
| EPI_ISL_3284353 | Lighthouse Laboratory Plymouth | Wellcome Sanger Institute for the COVID-19 Genomics UK (COG-UK) Consortium | Lighthouse Laboratory Plymouth and Alex Alderton, Roberto Amato, Jeffrey Barrett, Sonia Goncalves, Ewan Harrison, David K. Jackson, Ian Johnston, Dominic Kwiatkowski, Cordelia Langford, John Sillitoe on behalf of the Wellcome Sanger Institute COVID-19 Surveillance Team |
| EPI_ISL_3284354 | Berkshire and Surrey Pathology Services Lighthouse Laboratory | Wellcome Sanger Institute for the COVID-19 Genomics UK (COG-UK) Consortium | Berkshire and Surrey Pathology Services Lighthouse Laboratory and Alex Alderton, Roberto Amato, Jeffrey Barrett, Sonia Goncalves, Ewan Harrison, David K. Jackson, Ian Johnston, Dominic Kwiatkowski, Cordelia Langford, John Sillitoe on behalf of the Wellcome Sanger Institute COVID-19 Surveillance Team |
| EPI_ISL_3284355 | Lighthouse Laboratory Plymouth | Wellcome Sanger Institute for the COVID-19 Genomics UK (COG-UK) Consortium | Lighthouse Laboratory Plymouth and Alex Alderton, Roberto Amato, Jeffrey Barrett, Sonia Goncalves, Ewan Harrison, David K. Jackson, Ian Johnston, Dominic Kwiatkowski, Cordelia Langford, John Sillitoe on behalf of the Wellcome Sanger Institute COVID-19 Surveillance Team |
| EPI_ISL_3284356 | Berkshire and Surrey Pathology Services Lighthouse Laboratory | Wellcome Sanger Institute for the COVID-19 Genomics UK (COG-UK) Consortium | Berkshire and Surrey Pathology Services Lighthouse Laboratory and Alex Alderton, Roberto Amato, Jeffrey Barrett, Sonia Goncalves, Ewan Harrison, David K. Jackson, Ian Johnston, Dominic Kwiatkowski, Cordelia Langford, John Sillitoe on behalf of the Wellcome Sanger Institute COVID-19 Surveillance Team |
| EPI_ISL_3284357 | Lighthouse Laboratory Plymouth | Wellcome Sanger Institute for the COVID-19 Genomics UK (COG-UK) Consortium | Lighthouse Laboratory Plymouth and Alex Alderton, Roberto Amato, Jeffrey Barrett, Sonia Goncalves, Ewan Harrison, David K. Jackson, Ian Johnston, Dominic Kwiatkowski, Cordelia Langford, John Sillitoe on behalf of the Wellcome Sanger Institute COVID-19 Surveillance Team |
| EPI_ISL_3284358 | Berkshire and Surrey Pathology Services Lighthouse Laboratory | Wellcome Sanger Institute for the COVID-19 Genomics UK (COG-UK) Consortium | Berkshire and Surrey Pathology Services Lighthouse Laboratory and Alex Alderton, Roberto Amato, Jeffrey Barrett, Sonia Goncalves, Ewan Harrison, David K. Jackson, Ian Johnston, Dominic Kwiatkowski, Cordelia Langford, John Sillitoe on behalf of the Wellcome Sanger Institute COVID-19 Surveillance Team |
| EPI_ISL_3284359 | Lighthouse Laboratory Plymouth | Wellcome Sanger Institute for the COVID-19 Genomics UK (COG-UK) Consortium | Lighthouse Laboratory Plymouth and Alex Alderton, Roberto Amato, Jeffrey Barrett, Sonia Goncalves, Ewan Harrison, David K. Jackson, Ian Johnston, Dominic Kwiatkowski, Cordelia Langford, John Sillitoe on behalf of the Wellcome Sanger Institute COVID-19 Surveillance Team |
| EPI_ISL_3284360, EPI_ISL_3284361 | Berkshire and Surrey Pathology Services Lighthouse Laboratory | Wellcome Sanger Institute for the COVID-19 Genomics UK (COG-UK) Consortium | Berkshire and Surrey Pathology Services Lighthouse Laboratory and Alex Alderton, Roberto Amato, Jeffrey Barrett, Sonia Goncalves, Ewan Harrison, David K. Jackson, Ian Johnston, Dominic Kwiatkowski, Cordelia Langford, John Sillitoe on behalf of the Wellcome Sanger Institute COVID-19 Surveillance Team |
| EPI_ISL_3284362 | Lighthouse Laboratory Plymouth | Wellcome Sanger Institute for the COVID-19 Genomics UK (COG-UK) Consortium | Lighthouse Laboratory Plymouth and Alex Alderton, Roberto Amato, Jeffrey Barrett, Sonia Goncalves, Ewan Harrison, David K. Jackson, Ian Johnston, Dominic Kwiatkowski, Cordelia Langford, John Sillitoe on behalf of the Wellcome Sanger Institute COVID-19 Surveillance Team |
| EPI_ISL_3284363, EPI_ISL_3284364, EPI_ISL_3284365, EPI_ISL_3284366 | Berkshire and Surrey Pathology Services Lighthouse Laboratory | Wellcome Sanger Institute for the COVID-19 Genomics UK (COG-UK) Consortium | Berkshire and Surrey Pathology Services Lighthouse Laboratory and Alex Alderton, Roberto Amato, Jeffrey Barrett, Sonia Goncalves, Ewan Harrison, David K. Jackson, Ian Johnston, Dominic Kwiatkowski, Cordelia Langford, John Sillitoe on behalf of the Wellcome Sanger Institute COVID-19 Surveillance Team |
| EPI_ISL_3284367 | Lighthouse Laboratory Plymouth | Wellcome Sanger Institute for the COVID-19 Genomics UK (COG-UK) Consortium | Lighthouse Laboratory Plymouth and Alex Alderton, Roberto Amato, Jeffrey Barrett, Sonia Goncalves, Ewan Harrison, David K. Jackson, Ian Johnston, Dominic Kwiatkowski, Cordelia Langford, John Sillitoe on behalf of the Wellcome Sanger Institute COVID-19 Surveillance Team |
| EPI_ISL_3284368, EPI_ISL_3284369, EPI_ISL_3284370, EPI_ISL_3284371, EPI_ISL_3284372, EPI_ISL_3284373, EPI_ISL_3284374, EPI_ISL_3284375, EPI_ISL_3284376, EPI_ISL_3284377, EPI_ISL_3284378, EPI_ISL_3284379 | | | |
| see above | Berkshire and Surrey Pathology Services Lighthouse Laboratory | Wellcome Sanger Institute for the COVID-19 Genomics UK (COG-UK) Consortium | Berkshire and Surrey Pathology Services Lighthouse Laboratory and Alex Alderton, Roberto Amato, Jeffrey Barrett, Sonia Goncalves, Ewan Harrison, David K. Jackson, Ian Johnston, Dominic Kwiatkowski, Cordelia Langford, John Sillitoe on behalf of the Wellcome Sanger Institute COVID-19 Surveillance Team |
| EPI_ISL_3284380 | Lighthouse Lab in Alderley Park | Wellcome Sanger Institute for the COVID-19 Genomics UK (COG-UK) Consortium | Jacquelyn Wynn, Mairead Hyland, The Lighthouse Lab in Alderley Park and Alex Alderton, Roberto Amato, Jeffrey Barrett, Sonia Goncalves, Ewan Harrison, David K. Jackson, Ian Johnston, Dominic Kwiatkowski, Cordelia Langford, John Sillitoe on behalf of the Wellcome Sanger Institute COVID-19 Surveillance Team |
| EPI_ISL_3284381 | Lighthouse Lab in Glasgow | Wellcome Sanger Institute for the COVID-19 Genomics UK (COG-UK) Consortium | Harper VanSteenhouse, Yumi Kasai, David Gray, Carol Clugston, Anna Dominiczak and Alex Alderton, Roberto Amato, Jeffrey Barrett, Sonia Goncalves, Ewan Harrison, David K. Jackson, Ian Johnston, Dominic Kwiatkowski, Cordelia Langford, John Sillitoe on behalf of the Wellcome Sanger Institute COVID-19 Surveillance Team |
| EPI_ISL_3284382, EPI_ISL_3284383 | Lighthouse Lab in Alderley Park | Wellcome Sanger Institute for the COVID-19 Genomics UK (COG-UK) Consortium | Jacquelyn Wynn, Mairead Hyland, The Lighthouse Lab in Alderley Park and Alex Alderton, Roberto Amato, Jeffrey Barrett, Sonia Goncalves, Ewan Harrison, David K. Jackson, Ian Johnston, Dominic Kwiatkowski, Cordelia Langford, John Sillitoe on behalf of the Wellcome Sanger Institute COVID-19 Surveillance Team |
| EPI_ISL_3284384, EPI_ISL_3284385 | Lighthouse Lab in Glasgow | Wellcome Sanger Institute for the COVID-19 Genomics UK (COG-UK) Consortium | Harper VanSteenhouse, Yumi Kasai, David Gray, Carol Clugston, Anna Dominiczak and Alex Alderton, Roberto Amato, Jeffrey Barrett, Sonia Goncalves, Ewan Harrison, David K. Jackson, Ian Johnston, Dominic Kwiatkowski, Cordelia Langford, John Sillitoe on behalf of the Wellcome Sanger Institute COVID-19 Surveillance Team |
| EPI_ISL_3284386 | Lighthouse Lab in Alderley Park | Wellcome Sanger Institute for the COVID-19 Genomics UK (COG-UK) Consortium | Jacquelyn Wynn, Mairead Hyland, The Lighthouse Lab in Alderley Park and Alex Alderton, Roberto Amato, Jeffrey Barrett, Sonia Goncalves, Ewan Harrison, David K. Jackson, Ian Johnston, Dominic Kwiatkowski, Cordelia Langford, John Sillitoe on behalf of the Wellcome Sanger Institute COVID-19 Surveillance Team |
| EPI_ISL_3284387 | Lighthouse Lab in Glasgow | Wellcome Sanger Institute for the COVID-19 Genomics UK (COG-UK) Consortium | Harper VanSteenhouse, Yumi Kasai, David Gray, Carol Clugston, Anna Dominiczak and Alex Alderton, Roberto Amato, Jeffrey Barrett, Sonia Goncalves, Ewan Harrison, David K. Jackson, Ian Johnston, Dominic Kwiatkowski, Cordelia Langford, John Sillitoe on behalf of the Wellcome Sanger Institute COVID-19 Surveillance Team |
| EPI_ISL_3284388, EPI_ISL_3284389, EPI_ISL_3284390, EPI_ISL_3284391, EPI_ISL_3284392, EPI_ISL_3284393 | Lighthouse Lab in Alderley Park | Wellcome Sanger Institute for the COVID-19 Genomics UK (COG-UK) Consortium | Jacquelyn Wynn, Mairead Hyland, The Lighthouse Lab in Alderley Park and Alex Alderton, Roberto Amato, Jeffrey Barrett, Sonia Goncalves, Ewan Harrison, David K. Jackson, Ian Johnston, Dominic Kwiatkowski, Cordelia Langford, John Sillitoe on behalf of the Wellcome Sanger Institute COVID-19 Surveillance Team |
| EPI_ISL_3284394 | Lighthouse Lab in Glasgow | Wellcome Sanger Institute for the COVID-19 Genomics UK (COG-UK) Consortium | Harper VanSteenhouse, Yumi Kasai, David Gray, Carol Clugston, Anna Dominiczak and Alex Alderton, Roberto Amato, Jeffrey Barrett, Sonia Goncalves, Ewan Harrison, David K. Jackson, Ian Johnston, Dominic Kwiatkowski, Cordelia Langford, John Sillitoe on behalf of the Wellcome Sanger Institute COVID-19 Surveillance Team |
| EPI_ISL_3284395 | Lighthouse Lab in Alderley Park | Wellcome Sanger Institute for the COVID-19 Genomics UK (COG-UK) Consortium | Jacquelyn Wynn, Mairead Hyland, The Lighthouse Lab in Alderley Park and Alex Alderton, Roberto Amato, Jeffrey Barrett, Sonia Goncalves, Ewan Harrison, David K. Jackson, Ian Johnston, Dominic Kwiatkowski, Cordelia Langford, John Sillitoe on behalf of the Wellcome Sanger Institute COVID-19 Surveillance Team |
| EPI_ISL_3284396, EPI_ISL_3284397 | Lighthouse Lab in Glasgow | Wellcome Sanger Institute for the COVID-19 Genomics UK (COG-UK) Consortium | Harper VanSteenhouse, Yumi Kasai, David Gray, Carol Clugston, Anna Dominiczak and Alex Alderton, Roberto Amato, Jeffrey Barrett, Sonia Goncalves, Ewan Harrison, David K. Jackson, Ian Johnston, Dominic Kwiatkowski, Cordelia Langford, John Sillitoe on behalf of the Wellcome Sanger Institute COVID-19 Surveillance Team |
| EPI_ISL_3284398 | Lighthouse Lab in Alderley Park | Wellcome Sanger Institute for the COVID-19 Genomics UK (COG-UK) Consortium | Jacquelyn Wynn, Mairead Hyland, The Lighthouse Lab in Alderley Park and Alex Alderton, Roberto Amato, Jeffrey Barrett, Sonia Goncalves, Ewan Harrison, David K. Jackson, Ian Johnston, Dominic Kwiatkowski, Cordelia Langford, John Sillitoe on behalf of the Wellcome Sanger Institute COVID-19 Surveillance Team |
| EPI_ISL_3284399, EPI_ISL_3284400 | Lighthouse Lab in Glasgow | Wellcome Sanger Institute for the COVID-19 Genomics UK (COG-UK) Consortium | Harper VanSteenhouse, Yumi Kasai, David Gray, Carol Clugston, Anna Dominiczak and Alex Alderton, Roberto Amato, Jeffrey Barrett, Sonia Goncalves, Ewan Harrison, David K. Jackson, Ian Johnston, Dominic Kwiatkowski, Cordelia Langford, John Sillitoe on behalf of the Wellcome Sanger Institute COVID-19 Surveillance Team |
| EPI_ISL_3284401, EPI_ISL_3284402, EPI_ISL_3284403, EPI_ISL_3284404 | Lighthouse Lab in Alderley Park | Wellcome Sanger Institute for the COVID-19 Genomics UK (COG-UK) Consortium | Jacquelyn Wynn, Mairead Hyland, The Lighthouse Lab in Alderley Park and Alex Alderton, Roberto Amato, Jeffrey Barrett, Sonia Goncalves, Ewan Harrison, David K. Jackson, Ian Johnston, Dominic Kwiatkowski, Cordelia Langford, John Sillitoe on behalf of the Wellcome Sanger Institute COVID-19 Surveillance Team |
| EPI_ISL_3284405 | Lighthouse Lab in Glasgow | Wellcome Sanger Institute for the COVID-19 Genomics UK (COG-UK) Consortium | Harper VanSteenhouse, Yumi Kasai, David Gray, Carol Clugston, Anna Dominiczak and Alex Alderton, Roberto Amato, Jeffrey Barrett, Sonia Goncalves, Ewan Harrison, David K. Jackson, Ian Johnston, Dominic Kwiatkowski, Cordelia Langford, John Sillitoe on behalf of the Wellcome Sanger Institute COVID-19 Surveillance Team |
| EPI_ISL_3284406, EPI_ISL_3284407, EPI_ISL_3284408, EPI_ISL_3284409, EPI_ISL_3284410 | Lighthouse Lab in Alderley Park | Wellcome Sanger Institute for the COVID-19 Genomics UK (COG-UK) Consortium | Jacquelyn Wynn, Mairead Hyland, The Lighthouse Lab in Alderley Park and Alex Alderton, Roberto Amato, Jeffrey Barrett, Sonia Goncalves, Ewan Harrison, David K. Jackson, Ian Johnston, Dominic Kwiatkowski, Cordelia Langford, John Sillitoe on behalf of the Wellcome Sanger Institute COVID-19 Surveillance Team |
| EPI_ISL_3284411 | Lighthouse Lab in Glasgow | Wellcome Sanger Institute for the COVID-19 Genomics UK (COG-UK) Consortium | Harper VanSteenhouse, Yumi Kasai, David Gray, Carol Clugston, Anna Dominiczak and Alex Alderton, Roberto Amato, Jeffrey Barrett, Sonia Goncalves, Ewan Harrison, David K. Jackson, Ian Johnston, Dominic Kwiatkowski, Cordelia Langford, John Sillitoe on behalf of the Wellcome Sanger Institute COVID-19 Surveillance Team |
| EPI_ISL_3284412 | Lighthouse Lab in Alderley Park | Wellcome Sanger Institute for the COVID-19 Genomics UK (COG-UK) Consortium | Jacquelyn Wynn, Mairead Hyland, The Lighthouse Lab in Alderley Park and Alex Alderton, Roberto Amato, Jeffrey Barrett, Sonia Goncalves, Ewan Harrison, David K. Jackson, Ian Johnston, Dominic Kwiatkowski, Cordelia Langford, John Sillitoe on behalf of the Wellcome Sanger Institute COVID-19 Surveillance Team |
| EPI_ISL_3284413 | Lighthouse Lab in Glasgow | Wellcome Sanger Institute for the COVID-19 Genomics UK (COG-UK) Consortium | Harper VanSteenhouse, Yumi Kasai, David Gray, Carol Clugston, Anna Dominiczak and Alex Alderton, Roberto Amato, Jeffrey Barrett, Sonia Goncalves, Ewan Harrison, David K. Jackson, Ian Johnston, Dominic Kwiatkowski, Cordelia Langford, John Sillitoe on behalf of the Wellcome Sanger Institute COVID-19 Surveillance Team |
| EPI_ISL_3284414, EPI_ISL_3284415, EPI_ISL_3284416, | Lighthouse Lab in Alderley Park | Wellcome Sanger Institute for the COVID-19 Genomics UK (COG-UK) Consortium | Jacquelyn Wynn, Mairead Hyland, The Lighthouse Lab in Alderley Park and Alex Alderton, Roberto Amato, Jeffrey Barrett, Sonia Goncalves, Ewan Harrison, David K. Jackson, Ian Johnston, Dominic Kwiatkowski, Cordelia Langford, John Sillitoe on behalf of the Wellcome Sanger Institute COVID-19 Surveillance Team |

| | | | |
|---|---|---|---|
| EPI_ISL_3284417, EPI_ISL_3284418, EPI_ISL_3284419 | | | |
| EPI_ISL_3284420 | Lighthouse Lab in Glasgow | Wellcome Sanger Institute for the COVID-19 Genomics UK (COG-UK) Consortium | Harper VanSteenhouse, Yumi Kasai, David Gray, Carol Clugston, Anna Dominiczak and Alex Alderton, Roberto Amato, Jeffrey Barrett, Sonia Goncalves, Ewan Harrison, David K. Jackson, Ian Johnston, Dominic Kwiatkowski, Cordelia Langford, John Sillitoe on behalf of the Wellcome Sanger Institute COVID-19 Surveillance Team |
| EPI_ISL_3284421, EPI_ISL_3284422, EPI_ISL_3284423 | Lighthouse Lab in Alderley Park | Wellcome Sanger Institute for the COVID-19 Genomics UK (COG-UK) Consortium | Jacquelyn Wynn, Mairead Hyland, The Lighthouse Lab in Alderley Park and Alex Alderton, Roberto Amato, Jeffrey Barrett, Sonia Goncalves, Ewan Harrison, David K. Jackson, Ian Johnston, Dominic Kwiatkowski, Cordelia Langford, John Sillitoe on behalf of the Wellcome Sanger Institute COVID-19 Surveillance Team |
| EPI_ISL_3284424 | Lighthouse Lab in Glasgow | Wellcome Sanger Institute for the COVID-19 Genomics UK (COG-UK) Consortium | Harper VanSteenhouse, Yumi Kasai, David Gray, Carol Clugston, Anna Dominiczak and Alex Alderton, Roberto Amato, Jeffrey Barrett, Sonia Goncalves, Ewan Harrison, David K. Jackson, Ian Johnston, Dominic Kwiatkowski, Cordelia Langford, John Sillitoe on behalf of the Wellcome Sanger Institute COVID-19 Surveillance Team |
| EPI_ISL_3284425, EPI_ISL_3284426 | Lighthouse Lab in Alderley Park | Wellcome Sanger Institute for the COVID-19 Genomics UK (COG-UK) Consortium | Jacquelyn Wynn, Mairead Hyland, The Lighthouse Lab in Alderley Park and Alex Alderton, Roberto Amato, Jeffrey Barrett, Sonia Goncalves, Ewan Harrison, David K. Jackson, Ian Johnston, Dominic Kwiatkowski, Cordelia Langford, John Sillitoe on behalf of the Wellcome Sanger Institute COVID-19 Surveillance Team |
| EPI_ISL_3284427 | Lighthouse Lab in Glasgow | Wellcome Sanger Institute for the COVID-19 Genomics UK (COG-UK) Consortium | Harper VanSteenhouse, Yumi Kasai, David Gray, Carol Clugston, Anna Dominiczak and Alex Alderton, Roberto Amato, Jeffrey Barrett, Sonia Goncalves, Ewan Harrison, David K. Jackson, Ian Johnston, Dominic Kwiatkowski, Cordelia Langford, John Sillitoe on behalf of the Wellcome Sanger Institute COVID-19 Surveillance Team |
| EPI_ISL_3284428, EPI_ISL_3284429, EPI_ISL_3284430, EPI_ISL_3284431, EPI_ISL_3284432 | Lighthouse Lab in Alderley Park | Wellcome Sanger Institute for the COVID-19 Genomics UK (COG-UK) Consortium | Jacquelyn Wynn, Mairead Hyland, The Lighthouse Lab in Alderley Park and Alex Alderton, Roberto Amato, Jeffrey Barrett, Sonia Goncalves, Ewan Harrison, David K. Jackson, Ian Johnston, Dominic Kwiatkowski, Cordelia Langford, John Sillitoe on behalf of the Wellcome Sanger Institute COVID-19 Surveillance Team |
| EPI_ISL_3284433 | Lighthouse Lab in Glasgow | Wellcome Sanger Institute for the COVID-19 Genomics UK (COG-UK) Consortium | Harper VanSteenhouse, Yumi Kasai, David Gray, Carol Clugston, Anna Dominiczak and Alex Alderton, Roberto Amato, Jeffrey Barrett, Sonia Goncalves, Ewan Harrison, David K. Jackson, Ian Johnston, Dominic Kwiatkowski, Cordelia Langford, John Sillitoe on behalf of the Wellcome Sanger Institute COVID-19 Surveillance Team |
| EPI_ISL_3284434, EPI_ISL_3284435 | Lighthouse Lab in Alderley Park | Wellcome Sanger Institute for the COVID-19 Genomics UK (COG-UK) Consortium | Jacquelyn Wynn, Mairead Hyland, The Lighthouse Lab in Alderley Park and Alex Alderton, Roberto Amato, Jeffrey Barrett, Sonia Goncalves, Ewan Harrison, David K. Jackson, Ian Johnston, Dominic Kwiatkowski, Cordelia Langford, John Sillitoe on behalf of the Wellcome Sanger Institute COVID-19 Surveillance Team |
| EPI_ISL_3284436, EPI_ISL_3284437 | Lighthouse Lab in Glasgow | Wellcome Sanger Institute for the COVID-19 Genomics UK (COG-UK) Consortium | Harper VanSteenhouse, Yumi Kasai, David Gray, Carol Clugston, Anna Dominiczak and Alex Alderton, Roberto Amato, Jeffrey Barrett, Sonia Goncalves, Ewan Harrison, David K. Jackson, Ian Johnston, Dominic Kwiatkowski, Cordelia Langford, John Sillitoe on behalf of the Wellcome Sanger Institute COVID-19 Surveillance Team |
| EPI_ISL_3284438, EPI_ISL_3284439, EPI_ISL_3284440, EPI_ISL_3284441 | Lighthouse Lab in Alderley Park | Wellcome Sanger Institute for the COVID-19 Genomics UK (COG-UK) Consortium | Jacquelyn Wynn, Mairead Hyland, The Lighthouse Lab in Alderley Park and Alex Alderton, Roberto Amato, Jeffrey Barrett, Sonia Goncalves, Ewan Harrison, David K. Jackson, Ian Johnston, Dominic Kwiatkowski, Cordelia Langford, John Sillitoe on behalf of the Wellcome Sanger Institute COVID-19 Surveillance Team |
| EPI_ISL_3284442 | Lighthouse Lab in Glasgow | Wellcome Sanger Institute for the COVID-19 Genomics UK (COG-UK) Consortium | Harper VanSteenhouse, Yumi Kasai, David Gray, Carol Clugston, Anna Dominiczak and Alex Alderton, Roberto Amato, Jeffrey Barrett, Sonia Goncalves, Ewan Harrison, David K. Jackson, Ian Johnston, Dominic Kwiatkowski, Cordelia Langford, John Sillitoe on behalf of the Wellcome Sanger Institute COVID-19 Surveillance Team |
| EPI_ISL_3284443, EPI_ISL_3284444, EPI_ISL_3284445 | Lighthouse Lab in Alderley Park | Wellcome Sanger Institute for the COVID-19 Genomics UK (COG-UK) Consortium | Jacquelyn Wynn, Mairead Hyland, The Lighthouse Lab in Alderley Park and Alex Alderton, Roberto Amato, Jeffrey Barrett, Sonia Goncalves, Ewan Harrison, David K. Jackson, Ian Johnston, Dominic Kwiatkowski, Cordelia Langford, John Sillitoe on behalf of the Wellcome Sanger Institute COVID-19 Surveillance Team |
| EPI_ISL_3284446, EPI_ISL_3284447 | Lighthouse Lab in Glasgow | Wellcome Sanger Institute for the COVID-19 Genomics UK (COG-UK) Consortium | Harper VanSteenhouse, Yumi Kasai, David Gray, Carol Clugston, Anna Dominiczak and Alex Alderton, Roberto Amato, Jeffrey Barrett, Sonia Goncalves, Ewan Harrison, David K. Jackson, Ian Johnston, Dominic Kwiatkowski, Cordelia Langford, John Sillitoe on behalf of the Wellcome Sanger Institute COVID-19 Surveillance Team |
| EPI_ISL_3284448, EPI_ISL_3284449 | Lighthouse Lab in Alderley Park | Wellcome Sanger Institute for the COVID-19 Genomics UK (COG-UK) Consortium | Jacquelyn Wynn, Mairead Hyland, The Lighthouse Lab in Alderley Park and Alex Alderton, Roberto Amato, Jeffrey Barrett, Sonia Goncalves, Ewan Harrison, David K. Jackson, Ian Johnston, Dominic Kwiatkowski, Cordelia Langford, John Sillitoe on behalf of the Wellcome Sanger Institute COVID-19 Surveillance Team |
| EPI_ISL_3284450, EPI_ISL_3284451, EPI_ISL_3284452 | Lighthouse Lab in Glasgow | Wellcome Sanger Institute for the COVID-19 Genomics UK (COG-UK) Consortium | Harper VanSteenhouse, Yumi Kasai, David Gray, Carol Clugston, Anna Dominiczak and Alex Alderton, Roberto Amato, Jeffrey Barrett, Sonia Goncalves, Ewan Harrison, David K. Jackson, Ian Johnston, Dominic Kwiatkowski, Cordelia Langford, John Sillitoe on behalf of the Wellcome Sanger Institute COVID-19 Surveillance Team |
| EPI_ISL_3284453 | Lighthouse Lab in Alderley Park | Wellcome Sanger Institute for the COVID-19 Genomics UK (COG-UK) Consortium | Jacquelyn Wynn, Mairead Hyland, The Lighthouse Lab in Alderley Park and Alex Alderton, Roberto Amato, Jeffrey Barrett, Sonia Goncalves, Ewan Harrison, David K. Jackson, Ian Johnston, Dominic Kwiatkowski, Cordelia Langford, John Sillitoe on behalf of the Wellcome Sanger Institute COVID-19 Surveillance Team |
| EPI_ISL_3284454 | Lighthouse Lab in Glasgow | Wellcome Sanger Institute for the COVID-19 Genomics UK (COG-UK) Consortium | Harper VanSteenhouse, Yumi Kasai, David Gray, Carol Clugston, Anna Dominiczak and Alex Alderton, Roberto Amato, Jeffrey Barrett, Sonia Goncalves, Ewan Harrison, David K. Jackson, Ian Johnston, Dominic Kwiatkowski, Cordelia Langford, John Sillitoe on behalf of the Wellcome Sanger Institute COVID-19 Surveillance Team |
| EPI_ISL_3284455, EPI_ISL_3284456, EPI_ISL_3284457 | Lighthouse Lab in Alderley Park | Wellcome Sanger Institute for the COVID-19 Genomics UK (COG-UK) Consortium | Jacquelyn Wynn, Mairead Hyland, The Lighthouse Lab in Alderley Park and Alex Alderton, Roberto Amato, Jeffrey Barrett, Sonia Goncalves, Ewan Harrison, David K. Jackson, Ian Johnston, Dominic Kwiatkowski, Cordelia Langford, John Sillitoe on behalf of the Wellcome Sanger Institute COVID-19 Surveillance Team |
| EPI_ISL_3284458 | Lighthouse Lab in Glasgow | Wellcome Sanger Institute for the COVID-19 Genomics UK (COG-UK) Consortium | Harper VanSteenhouse, Yumi Kasai, David Gray, Carol Clugston, Anna Dominiczak and Alex Alderton, Roberto Amato, Jeffrey Barrett, Sonia Goncalves, Ewan Harrison, David K. Jackson, Ian Johnston, Dominic Kwiatkowski, Cordelia Langford, John Sillitoe on behalf of the Wellcome Sanger Institute COVID-19 Surveillance Team |
| EPI_ISL_3284459, EPI_ISL_3284460 | Lighthouse Lab in Alderley Park | Wellcome Sanger Institute for the COVID-19 Genomics UK (COG-UK) Consortium | Jacquelyn Wynn, Mairead Hyland, The Lighthouse Lab in Alderley Park and Alex Alderton, Roberto Amato, Jeffrey Barrett, Sonia Goncalves, Ewan Harrison, David K. Jackson, Ian Johnston, Dominic Kwiatkowski, Cordelia Langford, John Sillitoe on behalf of the Wellcome Sanger Institute COVID-19 Surveillance Team |
| EPI_ISL_3284461 | Lighthouse Lab in Glasgow | Wellcome Sanger Institute for the COVID-19 Genomics UK (COG-UK) Consortium | Harper VanSteenhouse, Yumi Kasai, David Gray, Carol Clugston, Anna Dominiczak and Alex Alderton, Roberto Amato, Jeffrey Barrett, Sonia Goncalves, Ewan Harrison, David K. Jackson, Ian Johnston, Dominic Kwiatkowski, Cordelia Langford, John Sillitoe on behalf of the Wellcome Sanger Institute COVID-19 Surveillance Team |
| EPI_ISL_3284462, EPI_ISL_3284463, EPI_ISL_3284464 | Lighthouse Lab in Alderley Park | Wellcome Sanger Institute for the COVID-19 Genomics UK (COG-UK) Consortium | Jacquelyn Wynn, Mairead Hyland, The Lighthouse Lab in Alderley Park and Alex Alderton, Roberto Amato, Jeffrey Barrett, Sonia Goncalves, Ewan Harrison, David K. Jackson, Ian Johnston, Dominic Kwiatkowski, Cordelia Langford, John Sillitoe on behalf of the Wellcome Sanger Institute COVID-19 Surveillance Team |
| EPI_ISL_3284465 | Lighthouse Lab in Glasgow | Wellcome Sanger Institute for the COVID-19 Genomics UK (COG-UK) Consortium | Harper VanSteenhouse, Yumi Kasai, David Gray, Carol Clugston, Anna Dominiczak and Alex Alderton, Roberto Amato, Jeffrey Barrett, Sonia Goncalves, Ewan Harrison, David K. Jackson, Ian Johnston, Dominic Kwiatkowski, Cordelia Langford, John Sillitoe on behalf of the Wellcome Sanger Institute COVID-19 Surveillance Team |
| EPI_ISL_3284466, EPI_ISL_3284467 | Lighthouse Lab in Alderley Park | Wellcome Sanger Institute for the COVID-19 Genomics UK (COG-UK) Consortium | Jacquelyn Wynn, Mairead Hyland, The Lighthouse Lab in Alderley Park and Alex Alderton, Roberto Amato, Jeffrey Barrett, Sonia Goncalves, Ewan Harrison, David K. Jackson, Ian Johnston, Dominic Kwiatkowski, Cordelia Langford, John Sillitoe on behalf of the Wellcome Sanger Institute COVID-19 Surveillance Team |
| EPI_ISL_3284468, EPI_ISL_3284469, EPI_ISL_3284470 | Lighthouse Lab in Glasgow | Wellcome Sanger Institute for the COVID-19 Genomics UK (COG-UK) Consortium | Harper VanSteenhouse, Yumi Kasai, David Gray, Carol Clugston, Anna Dominiczak and Alex Alderton, Roberto Amato, Jeffrey Barrett, Sonia Goncalves, Ewan Harrison, David K. Jackson, Ian Johnston, Dominic Kwiatkowski, Cordelia Langford, John Sillitoe on behalf of the Wellcome Sanger Institute COVID-19 Surveillance Team |
| EPI_ISL_3284471, EPI_ISL_3284472 | Lighthouse Lab in Alderley Park | Wellcome Sanger Institute for the COVID-19 Genomics UK (COG-UK) Consortium | Jacquelyn Wynn, Mairead Hyland, The Lighthouse Lab in Alderley Park and Alex Alderton, Roberto Amato, Jeffrey Barrett, Sonia Goncalves, Ewan Harrison, David K. Jackson, Ian Johnston, Dominic Kwiatkowski, Cordelia Langford, John Sillitoe on behalf of the Wellcome Sanger Institute COVID-19 Surveillance Team |
| EPI_ISL_3284473 | Lighthouse Lab in Glasgow | Wellcome Sanger Institute for the COVID-19 Genomics UK (COG-UK) Consortium | Harper VanSteenhouse, Yumi Kasai, David Gray, Carol Clugston, Anna Dominiczak and Alex Alderton, Roberto Amato, Jeffrey Barrett, Sonia Goncalves, Ewan Harrison, David K. Jackson, Ian Johnston, Dominic Kwiatkowski, Cordelia Langford, John Sillitoe on behalf of the Wellcome Sanger Institute COVID-19 Surveillance Team |
| EPI_ISL_3284474, EPI_ISL_3284475 | Lighthouse Lab in Alderley Park | Wellcome Sanger Institute for the COVID-19 Genomics UK (COG-UK) Consortium | Jacquelyn Wynn, Mairead Hyland, The Lighthouse Lab in Alderley Park and Alex Alderton, Roberto Amato, Jeffrey Barrett, Sonia Goncalves, Ewan Harrison, David K. Jackson, Ian Johnston, Dominic Kwiatkowski, Cordelia Langford, John Sillitoe on behalf of the Wellcome Sanger Institute COVID-19 Surveillance Team |
| EPI_ISL_3284476, EPI_ISL_3284477, EPI_ISL_3284478 | Lighthouse Lab in Glasgow | Wellcome Sanger Institute for the COVID-19 Genomics UK (COG-UK) Consortium | Harper VanSteenhouse, Yumi Kasai, David Gray, Carol Clugston, Anna Dominiczak and Alex Alderton, Roberto Amato, Jeffrey Barrett, Sonia Goncalves, Ewan Harrison, David K. Jackson, Ian Johnston, Dominic Kwiatkowski, Cordelia Langford, John Sillitoe on behalf of the Wellcome Sanger Institute COVID-19 Surveillance Team |
| EPI_ISL_3284479, EPI_ISL_3284480, EPI_ISL_3284481 | Lighthouse Lab in Alderley Park | Wellcome Sanger Institute for the COVID-19 Genomics UK (COG-UK) Consortium | Jacquelyn Wynn, Mairead Hyland, The Lighthouse Lab in Alderley Park and Alex Alderton, Roberto Amato, Jeffrey Barrett, Sonia Goncalves, Ewan Harrison, David K. Jackson, Ian Johnston, Dominic Kwiatkowski, Cordelia Langford, John Sillitoe on behalf of the Wellcome Sanger Institute COVID-19 Surveillance Team |
| EPI_ISL_3284482 | Lighthouse Lab in Glasgow | Wellcome Sanger Institute for the COVID-19 Genomics UK (COG-UK) Consortium | Harper VanSteenhouse, Yumi Kasai, David Gray, Carol Clugston, Anna Dominiczak and Alex Alderton, Roberto Amato, Jeffrey Barrett, Sonia Goncalves, Ewan Harrison, David K. Jackson, Ian Johnston, Dominic Kwiatkowski, Cordelia Langford, John Sillitoe on behalf of the Wellcome Sanger Institute COVID-19 Surveillance Team |
| EPI_ISL_3284483, EPI_ISL_3284484 | Lighthouse Lab in Alderley Park | Wellcome Sanger Institute for the COVID-19 Genomics UK (COG-UK) Consortium | Jacquelyn Wynn, Mairead Hyland, The Lighthouse Lab in Alderley Park and Alex Alderton, Roberto Amato, Jeffrey Barrett, Sonia Goncalves, Ewan Harrison, David K. Jackson, Ian Johnston, Dominic Kwiatkowski, Cordelia Langford, John Sillitoe on behalf of the Wellcome Sanger Institute COVID-19 Surveillance Team |
| EPI_ISL_3284485, EPI_ISL_3284486, EPI_ISL_3284487 | Lighthouse Lab in Glasgow | Wellcome Sanger Institute for the COVID-19 Genomics UK (COG-UK) Consortium | Harper VanSteenhouse, Yumi Kasai, David Gray, Carol Clugston, Anna Dominiczak and Alex Alderton, Roberto Amato, Jeffrey Barrett, Sonia Goncalves, Ewan Harrison, David K. Jackson, Ian Johnston, Dominic Kwiatkowski, Cordelia Langford, John Sillitoe on behalf of the Wellcome Sanger Institute COVID-19 Surveillance Team |
| EPI_ISL_3284488, EPI_ISL_3284489 | Lighthouse Lab in Alderley Park | Wellcome Sanger Institute for the COVID-19 Genomics UK (COG-UK) Consortium | Jacquelyn Wynn, Mairead Hyland, The Lighthouse Lab in Alderley Park and Alex Alderton, Roberto Amato, Jeffrey Barrett, Sonia Goncalves, Ewan Harrison, David K. Jackson, Ian Johnston, Dominic Kwiatkowski, Cordelia Langford, John Sillitoe on behalf of the Wellcome Sanger Institute COVID-19 Surveillance Team |
| EPI_ISL_3284490, EPI_ISL_3284491, EPI_ISL_3284492, EPI_ISL_3284493 | Lighthouse Lab in Glasgow | Wellcome Sanger Institute for the COVID-19 Genomics UK (COG-UK) Consortium | Harper VanSteenhouse, Yumi Kasai, David Gray, Carol Clugston, Anna Dominiczak and Alex Alderton, Roberto Amato, Jeffrey Barrett, Sonia Goncalves, Ewan Harrison, David K. Jackson, Ian Johnston, Dominic Kwiatkowski, Cordelia Langford, John Sillitoe on behalf of the Wellcome Sanger Institute COVID-19 Surveillance Team |
| EPI_ISL_3284494, EPI_ISL_3284495, EPI_ISL_3284496 | Lighthouse Lab in Alderley Park | Wellcome Sanger Institute for the COVID-19 Genomics UK (COG-UK) Consortium | Jacquelyn Wynn, Mairead Hyland, The Lighthouse Lab in Alderley Park and Alex Alderton, Roberto Amato, Jeffrey Barrett, Sonia Goncalves, Ewan Harrison, David K. Jackson, Ian Johnston, Dominic Kwiatkowski, Cordelia Langford, John Sillitoe on behalf of the Wellcome Sanger Institute COVID-19 Surveillance Team |
| EPI_ISL_3284497, EPI_ISL_3284498, | Lighthouse Lab in Glasgow | Wellcome Sanger Institute for the COVID-19 Genomics UK (COG-UK) Consortium | Harper VanSteenhouse, Yumi Kasai, David Gray, Carol Clugston, Anna Dominiczak and Alex Alderton, Roberto Amato, Jeffrey Barrett, Sonia Goncalves, Ewan Harrison, David K. Jackson, Ian Johnston, Dominic Kwiatkowski, Cordelia Langford, John Sillitoe on behalf of the Wellcome Sanger Institute COVID-19 Surveillance Team |

| | | | |
|---|---|---|---|
| EPI_ISL_3284499, EPI_ISL_3284500, EPI_ISL_3284501 | | | |
| EPI_ISL_3284502, EPI_ISL_3284503, EPI_ISL_3284504 | Lighthouse Lab in Alderley Park | Wellcome Sanger Institute for the COVID-19 Genomics UK (COG-UK) Consortium | Jacquelyn Wynn, Mairead Hyland, The Lighthouse Lab in Alderley Park and Alex Alderton, Roberto Amato, Jeffrey Barrett, Sonia Goncalves, Ewan Harrison, David K. Jackson, Ian Johnston, Dominic Kwiatkowski, Cordelia Langford, John Sillitoe on behalf of the Wellcome Sanger Institute COVID-19 Surveillance Team |
| EPI_ISL_3284505 | Lighthouse Lab in Glasgow | Wellcome Sanger Institute for the COVID-19 Genomics UK (COG-UK) Consortium | Harper VanSteenhouse, Yumi Kasai, David Gray, Carol Clugston, Anna Dominiczak and Alex Alderton, Roberto Amato, Jeffrey Barrett, Sonia Goncalves, Ewan Harrison, David K. Jackson, Ian Johnston, Dominic Kwiatkowski, Cordelia Langford, John Sillitoe on behalf of the Wellcome Sanger Institute COVID-19 Surveillance Team |
| EPI_ISL_3284506 | Lighthouse Lab in Alderley Park | Wellcome Sanger Institute for the COVID-19 Genomics UK (COG-UK) Consortium | Jacquelyn Wynn, Mairead Hyland, The Lighthouse Lab in Alderley Park and Alex Alderton, Roberto Amato, Jeffrey Barrett, Sonia Goncalves, Ewan Harrison, David K. Jackson, Ian Johnston, Dominic Kwiatkowski, Cordelia Langford, John Sillitoe on behalf of the Wellcome Sanger Institute COVID-19 Surveillance Team |
| EPI_ISL_3284507 | Lighthouse Lab in Glasgow | Wellcome Sanger Institute for the COVID-19 Genomics UK (COG-UK) Consortium | Harper VanSteenhouse, Yumi Kasai, David Gray, Carol Clugston, Anna Dominiczak and Alex Alderton, Roberto Amato, Jeffrey Barrett, Sonia Goncalves, Ewan Harrison, David K. Jackson, Ian Johnston, Dominic Kwiatkowski, Cordelia Langford, John Sillitoe on behalf of the Wellcome Sanger Institute COVID-19 Surveillance Team |
| EPI_ISL_3284508 | Lighthouse Lab in Alderley Park | Wellcome Sanger Institute for the COVID-19 Genomics UK (COG-UK) Consortium | Jacquelyn Wynn, Mairead Hyland, The Lighthouse Lab in Alderley Park and Alex Alderton, Roberto Amato, Jeffrey Barrett, Sonia Goncalves, Ewan Harrison, David K. Jackson, Ian Johnston, Dominic Kwiatkowski, Cordelia Langford, John Sillitoe on behalf of the Wellcome Sanger Institute COVID-19 Surveillance Team |
| EPI_ISL_3284509 | Lighthouse Lab in Glasgow | Wellcome Sanger Institute for the COVID-19 Genomics UK (COG-UK) Consortium | Harper VanSteenhouse, Yumi Kasai, David Gray, Carol Clugston, Anna Dominiczak and Alex Alderton, Roberto Amato, Jeffrey Barrett, Sonia Goncalves, Ewan Harrison, David K. Jackson, Ian Johnston, Dominic Kwiatkowski, Cordelia Langford, John Sillitoe on behalf of the Wellcome Sanger Institute COVID-19 Surveillance Team |
| EPI_ISL_3284510, EPI_ISL_3284511, EPI_ISL_3284512, EPI_ISL_3284513 | Lighthouse Lab in Alderley Park | Wellcome Sanger Institute for the COVID-19 Genomics UK (COG-UK) Consortium | Jacquelyn Wynn, Mairead Hyland, The Lighthouse Lab in Alderley Park and Alex Alderton, Roberto Amato, Jeffrey Barrett, Sonia Goncalves, Ewan Harrison, David K. Jackson, Ian Johnston, Dominic Kwiatkowski, Cordelia Langford, John Sillitoe on behalf of the Wellcome Sanger Institute COVID-19 Surveillance Team |
| EPI_ISL_3284514 | Lighthouse Lab in Glasgow | Wellcome Sanger Institute for the COVID-19 Genomics UK (COG-UK) Consortium | Harper VanSteenhouse, Yumi Kasai, David Gray, Carol Clugston, Anna Dominiczak and Alex Alderton, Roberto Amato, Jeffrey Barrett, Sonia Goncalves, Ewan Harrison, David K. Jackson, Ian Johnston, Dominic Kwiatkowski, Cordelia Langford, John Sillitoe on behalf of the Wellcome Sanger Institute COVID-19 Surveillance Team |
| EPI_ISL_3284515 | Lighthouse Lab in Alderley Park | Wellcome Sanger Institute for the COVID-19 Genomics UK (COG-UK) Consortium | Jacquelyn Wynn, Mairead Hyland, The Lighthouse Lab in Alderley Park and Alex Alderton, Roberto Amato, Jeffrey Barrett, Sonia Goncalves, Ewan Harrison, David K. Jackson, Ian Johnston, Dominic Kwiatkowski, Cordelia Langford, John Sillitoe on behalf of the Wellcome Sanger Institute COVID-19 Surveillance Team |
| EPI_ISL_3284516 | Lighthouse Lab in Glasgow | Wellcome Sanger Institute for the COVID-19 Genomics UK (COG-UK) Consortium | Harper VanSteenhouse, Yumi Kasai, David Gray, Carol Clugston, Anna Dominiczak and Alex Alderton, Roberto Amato, Jeffrey Barrett, Sonia Goncalves, Ewan Harrison, David K. Jackson, Ian Johnston, Dominic Kwiatkowski, Cordelia Langford, John Sillitoe on behalf of the Wellcome Sanger Institute COVID-19 Surveillance Team |
| EPI_ISL_3284517, EPI_ISL_3284518, EPI_ISL_3284519 | Lighthouse Lab in Alderley Park | Wellcome Sanger Institute for the COVID-19 Genomics UK (COG-UK) Consortium | Jacquelyn Wynn, Mairead Hyland, The Lighthouse Lab in Alderley Park and Alex Alderton, Roberto Amato, Jeffrey Barrett, Sonia Goncalves, Ewan Harrison, David K. Jackson, Ian Johnston, Dominic Kwiatkowski, Cordelia Langford, John Sillitoe on behalf of the Wellcome Sanger Institute COVID-19 Surveillance Team |
| EPI_ISL_3284520, EPI_ISL_3284521, EPI_ISL_3284522 | Lighthouse Lab in Glasgow | Wellcome Sanger Institute for the COVID-19 Genomics UK (COG-UK) Consortium | Harper VanSteenhouse, Yumi Kasai, David Gray, Carol Clugston, Anna Dominiczak and Alex Alderton, Roberto Amato, Jeffrey Barrett, Sonia Goncalves, Ewan Harrison, David K. Jackson, Ian Johnston, Dominic Kwiatkowski, Cordelia Langford, John Sillitoe on behalf of the Wellcome Sanger Institute COVID-19 Surveillance Team |
| EPI_ISL_3284523, EPI_ISL_3284524, EPI_ISL_3284525, EPI_ISL_3284526 | Lighthouse Lab in Alderley Park | Wellcome Sanger Institute for the COVID-19 Genomics UK (COG-UK) Consortium | Jacquelyn Wynn, Mairead Hyland, The Lighthouse Lab in Alderley Park and Alex Alderton, Roberto Amato, Jeffrey Barrett, Sonia Goncalves, Ewan Harrison, David K. Jackson, Ian Johnston, Dominic Kwiatkowski, Cordelia Langford, John Sillitoe on behalf of the Wellcome Sanger Institute COVID-19 Surveillance Team |
| EPI_ISL_3284527 | Lighthouse Lab in Glasgow | Wellcome Sanger Institute for the COVID-19 Genomics UK (COG-UK) Consortium | Harper VanSteenhouse, Yumi Kasai, David Gray, Carol Clugston, Anna Dominiczak and Alex Alderton, Roberto Amato, Jeffrey Barrett, Sonia Goncalves, Ewan Harrison, David K. Jackson, Ian Johnston, Dominic Kwiatkowski, Cordelia Langford, John Sillitoe on behalf of the Wellcome Sanger Institute COVID-19 Surveillance Team |
| EPI_ISL_3284528, EPI_ISL_3284529 | Lighthouse Lab in Alderley Park | Wellcome Sanger Institute for the COVID-19 Genomics UK (COG-UK) Consortium | Jacquelyn Wynn, Mairead Hyland, The Lighthouse Lab in Alderley Park and Alex Alderton, Roberto Amato, Jeffrey Barrett, Sonia Goncalves, Ewan Harrison, David K. Jackson, Ian Johnston, Dominic Kwiatkowski, Cordelia Langford, John Sillitoe on behalf of the Wellcome Sanger Institute COVID-19 Surveillance Team |
| EPI_ISL_3284530, EPI_ISL_3284531, EPI_ISL_3284532 | Lighthouse Lab in Glasgow | Wellcome Sanger Institute for the COVID-19 Genomics UK (COG-UK) Consortium | Harper VanSteenhouse, Yumi Kasai, David Gray, Carol Clugston, Anna Dominiczak and Alex Alderton, Roberto Amato, Jeffrey Barrett, Sonia Goncalves, Ewan Harrison, David K. Jackson, Ian Johnston, Dominic Kwiatkowski, Cordelia Langford, John Sillitoe on behalf of the Wellcome Sanger Institute COVID-19 Surveillance Team |
| EPI_ISL_3284533 | Lighthouse Lab in Alderley Park | Wellcome Sanger Institute for the COVID-19 Genomics UK (COG-UK) Consortium | Jacquelyn Wynn, Mairead Hyland, The Lighthouse Lab in Alderley Park and Alex Alderton, Roberto Amato, Jeffrey Barrett, Sonia Goncalves, Ewan Harrison, David K. Jackson, Ian Johnston, Dominic Kwiatkowski, Cordelia Langford, John Sillitoe on behalf of the Wellcome Sanger Institute COVID-19 Surveillance Team |
| EPI_ISL_3284534 | Lighthouse Lab in Glasgow | Wellcome Sanger Institute for the COVID-19 Genomics UK (COG-UK) Consortium | Harper VanSteenhouse, Yumi Kasai, David Gray, Carol Clugston, Anna Dominiczak and Alex Alderton, Roberto Amato, Jeffrey Barrett, Sonia Goncalves, Ewan Harrison, David K. Jackson, Ian Johnston, Dominic Kwiatkowski, Cordelia Langford, John Sillitoe on behalf of the Wellcome Sanger Institute COVID-19 Surveillance Team |
| EPI_ISL_3284535, EPI_ISL_3284536, EPI_ISL_3284537, EPI_ISL_3284538 | Lighthouse Lab in Alderley Park | Wellcome Sanger Institute for the COVID-19 Genomics UK (COG-UK) Consortium | Jacquelyn Wynn, Mairead Hyland, The Lighthouse Lab in Alderley Park and Alex Alderton, Roberto Amato, Jeffrey Barrett, Sonia Goncalves, Ewan Harrison, David K. Jackson, Ian Johnston, Dominic Kwiatkowski, Cordelia Langford, John Sillitoe on behalf of the Wellcome Sanger Institute COVID-19 Surveillance Team |
| EPI_ISL_3284539, EPI_ISL_3284540 | Lighthouse Lab in Glasgow | Wellcome Sanger Institute for the COVID-19 Genomics UK (COG-UK) Consortium | Harper VanSteenhouse, Yumi Kasai, David Gray, Carol Clugston, Anna Dominiczak and Alex Alderton, Roberto Amato, Jeffrey Barrett, Sonia Goncalves, Ewan Harrison, David K. Jackson, Ian Johnston, Dominic Kwiatkowski, Cordelia Langford, John Sillitoe on behalf of the Wellcome Sanger Institute COVID-19 Surveillance Team |
| EPI_ISL_3284541, EPI_ISL_3284542, EPI_ISL_3284543 | Lighthouse Lab in Alderley Park | Wellcome Sanger Institute for the COVID-19 Genomics UK (COG-UK) Consortium | Jacquelyn Wynn, Mairead Hyland, The Lighthouse Lab in Alderley Park and Alex Alderton, Roberto Amato, Jeffrey Barrett, Sonia Goncalves, Ewan Harrison, David K. Jackson, Ian Johnston, Dominic Kwiatkowski, Cordelia Langford, John Sillitoe on behalf of the Wellcome Sanger Institute COVID-19 Surveillance Team |
| EPI_ISL_3284544, EPI_ISL_3284545 | Lighthouse Lab in Glasgow | Wellcome Sanger Institute for the COVID-19 Genomics UK (COG-UK) Consortium | Harper VanSteenhouse, Yumi Kasai, David Gray, Carol Clugston, Anna Dominiczak and Alex Alderton, Roberto Amato, Jeffrey Barrett, Sonia Goncalves, Ewan Harrison, David K. Jackson, Ian Johnston, Dominic Kwiatkowski, Cordelia Langford, John Sillitoe on behalf of the Wellcome Sanger Institute COVID-19 Surveillance Team |
| EPI_ISL_3284546 | Lighthouse Lab in Alderley Park | Wellcome Sanger Institute for the COVID-19 Genomics UK (COG-UK) Consortium | Jacquelyn Wynn, Mairead Hyland, The Lighthouse Lab in Alderley Park and Alex Alderton, Roberto Amato, Jeffrey Barrett, Sonia Goncalves, Ewan Harrison, David K. Jackson, Ian Johnston, Dominic Kwiatkowski, Cordelia Langford, John Sillitoe on behalf of the Wellcome Sanger Institute COVID-19 Surveillance Team |
| EPI_ISL_3284547 | Lighthouse Lab in Glasgow | Wellcome Sanger Institute for the COVID-19 Genomics UK (COG-UK) Consortium | Harper VanSteenhouse, Yumi Kasai, David Gray, Carol Clugston, Anna Dominiczak and Alex Alderton, Roberto Amato, Jeffrey Barrett, Sonia Goncalves, Ewan Harrison, David K. Jackson, Ian Johnston, Dominic Kwiatkowski, Cordelia Langford, John Sillitoe on behalf of the Wellcome Sanger Institute COVID-19 Surveillance Team |
| EPI_ISL_3284548, EPI_ISL_3284549, EPI_ISL_3284550 | Lighthouse Lab in Alderley Park | Wellcome Sanger Institute for the COVID-19 Genomics UK (COG-UK) Consortium | Jacquelyn Wynn, Mairead Hyland, The Lighthouse Lab in Alderley Park and Alex Alderton, Roberto Amato, Jeffrey Barrett, Sonia Goncalves, Ewan Harrison, David K. Jackson, Ian Johnston, Dominic Kwiatkowski, Cordelia Langford, John Sillitoe on behalf of the Wellcome Sanger Institute COVID-19 Surveillance Team |
| EPI_ISL_3284551 | Lighthouse Lab in Glasgow | Wellcome Sanger Institute for the COVID-19 Genomics UK (COG-UK) Consortium | Harper VanSteenhouse, Yumi Kasai, David Gray, Carol Clugston, Anna Dominiczak and Alex Alderton, Roberto Amato, Jeffrey Barrett, Sonia Goncalves, Ewan Harrison, David K. Jackson, Ian Johnston, Dominic Kwiatkowski, Cordelia Langford, John Sillitoe on behalf of the Wellcome Sanger Institute COVID-19 Surveillance Team |
| EPI_ISL_3284552 | Lighthouse Lab in Alderley Park | Wellcome Sanger Institute for the COVID-19 Genomics UK (COG-UK) Consortium | Jacquelyn Wynn, Mairead Hyland, The Lighthouse Lab in Alderley Park and Alex Alderton, Roberto Amato, Jeffrey Barrett, Sonia Goncalves, Ewan Harrison, David K. Jackson, Ian Johnston, Dominic Kwiatkowski, Cordelia Langford, John Sillitoe on behalf of the Wellcome Sanger Institute COVID-19 Surveillance Team |
| EPI_ISL_3284553 | Lighthouse Lab in Glasgow | Wellcome Sanger Institute for the COVID-19 Genomics UK (COG-UK) Consortium | Harper VanSteenhouse, Yumi Kasai, David Gray, Carol Clugston, Anna Dominiczak and Alex Alderton, Roberto Amato, Jeffrey Barrett, Sonia Goncalves, Ewan Harrison, David K. Jackson, Ian Johnston, Dominic Kwiatkowski, Cordelia Langford, John Sillitoe on behalf of the Wellcome Sanger Institute COVID-19 Surveillance Team |
| EPI_ISL_3284554 | Lighthouse Lab in Alderley Park | Wellcome Sanger Institute for the COVID-19 Genomics UK (COG-UK) Consortium | Jacquelyn Wynn, Mairead Hyland, The Lighthouse Lab in Alderley Park and Alex Alderton, Roberto Amato, Jeffrey Barrett, Sonia Goncalves, Ewan Harrison, David K. Jackson, Ian Johnston, Dominic Kwiatkowski, Cordelia Langford, John Sillitoe on behalf of the Wellcome Sanger Institute COVID-19 Surveillance Team |
| EPI_ISL_3284555, EPI_ISL_3284556 | Lighthouse Lab in Glasgow | Wellcome Sanger Institute for the COVID-19 Genomics UK (COG-UK) Consortium | Harper VanSteenhouse, Yumi Kasai, David Gray, Carol Clugston, Anna Dominiczak and Alex Alderton, Roberto Amato, Jeffrey Barrett, Sonia Goncalves, Ewan Harrison, David K. Jackson, Ian Johnston, Dominic Kwiatkowski, Cordelia Langford, John Sillitoe on behalf of the Wellcome Sanger Institute COVID-19 Surveillance Team |
| EPI_ISL_3284557 | Lighthouse Lab in Alderley Park | Wellcome Sanger Institute for the COVID-19 Genomics UK (COG-UK) Consortium | Jacquelyn Wynn, Mairead Hyland, The Lighthouse Lab in Alderley Park and Alex Alderton, Roberto Amato, Jeffrey Barrett, Sonia Goncalves, Ewan Harrison, David K. Jackson, Ian Johnston, Dominic Kwiatkowski, Cordelia Langford, John Sillitoe on behalf of the Wellcome Sanger Institute COVID-19 Surveillance Team |
| EPI_ISL_3284558 | Lighthouse Lab in Glasgow | Wellcome Sanger Institute for the COVID-19 Genomics UK (COG-UK) Consortium | Harper VanSteenhouse, Yumi Kasai, David Gray, Carol Clugston, Anna Dominiczak and Alex Alderton, Roberto Amato, Jeffrey Barrett, Sonia Goncalves, Ewan Harrison, David K. Jackson, Ian Johnston, Dominic Kwiatkowski, Cordelia Langford, John Sillitoe on behalf of the Wellcome Sanger Institute COVID-19 Surveillance Team |
| EPI_ISL_3284559, EPI_ISL_3284560, EPI_ISL_3284561 | Lighthouse Lab in Alderley Park | Wellcome Sanger Institute for the COVID-19 Genomics UK (COG-UK) Consortium | Jacquelyn Wynn, Mairead Hyland, The Lighthouse Lab in Alderley Park and Alex Alderton, Roberto Amato, Jeffrey Barrett, Sonia Goncalves, Ewan Harrison, David K. Jackson, Ian Johnston, Dominic Kwiatkowski, Cordelia Langford, John Sillitoe on behalf of the Wellcome Sanger Institute COVID-19 Surveillance Team |
| EPI_ISL_3284562 | Lighthouse Lab in Glasgow | Wellcome Sanger Institute for the COVID-19 Genomics UK (COG-UK) Consortium | Harper VanSteenhouse, Yumi Kasai, David Gray, Carol Clugston, Anna Dominiczak and Alex Alderton, Roberto Amato, Jeffrey Barrett, Sonia Goncalves, Ewan Harrison, David K. Jackson, Ian Johnston, Dominic Kwiatkowski, Cordelia Langford, John Sillitoe on behalf of the Wellcome Sanger Institute COVID-19 Surveillance Team |
| EPI_ISL_3284563, EPI_ISL_3284564 | Lighthouse Lab in Alderley Park | Wellcome Sanger Institute for the COVID-19 Genomics UK (COG-UK) Consortium | Jacquelyn Wynn, Mairead Hyland, The Lighthouse Lab in Alderley Park and Alex Alderton, Roberto Amato, Jeffrey Barrett, Sonia Goncalves, Ewan Harrison, David K. Jackson, Ian Johnston, Dominic Kwiatkowski, Cordelia Langford, John Sillitoe on behalf of the Wellcome Sanger Institute COVID-19 Surveillance Team |
| EPI_ISL_3284565 | Lighthouse Lab in Glasgow | Wellcome Sanger Institute for the COVID-19 Genomics UK (COG-UK) Consortium | Harper VanSteenhouse, Yumi Kasai, David Gray, Carol Clugston, Anna Dominiczak and Alex Alderton, Roberto Amato, Jeffrey Barrett, Sonia Goncalves, Ewan Harrison, David K. Jackson, Ian Johnston, Dominic Kwiatkowski, Cordelia Langford, John Sillitoe on behalf of the Wellcome Sanger Institute COVID-19 Surveillance Team |
| EPI_ISL_3284566, EPI_ISL_3284567, EPI_ISL_3284568 | Lighthouse Lab in Alderley Park | Wellcome Sanger Institute for the COVID-19 Genomics UK (COG-UK) Consortium | Jacquelyn Wynn, Mairead Hyland, The Lighthouse Lab in Alderley Park and Alex Alderton, Roberto Amato, Jeffrey Barrett, Sonia Goncalves, Ewan Harrison, David K. Jackson, Ian Johnston, Dominic Kwiatkowski, Cordelia Langford, John Sillitoe on behalf of the Wellcome Sanger Institute COVID-19 Surveillance Team |
| EPI_ISL_3284569, EPI_ISL_3284570, | Lighthouse Lab in Glasgow | Wellcome Sanger Institute for the COVID-19 Genomics UK (COG-UK) Consortium | Harper VanSteenhouse, Yumi Kasai, David Gray, Carol Clugston, Anna Dominiczak and Alex Alderton, Roberto Amato, Jeffrey Barrett, Sonia Goncalves, Ewan Harrison, David K. Jackson, Ian Johnston, Dominic Kwiatkowski, Cordelia Langford, John Sillitoe on behalf of the Wellcome Sanger Institute COVID-19 Surveillance Team |

| | | | |
|---|---|---|---|
| EPI_ISL_3284571, EPI_ISL_3284572, EPI_ISL_3284573 | | | |
| EPI_ISL_3284574, EPI_ISL_3284575, EPI_ISL_3284576 | Lighthouse Lab in Alderley Park | Wellcome Sanger Institute for the COVID-19 Genomics UK (COG-UK) Consortium | Jacquelyn Wynn, Mairead Hyland, The Lighthouse Lab in Alderley Park and Alex Alderton, Roberto Amato, Jeffrey Barrett, Sonia Goncalves, Ewan Harrison, David K. Jackson, Ian Johnston, Dominic Kwiatkowski, Cordelia Langford, John Sillitoe on behalf of the Wellcome Sanger Institute COVID-19 Surveillance Team |
| EPI_ISL_3284577 | Lighthouse Lab in Glasgow | Wellcome Sanger Institute for the COVID-19 Genomics UK (COG-UK) Consortium | Harper VanSteenhouse, Yumi Kasai, David Gray, Carol Clugston, Anna Dominiczak and Alex Alderton, Roberto Amato, Jeffrey Barrett, Sonia Goncalves, Ewan Harrison, David K. Jackson, Ian Johnston, Dominic Kwiatkowski, Cordelia Langford, John Sillitoe on behalf of the Wellcome Sanger Institute COVID-19 Surveillance Team |
| EPI_ISL_3284578, EPI_ISL_3284579 | Lighthouse Lab in Alderley Park | Wellcome Sanger Institute for the COVID-19 Genomics UK (COG-UK) Consortium | Jacquelyn Wynn, Mairead Hyland, The Lighthouse Lab in Alderley Park and Alex Alderton, Roberto Amato, Jeffrey Barrett, Sonia Goncalves, Ewan Harrison, David K. Jackson, Ian Johnston, Dominic Kwiatkowski, Cordelia Langford, John Sillitoe on behalf of the Wellcome Sanger Institute COVID-19 Surveillance Team |
| EPI_ISL_3284580, EPI_ISL_3284581 | Lighthouse Lab in Glasgow | Wellcome Sanger Institute for the COVID-19 Genomics UK (COG-UK) Consortium | Harper VanSteenhouse, Yumi Kasai, David Gray, Carol Clugston, Anna Dominiczak and Alex Alderton, Roberto Amato, Jeffrey Barrett, Sonia Goncalves, Ewan Harrison, David K. Jackson, Ian Johnston, Dominic Kwiatkowski, Cordelia Langford, John Sillitoe on behalf of the Wellcome Sanger Institute COVID-19 Surveillance Team |
| EPI_ISL_3284582, EPI_ISL_3284583, EPI_ISL_3284584, EPI_ISL_3284585, EPI_ISL_3284586 | Lighthouse Lab in Alderley Park | Wellcome Sanger Institute for the COVID-19 Genomics UK (COG-UK) Consortium | Jacquelyn Wynn, Mairead Hyland, The Lighthouse Lab in Alderley Park and Alex Alderton, Roberto Amato, Jeffrey Barrett, Sonia Goncalves, Ewan Harrison, David K. Jackson, Ian Johnston, Dominic Kwiatkowski, Cordelia Langford, John Sillitoe on behalf of the Wellcome Sanger Institute COVID-19 Surveillance Team |
| EPI_ISL_3284587 | Lighthouse Lab in Glasgow | Wellcome Sanger Institute for the COVID-19 Genomics UK (COG-UK) Consortium | Harper VanSteenhouse, Yumi Kasai, David Gray, Carol Clugston, Anna Dominiczak and Alex Alderton, Roberto Amato, Jeffrey Barrett, Sonia Goncalves, Ewan Harrison, David K. Jackson, Ian Johnston, Dominic Kwiatkowski, Cordelia Langford, John Sillitoe on behalf of the Wellcome Sanger Institute COVID-19 Surveillance Team |
| EPI_ISL_3284588 | Lighthouse Lab in Alderley Park | Wellcome Sanger Institute for the COVID-19 Genomics UK (COG-UK) Consortium | Jacquelyn Wynn, Mairead Hyland, The Lighthouse Lab in Alderley Park and Alex Alderton, Roberto Amato, Jeffrey Barrett, Sonia Goncalves, Ewan Harrison, David K. Jackson, Ian Johnston, Dominic Kwiatkowski, Cordelia Langford, John Sillitoe on behalf of the Wellcome Sanger Institute COVID-19 Surveillance Team |
| EPI_ISL_3284589, EPI_ISL_3284590, EPI_ISL_3284591 | Lighthouse Lab in Glasgow | Wellcome Sanger Institute for the COVID-19 Genomics UK (COG-UK) Consortium | Harper VanSteenhouse, Yumi Kasai, David Gray, Carol Clugston, Anna Dominiczak and Alex Alderton, Roberto Amato, Jeffrey Barrett, Sonia Goncalves, Ewan Harrison, David K. Jackson, Ian Johnston, Dominic Kwiatkowski, Cordelia Langford, John Sillitoe on behalf of the Wellcome Sanger Institute COVID-19 Surveillance Team |
| EPI_ISL_3284592, EPI_ISL_3284593 | Lighthouse Lab in Alderley Park | Wellcome Sanger Institute for the COVID-19 Genomics UK (COG-UK) Consortium | Jacquelyn Wynn, Mairead Hyland, The Lighthouse Lab in Alderley Park and Alex Alderton, Roberto Amato, Jeffrey Barrett, Sonia Goncalves, Ewan Harrison, David K. Jackson, Ian Johnston, Dominic Kwiatkowski, Cordelia Langford, John Sillitoe on behalf of the Wellcome Sanger Institute COVID-19 Surveillance Team |
| EPI_ISL_3284594, EPI_ISL_3284595 | Lighthouse Lab in Glasgow | Wellcome Sanger Institute for the COVID-19 Genomics UK (COG-UK) Consortium | Harper VanSteenhouse, Yumi Kasai, David Gray, Carol Clugston, Anna Dominiczak and Alex Alderton, Roberto Amato, Jeffrey Barrett, Sonia Goncalves, Ewan Harrison, David K. Jackson, Ian Johnston, Dominic Kwiatkowski, Cordelia Langford, John Sillitoe on behalf of the Wellcome Sanger Institute COVID-19 Surveillance Team |
| EPI_ISL_3284596, EPI_ISL_3284597, EPI_ISL_3284598 | Lighthouse Lab in Alderley Park | Wellcome Sanger Institute for the COVID-19 Genomics UK (COG-UK) Consortium | Jacquelyn Wynn, Mairead Hyland, The Lighthouse Lab in Alderley Park and Alex Alderton, Roberto Amato, Jeffrey Barrett, Sonia Goncalves, Ewan Harrison, David K. Jackson, Ian Johnston, Dominic Kwiatkowski, Cordelia Langford, John Sillitoe on behalf of the Wellcome Sanger Institute COVID-19 Surveillance Team |
| EPI_ISL_3284599 | Lighthouse Lab in Glasgow | Wellcome Sanger Institute for the COVID-19 Genomics UK (COG-UK) Consortium | Harper VanSteenhouse, Yumi Kasai, David Gray, Carol Clugston, Anna Dominiczak and Alex Alderton, Roberto Amato, Jeffrey Barrett, Sonia Goncalves, Ewan Harrison, David K. Jackson, Ian Johnston, Dominic Kwiatkowski, Cordelia Langford, John Sillitoe on behalf of the Wellcome Sanger Institute COVID-19 Surveillance Team |
| EPI_ISL_3284600 | Lighthouse Lab in Alderley Park | Wellcome Sanger Institute for the COVID-19 Genomics UK (COG-UK) Consortium | Jacquelyn Wynn, Mairead Hyland, The Lighthouse Lab in Alderley Park and Alex Alderton, Roberto Amato, Jeffrey Barrett, Sonia Goncalves, Ewan Harrison, David K. Jackson, Ian Johnston, Dominic Kwiatkowski, Cordelia Langford, John Sillitoe on behalf of the Wellcome Sanger Institute COVID-19 Surveillance Team |
| EPI_ISL_3284601 | Lighthouse Lab in Glasgow | Wellcome Sanger Institute for the COVID-19 Genomics UK (COG-UK) Consortium | Harper VanSteenhouse, Yumi Kasai, David Gray, Carol Clugston, Anna Dominiczak and Alex Alderton, Roberto Amato, Jeffrey Barrett, Sonia Goncalves, Ewan Harrison, David K. Jackson, Ian Johnston, Dominic Kwiatkowski, Cordelia Langford, John Sillitoe on behalf of the Wellcome Sanger Institute COVID-19 Surveillance Team |
| EPI_ISL_3284602 | Lighthouse Lab in Alderley Park | Wellcome Sanger Institute for the COVID-19 Genomics UK (COG-UK) Consortium | Jacquelyn Wynn, Mairead Hyland, The Lighthouse Lab in Alderley Park and Alex Alderton, Roberto Amato, Jeffrey Barrett, Sonia Goncalves, Ewan Harrison, David K. Jackson, Ian Johnston, Dominic Kwiatkowski, Cordelia Langford, John Sillitoe on behalf of the Wellcome Sanger Institute COVID-19 Surveillance Team |
| EPI_ISL_3284603 | Lighthouse Lab in Glasgow | Wellcome Sanger Institute for the COVID-19 Genomics UK (COG-UK) Consortium | Harper VanSteenhouse, Yumi Kasai, David Gray, Carol Clugston, Anna Dominiczak and Alex Alderton, Roberto Amato, Jeffrey Barrett, Sonia Goncalves, Ewan Harrison, David K. Jackson, Ian Johnston, Dominic Kwiatkowski, Cordelia Langford, John Sillitoe on behalf of the Wellcome Sanger Institute COVID-19 Surveillance Team |
| EPI_ISL_3284604, EPI_ISL_3284605 | Lighthouse Lab in Alderley Park | Wellcome Sanger Institute for the COVID-19 Genomics UK (COG-UK) Consortium | Jacquelyn Wynn, Mairead Hyland, The Lighthouse Lab in Alderley Park and Alex Alderton, Roberto Amato, Jeffrey Barrett, Sonia Goncalves, Ewan Harrison, David K. Jackson, Ian Johnston, Dominic Kwiatkowski, Cordelia Langford, John Sillitoe on behalf of the Wellcome Sanger Institute COVID-19 Surveillance Team |
| EPI_ISL_3284606 | Lighthouse Lab in Glasgow | Wellcome Sanger Institute for the COVID-19 Genomics UK (COG-UK) Consortium | Harper VanSteenhouse, Yumi Kasai, David Gray, Carol Clugston, Anna Dominiczak and Alex Alderton, Roberto Amato, Jeffrey Barrett, Sonia Goncalves, Ewan Harrison, David K. Jackson, Ian Johnston, Dominic Kwiatkowski, Cordelia Langford, John Sillitoe on behalf of the Wellcome Sanger Institute COVID-19 Surveillance Team |
| EPI_ISL_3284607, EPI_ISL_3284608, EPI_ISL_3284609, EPI_ISL_3284610 | Lighthouse Lab in Alderley Park | Wellcome Sanger Institute for the COVID-19 Genomics UK (COG-UK) Consortium | Jacquelyn Wynn, Mairead Hyland, The Lighthouse Lab in Alderley Park and Alex Alderton, Roberto Amato, Jeffrey Barrett, Sonia Goncalves, Ewan Harrison, David K. Jackson, Ian Johnston, Dominic Kwiatkowski, Cordelia Langford, John Sillitoe on behalf of the Wellcome Sanger Institute COVID-19 Surveillance Team |
| EPI_ISL_3284611 | Lighthouse Lab in Glasgow | Wellcome Sanger Institute for the COVID-19 Genomics UK (COG-UK) Consortium | Harper VanSteenhouse, Yumi Kasai, David Gray, Carol Clugston, Anna Dominiczak and Alex Alderton, Roberto Amato, Jeffrey Barrett, Sonia Goncalves, Ewan Harrison, David K. Jackson, Ian Johnston, Dominic Kwiatkowski, Cordelia Langford, John Sillitoe on behalf of the Wellcome Sanger Institute COVID-19 Surveillance Team |
| EPI_ISL_3284612, EPI_ISL_3284613 | Lighthouse Lab in Alderley Park | Wellcome Sanger Institute for the COVID-19 Genomics UK (COG-UK) Consortium | Jacquelyn Wynn, Mairead Hyland, The Lighthouse Lab in Alderley Park and Alex Alderton, Roberto Amato, Jeffrey Barrett, Sonia Goncalves, Ewan Harrison, David K. Jackson, Ian Johnston, Dominic Kwiatkowski, Cordelia Langford, John Sillitoe on behalf of the Wellcome Sanger Institute COVID-19 Surveillance Team |
| EPI_ISL_3284614 | Lighthouse Lab in Glasgow | Wellcome Sanger Institute for the COVID-19 Genomics UK (COG-UK) Consortium | Harper VanSteenhouse, Yumi Kasai, David Gray, Carol Clugston, Anna Dominiczak and Alex Alderton, Roberto Amato, Jeffrey Barrett, Sonia Goncalves, Ewan Harrison, David K. Jackson, Ian Johnston, Dominic Kwiatkowski, Cordelia Langford, John Sillitoe on behalf of the Wellcome Sanger Institute COVID-19 Surveillance Team |
| EPI_ISL_3284615, EPI_ISL_3284616, EPI_ISL_3284617, EPI_ISL_3284618, EPI_ISL_3284619 | Lighthouse Lab in Alderley Park | Wellcome Sanger Institute for the COVID-19 Genomics UK (COG-UK) Consortium | Jacquelyn Wynn, Mairead Hyland, The Lighthouse Lab in Alderley Park and Alex Alderton, Roberto Amato, Jeffrey Barrett, Sonia Goncalves, Ewan Harrison, David K. Jackson, Ian Johnston, Dominic Kwiatkowski, Cordelia Langford, John Sillitoe on behalf of the Wellcome Sanger Institute COVID-19 Surveillance Team |
| EPI_ISL_3284620 | Lighthouse Lab in Glasgow | Wellcome Sanger Institute for the COVID-19 Genomics UK (COG-UK) Consortium | Harper VanSteenhouse, Yumi Kasai, David Gray, Carol Clugston, Anna Dominiczak and Alex Alderton, Roberto Amato, Jeffrey Barrett, Sonia Goncalves, Ewan Harrison, David K. Jackson, Ian Johnston, Dominic Kwiatkowski, Cordelia Langford, John Sillitoe on behalf of the Wellcome Sanger Institute COVID-19 Surveillance Team |
| EPI_ISL_3284621 | Lighthouse Lab in Alderley Park | Wellcome Sanger Institute for the COVID-19 Genomics UK (COG-UK) Consortium | Jacquelyn Wynn, Mairead Hyland, The Lighthouse Lab in Alderley Park and Alex Alderton, Roberto Amato, Jeffrey Barrett, Sonia Goncalves, Ewan Harrison, David K. Jackson, Ian Johnston, Dominic Kwiatkowski, Cordelia Langford, John Sillitoe on behalf of the Wellcome Sanger Institute COVID-19 Surveillance Team |
| EPI_ISL_3284622, EPI_ISL_3284623, EPI_ISL_3284624, EPI_ISL_3284625 | Lighthouse Lab in Glasgow | Wellcome Sanger Institute for the COVID-19 Genomics UK (COG-UK) Consortium | Harper VanSteenhouse, Yumi Kasai, David Gray, Carol Clugston, Anna Dominiczak and Alex Alderton, Roberto Amato, Jeffrey Barrett, Sonia Goncalves, Ewan Harrison, David K. Jackson, Ian Johnston, Dominic Kwiatkowski, Cordelia Langford, John Sillitoe on behalf of the Wellcome Sanger Institute COVID-19 Surveillance Team |
| EPI_ISL_3284626 | Lighthouse Lab in Alderley Park | Wellcome Sanger Institute for the COVID-19 Genomics UK (COG-UK) Consortium | Jacquelyn Wynn, Mairead Hyland, The Lighthouse Lab in Alderley Park and Alex Alderton, Roberto Amato, Jeffrey Barrett, Sonia Goncalves, Ewan Harrison, David K. Jackson, Ian Johnston, Dominic Kwiatkowski, Cordelia Langford, John Sillitoe on behalf of the Wellcome Sanger Institute COVID-19 Surveillance Team |
| EPI_ISL_3284627 | Lighthouse Lab in Glasgow | Wellcome Sanger Institute for the COVID-19 Genomics UK (COG-UK) Consortium | Harper VanSteenhouse, Yumi Kasai, David Gray, Carol Clugston, Anna Dominiczak and Alex Alderton, Roberto Amato, Jeffrey Barrett, Sonia Goncalves, Ewan Harrison, David K. Jackson, Ian Johnston, Dominic Kwiatkowski, Cordelia Langford, John Sillitoe on behalf of the Wellcome Sanger Institute COVID-19 Surveillance Team |
| EPI_ISL_3284628, EPI_ISL_3284629, EPI_ISL_3284630 | Lighthouse Lab in Alderley Park | Wellcome Sanger Institute for the COVID-19 Genomics UK (COG-UK) Consortium | Jacquelyn Wynn, Mairead Hyland, The Lighthouse Lab in Alderley Park and Alex Alderton, Roberto Amato, Jeffrey Barrett, Sonia Goncalves, Ewan Harrison, David K. Jackson, Ian Johnston, Dominic Kwiatkowski, Cordelia Langford, John Sillitoe on behalf of the Wellcome Sanger Institute COVID-19 Surveillance Team |
| EPI_ISL_3284631 | Lighthouse Lab in Glasgow | Wellcome Sanger Institute for the COVID-19 Genomics UK (COG-UK) Consortium | Harper VanSteenhouse, Yumi Kasai, David Gray, Carol Clugston, Anna Dominiczak and Alex Alderton, Roberto Amato, Jeffrey Barrett, Sonia Goncalves, Ewan Harrison, David K. Jackson, Ian Johnston, Dominic Kwiatkowski, Cordelia Langford, John Sillitoe on behalf of the Wellcome Sanger Institute COVID-19 Surveillance Team |
| EPI_ISL_3284632, EPI_ISL_3284633, EPI_ISL_3284634, EPI_ISL_3284635 | Lighthouse Lab in Alderley Park | Wellcome Sanger Institute for the COVID-19 Genomics UK (COG-UK) Consortium | Jacquelyn Wynn, Mairead Hyland, The Lighthouse Lab in Alderley Park and Alex Alderton, Roberto Amato, Jeffrey Barrett, Sonia Goncalves, Ewan Harrison, David K. Jackson, Ian Johnston, Dominic Kwiatkowski, Cordelia Langford, John Sillitoe on behalf of the Wellcome Sanger Institute COVID-19 Surveillance Team |
| EPI_ISL_3284636, EPI_ISL_3284637 | Lighthouse Lab in Glasgow | Wellcome Sanger Institute for the COVID-19 Genomics UK (COG-UK) Consortium | Harper VanSteenhouse, Yumi Kasai, David Gray, Carol Clugston, Anna Dominiczak and Alex Alderton, Roberto Amato, Jeffrey Barrett, Sonia Goncalves, Ewan Harrison, David K. Jackson, Ian Johnston, Dominic Kwiatkowski, Cordelia Langford, John Sillitoe on behalf of the Wellcome Sanger Institute COVID-19 Surveillance Team |
| EPI_ISL_3284638, EPI_ISL_3284639 | Lighthouse Lab in Alderley Park | Wellcome Sanger Institute for the COVID-19 Genomics UK (COG-UK) Consortium | Jacquelyn Wynn, Mairead Hyland, The Lighthouse Lab in Alderley Park and Alex Alderton, Roberto Amato, Jeffrey Barrett, Sonia Goncalves, Ewan Harrison, David K. Jackson, Ian Johnston, Dominic Kwiatkowski, Cordelia Langford, John Sillitoe on behalf of the Wellcome Sanger Institute COVID-19 Surveillance Team |
| EPI_ISL_3284640 | Lighthouse Lab in Glasgow | Wellcome Sanger Institute for the COVID-19 Genomics UK (COG-UK) Consortium | Harper VanSteenhouse, Yumi Kasai, David Gray, Carol Clugston, Anna Dominiczak and Alex Alderton, Roberto Amato, Jeffrey Barrett, Sonia Goncalves, Ewan Harrison, David K. Jackson, Ian Johnston, Dominic Kwiatkowski, Cordelia Langford, John Sillitoe on behalf of the Wellcome Sanger Institute COVID-19 Surveillance Team |
| EPI_ISL_3284641 | Lighthouse Lab in Alderley Park | Wellcome Sanger Institute for the COVID-19 Genomics UK (COG-UK) Consortium | Jacquelyn Wynn, Mairead Hyland, The Lighthouse Lab in Alderley Park and Alex Alderton, Roberto Amato, Jeffrey Barrett, Sonia Goncalves, Ewan Harrison, David K. Jackson, Ian Johnston, Dominic Kwiatkowski, Cordelia Langford, John Sillitoe on behalf of the Wellcome Sanger Institute COVID-19 Surveillance Team |
| EPI_ISL_3284642, EPI_ISL_3284643, EPI_ISL_3284644 | Lighthouse Lab in Glasgow | Wellcome Sanger Institute for the COVID-19 Genomics UK (COG-UK) Consortium | Harper VanSteenhouse, Yumi Kasai, David Gray, Carol Clugston, Anna Dominiczak and Alex Alderton, Roberto Amato, Jeffrey Barrett, Sonia Goncalves, Ewan Harrison, David K. Jackson, Ian Johnston, Dominic Kwiatkowski, Cordelia Langford, John Sillitoe on behalf of the Wellcome Sanger Institute COVID-19 Surveillance Team |
| EPI_ISL_3284645, | Lighthouse Lab in Alderley Park | Wellcome Sanger Institute for the COVID-19 Genomics UK (COG-UK) | Jacquelyn Wynn, Mairead Hyland, The Lighthouse Lab in Alderley Park and Alex Alderton, Roberto Amato, Jeffrey Barrett, Sonia Goncalves, Ewan Harrison, David K. Jackson, Ian Johnston, Dominic Kwiatkowski, Cordelia Langford, John Sillitoe on |

| | | | |
|---|---|---|---|
| EPI_ISL_3284646 | | Consortium | behalf of the Wellcome Sanger Institute COVID-19 Surveillance Team |
| EPI_ISL_3284647, EPI_ISL_3284648 | Lighthouse Lab in Glasgow | Wellcome Sanger Institute for the COVID-19 Genomics UK (COG-UK) Consortium | Harper VanSteenhouse, Yumi Kasai, David Gray, Carol Clugston, Anna Dominiczak and Alex Alderton, Roberto Amato, Jeffrey Barrett, Sonia Goncalves, Ewan Harrison, David K. Jackson, Ian Johnston, Dominic Kwiatkowski, Cordelia Langford, John Sillitoe on behalf of the Wellcome Sanger Institute COVID-19 Surveillance Team |
| EPI_ISL_3284649 | Lighthouse Lab in Alderley Park | Wellcome Sanger Institute for the COVID-19 Genomics UK (COG-UK) Consortium | Jacquelyn Wynn, Mairead Hyland, The Lighthouse Lab in Alderley Park and Alex Alderton, Roberto Amato, Jeffrey Barrett, Sonia Goncalves, Ewan Harrison, David K. Jackson, Ian Johnston, Dominic Kwiatkowski, Cordelia Langford, John Sillitoe on behalf of the Wellcome Sanger Institute COVID-19 Surveillance Team |
| EPI_ISL_3284650 | Lighthouse Lab in Glasgow | Wellcome Sanger Institute for the COVID-19 Genomics UK (COG-UK) Consortium | Harper VanSteenhouse, Yumi Kasai, David Gray, Carol Clugston, Anna Dominiczak and Alex Alderton, Roberto Amato, Jeffrey Barrett, Sonia Goncalves, Ewan Harrison, David K. Jackson, Ian Johnston, Dominic Kwiatkowski, Cordelia Langford, John Sillitoe on behalf of the Wellcome Sanger Institute COVID-19 Surveillance Team |
| EPI_ISL_3284651 | Lighthouse Lab in Alderley Park | Wellcome Sanger Institute for the COVID-19 Genomics UK (COG-UK) Consortium | Jacquelyn Wynn, Mairead Hyland, The Lighthouse Lab in Alderley Park and Alex Alderton, Roberto Amato, Jeffrey Barrett, Sonia Goncalves, Ewan Harrison, David K. Jackson, Ian Johnston, Dominic Kwiatkowski, Cordelia Langford, John Sillitoe on behalf of the Wellcome Sanger Institute COVID-19 Surveillance Team |
| EPI_ISL_3284652 | Lighthouse Lab in Glasgow | Wellcome Sanger Institute for the COVID-19 Genomics UK (COG-UK) Consortium | Harper VanSteenhouse, Yumi Kasai, David Gray, Carol Clugston, Anna Dominiczak and Alex Alderton, Roberto Amato, Jeffrey Barrett, Sonia Goncalves, Ewan Harrison, David K. Jackson, Ian Johnston, Dominic Kwiatkowski, Cordelia Langford, John Sillitoe on behalf of the Wellcome Sanger Institute COVID-19 Surveillance Team |
| EPI_ISL_3284653 | Lighthouse Lab in Alderley Park | Wellcome Sanger Institute for the COVID-19 Genomics UK (COG-UK) Consortium | Jacquelyn Wynn, Mairead Hyland, The Lighthouse Lab in Alderley Park and Alex Alderton, Roberto Amato, Jeffrey Barrett, Sonia Goncalves, Ewan Harrison, David K. Jackson, Ian Johnston, Dominic Kwiatkowski, Cordelia Langford, John Sillitoe on behalf of the Wellcome Sanger Institute COVID-19 Surveillance Team |
| EPI_ISL_3284654, EPI_ISL_3284655, EPI_ISL_3284656 | Lighthouse Lab in Glasgow | Wellcome Sanger Institute for the COVID-19 Genomics UK (COG-UK) Consortium | Harper VanSteenhouse, Yumi Kasai, David Gray, Carol Clugston, Anna Dominiczak and Alex Alderton, Roberto Amato, Jeffrey Barrett, Sonia Goncalves, Ewan Harrison, David K. Jackson, Ian Johnston, Dominic Kwiatkowski, Cordelia Langford, John Sillitoe on behalf of the Wellcome Sanger Institute COVID-19 Surveillance Team |
| EPI_ISL_3284657 | Health Services Laboratories | Wellcome Sanger Institute for the COVID-19 Genomics UK (COG-UK) Consortium | Health Services Laboratories and Alex Alderton, Roberto Amato, Jeffrey Barrett, Sonia Goncalves, Ewan Harrison, David K. Jackson, Ian Johnston, Dominic Kwiatkowski, Cordelia Langford, John Sillitoe on behalf of the Wellcome Sanger Institute COVID-19 Surveillance Team |
| EPI_ISL_3284658 | Lighthouse Lab in Milton Keynes | Wellcome Sanger Institute for the COVID-19 Genomics UK (COG-UK) Consortium | The Lighthouse Lab in Milton Keynes and Alex Alderton, Roberto Amato, Jeffrey Barrett, Sonia Goncalves, Ewan Harrison, David K. Jackson, Ian Johnston, Dominic Kwiatkowski, Cordelia Langford, John Sillitoe on behalf of the Wellcome Sanger Institute COVID-19 Surveillance Team |
| EPI_ISL_3284659 | Health Services Laboratories | Wellcome Sanger Institute for the COVID-19 Genomics UK (COG-UK) Consortium | Health Services Laboratories and Alex Alderton, Roberto Amato, Jeffrey Barrett, Sonia Goncalves, Ewan Harrison, David K. Jackson, Ian Johnston, Dominic Kwiatkowski, Cordelia Langford, John Sillitoe on behalf of the Wellcome Sanger Institute COVID-19 Surveillance Team |
| EPI_ISL_3284660, EPI_ISL_3284661 | Lighthouse Lab in Milton Keynes | Wellcome Sanger Institute for the COVID-19 Genomics UK (COG-UK) Consortium | The Lighthouse Lab in Milton Keynes and Alex Alderton, Roberto Amato, Jeffrey Barrett, Sonia Goncalves, Ewan Harrison, David K. Jackson, Ian Johnston, Dominic Kwiatkowski, Cordelia Langford, John Sillitoe on behalf of the Wellcome Sanger Institute COVID-19 Surveillance Team |
| EPI_ISL_3284662 | Health Services Laboratories | Wellcome Sanger Institute for the COVID-19 Genomics UK (COG-UK) Consortium | Health Services Laboratories and Alex Alderton, Roberto Amato, Jeffrey Barrett, Sonia Goncalves, Ewan Harrison, David K. Jackson, Ian Johnston, Dominic Kwiatkowski, Cordelia Langford, John Sillitoe on behalf of the Wellcome Sanger Institute COVID-19 Surveillance Team |
| EPI_ISL_3284663 | Lighthouse Lab in Milton Keynes | Wellcome Sanger Institute for the COVID-19 Genomics UK (COG-UK) Consortium | The Lighthouse Lab in Milton Keynes and Alex Alderton, Roberto Amato, Jeffrey Barrett, Sonia Goncalves, Ewan Harrison, David K. Jackson, Ian Johnston, Dominic Kwiatkowski, Cordelia Langford, John Sillitoe on behalf of the Wellcome Sanger Institute COVID-19 Surveillance Team |
| EPI_ISL_3284664, EPI_ISL_3284665 | Lighthouse Lab in Alderley Park | Wellcome Sanger Institute for the COVID-19 Genomics UK (COG-UK) Consortium | Jacquelyn Wynn, Mairead Hyland, The Lighthouse Lab in Alderley Park and Alex Alderton, Roberto Amato, Jeffrey Barrett, Sonia Goncalves, Ewan Harrison, David K. Jackson, Ian Johnston, Dominic Kwiatkowski, Cordelia Langford, John Sillitoe on behalf of the Wellcome Sanger Institute COVID-19 Surveillance Team |
| EPI_ISL_3284666 | Lighthouse Lab in Milton Keynes | Wellcome Sanger Institute for the COVID-19 Genomics UK (COG-UK) Consortium | The Lighthouse Lab in Milton Keynes and Alex Alderton, Roberto Amato, Jeffrey Barrett, Sonia Goncalves, Ewan Harrison, David K. Jackson, Ian Johnston, Dominic Kwiatkowski, Cordelia Langford, John Sillitoe on behalf of the Wellcome Sanger Institute COVID-19 Surveillance Team |
| EPI_ISL_3284667, EPI_ISL_3284668, EPI_ISL_3284669, EPI_ISL_3284670 | Health Services Laboratories | Wellcome Sanger Institute for the COVID-19 Genomics UK (COG-UK) Consortium | Health Services Laboratories and Alex Alderton, Roberto Amato, Jeffrey Barrett, Sonia Goncalves, Ewan Harrison, David K. Jackson, Ian Johnston, Dominic Kwiatkowski, Cordelia Langford, John Sillitoe on behalf of the Wellcome Sanger Institute COVID-19 Surveillance Team |
| EPI_ISL_3284671, EPI_ISL_3284672, EPI_ISL_3284673, EPI_ISL_3284674 | Lighthouse Lab in Milton Keynes | Wellcome Sanger Institute for the COVID-19 Genomics UK (COG-UK) Consortium | The Lighthouse Lab in Milton Keynes and Alex Alderton, Roberto Amato, Jeffrey Barrett, Sonia Goncalves, Ewan Harrison, David K. Jackson, Ian Johnston, Dominic Kwiatkowski, Cordelia Langford, John Sillitoe on behalf of the Wellcome Sanger Institute COVID-19 Surveillance Team |
| EPI_ISL_3284675 | Lighthouse Lab in Alderley Park | Wellcome Sanger Institute for the COVID-19 Genomics UK (COG-UK) Consortium | Jacquelyn Wynn, Mairead Hyland, The Lighthouse Lab in Alderley Park and Alex Alderton, Roberto Amato, Jeffrey Barrett, Sonia Goncalves, Ewan Harrison, David K. Jackson, Ian Johnston, Dominic Kwiatkowski, Cordelia Langford, John Sillitoe on behalf of the Wellcome Sanger Institute COVID-19 Surveillance Team |
| EPI_ISL_3284676 | Health Services Laboratories | Wellcome Sanger Institute for the COVID-19 Genomics UK (COG-UK) Consortium | Health Services Laboratories and Alex Alderton, Roberto Amato, Jeffrey Barrett, Sonia Goncalves, Ewan Harrison, David K. Jackson, Ian Johnston, Dominic Kwiatkowski, Cordelia Langford, John Sillitoe on behalf of the Wellcome Sanger Institute COVID-19 Surveillance Team |
| EPI_ISL_3284677 | Lighthouse Lab in Milton Keynes | Wellcome Sanger Institute for the COVID-19 Genomics UK (COG-UK) Consortium | The Lighthouse Lab in Milton Keynes and Alex Alderton, Roberto Amato, Jeffrey Barrett, Sonia Goncalves, Ewan Harrison, David K. Jackson, Ian Johnston, Dominic Kwiatkowski, Cordelia Langford, John Sillitoe on behalf of the Wellcome Sanger Institute COVID-19 Surveillance Team |
| EPI_ISL_3284678 | Lighthouse Lab in Alderley Park | Wellcome Sanger Institute for the COVID-19 Genomics UK (COG-UK) Consortium | Jacquelyn Wynn, Mairead Hyland, The Lighthouse Lab in Alderley Park and Alex Alderton, Roberto Amato, Jeffrey Barrett, Sonia Goncalves, Ewan Harrison, David K. Jackson, Ian Johnston, Dominic Kwiatkowski, Cordelia Langford, John Sillitoe on behalf of the Wellcome Sanger Institute COVID-19 Surveillance Team |
| EPI_ISL_3284679 | Health Services Laboratories | Wellcome Sanger Institute for the COVID-19 Genomics UK (COG-UK) Consortium | Health Services Laboratories and Alex Alderton, Roberto Amato, Jeffrey Barrett, Sonia Goncalves, Ewan Harrison, David K. Jackson, Ian Johnston, Dominic Kwiatkowski, Cordelia Langford, John Sillitoe on behalf of the Wellcome Sanger Institute COVID-19 Surveillance Team |
| EPI_ISL_3284680, EPI_ISL_3284681, EPI_ISL_3284682 | Lighthouse Lab in Alderley Park | Wellcome Sanger Institute for the COVID-19 Genomics UK (COG-UK) Consortium | Jacquelyn Wynn, Mairead Hyland, The Lighthouse Lab in Alderley Park and Alex Alderton, Roberto Amato, Jeffrey Barrett, Sonia Goncalves, Ewan Harrison, David K. Jackson, Ian Johnston, Dominic Kwiatkowski, Cordelia Langford, John Sillitoe on behalf of the Wellcome Sanger Institute COVID-19 Surveillance Team |
| EPI_ISL_3284683 | Health Services Laboratories | Wellcome Sanger Institute for the COVID-19 Genomics UK (COG-UK) Consortium | Health Services Laboratories and Alex Alderton, Roberto Amato, Jeffrey Barrett, Sonia Goncalves, Ewan Harrison, David K. Jackson, Ian Johnston, Dominic Kwiatkowski, Cordelia Langford, John Sillitoe on behalf of the Wellcome Sanger Institute COVID-19 Surveillance Team |
| EPI_ISL_3284684 | Lighthouse Lab in Milton Keynes | Wellcome Sanger Institute for the COVID-19 Genomics UK (COG-UK) Consortium | The Lighthouse Lab in Milton Keynes and Alex Alderton, Roberto Amato, Jeffrey Barrett, Sonia Goncalves, Ewan Harrison, David K. Jackson, Ian Johnston, Dominic Kwiatkowski, Cordelia Langford, John Sillitoe on behalf of the Wellcome Sanger Institute COVID-19 Surveillance Team |
| EPI_ISL_3284685, EPI_ISL_3284686, EPI_ISL_3284687, EPI_ISL_3284688 | Health Services Laboratories | Wellcome Sanger Institute for the COVID-19 Genomics UK (COG-UK) Consortium | Health Services Laboratories and Alex Alderton, Roberto Amato, Jeffrey Barrett, Sonia Goncalves, Ewan Harrison, David K. Jackson, Ian Johnston, Dominic Kwiatkowski, Cordelia Langford, John Sillitoe on behalf of the Wellcome Sanger Institute COVID-19 Surveillance Team |
| EPI_ISL_3284689 | Lighthouse Lab in Milton Keynes | Wellcome Sanger Institute for the COVID-19 Genomics UK (COG-UK) Consortium | The Lighthouse Lab in Milton Keynes and Alex Alderton, Roberto Amato, Jeffrey Barrett, Sonia Goncalves, Ewan Harrison, David K. Jackson, Ian Johnston, Dominic Kwiatkowski, Cordelia Langford, John Sillitoe on behalf of the Wellcome Sanger Institute COVID-19 Surveillance Team |
| EPI_ISL_3284690, EPI_ISL_3284691 | Health Services Laboratories | Wellcome Sanger Institute for the COVID-19 Genomics UK (COG-UK) Consortium | Health Services Laboratories and Alex Alderton, Roberto Amato, Jeffrey Barrett, Sonia Goncalves, Ewan Harrison, David K. Jackson, Ian Johnston, Dominic Kwiatkowski, Cordelia Langford, John Sillitoe on behalf of the Wellcome Sanger Institute COVID-19 Surveillance Team |
| EPI_ISL_3284692 | Lighthouse Lab in Alderley Park | Wellcome Sanger Institute for the COVID-19 Genomics UK (COG-UK) Consortium | Jacquelyn Wynn, Mairead Hyland, The Lighthouse Lab in Alderley Park and Alex Alderton, Roberto Amato, Jeffrey Barrett, Sonia Goncalves, Ewan Harrison, David K. Jackson, Ian Johnston, Dominic Kwiatkowski, Cordelia Langford, John Sillitoe on behalf of the Wellcome Sanger Institute COVID-19 Surveillance Team |
| EPI_ISL_3284693 | Health Services Laboratories | Wellcome Sanger Institute for the COVID-19 Genomics UK (COG-UK) Consortium | Health Services Laboratories and Alex Alderton, Roberto Amato, Jeffrey Barrett, Sonia Goncalves, Ewan Harrison, David K. Jackson, Ian Johnston, Dominic Kwiatkowski, Cordelia Langford, John Sillitoe on behalf of the Wellcome Sanger Institute COVID-19 Surveillance Team |
| EPI_ISL_3284694 | Lighthouse Lab in Alderley Park | Wellcome Sanger Institute for the COVID-19 Genomics UK (COG-UK) Consortium | Jacquelyn Wynn, Mairead Hyland, The Lighthouse Lab in Alderley Park and Alex Alderton, Roberto Amato, Jeffrey Barrett, Sonia Goncalves, Ewan Harrison, David K. Jackson, Ian Johnston, Dominic Kwiatkowski, Cordelia Langford, John Sillitoe on behalf of the Wellcome Sanger Institute COVID-19 Surveillance Team |
| EPI_ISL_3284695 | Lighthouse Lab in Milton Keynes | Wellcome Sanger Institute for the COVID-19 Genomics UK (COG-UK) Consortium | The Lighthouse Lab in Milton Keynes and Alex Alderton, Roberto Amato, Jeffrey Barrett, Sonia Goncalves, Ewan Harrison, David K. Jackson, Ian Johnston, Dominic Kwiatkowski, Cordelia Langford, John Sillitoe on behalf of the Wellcome Sanger Institute COVID-19 Surveillance Team |
| EPI_ISL_3284696, EPI_ISL_3284697 | Lighthouse Lab in Alderley Park | Wellcome Sanger Institute for the COVID-19 Genomics UK (COG-UK) Consortium | Jacquelyn Wynn, Mairead Hyland, The Lighthouse Lab in Alderley Park and Alex Alderton, Roberto Amato, Jeffrey Barrett, Sonia Goncalves, Ewan Harrison, David K. Jackson, Ian Johnston, Dominic Kwiatkowski, Cordelia Langford, John Sillitoe on behalf of the Wellcome Sanger Institute COVID-19 Surveillance Team |
| EPI_ISL_3284698 | Lighthouse Lab in Milton Keynes | Wellcome Sanger Institute for the COVID-19 Genomics UK (COG-UK) Consortium | The Lighthouse Lab in Milton Keynes and Alex Alderton, Roberto Amato, Jeffrey Barrett, Sonia Goncalves, Ewan Harrison, David K. Jackson, Ian Johnston, Dominic Kwiatkowski, Cordelia Langford, John Sillitoe on behalf of the Wellcome Sanger Institute COVID-19 Surveillance Team |
| EPI_ISL_3284699 | Lighthouse Lab in Alderley Park | Wellcome Sanger Institute for the COVID-19 Genomics UK (COG-UK) Consortium | Jacquelyn Wynn, Mairead Hyland, The Lighthouse Lab in Alderley Park and Alex Alderton, Roberto Amato, Jeffrey Barrett, Sonia Goncalves, Ewan Harrison, David K. Jackson, Ian Johnston, Dominic Kwiatkowski, Cordelia Langford, John Sillitoe on behalf of the Wellcome Sanger Institute COVID-19 Surveillance Team |
| EPI_ISL_3284700, EPI_ISL_3284701 | Lighthouse Lab in Milton Keynes | Wellcome Sanger Institute for the COVID-19 Genomics UK (COG-UK) Consortium | The Lighthouse Lab in Milton Keynes and Alex Alderton, Roberto Amato, Jeffrey Barrett, Sonia Goncalves, Ewan Harrison, David K. Jackson, Ian Johnston, Dominic Kwiatkowski, Cordelia Langford, John Sillitoe on behalf of the Wellcome Sanger Institute COVID-19 Surveillance Team |
| EPI_ISL_3284702 | Lighthouse Lab in Alderley Park | Wellcome Sanger Institute for the COVID-19 Genomics UK (COG-UK) Consortium | Jacquelyn Wynn, Mairead Hyland, The Lighthouse Lab in Alderley Park and Alex Alderton, Roberto Amato, Jeffrey Barrett, Sonia Goncalves, Ewan Harrison, David K. Jackson, Ian Johnston, Dominic Kwiatkowski, Cordelia Langford, John Sillitoe on behalf of the Wellcome Sanger Institute COVID-19 Surveillance Team |
| EPI_ISL_3284703, EPI_ISL_3284704 | Health Services Laboratories | Wellcome Sanger Institute for the COVID-19 Genomics UK (COG-UK) Consortium | Health Services Laboratories and Alex Alderton, Roberto Amato, Jeffrey Barrett, Sonia Goncalves, Ewan Harrison, David K. Jackson, Ian Johnston, Dominic Kwiatkowski, Cordelia Langford, John Sillitoe on behalf of the Wellcome Sanger Institute COVID-19 Surveillance Team |
| EPI_ISL_3284705 | Lighthouse Lab in Milton Keynes | Wellcome Sanger Institute for the COVID-19 Genomics UK (COG-UK) Consortium | The Lighthouse Lab in Milton Keynes and Alex Alderton, Roberto Amato, Jeffrey Barrett, Sonia Goncalves, Ewan Harrison, David K. Jackson, Ian Johnston, Dominic Kwiatkowski, Cordelia Langford, John Sillitoe on behalf of the Wellcome Sanger Institute COVID-19 Surveillance Team |
| EPI_ISL_3284706 | Lighthouse Lab in Alderley Park | Wellcome Sanger Institute for the COVID-19 Genomics UK (COG-UK) Consortium | Jacquelyn Wynn, Mairead Hyland, The Lighthouse Lab in Alderley Park and Alex Alderton, Roberto Amato, Jeffrey Barrett, Sonia Goncalves, Ewan Harrison, David K. Jackson, Ian Johnston, Dominic Kwiatkowski, Cordelia Langford, John Sillitoe on behalf of the Wellcome Sanger Institute COVID-19 Surveillance Team |
| EPI_ISL_3284707, EPI_ISL_3284708, EPI_ISL_3284709, | Lighthouse Lab in Milton Keynes | Wellcome Sanger Institute for the COVID-19 Genomics UK (COG-UK) Consortium | The Lighthouse Lab in Milton Keynes and Alex Alderton, Roberto Amato, Jeffrey Barrett, Sonia Goncalves, Ewan Harrison, David K. Jackson, Ian Johnston, Dominic Kwiatkowski, Cordelia Langford, John Sillitoe on behalf of the Wellcome Sanger Institute COVID-19 Surveillance Team |

| | | | |
|---|---|---|---|
| EPI_ISL_3284710 | | | |
| EPI_ISL_3284711 | Health Services Laboratories | Wellcome Sanger Institute for the COVID-19 Genomics UK (COG-UK) Consortium | Health Services Laboratories and Alex Alderton, Roberto Amato, Jeffrey Barrett, Sonia Goncalves, Ewan Harrison, David K. Jackson, Ian Johnston, Dominic Kwiatkowski, Cordelia Langford, John Sillitoe on behalf of the Wellcome Sanger Institute COVID-19 Surveillance Team |
| EPI_ISL_3284712 | Lighthouse Lab in Milton Keynes | Wellcome Sanger Institute for the COVID-19 Genomics UK (COG-UK) Consortium | The Lighthouse Lab in Milton Keynes and Alex Alderton, Roberto Amato, Jeffrey Barrett, Sonia Goncalves, Ewan Harrison, David K. Jackson, Ian Johnston, Dominic Kwiatkowski, Cordelia Langford, John Sillitoe on behalf of the Wellcome Sanger Institute COVID-19 Surveillance Team |
| EPI_ISL_3284713, EPI_ISL_3284714, EPI_ISL_3284715 | Lighthouse Lab in Alderley Park | Wellcome Sanger Institute for the COVID-19 Genomics UK (COG-UK) Consortium | Jacquelyn Wynn, Mairead Hyland, The Lighthouse Lab in Alderley Park and Alex Alderton, Roberto Amato, Jeffrey Barrett, Sonia Goncalves, Ewan Harrison, David K. Jackson, Ian Johnston, Dominic Kwiatkowski, Cordelia Langford, John Sillitoe on behalf of the Wellcome Sanger Institute COVID-19 Surveillance Team |
| EPI_ISL_3284716, EPI_ISL_3284717 | Lighthouse Lab in Milton Keynes | Wellcome Sanger Institute for the COVID-19 Genomics UK (COG-UK) Consortium | The Lighthouse Lab in Milton Keynes and Alex Alderton, Roberto Amato, Jeffrey Barrett, Sonia Goncalves, Ewan Harrison, David K. Jackson, Ian Johnston, Dominic Kwiatkowski, Cordelia Langford, John Sillitoe on behalf of the Wellcome Sanger Institute COVID-19 Surveillance Team |
| EPI_ISL_3284718 | Lighthouse Lab in Alderley Park | Wellcome Sanger Institute for the COVID-19 Genomics UK (COG-UK) Consortium | Jacquelyn Wynn, Mairead Hyland, The Lighthouse Lab in Alderley Park and Alex Alderton, Roberto Amato, Jeffrey Barrett, Sonia Goncalves, Ewan Harrison, David K. Jackson, Ian Johnston, Dominic Kwiatkowski, Cordelia Langford, John Sillitoe on behalf of the Wellcome Sanger Institute COVID-19 Surveillance Team |
| EPI_ISL_3284719, EPI_ISL_3284720 | Lighthouse Lab in Milton Keynes | Wellcome Sanger Institute for the COVID-19 Genomics UK (COG-UK) Consortium | The Lighthouse Lab in Milton Keynes and Alex Alderton, Roberto Amato, Jeffrey Barrett, Sonia Goncalves, Ewan Harrison, David K. Jackson, Ian Johnston, Dominic Kwiatkowski, Cordelia Langford, John Sillitoe on behalf of the Wellcome Sanger Institute COVID-19 Surveillance Team |
| EPI_ISL_3284721 | Lighthouse Lab in Alderley Park | Wellcome Sanger Institute for the COVID-19 Genomics UK (COG-UK) Consortium | Jacquelyn Wynn, Mairead Hyland, The Lighthouse Lab in Alderley Park and Alex Alderton, Roberto Amato, Jeffrey Barrett, Sonia Goncalves, Ewan Harrison, David K. Jackson, Ian Johnston, Dominic Kwiatkowski, Cordelia Langford, John Sillitoe on behalf of the Wellcome Sanger Institute COVID-19 Surveillance Team |
| EPI_ISL_3284722, EPI_ISL_3284723 | Lighthouse Lab in Milton Keynes | Wellcome Sanger Institute for the COVID-19 Genomics UK (COG-UK) Consortium | The Lighthouse Lab in Milton Keynes and Alex Alderton, Roberto Amato, Jeffrey Barrett, Sonia Goncalves, Ewan Harrison, David K. Jackson, Ian Johnston, Dominic Kwiatkowski, Cordelia Langford, John Sillitoe on behalf of the Wellcome Sanger Institute COVID-19 Surveillance Team |
| EPI_ISL_3284724 | Lighthouse Lab in Alderley Park | Wellcome Sanger Institute for the COVID-19 Genomics UK (COG-UK) Consortium | Jacquelyn Wynn, Mairead Hyland, The Lighthouse Lab in Alderley Park and Alex Alderton, Roberto Amato, Jeffrey Barrett, Sonia Goncalves, Ewan Harrison, David K. Jackson, Ian Johnston, Dominic Kwiatkowski, Cordelia Langford, John Sillitoe on behalf of the Wellcome Sanger Institute COVID-19 Surveillance Team |
| EPI_ISL_3284725 | Lighthouse Lab in Milton Keynes | Wellcome Sanger Institute for the COVID-19 Genomics UK (COG-UK) Consortium | The Lighthouse Lab in Milton Keynes and Alex Alderton, Roberto Amato, Jeffrey Barrett, Sonia Goncalves, Ewan Harrison, David K. Jackson, Ian Johnston, Dominic Kwiatkowski, Cordelia Langford, John Sillitoe on behalf of the Wellcome Sanger Institute COVID-19 Surveillance Team |
| EPI_ISL_3284726 | Lighthouse Lab in Alderley Park | Wellcome Sanger Institute for the COVID-19 Genomics UK (COG-UK) Consortium | Jacquelyn Wynn, Mairead Hyland, The Lighthouse Lab in Alderley Park and Alex Alderton, Roberto Amato, Jeffrey Barrett, Sonia Goncalves, Ewan Harrison, David K. Jackson, Ian Johnston, Dominic Kwiatkowski, Cordelia Langford, John Sillitoe on behalf of the Wellcome Sanger Institute COVID-19 Surveillance Team |
| EPI_ISL_3284727 | Lighthouse Lab in Milton Keynes | Wellcome Sanger Institute for the COVID-19 Genomics UK (COG-UK) Consortium | The Lighthouse Lab in Milton Keynes and Alex Alderton, Roberto Amato, Jeffrey Barrett, Sonia Goncalves, Ewan Harrison, David K. Jackson, Ian Johnston, Dominic Kwiatkowski, Cordelia Langford, John Sillitoe on behalf of the Wellcome Sanger Institute COVID-19 Surveillance Team |
| EPI_ISL_3284728 | Lighthouse Lab in Alderley Park | Wellcome Sanger Institute for the COVID-19 Genomics UK (COG-UK) Consortium | Jacquelyn Wynn, Mairead Hyland, The Lighthouse Lab in Alderley Park and Alex Alderton, Roberto Amato, Jeffrey Barrett, Sonia Goncalves, Ewan Harrison, David K. Jackson, Ian Johnston, Dominic Kwiatkowski, Cordelia Langford, John Sillitoe on behalf of the Wellcome Sanger Institute COVID-19 Surveillance Team |
| EPI_ISL_3284729 | Lighthouse Lab in Milton Keynes | Wellcome Sanger Institute for the COVID-19 Genomics UK (COG-UK) Consortium | The Lighthouse Lab in Milton Keynes and Alex Alderton, Roberto Amato, Jeffrey Barrett, Sonia Goncalves, Ewan Harrison, David K. Jackson, Ian Johnston, Dominic Kwiatkowski, Cordelia Langford, John Sillitoe on behalf of the Wellcome Sanger Institute COVID-19 Surveillance Team |
| EPI_ISL_3284730, EPI_ISL_3284731 | Lighthouse Lab in Alderley Park | Wellcome Sanger Institute for the COVID-19 Genomics UK (COG-UK) Consortium | Jacquelyn Wynn, Mairead Hyland, The Lighthouse Lab in Alderley Park and Alex Alderton, Roberto Amato, Jeffrey Barrett, Sonia Goncalves, Ewan Harrison, David K. Jackson, Ian Johnston, Dominic Kwiatkowski, Cordelia Langford, John Sillitoe on behalf of the Wellcome Sanger Institute COVID-19 Surveillance Team |
| EPI_ISL_3284732 | Lighthouse Lab in Milton Keynes | Wellcome Sanger Institute for the COVID-19 Genomics UK (COG-UK) Consortium | The Lighthouse Lab in Milton Keynes and Alex Alderton, Roberto Amato, Jeffrey Barrett, Sonia Goncalves, Ewan Harrison, David K. Jackson, Ian Johnston, Dominic Kwiatkowski, Cordelia Langford, John Sillitoe on behalf of the Wellcome Sanger Institute COVID-19 Surveillance Team |
| EPI_ISL_3284733 | Health Services Laboratories | Wellcome Sanger Institute for the COVID-19 Genomics UK (COG-UK) Consortium | Health Services Laboratories and Alex Alderton, Roberto Amato, Jeffrey Barrett, Sonia Goncalves, Ewan Harrison, David K. Jackson, Ian Johnston, Dominic Kwiatkowski, Cordelia Langford, John Sillitoe on behalf of the Wellcome Sanger Institute COVID-19 Surveillance Team |
| EPI_ISL_3284734, EPI_ISL_3284735 | Lighthouse Lab in Milton Keynes | Wellcome Sanger Institute for the COVID-19 Genomics UK (COG-UK) Consortium | The Lighthouse Lab in Milton Keynes and Alex Alderton, Roberto Amato, Jeffrey Barrett, Sonia Goncalves, Ewan Harrison, David K. Jackson, Ian Johnston, Dominic Kwiatkowski, Cordelia Langford, John Sillitoe on behalf of the Wellcome Sanger Institute COVID-19 Surveillance Team |
| EPI_ISL_3284736 | Lighthouse Lab in Alderley Park | Wellcome Sanger Institute for the COVID-19 Genomics UK (COG-UK) Consortium | Jacquelyn Wynn, Mairead Hyland, The Lighthouse Lab in Alderley Park and Alex Alderton, Roberto Amato, Jeffrey Barrett, Sonia Goncalves, Ewan Harrison, David K. Jackson, Ian Johnston, Dominic Kwiatkowski, Cordelia Langford, John Sillitoe on behalf of the Wellcome Sanger Institute COVID-19 Surveillance Team |
| EPI_ISL_3284737 | Health Services Laboratories | Wellcome Sanger Institute for the COVID-19 Genomics UK (COG-UK) Consortium | Health Services Laboratories and Alex Alderton, Roberto Amato, Jeffrey Barrett, Sonia Goncalves, Ewan Harrison, David K. Jackson, Ian Johnston, Dominic Kwiatkowski, Cordelia Langford, John Sillitoe on behalf of the Wellcome Sanger Institute COVID-19 Surveillance Team |
| EPI_ISL_3284738, EPI_ISL_3284739 | Lighthouse Lab in Milton Keynes | Wellcome Sanger Institute for the COVID-19 Genomics UK (COG-UK) Consortium | The Lighthouse Lab in Milton Keynes and Alex Alderton, Roberto Amato, Jeffrey Barrett, Sonia Goncalves, Ewan Harrison, David K. Jackson, Ian Johnston, Dominic Kwiatkowski, Cordelia Langford, John Sillitoe on behalf of the Wellcome Sanger Institute COVID-19 Surveillance Team |
| EPI_ISL_3284740, EPI_ISL_3284741 | Health Services Laboratories | Wellcome Sanger Institute for the COVID-19 Genomics UK (COG-UK) Consortium | Health Services Laboratories and Alex Alderton, Roberto Amato, Jeffrey Barrett, Sonia Goncalves, Ewan Harrison, David K. Jackson, Ian Johnston, Dominic Kwiatkowski, Cordelia Langford, John Sillitoe on behalf of the Wellcome Sanger Institute COVID-19 Surveillance Team |
| EPI_ISL_3284742 | Lighthouse Lab in Alderley Park | Wellcome Sanger Institute for the COVID-19 Genomics UK (COG-UK) Consortium | Jacquelyn Wynn, Mairead Hyland, The Lighthouse Lab in Alderley Park and Alex Alderton, Roberto Amato, Jeffrey Barrett, Sonia Goncalves, Ewan Harrison, David K. Jackson, Ian Johnston, Dominic Kwiatkowski, Cordelia Langford, John Sillitoe on behalf of the Wellcome Sanger Institute COVID-19 Surveillance Team |
| EPI_ISL_3284743 | Health Services Laboratories | Wellcome Sanger Institute for the COVID-19 Genomics UK (COG-UK) Consortium | Health Services Laboratories and Alex Alderton, Roberto Amato, Jeffrey Barrett, Sonia Goncalves, Ewan Harrison, David K. Jackson, Ian Johnston, Dominic Kwiatkowski, Cordelia Langford, John Sillitoe on behalf of the Wellcome Sanger Institute COVID-19 Surveillance Team |
| EPI_ISL_3284744, EPI_ISL_3284745 | Lighthouse Lab in Milton Keynes | Wellcome Sanger Institute for the COVID-19 Genomics UK (COG-UK) Consortium | The Lighthouse Lab in Milton Keynes and Alex Alderton, Roberto Amato, Jeffrey Barrett, Sonia Goncalves, Ewan Harrison, David K. Jackson, Ian Johnston, Dominic Kwiatkowski, Cordelia Langford, John Sillitoe on behalf of the Wellcome Sanger Institute COVID-19 Surveillance Team |
| EPI_ISL_3284746, EPI_ISL_3284747 | Health Services Laboratories | Wellcome Sanger Institute for the COVID-19 Genomics UK (COG-UK) Consortium | Health Services Laboratories and Alex Alderton, Roberto Amato, Jeffrey Barrett, Sonia Goncalves, Ewan Harrison, David K. Jackson, Ian Johnston, Dominic Kwiatkowski, Cordelia Langford, John Sillitoe on behalf of the Wellcome Sanger Institute COVID-19 Surveillance Team |
| EPI_ISL_3284748 | Lighthouse Lab in Alderley Park | Wellcome Sanger Institute for the COVID-19 Genomics UK (COG-UK) Consortium | Jacquelyn Wynn, Mairead Hyland, The Lighthouse Lab in Alderley Park and Alex Alderton, Roberto Amato, Jeffrey Barrett, Sonia Goncalves, Ewan Harrison, David K. Jackson, Ian Johnston, Dominic Kwiatkowski, Cordelia Langford, John Sillitoe on behalf of the Wellcome Sanger Institute COVID-19 Surveillance Team |
| EPI_ISL_3284749 | Health Services Laboratories | Wellcome Sanger Institute for the COVID-19 Genomics UK (COG-UK) Consortium | Health Services Laboratories and Alex Alderton, Roberto Amato, Jeffrey Barrett, Sonia Goncalves, Ewan Harrison, David K. Jackson, Ian Johnston, Dominic Kwiatkowski, Cordelia Langford, John Sillitoe on behalf of the Wellcome Sanger Institute COVID-19 Surveillance Team |
| EPI_ISL_3284750, EPI_ISL_3284751, EPI_ISL_3284752 | Lighthouse Lab in Milton Keynes | Wellcome Sanger Institute for the COVID-19 Genomics UK (COG-UK) Consortium | The Lighthouse Lab in Milton Keynes and Alex Alderton, Roberto Amato, Jeffrey Barrett, Sonia Goncalves, Ewan Harrison, David K. Jackson, Ian Johnston, Dominic Kwiatkowski, Cordelia Langford, John Sillitoe on behalf of the Wellcome Sanger Institute COVID-19 Surveillance Team |
| EPI_ISL_3284753 | Lighthouse Lab in Alderley Park | Wellcome Sanger Institute for the COVID-19 Genomics UK (COG-UK) Consortium | Jacquelyn Wynn, Mairead Hyland, The Lighthouse Lab in Alderley Park and Alex Alderton, Roberto Amato, Jeffrey Barrett, Sonia Goncalves, Ewan Harrison, David K. Jackson, Ian Johnston, Dominic Kwiatkowski, Cordelia Langford, John Sillitoe on behalf of the Wellcome Sanger Institute COVID-19 Surveillance Team |
| EPI_ISL_3284754, EPI_ISL_3284755, EPI_ISL_3284756 | Lighthouse Lab in Milton Keynes | Wellcome Sanger Institute for the COVID-19 Genomics UK (COG-UK) Consortium | The Lighthouse Lab in Milton Keynes and Alex Alderton, Roberto Amato, Jeffrey Barrett, Sonia Goncalves, Ewan Harrison, David K. Jackson, Ian Johnston, Dominic Kwiatkowski, Cordelia Langford, John Sillitoe on behalf of the Wellcome Sanger Institute COVID-19 Surveillance Team |
| EPI_ISL_3284757 | Health Services Laboratories | Wellcome Sanger Institute for the COVID-19 Genomics UK (COG-UK) Consortium | Health Services Laboratories and Alex Alderton, Roberto Amato, Jeffrey Barrett, Sonia Goncalves, Ewan Harrison, David K. Jackson, Ian Johnston, Dominic Kwiatkowski, Cordelia Langford, John Sillitoe on behalf of the Wellcome Sanger Institute COVID-19 Surveillance Team |
| EPI_ISL_3284758 | Lighthouse Lab in Milton Keynes | Wellcome Sanger Institute for the COVID-19 Genomics UK (COG-UK) Consortium | The Lighthouse Lab in Milton Keynes and Alex Alderton, Roberto Amato, Jeffrey Barrett, Sonia Goncalves, Ewan Harrison, David K. Jackson, Ian Johnston, Dominic Kwiatkowski, Cordelia Langford, John Sillitoe on behalf of the Wellcome Sanger Institute COVID-19 Surveillance Team |
| EPI_ISL_3284759 | Health Services Laboratories | Wellcome Sanger Institute for the COVID-19 Genomics UK (COG-UK) Consortium | Health Services Laboratories and Alex Alderton, Roberto Amato, Jeffrey Barrett, Sonia Goncalves, Ewan Harrison, David K. Jackson, Ian Johnston, Dominic Kwiatkowski, Cordelia Langford, John Sillitoe on behalf of the Wellcome Sanger Institute COVID-19 Surveillance Team |
| EPI_ISL_3284760 | Lighthouse Lab in Alderley Park | Wellcome Sanger Institute for the COVID-19 Genomics UK (COG-UK) Consortium | Jacquelyn Wynn, Mairead Hyland, The Lighthouse Lab in Alderley Park and Alex Alderton, Roberto Amato, Jeffrey Barrett, Sonia Goncalves, Ewan Harrison, David K. Jackson, Ian Johnston, Dominic Kwiatkowski, Cordelia Langford, John Sillitoe on behalf of the Wellcome Sanger Institute COVID-19 Surveillance Team |
| EPI_ISL_3284761, EPI_ISL_3284762, EPI_ISL_3284763, EPI_ISL_3284764 | Lighthouse Lab in Milton Keynes | Wellcome Sanger Institute for the COVID-19 Genomics UK (COG-UK) Consortium | The Lighthouse Lab in Milton Keynes and Alex Alderton, Roberto Amato, Jeffrey Barrett, Sonia Goncalves, Ewan Harrison, David K. Jackson, Ian Johnston, Dominic Kwiatkowski, Cordelia Langford, John Sillitoe on behalf of the Wellcome Sanger Institute COVID-19 Surveillance Team |
| EPI_ISL_3284765 | Lighthouse Lab in Alderley Park | Wellcome Sanger Institute for the COVID-19 Genomics UK (COG-UK) Consortium | Jacquelyn Wynn, Mairead Hyland, The Lighthouse Lab in Alderley Park and Alex Alderton, Roberto Amato, Jeffrey Barrett, Sonia Goncalves, Ewan Harrison, David K. Jackson, Ian Johnston, Dominic Kwiatkowski, Cordelia Langford, John Sillitoe on behalf of the Wellcome Sanger Institute COVID-19 Surveillance Team |
| EPI_ISL_3284766 | Health Services Laboratories | Wellcome Sanger Institute for the COVID-19 Genomics UK (COG-UK) Consortium | Health Services Laboratories and Alex Alderton, Roberto Amato, Jeffrey Barrett, Sonia Goncalves, Ewan Harrison, David K. Jackson, Ian Johnston, Dominic Kwiatkowski, Cordelia Langford, John Sillitoe on behalf of the Wellcome Sanger Institute COVID-19 Surveillance Team |
| EPI_ISL_3284767 | Lighthouse Lab in Milton Keynes | Wellcome Sanger Institute for the COVID-19 Genomics UK (COG-UK) Consortium | The Lighthouse Lab in Milton Keynes and Alex Alderton, Roberto Amato, Jeffrey Barrett, Sonia Goncalves, Ewan Harrison, David K. Jackson, Ian Johnston, Dominic Kwiatkowski, Cordelia Langford, John Sillitoe on behalf of the Wellcome Sanger Institute COVID-19 Surveillance Team |
| EPI_ISL_3284768, EPI_ISL_3284769 | Health Services Laboratories | Wellcome Sanger Institute for the COVID-19 Genomics UK (COG-UK) Consortium | Health Services Laboratories and Alex Alderton, Roberto Amato, Jeffrey Barrett, Sonia Goncalves, Ewan Harrison, David K. Jackson, Ian Johnston, Dominic Kwiatkowski, Cordelia Langford, John Sillitoe on behalf of the Wellcome Sanger Institute COVID-19 Surveillance Team |
| EPI_ISL_3284770, EPI_ISL_3284771, EPI_ISL_3284772, EPI_ISL_3284773 | Lighthouse Lab in Milton Keynes | Wellcome Sanger Institute for the COVID-19 Genomics UK (COG-UK) Consortium | The Lighthouse Lab in Milton Keynes and Alex Alderton, Roberto Amato, Jeffrey Barrett, Sonia Goncalves, Ewan Harrison, David K. Jackson, Ian Johnston, Dominic Kwiatkowski, Cordelia Langford, John Sillitoe on behalf of the Wellcome Sanger Institute COVID-19 Surveillance Team |
| EPI_ISL_3284774 | Health Services Laboratories | Wellcome Sanger Institute for the COVID-19 Genomics UK (COG-UK) | Health Services Laboratories and Alex Alderton, Roberto Amato, Jeffrey Barrett, Sonia Goncalves, Ewan Harrison, David K. Jackson, Ian Johnston, Dominic Kwiatkowski, Cordelia Langford, John Sillitoe on behalf of the Wellcome Sanger Institute |

| | | | |
|---|---|---|---|
| | | Consortium | COVID-19 Surveillance Team |
| EPI_ISL_3284775, EPI_ISL_3284776 | Lighthouse Lab in Milton Keynes | Wellcome Sanger Institute for the COVID-19 Genomics UK (COG-UK) Consortium | The Lighthouse Lab in Milton Keynes and Alex Alderton, Roberto Amato, Jeffrey Barrett, Sonia Goncalves, Ewan Harrison, David K. Jackson, Ian Johnston, Dominic Kwiatkowski, Cordelia Langford, John Sillitoe on behalf of the Wellcome Sanger Institute COVID-19 Surveillance Team |
| EPI_ISL_3284777, EPI_ISL_3284778 | Lighthouse Lab in Alderley Park | Wellcome Sanger Institute for the COVID-19 Genomics UK (COG-UK) Consortium | Jacquelyn Wynn, Mairead Hyland, The Lighthouse Lab in Alderley Park and Alex Alderton, Roberto Amato, Jeffrey Barrett, Sonia Goncalves, Ewan Harrison, David K. Jackson, Ian Johnston, Dominic Kwiatkowski, Cordelia Langford, John Sillitoe on behalf of the Wellcome Sanger Institute COVID-19 Surveillance Team |
| EPI_ISL_3284779, EPI_ISL_3284780, EPI_ISL_3284781 | Lighthouse Lab in Milton Keynes | Wellcome Sanger Institute for the COVID-19 Genomics UK (COG-UK) Consortium | The Lighthouse Lab in Milton Keynes and Alex Alderton, Roberto Amato, Jeffrey Barrett, Sonia Goncalves, Ewan Harrison, David K. Jackson, Ian Johnston, Dominic Kwiatkowski, Cordelia Langford, John Sillitoe on behalf of the Wellcome Sanger Institute COVID-19 Surveillance Team |
| EPI_ISL_3284782 | Lighthouse Lab in Alderley Park | Wellcome Sanger Institute for the COVID-19 Genomics UK (COG-UK) Consortium | Jacquelyn Wynn, Mairead Hyland, The Lighthouse Lab in Alderley Park and Alex Alderton, Roberto Amato, Jeffrey Barrett, Sonia Goncalves, Ewan Harrison, David K. Jackson, Ian Johnston, Dominic Kwiatkowski, Cordelia Langford, John Sillitoe on behalf of the Wellcome Sanger Institute COVID-19 Surveillance Team |
| EPI_ISL_3284783 | Lighthouse Lab in Milton Keynes | Wellcome Sanger Institute for the COVID-19 Genomics UK (COG-UK) Consortium | The Lighthouse Lab in Milton Keynes and Alex Alderton, Roberto Amato, Jeffrey Barrett, Sonia Goncalves, Ewan Harrison, David K. Jackson, Ian Johnston, Dominic Kwiatkowski, Cordelia Langford, John Sillitoe on behalf of the Wellcome Sanger Institute COVID-19 Surveillance Team |
| EPI_ISL_3284784 | Lighthouse Lab in Alderley Park | Wellcome Sanger Institute for the COVID-19 Genomics UK (COG-UK) Consortium | Jacquelyn Wynn, Mairead Hyland, The Lighthouse Lab in Alderley Park and Alex Alderton, Roberto Amato, Jeffrey Barrett, Sonia Goncalves, Ewan Harrison, David K. Jackson, Ian Johnston, Dominic Kwiatkowski, Cordelia Langford, John Sillitoe on behalf of the Wellcome Sanger Institute COVID-19 Surveillance Team |
| EPI_ISL_3284785, EPI_ISL_3284786, EPI_ISL_3284787 | Lighthouse Lab in Milton Keynes | Wellcome Sanger Institute for the COVID-19 Genomics UK (COG-UK) Consortium | The Lighthouse Lab in Milton Keynes and Alex Alderton, Roberto Amato, Jeffrey Barrett, Sonia Goncalves, Ewan Harrison, David K. Jackson, Ian Johnston, Dominic Kwiatkowski, Cordelia Langford, John Sillitoe on behalf of the Wellcome Sanger Institute COVID-19 Surveillance Team |
| EPI_ISL_3284788, EPI_ISL_3284789, EPI_ISL_3284790 | Lighthouse Lab in Alderley Park | Wellcome Sanger Institute for the COVID-19 Genomics UK (COG-UK) Consortium | Jacquelyn Wynn, Mairead Hyland, The Lighthouse Lab in Alderley Park and Alex Alderton, Roberto Amato, Jeffrey Barrett, Sonia Goncalves, Ewan Harrison, David K. Jackson, Ian Johnston, Dominic Kwiatkowski, Cordelia Langford, John Sillitoe on behalf of the Wellcome Sanger Institute COVID-19 Surveillance Team |
| EPI_ISL_3284791, EPI_ISL_3284792 | Lighthouse Lab in Milton Keynes | Wellcome Sanger Institute for the COVID-19 Genomics UK (COG-UK) Consortium | The Lighthouse Lab in Milton Keynes and Alex Alderton, Roberto Amato, Jeffrey Barrett, Sonia Goncalves, Ewan Harrison, David K. Jackson, Ian Johnston, Dominic Kwiatkowski, Cordelia Langford, John Sillitoe on behalf of the Wellcome Sanger Institute COVID-19 Surveillance Team |
| EPI_ISL_3284793 | Lighthouse Lab in Alderley Park | Wellcome Sanger Institute for the COVID-19 Genomics UK (COG-UK) Consortium | Jacquelyn Wynn, Mairead Hyland, The Lighthouse Lab in Alderley Park and Alex Alderton, Roberto Amato, Jeffrey Barrett, Sonia Goncalves, Ewan Harrison, David K. Jackson, Ian Johnston, Dominic Kwiatkowski, Cordelia Langford, John Sillitoe on behalf of the Wellcome Sanger Institute COVID-19 Surveillance Team |
| EPI_ISL_3284794 | Lighthouse Lab in Milton Keynes | Wellcome Sanger Institute for the COVID-19 Genomics UK (COG-UK) Consortium | The Lighthouse Lab in Milton Keynes and Alex Alderton, Roberto Amato, Jeffrey Barrett, Sonia Goncalves, Ewan Harrison, David K. Jackson, Ian Johnston, Dominic Kwiatkowski, Cordelia Langford, John Sillitoe on behalf of the Wellcome Sanger Institute COVID-19 Surveillance Team |
| EPI_ISL_3284795 | Lighthouse Lab in Alderley Park | Wellcome Sanger Institute for the COVID-19 Genomics UK (COG-UK) Consortium | Jacquelyn Wynn, Mairead Hyland, The Lighthouse Lab in Alderley Park and Alex Alderton, Roberto Amato, Jeffrey Barrett, Sonia Goncalves, Ewan Harrison, David K. Jackson, Ian Johnston, Dominic Kwiatkowski, Cordelia Langford, John Sillitoe on behalf of the Wellcome Sanger Institute COVID-19 Surveillance Team |
| EPI_ISL_3284796 | Lighthouse Lab in Milton Keynes | Wellcome Sanger Institute for the COVID-19 Genomics UK (COG-UK) Consortium | The Lighthouse Lab in Milton Keynes and Alex Alderton, Roberto Amato, Jeffrey Barrett, Sonia Goncalves, Ewan Harrison, David K. Jackson, Ian Johnston, Dominic Kwiatkowski, Cordelia Langford, John Sillitoe on behalf of the Wellcome Sanger Institute COVID-19 Surveillance Team |
| EPI_ISL_3284797 | Health Services Laboratories | Wellcome Sanger Institute for the COVID-19 Genomics UK (COG-UK) Consortium | Health Services Laboratories and Alex Alderton, Roberto Amato, Jeffrey Barrett, Sonia Goncalves, Ewan Harrison, David K. Jackson, Ian Johnston, Dominic Kwiatkowski, Cordelia Langford, John Sillitoe on behalf of the Wellcome Sanger Institute COVID-19 Surveillance Team |
| EPI_ISL_3284798 | Lighthouse Lab in Milton Keynes | Wellcome Sanger Institute for the COVID-19 Genomics UK (COG-UK) Consortium | The Lighthouse Lab in Milton Keynes and Alex Alderton, Roberto Amato, Jeffrey Barrett, Sonia Goncalves, Ewan Harrison, David K. Jackson, Ian Johnston, Dominic Kwiatkowski, Cordelia Langford, John Sillitoe on behalf of the Wellcome Sanger Institute COVID-19 Surveillance Team |
| EPI_ISL_3284799, EPI_ISL_3284800 | Lighthouse Lab in Alderley Park | Wellcome Sanger Institute for the COVID-19 Genomics UK (COG-UK) Consortium | Jacquelyn Wynn, Mairead Hyland, The Lighthouse Lab in Alderley Park and Alex Alderton, Roberto Amato, Jeffrey Barrett, Sonia Goncalves, Ewan Harrison, David K. Jackson, Ian Johnston, Dominic Kwiatkowski, Cordelia Langford, John Sillitoe on behalf of the Wellcome Sanger Institute COVID-19 Surveillance Team |
| EPI_ISL_3284801 | Lighthouse Lab in Milton Keynes | Wellcome Sanger Institute for the COVID-19 Genomics UK (COG-UK) Consortium | The Lighthouse Lab in Milton Keynes and Alex Alderton, Roberto Amato, Jeffrey Barrett, Sonia Goncalves, Ewan Harrison, David K. Jackson, Ian Johnston, Dominic Kwiatkowski, Cordelia Langford, John Sillitoe on behalf of the Wellcome Sanger Institute COVID-19 Surveillance Team |
| EPI_ISL_3284802 | Health Services Laboratories | Wellcome Sanger Institute for the COVID-19 Genomics UK (COG-UK) Consortium | Health Services Laboratories and Alex Alderton, Roberto Amato, Jeffrey Barrett, Sonia Goncalves, Ewan Harrison, David K. Jackson, Ian Johnston, Dominic Kwiatkowski, Cordelia Langford, John Sillitoe on behalf of the Wellcome Sanger Institute COVID-19 Surveillance Team |
| EPI_ISL_3284803 | Lighthouse Lab in Alderley Park | Wellcome Sanger Institute for the COVID-19 Genomics UK (COG-UK) Consortium | Jacquelyn Wynn, Mairead Hyland, The Lighthouse Lab in Alderley Park and Alex Alderton, Roberto Amato, Jeffrey Barrett, Sonia Goncalves, Ewan Harrison, David K. Jackson, Ian Johnston, Dominic Kwiatkowski, Cordelia Langford, John Sillitoe on behalf of the Wellcome Sanger Institute COVID-19 Surveillance Team |
| EPI_ISL_3284804, EPI_ISL_3284805 | Lighthouse Lab in Milton Keynes | Wellcome Sanger Institute for the COVID-19 Genomics UK (COG-UK) Consortium | The Lighthouse Lab in Milton Keynes and Alex Alderton, Roberto Amato, Jeffrey Barrett, Sonia Goncalves, Ewan Harrison, David K. Jackson, Ian Johnston, Dominic Kwiatkowski, Cordelia Langford, John Sillitoe on behalf of the Wellcome Sanger Institute COVID-19 Surveillance Team |
| EPI_ISL_3284806 | Lighthouse Lab in Alderley Park | Wellcome Sanger Institute for the COVID-19 Genomics UK (COG-UK) Consortium | Jacquelyn Wynn, Mairead Hyland, The Lighthouse Lab in Alderley Park and Alex Alderton, Roberto Amato, Jeffrey Barrett, Sonia Goncalves, Ewan Harrison, David K. Jackson, Ian Johnston, Dominic Kwiatkowski, Cordelia Langford, John Sillitoe on behalf of the Wellcome Sanger Institute COVID-19 Surveillance Team |
| EPI_ISL_3284807 | Lighthouse Lab in Milton Keynes | Wellcome Sanger Institute for the COVID-19 Genomics UK (COG-UK) Consortium | The Lighthouse Lab in Milton Keynes and Alex Alderton, Roberto Amato, Jeffrey Barrett, Sonia Goncalves, Ewan Harrison, David K. Jackson, Ian Johnston, Dominic Kwiatkowski, Cordelia Langford, John Sillitoe on behalf of the Wellcome Sanger Institute COVID-19 Surveillance Team |
| EPI_ISL_3284808 | Lighthouse Lab in Alderley Park | Wellcome Sanger Institute for the COVID-19 Genomics UK (COG-UK) Consortium | Jacquelyn Wynn, Mairead Hyland, The Lighthouse Lab in Alderley Park and Alex Alderton, Roberto Amato, Jeffrey Barrett, Sonia Goncalves, Ewan Harrison, David K. Jackson, Ian Johnston, Dominic Kwiatkowski, Cordelia Langford, John Sillitoe on behalf of the Wellcome Sanger Institute COVID-19 Surveillance Team |
| EPI_ISL_3284809, EPI_ISL_3284810 | Lighthouse Lab in Milton Keynes | Wellcome Sanger Institute for the COVID-19 Genomics UK (COG-UK) Consortium | The Lighthouse Lab in Milton Keynes and Alex Alderton, Roberto Amato, Jeffrey Barrett, Sonia Goncalves, Ewan Harrison, David K. Jackson, Ian Johnston, Dominic Kwiatkowski, Cordelia Langford, John Sillitoe on behalf of the Wellcome Sanger Institute COVID-19 Surveillance Team |
| EPI_ISL_3284811 | Health Services Laboratories | Wellcome Sanger Institute for the COVID-19 Genomics UK (COG-UK) Consortium | Health Services Laboratories and Alex Alderton, Roberto Amato, Jeffrey Barrett, Sonia Goncalves, Ewan Harrison, David K. Jackson, Ian Johnston, Dominic Kwiatkowski, Cordelia Langford, John Sillitoe on behalf of the Wellcome Sanger Institute COVID-19 Surveillance Team |
| EPI_ISL_3284812 | Lighthouse Lab in Alderley Park | Wellcome Sanger Institute for the COVID-19 Genomics UK (COG-UK) Consortium | Jacquelyn Wynn, Mairead Hyland, The Lighthouse Lab in Alderley Park and Alex Alderton, Roberto Amato, Jeffrey Barrett, Sonia Goncalves, Ewan Harrison, David K. Jackson, Ian Johnston, Dominic Kwiatkowski, Cordelia Langford, John Sillitoe on behalf of the Wellcome Sanger Institute COVID-19 Surveillance Team |
| EPI_ISL_3284813 | Lighthouse Lab in Milton Keynes | Wellcome Sanger Institute for the COVID-19 Genomics UK (COG-UK) Consortium | The Lighthouse Lab in Milton Keynes and Alex Alderton, Roberto Amato, Jeffrey Barrett, Sonia Goncalves, Ewan Harrison, David K. Jackson, Ian Johnston, Dominic Kwiatkowski, Cordelia Langford, John Sillitoe on behalf of the Wellcome Sanger Institute COVID-19 Surveillance Team |
| EPI_ISL_3284814 | Health Services Laboratories | Wellcome Sanger Institute for the COVID-19 Genomics UK (COG-UK) Consortium | Health Services Laboratories and Alex Alderton, Roberto Amato, Jeffrey Barrett, Sonia Goncalves, Ewan Harrison, David K. Jackson, Ian Johnston, Dominic Kwiatkowski, Cordelia Langford, John Sillitoe on behalf of the Wellcome Sanger Institute COVID-19 Surveillance Team |
| EPI_ISL_3284815 | Lighthouse Lab in Milton Keynes | Wellcome Sanger Institute for the COVID-19 Genomics UK (COG-UK) Consortium | The Lighthouse Lab in Milton Keynes and Alex Alderton, Roberto Amato, Jeffrey Barrett, Sonia Goncalves, Ewan Harrison, David K. Jackson, Ian Johnston, Dominic Kwiatkowski, Cordelia Langford, John Sillitoe on behalf of the Wellcome Sanger Institute COVID-19 Surveillance Team |
| EPI_ISL_3284816, EPI_ISL_3284817 | Health Services Laboratories | Wellcome Sanger Institute for the COVID-19 Genomics UK (COG-UK) Consortium | Health Services Laboratories and Alex Alderton, Roberto Amato, Jeffrey Barrett, Sonia Goncalves, Ewan Harrison, David K. Jackson, Ian Johnston, Dominic Kwiatkowski, Cordelia Langford, John Sillitoe on behalf of the Wellcome Sanger Institute COVID-19 Surveillance Team |
| EPI_ISL_3284818, EPI_ISL_3284819, EPI_ISL_3284820 | Lighthouse Lab in Milton Keynes | Wellcome Sanger Institute for the COVID-19 Genomics UK (COG-UK) Consortium | The Lighthouse Lab in Milton Keynes and Alex Alderton, Roberto Amato, Jeffrey Barrett, Sonia Goncalves, Ewan Harrison, David K. Jackson, Ian Johnston, Dominic Kwiatkowski, Cordelia Langford, John Sillitoe on behalf of the Wellcome Sanger Institute COVID-19 Surveillance Team |
| EPI_ISL_3284821 | Health Services Laboratories | Wellcome Sanger Institute for the COVID-19 Genomics UK (COG-UK) Consortium | Health Services Laboratories and Alex Alderton, Roberto Amato, Jeffrey Barrett, Sonia Goncalves, Ewan Harrison, David K. Jackson, Ian Johnston, Dominic Kwiatkowski, Cordelia Langford, John Sillitoe on behalf of the Wellcome Sanger Institute COVID-19 Surveillance Team |
| EPI_ISL_3284822, EPI_ISL_3284823, EPI_ISL_3284824 | Lighthouse Lab in Milton Keynes | Wellcome Sanger Institute for the COVID-19 Genomics UK (COG-UK) Consortium | The Lighthouse Lab in Milton Keynes and Alex Alderton, Roberto Amato, Jeffrey Barrett, Sonia Goncalves, Ewan Harrison, David K. Jackson, Ian Johnston, Dominic Kwiatkowski, Cordelia Langford, John Sillitoe on behalf of the Wellcome Sanger Institute COVID-19 Surveillance Team |
| EPI_ISL_3284825, EPI_ISL_3284826 | Lighthouse Lab in Alderley Park | Wellcome Sanger Institute for the COVID-19 Genomics UK (COG-UK) Consortium | Jacquelyn Wynn, Mairead Hyland, The Lighthouse Lab in Alderley Park and Alex Alderton, Roberto Amato, Jeffrey Barrett, Sonia Goncalves, Ewan Harrison, David K. Jackson, Ian Johnston, Dominic Kwiatkowski, Cordelia Langford, John Sillitoe on behalf of the Wellcome Sanger Institute COVID-19 Surveillance Team |
| EPI_ISL_3284827 | Health Services Laboratories | Wellcome Sanger Institute for the COVID-19 Genomics UK (COG-UK) Consortium | Health Services Laboratories and Alex Alderton, Roberto Amato, Jeffrey Barrett, Sonia Goncalves, Ewan Harrison, David K. Jackson, Ian Johnston, Dominic Kwiatkowski, Cordelia Langford, John Sillitoe on behalf of the Wellcome Sanger Institute COVID-19 Surveillance Team |
| EPI_ISL_3284828, EPI_ISL_3284829, EPI_ISL_3284830 | Lighthouse Lab in Milton Keynes | Wellcome Sanger Institute for the COVID-19 Genomics UK (COG-UK) Consortium | The Lighthouse Lab in Milton Keynes and Alex Alderton, Roberto Amato, Jeffrey Barrett, Sonia Goncalves, Ewan Harrison, David K. Jackson, Ian Johnston, Dominic Kwiatkowski, Cordelia Langford, John Sillitoe on behalf of the Wellcome Sanger Institute COVID-19 Surveillance Team |
| EPI_ISL_3284831 | Health Services Laboratories | Wellcome Sanger Institute for the COVID-19 Genomics UK (COG-UK) Consortium | Health Services Laboratories and Alex Alderton, Roberto Amato, Jeffrey Barrett, Sonia Goncalves, Ewan Harrison, David K. Jackson, Ian Johnston, Dominic Kwiatkowski, Cordelia Langford, John Sillitoe on behalf of the Wellcome Sanger Institute COVID-19 Surveillance Team |
| EPI_ISL_3284832 | Lighthouse Lab in Milton Keynes | Wellcome Sanger Institute for the COVID-19 Genomics UK (COG-UK) Consortium | The Lighthouse Lab in Milton Keynes and Alex Alderton, Roberto Amato, Jeffrey Barrett, Sonia Goncalves, Ewan Harrison, David K. Jackson, Ian Johnston, Dominic Kwiatkowski, Cordelia Langford, John Sillitoe on behalf of the Wellcome Sanger Institute COVID-19 Surveillance Team |
| EPI_ISL_3284833 | Lighthouse Lab in Alderley Park | Wellcome Sanger Institute for the COVID-19 Genomics UK (COG-UK) Consortium | Jacquelyn Wynn, Mairead Hyland, The Lighthouse Lab in Alderley Park and Alex Alderton, Roberto Amato, Jeffrey Barrett, Sonia Goncalves, Ewan Harrison, David K. Jackson, Ian Johnston, Dominic Kwiatkowski, Cordelia Langford, John Sillitoe on behalf of the Wellcome Sanger Institute COVID-19 Surveillance Team |
| EPI_ISL_3284834, EPI_ISL_3284835, EPI_ISL_3284836, EPI_ISL_3284837, EPI_ISL_3284838 | Lighthouse Lab in Milton Keynes | Wellcome Sanger Institute for the COVID-19 Genomics UK (COG-UK) Consortium | The Lighthouse Lab in Milton Keynes and Alex Alderton, Roberto Amato, Jeffrey Barrett, Sonia Goncalves, Ewan Harrison, David K. Jackson, Ian Johnston, Dominic Kwiatkowski, Cordelia Langford, John Sillitoe on behalf of the Wellcome Sanger Institute COVID-19 Surveillance Team |
| EPI_ISL_3284839 | Lighthouse Lab in Alderley Park | Wellcome Sanger Institute for the COVID-19 Genomics UK (COG-UK) Consortium | Jacquelyn Wynn, Mairead Hyland, The Lighthouse Lab in Alderley Park and Alex Alderton, Roberto Amato, Jeffrey Barrett, Sonia Goncalves, Ewan Harrison, David K. Jackson, Ian Johnston, Dominic Kwiatkowski, Cordelia Langford, John Sillitoe on behalf of the Wellcome Sanger Institute COVID-19 Surveillance Team |

| | | | |
|---|---|---|---|
| EPI_ISL_3284840, EPI_ISL_3284841 | Lighthouse Lab in Milton Keynes | Wellcome Sanger Institute for the COVID-19 Genomics UK (COG-UK) Consortium | The Lighthouse Lab in Milton Keynes and Alex Alderton, Roberto Amato, Jeffrey Barrett, Sonia Goncalves, Ewan Harrison, David K. Jackson, Ian Johnston, Dominic Kwiatkowski, Cordelia Langford, John Sillitoe on behalf of the Wellcome Sanger Institute COVID-19 Surveillance Team |
| EPI_ISL_3284842 | Health Services Laboratories | Wellcome Sanger Institute for the COVID-19 Genomics UK (COG-UK) Consortium | Health Services Laboratories and Alex Alderton, Roberto Amato, Jeffrey Barrett, Sonia Goncalves, Ewan Harrison, David K. Jackson, Ian Johnston, Dominic Kwiatkowski, Cordelia Langford, John Sillitoe on behalf of the Wellcome Sanger Institute COVID-19 Surveillance Team |
| EPI_ISL_3284843, EPI_ISL_3284844 | Lighthouse Lab in Milton Keynes | Wellcome Sanger Institute for the COVID-19 Genomics UK (COG-UK) Consortium | The Lighthouse Lab in Milton Keynes and Alex Alderton, Roberto Amato, Jeffrey Barrett, Sonia Goncalves, Ewan Harrison, David K. Jackson, Ian Johnston, Dominic Kwiatkowski, Cordelia Langford, John Sillitoe on behalf of the Wellcome Sanger Institute COVID-19 Surveillance Team |
| EPI_ISL_3284845 | Lighthouse Lab in Alderley Park | Wellcome Sanger Institute for the COVID-19 Genomics UK (COG-UK) Consortium | Jacquelyn Wynn, Mairead Hyland, The Lighthouse Lab in Alderley Park and Alex Alderton, Roberto Amato, Jeffrey Barrett, Sonia Goncalves, Ewan Harrison, David K. Jackson, Ian Johnston, Dominic Kwiatkowski, Cordelia Langford, John Sillitoe on behalf of the Wellcome Sanger Institute COVID-19 Surveillance Team |
| EPI_ISL_3284846 | Health Services Laboratories | Wellcome Sanger Institute for the COVID-19 Genomics UK (COG-UK) Consortium | Health Services Laboratories and Alex Alderton, Roberto Amato, Jeffrey Barrett, Sonia Goncalves, Ewan Harrison, David K. Jackson, Ian Johnston, Dominic Kwiatkowski, Cordelia Langford, John Sillitoe on behalf of the Wellcome Sanger Institute COVID-19 Surveillance Team |
| EPI_ISL_3284847 | Lighthouse Lab in Milton Keynes | Wellcome Sanger Institute for the COVID-19 Genomics UK (COG-UK) Consortium | The Lighthouse Lab in Milton Keynes and Alex Alderton, Roberto Amato, Jeffrey Barrett, Sonia Goncalves, Ewan Harrison, David K. Jackson, Ian Johnston, Dominic Kwiatkowski, Cordelia Langford, John Sillitoe on behalf of the Wellcome Sanger Institute COVID-19 Surveillance Team |
| EPI_ISL_3284848 | Health Services Laboratories | Wellcome Sanger Institute for the COVID-19 Genomics UK (COG-UK) Consortium | Health Services Laboratories and Alex Alderton, Roberto Amato, Jeffrey Barrett, Sonia Goncalves, Ewan Harrison, David K. Jackson, Ian Johnston, Dominic Kwiatkowski, Cordelia Langford, John Sillitoe on behalf of the Wellcome Sanger Institute COVID-19 Surveillance Team |
| EPI_ISL_3284849 | Lighthouse Lab in Alderley Park | Wellcome Sanger Institute for the COVID-19 Genomics UK (COG-UK) Consortium | Jacquelyn Wynn, Mairead Hyland, The Lighthouse Lab in Alderley Park and Alex Alderton, Roberto Amato, Jeffrey Barrett, Sonia Goncalves, Ewan Harrison, David K. Jackson, Ian Johnston, Dominic Kwiatkowski, Cordelia Langford, John Sillitoe on behalf of the Wellcome Sanger Institute COVID-19 Surveillance Team |
| EPI_ISL_3284850, EPI_ISL_3284851 | Health Services Laboratories | Wellcome Sanger Institute for the COVID-19 Genomics UK (COG-UK) Consortium | Health Services Laboratories and Alex Alderton, Roberto Amato, Jeffrey Barrett, Sonia Goncalves, Ewan Harrison, David K. Jackson, Ian Johnston, Dominic Kwiatkowski, Cordelia Langford, John Sillitoe on behalf of the Wellcome Sanger Institute COVID-19 Surveillance Team |
| EPI_ISL_3284852 | Lighthouse Lab in Milton Keynes | Wellcome Sanger Institute for the COVID-19 Genomics UK (COG-UK) Consortium | The Lighthouse Lab in Milton Keynes and Alex Alderton, Roberto Amato, Jeffrey Barrett, Sonia Goncalves, Ewan Harrison, David K. Jackson, Ian Johnston, Dominic Kwiatkowski, Cordelia Langford, John Sillitoe on behalf of the Wellcome Sanger Institute COVID-19 Surveillance Team |
| EPI_ISL_3284853 | Lighthouse Lab in Alderley Park | Wellcome Sanger Institute for the COVID-19 Genomics UK (COG-UK) Consortium | Jacquelyn Wynn, Mairead Hyland, The Lighthouse Lab in Alderley Park and Alex Alderton, Roberto Amato, Jeffrey Barrett, Sonia Goncalves, Ewan Harrison, David K. Jackson, Ian Johnston, Dominic Kwiatkowski, Cordelia Langford, John Sillitoe on behalf of the Wellcome Sanger Institute COVID-19 Surveillance Team |
| EPI_ISL_3284854, EPI_ISL_3284855, EPI_ISL_3284856, EPI_ISL_3284857, EPI_ISL_3284858 | Lighthouse Lab in Milton Keynes | Wellcome Sanger Institute for the COVID-19 Genomics UK (COG-UK) Consortium | The Lighthouse Lab in Milton Keynes and Alex Alderton, Roberto Amato, Jeffrey Barrett, Sonia Goncalves, Ewan Harrison, David K. Jackson, Ian Johnston, Dominic Kwiatkowski, Cordelia Langford, John Sillitoe on behalf of the Wellcome Sanger Institute COVID-19 Surveillance Team |
| EPI_ISL_3284859 | Health Services Laboratories | Wellcome Sanger Institute for the COVID-19 Genomics UK (COG-UK) Consortium | Health Services Laboratories and Alex Alderton, Roberto Amato, Jeffrey Barrett, Sonia Goncalves, Ewan Harrison, David K. Jackson, Ian Johnston, Dominic Kwiatkowski, Cordelia Langford, John Sillitoe on behalf of the Wellcome Sanger Institute COVID-19 Surveillance Team |
| EPI_ISL_3284860 | Lighthouse Lab in Alderley Park | Wellcome Sanger Institute for the COVID-19 Genomics UK (COG-UK) Consortium | Jacquelyn Wynn, Mairead Hyland, The Lighthouse Lab in Alderley Park and Alex Alderton, Roberto Amato, Jeffrey Barrett, Sonia Goncalves, Ewan Harrison, David K. Jackson, Ian Johnston, Dominic Kwiatkowski, Cordelia Langford, John Sillitoe on behalf of the Wellcome Sanger Institute COVID-19 Surveillance Team |
| EPI_ISL_3284861 | Health Services Laboratories | Wellcome Sanger Institute for the COVID-19 Genomics UK (COG-UK) Consortium | Health Services Laboratories and Alex Alderton, Roberto Amato, Jeffrey Barrett, Sonia Goncalves, Ewan Harrison, David K. Jackson, Ian Johnston, Dominic Kwiatkowski, Cordelia Langford, John Sillitoe on behalf of the Wellcome Sanger Institute COVID-19 Surveillance Team |
| EPI_ISL_3284862 | Lighthouse Lab in Alderley Park | Wellcome Sanger Institute for the COVID-19 Genomics UK (COG-UK) Consortium | Jacquelyn Wynn, Mairead Hyland, The Lighthouse Lab in Alderley Park and Alex Alderton, Roberto Amato, Jeffrey Barrett, Sonia Goncalves, Ewan Harrison, David K. Jackson, Ian Johnston, Dominic Kwiatkowski, Cordelia Langford, John Sillitoe on behalf of the Wellcome Sanger Institute COVID-19 Surveillance Team |
| EPI_ISL_3284863 | Health Services Laboratories | Wellcome Sanger Institute for the COVID-19 Genomics UK (COG-UK) Consortium | Health Services Laboratories and Alex Alderton, Roberto Amato, Jeffrey Barrett, Sonia Goncalves, Ewan Harrison, David K. Jackson, Ian Johnston, Dominic Kwiatkowski, Cordelia Langford, John Sillitoe on behalf of the Wellcome Sanger Institute COVID-19 Surveillance Team |
| EPI_ISL_3284864, EPI_ISL_3284865 | Lighthouse Lab in Milton Keynes | Wellcome Sanger Institute for the COVID-19 Genomics UK (COG-UK) Consortium | The Lighthouse Lab in Milton Keynes and Alex Alderton, Roberto Amato, Jeffrey Barrett, Sonia Goncalves, Ewan Harrison, David K. Jackson, Ian Johnston, Dominic Kwiatkowski, Cordelia Langford, John Sillitoe on behalf of the Wellcome Sanger Institute COVID-19 Surveillance Team |
| EPI_ISL_3284866 | Lighthouse Lab in Alderley Park | Wellcome Sanger Institute for the COVID-19 Genomics UK (COG-UK) Consortium | Jacquelyn Wynn, Mairead Hyland, The Lighthouse Lab in Alderley Park and Alex Alderton, Roberto Amato, Jeffrey Barrett, Sonia Goncalves, Ewan Harrison, David K. Jackson, Ian Johnston, Dominic Kwiatkowski, Cordelia Langford, John Sillitoe on behalf of the Wellcome Sanger Institute COVID-19 Surveillance Team |
| EPI_ISL_3284867 | Lighthouse Lab in Milton Keynes | Wellcome Sanger Institute for the COVID-19 Genomics UK (COG-UK) Consortium | The Lighthouse Lab in Milton Keynes and Alex Alderton, Roberto Amato, Jeffrey Barrett, Sonia Goncalves, Ewan Harrison, David K. Jackson, Ian Johnston, Dominic Kwiatkowski, Cordelia Langford, John Sillitoe on behalf of the Wellcome Sanger Institute COVID-19 Surveillance Team |
| EPI_ISL_3284868 | Lighthouse Lab in Alderley Park | Wellcome Sanger Institute for the COVID-19 Genomics UK (COG-UK) Consortium | Jacquelyn Wynn, Mairead Hyland, The Lighthouse Lab in Alderley Park and Alex Alderton, Roberto Amato, Jeffrey Barrett, Sonia Goncalves, Ewan Harrison, David K. Jackson, Ian Johnston, Dominic Kwiatkowski, Cordelia Langford, John Sillitoe on behalf of the Wellcome Sanger Institute COVID-19 Surveillance Team |
| EPI_ISL_3284869, EPI_ISL_3284870 | Health Services Laboratories | Wellcome Sanger Institute for the COVID-19 Genomics UK (COG-UK) Consortium | Health Services Laboratories and Alex Alderton, Roberto Amato, Jeffrey Barrett, Sonia Goncalves, Ewan Harrison, David K. Jackson, Ian Johnston, Dominic Kwiatkowski, Cordelia Langford, John Sillitoe on behalf of the Wellcome Sanger Institute COVID-19 Surveillance Team |
| EPI_ISL_3284871, EPI_ISL_3284872 | Lighthouse Lab in Milton Keynes | Wellcome Sanger Institute for the COVID-19 Genomics UK (COG-UK) Consortium | The Lighthouse Lab in Milton Keynes and Alex Alderton, Roberto Amato, Jeffrey Barrett, Sonia Goncalves, Ewan Harrison, David K. Jackson, Ian Johnston, Dominic Kwiatkowski, Cordelia Langford, John Sillitoe on behalf of the Wellcome Sanger Institute COVID-19 Surveillance Team |
| EPI_ISL_3284873 | Lighthouse Lab in Alderley Park | Wellcome Sanger Institute for the COVID-19 Genomics UK (COG-UK) Consortium | Jacquelyn Wynn, Mairead Hyland, The Lighthouse Lab in Alderley Park and Alex Alderton, Roberto Amato, Jeffrey Barrett, Sonia Goncalves, Ewan Harrison, David K. Jackson, Ian Johnston, Dominic Kwiatkowski, Cordelia Langford, John Sillitoe on behalf of the Wellcome Sanger Institute COVID-19 Surveillance Team |
| EPI_ISL_3284874 | Lighthouse Lab in Milton Keynes | Wellcome Sanger Institute for the COVID-19 Genomics UK (COG-UK) Consortium | The Lighthouse Lab in Milton Keynes and Alex Alderton, Roberto Amato, Jeffrey Barrett, Sonia Goncalves, Ewan Harrison, David K. Jackson, Ian Johnston, Dominic Kwiatkowski, Cordelia Langford, John Sillitoe on behalf of the Wellcome Sanger Institute COVID-19 Surveillance Team |
| EPI_ISL_3284875 | Lighthouse Lab in Alderley Park | Wellcome Sanger Institute for the COVID-19 Genomics UK (COG-UK) Consortium | Jacquelyn Wynn, Mairead Hyland, The Lighthouse Lab in Alderley Park and Alex Alderton, Roberto Amato, Jeffrey Barrett, Sonia Goncalves, Ewan Harrison, David K. Jackson, Ian Johnston, Dominic Kwiatkowski, Cordelia Langford, John Sillitoe on behalf of the Wellcome Sanger Institute COVID-19 Surveillance Team |
| EPI_ISL_3284876, EPI_ISL_3284877 | Lighthouse Lab in Milton Keynes | Wellcome Sanger Institute for the COVID-19 Genomics UK (COG-UK) Consortium | The Lighthouse Lab in Milton Keynes and Alex Alderton, Roberto Amato, Jeffrey Barrett, Sonia Goncalves, Ewan Harrison, David K. Jackson, Ian Johnston, Dominic Kwiatkowski, Cordelia Langford, John Sillitoe on behalf of the Wellcome Sanger Institute COVID-19 Surveillance Team |
| EPI_ISL_3284878 | Lighthouse Lab in Alderley Park | Wellcome Sanger Institute for the COVID-19 Genomics UK (COG-UK) Consortium | Jacquelyn Wynn, Mairead Hyland, The Lighthouse Lab in Alderley Park and Alex Alderton, Roberto Amato, Jeffrey Barrett, Sonia Goncalves, Ewan Harrison, David K. Jackson, Ian Johnston, Dominic Kwiatkowski, Cordelia Langford, John Sillitoe on behalf of the Wellcome Sanger Institute COVID-19 Surveillance Team |
| EPI_ISL_3284879 | Lighthouse Lab in Milton Keynes | Wellcome Sanger Institute for the COVID-19 Genomics UK (COG-UK) Consortium | The Lighthouse Lab in Milton Keynes and Alex Alderton, Roberto Amato, Jeffrey Barrett, Sonia Goncalves, Ewan Harrison, David K. Jackson, Ian Johnston, Dominic Kwiatkowski, Cordelia Langford, John Sillitoe on behalf of the Wellcome Sanger Institute COVID-19 Surveillance Team |
| EPI_ISL_3284880, EPI_ISL_3284881 | Health Services Laboratories | Wellcome Sanger Institute for the COVID-19 Genomics UK (COG-UK) Consortium | Health Services Laboratories and Alex Alderton, Roberto Amato, Jeffrey Barrett, Sonia Goncalves, Ewan Harrison, David K. Jackson, Ian Johnston, Dominic Kwiatkowski, Cordelia Langford, John Sillitoe on behalf of the Wellcome Sanger Institute COVID-19 Surveillance Team |
| EPI_ISL_3284882 | Lighthouse Lab in Milton Keynes | Wellcome Sanger Institute for the COVID-19 Genomics UK (COG-UK) Consortium | The Lighthouse Lab in Milton Keynes and Alex Alderton, Roberto Amato, Jeffrey Barrett, Sonia Goncalves, Ewan Harrison, David K. Jackson, Ian Johnston, Dominic Kwiatkowski, Cordelia Langford, John Sillitoe on behalf of the Wellcome Sanger Institute COVID-19 Surveillance Team |
| EPI_ISL_3284883, EPI_ISL_3284884 | Health Services Laboratories | Wellcome Sanger Institute for the COVID-19 Genomics UK (COG-UK) Consortium | Health Services Laboratories and Alex Alderton, Roberto Amato, Jeffrey Barrett, Sonia Goncalves, Ewan Harrison, David K. Jackson, Ian Johnston, Dominic Kwiatkowski, Cordelia Langford, John Sillitoe on behalf of the Wellcome Sanger Institute COVID-19 Surveillance Team |
| EPI_ISL_3284885 | Lighthouse Lab in Alderley Park | Wellcome Sanger Institute for the COVID-19 Genomics UK (COG-UK) Consortium | Jacquelyn Wynn, Mairead Hyland, The Lighthouse Lab in Alderley Park and Alex Alderton, Roberto Amato, Jeffrey Barrett, Sonia Goncalves, Ewan Harrison, David K. Jackson, Ian Johnston, Dominic Kwiatkowski, Cordelia Langford, John Sillitoe on behalf of the Wellcome Sanger Institute COVID-19 Surveillance Team |
| EPI_ISL_3284886, EPI_ISL_3284887 | Lighthouse Lab in Milton Keynes | Wellcome Sanger Institute for the COVID-19 Genomics UK (COG-UK) Consortium | The Lighthouse Lab in Milton Keynes and Alex Alderton, Roberto Amato, Jeffrey Barrett, Sonia Goncalves, Ewan Harrison, David K. Jackson, Ian Johnston, Dominic Kwiatkowski, Cordelia Langford, John Sillitoe on behalf of the Wellcome Sanger Institute COVID-19 Surveillance Team |
| EPI_ISL_3284888 | Health Services Laboratories | Wellcome Sanger Institute for the COVID-19 Genomics UK (COG-UK) Consortium | Health Services Laboratories and Alex Alderton, Roberto Amato, Jeffrey Barrett, Sonia Goncalves, Ewan Harrison, David K. Jackson, Ian Johnston, Dominic Kwiatkowski, Cordelia Langford, John Sillitoe on behalf of the Wellcome Sanger Institute COVID-19 Surveillance Team |
| EPI_ISL_3284889 | Lighthouse Lab in Milton Keynes | Wellcome Sanger Institute for the COVID-19 Genomics UK (COG-UK) Consortium | The Lighthouse Lab in Milton Keynes and Alex Alderton, Roberto Amato, Jeffrey Barrett, Sonia Goncalves, Ewan Harrison, David K. Jackson, Ian Johnston, Dominic Kwiatkowski, Cordelia Langford, John Sillitoe on behalf of the Wellcome Sanger Institute COVID-19 Surveillance Team |
| EPI_ISL_3284890 | Health Services Laboratories | Wellcome Sanger Institute for the COVID-19 Genomics UK (COG-UK) Consortium | Health Services Laboratories and Alex Alderton, Roberto Amato, Jeffrey Barrett, Sonia Goncalves, Ewan Harrison, David K. Jackson, Ian Johnston, Dominic Kwiatkowski, Cordelia Langford, John Sillitoe on behalf of the Wellcome Sanger Institute COVID-19 Surveillance Team |
| EPI_ISL_3284891 | Lighthouse Lab in Alderley Park | Wellcome Sanger Institute for the COVID-19 Genomics UK (COG-UK) Consortium | Jacquelyn Wynn, Mairead Hyland, The Lighthouse Lab in Alderley Park and Alex Alderton, Roberto Amato, Jeffrey Barrett, Sonia Goncalves, Ewan Harrison, David K. Jackson, Ian Johnston, Dominic Kwiatkowski, Cordelia Langford, John Sillitoe on behalf of the Wellcome Sanger Institute COVID-19 Surveillance Team |
| EPI_ISL_3284892, EPI_ISL_3284893 | Health Services Laboratories | Wellcome Sanger Institute for the COVID-19 Genomics UK (COG-UK) Consortium | Health Services Laboratories and Alex Alderton, Roberto Amato, Jeffrey Barrett, Sonia Goncalves, Ewan Harrison, David K. Jackson, Ian Johnston, Dominic Kwiatkowski, Cordelia Langford, John Sillitoe on behalf of the Wellcome Sanger Institute COVID-19 Surveillance Team |
| EPI_ISL_3284894 | Lighthouse Lab in Alderley Park | Wellcome Sanger Institute for the COVID-19 Genomics UK (COG-UK) Consortium | Jacquelyn Wynn, Mairead Hyland, The Lighthouse Lab in Alderley Park and Alex Alderton, Roberto Amato, Jeffrey Barrett, Sonia Goncalves, Ewan Harrison, David K. Jackson, Ian Johnston, Dominic Kwiatkowski, Cordelia Langford, John Sillitoe on behalf of the Wellcome Sanger Institute COVID-19 Surveillance Team |
| EPI_ISL_3284895, EPI_ISL_3284896 | Lighthouse Lab in Milton Keynes | Wellcome Sanger Institute for the COVID-19 Genomics UK (COG-UK) Consortium | The Lighthouse Lab in Milton Keynes and Alex Alderton, Roberto Amato, Jeffrey Barrett, Sonia Goncalves, Ewan Harrison, David K. Jackson, Ian Johnston, Dominic Kwiatkowski, Cordelia Langford, John Sillitoe on behalf of the Wellcome Sanger Institute COVID-19 Surveillance Team |
| EPI_ISL_3284897 | Health Services Laboratories | Wellcome Sanger Institute for the COVID-19 Genomics UK (COG-UK) Consortium | Health Services Laboratories and Alex Alderton, Roberto Amato, Jeffrey Barrett, Sonia Goncalves, Ewan Harrison, David K. Jackson, Ian Johnston, Dominic Kwiatkowski, Cordelia Langford, John Sillitoe on behalf of the Wellcome Sanger Institute COVID-19 Surveillance Team |
| EPI_ISL_3284898, EPI_ISL_3284899, EPI_ISL_3284900 | Lighthouse Lab in Milton Keynes | Wellcome Sanger Institute for the COVID-19 Genomics UK (COG-UK) Consortium | The Lighthouse Lab in Milton Keynes and Alex Alderton, Roberto Amato, Jeffrey Barrett, Sonia Goncalves, Ewan Harrison, David K. Jackson, Ian Johnston, Dominic Kwiatkowski, Cordelia Langford, John Sillitoe on behalf of the Wellcome Sanger Institute COVID-19 Surveillance Team |

| | | | |
|---|---|---|---|
| EPI_ISL_3284901, EPI_ISL_3284902 | Lighthouse Lab in Alderley Park | Wellcome Sanger Institute for the COVID-19 Genomics UK (COG-UK) Consortium | Jacquelyn Wynn, Mairead Hyland, The Lighthouse Lab in Alderley Park and Alex Alderton, Roberto Amato, Jeffrey Barrett, Sonia Goncalves, Ewan Harrison, David K. Jackson, Ian Johnston, Dominic Kwiatkowski, Cordelia Langford, John Sillitoe on behalf of the Wellcome Sanger Institute COVID-19 Surveillance Team |
| EPI_ISL_3284903 | Lighthouse Lab in Milton Keynes | Wellcome Sanger Institute for the COVID-19 Genomics UK (COG-UK) Consortium | The Lighthouse Lab in Milton Keynes and Alex Alderton, Roberto Amato, Jeffrey Barrett, Sonia Goncalves, Ewan Harrison, David K. Jackson, Ian Johnston, Dominic Kwiatkowski, Cordelia Langford, John Sillitoe on behalf of the Wellcome Sanger Institute COVID-19 Surveillance Team |
| EPI_ISL_3284904 | Health Services Laboratories | Wellcome Sanger Institute for the COVID-19 Genomics UK (COG-UK) Consortium | Health Services Laboratories and Alex Alderton, Roberto Amato, Jeffrey Barrett, Sonia Goncalves, Ewan Harrison, David K. Jackson, Ian Johnston, Dominic Kwiatkowski, Cordelia Langford, John Sillitoe on behalf of the Wellcome Sanger Institute COVID-19 Surveillance Team |
| EPI_ISL_3284905, EPI_ISL_3284906, EPI_ISL_3284907 | Lighthouse Lab in Milton Keynes | Wellcome Sanger Institute for the COVID-19 Genomics UK (COG-UK) Consortium | The Lighthouse Lab in Milton Keynes and Alex Alderton, Roberto Amato, Jeffrey Barrett, Sonia Goncalves, Ewan Harrison, David K. Jackson, Ian Johnston, Dominic Kwiatkowski, Cordelia Langford, John Sillitoe on behalf of the Wellcome Sanger Institute COVID-19 Surveillance Team |
| EPI_ISL_3284908 | Lighthouse Lab in Alderley Park | Wellcome Sanger Institute for the COVID-19 Genomics UK (COG-UK) Consortium | Jacquelyn Wynn, Mairead Hyland, The Lighthouse Lab in Alderley Park and Alex Alderton, Roberto Amato, Jeffrey Barrett, Sonia Goncalves, Ewan Harrison, David K. Jackson, Ian Johnston, Dominic Kwiatkowski, Cordelia Langford, John Sillitoe on behalf of the Wellcome Sanger Institute COVID-19 Surveillance Team |
| EPI_ISL_3284909, EPI_ISL_3284910, EPI_ISL_3284911 | Lighthouse Lab in Milton Keynes | Wellcome Sanger Institute for the COVID-19 Genomics UK (COG-UK) Consortium | The Lighthouse Lab in Milton Keynes and Alex Alderton, Roberto Amato, Jeffrey Barrett, Sonia Goncalves, Ewan Harrison, David K. Jackson, Ian Johnston, Dominic Kwiatkowski, Cordelia Langford, John Sillitoe on behalf of the Wellcome Sanger Institute COVID-19 Surveillance Team |
| EPI_ISL_3284912, EPI_ISL_3284913 | Health Services Laboratories | Wellcome Sanger Institute for the COVID-19 Genomics UK (COG-UK) Consortium | Health Services Laboratories and Alex Alderton, Roberto Amato, Jeffrey Barrett, Sonia Goncalves, Ewan Harrison, David K. Jackson, Ian Johnston, Dominic Kwiatkowski, Cordelia Langford, John Sillitoe on behalf of the Wellcome Sanger Institute COVID-19 Surveillance Team |
| EPI_ISL_3284914, EPI_ISL_3284915 | Lighthouse Lab in Alderley Park | Wellcome Sanger Institute for the COVID-19 Genomics UK (COG-UK) Consortium | Jacquelyn Wynn, Mairead Hyland, The Lighthouse Lab in Alderley Park and Alex Alderton, Roberto Amato, Jeffrey Barrett, Sonia Goncalves, Ewan Harrison, David K. Jackson, Ian Johnston, Dominic Kwiatkowski, Cordelia Langford, John Sillitoe on behalf of the Wellcome Sanger Institute COVID-19 Surveillance Team |
| EPI_ISL_3284916 | Lighthouse Lab in Milton Keynes | Wellcome Sanger Institute for the COVID-19 Genomics UK (COG-UK) Consortium | The Lighthouse Lab in Milton Keynes and Alex Alderton, Roberto Amato, Jeffrey Barrett, Sonia Goncalves, Ewan Harrison, David K. Jackson, Ian Johnston, Dominic Kwiatkowski, Cordelia Langford, John Sillitoe on behalf of the Wellcome Sanger Institute COVID-19 Surveillance Team |
| EPI_ISL_3284917 | Health Services Laboratories | Wellcome Sanger Institute for the COVID-19 Genomics UK (COG-UK) Consortium | Health Services Laboratories and Alex Alderton, Roberto Amato, Jeffrey Barrett, Sonia Goncalves, Ewan Harrison, David K. Jackson, Ian Johnston, Dominic Kwiatkowski, Cordelia Langford, John Sillitoe on behalf of the Wellcome Sanger Institute COVID-19 Surveillance Team |
| EPI_ISL_3284918 | Lighthouse Lab in Milton Keynes | Wellcome Sanger Institute for the COVID-19 Genomics UK (COG-UK) Consortium | The Lighthouse Lab in Milton Keynes and Alex Alderton, Roberto Amato, Jeffrey Barrett, Sonia Goncalves, Ewan Harrison, David K. Jackson, Ian Johnston, Dominic Kwiatkowski, Cordelia Langford, John Sillitoe on behalf of the Wellcome Sanger Institute COVID-19 Surveillance Team |
| EPI_ISL_3284919 | Health Services Laboratories | Wellcome Sanger Institute for the COVID-19 Genomics UK (COG-UK) Consortium | Health Services Laboratories and Alex Alderton, Roberto Amato, Jeffrey Barrett, Sonia Goncalves, Ewan Harrison, David K. Jackson, Ian Johnston, Dominic Kwiatkowski, Cordelia Langford, John Sillitoe on behalf of the Wellcome Sanger Institute COVID-19 Surveillance Team |
| EPI_ISL_3284920, EPI_ISL_3284921, EPI_ISL_3284922, EPI_ISL_3284923 | Lighthouse Lab in Milton Keynes | Wellcome Sanger Institute for the COVID-19 Genomics UK (COG-UK) Consortium | The Lighthouse Lab in Milton Keynes and Alex Alderton, Roberto Amato, Jeffrey Barrett, Sonia Goncalves, Ewan Harrison, David K. Jackson, Ian Johnston, Dominic Kwiatkowski, Cordelia Langford, John Sillitoe on behalf of the Wellcome Sanger Institute COVID-19 Surveillance Team |
| EPI_ISL_3284924 | Lighthouse Lab in Alderley Park | Wellcome Sanger Institute for the COVID-19 Genomics UK (COG-UK) Consortium | Jacquelyn Wynn, Mairead Hyland, The Lighthouse Lab in Alderley Park and Alex Alderton, Roberto Amato, Jeffrey Barrett, Sonia Goncalves, Ewan Harrison, David K. Jackson, Ian Johnston, Dominic Kwiatkowski, Cordelia Langford, John Sillitoe on behalf of the Wellcome Sanger Institute COVID-19 Surveillance Team |
| EPI_ISL_3284925 | Health Services Laboratories | Wellcome Sanger Institute for the COVID-19 Genomics UK (COG-UK) Consortium | Health Services Laboratories and Alex Alderton, Roberto Amato, Jeffrey Barrett, Sonia Goncalves, Ewan Harrison, David K. Jackson, Ian Johnston, Dominic Kwiatkowski, Cordelia Langford, John Sillitoe on behalf of the Wellcome Sanger Institute COVID-19 Surveillance Team |
| EPI_ISL_3284926, EPI_ISL_3284927 | Lighthouse Lab in Alderley Park | Wellcome Sanger Institute for the COVID-19 Genomics UK (COG-UK) Consortium | Jacquelyn Wynn, Mairead Hyland, The Lighthouse Lab in Alderley Park and Alex Alderton, Roberto Amato, Jeffrey Barrett, Sonia Goncalves, Ewan Harrison, David K. Jackson, Ian Johnston, Dominic Kwiatkowski, Cordelia Langford, John Sillitoe on behalf of the Wellcome Sanger Institute COVID-19 Surveillance Team |
| EPI_ISL_3284928 | Health Services Laboratories | Wellcome Sanger Institute for the COVID-19 Genomics UK (COG-UK) Consortium | Health Services Laboratories and Alex Alderton, Roberto Amato, Jeffrey Barrett, Sonia Goncalves, Ewan Harrison, David K. Jackson, Ian Johnston, Dominic Kwiatkowski, Cordelia Langford, John Sillitoe on behalf of the Wellcome Sanger Institute COVID-19 Surveillance Team |
| EPI_ISL_3284929 | Lighthouse Lab in Milton Keynes | Wellcome Sanger Institute for the COVID-19 Genomics UK (COG-UK) Consortium | The Lighthouse Lab in Milton Keynes and Alex Alderton, Roberto Amato, Jeffrey Barrett, Sonia Goncalves, Ewan Harrison, David K. Jackson, Ian Johnston, Dominic Kwiatkowski, Cordelia Langford, John Sillitoe on behalf of the Wellcome Sanger Institute COVID-19 Surveillance Team |
| EPI_ISL_3284930, EPI_ISL_3284931, EPI_ISL_3284932 | Health Services Laboratories | Wellcome Sanger Institute for the COVID-19 Genomics UK (COG-UK) Consortium | Health Services Laboratories and Alex Alderton, Roberto Amato, Jeffrey Barrett, Sonia Goncalves, Ewan Harrison, David K. Jackson, Ian Johnston, Dominic Kwiatkowski, Cordelia Langford, John Sillitoe on behalf of the Wellcome Sanger Institute COVID-19 Surveillance Team |
| EPI_ISL_3284933 | Lighthouse Lab in Milton Keynes | Wellcome Sanger Institute for the COVID-19 Genomics UK (COG-UK) Consortium | The Lighthouse Lab in Milton Keynes and Alex Alderton, Roberto Amato, Jeffrey Barrett, Sonia Goncalves, Ewan Harrison, David K. Jackson, Ian Johnston, Dominic Kwiatkowski, Cordelia Langford, John Sillitoe on behalf of the Wellcome Sanger Institute COVID-19 Surveillance Team |
| EPI_ISL_3284934 | Lighthouse Lab in Alderley Park | Wellcome Sanger Institute for the COVID-19 Genomics UK (COG-UK) Consortium | Jacquelyn Wynn, Mairead Hyland, The Lighthouse Lab in Alderley Park and Alex Alderton, Roberto Amato, Jeffrey Barrett, Sonia Goncalves, Ewan Harrison, David K. Jackson, Ian Johnston, Dominic Kwiatkowski, Cordelia Langford, John Sillitoe on behalf of the Wellcome Sanger Institute COVID-19 Surveillance Team |
| EPI_ISL_3284935 | Lighthouse Lab in Milton Keynes | Wellcome Sanger Institute for the COVID-19 Genomics UK (COG-UK) Consortium | The Lighthouse Lab in Milton Keynes and Alex Alderton, Roberto Amato, Jeffrey Barrett, Sonia Goncalves, Ewan Harrison, David K. Jackson, Ian Johnston, Dominic Kwiatkowski, Cordelia Langford, John Sillitoe on behalf of the Wellcome Sanger Institute COVID-19 Surveillance Team |
| EPI_ISL_3284936 | Lighthouse Lab in Alderley Park | Wellcome Sanger Institute for the COVID-19 Genomics UK (COG-UK) Consortium | Jacquelyn Wynn, Mairead Hyland, The Lighthouse Lab in Alderley Park and Alex Alderton, Roberto Amato, Jeffrey Barrett, Sonia Goncalves, Ewan Harrison, David K. Jackson, Ian Johnston, Dominic Kwiatkowski, Cordelia Langford, John Sillitoe on behalf of the Wellcome Sanger Institute COVID-19 Surveillance Team |
| EPI_ISL_3284937 | Lighthouse Lab in Milton Keynes | Wellcome Sanger Institute for the COVID-19 Genomics UK (COG-UK) Consortium | The Lighthouse Lab in Milton Keynes and Alex Alderton, Roberto Amato, Jeffrey Barrett, Sonia Goncalves, Ewan Harrison, David K. Jackson, Ian Johnston, Dominic Kwiatkowski, Cordelia Langford, John Sillitoe on behalf of the Wellcome Sanger Institute COVID-19 Surveillance Team |
| EPI_ISL_3284938 | Lighthouse Lab in Alderley Park | Wellcome Sanger Institute for the COVID-19 Genomics UK (COG-UK) Consortium | Jacquelyn Wynn, Mairead Hyland, The Lighthouse Lab in Alderley Park and Alex Alderton, Roberto Amato, Jeffrey Barrett, Sonia Goncalves, Ewan Harrison, David K. Jackson, Ian Johnston, Dominic Kwiatkowski, Cordelia Langford, John Sillitoe on behalf of the Wellcome Sanger Institute COVID-19 Surveillance Team |
| EPI_ISL_3284939, EPI_ISL_3284940 | Health Services Laboratories | Wellcome Sanger Institute for the COVID-19 Genomics UK (COG-UK) Consortium | Health Services Laboratories and Alex Alderton, Roberto Amato, Jeffrey Barrett, Sonia Goncalves, Ewan Harrison, David K. Jackson, Ian Johnston, Dominic Kwiatkowski, Cordelia Langford, John Sillitoe on behalf of the Wellcome Sanger Institute COVID-19 Surveillance Team |
| EPI_ISL_3284941 | Lighthouse Lab in Milton Keynes | Wellcome Sanger Institute for the COVID-19 Genomics UK (COG-UK) Consortium | The Lighthouse Lab in Milton Keynes and Alex Alderton, Roberto Amato, Jeffrey Barrett, Sonia Goncalves, Ewan Harrison, David K. Jackson, Ian Johnston, Dominic Kwiatkowski, Cordelia Langford, John Sillitoe on behalf of the Wellcome Sanger Institute COVID-19 Surveillance Team |
| EPI_ISL_3284942 | Health Services Laboratories | Wellcome Sanger Institute for the COVID-19 Genomics UK (COG-UK) Consortium | Health Services Laboratories and Alex Alderton, Roberto Amato, Jeffrey Barrett, Sonia Goncalves, Ewan Harrison, David K. Jackson, Ian Johnston, Dominic Kwiatkowski, Cordelia Langford, John Sillitoe on behalf of the Wellcome Sanger Institute COVID-19 Surveillance Team |
| EPI_ISL_3284943 | Lighthouse Lab in Milton Keynes | Wellcome Sanger Institute for the COVID-19 Genomics UK (COG-UK) Consortium | The Lighthouse Lab in Milton Keynes and Alex Alderton, Roberto Amato, Jeffrey Barrett, Sonia Goncalves, Ewan Harrison, David K. Jackson, Ian Johnston, Dominic Kwiatkowski, Cordelia Langford, John Sillitoe on behalf of the Wellcome Sanger Institute COVID-19 Surveillance Team |
| EPI_ISL_3284944 | Health Services Laboratories | Wellcome Sanger Institute for the COVID-19 Genomics UK (COG-UK) Consortium | Health Services Laboratories and Alex Alderton, Roberto Amato, Jeffrey Barrett, Sonia Goncalves, Ewan Harrison, David K. Jackson, Ian Johnston, Dominic Kwiatkowski, Cordelia Langford, John Sillitoe on behalf of the Wellcome Sanger Institute COVID-19 Surveillance Team |
| EPI_ISL_3284945, EPI_ISL_3284946, EPI_ISL_3284947 | Lighthouse Lab in Milton Keynes | Wellcome Sanger Institute for the COVID-19 Genomics UK (COG-UK) Consortium | The Lighthouse Lab in Milton Keynes and Alex Alderton, Roberto Amato, Jeffrey Barrett, Sonia Goncalves, Ewan Harrison, David K. Jackson, Ian Johnston, Dominic Kwiatkowski, Cordelia Langford, John Sillitoe on behalf of the Wellcome Sanger Institute COVID-19 Surveillance Team |
| EPI_ISL_3284948 | Berkshire and Surrey Pathology Services Lighthouse Laboratory | Wellcome Sanger Institute for the COVID-19 Genomics UK (COG-UK) Consortium | Berkshire and Surrey Pathology Services Lighthouse Laboratory and Alex Alderton, Roberto Amato, Jeffrey Barrett, Sonia Goncalves, Ewan Harrison, David K. Jackson, Ian Johnston, Dominic Kwiatkowski, Cordelia Langford, John Sillitoe on behalf of the Wellcome Sanger Institute COVID-19 Surveillance Team |
| EPI_ISL_3284949, EPI_ISL_3284950, EPI_ISL_3284951, EPI_ISL_3284952, EPI_ISL_3284953, EPI_ISL_3284954 | Lighthouse Lab in Milton Keynes | Wellcome Sanger Institute for the COVID-19 Genomics UK (COG-UK) Consortium | The Lighthouse Lab in Milton Keynes and Alex Alderton, Roberto Amato, Jeffrey Barrett, Sonia Goncalves, Ewan Harrison, David K. Jackson, Ian Johnston, Dominic Kwiatkowski, Cordelia Langford, John Sillitoe on behalf of the Wellcome Sanger Institute COVID-19 Surveillance Team |
| EPI_ISL_3284955 | Health Services Laboratories | Wellcome Sanger Institute for the COVID-19 Genomics UK (COG-UK) Consortium | Health Services Laboratories and Alex Alderton, Roberto Amato, Jeffrey Barrett, Sonia Goncalves, Ewan Harrison, David K. Jackson, Ian Johnston, Dominic Kwiatkowski, Cordelia Langford, John Sillitoe on behalf of the Wellcome Sanger Institute COVID-19 Surveillance Team |
| EPI_ISL_3284956, EPI_ISL_3284957 | Lighthouse Lab in Milton Keynes | Wellcome Sanger Institute for the COVID-19 Genomics UK (COG-UK) Consortium | The Lighthouse Lab in Milton Keynes and Alex Alderton, Roberto Amato, Jeffrey Barrett, Sonia Goncalves, Ewan Harrison, David K. Jackson, Ian Johnston, Dominic Kwiatkowski, Cordelia Langford, John Sillitoe on behalf of the Wellcome Sanger Institute COVID-19 Surveillance Team |
| EPI_ISL_3284958 | Lighthouse Lab in Alderley Park | Wellcome Sanger Institute for the COVID-19 Genomics UK (COG-UK) Consortium | Jacquelyn Wynn, Mairead Hyland, The Lighthouse Lab in Alderley Park and Alex Alderton, Roberto Amato, Jeffrey Barrett, Sonia Goncalves, Ewan Harrison, David K. Jackson, Ian Johnston, Dominic Kwiatkowski, Cordelia Langford, John Sillitoe on behalf of the Wellcome Sanger Institute COVID-19 Surveillance Team |
| EPI_ISL_3284959 | Lighthouse Lab in Milton Keynes | Wellcome Sanger Institute for the COVID-19 Genomics UK (COG-UK) Consortium | The Lighthouse Lab in Milton Keynes and Alex Alderton, Roberto Amato, Jeffrey Barrett, Sonia Goncalves, Ewan Harrison, David K. Jackson, Ian Johnston, Dominic Kwiatkowski, Cordelia Langford, John Sillitoe on behalf of the Wellcome Sanger Institute COVID-19 Surveillance Team |
| EPI_ISL_3284960 | Health Services Laboratories | Wellcome Sanger Institute for the COVID-19 Genomics UK (COG-UK) Consortium | Health Services Laboratories and Alex Alderton, Roberto Amato, Jeffrey Barrett, Sonia Goncalves, Ewan Harrison, David K. Jackson, Ian Johnston, Dominic Kwiatkowski, Cordelia Langford, John Sillitoe on behalf of the Wellcome Sanger Institute COVID-19 Surveillance Team |
| EPI_ISL_3284961, EPI_ISL_3284962 | Lighthouse Lab in Milton Keynes | Wellcome Sanger Institute for the COVID-19 Genomics UK (COG-UK) Consortium | The Lighthouse Lab in Milton Keynes and Alex Alderton, Roberto Amato, Jeffrey Barrett, Sonia Goncalves, Ewan Harrison, David K. Jackson, Ian Johnston, Dominic Kwiatkowski, Cordelia Langford, John Sillitoe on behalf of the Wellcome Sanger Institute COVID-19 Surveillance Team |
| EPI_ISL_3284963 | Lighthouse Lab in Alderley Park | Wellcome Sanger Institute for the COVID-19 Genomics UK (COG-UK) Consortium | Jacquelyn Wynn, Mairead Hyland, The Lighthouse Lab in Alderley Park and Alex Alderton, Roberto Amato, Jeffrey Barrett, Sonia Goncalves, Ewan Harrison, David K. Jackson, Ian Johnston, Dominic Kwiatkowski, Cordelia Langford, John Sillitoe on behalf of the Wellcome Sanger Institute COVID-19 Surveillance Team |
| EPI_ISL_3284964 | Health Services Laboratories | Wellcome Sanger Institute for the COVID-19 Genomics UK (COG-UK) Consortium | Health Services Laboratories and Alex Alderton, Roberto Amato, Jeffrey Barrett, Sonia Goncalves, Ewan Harrison, David K. Jackson, Ian Johnston, Dominic Kwiatkowski, Cordelia Langford, John Sillitoe on behalf of the Wellcome Sanger Institute COVID-19 Surveillance Team |

| | | | |
|---|---|---|---|
| EPI_ISL_3284965 | Berkshire and Surrey Pathology Services Lighthouse Laboratory | Wellcome Sanger Institute for the COVID-19 Genomics UK (COG-UK) Consortium | Berkshire and Surrey Pathology Services Lighthouse Laboratory and Alex Alderton, Roberto Amato, Jeffrey Barrett, Sonia Goncalves, Ewan Harrison, David K. Jackson, Ian Johnston, Dominic Kwiatkowski, Cordelia Langford, John Sillitoe on behalf of the Wellcome Sanger Institute COVID-19 Surveillance Team |
| EPI_ISL_3284966 | Lighthouse Lab in Milton Keynes | Wellcome Sanger Institute for the COVID-19 Genomics UK (COG-UK) Consortium | The Lighthouse Lab in Milton Keynes and Alex Alderton, Roberto Amato, Jeffrey Barrett, Sonia Goncalves, Ewan Harrison, David K. Jackson, Ian Johnston, Dominic Kwiatkowski, Cordelia Langford, John Sillitoe on behalf of the Wellcome Sanger Institute COVID-19 Surveillance Team |
| EPI_ISL_3284967 | Lighthouse Lab in Alderley Park | Wellcome Sanger Institute for the COVID-19 Genomics UK (COG-UK) Consortium | Jacquelyn Wynn, Mairead Hyland, The Lighthouse Lab in Alderley Park and Alex Alderton, Roberto Amato, Jeffrey Barrett, Sonia Goncalves, Ewan Harrison, David K. Jackson, Ian Johnston, Dominic Kwiatkowski, Cordelia Langford, John Sillitoe on behalf of the Wellcome Sanger Institute COVID-19 Surveillance Team |
| EPI_ISL_3284968 | Lighthouse Lab in Milton Keynes | Wellcome Sanger Institute for the COVID-19 Genomics UK (COG-UK) Consortium | The Lighthouse Lab in Milton Keynes and Alex Alderton, Roberto Amato, Jeffrey Barrett, Sonia Goncalves, Ewan Harrison, David K. Jackson, Ian Johnston, Dominic Kwiatkowski, Cordelia Langford, John Sillitoe on behalf of the Wellcome Sanger Institute COVID-19 Surveillance Team |
| EPI_ISL_3284969 | Health Services Laboratories | Wellcome Sanger Institute for the COVID-19 Genomics UK (COG-UK) Consortium | Health Services Laboratories and Alex Alderton, Roberto Amato, Jeffrey Barrett, Sonia Goncalves, Ewan Harrison, David K. Jackson, Ian Johnston, Dominic Kwiatkowski, Cordelia Langford, John Sillitoe on behalf of the Wellcome Sanger Institute COVID-19 Surveillance Team |
| EPI_ISL_3284970 | Lighthouse Lab in Alderley Park | Wellcome Sanger Institute for the COVID-19 Genomics UK (COG-UK) Consortium | Jacquelyn Wynn, Mairead Hyland, The Lighthouse Lab in Alderley Park and Alex Alderton, Roberto Amato, Jeffrey Barrett, Sonia Goncalves, Ewan Harrison, David K. Jackson, Ian Johnston, Dominic Kwiatkowski, Cordelia Langford, John Sillitoe on behalf of the Wellcome Sanger Institute COVID-19 Surveillance Team |
| EPI_ISL_3284971 | Lighthouse Lab in Milton Keynes | Wellcome Sanger Institute for the COVID-19 Genomics UK (COG-UK) Consortium | The Lighthouse Lab in Milton Keynes and Alex Alderton, Roberto Amato, Jeffrey Barrett, Sonia Goncalves, Ewan Harrison, David K. Jackson, Ian Johnston, Dominic Kwiatkowski, Cordelia Langford, John Sillitoe on behalf of the Wellcome Sanger Institute COVID-19 Surveillance Team |
| EPI_ISL_3284972 | Health Services Laboratories | Wellcome Sanger Institute for the COVID-19 Genomics UK (COG-UK) Consortium | Health Services Laboratories and Alex Alderton, Roberto Amato, Jeffrey Barrett, Sonia Goncalves, Ewan Harrison, David K. Jackson, Ian Johnston, Dominic Kwiatkowski, Cordelia Langford, John Sillitoe on behalf of the Wellcome Sanger Institute COVID-19 Surveillance Team |
| EPI_ISL_3284973 | Lighthouse Lab in Milton Keynes | Wellcome Sanger Institute for the COVID-19 Genomics UK (COG-UK) Consortium | The Lighthouse Lab in Milton Keynes and Alex Alderton, Roberto Amato, Jeffrey Barrett, Sonia Goncalves, Ewan Harrison, David K. Jackson, Ian Johnston, Dominic Kwiatkowski, Cordelia Langford, John Sillitoe on behalf of the Wellcome Sanger Institute COVID-19 Surveillance Team |
| EPI_ISL_3284974, EPI_ISL_3284975 | Health Services Laboratories | Wellcome Sanger Institute for the COVID-19 Genomics UK (COG-UK) Consortium | Health Services Laboratories and Alex Alderton, Roberto Amato, Jeffrey Barrett, Sonia Goncalves, Ewan Harrison, David K. Jackson, Ian Johnston, Dominic Kwiatkowski, Cordelia Langford, John Sillitoe on behalf of the Wellcome Sanger Institute COVID-19 Surveillance Team |
| EPI_ISL_3284976 | Lighthouse Lab in Milton Keynes | Wellcome Sanger Institute for the COVID-19 Genomics UK (COG-UK) Consortium | The Lighthouse Lab in Milton Keynes and Alex Alderton, Roberto Amato, Jeffrey Barrett, Sonia Goncalves, Ewan Harrison, David K. Jackson, Ian Johnston, Dominic Kwiatkowski, Cordelia Langford, John Sillitoe on behalf of the Wellcome Sanger Institute COVID-19 Surveillance Team |
| EPI_ISL_3284977, EPI_ISL_3284978, EPI_ISL_3284979 | Lighthouse Lab in Alderley Park | Wellcome Sanger Institute for the COVID-19 Genomics UK (COG-UK) Consortium | Jacquelyn Wynn, Mairead Hyland, The Lighthouse Lab in Alderley Park and Alex Alderton, Roberto Amato, Jeffrey Barrett, Sonia Goncalves, Ewan Harrison, David K. Jackson, Ian Johnston, Dominic Kwiatkowski, Cordelia Langford, John Sillitoe on behalf of the Wellcome Sanger Institute COVID-19 Surveillance Team |
| EPI_ISL_3284980, EPI_ISL_3284981 | Health Services Laboratories | Wellcome Sanger Institute for the COVID-19 Genomics UK (COG-UK) Consortium | Health Services Laboratories and Alex Alderton, Roberto Amato, Jeffrey Barrett, Sonia Goncalves, Ewan Harrison, David K. Jackson, Ian Johnston, Dominic Kwiatkowski, Cordelia Langford, John Sillitoe on behalf of the Wellcome Sanger Institute COVID-19 Surveillance Team |
| EPI_ISL_3284982 | Lighthouse Lab in Milton Keynes | Wellcome Sanger Institute for the COVID-19 Genomics UK (COG-UK) Consortium | The Lighthouse Lab in Milton Keynes and Alex Alderton, Roberto Amato, Jeffrey Barrett, Sonia Goncalves, Ewan Harrison, David K. Jackson, Ian Johnston, Dominic Kwiatkowski, Cordelia Langford, John Sillitoe on behalf of the Wellcome Sanger Institute COVID-19 Surveillance Team |
| EPI_ISL_3284983 | Lighthouse Lab in Alderley Park | Wellcome Sanger Institute for the COVID-19 Genomics UK (COG-UK) Consortium | Jacquelyn Wynn, Mairead Hyland, The Lighthouse Lab in Alderley Park and Alex Alderton, Roberto Amato, Jeffrey Barrett, Sonia Goncalves, Ewan Harrison, David K. Jackson, Ian Johnston, Dominic Kwiatkowski, Cordelia Langford, John Sillitoe on behalf of the Wellcome Sanger Institute COVID-19 Surveillance Team |
| EPI_ISL_3284984 | Health Services Laboratories | Wellcome Sanger Institute for the COVID-19 Genomics UK (COG-UK) Consortium | Health Services Laboratories and Alex Alderton, Roberto Amato, Jeffrey Barrett, Sonia Goncalves, Ewan Harrison, David K. Jackson, Ian Johnston, Dominic Kwiatkowski, Cordelia Langford, John Sillitoe on behalf of the Wellcome Sanger Institute COVID-19 Surveillance Team |
| EPI_ISL_3284985, EPI_ISL_3284986 | Lighthouse Lab in Milton Keynes | Wellcome Sanger Institute for the COVID-19 Genomics UK (COG-UK) Consortium | The Lighthouse Lab in Milton Keynes and Alex Alderton, Roberto Amato, Jeffrey Barrett, Sonia Goncalves, Ewan Harrison, David K. Jackson, Ian Johnston, Dominic Kwiatkowski, Cordelia Langford, John Sillitoe on behalf of the Wellcome Sanger Institute COVID-19 Surveillance Team |
| EPI_ISL_3284987, EPI_ISL_3284988, EPI_ISL_3284989 | Lighthouse Lab in Alderley Park | Wellcome Sanger Institute for the COVID-19 Genomics UK (COG-UK) Consortium | Jacquelyn Wynn, Mairead Hyland, The Lighthouse Lab in Alderley Park and Alex Alderton, Roberto Amato, Jeffrey Barrett, Sonia Goncalves, Ewan Harrison, David K. Jackson, Ian Johnston, Dominic Kwiatkowski, Cordelia Langford, John Sillitoe on behalf of the Wellcome Sanger Institute COVID-19 Surveillance Team |
| EPI_ISL_3284990 | Lighthouse Lab in Milton Keynes | Wellcome Sanger Institute for the COVID-19 Genomics UK (COG-UK) Consortium | The Lighthouse Lab in Milton Keynes and Alex Alderton, Roberto Amato, Jeffrey Barrett, Sonia Goncalves, Ewan Harrison, David K. Jackson, Ian Johnston, Dominic Kwiatkowski, Cordelia Langford, John Sillitoe on behalf of the Wellcome Sanger Institute COVID-19 Surveillance Team |
| EPI_ISL_3284991, EPI_ISL_3284992 | Lighthouse Lab in Alderley Park | Wellcome Sanger Institute for the COVID-19 Genomics UK (COG-UK) Consortium | Jacquelyn Wynn, Mairead Hyland, The Lighthouse Lab in Alderley Park and Alex Alderton, Roberto Amato, Jeffrey Barrett, Sonia Goncalves, Ewan Harrison, David K. Jackson, Ian Johnston, Dominic Kwiatkowski, Cordelia Langford, John Sillitoe on behalf of the Wellcome Sanger Institute COVID-19 Surveillance Team |
| EPI_ISL_3284993 | Berkshire and Surrey Pathology Services Lighthouse Laboratory | Wellcome Sanger Institute for the COVID-19 Genomics UK (COG-UK) Consortium | Berkshire and Surrey Pathology Services Lighthouse Laboratory and Alex Alderton, Roberto Amato, Jeffrey Barrett, Sonia Goncalves, Ewan Harrison, David K. Jackson, Ian Johnston, Dominic Kwiatkowski, Cordelia Langford, John Sillitoe on behalf of the Wellcome Sanger Institute COVID-19 Surveillance Team |
| EPI_ISL_3284994, EPI_ISL_3284995 | Lighthouse Lab in Alderley Park | Wellcome Sanger Institute for the COVID-19 Genomics UK (COG-UK) Consortium | Jacquelyn Wynn, Mairead Hyland, The Lighthouse Lab in Alderley Park and Alex Alderton, Roberto Amato, Jeffrey Barrett, Sonia Goncalves, Ewan Harrison, David K. Jackson, Ian Johnston, Dominic Kwiatkowski, Cordelia Langford, John Sillitoe on behalf of the Wellcome Sanger Institute COVID-19 Surveillance Team |
| EPI_ISL_3284996 | Lighthouse Lab in Milton Keynes | Wellcome Sanger Institute for the COVID-19 Genomics UK (COG-UK) Consortium | The Lighthouse Lab in Milton Keynes and Alex Alderton, Roberto Amato, Jeffrey Barrett, Sonia Goncalves, Ewan Harrison, David K. Jackson, Ian Johnston, Dominic Kwiatkowski, Cordelia Langford, John Sillitoe on behalf of the Wellcome Sanger Institute COVID-19 Surveillance Team |
| EPI_ISL_3284997, EPI_ISL_3284998, EPI_ISL_3284999, EPI_ISL_3285000 | Lighthouse Lab in Alderley Park | Wellcome Sanger Institute for the COVID-19 Genomics UK (COG-UK) Consortium | Jacquelyn Wynn, Mairead Hyland, The Lighthouse Lab in Alderley Park and Alex Alderton, Roberto Amato, Jeffrey Barrett, Sonia Goncalves, Ewan Harrison, David K. Jackson, Ian Johnston, Dominic Kwiatkowski, Cordelia Langford, John Sillitoe on behalf of the Wellcome Sanger Institute COVID-19 Surveillance Team |
| EPI_ISL_3285001, EPI_ISL_3285002 | Lighthouse Lab in Milton Keynes | Wellcome Sanger Institute for the COVID-19 Genomics UK (COG-UK) Consortium | The Lighthouse Lab in Milton Keynes and Alex Alderton, Roberto Amato, Jeffrey Barrett, Sonia Goncalves, Ewan Harrison, David K. Jackson, Ian Johnston, Dominic Kwiatkowski, Cordelia Langford, John Sillitoe on behalf of the Wellcome Sanger Institute COVID-19 Surveillance Team |
| EPI_ISL_3285003 | Lighthouse Lab in Alderley Park | Wellcome Sanger Institute for the COVID-19 Genomics UK (COG-UK) Consortium | Jacquelyn Wynn, Mairead Hyland, The Lighthouse Lab in Alderley Park and Alex Alderton, Roberto Amato, Jeffrey Barrett, Sonia Goncalves, Ewan Harrison, David K. Jackson, Ian Johnston, Dominic Kwiatkowski, Cordelia Langford, John Sillitoe on behalf of the Wellcome Sanger Institute COVID-19 Surveillance Team |
| EPI_ISL_3285004 | Lighthouse Lab in Milton Keynes | Wellcome Sanger Institute for the COVID-19 Genomics UK (COG-UK) Consortium | The Lighthouse Lab in Milton Keynes and Alex Alderton, Roberto Amato, Jeffrey Barrett, Sonia Goncalves, Ewan Harrison, David K. Jackson, Ian Johnston, Dominic Kwiatkowski, Cordelia Langford, John Sillitoe on behalf of the Wellcome Sanger Institute COVID-19 Surveillance Team |
| EPI_ISL_3285005, EPI_ISL_3285006 | Lighthouse Lab in Alderley Park | Wellcome Sanger Institute for the COVID-19 Genomics UK (COG-UK) Consortium | Jacquelyn Wynn, Mairead Hyland, The Lighthouse Lab in Alderley Park and Alex Alderton, Roberto Amato, Jeffrey Barrett, Sonia Goncalves, Ewan Harrison, David K. Jackson, Ian Johnston, Dominic Kwiatkowski, Cordelia Langford, John Sillitoe on behalf of the Wellcome Sanger Institute COVID-19 Surveillance Team |
| EPI_ISL_3285007 | Lighthouse Lab in Milton Keynes | Wellcome Sanger Institute for the COVID-19 Genomics UK (COG-UK) Consortium | The Lighthouse Lab in Milton Keynes and Alex Alderton, Roberto Amato, Jeffrey Barrett, Sonia Goncalves, Ewan Harrison, David K. Jackson, Ian Johnston, Dominic Kwiatkowski, Cordelia Langford, John Sillitoe on behalf of the Wellcome Sanger Institute COVID-19 Surveillance Team |
| EPI_ISL_3285008 | Lighthouse Lab in Alderley Park | Wellcome Sanger Institute for the COVID-19 Genomics UK (COG-UK) Consortium | Jacquelyn Wynn, Mairead Hyland, The Lighthouse Lab in Alderley Park and Alex Alderton, Roberto Amato, Jeffrey Barrett, Sonia Goncalves, Ewan Harrison, David K. Jackson, Ian Johnston, Dominic Kwiatkowski, Cordelia Langford, John Sillitoe on behalf of the Wellcome Sanger Institute COVID-19 Surveillance Team |
| EPI_ISL_3285009 | Berkshire and Surrey Pathology Services Lighthouse Laboratory | Wellcome Sanger Institute for the COVID-19 Genomics UK (COG-UK) Consortium | Berkshire and Surrey Pathology Services Lighthouse Laboratory and Alex Alderton, Roberto Amato, Jeffrey Barrett, Sonia Goncalves, Ewan Harrison, David K. Jackson, Ian Johnston, Dominic Kwiatkowski, Cordelia Langford, John Sillitoe on behalf of the Wellcome Sanger Institute COVID-19 Surveillance Team |
| EPI_ISL_3285010, EPI_ISL_3285011, EPI_ISL_3285012 | Lighthouse Lab in Alderley Park | Wellcome Sanger Institute for the COVID-19 Genomics UK (COG-UK) Consortium | Jacquelyn Wynn, Mairead Hyland, The Lighthouse Lab in Alderley Park and Alex Alderton, Roberto Amato, Jeffrey Barrett, Sonia Goncalves, Ewan Harrison, David K. Jackson, Ian Johnston, Dominic Kwiatkowski, Cordelia Langford, John Sillitoe on behalf of the Wellcome Sanger Institute COVID-19 Surveillance Team |
| EPI_ISL_3285013 | Lighthouse Lab in Milton Keynes | Wellcome Sanger Institute for the COVID-19 Genomics UK (COG-UK) Consortium | The Lighthouse Lab in Milton Keynes and Alex Alderton, Roberto Amato, Jeffrey Barrett, Sonia Goncalves, Ewan Harrison, David K. Jackson, Ian Johnston, Dominic Kwiatkowski, Cordelia Langford, John Sillitoe on behalf of the Wellcome Sanger Institute COVID-19 Surveillance Team |
| EPI_ISL_3285014, EPI_ISL_3285015 | Lighthouse Lab in Alderley Park | Wellcome Sanger Institute for the COVID-19 Genomics UK (COG-UK) Consortium | Jacquelyn Wynn, Mairead Hyland, The Lighthouse Lab in Alderley Park and Alex Alderton, Roberto Amato, Jeffrey Barrett, Sonia Goncalves, Ewan Harrison, David K. Jackson, Ian Johnston, Dominic Kwiatkowski, Cordelia Langford, John Sillitoe on behalf of the Wellcome Sanger Institute COVID-19 Surveillance Team |
| EPI_ISL_3285016 | Lighthouse Lab in Milton Keynes | Wellcome Sanger Institute for the COVID-19 Genomics UK (COG-UK) Consortium | The Lighthouse Lab in Milton Keynes and Alex Alderton, Roberto Amato, Jeffrey Barrett, Sonia Goncalves, Ewan Harrison, David K. Jackson, Ian Johnston, Dominic Kwiatkowski, Cordelia Langford, John Sillitoe on behalf of the Wellcome Sanger Institute COVID-19 Surveillance Team |
| EPI_ISL_3285017 | Berkshire and Surrey Pathology Services Lighthouse Laboratory | Wellcome Sanger Institute for the COVID-19 Genomics UK (COG-UK) Consortium | Berkshire and Surrey Pathology Services Lighthouse Laboratory and Alex Alderton, Roberto Amato, Jeffrey Barrett, Sonia Goncalves, Ewan Harrison, David K. Jackson, Ian Johnston, Dominic Kwiatkowski, Cordelia Langford, John Sillitoe on behalf of the Wellcome Sanger Institute COVID-19 Surveillance Team |
| EPI_ISL_3285018 | Lighthouse Lab in Alderley Park | Wellcome Sanger Institute for the COVID-19 Genomics UK (COG-UK) Consortium | Jacquelyn Wynn, Mairead Hyland, The Lighthouse Lab in Alderley Park and Alex Alderton, Roberto Amato, Jeffrey Barrett, Sonia Goncalves, Ewan Harrison, David K. Jackson, Ian Johnston, Dominic Kwiatkowski, Cordelia Langford, John Sillitoe on behalf of the Wellcome Sanger Institute COVID-19 Surveillance Team |
| EPI_ISL_3285019 | Berkshire and Surrey Pathology Services Lighthouse Laboratory | Wellcome Sanger Institute for the COVID-19 Genomics UK (COG-UK) Consortium | Berkshire and Surrey Pathology Services Lighthouse Laboratory and Alex Alderton, Roberto Amato, Jeffrey Barrett, Sonia Goncalves, Ewan Harrison, David K. Jackson, Ian Johnston, Dominic Kwiatkowski, Cordelia Langford, John Sillitoe on behalf of the Wellcome Sanger Institute COVID-19 Surveillance Team |
| EPI_ISL_3285020 | Lighthouse Lab in Alderley Park | Wellcome Sanger Institute for the COVID-19 Genomics UK (COG-UK) Consortium | Jacquelyn Wynn, Mairead Hyland, The Lighthouse Lab in Alderley Park and Alex Alderton, Roberto Amato, Jeffrey Barrett, Sonia Goncalves, Ewan Harrison, David K. Jackson, Ian Johnston, Dominic Kwiatkowski, Cordelia Langford, John Sillitoe on behalf of the Wellcome Sanger Institute COVID-19 Surveillance Team |
| EPI_ISL_3285021, EPI_ISL_3285022 | Lighthouse Lab in Milton Keynes | Wellcome Sanger Institute for the COVID-19 Genomics UK (COG-UK) Consortium | The Lighthouse Lab in Milton Keynes and Alex Alderton, Roberto Amato, Jeffrey Barrett, Sonia Goncalves, Ewan Harrison, David K. Jackson, Ian Johnston, Dominic Kwiatkowski, Cordelia Langford, John Sillitoe on behalf of the Wellcome Sanger Institute COVID-19 Surveillance Team |
| EPI_ISL_3285023, EPI_ISL_3285024 | Lighthouse Lab in Alderley Park | Wellcome Sanger Institute for the COVID-19 Genomics UK (COG-UK) Consortium | Jacquelyn Wynn, Mairead Hyland, The Lighthouse Lab in Alderley Park and Alex Alderton, Roberto Amato, Jeffrey Barrett, Sonia Goncalves, Ewan Harrison, David K. Jackson, Ian Johnston, Dominic Kwiatkowski, Cordelia Langford, John Sillitoe on behalf of the Wellcome Sanger Institute COVID-19 Surveillance Team |
| EPI_ISL_3285025, EPI_ISL_3285026, EPI_ISL_3285027, EPI_ISL_3285028, EPI_ISL_3285029 | Lighthouse Lab in Milton Keynes | Wellcome Sanger Institute for the COVID-19 Genomics UK (COG-UK) Consortium | The Lighthouse Lab in Milton Keynes and Alex Alderton, Roberto Amato, Jeffrey Barrett, Sonia Goncalves, Ewan Harrison, David K. Jackson, Ian Johnston, Dominic Kwiatkowski, Cordelia Langford, John Sillitoe on behalf of the Wellcome Sanger Institute COVID-19 Surveillance Team |

| | | | |
|---|---|---|---|
| EPI_ISL_3285030, EPI_ISL_3285031, EPI_ISL_3285032, EPI_ISL_3285033 | Lighthouse Lab in Alderley Park | Wellcome Sanger Institute for the COVID-19 Genomics UK (COG-UK) Consortium | Jacquelyn Wynn, Mairead Hyland, The Lighthouse Lab in Alderley Park and Alex Alderton, Roberto Amato, Jeffrey Barrett, Sonia Goncalves, Ewan Harrison, David K. Jackson, Ian Johnston, Dominic Kwiatkowski, Cordelia Langford, John Sillitoe on behalf of the Wellcome Sanger Institute COVID-19 Surveillance Team |
| EPI_ISL_3285034 | Berkshire and Surrey Pathology Services Lighthouse Laboratory | Wellcome Sanger Institute for the COVID-19 Genomics UK (COG-UK) Consortium | Berkshire and Surrey Pathology Services Lighthouse Laboratory and Alex Alderton, Roberto Amato, Jeffrey Barrett, Sonia Goncalves, Ewan Harrison, David K. Jackson, Ian Johnston, Dominic Kwiatkowski, Cordelia Langford, John Sillitoe on behalf of the Wellcome Sanger Institute COVID-19 Surveillance Team |
| EPI_ISL_3285035 | Lighthouse Lab in Alderley Park | Wellcome Sanger Institute for the COVID-19 Genomics UK (COG-UK) Consortium | Jacquelyn Wynn, Mairead Hyland, The Lighthouse Lab in Alderley Park and Alex Alderton, Roberto Amato, Jeffrey Barrett, Sonia Goncalves, Ewan Harrison, David K. Jackson, Ian Johnston, Dominic Kwiatkowski, Cordelia Langford, John Sillitoe on behalf of the Wellcome Sanger Institute COVID-19 Surveillance Team |
| EPI_ISL_3285036, EPI_ISL_3285037 | Lighthouse Lab in Milton Keynes | Wellcome Sanger Institute for the COVID-19 Genomics UK (COG-UK) Consortium | The Lighthouse Lab in Milton Keynes and Alex Alderton, Roberto Amato, Jeffrey Barrett, Sonia Goncalves, Ewan Harrison, David K. Jackson, Ian Johnston, Dominic Kwiatkowski, Cordelia Langford, John Sillitoe on behalf of the Wellcome Sanger Institute COVID-19 Surveillance Team |
| EPI_ISL_3285038 | Berkshire and Surrey Pathology Services Lighthouse Laboratory | Wellcome Sanger Institute for the COVID-19 Genomics UK (COG-UK) Consortium | Berkshire and Surrey Pathology Services Lighthouse Laboratory and Alex Alderton, Roberto Amato, Jeffrey Barrett, Sonia Goncalves, Ewan Harrison, David K. Jackson, Ian Johnston, Dominic Kwiatkowski, Cordelia Langford, John Sillitoe on behalf of the Wellcome Sanger Institute COVID-19 Surveillance Team |
| EPI_ISL_3285039 | Lighthouse Lab in Alderley Park | Wellcome Sanger Institute for the COVID-19 Genomics UK (COG-UK) Consortium | Jacquelyn Wynn, Mairead Hyland, The Lighthouse Lab in Alderley Park and Alex Alderton, Roberto Amato, Jeffrey Barrett, Sonia Goncalves, Ewan Harrison, David K. Jackson, Ian Johnston, Dominic Kwiatkowski, Cordelia Langford, John Sillitoe on behalf of the Wellcome Sanger Institute COVID-19 Surveillance Team |
| EPI_ISL_3285040 | Lighthouse Lab in Milton Keynes | Wellcome Sanger Institute for the COVID-19 Genomics UK (COG-UK) Consortium | The Lighthouse Lab in Milton Keynes and Alex Alderton, Roberto Amato, Jeffrey Barrett, Sonia Goncalves, Ewan Harrison, David K. Jackson, Ian Johnston, Dominic Kwiatkowski, Cordelia Langford, John Sillitoe on behalf of the Wellcome Sanger Institute COVID-19 Surveillance Team |
| EPI_ISL_3285041, EPI_ISL_3285042 | Berkshire and Surrey Pathology Services Lighthouse Laboratory | Wellcome Sanger Institute for the COVID-19 Genomics UK (COG-UK) Consortium | Berkshire and Surrey Pathology Services Lighthouse Laboratory and Alex Alderton, Roberto Amato, Jeffrey Barrett, Sonia Goncalves, Ewan Harrison, David K. Jackson, Ian Johnston, Dominic Kwiatkowski, Cordelia Langford, John Sillitoe on behalf of the Wellcome Sanger Institute COVID-19 Surveillance Team |
| EPI_ISL_3285043, EPI_ISL_3285044 | Lighthouse Lab in Alderley Park | Wellcome Sanger Institute for the COVID-19 Genomics UK (COG-UK) Consortium | Jacquelyn Wynn, Mairead Hyland, The Lighthouse Lab in Alderley Park and Alex Alderton, Roberto Amato, Jeffrey Barrett, Sonia Goncalves, Ewan Harrison, David K. Jackson, Ian Johnston, Dominic Kwiatkowski, Cordelia Langford, John Sillitoe on behalf of the Wellcome Sanger Institute COVID-19 Surveillance Team |
| EPI_ISL_3285045 | Lighthouse Lab in Milton Keynes | Wellcome Sanger Institute for the COVID-19 Genomics UK (COG-UK) Consortium | The Lighthouse Lab in Milton Keynes and Alex Alderton, Roberto Amato, Jeffrey Barrett, Sonia Goncalves, Ewan Harrison, David K. Jackson, Ian Johnston, Dominic Kwiatkowski, Cordelia Langford, John Sillitoe on behalf of the Wellcome Sanger Institute COVID-19 Surveillance Team |
| EPI_ISL_3285046, EPI_ISL_3285047, EPI_ISL_3285048, EPI_ISL_3285049, EPI_ISL_3285050 | Lighthouse Lab in Alderley Park | Wellcome Sanger Institute for the COVID-19 Genomics UK (COG-UK) Consortium | Jacquelyn Wynn, Mairead Hyland, The Lighthouse Lab in Alderley Park and Alex Alderton, Roberto Amato, Jeffrey Barrett, Sonia Goncalves, Ewan Harrison, David K. Jackson, Ian Johnston, Dominic Kwiatkowski, Cordelia Langford, John Sillitoe on behalf of the Wellcome Sanger Institute COVID-19 Surveillance Team |
| EPI_ISL_3285051 | Berkshire and Surrey Pathology Services Lighthouse Laboratory | Wellcome Sanger Institute for the COVID-19 Genomics UK (COG-UK) Consortium | Berkshire and Surrey Pathology Services Lighthouse Laboratory and Alex Alderton, Roberto Amato, Jeffrey Barrett, Sonia Goncalves, Ewan Harrison, David K. Jackson, Ian Johnston, Dominic Kwiatkowski, Cordelia Langford, John Sillitoe on behalf of the Wellcome Sanger Institute COVID-19 Surveillance Team |
| EPI_ISL_3285052 | Lighthouse Lab in Alderley Park | Wellcome Sanger Institute for the COVID-19 Genomics UK (COG-UK) Consortium | Jacquelyn Wynn, Mairead Hyland, The Lighthouse Lab in Alderley Park and Alex Alderton, Roberto Amato, Jeffrey Barrett, Sonia Goncalves, Ewan Harrison, David K. Jackson, Ian Johnston, Dominic Kwiatkowski, Cordelia Langford, John Sillitoe on behalf of the Wellcome Sanger Institute COVID-19 Surveillance Team |
| EPI_ISL_3285053 | Berkshire and Surrey Pathology Services Lighthouse Laboratory | Wellcome Sanger Institute for the COVID-19 Genomics UK (COG-UK) Consortium | Berkshire and Surrey Pathology Services Lighthouse Laboratory and Alex Alderton, Roberto Amato, Jeffrey Barrett, Sonia Goncalves, Ewan Harrison, David K. Jackson, Ian Johnston, Dominic Kwiatkowski, Cordelia Langford, John Sillitoe on behalf of the Wellcome Sanger Institute COVID-19 Surveillance Team |
| EPI_ISL_3285054 | Lighthouse Lab in Milton Keynes | Wellcome Sanger Institute for the COVID-19 Genomics UK (COG-UK) Consortium | The Lighthouse Lab in Milton Keynes and Alex Alderton, Roberto Amato, Jeffrey Barrett, Sonia Goncalves, Ewan Harrison, David K. Jackson, Ian Johnston, Dominic Kwiatkowski, Cordelia Langford, John Sillitoe on behalf of the Wellcome Sanger Institute COVID-19 Surveillance Team |
| EPI_ISL_3285055, EPI_ISL_3285056, EPI_ISL_3285057, EPI_ISL_3285058, EPI_ISL_3285059 | Lighthouse Lab in Alderley Park | Wellcome Sanger Institute for the COVID-19 Genomics UK (COG-UK) Consortium | Jacquelyn Wynn, Mairead Hyland, The Lighthouse Lab in Alderley Park and Alex Alderton, Roberto Amato, Jeffrey Barrett, Sonia Goncalves, Ewan Harrison, David K. Jackson, Ian Johnston, Dominic Kwiatkowski, Cordelia Langford, John Sillitoe on behalf of the Wellcome Sanger Institute COVID-19 Surveillance Team |
| EPI_ISL_3285060 | Lighthouse Lab in Milton Keynes | Wellcome Sanger Institute for the COVID-19 Genomics UK (COG-UK) Consortium | The Lighthouse Lab in Milton Keynes and Alex Alderton, Roberto Amato, Jeffrey Barrett, Sonia Goncalves, Ewan Harrison, David K. Jackson, Ian Johnston, Dominic Kwiatkowski, Cordelia Langford, John Sillitoe on behalf of the Wellcome Sanger Institute COVID-19 Surveillance Team |
| EPI_ISL_3285061 | Lighthouse Lab in Alderley Park | Wellcome Sanger Institute for the COVID-19 Genomics UK (COG-UK) Consortium | Jacquelyn Wynn, Mairead Hyland, The Lighthouse Lab in Alderley Park and Alex Alderton, Roberto Amato, Jeffrey Barrett, Sonia Goncalves, Ewan Harrison, David K. Jackson, Ian Johnston, Dominic Kwiatkowski, Cordelia Langford, John Sillitoe on behalf of the Wellcome Sanger Institute COVID-19 Surveillance Team |
| EPI_ISL_3285062 | Lighthouse Lab in Milton Keynes | Wellcome Sanger Institute for the COVID-19 Genomics UK (COG-UK) Consortium | The Lighthouse Lab in Milton Keynes and Alex Alderton, Roberto Amato, Jeffrey Barrett, Sonia Goncalves, Ewan Harrison, David K. Jackson, Ian Johnston, Dominic Kwiatkowski, Cordelia Langford, John Sillitoe on behalf of the Wellcome Sanger Institute COVID-19 Surveillance Team |
| EPI_ISL_3285063, EPI_ISL_3285064, EPI_ISL_3285065, EPI_ISL_3285066, EPI_ISL_3285067, EPI_ISL_3285068, EPI_ISL_3285069 | Lighthouse Lab in Alderley Park | Wellcome Sanger Institute for the COVID-19 Genomics UK (COG-UK) Consortium | Jacquelyn Wynn, Mairead Hyland, The Lighthouse Lab in Alderley Park and Alex Alderton, Roberto Amato, Jeffrey Barrett, Sonia Goncalves, Ewan Harrison, David K. Jackson, Ian Johnston, Dominic Kwiatkowski, Cordelia Langford, John Sillitoe on behalf of the Wellcome Sanger Institute COVID-19 Surveillance Team |
| EPI_ISL_3285070 | Berkshire and Surrey Pathology Services Lighthouse Laboratory | Wellcome Sanger Institute for the COVID-19 Genomics UK (COG-UK) Consortium | Berkshire and Surrey Pathology Services Lighthouse Laboratory and Alex Alderton, Roberto Amato, Jeffrey Barrett, Sonia Goncalves, Ewan Harrison, David K. Jackson, Ian Johnston, Dominic Kwiatkowski, Cordelia Langford, John Sillitoe on behalf of the Wellcome Sanger Institute COVID-19 Surveillance Team |
| EPI_ISL_3285071 | Lighthouse Lab in Alderley Park | Wellcome Sanger Institute for the COVID-19 Genomics UK (COG-UK) Consortium | Jacquelyn Wynn, Mairead Hyland, The Lighthouse Lab in Alderley Park and Alex Alderton, Roberto Amato, Jeffrey Barrett, Sonia Goncalves, Ewan Harrison, David K. Jackson, Ian Johnston, Dominic Kwiatkowski, Cordelia Langford, John Sillitoe on behalf of the Wellcome Sanger Institute COVID-19 Surveillance Team |
| EPI_ISL_3285072 | Lighthouse Lab in Milton Keynes | Wellcome Sanger Institute for the COVID-19 Genomics UK (COG-UK) Consortium | The Lighthouse Lab in Milton Keynes and Alex Alderton, Roberto Amato, Jeffrey Barrett, Sonia Goncalves, Ewan Harrison, David K. Jackson, Ian Johnston, Dominic Kwiatkowski, Cordelia Langford, John Sillitoe on behalf of the Wellcome Sanger Institute COVID-19 Surveillance Team |
| EPI_ISL_3285073 | Lighthouse Lab in Alderley Park | Wellcome Sanger Institute for the COVID-19 Genomics UK (COG-UK) Consortium | Jacquelyn Wynn, Mairead Hyland, The Lighthouse Lab in Alderley Park and Alex Alderton, Roberto Amato, Jeffrey Barrett, Sonia Goncalves, Ewan Harrison, David K. Jackson, Ian Johnston, Dominic Kwiatkowski, Cordelia Langford, John Sillitoe on behalf of the Wellcome Sanger Institute COVID-19 Surveillance Team |
| EPI_ISL_3285074 | Lighthouse Lab in Milton Keynes | Wellcome Sanger Institute for the COVID-19 Genomics UK (COG-UK) Consortium | The Lighthouse Lab in Milton Keynes and Alex Alderton, Roberto Amato, Jeffrey Barrett, Sonia Goncalves, Ewan Harrison, David K. Jackson, Ian Johnston, Dominic Kwiatkowski, Cordelia Langford, John Sillitoe on behalf of the Wellcome Sanger Institute COVID-19 Surveillance Team |
| EPI_ISL_3285075, EPI_ISL_3285076 | Lighthouse Lab in Alderley Park | Wellcome Sanger Institute for the COVID-19 Genomics UK (COG-UK) Consortium | Jacquelyn Wynn, Mairead Hyland, The Lighthouse Lab in Alderley Park and Alex Alderton, Roberto Amato, Jeffrey Barrett, Sonia Goncalves, Ewan Harrison, David K. Jackson, Ian Johnston, Dominic Kwiatkowski, Cordelia Langford, John Sillitoe on behalf of the Wellcome Sanger Institute COVID-19 Surveillance Team |
| EPI_ISL_3285077 | Berkshire and Surrey Pathology Services Lighthouse Laboratory | Wellcome Sanger Institute for the COVID-19 Genomics UK (COG-UK) Consortium | Berkshire and Surrey Pathology Services Lighthouse Laboratory and Alex Alderton, Roberto Amato, Jeffrey Barrett, Sonia Goncalves, Ewan Harrison, David K. Jackson, Ian Johnston, Dominic Kwiatkowski, Cordelia Langford, John Sillitoe on behalf of the Wellcome Sanger Institute COVID-19 Surveillance Team |
| EPI_ISL_3285078, EPI_ISL_3285079, EPI_ISL_3285080 | Lighthouse Lab in Alderley Park | Wellcome Sanger Institute for the COVID-19 Genomics UK (COG-UK) Consortium | Jacquelyn Wynn, Mairead Hyland, The Lighthouse Lab in Alderley Park and Alex Alderton, Roberto Amato, Jeffrey Barrett, Sonia Goncalves, Ewan Harrison, David K. Jackson, Ian Johnston, Dominic Kwiatkowski, Cordelia Langford, John Sillitoe on behalf of the Wellcome Sanger Institute COVID-19 Surveillance Team |
| EPI_ISL_3285081, EPI_ISL_3285082, EPI_ISL_3285083 | Berkshire and Surrey Pathology Services Lighthouse Laboratory | Wellcome Sanger Institute for the COVID-19 Genomics UK (COG-UK) Consortium | Berkshire and Surrey Pathology Services Lighthouse Laboratory and Alex Alderton, Roberto Amato, Jeffrey Barrett, Sonia Goncalves, Ewan Harrison, David K. Jackson, Ian Johnston, Dominic Kwiatkowski, Cordelia Langford, John Sillitoe on behalf of the Wellcome Sanger Institute COVID-19 Surveillance Team |
| EPI_ISL_3285084, EPI_ISL_3285085, EPI_ISL_3285086 | Lighthouse Lab in Alderley Park | Wellcome Sanger Institute for the COVID-19 Genomics UK (COG-UK) Consortium | Jacquelyn Wynn, Mairead Hyland, The Lighthouse Lab in Alderley Park and Alex Alderton, Roberto Amato, Jeffrey Barrett, Sonia Goncalves, Ewan Harrison, David K. Jackson, Ian Johnston, Dominic Kwiatkowski, Cordelia Langford, John Sillitoe on behalf of the Wellcome Sanger Institute COVID-19 Surveillance Team |
| EPI_ISL_3285087 | Lighthouse Lab in Milton Keynes | Wellcome Sanger Institute for the COVID-19 Genomics UK (COG-UK) Consortium | The Lighthouse Lab in Milton Keynes and Alex Alderton, Roberto Amato, Jeffrey Barrett, Sonia Goncalves, Ewan Harrison, David K. Jackson, Ian Johnston, Dominic Kwiatkowski, Cordelia Langford, John Sillitoe on behalf of the Wellcome Sanger Institute COVID-19 Surveillance Team |
| EPI_ISL_3285088, EPI_ISL_3285089 | Lighthouse Lab in Alderley Park | Wellcome Sanger Institute for the COVID-19 Genomics UK (COG-UK) Consortium | Jacquelyn Wynn, Mairead Hyland, The Lighthouse Lab in Alderley Park and Alex Alderton, Roberto Amato, Jeffrey Barrett, Sonia Goncalves, Ewan Harrison, David K. Jackson, Ian Johnston, Dominic Kwiatkowski, Cordelia Langford, John Sillitoe on behalf of the Wellcome Sanger Institute COVID-19 Surveillance Team |
| EPI_ISL_3285090 | Berkshire and Surrey Pathology Services Lighthouse Laboratory | Wellcome Sanger Institute for the COVID-19 Genomics UK (COG-UK) Consortium | Berkshire and Surrey Pathology Services Lighthouse Laboratory and Alex Alderton, Roberto Amato, Jeffrey Barrett, Sonia Goncalves, Ewan Harrison, David K. Jackson, Ian Johnston, Dominic Kwiatkowski, Cordelia Langford, John Sillitoe on behalf of the Wellcome Sanger Institute COVID-19 Surveillance Team |
| EPI_ISL_3285091 | Lighthouse Lab in Alderley Park | Wellcome Sanger Institute for the COVID-19 Genomics UK (COG-UK) Consortium | Jacquelyn Wynn, Mairead Hyland, The Lighthouse Lab in Alderley Park and Alex Alderton, Roberto Amato, Jeffrey Barrett, Sonia Goncalves, Ewan Harrison, David K. Jackson, Ian Johnston, Dominic Kwiatkowski, Cordelia Langford, John Sillitoe on behalf of the Wellcome Sanger Institute COVID-19 Surveillance Team |
| EPI_ISL_3285092 | Lighthouse Lab in Milton Keynes | Wellcome Sanger Institute for the COVID-19 Genomics UK (COG-UK) Consortium | The Lighthouse Lab in Milton Keynes and Alex Alderton, Roberto Amato, Jeffrey Barrett, Sonia Goncalves, Ewan Harrison, David K. Jackson, Ian Johnston, Dominic Kwiatkowski, Cordelia Langford, John Sillitoe on behalf of the Wellcome Sanger Institute COVID-19 Surveillance Team |
| EPI_ISL_3285093, EPI_ISL_3285094, EPI_ISL_3285095 | Lighthouse Lab in Alderley Park | Wellcome Sanger Institute for the COVID-19 Genomics UK (COG-UK) Consortium | Jacquelyn Wynn, Mairead Hyland, The Lighthouse Lab in Alderley Park and Alex Alderton, Roberto Amato, Jeffrey Barrett, Sonia Goncalves, Ewan Harrison, David K. Jackson, Ian Johnston, Dominic Kwiatkowski, Cordelia Langford, John Sillitoe on behalf of the Wellcome Sanger Institute COVID-19 Surveillance Team |
| EPI_ISL_3285096, EPI_ISL_3285097 | Berkshire and Surrey Pathology Services Lighthouse Laboratory | Wellcome Sanger Institute for the COVID-19 Genomics UK (COG-UK) Consortium | Berkshire and Surrey Pathology Services Lighthouse Laboratory and Alex Alderton, Roberto Amato, Jeffrey Barrett, Sonia Goncalves, Ewan Harrison, David K. Jackson, Ian Johnston, Dominic Kwiatkowski, Cordelia Langford, John Sillitoe on behalf of the Wellcome Sanger Institute COVID-19 Surveillance Team |
| EPI_ISL_3285098, EPI_ISL_3285099 | Lighthouse Lab in Alderley Park | Wellcome Sanger Institute for the COVID-19 Genomics UK (COG-UK) Consortium | Jacquelyn Wynn, Mairead Hyland, The Lighthouse Lab in Alderley Park and Alex Alderton, Roberto Amato, Jeffrey Barrett, Sonia Goncalves, Ewan Harrison, David K. Jackson, Ian Johnston, Dominic Kwiatkowski, Cordelia Langford, John Sillitoe on behalf of the Wellcome Sanger Institute COVID-19 Surveillance Team |

| | | | |
|---|---|---|---|
| EPI_ISL_3285100 | Lighthouse Lab in Milton Keynes | Wellcome Sanger Institute for the COVID-19 Genomics UK (COG-UK) Consortium | The Lighthouse Lab in Milton Keynes and Alex Alderton, Roberto Amato, Jeffrey Barrett, Sonia Goncalves, Ewan Harrison, David K. Jackson, Ian Johnston, Dominic Kwiatkowski, Cordelia Langford, John Sillitoe on behalf of the Wellcome Sanger Institute COVID-19 Surveillance Team |
| EPI_ISL_3285101 | Berkshire and Surrey Pathology Services Lighthouse Laboratory | Wellcome Sanger Institute for the COVID-19 Genomics UK (COG-UK) Consortium | Berkshire and Surrey Pathology Services Lighthouse Laboratory and Alex Alderton, Roberto Amato, Jeffrey Barrett, Sonia Goncalves, Ewan Harrison, David K. Jackson, Ian Johnston, Dominic Kwiatkowski, Cordelia Langford, John Sillitoe on behalf of the Wellcome Sanger Institute COVID-19 Surveillance Team |
| EPI_ISL_3285102, EPI_ISL_3285103, EPI_ISL_3285104 | Lighthouse Lab in Alderley Park | Wellcome Sanger Institute for the COVID-19 Genomics UK (COG-UK) Consortium | Jacquelyn Wynn, Mairead Hyland, The Lighthouse Lab in Alderley Park and Alex Alderton, Roberto Amato, Jeffrey Barrett, Sonia Goncalves, Ewan Harrison, David K. Jackson, Ian Johnston, Dominic Kwiatkowski, Cordelia Langford, John Sillitoe on behalf of the Wellcome Sanger Institute COVID-19 Surveillance Team |
| EPI_ISL_3285105 | Lighthouse Lab in Milton Keynes | Wellcome Sanger Institute for the COVID-19 Genomics UK (COG-UK) Consortium | The Lighthouse Lab in Milton Keynes and Alex Alderton, Roberto Amato, Jeffrey Barrett, Sonia Goncalves, Ewan Harrison, David K. Jackson, Ian Johnston, Dominic Kwiatkowski, Cordelia Langford, John Sillitoe on behalf of the Wellcome Sanger Institute COVID-19 Surveillance Team |
| EPI_ISL_3285106, EPI_ISL_3285107, EPI_ISL_3285108, EPI_ISL_3285109, EPI_ISL_3285110 | Lighthouse Lab in Alderley Park | Wellcome Sanger Institute for the COVID-19 Genomics UK (COG-UK) Consortium | Jacquelyn Wynn, Mairead Hyland, The Lighthouse Lab in Alderley Park and Alex Alderton, Roberto Amato, Jeffrey Barrett, Sonia Goncalves, Ewan Harrison, David K. Jackson, Ian Johnston, Dominic Kwiatkowski, Cordelia Langford, John Sillitoe on behalf of the Wellcome Sanger Institute COVID-19 Surveillance Team |
| EPI_ISL_3285111 | Lighthouse Lab in Milton Keynes | Wellcome Sanger Institute for the COVID-19 Genomics UK (COG-UK) Consortium | The Lighthouse Lab in Milton Keynes and Alex Alderton, Roberto Amato, Jeffrey Barrett, Sonia Goncalves, Ewan Harrison, David K. Jackson, Ian Johnston, Dominic Kwiatkowski, Cordelia Langford, John Sillitoe on behalf of the Wellcome Sanger Institute COVID-19 Surveillance Team |
| EPI_ISL_3285112 | Berkshire and Surrey Pathology Services Lighthouse Laboratory | Wellcome Sanger Institute for the COVID-19 Genomics UK (COG-UK) Consortium | Berkshire and Surrey Pathology Services Lighthouse Laboratory and Alex Alderton, Roberto Amato, Jeffrey Barrett, Sonia Goncalves, Ewan Harrison, David K. Jackson, Ian Johnston, Dominic Kwiatkowski, Cordelia Langford, John Sillitoe on behalf of the Wellcome Sanger Institute COVID-19 Surveillance Team |
| EPI_ISL_3285113 | Lighthouse Lab in Milton Keynes | Wellcome Sanger Institute for the COVID-19 Genomics UK (COG-UK) Consortium | The Lighthouse Lab in Milton Keynes and Alex Alderton, Roberto Amato, Jeffrey Barrett, Sonia Goncalves, Ewan Harrison, David K. Jackson, Ian Johnston, Dominic Kwiatkowski, Cordelia Langford, John Sillitoe on behalf of the Wellcome Sanger Institute COVID-19 Surveillance Team |
| EPI_ISL_3285114, EPI_ISL_3285115 | Lighthouse Lab in Alderley Park | Wellcome Sanger Institute for the COVID-19 Genomics UK (COG-UK) Consortium | Jacquelyn Wynn, Mairead Hyland, The Lighthouse Lab in Alderley Park and Alex Alderton, Roberto Amato, Jeffrey Barrett, Sonia Goncalves, Ewan Harrison, David K. Jackson, Ian Johnston, Dominic Kwiatkowski, Cordelia Langford, John Sillitoe on behalf of the Wellcome Sanger Institute COVID-19 Surveillance Team |
| EPI_ISL_3285116 | Berkshire and Surrey Pathology Services Lighthouse Laboratory | Wellcome Sanger Institute for the COVID-19 Genomics UK (COG-UK) Consortium | Berkshire and Surrey Pathology Services Lighthouse Laboratory and Alex Alderton, Roberto Amato, Jeffrey Barrett, Sonia Goncalves, Ewan Harrison, David K. Jackson, Ian Johnston, Dominic Kwiatkowski, Cordelia Langford, John Sillitoe on behalf of the Wellcome Sanger Institute COVID-19 Surveillance Team |
| EPI_ISL_3285117 | Lighthouse Lab in Milton Keynes | Wellcome Sanger Institute for the COVID-19 Genomics UK (COG-UK) Consortium | The Lighthouse Lab in Milton Keynes and Alex Alderton, Roberto Amato, Jeffrey Barrett, Sonia Goncalves, Ewan Harrison, David K. Jackson, Ian Johnston, Dominic Kwiatkowski, Cordelia Langford, John Sillitoe on behalf of the Wellcome Sanger Institute COVID-19 Surveillance Team |
| EPI_ISL_3285118 | Berkshire and Surrey Pathology Services Lighthouse Laboratory | Wellcome Sanger Institute for the COVID-19 Genomics UK (COG-UK) Consortium | Berkshire and Surrey Pathology Services Lighthouse Laboratory and Alex Alderton, Roberto Amato, Jeffrey Barrett, Sonia Goncalves, Ewan Harrison, David K. Jackson, Ian Johnston, Dominic Kwiatkowski, Cordelia Langford, John Sillitoe on behalf of the Wellcome Sanger Institute COVID-19 Surveillance Team |
| EPI_ISL_3285119 | Lighthouse Lab in Alderley Park | Wellcome Sanger Institute for the COVID-19 Genomics UK (COG-UK) Consortium | Jacquelyn Wynn, Mairead Hyland, The Lighthouse Lab in Alderley Park and Alex Alderton, Roberto Amato, Jeffrey Barrett, Sonia Goncalves, Ewan Harrison, David K. Jackson, Ian Johnston, Dominic Kwiatkowski, Cordelia Langford, John Sillitoe on behalf of the Wellcome Sanger Institute COVID-19 Surveillance Team |
| EPI_ISL_3285120 | Lighthouse Lab in Milton Keynes | Wellcome Sanger Institute for the COVID-19 Genomics UK (COG-UK) Consortium | The Lighthouse Lab in Milton Keynes and Alex Alderton, Roberto Amato, Jeffrey Barrett, Sonia Goncalves, Ewan Harrison, David K. Jackson, Ian Johnston, Dominic Kwiatkowski, Cordelia Langford, John Sillitoe on behalf of the Wellcome Sanger Institute COVID-19 Surveillance Team |
| EPI_ISL_3285121 | Lighthouse Lab in Alderley Park | Wellcome Sanger Institute for the COVID-19 Genomics UK (COG-UK) Consortium | Jacquelyn Wynn, Mairead Hyland, The Lighthouse Lab in Alderley Park and Alex Alderton, Roberto Amato, Jeffrey Barrett, Sonia Goncalves, Ewan Harrison, David K. Jackson, Ian Johnston, Dominic Kwiatkowski, Cordelia Langford, John Sillitoe on behalf of the Wellcome Sanger Institute COVID-19 Surveillance Team |
| EPI_ISL_3285122 | Lighthouse Lab in Milton Keynes | Wellcome Sanger Institute for the COVID-19 Genomics UK (COG-UK) Consortium | The Lighthouse Lab in Milton Keynes and Alex Alderton, Roberto Amato, Jeffrey Barrett, Sonia Goncalves, Ewan Harrison, David K. Jackson, Ian Johnston, Dominic Kwiatkowski, Cordelia Langford, John Sillitoe on behalf of the Wellcome Sanger Institute COVID-19 Surveillance Team |
| EPI_ISL_3285123 | Lighthouse Lab in Alderley Park | Wellcome Sanger Institute for the COVID-19 Genomics UK (COG-UK) Consortium | Jacquelyn Wynn, Mairead Hyland, The Lighthouse Lab in Alderley Park and Alex Alderton, Roberto Amato, Jeffrey Barrett, Sonia Goncalves, Ewan Harrison, David K. Jackson, Ian Johnston, Dominic Kwiatkowski, Cordelia Langford, John Sillitoe on behalf of the Wellcome Sanger Institute COVID-19 Surveillance Team |
| EPI_ISL_3285124 | Berkshire and Surrey Pathology Services Lighthouse Laboratory | Wellcome Sanger Institute for the COVID-19 Genomics UK (COG-UK) Consortium | Berkshire and Surrey Pathology Services Lighthouse Laboratory and Alex Alderton, Roberto Amato, Jeffrey Barrett, Sonia Goncalves, Ewan Harrison, David K. Jackson, Ian Johnston, Dominic Kwiatkowski, Cordelia Langford, John Sillitoe on behalf of the Wellcome Sanger Institute COVID-19 Surveillance Team |
| EPI_ISL_3285125 | Lighthouse Lab in Milton Keynes | Wellcome Sanger Institute for the COVID-19 Genomics UK (COG-UK) Consortium | The Lighthouse Lab in Milton Keynes and Alex Alderton, Roberto Amato, Jeffrey Barrett, Sonia Goncalves, Ewan Harrison, David K. Jackson, Ian Johnston, Dominic Kwiatkowski, Cordelia Langford, John Sillitoe on behalf of the Wellcome Sanger Institute COVID-19 Surveillance Team |
| EPI_ISL_3285126 | Lighthouse Lab in Alderley Park | Wellcome Sanger Institute for the COVID-19 Genomics UK (COG-UK) Consortium | Jacquelyn Wynn, Mairead Hyland, The Lighthouse Lab in Alderley Park and Alex Alderton, Roberto Amato, Jeffrey Barrett, Sonia Goncalves, Ewan Harrison, David K. Jackson, Ian Johnston, Dominic Kwiatkowski, Cordelia Langford, John Sillitoe on behalf of the Wellcome Sanger Institute COVID-19 Surveillance Team |
| EPI_ISL_3285127 | Berkshire and Surrey Pathology Services Lighthouse Laboratory | Wellcome Sanger Institute for the COVID-19 Genomics UK (COG-UK) Consortium | Berkshire and Surrey Pathology Services Lighthouse Laboratory and Alex Alderton, Roberto Amato, Jeffrey Barrett, Sonia Goncalves, Ewan Harrison, David K. Jackson, Ian Johnston, Dominic Kwiatkowski, Cordelia Langford, John Sillitoe on behalf of the Wellcome Sanger Institute COVID-19 Surveillance Team |
| EPI_ISL_3285128, EPI_ISL_3285129 | Lighthouse Lab in Alderley Park | Wellcome Sanger Institute for the COVID-19 Genomics UK (COG-UK) Consortium | Jacquelyn Wynn, Mairead Hyland, The Lighthouse Lab in Alderley Park and Alex Alderton, Roberto Amato, Jeffrey Barrett, Sonia Goncalves, Ewan Harrison, David K. Jackson, Ian Johnston, Dominic Kwiatkowski, Cordelia Langford, John Sillitoe on behalf of the Wellcome Sanger Institute COVID-19 Surveillance Team |
| EPI_ISL_3285130, EPI_ISL_3285131 | Berkshire and Surrey Pathology Services Lighthouse Laboratory | Wellcome Sanger Institute for the COVID-19 Genomics UK (COG-UK) Consortium | Berkshire and Surrey Pathology Services Lighthouse Laboratory and Alex Alderton, Roberto Amato, Jeffrey Barrett, Sonia Goncalves, Ewan Harrison, David K. Jackson, Ian Johnston, Dominic Kwiatkowski, Cordelia Langford, John Sillitoe on behalf of the Wellcome Sanger Institute COVID-19 Surveillance Team |
| EPI_ISL_3285132 | Lighthouse Lab in Alderley Park | Wellcome Sanger Institute for the COVID-19 Genomics UK (COG-UK) Consortium | Jacquelyn Wynn, Mairead Hyland, The Lighthouse Lab in Alderley Park and Alex Alderton, Roberto Amato, Jeffrey Barrett, Sonia Goncalves, Ewan Harrison, David K. Jackson, Ian Johnston, Dominic Kwiatkowski, Cordelia Langford, John Sillitoe on behalf of the Wellcome Sanger Institute COVID-19 Surveillance Team |
| EPI_ISL_3285133 | Lighthouse Lab in Milton Keynes | Wellcome Sanger Institute for the COVID-19 Genomics UK (COG-UK) Consortium | The Lighthouse Lab in Milton Keynes and Alex Alderton, Roberto Amato, Jeffrey Barrett, Sonia Goncalves, Ewan Harrison, David K. Jackson, Ian Johnston, Dominic Kwiatkowski, Cordelia Langford, John Sillitoe on behalf of the Wellcome Sanger Institute COVID-19 Surveillance Team |
| EPI_ISL_3285134, EPI_ISL_3285135, EPI_ISL_3285136 | Lighthouse Lab in Alderley Park | Wellcome Sanger Institute for the COVID-19 Genomics UK (COG-UK) Consortium | Jacquelyn Wynn, Mairead Hyland, The Lighthouse Lab in Alderley Park and Alex Alderton, Roberto Amato, Jeffrey Barrett, Sonia Goncalves, Ewan Harrison, David K. Jackson, Ian Johnston, Dominic Kwiatkowski, Cordelia Langford, John Sillitoe on behalf of the Wellcome Sanger Institute COVID-19 Surveillance Team |
| EPI_ISL_3285137 | Berkshire and Surrey Pathology Services Lighthouse Laboratory | Wellcome Sanger Institute for the COVID-19 Genomics UK (COG-UK) Consortium | Berkshire and Surrey Pathology Services Lighthouse Laboratory and Alex Alderton, Roberto Amato, Jeffrey Barrett, Sonia Goncalves, Ewan Harrison, David K. Jackson, Ian Johnston, Dominic Kwiatkowski, Cordelia Langford, John Sillitoe on behalf of the Wellcome Sanger Institute COVID-19 Surveillance Team |
| EPI_ISL_3285138 | Lighthouse Lab in Alderley Park | Wellcome Sanger Institute for the COVID-19 Genomics UK (COG-UK) Consortium | Jacquelyn Wynn, Mairead Hyland, The Lighthouse Lab in Alderley Park and Alex Alderton, Roberto Amato, Jeffrey Barrett, Sonia Goncalves, Ewan Harrison, David K. Jackson, Ian Johnston, Dominic Kwiatkowski, Cordelia Langford, John Sillitoe on behalf of the Wellcome Sanger Institute COVID-19 Surveillance Team |
| EPI_ISL_3285139, EPI_ISL_3285140, EPI_ISL_3285141 | Berkshire and Surrey Pathology Services Lighthouse Laboratory | Wellcome Sanger Institute for the COVID-19 Genomics UK (COG-UK) Consortium | Berkshire and Surrey Pathology Services Lighthouse Laboratory and Alex Alderton, Roberto Amato, Jeffrey Barrett, Sonia Goncalves, Ewan Harrison, David K. Jackson, Ian Johnston, Dominic Kwiatkowski, Cordelia Langford, John Sillitoe on behalf of the Wellcome Sanger Institute COVID-19 Surveillance Team |
| EPI_ISL_3285142, EPI_ISL_3285143 | Lighthouse Lab in Alderley Park | Wellcome Sanger Institute for the COVID-19 Genomics UK (COG-UK) Consortium | Jacquelyn Wynn, Mairead Hyland, The Lighthouse Lab in Alderley Park and Alex Alderton, Roberto Amato, Jeffrey Barrett, Sonia Goncalves, Ewan Harrison, David K. Jackson, Ian Johnston, Dominic Kwiatkowski, Cordelia Langford, John Sillitoe on behalf of the Wellcome Sanger Institute COVID-19 Surveillance Team |
| EPI_ISL_3285144 | Berkshire and Surrey Pathology Services Lighthouse Laboratory | Wellcome Sanger Institute for the COVID-19 Genomics UK (COG-UK) Consortium | Berkshire and Surrey Pathology Services Lighthouse Laboratory and Alex Alderton, Roberto Amato, Jeffrey Barrett, Sonia Goncalves, Ewan Harrison, David K. Jackson, Ian Johnston, Dominic Kwiatkowski, Cordelia Langford, John Sillitoe on behalf of the Wellcome Sanger Institute COVID-19 Surveillance Team |
| EPI_ISL_3285145 | Lighthouse Lab in Alderley Park | Wellcome Sanger Institute for the COVID-19 Genomics UK (COG-UK) Consortium | Jacquelyn Wynn, Mairead Hyland, The Lighthouse Lab in Alderley Park and Alex Alderton, Roberto Amato, Jeffrey Barrett, Sonia Goncalves, Ewan Harrison, David K. Jackson, Ian Johnston, Dominic Kwiatkowski, Cordelia Langford, John Sillitoe on behalf of the Wellcome Sanger Institute COVID-19 Surveillance Team |
| EPI_ISL_3285146 | Lighthouse Lab in Milton Keynes | Wellcome Sanger Institute for the COVID-19 Genomics UK (COG-UK) Consortium | The Lighthouse Lab in Milton Keynes and Alex Alderton, Roberto Amato, Jeffrey Barrett, Sonia Goncalves, Ewan Harrison, David K. Jackson, Ian Johnston, Dominic Kwiatkowski, Cordelia Langford, John Sillitoe on behalf of the Wellcome Sanger Institute COVID-19 Surveillance Team |
| EPI_ISL_3285147, EPI_ISL_3285148, EPI_ISL_3285149 | Lighthouse Lab in Alderley Park | Wellcome Sanger Institute for the COVID-19 Genomics UK (COG-UK) Consortium | Jacquelyn Wynn, Mairead Hyland, The Lighthouse Lab in Alderley Park and Alex Alderton, Roberto Amato, Jeffrey Barrett, Sonia Goncalves, Ewan Harrison, David K. Jackson, Ian Johnston, Dominic Kwiatkowski, Cordelia Langford, John Sillitoe on behalf of the Wellcome Sanger Institute COVID-19 Surveillance Team |
| EPI_ISL_3285150 | Berkshire and Surrey Pathology Services Lighthouse Laboratory | Wellcome Sanger Institute for the COVID-19 Genomics UK (COG-UK) Consortium | Berkshire and Surrey Pathology Services Lighthouse Laboratory and Alex Alderton, Roberto Amato, Jeffrey Barrett, Sonia Goncalves, Ewan Harrison, David K. Jackson, Ian Johnston, Dominic Kwiatkowski, Cordelia Langford, John Sillitoe on behalf of the Wellcome Sanger Institute COVID-19 Surveillance Team |
| EPI_ISL_3285151, EPI_ISL_3285152, EPI_ISL_3285153, EPI_ISL_3285154 | Lighthouse Lab in Alderley Park | Wellcome Sanger Institute for the COVID-19 Genomics UK (COG-UK) Consortium | Jacquelyn Wynn, Mairead Hyland, The Lighthouse Lab in Alderley Park and Alex Alderton, Roberto Amato, Jeffrey Barrett, Sonia Goncalves, Ewan Harrison, David K. Jackson, Ian Johnston, Dominic Kwiatkowski, Cordelia Langford, John Sillitoe on behalf of the Wellcome Sanger Institute COVID-19 Surveillance Team |
| EPI_ISL_3285155 | Lighthouse Lab in Milton Keynes | Wellcome Sanger Institute for the COVID-19 Genomics UK (COG-UK) Consortium | The Lighthouse Lab in Milton Keynes and Alex Alderton, Roberto Amato, Jeffrey Barrett, Sonia Goncalves, Ewan Harrison, David K. Jackson, Ian Johnston, Dominic Kwiatkowski, Cordelia Langford, John Sillitoe on behalf of the Wellcome Sanger Institute COVID-19 Surveillance Team |
| EPI_ISL_3285156 | Lighthouse Lab in Alderley Park | Wellcome Sanger Institute for the COVID-19 Genomics UK (COG-UK) Consortium | Jacquelyn Wynn, Mairead Hyland, The Lighthouse Lab in Alderley Park and Alex Alderton, Roberto Amato, Jeffrey Barrett, Sonia Goncalves, Ewan Harrison, David K. Jackson, Ian Johnston, Dominic Kwiatkowski, Cordelia Langford, John Sillitoe on behalf of the Wellcome Sanger Institute COVID-19 Surveillance Team |
| EPI_ISL_3285157 | Berkshire and Surrey Pathology Services Lighthouse Laboratory | Wellcome Sanger Institute for the COVID-19 Genomics UK (COG-UK) Consortium | Berkshire and Surrey Pathology Services Lighthouse Laboratory and Alex Alderton, Roberto Amato, Jeffrey Barrett, Sonia Goncalves, Ewan Harrison, David K. Jackson, Ian Johnston, Dominic Kwiatkowski, Cordelia Langford, John Sillitoe on behalf of the Wellcome Sanger Institute COVID-19 Surveillance Team |
| EPI_ISL_3285158, EPI_ISL_3285159, EPI_ISL_3285160 | Lighthouse Lab in Alderley Park | Wellcome Sanger Institute for the COVID-19 Genomics UK (COG-UK) Consortium | Jacquelyn Wynn, Mairead Hyland, The Lighthouse Lab in Alderley Park and Alex Alderton, Roberto Amato, Jeffrey Barrett, Sonia Goncalves, Ewan Harrison, David K. Jackson, Ian Johnston, Dominic Kwiatkowski, Cordelia Langford, John Sillitoe on behalf of the Wellcome Sanger Institute COVID-19 Surveillance Team |
| EPI_ISL_3285161, EPI_ISL_3285162 | Lighthouse Lab in Milton Keynes | Wellcome Sanger Institute for the COVID-19 Genomics UK (COG-UK) Consortium | The Lighthouse Lab in Milton Keynes and Alex Alderton, Roberto Amato, Jeffrey Barrett, Sonia Goncalves, Ewan Harrison, David K. Jackson, Ian Johnston, Dominic Kwiatkowski, Cordelia Langford, John Sillitoe on behalf of the Wellcome Sanger Institute COVID-19 Surveillance Team |

| | | | |
|---|---|---|---|
| EPI_ISL_3285163, EPI_ISL_3285164 | Lighthouse Lab in Alderley Park | Wellcome Sanger Institute for the COVID-19 Genomics UK (COG-UK) Consortium | Jacquelyn Wynn, Mairead Hyland, The Lighthouse Lab in Alderley Park and Alex Alderton, Roberto Amato, Jeffrey Barrett, Sonia Goncalves, Ewan Harrison, David K. Jackson, Ian Johnston, Dominic Kwiatkowski, Cordelia Langford, John Sillitoe on behalf of the Wellcome Sanger Institute COVID-19 Surveillance Team |
| EPI_ISL_3285165 | Berkshire and Surrey Pathology Services Lighthouse Laboratory | Wellcome Sanger Institute for the COVID-19 Genomics UK (COG-UK) Consortium | Berkshire and Surrey Pathology Services Lighthouse Laboratory and Alex Alderton, Roberto Amato, Jeffrey Barrett, Sonia Goncalves, Ewan Harrison, David K. Jackson, Ian Johnston, Dominic Kwiatkowski, Cordelia Langford, John Sillitoe on behalf of the Wellcome Sanger Institute COVID-19 Surveillance Team |
| EPI_ISL_3285166, EPI_ISL_3285167, EPI_ISL_3285168 | Lighthouse Lab in Alderley Park | Wellcome Sanger Institute for the COVID-19 Genomics UK (COG-UK) Consortium | Jacquelyn Wynn, Mairead Hyland, The Lighthouse Lab in Alderley Park and Alex Alderton, Roberto Amato, Jeffrey Barrett, Sonia Goncalves, Ewan Harrison, David K. Jackson, Ian Johnston, Dominic Kwiatkowski, Cordelia Langford, John Sillitoe on behalf of the Wellcome Sanger Institute COVID-19 Surveillance Team |
| EPI_ISL_3285169, EPI_ISL_3285170 | Berkshire and Surrey Pathology Services Lighthouse Laboratory | Wellcome Sanger Institute for the COVID-19 Genomics UK (COG-UK) Consortium | Berkshire and Surrey Pathology Services Lighthouse Laboratory and Alex Alderton, Roberto Amato, Jeffrey Barrett, Sonia Goncalves, Ewan Harrison, David K. Jackson, Ian Johnston, Dominic Kwiatkowski, Cordelia Langford, John Sillitoe on behalf of the Wellcome Sanger Institute COVID-19 Surveillance Team |
| EPI_ISL_3285171 | Lighthouse Lab in Alderley Park | Wellcome Sanger Institute for the COVID-19 Genomics UK (COG-UK) Consortium | Jacquelyn Wynn, Mairead Hyland, The Lighthouse Lab in Alderley Park and Alex Alderton, Roberto Amato, Jeffrey Barrett, Sonia Goncalves, Ewan Harrison, David K. Jackson, Ian Johnston, Dominic Kwiatkowski, Cordelia Langford, John Sillitoe on behalf of the Wellcome Sanger Institute COVID-19 Surveillance Team |
| EPI_ISL_3285172, EPI_ISL_3285173 | Lighthouse Lab in Milton Keynes | Wellcome Sanger Institute for the COVID-19 Genomics UK (COG-UK) Consortium | The Lighthouse Lab in Milton Keynes and Alex Alderton, Roberto Amato, Jeffrey Barrett, Sonia Goncalves, Ewan Harrison, David K. Jackson, Ian Johnston, Dominic Kwiatkowski, Cordelia Langford, John Sillitoe on behalf of the Wellcome Sanger Institute COVID-19 Surveillance Team |
| EPI_ISL_3285174, EPI_ISL_3285175, EPI_ISL_3285176, EPI_ISL_3285177, EPI_ISL_3285178 | Lighthouse Lab in Alderley Park | Wellcome Sanger Institute for the COVID-19 Genomics UK (COG-UK) Consortium | Jacquelyn Wynn, Mairead Hyland, The Lighthouse Lab in Alderley Park and Alex Alderton, Roberto Amato, Jeffrey Barrett, Sonia Goncalves, Ewan Harrison, David K. Jackson, Ian Johnston, Dominic Kwiatkowski, Cordelia Langford, John Sillitoe on behalf of the Wellcome Sanger Institute COVID-19 Surveillance Team |
| EPI_ISL_3285179, EPI_ISL_3285180, EPI_ISL_3285181 | Lighthouse Lab in Milton Keynes | Wellcome Sanger Institute for the COVID-19 Genomics UK (COG-UK) Consortium | The Lighthouse Lab in Milton Keynes and Alex Alderton, Roberto Amato, Jeffrey Barrett, Sonia Goncalves, Ewan Harrison, David K. Jackson, Ian Johnston, Dominic Kwiatkowski, Cordelia Langford, John Sillitoe on behalf of the Wellcome Sanger Institute COVID-19 Surveillance Team |
| EPI_ISL_3285182 | Berkshire and Surrey Pathology Services Lighthouse Laboratory | Wellcome Sanger Institute for the COVID-19 Genomics UK (COG-UK) Consortium | Berkshire and Surrey Pathology Services Lighthouse Laboratory and Alex Alderton, Roberto Amato, Jeffrey Barrett, Sonia Goncalves, Ewan Harrison, David K. Jackson, Ian Johnston, Dominic Kwiatkowski, Cordelia Langford, John Sillitoe on behalf of the Wellcome Sanger Institute COVID-19 Surveillance Team |
| EPI_ISL_3285183, EPI_ISL_3285184, EPI_ISL_3285185 | Lighthouse Lab in Alderley Park | Wellcome Sanger Institute for the COVID-19 Genomics UK (COG-UK) Consortium | Jacquelyn Wynn, Mairead Hyland, The Lighthouse Lab in Alderley Park and Alex Alderton, Roberto Amato, Jeffrey Barrett, Sonia Goncalves, Ewan Harrison, David K. Jackson, Ian Johnston, Dominic Kwiatkowski, Cordelia Langford, John Sillitoe on behalf of the Wellcome Sanger Institute COVID-19 Surveillance Team |
| EPI_ISL_3285186, EPI_ISL_3285187 | Lighthouse Lab in Milton Keynes | Wellcome Sanger Institute for the COVID-19 Genomics UK (COG-UK) Consortium | The Lighthouse Lab in Milton Keynes and Alex Alderton, Roberto Amato, Jeffrey Barrett, Sonia Goncalves, Ewan Harrison, David K. Jackson, Ian Johnston, Dominic Kwiatkowski, Cordelia Langford, John Sillitoe on behalf of the Wellcome Sanger Institute COVID-19 Surveillance Team |
| EPI_ISL_3285188, EPI_ISL_3285189, EPI_ISL_3285190, EPI_ISL_3285191, EPI_ISL_3285192 | Lighthouse Lab in Alderley Park | Wellcome Sanger Institute for the COVID-19 Genomics UK (COG-UK) Consortium | Jacquelyn Wynn, Mairead Hyland, The Lighthouse Lab in Alderley Park and Alex Alderton, Roberto Amato, Jeffrey Barrett, Sonia Goncalves, Ewan Harrison, David K. Jackson, Ian Johnston, Dominic Kwiatkowski, Cordelia Langford, John Sillitoe on behalf of the Wellcome Sanger Institute COVID-19 Surveillance Team |
| EPI_ISL_3285193 | Berkshire and Surrey Pathology Services Lighthouse Laboratory | Wellcome Sanger Institute for the COVID-19 Genomics UK (COG-UK) Consortium | Berkshire and Surrey Pathology Services Lighthouse Laboratory and Alex Alderton, Roberto Amato, Jeffrey Barrett, Sonia Goncalves, Ewan Harrison, David K. Jackson, Ian Johnston, Dominic Kwiatkowski, Cordelia Langford, John Sillitoe on behalf of the Wellcome Sanger Institute COVID-19 Surveillance Team |
| EPI_ISL_3285194 | Lighthouse Lab in Alderley Park | Wellcome Sanger Institute for the COVID-19 Genomics UK (COG-UK) Consortium | Jacquelyn Wynn, Mairead Hyland, The Lighthouse Lab in Alderley Park and Alex Alderton, Roberto Amato, Jeffrey Barrett, Sonia Goncalves, Ewan Harrison, David K. Jackson, Ian Johnston, Dominic Kwiatkowski, Cordelia Langford, John Sillitoe on behalf of the Wellcome Sanger Institute COVID-19 Surveillance Team |
| EPI_ISL_3285195 | Lighthouse Lab in Milton Keynes | Wellcome Sanger Institute for the COVID-19 Genomics UK (COG-UK) Consortium | The Lighthouse Lab in Milton Keynes and Alex Alderton, Roberto Amato, Jeffrey Barrett, Sonia Goncalves, Ewan Harrison, David K. Jackson, Ian Johnston, Dominic Kwiatkowski, Cordelia Langford, John Sillitoe on behalf of the Wellcome Sanger Institute COVID-19 Surveillance Team |
| EPI_ISL_3285196, EPI_ISL_3285197 | Lighthouse Lab in Alderley Park | Wellcome Sanger Institute for the COVID-19 Genomics UK (COG-UK) Consortium | Jacquelyn Wynn, Mairead Hyland, The Lighthouse Lab in Alderley Park and Alex Alderton, Roberto Amato, Jeffrey Barrett, Sonia Goncalves, Ewan Harrison, David K. Jackson, Ian Johnston, Dominic Kwiatkowski, Cordelia Langford, John Sillitoe on behalf of the Wellcome Sanger Institute COVID-19 Surveillance Team |
| EPI_ISL_3285198 | Lighthouse Lab in Milton Keynes | Wellcome Sanger Institute for the COVID-19 Genomics UK (COG-UK) Consortium | The Lighthouse Lab in Milton Keynes and Alex Alderton, Roberto Amato, Jeffrey Barrett, Sonia Goncalves, Ewan Harrison, David K. Jackson, Ian Johnston, Dominic Kwiatkowski, Cordelia Langford, John Sillitoe on behalf of the Wellcome Sanger Institute COVID-19 Surveillance Team |
| EPI_ISL_3285199, EPI_ISL_3285200 | Lighthouse Lab in Alderley Park | Wellcome Sanger Institute for the COVID-19 Genomics UK (COG-UK) Consortium | Jacquelyn Wynn, Mairead Hyland, The Lighthouse Lab in Alderley Park and Alex Alderton, Roberto Amato, Jeffrey Barrett, Sonia Goncalves, Ewan Harrison, David K. Jackson, Ian Johnston, Dominic Kwiatkowski, Cordelia Langford, John Sillitoe on behalf of the Wellcome Sanger Institute COVID-19 Surveillance Team |
| EPI_ISL_3285201 | Lighthouse Lab in Milton Keynes | Wellcome Sanger Institute for the COVID-19 Genomics UK (COG-UK) Consortium | The Lighthouse Lab in Milton Keynes and Alex Alderton, Roberto Amato, Jeffrey Barrett, Sonia Goncalves, Ewan Harrison, David K. Jackson, Ian Johnston, Dominic Kwiatkowski, Cordelia Langford, John Sillitoe on behalf of the Wellcome Sanger Institute COVID-19 Surveillance Team |
| EPI_ISL_3285202, EPI_ISL_3285203 | Lighthouse Lab in Alderley Park | Wellcome Sanger Institute for the COVID-19 Genomics UK (COG-UK) Consortium | Jacquelyn Wynn, Mairead Hyland, The Lighthouse Lab in Alderley Park and Alex Alderton, Roberto Amato, Jeffrey Barrett, Sonia Goncalves, Ewan Harrison, David K. Jackson, Ian Johnston, Dominic Kwiatkowski, Cordelia Langford, John Sillitoe on behalf of the Wellcome Sanger Institute COVID-19 Surveillance Team |
| EPI_ISL_3285204, EPI_ISL_3285205 | Berkshire and Surrey Pathology Services Lighthouse Laboratory | Wellcome Sanger Institute for the COVID-19 Genomics UK (COG-UK) Consortium | Berkshire and Surrey Pathology Services Lighthouse Laboratory and Alex Alderton, Roberto Amato, Jeffrey Barrett, Sonia Goncalves, Ewan Harrison, David K. Jackson, Ian Johnston, Dominic Kwiatkowski, Cordelia Langford, John Sillitoe on behalf of the Wellcome Sanger Institute COVID-19 Surveillance Team |
| EPI_ISL_3285206 | Lighthouse Lab in Milton Keynes | Wellcome Sanger Institute for the COVID-19 Genomics UK (COG-UK) Consortium | The Lighthouse Lab in Milton Keynes and Alex Alderton, Roberto Amato, Jeffrey Barrett, Sonia Goncalves, Ewan Harrison, David K. Jackson, Ian Johnston, Dominic Kwiatkowski, Cordelia Langford, John Sillitoe on behalf of the Wellcome Sanger Institute COVID-19 Surveillance Team |
| EPI_ISL_3285207 | Lighthouse Lab in Alderley Park | Wellcome Sanger Institute for the COVID-19 Genomics UK (COG-UK) Consortium | Jacquelyn Wynn, Mairead Hyland, The Lighthouse Lab in Alderley Park and Alex Alderton, Roberto Amato, Jeffrey Barrett, Sonia Goncalves, Ewan Harrison, David K. Jackson, Ian Johnston, Dominic Kwiatkowski, Cordelia Langford, John Sillitoe on behalf of the Wellcome Sanger Institute COVID-19 Surveillance Team |
| EPI_ISL_3285208, EPI_ISL_3285209 | Lighthouse Lab in Milton Keynes | Wellcome Sanger Institute for the COVID-19 Genomics UK (COG-UK) Consortium | The Lighthouse Lab in Milton Keynes and Alex Alderton, Roberto Amato, Jeffrey Barrett, Sonia Goncalves, Ewan Harrison, David K. Jackson, Ian Johnston, Dominic Kwiatkowski, Cordelia Langford, John Sillitoe on behalf of the Wellcome Sanger Institute COVID-19 Surveillance Team |
| EPI_ISL_3285210, EPI_ISL_3285211 | Lighthouse Lab in Alderley Park | Wellcome Sanger Institute for the COVID-19 Genomics UK (COG-UK) Consortium | Jacquelyn Wynn, Mairead Hyland, The Lighthouse Lab in Alderley Park and Alex Alderton, Roberto Amato, Jeffrey Barrett, Sonia Goncalves, Ewan Harrison, David K. Jackson, Ian Johnston, Dominic Kwiatkowski, Cordelia Langford, John Sillitoe on behalf of the Wellcome Sanger Institute COVID-19 Surveillance Team |
| EPI_ISL_3285212 | Lighthouse Lab in Milton Keynes | Wellcome Sanger Institute for the COVID-19 Genomics UK (COG-UK) Consortium | The Lighthouse Lab in Milton Keynes and Alex Alderton, Roberto Amato, Jeffrey Barrett, Sonia Goncalves, Ewan Harrison, David K. Jackson, Ian Johnston, Dominic Kwiatkowski, Cordelia Langford, John Sillitoe on behalf of the Wellcome Sanger Institute COVID-19 Surveillance Team |
| EPI_ISL_3285213 | Lighthouse Lab in Alderley Park | Wellcome Sanger Institute for the COVID-19 Genomics UK (COG-UK) Consortium | Jacquelyn Wynn, Mairead Hyland, The Lighthouse Lab in Alderley Park and Alex Alderton, Roberto Amato, Jeffrey Barrett, Sonia Goncalves, Ewan Harrison, David K. Jackson, Ian Johnston, Dominic Kwiatkowski, Cordelia Langford, John Sillitoe on behalf of the Wellcome Sanger Institute COVID-19 Surveillance Team |
| EPI_ISL_3285214, EPI_ISL_3285215, EPI_ISL_3285216 | Lighthouse Lab in Milton Keynes | Wellcome Sanger Institute for the COVID-19 Genomics UK (COG-UK) Consortium | The Lighthouse Lab in Milton Keynes and Alex Alderton, Roberto Amato, Jeffrey Barrett, Sonia Goncalves, Ewan Harrison, David K. Jackson, Ian Johnston, Dominic Kwiatkowski, Cordelia Langford, John Sillitoe on behalf of the Wellcome Sanger Institute COVID-19 Surveillance Team |
| EPI_ISL_3285217, EPI_ISL_3285218, EPI_ISL_3285219, EPI_ISL_3285220, EPI_ISL_3285221 | Lighthouse Lab in Alderley Park | Wellcome Sanger Institute for the COVID-19 Genomics UK (COG-UK) Consortium | Jacquelyn Wynn, Mairead Hyland, The Lighthouse Lab in Alderley Park and Alex Alderton, Roberto Amato, Jeffrey Barrett, Sonia Goncalves, Ewan Harrison, David K. Jackson, Ian Johnston, Dominic Kwiatkowski, Cordelia Langford, John Sillitoe on behalf of the Wellcome Sanger Institute COVID-19 Surveillance Team |
| EPI_ISL_3285222 | Lighthouse Lab in Milton Keynes | Wellcome Sanger Institute for the COVID-19 Genomics UK (COG-UK) Consortium | The Lighthouse Lab in Milton Keynes and Alex Alderton, Roberto Amato, Jeffrey Barrett, Sonia Goncalves, Ewan Harrison, David K. Jackson, Ian Johnston, Dominic Kwiatkowski, Cordelia Langford, John Sillitoe on behalf of the Wellcome Sanger Institute COVID-19 Surveillance Team |
| EPI_ISL_3285223 | Lighthouse Lab in Alderley Park | Wellcome Sanger Institute for the COVID-19 Genomics UK (COG-UK) Consortium | Jacquelyn Wynn, Mairead Hyland, The Lighthouse Lab in Alderley Park and Alex Alderton, Roberto Amato, Jeffrey Barrett, Sonia Goncalves, Ewan Harrison, David K. Jackson, Ian Johnston, Dominic Kwiatkowski, Cordelia Langford, John Sillitoe on behalf of the Wellcome Sanger Institute COVID-19 Surveillance Team |
| EPI_ISL_3285224, EPI_ISL_3285225 | Berkshire and Surrey Pathology Services Lighthouse Laboratory | Wellcome Sanger Institute for the COVID-19 Genomics UK (COG-UK) Consortium | Berkshire and Surrey Pathology Services Lighthouse Laboratory and Alex Alderton, Roberto Amato, Jeffrey Barrett, Sonia Goncalves, Ewan Harrison, David K. Jackson, Ian Johnston, Dominic Kwiatkowski, Cordelia Langford, John Sillitoe on behalf of the Wellcome Sanger Institute COVID-19 Surveillance Team |
| EPI_ISL_3285226, EPI_ISL_3285227, EPI_ISL_3285228 | Lighthouse Lab in Alderley Park | Wellcome Sanger Institute for the COVID-19 Genomics UK (COG-UK) Consortium | Jacquelyn Wynn, Mairead Hyland, The Lighthouse Lab in Alderley Park and Alex Alderton, Roberto Amato, Jeffrey Barrett, Sonia Goncalves, Ewan Harrison, David K. Jackson, Ian Johnston, Dominic Kwiatkowski, Cordelia Langford, John Sillitoe on behalf of the Wellcome Sanger Institute COVID-19 Surveillance Team |
| EPI_ISL_3285229 | Berkshire and Surrey Pathology Services Lighthouse Laboratory | Wellcome Sanger Institute for the COVID-19 Genomics UK (COG-UK) Consortium | Berkshire and Surrey Pathology Services Lighthouse Laboratory and Alex Alderton, Roberto Amato, Jeffrey Barrett, Sonia Goncalves, Ewan Harrison, David K. Jackson, Ian Johnston, Dominic Kwiatkowski, Cordelia Langford, John Sillitoe on behalf of the Wellcome Sanger Institute COVID-19 Surveillance Team |
| EPI_ISL_3285230, EPI_ISL_3285231 | Lighthouse Lab in Alderley Park | Wellcome Sanger Institute for the COVID-19 Genomics UK (COG-UK) Consortium | Jacquelyn Wynn, Mairead Hyland, The Lighthouse Lab in Alderley Park and Alex Alderton, Roberto Amato, Jeffrey Barrett, Sonia Goncalves, Ewan Harrison, David K. Jackson, Ian Johnston, Dominic Kwiatkowski, Cordelia Langford, John Sillitoe on behalf of the Wellcome Sanger Institute COVID-19 Surveillance Team |
| EPI_ISL_3285232, EPI_ISL_3285233, EPI_ISL_3285234 | Lighthouse Lab in Milton Keynes | Wellcome Sanger Institute for the COVID-19 Genomics UK (COG-UK) Consortium | The Lighthouse Lab in Milton Keynes and Alex Alderton, Roberto Amato, Jeffrey Barrett, Sonia Goncalves, Ewan Harrison, David K. Jackson, Ian Johnston, Dominic Kwiatkowski, Cordelia Langford, John Sillitoe on behalf of the Wellcome Sanger Institute COVID-19 Surveillance Team |
| EPI_ISL_3285235 | Berkshire and Surrey Pathology Services Lighthouse Laboratory | Wellcome Sanger Institute for the COVID-19 Genomics UK (COG-UK) Consortium | Berkshire and Surrey Pathology Services Lighthouse Laboratory and Alex Alderton, Roberto Amato, Jeffrey Barrett, Sonia Goncalves, Ewan Harrison, David K. Jackson, Ian Johnston, Dominic Kwiatkowski, Cordelia Langford, John Sillitoe on behalf of the Wellcome Sanger Institute COVID-19 Surveillance Team |
| EPI_ISL_3285236 | Lighthouse Lab in Alderley Park | Wellcome Sanger Institute for the COVID-19 Genomics UK (COG-UK) Consortium | Jacquelyn Wynn, Mairead Hyland, The Lighthouse Lab in Alderley Park and Alex Alderton, Roberto Amato, Jeffrey Barrett, Sonia Goncalves, Ewan Harrison, David K. Jackson, Ian Johnston, Dominic Kwiatkowski, Cordelia Langford, John Sillitoe on behalf of the Wellcome Sanger Institute COVID-19 Surveillance Team |
| EPI_ISL_3285237 | Lighthouse Lab in Milton Keynes | Wellcome Sanger Institute for the COVID-19 Genomics UK (COG-UK) Consortium | The Lighthouse Lab in Milton Keynes and Alex Alderton, Roberto Amato, Jeffrey Barrett, Sonia Goncalves, Ewan Harrison, David K. Jackson, Ian Johnston, Dominic Kwiatkowski, Cordelia Langford, John Sillitoe on behalf of the Wellcome Sanger Institute COVID-19 Surveillance Team |

| | | | |
|---|---|---|---|
| EPI_ISL_3285238, EPI_ISL_3285239, EPI_ISL_3285240 | Lighthouse Lab in Alderley Park | Wellcome Sanger Institute for the COVID-19 Genomics UK (COG-UK) Consortium | Jacquelyn Wynn, Mairead Hyland, The Lighthouse Lab in Alderley Park and Alex Alderton, Roberto Amato, Jeffrey Barrett, Sonia Goncalves, Ewan Harrison, David K. Jackson, Ian Johnston, Dominic Kwiatkowski, Cordelia Langford, John Sillitoe on behalf of the Wellcome Sanger Institute COVID-19 Surveillance Team |
| EPI_ISL_3285241 | Lighthouse Lab in Milton Keynes | Wellcome Sanger Institute for the COVID-19 Genomics UK (COG-UK) Consortium | The Lighthouse Lab in Milton Keynes and Alex Alderton, Roberto Amato, Jeffrey Barrett, Sonia Goncalves, Ewan Harrison, David K. Jackson, Ian Johnston, Dominic Kwiatkowski, Cordelia Langford, John Sillitoe on behalf of the Wellcome Sanger Institute COVID-19 Surveillance Team |
| EPI_ISL_3285242 | Berkshire and Surrey Pathology Services Lighthouse Laboratory | Wellcome Sanger Institute for the COVID-19 Genomics UK (COG-UK) Consortium | Berkshire and Surrey Pathology Services Lighthouse Laboratory and Alex Alderton, Roberto Amato, Jeffrey Barrett, Sonia Goncalves, Ewan Harrison, David K. Jackson, Ian Johnston, Dominic Kwiatkowski, Cordelia Langford, John Sillitoe on behalf of the Wellcome Sanger Institute COVID-19 Surveillance Team |
| EPI_ISL_3285243, EPI_ISL_3285244, EPI_ISL_3285245 | Lighthouse Lab in Alderley Park | Wellcome Sanger Institute for the COVID-19 Genomics UK (COG-UK) Consortium | Jacquelyn Wynn, Mairead Hyland, The Lighthouse Lab in Alderley Park and Alex Alderton, Roberto Amato, Jeffrey Barrett, Sonia Goncalves, Ewan Harrison, David K. Jackson, Ian Johnston, Dominic Kwiatkowski, Cordelia Langford, John Sillitoe on behalf of the Wellcome Sanger Institute COVID-19 Surveillance Team |
| EPI_ISL_3285246, EPI_ISL_3285247, EPI_ISL_3285248, EPI_ISL_3285249 | Lighthouse Lab in Milton Keynes | Wellcome Sanger Institute for the COVID-19 Genomics UK (COG-UK) Consortium | The Lighthouse Lab in Milton Keynes and Alex Alderton, Roberto Amato, Jeffrey Barrett, Sonia Goncalves, Ewan Harrison, David K. Jackson, Ian Johnston, Dominic Kwiatkowski, Cordelia Langford, John Sillitoe on behalf of the Wellcome Sanger Institute COVID-19 Surveillance Team |
| EPI_ISL_3285250, EPI_ISL_3285251, EPI_ISL_3285252, EPI_ISL_3285253 | Lighthouse Lab in Alderley Park | Wellcome Sanger Institute for the COVID-19 Genomics UK (COG-UK) Consortium | Jacquelyn Wynn, Mairead Hyland, The Lighthouse Lab in Alderley Park and Alex Alderton, Roberto Amato, Jeffrey Barrett, Sonia Goncalves, Ewan Harrison, David K. Jackson, Ian Johnston, Dominic Kwiatkowski, Cordelia Langford, John Sillitoe on behalf of the Wellcome Sanger Institute COVID-19 Surveillance Team |
| EPI_ISL_3285254, EPI_ISL_3285255, EPI_ISL_3285256, EPI_ISL_3285257 | Berkshire and Surrey Pathology Services Lighthouse Laboratory | Wellcome Sanger Institute for the COVID-19 Genomics UK (COG-UK) Consortium | Berkshire and Surrey Pathology Services Lighthouse Laboratory and Alex Alderton, Roberto Amato, Jeffrey Barrett, Sonia Goncalves, Ewan Harrison, David K. Jackson, Ian Johnston, Dominic Kwiatkowski, Cordelia Langford, John Sillitoe on behalf of the Wellcome Sanger Institute COVID-19 Surveillance Team |
| EPI_ISL_3285258 | Lighthouse Lab in Alderley Park | Wellcome Sanger Institute for the COVID-19 Genomics UK (COG-UK) Consortium | Jacquelyn Wynn, Mairead Hyland, The Lighthouse Lab in Alderley Park and Alex Alderton, Roberto Amato, Jeffrey Barrett, Sonia Goncalves, Ewan Harrison, David K. Jackson, Ian Johnston, Dominic Kwiatkowski, Cordelia Langford, John Sillitoe on behalf of the Wellcome Sanger Institute COVID-19 Surveillance Team |
| EPI_ISL_3285259 | Lighthouse Lab in Milton Keynes | Wellcome Sanger Institute for the COVID-19 Genomics UK (COG-UK) Consortium | The Lighthouse Lab in Milton Keynes and Alex Alderton, Roberto Amato, Jeffrey Barrett, Sonia Goncalves, Ewan Harrison, David K. Jackson, Ian Johnston, Dominic Kwiatkowski, Cordelia Langford, John Sillitoe on behalf of the Wellcome Sanger Institute COVID-19 Surveillance Team |
| EPI_ISL_3285260 | Berkshire and Surrey Pathology Services Lighthouse Laboratory | Wellcome Sanger Institute for the COVID-19 Genomics UK (COG-UK) Consortium | Berkshire and Surrey Pathology Services Lighthouse Laboratory and Alex Alderton, Roberto Amato, Jeffrey Barrett, Sonia Goncalves, Ewan Harrison, David K. Jackson, Ian Johnston, Dominic Kwiatkowski, Cordelia Langford, John Sillitoe on behalf of the Wellcome Sanger Institute COVID-19 Surveillance Team |
| EPI_ISL_3285261, EPI_ISL_3285262 | Lighthouse Lab in Alderley Park | Wellcome Sanger Institute for the COVID-19 Genomics UK (COG-UK) Consortium | Jacquelyn Wynn, Mairead Hyland, The Lighthouse Lab in Alderley Park and Alex Alderton, Roberto Amato, Jeffrey Barrett, Sonia Goncalves, Ewan Harrison, David K. Jackson, Ian Johnston, Dominic Kwiatkowski, Cordelia Langford, John Sillitoe on behalf of the Wellcome Sanger Institute COVID-19 Surveillance Team |
| EPI_ISL_3285263, EPI_ISL_3285264 | Berkshire and Surrey Pathology Services Lighthouse Laboratory | Wellcome Sanger Institute for the COVID-19 Genomics UK (COG-UK) Consortium | Berkshire and Surrey Pathology Services Lighthouse Laboratory and Alex Alderton, Roberto Amato, Jeffrey Barrett, Sonia Goncalves, Ewan Harrison, David K. Jackson, Ian Johnston, Dominic Kwiatkowski, Cordelia Langford, John Sillitoe on behalf of the Wellcome Sanger Institute COVID-19 Surveillance Team |
| EPI_ISL_3285265 | Lighthouse Lab in Milton Keynes | Wellcome Sanger Institute for the COVID-19 Genomics UK (COG-UK) Consortium | The Lighthouse Lab in Milton Keynes and Alex Alderton, Roberto Amato, Jeffrey Barrett, Sonia Goncalves, Ewan Harrison, David K. Jackson, Ian Johnston, Dominic Kwiatkowski, Cordelia Langford, John Sillitoe on behalf of the Wellcome Sanger Institute COVID-19 Surveillance Team |
| EPI_ISL_3285266, EPI_ISL_3285267, EPI_ISL_3285268 | Lighthouse Lab in Alderley Park | Wellcome Sanger Institute for the COVID-19 Genomics UK (COG-UK) Consortium | Jacquelyn Wynn, Mairead Hyland, The Lighthouse Lab in Alderley Park and Alex Alderton, Roberto Amato, Jeffrey Barrett, Sonia Goncalves, Ewan Harrison, David K. Jackson, Ian Johnston, Dominic Kwiatkowski, Cordelia Langford, John Sillitoe on behalf of the Wellcome Sanger Institute COVID-19 Surveillance Team |
| EPI_ISL_3285269, EPI_ISL_3285270, EPI_ISL_3285271 | Berkshire and Surrey Pathology Services Lighthouse Laboratory | Wellcome Sanger Institute for the COVID-19 Genomics UK (COG-UK) Consortium | Berkshire and Surrey Pathology Services Lighthouse Laboratory and Alex Alderton, Roberto Amato, Jeffrey Barrett, Sonia Goncalves, Ewan Harrison, David K. Jackson, Ian Johnston, Dominic Kwiatkowski, Cordelia Langford, John Sillitoe on behalf of the Wellcome Sanger Institute COVID-19 Surveillance Team |
| EPI_ISL_3285272 | Lighthouse Lab in Alderley Park | Wellcome Sanger Institute for the COVID-19 Genomics UK (COG-UK) Consortium | Jacquelyn Wynn, Mairead Hyland, The Lighthouse Lab in Alderley Park and Alex Alderton, Roberto Amato, Jeffrey Barrett, Sonia Goncalves, Ewan Harrison, David K. Jackson, Ian Johnston, Dominic Kwiatkowski, Cordelia Langford, John Sillitoe on behalf of the Wellcome Sanger Institute COVID-19 Surveillance Team |
| EPI_ISL_3285273, EPI_ISL_3285274 | Berkshire and Surrey Pathology Services Lighthouse Laboratory | Wellcome Sanger Institute for the COVID-19 Genomics UK (COG-UK) Consortium | Berkshire and Surrey Pathology Services Lighthouse Laboratory and Alex Alderton, Roberto Amato, Jeffrey Barrett, Sonia Goncalves, Ewan Harrison, David K. Jackson, Ian Johnston, Dominic Kwiatkowski, Cordelia Langford, John Sillitoe on behalf of the Wellcome Sanger Institute COVID-19 Surveillance Team |
| EPI_ISL_3285275, EPI_ISL_3285276, EPI_ISL_3285277, EPI_ISL_3285278 | Lighthouse Lab in Alderley Park | Wellcome Sanger Institute for the COVID-19 Genomics UK (COG-UK) Consortium | Jacquelyn Wynn, Mairead Hyland, The Lighthouse Lab in Alderley Park and Alex Alderton, Roberto Amato, Jeffrey Barrett, Sonia Goncalves, Ewan Harrison, David K. Jackson, Ian Johnston, Dominic Kwiatkowski, Cordelia Langford, John Sillitoe on behalf of the Wellcome Sanger Institute COVID-19 Surveillance Team |
| EPI_ISL_3285279 | Lighthouse Lab in Milton Keynes | Wellcome Sanger Institute for the COVID-19 Genomics UK (COG-UK) Consortium | The Lighthouse Lab in Milton Keynes and Alex Alderton, Roberto Amato, Jeffrey Barrett, Sonia Goncalves, Ewan Harrison, David K. Jackson, Ian Johnston, Dominic Kwiatkowski, Cordelia Langford, John Sillitoe on behalf of the Wellcome Sanger Institute COVID-19 Surveillance Team |
| EPI_ISL_3285280 | Lighthouse Lab in Alderley Park | Wellcome Sanger Institute for the COVID-19 Genomics UK (COG-UK) Consortium | Jacquelyn Wynn, Mairead Hyland, The Lighthouse Lab in Alderley Park and Alex Alderton, Roberto Amato, Jeffrey Barrett, Sonia Goncalves, Ewan Harrison, David K. Jackson, Ian Johnston, Dominic Kwiatkowski, Cordelia Langford, John Sillitoe on behalf of the Wellcome Sanger Institute COVID-19 Surveillance Team |
| EPI_ISL_3285281 | Lighthouse Lab in Milton Keynes | Wellcome Sanger Institute for the COVID-19 Genomics UK (COG-UK) Consortium | The Lighthouse Lab in Milton Keynes and Alex Alderton, Roberto Amato, Jeffrey Barrett, Sonia Goncalves, Ewan Harrison, David K. Jackson, Ian Johnston, Dominic Kwiatkowski, Cordelia Langford, John Sillitoe on behalf of the Wellcome Sanger Institute COVID-19 Surveillance Team |
| EPI_ISL_3285282 | Berkshire and Surrey Pathology Services Lighthouse Laboratory | Wellcome Sanger Institute for the COVID-19 Genomics UK (COG-UK) Consortium | Berkshire and Surrey Pathology Services Lighthouse Laboratory and Alex Alderton, Roberto Amato, Jeffrey Barrett, Sonia Goncalves, Ewan Harrison, David K. Jackson, Ian Johnston, Dominic Kwiatkowski, Cordelia Langford, John Sillitoe on behalf of the Wellcome Sanger Institute COVID-19 Surveillance Team |
| EPI_ISL_3285283, EPI_ISL_3285284, EPI_ISL_3285285 | Lighthouse Lab in Alderley Park | Wellcome Sanger Institute for the COVID-19 Genomics UK (COG-UK) Consortium | Jacquelyn Wynn, Mairead Hyland, The Lighthouse Lab in Alderley Park and Alex Alderton, Roberto Amato, Jeffrey Barrett, Sonia Goncalves, Ewan Harrison, David K. Jackson, Ian Johnston, Dominic Kwiatkowski, Cordelia Langford, John Sillitoe on behalf of the Wellcome Sanger Institute COVID-19 Surveillance Team |
| EPI_ISL_3285286 | Berkshire and Surrey Pathology Services Lighthouse Laboratory | Wellcome Sanger Institute for the COVID-19 Genomics UK (COG-UK) Consortium | Berkshire and Surrey Pathology Services Lighthouse Laboratory and Alex Alderton, Roberto Amato, Jeffrey Barrett, Sonia Goncalves, Ewan Harrison, David K. Jackson, Ian Johnston, Dominic Kwiatkowski, Cordelia Langford, John Sillitoe on behalf of the Wellcome Sanger Institute COVID-19 Surveillance Team |
| EPI_ISL_3285287, EPI_ISL_3285288 | Lighthouse Lab in Milton Keynes | Wellcome Sanger Institute for the COVID-19 Genomics UK (COG-UK) Consortium | The Lighthouse Lab in Milton Keynes and Alex Alderton, Roberto Amato, Jeffrey Barrett, Sonia Goncalves, Ewan Harrison, David K. Jackson, Ian Johnston, Dominic Kwiatkowski, Cordelia Langford, John Sillitoe on behalf of the Wellcome Sanger Institute COVID-19 Surveillance Team |
| EPI_ISL_3285289 | Lighthouse Lab in Alderley Park | Wellcome Sanger Institute for the COVID-19 Genomics UK (COG-UK) Consortium | Jacquelyn Wynn, Mairead Hyland, The Lighthouse Lab in Alderley Park and Alex Alderton, Roberto Amato, Jeffrey Barrett, Sonia Goncalves, Ewan Harrison, David K. Jackson, Ian Johnston, Dominic Kwiatkowski, Cordelia Langford, John Sillitoe on behalf of the Wellcome Sanger Institute COVID-19 Surveillance Team |
| EPI_ISL_3285290, EPI_ISL_3285291 | Lighthouse Lab in Milton Keynes | Wellcome Sanger Institute for the COVID-19 Genomics UK (COG-UK) Consortium | The Lighthouse Lab in Milton Keynes and Alex Alderton, Roberto Amato, Jeffrey Barrett, Sonia Goncalves, Ewan Harrison, David K. Jackson, Ian Johnston, Dominic Kwiatkowski, Cordelia Langford, John Sillitoe on behalf of the Wellcome Sanger Institute COVID-19 Surveillance Team |
| EPI_ISL_3285292 | Lighthouse Lab in Alderley Park | Wellcome Sanger Institute for the COVID-19 Genomics UK (COG-UK) Consortium | Jacquelyn Wynn, Mairead Hyland, The Lighthouse Lab in Alderley Park and Alex Alderton, Roberto Amato, Jeffrey Barrett, Sonia Goncalves, Ewan Harrison, David K. Jackson, Ian Johnston, Dominic Kwiatkowski, Cordelia Langford, John Sillitoe on behalf of the Wellcome Sanger Institute COVID-19 Surveillance Team |
| EPI_ISL_3285293, EPI_ISL_3285294, EPI_ISL_3285295, EPI_ISL_3285296 | Berkshire and Surrey Pathology Services Lighthouse Laboratory | Wellcome Sanger Institute for the COVID-19 Genomics UK (COG-UK) Consortium | Berkshire and Surrey Pathology Services Lighthouse Laboratory and Alex Alderton, Roberto Amato, Jeffrey Barrett, Sonia Goncalves, Ewan Harrison, David K. Jackson, Ian Johnston, Dominic Kwiatkowski, Cordelia Langford, John Sillitoe on behalf of the Wellcome Sanger Institute COVID-19 Surveillance Team |
| EPI_ISL_3285297 | Lighthouse Lab in Alderley Park | Wellcome Sanger Institute for the COVID-19 Genomics UK (COG-UK) Consortium | Jacquelyn Wynn, Mairead Hyland, The Lighthouse Lab in Alderley Park and Alex Alderton, Roberto Amato, Jeffrey Barrett, Sonia Goncalves, Ewan Harrison, David K. Jackson, Ian Johnston, Dominic Kwiatkowski, Cordelia Langford, John Sillitoe on behalf of the Wellcome Sanger Institute COVID-19 Surveillance Team |
| EPI_ISL_3285298 | Berkshire and Surrey Pathology Services Lighthouse Laboratory | Wellcome Sanger Institute for the COVID-19 Genomics UK (COG-UK) Consortium | Berkshire and Surrey Pathology Services Lighthouse Laboratory and Alex Alderton, Roberto Amato, Jeffrey Barrett, Sonia Goncalves, Ewan Harrison, David K. Jackson, Ian Johnston, Dominic Kwiatkowski, Cordelia Langford, John Sillitoe on behalf of the Wellcome Sanger Institute COVID-19 Surveillance Team |
| EPI_ISL_3285299 | Lighthouse Lab in Milton Keynes | Wellcome Sanger Institute for the COVID-19 Genomics UK (COG-UK) Consortium | The Lighthouse Lab in Milton Keynes and Alex Alderton, Roberto Amato, Jeffrey Barrett, Sonia Goncalves, Ewan Harrison, David K. Jackson, Ian Johnston, Dominic Kwiatkowski, Cordelia Langford, John Sillitoe on behalf of the Wellcome Sanger Institute COVID-19 Surveillance Team |
| EPI_ISL_3285300 | Lighthouse Lab in Alderley Park | Wellcome Sanger Institute for the COVID-19 Genomics UK (COG-UK) Consortium | Jacquelyn Wynn, Mairead Hyland, The Lighthouse Lab in Alderley Park and Alex Alderton, Roberto Amato, Jeffrey Barrett, Sonia Goncalves, Ewan Harrison, David K. Jackson, Ian Johnston, Dominic Kwiatkowski, Cordelia Langford, John Sillitoe on behalf of the Wellcome Sanger Institute COVID-19 Surveillance Team |
| EPI_ISL_3285301 | Lighthouse Lab in Milton Keynes | Wellcome Sanger Institute for the COVID-19 Genomics UK (COG-UK) Consortium | The Lighthouse Lab in Milton Keynes and Alex Alderton, Roberto Amato, Jeffrey Barrett, Sonia Goncalves, Ewan Harrison, David K. Jackson, Ian Johnston, Dominic Kwiatkowski, Cordelia Langford, John Sillitoe on behalf of the Wellcome Sanger Institute COVID-19 Surveillance Team |
| EPI_ISL_3285302 | Lighthouse Lab in Alderley Park | Wellcome Sanger Institute for the COVID-19 Genomics UK (COG-UK) Consortium | Jacquelyn Wynn, Mairead Hyland, The Lighthouse Lab in Alderley Park and Alex Alderton, Roberto Amato, Jeffrey Barrett, Sonia Goncalves, Ewan Harrison, David K. Jackson, Ian Johnston, Dominic Kwiatkowski, Cordelia Langford, John Sillitoe on behalf of the Wellcome Sanger Institute COVID-19 Surveillance Team |
| EPI_ISL_3285303, EPI_ISL_3285304 | Lighthouse Lab in Milton Keynes | Wellcome Sanger Institute for the COVID-19 Genomics UK (COG-UK) Consortium | The Lighthouse Lab in Milton Keynes and Alex Alderton, Roberto Amato, Jeffrey Barrett, Sonia Goncalves, Ewan Harrison, David K. Jackson, Ian Johnston, Dominic Kwiatkowski, Cordelia Langford, John Sillitoe on behalf of the Wellcome Sanger Institute COVID-19 Surveillance Team |
| EPI_ISL_3285305 | Lighthouse Lab in Alderley Park | Wellcome Sanger Institute for the COVID-19 Genomics UK (COG-UK) Consortium | Jacquelyn Wynn, Mairead Hyland, The Lighthouse Lab in Alderley Park and Alex Alderton, Roberto Amato, Jeffrey Barrett, Sonia Goncalves, Ewan Harrison, David K. Jackson, Ian Johnston, Dominic Kwiatkowski, Cordelia Langford, John Sillitoe on behalf of the Wellcome Sanger Institute COVID-19 Surveillance Team |
| EPI_ISL_3285306 | Lighthouse Lab in Glasgow | Wellcome Sanger Institute for the COVID-19 Genomics UK (COG-UK) Consortium | Harper VanSteenhouse, Yumi Kasai, David Gray, Carol Clugston, Anna Dominiczak and Alex Alderton, Roberto Amato, Jeffrey Barrett, Sonia Goncalves, Ewan Harrison, David K. Jackson, Ian Johnston, Dominic Kwiatkowski, Cordelia Langford, John Sillitoe on behalf of the Wellcome Sanger Institute COVID-19 Surveillance Team |
| EPI_ISL_3285307 | Berkshire and Surrey Pathology Services Lighthouse Laboratory | Wellcome Sanger Institute for the COVID-19 Genomics UK (COG-UK) Consortium | Berkshire and Surrey Pathology Services Lighthouse Laboratory and Alex Alderton, Roberto Amato, Jeffrey Barrett, Sonia Goncalves, Ewan Harrison, David K. Jackson, Ian Johnston, Dominic Kwiatkowski, Cordelia Langford, John Sillitoe on behalf of the Wellcome Sanger Institute COVID-19 Surveillance Team |

| | | | |
|---|---|---|---|
| EPI_ISL_3285308 | Lighthouse Lab in Milton Keynes | Wellcome Sanger Institute for the COVID-19 Genomics UK (COG-UK) Consortium | The Lighthouse Lab in Milton Keynes and Alex Alderton, Roberto Amato, Jeffrey Barrett, Sonia Goncalves, Ewan Harrison, David K. Jackson, Ian Johnston, Dominic Kwiatkowski, Cordelia Langford, John Sillitoe on behalf of the Wellcome Sanger Institute COVID-19 Surveillance Team |
| EPI_ISL_3285309 | Lighthouse Lab in Alderley Park | Wellcome Sanger Institute for the COVID-19 Genomics UK (COG-UK) Consortium | Jacquelyn Wynn, Mairead Hyland, The Lighthouse Lab in Alderley Park and Alex Alderton, Roberto Amato, Jeffrey Barrett, Sonia Goncalves, Ewan Harrison, David K. Jackson, Ian Johnston, Dominic Kwiatkowski, Cordelia Langford, John Sillitoe on behalf of the Wellcome Sanger Institute COVID-19 Surveillance Team |
| EPI_ISL_3285310 | Berkshire and Surrey Pathology Services Lighthouse Laboratory | Wellcome Sanger Institute for the COVID-19 Genomics UK (COG-UK) Consortium | Berkshire and Surrey Pathology Services Lighthouse Laboratory and Alex Alderton, Roberto Amato, Jeffrey Barrett, Sonia Goncalves, Ewan Harrison, David K. Jackson, Ian Johnston, Dominic Kwiatkowski, Cordelia Langford, John Sillitoe on behalf of the Wellcome Sanger Institute COVID-19 Surveillance Team |
| EPI_ISL_3285311 | Lighthouse Lab in Glasgow | Wellcome Sanger Institute for the COVID-19 Genomics UK (COG-UK) Consortium | Harper VanSteenhouse, Yumi Kasai, David Gray, Carol Clugston, Anna Dominiczak and Alex Alderton, Roberto Amato, Jeffrey Barrett, Sonia Goncalves, Ewan Harrison, David K. Jackson, Ian Johnston, Dominic Kwiatkowski, Cordelia Langford, John Sillitoe on behalf of the Wellcome Sanger Institute COVID-19 Surveillance Team |
| EPI_ISL_3285312 | Lighthouse Lab in Alderley Park | Wellcome Sanger Institute for the COVID-19 Genomics UK (COG-UK) Consortium | Jacquelyn Wynn, Mairead Hyland, The Lighthouse Lab in Alderley Park and Alex Alderton, Roberto Amato, Jeffrey Barrett, Sonia Goncalves, Ewan Harrison, David K. Jackson, Ian Johnston, Dominic Kwiatkowski, Cordelia Langford, John Sillitoe on behalf of the Wellcome Sanger Institute COVID-19 Surveillance Team |
| EPI_ISL_3285313 | Lighthouse Lab in Glasgow | Wellcome Sanger Institute for the COVID-19 Genomics UK (COG-UK) Consortium | Harper VanSteenhouse, Yumi Kasai, David Gray, Carol Clugston, Anna Dominiczak and Alex Alderton, Roberto Amato, Jeffrey Barrett, Sonia Goncalves, Ewan Harrison, David K. Jackson, Ian Johnston, Dominic Kwiatkowski, Cordelia Langford, John Sillitoe on behalf of the Wellcome Sanger Institute COVID-19 Surveillance Team |
| EPI_ISL_3285314, EPI_ISL_3285315, EPI_ISL_3285316 | Lighthouse Lab in Alderley Park | Wellcome Sanger Institute for the COVID-19 Genomics UK (COG-UK) Consortium | Jacquelyn Wynn, Mairead Hyland, The Lighthouse Lab in Alderley Park and Alex Alderton, Roberto Amato, Jeffrey Barrett, Sonia Goncalves, Ewan Harrison, David K. Jackson, Ian Johnston, Dominic Kwiatkowski, Cordelia Langford, John Sillitoe on behalf of the Wellcome Sanger Institute COVID-19 Surveillance Team |
| EPI_ISL_3285317 | Lighthouse Lab in Glasgow | Wellcome Sanger Institute for the COVID-19 Genomics UK (COG-UK) Consortium | Harper VanSteenhouse, Yumi Kasai, David Gray, Carol Clugston, Anna Dominiczak and Alex Alderton, Roberto Amato, Jeffrey Barrett, Sonia Goncalves, Ewan Harrison, David K. Jackson, Ian Johnston, Dominic Kwiatkowski, Cordelia Langford, John Sillitoe on behalf of the Wellcome Sanger Institute COVID-19 Surveillance Team |
| EPI_ISL_3285318, EPI_ISL_3285319, EPI_ISL_3285320 | Berkshire and Surrey Pathology Services Lighthouse Laboratory | Wellcome Sanger Institute for the COVID-19 Genomics UK (COG-UK) Consortium | Berkshire and Surrey Pathology Services Lighthouse Laboratory and Alex Alderton, Roberto Amato, Jeffrey Barrett, Sonia Goncalves, Ewan Harrison, David K. Jackson, Ian Johnston, Dominic Kwiatkowski, Cordelia Langford, John Sillitoe on behalf of the Wellcome Sanger Institute COVID-19 Surveillance Team |
| EPI_ISL_3285321 | Lighthouse Lab in Alderley Park | Wellcome Sanger Institute for the COVID-19 Genomics UK (COG-UK) Consortium | Jacquelyn Wynn, Mairead Hyland, The Lighthouse Lab in Alderley Park and Alex Alderton, Roberto Amato, Jeffrey Barrett, Sonia Goncalves, Ewan Harrison, David K. Jackson, Ian Johnston, Dominic Kwiatkowski, Cordelia Langford, John Sillitoe on behalf of the Wellcome Sanger Institute COVID-19 Surveillance Team |
| EPI_ISL_3285322 | Berkshire and Surrey Pathology Services Lighthouse Laboratory | Wellcome Sanger Institute for the COVID-19 Genomics UK (COG-UK) Consortium | Berkshire and Surrey Pathology Services Lighthouse Laboratory and Alex Alderton, Roberto Amato, Jeffrey Barrett, Sonia Goncalves, Ewan Harrison, David K. Jackson, Ian Johnston, Dominic Kwiatkowski, Cordelia Langford, John Sillitoe on behalf of the Wellcome Sanger Institute COVID-19 Surveillance Team |
| EPI_ISL_3285323, EPI_ISL_3285324 | Lighthouse Lab in Milton Keynes | Wellcome Sanger Institute for the COVID-19 Genomics UK (COG-UK) Consortium | The Lighthouse Lab in Milton Keynes and Alex Alderton, Roberto Amato, Jeffrey Barrett, Sonia Goncalves, Ewan Harrison, David K. Jackson, Ian Johnston, Dominic Kwiatkowski, Cordelia Langford, John Sillitoe on behalf of the Wellcome Sanger Institute COVID-19 Surveillance Team |
| EPI_ISL_3285325 | Lighthouse Lab in Glasgow | Wellcome Sanger Institute for the COVID-19 Genomics UK (COG-UK) Consortium | Harper VanSteenhouse, Yumi Kasai, David Gray, Carol Clugston, Anna Dominiczak and Alex Alderton, Roberto Amato, Jeffrey Barrett, Sonia Goncalves, Ewan Harrison, David K. Jackson, Ian Johnston, Dominic Kwiatkowski, Cordelia Langford, John Sillitoe on behalf of the Wellcome Sanger Institute COVID-19 Surveillance Team |
| EPI_ISL_3285326, EPI_ISL_3285327 | Lighthouse Lab in Milton Keynes | Wellcome Sanger Institute for the COVID-19 Genomics UK (COG-UK) Consortium | The Lighthouse Lab in Milton Keynes and Alex Alderton, Roberto Amato, Jeffrey Barrett, Sonia Goncalves, Ewan Harrison, David K. Jackson, Ian Johnston, Dominic Kwiatkowski, Cordelia Langford, John Sillitoe on behalf of the Wellcome Sanger Institute COVID-19 Surveillance Team |
| EPI_ISL_3285328 | Lighthouse Lab in Glasgow | Wellcome Sanger Institute for the COVID-19 Genomics UK (COG-UK) Consortium | Harper VanSteenhouse, Yumi Kasai, David Gray, Carol Clugston, Anna Dominiczak and Alex Alderton, Roberto Amato, Jeffrey Barrett, Sonia Goncalves, Ewan Harrison, David K. Jackson, Ian Johnston, Dominic Kwiatkowski, Cordelia Langford, John Sillitoe on behalf of the Wellcome Sanger Institute COVID-19 Surveillance Team |
| EPI_ISL_3285329 | Berkshire and Surrey Pathology Services Lighthouse Laboratory | Wellcome Sanger Institute for the COVID-19 Genomics UK (COG-UK) Consortium | Berkshire and Surrey Pathology Services Lighthouse Laboratory and Alex Alderton, Roberto Amato, Jeffrey Barrett, Sonia Goncalves, Ewan Harrison, David K. Jackson, Ian Johnston, Dominic Kwiatkowski, Cordelia Langford, John Sillitoe on behalf of the Wellcome Sanger Institute COVID-19 Surveillance Team |
| EPI_ISL_3285330 | Lighthouse Lab in Alderley Park | Wellcome Sanger Institute for the COVID-19 Genomics UK (COG-UK) Consortium | Jacquelyn Wynn, Mairead Hyland, The Lighthouse Lab in Alderley Park and Alex Alderton, Roberto Amato, Jeffrey Barrett, Sonia Goncalves, Ewan Harrison, David K. Jackson, Ian Johnston, Dominic Kwiatkowski, Cordelia Langford, John Sillitoe on behalf of the Wellcome Sanger Institute COVID-19 Surveillance Team |
| EPI_ISL_3285331 | Berkshire and Surrey Pathology Services Lighthouse Laboratory | Wellcome Sanger Institute for the COVID-19 Genomics UK (COG-UK) Consortium | Berkshire and Surrey Pathology Services Lighthouse Laboratory and Alex Alderton, Roberto Amato, Jeffrey Barrett, Sonia Goncalves, Ewan Harrison, David K. Jackson, Ian Johnston, Dominic Kwiatkowski, Cordelia Langford, John Sillitoe on behalf of the Wellcome Sanger Institute COVID-19 Surveillance Team |
| EPI_ISL_3285332 | Lighthouse Lab in Glasgow | Wellcome Sanger Institute for the COVID-19 Genomics UK (COG-UK) Consortium | Harper VanSteenhouse, Yumi Kasai, David Gray, Carol Clugston, Anna Dominiczak and Alex Alderton, Roberto Amato, Jeffrey Barrett, Sonia Goncalves, Ewan Harrison, David K. Jackson, Ian Johnston, Dominic Kwiatkowski, Cordelia Langford, John Sillitoe on behalf of the Wellcome Sanger Institute COVID-19 Surveillance Team |
| EPI_ISL_3285333 | Berkshire and Surrey Pathology Services Lighthouse Laboratory | Wellcome Sanger Institute for the COVID-19 Genomics UK (COG-UK) Consortium | Berkshire and Surrey Pathology Services Lighthouse Laboratory and Alex Alderton, Roberto Amato, Jeffrey Barrett, Sonia Goncalves, Ewan Harrison, David K. Jackson, Ian Johnston, Dominic Kwiatkowski, Cordelia Langford, John Sillitoe on behalf of the Wellcome Sanger Institute COVID-19 Surveillance Team |
| EPI_ISL_3285334 | Lighthouse Lab in Milton Keynes | Wellcome Sanger Institute for the COVID-19 Genomics UK (COG-UK) Consortium | The Lighthouse Lab in Milton Keynes and Alex Alderton, Roberto Amato, Jeffrey Barrett, Sonia Goncalves, Ewan Harrison, David K. Jackson, Ian Johnston, Dominic Kwiatkowski, Cordelia Langford, John Sillitoe on behalf of the Wellcome Sanger Institute COVID-19 Surveillance Team |
| EPI_ISL_3285335 | Lighthouse Lab in Alderley Park | Wellcome Sanger Institute for the COVID-19 Genomics UK (COG-UK) Consortium | Jacquelyn Wynn, Mairead Hyland, The Lighthouse Lab in Alderley Park and Alex Alderton, Roberto Amato, Jeffrey Barrett, Sonia Goncalves, Ewan Harrison, David K. Jackson, Ian Johnston, Dominic Kwiatkowski, Cordelia Langford, John Sillitoe on behalf of the Wellcome Sanger Institute COVID-19 Surveillance Team |
| EPI_ISL_3285336 | Lighthouse Lab in Glasgow | Wellcome Sanger Institute for the COVID-19 Genomics UK (COG-UK) Consortium | Harper VanSteenhouse, Yumi Kasai, David Gray, Carol Clugston, Anna Dominiczak and Alex Alderton, Roberto Amato, Jeffrey Barrett, Sonia Goncalves, Ewan Harrison, David K. Jackson, Ian Johnston, Dominic Kwiatkowski, Cordelia Langford, John Sillitoe on behalf of the Wellcome Sanger Institute COVID-19 Surveillance Team |
| EPI_ISL_3285337, EPI_ISL_3285338 | Lighthouse Lab in Alderley Park | Wellcome Sanger Institute for the COVID-19 Genomics UK (COG-UK) Consortium | Jacquelyn Wynn, Mairead Hyland, The Lighthouse Lab in Alderley Park and Alex Alderton, Roberto Amato, Jeffrey Barrett, Sonia Goncalves, Ewan Harrison, David K. Jackson, Ian Johnston, Dominic Kwiatkowski, Cordelia Langford, John Sillitoe on behalf of the Wellcome Sanger Institute COVID-19 Surveillance Team |
| EPI_ISL_3285339 | Berkshire and Surrey Pathology Services Lighthouse Laboratory | Wellcome Sanger Institute for the COVID-19 Genomics UK (COG-UK) Consortium | Berkshire and Surrey Pathology Services Lighthouse Laboratory and Alex Alderton, Roberto Amato, Jeffrey Barrett, Sonia Goncalves, Ewan Harrison, David K. Jackson, Ian Johnston, Dominic Kwiatkowski, Cordelia Langford, John Sillitoe on behalf of the Wellcome Sanger Institute COVID-19 Surveillance Team |
| EPI_ISL_3285340, EPI_ISL_3285341 | Lighthouse Lab in Milton Keynes | Wellcome Sanger Institute for the COVID-19 Genomics UK (COG-UK) Consortium | The Lighthouse Lab in Milton Keynes and Alex Alderton, Roberto Amato, Jeffrey Barrett, Sonia Goncalves, Ewan Harrison, David K. Jackson, Ian Johnston, Dominic Kwiatkowski, Cordelia Langford, John Sillitoe on behalf of the Wellcome Sanger Institute COVID-19 Surveillance Team |
| EPI_ISL_3285342 | Berkshire and Surrey Pathology Services Lighthouse Laboratory | Wellcome Sanger Institute for the COVID-19 Genomics UK (COG-UK) Consortium | Berkshire and Surrey Pathology Services Lighthouse Laboratory and Alex Alderton, Roberto Amato, Jeffrey Barrett, Sonia Goncalves, Ewan Harrison, David K. Jackson, Ian Johnston, Dominic Kwiatkowski, Cordelia Langford, John Sillitoe on behalf of the Wellcome Sanger Institute COVID-19 Surveillance Team |
| EPI_ISL_3285343 | Lighthouse Lab in Glasgow | Wellcome Sanger Institute for the COVID-19 Genomics UK (COG-UK) Consortium | Harper VanSteenhouse, Yumi Kasai, David Gray, Carol Clugston, Anna Dominiczak and Alex Alderton, Roberto Amato, Jeffrey Barrett, Sonia Goncalves, Ewan Harrison, David K. Jackson, Ian Johnston, Dominic Kwiatkowski, Cordelia Langford, John Sillitoe on behalf of the Wellcome Sanger Institute COVID-19 Surveillance Team |
| EPI_ISL_3285344, EPI_ISL_3285345 | Lighthouse Lab in Milton Keynes | Wellcome Sanger Institute for the COVID-19 Genomics UK (COG-UK) Consortium | The Lighthouse Lab in Milton Keynes and Alex Alderton, Roberto Amato, Jeffrey Barrett, Sonia Goncalves, Ewan Harrison, David K. Jackson, Ian Johnston, Dominic Kwiatkowski, Cordelia Langford, John Sillitoe on behalf of the Wellcome Sanger Institute COVID-19 Surveillance Team |
| EPI_ISL_3285346 | Berkshire and Surrey Pathology Services Lighthouse Laboratory | Wellcome Sanger Institute for the COVID-19 Genomics UK (COG-UK) Consortium | Berkshire and Surrey Pathology Services Lighthouse Laboratory and Alex Alderton, Roberto Amato, Jeffrey Barrett, Sonia Goncalves, Ewan Harrison, David K. Jackson, Ian Johnston, Dominic Kwiatkowski, Cordelia Langford, John Sillitoe on behalf of the Wellcome Sanger Institute COVID-19 Surveillance Team |
| EPI_ISL_3285347 | Lighthouse Lab in Glasgow | Wellcome Sanger Institute for the COVID-19 Genomics UK (COG-UK) Consortium | Harper VanSteenhouse, Yumi Kasai, David Gray, Carol Clugston, Anna Dominiczak and Alex Alderton, Roberto Amato, Jeffrey Barrett, Sonia Goncalves, Ewan Harrison, David K. Jackson, Ian Johnston, Dominic Kwiatkowski, Cordelia Langford, John Sillitoe on behalf of the Wellcome Sanger Institute COVID-19 Surveillance Team |
| EPI_ISL_3285348 | Lighthouse Lab in Alderley Park | Wellcome Sanger Institute for the COVID-19 Genomics UK (COG-UK) Consortium | Jacquelyn Wynn, Mairead Hyland, The Lighthouse Lab in Alderley Park and Alex Alderton, Roberto Amato, Jeffrey Barrett, Sonia Goncalves, Ewan Harrison, David K. Jackson, Ian Johnston, Dominic Kwiatkowski, Cordelia Langford, John Sillitoe on behalf of the Wellcome Sanger Institute COVID-19 Surveillance Team |
| EPI_ISL_3285349, EPI_ISL_3285350 | Lighthouse Lab in Milton Keynes | Wellcome Sanger Institute for the COVID-19 Genomics UK (COG-UK) Consortium | The Lighthouse Lab in Milton Keynes and Alex Alderton, Roberto Amato, Jeffrey Barrett, Sonia Goncalves, Ewan Harrison, David K. Jackson, Ian Johnston, Dominic Kwiatkowski, Cordelia Langford, John Sillitoe on behalf of the Wellcome Sanger Institute COVID-19 Surveillance Team |
| EPI_ISL_3285351 | Lighthouse Lab in Glasgow | Wellcome Sanger Institute for the COVID-19 Genomics UK (COG-UK) Consortium | Harper VanSteenhouse, Yumi Kasai, David Gray, Carol Clugston, Anna Dominiczak and Alex Alderton, Roberto Amato, Jeffrey Barrett, Sonia Goncalves, Ewan Harrison, David K. Jackson, Ian Johnston, Dominic Kwiatkowski, Cordelia Langford, John Sillitoe on behalf of the Wellcome Sanger Institute COVID-19 Surveillance Team |
| EPI_ISL_3285352 | Berkshire and Surrey Pathology Services Lighthouse Laboratory | Wellcome Sanger Institute for the COVID-19 Genomics UK (COG-UK) Consortium | Berkshire and Surrey Pathology Services Lighthouse Laboratory and Alex Alderton, Roberto Amato, Jeffrey Barrett, Sonia Goncalves, Ewan Harrison, David K. Jackson, Ian Johnston, Dominic Kwiatkowski, Cordelia Langford, John Sillitoe on behalf of the Wellcome Sanger Institute COVID-19 Surveillance Team |
| EPI_ISL_3285353 | Lighthouse Lab in Alderley Park | Wellcome Sanger Institute for the COVID-19 Genomics UK (COG-UK) Consortium | Jacquelyn Wynn, Mairead Hyland, The Lighthouse Lab in Alderley Park and Alex Alderton, Roberto Amato, Jeffrey Barrett, Sonia Goncalves, Ewan Harrison, David K. Jackson, Ian Johnston, Dominic Kwiatkowski, Cordelia Langford, John Sillitoe on behalf of the Wellcome Sanger Institute COVID-19 Surveillance Team |
| EPI_ISL_3285354, EPI_ISL_3285355 | Lighthouse Lab in Milton Keynes | Wellcome Sanger Institute for the COVID-19 Genomics UK (COG-UK) Consortium | The Lighthouse Lab in Milton Keynes and Alex Alderton, Roberto Amato, Jeffrey Barrett, Sonia Goncalves, Ewan Harrison, David K. Jackson, Ian Johnston, Dominic Kwiatkowski, Cordelia Langford, John Sillitoe on behalf of the Wellcome Sanger Institute COVID-19 Surveillance Team |
| EPI_ISL_3285356 | Lighthouse Lab in Alderley Park | Wellcome Sanger Institute for the COVID-19 Genomics UK (COG-UK) Consortium | Jacquelyn Wynn, Mairead Hyland, The Lighthouse Lab in Alderley Park and Alex Alderton, Roberto Amato, Jeffrey Barrett, Sonia Goncalves, Ewan Harrison, David K. Jackson, Ian Johnston, Dominic Kwiatkowski, Cordelia Langford, John Sillitoe on behalf of the Wellcome Sanger Institute COVID-19 Surveillance Team |
| EPI_ISL_3285357 | Lighthouse Lab in Milton Keynes | Wellcome Sanger Institute for the COVID-19 Genomics UK (COG-UK) Consortium | The Lighthouse Lab in Milton Keynes and Alex Alderton, Roberto Amato, Jeffrey Barrett, Sonia Goncalves, Ewan Harrison, David K. Jackson, Ian Johnston, Dominic Kwiatkowski, Cordelia Langford, John Sillitoe on behalf of the Wellcome Sanger Institute COVID-19 Surveillance Team |
| EPI_ISL_3285358 | Lighthouse Lab in Glasgow | Wellcome Sanger Institute for the COVID-19 Genomics UK (COG-UK) Consortium | Harper VanSteenhouse, Yumi Kasai, David Gray, Carol Clugston, Anna Dominiczak and Alex Alderton, Roberto Amato, Jeffrey Barrett, Sonia Goncalves, Ewan Harrison, David K. Jackson, Ian Johnston, Dominic Kwiatkowski, Cordelia Langford, John Sillitoe on behalf of the Wellcome Sanger Institute COVID-19 Surveillance Team |
| EPI_ISL_3285359, EPI_ISL_3285360 | Lighthouse Lab in Alderley Park | Wellcome Sanger Institute for the COVID-19 Genomics UK (COG-UK) Consortium | Jacquelyn Wynn, Mairead Hyland, The Lighthouse Lab in Alderley Park and Alex Alderton, Roberto Amato, Jeffrey Barrett, Sonia Goncalves, Ewan Harrison, David K. Jackson, Ian Johnston, Dominic Kwiatkowski, Cordelia Langford, John Sillitoe on behalf of the Wellcome Sanger Institute COVID-19 Surveillance Team |
| EPI_ISL_3285361, EPI_ISL_3285362 | Lighthouse Lab in Milton Keynes | Wellcome Sanger Institute for the COVID-19 Genomics UK (COG-UK) Consortium | The Lighthouse Lab in Milton Keynes and Alex Alderton, Roberto Amato, Jeffrey Barrett, Sonia Goncalves, Ewan Harrison, David K. Jackson, Ian Johnston, Dominic Kwiatkowski, Cordelia Langford, John Sillitoe on behalf of the Wellcome Sanger Institute COVID-19 Surveillance Team |
| EPI_ISL_3285363 | Lighthouse Lab in Glasgow | Wellcome Sanger Institute for the COVID-19 Genomics UK (COG-UK) Consortium | Harper VanSteenhouse, Yumi Kasai, David Gray, Carol Clugston, Anna Dominiczak and Alex Alderton, Roberto Amato, Jeffrey Barrett, Sonia Goncalves, Ewan Harrison, David K. Jackson, Ian Johnston, Dominic Kwiatkowski, Cordelia Langford, John Sillitoe on behalf of the Wellcome Sanger Institute COVID-19 Surveillance Team |
| EPI_ISL_3285364 | Lighthouse Lab in Alderley Park | Wellcome Sanger Institute for the COVID-19 Genomics UK (COG-UK) Consortium | Jacquelyn Wynn, Mairead Hyland, The Lighthouse Lab in Alderley Park and Alex Alderton, Roberto Amato, Jeffrey Barrett, Sonia Goncalves, Ewan Harrison, David K. Jackson, Ian Johnston, Dominic Kwiatkowski, Cordelia Langford, John Sillitoe on behalf of the Wellcome Sanger Institute COVID-19 Surveillance Team |

| | | | |
|---|---|---|---|
| EPI_ISL_3285365 | Lighthouse Lab in Glasgow | Wellcome Sanger Institute for the COVID-19 Genomics UK (COG-UK) Consortium | Harper VanSteenhouse, Yumi Kasai, David Gray, Carol Clugston, Anna Dominiczak and Alex Alderton, Roberto Amato, Jeffrey Barrett, Sonia Goncalves, Ewan Harrison, David K. Jackson, Ian Johnston, Dominic Kwiatkowski, Cordelia Langford, John Sillitoe on behalf of the Wellcome Sanger Institute COVID-19 Surveillance Team |
| EPI_ISL_3285366 | Lighthouse Lab in Milton Keynes | Wellcome Sanger Institute for the COVID-19 Genomics UK (COG-UK) Consortium | The Lighthouse Lab in Milton Keynes and Alex Alderton, Roberto Amato, Jeffrey Barrett, Sonia Goncalves, Ewan Harrison, David K. Jackson, Ian Johnston, Dominic Kwiatkowski, Cordelia Langford, John Sillitoe on behalf of the Wellcome Sanger Institute COVID-19 Surveillance Team |
| EPI_ISL_3285367, EPI_ISL_3285368 | Lighthouse Lab in Alderley Park | Wellcome Sanger Institute for the COVID-19 Genomics UK (COG-UK) Consortium | Jacquelyn Wynn, Mairead Hyland, The Lighthouse Lab in Alderley Park and Alex Alderton, Roberto Amato, Jeffrey Barrett, Sonia Goncalves, Ewan Harrison, David K. Jackson, Ian Johnston, Dominic Kwiatkowski, Cordelia Langford, John Sillitoe on behalf of the Wellcome Sanger Institute COVID-19 Surveillance Team |
| EPI_ISL_3285369 | Berkshire and Surrey Pathology Services Lighthouse Laboratory | Wellcome Sanger Institute for the COVID-19 Genomics UK (COG-UK) Consortium | Berkshire and Surrey Pathology Services Lighthouse Laboratory and Alex Alderton, Roberto Amato, Jeffrey Barrett, Sonia Goncalves, Ewan Harrison, David K. Jackson, Ian Johnston, Dominic Kwiatkowski, Cordelia Langford, John Sillitoe on behalf of the Wellcome Sanger Institute COVID-19 Surveillance Team |
| EPI_ISL_3285370 | Lighthouse Lab in Milton Keynes | Wellcome Sanger Institute for the COVID-19 Genomics UK (COG-UK) Consortium | The Lighthouse Lab in Milton Keynes and Alex Alderton, Roberto Amato, Jeffrey Barrett, Sonia Goncalves, Ewan Harrison, David K. Jackson, Ian Johnston, Dominic Kwiatkowski, Cordelia Langford, John Sillitoe on behalf of the Wellcome Sanger Institute COVID-19 Surveillance Team |
| EPI_ISL_3285371, EPI_ISL_3285372 | Lighthouse Lab in Glasgow | Wellcome Sanger Institute for the COVID-19 Genomics UK (COG-UK) Consortium | Harper VanSteenhouse, Yumi Kasai, David Gray, Carol Clugston, Anna Dominiczak and Alex Alderton, Roberto Amato, Jeffrey Barrett, Sonia Goncalves, Ewan Harrison, David K. Jackson, Ian Johnston, Dominic Kwiatkowski, Cordelia Langford, John Sillitoe on behalf of the Wellcome Sanger Institute COVID-19 Surveillance Team |
| EPI_ISL_3285373, EPI_ISL_3285374, EPI_ISL_3285375 | Berkshire and Surrey Pathology Services Lighthouse Laboratory | Wellcome Sanger Institute for the COVID-19 Genomics UK (COG-UK) Consortium | Berkshire and Surrey Pathology Services Lighthouse Laboratory and Alex Alderton, Roberto Amato, Jeffrey Barrett, Sonia Goncalves, Ewan Harrison, David K. Jackson, Ian Johnston, Dominic Kwiatkowski, Cordelia Langford, John Sillitoe on behalf of the Wellcome Sanger Institute COVID-19 Surveillance Team |
| EPI_ISL_3285376, EPI_ISL_3285377 | Lighthouse Lab in Milton Keynes | Wellcome Sanger Institute for the COVID-19 Genomics UK (COG-UK) Consortium | The Lighthouse Lab in Milton Keynes and Alex Alderton, Roberto Amato, Jeffrey Barrett, Sonia Goncalves, Ewan Harrison, David K. Jackson, Ian Johnston, Dominic Kwiatkowski, Cordelia Langford, John Sillitoe on behalf of the Wellcome Sanger Institute COVID-19 Surveillance Team |
| EPI_ISL_3285378 | Lighthouse Lab in Alderley Park | Wellcome Sanger Institute for the COVID-19 Genomics UK (COG-UK) Consortium | Jacquelyn Wynn, Mairead Hyland, The Lighthouse Lab in Alderley Park and Alex Alderton, Roberto Amato, Jeffrey Barrett, Sonia Goncalves, Ewan Harrison, David K. Jackson, Ian Johnston, Dominic Kwiatkowski, Cordelia Langford, John Sillitoe on behalf of the Wellcome Sanger Institute COVID-19 Surveillance Team |
| EPI_ISL_3285379 | Lighthouse Lab in Glasgow | Wellcome Sanger Institute for the COVID-19 Genomics UK (COG-UK) Consortium | Harper VanSteenhouse, Yumi Kasai, David Gray, Carol Clugston, Anna Dominiczak and Alex Alderton, Roberto Amato, Jeffrey Barrett, Sonia Goncalves, Ewan Harrison, David K. Jackson, Ian Johnston, Dominic Kwiatkowski, Cordelia Langford, John Sillitoe on behalf of the Wellcome Sanger Institute COVID-19 Surveillance Team |
| EPI_ISL_3285380 | Lighthouse Lab in Milton Keynes | Wellcome Sanger Institute for the COVID-19 Genomics UK (COG-UK) Consortium | The Lighthouse Lab in Milton Keynes and Alex Alderton, Roberto Amato, Jeffrey Barrett, Sonia Goncalves, Ewan Harrison, David K. Jackson, Ian Johnston, Dominic Kwiatkowski, Cordelia Langford, John Sillitoe on behalf of the Wellcome Sanger Institute COVID-19 Surveillance Team |
| EPI_ISL_3285381 | Lighthouse Lab in Glasgow | Wellcome Sanger Institute for the COVID-19 Genomics UK (COG-UK) Consortium | Harper VanSteenhouse, Yumi Kasai, David Gray, Carol Clugston, Anna Dominiczak and Alex Alderton, Roberto Amato, Jeffrey Barrett, Sonia Goncalves, Ewan Harrison, David K. Jackson, Ian Johnston, Dominic Kwiatkowski, Cordelia Langford, John Sillitoe on behalf of the Wellcome Sanger Institute COVID-19 Surveillance Team |
| EPI_ISL_3285382 | Lighthouse Lab in Alderley Park | Wellcome Sanger Institute for the COVID-19 Genomics UK (COG-UK) Consortium | Jacquelyn Wynn, Mairead Hyland, The Lighthouse Lab in Alderley Park and Alex Alderton, Roberto Amato, Jeffrey Barrett, Sonia Goncalves, Ewan Harrison, David K. Jackson, Ian Johnston, Dominic Kwiatkowski, Cordelia Langford, John Sillitoe on behalf of the Wellcome Sanger Institute COVID-19 Surveillance Team |
| EPI_ISL_3285383 | Lighthouse Lab in Milton Keynes | Wellcome Sanger Institute for the COVID-19 Genomics UK (COG-UK) Consortium | The Lighthouse Lab in Milton Keynes and Alex Alderton, Roberto Amato, Jeffrey Barrett, Sonia Goncalves, Ewan Harrison, David K. Jackson, Ian Johnston, Dominic Kwiatkowski, Cordelia Langford, John Sillitoe on behalf of the Wellcome Sanger Institute COVID-19 Surveillance Team |
| EPI_ISL_3285384, EPI_ISL_3285385, EPI_ISL_3285386, EPI_ISL_3285387, EPI_ISL_3285388, EPI_ISL_3285389 | Lighthouse Lab in Alderley Park | Wellcome Sanger Institute for the COVID-19 Genomics UK (COG-UK) Consortium | Jacquelyn Wynn, Mairead Hyland, The Lighthouse Lab in Alderley Park and Alex Alderton, Roberto Amato, Jeffrey Barrett, Sonia Goncalves, Ewan Harrison, David K. Jackson, Ian Johnston, Dominic Kwiatkowski, Cordelia Langford, John Sillitoe on behalf of the Wellcome Sanger Institute COVID-19 Surveillance Team |
| EPI_ISL_3285390 | Berkshire and Surrey Pathology Services Lighthouse Laboratory | Wellcome Sanger Institute for the COVID-19 Genomics UK (COG-UK) Consortium | Berkshire and Surrey Pathology Services Lighthouse Laboratory and Alex Alderton, Roberto Amato, Jeffrey Barrett, Sonia Goncalves, Ewan Harrison, David K. Jackson, Ian Johnston, Dominic Kwiatkowski, Cordelia Langford, John Sillitoe on behalf of the Wellcome Sanger Institute COVID-19 Surveillance Team |
| EPI_ISL_3285391, EPI_ISL_3285392 | Lighthouse Lab in Alderley Park | Wellcome Sanger Institute for the COVID-19 Genomics UK (COG-UK) Consortium | Jacquelyn Wynn, Mairead Hyland, The Lighthouse Lab in Alderley Park and Alex Alderton, Roberto Amato, Jeffrey Barrett, Sonia Goncalves, Ewan Harrison, David K. Jackson, Ian Johnston, Dominic Kwiatkowski, Cordelia Langford, John Sillitoe on behalf of the Wellcome Sanger Institute COVID-19 Surveillance Team |
| EPI_ISL_3285393 | Lighthouse Lab in Glasgow | Wellcome Sanger Institute for the COVID-19 Genomics UK (COG-UK) Consortium | Harper VanSteenhouse, Yumi Kasai, David Gray, Carol Clugston, Anna Dominiczak and Alex Alderton, Roberto Amato, Jeffrey Barrett, Sonia Goncalves, Ewan Harrison, David K. Jackson, Ian Johnston, Dominic Kwiatkowski, Cordelia Langford, John Sillitoe on behalf of the Wellcome Sanger Institute COVID-19 Surveillance Team |
| EPI_ISL_3285394 | Berkshire and Surrey Pathology Services Lighthouse Laboratory | Wellcome Sanger Institute for the COVID-19 Genomics UK (COG-UK) Consortium | Berkshire and Surrey Pathology Services Lighthouse Laboratory and Alex Alderton, Roberto Amato, Jeffrey Barrett, Sonia Goncalves, Ewan Harrison, David K. Jackson, Ian Johnston, Dominic Kwiatkowski, Cordelia Langford, John Sillitoe on behalf of the Wellcome Sanger Institute COVID-19 Surveillance Team |
| EPI_ISL_3285395 | Lighthouse Lab in Glasgow | Wellcome Sanger Institute for the COVID-19 Genomics UK (COG-UK) Consortium | Harper VanSteenhouse, Yumi Kasai, David Gray, Carol Clugston, Anna Dominiczak and Alex Alderton, Roberto Amato, Jeffrey Barrett, Sonia Goncalves, Ewan Harrison, David K. Jackson, Ian Johnston, Dominic Kwiatkowski, Cordelia Langford, John Sillitoe on behalf of the Wellcome Sanger Institute COVID-19 Surveillance Team |
| EPI_ISL_3285396, EPI_ISL_3285397 | Lighthouse Lab in Milton Keynes | Wellcome Sanger Institute for the COVID-19 Genomics UK (COG-UK) Consortium | The Lighthouse Lab in Milton Keynes and Alex Alderton, Roberto Amato, Jeffrey Barrett, Sonia Goncalves, Ewan Harrison, David K. Jackson, Ian Johnston, Dominic Kwiatkowski, Cordelia Langford, John Sillitoe on behalf of the Wellcome Sanger Institute COVID-19 Surveillance Team |
| EPI_ISL_3285398, EPI_ISL_3285399 | Lighthouse Lab in Alderley Park | Wellcome Sanger Institute for the COVID-19 Genomics UK (COG-UK) Consortium | Jacquelyn Wynn, Mairead Hyland, The Lighthouse Lab in Alderley Park and Alex Alderton, Roberto Amato, Jeffrey Barrett, Sonia Goncalves, Ewan Harrison, David K. Jackson, Ian Johnston, Dominic Kwiatkowski, Cordelia Langford, John Sillitoe on behalf of the Wellcome Sanger Institute COVID-19 Surveillance Team |
| EPI_ISL_3285400 | Lighthouse Lab in Glasgow | Wellcome Sanger Institute for the COVID-19 Genomics UK (COG-UK) Consortium | Harper VanSteenhouse, Yumi Kasai, David Gray, Carol Clugston, Anna Dominiczak and Alex Alderton, Roberto Amato, Jeffrey Barrett, Sonia Goncalves, Ewan Harrison, David K. Jackson, Ian Johnston, Dominic Kwiatkowski, Cordelia Langford, John Sillitoe on behalf of the Wellcome Sanger Institute COVID-19 Surveillance Team |
| EPI_ISL_3285401 | Lighthouse Lab in Milton Keynes | Wellcome Sanger Institute for the COVID-19 Genomics UK (COG-UK) Consortium | The Lighthouse Lab in Milton Keynes and Alex Alderton, Roberto Amato, Jeffrey Barrett, Sonia Goncalves, Ewan Harrison, David K. Jackson, Ian Johnston, Dominic Kwiatkowski, Cordelia Langford, John Sillitoe on behalf of the Wellcome Sanger Institute COVID-19 Surveillance Team |
| EPI_ISL_3285402 | Lighthouse Lab in Alderley Park | Wellcome Sanger Institute for the COVID-19 Genomics UK (COG-UK) Consortium | Jacquelyn Wynn, Mairead Hyland, The Lighthouse Lab in Alderley Park and Alex Alderton, Roberto Amato, Jeffrey Barrett, Sonia Goncalves, Ewan Harrison, David K. Jackson, Ian Johnston, Dominic Kwiatkowski, Cordelia Langford, John Sillitoe on behalf of the Wellcome Sanger Institute COVID-19 Surveillance Team |
| EPI_ISL_3285403 | Lighthouse Lab in Glasgow | Wellcome Sanger Institute for the COVID-19 Genomics UK (COG-UK) Consortium | Harper VanSteenhouse, Yumi Kasai, David Gray, Carol Clugston, Anna Dominiczak and Alex Alderton, Roberto Amato, Jeffrey Barrett, Sonia Goncalves, Ewan Harrison, David K. Jackson, Ian Johnston, Dominic Kwiatkowski, Cordelia Langford, John Sillitoe on behalf of the Wellcome Sanger Institute COVID-19 Surveillance Team |
| EPI_ISL_3285404 | Berkshire and Surrey Pathology Services Lighthouse Laboratory | Wellcome Sanger Institute for the COVID-19 Genomics UK (COG-UK) Consortium | Berkshire and Surrey Pathology Services Lighthouse Laboratory and Alex Alderton, Roberto Amato, Jeffrey Barrett, Sonia Goncalves, Ewan Harrison, David K. Jackson, Ian Johnston, Dominic Kwiatkowski, Cordelia Langford, John Sillitoe on behalf of the Wellcome Sanger Institute COVID-19 Surveillance Team |
| EPI_ISL_3285405 | Lighthouse Lab in Milton Keynes | Wellcome Sanger Institute for the COVID-19 Genomics UK (COG-UK) Consortium | The Lighthouse Lab in Milton Keynes and Alex Alderton, Roberto Amato, Jeffrey Barrett, Sonia Goncalves, Ewan Harrison, David K. Jackson, Ian Johnston, Dominic Kwiatkowski, Cordelia Langford, John Sillitoe on behalf of the Wellcome Sanger Institute COVID-19 Surveillance Team |
| EPI_ISL_3285406 | Lighthouse Lab in Alderley Park | Wellcome Sanger Institute for the COVID-19 Genomics UK (COG-UK) Consortium | Jacquelyn Wynn, Mairead Hyland, The Lighthouse Lab in Alderley Park and Alex Alderton, Roberto Amato, Jeffrey Barrett, Sonia Goncalves, Ewan Harrison, David K. Jackson, Ian Johnston, Dominic Kwiatkowski, Cordelia Langford, John Sillitoe on behalf of the Wellcome Sanger Institute COVID-19 Surveillance Team |
| EPI_ISL_3285407, EPI_ISL_3285408 | Berkshire and Surrey Pathology Services Lighthouse Laboratory | Wellcome Sanger Institute for the COVID-19 Genomics UK (COG-UK) Consortium | Berkshire and Surrey Pathology Services Lighthouse Laboratory and Alex Alderton, Roberto Amato, Jeffrey Barrett, Sonia Goncalves, Ewan Harrison, David K. Jackson, Ian Johnston, Dominic Kwiatkowski, Cordelia Langford, John Sillitoe on behalf of the Wellcome Sanger Institute COVID-19 Surveillance Team |
| EPI_ISL_3285409 | Lighthouse Lab in Glasgow | Wellcome Sanger Institute for the COVID-19 Genomics UK (COG-UK) Consortium | Harper VanSteenhouse, Yumi Kasai, David Gray, Carol Clugston, Anna Dominiczak and Alex Alderton, Roberto Amato, Jeffrey Barrett, Sonia Goncalves, Ewan Harrison, David K. Jackson, Ian Johnston, Dominic Kwiatkowski, Cordelia Langford, John Sillitoe on behalf of the Wellcome Sanger Institute COVID-19 Surveillance Team |
| EPI_ISL_3285410 | Lighthouse Lab in Alderley Park | Wellcome Sanger Institute for the COVID-19 Genomics UK (COG-UK) Consortium | Jacquelyn Wynn, Mairead Hyland, The Lighthouse Lab in Alderley Park and Alex Alderton, Roberto Amato, Jeffrey Barrett, Sonia Goncalves, Ewan Harrison, David K. Jackson, Ian Johnston, Dominic Kwiatkowski, Cordelia Langford, John Sillitoe on behalf of the Wellcome Sanger Institute COVID-19 Surveillance Team |
| EPI_ISL_3285411 | Lighthouse Lab in Glasgow | Wellcome Sanger Institute for the COVID-19 Genomics UK (COG-UK) Consortium | Harper VanSteenhouse, Yumi Kasai, David Gray, Carol Clugston, Anna Dominiczak and Alex Alderton, Roberto Amato, Jeffrey Barrett, Sonia Goncalves, Ewan Harrison, David K. Jackson, Ian Johnston, Dominic Kwiatkowski, Cordelia Langford, John Sillitoe on behalf of the Wellcome Sanger Institute COVID-19 Surveillance Team |
| EPI_ISL_3285412 | Lighthouse Lab in Milton Keynes | Wellcome Sanger Institute for the COVID-19 Genomics UK (COG-UK) Consortium | The Lighthouse Lab in Milton Keynes and Alex Alderton, Roberto Amato, Jeffrey Barrett, Sonia Goncalves, Ewan Harrison, David K. Jackson, Ian Johnston, Dominic Kwiatkowski, Cordelia Langford, John Sillitoe on behalf of the Wellcome Sanger Institute COVID-19 Surveillance Team |
| EPI_ISL_3285413 | Berkshire and Surrey Pathology Services Lighthouse Laboratory | Wellcome Sanger Institute for the COVID-19 Genomics UK (COG-UK) Consortium | Berkshire and Surrey Pathology Services Lighthouse Laboratory and Alex Alderton, Roberto Amato, Jeffrey Barrett, Sonia Goncalves, Ewan Harrison, David K. Jackson, Ian Johnston, Dominic Kwiatkowski, Cordelia Langford, John Sillitoe on behalf of the Wellcome Sanger Institute COVID-19 Surveillance Team |
| EPI_ISL_3285414 | Lighthouse Lab in Alderley Park | Wellcome Sanger Institute for the COVID-19 Genomics UK (COG-UK) Consortium | Jacquelyn Wynn, Mairead Hyland, The Lighthouse Lab in Alderley Park and Alex Alderton, Roberto Amato, Jeffrey Barrett, Sonia Goncalves, Ewan Harrison, David K. Jackson, Ian Johnston, Dominic Kwiatkowski, Cordelia Langford, John Sillitoe on behalf of the Wellcome Sanger Institute COVID-19 Surveillance Team |
| EPI_ISL_3285415, EPI_ISL_3285416 | Lighthouse Lab in Milton Keynes | Wellcome Sanger Institute for the COVID-19 Genomics UK (COG-UK) Consortium | The Lighthouse Lab in Milton Keynes and Alex Alderton, Roberto Amato, Jeffrey Barrett, Sonia Goncalves, Ewan Harrison, David K. Jackson, Ian Johnston, Dominic Kwiatkowski, Cordelia Langford, John Sillitoe on behalf of the Wellcome Sanger Institute COVID-19 Surveillance Team |
| EPI_ISL_3285417 | Lighthouse Lab in Alderley Park | Wellcome Sanger Institute for the COVID-19 Genomics UK (COG-UK) Consortium | Jacquelyn Wynn, Mairead Hyland, The Lighthouse Lab in Alderley Park and Alex Alderton, Roberto Amato, Jeffrey Barrett, Sonia Goncalves, Ewan Harrison, David K. Jackson, Ian Johnston, Dominic Kwiatkowski, Cordelia Langford, John Sillitoe on behalf of the Wellcome Sanger Institute COVID-19 Surveillance Team |
| EPI_ISL_3285418 | Lighthouse Lab in Glasgow | Wellcome Sanger Institute for the COVID-19 Genomics UK (COG-UK) Consortium | Harper VanSteenhouse, Yumi Kasai, David Gray, Carol Clugston, Anna Dominiczak and Alex Alderton, Roberto Amato, Jeffrey Barrett, Sonia Goncalves, Ewan Harrison, David K. Jackson, Ian Johnston, Dominic Kwiatkowski, Cordelia Langford, John Sillitoe on behalf of the Wellcome Sanger Institute COVID-19 Surveillance Team |
| EPI_ISL_3285419 | Lighthouse Lab in Alderley Park | Wellcome Sanger Institute for the COVID-19 Genomics UK (COG-UK) Consortium | Jacquelyn Wynn, Mairead Hyland, The Lighthouse Lab in Alderley Park and Alex Alderton, Roberto Amato, Jeffrey Barrett, Sonia Goncalves, Ewan Harrison, David K. Jackson, Ian Johnston, Dominic Kwiatkowski, Cordelia Langford, John Sillitoe on behalf of the Wellcome Sanger Institute COVID-19 Surveillance Team |
| EPI_ISL_3285420 | Lighthouse Lab in Glasgow | Wellcome Sanger Institute for the COVID-19 Genomics UK (COG-UK) Consortium | Harper VanSteenhouse, Yumi Kasai, David Gray, Carol Clugston, Anna Dominiczak and Alex Alderton, Roberto Amato, Jeffrey Barrett, Sonia Goncalves, Ewan Harrison, David K. Jackson, Ian Johnston, Dominic Kwiatkowski, Cordelia Langford, John Sillitoe on behalf of the Wellcome Sanger Institute COVID-19 Surveillance Team |
| EPI_ISL_3285421 | Lighthouse Lab in Alderley Park | Wellcome Sanger Institute for the COVID-19 Genomics UK (COG-UK) Consortium | Jacquelyn Wynn, Mairead Hyland, The Lighthouse Lab in Alderley Park and Alex Alderton, Roberto Amato, Jeffrey Barrett, Sonia Goncalves, Ewan Harrison, David K. Jackson, Ian Johnston, Dominic Kwiatkowski, Cordelia Langford, John Sillitoe on behalf of the Wellcome Sanger Institute COVID-19 Surveillance Team |
| EPI_ISL_3285422, EPI_ISL_3285423, EPI_ISL_3285424 | Lighthouse Lab in Milton Keynes | Wellcome Sanger Institute for the COVID-19 Genomics UK (COG-UK) Consortium | The Lighthouse Lab in Milton Keynes and Alex Alderton, Roberto Amato, Jeffrey Barrett, Sonia Goncalves, Ewan Harrison, David K. Jackson, Ian Johnston, Dominic Kwiatkowski, Cordelia Langford, John Sillitoe on behalf of the Wellcome Sanger Institute COVID-19 Surveillance Team |

| | | | |
|---|---|---|---|
| EPI_ISL_3285425, EPI_ISL_3285426 | Lighthouse Lab in Alderley Park | Wellcome Sanger Institute for the COVID-19 Genomics UK (COG-UK) Consortium | Jacquelyn Wynn, Mairead Hyland, The Lighthouse Lab in Alderley Park and Alex Alderton, Roberto Amato, Jeffrey Barrett, Sonia Goncalves, Ewan Harrison, David K. Jackson, Ian Johnston, Dominic Kwiatkowski, Cordelia Langford, John Sillitoe on behalf of the Wellcome Sanger Institute COVID-19 Surveillance Team |
| EPI_ISL_3285427 | Lighthouse Lab in Glasgow | Wellcome Sanger Institute for the COVID-19 Genomics UK (COG-UK) Consortium | Harper VanSteenhouse, Yumi Kasai, David Gray, Carol Clugston, Anna Dominiczak and Alex Alderton, Roberto Amato, Jeffrey Barrett, Sonia Goncalves, Ewan Harrison, David K. Jackson, Ian Johnston, Dominic Kwiatkowski, Cordelia Langford, John Sillitoe on behalf of the Wellcome Sanger Institute COVID-19 Surveillance Team |
| EPI_ISL_3285428 | Lighthouse Lab in Milton Keynes | Wellcome Sanger Institute for the COVID-19 Genomics UK (COG-UK) Consortium | The Lighthouse Lab in Milton Keynes and Alex Alderton, Roberto Amato, Jeffrey Barrett, Sonia Goncalves, Ewan Harrison, David K. Jackson, Ian Johnston, Dominic Kwiatkowski, Cordelia Langford, John Sillitoe on behalf of the Wellcome Sanger Institute COVID-19 Surveillance Team |
| EPI_ISL_3285429 | Lighthouse Lab in Glasgow | Wellcome Sanger Institute for the COVID-19 Genomics UK (COG-UK) Consortium | Harper VanSteenhouse, Yumi Kasai, David Gray, Carol Clugston, Anna Dominiczak and Alex Alderton, Roberto Amato, Jeffrey Barrett, Sonia Goncalves, Ewan Harrison, David K. Jackson, Ian Johnston, Dominic Kwiatkowski, Cordelia Langford, John Sillitoe on behalf of the Wellcome Sanger Institute COVID-19 Surveillance Team |
| EPI_ISL_3285430 | Lighthouse Lab in Alderley Park | Wellcome Sanger Institute for the COVID-19 Genomics UK (COG-UK) Consortium | Jacquelyn Wynn, Mairead Hyland, The Lighthouse Lab in Alderley Park and Alex Alderton, Roberto Amato, Jeffrey Barrett, Sonia Goncalves, Ewan Harrison, David K. Jackson, Ian Johnston, Dominic Kwiatkowski, Cordelia Langford, John Sillitoe on behalf of the Wellcome Sanger Institute COVID-19 Surveillance Team |
| EPI_ISL_3285431, EPI_ISL_3285432, EPI_ISL_3285433 | Lighthouse Lab in Glasgow | Wellcome Sanger Institute for the COVID-19 Genomics UK (COG-UK) Consortium | Harper VanSteenhouse, Yumi Kasai, David Gray, Carol Clugston, Anna Dominiczak and Alex Alderton, Roberto Amato, Jeffrey Barrett, Sonia Goncalves, Ewan Harrison, David K. Jackson, Ian Johnston, Dominic Kwiatkowski, Cordelia Langford, John Sillitoe on behalf of the Wellcome Sanger Institute COVID-19 Surveillance Team |
| EPI_ISL_3285434 | Lighthouse Lab in Milton Keynes | Wellcome Sanger Institute for the COVID-19 Genomics UK (COG-UK) Consortium | The Lighthouse Lab in Milton Keynes and Alex Alderton, Roberto Amato, Jeffrey Barrett, Sonia Goncalves, Ewan Harrison, David K. Jackson, Ian Johnston, Dominic Kwiatkowski, Cordelia Langford, John Sillitoe on behalf of the Wellcome Sanger Institute COVID-19 Surveillance Team |
| EPI_ISL_3285435, EPI_ISL_3285436 | Berkshire and Surrey Pathology Services Lighthouse Laboratory | Wellcome Sanger Institute for the COVID-19 Genomics UK (COG-UK) Consortium | Berkshire and Surrey Pathology Services Lighthouse Laboratory and Alex Alderton, Roberto Amato, Jeffrey Barrett, Sonia Goncalves, Ewan Harrison, David K. Jackson, Ian Johnston, Dominic Kwiatkowski, Cordelia Langford, John Sillitoe on behalf of the Wellcome Sanger Institute COVID-19 Surveillance Team |
| EPI_ISL_3285437 | Lighthouse Lab in Milton Keynes | Wellcome Sanger Institute for the COVID-19 Genomics UK (COG-UK) Consortium | The Lighthouse Lab in Milton Keynes and Alex Alderton, Roberto Amato, Jeffrey Barrett, Sonia Goncalves, Ewan Harrison, David K. Jackson, Ian Johnston, Dominic Kwiatkowski, Cordelia Langford, John Sillitoe on behalf of the Wellcome Sanger Institute COVID-19 Surveillance Team |
| EPI_ISL_3285438 | Lighthouse Lab in Glasgow | Wellcome Sanger Institute for the COVID-19 Genomics UK (COG-UK) Consortium | Harper VanSteenhouse, Yumi Kasai, David Gray, Carol Clugston, Anna Dominiczak and Alex Alderton, Roberto Amato, Jeffrey Barrett, Sonia Goncalves, Ewan Harrison, David K. Jackson, Ian Johnston, Dominic Kwiatkowski, Cordelia Langford, John Sillitoe on behalf of the Wellcome Sanger Institute COVID-19 Surveillance Team |
| EPI_ISL_3285439 | Lighthouse Lab in Alderley Park | Wellcome Sanger Institute for the COVID-19 Genomics UK (COG-UK) Consortium | Jacquelyn Wynn, Mairead Hyland, The Lighthouse Lab in Alderley Park and Alex Alderton, Roberto Amato, Jeffrey Barrett, Sonia Goncalves, Ewan Harrison, David K. Jackson, Ian Johnston, Dominic Kwiatkowski, Cordelia Langford, John Sillitoe on behalf of the Wellcome Sanger Institute COVID-19 Surveillance Team |
| EPI_ISL_3285440 | Lighthouse Lab in Milton Keynes | Wellcome Sanger Institute for the COVID-19 Genomics UK (COG-UK) Consortium | The Lighthouse Lab in Milton Keynes and Alex Alderton, Roberto Amato, Jeffrey Barrett, Sonia Goncalves, Ewan Harrison, David K. Jackson, Ian Johnston, Dominic Kwiatkowski, Cordelia Langford, John Sillitoe on behalf of the Wellcome Sanger Institute COVID-19 Surveillance Team |
| EPI_ISL_3285441 | Lighthouse Lab in Glasgow | Wellcome Sanger Institute for the COVID-19 Genomics UK (COG-UK) Consortium | Harper VanSteenhouse, Yumi Kasai, David Gray, Carol Clugston, Anna Dominiczak and Alex Alderton, Roberto Amato, Jeffrey Barrett, Sonia Goncalves, Ewan Harrison, David K. Jackson, Ian Johnston, Dominic Kwiatkowski, Cordelia Langford, John Sillitoe on behalf of the Wellcome Sanger Institute COVID-19 Surveillance Team |
| EPI_ISL_3285442 | Lighthouse Lab in Alderley Park | Wellcome Sanger Institute for the COVID-19 Genomics UK (COG-UK) Consortium | Jacquelyn Wynn, Mairead Hyland, The Lighthouse Lab in Alderley Park and Alex Alderton, Roberto Amato, Jeffrey Barrett, Sonia Goncalves, Ewan Harrison, David K. Jackson, Ian Johnston, Dominic Kwiatkowski, Cordelia Langford, John Sillitoe on behalf of the Wellcome Sanger Institute COVID-19 Surveillance Team |
| EPI_ISL_3285443 | Lighthouse Lab in Glasgow | Wellcome Sanger Institute for the COVID-19 Genomics UK (COG-UK) Consortium | Harper VanSteenhouse, Yumi Kasai, David Gray, Carol Clugston, Anna Dominiczak and Alex Alderton, Roberto Amato, Jeffrey Barrett, Sonia Goncalves, Ewan Harrison, David K. Jackson, Ian Johnston, Dominic Kwiatkowski, Cordelia Langford, John Sillitoe on behalf of the Wellcome Sanger Institute COVID-19 Surveillance Team |
| EPI_ISL_3285444 | Lighthouse Lab in Milton Keynes | Wellcome Sanger Institute for the COVID-19 Genomics UK (COG-UK) Consortium | The Lighthouse Lab in Milton Keynes and Alex Alderton, Roberto Amato, Jeffrey Barrett, Sonia Goncalves, Ewan Harrison, David K. Jackson, Ian Johnston, Dominic Kwiatkowski, Cordelia Langford, John Sillitoe on behalf of the Wellcome Sanger Institute COVID-19 Surveillance Team |
| EPI_ISL_3285445, EPI_ISL_3285446, EPI_ISL_3285447, EPI_ISL_3285448, EPI_ISL_3285449 | Berkshire and Surrey Pathology Services Lighthouse Laboratory | Wellcome Sanger Institute for the COVID-19 Genomics UK (COG-UK) Consortium | Berkshire and Surrey Pathology Services Lighthouse Laboratory and Alex Alderton, Roberto Amato, Jeffrey Barrett, Sonia Goncalves, Ewan Harrison, David K. Jackson, Ian Johnston, Dominic Kwiatkowski, Cordelia Langford, John Sillitoe on behalf of the Wellcome Sanger Institute COVID-19 Surveillance Team |
| EPI_ISL_3285450 | Lighthouse Lab in Alderley Park | Wellcome Sanger Institute for the COVID-19 Genomics UK (COG-UK) Consortium | Jacquelyn Wynn, Mairead Hyland, The Lighthouse Lab in Alderley Park and Alex Alderton, Roberto Amato, Jeffrey Barrett, Sonia Goncalves, Ewan Harrison, David K. Jackson, Ian Johnston, Dominic Kwiatkowski, Cordelia Langford, John Sillitoe on behalf of the Wellcome Sanger Institute COVID-19 Surveillance Team |
| EPI_ISL_3285451, EPI_ISL_3285452, EPI_ISL_3285453, EPI_ISL_3285454 | Lighthouse Lab in Milton Keynes | Wellcome Sanger Institute for the COVID-19 Genomics UK (COG-UK) Consortium | The Lighthouse Lab in Milton Keynes and Alex Alderton, Roberto Amato, Jeffrey Barrett, Sonia Goncalves, Ewan Harrison, David K. Jackson, Ian Johnston, Dominic Kwiatkowski, Cordelia Langford, John Sillitoe on behalf of the Wellcome Sanger Institute COVID-19 Surveillance Team |
| EPI_ISL_3285455, EPI_ISL_3285456 | Berkshire and Surrey Pathology Services Lighthouse Laboratory | Wellcome Sanger Institute for the COVID-19 Genomics UK (COG-UK) Consortium | Berkshire and Surrey Pathology Services Lighthouse Laboratory and Alex Alderton, Roberto Amato, Jeffrey Barrett, Sonia Goncalves, Ewan Harrison, David K. Jackson, Ian Johnston, Dominic Kwiatkowski, Cordelia Langford, John Sillitoe on behalf of the Wellcome Sanger Institute COVID-19 Surveillance Team |
| EPI_ISL_3285457, EPI_ISL_3285458 | Lighthouse Lab in Alderley Park | Wellcome Sanger Institute for the COVID-19 Genomics UK (COG-UK) Consortium | Jacquelyn Wynn, Mairead Hyland, The Lighthouse Lab in Alderley Park and Alex Alderton, Roberto Amato, Jeffrey Barrett, Sonia Goncalves, Ewan Harrison, David K. Jackson, Ian Johnston, Dominic Kwiatkowski, Cordelia Langford, John Sillitoe on behalf of the Wellcome Sanger Institute COVID-19 Surveillance Team |
| EPI_ISL_3285459 | Lighthouse Lab in Milton Keynes | Wellcome Sanger Institute for the COVID-19 Genomics UK (COG-UK) Consortium | The Lighthouse Lab in Milton Keynes and Alex Alderton, Roberto Amato, Jeffrey Barrett, Sonia Goncalves, Ewan Harrison, David K. Jackson, Ian Johnston, Dominic Kwiatkowski, Cordelia Langford, John Sillitoe on behalf of the Wellcome Sanger Institute COVID-19 Surveillance Team |
| EPI_ISL_3285460 | Berkshire and Surrey Pathology Services Lighthouse Laboratory | Wellcome Sanger Institute for the COVID-19 Genomics UK (COG-UK) Consortium | Berkshire and Surrey Pathology Services Lighthouse Laboratory and Alex Alderton, Roberto Amato, Jeffrey Barrett, Sonia Goncalves, Ewan Harrison, David K. Jackson, Ian Johnston, Dominic Kwiatkowski, Cordelia Langford, John Sillitoe on behalf of the Wellcome Sanger Institute COVID-19 Surveillance Team |
| EPI_ISL_3285461 | Lighthouse Lab in Alderley Park | Wellcome Sanger Institute for the COVID-19 Genomics UK (COG-UK) Consortium | Jacquelyn Wynn, Mairead Hyland, The Lighthouse Lab in Alderley Park and Alex Alderton, Roberto Amato, Jeffrey Barrett, Sonia Goncalves, Ewan Harrison, David K. Jackson, Ian Johnston, Dominic Kwiatkowski, Cordelia Langford, John Sillitoe on behalf of the Wellcome Sanger Institute COVID-19 Surveillance Team |
| EPI_ISL_3285462, EPI_ISL_3285463 | Berkshire and Surrey Pathology Services Lighthouse Laboratory | Wellcome Sanger Institute for the COVID-19 Genomics UK (COG-UK) Consortium | Berkshire and Surrey Pathology Services Lighthouse Laboratory and Alex Alderton, Roberto Amato, Jeffrey Barrett, Sonia Goncalves, Ewan Harrison, David K. Jackson, Ian Johnston, Dominic Kwiatkowski, Cordelia Langford, John Sillitoe on behalf of the Wellcome Sanger Institute COVID-19 Surveillance Team |
| EPI_ISL_3285464, EPI_ISL_3285465 | Lighthouse Lab in Glasgow | Wellcome Sanger Institute for the COVID-19 Genomics UK (COG-UK) Consortium | Harper VanSteenhouse, Yumi Kasai, David Gray, Carol Clugston, Anna Dominiczak and Alex Alderton, Roberto Amato, Jeffrey Barrett, Sonia Goncalves, Ewan Harrison, David K. Jackson, Ian Johnston, Dominic Kwiatkowski, Cordelia Langford, John Sillitoe on behalf of the Wellcome Sanger Institute COVID-19 Surveillance Team |
| EPI_ISL_3285466 | Berkshire and Surrey Pathology Services Lighthouse Laboratory | Wellcome Sanger Institute for the COVID-19 Genomics UK (COG-UK) Consortium | Berkshire and Surrey Pathology Services Lighthouse Laboratory and Alex Alderton, Roberto Amato, Jeffrey Barrett, Sonia Goncalves, Ewan Harrison, David K. Jackson, Ian Johnston, Dominic Kwiatkowski, Cordelia Langford, John Sillitoe on behalf of the Wellcome Sanger Institute COVID-19 Surveillance Team |
| EPI_ISL_3285467 | Lighthouse Lab in Alderley Park | Wellcome Sanger Institute for the COVID-19 Genomics UK (COG-UK) Consortium | Jacquelyn Wynn, Mairead Hyland, The Lighthouse Lab in Alderley Park and Alex Alderton, Roberto Amato, Jeffrey Barrett, Sonia Goncalves, Ewan Harrison, David K. Jackson, Ian Johnston, Dominic Kwiatkowski, Cordelia Langford, John Sillitoe on behalf of the Wellcome Sanger Institute COVID-19 Surveillance Team |
| EPI_ISL_3285468, EPI_ISL_3285469 | Lighthouse Lab in Glasgow | Wellcome Sanger Institute for the COVID-19 Genomics UK (COG-UK) Consortium | Harper VanSteenhouse, Yumi Kasai, David Gray, Carol Clugston, Anna Dominiczak and Alex Alderton, Roberto Amato, Jeffrey Barrett, Sonia Goncalves, Ewan Harrison, David K. Jackson, Ian Johnston, Dominic Kwiatkowski, Cordelia Langford, John Sillitoe on behalf of the Wellcome Sanger Institute COVID-19 Surveillance Team |
| EPI_ISL_3285470, EPI_ISL_3285471 | Lighthouse Lab in Alderley Park | Wellcome Sanger Institute for the COVID-19 Genomics UK (COG-UK) Consortium | Jacquelyn Wynn, Mairead Hyland, The Lighthouse Lab in Alderley Park and Alex Alderton, Roberto Amato, Jeffrey Barrett, Sonia Goncalves, Ewan Harrison, David K. Jackson, Ian Johnston, Dominic Kwiatkowski, Cordelia Langford, John Sillitoe on behalf of the Wellcome Sanger Institute COVID-19 Surveillance Team |
| EPI_ISL_3285472 | Lighthouse Lab in Glasgow | Wellcome Sanger Institute for the COVID-19 Genomics UK (COG-UK) Consortium | Harper VanSteenhouse, Yumi Kasai, David Gray, Carol Clugston, Anna Dominiczak and Alex Alderton, Roberto Amato, Jeffrey Barrett, Sonia Goncalves, Ewan Harrison, David K. Jackson, Ian Johnston, Dominic Kwiatkowski, Cordelia Langford, John Sillitoe on behalf of the Wellcome Sanger Institute COVID-19 Surveillance Team |
| EPI_ISL_3285473, EPI_ISL_3285474 | Berkshire and Surrey Pathology Services Lighthouse Laboratory | Wellcome Sanger Institute for the COVID-19 Genomics UK (COG-UK) Consortium | Berkshire and Surrey Pathology Services Lighthouse Laboratory and Alex Alderton, Roberto Amato, Jeffrey Barrett, Sonia Goncalves, Ewan Harrison, David K. Jackson, Ian Johnston, Dominic Kwiatkowski, Cordelia Langford, John Sillitoe on behalf of the Wellcome Sanger Institute COVID-19 Surveillance Team |
| EPI_ISL_3285475 | Lighthouse Lab in Glasgow | Wellcome Sanger Institute for the COVID-19 Genomics UK (COG-UK) Consortium | Harper VanSteenhouse, Yumi Kasai, David Gray, Carol Clugston, Anna Dominiczak and Alex Alderton, Roberto Amato, Jeffrey Barrett, Sonia Goncalves, Ewan Harrison, David K. Jackson, Ian Johnston, Dominic Kwiatkowski, Cordelia Langford, John Sillitoe on behalf of the Wellcome Sanger Institute COVID-19 Surveillance Team |
| EPI_ISL_3285476 | Lighthouse Lab in Milton Keynes | Wellcome Sanger Institute for the COVID-19 Genomics UK (COG-UK) Consortium | The Lighthouse Lab in Milton Keynes and Alex Alderton, Roberto Amato, Jeffrey Barrett, Sonia Goncalves, Ewan Harrison, David K. Jackson, Ian Johnston, Dominic Kwiatkowski, Cordelia Langford, John Sillitoe on behalf of the Wellcome Sanger Institute COVID-19 Surveillance Team |
| EPI_ISL_3285477, EPI_ISL_3285478, EPI_ISL_3285479 | Lighthouse Lab in Glasgow | Wellcome Sanger Institute for the COVID-19 Genomics UK (COG-UK) Consortium | Harper VanSteenhouse, Yumi Kasai, David Gray, Carol Clugston, Anna Dominiczak and Alex Alderton, Roberto Amato, Jeffrey Barrett, Sonia Goncalves, Ewan Harrison, David K. Jackson, Ian Johnston, Dominic Kwiatkowski, Cordelia Langford, John Sillitoe on behalf of the Wellcome Sanger Institute COVID-19 Surveillance Team |
| EPI_ISL_3285480, EPI_ISL_3285481 | Lighthouse Lab in Milton Keynes | Wellcome Sanger Institute for the COVID-19 Genomics UK (COG-UK) Consortium | The Lighthouse Lab in Milton Keynes and Alex Alderton, Roberto Amato, Jeffrey Barrett, Sonia Goncalves, Ewan Harrison, David K. Jackson, Ian Johnston, Dominic Kwiatkowski, Cordelia Langford, John Sillitoe on behalf of the Wellcome Sanger Institute COVID-19 Surveillance Team |
| EPI_ISL_3285482 | Lighthouse Lab in Alderley Park | Wellcome Sanger Institute for the COVID-19 Genomics UK (COG-UK) Consortium | Jacquelyn Wynn, Mairead Hyland, The Lighthouse Lab in Alderley Park and Alex Alderton, Roberto Amato, Jeffrey Barrett, Sonia Goncalves, Ewan Harrison, David K. Jackson, Ian Johnston, Dominic Kwiatkowski, Cordelia Langford, John Sillitoe on behalf of the Wellcome Sanger Institute COVID-19 Surveillance Team |
| EPI_ISL_3285483, EPI_ISL_3285484 | Lighthouse Lab in Glasgow | Wellcome Sanger Institute for the COVID-19 Genomics UK (COG-UK) Consortium | Harper VanSteenhouse, Yumi Kasai, David Gray, Carol Clugston, Anna Dominiczak and Alex Alderton, Roberto Amato, Jeffrey Barrett, Sonia Goncalves, Ewan Harrison, David K. Jackson, Ian Johnston, Dominic Kwiatkowski, Cordelia Langford, John Sillitoe on behalf of the Wellcome Sanger Institute COVID-19 Surveillance Team |
| EPI_ISL_3285485 | Berkshire and Surrey Pathology Services Lighthouse Laboratory | Wellcome Sanger Institute for the COVID-19 Genomics UK (COG-UK) Consortium | Berkshire and Surrey Pathology Services Lighthouse Laboratory and Alex Alderton, Roberto Amato, Jeffrey Barrett, Sonia Goncalves, Ewan Harrison, David K. Jackson, Ian Johnston, Dominic Kwiatkowski, Cordelia Langford, John Sillitoe on behalf of the Wellcome Sanger Institute COVID-19 Surveillance Team |
| EPI_ISL_3285486 | Lighthouse Lab in Milton Keynes | Wellcome Sanger Institute for the COVID-19 Genomics UK (COG-UK) Consortium | The Lighthouse Lab in Milton Keynes and Alex Alderton, Roberto Amato, Jeffrey Barrett, Sonia Goncalves, Ewan Harrison, David K. Jackson, Ian Johnston, Dominic Kwiatkowski, Cordelia Langford, John Sillitoe on behalf of the Wellcome Sanger Institute COVID-19 Surveillance Team |
| EPI_ISL_3285487 | Lighthouse Lab in Alderley Park | Wellcome Sanger Institute for the COVID-19 Genomics UK (COG-UK) Consortium | Jacquelyn Wynn, Mairead Hyland, The Lighthouse Lab in Alderley Park and Alex Alderton, Roberto Amato, Jeffrey Barrett, Sonia Goncalves, Ewan Harrison, David K. Jackson, Ian Johnston, Dominic Kwiatkowski, Cordelia Langford, John Sillitoe on behalf of the Wellcome Sanger Institute COVID-19 Surveillance Team |
| EPI_ISL_3285488, EPI_ISL_3285489 | Berkshire and Surrey Pathology Services Lighthouse Laboratory | Wellcome Sanger Institute for the COVID-19 Genomics UK (COG-UK) Consortium | Berkshire and Surrey Pathology Services Lighthouse Laboratory and Alex Alderton, Roberto Amato, Jeffrey Barrett, Sonia Goncalves, Ewan Harrison, David K. Jackson, Ian Johnston, Dominic Kwiatkowski, Cordelia Langford, John Sillitoe on behalf of the Wellcome Sanger Institute COVID-19 Surveillance Team |

| | | | |
|---|---|---|---|
| EPI_ISL_3285490 | Lighthouse Lab in Alderley Park | Wellcome Sanger Institute for the COVID-19 Genomics UK (COG-UK) Consortium | Jacquelyn Wynn, Mairead Hyland, The Lighthouse Lab in Alderley Park and Alex Alderton, Roberto Amato, Jeffrey Barrett, Sonia Goncalves, Ewan Harrison, David K. Jackson, Ian Johnston, Dominic Kwiatkowski, Cordelia Langford, John Sillitoe on behalf of the Wellcome Sanger Institute COVID-19 Surveillance Team |
| EPI_ISL_3285491 | Lighthouse Lab in Milton Keynes | Wellcome Sanger Institute for the COVID-19 Genomics UK (COG-UK) Consortium | The Lighthouse Lab in Milton Keynes and Alex Alderton, Roberto Amato, Jeffrey Barrett, Sonia Goncalves, Ewan Harrison, David K. Jackson, Ian Johnston, Dominic Kwiatkowski, Cordelia Langford, John Sillitoe on behalf of the Wellcome Sanger Institute COVID-19 Surveillance Team |
| EPI_ISL_3285492 | Berkshire and Surrey Pathology Services Lighthouse Laboratory | Wellcome Sanger Institute for the COVID-19 Genomics UK (COG-UK) Consortium | Berkshire and Surrey Pathology Services Lighthouse Laboratory and Alex Alderton, Roberto Amato, Jeffrey Barrett, Sonia Goncalves, Ewan Harrison, David K. Jackson, Ian Johnston, Dominic Kwiatkowski, Cordelia Langford, John Sillitoe on behalf of the Wellcome Sanger Institute COVID-19 Surveillance Team |
| EPI_ISL_3285493 | Lighthouse Lab in Glasgow | Wellcome Sanger Institute for the COVID-19 Genomics UK (COG-UK) Consortium | Harper VanSteenhouse, Yumi Kasai, David Gray, Carol Clugston, Anna Dominiczak and Alex Alderton, Roberto Amato, Jeffrey Barrett, Sonia Goncalves, Ewan Harrison, David K. Jackson, Ian Johnston, Dominic Kwiatkowski, Cordelia Langford, John Sillitoe on behalf of the Wellcome Sanger Institute COVID-19 Surveillance Team |
| EPI_ISL_3285494 | Berkshire and Surrey Pathology Services Lighthouse Laboratory | Wellcome Sanger Institute for the COVID-19 Genomics UK (COG-UK) Consortium | Berkshire and Surrey Pathology Services Lighthouse Laboratory and Alex Alderton, Roberto Amato, Jeffrey Barrett, Sonia Goncalves, Ewan Harrison, David K. Jackson, Ian Johnston, Dominic Kwiatkowski, Cordelia Langford, John Sillitoe on behalf of the Wellcome Sanger Institute COVID-19 Surveillance Team |
| EPI_ISL_3285495 | Lighthouse Lab in Alderley Park | Wellcome Sanger Institute for the COVID-19 Genomics UK (COG-UK) Consortium | Jacquelyn Wynn, Mairead Hyland, The Lighthouse Lab in Alderley Park and Alex Alderton, Roberto Amato, Jeffrey Barrett, Sonia Goncalves, Ewan Harrison, David K. Jackson, Ian Johnston, Dominic Kwiatkowski, Cordelia Langford, John Sillitoe on behalf of the Wellcome Sanger Institute COVID-19 Surveillance Team |
| EPI_ISL_3285496, EPI_ISL_3285497, EPI_ISL_3285498 | Lighthouse Lab in Milton Keynes | Wellcome Sanger Institute for the COVID-19 Genomics UK (COG-UK) Consortium | The Lighthouse Lab in Milton Keynes and Alex Alderton, Roberto Amato, Jeffrey Barrett, Sonia Goncalves, Ewan Harrison, David K. Jackson, Ian Johnston, Dominic Kwiatkowski, Cordelia Langford, John Sillitoe on behalf of the Wellcome Sanger Institute COVID-19 Surveillance Team |
| EPI_ISL_3285499 | Berkshire and Surrey Pathology Services Lighthouse Laboratory | Wellcome Sanger Institute for the COVID-19 Genomics UK (COG-UK) Consortium | Berkshire and Surrey Pathology Services Lighthouse Laboratory and Alex Alderton, Roberto Amato, Jeffrey Barrett, Sonia Goncalves, Ewan Harrison, David K. Jackson, Ian Johnston, Dominic Kwiatkowski, Cordelia Langford, John Sillitoe on behalf of the Wellcome Sanger Institute COVID-19 Surveillance Team |
| EPI_ISL_3285500 | Lighthouse Lab in Glasgow | Wellcome Sanger Institute for the COVID-19 Genomics UK (COG-UK) Consortium | Harper VanSteenhouse, Yumi Kasai, David Gray, Carol Clugston, Anna Dominiczak and Alex Alderton, Roberto Amato, Jeffrey Barrett, Sonia Goncalves, Ewan Harrison, David K. Jackson, Ian Johnston, Dominic Kwiatkowski, Cordelia Langford, John Sillitoe on behalf of the Wellcome Sanger Institute COVID-19 Surveillance Team |
| EPI_ISL_3285501, EPI_ISL_3285502 | Lighthouse Lab in Milton Keynes | Wellcome Sanger Institute for the COVID-19 Genomics UK (COG-UK) Consortium | The Lighthouse Lab in Milton Keynes and Alex Alderton, Roberto Amato, Jeffrey Barrett, Sonia Goncalves, Ewan Harrison, David K. Jackson, Ian Johnston, Dominic Kwiatkowski, Cordelia Langford, John Sillitoe on behalf of the Wellcome Sanger Institute COVID-19 Surveillance Team |
| EPI_ISL_3285503 | Lighthouse Lab in Alderley Park | Wellcome Sanger Institute for the COVID-19 Genomics UK (COG-UK) Consortium | Jacquelyn Wynn, Mairead Hyland, The Lighthouse Lab in Alderley Park and Alex Alderton, Roberto Amato, Jeffrey Barrett, Sonia Goncalves, Ewan Harrison, David K. Jackson, Ian Johnston, Dominic Kwiatkowski, Cordelia Langford, John Sillitoe on behalf of the Wellcome Sanger Institute COVID-19 Surveillance Team |
| EPI_ISL_3285504, EPI_ISL_3285505 | Berkshire and Surrey Pathology Services Lighthouse Laboratory | Wellcome Sanger Institute for the COVID-19 Genomics UK (COG-UK) Consortium | Berkshire and Surrey Pathology Services Lighthouse Laboratory and Alex Alderton, Roberto Amato, Jeffrey Barrett, Sonia Goncalves, Ewan Harrison, David K. Jackson, Ian Johnston, Dominic Kwiatkowski, Cordelia Langford, John Sillitoe on behalf of the Wellcome Sanger Institute COVID-19 Surveillance Team |
| EPI_ISL_3285506, EPI_ISL_3285507 | Lighthouse Lab in Glasgow | Wellcome Sanger Institute for the COVID-19 Genomics UK (COG-UK) Consortium | Harper VanSteenhouse, Yumi Kasai, David Gray, Carol Clugston, Anna Dominiczak and Alex Alderton, Roberto Amato, Jeffrey Barrett, Sonia Goncalves, Ewan Harrison, David K. Jackson, Ian Johnston, Dominic Kwiatkowski, Cordelia Langford, John Sillitoe on behalf of the Wellcome Sanger Institute COVID-19 Surveillance Team |
| EPI_ISL_3285508 | Lighthouse Lab in Alderley Park | Wellcome Sanger Institute for the COVID-19 Genomics UK (COG-UK) Consortium | Jacquelyn Wynn, Mairead Hyland, The Lighthouse Lab in Alderley Park and Alex Alderton, Roberto Amato, Jeffrey Barrett, Sonia Goncalves, Ewan Harrison, David K. Jackson, Ian Johnston, Dominic Kwiatkowski, Cordelia Langford, John Sillitoe on behalf of the Wellcome Sanger Institute COVID-19 Surveillance Team |
| EPI_ISL_3285509 | Berkshire and Surrey Pathology Services Lighthouse Laboratory | Wellcome Sanger Institute for the COVID-19 Genomics UK (COG-UK) Consortium | Berkshire and Surrey Pathology Services Lighthouse Laboratory and Alex Alderton, Roberto Amato, Jeffrey Barrett, Sonia Goncalves, Ewan Harrison, David K. Jackson, Ian Johnston, Dominic Kwiatkowski, Cordelia Langford, John Sillitoe on behalf of the Wellcome Sanger Institute COVID-19 Surveillance Team |
| EPI_ISL_3285510 | Lighthouse Lab in Alderley Park | Wellcome Sanger Institute for the COVID-19 Genomics UK (COG-UK) Consortium | Jacquelyn Wynn, Mairead Hyland, The Lighthouse Lab in Alderley Park and Alex Alderton, Roberto Amato, Jeffrey Barrett, Sonia Goncalves, Ewan Harrison, David K. Jackson, Ian Johnston, Dominic Kwiatkowski, Cordelia Langford, John Sillitoe on behalf of the Wellcome Sanger Institute COVID-19 Surveillance Team |
| EPI_ISL_3285511 | Lighthouse Lab in Glasgow | Wellcome Sanger Institute for the COVID-19 Genomics UK (COG-UK) Consortium | Harper VanSteenhouse, Yumi Kasai, David Gray, Carol Clugston, Anna Dominiczak and Alex Alderton, Roberto Amato, Jeffrey Barrett, Sonia Goncalves, Ewan Harrison, David K. Jackson, Ian Johnston, Dominic Kwiatkowski, Cordelia Langford, John Sillitoe on behalf of the Wellcome Sanger Institute COVID-19 Surveillance Team |
| EPI_ISL_3285512 | Lighthouse Lab in Alderley Park | Wellcome Sanger Institute for the COVID-19 Genomics UK (COG-UK) Consortium | Jacquelyn Wynn, Mairead Hyland, The Lighthouse Lab in Alderley Park and Alex Alderton, Roberto Amato, Jeffrey Barrett, Sonia Goncalves, Ewan Harrison, David K. Jackson, Ian Johnston, Dominic Kwiatkowski, Cordelia Langford, John Sillitoe on behalf of the Wellcome Sanger Institute COVID-19 Surveillance Team |
| EPI_ISL_3285513 | Berkshire and Surrey Pathology Services Lighthouse Laboratory | Wellcome Sanger Institute for the COVID-19 Genomics UK (COG-UK) Consortium | Berkshire and Surrey Pathology Services Lighthouse Laboratory and Alex Alderton, Roberto Amato, Jeffrey Barrett, Sonia Goncalves, Ewan Harrison, David K. Jackson, Ian Johnston, Dominic Kwiatkowski, Cordelia Langford, John Sillitoe on behalf of the Wellcome Sanger Institute COVID-19 Surveillance Team |
| EPI_ISL_3285514 | Lighthouse Lab in Milton Keynes | Wellcome Sanger Institute for the COVID-19 Genomics UK (COG-UK) Consortium | The Lighthouse Lab in Milton Keynes and Alex Alderton, Roberto Amato, Jeffrey Barrett, Sonia Goncalves, Ewan Harrison, David K. Jackson, Ian Johnston, Dominic Kwiatkowski, Cordelia Langford, John Sillitoe on behalf of the Wellcome Sanger Institute COVID-19 Surveillance Team |
| EPI_ISL_3285515 | Berkshire and Surrey Pathology Services Lighthouse Laboratory | Wellcome Sanger Institute for the COVID-19 Genomics UK (COG-UK) Consortium | Berkshire and Surrey Pathology Services Lighthouse Laboratory and Alex Alderton, Roberto Amato, Jeffrey Barrett, Sonia Goncalves, Ewan Harrison, David K. Jackson, Ian Johnston, Dominic Kwiatkowski, Cordelia Langford, John Sillitoe on behalf of the Wellcome Sanger Institute COVID-19 Surveillance Team |
| EPI_ISL_3285516, EPI_ISL_3285517 | Lighthouse Lab in Milton Keynes | Wellcome Sanger Institute for the COVID-19 Genomics UK (COG-UK) Consortium | The Lighthouse Lab in Milton Keynes and Alex Alderton, Roberto Amato, Jeffrey Barrett, Sonia Goncalves, Ewan Harrison, David K. Jackson, Ian Johnston, Dominic Kwiatkowski, Cordelia Langford, John Sillitoe on behalf of the Wellcome Sanger Institute COVID-19 Surveillance Team |
| EPI_ISL_3285518, EPI_ISL_3285519 | Lighthouse Lab in Alderley Park | Wellcome Sanger Institute for the COVID-19 Genomics UK (COG-UK) Consortium | Jacquelyn Wynn, Mairead Hyland, The Lighthouse Lab in Alderley Park and Alex Alderton, Roberto Amato, Jeffrey Barrett, Sonia Goncalves, Ewan Harrison, David K. Jackson, Ian Johnston, Dominic Kwiatkowski, Cordelia Langford, John Sillitoe on behalf of the Wellcome Sanger Institute COVID-19 Surveillance Team |
| EPI_ISL_3285520 | Lighthouse Lab in Milton Keynes | Wellcome Sanger Institute for the COVID-19 Genomics UK (COG-UK) Consortium | The Lighthouse Lab in Milton Keynes and Alex Alderton, Roberto Amato, Jeffrey Barrett, Sonia Goncalves, Ewan Harrison, David K. Jackson, Ian Johnston, Dominic Kwiatkowski, Cordelia Langford, John Sillitoe on behalf of the Wellcome Sanger Institute COVID-19 Surveillance Team |
| EPI_ISL_3285521, EPI_ISL_3285522 | Lighthouse Lab in Glasgow | Wellcome Sanger Institute for the COVID-19 Genomics UK (COG-UK) Consortium | Harper VanSteenhouse, Yumi Kasai, David Gray, Carol Clugston, Anna Dominiczak and Alex Alderton, Roberto Amato, Jeffrey Barrett, Sonia Goncalves, Ewan Harrison, David K. Jackson, Ian Johnston, Dominic Kwiatkowski, Cordelia Langford, John Sillitoe on behalf of the Wellcome Sanger Institute COVID-19 Surveillance Team |
| EPI_ISL_3285523 | Berkshire and Surrey Pathology Services Lighthouse Laboratory | Wellcome Sanger Institute for the COVID-19 Genomics UK (COG-UK) Consortium | Berkshire and Surrey Pathology Services Lighthouse Laboratory and Alex Alderton, Roberto Amato, Jeffrey Barrett, Sonia Goncalves, Ewan Harrison, David K. Jackson, Ian Johnston, Dominic Kwiatkowski, Cordelia Langford, John Sillitoe on behalf of the Wellcome Sanger Institute COVID-19 Surveillance Team |
| EPI_ISL_3285524 | Lighthouse Lab in Milton Keynes | Wellcome Sanger Institute for the COVID-19 Genomics UK (COG-UK) Consortium | The Lighthouse Lab in Milton Keynes and Alex Alderton, Roberto Amato, Jeffrey Barrett, Sonia Goncalves, Ewan Harrison, David K. Jackson, Ian Johnston, Dominic Kwiatkowski, Cordelia Langford, John Sillitoe on behalf of the Wellcome Sanger Institute COVID-19 Surveillance Team |
| EPI_ISL_3285525, EPI_ISL_3285526 | Berkshire and Surrey Pathology Services Lighthouse Laboratory | Wellcome Sanger Institute for the COVID-19 Genomics UK (COG-UK) Consortium | Berkshire and Surrey Pathology Services Lighthouse Laboratory and Alex Alderton, Roberto Amato, Jeffrey Barrett, Sonia Goncalves, Ewan Harrison, David K. Jackson, Ian Johnston, Dominic Kwiatkowski, Cordelia Langford, John Sillitoe on behalf of the Wellcome Sanger Institute COVID-19 Surveillance Team |
| EPI_ISL_3285527 | Lighthouse Lab in Milton Keynes | Wellcome Sanger Institute for the COVID-19 Genomics UK (COG-UK) Consortium | The Lighthouse Lab in Milton Keynes and Alex Alderton, Roberto Amato, Jeffrey Barrett, Sonia Goncalves, Ewan Harrison, David K. Jackson, Ian Johnston, Dominic Kwiatkowski, Cordelia Langford, John Sillitoe on behalf of the Wellcome Sanger Institute COVID-19 Surveillance Team |
| EPI_ISL_3285528 | Lighthouse Lab in Glasgow | Wellcome Sanger Institute for the COVID-19 Genomics UK (COG-UK) Consortium | Harper VanSteenhouse, Yumi Kasai, David Gray, Carol Clugston, Anna Dominiczak and Alex Alderton, Roberto Amato, Jeffrey Barrett, Sonia Goncalves, Ewan Harrison, David K. Jackson, Ian Johnston, Dominic Kwiatkowski, Cordelia Langford, John Sillitoe on behalf of the Wellcome Sanger Institute COVID-19 Surveillance Team |
| EPI_ISL_3285529, EPI_ISL_3285530, EPI_ISL_3285531 | Lighthouse Lab in Milton Keynes | Wellcome Sanger Institute for the COVID-19 Genomics UK (COG-UK) Consortium | The Lighthouse Lab in Milton Keynes and Alex Alderton, Roberto Amato, Jeffrey Barrett, Sonia Goncalves, Ewan Harrison, David K. Jackson, Ian Johnston, Dominic Kwiatkowski, Cordelia Langford, John Sillitoe on behalf of the Wellcome Sanger Institute COVID-19 Surveillance Team |
| EPI_ISL_3285532, EPI_ISL_3285533 | Lighthouse Lab in Alderley Park | Wellcome Sanger Institute for the COVID-19 Genomics UK (COG-UK) Consortium | Jacquelyn Wynn, Mairead Hyland, The Lighthouse Lab in Alderley Park and Alex Alderton, Roberto Amato, Jeffrey Barrett, Sonia Goncalves, Ewan Harrison, David K. Jackson, Ian Johnston, Dominic Kwiatkowski, Cordelia Langford, John Sillitoe on behalf of the Wellcome Sanger Institute COVID-19 Surveillance Team |
| EPI_ISL_3285534 | Lighthouse Lab in Glasgow | Wellcome Sanger Institute for the COVID-19 Genomics UK (COG-UK) Consortium | Harper VanSteenhouse, Yumi Kasai, David Gray, Carol Clugston, Anna Dominiczak and Alex Alderton, Roberto Amato, Jeffrey Barrett, Sonia Goncalves, Ewan Harrison, David K. Jackson, Ian Johnston, Dominic Kwiatkowski, Cordelia Langford, John Sillitoe on behalf of the Wellcome Sanger Institute COVID-19 Surveillance Team |
| EPI_ISL_3285535 | Lighthouse Lab in Alderley Park | Wellcome Sanger Institute for the COVID-19 Genomics UK (COG-UK) Consortium | Jacquelyn Wynn, Mairead Hyland, The Lighthouse Lab in Alderley Park and Alex Alderton, Roberto Amato, Jeffrey Barrett, Sonia Goncalves, Ewan Harrison, David K. Jackson, Ian Johnston, Dominic Kwiatkowski, Cordelia Langford, John Sillitoe on behalf of the Wellcome Sanger Institute COVID-19 Surveillance Team |
| EPI_ISL_3285536 | Berkshire and Surrey Pathology Services Lighthouse Laboratory | Wellcome Sanger Institute for the COVID-19 Genomics UK (COG-UK) Consortium | Berkshire and Surrey Pathology Services Lighthouse Laboratory and Alex Alderton, Roberto Amato, Jeffrey Barrett, Sonia Goncalves, Ewan Harrison, David K. Jackson, Ian Johnston, Dominic Kwiatkowski, Cordelia Langford, John Sillitoe on behalf of the Wellcome Sanger Institute COVID-19 Surveillance Team |
| EPI_ISL_3285537, EPI_ISL_3285538 | Lighthouse Lab in Glasgow | Wellcome Sanger Institute for the COVID-19 Genomics UK (COG-UK) Consortium | Harper VanSteenhouse, Yumi Kasai, David Gray, Carol Clugston, Anna Dominiczak and Alex Alderton, Roberto Amato, Jeffrey Barrett, Sonia Goncalves, Ewan Harrison, David K. Jackson, Ian Johnston, Dominic Kwiatkowski, Cordelia Langford, John Sillitoe on behalf of the Wellcome Sanger Institute COVID-19 Surveillance Team |
| EPI_ISL_3285539 | Lighthouse Lab in Milton Keynes | Wellcome Sanger Institute for the COVID-19 Genomics UK (COG-UK) Consortium | The Lighthouse Lab in Milton Keynes and Alex Alderton, Roberto Amato, Jeffrey Barrett, Sonia Goncalves, Ewan Harrison, David K. Jackson, Ian Johnston, Dominic Kwiatkowski, Cordelia Langford, John Sillitoe on behalf of the Wellcome Sanger Institute COVID-19 Surveillance Team |
| EPI_ISL_3285540, EPI_ISL_3285541 | Berkshire and Surrey Pathology Services Lighthouse Laboratory | Wellcome Sanger Institute for the COVID-19 Genomics UK (COG-UK) Consortium | Berkshire and Surrey Pathology Services Lighthouse Laboratory and Alex Alderton, Roberto Amato, Jeffrey Barrett, Sonia Goncalves, Ewan Harrison, David K. Jackson, Ian Johnston, Dominic Kwiatkowski, Cordelia Langford, John Sillitoe on behalf of the Wellcome Sanger Institute COVID-19 Surveillance Team |
| EPI_ISL_3285542 | Lighthouse Lab in Milton Keynes | Wellcome Sanger Institute for the COVID-19 Genomics UK (COG-UK) Consortium | The Lighthouse Lab in Milton Keynes and Alex Alderton, Roberto Amato, Jeffrey Barrett, Sonia Goncalves, Ewan Harrison, David K. Jackson, Ian Johnston, Dominic Kwiatkowski, Cordelia Langford, John Sillitoe on behalf of the Wellcome Sanger Institute COVID-19 Surveillance Team |
| EPI_ISL_3285543 | Lighthouse Lab in Glasgow | Wellcome Sanger Institute for the COVID-19 Genomics UK (COG-UK) Consortium | Harper VanSteenhouse, Yumi Kasai, David Gray, Carol Clugston, Anna Dominiczak and Alex Alderton, Roberto Amato, Jeffrey Barrett, Sonia Goncalves, Ewan Harrison, David K. Jackson, Ian Johnston, Dominic Kwiatkowski, Cordelia Langford, John Sillitoe on behalf of the Wellcome Sanger Institute COVID-19 Surveillance Team |
| EPI_ISL_3285544 | Berkshire and Surrey Pathology Services Lighthouse Laboratory | Wellcome Sanger Institute for the COVID-19 Genomics UK (COG-UK) Consortium | Berkshire and Surrey Pathology Services Lighthouse Laboratory and Alex Alderton, Roberto Amato, Jeffrey Barrett, Sonia Goncalves, Ewan Harrison, David K. Jackson, Ian Johnston, Dominic Kwiatkowski, Cordelia Langford, John Sillitoe on behalf of the Wellcome Sanger Institute COVID-19 Surveillance Team |
| EPI_ISL_3285545 | Lighthouse Lab in Glasgow | Wellcome Sanger Institute for the COVID-19 Genomics UK (COG-UK) Consortium | Harper VanSteenhouse, Yumi Kasai, David Gray, Carol Clugston, Anna Dominiczak and Alex Alderton, Roberto Amato, Jeffrey Barrett, Sonia Goncalves, Ewan Harrison, David K. Jackson, Ian Johnston, Dominic Kwiatkowski, Cordelia Langford, John Sillitoe on behalf of the Wellcome Sanger Institute COVID-19 Surveillance Team |
| EPI_ISL_3285546 | Berkshire and Surrey Pathology Services Lighthouse Laboratory | Wellcome Sanger Institute for the COVID-19 Genomics UK (COG-UK) Consortium | Berkshire and Surrey Pathology Services Lighthouse Laboratory and Alex Alderton, Roberto Amato, Jeffrey Barrett, Sonia Goncalves, Ewan Harrison, David K. Jackson, Ian Johnston, Dominic Kwiatkowski, Cordelia Langford, John Sillitoe on behalf of the Wellcome Sanger Institute COVID-19 Surveillance Team |
| EPI_ISL_3285547 | Lighthouse Lab in Milton Keynes | Wellcome Sanger Institute for the COVID-19 Genomics UK (COG-UK) Consortium | The Lighthouse Lab in Milton Keynes and Alex Alderton, Roberto Amato, Jeffrey Barrett, Sonia Goncalves, Ewan Harrison, David K. Jackson, Ian Johnston, Dominic Kwiatkowski, Cordelia Langford, John Sillitoe on behalf of the Wellcome Sanger Institute COVID-19 Surveillance Team |

| | | | |
|---|---|---|---|
| EPI_ISL_3285548 | Lighthouse Lab in Alderley Park | Wellcome Sanger Institute for the COVID-19 Genomics UK (COG-UK) Consortium | Jacquelyn Wynn, Mairead Hyland, The Lighthouse Lab in Alderley Park and Alex Alderton, Roberto Amato, Jeffrey Barrett, Sonia Goncalves, Ewan Harrison, David K. Jackson, Ian Johnston, Dominic Kwiatkowski, Cordelia Langford, John Sillitoe on behalf of the Wellcome Sanger Institute COVID-19 Surveillance Team |
| EPI_ISL_3285549, EPI_ISL_3285550, EPI_ISL_3285551 | Berkshire and Surrey Pathology Services Lighthouse Laboratory | Wellcome Sanger Institute for the COVID-19 Genomics UK (COG-UK) Consortium | Berkshire and Surrey Pathology Services Lighthouse Laboratory and Alex Alderton, Roberto Amato, Jeffrey Barrett, Sonia Goncalves, Ewan Harrison, David K. Jackson, Ian Johnston, Dominic Kwiatkowski, Cordelia Langford, John Sillitoe on behalf of the Wellcome Sanger Institute COVID-19 Surveillance Team |
| EPI_ISL_3285552 | Lighthouse Lab in Alderley Park | Wellcome Sanger Institute for the COVID-19 Genomics UK (COG-UK) Consortium | Jacquelyn Wynn, Mairead Hyland, The Lighthouse Lab in Alderley Park and Alex Alderton, Roberto Amato, Jeffrey Barrett, Sonia Goncalves, Ewan Harrison, David K. Jackson, Ian Johnston, Dominic Kwiatkowski, Cordelia Langford, John Sillitoe on behalf of the Wellcome Sanger Institute COVID-19 Surveillance Team |
| EPI_ISL_3285553 | Lighthouse Lab in Milton Keynes | Wellcome Sanger Institute for the COVID-19 Genomics UK (COG-UK) Consortium | The Lighthouse Lab in Milton Keynes and Alex Alderton, Roberto Amato, Jeffrey Barrett, Sonia Goncalves, Ewan Harrison, David K. Jackson, Ian Johnston, Dominic Kwiatkowski, Cordelia Langford, John Sillitoe on behalf of the Wellcome Sanger Institute COVID-19 Surveillance Team |
| EPI_ISL_3285554 | Lighthouse Lab in Alderley Park | Wellcome Sanger Institute for the COVID-19 Genomics UK (COG-UK) Consortium | Jacquelyn Wynn, Mairead Hyland, The Lighthouse Lab in Alderley Park and Alex Alderton, Roberto Amato, Jeffrey Barrett, Sonia Goncalves, Ewan Harrison, David K. Jackson, Ian Johnston, Dominic Kwiatkowski, Cordelia Langford, John Sillitoe on behalf of the Wellcome Sanger Institute COVID-19 Surveillance Team |
| EPI_ISL_3285555 | Lighthouse Lab in Milton Keynes | Wellcome Sanger Institute for the COVID-19 Genomics UK (COG-UK) Consortium | The Lighthouse Lab in Milton Keynes and Alex Alderton, Roberto Amato, Jeffrey Barrett, Sonia Goncalves, Ewan Harrison, David K. Jackson, Ian Johnston, Dominic Kwiatkowski, Cordelia Langford, John Sillitoe on behalf of the Wellcome Sanger Institute COVID-19 Surveillance Team |
| EPI_ISL_3285556 | Lighthouse Lab in Alderley Park | Wellcome Sanger Institute for the COVID-19 Genomics UK (COG-UK) Consortium | Jacquelyn Wynn, Mairead Hyland, The Lighthouse Lab in Alderley Park and Alex Alderton, Roberto Amato, Jeffrey Barrett, Sonia Goncalves, Ewan Harrison, David K. Jackson, Ian Johnston, Dominic Kwiatkowski, Cordelia Langford, John Sillitoe on behalf of the Wellcome Sanger Institute COVID-19 Surveillance Team |
| EPI_ISL_3285557, EPI_ISL_3285558 | Lighthouse Lab in Glasgow | Wellcome Sanger Institute for the COVID-19 Genomics UK (COG-UK) Consortium | Harper VanSteenhouse, Yumi Kasai, David Gray, Carol Clugston, Anna Dominiczak and Alex Alderton, Roberto Amato, Jeffrey Barrett, Sonia Goncalves, Ewan Harrison, David K. Jackson, Ian Johnston, Dominic Kwiatkowski, Cordelia Langford, John Sillitoe on behalf of the Wellcome Sanger Institute COVID-19 Surveillance Team |
| EPI_ISL_3285559 | Lighthouse Lab in Milton Keynes | Wellcome Sanger Institute for the COVID-19 Genomics UK (COG-UK) Consortium | The Lighthouse Lab in Milton Keynes and Alex Alderton, Roberto Amato, Jeffrey Barrett, Sonia Goncalves, Ewan Harrison, David K. Jackson, Ian Johnston, Dominic Kwiatkowski, Cordelia Langford, John Sillitoe on behalf of the Wellcome Sanger Institute COVID-19 Surveillance Team |
| EPI_ISL_3285560 | Berkshire and Surrey Pathology Services Lighthouse Laboratory | Wellcome Sanger Institute for the COVID-19 Genomics UK (COG-UK) Consortium | Berkshire and Surrey Pathology Services Lighthouse Laboratory and Alex Alderton, Roberto Amato, Jeffrey Barrett, Sonia Goncalves, Ewan Harrison, David K. Jackson, Ian Johnston, Dominic Kwiatkowski, Cordelia Langford, John Sillitoe on behalf of the Wellcome Sanger Institute COVID-19 Surveillance Team |
| EPI_ISL_3285561 | Lighthouse Lab in Alderley Park | Wellcome Sanger Institute for the COVID-19 Genomics UK (COG-UK) Consortium | Jacquelyn Wynn, Mairead Hyland, The Lighthouse Lab in Alderley Park and Alex Alderton, Roberto Amato, Jeffrey Barrett, Sonia Goncalves, Ewan Harrison, David K. Jackson, Ian Johnston, Dominic Kwiatkowski, Cordelia Langford, John Sillitoe on behalf of the Wellcome Sanger Institute COVID-19 Surveillance Team |
| EPI_ISL_3285562 | Berkshire and Surrey Pathology Services Lighthouse Laboratory | Wellcome Sanger Institute for the COVID-19 Genomics UK (COG-UK) Consortium | Berkshire and Surrey Pathology Services Lighthouse Laboratory and Alex Alderton, Roberto Amato, Jeffrey Barrett, Sonia Goncalves, Ewan Harrison, David K. Jackson, Ian Johnston, Dominic Kwiatkowski, Cordelia Langford, John Sillitoe on behalf of the Wellcome Sanger Institute COVID-19 Surveillance Team |
| EPI_ISL_3285563 | Lighthouse Lab in Alderley Park | Wellcome Sanger Institute for the COVID-19 Genomics UK (COG-UK) Consortium | Jacquelyn Wynn, Mairead Hyland, The Lighthouse Lab in Alderley Park and Alex Alderton, Roberto Amato, Jeffrey Barrett, Sonia Goncalves, Ewan Harrison, David K. Jackson, Ian Johnston, Dominic Kwiatkowski, Cordelia Langford, John Sillitoe on behalf of the Wellcome Sanger Institute COVID-19 Surveillance Team |
| EPI_ISL_3285564 | Lighthouse Lab in Glasgow | Wellcome Sanger Institute for the COVID-19 Genomics UK (COG-UK) Consortium | Harper VanSteenhouse, Yumi Kasai, David Gray, Carol Clugston, Anna Dominiczak and Alex Alderton, Roberto Amato, Jeffrey Barrett, Sonia Goncalves, Ewan Harrison, David K. Jackson, Ian Johnston, Dominic Kwiatkowski, Cordelia Langford, John Sillitoe on behalf of the Wellcome Sanger Institute COVID-19 Surveillance Team |
| EPI_ISL_3285565, EPI_ISL_3285566 | Berkshire and Surrey Pathology Services Lighthouse Laboratory | Wellcome Sanger Institute for the COVID-19 Genomics UK (COG-UK) Consortium | Berkshire and Surrey Pathology Services Lighthouse Laboratory and Alex Alderton, Roberto Amato, Jeffrey Barrett, Sonia Goncalves, Ewan Harrison, David K. Jackson, Ian Johnston, Dominic Kwiatkowski, Cordelia Langford, John Sillitoe on behalf of the Wellcome Sanger Institute COVID-19 Surveillance Team |
| EPI_ISL_3285567, EPI_ISL_3285568 | Lighthouse Lab in Glasgow | Wellcome Sanger Institute for the COVID-19 Genomics UK (COG-UK) Consortium | Harper VanSteenhouse, Yumi Kasai, David Gray, Carol Clugston, Anna Dominiczak and Alex Alderton, Roberto Amato, Jeffrey Barrett, Sonia Goncalves, Ewan Harrison, David K. Jackson, Ian Johnston, Dominic Kwiatkowski, Cordelia Langford, John Sillitoe on behalf of the Wellcome Sanger Institute COVID-19 Surveillance Team |
| EPI_ISL_3285569 | Lighthouse Lab in Alderley Park | Wellcome Sanger Institute for the COVID-19 Genomics UK (COG-UK) Consortium | Jacquelyn Wynn, Mairead Hyland, The Lighthouse Lab in Alderley Park and Alex Alderton, Roberto Amato, Jeffrey Barrett, Sonia Goncalves, Ewan Harrison, David K. Jackson, Ian Johnston, Dominic Kwiatkowski, Cordelia Langford, John Sillitoe on behalf of the Wellcome Sanger Institute COVID-19 Surveillance Team |
| EPI_ISL_3285570 | Berkshire and Surrey Pathology Services Lighthouse Laboratory | Wellcome Sanger Institute for the COVID-19 Genomics UK (COG-UK) Consortium | Berkshire and Surrey Pathology Services Lighthouse Laboratory and Alex Alderton, Roberto Amato, Jeffrey Barrett, Sonia Goncalves, Ewan Harrison, David K. Jackson, Ian Johnston, Dominic Kwiatkowski, Cordelia Langford, John Sillitoe on behalf of the Wellcome Sanger Institute COVID-19 Surveillance Team |
| EPI_ISL_3285571 | Lighthouse Lab in Milton Keynes | Wellcome Sanger Institute for the COVID-19 Genomics UK (COG-UK) Consortium | The Lighthouse Lab in Milton Keynes and Alex Alderton, Roberto Amato, Jeffrey Barrett, Sonia Goncalves, Ewan Harrison, David K. Jackson, Ian Johnston, Dominic Kwiatkowski, Cordelia Langford, John Sillitoe on behalf of the Wellcome Sanger Institute COVID-19 Surveillance Team |
| EPI_ISL_3285572, EPI_ISL_3285573 | Lighthouse Lab in Glasgow | Wellcome Sanger Institute for the COVID-19 Genomics UK (COG-UK) Consortium | Harper VanSteenhouse, Yumi Kasai, David Gray, Carol Clugston, Anna Dominiczak and Alex Alderton, Roberto Amato, Jeffrey Barrett, Sonia Goncalves, Ewan Harrison, David K. Jackson, Ian Johnston, Dominic Kwiatkowski, Cordelia Langford, John Sillitoe on behalf of the Wellcome Sanger Institute COVID-19 Surveillance Team |
| EPI_ISL_3285574, EPI_ISL_3285575 | Lighthouse Lab in Alderley Park | Wellcome Sanger Institute for the COVID-19 Genomics UK (COG-UK) Consortium | Jacquelyn Wynn, Mairead Hyland, The Lighthouse Lab in Alderley Park and Alex Alderton, Roberto Amato, Jeffrey Barrett, Sonia Goncalves, Ewan Harrison, David K. Jackson, Ian Johnston, Dominic Kwiatkowski, Cordelia Langford, John Sillitoe on behalf of the Wellcome Sanger Institute COVID-19 Surveillance Team |
| EPI_ISL_3285576, EPI_ISL_3285577, EPI_ISL_3285578 | Lighthouse Lab in Glasgow | Wellcome Sanger Institute for the COVID-19 Genomics UK (COG-UK) Consortium | Harper VanSteenhouse, Yumi Kasai, David Gray, Carol Clugston, Anna Dominiczak and Alex Alderton, Roberto Amato, Jeffrey Barrett, Sonia Goncalves, Ewan Harrison, David K. Jackson, Ian Johnston, Dominic Kwiatkowski, Cordelia Langford, John Sillitoe on behalf of the Wellcome Sanger Institute COVID-19 Surveillance Team |
| EPI_ISL_3285579 | Lighthouse Lab in Milton Keynes | Wellcome Sanger Institute for the COVID-19 Genomics UK (COG-UK) Consortium | The Lighthouse Lab in Milton Keynes and Alex Alderton, Roberto Amato, Jeffrey Barrett, Sonia Goncalves, Ewan Harrison, David K. Jackson, Ian Johnston, Dominic Kwiatkowski, Cordelia Langford, John Sillitoe on behalf of the Wellcome Sanger Institute COVID-19 Surveillance Team |
| EPI_ISL_3285580, EPI_ISL_3285581, EPI_ISL_3285582 | Lighthouse Lab in Glasgow | Wellcome Sanger Institute for the COVID-19 Genomics UK (COG-UK) Consortium | Harper VanSteenhouse, Yumi Kasai, David Gray, Carol Clugston, Anna Dominiczak and Alex Alderton, Roberto Amato, Jeffrey Barrett, Sonia Goncalves, Ewan Harrison, David K. Jackson, Ian Johnston, Dominic Kwiatkowski, Cordelia Langford, John Sillitoe on behalf of the Wellcome Sanger Institute COVID-19 Surveillance Team |
| EPI_ISL_3285583 | Lighthouse Lab in Alderley Park | Wellcome Sanger Institute for the COVID-19 Genomics UK (COG-UK) Consortium | Jacquelyn Wynn, Mairead Hyland, The Lighthouse Lab in Alderley Park and Alex Alderton, Roberto Amato, Jeffrey Barrett, Sonia Goncalves, Ewan Harrison, David K. Jackson, Ian Johnston, Dominic Kwiatkowski, Cordelia Langford, John Sillitoe on behalf of the Wellcome Sanger Institute COVID-19 Surveillance Team |
| EPI_ISL_3285584 | Lighthouse Lab in Glasgow | Wellcome Sanger Institute for the COVID-19 Genomics UK (COG-UK) Consortium | Harper VanSteenhouse, Yumi Kasai, David Gray, Carol Clugston, Anna Dominiczak and Alex Alderton, Roberto Amato, Jeffrey Barrett, Sonia Goncalves, Ewan Harrison, David K. Jackson, Ian Johnston, Dominic Kwiatkowski, Cordelia Langford, John Sillitoe on behalf of the Wellcome Sanger Institute COVID-19 Surveillance Team |
| EPI_ISL_3285585, EPI_ISL_3285586, EPI_ISL_3285587, EPI_ISL_3285588, EPI_ISL_3285589 | Lighthouse Lab in Alderley Park | Wellcome Sanger Institute for the COVID-19 Genomics UK (COG-UK) Consortium | Jacquelyn Wynn, Mairead Hyland, The Lighthouse Lab in Alderley Park and Alex Alderton, Roberto Amato, Jeffrey Barrett, Sonia Goncalves, Ewan Harrison, David K. Jackson, Ian Johnston, Dominic Kwiatkowski, Cordelia Langford, John Sillitoe on behalf of the Wellcome Sanger Institute COVID-19 Surveillance Team |
| EPI_ISL_3285590 | Lighthouse Lab in Glasgow | Wellcome Sanger Institute for the COVID-19 Genomics UK (COG-UK) Consortium | Harper VanSteenhouse, Yumi Kasai, David Gray, Carol Clugston, Anna Dominiczak and Alex Alderton, Roberto Amato, Jeffrey Barrett, Sonia Goncalves, Ewan Harrison, David K. Jackson, Ian Johnston, Dominic Kwiatkowski, Cordelia Langford, John Sillitoe on behalf of the Wellcome Sanger Institute COVID-19 Surveillance Team |
| EPI_ISL_3285591 | Lighthouse Lab in Milton Keynes | Wellcome Sanger Institute for the COVID-19 Genomics UK (COG-UK) Consortium | The Lighthouse Lab in Milton Keynes and Alex Alderton, Roberto Amato, Jeffrey Barrett, Sonia Goncalves, Ewan Harrison, David K. Jackson, Ian Johnston, Dominic Kwiatkowski, Cordelia Langford, John Sillitoe on behalf of the Wellcome Sanger Institute COVID-19 Surveillance Team |
| EPI_ISL_3285592 | Lighthouse Lab in Alderley Park | Wellcome Sanger Institute for the COVID-19 Genomics UK (COG-UK) Consortium | Jacquelyn Wynn, Mairead Hyland, The Lighthouse Lab in Alderley Park and Alex Alderton, Roberto Amato, Jeffrey Barrett, Sonia Goncalves, Ewan Harrison, David K. Jackson, Ian Johnston, Dominic Kwiatkowski, Cordelia Langford, John Sillitoe on behalf of the Wellcome Sanger Institute COVID-19 Surveillance Team |
| EPI_ISL_3285593 | Lighthouse Lab in Milton Keynes | Wellcome Sanger Institute for the COVID-19 Genomics UK (COG-UK) Consortium | The Lighthouse Lab in Milton Keynes and Alex Alderton, Roberto Amato, Jeffrey Barrett, Sonia Goncalves, Ewan Harrison, David K. Jackson, Ian Johnston, Dominic Kwiatkowski, Cordelia Langford, John Sillitoe on behalf of the Wellcome Sanger Institute COVID-19 Surveillance Team |
| EPI_ISL_3285594 | Berkshire and Surrey Pathology Services Lighthouse Laboratory | Wellcome Sanger Institute for the COVID-19 Genomics UK (COG-UK) Consortium | Berkshire and Surrey Pathology Services Lighthouse Laboratory and Alex Alderton, Roberto Amato, Jeffrey Barrett, Sonia Goncalves, Ewan Harrison, David K. Jackson, Ian Johnston, Dominic Kwiatkowski, Cordelia Langford, John Sillitoe on behalf of the Wellcome Sanger Institute COVID-19 Surveillance Team |
| EPI_ISL_3285595, EPI_ISL_3285596 | Lighthouse Lab in Milton Keynes | Wellcome Sanger Institute for the COVID-19 Genomics UK (COG-UK) Consortium | The Lighthouse Lab in Milton Keynes and Alex Alderton, Roberto Amato, Jeffrey Barrett, Sonia Goncalves, Ewan Harrison, David K. Jackson, Ian Johnston, Dominic Kwiatkowski, Cordelia Langford, John Sillitoe on behalf of the Wellcome Sanger Institute COVID-19 Surveillance Team |
| EPI_ISL_3285597 | Berkshire and Surrey Pathology Services Lighthouse Laboratory | Wellcome Sanger Institute for the COVID-19 Genomics UK (COG-UK) Consortium | Berkshire and Surrey Pathology Services Lighthouse Laboratory and Alex Alderton, Roberto Amato, Jeffrey Barrett, Sonia Goncalves, Ewan Harrison, David K. Jackson, Ian Johnston, Dominic Kwiatkowski, Cordelia Langford, John Sillitoe on behalf of the Wellcome Sanger Institute COVID-19 Surveillance Team |
| EPI_ISL_3285598, EPI_ISL_3285599 | Lighthouse Lab in Alderley Park | Wellcome Sanger Institute for the COVID-19 Genomics UK (COG-UK) Consortium | Jacquelyn Wynn, Mairead Hyland, The Lighthouse Lab in Alderley Park and Alex Alderton, Roberto Amato, Jeffrey Barrett, Sonia Goncalves, Ewan Harrison, David K. Jackson, Ian Johnston, Dominic Kwiatkowski, Cordelia Langford, John Sillitoe on behalf of the Wellcome Sanger Institute COVID-19 Surveillance Team |
| EPI_ISL_3285600 | Berkshire and Surrey Pathology Services Lighthouse Laboratory | Wellcome Sanger Institute for the COVID-19 Genomics UK (COG-UK) Consortium | Berkshire and Surrey Pathology Services Lighthouse Laboratory and Alex Alderton, Roberto Amato, Jeffrey Barrett, Sonia Goncalves, Ewan Harrison, David K. Jackson, Ian Johnston, Dominic Kwiatkowski, Cordelia Langford, John Sillitoe on behalf of the Wellcome Sanger Institute COVID-19 Surveillance Team |
| EPI_ISL_3285601 | Lighthouse Lab in Alderley Park | Wellcome Sanger Institute for the COVID-19 Genomics UK (COG-UK) Consortium | Jacquelyn Wynn, Mairead Hyland, The Lighthouse Lab in Alderley Park and Alex Alderton, Roberto Amato, Jeffrey Barrett, Sonia Goncalves, Ewan Harrison, David K. Jackson, Ian Johnston, Dominic Kwiatkowski, Cordelia Langford, John Sillitoe on behalf of the Wellcome Sanger Institute COVID-19 Surveillance Team |
| EPI_ISL_3285602 | Lighthouse Lab in Milton Keynes | Wellcome Sanger Institute for the COVID-19 Genomics UK (COG-UK) Consortium | The Lighthouse Lab in Milton Keynes and Alex Alderton, Roberto Amato, Jeffrey Barrett, Sonia Goncalves, Ewan Harrison, David K. Jackson, Ian Johnston, Dominic Kwiatkowski, Cordelia Langford, John Sillitoe on behalf of the Wellcome Sanger Institute COVID-19 Surveillance Team |
| EPI_ISL_3285603, EPI_ISL_3285604 | Lighthouse Lab in Glasgow | Wellcome Sanger Institute for the COVID-19 Genomics UK (COG-UK) Consortium | Harper VanSteenhouse, Yumi Kasai, David Gray, Carol Clugston, Anna Dominiczak and Alex Alderton, Roberto Amato, Jeffrey Barrett, Sonia Goncalves, Ewan Harrison, David K. Jackson, Ian Johnston, Dominic Kwiatkowski, Cordelia Langford, John Sillitoe on behalf of the Wellcome Sanger Institute COVID-19 Surveillance Team |
| EPI_ISL_3285605, EPI_ISL_3285606 | Berkshire and Surrey Pathology Services Lighthouse Laboratory | Wellcome Sanger Institute for the COVID-19 Genomics UK (COG-UK) Consortium | Berkshire and Surrey Pathology Services Lighthouse Laboratory and Alex Alderton, Roberto Amato, Jeffrey Barrett, Sonia Goncalves, Ewan Harrison, David K. Jackson, Ian Johnston, Dominic Kwiatkowski, Cordelia Langford, John Sillitoe on behalf of the Wellcome Sanger Institute COVID-19 Surveillance Team |
| EPI_ISL_3285607 | Lighthouse Lab in Glasgow | Wellcome Sanger Institute for the COVID-19 Genomics UK (COG-UK) Consortium | Harper VanSteenhouse, Yumi Kasai, David Gray, Carol Clugston, Anna Dominiczak and Alex Alderton, Roberto Amato, Jeffrey Barrett, Sonia Goncalves, Ewan Harrison, David K. Jackson, Ian Johnston, Dominic Kwiatkowski, Cordelia Langford, John Sillitoe on behalf of the Wellcome Sanger Institute COVID-19 Surveillance Team |
| EPI_ISL_3285608 | Lighthouse Lab in Alderley Park | Wellcome Sanger Institute for the COVID-19 Genomics UK (COG-UK) Consortium | Jacquelyn Wynn, Mairead Hyland, The Lighthouse Lab in Alderley Park and Alex Alderton, Roberto Amato, Jeffrey Barrett, Sonia Goncalves, Ewan Harrison, David K. Jackson, Ian Johnston, Dominic Kwiatkowski, Cordelia Langford, John Sillitoe on behalf of the Wellcome Sanger Institute COVID-19 Surveillance Team |

| | | | |
|---|---|---|---|
| EPI_ISL_3285609 | Berkshire and Surrey Pathology Services Lighthouse Laboratory | Wellcome Sanger Institute for the COVID-19 Genomics UK (COG-UK) Consortium | Berkshire and Surrey Pathology Services Lighthouse Laboratory and Alex Alderton, Roberto Amato, Jeffrey Barrett, Sonia Goncalves, Ewan Harrison, David K. Jackson, Ian Johnston, Dominic Kwiatkowski, Cordelia Langford, John Sillitoe on behalf of the Wellcome Sanger Institute COVID-19 Surveillance Team |
| EPI_ISL_3285610, EPI_ISL_3285611 | Lighthouse Lab in Glasgow | Wellcome Sanger Institute for the COVID-19 Genomics UK (COG-UK) Consortium | Harper VanSteenhouse, Yumi Kasai, David Gray, Carol Clugston, Anna Dominiczak and Alex Alderton, Roberto Amato, Jeffrey Barrett, Sonia Goncalves, Ewan Harrison, David K. Jackson, Ian Johnston, Dominic Kwiatkowski, Cordelia Langford, John Sillitoe on behalf of the Wellcome Sanger Institute COVID-19 Surveillance Team |
| EPI_ISL_3285612 | Lighthouse Lab in Alderley Park | Wellcome Sanger Institute for the COVID-19 Genomics UK (COG-UK) Consortium | Jacquelyn Wynn, Mairead Hyland, The Lighthouse Lab in Alderley Park and Alex Alderton, Roberto Amato, Jeffrey Barrett, Sonia Goncalves, Ewan Harrison, David K. Jackson, Ian Johnston, Dominic Kwiatkowski, Cordelia Langford, John Sillitoe on behalf of the Wellcome Sanger Institute COVID-19 Surveillance Team |
| EPI_ISL_3285613 | Berkshire and Surrey Pathology Services Lighthouse Laboratory | Wellcome Sanger Institute for the COVID-19 Genomics UK (COG-UK) Consortium | Berkshire and Surrey Pathology Services Lighthouse Laboratory and Alex Alderton, Roberto Amato, Jeffrey Barrett, Sonia Goncalves, Ewan Harrison, David K. Jackson, Ian Johnston, Dominic Kwiatkowski, Cordelia Langford, John Sillitoe on behalf of the Wellcome Sanger Institute COVID-19 Surveillance Team |
| EPI_ISL_3285614, EPI_ISL_3285615 | Lighthouse Lab in Glasgow | Wellcome Sanger Institute for the COVID-19 Genomics UK (COG-UK) Consortium | Harper VanSteenhouse, Yumi Kasai, David Gray, Carol Clugston, Anna Dominiczak and Alex Alderton, Roberto Amato, Jeffrey Barrett, Sonia Goncalves, Ewan Harrison, David K. Jackson, Ian Johnston, Dominic Kwiatkowski, Cordelia Langford, John Sillitoe on behalf of the Wellcome Sanger Institute COVID-19 Surveillance Team |
| EPI_ISL_3285616 | Lighthouse Lab in Milton Keynes | Wellcome Sanger Institute for the COVID-19 Genomics UK (COG-UK) Consortium | The Lighthouse Lab in Milton Keynes and Alex Alderton, Roberto Amato, Jeffrey Barrett, Sonia Goncalves, Ewan Harrison, David K. Jackson, Ian Johnston, Dominic Kwiatkowski, Cordelia Langford, John Sillitoe on behalf of the Wellcome Sanger Institute COVID-19 Surveillance Team |
| EPI_ISL_3285617 | Berkshire and Surrey Pathology Services Lighthouse Laboratory | Wellcome Sanger Institute for the COVID-19 Genomics UK (COG-UK) Consortium | Berkshire and Surrey Pathology Services Lighthouse Laboratory and Alex Alderton, Roberto Amato, Jeffrey Barrett, Sonia Goncalves, Ewan Harrison, David K. Jackson, Ian Johnston, Dominic Kwiatkowski, Cordelia Langford, John Sillitoe on behalf of the Wellcome Sanger Institute COVID-19 Surveillance Team |
| EPI_ISL_3285618 | Lighthouse Lab in Milton Keynes | Wellcome Sanger Institute for the COVID-19 Genomics UK (COG-UK) Consortium | The Lighthouse Lab in Milton Keynes and Alex Alderton, Roberto Amato, Jeffrey Barrett, Sonia Goncalves, Ewan Harrison, David K. Jackson, Ian Johnston, Dominic Kwiatkowski, Cordelia Langford, John Sillitoe on behalf of the Wellcome Sanger Institute COVID-19 Surveillance Team |
| EPI_ISL_3285619 | Lighthouse Lab in Glasgow | Wellcome Sanger Institute for the COVID-19 Genomics UK (COG-UK) Consortium | Harper VanSteenhouse, Yumi Kasai, David Gray, Carol Clugston, Anna Dominiczak and Alex Alderton, Roberto Amato, Jeffrey Barrett, Sonia Goncalves, Ewan Harrison, David K. Jackson, Ian Johnston, Dominic Kwiatkowski, Cordelia Langford, John Sillitoe on behalf of the Wellcome Sanger Institute COVID-19 Surveillance Team |
| EPI_ISL_3285620, EPI_ISL_3285621 | Lighthouse Lab in Milton Keynes | Wellcome Sanger Institute for the COVID-19 Genomics UK (COG-UK) Consortium | The Lighthouse Lab in Milton Keynes and Alex Alderton, Roberto Amato, Jeffrey Barrett, Sonia Goncalves, Ewan Harrison, David K. Jackson, Ian Johnston, Dominic Kwiatkowski, Cordelia Langford, John Sillitoe on behalf of the Wellcome Sanger Institute COVID-19 Surveillance Team |
| EPI_ISL_3285622 | Berkshire and Surrey Pathology Services Lighthouse Laboratory | Wellcome Sanger Institute for the COVID-19 Genomics UK (COG-UK) Consortium | Berkshire and Surrey Pathology Services Lighthouse Laboratory and Alex Alderton, Roberto Amato, Jeffrey Barrett, Sonia Goncalves, Ewan Harrison, David K. Jackson, Ian Johnston, Dominic Kwiatkowski, Cordelia Langford, John Sillitoe on behalf of the Wellcome Sanger Institute COVID-19 Surveillance Team |
| EPI_ISL_3285623 | Lighthouse Lab in Milton Keynes | Wellcome Sanger Institute for the COVID-19 Genomics UK (COG-UK) Consortium | The Lighthouse Lab in Milton Keynes and Alex Alderton, Roberto Amato, Jeffrey Barrett, Sonia Goncalves, Ewan Harrison, David K. Jackson, Ian Johnston, Dominic Kwiatkowski, Cordelia Langford, John Sillitoe on behalf of the Wellcome Sanger Institute COVID-19 Surveillance Team |
| EPI_ISL_3285624 | Lighthouse Lab in Glasgow | Wellcome Sanger Institute for the COVID-19 Genomics UK (COG-UK) Consortium | Harper VanSteenhouse, Yumi Kasai, David Gray, Carol Clugston, Anna Dominiczak and Alex Alderton, Roberto Amato, Jeffrey Barrett, Sonia Goncalves, Ewan Harrison, David K. Jackson, Ian Johnston, Dominic Kwiatkowski, Cordelia Langford, John Sillitoe on behalf of the Wellcome Sanger Institute COVID-19 Surveillance Team |
| EPI_ISL_3285625, EPI_ISL_3285626 | Lighthouse Lab in Milton Keynes | Wellcome Sanger Institute for the COVID-19 Genomics UK (COG-UK) Consortium | The Lighthouse Lab in Milton Keynes and Alex Alderton, Roberto Amato, Jeffrey Barrett, Sonia Goncalves, Ewan Harrison, David K. Jackson, Ian Johnston, Dominic Kwiatkowski, Cordelia Langford, John Sillitoe on behalf of the Wellcome Sanger Institute COVID-19 Surveillance Team |
| EPI_ISL_3285627 | Berkshire and Surrey Pathology Services Lighthouse Laboratory | Wellcome Sanger Institute for the COVID-19 Genomics UK (COG-UK) Consortium | Berkshire and Surrey Pathology Services Lighthouse Laboratory and Alex Alderton, Roberto Amato, Jeffrey Barrett, Sonia Goncalves, Ewan Harrison, David K. Jackson, Ian Johnston, Dominic Kwiatkowski, Cordelia Langford, John Sillitoe on behalf of the Wellcome Sanger Institute COVID-19 Surveillance Team |
| EPI_ISL_3285628 | Lighthouse Lab in Glasgow | Wellcome Sanger Institute for the COVID-19 Genomics UK (COG-UK) Consortium | Harper VanSteenhouse, Yumi Kasai, David Gray, Carol Clugston, Anna Dominiczak and Alex Alderton, Roberto Amato, Jeffrey Barrett, Sonia Goncalves, Ewan Harrison, David K. Jackson, Ian Johnston, Dominic Kwiatkowski, Cordelia Langford, John Sillitoe on behalf of the Wellcome Sanger Institute COVID-19 Surveillance Team |
| EPI_ISL_3285629 | Berkshire and Surrey Pathology Services Lighthouse Laboratory | Wellcome Sanger Institute for the COVID-19 Genomics UK (COG-UK) Consortium | Berkshire and Surrey Pathology Services Lighthouse Laboratory and Alex Alderton, Roberto Amato, Jeffrey Barrett, Sonia Goncalves, Ewan Harrison, David K. Jackson, Ian Johnston, Dominic Kwiatkowski, Cordelia Langford, John Sillitoe on behalf of the Wellcome Sanger Institute COVID-19 Surveillance Team |
| EPI_ISL_3285630 | Lighthouse Lab in Milton Keynes | Wellcome Sanger Institute for the COVID-19 Genomics UK (COG-UK) Consortium | The Lighthouse Lab in Milton Keynes and Alex Alderton, Roberto Amato, Jeffrey Barrett, Sonia Goncalves, Ewan Harrison, David K. Jackson, Ian Johnston, Dominic Kwiatkowski, Cordelia Langford, John Sillitoe on behalf of the Wellcome Sanger Institute COVID-19 Surveillance Team |
| EPI_ISL_3285631 | Berkshire and Surrey Pathology Services Lighthouse Laboratory | Wellcome Sanger Institute for the COVID-19 Genomics UK (COG-UK) Consortium | Berkshire and Surrey Pathology Services Lighthouse Laboratory and Alex Alderton, Roberto Amato, Jeffrey Barrett, Sonia Goncalves, Ewan Harrison, David K. Jackson, Ian Johnston, Dominic Kwiatkowski, Cordelia Langford, John Sillitoe on behalf of the Wellcome Sanger Institute COVID-19 Surveillance Team |
| EPI_ISL_3285632, EPI_ISL_3285633 | Lighthouse Lab in Glasgow | Wellcome Sanger Institute for the COVID-19 Genomics UK (COG-UK) Consortium | Harper VanSteenhouse, Yumi Kasai, David Gray, Carol Clugston, Anna Dominiczak and Alex Alderton, Roberto Amato, Jeffrey Barrett, Sonia Goncalves, Ewan Harrison, David K. Jackson, Ian Johnston, Dominic Kwiatkowski, Cordelia Langford, John Sillitoe on behalf of the Wellcome Sanger Institute COVID-19 Surveillance Team |
| EPI_ISL_3285634, EPI_ISL_3285635, EPI_ISL_3285636, EPI_ISL_3285637 | Lighthouse Lab in Milton Keynes | Wellcome Sanger Institute for the COVID-19 Genomics UK (COG-UK) Consortium | The Lighthouse Lab in Milton Keynes and Alex Alderton, Roberto Amato, Jeffrey Barrett, Sonia Goncalves, Ewan Harrison, David K. Jackson, Ian Johnston, Dominic Kwiatkowski, Cordelia Langford, John Sillitoe on behalf of the Wellcome Sanger Institute COVID-19 Surveillance Team |
| EPI_ISL_3285638 | Berkshire and Surrey Pathology Services Lighthouse Laboratory | Wellcome Sanger Institute for the COVID-19 Genomics UK (COG-UK) Consortium | Berkshire and Surrey Pathology Services Lighthouse Laboratory and Alex Alderton, Roberto Amato, Jeffrey Barrett, Sonia Goncalves, Ewan Harrison, David K. Jackson, Ian Johnston, Dominic Kwiatkowski, Cordelia Langford, John Sillitoe on behalf of the Wellcome Sanger Institute COVID-19 Surveillance Team |
| EPI_ISL_3285639, EPI_ISL_3285640 | Lighthouse Lab in Milton Keynes | Wellcome Sanger Institute for the COVID-19 Genomics UK (COG-UK) Consortium | The Lighthouse Lab in Milton Keynes and Alex Alderton, Roberto Amato, Jeffrey Barrett, Sonia Goncalves, Ewan Harrison, David K. Jackson, Ian Johnston, Dominic Kwiatkowski, Cordelia Langford, John Sillitoe on behalf of the Wellcome Sanger Institute COVID-19 Surveillance Team |
| EPI_ISL_3285641 | Berkshire and Surrey Pathology Services Lighthouse Laboratory | Wellcome Sanger Institute for the COVID-19 Genomics UK (COG-UK) Consortium | Berkshire and Surrey Pathology Services Lighthouse Laboratory and Alex Alderton, Roberto Amato, Jeffrey Barrett, Sonia Goncalves, Ewan Harrison, David K. Jackson, Ian Johnston, Dominic Kwiatkowski, Cordelia Langford, John Sillitoe on behalf of the Wellcome Sanger Institute COVID-19 Surveillance Team |
| EPI_ISL_3285642, EPI_ISL_3285643, EPI_ISL_3285644 | Lighthouse Lab in Milton Keynes | Wellcome Sanger Institute for the COVID-19 Genomics UK (COG-UK) Consortium | The Lighthouse Lab in Milton Keynes and Alex Alderton, Roberto Amato, Jeffrey Barrett, Sonia Goncalves, Ewan Harrison, David K. Jackson, Ian Johnston, Dominic Kwiatkowski, Cordelia Langford, John Sillitoe on behalf of the Wellcome Sanger Institute COVID-19 Surveillance Team |
| EPI_ISL_3285645, EPI_ISL_3285646, EPI_ISL_3285647 | Lighthouse Lab in Glasgow | Wellcome Sanger Institute for the COVID-19 Genomics UK (COG-UK) Consortium | Harper VanSteenhouse, Yumi Kasai, David Gray, Carol Clugston, Anna Dominiczak and Alex Alderton, Roberto Amato, Jeffrey Barrett, Sonia Goncalves, Ewan Harrison, David K. Jackson, Ian Johnston, Dominic Kwiatkowski, Cordelia Langford, John Sillitoe on behalf of the Wellcome Sanger Institute COVID-19 Surveillance Team |
| EPI_ISL_3285648, EPI_ISL_3285649, EPI_ISL_3285650 | Lighthouse Lab in Milton Keynes | Wellcome Sanger Institute for the COVID-19 Genomics UK (COG-UK) Consortium | The Lighthouse Lab in Milton Keynes and Alex Alderton, Roberto Amato, Jeffrey Barrett, Sonia Goncalves, Ewan Harrison, David K. Jackson, Ian Johnston, Dominic Kwiatkowski, Cordelia Langford, John Sillitoe on behalf of the Wellcome Sanger Institute COVID-19 Surveillance Team |
| EPI_ISL_3285651 | Berkshire and Surrey Pathology Services Lighthouse Laboratory | Wellcome Sanger Institute for the COVID-19 Genomics UK (COG-UK) Consortium | Berkshire and Surrey Pathology Services Lighthouse Laboratory and Alex Alderton, Roberto Amato, Jeffrey Barrett, Sonia Goncalves, Ewan Harrison, David K. Jackson, Ian Johnston, Dominic Kwiatkowski, Cordelia Langford, John Sillitoe on behalf of the Wellcome Sanger Institute COVID-19 Surveillance Team |
| EPI_ISL_3285652 | Lighthouse Lab in Glasgow | Wellcome Sanger Institute for the COVID-19 Genomics UK (COG-UK) Consortium | Harper VanSteenhouse, Yumi Kasai, David Gray, Carol Clugston, Anna Dominiczak and Alex Alderton, Roberto Amato, Jeffrey Barrett, Sonia Goncalves, Ewan Harrison, David K. Jackson, Ian Johnston, Dominic Kwiatkowski, Cordelia Langford, John Sillitoe on behalf of the Wellcome Sanger Institute COVID-19 Surveillance Team |
| EPI_ISL_3285653, EPI_ISL_3285654, EPI_ISL_3285655, EPI_ISL_3285656, EPI_ISL_3285657, EPI_ISL_3285658 | Lighthouse Lab in Milton Keynes | Wellcome Sanger Institute for the COVID-19 Genomics UK (COG-UK) Consortium | The Lighthouse Lab in Milton Keynes and Alex Alderton, Roberto Amato, Jeffrey Barrett, Sonia Goncalves, Ewan Harrison, David K. Jackson, Ian Johnston, Dominic Kwiatkowski, Cordelia Langford, John Sillitoe on behalf of the Wellcome Sanger Institute COVID-19 Surveillance Team |
| EPI_ISL_3285659 | Berkshire and Surrey Pathology Services Lighthouse Laboratory | Wellcome Sanger Institute for the COVID-19 Genomics UK (COG-UK) Consortium | Berkshire and Surrey Pathology Services Lighthouse Laboratory and Alex Alderton, Roberto Amato, Jeffrey Barrett, Sonia Goncalves, Ewan Harrison, David K. Jackson, Ian Johnston, Dominic Kwiatkowski, Cordelia Langford, John Sillitoe on behalf of the Wellcome Sanger Institute COVID-19 Surveillance Team |
| EPI_ISL_3285660 | Lighthouse Lab in Glasgow | Wellcome Sanger Institute for the COVID-19 Genomics UK (COG-UK) Consortium | Harper VanSteenhouse, Yumi Kasai, David Gray, Carol Clugston, Anna Dominiczak and Alex Alderton, Roberto Amato, Jeffrey Barrett, Sonia Goncalves, Ewan Harrison, David K. Jackson, Ian Johnston, Dominic Kwiatkowski, Cordelia Langford, John Sillitoe on behalf of the Wellcome Sanger Institute COVID-19 Surveillance Team |
| EPI_ISL_3285661 | Lighthouse Lab in Milton Keynes | Wellcome Sanger Institute for the COVID-19 Genomics UK (COG-UK) Consortium | The Lighthouse Lab in Milton Keynes and Alex Alderton, Roberto Amato, Jeffrey Barrett, Sonia Goncalves, Ewan Harrison, David K. Jackson, Ian Johnston, Dominic Kwiatkowski, Cordelia Langford, John Sillitoe on behalf of the Wellcome Sanger Institute COVID-19 Surveillance Team |
| EPI_ISL_3285662 | Lighthouse Lab in Glasgow | Wellcome Sanger Institute for the COVID-19 Genomics UK (COG-UK) Consortium | Harper VanSteenhouse, Yumi Kasai, David Gray, Carol Clugston, Anna Dominiczak and Alex Alderton, Roberto Amato, Jeffrey Barrett, Sonia Goncalves, Ewan Harrison, David K. Jackson, Ian Johnston, Dominic Kwiatkowski, Cordelia Langford, John Sillitoe on behalf of the Wellcome Sanger Institute COVID-19 Surveillance Team |
| EPI_ISL_3285663 | Lighthouse Lab in Milton Keynes | Wellcome Sanger Institute for the COVID-19 Genomics UK (COG-UK) Consortium | The Lighthouse Lab in Milton Keynes and Alex Alderton, Roberto Amato, Jeffrey Barrett, Sonia Goncalves, Ewan Harrison, David K. Jackson, Ian Johnston, Dominic Kwiatkowski, Cordelia Langford, John Sillitoe on behalf of the Wellcome Sanger Institute COVID-19 Surveillance Team |
| EPI_ISL_3285664 | Berkshire and Surrey Pathology Services Lighthouse Laboratory | Wellcome Sanger Institute for the COVID-19 Genomics UK (COG-UK) Consortium | Berkshire and Surrey Pathology Services Lighthouse Laboratory and Alex Alderton, Roberto Amato, Jeffrey Barrett, Sonia Goncalves, Ewan Harrison, David K. Jackson, Ian Johnston, Dominic Kwiatkowski, Cordelia Langford, John Sillitoe on behalf of the Wellcome Sanger Institute COVID-19 Surveillance Team |
| EPI_ISL_3285665, EPI_ISL_3285666 | Lighthouse Lab in Milton Keynes | Wellcome Sanger Institute for the COVID-19 Genomics UK (COG-UK) Consortium | The Lighthouse Lab in Milton Keynes and Alex Alderton, Roberto Amato, Jeffrey Barrett, Sonia Goncalves, Ewan Harrison, David K. Jackson, Ian Johnston, Dominic Kwiatkowski, Cordelia Langford, John Sillitoe on behalf of the Wellcome Sanger Institute COVID-19 Surveillance Team |
| EPI_ISL_3285667, EPI_ISL_3285668, EPI_ISL_3285669, EPI_ISL_3285670 | Lighthouse Lab in Glasgow | Wellcome Sanger Institute for the COVID-19 Genomics UK (COG-UK) Consortium | Harper VanSteenhouse, Yumi Kasai, David Gray, Carol Clugston, Anna Dominiczak and Alex Alderton, Roberto Amato, Jeffrey Barrett, Sonia Goncalves, Ewan Harrison, David K. Jackson, Ian Johnston, Dominic Kwiatkowski, Cordelia Langford, John Sillitoe on behalf of the Wellcome Sanger Institute COVID-19 Surveillance Team |
| EPI_ISL_3285671 | Lighthouse Lab in Milton Keynes | Wellcome Sanger Institute for the COVID-19 Genomics UK (COG-UK) Consortium | The Lighthouse Lab in Milton Keynes and Alex Alderton, Roberto Amato, Jeffrey Barrett, Sonia Goncalves, Ewan Harrison, David K. Jackson, Ian Johnston, Dominic Kwiatkowski, Cordelia Langford, John Sillitoe on behalf of the Wellcome Sanger Institute COVID-19 Surveillance Team |
| EPI_ISL_3285672 | Berkshire and Surrey Pathology Services Lighthouse Laboratory | Wellcome Sanger Institute for the COVID-19 Genomics UK (COG-UK) Consortium | Berkshire and Surrey Pathology Services Lighthouse Laboratory and Alex Alderton, Roberto Amato, Jeffrey Barrett, Sonia Goncalves, Ewan Harrison, David K. Jackson, Ian Johnston, Dominic Kwiatkowski, Cordelia Langford, John Sillitoe on behalf of the Wellcome Sanger Institute COVID-19 Surveillance Team |
| EPI_ISL_3285673, EPI_ISL_3285674, | Lighthouse Lab in Milton Keynes | Wellcome Sanger Institute for the COVID-19 Genomics UK (COG-UK) Consortium | The Lighthouse Lab in Milton Keynes and Alex Alderton, Roberto Amato, Jeffrey Barrett, Sonia Goncalves, Ewan Harrison, David K. Jackson, Ian Johnston, Dominic Kwiatkowski, Cordelia Langford, John Sillitoe on behalf of the Wellcome Sanger Institute COVID-19 Surveillance Team |

| | | | |
|---|---|---|---|
| EPI_ISL_3285675 | | | |
| EPI_ISL_3285676, EPI_ISL_3285677, EPI_ISL_3285678, EPI_ISL_3285679, EPI_ISL_3285680 | Lighthouse Lab in Glasgow | Wellcome Sanger Institute for the COVID-19 Genomics UK (COG-UK) Consortium | Harper VanSteenhouse, Yumi Kasai, David Gray, Carol Clugston, Anna Dominiczak and Alex Alderton, Roberto Amato, Jeffrey Barrett, Sonia Goncalves, Ewan Harrison, David K. Jackson, Ian Johnston, Dominic Kwiatkowski, Cordelia Langford, John Sillitoe on behalf of the Wellcome Sanger Institute COVID-19 Surveillance Team |
| EPI_ISL_3285681, EPI_ISL_3285682, EPI_ISL_3285683, EPI_ISL_3285684, EPI_ISL_3285685 | Berkshire and Surrey Pathology Services Lighthouse Laboratory | Wellcome Sanger Institute for the COVID-19 Genomics UK (COG-UK) Consortium | Berkshire and Surrey Pathology Services Lighthouse Laboratory and Alex Alderton, Roberto Amato, Jeffrey Barrett, Sonia Goncalves, Ewan Harrison, David K. Jackson, Ian Johnston, Dominic Kwiatkowski, Cordelia Langford, John Sillitoe on behalf of the Wellcome Sanger Institute COVID-19 Surveillance Team |
| EPI_ISL_3285686, EPI_ISL_3285687 | Lighthouse Lab in Milton Keynes | Wellcome Sanger Institute for the COVID-19 Genomics UK (COG-UK) Consortium | The Lighthouse Lab in Milton Keynes and Alex Alderton, Roberto Amato, Jeffrey Barrett, Sonia Goncalves, Ewan Harrison, David K. Jackson, Ian Johnston, Dominic Kwiatkowski, Cordelia Langford, John Sillitoe on behalf of the Wellcome Sanger Institute COVID-19 Surveillance Team |
| EPI_ISL_3285688 | Berkshire and Surrey Pathology Services Lighthouse Laboratory | Wellcome Sanger Institute for the COVID-19 Genomics UK (COG-UK) Consortium | Berkshire and Surrey Pathology Services Lighthouse Laboratory and Alex Alderton, Roberto Amato, Jeffrey Barrett, Sonia Goncalves, Ewan Harrison, David K. Jackson, Ian Johnston, Dominic Kwiatkowski, Cordelia Langford, John Sillitoe on behalf of the Wellcome Sanger Institute COVID-19 Surveillance Team |
| EPI_ISL_3285689, EPI_ISL_3285690 | Lighthouse Lab in Milton Keynes | Wellcome Sanger Institute for the COVID-19 Genomics UK (COG-UK) Consortium | The Lighthouse Lab in Milton Keynes and Alex Alderton, Roberto Amato, Jeffrey Barrett, Sonia Goncalves, Ewan Harrison, David K. Jackson, Ian Johnston, Dominic Kwiatkowski, Cordelia Langford, John Sillitoe on behalf of the Wellcome Sanger Institute COVID-19 Surveillance Team |
| EPI_ISL_3285691 | Lighthouse Lab in Glasgow | Wellcome Sanger Institute for the COVID-19 Genomics UK (COG-UK) Consortium | Harper VanSteenhouse, Yumi Kasai, David Gray, Carol Clugston, Anna Dominiczak and Alex Alderton, Roberto Amato, Jeffrey Barrett, Sonia Goncalves, Ewan Harrison, David K. Jackson, Ian Johnston, Dominic Kwiatkowski, Cordelia Langford, John Sillitoe on behalf of the Wellcome Sanger Institute COVID-19 Surveillance Team |
| EPI_ISL_3285692, EPI_ISL_3285693 | Lighthouse Lab in Milton Keynes | Wellcome Sanger Institute for the COVID-19 Genomics UK (COG-UK) Consortium | The Lighthouse Lab in Milton Keynes and Alex Alderton, Roberto Amato, Jeffrey Barrett, Sonia Goncalves, Ewan Harrison, David K. Jackson, Ian Johnston, Dominic Kwiatkowski, Cordelia Langford, John Sillitoe on behalf of the Wellcome Sanger Institute COVID-19 Surveillance Team |
| EPI_ISL_3285694 | Lighthouse Lab in Glasgow | Wellcome Sanger Institute for the COVID-19 Genomics UK (COG-UK) Consortium | Harper VanSteenhouse, Yumi Kasai, David Gray, Carol Clugston, Anna Dominiczak and Alex Alderton, Roberto Amato, Jeffrey Barrett, Sonia Goncalves, Ewan Harrison, David K. Jackson, Ian Johnston, Dominic Kwiatkowski, Cordelia Langford, John Sillitoe on behalf of the Wellcome Sanger Institute COVID-19 Surveillance Team |
| EPI_ISL_3285695, EPI_ISL_3285696 | Lighthouse Lab in Milton Keynes | Wellcome Sanger Institute for the COVID-19 Genomics UK (COG-UK) Consortium | The Lighthouse Lab in Milton Keynes and Alex Alderton, Roberto Amato, Jeffrey Barrett, Sonia Goncalves, Ewan Harrison, David K. Jackson, Ian Johnston, Dominic Kwiatkowski, Cordelia Langford, John Sillitoe on behalf of the Wellcome Sanger Institute COVID-19 Surveillance Team |
| EPI_ISL_3285697, EPI_ISL_3285698, EPI_ISL_3285699, EPI_ISL_3285700 | Berkshire and Surrey Pathology Services Lighthouse Laboratory | Wellcome Sanger Institute for the COVID-19 Genomics UK (COG-UK) Consortium | Berkshire and Surrey Pathology Services Lighthouse Laboratory and Alex Alderton, Roberto Amato, Jeffrey Barrett, Sonia Goncalves, Ewan Harrison, David K. Jackson, Ian Johnston, Dominic Kwiatkowski, Cordelia Langford, John Sillitoe on behalf of the Wellcome Sanger Institute COVID-19 Surveillance Team |
| EPI_ISL_3285701, EPI_ISL_3285702, EPI_ISL_3285703 | Lighthouse Lab in Glasgow | Wellcome Sanger Institute for the COVID-19 Genomics UK (COG-UK) Consortium | Harper VanSteenhouse, Yumi Kasai, David Gray, Carol Clugston, Anna Dominiczak and Alex Alderton, Roberto Amato, Jeffrey Barrett, Sonia Goncalves, Ewan Harrison, David K. Jackson, Ian Johnston, Dominic Kwiatkowski, Cordelia Langford, John Sillitoe on behalf of the Wellcome Sanger Institute COVID-19 Surveillance Team |
| EPI_ISL_3285704 | Lighthouse Lab in Milton Keynes | Wellcome Sanger Institute for the COVID-19 Genomics UK (COG-UK) Consortium | The Lighthouse Lab in Milton Keynes and Alex Alderton, Roberto Amato, Jeffrey Barrett, Sonia Goncalves, Ewan Harrison, David K. Jackson, Ian Johnston, Dominic Kwiatkowski, Cordelia Langford, John Sillitoe on behalf of the Wellcome Sanger Institute COVID-19 Surveillance Team |
| EPI_ISL_3285705 | Lighthouse Lab in Glasgow | Wellcome Sanger Institute for the COVID-19 Genomics UK (COG-UK) Consortium | Harper VanSteenhouse, Yumi Kasai, David Gray, Carol Clugston, Anna Dominiczak and Alex Alderton, Roberto Amato, Jeffrey Barrett, Sonia Goncalves, Ewan Harrison, David K. Jackson, Ian Johnston, Dominic Kwiatkowski, Cordelia Langford, John Sillitoe on behalf of the Wellcome Sanger Institute COVID-19 Surveillance Team |
| EPI_ISL_3285706 | Lighthouse Lab in Milton Keynes | Wellcome Sanger Institute for the COVID-19 Genomics UK (COG-UK) Consortium | The Lighthouse Lab in Milton Keynes and Alex Alderton, Roberto Amato, Jeffrey Barrett, Sonia Goncalves, Ewan Harrison, David K. Jackson, Ian Johnston, Dominic Kwiatkowski, Cordelia Langford, John Sillitoe on behalf of the Wellcome Sanger Institute COVID-19 Surveillance Team |
| EPI_ISL_3285707 | Lighthouse Lab in Glasgow | Wellcome Sanger Institute for the COVID-19 Genomics UK (COG-UK) Consortium | Harper VanSteenhouse, Yumi Kasai, David Gray, Carol Clugston, Anna Dominiczak and Alex Alderton, Roberto Amato, Jeffrey Barrett, Sonia Goncalves, Ewan Harrison, David K. Jackson, Ian Johnston, Dominic Kwiatkowski, Cordelia Langford, John Sillitoe on behalf of the Wellcome Sanger Institute COVID-19 Surveillance Team |
| EPI_ISL_3285708 | Lighthouse Lab in Milton Keynes | Wellcome Sanger Institute for the COVID-19 Genomics UK (COG-UK) Consortium | The Lighthouse Lab in Milton Keynes and Alex Alderton, Roberto Amato, Jeffrey Barrett, Sonia Goncalves, Ewan Harrison, David K. Jackson, Ian Johnston, Dominic Kwiatkowski, Cordelia Langford, John Sillitoe on behalf of the Wellcome Sanger Institute COVID-19 Surveillance Team |
| EPI_ISL_3285709 | Lighthouse Lab in Glasgow | Wellcome Sanger Institute for the COVID-19 Genomics UK (COG-UK) Consortium | Harper VanSteenhouse, Yumi Kasai, David Gray, Carol Clugston, Anna Dominiczak and Alex Alderton, Roberto Amato, Jeffrey Barrett, Sonia Goncalves, Ewan Harrison, David K. Jackson, Ian Johnston, Dominic Kwiatkowski, Cordelia Langford, John Sillitoe on behalf of the Wellcome Sanger Institute COVID-19 Surveillance Team |
| EPI_ISL_3285710 | Berkshire and Surrey Pathology Services Lighthouse Laboratory | Wellcome Sanger Institute for the COVID-19 Genomics UK (COG-UK) Consortium | Berkshire and Surrey Pathology Services Lighthouse Laboratory and Alex Alderton, Roberto Amato, Jeffrey Barrett, Sonia Goncalves, Ewan Harrison, David K. Jackson, Ian Johnston, Dominic Kwiatkowski, Cordelia Langford, John Sillitoe on behalf of the Wellcome Sanger Institute COVID-19 Surveillance Team |
| EPI_ISL_3285711, EPI_ISL_3285712, EPI_ISL_3285713, EPI_ISL_3285714 | Lighthouse Lab in Milton Keynes | Wellcome Sanger Institute for the COVID-19 Genomics UK (COG-UK) Consortium | The Lighthouse Lab in Milton Keynes and Alex Alderton, Roberto Amato, Jeffrey Barrett, Sonia Goncalves, Ewan Harrison, David K. Jackson, Ian Johnston, Dominic Kwiatkowski, Cordelia Langford, John Sillitoe on behalf of the Wellcome Sanger Institute COVID-19 Surveillance Team |
| EPI_ISL_3285715 | Lighthouse Lab in Glasgow | Wellcome Sanger Institute for the COVID-19 Genomics UK (COG-UK) Consortium | Harper VanSteenhouse, Yumi Kasai, David Gray, Carol Clugston, Anna Dominiczak and Alex Alderton, Roberto Amato, Jeffrey Barrett, Sonia Goncalves, Ewan Harrison, David K. Jackson, Ian Johnston, Dominic Kwiatkowski, Cordelia Langford, John Sillitoe on behalf of the Wellcome Sanger Institute COVID-19 Surveillance Team |
| EPI_ISL_3285716 | Lighthouse Lab in Milton Keynes | Wellcome Sanger Institute for the COVID-19 Genomics UK (COG-UK) Consortium | The Lighthouse Lab in Milton Keynes and Alex Alderton, Roberto Amato, Jeffrey Barrett, Sonia Goncalves, Ewan Harrison, David K. Jackson, Ian Johnston, Dominic Kwiatkowski, Cordelia Langford, John Sillitoe on behalf of the Wellcome Sanger Institute COVID-19 Surveillance Team |
| EPI_ISL_3285717, EPI_ISL_3285718 | Lighthouse Lab in Glasgow | Wellcome Sanger Institute for the COVID-19 Genomics UK (COG-UK) Consortium | Harper VanSteenhouse, Yumi Kasai, David Gray, Carol Clugston, Anna Dominiczak and Alex Alderton, Roberto Amato, Jeffrey Barrett, Sonia Goncalves, Ewan Harrison, David K. Jackson, Ian Johnston, Dominic Kwiatkowski, Cordelia Langford, John Sillitoe on behalf of the Wellcome Sanger Institute COVID-19 Surveillance Team |
| EPI_ISL_3285719, EPI_ISL_3285720 | Lighthouse Lab in Milton Keynes | Wellcome Sanger Institute for the COVID-19 Genomics UK (COG-UK) Consortium | The Lighthouse Lab in Milton Keynes and Alex Alderton, Roberto Amato, Jeffrey Barrett, Sonia Goncalves, Ewan Harrison, David K. Jackson, Ian Johnston, Dominic Kwiatkowski, Cordelia Langford, John Sillitoe on behalf of the Wellcome Sanger Institute COVID-19 Surveillance Team |
| EPI_ISL_3285721 | Berkshire and Surrey Pathology Services Lighthouse Laboratory | Wellcome Sanger Institute for the COVID-19 Genomics UK (COG-UK) Consortium | Berkshire and Surrey Pathology Services Lighthouse Laboratory and Alex Alderton, Roberto Amato, Jeffrey Barrett, Sonia Goncalves, Ewan Harrison, David K. Jackson, Ian Johnston, Dominic Kwiatkowski, Cordelia Langford, John Sillitoe on behalf of the Wellcome Sanger Institute COVID-19 Surveillance Team |
| EPI_ISL_3285722 | Lighthouse Lab in Milton Keynes | Wellcome Sanger Institute for the COVID-19 Genomics UK (COG-UK) Consortium | The Lighthouse Lab in Milton Keynes and Alex Alderton, Roberto Amato, Jeffrey Barrett, Sonia Goncalves, Ewan Harrison, David K. Jackson, Ian Johnston, Dominic Kwiatkowski, Cordelia Langford, John Sillitoe on behalf of the Wellcome Sanger Institute COVID-19 Surveillance Team |
| EPI_ISL_3285723 | Lighthouse Lab in Glasgow | Wellcome Sanger Institute for the COVID-19 Genomics UK (COG-UK) Consortium | Harper VanSteenhouse, Yumi Kasai, David Gray, Carol Clugston, Anna Dominiczak and Alex Alderton, Roberto Amato, Jeffrey Barrett, Sonia Goncalves, Ewan Harrison, David K. Jackson, Ian Johnston, Dominic Kwiatkowski, Cordelia Langford, John Sillitoe on behalf of the Wellcome Sanger Institute COVID-19 Surveillance Team |
| EPI_ISL_3285724, EPI_ISL_3285725 | Lighthouse Lab in Milton Keynes | Wellcome Sanger Institute for the COVID-19 Genomics UK (COG-UK) Consortium | The Lighthouse Lab in Milton Keynes and Alex Alderton, Roberto Amato, Jeffrey Barrett, Sonia Goncalves, Ewan Harrison, David K. Jackson, Ian Johnston, Dominic Kwiatkowski, Cordelia Langford, John Sillitoe on behalf of the Wellcome Sanger Institute COVID-19 Surveillance Team |
| EPI_ISL_3285726, EPI_ISL_3285727 | Berkshire and Surrey Pathology Services Lighthouse Laboratory | Wellcome Sanger Institute for the COVID-19 Genomics UK (COG-UK) Consortium | Berkshire and Surrey Pathology Services Lighthouse Laboratory and Alex Alderton, Roberto Amato, Jeffrey Barrett, Sonia Goncalves, Ewan Harrison, David K. Jackson, Ian Johnston, Dominic Kwiatkowski, Cordelia Langford, John Sillitoe on behalf of the Wellcome Sanger Institute COVID-19 Surveillance Team |
| EPI_ISL_3285728 | Lighthouse Lab in Glasgow | Wellcome Sanger Institute for the COVID-19 Genomics UK (COG-UK) Consortium | Harper VanSteenhouse, Yumi Kasai, David Gray, Carol Clugston, Anna Dominiczak and Alex Alderton, Roberto Amato, Jeffrey Barrett, Sonia Goncalves, Ewan Harrison, David K. Jackson, Ian Johnston, Dominic Kwiatkowski, Cordelia Langford, John Sillitoe on behalf of the Wellcome Sanger Institute COVID-19 Surveillance Team |
| EPI_ISL_3285729 | Berkshire and Surrey Pathology Services Lighthouse Laboratory | Wellcome Sanger Institute for the COVID-19 Genomics UK (COG-UK) Consortium | Berkshire and Surrey Pathology Services Lighthouse Laboratory and Alex Alderton, Roberto Amato, Jeffrey Barrett, Sonia Goncalves, Ewan Harrison, David K. Jackson, Ian Johnston, Dominic Kwiatkowski, Cordelia Langford, John Sillitoe on behalf of the Wellcome Sanger Institute COVID-19 Surveillance Team |
| EPI_ISL_3285730 | Lighthouse Lab in Milton Keynes | Wellcome Sanger Institute for the COVID-19 Genomics UK (COG-UK) Consortium | The Lighthouse Lab in Milton Keynes and Alex Alderton, Roberto Amato, Jeffrey Barrett, Sonia Goncalves, Ewan Harrison, David K. Jackson, Ian Johnston, Dominic Kwiatkowski, Cordelia Langford, John Sillitoe on behalf of the Wellcome Sanger Institute COVID-19 Surveillance Team |
| EPI_ISL_3285731, EPI_ISL_3285732 | Lighthouse Lab in Glasgow | Wellcome Sanger Institute for the COVID-19 Genomics UK (COG-UK) Consortium | Harper VanSteenhouse, Yumi Kasai, David Gray, Carol Clugston, Anna Dominiczak and Alex Alderton, Roberto Amato, Jeffrey Barrett, Sonia Goncalves, Ewan Harrison, David K. Jackson, Ian Johnston, Dominic Kwiatkowski, Cordelia Langford, John Sillitoe on behalf of the Wellcome Sanger Institute COVID-19 Surveillance Team |
| EPI_ISL_3285733 | Berkshire and Surrey Pathology Services Lighthouse Laboratory | Wellcome Sanger Institute for the COVID-19 Genomics UK (COG-UK) Consortium | Berkshire and Surrey Pathology Services Lighthouse Laboratory and Alex Alderton, Roberto Amato, Jeffrey Barrett, Sonia Goncalves, Ewan Harrison, David K. Jackson, Ian Johnston, Dominic Kwiatkowski, Cordelia Langford, John Sillitoe on behalf of the Wellcome Sanger Institute COVID-19 Surveillance Team |
| EPI_ISL_3285734, EPI_ISL_3285735, EPI_ISL_3285736 | Lighthouse Lab in Glasgow | Wellcome Sanger Institute for the COVID-19 Genomics UK (COG-UK) Consortium | Harper VanSteenhouse, Yumi Kasai, David Gray, Carol Clugston, Anna Dominiczak and Alex Alderton, Roberto Amato, Jeffrey Barrett, Sonia Goncalves, Ewan Harrison, David K. Jackson, Ian Johnston, Dominic Kwiatkowski, Cordelia Langford, John Sillitoe on behalf of the Wellcome Sanger Institute COVID-19 Surveillance Team |
| EPI_ISL_3285737, EPI_ISL_3285738, EPI_ISL_3285739 | Lighthouse Lab in Milton Keynes | Wellcome Sanger Institute for the COVID-19 Genomics UK (COG-UK) Consortium | The Lighthouse Lab in Milton Keynes and Alex Alderton, Roberto Amato, Jeffrey Barrett, Sonia Goncalves, Ewan Harrison, David K. Jackson, Ian Johnston, Dominic Kwiatkowski, Cordelia Langford, John Sillitoe on behalf of the Wellcome Sanger Institute COVID-19 Surveillance Team |
| EPI_ISL_3285740 | Berkshire and Surrey Pathology Services Lighthouse Laboratory | Wellcome Sanger Institute for the COVID-19 Genomics UK (COG-UK) Consortium | Berkshire and Surrey Pathology Services Lighthouse Laboratory and Alex Alderton, Roberto Amato, Jeffrey Barrett, Sonia Goncalves, Ewan Harrison, David K. Jackson, Ian Johnston, Dominic Kwiatkowski, Cordelia Langford, John Sillitoe on behalf of the Wellcome Sanger Institute COVID-19 Surveillance Team |
| EPI_ISL_3285741, EPI_ISL_3285742, EPI_ISL_3285743 | Lighthouse Lab in Milton Keynes | Wellcome Sanger Institute for the COVID-19 Genomics UK (COG-UK) Consortium | The Lighthouse Lab in Milton Keynes and Alex Alderton, Roberto Amato, Jeffrey Barrett, Sonia Goncalves, Ewan Harrison, David K. Jackson, Ian Johnston, Dominic Kwiatkowski, Cordelia Langford, John Sillitoe on behalf of the Wellcome Sanger Institute COVID-19 Surveillance Team |
| EPI_ISL_3285744 | Berkshire and Surrey Pathology Services Lighthouse Laboratory | Wellcome Sanger Institute for the COVID-19 Genomics UK (COG-UK) Consortium | Berkshire and Surrey Pathology Services Lighthouse Laboratory and Alex Alderton, Roberto Amato, Jeffrey Barrett, Sonia Goncalves, Ewan Harrison, David K. Jackson, Ian Johnston, Dominic Kwiatkowski, Cordelia Langford, John Sillitoe on behalf of the Wellcome Sanger Institute COVID-19 Surveillance Team |
| EPI_ISL_3285745 | Lighthouse Lab in Glasgow | Wellcome Sanger Institute for the COVID-19 Genomics UK (COG-UK) | Harper VanSteenhouse, Yumi Kasai, David Gray, Carol Clugston, Anna Dominiczak and Alex Alderton, Roberto Amato, Jeffrey Barrett, Sonia Goncalves, Ewan Harrison, David K. Jackson, Ian Johnston, Dominic Kwiatkowski, Cordelia Langford, |

| | | | |
|---|---|---|---|
| | | Consortium | John Sillitoe on behalf of the Wellcome Sanger Institute COVID-19 Surveillance Team |
| EPI_ISL_3285746 | Lighthouse Lab in Milton Keynes | Wellcome Sanger Institute for the COVID-19 Genomics UK (COG-UK) Consortium | The Lighthouse Lab in Milton Keynes and Alex Alderton, Roberto Amato, Jeffrey Barrett, Sonia Goncalves, Ewan Harrison, David K. Jackson, Ian Johnston, Dominic Kwiatkowski, Cordelia Langford, John Sillitoe on behalf of the Wellcome Sanger Institute COVID-19 Surveillance Team |
| EPI_ISL_3285747, EPI_ISL_3285748, EPI_ISL_3285749 | Berkshire and Surrey Pathology Services Lighthouse Laboratory | Wellcome Sanger Institute for the COVID-19 Genomics UK (COG-UK) Consortium | Berkshire and Surrey Pathology Services Lighthouse Laboratory and Alex Alderton, Roberto Amato, Jeffrey Barrett, Sonia Goncalves, Ewan Harrison, David K. Jackson, Ian Johnston, Dominic Kwiatkowski, Cordelia Langford, John Sillitoe on behalf of the Wellcome Sanger Institute COVID-19 Surveillance Team |
| EPI_ISL_3285750 | Lighthouse Lab in Milton Keynes | Wellcome Sanger Institute for the COVID-19 Genomics UK (COG-UK) Consortium | The Lighthouse Lab in Milton Keynes and Alex Alderton, Roberto Amato, Jeffrey Barrett, Sonia Goncalves, Ewan Harrison, David K. Jackson, Ian Johnston, Dominic Kwiatkowski, Cordelia Langford, John Sillitoe on behalf of the Wellcome Sanger Institute COVID-19 Surveillance Team |
| EPI_ISL_3285751, EPI_ISL_3285752, EPI_ISL_3285753 | Lighthouse Lab in Glasgow | Wellcome Sanger Institute for the COVID-19 Genomics UK (COG-UK) Consortium | Harper VanSteenhouse, Yumi Kasai, David Gray, Carol Clugston, Anna Dominiczak and Alex Alderton, Roberto Amato, Jeffrey Barrett, Sonia Goncalves, Ewan Harrison, David K. Jackson, Ian Johnston, Dominic Kwiatkowski, Cordelia Langford, John Sillitoe on behalf of the Wellcome Sanger Institute COVID-19 Surveillance Team |
| EPI_ISL_3285754 | Lighthouse Lab in Milton Keynes | Wellcome Sanger Institute for the COVID-19 Genomics UK (COG-UK) Consortium | The Lighthouse Lab in Milton Keynes and Alex Alderton, Roberto Amato, Jeffrey Barrett, Sonia Goncalves, Ewan Harrison, David K. Jackson, Ian Johnston, Dominic Kwiatkowski, Cordelia Langford, John Sillitoe on behalf of the Wellcome Sanger Institute COVID-19 Surveillance Team |
| EPI_ISL_3285755 | Lighthouse Lab in Glasgow | Wellcome Sanger Institute for the COVID-19 Genomics UK (COG-UK) Consortium | Harper VanSteenhouse, Yumi Kasai, David Gray, Carol Clugston, Anna Dominiczak and Alex Alderton, Roberto Amato, Jeffrey Barrett, Sonia Goncalves, Ewan Harrison, David K. Jackson, Ian Johnston, Dominic Kwiatkowski, Cordelia Langford, John Sillitoe on behalf of the Wellcome Sanger Institute COVID-19 Surveillance Team |
| EPI_ISL_3285756, EPI_ISL_3285757 | Lighthouse Lab in Milton Keynes | Wellcome Sanger Institute for the COVID-19 Genomics UK (COG-UK) Consortium | The Lighthouse Lab in Milton Keynes and Alex Alderton, Roberto Amato, Jeffrey Barrett, Sonia Goncalves, Ewan Harrison, David K. Jackson, Ian Johnston, Dominic Kwiatkowski, Cordelia Langford, John Sillitoe on behalf of the Wellcome Sanger Institute COVID-19 Surveillance Team |
| EPI_ISL_3285758 | Berkshire and Surrey Pathology Services Lighthouse Laboratory | Wellcome Sanger Institute for the COVID-19 Genomics UK (COG-UK) Consortium | Berkshire and Surrey Pathology Services Lighthouse Laboratory and Alex Alderton, Roberto Amato, Jeffrey Barrett, Sonia Goncalves, Ewan Harrison, David K. Jackson, Ian Johnston, Dominic Kwiatkowski, Cordelia Langford, John Sillitoe on behalf of the Wellcome Sanger Institute COVID-19 Surveillance Team |
| EPI_ISL_3285759, EPI_ISL_3285760 | Lighthouse Lab in Milton Keynes | Wellcome Sanger Institute for the COVID-19 Genomics UK (COG-UK) Consortium | The Lighthouse Lab in Milton Keynes and Alex Alderton, Roberto Amato, Jeffrey Barrett, Sonia Goncalves, Ewan Harrison, David K. Jackson, Ian Johnston, Dominic Kwiatkowski, Cordelia Langford, John Sillitoe on behalf of the Wellcome Sanger Institute COVID-19 Surveillance Team |
| EPI_ISL_3285761 | Lighthouse Lab in Glasgow | Wellcome Sanger Institute for the COVID-19 Genomics UK (COG-UK) Consortium | Harper VanSteenhouse, Yumi Kasai, David Gray, Carol Clugston, Anna Dominiczak and Alex Alderton, Roberto Amato, Jeffrey Barrett, Sonia Goncalves, Ewan Harrison, David K. Jackson, Ian Johnston, Dominic Kwiatkowski, Cordelia Langford, John Sillitoe on behalf of the Wellcome Sanger Institute COVID-19 Surveillance Team |
| EPI_ISL_3285762 | Lighthouse Lab in Milton Keynes | Wellcome Sanger Institute for the COVID-19 Genomics UK (COG-UK) Consortium | The Lighthouse Lab in Milton Keynes and Alex Alderton, Roberto Amato, Jeffrey Barrett, Sonia Goncalves, Ewan Harrison, David K. Jackson, Ian Johnston, Dominic Kwiatkowski, Cordelia Langford, John Sillitoe on behalf of the Wellcome Sanger Institute COVID-19 Surveillance Team |
| EPI_ISL_3285763 | Lighthouse Lab in Glasgow | Wellcome Sanger Institute for the COVID-19 Genomics UK (COG-UK) Consortium | Harper VanSteenhouse, Yumi Kasai, David Gray, Carol Clugston, Anna Dominiczak and Alex Alderton, Roberto Amato, Jeffrey Barrett, Sonia Goncalves, Ewan Harrison, David K. Jackson, Ian Johnston, Dominic Kwiatkowski, Cordelia Langford, John Sillitoe on behalf of the Wellcome Sanger Institute COVID-19 Surveillance Team |
| EPI_ISL_3285764 | Lighthouse Lab in Milton Keynes | Wellcome Sanger Institute for the COVID-19 Genomics UK (COG-UK) Consortium | The Lighthouse Lab in Milton Keynes and Alex Alderton, Roberto Amato, Jeffrey Barrett, Sonia Goncalves, Ewan Harrison, David K. Jackson, Ian Johnston, Dominic Kwiatkowski, Cordelia Langford, John Sillitoe on behalf of the Wellcome Sanger Institute COVID-19 Surveillance Team |
| EPI_ISL_3285765 | Lighthouse Lab in Glasgow | Wellcome Sanger Institute for the COVID-19 Genomics UK (COG-UK) Consortium | Harper VanSteenhouse, Yumi Kasai, David Gray, Carol Clugston, Anna Dominiczak and Alex Alderton, Roberto Amato, Jeffrey Barrett, Sonia Goncalves, Ewan Harrison, David K. Jackson, Ian Johnston, Dominic Kwiatkowski, Cordelia Langford, John Sillitoe on behalf of the Wellcome Sanger Institute COVID-19 Surveillance Team |
| EPI_ISL_3285766, EPI_ISL_3285767 | Lighthouse Lab in Milton Keynes | Wellcome Sanger Institute for the COVID-19 Genomics UK (COG-UK) Consortium | The Lighthouse Lab in Milton Keynes and Alex Alderton, Roberto Amato, Jeffrey Barrett, Sonia Goncalves, Ewan Harrison, David K. Jackson, Ian Johnston, Dominic Kwiatkowski, Cordelia Langford, John Sillitoe on behalf of the Wellcome Sanger Institute COVID-19 Surveillance Team |
| EPI_ISL_3285768 | Lighthouse Lab in Glasgow | Wellcome Sanger Institute for the COVID-19 Genomics UK (COG-UK) Consortium | Harper VanSteenhouse, Yumi Kasai, David Gray, Carol Clugston, Anna Dominiczak and Alex Alderton, Roberto Amato, Jeffrey Barrett, Sonia Goncalves, Ewan Harrison, David K. Jackson, Ian Johnston, Dominic Kwiatkowski, Cordelia Langford, John Sillitoe on behalf of the Wellcome Sanger Institute COVID-19 Surveillance Team |
| EPI_ISL_3285769 | Berkshire and Surrey Pathology Services Lighthouse Laboratory | Wellcome Sanger Institute for the COVID-19 Genomics UK (COG-UK) Consortium | Berkshire and Surrey Pathology Services Lighthouse Laboratory and Alex Alderton, Roberto Amato, Jeffrey Barrett, Sonia Goncalves, Ewan Harrison, David K. Jackson, Ian Johnston, Dominic Kwiatkowski, Cordelia Langford, John Sillitoe on behalf of the Wellcome Sanger Institute COVID-19 Surveillance Team |
| EPI_ISL_3285770, EPI_ISL_3285771, EPI_ISL_3285772, EPI_ISL_3285773 | Lighthouse Lab in Milton Keynes | Wellcome Sanger Institute for the COVID-19 Genomics UK (COG-UK) Consortium | The Lighthouse Lab in Milton Keynes and Alex Alderton, Roberto Amato, Jeffrey Barrett, Sonia Goncalves, Ewan Harrison, David K. Jackson, Ian Johnston, Dominic Kwiatkowski, Cordelia Langford, John Sillitoe on behalf of the Wellcome Sanger Institute COVID-19 Surveillance Team |
| EPI_ISL_3285774, EPI_ISL_3285775, EPI_ISL_3285776 | Berkshire and Surrey Pathology Services Lighthouse Laboratory | Wellcome Sanger Institute for the COVID-19 Genomics UK (COG-UK) Consortium | Berkshire and Surrey Pathology Services Lighthouse Laboratory and Alex Alderton, Roberto Amato, Jeffrey Barrett, Sonia Goncalves, Ewan Harrison, David K. Jackson, Ian Johnston, Dominic Kwiatkowski, Cordelia Langford, John Sillitoe on behalf of the Wellcome Sanger Institute COVID-19 Surveillance Team |
| EPI_ISL_3285777 | Lighthouse Lab in Milton Keynes | Wellcome Sanger Institute for the COVID-19 Genomics UK (COG-UK) Consortium | The Lighthouse Lab in Milton Keynes and Alex Alderton, Roberto Amato, Jeffrey Barrett, Sonia Goncalves, Ewan Harrison, David K. Jackson, Ian Johnston, Dominic Kwiatkowski, Cordelia Langford, John Sillitoe on behalf of the Wellcome Sanger Institute COVID-19 Surveillance Team |
| EPI_ISL_3285778, EPI_ISL_3285779 | Lighthouse Lab in Glasgow | Wellcome Sanger Institute for the COVID-19 Genomics UK (COG-UK) Consortium | Harper VanSteenhouse, Yumi Kasai, David Gray, Carol Clugston, Anna Dominiczak and Alex Alderton, Roberto Amato, Jeffrey Barrett, Sonia Goncalves, Ewan Harrison, David K. Jackson, Ian Johnston, Dominic Kwiatkowski, Cordelia Langford, John Sillitoe on behalf of the Wellcome Sanger Institute COVID-19 Surveillance Team |
| EPI_ISL_3285780 | Berkshire and Surrey Pathology Services Lighthouse Laboratory | Wellcome Sanger Institute for the COVID-19 Genomics UK (COG-UK) Consortium | Berkshire and Surrey Pathology Services Lighthouse Laboratory and Alex Alderton, Roberto Amato, Jeffrey Barrett, Sonia Goncalves, Ewan Harrison, David K. Jackson, Ian Johnston, Dominic Kwiatkowski, Cordelia Langford, John Sillitoe on behalf of the Wellcome Sanger Institute COVID-19 Surveillance Team |
| EPI_ISL_3285781 | Lighthouse Lab in Milton Keynes | Wellcome Sanger Institute for the COVID-19 Genomics UK (COG-UK) Consortium | The Lighthouse Lab in Milton Keynes and Alex Alderton, Roberto Amato, Jeffrey Barrett, Sonia Goncalves, Ewan Harrison, David K. Jackson, Ian Johnston, Dominic Kwiatkowski, Cordelia Langford, John Sillitoe on behalf of the Wellcome Sanger Institute COVID-19 Surveillance Team |
| EPI_ISL_3285782 | Berkshire and Surrey Pathology Services Lighthouse Laboratory | Wellcome Sanger Institute for the COVID-19 Genomics UK (COG-UK) Consortium | Berkshire and Surrey Pathology Services Lighthouse Laboratory and Alex Alderton, Roberto Amato, Jeffrey Barrett, Sonia Goncalves, Ewan Harrison, David K. Jackson, Ian Johnston, Dominic Kwiatkowski, Cordelia Langford, John Sillitoe on behalf of the Wellcome Sanger Institute COVID-19 Surveillance Team |
| EPI_ISL_3285783, EPI_ISL_3285784, EPI_ISL_3285785 | Lighthouse Lab in Milton Keynes | Wellcome Sanger Institute for the COVID-19 Genomics UK (COG-UK) Consortium | The Lighthouse Lab in Milton Keynes and Alex Alderton, Roberto Amato, Jeffrey Barrett, Sonia Goncalves, Ewan Harrison, David K. Jackson, Ian Johnston, Dominic Kwiatkowski, Cordelia Langford, John Sillitoe on behalf of the Wellcome Sanger Institute COVID-19 Surveillance Team |
| EPI_ISL_3285786 | Lighthouse Lab in Glasgow | Wellcome Sanger Institute for the COVID-19 Genomics UK (COG-UK) Consortium | Harper VanSteenhouse, Yumi Kasai, David Gray, Carol Clugston, Anna Dominiczak and Alex Alderton, Roberto Amato, Jeffrey Barrett, Sonia Goncalves, Ewan Harrison, David K. Jackson, Ian Johnston, Dominic Kwiatkowski, Cordelia Langford, John Sillitoe on behalf of the Wellcome Sanger Institute COVID-19 Surveillance Team |
| EPI_ISL_3285787, EPI_ISL_3285788 | Lighthouse Lab in Milton Keynes | Wellcome Sanger Institute for the COVID-19 Genomics UK (COG-UK) Consortium | The Lighthouse Lab in Milton Keynes and Alex Alderton, Roberto Amato, Jeffrey Barrett, Sonia Goncalves, Ewan Harrison, David K. Jackson, Ian Johnston, Dominic Kwiatkowski, Cordelia Langford, John Sillitoe on behalf of the Wellcome Sanger Institute COVID-19 Surveillance Team |
| EPI_ISL_3285789, EPI_ISL_3285790, EPI_ISL_3285791 | Berkshire and Surrey Pathology Services Lighthouse Laboratory | Wellcome Sanger Institute for the COVID-19 Genomics UK (COG-UK) Consortium | Berkshire and Surrey Pathology Services Lighthouse Laboratory and Alex Alderton, Roberto Amato, Jeffrey Barrett, Sonia Goncalves, Ewan Harrison, David K. Jackson, Ian Johnston, Dominic Kwiatkowski, Cordelia Langford, John Sillitoe on behalf of the Wellcome Sanger Institute COVID-19 Surveillance Team |
| EPI_ISL_3285792, EPI_ISL_3285793 | Lighthouse Lab in Glasgow | Wellcome Sanger Institute for the COVID-19 Genomics UK (COG-UK) Consortium | Harper VanSteenhouse, Yumi Kasai, David Gray, Carol Clugston, Anna Dominiczak and Alex Alderton, Roberto Amato, Jeffrey Barrett, Sonia Goncalves, Ewan Harrison, David K. Jackson, Ian Johnston, Dominic Kwiatkowski, Cordelia Langford, John Sillitoe on behalf of the Wellcome Sanger Institute COVID-19 Surveillance Team |
| EPI_ISL_3285794 | Lighthouse Lab in Milton Keynes | Wellcome Sanger Institute for the COVID-19 Genomics UK (COG-UK) Consortium | The Lighthouse Lab in Milton Keynes and Alex Alderton, Roberto Amato, Jeffrey Barrett, Sonia Goncalves, Ewan Harrison, David K. Jackson, Ian Johnston, Dominic Kwiatkowski, Cordelia Langford, John Sillitoe on behalf of the Wellcome Sanger Institute COVID-19 Surveillance Team |
| EPI_ISL_3285795 | Berkshire and Surrey Pathology Services Lighthouse Laboratory | Wellcome Sanger Institute for the COVID-19 Genomics UK (COG-UK) Consortium | Berkshire and Surrey Pathology Services Lighthouse Laboratory and Alex Alderton, Roberto Amato, Jeffrey Barrett, Sonia Goncalves, Ewan Harrison, David K. Jackson, Ian Johnston, Dominic Kwiatkowski, Cordelia Langford, John Sillitoe on behalf of the Wellcome Sanger Institute COVID-19 Surveillance Team |
| EPI_ISL_3285796, EPI_ISL_3285797, EPI_ISL_3285798, EPI_ISL_3285799 | Lighthouse Lab in Milton Keynes | Wellcome Sanger Institute for the COVID-19 Genomics UK (COG-UK) Consortium | The Lighthouse Lab in Milton Keynes and Alex Alderton, Roberto Amato, Jeffrey Barrett, Sonia Goncalves, Ewan Harrison, David K. Jackson, Ian Johnston, Dominic Kwiatkowski, Cordelia Langford, John Sillitoe on behalf of the Wellcome Sanger Institute COVID-19 Surveillance Team |
| EPI_ISL_3285800 | Lighthouse Lab in Glasgow | Wellcome Sanger Institute for the COVID-19 Genomics UK (COG-UK) Consortium | Harper VanSteenhouse, Yumi Kasai, David Gray, Carol Clugston, Anna Dominiczak and Alex Alderton, Roberto Amato, Jeffrey Barrett, Sonia Goncalves, Ewan Harrison, David K. Jackson, Ian Johnston, Dominic Kwiatkowski, Cordelia Langford, John Sillitoe on behalf of the Wellcome Sanger Institute COVID-19 Surveillance Team |
| EPI_ISL_3285801 | Lighthouse Lab in Milton Keynes | Wellcome Sanger Institute for the COVID-19 Genomics UK (COG-UK) Consortium | The Lighthouse Lab in Milton Keynes and Alex Alderton, Roberto Amato, Jeffrey Barrett, Sonia Goncalves, Ewan Harrison, David K. Jackson, Ian Johnston, Dominic Kwiatkowski, Cordelia Langford, John Sillitoe on behalf of the Wellcome Sanger Institute COVID-19 Surveillance Team |
| EPI_ISL_3285802 | Lighthouse Lab in Glasgow | Wellcome Sanger Institute for the COVID-19 Genomics UK (COG-UK) Consortium | Harper VanSteenhouse, Yumi Kasai, David Gray, Carol Clugston, Anna Dominiczak and Alex Alderton, Roberto Amato, Jeffrey Barrett, Sonia Goncalves, Ewan Harrison, David K. Jackson, Ian Johnston, Dominic Kwiatkowski, Cordelia Langford, John Sillitoe on behalf of the Wellcome Sanger Institute COVID-19 Surveillance Team |
| EPI_ISL_3285803, EPI_ISL_3285804 | Lighthouse Lab in Milton Keynes | Wellcome Sanger Institute for the COVID-19 Genomics UK (COG-UK) Consortium | The Lighthouse Lab in Milton Keynes and Alex Alderton, Roberto Amato, Jeffrey Barrett, Sonia Goncalves, Ewan Harrison, David K. Jackson, Ian Johnston, Dominic Kwiatkowski, Cordelia Langford, John Sillitoe on behalf of the Wellcome Sanger Institute COVID-19 Surveillance Team |
| EPI_ISL_3285805 | Berkshire and Surrey Pathology Services Lighthouse Laboratory | Wellcome Sanger Institute for the COVID-19 Genomics UK (COG-UK) Consortium | Berkshire and Surrey Pathology Services Lighthouse Laboratory and Alex Alderton, Roberto Amato, Jeffrey Barrett, Sonia Goncalves, Ewan Harrison, David K. Jackson, Ian Johnston, Dominic Kwiatkowski, Cordelia Langford, John Sillitoe on behalf of the Wellcome Sanger Institute COVID-19 Surveillance Team |
| EPI_ISL_3285806, EPI_ISL_3285807, EPI_ISL_3285808 | Lighthouse Lab in Milton Keynes | Wellcome Sanger Institute for the COVID-19 Genomics UK (COG-UK) Consortium | The Lighthouse Lab in Milton Keynes and Alex Alderton, Roberto Amato, Jeffrey Barrett, Sonia Goncalves, Ewan Harrison, David K. Jackson, Ian Johnston, Dominic Kwiatkowski, Cordelia Langford, John Sillitoe on behalf of the Wellcome Sanger Institute COVID-19 Surveillance Team |
| EPI_ISL_3285809 | Lighthouse Lab in Glasgow | Wellcome Sanger Institute for the COVID-19 Genomics UK (COG-UK) Consortium | Harper VanSteenhouse, Yumi Kasai, David Gray, Carol Clugston, Anna Dominiczak and Alex Alderton, Roberto Amato, Jeffrey Barrett, Sonia Goncalves, Ewan Harrison, David K. Jackson, Ian Johnston, Dominic Kwiatkowski, Cordelia Langford, John Sillitoe on behalf of the Wellcome Sanger Institute COVID-19 Surveillance Team |
| EPI_ISL_3285810 | Berkshire and Surrey Pathology Services Lighthouse Laboratory | Wellcome Sanger Institute for the COVID-19 Genomics UK (COG-UK) Consortium | Berkshire and Surrey Pathology Services Lighthouse Laboratory and Alex Alderton, Roberto Amato, Jeffrey Barrett, Sonia Goncalves, Ewan Harrison, David K. Jackson, Ian Johnston, Dominic Kwiatkowski, Cordelia Langford, John Sillitoe on behalf of the Wellcome Sanger Institute COVID-19 Surveillance Team |
| EPI_ISL_3285811, | Lighthouse Lab in Milton Keynes | Wellcome Sanger Institute for the COVID-19 Genomics UK (COG-UK) | The Lighthouse Lab in Milton Keynes and Alex Alderton, Roberto Amato, Jeffrey Barrett, Sonia Goncalves, Ewan Harrison, David K. Jackson, Ian Johnston, Dominic Kwiatkowski, Cordelia Langford, John Sillitoe on behalf of the Wellcome Sanger |

| | | | |
|---|---|---|---|
| EPI_ISL_3285812 | | Consortium | Institute COVID-19 Surveillance Team |
| EPI_ISL_3285813 | Lighthouse Lab in Glasgow | Wellcome Sanger Institute for the COVID-19 Genomics UK (COG-UK) Consortium | Harper VanSteenhouse, Yumi Kasai, David Gray, Carol Clugston, Anna Dominiczak and Alex Alderton, Roberto Amato, Jeffrey Barrett, Sonia Goncalves, Ewan Harrison, David K. Jackson, Ian Johnston, Dominic Kwiatkowski, Cordelia Langford, John Sillitoe on behalf of the Wellcome Sanger Institute COVID-19 Surveillance Team |
| EPI_ISL_3285814, EPI_ISL_3285815 | Lighthouse Lab in Milton Keynes | Wellcome Sanger Institute for the COVID-19 Genomics UK (COG-UK) Consortium | The Lighthouse Lab in Milton Keynes and Alex Alderton, Roberto Amato, Jeffrey Barrett, Sonia Goncalves, Ewan Harrison, David K. Jackson, Ian Johnston, Dominic Kwiatkowski, Cordelia Langford, John Sillitoe on behalf of the Wellcome Sanger Institute COVID-19 Surveillance Team |
| EPI_ISL_3285816 | Lighthouse Lab in Glasgow | Wellcome Sanger Institute for the COVID-19 Genomics UK (COG-UK) Consortium | Harper VanSteenhouse, Yumi Kasai, David Gray, Carol Clugston, Anna Dominiczak and Alex Alderton, Roberto Amato, Jeffrey Barrett, Sonia Goncalves, Ewan Harrison, David K. Jackson, Ian Johnston, Dominic Kwiatkowski, Cordelia Langford, John Sillitoe on behalf of the Wellcome Sanger Institute COVID-19 Surveillance Team |
| EPI_ISL_3285817 | Lighthouse Lab in Milton Keynes | Wellcome Sanger Institute for the COVID-19 Genomics UK (COG-UK) Consortium | The Lighthouse Lab in Milton Keynes and Alex Alderton, Roberto Amato, Jeffrey Barrett, Sonia Goncalves, Ewan Harrison, David K. Jackson, Ian Johnston, Dominic Kwiatkowski, Cordelia Langford, John Sillitoe on behalf of the Wellcome Sanger Institute COVID-19 Surveillance Team |
| EPI_ISL_3285818 | Lighthouse Lab in Glasgow | Wellcome Sanger Institute for the COVID-19 Genomics UK (COG-UK) Consortium | Harper VanSteenhouse, Yumi Kasai, David Gray, Carol Clugston, Anna Dominiczak and Alex Alderton, Roberto Amato, Jeffrey Barrett, Sonia Goncalves, Ewan Harrison, David K. Jackson, Ian Johnston, Dominic Kwiatkowski, Cordelia Langford, John Sillitoe on behalf of the Wellcome Sanger Institute COVID-19 Surveillance Team |
| EPI_ISL_3285819 | Lighthouse Lab in Milton Keynes | Wellcome Sanger Institute for the COVID-19 Genomics UK (COG-UK) Consortium | The Lighthouse Lab in Milton Keynes and Alex Alderton, Roberto Amato, Jeffrey Barrett, Sonia Goncalves, Ewan Harrison, David K. Jackson, Ian Johnston, Dominic Kwiatkowski, Cordelia Langford, John Sillitoe on behalf of the Wellcome Sanger Institute COVID-19 Surveillance Team |
| EPI_ISL_3285820 | Lighthouse Lab in Glasgow | Wellcome Sanger Institute for the COVID-19 Genomics UK (COG-UK) Consortium | Harper VanSteenhouse, Yumi Kasai, David Gray, Carol Clugston, Anna Dominiczak and Alex Alderton, Roberto Amato, Jeffrey Barrett, Sonia Goncalves, Ewan Harrison, David K. Jackson, Ian Johnston, Dominic Kwiatkowski, Cordelia Langford, John Sillitoe on behalf of the Wellcome Sanger Institute COVID-19 Surveillance Team |
| EPI_ISL_3285821 | Lighthouse Lab in Milton Keynes | Wellcome Sanger Institute for the COVID-19 Genomics UK (COG-UK) Consortium | The Lighthouse Lab in Milton Keynes and Alex Alderton, Roberto Amato, Jeffrey Barrett, Sonia Goncalves, Ewan Harrison, David K. Jackson, Ian Johnston, Dominic Kwiatkowski, Cordelia Langford, John Sillitoe on behalf of the Wellcome Sanger Institute COVID-19 Surveillance Team |
| EPI_ISL_3285822 | Lighthouse Lab in Glasgow | Wellcome Sanger Institute for the COVID-19 Genomics UK (COG-UK) Consortium | Harper VanSteenhouse, Yumi Kasai, David Gray, Carol Clugston, Anna Dominiczak and Alex Alderton, Roberto Amato, Jeffrey Barrett, Sonia Goncalves, Ewan Harrison, David K. Jackson, Ian Johnston, Dominic Kwiatkowski, Cordelia Langford, John Sillitoe on behalf of the Wellcome Sanger Institute COVID-19 Surveillance Team |
| EPI_ISL_3285823 | Lighthouse Lab in Milton Keynes | Wellcome Sanger Institute for the COVID-19 Genomics UK (COG-UK) Consortium | The Lighthouse Lab in Milton Keynes and Alex Alderton, Roberto Amato, Jeffrey Barrett, Sonia Goncalves, Ewan Harrison, David K. Jackson, Ian Johnston, Dominic Kwiatkowski, Cordelia Langford, John Sillitoe on behalf of the Wellcome Sanger Institute COVID-19 Surveillance Team |
| EPI_ISL_3285824 | Lighthouse Lab in Glasgow | Wellcome Sanger Institute for the COVID-19 Genomics UK (COG-UK) Consortium | Harper VanSteenhouse, Yumi Kasai, David Gray, Carol Clugston, Anna Dominiczak and Alex Alderton, Roberto Amato, Jeffrey Barrett, Sonia Goncalves, Ewan Harrison, David K. Jackson, Ian Johnston, Dominic Kwiatkowski, Cordelia Langford, John Sillitoe on behalf of the Wellcome Sanger Institute COVID-19 Surveillance Team |
| EPI_ISL_3285825 | Lighthouse Lab in Milton Keynes | Wellcome Sanger Institute for the COVID-19 Genomics UK (COG-UK) Consortium | The Lighthouse Lab in Milton Keynes and Alex Alderton, Roberto Amato, Jeffrey Barrett, Sonia Goncalves, Ewan Harrison, David K. Jackson, Ian Johnston, Dominic Kwiatkowski, Cordelia Langford, John Sillitoe on behalf of the Wellcome Sanger Institute COVID-19 Surveillance Team |
| EPI_ISL_3285826 | Berkshire and Surrey Pathology Services Lighthouse Laboratory | Wellcome Sanger Institute for the COVID-19 Genomics UK (COG-UK) Consortium | Berkshire and Surrey Pathology Services Lighthouse Laboratory and Alex Alderton, Roberto Amato, Jeffrey Barrett, Sonia Goncalves, Ewan Harrison, David K. Jackson, Ian Johnston, Dominic Kwiatkowski, Cordelia Langford, John Sillitoe on behalf of the Wellcome Sanger Institute COVID-19 Surveillance Team |
| EPI_ISL_3285827 | Lighthouse Lab in Glasgow | Wellcome Sanger Institute for the COVID-19 Genomics UK (COG-UK) Consortium | Harper VanSteenhouse, Yumi Kasai, David Gray, Carol Clugston, Anna Dominiczak and Alex Alderton, Roberto Amato, Jeffrey Barrett, Sonia Goncalves, Ewan Harrison, David K. Jackson, Ian Johnston, Dominic Kwiatkowski, Cordelia Langford, John Sillitoe on behalf of the Wellcome Sanger Institute COVID-19 Surveillance Team |
| EPI_ISL_3285828 | Lighthouse Lab in Milton Keynes | Wellcome Sanger Institute for the COVID-19 Genomics UK (COG-UK) Consortium | The Lighthouse Lab in Milton Keynes and Alex Alderton, Roberto Amato, Jeffrey Barrett, Sonia Goncalves, Ewan Harrison, David K. Jackson, Ian Johnston, Dominic Kwiatkowski, Cordelia Langford, John Sillitoe on behalf of the Wellcome Sanger Institute COVID-19 Surveillance Team |
| EPI_ISL_3285829 | Lighthouse Lab in Glasgow | Wellcome Sanger Institute for the COVID-19 Genomics UK (COG-UK) Consortium | Harper VanSteenhouse, Yumi Kasai, David Gray, Carol Clugston, Anna Dominiczak and Alex Alderton, Roberto Amato, Jeffrey Barrett, Sonia Goncalves, Ewan Harrison, David K. Jackson, Ian Johnston, Dominic Kwiatkowski, Cordelia Langford, John Sillitoe on behalf of the Wellcome Sanger Institute COVID-19 Surveillance Team |
| EPI_ISL_3285830 | Lighthouse Lab in Milton Keynes | Wellcome Sanger Institute for the COVID-19 Genomics UK (COG-UK) Consortium | The Lighthouse Lab in Milton Keynes and Alex Alderton, Roberto Amato, Jeffrey Barrett, Sonia Goncalves, Ewan Harrison, David K. Jackson, Ian Johnston, Dominic Kwiatkowski, Cordelia Langford, John Sillitoe on behalf of the Wellcome Sanger Institute COVID-19 Surveillance Team |
| EPI_ISL_3285831 | Lighthouse Lab in Glasgow | Wellcome Sanger Institute for the COVID-19 Genomics UK (COG-UK) Consortium | Harper VanSteenhouse, Yumi Kasai, David Gray, Carol Clugston, Anna Dominiczak and Alex Alderton, Roberto Amato, Jeffrey Barrett, Sonia Goncalves, Ewan Harrison, David K. Jackson, Ian Johnston, Dominic Kwiatkowski, Cordelia Langford, John Sillitoe on behalf of the Wellcome Sanger Institute COVID-19 Surveillance Team |
| EPI_ISL_3285832 | Lighthouse Lab in Milton Keynes | Wellcome Sanger Institute for the COVID-19 Genomics UK (COG-UK) Consortium | The Lighthouse Lab in Milton Keynes and Alex Alderton, Roberto Amato, Jeffrey Barrett, Sonia Goncalves, Ewan Harrison, David K. Jackson, Ian Johnston, Dominic Kwiatkowski, Cordelia Langford, John Sillitoe on behalf of the Wellcome Sanger Institute COVID-19 Surveillance Team |
| EPI_ISL_3285833 | Lighthouse Lab in Glasgow | Wellcome Sanger Institute for the COVID-19 Genomics UK (COG-UK) Consortium | Harper VanSteenhouse, Yumi Kasai, David Gray, Carol Clugston, Anna Dominiczak and Alex Alderton, Roberto Amato, Jeffrey Barrett, Sonia Goncalves, Ewan Harrison, David K. Jackson, Ian Johnston, Dominic Kwiatkowski, Cordelia Langford, John Sillitoe on behalf of the Wellcome Sanger Institute COVID-19 Surveillance Team |
| EPI_ISL_3285834 | Lighthouse Lab in Milton Keynes | Wellcome Sanger Institute for the COVID-19 Genomics UK (COG-UK) Consortium | The Lighthouse Lab in Milton Keynes and Alex Alderton, Roberto Amato, Jeffrey Barrett, Sonia Goncalves, Ewan Harrison, David K. Jackson, Ian Johnston, Dominic Kwiatkowski, Cordelia Langford, John Sillitoe on behalf of the Wellcome Sanger Institute COVID-19 Surveillance Team |
| EPI_ISL_3285835 | Berkshire and Surrey Pathology Services Lighthouse Laboratory | Wellcome Sanger Institute for the COVID-19 Genomics UK (COG-UK) Consortium | Berkshire and Surrey Pathology Services Lighthouse Laboratory and Alex Alderton, Roberto Amato, Jeffrey Barrett, Sonia Goncalves, Ewan Harrison, David K. Jackson, Ian Johnston, Dominic Kwiatkowski, Cordelia Langford, John Sillitoe on behalf of the Wellcome Sanger Institute COVID-19 Surveillance Team |
| EPI_ISL_3285836 | Lighthouse Lab in Milton Keynes | Wellcome Sanger Institute for the COVID-19 Genomics UK (COG-UK) Consortium | The Lighthouse Lab in Milton Keynes and Alex Alderton, Roberto Amato, Jeffrey Barrett, Sonia Goncalves, Ewan Harrison, David K. Jackson, Ian Johnston, Dominic Kwiatkowski, Cordelia Langford, John Sillitoe on behalf of the Wellcome Sanger Institute COVID-19 Surveillance Team |
| EPI_ISL_3285837 | Berkshire and Surrey Pathology Services Lighthouse Laboratory | Wellcome Sanger Institute for the COVID-19 Genomics UK (COG-UK) Consortium | Berkshire and Surrey Pathology Services Lighthouse Laboratory and Alex Alderton, Roberto Amato, Jeffrey Barrett, Sonia Goncalves, Ewan Harrison, David K. Jackson, Ian Johnston, Dominic Kwiatkowski, Cordelia Langford, John Sillitoe on behalf of the Wellcome Sanger Institute COVID-19 Surveillance Team |
| EPI_ISL_3285838 | Lighthouse Lab in Milton Keynes | Wellcome Sanger Institute for the COVID-19 Genomics UK (COG-UK) Consortium | The Lighthouse Lab in Milton Keynes and Alex Alderton, Roberto Amato, Jeffrey Barrett, Sonia Goncalves, Ewan Harrison, David K. Jackson, Ian Johnston, Dominic Kwiatkowski, Cordelia Langford, John Sillitoe on behalf of the Wellcome Sanger Institute COVID-19 Surveillance Team |
| EPI_ISL_3285839, EPI_ISL_3285840 | Berkshire and Surrey Pathology Services Lighthouse Laboratory | Wellcome Sanger Institute for the COVID-19 Genomics UK (COG-UK) Consortium | Berkshire and Surrey Pathology Services Lighthouse Laboratory and Alex Alderton, Roberto Amato, Jeffrey Barrett, Sonia Goncalves, Ewan Harrison, David K. Jackson, Ian Johnston, Dominic Kwiatkowski, Cordelia Langford, John Sillitoe on behalf of the Wellcome Sanger Institute COVID-19 Surveillance Team |
| EPI_ISL_3285841 | Lighthouse Lab in Milton Keynes | Wellcome Sanger Institute for the COVID-19 Genomics UK (COG-UK) Consortium | The Lighthouse Lab in Milton Keynes and Alex Alderton, Roberto Amato, Jeffrey Barrett, Sonia Goncalves, Ewan Harrison, David K. Jackson, Ian Johnston, Dominic Kwiatkowski, Cordelia Langford, John Sillitoe on behalf of the Wellcome Sanger Institute COVID-19 Surveillance Team |
| EPI_ISL_3285842 | Berkshire and Surrey Pathology Services Lighthouse Laboratory | Wellcome Sanger Institute for the COVID-19 Genomics UK (COG-UK) Consortium | Berkshire and Surrey Pathology Services Lighthouse Laboratory and Alex Alderton, Roberto Amato, Jeffrey Barrett, Sonia Goncalves, Ewan Harrison, David K. Jackson, Ian Johnston, Dominic Kwiatkowski, Cordelia Langford, John Sillitoe on behalf of the Wellcome Sanger Institute COVID-19 Surveillance Team |
| EPI_ISL_3285843 | Lighthouse Lab in Milton Keynes | Wellcome Sanger Institute for the COVID-19 Genomics UK (COG-UK) Consortium | The Lighthouse Lab in Milton Keynes and Alex Alderton, Roberto Amato, Jeffrey Barrett, Sonia Goncalves, Ewan Harrison, David K. Jackson, Ian Johnston, Dominic Kwiatkowski, Cordelia Langford, John Sillitoe on behalf of the Wellcome Sanger Institute COVID-19 Surveillance Team |
| EPI_ISL_3285844 | Berkshire and Surrey Pathology Services Lighthouse Laboratory | Wellcome Sanger Institute for the COVID-19 Genomics UK (COG-UK) Consortium | Berkshire and Surrey Pathology Services Lighthouse Laboratory and Alex Alderton, Roberto Amato, Jeffrey Barrett, Sonia Goncalves, Ewan Harrison, David K. Jackson, Ian Johnston, Dominic Kwiatkowski, Cordelia Langford, John Sillitoe on behalf of the Wellcome Sanger Institute COVID-19 Surveillance Team |
| EPI_ISL_3285845 | Lighthouse Lab in Milton Keynes | Wellcome Sanger Institute for the COVID-19 Genomics UK (COG-UK) Consortium | The Lighthouse Lab in Milton Keynes and Alex Alderton, Roberto Amato, Jeffrey Barrett, Sonia Goncalves, Ewan Harrison, David K. Jackson, Ian Johnston, Dominic Kwiatkowski, Cordelia Langford, John Sillitoe on behalf of the Wellcome Sanger Institute COVID-19 Surveillance Team |
| EPI_ISL_3285846 | Lighthouse Lab in Glasgow | Wellcome Sanger Institute for the COVID-19 Genomics UK (COG-UK) Consortium | Harper VanSteenhouse, Yumi Kasai, David Gray, Carol Clugston, Anna Dominiczak and Alex Alderton, Roberto Amato, Jeffrey Barrett, Sonia Goncalves, Ewan Harrison, David K. Jackson, Ian Johnston, Dominic Kwiatkowski, Cordelia Langford, John Sillitoe on behalf of the Wellcome Sanger Institute COVID-19 Surveillance Team |
| EPI_ISL_3285847 | Berkshire and Surrey Pathology Services Lighthouse Laboratory | Wellcome Sanger Institute for the COVID-19 Genomics UK (COG-UK) Consortium | Berkshire and Surrey Pathology Services Lighthouse Laboratory and Alex Alderton, Roberto Amato, Jeffrey Barrett, Sonia Goncalves, Ewan Harrison, David K. Jackson, Ian Johnston, Dominic Kwiatkowski, Cordelia Langford, John Sillitoe on behalf of the Wellcome Sanger Institute COVID-19 Surveillance Team |
| EPI_ISL_3285848, EPI_ISL_3285849 | Lighthouse Lab in Milton Keynes | Wellcome Sanger Institute for the COVID-19 Genomics UK (COG-UK) Consortium | The Lighthouse Lab in Milton Keynes and Alex Alderton, Roberto Amato, Jeffrey Barrett, Sonia Goncalves, Ewan Harrison, David K. Jackson, Ian Johnston, Dominic Kwiatkowski, Cordelia Langford, John Sillitoe on behalf of the Wellcome Sanger Institute COVID-19 Surveillance Team |
| EPI_ISL_3285850, EPI_ISL_3285851, EPI_ISL_3285852, EPI_ISL_3285853 | Berkshire and Surrey Pathology Services Lighthouse Laboratory | Wellcome Sanger Institute for the COVID-19 Genomics UK (COG-UK) Consortium | Berkshire and Surrey Pathology Services Lighthouse Laboratory and Alex Alderton, Roberto Amato, Jeffrey Barrett, Sonia Goncalves, Ewan Harrison, David K. Jackson, Ian Johnston, Dominic Kwiatkowski, Cordelia Langford, John Sillitoe on behalf of the Wellcome Sanger Institute COVID-19 Surveillance Team |
| EPI_ISL_3285854, EPI_ISL_3285855, EPI_ISL_3285856, EPI_ISL_3285857, EPI_ISL_3285858 | Lighthouse Lab in Milton Keynes | Wellcome Sanger Institute for the COVID-19 Genomics UK (COG-UK) Consortium | The Lighthouse Lab in Milton Keynes and Alex Alderton, Roberto Amato, Jeffrey Barrett, Sonia Goncalves, Ewan Harrison, David K. Jackson, Ian Johnston, Dominic Kwiatkowski, Cordelia Langford, John Sillitoe on behalf of the Wellcome Sanger Institute COVID-19 Surveillance Team |
| EPI_ISL_3285859 | Lighthouse Lab in Glasgow | Wellcome Sanger Institute for the COVID-19 Genomics UK (COG-UK) Consortium | Harper VanSteenhouse, Yumi Kasai, David Gray, Carol Clugston, Anna Dominiczak and Alex Alderton, Roberto Amato, Jeffrey Barrett, Sonia Goncalves, Ewan Harrison, David K. Jackson, Ian Johnston, Dominic Kwiatkowski, Cordelia Langford, John Sillitoe on behalf of the Wellcome Sanger Institute COVID-19 Surveillance Team |
| EPI_ISL_3285860 | Berkshire and Surrey Pathology Services Lighthouse Laboratory | Wellcome Sanger Institute for the COVID-19 Genomics UK (COG-UK) Consortium | Berkshire and Surrey Pathology Services Lighthouse Laboratory and Alex Alderton, Roberto Amato, Jeffrey Barrett, Sonia Goncalves, Ewan Harrison, David K. Jackson, Ian Johnston, Dominic Kwiatkowski, Cordelia Langford, John Sillitoe on behalf of the Wellcome Sanger Institute COVID-19 Surveillance Team |
| EPI_ISL_3285861, EPI_ISL_3285862, EPI_ISL_3285863, EPI_ISL_3285864, EPI_ISL_3285865, EPI_ISL_3285866 | Lighthouse Lab in Milton Keynes | Wellcome Sanger Institute for the COVID-19 Genomics UK (COG-UK) Consortium | The Lighthouse Lab in Milton Keynes and Alex Alderton, Roberto Amato, Jeffrey Barrett, Sonia Goncalves, Ewan Harrison, David K. Jackson, Ian Johnston, Dominic Kwiatkowski, Cordelia Langford, John Sillitoe on behalf of the Wellcome Sanger Institute COVID-19 Surveillance Team |
| EPI_ISL_3285867, EPI_ISL_3285868 | Berkshire and Surrey Pathology Services Lighthouse Laboratory | Wellcome Sanger Institute for the COVID-19 Genomics UK (COG-UK) Consortium | Berkshire and Surrey Pathology Services Lighthouse Laboratory and Alex Alderton, Roberto Amato, Jeffrey Barrett, Sonia Goncalves, Ewan Harrison, David K. Jackson, Ian Johnston, Dominic Kwiatkowski, Cordelia Langford, John Sillitoe on behalf of the Wellcome Sanger Institute COVID-19 Surveillance Team |
| EPI_ISL_3285869, | Lighthouse Lab in Glasgow | Wellcome Sanger Institute for the COVID-19 Genomics UK (COG-UK) | Harper VanSteenhouse, Yumi Kasai, David Gray, Carol Clugston, Anna Dominiczak and Alex Alderton, Roberto Amato, Jeffrey Barrett, Sonia Goncalves, Ewan Harrison, David K. Jackson, Ian Johnston, Dominic Kwiatkowski, Cordelia Langford, |

| | | | |
|---|---|---|---|
| EPI_ISL_3285870 | | Consortium | John Sillitoe on behalf of the Wellcome Sanger Institute COVID-19 Surveillance Team |
| EPI_ISL_3285871 | Lighthouse Lab in Milton Keynes | Wellcome Sanger Institute for the COVID-19 Genomics UK (COG-UK) Consortium | The Lighthouse Lab in Milton Keynes and Alex Alderton, Roberto Amato, Jeffrey Barrett, Sonia Goncalves, Ewan Harrison, David K. Jackson, Ian Johnston, Dominic Kwiatkowski, Cordelia Langford, John Sillitoe on behalf of the Wellcome Sanger Institute COVID-19 Surveillance Team |
| EPI_ISL_3285872 | Berkshire and Surrey Pathology Services Lighthouse Laboratory | Wellcome Sanger Institute for the COVID-19 Genomics UK (COG-UK) Consortium | Berkshire and Surrey Pathology Services Lighthouse Laboratory and Alex Alderton, Roberto Amato, Jeffrey Barrett, Sonia Goncalves, Ewan Harrison, David K. Jackson, Ian Johnston, Dominic Kwiatkowski, Cordelia Langford, John Sillitoe on behalf of the Wellcome Sanger Institute COVID-19 Surveillance Team |
| EPI_ISL_3285873 | Lighthouse Lab in Glasgow | Wellcome Sanger Institute for the COVID-19 Genomics UK (COG-UK) Consortium | Harper VanSteenhouse, Yumi Kasai, David Gray, Carol Clugston, Anna Dominiczak and Alex Alderton, Roberto Amato, Jeffrey Barrett, Sonia Goncalves, Ewan Harrison, David K. Jackson, Ian Johnston, Dominic Kwiatkowski, Cordelia Langford, John Sillitoe on behalf of the Wellcome Sanger Institute COVID-19 Surveillance Team |
| EPI_ISL_3285874, EPI_ISL_3285875, EPI_ISL_3285876 | Lighthouse Lab in Milton Keynes | Wellcome Sanger Institute for the COVID-19 Genomics UK (COG-UK) Consortium | The Lighthouse Lab in Milton Keynes and Alex Alderton, Roberto Amato, Jeffrey Barrett, Sonia Goncalves, Ewan Harrison, David K. Jackson, Ian Johnston, Dominic Kwiatkowski, Cordelia Langford, John Sillitoe on behalf of the Wellcome Sanger Institute COVID-19 Surveillance Team |
| EPI_ISL_3285877, EPI_ISL_3285878 | Lighthouse Lab in Glasgow | Wellcome Sanger Institute for the COVID-19 Genomics UK (COG-UK) Consortium | Harper VanSteenhouse, Yumi Kasai, David Gray, Carol Clugston, Anna Dominiczak and Alex Alderton, Roberto Amato, Jeffrey Barrett, Sonia Goncalves, Ewan Harrison, David K. Jackson, Ian Johnston, Dominic Kwiatkowski, Cordelia Langford, John Sillitoe on behalf of the Wellcome Sanger Institute COVID-19 Surveillance Team |
| EPI_ISL_3285879 | Berkshire and Surrey Pathology Services Lighthouse Laboratory | Wellcome Sanger Institute for the COVID-19 Genomics UK (COG-UK) Consortium | Berkshire and Surrey Pathology Services Lighthouse Laboratory and Alex Alderton, Roberto Amato, Jeffrey Barrett, Sonia Goncalves, Ewan Harrison, David K. Jackson, Ian Johnston, Dominic Kwiatkowski, Cordelia Langford, John Sillitoe on behalf of the Wellcome Sanger Institute COVID-19 Surveillance Team |
| EPI_ISL_3285880 | Lighthouse Lab in Milton Keynes | Wellcome Sanger Institute for the COVID-19 Genomics UK (COG-UK) Consortium | The Lighthouse Lab in Milton Keynes and Alex Alderton, Roberto Amato, Jeffrey Barrett, Sonia Goncalves, Ewan Harrison, David K. Jackson, Ian Johnston, Dominic Kwiatkowski, Cordelia Langford, John Sillitoe on behalf of the Wellcome Sanger Institute COVID-19 Surveillance Team |
| EPI_ISL_3285881 | Berkshire and Surrey Pathology Services Lighthouse Laboratory | Wellcome Sanger Institute for the COVID-19 Genomics UK (COG-UK) Consortium | Berkshire and Surrey Pathology Services Lighthouse Laboratory and Alex Alderton, Roberto Amato, Jeffrey Barrett, Sonia Goncalves, Ewan Harrison, David K. Jackson, Ian Johnston, Dominic Kwiatkowski, Cordelia Langford, John Sillitoe on behalf of the Wellcome Sanger Institute COVID-19 Surveillance Team |
| EPI_ISL_3285882, EPI_ISL_3285883 | Lighthouse Lab in Glasgow | Wellcome Sanger Institute for the COVID-19 Genomics UK (COG-UK) Consortium | Harper VanSteenhouse, Yumi Kasai, David Gray, Carol Clugston, Anna Dominiczak and Alex Alderton, Roberto Amato, Jeffrey Barrett, Sonia Goncalves, Ewan Harrison, David K. Jackson, Ian Johnston, Dominic Kwiatkowski, Cordelia Langford, John Sillitoe on behalf of the Wellcome Sanger Institute COVID-19 Surveillance Team |
| EPI_ISL_3285884, EPI_ISL_3285885, EPI_ISL_3285886, EPI_ISL_3285887 | Lighthouse Lab in Milton Keynes | Wellcome Sanger Institute for the COVID-19 Genomics UK (COG-UK) Consortium | The Lighthouse Lab in Milton Keynes and Alex Alderton, Roberto Amato, Jeffrey Barrett, Sonia Goncalves, Ewan Harrison, David K. Jackson, Ian Johnston, Dominic Kwiatkowski, Cordelia Langford, John Sillitoe on behalf of the Wellcome Sanger Institute COVID-19 Surveillance Team |
| EPI_ISL_3285888, EPI_ISL_3285889 | Lighthouse Lab in Glasgow | Wellcome Sanger Institute for the COVID-19 Genomics UK (COG-UK) Consortium | Harper VanSteenhouse, Yumi Kasai, David Gray, Carol Clugston, Anna Dominiczak and Alex Alderton, Roberto Amato, Jeffrey Barrett, Sonia Goncalves, Ewan Harrison, David K. Jackson, Ian Johnston, Dominic Kwiatkowski, Cordelia Langford, John Sillitoe on behalf of the Wellcome Sanger Institute COVID-19 Surveillance Team |
| EPI_ISL_3285890, EPI_ISL_3285891, EPI_ISL_3285892, EPI_ISL_3285893, EPI_ISL_3285894, EPI_ISL_3285895 | Lighthouse Lab in Milton Keynes | Wellcome Sanger Institute for the COVID-19 Genomics UK (COG-UK) Consortium | The Lighthouse Lab in Milton Keynes and Alex Alderton, Roberto Amato, Jeffrey Barrett, Sonia Goncalves, Ewan Harrison, David K. Jackson, Ian Johnston, Dominic Kwiatkowski, Cordelia Langford, John Sillitoe on behalf of the Wellcome Sanger Institute COVID-19 Surveillance Team |
| EPI_ISL_3285896 | Lighthouse Lab in Glasgow | Wellcome Sanger Institute for the COVID-19 Genomics UK (COG-UK) Consortium | Harper VanSteenhouse, Yumi Kasai, David Gray, Carol Clugston, Anna Dominiczak and Alex Alderton, Roberto Amato, Jeffrey Barrett, Sonia Goncalves, Ewan Harrison, David K. Jackson, Ian Johnston, Dominic Kwiatkowski, Cordelia Langford, John Sillitoe on behalf of the Wellcome Sanger Institute COVID-19 Surveillance Team |
| EPI_ISL_3285897 | Lighthouse Lab in Milton Keynes | Wellcome Sanger Institute for the COVID-19 Genomics UK (COG-UK) Consortium | The Lighthouse Lab in Milton Keynes and Alex Alderton, Roberto Amato, Jeffrey Barrett, Sonia Goncalves, Ewan Harrison, David K. Jackson, Ian Johnston, Dominic Kwiatkowski, Cordelia Langford, John Sillitoe on behalf of the Wellcome Sanger Institute COVID-19 Surveillance Team |
| EPI_ISL_3285898 | Berkshire and Surrey Pathology Services Lighthouse Laboratory | Wellcome Sanger Institute for the COVID-19 Genomics UK (COG-UK) Consortium | Berkshire and Surrey Pathology Services Lighthouse Laboratory and Alex Alderton, Roberto Amato, Jeffrey Barrett, Sonia Goncalves, Ewan Harrison, David K. Jackson, Ian Johnston, Dominic Kwiatkowski, Cordelia Langford, John Sillitoe on behalf of the Wellcome Sanger Institute COVID-19 Surveillance Team |
| EPI_ISL_3285899 | Lighthouse Lab in Glasgow | Wellcome Sanger Institute for the COVID-19 Genomics UK (COG-UK) Consortium | Harper VanSteenhouse, Yumi Kasai, David Gray, Carol Clugston, Anna Dominiczak and Alex Alderton, Roberto Amato, Jeffrey Barrett, Sonia Goncalves, Ewan Harrison, David K. Jackson, Ian Johnston, Dominic Kwiatkowski, Cordelia Langford, John Sillitoe on behalf of the Wellcome Sanger Institute COVID-19 Surveillance Team |
| EPI_ISL_3285900 | Berkshire and Surrey Pathology Services Lighthouse Laboratory | Wellcome Sanger Institute for the COVID-19 Genomics UK (COG-UK) Consortium | Berkshire and Surrey Pathology Services Lighthouse Laboratory and Alex Alderton, Roberto Amato, Jeffrey Barrett, Sonia Goncalves, Ewan Harrison, David K. Jackson, Ian Johnston, Dominic Kwiatkowski, Cordelia Langford, John Sillitoe on behalf of the Wellcome Sanger Institute COVID-19 Surveillance Team |
| EPI_ISL_3285901, EPI_ISL_3285902, EPI_ISL_3285903, EPI_ISL_3285904, EPI_ISL_3285905, EPI_ISL_3285906, EPI_ISL_3285907, EPI_ISL_3285908, EPI_ISL_3285909 | Lighthouse Lab in Milton Keynes | Wellcome Sanger Institute for the COVID-19 Genomics UK (COG-UK) Consortium | The Lighthouse Lab in Milton Keynes and Alex Alderton, Roberto Amato, Jeffrey Barrett, Sonia Goncalves, Ewan Harrison, David K. Jackson, Ian Johnston, Dominic Kwiatkowski, Cordelia Langford, John Sillitoe on behalf of the Wellcome Sanger Institute COVID-19 Surveillance Team |
| EPI_ISL_3285910 | Lighthouse Lab in Glasgow | Wellcome Sanger Institute for the COVID-19 Genomics UK (COG-UK) Consortium | Harper VanSteenhouse, Yumi Kasai, David Gray, Carol Clugston, Anna Dominiczak and Alex Alderton, Roberto Amato, Jeffrey Barrett, Sonia Goncalves, Ewan Harrison, David K. Jackson, Ian Johnston, Dominic Kwiatkowski, Cordelia Langford, John Sillitoe on behalf of the Wellcome Sanger Institute COVID-19 Surveillance Team |
| EPI_ISL_3285911 | Lighthouse Lab in Milton Keynes | Wellcome Sanger Institute for the COVID-19 Genomics UK (COG-UK) Consortium | The Lighthouse Lab in Milton Keynes and Alex Alderton, Roberto Amato, Jeffrey Barrett, Sonia Goncalves, Ewan Harrison, David K. Jackson, Ian Johnston, Dominic Kwiatkowski, Cordelia Langford, John Sillitoe on behalf of the Wellcome Sanger Institute COVID-19 Surveillance Team |
| EPI_ISL_3285912, EPI_ISL_3285913, EPI_ISL_3285914, EPI_ISL_3285915 | Berkshire and Surrey Pathology Services Lighthouse Laboratory | Wellcome Sanger Institute for the COVID-19 Genomics UK (COG-UK) Consortium | Berkshire and Surrey Pathology Services Lighthouse Laboratory and Alex Alderton, Roberto Amato, Jeffrey Barrett, Sonia Goncalves, Ewan Harrison, David K. Jackson, Ian Johnston, Dominic Kwiatkowski, Cordelia Langford, John Sillitoe on behalf of the Wellcome Sanger Institute COVID-19 Surveillance Team |
| EPI_ISL_3285916 | Lighthouse Lab in Milton Keynes | Wellcome Sanger Institute for the COVID-19 Genomics UK (COG-UK) Consortium | The Lighthouse Lab in Milton Keynes and Alex Alderton, Roberto Amato, Jeffrey Barrett, Sonia Goncalves, Ewan Harrison, David K. Jackson, Ian Johnston, Dominic Kwiatkowski, Cordelia Langford, John Sillitoe on behalf of the Wellcome Sanger Institute COVID-19 Surveillance Team |
| EPI_ISL_3285917 | Berkshire and Surrey Pathology Services Lighthouse Laboratory | Wellcome Sanger Institute for the COVID-19 Genomics UK (COG-UK) Consortium | Berkshire and Surrey Pathology Services Lighthouse Laboratory and Alex Alderton, Roberto Amato, Jeffrey Barrett, Sonia Goncalves, Ewan Harrison, David K. Jackson, Ian Johnston, Dominic Kwiatkowski, Cordelia Langford, John Sillitoe on behalf of the Wellcome Sanger Institute COVID-19 Surveillance Team |
| EPI_ISL_3285918 | Lighthouse Lab in Glasgow | Wellcome Sanger Institute for the COVID-19 Genomics UK (COG-UK) Consortium | Harper VanSteenhouse, Yumi Kasai, David Gray, Carol Clugston, Anna Dominiczak and Alex Alderton, Roberto Amato, Jeffrey Barrett, Sonia Goncalves, Ewan Harrison, David K. Jackson, Ian Johnston, Dominic Kwiatkowski, Cordelia Langford, John Sillitoe on behalf of the Wellcome Sanger Institute COVID-19 Surveillance Team |
| EPI_ISL_3285919, EPI_ISL_3285920 | Lighthouse Lab in Milton Keynes | Wellcome Sanger Institute for the COVID-19 Genomics UK (COG-UK) Consortium | The Lighthouse Lab in Milton Keynes and Alex Alderton, Roberto Amato, Jeffrey Barrett, Sonia Goncalves, Ewan Harrison, David K. Jackson, Ian Johnston, Dominic Kwiatkowski, Cordelia Langford, John Sillitoe on behalf of the Wellcome Sanger Institute COVID-19 Surveillance Team |
| EPI_ISL_3285921 | Lighthouse Lab in Alderley Park | Wellcome Sanger Institute for the COVID-19 Genomics UK (COG-UK) Consortium | Jacquelyn Wynn, Mairead Hyland, The Lighthouse Lab in Alderley Park and Alex Alderton, Roberto Amato, Jeffrey Barrett, Sonia Goncalves, Ewan Harrison, David K. Jackson, Ian Johnston, Dominic Kwiatkowski, Cordelia Langford, John Sillitoe on behalf of the Wellcome Sanger Institute COVID-19 Surveillance Team |
| EPI_ISL_3285922 | Randox Laboratories | Wellcome Sanger Institute for the COVID-19 Genomics UK (COG-UK) Consortium | Randox Laboratories and Alex Alderton, Roberto Amato, Jeffrey Barrett, Sonia Goncalves, Ewan Harrison, David K. Jackson, Ian Johnston, Dominic Kwiatkowski, Cordelia Langford, John Sillitoe on behalf of the Wellcome Sanger Institute COVID-19 Surveillance Team |
| EPI_ISL_3285923 | Lighthouse Lab in Alderley Park | Wellcome Sanger Institute for the COVID-19 Genomics UK (COG-UK) Consortium | Jacquelyn Wynn, Mairead Hyland, The Lighthouse Lab in Alderley Park and Alex Alderton, Roberto Amato, Jeffrey Barrett, Sonia Goncalves, Ewan Harrison, David K. Jackson, Ian Johnston, Dominic Kwiatkowski, Cordelia Langford, John Sillitoe on behalf of the Wellcome Sanger Institute COVID-19 Surveillance Team |
| EPI_ISL_3285924 | Randox Laboratories | Wellcome Sanger Institute for the COVID-19 Genomics UK (COG-UK) Consortium | Randox Laboratories and Alex Alderton, Roberto Amato, Jeffrey Barrett, Sonia Goncalves, Ewan Harrison, David K. Jackson, Ian Johnston, Dominic Kwiatkowski, Cordelia Langford, John Sillitoe on behalf of the Wellcome Sanger Institute COVID-19 Surveillance Team |
| EPI_ISL_3285925, EPI_ISL_3285926, EPI_ISL_3285927, EPI_ISL_3285928, EPI_ISL_3285929 | Lighthouse Lab in Alderley Park | Wellcome Sanger Institute for the COVID-19 Genomics UK (COG-UK) Consortium | Jacquelyn Wynn, Mairead Hyland, The Lighthouse Lab in Alderley Park and Alex Alderton, Roberto Amato, Jeffrey Barrett, Sonia Goncalves, Ewan Harrison, David K. Jackson, Ian Johnston, Dominic Kwiatkowski, Cordelia Langford, John Sillitoe on behalf of the Wellcome Sanger Institute COVID-19 Surveillance Team |
| EPI_ISL_3285930 | Lighthouse Lab in Milton Keynes | Wellcome Sanger Institute for the COVID-19 Genomics UK (COG-UK) Consortium | The Lighthouse Lab in Milton Keynes and Alex Alderton, Roberto Amato, Jeffrey Barrett, Sonia Goncalves, Ewan Harrison, David K. Jackson, Ian Johnston, Dominic Kwiatkowski, Cordelia Langford, John Sillitoe on behalf of the Wellcome Sanger Institute COVID-19 Surveillance Team |
| EPI_ISL_3285931 | Randox Laboratories | Wellcome Sanger Institute for the COVID-19 Genomics UK (COG-UK) Consortium | Randox Laboratories and Alex Alderton, Roberto Amato, Jeffrey Barrett, Sonia Goncalves, Ewan Harrison, David K. Jackson, Ian Johnston, Dominic Kwiatkowski, Cordelia Langford, John Sillitoe on behalf of the Wellcome Sanger Institute COVID-19 Surveillance Team |
| EPI_ISL_3285932, EPI_ISL_3285933 | Lighthouse Lab in Alderley Park | Wellcome Sanger Institute for the COVID-19 Genomics UK (COG-UK) Consortium | Jacquelyn Wynn, Mairead Hyland, The Lighthouse Lab in Alderley Park and Alex Alderton, Roberto Amato, Jeffrey Barrett, Sonia Goncalves, Ewan Harrison, David K. Jackson, Ian Johnston, Dominic Kwiatkowski, Cordelia Langford, John Sillitoe on behalf of the Wellcome Sanger Institute COVID-19 Surveillance Team |
| EPI_ISL_3285934 | Randox Laboratories | Wellcome Sanger Institute for the COVID-19 Genomics UK (COG-UK) Consortium | Randox Laboratories and Alex Alderton, Roberto Amato, Jeffrey Barrett, Sonia Goncalves, Ewan Harrison, David K. Jackson, Ian Johnston, Dominic Kwiatkowski, Cordelia Langford, John Sillitoe on behalf of the Wellcome Sanger Institute COVID-19 Surveillance Team |
| EPI_ISL_3285935 | Lighthouse Lab in Alderley Park | Wellcome Sanger Institute for the COVID-19 Genomics UK (COG-UK) Consortium | Jacquelyn Wynn, Mairead Hyland, The Lighthouse Lab in Alderley Park and Alex Alderton, Roberto Amato, Jeffrey Barrett, Sonia Goncalves, Ewan Harrison, David K. Jackson, Ian Johnston, Dominic Kwiatkowski, Cordelia Langford, John Sillitoe on behalf of the Wellcome Sanger Institute COVID-19 Surveillance Team |
| EPI_ISL_3285936 | Randox Laboratories | Wellcome Sanger Institute for the COVID-19 Genomics UK (COG-UK) Consortium | Randox Laboratories and Alex Alderton, Roberto Amato, Jeffrey Barrett, Sonia Goncalves, Ewan Harrison, David K. Jackson, Ian Johnston, Dominic Kwiatkowski, Cordelia Langford, John Sillitoe on behalf of the Wellcome Sanger Institute COVID-19 Surveillance Team |
| EPI_ISL_3285937 | Lighthouse Lab in Alderley Park | Wellcome Sanger Institute for the COVID-19 Genomics UK (COG-UK) | Jacquelyn Wynn, Mairead Hyland, The Lighthouse Lab in Alderley Park and Alex Alderton, Roberto Amato, Jeffrey Barrett, Sonia Goncalves, Ewan Harrison, David K. Jackson, Ian Johnston, Dominic Kwiatkowski, Cordelia Langford, John Sillitoe on |

| | | | |
|---|---|---|---|
| | | Consortium | behalf of the Wellcome Sanger Institute COVID-19 Surveillance Team |
| EPI_ISL_3285938 | Lighthouse Lab in Milton Keynes | Wellcome Sanger Institute for the COVID-19 Genomics UK (COG-UK) Consortium | The Lighthouse Lab in Milton Keynes and Alex Alderton, Roberto Amato, Jeffrey Barrett, Sonia Goncalves, Ewan Harrison, David K. Jackson, Ian Johnston, Dominic Kwiatkowski, Cordelia Langford, John Sillitoe on behalf of the Wellcome Sanger Institute COVID-19 Surveillance Team |
| EPI_ISL_3285939 | Randox Laboratories | Wellcome Sanger Institute for the COVID-19 Genomics UK (COG-UK) Consortium | Randox Laboratories and Alex Alderton, Roberto Amato, Jeffrey Barrett, Sonia Goncalves, Ewan Harrison, David K. Jackson, Ian Johnston, Dominic Kwiatkowski, Cordelia Langford, John Sillitoe on behalf of the Wellcome Sanger Institute COVID-19 Surveillance Team |
| EPI_ISL_3285940, EPI_ISL_3285941, EPI_ISL_3285942, EPI_ISL_3285943, EPI_ISL_3285944, EPI_ISL_3285945 | Lighthouse Lab in Alderley Park | Wellcome Sanger Institute for the COVID-19 Genomics UK (COG-UK) Consortium | Jacquelyn Wynn, Mairead Hyland, The Lighthouse Lab in Alderley Park and Alex Alderton, Roberto Amato, Jeffrey Barrett, Sonia Goncalves, Ewan Harrison, David K. Jackson, Ian Johnston, Dominic Kwiatkowski, Cordelia Langford, John Sillitoe on behalf of the Wellcome Sanger Institute COVID-19 Surveillance Team |
| EPI_ISL_3285946 | Lighthouse Lab in Milton Keynes | Wellcome Sanger Institute for the COVID-19 Genomics UK (COG-UK) Consortium | The Lighthouse Lab in Milton Keynes and Alex Alderton, Roberto Amato, Jeffrey Barrett, Sonia Goncalves, Ewan Harrison, David K. Jackson, Ian Johnston, Dominic Kwiatkowski, Cordelia Langford, John Sillitoe on behalf of the Wellcome Sanger Institute COVID-19 Surveillance Team |
| EPI_ISL_3285947, EPI_ISL_3285948 | Lighthouse Lab in Alderley Park | Wellcome Sanger Institute for the COVID-19 Genomics UK (COG-UK) Consortium | Jacquelyn Wynn, Mairead Hyland, The Lighthouse Lab in Alderley Park and Alex Alderton, Roberto Amato, Jeffrey Barrett, Sonia Goncalves, Ewan Harrison, David K. Jackson, Ian Johnston, Dominic Kwiatkowski, Cordelia Langford, John Sillitoe on behalf of the Wellcome Sanger Institute COVID-19 Surveillance Team |
| EPI_ISL_3285949 | Lighthouse Lab in Milton Keynes | Wellcome Sanger Institute for the COVID-19 Genomics UK (COG-UK) Consortium | The Lighthouse Lab in Milton Keynes and Alex Alderton, Roberto Amato, Jeffrey Barrett, Sonia Goncalves, Ewan Harrison, David K. Jackson, Ian Johnston, Dominic Kwiatkowski, Cordelia Langford, John Sillitoe on behalf of the Wellcome Sanger Institute COVID-19 Surveillance Team |
| EPI_ISL_3285950, EPI_ISL_3285951, EPI_ISL_3285952 | Lighthouse Lab in Alderley Park | Wellcome Sanger Institute for the COVID-19 Genomics UK (COG-UK) Consortium | Jacquelyn Wynn, Mairead Hyland, The Lighthouse Lab in Alderley Park and Alex Alderton, Roberto Amato, Jeffrey Barrett, Sonia Goncalves, Ewan Harrison, David K. Jackson, Ian Johnston, Dominic Kwiatkowski, Cordelia Langford, John Sillitoe on behalf of the Wellcome Sanger Institute COVID-19 Surveillance Team |
| EPI_ISL_3285953, EPI_ISL_3285954 | Lighthouse Lab in Milton Keynes | Wellcome Sanger Institute for the COVID-19 Genomics UK (COG-UK) Consortium | The Lighthouse Lab in Milton Keynes and Alex Alderton, Roberto Amato, Jeffrey Barrett, Sonia Goncalves, Ewan Harrison, David K. Jackson, Ian Johnston, Dominic Kwiatkowski, Cordelia Langford, John Sillitoe on behalf of the Wellcome Sanger Institute COVID-19 Surveillance Team |
| EPI_ISL_3285955 | Lighthouse Lab in Alderley Park | Wellcome Sanger Institute for the COVID-19 Genomics UK (COG-UK) Consortium | Jacquelyn Wynn, Mairead Hyland, The Lighthouse Lab in Alderley Park and Alex Alderton, Roberto Amato, Jeffrey Barrett, Sonia Goncalves, Ewan Harrison, David K. Jackson, Ian Johnston, Dominic Kwiatkowski, Cordelia Langford, John Sillitoe on behalf of the Wellcome Sanger Institute COVID-19 Surveillance Team |
| EPI_ISL_3285956 | Lighthouse Lab in Milton Keynes | Wellcome Sanger Institute for the COVID-19 Genomics UK (COG-UK) Consortium | The Lighthouse Lab in Milton Keynes and Alex Alderton, Roberto Amato, Jeffrey Barrett, Sonia Goncalves, Ewan Harrison, David K. Jackson, Ian Johnston, Dominic Kwiatkowski, Cordelia Langford, John Sillitoe on behalf of the Wellcome Sanger Institute COVID-19 Surveillance Team |
| EPI_ISL_3285957 | Lighthouse Lab in Alderley Park | Wellcome Sanger Institute for the COVID-19 Genomics UK (COG-UK) Consortium | Jacquelyn Wynn, Mairead Hyland, The Lighthouse Lab in Alderley Park and Alex Alderton, Roberto Amato, Jeffrey Barrett, Sonia Goncalves, Ewan Harrison, David K. Jackson, Ian Johnston, Dominic Kwiatkowski, Cordelia Langford, John Sillitoe on behalf of the Wellcome Sanger Institute COVID-19 Surveillance Team |
| EPI_ISL_3285958 | Lighthouse Lab in Milton Keynes | Wellcome Sanger Institute for the COVID-19 Genomics UK (COG-UK) Consortium | The Lighthouse Lab in Milton Keynes and Alex Alderton, Roberto Amato, Jeffrey Barrett, Sonia Goncalves, Ewan Harrison, David K. Jackson, Ian Johnston, Dominic Kwiatkowski, Cordelia Langford, John Sillitoe on behalf of the Wellcome Sanger Institute COVID-19 Surveillance Team |
| EPI_ISL_3285959 | Randox Laboratories | Wellcome Sanger Institute for the COVID-19 Genomics UK (COG-UK) Consortium | Randox Laboratories and Alex Alderton, Roberto Amato, Jeffrey Barrett, Sonia Goncalves, Ewan Harrison, David K. Jackson, Ian Johnston, Dominic Kwiatkowski, Cordelia Langford, John Sillitoe on behalf of the Wellcome Sanger Institute COVID-19 Surveillance Team |
| EPI_ISL_3285960 | Lighthouse Lab in Milton Keynes | Wellcome Sanger Institute for the COVID-19 Genomics UK (COG-UK) Consortium | The Lighthouse Lab in Milton Keynes and Alex Alderton, Roberto Amato, Jeffrey Barrett, Sonia Goncalves, Ewan Harrison, David K. Jackson, Ian Johnston, Dominic Kwiatkowski, Cordelia Langford, John Sillitoe on behalf of the Wellcome Sanger Institute COVID-19 Surveillance Team |
| EPI_ISL_3285961 | Lighthouse Lab in Alderley Park | Wellcome Sanger Institute for the COVID-19 Genomics UK (COG-UK) Consortium | Jacquelyn Wynn, Mairead Hyland, The Lighthouse Lab in Alderley Park and Alex Alderton, Roberto Amato, Jeffrey Barrett, Sonia Goncalves, Ewan Harrison, David K. Jackson, Ian Johnston, Dominic Kwiatkowski, Cordelia Langford, John Sillitoe on behalf of the Wellcome Sanger Institute COVID-19 Surveillance Team |
| EPI_ISL_3285962, EPI_ISL_3285963 | Lighthouse Lab in Milton Keynes | Wellcome Sanger Institute for the COVID-19 Genomics UK (COG-UK) Consortium | The Lighthouse Lab in Milton Keynes and Alex Alderton, Roberto Amato, Jeffrey Barrett, Sonia Goncalves, Ewan Harrison, David K. Jackson, Ian Johnston, Dominic Kwiatkowski, Cordelia Langford, John Sillitoe on behalf of the Wellcome Sanger Institute COVID-19 Surveillance Team |
| EPI_ISL_3285964 | Lighthouse Lab in Alderley Park | Wellcome Sanger Institute for the COVID-19 Genomics UK (COG-UK) Consortium | Jacquelyn Wynn, Mairead Hyland, The Lighthouse Lab in Alderley Park and Alex Alderton, Roberto Amato, Jeffrey Barrett, Sonia Goncalves, Ewan Harrison, David K. Jackson, Ian Johnston, Dominic Kwiatkowski, Cordelia Langford, John Sillitoe on behalf of the Wellcome Sanger Institute COVID-19 Surveillance Team |
| EPI_ISL_3285965 | Lighthouse Lab in Milton Keynes | Wellcome Sanger Institute for the COVID-19 Genomics UK (COG-UK) Consortium | The Lighthouse Lab in Milton Keynes and Alex Alderton, Roberto Amato, Jeffrey Barrett, Sonia Goncalves, Ewan Harrison, David K. Jackson, Ian Johnston, Dominic Kwiatkowski, Cordelia Langford, John Sillitoe on behalf of the Wellcome Sanger Institute COVID-19 Surveillance Team |
| EPI_ISL_3285966, EPI_ISL_3285967 | Lighthouse Lab in Alderley Park | Wellcome Sanger Institute for the COVID-19 Genomics UK (COG-UK) Consortium | Jacquelyn Wynn, Mairead Hyland, The Lighthouse Lab in Alderley Park and Alex Alderton, Roberto Amato, Jeffrey Barrett, Sonia Goncalves, Ewan Harrison, David K. Jackson, Ian Johnston, Dominic Kwiatkowski, Cordelia Langford, John Sillitoe on behalf of the Wellcome Sanger Institute COVID-19 Surveillance Team |
| EPI_ISL_3285968 | Randox Laboratories | Wellcome Sanger Institute for the COVID-19 Genomics UK (COG-UK) Consortium | Randox Laboratories and Alex Alderton, Roberto Amato, Jeffrey Barrett, Sonia Goncalves, Ewan Harrison, David K. Jackson, Ian Johnston, Dominic Kwiatkowski, Cordelia Langford, John Sillitoe on behalf of the Wellcome Sanger Institute COVID-19 Surveillance Team |
| EPI_ISL_3285969, EPI_ISL_3285970, EPI_ISL_3285971, EPI_ISL_3285972 | Lighthouse Lab in Alderley Park | Wellcome Sanger Institute for the COVID-19 Genomics UK (COG-UK) Consortium | Jacquelyn Wynn, Mairead Hyland, The Lighthouse Lab in Alderley Park and Alex Alderton, Roberto Amato, Jeffrey Barrett, Sonia Goncalves, Ewan Harrison, David K. Jackson, Ian Johnston, Dominic Kwiatkowski, Cordelia Langford, John Sillitoe on behalf of the Wellcome Sanger Institute COVID-19 Surveillance Team |
| EPI_ISL_3285973 | Lighthouse Lab in Milton Keynes | Wellcome Sanger Institute for the COVID-19 Genomics UK (COG-UK) Consortium | The Lighthouse Lab in Milton Keynes and Alex Alderton, Roberto Amato, Jeffrey Barrett, Sonia Goncalves, Ewan Harrison, David K. Jackson, Ian Johnston, Dominic Kwiatkowski, Cordelia Langford, John Sillitoe on behalf of the Wellcome Sanger Institute COVID-19 Surveillance Team |
| EPI_ISL_3285974 | Lighthouse Lab in Alderley Park | Wellcome Sanger Institute for the COVID-19 Genomics UK (COG-UK) Consortium | Jacquelyn Wynn, Mairead Hyland, The Lighthouse Lab in Alderley Park and Alex Alderton, Roberto Amato, Jeffrey Barrett, Sonia Goncalves, Ewan Harrison, David K. Jackson, Ian Johnston, Dominic Kwiatkowski, Cordelia Langford, John Sillitoe on behalf of the Wellcome Sanger Institute COVID-19 Surveillance Team |
| EPI_ISL_3285975 | Randox Laboratories | Wellcome Sanger Institute for the COVID-19 Genomics UK (COG-UK) Consortium | Randox Laboratories and Alex Alderton, Roberto Amato, Jeffrey Barrett, Sonia Goncalves, Ewan Harrison, David K. Jackson, Ian Johnston, Dominic Kwiatkowski, Cordelia Langford, John Sillitoe on behalf of the Wellcome Sanger Institute COVID-19 Surveillance Team |
| EPI_ISL_3285976 | Lighthouse Lab in Alderley Park | Wellcome Sanger Institute for the COVID-19 Genomics UK (COG-UK) Consortium | Jacquelyn Wynn, Mairead Hyland, The Lighthouse Lab in Alderley Park and Alex Alderton, Roberto Amato, Jeffrey Barrett, Sonia Goncalves, Ewan Harrison, David K. Jackson, Ian Johnston, Dominic Kwiatkowski, Cordelia Langford, John Sillitoe on behalf of the Wellcome Sanger Institute COVID-19 Surveillance Team |
| EPI_ISL_3285977, EPI_ISL_3285978 | Lighthouse Lab in Milton Keynes | Wellcome Sanger Institute for the COVID-19 Genomics UK (COG-UK) Consortium | The Lighthouse Lab in Milton Keynes and Alex Alderton, Roberto Amato, Jeffrey Barrett, Sonia Goncalves, Ewan Harrison, David K. Jackson, Ian Johnston, Dominic Kwiatkowski, Cordelia Langford, John Sillitoe on behalf of the Wellcome Sanger Institute COVID-19 Surveillance Team |
| EPI_ISL_3285979 | Randox Laboratories | Wellcome Sanger Institute for the COVID-19 Genomics UK (COG-UK) Consortium | Randox Laboratories and Alex Alderton, Roberto Amato, Jeffrey Barrett, Sonia Goncalves, Ewan Harrison, David K. Jackson, Ian Johnston, Dominic Kwiatkowski, Cordelia Langford, John Sillitoe on behalf of the Wellcome Sanger Institute COVID-19 Surveillance Team |
| EPI_ISL_3285980 | Lighthouse Lab in Milton Keynes | Wellcome Sanger Institute for the COVID-19 Genomics UK (COG-UK) Consortium | The Lighthouse Lab in Milton Keynes and Alex Alderton, Roberto Amato, Jeffrey Barrett, Sonia Goncalves, Ewan Harrison, David K. Jackson, Ian Johnston, Dominic Kwiatkowski, Cordelia Langford, John Sillitoe on behalf of the Wellcome Sanger Institute COVID-19 Surveillance Team |
| EPI_ISL_3285981 | Randox Laboratories | Wellcome Sanger Institute for the COVID-19 Genomics UK (COG-UK) Consortium | Randox Laboratories and Alex Alderton, Roberto Amato, Jeffrey Barrett, Sonia Goncalves, Ewan Harrison, David K. Jackson, Ian Johnston, Dominic Kwiatkowski, Cordelia Langford, John Sillitoe on behalf of the Wellcome Sanger Institute COVID-19 Surveillance Team |
| EPI_ISL_3285982, EPI_ISL_3285983, EPI_ISL_3285984, EPI_ISL_3285985, EPI_ISL_3285986, EPI_ISL_3285987, EPI_ISL_3285988, EPI_ISL_3285989 | Lighthouse Lab in Alderley Park | Wellcome Sanger Institute for the COVID-19 Genomics UK (COG-UK) Consortium | Jacquelyn Wynn, Mairead Hyland, The Lighthouse Lab in Alderley Park and Alex Alderton, Roberto Amato, Jeffrey Barrett, Sonia Goncalves, Ewan Harrison, David K. Jackson, Ian Johnston, Dominic Kwiatkowski, Cordelia Langford, John Sillitoe on behalf of the Wellcome Sanger Institute COVID-19 Surveillance Team |
| EPI_ISL_3285990 | Randox Laboratories | Wellcome Sanger Institute for the COVID-19 Genomics UK (COG-UK) Consortium | Randox Laboratories and Alex Alderton, Roberto Amato, Jeffrey Barrett, Sonia Goncalves, Ewan Harrison, David K. Jackson, Ian Johnston, Dominic Kwiatkowski, Cordelia Langford, John Sillitoe on behalf of the Wellcome Sanger Institute COVID-19 Surveillance Team |
| EPI_ISL_3285991 | Lighthouse Lab in Milton Keynes | Wellcome Sanger Institute for the COVID-19 Genomics UK (COG-UK) Consortium | The Lighthouse Lab in Milton Keynes and Alex Alderton, Roberto Amato, Jeffrey Barrett, Sonia Goncalves, Ewan Harrison, David K. Jackson, Ian Johnston, Dominic Kwiatkowski, Cordelia Langford, John Sillitoe on behalf of the Wellcome Sanger Institute COVID-19 Surveillance Team |
| EPI_ISL_3285992, EPI_ISL_3285993, EPI_ISL_3285994, EPI_ISL_3285995 | Lighthouse Lab in Alderley Park | Wellcome Sanger Institute for the COVID-19 Genomics UK (COG-UK) Consortium | Jacquelyn Wynn, Mairead Hyland, The Lighthouse Lab in Alderley Park and Alex Alderton, Roberto Amato, Jeffrey Barrett, Sonia Goncalves, Ewan Harrison, David K. Jackson, Ian Johnston, Dominic Kwiatkowski, Cordelia Langford, John Sillitoe on behalf of the Wellcome Sanger Institute COVID-19 Surveillance Team |
| EPI_ISL_3285996, EPI_ISL_3285997 | Lighthouse Lab in Milton Keynes | Wellcome Sanger Institute for the COVID-19 Genomics UK (COG-UK) Consortium | The Lighthouse Lab in Milton Keynes and Alex Alderton, Roberto Amato, Jeffrey Barrett, Sonia Goncalves, Ewan Harrison, David K. Jackson, Ian Johnston, Dominic Kwiatkowski, Cordelia Langford, John Sillitoe on behalf of the Wellcome Sanger Institute COVID-19 Surveillance Team |
| EPI_ISL_3285998, EPI_ISL_3285999, EPI_ISL_3286000, EPI_ISL_3286001, EPI_ISL_3286002, EPI_ISL_3286003 | Lighthouse Lab in Alderley Park | Wellcome Sanger Institute for the COVID-19 Genomics UK (COG-UK) Consortium | Jacquelyn Wynn, Mairead Hyland, The Lighthouse Lab in Alderley Park and Alex Alderton, Roberto Amato, Jeffrey Barrett, Sonia Goncalves, Ewan Harrison, David K. Jackson, Ian Johnston, Dominic Kwiatkowski, Cordelia Langford, John Sillitoe on behalf of the Wellcome Sanger Institute COVID-19 Surveillance Team |
| EPI_ISL_3286004, EPI_ISL_3286005 | Randox Laboratories | Wellcome Sanger Institute for the COVID-19 Genomics UK (COG-UK) Consortium | Randox Laboratories and Alex Alderton, Roberto Amato, Jeffrey Barrett, Sonia Goncalves, Ewan Harrison, David K. Jackson, Ian Johnston, Dominic Kwiatkowski, Cordelia Langford, John Sillitoe on behalf of the Wellcome Sanger Institute COVID-19 Surveillance Team |
| EPI_ISL_3286006, EPI_ISL_3286007, | Lighthouse Lab in Alderley Park | Wellcome Sanger Institute for the COVID-19 Genomics UK (COG-UK) Consortium | Jacquelyn Wynn, Mairead Hyland, The Lighthouse Lab in Alderley Park and Alex Alderton, Roberto Amato, Jeffrey Barrett, Sonia Goncalves, Ewan Harrison, David K. Jackson, Ian Johnston, Dominic Kwiatkowski, Cordelia Langford, John Sillitoe on behalf of the Wellcome Sanger Institute COVID-19 Surveillance Team |

| | | | |
|---|---|---|---|
| EPI_ISL_3286008, EPI_ISL_3286009, EPI_ISL_3286010 | | | |
| EPI_ISL_3286011 | Lighthouse Lab in Milton Keynes | Wellcome Sanger Institute for the COVID-19 Genomics UK (COG-UK) Consortium | The Lighthouse Lab in Milton Keynes and Alex Alderton, Roberto Amato, Jeffrey Barrett, Sonia Goncalves, Ewan Harrison, David K. Jackson, Ian Johnston, Dominic Kwiatkowski, Cordelia Langford, John Sillitoe on behalf of the Wellcome Sanger Institute COVID-19 Surveillance Team |
| EPI_ISL_3286012, EPI_ISL_3286013, EPI_ISL_3286014, EPI_ISL_3286015, EPI_ISL_3286016, EPI_ISL_3286017, EPI_ISL_3286018 | Lighthouse Lab in Alderley Park | Wellcome Sanger Institute for the COVID-19 Genomics UK (COG-UK) Consortium | Jacquelyn Wynn, Mairead Hyland, The Lighthouse Lab in Alderley Park and Alex Alderton, Roberto Amato, Jeffrey Barrett, Sonia Goncalves, Ewan Harrison, David K. Jackson, Ian Johnston, Dominic Kwiatkowski, Cordelia Langford, John Sillitoe on behalf of the Wellcome Sanger Institute COVID-19 Surveillance Team |
| EPI_ISL_3286019, EPI_ISL_3286020, EPI_ISL_3286021 | Lighthouse Lab in Milton Keynes | Wellcome Sanger Institute for the COVID-19 Genomics UK (COG-UK) Consortium | The Lighthouse Lab in Milton Keynes and Alex Alderton, Roberto Amato, Jeffrey Barrett, Sonia Goncalves, Ewan Harrison, David K. Jackson, Ian Johnston, Dominic Kwiatkowski, Cordelia Langford, John Sillitoe on behalf of the Wellcome Sanger Institute COVID-19 Surveillance Team |
| EPI_ISL_3286022, EPI_ISL_3286023 | Lighthouse Lab in Alderley Park | Wellcome Sanger Institute for the COVID-19 Genomics UK (COG-UK) Consortium | Jacquelyn Wynn, Mairead Hyland, The Lighthouse Lab in Alderley Park and Alex Alderton, Roberto Amato, Jeffrey Barrett, Sonia Goncalves, Ewan Harrison, David K. Jackson, Ian Johnston, Dominic Kwiatkowski, Cordelia Langford, John Sillitoe on behalf of the Wellcome Sanger Institute COVID-19 Surveillance Team |
| EPI_ISL_3286024, EPI_ISL_3286025 | Lighthouse Lab in Milton Keynes | Wellcome Sanger Institute for the COVID-19 Genomics UK (COG-UK) Consortium | The Lighthouse Lab in Milton Keynes and Alex Alderton, Roberto Amato, Jeffrey Barrett, Sonia Goncalves, Ewan Harrison, David K. Jackson, Ian Johnston, Dominic Kwiatkowski, Cordelia Langford, John Sillitoe on behalf of the Wellcome Sanger Institute COVID-19 Surveillance Team |
| EPI_ISL_3286026 | Lighthouse Lab in Alderley Park | Wellcome Sanger Institute for the COVID-19 Genomics UK (COG-UK) Consortium | Jacquelyn Wynn, Mairead Hyland, The Lighthouse Lab in Alderley Park and Alex Alderton, Roberto Amato, Jeffrey Barrett, Sonia Goncalves, Ewan Harrison, David K. Jackson, Ian Johnston, Dominic Kwiatkowski, Cordelia Langford, John Sillitoe on behalf of the Wellcome Sanger Institute COVID-19 Surveillance Team |
| EPI_ISL_3286027 | Lighthouse Lab in Milton Keynes | Wellcome Sanger Institute for the COVID-19 Genomics UK (COG-UK) Consortium | The Lighthouse Lab in Milton Keynes and Alex Alderton, Roberto Amato, Jeffrey Barrett, Sonia Goncalves, Ewan Harrison, David K. Jackson, Ian Johnston, Dominic Kwiatkowski, Cordelia Langford, John Sillitoe on behalf of the Wellcome Sanger Institute COVID-19 Surveillance Team |
| EPI_ISL_3286028 | Lighthouse Lab in Alderley Park | Wellcome Sanger Institute for the COVID-19 Genomics UK (COG-UK) Consortium | Jacquelyn Wynn, Mairead Hyland, The Lighthouse Lab in Alderley Park and Alex Alderton, Roberto Amato, Jeffrey Barrett, Sonia Goncalves, Ewan Harrison, David K. Jackson, Ian Johnston, Dominic Kwiatkowski, Cordelia Langford, John Sillitoe on behalf of the Wellcome Sanger Institute COVID-19 Surveillance Team |
| EPI_ISL_3286029 | Lighthouse Lab in Milton Keynes | Wellcome Sanger Institute for the COVID-19 Genomics UK (COG-UK) Consortium | The Lighthouse Lab in Milton Keynes and Alex Alderton, Roberto Amato, Jeffrey Barrett, Sonia Goncalves, Ewan Harrison, David K. Jackson, Ian Johnston, Dominic Kwiatkowski, Cordelia Langford, John Sillitoe on behalf of the Wellcome Sanger Institute COVID-19 Surveillance Team |
| EPI_ISL_3286030, EPI_ISL_3286031 | Lighthouse Lab in Alderley Park | Wellcome Sanger Institute for the COVID-19 Genomics UK (COG-UK) Consortium | Jacquelyn Wynn, Mairead Hyland, The Lighthouse Lab in Alderley Park and Alex Alderton, Roberto Amato, Jeffrey Barrett, Sonia Goncalves, Ewan Harrison, David K. Jackson, Ian Johnston, Dominic Kwiatkowski, Cordelia Langford, John Sillitoe on behalf of the Wellcome Sanger Institute COVID-19 Surveillance Team |
| EPI_ISL_3286032 | Lighthouse Lab in Milton Keynes | Wellcome Sanger Institute for the COVID-19 Genomics UK (COG-UK) Consortium | The Lighthouse Lab in Milton Keynes and Alex Alderton, Roberto Amato, Jeffrey Barrett, Sonia Goncalves, Ewan Harrison, David K. Jackson, Ian Johnston, Dominic Kwiatkowski, Cordelia Langford, John Sillitoe on behalf of the Wellcome Sanger Institute COVID-19 Surveillance Team |
| EPI_ISL_3286033, EPI_ISL_3286034, EPI_ISL_3286035 | Lighthouse Lab in Alderley Park | Wellcome Sanger Institute for the COVID-19 Genomics UK (COG-UK) Consortium | Jacquelyn Wynn, Mairead Hyland, The Lighthouse Lab in Alderley Park and Alex Alderton, Roberto Amato, Jeffrey Barrett, Sonia Goncalves, Ewan Harrison, David K. Jackson, Ian Johnston, Dominic Kwiatkowski, Cordelia Langford, John Sillitoe on behalf of the Wellcome Sanger Institute COVID-19 Surveillance Team |
| EPI_ISL_3286036, EPI_ISL_3286037 | Lighthouse Lab in Milton Keynes | Wellcome Sanger Institute for the COVID-19 Genomics UK (COG-UK) Consortium | The Lighthouse Lab in Milton Keynes and Alex Alderton, Roberto Amato, Jeffrey Barrett, Sonia Goncalves, Ewan Harrison, David K. Jackson, Ian Johnston, Dominic Kwiatkowski, Cordelia Langford, John Sillitoe on behalf of the Wellcome Sanger Institute COVID-19 Surveillance Team |
| EPI_ISL_3286038, EPI_ISL_3286039, EPI_ISL_3286040, EPI_ISL_3286041, EPI_ISL_3286042 | Lighthouse Lab in Alderley Park | Wellcome Sanger Institute for the COVID-19 Genomics UK (COG-UK) Consortium | Jacquelyn Wynn, Mairead Hyland, The Lighthouse Lab in Alderley Park and Alex Alderton, Roberto Amato, Jeffrey Barrett, Sonia Goncalves, Ewan Harrison, David K. Jackson, Ian Johnston, Dominic Kwiatkowski, Cordelia Langford, John Sillitoe on behalf of the Wellcome Sanger Institute COVID-19 Surveillance Team |
| EPI_ISL_3286043 | Lighthouse Lab in Milton Keynes | Wellcome Sanger Institute for the COVID-19 Genomics UK (COG-UK) Consortium | The Lighthouse Lab in Milton Keynes and Alex Alderton, Roberto Amato, Jeffrey Barrett, Sonia Goncalves, Ewan Harrison, David K. Jackson, Ian Johnston, Dominic Kwiatkowski, Cordelia Langford, John Sillitoe on behalf of the Wellcome Sanger Institute COVID-19 Surveillance Team |
| EPI_ISL_3286044 | Lighthouse Lab in Alderley Park | Wellcome Sanger Institute for the COVID-19 Genomics UK (COG-UK) Consortium | Jacquelyn Wynn, Mairead Hyland, The Lighthouse Lab in Alderley Park and Alex Alderton, Roberto Amato, Jeffrey Barrett, Sonia Goncalves, Ewan Harrison, David K. Jackson, Ian Johnston, Dominic Kwiatkowski, Cordelia Langford, John Sillitoe on behalf of the Wellcome Sanger Institute COVID-19 Surveillance Team |
| EPI_ISL_3286045, EPI_ISL_3286046 | Lighthouse Lab in Milton Keynes | Wellcome Sanger Institute for the COVID-19 Genomics UK (COG-UK) Consortium | The Lighthouse Lab in Milton Keynes and Alex Alderton, Roberto Amato, Jeffrey Barrett, Sonia Goncalves, Ewan Harrison, David K. Jackson, Ian Johnston, Dominic Kwiatkowski, Cordelia Langford, John Sillitoe on behalf of the Wellcome Sanger Institute COVID-19 Surveillance Team |
| EPI_ISL_3286047 | Randox Laboratories | Wellcome Sanger Institute for the COVID-19 Genomics UK (COG-UK) Consortium | Randox Laboratories and Alex Alderton, Roberto Amato, Jeffrey Barrett, Sonia Goncalves, Ewan Harrison, David K. Jackson, Ian Johnston, Dominic Kwiatkowski, Cordelia Langford, John Sillitoe on behalf of the Wellcome Sanger Institute COVID-19 Surveillance Team |
| EPI_ISL_3286048 | Lighthouse Lab in Milton Keynes | Wellcome Sanger Institute for the COVID-19 Genomics UK (COG-UK) Consortium | The Lighthouse Lab in Milton Keynes and Alex Alderton, Roberto Amato, Jeffrey Barrett, Sonia Goncalves, Ewan Harrison, David K. Jackson, Ian Johnston, Dominic Kwiatkowski, Cordelia Langford, John Sillitoe on behalf of the Wellcome Sanger Institute COVID-19 Surveillance Team |
| EPI_ISL_3286049 | Randox Laboratories | Wellcome Sanger Institute for the COVID-19 Genomics UK (COG-UK) Consortium | Randox Laboratories and Alex Alderton, Roberto Amato, Jeffrey Barrett, Sonia Goncalves, Ewan Harrison, David K. Jackson, Ian Johnston, Dominic Kwiatkowski, Cordelia Langford, John Sillitoe on behalf of the Wellcome Sanger Institute COVID-19 Surveillance Team |
| EPI_ISL_3286050, EPI_ISL_3286051, EPI_ISL_3286052 | Lighthouse Lab in Alderley Park | Wellcome Sanger Institute for the COVID-19 Genomics UK (COG-UK) Consortium | Jacquelyn Wynn, Mairead Hyland, The Lighthouse Lab in Alderley Park and Alex Alderton, Roberto Amato, Jeffrey Barrett, Sonia Goncalves, Ewan Harrison, David K. Jackson, Ian Johnston, Dominic Kwiatkowski, Cordelia Langford, John Sillitoe on behalf of the Wellcome Sanger Institute COVID-19 Surveillance Team |
| EPI_ISL_3286053, EPI_ISL_3286054 | Randox Laboratories | Wellcome Sanger Institute for the COVID-19 Genomics UK (COG-UK) Consortium | Randox Laboratories and Alex Alderton, Roberto Amato, Jeffrey Barrett, Sonia Goncalves, Ewan Harrison, David K. Jackson, Ian Johnston, Dominic Kwiatkowski, Cordelia Langford, John Sillitoe on behalf of the Wellcome Sanger Institute COVID-19 Surveillance Team |
| EPI_ISL_3286055 | Lighthouse Lab in Alderley Park | Wellcome Sanger Institute for the COVID-19 Genomics UK (COG-UK) Consortium | Jacquelyn Wynn, Mairead Hyland, The Lighthouse Lab in Alderley Park and Alex Alderton, Roberto Amato, Jeffrey Barrett, Sonia Goncalves, Ewan Harrison, David K. Jackson, Ian Johnston, Dominic Kwiatkowski, Cordelia Langford, John Sillitoe on behalf of the Wellcome Sanger Institute COVID-19 Surveillance Team |
| EPI_ISL_3286056, EPI_ISL_3286057 | Lighthouse Lab in Milton Keynes | Wellcome Sanger Institute for the COVID-19 Genomics UK (COG-UK) Consortium | The Lighthouse Lab in Milton Keynes and Alex Alderton, Roberto Amato, Jeffrey Barrett, Sonia Goncalves, Ewan Harrison, David K. Jackson, Ian Johnston, Dominic Kwiatkowski, Cordelia Langford, John Sillitoe on behalf of the Wellcome Sanger Institute COVID-19 Surveillance Team |
| EPI_ISL_3286058, EPI_ISL_3286059 | Randox Laboratories | Wellcome Sanger Institute for the COVID-19 Genomics UK (COG-UK) Consortium | Randox Laboratories and Alex Alderton, Roberto Amato, Jeffrey Barrett, Sonia Goncalves, Ewan Harrison, David K. Jackson, Ian Johnston, Dominic Kwiatkowski, Cordelia Langford, John Sillitoe on behalf of the Wellcome Sanger Institute COVID-19 Surveillance Team |
| EPI_ISL_3286060, EPI_ISL_3286061 | Lighthouse Lab in Milton Keynes | Wellcome Sanger Institute for the COVID-19 Genomics UK (COG-UK) Consortium | The Lighthouse Lab in Milton Keynes and Alex Alderton, Roberto Amato, Jeffrey Barrett, Sonia Goncalves, Ewan Harrison, David K. Jackson, Ian Johnston, Dominic Kwiatkowski, Cordelia Langford, John Sillitoe on behalf of the Wellcome Sanger Institute COVID-19 Surveillance Team |
| EPI_ISL_3286062, EPI_ISL_3286063, EPI_ISL_3286064, EPI_ISL_3286065 | Lighthouse Lab in Alderley Park | Wellcome Sanger Institute for the COVID-19 Genomics UK (COG-UK) Consortium | Jacquelyn Wynn, Mairead Hyland, The Lighthouse Lab in Alderley Park and Alex Alderton, Roberto Amato, Jeffrey Barrett, Sonia Goncalves, Ewan Harrison, David K. Jackson, Ian Johnston, Dominic Kwiatkowski, Cordelia Langford, John Sillitoe on behalf of the Wellcome Sanger Institute COVID-19 Surveillance Team |
| EPI_ISL_3286066, EPI_ISL_3286067 | Lighthouse Lab in Milton Keynes | Wellcome Sanger Institute for the COVID-19 Genomics UK (COG-UK) Consortium | The Lighthouse Lab in Milton Keynes and Alex Alderton, Roberto Amato, Jeffrey Barrett, Sonia Goncalves, Ewan Harrison, David K. Jackson, Ian Johnston, Dominic Kwiatkowski, Cordelia Langford, John Sillitoe on behalf of the Wellcome Sanger Institute COVID-19 Surveillance Team |
| EPI_ISL_3286068, EPI_ISL_3286069, EPI_ISL_3286070 | Lighthouse Lab in Alderley Park | Wellcome Sanger Institute for the COVID-19 Genomics UK (COG-UK) Consortium | Jacquelyn Wynn, Mairead Hyland, The Lighthouse Lab in Alderley Park and Alex Alderton, Roberto Amato, Jeffrey Barrett, Sonia Goncalves, Ewan Harrison, David K. Jackson, Ian Johnston, Dominic Kwiatkowski, Cordelia Langford, John Sillitoe on behalf of the Wellcome Sanger Institute COVID-19 Surveillance Team |
| EPI_ISL_3286071 | Lighthouse Lab in Milton Keynes | Wellcome Sanger Institute for the COVID-19 Genomics UK (COG-UK) Consortium | The Lighthouse Lab in Milton Keynes and Alex Alderton, Roberto Amato, Jeffrey Barrett, Sonia Goncalves, Ewan Harrison, David K. Jackson, Ian Johnston, Dominic Kwiatkowski, Cordelia Langford, John Sillitoe on behalf of the Wellcome Sanger Institute COVID-19 Surveillance Team |
| EPI_ISL_3286072, EPI_ISL_3286073 | Lighthouse Lab in Alderley Park | Wellcome Sanger Institute for the COVID-19 Genomics UK (COG-UK) Consortium | Jacquelyn Wynn, Mairead Hyland, The Lighthouse Lab in Alderley Park and Alex Alderton, Roberto Amato, Jeffrey Barrett, Sonia Goncalves, Ewan Harrison, David K. Jackson, Ian Johnston, Dominic Kwiatkowski, Cordelia Langford, John Sillitoe on behalf of the Wellcome Sanger Institute COVID-19 Surveillance Team |
| EPI_ISL_3286074, EPI_ISL_3286075 | Randox Laboratories | Wellcome Sanger Institute for the COVID-19 Genomics UK (COG-UK) Consortium | Randox Laboratories and Alex Alderton, Roberto Amato, Jeffrey Barrett, Sonia Goncalves, Ewan Harrison, David K. Jackson, Ian Johnston, Dominic Kwiatkowski, Cordelia Langford, John Sillitoe on behalf of the Wellcome Sanger Institute COVID-19 Surveillance Team |
| EPI_ISL_3286076, EPI_ISL_3286077 | Lighthouse Lab in Alderley Park | Wellcome Sanger Institute for the COVID-19 Genomics UK (COG-UK) Consortium | Jacquelyn Wynn, Mairead Hyland, The Lighthouse Lab in Alderley Park and Alex Alderton, Roberto Amato, Jeffrey Barrett, Sonia Goncalves, Ewan Harrison, David K. Jackson, Ian Johnston, Dominic Kwiatkowski, Cordelia Langford, John Sillitoe on behalf of the Wellcome Sanger Institute COVID-19 Surveillance Team |
| EPI_ISL_3286078 | Lighthouse Lab in Milton Keynes | Wellcome Sanger Institute for the COVID-19 Genomics UK (COG-UK) Consortium | The Lighthouse Lab in Milton Keynes and Alex Alderton, Roberto Amato, Jeffrey Barrett, Sonia Goncalves, Ewan Harrison, David K. Jackson, Ian Johnston, Dominic Kwiatkowski, Cordelia Langford, John Sillitoe on behalf of the Wellcome Sanger Institute COVID-19 Surveillance Team |
| EPI_ISL_3286079, EPI_ISL_3286080, EPI_ISL_3286081, EPI_ISL_3286082, EPI_ISL_3286083, EPI_ISL_3286084 | Lighthouse Lab in Alderley Park | Wellcome Sanger Institute for the COVID-19 Genomics UK (COG-UK) Consortium | Jacquelyn Wynn, Mairead Hyland, The Lighthouse Lab in Alderley Park and Alex Alderton, Roberto Amato, Jeffrey Barrett, Sonia Goncalves, Ewan Harrison, David K. Jackson, Ian Johnston, Dominic Kwiatkowski, Cordelia Langford, John Sillitoe on behalf of the Wellcome Sanger Institute COVID-19 Surveillance Team |
| EPI_ISL_3286085 | Lighthouse Lab in Milton Keynes | Wellcome Sanger Institute for the COVID-19 Genomics UK (COG-UK) Consortium | The Lighthouse Lab in Milton Keynes and Alex Alderton, Roberto Amato, Jeffrey Barrett, Sonia Goncalves, Ewan Harrison, David K. Jackson, Ian Johnston, Dominic Kwiatkowski, Cordelia Langford, John Sillitoe on behalf of the Wellcome Sanger Institute COVID-19 Surveillance Team |

| | | | |
|---|---|---|---|
| EPI_ISL_3286086 | Lighthouse Lab in Alderley Park | Wellcome Sanger Institute for the COVID-19 Genomics UK (COG-UK) Consortium | Jacquelyn Wynn, Mairead Hyland, The Lighthouse Lab in Alderley Park and Alex Alderton, Roberto Amato, Jeffrey Barrett, Sonia Goncalves, Ewan Harrison, David K. Jackson, Ian Johnston, Dominic Kwiatkowski, Cordelia Langford, John Sillitoe on behalf of the Wellcome Sanger Institute COVID-19 Surveillance Team |
| EPI_ISL_3286087 | Randox Laboratories | Wellcome Sanger Institute for the COVID-19 Genomics UK (COG-UK) Consortium | Randox Laboratories and Alex Alderton, Roberto Amato, Jeffrey Barrett, Sonia Goncalves, Ewan Harrison, David K. Jackson, Ian Johnston, Dominic Kwiatkowski, Cordelia Langford, John Sillitoe on behalf of the Wellcome Sanger Institute COVID-19 Surveillance Team |
| EPI_ISL_3286088, EPI_ISL_3286089, EPI_ISL_3286090 | Lighthouse Lab in Alderley Park | Wellcome Sanger Institute for the COVID-19 Genomics UK (COG-UK) Consortium | Jacquelyn Wynn, Mairead Hyland, The Lighthouse Lab in Alderley Park and Alex Alderton, Roberto Amato, Jeffrey Barrett, Sonia Goncalves, Ewan Harrison, David K. Jackson, Ian Johnston, Dominic Kwiatkowski, Cordelia Langford, John Sillitoe on behalf of the Wellcome Sanger Institute COVID-19 Surveillance Team |
| EPI_ISL_3286091 | Randox Laboratories | Wellcome Sanger Institute for the COVID-19 Genomics UK (COG-UK) Consortium | Randox Laboratories and Alex Alderton, Roberto Amato, Jeffrey Barrett, Sonia Goncalves, Ewan Harrison, David K. Jackson, Ian Johnston, Dominic Kwiatkowski, Cordelia Langford, John Sillitoe on behalf of the Wellcome Sanger Institute COVID-19 Surveillance Team |
| EPI_ISL_3286092 | Lighthouse Lab in Alderley Park | Wellcome Sanger Institute for the COVID-19 Genomics UK (COG-UK) Consortium | Jacquelyn Wynn, Mairead Hyland, The Lighthouse Lab in Alderley Park and Alex Alderton, Roberto Amato, Jeffrey Barrett, Sonia Goncalves, Ewan Harrison, David K. Jackson, Ian Johnston, Dominic Kwiatkowski, Cordelia Langford, John Sillitoe on behalf of the Wellcome Sanger Institute COVID-19 Surveillance Team |
| EPI_ISL_3286093 | Randox Laboratories | Wellcome Sanger Institute for the COVID-19 Genomics UK (COG-UK) Consortium | Randox Laboratories and Alex Alderton, Roberto Amato, Jeffrey Barrett, Sonia Goncalves, Ewan Harrison, David K. Jackson, Ian Johnston, Dominic Kwiatkowski, Cordelia Langford, John Sillitoe on behalf of the Wellcome Sanger Institute COVID-19 Surveillance Team |
| EPI_ISL_3286094, EPI_ISL_3286095 | Lighthouse Lab in Milton Keynes | Wellcome Sanger Institute for the COVID-19 Genomics UK (COG-UK) Consortium | The Lighthouse Lab in Milton Keynes and Alex Alderton, Roberto Amato, Jeffrey Barrett, Sonia Goncalves, Ewan Harrison, David K. Jackson, Ian Johnston, Dominic Kwiatkowski, Cordelia Langford, John Sillitoe on behalf of the Wellcome Sanger Institute COVID-19 Surveillance Team |
| EPI_ISL_3286096 | Randox Laboratories | Wellcome Sanger Institute for the COVID-19 Genomics UK (COG-UK) Consortium | Randox Laboratories and Alex Alderton, Roberto Amato, Jeffrey Barrett, Sonia Goncalves, Ewan Harrison, David K. Jackson, Ian Johnston, Dominic Kwiatkowski, Cordelia Langford, John Sillitoe on behalf of the Wellcome Sanger Institute COVID-19 Surveillance Team |
| EPI_ISL_3286097, EPI_ISL_3286098 | Lighthouse Lab in Alderley Park | Wellcome Sanger Institute for the COVID-19 Genomics UK (COG-UK) Consortium | Jacquelyn Wynn, Mairead Hyland, The Lighthouse Lab in Alderley Park and Alex Alderton, Roberto Amato, Jeffrey Barrett, Sonia Goncalves, Ewan Harrison, David K. Jackson, Ian Johnston, Dominic Kwiatkowski, Cordelia Langford, John Sillitoe on behalf of the Wellcome Sanger Institute COVID-19 Surveillance Team |
| EPI_ISL_3286099 | Randox Laboratories | Wellcome Sanger Institute for the COVID-19 Genomics UK (COG-UK) Consortium | Randox Laboratories and Alex Alderton, Roberto Amato, Jeffrey Barrett, Sonia Goncalves, Ewan Harrison, David K. Jackson, Ian Johnston, Dominic Kwiatkowski, Cordelia Langford, John Sillitoe on behalf of the Wellcome Sanger Institute COVID-19 Surveillance Team |
| EPI_ISL_3286100 | Lighthouse Lab in Alderley Park | Wellcome Sanger Institute for the COVID-19 Genomics UK (COG-UK) Consortium | Jacquelyn Wynn, Mairead Hyland, The Lighthouse Lab in Alderley Park and Alex Alderton, Roberto Amato, Jeffrey Barrett, Sonia Goncalves, Ewan Harrison, David K. Jackson, Ian Johnston, Dominic Kwiatkowski, Cordelia Langford, John Sillitoe on behalf of the Wellcome Sanger Institute COVID-19 Surveillance Team |
| EPI_ISL_3286101 | Lighthouse Lab in Milton Keynes | Wellcome Sanger Institute for the COVID-19 Genomics UK (COG-UK) Consortium | The Lighthouse Lab in Milton Keynes and Alex Alderton, Roberto Amato, Jeffrey Barrett, Sonia Goncalves, Ewan Harrison, David K. Jackson, Ian Johnston, Dominic Kwiatkowski, Cordelia Langford, John Sillitoe on behalf of the Wellcome Sanger Institute COVID-19 Surveillance Team |
| EPI_ISL_3286102 | Lighthouse Lab in Alderley Park | Wellcome Sanger Institute for the COVID-19 Genomics UK (COG-UK) Consortium | Jacquelyn Wynn, Mairead Hyland, The Lighthouse Lab in Alderley Park and Alex Alderton, Roberto Amato, Jeffrey Barrett, Sonia Goncalves, Ewan Harrison, David K. Jackson, Ian Johnston, Dominic Kwiatkowski, Cordelia Langford, John Sillitoe on behalf of the Wellcome Sanger Institute COVID-19 Surveillance Team |
| EPI_ISL_3286103 | Lighthouse Lab in Milton Keynes | Wellcome Sanger Institute for the COVID-19 Genomics UK (COG-UK) Consortium | The Lighthouse Lab in Milton Keynes and Alex Alderton, Roberto Amato, Jeffrey Barrett, Sonia Goncalves, Ewan Harrison, David K. Jackson, Ian Johnston, Dominic Kwiatkowski, Cordelia Langford, John Sillitoe on behalf of the Wellcome Sanger Institute COVID-19 Surveillance Team |
| EPI_ISL_3286104, EPI_ISL_3286105, EPI_ISL_3286106 | Lighthouse Lab in Alderley Park | Wellcome Sanger Institute for the COVID-19 Genomics UK (COG-UK) Consortium | Jacquelyn Wynn, Mairead Hyland, The Lighthouse Lab in Alderley Park and Alex Alderton, Roberto Amato, Jeffrey Barrett, Sonia Goncalves, Ewan Harrison, David K. Jackson, Ian Johnston, Dominic Kwiatkowski, Cordelia Langford, John Sillitoe on behalf of the Wellcome Sanger Institute COVID-19 Surveillance Team |
| EPI_ISL_3286107 | Randox Laboratories | Wellcome Sanger Institute for the COVID-19 Genomics UK (COG-UK) Consortium | Randox Laboratories and Alex Alderton, Roberto Amato, Jeffrey Barrett, Sonia Goncalves, Ewan Harrison, David K. Jackson, Ian Johnston, Dominic Kwiatkowski, Cordelia Langford, John Sillitoe on behalf of the Wellcome Sanger Institute COVID-19 Surveillance Team |
| EPI_ISL_3286108 | Lighthouse Lab in Milton Keynes | Wellcome Sanger Institute for the COVID-19 Genomics UK (COG-UK) Consortium | The Lighthouse Lab in Milton Keynes and Alex Alderton, Roberto Amato, Jeffrey Barrett, Sonia Goncalves, Ewan Harrison, David K. Jackson, Ian Johnston, Dominic Kwiatkowski, Cordelia Langford, John Sillitoe on behalf of the Wellcome Sanger Institute COVID-19 Surveillance Team |
| EPI_ISL_3286109, EPI_ISL_3286110 | Lighthouse Lab in Alderley Park | Wellcome Sanger Institute for the COVID-19 Genomics UK (COG-UK) Consortium | Jacquelyn Wynn, Mairead Hyland, The Lighthouse Lab in Alderley Park and Alex Alderton, Roberto Amato, Jeffrey Barrett, Sonia Goncalves, Ewan Harrison, David K. Jackson, Ian Johnston, Dominic Kwiatkowski, Cordelia Langford, John Sillitoe on behalf of the Wellcome Sanger Institute COVID-19 Surveillance Team |
| EPI_ISL_3286111, EPI_ISL_3286112, EPI_ISL_3286113 | Lighthouse Lab in Milton Keynes | Wellcome Sanger Institute for the COVID-19 Genomics UK (COG-UK) Consortium | The Lighthouse Lab in Milton Keynes and Alex Alderton, Roberto Amato, Jeffrey Barrett, Sonia Goncalves, Ewan Harrison, David K. Jackson, Ian Johnston, Dominic Kwiatkowski, Cordelia Langford, John Sillitoe on behalf of the Wellcome Sanger Institute COVID-19 Surveillance Team |
| EPI_ISL_3286114, EPI_ISL_3286115, EPI_ISL_3286116, EPI_ISL_3286117, EPI_ISL_3286118 | Lighthouse Lab in Alderley Park | Wellcome Sanger Institute for the COVID-19 Genomics UK (COG-UK) Consortium | Jacquelyn Wynn, Mairead Hyland, The Lighthouse Lab in Alderley Park and Alex Alderton, Roberto Amato, Jeffrey Barrett, Sonia Goncalves, Ewan Harrison, David K. Jackson, Ian Johnston, Dominic Kwiatkowski, Cordelia Langford, John Sillitoe on behalf of the Wellcome Sanger Institute COVID-19 Surveillance Team |
| EPI_ISL_3286119 | Randox Laboratories | Wellcome Sanger Institute for the COVID-19 Genomics UK (COG-UK) Consortium | Randox Laboratories and Alex Alderton, Roberto Amato, Jeffrey Barrett, Sonia Goncalves, Ewan Harrison, David K. Jackson, Ian Johnston, Dominic Kwiatkowski, Cordelia Langford, John Sillitoe on behalf of the Wellcome Sanger Institute COVID-19 Surveillance Team |
| EPI_ISL_3286120, EPI_ISL_3286121, EPI_ISL_3286122 | Lighthouse Lab in Milton Keynes | Wellcome Sanger Institute for the COVID-19 Genomics UK (COG-UK) Consortium | The Lighthouse Lab in Milton Keynes and Alex Alderton, Roberto Amato, Jeffrey Barrett, Sonia Goncalves, Ewan Harrison, David K. Jackson, Ian Johnston, Dominic Kwiatkowski, Cordelia Langford, John Sillitoe on behalf of the Wellcome Sanger Institute COVID-19 Surveillance Team |
| EPI_ISL_3286123, EPI_ISL_3286124, EPI_ISL_3286125 | Lighthouse Lab in Alderley Park | Wellcome Sanger Institute for the COVID-19 Genomics UK (COG-UK) Consortium | Jacquelyn Wynn, Mairead Hyland, The Lighthouse Lab in Alderley Park and Alex Alderton, Roberto Amato, Jeffrey Barrett, Sonia Goncalves, Ewan Harrison, David K. Jackson, Ian Johnston, Dominic Kwiatkowski, Cordelia Langford, John Sillitoe on behalf of the Wellcome Sanger Institute COVID-19 Surveillance Team |
| EPI_ISL_3286126, EPI_ISL_3286127, EPI_ISL_3286128, EPI_ISL_3286129 | Lighthouse Lab in Milton Keynes | Wellcome Sanger Institute for the COVID-19 Genomics UK (COG-UK) Consortium | The Lighthouse Lab in Milton Keynes and Alex Alderton, Roberto Amato, Jeffrey Barrett, Sonia Goncalves, Ewan Harrison, David K. Jackson, Ian Johnston, Dominic Kwiatkowski, Cordelia Langford, John Sillitoe on behalf of the Wellcome Sanger Institute COVID-19 Surveillance Team |
| EPI_ISL_3286130, EPI_ISL_3286131 | Lighthouse Lab in Alderley Park | Wellcome Sanger Institute for the COVID-19 Genomics UK (COG-UK) Consortium | Jacquelyn Wynn, Mairead Hyland, The Lighthouse Lab in Alderley Park and Alex Alderton, Roberto Amato, Jeffrey Barrett, Sonia Goncalves, Ewan Harrison, David K. Jackson, Ian Johnston, Dominic Kwiatkowski, Cordelia Langford, John Sillitoe on behalf of the Wellcome Sanger Institute COVID-19 Surveillance Team |
| EPI_ISL_3286132 | Lighthouse Lab in Milton Keynes | Wellcome Sanger Institute for the COVID-19 Genomics UK (COG-UK) Consortium | The Lighthouse Lab in Milton Keynes and Alex Alderton, Roberto Amato, Jeffrey Barrett, Sonia Goncalves, Ewan Harrison, David K. Jackson, Ian Johnston, Dominic Kwiatkowski, Cordelia Langford, John Sillitoe on behalf of the Wellcome Sanger Institute COVID-19 Surveillance Team |
| EPI_ISL_3286133, EPI_ISL_3286134, EPI_ISL_3286135 | Lighthouse Lab in Alderley Park | Wellcome Sanger Institute for the COVID-19 Genomics UK (COG-UK) Consortium | Jacquelyn Wynn, Mairead Hyland, The Lighthouse Lab in Alderley Park and Alex Alderton, Roberto Amato, Jeffrey Barrett, Sonia Goncalves, Ewan Harrison, David K. Jackson, Ian Johnston, Dominic Kwiatkowski, Cordelia Langford, John Sillitoe on behalf of the Wellcome Sanger Institute COVID-19 Surveillance Team |
| EPI_ISL_3286136 | Lighthouse Lab in Milton Keynes | Wellcome Sanger Institute for the COVID-19 Genomics UK (COG-UK) Consortium | The Lighthouse Lab in Milton Keynes and Alex Alderton, Roberto Amato, Jeffrey Barrett, Sonia Goncalves, Ewan Harrison, David K. Jackson, Ian Johnston, Dominic Kwiatkowski, Cordelia Langford, John Sillitoe on behalf of the Wellcome Sanger Institute COVID-19 Surveillance Team |
| EPI_ISL_3286137 | Lighthouse Lab in Alderley Park | Wellcome Sanger Institute for the COVID-19 Genomics UK (COG-UK) Consortium | Jacquelyn Wynn, Mairead Hyland, The Lighthouse Lab in Alderley Park and Alex Alderton, Roberto Amato, Jeffrey Barrett, Sonia Goncalves, Ewan Harrison, David K. Jackson, Ian Johnston, Dominic Kwiatkowski, Cordelia Langford, John Sillitoe on behalf of the Wellcome Sanger Institute COVID-19 Surveillance Team |
| EPI_ISL_3286138 | Lighthouse Lab in Milton Keynes | Wellcome Sanger Institute for the COVID-19 Genomics UK (COG-UK) Consortium | The Lighthouse Lab in Milton Keynes and Alex Alderton, Roberto Amato, Jeffrey Barrett, Sonia Goncalves, Ewan Harrison, David K. Jackson, Ian Johnston, Dominic Kwiatkowski, Cordelia Langford, John Sillitoe on behalf of the Wellcome Sanger Institute COVID-19 Surveillance Team |
| EPI_ISL_3286139 | Lighthouse Lab in Alderley Park | Wellcome Sanger Institute for the COVID-19 Genomics UK (COG-UK) Consortium | Jacquelyn Wynn, Mairead Hyland, The Lighthouse Lab in Alderley Park and Alex Alderton, Roberto Amato, Jeffrey Barrett, Sonia Goncalves, Ewan Harrison, David K. Jackson, Ian Johnston, Dominic Kwiatkowski, Cordelia Langford, John Sillitoe on behalf of the Wellcome Sanger Institute COVID-19 Surveillance Team |
| EPI_ISL_3286140 | Randox Laboratories | Wellcome Sanger Institute for the COVID-19 Genomics UK (COG-UK) Consortium | Randox Laboratories and Alex Alderton, Roberto Amato, Jeffrey Barrett, Sonia Goncalves, Ewan Harrison, David K. Jackson, Ian Johnston, Dominic Kwiatkowski, Cordelia Langford, John Sillitoe on behalf of the Wellcome Sanger Institute COVID-19 Surveillance Team |
| EPI_ISL_3286141, EPI_ISL_3286142 | Lighthouse Lab in Milton Keynes | Wellcome Sanger Institute for the COVID-19 Genomics UK (COG-UK) Consortium | The Lighthouse Lab in Milton Keynes and Alex Alderton, Roberto Amato, Jeffrey Barrett, Sonia Goncalves, Ewan Harrison, David K. Jackson, Ian Johnston, Dominic Kwiatkowski, Cordelia Langford, John Sillitoe on behalf of the Wellcome Sanger Institute COVID-19 Surveillance Team |
| EPI_ISL_3286143 | Lighthouse Lab in Alderley Park | Wellcome Sanger Institute for the COVID-19 Genomics UK (COG-UK) Consortium | Jacquelyn Wynn, Mairead Hyland, The Lighthouse Lab in Alderley Park and Alex Alderton, Roberto Amato, Jeffrey Barrett, Sonia Goncalves, Ewan Harrison, David K. Jackson, Ian Johnston, Dominic Kwiatkowski, Cordelia Langford, John Sillitoe on behalf of the Wellcome Sanger Institute COVID-19 Surveillance Team |
| EPI_ISL_3286144 | Lighthouse Lab in Milton Keynes | Wellcome Sanger Institute for the COVID-19 Genomics UK (COG-UK) Consortium | The Lighthouse Lab in Milton Keynes and Alex Alderton, Roberto Amato, Jeffrey Barrett, Sonia Goncalves, Ewan Harrison, David K. Jackson, Ian Johnston, Dominic Kwiatkowski, Cordelia Langford, John Sillitoe on behalf of the Wellcome Sanger Institute COVID-19 Surveillance Team |
| EPI_ISL_3286145, EPI_ISL_3286146 | Lighthouse Lab in Alderley Park | Wellcome Sanger Institute for the COVID-19 Genomics UK (COG-UK) Consortium | Jacquelyn Wynn, Mairead Hyland, The Lighthouse Lab in Alderley Park and Alex Alderton, Roberto Amato, Jeffrey Barrett, Sonia Goncalves, Ewan Harrison, David K. Jackson, Ian Johnston, Dominic Kwiatkowski, Cordelia Langford, John Sillitoe on behalf of the Wellcome Sanger Institute COVID-19 Surveillance Team |
| EPI_ISL_3286147 | Lighthouse Lab in Milton Keynes | Wellcome Sanger Institute for the COVID-19 Genomics UK (COG-UK) Consortium | The Lighthouse Lab in Milton Keynes and Alex Alderton, Roberto Amato, Jeffrey Barrett, Sonia Goncalves, Ewan Harrison, David K. Jackson, Ian Johnston, Dominic Kwiatkowski, Cordelia Langford, John Sillitoe on behalf of the Wellcome Sanger Institute COVID-19 Surveillance Team |
| EPI_ISL_3286148 | Randox Laboratories | Wellcome Sanger Institute for the COVID-19 Genomics UK (COG-UK) Consortium | Randox Laboratories and Alex Alderton, Roberto Amato, Jeffrey Barrett, Sonia Goncalves, Ewan Harrison, David K. Jackson, Ian Johnston, Dominic Kwiatkowski, Cordelia Langford, John Sillitoe on behalf of the Wellcome Sanger Institute COVID-19 Surveillance Team |
| EPI_ISL_3286149, EPI_ISL_3286150, EPI_ISL_3286151 | Lighthouse Lab in Milton Keynes | Wellcome Sanger Institute for the COVID-19 Genomics UK (COG-UK) Consortium | The Lighthouse Lab in Milton Keynes and Alex Alderton, Roberto Amato, Jeffrey Barrett, Sonia Goncalves, Ewan Harrison, David K. Jackson, Ian Johnston, Dominic Kwiatkowski, Cordelia Langford, John Sillitoe on behalf of the Wellcome Sanger Institute COVID-19 Surveillance Team |
| EPI_ISL_3286152, EPI_ISL_3286153 | Lighthouse Lab in Alderley Park | Wellcome Sanger Institute for the COVID-19 Genomics UK (COG-UK) Consortium | Jacquelyn Wynn, Mairead Hyland, The Lighthouse Lab in Alderley Park and Alex Alderton, Roberto Amato, Jeffrey Barrett, Sonia Goncalves, Ewan Harrison, David K. Jackson, Ian Johnston, Dominic Kwiatkowski, Cordelia Langford, John Sillitoe on behalf of the Wellcome Sanger Institute COVID-19 Surveillance Team |

| | | | |
|---|---|---|---|
| EPI_ISL_3286154 | Lighthouse Lab in Milton Keynes | Wellcome Sanger Institute for the COVID-19 Genomics UK (COG-UK) Consortium | The Lighthouse Lab in Milton Keynes and Alex Alderton, Roberto Amato, Jeffrey Barrett, Sonia Goncalves, Ewan Harrison, David K. Jackson, Ian Johnston, Dominic Kwiatkowski, Cordelia Langford, John Sillitoe on behalf of the Wellcome Sanger Institute COVID-19 Surveillance Team |
| EPI_ISL_3286155 | Lighthouse Lab in Alderley Park | Wellcome Sanger Institute for the COVID-19 Genomics UK (COG-UK) Consortium | Jacquelyn Wynn, Mairead Hyland, The Lighthouse Lab in Alderley Park and Alex Alderton, Roberto Amato, Jeffrey Barrett, Sonia Goncalves, Ewan Harrison, David K. Jackson, Ian Johnston, Dominic Kwiatkowski, Cordelia Langford, John Sillitoe on behalf of the Wellcome Sanger Institute COVID-19 Surveillance Team |
| EPI_ISL_3286156 | Lighthouse Lab in Milton Keynes | Wellcome Sanger Institute for the COVID-19 Genomics UK (COG-UK) Consortium | The Lighthouse Lab in Milton Keynes and Alex Alderton, Roberto Amato, Jeffrey Barrett, Sonia Goncalves, Ewan Harrison, David K. Jackson, Ian Johnston, Dominic Kwiatkowski, Cordelia Langford, John Sillitoe on behalf of the Wellcome Sanger Institute COVID-19 Surveillance Team |
| EPI_ISL_3286157, EPI_ISL_3286158, EPI_ISL_3286159 | Lighthouse Lab in Alderley Park | Wellcome Sanger Institute for the COVID-19 Genomics UK (COG-UK) Consortium | Jacquelyn Wynn, Mairead Hyland, The Lighthouse Lab in Alderley Park and Alex Alderton, Roberto Amato, Jeffrey Barrett, Sonia Goncalves, Ewan Harrison, David K. Jackson, Ian Johnston, Dominic Kwiatkowski, Cordelia Langford, John Sillitoe on behalf of the Wellcome Sanger Institute COVID-19 Surveillance Team |
| EPI_ISL_3286160 | Lighthouse Lab in Milton Keynes | Wellcome Sanger Institute for the COVID-19 Genomics UK (COG-UK) Consortium | The Lighthouse Lab in Milton Keynes and Alex Alderton, Roberto Amato, Jeffrey Barrett, Sonia Goncalves, Ewan Harrison, David K. Jackson, Ian Johnston, Dominic Kwiatkowski, Cordelia Langford, John Sillitoe on behalf of the Wellcome Sanger Institute COVID-19 Surveillance Team |
| EPI_ISL_3286161 | Randox Laboratories | Wellcome Sanger Institute for the COVID-19 Genomics UK (COG-UK) Consortium | Randox Laboratories and Alex Alderton, Roberto Amato, Jeffrey Barrett, Sonia Goncalves, Ewan Harrison, David K. Jackson, Ian Johnston, Dominic Kwiatkowski, Cordelia Langford, John Sillitoe on behalf of the Wellcome Sanger Institute COVID-19 Surveillance Team |
| EPI_ISL_3286162 | Lighthouse Lab in Milton Keynes | Wellcome Sanger Institute for the COVID-19 Genomics UK (COG-UK) Consortium | The Lighthouse Lab in Milton Keynes and Alex Alderton, Roberto Amato, Jeffrey Barrett, Sonia Goncalves, Ewan Harrison, David K. Jackson, Ian Johnston, Dominic Kwiatkowski, Cordelia Langford, John Sillitoe on behalf of the Wellcome Sanger Institute COVID-19 Surveillance Team |
| EPI_ISL_3286163, EPI_ISL_3286164 | Lighthouse Lab in Alderley Park | Wellcome Sanger Institute for the COVID-19 Genomics UK (COG-UK) Consortium | Jacquelyn Wynn, Mairead Hyland, The Lighthouse Lab in Alderley Park and Alex Alderton, Roberto Amato, Jeffrey Barrett, Sonia Goncalves, Ewan Harrison, David K. Jackson, Ian Johnston, Dominic Kwiatkowski, Cordelia Langford, John Sillitoe on behalf of the Wellcome Sanger Institute COVID-19 Surveillance Team |
| EPI_ISL_3286165 | Lighthouse Lab in Milton Keynes | Wellcome Sanger Institute for the COVID-19 Genomics UK (COG-UK) Consortium | The Lighthouse Lab in Milton Keynes and Alex Alderton, Roberto Amato, Jeffrey Barrett, Sonia Goncalves, Ewan Harrison, David K. Jackson, Ian Johnston, Dominic Kwiatkowski, Cordelia Langford, John Sillitoe on behalf of the Wellcome Sanger Institute COVID-19 Surveillance Team |
| EPI_ISL_3286166, EPI_ISL_3286167, EPI_ISL_3286168 | Lighthouse Lab in Alderley Park | Wellcome Sanger Institute for the COVID-19 Genomics UK (COG-UK) Consortium | Jacquelyn Wynn, Mairead Hyland, The Lighthouse Lab in Alderley Park and Alex Alderton, Roberto Amato, Jeffrey Barrett, Sonia Goncalves, Ewan Harrison, David K. Jackson, Ian Johnston, Dominic Kwiatkowski, Cordelia Langford, John Sillitoe on behalf of the Wellcome Sanger Institute COVID-19 Surveillance Team |
| EPI_ISL_3286169 | Lighthouse Lab in Milton Keynes | Wellcome Sanger Institute for the COVID-19 Genomics UK (COG-UK) Consortium | The Lighthouse Lab in Milton Keynes and Alex Alderton, Roberto Amato, Jeffrey Barrett, Sonia Goncalves, Ewan Harrison, David K. Jackson, Ian Johnston, Dominic Kwiatkowski, Cordelia Langford, John Sillitoe on behalf of the Wellcome Sanger Institute COVID-19 Surveillance Team |
| EPI_ISL_3286170, EPI_ISL_3286171, EPI_ISL_3286172 | Lighthouse Lab in Alderley Park | Wellcome Sanger Institute for the COVID-19 Genomics UK (COG-UK) Consortium | Jacquelyn Wynn, Mairead Hyland, The Lighthouse Lab in Alderley Park and Alex Alderton, Roberto Amato, Jeffrey Barrett, Sonia Goncalves, Ewan Harrison, David K. Jackson, Ian Johnston, Dominic Kwiatkowski, Cordelia Langford, John Sillitoe on behalf of the Wellcome Sanger Institute COVID-19 Surveillance Team |
| EPI_ISL_3286173 | Lighthouse Lab in Milton Keynes | Wellcome Sanger Institute for the COVID-19 Genomics UK (COG-UK) Consortium | The Lighthouse Lab in Milton Keynes and Alex Alderton, Roberto Amato, Jeffrey Barrett, Sonia Goncalves, Ewan Harrison, David K. Jackson, Ian Johnston, Dominic Kwiatkowski, Cordelia Langford, John Sillitoe on behalf of the Wellcome Sanger Institute COVID-19 Surveillance Team |
| EPI_ISL_3286174 | Lighthouse Lab in Alderley Park | Wellcome Sanger Institute for the COVID-19 Genomics UK (COG-UK) Consortium | Jacquelyn Wynn, Mairead Hyland, The Lighthouse Lab in Alderley Park and Alex Alderton, Roberto Amato, Jeffrey Barrett, Sonia Goncalves, Ewan Harrison, David K. Jackson, Ian Johnston, Dominic Kwiatkowski, Cordelia Langford, John Sillitoe on behalf of the Wellcome Sanger Institute COVID-19 Surveillance Team |
| EPI_ISL_3286175 | Randox Laboratories | Wellcome Sanger Institute for the COVID-19 Genomics UK (COG-UK) Consortium | Randox Laboratories and Alex Alderton, Roberto Amato, Jeffrey Barrett, Sonia Goncalves, Ewan Harrison, David K. Jackson, Ian Johnston, Dominic Kwiatkowski, Cordelia Langford, John Sillitoe on behalf of the Wellcome Sanger Institute COVID-19 Surveillance Team |
| EPI_ISL_3286176 | Lighthouse Lab in Milton Keynes | Wellcome Sanger Institute for the COVID-19 Genomics UK (COG-UK) Consortium | The Lighthouse Lab in Milton Keynes and Alex Alderton, Roberto Amato, Jeffrey Barrett, Sonia Goncalves, Ewan Harrison, David K. Jackson, Ian Johnston, Dominic Kwiatkowski, Cordelia Langford, John Sillitoe on behalf of the Wellcome Sanger Institute COVID-19 Surveillance Team |
| EPI_ISL_3286177, EPI_ISL_3286178 | Lighthouse Lab in Alderley Park | Wellcome Sanger Institute for the COVID-19 Genomics UK (COG-UK) Consortium | Jacquelyn Wynn, Mairead Hyland, The Lighthouse Lab in Alderley Park and Alex Alderton, Roberto Amato, Jeffrey Barrett, Sonia Goncalves, Ewan Harrison, David K. Jackson, Ian Johnston, Dominic Kwiatkowski, Cordelia Langford, John Sillitoe on behalf of the Wellcome Sanger Institute COVID-19 Surveillance Team |
| EPI_ISL_3286179, EPI_ISL_3286180 | Lighthouse Lab in Milton Keynes | Wellcome Sanger Institute for the COVID-19 Genomics UK (COG-UK) Consortium | The Lighthouse Lab in Milton Keynes and Alex Alderton, Roberto Amato, Jeffrey Barrett, Sonia Goncalves, Ewan Harrison, David K. Jackson, Ian Johnston, Dominic Kwiatkowski, Cordelia Langford, John Sillitoe on behalf of the Wellcome Sanger Institute COVID-19 Surveillance Team |
| EPI_ISL_3286181, EPI_ISL_3286182, EPI_ISL_3286183, EPI_ISL_3286184 | Lighthouse Lab in Alderley Park | Wellcome Sanger Institute for the COVID-19 Genomics UK (COG-UK) Consortium | Jacquelyn Wynn, Mairead Hyland, The Lighthouse Lab in Alderley Park and Alex Alderton, Roberto Amato, Jeffrey Barrett, Sonia Goncalves, Ewan Harrison, David K. Jackson, Ian Johnston, Dominic Kwiatkowski, Cordelia Langford, John Sillitoe on behalf of the Wellcome Sanger Institute COVID-19 Surveillance Team |
| EPI_ISL_3286185 | Lighthouse Lab in Milton Keynes | Wellcome Sanger Institute for the COVID-19 Genomics UK (COG-UK) Consortium | The Lighthouse Lab in Milton Keynes and Alex Alderton, Roberto Amato, Jeffrey Barrett, Sonia Goncalves, Ewan Harrison, David K. Jackson, Ian Johnston, Dominic Kwiatkowski, Cordelia Langford, John Sillitoe on behalf of the Wellcome Sanger Institute COVID-19 Surveillance Team |
| EPI_ISL_3286186, EPI_ISL_3286187 | Lighthouse Lab in Alderley Park | Wellcome Sanger Institute for the COVID-19 Genomics UK (COG-UK) Consortium | Jacquelyn Wynn, Mairead Hyland, The Lighthouse Lab in Alderley Park and Alex Alderton, Roberto Amato, Jeffrey Barrett, Sonia Goncalves, Ewan Harrison, David K. Jackson, Ian Johnston, Dominic Kwiatkowski, Cordelia Langford, John Sillitoe on behalf of the Wellcome Sanger Institute COVID-19 Surveillance Team |
| EPI_ISL_3286188 | Randox Laboratories | Wellcome Sanger Institute for the COVID-19 Genomics UK (COG-UK) Consortium | Randox Laboratories and Alex Alderton, Roberto Amato, Jeffrey Barrett, Sonia Goncalves, Ewan Harrison, David K. Jackson, Ian Johnston, Dominic Kwiatkowski, Cordelia Langford, John Sillitoe on behalf of the Wellcome Sanger Institute COVID-19 Surveillance Team |
| EPI_ISL_3286189 | Lighthouse Lab in Alderley Park | Wellcome Sanger Institute for the COVID-19 Genomics UK (COG-UK) Consortium | Jacquelyn Wynn, Mairead Hyland, The Lighthouse Lab in Alderley Park and Alex Alderton, Roberto Amato, Jeffrey Barrett, Sonia Goncalves, Ewan Harrison, David K. Jackson, Ian Johnston, Dominic Kwiatkowski, Cordelia Langford, John Sillitoe on behalf of the Wellcome Sanger Institute COVID-19 Surveillance Team |
| EPI_ISL_3286190 | Randox Laboratories | Wellcome Sanger Institute for the COVID-19 Genomics UK (COG-UK) Consortium | Randox Laboratories and Alex Alderton, Roberto Amato, Jeffrey Barrett, Sonia Goncalves, Ewan Harrison, David K. Jackson, Ian Johnston, Dominic Kwiatkowski, Cordelia Langford, John Sillitoe on behalf of the Wellcome Sanger Institute COVID-19 Surveillance Team |
| EPI_ISL_3286191 | Lighthouse Lab in Milton Keynes | Wellcome Sanger Institute for the COVID-19 Genomics UK (COG-UK) Consortium | The Lighthouse Lab in Milton Keynes and Alex Alderton, Roberto Amato, Jeffrey Barrett, Sonia Goncalves, Ewan Harrison, David K. Jackson, Ian Johnston, Dominic Kwiatkowski, Cordelia Langford, John Sillitoe on behalf of the Wellcome Sanger Institute COVID-19 Surveillance Team |
| EPI_ISL_3286192 | Lighthouse Lab in Alderley Park | Wellcome Sanger Institute for the COVID-19 Genomics UK (COG-UK) Consortium | Jacquelyn Wynn, Mairead Hyland, The Lighthouse Lab in Alderley Park and Alex Alderton, Roberto Amato, Jeffrey Barrett, Sonia Goncalves, Ewan Harrison, David K. Jackson, Ian Johnston, Dominic Kwiatkowski, Cordelia Langford, John Sillitoe on behalf of the Wellcome Sanger Institute COVID-19 Surveillance Team |
| EPI_ISL_3286193 | Lighthouse Lab in Milton Keynes | Wellcome Sanger Institute for the COVID-19 Genomics UK (COG-UK) Consortium | The Lighthouse Lab in Milton Keynes and Alex Alderton, Roberto Amato, Jeffrey Barrett, Sonia Goncalves, Ewan Harrison, David K. Jackson, Ian Johnston, Dominic Kwiatkowski, Cordelia Langford, John Sillitoe on behalf of the Wellcome Sanger Institute COVID-19 Surveillance Team |
| EPI_ISL_3286194 | Randox Laboratories | Wellcome Sanger Institute for the COVID-19 Genomics UK (COG-UK) Consortium | Randox Laboratories and Alex Alderton, Roberto Amato, Jeffrey Barrett, Sonia Goncalves, Ewan Harrison, David K. Jackson, Ian Johnston, Dominic Kwiatkowski, Cordelia Langford, John Sillitoe on behalf of the Wellcome Sanger Institute COVID-19 Surveillance Team |
| EPI_ISL_3286195, EPI_ISL_3286196, EPI_ISL_3286197 | Lighthouse Lab in Alderley Park | Wellcome Sanger Institute for the COVID-19 Genomics UK (COG-UK) Consortium | Jacquelyn Wynn, Mairead Hyland, The Lighthouse Lab in Alderley Park and Alex Alderton, Roberto Amato, Jeffrey Barrett, Sonia Goncalves, Ewan Harrison, David K. Jackson, Ian Johnston, Dominic Kwiatkowski, Cordelia Langford, John Sillitoe on behalf of the Wellcome Sanger Institute COVID-19 Surveillance Team |
| EPI_ISL_3286198, EPI_ISL_3286199 | Lighthouse Lab in Milton Keynes | Wellcome Sanger Institute for the COVID-19 Genomics UK (COG-UK) Consortium | The Lighthouse Lab in Milton Keynes and Alex Alderton, Roberto Amato, Jeffrey Barrett, Sonia Goncalves, Ewan Harrison, David K. Jackson, Ian Johnston, Dominic Kwiatkowski, Cordelia Langford, John Sillitoe on behalf of the Wellcome Sanger Institute COVID-19 Surveillance Team |
| EPI_ISL_3286200 | Lighthouse Lab in Alderley Park | Wellcome Sanger Institute for the COVID-19 Genomics UK (COG-UK) Consortium | Jacquelyn Wynn, Mairead Hyland, The Lighthouse Lab in Alderley Park and Alex Alderton, Roberto Amato, Jeffrey Barrett, Sonia Goncalves, Ewan Harrison, David K. Jackson, Ian Johnston, Dominic Kwiatkowski, Cordelia Langford, John Sillitoe on behalf of the Wellcome Sanger Institute COVID-19 Surveillance Team |
| EPI_ISL_3286201 | Lighthouse Lab in Milton Keynes | Wellcome Sanger Institute for the COVID-19 Genomics UK (COG-UK) Consortium | The Lighthouse Lab in Milton Keynes and Alex Alderton, Roberto Amato, Jeffrey Barrett, Sonia Goncalves, Ewan Harrison, David K. Jackson, Ian Johnston, Dominic Kwiatkowski, Cordelia Langford, John Sillitoe on behalf of the Wellcome Sanger Institute COVID-19 Surveillance Team |
| EPI_ISL_3286202, EPI_ISL_3286203 | Lighthouse Lab in Alderley Park | Wellcome Sanger Institute for the COVID-19 Genomics UK (COG-UK) Consortium | Jacquelyn Wynn, Mairead Hyland, The Lighthouse Lab in Alderley Park and Alex Alderton, Roberto Amato, Jeffrey Barrett, Sonia Goncalves, Ewan Harrison, David K. Jackson, Ian Johnston, Dominic Kwiatkowski, Cordelia Langford, John Sillitoe on behalf of the Wellcome Sanger Institute COVID-19 Surveillance Team |
| EPI_ISL_3286204 | Randox Laboratories | Wellcome Sanger Institute for the COVID-19 Genomics UK (COG-UK) Consortium | Randox Laboratories and Alex Alderton, Roberto Amato, Jeffrey Barrett, Sonia Goncalves, Ewan Harrison, David K. Jackson, Ian Johnston, Dominic Kwiatkowski, Cordelia Langford, John Sillitoe on behalf of the Wellcome Sanger Institute COVID-19 Surveillance Team |
| EPI_ISL_3286205 | Lighthouse Lab in Alderley Park | Wellcome Sanger Institute for the COVID-19 Genomics UK (COG-UK) Consortium | Jacquelyn Wynn, Mairead Hyland, The Lighthouse Lab in Alderley Park and Alex Alderton, Roberto Amato, Jeffrey Barrett, Sonia Goncalves, Ewan Harrison, David K. Jackson, Ian Johnston, Dominic Kwiatkowski, Cordelia Langford, John Sillitoe on behalf of the Wellcome Sanger Institute COVID-19 Surveillance Team |
| EPI_ISL_3286206 | Lighthouse Lab in Milton Keynes | Wellcome Sanger Institute for the COVID-19 Genomics UK (COG-UK) Consortium | The Lighthouse Lab in Milton Keynes and Alex Alderton, Roberto Amato, Jeffrey Barrett, Sonia Goncalves, Ewan Harrison, David K. Jackson, Ian Johnston, Dominic Kwiatkowski, Cordelia Langford, John Sillitoe on behalf of the Wellcome Sanger Institute COVID-19 Surveillance Team |
| EPI_ISL_3286207 | Randox Laboratories | Wellcome Sanger Institute for the COVID-19 Genomics UK (COG-UK) Consortium | Randox Laboratories and Alex Alderton, Roberto Amato, Jeffrey Barrett, Sonia Goncalves, Ewan Harrison, David K. Jackson, Ian Johnston, Dominic Kwiatkowski, Cordelia Langford, John Sillitoe on behalf of the Wellcome Sanger Institute COVID-19 Surveillance Team |
| EPI_ISL_3286208 | Lighthouse Lab in Milton Keynes | Wellcome Sanger Institute for the COVID-19 Genomics UK (COG-UK) Consortium | The Lighthouse Lab in Milton Keynes and Alex Alderton, Roberto Amato, Jeffrey Barrett, Sonia Goncalves, Ewan Harrison, David K. Jackson, Ian Johnston, Dominic Kwiatkowski, Cordelia Langford, John Sillitoe on behalf of the Wellcome Sanger Institute COVID-19 Surveillance Team |
| EPI_ISL_3286209, EPI_ISL_3286210, EPI_ISL_3286211 | Lighthouse Lab in Alderley Park | Wellcome Sanger Institute for the COVID-19 Genomics UK (COG-UK) Consortium | Jacquelyn Wynn, Mairead Hyland, The Lighthouse Lab in Alderley Park and Alex Alderton, Roberto Amato, Jeffrey Barrett, Sonia Goncalves, Ewan Harrison, David K. Jackson, Ian Johnston, Dominic Kwiatkowski, Cordelia Langford, John Sillitoe on behalf of the Wellcome Sanger Institute COVID-19 Surveillance Team |
| EPI_ISL_3286212, EPI_ISL_3286213 | Randox Laboratories | Wellcome Sanger Institute for the COVID-19 Genomics UK (COG-UK) Consortium | Randox Laboratories and Alex Alderton, Roberto Amato, Jeffrey Barrett, Sonia Goncalves, Ewan Harrison, David K. Jackson, Ian Johnston, Dominic Kwiatkowski, Cordelia Langford, John Sillitoe on behalf of the Wellcome Sanger Institute COVID-19 Surveillance Team |
| EPI_ISL_3286214, EPI_ISL_3286215 | Lighthouse Lab in Milton Keynes | Wellcome Sanger Institute for the COVID-19 Genomics UK (COG-UK) Consortium | The Lighthouse Lab in Milton Keynes and Alex Alderton, Roberto Amato, Jeffrey Barrett, Sonia Goncalves, Ewan Harrison, David K. Jackson, Ian Johnston, Dominic Kwiatkowski, Cordelia Langford, John Sillitoe on behalf of the Wellcome Sanger Institute COVID-19 Surveillance Team |
| EPI_ISL_3286216, EPI_ISL_3286217, EPI_ISL_3286218, | Lighthouse Lab in Alderley Park | Wellcome Sanger Institute for the COVID-19 Genomics UK (COG-UK) Consortium | Jacquelyn Wynn, Mairead Hyland, The Lighthouse Lab in Alderley Park and Alex Alderton, Roberto Amato, Jeffrey Barrett, Sonia Goncalves, Ewan Harrison, David K. Jackson, Ian Johnston, Dominic Kwiatkowski, Cordelia Langford, John Sillitoe on behalf of the Wellcome Sanger Institute COVID-19 Surveillance Team |

| | | | |
|---|---|---|---|
| EPI_ISL_3286219, EPI_ISL_3286220, EPI_ISL_3286221, EPI_ISL_3286222, EPI_ISL_3286223, EPI_ISL_3286224 | | | |
| EPI_ISL_3286225, EPI_ISL_3286226 | Lighthouse Lab in Milton Keynes | Wellcome Sanger Institute for the COVID-19 Genomics UK (COG-UK) Consortium | The Lighthouse Lab in Milton Keynes and Alex Alderton, Roberto Amato, Jeffrey Barrett, Sonia Goncalves, Ewan Harrison, David K. Jackson, Ian Johnston, Dominic Kwiatkowski, Cordelia Langford, John Sillitoe on behalf of the Wellcome Sanger Institute COVID-19 Surveillance Team |
| EPI_ISL_3286227, EPI_ISL_3286228, EPI_ISL_3286229 | Lighthouse Lab in Alderley Park | Wellcome Sanger Institute for the COVID-19 Genomics UK (COG-UK) Consortium | Jacquelyn Wynn, Mairead Hyland, The Lighthouse Lab in Alderley Park and Alex Alderton, Roberto Amato, Jeffrey Barrett, Sonia Goncalves, Ewan Harrison, David K. Jackson, Ian Johnston, Dominic Kwiatkowski, Cordelia Langford, John Sillitoe on behalf of the Wellcome Sanger Institute COVID-19 Surveillance Team |
| EPI_ISL_3286230, EPI_ISL_3286231, EPI_ISL_3286232 | Lighthouse Lab in Milton Keynes | Wellcome Sanger Institute for the COVID-19 Genomics UK (COG-UK) Consortium | The Lighthouse Lab in Milton Keynes and Alex Alderton, Roberto Amato, Jeffrey Barrett, Sonia Goncalves, Ewan Harrison, David K. Jackson, Ian Johnston, Dominic Kwiatkowski, Cordelia Langford, John Sillitoe on behalf of the Wellcome Sanger Institute COVID-19 Surveillance Team |
| EPI_ISL_3286233 | Lighthouse Lab in Alderley Park | Wellcome Sanger Institute for the COVID-19 Genomics UK (COG-UK) Consortium | Jacquelyn Wynn, Mairead Hyland, The Lighthouse Lab in Alderley Park and Alex Alderton, Roberto Amato, Jeffrey Barrett, Sonia Goncalves, Ewan Harrison, David K. Jackson, Ian Johnston, Dominic Kwiatkowski, Cordelia Langford, John Sillitoe on behalf of the Wellcome Sanger Institute COVID-19 Surveillance Team |
| EPI_ISL_3286234 | Lighthouse Lab in Milton Keynes | Wellcome Sanger Institute for the COVID-19 Genomics UK (COG-UK) Consortium | The Lighthouse Lab in Milton Keynes and Alex Alderton, Roberto Amato, Jeffrey Barrett, Sonia Goncalves, Ewan Harrison, David K. Jackson, Ian Johnston, Dominic Kwiatkowski, Cordelia Langford, John Sillitoe on behalf of the Wellcome Sanger Institute COVID-19 Surveillance Team |
| EPI_ISL_3286235, EPI_ISL_3286236 | Lighthouse Lab in Alderley Park | Wellcome Sanger Institute for the COVID-19 Genomics UK (COG-UK) Consortium | Jacquelyn Wynn, Mairead Hyland, The Lighthouse Lab in Alderley Park and Alex Alderton, Roberto Amato, Jeffrey Barrett, Sonia Goncalves, Ewan Harrison, David K. Jackson, Ian Johnston, Dominic Kwiatkowski, Cordelia Langford, John Sillitoe on behalf of the Wellcome Sanger Institute COVID-19 Surveillance Team |
| EPI_ISL_3286237, EPI_ISL_3286238 | Lighthouse Lab in Milton Keynes | Wellcome Sanger Institute for the COVID-19 Genomics UK (COG-UK) Consortium | The Lighthouse Lab in Milton Keynes and Alex Alderton, Roberto Amato, Jeffrey Barrett, Sonia Goncalves, Ewan Harrison, David K. Jackson, Ian Johnston, Dominic Kwiatkowski, Cordelia Langford, John Sillitoe on behalf of the Wellcome Sanger Institute COVID-19 Surveillance Team |
| EPI_ISL_3286239 | Lighthouse Lab in Alderley Park | Wellcome Sanger Institute for the COVID-19 Genomics UK (COG-UK) Consortium | Jacquelyn Wynn, Mairead Hyland, The Lighthouse Lab in Alderley Park and Alex Alderton, Roberto Amato, Jeffrey Barrett, Sonia Goncalves, Ewan Harrison, David K. Jackson, Ian Johnston, Dominic Kwiatkowski, Cordelia Langford, John Sillitoe on behalf of the Wellcome Sanger Institute COVID-19 Surveillance Team |
| EPI_ISL_3286240 | Randox Laboratories | Wellcome Sanger Institute for the COVID-19 Genomics UK (COG-UK) Consortium | Randox Laboratories and Alex Alderton, Roberto Amato, Jeffrey Barrett, Sonia Goncalves, Ewan Harrison, David K. Jackson, Ian Johnston, Dominic Kwiatkowski, Cordelia Langford, John Sillitoe on behalf of the Wellcome Sanger Institute COVID-19 Surveillance Team |
| EPI_ISL_3286241 | Lighthouse Lab in Alderley Park | Wellcome Sanger Institute for the COVID-19 Genomics UK (COG-UK) Consortium | Jacquelyn Wynn, Mairead Hyland, The Lighthouse Lab in Alderley Park and Alex Alderton, Roberto Amato, Jeffrey Barrett, Sonia Goncalves, Ewan Harrison, David K. Jackson, Ian Johnston, Dominic Kwiatkowski, Cordelia Langford, John Sillitoe on behalf of the Wellcome Sanger Institute COVID-19 Surveillance Team |
| EPI_ISL_3286242 | Lighthouse Lab in Milton Keynes | Wellcome Sanger Institute for the COVID-19 Genomics UK (COG-UK) Consortium | The Lighthouse Lab in Milton Keynes and Alex Alderton, Roberto Amato, Jeffrey Barrett, Sonia Goncalves, Ewan Harrison, David K. Jackson, Ian Johnston, Dominic Kwiatkowski, Cordelia Langford, John Sillitoe on behalf of the Wellcome Sanger Institute COVID-19 Surveillance Team |
| EPI_ISL_3286243 | Lighthouse Lab in Alderley Park | Wellcome Sanger Institute for the COVID-19 Genomics UK (COG-UK) Consortium | Jacquelyn Wynn, Mairead Hyland, The Lighthouse Lab in Alderley Park and Alex Alderton, Roberto Amato, Jeffrey Barrett, Sonia Goncalves, Ewan Harrison, David K. Jackson, Ian Johnston, Dominic Kwiatkowski, Cordelia Langford, John Sillitoe on behalf of the Wellcome Sanger Institute COVID-19 Surveillance Team |
| EPI_ISL_3286244 | Randox Laboratories | Wellcome Sanger Institute for the COVID-19 Genomics UK (COG-UK) Consortium | Randox Laboratories and Alex Alderton, Roberto Amato, Jeffrey Barrett, Sonia Goncalves, Ewan Harrison, David K. Jackson, Ian Johnston, Dominic Kwiatkowski, Cordelia Langford, John Sillitoe on behalf of the Wellcome Sanger Institute COVID-19 Surveillance Team |
| EPI_ISL_3286245 | Lighthouse Lab in Alderley Park | Wellcome Sanger Institute for the COVID-19 Genomics UK (COG-UK) Consortium | Jacquelyn Wynn, Mairead Hyland, The Lighthouse Lab in Alderley Park and Alex Alderton, Roberto Amato, Jeffrey Barrett, Sonia Goncalves, Ewan Harrison, David K. Jackson, Ian Johnston, Dominic Kwiatkowski, Cordelia Langford, John Sillitoe on behalf of the Wellcome Sanger Institute COVID-19 Surveillance Team |
| EPI_ISL_3286246 | Lighthouse Lab in Milton Keynes | Wellcome Sanger Institute for the COVID-19 Genomics UK (COG-UK) Consortium | The Lighthouse Lab in Milton Keynes and Alex Alderton, Roberto Amato, Jeffrey Barrett, Sonia Goncalves, Ewan Harrison, David K. Jackson, Ian Johnston, Dominic Kwiatkowski, Cordelia Langford, John Sillitoe on behalf of the Wellcome Sanger Institute COVID-19 Surveillance Team |
| EPI_ISL_3286247, EPI_ISL_3286248 | Lighthouse Lab in Alderley Park | Wellcome Sanger Institute for the COVID-19 Genomics UK (COG-UK) Consortium | Jacquelyn Wynn, Mairead Hyland, The Lighthouse Lab in Alderley Park and Alex Alderton, Roberto Amato, Jeffrey Barrett, Sonia Goncalves, Ewan Harrison, David K. Jackson, Ian Johnston, Dominic Kwiatkowski, Cordelia Langford, John Sillitoe on behalf of the Wellcome Sanger Institute COVID-19 Surveillance Team |
| EPI_ISL_3286249 | Randox Laboratories | Wellcome Sanger Institute for the COVID-19 Genomics UK (COG-UK) Consortium | Randox Laboratories and Alex Alderton, Roberto Amato, Jeffrey Barrett, Sonia Goncalves, Ewan Harrison, David K. Jackson, Ian Johnston, Dominic Kwiatkowski, Cordelia Langford, John Sillitoe on behalf of the Wellcome Sanger Institute COVID-19 Surveillance Team |
| EPI_ISL_3286250 | Lighthouse Lab in Milton Keynes | Wellcome Sanger Institute for the COVID-19 Genomics UK (COG-UK) Consortium | The Lighthouse Lab in Milton Keynes and Alex Alderton, Roberto Amato, Jeffrey Barrett, Sonia Goncalves, Ewan Harrison, David K. Jackson, Ian Johnston, Dominic Kwiatkowski, Cordelia Langford, John Sillitoe on behalf of the Wellcome Sanger Institute COVID-19 Surveillance Team |
| EPI_ISL_3286251 | Randox Laboratories | Wellcome Sanger Institute for the COVID-19 Genomics UK (COG-UK) Consortium | Randox Laboratories and Alex Alderton, Roberto Amato, Jeffrey Barrett, Sonia Goncalves, Ewan Harrison, David K. Jackson, Ian Johnston, Dominic Kwiatkowski, Cordelia Langford, John Sillitoe on behalf of the Wellcome Sanger Institute COVID-19 Surveillance Team |
| EPI_ISL_3286252, EPI_ISL_3286253 | Lighthouse Lab in Alderley Park | Wellcome Sanger Institute for the COVID-19 Genomics UK (COG-UK) Consortium | Jacquelyn Wynn, Mairead Hyland, The Lighthouse Lab in Alderley Park and Alex Alderton, Roberto Amato, Jeffrey Barrett, Sonia Goncalves, Ewan Harrison, David K. Jackson, Ian Johnston, Dominic Kwiatkowski, Cordelia Langford, John Sillitoe on behalf of the Wellcome Sanger Institute COVID-19 Surveillance Team |
| EPI_ISL_3286254 | Lighthouse Lab in Milton Keynes | Wellcome Sanger Institute for the COVID-19 Genomics UK (COG-UK) Consortium | The Lighthouse Lab in Milton Keynes and Alex Alderton, Roberto Amato, Jeffrey Barrett, Sonia Goncalves, Ewan Harrison, David K. Jackson, Ian Johnston, Dominic Kwiatkowski, Cordelia Langford, John Sillitoe on behalf of the Wellcome Sanger Institute COVID-19 Surveillance Team |
| EPI_ISL_3286255, EPI_ISL_3286256, EPI_ISL_3286257, EPI_ISL_3286258, EPI_ISL_3286259 | Lighthouse Lab in Alderley Park | Wellcome Sanger Institute for the COVID-19 Genomics UK (COG-UK) Consortium | Jacquelyn Wynn, Mairead Hyland, The Lighthouse Lab in Alderley Park and Alex Alderton, Roberto Amato, Jeffrey Barrett, Sonia Goncalves, Ewan Harrison, David K. Jackson, Ian Johnston, Dominic Kwiatkowski, Cordelia Langford, John Sillitoe on behalf of the Wellcome Sanger Institute COVID-19 Surveillance Team |
| EPI_ISL_3286260, EPI_ISL_3286261 | Lighthouse Lab in Milton Keynes | Wellcome Sanger Institute for the COVID-19 Genomics UK (COG-UK) Consortium | The Lighthouse Lab in Milton Keynes and Alex Alderton, Roberto Amato, Jeffrey Barrett, Sonia Goncalves, Ewan Harrison, David K. Jackson, Ian Johnston, Dominic Kwiatkowski, Cordelia Langford, John Sillitoe on behalf of the Wellcome Sanger Institute COVID-19 Surveillance Team |
| EPI_ISL_3286262 | Lighthouse Lab in Alderley Park | Wellcome Sanger Institute for the COVID-19 Genomics UK (COG-UK) Consortium | Jacquelyn Wynn, Mairead Hyland, The Lighthouse Lab in Alderley Park and Alex Alderton, Roberto Amato, Jeffrey Barrett, Sonia Goncalves, Ewan Harrison, David K. Jackson, Ian Johnston, Dominic Kwiatkowski, Cordelia Langford, John Sillitoe on behalf of the Wellcome Sanger Institute COVID-19 Surveillance Team |
| EPI_ISL_3286263, EPI_ISL_3286264, EPI_ISL_3286265 | Lighthouse Lab in Milton Keynes | Wellcome Sanger Institute for the COVID-19 Genomics UK (COG-UK) Consortium | The Lighthouse Lab in Milton Keynes and Alex Alderton, Roberto Amato, Jeffrey Barrett, Sonia Goncalves, Ewan Harrison, David K. Jackson, Ian Johnston, Dominic Kwiatkowski, Cordelia Langford, John Sillitoe on behalf of the Wellcome Sanger Institute COVID-19 Surveillance Team |
| EPI_ISL_3286266 | Randox Laboratories | Wellcome Sanger Institute for the COVID-19 Genomics UK (COG-UK) Consortium | Randox Laboratories and Alex Alderton, Roberto Amato, Jeffrey Barrett, Sonia Goncalves, Ewan Harrison, David K. Jackson, Ian Johnston, Dominic Kwiatkowski, Cordelia Langford, John Sillitoe on behalf of the Wellcome Sanger Institute COVID-19 Surveillance Team |
| EPI_ISL_3286267 | Lighthouse Lab in Milton Keynes | Wellcome Sanger Institute for the COVID-19 Genomics UK (COG-UK) Consortium | The Lighthouse Lab in Milton Keynes and Alex Alderton, Roberto Amato, Jeffrey Barrett, Sonia Goncalves, Ewan Harrison, David K. Jackson, Ian Johnston, Dominic Kwiatkowski, Cordelia Langford, John Sillitoe on behalf of the Wellcome Sanger Institute COVID-19 Surveillance Team |
| EPI_ISL_3286268, EPI_ISL_3286269, EPI_ISL_3286270, EPI_ISL_3286271 | Lighthouse Lab in Alderley Park | Wellcome Sanger Institute for the COVID-19 Genomics UK (COG-UK) Consortium | Jacquelyn Wynn, Mairead Hyland, The Lighthouse Lab in Alderley Park and Alex Alderton, Roberto Amato, Jeffrey Barrett, Sonia Goncalves, Ewan Harrison, David K. Jackson, Ian Johnston, Dominic Kwiatkowski, Cordelia Langford, John Sillitoe on behalf of the Wellcome Sanger Institute COVID-19 Surveillance Team |
| EPI_ISL_3286272 | Lighthouse Lab in Milton Keynes | Wellcome Sanger Institute for the COVID-19 Genomics UK (COG-UK) Consortium | The Lighthouse Lab in Milton Keynes and Alex Alderton, Roberto Amato, Jeffrey Barrett, Sonia Goncalves, Ewan Harrison, David K. Jackson, Ian Johnston, Dominic Kwiatkowski, Cordelia Langford, John Sillitoe on behalf of the Wellcome Sanger Institute COVID-19 Surveillance Team |
| EPI_ISL_3286273, EPI_ISL_3286274, EPI_ISL_3286275 | Lighthouse Lab in Alderley Park | Wellcome Sanger Institute for the COVID-19 Genomics UK (COG-UK) Consortium | Jacquelyn Wynn, Mairead Hyland, The Lighthouse Lab in Alderley Park and Alex Alderton, Roberto Amato, Jeffrey Barrett, Sonia Goncalves, Ewan Harrison, David K. Jackson, Ian Johnston, Dominic Kwiatkowski, Cordelia Langford, John Sillitoe on behalf of the Wellcome Sanger Institute COVID-19 Surveillance Team |
| EPI_ISL_3286276 | Randox Laboratories | Wellcome Sanger Institute for the COVID-19 Genomics UK (COG-UK) Consortium | Randox Laboratories and Alex Alderton, Roberto Amato, Jeffrey Barrett, Sonia Goncalves, Ewan Harrison, David K. Jackson, Ian Johnston, Dominic Kwiatkowski, Cordelia Langford, John Sillitoe on behalf of the Wellcome Sanger Institute COVID-19 Surveillance Team |
| EPI_ISL_3286277 | Lighthouse Lab in Alderley Park | Wellcome Sanger Institute for the COVID-19 Genomics UK (COG-UK) Consortium | Jacquelyn Wynn, Mairead Hyland, The Lighthouse Lab in Alderley Park and Alex Alderton, Roberto Amato, Jeffrey Barrett, Sonia Goncalves, Ewan Harrison, David K. Jackson, Ian Johnston, Dominic Kwiatkowski, Cordelia Langford, John Sillitoe on behalf of the Wellcome Sanger Institute COVID-19 Surveillance Team |
| EPI_ISL_3286278 | Randox Laboratories | Wellcome Sanger Institute for the COVID-19 Genomics UK (COG-UK) Consortium | Randox Laboratories and Alex Alderton, Roberto Amato, Jeffrey Barrett, Sonia Goncalves, Ewan Harrison, David K. Jackson, Ian Johnston, Dominic Kwiatkowski, Cordelia Langford, John Sillitoe on behalf of the Wellcome Sanger Institute COVID-19 Surveillance Team |
| EPI_ISL_3286279, EPI_ISL_3286280 | Lighthouse Lab in Alderley Park | Wellcome Sanger Institute for the COVID-19 Genomics UK (COG-UK) Consortium | Jacquelyn Wynn, Mairead Hyland, The Lighthouse Lab in Alderley Park and Alex Alderton, Roberto Amato, Jeffrey Barrett, Sonia Goncalves, Ewan Harrison, David K. Jackson, Ian Johnston, Dominic Kwiatkowski, Cordelia Langford, John Sillitoe on behalf of the Wellcome Sanger Institute COVID-19 Surveillance Team |
| EPI_ISL_3286281 | Lighthouse Lab in Milton Keynes | Wellcome Sanger Institute for the COVID-19 Genomics UK (COG-UK) Consortium | The Lighthouse Lab in Milton Keynes and Alex Alderton, Roberto Amato, Jeffrey Barrett, Sonia Goncalves, Ewan Harrison, David K. Jackson, Ian Johnston, Dominic Kwiatkowski, Cordelia Langford, John Sillitoe on behalf of the Wellcome Sanger Institute COVID-19 Surveillance Team |
| EPI_ISL_3286282, EPI_ISL_3286283 | Lighthouse Lab in Alderley Park | Wellcome Sanger Institute for the COVID-19 Genomics UK (COG-UK) Consortium | Jacquelyn Wynn, Mairead Hyland, The Lighthouse Lab in Alderley Park and Alex Alderton, Roberto Amato, Jeffrey Barrett, Sonia Goncalves, Ewan Harrison, David K. Jackson, Ian Johnston, Dominic Kwiatkowski, Cordelia Langford, John Sillitoe on behalf of the Wellcome Sanger Institute COVID-19 Surveillance Team |
| EPI_ISL_3286284 | Lighthouse Lab in Milton Keynes | Wellcome Sanger Institute for the COVID-19 Genomics UK (COG-UK) Consortium | The Lighthouse Lab in Milton Keynes and Alex Alderton, Roberto Amato, Jeffrey Barrett, Sonia Goncalves, Ewan Harrison, David K. Jackson, Ian Johnston, Dominic Kwiatkowski, Cordelia Langford, John Sillitoe on behalf of the Wellcome Sanger Institute COVID-19 Surveillance Team |
| EPI_ISL_3286285 | Lighthouse Lab in Alderley Park | Wellcome Sanger Institute for the COVID-19 Genomics UK (COG-UK) Consortium | Jacquelyn Wynn, Mairead Hyland, The Lighthouse Lab in Alderley Park and Alex Alderton, Roberto Amato, Jeffrey Barrett, Sonia Goncalves, Ewan Harrison, David K. Jackson, Ian Johnston, Dominic Kwiatkowski, Cordelia Langford, John Sillitoe on behalf of the Wellcome Sanger Institute COVID-19 Surveillance Team |
| EPI_ISL_3286286, | Lighthouse Lab in Milton Keynes | Wellcome Sanger Institute for the COVID-19 Genomics UK (COG-UK) | The Lighthouse Lab in Milton Keynes and Alex Alderton, Roberto Amato, Jeffrey Barrett, Sonia Goncalves, Ewan Harrison, David K. Jackson, Ian Johnston, Dominic Kwiatkowski, Cordelia Langford, John Sillitoe on behalf of the Wellcome Sanger |

| | | | |
|---|---|---|---|
| EPI_ISL_3286287, EPI_ISL_3286288 | | Consortium | Institute COVID-19 Surveillance Team |
| EPI_ISL_3286289 | Lighthouse Lab in Alderley Park | Wellcome Sanger Institute for the COVID-19 Genomics UK (COG-UK) Consortium | Jacquelyn Wynn, Mairead Hyland, The Lighthouse Lab in Alderley Park and Alex Alderton, Roberto Amato, Jeffrey Barrett, Sonia Goncalves, Ewan Harrison, David K. Jackson, Ian Johnston, Dominic Kwiatkowski, Cordelia Langford, John Sillitoe on behalf of the Wellcome Sanger Institute COVID-19 Surveillance Team |
| EPI_ISL_3286290 | Randox Laboratories | Wellcome Sanger Institute for the COVID-19 Genomics UK (COG-UK) Consortium | Randox Laboratories and Alex Alderton, Roberto Amato, Jeffrey Barrett, Sonia Goncalves, Ewan Harrison, David K. Jackson, Ian Johnston, Dominic Kwiatkowski, Cordelia Langford, John Sillitoe on behalf of the Wellcome Sanger Institute COVID-19 Surveillance Team |
| EPI_ISL_3286291, EPI_ISL_3286292, EPI_ISL_3286293, EPI_ISL_3286294, EPI_ISL_3286295, EPI_ISL_3286296 | Lighthouse Lab in Alderley Park | Wellcome Sanger Institute for the COVID-19 Genomics UK (COG-UK) Consortium | Jacquelyn Wynn, Mairead Hyland, The Lighthouse Lab in Alderley Park and Alex Alderton, Roberto Amato, Jeffrey Barrett, Sonia Goncalves, Ewan Harrison, David K. Jackson, Ian Johnston, Dominic Kwiatkowski, Cordelia Langford, John Sillitoe on behalf of the Wellcome Sanger Institute COVID-19 Surveillance Team |
| EPI_ISL_3286297 | Lighthouse Lab in Milton Keynes | Wellcome Sanger Institute for the COVID-19 Genomics UK (COG-UK) Consortium | The Lighthouse Lab in Milton Keynes and Alex Alderton, Roberto Amato, Jeffrey Barrett, Sonia Goncalves, Ewan Harrison, David K. Jackson, Ian Johnston, Dominic Kwiatkowski, Cordelia Langford, John Sillitoe on behalf of the Wellcome Sanger Institute COVID-19 Surveillance Team |
| EPI_ISL_3286298 | Lighthouse Lab in Alderley Park | Wellcome Sanger Institute for the COVID-19 Genomics UK (COG-UK) Consortium | Jacquelyn Wynn, Mairead Hyland, The Lighthouse Lab in Alderley Park and Alex Alderton, Roberto Amato, Jeffrey Barrett, Sonia Goncalves, Ewan Harrison, David K. Jackson, Ian Johnston, Dominic Kwiatkowski, Cordelia Langford, John Sillitoe on behalf of the Wellcome Sanger Institute COVID-19 Surveillance Team |
| EPI_ISL_3286299 | Lighthouse Lab in Milton Keynes | Wellcome Sanger Institute for the COVID-19 Genomics UK (COG-UK) Consortium | The Lighthouse Lab in Milton Keynes and Alex Alderton, Roberto Amato, Jeffrey Barrett, Sonia Goncalves, Ewan Harrison, David K. Jackson, Ian Johnston, Dominic Kwiatkowski, Cordelia Langford, John Sillitoe on behalf of the Wellcome Sanger Institute COVID-19 Surveillance Team |
| EPI_ISL_3286300, EPI_ISL_3286301, EPI_ISL_3286302, EPI_ISL_3286303, EPI_ISL_3286304, EPI_ISL_3286305 | Lighthouse Lab in Alderley Park | Wellcome Sanger Institute for the COVID-19 Genomics UK (COG-UK) Consortium | Jacquelyn Wynn, Mairead Hyland, The Lighthouse Lab in Alderley Park and Alex Alderton, Roberto Amato, Jeffrey Barrett, Sonia Goncalves, Ewan Harrison, David K. Jackson, Ian Johnston, Dominic Kwiatkowski, Cordelia Langford, John Sillitoe on behalf of the Wellcome Sanger Institute COVID-19 Surveillance Team |
| EPI_ISL_3286306 | Lighthouse Lab in Milton Keynes | Wellcome Sanger Institute for the COVID-19 Genomics UK (COG-UK) Consortium | The Lighthouse Lab in Milton Keynes and Alex Alderton, Roberto Amato, Jeffrey Barrett, Sonia Goncalves, Ewan Harrison, David K. Jackson, Ian Johnston, Dominic Kwiatkowski, Cordelia Langford, John Sillitoe on behalf of the Wellcome Sanger Institute COVID-19 Surveillance Team |
| EPI_ISL_3286307 | Lighthouse Lab in Alderley Park | Wellcome Sanger Institute for the COVID-19 Genomics UK (COG-UK) Consortium | Jacquelyn Wynn, Mairead Hyland, The Lighthouse Lab in Alderley Park and Alex Alderton, Roberto Amato, Jeffrey Barrett, Sonia Goncalves, Ewan Harrison, David K. Jackson, Ian Johnston, Dominic Kwiatkowski, Cordelia Langford, John Sillitoe on behalf of the Wellcome Sanger Institute COVID-19 Surveillance Team |
| EPI_ISL_3286308 | Lighthouse Lab in Milton Keynes | Wellcome Sanger Institute for the COVID-19 Genomics UK (COG-UK) Consortium | The Lighthouse Lab in Milton Keynes and Alex Alderton, Roberto Amato, Jeffrey Barrett, Sonia Goncalves, Ewan Harrison, David K. Jackson, Ian Johnston, Dominic Kwiatkowski, Cordelia Langford, John Sillitoe on behalf of the Wellcome Sanger Institute COVID-19 Surveillance Team |
| EPI_ISL_3286309, EPI_ISL_3286310 | Lighthouse Lab in Alderley Park | Wellcome Sanger Institute for the COVID-19 Genomics UK (COG-UK) Consortium | Jacquelyn Wynn, Mairead Hyland, The Lighthouse Lab in Alderley Park and Alex Alderton, Roberto Amato, Jeffrey Barrett, Sonia Goncalves, Ewan Harrison, David K. Jackson, Ian Johnston, Dominic Kwiatkowski, Cordelia Langford, John Sillitoe on behalf of the Wellcome Sanger Institute COVID-19 Surveillance Team |
| EPI_ISL_3286311 | Randox Laboratories | Wellcome Sanger Institute for the COVID-19 Genomics UK (COG-UK) Consortium | Randox Laboratories and Alex Alderton, Roberto Amato, Jeffrey Barrett, Sonia Goncalves, Ewan Harrison, David K. Jackson, Ian Johnston, Dominic Kwiatkowski, Cordelia Langford, John Sillitoe on behalf of the Wellcome Sanger Institute COVID-19 Surveillance Team |
| EPI_ISL_3286312, EPI_ISL_3286313, EPI_ISL_3286314, EPI_ISL_3286315, EPI_ISL_3286316 | Lighthouse Lab in Alderley Park | Wellcome Sanger Institute for the COVID-19 Genomics UK (COG-UK) Consortium | Jacquelyn Wynn, Mairead Hyland, The Lighthouse Lab in Alderley Park and Alex Alderton, Roberto Amato, Jeffrey Barrett, Sonia Goncalves, Ewan Harrison, David K. Jackson, Ian Johnston, Dominic Kwiatkowski, Cordelia Langford, John Sillitoe on behalf of the Wellcome Sanger Institute COVID-19 Surveillance Team |
| EPI_ISL_3286317 | Randox Laboratories | Wellcome Sanger Institute for the COVID-19 Genomics UK (COG-UK) Consortium | Randox Laboratories and Alex Alderton, Roberto Amato, Jeffrey Barrett, Sonia Goncalves, Ewan Harrison, David K. Jackson, Ian Johnston, Dominic Kwiatkowski, Cordelia Langford, John Sillitoe on behalf of the Wellcome Sanger Institute COVID-19 Surveillance Team |
| EPI_ISL_3286318, EPI_ISL_3286319 | Lighthouse Lab in Alderley Park | Wellcome Sanger Institute for the COVID-19 Genomics UK (COG-UK) Consortium | Jacquelyn Wynn, Mairead Hyland, The Lighthouse Lab in Alderley Park and Alex Alderton, Roberto Amato, Jeffrey Barrett, Sonia Goncalves, Ewan Harrison, David K. Jackson, Ian Johnston, Dominic Kwiatkowski, Cordelia Langford, John Sillitoe on behalf of the Wellcome Sanger Institute COVID-19 Surveillance Team |
| EPI_ISL_3286320 | Lighthouse Lab in Milton Keynes | Wellcome Sanger Institute for the COVID-19 Genomics UK (COG-UK) Consortium | The Lighthouse Lab in Milton Keynes and Alex Alderton, Roberto Amato, Jeffrey Barrett, Sonia Goncalves, Ewan Harrison, David K. Jackson, Ian Johnston, Dominic Kwiatkowski, Cordelia Langford, John Sillitoe on behalf of the Wellcome Sanger Institute COVID-19 Surveillance Team |
| EPI_ISL_3286321, EPI_ISL_3286322, EPI_ISL_3286323, EPI_ISL_3286324 | Lighthouse Lab in Alderley Park | Wellcome Sanger Institute for the COVID-19 Genomics UK (COG-UK) Consortium | Jacquelyn Wynn, Mairead Hyland, The Lighthouse Lab in Alderley Park and Alex Alderton, Roberto Amato, Jeffrey Barrett, Sonia Goncalves, Ewan Harrison, David K. Jackson, Ian Johnston, Dominic Kwiatkowski, Cordelia Langford, John Sillitoe on behalf of the Wellcome Sanger Institute COVID-19 Surveillance Team |
| EPI_ISL_3286325 | Lighthouse Lab in Milton Keynes | Wellcome Sanger Institute for the COVID-19 Genomics UK (COG-UK) Consortium | The Lighthouse Lab in Milton Keynes and Alex Alderton, Roberto Amato, Jeffrey Barrett, Sonia Goncalves, Ewan Harrison, David K. Jackson, Ian Johnston, Dominic Kwiatkowski, Cordelia Langford, John Sillitoe on behalf of the Wellcome Sanger Institute COVID-19 Surveillance Team |
| EPI_ISL_3286326, EPI_ISL_3286327, EPI_ISL_3286328, EPI_ISL_3286329 | Lighthouse Lab in Alderley Park | Wellcome Sanger Institute for the COVID-19 Genomics UK (COG-UK) Consortium | Jacquelyn Wynn, Mairead Hyland, The Lighthouse Lab in Alderley Park and Alex Alderton, Roberto Amato, Jeffrey Barrett, Sonia Goncalves, Ewan Harrison, David K. Jackson, Ian Johnston, Dominic Kwiatkowski, Cordelia Langford, John Sillitoe on behalf of the Wellcome Sanger Institute COVID-19 Surveillance Team |
| EPI_ISL_3286330, EPI_ISL_3286331 | Randox Laboratories | Wellcome Sanger Institute for the COVID-19 Genomics UK (COG-UK) Consortium | Randox Laboratories and Alex Alderton, Roberto Amato, Jeffrey Barrett, Sonia Goncalves, Ewan Harrison, David K. Jackson, Ian Johnston, Dominic Kwiatkowski, Cordelia Langford, John Sillitoe on behalf of the Wellcome Sanger Institute COVID-19 Surveillance Team |
| EPI_ISL_3286332 | Lighthouse Lab in Alderley Park | Wellcome Sanger Institute for the COVID-19 Genomics UK (COG-UK) Consortium | Jacquelyn Wynn, Mairead Hyland, The Lighthouse Lab in Alderley Park and Alex Alderton, Roberto Amato, Jeffrey Barrett, Sonia Goncalves, Ewan Harrison, David K. Jackson, Ian Johnston, Dominic Kwiatkowski, Cordelia Langford, John Sillitoe on behalf of the Wellcome Sanger Institute COVID-19 Surveillance Team |
| EPI_ISL_3286333 | Randox Laboratories | Wellcome Sanger Institute for the COVID-19 Genomics UK (COG-UK) Consortium | Randox Laboratories and Alex Alderton, Roberto Amato, Jeffrey Barrett, Sonia Goncalves, Ewan Harrison, David K. Jackson, Ian Johnston, Dominic Kwiatkowski, Cordelia Langford, John Sillitoe on behalf of the Wellcome Sanger Institute COVID-19 Surveillance Team |
| EPI_ISL_3286334, EPI_ISL_3286335, EPI_ISL_3286336, EPI_ISL_3286337, EPI_ISL_3286338, EPI_ISL_3286339 | Lighthouse Lab in Alderley Park | Wellcome Sanger Institute for the COVID-19 Genomics UK (COG-UK) Consortium | Jacquelyn Wynn, Mairead Hyland, The Lighthouse Lab in Alderley Park and Alex Alderton, Roberto Amato, Jeffrey Barrett, Sonia Goncalves, Ewan Harrison, David K. Jackson, Ian Johnston, Dominic Kwiatkowski, Cordelia Langford, John Sillitoe on behalf of the Wellcome Sanger Institute COVID-19 Surveillance Team |
| EPI_ISL_3286340 | Lighthouse Lab in Milton Keynes | Wellcome Sanger Institute for the COVID-19 Genomics UK (COG-UK) Consortium | The Lighthouse Lab in Milton Keynes and Alex Alderton, Roberto Amato, Jeffrey Barrett, Sonia Goncalves, Ewan Harrison, David K. Jackson, Ian Johnston, Dominic Kwiatkowski, Cordelia Langford, John Sillitoe on behalf of the Wellcome Sanger Institute COVID-19 Surveillance Team |
| EPI_ISL_3286341, EPI_ISL_3286342, EPI_ISL_3286343, EPI_ISL_3286344, EPI_ISL_3286345, EPI_ISL_3286346, EPI_ISL_3286347, EPI_ISL_3286348 | Lighthouse Lab in Alderley Park | Wellcome Sanger Institute for the COVID-19 Genomics UK (COG-UK) Consortium | Jacquelyn Wynn, Mairead Hyland, The Lighthouse Lab in Alderley Park and Alex Alderton, Roberto Amato, Jeffrey Barrett, Sonia Goncalves, Ewan Harrison, David K. Jackson, Ian Johnston, Dominic Kwiatkowski, Cordelia Langford, John Sillitoe on behalf of the Wellcome Sanger Institute COVID-19 Surveillance Team |
| EPI_ISL_3286349, EPI_ISL_3286350, EPI_ISL_3286351, EPI_ISL_3286352, EPI_ISL_3286353 | Lighthouse Lab in Milton Keynes | Wellcome Sanger Institute for the COVID-19 Genomics UK (COG-UK) Consortium | The Lighthouse Lab in Milton Keynes and Alex Alderton, Roberto Amato, Jeffrey Barrett, Sonia Goncalves, Ewan Harrison, David K. Jackson, Ian Johnston, Dominic Kwiatkowski, Cordelia Langford, John Sillitoe on behalf of the Wellcome Sanger Institute COVID-19 Surveillance Team |
| EPI_ISL_3286354, EPI_ISL_3286355, EPI_ISL_3286356 | Lighthouse Lab in Alderley Park | Wellcome Sanger Institute for the COVID-19 Genomics UK (COG-UK) Consortium | Jacquelyn Wynn, Mairead Hyland, The Lighthouse Lab in Alderley Park and Alex Alderton, Roberto Amato, Jeffrey Barrett, Sonia Goncalves, Ewan Harrison, David K. Jackson, Ian Johnston, Dominic Kwiatkowski, Cordelia Langford, John Sillitoe on behalf of the Wellcome Sanger Institute COVID-19 Surveillance Team |
| EPI_ISL_3286357 | Lighthouse Lab in Milton Keynes | Wellcome Sanger Institute for the COVID-19 Genomics UK (COG-UK) Consortium | The Lighthouse Lab in Milton Keynes and Alex Alderton, Roberto Amato, Jeffrey Barrett, Sonia Goncalves, Ewan Harrison, David K. Jackson, Ian Johnston, Dominic Kwiatkowski, Cordelia Langford, John Sillitoe on behalf of the Wellcome Sanger Institute COVID-19 Surveillance Team |
| EPI_ISL_3286358 | Randox Laboratories | Wellcome Sanger Institute for the COVID-19 Genomics UK (COG-UK) Consortium | Randox Laboratories and Alex Alderton, Roberto Amato, Jeffrey Barrett, Sonia Goncalves, Ewan Harrison, David K. Jackson, Ian Johnston, Dominic Kwiatkowski, Cordelia Langford, John Sillitoe on behalf of the Wellcome Sanger Institute COVID-19 Surveillance Team |
| EPI_ISL_3286359, EPI_ISL_3286360, EPI_ISL_3286361, EPI_ISL_3286362, EPI_ISL_3286363, EPI_ISL_3286364, EPI_ISL_3286365, EPI_ISL_3286366, EPI_ISL_3286367, EPI_ISL_3286368, EPI_ISL_3286369 | | | |
| see above | Lighthouse Lab in Alderley Park | Wellcome Sanger Institute for the COVID-19 Genomics UK (COG-UK) Consortium | Jacquelyn Wynn, Mairead Hyland, The Lighthouse Lab in Alderley Park and Alex Alderton, Roberto Amato, Jeffrey Barrett, Sonia Goncalves, Ewan Harrison, David K. Jackson, Ian Johnston, Dominic Kwiatkowski, Cordelia Langford, John Sillitoe on behalf of the Wellcome Sanger Institute COVID-19 Surveillance Team |
| EPI_ISL_3286370 | Randox Laboratories | Wellcome Sanger Institute for the COVID-19 Genomics UK (COG-UK) Consortium | Randox Laboratories and Alex Alderton, Roberto Amato, Jeffrey Barrett, Sonia Goncalves, Ewan Harrison, David K. Jackson, Ian Johnston, Dominic Kwiatkowski, Cordelia Langford, John Sillitoe on behalf of the Wellcome Sanger Institute COVID-19 Surveillance Team |
| EPI_ISL_3286371, EPI_ISL_3286372, | Lighthouse Lab in Alderley Park | Wellcome Sanger Institute for the COVID-19 Genomics UK (COG-UK) Consortium | Jacquelyn Wynn, Mairead Hyland, The Lighthouse Lab in Alderley Park and Alex Alderton, Roberto Amato, Jeffrey Barrett, Sonia Goncalves, Ewan Harrison, David K. Jackson, Ian Johnston, Dominic Kwiatkowski, Cordelia Langford, John Sillitoe on behalf of the Wellcome Sanger Institute COVID-19 Surveillance Team |

| | | | |
|---|---|---|---|
| EPI_ISL_3286373 | | | |
| EPI_ISL_3286374 | Lighthouse Lab in Milton Keynes | Wellcome Sanger Institute for the COVID-19 Genomics UK (COG-UK) Consortium | The Lighthouse Lab in Milton Keynes and Alex Alderton, Roberto Amato, Jeffrey Barrett, Sonia Goncalves, Ewan Harrison, David K. Jackson, Ian Johnston, Dominic Kwiatkowski, Cordelia Langford, John Sillitoe on behalf of the Wellcome Sanger Institute COVID-19 Surveillance Team |
| EPI_ISL_3286375, EPI_ISL_3286376, EPI_ISL_3286377, EPI_ISL_3286378 | Lighthouse Lab in Alderley Park | Wellcome Sanger Institute for the COVID-19 Genomics UK (COG-UK) Consortium | Jacquelyn Wynn, Mairead Hyland, The Lighthouse Lab in Alderley Park and Alex Alderton, Roberto Amato, Jeffrey Barrett, Sonia Goncalves, Ewan Harrison, David K. Jackson, Ian Johnston, Dominic Kwiatkowski, Cordelia Langford, John Sillitoe on behalf of the Wellcome Sanger Institute COVID-19 Surveillance Team |
| EPI_ISL_3286379 | Randox Laboratories | Wellcome Sanger Institute for the COVID-19 Genomics UK (COG-UK) Consortium | Randox Laboratories and Alex Alderton, Roberto Amato, Jeffrey Barrett, Sonia Goncalves, Ewan Harrison, David K. Jackson, Ian Johnston, Dominic Kwiatkowski, Cordelia Langford, John Sillitoe on behalf of the Wellcome Sanger Institute COVID-19 Surveillance Team |
| EPI_ISL_3286380, EPI_ISL_3286381, EPI_ISL_3286382 | Lighthouse Lab in Milton Keynes | Wellcome Sanger Institute for the COVID-19 Genomics UK (COG-UK) Consortium | The Lighthouse Lab in Milton Keynes and Alex Alderton, Roberto Amato, Jeffrey Barrett, Sonia Goncalves, Ewan Harrison, David K. Jackson, Ian Johnston, Dominic Kwiatkowski, Cordelia Langford, John Sillitoe on behalf of the Wellcome Sanger Institute COVID-19 Surveillance Team |
| EPI_ISL_3286383, EPI_ISL_3286384 | Lighthouse Lab in Alderley Park | Wellcome Sanger Institute for the COVID-19 Genomics UK (COG-UK) Consortium | Jacquelyn Wynn, Mairead Hyland, The Lighthouse Lab in Alderley Park and Alex Alderton, Roberto Amato, Jeffrey Barrett, Sonia Goncalves, Ewan Harrison, David K. Jackson, Ian Johnston, Dominic Kwiatkowski, Cordelia Langford, John Sillitoe on behalf of the Wellcome Sanger Institute COVID-19 Surveillance Team |
| EPI_ISL_3286385, EPI_ISL_3286386 | Randox Laboratories | Wellcome Sanger Institute for the COVID-19 Genomics UK (COG-UK) Consortium | Randox Laboratories and Alex Alderton, Roberto Amato, Jeffrey Barrett, Sonia Goncalves, Ewan Harrison, David K. Jackson, Ian Johnston, Dominic Kwiatkowski, Cordelia Langford, John Sillitoe on behalf of the Wellcome Sanger Institute COVID-19 Surveillance Team |
| EPI_ISL_3286387, EPI_ISL_3286388 | Lighthouse Lab in Alderley Park | Wellcome Sanger Institute for the COVID-19 Genomics UK (COG-UK) Consortium | Jacquelyn Wynn, Mairead Hyland, The Lighthouse Lab in Alderley Park and Alex Alderton, Roberto Amato, Jeffrey Barrett, Sonia Goncalves, Ewan Harrison, David K. Jackson, Ian Johnston, Dominic Kwiatkowski, Cordelia Langford, John Sillitoe on behalf of the Wellcome Sanger Institute COVID-19 Surveillance Team |
| EPI_ISL_3286389, EPI_ISL_3286390 | Lighthouse Lab in Milton Keynes | Wellcome Sanger Institute for the COVID-19 Genomics UK (COG-UK) Consortium | The Lighthouse Lab in Milton Keynes and Alex Alderton, Roberto Amato, Jeffrey Barrett, Sonia Goncalves, Ewan Harrison, David K. Jackson, Ian Johnston, Dominic Kwiatkowski, Cordelia Langford, John Sillitoe on behalf of the Wellcome Sanger Institute COVID-19 Surveillance Team |
| EPI_ISL_3286391, EPI_ISL_3286392, EPI_ISL_3286393, EPI_ISL_3286394, EPI_ISL_3286395, EPI_ISL_3286396, EPI_ISL_3286397, EPI_ISL_3286398 | Lighthouse Lab in Alderley Park | Wellcome Sanger Institute for the COVID-19 Genomics UK (COG-UK) Consortium | Jacquelyn Wynn, Mairead Hyland, The Lighthouse Lab in Alderley Park and Alex Alderton, Roberto Amato, Jeffrey Barrett, Sonia Goncalves, Ewan Harrison, David K. Jackson, Ian Johnston, Dominic Kwiatkowski, Cordelia Langford, John Sillitoe on behalf of the Wellcome Sanger Institute COVID-19 Surveillance Team |
| EPI_ISL_3286399, EPI_ISL_3286400 | Lighthouse Lab in Milton Keynes | Wellcome Sanger Institute for the COVID-19 Genomics UK (COG-UK) Consortium | The Lighthouse Lab in Milton Keynes and Alex Alderton, Roberto Amato, Jeffrey Barrett, Sonia Goncalves, Ewan Harrison, David K. Jackson, Ian Johnston, Dominic Kwiatkowski, Cordelia Langford, John Sillitoe on behalf of the Wellcome Sanger Institute COVID-19 Surveillance Team |
| EPI_ISL_3286401 | Lighthouse Lab in Alderley Park | Wellcome Sanger Institute for the COVID-19 Genomics UK (COG-UK) Consortium | Jacquelyn Wynn, Mairead Hyland, The Lighthouse Lab in Alderley Park and Alex Alderton, Roberto Amato, Jeffrey Barrett, Sonia Goncalves, Ewan Harrison, David K. Jackson, Ian Johnston, Dominic Kwiatkowski, Cordelia Langford, John Sillitoe on behalf of the Wellcome Sanger Institute COVID-19 Surveillance Team |
| EPI_ISL_3286402, EPI_ISL_3286403 | Lighthouse Lab in Milton Keynes | Wellcome Sanger Institute for the COVID-19 Genomics UK (COG-UK) Consortium | The Lighthouse Lab in Milton Keynes and Alex Alderton, Roberto Amato, Jeffrey Barrett, Sonia Goncalves, Ewan Harrison, David K. Jackson, Ian Johnston, Dominic Kwiatkowski, Cordelia Langford, John Sillitoe on behalf of the Wellcome Sanger Institute COVID-19 Surveillance Team |
| EPI_ISL_3286404, EPI_ISL_3286405, EPI_ISL_3286406, EPI_ISL_3286407, EPI_ISL_3286408 | Lighthouse Lab in Alderley Park | Wellcome Sanger Institute for the COVID-19 Genomics UK (COG-UK) Consortium | Jacquelyn Wynn, Mairead Hyland, The Lighthouse Lab in Alderley Park and Alex Alderton, Roberto Amato, Jeffrey Barrett, Sonia Goncalves, Ewan Harrison, David K. Jackson, Ian Johnston, Dominic Kwiatkowski, Cordelia Langford, John Sillitoe on behalf of the Wellcome Sanger Institute COVID-19 Surveillance Team |
| EPI_ISL_3286409 | Lighthouse Lab in Milton Keynes | Wellcome Sanger Institute for the COVID-19 Genomics UK (COG-UK) Consortium | The Lighthouse Lab in Milton Keynes and Alex Alderton, Roberto Amato, Jeffrey Barrett, Sonia Goncalves, Ewan Harrison, David K. Jackson, Ian Johnston, Dominic Kwiatkowski, Cordelia Langford, John Sillitoe on behalf of the Wellcome Sanger Institute COVID-19 Surveillance Team |
| EPI_ISL_3286410, EPI_ISL_3286411 | Lighthouse Lab in Alderley Park | Wellcome Sanger Institute for the COVID-19 Genomics UK (COG-UK) Consortium | Jacquelyn Wynn, Mairead Hyland, The Lighthouse Lab in Alderley Park and Alex Alderton, Roberto Amato, Jeffrey Barrett, Sonia Goncalves, Ewan Harrison, David K. Jackson, Ian Johnston, Dominic Kwiatkowski, Cordelia Langford, John Sillitoe on behalf of the Wellcome Sanger Institute COVID-19 Surveillance Team |
| EPI_ISL_3286412 | Lighthouse Lab in Milton Keynes | Wellcome Sanger Institute for the COVID-19 Genomics UK (COG-UK) Consortium | The Lighthouse Lab in Milton Keynes and Alex Alderton, Roberto Amato, Jeffrey Barrett, Sonia Goncalves, Ewan Harrison, David K. Jackson, Ian Johnston, Dominic Kwiatkowski, Cordelia Langford, John Sillitoe on behalf of the Wellcome Sanger Institute COVID-19 Surveillance Team |
| EPI_ISL_3286413 | Lighthouse Lab in Alderley Park | Wellcome Sanger Institute for the COVID-19 Genomics UK (COG-UK) Consortium | Jacquelyn Wynn, Mairead Hyland, The Lighthouse Lab in Alderley Park and Alex Alderton, Roberto Amato, Jeffrey Barrett, Sonia Goncalves, Ewan Harrison, David K. Jackson, Ian Johnston, Dominic Kwiatkowski, Cordelia Langford, John Sillitoe on behalf of the Wellcome Sanger Institute COVID-19 Surveillance Team |
| EPI_ISL_3286414, EPI_ISL_3286415, EPI_ISL_3286416, EPI_ISL_3286417 | Lighthouse Lab in Milton Keynes | Wellcome Sanger Institute for the COVID-19 Genomics UK (COG-UK) Consortium | The Lighthouse Lab in Milton Keynes and Alex Alderton, Roberto Amato, Jeffrey Barrett, Sonia Goncalves, Ewan Harrison, David K. Jackson, Ian Johnston, Dominic Kwiatkowski, Cordelia Langford, John Sillitoe on behalf of the Wellcome Sanger Institute COVID-19 Surveillance Team |
| EPI_ISL_3286418, EPI_ISL_3286419 | Lighthouse Lab in Alderley Park | Wellcome Sanger Institute for the COVID-19 Genomics UK (COG-UK) Consortium | Jacquelyn Wynn, Mairead Hyland, The Lighthouse Lab in Alderley Park and Alex Alderton, Roberto Amato, Jeffrey Barrett, Sonia Goncalves, Ewan Harrison, David K. Jackson, Ian Johnston, Dominic Kwiatkowski, Cordelia Langford, John Sillitoe on behalf of the Wellcome Sanger Institute COVID-19 Surveillance Team |
| EPI_ISL_3286420 | Randox Laboratories | Wellcome Sanger Institute for the COVID-19 Genomics UK (COG-UK) Consortium | Randox Laboratories and Alex Alderton, Roberto Amato, Jeffrey Barrett, Sonia Goncalves, Ewan Harrison, David K. Jackson, Ian Johnston, Dominic Kwiatkowski, Cordelia Langford, John Sillitoe on behalf of the Wellcome Sanger Institute COVID-19 Surveillance Team |
| EPI_ISL_3286421, EPI_ISL_3286422 | Lighthouse Lab in Alderley Park | Wellcome Sanger Institute for the COVID-19 Genomics UK (COG-UK) Consortium | Jacquelyn Wynn, Mairead Hyland, The Lighthouse Lab in Alderley Park and Alex Alderton, Roberto Amato, Jeffrey Barrett, Sonia Goncalves, Ewan Harrison, David K. Jackson, Ian Johnston, Dominic Kwiatkowski, Cordelia Langford, John Sillitoe on behalf of the Wellcome Sanger Institute COVID-19 Surveillance Team |
| EPI_ISL_3286423, EPI_ISL_3286424, EPI_ISL_3286425, EPI_ISL_3286426, EPI_ISL_3286427, EPI_ISL_3286428 | Lighthouse Lab in Milton Keynes | Wellcome Sanger Institute for the COVID-19 Genomics UK (COG-UK) Consortium | The Lighthouse Lab in Milton Keynes and Alex Alderton, Roberto Amato, Jeffrey Barrett, Sonia Goncalves, Ewan Harrison, David K. Jackson, Ian Johnston, Dominic Kwiatkowski, Cordelia Langford, John Sillitoe on behalf of the Wellcome Sanger Institute COVID-19 Surveillance Team |
| EPI_ISL_3286429, EPI_ISL_3286430, EPI_ISL_3286431, EPI_ISL_3286432, EPI_ISL_3286433 | Lighthouse Lab in Alderley Park | Wellcome Sanger Institute for the COVID-19 Genomics UK (COG-UK) Consortium | Jacquelyn Wynn, Mairead Hyland, The Lighthouse Lab in Alderley Park and Alex Alderton, Roberto Amato, Jeffrey Barrett, Sonia Goncalves, Ewan Harrison, David K. Jackson, Ian Johnston, Dominic Kwiatkowski, Cordelia Langford, John Sillitoe on behalf of the Wellcome Sanger Institute COVID-19 Surveillance Team |
| EPI_ISL_3286434, EPI_ISL_3286435, EPI_ISL_3286436 | Lighthouse Lab in Milton Keynes | Wellcome Sanger Institute for the COVID-19 Genomics UK (COG-UK) Consortium | The Lighthouse Lab in Milton Keynes and Alex Alderton, Roberto Amato, Jeffrey Barrett, Sonia Goncalves, Ewan Harrison, David K. Jackson, Ian Johnston, Dominic Kwiatkowski, Cordelia Langford, John Sillitoe on behalf of the Wellcome Sanger Institute COVID-19 Surveillance Team |
| EPI_ISL_3286437, EPI_ISL_3286438, EPI_ISL_3286439, EPI_ISL_3286440, EPI_ISL_3286441, EPI_ISL_3286442, EPI_ISL_3286443 | Lighthouse Lab in Alderley Park | Wellcome Sanger Institute for the COVID-19 Genomics UK (COG-UK) Consortium | Jacquelyn Wynn, Mairead Hyland, The Lighthouse Lab in Alderley Park and Alex Alderton, Roberto Amato, Jeffrey Barrett, Sonia Goncalves, Ewan Harrison, David K. Jackson, Ian Johnston, Dominic Kwiatkowski, Cordelia Langford, John Sillitoe on behalf of the Wellcome Sanger Institute COVID-19 Surveillance Team |
| EPI_ISL_3286444 | Lighthouse Lab in Milton Keynes | Wellcome Sanger Institute for the COVID-19 Genomics UK (COG-UK) Consortium | The Lighthouse Lab in Milton Keynes and Alex Alderton, Roberto Amato, Jeffrey Barrett, Sonia Goncalves, Ewan Harrison, David K. Jackson, Ian Johnston, Dominic Kwiatkowski, Cordelia Langford, John Sillitoe on behalf of the Wellcome Sanger Institute COVID-19 Surveillance Team |
| EPI_ISL_3286445, EPI_ISL_3286446, EPI_ISL_3286447, EPI_ISL_3286448, EPI_ISL_3286449 | Lighthouse Lab in Alderley Park | Wellcome Sanger Institute for the COVID-19 Genomics UK (COG-UK) Consortium | Jacquelyn Wynn, Mairead Hyland, The Lighthouse Lab in Alderley Park and Alex Alderton, Roberto Amato, Jeffrey Barrett, Sonia Goncalves, Ewan Harrison, David K. Jackson, Ian Johnston, Dominic Kwiatkowski, Cordelia Langford, John Sillitoe on behalf of the Wellcome Sanger Institute COVID-19 Surveillance Team |
| EPI_ISL_3286450 | Lighthouse Lab in Milton Keynes | Wellcome Sanger Institute for the COVID-19 Genomics UK (COG-UK) Consortium | The Lighthouse Lab in Milton Keynes and Alex Alderton, Roberto Amato, Jeffrey Barrett, Sonia Goncalves, Ewan Harrison, David K. Jackson, Ian Johnston, Dominic Kwiatkowski, Cordelia Langford, John Sillitoe on behalf of the Wellcome Sanger Institute COVID-19 Surveillance Team |
| EPI_ISL_3286451, EPI_ISL_3286452 | Lighthouse Lab in Alderley Park | Wellcome Sanger Institute for the COVID-19 Genomics UK (COG-UK) Consortium | Jacquelyn Wynn, Mairead Hyland, The Lighthouse Lab in Alderley Park and Alex Alderton, Roberto Amato, Jeffrey Barrett, Sonia Goncalves, Ewan Harrison, David K. Jackson, Ian Johnston, Dominic Kwiatkowski, Cordelia Langford, John Sillitoe on behalf of the Wellcome Sanger Institute COVID-19 Surveillance Team |
| EPI_ISL_3286453 | Lighthouse Lab in Milton Keynes | Wellcome Sanger Institute for the COVID-19 Genomics UK (COG-UK) Consortium | The Lighthouse Lab in Milton Keynes and Alex Alderton, Roberto Amato, Jeffrey Barrett, Sonia Goncalves, Ewan Harrison, David K. Jackson, Ian Johnston, Dominic Kwiatkowski, Cordelia Langford, John Sillitoe on behalf of the Wellcome Sanger Institute COVID-19 Surveillance Team |
| EPI_ISL_3286454 | Lighthouse Lab in Alderley Park | Wellcome Sanger Institute for the COVID-19 Genomics UK (COG-UK) Consortium | Jacquelyn Wynn, Mairead Hyland, The Lighthouse Lab in Alderley Park and Alex Alderton, Roberto Amato, Jeffrey Barrett, Sonia Goncalves, Ewan Harrison, David K. Jackson, Ian Johnston, Dominic Kwiatkowski, Cordelia Langford, John Sillitoe on behalf of the Wellcome Sanger Institute COVID-19 Surveillance Team |
| EPI_ISL_3286455 | Lighthouse Lab in Milton Keynes | Wellcome Sanger Institute for the COVID-19 Genomics UK (COG-UK) Consortium | The Lighthouse Lab in Milton Keynes and Alex Alderton, Roberto Amato, Jeffrey Barrett, Sonia Goncalves, Ewan Harrison, David K. Jackson, Ian Johnston, Dominic Kwiatkowski, Cordelia Langford, John Sillitoe on behalf of the Wellcome Sanger Institute COVID-19 Surveillance Team |
| EPI_ISL_3286456, | Lighthouse Lab in Alderley Park | Wellcome Sanger Institute for the COVID-19 Genomics UK (COG-UK) | Jacquelyn Wynn, Mairead Hyland, The Lighthouse Lab in Alderley Park and Alex Alderton, Roberto Amato, Jeffrey Barrett, Sonia Goncalves, Ewan Harrison, David K. Jackson, Ian Johnston, Dominic Kwiatkowski, Cordelia Langford, John Sillitoe on |

| | | | |
|---|---|---|---|
| EPI_ISL_3286457, EPI_ISL_3286458 | | Consortium | behalf of the Wellcome Sanger Institute COVID-19 Surveillance Team |
| EPI_ISL_3286459 | Lighthouse Lab in Milton Keynes | Wellcome Sanger Institute for the COVID-19 Genomics UK (COG-UK) Consortium | The Lighthouse Lab in Milton Keynes and Alex Alderton, Roberto Amato, Jeffrey Barrett, Sonia Goncalves, Ewan Harrison, David K. Jackson, Ian Johnston, Dominic Kwiatkowski, Cordelia Langford, John Sillitoe on behalf of the Wellcome Sanger Institute COVID-19 Surveillance Team |
| EPI_ISL_3286460, EPI_ISL_3286461 | Lighthouse Lab in Alderley Park | Wellcome Sanger Institute for the COVID-19 Genomics UK (COG-UK) Consortium | Jacquelyn Wynn, Mairead Hyland, The Lighthouse Lab in Alderley Park and Alex Alderton, Roberto Amato, Jeffrey Barrett, Sonia Goncalves, Ewan Harrison, David K. Jackson, Ian Johnston, Dominic Kwiatkowski, Cordelia Langford, John Sillitoe on behalf of the Wellcome Sanger Institute COVID-19 Surveillance Team |
| EPI_ISL_3286462 | Lighthouse Lab in Milton Keynes | Wellcome Sanger Institute for the COVID-19 Genomics UK (COG-UK) Consortium | The Lighthouse Lab in Milton Keynes and Alex Alderton, Roberto Amato, Jeffrey Barrett, Sonia Goncalves, Ewan Harrison, David K. Jackson, Ian Johnston, Dominic Kwiatkowski, Cordelia Langford, John Sillitoe on behalf of the Wellcome Sanger Institute COVID-19 Surveillance Team |
| EPI_ISL_3286463 | Randox Laboratories | Wellcome Sanger Institute for the COVID-19 Genomics UK (COG-UK) Consortium | Randox Laboratories and Alex Alderton, Roberto Amato, Jeffrey Barrett, Sonia Goncalves, Ewan Harrison, David K. Jackson, Ian Johnston, Dominic Kwiatkowski, Cordelia Langford, John Sillitoe on behalf of the Wellcome Sanger Institute COVID-19 Surveillance Team |
| EPI_ISL_3286464 | Lighthouse Lab in Milton Keynes | Wellcome Sanger Institute for the COVID-19 Genomics UK (COG-UK) Consortium | The Lighthouse Lab in Milton Keynes and Alex Alderton, Roberto Amato, Jeffrey Barrett, Sonia Goncalves, Ewan Harrison, David K. Jackson, Ian Johnston, Dominic Kwiatkowski, Cordelia Langford, John Sillitoe on behalf of the Wellcome Sanger Institute COVID-19 Surveillance Team |
| EPI_ISL_3286465 | Lighthouse Lab in Alderley Park | Wellcome Sanger Institute for the COVID-19 Genomics UK (COG-UK) Consortium | Jacquelyn Wynn, Mairead Hyland, The Lighthouse Lab in Alderley Park and Alex Alderton, Roberto Amato, Jeffrey Barrett, Sonia Goncalves, Ewan Harrison, David K. Jackson, Ian Johnston, Dominic Kwiatkowski, Cordelia Langford, John Sillitoe on behalf of the Wellcome Sanger Institute COVID-19 Surveillance Team |
| EPI_ISL_3286466 | Lighthouse Lab in Milton Keynes | Wellcome Sanger Institute for the COVID-19 Genomics UK (COG-UK) Consortium | The Lighthouse Lab in Milton Keynes and Alex Alderton, Roberto Amato, Jeffrey Barrett, Sonia Goncalves, Ewan Harrison, David K. Jackson, Ian Johnston, Dominic Kwiatkowski, Cordelia Langford, John Sillitoe on behalf of the Wellcome Sanger Institute COVID-19 Surveillance Team |
| EPI_ISL_3286467 | Randox Laboratories | Wellcome Sanger Institute for the COVID-19 Genomics UK (COG-UK) Consortium | Randox Laboratories and Alex Alderton, Roberto Amato, Jeffrey Barrett, Sonia Goncalves, Ewan Harrison, David K. Jackson, Ian Johnston, Dominic Kwiatkowski, Cordelia Langford, John Sillitoe on behalf of the Wellcome Sanger Institute COVID-19 Surveillance Team |
| EPI_ISL_3286468, EPI_ISL_3286469 | Lighthouse Lab in Milton Keynes | Wellcome Sanger Institute for the COVID-19 Genomics UK (COG-UK) Consortium | The Lighthouse Lab in Milton Keynes and Alex Alderton, Roberto Amato, Jeffrey Barrett, Sonia Goncalves, Ewan Harrison, David K. Jackson, Ian Johnston, Dominic Kwiatkowski, Cordelia Langford, John Sillitoe on behalf of the Wellcome Sanger Institute COVID-19 Surveillance Team |
| EPI_ISL_3286470, EPI_ISL_3286471, EPI_ISL_3286472, EPI_ISL_3286473, EPI_ISL_3286474 | Lighthouse Lab in Alderley Park | Wellcome Sanger Institute for the COVID-19 Genomics UK (COG-UK) Consortium | Jacquelyn Wynn, Mairead Hyland, The Lighthouse Lab in Alderley Park and Alex Alderton, Roberto Amato, Jeffrey Barrett, Sonia Goncalves, Ewan Harrison, David K. Jackson, Ian Johnston, Dominic Kwiatkowski, Cordelia Langford, John Sillitoe on behalf of the Wellcome Sanger Institute COVID-19 Surveillance Team |
| EPI_ISL_3286475 | Randox Laboratories | Wellcome Sanger Institute for the COVID-19 Genomics UK (COG-UK) Consortium | Randox Laboratories and Alex Alderton, Roberto Amato, Jeffrey Barrett, Sonia Goncalves, Ewan Harrison, David K. Jackson, Ian Johnston, Dominic Kwiatkowski, Cordelia Langford, John Sillitoe on behalf of the Wellcome Sanger Institute COVID-19 Surveillance Team |
| EPI_ISL_3286476 | Lighthouse Lab in Milton Keynes | Wellcome Sanger Institute for the COVID-19 Genomics UK (COG-UK) Consortium | The Lighthouse Lab in Milton Keynes and Alex Alderton, Roberto Amato, Jeffrey Barrett, Sonia Goncalves, Ewan Harrison, David K. Jackson, Ian Johnston, Dominic Kwiatkowski, Cordelia Langford, John Sillitoe on behalf of the Wellcome Sanger Institute COVID-19 Surveillance Team |
| EPI_ISL_3286477 | Lighthouse Lab in Alderley Park | Wellcome Sanger Institute for the COVID-19 Genomics UK (COG-UK) Consortium | Jacquelyn Wynn, Mairead Hyland, The Lighthouse Lab in Alderley Park and Alex Alderton, Roberto Amato, Jeffrey Barrett, Sonia Goncalves, Ewan Harrison, David K. Jackson, Ian Johnston, Dominic Kwiatkowski, Cordelia Langford, John Sillitoe on behalf of the Wellcome Sanger Institute COVID-19 Surveillance Team |
| EPI_ISL_3286478, EPI_ISL_3286479 | Lighthouse Lab in Milton Keynes | Wellcome Sanger Institute for the COVID-19 Genomics UK (COG-UK) Consortium | The Lighthouse Lab in Milton Keynes and Alex Alderton, Roberto Amato, Jeffrey Barrett, Sonia Goncalves, Ewan Harrison, David K. Jackson, Ian Johnston, Dominic Kwiatkowski, Cordelia Langford, John Sillitoe on behalf of the Wellcome Sanger Institute COVID-19 Surveillance Team |
| EPI_ISL_3286480, EPI_ISL_3286481 | Randox Laboratories | Wellcome Sanger Institute for the COVID-19 Genomics UK (COG-UK) Consortium | Randox Laboratories and Alex Alderton, Roberto Amato, Jeffrey Barrett, Sonia Goncalves, Ewan Harrison, David K. Jackson, Ian Johnston, Dominic Kwiatkowski, Cordelia Langford, John Sillitoe on behalf of the Wellcome Sanger Institute COVID-19 Surveillance Team |
| EPI_ISL_3286482 | Lighthouse Lab in Alderley Park | Wellcome Sanger Institute for the COVID-19 Genomics UK (COG-UK) Consortium | Jacquelyn Wynn, Mairead Hyland, The Lighthouse Lab in Alderley Park and Alex Alderton, Roberto Amato, Jeffrey Barrett, Sonia Goncalves, Ewan Harrison, David K. Jackson, Ian Johnston, Dominic Kwiatkowski, Cordelia Langford, John Sillitoe on behalf of the Wellcome Sanger Institute COVID-19 Surveillance Team |
| EPI_ISL_3286483 | Randox Laboratories | Wellcome Sanger Institute for the COVID-19 Genomics UK (COG-UK) Consortium | Randox Laboratories and Alex Alderton, Roberto Amato, Jeffrey Barrett, Sonia Goncalves, Ewan Harrison, David K. Jackson, Ian Johnston, Dominic Kwiatkowski, Cordelia Langford, John Sillitoe on behalf of the Wellcome Sanger Institute COVID-19 Surveillance Team |
| EPI_ISL_3286484, EPI_ISL_3286485 | Lighthouse Lab in Milton Keynes | Wellcome Sanger Institute for the COVID-19 Genomics UK (COG-UK) Consortium | The Lighthouse Lab in Milton Keynes and Alex Alderton, Roberto Amato, Jeffrey Barrett, Sonia Goncalves, Ewan Harrison, David K. Jackson, Ian Johnston, Dominic Kwiatkowski, Cordelia Langford, John Sillitoe on behalf of the Wellcome Sanger Institute COVID-19 Surveillance Team |
| EPI_ISL_3286486, EPI_ISL_3286487 | Lighthouse Lab in Alderley Park | Wellcome Sanger Institute for the COVID-19 Genomics UK (COG-UK) Consortium | Jacquelyn Wynn, Mairead Hyland, The Lighthouse Lab in Alderley Park and Alex Alderton, Roberto Amato, Jeffrey Barrett, Sonia Goncalves, Ewan Harrison, David K. Jackson, Ian Johnston, Dominic Kwiatkowski, Cordelia Langford, John Sillitoe on behalf of the Wellcome Sanger Institute COVID-19 Surveillance Team |
| EPI_ISL_3286488 | Lighthouse Lab in Milton Keynes | Wellcome Sanger Institute for the COVID-19 Genomics UK (COG-UK) Consortium | The Lighthouse Lab in Milton Keynes and Alex Alderton, Roberto Amato, Jeffrey Barrett, Sonia Goncalves, Ewan Harrison, David K. Jackson, Ian Johnston, Dominic Kwiatkowski, Cordelia Langford, John Sillitoe on behalf of the Wellcome Sanger Institute COVID-19 Surveillance Team |
| EPI_ISL_3286489, EPI_ISL_3286490, EPI_ISL_3286491 | Lighthouse Lab in Alderley Park | Wellcome Sanger Institute for the COVID-19 Genomics UK (COG-UK) Consortium | Jacquelyn Wynn, Mairead Hyland, The Lighthouse Lab in Alderley Park and Alex Alderton, Roberto Amato, Jeffrey Barrett, Sonia Goncalves, Ewan Harrison, David K. Jackson, Ian Johnston, Dominic Kwiatkowski, Cordelia Langford, John Sillitoe on behalf of the Wellcome Sanger Institute COVID-19 Surveillance Team |
| EPI_ISL_3286492 | Lighthouse Lab in Milton Keynes | Wellcome Sanger Institute for the COVID-19 Genomics UK (COG-UK) Consortium | The Lighthouse Lab in Milton Keynes and Alex Alderton, Roberto Amato, Jeffrey Barrett, Sonia Goncalves, Ewan Harrison, David K. Jackson, Ian Johnston, Dominic Kwiatkowski, Cordelia Langford, John Sillitoe on behalf of the Wellcome Sanger Institute COVID-19 Surveillance Team |
| EPI_ISL_3286493 | Lighthouse Lab in Alderley Park | Wellcome Sanger Institute for the COVID-19 Genomics UK (COG-UK) Consortium | Jacquelyn Wynn, Mairead Hyland, The Lighthouse Lab in Alderley Park and Alex Alderton, Roberto Amato, Jeffrey Barrett, Sonia Goncalves, Ewan Harrison, David K. Jackson, Ian Johnston, Dominic Kwiatkowski, Cordelia Langford, John Sillitoe on behalf of the Wellcome Sanger Institute COVID-19 Surveillance Team |
| EPI_ISL_3286494 | Lighthouse Lab in Milton Keynes | Wellcome Sanger Institute for the COVID-19 Genomics UK (COG-UK) Consortium | The Lighthouse Lab in Milton Keynes and Alex Alderton, Roberto Amato, Jeffrey Barrett, Sonia Goncalves, Ewan Harrison, David K. Jackson, Ian Johnston, Dominic Kwiatkowski, Cordelia Langford, John Sillitoe on behalf of the Wellcome Sanger Institute COVID-19 Surveillance Team |
| EPI_ISL_3286495 | Lighthouse Lab in Alderley Park | Wellcome Sanger Institute for the COVID-19 Genomics UK (COG-UK) Consortium | Jacquelyn Wynn, Mairead Hyland, The Lighthouse Lab in Alderley Park and Alex Alderton, Roberto Amato, Jeffrey Barrett, Sonia Goncalves, Ewan Harrison, David K. Jackson, Ian Johnston, Dominic Kwiatkowski, Cordelia Langford, John Sillitoe on behalf of the Wellcome Sanger Institute COVID-19 Surveillance Team |
| EPI_ISL_3286496 | Lighthouse Lab in Milton Keynes | Wellcome Sanger Institute for the COVID-19 Genomics UK (COG-UK) Consortium | The Lighthouse Lab in Milton Keynes and Alex Alderton, Roberto Amato, Jeffrey Barrett, Sonia Goncalves, Ewan Harrison, David K. Jackson, Ian Johnston, Dominic Kwiatkowski, Cordelia Langford, John Sillitoe on behalf of the Wellcome Sanger Institute COVID-19 Surveillance Team |
| EPI_ISL_3286497, EPI_ISL_3286498, EPI_ISL_3286499, EPI_ISL_3286500 | Lighthouse Lab in Alderley Park | Wellcome Sanger Institute for the COVID-19 Genomics UK (COG-UK) Consortium | Jacquelyn Wynn, Mairead Hyland, The Lighthouse Lab in Alderley Park and Alex Alderton, Roberto Amato, Jeffrey Barrett, Sonia Goncalves, Ewan Harrison, David K. Jackson, Ian Johnston, Dominic Kwiatkowski, Cordelia Langford, John Sillitoe on behalf of the Wellcome Sanger Institute COVID-19 Surveillance Team |
| EPI_ISL_3286501, EPI_ISL_3286502, EPI_ISL_3286503, EPI_ISL_3286504 | Lighthouse Lab in Milton Keynes | Wellcome Sanger Institute for the COVID-19 Genomics UK (COG-UK) Consortium | The Lighthouse Lab in Milton Keynes and Alex Alderton, Roberto Amato, Jeffrey Barrett, Sonia Goncalves, Ewan Harrison, David K. Jackson, Ian Johnston, Dominic Kwiatkowski, Cordelia Langford, John Sillitoe on behalf of the Wellcome Sanger Institute COVID-19 Surveillance Team |
| EPI_ISL_3286505, EPI_ISL_3286506, EPI_ISL_3286507, EPI_ISL_3286508, EPI_ISL_3286509, EPI_ISL_3286510, EPI_ISL_3286511, EPI_ISL_3286512 | Lighthouse Lab in Alderley Park | Wellcome Sanger Institute for the COVID-19 Genomics UK (COG-UK) Consortium | Jacquelyn Wynn, Mairead Hyland, The Lighthouse Lab in Alderley Park and Alex Alderton, Roberto Amato, Jeffrey Barrett, Sonia Goncalves, Ewan Harrison, David K. Jackson, Ian Johnston, Dominic Kwiatkowski, Cordelia Langford, John Sillitoe on behalf of the Wellcome Sanger Institute COVID-19 Surveillance Team |
| EPI_ISL_3286513 | Lighthouse Lab in Milton Keynes | Wellcome Sanger Institute for the COVID-19 Genomics UK (COG-UK) Consortium | The Lighthouse Lab in Milton Keynes and Alex Alderton, Roberto Amato, Jeffrey Barrett, Sonia Goncalves, Ewan Harrison, David K. Jackson, Ian Johnston, Dominic Kwiatkowski, Cordelia Langford, John Sillitoe on behalf of the Wellcome Sanger Institute COVID-19 Surveillance Team |
| EPI_ISL_3286514 | Randox Laboratories | Wellcome Sanger Institute for the COVID-19 Genomics UK (COG-UK) Consortium | Randox Laboratories and Alex Alderton, Roberto Amato, Jeffrey Barrett, Sonia Goncalves, Ewan Harrison, David K. Jackson, Ian Johnston, Dominic Kwiatkowski, Cordelia Langford, John Sillitoe on behalf of the Wellcome Sanger Institute COVID-19 Surveillance Team |
| EPI_ISL_3286515 | Lighthouse Lab in Alderley Park | Wellcome Sanger Institute for the COVID-19 Genomics UK (COG-UK) Consortium | Jacquelyn Wynn, Mairead Hyland, The Lighthouse Lab in Alderley Park and Alex Alderton, Roberto Amato, Jeffrey Barrett, Sonia Goncalves, Ewan Harrison, David K. Jackson, Ian Johnston, Dominic Kwiatkowski, Cordelia Langford, John Sillitoe on behalf of the Wellcome Sanger Institute COVID-19 Surveillance Team |
| EPI_ISL_3286516, EPI_ISL_3286517, EPI_ISL_3286518 | Lighthouse Lab in Milton Keynes | Wellcome Sanger Institute for the COVID-19 Genomics UK (COG-UK) Consortium | The Lighthouse Lab in Milton Keynes and Alex Alderton, Roberto Amato, Jeffrey Barrett, Sonia Goncalves, Ewan Harrison, David K. Jackson, Ian Johnston, Dominic Kwiatkowski, Cordelia Langford, John Sillitoe on behalf of the Wellcome Sanger Institute COVID-19 Surveillance Team |
| EPI_ISL_3286519, EPI_ISL_3286520, EPI_ISL_3286521 | Lighthouse Lab in Alderley Park | Wellcome Sanger Institute for the COVID-19 Genomics UK (COG-UK) Consortium | Jacquelyn Wynn, Mairead Hyland, The Lighthouse Lab in Alderley Park and Alex Alderton, Roberto Amato, Jeffrey Barrett, Sonia Goncalves, Ewan Harrison, David K. Jackson, Ian Johnston, Dominic Kwiatkowski, Cordelia Langford, John Sillitoe on behalf of the Wellcome Sanger Institute COVID-19 Surveillance Team |
| EPI_ISL_3286522, EPI_ISL_3286523 | Lighthouse Lab in Milton Keynes | Wellcome Sanger Institute for the COVID-19 Genomics UK (COG-UK) Consortium | The Lighthouse Lab in Milton Keynes and Alex Alderton, Roberto Amato, Jeffrey Barrett, Sonia Goncalves, Ewan Harrison, David K. Jackson, Ian Johnston, Dominic Kwiatkowski, Cordelia Langford, John Sillitoe on behalf of the Wellcome Sanger Institute COVID-19 Surveillance Team |
| EPI_ISL_3286524 | Lighthouse Lab in Alderley Park | Wellcome Sanger Institute for the COVID-19 Genomics UK (COG-UK) Consortium | Jacquelyn Wynn, Mairead Hyland, The Lighthouse Lab in Alderley Park and Alex Alderton, Roberto Amato, Jeffrey Barrett, Sonia Goncalves, Ewan Harrison, David K. Jackson, Ian Johnston, Dominic Kwiatkowski, Cordelia Langford, John Sillitoe on behalf of the Wellcome Sanger Institute COVID-19 Surveillance Team |
| EPI_ISL_3286525 | Lighthouse Lab in Milton Keynes | Wellcome Sanger Institute for the COVID-19 Genomics UK (COG-UK) Consortium | The Lighthouse Lab in Milton Keynes and Alex Alderton, Roberto Amato, Jeffrey Barrett, Sonia Goncalves, Ewan Harrison, David K. Jackson, Ian Johnston, Dominic Kwiatkowski, Cordelia Langford, John Sillitoe on behalf of the Wellcome Sanger Institute COVID-19 Surveillance Team |
| EPI_ISL_3286526 | Lighthouse Lab in Alderley Park | Wellcome Sanger Institute for the COVID-19 Genomics UK (COG-UK) | Jacquelyn Wynn, Mairead Hyland, The Lighthouse Lab in Alderley Park and Alex Alderton, Roberto Amato, Jeffrey Barrett, Sonia Goncalves, Ewan Harrison, David K. Jackson, Ian Johnston, Dominic Kwiatkowski, Cordelia Langford, John Sillitoe on |

| | | | |
|---|---|---|---|
| | | Consortium | behalf of the Wellcome Sanger Institute COVID-19 Surveillance Team |
| EPI_ISL_3286527 | Lighthouse Lab in Milton Keynes | Wellcome Sanger Institute for the COVID-19 Genomics UK (COG-UK) Consortium | The Lighthouse Lab in Milton Keynes and Alex Alderton, Roberto Amato, Jeffrey Barrett, Sonia Goncalves, Ewan Harrison, David K. Jackson, Ian Johnston, Dominic Kwiatkowski, Cordelia Langford, John Sillitoe on behalf of the Wellcome Sanger Institute COVID-19 Surveillance Team |
| EPI_ISL_3286528 | Lighthouse Lab in Alderley Park | Wellcome Sanger Institute for the COVID-19 Genomics UK (COG-UK) Consortium | Jacquelyn Wynn, Mairead Hyland, The Lighthouse Lab in Alderley Park and Alex Alderton, Roberto Amato, Jeffrey Barrett, Sonia Goncalves, Ewan Harrison, David K. Jackson, Ian Johnston, Dominic Kwiatkowski, Cordelia Langford, John Sillitoe on behalf of the Wellcome Sanger Institute COVID-19 Surveillance Team |
| EPI_ISL_3286529, EPI_ISL_3286530, EPI_ISL_3286531, EPI_ISL_3286532, EPI_ISL_3286533, EPI_ISL_3286534, EPI_ISL_3286535, EPI_ISL_3286536, EPI_ISL_3286537, EPI_ISL_3286538, EPI_ISL_3286539, EPI_ISL_3286540, EPI_ISL_3286541 | | | |
| see above | Lighthouse Lab in Milton Keynes | Wellcome Sanger Institute for the COVID-19 Genomics UK (COG-UK) Consortium | The Lighthouse Lab in Milton Keynes and Alex Alderton, Roberto Amato, Jeffrey Barrett, Sonia Goncalves, Ewan Harrison, David K. Jackson, Ian Johnston, Dominic Kwiatkowski, Cordelia Langford, John Sillitoe on behalf of the Wellcome Sanger Institute COVID-19 Surveillance Team |
| EPI_ISL_3286542, EPI_ISL_3286543, EPI_ISL_3286544, EPI_ISL_3286545 | Lighthouse Lab in Alderley Park | Wellcome Sanger Institute for the COVID-19 Genomics UK (COG-UK) Consortium | Jacquelyn Wynn, Mairead Hyland, The Lighthouse Lab in Alderley Park and Alex Alderton, Roberto Amato, Jeffrey Barrett, Sonia Goncalves, Ewan Harrison, David K. Jackson, Ian Johnston, Dominic Kwiatkowski, Cordelia Langford, John Sillitoe on behalf of the Wellcome Sanger Institute COVID-19 Surveillance Team |
| EPI_ISL_3286546 | Lighthouse Lab in Milton Keynes | Wellcome Sanger Institute for the COVID-19 Genomics UK (COG-UK) Consortium | The Lighthouse Lab in Milton Keynes and Alex Alderton, Roberto Amato, Jeffrey Barrett, Sonia Goncalves, Ewan Harrison, David K. Jackson, Ian Johnston, Dominic Kwiatkowski, Cordelia Langford, John Sillitoe on behalf of the Wellcome Sanger Institute COVID-19 Surveillance Team |
| EPI_ISL_3286547 | Lighthouse Lab in Alderley Park | Wellcome Sanger Institute for the COVID-19 Genomics UK (COG-UK) Consortium | Jacquelyn Wynn, Mairead Hyland, The Lighthouse Lab in Alderley Park and Alex Alderton, Roberto Amato, Jeffrey Barrett, Sonia Goncalves, Ewan Harrison, David K. Jackson, Ian Johnston, Dominic Kwiatkowski, Cordelia Langford, John Sillitoe on behalf of the Wellcome Sanger Institute COVID-19 Surveillance Team |
| EPI_ISL_3286548 | Lighthouse Lab in Milton Keynes | Wellcome Sanger Institute for the COVID-19 Genomics UK (COG-UK) Consortium | The Lighthouse Lab in Milton Keynes and Alex Alderton, Roberto Amato, Jeffrey Barrett, Sonia Goncalves, Ewan Harrison, David K. Jackson, Ian Johnston, Dominic Kwiatkowski, Cordelia Langford, John Sillitoe on behalf of the Wellcome Sanger Institute COVID-19 Surveillance Team |
| EPI_ISL_3286549, EPI_ISL_3286550 | Lighthouse Lab in Alderley Park | Wellcome Sanger Institute for the COVID-19 Genomics UK (COG-UK) Consortium | Jacquelyn Wynn, Mairead Hyland, The Lighthouse Lab in Alderley Park and Alex Alderton, Roberto Amato, Jeffrey Barrett, Sonia Goncalves, Ewan Harrison, David K. Jackson, Ian Johnston, Dominic Kwiatkowski, Cordelia Langford, John Sillitoe on behalf of the Wellcome Sanger Institute COVID-19 Surveillance Team |
| EPI_ISL_3286551 | Lighthouse Lab in Milton Keynes | Wellcome Sanger Institute for the COVID-19 Genomics UK (COG-UK) Consortium | The Lighthouse Lab in Milton Keynes and Alex Alderton, Roberto Amato, Jeffrey Barrett, Sonia Goncalves, Ewan Harrison, David K. Jackson, Ian Johnston, Dominic Kwiatkowski, Cordelia Langford, John Sillitoe on behalf of the Wellcome Sanger Institute COVID-19 Surveillance Team |
| EPI_ISL_3286552, EPI_ISL_3286553, EPI_ISL_3286554, EPI_ISL_3286555 | Lighthouse Lab in Alderley Park | Wellcome Sanger Institute for the COVID-19 Genomics UK (COG-UK) Consortium | Jacquelyn Wynn, Mairead Hyland, The Lighthouse Lab in Alderley Park and Alex Alderton, Roberto Amato, Jeffrey Barrett, Sonia Goncalves, Ewan Harrison, David K. Jackson, Ian Johnston, Dominic Kwiatkowski, Cordelia Langford, John Sillitoe on behalf of the Wellcome Sanger Institute COVID-19 Surveillance Team |
| EPI_ISL_3286556, EPI_ISL_3286557, EPI_ISL_3286558 | Lighthouse Lab in Milton Keynes | Wellcome Sanger Institute for the COVID-19 Genomics UK (COG-UK) Consortium | The Lighthouse Lab in Milton Keynes and Alex Alderton, Roberto Amato, Jeffrey Barrett, Sonia Goncalves, Ewan Harrison, David K. Jackson, Ian Johnston, Dominic Kwiatkowski, Cordelia Langford, John Sillitoe on behalf of the Wellcome Sanger Institute COVID-19 Surveillance Team |
| EPI_ISL_3286559, EPI_ISL_3286560 | Lighthouse Lab in Alderley Park | Wellcome Sanger Institute for the COVID-19 Genomics UK (COG-UK) Consortium | Jacquelyn Wynn, Mairead Hyland, The Lighthouse Lab in Alderley Park and Alex Alderton, Roberto Amato, Jeffrey Barrett, Sonia Goncalves, Ewan Harrison, David K. Jackson, Ian Johnston, Dominic Kwiatkowski, Cordelia Langford, John Sillitoe on behalf of the Wellcome Sanger Institute COVID-19 Surveillance Team |
| EPI_ISL_3286561, EPI_ISL_3286562, EPI_ISL_3286563, EPI_ISL_3286564 | Lighthouse Lab in Milton Keynes | Wellcome Sanger Institute for the COVID-19 Genomics UK (COG-UK) Consortium | The Lighthouse Lab in Milton Keynes and Alex Alderton, Roberto Amato, Jeffrey Barrett, Sonia Goncalves, Ewan Harrison, David K. Jackson, Ian Johnston, Dominic Kwiatkowski, Cordelia Langford, John Sillitoe on behalf of the Wellcome Sanger Institute COVID-19 Surveillance Team |
| EPI_ISL_3286565 | Lighthouse Lab in Alderley Park | Wellcome Sanger Institute for the COVID-19 Genomics UK (COG-UK) Consortium | Jacquelyn Wynn, Mairead Hyland, The Lighthouse Lab in Alderley Park and Alex Alderton, Roberto Amato, Jeffrey Barrett, Sonia Goncalves, Ewan Harrison, David K. Jackson, Ian Johnston, Dominic Kwiatkowski, Cordelia Langford, John Sillitoe on behalf of the Wellcome Sanger Institute COVID-19 Surveillance Team |
| EPI_ISL_3286566 | Lighthouse Lab in Milton Keynes | Wellcome Sanger Institute for the COVID-19 Genomics UK (COG-UK) Consortium | The Lighthouse Lab in Milton Keynes and Alex Alderton, Roberto Amato, Jeffrey Barrett, Sonia Goncalves, Ewan Harrison, David K. Jackson, Ian Johnston, Dominic Kwiatkowski, Cordelia Langford, John Sillitoe on behalf of the Wellcome Sanger Institute COVID-19 Surveillance Team |
| EPI_ISL_3286567 | Lighthouse Lab in Alderley Park | Wellcome Sanger Institute for the COVID-19 Genomics UK (COG-UK) Consortium | Jacquelyn Wynn, Mairead Hyland, The Lighthouse Lab in Alderley Park and Alex Alderton, Roberto Amato, Jeffrey Barrett, Sonia Goncalves, Ewan Harrison, David K. Jackson, Ian Johnston, Dominic Kwiatkowski, Cordelia Langford, John Sillitoe on behalf of the Wellcome Sanger Institute COVID-19 Surveillance Team |
| EPI_ISL_3286568, EPI_ISL_3286569 | Lighthouse Lab in Milton Keynes | Wellcome Sanger Institute for the COVID-19 Genomics UK (COG-UK) Consortium | The Lighthouse Lab in Milton Keynes and Alex Alderton, Roberto Amato, Jeffrey Barrett, Sonia Goncalves, Ewan Harrison, David K. Jackson, Ian Johnston, Dominic Kwiatkowski, Cordelia Langford, John Sillitoe on behalf of the Wellcome Sanger Institute COVID-19 Surveillance Team |
| EPI_ISL_3286570 | Lighthouse Lab in Alderley Park | Wellcome Sanger Institute for the COVID-19 Genomics UK (COG-UK) Consortium | Jacquelyn Wynn, Mairead Hyland, The Lighthouse Lab in Alderley Park and Alex Alderton, Roberto Amato, Jeffrey Barrett, Sonia Goncalves, Ewan Harrison, David K. Jackson, Ian Johnston, Dominic Kwiatkowski, Cordelia Langford, John Sillitoe on behalf of the Wellcome Sanger Institute COVID-19 Surveillance Team |
| EPI_ISL_3286571, EPI_ISL_3286572 | Lighthouse Lab in Milton Keynes | Wellcome Sanger Institute for the COVID-19 Genomics UK (COG-UK) Consortium | The Lighthouse Lab in Milton Keynes and Alex Alderton, Roberto Amato, Jeffrey Barrett, Sonia Goncalves, Ewan Harrison, David K. Jackson, Ian Johnston, Dominic Kwiatkowski, Cordelia Langford, John Sillitoe on behalf of the Wellcome Sanger Institute COVID-19 Surveillance Team |
| EPI_ISL_3286573, EPI_ISL_3286574 | Lighthouse Lab in Alderley Park | Wellcome Sanger Institute for the COVID-19 Genomics UK (COG-UK) Consortium | Jacquelyn Wynn, Mairead Hyland, The Lighthouse Lab in Alderley Park and Alex Alderton, Roberto Amato, Jeffrey Barrett, Sonia Goncalves, Ewan Harrison, David K. Jackson, Ian Johnston, Dominic Kwiatkowski, Cordelia Langford, John Sillitoe on behalf of the Wellcome Sanger Institute COVID-19 Surveillance Team |
| EPI_ISL_3286575 | Lighthouse Lab in Milton Keynes | Wellcome Sanger Institute for the COVID-19 Genomics UK (COG-UK) Consortium | The Lighthouse Lab in Milton Keynes and Alex Alderton, Roberto Amato, Jeffrey Barrett, Sonia Goncalves, Ewan Harrison, David K. Jackson, Ian Johnston, Dominic Kwiatkowski, Cordelia Langford, John Sillitoe on behalf of the Wellcome Sanger Institute COVID-19 Surveillance Team |
| EPI_ISL_3286576 | Lighthouse Lab in Alderley Park | Wellcome Sanger Institute for the COVID-19 Genomics UK (COG-UK) Consortium | Jacquelyn Wynn, Mairead Hyland, The Lighthouse Lab in Alderley Park and Alex Alderton, Roberto Amato, Jeffrey Barrett, Sonia Goncalves, Ewan Harrison, David K. Jackson, Ian Johnston, Dominic Kwiatkowski, Cordelia Langford, John Sillitoe on behalf of the Wellcome Sanger Institute COVID-19 Surveillance Team |
| EPI_ISL_3286577 | Lighthouse Lab in Milton Keynes | Wellcome Sanger Institute for the COVID-19 Genomics UK (COG-UK) Consortium | The Lighthouse Lab in Milton Keynes and Alex Alderton, Roberto Amato, Jeffrey Barrett, Sonia Goncalves, Ewan Harrison, David K. Jackson, Ian Johnston, Dominic Kwiatkowski, Cordelia Langford, John Sillitoe on behalf of the Wellcome Sanger Institute COVID-19 Surveillance Team |
| EPI_ISL_3286578 | Lighthouse Lab in Alderley Park | Wellcome Sanger Institute for the COVID-19 Genomics UK (COG-UK) Consortium | Jacquelyn Wynn, Mairead Hyland, The Lighthouse Lab in Alderley Park and Alex Alderton, Roberto Amato, Jeffrey Barrett, Sonia Goncalves, Ewan Harrison, David K. Jackson, Ian Johnston, Dominic Kwiatkowski, Cordelia Langford, John Sillitoe on behalf of the Wellcome Sanger Institute COVID-19 Surveillance Team |
| EPI_ISL_3286579, EPI_ISL_3286580 | Lighthouse Lab in Milton Keynes | Wellcome Sanger Institute for the COVID-19 Genomics UK (COG-UK) Consortium | The Lighthouse Lab in Milton Keynes and Alex Alderton, Roberto Amato, Jeffrey Barrett, Sonia Goncalves, Ewan Harrison, David K. Jackson, Ian Johnston, Dominic Kwiatkowski, Cordelia Langford, John Sillitoe on behalf of the Wellcome Sanger Institute COVID-19 Surveillance Team |
| EPI_ISL_3286581 | Lighthouse Lab in Alderley Park | Wellcome Sanger Institute for the COVID-19 Genomics UK (COG-UK) Consortium | Jacquelyn Wynn, Mairead Hyland, The Lighthouse Lab in Alderley Park and Alex Alderton, Roberto Amato, Jeffrey Barrett, Sonia Goncalves, Ewan Harrison, David K. Jackson, Ian Johnston, Dominic Kwiatkowski, Cordelia Langford, John Sillitoe on behalf of the Wellcome Sanger Institute COVID-19 Surveillance Team |
| EPI_ISL_3286582, EPI_ISL_3286583 | Lighthouse Lab in Milton Keynes | Wellcome Sanger Institute for the COVID-19 Genomics UK (COG-UK) Consortium | The Lighthouse Lab in Milton Keynes and Alex Alderton, Roberto Amato, Jeffrey Barrett, Sonia Goncalves, Ewan Harrison, David K. Jackson, Ian Johnston, Dominic Kwiatkowski, Cordelia Langford, John Sillitoe on behalf of the Wellcome Sanger Institute COVID-19 Surveillance Team |
| EPI_ISL_3286584 | Lighthouse Lab in Alderley Park | Wellcome Sanger Institute for the COVID-19 Genomics UK (COG-UK) Consortium | Jacquelyn Wynn, Mairead Hyland, The Lighthouse Lab in Alderley Park and Alex Alderton, Roberto Amato, Jeffrey Barrett, Sonia Goncalves, Ewan Harrison, David K. Jackson, Ian Johnston, Dominic Kwiatkowski, Cordelia Langford, John Sillitoe on behalf of the Wellcome Sanger Institute COVID-19 Surveillance Team |
| EPI_ISL_3286585, EPI_ISL_3286586 | Lighthouse Lab in Milton Keynes | Wellcome Sanger Institute for the COVID-19 Genomics UK (COG-UK) Consortium | The Lighthouse Lab in Milton Keynes and Alex Alderton, Roberto Amato, Jeffrey Barrett, Sonia Goncalves, Ewan Harrison, David K. Jackson, Ian Johnston, Dominic Kwiatkowski, Cordelia Langford, John Sillitoe on behalf of the Wellcome Sanger Institute COVID-19 Surveillance Team |
| EPI_ISL_3286587, EPI_ISL_3286588, EPI_ISL_3286589, EPI_ISL_3286590 | Lighthouse Lab in Alderley Park | Wellcome Sanger Institute for the COVID-19 Genomics UK (COG-UK) Consortium | Jacquelyn Wynn, Mairead Hyland, The Lighthouse Lab in Alderley Park and Alex Alderton, Roberto Amato, Jeffrey Barrett, Sonia Goncalves, Ewan Harrison, David K. Jackson, Ian Johnston, Dominic Kwiatkowski, Cordelia Langford, John Sillitoe on behalf of the Wellcome Sanger Institute COVID-19 Surveillance Team |
| EPI_ISL_3286591, EPI_ISL_3286592 | Lighthouse Lab in Milton Keynes | Wellcome Sanger Institute for the COVID-19 Genomics UK (COG-UK) Consortium | The Lighthouse Lab in Milton Keynes and Alex Alderton, Roberto Amato, Jeffrey Barrett, Sonia Goncalves, Ewan Harrison, David K. Jackson, Ian Johnston, Dominic Kwiatkowski, Cordelia Langford, John Sillitoe on behalf of the Wellcome Sanger Institute COVID-19 Surveillance Team |
| EPI_ISL_3286593, EPI_ISL_3286594, EPI_ISL_3286595, EPI_ISL_3286596, EPI_ISL_3286597, EPI_ISL_3286598, EPI_ISL_3286599, EPI_ISL_3286600 | Lighthouse Lab in Alderley Park | Wellcome Sanger Institute for the COVID-19 Genomics UK (COG-UK) Consortium | Jacquelyn Wynn, Mairead Hyland, The Lighthouse Lab in Alderley Park and Alex Alderton, Roberto Amato, Jeffrey Barrett, Sonia Goncalves, Ewan Harrison, David K. Jackson, Ian Johnston, Dominic Kwiatkowski, Cordelia Langford, John Sillitoe on behalf of the Wellcome Sanger Institute COVID-19 Surveillance Team |
| EPI_ISL_3286601 | Lighthouse Lab in Milton Keynes | Wellcome Sanger Institute for the COVID-19 Genomics UK (COG-UK) Consortium | The Lighthouse Lab in Milton Keynes and Alex Alderton, Roberto Amato, Jeffrey Barrett, Sonia Goncalves, Ewan Harrison, David K. Jackson, Ian Johnston, Dominic Kwiatkowski, Cordelia Langford, John Sillitoe on behalf of the Wellcome Sanger Institute COVID-19 Surveillance Team |
| EPI_ISL_3286602 | Lighthouse Lab in Alderley Park | Wellcome Sanger Institute for the COVID-19 Genomics UK (COG-UK) Consortium | Jacquelyn Wynn, Mairead Hyland, The Lighthouse Lab in Alderley Park and Alex Alderton, Roberto Amato, Jeffrey Barrett, Sonia Goncalves, Ewan Harrison, David K. Jackson, Ian Johnston, Dominic Kwiatkowski, Cordelia Langford, John Sillitoe on behalf of the Wellcome Sanger Institute COVID-19 Surveillance Team |
| EPI_ISL_3286603 | Lighthouse Lab in Milton Keynes | Wellcome Sanger Institute for the COVID-19 Genomics UK (COG-UK) Consortium | The Lighthouse Lab in Milton Keynes and Alex Alderton, Roberto Amato, Jeffrey Barrett, Sonia Goncalves, Ewan Harrison, David K. Jackson, Ian Johnston, Dominic Kwiatkowski, Cordelia Langford, John Sillitoe on behalf of the Wellcome Sanger Institute COVID-19 Surveillance Team |
| EPI_ISL_3286604 | Lighthouse Lab in Alderley Park | Wellcome Sanger Institute for the COVID-19 Genomics UK (COG-UK) Consortium | Jacquelyn Wynn, Mairead Hyland, The Lighthouse Lab in Alderley Park and Alex Alderton, Roberto Amato, Jeffrey Barrett, Sonia Goncalves, Ewan Harrison, David K. Jackson, Ian Johnston, Dominic Kwiatkowski, Cordelia Langford, John Sillitoe on behalf of the Wellcome Sanger Institute COVID-19 Surveillance Team |
| EPI_ISL_3286605, EPI_ISL_3286606 | Lighthouse Lab in Milton Keynes | Wellcome Sanger Institute for the COVID-19 Genomics UK (COG-UK) Consortium | The Lighthouse Lab in Milton Keynes and Alex Alderton, Roberto Amato, Jeffrey Barrett, Sonia Goncalves, Ewan Harrison, David K. Jackson, Ian Johnston, Dominic Kwiatkowski, Cordelia Langford, John Sillitoe on behalf of the Wellcome Sanger Institute COVID-19 Surveillance Team |
| EPI_ISL_3286607, | Lighthouse Lab in Alderley Park | Wellcome Sanger Institute for the COVID-19 Genomics UK (COG-UK) | Jacquelyn Wynn, Mairead Hyland, The Lighthouse Lab in Alderley Park and Alex Alderton, Roberto Amato, Jeffrey Barrett, Sonia Goncalves, Ewan Harrison, David K. Jackson, Ian Johnston, Dominic Kwiatkowski, Cordelia Langford, John Sillitoe on |

| | | | |
|---|---|---|---|
| EPI_ISL_3286608, EPI_ISL_3286609 | | Consortium | behalf of the Wellcome Sanger Institute COVID-19 Surveillance Team |
| EPI_ISL_3286610 | Lighthouse Lab in Milton Keynes | Wellcome Sanger Institute for the COVID-19 Genomics UK (COG-UK) Consortium | The Lighthouse Lab in Milton Keynes and Alex Alderton, Roberto Amato, Jeffrey Barrett, Sonia Goncalves, Ewan Harrison, David K. Jackson, Ian Johnston, Dominic Kwiatkowski, Cordelia Langford, John Sillitoe on behalf of the Wellcome Sanger Institute COVID-19 Surveillance Team |
| EPI_ISL_3286611 | Lighthouse Lab in Alderley Park | Wellcome Sanger Institute for the COVID-19 Genomics UK (COG-UK) Consortium | Jacquelyn Wynn, Mairead Hyland, The Lighthouse Lab in Alderley Park and Alex Alderton, Roberto Amato, Jeffrey Barrett, Sonia Goncalves, Ewan Harrison, David K. Jackson, Ian Johnston, Dominic Kwiatkowski, Cordelia Langford, John Sillitoe on behalf of the Wellcome Sanger Institute COVID-19 Surveillance Team |
| EPI_ISL_3286612 | Lighthouse Lab in Milton Keynes | Wellcome Sanger Institute for the COVID-19 Genomics UK (COG-UK) Consortium | The Lighthouse Lab in Milton Keynes and Alex Alderton, Roberto Amato, Jeffrey Barrett, Sonia Goncalves, Ewan Harrison, David K. Jackson, Ian Johnston, Dominic Kwiatkowski, Cordelia Langford, John Sillitoe on behalf of the Wellcome Sanger Institute COVID-19 Surveillance Team |
| EPI_ISL_3286613 | Lighthouse Lab in Alderley Park | Wellcome Sanger Institute for the COVID-19 Genomics UK (COG-UK) Consortium | Jacquelyn Wynn, Mairead Hyland, The Lighthouse Lab in Alderley Park and Alex Alderton, Roberto Amato, Jeffrey Barrett, Sonia Goncalves, Ewan Harrison, David K. Jackson, Ian Johnston, Dominic Kwiatkowski, Cordelia Langford, John Sillitoe on behalf of the Wellcome Sanger Institute COVID-19 Surveillance Team |
| EPI_ISL_3286614 | Lighthouse Lab in Milton Keynes | Wellcome Sanger Institute for the COVID-19 Genomics UK (COG-UK) Consortium | The Lighthouse Lab in Milton Keynes and Alex Alderton, Roberto Amato, Jeffrey Barrett, Sonia Goncalves, Ewan Harrison, David K. Jackson, Ian Johnston, Dominic Kwiatkowski, Cordelia Langford, John Sillitoe on behalf of the Wellcome Sanger Institute COVID-19 Surveillance Team |
| EPI_ISL_3286615 | Lighthouse Lab in Alderley Park | Wellcome Sanger Institute for the COVID-19 Genomics UK (COG-UK) Consortium | Jacquelyn Wynn, Mairead Hyland, The Lighthouse Lab in Alderley Park and Alex Alderton, Roberto Amato, Jeffrey Barrett, Sonia Goncalves, Ewan Harrison, David K. Jackson, Ian Johnston, Dominic Kwiatkowski, Cordelia Langford, John Sillitoe on behalf of the Wellcome Sanger Institute COVID-19 Surveillance Team |
| EPI_ISL_3286616 | Lighthouse Lab in Milton Keynes | Wellcome Sanger Institute for the COVID-19 Genomics UK (COG-UK) Consortium | The Lighthouse Lab in Milton Keynes and Alex Alderton, Roberto Amato, Jeffrey Barrett, Sonia Goncalves, Ewan Harrison, David K. Jackson, Ian Johnston, Dominic Kwiatkowski, Cordelia Langford, John Sillitoe on behalf of the Wellcome Sanger Institute COVID-19 Surveillance Team |
| EPI_ISL_3286617, EPI_ISL_3286618, EPI_ISL_3286619 | Lighthouse Lab in Alderley Park | Wellcome Sanger Institute for the COVID-19 Genomics UK (COG-UK) Consortium | Jacquelyn Wynn, Mairead Hyland, The Lighthouse Lab in Alderley Park and Alex Alderton, Roberto Amato, Jeffrey Barrett, Sonia Goncalves, Ewan Harrison, David K. Jackson, Ian Johnston, Dominic Kwiatkowski, Cordelia Langford, John Sillitoe on behalf of the Wellcome Sanger Institute COVID-19 Surveillance Team |
| EPI_ISL_3286620, EPI_ISL_3286621 | Lighthouse Lab in Milton Keynes | Wellcome Sanger Institute for the COVID-19 Genomics UK (COG-UK) Consortium | The Lighthouse Lab in Milton Keynes and Alex Alderton, Roberto Amato, Jeffrey Barrett, Sonia Goncalves, Ewan Harrison, David K. Jackson, Ian Johnston, Dominic Kwiatkowski, Cordelia Langford, John Sillitoe on behalf of the Wellcome Sanger Institute COVID-19 Surveillance Team |
| EPI_ISL_3286622 | Lighthouse Lab in Alderley Park | Wellcome Sanger Institute for the COVID-19 Genomics UK (COG-UK) Consortium | Jacquelyn Wynn, Mairead Hyland, The Lighthouse Lab in Alderley Park and Alex Alderton, Roberto Amato, Jeffrey Barrett, Sonia Goncalves, Ewan Harrison, David K. Jackson, Ian Johnston, Dominic Kwiatkowski, Cordelia Langford, John Sillitoe on behalf of the Wellcome Sanger Institute COVID-19 Surveillance Team |
| EPI_ISL_3286623, EPI_ISL_3286624, EPI_ISL_3286625 | Lighthouse Lab in Milton Keynes | Wellcome Sanger Institute for the COVID-19 Genomics UK (COG-UK) Consortium | The Lighthouse Lab in Milton Keynes and Alex Alderton, Roberto Amato, Jeffrey Barrett, Sonia Goncalves, Ewan Harrison, David K. Jackson, Ian Johnston, Dominic Kwiatkowski, Cordelia Langford, John Sillitoe on behalf of the Wellcome Sanger Institute COVID-19 Surveillance Team |
| EPI_ISL_3286626, EPI_ISL_3286627, EPI_ISL_3286628 | Lighthouse Lab in Alderley Park | Wellcome Sanger Institute for the COVID-19 Genomics UK (COG-UK) Consortium | Jacquelyn Wynn, Mairead Hyland, The Lighthouse Lab in Alderley Park and Alex Alderton, Roberto Amato, Jeffrey Barrett, Sonia Goncalves, Ewan Harrison, David K. Jackson, Ian Johnston, Dominic Kwiatkowski, Cordelia Langford, John Sillitoe on behalf of the Wellcome Sanger Institute COVID-19 Surveillance Team |
| EPI_ISL_3286629, EPI_ISL_3286630, EPI_ISL_3286631 | Lighthouse Lab in Milton Keynes | Wellcome Sanger Institute for the COVID-19 Genomics UK (COG-UK) Consortium | The Lighthouse Lab in Milton Keynes and Alex Alderton, Roberto Amato, Jeffrey Barrett, Sonia Goncalves, Ewan Harrison, David K. Jackson, Ian Johnston, Dominic Kwiatkowski, Cordelia Langford, John Sillitoe on behalf of the Wellcome Sanger Institute COVID-19 Surveillance Team |
| EPI_ISL_3286632 | Lighthouse Lab in Alderley Park | Wellcome Sanger Institute for the COVID-19 Genomics UK (COG-UK) Consortium | Jacquelyn Wynn, Mairead Hyland, The Lighthouse Lab in Alderley Park and Alex Alderton, Roberto Amato, Jeffrey Barrett, Sonia Goncalves, Ewan Harrison, David K. Jackson, Ian Johnston, Dominic Kwiatkowski, Cordelia Langford, John Sillitoe on behalf of the Wellcome Sanger Institute COVID-19 Surveillance Team |
| EPI_ISL_3286633 | Lighthouse Lab in Milton Keynes | Wellcome Sanger Institute for the COVID-19 Genomics UK (COG-UK) Consortium | The Lighthouse Lab in Milton Keynes and Alex Alderton, Roberto Amato, Jeffrey Barrett, Sonia Goncalves, Ewan Harrison, David K. Jackson, Ian Johnston, Dominic Kwiatkowski, Cordelia Langford, John Sillitoe on behalf of the Wellcome Sanger Institute COVID-19 Surveillance Team |
| EPI_ISL_3286634 | Lighthouse Lab in Alderley Park | Wellcome Sanger Institute for the COVID-19 Genomics UK (COG-UK) Consortium | Jacquelyn Wynn, Mairead Hyland, The Lighthouse Lab in Alderley Park and Alex Alderton, Roberto Amato, Jeffrey Barrett, Sonia Goncalves, Ewan Harrison, David K. Jackson, Ian Johnston, Dominic Kwiatkowski, Cordelia Langford, John Sillitoe on behalf of the Wellcome Sanger Institute COVID-19 Surveillance Team |
| EPI_ISL_3286635 | Lighthouse Lab in Milton Keynes | Wellcome Sanger Institute for the COVID-19 Genomics UK (COG-UK) Consortium | The Lighthouse Lab in Milton Keynes and Alex Alderton, Roberto Amato, Jeffrey Barrett, Sonia Goncalves, Ewan Harrison, David K. Jackson, Ian Johnston, Dominic Kwiatkowski, Cordelia Langford, John Sillitoe on behalf of the Wellcome Sanger Institute COVID-19 Surveillance Team |
| EPI_ISL_3286636 | Lighthouse Lab in Alderley Park | Wellcome Sanger Institute for the COVID-19 Genomics UK (COG-UK) Consortium | Jacquelyn Wynn, Mairead Hyland, The Lighthouse Lab in Alderley Park and Alex Alderton, Roberto Amato, Jeffrey Barrett, Sonia Goncalves, Ewan Harrison, David K. Jackson, Ian Johnston, Dominic Kwiatkowski, Cordelia Langford, John Sillitoe on behalf of the Wellcome Sanger Institute COVID-19 Surveillance Team |
| EPI_ISL_3286637, EPI_ISL_3286638, EPI_ISL_3286639 | Lighthouse Lab in Milton Keynes | Wellcome Sanger Institute for the COVID-19 Genomics UK (COG-UK) Consortium | The Lighthouse Lab in Milton Keynes and Alex Alderton, Roberto Amato, Jeffrey Barrett, Sonia Goncalves, Ewan Harrison, David K. Jackson, Ian Johnston, Dominic Kwiatkowski, Cordelia Langford, John Sillitoe on behalf of the Wellcome Sanger Institute COVID-19 Surveillance Team |
| EPI_ISL_3286640, EPI_ISL_3286641, EPI_ISL_3286642, EPI_ISL_3286643 | Lighthouse Lab in Alderley Park | Wellcome Sanger Institute for the COVID-19 Genomics UK (COG-UK) Consortium | Jacquelyn Wynn, Mairead Hyland, The Lighthouse Lab in Alderley Park and Alex Alderton, Roberto Amato, Jeffrey Barrett, Sonia Goncalves, Ewan Harrison, David K. Jackson, Ian Johnston, Dominic Kwiatkowski, Cordelia Langford, John Sillitoe on behalf of the Wellcome Sanger Institute COVID-19 Surveillance Team |
| EPI_ISL_3286644, EPI_ISL_3286645 | Lighthouse Lab in Milton Keynes | Wellcome Sanger Institute for the COVID-19 Genomics UK (COG-UK) Consortium | The Lighthouse Lab in Milton Keynes and Alex Alderton, Roberto Amato, Jeffrey Barrett, Sonia Goncalves, Ewan Harrison, David K. Jackson, Ian Johnston, Dominic Kwiatkowski, Cordelia Langford, John Sillitoe on behalf of the Wellcome Sanger Institute COVID-19 Surveillance Team |
| EPI_ISL_3286646 | Lighthouse Lab in Alderley Park | Wellcome Sanger Institute for the COVID-19 Genomics UK (COG-UK) Consortium | Jacquelyn Wynn, Mairead Hyland, The Lighthouse Lab in Alderley Park and Alex Alderton, Roberto Amato, Jeffrey Barrett, Sonia Goncalves, Ewan Harrison, David K. Jackson, Ian Johnston, Dominic Kwiatkowski, Cordelia Langford, John Sillitoe on behalf of the Wellcome Sanger Institute COVID-19 Surveillance Team |
| EPI_ISL_3286647, EPI_ISL_3286648 | Lighthouse Lab in Milton Keynes | Wellcome Sanger Institute for the COVID-19 Genomics UK (COG-UK) Consortium | The Lighthouse Lab in Milton Keynes and Alex Alderton, Roberto Amato, Jeffrey Barrett, Sonia Goncalves, Ewan Harrison, David K. Jackson, Ian Johnston, Dominic Kwiatkowski, Cordelia Langford, John Sillitoe on behalf of the Wellcome Sanger Institute COVID-19 Surveillance Team |
| EPI_ISL_3286649 | Lighthouse Lab in Alderley Park | Wellcome Sanger Institute for the COVID-19 Genomics UK (COG-UK) Consortium | Jacquelyn Wynn, Mairead Hyland, The Lighthouse Lab in Alderley Park and Alex Alderton, Roberto Amato, Jeffrey Barrett, Sonia Goncalves, Ewan Harrison, David K. Jackson, Ian Johnston, Dominic Kwiatkowski, Cordelia Langford, John Sillitoe on behalf of the Wellcome Sanger Institute COVID-19 Surveillance Team |
| EPI_ISL_3286650, EPI_ISL_3286651 | Lighthouse Lab in Milton Keynes | Wellcome Sanger Institute for the COVID-19 Genomics UK (COG-UK) Consortium | The Lighthouse Lab in Milton Keynes and Alex Alderton, Roberto Amato, Jeffrey Barrett, Sonia Goncalves, Ewan Harrison, David K. Jackson, Ian Johnston, Dominic Kwiatkowski, Cordelia Langford, John Sillitoe on behalf of the Wellcome Sanger Institute COVID-19 Surveillance Team |
| EPI_ISL_3286652 | Lighthouse Lab in Alderley Park | Wellcome Sanger Institute for the COVID-19 Genomics UK (COG-UK) Consortium | Jacquelyn Wynn, Mairead Hyland, The Lighthouse Lab in Alderley Park and Alex Alderton, Roberto Amato, Jeffrey Barrett, Sonia Goncalves, Ewan Harrison, David K. Jackson, Ian Johnston, Dominic Kwiatkowski, Cordelia Langford, John Sillitoe on behalf of the Wellcome Sanger Institute COVID-19 Surveillance Team |
| EPI_ISL_3286653, EPI_ISL_3286654, EPI_ISL_3286655 | Lighthouse Lab in Milton Keynes | Wellcome Sanger Institute for the COVID-19 Genomics UK (COG-UK) Consortium | The Lighthouse Lab in Milton Keynes and Alex Alderton, Roberto Amato, Jeffrey Barrett, Sonia Goncalves, Ewan Harrison, David K. Jackson, Ian Johnston, Dominic Kwiatkowski, Cordelia Langford, John Sillitoe on behalf of the Wellcome Sanger Institute COVID-19 Surveillance Team |
| EPI_ISL_3286656, EPI_ISL_3286657, EPI_ISL_3286658, EPI_ISL_3286659 | Lighthouse Lab in Alderley Park | Wellcome Sanger Institute for the COVID-19 Genomics UK (COG-UK) Consortium | Jacquelyn Wynn, Mairead Hyland, The Lighthouse Lab in Alderley Park and Alex Alderton, Roberto Amato, Jeffrey Barrett, Sonia Goncalves, Ewan Harrison, David K. Jackson, Ian Johnston, Dominic Kwiatkowski, Cordelia Langford, John Sillitoe on behalf of the Wellcome Sanger Institute COVID-19 Surveillance Team |
| EPI_ISL_3286660 | Lighthouse Lab in Milton Keynes | Wellcome Sanger Institute for the COVID-19 Genomics UK (COG-UK) Consortium | The Lighthouse Lab in Milton Keynes and Alex Alderton, Roberto Amato, Jeffrey Barrett, Sonia Goncalves, Ewan Harrison, David K. Jackson, Ian Johnston, Dominic Kwiatkowski, Cordelia Langford, John Sillitoe on behalf of the Wellcome Sanger Institute COVID-19 Surveillance Team |
| EPI_ISL_3286661, EPI_ISL_3286662 | Lighthouse Lab in Alderley Park | Wellcome Sanger Institute for the COVID-19 Genomics UK (COG-UK) Consortium | Jacquelyn Wynn, Mairead Hyland, The Lighthouse Lab in Alderley Park and Alex Alderton, Roberto Amato, Jeffrey Barrett, Sonia Goncalves, Ewan Harrison, David K. Jackson, Ian Johnston, Dominic Kwiatkowski, Cordelia Langford, John Sillitoe on behalf of the Wellcome Sanger Institute COVID-19 Surveillance Team |
| EPI_ISL_3286663 | Lighthouse Lab in Milton Keynes | Wellcome Sanger Institute for the COVID-19 Genomics UK (COG-UK) Consortium | The Lighthouse Lab in Milton Keynes and Alex Alderton, Roberto Amato, Jeffrey Barrett, Sonia Goncalves, Ewan Harrison, David K. Jackson, Ian Johnston, Dominic Kwiatkowski, Cordelia Langford, John Sillitoe on behalf of the Wellcome Sanger Institute COVID-19 Surveillance Team |
| EPI_ISL_3286664, EPI_ISL_3286665 | Lighthouse Lab in Alderley Park | Wellcome Sanger Institute for the COVID-19 Genomics UK (COG-UK) Consortium | Jacquelyn Wynn, Mairead Hyland, The Lighthouse Lab in Alderley Park and Alex Alderton, Roberto Amato, Jeffrey Barrett, Sonia Goncalves, Ewan Harrison, David K. Jackson, Ian Johnston, Dominic Kwiatkowski, Cordelia Langford, John Sillitoe on behalf of the Wellcome Sanger Institute COVID-19 Surveillance Team |
| EPI_ISL_3286666 | Lighthouse Lab in Milton Keynes | Wellcome Sanger Institute for the COVID-19 Genomics UK (COG-UK) Consortium | The Lighthouse Lab in Milton Keynes and Alex Alderton, Roberto Amato, Jeffrey Barrett, Sonia Goncalves, Ewan Harrison, David K. Jackson, Ian Johnston, Dominic Kwiatkowski, Cordelia Langford, John Sillitoe on behalf of the Wellcome Sanger Institute COVID-19 Surveillance Team |
| EPI_ISL_3286667 | Lighthouse Lab in Alderley Park | Wellcome Sanger Institute for the COVID-19 Genomics UK (COG-UK) Consortium | Jacquelyn Wynn, Mairead Hyland, The Lighthouse Lab in Alderley Park and Alex Alderton, Roberto Amato, Jeffrey Barrett, Sonia Goncalves, Ewan Harrison, David K. Jackson, Ian Johnston, Dominic Kwiatkowski, Cordelia Langford, John Sillitoe on behalf of the Wellcome Sanger Institute COVID-19 Surveillance Team |
| EPI_ISL_3286668 | Lighthouse Lab in Milton Keynes | Wellcome Sanger Institute for the COVID-19 Genomics UK (COG-UK) Consortium | The Lighthouse Lab in Milton Keynes and Alex Alderton, Roberto Amato, Jeffrey Barrett, Sonia Goncalves, Ewan Harrison, David K. Jackson, Ian Johnston, Dominic Kwiatkowski, Cordelia Langford, John Sillitoe on behalf of the Wellcome Sanger Institute COVID-19 Surveillance Team |
| EPI_ISL_3286669 | Lighthouse Lab in Alderley Park | Wellcome Sanger Institute for the COVID-19 Genomics UK (COG-UK) Consortium | Jacquelyn Wynn, Mairead Hyland, The Lighthouse Lab in Alderley Park and Alex Alderton, Roberto Amato, Jeffrey Barrett, Sonia Goncalves, Ewan Harrison, David K. Jackson, Ian Johnston, Dominic Kwiatkowski, Cordelia Langford, John Sillitoe on behalf of the Wellcome Sanger Institute COVID-19 Surveillance Team |
| EPI_ISL_3286670 | Lighthouse Lab in Milton Keynes | Wellcome Sanger Institute for the COVID-19 Genomics UK (COG-UK) Consortium | The Lighthouse Lab in Milton Keynes and Alex Alderton, Roberto Amato, Jeffrey Barrett, Sonia Goncalves, Ewan Harrison, David K. Jackson, Ian Johnston, Dominic Kwiatkowski, Cordelia Langford, John Sillitoe on behalf of the Wellcome Sanger Institute COVID-19 Surveillance Team |
| EPI_ISL_3286671 | Lighthouse Lab in Alderley Park | Wellcome Sanger Institute for the COVID-19 Genomics UK (COG-UK) Consortium | Jacquelyn Wynn, Mairead Hyland, The Lighthouse Lab in Alderley Park and Alex Alderton, Roberto Amato, Jeffrey Barrett, Sonia Goncalves, Ewan Harrison, David K. Jackson, Ian Johnston, Dominic Kwiatkowski, Cordelia Langford, John Sillitoe on behalf of the Wellcome Sanger Institute COVID-19 Surveillance Team |
| EPI_ISL_3286672 | Lighthouse Lab in Milton Keynes | Wellcome Sanger Institute for the COVID-19 Genomics UK (COG-UK) Consortium | The Lighthouse Lab in Milton Keynes and Alex Alderton, Roberto Amato, Jeffrey Barrett, Sonia Goncalves, Ewan Harrison, David K. Jackson, Ian Johnston, Dominic Kwiatkowski, Cordelia Langford, John Sillitoe on behalf of the Wellcome Sanger Institute COVID-19 Surveillance Team |
| EPI_ISL_3286673, | Lighthouse Lab in Alderley Park | Wellcome Sanger Institute for the COVID-19 Genomics UK (COG-UK) | Jacquelyn Wynn, Mairead Hyland, The Lighthouse Lab in Alderley Park and Alex Alderton, Roberto Amato, Jeffrey Barrett, Sonia Goncalves, Ewan Harrison, David K. Jackson, Ian Johnston, Dominic Kwiatkowski, Cordelia Langford, John Sillitoe on |

| | | | |
|---|---|---|---|
| EPI_ISL_3286674 | | Consortium | behalf of the Wellcome Sanger Institute COVID-19 Surveillance Team |
| EPI_ISL_3286675 | Lighthouse Lab in Milton Keynes | Wellcome Sanger Institute for the COVID-19 Genomics UK (COG-UK) Consortium | The Lighthouse Lab in Milton Keynes and Alex Alderton, Roberto Amato, Jeffrey Barrett, Sonia Goncalves, Ewan Harrison, David K. Jackson, Ian Johnston, Dominic Kwiatkowski, Cordelia Langford, John Sillitoe on behalf of the Wellcome Sanger Institute COVID-19 Surveillance Team |
| EPI_ISL_3286676, EPI_ISL_3286677, EPI_ISL_3286678 | Lighthouse Lab in Alderley Park | Wellcome Sanger Institute for the COVID-19 Genomics UK (COG-UK) Consortium | Jacquelyn Wynn, Mairead Hyland, The Lighthouse Lab in Alderley Park and Alex Alderton, Roberto Amato, Jeffrey Barrett, Sonia Goncalves, Ewan Harrison, David K. Jackson, Ian Johnston, Dominic Kwiatkowski, Cordelia Langford, John Sillitoe on behalf of the Wellcome Sanger Institute COVID-19 Surveillance Team |
| EPI_ISL_3286679 | Lighthouse Lab in Milton Keynes | Wellcome Sanger Institute for the COVID-19 Genomics UK (COG-UK) Consortium | The Lighthouse Lab in Milton Keynes and Alex Alderton, Roberto Amato, Jeffrey Barrett, Sonia Goncalves, Ewan Harrison, David K. Jackson, Ian Johnston, Dominic Kwiatkowski, Cordelia Langford, John Sillitoe on behalf of the Wellcome Sanger Institute COVID-19 Surveillance Team |
| EPI_ISL_3286680, EPI_ISL_3286681, EPI_ISL_3286682 | Lighthouse Lab in Alderley Park | Wellcome Sanger Institute for the COVID-19 Genomics UK (COG-UK) Consortium | Jacquelyn Wynn, Mairead Hyland, The Lighthouse Lab in Alderley Park and Alex Alderton, Roberto Amato, Jeffrey Barrett, Sonia Goncalves, Ewan Harrison, David K. Jackson, Ian Johnston, Dominic Kwiatkowski, Cordelia Langford, John Sillitoe on behalf of the Wellcome Sanger Institute COVID-19 Surveillance Team |
| EPI_ISL_3286683 | Lighthouse Lab in Milton Keynes | Wellcome Sanger Institute for the COVID-19 Genomics UK (COG-UK) Consortium | The Lighthouse Lab in Milton Keynes and Alex Alderton, Roberto Amato, Jeffrey Barrett, Sonia Goncalves, Ewan Harrison, David K. Jackson, Ian Johnston, Dominic Kwiatkowski, Cordelia Langford, John Sillitoe on behalf of the Wellcome Sanger Institute COVID-19 Surveillance Team |
| EPI_ISL_3286684 | Lighthouse Lab in Alderley Park | Wellcome Sanger Institute for the COVID-19 Genomics UK (COG-UK) Consortium | Jacquelyn Wynn, Mairead Hyland, The Lighthouse Lab in Alderley Park and Alex Alderton, Roberto Amato, Jeffrey Barrett, Sonia Goncalves, Ewan Harrison, David K. Jackson, Ian Johnston, Dominic Kwiatkowski, Cordelia Langford, John Sillitoe on behalf of the Wellcome Sanger Institute COVID-19 Surveillance Team |
| EPI_ISL_3286685 | Lighthouse Lab in Milton Keynes | Wellcome Sanger Institute for the COVID-19 Genomics UK (COG-UK) Consortium | The Lighthouse Lab in Milton Keynes and Alex Alderton, Roberto Amato, Jeffrey Barrett, Sonia Goncalves, Ewan Harrison, David K. Jackson, Ian Johnston, Dominic Kwiatkowski, Cordelia Langford, John Sillitoe on behalf of the Wellcome Sanger Institute COVID-19 Surveillance Team |
| EPI_ISL_3286686 | Lighthouse Lab in Alderley Park | Wellcome Sanger Institute for the COVID-19 Genomics UK (COG-UK) Consortium | Jacquelyn Wynn, Mairead Hyland, The Lighthouse Lab in Alderley Park and Alex Alderton, Roberto Amato, Jeffrey Barrett, Sonia Goncalves, Ewan Harrison, David K. Jackson, Ian Johnston, Dominic Kwiatkowski, Cordelia Langford, John Sillitoe on behalf of the Wellcome Sanger Institute COVID-19 Surveillance Team |
| EPI_ISL_3286687 | Lighthouse Lab in Milton Keynes | Wellcome Sanger Institute for the COVID-19 Genomics UK (COG-UK) Consortium | The Lighthouse Lab in Milton Keynes and Alex Alderton, Roberto Amato, Jeffrey Barrett, Sonia Goncalves, Ewan Harrison, David K. Jackson, Ian Johnston, Dominic Kwiatkowski, Cordelia Langford, John Sillitoe on behalf of the Wellcome Sanger Institute COVID-19 Surveillance Team |
| EPI_ISL_3286688 | Lighthouse Lab in Alderley Park | Wellcome Sanger Institute for the COVID-19 Genomics UK (COG-UK) Consortium | Jacquelyn Wynn, Mairead Hyland, The Lighthouse Lab in Alderley Park and Alex Alderton, Roberto Amato, Jeffrey Barrett, Sonia Goncalves, Ewan Harrison, David K. Jackson, Ian Johnston, Dominic Kwiatkowski, Cordelia Langford, John Sillitoe on behalf of the Wellcome Sanger Institute COVID-19 Surveillance Team |
| EPI_ISL_3286689, EPI_ISL_3286690 | Lighthouse Lab in Milton Keynes | Wellcome Sanger Institute for the COVID-19 Genomics UK (COG-UK) Consortium | The Lighthouse Lab in Milton Keynes and Alex Alderton, Roberto Amato, Jeffrey Barrett, Sonia Goncalves, Ewan Harrison, David K. Jackson, Ian Johnston, Dominic Kwiatkowski, Cordelia Langford, John Sillitoe on behalf of the Wellcome Sanger Institute COVID-19 Surveillance Team |
| EPI_ISL_3286691, EPI_ISL_3286692 | Lighthouse Lab in Alderley Park | Wellcome Sanger Institute for the COVID-19 Genomics UK (COG-UK) Consortium | Jacquelyn Wynn, Mairead Hyland, The Lighthouse Lab in Alderley Park and Alex Alderton, Roberto Amato, Jeffrey Barrett, Sonia Goncalves, Ewan Harrison, David K. Jackson, Ian Johnston, Dominic Kwiatkowski, Cordelia Langford, John Sillitoe on behalf of the Wellcome Sanger Institute COVID-19 Surveillance Team |
| EPI_ISL_3286693 | Lighthouse Lab in Milton Keynes | Wellcome Sanger Institute for the COVID-19 Genomics UK (COG-UK) Consortium | The Lighthouse Lab in Milton Keynes and Alex Alderton, Roberto Amato, Jeffrey Barrett, Sonia Goncalves, Ewan Harrison, David K. Jackson, Ian Johnston, Dominic Kwiatkowski, Cordelia Langford, John Sillitoe on behalf of the Wellcome Sanger Institute COVID-19 Surveillance Team |
| EPI_ISL_3286694, EPI_ISL_3286695 | Lighthouse Lab in Alderley Park | Wellcome Sanger Institute for the COVID-19 Genomics UK (COG-UK) Consortium | Jacquelyn Wynn, Mairead Hyland, The Lighthouse Lab in Alderley Park and Alex Alderton, Roberto Amato, Jeffrey Barrett, Sonia Goncalves, Ewan Harrison, David K. Jackson, Ian Johnston, Dominic Kwiatkowski, Cordelia Langford, John Sillitoe on behalf of the Wellcome Sanger Institute COVID-19 Surveillance Team |
| EPI_ISL_3286696, EPI_ISL_3286697 | Lighthouse Lab in Milton Keynes | Wellcome Sanger Institute for the COVID-19 Genomics UK (COG-UK) Consortium | The Lighthouse Lab in Milton Keynes and Alex Alderton, Roberto Amato, Jeffrey Barrett, Sonia Goncalves, Ewan Harrison, David K. Jackson, Ian Johnston, Dominic Kwiatkowski, Cordelia Langford, John Sillitoe on behalf of the Wellcome Sanger Institute COVID-19 Surveillance Team |
| EPI_ISL_3286698, EPI_ISL_3286699 | Lighthouse Lab in Alderley Park | Wellcome Sanger Institute for the COVID-19 Genomics UK (COG-UK) Consortium | Jacquelyn Wynn, Mairead Hyland, The Lighthouse Lab in Alderley Park and Alex Alderton, Roberto Amato, Jeffrey Barrett, Sonia Goncalves, Ewan Harrison, David K. Jackson, Ian Johnston, Dominic Kwiatkowski, Cordelia Langford, John Sillitoe on behalf of the Wellcome Sanger Institute COVID-19 Surveillance Team |
| EPI_ISL_3286700 | Lighthouse Lab in Milton Keynes | Wellcome Sanger Institute for the COVID-19 Genomics UK (COG-UK) Consortium | The Lighthouse Lab in Milton Keynes and Alex Alderton, Roberto Amato, Jeffrey Barrett, Sonia Goncalves, Ewan Harrison, David K. Jackson, Ian Johnston, Dominic Kwiatkowski, Cordelia Langford, John Sillitoe on behalf of the Wellcome Sanger Institute COVID-19 Surveillance Team |
| EPI_ISL_3286701, EPI_ISL_3286702 | Lighthouse Lab in Alderley Park | Wellcome Sanger Institute for the COVID-19 Genomics UK (COG-UK) Consortium | Jacquelyn Wynn, Mairead Hyland, The Lighthouse Lab in Alderley Park and Alex Alderton, Roberto Amato, Jeffrey Barrett, Sonia Goncalves, Ewan Harrison, David K. Jackson, Ian Johnston, Dominic Kwiatkowski, Cordelia Langford, John Sillitoe on behalf of the Wellcome Sanger Institute COVID-19 Surveillance Team |
| EPI_ISL_3286703, EPI_ISL_3286704, EPI_ISL_3286705 | Lighthouse Lab in Milton Keynes | Wellcome Sanger Institute for the COVID-19 Genomics UK (COG-UK) Consortium | The Lighthouse Lab in Milton Keynes and Alex Alderton, Roberto Amato, Jeffrey Barrett, Sonia Goncalves, Ewan Harrison, David K. Jackson, Ian Johnston, Dominic Kwiatkowski, Cordelia Langford, John Sillitoe on behalf of the Wellcome Sanger Institute COVID-19 Surveillance Team |
| EPI_ISL_3286706, EPI_ISL_3286707, EPI_ISL_3286708, EPI_ISL_3286709, EPI_ISL_3286710 | Lighthouse Lab in Alderley Park | Wellcome Sanger Institute for the COVID-19 Genomics UK (COG-UK) Consortium | Jacquelyn Wynn, Mairead Hyland, The Lighthouse Lab in Alderley Park and Alex Alderton, Roberto Amato, Jeffrey Barrett, Sonia Goncalves, Ewan Harrison, David K. Jackson, Ian Johnston, Dominic Kwiatkowski, Cordelia Langford, John Sillitoe on behalf of the Wellcome Sanger Institute COVID-19 Surveillance Team |
| EPI_ISL_3286711, EPI_ISL_3286712 | Lighthouse Lab in Milton Keynes | Wellcome Sanger Institute for the COVID-19 Genomics UK (COG-UK) Consortium | The Lighthouse Lab in Milton Keynes and Alex Alderton, Roberto Amato, Jeffrey Barrett, Sonia Goncalves, Ewan Harrison, David K. Jackson, Ian Johnston, Dominic Kwiatkowski, Cordelia Langford, John Sillitoe on behalf of the Wellcome Sanger Institute COVID-19 Surveillance Team |
| EPI_ISL_3286713, EPI_ISL_3286714, EPI_ISL_3286715 | Lighthouse Lab in Alderley Park | Wellcome Sanger Institute for the COVID-19 Genomics UK (COG-UK) Consortium | Jacquelyn Wynn, Mairead Hyland, The Lighthouse Lab in Alderley Park and Alex Alderton, Roberto Amato, Jeffrey Barrett, Sonia Goncalves, Ewan Harrison, David K. Jackson, Ian Johnston, Dominic Kwiatkowski, Cordelia Langford, John Sillitoe on behalf of the Wellcome Sanger Institute COVID-19 Surveillance Team |
| EPI_ISL_3286716, EPI_ISL_3286717 | Lighthouse Lab in Milton Keynes | Wellcome Sanger Institute for the COVID-19 Genomics UK (COG-UK) Consortium | The Lighthouse Lab in Milton Keynes and Alex Alderton, Roberto Amato, Jeffrey Barrett, Sonia Goncalves, Ewan Harrison, David K. Jackson, Ian Johnston, Dominic Kwiatkowski, Cordelia Langford, John Sillitoe on behalf of the Wellcome Sanger Institute COVID-19 Surveillance Team |
| EPI_ISL_3286718 | Lighthouse Lab in Alderley Park | Wellcome Sanger Institute for the COVID-19 Genomics UK (COG-UK) Consortium | Jacquelyn Wynn, Mairead Hyland, The Lighthouse Lab in Alderley Park and Alex Alderton, Roberto Amato, Jeffrey Barrett, Sonia Goncalves, Ewan Harrison, David K. Jackson, Ian Johnston, Dominic Kwiatkowski, Cordelia Langford, John Sillitoe on behalf of the Wellcome Sanger Institute COVID-19 Surveillance Team |
| EPI_ISL_3286719, EPI_ISL_3286720, EPI_ISL_3286721, EPI_ISL_3286722, EPI_ISL_3286723, EPI_ISL_3286724, EPI_ISL_3286725, EPI_ISL_3286726 | Lighthouse Lab in Milton Keynes | Wellcome Sanger Institute for the COVID-19 Genomics UK (COG-UK) Consortium | The Lighthouse Lab in Milton Keynes and Alex Alderton, Roberto Amato, Jeffrey Barrett, Sonia Goncalves, Ewan Harrison, David K. Jackson, Ian Johnston, Dominic Kwiatkowski, Cordelia Langford, John Sillitoe on behalf of the Wellcome Sanger Institute COVID-19 Surveillance Team |
| EPI_ISL_3286727 | Lighthouse Lab in Alderley Park | Wellcome Sanger Institute for the COVID-19 Genomics UK (COG-UK) Consortium | Jacquelyn Wynn, Mairead Hyland, The Lighthouse Lab in Alderley Park and Alex Alderton, Roberto Amato, Jeffrey Barrett, Sonia Goncalves, Ewan Harrison, David K. Jackson, Ian Johnston, Dominic Kwiatkowski, Cordelia Langford, John Sillitoe on behalf of the Wellcome Sanger Institute COVID-19 Surveillance Team |
| EPI_ISL_3286728 | Lighthouse Lab in Milton Keynes | Wellcome Sanger Institute for the COVID-19 Genomics UK (COG-UK) Consortium | The Lighthouse Lab in Milton Keynes and Alex Alderton, Roberto Amato, Jeffrey Barrett, Sonia Goncalves, Ewan Harrison, David K. Jackson, Ian Johnston, Dominic Kwiatkowski, Cordelia Langford, John Sillitoe on behalf of the Wellcome Sanger Institute COVID-19 Surveillance Team |
| EPI_ISL_3286729 | Lighthouse Lab in Alderley Park | Wellcome Sanger Institute for the COVID-19 Genomics UK (COG-UK) Consortium | Jacquelyn Wynn, Mairead Hyland, The Lighthouse Lab in Alderley Park and Alex Alderton, Roberto Amato, Jeffrey Barrett, Sonia Goncalves, Ewan Harrison, David K. Jackson, Ian Johnston, Dominic Kwiatkowski, Cordelia Langford, John Sillitoe on behalf of the Wellcome Sanger Institute COVID-19 Surveillance Team |
| EPI_ISL_3286730 | Lighthouse Lab in Milton Keynes | Wellcome Sanger Institute for the COVID-19 Genomics UK (COG-UK) Consortium | The Lighthouse Lab in Milton Keynes and Alex Alderton, Roberto Amato, Jeffrey Barrett, Sonia Goncalves, Ewan Harrison, David K. Jackson, Ian Johnston, Dominic Kwiatkowski, Cordelia Langford, John Sillitoe on behalf of the Wellcome Sanger Institute COVID-19 Surveillance Team |
| EPI_ISL_3286731 | Lighthouse Lab in Alderley Park | Wellcome Sanger Institute for the COVID-19 Genomics UK (COG-UK) Consortium | Jacquelyn Wynn, Mairead Hyland, The Lighthouse Lab in Alderley Park and Alex Alderton, Roberto Amato, Jeffrey Barrett, Sonia Goncalves, Ewan Harrison, David K. Jackson, Ian Johnston, Dominic Kwiatkowski, Cordelia Langford, John Sillitoe on behalf of the Wellcome Sanger Institute COVID-19 Surveillance Team |
| EPI_ISL_3286732, EPI_ISL_3286733 | Lighthouse Lab in Milton Keynes | Wellcome Sanger Institute for the COVID-19 Genomics UK (COG-UK) Consortium | The Lighthouse Lab in Milton Keynes and Alex Alderton, Roberto Amato, Jeffrey Barrett, Sonia Goncalves, Ewan Harrison, David K. Jackson, Ian Johnston, Dominic Kwiatkowski, Cordelia Langford, John Sillitoe on behalf of the Wellcome Sanger Institute COVID-19 Surveillance Team |
| EPI_ISL_3286734 | Lighthouse Lab in Alderley Park | Wellcome Sanger Institute for the COVID-19 Genomics UK (COG-UK) Consortium | Jacquelyn Wynn, Mairead Hyland, The Lighthouse Lab in Alderley Park and Alex Alderton, Roberto Amato, Jeffrey Barrett, Sonia Goncalves, Ewan Harrison, David K. Jackson, Ian Johnston, Dominic Kwiatkowski, Cordelia Langford, John Sillitoe on behalf of the Wellcome Sanger Institute COVID-19 Surveillance Team |
| EPI_ISL_3286735, EPI_ISL_3286736, EPI_ISL_3286737 | Lighthouse Lab in Milton Keynes | Wellcome Sanger Institute for the COVID-19 Genomics UK (COG-UK) Consortium | The Lighthouse Lab in Milton Keynes and Alex Alderton, Roberto Amato, Jeffrey Barrett, Sonia Goncalves, Ewan Harrison, David K. Jackson, Ian Johnston, Dominic Kwiatkowski, Cordelia Langford, John Sillitoe on behalf of the Wellcome Sanger Institute COVID-19 Surveillance Team |
| EPI_ISL_3286738, EPI_ISL_3286739, EPI_ISL_3286740 | Lighthouse Lab in Alderley Park | Wellcome Sanger Institute for the COVID-19 Genomics UK (COG-UK) Consortium | Jacquelyn Wynn, Mairead Hyland, The Lighthouse Lab in Alderley Park and Alex Alderton, Roberto Amato, Jeffrey Barrett, Sonia Goncalves, Ewan Harrison, David K. Jackson, Ian Johnston, Dominic Kwiatkowski, Cordelia Langford, John Sillitoe on behalf of the Wellcome Sanger Institute COVID-19 Surveillance Team |
| EPI_ISL_3286741 | Lighthouse Lab in Milton Keynes | Wellcome Sanger Institute for the COVID-19 Genomics UK (COG-UK) Consortium | The Lighthouse Lab in Milton Keynes and Alex Alderton, Roberto Amato, Jeffrey Barrett, Sonia Goncalves, Ewan Harrison, David K. Jackson, Ian Johnston, Dominic Kwiatkowski, Cordelia Langford, John Sillitoe on behalf of the Wellcome Sanger Institute COVID-19 Surveillance Team |
| EPI_ISL_3286742, EPI_ISL_3286743, EPI_ISL_3286744 | Lighthouse Lab in Alderley Park | Wellcome Sanger Institute for the COVID-19 Genomics UK (COG-UK) Consortium | Jacquelyn Wynn, Mairead Hyland, The Lighthouse Lab in Alderley Park and Alex Alderton, Roberto Amato, Jeffrey Barrett, Sonia Goncalves, Ewan Harrison, David K. Jackson, Ian Johnston, Dominic Kwiatkowski, Cordelia Langford, John Sillitoe on behalf of the Wellcome Sanger Institute COVID-19 Surveillance Team |
| EPI_ISL_3286745, EPI_ISL_3286746, EPI_ISL_3286747 | Lighthouse Lab in Milton Keynes | Wellcome Sanger Institute for the COVID-19 Genomics UK (COG-UK) Consortium | The Lighthouse Lab in Milton Keynes and Alex Alderton, Roberto Amato, Jeffrey Barrett, Sonia Goncalves, Ewan Harrison, David K. Jackson, Ian Johnston, Dominic Kwiatkowski, Cordelia Langford, John Sillitoe on behalf of the Wellcome Sanger Institute COVID-19 Surveillance Team |
| EPI_ISL_3286748 | Lighthouse Lab in Alderley Park | Wellcome Sanger Institute for the COVID-19 Genomics UK (COG-UK) | Jacquelyn Wynn, Mairead Hyland, The Lighthouse Lab in Alderley Park and Alex Alderton, Roberto Amato, Jeffrey Barrett, Sonia Goncalves, Ewan Harrison, David K. Jackson, Ian Johnston, Dominic Kwiatkowski, Cordelia Langford, John Sillitoe on |

| | | | |
|---|---|---|---|
| | | Consortium | behalf of the Wellcome Sanger Institute COVID-19 Surveillance Team |
| EPI_ISL_3286749 | Lighthouse Lab in Milton Keynes | Wellcome Sanger Institute for the COVID-19 Genomics UK (COG-UK) Consortium | The Lighthouse Lab in Milton Keynes and Alex Alderton, Roberto Amato, Jeffrey Barrett, Sonia Goncalves, Ewan Harrison, David K. Jackson, Ian Johnston, Dominic Kwiatkowski, Cordelia Langford, John Sillitoe on behalf of the Wellcome Sanger Institute COVID-19 Surveillance Team |
| EPI_ISL_3286750, EPI_ISL_3286751, EPI_ISL_3286752, EPI_ISL_3286753, EPI_ISL_3286754 | Lighthouse Lab in Alderley Park | Wellcome Sanger Institute for the COVID-19 Genomics UK (COG-UK) Consortium | Jacquelyn Wynn, Mairead Hyland, The Lighthouse Lab in Alderley Park and Alex Alderton, Roberto Amato, Jeffrey Barrett, Sonia Goncalves, Ewan Harrison, David K. Jackson, Ian Johnston, Dominic Kwiatkowski, Cordelia Langford, John Sillitoe on behalf of the Wellcome Sanger Institute COVID-19 Surveillance Team |
| EPI_ISL_3286755, EPI_ISL_3286756 | Lighthouse Lab in Milton Keynes | Wellcome Sanger Institute for the COVID-19 Genomics UK (COG-UK) Consortium | The Lighthouse Lab in Milton Keynes and Alex Alderton, Roberto Amato, Jeffrey Barrett, Sonia Goncalves, Ewan Harrison, David K. Jackson, Ian Johnston, Dominic Kwiatkowski, Cordelia Langford, John Sillitoe on behalf of the Wellcome Sanger Institute COVID-19 Surveillance Team |
| EPI_ISL_3286757, EPI_ISL_3286758 | Lighthouse Lab in Alderley Park | Wellcome Sanger Institute for the COVID-19 Genomics UK (COG-UK) Consortium | Jacquelyn Wynn, Mairead Hyland, The Lighthouse Lab in Alderley Park and Alex Alderton, Roberto Amato, Jeffrey Barrett, Sonia Goncalves, Ewan Harrison, David K. Jackson, Ian Johnston, Dominic Kwiatkowski, Cordelia Langford, John Sillitoe on behalf of the Wellcome Sanger Institute COVID-19 Surveillance Team |
| EPI_ISL_3286759 | Lighthouse Lab in Milton Keynes | Wellcome Sanger Institute for the COVID-19 Genomics UK (COG-UK) Consortium | The Lighthouse Lab in Milton Keynes and Alex Alderton, Roberto Amato, Jeffrey Barrett, Sonia Goncalves, Ewan Harrison, David K. Jackson, Ian Johnston, Dominic Kwiatkowski, Cordelia Langford, John Sillitoe on behalf of the Wellcome Sanger Institute COVID-19 Surveillance Team |
| EPI_ISL_3286760 | Lighthouse Lab in Alderley Park | Wellcome Sanger Institute for the COVID-19 Genomics UK (COG-UK) Consortium | Jacquelyn Wynn, Mairead Hyland, The Lighthouse Lab in Alderley Park and Alex Alderton, Roberto Amato, Jeffrey Barrett, Sonia Goncalves, Ewan Harrison, David K. Jackson, Ian Johnston, Dominic Kwiatkowski, Cordelia Langford, John Sillitoe on behalf of the Wellcome Sanger Institute COVID-19 Surveillance Team |
| EPI_ISL_3286761 | Lighthouse Lab in Milton Keynes | Wellcome Sanger Institute for the COVID-19 Genomics UK (COG-UK) Consortium | The Lighthouse Lab in Milton Keynes and Alex Alderton, Roberto Amato, Jeffrey Barrett, Sonia Goncalves, Ewan Harrison, David K. Jackson, Ian Johnston, Dominic Kwiatkowski, Cordelia Langford, John Sillitoe on behalf of the Wellcome Sanger Institute COVID-19 Surveillance Team |
| EPI_ISL_3286762, EPI_ISL_3286763, EPI_ISL_3286764 | Lighthouse Lab in Alderley Park | Wellcome Sanger Institute for the COVID-19 Genomics UK (COG-UK) Consortium | Jacquelyn Wynn, Mairead Hyland, The Lighthouse Lab in Alderley Park and Alex Alderton, Roberto Amato, Jeffrey Barrett, Sonia Goncalves, Ewan Harrison, David K. Jackson, Ian Johnston, Dominic Kwiatkowski, Cordelia Langford, John Sillitoe on behalf of the Wellcome Sanger Institute COVID-19 Surveillance Team |
| EPI_ISL_3286765, EPI_ISL_3286766, EPI_ISL_3286767 | Lighthouse Lab in Milton Keynes | Wellcome Sanger Institute for the COVID-19 Genomics UK (COG-UK) Consortium | The Lighthouse Lab in Milton Keynes and Alex Alderton, Roberto Amato, Jeffrey Barrett, Sonia Goncalves, Ewan Harrison, David K. Jackson, Ian Johnston, Dominic Kwiatkowski, Cordelia Langford, John Sillitoe on behalf of the Wellcome Sanger Institute COVID-19 Surveillance Team |
| EPI_ISL_3286768 | Lighthouse Lab in Alderley Park | Wellcome Sanger Institute for the COVID-19 Genomics UK (COG-UK) Consortium | Jacquelyn Wynn, Mairead Hyland, The Lighthouse Lab in Alderley Park and Alex Alderton, Roberto Amato, Jeffrey Barrett, Sonia Goncalves, Ewan Harrison, David K. Jackson, Ian Johnston, Dominic Kwiatkowski, Cordelia Langford, John Sillitoe on behalf of the Wellcome Sanger Institute COVID-19 Surveillance Team |
| EPI_ISL_3286769, EPI_ISL_3286770, EPI_ISL_3286771 | Lighthouse Lab in Milton Keynes | Wellcome Sanger Institute for the COVID-19 Genomics UK (COG-UK) Consortium | The Lighthouse Lab in Milton Keynes and Alex Alderton, Roberto Amato, Jeffrey Barrett, Sonia Goncalves, Ewan Harrison, David K. Jackson, Ian Johnston, Dominic Kwiatkowski, Cordelia Langford, John Sillitoe on behalf of the Wellcome Sanger Institute COVID-19 Surveillance Team |
| EPI_ISL_3286772 | Lighthouse Lab in Alderley Park | Wellcome Sanger Institute for the COVID-19 Genomics UK (COG-UK) Consortium | Jacquelyn Wynn, Mairead Hyland, The Lighthouse Lab in Alderley Park and Alex Alderton, Roberto Amato, Jeffrey Barrett, Sonia Goncalves, Ewan Harrison, David K. Jackson, Ian Johnston, Dominic Kwiatkowski, Cordelia Langford, John Sillitoe on behalf of the Wellcome Sanger Institute COVID-19 Surveillance Team |
| EPI_ISL_3286773 | Lighthouse Lab in Milton Keynes | Wellcome Sanger Institute for the COVID-19 Genomics UK (COG-UK) Consortium | The Lighthouse Lab in Milton Keynes and Alex Alderton, Roberto Amato, Jeffrey Barrett, Sonia Goncalves, Ewan Harrison, David K. Jackson, Ian Johnston, Dominic Kwiatkowski, Cordelia Langford, John Sillitoe on behalf of the Wellcome Sanger Institute COVID-19 Surveillance Team |
| EPI_ISL_3286774 | Lighthouse Lab in Alderley Park | Wellcome Sanger Institute for the COVID-19 Genomics UK (COG-UK) Consortium | Jacquelyn Wynn, Mairead Hyland, The Lighthouse Lab in Alderley Park and Alex Alderton, Roberto Amato, Jeffrey Barrett, Sonia Goncalves, Ewan Harrison, David K. Jackson, Ian Johnston, Dominic Kwiatkowski, Cordelia Langford, John Sillitoe on behalf of the Wellcome Sanger Institute COVID-19 Surveillance Team |
| EPI_ISL_3286775 | Lighthouse Lab in Milton Keynes | Wellcome Sanger Institute for the COVID-19 Genomics UK (COG-UK) Consortium | The Lighthouse Lab in Milton Keynes and Alex Alderton, Roberto Amato, Jeffrey Barrett, Sonia Goncalves, Ewan Harrison, David K. Jackson, Ian Johnston, Dominic Kwiatkowski, Cordelia Langford, John Sillitoe on behalf of the Wellcome Sanger Institute COVID-19 Surveillance Team |
| EPI_ISL_3286776 | Lighthouse Lab in Alderley Park | Wellcome Sanger Institute for the COVID-19 Genomics UK (COG-UK) Consortium | Jacquelyn Wynn, Mairead Hyland, The Lighthouse Lab in Alderley Park and Alex Alderton, Roberto Amato, Jeffrey Barrett, Sonia Goncalves, Ewan Harrison, David K. Jackson, Ian Johnston, Dominic Kwiatkowski, Cordelia Langford, John Sillitoe on behalf of the Wellcome Sanger Institute COVID-19 Surveillance Team |
| EPI_ISL_3286777, EPI_ISL_3286778, EPI_ISL_3286779, EPI_ISL_3286780, EPI_ISL_3286781, EPI_ISL_3286782, EPI_ISL_3286783 | Lighthouse Lab in Milton Keynes | Wellcome Sanger Institute for the COVID-19 Genomics UK (COG-UK) Consortium | The Lighthouse Lab in Milton Keynes and Alex Alderton, Roberto Amato, Jeffrey Barrett, Sonia Goncalves, Ewan Harrison, David K. Jackson, Ian Johnston, Dominic Kwiatkowski, Cordelia Langford, John Sillitoe on behalf of the Wellcome Sanger Institute COVID-19 Surveillance Team |
| EPI_ISL_3286784 | Lighthouse Lab in Alderley Park | Wellcome Sanger Institute for the COVID-19 Genomics UK (COG-UK) Consortium | Jacquelyn Wynn, Mairead Hyland, The Lighthouse Lab in Alderley Park and Alex Alderton, Roberto Amato, Jeffrey Barrett, Sonia Goncalves, Ewan Harrison, David K. Jackson, Ian Johnston, Dominic Kwiatkowski, Cordelia Langford, John Sillitoe on behalf of the Wellcome Sanger Institute COVID-19 Surveillance Team |
| EPI_ISL_3286785 | Lighthouse Lab in Milton Keynes | Wellcome Sanger Institute for the COVID-19 Genomics UK (COG-UK) Consortium | The Lighthouse Lab in Milton Keynes and Alex Alderton, Roberto Amato, Jeffrey Barrett, Sonia Goncalves, Ewan Harrison, David K. Jackson, Ian Johnston, Dominic Kwiatkowski, Cordelia Langford, John Sillitoe on behalf of the Wellcome Sanger Institute COVID-19 Surveillance Team |
| EPI_ISL_3286786 | Lighthouse Lab in Alderley Park | Wellcome Sanger Institute for the COVID-19 Genomics UK (COG-UK) Consortium | Jacquelyn Wynn, Mairead Hyland, The Lighthouse Lab in Alderley Park and Alex Alderton, Roberto Amato, Jeffrey Barrett, Sonia Goncalves, Ewan Harrison, David K. Jackson, Ian Johnston, Dominic Kwiatkowski, Cordelia Langford, John Sillitoe on behalf of the Wellcome Sanger Institute COVID-19 Surveillance Team |
| EPI_ISL_3286787 | Lighthouse Lab in Milton Keynes | Wellcome Sanger Institute for the COVID-19 Genomics UK (COG-UK) Consortium | The Lighthouse Lab in Milton Keynes and Alex Alderton, Roberto Amato, Jeffrey Barrett, Sonia Goncalves, Ewan Harrison, David K. Jackson, Ian Johnston, Dominic Kwiatkowski, Cordelia Langford, John Sillitoe on behalf of the Wellcome Sanger Institute COVID-19 Surveillance Team |
| EPI_ISL_3286788, EPI_ISL_3286789, EPI_ISL_3286790, EPI_ISL_3286791 | Lighthouse Lab in Alderley Park | Wellcome Sanger Institute for the COVID-19 Genomics UK (COG-UK) Consortium | Jacquelyn Wynn, Mairead Hyland, The Lighthouse Lab in Alderley Park and Alex Alderton, Roberto Amato, Jeffrey Barrett, Sonia Goncalves, Ewan Harrison, David K. Jackson, Ian Johnston, Dominic Kwiatkowski, Cordelia Langford, John Sillitoe on behalf of the Wellcome Sanger Institute COVID-19 Surveillance Team |
| EPI_ISL_3286792 | Lighthouse Lab in Milton Keynes | Wellcome Sanger Institute for the COVID-19 Genomics UK (COG-UK) Consortium | The Lighthouse Lab in Milton Keynes and Alex Alderton, Roberto Amato, Jeffrey Barrett, Sonia Goncalves, Ewan Harrison, David K. Jackson, Ian Johnston, Dominic Kwiatkowski, Cordelia Langford, John Sillitoe on behalf of the Wellcome Sanger Institute COVID-19 Surveillance Team |
| EPI_ISL_3286793, EPI_ISL_3286794 | Lighthouse Lab in Alderley Park | Wellcome Sanger Institute for the COVID-19 Genomics UK (COG-UK) Consortium | Jacquelyn Wynn, Mairead Hyland, The Lighthouse Lab in Alderley Park and Alex Alderton, Roberto Amato, Jeffrey Barrett, Sonia Goncalves, Ewan Harrison, David K. Jackson, Ian Johnston, Dominic Kwiatkowski, Cordelia Langford, John Sillitoe on behalf of the Wellcome Sanger Institute COVID-19 Surveillance Team |
| EPI_ISL_3286795 | Lighthouse Lab in Milton Keynes | Wellcome Sanger Institute for the COVID-19 Genomics UK (COG-UK) Consortium | The Lighthouse Lab in Milton Keynes and Alex Alderton, Roberto Amato, Jeffrey Barrett, Sonia Goncalves, Ewan Harrison, David K. Jackson, Ian Johnston, Dominic Kwiatkowski, Cordelia Langford, John Sillitoe on behalf of the Wellcome Sanger Institute COVID-19 Surveillance Team |
| EPI_ISL_3286796, EPI_ISL_3286797, EPI_ISL_3286798, EPI_ISL_3286799 | Lighthouse Lab in Alderley Park | Wellcome Sanger Institute for the COVID-19 Genomics UK (COG-UK) Consortium | Jacquelyn Wynn, Mairead Hyland, The Lighthouse Lab in Alderley Park and Alex Alderton, Roberto Amato, Jeffrey Barrett, Sonia Goncalves, Ewan Harrison, David K. Jackson, Ian Johnston, Dominic Kwiatkowski, Cordelia Langford, John Sillitoe on behalf of the Wellcome Sanger Institute COVID-19 Surveillance Team |
| EPI_ISL_3286800 | Lighthouse Lab in Milton Keynes | Wellcome Sanger Institute for the COVID-19 Genomics UK (COG-UK) Consortium | The Lighthouse Lab in Milton Keynes and Alex Alderton, Roberto Amato, Jeffrey Barrett, Sonia Goncalves, Ewan Harrison, David K. Jackson, Ian Johnston, Dominic Kwiatkowski, Cordelia Langford, John Sillitoe on behalf of the Wellcome Sanger Institute COVID-19 Surveillance Team |
| EPI_ISL_3286801, EPI_ISL_3286802, EPI_ISL_3286803, EPI_ISL_3286804 | Lighthouse Lab in Alderley Park | Wellcome Sanger Institute for the COVID-19 Genomics UK (COG-UK) Consortium | Jacquelyn Wynn, Mairead Hyland, The Lighthouse Lab in Alderley Park and Alex Alderton, Roberto Amato, Jeffrey Barrett, Sonia Goncalves, Ewan Harrison, David K. Jackson, Ian Johnston, Dominic Kwiatkowski, Cordelia Langford, John Sillitoe on behalf of the Wellcome Sanger Institute COVID-19 Surveillance Team |
| EPI_ISL_3286805 | Lighthouse Lab in Milton Keynes | Wellcome Sanger Institute for the COVID-19 Genomics UK (COG-UK) Consortium | The Lighthouse Lab in Milton Keynes and Alex Alderton, Roberto Amato, Jeffrey Barrett, Sonia Goncalves, Ewan Harrison, David K. Jackson, Ian Johnston, Dominic Kwiatkowski, Cordelia Langford, John Sillitoe on behalf of the Wellcome Sanger Institute COVID-19 Surveillance Team |
| EPI_ISL_3286806, EPI_ISL_3286807, EPI_ISL_3286808, EPI_ISL_3286809, EPI_ISL_3286810, EPI_ISL_3286811 | Lighthouse Lab in Alderley Park | Wellcome Sanger Institute for the COVID-19 Genomics UK (COG-UK) Consortium | Jacquelyn Wynn, Mairead Hyland, The Lighthouse Lab in Alderley Park and Alex Alderton, Roberto Amato, Jeffrey Barrett, Sonia Goncalves, Ewan Harrison, David K. Jackson, Ian Johnston, Dominic Kwiatkowski, Cordelia Langford, John Sillitoe on behalf of the Wellcome Sanger Institute COVID-19 Surveillance Team |
| EPI_ISL_3286812, EPI_ISL_3286813, EPI_ISL_3286814, EPI_ISL_3286815, EPI_ISL_3286816, EPI_ISL_3286817 | Lighthouse Lab in Milton Keynes | Wellcome Sanger Institute for the COVID-19 Genomics UK (COG-UK) Consortium | The Lighthouse Lab in Milton Keynes and Alex Alderton, Roberto Amato, Jeffrey Barrett, Sonia Goncalves, Ewan Harrison, David K. Jackson, Ian Johnston, Dominic Kwiatkowski, Cordelia Langford, John Sillitoe on behalf of the Wellcome Sanger Institute COVID-19 Surveillance Team |
| EPI_ISL_3286818, EPI_ISL_3286819, EPI_ISL_3286820, EPI_ISL_3286821 | Lighthouse Lab in Alderley Park | Wellcome Sanger Institute for the COVID-19 Genomics UK (COG-UK) Consortium | Jacquelyn Wynn, Mairead Hyland, The Lighthouse Lab in Alderley Park and Alex Alderton, Roberto Amato, Jeffrey Barrett, Sonia Goncalves, Ewan Harrison, David K. Jackson, Ian Johnston, Dominic Kwiatkowski, Cordelia Langford, John Sillitoe on behalf of the Wellcome Sanger Institute COVID-19 Surveillance Team |
| EPI_ISL_3286822, EPI_ISL_3286823, EPI_ISL_3286824, EPI_ISL_3286825, | Lighthouse Lab in Milton Keynes | Wellcome Sanger Institute for the COVID-19 Genomics UK (COG-UK) Consortium | The Lighthouse Lab in Milton Keynes and Alex Alderton, Roberto Amato, Jeffrey Barrett, Sonia Goncalves, Ewan Harrison, David K. Jackson, Ian Johnston, Dominic Kwiatkowski, Cordelia Langford, John Sillitoe on behalf of the Wellcome Sanger Institute COVID-19 Surveillance Team |

| | | | |
|---|---|---|---|
| EPI_ISL_3286826, EPI_ISL_3286827, EPI_ISL_3286828, EPI_ISL_3286829, EPI_ISL_3286830 | | | |
| EPI_ISL_3286831 | Lighthouse Lab in Alderley Park | Wellcome Sanger Institute for the COVID-19 Genomics UK (COG-UK) Consortium | Jacquelyn Wynn, Mairead Hyland, The Lighthouse Lab in Alderley Park and Alex Alderton, Roberto Amato, Jeffrey Barrett, Sonia Goncalves, Ewan Harrison, David K. Jackson, Ian Johnston, Dominic Kwiatkowski, Cordelia Langford, John Sillitoe on behalf of the Wellcome Sanger Institute COVID-19 Surveillance Team |
| EPI_ISL_3286832, EPI_ISL_3286833, EPI_ISL_3286834, EPI_ISL_3286835 | Lighthouse Lab in Milton Keynes | Wellcome Sanger Institute for the COVID-19 Genomics UK (COG-UK) Consortium | The Lighthouse Lab in Milton Keynes and Alex Alderton, Roberto Amato, Jeffrey Barrett, Sonia Goncalves, Ewan Harrison, David K. Jackson, Ian Johnston, Dominic Kwiatkowski, Cordelia Langford, John Sillitoe on behalf of the Wellcome Sanger Institute COVID-19 Surveillance Team |
| EPI_ISL_3286836, EPI_ISL_3286837 | Lighthouse Lab in Alderley Park | Wellcome Sanger Institute for the COVID-19 Genomics UK (COG-UK) Consortium | Jacquelyn Wynn, Mairead Hyland, The Lighthouse Lab in Alderley Park and Alex Alderton, Roberto Amato, Jeffrey Barrett, Sonia Goncalves, Ewan Harrison, David K. Jackson, Ian Johnston, Dominic Kwiatkowski, Cordelia Langford, John Sillitoe on behalf of the Wellcome Sanger Institute COVID-19 Surveillance Team |
| EPI_ISL_3286838, EPI_ISL_3286839, EPI_ISL_3286840 | Lighthouse Lab in Milton Keynes | Wellcome Sanger Institute for the COVID-19 Genomics UK (COG-UK) Consortium | The Lighthouse Lab in Milton Keynes and Alex Alderton, Roberto Amato, Jeffrey Barrett, Sonia Goncalves, Ewan Harrison, David K. Jackson, Ian Johnston, Dominic Kwiatkowski, Cordelia Langford, John Sillitoe on behalf of the Wellcome Sanger Institute COVID-19 Surveillance Team |
| EPI_ISL_3286841 | Lighthouse Lab in Alderley Park | Wellcome Sanger Institute for the COVID-19 Genomics UK (COG-UK) Consortium | Jacquelyn Wynn, Mairead Hyland, The Lighthouse Lab in Alderley Park and Alex Alderton, Roberto Amato, Jeffrey Barrett, Sonia Goncalves, Ewan Harrison, David K. Jackson, Ian Johnston, Dominic Kwiatkowski, Cordelia Langford, John Sillitoe on behalf of the Wellcome Sanger Institute COVID-19 Surveillance Team |
| EPI_ISL_3286842, EPI_ISL_3286843, EPI_ISL_3286844 | Lighthouse Lab in Milton Keynes | Wellcome Sanger Institute for the COVID-19 Genomics UK (COG-UK) Consortium | The Lighthouse Lab in Milton Keynes and Alex Alderton, Roberto Amato, Jeffrey Barrett, Sonia Goncalves, Ewan Harrison, David K. Jackson, Ian Johnston, Dominic Kwiatkowski, Cordelia Langford, John Sillitoe on behalf of the Wellcome Sanger Institute COVID-19 Surveillance Team |
| EPI_ISL_3286845, EPI_ISL_3286846, EPI_ISL_3286847, EPI_ISL_3286848, EPI_ISL_3286849, EPI_ISL_3286850, EPI_ISL_3286851 | Lighthouse Lab in Alderley Park | Wellcome Sanger Institute for the COVID-19 Genomics UK (COG-UK) Consortium | Jacquelyn Wynn, Mairead Hyland, The Lighthouse Lab in Alderley Park and Alex Alderton, Roberto Amato, Jeffrey Barrett, Sonia Goncalves, Ewan Harrison, David K. Jackson, Ian Johnston, Dominic Kwiatkowski, Cordelia Langford, John Sillitoe on behalf of the Wellcome Sanger Institute COVID-19 Surveillance Team |
| EPI_ISL_3286852 | Lighthouse Lab in Milton Keynes | Wellcome Sanger Institute for the COVID-19 Genomics UK (COG-UK) Consortium | The Lighthouse Lab in Milton Keynes and Alex Alderton, Roberto Amato, Jeffrey Barrett, Sonia Goncalves, Ewan Harrison, David K. Jackson, Ian Johnston, Dominic Kwiatkowski, Cordelia Langford, John Sillitoe on behalf of the Wellcome Sanger Institute COVID-19 Surveillance Team |
| EPI_ISL_3286853, EPI_ISL_3286854 | Lighthouse Lab in Alderley Park | Wellcome Sanger Institute for the COVID-19 Genomics UK (COG-UK) Consortium | Jacquelyn Wynn, Mairead Hyland, The Lighthouse Lab in Alderley Park and Alex Alderton, Roberto Amato, Jeffrey Barrett, Sonia Goncalves, Ewan Harrison, David K. Jackson, Ian Johnston, Dominic Kwiatkowski, Cordelia Langford, John Sillitoe on behalf of the Wellcome Sanger Institute COVID-19 Surveillance Team |
| EPI_ISL_3286855, EPI_ISL_3286856, EPI_ISL_3286857 | Lighthouse Lab in Milton Keynes | Wellcome Sanger Institute for the COVID-19 Genomics UK (COG-UK) Consortium | The Lighthouse Lab in Milton Keynes and Alex Alderton, Roberto Amato, Jeffrey Barrett, Sonia Goncalves, Ewan Harrison, David K. Jackson, Ian Johnston, Dominic Kwiatkowski, Cordelia Langford, John Sillitoe on behalf of the Wellcome Sanger Institute COVID-19 Surveillance Team |
| EPI_ISL_3286858 | Lighthouse Lab in Alderley Park | Wellcome Sanger Institute for the COVID-19 Genomics UK (COG-UK) Consortium | Jacquelyn Wynn, Mairead Hyland, The Lighthouse Lab in Alderley Park and Alex Alderton, Roberto Amato, Jeffrey Barrett, Sonia Goncalves, Ewan Harrison, David K. Jackson, Ian Johnston, Dominic Kwiatkowski, Cordelia Langford, John Sillitoe on behalf of the Wellcome Sanger Institute COVID-19 Surveillance Team |
| EPI_ISL_3286859 | Lighthouse Lab in Glasgow | Wellcome Sanger Institute for the COVID-19 Genomics UK (COG-UK) Consortium | Harper VanSteenhouse, Yumi Kasai, David Gray, Carol Clugston, Anna Dominiczak and Alex Alderton, Roberto Amato, Jeffrey Barrett, Sonia Goncalves, Ewan Harrison, David K. Jackson, Ian Johnston, Dominic Kwiatkowski, Cordelia Langford, John Sillitoe on behalf of the Wellcome Sanger Institute COVID-19 Surveillance Team |
| EPI_ISL_3286860 | Lighthouse Lab in Milton Keynes | Wellcome Sanger Institute for the COVID-19 Genomics UK (COG-UK) Consortium | The Lighthouse Lab in Milton Keynes and Alex Alderton, Roberto Amato, Jeffrey Barrett, Sonia Goncalves, Ewan Harrison, David K. Jackson, Ian Johnston, Dominic Kwiatkowski, Cordelia Langford, John Sillitoe on behalf of the Wellcome Sanger Institute COVID-19 Surveillance Team |
| EPI_ISL_3286861 | Lighthouse Lab in Alderley Park | Wellcome Sanger Institute for the COVID-19 Genomics UK (COG-UK) Consortium | Jacquelyn Wynn, Mairead Hyland, The Lighthouse Lab in Alderley Park and Alex Alderton, Roberto Amato, Jeffrey Barrett, Sonia Goncalves, Ewan Harrison, David K. Jackson, Ian Johnston, Dominic Kwiatkowski, Cordelia Langford, John Sillitoe on behalf of the Wellcome Sanger Institute COVID-19 Surveillance Team |
| EPI_ISL_3286862 | Lighthouse Lab in Milton Keynes | Wellcome Sanger Institute for the COVID-19 Genomics UK (COG-UK) Consortium | The Lighthouse Lab in Milton Keynes and Alex Alderton, Roberto Amato, Jeffrey Barrett, Sonia Goncalves, Ewan Harrison, David K. Jackson, Ian Johnston, Dominic Kwiatkowski, Cordelia Langford, John Sillitoe on behalf of the Wellcome Sanger Institute COVID-19 Surveillance Team |
| EPI_ISL_3286863, EPI_ISL_3286864 | Lighthouse Laboratory Plymouth | Wellcome Sanger Institute for the COVID-19 Genomics UK (COG-UK) Consortium | Lighthouse Laboratory Plymouth and Alex Alderton, Roberto Amato, Jeffrey Barrett, Sonia Goncalves, Ewan Harrison, David K. Jackson, Ian Johnston, Dominic Kwiatkowski, Cordelia Langford, John Sillitoe on behalf of the Wellcome Sanger Institute COVID-19 Surveillance Team |
| EPI_ISL_3286865 | Lighthouse Lab in Glasgow | Wellcome Sanger Institute for the COVID-19 Genomics UK (COG-UK) Consortium | Harper VanSteenhouse, Yumi Kasai, David Gray, Carol Clugston, Anna Dominiczak and Alex Alderton, Roberto Amato, Jeffrey Barrett, Sonia Goncalves, Ewan Harrison, David K. Jackson, Ian Johnston, Dominic Kwiatkowski, Cordelia Langford, John Sillitoe on behalf of the Wellcome Sanger Institute COVID-19 Surveillance Team |
| EPI_ISL_3286866 | Lighthouse Lab in Alderley Park | Wellcome Sanger Institute for the COVID-19 Genomics UK (COG-UK) Consortium | Jacquelyn Wynn, Mairead Hyland, The Lighthouse Lab in Alderley Park and Alex Alderton, Roberto Amato, Jeffrey Barrett, Sonia Goncalves, Ewan Harrison, David K. Jackson, Ian Johnston, Dominic Kwiatkowski, Cordelia Langford, John Sillitoe on behalf of the Wellcome Sanger Institute COVID-19 Surveillance Team |
| EPI_ISL_3286867 | Lighthouse Lab in Milton Keynes | Wellcome Sanger Institute for the COVID-19 Genomics UK (COG-UK) Consortium | The Lighthouse Lab in Milton Keynes and Alex Alderton, Roberto Amato, Jeffrey Barrett, Sonia Goncalves, Ewan Harrison, David K. Jackson, Ian Johnston, Dominic Kwiatkowski, Cordelia Langford, John Sillitoe on behalf of the Wellcome Sanger Institute COVID-19 Surveillance Team |
| EPI_ISL_3286868, EPI_ISL_3286869 | Lighthouse Lab in Alderley Park | Wellcome Sanger Institute for the COVID-19 Genomics UK (COG-UK) Consortium | Jacquelyn Wynn, Mairead Hyland, The Lighthouse Lab in Alderley Park and Alex Alderton, Roberto Amato, Jeffrey Barrett, Sonia Goncalves, Ewan Harrison, David K. Jackson, Ian Johnston, Dominic Kwiatkowski, Cordelia Langford, John Sillitoe on behalf of the Wellcome Sanger Institute COVID-19 Surveillance Team |
| EPI_ISL_3286870 | Lighthouse Lab in Milton Keynes | Wellcome Sanger Institute for the COVID-19 Genomics UK (COG-UK) Consortium | The Lighthouse Lab in Milton Keynes and Alex Alderton, Roberto Amato, Jeffrey Barrett, Sonia Goncalves, Ewan Harrison, David K. Jackson, Ian Johnston, Dominic Kwiatkowski, Cordelia Langford, John Sillitoe on behalf of the Wellcome Sanger Institute COVID-19 Surveillance Team |
| EPI_ISL_3286871 | Lighthouse Lab in Glasgow | Wellcome Sanger Institute for the COVID-19 Genomics UK (COG-UK) Consortium | Harper VanSteenhouse, Yumi Kasai, David Gray, Carol Clugston, Anna Dominiczak and Alex Alderton, Roberto Amato, Jeffrey Barrett, Sonia Goncalves, Ewan Harrison, David K. Jackson, Ian Johnston, Dominic Kwiatkowski, Cordelia Langford, John Sillitoe on behalf of the Wellcome Sanger Institute COVID-19 Surveillance Team |
| EPI_ISL_3286872 | Lighthouse Lab in Milton Keynes | Wellcome Sanger Institute for the COVID-19 Genomics UK (COG-UK) Consortium | The Lighthouse Lab in Milton Keynes and Alex Alderton, Roberto Amato, Jeffrey Barrett, Sonia Goncalves, Ewan Harrison, David K. Jackson, Ian Johnston, Dominic Kwiatkowski, Cordelia Langford, John Sillitoe on behalf of the Wellcome Sanger Institute COVID-19 Surveillance Team |
| EPI_ISL_3286873 | Lighthouse Laboratory Plymouth | Wellcome Sanger Institute for the COVID-19 Genomics UK (COG-UK) Consortium | Lighthouse Laboratory Plymouth and Alex Alderton, Roberto Amato, Jeffrey Barrett, Sonia Goncalves, Ewan Harrison, David K. Jackson, Ian Johnston, Dominic Kwiatkowski, Cordelia Langford, John Sillitoe on behalf of the Wellcome Sanger Institute COVID-19 Surveillance Team |
| EPI_ISL_3286874 | Lighthouse Lab in Alderley Park | Wellcome Sanger Institute for the COVID-19 Genomics UK (COG-UK) Consortium | Jacquelyn Wynn, Mairead Hyland, The Lighthouse Lab in Alderley Park and Alex Alderton, Roberto Amato, Jeffrey Barrett, Sonia Goncalves, Ewan Harrison, David K. Jackson, Ian Johnston, Dominic Kwiatkowski, Cordelia Langford, John Sillitoe on behalf of the Wellcome Sanger Institute COVID-19 Surveillance Team |
| EPI_ISL_3286875, EPI_ISL_3286876, EPI_ISL_3286877 | Lighthouse Laboratory Plymouth | Wellcome Sanger Institute for the COVID-19 Genomics UK (COG-UK) Consortium | Lighthouse Laboratory Plymouth and Alex Alderton, Roberto Amato, Jeffrey Barrett, Sonia Goncalves, Ewan Harrison, David K. Jackson, Ian Johnston, Dominic Kwiatkowski, Cordelia Langford, John Sillitoe on behalf of the Wellcome Sanger Institute COVID-19 Surveillance Team |
| EPI_ISL_3286878, EPI_ISL_3286879 | Lighthouse Lab in Milton Keynes | Wellcome Sanger Institute for the COVID-19 Genomics UK (COG-UK) Consortium | The Lighthouse Lab in Milton Keynes and Alex Alderton, Roberto Amato, Jeffrey Barrett, Sonia Goncalves, Ewan Harrison, David K. Jackson, Ian Johnston, Dominic Kwiatkowski, Cordelia Langford, John Sillitoe on behalf of the Wellcome Sanger Institute COVID-19 Surveillance Team |
| EPI_ISL_3286880 | Lighthouse Laboratory Plymouth | Wellcome Sanger Institute for the COVID-19 Genomics UK (COG-UK) Consortium | Lighthouse Laboratory Plymouth and Alex Alderton, Roberto Amato, Jeffrey Barrett, Sonia Goncalves, Ewan Harrison, David K. Jackson, Ian Johnston, Dominic Kwiatkowski, Cordelia Langford, John Sillitoe on behalf of the Wellcome Sanger Institute COVID-19 Surveillance Team |
| EPI_ISL_3286881 | Lighthouse Lab in Milton Keynes | Wellcome Sanger Institute for the COVID-19 Genomics UK (COG-UK) Consortium | The Lighthouse Lab in Milton Keynes and Alex Alderton, Roberto Amato, Jeffrey Barrett, Sonia Goncalves, Ewan Harrison, David K. Jackson, Ian Johnston, Dominic Kwiatkowski, Cordelia Langford, John Sillitoe on behalf of the Wellcome Sanger Institute COVID-19 Surveillance Team |
| EPI_ISL_3286882, EPI_ISL_3286883 | Lighthouse Lab in Glasgow | Wellcome Sanger Institute for the COVID-19 Genomics UK (COG-UK) Consortium | Harper VanSteenhouse, Yumi Kasai, David Gray, Carol Clugston, Anna Dominiczak and Alex Alderton, Roberto Amato, Jeffrey Barrett, Sonia Goncalves, Ewan Harrison, David K. Jackson, Ian Johnston, Dominic Kwiatkowski, Cordelia Langford, John Sillitoe on behalf of the Wellcome Sanger Institute COVID-19 Surveillance Team |
| EPI_ISL_3286884 | Lighthouse Lab in Alderley Park | Wellcome Sanger Institute for the COVID-19 Genomics UK (COG-UK) Consortium | Jacquelyn Wynn, Mairead Hyland, The Lighthouse Lab in Alderley Park and Alex Alderton, Roberto Amato, Jeffrey Barrett, Sonia Goncalves, Ewan Harrison, David K. Jackson, Ian Johnston, Dominic Kwiatkowski, Cordelia Langford, John Sillitoe on behalf of the Wellcome Sanger Institute COVID-19 Surveillance Team |
| EPI_ISL_3286885, EPI_ISL_3286886 | Lighthouse Laboratory Plymouth | Wellcome Sanger Institute for the COVID-19 Genomics UK (COG-UK) Consortium | Lighthouse Laboratory Plymouth and Alex Alderton, Roberto Amato, Jeffrey Barrett, Sonia Goncalves, Ewan Harrison, David K. Jackson, Ian Johnston, Dominic Kwiatkowski, Cordelia Langford, John Sillitoe on behalf of the Wellcome Sanger Institute COVID-19 Surveillance Team |
| EPI_ISL_3286887 | Lighthouse Lab in Alderley Park | Wellcome Sanger Institute for the COVID-19 Genomics UK (COG-UK) Consortium | Jacquelyn Wynn, Mairead Hyland, The Lighthouse Lab in Alderley Park and Alex Alderton, Roberto Amato, Jeffrey Barrett, Sonia Goncalves, Ewan Harrison, David K. Jackson, Ian Johnston, Dominic Kwiatkowski, Cordelia Langford, John Sillitoe on behalf of the Wellcome Sanger Institute COVID-19 Surveillance Team |
| EPI_ISL_3286888 | Lighthouse Laboratory Plymouth | Wellcome Sanger Institute for the COVID-19 Genomics UK (COG-UK) Consortium | Lighthouse Laboratory Plymouth and Alex Alderton, Roberto Amato, Jeffrey Barrett, Sonia Goncalves, Ewan Harrison, David K. Jackson, Ian Johnston, Dominic Kwiatkowski, Cordelia Langford, John Sillitoe on behalf of the Wellcome Sanger Institute COVID-19 Surveillance Team |
| EPI_ISL_3286889 | Lighthouse Lab in Glasgow | Wellcome Sanger Institute for the COVID-19 Genomics UK (COG-UK) Consortium | Harper VanSteenhouse, Yumi Kasai, David Gray, Carol Clugston, Anna Dominiczak and Alex Alderton, Roberto Amato, Jeffrey Barrett, Sonia Goncalves, Ewan Harrison, David K. Jackson, Ian Johnston, Dominic Kwiatkowski, Cordelia Langford, John Sillitoe on behalf of the Wellcome Sanger Institute COVID-19 Surveillance Team |
| EPI_ISL_3286890 | Lighthouse Laboratory Plymouth | Wellcome Sanger Institute for the COVID-19 Genomics UK (COG-UK) Consortium | Lighthouse Laboratory Plymouth and Alex Alderton, Roberto Amato, Jeffrey Barrett, Sonia Goncalves, Ewan Harrison, David K. Jackson, Ian Johnston, Dominic Kwiatkowski, Cordelia Langford, John Sillitoe on behalf of the Wellcome Sanger Institute COVID-19 Surveillance Team |
| EPI_ISL_3286891 | Lighthouse Lab in Milton Keynes | Wellcome Sanger Institute for the COVID-19 Genomics UK (COG-UK) Consortium | The Lighthouse Lab in Milton Keynes and Alex Alderton, Roberto Amato, Jeffrey Barrett, Sonia Goncalves, Ewan Harrison, David K. Jackson, Ian Johnston, Dominic Kwiatkowski, Cordelia Langford, John Sillitoe on behalf of the Wellcome Sanger Institute COVID-19 Surveillance Team |
| EPI_ISL_3286892 | Lighthouse Lab in Alderley Park | Wellcome Sanger Institute for the COVID-19 Genomics UK (COG-UK) Consortium | Jacquelyn Wynn, Mairead Hyland, The Lighthouse Lab in Alderley Park and Alex Alderton, Roberto Amato, Jeffrey Barrett, Sonia Goncalves, Ewan Harrison, David K. Jackson, Ian Johnston, Dominic Kwiatkowski, Cordelia Langford, John Sillitoe on behalf of the Wellcome Sanger Institute COVID-19 Surveillance Team |
| EPI_ISL_3286893 | Lighthouse Lab in Milton Keynes | Wellcome Sanger Institute for the COVID-19 Genomics UK (COG-UK) | The Lighthouse Lab in Milton Keynes and Alex Alderton, Roberto Amato, Jeffrey Barrett, Sonia Goncalves, Ewan Harrison, David K. Jackson, Ian Johnston, Dominic Kwiatkowski, Cordelia Langford, John Sillitoe on behalf of the Wellcome Sanger |

| | | | |
|---|---|---|---|
| | | Consortium | Institute COVID-19 Surveillance Team |
| EPI_ISL_3286894 | Lighthouse Lab in Glasgow | Wellcome Sanger Institute for the COVID-19 Genomics UK (COG-UK) Consortium | Harper VanSteenhouse, Yumi Kasai, David Gray, Carol Clugston, Anna Dominiczak and Alex Alderton, Roberto Amato, Jeffrey Barrett, Sonia Goncalves, Ewan Harrison, David K. Jackson, Ian Johnston, Dominic Kwiatkowski, Cordelia Langford, John Sillitoe on behalf of the Wellcome Sanger Institute COVID-19 Surveillance Team |
| EPI_ISL_3286895 | Lighthouse Laboratory Plymouth | Wellcome Sanger Institute for the COVID-19 Genomics UK (COG-UK) Consortium | Lighthouse Laboratory Plymouth and Alex Alderton, Roberto Amato, Jeffrey Barrett, Sonia Goncalves, Ewan Harrison, David K. Jackson, Ian Johnston, Dominic Kwiatkowski, Cordelia Langford, John Sillitoe on behalf of the Wellcome Sanger Institute COVID-19 Surveillance Team |
| EPI_ISL_3286896 | Lighthouse Lab in Alderley Park | Wellcome Sanger Institute for the COVID-19 Genomics UK (COG-UK) Consortium | Jacquelyn Wynn, Mairead Hyland, The Lighthouse Lab in Alderley Park and Alex Alderton, Roberto Amato, Jeffrey Barrett, Sonia Goncalves, Ewan Harrison, David K. Jackson, Ian Johnston, Dominic Kwiatkowski, Cordelia Langford, John Sillitoe on behalf of the Wellcome Sanger Institute COVID-19 Surveillance Team |
| EPI_ISL_3286897 | Lighthouse Laboratory Plymouth | Wellcome Sanger Institute for the COVID-19 Genomics UK (COG-UK) Consortium | Lighthouse Laboratory Plymouth and Alex Alderton, Roberto Amato, Jeffrey Barrett, Sonia Goncalves, Ewan Harrison, David K. Jackson, Ian Johnston, Dominic Kwiatkowski, Cordelia Langford, John Sillitoe on behalf of the Wellcome Sanger Institute COVID-19 Surveillance Team |
| EPI_ISL_3286898 | Lighthouse Lab in Glasgow | Wellcome Sanger Institute for the COVID-19 Genomics UK (COG-UK) Consortium | Harper VanSteenhouse, Yumi Kasai, David Gray, Carol Clugston, Anna Dominiczak and Alex Alderton, Roberto Amato, Jeffrey Barrett, Sonia Goncalves, Ewan Harrison, David K. Jackson, Ian Johnston, Dominic Kwiatkowski, Cordelia Langford, John Sillitoe on behalf of the Wellcome Sanger Institute COVID-19 Surveillance Team |
| EPI_ISL_3286899, EPI_ISL_3286900 | Lighthouse Lab in Alderley Park | Wellcome Sanger Institute for the COVID-19 Genomics UK (COG-UK) Consortium | Jacquelyn Wynn, Mairead Hyland, The Lighthouse Lab in Alderley Park and Alex Alderton, Roberto Amato, Jeffrey Barrett, Sonia Goncalves, Ewan Harrison, David K. Jackson, Ian Johnston, Dominic Kwiatkowski, Cordelia Langford, John Sillitoe on behalf of the Wellcome Sanger Institute COVID-19 Surveillance Team |
| EPI_ISL_3286901 | Lighthouse Lab in Glasgow | Wellcome Sanger Institute for the COVID-19 Genomics UK (COG-UK) Consortium | Harper VanSteenhouse, Yumi Kasai, David Gray, Carol Clugston, Anna Dominiczak and Alex Alderton, Roberto Amato, Jeffrey Barrett, Sonia Goncalves, Ewan Harrison, David K. Jackson, Ian Johnston, Dominic Kwiatkowski, Cordelia Langford, John Sillitoe on behalf of the Wellcome Sanger Institute COVID-19 Surveillance Team |
| EPI_ISL_3286902 | Lighthouse Lab in Milton Keynes | Wellcome Sanger Institute for the COVID-19 Genomics UK (COG-UK) Consortium | The Lighthouse Lab in Milton Keynes and Alex Alderton, Roberto Amato, Jeffrey Barrett, Sonia Goncalves, Ewan Harrison, David K. Jackson, Ian Johnston, Dominic Kwiatkowski, Cordelia Langford, John Sillitoe on behalf of the Wellcome Sanger Institute COVID-19 Surveillance Team |
| EPI_ISL_3286903 | Lighthouse Lab in Glasgow | Wellcome Sanger Institute for the COVID-19 Genomics UK (COG-UK) Consortium | Harper VanSteenhouse, Yumi Kasai, David Gray, Carol Clugston, Anna Dominiczak and Alex Alderton, Roberto Amato, Jeffrey Barrett, Sonia Goncalves, Ewan Harrison, David K. Jackson, Ian Johnston, Dominic Kwiatkowski, Cordelia Langford, John Sillitoe on behalf of the Wellcome Sanger Institute COVID-19 Surveillance Team |
| EPI_ISL_3286904 | Lighthouse Lab in Alderley Park | Wellcome Sanger Institute for the COVID-19 Genomics UK (COG-UK) Consortium | Jacquelyn Wynn, Mairead Hyland, The Lighthouse Lab in Alderley Park and Alex Alderton, Roberto Amato, Jeffrey Barrett, Sonia Goncalves, Ewan Harrison, David K. Jackson, Ian Johnston, Dominic Kwiatkowski, Cordelia Langford, John Sillitoe on behalf of the Wellcome Sanger Institute COVID-19 Surveillance Team |
| EPI_ISL_3286905, EPI_ISL_3286906, EPI_ISL_3286907 | Lighthouse Lab in Glasgow | Wellcome Sanger Institute for the COVID-19 Genomics UK (COG-UK) Consortium | Harper VanSteenhouse, Yumi Kasai, David Gray, Carol Clugston, Anna Dominiczak and Alex Alderton, Roberto Amato, Jeffrey Barrett, Sonia Goncalves, Ewan Harrison, David K. Jackson, Ian Johnston, Dominic Kwiatkowski, Cordelia Langford, John Sillitoe on behalf of the Wellcome Sanger Institute COVID-19 Surveillance Team |
| EPI_ISL_3286908 | Lighthouse Laboratory Plymouth | Wellcome Sanger Institute for the COVID-19 Genomics UK (COG-UK) Consortium | Lighthouse Laboratory Plymouth and Alex Alderton, Roberto Amato, Jeffrey Barrett, Sonia Goncalves, Ewan Harrison, David K. Jackson, Ian Johnston, Dominic Kwiatkowski, Cordelia Langford, John Sillitoe on behalf of the Wellcome Sanger Institute COVID-19 Surveillance Team |
| EPI_ISL_3286909 | Lighthouse Lab in Alderley Park | Wellcome Sanger Institute for the COVID-19 Genomics UK (COG-UK) Consortium | Jacquelyn Wynn, Mairead Hyland, The Lighthouse Lab in Alderley Park and Alex Alderton, Roberto Amato, Jeffrey Barrett, Sonia Goncalves, Ewan Harrison, David K. Jackson, Ian Johnston, Dominic Kwiatkowski, Cordelia Langford, John Sillitoe on behalf of the Wellcome Sanger Institute COVID-19 Surveillance Team |
| EPI_ISL_3286910 | Lighthouse Lab in Milton Keynes | Wellcome Sanger Institute for the COVID-19 Genomics UK (COG-UK) Consortium | The Lighthouse Lab in Milton Keynes and Alex Alderton, Roberto Amato, Jeffrey Barrett, Sonia Goncalves, Ewan Harrison, David K. Jackson, Ian Johnston, Dominic Kwiatkowski, Cordelia Langford, John Sillitoe on behalf of the Wellcome Sanger Institute COVID-19 Surveillance Team |
| EPI_ISL_3286911 | Lighthouse Lab in Alderley Park | Wellcome Sanger Institute for the COVID-19 Genomics UK (COG-UK) Consortium | Jacquelyn Wynn, Mairead Hyland, The Lighthouse Lab in Alderley Park and Alex Alderton, Roberto Amato, Jeffrey Barrett, Sonia Goncalves, Ewan Harrison, David K. Jackson, Ian Johnston, Dominic Kwiatkowski, Cordelia Langford, John Sillitoe on behalf of the Wellcome Sanger Institute COVID-19 Surveillance Team |
| EPI_ISL_3286912 | Lighthouse Lab in Glasgow | Wellcome Sanger Institute for the COVID-19 Genomics UK (COG-UK) Consortium | Harper VanSteenhouse, Yumi Kasai, David Gray, Carol Clugston, Anna Dominiczak and Alex Alderton, Roberto Amato, Jeffrey Barrett, Sonia Goncalves, Ewan Harrison, David K. Jackson, Ian Johnston, Dominic Kwiatkowski, Cordelia Langford, John Sillitoe on behalf of the Wellcome Sanger Institute COVID-19 Surveillance Team |
| EPI_ISL_3286913, EPI_ISL_3286914, EPI_ISL_3286915, EPI_ISL_3286916 | Lighthouse Lab in Milton Keynes | Wellcome Sanger Institute for the COVID-19 Genomics UK (COG-UK) Consortium | The Lighthouse Lab in Milton Keynes and Alex Alderton, Roberto Amato, Jeffrey Barrett, Sonia Goncalves, Ewan Harrison, David K. Jackson, Ian Johnston, Dominic Kwiatkowski, Cordelia Langford, John Sillitoe on behalf of the Wellcome Sanger Institute COVID-19 Surveillance Team |
| EPI_ISL_3286917 | Lighthouse Laboratory Plymouth | Wellcome Sanger Institute for the COVID-19 Genomics UK (COG-UK) Consortium | Lighthouse Laboratory Plymouth and Alex Alderton, Roberto Amato, Jeffrey Barrett, Sonia Goncalves, Ewan Harrison, David K. Jackson, Ian Johnston, Dominic Kwiatkowski, Cordelia Langford, John Sillitoe on behalf of the Wellcome Sanger Institute COVID-19 Surveillance Team |
| EPI_ISL_3286918 | Lighthouse Lab in Milton Keynes | Wellcome Sanger Institute for the COVID-19 Genomics UK (COG-UK) Consortium | The Lighthouse Lab in Milton Keynes and Alex Alderton, Roberto Amato, Jeffrey Barrett, Sonia Goncalves, Ewan Harrison, David K. Jackson, Ian Johnston, Dominic Kwiatkowski, Cordelia Langford, John Sillitoe on behalf of the Wellcome Sanger Institute COVID-19 Surveillance Team |
| EPI_ISL_3286919 | Lighthouse Lab in Alderley Park | Wellcome Sanger Institute for the COVID-19 Genomics UK (COG-UK) Consortium | Jacquelyn Wynn, Mairead Hyland, The Lighthouse Lab in Alderley Park and Alex Alderton, Roberto Amato, Jeffrey Barrett, Sonia Goncalves, Ewan Harrison, David K. Jackson, Ian Johnston, Dominic Kwiatkowski, Cordelia Langford, John Sillitoe on behalf of the Wellcome Sanger Institute COVID-19 Surveillance Team |
| EPI_ISL_3286920, EPI_ISL_3286921 | Lighthouse Laboratory Plymouth | Wellcome Sanger Institute for the COVID-19 Genomics UK (COG-UK) Consortium | Lighthouse Laboratory Plymouth and Alex Alderton, Roberto Amato, Jeffrey Barrett, Sonia Goncalves, Ewan Harrison, David K. Jackson, Ian Johnston, Dominic Kwiatkowski, Cordelia Langford, John Sillitoe on behalf of the Wellcome Sanger Institute COVID-19 Surveillance Team |
| EPI_ISL_3286922, EPI_ISL_3286923, EPI_ISL_3286924 | Lighthouse Lab in Milton Keynes | Wellcome Sanger Institute for the COVID-19 Genomics UK (COG-UK) Consortium | The Lighthouse Lab in Milton Keynes and Alex Alderton, Roberto Amato, Jeffrey Barrett, Sonia Goncalves, Ewan Harrison, David K. Jackson, Ian Johnston, Dominic Kwiatkowski, Cordelia Langford, John Sillitoe on behalf of the Wellcome Sanger Institute COVID-19 Surveillance Team |
| EPI_ISL_3286925 | Lighthouse Laboratory Plymouth | Wellcome Sanger Institute for the COVID-19 Genomics UK (COG-UK) Consortium | Lighthouse Laboratory Plymouth and Alex Alderton, Roberto Amato, Jeffrey Barrett, Sonia Goncalves, Ewan Harrison, David K. Jackson, Ian Johnston, Dominic Kwiatkowski, Cordelia Langford, John Sillitoe on behalf of the Wellcome Sanger Institute COVID-19 Surveillance Team |
| EPI_ISL_3286926 | Lighthouse Lab in Glasgow | Wellcome Sanger Institute for the COVID-19 Genomics UK (COG-UK) Consortium | Harper VanSteenhouse, Yumi Kasai, David Gray, Carol Clugston, Anna Dominiczak and Alex Alderton, Roberto Amato, Jeffrey Barrett, Sonia Goncalves, Ewan Harrison, David K. Jackson, Ian Johnston, Dominic Kwiatkowski, Cordelia Langford, John Sillitoe on behalf of the Wellcome Sanger Institute COVID-19 Surveillance Team |
| EPI_ISL_3286927 | Lighthouse Lab in Alderley Park | Wellcome Sanger Institute for the COVID-19 Genomics UK (COG-UK) Consortium | Jacquelyn Wynn, Mairead Hyland, The Lighthouse Lab in Alderley Park and Alex Alderton, Roberto Amato, Jeffrey Barrett, Sonia Goncalves, Ewan Harrison, David K. Jackson, Ian Johnston, Dominic Kwiatkowski, Cordelia Langford, John Sillitoe on behalf of the Wellcome Sanger Institute COVID-19 Surveillance Team |
| EPI_ISL_3286928 | Lighthouse Lab in Glasgow | Wellcome Sanger Institute for the COVID-19 Genomics UK (COG-UK) Consortium | Harper VanSteenhouse, Yumi Kasai, David Gray, Carol Clugston, Anna Dominiczak and Alex Alderton, Roberto Amato, Jeffrey Barrett, Sonia Goncalves, Ewan Harrison, David K. Jackson, Ian Johnston, Dominic Kwiatkowski, Cordelia Langford, John Sillitoe on behalf of the Wellcome Sanger Institute COVID-19 Surveillance Team |
| EPI_ISL_3286929 | Lighthouse Lab in Alderley Park | Wellcome Sanger Institute for the COVID-19 Genomics UK (COG-UK) Consortium | Jacquelyn Wynn, Mairead Hyland, The Lighthouse Lab in Alderley Park and Alex Alderton, Roberto Amato, Jeffrey Barrett, Sonia Goncalves, Ewan Harrison, David K. Jackson, Ian Johnston, Dominic Kwiatkowski, Cordelia Langford, John Sillitoe on behalf of the Wellcome Sanger Institute COVID-19 Surveillance Team |
| EPI_ISL_3286930 | Lighthouse Lab in Milton Keynes | Wellcome Sanger Institute for the COVID-19 Genomics UK (COG-UK) Consortium | The Lighthouse Lab in Milton Keynes and Alex Alderton, Roberto Amato, Jeffrey Barrett, Sonia Goncalves, Ewan Harrison, David K. Jackson, Ian Johnston, Dominic Kwiatkowski, Cordelia Langford, John Sillitoe on behalf of the Wellcome Sanger Institute COVID-19 Surveillance Team |
| EPI_ISL_3286931 | Lighthouse Lab in Glasgow | Wellcome Sanger Institute for the COVID-19 Genomics UK (COG-UK) Consortium | Harper VanSteenhouse, Yumi Kasai, David Gray, Carol Clugston, Anna Dominiczak and Alex Alderton, Roberto Amato, Jeffrey Barrett, Sonia Goncalves, Ewan Harrison, David K. Jackson, Ian Johnston, Dominic Kwiatkowski, Cordelia Langford, John Sillitoe on behalf of the Wellcome Sanger Institute COVID-19 Surveillance Team |
| EPI_ISL_3286932 | Lighthouse Lab in Milton Keynes | Wellcome Sanger Institute for the COVID-19 Genomics UK (COG-UK) Consortium | The Lighthouse Lab in Milton Keynes and Alex Alderton, Roberto Amato, Jeffrey Barrett, Sonia Goncalves, Ewan Harrison, David K. Jackson, Ian Johnston, Dominic Kwiatkowski, Cordelia Langford, John Sillitoe on behalf of the Wellcome Sanger Institute COVID-19 Surveillance Team |
| EPI_ISL_3286933 | Lighthouse Laboratory Plymouth | Wellcome Sanger Institute for the COVID-19 Genomics UK (COG-UK) Consortium | Lighthouse Laboratory Plymouth and Alex Alderton, Roberto Amato, Jeffrey Barrett, Sonia Goncalves, Ewan Harrison, David K. Jackson, Ian Johnston, Dominic Kwiatkowski, Cordelia Langford, John Sillitoe on behalf of the Wellcome Sanger Institute COVID-19 Surveillance Team |
| EPI_ISL_3286934 | Lighthouse Lab in Alderley Park | Wellcome Sanger Institute for the COVID-19 Genomics UK (COG-UK) Consortium | Jacquelyn Wynn, Mairead Hyland, The Lighthouse Lab in Alderley Park and Alex Alderton, Roberto Amato, Jeffrey Barrett, Sonia Goncalves, Ewan Harrison, David K. Jackson, Ian Johnston, Dominic Kwiatkowski, Cordelia Langford, John Sillitoe on behalf of the Wellcome Sanger Institute COVID-19 Surveillance Team |
| EPI_ISL_3286935 | Lighthouse Lab in Milton Keynes | Wellcome Sanger Institute for the COVID-19 Genomics UK (COG-UK) Consortium | The Lighthouse Lab in Milton Keynes and Alex Alderton, Roberto Amato, Jeffrey Barrett, Sonia Goncalves, Ewan Harrison, David K. Jackson, Ian Johnston, Dominic Kwiatkowski, Cordelia Langford, John Sillitoe on behalf of the Wellcome Sanger Institute COVID-19 Surveillance Team |
| EPI_ISL_3286936 | Lighthouse Laboratory Plymouth | Wellcome Sanger Institute for the COVID-19 Genomics UK (COG-UK) Consortium | Lighthouse Laboratory Plymouth and Alex Alderton, Roberto Amato, Jeffrey Barrett, Sonia Goncalves, Ewan Harrison, David K. Jackson, Ian Johnston, Dominic Kwiatkowski, Cordelia Langford, John Sillitoe on behalf of the Wellcome Sanger Institute COVID-19 Surveillance Team |
| EPI_ISL_3286937 | Lighthouse Lab in Milton Keynes | Wellcome Sanger Institute for the COVID-19 Genomics UK (COG-UK) Consortium | The Lighthouse Lab in Milton Keynes and Alex Alderton, Roberto Amato, Jeffrey Barrett, Sonia Goncalves, Ewan Harrison, David K. Jackson, Ian Johnston, Dominic Kwiatkowski, Cordelia Langford, John Sillitoe on behalf of the Wellcome Sanger Institute COVID-19 Surveillance Team |
| EPI_ISL_3286938 | Lighthouse Laboratory Plymouth | Wellcome Sanger Institute for the COVID-19 Genomics UK (COG-UK) Consortium | Lighthouse Laboratory Plymouth and Alex Alderton, Roberto Amato, Jeffrey Barrett, Sonia Goncalves, Ewan Harrison, David K. Jackson, Ian Johnston, Dominic Kwiatkowski, Cordelia Langford, John Sillitoe on behalf of the Wellcome Sanger Institute COVID-19 Surveillance Team |
| EPI_ISL_3286939 | Lighthouse Lab in Milton Keynes | Wellcome Sanger Institute for the COVID-19 Genomics UK (COG-UK) Consortium | The Lighthouse Lab in Milton Keynes and Alex Alderton, Roberto Amato, Jeffrey Barrett, Sonia Goncalves, Ewan Harrison, David K. Jackson, Ian Johnston, Dominic Kwiatkowski, Cordelia Langford, John Sillitoe on behalf of the Wellcome Sanger Institute COVID-19 Surveillance Team |
| EPI_ISL_3286940 | Lighthouse Lab in Glasgow | Wellcome Sanger Institute for the COVID-19 Genomics UK (COG-UK) Consortium | Harper VanSteenhouse, Yumi Kasai, David Gray, Carol Clugston, Anna Dominiczak and Alex Alderton, Roberto Amato, Jeffrey Barrett, Sonia Goncalves, Ewan Harrison, David K. Jackson, Ian Johnston, Dominic Kwiatkowski, Cordelia Langford, John Sillitoe on behalf of the Wellcome Sanger Institute COVID-19 Surveillance Team |
| EPI_ISL_3286941 | Lighthouse Lab in Milton Keynes | Wellcome Sanger Institute for the COVID-19 Genomics UK (COG-UK) Consortium | The Lighthouse Lab in Milton Keynes and Alex Alderton, Roberto Amato, Jeffrey Barrett, Sonia Goncalves, Ewan Harrison, David K. Jackson, Ian Johnston, Dominic Kwiatkowski, Cordelia Langford, John Sillitoe on behalf of the Wellcome Sanger Institute COVID-19 Surveillance Team |
| EPI_ISL_3286942, EPI_ISL_3286943, EPI_ISL_3286944 | Lighthouse Lab in Glasgow | Wellcome Sanger Institute for the COVID-19 Genomics UK (COG-UK) Consortium | Harper VanSteenhouse, Yumi Kasai, David Gray, Carol Clugston, Anna Dominiczak and Alex Alderton, Roberto Amato, Jeffrey Barrett, Sonia Goncalves, Ewan Harrison, David K. Jackson, Ian Johnston, Dominic Kwiatkowski, Cordelia Langford, John Sillitoe on behalf of the Wellcome Sanger Institute COVID-19 Surveillance Team |
| EPI_ISL_3286945 | Lighthouse Laboratory Plymouth | Wellcome Sanger Institute for the COVID-19 Genomics UK (COG-UK) Consortium | Lighthouse Laboratory Plymouth and Alex Alderton, Roberto Amato, Jeffrey Barrett, Sonia Goncalves, Ewan Harrison, David K. Jackson, Ian Johnston, Dominic Kwiatkowski, Cordelia Langford, John Sillitoe on behalf of the Wellcome Sanger Institute COVID-19 Surveillance Team |
| EPI_ISL_3286946 | Lighthouse Lab in Glasgow | Wellcome Sanger Institute for the COVID-19 Genomics UK (COG-UK) Consortium | Harper VanSteenhouse, Yumi Kasai, David Gray, Carol Clugston, Anna Dominiczak and Alex Alderton, Roberto Amato, Jeffrey Barrett, Sonia Goncalves, Ewan Harrison, David K. Jackson, Ian Johnston, Dominic Kwiatkowski, Cordelia Langford, John Sillitoe on behalf of the Wellcome Sanger Institute COVID-19 Surveillance Team |
| EPI_ISL_3286947 | Lighthouse Lab in Alderley Park | Wellcome Sanger Institute for the COVID-19 Genomics UK (COG-UK) | Jacquelyn Wynn, Mairead Hyland, The Lighthouse Lab in Alderley Park and Alex Alderton, Roberto Amato, Jeffrey Barrett, Sonia Goncalves, Ewan Harrison, David K. Jackson, Ian Johnston, Dominic Kwiatkowski, Cordelia Langford, John Sillitoe on |

| | | | |
|---|---|---|---|
| | | Consortium | behalf of the Wellcome Sanger Institute COVID-19 Surveillance Team |
| EPI_ISL_3286948 | Lighthouse Lab in Glasgow | Wellcome Sanger Institute for the COVID-19 Genomics UK (COG-UK) Consortium | Harper VanSteenhouse, Yumi Kasai, David Gray, Carol Clugston, Anna Dominiczak and Alex Alderton, Roberto Amato, Jeffrey Barrett, Sonia Goncalves, Ewan Harrison, David K. Jackson, Ian Johnston, Dominic Kwiatkowski, Cordelia Langford, John Sillitoe on behalf of the Wellcome Sanger Institute COVID-19 Surveillance Team |
| EPI_ISL_3286949 | Lighthouse Lab in Alderley Park | Wellcome Sanger Institute for the COVID-19 Genomics UK (COG-UK) Consortium | Jacquelyn Wynn, Mairead Hyland, The Lighthouse Lab in Alderley Park and Alex Alderton, Roberto Amato, Jeffrey Barrett, Sonia Goncalves, Ewan Harrison, David K. Jackson, Ian Johnston, Dominic Kwiatkowski, Cordelia Langford, John Sillitoe on behalf of the Wellcome Sanger Institute COVID-19 Surveillance Team |
| EPI_ISL_3286950, EPI_ISL_3286951 | Lighthouse Laboratory Plymouth | Wellcome Sanger Institute for the COVID-19 Genomics UK (COG-UK) Consortium | Lighthouse Laboratory Plymouth and Alex Alderton, Roberto Amato, Jeffrey Barrett, Sonia Goncalves, Ewan Harrison, David K. Jackson, Ian Johnston, Dominic Kwiatkowski, Cordelia Langford, John Sillitoe on behalf of the Wellcome Sanger Institute COVID-19 Surveillance Team |
| EPI_ISL_3286952 | Lighthouse Lab in Alderley Park | Wellcome Sanger Institute for the COVID-19 Genomics UK (COG-UK) Consortium | Jacquelyn Wynn, Mairead Hyland, The Lighthouse Lab in Alderley Park and Alex Alderton, Roberto Amato, Jeffrey Barrett, Sonia Goncalves, Ewan Harrison, David K. Jackson, Ian Johnston, Dominic Kwiatkowski, Cordelia Langford, John Sillitoe on behalf of the Wellcome Sanger Institute COVID-19 Surveillance Team |
| EPI_ISL_3286953 | Lighthouse Lab in Glasgow | Wellcome Sanger Institute for the COVID-19 Genomics UK (COG-UK) Consortium | Harper VanSteenhouse, Yumi Kasai, David Gray, Carol Clugston, Anna Dominiczak and Alex Alderton, Roberto Amato, Jeffrey Barrett, Sonia Goncalves, Ewan Harrison, David K. Jackson, Ian Johnston, Dominic Kwiatkowski, Cordelia Langford, John Sillitoe on behalf of the Wellcome Sanger Institute COVID-19 Surveillance Team |
| EPI_ISL_3286954 | Lighthouse Lab in Milton Keynes | Wellcome Sanger Institute for the COVID-19 Genomics UK (COG-UK) Consortium | The Lighthouse Lab in Milton Keynes and Alex Alderton, Roberto Amato, Jeffrey Barrett, Sonia Goncalves, Ewan Harrison, David K. Jackson, Ian Johnston, Dominic Kwiatkowski, Cordelia Langford, John Sillitoe on behalf of the Wellcome Sanger Institute COVID-19 Surveillance Team |
| EPI_ISL_3286955 | Lighthouse Lab in Alderley Park | Wellcome Sanger Institute for the COVID-19 Genomics UK (COG-UK) Consortium | Jacquelyn Wynn, Mairead Hyland, The Lighthouse Lab in Alderley Park and Alex Alderton, Roberto Amato, Jeffrey Barrett, Sonia Goncalves, Ewan Harrison, David K. Jackson, Ian Johnston, Dominic Kwiatkowski, Cordelia Langford, John Sillitoe on behalf of the Wellcome Sanger Institute COVID-19 Surveillance Team |
| EPI_ISL_3286956, EPI_ISL_3286957 | Lighthouse Lab in Glasgow | Wellcome Sanger Institute for the COVID-19 Genomics UK (COG-UK) Consortium | Harper VanSteenhouse, Yumi Kasai, David Gray, Carol Clugston, Anna Dominiczak and Alex Alderton, Roberto Amato, Jeffrey Barrett, Sonia Goncalves, Ewan Harrison, David K. Jackson, Ian Johnston, Dominic Kwiatkowski, Cordelia Langford, John Sillitoe on behalf of the Wellcome Sanger Institute COVID-19 Surveillance Team |
| EPI_ISL_3286958 | Lighthouse Lab in Milton Keynes | Wellcome Sanger Institute for the COVID-19 Genomics UK (COG-UK) Consortium | The Lighthouse Lab in Milton Keynes and Alex Alderton, Roberto Amato, Jeffrey Barrett, Sonia Goncalves, Ewan Harrison, David K. Jackson, Ian Johnston, Dominic Kwiatkowski, Cordelia Langford, John Sillitoe on behalf of the Wellcome Sanger Institute COVID-19 Surveillance Team |
| EPI_ISL_3286959, EPI_ISL_3286960 | Lighthouse Laboratory Plymouth | Wellcome Sanger Institute for the COVID-19 Genomics UK (COG-UK) Consortium | Lighthouse Laboratory Plymouth and Alex Alderton, Roberto Amato, Jeffrey Barrett, Sonia Goncalves, Ewan Harrison, David K. Jackson, Ian Johnston, Dominic Kwiatkowski, Cordelia Langford, John Sillitoe on behalf of the Wellcome Sanger Institute COVID-19 Surveillance Team |
| EPI_ISL_3286961 | Lighthouse Lab in Milton Keynes | Wellcome Sanger Institute for the COVID-19 Genomics UK (COG-UK) Consortium | The Lighthouse Lab in Milton Keynes and Alex Alderton, Roberto Amato, Jeffrey Barrett, Sonia Goncalves, Ewan Harrison, David K. Jackson, Ian Johnston, Dominic Kwiatkowski, Cordelia Langford, John Sillitoe on behalf of the Wellcome Sanger Institute COVID-19 Surveillance Team |
| EPI_ISL_3286962 | Lighthouse Lab in Glasgow | Wellcome Sanger Institute for the COVID-19 Genomics UK (COG-UK) Consortium | Harper VanSteenhouse, Yumi Kasai, David Gray, Carol Clugston, Anna Dominiczak and Alex Alderton, Roberto Amato, Jeffrey Barrett, Sonia Goncalves, Ewan Harrison, David K. Jackson, Ian Johnston, Dominic Kwiatkowski, Cordelia Langford, John Sillitoe on behalf of the Wellcome Sanger Institute COVID-19 Surveillance Team |
| EPI_ISL_3286963 | Lighthouse Lab in Milton Keynes | Wellcome Sanger Institute for the COVID-19 Genomics UK (COG-UK) Consortium | The Lighthouse Lab in Milton Keynes and Alex Alderton, Roberto Amato, Jeffrey Barrett, Sonia Goncalves, Ewan Harrison, David K. Jackson, Ian Johnston, Dominic Kwiatkowski, Cordelia Langford, John Sillitoe on behalf of the Wellcome Sanger Institute COVID-19 Surveillance Team |
| EPI_ISL_3286964, EPI_ISL_3286965 | Lighthouse Laboratory Plymouth | Wellcome Sanger Institute for the COVID-19 Genomics UK (COG-UK) Consortium | Lighthouse Laboratory Plymouth and Alex Alderton, Roberto Amato, Jeffrey Barrett, Sonia Goncalves, Ewan Harrison, David K. Jackson, Ian Johnston, Dominic Kwiatkowski, Cordelia Langford, John Sillitoe on behalf of the Wellcome Sanger Institute COVID-19 Surveillance Team |
| EPI_ISL_3286966, EPI_ISL_3286967 | Lighthouse Lab in Alderley Park | Wellcome Sanger Institute for the COVID-19 Genomics UK (COG-UK) Consortium | Jacquelyn Wynn, Mairead Hyland, The Lighthouse Lab in Alderley Park and Alex Alderton, Roberto Amato, Jeffrey Barrett, Sonia Goncalves, Ewan Harrison, David K. Jackson, Ian Johnston, Dominic Kwiatkowski, Cordelia Langford, John Sillitoe on behalf of the Wellcome Sanger Institute COVID-19 Surveillance Team |
| EPI_ISL_3286968, EPI_ISL_3286969 | Lighthouse Lab in Glasgow | Wellcome Sanger Institute for the COVID-19 Genomics UK (COG-UK) Consortium | Harper VanSteenhouse, Yumi Kasai, David Gray, Carol Clugston, Anna Dominiczak and Alex Alderton, Roberto Amato, Jeffrey Barrett, Sonia Goncalves, Ewan Harrison, David K. Jackson, Ian Johnston, Dominic Kwiatkowski, Cordelia Langford, John Sillitoe on behalf of the Wellcome Sanger Institute COVID-19 Surveillance Team |
| EPI_ISL_3286970, EPI_ISL_3286971 | Lighthouse Laboratory Plymouth | Wellcome Sanger Institute for the COVID-19 Genomics UK (COG-UK) Consortium | Lighthouse Laboratory Plymouth and Alex Alderton, Roberto Amato, Jeffrey Barrett, Sonia Goncalves, Ewan Harrison, David K. Jackson, Ian Johnston, Dominic Kwiatkowski, Cordelia Langford, John Sillitoe on behalf of the Wellcome Sanger Institute COVID-19 Surveillance Team |
| EPI_ISL_3286972, EPI_ISL_3286973, EPI_ISL_3286974 | Lighthouse Lab in Alderley Park | Wellcome Sanger Institute for the COVID-19 Genomics UK (COG-UK) Consortium | Jacquelyn Wynn, Mairead Hyland, The Lighthouse Lab in Alderley Park and Alex Alderton, Roberto Amato, Jeffrey Barrett, Sonia Goncalves, Ewan Harrison, David K. Jackson, Ian Johnston, Dominic Kwiatkowski, Cordelia Langford, John Sillitoe on behalf of the Wellcome Sanger Institute COVID-19 Surveillance Team |
| EPI_ISL_3286975, EPI_ISL_3286976 | Lighthouse Lab in Glasgow | Wellcome Sanger Institute for the COVID-19 Genomics UK (COG-UK) Consortium | Harper VanSteenhouse, Yumi Kasai, David Gray, Carol Clugston, Anna Dominiczak and Alex Alderton, Roberto Amato, Jeffrey Barrett, Sonia Goncalves, Ewan Harrison, David K. Jackson, Ian Johnston, Dominic Kwiatkowski, Cordelia Langford, John Sillitoe on behalf of the Wellcome Sanger Institute COVID-19 Surveillance Team |
| EPI_ISL_3286977 | Lighthouse Lab in Alderley Park | Wellcome Sanger Institute for the COVID-19 Genomics UK (COG-UK) Consortium | Jacquelyn Wynn, Mairead Hyland, The Lighthouse Lab in Alderley Park and Alex Alderton, Roberto Amato, Jeffrey Barrett, Sonia Goncalves, Ewan Harrison, David K. Jackson, Ian Johnston, Dominic Kwiatkowski, Cordelia Langford, John Sillitoe on behalf of the Wellcome Sanger Institute COVID-19 Surveillance Team |
| EPI_ISL_3286978 | Lighthouse Lab in Glasgow | Wellcome Sanger Institute for the COVID-19 Genomics UK (COG-UK) Consortium | Harper VanSteenhouse, Yumi Kasai, David Gray, Carol Clugston, Anna Dominiczak and Alex Alderton, Roberto Amato, Jeffrey Barrett, Sonia Goncalves, Ewan Harrison, David K. Jackson, Ian Johnston, Dominic Kwiatkowski, Cordelia Langford, John Sillitoe on behalf of the Wellcome Sanger Institute COVID-19 Surveillance Team |
| EPI_ISL_3286979 | Lighthouse Lab in Alderley Park | Wellcome Sanger Institute for the COVID-19 Genomics UK (COG-UK) Consortium | Jacquelyn Wynn, Mairead Hyland, The Lighthouse Lab in Alderley Park and Alex Alderton, Roberto Amato, Jeffrey Barrett, Sonia Goncalves, Ewan Harrison, David K. Jackson, Ian Johnston, Dominic Kwiatkowski, Cordelia Langford, John Sillitoe on behalf of the Wellcome Sanger Institute COVID-19 Surveillance Team |
| EPI_ISL_3286980 | Lighthouse Laboratory Plymouth | Wellcome Sanger Institute for the COVID-19 Genomics UK (COG-UK) Consortium | Lighthouse Laboratory Plymouth and Alex Alderton, Roberto Amato, Jeffrey Barrett, Sonia Goncalves, Ewan Harrison, David K. Jackson, Ian Johnston, Dominic Kwiatkowski, Cordelia Langford, John Sillitoe on behalf of the Wellcome Sanger Institute COVID-19 Surveillance Team |
| EPI_ISL_3286981 | Lighthouse Lab in Alderley Park | Wellcome Sanger Institute for the COVID-19 Genomics UK (COG-UK) Consortium | Jacquelyn Wynn, Mairead Hyland, The Lighthouse Lab in Alderley Park and Alex Alderton, Roberto Amato, Jeffrey Barrett, Sonia Goncalves, Ewan Harrison, David K. Jackson, Ian Johnston, Dominic Kwiatkowski, Cordelia Langford, John Sillitoe on behalf of the Wellcome Sanger Institute COVID-19 Surveillance Team |
| EPI_ISL_3286982 | Lighthouse Laboratory Plymouth | Wellcome Sanger Institute for the COVID-19 Genomics UK (COG-UK) Consortium | Lighthouse Laboratory Plymouth and Alex Alderton, Roberto Amato, Jeffrey Barrett, Sonia Goncalves, Ewan Harrison, David K. Jackson, Ian Johnston, Dominic Kwiatkowski, Cordelia Langford, John Sillitoe on behalf of the Wellcome Sanger Institute COVID-19 Surveillance Team |
| EPI_ISL_3286983 | Lighthouse Lab in Glasgow | Wellcome Sanger Institute for the COVID-19 Genomics UK (COG-UK) Consortium | Harper VanSteenhouse, Yumi Kasai, David Gray, Carol Clugston, Anna Dominiczak and Alex Alderton, Roberto Amato, Jeffrey Barrett, Sonia Goncalves, Ewan Harrison, David K. Jackson, Ian Johnston, Dominic Kwiatkowski, Cordelia Langford, John Sillitoe on behalf of the Wellcome Sanger Institute COVID-19 Surveillance Team |
| EPI_ISL_3286984, EPI_ISL_3286985, EPI_ISL_3286986, EPI_ISL_3286987 | Lighthouse Laboratory Plymouth | Wellcome Sanger Institute for the COVID-19 Genomics UK (COG-UK) Consortium | Lighthouse Laboratory Plymouth and Alex Alderton, Roberto Amato, Jeffrey Barrett, Sonia Goncalves, Ewan Harrison, David K. Jackson, Ian Johnston, Dominic Kwiatkowski, Cordelia Langford, John Sillitoe on behalf of the Wellcome Sanger Institute COVID-19 Surveillance Team |
| EPI_ISL_3286988 | Lighthouse Lab in Milton Keynes | Wellcome Sanger Institute for the COVID-19 Genomics UK (COG-UK) Consortium | The Lighthouse Lab in Milton Keynes and Alex Alderton, Roberto Amato, Jeffrey Barrett, Sonia Goncalves, Ewan Harrison, David K. Jackson, Ian Johnston, Dominic Kwiatkowski, Cordelia Langford, John Sillitoe on behalf of the Wellcome Sanger Institute COVID-19 Surveillance Team |
| EPI_ISL_3286989, EPI_ISL_3286990 | Lighthouse Lab in Glasgow | Wellcome Sanger Institute for the COVID-19 Genomics UK (COG-UK) Consortium | Harper VanSteenhouse, Yumi Kasai, David Gray, Carol Clugston, Anna Dominiczak and Alex Alderton, Roberto Amato, Jeffrey Barrett, Sonia Goncalves, Ewan Harrison, David K. Jackson, Ian Johnston, Dominic Kwiatkowski, Cordelia Langford, John Sillitoe on behalf of the Wellcome Sanger Institute COVID-19 Surveillance Team |
| EPI_ISL_3286991 | Lighthouse Laboratory Plymouth | Wellcome Sanger Institute for the COVID-19 Genomics UK (COG-UK) Consortium | Lighthouse Laboratory Plymouth and Alex Alderton, Roberto Amato, Jeffrey Barrett, Sonia Goncalves, Ewan Harrison, David K. Jackson, Ian Johnston, Dominic Kwiatkowski, Cordelia Langford, John Sillitoe on behalf of the Wellcome Sanger Institute COVID-19 Surveillance Team |
| EPI_ISL_3286992, EPI_ISL_3286993 | Lighthouse Lab in Milton Keynes | Wellcome Sanger Institute for the COVID-19 Genomics UK (COG-UK) Consortium | The Lighthouse Lab in Milton Keynes and Alex Alderton, Roberto Amato, Jeffrey Barrett, Sonia Goncalves, Ewan Harrison, David K. Jackson, Ian Johnston, Dominic Kwiatkowski, Cordelia Langford, John Sillitoe on behalf of the Wellcome Sanger Institute COVID-19 Surveillance Team |
| EPI_ISL_3286994, EPI_ISL_3286995 | Lighthouse Laboratory Plymouth | Wellcome Sanger Institute for the COVID-19 Genomics UK (COG-UK) Consortium | Lighthouse Laboratory Plymouth and Alex Alderton, Roberto Amato, Jeffrey Barrett, Sonia Goncalves, Ewan Harrison, David K. Jackson, Ian Johnston, Dominic Kwiatkowski, Cordelia Langford, John Sillitoe on behalf of the Wellcome Sanger Institute COVID-19 Surveillance Team |
| EPI_ISL_3286996 | Lighthouse Lab in Milton Keynes | Wellcome Sanger Institute for the COVID-19 Genomics UK (COG-UK) Consortium | The Lighthouse Lab in Milton Keynes and Alex Alderton, Roberto Amato, Jeffrey Barrett, Sonia Goncalves, Ewan Harrison, David K. Jackson, Ian Johnston, Dominic Kwiatkowski, Cordelia Langford, John Sillitoe on behalf of the Wellcome Sanger Institute COVID-19 Surveillance Team |
| EPI_ISL_3286997, EPI_ISL_3286998, EPI_ISL_3286999 | Lighthouse Lab in Alderley Park | Wellcome Sanger Institute for the COVID-19 Genomics UK (COG-UK) Consortium | Jacquelyn Wynn, Mairead Hyland, The Lighthouse Lab in Alderley Park and Alex Alderton, Roberto Amato, Jeffrey Barrett, Sonia Goncalves, Ewan Harrison, David K. Jackson, Ian Johnston, Dominic Kwiatkowski, Cordelia Langford, John Sillitoe on behalf of the Wellcome Sanger Institute COVID-19 Surveillance Team |
| EPI_ISL_3287001 | Lighthouse Lab in Milton Keynes | Wellcome Sanger Institute for the COVID-19 Genomics UK (COG-UK) Consortium | The Lighthouse Lab in Milton Keynes and Alex Alderton, Roberto Amato, Jeffrey Barrett, Sonia Goncalves, Ewan Harrison, David K. Jackson, Ian Johnston, Dominic Kwiatkowski, Cordelia Langford, John Sillitoe on behalf of the Wellcome Sanger Institute COVID-19 Surveillance Team |
| EPI_ISL_3287002 | Lighthouse Lab in Alderley Park | Wellcome Sanger Institute for the COVID-19 Genomics UK (COG-UK) Consortium | Jacquelyn Wynn, Mairead Hyland, The Lighthouse Lab in Alderley Park and Alex Alderton, Roberto Amato, Jeffrey Barrett, Sonia Goncalves, Ewan Harrison, David K. Jackson, Ian Johnston, Dominic Kwiatkowski, Cordelia Langford, John Sillitoe on behalf of the Wellcome Sanger Institute COVID-19 Surveillance Team |
| EPI_ISL_3287003 | Lighthouse Lab in Milton Keynes | Wellcome Sanger Institute for the COVID-19 Genomics UK (COG-UK) Consortium | The Lighthouse Lab in Milton Keynes and Alex Alderton, Roberto Amato, Jeffrey Barrett, Sonia Goncalves, Ewan Harrison, David K. Jackson, Ian Johnston, Dominic Kwiatkowski, Cordelia Langford, John Sillitoe on behalf of the Wellcome Sanger Institute COVID-19 Surveillance Team |
| EPI_ISL_3287004 | Lighthouse Laboratory Plymouth | Wellcome Sanger Institute for the COVID-19 Genomics UK (COG-UK) Consortium | Lighthouse Laboratory Plymouth and Alex Alderton, Roberto Amato, Jeffrey Barrett, Sonia Goncalves, Ewan Harrison, David K. Jackson, Ian Johnston, Dominic Kwiatkowski, Cordelia Langford, John Sillitoe on behalf of the Wellcome Sanger Institute COVID-19 Surveillance Team |
| EPI_ISL_3287005 | Lighthouse Lab in Milton Keynes | Wellcome Sanger Institute for the COVID-19 Genomics UK (COG-UK) Consortium | The Lighthouse Lab in Milton Keynes and Alex Alderton, Roberto Amato, Jeffrey Barrett, Sonia Goncalves, Ewan Harrison, David K. Jackson, Ian Johnston, Dominic Kwiatkowski, Cordelia Langford, John Sillitoe on behalf of the Wellcome Sanger Institute COVID-19 Surveillance Team |
| EPI_ISL_3287006 | Lighthouse Lab in Alderley Park | Wellcome Sanger Institute for the COVID-19 Genomics UK (COG-UK) Consortium | Jacquelyn Wynn, Mairead Hyland, The Lighthouse Lab in Alderley Park and Alex Alderton, Roberto Amato, Jeffrey Barrett, Sonia Goncalves, Ewan Harrison, David K. Jackson, Ian Johnston, Dominic Kwiatkowski, Cordelia Langford, John Sillitoe on behalf of the Wellcome Sanger Institute COVID-19 Surveillance Team |
| EPI_ISL_3287007 | Lighthouse Laboratory Plymouth | Wellcome Sanger Institute for the COVID-19 Genomics UK (COG-UK) Consortium | Lighthouse Laboratory Plymouth and Alex Alderton, Roberto Amato, Jeffrey Barrett, Sonia Goncalves, Ewan Harrison, David K. Jackson, Ian Johnston, Dominic Kwiatkowski, Cordelia Langford, John Sillitoe on behalf of the Wellcome Sanger Institute COVID-19 Surveillance Team |
| EPI_ISL_3287008 | Lighthouse Lab in Glasgow | Wellcome Sanger Institute for the COVID-19 Genomics UK (COG-UK) Consortium | Harper VanSteenhouse, Yumi Kasai, David Gray, Carol Clugston, Anna Dominiczak and Alex Alderton, Roberto Amato, Jeffrey Barrett, Sonia Goncalves, Ewan Harrison, David K. Jackson, Ian Johnston, Dominic Kwiatkowski, Cordelia Langford, John Sillitoe on behalf of the Wellcome Sanger Institute COVID-19 Surveillance Team |
| EPI_ISL_3287009 | Lighthouse Lab in Milton Keynes | Wellcome Sanger Institute for the COVID-19 Genomics UK (COG-UK) Consortium | The Lighthouse Lab in Milton Keynes and Alex Alderton, Roberto Amato, Jeffrey Barrett, Sonia Goncalves, Ewan Harrison, David K. Jackson, Ian Johnston, Dominic Kwiatkowski, Cordelia Langford, John Sillitoe on behalf of the Wellcome Sanger Institute COVID-19 Surveillance Team |

| | | | |
|---|---|---|---|
| EPI_ISL_3287010, EPI_ISL_3287011 | Lighthouse Lab in Alderley Park | Wellcome Sanger Institute for the COVID-19 Genomics UK (COG-UK) Consortium | Jacquelyn Wynn, Mairead Hyland, The Lighthouse Lab in Alderley Park and Alex Alderton, Roberto Amato, Jeffrey Barrett, Sonia Goncalves, Ewan Harrison, David K. Jackson, Ian Johnston, Dominic Kwiatkowski, Cordelia Langford, John Sillitoe on behalf of the Wellcome Sanger Institute COVID-19 Surveillance Team |
| EPI_ISL_3287012 | Lighthouse Laboratory Plymouth | Wellcome Sanger Institute for the COVID-19 Genomics UK (COG-UK) Consortium | Lighthouse Laboratory Plymouth and Alex Alderton, Roberto Amato, Jeffrey Barrett, Sonia Goncalves, Ewan Harrison, David K. Jackson, Ian Johnston, Dominic Kwiatkowski, Cordelia Langford, John Sillitoe on behalf of the Wellcome Sanger Institute COVID-19 Surveillance Team |
| EPI_ISL_3287013, EPI_ISL_3287014 | Lighthouse Lab in Milton Keynes | Wellcome Sanger Institute for the COVID-19 Genomics UK (COG-UK) Consortium | The Lighthouse Lab in Milton Keynes and Alex Alderton, Roberto Amato, Jeffrey Barrett, Sonia Goncalves, Ewan Harrison, David K. Jackson, Ian Johnston, Dominic Kwiatkowski, Cordelia Langford, John Sillitoe on behalf of the Wellcome Sanger Institute COVID-19 Surveillance Team |
| EPI_ISL_3287015 | Lighthouse Lab in Glasgow | Wellcome Sanger Institute for the COVID-19 Genomics UK (COG-UK) Consortium | Harper VanSteenhouse, Yumi Kasai, David Gray, Carol Clugston, Anna Dominiczak and Alex Alderton, Roberto Amato, Jeffrey Barrett, Sonia Goncalves, Ewan Harrison, David K. Jackson, Ian Johnston, Dominic Kwiatkowski, Cordelia Langford, John Sillitoe on behalf of the Wellcome Sanger Institute COVID-19 Surveillance Team |
| EPI_ISL_3287016 | Lighthouse Laboratory Plymouth | Wellcome Sanger Institute for the COVID-19 Genomics UK (COG-UK) Consortium | Lighthouse Laboratory Plymouth and Alex Alderton, Roberto Amato, Jeffrey Barrett, Sonia Goncalves, Ewan Harrison, David K. Jackson, Ian Johnston, Dominic Kwiatkowski, Cordelia Langford, John Sillitoe on behalf of the Wellcome Sanger Institute COVID-19 Surveillance Team |
| EPI_ISL_3287017, EPI_ISL_3287018 | Lighthouse Lab in Milton Keynes | Wellcome Sanger Institute for the COVID-19 Genomics UK (COG-UK) Consortium | The Lighthouse Lab in Milton Keynes and Alex Alderton, Roberto Amato, Jeffrey Barrett, Sonia Goncalves, Ewan Harrison, David K. Jackson, Ian Johnston, Dominic Kwiatkowski, Cordelia Langford, John Sillitoe on behalf of the Wellcome Sanger Institute COVID-19 Surveillance Team |
| EPI_ISL_3287019 | Lighthouse Lab in Alderley Park | Wellcome Sanger Institute for the COVID-19 Genomics UK (COG-UK) Consortium | Jacquelyn Wynn, Mairead Hyland, The Lighthouse Lab in Alderley Park and Alex Alderton, Roberto Amato, Jeffrey Barrett, Sonia Goncalves, Ewan Harrison, David K. Jackson, Ian Johnston, Dominic Kwiatkowski, Cordelia Langford, John Sillitoe on behalf of the Wellcome Sanger Institute COVID-19 Surveillance Team |
| EPI_ISL_3287020 | Lighthouse Laboratory Plymouth | Wellcome Sanger Institute for the COVID-19 Genomics UK (COG-UK) Consortium | Lighthouse Laboratory Plymouth and Alex Alderton, Roberto Amato, Jeffrey Barrett, Sonia Goncalves, Ewan Harrison, David K. Jackson, Ian Johnston, Dominic Kwiatkowski, Cordelia Langford, John Sillitoe on behalf of the Wellcome Sanger Institute COVID-19 Surveillance Team |
| EPI_ISL_3287021 | Lighthouse Lab in Alderley Park | Wellcome Sanger Institute for the COVID-19 Genomics UK (COG-UK) Consortium | Jacquelyn Wynn, Mairead Hyland, The Lighthouse Lab in Alderley Park and Alex Alderton, Roberto Amato, Jeffrey Barrett, Sonia Goncalves, Ewan Harrison, David K. Jackson, Ian Johnston, Dominic Kwiatkowski, Cordelia Langford, John Sillitoe on behalf of the Wellcome Sanger Institute COVID-19 Surveillance Team |
| EPI_ISL_3287022, EPI_ISL_3287023 | Lighthouse Laboratory Plymouth | Wellcome Sanger Institute for the COVID-19 Genomics UK (COG-UK) Consortium | Lighthouse Laboratory Plymouth and Alex Alderton, Roberto Amato, Jeffrey Barrett, Sonia Goncalves, Ewan Harrison, David K. Jackson, Ian Johnston, Dominic Kwiatkowski, Cordelia Langford, John Sillitoe on behalf of the Wellcome Sanger Institute COVID-19 Surveillance Team |
| EPI_ISL_3287024 | Lighthouse Lab in Glasgow | Wellcome Sanger Institute for the COVID-19 Genomics UK (COG-UK) Consortium | Harper VanSteenhouse, Yumi Kasai, David Gray, Carol Clugston, Anna Dominiczak and Alex Alderton, Roberto Amato, Jeffrey Barrett, Sonia Goncalves, Ewan Harrison, David K. Jackson, Ian Johnston, Dominic Kwiatkowski, Cordelia Langford, John Sillitoe on behalf of the Wellcome Sanger Institute COVID-19 Surveillance Team |
| EPI_ISL_3287025, EPI_ISL_3287026 | Lighthouse Lab in Milton Keynes | Wellcome Sanger Institute for the COVID-19 Genomics UK (COG-UK) Consortium | The Lighthouse Lab in Milton Keynes and Alex Alderton, Roberto Amato, Jeffrey Barrett, Sonia Goncalves, Ewan Harrison, David K. Jackson, Ian Johnston, Dominic Kwiatkowski, Cordelia Langford, John Sillitoe on behalf of the Wellcome Sanger Institute COVID-19 Surveillance Team |
| EPI_ISL_3287027 | Lighthouse Lab in Glasgow | Wellcome Sanger Institute for the COVID-19 Genomics UK (COG-UK) Consortium | Harper VanSteenhouse, Yumi Kasai, David Gray, Carol Clugston, Anna Dominiczak and Alex Alderton, Roberto Amato, Jeffrey Barrett, Sonia Goncalves, Ewan Harrison, David K. Jackson, Ian Johnston, Dominic Kwiatkowski, Cordelia Langford, John Sillitoe on behalf of the Wellcome Sanger Institute COVID-19 Surveillance Team |
| EPI_ISL_3287028, EPI_ISL_3287029 | Lighthouse Lab in Alderley Park | Wellcome Sanger Institute for the COVID-19 Genomics UK (COG-UK) Consortium | Jacquelyn Wynn, Mairead Hyland, The Lighthouse Lab in Alderley Park and Alex Alderton, Roberto Amato, Jeffrey Barrett, Sonia Goncalves, Ewan Harrison, David K. Jackson, Ian Johnston, Dominic Kwiatkowski, Cordelia Langford, John Sillitoe on behalf of the Wellcome Sanger Institute COVID-19 Surveillance Team |
| EPI_ISL_3287030, EPI_ISL_3287031 | Lighthouse Lab in Glasgow | Wellcome Sanger Institute for the COVID-19 Genomics UK (COG-UK) Consortium | Harper VanSteenhouse, Yumi Kasai, David Gray, Carol Clugston, Anna Dominiczak and Alex Alderton, Roberto Amato, Jeffrey Barrett, Sonia Goncalves, Ewan Harrison, David K. Jackson, Ian Johnston, Dominic Kwiatkowski, Cordelia Langford, John Sillitoe on behalf of the Wellcome Sanger Institute COVID-19 Surveillance Team |
| EPI_ISL_3287032, EPI_ISL_3287033, EPI_ISL_3287034, EPI_ISL_3287035 | Lighthouse Laboratory Plymouth | Wellcome Sanger Institute for the COVID-19 Genomics UK (COG-UK) Consortium | Lighthouse Laboratory Plymouth and Alex Alderton, Roberto Amato, Jeffrey Barrett, Sonia Goncalves, Ewan Harrison, David K. Jackson, Ian Johnston, Dominic Kwiatkowski, Cordelia Langford, John Sillitoe on behalf of the Wellcome Sanger Institute COVID-19 Surveillance Team |
| EPI_ISL_3287036 | Lighthouse Lab in Milton Keynes | Wellcome Sanger Institute for the COVID-19 Genomics UK (COG-UK) Consortium | The Lighthouse Lab in Milton Keynes and Alex Alderton, Roberto Amato, Jeffrey Barrett, Sonia Goncalves, Ewan Harrison, David K. Jackson, Ian Johnston, Dominic Kwiatkowski, Cordelia Langford, John Sillitoe on behalf of the Wellcome Sanger Institute COVID-19 Surveillance Team |
| EPI_ISL_3287037, EPI_ISL_3287038 | Lighthouse Laboratory Plymouth | Wellcome Sanger Institute for the COVID-19 Genomics UK (COG-UK) Consortium | Lighthouse Laboratory Plymouth and Alex Alderton, Roberto Amato, Jeffrey Barrett, Sonia Goncalves, Ewan Harrison, David K. Jackson, Ian Johnston, Dominic Kwiatkowski, Cordelia Langford, John Sillitoe on behalf of the Wellcome Sanger Institute COVID-19 Surveillance Team |
| EPI_ISL_3287039 | Lighthouse Lab in Alderley Park | Wellcome Sanger Institute for the COVID-19 Genomics UK (COG-UK) Consortium | Jacquelyn Wynn, Mairead Hyland, The Lighthouse Lab in Alderley Park and Alex Alderton, Roberto Amato, Jeffrey Barrett, Sonia Goncalves, Ewan Harrison, David K. Jackson, Ian Johnston, Dominic Kwiatkowski, Cordelia Langford, John Sillitoe on behalf of the Wellcome Sanger Institute COVID-19 Surveillance Team |
| EPI_ISL_3287040, EPI_ISL_3287041 | Lighthouse Lab in Milton Keynes | Wellcome Sanger Institute for the COVID-19 Genomics UK (COG-UK) Consortium | The Lighthouse Lab in Milton Keynes and Alex Alderton, Roberto Amato, Jeffrey Barrett, Sonia Goncalves, Ewan Harrison, David K. Jackson, Ian Johnston, Dominic Kwiatkowski, Cordelia Langford, John Sillitoe on behalf of the Wellcome Sanger Institute COVID-19 Surveillance Team |
| EPI_ISL_3287042 | Lighthouse Lab in Alderley Park | Wellcome Sanger Institute for the COVID-19 Genomics UK (COG-UK) Consortium | Jacquelyn Wynn, Mairead Hyland, The Lighthouse Lab in Alderley Park and Alex Alderton, Roberto Amato, Jeffrey Barrett, Sonia Goncalves, Ewan Harrison, David K. Jackson, Ian Johnston, Dominic Kwiatkowski, Cordelia Langford, John Sillitoe on behalf of the Wellcome Sanger Institute COVID-19 Surveillance Team |
| EPI_ISL_3287043 | Lighthouse Lab in Milton Keynes | Wellcome Sanger Institute for the COVID-19 Genomics UK (COG-UK) Consortium | The Lighthouse Lab in Milton Keynes and Alex Alderton, Roberto Amato, Jeffrey Barrett, Sonia Goncalves, Ewan Harrison, David K. Jackson, Ian Johnston, Dominic Kwiatkowski, Cordelia Langford, John Sillitoe on behalf of the Wellcome Sanger Institute COVID-19 Surveillance Team |
| EPI_ISL_3287044 | Lighthouse Lab in Glasgow | Wellcome Sanger Institute for the COVID-19 Genomics UK (COG-UK) Consortium | Harper VanSteenhouse, Yumi Kasai, David Gray, Carol Clugston, Anna Dominiczak and Alex Alderton, Roberto Amato, Jeffrey Barrett, Sonia Goncalves, Ewan Harrison, David K. Jackson, Ian Johnston, Dominic Kwiatkowski, Cordelia Langford, John Sillitoe on behalf of the Wellcome Sanger Institute COVID-19 Surveillance Team |
| EPI_ISL_3287045 | Lighthouse Laboratory Plymouth | Wellcome Sanger Institute for the COVID-19 Genomics UK (COG-UK) Consortium | Lighthouse Laboratory Plymouth and Alex Alderton, Roberto Amato, Jeffrey Barrett, Sonia Goncalves, Ewan Harrison, David K. Jackson, Ian Johnston, Dominic Kwiatkowski, Cordelia Langford, John Sillitoe on behalf of the Wellcome Sanger Institute COVID-19 Surveillance Team |
| EPI_ISL_3287046 | Lighthouse Lab in Alderley Park | Wellcome Sanger Institute for the COVID-19 Genomics UK (COG-UK) Consortium | Jacquelyn Wynn, Mairead Hyland, The Lighthouse Lab in Alderley Park and Alex Alderton, Roberto Amato, Jeffrey Barrett, Sonia Goncalves, Ewan Harrison, David K. Jackson, Ian Johnston, Dominic Kwiatkowski, Cordelia Langford, John Sillitoe on behalf of the Wellcome Sanger Institute COVID-19 Surveillance Team |
| EPI_ISL_3287047 | Lighthouse Lab in Glasgow | Wellcome Sanger Institute for the COVID-19 Genomics UK (COG-UK) Consortium | Harper VanSteenhouse, Yumi Kasai, David Gray, Carol Clugston, Anna Dominiczak and Alex Alderton, Roberto Amato, Jeffrey Barrett, Sonia Goncalves, Ewan Harrison, David K. Jackson, Ian Johnston, Dominic Kwiatkowski, Cordelia Langford, John Sillitoe on behalf of the Wellcome Sanger Institute COVID-19 Surveillance Team |
| EPI_ISL_3287048, EPI_ISL_3287049, EPI_ISL_3287050 | Lighthouse Lab in Milton Keynes | Wellcome Sanger Institute for the COVID-19 Genomics UK (COG-UK) Consortium | The Lighthouse Lab in Milton Keynes and Alex Alderton, Roberto Amato, Jeffrey Barrett, Sonia Goncalves, Ewan Harrison, David K. Jackson, Ian Johnston, Dominic Kwiatkowski, Cordelia Langford, John Sillitoe on behalf of the Wellcome Sanger Institute COVID-19 Surveillance Team |
| EPI_ISL_3287051, EPI_ISL_3287052 | Lighthouse Laboratory Plymouth | Wellcome Sanger Institute for the COVID-19 Genomics UK (COG-UK) Consortium | Lighthouse Laboratory Plymouth and Alex Alderton, Roberto Amato, Jeffrey Barrett, Sonia Goncalves, Ewan Harrison, David K. Jackson, Ian Johnston, Dominic Kwiatkowski, Cordelia Langford, John Sillitoe on behalf of the Wellcome Sanger Institute COVID-19 Surveillance Team |
| EPI_ISL_3287053, EPI_ISL_3287054 | Lighthouse Lab in Alderley Park | Wellcome Sanger Institute for the COVID-19 Genomics UK (COG-UK) Consortium | Jacquelyn Wynn, Mairead Hyland, The Lighthouse Lab in Alderley Park and Alex Alderton, Roberto Amato, Jeffrey Barrett, Sonia Goncalves, Ewan Harrison, David K. Jackson, Ian Johnston, Dominic Kwiatkowski, Cordelia Langford, John Sillitoe on behalf of the Wellcome Sanger Institute COVID-19 Surveillance Team |
| EPI_ISL_3287055 | Lighthouse Lab in Glasgow | Wellcome Sanger Institute for the COVID-19 Genomics UK (COG-UK) Consortium | Harper VanSteenhouse, Yumi Kasai, David Gray, Carol Clugston, Anna Dominiczak and Alex Alderton, Roberto Amato, Jeffrey Barrett, Sonia Goncalves, Ewan Harrison, David K. Jackson, Ian Johnston, Dominic Kwiatkowski, Cordelia Langford, John Sillitoe on behalf of the Wellcome Sanger Institute COVID-19 Surveillance Team |
| EPI_ISL_3287056 | Lighthouse Lab in Milton Keynes | Wellcome Sanger Institute for the COVID-19 Genomics UK (COG-UK) Consortium | The Lighthouse Lab in Milton Keynes and Alex Alderton, Roberto Amato, Jeffrey Barrett, Sonia Goncalves, Ewan Harrison, David K. Jackson, Ian Johnston, Dominic Kwiatkowski, Cordelia Langford, John Sillitoe on behalf of the Wellcome Sanger Institute COVID-19 Surveillance Team |
| EPI_ISL_3287057 | Lighthouse Lab in Alderley Park | Wellcome Sanger Institute for the COVID-19 Genomics UK (COG-UK) Consortium | Jacquelyn Wynn, Mairead Hyland, The Lighthouse Lab in Alderley Park and Alex Alderton, Roberto Amato, Jeffrey Barrett, Sonia Goncalves, Ewan Harrison, David K. Jackson, Ian Johnston, Dominic Kwiatkowski, Cordelia Langford, John Sillitoe on behalf of the Wellcome Sanger Institute COVID-19 Surveillance Team |
| EPI_ISL_3287058, EPI_ISL_3287059, EPI_ISL_3287060 | Lighthouse Lab in Glasgow | Wellcome Sanger Institute for the COVID-19 Genomics UK (COG-UK) Consortium | Harper VanSteenhouse, Yumi Kasai, David Gray, Carol Clugston, Anna Dominiczak and Alex Alderton, Roberto Amato, Jeffrey Barrett, Sonia Goncalves, Ewan Harrison, David K. Jackson, Ian Johnston, Dominic Kwiatkowski, Cordelia Langford, John Sillitoe on behalf of the Wellcome Sanger Institute COVID-19 Surveillance Team |
| EPI_ISL_3287061 | Lighthouse Lab in Alderley Park | Wellcome Sanger Institute for the COVID-19 Genomics UK (COG-UK) Consortium | Jacquelyn Wynn, Mairead Hyland, The Lighthouse Lab in Alderley Park and Alex Alderton, Roberto Amato, Jeffrey Barrett, Sonia Goncalves, Ewan Harrison, David K. Jackson, Ian Johnston, Dominic Kwiatkowski, Cordelia Langford, John Sillitoe on behalf of the Wellcome Sanger Institute COVID-19 Surveillance Team |
| EPI_ISL_3287062 | Lighthouse Laboratory Plymouth | Wellcome Sanger Institute for the COVID-19 Genomics UK (COG-UK) Consortium | Lighthouse Laboratory Plymouth and Alex Alderton, Roberto Amato, Jeffrey Barrett, Sonia Goncalves, Ewan Harrison, David K. Jackson, Ian Johnston, Dominic Kwiatkowski, Cordelia Langford, John Sillitoe on behalf of the Wellcome Sanger Institute COVID-19 Surveillance Team |
| EPI_ISL_3287063 | Lighthouse Lab in Alderley Park | Wellcome Sanger Institute for the COVID-19 Genomics UK (COG-UK) Consortium | Jacquelyn Wynn, Mairead Hyland, The Lighthouse Lab in Alderley Park and Alex Alderton, Roberto Amato, Jeffrey Barrett, Sonia Goncalves, Ewan Harrison, David K. Jackson, Ian Johnston, Dominic Kwiatkowski, Cordelia Langford, John Sillitoe on behalf of the Wellcome Sanger Institute COVID-19 Surveillance Team |
| EPI_ISL_3287064 | Lighthouse Laboratory Plymouth | Wellcome Sanger Institute for the COVID-19 Genomics UK (COG-UK) Consortium | Lighthouse Laboratory Plymouth and Alex Alderton, Roberto Amato, Jeffrey Barrett, Sonia Goncalves, Ewan Harrison, David K. Jackson, Ian Johnston, Dominic Kwiatkowski, Cordelia Langford, John Sillitoe on behalf of the Wellcome Sanger Institute COVID-19 Surveillance Team |
| EPI_ISL_3287065 | Lighthouse Lab in Glasgow | Wellcome Sanger Institute for the COVID-19 Genomics UK (COG-UK) Consortium | Harper VanSteenhouse, Yumi Kasai, David Gray, Carol Clugston, Anna Dominiczak and Alex Alderton, Roberto Amato, Jeffrey Barrett, Sonia Goncalves, Ewan Harrison, David K. Jackson, Ian Johnston, Dominic Kwiatkowski, Cordelia Langford, John Sillitoe on behalf of the Wellcome Sanger Institute COVID-19 Surveillance Team |
| EPI_ISL_3287066 | Lighthouse Laboratory Plymouth | Wellcome Sanger Institute for the COVID-19 Genomics UK (COG-UK) Consortium | Lighthouse Laboratory Plymouth and Alex Alderton, Roberto Amato, Jeffrey Barrett, Sonia Goncalves, Ewan Harrison, David K. Jackson, Ian Johnston, Dominic Kwiatkowski, Cordelia Langford, John Sillitoe on behalf of the Wellcome Sanger Institute COVID-19 Surveillance Team |
| EPI_ISL_3287067 | Lighthouse Lab in Alderley Park | Wellcome Sanger Institute for the COVID-19 Genomics UK (COG-UK) Consortium | Jacquelyn Wynn, Mairead Hyland, The Lighthouse Lab in Alderley Park and Alex Alderton, Roberto Amato, Jeffrey Barrett, Sonia Goncalves, Ewan Harrison, David K. Jackson, Ian Johnston, Dominic Kwiatkowski, Cordelia Langford, John Sillitoe on behalf of the Wellcome Sanger Institute COVID-19 Surveillance Team |
| EPI_ISL_3287068 | Lighthouse Lab in Milton Keynes | Wellcome Sanger Institute for the COVID-19 Genomics UK (COG-UK) Consortium | The Lighthouse Lab in Milton Keynes and Alex Alderton, Roberto Amato, Jeffrey Barrett, Sonia Goncalves, Ewan Harrison, David K. Jackson, Ian Johnston, Dominic Kwiatkowski, Cordelia Langford, John Sillitoe on behalf of the Wellcome Sanger Institute COVID-19 Surveillance Team |
| EPI_ISL_3287069 | Lighthouse Lab in Glasgow | Wellcome Sanger Institute for the COVID-19 Genomics UK (COG-UK) Consortium | Harper VanSteenhouse, Yumi Kasai, David Gray, Carol Clugston, Anna Dominiczak and Alex Alderton, Roberto Amato, Jeffrey Barrett, Sonia Goncalves, Ewan Harrison, David K. Jackson, Ian Johnston, Dominic Kwiatkowski, Cordelia Langford, John Sillitoe on behalf of the Wellcome Sanger Institute COVID-19 Surveillance Team |
| EPI_ISL_3287070, EPI_ISL_3287071 | Lighthouse Lab in Alderley Park | Wellcome Sanger Institute for the COVID-19 Genomics UK (COG-UK) Consortium | Jacquelyn Wynn, Mairead Hyland, The Lighthouse Lab in Alderley Park and Alex Alderton, Roberto Amato, Jeffrey Barrett, Sonia Goncalves, Ewan Harrison, David K. Jackson, Ian Johnston, Dominic Kwiatkowski, Cordelia Langford, John Sillitoe on behalf of the Wellcome Sanger Institute COVID-19 Surveillance Team |
| EPI_ISL_3287072, | Lighthouse Laboratory Plymouth | Wellcome Sanger Institute for the COVID-19 Genomics UK (COG-UK) | Lighthouse Laboratory Plymouth and Alex Alderton, Roberto Amato, Jeffrey Barrett, Sonia Goncalves, Ewan Harrison, David K. Jackson, Ian Johnston, Dominic Kwiatkowski, Cordelia Langford, John Sillitoe on behalf of the Wellcome Sanger |

| | | | |
|---|---|---|---|
| EPI_ISL_3287073 | | Consortium | Institute COVID-19 Surveillance Team |
| EPI_ISL_3287074 | Lighthouse Lab in Milton Keynes | Wellcome Sanger Institute for the COVID-19 Genomics UK (COG-UK) Consortium | The Lighthouse Lab in Milton Keynes and Alex Alderton, Roberto Amato, Jeffrey Barrett, Sonia Goncalves, Ewan Harrison, David K. Jackson, Ian Johnston, Dominic Kwiatkowski, Cordelia Langford, John Sillitoe on behalf of the Wellcome Sanger Institute COVID-19 Surveillance Team |
| EPI_ISL_3287075, EPI_ISL_3287076 | Lighthouse Lab in Alderley Park | Wellcome Sanger Institute for the COVID-19 Genomics UK (COG-UK) Consortium | Jacquelyn Wynn, Mairead Hyland, The Lighthouse Lab in Alderley Park and Alex Alderton, Roberto Amato, Jeffrey Barrett, Sonia Goncalves, Ewan Harrison, David K. Jackson, Ian Johnston, Dominic Kwiatkowski, Cordelia Langford, John Sillitoe on behalf of the Wellcome Sanger Institute COVID-19 Surveillance Team |
| EPI_ISL_3287077, EPI_ISL_3287078 | Lighthouse Laboratory Plymouth | Wellcome Sanger Institute for the COVID-19 Genomics UK (COG-UK) Consortium | Lighthouse Laboratory Plymouth and Alex Alderton, Roberto Amato, Jeffrey Barrett, Sonia Goncalves, Ewan Harrison, David K. Jackson, Ian Johnston, Dominic Kwiatkowski, Cordelia Langford, John Sillitoe on behalf of the Wellcome Sanger Institute COVID-19 Surveillance Team |
| EPI_ISL_3287079 | Lighthouse Lab in Milton Keynes | Wellcome Sanger Institute for the COVID-19 Genomics UK (COG-UK) Consortium | The Lighthouse Lab in Milton Keynes and Alex Alderton, Roberto Amato, Jeffrey Barrett, Sonia Goncalves, Ewan Harrison, David K. Jackson, Ian Johnston, Dominic Kwiatkowski, Cordelia Langford, John Sillitoe on behalf of the Wellcome Sanger Institute COVID-19 Surveillance Team |
| EPI_ISL_3287080, EPI_ISL_3287081 | Lighthouse Lab in Glasgow | Wellcome Sanger Institute for the COVID-19 Genomics UK (COG-UK) Consortium | Harper VanSteenhouse, Yumi Kasai, David Gray, Carol Clugston, Anna Dominiczak and Alex Alderton, Roberto Amato, Jeffrey Barrett, Sonia Goncalves, Ewan Harrison, David K. Jackson, Ian Johnston, Dominic Kwiatkowski, Cordelia Langford, John Sillitoe on behalf of the Wellcome Sanger Institute COVID-19 Surveillance Team |
| EPI_ISL_3287082 | Lighthouse Lab in Milton Keynes | Wellcome Sanger Institute for the COVID-19 Genomics UK (COG-UK) Consortium | The Lighthouse Lab in Milton Keynes and Alex Alderton, Roberto Amato, Jeffrey Barrett, Sonia Goncalves, Ewan Harrison, David K. Jackson, Ian Johnston, Dominic Kwiatkowski, Cordelia Langford, John Sillitoe on behalf of the Wellcome Sanger Institute COVID-19 Surveillance Team |
| EPI_ISL_3287083 | Lighthouse Laboratory Plymouth | Wellcome Sanger Institute for the COVID-19 Genomics UK (COG-UK) Consortium | Lighthouse Laboratory Plymouth and Alex Alderton, Roberto Amato, Jeffrey Barrett, Sonia Goncalves, Ewan Harrison, David K. Jackson, Ian Johnston, Dominic Kwiatkowski, Cordelia Langford, John Sillitoe on behalf of the Wellcome Sanger Institute COVID-19 Surveillance Team |
| EPI_ISL_3287084 | Lighthouse Lab in Glasgow | Wellcome Sanger Institute for the COVID-19 Genomics UK (COG-UK) Consortium | Harper VanSteenhouse, Yumi Kasai, David Gray, Carol Clugston, Anna Dominiczak and Alex Alderton, Roberto Amato, Jeffrey Barrett, Sonia Goncalves, Ewan Harrison, David K. Jackson, Ian Johnston, Dominic Kwiatkowski, Cordelia Langford, John Sillitoe on behalf of the Wellcome Sanger Institute COVID-19 Surveillance Team |
| EPI_ISL_3287085, EPI_ISL_3287086 | Lighthouse Laboratory Plymouth | Wellcome Sanger Institute for the COVID-19 Genomics UK (COG-UK) Consortium | Lighthouse Laboratory Plymouth and Alex Alderton, Roberto Amato, Jeffrey Barrett, Sonia Goncalves, Ewan Harrison, David K. Jackson, Ian Johnston, Dominic Kwiatkowski, Cordelia Langford, John Sillitoe on behalf of the Wellcome Sanger Institute COVID-19 Surveillance Team |
| EPI_ISL_3287087 | Lighthouse Lab in Milton Keynes | Wellcome Sanger Institute for the COVID-19 Genomics UK (COG-UK) Consortium | The Lighthouse Lab in Milton Keynes and Alex Alderton, Roberto Amato, Jeffrey Barrett, Sonia Goncalves, Ewan Harrison, David K. Jackson, Ian Johnston, Dominic Kwiatkowski, Cordelia Langford, John Sillitoe on behalf of the Wellcome Sanger Institute COVID-19 Surveillance Team |
| EPI_ISL_3287088 | Lighthouse Lab in Alderley Park | Wellcome Sanger Institute for the COVID-19 Genomics UK (COG-UK) Consortium | Jacquelyn Wynn, Mairead Hyland, The Lighthouse Lab in Alderley Park and Alex Alderton, Roberto Amato, Jeffrey Barrett, Sonia Goncalves, Ewan Harrison, David K. Jackson, Ian Johnston, Dominic Kwiatkowski, Cordelia Langford, John Sillitoe on behalf of the Wellcome Sanger Institute COVID-19 Surveillance Team |
| EPI_ISL_3287089 | Lighthouse Lab in Glasgow | Wellcome Sanger Institute for the COVID-19 Genomics UK (COG-UK) Consortium | Harper VanSteenhouse, Yumi Kasai, David Gray, Carol Clugston, Anna Dominiczak and Alex Alderton, Roberto Amato, Jeffrey Barrett, Sonia Goncalves, Ewan Harrison, David K. Jackson, Ian Johnston, Dominic Kwiatkowski, Cordelia Langford, John Sillitoe on behalf of the Wellcome Sanger Institute COVID-19 Surveillance Team |
| EPI_ISL_3287090, EPI_ISL_3287091 | Lighthouse Laboratory Plymouth | Wellcome Sanger Institute for the COVID-19 Genomics UK (COG-UK) Consortium | Lighthouse Laboratory Plymouth and Alex Alderton, Roberto Amato, Jeffrey Barrett, Sonia Goncalves, Ewan Harrison, David K. Jackson, Ian Johnston, Dominic Kwiatkowski, Cordelia Langford, John Sillitoe on behalf of the Wellcome Sanger Institute COVID-19 Surveillance Team |
| EPI_ISL_3287092 | Lighthouse Lab in Milton Keynes | Wellcome Sanger Institute for the COVID-19 Genomics UK (COG-UK) Consortium | The Lighthouse Lab in Milton Keynes and Alex Alderton, Roberto Amato, Jeffrey Barrett, Sonia Goncalves, Ewan Harrison, David K. Jackson, Ian Johnston, Dominic Kwiatkowski, Cordelia Langford, John Sillitoe on behalf of the Wellcome Sanger Institute COVID-19 Surveillance Team |
| EPI_ISL_3287093, EPI_ISL_3287094 | Lighthouse Laboratory Plymouth | Wellcome Sanger Institute for the COVID-19 Genomics UK (COG-UK) Consortium | Lighthouse Laboratory Plymouth and Alex Alderton, Roberto Amato, Jeffrey Barrett, Sonia Goncalves, Ewan Harrison, David K. Jackson, Ian Johnston, Dominic Kwiatkowski, Cordelia Langford, John Sillitoe on behalf of the Wellcome Sanger Institute COVID-19 Surveillance Team |
| EPI_ISL_3287095 | Lighthouse Lab in Alderley Park | Wellcome Sanger Institute for the COVID-19 Genomics UK (COG-UK) Consortium | Jacquelyn Wynn, Mairead Hyland, The Lighthouse Lab in Alderley Park and Alex Alderton, Roberto Amato, Jeffrey Barrett, Sonia Goncalves, Ewan Harrison, David K. Jackson, Ian Johnston, Dominic Kwiatkowski, Cordelia Langford, John Sillitoe on behalf of the Wellcome Sanger Institute COVID-19 Surveillance Team |
| EPI_ISL_3287096, EPI_ISL_3287097 | Lighthouse Lab in Milton Keynes | Wellcome Sanger Institute for the COVID-19 Genomics UK (COG-UK) Consortium | The Lighthouse Lab in Milton Keynes and Alex Alderton, Roberto Amato, Jeffrey Barrett, Sonia Goncalves, Ewan Harrison, David K. Jackson, Ian Johnston, Dominic Kwiatkowski, Cordelia Langford, John Sillitoe on behalf of the Wellcome Sanger Institute COVID-19 Surveillance Team |
| EPI_ISL_3287098 | Lighthouse Lab in Alderley Park | Wellcome Sanger Institute for the COVID-19 Genomics UK (COG-UK) Consortium | Jacquelyn Wynn, Mairead Hyland, The Lighthouse Lab in Alderley Park and Alex Alderton, Roberto Amato, Jeffrey Barrett, Sonia Goncalves, Ewan Harrison, David K. Jackson, Ian Johnston, Dominic Kwiatkowski, Cordelia Langford, John Sillitoe on behalf of the Wellcome Sanger Institute COVID-19 Surveillance Team |
| EPI_ISL_3287099 | Lighthouse Laboratory Plymouth | Wellcome Sanger Institute for the COVID-19 Genomics UK (COG-UK) Consortium | Lighthouse Laboratory Plymouth and Alex Alderton, Roberto Amato, Jeffrey Barrett, Sonia Goncalves, Ewan Harrison, David K. Jackson, Ian Johnston, Dominic Kwiatkowski, Cordelia Langford, John Sillitoe on behalf of the Wellcome Sanger Institute COVID-19 Surveillance Team |
| EPI_ISL_3287100 | Lighthouse Lab in Alderley Park | Wellcome Sanger Institute for the COVID-19 Genomics UK (COG-UK) Consortium | Jacquelyn Wynn, Mairead Hyland, The Lighthouse Lab in Alderley Park and Alex Alderton, Roberto Amato, Jeffrey Barrett, Sonia Goncalves, Ewan Harrison, David K. Jackson, Ian Johnston, Dominic Kwiatkowski, Cordelia Langford, John Sillitoe on behalf of the Wellcome Sanger Institute COVID-19 Surveillance Team |
| EPI_ISL_3287101 | Lighthouse Laboratory Plymouth | Wellcome Sanger Institute for the COVID-19 Genomics UK (COG-UK) Consortium | Lighthouse Laboratory Plymouth and Alex Alderton, Roberto Amato, Jeffrey Barrett, Sonia Goncalves, Ewan Harrison, David K. Jackson, Ian Johnston, Dominic Kwiatkowski, Cordelia Langford, John Sillitoe on behalf of the Wellcome Sanger Institute COVID-19 Surveillance Team |
| EPI_ISL_3287102 | Lighthouse Lab in Alderley Park | Wellcome Sanger Institute for the COVID-19 Genomics UK (COG-UK) Consortium | Jacquelyn Wynn, Mairead Hyland, The Lighthouse Lab in Alderley Park and Alex Alderton, Roberto Amato, Jeffrey Barrett, Sonia Goncalves, Ewan Harrison, David K. Jackson, Ian Johnston, Dominic Kwiatkowski, Cordelia Langford, John Sillitoe on behalf of the Wellcome Sanger Institute COVID-19 Surveillance Team |
| EPI_ISL_3287103 | Lighthouse Lab in Glasgow | Wellcome Sanger Institute for the COVID-19 Genomics UK (COG-UK) Consortium | Harper VanSteenhouse, Yumi Kasai, David Gray, Carol Clugston, Anna Dominiczak and Alex Alderton, Roberto Amato, Jeffrey Barrett, Sonia Goncalves, Ewan Harrison, David K. Jackson, Ian Johnston, Dominic Kwiatkowski, Cordelia Langford, John Sillitoe on behalf of the Wellcome Sanger Institute COVID-19 Surveillance Team |
| EPI_ISL_3287104 | Lighthouse Lab in Milton Keynes | Wellcome Sanger Institute for the COVID-19 Genomics UK (COG-UK) Consortium | The Lighthouse Lab in Milton Keynes and Alex Alderton, Roberto Amato, Jeffrey Barrett, Sonia Goncalves, Ewan Harrison, David K. Jackson, Ian Johnston, Dominic Kwiatkowski, Cordelia Langford, John Sillitoe on behalf of the Wellcome Sanger Institute COVID-19 Surveillance Team |
| EPI_ISL_3287105 | Lighthouse Laboratory Plymouth | Wellcome Sanger Institute for the COVID-19 Genomics UK (COG-UK) Consortium | Lighthouse Laboratory Plymouth and Alex Alderton, Roberto Amato, Jeffrey Barrett, Sonia Goncalves, Ewan Harrison, David K. Jackson, Ian Johnston, Dominic Kwiatkowski, Cordelia Langford, John Sillitoe on behalf of the Wellcome Sanger Institute COVID-19 Surveillance Team |
| EPI_ISL_3287106 | Lighthouse Lab in Milton Keynes | Wellcome Sanger Institute for the COVID-19 Genomics UK (COG-UK) Consortium | The Lighthouse Lab in Milton Keynes and Alex Alderton, Roberto Amato, Jeffrey Barrett, Sonia Goncalves, Ewan Harrison, David K. Jackson, Ian Johnston, Dominic Kwiatkowski, Cordelia Langford, John Sillitoe on behalf of the Wellcome Sanger Institute COVID-19 Surveillance Team |
| EPI_ISL_3287107 | Lighthouse Laboratory Plymouth | Wellcome Sanger Institute for the COVID-19 Genomics UK (COG-UK) Consortium | Lighthouse Laboratory Plymouth and Alex Alderton, Roberto Amato, Jeffrey Barrett, Sonia Goncalves, Ewan Harrison, David K. Jackson, Ian Johnston, Dominic Kwiatkowski, Cordelia Langford, John Sillitoe on behalf of the Wellcome Sanger Institute COVID-19 Surveillance Team |
| EPI_ISL_3287108 | Lighthouse Lab in Glasgow | Wellcome Sanger Institute for the COVID-19 Genomics UK (COG-UK) Consortium | Harper VanSteenhouse, Yumi Kasai, David Gray, Carol Clugston, Anna Dominiczak and Alex Alderton, Roberto Amato, Jeffrey Barrett, Sonia Goncalves, Ewan Harrison, David K. Jackson, Ian Johnston, Dominic Kwiatkowski, Cordelia Langford, John Sillitoe on behalf of the Wellcome Sanger Institute COVID-19 Surveillance Team |
| EPI_ISL_3287109 | Lighthouse Lab in Milton Keynes | Wellcome Sanger Institute for the COVID-19 Genomics UK (COG-UK) Consortium | The Lighthouse Lab in Milton Keynes and Alex Alderton, Roberto Amato, Jeffrey Barrett, Sonia Goncalves, Ewan Harrison, David K. Jackson, Ian Johnston, Dominic Kwiatkowski, Cordelia Langford, John Sillitoe on behalf of the Wellcome Sanger Institute COVID-19 Surveillance Team |
| EPI_ISL_3287110 | Lighthouse Lab in Alderley Park | Wellcome Sanger Institute for the COVID-19 Genomics UK (COG-UK) Consortium | Jacquelyn Wynn, Mairead Hyland, The Lighthouse Lab in Alderley Park and Alex Alderton, Roberto Amato, Jeffrey Barrett, Sonia Goncalves, Ewan Harrison, David K. Jackson, Ian Johnston, Dominic Kwiatkowski, Cordelia Langford, John Sillitoe on behalf of the Wellcome Sanger Institute COVID-19 Surveillance Team |
| EPI_ISL_3287111 | Lighthouse Lab in Milton Keynes | Wellcome Sanger Institute for the COVID-19 Genomics UK (COG-UK) Consortium | The Lighthouse Lab in Milton Keynes and Alex Alderton, Roberto Amato, Jeffrey Barrett, Sonia Goncalves, Ewan Harrison, David K. Jackson, Ian Johnston, Dominic Kwiatkowski, Cordelia Langford, John Sillitoe on behalf of the Wellcome Sanger Institute COVID-19 Surveillance Team |
| EPI_ISL_3287112 | Lighthouse Lab in Alderley Park | Wellcome Sanger Institute for the COVID-19 Genomics UK (COG-UK) Consortium | Jacquelyn Wynn, Mairead Hyland, The Lighthouse Lab in Alderley Park and Alex Alderton, Roberto Amato, Jeffrey Barrett, Sonia Goncalves, Ewan Harrison, David K. Jackson, Ian Johnston, Dominic Kwiatkowski, Cordelia Langford, John Sillitoe on behalf of the Wellcome Sanger Institute COVID-19 Surveillance Team |
| EPI_ISL_3287113 | Lighthouse Lab in Milton Keynes | Wellcome Sanger Institute for the COVID-19 Genomics UK (COG-UK) Consortium | The Lighthouse Lab in Milton Keynes and Alex Alderton, Roberto Amato, Jeffrey Barrett, Sonia Goncalves, Ewan Harrison, David K. Jackson, Ian Johnston, Dominic Kwiatkowski, Cordelia Langford, John Sillitoe on behalf of the Wellcome Sanger Institute COVID-19 Surveillance Team |
| EPI_ISL_3287114 | Lighthouse Lab in Glasgow | Wellcome Sanger Institute for the COVID-19 Genomics UK (COG-UK) Consortium | Harper VanSteenhouse, Yumi Kasai, David Gray, Carol Clugston, Anna Dominiczak and Alex Alderton, Roberto Amato, Jeffrey Barrett, Sonia Goncalves, Ewan Harrison, David K. Jackson, Ian Johnston, Dominic Kwiatkowski, Cordelia Langford, John Sillitoe on behalf of the Wellcome Sanger Institute COVID-19 Surveillance Team |
| EPI_ISL_3287115 | Lighthouse Lab in Milton Keynes | Wellcome Sanger Institute for the COVID-19 Genomics UK (COG-UK) Consortium | The Lighthouse Lab in Milton Keynes and Alex Alderton, Roberto Amato, Jeffrey Barrett, Sonia Goncalves, Ewan Harrison, David K. Jackson, Ian Johnston, Dominic Kwiatkowski, Cordelia Langford, John Sillitoe on behalf of the Wellcome Sanger Institute COVID-19 Surveillance Team |
| EPI_ISL_3287116 | Lighthouse Lab in Alderley Park | Wellcome Sanger Institute for the COVID-19 Genomics UK (COG-UK) Consortium | Jacquelyn Wynn, Mairead Hyland, The Lighthouse Lab in Alderley Park and Alex Alderton, Roberto Amato, Jeffrey Barrett, Sonia Goncalves, Ewan Harrison, David K. Jackson, Ian Johnston, Dominic Kwiatkowski, Cordelia Langford, John Sillitoe on behalf of the Wellcome Sanger Institute COVID-19 Surveillance Team |
| EPI_ISL_3287117, EPI_ISL_3287118 | Lighthouse Lab in Milton Keynes | Wellcome Sanger Institute for the COVID-19 Genomics UK (COG-UK) Consortium | The Lighthouse Lab in Milton Keynes and Alex Alderton, Roberto Amato, Jeffrey Barrett, Sonia Goncalves, Ewan Harrison, David K. Jackson, Ian Johnston, Dominic Kwiatkowski, Cordelia Langford, John Sillitoe on behalf of the Wellcome Sanger Institute COVID-19 Surveillance Team |
| EPI_ISL_3287119 | Lighthouse Laboratory Plymouth | Wellcome Sanger Institute for the COVID-19 Genomics UK (COG-UK) Consortium | Lighthouse Laboratory Plymouth and Alex Alderton, Roberto Amato, Jeffrey Barrett, Sonia Goncalves, Ewan Harrison, David K. Jackson, Ian Johnston, Dominic Kwiatkowski, Cordelia Langford, John Sillitoe on behalf of the Wellcome Sanger Institute COVID-19 Surveillance Team |
| EPI_ISL_3287120 | Lighthouse Lab in Alderley Park | Wellcome Sanger Institute for the COVID-19 Genomics UK (COG-UK) Consortium | Jacquelyn Wynn, Mairead Hyland, The Lighthouse Lab in Alderley Park and Alex Alderton, Roberto Amato, Jeffrey Barrett, Sonia Goncalves, Ewan Harrison, David K. Jackson, Ian Johnston, Dominic Kwiatkowski, Cordelia Langford, John Sillitoe on behalf of the Wellcome Sanger Institute COVID-19 Surveillance Team |
| EPI_ISL_3287121 | Lighthouse Lab in Milton Keynes | Wellcome Sanger Institute for the COVID-19 Genomics UK (COG-UK) Consortium | The Lighthouse Lab in Milton Keynes and Alex Alderton, Roberto Amato, Jeffrey Barrett, Sonia Goncalves, Ewan Harrison, David K. Jackson, Ian Johnston, Dominic Kwiatkowski, Cordelia Langford, John Sillitoe on behalf of the Wellcome Sanger Institute COVID-19 Surveillance Team |
| EPI_ISL_3287122, EPI_ISL_3287123 | Lighthouse Lab in Alderley Park | Wellcome Sanger Institute for the COVID-19 Genomics UK (COG-UK) Consortium | Jacquelyn Wynn, Mairead Hyland, The Lighthouse Lab in Alderley Park and Alex Alderton, Roberto Amato, Jeffrey Barrett, Sonia Goncalves, Ewan Harrison, David K. Jackson, Ian Johnston, Dominic Kwiatkowski, Cordelia Langford, John Sillitoe on behalf of the Wellcome Sanger Institute COVID-19 Surveillance Team |
| EPI_ISL_3287124 | Lighthouse Lab in Milton Keynes | Wellcome Sanger Institute for the COVID-19 Genomics UK (COG-UK) Consortium | The Lighthouse Lab in Milton Keynes and Alex Alderton, Roberto Amato, Jeffrey Barrett, Sonia Goncalves, Ewan Harrison, David K. Jackson, Ian Johnston, Dominic Kwiatkowski, Cordelia Langford, John Sillitoe on behalf of the Wellcome Sanger Institute COVID-19 Surveillance Team |
| EPI_ISL_3287125 | Lighthouse Laboratory Plymouth | Wellcome Sanger Institute for the COVID-19 Genomics UK (COG-UK) Consortium | Lighthouse Laboratory Plymouth and Alex Alderton, Roberto Amato, Jeffrey Barrett, Sonia Goncalves, Ewan Harrison, David K. Jackson, Ian Johnston, Dominic Kwiatkowski, Cordelia Langford, John Sillitoe on behalf of the Wellcome Sanger Institute COVID-19 Surveillance Team |
| EPI_ISL_3287126, EPI_ISL_3287127 | Lighthouse Lab in Glasgow | Wellcome Sanger Institute for the COVID-19 Genomics UK (COG-UK) Consortium | Harper VanSteenhouse, Yumi Kasai, David Gray, Carol Clugston, Anna Dominiczak and Alex Alderton, Roberto Amato, Jeffrey Barrett, Sonia Goncalves, Ewan Harrison, David K. Jackson, Ian Johnston, Dominic Kwiatkowski, Cordelia Langford, John Sillitoe on behalf of the Wellcome Sanger Institute COVID-19 Surveillance Team |
| EPI_ISL_3287128 | Lighthouse Laboratory Plymouth | Wellcome Sanger Institute for the COVID-19 Genomics UK (COG-UK) | Lighthouse Laboratory Plymouth and Alex Alderton, Roberto Amato, Jeffrey Barrett, Sonia Goncalves, Ewan Harrison, David K. Jackson, Ian Johnston, Dominic Kwiatkowski, Cordelia Langford, John Sillitoe on behalf of the Wellcome Sanger |

| | | | |
|---|---|---|---|
| | | Consortium | Institute COVID-19 Surveillance Team |
| EPI_ISL_3287129 | Lighthouse Lab in Glasgow | Wellcome Sanger Institute for the COVID-19 Genomics UK (COG-UK) Consortium | Harper VanSteenhouse, Yumi Kasai, David Gray, Carol Clugston, Anna Dominiczak and Alex Alderton, Roberto Amato, Jeffrey Barrett, Sonia Goncalves, Ewan Harrison, David K. Jackson, Ian Johnston, Dominic Kwiatkowski, Cordelia Langford, John Sillitoe on behalf of the Wellcome Sanger Institute COVID-19 Surveillance Team |
| EPI_ISL_3287130, EPI_ISL_3287131, EPI_ISL_3287132 | Lighthouse Lab in Milton Keynes | Wellcome Sanger Institute for the COVID-19 Genomics UK (COG-UK) Consortium | The Lighthouse Lab in Milton Keynes and Alex Alderton, Roberto Amato, Jeffrey Barrett, Sonia Goncalves, Ewan Harrison, David K. Jackson, Ian Johnston, Dominic Kwiatkowski, Cordelia Langford, John Sillitoe on behalf of the Wellcome Sanger Institute COVID-19 Surveillance Team |
| EPI_ISL_3287133 | Lighthouse Laboratory Plymouth | Wellcome Sanger Institute for the COVID-19 Genomics UK (COG-UK) Consortium | Lighthouse Laboratory Plymouth and Alex Alderton, Roberto Amato, Jeffrey Barrett, Sonia Goncalves, Ewan Harrison, David K. Jackson, Ian Johnston, Dominic Kwiatkowski, Cordelia Langford, John Sillitoe on behalf of the Wellcome Sanger Institute COVID-19 Surveillance Team |
| EPI_ISL_3287134 | Lighthouse Lab in Alderley Park | Wellcome Sanger Institute for the COVID-19 Genomics UK (COG-UK) Consortium | Jacquelyn Wynn, Mairead Hyland, The Lighthouse Lab in Alderley Park and Alex Alderton, Roberto Amato, Jeffrey Barrett, Sonia Goncalves, Ewan Harrison, David K. Jackson, Ian Johnston, Dominic Kwiatkowski, Cordelia Langford, John Sillitoe on behalf of the Wellcome Sanger Institute COVID-19 Surveillance Team |
| EPI_ISL_3287135 | Lighthouse Laboratory Plymouth | Wellcome Sanger Institute for the COVID-19 Genomics UK (COG-UK) Consortium | Lighthouse Laboratory Plymouth and Alex Alderton, Roberto Amato, Jeffrey Barrett, Sonia Goncalves, Ewan Harrison, David K. Jackson, Ian Johnston, Dominic Kwiatkowski, Cordelia Langford, John Sillitoe on behalf of the Wellcome Sanger Institute COVID-19 Surveillance Team |
| EPI_ISL_3287136, EPI_ISL_3287137 | Lighthouse Lab in Milton Keynes | Wellcome Sanger Institute for the COVID-19 Genomics UK (COG-UK) Consortium | The Lighthouse Lab in Milton Keynes and Alex Alderton, Roberto Amato, Jeffrey Barrett, Sonia Goncalves, Ewan Harrison, David K. Jackson, Ian Johnston, Dominic Kwiatkowski, Cordelia Langford, John Sillitoe on behalf of the Wellcome Sanger Institute COVID-19 Surveillance Team |
| EPI_ISL_3287138 | Lighthouse Lab in Alderley Park | Wellcome Sanger Institute for the COVID-19 Genomics UK (COG-UK) Consortium | Jacquelyn Wynn, Mairead Hyland, The Lighthouse Lab in Alderley Park and Alex Alderton, Roberto Amato, Jeffrey Barrett, Sonia Goncalves, Ewan Harrison, David K. Jackson, Ian Johnston, Dominic Kwiatkowski, Cordelia Langford, John Sillitoe on behalf of the Wellcome Sanger Institute COVID-19 Surveillance Team |
| EPI_ISL_3287139 | Lighthouse Lab in Glasgow | Wellcome Sanger Institute for the COVID-19 Genomics UK (COG-UK) Consortium | Harper VanSteenhouse, Yumi Kasai, David Gray, Carol Clugston, Anna Dominiczak and Alex Alderton, Roberto Amato, Jeffrey Barrett, Sonia Goncalves, Ewan Harrison, David K. Jackson, Ian Johnston, Dominic Kwiatkowski, Cordelia Langford, John Sillitoe on behalf of the Wellcome Sanger Institute COVID-19 Surveillance Team |
| EPI_ISL_3287140 | Lighthouse Lab in Milton Keynes | Wellcome Sanger Institute for the COVID-19 Genomics UK (COG-UK) Consortium | The Lighthouse Lab in Milton Keynes and Alex Alderton, Roberto Amato, Jeffrey Barrett, Sonia Goncalves, Ewan Harrison, David K. Jackson, Ian Johnston, Dominic Kwiatkowski, Cordelia Langford, John Sillitoe on behalf of the Wellcome Sanger Institute COVID-19 Surveillance Team |
| EPI_ISL_3287141 | Lighthouse Lab in Alderley Park | Wellcome Sanger Institute for the COVID-19 Genomics UK (COG-UK) Consortium | Jacquelyn Wynn, Mairead Hyland, The Lighthouse Lab in Alderley Park and Alex Alderton, Roberto Amato, Jeffrey Barrett, Sonia Goncalves, Ewan Harrison, David K. Jackson, Ian Johnston, Dominic Kwiatkowski, Cordelia Langford, John Sillitoe on behalf of the Wellcome Sanger Institute COVID-19 Surveillance Team |
| EPI_ISL_3287142, EPI_ISL_3287143 | Lighthouse Lab in Glasgow | Wellcome Sanger Institute for the COVID-19 Genomics UK (COG-UK) Consortium | Harper VanSteenhouse, Yumi Kasai, David Gray, Carol Clugston, Anna Dominiczak and Alex Alderton, Roberto Amato, Jeffrey Barrett, Sonia Goncalves, Ewan Harrison, David K. Jackson, Ian Johnston, Dominic Kwiatkowski, Cordelia Langford, John Sillitoe on behalf of the Wellcome Sanger Institute COVID-19 Surveillance Team |
| EPI_ISL_3287144 | Lighthouse Lab in Milton Keynes | Wellcome Sanger Institute for the COVID-19 Genomics UK (COG-UK) Consortium | The Lighthouse Lab in Milton Keynes and Alex Alderton, Roberto Amato, Jeffrey Barrett, Sonia Goncalves, Ewan Harrison, David K. Jackson, Ian Johnston, Dominic Kwiatkowski, Cordelia Langford, John Sillitoe on behalf of the Wellcome Sanger Institute COVID-19 Surveillance Team |
| EPI_ISL_3287145, EPI_ISL_3287146, EPI_ISL_3287147 | Lighthouse Lab in Alderley Park | Wellcome Sanger Institute for the COVID-19 Genomics UK (COG-UK) Consortium | Jacquelyn Wynn, Mairead Hyland, The Lighthouse Lab in Alderley Park and Alex Alderton, Roberto Amato, Jeffrey Barrett, Sonia Goncalves, Ewan Harrison, David K. Jackson, Ian Johnston, Dominic Kwiatkowski, Cordelia Langford, John Sillitoe on behalf of the Wellcome Sanger Institute COVID-19 Surveillance Team |
| EPI_ISL_3287148 | Lighthouse Lab in Glasgow | Wellcome Sanger Institute for the COVID-19 Genomics UK (COG-UK) Consortium | Harper VanSteenhouse, Yumi Kasai, David Gray, Carol Clugston, Anna Dominiczak and Alex Alderton, Roberto Amato, Jeffrey Barrett, Sonia Goncalves, Ewan Harrison, David K. Jackson, Ian Johnston, Dominic Kwiatkowski, Cordelia Langford, John Sillitoe on behalf of the Wellcome Sanger Institute COVID-19 Surveillance Team |
| EPI_ISL_3287149, EPI_ISL_3287150 | Lighthouse Lab in Milton Keynes | Wellcome Sanger Institute for the COVID-19 Genomics UK (COG-UK) Consortium | The Lighthouse Lab in Milton Keynes and Alex Alderton, Roberto Amato, Jeffrey Barrett, Sonia Goncalves, Ewan Harrison, David K. Jackson, Ian Johnston, Dominic Kwiatkowski, Cordelia Langford, John Sillitoe on behalf of the Wellcome Sanger Institute COVID-19 Surveillance Team |
| EPI_ISL_3287151 | Lighthouse Lab in Alderley Park | Wellcome Sanger Institute for the COVID-19 Genomics UK (COG-UK) Consortium | Jacquelyn Wynn, Mairead Hyland, The Lighthouse Lab in Alderley Park and Alex Alderton, Roberto Amato, Jeffrey Barrett, Sonia Goncalves, Ewan Harrison, David K. Jackson, Ian Johnston, Dominic Kwiatkowski, Cordelia Langford, John Sillitoe on behalf of the Wellcome Sanger Institute COVID-19 Surveillance Team |
| EPI_ISL_3287152 | Lighthouse Laboratory Plymouth | Wellcome Sanger Institute for the COVID-19 Genomics UK (COG-UK) Consortium | Lighthouse Laboratory Plymouth and Alex Alderton, Roberto Amato, Jeffrey Barrett, Sonia Goncalves, Ewan Harrison, David K. Jackson, Ian Johnston, Dominic Kwiatkowski, Cordelia Langford, John Sillitoe on behalf of the Wellcome Sanger Institute COVID-19 Surveillance Team |
| EPI_ISL_3287153 | Lighthouse Lab in Glasgow | Wellcome Sanger Institute for the COVID-19 Genomics UK (COG-UK) Consortium | Harper VanSteenhouse, Yumi Kasai, David Gray, Carol Clugston, Anna Dominiczak and Alex Alderton, Roberto Amato, Jeffrey Barrett, Sonia Goncalves, Ewan Harrison, David K. Jackson, Ian Johnston, Dominic Kwiatkowski, Cordelia Langford, John Sillitoe on behalf of the Wellcome Sanger Institute COVID-19 Surveillance Team |
| EPI_ISL_3287154, EPI_ISL_3287155 | Lighthouse Lab in Alderley Park | Wellcome Sanger Institute for the COVID-19 Genomics UK (COG-UK) Consortium | Jacquelyn Wynn, Mairead Hyland, The Lighthouse Lab in Alderley Park and Alex Alderton, Roberto Amato, Jeffrey Barrett, Sonia Goncalves, Ewan Harrison, David K. Jackson, Ian Johnston, Dominic Kwiatkowski, Cordelia Langford, John Sillitoe on behalf of the Wellcome Sanger Institute COVID-19 Surveillance Team |
| EPI_ISL_3287156 | Lighthouse Laboratory Plymouth | Wellcome Sanger Institute for the COVID-19 Genomics UK (COG-UK) Consortium | Lighthouse Laboratory Plymouth and Alex Alderton, Roberto Amato, Jeffrey Barrett, Sonia Goncalves, Ewan Harrison, David K. Jackson, Ian Johnston, Dominic Kwiatkowski, Cordelia Langford, John Sillitoe on behalf of the Wellcome Sanger Institute COVID-19 Surveillance Team |
| EPI_ISL_3287157, EPI_ISL_3287158, EPI_ISL_3287159, EPI_ISL_3287160 | Lighthouse Lab in Milton Keynes | Wellcome Sanger Institute for the COVID-19 Genomics UK (COG-UK) Consortium | The Lighthouse Lab in Milton Keynes and Alex Alderton, Roberto Amato, Jeffrey Barrett, Sonia Goncalves, Ewan Harrison, David K. Jackson, Ian Johnston, Dominic Kwiatkowski, Cordelia Langford, John Sillitoe on behalf of the Wellcome Sanger Institute COVID-19 Surveillance Team |
| EPI_ISL_3287161 | Lighthouse Laboratory Plymouth | Wellcome Sanger Institute for the COVID-19 Genomics UK (COG-UK) Consortium | Lighthouse Laboratory Plymouth and Alex Alderton, Roberto Amato, Jeffrey Barrett, Sonia Goncalves, Ewan Harrison, David K. Jackson, Ian Johnston, Dominic Kwiatkowski, Cordelia Langford, John Sillitoe on behalf of the Wellcome Sanger Institute COVID-19 Surveillance Team |
| EPI_ISL_3287162 | Lighthouse Lab in Glasgow | Wellcome Sanger Institute for the COVID-19 Genomics UK (COG-UK) Consortium | Harper VanSteenhouse, Yumi Kasai, David Gray, Carol Clugston, Anna Dominiczak and Alex Alderton, Roberto Amato, Jeffrey Barrett, Sonia Goncalves, Ewan Harrison, David K. Jackson, Ian Johnston, Dominic Kwiatkowski, Cordelia Langford, John Sillitoe on behalf of the Wellcome Sanger Institute COVID-19 Surveillance Team |
| EPI_ISL_3287163 | Lighthouse Lab in Alderley Park | Wellcome Sanger Institute for the COVID-19 Genomics UK (COG-UK) Consortium | Jacquelyn Wynn, Mairead Hyland, The Lighthouse Lab in Alderley Park and Alex Alderton, Roberto Amato, Jeffrey Barrett, Sonia Goncalves, Ewan Harrison, David K. Jackson, Ian Johnston, Dominic Kwiatkowski, Cordelia Langford, John Sillitoe on behalf of the Wellcome Sanger Institute COVID-19 Surveillance Team |
| EPI_ISL_3287164, EPI_ISL_3287165 | Lighthouse Laboratory Plymouth | Wellcome Sanger Institute for the COVID-19 Genomics UK (COG-UK) Consortium | Lighthouse Laboratory Plymouth and Alex Alderton, Roberto Amato, Jeffrey Barrett, Sonia Goncalves, Ewan Harrison, David K. Jackson, Ian Johnston, Dominic Kwiatkowski, Cordelia Langford, John Sillitoe on behalf of the Wellcome Sanger Institute COVID-19 Surveillance Team |
| EPI_ISL_3287166, EPI_ISL_3287167 | Lighthouse Lab in Alderley Park | Wellcome Sanger Institute for the COVID-19 Genomics UK (COG-UK) Consortium | Jacquelyn Wynn, Mairead Hyland, The Lighthouse Lab in Alderley Park and Alex Alderton, Roberto Amato, Jeffrey Barrett, Sonia Goncalves, Ewan Harrison, David K. Jackson, Ian Johnston, Dominic Kwiatkowski, Cordelia Langford, John Sillitoe on behalf of the Wellcome Sanger Institute COVID-19 Surveillance Team |
| EPI_ISL_3287168 | Lighthouse Laboratory Plymouth | Wellcome Sanger Institute for the COVID-19 Genomics UK (COG-UK) Consortium | Lighthouse Laboratory Plymouth and Alex Alderton, Roberto Amato, Jeffrey Barrett, Sonia Goncalves, Ewan Harrison, David K. Jackson, Ian Johnston, Dominic Kwiatkowski, Cordelia Langford, John Sillitoe on behalf of the Wellcome Sanger Institute COVID-19 Surveillance Team |
| EPI_ISL_3287169, EPI_ISL_3287170, EPI_ISL_3287171 | Lighthouse Lab in Alderley Park | Wellcome Sanger Institute for the COVID-19 Genomics UK (COG-UK) Consortium | Jacquelyn Wynn, Mairead Hyland, The Lighthouse Lab in Alderley Park and Alex Alderton, Roberto Amato, Jeffrey Barrett, Sonia Goncalves, Ewan Harrison, David K. Jackson, Ian Johnston, Dominic Kwiatkowski, Cordelia Langford, John Sillitoe on behalf of the Wellcome Sanger Institute COVID-19 Surveillance Team |
| EPI_ISL_3287172 | Lighthouse Lab in Glasgow | Wellcome Sanger Institute for the COVID-19 Genomics UK (COG-UK) Consortium | Harper VanSteenhouse, Yumi Kasai, David Gray, Carol Clugston, Anna Dominiczak and Alex Alderton, Roberto Amato, Jeffrey Barrett, Sonia Goncalves, Ewan Harrison, David K. Jackson, Ian Johnston, Dominic Kwiatkowski, Cordelia Langford, John Sillitoe on behalf of the Wellcome Sanger Institute COVID-19 Surveillance Team |
| EPI_ISL_3287173 | Lighthouse Lab in Alderley Park | Wellcome Sanger Institute for the COVID-19 Genomics UK (COG-UK) Consortium | Jacquelyn Wynn, Mairead Hyland, The Lighthouse Lab in Alderley Park and Alex Alderton, Roberto Amato, Jeffrey Barrett, Sonia Goncalves, Ewan Harrison, David K. Jackson, Ian Johnston, Dominic Kwiatkowski, Cordelia Langford, John Sillitoe on behalf of the Wellcome Sanger Institute COVID-19 Surveillance Team |
| EPI_ISL_3287174 | Lighthouse Lab in Glasgow | Wellcome Sanger Institute for the COVID-19 Genomics UK (COG-UK) Consortium | Harper VanSteenhouse, Yumi Kasai, David Gray, Carol Clugston, Anna Dominiczak and Alex Alderton, Roberto Amato, Jeffrey Barrett, Sonia Goncalves, Ewan Harrison, David K. Jackson, Ian Johnston, Dominic Kwiatkowski, Cordelia Langford, John Sillitoe on behalf of the Wellcome Sanger Institute COVID-19 Surveillance Team |
| EPI_ISL_3287175 | Lighthouse Lab in Alderley Park | Wellcome Sanger Institute for the COVID-19 Genomics UK (COG-UK) Consortium | Jacquelyn Wynn, Mairead Hyland, The Lighthouse Lab in Alderley Park and Alex Alderton, Roberto Amato, Jeffrey Barrett, Sonia Goncalves, Ewan Harrison, David K. Jackson, Ian Johnston, Dominic Kwiatkowski, Cordelia Langford, John Sillitoe on behalf of the Wellcome Sanger Institute COVID-19 Surveillance Team |
| EPI_ISL_3287176 | Lighthouse Lab in Milton Keynes | Wellcome Sanger Institute for the COVID-19 Genomics UK (COG-UK) Consortium | The Lighthouse Lab in Milton Keynes and Alex Alderton, Roberto Amato, Jeffrey Barrett, Sonia Goncalves, Ewan Harrison, David K. Jackson, Ian Johnston, Dominic Kwiatkowski, Cordelia Langford, John Sillitoe on behalf of the Wellcome Sanger Institute COVID-19 Surveillance Team |
| EPI_ISL_3287177 | Lighthouse Lab in Glasgow | Wellcome Sanger Institute for the COVID-19 Genomics UK (COG-UK) Consortium | Harper VanSteenhouse, Yumi Kasai, David Gray, Carol Clugston, Anna Dominiczak and Alex Alderton, Roberto Amato, Jeffrey Barrett, Sonia Goncalves, Ewan Harrison, David K. Jackson, Ian Johnston, Dominic Kwiatkowski, Cordelia Langford, John Sillitoe on behalf of the Wellcome Sanger Institute COVID-19 Surveillance Team |
| EPI_ISL_3287178 | Lighthouse Lab in Milton Keynes | Wellcome Sanger Institute for the COVID-19 Genomics UK (COG-UK) Consortium | The Lighthouse Lab in Milton Keynes and Alex Alderton, Roberto Amato, Jeffrey Barrett, Sonia Goncalves, Ewan Harrison, David K. Jackson, Ian Johnston, Dominic Kwiatkowski, Cordelia Langford, John Sillitoe on behalf of the Wellcome Sanger Institute COVID-19 Surveillance Team |
| EPI_ISL_3287179, EPI_ISL_3287180 | Lighthouse Lab in Alderley Park | Wellcome Sanger Institute for the COVID-19 Genomics UK (COG-UK) Consortium | Jacquelyn Wynn, Mairead Hyland, The Lighthouse Lab in Alderley Park and Alex Alderton, Roberto Amato, Jeffrey Barrett, Sonia Goncalves, Ewan Harrison, David K. Jackson, Ian Johnston, Dominic Kwiatkowski, Cordelia Langford, John Sillitoe on behalf of the Wellcome Sanger Institute COVID-19 Surveillance Team |
| EPI_ISL_3287181, EPI_ISL_3287182, EPI_ISL_3287183, EPI_ISL_3287184, EPI_ISL_3287185, EPI_ISL_3287186, EPI_ISL_3287187, EPI_ISL_3287188, EPI_ISL_3287189, EPI_ISL_3287190, EPI_ISL_3287191, EPI_ISL_3287192, EPI_ISL_3287193, EPI_ISL_3287194, EPI_ISL_3287195, EPI_ISL_3287196, EPI_ISL_3287197, EPI_ISL_3287198, EPI_ISL_3287199, EPI_ISL_3287200, EPI_ISL_3287201, EPI_ISL_3287202, EPI_ISL_3287203, EPI_ISL_3287204, EPI_ISL_3287205, EPI_ISL_3287206, EPI_ISL_3287207, EPI_ISL_3287208, EPI_ISL_3287209, EPI_ISL_3287210, EPI_ISL_3287211, EPI_ISL_3287212, EPI_ISL_3287213, EPI_ISL_3287214, EPI_ISL_3287215, EPI_ISL_3287216, EPI_ISL_3287217, EPI_ISL_3287218, EPI_ISL_3287219, EPI_ISL_3287220, EPI_ISL_3287221, EPI_ISL_3287222, EPI_ISL_3287223, EPI_ISL_3287224, EPI_ISL_3287225, EPI_ISL_3287226, EPI_ISL_3287227, EPI_ISL_3287228, EPI_ISL_3287229, EPI_ISL_3287230, EPI_ISL_3287231, EPI_ISL_3287232, EPI_ISL_3287233, EPI_ISL_3287234, EPI_ISL_3287235, EPI_ISL_3287236, EPI_ISL_3287237, EPI_ISL_3287238, EPI_ISL_3287239, EPI_ISL_3287240, EPI_ISL_3287241, EPI_ISL_3287242 | | | |
| see above | Lighthouse Lab in Glasgow | Wellcome Sanger Institute for the COVID-19 Genomics UK (COG-UK) Consortium | Harper VanSteenhouse, Yumi Kasai, David Gray, Carol Clugston, Anna Dominiczak and Alex Alderton, Roberto Amato, Jeffrey Barrett, Sonia Goncalves, Ewan Harrison, David K. Jackson, Ian Johnston, Dominic Kwiatkowski, Cordelia Langford, John Sillitoe on behalf of the Wellcome Sanger Institute COVID-19 Surveillance Team |
| EPI_ISL_3287243 | Lighthouse Lab in Alderley Park | Wellcome Sanger Institute for the COVID-19 Genomics UK (COG-UK) Consortium | Jacquelyn Wynn, Mairead Hyland, The Lighthouse Lab in Alderley Park and Alex Alderton, Roberto Amato, Jeffrey Barrett, Sonia Goncalves, Ewan Harrison, David K. Jackson, Ian Johnston, Dominic Kwiatkowski, Cordelia Langford, John Sillitoe on behalf of the Wellcome Sanger Institute COVID-19 Surveillance Team |
| EPI_ISL_3287244, EPI_ISL_3287245, EPI_ISL_3287246, EPI_ISL_3287247, EPI_ISL_3287248, EPI_ISL_3287249, EPI_ISL_3287250, EPI_ISL_3287251, EPI_ISL_3287252, EPI_ISL_3287253, EPI_ISL_3287254, EPI_ISL_3287255, EPI_ISL_3287256, EPI_ISL_3287257, EPI_ISL_3287258, EPI_ISL_3287259, EPI_ISL_3287260, EPI_ISL_3287261, EPI_ISL_3287262, EPI_ISL_3287263, EPI_ISL_3287264, EPI_ISL_3287265, EPI_ISL_3287266, EPI_ISL_3287267, EPI_ISL_3287268, EPI_ISL_3287269, EPI_ISL_3287270, EPI_ISL_3287271, EPI_ISL_3287272, EPI_ISL_3287273, EPI_ISL_3287274, EPI_ISL_3287275, EPI_ISL_3287276, EPI_ISL_3287277, EPI_ISL_3287278, EPI_ISL_3287279, EPI_ISL_3287280, EPI_ISL_3287281, EPI_ISL_3287282, EPI_ISL_3287283, EPI_ISL_3287284, EPI_ISL_3287285, EPI_ISL_3287286, EPI_ISL_3287287, EPI_ISL_3287288, EPI_ISL_3287289, EPI_ISL_3287290, EPI_ISL_3287291, EPI_ISL_3287292, EPI_ISL_3287293, EPI_ISL_3287294, EPI_ISL_3287295, EPI_ISL_3287296, EPI_ISL_3287297, EPI_ISL_3287298, EPI_ISL_3287299, EPI_ISL_3287300, EPI_ISL_3287301, EPI_ISL_3287302, EPI_ISL_3287303, EPI_ISL_3287304, EPI_ISL_3287305, EPI_ISL_3287306, EPI_ISL_3287307, EPI_ISL_3287308, EPI_ISL_3287309, EPI_ISL_3287310, EPI_ISL_3287311, EPI_ISL_3287312, EPI_ISL_3287313, EPI_ISL_3287314, EPI_ISL_3287315, EPI_ISL_3287316, EPI_ISL_3287317, EPI_ISL_3287318, EPI_ISL_3287319, EPI_ISL_3287320, EPI_ISL_3287321, EPI_ISL_3287322, EPI_ISL_3287323, EPI_ISL_3287324, EPI_ISL_3287325, EPI_ISL_3287326, EPI_ISL_3287327, EPI_ISL_3287328, EPI_ISL_3287329, EPI_ISL_3287330, EPI_ISL_3287331, EPI_ISL_3287332, EPI_ISL_3287333, EPI_ISL_3287334, EPI_ISL_3287335, EPI_ISL_3287336 | | | |
| see above | Lighthouse Lab in Glasgow | Wellcome Sanger Institute for the COVID-19 Genomics UK (COG-UK) | Harper VanSteenhouse, Yumi Kasai, David Gray, Carol Clugston, Anna Dominiczak and Alex Alderton, Roberto Amato, Jeffrey Barrett, Sonia Goncalves, Ewan Harrison, David K. Jackson, Ian Johnston, Dominic Kwiatkowski, Cordelia Langford, |

| | | | |
|---|---|---|---|
| | | Consortium | John Sillitoe on behalf of the Wellcome Sanger Institute COVID-19 Surveillance Team |
| EPI_ISL_3287337 | Lighthouse Laboratory Plymouth | Wellcome Sanger Institute for the COVID-19 Genomics UK (COG-UK) Consortium | Lighthouse Laboratory Plymouth and Alex Alderton, Roberto Amato, Jeffrey Barrett, Sonia Goncalves, Ewan Harrison, David K. Jackson, Ian Johnston, Dominic Kwiatkowski, Cordelia Langford, John Sillitoe on behalf of the Wellcome Sanger Institute COVID-19 Surveillance Team |
| EPI_ISL_3287338, EPI_ISL_3287339, EPI_ISL_3287340, EPI_ISL_3287341, EPI_ISL_3287342, EPI_ISL_3287343, EPI_ISL_3287344, EPI_ISL_3287345, EPI_ISL_3287346, EPI_ISL_3287347, EPI_ISL_3287348 | | | |
| see above | Lighthouse Lab in Glasgow | Wellcome Sanger Institute for the COVID-19 Genomics UK (COG-UK) Consortium | Harper VanSteenhouse, Yumi Kasai, David Gray, Carol Clugston, Anna Dominiczak and Alex Alderton, Roberto Amato, Jeffrey Barrett, Sonia Goncalves, Ewan Harrison, David K. Jackson, Ian Johnston, Dominic Kwiatkowski, Cordelia Langford, John Sillitoe on behalf of the Wellcome Sanger Institute COVID-19 Surveillance Team |
| EPI_ISL_3287349 | Lighthouse Laboratory Plymouth | Wellcome Sanger Institute for the COVID-19 Genomics UK (COG-UK) Consortium | Lighthouse Laboratory Plymouth and Alex Alderton, Roberto Amato, Jeffrey Barrett, Sonia Goncalves, Ewan Harrison, David K. Jackson, Ian Johnston, Dominic Kwiatkowski, Cordelia Langford, John Sillitoe on behalf of the Wellcome Sanger Institute COVID-19 Surveillance Team |
| EPI_ISL_3287350, EPI_ISL_3287351, EPI_ISL_3287352, EPI_ISL_3287353, EPI_ISL_3287354, EPI_ISL_3287355, EPI_ISL_3287356, EPI_ISL_3287357, EPI_ISL_3287358, EPI_ISL_3287359, EPI_ISL_3287360, EPI_ISL_3287361, EPI_ISL_3287362, EPI_ISL_3287363, EPI_ISL_3287364, EPI_ISL_3287365, EPI_ISL_3287366, EPI_ISL_3287367, EPI_ISL_3287368, EPI_ISL_3287369, EPI_ISL_3287370, EPI_ISL_3287371, EPI_ISL_3287372, EPI_ISL_3287373, EPI_ISL_3287374, EPI_ISL_3287375, EPI_ISL_3287376, EPI_ISL_3287377, EPI_ISL_3287378, EPI_ISL_3287379, EPI_ISL_3287380, EPI_ISL_3287381, EPI_ISL_3287382, EPI_ISL_3287383, EPI_ISL_3287384, EPI_ISL_3287385, EPI_ISL_3287386, EPI_ISL_3287387, EPI_ISL_3287388, EPI_ISL_3287389, EPI_ISL_3287390, EPI_ISL_3287391, EPI_ISL_3287392, EPI_ISL_3287393, EPI_ISL_3287394, EPI_ISL_3287395, EPI_ISL_3287396, EPI_ISL_3287397, EPI_ISL_3287398, EPI_ISL_3287399, EPI_ISL_3287400, EPI_ISL_3287401, EPI_ISL_3287402, EPI_ISL_3287403, EPI_ISL_3287404, EPI_ISL_3287405 | | | |
| see above | Lighthouse Lab in Glasgow | Wellcome Sanger Institute for the COVID-19 Genomics UK (COG-UK) Consortium | Harper VanSteenhouse, Yumi Kasai, David Gray, Carol Clugston, Anna Dominiczak and Alex Alderton, Roberto Amato, Jeffrey Barrett, Sonia Goncalves, Ewan Harrison, David K. Jackson, Ian Johnston, Dominic Kwiatkowski, Cordelia Langford, John Sillitoe on behalf of the Wellcome Sanger Institute COVID-19 Surveillance Team |
| EPI_ISL_3287406 | Lighthouse Lab in Alderley Park | Wellcome Sanger Institute for the COVID-19 Genomics UK (COG-UK) Consortium | Jacquelyn Wynn, Mairead Hyland, The Lighthouse Lab in Alderley Park and Alex Alderton, Roberto Amato, Jeffrey Barrett, Sonia Goncalves, Ewan Harrison, David K. Jackson, Ian Johnston, Dominic Kwiatkowski, Cordelia Langford, John Sillitoe on behalf of the Wellcome Sanger Institute COVID-19 Surveillance Team |
| EPI_ISL_3287407, EPI_ISL_3287408, EPI_ISL_3287409, EPI_ISL_3287410, EPI_ISL_3287411, EPI_ISL_3287412, EPI_ISL_3287413, EPI_ISL_3287414, EPI_ISL_3287415, EPI_ISL_3287416, EPI_ISL_3287417, EPI_ISL_3287418, EPI_ISL_3287419, EPI_ISL_3287420, EPI_ISL_3287421, EPI_ISL_3287422, EPI_ISL_3287423, EPI_ISL_3287424, EPI_ISL_3287425, EPI_ISL_3287426, EPI_ISL_3287427, EPI_ISL_3287428, EPI_ISL_3287429, EPI_ISL_3287430, EPI_ISL_3287431, EPI_ISL_3287432, EPI_ISL_3287433, EPI_ISL_3287434, EPI_ISL_3287435, EPI_ISL_3287436, EPI_ISL_3287437, EPI_ISL_3287438, EPI_ISL_3287439, EPI_ISL_3287440, EPI_ISL_3287441, EPI_ISL_3287442, EPI_ISL_3287443, EPI_ISL_3287444, EPI_ISL_3287445, EPI_ISL_3287446, EPI_ISL_3287447, EPI_ISL_3287448, EPI_ISL_3287449, EPI_ISL_3287450, EPI_ISL_3287451, EPI_ISL_3287452, EPI_ISL_3287453, EPI_ISL_3287454, EPI_ISL_3287455, EPI_ISL_3287456 | | | |
| see above | Lighthouse Lab in Glasgow | Wellcome Sanger Institute for the COVID-19 Genomics UK (COG-UK) Consortium | Harper VanSteenhouse, Yumi Kasai, David Gray, Carol Clugston, Anna Dominiczak and Alex Alderton, Roberto Amato, Jeffrey Barrett, Sonia Goncalves, Ewan Harrison, David K. Jackson, Ian Johnston, Dominic Kwiatkowski, Cordelia Langford, John Sillitoe on behalf of the Wellcome Sanger Institute COVID-19 Surveillance Team |
| EPI_ISL_3287457 | Health Services Laboratories | Wellcome Sanger Institute for the COVID-19 Genomics UK (COG-UK) Consortium | Health Services Laboratories and Alex Alderton, Roberto Amato, Jeffrey Barrett, Sonia Goncalves, Ewan Harrison, David K. Jackson, Ian Johnston, Dominic Kwiatkowski, Cordelia Langford, John Sillitoe on behalf of the Wellcome Sanger Institute COVID-19 Surveillance Team |
| EPI_ISL_3287458, EPI_ISL_3287459 | Lighthouse Lab in Glasgow | Wellcome Sanger Institute for the COVID-19 Genomics UK (COG-UK) Consortium | Harper VanSteenhouse, Yumi Kasai, David Gray, Carol Clugston, Anna Dominiczak and Alex Alderton, Roberto Amato, Jeffrey Barrett, Sonia Goncalves, Ewan Harrison, David K. Jackson, Ian Johnston, Dominic Kwiatkowski, Cordelia Langford, John Sillitoe on behalf of the Wellcome Sanger Institute COVID-19 Surveillance Team |
| EPI_ISL_3287460 | Health Services Laboratories | Wellcome Sanger Institute for the COVID-19 Genomics UK (COG-UK) Consortium | Health Services Laboratories and Alex Alderton, Roberto Amato, Jeffrey Barrett, Sonia Goncalves, Ewan Harrison, David K. Jackson, Ian Johnston, Dominic Kwiatkowski, Cordelia Langford, John Sillitoe on behalf of the Wellcome Sanger Institute COVID-19 Surveillance Team |
| EPI_ISL_3287461, EPI_ISL_3287462 | Lighthouse Lab in Milton Keynes | Wellcome Sanger Institute for the COVID-19 Genomics UK (COG-UK) Consortium | The Lighthouse Lab in Milton Keynes and Alex Alderton, Roberto Amato, Jeffrey Barrett, Sonia Goncalves, Ewan Harrison, David K. Jackson, Ian Johnston, Dominic Kwiatkowski, Cordelia Langford, John Sillitoe on behalf of the Wellcome Sanger Institute COVID-19 Surveillance Team |
| EPI_ISL_3287463, EPI_ISL_3287464 | Health Services Laboratories | Wellcome Sanger Institute for the COVID-19 Genomics UK (COG-UK) Consortium | Health Services Laboratories and Alex Alderton, Roberto Amato, Jeffrey Barrett, Sonia Goncalves, Ewan Harrison, David K. Jackson, Ian Johnston, Dominic Kwiatkowski, Cordelia Langford, John Sillitoe on behalf of the Wellcome Sanger Institute COVID-19 Surveillance Team |
| EPI_ISL_3287465 | Lighthouse Lab in Milton Keynes | Wellcome Sanger Institute for the COVID-19 Genomics UK (COG-UK) Consortium | The Lighthouse Lab in Milton Keynes and Alex Alderton, Roberto Amato, Jeffrey Barrett, Sonia Goncalves, Ewan Harrison, David K. Jackson, Ian Johnston, Dominic Kwiatkowski, Cordelia Langford, John Sillitoe on behalf of the Wellcome Sanger Institute COVID-19 Surveillance Team |
| EPI_ISL_3287466 | Health Services Laboratories | Wellcome Sanger Institute for the COVID-19 Genomics UK (COG-UK) Consortium | Health Services Laboratories and Alex Alderton, Roberto Amato, Jeffrey Barrett, Sonia Goncalves, Ewan Harrison, David K. Jackson, Ian Johnston, Dominic Kwiatkowski, Cordelia Langford, John Sillitoe on behalf of the Wellcome Sanger Institute COVID-19 Surveillance Team |
| EPI_ISL_3287467, EPI_ISL_3287468, EPI_ISL_3287469 | Lighthouse Lab in Glasgow | Wellcome Sanger Institute for the COVID-19 Genomics UK (COG-UK) Consortium | Harper VanSteenhouse, Yumi Kasai, David Gray, Carol Clugston, Anna Dominiczak and Alex Alderton, Roberto Amato, Jeffrey Barrett, Sonia Goncalves, Ewan Harrison, David K. Jackson, Ian Johnston, Dominic Kwiatkowski, Cordelia Langford, John Sillitoe on behalf of the Wellcome Sanger Institute COVID-19 Surveillance Team |
| EPI_ISL_3287470, EPI_ISL_3287471, EPI_ISL_3287472 | Lighthouse Lab in Milton Keynes | Wellcome Sanger Institute for the COVID-19 Genomics UK (COG-UK) Consortium | The Lighthouse Lab in Milton Keynes and Alex Alderton, Roberto Amato, Jeffrey Barrett, Sonia Goncalves, Ewan Harrison, David K. Jackson, Ian Johnston, Dominic Kwiatkowski, Cordelia Langford, John Sillitoe on behalf of the Wellcome Sanger Institute COVID-19 Surveillance Team |
| EPI_ISL_3287473 | Lighthouse Lab in Glasgow | Wellcome Sanger Institute for the COVID-19 Genomics UK (COG-UK) Consortium | Harper VanSteenhouse, Yumi Kasai, David Gray, Carol Clugston, Anna Dominiczak and Alex Alderton, Roberto Amato, Jeffrey Barrett, Sonia Goncalves, Ewan Harrison, David K. Jackson, Ian Johnston, Dominic Kwiatkowski, Cordelia Langford, John Sillitoe on behalf of the Wellcome Sanger Institute COVID-19 Surveillance Team |
| EPI_ISL_3287474 | Lighthouse Lab in Milton Keynes | Wellcome Sanger Institute for the COVID-19 Genomics UK (COG-UK) Consortium | The Lighthouse Lab in Milton Keynes and Alex Alderton, Roberto Amato, Jeffrey Barrett, Sonia Goncalves, Ewan Harrison, David K. Jackson, Ian Johnston, Dominic Kwiatkowski, Cordelia Langford, John Sillitoe on behalf of the Wellcome Sanger Institute COVID-19 Surveillance Team |
| EPI_ISL_3287475 | Health Services Laboratories | Wellcome Sanger Institute for the COVID-19 Genomics UK (COG-UK) Consortium | Health Services Laboratories and Alex Alderton, Roberto Amato, Jeffrey Barrett, Sonia Goncalves, Ewan Harrison, David K. Jackson, Ian Johnston, Dominic Kwiatkowski, Cordelia Langford, John Sillitoe on behalf of the Wellcome Sanger Institute COVID-19 Surveillance Team |
| EPI_ISL_3287476 | Lighthouse Lab in Glasgow | Wellcome Sanger Institute for the COVID-19 Genomics UK (COG-UK) Consortium | Harper VanSteenhouse, Yumi Kasai, David Gray, Carol Clugston, Anna Dominiczak and Alex Alderton, Roberto Amato, Jeffrey Barrett, Sonia Goncalves, Ewan Harrison, David K. Jackson, Ian Johnston, Dominic Kwiatkowski, Cordelia Langford, John Sillitoe on behalf of the Wellcome Sanger Institute COVID-19 Surveillance Team |
| EPI_ISL_3287477 | Health Services Laboratories | Wellcome Sanger Institute for the COVID-19 Genomics UK (COG-UK) Consortium | Health Services Laboratories and Alex Alderton, Roberto Amato, Jeffrey Barrett, Sonia Goncalves, Ewan Harrison, David K. Jackson, Ian Johnston, Dominic Kwiatkowski, Cordelia Langford, John Sillitoe on behalf of the Wellcome Sanger Institute COVID-19 Surveillance Team |
| EPI_ISL_3287478, EPI_ISL_3287479 | Lighthouse Lab in Glasgow | Wellcome Sanger Institute for the COVID-19 Genomics UK (COG-UK) Consortium | Harper VanSteenhouse, Yumi Kasai, David Gray, Carol Clugston, Anna Dominiczak and Alex Alderton, Roberto Amato, Jeffrey Barrett, Sonia Goncalves, Ewan Harrison, David K. Jackson, Ian Johnston, Dominic Kwiatkowski, Cordelia Langford, John Sillitoe on behalf of the Wellcome Sanger Institute COVID-19 Surveillance Team |
| EPI_ISL_3287480 | Health Services Laboratories | Wellcome Sanger Institute for the COVID-19 Genomics UK (COG-UK) Consortium | Health Services Laboratories and Alex Alderton, Roberto Amato, Jeffrey Barrett, Sonia Goncalves, Ewan Harrison, David K. Jackson, Ian Johnston, Dominic Kwiatkowski, Cordelia Langford, John Sillitoe on behalf of the Wellcome Sanger Institute COVID-19 Surveillance Team |
| EPI_ISL_3287481, EPI_ISL_3287482 | Lighthouse Lab in Milton Keynes | Wellcome Sanger Institute for the COVID-19 Genomics UK (COG-UK) Consortium | The Lighthouse Lab in Milton Keynes and Alex Alderton, Roberto Amato, Jeffrey Barrett, Sonia Goncalves, Ewan Harrison, David K. Jackson, Ian Johnston, Dominic Kwiatkowski, Cordelia Langford, John Sillitoe on behalf of the Wellcome Sanger Institute COVID-19 Surveillance Team |
| EPI_ISL_3287483, EPI_ISL_3287484, EPI_ISL_3287485 | Health Services Laboratories | Wellcome Sanger Institute for the COVID-19 Genomics UK (COG-UK) Consortium | Health Services Laboratories and Alex Alderton, Roberto Amato, Jeffrey Barrett, Sonia Goncalves, Ewan Harrison, David K. Jackson, Ian Johnston, Dominic Kwiatkowski, Cordelia Langford, John Sillitoe on behalf of the Wellcome Sanger Institute COVID-19 Surveillance Team |
| EPI_ISL_3287486 | Lighthouse Lab in Glasgow | Wellcome Sanger Institute for the COVID-19 Genomics UK (COG-UK) Consortium | Harper VanSteenhouse, Yumi Kasai, David Gray, Carol Clugston, Anna Dominiczak and Alex Alderton, Roberto Amato, Jeffrey Barrett, Sonia Goncalves, Ewan Harrison, David K. Jackson, Ian Johnston, Dominic Kwiatkowski, Cordelia Langford, John Sillitoe on behalf of the Wellcome Sanger Institute COVID-19 Surveillance Team |
| EPI_ISL_3287487, EPI_ISL_3287488 | Lighthouse Lab in Milton Keynes | Wellcome Sanger Institute for the COVID-19 Genomics UK (COG-UK) Consortium | The Lighthouse Lab in Milton Keynes and Alex Alderton, Roberto Amato, Jeffrey Barrett, Sonia Goncalves, Ewan Harrison, David K. Jackson, Ian Johnston, Dominic Kwiatkowski, Cordelia Langford, John Sillitoe on behalf of the Wellcome Sanger Institute COVID-19 Surveillance Team |
| EPI_ISL_3287489 | Health Services Laboratories | Wellcome Sanger Institute for the COVID-19 Genomics UK (COG-UK) Consortium | Health Services Laboratories and Alex Alderton, Roberto Amato, Jeffrey Barrett, Sonia Goncalves, Ewan Harrison, David K. Jackson, Ian Johnston, Dominic Kwiatkowski, Cordelia Langford, John Sillitoe on behalf of the Wellcome Sanger Institute COVID-19 Surveillance Team |
| EPI_ISL_3287490 | Lighthouse Lab in Milton Keynes | Wellcome Sanger Institute for the COVID-19 Genomics UK (COG-UK) Consortium | The Lighthouse Lab in Milton Keynes and Alex Alderton, Roberto Amato, Jeffrey Barrett, Sonia Goncalves, Ewan Harrison, David K. Jackson, Ian Johnston, Dominic Kwiatkowski, Cordelia Langford, John Sillitoe on behalf of the Wellcome Sanger Institute COVID-19 Surveillance Team |
| EPI_ISL_3287491 | Lighthouse Lab in Glasgow | Wellcome Sanger Institute for the COVID-19 Genomics UK (COG-UK) Consortium | Harper VanSteenhouse, Yumi Kasai, David Gray, Carol Clugston, Anna Dominiczak and Alex Alderton, Roberto Amato, Jeffrey Barrett, Sonia Goncalves, Ewan Harrison, David K. Jackson, Ian Johnston, Dominic Kwiatkowski, Cordelia Langford, John Sillitoe on behalf of the Wellcome Sanger Institute COVID-19 Surveillance Team |
| EPI_ISL_3287492 | Health Services Laboratories | Wellcome Sanger Institute for the COVID-19 Genomics UK (COG-UK) Consortium | Health Services Laboratories and Alex Alderton, Roberto Amato, Jeffrey Barrett, Sonia Goncalves, Ewan Harrison, David K. Jackson, Ian Johnston, Dominic Kwiatkowski, Cordelia Langford, John Sillitoe on behalf of the Wellcome Sanger Institute COVID-19 Surveillance Team |
| EPI_ISL_3287493 | Lighthouse Lab in Milton Keynes | Wellcome Sanger Institute for the COVID-19 Genomics UK (COG-UK) Consortium | The Lighthouse Lab in Milton Keynes and Alex Alderton, Roberto Amato, Jeffrey Barrett, Sonia Goncalves, Ewan Harrison, David K. Jackson, Ian Johnston, Dominic Kwiatkowski, Cordelia Langford, John Sillitoe on behalf of the Wellcome Sanger Institute COVID-19 Surveillance Team |
| EPI_ISL_3287494 | Lighthouse Lab in Glasgow | Wellcome Sanger Institute for the COVID-19 Genomics UK (COG-UK) Consortium | Harper VanSteenhouse, Yumi Kasai, David Gray, Carol Clugston, Anna Dominiczak and Alex Alderton, Roberto Amato, Jeffrey Barrett, Sonia Goncalves, Ewan Harrison, David K. Jackson, Ian Johnston, Dominic Kwiatkowski, Cordelia Langford, John Sillitoe on behalf of the Wellcome Sanger Institute COVID-19 Surveillance Team |
| EPI_ISL_3287495, EPI_ISL_3287496 | Lighthouse Lab in Milton Keynes | Wellcome Sanger Institute for the COVID-19 Genomics UK (COG-UK) Consortium | The Lighthouse Lab in Milton Keynes and Alex Alderton, Roberto Amato, Jeffrey Barrett, Sonia Goncalves, Ewan Harrison, David K. Jackson, Ian Johnston, Dominic Kwiatkowski, Cordelia Langford, John Sillitoe on behalf of the Wellcome Sanger Institute COVID-19 Surveillance Team |
| EPI_ISL_3287497, EPI_ISL_3287498, EPI_ISL_3287499 | Health Services Laboratories | Wellcome Sanger Institute for the COVID-19 Genomics UK (COG-UK) Consortium | Health Services Laboratories and Alex Alderton, Roberto Amato, Jeffrey Barrett, Sonia Goncalves, Ewan Harrison, David K. Jackson, Ian Johnston, Dominic Kwiatkowski, Cordelia Langford, John Sillitoe on behalf of the Wellcome Sanger Institute COVID-19 Surveillance Team |
| EPI_ISL_3287500, EPI_ISL_3287501, EPI_ISL_3287502 | Lighthouse Lab in Milton Keynes | Wellcome Sanger Institute for the COVID-19 Genomics UK (COG-UK) Consortium | The Lighthouse Lab in Milton Keynes and Alex Alderton, Roberto Amato, Jeffrey Barrett, Sonia Goncalves, Ewan Harrison, David K. Jackson, Ian Johnston, Dominic Kwiatkowski, Cordelia Langford, John Sillitoe on behalf of the Wellcome Sanger Institute COVID-19 Surveillance Team |
| EPI_ISL_3287503 | Lighthouse Lab in Glasgow | Wellcome Sanger Institute for the COVID-19 Genomics UK (COG-UK) Consortium | Harper VanSteenhouse, Yumi Kasai, David Gray, Carol Clugston, Anna Dominiczak and Alex Alderton, Roberto Amato, Jeffrey Barrett, Sonia Goncalves, Ewan Harrison, David K. Jackson, Ian Johnston, Dominic Kwiatkowski, Cordelia Langford, John Sillitoe on behalf of the Wellcome Sanger Institute COVID-19 Surveillance Team |
| EPI_ISL_3287504 | Health Services Laboratories | Wellcome Sanger Institute for the COVID-19 Genomics UK (COG-UK) Consortium | Health Services Laboratories and Alex Alderton, Roberto Amato, Jeffrey Barrett, Sonia Goncalves, Ewan Harrison, David K. Jackson, Ian Johnston, Dominic Kwiatkowski, Cordelia Langford, John Sillitoe on behalf of the Wellcome Sanger Institute COVID-19 Surveillance Team |
| EPI_ISL_3287505 | Lighthouse Lab in Milton Keynes | Wellcome Sanger Institute for the COVID-19 Genomics UK (COG-UK) Consortium | The Lighthouse Lab in Milton Keynes and Alex Alderton, Roberto Amato, Jeffrey Barrett, Sonia Goncalves, Ewan Harrison, David K. Jackson, Ian Johnston, Dominic Kwiatkowski, Cordelia Langford, John Sillitoe on behalf of the Wellcome Sanger Institute COVID-19 Surveillance Team |
| EPI_ISL_3287506 | Lighthouse Lab in Glasgow | Wellcome Sanger Institute for the COVID-19 Genomics UK (COG-UK) Consortium | Harper VanSteenhouse, Yumi Kasai, David Gray, Carol Clugston, Anna Dominiczak and Alex Alderton, Roberto Amato, Jeffrey Barrett, Sonia Goncalves, Ewan Harrison, David K. Jackson, Ian Johnston, Dominic Kwiatkowski, Cordelia Langford, John Sillitoe on behalf of the Wellcome Sanger Institute COVID-19 Surveillance Team |

| | | | |
|---|---|---|---|
| EPI_ISL_3287507, EPI_ISL_3287508, EPI_ISL_3287509, EPI_ISL_3287510 | Health Services Laboratories | Wellcome Sanger Institute for the COVID-19 Genomics UK (COG-UK) Consortium | Health Services Laboratories and Alex Alderton, Roberto Amato, Jeffrey Barrett, Sonia Goncalves, Ewan Harrison, David K. Jackson, Ian Johnston, Dominic Kwiatkowski, Cordelia Langford, John Sillitoe on behalf of the Wellcome Sanger Institute COVID-19 Surveillance Team |
| EPI_ISL_3287511 | Lighthouse Lab in Milton Keynes | Wellcome Sanger Institute for the COVID-19 Genomics UK (COG-UK) Consortium | The Lighthouse Lab in Milton Keynes and Alex Alderton, Roberto Amato, Jeffrey Barrett, Sonia Goncalves, Ewan Harrison, David K. Jackson, Ian Johnston, Dominic Kwiatkowski, Cordelia Langford, John Sillitoe on behalf of the Wellcome Sanger Institute COVID-19 Surveillance Team |
| EPI_ISL_3287512 | Lighthouse Lab in Glasgow | Wellcome Sanger Institute for the COVID-19 Genomics UK (COG-UK) Consortium | Harper VanSteenhouse, Yumi Kasai, David Gray, Carol Clugston, Anna Dominiczak and Alex Alderton, Roberto Amato, Jeffrey Barrett, Sonia Goncalves, Ewan Harrison, David K. Jackson, Ian Johnston, Dominic Kwiatkowski, Cordelia Langford, John Sillitoe on behalf of the Wellcome Sanger Institute COVID-19 Surveillance Team |
| EPI_ISL_3287513 | Health Services Laboratories | Wellcome Sanger Institute for the COVID-19 Genomics UK (COG-UK) Consortium | Health Services Laboratories and Alex Alderton, Roberto Amato, Jeffrey Barrett, Sonia Goncalves, Ewan Harrison, David K. Jackson, Ian Johnston, Dominic Kwiatkowski, Cordelia Langford, John Sillitoe on behalf of the Wellcome Sanger Institute COVID-19 Surveillance Team |
| EPI_ISL_3287514 | Lighthouse Lab in Milton Keynes | Wellcome Sanger Institute for the COVID-19 Genomics UK (COG-UK) Consortium | The Lighthouse Lab in Milton Keynes and Alex Alderton, Roberto Amato, Jeffrey Barrett, Sonia Goncalves, Ewan Harrison, David K. Jackson, Ian Johnston, Dominic Kwiatkowski, Cordelia Langford, John Sillitoe on behalf of the Wellcome Sanger Institute COVID-19 Surveillance Team |
| EPI_ISL_3287515 | Lighthouse Lab in Glasgow | Wellcome Sanger Institute for the COVID-19 Genomics UK (COG-UK) Consortium | Harper VanSteenhouse, Yumi Kasai, David Gray, Carol Clugston, Anna Dominiczak and Alex Alderton, Roberto Amato, Jeffrey Barrett, Sonia Goncalves, Ewan Harrison, David K. Jackson, Ian Johnston, Dominic Kwiatkowski, Cordelia Langford, John Sillitoe on behalf of the Wellcome Sanger Institute COVID-19 Surveillance Team |
| EPI_ISL_3287516 | Health Services Laboratories | Wellcome Sanger Institute for the COVID-19 Genomics UK (COG-UK) Consortium | Health Services Laboratories and Alex Alderton, Roberto Amato, Jeffrey Barrett, Sonia Goncalves, Ewan Harrison, David K. Jackson, Ian Johnston, Dominic Kwiatkowski, Cordelia Langford, John Sillitoe on behalf of the Wellcome Sanger Institute COVID-19 Surveillance Team |
| EPI_ISL_3287517 | Lighthouse Lab in Milton Keynes | Wellcome Sanger Institute for the COVID-19 Genomics UK (COG-UK) Consortium | The Lighthouse Lab in Milton Keynes and Alex Alderton, Roberto Amato, Jeffrey Barrett, Sonia Goncalves, Ewan Harrison, David K. Jackson, Ian Johnston, Dominic Kwiatkowski, Cordelia Langford, John Sillitoe on behalf of the Wellcome Sanger Institute COVID-19 Surveillance Team |
| EPI_ISL_3287518 | Health Services Laboratories | Wellcome Sanger Institute for the COVID-19 Genomics UK (COG-UK) Consortium | Health Services Laboratories and Alex Alderton, Roberto Amato, Jeffrey Barrett, Sonia Goncalves, Ewan Harrison, David K. Jackson, Ian Johnston, Dominic Kwiatkowski, Cordelia Langford, John Sillitoe on behalf of the Wellcome Sanger Institute COVID-19 Surveillance Team |
| EPI_ISL_3287519 | Lighthouse Lab in Milton Keynes | Wellcome Sanger Institute for the COVID-19 Genomics UK (COG-UK) Consortium | The Lighthouse Lab in Milton Keynes and Alex Alderton, Roberto Amato, Jeffrey Barrett, Sonia Goncalves, Ewan Harrison, David K. Jackson, Ian Johnston, Dominic Kwiatkowski, Cordelia Langford, John Sillitoe on behalf of the Wellcome Sanger Institute COVID-19 Surveillance Team |
| EPI_ISL_3287520, EPI_ISL_3287521 | Lighthouse Lab in Glasgow | Wellcome Sanger Institute for the COVID-19 Genomics UK (COG-UK) Consortium | Harper VanSteenhouse, Yumi Kasai, David Gray, Carol Clugston, Anna Dominiczak and Alex Alderton, Roberto Amato, Jeffrey Barrett, Sonia Goncalves, Ewan Harrison, David K. Jackson, Ian Johnston, Dominic Kwiatkowski, Cordelia Langford, John Sillitoe on behalf of the Wellcome Sanger Institute COVID-19 Surveillance Team |
| EPI_ISL_3287522 | Health Services Laboratories | Wellcome Sanger Institute for the COVID-19 Genomics UK (COG-UK) Consortium | Health Services Laboratories and Alex Alderton, Roberto Amato, Jeffrey Barrett, Sonia Goncalves, Ewan Harrison, David K. Jackson, Ian Johnston, Dominic Kwiatkowski, Cordelia Langford, John Sillitoe on behalf of the Wellcome Sanger Institute COVID-19 Surveillance Team |
| EPI_ISL_3287523, EPI_ISL_3287524 | Lighthouse Lab in Glasgow | Wellcome Sanger Institute for the COVID-19 Genomics UK (COG-UK) Consortium | Harper VanSteenhouse, Yumi Kasai, David Gray, Carol Clugston, Anna Dominiczak and Alex Alderton, Roberto Amato, Jeffrey Barrett, Sonia Goncalves, Ewan Harrison, David K. Jackson, Ian Johnston, Dominic Kwiatkowski, Cordelia Langford, John Sillitoe on behalf of the Wellcome Sanger Institute COVID-19 Surveillance Team |
| EPI_ISL_3287525, EPI_ISL_3287526, EPI_ISL_3287527 | Health Services Laboratories | Wellcome Sanger Institute for the COVID-19 Genomics UK (COG-UK) Consortium | Health Services Laboratories and Alex Alderton, Roberto Amato, Jeffrey Barrett, Sonia Goncalves, Ewan Harrison, David K. Jackson, Ian Johnston, Dominic Kwiatkowski, Cordelia Langford, John Sillitoe on behalf of the Wellcome Sanger Institute COVID-19 Surveillance Team |
| EPI_ISL_3287528 | Lighthouse Lab in Milton Keynes | Wellcome Sanger Institute for the COVID-19 Genomics UK (COG-UK) Consortium | The Lighthouse Lab in Milton Keynes and Alex Alderton, Roberto Amato, Jeffrey Barrett, Sonia Goncalves, Ewan Harrison, David K. Jackson, Ian Johnston, Dominic Kwiatkowski, Cordelia Langford, John Sillitoe on behalf of the Wellcome Sanger Institute COVID-19 Surveillance Team |
| EPI_ISL_3287529 | Lighthouse Lab in Glasgow | Wellcome Sanger Institute for the COVID-19 Genomics UK (COG-UK) Consortium | Harper VanSteenhouse, Yumi Kasai, David Gray, Carol Clugston, Anna Dominiczak and Alex Alderton, Roberto Amato, Jeffrey Barrett, Sonia Goncalves, Ewan Harrison, David K. Jackson, Ian Johnston, Dominic Kwiatkowski, Cordelia Langford, John Sillitoe on behalf of the Wellcome Sanger Institute COVID-19 Surveillance Team |
| EPI_ISL_3287530 | Lighthouse Lab in Milton Keynes | Wellcome Sanger Institute for the COVID-19 Genomics UK (COG-UK) Consortium | The Lighthouse Lab in Milton Keynes and Alex Alderton, Roberto Amato, Jeffrey Barrett, Sonia Goncalves, Ewan Harrison, David K. Jackson, Ian Johnston, Dominic Kwiatkowski, Cordelia Langford, John Sillitoe on behalf of the Wellcome Sanger Institute COVID-19 Surveillance Team |
| EPI_ISL_3287531 | Health Services Laboratories | Wellcome Sanger Institute for the COVID-19 Genomics UK (COG-UK) Consortium | Health Services Laboratories and Alex Alderton, Roberto Amato, Jeffrey Barrett, Sonia Goncalves, Ewan Harrison, David K. Jackson, Ian Johnston, Dominic Kwiatkowski, Cordelia Langford, John Sillitoe on behalf of the Wellcome Sanger Institute COVID-19 Surveillance Team |
| EPI_ISL_3287532 | Lighthouse Lab in Glasgow | Wellcome Sanger Institute for the COVID-19 Genomics UK (COG-UK) Consortium | Harper VanSteenhouse, Yumi Kasai, David Gray, Carol Clugston, Anna Dominiczak and Alex Alderton, Roberto Amato, Jeffrey Barrett, Sonia Goncalves, Ewan Harrison, David K. Jackson, Ian Johnston, Dominic Kwiatkowski, Cordelia Langford, John Sillitoe on behalf of the Wellcome Sanger Institute COVID-19 Surveillance Team |
| EPI_ISL_3287533, EPI_ISL_3287534 | Health Services Laboratories | Wellcome Sanger Institute for the COVID-19 Genomics UK (COG-UK) Consortium | Health Services Laboratories and Alex Alderton, Roberto Amato, Jeffrey Barrett, Sonia Goncalves, Ewan Harrison, David K. Jackson, Ian Johnston, Dominic Kwiatkowski, Cordelia Langford, John Sillitoe on behalf of the Wellcome Sanger Institute COVID-19 Surveillance Team |
| EPI_ISL_3287535 | Lighthouse Lab in Glasgow | Wellcome Sanger Institute for the COVID-19 Genomics UK (COG-UK) Consortium | Harper VanSteenhouse, Yumi Kasai, David Gray, Carol Clugston, Anna Dominiczak and Alex Alderton, Roberto Amato, Jeffrey Barrett, Sonia Goncalves, Ewan Harrison, David K. Jackson, Ian Johnston, Dominic Kwiatkowski, Cordelia Langford, John Sillitoe on behalf of the Wellcome Sanger Institute COVID-19 Surveillance Team |
| EPI_ISL_3287536, EPI_ISL_3287537 | Lighthouse Lab in Milton Keynes | Wellcome Sanger Institute for the COVID-19 Genomics UK (COG-UK) Consortium | The Lighthouse Lab in Milton Keynes and Alex Alderton, Roberto Amato, Jeffrey Barrett, Sonia Goncalves, Ewan Harrison, David K. Jackson, Ian Johnston, Dominic Kwiatkowski, Cordelia Langford, John Sillitoe on behalf of the Wellcome Sanger Institute COVID-19 Surveillance Team |
| EPI_ISL_3287538, EPI_ISL_3287539 | Health Services Laboratories | Wellcome Sanger Institute for the COVID-19 Genomics UK (COG-UK) Consortium | Health Services Laboratories and Alex Alderton, Roberto Amato, Jeffrey Barrett, Sonia Goncalves, Ewan Harrison, David K. Jackson, Ian Johnston, Dominic Kwiatkowski, Cordelia Langford, John Sillitoe on behalf of the Wellcome Sanger Institute COVID-19 Surveillance Team |
| EPI_ISL_3287540, EPI_ISL_3287541 | Lighthouse Lab in Glasgow | Wellcome Sanger Institute for the COVID-19 Genomics UK (COG-UK) Consortium | Harper VanSteenhouse, Yumi Kasai, David Gray, Carol Clugston, Anna Dominiczak and Alex Alderton, Roberto Amato, Jeffrey Barrett, Sonia Goncalves, Ewan Harrison, David K. Jackson, Ian Johnston, Dominic Kwiatkowski, Cordelia Langford, John Sillitoe on behalf of the Wellcome Sanger Institute COVID-19 Surveillance Team |
| EPI_ISL_3287542 | Lighthouse Lab in Milton Keynes | Wellcome Sanger Institute for the COVID-19 Genomics UK (COG-UK) Consortium | The Lighthouse Lab in Milton Keynes and Alex Alderton, Roberto Amato, Jeffrey Barrett, Sonia Goncalves, Ewan Harrison, David K. Jackson, Ian Johnston, Dominic Kwiatkowski, Cordelia Langford, John Sillitoe on behalf of the Wellcome Sanger Institute COVID-19 Surveillance Team |
| EPI_ISL_3287543, EPI_ISL_3287544 | Lighthouse Lab in Glasgow | Wellcome Sanger Institute for the COVID-19 Genomics UK (COG-UK) Consortium | Harper VanSteenhouse, Yumi Kasai, David Gray, Carol Clugston, Anna Dominiczak and Alex Alderton, Roberto Amato, Jeffrey Barrett, Sonia Goncalves, Ewan Harrison, David K. Jackson, Ian Johnston, Dominic Kwiatkowski, Cordelia Langford, John Sillitoe on behalf of the Wellcome Sanger Institute COVID-19 Surveillance Team |
| EPI_ISL_3287545 | Health Services Laboratories | Wellcome Sanger Institute for the COVID-19 Genomics UK (COG-UK) Consortium | Health Services Laboratories and Alex Alderton, Roberto Amato, Jeffrey Barrett, Sonia Goncalves, Ewan Harrison, David K. Jackson, Ian Johnston, Dominic Kwiatkowski, Cordelia Langford, John Sillitoe on behalf of the Wellcome Sanger Institute COVID-19 Surveillance Team |
| EPI_ISL_3287546 | Lighthouse Lab in Glasgow | Wellcome Sanger Institute for the COVID-19 Genomics UK (COG-UK) Consortium | Harper VanSteenhouse, Yumi Kasai, David Gray, Carol Clugston, Anna Dominiczak and Alex Alderton, Roberto Amato, Jeffrey Barrett, Sonia Goncalves, Ewan Harrison, David K. Jackson, Ian Johnston, Dominic Kwiatkowski, Cordelia Langford, John Sillitoe on behalf of the Wellcome Sanger Institute COVID-19 Surveillance Team |
| EPI_ISL_3287547 | Lighthouse Lab in Milton Keynes | Wellcome Sanger Institute for the COVID-19 Genomics UK (COG-UK) Consortium | The Lighthouse Lab in Milton Keynes and Alex Alderton, Roberto Amato, Jeffrey Barrett, Sonia Goncalves, Ewan Harrison, David K. Jackson, Ian Johnston, Dominic Kwiatkowski, Cordelia Langford, John Sillitoe on behalf of the Wellcome Sanger Institute COVID-19 Surveillance Team |
| EPI_ISL_3287548 | Lighthouse Lab in Glasgow | Wellcome Sanger Institute for the COVID-19 Genomics UK (COG-UK) Consortium | Harper VanSteenhouse, Yumi Kasai, David Gray, Carol Clugston, Anna Dominiczak and Alex Alderton, Roberto Amato, Jeffrey Barrett, Sonia Goncalves, Ewan Harrison, David K. Jackson, Ian Johnston, Dominic Kwiatkowski, Cordelia Langford, John Sillitoe on behalf of the Wellcome Sanger Institute COVID-19 Surveillance Team |
| EPI_ISL_3287549 | Health Services Laboratories | Wellcome Sanger Institute for the COVID-19 Genomics UK (COG-UK) Consortium | Health Services Laboratories and Alex Alderton, Roberto Amato, Jeffrey Barrett, Sonia Goncalves, Ewan Harrison, David K. Jackson, Ian Johnston, Dominic Kwiatkowski, Cordelia Langford, John Sillitoe on behalf of the Wellcome Sanger Institute COVID-19 Surveillance Team |
| EPI_ISL_3287550 | Lighthouse Lab in Glasgow | Wellcome Sanger Institute for the COVID-19 Genomics UK (COG-UK) Consortium | Harper VanSteenhouse, Yumi Kasai, David Gray, Carol Clugston, Anna Dominiczak and Alex Alderton, Roberto Amato, Jeffrey Barrett, Sonia Goncalves, Ewan Harrison, David K. Jackson, Ian Johnston, Dominic Kwiatkowski, Cordelia Langford, John Sillitoe on behalf of the Wellcome Sanger Institute COVID-19 Surveillance Team |
| EPI_ISL_3287551, EPI_ISL_3287552, EPI_ISL_3287553 | Lighthouse Lab in Milton Keynes | Wellcome Sanger Institute for the COVID-19 Genomics UK (COG-UK) Consortium | The Lighthouse Lab in Milton Keynes and Alex Alderton, Roberto Amato, Jeffrey Barrett, Sonia Goncalves, Ewan Harrison, David K. Jackson, Ian Johnston, Dominic Kwiatkowski, Cordelia Langford, John Sillitoe on behalf of the Wellcome Sanger Institute COVID-19 Surveillance Team |
| EPI_ISL_3287554 | Health Services Laboratories | Wellcome Sanger Institute for the COVID-19 Genomics UK (COG-UK) Consortium | Health Services Laboratories and Alex Alderton, Roberto Amato, Jeffrey Barrett, Sonia Goncalves, Ewan Harrison, David K. Jackson, Ian Johnston, Dominic Kwiatkowski, Cordelia Langford, John Sillitoe on behalf of the Wellcome Sanger Institute COVID-19 Surveillance Team |
| EPI_ISL_3287555 | Lighthouse Lab in Glasgow | Wellcome Sanger Institute for the COVID-19 Genomics UK (COG-UK) Consortium | Harper VanSteenhouse, Yumi Kasai, David Gray, Carol Clugston, Anna Dominiczak and Alex Alderton, Roberto Amato, Jeffrey Barrett, Sonia Goncalves, Ewan Harrison, David K. Jackson, Ian Johnston, Dominic Kwiatkowski, Cordelia Langford, John Sillitoe on behalf of the Wellcome Sanger Institute COVID-19 Surveillance Team |
| EPI_ISL_3287556, EPI_ISL_3287557 | Health Services Laboratories | Wellcome Sanger Institute for the COVID-19 Genomics UK (COG-UK) Consortium | Health Services Laboratories and Alex Alderton, Roberto Amato, Jeffrey Barrett, Sonia Goncalves, Ewan Harrison, David K. Jackson, Ian Johnston, Dominic Kwiatkowski, Cordelia Langford, John Sillitoe on behalf of the Wellcome Sanger Institute COVID-19 Surveillance Team |
| EPI_ISL_3287558 | Lighthouse Lab in Glasgow | Wellcome Sanger Institute for the COVID-19 Genomics UK (COG-UK) Consortium | Harper VanSteenhouse, Yumi Kasai, David Gray, Carol Clugston, Anna Dominiczak and Alex Alderton, Roberto Amato, Jeffrey Barrett, Sonia Goncalves, Ewan Harrison, David K. Jackson, Ian Johnston, Dominic Kwiatkowski, Cordelia Langford, John Sillitoe on behalf of the Wellcome Sanger Institute COVID-19 Surveillance Team |
| EPI_ISL_3287559, EPI_ISL_3287560 | Lighthouse Lab in Milton Keynes | Wellcome Sanger Institute for the COVID-19 Genomics UK (COG-UK) Consortium | The Lighthouse Lab in Milton Keynes and Alex Alderton, Roberto Amato, Jeffrey Barrett, Sonia Goncalves, Ewan Harrison, David K. Jackson, Ian Johnston, Dominic Kwiatkowski, Cordelia Langford, John Sillitoe on behalf of the Wellcome Sanger Institute COVID-19 Surveillance Team |
| EPI_ISL_3287561 | Lighthouse Lab in Glasgow | Wellcome Sanger Institute for the COVID-19 Genomics UK (COG-UK) Consortium | Harper VanSteenhouse, Yumi Kasai, David Gray, Carol Clugston, Anna Dominiczak and Alex Alderton, Roberto Amato, Jeffrey Barrett, Sonia Goncalves, Ewan Harrison, David K. Jackson, Ian Johnston, Dominic Kwiatkowski, Cordelia Langford, John Sillitoe on behalf of the Wellcome Sanger Institute COVID-19 Surveillance Team |
| EPI_ISL_3287562, EPI_ISL_3287563, EPI_ISL_3287564 | Health Services Laboratories | Wellcome Sanger Institute for the COVID-19 Genomics UK (COG-UK) Consortium | Health Services Laboratories and Alex Alderton, Roberto Amato, Jeffrey Barrett, Sonia Goncalves, Ewan Harrison, David K. Jackson, Ian Johnston, Dominic Kwiatkowski, Cordelia Langford, John Sillitoe on behalf of the Wellcome Sanger Institute COVID-19 Surveillance Team |
| EPI_ISL_3287565 | Lighthouse Lab in Milton Keynes | Wellcome Sanger Institute for the COVID-19 Genomics UK (COG-UK) Consortium | The Lighthouse Lab in Milton Keynes and Alex Alderton, Roberto Amato, Jeffrey Barrett, Sonia Goncalves, Ewan Harrison, David K. Jackson, Ian Johnston, Dominic Kwiatkowski, Cordelia Langford, John Sillitoe on behalf of the Wellcome Sanger Institute COVID-19 Surveillance Team |
| EPI_ISL_3287566, EPI_ISL_3287567, EPI_ISL_3287568, EPI_ISL_3287569, EPI_ISL_3287570 | Lighthouse Lab in Glasgow | Wellcome Sanger Institute for the COVID-19 Genomics UK (COG-UK) Consortium | Harper VanSteenhouse, Yumi Kasai, David Gray, Carol Clugston, Anna Dominiczak and Alex Alderton, Roberto Amato, Jeffrey Barrett, Sonia Goncalves, Ewan Harrison, David K. Jackson, Ian Johnston, Dominic Kwiatkowski, Cordelia Langford, John Sillitoe on behalf of the Wellcome Sanger Institute COVID-19 Surveillance Team |

| | | | |
|---|---|---|---|
| EPI_ISL_3287571 | Lighthouse Lab in Milton Keynes | Wellcome Sanger Institute for the COVID-19 Genomics UK (COG-UK) Consortium | The Lighthouse Lab in Milton Keynes and Alex Alderton, Roberto Amato, Jeffrey Barrett, Sonia Goncalves, Ewan Harrison, David K. Jackson, Ian Johnston, Dominic Kwiatkowski, Cordelia Langford, John Sillitoe on behalf of the Wellcome Sanger Institute COVID-19 Surveillance Team |
| EPI_ISL_3287572, EPI_ISL_3287573, EPI_ISL_3287574, EPI_ISL_3287575 | Health Services Laboratories | Wellcome Sanger Institute for the COVID-19 Genomics UK (COG-UK) Consortium | Health Services Laboratories and Alex Alderton, Roberto Amato, Jeffrey Barrett, Sonia Goncalves, Ewan Harrison, David K. Jackson, Ian Johnston, Dominic Kwiatkowski, Cordelia Langford, John Sillitoe on behalf of the Wellcome Sanger Institute COVID-19 Surveillance Team |
| EPI_ISL_3287576, EPI_ISL_3287577 | Lighthouse Lab in Milton Keynes | Wellcome Sanger Institute for the COVID-19 Genomics UK (COG-UK) Consortium | The Lighthouse Lab in Milton Keynes and Alex Alderton, Roberto Amato, Jeffrey Barrett, Sonia Goncalves, Ewan Harrison, David K. Jackson, Ian Johnston, Dominic Kwiatkowski, Cordelia Langford, John Sillitoe on behalf of the Wellcome Sanger Institute COVID-19 Surveillance Team |
| EPI_ISL_3287578 | Health Services Laboratories | Wellcome Sanger Institute for the COVID-19 Genomics UK (COG-UK) Consortium | Health Services Laboratories and Alex Alderton, Roberto Amato, Jeffrey Barrett, Sonia Goncalves, Ewan Harrison, David K. Jackson, Ian Johnston, Dominic Kwiatkowski, Cordelia Langford, John Sillitoe on behalf of the Wellcome Sanger Institute COVID-19 Surveillance Team |
| EPI_ISL_3287579 | Lighthouse Lab in Milton Keynes | Wellcome Sanger Institute for the COVID-19 Genomics UK (COG-UK) Consortium | The Lighthouse Lab in Milton Keynes and Alex Alderton, Roberto Amato, Jeffrey Barrett, Sonia Goncalves, Ewan Harrison, David K. Jackson, Ian Johnston, Dominic Kwiatkowski, Cordelia Langford, John Sillitoe on behalf of the Wellcome Sanger Institute COVID-19 Surveillance Team |
| EPI_ISL_3287580 | Lighthouse Lab in Glasgow | Wellcome Sanger Institute for the COVID-19 Genomics UK (COG-UK) Consortium | Harper VanSteenhouse, Yumi Kasai, David Gray, Carol Clugston, Anna Dominiczak and Alex Alderton, Roberto Amato, Jeffrey Barrett, Sonia Goncalves, Ewan Harrison, David K. Jackson, Ian Johnston, Dominic Kwiatkowski, Cordelia Langford, John Sillitoe on behalf of the Wellcome Sanger Institute COVID-19 Surveillance Team |
| EPI_ISL_3287581, EPI_ISL_3287582, EPI_ISL_3287583 | Health Services Laboratories | Wellcome Sanger Institute for the COVID-19 Genomics UK (COG-UK) Consortium | Health Services Laboratories and Alex Alderton, Roberto Amato, Jeffrey Barrett, Sonia Goncalves, Ewan Harrison, David K. Jackson, Ian Johnston, Dominic Kwiatkowski, Cordelia Langford, John Sillitoe on behalf of the Wellcome Sanger Institute COVID-19 Surveillance Team |
| EPI_ISL_3287584 | Lighthouse Lab in Milton Keynes | Wellcome Sanger Institute for the COVID-19 Genomics UK (COG-UK) Consortium | The Lighthouse Lab in Milton Keynes and Alex Alderton, Roberto Amato, Jeffrey Barrett, Sonia Goncalves, Ewan Harrison, David K. Jackson, Ian Johnston, Dominic Kwiatkowski, Cordelia Langford, John Sillitoe on behalf of the Wellcome Sanger Institute COVID-19 Surveillance Team |
| EPI_ISL_3287585, EPI_ISL_3287586 | Health Services Laboratories | Wellcome Sanger Institute for the COVID-19 Genomics UK (COG-UK) Consortium | Health Services Laboratories and Alex Alderton, Roberto Amato, Jeffrey Barrett, Sonia Goncalves, Ewan Harrison, David K. Jackson, Ian Johnston, Dominic Kwiatkowski, Cordelia Langford, John Sillitoe on behalf of the Wellcome Sanger Institute COVID-19 Surveillance Team |
| EPI_ISL_3287587 | Lighthouse Lab in Glasgow | Wellcome Sanger Institute for the COVID-19 Genomics UK (COG-UK) Consortium | Harper VanSteenhouse, Yumi Kasai, David Gray, Carol Clugston, Anna Dominiczak and Alex Alderton, Roberto Amato, Jeffrey Barrett, Sonia Goncalves, Ewan Harrison, David K. Jackson, Ian Johnston, Dominic Kwiatkowski, Cordelia Langford, John Sillitoe on behalf of the Wellcome Sanger Institute COVID-19 Surveillance Team |
| EPI_ISL_3287588 | Health Services Laboratories | Wellcome Sanger Institute for the COVID-19 Genomics UK (COG-UK) Consortium | Health Services Laboratories and Alex Alderton, Roberto Amato, Jeffrey Barrett, Sonia Goncalves, Ewan Harrison, David K. Jackson, Ian Johnston, Dominic Kwiatkowski, Cordelia Langford, John Sillitoe on behalf of the Wellcome Sanger Institute COVID-19 Surveillance Team |
| EPI_ISL_3287589 | Lighthouse Lab in Milton Keynes | Wellcome Sanger Institute for the COVID-19 Genomics UK (COG-UK) Consortium | The Lighthouse Lab in Milton Keynes and Alex Alderton, Roberto Amato, Jeffrey Barrett, Sonia Goncalves, Ewan Harrison, David K. Jackson, Ian Johnston, Dominic Kwiatkowski, Cordelia Langford, John Sillitoe on behalf of the Wellcome Sanger Institute COVID-19 Surveillance Team |
| EPI_ISL_3287590, EPI_ISL_3287591, EPI_ISL_3287592 | Lighthouse Lab in Glasgow | Wellcome Sanger Institute for the COVID-19 Genomics UK (COG-UK) Consortium | Harper VanSteenhouse, Yumi Kasai, David Gray, Carol Clugston, Anna Dominiczak and Alex Alderton, Roberto Amato, Jeffrey Barrett, Sonia Goncalves, Ewan Harrison, David K. Jackson, Ian Johnston, Dominic Kwiatkowski, Cordelia Langford, John Sillitoe on behalf of the Wellcome Sanger Institute COVID-19 Surveillance Team |
| EPI_ISL_3287593 | Lighthouse Lab in Milton Keynes | Wellcome Sanger Institute for the COVID-19 Genomics UK (COG-UK) Consortium | The Lighthouse Lab in Milton Keynes and Alex Alderton, Roberto Amato, Jeffrey Barrett, Sonia Goncalves, Ewan Harrison, David K. Jackson, Ian Johnston, Dominic Kwiatkowski, Cordelia Langford, John Sillitoe on behalf of the Wellcome Sanger Institute COVID-19 Surveillance Team |
| EPI_ISL_3287594 | Lighthouse Lab in Glasgow | Wellcome Sanger Institute for the COVID-19 Genomics UK (COG-UK) Consortium | Harper VanSteenhouse, Yumi Kasai, David Gray, Carol Clugston, Anna Dominiczak and Alex Alderton, Roberto Amato, Jeffrey Barrett, Sonia Goncalves, Ewan Harrison, David K. Jackson, Ian Johnston, Dominic Kwiatkowski, Cordelia Langford, John Sillitoe on behalf of the Wellcome Sanger Institute COVID-19 Surveillance Team |
| EPI_ISL_3287595, EPI_ISL_3287596, EPI_ISL_3287597 | Lighthouse Lab in Milton Keynes | Wellcome Sanger Institute for the COVID-19 Genomics UK (COG-UK) Consortium | The Lighthouse Lab in Milton Keynes and Alex Alderton, Roberto Amato, Jeffrey Barrett, Sonia Goncalves, Ewan Harrison, David K. Jackson, Ian Johnston, Dominic Kwiatkowski, Cordelia Langford, John Sillitoe on behalf of the Wellcome Sanger Institute COVID-19 Surveillance Team |
| EPI_ISL_3287598, EPI_ISL_3287599 | Lighthouse Lab in Glasgow | Wellcome Sanger Institute for the COVID-19 Genomics UK (COG-UK) Consortium | Harper VanSteenhouse, Yumi Kasai, David Gray, Carol Clugston, Anna Dominiczak and Alex Alderton, Roberto Amato, Jeffrey Barrett, Sonia Goncalves, Ewan Harrison, David K. Jackson, Ian Johnston, Dominic Kwiatkowski, Cordelia Langford, John Sillitoe on behalf of the Wellcome Sanger Institute COVID-19 Surveillance Team |
| EPI_ISL_3287600 | Lighthouse Lab in Milton Keynes | Wellcome Sanger Institute for the COVID-19 Genomics UK (COG-UK) Consortium | The Lighthouse Lab in Milton Keynes and Alex Alderton, Roberto Amato, Jeffrey Barrett, Sonia Goncalves, Ewan Harrison, David K. Jackson, Ian Johnston, Dominic Kwiatkowski, Cordelia Langford, John Sillitoe on behalf of the Wellcome Sanger Institute COVID-19 Surveillance Team |
| EPI_ISL_3287601 | Lighthouse Lab in Glasgow | Wellcome Sanger Institute for the COVID-19 Genomics UK (COG-UK) Consortium | Harper VanSteenhouse, Yumi Kasai, David Gray, Carol Clugston, Anna Dominiczak and Alex Alderton, Roberto Amato, Jeffrey Barrett, Sonia Goncalves, Ewan Harrison, David K. Jackson, Ian Johnston, Dominic Kwiatkowski, Cordelia Langford, John Sillitoe on behalf of the Wellcome Sanger Institute COVID-19 Surveillance Team |
| EPI_ISL_3287602 | Lighthouse Lab in Milton Keynes | Wellcome Sanger Institute for the COVID-19 Genomics UK (COG-UK) Consortium | The Lighthouse Lab in Milton Keynes and Alex Alderton, Roberto Amato, Jeffrey Barrett, Sonia Goncalves, Ewan Harrison, David K. Jackson, Ian Johnston, Dominic Kwiatkowski, Cordelia Langford, John Sillitoe on behalf of the Wellcome Sanger Institute COVID-19 Surveillance Team |
| EPI_ISL_3287603, EPI_ISL_3287604 | Lighthouse Lab in Glasgow | Wellcome Sanger Institute for the COVID-19 Genomics UK (COG-UK) Consortium | Harper VanSteenhouse, Yumi Kasai, David Gray, Carol Clugston, Anna Dominiczak and Alex Alderton, Roberto Amato, Jeffrey Barrett, Sonia Goncalves, Ewan Harrison, David K. Jackson, Ian Johnston, Dominic Kwiatkowski, Cordelia Langford, John Sillitoe on behalf of the Wellcome Sanger Institute COVID-19 Surveillance Team |
| EPI_ISL_3287605 | Lighthouse Lab in Milton Keynes | Wellcome Sanger Institute for the COVID-19 Genomics UK (COG-UK) Consortium | The Lighthouse Lab in Milton Keynes and Alex Alderton, Roberto Amato, Jeffrey Barrett, Sonia Goncalves, Ewan Harrison, David K. Jackson, Ian Johnston, Dominic Kwiatkowski, Cordelia Langford, John Sillitoe on behalf of the Wellcome Sanger Institute COVID-19 Surveillance Team |
| EPI_ISL_3287606, EPI_ISL_3287607, EPI_ISL_3287608 | Lighthouse Lab in Glasgow | Wellcome Sanger Institute for the COVID-19 Genomics UK (COG-UK) Consortium | Harper VanSteenhouse, Yumi Kasai, David Gray, Carol Clugston, Anna Dominiczak and Alex Alderton, Roberto Amato, Jeffrey Barrett, Sonia Goncalves, Ewan Harrison, David K. Jackson, Ian Johnston, Dominic Kwiatkowski, Cordelia Langford, John Sillitoe on behalf of the Wellcome Sanger Institute COVID-19 Surveillance Team |
| EPI_ISL_3287609 | Lighthouse Lab in Milton Keynes | Wellcome Sanger Institute for the COVID-19 Genomics UK (COG-UK) Consortium | The Lighthouse Lab in Milton Keynes and Alex Alderton, Roberto Amato, Jeffrey Barrett, Sonia Goncalves, Ewan Harrison, David K. Jackson, Ian Johnston, Dominic Kwiatkowski, Cordelia Langford, John Sillitoe on behalf of the Wellcome Sanger Institute COVID-19 Surveillance Team |
| EPI_ISL_3287610 | Lighthouse Lab in Glasgow | Wellcome Sanger Institute for the COVID-19 Genomics UK (COG-UK) Consortium | Harper VanSteenhouse, Yumi Kasai, David Gray, Carol Clugston, Anna Dominiczak and Alex Alderton, Roberto Amato, Jeffrey Barrett, Sonia Goncalves, Ewan Harrison, David K. Jackson, Ian Johnston, Dominic Kwiatkowski, Cordelia Langford, John Sillitoe on behalf of the Wellcome Sanger Institute COVID-19 Surveillance Team |
| EPI_ISL_3287611 | Lighthouse Lab in Milton Keynes | Wellcome Sanger Institute for the COVID-19 Genomics UK (COG-UK) Consortium | The Lighthouse Lab in Milton Keynes and Alex Alderton, Roberto Amato, Jeffrey Barrett, Sonia Goncalves, Ewan Harrison, David K. Jackson, Ian Johnston, Dominic Kwiatkowski, Cordelia Langford, John Sillitoe on behalf of the Wellcome Sanger Institute COVID-19 Surveillance Team |
| EPI_ISL_3287612 | Lighthouse Lab in Glasgow | Wellcome Sanger Institute for the COVID-19 Genomics UK (COG-UK) Consortium | Harper VanSteenhouse, Yumi Kasai, David Gray, Carol Clugston, Anna Dominiczak and Alex Alderton, Roberto Amato, Jeffrey Barrett, Sonia Goncalves, Ewan Harrison, David K. Jackson, Ian Johnston, Dominic Kwiatkowski, Cordelia Langford, John Sillitoe on behalf of the Wellcome Sanger Institute COVID-19 Surveillance Team |
| EPI_ISL_3287613 | Lighthouse Lab in Milton Keynes | Wellcome Sanger Institute for the COVID-19 Genomics UK (COG-UK) Consortium | The Lighthouse Lab in Milton Keynes and Alex Alderton, Roberto Amato, Jeffrey Barrett, Sonia Goncalves, Ewan Harrison, David K. Jackson, Ian Johnston, Dominic Kwiatkowski, Cordelia Langford, John Sillitoe on behalf of the Wellcome Sanger Institute COVID-19 Surveillance Team |
| EPI_ISL_3287614 | Lighthouse Lab in Glasgow | Wellcome Sanger Institute for the COVID-19 Genomics UK (COG-UK) Consortium | Harper VanSteenhouse, Yumi Kasai, David Gray, Carol Clugston, Anna Dominiczak and Alex Alderton, Roberto Amato, Jeffrey Barrett, Sonia Goncalves, Ewan Harrison, David K. Jackson, Ian Johnston, Dominic Kwiatkowski, Cordelia Langford, John Sillitoe on behalf of the Wellcome Sanger Institute COVID-19 Surveillance Team |
| EPI_ISL_3287615, EPI_ISL_3287616 | Lighthouse Lab in Milton Keynes | Wellcome Sanger Institute for the COVID-19 Genomics UK (COG-UK) Consortium | The Lighthouse Lab in Milton Keynes and Alex Alderton, Roberto Amato, Jeffrey Barrett, Sonia Goncalves, Ewan Harrison, David K. Jackson, Ian Johnston, Dominic Kwiatkowski, Cordelia Langford, John Sillitoe on behalf of the Wellcome Sanger Institute COVID-19 Surveillance Team |
| EPI_ISL_3287617 | Lighthouse Lab in Glasgow | Wellcome Sanger Institute for the COVID-19 Genomics UK (COG-UK) Consortium | Harper VanSteenhouse, Yumi Kasai, David Gray, Carol Clugston, Anna Dominiczak and Alex Alderton, Roberto Amato, Jeffrey Barrett, Sonia Goncalves, Ewan Harrison, David K. Jackson, Ian Johnston, Dominic Kwiatkowski, Cordelia Langford, John Sillitoe on behalf of the Wellcome Sanger Institute COVID-19 Surveillance Team |
| EPI_ISL_3287618 | Lighthouse Lab in Milton Keynes | Wellcome Sanger Institute for the COVID-19 Genomics UK (COG-UK) Consortium | The Lighthouse Lab in Milton Keynes and Alex Alderton, Roberto Amato, Jeffrey Barrett, Sonia Goncalves, Ewan Harrison, David K. Jackson, Ian Johnston, Dominic Kwiatkowski, Cordelia Langford, John Sillitoe on behalf of the Wellcome Sanger Institute COVID-19 Surveillance Team |
| EPI_ISL_3287619 | Lighthouse Lab in Glasgow | Wellcome Sanger Institute for the COVID-19 Genomics UK (COG-UK) Consortium | Harper VanSteenhouse, Yumi Kasai, David Gray, Carol Clugston, Anna Dominiczak and Alex Alderton, Roberto Amato, Jeffrey Barrett, Sonia Goncalves, Ewan Harrison, David K. Jackson, Ian Johnston, Dominic Kwiatkowski, Cordelia Langford, John Sillitoe on behalf of the Wellcome Sanger Institute COVID-19 Surveillance Team |
| EPI_ISL_3287620, EPI_ISL_3287621 | Lighthouse Lab in Milton Keynes | Wellcome Sanger Institute for the COVID-19 Genomics UK (COG-UK) Consortium | The Lighthouse Lab in Milton Keynes and Alex Alderton, Roberto Amato, Jeffrey Barrett, Sonia Goncalves, Ewan Harrison, David K. Jackson, Ian Johnston, Dominic Kwiatkowski, Cordelia Langford, John Sillitoe on behalf of the Wellcome Sanger Institute COVID-19 Surveillance Team |
| EPI_ISL_3287622, EPI_ISL_3287623, EPI_ISL_3287624, EPI_ISL_3287625 | Lighthouse Lab in Glasgow | Wellcome Sanger Institute for the COVID-19 Genomics UK (COG-UK) Consortium | Harper VanSteenhouse, Yumi Kasai, David Gray, Carol Clugston, Anna Dominiczak and Alex Alderton, Roberto Amato, Jeffrey Barrett, Sonia Goncalves, Ewan Harrison, David K. Jackson, Ian Johnston, Dominic Kwiatkowski, Cordelia Langford, John Sillitoe on behalf of the Wellcome Sanger Institute COVID-19 Surveillance Team |
| EPI_ISL_3287626 | Health Services Laboratories | Wellcome Sanger Institute for the COVID-19 Genomics UK (COG-UK) Consortium | Health Services Laboratories and Alex Alderton, Roberto Amato, Jeffrey Barrett, Sonia Goncalves, Ewan Harrison, David K. Jackson, Ian Johnston, Dominic Kwiatkowski, Cordelia Langford, John Sillitoe on behalf of the Wellcome Sanger Institute COVID-19 Surveillance Team |
| EPI_ISL_3287627, EPI_ISL_3287628, EPI_ISL_3287629 | Lighthouse Lab in Milton Keynes | Wellcome Sanger Institute for the COVID-19 Genomics UK (COG-UK) Consortium | The Lighthouse Lab in Milton Keynes and Alex Alderton, Roberto Amato, Jeffrey Barrett, Sonia Goncalves, Ewan Harrison, David K. Jackson, Ian Johnston, Dominic Kwiatkowski, Cordelia Langford, John Sillitoe on behalf of the Wellcome Sanger Institute COVID-19 Surveillance Team |
| EPI_ISL_3287630 | Lighthouse Lab in Glasgow | Wellcome Sanger Institute for the COVID-19 Genomics UK (COG-UK) Consortium | Harper VanSteenhouse, Yumi Kasai, David Gray, Carol Clugston, Anna Dominiczak and Alex Alderton, Roberto Amato, Jeffrey Barrett, Sonia Goncalves, Ewan Harrison, David K. Jackson, Ian Johnston, Dominic Kwiatkowski, Cordelia Langford, John Sillitoe on behalf of the Wellcome Sanger Institute COVID-19 Surveillance Team |
| EPI_ISL_3287631 | Lighthouse Lab in Milton Keynes | Wellcome Sanger Institute for the COVID-19 Genomics UK (COG-UK) Consortium | The Lighthouse Lab in Milton Keynes and Alex Alderton, Roberto Amato, Jeffrey Barrett, Sonia Goncalves, Ewan Harrison, David K. Jackson, Ian Johnston, Dominic Kwiatkowski, Cordelia Langford, John Sillitoe on behalf of the Wellcome Sanger Institute COVID-19 Surveillance Team |
| EPI_ISL_3287632 | Health Services Laboratories | Wellcome Sanger Institute for the COVID-19 Genomics UK (COG-UK) Consortium | Health Services Laboratories and Alex Alderton, Roberto Amato, Jeffrey Barrett, Sonia Goncalves, Ewan Harrison, David K. Jackson, Ian Johnston, Dominic Kwiatkowski, Cordelia Langford, John Sillitoe on behalf of the Wellcome Sanger Institute COVID-19 Surveillance Team |
| EPI_ISL_3287633, EPI_ISL_3287634, EPI_ISL_3287635 | Lighthouse Lab in Milton Keynes | Wellcome Sanger Institute for the COVID-19 Genomics UK (COG-UK) Consortium | The Lighthouse Lab in Milton Keynes and Alex Alderton, Roberto Amato, Jeffrey Barrett, Sonia Goncalves, Ewan Harrison, David K. Jackson, Ian Johnston, Dominic Kwiatkowski, Cordelia Langford, John Sillitoe on behalf of the Wellcome Sanger Institute COVID-19 Surveillance Team |

| | | | |
|---|---|---|---|
| EPI_ISL_3287636 | Health Services Laboratories | Wellcome Sanger Institute for the COVID-19 Genomics UK (COG-UK) Consortium | Health Services Laboratories and Alex Alderton, Roberto Amato, Jeffrey Barrett, Sonia Goncalves, Ewan Harrison, David K. Jackson, Ian Johnston, Dominic Kwiatkowski, Cordelia Langford, John Sillitoe on behalf of the Wellcome Sanger Institute COVID-19 Surveillance Team |
| EPI_ISL_3287637 | Lighthouse Lab in Milton Keynes | Wellcome Sanger Institute for the COVID-19 Genomics UK (COG-UK) Consortium | The Lighthouse Lab in Milton Keynes and Alex Alderton, Roberto Amato, Jeffrey Barrett, Sonia Goncalves, Ewan Harrison, David K. Jackson, Ian Johnston, Dominic Kwiatkowski, Cordelia Langford, John Sillitoe on behalf of the Wellcome Sanger Institute COVID-19 Surveillance Team |
| EPI_ISL_3287638 | Lighthouse Lab in Glasgow | Wellcome Sanger Institute for the COVID-19 Genomics UK (COG-UK) Consortium | Harper VanSteenhouse, Yumi Kasai, David Gray, Carol Clugston, Anna Dominiczak and Alex Alderton, Roberto Amato, Jeffrey Barrett, Sonia Goncalves, Ewan Harrison, David K. Jackson, Ian Johnston, Dominic Kwiatkowski, Cordelia Langford, John Sillitoe on behalf of the Wellcome Sanger Institute COVID-19 Surveillance Team |
| EPI_ISL_3287639 | Lighthouse Lab in Milton Keynes | Wellcome Sanger Institute for the COVID-19 Genomics UK (COG-UK) Consortium | The Lighthouse Lab in Milton Keynes and Alex Alderton, Roberto Amato, Jeffrey Barrett, Sonia Goncalves, Ewan Harrison, David K. Jackson, Ian Johnston, Dominic Kwiatkowski, Cordelia Langford, John Sillitoe on behalf of the Wellcome Sanger Institute COVID-19 Surveillance Team |
| EPI_ISL_3287640, EPI_ISL_3287641 | Lighthouse Lab in Glasgow | Wellcome Sanger Institute for the COVID-19 Genomics UK (COG-UK) Consortium | Harper VanSteenhouse, Yumi Kasai, David Gray, Carol Clugston, Anna Dominiczak and Alex Alderton, Roberto Amato, Jeffrey Barrett, Sonia Goncalves, Ewan Harrison, David K. Jackson, Ian Johnston, Dominic Kwiatkowski, Cordelia Langford, John Sillitoe on behalf of the Wellcome Sanger Institute COVID-19 Surveillance Team |
| EPI_ISL_3287642 | Lighthouse Lab in Milton Keynes | Wellcome Sanger Institute for the COVID-19 Genomics UK (COG-UK) Consortium | The Lighthouse Lab in Milton Keynes and Alex Alderton, Roberto Amato, Jeffrey Barrett, Sonia Goncalves, Ewan Harrison, David K. Jackson, Ian Johnston, Dominic Kwiatkowski, Cordelia Langford, John Sillitoe on behalf of the Wellcome Sanger Institute COVID-19 Surveillance Team |
| EPI_ISL_3287643, EPI_ISL_3287644, EPI_ISL_3287645, EPI_ISL_3287646 | Lighthouse Lab in Glasgow | Wellcome Sanger Institute for the COVID-19 Genomics UK (COG-UK) Consortium | Harper VanSteenhouse, Yumi Kasai, David Gray, Carol Clugston, Anna Dominiczak and Alex Alderton, Roberto Amato, Jeffrey Barrett, Sonia Goncalves, Ewan Harrison, David K. Jackson, Ian Johnston, Dominic Kwiatkowski, Cordelia Langford, John Sillitoe on behalf of the Wellcome Sanger Institute COVID-19 Surveillance Team |
| EPI_ISL_3287647 | Health Services Laboratories | Wellcome Sanger Institute for the COVID-19 Genomics UK (COG-UK) Consortium | Health Services Laboratories and Alex Alderton, Roberto Amato, Jeffrey Barrett, Sonia Goncalves, Ewan Harrison, David K. Jackson, Ian Johnston, Dominic Kwiatkowski, Cordelia Langford, John Sillitoe on behalf of the Wellcome Sanger Institute COVID-19 Surveillance Team |
| EPI_ISL_3287648 | Lighthouse Lab in Milton Keynes | Wellcome Sanger Institute for the COVID-19 Genomics UK (COG-UK) Consortium | The Lighthouse Lab in Milton Keynes and Alex Alderton, Roberto Amato, Jeffrey Barrett, Sonia Goncalves, Ewan Harrison, David K. Jackson, Ian Johnston, Dominic Kwiatkowski, Cordelia Langford, John Sillitoe on behalf of the Wellcome Sanger Institute COVID-19 Surveillance Team |
| EPI_ISL_3287649 | Lighthouse Lab in Glasgow | Wellcome Sanger Institute for the COVID-19 Genomics UK (COG-UK) Consortium | Harper VanSteenhouse, Yumi Kasai, David Gray, Carol Clugston, Anna Dominiczak and Alex Alderton, Roberto Amato, Jeffrey Barrett, Sonia Goncalves, Ewan Harrison, David K. Jackson, Ian Johnston, Dominic Kwiatkowski, Cordelia Langford, John Sillitoe on behalf of the Wellcome Sanger Institute COVID-19 Surveillance Team |
| EPI_ISL_3287650, EPI_ISL_3287651, EPI_ISL_3287652 | Health Services Laboratories | Wellcome Sanger Institute for the COVID-19 Genomics UK (COG-UK) Consortium | Health Services Laboratories and Alex Alderton, Roberto Amato, Jeffrey Barrett, Sonia Goncalves, Ewan Harrison, David K. Jackson, Ian Johnston, Dominic Kwiatkowski, Cordelia Langford, John Sillitoe on behalf of the Wellcome Sanger Institute COVID-19 Surveillance Team |
| EPI_ISL_3287653 | Lighthouse Lab in Milton Keynes | Wellcome Sanger Institute for the COVID-19 Genomics UK (COG-UK) Consortium | The Lighthouse Lab in Milton Keynes and Alex Alderton, Roberto Amato, Jeffrey Barrett, Sonia Goncalves, Ewan Harrison, David K. Jackson, Ian Johnston, Dominic Kwiatkowski, Cordelia Langford, John Sillitoe on behalf of the Wellcome Sanger Institute COVID-19 Surveillance Team |
| EPI_ISL_3287654 | Lighthouse Lab in Glasgow | Wellcome Sanger Institute for the COVID-19 Genomics UK (COG-UK) Consortium | Harper VanSteenhouse, Yumi Kasai, David Gray, Carol Clugston, Anna Dominiczak and Alex Alderton, Roberto Amato, Jeffrey Barrett, Sonia Goncalves, Ewan Harrison, David K. Jackson, Ian Johnston, Dominic Kwiatkowski, Cordelia Langford, John Sillitoe on behalf of the Wellcome Sanger Institute COVID-19 Surveillance Team |
| EPI_ISL_3287655 | Lighthouse Lab in Milton Keynes | Wellcome Sanger Institute for the COVID-19 Genomics UK (COG-UK) Consortium | The Lighthouse Lab in Milton Keynes and Alex Alderton, Roberto Amato, Jeffrey Barrett, Sonia Goncalves, Ewan Harrison, David K. Jackson, Ian Johnston, Dominic Kwiatkowski, Cordelia Langford, John Sillitoe on behalf of the Wellcome Sanger Institute COVID-19 Surveillance Team |
| EPI_ISL_3287656 | Health Services Laboratories | Wellcome Sanger Institute for the COVID-19 Genomics UK (COG-UK) Consortium | Health Services Laboratories and Alex Alderton, Roberto Amato, Jeffrey Barrett, Sonia Goncalves, Ewan Harrison, David K. Jackson, Ian Johnston, Dominic Kwiatkowski, Cordelia Langford, John Sillitoe on behalf of the Wellcome Sanger Institute COVID-19 Surveillance Team |
| EPI_ISL_3287657 | Lighthouse Lab in Glasgow | Wellcome Sanger Institute for the COVID-19 Genomics UK (COG-UK) Consortium | Harper VanSteenhouse, Yumi Kasai, David Gray, Carol Clugston, Anna Dominiczak and Alex Alderton, Roberto Amato, Jeffrey Barrett, Sonia Goncalves, Ewan Harrison, David K. Jackson, Ian Johnston, Dominic Kwiatkowski, Cordelia Langford, John Sillitoe on behalf of the Wellcome Sanger Institute COVID-19 Surveillance Team |
| EPI_ISL_3287658 | Health Services Laboratories | Wellcome Sanger Institute for the COVID-19 Genomics UK (COG-UK) Consortium | Health Services Laboratories and Alex Alderton, Roberto Amato, Jeffrey Barrett, Sonia Goncalves, Ewan Harrison, David K. Jackson, Ian Johnston, Dominic Kwiatkowski, Cordelia Langford, John Sillitoe on behalf of the Wellcome Sanger Institute COVID-19 Surveillance Team |
| EPI_ISL_3287659 | Lighthouse Lab in Milton Keynes | Wellcome Sanger Institute for the COVID-19 Genomics UK (COG-UK) Consortium | The Lighthouse Lab in Milton Keynes and Alex Alderton, Roberto Amato, Jeffrey Barrett, Sonia Goncalves, Ewan Harrison, David K. Jackson, Ian Johnston, Dominic Kwiatkowski, Cordelia Langford, John Sillitoe on behalf of the Wellcome Sanger Institute COVID-19 Surveillance Team |
| EPI_ISL_3287660 | Lighthouse Lab in Glasgow | Wellcome Sanger Institute for the COVID-19 Genomics UK (COG-UK) Consortium | Harper VanSteenhouse, Yumi Kasai, David Gray, Carol Clugston, Anna Dominiczak and Alex Alderton, Roberto Amato, Jeffrey Barrett, Sonia Goncalves, Ewan Harrison, David K. Jackson, Ian Johnston, Dominic Kwiatkowski, Cordelia Langford, John Sillitoe on behalf of the Wellcome Sanger Institute COVID-19 Surveillance Team |
| EPI_ISL_3287661, EPI_ISL_3287662 | Lighthouse Lab in Milton Keynes | Wellcome Sanger Institute for the COVID-19 Genomics UK (COG-UK) Consortium | The Lighthouse Lab in Milton Keynes and Alex Alderton, Roberto Amato, Jeffrey Barrett, Sonia Goncalves, Ewan Harrison, David K. Jackson, Ian Johnston, Dominic Kwiatkowski, Cordelia Langford, John Sillitoe on behalf of the Wellcome Sanger Institute COVID-19 Surveillance Team |
| EPI_ISL_3287663 | Health Services Laboratories | Wellcome Sanger Institute for the COVID-19 Genomics UK (COG-UK) Consortium | Health Services Laboratories and Alex Alderton, Roberto Amato, Jeffrey Barrett, Sonia Goncalves, Ewan Harrison, David K. Jackson, Ian Johnston, Dominic Kwiatkowski, Cordelia Langford, John Sillitoe on behalf of the Wellcome Sanger Institute COVID-19 Surveillance Team |
| EPI_ISL_3287664 | Lighthouse Lab in Glasgow | Wellcome Sanger Institute for the COVID-19 Genomics UK (COG-UK) Consortium | Harper VanSteenhouse, Yumi Kasai, David Gray, Carol Clugston, Anna Dominiczak and Alex Alderton, Roberto Amato, Jeffrey Barrett, Sonia Goncalves, Ewan Harrison, David K. Jackson, Ian Johnston, Dominic Kwiatkowski, Cordelia Langford, John Sillitoe on behalf of the Wellcome Sanger Institute COVID-19 Surveillance Team |
| EPI_ISL_3287665, EPI_ISL_3287666, EPI_ISL_3287667, EPI_ISL_3287668 | Health Services Laboratories | Wellcome Sanger Institute for the COVID-19 Genomics UK (COG-UK) Consortium | Health Services Laboratories and Alex Alderton, Roberto Amato, Jeffrey Barrett, Sonia Goncalves, Ewan Harrison, David K. Jackson, Ian Johnston, Dominic Kwiatkowski, Cordelia Langford, John Sillitoe on behalf of the Wellcome Sanger Institute COVID-19 Surveillance Team |
| EPI_ISL_3287669 | Lighthouse Lab in Milton Keynes | Wellcome Sanger Institute for the COVID-19 Genomics UK (COG-UK) Consortium | The Lighthouse Lab in Milton Keynes and Alex Alderton, Roberto Amato, Jeffrey Barrett, Sonia Goncalves, Ewan Harrison, David K. Jackson, Ian Johnston, Dominic Kwiatkowski, Cordelia Langford, John Sillitoe on behalf of the Wellcome Sanger Institute COVID-19 Surveillance Team |
| EPI_ISL_3287670 | Lighthouse Lab in Glasgow | Wellcome Sanger Institute for the COVID-19 Genomics UK (COG-UK) Consortium | Harper VanSteenhouse, Yumi Kasai, David Gray, Carol Clugston, Anna Dominiczak and Alex Alderton, Roberto Amato, Jeffrey Barrett, Sonia Goncalves, Ewan Harrison, David K. Jackson, Ian Johnston, Dominic Kwiatkowski, Cordelia Langford, John Sillitoe on behalf of the Wellcome Sanger Institute COVID-19 Surveillance Team |
| EPI_ISL_3287671 | Health Services Laboratories | Wellcome Sanger Institute for the COVID-19 Genomics UK (COG-UK) Consortium | Health Services Laboratories and Alex Alderton, Roberto Amato, Jeffrey Barrett, Sonia Goncalves, Ewan Harrison, David K. Jackson, Ian Johnston, Dominic Kwiatkowski, Cordelia Langford, John Sillitoe on behalf of the Wellcome Sanger Institute COVID-19 Surveillance Team |
| EPI_ISL_3287672 | Lighthouse Lab in Glasgow | Wellcome Sanger Institute for the COVID-19 Genomics UK (COG-UK) Consortium | Harper VanSteenhouse, Yumi Kasai, David Gray, Carol Clugston, Anna Dominiczak and Alex Alderton, Roberto Amato, Jeffrey Barrett, Sonia Goncalves, Ewan Harrison, David K. Jackson, Ian Johnston, Dominic Kwiatkowski, Cordelia Langford, John Sillitoe on behalf of the Wellcome Sanger Institute COVID-19 Surveillance Team |
| EPI_ISL_3287673 | Lighthouse Lab in Milton Keynes | Wellcome Sanger Institute for the COVID-19 Genomics UK (COG-UK) Consortium | The Lighthouse Lab in Milton Keynes and Alex Alderton, Roberto Amato, Jeffrey Barrett, Sonia Goncalves, Ewan Harrison, David K. Jackson, Ian Johnston, Dominic Kwiatkowski, Cordelia Langford, John Sillitoe on behalf of the Wellcome Sanger Institute COVID-19 Surveillance Team |
| EPI_ISL_3287674, EPI_ISL_3287675 | Health Services Laboratories | Wellcome Sanger Institute for the COVID-19 Genomics UK (COG-UK) Consortium | Health Services Laboratories and Alex Alderton, Roberto Amato, Jeffrey Barrett, Sonia Goncalves, Ewan Harrison, David K. Jackson, Ian Johnston, Dominic Kwiatkowski, Cordelia Langford, John Sillitoe on behalf of the Wellcome Sanger Institute COVID-19 Surveillance Team |
| EPI_ISL_3287676, EPI_ISL_3287677 | Lighthouse Lab in Milton Keynes | Wellcome Sanger Institute for the COVID-19 Genomics UK (COG-UK) Consortium | The Lighthouse Lab in Milton Keynes and Alex Alderton, Roberto Amato, Jeffrey Barrett, Sonia Goncalves, Ewan Harrison, David K. Jackson, Ian Johnston, Dominic Kwiatkowski, Cordelia Langford, John Sillitoe on behalf of the Wellcome Sanger Institute COVID-19 Surveillance Team |
| EPI_ISL_3287678, EPI_ISL_3287679, EPI_ISL_3287680, EPI_ISL_3287681, EPI_ISL_3287682 | Health Services Laboratories | Wellcome Sanger Institute for the COVID-19 Genomics UK (COG-UK) Consortium | Health Services Laboratories and Alex Alderton, Roberto Amato, Jeffrey Barrett, Sonia Goncalves, Ewan Harrison, David K. Jackson, Ian Johnston, Dominic Kwiatkowski, Cordelia Langford, John Sillitoe on behalf of the Wellcome Sanger Institute COVID-19 Surveillance Team |
| EPI_ISL_3287683 | Lighthouse Lab in Glasgow | Wellcome Sanger Institute for the COVID-19 Genomics UK (COG-UK) Consortium | Harper VanSteenhouse, Yumi Kasai, David Gray, Carol Clugston, Anna Dominiczak and Alex Alderton, Roberto Amato, Jeffrey Barrett, Sonia Goncalves, Ewan Harrison, David K. Jackson, Ian Johnston, Dominic Kwiatkowski, Cordelia Langford, John Sillitoe on behalf of the Wellcome Sanger Institute COVID-19 Surveillance Team |
| EPI_ISL_3287684 | Health Services Laboratories | Wellcome Sanger Institute for the COVID-19 Genomics UK (COG-UK) Consortium | Health Services Laboratories and Alex Alderton, Roberto Amato, Jeffrey Barrett, Sonia Goncalves, Ewan Harrison, David K. Jackson, Ian Johnston, Dominic Kwiatkowski, Cordelia Langford, John Sillitoe on behalf of the Wellcome Sanger Institute COVID-19 Surveillance Team |
| EPI_ISL_3287685, EPI_ISL_3287686, EPI_ISL_3287687 | Lighthouse Lab in Milton Keynes | Wellcome Sanger Institute for the COVID-19 Genomics UK (COG-UK) Consortium | The Lighthouse Lab in Milton Keynes and Alex Alderton, Roberto Amato, Jeffrey Barrett, Sonia Goncalves, Ewan Harrison, David K. Jackson, Ian Johnston, Dominic Kwiatkowski, Cordelia Langford, John Sillitoe on behalf of the Wellcome Sanger Institute COVID-19 Surveillance Team |
| EPI_ISL_3287688, EPI_ISL_3287689 | Lighthouse Lab in Glasgow | Wellcome Sanger Institute for the COVID-19 Genomics UK (COG-UK) Consortium | Harper VanSteenhouse, Yumi Kasai, David Gray, Carol Clugston, Anna Dominiczak and Alex Alderton, Roberto Amato, Jeffrey Barrett, Sonia Goncalves, Ewan Harrison, David K. Jackson, Ian Johnston, Dominic Kwiatkowski, Cordelia Langford, John Sillitoe on behalf of the Wellcome Sanger Institute COVID-19 Surveillance Team |
| EPI_ISL_3287690, EPI_ISL_3287691 | Lighthouse Lab in Milton Keynes | Wellcome Sanger Institute for the COVID-19 Genomics UK (COG-UK) Consortium | The Lighthouse Lab in Milton Keynes and Alex Alderton, Roberto Amato, Jeffrey Barrett, Sonia Goncalves, Ewan Harrison, David K. Jackson, Ian Johnston, Dominic Kwiatkowski, Cordelia Langford, John Sillitoe on behalf of the Wellcome Sanger Institute COVID-19 Surveillance Team |
| EPI_ISL_3287692 | Lighthouse Lab in Glasgow | Wellcome Sanger Institute for the COVID-19 Genomics UK (COG-UK) Consortium | Harper VanSteenhouse, Yumi Kasai, David Gray, Carol Clugston, Anna Dominiczak and Alex Alderton, Roberto Amato, Jeffrey Barrett, Sonia Goncalves, Ewan Harrison, David K. Jackson, Ian Johnston, Dominic Kwiatkowski, Cordelia Langford, John Sillitoe on behalf of the Wellcome Sanger Institute COVID-19 Surveillance Team |
| EPI_ISL_3287693 | Lighthouse Lab in Milton Keynes | Wellcome Sanger Institute for the COVID-19 Genomics UK (COG-UK) Consortium | The Lighthouse Lab in Milton Keynes and Alex Alderton, Roberto Amato, Jeffrey Barrett, Sonia Goncalves, Ewan Harrison, David K. Jackson, Ian Johnston, Dominic Kwiatkowski, Cordelia Langford, John Sillitoe on behalf of the Wellcome Sanger Institute COVID-19 Surveillance Team |
| EPI_ISL_3287694, EPI_ISL_3287695 | Lighthouse Lab in Glasgow | Wellcome Sanger Institute for the COVID-19 Genomics UK (COG-UK) Consortium | Harper VanSteenhouse, Yumi Kasai, David Gray, Carol Clugston, Anna Dominiczak and Alex Alderton, Roberto Amato, Jeffrey Barrett, Sonia Goncalves, Ewan Harrison, David K. Jackson, Ian Johnston, Dominic Kwiatkowski, Cordelia Langford, John Sillitoe on behalf of the Wellcome Sanger Institute COVID-19 Surveillance Team |
| EPI_ISL_3287696 | Health Services Laboratories | Wellcome Sanger Institute for the COVID-19 Genomics UK (COG-UK) Consortium | Health Services Laboratories and Alex Alderton, Roberto Amato, Jeffrey Barrett, Sonia Goncalves, Ewan Harrison, David K. Jackson, Ian Johnston, Dominic Kwiatkowski, Cordelia Langford, John Sillitoe on behalf of the Wellcome Sanger Institute COVID-19 Surveillance Team |
| EPI_ISL_3287697, EPI_ISL_3287698 | Lighthouse Lab in Glasgow | Wellcome Sanger Institute for the COVID-19 Genomics UK (COG-UK) Consortium | Harper VanSteenhouse, Yumi Kasai, David Gray, Carol Clugston, Anna Dominiczak and Alex Alderton, Roberto Amato, Jeffrey Barrett, Sonia Goncalves, Ewan Harrison, David K. Jackson, Ian Johnston, Dominic Kwiatkowski, Cordelia Langford, John Sillitoe on behalf of the Wellcome Sanger Institute COVID-19 Surveillance Team |
| EPI_ISL_3287699, | Health Services Laboratories | Wellcome Sanger Institute for the COVID-19 Genomics UK (COG-UK) | Health Services Laboratories and Alex Alderton, Roberto Amato, Jeffrey Barrett, Sonia Goncalves, Ewan Harrison, David K. Jackson, Ian Johnston, Dominic Kwiatkowski, Cordelia Langford, John Sillitoe on behalf of the Wellcome Sanger Institute |

| | | | |
|---|---|---|---|
| EPI_ISL_3287700, EPI_ISL_3287701 | | Consortium | COVID-19 Surveillance Team |
| EPI_ISL_3287702 | Lighthouse Lab in Glasgow | Wellcome Sanger Institute for the COVID-19 Genomics UK (COG-UK) Consortium | Harper VanSteenhouse, Yumi Kasai, David Gray, Carol Clugston, Anna Dominiczak and Alex Alderton, Roberto Amato, Jeffrey Barrett, Sonia Goncalves, Ewan Harrison, David K. Jackson, Ian Johnston, Dominic Kwiatkowski, Cordelia Langford, John Sillitoe on behalf of the Wellcome Sanger Institute COVID-19 Surveillance Team |
| EPI_ISL_3287703 | Lighthouse Lab in Milton Keynes | Wellcome Sanger Institute for the COVID-19 Genomics UK (COG-UK) Consortium | The Lighthouse Lab in Milton Keynes and Alex Alderton, Roberto Amato, Jeffrey Barrett, Sonia Goncalves, Ewan Harrison, David K. Jackson, Ian Johnston, Dominic Kwiatkowski, Cordelia Langford, John Sillitoe on behalf of the Wellcome Sanger Institute COVID-19 Surveillance Team |
| EPI_ISL_3287704 | Lighthouse Lab in Glasgow | Wellcome Sanger Institute for the COVID-19 Genomics UK (COG-UK) Consortium | Harper VanSteenhouse, Yumi Kasai, David Gray, Carol Clugston, Anna Dominiczak and Alex Alderton, Roberto Amato, Jeffrey Barrett, Sonia Goncalves, Ewan Harrison, David K. Jackson, Ian Johnston, Dominic Kwiatkowski, Cordelia Langford, John Sillitoe on behalf of the Wellcome Sanger Institute COVID-19 Surveillance Team |
| EPI_ISL_3287705 | Health Services Laboratories | Wellcome Sanger Institute for the COVID-19 Genomics UK (COG-UK) Consortium | Health Services Laboratories and Alex Alderton, Roberto Amato, Jeffrey Barrett, Sonia Goncalves, Ewan Harrison, David K. Jackson, Ian Johnston, Dominic Kwiatkowski, Cordelia Langford, John Sillitoe on behalf of the Wellcome Sanger Institute COVID-19 Surveillance Team |
| EPI_ISL_3287706, EPI_ISL_3287707, EPI_ISL_3287708, EPI_ISL_3287709, EPI_ISL_3287710 | Lighthouse Lab in Glasgow | Wellcome Sanger Institute for the COVID-19 Genomics UK (COG-UK) Consortium | Harper VanSteenhouse, Yumi Kasai, David Gray, Carol Clugston, Anna Dominiczak and Alex Alderton, Roberto Amato, Jeffrey Barrett, Sonia Goncalves, Ewan Harrison, David K. Jackson, Ian Johnston, Dominic Kwiatkowski, Cordelia Langford, John Sillitoe on behalf of the Wellcome Sanger Institute COVID-19 Surveillance Team |
| EPI_ISL_3287711 | Lighthouse Lab in Milton Keynes | Wellcome Sanger Institute for the COVID-19 Genomics UK (COG-UK) Consortium | The Lighthouse Lab in Milton Keynes and Alex Alderton, Roberto Amato, Jeffrey Barrett, Sonia Goncalves, Ewan Harrison, David K. Jackson, Ian Johnston, Dominic Kwiatkowski, Cordelia Langford, John Sillitoe on behalf of the Wellcome Sanger Institute COVID-19 Surveillance Team |
| EPI_ISL_3287713 | Lighthouse Lab in Glasgow | Wellcome Sanger Institute for the COVID-19 Genomics UK (COG-UK) Consortium | Harper VanSteenhouse, Yumi Kasai, David Gray, Carol Clugston, Anna Dominiczak and Alex Alderton, Roberto Amato, Jeffrey Barrett, Sonia Goncalves, Ewan Harrison, David K. Jackson, Ian Johnston, Dominic Kwiatkowski, Cordelia Langford, John Sillitoe on behalf of the Wellcome Sanger Institute COVID-19 Surveillance Team |
| EPI_ISL_3287714 | Health Services Laboratories | Wellcome Sanger Institute for the COVID-19 Genomics UK (COG-UK) Consortium | Health Services Laboratories and Alex Alderton, Roberto Amato, Jeffrey Barrett, Sonia Goncalves, Ewan Harrison, David K. Jackson, Ian Johnston, Dominic Kwiatkowski, Cordelia Langford, John Sillitoe on behalf of the Wellcome Sanger Institute COVID-19 Surveillance Team |
| EPI_ISL_3287715 | Lighthouse Lab in Milton Keynes | Wellcome Sanger Institute for the COVID-19 Genomics UK (COG-UK) Consortium | The Lighthouse Lab in Milton Keynes and Alex Alderton, Roberto Amato, Jeffrey Barrett, Sonia Goncalves, Ewan Harrison, David K. Jackson, Ian Johnston, Dominic Kwiatkowski, Cordelia Langford, John Sillitoe on behalf of the Wellcome Sanger Institute COVID-19 Surveillance Team |
| EPI_ISL_3287716 | Health Services Laboratories | Wellcome Sanger Institute for the COVID-19 Genomics UK (COG-UK) Consortium | Health Services Laboratories and Alex Alderton, Roberto Amato, Jeffrey Barrett, Sonia Goncalves, Ewan Harrison, David K. Jackson, Ian Johnston, Dominic Kwiatkowski, Cordelia Langford, John Sillitoe on behalf of the Wellcome Sanger Institute COVID-19 Surveillance Team |
| EPI_ISL_3287717 | Lighthouse Lab in Glasgow | Wellcome Sanger Institute for the COVID-19 Genomics UK (COG-UK) Consortium | Harper VanSteenhouse, Yumi Kasai, David Gray, Carol Clugston, Anna Dominiczak and Alex Alderton, Roberto Amato, Jeffrey Barrett, Sonia Goncalves, Ewan Harrison, David K. Jackson, Ian Johnston, Dominic Kwiatkowski, Cordelia Langford, John Sillitoe on behalf of the Wellcome Sanger Institute COVID-19 Surveillance Team |
| EPI_ISL_3287718 | Health Services Laboratories | Wellcome Sanger Institute for the COVID-19 Genomics UK (COG-UK) Consortium | Health Services Laboratories and Alex Alderton, Roberto Amato, Jeffrey Barrett, Sonia Goncalves, Ewan Harrison, David K. Jackson, Ian Johnston, Dominic Kwiatkowski, Cordelia Langford, John Sillitoe on behalf of the Wellcome Sanger Institute COVID-19 Surveillance Team |
| EPI_ISL_3287719, EPI_ISL_3287720, EPI_ISL_3287721 | Lighthouse Lab in Glasgow | Wellcome Sanger Institute for the COVID-19 Genomics UK (COG-UK) Consortium | Harper VanSteenhouse, Yumi Kasai, David Gray, Carol Clugston, Anna Dominiczak and Alex Alderton, Roberto Amato, Jeffrey Barrett, Sonia Goncalves, Ewan Harrison, David K. Jackson, Ian Johnston, Dominic Kwiatkowski, Cordelia Langford, John Sillitoe on behalf of the Wellcome Sanger Institute COVID-19 Surveillance Team |
| EPI_ISL_3287722, EPI_ISL_3287723 | Lighthouse Lab in Milton Keynes | Wellcome Sanger Institute for the COVID-19 Genomics UK (COG-UK) Consortium | The Lighthouse Lab in Milton Keynes and Alex Alderton, Roberto Amato, Jeffrey Barrett, Sonia Goncalves, Ewan Harrison, David K. Jackson, Ian Johnston, Dominic Kwiatkowski, Cordelia Langford, John Sillitoe on behalf of the Wellcome Sanger Institute COVID-19 Surveillance Team |
| EPI_ISL_3287724 | Lighthouse Lab in Glasgow | Wellcome Sanger Institute for the COVID-19 Genomics UK (COG-UK) Consortium | Harper VanSteenhouse, Yumi Kasai, David Gray, Carol Clugston, Anna Dominiczak and Alex Alderton, Roberto Amato, Jeffrey Barrett, Sonia Goncalves, Ewan Harrison, David K. Jackson, Ian Johnston, Dominic Kwiatkowski, Cordelia Langford, John Sillitoe on behalf of the Wellcome Sanger Institute COVID-19 Surveillance Team |
| EPI_ISL_3287725, EPI_ISL_3287726, EPI_ISL_3287727, EPI_ISL_3287728, EPI_ISL_3287729, EPI_ISL_3287730, EPI_ISL_3287731, EPI_ISL_3287732, EPI_ISL_3287733, EPI_ISL_3287734, EPI_ISL_3287735, EPI_ISL_3287736, EPI_ISL_3287737, EPI_ISL_3287738, EPI_ISL_3287739, EPI_ISL_3287740, EPI_ISL_3287741, EPI_ISL_3287742, EPI_ISL_3287743, EPI_ISL_3287744, EPI_ISL_3287745, EPI_ISL_3287746, EPI_ISL_3287747, EPI_ISL_3287748, EPI_ISL_3287749, EPI_ISL_3287750, EPI_ISL_3287751, EPI_ISL_3287752, EPI_ISL_3287753, EPI_ISL_3287754, EPI_ISL_3287755, EPI_ISL_3287757, EPI_ISL_3287758, EPI_ISL_3287759, EPI_ISL_3287760, EPI_ISL_3287761, EPI_ISL_3287762, EPI_ISL_3287763, EPI_ISL_3287764, EPI_ISL_3287765, EPI_ISL_3287766, EPI_ISL_3287767, EPI_ISL_3287768, EPI_ISL_3287769, EPI_ISL_3287770, EPI_ISL_3287771, EPI_ISL_3287772, EPI_ISL_3287773, EPI_ISL_3287774, EPI_ISL_3287776, EPI_ISL_3287777, EPI_ISL_3287778, EPI_ISL_3287779, EPI_ISL_3287780, EPI_ISL_3287781, EPI_ISL_3287782, EPI_ISL_3287783, EPI_ISL_3287784, EPI_ISL_3287785, EPI_ISL_3287786, EPI_ISL_3287787, EPI_ISL_3287788, EPI_ISL_3287789, EPI_ISL_3287790, EPI_ISL_3287791, EPI_ISL_3287792, EPI_ISL_3287793, EPI_ISL_3287794, EPI_ISL_3287795, EPI_ISL_3287796, EPI_ISL_3287797, EPI_ISL_3287798, EPI_ISL_3287799, EPI_ISL_3287800, EPI_ISL_3287801, EPI_ISL_3287802, EPI_ISL_3287803, EPI_ISL_3287804, EPI_ISL_3287805, EPI_ISL_3287806, EPI_ISL_3287807, EPI_ISL_3287808, EPI_ISL_3287809, EPI_ISL_3287810, EPI_ISL_3287812, EPI_ISL_3287813, EPI_ISL_3287814, EPI_ISL_3287815, EPI_ISL_3287816, EPI_ISL_3287817, EPI_ISL_3287818, EPI_ISL_3287819, EPI_ISL_3287820, EPI_ISL_3287821, EPI_ISL_3287822, EPI_ISL_3287823, EPI_ISL_3287824, EPI_ISL_3287825, EPI_ISL_3287826, EPI_ISL_3287827, EPI_ISL_3287828, EPI_ISL_3287829, EPI_ISL_3287830, EPI_ISL_3287831, EPI_ISL_3287832, EPI_ISL_3287833, EPI_ISL_3287834, EPI_ISL_3287835, EPI_ISL_3287836, EPI_ISL_3287837, EPI_ISL_3287838, EPI_ISL_3287839, EPI_ISL_3287840, EPI_ISL_3287841, EPI_ISL_3287842, EPI_ISL_3287843, EPI_ISL_3287844, EPI_ISL_3287845, EPI_ISL_3287847, EPI_ISL_3287848, EPI_ISL_3287849, EPI_ISL_3287850, EPI_ISL_3287851, EPI_ISL_3287852, EPI_ISL_3287853, EPI_ISL_3287854, EPI_ISL_3287855, EPI_ISL_3287856, EPI_ISL_3287857, EPI_ISL_3287858, EPI_ISL_3287859, EPI_ISL_3287860, EPI_ISL_3287861, EPI_ISL_3287862, EPI_ISL_3287863, EPI_ISL_3287864, EPI_ISL_3287865, EPI_ISL_3287866, EPI_ISL_3287867, EPI_ISL_3287868, EPI_ISL_3287869, EPI_ISL_3287870, EPI_ISL_3287871, EPI_ISL_3287872, EPI_ISL_3287873, EPI_ISL_3287874, EPI_ISL_3287875, EPI_ISL_3287876, EPI_ISL_3287877, EPI_ISL_3287878, EPI_ISL_3287879, EPI_ISL_3287880, EPI_ISL_3287881, EPI_ISL_3287882, EPI_ISL_3287883, EPI_ISL_3287884, EPI_ISL_3287885, EPI_ISL_3287886, EPI_ISL_3287887, EPI_ISL_3287888, EPI_ISL_3287889, EPI_ISL_3287890, EPI_ISL_3287891, EPI_ISL_3287892, EPI_ISL_3287893, EPI_ISL_3287894, EPI_ISL_3287895, EPI_ISL_3287896, EPI_ISL_3287897, EPI_ISL_3287898, EPI_ISL_3287899, EPI_ISL_3287900, EPI_ISL_3287902, EPI_ISL_3287903, EPI_ISL_3287904, EPI_ISL_3287905, EPI_ISL_3287906, EPI_ISL_3287907, EPI_ISL_3287908, EPI_ISL_3287909, EPI_ISL_3287910, EPI_ISL_3287911, EPI_ISL_3287912, EPI_ISL_3287913, EPI_ISL_3287914, EPI_ISL_3287915, EPI_ISL_3287916, EPI_ISL_3287917, EPI_ISL_3287918, EPI_ISL_3287919, EPI_ISL_3287920, EPI_ISL_3287921, EPI_ISL_3287922, EPI_ISL_3287923, EPI_ISL_3287924, EPI_ISL_3287925, EPI_ISL_3287926, EPI_ISL_3287927, EPI_ISL_3287928, EPI_ISL_3287931, EPI_ISL_3287932, EPI_ISL_3287933, EPI_ISL_3287934, EPI_ISL_3287935, EPI_ISL_3287936, EPI_ISL_3287937, EPI_ISL_3287938, EPI_ISL_3287939, EPI_ISL_3287940, EPI_ISL_3287941, EPI_ISL_3287942, EPI_ISL_3287943, EPI_ISL_3287944, EPI_ISL_3287945, EPI_ISL_3287946, EPI_ISL_3287947, EPI_ISL_3287948, EPI_ISL_3287949, EPI_ISL_3287950, EPI_ISL_3287951, EPI_ISL_3287952, EPI_ISL_3287953, EPI_ISL_3287954, EPI_ISL_3287955, EPI_ISL_3287956, EPI_ISL_3287957 | | | |
| see above | Virology Department, Royal Infirmary of Edinburgh, NHS Lothian / School of Biological Sciences, University of Edinburgh | COVID-19 Genomics UK (COG-UK) Consortium | McHugh M, Dewar R, Cotton S, Gallagher A, Maloney D, Fernandez G, O'Toole Á, Scher E, Hill V, McCrone JT, Colquhoun R, Yu X, Jackson B, Rambaut A, Templeton K |
| EPI_ISL_3288031, EPI_ISL_3288032, EPI_ISL_3288033, EPI_ISL_3288034, EPI_ISL_3288035, EPI_ISL_3288036, EPI_ISL_3288037, EPI_ISL_3288038, EPI_ISL_3288039, EPI_ISL_3288040, EPI_ISL_3288041, EPI_ISL_3288042, EPI_ISL_3288043, EPI_ISL_3288044, EPI_ISL_3288045, EPI_ISL_3288046, EPI_ISL_3288047, EPI_ISL_3288048, EPI_ISL_3288049, EPI_ISL_3288050, EPI_ISL_3288051, EPI_ISL_3288052, EPI_ISL_3288053, EPI_ISL_3288054, EPI_ISL_3288055, EPI_ISL_3288056, EPI_ISL_3288057, EPI_ISL_3288058, EPI_ISL_3288059, EPI_ISL_3288060, EPI_ISL_3288061, EPI_ISL_3288062, EPI_ISL_3288063, EPI_ISL_3288064, EPI_ISL_3288065, EPI_ISL_3288066, EPI_ISL_3288067, EPI_ISL_3288068, EPI_ISL_3288069, EPI_ISL_3288070, EPI_ISL_3288071, EPI_ISL_3288072, EPI_ISL_3288073, EPI_ISL_3288074, EPI_ISL_3288075, EPI_ISL_3288076, EPI_ISL_3288077, EPI_ISL_3288078, EPI_ISL_3288079, EPI_ISL_3288080, EPI_ISL_3288081, EPI_ISL_3288082, EPI_ISL_3288083, EPI_ISL_3288084, EPI_ISL_3288085, EPI_ISL_3288086, EPI_ISL_3288087, EPI_ISL_3288088, EPI_ISL_3288089, EPI_ISL_3288090, EPI_ISL_3288091, EPI_ISL_3288092, EPI_ISL_3288093, EPI_ISL_3288094, EPI_ISL_3288095, EPI_ISL_3288096, EPI_ISL_3288097, EPI_ISL_3288098, EPI_ISL_3288099, EPI_ISL_3288100, EPI_ISL_3288101, EPI_ISL_3288102, EPI_ISL_3288103, EPI_ISL_3288104, EPI_ISL_3288110, EPI_ISL_3288112, EPI_ISL_3288113, EPI_ISL_3288114, EPI_ISL_3288115, EPI_ISL_3288116, EPI_ISL_3288117, EPI_ISL_3288118, EPI_ISL_3288119, EPI_ISL_3288121, EPI_ISL_3288129, EPI_ISL_3288130, EPI_ISL_3288132, EPI_ISL_3288143, EPI_ISL_3288149, EPI_ISL_3288150, EPI_ISL_3288151, EPI_ISL_3288152, EPI_ISL_3288153, EPI_ISL_3288154, EPI_ISL_3288155, EPI_ISL_3288156, EPI_ISL_3288157, EPI_ISL_3288158, EPI_ISL_3288159, EPI_ISL_3288160, EPI_ISL_3288161, EPI_ISL_3288162, EPI_ISL_3288163, EPI_ISL_3288164, EPI_ISL_3288165, EPI_ISL_3288166, EPI_ISL_3288167, EPI_ISL_3288168, EPI_ISL_3288169, EPI_ISL_3288170, EPI_ISL_3288171, EPI_ISL_3288172, EPI_ISL_3288173, EPI_ISL_3288174, EPI_ISL_3288175, EPI_ISL_3288176, EPI_ISL_3288177, EPI_ISL_3288178, EPI_ISL_3288179, EPI_ISL_3288180, EPI_ISL_3288181, EPI_ISL_3288182, EPI_ISL_3288183, EPI_ISL_3288184, EPI_ISL_3288185, EPI_ISL_3288186, EPI_ISL_3288187, EPI_ISL_3288188, EPI_ISL_3288189, EPI_ISL_3288190, EPI_ISL_3288191, EPI_ISL_3288192, EPI_ISL_3288193, EPI_ISL_3288194, EPI_ISL_3288195, EPI_ISL_3288196, EPI_ISL_3288197, EPI_ISL_3288198, EPI_ISL_3288199, EPI_ISL_3288200, EPI_ISL_3288201, EPI_ISL_3288202 | | | |
| see above | University College London, Great Ormond Street Hospital for Children NHS Foundation Trust, Imperial College Healthcare NHS Trust | COVID-19 Genomics UK (COG-UK) Consortium | Sergi Castellano, Rachel Williams, Mark Kristiansen, Sunando Roy, Charlotte Williams, Nadua Bayzid, Marius Cotic, Helena Tutill, Patricia Dyal, Judith Breuer |
| EPI_ISL_3288203 | U.O. Microbiologia Laboratorio Unico Centro Servizi - AUSL della Romagna | U.O. Microbiologia, Laboratorio Unico Centro Servizi - AUSL della Romagna | Giorgio Dirani |
| EPI_ISL_3288204 | University College London, Great Ormond Street Hospital for Children NHS Foundation Trust, Imperial College Healthcare NHS Trust | COVID-19 Genomics UK (COG-UK) Consortium | Sergi Castellano, Rachel Williams, Mark Kristiansen, Sunando Roy, Charlotte Williams, Nadua Bayzid, Marius Cotic, Helena Tutill, Patricia Dyal, Judith Breuer |
| EPI_ISL_3288205 | U.O. Microbiologia Laboratorio Unico Centro Servizi - AUSL della Romagna | U.O. Microbiologia, Laboratorio Unico Centro Servizi - AUSL della Romagna | Giorgio Dirani |
| EPI_ISL_3288206 | University College London, Great Ormond Street Hospital for Children NHS Foundation Trust, Imperial College Healthcare NHS Trust | COVID-19 Genomics UK (COG-UK) Consortium | Sergi Castellano, Rachel Williams, Mark Kristiansen, Sunando Roy, Charlotte Williams, Nadua Bayzid, Marius Cotic, Helena Tutill, Patricia Dyal, Judith Breuer |
| EPI_ISL_3288207 | U.O. Microbiologia Laboratorio Unico Centro Servizi - AUSL della Romagna | U.O. Microbiologia, Laboratorio Unico Centro Servizi - AUSL della Romagna | Giorgio Dirani |
| EPI_ISL_3288208, EPI_ISL_3288209 | University College London, Great Ormond Street Hospital for Children NHS Foundation Trust, Imperial College Healthcare NHS Trust | COVID-19 Genomics UK (COG-UK) Consortium | Sergi Castellano, Rachel Williams, Mark Kristiansen, Sunando Roy, Charlotte Williams, Nadua Bayzid, Marius Cotic, Helena Tutill, Patricia Dyal, Judith Breuer |
| EPI_ISL_3288210 | U.O. Microbiologia Laboratorio Unico Centro Servizi - AUSL della Romagna | U.O. Microbiologia, Laboratorio Unico Centro Servizi - AUSL della Romagna | Giorgio Dirani |
| EPI_ISL_3288211 | University College London, Great Ormond Street Hospital for Children NHS Foundation Trust, Imperial College Healthcare NHS Trust | COVID-19 Genomics UK (COG-UK) Consortium | Sergi Castellano, Rachel Williams, Mark Kristiansen, Sunando Roy, Charlotte Williams, Nadua Bayzid, Marius Cotic, Helena Tutill, Patricia Dyal, Judith Breuer |
| EPI_ISL_3288212 | U.O. Microbiologia Laboratorio Unico Centro Servizi - AUSL della Romagna | U.O. Microbiologia, Laboratorio Unico Centro Servizi - AUSL della Romagna | Giorgio Dirani |
| EPI_ISL_3288213 | University College London, Great Ormond Street Hospital for Children NHS Foundation Trust, Imperial College Healthcare NHS Trust | COVID-19 Genomics UK (COG-UK) Consortium | Sergi Castellano, Rachel Williams, Mark Kristiansen, Sunando Roy, Charlotte Williams, Nadua Bayzid, Marius Cotic, Helena Tutill, Patricia Dyal, Judith Breuer |
| EPI_ISL_3288214 | U.O. Microbiologia Laboratorio Unico Centro Servizi - AUSL della Romagna | U.O. Microbiologia, Laboratorio Unico Centro Servizi - AUSL della Romagna | Giorgio Dirani |
| EPI_ISL_3288215 | University College London, Great Ormond Street Hospital for Children NHS Foundation Trust, Imperial College Healthcare NHS Trust | COVID-19 Genomics UK (COG-UK) Consortium | Sergi Castellano, Rachel Williams, Mark Kristiansen, Sunando Roy, Charlotte Williams, Nadua Bayzid, Marius Cotic, Helena Tutill, Patricia Dyal, Judith Breuer |
| EPI_ISL_3288216 | U.O. Microbiologia Laboratorio Unico Centro Servizi - AUSL della Romagna | U.O. Microbiologia, Laboratorio Unico Centro Servizi - AUSL della Romagna | Giorgio Dirani |
| EPI_ISL_3288217 | University College London, Great Ormond Street Hospital for Children NHS Foundation Trust, Imperial College Healthcare NHS Trust | COVID-19 Genomics UK (COG-UK) Consortium | Sergi Castellano, Rachel Williams, Mark Kristiansen, Sunando Roy, Charlotte Williams, Nadua Bayzid, Marius Cotic, Helena Tutill, Patricia Dyal, Judith Breuer |
| EPI_ISL_3288218 | U.O. Microbiologia Laboratorio Unico Centro Servizi - AUSL della | U.O. Microbiologia, Laboratorio Unico Centro Servizi - AUSL della | Giorgio Dirani |

| | | | |
|---|---|---|---|
| | Romagna | Romagna | |
| EPI_ISL_3288219 | University College London, Great Ormond Street Hospital for Children NHS Foundation Trust, Imperial College Healthcare NHS Trust | COVID-19 Genomics UK (COG-UK) Consortium | Sergi Castellano, Rachel Williams, Mark Kristiansen, Sunando Roy, Charlotte Williams, Nadua Bayzid, Marius Cotic, Helena Tutill, Patricia Dyal, Judith Breuer |
| EPI_ISL_3288220 | U.O. Microbiologia Laboratorio Unico Centro Servizi - AUSL della Romagna | U.O. Microbiologia, Laboratorio Unico Centro Servizi - AUSL della Romagna | Giorgio Dirani |
| EPI_ISL_3288221, EPI_ISL_3288222, EPI_ISL_3288224 | University College London, Great Ormond Street Hospital for Children NHS Foundation Trust, Imperial College Healthcare NHS Trust | COVID-19 Genomics UK (COG-UK) Consortium | Sergi Castellano, Rachel Williams, Mark Kristiansen, Sunando Roy, Charlotte Williams, Nadua Bayzid, Marius Cotic, Helena Tutill, Patricia Dyal, Judith Breuer |
| EPI_ISL_3288225 | U.O. Microbiologia Laboratorio Unico Centro Servizi - AUSL della Romagna | U.O. Microbiologia, Laboratorio Unico Centro Servizi - AUSL della Romagna | Giorgio Dirani |
| EPI_ISL_3288226 | University College London, Great Ormond Street Hospital for Children NHS Foundation Trust, Imperial College Healthcare NHS Trust | COVID-19 Genomics UK (COG-UK) Consortium | Sergi Castellano, Rachel Williams, Mark Kristiansen, Sunando Roy, Charlotte Williams, Nadua Bayzid, Marius Cotic, Helena Tutill, Patricia Dyal, Judith Breuer |
| EPI_ISL_3288227 | U.O. Microbiologia Laboratorio Unico Centro Servizi - AUSL della Romagna | U.O. Microbiologia, Laboratorio Unico Centro Servizi - AUSL della Romagna | Giorgio Dirani |
| EPI_ISL_3288228 | University College London, Great Ormond Street Hospital for Children NHS Foundation Trust, Imperial College Healthcare NHS Trust | COVID-19 Genomics UK (COG-UK) Consortium | Sergi Castellano, Rachel Williams, Mark Kristiansen, Sunando Roy, Charlotte Williams, Nadua Bayzid, Marius Cotic, Helena Tutill, Patricia Dyal, Judith Breuer |
| EPI_ISL_3288229 | U.O. Microbiologia Laboratorio Unico Centro Servizi - AUSL della Romagna | U.O. Microbiologia, Laboratorio Unico Centro Servizi - AUSL della Romagna | Giorgio Dirani |
| EPI_ISL_3288230 | University College London, Great Ormond Street Hospital for Children NHS Foundation Trust, Imperial College Healthcare NHS Trust | COVID-19 Genomics UK (COG-UK) Consortium | Sergi Castellano, Rachel Williams, Mark Kristiansen, Sunando Roy, Charlotte Williams, Nadua Bayzid, Marius Cotic, Helena Tutill, Patricia Dyal, Judith Breuer |
| EPI_ISL_3288231 | U.O. Microbiologia Laboratorio Unico Centro Servizi - AUSL della Romagna | U.O. Microbiologia, Laboratorio Unico Centro Servizi - AUSL della Romagna | Giorgio Dirani |
| EPI_ISL_3288232 | University College London, Great Ormond Street Hospital for Children NHS Foundation Trust, Imperial College Healthcare NHS Trust | COVID-19 Genomics UK (COG-UK) Consortium | Sergi Castellano, Rachel Williams, Mark Kristiansen, Sunando Roy, Charlotte Williams, Nadua Bayzid, Marius Cotic, Helena Tutill, Patricia Dyal, Judith Breuer |
| EPI_ISL_3288233 | U.O. Microbiologia Laboratorio Unico Centro Servizi - AUSL della Romagna | U.O. Microbiologia, Laboratorio Unico Centro Servizi - AUSL della Romagna | Giorgio Dirani |
| EPI_ISL_3288234 | University College London, Great Ormond Street Hospital for Children NHS Foundation Trust, Imperial College Healthcare NHS Trust | COVID-19 Genomics UK (COG-UK) Consortium | Sergi Castellano, Rachel Williams, Mark Kristiansen, Sunando Roy, Charlotte Williams, Nadua Bayzid, Marius Cotic, Helena Tutill, Patricia Dyal, Judith Breuer |
| EPI_ISL_3288235 | U.O. Microbiologia Laboratorio Unico Centro Servizi - AUSL della Romagna | U.O. Microbiologia, Laboratorio Unico Centro Servizi - AUSL della Romagna | Giorgio Dirani |
| EPI_ISL_3288236 | University College London, Great Ormond Street Hospital for Children NHS Foundation Trust, Imperial College Healthcare NHS Trust | COVID-19 Genomics UK (COG-UK) Consortium | Sergi Castellano, Rachel Williams, Mark Kristiansen, Sunando Roy, Charlotte Williams, Nadua Bayzid, Marius Cotic, Helena Tutill, Patricia Dyal, Judith Breuer |
| EPI_ISL_3288237 | U.O. Microbiologia Laboratorio Unico Centro Servizi - AUSL della Romagna | U.O. Microbiologia, Laboratorio Unico Centro Servizi - AUSL della Romagna | Giorgio Dirani |
| EPI_ISL_3288238 | University College London, Great Ormond Street Hospital for Children NHS Foundation Trust, Imperial College Healthcare NHS Trust | COVID-19 Genomics UK (COG-UK) Consortium | Sergi Castellano, Rachel Williams, Mark Kristiansen, Sunando Roy, Charlotte Williams, Nadua Bayzid, Marius Cotic, Helena Tutill, Patricia Dyal, Judith Breuer |
| EPI_ISL_3288239 | U.O. Microbiologia Laboratorio Unico Centro Servizi - AUSL della Romagna | U.O. Microbiologia, Laboratorio Unico Centro Servizi - AUSL della Romagna | Giorgio Dirani |
| EPI_ISL_3288240 | University College London, Great Ormond Street Hospital for Children NHS Foundation Trust, Imperial College Healthcare NHS Trust | COVID-19 Genomics UK (COG-UK) Consortium | Sergi Castellano, Rachel Williams, Mark Kristiansen, Sunando Roy, Charlotte Williams, Nadua Bayzid, Marius Cotic, Helena Tutill, Patricia Dyal, Judith Breuer |
| EPI_ISL_3288241 | U.O. Microbiologia Laboratorio Unico Centro Servizi - AUSL della Romagna | U.O. Microbiologia, Laboratorio Unico Centro Servizi - AUSL della Romagna | Giorgio Dirani |
| EPI_ISL_3288242 | University College London, Great Ormond Street Hospital for Children NHS Foundation Trust, Imperial College Healthcare NHS Trust | COVID-19 Genomics UK (COG-UK) Consortium | Sergi Castellano, Rachel Williams, Mark Kristiansen, Sunando Roy, Charlotte Williams, Nadua Bayzid, Marius Cotic, Helena Tutill, Patricia Dyal, Judith Breuer |
| EPI_ISL_3288243 | U.O. Microbiologia Laboratorio Unico Centro Servizi - AUSL della Romagna | U.O. Microbiologia, Laboratorio Unico Centro Servizi - AUSL della Romagna | Giorgio Dirani |
| EPI_ISL_3288244, EPI_ISL_3288245 | University College London, Great Ormond Street Hospital for Children NHS Foundation Trust, Imperial College Healthcare NHS Trust | COVID-19 Genomics UK (COG-UK) Consortium | Sergi Castellano, Rachel Williams, Mark Kristiansen, Sunando Roy, Charlotte Williams, Nadua Bayzid, Marius Cotic, Helena Tutill, Patricia Dyal, Judith Breuer |
| EPI_ISL_3288246 | U.O. Microbiologia Laboratorio Unico Centro Servizi - AUSL della Romagna | U.O. Microbiologia, Laboratorio Unico Centro Servizi - AUSL della Romagna | Giorgio Dirani |
| EPI_ISL_3288247 | University College London, Great Ormond Street Hospital for Children NHS Foundation Trust, Imperial College Healthcare NHS Trust | COVID-19 Genomics UK (COG-UK) Consortium | Sergi Castellano, Rachel Williams, Mark Kristiansen, Sunando Roy, Charlotte Williams, Nadua Bayzid, Marius Cotic, Helena Tutill, Patricia Dyal, Judith Breuer |
| EPI_ISL_3288248 | U.O. Microbiologia Laboratorio Unico Centro Servizi - AUSL della Romagna | U.O. Microbiologia, Laboratorio Unico Centro Servizi - AUSL della Romagna | Giorgio Dirani |
| EPI_ISL_3288249 | University College London, Great Ormond Street Hospital for Children NHS Foundation Trust, Imperial College Healthcare NHS Trust | COVID-19 Genomics UK (COG-UK) Consortium | Sergi Castellano, Rachel Williams, Mark Kristiansen, Sunando Roy, Charlotte Williams, Nadua Bayzid, Marius Cotic, Helena Tutill, Patricia Dyal, Judith Breuer |
| EPI_ISL_3288250 | U.O. Microbiologia Laboratorio Unico Centro Servizi - AUSL della Romagna | U.O. Microbiologia, Laboratorio Unico Centro Servizi - AUSL della Romagna | Giorgio Dirani |
| EPI_ISL_3288251 | University College London, Great Ormond Street Hospital for Children NHS Foundation Trust, Imperial College Healthcare NHS Trust | COVID-19 Genomics UK (COG-UK) Consortium | Sergi Castellano, Rachel Williams, Mark Kristiansen, Sunando Roy, Charlotte Williams, Nadua Bayzid, Marius Cotic, Helena Tutill, Patricia Dyal, Judith Breuer |
| EPI_ISL_3288252 | U.O. Microbiologia Laboratorio Unico Centro Servizi - AUSL della Romagna | U.O. Microbiologia, Laboratorio Unico Centro Servizi - AUSL della Romagna | Giorgio Dirani |
| EPI_ISL_3288253 | University College London, Great Ormond Street Hospital for Children NHS Foundation Trust, Imperial College Healthcare NHS Trust | COVID-19 Genomics UK (COG-UK) Consortium | Sergi Castellano, Rachel Williams, Mark Kristiansen, Sunando Roy, Charlotte Williams, Nadua Bayzid, Marius Cotic, Helena Tutill, Patricia Dyal, Judith Breuer |
| EPI_ISL_3288254 | U.O. Microbiologia Laboratorio Unico Centro Servizi - AUSL della Romagna | U.O. Microbiologia, Laboratorio Unico Centro Servizi - AUSL della Romagna | Giorgio Dirani |
| EPI_ISL_3288255 | University College London, Great Ormond Street Hospital for Children NHS Foundation Trust, Imperial College Healthcare NHS Trust | COVID-19 Genomics UK (COG-UK) Consortium | Sergi Castellano, Rachel Williams, Mark Kristiansen, Sunando Roy, Charlotte Williams, Nadua Bayzid, Marius Cotic, Helena Tutill, Patricia Dyal, Judith Breuer |
| EPI_ISL_3288256 | U.O. Microbiologia Laboratorio Unico Centro Servizi - AUSL della Romagna | U.O. Microbiologia, Laboratorio Unico Centro Servizi - AUSL della Romagna | Giorgio Dirani |
| EPI_ISL_3288257 | University College London, Great Ormond Street Hospital for Children NHS Foundation Trust, Imperial College Healthcare NHS Trust | COVID-19 Genomics UK (COG-UK) Consortium | Sergi Castellano, Rachel Williams, Mark Kristiansen, Sunando Roy, Charlotte Williams, Nadua Bayzid, Marius Cotic, Helena Tutill, Patricia Dyal, Judith Breuer |
| EPI_ISL_3288258 | U.O. Microbiologia Laboratorio Unico Centro Servizi - AUSL della Romagna | U.O. Microbiologia, Laboratorio Unico Centro Servizi - AUSL della Romagna | Giorgio Dirani |
| EPI_ISL_3288259 | University College London, Great Ormond Street Hospital for Children NHS Foundation Trust, Imperial College Healthcare NHS Trust | COVID-19 Genomics UK (COG-UK) Consortium | Sergi Castellano, Rachel Williams, Mark Kristiansen, Sunando Roy, Charlotte Williams, Nadua Bayzid, Marius Cotic, Helena Tutill, Patricia Dyal, Judith Breuer |

| | | | |
|---|---|---|---|
| EPI_ISL_3288260 | U.O. Microbiologia Laboratorio Unico Centro Servizi - AUSL della Romagna | U.O. Microbiologia, Laboratorio Unico Centro Servizi - AUSL della Romagna | Giorgio Dirani |
| EPI_ISL_3288261 | University College London, Great Ormond Street Hospital for Children NHS Foundation Trust, Imperial College Healthcare NHS Trust | COVID-19 Genomics UK (COG-UK) Consortium | Sergi Castellano, Rachel Williams, Mark Kristiansen, Sunando Roy, Charlotte Williams, Nadua Bayzid, Marius Cotic, Helena Tutill, Patricia Dyal, Judith Breuer |
| EPI_ISL_3288262 | U.O. Microbiologia Laboratorio Unico Centro Servizi - AUSL della Romagna | U.O. Microbiologia, Laboratorio Unico Centro Servizi - AUSL della Romagna | Giorgio Dirani |
| EPI_ISL_3288263 | University College London, Great Ormond Street Hospital for Children NHS Foundation Trust, Imperial College Healthcare NHS Trust | COVID-19 Genomics UK (COG-UK) Consortium | Sergi Castellano, Rachel Williams, Mark Kristiansen, Sunando Roy, Charlotte Williams, Nadua Bayzid, Marius Cotic, Helena Tutill, Patricia Dyal, Judith Breuer |
| EPI_ISL_3288264 | U.O. Microbiologia Laboratorio Unico Centro Servizi - AUSL della Romagna | U.O. Microbiologia, Laboratorio Unico Centro Servizi - AUSL della Romagna | Giorgio Dirani |
| EPI_ISL_3288265 | University College London, Great Ormond Street Hospital for Children NHS Foundation Trust, Imperial College Healthcare NHS Trust | COVID-19 Genomics UK (COG-UK) Consortium | Sergi Castellano, Rachel Williams, Mark Kristiansen, Sunando Roy, Charlotte Williams, Nadua Bayzid, Marius Cotic, Helena Tutill, Patricia Dyal, Judith Breuer |
| EPI_ISL_3288266 | U.O. Microbiologia Laboratorio Unico Centro Servizi - AUSL della Romagna | U.O. Microbiologia, Laboratorio Unico Centro Servizi - AUSL della Romagna | Giorgio Dirani |
| EPI_ISL_3288267 | University College London, Great Ormond Street Hospital for Children NHS Foundation Trust, Imperial College Healthcare NHS Trust | COVID-19 Genomics UK (COG-UK) Consortium | Sergi Castellano, Rachel Williams, Mark Kristiansen, Sunando Roy, Charlotte Williams, Nadua Bayzid, Marius Cotic, Helena Tutill, Patricia Dyal, Judith Breuer |
| EPI_ISL_3288268 | U.O. Microbiologia Laboratorio Unico Centro Servizi - AUSL della Romagna | U.O. Microbiologia, Laboratorio Unico Centro Servizi - AUSL della Romagna | Giorgio Dirani |
| EPI_ISL_3288269 | University College London, Great Ormond Street Hospital for Children NHS Foundation Trust, Imperial College Healthcare NHS Trust | COVID-19 Genomics UK (COG-UK) Consortium | Sergi Castellano, Rachel Williams, Mark Kristiansen, Sunando Roy, Charlotte Williams, Nadua Bayzid, Marius Cotic, Helena Tutill, Patricia Dyal, Judith Breuer |
| EPI_ISL_3288270 | U.O. Microbiologia Laboratorio Unico Centro Servizi - AUSL della Romagna | U.O. Microbiologia, Laboratorio Unico Centro Servizi - AUSL della Romagna | Giorgio Dirani |
| EPI_ISL_3288271 | University College London, Great Ormond Street Hospital for Children NHS Foundation Trust, Imperial College Healthcare NHS Trust | COVID-19 Genomics UK (COG-UK) Consortium | Sergi Castellano, Rachel Williams, Mark Kristiansen, Sunando Roy, Charlotte Williams, Nadua Bayzid, Marius Cotic, Helena Tutill, Patricia Dyal, Judith Breuer |
| EPI_ISL_3288272 | U.O. Microbiologia Laboratorio Unico Centro Servizi - AUSL della Romagna | U.O. Microbiologia, Laboratorio Unico Centro Servizi - AUSL della Romagna | Giorgio Dirani |
| EPI_ISL_3288273 | University College London, Great Ormond Street Hospital for Children NHS Foundation Trust, Imperial College Healthcare NHS Trust | COVID-19 Genomics UK (COG-UK) Consortium | Sergi Castellano, Rachel Williams, Mark Kristiansen, Sunando Roy, Charlotte Williams, Nadua Bayzid, Marius Cotic, Helena Tutill, Patricia Dyal, Judith Breuer |
| EPI_ISL_3288274 | U.O. Microbiologia Laboratorio Unico Centro Servizi - AUSL della Romagna | U.O. Microbiologia, Laboratorio Unico Centro Servizi - AUSL della Romagna | Giorgio Dirani |
| EPI_ISL_3288275 | University College London, Great Ormond Street Hospital for Children NHS Foundation Trust, Imperial College Healthcare NHS Trust | COVID-19 Genomics UK (COG-UK) Consortium | Sergi Castellano, Rachel Williams, Mark Kristiansen, Sunando Roy, Charlotte Williams, Nadua Bayzid, Marius Cotic, Helena Tutill, Patricia Dyal, Judith Breuer |
| EPI_ISL_3288276 | U.O. Microbiologia Laboratorio Unico Centro Servizi - AUSL della Romagna | U.O. Microbiologia, Laboratorio Unico Centro Servizi - AUSL della Romagna | Giorgio Dirani |
| EPI_ISL_3288277 | University College London, Great Ormond Street Hospital for Children NHS Foundation Trust, Imperial College Healthcare NHS Trust | COVID-19 Genomics UK (COG-UK) Consortium | Sergi Castellano, Rachel Williams, Mark Kristiansen, Sunando Roy, Charlotte Williams, Nadua Bayzid, Marius Cotic, Helena Tutill, Patricia Dyal, Judith Breuer |
| EPI_ISL_3288278 | U.O. Microbiologia Laboratorio Unico Centro Servizi - AUSL della Romagna | U.O. Microbiologia, Laboratorio Unico Centro Servizi - AUSL della Romagna | Giorgio Dirani |
| EPI_ISL_3288279 | University College London, Great Ormond Street Hospital for Children NHS Foundation Trust, Imperial College Healthcare NHS Trust | COVID-19 Genomics UK (COG-UK) Consortium | Sergi Castellano, Rachel Williams, Mark Kristiansen, Sunando Roy, Charlotte Williams, Nadua Bayzid, Marius Cotic, Helena Tutill, Patricia Dyal, Judith Breuer |
| EPI_ISL_3288280 | U.O. Microbiologia Laboratorio Unico Centro Servizi - AUSL della Romagna | U.O. Microbiologia, Laboratorio Unico Centro Servizi - AUSL della Romagna | Giorgio Dirani |
| EPI_ISL_3288281, EPI_ISL_3288282, EPI_ISL_3288283, EPI_ISL_3288284, EPI_ISL_3288285, EPI_ISL_3288286, EPI_ISL_3288287, EPI_ISL_3288288, EPI_ISL_3288289, EPI_ISL_3288290, EPI_ISL_3288291, EPI_ISL_3288292, EPI_ISL_3288293, EPI_ISL_3288294, EPI_ISL_3288295, EPI_ISL_3288296, EPI_ISL_3288297, EPI_ISL_3288298, EPI_ISL_3288299, EPI_ISL_3288300, EPI_ISL_3288301, EPI_ISL_3288302, EPI_ISL_3288303, EPI_ISL_3288304, EPI_ISL_3288305, EPI_ISL_3288306, EPI_ISL_3288307, EPI_ISL_3288308, EPI_ISL_3288309, EPI_ISL_3288310, EPI_ISL_3288311, EPI_ISL_3288312, EPI_ISL_3288313, EPI_ISL_3288314, EPI_ISL_3288315, EPI_ISL_3288318, EPI_ISL_3288319, EPI_ISL_3288320, EPI_ISL_3288321, EPI_ISL_3288322, EPI_ISL_3288323, EPI_ISL_3288324, EPI_ISL_3288325, EPI_ISL_3288326, EPI_ISL_3288327, EPI_ISL_3288328, EPI_ISL_3288329, EPI_ISL_3288330, EPI_ISL_3288331, EPI_ISL_3288332, EPI_ISL_3288333, EPI_ISL_3288334, EPI_ISL_3288342, EPI_ISL_3288350, EPI_ISL_3288351, EPI_ISL_3288352, EPI_ISL_3288353, EPI_ISL_3288354, EPI_ISL_3288355, EPI_ISL_3288356, EPI_ISL_3288357, EPI_ISL_3288358, EPI_ISL_3288359, EPI_ISL_3288360, EPI_ISL_3288361, EPI_ISL_3288362, EPI_ISL_3288363, EPI_ISL_3288364, EPI_ISL_3288365, EPI_ISL_3288366, EPI_ISL_3288367, EPI_ISL_3288368, EPI_ISL_3288369, EPI_ISL_3288370, EPI_ISL_3288371, EPI_ISL_3288372, EPI_ISL_3288373, EPI_ISL_3288374, EPI_ISL_3288375, EPI_ISL_3288376, EPI_ISL_3288377, EPI_ISL_3288378, EPI_ISL_3288379, EPI_ISL_3288380, EPI_ISL_3288381, EPI_ISL_3288382, EPI_ISL_3288383, EPI_ISL_3288384, EPI_ISL_3288385, EPI_ISL_3288386, EPI_ISL_3288387, EPI_ISL_3288388, EPI_ISL_3288389, EPI_ISL_3288390, EPI_ISL_3288391, EPI_ISL_3288392, EPI_ISL_3288393, EPI_ISL_3288394, EPI_ISL_3288395, EPI_ISL_3288396, EPI_ISL_3288397, EPI_ISL_3288398, EPI_ISL_3288399, EPI_ISL_3288400, EPI_ISL_3288401, EPI_ISL_3288428 | | | |
| see above | University College London, Great Ormond Street Hospital for Children NHS Foundation Trust, Imperial College Healthcare NHS Trust | COVID-19 Genomics UK (COG-UK) Consortium | Sergi Castellano, Rachel Williams, Mark Kristiansen, Sunando Roy, Charlotte Williams, Nadua Bayzid, Marius Cotic, Helena Tutill, Patricia Dyal, Judith Breuer |
| EPI_ISL_3288429, EPI_ISL_3288430, EPI_ISL_3288431, EPI_ISL_3288432, EPI_ISL_3288433, EPI_ISL_3288434, EPI_ISL_3288435, EPI_ISL_3288436, EPI_ISL_3288437, EPI_ISL_3288438, EPI_ISL_3288439, EPI_ISL_3288440, EPI_ISL_3288441, EPI_ISL_3288442, EPI_ISL_3288443, EPI_ISL_3288444, EPI_ISL_3288445, EPI_ISL_3288446, EPI_ISL_3288447, EPI_ISL_3288448, EPI_ISL_3288449, EPI_ISL_3288450, EPI_ISL_3288451, EPI_ISL_3288452, EPI_ISL_3288453, EPI_ISL_3288454, EPI_ISL_3288455, EPI_ISL_3288456, EPI_ISL_3288457, EPI_ISL_3288458, EPI_ISL_3288459, EPI_ISL_3288460, EPI_ISL_3288461, EPI_ISL_3288462, EPI_ISL_3288463, EPI_ISL_3288464, EPI_ISL_3288465, EPI_ISL_3288466, EPI_ISL_3288467, EPI_ISL_3288468, EPI_ISL_3288469, EPI_ISL_3288470, EPI_ISL_3288471, EPI_ISL_3288472, EPI_ISL_3288473, EPI_ISL_3288474, EPI_ISL_3288475, EPI_ISL_3288476, EPI_ISL_3288477, EPI_ISL_3288478, EPI_ISL_3288479, EPI_ISL_3288480, EPI_ISL_3288481, EPI_ISL_3288482, EPI_ISL_3288483, EPI_ISL_3288484, EPI_ISL_3288485, EPI_ISL_3288486, EPI_ISL_3288487, EPI_ISL_3288488, EPI_ISL_3288489, EPI_ISL_3288490, EPI_ISL_3288491, EPI_ISL_3288492, EPI_ISL_3288493, EPI_ISL_3288494, EPI_ISL_3288495, EPI_ISL_3288496, EPI_ISL_3288497, EPI_ISL_3288498, EPI_ISL_3288499, EPI_ISL_3288500, EPI_ISL_3288501, EPI_ISL_3288502, EPI_ISL_3288503, EPI_ISL_3288504, EPI_ISL_3288505, EPI_ISL_3288506, EPI_ISL_3288507, EPI_ISL_3288508, EPI_ISL_3288509, EPI_ISL_3288510, EPI_ISL_3288511, EPI_ISL_3288512, EPI_ISL_3288513, EPI_ISL_3288514, EPI_ISL_3288515, EPI_ISL_3288516, EPI_ISL_3288517, EPI_ISL_3288518, EPI_ISL_3288519, EPI_ISL_3288520, EPI_ISL_3288521, EPI_ISL_3288522, EPI_ISL_3288523, EPI_ISL_3288524, EPI_ISL_3288525, EPI_ISL_3288526, EPI_ISL_3288527, EPI_ISL_3288528, EPI_ISL_3288529, EPI_ISL_3288530, EPI_ISL_3288531, EPI_ISL_3288532, EPI_ISL_3288533, EPI_ISL_3288534, EPI_ISL_3288535, EPI_ISL_3288536, EPI_ISL_3288537, EPI_ISL_3288538, EPI_ISL_3288539, EPI_ISL_3288540, EPI_ISL_3288541, EPI_ISL_3288542, EPI_ISL_3288543, EPI_ISL_3288544, EPI_ISL_3288545, EPI_ISL_3288546, EPI_ISL_3288547, EPI_ISL_3288548, EPI_ISL_3288549, EPI_ISL_3288550, EPI_ISL_3288551, EPI_ISL_3288552, EPI_ISL_3288553, EPI_ISL_3288554, EPI_ISL_3288555, EPI_ISL_3288556, EPI_ISL_3288557, EPI_ISL_3288558, EPI_ISL_3288559, EPI_ISL_3288560, EPI_ISL_3288561, EPI_ISL_3288562, EPI_ISL_3288563, EPI_ISL_3288564, EPI_ISL_3288565, EPI_ISL_3288566, EPI_ISL_3288567, EPI_ISL_3288568, EPI_ISL_3288569, EPI_ISL_3288570, EPI_ISL_3288571, EPI_ISL_3288572, EPI_ISL_3288573, EPI_ISL_3288574, EPI_ISL_3288575, EPI_ISL_3288576, EPI_ISL_3288577, EPI_ISL_3288578, EPI_ISL_3288579, EPI_ISL_3288580, EPI_ISL_3288581, EPI_ISL_3288582, EPI_ISL_3288583, EPI_ISL_3288584, EPI_ISL_3288585, EPI_ISL_3288586, EPI_ISL_3288587, EPI_ISL_3288588, EPI_ISL_3288589, EPI_ISL_3288590, EPI_ISL_3288591, EPI_ISL_3288592, EPI_ISL_3288593 | | | |
| see above | Northumbria University / South Tees Hospitals NHS Foundation Trust / North Cumbria Integrated Care NHS Foundation Trust / North Tees and Hartlepool NHS Foundation Trust / Newcastle Hospitals NHS Foundation Trust | COVID-19 Genomics UK (COG-UK) Consortium | Darren L Smith,Andrew Nelson,Matthew Bashton,Greg R Young,Joshua Loh,John Allan,Mohammad A Tariq,Giles S Holt,Gary Black,Wen C Yew,Lynn Dover,Paul Baker,Steve Liggett,Sarah Essex,Jane Greenaway,Debra Padgett,Clive Graham,Garren Scott,Edward Barton,Emma Swindells,Brendan Payne,Jennifer Collins,Yusri Taha,Gary Eltringham |
| EPI_ISL_3288594 | Pathology and Laboratory Medicine Institute, Cleveland Clinic, Ohio, USA | Pathology and Laboratory Medicine Institute, Cleveland Clinic, Ohio, USA | Jessica Spildener, Concetta Peck, Jennifer Starbuck, Joy Nakitandwe, Kristen McDonnell, David Plunkett, Zheng Jin Tu, Jay Brock, Yu-Wei Cheng, Gary Procop, Daniel Rhoads, Daniel H. Farkas, David Bosler |
| EPI_ISL_3288595, EPI_ISL_3288596 | Northumbria University / South Tees Hospitals NHS Foundation Trust / North Cumbria Integrated Care NHS Foundation Trust / North Tees and Hartlepool NHS Foundation Trust / Newcastle Hospitals NHS Foundation Trust | COVID-19 Genomics UK (COG-UK) Consortium | Darren L Smith,Andrew Nelson,Matthew Bashton,Greg R Young,Joshua Loh,John Allan,Mohammad A Tariq,Giles S Holt,Gary Black,Wen C Yew,Lynn Dover,Paul Baker,Steve Liggett,Sarah Essex,Jane Greenaway,Debra Padgett,Clive Graham,Garren Scott,Edward Barton,Emma Swindells,Brendan Payne,Jennifer Collins,Yusri Taha,Gary Eltringham |
| EPI_ISL_3288597 | Pathology and Laboratory Medicine Institute, Cleveland Clinic, Ohio, USA | Pathology and Laboratory Medicine Institute, Cleveland Clinic, Ohio, USA | Jessica Spildener, Concetta Peck, Jennifer Starbuck, Joy Nakitandwe, Kristen McDonnell, David Plunkett, Zheng Jin Tu, Jay Brock, Yu-Wei Cheng, Gary Procop, Daniel Rhoads, Daniel H. Farkas, David Bosler |
| EPI_ISL_3288598 | Northumbria University / South Tees Hospitals NHS Foundation Trust / North Cumbria Integrated Care NHS Foundation Trust / North Tees and Hartlepool NHS Foundation Trust / Newcastle Hospitals NHS Foundation Trust | COVID-19 Genomics UK (COG-UK) Consortium | Darren L Smith,Andrew Nelson,Matthew Bashton,Greg R Young,Joshua Loh,John Allan,Mohammad A Tariq,Giles S Holt,Gary Black,Wen C Yew,Lynn Dover,Paul Baker,Steve Liggett,Sarah Essex,Jane Greenaway,Debra Padgett,Clive Graham,Garren Scott,Edward Barton,Emma Swindells,Brendan Payne,Jennifer Collins,Yusri Taha,Gary Eltringham |
| EPI_ISL_3288599 | Pathology and Laboratory Medicine Institute, Cleveland Clinic, Ohio, USA | Pathology and Laboratory Medicine Institute, Cleveland Clinic, Ohio, USA | Jessica Spildener, Concetta Peck, Jennifer Starbuck, Joy Nakitandwe, Kristen McDonnell, David Plunkett, Zheng Jin Tu, Jay Brock, Yu-Wei Cheng, Gary Procop, Daniel Rhoads, Daniel H. Farkas, David Bosler |
| EPI_ISL_3288600 | Northumbria University / South Tees Hospitals NHS Foundation Trust / North Cumbria Integrated Care NHS Foundation Trust / North Tees and Hartlepool NHS Foundation Trust / Newcastle Hospitals NHS Foundation Trust | COVID-19 Genomics UK (COG-UK) Consortium | Darren L Smith,Andrew Nelson,Matthew Bashton,Greg R Young,Joshua Loh,John Allan,Mohammad A Tariq,Giles S Holt,Gary Black,Wen C Yew,Lynn Dover,Paul Baker,Steve Liggett,Sarah Essex,Jane Greenaway,Debra Padgett,Clive Graham,Garren Scott,Edward Barton,Emma Swindells,Brendan Payne,Jennifer Collins,Yusri Taha,Gary Eltringham |
| EPI_ISL_3288601 | Pathology and Laboratory Medicine Institute, Cleveland Clinic, Ohio, USA | Pathology and Laboratory Medicine Institute, Cleveland Clinic, Ohio, USA | Jessica Spildener, Concetta Peck, Jennifer Starbuck, Joy Nakitandwe, Kristen McDonnell, David Plunkett, Zheng Jin Tu, Jay Brock, Yu-Wei Cheng, Gary Procop, Daniel Rhoads, Daniel H. Farkas, David Bosler |
| EPI_ISL_3288602, EPI_ISL_3288603 | Northumbria University / South Tees Hospitals NHS Foundation Trust / North Cumbria Integrated Care NHS Foundation Trust / North Tees and Hartlepool NHS Foundation Trust / Newcastle | COVID-19 Genomics UK (COG-UK) Consortium | Darren L Smith,Andrew Nelson,Matthew Bashton,Greg R Young,Joshua Loh,John Allan,Mohammad A Tariq,Giles S Holt,Gary Black,Wen C Yew,Lynn Dover,Paul Baker,Steve Liggett,Sarah Essex,Jane Greenaway,Debra Padgett,Clive Graham,Garren Scott,Edward Barton,Emma Swindells,Brendan Payne,Jennifer Collins,Yusri Taha,Gary Eltringham |

| | | | |
|---|---|---|---|
| | Hospitals NHS Foundation Trust | | |
| EPI_ISL_3288604 | Pathology and Laboratory Medicine Institute, Cleveland Clinic, Ohio, USA | Pathology and Laboratory Medicine Institute, Cleveland Clinic, Ohio, USA | Jessica Spildener, Concetta Peck, Jennifer Starbuck, Joy Nakitandwe, Kristen McDonnell, David Plunkett, Zheng Jin Tu, Jay Brock, Yu-Wei Cheng, Gary Procop, Daniel Rhoads, Daniel H. Farkas, David Bosler |
| EPI_ISL_3288605 | Northumbria University / South Tees Hospitals NHS Foundation Trust / North Cumbria Integrated Care NHS Foundation Trust / North Tees and Hartlepool NHS Foundation Trust / Newcastle Hospitals NHS Foundation Trust | COVID-19 Genomics UK (COG-UK) Consortium | Darren L Smith,Andrew Nelson,Matthew Bashton,Greg R Young,Joshua Loh,John Allan,Mohammad A Tariq,Giles S Holt,Gary Black,Wen C Yew,Lynn Dover,Paul Baker,Steve Liggett,Sarah Essex,Jane Greenaway,Debra Padgett,Clive Graham,Garren Scott,Edward Barton,Emma Swindells,Brendan Payne,Jennifer Collins,Yusri Taha,Gary Eltringham |
| EPI_ISL_3288606 | Pathology and Laboratory Medicine Institute, Cleveland Clinic, Ohio, USA | Pathology and Laboratory Medicine Institute, Cleveland Clinic, Ohio, USA | Jessica Spildener, Concetta Peck, Jennifer Starbuck, Joy Nakitandwe, Kristen McDonnell, David Plunkett, Zheng Jin Tu, Jay Brock, Yu-Wei Cheng, Gary Procop, Daniel Rhoads, Daniel H. Farkas, David Bosler |
| EPI_ISL_3288607 | Northumbria University / South Tees Hospitals NHS Foundation Trust / North Cumbria Integrated Care NHS Foundation Trust / North Tees and Hartlepool NHS Foundation Trust / Newcastle Hospitals NHS Foundation Trust | COVID-19 Genomics UK (COG-UK) Consortium | Darren L Smith,Andrew Nelson,Matthew Bashton,Greg R Young,Joshua Loh,John Allan,Mohammad A Tariq,Giles S Holt,Gary Black,Wen C Yew,Lynn Dover,Paul Baker,Steve Liggett,Sarah Essex,Jane Greenaway,Debra Padgett,Clive Graham,Garren Scott,Edward Barton,Emma Swindells,Brendan Payne,Jennifer Collins,Yusri Taha,Gary Eltringham |
| EPI_ISL_3288608 | Pathology and Laboratory Medicine Institute, Cleveland Clinic, Ohio, USA | Pathology and Laboratory Medicine Institute, Cleveland Clinic, Ohio, USA | Jessica Spildener, Concetta Peck, Jennifer Starbuck, Joy Nakitandwe, Kristen McDonnell, David Plunkett, Zheng Jin Tu, Jay Brock, Yu-Wei Cheng, Gary Procop, Daniel Rhoads, Daniel H. Farkas, David Bosler |
| EPI_ISL_3288609 | Northumbria University / South Tees Hospitals NHS Foundation Trust / North Cumbria Integrated Care NHS Foundation Trust / North Tees and Hartlepool NHS Foundation Trust / Newcastle Hospitals NHS Foundation Trust | COVID-19 Genomics UK (COG-UK) Consortium | Darren L Smith,Andrew Nelson,Matthew Bashton,Greg R Young,Joshua Loh,John Allan,Mohammad A Tariq,Giles S Holt,Gary Black,Wen C Yew,Lynn Dover,Paul Baker,Steve Liggett,Sarah Essex,Jane Greenaway,Debra Padgett,Clive Graham,Garren Scott,Edward Barton,Emma Swindells,Brendan Payne,Jennifer Collins,Yusri Taha,Gary Eltringham |
| EPI_ISL_3288610 | Pathology and Laboratory Medicine Institute, Cleveland Clinic, Ohio, USA | Pathology and Laboratory Medicine Institute, Cleveland Clinic, Ohio, USA | Jessica Spildener, Concetta Peck, Jennifer Starbuck, Joy Nakitandwe, Kristen McDonnell, David Plunkett, Zheng Jin Tu, Jay Brock, Yu-Wei Cheng, Gary Procop, Daniel Rhoads, Daniel H. Farkas, David Bosler |

EPI_ISL_3288611, EPI_ISL_3288612, EPI_ISL_3288613, EPI_ISL_3288614, EPI_ISL_3288615, EPI_ISL_3288616, EPI_ISL_3288617, EPI_ISL_3288618, EPI_ISL_3288619, EPI_ISL_3288620, EPI_ISL_3288621, EPI_ISL_3288622, EPI_ISL_3288623, EPI_ISL_3288624, EPI_ISL_3288625, EPI_ISL_3288626, EPI_ISL_3288627, EPI_ISL_3288628, EPI_ISL_3288629, EPI_ISL_3288630, EPI_ISL_3288631, EPI_ISL_3288632,
EPI_ISL_3288633, EPI_ISL_3288634, EPI_ISL_3288635, EPI_ISL_3288636, EPI_ISL_3288637, EPI_ISL_3288638, EPI_ISL_3288639, EPI_ISL_3288640, EPI_ISL_3288641, EPI_ISL_3288642, EPI_ISL_3288643, EPI_ISL_3288644, EPI_ISL_3288645, EPI_ISL_3288646, EPI_ISL_3288647, EPI_ISL_3288648, EPI_ISL_3288649, EPI_ISL_3288650, EPI_ISL_3288651, EPI_ISL_3288652, EPI_ISL_3288653, EPI_ISL_3288654,
EPI_ISL_3288655, EPI_ISL_3288656, EPI_ISL_3288657, EPI_ISL_3288658, EPI_ISL_3288659, EPI_ISL_3288660, EPI_ISL_3288661, EPI_ISL_3288662, EPI_ISL_3288663, EPI_ISL_3288664, EPI_ISL_3288665, EPI_ISL_3288666, EPI_ISL_3288667, EPI_ISL_3288668, EPI_ISL_3288669, EPI_ISL_3288670, EPI_ISL_3288671, EPI_ISL_3288672, EPI_ISL_3288673, EPI_ISL_3288674, EPI_ISL_3288675, EPI_ISL_3288676,
EPI_ISL_3288677, EPI_ISL_3288678, EPI_ISL_3288679, EPI_ISL_3288680, EPI_ISL_3288681, EPI_ISL_3288682, EPI_ISL_3288683, EPI_ISL_3288684, EPI_ISL_3288685, EPI_ISL_3288686, EPI_ISL_3288687, EPI_ISL_3288688, EPI_ISL_3288689, EPI_ISL_3288690, EPI_ISL_3288691, EPI_ISL_3288692, EPI_ISL_3288693, EPI_ISL_3288694, EPI_ISL_3288695, EPI_ISL_3288696, EPI_ISL_3288697, EPI_ISL_3288698,
EPI_ISL_3288699, EPI_ISL_3288700, EPI_ISL_3288701, EPI_ISL_3288702, EPI_ISL_3288703, EPI_ISL_3288704, EPI_ISL_3288705, EPI_ISL_3288706, EPI_ISL_3288707, EPI_ISL_3288708, EPI_ISL_3288709, EPI_ISL_3288710, EPI_ISL_3288711, EPI_ISL_3288712, EPI_ISL_3288713, EPI_ISL_3288714, EPI_ISL_3288715, EPI_ISL_3288716, EPI_ISL_3288717, EPI_ISL_3288718, EPI_ISL_3288719, EPI_ISL_3288720,
EPI_ISL_3288721, EPI_ISL_3288722, EPI_ISL_3288723, EPI_ISL_3288724, EPI_ISL_3288725, EPI_ISL_3288726, EPI_ISL_3288727, EPI_ISL_3288728, EPI_ISL_3288729, EPI_ISL_3288730, EPI_ISL_3288731, EPI_ISL_3288732, EPI_ISL_3288733, EPI_ISL_3288734, EPI_ISL_3288735, EPI_ISL_3288736, EPI_ISL_3288737, EPI_ISL_3288738, EPI_ISL_3288739, EPI_ISL_3288740, EPI_ISL_3288741, EPI_ISL_3288742,
EPI_ISL_3288743, EPI_ISL_3288744, EPI_ISL_3288745, EPI_ISL_3288746, EPI_ISL_3288747, EPI_ISL_3288748, EPI_ISL_3288749, EPI_ISL_3288750, EPI_ISL_3288751, EPI_ISL_3288752, EPI_ISL_3288753, EPI_ISL_3288754, EPI_ISL_3288755, EPI_ISL_3288756, EPI_ISL_3288757, EPI_ISL_3288758, EPI_ISL_3288759, EPI_ISL_3288760, EPI_ISL_3288761, EPI_ISL_3288762, EPI_ISL_3288763, EPI_ISL_3288764,
EPI_ISL_3288765, EPI_ISL_3288766, EPI_ISL_3288767, EPI_ISL_3288768, EPI_ISL_3288769, EPI_ISL_3288770, EPI_ISL_3288771, EPI_ISL_3288772, EPI_ISL_3288773, EPI_ISL_3288774, EPI_ISL_3288775, EPI_ISL_3288776, EPI_ISL_3288777, EPI_ISL_3288778, EPI_ISL_3288779, EPI_ISL_3288780, EPI_ISL_3288781, EPI_ISL_3288782, EPI_ISL_3288783, EPI_ISL_3288784, EPI_ISL_3288785, EPI_ISL_3288786,
EPI_ISL_3288787, EPI_ISL_3288788, EPI_ISL_3288789, EPI_ISL_3288790, EPI_ISL_3288791, EPI_ISL_3288792, EPI_ISL_3288793, EPI_ISL_3288794, EPI_ISL_3288795, EPI_ISL_3288796, EPI_ISL_3288797, EPI_ISL_3288798, EPI_ISL_3288799, EPI_ISL_3288800, EPI_ISL_3288801, EPI_ISL_3288802, EPI_ISL_3288803, EPI_ISL_3288804, EPI_ISL_3288805, EPI_ISL_3288806, EPI_ISL_3288807, EPI_ISL_3288808,
EPI_ISL_3288809, EPI_ISL_3288810, EPI_ISL_3288811, EPI_ISL_3288812, EPI_ISL_3288813, EPI_ISL_3288814, EPI_ISL_3288815, EPI_ISL_3288816, EPI_ISL_3288817, EPI_ISL_3288818, EPI_ISL_3288819, EPI_ISL_3288820, EPI_ISL_3288821, EPI_ISL_3288822, EPI_ISL_3288823, EPI_ISL_3288824, EPI_ISL_3288825, EPI_ISL_3288826, EPI_ISL_3288827, EPI_ISL_3288828, EPI_ISL_3288829, EPI_ISL_3288830,
EPI_ISL_3288831, EPI_ISL_3288832, EPI_ISL_3288833, EPI_ISL_3288834, EPI_ISL_3288835, EPI_ISL_3288836, EPI_ISL_3288837, EPI_ISL_3288838, EPI_ISL_3288839, EPI_ISL_3288840, EPI_ISL_3288841, EPI_ISL_3288842, EPI_ISL_3288843, EPI_ISL_3288844, EPI_ISL_3288845, EPI_ISL_3288846, EPI_ISL_3288847, EPI_ISL_3288848, EPI_ISL_3288849, EPI_ISL_3288850, EPI_ISL_3288851, EPI_ISL_3288852,
EPI_ISL_3288853, EPI_ISL_3288854, EPI_ISL_3288855, EPI_ISL_3288856, EPI_ISL_3288857, EPI_ISL_3288858, EPI_ISL_3288859, EPI_ISL_3288860, EPI_ISL_3288861, EPI_ISL_3288862, EPI_ISL_3288863, EPI_ISL_3288864, EPI_ISL_3288865, EPI_ISL_3288866, EPI_ISL_3288867, EPI_ISL_3288868, EPI_ISL_3288869, EPI_ISL_3288870, EPI_ISL_3288871, EPI_ISL_3288872, EPI_ISL_3288873, EPI_ISL_3288874,
EPI_ISL_3288875, EPI_ISL_3288876, EPI_ISL_3288877, EPI_ISL_3288878, EPI_ISL_3288879, EPI_ISL_3288880, EPI_ISL_3288881, EPI_ISL_3288882, EPI_ISL_3288883, EPI_ISL_3288884, EPI_ISL_3288885, EPI_ISL_3288886, EPI_ISL_3288887, EPI_ISL_3288888, EPI_ISL_3288889, EPI_ISL_3288890, EPI_ISL_3288891, EPI_ISL_3288892, EPI_ISL_3288893, EPI_ISL_3288894, EPI_ISL_3288895, EPI_ISL_3288896,
EPI_ISL_3288897, EPI_ISL_3288898, EPI_ISL_3288899, EPI_ISL_3288900, EPI_ISL_3288901, EPI_ISL_3288902, EPI_ISL_3288903, EPI_ISL_3288904, EPI_ISL_3288905, EPI_ISL_3288906, EPI_ISL_3288907, EPI_ISL_3288908, EPI_ISL_3288909, EPI_ISL_3288910, EPI_ISL_3288911, EPI_ISL_3288912, EPI_ISL_3288913, EPI_ISL_3288914, EPI_ISL_3288915, EPI_ISL_3288916, EPI_ISL_3288917, EPI_ISL_3288918,
EPI_ISL_3288919, EPI_ISL_3288920, EPI_ISL_3288921, EPI_ISL_3288922, EPI_ISL_3288923, EPI_ISL_3288924, EPI_ISL_3288925, EPI_ISL_3288926, EPI_ISL_3288927, EPI_ISL_3288928, EPI_ISL_3288929, EPI_ISL_3288930, EPI_ISL_3288931, EPI_ISL_3288932, EPI_ISL_3288933, EPI_ISL_3288934, EPI_ISL_3288935, EPI_ISL_3288936, EPI_ISL_3288937, EPI_ISL_3288938, EPI_ISL_3288939, EPI_ISL_3288940,
EPI_ISL_3288941, EPI_ISL_3288942, EPI_ISL_3288943, EPI_ISL_3288944, EPI_ISL_3288945, EPI_ISL_3288946, EPI_ISL_3288947, EPI_ISL_3288948, EPI_ISL_3288949, EPI_ISL_3288950, EPI_ISL_3288951

| | | | |
|---|---|---|---|
| see above | Northumbria University / South Tees Hospitals NHS Foundation Trust / North Cumbria Integrated Care NHS Foundation Trust / North Tees and Hartlepool NHS Foundation Trust / Newcastle Hospitals NHS Foundation Trust | COVID-19 Genomics UK (COG-UK) Consortium | Darren L Smith,Andrew Nelson,Matthew Bashton,Greg R Young,Joshua Loh,John Allan,Mohammad A Tariq,Giles S Holt,Gary Black,Wen C Yew,Lynn Dover,Paul Baker,Steve Liggett,Sarah Essex,Jane Greenaway,Debra Padgett,Clive Graham,Garren Scott,Edward Barton,Emma Swindells,Brendan Payne,Jennifer Collins,Yusri Taha,Gary Eltringham |

EPI_ISL_3288952, EPI_ISL_3288953, EPI_ISL_3288954, EPI_ISL_3288955, EPI_ISL_3288956, EPI_ISL_3288957, EPI_ISL_3288958, EPI_ISL_3288959, EPI_ISL_3288960, EPI_ISL_3288961, EPI_ISL_3288962, EPI_ISL_3288963, EPI_ISL_3288964, EPI_ISL_3288965, EPI_ISL_3288966, EPI_ISL_3288967, EPI_ISL_3288968, EPI_ISL_3288969, EPI_ISL_3288970, EPI_ISL_3288971, EPI_ISL_3288972, EPI_ISL_3288973,
EPI_ISL_3288974, EPI_ISL_3288975, EPI_ISL_3288976, EPI_ISL_3288977, EPI_ISL_3288978, EPI_ISL_3288979, EPI_ISL_3288980, EPI_ISL_3288981, EPI_ISL_3288982, EPI_ISL_3288983, EPI_ISL_3288984, EPI_ISL_3288985, EPI_ISL_3288986, EPI_ISL_3288987, EPI_ISL_3288988, EPI_ISL_3288989, EPI_ISL_3288990, EPI_ISL_3288991, EPI_ISL_3288992, EPI_ISL_3288993, EPI_ISL_3288994, EPI_ISL_3288995,
EPI_ISL_3288996, EPI_ISL_3288997, EPI_ISL_3288998, EPI_ISL_3288999, EPI_ISL_3289000, EPI_ISL_3289001, EPI_ISL_3289002, EPI_ISL_3289003, EPI_ISL_3289004, EPI_ISL_3289005, EPI_ISL_3289006, EPI_ISL_3289007, EPI_ISL_3289008, EPI_ISL_3289009, EPI_ISL_3289010, EPI_ISL_3289011, EPI_ISL_3289012, EPI_ISL_3289013, EPI_ISL_3289014, EPI_ISL_3289015, EPI_ISL_3289016, EPI_ISL_3289017,
EPI_ISL_3289018, EPI_ISL_3289019, EPI_ISL_3289020, EPI_ISL_3289021, EPI_ISL_3289022, EPI_ISL_3289023, EPI_ISL_3289024, EPI_ISL_3289025, EPI_ISL_3289026, EPI_ISL_3289027, EPI_ISL_3289028, EPI_ISL_3289029, EPI_ISL_3289030, EPI_ISL_3289031, EPI_ISL_3289032, EPI_ISL_3289033, EPI_ISL_3289034, EPI_ISL_3289035, EPI_ISL_3289036, EPI_ISL_3289037, EPI_ISL_3289038, EPI_ISL_3289039,
EPI_ISL_3289040, EPI_ISL_3289041, EPI_ISL_3289042, EPI_ISL_3289043, EPI_ISL_3289044, EPI_ISL_3289045, EPI_ISL_3289046, EPI_ISL_3289047, EPI_ISL_3289048, EPI_ISL_3289049, EPI_ISL_3289050, EPI_ISL_3289051, EPI_ISL_3289052, EPI_ISL_3289053, EPI_ISL_3289054, EPI_ISL_3289055, EPI_ISL_3289056, EPI_ISL_3289057, EPI_ISL_3289058, EPI_ISL_3289059, EPI_ISL_3289060, EPI_ISL_3289061,
EPI_ISL_3289062, EPI_ISL_3289063, EPI_ISL_3289064, EPI_ISL_3289065, EPI_ISL_3289066, EPI_ISL_3289067, EPI_ISL_3289068, EPI_ISL_3289070, EPI_ISL_3289071, EPI_ISL_3289072, EPI_ISL_3289073, EPI_ISL_3289074, EPI_ISL_3289075, EPI_ISL_3289076, EPI_ISL_3289077, EPI_ISL_3289078, EPI_ISL_3289079, EPI_ISL_3289080, EPI_ISL_3289081, EPI_ISL_3289082, EPI_ISL_3289083, EPI_ISL_3289084,
EPI_ISL_3289085, EPI_ISL_3289086, EPI_ISL_3289087, EPI_ISL_3289088, EPI_ISL_3289089, EPI_ISL_3289090, EPI_ISL_3289091, EPI_ISL_3289092, EPI_ISL_3289093, EPI_ISL_3289094, EPI_ISL_3289095, EPI_ISL_3289096, EPI_ISL_3289097, EPI_ISL_3289098, EPI_ISL_3289099, EPI_ISL_3289100, EPI_ISL_3289101, EPI_ISL_3289102, EPI_ISL_3289103, EPI_ISL_3289104, EPI_ISL_3289105, EPI_ISL_3289106,
EPI_ISL_3289107, EPI_ISL_3289108, EPI_ISL_3289109, EPI_ISL_3289110, EPI_ISL_3289111, EPI_ISL_3289112, EPI_ISL_3289113, EPI_ISL_3289114, EPI_ISL_3289115, EPI_ISL_3289116, EPI_ISL_3289117, EPI_ISL_3289118, EPI_ISL_3289119, EPI_ISL_3289120, EPI_ISL_3289121, EPI_ISL_3289122, EPI_ISL_3289123, EPI_ISL_3289124, EPI_ISL_3289125, EPI_ISL_3289126, EPI_ISL_3289127, EPI_ISL_3289128,
EPI_ISL_3289129, EPI_ISL_3289130, EPI_ISL_3289131, EPI_ISL_3289132, EPI_ISL_3289133, EPI_ISL_3289134, EPI_ISL_3289135, EPI_ISL_3289136, EPI_ISL_3289137, EPI_ISL_3289138, EPI_ISL_3289139, EPI_ISL_3289140, EPI_ISL_3289141, EPI_ISL_3289142, EPI_ISL_3289143, EPI_ISL_3289144, EPI_ISL_3289145, EPI_ISL_3289146, EPI_ISL_3289147, EPI_ISL_3289148, EPI_ISL_3289149, EPI_ISL_3289150,
EPI_ISL_3289151, EPI_ISL_3289152, EPI_ISL_3289153, EPI_ISL_3289154, EPI_ISL_3289155, EPI_ISL_3289156, EPI_ISL_3289157, EPI_ISL_3289158, EPI_ISL_3289159, EPI_ISL_3289160, EPI_ISL_3289161, EPI_ISL_3289162, EPI_ISL_3289163, EPI_ISL_3289164, EPI_ISL_3289165, EPI_ISL_3289166, EPI_ISL_3289167, EPI_ISL_3289168, EPI_ISL_3289169, EPI_ISL_3289170, EPI_ISL_3289171, EPI_ISL_3289172,
EPI_ISL_3289173, EPI_ISL_3289174, EPI_ISL_3289175, EPI_ISL_3289176, EPI_ISL_3289177, EPI_ISL_3289178, EPI_ISL_3289179, EPI_ISL_3289180, EPI_ISL_3289181, EPI_ISL_3289182, EPI_ISL_3289183, EPI_ISL_3289184, EPI_ISL_3289185, EPI_ISL_3289186, EPI_ISL_3289187, EPI_ISL_3289188, EPI_ISL_3289189, EPI_ISL_3289190, EPI_ISL_3289191, EPI_ISL_3289192, EPI_ISL_3289193, EPI_ISL_3289194,
EPI_ISL_3289195, EPI_ISL_3289196, EPI_ISL_3289197, EPI_ISL_3289198, EPI_ISL_3289199, EPI_ISL_3289200, EPI_ISL_3289201, EPI_ISL_3289202, EPI_ISL_3289203, EPI_ISL_3289204, EPI_ISL_3289205, EPI_ISL_3289206, EPI_ISL_3289207, EPI_ISL_3289208, EPI_ISL_3289209, EPI_ISL_3289210, EPI_ISL_3289211, EPI_ISL_3289212, EPI_ISL_3289213, EPI_ISL_3289214, EPI_ISL_3289215, EPI_ISL_3289216,
EPI_ISL_3289217, EPI_ISL_3289218, EPI_ISL_3289219, EPI_ISL_3289220, EPI_ISL_3289221, EPI_ISL_3289222, EPI_ISL_3289223, EPI_ISL_3289224, EPI_ISL_3289225, EPI_ISL_3289226, EPI_ISL_3289227, EPI_ISL_3289228, EPI_ISL_3289229, EPI_ISL_3289230, EPI_ISL_3289231, EPI_ISL_3289232, EPI_ISL_3289233, EPI_ISL_3289234, EPI_ISL_3289235, EPI_ISL_3289236, EPI_ISL_3289237, EPI_ISL_3289238,
EPI_ISL_3289239, EPI_ISL_3289240, EPI_ISL_3289241, EPI_ISL_3289242, EPI_ISL_3289243, EPI_ISL_3289244, EPI_ISL_3289245, EPI_ISL_3289246, EPI_ISL_3289247, EPI_ISL_3289248, EPI_ISL_3289249, EPI_ISL_3289250, EPI_ISL_3289251, EPI_ISL_3289252, EPI_ISL_3289253, EPI_ISL_3289254, EPI_ISL_3289255, EPI_ISL_3289256, EPI_ISL_3289257, EPI_ISL_3289258, EPI_ISL_3289259, EPI_ISL_3289260,
EPI_ISL_3289261, EPI_ISL_3289262, EPI_ISL_3289263, EPI_ISL_3289264, EPI_ISL_3289265, EPI_ISL_3289266, EPI_ISL_3289267, EPI_ISL_3289268, EPI_ISL_3289269, EPI_ISL_3289270, EPI_ISL_3289271, EPI_ISL_3289272, EPI_ISL_3289273, EPI_ISL_3289274, EPI_ISL_3289275, EPI_ISL_3289276, EPI_ISL_3289277, EPI_ISL_3289278, EPI_ISL_3289279, EPI_ISL_3289280, EPI_ISL_3289281, EPI_ISL_3289282,
EPI_ISL_3289283, EPI_ISL_3289284, EPI_ISL_3289285, EPI_ISL_3289286, EPI_ISL_3289287, EPI_ISL_3289288, EPI_ISL_3289289, EPI_ISL_3289290, EPI_ISL_3289291, EPI_ISL_3289292, EPI_ISL_3289293, EPI_ISL_3289294, EPI_ISL_3289295, EPI_ISL_3289296, EPI_ISL_3289297, EPI_ISL_3289298, EPI_ISL_3289299, EPI_ISL_3289300, EPI_ISL_3289301, EPI_ISL_3289302, EPI_ISL_3289303, EPI_ISL_3289304,
EPI_ISL_3289305, EPI_ISL_3289306, EPI_ISL_3289307, EPI_ISL_3289308, EPI_ISL_3289309, EPI_ISL_3289310, EPI_ISL_3289311, EPI_ISL_3289312, EPI_ISL_3289313, EPI_ISL_3289314, EPI_ISL_3289315, EPI_ISL_3289316, EPI_ISL_3289317, EPI_ISL_3289318, EPI_ISL_3289319, EPI_ISL_3289320, EPI_ISL_3289321, EPI_ISL_3289322, EPI_ISL_3289323, EPI_ISL_3289324, EPI_ISL_3289325, EPI_ISL_3289326,
EPI_ISL_3289327, EPI_ISL_3289328, EPI_ISL_3289329, EPI_ISL_3289330, EPI_ISL_3289331, EPI_ISL_3289332, EPI_ISL_3289333, EPI_ISL_3289334, EPI_ISL_3289335, EPI_ISL_3289336, EPI_ISL_3289337, EPI_ISL_3289338, EPI_ISL_3289339, EPI_ISL_3289340, EPI_ISL_3289341, EPI_ISL_3289342, EPI_ISL_3289343, EPI_ISL_3289344, EPI_ISL_3289345, EPI_ISL_3289346, EPI_ISL_3289347, EPI_ISL_3289348,
EPI_ISL_3289349, EPI_ISL_3289350, EPI_ISL_3289351, EPI_ISL_3289352, EPI_ISL_3289353, EPI_ISL_3289354, EPI_ISL_3289355, EPI_ISL_3289356, EPI_ISL_3289357, EPI_ISL_3289358, EPI_ISL_3289359, EPI_ISL_3289360, EPI_ISL_3289361, EPI_ISL_3289362, EPI_ISL_3289363, EPI_ISL_3289364, EPI_ISL_3289365, EPI_ISL_3289366, EPI_ISL_3289367, EPI_ISL_3289368, EPI_ISL_3289369, EPI_ISL_3289370,
EPI_ISL_3289371, EPI_ISL_3289372, EPI_ISL_3289373, EPI_ISL_3289374, EPI_ISL_3289375, EPI_ISL_3289376, EPI_ISL_3289377, EPI_ISL_3289378, EPI_ISL_3289379, EPI_ISL_3289380, EPI_ISL_3289381, EPI_ISL_3289382, EPI_ISL_3289383, EPI_ISL_3289384, EPI_ISL_3289385, EPI_ISL_3289386, EPI_ISL_3289387, EPI_ISL_3289388, EPI_ISL_3289389, EPI_ISL_3289390, EPI_ISL_3289391, EPI_ISL_3289392,
EPI_ISL_3289393, EPI_ISL_3289394, EPI_ISL_3289395, EPI_ISL_3289396, EPI_ISL_3289397, EPI_ISL_3289398, EPI_ISL_3289399, EPI_ISL_3289400, EPI_ISL_3289401, EPI_ISL_3289402, EPI_ISL_3289403, EPI_ISL_3289404, EPI_ISL_3289405, EPI_ISL_3289406, EPI_ISL_3289407, EPI_ISL_3289408, EPI_ISL_3289409, EPI_ISL_3289410, EPI_ISL_3289411, EPI_ISL_3289412, EPI_ISL_3289413, EPI_ISL_3289414,
EPI_ISL_3289415, EPI_ISL_3289416, EPI_ISL_3289417, EPI_ISL_3289418, EPI_ISL_3289419, EPI_ISL_3289420, EPI_ISL_3289421, EPI_ISL_3289422, EPI_ISL_3289423, EPI_ISL_3289424, EPI_ISL_3289425, EPI_ISL_3289426, EPI_ISL_3289427, EPI_ISL_3289428, EPI_ISL_3289429, EPI_ISL_3289430, EPI_ISL_3289431, EPI_ISL_3289432, EPI_ISL_3289433, EPI_ISL_3289434, EPI_ISL_3289435, EPI_ISL_3289436,
EPI_ISL_3289437, EPI_ISL_3289438, EPI_ISL_3289439, EPI_ISL_3289440, EPI_ISL_3289441, EPI_ISL_3289442, EPI_ISL_3289443, EPI_ISL_3289444, EPI_ISL_3289445, EPI_ISL_3289446, EPI_ISL_3289447, EPI_ISL_3289448, EPI_ISL_3289449, EPI_ISL_3289450, EPI_ISL_3289451, EPI_ISL_3289452, EPI_ISL_3289453, EPI_ISL_3289454, EPI_ISL_3289455, EPI_ISL_3289456, EPI_ISL_3289457, EPI_ISL_3289458,
EPI_ISL_3289459, EPI_ISL_3289460, EPI_ISL_3289461, EPI_ISL_3289462, EPI_ISL_3289463, EPI_ISL_3289464, EPI_ISL_3289465, EPI_ISL_3289466, EPI_ISL_3289467, EPI_ISL_3289468, EPI_ISL_3289469, EPI_ISL_3289470, EPI_ISL_3289471, EPI_ISL_3289472, EPI_ISL_3289473, EPI_ISL_3289474, EPI_ISL_3289475, EPI_ISL_3289476, EPI_ISL_3289477, EPI_ISL_3289478, EPI_ISL_3289479, EPI_ISL_3289480,
EPI_ISL_3289481, EPI_ISL_3289482, EPI_ISL_3289483, EPI_ISL_3289484, EPI_ISL_3289485, EPI_ISL_3289486, EPI_ISL_3289487, EPI_ISL_3289488, EPI_ISL_3289489, EPI_ISL_3289490, EPI_ISL_3289491, EPI_ISL_3289492, EPI_ISL_3289493, EPI_ISL_3289494, EPI_ISL_3289495, EPI_ISL_3289496, EPI_ISL_3289497, EPI_ISL_3289498, EPI_ISL_3289499, EPI_ISL_3289500, EPI_ISL_3289501, EPI_ISL_3289502,
EPI_ISL_3289503, EPI_ISL_3289504, EPI_ISL_3289505, EPI_ISL_3289506, EPI_ISL_3289507, EPI_ISL_3289508, EPI_ISL_3289509, EPI_ISL_3289510, EPI_ISL_3289511, EPI_ISL_3289512, EPI_ISL_3289513, EPI_ISL_3289514, EPI_ISL_3289515, EPI_ISL_3289516, EPI_ISL_3289517, EPI_ISL_3289518, EPI_ISL_3289519, EPI_ISL_3289520, EPI_ISL_3289521, EPI_ISL_3289522, EPI_ISL_3289523, EPI_ISL_3289524,
EPI_ISL_3289525, EPI_ISL_3289526, EPI_ISL_3289527, EPI_ISL_3289528, EPI_ISL_3289529, EPI_ISL_3289530, EPI_ISL_3289531, EPI_ISL_3289532, EPI_ISL_3289533, EPI_ISL_3289534, EPI_ISL_3289535, EPI_ISL_3289536, EPI_ISL_3289537, EPI_ISL_3289538, EPI_ISL_3289539, EPI_ISL_3289540, EPI_ISL_3289541, EPI_ISL_3289542, EPI_ISL_3289543, EPI_ISL_3289544, EPI_ISL_3289545, EPI_ISL_3289546,
EPI_ISL_3289547, EPI_ISL_3289548, EPI_ISL_3289549, EPI_ISL_3289550, EPI_ISL_3289551, EPI_ISL_3289552, EPI_ISL_3289553, EPI_ISL_3289554, EPI_ISL_3289555, EPI_ISL_3289556, EPI_ISL_3289557, EPI_ISL_3289558, EPI_ISL_3289559, EPI_ISL_3289560, EPI_ISL_3289561, EPI_ISL_3289562, EPI_ISL_3289563, EPI_ISL_3289564, EPI_ISL_3289565, EPI_ISL_3289566, EPI_ISL_3289567, EPI_ISL_3289568,
EPI_ISL_3289569, EPI_ISL_3289570, EPI_ISL_3289571, EPI_ISL_3289572, EPI_ISL_3289573, EPI_ISL_3289574, EPI_ISL_3289575, EPI_ISL_3289576, EPI_ISL_3289577, EPI_ISL_3289578, EPI_ISL_3289579, EPI_ISL_3289580, EPI_ISL_3289581, EPI_ISL_3289582, EPI_ISL_3289583, EPI_ISL_3289584, EPI_ISL_3289585, EPI_ISL_3289586, EPI_ISL_3289587, EPI_ISL_3289588, EPI_ISL_3289589

| | | | |
|---|---|---|---|
| see above | Quadram Institute Bioscience | COVID-19 Genomics UK (COG-UK) Consortium | Dave J. Baker, Gemma L. Kay, Alp Aydin, Thanh Le-Viet, Steven Rudder, Ana P. Tedim, Anastasia Kolyva, Maria Diaz, Leonardo de Oliveira Martins, Nabil-Fareed Alikhan, Lizzie Meadows, Rachael Stanley, Rose K Davidson, Ngozi Elumogo, Muhammed Yasir, Sophie J. Prosolek, Ebenezer Foster-Nyarko, Martin Lott, Christopher Jeanes, Lewis G. Spurgin, Nicholas M. Thomson, Alexander J Trotter, Rachel Gilroy, Luke Bedford, Samuel Bloomfield, Claire Stuart, Andrew Bell, Reenesh Prakash, Lindsay Coupland, Samir Dervisevic, Alison E. Mather, John Wain, Mark Webber, Andrew J. Page, Justin O'Grady |

EPI_ISL_3289590, EPI_ISL_3289591, EPI_ISL_3289592, EPI_ISL_3289593, EPI_ISL_3289594, EPI_ISL_3289595, EPI_ISL_3289596, EPI_ISL_3289597, EPI_ISL_3289599, EPI_ISL_3289600, EPI_ISL_3289601, EPI_ISL_3289602, EPI_ISL_3289603, EPI_ISL_3289604, EPI_ISL_3289605, EPI_ISL_3289606, EPI_ISL_3289607, EPI_ISL_3289608, EPI_ISL_3289609, EPI_ISL_3289610, EPI_ISL_3289611, EPI_ISL_3289612,
EPI_ISL_3289613, EPI_ISL_3289614, EPI_ISL_3289615, EPI_ISL_3289616, EPI_ISL_3289617, EPI_ISL_3289618, EPI_ISL_3289619, EPI_ISL_3289620, EPI_ISL_3289621, EPI_ISL_3289622, EPI_ISL_3289623, EPI_ISL_3289624, EPI_ISL_3289625, EPI_ISL_3289626, EPI_ISL_3289627, EPI_ISL_3289628, EPI_ISL_3289629, EPI_ISL_3289630, EPI_ISL_3289631, EPI_ISL_3289632, EPI_ISL_3289633, EPI_ISL_3289634,
EPI_ISL_3289635, EPI_ISL_3289636, EPI_ISL_3289637, EPI_ISL_3289638, EPI_ISL_3289639, EPI_ISL_3289640, EPI_ISL_3289641, EPI_ISL_3289642, EPI_ISL_3289643, EPI_ISL_3289644, EPI_ISL_3289645, EPI_ISL_3289647, EPI_ISL_3289648, EPI_ISL_3289649, EPI_ISL_3289650, EPI_ISL_3289651, EPI_ISL_3289652, EPI_ISL_3289653, EPI_ISL_3289654, EPI_ISL_3289655, EPI_ISL_3289656, EPI_ISL_3289657,
EPI_ISL_3289658, EPI_ISL_3289659, EPI_ISL_3289660, EPI_ISL_3289661, EPI_ISL_3289662, EPI_ISL_3289663, EPI_ISL_3289664, EPI_ISL_3289665, EPI_ISL_3289666, EPI_ISL_3289667, EPI_ISL_3289668, EPI_ISL_3289669, EPI_ISL_3289670, EPI_ISL_3289671, EPI_ISL_3289672, EPI_ISL_3289673, EPI_ISL_3289674, EPI_ISL_3289675, EPI_ISL_3289676, EPI_ISL_3289677, EPI_ISL_3289678, EPI_ISL_3289679,
EPI_ISL_3289680, EPI_ISL_3289681, EPI_ISL_3289682, EPI_ISL_3289683, EPI_ISL_3289684, EPI_ISL_3289685, EPI_ISL_3289686, EPI_ISL_3289687, EPI_ISL_3289688, EPI_ISL_3289689, EPI_ISL_3289690, EPI_ISL_3289691, EPI_ISL_3289692, EPI_ISL_3289693, EPI_ISL_3289694, EPI_ISL_3289695, EPI_ISL_3289696, EPI_ISL_3289697, EPI_ISL_3289698, EPI_ISL_3289699, EPI_ISL_3289700, EPI_ISL_3289701,
EPI_ISL_3289702, EPI_ISL_3289703, EPI_ISL_3289704, EPI_ISL_3289705, EPI_ISL_3289706, EPI_ISL_3289707, EPI_ISL_3289708, EPI_ISL_3289710, EPI_ISL_3289711, EPI_ISL_3289712, EPI_ISL_3289713, EPI_ISL_3289714, EPI_ISL_3289715, EPI_ISL_3289716, EPI_ISL_3289717, EPI_ISL_3289718, EPI_ISL_3289719, EPI_ISL_3289720, EPI_ISL_3289721, EPI_ISL_3289722, EPI_ISL_3289723, EPI_ISL_3289724,
EPI_ISL_3289725, EPI_ISL_3289726, EPI_ISL_3289727, EPI_ISL_3289728, EPI_ISL_3289729, EPI_ISL_3289730, EPI_ISL_3289731, EPI_ISL_3289732, EPI_ISL_3289733, EPI_ISL_3289734, EPI_ISL_3289735, EPI_ISL_3289736, EPI_ISL_3289737, EPI_ISL_3289738, EPI_ISL_3289739, EPI_ISL_3289740, EPI_ISL_3289741, EPI_ISL_3289742, EPI_ISL_3289743, EPI_ISL_3289744, EPI_ISL_3289745, EPI_ISL_3289746,
EPI_ISL_3289747, EPI_ISL_3289748, EPI_ISL_3289749, EPI_ISL_3289750, EPI_ISL_3289751, EPI_ISL_3289752, EPI_ISL_3289753, EPI_ISL_3289754, EPI_ISL_3289755, EPI_ISL_3289756, EPI_ISL_3289757, EPI_ISL_3289758, EPI_ISL_3289759, EPI_ISL_3289760, EPI_ISL_3289761, EPI_ISL_3289762, EPI_ISL_3289763, EPI_ISL_3289764, EPI_ISL_3289765, EPI_ISL_3289766, EPI_ISL_3289767, EPI_ISL_3289768,
EPI_ISL_3289769, EPI_ISL_3289770, EPI_ISL_3289771, EPI_ISL_3289772, EPI_ISL_3289773, EPI_ISL_3289774, EPI_ISL_3289775, EPI_ISL_3289776, EPI_ISL_3289777, EPI_ISL_3289778, EPI_ISL_3289779, EPI_ISL_3289780, EPI_ISL_3289781, EPI_ISL_3289782, EPI_ISL_3289783, EPI_ISL_3289784, EPI_ISL_3289785, EPI_ISL_3289786, EPI_ISL_3289787, EPI_ISL_3289788, EPI_ISL_3289789, EPI_ISL_3289790,
EPI_ISL_3289791, EPI_ISL_3289792, EPI_ISL_3289793, EPI_ISL_3289794, EPI_ISL_3289795, EPI_ISL_3289796, EPI_ISL_3289797, EPI_ISL_3289798, EPI_ISL_3289799, EPI_ISL_3289800, EPI_ISL_3289801, EPI_ISL_3289802, EPI_ISL_3289803, EPI_ISL_3289804, EPI_ISL_3289805, EPI_ISL_3289806, EPI_ISL_3289807, EPI_ISL_3289808, EPI_ISL_3289809, EPI_ISL_3289810, EPI_ISL_3289811, EPI_ISL_3289812,
EPI_ISL_3289813, EPI_ISL_3289814, EPI_ISL_3289815, EPI_ISL_3289816, EPI_ISL_3289817, EPI_ISL_3289818, EPI_ISL_3289819, EPI_ISL_3289820, EPI_ISL_3289821, EPI_ISL_3289822, EPI_ISL_3289823, EPI_ISL_3289824, EPI_ISL_3289825, EPI_ISL_3289826, EPI_ISL_3289827, EPI_ISL_3289828, EPI_ISL_3289829, EPI_ISL_3289830, EPI_ISL_3289831, EPI_ISL_3289832, EPI_ISL_3289833, EPI_ISL_3289834,
EPI_ISL_3289835, EPI_ISL_3289836, EPI_ISL_3289837, EPI_ISL_3289838, EPI_ISL_3289839, EPI_ISL_3289840, EPI_ISL_3289845, EPI_ISL_3289846, EPI_ISL_3289847, EPI_ISL_3289848, EPI_ISL_3289849, EPI_ISL_3289850, EPI_ISL_3289851, EPI_ISL_3289852, EPI_ISL_3289853, EPI_ISL_3289854, EPI_ISL_3289855, EPI_ISL_3289856, EPI_ISL_3289857, EPI_ISL_3289858, EPI_ISL_3289859, EPI_ISL_3289860,
EPI_ISL_3289861, EPI_ISL_3289862, EPI_ISL_3289863, EPI_ISL_3289864, EPI_ISL_3289865, EPI_ISL_3289866, EPI_ISL_3289867, EPI_ISL_3289868, EPI_ISL_3289869, EPI_ISL_3289870, EPI_ISL_3289871, EPI_ISL_3289872, EPI_ISL_3289873, EPI_ISL_3289874, EPI_ISL_3289875, EPI_ISL_3289876, EPI_ISL_3289877, EPI_ISL_3289878, EPI_ISL_3289879, EPI_ISL_3289880, EPI_ISL_3289881, EPI_ISL_3289882,
EPI_ISL_3289883, EPI_ISL_3289884, EPI_ISL_3289885, EPI_ISL_3289886, EPI_ISL_3289887, EPI_ISL_3289888, EPI_ISL_3289890, EPI_ISL_3289891, EPI_ISL_3289892, EPI_ISL_3289893, EPI_ISL_3289894, EPI_ISL_3289895, EPI_ISL_3289896, EPI_ISL_3289897, EPI_ISL_3289898, EPI_ISL_3289899, EPI_ISL_3289900, EPI_ISL_3289901, EPI_ISL_3289902, EPI_ISL_3289903, EPI_ISL_3289904, EPI_ISL_3289905,
EPI_ISL_3289906, EPI_ISL_3289907, EPI_ISL_3289908, EPI_ISL_3289909, EPI_ISL_3289910, EPI_ISL_3289911, EPI_ISL_3289912, EPI_ISL_3289913, EPI_ISL_3289914, EPI_ISL_3289915, EPI_ISL_3289916, EPI_ISL_3289917, EPI_ISL_3289918, EPI_ISL_3289919, EPI_ISL_3289920, EPI_ISL_3289921, EPI_ISL_3289922, EPI_ISL_3289923, EPI_ISL_3289924, EPI_ISL_3289925, EPI_ISL_3289926, EPI_ISL_3289927,
EPI_ISL_3289928, EPI_ISL_3289929, EPI_ISL_3289930, EPI_ISL_3289931, EPI_ISL_3289932, EPI_ISL_3289933, EPI_ISL_3289934, EPI_ISL_3289935, EPI_ISL_3289936, EPI_ISL_3289937, EPI_ISL_3289938, EPI_ISL_3289939, EPI_ISL_3289940, EPI_ISL_3289941, EPI_ISL_3289942, EPI_ISL_3289943, EPI_ISL_3289944, EPI_ISL_3289945, EPI_ISL_3289946, EPI_ISL_3289947, EPI_ISL_3289948, EPI_ISL_3289949,
EPI_ISL_3289950, EPI_ISL_3289951, EPI_ISL_3289952, EPI_ISL_3289953, EPI_ISL_3289954, EPI_ISL_3289955, EPI_ISL_3289956, EPI_ISL_3289957, EPI_ISL_3289958, EPI_ISL_3289959, EPI_ISL_3289960, EPI_ISL_3289961, EPI_ISL_3289962, EPI_ISL_3289963, EPI_ISL_3289964, EPI_ISL_3289965, EPI_ISL_3289966, EPI_ISL_3289967, EPI_ISL_3289968, EPI_ISL_3289969, EPI_ISL_3289970, EPI_ISL_3289971,
EPI_ISL_3289972, EPI_ISL_3289973, EPI_ISL_3289974, EPI_ISL_3289975, EPI_ISL_3289976, EPI_ISL_3289977, EPI_ISL_3289978, EPI_ISL_3289979, EPI_ISL_3289980, EPI_ISL_3289981, EPI_ISL_3289982, EPI_ISL_3289983, EPI_ISL_3289985, EPI_ISL_3289986, EPI_ISL_3289987, EPI_ISL_3289988, EPI_ISL_3289989, EPI_ISL_3289990, EPI_ISL_3289991, EPI_ISL_3289992, EPI_ISL_3289993, EPI_ISL_3289994,
EPI_ISL_3289995, EPI_ISL_3289996, EPI_ISL_3289997, EPI_ISL_3289998, EPI_ISL_3289999, EPI_ISL_3290000, EPI_ISL_3290001, EPI_ISL_3290002, EPI_ISL_3290003, EPI_ISL_3290004, EPI_ISL_3290005, EPI_ISL_3290006, EPI_ISL_3290007, EPI_ISL_3290008, EPI_ISL_3290009, EPI_ISL_3290010, EPI_ISL_3290011, EPI_ISL_3290012, EPI_ISL_3290013, EPI_ISL_3290014, EPI_ISL_3290015, EPI_ISL_3290016,
EPI_ISL_3290017, EPI_ISL_3290018, EPI_ISL_3290019, EPI_ISL_3290020, EPI_ISL_3290021, EPI_ISL_3290022, EPI_ISL_3290023, EPI_ISL_3290024, EPI_ISL_3290025, EPI_ISL_3290026, EPI_ISL_3290027, EPI_ISL_3290028, EPI_ISL_3290029, EPI_ISL_3290030, EPI_ISL_3290031, EPI_ISL_3290032, EPI_ISL_3290033, EPI_ISL_3290034, EPI_ISL_3290035, EPI_ISL_3290036, EPI_ISL_3290037, EPI_ISL_3290038,
EPI_ISL_3290039, EPI_ISL_3290040, EPI_ISL_3290041, EPI_ISL_3290042, EPI_ISL_3290043, EPI_ISL_3290044, EPI_ISL_3290045, EPI_ISL_3290046, EPI_ISL_3290047, EPI_ISL_3290048, EPI_ISL_3290049, EPI_ISL_3290050, EPI_ISL_3290051, EPI_ISL_3290052, EPI_ISL_3290053, EPI_ISL_3290054, EPI_ISL_3290055, EPI_ISL_3290056, EPI_ISL_3290057, EPI_ISL_3290058, EPI_ISL_3290059, EPI_ISL_3290060,
EPI_ISL_3290061, EPI_ISL_3290062, EPI_ISL_3290063, EPI_ISL_3290064, EPI_ISL_3290065, EPI_ISL_3290066, EPI_ISL_3290067, EPI_ISL_3290068, EPI_ISL_3290069, EPI_ISL_3290070, EPI_ISL_3290071, EPI_ISL_3290072, EPI_ISL_3290073, EPI_ISL_3290074, EPI_ISL_3290075, EPI_ISL_3290076, EPI_ISL_3290078, EPI_ISL_3290079, EPI_ISL_3290080, EPI_ISL_3290081, EPI_ISL_3290082, EPI_ISL_3290083,
EPI_ISL_3290084, EPI_ISL_3290085, EPI_ISL_3290086, EPI_ISL_3290087, EPI_ISL_3290088, EPI_ISL_3290089, EPI_ISL_3290090, EPI_ISL_3290091, EPI_ISL_3290092, EPI_ISL_3290093, EPI_ISL_3290095, EPI_ISL_3290096, EPI_ISL_3290097, EPI_ISL_3290098, EPI_ISL_3290099, EPI_ISL_3290100, EPI_ISL_3290101, EPI_ISL_3290102, EPI_ISL_3290103, EPI_ISL_3290104, EPI_ISL_3290105, EPI_ISL_3290106,
EPI_ISL_3290107, EPI_ISL_3290108, EPI_ISL_3290109, EPI_ISL_3290110, EPI_ISL_3290111, EPI_ISL_3290112, EPI_ISL_3290113, EPI_ISL_3290114, EPI_ISL_3290115, EPI_ISL_3290116, EPI_ISL_3290117, EPI_ISL_3290118, EPI_ISL_3290119, EPI_ISL_3290120, EPI_ISL_3290121, EPI_ISL_3290122, EPI_ISL_3290123, EPI_ISL_3290124, EPI_ISL_3290125, EPI_ISL_3290126, EPI_ISL_3290127, EPI_ISL_3290128,
EPI_ISL_3290129, EPI_ISL_3290130, EPI_ISL_3290131, EPI_ISL_3290132, EPI_ISL_3290133, EPI_ISL_3290134, EPI_ISL_3290135, EPI_ISL_3290136, EPI_ISL_3290137, EPI_ISL_3290138, EPI_ISL_3290139, EPI_ISL_3290140, EPI_ISL_3290141, EPI_ISL_3290142, EPI_ISL_3290143, EPI_ISL_3290144, EPI_ISL_3290145, EPI_ISL_3290146, EPI_ISL_3290147, EPI_ISL_3290148, EPI_ISL_3290149, EPI_ISL_3290150,
EPI_ISL_3290151, EPI_ISL_3290152, EPI_ISL_3290153, EPI_ISL_3290154, EPI_ISL_3290155, EPI_ISL_3290156, EPI_ISL_3290157, EPI_ISL_3290158, EPI_ISL_3290159, EPI_ISL_3290160, EPI_ISL_3290161, EPI_ISL_3290162, EPI_ISL_3290163, EPI_ISL_3290164, EPI_ISL_3290165, EPI_ISL_3290166, EPI_ISL_3290167, EPI_ISL_3290168, EPI_ISL_3290169, EPI_ISL_3290170, EPI_ISL_3290171, EPI_ISL_3290172,
EPI_ISL_3290173, EPI_ISL_3290174, EPI_ISL_3290175, EPI_ISL_3290176, EPI_ISL_3290177, EPI_ISL_3290178, EPI_ISL_3290179, EPI_ISL_3290180, EPI_ISL_3290181, EPI_ISL_3290182, EPI_ISL_3290185, EPI_ISL_3290186, EPI_ISL_3290187, EPI_ISL_3290188, EPI_ISL_3290189, EPI_ISL_3290190, EPI_ISL_3290191, EPI_ISL_3290192, EPI_ISL_3290193, EPI_ISL_3290194, EPI_ISL_3290195, EPI_ISL_3290196,
EPI_ISL_3290197, EPI_ISL_3290198, EPI_ISL_3290199, EPI_ISL_3290200, EPI_ISL_3290201, EPI_ISL_3290202, EPI_ISL_3290203, EPI_ISL_3290204, EPI_ISL_3290205, EPI_ISL_3290206, EPI_ISL_3290207, EPI_ISL_3290208, EPI_ISL_3290209, EPI_ISL_3290210, EPI_ISL_3290211, EPI_ISL_3290212, EPI_ISL_3290213, EPI_ISL_3290214, EPI_ISL_3290215, EPI_ISL_3290216, EPI_ISL_3290217, EPI_ISL_3290218,
EPI_ISL_3290219, EPI_ISL_3290220, EPI_ISL_3290221, EPI_ISL_3290222, EPI_ISL_3290223, EPI_ISL_3290224, EPI_ISL_3290225, EPI_ISL_3290226, EPI_ISL_3290227, EPI_ISL_3290228, EPI_ISL_3290229, EPI_ISL_3290230, EPI_ISL_3290231, EPI_ISL_3290232, EPI_ISL_3290233, EPI_ISL_3290234, EPI_ISL_3290235, EPI_ISL_3290236, EPI_ISL_3290237, EPI_ISL_3290238, EPI_ISL_3290239, EPI_ISL_3290240,
EPI_ISL_3290241, EPI_ISL_3290242, EPI_ISL_3290243, EPI_ISL_3290244, EPI_ISL_3290245, EPI_ISL_3290246, EPI_ISL_3290247, EPI_ISL_3290248, EPI_ISL_3290249, EPI_ISL_3290250, EPI_ISL_3290251, EPI_ISL_3290252, EPI_ISL_3290253, EPI_ISL_3290254, EPI_ISL_3290255, EPI_ISL_3290256, EPI_ISL_3290257, EPI_ISL_3290258, EPI_ISL_3290259, EPI_ISL_3290260, EPI_ISL_3290261, EPI_ISL_3290262,
EPI_ISL_3290263, EPI_ISL_3290264, EPI_ISL_3290265, EPI_ISL_3290266, EPI_ISL_3290267, EPI_ISL_3290268, EPI_ISL_3290269, EPI_ISL_3290270, EPI_ISL_3290271, EPI_ISL_3290272, EPI_ISL_3290273, EPI_ISL_3290274, EPI_ISL_3290275, EPI_ISL_3290276, EPI_ISL_3290277, EPI_ISL_3290278, EPI_ISL_3290279, EPI_ISL_3290280, EPI_ISL_3290281, EPI_ISL_3290282, EPI_ISL_3290283, EPI_ISL_3290284,
EPI_ISL_3290285, EPI_ISL_3290286, EPI_ISL_3290287, EPI_ISL_3290288, EPI_ISL_3290289, EPI_ISL_3290290, EPI_ISL_3290291, EPI_ISL_3290292, EPI_ISL_3290293, EPI_ISL_3290294, EPI_ISL_3290295, EPI_ISL_3290296, EPI_ISL_3290297, EPI_ISL_3290298, EPI_ISL_3290299, EPI_ISL_3290300, EPI_ISL_3290301, EPI_ISL_3290302, EPI_ISL_3290303, EPI_ISL_3290304, EPI_ISL_3290305, EPI_ISL_3290306,

EPI_ISL_3290307, EPI_ISL_3290308, EPI_ISL_3290309, EPI_ISL_3290310, EPI_ISL_3290311, EPI_ISL_3290312, EPI_ISL_3290313, EPI_ISL_3290314, EPI_ISL_3290315, EPI_ISL_3290316, EPI_ISL_3290317, EPI_ISL_3290318, EPI_ISL_3290320, EPI_ISL_3290321, EPI_ISL_3290322, EPI_ISL_3290323, EPI_ISL_3290324, EPI_ISL_3290325, EPI_ISL_3290326, EPI_ISL_3290327, EPI_ISL_3290328, EPI_ISL_3290329, EPI_ISL_3290330, EPI_ISL_3290331, EPI_ISL_3290332, EPI_ISL_3290333, EPI_ISL_3290334, EPI_ISL_3290335, EPI_ISL_3290336, EPI_ISL_3290337, EPI_ISL_3290338, EPI_ISL_3290339, EPI_ISL_3290340, EPI_ISL_3290341, EPI_ISL_3290342, EPI_ISL_3290343, EPI_ISL_3290344, EPI_ISL_3290345, EPI_ISL_3290346, EPI_ISL_3290347, EPI_ISL_3290349, EPI_ISL_3290350, EPI_ISL_3290351, EPI_ISL_3290352, EPI_ISL_3290353, EPI_ISL_3290354, EPI_ISL_3290355, EPI_ISL_3290356, EPI_ISL_3290357, EPI_ISL_3290358, EPI_ISL_3290359, EPI_ISL_3290360, EPI_ISL_3290361, EPI_ISL_3290362, EPI_ISL_3290363, EPI_ISL_3290364, EPI_ISL_3290365, EPI_ISL_3290366, EPI_ISL_3290367, EPI_ISL_3290368, EPI_ISL_3290369, EPI_ISL_3290370, EPI_ISL_3290371, EPI_ISL_3290372, EPI_ISL_3290373, EPI_ISL_3290374, EPI_ISL_3290375, EPI_ISL_3290376, EPI_ISL_3290377, EPI_ISL_3290378, EPI_ISL_3290379, EPI_ISL_3290380, EPI_ISL_3290381, EPI_ISL_3290382, EPI_ISL_3290383, EPI_ISL_3290384, EPI_ISL_3290385, EPI_ISL_3290386, EPI_ISL_3290387, EPI_ISL_3290388, EPI_ISL_3290389, EPI_ISL_3290390, EPI_ISL_3290391, EPI_ISL_3290392, EPI_ISL_3290393, EPI_ISL_3290394, EPI_ISL_3290395, EPI_ISL_3290396, EPI_ISL_3290397, EPI_ISL_3290398, EPI_ISL_3290399, EPI_ISL_3290400, EPI_ISL_3290401, EPI_ISL_3290402, EPI_ISL_3290403, EPI_ISL_3290404, EPI_ISL_3290405, EPI_ISL_3290406, EPI_ISL_3290407, EPI_ISL_3290408, EPI_ISL_3290409, EPI_ISL_3290410, EPI_ISL_3290411, EPI_ISL_3290412, EPI_ISL_3290413, EPI_ISL_3290414, EPI_ISL_3290415, EPI_ISL_3290416, EPI_ISL_3290417, EPI_ISL_3290418, EPI_ISL_3290419, EPI_ISL_3290420, EPI_ISL_3290421, EPI_ISL_3290422, EPI_ISL_3290423, EPI_ISL_3290424, EPI_ISL_3290425, EPI_ISL_3290426, EPI_ISL_3290427, EPI_ISL_3290428, EPI_ISL_3290429, EPI_ISL_3290430, EPI_ISL_3290431, EPI_ISL_3290432, EPI_ISL_3290433, EPI_ISL_3290434, EPI_ISL_3290435, EPI_ISL_3290436, EPI_ISL_3290437, EPI_ISL_3290438, EPI_ISL_3290439, EPI_ISL_3290440, EPI_ISL_3290441, EPI_ISL_3290442, EPI_ISL_3290443, EPI_ISL_3290444, EPI_ISL_3290445, EPI_ISL_3290446, EPI_ISL_3290447, EPI_ISL_3290448, EPI_ISL_3290449, EPI_ISL_3290450, EPI_ISL_3290451, EPI_ISL_3290452, EPI_ISL_3290453, EPI_ISL_3290454, EPI_ISL_3290455, EPI_ISL_3290456, EPI_ISL_3290457, EPI_ISL_3290458, EPI_ISL_3290459, EPI_ISL_3290460, EPI_ISL_3290461, EPI_ISL_3290462, EPI_ISL_3290463, EPI_ISL_3290464, EPI_ISL_3290465, EPI_ISL_3290466, EPI_ISL_3290467, EPI_ISL_3290468, EPI_ISL_3290469, EPI_ISL_3290470, EPI_ISL_3290471, EPI_ISL_3290472, EPI_ISL_3290473, EPI_ISL_3290474, EPI_ISL_3290475, EPI_ISL_3290476, EPI_ISL_3290477, EPI_ISL_3290478, EPI_ISL_3290479, EPI_ISL_3290480, EPI_ISL_3290481, EPI_ISL_3290482, EPI_ISL_3290483, EPI_ISL_3290484, EPI_ISL_3290485, EPI_ISL_3290486, EPI_ISL_3290487, EPI_ISL_3290488, EPI_ISL_3290489, EPI_ISL_3290490, EPI_ISL_3290491, EPI_ISL_3290492, EPI_ISL_3290493, EPI_ISL_3290494, EPI_ISL_3290495, EPI_ISL_3290496, EPI_ISL_3290497, EPI_ISL_3290498, EPI_ISL_3290499, EPI_ISL_3290500, EPI_ISL_3290501, EPI_ISL_3290502, EPI_ISL_3290503, EPI_ISL_3290504, EPI_ISL_3290505, EPI_ISL_3290506, EPI_ISL_3290507, EPI_ISL_3290508, EPI_ISL_3290509, EPI_ISL_3290510, EPI_ISL_3290511, EPI_ISL_3290512, EPI_ISL_3290513, EPI_ISL_3290514, EPI_ISL_3290515, EPI_ISL_3290516, EPI_ISL_3290517, EPI_ISL_3290518, EPI_ISL_3290519, EPI_ISL_3290520, EPI_ISL_3290521, EPI_ISL_3290522, EPI_ISL_3290523, EPI_ISL_3290524, EPI_ISL_3290525, EPI_ISL_3290526, EPI_ISL_3290527, EPI_ISL_3290528, EPI_ISL_3290529, EPI_ISL_3290530, EPI_ISL_3290531, EPI_ISL_3290532, EPI_ISL_3290533, EPI_ISL_3290534, EPI_ISL_3290535, EPI_ISL_3290536, EPI_ISL_3290537, EPI_ISL_3290538, EPI_ISL_3290539, EPI_ISL_3290540, EPI_ISL_3290541, EPI_ISL_3290542, EPI_ISL_3290543, EPI_ISL_3290544, EPI_ISL_3290545, EPI_ISL_3290546, EPI_ISL_3290547, EPI_ISL_3290548, EPI_ISL_3290549, EPI_ISL_3290550, EPI_ISL_3290551, EPI_ISL_3290552, EPI_ISL_3290553, EPI_ISL_3290554, EPI_ISL_3290555, EPI_ISL_3290556, EPI_ISL_3290557, EPI_ISL_3290558, EPI_ISL_3290559, EPI_ISL_3290560, EPI_ISL_3290561, EPI_ISL_3290562, EPI_ISL_3290563, EPI_ISL_3290564, EPI_ISL_3290565, EPI_ISL_3290566, EPI_ISL_3290567, EPI_ISL_3290568, EPI_ISL_3290569, EPI_ISL_3290570, EPI_ISL_3290571, EPI_ISL_3290572, EPI_ISL_3290573, EPI_ISL_3290574, EPI_ISL_3290575, EPI_ISL_3290576, EPI_ISL_3290577, EPI_ISL_3290578, EPI_ISL_3290579, EPI_ISL_3290580, EPI_ISL_3290581, EPI_ISL_3290582, EPI_ISL_3290583, EPI_ISL_3290584, EPI_ISL_3290585, EPI_ISL_3290586, EPI_ISL_3290587, EPI_ISL_3290588, EPI_ISL_3290589, EPI_ISL_3290590, EPI_ISL_3290591, EPI_ISL_3290592, EPI_ISL_3290593, EPI_ISL_3290594, EPI_ISL_3290595, EPI_ISL_3290596, EPI_ISL_3290597, EPI_ISL_3290598, EPI_ISL_3290599, EPI_ISL_3290600, EPI_ISL_3290601, EPI_ISL_3290602, EPI_ISL_3290603, EPI_ISL_3290604, EPI_ISL_3290606, EPI_ISL_3290607, EPI_ISL_3290608, EPI_ISL_3290609, EPI_ISL_3290610, EPI_ISL_3290611, EPI_ISL_3290612, EPI_ISL_3290613, EPI_ISL_3290614, EPI_ISL_3290615, EPI_ISL_3290616, EPI_ISL_3290617, EPI_ISL_3290618, EPI_ISL_3290619, EPI_ISL_3290620, EPI_ISL_3290621, EPI_ISL_3290622, EPI_ISL_3290623, EPI_ISL_3290624, EPI_ISL_3290625, EPI_ISL_3290626, EPI_ISL_3290627, EPI_ISL_3290628, EPI_ISL_3290629, EPI_ISL_3290630, EPI_ISL_3290631, EPI_ISL_3290632, EPI_ISL_3290633, EPI_ISL_3290634, EPI_ISL_3290635, EPI_ISL_3290636, EPI_ISL_3290637, EPI_ISL_3290638, EPI_ISL_3290639, EPI_ISL_3290640, EPI_ISL_3290641, EPI_ISL_3290642, EPI_ISL_3290643, EPI_ISL_3290644, EPI_ISL_3290645, EPI_ISL_3290646, EPI_ISL_3290647, EPI_ISL_3290648, EPI_ISL_3290649, EPI_ISL_3290650, EPI_ISL_3290651, EPI_ISL_3290652, EPI_ISL_3290653, EPI_ISL_3290654, EPI_ISL_3290655, EPI_ISL_3290656, EPI_ISL_3290657, EPI_ISL_3290658, EPI_ISL_3290659, EPI_ISL_3290660, EPI_ISL_3290661, EPI_ISL_3290662, EPI_ISL_3290663, EPI_ISL_3290664, EPI_ISL_3290665, EPI_ISL_3290666, EPI_ISL_3290667, EPI_ISL_3290668, EPI_ISL_3290669, EPI_ISL_3290670, EPI_ISL_3290671, EPI_ISL_3290672, EPI_ISL_3290673, EPI_ISL_3290674, EPI_ISL_3290675, EPI_ISL_3290676, EPI_ISL_3290677, EPI_ISL_3290678, EPI_ISL_3290679, EPI_ISL_3290680, EPI_ISL_3290681, EPI_ISL_3290682, EPI_ISL_3290684, EPI_ISL_3290685, EPI_ISL_3290686, EPI_ISL_3290687, EPI_ISL_3290688, EPI_ISL_3290689, EPI_ISL_3290690, EPI_ISL_3290691, EPI_ISL_3290692, EPI_ISL_3290693, EPI_ISL_3290695, EPI_ISL_3290697, EPI_ISL_3290698, EPI_ISL_3290699, EPI_ISL_3290700, EPI_ISL_3290702, EPI_ISL_3290703, EPI_ISL_3290704, EPI_ISL_3290705, EPI_ISL_3290706, EPI_ISL_3290707, EPI_ISL_3290708, EPI_ISL_3290709, EPI_ISL_3290710, EPI_ISL_3290711, EPI_ISL_3290712, EPI_ISL_3290713, EPI_ISL_3290714, EPI_ISL_3290715, EPI_ISL_3290716, EPI_ISL_3290717, EPI_ISL_3290718, EPI_ISL_3290719, EPI_ISL_3290720, EPI_ISL_3290721, EPI_ISL_3290722, EPI_ISL_3290723, EPI_ISL_3290724, EPI_ISL_3290725, EPI_ISL_3290726, EPI_ISL_3290727, EPI_ISL_3290728, EPI_ISL_3290729, EPI_ISL_3290730, EPI_ISL_3290731, EPI_ISL_3290732, EPI_ISL_3290733, EPI_ISL_3290734, EPI_ISL_3290735, EPI_ISL_3290736, EPI_ISL_3290737, EPI_ISL_3290738, EPI_ISL_3290739, EPI_ISL_3290740, EPI_ISL_3290741, EPI_ISL_3290742, EPI_ISL_3290743, EPI_ISL_3290744, EPI_ISL_3290745, EPI_ISL_3290746, EPI_ISL_3290747, EPI_ISL_3290748, EPI_ISL_3290749, EPI_ISL_3290750, EPI_ISL_3290751, EPI_ISL_3290752, EPI_ISL_3290753, EPI_ISL_3290754, EPI_ISL_3290755, EPI_ISL_3290756, EPI_ISL_3290757, EPI_ISL_3290758, EPI_ISL_3290759, EPI_ISL_3290760, EPI_ISL_3290761, EPI_ISL_3290762, EPI_ISL_3290763, EPI_ISL_3290764, EPI_ISL_3290765, EPI_ISL_3290766, EPI_ISL_3290767, EPI_ISL_3290768, EPI_ISL_3290769, EPI_ISL_3290770, EPI_ISL_3290771, EPI_ISL_3290772, EPI_ISL_3290773, EPI_ISL_3290774, EPI_ISL_3290775, EPI_ISL_3290776, EPI_ISL_3290777, EPI_ISL_3290778, EPI_ISL_3290779, EPI_ISL_3290780, EPI_ISL_3290781, EPI_ISL_3290782, EPI_ISL_3290783, EPI_ISL_3290784, EPI_ISL_3290785, EPI_ISL_3290786, EPI_ISL_3290787, EPI_ISL_3290788, EPI_ISL_3290789, EPI_ISL_3290790, EPI_ISL_3290791, EPI_ISL_3290792, EPI_ISL_3290793, EPI_ISL_3290794, EPI_ISL_3290795, EPI_ISL_3290796, EPI_ISL_3290797, EPI_ISL_3290798, EPI_ISL_3290799, EPI_ISL_3290800, EPI_ISL_3290801, EPI_ISL_3290802, EPI_ISL_3290803, EPI_ISL_3290804, EPI_ISL_3290806, EPI_ISL_3290807, EPI_ISL_3290808, EPI_ISL_3290809, EPI_ISL_3290810, EPI_ISL_3290811, EPI_ISL_3290812, EPI_ISL_3290813, EPI_ISL_3290814, EPI_ISL_3290815, EPI_ISL_3290816, EPI_ISL_3290817, EPI_ISL_3290818, EPI_ISL_3290819, EPI_ISL_3290820, EPI_ISL_3290821, EPI_ISL_3290822, EPI_ISL_3290823, EPI_ISL_3290824, EPI_ISL_3290825, EPI_ISL_3290826, EPI_ISL_3290827, EPI_ISL_3290828, EPI_ISL_3290829, EPI_ISL_3290830, EPI_ISL_3290831, EPI_ISL_3290832, EPI_ISL_3290833, EPI_ISL_3290834, EPI_ISL_3290835, EPI_ISL_3290836, EPI_ISL_3290837, EPI_ISL_3290838, EPI_ISL_3290839, EPI_ISL_3290840, EPI_ISL_3290841, EPI_ISL_3290842, EPI_ISL_3290843, EPI_ISL_3290844, EPI_ISL_3290845, EPI_ISL_3290846, EPI_ISL_3290847, EPI_ISL_3290848, EPI_ISL_3290849, EPI_ISL_3290850, EPI_ISL_3290851, EPI_ISL_3290852, EPI_ISL_3290853, EPI_ISL_3290854, EPI_ISL_3290855, EPI_ISL_3290856, EPI_ISL_3290857, EPI_ISL_3290858, EPI_ISL_3290859, EPI_ISL_3290860, EPI_ISL_3290861, EPI_ISL_3290862, EPI_ISL_3290863, EPI_ISL_3290864, EPI_ISL_3290865, EPI_ISL_3290866, EPI_ISL_3290867, EPI_ISL_3290868, EPI_ISL_3290869, EPI_ISL_3290871, EPI_ISL_3290872, EPI_ISL_3290873, EPI_ISL_3290874, EPI_ISL_3290875, EPI_ISL_3290876, EPI_ISL_3290877, EPI_ISL_3290878, EPI_ISL_3290880, EPI_ISL_3290881, EPI_ISL_3290882, EPI_ISL_3290883, EPI_ISL_3290884, EPI_ISL_3290885, EPI_ISL_3290886, EPI_ISL_3290887, EPI_ISL_3290888, EPI_ISL_3290889, EPI_ISL_3290890, EPI_ISL_3290891, EPI_ISL_3290892, EPI_ISL_3290893, EPI_ISL_3290894, EPI_ISL_3290895, EPI_ISL_3290896, EPI_ISL_3290897, EPI_ISL_3290898, EPI_ISL_3290899, EPI_ISL_3290900, EPI_ISL_3290901, EPI_ISL_3290902, EPI_ISL_3290903, EPI_ISL_3290904, EPI_ISL_3290905, EPI_ISL_3290906, EPI_ISL_3290907, EPI_ISL_3290908, EPI_ISL_3290909, EPI_ISL_3290910, EPI_ISL_3290911, EPI_ISL_3290912, EPI_ISL_3290913, EPI_ISL_3290914, EPI_ISL_3290915, EPI_ISL_3290916, EPI_ISL_3290917, EPI_ISL_3290918, EPI_ISL_3290919, EPI_ISL_3290920, EPI_ISL_3290921, EPI_ISL_3290922, EPI_ISL_3290923, EPI_ISL_3290924, EPI_ISL_3290925, EPI_ISL_3290926, EPI_ISL_3290927, EPI_ISL_3290928, EPI_ISL_3290929, EPI_ISL_3290930, EPI_ISL_3290931, EPI_ISL_3290932, EPI_ISL_3290933, EPI_ISL_3290934, EPI_ISL_3290935, EPI_ISL_3290936, EPI_ISL_3290938, EPI_ISL_3290939, EPI_ISL_3290940, EPI_ISL_3290941, EPI_ISL_3290942, EPI_ISL_3290943, EPI_ISL_3290944, EPI_ISL_3290945, EPI_ISL_3290946, EPI_ISL_3290947, EPI_ISL_3290948, EPI_ISL_3290949, EPI_ISL_3290950, EPI_ISL_3290951, EPI_ISL_3290952, EPI_ISL_3290953, EPI_ISL_3290954, EPI_ISL_3290955, EPI_ISL_3290956, EPI_ISL_3290957, EPI_ISL_3290958, EPI_ISL_3290959, EPI_ISL_3290960, EPI_ISL_3290961, EPI_ISL_3290962, EPI_ISL_3290963, EPI_ISL_3290964, EPI_ISL_3290965, EPI_ISL_3290966, EPI_ISL_3290967, EPI_ISL_3290968, EPI_ISL_3290969, EPI_ISL_3290970, EPI_ISL_3290971, EPI_ISL_3290972, EPI_ISL_3290973, EPI_ISL_3290974, EPI_ISL_3290975, EPI_ISL_3290976, EPI_ISL_3290977, EPI_ISL_3290978, EPI_ISL_3290979, EPI_ISL_3290980, EPI_ISL_3290981, EPI_ISL_3290982, EPI_ISL_3290983, EPI_ISL_3290984, EPI_ISL_3290986, EPI_ISL_3290987, EPI_ISL_3290988, EPI_ISL_3290989, EPI_ISL_3290990, EPI_ISL_3290991, EPI_ISL_3290992, EPI_ISL_3290993, EPI_ISL_3290994, EPI_ISL_3290995, EPI_ISL_3290996, EPI_ISL_3290997, EPI_ISL_3290998, EPI_ISL_3290999, EPI_ISL_3291000, EPI_ISL_3291001, EPI_ISL_3291002, EPI_ISL_3291003, EPI_ISL_3291004, EPI_ISL_3291005, EPI_ISL_3291006, EPI_ISL_3291007, EPI_ISL_3291008, EPI_ISL_3291009, EPI_ISL_3291010, EPI_ISL_3291011, EPI_ISL_3291012, EPI_ISL_3291013, EPI_ISL_3291014, EPI_ISL_3291015, EPI_ISL_3291016, EPI_ISL_3291017, EPI_ISL_3291018, EPI_ISL_3291019, EPI_ISL_3291020, EPI_ISL_3291021, EPI_ISL_3291022, EPI_ISL_3291023, EPI_ISL_3291024, EPI_ISL_3291025, EPI_ISL_3291026, EPI_ISL_3291027, EPI_ISL_3291028, EPI_ISL_3291029, EPI_ISL_3291030, EPI_ISL_3291031, EPI_ISL_3291032, EPI_ISL_3291033, EPI_ISL_3291034, EPI_ISL_3291035, EPI_ISL_3291036, EPI_ISL_3291037, EPI_ISL_3291038, EPI_ISL_3291039, EPI_ISL_3291040, EPI_ISL_3291041, EPI_ISL_3291042, EPI_ISL_3291043, EPI_ISL_3291044, EPI_ISL_3291045, EPI_ISL_3291046, EPI_ISL_3291047, EPI_ISL_3291048, EPI_ISL_3291049, EPI_ISL_3291050, EPI_ISL_3291051, EPI_ISL_3291052, EPI_ISL_3291053, EPI_ISL_3291054, EPI_ISL_3291055, EPI_ISL_3291056, EPI_ISL_3291057, EPI_ISL_3291058, EPI_ISL_3291059, EPI_ISL_3291060, EPI_ISL_3291061, EPI_ISL_3291062, EPI_ISL_3291063, EPI_ISL_3291064, EPI_ISL_3291065, EPI_ISL_3291066, EPI_ISL_3291067, EPI_ISL_3291068, EPI_ISL_3291069, EPI_ISL_3291070, EPI_ISL_3291072, EPI_ISL_3291073, EPI_ISL_3291074, EPI_ISL_3291075, EPI_ISL_3291076, EPI_ISL_3291077, EPI_ISL_3291078, EPI_ISL_3291079, EPI_ISL_3291080, EPI_ISL_3291081, EPI_ISL_3291082, EPI_ISL_3291083, EPI_ISL_3291084, EPI_ISL_3291085, EPI_ISL_3291086, EPI_ISL_3291087, EPI_ISL_3291088, EPI_ISL_3291089, EPI_ISL_3291090, EPI_ISL_3291091, EPI_ISL_3291092, EPI_ISL_3291093, EPI_ISL_3291094, EPI_ISL_3291095, EPI_ISL_3291096, EPI_ISL_3291097, EPI_ISL_3291098, EPI_ISL_3291099, EPI_ISL_3291100, EPI_ISL_3291102, EPI_ISL_3291103, EPI_ISL_3291104, EPI_ISL_3291105, EPI_ISL_3291106, EPI_ISL_3291107, EPI_ISL_3291108, EPI_ISL_3291109, EPI_ISL_3291110, EPI_ISL_3291111, EPI_ISL_3291112, EPI_ISL_3291113, EPI_ISL_3291114, EPI_ISL_3291115, EPI_ISL_3291116, EPI_ISL_3291117, EPI_ISL_3291118, EPI_ISL_3291120, EPI_ISL_3291122, EPI_ISL_3291123, EPI_ISL_3291124, EPI_ISL_3291125, EPI_ISL_3291126, EPI_ISL_3291128, EPI_ISL_3291130, EPI_ISL_3291131, EPI_ISL_3291132, EPI_ISL_3291133, EPI_ISL_3291134, EPI_ISL_3291135, EPI_ISL_3291136, EPI_ISL_3291137, EPI_ISL_3291138, EPI_ISL_3291139, EPI_ISL_3291140, EPI_ISL_3291141, EPI_ISL_3291142, EPI_ISL_3291143, EPI_ISL_3291144, EPI_ISL_3291145, EPI_ISL_3291146, EPI_ISL_3291147, EPI_ISL_3291148, EPI_ISL_3291149, EPI_ISL_3291150, EPI_ISL_3291151, EPI_ISL_3291152, EPI_ISL_3291153, EPI_ISL_3291155, EPI_ISL_3291156, EPI_ISL_3291157, EPI_ISL_3291158, EPI_ISL_3291159, EPI_ISL_3291160, EPI_ISL_3291161, EPI_ISL_3291162, EPI_ISL_3291163, EPI_ISL_3291164, EPI_ISL_3291165, EPI_ISL_3291166, EPI_ISL_3291167, EPI_ISL_3291168, EPI_ISL_3291169, EPI_ISL_3291170, EPI_ISL_3291172, EPI_ISL_3291173, EPI_ISL_3291174, EPI_ISL_3291175, EPI_ISL_3291176, EPI_ISL_3291177, EPI_ISL_3291178, EPI_ISL_3291179, EPI_ISL_3291180, EPI_ISL_3291181, EPI_ISL_3291182, EPI_ISL_3291183, EPI_ISL_3291184, EPI_ISL_3291185, EPI_ISL_3291186, EPI_ISL_3291187, EPI_ISL_3291188, EPI_ISL_3291189, EPI_ISL_3291190, EPI_ISL_3291191, EPI_ISL_3291192, EPI_ISL_3291193, EPI_ISL_3291194, EPI_ISL_3291195, EPI_ISL_3291197, EPI_ISL_3291198, EPI_ISL_3291199, EPI_ISL_3291200, EPI_ISL_3291201, EPI_ISL_3291202, EPI_ISL_3291203, EPI_ISL_3291204, EPI_ISL_3291206, EPI_ISL_3291207, EPI_ISL_3291208, EPI_ISL_3291210, EPI_ISL_3291211, EPI_ISL_3291212, EPI_ISL_3291213, EPI_ISL_3291214, EPI_ISL_3291215, EPI_ISL_3291216, EPI_ISL_3291218, EPI_ISL_3291219, EPI_ISL_3291228, EPI_ISL_3291233, EPI_ISL_3291241, EPI_ISL_3291242, EPI_ISL_3291243, EPI_ISL_3291245, EPI_ISL_3291246, EPI_ISL_3291247, EPI_ISL_3291248, EPI_ISL_3291249, EPI_ISL_3291250, EPI_ISL_3291251

| | | | |
|---|---|---|---|
| see above | Respiratory Virus Unit, Microbiology Services Colindale, Public Health England | COVID-19 Genomics UK (COG-UK) Consortium | PHE Covid Sequencing Team |
| EPI_ISL_3291252 | U.O. Microbiologia Laboratorio Unico Centro Servizi - AUSL della Romagna | U.O. Microbiologia, Laboratorio Unico Centro Servizi - AUSL della Romagna | Giorgio Dirani |
| EPI_ISL_3291253 | Respiratory Virus Unit, Microbiology Services Colindale, Public Health England | COVID-19 Genomics UK (COG-UK) Consortium | PHE Covid Sequencing Team |
| EPI_ISL_3291254, EPI_ISL_3291256, EPI_ISL_3291258 | U.O. Microbiologia Laboratorio Unico Centro Servizi - AUSL della Romagna | U.O. Microbiologia, Laboratorio Unico Centro Servizi - AUSL della Romagna | Giorgio Dirani |
| EPI_ISL_3291259 | Respiratory Virus Unit, Microbiology Services Colindale, Public Health England | COVID-19 Genomics UK (COG-UK) Consortium | PHE Covid Sequencing Team |
| EPI_ISL_3291260, EPI_ISL_3291261 | U.O. Microbiologia Laboratorio Unico Centro Servizi - AUSL della Romagna | U.O. Microbiologia, Laboratorio Unico Centro Servizi - AUSL della Romagna | Giorgio Dirani |
| EPI_ISL_3291262, EPI_ISL_3291263 | Respiratory Virus Unit, Microbiology Services Colindale, Public Health England | COVID-19 Genomics UK (COG-UK) Consortium | PHE Covid Sequencing Team |
| EPI_ISL_3291264 | U.O. Microbiologia Laboratorio Unico Centro Servizi - AUSL della Romagna | U.O. Microbiologia, Laboratorio Unico Centro Servizi - AUSL della Romagna | Giorgio Dirani |
| EPI_ISL_3291265 | Respiratory Virus Unit, Microbiology Services Colindale, Public Health England | COVID-19 Genomics UK (COG-UK) Consortium | PHE Covid Sequencing Team |
| EPI_ISL_3291266 | U.O. Microbiologia Laboratorio Unico Centro Servizi - AUSL della Romagna | U.O. Microbiologia, Laboratorio Unico Centro Servizi - AUSL della Romagna | Giorgio Dirani |
| EPI_ISL_3291267 | Respiratory Virus Unit, Microbiology Services Colindale, Public Health England | COVID-19 Genomics UK (COG-UK) Consortium | PHE Covid Sequencing Team |
| EPI_ISL_3291268 | U.O. Microbiologia Laboratorio Unico Centro Servizi - AUSL della Romagna | U.O. Microbiologia, Laboratorio Unico Centro Servizi - AUSL della Romagna | Giorgio Dirani |
| EPI_ISL_3291269 | Respiratory Virus Unit, Microbiology Services Colindale, Public Health England | COVID-19 Genomics UK (COG-UK) Consortium | PHE Covid Sequencing Team |
| EPI_ISL_3291270, EPI_ISL_3291272 | U.O. Microbiologia Laboratorio Unico Centro Servizi - AUSL della Romagna | U.O. Microbiologia, Laboratorio Unico Centro Servizi - AUSL della Romagna | Giorgio Dirani |
| EPI_ISL_3291273 | Respiratory Virus Unit, Microbiology Services Colindale, Public Health England | COVID-19 Genomics UK (COG-UK) Consortium | PHE Covid Sequencing Team |
| EPI_ISL_3291274, EPI_ISL_3291276 | U.O. Microbiologia Laboratorio Unico Centro Servizi - AUSL della Romagna | U.O. Microbiologia, Laboratorio Unico Centro Servizi - AUSL della Romagna | Giorgio Dirani |
| EPI_ISL_3291277 | Respiratory Virus Unit, Microbiology Services Colindale, Public Health England | COVID-19 Genomics UK (COG-UK) Consortium | PHE Covid Sequencing Team |
| EPI_ISL_3291278 | U.O. Microbiologia Laboratorio Unico Centro Servizi - AUSL della Romagna | U.O. Microbiologia, Laboratorio Unico Centro Servizi - AUSL della Romagna | Giorgio Dirani |
| EPI_ISL_3291279 | Respiratory Virus Unit, Microbiology Services Colindale, Public Health England | COVID-19 Genomics UK (COG-UK) Consortium | PHE Covid Sequencing Team |
| EPI_ISL_3291280 | U.O. Microbiologia Laboratorio Unico Centro Servizi - AUSL della Romagna | U.O. Microbiologia, Laboratorio Unico Centro Servizi - AUSL della Romagna | Giorgio Dirani |
| EPI_ISL_3291281 | Respiratory Virus Unit, Microbiology Services Colindale, Public Health England | COVID-19 Genomics UK (COG-UK) Consortium | PHE Covid Sequencing Team |
| EPI_ISL_3291282 | U.O. Microbiologia Laboratorio Unico Centro Servizi - AUSL della Romagna | U.O. Microbiologia, Laboratorio Unico Centro Servizi - AUSL della Romagna | Giorgio Dirani |
| EPI_ISL_3291283 | Respiratory Virus Unit, Microbiology Services Colindale, Public Health England | COVID-19 Genomics UK (COG-UK) Consortium | PHE Covid Sequencing Team |
| EPI_ISL_3291284 | U.O. Microbiologia Laboratorio Unico Centro Servizi - AUSL della Romagna | U.O. Microbiologia, Laboratorio Unico Centro Servizi - AUSL della Romagna | Giorgio Dirani |
| EPI_ISL_3291285 | Respiratory Virus Unit, Microbiology Services Colindale, Public Health England | COVID-19 Genomics UK (COG-UK) Consortium | PHE Covid Sequencing Team |
| EPI_ISL_3291286 | U.O. Microbiologia Laboratorio Unico Centro Servizi - AUSL della Romagna | U.O. Microbiologia, Laboratorio Unico Centro Servizi - AUSL della Romagna | Giorgio Dirani |
| EPI_ISL_3291287 | Respiratory Virus Unit, Microbiology Services Colindale, Public Health England | COVID-19 Genomics UK (COG-UK) Consortium | PHE Covid Sequencing Team |
| EPI_ISL_3291288 | U.O. Microbiologia Laboratorio Unico Centro Servizi - AUSL della | U.O. Microbiologia, Laboratorio Unico Centro Servizi - AUSL della | Giorgio Dirani |

| | | | |
|---|---|---|---|
| | Romagna | Romagna | |
| EPI_ISL_3291289 | Respiratory Virus Unit, Microbiology Services Colindale, Public Health England | COVID-19 Genomics UK (COG-UK) Consortium | PHE Covid Sequencing Team |
| EPI_ISL_3291290 | U.O. Microbiologia Laboratorio Unico Centro Servizi - AUSL della Romagna | U.O. Microbiologia, Laboratorio Unico Centro Servizi - AUSL della Romagna | Giorgio Dirani |
| EPI_ISL_3291291 | Respiratory Virus Unit, Microbiology Services Colindale, Public Health England | COVID-19 Genomics UK (COG-UK) Consortium | PHE Covid Sequencing Team |
| EPI_ISL_3291292 | U.O. Microbiologia Laboratorio Unico Centro Servizi - AUSL della Romagna | U.O. Microbiologia, Laboratorio Unico Centro Servizi - AUSL della Romagna | Giorgio Dirani |
| EPI_ISL_3291293 | Respiratory Virus Unit, Microbiology Services Colindale, Public Health England | COVID-19 Genomics UK (COG-UK) Consortium | PHE Covid Sequencing Team |
| EPI_ISL_3291294, EPI_ISL_3291296, EPI_ISL_3291298 | U.O. Microbiologia Laboratorio Unico Centro Servizi - AUSL della Romagna | U.O. Microbiologia, Laboratorio Unico Centro Servizi - AUSL della Romagna | Giorgio Dirani |
| EPI_ISL_3291299 | Respiratory Virus Unit, Microbiology Services Colindale, Public Health England | COVID-19 Genomics UK (COG-UK) Consortium | PHE Covid Sequencing Team |
| EPI_ISL_3291300 | U.O. Microbiologia Laboratorio Unico Centro Servizi - AUSL della Romagna | U.O. Microbiologia, Laboratorio Unico Centro Servizi - AUSL della Romagna | Giorgio Dirani |
| EPI_ISL_3291301 | Respiratory Virus Unit, Microbiology Services Colindale, Public Health England | COVID-19 Genomics UK (COG-UK) Consortium | PHE Covid Sequencing Team |
| EPI_ISL_3291302 | U.O. Microbiologia Laboratorio Unico Centro Servizi - AUSL della Romagna | U.O. Microbiologia, Laboratorio Unico Centro Servizi - AUSL della Romagna | Giorgio Dirani |
| EPI_ISL_3291303 | Respiratory Virus Unit, Microbiology Services Colindale, Public Health England | COVID-19 Genomics UK (COG-UK) Consortium | PHE Covid Sequencing Team |
| EPI_ISL_3291304 | U.O. Microbiologia Laboratorio Unico Centro Servizi - AUSL della Romagna | U.O. Microbiologia, Laboratorio Unico Centro Servizi - AUSL della Romagna | Giorgio Dirani |
| EPI_ISL_3291305, EPI_ISL_3291307 | Respiratory Virus Unit, Microbiology Services Colindale, Public Health England | COVID-19 Genomics UK (COG-UK) Consortium | PHE Covid Sequencing Team |
| EPI_ISL_3291308 | U.O. Microbiologia Laboratorio Unico Centro Servizi - AUSL della Romagna | U.O. Microbiologia, Laboratorio Unico Centro Servizi - AUSL della Romagna | Giorgio Dirani |
| EPI_ISL_3291309 | Respiratory Virus Unit, Microbiology Services Colindale, Public Health England | COVID-19 Genomics UK (COG-UK) Consortium | PHE Covid Sequencing Team |
| EPI_ISL_3291310 | U.O. Microbiologia Laboratorio Unico Centro Servizi - AUSL della Romagna | U.O. Microbiologia, Laboratorio Unico Centro Servizi - AUSL della Romagna | Giorgio Dirani |
| EPI_ISL_3291311, EPI_ISL_3291312, EPI_ISL_3291314 | Respiratory Virus Unit, Microbiology Services Colindale, Public Health England | COVID-19 Genomics UK (COG-UK) Consortium | PHE Covid Sequencing Team |
| EPI_ISL_3291315 | U.O. Microbiologia Laboratorio Unico Centro Servizi - AUSL della Romagna | U.O. Microbiologia, Laboratorio Unico Centro Servizi - AUSL della Romagna | Giorgio Dirani |
| EPI_ISL_3291316 | Respiratory Virus Unit, Microbiology Services Colindale, Public Health England | COVID-19 Genomics UK (COG-UK) Consortium | PHE Covid Sequencing Team |
| EPI_ISL_3291317 | U.O. Microbiologia Laboratorio Unico Centro Servizi - AUSL della Romagna | U.O. Microbiologia, Laboratorio Unico Centro Servizi - AUSL della Romagna | Giorgio Dirani |
| EPI_ISL_3291318, EPI_ISL_3291320 | Respiratory Virus Unit, Microbiology Services Colindale, Public Health England | COVID-19 Genomics UK (COG-UK) Consortium | PHE Covid Sequencing Team |
| EPI_ISL_3291321 | U.O. Microbiologia Laboratorio Unico Centro Servizi - AUSL della Romagna | U.O. Microbiologia, Laboratorio Unico Centro Servizi - AUSL della Romagna | Giorgio Dirani |
| EPI_ISL_3291322 | Respiratory Virus Unit, Microbiology Services Colindale, Public Health England | COVID-19 Genomics UK (COG-UK) Consortium | PHE Covid Sequencing Team |
| EPI_ISL_3291323 | U.O. Microbiologia Laboratorio Unico Centro Servizi - AUSL della Romagna | U.O. Microbiologia, Laboratorio Unico Centro Servizi - AUSL della Romagna | Giorgio Dirani |
| EPI_ISL_3291324 | Respiratory Virus Unit, Microbiology Services Colindale, Public Health England | COVID-19 Genomics UK (COG-UK) Consortium | PHE Covid Sequencing Team |
| EPI_ISL_3291325 | U.O. Microbiologia Laboratorio Unico Centro Servizi - AUSL della Romagna | U.O. Microbiologia, Laboratorio Unico Centro Servizi - AUSL della Romagna | Giorgio Dirani |

EPI_ISL_3291327, EPI_ISL_3291328, EPI_ISL_3291329, EPI_ISL_3291330, EPI_ISL_3291331, EPI_ISL_3291332, EPI_ISL_3291333, EPI_ISL_3291335, EPI_ISL_3291336, EPI_ISL_3291337, EPI_ISL_3291338, EPI_ISL_3291339, EPI_ISL_3291340, EPI_ISL_3291342, EPI_ISL_3291343, EPI_ISL_3291344, EPI_ISL_3291345, EPI_ISL_3291346, EPI_ISL_3291347, EPI_ISL_3291348, EPI_ISL_3291349, EPI_ISL_3291350, EPI_ISL_3291351, EPI_ISL_3291352, EPI_ISL_3291353, EPI_ISL_3291355, EPI_ISL_3291356, EPI_ISL_3291357, EPI_ISL_3291358, EPI_ISL_3291359, EPI_ISL_3291360, EPI_ISL_3291361, EPI_ISL_3291362, EPI_ISL_3291364, EPI_ISL_3291365, EPI_ISL_3291366, EPI_ISL_3291367, EPI_ISL_3291368, EPI_ISL_3291369, EPI_ISL_3291370, EPI_ISL_3291372, EPI_ISL_3291373, EPI_ISL_3291374, EPI_ISL_3291375, EPI_ISL_3291376, EPI_ISL_3291377, EPI_ISL_3291378, EPI_ISL_3291380, EPI_ISL_3291381, EPI_ISL_3291382, EPI_ISL_3291383, EPI_ISL_3291384, EPI_ISL_3291385, EPI_ISL_3291386, EPI_ISL_3291387, EPI_ISL_3291388, EPI_ISL_3291389, EPI_ISL_3291390, EPI_ISL_3291391, EPI_ISL_3291392, EPI_ISL_3291393, EPI_ISL_3291396, EPI_ISL_3291398, EPI_ISL_3291399, EPI_ISL_3291400, EPI_ISL_3291401, EPI_ISL_3291402, EPI_ISL_3291403, EPI_ISL_3291407, EPI_ISL_3291408, EPI_ISL_3291409, EPI_ISL_3291410, EPI_ISL_3291411, EPI_ISL_3291412, EPI_ISL_3291414, EPI_ISL_3291415, EPI_ISL_3291416, EPI_ISL_3291418, EPI_ISL_3291419, EPI_ISL_3291420, EPI_ISL_3291421, EPI_ISL_3291422, EPI_ISL_3291423, EPI_ISL_3291424, EPI_ISL_3291425, EPI_ISL_3291426, EPI_ISL_3291427, EPI_ISL_3291428, EPI_ISL_3291429, EPI_ISL_3291430, EPI_ISL_3291431, EPI_ISL_3291432, EPI_ISL_3291433, EPI_ISL_3291434, EPI_ISL_3291435, EPI_ISL_3291436, EPI_ISL_3291437, EPI_ISL_3291438, EPI_ISL_3291439, EPI_ISL_3291440, EPI_ISL_3291441, EPI_ISL_3291442, EPI_ISL_3291443, EPI_ISL_3291444, EPI_ISL_3291445, EPI_ISL_3291446, EPI_ISL_3291447, EPI_ISL_3291448, EPI_ISL_3291449, EPI_ISL_3291450, EPI_ISL_3291451, EPI_ISL_3291452, EPI_ISL_3291453, EPI_ISL_3291454, EPI_ISL_3291455, EPI_ISL_3291456, EPI_ISL_3291457, EPI_ISL_3291458, EPI_ISL_3291459, EPI_ISL_3291460, EPI_ISL_3291461, EPI_ISL_3291462, EPI_ISL_3291463, EPI_ISL_3291464, EPI_ISL_3291466, EPI_ISL_3291468, EPI_ISL_3291469, EPI_ISL_3291470, EPI_ISL_3291471, EPI_ISL_3291472, EPI_ISL_3291473, EPI_ISL_3291474, EPI_ISL_3291475, EPI_ISL_3291476, EPI_ISL_3291477, EPI_ISL_3291478, EPI_ISL_3291479, EPI_ISL_3291480, EPI_ISL_3291481, EPI_ISL_3291482, EPI_ISL_3291483, EPI_ISL_3291484, EPI_ISL_3291485, EPI_ISL_3291486, EPI_ISL_3291487, EPI_ISL_3291488, EPI_ISL_3291489, EPI_ISL_3291490, EPI_ISL_3291491, EPI_ISL_3291492, EPI_ISL_3291493, EPI_ISL_3291494, EPI_ISL_3291495, EPI_ISL_3291496, EPI_ISL_3291497, EPI_ISL_3291498, EPI_ISL_3291499, EPI_ISL_3291500, EPI_ISL_3291501, EPI_ISL_3291503, EPI_ISL_3291504, EPI_ISL_3291505, EPI_ISL_3291506, EPI_ISL_3291510, EPI_ISL_3291511, EPI_ISL_3291512, EPI_ISL_3291514, EPI_ISL_3291515, EPI_ISL_3291516, EPI_ISL_3291517, EPI_ISL_3291518, EPI_ISL_3291519, EPI_ISL_3291520, EPI_ISL_3291521, EPI_ISL_3291522, EPI_ISL_3291523, EPI_ISL_3291524, EPI_ISL_3291525, EPI_ISL_3291526, EPI_ISL_3291527, EPI_ISL_3291528, EPI_ISL_3291530, EPI_ISL_3291531, EPI_ISL_3291532, EPI_ISL_3291533, EPI_ISL_3291535, EPI_ISL_3291536, EPI_ISL_3291537, EPI_ISL_3291538, EPI_ISL_3291539, EPI_ISL_3291540, EPI_ISL_3291541, EPI_ISL_3291542, EPI_ISL_3291543, EPI_ISL_3291544, EPI_ISL_3291546, EPI_ISL_3291547, EPI_ISL_3291548, EPI_ISL_3291549, EPI_ISL_3291550, EPI_ISL_3291552, EPI_ISL_3291553, EPI_ISL_3291554, EPI_ISL_3291555, EPI_ISL_3291556, EPI_ISL_3291557, EPI_ISL_3291558, EPI_ISL_3291559, EPI_ISL_3291560, EPI_ISL_3291561, EPI_ISL_3291562, EPI_ISL_3291563, EPI_ISL_3291564, EPI_ISL_3291565, EPI_ISL_3291566, EPI_ISL_3291567, EPI_ISL_3291568, EPI_ISL_3291569, EPI_ISL_3291570, EPI_ISL_3291571, EPI_ISL_3291572, EPI_ISL_3291573, EPI_ISL_3291574, EPI_ISL_3291575, EPI_ISL_3291576, EPI_ISL_3291577, EPI_ISL_3291578, EPI_ISL_3291579, EPI_ISL_3291580, EPI_ISL_3291581, EPI_ISL_3291582, EPI_ISL_3291583, EPI_ISL_3291584, EPI_ISL_3291585, EPI_ISL_3291586, EPI_ISL_3291587, EPI_ISL_3291588, EPI_ISL_3291589, EPI_ISL_3291590, EPI_ISL_3291591, EPI_ISL_3291592, EPI_ISL_3291593, EPI_ISL_3291594, EPI_ISL_3291595, EPI_ISL_3291597, EPI_ISL_3291598, EPI_ISL_3291599, EPI_ISL_3291600, EPI_ISL_3291601, EPI_ISL_3291602, EPI_ISL_3291603, EPI_ISL_3291604, EPI_ISL_3291605, EPI_ISL_3291606, EPI_ISL_3291607, EPI_ISL_3291608, EPI_ISL_3291609, EPI_ISL_3291610, EPI_ISL_3291611, EPI_ISL_3291612, EPI_ISL_3291613, EPI_ISL_3291614, EPI_ISL_3291615, EPI_ISL_3291616, EPI_ISL_3291617, EPI_ISL_3291618, EPI_ISL_3291619, EPI_ISL_3291620, EPI_ISL_3291621, EPI_ISL_3291622, EPI_ISL_3291623, EPI_ISL_3291624, EPI_ISL_3291625, EPI_ISL_3291626, EPI_ISL_3291627, EPI_ISL_3291628, EPI_ISL_3291629, EPI_ISL_3291630, EPI_ISL_3291631, EPI_ISL_3291632, EPI_ISL_3291633, EPI_ISL_3291634, EPI_ISL_3291635, EPI_ISL_3291636, EPI_ISL_3291637, EPI_ISL_3291638, EPI_ISL_3291639, EPI_ISL_3291640, EPI_ISL_3291641, EPI_ISL_3291642, EPI_ISL_3291643, EPI_ISL_3291644, EPI_ISL_3291645, EPI_ISL_3291647, EPI_ISL_3291648, EPI_ISL_3291649, EPI_ISL_3291650, EPI_ISL_3291651, EPI_ISL_3291652, EPI_ISL_3291653, EPI_ISL_3291654, EPI_ISL_3291655, EPI_ISL_3291656, EPI_ISL_3291657, EPI_ISL_3291658, EPI_ISL_3291659, EPI_ISL_3291660, EPI_ISL_3291661, EPI_ISL_3291662, EPI_ISL_3291663, EPI_ISL_3291664, EPI_ISL_3291665, EPI_ISL_3291666, EPI_ISL_3291667, EPI_ISL_3291668, EPI_ISL_3291669, EPI_ISL_3291670, EPI_ISL_3291671, EPI_ISL_3291672, EPI_ISL_3291673, EPI_ISL_3291674, EPI_ISL_3291675, EPI_ISL_3291676, EPI_ISL_3291677, EPI_ISL_3291678, EPI_ISL_3291679, EPI_ISL_3291680, EPI_ISL_3291681, EPI_ISL_3291682, EPI_ISL_3291683, EPI_ISL_3291684, EPI_ISL_3291685, EPI_ISL_3291686, EPI_ISL_3291687, EPI_ISL_3291689, EPI_ISL_3291690, EPI_ISL_3291691, EPI_ISL_3291692, EPI_ISL_3291693, EPI_ISL_3291694, EPI_ISL_3291695, EPI_ISL_3291696, EPI_ISL_3291698, EPI_ISL_3291699, EPI_ISL_3291700, EPI_ISL_3291701, EPI_ISL_3291702, EPI_ISL_3291703, EPI_ISL_3291704, EPI_ISL_3291705, EPI_ISL_3291706, EPI_ISL_3291707, EPI_ISL_3291708, EPI_ISL_3291709, EPI_ISL_3291710, EPI_ISL_3291711, EPI_ISL_3291712, EPI_ISL_3291713, EPI_ISL_3291714, EPI_ISL_3291715, EPI_ISL_3291716, EPI_ISL_3291717, EPI_ISL_3291718, EPI_ISL_3291719, EPI_ISL_3291720, EPI_ISL_3291721, EPI_ISL_3291722, EPI_ISL_3291723, EPI_ISL_3291724, EPI_ISL_3291725, EPI_ISL_3291726, EPI_ISL_3291727, EPI_ISL_3291728, EPI_ISL_3291729, EPI_ISL_3291730, EPI_ISL_3291731, EPI_ISL_3291732, EPI_ISL_3291733, EPI_ISL_3291734, EPI_ISL_3291735, EPI_ISL_3291737, EPI_ISL_3291738, EPI_ISL_3291739, EPI_ISL_3291740, EPI_ISL_3291741, EPI_ISL_3291742, EPI_ISL_3291743, EPI_ISL_3291744, EPI_ISL_3291745, EPI_ISL_3291746, EPI_ISL_3291747, EPI_ISL_3291748, EPI_ISL_3291749, EPI_ISL_3291750, EPI_ISL_3291751, EPI_ISL_3291752, EPI_ISL_3291754, EPI_ISL_3291755, EPI_ISL_3291756, EPI_ISL_3291758, EPI_ISL_3291759, EPI_ISL_3291760, EPI_ISL_3291761, EPI_ISL_3291762, EPI_ISL_3291763, EPI_ISL_3291764, EPI_ISL_3291765, EPI_ISL_3291766, EPI_ISL_3291767, EPI_ISL_3291768, EPI_ISL_3291769, EPI_ISL_3291771, EPI_ISL_3291773, EPI_ISL_3291774, EPI_ISL_3291775, EPI_ISL_3291776, EPI_ISL_3291777, EPI_ISL_3291778, EPI_ISL_3291779, EPI_ISL_3291780, EPI_ISL_3291781, EPI_ISL_3291782, EPI_ISL_3291783, EPI_ISL_3291784, EPI_ISL_3291785, EPI_ISL_3291786, EPI_ISL_3291787, EPI_ISL_3291788, EPI_ISL_3291789, EPI_ISL_3291790, EPI_ISL_3291791, EPI_ISL_3291792, EPI_ISL_3291793, EPI_ISL_3291794, EPI_ISL_3291795, EPI_ISL_3291796, EPI_ISL_3291797, EPI_ISL_3291798, EPI_ISL_3291799, EPI_ISL_3291800, EPI_ISL_3291801, EPI_ISL_3291802, EPI_ISL_3291803, EPI_ISL_3291804, EPI_ISL_3291805, EPI_ISL_3291806, EPI_ISL_3291808, EPI_ISL_3291809, EPI_ISL_3291810, EPI_ISL_3291811, EPI_ISL_3291812, EPI_ISL_3291813, EPI_ISL_3291814, EPI_ISL_3291815, EPI_ISL_3291816, EPI_ISL_3291817, EPI_ISL_3291818, EPI_ISL_3291819, EPI_ISL_3291820, EPI_ISL_3291821, EPI_ISL_3291822, EPI_ISL_3291823, EPI_ISL_3291824, EPI_ISL_3291825, EPI_ISL_3291826

| | | | |
|---|---|---|---|
| see above | Respiratory Virus Unit, Microbiology Services Colindale, Public Health England | COVID-19 Genomics UK (COG-UK) Consortium | PHE Covid Sequencing Team |
| EPI_ISL_3291827 | U.O. Microbiologia Laboratorio Unico Centro Servizi - AUSL della Romagna | U.O. Microbiologia, Laboratorio Unico Centro Servizi - AUSL della Romagna | Giorgio Dirani |
| EPI_ISL_3291828 | Respiratory Virus Unit, Microbiology Services Colindale, Public Health England | COVID-19 Genomics UK (COG-UK) Consortium | PHE Covid Sequencing Team |
| EPI_ISL_3291829 | U.O. Microbiologia Laboratorio Unico Centro Servizi - AUSL della Romagna | U.O. Microbiologia, Laboratorio Unico Centro Servizi - AUSL della Romagna | Giorgio Dirani |
| EPI_ISL_3291830 | Respiratory Virus Unit, Microbiology Services Colindale, Public Health England | COVID-19 Genomics UK (COG-UK) Consortium | PHE Covid Sequencing Team |
| EPI_ISL_3291831 | U.O. Microbiologia Laboratorio Unico Centro Servizi - AUSL della Romagna | U.O. Microbiologia, Laboratorio Unico Centro Servizi - AUSL della Romagna | Giorgio Dirani |
| EPI_ISL_3291832 | Respiratory Virus Unit, Microbiology Services Colindale, Public Health England | COVID-19 Genomics UK (COG-UK) Consortium | PHE Covid Sequencing Team |
| EPI_ISL_3291833 | U.O. Microbiologia Laboratorio Unico Centro Servizi - AUSL della Romagna | U.O. Microbiologia, Laboratorio Unico Centro Servizi - AUSL della Romagna | Giorgio Dirani |
| EPI_ISL_3291834 | Respiratory Virus Unit, Microbiology Services Colindale, Public | COVID-19 Genomics UK (COG-UK) Consortium | PHE Covid Sequencing Team |

| | | | |
|---|---|---|---|
| | Health England | | |
| EPI_ISL_3291835 | U.O. Microbiologia Laboratorio Unico Centro Servizi - AUSL della Romagna | U.O. Microbiologia, Laboratorio Unico Centro Servizi - AUSL della Romagna | Giorgio Dirani |
| EPI_ISL_3291836 | Respiratory Virus Unit, Microbiology Services Colindale, Public Health England | COVID-19 Genomics UK (COG-UK) Consortium | PHE Covid Sequencing Team |
| EPI_ISL_3291837 | U.O. Microbiologia Laboratorio Unico Centro Servizi - AUSL della Romagna | U.O. Microbiologia, Laboratorio Unico Centro Servizi - AUSL della Romagna | Giorgio Dirani |
| EPI_ISL_3291838, EPI_ISL_3291839 | Respiratory Virus Unit, Microbiology Services Colindale, Public Health England | COVID-19 Genomics UK (COG-UK) Consortium | PHE Covid Sequencing Team |
| EPI_ISL_3291840, EPI_ISL_3291842 | U.O. Microbiologia Laboratorio Unico Centro Servizi - AUSL della Romagna | U.O. Microbiologia, Laboratorio Unico Centro Servizi - AUSL della Romagna | Giorgio Dirani |
| EPI_ISL_3291843 | Respiratory Virus Unit, Microbiology Services Colindale, Public Health England | COVID-19 Genomics UK (COG-UK) Consortium | PHE Covid Sequencing Team |
| EPI_ISL_3291844 | U.O. Microbiologia Laboratorio Unico Centro Servizi - AUSL della Romagna | U.O. Microbiologia, Laboratorio Unico Centro Servizi - AUSL della Romagna | Giorgio Dirani |
| EPI_ISL_3291845 | Respiratory Virus Unit, Microbiology Services Colindale, Public Health England | COVID-19 Genomics UK (COG-UK) Consortium | PHE Covid Sequencing Team |
| EPI_ISL_3291846 | U.O. Microbiologia Laboratorio Unico Centro Servizi - AUSL della Romagna | U.O. Microbiologia, Laboratorio Unico Centro Servizi - AUSL della Romagna | Giorgio Dirani |
| EPI_ISL_3291847 | Respiratory Virus Unit, Microbiology Services Colindale, Public Health England | COVID-19 Genomics UK (COG-UK) Consortium | PHE Covid Sequencing Team |
| EPI_ISL_3291848 | U.O. Microbiologia Laboratorio Unico Centro Servizi - AUSL della Romagna | U.O. Microbiologia, Laboratorio Unico Centro Servizi - AUSL della Romagna | Giorgio Dirani |
| EPI_ISL_3291849 | Respiratory Virus Unit, Microbiology Services Colindale, Public Health England | COVID-19 Genomics UK (COG-UK) Consortium | PHE Covid Sequencing Team |
| EPI_ISL_3291850, EPI_ISL_3291852 | U.O. Microbiologia Laboratorio Unico Centro Servizi - AUSL della Romagna | U.O. Microbiologia, Laboratorio Unico Centro Servizi - AUSL della Romagna | Giorgio Dirani |
| EPI_ISL_3291853 | Respiratory Virus Unit, Microbiology Services Colindale, Public Health England | COVID-19 Genomics UK (COG-UK) Consortium | PHE Covid Sequencing Team |
| EPI_ISL_3291854 | U.O. Microbiologia Laboratorio Unico Centro Servizi - AUSL della Romagna | U.O. Microbiologia, Laboratorio Unico Centro Servizi - AUSL della Romagna | Giorgio Dirani |
| EPI_ISL_3291855 | Respiratory Virus Unit, Microbiology Services Colindale, Public Health England | COVID-19 Genomics UK (COG-UK) Consortium | PHE Covid Sequencing Team |
| EPI_ISL_3291856 | U.O. Microbiologia Laboratorio Unico Centro Servizi - AUSL della Romagna | U.O. Microbiologia, Laboratorio Unico Centro Servizi - AUSL della Romagna | Giorgio Dirani |
| EPI_ISL_3291857 | Respiratory Virus Unit, Microbiology Services Colindale, Public Health England | COVID-19 Genomics UK (COG-UK) Consortium | PHE Covid Sequencing Team |
| EPI_ISL_3291858 | U.O. Microbiologia Laboratorio Unico Centro Servizi - AUSL della Romagna | U.O. Microbiologia, Laboratorio Unico Centro Servizi - AUSL della Romagna | Giorgio Dirani |
| EPI_ISL_3291859 | Respiratory Virus Unit, Microbiology Services Colindale, Public Health England | COVID-19 Genomics UK (COG-UK) Consortium | PHE Covid Sequencing Team |
| EPI_ISL_3291860 | U.O. Microbiologia Laboratorio Unico Centro Servizi - AUSL della Romagna | U.O. Microbiologia, Laboratorio Unico Centro Servizi - AUSL della Romagna | Giorgio Dirani |
| EPI_ISL_3291861 | Respiratory Virus Unit, Microbiology Services Colindale, Public Health England | COVID-19 Genomics UK (COG-UK) Consortium | PHE Covid Sequencing Team |
| EPI_ISL_3291862 | U.O. Microbiologia Laboratorio Unico Centro Servizi - AUSL della Romagna | U.O. Microbiologia, Laboratorio Unico Centro Servizi - AUSL della Romagna | Giorgio Dirani |
| EPI_ISL_3291863 | Respiratory Virus Unit, Microbiology Services Colindale, Public Health England | COVID-19 Genomics UK (COG-UK) Consortium | PHE Covid Sequencing Team |
| EPI_ISL_3291864 | U.O. Microbiologia Laboratorio Unico Centro Servizi - AUSL della Romagna | U.O. Microbiologia, Laboratorio Unico Centro Servizi - AUSL della Romagna | Giorgio Dirani |
| EPI_ISL_3291865 | Respiratory Virus Unit, Microbiology Services Colindale, Public Health England | COVID-19 Genomics UK (COG-UK) Consortium | PHE Covid Sequencing Team |
| EPI_ISL_3291866 | U.O. Microbiologia Laboratorio Unico Centro Servizi - AUSL della Romagna | U.O. Microbiologia, Laboratorio Unico Centro Servizi - AUSL della Romagna | Giorgio Dirani |
| EPI_ISL_3291867 | Respiratory Virus Unit, Microbiology Services Colindale, Public Health England | COVID-19 Genomics UK (COG-UK) Consortium | PHE Covid Sequencing Team |
| EPI_ISL_3291868 | U.O. Microbiologia Laboratorio Unico Centro Servizi - AUSL della Romagna | U.O. Microbiologia, Laboratorio Unico Centro Servizi - AUSL della Romagna | Giorgio Dirani |
| EPI_ISL_3291869 | Respiratory Virus Unit, Microbiology Services Colindale, Public Health England | COVID-19 Genomics UK (COG-UK) Consortium | PHE Covid Sequencing Team |
| EPI_ISL_3291870 | U.O. Microbiologia Laboratorio Unico Centro Servizi - AUSL della Romagna | U.O. Microbiologia, Laboratorio Unico Centro Servizi - AUSL della Romagna | Giorgio Dirani |
| EPI_ISL_3291871 | Respiratory Virus Unit, Microbiology Services Colindale, Public Health England | COVID-19 Genomics UK (COG-UK) Consortium | PHE Covid Sequencing Team |
| EPI_ISL_3291872 | U.O. Microbiologia Laboratorio Unico Centro Servizi - AUSL della Romagna | U.O. Microbiologia, Laboratorio Unico Centro Servizi - AUSL della Romagna | Giorgio Dirani |
| EPI_ISL_3291873 | Respiratory Virus Unit, Microbiology Services Colindale, Public Health England | COVID-19 Genomics UK (COG-UK) Consortium | PHE Covid Sequencing Team |
| EPI_ISL_3291874 | U.O. Microbiologia Laboratorio Unico Centro Servizi - AUSL della Romagna | U.O. Microbiologia, Laboratorio Unico Centro Servizi - AUSL della Romagna | Giorgio Dirani |
| EPI_ISL_3291875 | Respiratory Virus Unit, Microbiology Services Colindale, Public Health England | COVID-19 Genomics UK (COG-UK) Consortium | PHE Covid Sequencing Team |
| EPI_ISL_3291876, EPI_ISL_3291877 | U.O. Microbiologia Laboratorio Unico Centro Servizi - AUSL della Romagna | U.O. Microbiologia, Laboratorio Unico Centro Servizi - AUSL della Romagna | Giorgio Dirani |
| EPI_ISL_3291878 | Respiratory Virus Unit, Microbiology Services Colindale, Public Health England | COVID-19 Genomics UK (COG-UK) Consortium | PHE Covid Sequencing Team |
| EPI_ISL_3291879 | U.O. Microbiologia Laboratorio Unico Centro Servizi - AUSL della Romagna | U.O. Microbiologia, Laboratorio Unico Centro Servizi - AUSL della Romagna | Giorgio Dirani |
| EPI_ISL_3291880 | Respiratory Virus Unit, Microbiology Services Colindale, Public Health England | COVID-19 Genomics UK (COG-UK) Consortium | PHE Covid Sequencing Team |
| EPI_ISL_3291881 | U.O. Microbiologia Laboratorio Unico Centro Servizi - AUSL della Romagna | U.O. Microbiologia, Laboratorio Unico Centro Servizi - AUSL della Romagna | Giorgio Dirani |
| EPI_ISL_3291882, EPI_ISL_3291883 | Respiratory Virus Unit, Microbiology Services Colindale, Public Health England | COVID-19 Genomics UK (COG-UK) Consortium | PHE Covid Sequencing Team |
| EPI_ISL_3291884 | U.O. Microbiologia Laboratorio Unico Centro Servizi - AUSL della Romagna | U.O. Microbiologia, Laboratorio Unico Centro Servizi - AUSL della Romagna | Giorgio Dirani |
| EPI_ISL_3291885 | Respiratory Virus Unit, Microbiology Services Colindale, Public Health England | COVID-19 Genomics UK (COG-UK) Consortium | PHE Covid Sequencing Team |
| EPI_ISL_3291886, | U.O. Microbiologia Laboratorio Unico Centro Servizi - AUSL della | U.O. Microbiologia, Laboratorio Unico Centro Servizi - AUSL della | Giorgio Dirani |

| | | | |
|---|---|---|---|
| | Romagna | Romagna | |
| EPI_ISL_3291888 | Respiratory Virus Unit, Microbiology Services Colindale, Public Health England | COVID-19 Genomics UK (COG-UK) Consortium | PHE Covid Sequencing Team |
| EPI_ISL_3291889 | U.O. Microbiologia Laboratorio Unico Centro Servizi - AUSL della Romagna | U.O. Microbiologia, Laboratorio Unico Centro Servizi - AUSL della Romagna | Giorgio Dirani |
| EPI_ISL_3291890, EPI_ISL_3291891 | Respiratory Virus Unit, Microbiology Services Colindale, Public Health England | COVID-19 Genomics UK (COG-UK) Consortium | PHE Covid Sequencing Team |
| EPI_ISL_3291892 | U.O. Microbiologia Laboratorio Unico Centro Servizi - AUSL della Romagna | U.O. Microbiologia, Laboratorio Unico Centro Servizi - AUSL della Romagna | Giorgio Dirani |
| EPI_ISL_3291893 | Respiratory Virus Unit, Microbiology Services Colindale, Public Health England | COVID-19 Genomics UK (COG-UK) Consortium | PHE Covid Sequencing Team |
| EPI_ISL_3291894 | U.O. Microbiologia Laboratorio Unico Centro Servizi - AUSL della Romagna | U.O. Microbiologia, Laboratorio Unico Centro Servizi - AUSL della Romagna | Giorgio Dirani |
| EPI_ISL_3291895 | Respiratory Virus Unit, Microbiology Services Colindale, Public Health England | COVID-19 Genomics UK (COG-UK) Consortium | PHE Covid Sequencing Team |
| EPI_ISL_3291896 | U.O. Microbiologia Laboratorio Unico Centro Servizi - AUSL della Romagna | U.O. Microbiologia, Laboratorio Unico Centro Servizi - AUSL della Romagna | Giorgio Dirani |
| EPI_ISL_3291897 | Respiratory Virus Unit, Microbiology Services Colindale, Public Health England | COVID-19 Genomics UK (COG-UK) Consortium | PHE Covid Sequencing Team |
| EPI_ISL_3291898, EPI_ISL_3291899 | U.O. Microbiologia Laboratorio Unico Centro Servizi - AUSL della Romagna | U.O. Microbiologia, Laboratorio Unico Centro Servizi - AUSL della Romagna | Giorgio Dirani |
| EPI_ISL_3291900 | Respiratory Virus Unit, Microbiology Services Colindale, Public Health England | COVID-19 Genomics UK (COG-UK) Consortium | PHE Covid Sequencing Team |
| EPI_ISL_3291901 | U.O. Microbiologia Laboratorio Unico Centro Servizi - AUSL della Romagna | U.O. Microbiologia, Laboratorio Unico Centro Servizi - AUSL della Romagna | Giorgio Dirani |
| EPI_ISL_3291902 | Respiratory Virus Unit, Microbiology Services Colindale, Public Health England | COVID-19 Genomics UK (COG-UK) Consortium | PHE Covid Sequencing Team |
| EPI_ISL_3291903 | U.O. Microbiologia Laboratorio Unico Centro Servizi - AUSL della Romagna | U.O. Microbiologia, Laboratorio Unico Centro Servizi - AUSL della Romagna | Giorgio Dirani |

EPI_ISL_3291904, EPI_ISL_3291905, EPI_ISL_3291907, EPI_ISL_3291908, EPI_ISL_3291909, EPI_ISL_3291910, EPI_ISL_3291911, EPI_ISL_3291912, EPI_ISL_3291913, EPI_ISL_3291914, EPI_ISL_3291915, EPI_ISL_3291916, EPI_ISL_3291917, EPI_ISL_3291918, EPI_ISL_3291919, EPI_ISL_3291920, EPI_ISL_3291921, EPI_ISL_3291922, EPI_ISL_3291923, EPI_ISL_3291924, EPI_ISL_3291925, EPI_ISL_3291926, EPI_ISL_3291927, EPI_ISL_3291928, EPI_ISL_3291929, EPI_ISL_3291930, EPI_ISL_3291931, EPI_ISL_3291932, EPI_ISL_3291933, EPI_ISL_3291934, EPI_ISL_3291935, EPI_ISL_3291936, EPI_ISL_3291938, EPI_ISL_3291939, EPI_ISL_3291940, EPI_ISL_3291941, EPI_ISL_3291942, EPI_ISL_3291943, EPI_ISL_3291944, EPI_ISL_3291946, EPI_ISL_3291947, EPI_ISL_3291948, EPI_ISL_3291950, EPI_ISL_3291951, EPI_ISL_3291952, EPI_ISL_3291953, EPI_ISL_3291954, EPI_ISL_3291955, EPI_ISL_3291956, EPI_ISL_3291957, EPI_ISL_3291958, EPI_ISL_3291959, EPI_ISL_3291960, EPI_ISL_3291961, EPI_ISL_3291962, EPI_ISL_3291963, EPI_ISL_3291964, EPI_ISL_3291965, EPI_ISL_3291966, EPI_ISL_3291967, EPI_ISL_3291968, EPI_ISL_3291969, EPI_ISL_3291970, EPI_ISL_3291971, EPI_ISL_3291972, EPI_ISL_3291973, EPI_ISL_3291974, EPI_ISL_3291975, EPI_ISL_3291976, EPI_ISL_3291977, EPI_ISL_3291978, EPI_ISL_3291979, EPI_ISL_3291980, EPI_ISL_3291981, EPI_ISL_3291982, EPI_ISL_3291983, EPI_ISL_3291984, EPI_ISL_3291985, EPI_ISL_3291986, EPI_ISL_3291987, EPI_ISL_3291988, EPI_ISL_3291989, EPI_ISL_3291990, EPI_ISL_3291991, EPI_ISL_3291992, EPI_ISL_3291993, EPI_ISL_3291994, EPI_ISL_3291995, EPI_ISL_3291996, EPI_ISL_3291997, EPI_ISL_3291998, EPI_ISL_3291999, EPI_ISL_3292000, EPI_ISL_3292001, EPI_ISL_3292003, EPI_ISL_3292004, EPI_ISL_3292005, EPI_ISL_3292006, EPI_ISL_3292007, EPI_ISL_3292008, EPI_ISL_3292009, EPI_ISL_3292010, EPI_ISL_3292011, EPI_ISL_3292012, EPI_ISL_3292013, EPI_ISL_3292014, EPI_ISL_3292015, EPI_ISL_3292016, EPI_ISL_3292017, EPI_ISL_3292018, EPI_ISL_3292019, EPI_ISL_3292020, EPI_ISL_3292021, EPI_ISL_3292022, EPI_ISL_3292023, EPI_ISL_3292024, EPI_ISL_3292025, EPI_ISL_3292026, EPI_ISL_3292027, EPI_ISL_3292028, EPI_ISL_3292029, EPI_ISL_3292030, EPI_ISL_3292031, EPI_ISL_3292032, EPI_ISL_3292033, EPI_ISL_3292034, EPI_ISL_3292035, EPI_ISL_3292036, EPI_ISL_3292037, EPI_ISL_3292038, EPI_ISL_3292039, EPI_ISL_3292040, EPI_ISL_3292041, EPI_ISL_3292042, EPI_ISL_3292043, EPI_ISL_3292044, EPI_ISL_3292045, EPI_ISL_3292046, EPI_ISL_3292047, EPI_ISL_3292048, EPI_ISL_3292049, EPI_ISL_3292050, EPI_ISL_3292051, EPI_ISL_3292053, EPI_ISL_3292054, EPI_ISL_3292055, EPI_ISL_3292056, EPI_ISL_3292057, EPI_ISL_3292058, EPI_ISL_3292059, EPI_ISL_3292060, EPI_ISL_3292061, EPI_ISL_3292062, EPI_ISL_3292063, EPI_ISL_3292064, EPI_ISL_3292065, EPI_ISL_3292066, EPI_ISL_3292068, EPI_ISL_3292069, EPI_ISL_3292070, EPI_ISL_3292071, EPI_ISL_3292072, EPI_ISL_3292073, EPI_ISL_3292074, EPI_ISL_3292075, EPI_ISL_3292076, EPI_ISL_3292077, EPI_ISL_3292078, EPI_ISL_3292079, EPI_ISL_3292080, EPI_ISL_3292082, EPI_ISL_3292083, EPI_ISL_3292084, EPI_ISL_3292085, EPI_ISL_3292086, EPI_ISL_3292087, EPI_ISL_3292088, EPI_ISL_3292089, EPI_ISL_3292090, EPI_ISL_3292091, EPI_ISL_3292092, EPI_ISL_3292093, EPI_ISL_3292094, EPI_ISL_3292095, EPI_ISL_3292096, EPI_ISL_3292097, EPI_ISL_3292098, EPI_ISL_3292099, EPI_ISL_3292100, EPI_ISL_3292101, EPI_ISL_3292102, EPI_ISL_3292103, EPI_ISL_3292104, EPI_ISL_3292105, EPI_ISL_3292106, EPI_ISL_3292107, EPI_ISL_3292108, EPI_ISL_3292109, EPI_ISL_3292110, EPI_ISL_3292111, EPI_ISL_3292113, EPI_ISL_3292114, EPI_ISL_3292115, EPI_ISL_3292116, EPI_ISL_3292117, EPI_ISL_3292118, EPI_ISL_3292119, EPI_ISL_3292120, EPI_ISL_3292121, EPI_ISL_3292122, EPI_ISL_3292123, EPI_ISL_3292124, EPI_ISL_3292125, EPI_ISL_3292126, EPI_ISL_3292127, EPI_ISL_3292128, EPI_ISL_3292129, EPI_ISL_3292130, EPI_ISL_3292131, EPI_ISL_3292132, EPI_ISL_3292133, EPI_ISL_3292134, EPI_ISL_3292135, EPI_ISL_3292136, EPI_ISL_3292137, EPI_ISL_3292139, EPI_ISL_3292140, EPI_ISL_3292141, EPI_ISL_3292142, EPI_ISL_3292143, EPI_ISL_3292144, EPI_ISL_3292145, EPI_ISL_3292146, EPI_ISL_3292147, EPI_ISL_3292148, EPI_ISL_3292149, EPI_ISL_3292150, EPI_ISL_3292151, EPI_ISL_3292152, EPI_ISL_3292153, EPI_ISL_3292154, EPI_ISL_3292155, EPI_ISL_3292156, EPI_ISL_3292157, EPI_ISL_3292158, EPI_ISL_3292159, EPI_ISL_3292160, EPI_ISL_3292161, EPI_ISL_3292162, EPI_ISL_3292163, EPI_ISL_3292164, EPI_ISL_3292165, EPI_ISL_3292166, EPI_ISL_3292167, EPI_ISL_3292168, EPI_ISL_3292169, EPI_ISL_3292170, EPI_ISL_3292171, EPI_ISL_3292172, EPI_ISL_3292173, EPI_ISL_3292174, EPI_ISL_3292175, EPI_ISL_3292176, EPI_ISL_3292177, EPI_ISL_3292178, EPI_ISL_3292179, EPI_ISL_3292180, EPI_ISL_3292181, EPI_ISL_3292182, EPI_ISL_3292183, EPI_ISL_3292184, EPI_ISL_3292185, EPI_ISL_3292186, EPI_ISL_3292187, EPI_ISL_3292188, EPI_ISL_3292189, EPI_ISL_3292190, EPI_ISL_3292191, EPI_ISL_3292192, EPI_ISL_3292193, EPI_ISL_3292194, EPI_ISL_3292195, EPI_ISL_3292196, EPI_ISL_3292197, EPI_ISL_3292198, EPI_ISL_3292199, EPI_ISL_3292200, EPI_ISL_3292201, EPI_ISL_3292202, EPI_ISL_3292203, EPI_ISL_3292204, EPI_ISL_3292205, EPI_ISL_3292206, EPI_ISL_3292207, EPI_ISL_3292208, EPI_ISL_3292209, EPI_ISL_3292210, EPI_ISL_3292211, EPI_ISL_3292212, EPI_ISL_3292213, EPI_ISL_3292214, EPI_ISL_3292216, EPI_ISL_3292217, EPI_ISL_3292218, EPI_ISL_3292219, EPI_ISL_3292220, EPI_ISL_3292221, EPI_ISL_3292222, EPI_ISL_3292223, EPI_ISL_3292224, EPI_ISL_3292225, EPI_ISL_3292226, EPI_ISL_3292227, EPI_ISL_3292228, EPI_ISL_3292229, EPI_ISL_3292230, EPI_ISL_3292231, EPI_ISL_3292232, EPI_ISL_3292233, EPI_ISL_3292234, EPI_ISL_3292235, EPI_ISL_3292236, EPI_ISL_3292237, EPI_ISL_3292238, EPI_ISL_3292239, EPI_ISL_3292240, EPI_ISL_3292241, EPI_ISL_3292242, EPI_ISL_3292243, EPI_ISL_3292244, EPI_ISL_3292245, EPI_ISL_3292246, EPI_ISL_3292247, EPI_ISL_3292248, EPI_ISL_3292249, EPI_ISL_3292250, EPI_ISL_3292251, EPI_ISL_3292252, EPI_ISL_3292253, EPI_ISL_3292254, EPI_ISL_3292255, EPI_ISL_3292256, EPI_ISL_3292257, EPI_ISL_3292258, EPI_ISL_3292259, EPI_ISL_3292260, EPI_ISL_3292261, EPI_ISL_3292262, EPI_ISL_3292263, EPI_ISL_3292264, EPI_ISL_3292265, EPI_ISL_3292266, EPI_ISL_3292267, EPI_ISL_3292268, EPI_ISL_3292269, EPI_ISL_3292270, EPI_ISL_3292271, EPI_ISL_3292272, EPI_ISL_3292273, EPI_ISL_3292274, EPI_ISL_3292275, EPI_ISL_3292276, EPI_ISL_3292277, EPI_ISL_3292278, EPI_ISL_3292279, EPI_ISL_3292280, EPI_ISL_3292281, EPI_ISL_3292282, EPI_ISL_3292283, EPI_ISL_3292284, EPI_ISL_3292285, EPI_ISL_3292286, EPI_ISL_3292287, EPI_ISL_3292289, EPI_ISL_3292290, EPI_ISL_3292291, EPI_ISL_3292292, EPI_ISL_3292293, EPI_ISL_3292294, EPI_ISL_3292295, EPI_ISL_3292296, EPI_ISL_3292297, EPI_ISL_3292298, EPI_ISL_3292299, EPI_ISL_3292300, EPI_ISL_3292301, EPI_ISL_3292302, EPI_ISL_3292303, EPI_ISL_3292304, EPI_ISL_3292305, EPI_ISL_3292306, EPI_ISL_3292307, EPI_ISL_3292308, EPI_ISL_3292309, EPI_ISL_3292310, EPI_ISL_3292311, EPI_ISL_3292312, EPI_ISL_3292313, EPI_ISL_3292314, EPI_ISL_3292315, EPI_ISL_3292316, EPI_ISL_3292317, EPI_ISL_3292318, EPI_ISL_3292320, EPI_ISL_3292321, EPI_ISL_3292322, EPI_ISL_3292323, EPI_ISL_3292324, EPI_ISL_3292325, EPI_ISL_3292326, EPI_ISL_3292327, EPI_ISL_3292328, EPI_ISL_3292329, EPI_ISL_3292330, EPI_ISL_3292331, EPI_ISL_3292332, EPI_ISL_3292333, EPI_ISL_3292334, EPI_ISL_3292335, EPI_ISL_3292336, EPI_ISL_3292338, EPI_ISL_3292339, EPI_ISL_3292340, EPI_ISL_3292341, EPI_ISL_3292342, EPI_ISL_3292343, EPI_ISL_3292344, EPI_ISL_3292345, EPI_ISL_3292346, EPI_ISL_3292347, EPI_ISL_3292348, EPI_ISL_3292349, EPI_ISL_3292350, EPI_ISL_3292351, EPI_ISL_3292352, EPI_ISL_3292353, EPI_ISL_3292354, EPI_ISL_3292355, EPI_ISL_3292356, EPI_ISL_3292357, EPI_ISL_3292358, EPI_ISL_3292359, EPI_ISL_3292360, EPI_ISL_3292361, EPI_ISL_3292362, EPI_ISL_3292363, EPI_ISL_3292364, EPI_ISL_3292365, EPI_ISL_3292366, EPI_ISL_3292367, EPI_ISL_3292368, EPI_ISL_3292369, EPI_ISL_3292370, EPI_ISL_3292371, EPI_ISL_3292372, EPI_ISL_3292373, EPI_ISL_3292375, EPI_ISL_3292376, EPI_ISL_3292377, EPI_ISL_3292378, EPI_ISL_3292379, EPI_ISL_3292380, EPI_ISL_3292381, EPI_ISL_3292382, EPI_ISL_3292383, EPI_ISL_3292384, EPI_ISL_3292385, EPI_ISL_3292386, EPI_ISL_3292387, EPI_ISL_3292388, EPI_ISL_3292389, EPI_ISL_3292391, EPI_ISL_3292392, EPI_ISL_3292393, EPI_ISL_3292394, EPI_ISL_3292395, EPI_ISL_3292396, EPI_ISL_3292397, EPI_ISL_3292399, EPI_ISL_3292400, EPI_ISL_3292401, EPI_ISL_3292402, EPI_ISL_3292403, EPI_ISL_3292404, EPI_ISL_3292405, EPI_ISL_3292406, EPI_ISL_3292407, EPI_ISL_3292408, EPI_ISL_3292409, EPI_ISL_3292410, EPI_ISL_3292411, EPI_ISL_3292412, EPI_ISL_3292413, EPI_ISL_3292414, EPI_ISL_3292415, EPI_ISL_3292416, EPI_ISL_3292417, EPI_ISL_3292419, EPI_ISL_3292420, EPI_ISL_3292421, EPI_ISL_3292422, EPI_ISL_3292423, EPI_ISL_3292424, EPI_ISL_3292425, EPI_ISL_3292426, EPI_ISL_3292427, EPI_ISL_3292428, EPI_ISL_3292429, EPI_ISL_3292430, EPI_ISL_3292431, EPI_ISL_3292432, EPI_ISL_3292433, EPI_ISL_3292434, EPI_ISL_3292435, EPI_ISL_3292436, EPI_ISL_3292437, EPI_ISL_3292438, EPI_ISL_3292439, EPI_ISL_3292441, EPI_ISL_3292442, EPI_ISL_3292443, EPI_ISL_3292444, EPI_ISL_3292445, EPI_ISL_3292446, EPI_ISL_3292447, EPI_ISL_3292448, EPI_ISL_3292449, EPI_ISL_3292450, EPI_ISL_3292451, EPI_ISL_3292452, EPI_ISL_3292453, EPI_ISL_3292454, EPI_ISL_3292455, EPI_ISL_3292456, EPI_ISL_3292457, EPI_ISL_3292458, EPI_ISL_3292459, EPI_ISL_3292460, EPI_ISL_3292461, EPI_ISL_3292462, EPI_ISL_3292463, EPI_ISL_3292464, EPI_ISL_3292465, EPI_ISL_3292466, EPI_ISL_3292467, EPI_ISL_3292468, EPI_ISL_3292469, EPI_ISL_3292470, EPI_ISL_3292471, EPI_ISL_3292472, EPI_ISL_3292473, EPI_ISL_3292474, EPI_ISL_3292475, EPI_ISL_3292476, EPI_ISL_3292477, EPI_ISL_3292478, EPI_ISL_3292479, EPI_ISL_3292480, EPI_ISL_3292481, EPI_ISL_3292482, EPI_ISL_3292483, EPI_ISL_3292484, EPI_ISL_3292485, EPI_ISL_3292486, EPI_ISL_3292487, EPI_ISL_3292488, EPI_ISL_3292489, EPI_ISL_3292490, EPI_ISL_3292491, EPI_ISL_3292492, EPI_ISL_3292493, EPI_ISL_3292494, EPI_ISL_3292495, EPI_ISL_3292496, EPI_ISL_3292497, EPI_ISL_3292498, EPI_ISL_3292499, EPI_ISL_3292500, EPI_ISL_3292501, EPI_ISL_3292502, EPI_ISL_3292503, EPI_ISL_3292504, EPI_ISL_3292505, EPI_ISL_3292506, EPI_ISL_3292507, EPI_ISL_3292508, EPI_ISL_3292510, EPI_ISL_3292511, EPI_ISL_3292512, EPI_ISL_3292513, EPI_ISL_3292514, EPI_ISL_3292515, EPI_ISL_3292516, EPI_ISL_3292517, EPI_ISL_3292518, EPI_ISL_3292519, EPI_ISL_3292520, EPI_ISL_3292521, EPI_ISL_3292522, EPI_ISL_3292523, EPI_ISL_3292524, EPI_ISL_3292525, EPI_ISL_3292526, EPI_ISL_3292527, EPI_ISL_3292528, EPI_ISL_3292529, EPI_ISL_3292530, EPI_ISL_3292531, EPI_ISL_3292532, EPI_ISL_3292533, EPI_ISL_3292534, EPI_ISL_3292535, EPI_ISL_3292536, EPI_ISL_3292537, EPI_ISL_3292538, EPI_ISL_3292539, EPI_ISL_3292540, EPI_ISL_3292541, EPI_ISL_3292542, EPI_ISL_3292543, EPI_ISL_3292544, EPI_ISL_3292545, EPI_ISL_3292547, EPI_ISL_3292548, EPI_ISL_3292549, EPI_ISL_3292550, EPI_ISL_3292551, EPI_ISL_3292552, EPI_ISL_3292553, EPI_ISL_3292554, EPI_ISL_3292555, EPI_ISL_3292556, EPI_ISL_3292557, EPI_ISL_3292558, EPI_ISL_3292559, EPI_ISL_3292560, EPI_ISL_3292561, EPI_ISL_3292562, EPI_ISL_3292563, EPI_ISL_3292564, EPI_ISL_3292565, EPI_ISL_3292566, EPI_ISL_3292567, EPI_ISL_3292569, EPI_ISL_3292570, EPI_ISL_3292571, EPI_ISL_3292572, EPI_ISL_3292573, EPI_ISL_3292574, EPI_ISL_3292575, EPI_ISL_3292576, EPI_ISL_3292577, EPI_ISL_3292578, EPI_ISL_3292579, EPI_ISL_3292581, EPI_ISL_3292582, EPI_ISL_3292583, EPI_ISL_3292584, EPI_ISL_3292585, EPI_ISL_3292586, EPI_ISL_3292587, EPI_ISL_3292588, EPI_ISL_3292589, EPI_ISL_3292590, EPI_ISL_3292591, EPI_ISL_3292592, EPI_ISL_3292593, EPI_ISL_3292594, EPI_ISL_3292595, EPI_ISL_3292596, EPI_ISL_3292597, EPI_ISL_3292598, EPI_ISL_3292599, EPI_ISL_3292600, EPI_ISL_3292602, EPI_ISL_3292603, EPI_ISL_3292604, EPI_ISL_3292605, EPI_ISL_3292606, EPI_ISL_3292607, EPI_ISL_3292608, EPI_ISL_3292609, EPI_ISL_3292610, EPI_ISL_3292611, EPI_ISL_3292612, EPI_ISL_3292613, EPI_ISL_3292614, EPI_ISL_3292615, EPI_ISL_3292616, EPI_ISL_3292617, EPI_ISL_3292618, EPI_ISL_3292619, EPI_ISL_3292620, EPI_ISL_3292621, EPI_ISL_3292622, EPI_ISL_3292623, EPI_ISL_3292624, EPI_ISL_3292625, EPI_ISL_3292626, EPI_ISL_3292627, EPI_ISL_3292628, EPI_ISL_3292629, EPI_ISL_3292630, EPI_ISL_3292631, EPI_ISL_3292632, EPI_ISL_3292633, EPI_ISL_3292634, EPI_ISL_3292635, EPI_ISL_3292636, EPI_ISL_3292637, EPI_ISL_3292638, EPI_ISL_3292639, EPI_ISL_3292640, EPI_ISL_3292641, EPI_ISL_3292642, EPI_ISL_3292643, EPI_ISL_3292644, EPI_ISL_3292645, EPI_ISL_3292646, EPI_ISL_3292647, EPI_ISL_3292648, EPI_ISL_3292649, EPI_ISL_3292650, EPI_ISL_3292651, EPI_ISL_3292652, EPI_ISL_3292653, EPI_ISL_3292654, EPI_ISL_3292655, EPI_ISL_3292656, EPI_ISL_3292657, EPI_ISL_3292658, EPI_ISL_3292659, EPI_ISL_3292660, EPI_ISL_3292661, EPI_ISL_3292662, EPI_ISL_3292663, EPI_ISL_3292664, EPI_ISL_3292665, EPI_ISL_3292666, EPI_ISL_3292667, EPI_ISL_3292668, EPI_ISL_3292669, EPI_ISL_3292670, EPI_ISL_3292671, EPI_ISL_3292672, EPI_ISL_3292673, EPI_ISL_3292674, EPI_ISL_3292675, EPI_ISL_3292676, EPI_ISL_3292677, EPI_ISL_3292678, EPI_ISL_3292679, EPI_ISL_3292680, EPI_ISL_3292681, EPI_ISL_3292682, EPI_ISL_3292683, EPI_ISL_3292684, EPI_ISL_3292685, EPI_ISL_3292686, EPI_ISL_3292687, EPI_ISL_3292688, EPI_ISL_3292689, EPI_ISL_3292690, EPI_ISL_3292691, EPI_ISL_3292692, EPI_ISL_3292693, EPI_ISL_3292694, EPI_ISL_3292695, EPI_ISL_3292696, EPI_ISL_3292697, EPI_ISL_3292698, EPI_ISL_3292699, EPI_ISL_3292700, EPI_ISL_3292701, EPI_ISL_3292702, EPI_ISL_3292703, EPI_ISL_3292704, EPI_ISL_3292705, EPI_ISL_3292706, EPI_ISL_3292707, EPI_ISL_3292708, EPI_ISL_3292709, EPI_ISL_3292710, EPI_ISL_3292711, EPI_ISL_3292712, EPI_ISL_3292713, EPI_ISL_3292714, EPI_ISL_3292715, EPI_ISL_3292716, EPI_ISL_3292717, EPI_ISL_3292718, EPI_ISL_3292719, EPI_ISL_3292720, EPI_ISL_3292721, EPI_ISL_3292722, EPI_ISL_3292723, EPI_ISL_3292724, EPI_ISL_3292725, EPI_ISL_3292726, EPI_ISL_3292727, EPI_ISL_3292728, EPI_ISL_3292729, EPI_ISL_3292730, EPI_ISL_3292731, EPI_ISL_3292732, EPI_ISL_3292733, EPI_ISL_3292734, EPI_ISL_3292735, EPI_ISL_3292736, EPI_ISL_3292737, EPI_ISL_3292738, EPI_ISL_3292739, EPI_ISL_3292740, EPI_ISL_3292741, EPI_ISL_3292742, EPI_ISL_3292743, EPI_ISL_3292744, EPI_ISL_3292745, EPI_ISL_3292746, EPI_ISL_3292747, EPI_ISL_3292748, EPI_ISL_3292749, EPI_ISL_3292750, EPI_ISL_3292751, EPI_ISL_3292752, EPI_ISL_3292753, EPI_ISL_3292754, EPI_ISL_3292755, EPI_ISL_3292756, EPI_ISL_3292757, EPI_ISL_3292758, EPI_ISL_3292759, EPI_ISL_3292760, EPI_ISL_3292761, EPI_ISL_3292762, EPI_ISL_3292763, EPI_ISL_3292764, EPI_ISL_3292765, EPI_ISL_3292766, EPI_ISL_3292767, EPI_ISL_3292768, EPI_ISL_3292769, EPI_ISL_3292770, EPI_ISL_3292771, EPI_ISL_3292772, EPI_ISL_3292773, EPI_ISL_3292774, EPI_ISL_3292775, EPI_ISL_3292776, EPI_ISL_3292777, EPI_ISL_3292778, EPI_ISL_3292779, EPI_ISL_3292780, EPI_ISL_3292781, EPI_ISL_3292782, EPI_ISL_3292783, EPI_ISL_3292784, EPI_ISL_3292785, EPI_ISL_3292786, EPI_ISL_3292787, EPI_ISL_3292788, EPI_ISL_3292789, EPI_ISL_3292790, EPI_ISL_3292791, EPI_ISL_3292792, EPI_ISL_3292793, EPI_ISL_3292794, EPI_ISL_3292795, EPI_ISL_3292796, EPI_ISL_3292797, EPI_ISL_3292798, EPI_ISL_3292799, EPI_ISL_3292800, EPI_ISL_3292801, EPI_ISL_3292802, EPI_ISL_3292803, EPI_ISL_3292804, EPI_ISL_3292806, EPI_ISL_3292807, EPI_ISL_3292808, EPI_ISL_3292809, EPI_ISL_3292810, EPI_ISL_3292811, EPI_ISL_3292812, EPI_ISL_3292813, EPI_ISL_3292814, EPI_ISL_3292815, EPI_ISL_3292816, EPI_ISL_3292817, EPI_ISL_3292818, EPI_ISL_3292819, EPI_ISL_3292820, EPI_ISL_3292821, EPI_ISL_3292822, EPI_ISL_3292823, EPI_ISL_3292824, EPI_ISL_3292825, EPI_ISL_3292826, EPI_ISL_3292827, EPI_ISL_3292828, EPI_ISL_3292829, EPI_ISL_3292830, EPI_ISL_3292831, EPI_ISL_3292832, EPI_ISL_3292833, EPI_ISL_3292834, EPI_ISL_3292835, EPI_ISL_3292836, EPI_ISL_3292837, EPI_ISL_3292838, EPI_ISL_3292839, EPI_ISL_3292840, EPI_ISL_3292841, EPI_ISL_3292842, EPI_ISL_3292843, EPI_ISL_3292844, EPI_ISL_3292845, EPI_ISL_3292846, EPI_ISL_3292847, EPI_ISL_3292848, EPI_ISL_3292849, EPI_ISL_3292850, EPI_ISL_3292851, EPI_ISL_3292852, EPI_ISL_3292853, EPI_ISL_3292854, EPI_ISL_3292855, EPI_ISL_3292856, EPI_ISL_3292857, EPI_ISL_3292858, EPI_ISL_3292859, EPI_ISL_3292860, EPI_ISL_3292861, EPI_ISL_3292862, EPI_ISL_3292863, EPI_ISL_3292864, EPI_ISL_3292865, EPI_ISL_3292866, EPI_ISL_3292867, EPI_ISL_3292868, EPI_ISL_3292869, EPI_ISL_3292870, EPI_ISL_3292871, EPI_ISL_3292872, EPI_ISL_3292873, EPI_ISL_3292874, EPI_ISL_3292875, EPI_ISL_3292876, EPI_ISL_3292877, EPI_ISL_3292878, EPI_ISL_3292879, EPI_ISL_3292880, EPI_ISL_3292881, EPI_ISL_3292882, EPI_ISL_3292883, EPI_ISL_3292884, EPI_ISL_3292885, EPI_ISL_3292886, EPI_ISL_3292887, EPI_ISL_3292888, EPI_ISL_3292889, EPI_ISL_3292890, EPI_ISL_3292891, EPI_ISL_3292892, EPI_ISL_3292893, EPI_ISL_3292894, EPI_ISL_3292895, EPI_ISL_3292897, EPI_ISL_3292899, EPI_ISL_3292900, EPI_ISL_3292901, EPI_ISL_3292902, EPI_ISL_3292903, EPI_ISL_3292904, EPI_ISL_3292905, EPI_ISL_3292906, EPI_ISL_3292907, EPI_ISL_3292909, EPI_ISL_3292910, EPI_ISL_3292911, EPI_ISL_3292914, EPI_ISL_3292915, EPI_ISL_3292916, EPI_ISL_3292917, EPI_ISL_3292918, EPI_ISL_3292919, EPI_ISL_3292922, EPI_ISL_3292923, EPI_ISL_3292924, EPI_ISL_3292925, EPI_ISL_3292926, EPI_ISL_3292927, EPI_ISL_3292928, EPI_ISL_3292929, EPI_ISL_3292930, EPI_ISL_3292931, EPI_ISL_3292932, EPI_ISL_3292933, EPI_ISL_3292934, EPI_ISL_3292935, EPI_ISL_3292938, EPI_ISL_3292939, EPI_ISL_3292940, EPI_ISL_3292941, EPI_ISL_3292942, EPI_ISL_3292943, EPI_ISL_3292944, EPI_ISL_3292947, EPI_ISL_3292948, EPI_ISL_3292949, EPI_ISL_3292950, EPI_ISL_3292952, EPI_ISL_3292954, EPI_ISL_3292955, EPI_ISL_3292956, EPI_ISL_3292957, EPI_ISL_3292958, EPI_ISL_3292959, EPI_ISL_3292960, EPI_ISL_3292961, EPI_ISL_3292962, EPI_ISL_3292963, EPI_ISL_3292964, EPI_ISL_3292965, EPI_ISL_3292966, EPI_ISL_3292967, EPI_ISL_3292968, EPI_ISL_3292969, EPI_ISL_3292970, EPI_ISL_3292971, EPI_ISL_3292972, EPI_ISL_3292973, EPI_ISL_3292974, EPI_ISL_3292975, EPI_ISL_3292976, EPI_ISL_3292977, EPI_ISL_3292978, EPI_ISL_3292979, EPI_ISL_3292980, EPI_ISL_3292981, EPI_ISL_3292982, EPI_ISL_3292983, EPI_ISL_3292984, EPI_ISL_3292985, EPI_ISL_3292986, EPI_ISL_3292987, EPI_ISL_3292988, EPI_ISL_3292989, EPI_ISL_3292990, EPI_ISL_3292991, EPI_ISL_3292992, EPI_ISL_3292993, EPI_ISL_3292994, EPI_ISL_3292995, EPI_ISL_3292996, EPI_ISL_3292997, EPI_ISL_3292998, EPI_ISL_3293000, EPI_ISL_3293001, EPI_ISL_3293002, EPI_ISL_3293003, EPI_ISL_3293004, EPI_ISL_3293005, EPI_ISL_3293006, EPI_ISL_3293007, EPI_ISL_3293008, EPI_ISL_3293009, EPI_ISL_3293010, EPI_ISL_3293011, EPI_ISL_3293012, EPI_ISL_3293013, EPI_ISL_3293014, EPI_ISL_3293015, EPI_ISL_3293016, EPI_ISL_3293017, EPI_ISL_3293018, EPI_ISL_3293019, EPI_ISL_3293020, EPI_ISL_3293022, EPI_ISL_3293023, EPI_ISL_3293024, EPI_ISL_3293025, EPI_ISL_3293026, EPI_ISL_3293027, EPI_ISL_3293028, EPI_ISL_3293029, EPI_ISL_3293030, EPI_ISL_3293031, EPI_ISL_3293032, EPI_ISL_3293033, EPI_ISL_3293034, EPI_ISL_3293035, EPI_ISL_3293036, EPI_ISL_3293037, EPI_ISL_3293038, EPI_ISL_3293039, EPI_ISL_3293040, EPI_ISL_3293041, EPI_ISL_3293042, EPI_ISL_3293043, EPI_ISL_3293044, EPI_ISL_3293046, EPI_ISL_3293047, EPI_ISL_3293048, EPI_ISL_3293049, EPI_ISL_3293050, EPI_ISL_3293051, EPI_ISL_3293052, EPI_ISL_3293053, EPI_ISL_3293054, EPI_ISL_3293055, EPI_ISL_3293056, EPI_ISL_3293058, EPI_ISL_3293059, EPI_ISL_3293060, EPI_ISL_3293061, EPI_ISL_3293062, EPI_ISL_3293063, EPI_ISL_3293064, EPI_ISL_3293065, EPI_ISL_3293066, EPI_ISL_3293067, EPI_ISL_3293068, EPI_ISL_3293069, EPI_ISL_3293070, EPI_ISL_3293071, EPI_ISL_3293072, EPI_ISL_3293073, EPI_ISL_3293074, EPI_ISL_3293075, EPI_ISL_3293076, EPI_ISL_3293077, EPI_ISL_3293078, EPI_ISL_3293079, EPI_ISL_3293080, EPI_ISL_3293082, EPI_ISL_3293083, EPI_ISL_3293084, EPI_ISL_3293085, EPI_ISL_3293086, EPI_ISL_3293087, EPI_ISL_3293088, EPI_ISL_3293089, EPI_ISL_3293090, EPI_ISL_3293091, EPI_ISL_3293092, EPI_ISL_3293093, EPI_ISL_3293094, EPI_ISL_3293095, EPI_ISL_3293096, EPI_ISL_3293097, EPI_ISL_3293098, EPI_ISL_3293099, EPI_ISL_3293100, EPI_ISL_3293101, EPI_ISL_3293102, EPI_ISL_3293104, EPI_ISL_3293105, EPI_ISL_3293106, EPI_ISL_3293107, EPI_ISL_3293108, EPI_ISL_3293109, EPI_ISL_3293110, EPI_ISL_3293111, EPI_ISL_3293112, EPI_ISL_3293113, EPI_ISL_3293114, EPI_ISL_3293115, EPI_ISL_3293116, EPI_ISL_3293117, EPI_ISL_3293118, EPI_ISL_3293119, EPI_ISL_3293120, EPI_ISL_3293121, EPI_ISL_3293122, EPI_ISL_3293123, EPI_ISL_3293124, EPI_ISL_3293125, EPI_ISL_3293126, EPI_ISL_3293127, EPI_ISL_3293128, EPI_ISL_3293129, EPI_ISL_3293130, EPI_ISL_3293131, EPI_ISL_3293132, EPI_ISL_3293133, EPI_ISL_3293134, EPI_ISL_3293135, EPI_ISL_3293136, EPI_ISL_3293137, EPI_ISL_3293138, EPI_ISL_3293139, EPI_ISL_3293140, EPI_ISL_3293141, EPI_ISL_3293142, EPI_ISL_3293143, EPI_ISL_3293144, EPI_ISL_3293145, EPI_ISL_3293146, EPI_ISL_3293147, EPI_ISL_3293148, EPI_ISL_3293149, EPI_ISL_3293150, EPI_ISL_3293151, EPI_ISL_3293152, EPI_ISL_3293153, EPI_ISL_3293154, EPI_ISL_3293155, EPI_ISL_3293156, EPI_ISL_3293157, EPI_ISL_3293158, EPI_ISL_3293159, EPI_ISL_3293160, EPI_ISL_3293161, EPI_ISL_3293162, EPI_ISL_3293163, EPI_ISL_3293164, EPI_ISL_3293165, EPI_ISL_3293166, EPI_ISL_3293167, EPI_ISL_3293168, EPI_ISL_3293169, EPI_ISL_3293170, EPI_ISL_3293171, EPI_ISL_3293172, EPI_ISL_3293173, EPI_ISL_3293174, EPI_ISL_3293175, EPI_ISL_3293176, EPI_ISL_3293177, EPI_ISL_3293179, EPI_ISL_3293180, EPI_ISL_3293181, EPI_ISL_3293182, EPI_ISL_3293183, EPI_ISL_3293184, EPI_ISL_3293185, EPI_ISL_3293186, EPI_ISL_3293187, EPI_ISL_3293188, EPI_ISL_3293189, EPI_ISL_3293190, EPI_ISL_3293191, EPI_ISL_3293192, EPI_ISL_3293193, EPI_ISL_3293194, EPI_ISL_3293195, EPI_ISL_3293196, EPI_ISL_3293197, EPI_ISL_3293198, EPI_ISL_3293199, EPI_ISL_3293200, EPI_ISL_3293201, EPI_ISL_3293202, EPI_ISL_3293203, EPI_ISL_3293204, EPI_ISL_3293205, EPI_ISL_3293206, EPI_ISL_3293207, EPI_ISL_3293208, EPI_ISL_3293209, EPI_ISL_3293210, EPI_ISL_3293211, EPI_ISL_3293212, EPI_ISL_3293213, EPI_ISL_3293214, EPI_ISL_3293215, EPI_ISL_3293216, EPI_ISL_3293217, EPI_ISL_3293218, EPI_ISL_3293219, EPI_ISL_3293220, EPI_ISL_3293221, EPI_ISL_3293222, EPI_ISL_3293223, EPI_ISL_3293224, EPI_ISL_3293225, EPI_ISL_3293226, EPI_ISL_3293228, EPI_ISL_3293229, EPI_ISL_3293230, EPI_ISL_3293231, EPI_ISL_3293232, EPI_ISL_3293233, EPI_ISL_3293234, EPI_ISL_3293235, EPI_ISL_3293236, EPI_ISL_3293237, EPI_ISL_3293238, EPI_ISL_3293239, EPI_ISL_3293240, EPI_ISL_3293241, EPI_ISL_3293242, EPI_ISL_3293243, EPI_ISL_3293244, EPI_ISL_3293245, EPI_ISL_3293247, EPI_ISL_3293248, EPI_ISL_3293249, EPI_ISL_3293250, EPI_ISL_3293251, EPI_ISL_3293252, EPI_ISL_3293253, EPI_ISL_3293254, EPI_ISL_3293255, EPI_ISL_3293256, EPI_ISL_3293257, EPI_ISL_3293258, EPI_ISL_3293259, EPI_ISL_3293260, EPI_ISL_3293261, EPI_ISL_3293262, EPI_ISL_3293263, EPI_ISL_3293264, EPI_ISL_3293265, EPI_ISL_3293266, EPI_ISL_3293267, EPI_ISL_3293268, EPI_ISL_3293269, EPI_ISL_3293270, EPI_ISL_3293271, EPI_ISL_3293272, EPI_ISL_3293273, EPI_ISL_3293274, EPI_ISL_3293275, EPI_ISL_3293276, EPI_ISL_3293277, EPI_ISL_3293278, EPI_ISL_3293279, EPI_ISL_3293281, EPI_ISL_3293282, EPI_ISL_3293283, EPI_ISL_3293284, EPI_ISL_3293285, EPI_ISL_3293286, EPI_ISL_3293287, EPI_ISL_3293288, EPI_ISL_3293289, EPI_ISL_3293290, EPI_ISL_3293291, EPI_ISL_3293292, EPI_ISL_3293293, EPI_ISL_3293294, EPI_ISL_3293295, EPI_ISL_3293296, EPI_ISL_3293298, EPI_ISL_3293299, EPI_ISL_3293300, EPI_ISL_3293301, EPI_ISL_3293302, EPI_ISL_3293303, EPI_ISL_3293304, EPI_ISL_3293305, EPI_ISL_3293306, EPI_ISL_3293307, EPI_ISL_3293308, EPI_ISL_3293309, EPI_ISL_3293310, EPI_ISL_3293311, EPI_ISL_3293312, EPI_ISL_3293313, EPI_ISL_3293314, EPI_ISL_3293315, EPI_ISL_3293316, EPI_ISL_3293317, EPI_ISL_3293318, EPI_ISL_3293319, EPI_ISL_3293320, EPI_ISL_3293321, EPI_ISL_3293322, EPI_ISL_3293323, EPI_ISL_3293324, EPI_ISL_3293325, EPI_ISL_3293326, EPI_ISL_3293327, EPI_ISL_3293328, EPI_ISL_3293329, EPI_ISL_3293330, EPI_ISL_3293331, EPI_ISL_3293332, EPI_ISL_3293333, EPI_ISL_3293334, EPI_ISL_3293335, EPI_ISL_3293336, EPI_ISL_3293337, EPI_ISL_3293338, EPI_ISL_3293339, EPI_ISL_3293340, EPI_ISL_3293341, EPI_ISL_3293342, EPI_ISL_3293343, EPI_ISL_3293344, EPI_ISL_3293345, EPI_ISL_3293346, EPI_ISL_3293347, EPI_ISL_3293348, EPI_ISL_3293349, EPI_ISL_3293350, EPI_ISL_3293351, EPI_ISL_3293352, EPI_ISL_3293353, EPI_ISL_3293354, EPI_ISL_3293355, EPI_ISL_3293356, EPI_ISL_3293357, EPI_ISL_3293358, EPI_ISL_3293359, EPI_ISL_3293360, EPI_ISL_3293361, EPI_ISL_3293362, EPI_ISL_3293363, EPI_ISL_3293365, EPI_ISL_3293366, EPI_ISL_3293367, EPI_ISL_3293368, EPI_ISL_3293369, EPI_ISL_3293370, EPI_ISL_3293371, EPI_ISL_3293372, EPI_ISL_3293373, EPI_ISL_3293374, EPI_ISL_3293375, EPI_ISL_3293376, EPI_ISL_3293377, EPI_ISL_3293378, EPI_ISL_3293379, EPI_ISL_3293380, EPI_ISL_3293381, EPI_ISL_3293382, EPI_ISL_3293383, EPI_ISL_3293384, EPI_ISL_3293385, EPI_ISL_3293386, EPI_ISL_3293387, EPI_ISL_3293388, EPI_ISL_3293389, EPI_ISL_3293390, EPI_ISL_3293391, EPI_ISL_3293392, EPI_ISL_3293393, EPI_ISL_3293394, EPI_ISL_3293395, EPI_ISL_3293396, EPI_ISL_3293397, EPI_ISL_3293398, EPI_ISL_3293399, EPI_ISL_3293400, EPI_ISL_3293401, EPI_ISL_3293402, EPI_ISL_3293403, EPI_ISL_3293404, EPI_ISL_3293405, EPI_ISL_3293406, EPI_ISL_3293407, EPI_ISL_3293408, EPI_ISL_3293409, EPI_ISL_3293410, EPI_ISL_3293411, EPI_ISL_3293412, EPI_ISL_3293413, EPI_ISL_3293414, EPI_ISL_3293415, EPI_ISL_3293416, EPI_ISL_3293417, EPI_ISL_3293418, EPI_ISL_3293419, EPI_ISL_3293420, EPI_ISL_3293421, EPI_ISL_3293422, EPI_ISL_3293423, EPI_ISL_3293424, EPI_ISL_3293425, EPI_ISL_3293426, EPI_ISL_3293427, EPI_ISL_3293428, EPI_ISL_3293429, EPI_ISL_3293430, EPI_ISL_3293431, EPI_ISL_3293432, EPI_ISL_3293433, EPI_ISL_3293434, EPI_ISL_3293435, EPI_ISL_3293436, EPI_ISL_3293437, EPI_ISL_3293438, EPI_ISL_3293439, EPI_ISL_3293440, EPI_ISL_3293441, EPI_ISL_3293442, EPI_ISL_3293443, EPI_ISL_3293444, EPI_ISL_3293445, EPI_ISL_3293446, EPI_ISL_3293447, EPI_ISL_3293448, EPI_ISL_3293449, EPI_ISL_3293450, EPI_ISL_3293451, EPI_ISL_3293452, EPI_ISL_3293453, EPI_ISL_3293454

| | | | |
|---|---|---|---|
| see above | Respiratory Virus Unit, Microbiology Services Colindale, Public Health England | COVID-19 Genomics UK (COG-UK) Consortium | PHE Covid Sequencing Team |

EPI_ISL_3293455, EPI_ISL_3293456, EPI_ISL_3293457, EPI_ISL_3293458, EPI_ISL_3293459, EPI_ISL_3293460, EPI_ISL_3293461, EPI_ISL_3293462, EPI_ISL_3293463, EPI_ISL_3293464, EPI_ISL_3293465, EPI_ISL_3293466, EPI_ISL_3293467, EPI_ISL_3293468, EPI_ISL_3293469, EPI_ISL_3293470, EPI_ISL_3293471, EPI_ISL_3293472, EPI_ISL_3293473, EPI_ISL_3293474, EPI_ISL_3293475, EPI_ISL_3293476, EPI_ISL_3293477, EPI_ISL_3293478, EPI_ISL_3293479, EPI_ISL_3293480, EPI_ISL_3293481, EPI_ISL_3293482, EPI_ISL_3293483, EPI_ISL_3293484, EPI_ISL_3293485, EPI_ISL_3293486, EPI_ISL_3293487, EPI_ISL_3293488, EPI_ISL_3293489, EPI_ISL_3293490, EPI_ISL_3293491, EPI_ISL_3293492, EPI_ISL_3293493, EPI_ISL_3293494, EPI_ISL_3293495, EPI_ISL_3293496, EPI_ISL_3293497, EPI_ISL_3293498, EPI_ISL_3293499, EPI_ISL_3293500, EPI_ISL_3293501, EPI_ISL_3293502, EPI_ISL_3293503, EPI_ISL_3293504, EPI_ISL_3293505, EPI_ISL_3293506, EPI_ISL_3293507, EPI_ISL_3293508, EPI_ISL_3293509, EPI_ISL_3293510, EPI_ISL_3293511, EPI_ISL_3293512, EPI_ISL_3293513, EPI_ISL_3293514, EPI_ISL_3293515, EPI_ISL_3293516, EPI_ISL_3293517, EPI_ISL_3293518, EPI_ISL_3293519, EPI_ISL_3293520,

EPI_ISL_3293521, EPI_ISL_3293522, EPI_ISL_3293523, EPI_ISL_3293524, EPI_ISL_3293525, EPI_ISL_3293526, EPI_ISL_3293527, EPI_ISL_3293528, EPI_ISL_3293529, EPI_ISL_3293530, EPI_ISL_3293531, EPI_ISL_3293532, EPI_ISL_3293533, EPI_ISL_3293534, EPI_ISL_3293535, EPI_ISL_3293536, EPI_ISL_3293537, EPI_ISL_3293538, EPI_ISL_3293539, EPI_ISL_3293540, EPI_ISL_3293541, EPI_ISL_3293542,
EPI_ISL_3293543, EPI_ISL_3293544, EPI_ISL_3293545, EPI_ISL_3293546, EPI_ISL_3293547, EPI_ISL_3293548, EPI_ISL_3293549, EPI_ISL_3293550, EPI_ISL_3293551, EPI_ISL_3293552, EPI_ISL_3293553, EPI_ISL_3293554, EPI_ISL_3293555, EPI_ISL_3293556, EPI_ISL_3293557, EPI_ISL_3293558, EPI_ISL_3293559, EPI_ISL_3293560, EPI_ISL_3293561, EPI_ISL_3293562, EPI_ISL_3293563, EPI_ISL_3293564,
EPI_ISL_3293565, EPI_ISL_3293566, EPI_ISL_3293567, EPI_ISL_3293568, EPI_ISL_3293569, EPI_ISL_3293570, EPI_ISL_3293571, EPI_ISL_3293572, EPI_ISL_3293573, EPI_ISL_3293574, EPI_ISL_3293575, EPI_ISL_3293576, EPI_ISL_3293577, EPI_ISL_3293578, EPI_ISL_3293579, EPI_ISL_3293580, EPI_ISL_3293581, EPI_ISL_3293582, EPI_ISL_3293583, EPI_ISL_3293584, EPI_ISL_3293585, EPI_ISL_3293586,
EPI_ISL_3293587, EPI_ISL_3293588, EPI_ISL_3293589, EPI_ISL_3293590, EPI_ISL_3293591, EPI_ISL_3293592, EPI_ISL_3293593, EPI_ISL_3293594, EPI_ISL_3293595, EPI_ISL_3293596, EPI_ISL_3293597, EPI_ISL_3293598, EPI_ISL_3293599, EPI_ISL_3293600, EPI_ISL_3293601, EPI_ISL_3293602, EPI_ISL_3293603, EPI_ISL_3293604, EPI_ISL_3293605, EPI_ISL_3293606, EPI_ISL_3293607, EPI_ISL_3293608,
EPI_ISL_3293609, EPI_ISL_3293610, EPI_ISL_3293611, EPI_ISL_3293612, EPI_ISL_3293613, EPI_ISL_3293614, EPI_ISL_3293615, EPI_ISL_3293616, EPI_ISL_3293617, EPI_ISL_3293618, EPI_ISL_3293619, EPI_ISL_3293620, EPI_ISL_3293621, EPI_ISL_3293622, EPI_ISL_3293623, EPI_ISL_3293624, EPI_ISL_3293625, EPI_ISL_3293626, EPI_ISL_3293627, EPI_ISL_3293628, EPI_ISL_3293629, EPI_ISL_3293630,
EPI_ISL_3293631, EPI_ISL_3293632, EPI_ISL_3293633, EPI_ISL_3293634, EPI_ISL_3293635, EPI_ISL_3293636, EPI_ISL_3293637, EPI_ISL_3293638, EPI_ISL_3293639, EPI_ISL_3293640, EPI_ISL_3293641, EPI_ISL_3293642, EPI_ISL_3293643, EPI_ISL_3293644, EPI_ISL_3293645, EPI_ISL_3293646, EPI_ISL_3293647, EPI_ISL_3293648, EPI_ISL_3293649, EPI_ISL_3293650, EPI_ISL_3293651, EPI_ISL_3293652,
EPI_ISL_3293653, EPI_ISL_3293654, EPI_ISL_3293655, EPI_ISL_3293656, EPI_ISL_3293657, EPI_ISL_3293658, EPI_ISL_3293659, EPI_ISL_3293660, EPI_ISL_3293661, EPI_ISL_3293662, EPI_ISL_3293663, EPI_ISL_3293664, EPI_ISL_3293665, EPI_ISL_3293666, EPI_ISL_3293667, EPI_ISL_3293668, EPI_ISL_3293669, EPI_ISL_3293670, EPI_ISL_3293671, EPI_ISL_3293672, EPI_ISL_3293673, EPI_ISL_3293674,
EPI_ISL_3293675, EPI_ISL_3293676, EPI_ISL_3293677, EPI_ISL_3293678, EPI_ISL_3293679, EPI_ISL_3293680, EPI_ISL_3293681, EPI_ISL_3293682, EPI_ISL_3293683, EPI_ISL_3293684, EPI_ISL_3293685, EPI_ISL_3293686, EPI_ISL_3293687, EPI_ISL_3293688, EPI_ISL_3293689, EPI_ISL_3293690, EPI_ISL_3293691, EPI_ISL_3293692, EPI_ISL_3293693, EPI_ISL_3293694, EPI_ISL_3293695, EPI_ISL_3293696,
EPI_ISL_3293697, EPI_ISL_3293698, EPI_ISL_3293699, EPI_ISL_3293700, EPI_ISL_3293701, EPI_ISL_3293702, EPI_ISL_3293703, EPI_ISL_3293704, EPI_ISL_3293705, EPI_ISL_3293706, EPI_ISL_3293707, EPI_ISL_3293708, EPI_ISL_3293709, EPI_ISL_3293710, EPI_ISL_3293711, EPI_ISL_3293712, EPI_ISL_3293713, EPI_ISL_3293714, EPI_ISL_3293715, EPI_ISL_3293716, EPI_ISL_3293717, EPI_ISL_3293718,
EPI_ISL_3293719, EPI_ISL_3293720, EPI_ISL_3293721, EPI_ISL_3293722, EPI_ISL_3293723, EPI_ISL_3293724, EPI_ISL_3293725, EPI_ISL_3293726, EPI_ISL_3293727, EPI_ISL_3293728, EPI_ISL_3293729, EPI_ISL_3293730, EPI_ISL_3293731, EPI_ISL_3293732, EPI_ISL_3293733, EPI_ISL_3293734, EPI_ISL_3293735, EPI_ISL_3293736, EPI_ISL_3293737, EPI_ISL_3293738, EPI_ISL_3293739, EPI_ISL_3293740,
EPI_ISL_3293741, EPI_ISL_3293742, EPI_ISL_3293743, EPI_ISL_3293744, EPI_ISL_3293745, EPI_ISL_3293746, EPI_ISL_3293747, EPI_ISL_3293748, EPI_ISL_3293749, EPI_ISL_3293750, EPI_ISL_3293751, EPI_ISL_3293752, EPI_ISL_3293753, EPI_ISL_3293754, EPI_ISL_3293755, EPI_ISL_3293756, EPI_ISL_3293757, EPI_ISL_3293758, EPI_ISL_3293759, EPI_ISL_3293760, EPI_ISL_3293761, EPI_ISL_3293762,
EPI_ISL_3293763, EPI_ISL_3293764, EPI_ISL_3293765, EPI_ISL_3293766, EPI_ISL_3293767, EPI_ISL_3293768, EPI_ISL_3293769, EPI_ISL_3293770, EPI_ISL_3293771, EPI_ISL_3293772, EPI_ISL_3293773, EPI_ISL_3293774, EPI_ISL_3293775, EPI_ISL_3293776, EPI_ISL_3293777, EPI_ISL_3293778, EPI_ISL_3293779, EPI_ISL_3293780, EPI_ISL_3293781, EPI_ISL_3293782, EPI_ISL_3293783, EPI_ISL_3293784,
EPI_ISL_3293785, EPI_ISL_3293786, EPI_ISL_3293787, EPI_ISL_3293788, EPI_ISL_3293789, EPI_ISL_3293790, EPI_ISL_3293791, EPI_ISL_3293792, EPI_ISL_3293793, EPI_ISL_3293794, EPI_ISL_3293795, EPI_ISL_3293796, EPI_ISL_3293797, EPI_ISL_3293798, EPI_ISL_3293799, EPI_ISL_3293800, EPI_ISL_3293801, EPI_ISL_3293802, EPI_ISL_3293803, EPI_ISL_3293804, EPI_ISL_3293805, EPI_ISL_3293806,
EPI_ISL_3293807, EPI_ISL_3293808, EPI_ISL_3293809, EPI_ISL_3293810, EPI_ISL_3293811, EPI_ISL_3293812, EPI_ISL_3293813, EPI_ISL_3293814, EPI_ISL_3293815, EPI_ISL_3293816, EPI_ISL_3293817, EPI_ISL_3293818, EPI_ISL_3293819, EPI_ISL_3293820, EPI_ISL_3293821, EPI_ISL_3293822, EPI_ISL_3293823, EPI_ISL_3293824, EPI_ISL_3293825, EPI_ISL_3293826, EPI_ISL_3293827, EPI_ISL_3293828,
EPI_ISL_3293829, EPI_ISL_3293830, EPI_ISL_3293831, EPI_ISL_3293832, EPI_ISL_3293833, EPI_ISL_3293834, EPI_ISL_3293835, EPI_ISL_3293836, EPI_ISL_3293837, EPI_ISL_3293838, EPI_ISL_3293839, EPI_ISL_3293840, EPI_ISL_3293841, EPI_ISL_3293842, EPI_ISL_3293843, EPI_ISL_3293844, EPI_ISL_3293845, EPI_ISL_3293846, EPI_ISL_3293847, EPI_ISL_3293848, EPI_ISL_3293849, EPI_ISL_3293850,
EPI_ISL_3293851, EPI_ISL_3293852, EPI_ISL_3293853, EPI_ISL_3293854, EPI_ISL_3293855, EPI_ISL_3293856, EPI_ISL_3293857, EPI_ISL_3293858, EPI_ISL_3293859, EPI_ISL_3293860, EPI_ISL_3293861, EPI_ISL_3293862, EPI_ISL_3293863, EPI_ISL_3293864, EPI_ISL_3293865, EPI_ISL_3293866, EPI_ISL_3293867, EPI_ISL_3293868, EPI_ISL_3293869, EPI_ISL_3293870, EPI_ISL_3293871, EPI_ISL_3293872,
EPI_ISL_3293873, EPI_ISL_3293874, EPI_ISL_3293875, EPI_ISL_3293876, EPI_ISL_3293877, EPI_ISL_3293878, EPI_ISL_3293879, EPI_ISL_3293880, EPI_ISL_3293881, EPI_ISL_3293882, EPI_ISL_3293883, EPI_ISL_3293884, EPI_ISL_3293885, EPI_ISL_3293886, EPI_ISL_3293887, EPI_ISL_3293888, EPI_ISL_3293889, EPI_ISL_3293890, EPI_ISL_3293891, EPI_ISL_3293892, EPI_ISL_3293893, EPI_ISL_3293894,
EPI_ISL_3293895, EPI_ISL_3293896, EPI_ISL_3293897, EPI_ISL_3293898, EPI_ISL_3293899, EPI_ISL_3293900, EPI_ISL_3293901, EPI_ISL_3293902, EPI_ISL_3293903, EPI_ISL_3293904, EPI_ISL_3293905, EPI_ISL_3293906, EPI_ISL_3293907, EPI_ISL_3293908, EPI_ISL_3293909, EPI_ISL_3293910, EPI_ISL_3293911, EPI_ISL_3293912, EPI_ISL_3293913, EPI_ISL_3293914, EPI_ISL_3293915, EPI_ISL_3293916,
EPI_ISL_3293917, EPI_ISL_3293918, EPI_ISL_3293919, EPI_ISL_3293920, EPI_ISL_3293921, EPI_ISL_3293922, EPI_ISL_3293923, EPI_ISL_3293924, EPI_ISL_3293925, EPI_ISL_3293926, EPI_ISL_3293927, EPI_ISL_3293928, EPI_ISL_3293929, EPI_ISL_3293930, EPI_ISL_3293931, EPI_ISL_3293932, EPI_ISL_3293933, EPI_ISL_3293934, EPI_ISL_3293935, EPI_ISL_3293936, EPI_ISL_3293937, EPI_ISL_3293938,
EPI_ISL_3293939, EPI_ISL_3293940, EPI_ISL_3293941, EPI_ISL_3293942, EPI_ISL_3293943, EPI_ISL_3293944, EPI_ISL_3293945, EPI_ISL_3293946, EPI_ISL_3293947, EPI_ISL_3293948, EPI_ISL_3293949, EPI_ISL_3293950, EPI_ISL_3293951, EPI_ISL_3293952, EPI_ISL_3293953, EPI_ISL_3293954, EPI_ISL_3293955, EPI_ISL_3293956, EPI_ISL_3293957, EPI_ISL_3293958, EPI_ISL_3293959, EPI_ISL_3293960,
EPI_ISL_3293961, EPI_ISL_3293962, EPI_ISL_3293963, EPI_ISL_3293964, EPI_ISL_3293965, EPI_ISL_3293966, EPI_ISL_3293967, EPI_ISL_3293968, EPI_ISL_3293969, EPI_ISL_3293970, EPI_ISL_3293971, EPI_ISL_3293972, EPI_ISL_3293973, EPI_ISL_3293974, EPI_ISL_3293975, EPI_ISL_3293976, EPI_ISL_3293977, EPI_ISL_3293978, EPI_ISL_3293979, EPI_ISL_3293980, EPI_ISL_3293981, EPI_ISL_3293982,
EPI_ISL_3293983, EPI_ISL_3293984, EPI_ISL_3293985, EPI_ISL_3293986, EPI_ISL_3293987, EPI_ISL_3293988, EPI_ISL_3293989, EPI_ISL_3293990, EPI_ISL_3293991, EPI_ISL_3293992, EPI_ISL_3293993, EPI_ISL_3293994, EPI_ISL_3293995, EPI_ISL_3293996, EPI_ISL_3293997, EPI_ISL_3293998, EPI_ISL_3293999, EPI_ISL_3294000, EPI_ISL_3294001, EPI_ISL_3294002, EPI_ISL_3294003, EPI_ISL_3294004,
EPI_ISL_3294005, EPI_ISL_3294006, EPI_ISL_3294007, EPI_ISL_3294008, EPI_ISL_3294009, EPI_ISL_3294010, EPI_ISL_3294011, EPI_ISL_3294012, EPI_ISL_3294013, EPI_ISL_3294014, EPI_ISL_3294015, EPI_ISL_3294016, EPI_ISL_3294017, EPI_ISL_3294018, EPI_ISL_3294019, EPI_ISL_3294020, EPI_ISL_3294021, EPI_ISL_3294022, EPI_ISL_3294023, EPI_ISL_3294024, EPI_ISL_3294025, EPI_ISL_3294026,
EPI_ISL_3294027, EPI_ISL_3294028, EPI_ISL_3294029, EPI_ISL_3294030, EPI_ISL_3294031, EPI_ISL_3294032, EPI_ISL_3294033, EPI_ISL_3294034, EPI_ISL_3294035, EPI_ISL_3294036, EPI_ISL_3294037, EPI_ISL_3294038, EPI_ISL_3294039, EPI_ISL_3294040, EPI_ISL_3294041, EPI_ISL_3294042, EPI_ISL_3294043, EPI_ISL_3294044, EPI_ISL_3294045, EPI_ISL_3294046, EPI_ISL_3294047, EPI_ISL_3294048,
EPI_ISL_3294049, EPI_ISL_3294050, EPI_ISL_3294051, EPI_ISL_3294052, EPI_ISL_3294053, EPI_ISL_3294054, EPI_ISL_3294055, EPI_ISL_3294056, EPI_ISL_3294057, EPI_ISL_3294058, EPI_ISL_3294059, EPI_ISL_3294060, EPI_ISL_3294061, EPI_ISL_3294062, EPI_ISL_3294063, EPI_ISL_3294064, EPI_ISL_3294065, EPI_ISL_3294066, EPI_ISL_3294067, EPI_ISL_3294068, EPI_ISL_3294069, EPI_ISL_3294070,
EPI_ISL_3294071, EPI_ISL_3294072, EPI_ISL_3294073, EPI_ISL_3294074, EPI_ISL_3294075, EPI_ISL_3294076, EPI_ISL_3294077, EPI_ISL_3294078, EPI_ISL_3294079, EPI_ISL_3294080, EPI_ISL_3294081, EPI_ISL_3294082, EPI_ISL_3294083, EPI_ISL_3294084, EPI_ISL_3294085, EPI_ISL_3294086, EPI_ISL_3294087, EPI_ISL_3294088, EPI_ISL_3294089, EPI_ISL_3294090, EPI_ISL_3294091, EPI_ISL_3294092,
EPI_ISL_3294093, EPI_ISL_3294094, EPI_ISL_3294095, EPI_ISL_3294096, EPI_ISL_3294097, EPI_ISL_3294098, EPI_ISL_3294099, EPI_ISL_3294100, EPI_ISL_3294101, EPI_ISL_3294102, EPI_ISL_3294103, EPI_ISL_3294104, EPI_ISL_3294105, EPI_ISL_3294106, EPI_ISL_3294107, EPI_ISL_3294108, EPI_ISL_3294109, EPI_ISL_3294110, EPI_ISL_3294111, EPI_ISL_3294112, EPI_ISL_3294113, EPI_ISL_3294114,
EPI_ISL_3294115, EPI_ISL_3294116, EPI_ISL_3294117, EPI_ISL_3294118, EPI_ISL_3294119, EPI_ISL_3294120, EPI_ISL_3294121, EPI_ISL_3294122, EPI_ISL_3294123, EPI_ISL_3294124, EPI_ISL_3294125, EPI_ISL_3294126, EPI_ISL_3294127, EPI_ISL_3294128, EPI_ISL_3294129, EPI_ISL_3294130, EPI_ISL_3294131, EPI_ISL_3294132, EPI_ISL_3294133, EPI_ISL_3294134, EPI_ISL_3294135, EPI_ISL_3294136,
EPI_ISL_3294137, EPI_ISL_3294138, EPI_ISL_3294139, EPI_ISL_3294140, EPI_ISL_3294141, EPI_ISL_3294142, EPI_ISL_3294143, EPI_ISL_3294144, EPI_ISL_3294145, EPI_ISL_3294146, EPI_ISL_3294147, EPI_ISL_3294148, EPI_ISL_3294149, EPI_ISL_3294150, EPI_ISL_3294151, EPI_ISL_3294152, EPI_ISL_3294153, EPI_ISL_3294154, EPI_ISL_3294155, EPI_ISL_3294156, EPI_ISL_3294157, EPI_ISL_3294158,
EPI_ISL_3294159, EPI_ISL_3294160, EPI_ISL_3294161, EPI_ISL_3294162, EPI_ISL_3294163, EPI_ISL_3294164, EPI_ISL_3294165, EPI_ISL_3294166, EPI_ISL_3294167, EPI_ISL_3294168, EPI_ISL_3294169, EPI_ISL_3294170, EPI_ISL_3294171, EPI_ISL_3294172, EPI_ISL_3294173, EPI_ISL_3294174, EPI_ISL_3294175, EPI_ISL_3294176, EPI_ISL_3294177, EPI_ISL_3294178, EPI_ISL_3294179, EPI_ISL_3294180,
EPI_ISL_3294181, EPI_ISL_3294182, EPI_ISL_3294183, EPI_ISL_3294184, EPI_ISL_3294185, EPI_ISL_3294186, EPI_ISL_3294187, EPI_ISL_3294188, EPI_ISL_3294189, EPI_ISL_3294190, EPI_ISL_3294191, EPI_ISL_3294192, EPI_ISL_3294193, EPI_ISL_3294194, EPI_ISL_3294195, EPI_ISL_3294196, EPI_ISL_3294197, EPI_ISL_3294198, EPI_ISL_3294199, EPI_ISL_3294200, EPI_ISL_3294201, EPI_ISL_3294202,
EPI_ISL_3294203, EPI_ISL_3294204, EPI_ISL_3294205, EPI_ISL_3294206, EPI_ISL_3294207, EPI_ISL_3294208, EPI_ISL_3294209, EPI_ISL_3294210, EPI_ISL_3294211, EPI_ISL_3294212, EPI_ISL_3294213, EPI_ISL_3294214, EPI_ISL_3294215, EPI_ISL_3294216, EPI_ISL_3294217, EPI_ISL_3294218, EPI_ISL_3294219, EPI_ISL_3294220, EPI_ISL_3294221, EPI_ISL_3294222, EPI_ISL_3294223, EPI_ISL_3294224,
EPI_ISL_3294225, EPI_ISL_3294226, EPI_ISL_3294227, EPI_ISL_3294228, EPI_ISL_3294229, EPI_ISL_3294230, EPI_ISL_3294231, EPI_ISL_3294232, EPI_ISL_3294233, EPI_ISL_3294234, EPI_ISL_3294235, EPI_ISL_3294236, EPI_ISL_3294237, EPI_ISL_3294238, EPI_ISL_3294239, EPI_ISL_3294240, EPI_ISL_3294241, EPI_ISL_3294242, EPI_ISL_3294243, EPI_ISL_3294244, EPI_ISL_3294245, EPI_ISL_3294246,
EPI_ISL_3294247, EPI_ISL_3294248, EPI_ISL_3294249, EPI_ISL_3294250, EPI_ISL_3294251, EPI_ISL_3294252, EPI_ISL_3294253, EPI_ISL_3294254, EPI_ISL_3294255, EPI_ISL_3294256, EPI_ISL_3294257, EPI_ISL_3294258, EPI_ISL_3294259, EPI_ISL_3294260, EPI_ISL_3294261, EPI_ISL_3294262, EPI_ISL_3294263, EPI_ISL_3294264, EPI_ISL_3294265, EPI_ISL_3294266, EPI_ISL_3294267, EPI_ISL_3294268,
EPI_ISL_3294269, EPI_ISL_3294270, EPI_ISL_3294271, EPI_ISL_3294272, EPI_ISL_3294273, EPI_ISL_3294274, EPI_ISL_3294275, EPI_ISL_3294276, EPI_ISL_3294277, EPI_ISL_3294278, EPI_ISL_3294279, EPI_ISL_3294280, EPI_ISL_3294281, EPI_ISL_3294282, EPI_ISL_3294283, EPI_ISL_3294284, EPI_ISL_3294285, EPI_ISL_3294286, EPI_ISL_3294287, EPI_ISL_3294288, EPI_ISL_3294289, EPI_ISL_3294290,
EPI_ISL_3294291, EPI_ISL_3294292, EPI_ISL_3294293, EPI_ISL_3294294, EPI_ISL_3294295, EPI_ISL_3294296, EPI_ISL_3294297, EPI_ISL_3294298, EPI_ISL_3294299, EPI_ISL_3294300, EPI_ISL_3294301, EPI_ISL_3294302, EPI_ISL_3294303, EPI_ISL_3294304, EPI_ISL_3294305, EPI_ISL_3294306, EPI_ISL_3294307, EPI_ISL_3294308, EPI_ISL_3294309, EPI_ISL_3294310, EPI_ISL_3294311, EPI_ISL_3294312,
EPI_ISL_3294313, EPI_ISL_3294314, EPI_ISL_3294315, EPI_ISL_3294316, EPI_ISL_3294317, EPI_ISL_3294318, EPI_ISL_3294319, EPI_ISL_3294320, EPI_ISL_3294321, EPI_ISL_3294322, EPI_ISL_3294323, EPI_ISL_3294324, EPI_ISL_3294325, EPI_ISL_3294326, EPI_ISL_3294327, EPI_ISL_3294328, EPI_ISL_3294329, EPI_ISL_3294330, EPI_ISL_3294331, EPI_ISL_3294332, EPI_ISL_3294333, EPI_ISL_3294334,
EPI_ISL_3294335, EPI_ISL_3294336, EPI_ISL_3294337, EPI_ISL_3294338, EPI_ISL_3294339, EPI_ISL_3294340, EPI_ISL_3294341, EPI_ISL_3294342, EPI_ISL_3294343, EPI_ISL_3294344, EPI_ISL_3294345, EPI_ISL_3294346, EPI_ISL_3294347, EPI_ISL_3294348, EPI_ISL_3294349, EPI_ISL_3294350, EPI_ISL_3294351, EPI_ISL_3294352, EPI_ISL_3294353, EPI_ISL_3294354, EPI_ISL_3294355, EPI_ISL_3294356,
EPI_ISL_3294357, EPI_ISL_3294358, EPI_ISL_3294359, EPI_ISL_3294360, EPI_ISL_3294361, EPI_ISL_3294362, EPI_ISL_3294363, EPI_ISL_3294364, EPI_ISL_3294365, EPI_ISL_3294366, EPI_ISL_3294367, EPI_ISL_3294368, EPI_ISL_3294369, EPI_ISL_3294370, EPI_ISL_3294371, EPI_ISL_3294372, EPI_ISL_3294373, EPI_ISL_3294374, EPI_ISL_3294375, EPI_ISL_3294376, EPI_ISL_3294377, EPI_ISL_3294378,
EPI_ISL_3294379, EPI_ISL_3294380, EPI_ISL_3294381, EPI_ISL_3294382, EPI_ISL_3294383, EPI_ISL_3294384, EPI_ISL_3294385, EPI_ISL_3294386, EPI_ISL_3294387, EPI_ISL_3294388, EPI_ISL_3294389, EPI_ISL_3294390, EPI_ISL_3294391, EPI_ISL_3294392, EPI_ISL_3294393, EPI_ISL_3294394, EPI_ISL_3294395, EPI_ISL_3294396, EPI_ISL_3294397, EPI_ISL_3294398, EPI_ISL_3294399, EPI_ISL_3294400,
EPI_ISL_3294401, EPI_ISL_3294402, EPI_ISL_3294403, EPI_ISL_3294404, EPI_ISL_3294405, EPI_ISL_3294406, EPI_ISL_3294407, EPI_ISL_3294408, EPI_ISL_3294409, EPI_ISL_3294410, EPI_ISL_3294411, EPI_ISL_3294412, EPI_ISL_3294413, EPI_ISL_3294414, EPI_ISL_3294415, EPI_ISL_3294416, EPI_ISL_3294417, EPI_ISL_3294418, EPI_ISL_3294419, EPI_ISL_3294420, EPI_ISL_3294421, EPI_ISL_3294422,
EPI_ISL_3294423, EPI_ISL_3294424, EPI_ISL_3294425, EPI_ISL_3294426, EPI_ISL_3294427, EPI_ISL_3294428, EPI_ISL_3294429, EPI_ISL_3294430, EPI_ISL_3294431, EPI_ISL_3294432, EPI_ISL_3294433, EPI_ISL_3294434, EPI_ISL_3294435, EPI_ISL_3294436, EPI_ISL_3294437, EPI_ISL_3294438, EPI_ISL_3294439, EPI_ISL_3294440, EPI_ISL_3294441, EPI_ISL_3294442, EPI_ISL_3294443, EPI_ISL_3294444,
EPI_ISL_3294445, EPI_ISL_3294446, EPI_ISL_3294447, EPI_ISL_3294448, EPI_ISL_3294449, EPI_ISL_3294450, EPI_ISL_3294451, EPI_ISL_3294452, EPI_ISL_3294453, EPI_ISL_3294454, EPI_ISL_3294455, EPI_ISL_3294456, EPI_ISL_3294457, EPI_ISL_3294458, EPI_ISL_3294459, EPI_ISL_3294460, EPI_ISL_3294461, EPI_ISL_3294462, EPI_ISL_3294463, EPI_ISL_3294464, EPI_ISL_3294465, EPI_ISL_3294466,
EPI_ISL_3294467, EPI_ISL_3294468, EPI_ISL_3294469, EPI_ISL_3294470, EPI_ISL_3294471, EPI_ISL_3294472, EPI_ISL_3294473, EPI_ISL_3294474, EPI_ISL_3294475, EPI_ISL_3294476, EPI_ISL_3294477, EPI_ISL_3294478, EPI_ISL_3294479, EPI_ISL_3294480, EPI_ISL_3294481, EPI_ISL_3294482, EPI_ISL_3294483, EPI_ISL_3294484, EPI_ISL_3294485, EPI_ISL_3294486, EPI_ISL_3294487, EPI_ISL_3294488,
EPI_ISL_3294489, EPI_ISL_3294490, EPI_ISL_3294491, EPI_ISL_3294492, EPI_ISL_3294493, EPI_ISL_3294494, EPI_ISL_3294495, EPI_ISL_3294496, EPI_ISL_3294497, EPI_ISL_3294498, EPI_ISL_3294499, EPI_ISL_3294500, EPI_ISL_3294501, EPI_ISL_3294502, EPI_ISL_3294503, EPI_ISL_3294504, EPI_ISL_3294505, EPI_ISL_3294506, EPI_ISL_3294507, EPI_ISL_3294508, EPI_ISL_3294509, EPI_ISL_3294510,
EPI_ISL_3294511, EPI_ISL_3294512, EPI_ISL_3294513, EPI_ISL_3294514, EPI_ISL_3294515, EPI_ISL_3294516, EPI_ISL_3294517, EPI_ISL_3294518, EPI_ISL_3294519, EPI_ISL_3294520, EPI_ISL_3294521, EPI_ISL_3294522, EPI_ISL_3294523, EPI_ISL_3294524, EPI_ISL_3294525, EPI_ISL_3294526, EPI_ISL_3294527, EPI_ISL_3294528, EPI_ISL_3294529, EPI_ISL_3294530, EPI_ISL_3294531, EPI_ISL_3294532,
EPI_ISL_3294533, EPI_ISL_3294534, EPI_ISL_3294535, EPI_ISL_3294536, EPI_ISL_3294537, EPI_ISL_3294538, EPI_ISL_3294539, EPI_ISL_3294540, EPI_ISL_3294541, EPI_ISL_3294542, EPI_ISL_3294543, EPI_ISL_3294544, EPI_ISL_3294545, EPI_ISL_3294546, EPI_ISL_3294547, EPI_ISL_3294548, EPI_ISL_3294549, EPI_ISL_3294550, EPI_ISL_3294551, EPI_ISL_3294552, EPI_ISL_3294553, EPI_ISL_3294554,
EPI_ISL_3294555, EPI_ISL_3294556, EPI_ISL_3294557, EPI_ISL_3294558, EPI_ISL_3294559, EPI_ISL_3294560, EPI_ISL_3294561, EPI_ISL_3294562, EPI_ISL_3294563, EPI_ISL_3294564, EPI_ISL_3294565, EPI_ISL_3294566, EPI_ISL_3294567, EPI_ISL_3294568, EPI_ISL_3294569, EPI_ISL_3294570, EPI_ISL_3294571, EPI_ISL_3294572, EPI_ISL_3294573, EPI_ISL_3294574, EPI_ISL_3294575, EPI_ISL_3294576,
EPI_ISL_3294577, EPI_ISL_3294578, EPI_ISL_3294579, EPI_ISL_3294580, EPI_ISL_3294581, EPI_ISL_3294582, EPI_ISL_3294583, EPI_ISL_3294584, EPI_ISL_3294585, EPI_ISL_3294586, EPI_ISL_3294587, EPI_ISL_3294588, EPI_ISL_3294589, EPI_ISL_3294590, EPI_ISL_3294591, EPI_ISL_3294592, EPI_ISL_3294593, EPI_ISL_3294594, EPI_ISL_3294595, EPI_ISL_3294596, EPI_ISL_3294597, EPI_ISL_3294598, EPI_ISL_3294599,
EPI_ISL_3294600, EPI_ISL_3294601, EPI_ISL_3294602, EPI_ISL_3294603, EPI_ISL_3294604, EPI_ISL_3294605, EPI_ISL_3294606, EPI_ISL_3294607, EPI_ISL_3294608, EPI_ISL_3294609, EPI_ISL_3294610, EPI_ISL_3294611, EPI_ISL_3294612, EPI_ISL_3294613, EPI_ISL_3294614, EPI_ISL_3294615, EPI_ISL_3294616, EPI_ISL_3294617, EPI_ISL_3294618, EPI_ISL_3294619, EPI_ISL_3294620, EPI_ISL_3294621,
EPI_ISL_3294622, EPI_ISL_3294623, EPI_ISL_3294624, EPI_ISL_3294625, EPI_ISL_3294626, EPI_ISL_3294627, EPI_ISL_3294628, EPI_ISL_3294629, EPI_ISL_3294630, EPI_ISL_3294631, EPI_ISL_3294632, EPI_ISL_3294633, EPI_ISL_3294634, EPI_ISL_3294635, EPI_ISL_3294636, EPI_ISL_3294637, EPI_ISL_3294638, EPI_ISL_3294639, EPI_ISL_3294640, EPI_ISL_3294641, EPI_ISL_3294642, EPI_ISL_3294643,
EPI_ISL_3294644, EPI_ISL_3294645, EPI_ISL_3294646, EPI_ISL_3294647, EPI_ISL_3294648, EPI_ISL_3294649, EPI_ISL_3294650, EPI_ISL_3294651, EPI_ISL_3294652, EPI_ISL_3294653, EPI_ISL_3294654, EPI_ISL_3294655, EPI_ISL_3294656, EPI_ISL_3294657, EPI_ISL_3294658, EPI_ISL_3294659, EPI_ISL_3294660, EPI_ISL_3294661, EPI_ISL_3294662, EPI_ISL_3294663, EPI_ISL_3294664, EPI_ISL_3294665,
EPI_ISL_3294666, EPI_ISL_3294667, EPI_ISL_3294668, EPI_ISL_3294669, EPI_ISL_3294670, EPI_ISL_3294671, EPI_ISL_3294672, EPI_ISL_3294673, EPI_ISL_3294674, EPI_ISL_3294675, EPI_ISL_3294676, EPI_ISL_3294677, EPI_ISL_3294678, EPI_ISL_3294679, EPI_ISL_3294680, EPI_ISL_3294681, EPI_ISL_3294682, EPI_ISL_3294683, EPI_ISL_3294684, EPI_ISL_3294685, EPI_ISL_3294686, EPI_ISL_3294687,
EPI_ISL_3294688, EPI_ISL_3294689, EPI_ISL_3294690, EPI_ISL_3294691, EPI_ISL_3294692, EPI_ISL_3294693, EPI_ISL_3294694, EPI_ISL_3294695, EPI_ISL_3294696, EPI_ISL_3294697, EPI_ISL_3294698, EPI_ISL_3294699, EPI_ISL_3294700, EPI_ISL_3294701, EPI_ISL_3294702, EPI_ISL_3294703, EPI_ISL_3294704, EPI_ISL_3294705, EPI_ISL_3294706, EPI_ISL_3294707, EPI_ISL_3294708, EPI_ISL_3294709,
EPI_ISL_3294710, EPI_ISL_3294711, EPI_ISL_3294712, EPI_ISL_3294713, EPI_ISL_3294714, EPI_ISL_3294715, EPI_ISL_3294716, EPI_ISL_3294717, EPI_ISL_3294718, EPI_ISL_3294719, EPI_ISL_3294720, EPI_ISL_3294721, EPI_ISL_3294722, EPI_ISL_3294723, EPI_ISL_3294724, EPI_ISL_3294725, EPI_ISL_3294726, EPI_ISL_3294727, EPI_ISL_3294728, EPI_ISL_3294729, EPI_ISL_3294730, EPI_ISL_3294731,
EPI_ISL_3294732, EPI_ISL_3294733, EPI_ISL_3294734, EPI_ISL_3294735, EPI_ISL_3294736, EPI_ISL_3294737, EPI_ISL_3294738, EPI_ISL_3294739, EPI_ISL_3294740, EPI_ISL_3294741, EPI_ISL_3294742, EPI_ISL_3294743, EPI_ISL_3294744, EPI_ISL_3294745, EPI_ISL_3294746, EPI_ISL_3294747, EPI_ISL_3294748, EPI_ISL_3294749, EPI_ISL_3294750, EPI_ISL_3294751, EPI_ISL_3294752, EPI_ISL_3294753,
EPI_ISL_3294754, EPI_ISL_3294755, EPI_ISL_3294756, EPI_ISL_3294757, EPI_ISL_3294758, EPI_ISL_3294759, EPI_ISL_3294760, EPI_ISL_3294761, EPI_ISL_3294762, EPI_ISL_3294763, EPI_ISL_3294764, EPI_ISL_3294765, EPI_ISL_3294766, EPI_ISL_3294767, EPI_ISL_3294768, EPI_ISL_3294769, EPI_ISL_3294770, EPI_ISL_3294771, EPI_ISL_3294772, EPI_ISL_3294773, EPI_ISL_3294774, EPI_ISL_3294775,
EPI_ISL_3294776, EPI_ISL_3294777, EPI_ISL_3294778, EPI_ISL_3294779, EPI_ISL_3294780, EPI_ISL_3294781, EPI_ISL_3294782, EPI_ISL_3294783, EPI_ISL_3294784, EPI_ISL_3294785, EPI_ISL_3294786, EPI_ISL_3294787, EPI_ISL_3294788, EPI_ISL_3294789, EPI_ISL_3294790, EPI_ISL_3294791, EPI_ISL_3294792, EPI_ISL_3294793, EPI_ISL_3294794, EPI_ISL_3294795, EPI_ISL_3294796, EPI_ISL_3294797,
EPI_ISL_3294798, EPI_ISL_3294799, EPI_ISL_3294800, EPI_ISL_3294801, EPI_ISL_3294802, EPI_ISL_3294803, EPI_ISL_3294804, EPI_ISL_3294805, EPI_ISL_3294806, EPI_ISL_3294807, EPI_ISL_3294808, EPI_ISL_3294809, EPI_ISL_3294810, EPI_ISL_3294811, EPI_ISL_3294812, EPI_ISL_3294813, EPI_ISL_3294814, EPI_ISL_3294815, EPI_ISL_3294816, EPI_ISL_3294817, EPI_ISL_3294818, EPI_ISL_3294819,
EPI_ISL_3294820, EPI_ISL_3294821, EPI_ISL_3294822, EPI_ISL_3294823, EPI_ISL_3294824, EPI_ISL_3294825, EPI_ISL_3294826, EPI_ISL_3294827, EPI_ISL_3294828, EPI_ISL_3294829, EPI_ISL_3294830, EPI_ISL_3294831, EPI_ISL_3294832, EPI_ISL_3294833, EPI_ISL_3294834, EPI_ISL_3294835, EPI_ISL_3294836, EPI_ISL_3294837, EPI_ISL_3294838, EPI_ISL_3294839, EPI_ISL_3294840, EPI_ISL_3294841,
EPI_ISL_3294842, EPI_ISL_3294843, EPI_ISL_3294844, EPI_ISL_3294845, EPI_ISL_3294846, EPI_ISL_3294847, EPI_ISL_3294848, EPI_ISL_3294849, EPI_ISL_3294850, EPI_ISL_3294851, EPI_ISL_3294852, EPI_ISL_3294853, EPI_ISL_3294854, EPI_ISL_3294855, EPI_ISL_3294856, EPI_ISL_3294857, EPI_ISL_3294858, EPI_ISL_3294859, EPI_ISL_3294860, EPI_ISL_3294861, EPI_ISL_3294862, EPI_ISL_3294863,
EPI_ISL_3294864, EPI_ISL_3294865, EPI_ISL_3294866, EPI_ISL_3294867, EPI_ISL_3294868, EPI_ISL_3294869, EPI_ISL_3294870, EPI_ISL_3294871, EPI_ISL_3294872, EPI_ISL_3294873, EPI_ISL_3294874, EPI_ISL_3294875, EPI_ISL_3294876, EPI_ISL_3294877, EPI_ISL_3294878, EPI_ISL_3294879, EPI_ISL_3294880, EPI_ISL_3294881, EPI_ISL_3294882, EPI_ISL_3294883, EPI_ISL_3294884, EPI_ISL_3294885,
EPI_ISL_3294886, EPI_ISL_3294887, EPI_ISL_3294888, EPI_ISL_3294889, EPI_ISL_3294890, EPI_ISL_3294891, EPI_ISL_3294892, EPI_ISL_3294893, EPI_ISL_3294894, EPI_ISL_3294895, EPI_ISL_3294896, EPI_ISL_3294897, EPI_ISL_3294898, EPI_ISL_3294899, EPI_ISL_3294900, EPI_ISL_3294901, EPI_ISL_3294902, EPI_ISL_3294903, EPI_ISL_3294904, EPI_ISL_3294905, EPI_ISL_3294906, EPI_ISL_3294907,
EPI_ISL_3294908, EPI_ISL_3294909, EPI_ISL_3294910, EPI_ISL_3294911, EPI_ISL_3294912, EPI_ISL_3294913, EPI_ISL_3294914, EPI_ISL_3294915, EPI_ISL_3294916, EPI_ISL_3294917, EPI_ISL_3294918, EPI_ISL_3294919, EPI_ISL_3294920, EPI_ISL_3294921, EPI_ISL_3294922, EPI_ISL_3294923, EPI_ISL_3294924, EPI_ISL_3294925, EPI_ISL_3294926, EPI_ISL_3294927, EPI_ISL_3294928, EPI_ISL_3294929,
EPI_ISL_3294930, EPI_ISL_3294931, EPI_ISL_3294932, EPI_ISL_3294933, EPI_ISL_3294934, EPI_ISL_3294935, EPI_ISL_3294936, EPI_ISL_3294937, EPI_ISL_3294938, EPI_ISL_3294939, EPI_ISL_3294940, EPI_ISL_3294941, EPI_ISL_3294942, EPI_ISL_3294943, EPI_ISL_3294944, EPI_ISL_3294945, EPI_ISL_3294946, EPI_ISL_3294947, EPI_ISL_3294948, EPI_ISL_3294949, EPI_ISL_3294950, EPI_ISL_3294951,
EPI_ISL_3294952, EPI_ISL_3294953, EPI_ISL_3294954, EPI_ISL_3294955, EPI_ISL_3294956, EPI_ISL_3294957, EPI_ISL_3294958, EPI_ISL_3294959, EPI_ISL_3294960, EPI_ISL_3294961, EPI_ISL_3294962, EPI_ISL_3294963, EPI_ISL_3294964, EPI_ISL_3294965, EPI_ISL_3294966, EPI_ISL_3294967, EPI_ISL_3294968, EPI_ISL_3294969, EPI_ISL_3294970, EPI_ISL_3294971, EPI_ISL_3294972, EPI_ISL_3294973,
EPI_ISL_3294974, EPI_ISL_3294975, EPI_ISL_3294976, EPI_ISL_3294977, EPI_ISL_3294978, EPI_ISL_3294979, EPI_ISL_3294980, EPI_ISL_3294981, EPI_ISL_3294982, EPI_ISL_3294983, EPI_ISL_3294984, EPI_ISL_3294985, EPI_ISL_3294986, EPI_ISL_3294987, EPI_ISL_3294988, EPI_ISL_3294989, EPI_ISL_3294990, EPI_ISL_3294991, EPI_ISL_3294992, EPI_ISL_3294993, EPI_ISL_3294994, EPI_ISL_3294995,
EPI_ISL_3294996, EPI_ISL_3294997, EPI_ISL_3294998, EPI_ISL_3294999, EPI_ISL_3295000, EPI_ISL_3295001, EPI_ISL_3295002, EPI_ISL_3295003, EPI_ISL_3295004, EPI_ISL_3295005, EPI_ISL_3295006, EPI_ISL_3295007, EPI_ISL_3295008, EPI_ISL_3295009, EPI_ISL_3295010, EPI_ISL_3295011, EPI_ISL_3295012, EPI_ISL_3295013, EPI_ISL_3295014, EPI_ISL_3295015, EPI_ISL_3295016, EPI_ISL_3295017,
EPI_ISL_3295018, EPI_ISL_3295019, EPI_ISL_3295020, EPI_ISL_3295021, EPI_ISL_3295022, EPI_ISL_3295023, EPI_ISL_3295024, EPI_ISL_3295025, EPI_ISL_3295026, EPI_ISL_3295027, EPI_ISL_3295028, EPI_ISL_3295029, EPI_ISL_3295030, EPI_ISL_3295031, EPI_ISL_3295032, EPI_ISL_3295033, EPI_ISL_3295034, EPI_ISL_3295035, EPI_ISL_3295036, EPI_ISL_3295037, EPI_ISL_3295038, EPI_ISL_3295039,
EPI_ISL_3295040, EPI_ISL_3295041, EPI_ISL_3295042, EPI_ISL_3295043, EPI_ISL_3295044, EPI_ISL_3295045, EPI_ISL_3295046, EPI_ISL_3295047, EPI_ISL_3295048, EPI_ISL_3295049, EPI_ISL_3295050, EPI_ISL_3295051, EPI_ISL_3295052, EPI_ISL_3295053, EPI_ISL_3295054, EPI_ISL_3295055, EPI_ISL_3295056, EPI_ISL_3295057, EPI_ISL_3295058, EPI_ISL_3295059, EPI_ISL_3295060, EPI_ISL_3295061,
EPI_ISL_3295062, EPI_ISL_3295063, EPI_ISL_3295064, EPI_ISL_3295065, EPI_ISL_3295066, EPI_ISL_3295067, EPI_ISL_3295068, EPI_ISL_3295069, EPI_ISL_3295070, EPI_ISL_3295071, EPI_ISL_3295072, EPI_ISL_3295073, EPI_ISL_3295074, EPI_ISL_3295075, EPI_ISL_3295076, EPI_ISL_3295077, EPI_ISL_3295078, EPI_ISL_3295079, EPI_ISL_3295080, EPI_ISL_3295081, EPI_ISL_3295082, EPI_ISL_3295083,
EPI_ISL_3295084, EPI_ISL_3295085, EPI_ISL_3295086, EPI_ISL_3295087, EPI_ISL_3295088, EPI_ISL_3295089, EPI_ISL_3295090, EPI_ISL_3295091, EPI_ISL_3295092, EPI_ISL_3295093, EPI_ISL_3295094, EPI_ISL_3295095, EPI_ISL_3295096, EPI_ISL_3295097, EPI_ISL_3295098, EPI_ISL_3295099, EPI_ISL_3295100, EPI_ISL_3295101, EPI_ISL_3295102, EPI_ISL_3295103, EPI_ISL_3295104, EPI_ISL_3295105,
EPI_ISL_3295106, EPI_ISL_3295107, EPI_ISL_3295108, EPI_ISL_3295109, EPI_ISL_3295110, EPI_ISL_3295111, EPI_ISL_3295112, EPI_ISL_3295113, EPI_ISL_3295114, EPI_ISL_3295115, EPI_ISL_3295116, EPI_ISL_3295117, EPI_ISL_3295118, EPI_ISL_3295119, EPI_ISL_3295120, EPI_ISL_3295121, EPI_ISL_3295122, EPI_ISL_3295123, EPI_ISL_3295124, EPI_ISL_3295125, EPI_ISL_3295126, EPI_ISL_3295127,
EPI_ISL_3295128, EPI_ISL_3295129, EPI_ISL_3295130, EPI_ISL_3295131, EPI_ISL_3295132, EPI_ISL_3295133, EPI_ISL_3295134, EPI_ISL_3295135, EPI_ISL_3295136, EPI_ISL_3295137, EPI_ISL_3295138, EPI_ISL_3295139, EPI_ISL_3295140, EPI_ISL_3295141, EPI_ISL_3295142, EPI_ISL_3295143, EPI_ISL_3295144, EPI_ISL_3295145, EPI_ISL_3295146, EPI_ISL_3295147, EPI_ISL_3295148, EPI_ISL_3295149,
EPI_ISL_3295150, EPI_ISL_3295151, EPI_ISL_3295152, EPI_ISL_3295153, EPI_ISL_3295154, EPI_ISL_3295155, EPI_ISL_3295156, EPI_ISL_3295157, EPI_ISL_3295158, EPI_ISL_3295159, EPI_ISL_3295160, EPI_ISL_3295161, EPI_ISL_3295162, EPI_ISL_3295163, EPI_ISL_3295164, EPI_ISL_3295165, EPI_ISL_3295166, EPI_ISL_3295167, EPI_ISL_3295168, EPI_ISL_3295169, EPI_ISL_3295170, EPI_ISL_3295171,
EPI_ISL_3295172, EPI_ISL_3295173, EPI_ISL_3295174, EPI_ISL_3295175, EPI_ISL_3295176, EPI_ISL_3295177, EPI_ISL_3295178, EPI_ISL_3295179, EPI_ISL_3295180, EPI_ISL_3295181, EPI_ISL_3295182, EPI_ISL_3295183, EPI_ISL_3295184, EPI_ISL_3295185, EPI_ISL_3295186, EPI_ISL_3295187, EPI_ISL_3295188, EPI_ISL_3295189, EPI_ISL_3295190, EPI_ISL_3295191, EPI_ISL_3295192, EPI_ISL_3295193,
EPI_ISL_3295194, EPI_ISL_3295195, EPI_ISL_3295196, EPI_ISL_3295197, EPI_ISL_3295198, EPI_ISL_3295199, EPI_ISL_3295200, EPI_ISL_3295201, EPI_ISL_3295202, EPI_ISL_3295203, EPI_ISL_3295204, EPI_ISL_3295205, EPI_ISL_3295206, EPI_ISL_3295207, EPI_ISL_3295208, EPI_ISL_3295209, EPI_ISL_3295210, EPI_ISL_3295211, EPI_ISL_3295212, EPI_ISL_3295213, EPI_ISL_3295214, EPI_ISL_3295215,
EPI_ISL_3295216, EPI_ISL_3295217, EPI_ISL_3295218, EPI_ISL_3295219, EPI_ISL_3295220, EPI_ISL_3295221, EPI_ISL_3295222, EPI_ISL_3295223, EPI_ISL_3295224, EPI_ISL_3295225, EPI_ISL_3295226, EPI_ISL_3295227, EPI_ISL_3295228, EPI_ISL_3295229, EPI_ISL_3295230, EPI_ISL_3295231, EPI_ISL_3295232, EPI_ISL_3295233, EPI_ISL_3295234, EPI_ISL_3295235, EPI_ISL_3295236, EPI_ISL_3295237,
EPI_ISL_3295238, EPI_ISL_3295239, EPI_ISL_3295240, EPI_ISL_3295241, EPI_ISL_3295242, EPI_ISL_3295243, EPI_ISL_3295244, EPI_ISL_3295245, EPI_ISL_3295246, EPI_ISL_3295247, EPI_ISL_3295248, EPI_ISL_3295249, EPI_ISL_3295250, EPI_ISL_3295251, EPI_ISL_3295252, EPI_ISL_3295253, EPI_ISL_3295254, EPI_ISL_3295255, EPI_ISL_3295256, EPI_ISL_3295257, EPI_ISL_3295258, EPI_ISL_3295259,
EPI_ISL_3295260, EPI_ISL_3295261, EPI_ISL_3295262, EPI_ISL_3295263, EPI_ISL_3295264, EPI_ISL_3295265, EPI_ISL_3295266, EPI_ISL_3295267, EPI_ISL_3295268, EPI_ISL_3295269, EPI_ISL_3295270, EPI_ISL_3295271, EPI_ISL_3295272, EPI_ISL_3295273, EPI_ISL_3295274, EPI_ISL_3295275, EPI_ISL_3295276, EPI_ISL_3295277, EPI_ISL_3295278, EPI_ISL_3295279, EPI_ISL_3295280, EPI_ISL_3295281,
EPI_ISL_3295282, EPI_ISL_3295283, EPI_ISL_3295284, EPI_ISL_3295285, EPI_ISL_3295286, EPI_ISL_3295287, EPI_ISL_3295288, EPI_ISL_3295289, EPI_ISL_3295290, EPI_ISL_3295291, EPI_ISL_3295292, EPI_ISL_3295293, EPI_ISL_3295294, EPI_ISL_3295295, EPI_ISL_3295296, EPI_ISL_3295297, EPI_ISL_3295298, EPI_ISL_3295299, EPI_ISL_3295300, EPI_ISL_3295301, EPI_ISL_3295302, EPI_ISL_3295303,
EPI_ISL_3295304, EPI_ISL_3295305, EPI_ISL_3295306, EPI_ISL_3295307, EPI_ISL_3295308, EPI_ISL_3295309, EPI_ISL_3295310, EPI_ISL_3295311, EPI_ISL_3295312, EPI_ISL_3295313, EPI_ISL_3295314, EPI_ISL_3295315, EPI_ISL_3295316, EPI_ISL_3295317, EPI_ISL_3295318, EPI_ISL_3295319, EPI_ISL_3295320, EPI_ISL_3295321, EPI_ISL_3295322, EPI_ISL_3295323, EPI_ISL_3295324, EPI_ISL_3295325,
EPI_ISL_3295326, EPI_ISL_3295327, EPI_ISL_3295328, EPI_ISL_3295329, EPI_ISL_3295330, EPI_ISL_3295331, EPI_ISL_3295332, EPI_ISL_3295333, EPI_ISL_3295334, EPI_ISL_3295335, EPI_ISL_3295336, EPI_ISL_3295337, EPI_ISL_3295338, EPI_ISL_3295339, EPI_ISL_3295340, EPI_ISL_3295341, EPI_ISL_3295342, EPI_ISL_3295343, EPI_ISL_3295344, EPI_ISL_3295345, EPI_ISL_3295346, EPI_ISL_3295347,
EPI_ISL_3295348, EPI_ISL_3295349, EPI_ISL_3295350, EPI_ISL_3295351, EPI_ISL_3295352, EPI_ISL_3295353, EPI_ISL_3295354, EPI_ISL_3295355, EPI_ISL_3295356, EPI_ISL_3295357, EPI_ISL_3295358, EPI_ISL_3295359, EPI_ISL_3295360, EPI_ISL_3295361, EPI_ISL_3295362, EPI_ISL_3295363, EPI_ISL_3295364, EPI_ISL_3295365, EPI_ISL_3295366, EPI_ISL_3295367, EPI_ISL_3295368, EPI_ISL_3295369,
EPI_ISL_3295370, EPI_ISL_3295371, EPI_ISL_3295372, EPI_ISL_3295373, EPI_ISL_3295374, EPI_ISL_3295375, EPI_ISL_3295376, EPI_ISL_3295377, EPI_ISL_3295378, EPI_ISL_3295379, EPI_ISL_3295380, EPI_ISL_3295381, EPI_ISL_3295382, EPI_ISL_3295383, EPI_ISL_3295384, EPI_ISL_3295385, EPI_ISL_3295386, EPI_ISL_3295387, EPI_ISL_3295388, EPI_ISL_3295389, EPI_ISL_3295390, EPI_ISL_3295391,
EPI_ISL_3295392, EPI_ISL_3295393, EPI_ISL_3295394, EPI_ISL_3295395, EPI_ISL_3295396, EPI_ISL_3295397, EPI_ISL_3295398, EPI_ISL_3295399, EPI_ISL_3295400, EPI_ISL_3295401, EPI_ISL_3295402, EPI_ISL_3295403, EPI_ISL_3295404, EPI_ISL_3295405, EPI_ISL_3295406, EPI_ISL_3295407, EPI_ISL_3295408, EPI_ISL_3295409, EPI_ISL_3295410, EPI_ISL_3295411, EPI_ISL_3295412, EPI_ISL_3295413,
EPI_ISL_3295414, EPI_ISL_3295415, EPI_ISL_3295416, EPI_ISL_3295417, EPI_ISL_3295418, EPI_ISL_3295419, EPI_ISL_3295420, EPI_ISL_3295421, EPI_ISL_3295422, EPI_ISL_3295423, EPI_ISL_3295424, EPI_ISL_3295425, EPI_ISL_3295426, EPI_ISL_3295427, EPI_ISL_3295428, EPI_ISL_3295429, EPI_ISL_3295430, EPI_ISL_3295431, EPI_ISL_3295432, EPI_ISL_3295433, EPI_ISL_3295434, EPI_ISL_3295435,
EPI_ISL_3295436, EPI_ISL_3295437, EPI_ISL_3295438, EPI_ISL_3295439, EPI_ISL_3295440, EPI_ISL_3295441, EPI_ISL_3295442, EPI_ISL_3295443, EPI_ISL_3295444, EPI_ISL_3295445, EPI_ISL_3295446, EPI_ISL_3295447, EPI_ISL_3295448, EPI_ISL_3295449, EPI_ISL_3295450, EPI_ISL_3295451, EPI_ISL_3295452, EPI_ISL_3295453, EPI_ISL_3295454, EPI_ISL_3295455, EPI_ISL_3295456, EPI_ISL_3295457,
EPI_ISL_3295458, EPI_ISL_3295459, EPI_ISL_3295460, EPI_ISL_3295461, EPI_ISL_3295462, EPI_ISL_3295463, EPI_ISL_3295464, EPI_ISL_3295465, EPI_ISL_3295466, EPI_ISL_3295467, EPI_ISL_3295468, EPI_ISL_3295469, EPI_ISL_3295470, EPI_ISL_3295471, EPI_ISL_3295472, EPI_ISL_3295473, EPI_ISL_3295474, EPI_ISL_3295475, EPI_ISL_3295476, EPI_ISL_3295477, EPI_ISL_3295478, EPI_ISL_3295479,
EPI_ISL_3295480, EPI_ISL_3295481, EPI_ISL_3295482, EPI_ISL_3295483, EPI_ISL_3295484, EPI_ISL_3295485, EPI_ISL_3295486, EPI_ISL_3295487, EPI_ISL_3295488, EPI_ISL_3295489, EPI_ISL_3295490, EPI_ISL_3295491, EPI_ISL_3295492, EPI_ISL_3295493, EPI_ISL_3295494, EPI_ISL_3295495, EPI_ISL_3295496, EPI_ISL_3295497, EPI_ISL_3295498, EPI_ISL_3295499, EPI_ISL_3295500, EPI_ISL_3295501,
EPI_ISL_3295502, EPI_ISL_3295503, EPI_ISL_3295504, EPI_ISL_3295505

| | | | |
|---|---|---|---|
| see above | Wales Specialist Virology Centre Sequencing lab: Pathogen Genomics Unit | Public Health Wales Microbiology Cardiff Wales Specialist Virology Centre | Catherine Moore, Johnathan Evans, Laura Gifford, Malorie Perry, Simon Cottrell, Angela Marchbank, Alec Birchley, Alexander Adams, Amy Gaskin, Bree Gatica-Wilcox, Jason Coombes, Joel Southgate, Lauren Gilbert, Lee Graham, Nicole Pacchiarini, Sara Kumziene-Summerhayes, Sarah Taylor, Sophie Jones, Sara Rey, Matthew Bull, Joanne Watkins, Sally Corden, Tom Connor |
| EPI_ISL_3295506 | Viollier AG | Department of Biosystems Science and Engineering, ETH Zürich | Christian Beisel, Sarah Nadeau, Chaoran Chen, Ivan Topolsky, Kim Philipp Jablonski, Lara Fuhrmann, Rebecca Denes, Mirjam Feldkamp, Ina Nissen, Natascha Santacroce, Elodie Burcklen, Christiane Beckmann, Maurice Redondo, Olivier Kobel, Christoph Noppen, Niko Beerenwinkel, Tanja Stadler |
| EPI_ISL_3295507 | Wales Specialist Virology Centre Sequencing lab: Pathogen Genomics Unit | Public Health Wales Microbiology Cardiff Wales Specialist Virology Centre | Catherine Moore, Johnathan Evans, Laura Gifford, Malorie Perry, Simon Cottrell, Angela Marchbank, Alec Birchley, Alexander Adams, Amy Gaskin, Bree Gatica-Wilcox, Jason Coombes, Joel Southgate, Lauren Gilbert, Lee Graham, Nicole Pacchiarini, Sara Kumziene-Summerhayes, Sarah Taylor, Sophie Jones, Sara Rey, Matthew Bull, Joanne Watkins, Sally Corden, Tom Connor |
| EPI_ISL_3295508 | Viollier AG | Department of Biosystems Science and Engineering, ETH Zürich | Christian Beisel, Sarah Nadeau, Chaoran Chen, Ivan Topolsky, Kim Philipp Jablonski, Lara Fuhrmann, Rebecca Denes, Mirjam Feldkamp, Ina Nissen, Natascha Santacroce, Elodie Burcklen, Christiane Beckmann, Maurice Redondo, Olivier Kobel, Christoph Noppen, Niko Beerenwinkel, Tanja Stadler |
| EPI_ISL_3295509 | Wales Specialist Virology Centre Sequencing lab: Pathogen Genomics Unit | Public Health Wales Microbiology Cardiff Wales Specialist Virology Centre | Catherine Moore, Johnathan Evans, Laura Gifford, Malorie Perry, Simon Cottrell, Angela Marchbank, Alec Birchley, Alexander Adams, Amy Gaskin, Bree Gatica-Wilcox, Jason Coombes, Joel Southgate, Lauren Gilbert, Lee Graham, Nicole Pacchiarini, Sara Kumziene-Summerhayes, Sarah Taylor, Sophie Jones, Sara Rey, Matthew Bull, Joanne Watkins, Sally Corden, Tom Connor |
| EPI_ISL_3295510 | Viollier AG | Department of Biosystems Science and Engineering, ETH Zürich | Christian Beisel, Sarah Nadeau, Chaoran Chen, Ivan Topolsky, Kim Philipp Jablonski, Lara Fuhrmann, Rebecca Denes, Mirjam Feldkamp, Ina Nissen, Natascha Santacroce, Elodie Burcklen, Christiane Beckmann, Maurice Redondo, Olivier Kobel, Christoph Noppen, Niko Beerenwinkel, Tanja Stadler |
| EPI_ISL_3295511 | Wales Specialist Virology Centre Sequencing lab: Pathogen Genomics Unit | Public Health Wales Microbiology Cardiff Wales Specialist Virology Centre | Catherine Moore, Johnathan Evans, Laura Gifford, Malorie Perry, Simon Cottrell, Angela Marchbank, Alec Birchley, Alexander Adams, Amy Gaskin, Bree Gatica-Wilcox, Jason Coombes, Joel Southgate, Lauren Gilbert, Lee Graham, Nicole Pacchiarini, Sara Kumziene-Summerhayes, Sarah Taylor, Sophie Jones, Sara Rey, Matthew Bull, Joanne Watkins, Sally Corden, Tom Connor |
| EPI_ISL_3295512 | Viollier AG | Department of Biosystems Science and Engineering, ETH Zürich | Christian Beisel, Sarah Nadeau, Chaoran Chen, Ivan Topolsky, Kim Philipp Jablonski, Lara Fuhrmann, Rebecca Denes, Mirjam Feldkamp, Ina Nissen, Natascha Santacroce, Elodie Burcklen, Christiane Beckmann, Maurice Redondo, Olivier Kobel, |

| | | | |
|---|---|---|---|
| | | | Christoph Noppen, Niko Beerenwinkel, Tanja Stadler |
| EPI_ISL_3295513 | Wales Specialist Virology Centre Sequencing lab: Pathogen Genomics Unit | Public Health Wales Microbiology Cardiff Wales Specialist Virology Centre | Catherine Moore, Johnathan Evans, Laura Gifford, Malorie Perry, Simon Cottrell, Angela Marchbank, Alec Birchley, Alexander Adams, Amy Gaskin, Bree Gatica-Wilcox, Jason Coombes, Joel Southgate, Lauren Gilbert, Lee Graham, Nicole Pacchiarini, Sara Kumziene-Summerhayes, Sarah Taylor, Sophie Jones, Sara Rey, Matthew Bull, Joanne Watkins, Sally Corden, Tom Connor |
| EPI_ISL_3295514 | Viollier AG | Department of Biosystems Science and Engineering, ETH Zürich | Christian Beisel, Sarah Nadeau, Chaoran Chen, Ivan Topolsky, Kim Philipp Jablonski, Lara Fuhrmann, Rebecca Denes, Mirjam Feldkamp, Ina Nissen, Natascha Santacroce, Elodie Burcklen, Christiane Beckmann, Maurice Redondo, Olivier Kobel, Christoph Noppen, Niko Beerenwinkel, Tanja Stadler |
| EPI_ISL_3295515 | Wales Specialist Virology Centre Sequencing lab: Pathogen Genomics Unit | Public Health Wales Microbiology Cardiff Wales Specialist Virology Centre | Catherine Moore, Johnathan Evans, Laura Gifford, Malorie Perry, Simon Cottrell, Angela Marchbank, Alec Birchley, Alexander Adams, Amy Gaskin, Bree Gatica-Wilcox, Jason Coombes, Joel Southgate, Lauren Gilbert, Lee Graham, Nicole Pacchiarini, Sara Kumziene-Summerhayes, Sarah Taylor, Sophie Jones, Sara Rey, Matthew Bull, Joanne Watkins, Sally Corden, Tom Connor |
| EPI_ISL_3295516 | Viollier AG | Department of Biosystems Science and Engineering, ETH Zürich | Christian Beisel, Sarah Nadeau, Chaoran Chen, Ivan Topolsky, Kim Philipp Jablonski, Lara Fuhrmann, Rebecca Denes, Mirjam Feldkamp, Ina Nissen, Natascha Santacroce, Elodie Burcklen, Christiane Beckmann, Maurice Redondo, Olivier Kobel, Christoph Noppen, Niko Beerenwinkel, Tanja Stadler |
| EPI_ISL_3295517 | Wales Specialist Virology Centre Sequencing lab: Pathogen Genomics Unit | Public Health Wales Microbiology Cardiff Wales Specialist Virology Centre | Catherine Moore, Johnathan Evans, Laura Gifford, Malorie Perry, Simon Cottrell, Angela Marchbank, Alec Birchley, Alexander Adams, Amy Gaskin, Bree Gatica-Wilcox, Jason Coombes, Joel Southgate, Lauren Gilbert, Lee Graham, Nicole Pacchiarini, Sara Kumziene-Summerhayes, Sarah Taylor, Sophie Jones, Sara Rey, Matthew Bull, Joanne Watkins, Sally Corden, Tom Connor |
| EPI_ISL_3295518 | Viollier AG | Department of Biosystems Science and Engineering, ETH Zürich | Christian Beisel, Sarah Nadeau, Chaoran Chen, Ivan Topolsky, Kim Philipp Jablonski, Lara Fuhrmann, Rebecca Denes, Mirjam Feldkamp, Ina Nissen, Natascha Santacroce, Elodie Burcklen, Christiane Beckmann, Maurice Redondo, Olivier Kobel, Christoph Noppen, Niko Beerenwinkel, Tanja Stadler |
| EPI_ISL_3295519, EPI_ISL_3295520 | Wales Specialist Virology Centre Sequencing lab: Pathogen Genomics Unit | Public Health Wales Microbiology Cardiff Wales Specialist Virology Centre | Catherine Moore, Johnathan Evans, Laura Gifford, Malorie Perry, Simon Cottrell, Angela Marchbank, Alec Birchley, Alexander Adams, Amy Gaskin, Bree Gatica-Wilcox, Jason Coombes, Joel Southgate, Lauren Gilbert, Lee Graham, Nicole Pacchiarini, Sara Kumziene-Summerhayes, Sarah Taylor, Sophie Jones, Sara Rey, Matthew Bull, Joanne Watkins, Sally Corden, Tom Connor |
| EPI_ISL_3295521 | Viollier AG | Department of Biosystems Science and Engineering, ETH Zürich | Christian Beisel, Sarah Nadeau, Chaoran Chen, Ivan Topolsky, Kim Philipp Jablonski, Lara Fuhrmann, Rebecca Denes, Mirjam Feldkamp, Ina Nissen, Natascha Santacroce, Elodie Burcklen, Christiane Beckmann, Maurice Redondo, Olivier Kobel, Christoph Noppen, Niko Beerenwinkel, Tanja Stadler |
| EPI_ISL_3295522 | Wales Specialist Virology Centre Sequencing lab: Pathogen Genomics Unit | Public Health Wales Microbiology Cardiff Wales Specialist Virology Centre | Catherine Moore, Johnathan Evans, Laura Gifford, Malorie Perry, Simon Cottrell, Angela Marchbank, Alec Birchley, Alexander Adams, Amy Gaskin, Bree Gatica-Wilcox, Jason Coombes, Joel Southgate, Lauren Gilbert, Lee Graham, Nicole Pacchiarini, Sara Kumziene-Summerhayes, Sarah Taylor, Sophie Jones, Sara Rey, Matthew Bull, Joanne Watkins, Sally Corden, Tom Connor |
| EPI_ISL_3295523 | Viollier AG | Department of Biosystems Science and Engineering, ETH Zürich | Christian Beisel, Sarah Nadeau, Chaoran Chen, Ivan Topolsky, Kim Philipp Jablonski, Lara Fuhrmann, Rebecca Denes, Mirjam Feldkamp, Ina Nissen, Natascha Santacroce, Elodie Burcklen, Christiane Beckmann, Maurice Redondo, Olivier Kobel, Christoph Noppen, Niko Beerenwinkel, Tanja Stadler |
| EPI_ISL_3295524 | Wales Specialist Virology Centre Sequencing lab: Pathogen Genomics Unit | Public Health Wales Microbiology Cardiff Wales Specialist Virology Centre | Catherine Moore, Johnathan Evans, Laura Gifford, Malorie Perry, Simon Cottrell, Angela Marchbank, Alec Birchley, Alexander Adams, Amy Gaskin, Bree Gatica-Wilcox, Jason Coombes, Joel Southgate, Lauren Gilbert, Lee Graham, Nicole Pacchiarini, Sara Kumziene-Summerhayes, Sarah Taylor, Sophie Jones, Sara Rey, Matthew Bull, Joanne Watkins, Sally Corden, Tom Connor |
| EPI_ISL_3295525 | Viollier AG | Department of Biosystems Science and Engineering, ETH Zürich | Christian Beisel, Sarah Nadeau, Chaoran Chen, Ivan Topolsky, Kim Philipp Jablonski, Lara Fuhrmann, Rebecca Denes, Mirjam Feldkamp, Ina Nissen, Natascha Santacroce, Elodie Burcklen, Christiane Beckmann, Maurice Redondo, Olivier Kobel, Christoph Noppen, Niko Beerenwinkel, Tanja Stadler |
| EPI_ISL_3295526 | Wales Specialist Virology Centre Sequencing lab: Pathogen Genomics Unit | Public Health Wales Microbiology Cardiff Wales Specialist Virology Centre | Catherine Moore, Johnathan Evans, Laura Gifford, Malorie Perry, Simon Cottrell, Angela Marchbank, Alec Birchley, Alexander Adams, Amy Gaskin, Bree Gatica-Wilcox, Jason Coombes, Joel Southgate, Lauren Gilbert, Lee Graham, Nicole Pacchiarini, Sara Kumziene-Summerhayes, Sarah Taylor, Sophie Jones, Sara Rey, Matthew Bull, Joanne Watkins, Sally Corden, Tom Connor |
| EPI_ISL_3295527 | Viollier AG | Department of Biosystems Science and Engineering, ETH Zürich | Christian Beisel, Sarah Nadeau, Chaoran Chen, Ivan Topolsky, Kim Philipp Jablonski, Lara Fuhrmann, Rebecca Denes, Mirjam Feldkamp, Ina Nissen, Natascha Santacroce, Elodie Burcklen, Christiane Beckmann, Maurice Redondo, Olivier Kobel, Christoph Noppen, Niko Beerenwinkel, Tanja Stadler |
| EPI_ISL_3295528 | Wales Specialist Virology Centre Sequencing lab: Pathogen Genomics Unit | Public Health Wales Microbiology Cardiff Wales Specialist Virology Centre | Catherine Moore, Johnathan Evans, Laura Gifford, Malorie Perry, Simon Cottrell, Angela Marchbank, Alec Birchley, Alexander Adams, Amy Gaskin, Bree Gatica-Wilcox, Jason Coombes, Joel Southgate, Lauren Gilbert, Lee Graham, Nicole Pacchiarini, Sara Kumziene-Summerhayes, Sarah Taylor, Sophie Jones, Sara Rey, Matthew Bull, Joanne Watkins, Sally Corden, Tom Connor |
| EPI_ISL_3295529 | Viollier AG | Department of Biosystems Science and Engineering, ETH Zürich | Christian Beisel, Sarah Nadeau, Chaoran Chen, Ivan Topolsky, Kim Philipp Jablonski, Lara Fuhrmann, Rebecca Denes, Mirjam Feldkamp, Ina Nissen, Natascha Santacroce, Elodie Burcklen, Christiane Beckmann, Maurice Redondo, Olivier Kobel, Christoph Noppen, Niko Beerenwinkel, Tanja Stadler |
| EPI_ISL_3295530 | Wales Specialist Virology Centre Sequencing lab: Pathogen Genomics Unit | Public Health Wales Microbiology Cardiff Wales Specialist Virology Centre | Catherine Moore, Johnathan Evans, Laura Gifford, Malorie Perry, Simon Cottrell, Angela Marchbank, Alec Birchley, Alexander Adams, Amy Gaskin, Bree Gatica-Wilcox, Jason Coombes, Joel Southgate, Lauren Gilbert, Lee Graham, Nicole Pacchiarini, Sara Kumziene-Summerhayes, Sarah Taylor, Sophie Jones, Sara Rey, Matthew Bull, Joanne Watkins, Sally Corden, Tom Connor |
| EPI_ISL_3295531 | Viollier AG | Department of Biosystems Science and Engineering, ETH Zürich | Christian Beisel, Sarah Nadeau, Chaoran Chen, Ivan Topolsky, Kim Philipp Jablonski, Lara Fuhrmann, Rebecca Denes, Mirjam Feldkamp, Ina Nissen, Natascha Santacroce, Elodie Burcklen, Christiane Beckmann, Maurice Redondo, Olivier Kobel, Christoph Noppen, Niko Beerenwinkel, Tanja Stadler |
| EPI_ISL_3295532 | Wales Specialist Virology Centre Sequencing lab: Pathogen Genomics Unit | Public Health Wales Microbiology Cardiff Wales Specialist Virology Centre | Catherine Moore, Johnathan Evans, Laura Gifford, Malorie Perry, Simon Cottrell, Angela Marchbank, Alec Birchley, Alexander Adams, Amy Gaskin, Bree Gatica-Wilcox, Jason Coombes, Joel Southgate, Lauren Gilbert, Lee Graham, Nicole Pacchiarini, Sara Kumziene-Summerhayes, Sarah Taylor, Sophie Jones, Sara Rey, Matthew Bull, Joanne Watkins, Sally Corden, Tom Connor |
| EPI_ISL_3295533 | Viollier AG | Department of Biosystems Science and Engineering, ETH Zürich | Christian Beisel, Sarah Nadeau, Chaoran Chen, Ivan Topolsky, Kim Philipp Jablonski, Lara Fuhrmann, Rebecca Denes, Mirjam Feldkamp, Ina Nissen, Natascha Santacroce, Elodie Burcklen, Christiane Beckmann, Maurice Redondo, Olivier Kobel, Christoph Noppen, Niko Beerenwinkel, Tanja Stadler |
| EPI_ISL_3295534 | Wales Specialist Virology Centre Sequencing lab: Pathogen Genomics Unit | Public Health Wales Microbiology Cardiff Wales Specialist Virology Centre | Catherine Moore, Johnathan Evans, Laura Gifford, Malorie Perry, Simon Cottrell, Angela Marchbank, Alec Birchley, Alexander Adams, Amy Gaskin, Bree Gatica-Wilcox, Jason Coombes, Joel Southgate, Lauren Gilbert, Lee Graham, Nicole Pacchiarini, Sara Kumziene-Summerhayes, Sarah Taylor, Sophie Jones, Sara Rey, Matthew Bull, Joanne Watkins, Sally Corden, Tom Connor |
| EPI_ISL_3295535 | Viollier AG | Department of Biosystems Science and Engineering, ETH Zürich | Christian Beisel, Sarah Nadeau, Chaoran Chen, Ivan Topolsky, Kim Philipp Jablonski, Lara Fuhrmann, Rebecca Denes, Mirjam Feldkamp, Ina Nissen, Natascha Santacroce, Elodie Burcklen, Christiane Beckmann, Maurice Redondo, Olivier Kobel, Christoph Noppen, Niko Beerenwinkel, Tanja Stadler |
| EPI_ISL_3295536 | Wales Specialist Virology Centre Sequencing lab: Pathogen Genomics Unit | Public Health Wales Microbiology Cardiff Wales Specialist Virology Centre | Catherine Moore, Johnathan Evans, Laura Gifford, Malorie Perry, Simon Cottrell, Angela Marchbank, Alec Birchley, Alexander Adams, Amy Gaskin, Bree Gatica-Wilcox, Jason Coombes, Joel Southgate, Lauren Gilbert, Lee Graham, Nicole Pacchiarini, Sara Kumziene-Summerhayes, Sarah Taylor, Sophie Jones, Sara Rey, Matthew Bull, Joanne Watkins, Sally Corden, Tom Connor |
| EPI_ISL_3295537 | Viollier AG | Department of Biosystems Science and Engineering, ETH Zürich | Christian Beisel, Sarah Nadeau, Chaoran Chen, Ivan Topolsky, Kim Philipp Jablonski, Lara Fuhrmann, Rebecca Denes, Mirjam Feldkamp, Ina Nissen, Natascha Santacroce, Elodie Burcklen, Christiane Beckmann, Maurice Redondo, Olivier Kobel, Christoph Noppen, Niko Beerenwinkel, Tanja Stadler |
| EPI_ISL_3295538 | Wales Specialist Virology Centre Sequencing lab: Pathogen Genomics Unit | Public Health Wales Microbiology Cardiff Wales Specialist Virology Centre | Catherine Moore, Johnathan Evans, Laura Gifford, Malorie Perry, Simon Cottrell, Angela Marchbank, Alec Birchley, Alexander Adams, Amy Gaskin, Bree Gatica-Wilcox, Jason Coombes, Joel Southgate, Lauren Gilbert, Lee Graham, Nicole Pacchiarini, Sara Kumziene-Summerhayes, Sarah Taylor, Sophie Jones, Sara Rey, Matthew Bull, Joanne Watkins, Sally Corden, Tom Connor |
| EPI_ISL_3295539 | Viollier AG | Department of Biosystems Science and Engineering, ETH Zürich | Christian Beisel, Sarah Nadeau, Chaoran Chen, Ivan Topolsky, Kim Philipp Jablonski, Lara Fuhrmann, Rebecca Denes, Mirjam Feldkamp, Ina Nissen, Natascha Santacroce, Elodie Burcklen, Christiane Beckmann, Maurice Redondo, Olivier Kobel, Christoph Noppen, Niko Beerenwinkel, Tanja Stadler |
| EPI_ISL_3295540 | Wales Specialist Virology Centre Sequencing lab: Pathogen Genomics Unit | Public Health Wales Microbiology Cardiff Wales Specialist Virology Centre | Catherine Moore, Johnathan Evans, Laura Gifford, Malorie Perry, Simon Cottrell, Angela Marchbank, Alec Birchley, Alexander Adams, Amy Gaskin, Bree Gatica-Wilcox, Jason Coombes, Joel Southgate, Lauren Gilbert, Lee Graham, Nicole Pacchiarini, Sara Kumziene-Summerhayes, Sarah Taylor, Sophie Jones, Sara Rey, Matthew Bull, Joanne Watkins, Sally Corden, Tom Connor |
| EPI_ISL_3295541 | Viollier AG | Department of Biosystems Science and Engineering, ETH Zürich | Christian Beisel, Sarah Nadeau, Chaoran Chen, Ivan Topolsky, Kim Philipp Jablonski, Lara Fuhrmann, Rebecca Denes, Mirjam Feldkamp, Ina Nissen, Natascha Santacroce, Elodie Burcklen, Christiane Beckmann, Maurice Redondo, Olivier Kobel, Christoph Noppen, Niko Beerenwinkel, Tanja Stadler |
| EPI_ISL_3295542, EPI_ISL_3295543 | Wales Specialist Virology Centre Sequencing lab: Pathogen Genomics Unit | Public Health Wales Microbiology Cardiff Wales Specialist Virology Centre | Catherine Moore, Johnathan Evans, Laura Gifford, Malorie Perry, Simon Cottrell, Angela Marchbank, Alec Birchley, Alexander Adams, Amy Gaskin, Bree Gatica-Wilcox, Jason Coombes, Joel Southgate, Lauren Gilbert, Lee Graham, Nicole Pacchiarini, Sara Kumziene-Summerhayes, Sarah Taylor, Sophie Jones, Sara Rey, Matthew Bull, Joanne Watkins, Sally Corden, Tom Connor |
| EPI_ISL_3295544 | Viollier AG | Department of Biosystems Science and Engineering, ETH Zürich | Christian Beisel, Sarah Nadeau, Chaoran Chen, Ivan Topolsky, Kim Philipp Jablonski, Lara Fuhrmann, Rebecca Denes, Mirjam Feldkamp, Ina Nissen, Natascha Santacroce, Elodie Burcklen, Christiane Beckmann, Maurice Redondo, Olivier Kobel, Christoph Noppen, Niko Beerenwinkel, Tanja Stadler |
| EPI_ISL_3295545 | Wales Specialist Virology Centre Sequencing lab: Pathogen Genomics Unit | Public Health Wales Microbiology Cardiff Wales Specialist Virology Centre | Catherine Moore, Johnathan Evans, Laura Gifford, Malorie Perry, Simon Cottrell, Angela Marchbank, Alec Birchley, Alexander Adams, Amy Gaskin, Bree Gatica-Wilcox, Jason Coombes, Joel Southgate, Lauren Gilbert, Lee Graham, Nicole Pacchiarini, Sara Kumziene-Summerhayes, Sarah Taylor, Sophie Jones, Sara Rey, Matthew Bull, Joanne Watkins, Sally Corden, Tom Connor |
| EPI_ISL_3295546 | Viollier AG | Department of Biosystems Science and Engineering, ETH Zürich | Christian Beisel, Sarah Nadeau, Chaoran Chen, Ivan Topolsky, Kim Philipp Jablonski, Lara Fuhrmann, Rebecca Denes, Mirjam Feldkamp, Ina Nissen, Natascha Santacroce, Elodie Burcklen, Christiane Beckmann, Maurice Redondo, Olivier Kobel, Christoph Noppen, Niko Beerenwinkel, Tanja Stadler |
| EPI_ISL_3295547 | Wales Specialist Virology Centre Sequencing lab: Pathogen Genomics Unit | Public Health Wales Microbiology Cardiff Wales Specialist Virology Centre | Catherine Moore, Johnathan Evans, Laura Gifford, Malorie Perry, Simon Cottrell, Angela Marchbank, Alec Birchley, Alexander Adams, Amy Gaskin, Bree Gatica-Wilcox, Jason Coombes, Joel Southgate, Lauren Gilbert, Lee Graham, Nicole Pacchiarini, Sara Kumziene-Summerhayes, Sarah Taylor, Sophie Jones, Sara Rey, Matthew Bull, Joanne Watkins, Sally Corden, Tom Connor |
| EPI_ISL_3295548 | Viollier AG | Department of Biosystems Science and Engineering, ETH Zürich | Christian Beisel, Sarah Nadeau, Chaoran Chen, Ivan Topolsky, Kim Philipp Jablonski, Lara Fuhrmann, Rebecca Denes, Mirjam Feldkamp, Ina Nissen, Natascha Santacroce, Elodie Burcklen, Christiane Beckmann, Maurice Redondo, Olivier Kobel, Christoph Noppen, Niko Beerenwinkel, Tanja Stadler |
| EPI_ISL_3295549 | Wales Specialist Virology Centre Sequencing lab: Pathogen Genomics Unit | Public Health Wales Microbiology Cardiff Wales Specialist Virology Centre | Catherine Moore, Johnathan Evans, Laura Gifford, Malorie Perry, Simon Cottrell, Angela Marchbank, Alec Birchley, Alexander Adams, Amy Gaskin, Bree Gatica-Wilcox, Jason Coombes, Joel Southgate, Lauren Gilbert, Lee Graham, Nicole Pacchiarini, Sara Kumziene-Summerhayes, Sarah Taylor, Sophie Jones, Sara Rey, Matthew Bull, Joanne Watkins, Sally Corden, Tom Connor |
| EPI_ISL_3295550 | Viollier AG | Department of Biosystems Science and Engineering, ETH Zürich | Christian Beisel, Sarah Nadeau, Chaoran Chen, Ivan Topolsky, Kim Philipp Jablonski, Lara Fuhrmann, Rebecca Denes, Mirjam Feldkamp, Ina Nissen, Natascha Santacroce, Elodie Burcklen, Christiane Beckmann, Maurice Redondo, Olivier Kobel, Christoph Noppen, Niko Beerenwinkel, Tanja Stadler |
| EPI_ISL_3295551 | Wales Specialist Virology Centre Sequencing lab: Pathogen Genomics Unit | Public Health Wales Microbiology Cardiff Wales Specialist Virology Centre | Catherine Moore, Johnathan Evans, Laura Gifford, Malorie Perry, Simon Cottrell, Angela Marchbank, Alec Birchley, Alexander Adams, Amy Gaskin, Bree Gatica-Wilcox, Jason Coombes, Joel Southgate, Lauren Gilbert, Lee Graham, Nicole Pacchiarini, Sara Kumziene-Summerhayes, Sarah Taylor, Sophie Jones, Sara Rey, Matthew Bull, Joanne Watkins, Sally Corden, Tom Connor |
| EPI_ISL_3295552 | Viollier AG | Department of Biosystems Science and Engineering, ETH Zürich | Christian Beisel, Sarah Nadeau, Chaoran Chen, Ivan Topolsky, Kim Philipp Jablonski, Lara Fuhrmann, Rebecca Denes, Mirjam Feldkamp, Ina Nissen, Natascha Santacroce, Elodie Burcklen, Christiane Beckmann, Maurice Redondo, Olivier Kobel, Christoph Noppen, Niko Beerenwinkel, Tanja Stadler |
| EPI_ISL_3295553 | Wales Specialist Virology Centre Sequencing lab: Pathogen Genomics Unit | Public Health Wales Microbiology Cardiff Wales Specialist Virology Centre | Catherine Moore, Johnathan Evans, Laura Gifford, Malorie Perry, Simon Cottrell, Angela Marchbank, Alec Birchley, Alexander Adams, Amy Gaskin, Bree Gatica-Wilcox, Jason Coombes, Joel Southgate, Lauren Gilbert, Lee Graham, Nicole Pacchiarini, Sara Kumziene-Summerhayes, Sarah Taylor, Sophie Jones, Sara Rey, Matthew Bull, Joanne Watkins, Sally Corden, Tom Connor |
| EPI_ISL_3295554 | Viollier AG | Department of Biosystems Science and Engineering, ETH Zürich | Christian Beisel, Sarah Nadeau, Chaoran Chen, Ivan Topolsky, Kim Philipp Jablonski, Lara Fuhrmann, Rebecca Denes, Mirjam Feldkamp, Ina Nissen, Natascha Santacroce, Elodie Burcklen, Christiane Beckmann, Maurice Redondo, Olivier Kobel, Christoph Noppen, Niko Beerenwinkel, Tanja Stadler |
| EPI_ISL_3295555, EPI_ISL_3295557 | Wales Specialist Virology Centre Sequencing lab: Pathogen Genomics Unit | Public Health Wales Microbiology Cardiff Wales Specialist Virology Centre | Catherine Moore, Johnathan Evans, Laura Gifford, Malorie Perry, Simon Cottrell, Angela Marchbank, Alec Birchley, Alexander Adams, Amy Gaskin, Bree Gatica-Wilcox, Jason Coombes, Joel Southgate, Lauren Gilbert, Lee Graham, Nicole Pacchiarini, Sara Kumziene-Summerhayes, Sarah Taylor, Sophie Jones, Sara Rey, Matthew Bull, Joanne Watkins, Sally Corden, Tom Connor |
| EPI_ISL_3295558 | Viollier AG | Department of Biosystems Science and Engineering, ETH Zürich | Christian Beisel, Sarah Nadeau, Chaoran Chen, Ivan Topolsky, Kim Philipp Jablonski, Lara Fuhrmann, Rebecca Denes, Mirjam Feldkamp, Ina Nissen, Natascha Santacroce, Elodie Burcklen, Christiane Beckmann, Maurice Redondo, Olivier Kobel, Christoph Noppen, Niko Beerenwinkel, Tanja Stadler |
| EPI_ISL_3295559 | Wales Specialist Virology Centre Sequencing lab: Pathogen Genomics Unit | Public Health Wales Microbiology Cardiff Wales Specialist Virology Centre | Catherine Moore, Johnathan Evans, Laura Gifford, Malorie Perry, Simon Cottrell, Angela Marchbank, Alec Birchley, Alexander Adams, Amy Gaskin, Bree Gatica-Wilcox, Jason Coombes, Joel Southgate, Lauren Gilbert, Lee Graham, Nicole Pacchiarini, Sara Kumziene-Summerhayes, Sarah Taylor, Sophie Jones, Sara Rey, Matthew Bull, Joanne Watkins, Sally Corden, Tom Connor |
| EPI_ISL_3295560 | Viollier AG | Department of Biosystems Science and Engineering, ETH Zürich | Christian Beisel, Sarah Nadeau, Chaoran Chen, Ivan Topolsky, Kim Philipp Jablonski, Lara Fuhrmann, Rebecca Denes, Mirjam Feldkamp, Ina Nissen, Natascha Santacroce, Elodie Burcklen, Christiane Beckmann, Maurice Redondo, Olivier Kobel, Christoph Noppen, Niko Beerenwinkel, Tanja Stadler |
| EPI_ISL_3295561 | Wales Specialist Virology Centre Sequencing lab: Pathogen | Public Health Wales Microbiology Cardiff Wales Specialist Virology | Catherine Moore, Johnathan Evans, Laura Gifford, Malorie Perry, Simon Cottrell, Angela Marchbank, Alec Birchley, Alexander Adams, Amy Gaskin, Bree Gatica-Wilcox, Jason Coombes, Joel Southgate, Lauren Gilbert, Lee Graham, Nicole |

| | | | |
|---|---|---|---|
| | Genomics Unit | Centre | Pacchiarini, Sara Kumziene-Summerhayes, Sarah Taylor, Sophie Jones, Sara Rey, Matthew Bull, Joanne Watkins, Sally Corden, Tom Connor |
| EPI_ISL_3295562 | Viollier AG | Department of Biosystems Science and Engineering, ETH Zürich | Christian Beisel, Sarah Nadeau, Chaoran Chen, Ivan Topolsky, Kim Philipp Jablonski, Lara Fuhrmann, Rebecca Denes, Mirjam Feldkamp, Ina Nissen, Natascha Santacroce, Elodie Burcklen, Christiane Beckmann, Maurice Redondo, Olivier Kobel, Christoph Noppen, Niko Beerenwinkel, Tanja Stadler |
| EPI_ISL_3295563 | Wales Specialist Virology Centre Sequencing lab: Pathogen Genomics Unit | Public Health Wales Microbiology Cardiff Wales Specialist Virology Centre | Catherine Moore, Johnathan Evans, Laura Gifford, Malorie Perry, Simon Cottrell, Angela Marchbank, Alec Birchley, Alexander Adams, Amy Gaskin, Bree Gatica-Wilcox, Jason Coombes, Joel Southgate, Lauren Gilbert, Lee Graham, Nicole Pacchiarini, Sara Kumziene-Summerhayes, Sarah Taylor, Sophie Jones, Sara Rey, Matthew Bull, Joanne Watkins, Sally Corden, Tom Connor |
| EPI_ISL_3295564 | Viollier AG | Department of Biosystems Science and Engineering, ETH Zürich | Christian Beisel, Sarah Nadeau, Chaoran Chen, Ivan Topolsky, Kim Philipp Jablonski, Lara Fuhrmann, Rebecca Denes, Mirjam Feldkamp, Ina Nissen, Natascha Santacroce, Elodie Burcklen, Christiane Beckmann, Maurice Redondo, Olivier Kobel, Christoph Noppen, Niko Beerenwinkel, Tanja Stadler |
| EPI_ISL_3295565 | Wales Specialist Virology Centre Sequencing lab: Pathogen Genomics Unit | Public Health Wales Microbiology Cardiff Wales Specialist Virology Centre | Catherine Moore, Johnathan Evans, Laura Gifford, Malorie Perry, Simon Cottrell, Angela Marchbank, Alec Birchley, Alexander Adams, Amy Gaskin, Bree Gatica-Wilcox, Jason Coombes, Joel Southgate, Lauren Gilbert, Lee Graham, Nicole Pacchiarini, Sara Kumziene-Summerhayes, Sarah Taylor, Sophie Jones, Sara Rey, Matthew Bull, Joanne Watkins, Sally Corden, Tom Connor |
| EPI_ISL_3295566 | Viollier AG | Department of Biosystems Science and Engineering, ETH Zürich | Christian Beisel, Sarah Nadeau, Chaoran Chen, Ivan Topolsky, Kim Philipp Jablonski, Lara Fuhrmann, Rebecca Denes, Mirjam Feldkamp, Ina Nissen, Natascha Santacroce, Elodie Burcklen, Christiane Beckmann, Maurice Redondo, Olivier Kobel, Christoph Noppen, Niko Beerenwinkel, Tanja Stadler |
| EPI_ISL_3295567 | Wales Specialist Virology Centre Sequencing lab: Pathogen Genomics Unit | Public Health Wales Microbiology Cardiff Wales Specialist Virology Centre | Catherine Moore, Johnathan Evans, Laura Gifford, Malorie Perry, Simon Cottrell, Angela Marchbank, Alec Birchley, Alexander Adams, Amy Gaskin, Bree Gatica-Wilcox, Jason Coombes, Joel Southgate, Lauren Gilbert, Lee Graham, Nicole Pacchiarini, Sara Kumziene-Summerhayes, Sarah Taylor, Sophie Jones, Sara Rey, Matthew Bull, Joanne Watkins, Sally Corden, Tom Connor |
| EPI_ISL_3295568 | Viollier AG | Department of Biosystems Science and Engineering, ETH Zürich | Christian Beisel, Sarah Nadeau, Chaoran Chen, Ivan Topolsky, Kim Philipp Jablonski, Lara Fuhrmann, Rebecca Denes, Mirjam Feldkamp, Ina Nissen, Natascha Santacroce, Elodie Burcklen, Christiane Beckmann, Maurice Redondo, Olivier Kobel, Christoph Noppen, Niko Beerenwinkel, Tanja Stadler |
| EPI_ISL_3295569 | Wales Specialist Virology Centre Sequencing lab: Pathogen Genomics Unit | Public Health Wales Microbiology Cardiff Wales Specialist Virology Centre | Catherine Moore, Johnathan Evans, Laura Gifford, Malorie Perry, Simon Cottrell, Angela Marchbank, Alec Birchley, Alexander Adams, Amy Gaskin, Bree Gatica-Wilcox, Jason Coombes, Joel Southgate, Lauren Gilbert, Lee Graham, Nicole Pacchiarini, Sara Kumziene-Summerhayes, Sarah Taylor, Sophie Jones, Sara Rey, Matthew Bull, Joanne Watkins, Sally Corden, Tom Connor |
| EPI_ISL_3295570 | Viollier AG | Department of Biosystems Science and Engineering, ETH Zürich | Christian Beisel, Sarah Nadeau, Chaoran Chen, Ivan Topolsky, Kim Philipp Jablonski, Lara Fuhrmann, Rebecca Denes, Mirjam Feldkamp, Ina Nissen, Natascha Santacroce, Elodie Burcklen, Christiane Beckmann, Maurice Redondo, Olivier Kobel, Christoph Noppen, Niko Beerenwinkel, Tanja Stadler |
| EPI_ISL_3295571, EPI_ISL_3295572 | Wales Specialist Virology Centre Sequencing lab: Pathogen Genomics Unit | Public Health Wales Microbiology Cardiff Wales Specialist Virology Centre | Catherine Moore, Johnathan Evans, Laura Gifford, Malorie Perry, Simon Cottrell, Angela Marchbank, Alec Birchley, Alexander Adams, Amy Gaskin, Bree Gatica-Wilcox, Jason Coombes, Joel Southgate, Lauren Gilbert, Lee Graham, Nicole Pacchiarini, Sara Kumziene-Summerhayes, Sarah Taylor, Sophie Jones, Sara Rey, Matthew Bull, Joanne Watkins, Sally Corden, Tom Connor |
| EPI_ISL_3295573 | Viollier AG | Department of Biosystems Science and Engineering, ETH Zürich | Christian Beisel, Sarah Nadeau, Chaoran Chen, Ivan Topolsky, Kim Philipp Jablonski, Lara Fuhrmann, Rebecca Denes, Mirjam Feldkamp, Ina Nissen, Natascha Santacroce, Elodie Burcklen, Christiane Beckmann, Maurice Redondo, Olivier Kobel, Christoph Noppen, Niko Beerenwinkel, Tanja Stadler |
| EPI_ISL_3295574 | Wales Specialist Virology Centre Sequencing lab: Pathogen Genomics Unit | Public Health Wales Microbiology Cardiff Wales Specialist Virology Centre | Catherine Moore, Johnathan Evans, Laura Gifford, Malorie Perry, Simon Cottrell, Angela Marchbank, Alec Birchley, Alexander Adams, Amy Gaskin, Bree Gatica-Wilcox, Jason Coombes, Joel Southgate, Lauren Gilbert, Lee Graham, Nicole Pacchiarini, Sara Kumziene-Summerhayes, Sarah Taylor, Sophie Jones, Sara Rey, Matthew Bull, Joanne Watkins, Sally Corden, Tom Connor |
| EPI_ISL_3295575 | Viollier AG | Department of Biosystems Science and Engineering, ETH Zürich | Christian Beisel, Sarah Nadeau, Chaoran Chen, Ivan Topolsky, Kim Philipp Jablonski, Lara Fuhrmann, Rebecca Denes, Mirjam Feldkamp, Ina Nissen, Natascha Santacroce, Elodie Burcklen, Christiane Beckmann, Maurice Redondo, Olivier Kobel, Christoph Noppen, Niko Beerenwinkel, Tanja Stadler |
| EPI_ISL_3295576 | Wales Specialist Virology Centre Sequencing lab: Pathogen Genomics Unit | Public Health Wales Microbiology Cardiff Wales Specialist Virology Centre | Catherine Moore, Johnathan Evans, Laura Gifford, Malorie Perry, Simon Cottrell, Angela Marchbank, Alec Birchley, Alexander Adams, Amy Gaskin, Bree Gatica-Wilcox, Jason Coombes, Joel Southgate, Lauren Gilbert, Lee Graham, Nicole Pacchiarini, Sara Kumziene-Summerhayes, Sarah Taylor, Sophie Jones, Sara Rey, Matthew Bull, Joanne Watkins, Sally Corden, Tom Connor |
| EPI_ISL_3295577 | Viollier AG | Department of Biosystems Science and Engineering, ETH Zürich | Christian Beisel, Sarah Nadeau, Chaoran Chen, Ivan Topolsky, Kim Philipp Jablonski, Lara Fuhrmann, Rebecca Denes, Mirjam Feldkamp, Ina Nissen, Natascha Santacroce, Elodie Burcklen, Christiane Beckmann, Maurice Redondo, Olivier Kobel, Christoph Noppen, Niko Beerenwinkel, Tanja Stadler |
| EPI_ISL_3295578 | Wales Specialist Virology Centre Sequencing lab: Pathogen Genomics Unit | Public Health Wales Microbiology Cardiff Wales Specialist Virology Centre | Catherine Moore, Johnathan Evans, Laura Gifford, Malorie Perry, Simon Cottrell, Angela Marchbank, Alec Birchley, Alexander Adams, Amy Gaskin, Bree Gatica-Wilcox, Jason Coombes, Joel Southgate, Lauren Gilbert, Lee Graham, Nicole Pacchiarini, Sara Kumziene-Summerhayes, Sarah Taylor, Sophie Jones, Sara Rey, Matthew Bull, Joanne Watkins, Sally Corden, Tom Connor |
| EPI_ISL_3295579 | Viollier AG | Department of Biosystems Science and Engineering, ETH Zürich | Christian Beisel, Sarah Nadeau, Chaoran Chen, Ivan Topolsky, Kim Philipp Jablonski, Lara Fuhrmann, Rebecca Denes, Mirjam Feldkamp, Ina Nissen, Natascha Santacroce, Elodie Burcklen, Christiane Beckmann, Maurice Redondo, Olivier Kobel, Christoph Noppen, Niko Beerenwinkel, Tanja Stadler |
| EPI_ISL_3295580, EPI_ISL_3295582 | Wales Specialist Virology Centre Sequencing lab: Pathogen Genomics Unit | Public Health Wales Microbiology Cardiff Wales Specialist Virology Centre | Catherine Moore, Johnathan Evans, Laura Gifford, Malorie Perry, Simon Cottrell, Angela Marchbank, Alec Birchley, Alexander Adams, Amy Gaskin, Bree Gatica-Wilcox, Jason Coombes, Joel Southgate, Lauren Gilbert, Lee Graham, Nicole Pacchiarini, Sara Kumziene-Summerhayes, Sarah Taylor, Sophie Jones, Sara Rey, Matthew Bull, Joanne Watkins, Sally Corden, Tom Connor |
| EPI_ISL_3295583 | Viollier AG | Department of Biosystems Science and Engineering, ETH Zürich | Christian Beisel, Sarah Nadeau, Chaoran Chen, Ivan Topolsky, Kim Philipp Jablonski, Lara Fuhrmann, Rebecca Denes, Mirjam Feldkamp, Ina Nissen, Natascha Santacroce, Elodie Burcklen, Christiane Beckmann, Maurice Redondo, Olivier Kobel, Christoph Noppen, Niko Beerenwinkel, Tanja Stadler |
| EPI_ISL_3295584 | Wales Specialist Virology Centre Sequencing lab: Pathogen Genomics Unit | Public Health Wales Microbiology Cardiff Wales Specialist Virology Centre | Catherine Moore, Johnathan Evans, Laura Gifford, Malorie Perry, Simon Cottrell, Angela Marchbank, Alec Birchley, Alexander Adams, Amy Gaskin, Bree Gatica-Wilcox, Jason Coombes, Joel Southgate, Lauren Gilbert, Lee Graham, Nicole Pacchiarini, Sara Kumziene-Summerhayes, Sarah Taylor, Sophie Jones, Sara Rey, Matthew Bull, Joanne Watkins, Sally Corden, Tom Connor |
| EPI_ISL_3295585 | Viollier AG | Department of Biosystems Science and Engineering, ETH Zürich | Christian Beisel, Sarah Nadeau, Chaoran Chen, Ivan Topolsky, Kim Philipp Jablonski, Lara Fuhrmann, Rebecca Denes, Mirjam Feldkamp, Ina Nissen, Natascha Santacroce, Elodie Burcklen, Christiane Beckmann, Maurice Redondo, Olivier Kobel, Christoph Noppen, Niko Beerenwinkel, Tanja Stadler |
| EPI_ISL_3295586, EPI_ISL_3295588 | Wales Specialist Virology Centre Sequencing lab: Pathogen Genomics Unit | Public Health Wales Microbiology Cardiff Wales Specialist Virology Centre | Catherine Moore, Johnathan Evans, Laura Gifford, Malorie Perry, Simon Cottrell, Angela Marchbank, Alec Birchley, Alexander Adams, Amy Gaskin, Bree Gatica-Wilcox, Jason Coombes, Joel Southgate, Lauren Gilbert, Lee Graham, Nicole Pacchiarini, Sara Kumziene-Summerhayes, Sarah Taylor, Sophie Jones, Sara Rey, Matthew Bull, Joanne Watkins, Sally Corden, Tom Connor |
| EPI_ISL_3295589 | Viollier AG | Department of Biosystems Science and Engineering, ETH Zürich | Christian Beisel, Sarah Nadeau, Chaoran Chen, Ivan Topolsky, Kim Philipp Jablonski, Lara Fuhrmann, Rebecca Denes, Mirjam Feldkamp, Ina Nissen, Natascha Santacroce, Elodie Burcklen, Christiane Beckmann, Maurice Redondo, Olivier Kobel, Christoph Noppen, Niko Beerenwinkel, Tanja Stadler |
| EPI_ISL_3295590 | Wales Specialist Virology Centre Sequencing lab: Pathogen Genomics Unit | Public Health Wales Microbiology Cardiff Wales Specialist Virology Centre | Catherine Moore, Johnathan Evans, Laura Gifford, Malorie Perry, Simon Cottrell, Angela Marchbank, Alec Birchley, Alexander Adams, Amy Gaskin, Bree Gatica-Wilcox, Jason Coombes, Joel Southgate, Lauren Gilbert, Lee Graham, Nicole Pacchiarini, Sara Kumziene-Summerhayes, Sarah Taylor, Sophie Jones, Sara Rey, Matthew Bull, Joanne Watkins, Sally Corden, Tom Connor |
| EPI_ISL_3295591 | Viollier AG | Department of Biosystems Science and Engineering, ETH Zürich | Christian Beisel, Sarah Nadeau, Chaoran Chen, Ivan Topolsky, Kim Philipp Jablonski, Lara Fuhrmann, Rebecca Denes, Mirjam Feldkamp, Ina Nissen, Natascha Santacroce, Elodie Burcklen, Christiane Beckmann, Maurice Redondo, Olivier Kobel, Christoph Noppen, Niko Beerenwinkel, Tanja Stadler |
| EPI_ISL_3295592 | Wales Specialist Virology Centre Sequencing lab: Pathogen Genomics Unit | Public Health Wales Microbiology Cardiff Wales Specialist Virology Centre | Catherine Moore, Johnathan Evans, Laura Gifford, Malorie Perry, Simon Cottrell, Angela Marchbank, Alec Birchley, Alexander Adams, Amy Gaskin, Bree Gatica-Wilcox, Jason Coombes, Joel Southgate, Lauren Gilbert, Lee Graham, Nicole Pacchiarini, Sara Kumziene-Summerhayes, Sarah Taylor, Sophie Jones, Sara Rey, Matthew Bull, Joanne Watkins, Sally Corden, Tom Connor |
| EPI_ISL_3295593 | Viollier AG | Department of Biosystems Science and Engineering, ETH Zürich | Christian Beisel, Sarah Nadeau, Chaoran Chen, Ivan Topolsky, Kim Philipp Jablonski, Lara Fuhrmann, Rebecca Denes, Mirjam Feldkamp, Ina Nissen, Natascha Santacroce, Elodie Burcklen, Christiane Beckmann, Maurice Redondo, Olivier Kobel, Christoph Noppen, Niko Beerenwinkel, Tanja Stadler |
| EPI_ISL_3295594 | Wales Specialist Virology Centre Sequencing lab: Pathogen Genomics Unit | Public Health Wales Microbiology Cardiff Wales Specialist Virology Centre | Catherine Moore, Johnathan Evans, Laura Gifford, Malorie Perry, Simon Cottrell, Angela Marchbank, Alec Birchley, Alexander Adams, Amy Gaskin, Bree Gatica-Wilcox, Jason Coombes, Joel Southgate, Lauren Gilbert, Lee Graham, Nicole Pacchiarini, Sara Kumziene-Summerhayes, Sarah Taylor, Sophie Jones, Sara Rey, Matthew Bull, Joanne Watkins, Sally Corden, Tom Connor |
| EPI_ISL_3295595 | Viollier AG | Department of Biosystems Science and Engineering, ETH Zürich | Christian Beisel, Sarah Nadeau, Chaoran Chen, Ivan Topolsky, Kim Philipp Jablonski, Lara Fuhrmann, Rebecca Denes, Mirjam Feldkamp, Ina Nissen, Natascha Santacroce, Elodie Burcklen, Christiane Beckmann, Maurice Redondo, Olivier Kobel, Christoph Noppen, Niko Beerenwinkel, Tanja Stadler |
| EPI_ISL_3295596 | Wales Specialist Virology Centre Sequencing lab: Pathogen Genomics Unit | Public Health Wales Microbiology Cardiff Wales Specialist Virology Centre | Catherine Moore, Johnathan Evans, Laura Gifford, Malorie Perry, Simon Cottrell, Angela Marchbank, Alec Birchley, Alexander Adams, Amy Gaskin, Bree Gatica-Wilcox, Jason Coombes, Joel Southgate, Lauren Gilbert, Lee Graham, Nicole Pacchiarini, Sara Kumziene-Summerhayes, Sarah Taylor, Sophie Jones, Sara Rey, Matthew Bull, Joanne Watkins, Sally Corden, Tom Connor |
| EPI_ISL_3295597 | Viollier AG | Department of Biosystems Science and Engineering, ETH Zürich | Christian Beisel, Sarah Nadeau, Chaoran Chen, Ivan Topolsky, Kim Philipp Jablonski, Lara Fuhrmann, Rebecca Denes, Mirjam Feldkamp, Ina Nissen, Natascha Santacroce, Elodie Burcklen, Christiane Beckmann, Maurice Redondo, Olivier Kobel, Christoph Noppen, Niko Beerenwinkel, Tanja Stadler |
| EPI_ISL_3295598 | Wales Specialist Virology Centre Sequencing lab: Pathogen Genomics Unit | Public Health Wales Microbiology Cardiff Wales Specialist Virology Centre | Catherine Moore, Johnathan Evans, Laura Gifford, Malorie Perry, Simon Cottrell, Angela Marchbank, Alec Birchley, Alexander Adams, Amy Gaskin, Bree Gatica-Wilcox, Jason Coombes, Joel Southgate, Lauren Gilbert, Lee Graham, Nicole Pacchiarini, Sara Kumziene-Summerhayes, Sarah Taylor, Sophie Jones, Sara Rey, Matthew Bull, Joanne Watkins, Sally Corden, Tom Connor |
| EPI_ISL_3295599 | Viollier AG | Department of Biosystems Science and Engineering, ETH Zürich | Christian Beisel, Sarah Nadeau, Chaoran Chen, Ivan Topolsky, Kim Philipp Jablonski, Lara Fuhrmann, Rebecca Denes, Mirjam Feldkamp, Ina Nissen, Natascha Santacroce, Elodie Burcklen, Christiane Beckmann, Maurice Redondo, Olivier Kobel, Christoph Noppen, Niko Beerenwinkel, Tanja Stadler |
| EPI_ISL_3295600 | Wales Specialist Virology Centre Sequencing lab: Pathogen Genomics Unit | Public Health Wales Microbiology Cardiff Wales Specialist Virology Centre | Catherine Moore, Johnathan Evans, Laura Gifford, Malorie Perry, Simon Cottrell, Angela Marchbank, Alec Birchley, Alexander Adams, Amy Gaskin, Bree Gatica-Wilcox, Jason Coombes, Joel Southgate, Lauren Gilbert, Lee Graham, Nicole Pacchiarini, Sara Kumziene-Summerhayes, Sarah Taylor, Sophie Jones, Sara Rey, Matthew Bull, Joanne Watkins, Sally Corden, Tom Connor |
| EPI_ISL_3295601 | Viollier AG | Department of Biosystems Science and Engineering, ETH Zürich | Christian Beisel, Sarah Nadeau, Chaoran Chen, Ivan Topolsky, Kim Philipp Jablonski, Lara Fuhrmann, Rebecca Denes, Mirjam Feldkamp, Ina Nissen, Natascha Santacroce, Elodie Burcklen, Christiane Beckmann, Maurice Redondo, Olivier Kobel, Christoph Noppen, Niko Beerenwinkel, Tanja Stadler |
| EPI_ISL_3295602 | Wales Specialist Virology Centre Sequencing lab: Pathogen Genomics Unit | Public Health Wales Microbiology Cardiff Wales Specialist Virology Centre | Catherine Moore, Johnathan Evans, Laura Gifford, Malorie Perry, Simon Cottrell, Angela Marchbank, Alec Birchley, Alexander Adams, Amy Gaskin, Bree Gatica-Wilcox, Jason Coombes, Joel Southgate, Lauren Gilbert, Lee Graham, Nicole Pacchiarini, Sara Kumziene-Summerhayes, Sarah Taylor, Sophie Jones, Sara Rey, Matthew Bull, Joanne Watkins, Sally Corden, Tom Connor |
| EPI_ISL_3295603 | Viollier AG | Department of Biosystems Science and Engineering, ETH Zürich | Christian Beisel, Sarah Nadeau, Chaoran Chen, Ivan Topolsky, Kim Philipp Jablonski, Lara Fuhrmann, Rebecca Denes, Mirjam Feldkamp, Ina Nissen, Natascha Santacroce, Elodie Burcklen, Christiane Beckmann, Maurice Redondo, Olivier Kobel, Christoph Noppen, Niko Beerenwinkel, Tanja Stadler |
| EPI_ISL_3295604, EPI_ISL_3295605 | Wales Specialist Virology Centre Sequencing lab: Pathogen Genomics Unit | Public Health Wales Microbiology Cardiff Wales Specialist Virology Centre | Catherine Moore, Johnathan Evans, Laura Gifford, Malorie Perry, Simon Cottrell, Angela Marchbank, Alec Birchley, Alexander Adams, Amy Gaskin, Bree Gatica-Wilcox, Jason Coombes, Joel Southgate, Lauren Gilbert, Lee Graham, Nicole Pacchiarini, Sara Kumziene-Summerhayes, Sarah Taylor, Sophie Jones, Sara Rey, Matthew Bull, Joanne Watkins, Sally Corden, Tom Connor |
| EPI_ISL_3295606 | Viollier AG | Department of Biosystems Science and Engineering, ETH Zürich | Christian Beisel, Sarah Nadeau, Chaoran Chen, Ivan Topolsky, Kim Philipp Jablonski, Lara Fuhrmann, Rebecca Denes, Mirjam Feldkamp, Ina Nissen, Natascha Santacroce, Elodie Burcklen, Christiane Beckmann, Maurice Redondo, Olivier Kobel, Christoph Noppen, Niko Beerenwinkel, Tanja Stadler |
| EPI_ISL_3295607 | Wales Specialist Virology Centre Sequencing lab: Pathogen Genomics Unit | Public Health Wales Microbiology Cardiff Wales Specialist Virology Centre | Catherine Moore, Johnathan Evans, Laura Gifford, Malorie Perry, Simon Cottrell, Angela Marchbank, Alec Birchley, Alexander Adams, Amy Gaskin, Bree Gatica-Wilcox, Jason Coombes, Joel Southgate, Lauren Gilbert, Lee Graham, Nicole Pacchiarini, Sara Kumziene-Summerhayes, Sarah Taylor, Sophie Jones, Sara Rey, Matthew Bull, Joanne Watkins, Sally Corden, Tom Connor |
| EPI_ISL_3295608 | Viollier AG | Department of Biosystems Science and Engineering, ETH Zürich | Christian Beisel, Sarah Nadeau, Chaoran Chen, Ivan Topolsky, Kim Philipp Jablonski, Lara Fuhrmann, Rebecca Denes, Mirjam Feldkamp, Ina Nissen, Natascha Santacroce, Elodie Burcklen, Christiane Beckmann, Maurice Redondo, Olivier Kobel, Christoph Noppen, Niko Beerenwinkel, Tanja Stadler |
| EPI_ISL_3295609 | Wales Specialist Virology Centre Sequencing lab: Pathogen Genomics Unit | Public Health Wales Microbiology Cardiff Wales Specialist Virology Centre | Catherine Moore, Johnathan Evans, Laura Gifford, Malorie Perry, Simon Cottrell, Angela Marchbank, Alec Birchley, Alexander Adams, Amy Gaskin, Bree Gatica-Wilcox, Jason Coombes, Joel Southgate, Lauren Gilbert, Lee Graham, Nicole Pacchiarini, Sara Kumziene-Summerhayes, Sarah Taylor, Sophie Jones, Sara Rey, Matthew Bull, Joanne Watkins, Sally Corden, Tom Connor |
| EPI_ISL_3295610 | Viollier AG | Department of Biosystems Science and Engineering, ETH Zürich | Christian Beisel, Sarah Nadeau, Chaoran Chen, Ivan Topolsky, Kim Philipp Jablonski, Lara Fuhrmann, Rebecca Denes, Mirjam Feldkamp, Ina Nissen, Natascha Santacroce, Elodie Burcklen, Christiane Beckmann, Maurice Redondo, Olivier Kobel, Christoph Noppen, Niko Beerenwinkel, Tanja Stadler |
| EPI_ISL_3295611 | Wales Specialist Virology Centre Sequencing lab: Pathogen Genomics Unit | Public Health Wales Microbiology Cardiff Wales Specialist Virology Centre | Catherine Moore, Johnathan Evans, Laura Gifford, Malorie Perry, Simon Cottrell, Angela Marchbank, Alec Birchley, Alexander Adams, Amy Gaskin, Bree Gatica-Wilcox, Jason Coombes, Joel Southgate, Lauren Gilbert, Lee Graham, Nicole Pacchiarini, Sara Kumziene-Summerhayes, Sarah Taylor, Sophie Jones, Sara Rey, Matthew Bull, Joanne Watkins, Sally Corden, Tom Connor |
| EPI_ISL_3295612 | Viollier AG | Department of Biosystems Science and Engineering, ETH Zürich | Christian Beisel, Sarah Nadeau, Chaoran Chen, Ivan Topolsky, Kim Philipp Jablonski, Lara Fuhrmann, Rebecca Denes, Mirjam Feldkamp, Ina Nissen, Natascha Santacroce, Elodie Burcklen, Christiane Beckmann, Maurice Redondo, Olivier Kobel, |

| | | | |
|---|---|---|---|
| | | | Christoph Noppen, Niko Beerenwinkel, Tanja Stadler |
| EPI_ISL_3295613 | Wales Specialist Virology Centre Sequencing lab: Pathogen Genomics Unit | Public Health Wales Microbiology Cardiff Wales Specialist Virology Centre | Catherine Moore, Johnathan Evans, Laura Gifford, Malorie Perry, Simon Cottrell, Angela Marchbank, Alec Birchley, Alexander Adams, Amy Gaskin, Bree Gatica-Wilcox, Jason Coombes, Joel Southgate, Lauren Gilbert, Lee Graham, Nicole Pacchiarini, Sara Kumziene-Summerhayes, Sarah Taylor, Sophie Jones, Sara Rey, Matthew Bull, Joanne Watkins, Sally Corden, Tom Connor |
| EPI_ISL_3295614 | Viollier AG | Department of Biosystems Science and Engineering, ETH Zürich | Christian Beisel, Sarah Nadeau, Chaoran Chen, Ivan Topolsky, Kim Philipp Jablonski, Lara Fuhrmann, Rebecca Denes, Mirjam Feldkamp, Ina Nissen, Natascha Santacroce, Elodie Burcklen, Christiane Beckmann, Maurice Redondo, Olivier Kobel, Christoph Noppen, Niko Beerenwinkel, Tanja Stadler |
| EPI_ISL_3295615 | Wales Specialist Virology Centre Sequencing lab: Pathogen Genomics Unit | Public Health Wales Microbiology Cardiff Wales Specialist Virology Centre | Catherine Moore, Johnathan Evans, Laura Gifford, Malorie Perry, Simon Cottrell, Angela Marchbank, Alec Birchley, Alexander Adams, Amy Gaskin, Bree Gatica-Wilcox, Jason Coombes, Joel Southgate, Lauren Gilbert, Lee Graham, Nicole Pacchiarini, Sara Kumziene-Summerhayes, Sarah Taylor, Sophie Jones, Sara Rey, Matthew Bull, Joanne Watkins, Sally Corden, Tom Connor |
| EPI_ISL_3295616 | Viollier AG | Department of Biosystems Science and Engineering, ETH Zürich | Christian Beisel, Sarah Nadeau, Chaoran Chen, Ivan Topolsky, Kim Philipp Jablonski, Lara Fuhrmann, Rebecca Denes, Mirjam Feldkamp, Ina Nissen, Natascha Santacroce, Elodie Burcklen, Christiane Beckmann, Maurice Redondo, Olivier Kobel, Christoph Noppen, Niko Beerenwinkel, Tanja Stadler |
| EPI_ISL_3295617 | Wales Specialist Virology Centre Sequencing lab: Pathogen Genomics Unit | Public Health Wales Microbiology Cardiff Wales Specialist Virology Centre | Catherine Moore, Johnathan Evans, Laura Gifford, Malorie Perry, Simon Cottrell, Angela Marchbank, Alec Birchley, Alexander Adams, Amy Gaskin, Bree Gatica-Wilcox, Jason Coombes, Joel Southgate, Lauren Gilbert, Lee Graham, Nicole Pacchiarini, Sara Kumziene-Summerhayes, Sarah Taylor, Sophie Jones, Sara Rey, Matthew Bull, Joanne Watkins, Sally Corden, Tom Connor |
| EPI_ISL_3295618 | Viollier AG | Department of Biosystems Science and Engineering, ETH Zürich | Christian Beisel, Sarah Nadeau, Chaoran Chen, Ivan Topolsky, Kim Philipp Jablonski, Lara Fuhrmann, Rebecca Denes, Mirjam Feldkamp, Ina Nissen, Natascha Santacroce, Elodie Burcklen, Christiane Beckmann, Maurice Redondo, Olivier Kobel, Christoph Noppen, Niko Beerenwinkel, Tanja Stadler |
| EPI_ISL_3295619 | Wales Specialist Virology Centre Sequencing lab: Pathogen Genomics Unit | Public Health Wales Microbiology Cardiff Wales Specialist Virology Centre | Catherine Moore, Johnathan Evans, Laura Gifford, Malorie Perry, Simon Cottrell, Angela Marchbank, Alec Birchley, Alexander Adams, Amy Gaskin, Bree Gatica-Wilcox, Jason Coombes, Joel Southgate, Lauren Gilbert, Lee Graham, Nicole Pacchiarini, Sara Kumziene-Summerhayes, Sarah Taylor, Sophie Jones, Sara Rey, Matthew Bull, Joanne Watkins, Sally Corden, Tom Connor |
| EPI_ISL_3295620 | Viollier AG | Department of Biosystems Science and Engineering, ETH Zürich | Christian Beisel, Sarah Nadeau, Chaoran Chen, Ivan Topolsky, Kim Philipp Jablonski, Lara Fuhrmann, Rebecca Denes, Mirjam Feldkamp, Ina Nissen, Natascha Santacroce, Elodie Burcklen, Christiane Beckmann, Maurice Redondo, Olivier Kobel, Christoph Noppen, Niko Beerenwinkel, Tanja Stadler |
| EPI_ISL_3295621, EPI_ISL_3295622 | Wales Specialist Virology Centre Sequencing lab: Pathogen Genomics Unit | Public Health Wales Microbiology Cardiff Wales Specialist Virology Centre | Catherine Moore, Johnathan Evans, Laura Gifford, Malorie Perry, Simon Cottrell, Angela Marchbank, Alec Birchley, Alexander Adams, Amy Gaskin, Bree Gatica-Wilcox, Jason Coombes, Joel Southgate, Lauren Gilbert, Lee Graham, Nicole Pacchiarini, Sara Kumziene-Summerhayes, Sarah Taylor, Sophie Jones, Sara Rey, Matthew Bull, Joanne Watkins, Sally Corden, Tom Connor |
| EPI_ISL_3295623, EPI_ISL_3295624 | Viollier AG | Department of Biosystems Science and Engineering, ETH Zürich | Christian Beisel, Sarah Nadeau, Chaoran Chen, Ivan Topolsky, Kim Philipp Jablonski, Lara Fuhrmann, Rebecca Denes, Mirjam Feldkamp, Ina Nissen, Natascha Santacroce, Elodie Burcklen, Christiane Beckmann, Maurice Redondo, Olivier Kobel, Christoph Noppen, Niko Beerenwinkel, Tanja Stadler |
| EPI_ISL_3295625 | Wales Specialist Virology Centre Sequencing lab: Pathogen Genomics Unit | Public Health Wales Microbiology Cardiff Wales Specialist Virology Centre | Catherine Moore, Johnathan Evans, Laura Gifford, Malorie Perry, Simon Cottrell, Angela Marchbank, Alec Birchley, Alexander Adams, Amy Gaskin, Bree Gatica-Wilcox, Jason Coombes, Joel Southgate, Lauren Gilbert, Lee Graham, Nicole Pacchiarini, Sara Kumziene-Summerhayes, Sarah Taylor, Sophie Jones, Sara Rey, Matthew Bull, Joanne Watkins, Sally Corden, Tom Connor |
| EPI_ISL_3295626 | Viollier AG | Department of Biosystems Science and Engineering, ETH Zürich | Christian Beisel, Sarah Nadeau, Chaoran Chen, Ivan Topolsky, Kim Philipp Jablonski, Lara Fuhrmann, Rebecca Denes, Mirjam Feldkamp, Ina Nissen, Natascha Santacroce, Elodie Burcklen, Christiane Beckmann, Maurice Redondo, Olivier Kobel, Christoph Noppen, Niko Beerenwinkel, Tanja Stadler |
| EPI_ISL_3295627 | Wales Specialist Virology Centre Sequencing lab: Pathogen Genomics Unit | Public Health Wales Microbiology Cardiff Wales Specialist Virology Centre | Catherine Moore, Johnathan Evans, Laura Gifford, Malorie Perry, Simon Cottrell, Angela Marchbank, Alec Birchley, Alexander Adams, Amy Gaskin, Bree Gatica-Wilcox, Jason Coombes, Joel Southgate, Lauren Gilbert, Lee Graham, Nicole Pacchiarini, Sara Kumziene-Summerhayes, Sarah Taylor, Sophie Jones, Sara Rey, Matthew Bull, Joanne Watkins, Sally Corden, Tom Connor |
| EPI_ISL_3295628 | Viollier AG | Department of Biosystems Science and Engineering, ETH Zürich | Christian Beisel, Sarah Nadeau, Chaoran Chen, Ivan Topolsky, Kim Philipp Jablonski, Lara Fuhrmann, Rebecca Denes, Mirjam Feldkamp, Ina Nissen, Natascha Santacroce, Elodie Burcklen, Christiane Beckmann, Maurice Redondo, Olivier Kobel, Christoph Noppen, Niko Beerenwinkel, Tanja Stadler |
| EPI_ISL_3295629, EPI_ISL_3295631, EPI_ISL_3295632 | Wales Specialist Virology Centre Sequencing lab: Pathogen Genomics Unit | Public Health Wales Microbiology Cardiff Wales Specialist Virology Centre | Catherine Moore, Johnathan Evans, Laura Gifford, Malorie Perry, Simon Cottrell, Angela Marchbank, Alec Birchley, Alexander Adams, Amy Gaskin, Bree Gatica-Wilcox, Jason Coombes, Joel Southgate, Lauren Gilbert, Lee Graham, Nicole Pacchiarini, Sara Kumziene-Summerhayes, Sarah Taylor, Sophie Jones, Sara Rey, Matthew Bull, Joanne Watkins, Sally Corden, Tom Connor |
| EPI_ISL_3295633 | Viollier AG | Department of Biosystems Science and Engineering, ETH Zürich | Christian Beisel, Sarah Nadeau, Chaoran Chen, Ivan Topolsky, Kim Philipp Jablonski, Lara Fuhrmann, Rebecca Denes, Mirjam Feldkamp, Ina Nissen, Natascha Santacroce, Elodie Burcklen, Christiane Beckmann, Maurice Redondo, Olivier Kobel, Christoph Noppen, Niko Beerenwinkel, Tanja Stadler |
| EPI_ISL_3295634 | Wales Specialist Virology Centre Sequencing lab: Pathogen Genomics Unit | Public Health Wales Microbiology Cardiff Wales Specialist Virology Centre | Catherine Moore, Johnathan Evans, Laura Gifford, Malorie Perry, Simon Cottrell, Angela Marchbank, Alec Birchley, Alexander Adams, Amy Gaskin, Bree Gatica-Wilcox, Jason Coombes, Joel Southgate, Lauren Gilbert, Lee Graham, Nicole Pacchiarini, Sara Kumziene-Summerhayes, Sarah Taylor, Sophie Jones, Sara Rey, Matthew Bull, Joanne Watkins, Sally Corden, Tom Connor |
| EPI_ISL_3295635 | Viollier AG | Department of Biosystems Science and Engineering, ETH Zürich | Christian Beisel, Sarah Nadeau, Chaoran Chen, Ivan Topolsky, Kim Philipp Jablonski, Lara Fuhrmann, Rebecca Denes, Mirjam Feldkamp, Ina Nissen, Natascha Santacroce, Elodie Burcklen, Christiane Beckmann, Maurice Redondo, Olivier Kobel, Christoph Noppen, Niko Beerenwinkel, Tanja Stadler |
| EPI_ISL_3295636 | Wales Specialist Virology Centre Sequencing lab: Pathogen Genomics Unit | Public Health Wales Microbiology Cardiff Wales Specialist Virology Centre | Catherine Moore, Johnathan Evans, Laura Gifford, Malorie Perry, Simon Cottrell, Angela Marchbank, Alec Birchley, Alexander Adams, Amy Gaskin, Bree Gatica-Wilcox, Jason Coombes, Joel Southgate, Lauren Gilbert, Lee Graham, Nicole Pacchiarini, Sara Kumziene-Summerhayes, Sarah Taylor, Sophie Jones, Sara Rey, Matthew Bull, Joanne Watkins, Sally Corden, Tom Connor |
| EPI_ISL_3295637 | Viollier AG | Department of Biosystems Science and Engineering, ETH Zürich | Christian Beisel, Sarah Nadeau, Chaoran Chen, Ivan Topolsky, Kim Philipp Jablonski, Lara Fuhrmann, Rebecca Denes, Mirjam Feldkamp, Ina Nissen, Natascha Santacroce, Elodie Burcklen, Christiane Beckmann, Maurice Redondo, Olivier Kobel, Christoph Noppen, Niko Beerenwinkel, Tanja Stadler |
| EPI_ISL_3295638 | Wales Specialist Virology Centre Sequencing lab: Pathogen Genomics Unit | Public Health Wales Microbiology Cardiff Wales Specialist Virology Centre | Catherine Moore, Johnathan Evans, Laura Gifford, Malorie Perry, Simon Cottrell, Angela Marchbank, Alec Birchley, Alexander Adams, Amy Gaskin, Bree Gatica-Wilcox, Jason Coombes, Joel Southgate, Lauren Gilbert, Lee Graham, Nicole Pacchiarini, Sara Kumziene-Summerhayes, Sarah Taylor, Sophie Jones, Sara Rey, Matthew Bull, Joanne Watkins, Sally Corden, Tom Connor |
| EPI_ISL_3295639 | Viollier AG | Department of Biosystems Science and Engineering, ETH Zürich | Christian Beisel, Sarah Nadeau, Chaoran Chen, Ivan Topolsky, Kim Philipp Jablonski, Lara Fuhrmann, Rebecca Denes, Mirjam Feldkamp, Ina Nissen, Natascha Santacroce, Elodie Burcklen, Christiane Beckmann, Maurice Redondo, Olivier Kobel, Christoph Noppen, Niko Beerenwinkel, Tanja Stadler |
| EPI_ISL_3295640 | Wales Specialist Virology Centre Sequencing lab: Pathogen Genomics Unit | Public Health Wales Microbiology Cardiff Wales Specialist Virology Centre | Catherine Moore, Johnathan Evans, Laura Gifford, Malorie Perry, Simon Cottrell, Angela Marchbank, Alec Birchley, Alexander Adams, Amy Gaskin, Bree Gatica-Wilcox, Jason Coombes, Joel Southgate, Lauren Gilbert, Lee Graham, Nicole Pacchiarini, Sara Kumziene-Summerhayes, Sarah Taylor, Sophie Jones, Sara Rey, Matthew Bull, Joanne Watkins, Sally Corden, Tom Connor |
| EPI_ISL_3295641 | Viollier AG | Department of Biosystems Science and Engineering, ETH Zürich | Christian Beisel, Sarah Nadeau, Chaoran Chen, Ivan Topolsky, Kim Philipp Jablonski, Lara Fuhrmann, Rebecca Denes, Mirjam Feldkamp, Ina Nissen, Natascha Santacroce, Elodie Burcklen, Christiane Beckmann, Maurice Redondo, Olivier Kobel, Christoph Noppen, Niko Beerenwinkel, Tanja Stadler |
| EPI_ISL_3295642, EPI_ISL_3295643 | Wales Specialist Virology Centre Sequencing lab: Pathogen Genomics Unit | Public Health Wales Microbiology Cardiff Wales Specialist Virology Centre | Catherine Moore, Johnathan Evans, Laura Gifford, Malorie Perry, Simon Cottrell, Angela Marchbank, Alec Birchley, Alexander Adams, Amy Gaskin, Bree Gatica-Wilcox, Jason Coombes, Joel Southgate, Lauren Gilbert, Lee Graham, Nicole Pacchiarini, Sara Kumziene-Summerhayes, Sarah Taylor, Sophie Jones, Sara Rey, Matthew Bull, Joanne Watkins, Sally Corden, Tom Connor |
| EPI_ISL_3295644 | Viollier AG | Department of Biosystems Science and Engineering, ETH Zürich | Christian Beisel, Sarah Nadeau, Chaoran Chen, Ivan Topolsky, Kim Philipp Jablonski, Lara Fuhrmann, Rebecca Denes, Mirjam Feldkamp, Ina Nissen, Natascha Santacroce, Elodie Burcklen, Christiane Beckmann, Maurice Redondo, Olivier Kobel, Christoph Noppen, Niko Beerenwinkel, Tanja Stadler |
| EPI_ISL_3295645 | Wales Specialist Virology Centre Sequencing lab: Pathogen Genomics Unit | Public Health Wales Microbiology Cardiff Wales Specialist Virology Centre | Catherine Moore, Johnathan Evans, Laura Gifford, Malorie Perry, Simon Cottrell, Angela Marchbank, Alec Birchley, Alexander Adams, Amy Gaskin, Bree Gatica-Wilcox, Jason Coombes, Joel Southgate, Lauren Gilbert, Lee Graham, Nicole Pacchiarini, Sara Kumziene-Summerhayes, Sarah Taylor, Sophie Jones, Sara Rey, Matthew Bull, Joanne Watkins, Sally Corden, Tom Connor |
| EPI_ISL_3295646 | Viollier AG | Department of Biosystems Science and Engineering, ETH Zürich | Christian Beisel, Sarah Nadeau, Chaoran Chen, Ivan Topolsky, Kim Philipp Jablonski, Lara Fuhrmann, Rebecca Denes, Mirjam Feldkamp, Ina Nissen, Natascha Santacroce, Elodie Burcklen, Christiane Beckmann, Maurice Redondo, Olivier Kobel, Christoph Noppen, Niko Beerenwinkel, Tanja Stadler |
| EPI_ISL_3295647 | Wales Specialist Virology Centre Sequencing lab: Pathogen Genomics Unit | Public Health Wales Microbiology Cardiff Wales Specialist Virology Centre | Catherine Moore, Johnathan Evans, Laura Gifford, Malorie Perry, Simon Cottrell, Angela Marchbank, Alec Birchley, Alexander Adams, Amy Gaskin, Bree Gatica-Wilcox, Jason Coombes, Joel Southgate, Lauren Gilbert, Lee Graham, Nicole Pacchiarini, Sara Kumziene-Summerhayes, Sarah Taylor, Sophie Jones, Sara Rey, Matthew Bull, Joanne Watkins, Sally Corden, Tom Connor |
| EPI_ISL_3295648 | Viollier AG | Department of Biosystems Science and Engineering, ETH Zürich | Christian Beisel, Sarah Nadeau, Chaoran Chen, Ivan Topolsky, Kim Philipp Jablonski, Lara Fuhrmann, Rebecca Denes, Mirjam Feldkamp, Ina Nissen, Natascha Santacroce, Elodie Burcklen, Christiane Beckmann, Maurice Redondo, Olivier Kobel, Christoph Noppen, Niko Beerenwinkel, Tanja Stadler |
| EPI_ISL_3295649 | Wales Specialist Virology Centre Sequencing lab: Pathogen Genomics Unit | Public Health Wales Microbiology Cardiff Wales Specialist Virology Centre | Catherine Moore, Johnathan Evans, Laura Gifford, Malorie Perry, Simon Cottrell, Angela Marchbank, Alec Birchley, Alexander Adams, Amy Gaskin, Bree Gatica-Wilcox, Jason Coombes, Joel Southgate, Lauren Gilbert, Lee Graham, Nicole Pacchiarini, Sara Kumziene-Summerhayes, Sarah Taylor, Sophie Jones, Sara Rey, Matthew Bull, Joanne Watkins, Sally Corden, Tom Connor |
| EPI_ISL_3295650 | Viollier AG | Department of Biosystems Science and Engineering, ETH Zürich | Christian Beisel, Sarah Nadeau, Chaoran Chen, Ivan Topolsky, Kim Philipp Jablonski, Lara Fuhrmann, Rebecca Denes, Mirjam Feldkamp, Ina Nissen, Natascha Santacroce, Elodie Burcklen, Christiane Beckmann, Maurice Redondo, Olivier Kobel, Christoph Noppen, Niko Beerenwinkel, Tanja Stadler |
| EPI_ISL_3295651 | Wales Specialist Virology Centre Sequencing lab: Pathogen Genomics Unit | Public Health Wales Microbiology Cardiff Wales Specialist Virology Centre | Catherine Moore, Johnathan Evans, Laura Gifford, Malorie Perry, Simon Cottrell, Angela Marchbank, Alec Birchley, Alexander Adams, Amy Gaskin, Bree Gatica-Wilcox, Jason Coombes, Joel Southgate, Lauren Gilbert, Lee Graham, Nicole Pacchiarini, Sara Kumziene-Summerhayes, Sarah Taylor, Sophie Jones, Sara Rey, Matthew Bull, Joanne Watkins, Sally Corden, Tom Connor |
| EPI_ISL_3295652 | Viollier AG | Department of Biosystems Science and Engineering, ETH Zürich | Christian Beisel, Sarah Nadeau, Chaoran Chen, Ivan Topolsky, Kim Philipp Jablonski, Lara Fuhrmann, Rebecca Denes, Mirjam Feldkamp, Ina Nissen, Natascha Santacroce, Elodie Burcklen, Christiane Beckmann, Maurice Redondo, Olivier Kobel, Christoph Noppen, Niko Beerenwinkel, Tanja Stadler |
| EPI_ISL_3295653 | Wales Specialist Virology Centre Sequencing lab: Pathogen Genomics Unit | Public Health Wales Microbiology Cardiff Wales Specialist Virology Centre | Catherine Moore, Johnathan Evans, Laura Gifford, Malorie Perry, Simon Cottrell, Angela Marchbank, Alec Birchley, Alexander Adams, Amy Gaskin, Bree Gatica-Wilcox, Jason Coombes, Joel Southgate, Lauren Gilbert, Lee Graham, Nicole Pacchiarini, Sara Kumziene-Summerhayes, Sarah Taylor, Sophie Jones, Sara Rey, Matthew Bull, Joanne Watkins, Sally Corden, Tom Connor |
| EPI_ISL_3295654 | Viollier AG | Department of Biosystems Science and Engineering, ETH Zürich | Christian Beisel, Sarah Nadeau, Chaoran Chen, Ivan Topolsky, Kim Philipp Jablonski, Lara Fuhrmann, Rebecca Denes, Mirjam Feldkamp, Ina Nissen, Natascha Santacroce, Elodie Burcklen, Christiane Beckmann, Maurice Redondo, Olivier Kobel, Christoph Noppen, Niko Beerenwinkel, Tanja Stadler |
| EPI_ISL_3295655 | Wales Specialist Virology Centre Sequencing lab: Pathogen Genomics Unit | Public Health Wales Microbiology Cardiff Wales Specialist Virology Centre | Catherine Moore, Johnathan Evans, Laura Gifford, Malorie Perry, Simon Cottrell, Angela Marchbank, Alec Birchley, Alexander Adams, Amy Gaskin, Bree Gatica-Wilcox, Jason Coombes, Joel Southgate, Lauren Gilbert, Lee Graham, Nicole Pacchiarini, Sara Kumziene-Summerhayes, Sarah Taylor, Sophie Jones, Sara Rey, Matthew Bull, Joanne Watkins, Sally Corden, Tom Connor |
| EPI_ISL_3295656 | Viollier AG | Department of Biosystems Science and Engineering, ETH Zürich | Christian Beisel, Sarah Nadeau, Chaoran Chen, Ivan Topolsky, Kim Philipp Jablonski, Lara Fuhrmann, Rebecca Denes, Mirjam Feldkamp, Ina Nissen, Natascha Santacroce, Elodie Burcklen, Christiane Beckmann, Maurice Redondo, Olivier Kobel, Christoph Noppen, Niko Beerenwinkel, Tanja Stadler |
| EPI_ISL_3295657 | Wales Specialist Virology Centre Sequencing lab: Pathogen Genomics Unit | Public Health Wales Microbiology Cardiff Wales Specialist Virology Centre | Catherine Moore, Johnathan Evans, Laura Gifford, Malorie Perry, Simon Cottrell, Angela Marchbank, Alec Birchley, Alexander Adams, Amy Gaskin, Bree Gatica-Wilcox, Jason Coombes, Joel Southgate, Lauren Gilbert, Lee Graham, Nicole Pacchiarini, Sara Kumziene-Summerhayes, Sarah Taylor, Sophie Jones, Sara Rey, Matthew Bull, Joanne Watkins, Sally Corden, Tom Connor |
| EPI_ISL_3295658 | Viollier AG | Department of Biosystems Science and Engineering, ETH Zürich | Christian Beisel, Sarah Nadeau, Chaoran Chen, Ivan Topolsky, Kim Philipp Jablonski, Lara Fuhrmann, Rebecca Denes, Mirjam Feldkamp, Ina Nissen, Natascha Santacroce, Elodie Burcklen, Christiane Beckmann, Maurice Redondo, Olivier Kobel, Christoph Noppen, Niko Beerenwinkel, Tanja Stadler |
| EPI_ISL_3295659 | Wales Specialist Virology Centre Sequencing lab: Pathogen Genomics Unit | Public Health Wales Microbiology Cardiff Wales Specialist Virology Centre | Catherine Moore, Johnathan Evans, Laura Gifford, Malorie Perry, Simon Cottrell, Angela Marchbank, Alec Birchley, Alexander Adams, Amy Gaskin, Bree Gatica-Wilcox, Jason Coombes, Joel Southgate, Lauren Gilbert, Lee Graham, Nicole Pacchiarini, Sara Kumziene-Summerhayes, Sarah Taylor, Sophie Jones, Sara Rey, Matthew Bull, Joanne Watkins, Sally Corden, Tom Connor |
| EPI_ISL_3295660 | Viollier AG | Department of Biosystems Science and Engineering, ETH Zürich | Christian Beisel, Sarah Nadeau, Chaoran Chen, Ivan Topolsky, Kim Philipp Jablonski, Lara Fuhrmann, Rebecca Denes, Mirjam Feldkamp, Ina Nissen, Natascha Santacroce, Elodie Burcklen, Christiane Beckmann, Maurice Redondo, Olivier Kobel, Christoph Noppen, Niko Beerenwinkel, Tanja Stadler |
| EPI_ISL_3295661 | Wales Specialist Virology Centre Sequencing lab: Pathogen Genomics Unit | Public Health Wales Microbiology Cardiff Wales Specialist Virology Centre | Catherine Moore, Johnathan Evans, Laura Gifford, Malorie Perry, Simon Cottrell, Angela Marchbank, Alec Birchley, Alexander Adams, Amy Gaskin, Bree Gatica-Wilcox, Jason Coombes, Joel Southgate, Lauren Gilbert, Lee Graham, Nicole Pacchiarini, Sara Kumziene-Summerhayes, Sarah Taylor, Sophie Jones, Sara Rey, Matthew Bull, Joanne Watkins, Sally Corden, Tom Connor |
| EPI_ISL_3295662 | Viollier AG | Department of Biosystems Science and Engineering, ETH Zürich | Christian Beisel, Sarah Nadeau, Chaoran Chen, Ivan Topolsky, Kim Philipp Jablonski, Lara Fuhrmann, Rebecca Denes, Mirjam Feldkamp, Ina Nissen, Natascha Santacroce, Elodie Burcklen, Christiane Beckmann, Maurice Redondo, Olivier Kobel, Christoph Noppen, Niko Beerenwinkel, Tanja Stadler |

| | | | |
|---|---|---|---|
| EPI_ISL_3295663 | Wales Specialist Virology Centre Sequencing lab: Pathogen Genomics Unit | Public Health Wales Microbiology Cardiff Wales Specialist Virology Centre | Catherine Moore, Johnathan Evans, Laura Gifford, Malorie Perry, Simon Cottrell, Angela Marchbank, Alec Birchley, Alexander Adams, Amy Gaskin, Bree Gatica-Wilcox, Jason Coombes, Joel Southgate, Lauren Gilbert, Lee Graham, Nicole Pacchiarini, Sara Kumziene-Summerhayes, Sarah Taylor, Sophie Jones, Sara Rey, Matthew Bull, Joanne Watkins, Sally Corden, Tom Connor |
| EPI_ISL_3295664 | Viollier AG | Department of Biosystems Science and Engineering, ETH Zürich | Christian Beisel, Sarah Nadeau, Chaoran Chen, Ivan Topolsky, Kim Philipp Jablonski, Lara Fuhrmann, Rebecca Denes, Mirjam Feldkamp, Ina Nissen, Natascha Santacroce, Elodie Burcklen, Christiane Beckmann, Maurice Redondo, Olivier Kobel, Christoph Noppen, Niko Beerenwinkel, Tanja Stadler |
| EPI_ISL_3295665 | Wales Specialist Virology Centre Sequencing lab: Pathogen Genomics Unit | Public Health Wales Microbiology Cardiff Wales Specialist Virology Centre | Catherine Moore, Johnathan Evans, Laura Gifford, Malorie Perry, Simon Cottrell, Angela Marchbank, Alec Birchley, Alexander Adams, Amy Gaskin, Bree Gatica-Wilcox, Jason Coombes, Joel Southgate, Lauren Gilbert, Lee Graham, Nicole Pacchiarini, Sara Kumziene-Summerhayes, Sarah Taylor, Sophie Jones, Sara Rey, Matthew Bull, Joanne Watkins, Sally Corden, Tom Connor |
| EPI_ISL_3295666, EPI_ISL_3295667, EPI_ISL_3295668 | Viollier AG | Department of Biosystems Science and Engineering, ETH Zürich | Christian Beisel, Sarah Nadeau, Chaoran Chen, Ivan Topolsky, Kim Philipp Jablonski, Lara Fuhrmann, Rebecca Denes, Mirjam Feldkamp, Ina Nissen, Natascha Santacroce, Elodie Burcklen, Christiane Beckmann, Maurice Redondo, Olivier Kobel, Christoph Noppen, Niko Beerenwinkel, Tanja Stadler |
| EPI_ISL_3295669 | Wales Specialist Virology Centre Sequencing lab: Pathogen Genomics Unit | Public Health Wales Microbiology Cardiff Wales Specialist Virology Centre | Catherine Moore, Johnathan Evans, Laura Gifford, Malorie Perry, Simon Cottrell, Angela Marchbank, Alec Birchley, Alexander Adams, Amy Gaskin, Bree Gatica-Wilcox, Jason Coombes, Joel Southgate, Lauren Gilbert, Lee Graham, Nicole Pacchiarini, Sara Kumziene-Summerhayes, Sarah Taylor, Sophie Jones, Sara Rey, Matthew Bull, Joanne Watkins, Sally Corden, Tom Connor |
| EPI_ISL_3295670 | Viollier AG | Department of Biosystems Science and Engineering, ETH Zürich | Christian Beisel, Sarah Nadeau, Chaoran Chen, Ivan Topolsky, Kim Philipp Jablonski, Lara Fuhrmann, Rebecca Denes, Mirjam Feldkamp, Ina Nissen, Natascha Santacroce, Elodie Burcklen, Christiane Beckmann, Maurice Redondo, Olivier Kobel, Christoph Noppen, Niko Beerenwinkel, Tanja Stadler |
| EPI_ISL_3295671 | Wales Specialist Virology Centre Sequencing lab: Pathogen Genomics Unit | Public Health Wales Microbiology Cardiff Wales Specialist Virology Centre | Catherine Moore, Johnathan Evans, Laura Gifford, Malorie Perry, Simon Cottrell, Angela Marchbank, Alec Birchley, Alexander Adams, Amy Gaskin, Bree Gatica-Wilcox, Jason Coombes, Joel Southgate, Lauren Gilbert, Lee Graham, Nicole Pacchiarini, Sara Kumziene-Summerhayes, Sarah Taylor, Sophie Jones, Sara Rey, Matthew Bull, Joanne Watkins, Sally Corden, Tom Connor |
| EPI_ISL_3295672 | Viollier AG | Department of Biosystems Science and Engineering, ETH Zürich | Christian Beisel, Sarah Nadeau, Chaoran Chen, Ivan Topolsky, Kim Philipp Jablonski, Lara Fuhrmann, Rebecca Denes, Mirjam Feldkamp, Ina Nissen, Natascha Santacroce, Elodie Burcklen, Christiane Beckmann, Maurice Redondo, Olivier Kobel, Christoph Noppen, Niko Beerenwinkel, Tanja Stadler |
| EPI_ISL_3295673 | Wales Specialist Virology Centre Sequencing lab: Pathogen Genomics Unit | Public Health Wales Microbiology Cardiff Wales Specialist Virology Centre | Catherine Moore, Johnathan Evans, Laura Gifford, Malorie Perry, Simon Cottrell, Angela Marchbank, Alec Birchley, Alexander Adams, Amy Gaskin, Bree Gatica-Wilcox, Jason Coombes, Joel Southgate, Lauren Gilbert, Lee Graham, Nicole Pacchiarini, Sara Kumziene-Summerhayes, Sarah Taylor, Sophie Jones, Sara Rey, Matthew Bull, Joanne Watkins, Sally Corden, Tom Connor |
| EPI_ISL_3295674 | Viollier AG | Department of Biosystems Science and Engineering, ETH Zürich | Christian Beisel, Sarah Nadeau, Chaoran Chen, Ivan Topolsky, Kim Philipp Jablonski, Lara Fuhrmann, Rebecca Denes, Mirjam Feldkamp, Ina Nissen, Natascha Santacroce, Elodie Burcklen, Christiane Beckmann, Maurice Redondo, Olivier Kobel, Christoph Noppen, Niko Beerenwinkel, Tanja Stadler |
| EPI_ISL_3295675 | Wales Specialist Virology Centre Sequencing lab: Pathogen Genomics Unit | Public Health Wales Microbiology Cardiff Wales Specialist Virology Centre | Catherine Moore, Johnathan Evans, Laura Gifford, Malorie Perry, Simon Cottrell, Angela Marchbank, Alec Birchley, Alexander Adams, Amy Gaskin, Bree Gatica-Wilcox, Jason Coombes, Joel Southgate, Lauren Gilbert, Lee Graham, Nicole Pacchiarini, Sara Kumziene-Summerhayes, Sarah Taylor, Sophie Jones, Sara Rey, Matthew Bull, Joanne Watkins, Sally Corden, Tom Connor |
| EPI_ISL_3295676 | Viollier AG | Department of Biosystems Science and Engineering, ETH Zürich | Christian Beisel, Sarah Nadeau, Chaoran Chen, Ivan Topolsky, Kim Philipp Jablonski, Lara Fuhrmann, Rebecca Denes, Mirjam Feldkamp, Ina Nissen, Natascha Santacroce, Elodie Burcklen, Christiane Beckmann, Maurice Redondo, Olivier Kobel, Christoph Noppen, Niko Beerenwinkel, Tanja Stadler |
| EPI_ISL_3295677 | Wales Specialist Virology Centre Sequencing lab: Pathogen Genomics Unit | Public Health Wales Microbiology Cardiff Wales Specialist Virology Centre | Catherine Moore, Johnathan Evans, Laura Gifford, Malorie Perry, Simon Cottrell, Angela Marchbank, Alec Birchley, Alexander Adams, Amy Gaskin, Bree Gatica-Wilcox, Jason Coombes, Joel Southgate, Lauren Gilbert, Lee Graham, Nicole Pacchiarini, Sara Kumziene-Summerhayes, Sarah Taylor, Sophie Jones, Sara Rey, Matthew Bull, Joanne Watkins, Sally Corden, Tom Connor |
| EPI_ISL_3295678 | Viollier AG | Department of Biosystems Science and Engineering, ETH Zürich | Christian Beisel, Sarah Nadeau, Chaoran Chen, Ivan Topolsky, Kim Philipp Jablonski, Lara Fuhrmann, Rebecca Denes, Mirjam Feldkamp, Ina Nissen, Natascha Santacroce, Elodie Burcklen, Christiane Beckmann, Maurice Redondo, Olivier Kobel, Christoph Noppen, Niko Beerenwinkel, Tanja Stadler |
| EPI_ISL_3295679 | Wales Specialist Virology Centre Sequencing lab: Pathogen Genomics Unit | Public Health Wales Microbiology Cardiff Wales Specialist Virology Centre | Catherine Moore, Johnathan Evans, Laura Gifford, Malorie Perry, Simon Cottrell, Angela Marchbank, Alec Birchley, Alexander Adams, Amy Gaskin, Bree Gatica-Wilcox, Jason Coombes, Joel Southgate, Lauren Gilbert, Lee Graham, Nicole Pacchiarini, Sara Kumziene-Summerhayes, Sarah Taylor, Sophie Jones, Sara Rey, Matthew Bull, Joanne Watkins, Sally Corden, Tom Connor |
| EPI_ISL_3295680 | Viollier AG | Department of Biosystems Science and Engineering, ETH Zürich | Christian Beisel, Sarah Nadeau, Chaoran Chen, Ivan Topolsky, Kim Philipp Jablonski, Lara Fuhrmann, Rebecca Denes, Mirjam Feldkamp, Ina Nissen, Natascha Santacroce, Elodie Burcklen, Christiane Beckmann, Maurice Redondo, Olivier Kobel, Christoph Noppen, Niko Beerenwinkel, Tanja Stadler |
| EPI_ISL_3295681 | Wales Specialist Virology Centre Sequencing lab: Pathogen Genomics Unit | Public Health Wales Microbiology Cardiff Wales Specialist Virology Centre | Catherine Moore, Johnathan Evans, Laura Gifford, Malorie Perry, Simon Cottrell, Angela Marchbank, Alec Birchley, Alexander Adams, Amy Gaskin, Bree Gatica-Wilcox, Jason Coombes, Joel Southgate, Lauren Gilbert, Lee Graham, Nicole Pacchiarini, Sara Kumziene-Summerhayes, Sarah Taylor, Sophie Jones, Sara Rey, Matthew Bull, Joanne Watkins, Sally Corden, Tom Connor |
| EPI_ISL_3295682 | Viollier AG | Department of Biosystems Science and Engineering, ETH Zürich | Christian Beisel, Sarah Nadeau, Chaoran Chen, Ivan Topolsky, Kim Philipp Jablonski, Lara Fuhrmann, Rebecca Denes, Mirjam Feldkamp, Ina Nissen, Natascha Santacroce, Elodie Burcklen, Christiane Beckmann, Maurice Redondo, Olivier Kobel, Christoph Noppen, Niko Beerenwinkel, Tanja Stadler |
| EPI_ISL_3295683 | Wales Specialist Virology Centre Sequencing lab: Pathogen Genomics Unit | Public Health Wales Microbiology Cardiff Wales Specialist Virology Centre | Catherine Moore, Johnathan Evans, Laura Gifford, Malorie Perry, Simon Cottrell, Angela Marchbank, Alec Birchley, Alexander Adams, Amy Gaskin, Bree Gatica-Wilcox, Jason Coombes, Joel Southgate, Lauren Gilbert, Lee Graham, Nicole Pacchiarini, Sara Kumziene-Summerhayes, Sarah Taylor, Sophie Jones, Sara Rey, Matthew Bull, Joanne Watkins, Sally Corden, Tom Connor |
| EPI_ISL_3295684 | Viollier AG | Department of Biosystems Science and Engineering, ETH Zürich | Christian Beisel, Sarah Nadeau, Chaoran Chen, Ivan Topolsky, Kim Philipp Jablonski, Lara Fuhrmann, Rebecca Denes, Mirjam Feldkamp, Ina Nissen, Natascha Santacroce, Elodie Burcklen, Christiane Beckmann, Maurice Redondo, Olivier Kobel, Christoph Noppen, Niko Beerenwinkel, Tanja Stadler |
| EPI_ISL_3295685, EPI_ISL_3295686 | Wales Specialist Virology Centre Sequencing lab: Pathogen Genomics Unit | Public Health Wales Microbiology Cardiff Wales Specialist Virology Centre | Catherine Moore, Johnathan Evans, Laura Gifford, Malorie Perry, Simon Cottrell, Angela Marchbank, Alec Birchley, Alexander Adams, Amy Gaskin, Bree Gatica-Wilcox, Jason Coombes, Joel Southgate, Lauren Gilbert, Lee Graham, Nicole Pacchiarini, Sara Kumziene-Summerhayes, Sarah Taylor, Sophie Jones, Sara Rey, Matthew Bull, Joanne Watkins, Sally Corden, Tom Connor |
| EPI_ISL_3295687 | Viollier AG | Department of Biosystems Science and Engineering, ETH Zürich | Christian Beisel, Sarah Nadeau, Chaoran Chen, Ivan Topolsky, Kim Philipp Jablonski, Lara Fuhrmann, Rebecca Denes, Mirjam Feldkamp, Ina Nissen, Natascha Santacroce, Elodie Burcklen, Christiane Beckmann, Maurice Redondo, Olivier Kobel, Christoph Noppen, Niko Beerenwinkel, Tanja Stadler |
| EPI_ISL_3295688 | Wales Specialist Virology Centre Sequencing lab: Pathogen Genomics Unit | Public Health Wales Microbiology Cardiff Wales Specialist Virology Centre | Catherine Moore, Johnathan Evans, Laura Gifford, Malorie Perry, Simon Cottrell, Angela Marchbank, Alec Birchley, Alexander Adams, Amy Gaskin, Bree Gatica-Wilcox, Jason Coombes, Joel Southgate, Lauren Gilbert, Lee Graham, Nicole Pacchiarini, Sara Kumziene-Summerhayes, Sarah Taylor, Sophie Jones, Sara Rey, Matthew Bull, Joanne Watkins, Sally Corden, Tom Connor |
| EPI_ISL_3295689 | Viollier AG | Department of Biosystems Science and Engineering, ETH Zürich | Christian Beisel, Sarah Nadeau, Chaoran Chen, Ivan Topolsky, Kim Philipp Jablonski, Lara Fuhrmann, Rebecca Denes, Mirjam Feldkamp, Ina Nissen, Natascha Santacroce, Elodie Burcklen, Christiane Beckmann, Maurice Redondo, Olivier Kobel, Christoph Noppen, Niko Beerenwinkel, Tanja Stadler |
| EPI_ISL_3295690 | Wales Specialist Virology Centre Sequencing lab: Pathogen Genomics Unit | Public Health Wales Microbiology Cardiff Wales Specialist Virology Centre | Catherine Moore, Johnathan Evans, Laura Gifford, Malorie Perry, Simon Cottrell, Angela Marchbank, Alec Birchley, Alexander Adams, Amy Gaskin, Bree Gatica-Wilcox, Jason Coombes, Joel Southgate, Lauren Gilbert, Lee Graham, Nicole Pacchiarini, Sara Kumziene-Summerhayes, Sarah Taylor, Sophie Jones, Sara Rey, Matthew Bull, Joanne Watkins, Sally Corden, Tom Connor |
| EPI_ISL_3295691 | Viollier AG | Department of Biosystems Science and Engineering, ETH Zürich | Christian Beisel, Sarah Nadeau, Chaoran Chen, Ivan Topolsky, Kim Philipp Jablonski, Lara Fuhrmann, Rebecca Denes, Mirjam Feldkamp, Ina Nissen, Natascha Santacroce, Elodie Burcklen, Christiane Beckmann, Maurice Redondo, Olivier Kobel, Christoph Noppen, Niko Beerenwinkel, Tanja Stadler |
| EPI_ISL_3295692 | Wales Specialist Virology Centre Sequencing lab: Pathogen Genomics Unit | Public Health Wales Microbiology Cardiff Wales Specialist Virology Centre | Catherine Moore, Johnathan Evans, Laura Gifford, Malorie Perry, Simon Cottrell, Angela Marchbank, Alec Birchley, Alexander Adams, Amy Gaskin, Bree Gatica-Wilcox, Jason Coombes, Joel Southgate, Lauren Gilbert, Lee Graham, Nicole Pacchiarini, Sara Kumziene-Summerhayes, Sarah Taylor, Sophie Jones, Sara Rey, Matthew Bull, Joanne Watkins, Sally Corden, Tom Connor |
| EPI_ISL_3295693 | Viollier AG | Department of Biosystems Science and Engineering, ETH Zürich | Christian Beisel, Sarah Nadeau, Chaoran Chen, Ivan Topolsky, Kim Philipp Jablonski, Lara Fuhrmann, Rebecca Denes, Mirjam Feldkamp, Ina Nissen, Natascha Santacroce, Elodie Burcklen, Christiane Beckmann, Maurice Redondo, Olivier Kobel, Christoph Noppen, Niko Beerenwinkel, Tanja Stadler |
| EPI_ISL_3295694, EPI_ISL_3295696 | Wales Specialist Virology Centre Sequencing lab: Pathogen Genomics Unit | Public Health Wales Microbiology Cardiff Wales Specialist Virology Centre | Catherine Moore, Johnathan Evans, Laura Gifford, Malorie Perry, Simon Cottrell, Angela Marchbank, Alec Birchley, Alexander Adams, Amy Gaskin, Bree Gatica-Wilcox, Jason Coombes, Joel Southgate, Lauren Gilbert, Lee Graham, Nicole Pacchiarini, Sara Kumziene-Summerhayes, Sarah Taylor, Sophie Jones, Sara Rey, Matthew Bull, Joanne Watkins, Sally Corden, Tom Connor |
| EPI_ISL_3295697 | Viollier AG | Department of Biosystems Science and Engineering, ETH Zürich | Christian Beisel, Sarah Nadeau, Chaoran Chen, Ivan Topolsky, Kim Philipp Jablonski, Lara Fuhrmann, Rebecca Denes, Mirjam Feldkamp, Ina Nissen, Natascha Santacroce, Elodie Burcklen, Christiane Beckmann, Maurice Redondo, Olivier Kobel, Christoph Noppen, Niko Beerenwinkel, Tanja Stadler |
| EPI_ISL_3295698 | Wales Specialist Virology Centre Sequencing lab: Pathogen Genomics Unit | Public Health Wales Microbiology Cardiff Wales Specialist Virology Centre | Catherine Moore, Johnathan Evans, Laura Gifford, Malorie Perry, Simon Cottrell, Angela Marchbank, Alec Birchley, Alexander Adams, Amy Gaskin, Bree Gatica-Wilcox, Jason Coombes, Joel Southgate, Lauren Gilbert, Lee Graham, Nicole Pacchiarini, Sara Kumziene-Summerhayes, Sarah Taylor, Sophie Jones, Sara Rey, Matthew Bull, Joanne Watkins, Sally Corden, Tom Connor |
| EPI_ISL_3295699 | Viollier AG | Department of Biosystems Science and Engineering, ETH Zürich | Christian Beisel, Sarah Nadeau, Chaoran Chen, Ivan Topolsky, Kim Philipp Jablonski, Lara Fuhrmann, Rebecca Denes, Mirjam Feldkamp, Ina Nissen, Natascha Santacroce, Elodie Burcklen, Christiane Beckmann, Maurice Redondo, Olivier Kobel, Christoph Noppen, Niko Beerenwinkel, Tanja Stadler |
| EPI_ISL_3295700 | Wales Specialist Virology Centre Sequencing lab: Pathogen Genomics Unit | Public Health Wales Microbiology Cardiff Wales Specialist Virology Centre | Catherine Moore, Johnathan Evans, Laura Gifford, Malorie Perry, Simon Cottrell, Angela Marchbank, Alec Birchley, Alexander Adams, Amy Gaskin, Bree Gatica-Wilcox, Jason Coombes, Joel Southgate, Lauren Gilbert, Lee Graham, Nicole Pacchiarini, Sara Kumziene-Summerhayes, Sarah Taylor, Sophie Jones, Sara Rey, Matthew Bull, Joanne Watkins, Sally Corden, Tom Connor |
| EPI_ISL_3295701 | Viollier AG | Department of Biosystems Science and Engineering, ETH Zürich | Christian Beisel, Sarah Nadeau, Chaoran Chen, Ivan Topolsky, Kim Philipp Jablonski, Lara Fuhrmann, Rebecca Denes, Mirjam Feldkamp, Ina Nissen, Natascha Santacroce, Elodie Burcklen, Christiane Beckmann, Maurice Redondo, Olivier Kobel, Christoph Noppen, Niko Beerenwinkel, Tanja Stadler |
| EPI_ISL_3295702 | Wales Specialist Virology Centre Sequencing lab: Pathogen Genomics Unit | Public Health Wales Microbiology Cardiff Wales Specialist Virology Centre | Catherine Moore, Johnathan Evans, Laura Gifford, Malorie Perry, Simon Cottrell, Angela Marchbank, Alec Birchley, Alexander Adams, Amy Gaskin, Bree Gatica-Wilcox, Jason Coombes, Joel Southgate, Lauren Gilbert, Lee Graham, Nicole Pacchiarini, Sara Kumziene-Summerhayes, Sarah Taylor, Sophie Jones, Sara Rey, Matthew Bull, Joanne Watkins, Sally Corden, Tom Connor |
| EPI_ISL_3295703 | Viollier AG | Department of Biosystems Science and Engineering, ETH Zürich | Christian Beisel, Sarah Nadeau, Chaoran Chen, Ivan Topolsky, Kim Philipp Jablonski, Lara Fuhrmann, Rebecca Denes, Mirjam Feldkamp, Ina Nissen, Natascha Santacroce, Elodie Burcklen, Christiane Beckmann, Maurice Redondo, Olivier Kobel, Christoph Noppen, Niko Beerenwinkel, Tanja Stadler |
| EPI_ISL_3295704, EPI_ISL_3295705 | Wales Specialist Virology Centre Sequencing lab: Pathogen Genomics Unit | Public Health Wales Microbiology Cardiff Wales Specialist Virology Centre | Catherine Moore, Johnathan Evans, Laura Gifford, Malorie Perry, Simon Cottrell, Angela Marchbank, Alec Birchley, Alexander Adams, Amy Gaskin, Bree Gatica-Wilcox, Jason Coombes, Joel Southgate, Lauren Gilbert, Lee Graham, Nicole Pacchiarini, Sara Kumziene-Summerhayes, Sarah Taylor, Sophie Jones, Sara Rey, Matthew Bull, Joanne Watkins, Sally Corden, Tom Connor |
| EPI_ISL_3295706 | Viollier AG | Department of Biosystems Science and Engineering, ETH Zürich | Christian Beisel, Sarah Nadeau, Chaoran Chen, Ivan Topolsky, Kim Philipp Jablonski, Lara Fuhrmann, Rebecca Denes, Mirjam Feldkamp, Ina Nissen, Natascha Santacroce, Elodie Burcklen, Christiane Beckmann, Maurice Redondo, Olivier Kobel, Christoph Noppen, Niko Beerenwinkel, Tanja Stadler |
| EPI_ISL_3295707, EPI_ISL_3295709 | Wales Specialist Virology Centre Sequencing lab: Pathogen Genomics Unit | Public Health Wales Microbiology Cardiff Wales Specialist Virology Centre | Catherine Moore, Johnathan Evans, Laura Gifford, Malorie Perry, Simon Cottrell, Angela Marchbank, Alec Birchley, Alexander Adams, Amy Gaskin, Bree Gatica-Wilcox, Jason Coombes, Joel Southgate, Lauren Gilbert, Lee Graham, Nicole Pacchiarini, Sara Kumziene-Summerhayes, Sarah Taylor, Sophie Jones, Sara Rey, Matthew Bull, Joanne Watkins, Sally Corden, Tom Connor |
| EPI_ISL_3295710 | Viollier AG | Department of Biosystems Science and Engineering, ETH Zürich | Christian Beisel, Sarah Nadeau, Chaoran Chen, Ivan Topolsky, Kim Philipp Jablonski, Lara Fuhrmann, Rebecca Denes, Mirjam Feldkamp, Ina Nissen, Natascha Santacroce, Elodie Burcklen, Christiane Beckmann, Maurice Redondo, Olivier Kobel, Christoph Noppen, Niko Beerenwinkel, Tanja Stadler |
| EPI_ISL_3295711 | Wales Specialist Virology Centre Sequencing lab: Pathogen Genomics Unit | Public Health Wales Microbiology Cardiff Wales Specialist Virology Centre | Catherine Moore, Johnathan Evans, Laura Gifford, Malorie Perry, Simon Cottrell, Angela Marchbank, Alec Birchley, Alexander Adams, Amy Gaskin, Bree Gatica-Wilcox, Jason Coombes, Joel Southgate, Lauren Gilbert, Lee Graham, Nicole Pacchiarini, Sara Kumziene-Summerhayes, Sarah Taylor, Sophie Jones, Sara Rey, Matthew Bull, Joanne Watkins, Sally Corden, Tom Connor |
| EPI_ISL_3295712 | Viollier AG | Department of Biosystems Science and Engineering, ETH Zürich | Christian Beisel, Sarah Nadeau, Chaoran Chen, Ivan Topolsky, Kim Philipp Jablonski, Lara Fuhrmann, Rebecca Denes, Mirjam Feldkamp, Ina Nissen, Natascha Santacroce, Elodie Burcklen, Christiane Beckmann, Maurice Redondo, Olivier Kobel, Christoph Noppen, Niko Beerenwinkel, Tanja Stadler |
| EPI_ISL_3295713 | Wales Specialist Virology Centre Sequencing lab: Pathogen Genomics Unit | Public Health Wales Microbiology Cardiff Wales Specialist Virology Centre | Catherine Moore, Johnathan Evans, Laura Gifford, Malorie Perry, Simon Cottrell, Angela Marchbank, Alec Birchley, Alexander Adams, Amy Gaskin, Bree Gatica-Wilcox, Jason Coombes, Joel Southgate, Lauren Gilbert, Lee Graham, Nicole Pacchiarini, Sara Kumziene-Summerhayes, Sarah Taylor, Sophie Jones, Sara Rey, Matthew Bull, Joanne Watkins, Sally Corden, Tom Connor |
| EPI_ISL_3295714 | Viollier AG | Department of Biosystems Science and Engineering, ETH Zürich | Christian Beisel, Sarah Nadeau, Chaoran Chen, Ivan Topolsky, Kim Philipp Jablonski, Lara Fuhrmann, Rebecca Denes, Mirjam Feldkamp, Ina Nissen, Natascha Santacroce, Elodie Burcklen, Christiane Beckmann, Maurice Redondo, Olivier Kobel, Christoph Noppen, Niko Beerenwinkel, Tanja Stadler |
| EPI_ISL_3295715 | Wales Specialist Virology Centre Sequencing lab: Pathogen Genomics Unit | Public Health Wales Microbiology Cardiff Wales Specialist Virology Centre | Catherine Moore, Johnathan Evans, Laura Gifford, Malorie Perry, Simon Cottrell, Angela Marchbank, Alec Birchley, Alexander Adams, Amy Gaskin, Bree Gatica-Wilcox, Jason Coombes, Joel Southgate, Lauren Gilbert, Lee Graham, Nicole Pacchiarini, Sara Kumziene-Summerhayes, Sarah Taylor, Sophie Jones, Sara Rey, Matthew Bull, Joanne Watkins, Sally Corden, Tom Connor |

| | | | |
|---|---|---|---|
| EPI_ISL_3295716 | Viollier AG | Department of Biosystems Science and Engineering, ETH Zürich | Christian Beisel, Sarah Nadeau, Chaoran Chen, Ivan Topolsky, Kim Philipp Jablonski, Lara Fuhrmann, Rebecca Denes, Mirjam Feldkamp, Ina Nissen, Natascha Santacroce, Elodie Burcklen, Christiane Beckmann, Maurice Redondo, Olivier Kobel, Christoph Noppen, Niko Beerenwinkel, Tanja Stadler |
| EPI_ISL_3295717, EPI_ISL_3295719 | Wales Specialist Virology Centre Sequencing lab: Pathogen Genomics Unit | Public Health Wales Microbiology Cardiff Wales Specialist Virology Centre | Catherine Moore, Johnathan Evans, Laura Gifford, Malorie Perry, Simon Cottrell, Angela Marchbank, Alec Birchley, Alexander Adams, Amy Gaskin, Bree Gatica-Wilcox, Jason Coombes, Joel Southgate, Lauren Gilbert, Lee Graham, Nicole Pacchiarini, Sara Kumziene-Summerhayes, Sarah Taylor, Sophie Jones, Sara Rey, Matthew Bull, Joanne Watkins, Sally Corden, Tom Connor |
| EPI_ISL_3295720 | Viollier AG | Department of Biosystems Science and Engineering, ETH Zürich | Christian Beisel, Sarah Nadeau, Chaoran Chen, Ivan Topolsky, Kim Philipp Jablonski, Lara Fuhrmann, Rebecca Denes, Mirjam Feldkamp, Ina Nissen, Natascha Santacroce, Elodie Burcklen, Christiane Beckmann, Maurice Redondo, Olivier Kobel, Christoph Noppen, Niko Beerenwinkel, Tanja Stadler |
| EPI_ISL_3295721 | Wales Specialist Virology Centre Sequencing lab: Pathogen Genomics Unit | Public Health Wales Microbiology Cardiff Wales Specialist Virology Centre | Catherine Moore, Johnathan Evans, Laura Gifford, Malorie Perry, Simon Cottrell, Angela Marchbank, Alec Birchley, Alexander Adams, Amy Gaskin, Bree Gatica-Wilcox, Jason Coombes, Joel Southgate, Lauren Gilbert, Lee Graham, Nicole Pacchiarini, Sara Kumziene-Summerhayes, Sarah Taylor, Sophie Jones, Sara Rey, Matthew Bull, Joanne Watkins, Sally Corden, Tom Connor |
| EPI_ISL_3295722 | Viollier AG | Department of Biosystems Science and Engineering, ETH Zürich | Christian Beisel, Sarah Nadeau, Chaoran Chen, Ivan Topolsky, Kim Philipp Jablonski, Lara Fuhrmann, Rebecca Denes, Mirjam Feldkamp, Ina Nissen, Natascha Santacroce, Elodie Burcklen, Christiane Beckmann, Maurice Redondo, Olivier Kobel, Christoph Noppen, Niko Beerenwinkel, Tanja Stadler |
| EPI_ISL_3295723 | Wales Specialist Virology Centre Sequencing lab: Pathogen Genomics Unit | Public Health Wales Microbiology Cardiff Wales Specialist Virology Centre | Catherine Moore, Johnathan Evans, Laura Gifford, Malorie Perry, Simon Cottrell, Angela Marchbank, Alec Birchley, Alexander Adams, Amy Gaskin, Bree Gatica-Wilcox, Jason Coombes, Joel Southgate, Lauren Gilbert, Lee Graham, Nicole Pacchiarini, Sara Kumziene-Summerhayes, Sarah Taylor, Sophie Jones, Sara Rey, Matthew Bull, Joanne Watkins, Sally Corden, Tom Connor |
| EPI_ISL_3295724 | Viollier AG | Department of Biosystems Science and Engineering, ETH Zürich | Christian Beisel, Sarah Nadeau, Chaoran Chen, Ivan Topolsky, Kim Philipp Jablonski, Lara Fuhrmann, Rebecca Denes, Mirjam Feldkamp, Ina Nissen, Natascha Santacroce, Elodie Burcklen, Christiane Beckmann, Maurice Redondo, Olivier Kobel, Christoph Noppen, Niko Beerenwinkel, Tanja Stadler |
| EPI_ISL_3295725 | Wales Specialist Virology Centre Sequencing lab: Pathogen Genomics Unit | Public Health Wales Microbiology Cardiff Wales Specialist Virology Centre | Catherine Moore, Johnathan Evans, Laura Gifford, Malorie Perry, Simon Cottrell, Angela Marchbank, Alec Birchley, Alexander Adams, Amy Gaskin, Bree Gatica-Wilcox, Jason Coombes, Joel Southgate, Lauren Gilbert, Lee Graham, Nicole Pacchiarini, Sara Kumziene-Summerhayes, Sarah Taylor, Sophie Jones, Sara Rey, Matthew Bull, Joanne Watkins, Sally Corden, Tom Connor |
| EPI_ISL_3295726 | Viollier AG | Department of Biosystems Science and Engineering, ETH Zürich | Christian Beisel, Sarah Nadeau, Chaoran Chen, Ivan Topolsky, Kim Philipp Jablonski, Lara Fuhrmann, Rebecca Denes, Mirjam Feldkamp, Ina Nissen, Natascha Santacroce, Elodie Burcklen, Christiane Beckmann, Maurice Redondo, Olivier Kobel, Christoph Noppen, Niko Beerenwinkel, Tanja Stadler |
| EPI_ISL_3295727 | Wales Specialist Virology Centre Sequencing lab: Pathogen Genomics Unit | Public Health Wales Microbiology Cardiff Wales Specialist Virology Centre | Catherine Moore, Johnathan Evans, Laura Gifford, Malorie Perry, Simon Cottrell, Angela Marchbank, Alec Birchley, Alexander Adams, Amy Gaskin, Bree Gatica-Wilcox, Jason Coombes, Joel Southgate, Lauren Gilbert, Lee Graham, Nicole Pacchiarini, Sara Kumziene-Summerhayes, Sarah Taylor, Sophie Jones, Sara Rey, Matthew Bull, Joanne Watkins, Sally Corden, Tom Connor |
| EPI_ISL_3295728 | Viollier AG | Department of Biosystems Science and Engineering, ETH Zürich | Christian Beisel, Sarah Nadeau, Chaoran Chen, Ivan Topolsky, Kim Philipp Jablonski, Lara Fuhrmann, Rebecca Denes, Mirjam Feldkamp, Ina Nissen, Natascha Santacroce, Elodie Burcklen, Christiane Beckmann, Maurice Redondo, Olivier Kobel, Christoph Noppen, Niko Beerenwinkel, Tanja Stadler |
| EPI_ISL_3295729 | Wales Specialist Virology Centre Sequencing lab: Pathogen Genomics Unit | Public Health Wales Microbiology Cardiff Wales Specialist Virology Centre | Catherine Moore, Johnathan Evans, Laura Gifford, Malorie Perry, Simon Cottrell, Angela Marchbank, Alec Birchley, Alexander Adams, Amy Gaskin, Bree Gatica-Wilcox, Jason Coombes, Joel Southgate, Lauren Gilbert, Lee Graham, Nicole Pacchiarini, Sara Kumziene-Summerhayes, Sarah Taylor, Sophie Jones, Sara Rey, Matthew Bull, Joanne Watkins, Sally Corden, Tom Connor |
| EPI_ISL_3295730 | Viollier AG | Department of Biosystems Science and Engineering, ETH Zürich | Christian Beisel, Sarah Nadeau, Chaoran Chen, Ivan Topolsky, Kim Philipp Jablonski, Lara Fuhrmann, Rebecca Denes, Mirjam Feldkamp, Ina Nissen, Natascha Santacroce, Elodie Burcklen, Christiane Beckmann, Maurice Redondo, Olivier Kobel, Christoph Noppen, Niko Beerenwinkel, Tanja Stadler |
| EPI_ISL_3295731 | Wales Specialist Virology Centre Sequencing lab: Pathogen Genomics Unit | Public Health Wales Microbiology Cardiff Wales Specialist Virology Centre | Catherine Moore, Johnathan Evans, Laura Gifford, Malorie Perry, Simon Cottrell, Angela Marchbank, Alec Birchley, Alexander Adams, Amy Gaskin, Bree Gatica-Wilcox, Jason Coombes, Joel Southgate, Lauren Gilbert, Lee Graham, Nicole Pacchiarini, Sara Kumziene-Summerhayes, Sarah Taylor, Sophie Jones, Sara Rey, Matthew Bull, Joanne Watkins, Sally Corden, Tom Connor |
| EPI_ISL_3295732 | Viollier AG | Department of Biosystems Science and Engineering, ETH Zürich | Christian Beisel, Sarah Nadeau, Chaoran Chen, Ivan Topolsky, Kim Philipp Jablonski, Lara Fuhrmann, Rebecca Denes, Mirjam Feldkamp, Ina Nissen, Natascha Santacroce, Elodie Burcklen, Christiane Beckmann, Maurice Redondo, Olivier Kobel, Christoph Noppen, Niko Beerenwinkel, Tanja Stadler |
| EPI_ISL_3295733 | Wales Specialist Virology Centre Sequencing lab: Pathogen Genomics Unit | Public Health Wales Microbiology Cardiff Wales Specialist Virology Centre | Catherine Moore, Johnathan Evans, Laura Gifford, Malorie Perry, Simon Cottrell, Angela Marchbank, Alec Birchley, Alexander Adams, Amy Gaskin, Bree Gatica-Wilcox, Jason Coombes, Joel Southgate, Lauren Gilbert, Lee Graham, Nicole Pacchiarini, Sara Kumziene-Summerhayes, Sarah Taylor, Sophie Jones, Sara Rey, Matthew Bull, Joanne Watkins, Sally Corden, Tom Connor |
| EPI_ISL_3295734 | Viollier AG | Department of Biosystems Science and Engineering, ETH Zürich | Christian Beisel, Sarah Nadeau, Chaoran Chen, Ivan Topolsky, Kim Philipp Jablonski, Lara Fuhrmann, Rebecca Denes, Mirjam Feldkamp, Ina Nissen, Natascha Santacroce, Elodie Burcklen, Christiane Beckmann, Maurice Redondo, Olivier Kobel, Christoph Noppen, Niko Beerenwinkel, Tanja Stadler |
| EPI_ISL_3295735, EPI_ISL_3295737, EPI_ISL_3295738 | Wales Specialist Virology Centre Sequencing lab: Pathogen Genomics Unit | Public Health Wales Microbiology Cardiff Wales Specialist Virology Centre | Catherine Moore, Johnathan Evans, Laura Gifford, Malorie Perry, Simon Cottrell, Angela Marchbank, Alec Birchley, Alexander Adams, Amy Gaskin, Bree Gatica-Wilcox, Jason Coombes, Joel Southgate, Lauren Gilbert, Lee Graham, Nicole Pacchiarini, Sara Kumziene-Summerhayes, Sarah Taylor, Sophie Jones, Sara Rey, Matthew Bull, Joanne Watkins, Sally Corden, Tom Connor |
| EPI_ISL_3295739 | Viollier AG | Department of Biosystems Science and Engineering, ETH Zürich | Christian Beisel, Sarah Nadeau, Chaoran Chen, Ivan Topolsky, Kim Philipp Jablonski, Lara Fuhrmann, Rebecca Denes, Mirjam Feldkamp, Ina Nissen, Natascha Santacroce, Elodie Burcklen, Christiane Beckmann, Maurice Redondo, Olivier Kobel, Christoph Noppen, Niko Beerenwinkel, Tanja Stadler |
| EPI_ISL_3295740 | Wales Specialist Virology Centre Sequencing lab: Pathogen Genomics Unit | Public Health Wales Microbiology Cardiff Wales Specialist Virology Centre | Catherine Moore, Johnathan Evans, Laura Gifford, Malorie Perry, Simon Cottrell, Angela Marchbank, Alec Birchley, Alexander Adams, Amy Gaskin, Bree Gatica-Wilcox, Jason Coombes, Joel Southgate, Lauren Gilbert, Lee Graham, Nicole Pacchiarini, Sara Kumziene-Summerhayes, Sarah Taylor, Sophie Jones, Sara Rey, Matthew Bull, Joanne Watkins, Sally Corden, Tom Connor |
| EPI_ISL_3295741 | Viollier AG | Department of Biosystems Science and Engineering, ETH Zürich | Christian Beisel, Sarah Nadeau, Chaoran Chen, Ivan Topolsky, Kim Philipp Jablonski, Lara Fuhrmann, Rebecca Denes, Mirjam Feldkamp, Ina Nissen, Natascha Santacroce, Elodie Burcklen, Christiane Beckmann, Maurice Redondo, Olivier Kobel, Christoph Noppen, Niko Beerenwinkel, Tanja Stadler |
| EPI_ISL_3295742 | Wales Specialist Virology Centre Sequencing lab: Pathogen Genomics Unit | Public Health Wales Microbiology Cardiff Wales Specialist Virology Centre | Catherine Moore, Johnathan Evans, Laura Gifford, Malorie Perry, Simon Cottrell, Angela Marchbank, Alec Birchley, Alexander Adams, Amy Gaskin, Bree Gatica-Wilcox, Jason Coombes, Joel Southgate, Lauren Gilbert, Lee Graham, Nicole Pacchiarini, Sara Kumziene-Summerhayes, Sarah Taylor, Sophie Jones, Sara Rey, Matthew Bull, Joanne Watkins, Sally Corden, Tom Connor |
| EPI_ISL_3295743 | Viollier AG | Department of Biosystems Science and Engineering, ETH Zürich | Christian Beisel, Sarah Nadeau, Chaoran Chen, Ivan Topolsky, Kim Philipp Jablonski, Lara Fuhrmann, Rebecca Denes, Mirjam Feldkamp, Ina Nissen, Natascha Santacroce, Elodie Burcklen, Christiane Beckmann, Maurice Redondo, Olivier Kobel, Christoph Noppen, Niko Beerenwinkel, Tanja Stadler |
| EPI_ISL_3295744 | Wales Specialist Virology Centre Sequencing lab: Pathogen Genomics Unit | Public Health Wales Microbiology Cardiff Wales Specialist Virology Centre | Catherine Moore, Johnathan Evans, Laura Gifford, Malorie Perry, Simon Cottrell, Angela Marchbank, Alec Birchley, Alexander Adams, Amy Gaskin, Bree Gatica-Wilcox, Jason Coombes, Joel Southgate, Lauren Gilbert, Lee Graham, Nicole Pacchiarini, Sara Kumziene-Summerhayes, Sarah Taylor, Sophie Jones, Sara Rey, Matthew Bull, Joanne Watkins, Sally Corden, Tom Connor |
| EPI_ISL_3295745 | Viollier AG | Department of Biosystems Science and Engineering, ETH Zürich | Christian Beisel, Sarah Nadeau, Chaoran Chen, Ivan Topolsky, Kim Philipp Jablonski, Lara Fuhrmann, Rebecca Denes, Mirjam Feldkamp, Ina Nissen, Natascha Santacroce, Elodie Burcklen, Christiane Beckmann, Maurice Redondo, Olivier Kobel, Christoph Noppen, Niko Beerenwinkel, Tanja Stadler |
| EPI_ISL_3295746 | Wales Specialist Virology Centre Sequencing lab: Pathogen Genomics Unit | Public Health Wales Microbiology Cardiff Wales Specialist Virology Centre | Catherine Moore, Johnathan Evans, Laura Gifford, Malorie Perry, Simon Cottrell, Angela Marchbank, Alec Birchley, Alexander Adams, Amy Gaskin, Bree Gatica-Wilcox, Jason Coombes, Joel Southgate, Lauren Gilbert, Lee Graham, Nicole Pacchiarini, Sara Kumziene-Summerhayes, Sarah Taylor, Sophie Jones, Sara Rey, Matthew Bull, Joanne Watkins, Sally Corden, Tom Connor |
| EPI_ISL_3295747 | Viollier AG | Department of Biosystems Science and Engineering, ETH Zürich | Christian Beisel, Sarah Nadeau, Chaoran Chen, Ivan Topolsky, Kim Philipp Jablonski, Lara Fuhrmann, Rebecca Denes, Mirjam Feldkamp, Ina Nissen, Natascha Santacroce, Elodie Burcklen, Christiane Beckmann, Maurice Redondo, Olivier Kobel, Christoph Noppen, Niko Beerenwinkel, Tanja Stadler |
| EPI_ISL_3295748 | Wales Specialist Virology Centre Sequencing lab: Pathogen Genomics Unit | Public Health Wales Microbiology Cardiff Wales Specialist Virology Centre | Catherine Moore, Johnathan Evans, Laura Gifford, Malorie Perry, Simon Cottrell, Angela Marchbank, Alec Birchley, Alexander Adams, Amy Gaskin, Bree Gatica-Wilcox, Jason Coombes, Joel Southgate, Lauren Gilbert, Lee Graham, Nicole Pacchiarini, Sara Kumziene-Summerhayes, Sarah Taylor, Sophie Jones, Sara Rey, Matthew Bull, Joanne Watkins, Sally Corden, Tom Connor |
| EPI_ISL_3295749 | Viollier AG | Department of Biosystems Science and Engineering, ETH Zürich | Christian Beisel, Sarah Nadeau, Chaoran Chen, Ivan Topolsky, Kim Philipp Jablonski, Lara Fuhrmann, Rebecca Denes, Mirjam Feldkamp, Ina Nissen, Natascha Santacroce, Elodie Burcklen, Christiane Beckmann, Maurice Redondo, Olivier Kobel, Christoph Noppen, Niko Beerenwinkel, Tanja Stadler |
| EPI_ISL_3295750 | Wales Specialist Virology Centre Sequencing lab: Pathogen Genomics Unit | Public Health Wales Microbiology Cardiff Wales Specialist Virology Centre | Catherine Moore, Johnathan Evans, Laura Gifford, Malorie Perry, Simon Cottrell, Angela Marchbank, Alec Birchley, Alexander Adams, Amy Gaskin, Bree Gatica-Wilcox, Jason Coombes, Joel Southgate, Lauren Gilbert, Lee Graham, Nicole Pacchiarini, Sara Kumziene-Summerhayes, Sarah Taylor, Sophie Jones, Sara Rey, Matthew Bull, Joanne Watkins, Sally Corden, Tom Connor |
| EPI_ISL_3295751 | Viollier AG | Department of Biosystems Science and Engineering, ETH Zürich | Christian Beisel, Sarah Nadeau, Chaoran Chen, Ivan Topolsky, Kim Philipp Jablonski, Lara Fuhrmann, Rebecca Denes, Mirjam Feldkamp, Ina Nissen, Natascha Santacroce, Elodie Burcklen, Christiane Beckmann, Maurice Redondo, Olivier Kobel, Christoph Noppen, Niko Beerenwinkel, Tanja Stadler |
| EPI_ISL_3295752 | Wales Specialist Virology Centre Sequencing lab: Pathogen Genomics Unit | Public Health Wales Microbiology Cardiff Wales Specialist Virology Centre | Catherine Moore, Johnathan Evans, Laura Gifford, Malorie Perry, Simon Cottrell, Angela Marchbank, Alec Birchley, Alexander Adams, Amy Gaskin, Bree Gatica-Wilcox, Jason Coombes, Joel Southgate, Lauren Gilbert, Lee Graham, Nicole Pacchiarini, Sara Kumziene-Summerhayes, Sarah Taylor, Sophie Jones, Sara Rey, Matthew Bull, Joanne Watkins, Sally Corden, Tom Connor |
| EPI_ISL_3295753 | Viollier AG | Department of Biosystems Science and Engineering, ETH Zürich | Christian Beisel, Sarah Nadeau, Chaoran Chen, Ivan Topolsky, Kim Philipp Jablonski, Lara Fuhrmann, Rebecca Denes, Mirjam Feldkamp, Ina Nissen, Natascha Santacroce, Elodie Burcklen, Christiane Beckmann, Maurice Redondo, Olivier Kobel, Christoph Noppen, Niko Beerenwinkel, Tanja Stadler |
| EPI_ISL_3295754 | Wales Specialist Virology Centre Sequencing lab: Pathogen Genomics Unit | Public Health Wales Microbiology Cardiff Wales Specialist Virology Centre | Catherine Moore, Johnathan Evans, Laura Gifford, Malorie Perry, Simon Cottrell, Angela Marchbank, Alec Birchley, Alexander Adams, Amy Gaskin, Bree Gatica-Wilcox, Jason Coombes, Joel Southgate, Lauren Gilbert, Lee Graham, Nicole Pacchiarini, Sara Kumziene-Summerhayes, Sarah Taylor, Sophie Jones, Sara Rey, Matthew Bull, Joanne Watkins, Sally Corden, Tom Connor |
| EPI_ISL_3295755 | Viollier AG | Department of Biosystems Science and Engineering, ETH Zürich | Christian Beisel, Sarah Nadeau, Chaoran Chen, Ivan Topolsky, Kim Philipp Jablonski, Lara Fuhrmann, Rebecca Denes, Mirjam Feldkamp, Ina Nissen, Natascha Santacroce, Elodie Burcklen, Christiane Beckmann, Maurice Redondo, Olivier Kobel, Christoph Noppen, Niko Beerenwinkel, Tanja Stadler |
| EPI_ISL_3295756 | Wales Specialist Virology Centre Sequencing lab: Pathogen Genomics Unit | Public Health Wales Microbiology Cardiff Wales Specialist Virology Centre | Catherine Moore, Johnathan Evans, Laura Gifford, Malorie Perry, Simon Cottrell, Angela Marchbank, Alec Birchley, Alexander Adams, Amy Gaskin, Bree Gatica-Wilcox, Jason Coombes, Joel Southgate, Lauren Gilbert, Lee Graham, Nicole Pacchiarini, Sara Kumziene-Summerhayes, Sarah Taylor, Sophie Jones, Sara Rey, Matthew Bull, Joanne Watkins, Sally Corden, Tom Connor |
| EPI_ISL_3295757 | Viollier AG | Department of Biosystems Science and Engineering, ETH Zürich | Christian Beisel, Sarah Nadeau, Chaoran Chen, Ivan Topolsky, Kim Philipp Jablonski, Lara Fuhrmann, Rebecca Denes, Mirjam Feldkamp, Ina Nissen, Natascha Santacroce, Elodie Burcklen, Christiane Beckmann, Maurice Redondo, Olivier Kobel, Christoph Noppen, Niko Beerenwinkel, Tanja Stadler |
| EPI_ISL_3295758, EPI_ISL_3295759 | Wales Specialist Virology Centre Sequencing lab: Pathogen Genomics Unit | Public Health Wales Microbiology Cardiff Wales Specialist Virology Centre | Catherine Moore, Johnathan Evans, Laura Gifford, Malorie Perry, Simon Cottrell, Angela Marchbank, Alec Birchley, Alexander Adams, Amy Gaskin, Bree Gatica-Wilcox, Jason Coombes, Joel Southgate, Lauren Gilbert, Lee Graham, Nicole Pacchiarini, Sara Kumziene-Summerhayes, Sarah Taylor, Sophie Jones, Sara Rey, Matthew Bull, Joanne Watkins, Sally Corden, Tom Connor |
| EPI_ISL_3295760 | Viollier AG | Department of Biosystems Science and Engineering, ETH Zürich | Christian Beisel, Sarah Nadeau, Chaoran Chen, Ivan Topolsky, Kim Philipp Jablonski, Lara Fuhrmann, Rebecca Denes, Mirjam Feldkamp, Ina Nissen, Natascha Santacroce, Elodie Burcklen, Christiane Beckmann, Maurice Redondo, Olivier Kobel, Christoph Noppen, Niko Beerenwinkel, Tanja Stadler |
| EPI_ISL_3295761 | Wales Specialist Virology Centre Sequencing lab: Pathogen Genomics Unit | Public Health Wales Microbiology Cardiff Wales Specialist Virology Centre | Catherine Moore, Johnathan Evans, Laura Gifford, Malorie Perry, Simon Cottrell, Angela Marchbank, Alec Birchley, Alexander Adams, Amy Gaskin, Bree Gatica-Wilcox, Jason Coombes, Joel Southgate, Lauren Gilbert, Lee Graham, Nicole Pacchiarini, Sara Kumziene-Summerhayes, Sarah Taylor, Sophie Jones, Sara Rey, Matthew Bull, Joanne Watkins, Sally Corden, Tom Connor |
| EPI_ISL_3295762 | Viollier AG | Department of Biosystems Science and Engineering, ETH Zürich | Christian Beisel, Sarah Nadeau, Chaoran Chen, Ivan Topolsky, Kim Philipp Jablonski, Lara Fuhrmann, Rebecca Denes, Mirjam Feldkamp, Ina Nissen, Natascha Santacroce, Elodie Burcklen, Christiane Beckmann, Maurice Redondo, Olivier Kobel, Christoph Noppen, Niko Beerenwinkel, Tanja Stadler |
| EPI_ISL_3295763 | Wales Specialist Virology Centre Sequencing lab: Pathogen Genomics Unit | Public Health Wales Microbiology Cardiff Wales Specialist Virology Centre | Catherine Moore, Johnathan Evans, Laura Gifford, Malorie Perry, Simon Cottrell, Angela Marchbank, Alec Birchley, Alexander Adams, Amy Gaskin, Bree Gatica-Wilcox, Jason Coombes, Joel Southgate, Lauren Gilbert, Lee Graham, Nicole Pacchiarini, Sara Kumziene-Summerhayes, Sarah Taylor, Sophie Jones, Sara Rey, Matthew Bull, Joanne Watkins, Sally Corden, Tom Connor |
| EPI_ISL_3295764 | Viollier AG | Department of Biosystems Science and Engineering, ETH Zürich | Christian Beisel, Sarah Nadeau, Chaoran Chen, Ivan Topolsky, Kim Philipp Jablonski, Lara Fuhrmann, Rebecca Denes, Mirjam Feldkamp, Ina Nissen, Natascha Santacroce, Elodie Burcklen, Christiane Beckmann, Maurice Redondo, Olivier Kobel, Christoph Noppen, Niko Beerenwinkel, Tanja Stadler |
| EPI_ISL_3295765 | Wales Specialist Virology Centre Sequencing lab: Pathogen Genomics Unit | Public Health Wales Microbiology Cardiff Wales Specialist Virology Centre | Catherine Moore, Johnathan Evans, Laura Gifford, Malorie Perry, Simon Cottrell, Angela Marchbank, Alec Birchley, Alexander Adams, Amy Gaskin, Bree Gatica-Wilcox, Jason Coombes, Joel Southgate, Lauren Gilbert, Lee Graham, Nicole Pacchiarini, Sara Kumziene-Summerhayes, Sarah Taylor, Sophie Jones, Sara Rey, Matthew Bull, Joanne Watkins, Sally Corden, Tom Connor |
| EPI_ISL_3295766 | Viollier AG | Department of Biosystems Science and Engineering, ETH Zürich | Christian Beisel, Sarah Nadeau, Chaoran Chen, Ivan Topolsky, Kim Philipp Jablonski, Lara Fuhrmann, Rebecca Denes, Mirjam Feldkamp, Ina Nissen, Natascha Santacroce, Elodie Burcklen, Christiane Beckmann, Maurice Redondo, Olivier Kobel, |

| | | | |
|---|---|---|---|
| | | | Christoph Noppen, Niko Beerenwinkel, Tanja Stadler |
| EPI_ISL_3295767 | Wales Specialist Virology Centre Sequencing lab: Pathogen Genomics Unit | Public Health Wales Microbiology Cardiff Wales Specialist Virology Centre | Catherine Moore, Johnathan Evans, Laura Gifford, Malorie Perry, Simon Cottrell, Angela Marchbank, Alec Birchley, Alexander Adams, Amy Gaskin, Bree Gatica-Wilcox, Jason Coombes, Joel Southgate, Lauren Gilbert, Lee Graham, Nicole Pacchiarini, Sara Kumziene-Summerhayes, Sarah Taylor, Sophie Jones, Sara Rey, Matthew Bull, Joanne Watkins, Sally Corden, Tom Connor |
| EPI_ISL_3295768 | Viollier AG | Department of Biosystems Science and Engineering, ETH Zürich | Christian Beisel, Sarah Nadeau, Chaoran Chen, Ivan Topolsky, Kim Philipp Jablonski, Lara Fuhrmann, Rebecca Denes, Mirjam Feldkamp, Ina Nissen, Natascha Santacroce, Elodie Burcklen, Christiane Beckmann, Maurice Redondo, Olivier Kobel, Christoph Noppen, Niko Beerenwinkel, Tanja Stadler |
| EPI_ISL_3295769, EPI_ISL_3295771 | Wales Specialist Virology Centre Sequencing lab: Pathogen Genomics Unit | Public Health Wales Microbiology Cardiff Wales Specialist Virology Centre | Catherine Moore, Johnathan Evans, Laura Gifford, Malorie Perry, Simon Cottrell, Angela Marchbank, Alec Birchley, Alexander Adams, Amy Gaskin, Bree Gatica-Wilcox, Jason Coombes, Joel Southgate, Lauren Gilbert, Lee Graham, Nicole Pacchiarini, Sara Kumziene-Summerhayes, Sarah Taylor, Sophie Jones, Sara Rey, Matthew Bull, Joanne Watkins, Sally Corden, Tom Connor |
| EPI_ISL_3295772 | Viollier AG | Department of Biosystems Science and Engineering, ETH Zürich | Christian Beisel, Sarah Nadeau, Chaoran Chen, Ivan Topolsky, Kim Philipp Jablonski, Lara Fuhrmann, Rebecca Denes, Mirjam Feldkamp, Ina Nissen, Natascha Santacroce, Elodie Burcklen, Christiane Beckmann, Maurice Redondo, Olivier Kobel, Christoph Noppen, Niko Beerenwinkel, Tanja Stadler |
| EPI_ISL_3295773 | Wales Specialist Virology Centre Sequencing lab: Pathogen Genomics Unit | Public Health Wales Microbiology Cardiff Wales Specialist Virology Centre | Catherine Moore, Johnathan Evans, Laura Gifford, Malorie Perry, Simon Cottrell, Angela Marchbank, Alec Birchley, Alexander Adams, Amy Gaskin, Bree Gatica-Wilcox, Jason Coombes, Joel Southgate, Lauren Gilbert, Lee Graham, Nicole Pacchiarini, Sara Kumziene-Summerhayes, Sarah Taylor, Sophie Jones, Sara Rey, Matthew Bull, Joanne Watkins, Sally Corden, Tom Connor |
| EPI_ISL_3295774 | Viollier AG | Department of Biosystems Science and Engineering, ETH Zürich | Christian Beisel, Sarah Nadeau, Chaoran Chen, Ivan Topolsky, Kim Philipp Jablonski, Lara Fuhrmann, Rebecca Denes, Mirjam Feldkamp, Ina Nissen, Natascha Santacroce, Elodie Burcklen, Christiane Beckmann, Maurice Redondo, Olivier Kobel, Christoph Noppen, Niko Beerenwinkel, Tanja Stadler |
| EPI_ISL_3295775 | Wales Specialist Virology Centre Sequencing lab: Pathogen Genomics Unit | Public Health Wales Microbiology Cardiff Wales Specialist Virology Centre | Catherine Moore, Johnathan Evans, Laura Gifford, Malorie Perry, Simon Cottrell, Angela Marchbank, Alec Birchley, Alexander Adams, Amy Gaskin, Bree Gatica-Wilcox, Jason Coombes, Joel Southgate, Lauren Gilbert, Lee Graham, Nicole Pacchiarini, Sara Kumziene-Summerhayes, Sarah Taylor, Sophie Jones, Sara Rey, Matthew Bull, Joanne Watkins, Sally Corden, Tom Connor |
| EPI_ISL_3295776 | Viollier AG | Department of Biosystems Science and Engineering, ETH Zürich | Christian Beisel, Sarah Nadeau, Chaoran Chen, Ivan Topolsky, Kim Philipp Jablonski, Lara Fuhrmann, Rebecca Denes, Mirjam Feldkamp, Ina Nissen, Natascha Santacroce, Elodie Burcklen, Christiane Beckmann, Maurice Redondo, Olivier Kobel, Christoph Noppen, Niko Beerenwinkel, Tanja Stadler |
| EPI_ISL_3295777 | Wales Specialist Virology Centre Sequencing lab: Pathogen Genomics Unit | Public Health Wales Microbiology Cardiff Wales Specialist Virology Centre | Catherine Moore, Johnathan Evans, Laura Gifford, Malorie Perry, Simon Cottrell, Angela Marchbank, Alec Birchley, Alexander Adams, Amy Gaskin, Bree Gatica-Wilcox, Jason Coombes, Joel Southgate, Lauren Gilbert, Lee Graham, Nicole Pacchiarini, Sara Kumziene-Summerhayes, Sarah Taylor, Sophie Jones, Sara Rey, Matthew Bull, Joanne Watkins, Sally Corden, Tom Connor |
| EPI_ISL_3295778 | Viollier AG | Department of Biosystems Science and Engineering, ETH Zürich | Christian Beisel, Sarah Nadeau, Chaoran Chen, Ivan Topolsky, Kim Philipp Jablonski, Lara Fuhrmann, Rebecca Denes, Mirjam Feldkamp, Ina Nissen, Natascha Santacroce, Elodie Burcklen, Christiane Beckmann, Maurice Redondo, Olivier Kobel, Christoph Noppen, Niko Beerenwinkel, Tanja Stadler |
| EPI_ISL_3295779 | Wales Specialist Virology Centre Sequencing lab: Pathogen Genomics Unit | Public Health Wales Microbiology Cardiff Wales Specialist Virology Centre | Catherine Moore, Johnathan Evans, Laura Gifford, Malorie Perry, Simon Cottrell, Angela Marchbank, Alec Birchley, Alexander Adams, Amy Gaskin, Bree Gatica-Wilcox, Jason Coombes, Joel Southgate, Lauren Gilbert, Lee Graham, Nicole Pacchiarini, Sara Kumziene-Summerhayes, Sarah Taylor, Sophie Jones, Sara Rey, Matthew Bull, Joanne Watkins, Sally Corden, Tom Connor |
| EPI_ISL_3295780 | Viollier AG | Department of Biosystems Science and Engineering, ETH Zürich | Christian Beisel, Sarah Nadeau, Chaoran Chen, Ivan Topolsky, Kim Philipp Jablonski, Lara Fuhrmann, Rebecca Denes, Mirjam Feldkamp, Ina Nissen, Natascha Santacroce, Elodie Burcklen, Christiane Beckmann, Maurice Redondo, Olivier Kobel, Christoph Noppen, Niko Beerenwinkel, Tanja Stadler |
| EPI_ISL_3295781 | Wales Specialist Virology Centre Sequencing lab: Pathogen Genomics Unit | Public Health Wales Microbiology Cardiff Wales Specialist Virology Centre | Catherine Moore, Johnathan Evans, Laura Gifford, Malorie Perry, Simon Cottrell, Angela Marchbank, Alec Birchley, Alexander Adams, Amy Gaskin, Bree Gatica-Wilcox, Jason Coombes, Joel Southgate, Lauren Gilbert, Lee Graham, Nicole Pacchiarini, Sara Kumziene-Summerhayes, Sarah Taylor, Sophie Jones, Sara Rey, Matthew Bull, Joanne Watkins, Sally Corden, Tom Connor |
| EPI_ISL_3295782 | Viollier AG | Department of Biosystems Science and Engineering, ETH Zürich | Christian Beisel, Sarah Nadeau, Chaoran Chen, Ivan Topolsky, Kim Philipp Jablonski, Lara Fuhrmann, Rebecca Denes, Mirjam Feldkamp, Ina Nissen, Natascha Santacroce, Elodie Burcklen, Christiane Beckmann, Maurice Redondo, Olivier Kobel, Christoph Noppen, Niko Beerenwinkel, Tanja Stadler |
| EPI_ISL_3295783 | Wales Specialist Virology Centre Sequencing lab: Pathogen Genomics Unit | Public Health Wales Microbiology Cardiff Wales Specialist Virology Centre | Catherine Moore, Johnathan Evans, Laura Gifford, Malorie Perry, Simon Cottrell, Angela Marchbank, Alec Birchley, Alexander Adams, Amy Gaskin, Bree Gatica-Wilcox, Jason Coombes, Joel Southgate, Lauren Gilbert, Lee Graham, Nicole Pacchiarini, Sara Kumziene-Summerhayes, Sarah Taylor, Sophie Jones, Sara Rey, Matthew Bull, Joanne Watkins, Sally Corden, Tom Connor |
| EPI_ISL_3295784 | Viollier AG | Department of Biosystems Science and Engineering, ETH Zürich | Christian Beisel, Sarah Nadeau, Chaoran Chen, Ivan Topolsky, Kim Philipp Jablonski, Lara Fuhrmann, Rebecca Denes, Mirjam Feldkamp, Ina Nissen, Natascha Santacroce, Elodie Burcklen, Christiane Beckmann, Maurice Redondo, Olivier Kobel, Christoph Noppen, Niko Beerenwinkel, Tanja Stadler |
| EPI_ISL_3295785 | Wales Specialist Virology Centre Sequencing lab: Pathogen Genomics Unit | Public Health Wales Microbiology Cardiff Wales Specialist Virology Centre | Catherine Moore, Johnathan Evans, Laura Gifford, Malorie Perry, Simon Cottrell, Angela Marchbank, Alec Birchley, Alexander Adams, Amy Gaskin, Bree Gatica-Wilcox, Jason Coombes, Joel Southgate, Lauren Gilbert, Lee Graham, Nicole Pacchiarini, Sara Kumziene-Summerhayes, Sarah Taylor, Sophie Jones, Sara Rey, Matthew Bull, Joanne Watkins, Sally Corden, Tom Connor |
| EPI_ISL_3295786 | Viollier AG | Department of Biosystems Science and Engineering, ETH Zürich | Christian Beisel, Sarah Nadeau, Chaoran Chen, Ivan Topolsky, Kim Philipp Jablonski, Lara Fuhrmann, Rebecca Denes, Mirjam Feldkamp, Ina Nissen, Natascha Santacroce, Elodie Burcklen, Christiane Beckmann, Maurice Redondo, Olivier Kobel, Christoph Noppen, Niko Beerenwinkel, Tanja Stadler |
| EPI_ISL_3295787, EPI_ISL_3295788 | Wales Specialist Virology Centre Sequencing lab: Pathogen Genomics Unit | Public Health Wales Microbiology Cardiff Wales Specialist Virology Centre | Catherine Moore, Johnathan Evans, Laura Gifford, Malorie Perry, Simon Cottrell, Angela Marchbank, Alec Birchley, Alexander Adams, Amy Gaskin, Bree Gatica-Wilcox, Jason Coombes, Joel Southgate, Lauren Gilbert, Lee Graham, Nicole Pacchiarini, Sara Kumziene-Summerhayes, Sarah Taylor, Sophie Jones, Sara Rey, Matthew Bull, Joanne Watkins, Sally Corden, Tom Connor |
| EPI_ISL_3295789 | Viollier AG | Department of Biosystems Science and Engineering, ETH Zürich | Christian Beisel, Sarah Nadeau, Chaoran Chen, Ivan Topolsky, Kim Philipp Jablonski, Lara Fuhrmann, Rebecca Denes, Mirjam Feldkamp, Ina Nissen, Natascha Santacroce, Elodie Burcklen, Christiane Beckmann, Maurice Redondo, Olivier Kobel, Christoph Noppen, Niko Beerenwinkel, Tanja Stadler |
| EPI_ISL_3295790 | Wales Specialist Virology Centre Sequencing lab: Pathogen Genomics Unit | Public Health Wales Microbiology Cardiff Wales Specialist Virology Centre | Catherine Moore, Johnathan Evans, Laura Gifford, Malorie Perry, Simon Cottrell, Angela Marchbank, Alec Birchley, Alexander Adams, Amy Gaskin, Bree Gatica-Wilcox, Jason Coombes, Joel Southgate, Lauren Gilbert, Lee Graham, Nicole Pacchiarini, Sara Kumziene-Summerhayes, Sarah Taylor, Sophie Jones, Sara Rey, Matthew Bull, Joanne Watkins, Sally Corden, Tom Connor |
| EPI_ISL_3295791 | Viollier AG | Department of Biosystems Science and Engineering, ETH Zürich | Christian Beisel, Sarah Nadeau, Chaoran Chen, Ivan Topolsky, Kim Philipp Jablonski, Lara Fuhrmann, Rebecca Denes, Mirjam Feldkamp, Ina Nissen, Natascha Santacroce, Elodie Burcklen, Christiane Beckmann, Maurice Redondo, Olivier Kobel, Christoph Noppen, Niko Beerenwinkel, Tanja Stadler |
| EPI_ISL_3295792 | Wales Specialist Virology Centre Sequencing lab: Pathogen Genomics Unit | Public Health Wales Microbiology Cardiff Wales Specialist Virology Centre | Catherine Moore, Johnathan Evans, Laura Gifford, Malorie Perry, Simon Cottrell, Angela Marchbank, Alec Birchley, Alexander Adams, Amy Gaskin, Bree Gatica-Wilcox, Jason Coombes, Joel Southgate, Lauren Gilbert, Lee Graham, Nicole Pacchiarini, Sara Kumziene-Summerhayes, Sarah Taylor, Sophie Jones, Sara Rey, Matthew Bull, Joanne Watkins, Sally Corden, Tom Connor |
| EPI_ISL_3295793 | Viollier AG | Department of Biosystems Science and Engineering, ETH Zürich | Christian Beisel, Sarah Nadeau, Chaoran Chen, Ivan Topolsky, Kim Philipp Jablonski, Lara Fuhrmann, Rebecca Denes, Mirjam Feldkamp, Ina Nissen, Natascha Santacroce, Elodie Burcklen, Christiane Beckmann, Maurice Redondo, Olivier Kobel, Christoph Noppen, Niko Beerenwinkel, Tanja Stadler |
| EPI_ISL_3295794 | Wales Specialist Virology Centre Sequencing lab: Pathogen Genomics Unit | Public Health Wales Microbiology Cardiff Wales Specialist Virology Centre | Catherine Moore, Johnathan Evans, Laura Gifford, Malorie Perry, Simon Cottrell, Angela Marchbank, Alec Birchley, Alexander Adams, Amy Gaskin, Bree Gatica-Wilcox, Jason Coombes, Joel Southgate, Lauren Gilbert, Lee Graham, Nicole Pacchiarini, Sara Kumziene-Summerhayes, Sarah Taylor, Sophie Jones, Sara Rey, Matthew Bull, Joanne Watkins, Sally Corden, Tom Connor |
| EPI_ISL_3295795 | Viollier AG | Department of Biosystems Science and Engineering, ETH Zürich | Christian Beisel, Sarah Nadeau, Chaoran Chen, Ivan Topolsky, Kim Philipp Jablonski, Lara Fuhrmann, Rebecca Denes, Mirjam Feldkamp, Ina Nissen, Natascha Santacroce, Elodie Burcklen, Christiane Beckmann, Maurice Redondo, Olivier Kobel, Christoph Noppen, Niko Beerenwinkel, Tanja Stadler |
| EPI_ISL_3295796 | Wales Specialist Virology Centre Sequencing lab: Pathogen Genomics Unit | Public Health Wales Microbiology Cardiff Wales Specialist Virology Centre | Catherine Moore, Johnathan Evans, Laura Gifford, Malorie Perry, Simon Cottrell, Angela Marchbank, Alec Birchley, Alexander Adams, Amy Gaskin, Bree Gatica-Wilcox, Jason Coombes, Joel Southgate, Lauren Gilbert, Lee Graham, Nicole Pacchiarini, Sara Kumziene-Summerhayes, Sarah Taylor, Sophie Jones, Sara Rey, Matthew Bull, Joanne Watkins, Sally Corden, Tom Connor |
| EPI_ISL_3295797 | Viollier AG | Department of Biosystems Science and Engineering, ETH Zürich | Christian Beisel, Sarah Nadeau, Chaoran Chen, Ivan Topolsky, Kim Philipp Jablonski, Lara Fuhrmann, Rebecca Denes, Mirjam Feldkamp, Ina Nissen, Natascha Santacroce, Elodie Burcklen, Christiane Beckmann, Maurice Redondo, Olivier Kobel, Christoph Noppen, Niko Beerenwinkel, Tanja Stadler |
| EPI_ISL_3295798 | Wales Specialist Virology Centre Sequencing lab: Pathogen Genomics Unit | Public Health Wales Microbiology Cardiff Wales Specialist Virology Centre | Catherine Moore, Johnathan Evans, Laura Gifford, Malorie Perry, Simon Cottrell, Angela Marchbank, Alec Birchley, Alexander Adams, Amy Gaskin, Bree Gatica-Wilcox, Jason Coombes, Joel Southgate, Lauren Gilbert, Lee Graham, Nicole Pacchiarini, Sara Kumziene-Summerhayes, Sarah Taylor, Sophie Jones, Sara Rey, Matthew Bull, Joanne Watkins, Sally Corden, Tom Connor |
| EPI_ISL_3295799 | Viollier AG | Department of Biosystems Science and Engineering, ETH Zürich | Christian Beisel, Sarah Nadeau, Chaoran Chen, Ivan Topolsky, Kim Philipp Jablonski, Lara Fuhrmann, Rebecca Denes, Mirjam Feldkamp, Ina Nissen, Natascha Santacroce, Elodie Burcklen, Christiane Beckmann, Maurice Redondo, Olivier Kobel, Christoph Noppen, Niko Beerenwinkel, Tanja Stadler |
| EPI_ISL_3295800 | Wales Specialist Virology Centre Sequencing lab: Pathogen Genomics Unit | Public Health Wales Microbiology Cardiff Wales Specialist Virology Centre | Catherine Moore, Johnathan Evans, Laura Gifford, Malorie Perry, Simon Cottrell, Angela Marchbank, Alec Birchley, Alexander Adams, Amy Gaskin, Bree Gatica-Wilcox, Jason Coombes, Joel Southgate, Lauren Gilbert, Lee Graham, Nicole Pacchiarini, Sara Kumziene-Summerhayes, Sarah Taylor, Sophie Jones, Sara Rey, Matthew Bull, Joanne Watkins, Sally Corden, Tom Connor |
| EPI_ISL_3295801 | Viollier AG | Department of Biosystems Science and Engineering, ETH Zürich | Christian Beisel, Sarah Nadeau, Chaoran Chen, Ivan Topolsky, Kim Philipp Jablonski, Lara Fuhrmann, Rebecca Denes, Mirjam Feldkamp, Ina Nissen, Natascha Santacroce, Elodie Burcklen, Christiane Beckmann, Maurice Redondo, Olivier Kobel, Christoph Noppen, Niko Beerenwinkel, Tanja Stadler |
| EPI_ISL_3295802 | Wales Specialist Virology Centre Sequencing lab: Pathogen Genomics Unit | Public Health Wales Microbiology Cardiff Wales Specialist Virology Centre | Catherine Moore, Johnathan Evans, Laura Gifford, Malorie Perry, Simon Cottrell, Angela Marchbank, Alec Birchley, Alexander Adams, Amy Gaskin, Bree Gatica-Wilcox, Jason Coombes, Joel Southgate, Lauren Gilbert, Lee Graham, Nicole Pacchiarini, Sara Kumziene-Summerhayes, Sarah Taylor, Sophie Jones, Sara Rey, Matthew Bull, Joanne Watkins, Sally Corden, Tom Connor |
| EPI_ISL_3295803 | Viollier AG | Department of Biosystems Science and Engineering, ETH Zürich | Christian Beisel, Sarah Nadeau, Chaoran Chen, Ivan Topolsky, Kim Philipp Jablonski, Lara Fuhrmann, Rebecca Denes, Mirjam Feldkamp, Ina Nissen, Natascha Santacroce, Elodie Burcklen, Christiane Beckmann, Maurice Redondo, Olivier Kobel, Christoph Noppen, Niko Beerenwinkel, Tanja Stadler |
| EPI_ISL_3295804 | Wales Specialist Virology Centre Sequencing lab: Pathogen Genomics Unit | Public Health Wales Microbiology Cardiff Wales Specialist Virology Centre | Catherine Moore, Johnathan Evans, Laura Gifford, Malorie Perry, Simon Cottrell, Angela Marchbank, Alec Birchley, Alexander Adams, Amy Gaskin, Bree Gatica-Wilcox, Jason Coombes, Joel Southgate, Lauren Gilbert, Lee Graham, Nicole Pacchiarini, Sara Kumziene-Summerhayes, Sarah Taylor, Sophie Jones, Sara Rey, Matthew Bull, Joanne Watkins, Sally Corden, Tom Connor |
| EPI_ISL_3295805 | Viollier AG | Department of Biosystems Science and Engineering, ETH Zürich | Christian Beisel, Sarah Nadeau, Chaoran Chen, Ivan Topolsky, Kim Philipp Jablonski, Lara Fuhrmann, Rebecca Denes, Mirjam Feldkamp, Ina Nissen, Natascha Santacroce, Elodie Burcklen, Christiane Beckmann, Maurice Redondo, Olivier Kobel, Christoph Noppen, Niko Beerenwinkel, Tanja Stadler |
| EPI_ISL_3295806 | Wales Specialist Virology Centre Sequencing lab: Pathogen Genomics Unit | Public Health Wales Microbiology Cardiff Wales Specialist Virology Centre | Catherine Moore, Johnathan Evans, Laura Gifford, Malorie Perry, Simon Cottrell, Angela Marchbank, Alec Birchley, Alexander Adams, Amy Gaskin, Bree Gatica-Wilcox, Jason Coombes, Joel Southgate, Lauren Gilbert, Lee Graham, Nicole Pacchiarini, Sara Kumziene-Summerhayes, Sarah Taylor, Sophie Jones, Sara Rey, Matthew Bull, Joanne Watkins, Sally Corden, Tom Connor |
| EPI_ISL_3295807 | Viollier AG | Department of Biosystems Science and Engineering, ETH Zürich | Christian Beisel, Sarah Nadeau, Chaoran Chen, Ivan Topolsky, Kim Philipp Jablonski, Lara Fuhrmann, Rebecca Denes, Mirjam Feldkamp, Ina Nissen, Natascha Santacroce, Elodie Burcklen, Christiane Beckmann, Maurice Redondo, Olivier Kobel, Christoph Noppen, Niko Beerenwinkel, Tanja Stadler |
| EPI_ISL_3295808 | Wales Specialist Virology Centre Sequencing lab: Pathogen Genomics Unit | Public Health Wales Microbiology Cardiff Wales Specialist Virology Centre | Catherine Moore, Johnathan Evans, Laura Gifford, Malorie Perry, Simon Cottrell, Angela Marchbank, Alec Birchley, Alexander Adams, Amy Gaskin, Bree Gatica-Wilcox, Jason Coombes, Joel Southgate, Lauren Gilbert, Lee Graham, Nicole Pacchiarini, Sara Kumziene-Summerhayes, Sarah Taylor, Sophie Jones, Sara Rey, Matthew Bull, Joanne Watkins, Sally Corden, Tom Connor |
| EPI_ISL_3295809 | Viollier AG | Department of Biosystems Science and Engineering, ETH Zürich | Christian Beisel, Sarah Nadeau, Chaoran Chen, Ivan Topolsky, Kim Philipp Jablonski, Lara Fuhrmann, Rebecca Denes, Mirjam Feldkamp, Ina Nissen, Natascha Santacroce, Elodie Burcklen, Christiane Beckmann, Maurice Redondo, Olivier Kobel, Christoph Noppen, Niko Beerenwinkel, Tanja Stadler |
| EPI_ISL_3295810 | Wales Specialist Virology Centre Sequencing lab: Pathogen Genomics Unit | Public Health Wales Microbiology Cardiff Wales Specialist Virology Centre | Catherine Moore, Johnathan Evans, Laura Gifford, Malorie Perry, Simon Cottrell, Angela Marchbank, Alec Birchley, Alexander Adams, Amy Gaskin, Bree Gatica-Wilcox, Jason Coombes, Joel Southgate, Lauren Gilbert, Lee Graham, Nicole Pacchiarini, Sara Kumziene-Summerhayes, Sarah Taylor, Sophie Jones, Sara Rey, Matthew Bull, Joanne Watkins, Sally Corden, Tom Connor |
| EPI_ISL_3295811 | Viollier AG | Department of Biosystems Science and Engineering, ETH Zürich | Christian Beisel, Sarah Nadeau, Chaoran Chen, Ivan Topolsky, Kim Philipp Jablonski, Lara Fuhrmann, Rebecca Denes, Mirjam Feldkamp, Ina Nissen, Natascha Santacroce, Elodie Burcklen, Christiane Beckmann, Maurice Redondo, Olivier Kobel, Christoph Noppen, Niko Beerenwinkel, Tanja Stadler |
| EPI_ISL_3295812 | Wales Specialist Virology Centre Sequencing lab: Pathogen Genomics Unit | Public Health Wales Microbiology Cardiff Wales Specialist Virology Centre | Catherine Moore, Johnathan Evans, Laura Gifford, Malorie Perry, Simon Cottrell, Angela Marchbank, Alec Birchley, Alexander Adams, Amy Gaskin, Bree Gatica-Wilcox, Jason Coombes, Joel Southgate, Lauren Gilbert, Lee Graham, Nicole Pacchiarini, Sara Kumziene-Summerhayes, Sarah Taylor, Sophie Jones, Sara Rey, Matthew Bull, Joanne Watkins, Sally Corden, Tom Connor |
| EPI_ISL_3295813 | Viollier AG | Department of Biosystems Science and Engineering, ETH Zürich | Christian Beisel, Sarah Nadeau, Chaoran Chen, Ivan Topolsky, Kim Philipp Jablonski, Lara Fuhrmann, Rebecca Denes, Mirjam Feldkamp, Ina Nissen, Natascha Santacroce, Elodie Burcklen, Christiane Beckmann, Maurice Redondo, Olivier Kobel, Christoph Noppen, Niko Beerenwinkel, Tanja Stadler |
| EPI_ISL_3295814 | Wales Specialist Virology Centre Sequencing lab: Pathogen | Public Health Wales Microbiology Cardiff Wales Specialist Virology | Catherine Moore, Johnathan Evans, Laura Gifford, Malorie Perry, Simon Cottrell, Angela Marchbank, Alec Birchley, Alexander Adams, Amy Gaskin, Bree Gatica-Wilcox, Jason Coombes, Joel Southgate, Lauren Gilbert, Lee Graham, Nicole |

| | | | |
|---|---|---|---|
| | Genomics Unit | Centre | Pacchiarini, Sara Kumziene-Summerhayes, Sarah Taylor, Sophie Jones, Sara Rey, Matthew Bull, Joanne Watkins, Sally Corden, Tom Connor |
| EPI_ISL_3295815 | Viollier AG | Department of Biosystems Science and Engineering, ETH Zürich | Christian Beisel, Sarah Nadeau, Chaoran Chen, Ivan Topolsky, Kim Philipp Jablonski, Lara Fuhrmann, Rebecca Denes, Mirjam Feldkamp, Ina Nissen, Natascha Santacroce, Elodie Burcklen, Christiane Beckmann, Maurice Redondo, Olivier Kobel, Christoph Noppen, Niko Beerenwinkel, Tanja Stadler |
| EPI_ISL_3295816, EPI_ISL_3295817 | Wales Specialist Virology Centre Sequencing lab: Pathogen Genomics Unit | Public Health Wales Microbiology Cardiff Wales Specialist Virology Centre | Catherine Moore, Johnathan Evans, Laura Gifford, Malorie Perry, Simon Cottrell, Angela Marchbank, Alec Birchley, Alexander Adams, Amy Gaskin, Bree Gatica-Wilcox, Jason Coombes, Joel Southgate, Lauren Gilbert, Lee Graham, Nicole Pacchiarini, Sara Kumziene-Summerhayes, Sarah Taylor, Sophie Jones, Sara Rey, Matthew Bull, Joanne Watkins, Sally Corden, Tom Connor |
| EPI_ISL_3295818 | Viollier AG | Department of Biosystems Science and Engineering, ETH Zürich | Christian Beisel, Sarah Nadeau, Chaoran Chen, Ivan Topolsky, Kim Philipp Jablonski, Lara Fuhrmann, Rebecca Denes, Mirjam Feldkamp, Ina Nissen, Natascha Santacroce, Elodie Burcklen, Christiane Beckmann, Maurice Redondo, Olivier Kobel, Christoph Noppen, Niko Beerenwinkel, Tanja Stadler |
| EPI_ISL_3295819 | Wales Specialist Virology Centre Sequencing lab: Pathogen Genomics Unit | Public Health Wales Microbiology Cardiff Wales Specialist Virology Centre | Catherine Moore, Johnathan Evans, Laura Gifford, Malorie Perry, Simon Cottrell, Angela Marchbank, Alec Birchley, Alexander Adams, Amy Gaskin, Bree Gatica-Wilcox, Jason Coombes, Joel Southgate, Lauren Gilbert, Lee Graham, Nicole Pacchiarini, Sara Kumziene-Summerhayes, Sarah Taylor, Sophie Jones, Sara Rey, Matthew Bull, Joanne Watkins, Sally Corden, Tom Connor |
| EPI_ISL_3295820, EPI_ISL_3295821 | Viollier AG | Department of Biosystems Science and Engineering, ETH Zürich | Christian Beisel, Sarah Nadeau, Chaoran Chen, Ivan Topolsky, Kim Philipp Jablonski, Lara Fuhrmann, Rebecca Denes, Mirjam Feldkamp, Ina Nissen, Natascha Santacroce, Elodie Burcklen, Christiane Beckmann, Maurice Redondo, Olivier Kobel, Christoph Noppen, Niko Beerenwinkel, Tanja Stadler |
| EPI_ISL_3295822, EPI_ISL_3295823 | Wales Specialist Virology Centre Sequencing lab: Pathogen Genomics Unit | Public Health Wales Microbiology Cardiff Wales Specialist Virology Centre | Catherine Moore, Johnathan Evans, Laura Gifford, Malorie Perry, Simon Cottrell, Angela Marchbank, Alec Birchley, Alexander Adams, Amy Gaskin, Bree Gatica-Wilcox, Jason Coombes, Joel Southgate, Lauren Gilbert, Lee Graham, Nicole Pacchiarini, Sara Kumziene-Summerhayes, Sarah Taylor, Sophie Jones, Sara Rey, Matthew Bull, Joanne Watkins, Sally Corden, Tom Connor |
| EPI_ISL_3295824 | Viollier AG | Department of Biosystems Science and Engineering, ETH Zürich | Christian Beisel, Sarah Nadeau, Chaoran Chen, Ivan Topolsky, Kim Philipp Jablonski, Lara Fuhrmann, Rebecca Denes, Mirjam Feldkamp, Ina Nissen, Natascha Santacroce, Elodie Burcklen, Christiane Beckmann, Maurice Redondo, Olivier Kobel, Christoph Noppen, Niko Beerenwinkel, Tanja Stadler |
| EPI_ISL_3295825 | Wales Specialist Virology Centre Sequencing lab: Pathogen Genomics Unit | Public Health Wales Microbiology Cardiff Wales Specialist Virology Centre | Catherine Moore, Johnathan Evans, Laura Gifford, Malorie Perry, Simon Cottrell, Angela Marchbank, Alec Birchley, Alexander Adams, Amy Gaskin, Bree Gatica-Wilcox, Jason Coombes, Joel Southgate, Lauren Gilbert, Lee Graham, Nicole Pacchiarini, Sara Kumziene-Summerhayes, Sarah Taylor, Sophie Jones, Sara Rey, Matthew Bull, Joanne Watkins, Sally Corden, Tom Connor |
| EPI_ISL_3295826 | Viollier AG | Department of Biosystems Science and Engineering, ETH Zürich | Christian Beisel, Sarah Nadeau, Chaoran Chen, Ivan Topolsky, Kim Philipp Jablonski, Lara Fuhrmann, Rebecca Denes, Mirjam Feldkamp, Ina Nissen, Natascha Santacroce, Elodie Burcklen, Christiane Beckmann, Maurice Redondo, Olivier Kobel, Christoph Noppen, Niko Beerenwinkel, Tanja Stadler |
| EPI_ISL_3295827 | Wales Specialist Virology Centre Sequencing lab: Pathogen Genomics Unit | Public Health Wales Microbiology Cardiff Wales Specialist Virology Centre | Catherine Moore, Johnathan Evans, Laura Gifford, Malorie Perry, Simon Cottrell, Angela Marchbank, Alec Birchley, Alexander Adams, Amy Gaskin, Bree Gatica-Wilcox, Jason Coombes, Joel Southgate, Lauren Gilbert, Lee Graham, Nicole Pacchiarini, Sara Kumziene-Summerhayes, Sarah Taylor, Sophie Jones, Sara Rey, Matthew Bull, Joanne Watkins, Sally Corden, Tom Connor |
| EPI_ISL_3295828 | Viollier AG | Department of Biosystems Science and Engineering, ETH Zürich | Christian Beisel, Sarah Nadeau, Chaoran Chen, Ivan Topolsky, Kim Philipp Jablonski, Lara Fuhrmann, Rebecca Denes, Mirjam Feldkamp, Ina Nissen, Natascha Santacroce, Elodie Burcklen, Christiane Beckmann, Maurice Redondo, Olivier Kobel, Christoph Noppen, Niko Beerenwinkel, Tanja Stadler |
| EPI_ISL_3295829 | Wales Specialist Virology Centre Sequencing lab: Pathogen Genomics Unit | Public Health Wales Microbiology Cardiff Wales Specialist Virology Centre | Catherine Moore, Johnathan Evans, Laura Gifford, Malorie Perry, Simon Cottrell, Angela Marchbank, Alec Birchley, Alexander Adams, Amy Gaskin, Bree Gatica-Wilcox, Jason Coombes, Joel Southgate, Lauren Gilbert, Lee Graham, Nicole Pacchiarini, Sara Kumziene-Summerhayes, Sarah Taylor, Sophie Jones, Sara Rey, Matthew Bull, Joanne Watkins, Sally Corden, Tom Connor |
| EPI_ISL_3295830 | Viollier AG | Department of Biosystems Science and Engineering, ETH Zürich | Christian Beisel, Sarah Nadeau, Chaoran Chen, Ivan Topolsky, Kim Philipp Jablonski, Lara Fuhrmann, Rebecca Denes, Mirjam Feldkamp, Ina Nissen, Natascha Santacroce, Elodie Burcklen, Christiane Beckmann, Maurice Redondo, Olivier Kobel, Christoph Noppen, Niko Beerenwinkel, Tanja Stadler |
| EPI_ISL_3295831 | Wales Specialist Virology Centre Sequencing lab: Pathogen Genomics Unit | Public Health Wales Microbiology Cardiff Wales Specialist Virology Centre | Catherine Moore, Johnathan Evans, Laura Gifford, Malorie Perry, Simon Cottrell, Angela Marchbank, Alec Birchley, Alexander Adams, Amy Gaskin, Bree Gatica-Wilcox, Jason Coombes, Joel Southgate, Lauren Gilbert, Lee Graham, Nicole Pacchiarini, Sara Kumziene-Summerhayes, Sarah Taylor, Sophie Jones, Sara Rey, Matthew Bull, Joanne Watkins, Sally Corden, Tom Connor |
| EPI_ISL_3295832 | Viollier AG | Department of Biosystems Science and Engineering, ETH Zürich | Christian Beisel, Sarah Nadeau, Chaoran Chen, Ivan Topolsky, Kim Philipp Jablonski, Lara Fuhrmann, Rebecca Denes, Mirjam Feldkamp, Ina Nissen, Natascha Santacroce, Elodie Burcklen, Christiane Beckmann, Maurice Redondo, Olivier Kobel, Christoph Noppen, Niko Beerenwinkel, Tanja Stadler |
| EPI_ISL_3295833 | Wales Specialist Virology Centre Sequencing lab: Pathogen Genomics Unit | Public Health Wales Microbiology Cardiff Wales Specialist Virology Centre | Catherine Moore, Johnathan Evans, Laura Gifford, Malorie Perry, Simon Cottrell, Angela Marchbank, Alec Birchley, Alexander Adams, Amy Gaskin, Bree Gatica-Wilcox, Jason Coombes, Joel Southgate, Lauren Gilbert, Lee Graham, Nicole Pacchiarini, Sara Kumziene-Summerhayes, Sarah Taylor, Sophie Jones, Sara Rey, Matthew Bull, Joanne Watkins, Sally Corden, Tom Connor |
| EPI_ISL_3295834 | Viollier AG | Department of Biosystems Science and Engineering, ETH Zürich | Christian Beisel, Sarah Nadeau, Chaoran Chen, Ivan Topolsky, Kim Philipp Jablonski, Lara Fuhrmann, Rebecca Denes, Mirjam Feldkamp, Ina Nissen, Natascha Santacroce, Elodie Burcklen, Christiane Beckmann, Maurice Redondo, Olivier Kobel, Christoph Noppen, Niko Beerenwinkel, Tanja Stadler |
| EPI_ISL_3295835 | Wales Specialist Virology Centre Sequencing lab: Pathogen Genomics Unit | Public Health Wales Microbiology Cardiff Wales Specialist Virology Centre | Catherine Moore, Johnathan Evans, Laura Gifford, Malorie Perry, Simon Cottrell, Angela Marchbank, Alec Birchley, Alexander Adams, Amy Gaskin, Bree Gatica-Wilcox, Jason Coombes, Joel Southgate, Lauren Gilbert, Lee Graham, Nicole Pacchiarini, Sara Kumziene-Summerhayes, Sarah Taylor, Sophie Jones, Sara Rey, Matthew Bull, Joanne Watkins, Sally Corden, Tom Connor |
| EPI_ISL_3295836 | Viollier AG | Department of Biosystems Science and Engineering, ETH Zürich | Christian Beisel, Sarah Nadeau, Chaoran Chen, Ivan Topolsky, Kim Philipp Jablonski, Lara Fuhrmann, Rebecca Denes, Mirjam Feldkamp, Ina Nissen, Natascha Santacroce, Elodie Burcklen, Christiane Beckmann, Maurice Redondo, Olivier Kobel, Christoph Noppen, Niko Beerenwinkel, Tanja Stadler |
| EPI_ISL_3295837 | Wales Specialist Virology Centre Sequencing lab: Pathogen Genomics Unit | Public Health Wales Microbiology Cardiff Wales Specialist Virology Centre | Catherine Moore, Johnathan Evans, Laura Gifford, Malorie Perry, Simon Cottrell, Angela Marchbank, Alec Birchley, Alexander Adams, Amy Gaskin, Bree Gatica-Wilcox, Jason Coombes, Joel Southgate, Lauren Gilbert, Lee Graham, Nicole Pacchiarini, Sara Kumziene-Summerhayes, Sarah Taylor, Sophie Jones, Sara Rey, Matthew Bull, Joanne Watkins, Sally Corden, Tom Connor |
| EPI_ISL_3295838 | Viollier AG | Department of Biosystems Science and Engineering, ETH Zürich | Christian Beisel, Sarah Nadeau, Chaoran Chen, Ivan Topolsky, Kim Philipp Jablonski, Lara Fuhrmann, Rebecca Denes, Mirjam Feldkamp, Ina Nissen, Natascha Santacroce, Elodie Burcklen, Christiane Beckmann, Maurice Redondo, Olivier Kobel, Christoph Noppen, Niko Beerenwinkel, Tanja Stadler |
| EPI_ISL_3295839 | Wales Specialist Virology Centre Sequencing lab: Pathogen Genomics Unit | Public Health Wales Microbiology Cardiff Wales Specialist Virology Centre | Catherine Moore, Johnathan Evans, Laura Gifford, Malorie Perry, Simon Cottrell, Angela Marchbank, Alec Birchley, Alexander Adams, Amy Gaskin, Bree Gatica-Wilcox, Jason Coombes, Joel Southgate, Lauren Gilbert, Lee Graham, Nicole Pacchiarini, Sara Kumziene-Summerhayes, Sarah Taylor, Sophie Jones, Sara Rey, Matthew Bull, Joanne Watkins, Sally Corden, Tom Connor |
| EPI_ISL_3295840 | Viollier AG | Department of Biosystems Science and Engineering, ETH Zürich | Christian Beisel, Sarah Nadeau, Chaoran Chen, Ivan Topolsky, Kim Philipp Jablonski, Lara Fuhrmann, Rebecca Denes, Mirjam Feldkamp, Ina Nissen, Natascha Santacroce, Elodie Burcklen, Christiane Beckmann, Maurice Redondo, Olivier Kobel, Christoph Noppen, Niko Beerenwinkel, Tanja Stadler |
| EPI_ISL_3295841 | Wales Specialist Virology Centre Sequencing lab: Pathogen Genomics Unit | Public Health Wales Microbiology Cardiff Wales Specialist Virology Centre | Catherine Moore, Johnathan Evans, Laura Gifford, Malorie Perry, Simon Cottrell, Angela Marchbank, Alec Birchley, Alexander Adams, Amy Gaskin, Bree Gatica-Wilcox, Jason Coombes, Joel Southgate, Lauren Gilbert, Lee Graham, Nicole Pacchiarini, Sara Kumziene-Summerhayes, Sarah Taylor, Sophie Jones, Sara Rey, Matthew Bull, Joanne Watkins, Sally Corden, Tom Connor |
| EPI_ISL_3295842 | Viollier AG | Department of Biosystems Science and Engineering, ETH Zürich | Christian Beisel, Sarah Nadeau, Chaoran Chen, Ivan Topolsky, Kim Philipp Jablonski, Lara Fuhrmann, Rebecca Denes, Mirjam Feldkamp, Ina Nissen, Natascha Santacroce, Elodie Burcklen, Christiane Beckmann, Maurice Redondo, Olivier Kobel, Christoph Noppen, Niko Beerenwinkel, Tanja Stadler |
| EPI_ISL_3295844 | Wales Specialist Virology Centre Sequencing lab: Pathogen Genomics Unit | Public Health Wales Microbiology Cardiff Wales Specialist Virology Centre | Catherine Moore, Johnathan Evans, Laura Gifford, Malorie Perry, Simon Cottrell, Angela Marchbank, Alec Birchley, Alexander Adams, Amy Gaskin, Bree Gatica-Wilcox, Jason Coombes, Joel Southgate, Lauren Gilbert, Lee Graham, Nicole Pacchiarini, Sara Kumziene-Summerhayes, Sarah Taylor, Sophie Jones, Sara Rey, Matthew Bull, Joanne Watkins, Sally Corden, Tom Connor |
| EPI_ISL_3295845, EPI_ISL_3295846 | Viollier AG | Department of Biosystems Science and Engineering, ETH Zürich | Christian Beisel, Sarah Nadeau, Chaoran Chen, Ivan Topolsky, Kim Philipp Jablonski, Lara Fuhrmann, Rebecca Denes, Mirjam Feldkamp, Ina Nissen, Natascha Santacroce, Elodie Burcklen, Christiane Beckmann, Maurice Redondo, Olivier Kobel, Christoph Noppen, Niko Beerenwinkel, Tanja Stadler |
| EPI_ISL_3295847, EPI_ISL_3295848 | Wales Specialist Virology Centre Sequencing lab: Pathogen Genomics Unit | Public Health Wales Microbiology Cardiff Wales Specialist Virology Centre | Catherine Moore, Johnathan Evans, Laura Gifford, Malorie Perry, Simon Cottrell, Angela Marchbank, Alec Birchley, Alexander Adams, Amy Gaskin, Bree Gatica-Wilcox, Jason Coombes, Joel Southgate, Lauren Gilbert, Lee Graham, Nicole Pacchiarini, Sara Kumziene-Summerhayes, Sarah Taylor, Sophie Jones, Sara Rey, Matthew Bull, Joanne Watkins, Sally Corden, Tom Connor |
| EPI_ISL_3295849 | Viollier AG | Department of Biosystems Science and Engineering, ETH Zürich | Christian Beisel, Sarah Nadeau, Chaoran Chen, Ivan Topolsky, Kim Philipp Jablonski, Lara Fuhrmann, Rebecca Denes, Mirjam Feldkamp, Ina Nissen, Natascha Santacroce, Elodie Burcklen, Christiane Beckmann, Maurice Redondo, Olivier Kobel, Christoph Noppen, Niko Beerenwinkel, Tanja Stadler |
| EPI_ISL_3295850 | Wales Specialist Virology Centre Sequencing lab: Pathogen Genomics Unit | Public Health Wales Microbiology Cardiff Wales Specialist Virology Centre | Catherine Moore, Johnathan Evans, Laura Gifford, Malorie Perry, Simon Cottrell, Angela Marchbank, Alec Birchley, Alexander Adams, Amy Gaskin, Bree Gatica-Wilcox, Jason Coombes, Joel Southgate, Lauren Gilbert, Lee Graham, Nicole Pacchiarini, Sara Kumziene-Summerhayes, Sarah Taylor, Sophie Jones, Sara Rey, Matthew Bull, Joanne Watkins, Sally Corden, Tom Connor |
| EPI_ISL_3295851 | Viollier AG | Department of Biosystems Science and Engineering, ETH Zürich | Christian Beisel, Sarah Nadeau, Chaoran Chen, Ivan Topolsky, Kim Philipp Jablonski, Lara Fuhrmann, Rebecca Denes, Mirjam Feldkamp, Ina Nissen, Natascha Santacroce, Elodie Burcklen, Christiane Beckmann, Maurice Redondo, Olivier Kobel, Christoph Noppen, Niko Beerenwinkel, Tanja Stadler |
| EPI_ISL_3295852 | Wales Specialist Virology Centre Sequencing lab: Pathogen Genomics Unit | Public Health Wales Microbiology Cardiff Wales Specialist Virology Centre | Catherine Moore, Johnathan Evans, Laura Gifford, Malorie Perry, Simon Cottrell, Angela Marchbank, Alec Birchley, Alexander Adams, Amy Gaskin, Bree Gatica-Wilcox, Jason Coombes, Joel Southgate, Lauren Gilbert, Lee Graham, Nicole Pacchiarini, Sara Kumziene-Summerhayes, Sarah Taylor, Sophie Jones, Sara Rey, Matthew Bull, Joanne Watkins, Sally Corden, Tom Connor |
| EPI_ISL_3295853 | Viollier AG | Department of Biosystems Science and Engineering, ETH Zürich | Christian Beisel, Sarah Nadeau, Chaoran Chen, Ivan Topolsky, Kim Philipp Jablonski, Lara Fuhrmann, Rebecca Denes, Mirjam Feldkamp, Ina Nissen, Natascha Santacroce, Elodie Burcklen, Christiane Beckmann, Maurice Redondo, Olivier Kobel, Christoph Noppen, Niko Beerenwinkel, Tanja Stadler |
| EPI_ISL_3295854 | Wales Specialist Virology Centre Sequencing lab: Pathogen Genomics Unit | Public Health Wales Microbiology Cardiff Wales Specialist Virology Centre | Catherine Moore, Johnathan Evans, Laura Gifford, Malorie Perry, Simon Cottrell, Angela Marchbank, Alec Birchley, Alexander Adams, Amy Gaskin, Bree Gatica-Wilcox, Jason Coombes, Joel Southgate, Lauren Gilbert, Lee Graham, Nicole Pacchiarini, Sara Kumziene-Summerhayes, Sarah Taylor, Sophie Jones, Sara Rey, Matthew Bull, Joanne Watkins, Sally Corden, Tom Connor |
| EPI_ISL_3295855 | Viollier AG | Department of Biosystems Science and Engineering, ETH Zürich | Christian Beisel, Sarah Nadeau, Chaoran Chen, Ivan Topolsky, Kim Philipp Jablonski, Lara Fuhrmann, Rebecca Denes, Mirjam Feldkamp, Ina Nissen, Natascha Santacroce, Elodie Burcklen, Christiane Beckmann, Maurice Redondo, Olivier Kobel, Christoph Noppen, Niko Beerenwinkel, Tanja Stadler |
| EPI_ISL_3295856 | Wales Specialist Virology Centre Sequencing lab: Pathogen Genomics Unit | Public Health Wales Microbiology Cardiff Wales Specialist Virology Centre | Catherine Moore, Johnathan Evans, Laura Gifford, Malorie Perry, Simon Cottrell, Angela Marchbank, Alec Birchley, Alexander Adams, Amy Gaskin, Bree Gatica-Wilcox, Jason Coombes, Joel Southgate, Lauren Gilbert, Lee Graham, Nicole Pacchiarini, Sara Kumziene-Summerhayes, Sarah Taylor, Sophie Jones, Sara Rey, Matthew Bull, Joanne Watkins, Sally Corden, Tom Connor |
| EPI_ISL_3295857 | Viollier AG | Department of Biosystems Science and Engineering, ETH Zürich | Christian Beisel, Sarah Nadeau, Chaoran Chen, Ivan Topolsky, Kim Philipp Jablonski, Lara Fuhrmann, Rebecca Denes, Mirjam Feldkamp, Ina Nissen, Natascha Santacroce, Elodie Burcklen, Christiane Beckmann, Maurice Redondo, Olivier Kobel, Christoph Noppen, Niko Beerenwinkel, Tanja Stadler |
| EPI_ISL_3295858 | Wales Specialist Virology Centre Sequencing lab: Pathogen Genomics Unit | Public Health Wales Microbiology Cardiff Wales Specialist Virology Centre | Catherine Moore, Johnathan Evans, Laura Gifford, Malorie Perry, Simon Cottrell, Angela Marchbank, Alec Birchley, Alexander Adams, Amy Gaskin, Bree Gatica-Wilcox, Jason Coombes, Joel Southgate, Lauren Gilbert, Lee Graham, Nicole Pacchiarini, Sara Kumziene-Summerhayes, Sarah Taylor, Sophie Jones, Sara Rey, Matthew Bull, Joanne Watkins, Sally Corden, Tom Connor |
| EPI_ISL_3295859 | Viollier AG | Department of Biosystems Science and Engineering, ETH Zürich | Christian Beisel, Sarah Nadeau, Chaoran Chen, Ivan Topolsky, Kim Philipp Jablonski, Lara Fuhrmann, Rebecca Denes, Mirjam Feldkamp, Ina Nissen, Natascha Santacroce, Elodie Burcklen, Christiane Beckmann, Maurice Redondo, Olivier Kobel, Christoph Noppen, Niko Beerenwinkel, Tanja Stadler |
| EPI_ISL_3295860, EPI_ISL_3295862 | Wales Specialist Virology Centre Sequencing lab: Pathogen Genomics Unit | Public Health Wales Microbiology Cardiff Wales Specialist Virology Centre | Catherine Moore, Johnathan Evans, Laura Gifford, Malorie Perry, Simon Cottrell, Angela Marchbank, Alec Birchley, Alexander Adams, Amy Gaskin, Bree Gatica-Wilcox, Jason Coombes, Joel Southgate, Lauren Gilbert, Lee Graham, Nicole Pacchiarini, Sara Kumziene-Summerhayes, Sarah Taylor, Sophie Jones, Sara Rey, Matthew Bull, Joanne Watkins, Sally Corden, Tom Connor |
| EPI_ISL_3295863 | Viollier AG | Department of Biosystems Science and Engineering, ETH Zürich | Christian Beisel, Sarah Nadeau, Chaoran Chen, Ivan Topolsky, Kim Philipp Jablonski, Lara Fuhrmann, Rebecca Denes, Mirjam Feldkamp, Ina Nissen, Natascha Santacroce, Elodie Burcklen, Christiane Beckmann, Maurice Redondo, Olivier Kobel, Christoph Noppen, Niko Beerenwinkel, Tanja Stadler |
| EPI_ISL_3295864 | Wales Specialist Virology Centre Sequencing lab: Pathogen Genomics Unit | Public Health Wales Microbiology Cardiff Wales Specialist Virology Centre | Catherine Moore, Johnathan Evans, Laura Gifford, Malorie Perry, Simon Cottrell, Angela Marchbank, Alec Birchley, Alexander Adams, Amy Gaskin, Bree Gatica-Wilcox, Jason Coombes, Joel Southgate, Lauren Gilbert, Lee Graham, Nicole Pacchiarini, Sara Kumziene-Summerhayes, Sarah Taylor, Sophie Jones, Sara Rey, Matthew Bull, Joanne Watkins, Sally Corden, Tom Connor |
| EPI_ISL_3295865 | Viollier AG | Department of Biosystems Science and Engineering, ETH Zürich | Christian Beisel, Sarah Nadeau, Chaoran Chen, Ivan Topolsky, Kim Philipp Jablonski, Lara Fuhrmann, Rebecca Denes, Mirjam Feldkamp, Ina Nissen, Natascha Santacroce, Elodie Burcklen, Christiane Beckmann, Maurice Redondo, Olivier Kobel, Christoph Noppen, Niko Beerenwinkel, Tanja Stadler |
| EPI_ISL_3295866, EPI_ISL_3295867 | Wales Specialist Virology Centre Sequencing lab: Pathogen Genomics Unit | Public Health Wales Microbiology Cardiff Wales Specialist Virology Centre | Catherine Moore, Johnathan Evans, Laura Gifford, Malorie Perry, Simon Cottrell, Angela Marchbank, Alec Birchley, Alexander Adams, Amy Gaskin, Bree Gatica-Wilcox, Jason Coombes, Joel Southgate, Lauren Gilbert, Lee Graham, Nicole Pacchiarini, Sara Kumziene-Summerhayes, Sarah Taylor, Sophie Jones, Sara Rey, Matthew Bull, Joanne Watkins, Sally Corden, Tom Connor |
| EPI_ISL_3295868 | Viollier AG | Department of Biosystems Science and Engineering, ETH Zürich | Christian Beisel, Sarah Nadeau, Chaoran Chen, Ivan Topolsky, Kim Philipp Jablonski, Lara Fuhrmann, Rebecca Denes, Mirjam Feldkamp, Ina Nissen, Natascha Santacroce, Elodie Burcklen, Christiane Beckmann, Maurice Redondo, Olivier Kobel, |

| | | | |
|---|---|---|---|
| | | | Christoph Noppen, Niko Beerenwinkel, Tanja Stadler |
| EPI_ISL_3295869 | Wales Specialist Virology Centre Sequencing lab: Pathogen Genomics Unit | Public Health Wales Microbiology Cardiff Wales Specialist Virology Centre | Catherine Moore, Johnathan Evans, Laura Gifford, Malorie Perry, Simon Cottrell, Angela Marchbank, Alec Birchley, Alexander Adams, Amy Gaskin, Bree Gatica-Wilcox, Jason Coombes, Joel Southgate, Lauren Gilbert, Lee Graham, Nicole Pacchiarini, Sara Kumziene-Summerhayes, Sarah Taylor, Sophie Jones, Sara Rey, Matthew Bull, Joanne Watkins, Sally Corden, Tom Connor |
| EPI_ISL_3295870 | Viollier AG | Department of Biosystems Science and Engineering, ETH Zürich | Christian Beisel, Sarah Nadeau, Chaoran Chen, Ivan Topolsky, Kim Philipp Jablonski, Lara Fuhrmann, Rebecca Denes, Mirjam Feldkamp, Ina Nissen, Natascha Santacroce, Elodie Burcklen, Christiane Beckmann, Maurice Redondo, Olivier Kobel, Christoph Noppen, Niko Beerenwinkel, Tanja Stadler |
| EPI_ISL_3295871 | Wales Specialist Virology Centre Sequencing lab: Pathogen Genomics Unit | Public Health Wales Microbiology Cardiff Wales Specialist Virology Centre | Catherine Moore, Johnathan Evans, Laura Gifford, Malorie Perry, Simon Cottrell, Angela Marchbank, Alec Birchley, Alexander Adams, Amy Gaskin, Bree Gatica-Wilcox, Jason Coombes, Joel Southgate, Lauren Gilbert, Lee Graham, Nicole Pacchiarini, Sara Kumziene-Summerhayes, Sarah Taylor, Sophie Jones, Sara Rey, Matthew Bull, Joanne Watkins, Sally Corden, Tom Connor |
| EPI_ISL_3295872 | Viollier AG | Department of Biosystems Science and Engineering, ETH Zürich | Christian Beisel, Sarah Nadeau, Chaoran Chen, Ivan Topolsky, Kim Philipp Jablonski, Lara Fuhrmann, Rebecca Denes, Mirjam Feldkamp, Ina Nissen, Natascha Santacroce, Elodie Burcklen, Christiane Beckmann, Maurice Redondo, Olivier Kobel, Christoph Noppen, Niko Beerenwinkel, Tanja Stadler |
| EPI_ISL_3295873 | Wales Specialist Virology Centre Sequencing lab: Pathogen Genomics Unit | Public Health Wales Microbiology Cardiff Wales Specialist Virology Centre | Catherine Moore, Johnathan Evans, Laura Gifford, Malorie Perry, Simon Cottrell, Angela Marchbank, Alec Birchley, Alexander Adams, Amy Gaskin, Bree Gatica-Wilcox, Jason Coombes, Joel Southgate, Lauren Gilbert, Lee Graham, Nicole Pacchiarini, Sara Kumziene-Summerhayes, Sarah Taylor, Sophie Jones, Sara Rey, Matthew Bull, Joanne Watkins, Sally Corden, Tom Connor |
| EPI_ISL_3295874 | Viollier AG | Department of Biosystems Science and Engineering, ETH Zürich | Christian Beisel, Sarah Nadeau, Chaoran Chen, Ivan Topolsky, Kim Philipp Jablonski, Lara Fuhrmann, Rebecca Denes, Mirjam Feldkamp, Ina Nissen, Natascha Santacroce, Elodie Burcklen, Christiane Beckmann, Maurice Redondo, Olivier Kobel, Christoph Noppen, Niko Beerenwinkel, Tanja Stadler |
| EPI_ISL_3295875 | Wales Specialist Virology Centre Sequencing lab: Pathogen Genomics Unit | Public Health Wales Microbiology Cardiff Wales Specialist Virology Centre | Catherine Moore, Johnathan Evans, Laura Gifford, Malorie Perry, Simon Cottrell, Angela Marchbank, Alec Birchley, Alexander Adams, Amy Gaskin, Bree Gatica-Wilcox, Jason Coombes, Joel Southgate, Lauren Gilbert, Lee Graham, Nicole Pacchiarini, Sara Kumziene-Summerhayes, Sarah Taylor, Sophie Jones, Sara Rey, Matthew Bull, Joanne Watkins, Sally Corden, Tom Connor |
| EPI_ISL_3295876 | Viollier AG | Department of Biosystems Science and Engineering, ETH Zürich | Christian Beisel, Sarah Nadeau, Chaoran Chen, Ivan Topolsky, Kim Philipp Jablonski, Lara Fuhrmann, Rebecca Denes, Mirjam Feldkamp, Ina Nissen, Natascha Santacroce, Elodie Burcklen, Christiane Beckmann, Maurice Redondo, Olivier Kobel, Christoph Noppen, Niko Beerenwinkel, Tanja Stadler |
| EPI_ISL_3295877 | Wales Specialist Virology Centre Sequencing lab: Pathogen Genomics Unit | Public Health Wales Microbiology Cardiff Wales Specialist Virology Centre | Catherine Moore, Johnathan Evans, Laura Gifford, Malorie Perry, Simon Cottrell, Angela Marchbank, Alec Birchley, Alexander Adams, Amy Gaskin, Bree Gatica-Wilcox, Jason Coombes, Joel Southgate, Lauren Gilbert, Lee Graham, Nicole Pacchiarini, Sara Kumziene-Summerhayes, Sarah Taylor, Sophie Jones, Sara Rey, Matthew Bull, Joanne Watkins, Sally Corden, Tom Connor |
| EPI_ISL_3295878 | Viollier AG | Department of Biosystems Science and Engineering, ETH Zürich | Christian Beisel, Sarah Nadeau, Chaoran Chen, Ivan Topolsky, Kim Philipp Jablonski, Lara Fuhrmann, Rebecca Denes, Mirjam Feldkamp, Ina Nissen, Natascha Santacroce, Elodie Burcklen, Christiane Beckmann, Maurice Redondo, Olivier Kobel, Christoph Noppen, Niko Beerenwinkel, Tanja Stadler |
| EPI_ISL_3295879 | Wales Specialist Virology Centre Sequencing lab: Pathogen Genomics Unit | Public Health Wales Microbiology Cardiff Wales Specialist Virology Centre | Catherine Moore, Johnathan Evans, Laura Gifford, Malorie Perry, Simon Cottrell, Angela Marchbank, Alec Birchley, Alexander Adams, Amy Gaskin, Bree Gatica-Wilcox, Jason Coombes, Joel Southgate, Lauren Gilbert, Lee Graham, Nicole Pacchiarini, Sara Kumziene-Summerhayes, Sarah Taylor, Sophie Jones, Sara Rey, Matthew Bull, Joanne Watkins, Sally Corden, Tom Connor |
| EPI_ISL_3295880 | Viollier AG | Department of Biosystems Science and Engineering, ETH Zürich | Christian Beisel, Sarah Nadeau, Chaoran Chen, Ivan Topolsky, Kim Philipp Jablonski, Lara Fuhrmann, Rebecca Denes, Mirjam Feldkamp, Ina Nissen, Natascha Santacroce, Elodie Burcklen, Christiane Beckmann, Maurice Redondo, Olivier Kobel, Christoph Noppen, Niko Beerenwinkel, Tanja Stadler |
| EPI_ISL_3295881 | Wales Specialist Virology Centre Sequencing lab: Pathogen Genomics Unit | Public Health Wales Microbiology Cardiff Wales Specialist Virology Centre | Catherine Moore, Johnathan Evans, Laura Gifford, Malorie Perry, Simon Cottrell, Angela Marchbank, Alec Birchley, Alexander Adams, Amy Gaskin, Bree Gatica-Wilcox, Jason Coombes, Joel Southgate, Lauren Gilbert, Lee Graham, Nicole Pacchiarini, Sara Kumziene-Summerhayes, Sarah Taylor, Sophie Jones, Sara Rey, Matthew Bull, Joanne Watkins, Sally Corden, Tom Connor |
| EPI_ISL_3295882 | Viollier AG | Department of Biosystems Science and Engineering, ETH Zürich | Christian Beisel, Sarah Nadeau, Chaoran Chen, Ivan Topolsky, Kim Philipp Jablonski, Lara Fuhrmann, Rebecca Denes, Mirjam Feldkamp, Ina Nissen, Natascha Santacroce, Elodie Burcklen, Christiane Beckmann, Maurice Redondo, Olivier Kobel, Christoph Noppen, Niko Beerenwinkel, Tanja Stadler |
| EPI_ISL_3295883, EPI_ISL_3295884 | Wales Specialist Virology Centre Sequencing lab: Pathogen Genomics Unit | Public Health Wales Microbiology Cardiff Wales Specialist Virology Centre | Catherine Moore, Johnathan Evans, Laura Gifford, Malorie Perry, Simon Cottrell, Angela Marchbank, Alec Birchley, Alexander Adams, Amy Gaskin, Bree Gatica-Wilcox, Jason Coombes, Joel Southgate, Lauren Gilbert, Lee Graham, Nicole Pacchiarini, Sara Kumziene-Summerhayes, Sarah Taylor, Sophie Jones, Sara Rey, Matthew Bull, Joanne Watkins, Sally Corden, Tom Connor |
| EPI_ISL_3295885 | Viollier AG | Department of Biosystems Science and Engineering, ETH Zürich | Christian Beisel, Sarah Nadeau, Chaoran Chen, Ivan Topolsky, Kim Philipp Jablonski, Lara Fuhrmann, Rebecca Denes, Mirjam Feldkamp, Ina Nissen, Natascha Santacroce, Elodie Burcklen, Christiane Beckmann, Maurice Redondo, Olivier Kobel, Christoph Noppen, Niko Beerenwinkel, Tanja Stadler |
| EPI_ISL_3295886 | Wales Specialist Virology Centre Sequencing lab: Pathogen Genomics Unit | Public Health Wales Microbiology Cardiff Wales Specialist Virology Centre | Catherine Moore, Johnathan Evans, Laura Gifford, Malorie Perry, Simon Cottrell, Angela Marchbank, Alec Birchley, Alexander Adams, Amy Gaskin, Bree Gatica-Wilcox, Jason Coombes, Joel Southgate, Lauren Gilbert, Lee Graham, Nicole Pacchiarini, Sara Kumziene-Summerhayes, Sarah Taylor, Sophie Jones, Sara Rey, Matthew Bull, Joanne Watkins, Sally Corden, Tom Connor |
| EPI_ISL_3295887 | Viollier AG | Department of Biosystems Science and Engineering, ETH Zürich | Christian Beisel, Sarah Nadeau, Chaoran Chen, Ivan Topolsky, Kim Philipp Jablonski, Lara Fuhrmann, Rebecca Denes, Mirjam Feldkamp, Ina Nissen, Natascha Santacroce, Elodie Burcklen, Christiane Beckmann, Maurice Redondo, Olivier Kobel, Christoph Noppen, Niko Beerenwinkel, Tanja Stadler |
| EPI_ISL_3295888 | Wales Specialist Virology Centre Sequencing lab: Pathogen Genomics Unit | Public Health Wales Microbiology Cardiff Wales Specialist Virology Centre | Catherine Moore, Johnathan Evans, Laura Gifford, Malorie Perry, Simon Cottrell, Angela Marchbank, Alec Birchley, Alexander Adams, Amy Gaskin, Bree Gatica-Wilcox, Jason Coombes, Joel Southgate, Lauren Gilbert, Lee Graham, Nicole Pacchiarini, Sara Kumziene-Summerhayes, Sarah Taylor, Sophie Jones, Sara Rey, Matthew Bull, Joanne Watkins, Sally Corden, Tom Connor |
| EPI_ISL_3295889 | Viollier AG | Department of Biosystems Science and Engineering, ETH Zürich | Christian Beisel, Sarah Nadeau, Chaoran Chen, Ivan Topolsky, Kim Philipp Jablonski, Lara Fuhrmann, Rebecca Denes, Mirjam Feldkamp, Ina Nissen, Natascha Santacroce, Elodie Burcklen, Christiane Beckmann, Maurice Redondo, Olivier Kobel, Christoph Noppen, Niko Beerenwinkel, Tanja Stadler |
| EPI_ISL_3295890 | Wales Specialist Virology Centre Sequencing lab: Pathogen Genomics Unit | Public Health Wales Microbiology Cardiff Wales Specialist Virology Centre | Catherine Moore, Johnathan Evans, Laura Gifford, Malorie Perry, Simon Cottrell, Angela Marchbank, Alec Birchley, Alexander Adams, Amy Gaskin, Bree Gatica-Wilcox, Jason Coombes, Joel Southgate, Lauren Gilbert, Lee Graham, Nicole Pacchiarini, Sara Kumziene-Summerhayes, Sarah Taylor, Sophie Jones, Sara Rey, Matthew Bull, Joanne Watkins, Sally Corden, Tom Connor |
| EPI_ISL_3295891 | Viollier AG | Department of Biosystems Science and Engineering, ETH Zürich | Christian Beisel, Sarah Nadeau, Chaoran Chen, Ivan Topolsky, Kim Philipp Jablonski, Lara Fuhrmann, Rebecca Denes, Mirjam Feldkamp, Ina Nissen, Natascha Santacroce, Elodie Burcklen, Christiane Beckmann, Maurice Redondo, Olivier Kobel, Christoph Noppen, Niko Beerenwinkel, Tanja Stadler |
| EPI_ISL_3295892 | Wales Specialist Virology Centre Sequencing lab: Pathogen Genomics Unit | Public Health Wales Microbiology Cardiff Wales Specialist Virology Centre | Catherine Moore, Johnathan Evans, Laura Gifford, Malorie Perry, Simon Cottrell, Angela Marchbank, Alec Birchley, Alexander Adams, Amy Gaskin, Bree Gatica-Wilcox, Jason Coombes, Joel Southgate, Lauren Gilbert, Lee Graham, Nicole Pacchiarini, Sara Kumziene-Summerhayes, Sarah Taylor, Sophie Jones, Sara Rey, Matthew Bull, Joanne Watkins, Sally Corden, Tom Connor |
| EPI_ISL_3295893 | Viollier AG | Department of Biosystems Science and Engineering, ETH Zürich | Christian Beisel, Sarah Nadeau, Chaoran Chen, Ivan Topolsky, Kim Philipp Jablonski, Lara Fuhrmann, Rebecca Denes, Mirjam Feldkamp, Ina Nissen, Natascha Santacroce, Elodie Burcklen, Christiane Beckmann, Maurice Redondo, Olivier Kobel, Christoph Noppen, Niko Beerenwinkel, Tanja Stadler |
| EPI_ISL_3295894 | Wales Specialist Virology Centre Sequencing lab: Pathogen Genomics Unit | Public Health Wales Microbiology Cardiff Wales Specialist Virology Centre | Catherine Moore, Johnathan Evans, Laura Gifford, Malorie Perry, Simon Cottrell, Angela Marchbank, Alec Birchley, Alexander Adams, Amy Gaskin, Bree Gatica-Wilcox, Jason Coombes, Joel Southgate, Lauren Gilbert, Lee Graham, Nicole Pacchiarini, Sara Kumziene-Summerhayes, Sarah Taylor, Sophie Jones, Sara Rey, Matthew Bull, Joanne Watkins, Sally Corden, Tom Connor |
| EPI_ISL_3295895 | Viollier AG | Department of Biosystems Science and Engineering, ETH Zürich | Christian Beisel, Sarah Nadeau, Chaoran Chen, Ivan Topolsky, Kim Philipp Jablonski, Lara Fuhrmann, Rebecca Denes, Mirjam Feldkamp, Ina Nissen, Natascha Santacroce, Elodie Burcklen, Christiane Beckmann, Maurice Redondo, Olivier Kobel, Christoph Noppen, Niko Beerenwinkel, Tanja Stadler |
| EPI_ISL_3295896 | Wales Specialist Virology Centre Sequencing lab: Pathogen Genomics Unit | Public Health Wales Microbiology Cardiff Wales Specialist Virology Centre | Catherine Moore, Johnathan Evans, Laura Gifford, Malorie Perry, Simon Cottrell, Angela Marchbank, Alec Birchley, Alexander Adams, Amy Gaskin, Bree Gatica-Wilcox, Jason Coombes, Joel Southgate, Lauren Gilbert, Lee Graham, Nicole Pacchiarini, Sara Kumziene-Summerhayes, Sarah Taylor, Sophie Jones, Sara Rey, Matthew Bull, Joanne Watkins, Sally Corden, Tom Connor |
| EPI_ISL_3295897 | Viollier AG | Department of Biosystems Science and Engineering, ETH Zürich | Christian Beisel, Sarah Nadeau, Chaoran Chen, Ivan Topolsky, Kim Philipp Jablonski, Lara Fuhrmann, Rebecca Denes, Mirjam Feldkamp, Ina Nissen, Natascha Santacroce, Elodie Burcklen, Christiane Beckmann, Maurice Redondo, Olivier Kobel, Christoph Noppen, Niko Beerenwinkel, Tanja Stadler |
| EPI_ISL_3295898 | Wales Specialist Virology Centre Sequencing lab: Pathogen Genomics Unit | Public Health Wales Microbiology Cardiff Wales Specialist Virology Centre | Catherine Moore, Johnathan Evans, Laura Gifford, Malorie Perry, Simon Cottrell, Angela Marchbank, Alec Birchley, Alexander Adams, Amy Gaskin, Bree Gatica-Wilcox, Jason Coombes, Joel Southgate, Lauren Gilbert, Lee Graham, Nicole Pacchiarini, Sara Kumziene-Summerhayes, Sarah Taylor, Sophie Jones, Sara Rey, Matthew Bull, Joanne Watkins, Sally Corden, Tom Connor |
| EPI_ISL_3295899 | Viollier AG | Department of Biosystems Science and Engineering, ETH Zürich | Christian Beisel, Sarah Nadeau, Chaoran Chen, Ivan Topolsky, Kim Philipp Jablonski, Lara Fuhrmann, Rebecca Denes, Mirjam Feldkamp, Ina Nissen, Natascha Santacroce, Elodie Burcklen, Christiane Beckmann, Maurice Redondo, Olivier Kobel, Christoph Noppen, Niko Beerenwinkel, Tanja Stadler |
| EPI_ISL_3295900, EPI_ISL_3295901 | Wales Specialist Virology Centre Sequencing lab: Pathogen Genomics Unit | Public Health Wales Microbiology Cardiff Wales Specialist Virology Centre | Catherine Moore, Johnathan Evans, Laura Gifford, Malorie Perry, Simon Cottrell, Angela Marchbank, Alec Birchley, Alexander Adams, Amy Gaskin, Bree Gatica-Wilcox, Jason Coombes, Joel Southgate, Lauren Gilbert, Lee Graham, Nicole Pacchiarini, Sara Kumziene-Summerhayes, Sarah Taylor, Sophie Jones, Sara Rey, Matthew Bull, Joanne Watkins, Sally Corden, Tom Connor |
| EPI_ISL_3295902 | Viollier AG | Department of Biosystems Science and Engineering, ETH Zürich | Christian Beisel, Sarah Nadeau, Chaoran Chen, Ivan Topolsky, Kim Philipp Jablonski, Lara Fuhrmann, Rebecca Denes, Mirjam Feldkamp, Ina Nissen, Natascha Santacroce, Elodie Burcklen, Christiane Beckmann, Maurice Redondo, Olivier Kobel, Christoph Noppen, Niko Beerenwinkel, Tanja Stadler |
| EPI_ISL_3295903 | Wales Specialist Virology Centre Sequencing lab: Pathogen Genomics Unit | Public Health Wales Microbiology Cardiff Wales Specialist Virology Centre | Catherine Moore, Johnathan Evans, Laura Gifford, Malorie Perry, Simon Cottrell, Angela Marchbank, Alec Birchley, Alexander Adams, Amy Gaskin, Bree Gatica-Wilcox, Jason Coombes, Joel Southgate, Lauren Gilbert, Lee Graham, Nicole Pacchiarini, Sara Kumziene-Summerhayes, Sarah Taylor, Sophie Jones, Sara Rey, Matthew Bull, Joanne Watkins, Sally Corden, Tom Connor |
| EPI_ISL_3295904 | Viollier AG | Department of Biosystems Science and Engineering, ETH Zürich | Christian Beisel, Sarah Nadeau, Chaoran Chen, Ivan Topolsky, Kim Philipp Jablonski, Lara Fuhrmann, Rebecca Denes, Mirjam Feldkamp, Ina Nissen, Natascha Santacroce, Elodie Burcklen, Christiane Beckmann, Maurice Redondo, Olivier Kobel, Christoph Noppen, Niko Beerenwinkel, Tanja Stadler |
| EPI_ISL_3295905 | Wales Specialist Virology Centre Sequencing lab: Pathogen Genomics Unit | Public Health Wales Microbiology Cardiff Wales Specialist Virology Centre | Catherine Moore, Johnathan Evans, Laura Gifford, Malorie Perry, Simon Cottrell, Angela Marchbank, Alec Birchley, Alexander Adams, Amy Gaskin, Bree Gatica-Wilcox, Jason Coombes, Joel Southgate, Lauren Gilbert, Lee Graham, Nicole Pacchiarini, Sara Kumziene-Summerhayes, Sarah Taylor, Sophie Jones, Sara Rey, Matthew Bull, Joanne Watkins, Sally Corden, Tom Connor |
| EPI_ISL_3295906 | Viollier AG | Department of Biosystems Science and Engineering, ETH Zürich | Christian Beisel, Sarah Nadeau, Chaoran Chen, Ivan Topolsky, Kim Philipp Jablonski, Lara Fuhrmann, Rebecca Denes, Mirjam Feldkamp, Ina Nissen, Natascha Santacroce, Elodie Burcklen, Christiane Beckmann, Maurice Redondo, Olivier Kobel, Christoph Noppen, Niko Beerenwinkel, Tanja Stadler |
| EPI_ISL_3295907 | Wales Specialist Virology Centre Sequencing lab: Pathogen Genomics Unit | Public Health Wales Microbiology Cardiff Wales Specialist Virology Centre | Catherine Moore, Johnathan Evans, Laura Gifford, Malorie Perry, Simon Cottrell, Angela Marchbank, Alec Birchley, Alexander Adams, Amy Gaskin, Bree Gatica-Wilcox, Jason Coombes, Joel Southgate, Lauren Gilbert, Lee Graham, Nicole Pacchiarini, Sara Kumziene-Summerhayes, Sarah Taylor, Sophie Jones, Sara Rey, Matthew Bull, Joanne Watkins, Sally Corden, Tom Connor |
| EPI_ISL_3295908 | Viollier AG | Department of Biosystems Science and Engineering, ETH Zürich | Christian Beisel, Sarah Nadeau, Chaoran Chen, Ivan Topolsky, Kim Philipp Jablonski, Lara Fuhrmann, Rebecca Denes, Mirjam Feldkamp, Ina Nissen, Natascha Santacroce, Elodie Burcklen, Christiane Beckmann, Maurice Redondo, Olivier Kobel, Christoph Noppen, Niko Beerenwinkel, Tanja Stadler |
| EPI_ISL_3295909 | Wales Specialist Virology Centre Sequencing lab: Pathogen Genomics Unit | Public Health Wales Microbiology Cardiff Wales Specialist Virology Centre | Catherine Moore, Johnathan Evans, Laura Gifford, Malorie Perry, Simon Cottrell, Angela Marchbank, Alec Birchley, Alexander Adams, Amy Gaskin, Bree Gatica-Wilcox, Jason Coombes, Joel Southgate, Lauren Gilbert, Lee Graham, Nicole Pacchiarini, Sara Kumziene-Summerhayes, Sarah Taylor, Sophie Jones, Sara Rey, Matthew Bull, Joanne Watkins, Sally Corden, Tom Connor |
| EPI_ISL_3295910 | Viollier AG | Department of Biosystems Science and Engineering, ETH Zürich | Christian Beisel, Sarah Nadeau, Chaoran Chen, Ivan Topolsky, Kim Philipp Jablonski, Lara Fuhrmann, Rebecca Denes, Mirjam Feldkamp, Ina Nissen, Natascha Santacroce, Elodie Burcklen, Christiane Beckmann, Maurice Redondo, Olivier Kobel, Christoph Noppen, Niko Beerenwinkel, Tanja Stadler |
| EPI_ISL_3295911 | Wales Specialist Virology Centre Sequencing lab: Pathogen Genomics Unit | Public Health Wales Microbiology Cardiff Wales Specialist Virology Centre | Catherine Moore, Johnathan Evans, Laura Gifford, Malorie Perry, Simon Cottrell, Angela Marchbank, Alec Birchley, Alexander Adams, Amy Gaskin, Bree Gatica-Wilcox, Jason Coombes, Joel Southgate, Lauren Gilbert, Lee Graham, Nicole Pacchiarini, Sara Kumziene-Summerhayes, Sarah Taylor, Sophie Jones, Sara Rey, Matthew Bull, Joanne Watkins, Sally Corden, Tom Connor |
| EPI_ISL_3295912 | Viollier AG | Department of Biosystems Science and Engineering, ETH Zürich | Christian Beisel, Sarah Nadeau, Chaoran Chen, Ivan Topolsky, Kim Philipp Jablonski, Lara Fuhrmann, Rebecca Denes, Mirjam Feldkamp, Ina Nissen, Natascha Santacroce, Elodie Burcklen, Christiane Beckmann, Maurice Redondo, Olivier Kobel, Christoph Noppen, Niko Beerenwinkel, Tanja Stadler |
| EPI_ISL_3295913 | Wales Specialist Virology Centre Sequencing lab: Pathogen Genomics Unit | Public Health Wales Microbiology Cardiff Wales Specialist Virology Centre | Catherine Moore, Johnathan Evans, Laura Gifford, Malorie Perry, Simon Cottrell, Angela Marchbank, Alec Birchley, Alexander Adams, Amy Gaskin, Bree Gatica-Wilcox, Jason Coombes, Joel Southgate, Lauren Gilbert, Lee Graham, Nicole Pacchiarini, Sara Kumziene-Summerhayes, Sarah Taylor, Sophie Jones, Sara Rey, Matthew Bull, Joanne Watkins, Sally Corden, Tom Connor |
| EPI_ISL_3295914 | Viollier AG | Department of Biosystems Science and Engineering, ETH Zürich | Christian Beisel, Sarah Nadeau, Chaoran Chen, Ivan Topolsky, Kim Philipp Jablonski, Lara Fuhrmann, Rebecca Denes, Mirjam Feldkamp, Ina Nissen, Natascha Santacroce, Elodie Burcklen, Christiane Beckmann, Maurice Redondo, Olivier Kobel, Christoph Noppen, Niko Beerenwinkel, Tanja Stadler |
| EPI_ISL_3295915 | Wales Specialist Virology Centre Sequencing lab: Pathogen | Public Health Wales Microbiology Cardiff Wales Specialist Virology | Catherine Moore, Johnathan Evans, Laura Gifford, Malorie Perry, Simon Cottrell, Angela Marchbank, Alec Birchley, Alexander Adams, Amy Gaskin, Bree Gatica-Wilcox, Jason Coombes, Joel Southgate, Lauren Gilbert, Lee Graham, Nicole |

| | | | |
|---|---|---|---|
| | Genomics Unit | Centre | Pacchiarini, Sara Kumziene-Summerhayes, Sarah Taylor, Sophie Jones, Sara Rey, Matthew Bull, Joanne Watkins, Sally Corden, Tom Connor |
| EPI_ISL_3295916 | Viollier AG | Department of Biosystems Science and Engineering, ETH Zürich | Christian Beisel, Sarah Nadeau, Chaoran Chen, Ivan Topolsky, Kim Philipp Jablonski, Lara Fuhrmann, Rebecca Denes, Mirjam Feldkamp, Ina Nissen, Natascha Santacroce, Elodie Burcklen, Christiane Beckmann, Maurice Redondo, Olivier Kobel, Christoph Noppen, Niko Beerenwinkel, Tanja Stadler |
| EPI_ISL_3295917 | Wales Specialist Virology Centre Sequencing lab: Pathogen Genomics Unit | Public Health Wales Microbiology Cardiff Wales Specialist Virology Centre | Catherine Moore, Johnathan Evans, Laura Gifford, Malorie Perry, Simon Cottrell, Angela Marchbank, Alec Birchley, Alexander Adams, Amy Gaskin, Bree Gatica-Wilcox, Jason Coombes, Joel Southgate, Lauren Gilbert, Lee Graham, Nicole Pacchiarini, Sara Kumziene-Summerhayes, Sarah Taylor, Sophie Jones, Sara Rey, Matthew Bull, Joanne Watkins, Sally Corden, Tom Connor |
| EPI_ISL_3295918 | Viollier AG | Department of Biosystems Science and Engineering, ETH Zürich | Christian Beisel, Sarah Nadeau, Chaoran Chen, Ivan Topolsky, Kim Philipp Jablonski, Lara Fuhrmann, Rebecca Denes, Mirjam Feldkamp, Ina Nissen, Natascha Santacroce, Elodie Burcklen, Christiane Beckmann, Maurice Redondo, Olivier Kobel, Christoph Noppen, Niko Beerenwinkel, Tanja Stadler |
| EPI_ISL_3295919 | Wales Specialist Virology Centre Sequencing lab: Pathogen Genomics Unit | Public Health Wales Microbiology Cardiff Wales Specialist Virology Centre | Catherine Moore, Johnathan Evans, Laura Gifford, Malorie Perry, Simon Cottrell, Angela Marchbank, Alec Birchley, Alexander Adams, Amy Gaskin, Bree Gatica-Wilcox, Jason Coombes, Joel Southgate, Lauren Gilbert, Lee Graham, Nicole Pacchiarini, Sara Kumziene-Summerhayes, Sarah Taylor, Sophie Jones, Sara Rey, Matthew Bull, Joanne Watkins, Sally Corden, Tom Connor |
| EPI_ISL_3295920 | Viollier AG | Department of Biosystems Science and Engineering, ETH Zürich | Christian Beisel, Sarah Nadeau, Chaoran Chen, Ivan Topolsky, Kim Philipp Jablonski, Lara Fuhrmann, Rebecca Denes, Mirjam Feldkamp, Ina Nissen, Natascha Santacroce, Elodie Burcklen, Christiane Beckmann, Maurice Redondo, Olivier Kobel, Christoph Noppen, Niko Beerenwinkel, Tanja Stadler |
| EPI_ISL_3295921 | Wales Specialist Virology Centre Sequencing lab: Pathogen Genomics Unit | Public Health Wales Microbiology Cardiff Wales Specialist Virology Centre | Catherine Moore, Johnathan Evans, Laura Gifford, Malorie Perry, Simon Cottrell, Angela Marchbank, Alec Birchley, Alexander Adams, Amy Gaskin, Bree Gatica-Wilcox, Jason Coombes, Joel Southgate, Lauren Gilbert, Lee Graham, Nicole Pacchiarini, Sara Kumziene-Summerhayes, Sarah Taylor, Sophie Jones, Sara Rey, Matthew Bull, Joanne Watkins, Sally Corden, Tom Connor |
| EPI_ISL_3295922 | Viollier AG | Department of Biosystems Science and Engineering, ETH Zürich | Christian Beisel, Sarah Nadeau, Chaoran Chen, Ivan Topolsky, Kim Philipp Jablonski, Lara Fuhrmann, Rebecca Denes, Mirjam Feldkamp, Ina Nissen, Natascha Santacroce, Elodie Burcklen, Christiane Beckmann, Maurice Redondo, Olivier Kobel, Christoph Noppen, Niko Beerenwinkel, Tanja Stadler |
| EPI_ISL_3295923 | Wales Specialist Virology Centre Sequencing lab: Pathogen Genomics Unit | Public Health Wales Microbiology Cardiff Wales Specialist Virology Centre | Catherine Moore, Johnathan Evans, Laura Gifford, Malorie Perry, Simon Cottrell, Angela Marchbank, Alec Birchley, Alexander Adams, Amy Gaskin, Bree Gatica-Wilcox, Jason Coombes, Joel Southgate, Lauren Gilbert, Lee Graham, Nicole Pacchiarini, Sara Kumziene-Summerhayes, Sarah Taylor, Sophie Jones, Sara Rey, Matthew Bull, Joanne Watkins, Sally Corden, Tom Connor |
| EPI_ISL_3295924 | Viollier AG | Department of Biosystems Science and Engineering, ETH Zürich | Christian Beisel, Sarah Nadeau, Chaoran Chen, Ivan Topolsky, Kim Philipp Jablonski, Lara Fuhrmann, Rebecca Denes, Mirjam Feldkamp, Ina Nissen, Natascha Santacroce, Elodie Burcklen, Christiane Beckmann, Maurice Redondo, Olivier Kobel, Christoph Noppen, Niko Beerenwinkel, Tanja Stadler |
| EPI_ISL_3295925, EPI_ISL_3295926 | Wales Specialist Virology Centre Sequencing lab: Pathogen Genomics Unit | Public Health Wales Microbiology Cardiff Wales Specialist Virology Centre | Catherine Moore, Johnathan Evans, Laura Gifford, Malorie Perry, Simon Cottrell, Angela Marchbank, Alec Birchley, Alexander Adams, Amy Gaskin, Bree Gatica-Wilcox, Jason Coombes, Joel Southgate, Lauren Gilbert, Lee Graham, Nicole Pacchiarini, Sara Kumziene-Summerhayes, Sarah Taylor, Sophie Jones, Sara Rey, Matthew Bull, Joanne Watkins, Sally Corden, Tom Connor |
| EPI_ISL_3295927 | Viollier AG | Department of Biosystems Science and Engineering, ETH Zürich | Christian Beisel, Sarah Nadeau, Chaoran Chen, Ivan Topolsky, Kim Philipp Jablonski, Lara Fuhrmann, Rebecca Denes, Mirjam Feldkamp, Ina Nissen, Natascha Santacroce, Elodie Burcklen, Christiane Beckmann, Maurice Redondo, Olivier Kobel, Christoph Noppen, Niko Beerenwinkel, Tanja Stadler |
| EPI_ISL_3295928 | Wales Specialist Virology Centre Sequencing lab: Pathogen Genomics Unit | Public Health Wales Microbiology Cardiff Wales Specialist Virology Centre | Catherine Moore, Johnathan Evans, Laura Gifford, Malorie Perry, Simon Cottrell, Angela Marchbank, Alec Birchley, Alexander Adams, Amy Gaskin, Bree Gatica-Wilcox, Jason Coombes, Joel Southgate, Lauren Gilbert, Lee Graham, Nicole Pacchiarini, Sara Kumziene-Summerhayes, Sarah Taylor, Sophie Jones, Sara Rey, Matthew Bull, Joanne Watkins, Sally Corden, Tom Connor |
| EPI_ISL_3295929 | Viollier AG | Department of Biosystems Science and Engineering, ETH Zürich | Christian Beisel, Sarah Nadeau, Chaoran Chen, Ivan Topolsky, Kim Philipp Jablonski, Lara Fuhrmann, Rebecca Denes, Mirjam Feldkamp, Ina Nissen, Natascha Santacroce, Elodie Burcklen, Christiane Beckmann, Maurice Redondo, Olivier Kobel, Christoph Noppen, Niko Beerenwinkel, Tanja Stadler |
| EPI_ISL_3295930 | Wales Specialist Virology Centre Sequencing lab: Pathogen Genomics Unit | Public Health Wales Microbiology Cardiff Wales Specialist Virology Centre | Catherine Moore, Johnathan Evans, Laura Gifford, Malorie Perry, Simon Cottrell, Angela Marchbank, Alec Birchley, Alexander Adams, Amy Gaskin, Bree Gatica-Wilcox, Jason Coombes, Joel Southgate, Lauren Gilbert, Lee Graham, Nicole Pacchiarini, Sara Kumziene-Summerhayes, Sarah Taylor, Sophie Jones, Sara Rey, Matthew Bull, Joanne Watkins, Sally Corden, Tom Connor |
| EPI_ISL_3295931 | Viollier AG | Department of Biosystems Science and Engineering, ETH Zürich | Christian Beisel, Sarah Nadeau, Chaoran Chen, Ivan Topolsky, Kim Philipp Jablonski, Lara Fuhrmann, Rebecca Denes, Mirjam Feldkamp, Ina Nissen, Natascha Santacroce, Elodie Burcklen, Christiane Beckmann, Maurice Redondo, Olivier Kobel, Christoph Noppen, Niko Beerenwinkel, Tanja Stadler |
| EPI_ISL_3295932 | Wales Specialist Virology Centre Sequencing lab: Pathogen Genomics Unit | Public Health Wales Microbiology Cardiff Wales Specialist Virology Centre | Catherine Moore, Johnathan Evans, Laura Gifford, Malorie Perry, Simon Cottrell, Angela Marchbank, Alec Birchley, Alexander Adams, Amy Gaskin, Bree Gatica-Wilcox, Jason Coombes, Joel Southgate, Lauren Gilbert, Lee Graham, Nicole Pacchiarini, Sara Kumziene-Summerhayes, Sarah Taylor, Sophie Jones, Sara Rey, Matthew Bull, Joanne Watkins, Sally Corden, Tom Connor |
| EPI_ISL_3295933 | Viollier AG | Department of Biosystems Science and Engineering, ETH Zürich | Christian Beisel, Sarah Nadeau, Chaoran Chen, Ivan Topolsky, Kim Philipp Jablonski, Lara Fuhrmann, Rebecca Denes, Mirjam Feldkamp, Ina Nissen, Natascha Santacroce, Elodie Burcklen, Christiane Beckmann, Maurice Redondo, Olivier Kobel, Christoph Noppen, Niko Beerenwinkel, Tanja Stadler |
| EPI_ISL_3295934 | Wales Specialist Virology Centre Sequencing lab: Pathogen Genomics Unit | Public Health Wales Microbiology Cardiff Wales Specialist Virology Centre | Catherine Moore, Johnathan Evans, Laura Gifford, Malorie Perry, Simon Cottrell, Angela Marchbank, Alec Birchley, Alexander Adams, Amy Gaskin, Bree Gatica-Wilcox, Jason Coombes, Joel Southgate, Lauren Gilbert, Lee Graham, Nicole Pacchiarini, Sara Kumziene-Summerhayes, Sarah Taylor, Sophie Jones, Sara Rey, Matthew Bull, Joanne Watkins, Sally Corden, Tom Connor |
| EPI_ISL_3295935 | Viollier AG | Department of Biosystems Science and Engineering, ETH Zürich | Christian Beisel, Sarah Nadeau, Chaoran Chen, Ivan Topolsky, Kim Philipp Jablonski, Lara Fuhrmann, Rebecca Denes, Mirjam Feldkamp, Ina Nissen, Natascha Santacroce, Elodie Burcklen, Christiane Beckmann, Maurice Redondo, Olivier Kobel, Christoph Noppen, Niko Beerenwinkel, Tanja Stadler |
| EPI_ISL_3295936 | Wales Specialist Virology Centre Sequencing lab: Pathogen Genomics Unit | Public Health Wales Microbiology Cardiff Wales Specialist Virology Centre | Catherine Moore, Johnathan Evans, Laura Gifford, Malorie Perry, Simon Cottrell, Angela Marchbank, Alec Birchley, Alexander Adams, Amy Gaskin, Bree Gatica-Wilcox, Jason Coombes, Joel Southgate, Lauren Gilbert, Lee Graham, Nicole Pacchiarini, Sara Kumziene-Summerhayes, Sarah Taylor, Sophie Jones, Sara Rey, Matthew Bull, Joanne Watkins, Sally Corden, Tom Connor |
| EPI_ISL_3295937 | Viollier AG | Department of Biosystems Science and Engineering, ETH Zürich | Christian Beisel, Sarah Nadeau, Chaoran Chen, Ivan Topolsky, Kim Philipp Jablonski, Lara Fuhrmann, Rebecca Denes, Mirjam Feldkamp, Ina Nissen, Natascha Santacroce, Elodie Burcklen, Christiane Beckmann, Maurice Redondo, Olivier Kobel, Christoph Noppen, Niko Beerenwinkel, Tanja Stadler |
| EPI_ISL_3295938, EPI_ISL_3295939 | Wales Specialist Virology Centre Sequencing lab: Pathogen Genomics Unit | Public Health Wales Microbiology Cardiff Wales Specialist Virology Centre | Catherine Moore, Johnathan Evans, Laura Gifford, Malorie Perry, Simon Cottrell, Angela Marchbank, Alec Birchley, Alexander Adams, Amy Gaskin, Bree Gatica-Wilcox, Jason Coombes, Joel Southgate, Lauren Gilbert, Lee Graham, Nicole Pacchiarini, Sara Kumziene-Summerhayes, Sarah Taylor, Sophie Jones, Sara Rey, Matthew Bull, Joanne Watkins, Sally Corden, Tom Connor |
| EPI_ISL_3295940 | Viollier AG | Department of Biosystems Science and Engineering, ETH Zürich | Christian Beisel, Sarah Nadeau, Chaoran Chen, Ivan Topolsky, Kim Philipp Jablonski, Lara Fuhrmann, Rebecca Denes, Mirjam Feldkamp, Ina Nissen, Natascha Santacroce, Elodie Burcklen, Christiane Beckmann, Maurice Redondo, Olivier Kobel, Christoph Noppen, Niko Beerenwinkel, Tanja Stadler |
| EPI_ISL_3295941 | Wales Specialist Virology Centre Sequencing lab: Pathogen Genomics Unit | Public Health Wales Microbiology Cardiff Wales Specialist Virology Centre | Catherine Moore, Johnathan Evans, Laura Gifford, Malorie Perry, Simon Cottrell, Angela Marchbank, Alec Birchley, Alexander Adams, Amy Gaskin, Bree Gatica-Wilcox, Jason Coombes, Joel Southgate, Lauren Gilbert, Lee Graham, Nicole Pacchiarini, Sara Kumziene-Summerhayes, Sarah Taylor, Sophie Jones, Sara Rey, Matthew Bull, Joanne Watkins, Sally Corden, Tom Connor |
| EPI_ISL_3295942 | Viollier AG | Department of Biosystems Science and Engineering, ETH Zürich | Christian Beisel, Sarah Nadeau, Chaoran Chen, Ivan Topolsky, Kim Philipp Jablonski, Lara Fuhrmann, Rebecca Denes, Mirjam Feldkamp, Ina Nissen, Natascha Santacroce, Elodie Burcklen, Christiane Beckmann, Maurice Redondo, Olivier Kobel, Christoph Noppen, Niko Beerenwinkel, Tanja Stadler |
| EPI_ISL_3295943 | Wales Specialist Virology Centre Sequencing lab: Pathogen Genomics Unit | Public Health Wales Microbiology Cardiff Wales Specialist Virology Centre | Catherine Moore, Johnathan Evans, Laura Gifford, Malorie Perry, Simon Cottrell, Angela Marchbank, Alec Birchley, Alexander Adams, Amy Gaskin, Bree Gatica-Wilcox, Jason Coombes, Joel Southgate, Lauren Gilbert, Lee Graham, Nicole Pacchiarini, Sara Kumziene-Summerhayes, Sarah Taylor, Sophie Jones, Sara Rey, Matthew Bull, Joanne Watkins, Sally Corden, Tom Connor |
| EPI_ISL_3295944, EPI_ISL_3295945 | Viollier AG | Department of Biosystems Science and Engineering, ETH Zürich | Christian Beisel, Sarah Nadeau, Chaoran Chen, Ivan Topolsky, Kim Philipp Jablonski, Lara Fuhrmann, Rebecca Denes, Mirjam Feldkamp, Ina Nissen, Natascha Santacroce, Elodie Burcklen, Christiane Beckmann, Maurice Redondo, Olivier Kobel, Christoph Noppen, Niko Beerenwinkel, Tanja Stadler |
| EPI_ISL_3295946, EPI_ISL_3295947 | Wales Specialist Virology Centre Sequencing lab: Pathogen Genomics Unit | Public Health Wales Microbiology Cardiff Wales Specialist Virology Centre | Catherine Moore, Johnathan Evans, Laura Gifford, Malorie Perry, Simon Cottrell, Angela Marchbank, Alec Birchley, Alexander Adams, Amy Gaskin, Bree Gatica-Wilcox, Jason Coombes, Joel Southgate, Lauren Gilbert, Lee Graham, Nicole Pacchiarini, Sara Kumziene-Summerhayes, Sarah Taylor, Sophie Jones, Sara Rey, Matthew Bull, Joanne Watkins, Sally Corden, Tom Connor |
| EPI_ISL_3295948 | Viollier AG | Department of Biosystems Science and Engineering, ETH Zürich | Christian Beisel, Sarah Nadeau, Chaoran Chen, Ivan Topolsky, Kim Philipp Jablonski, Lara Fuhrmann, Rebecca Denes, Mirjam Feldkamp, Ina Nissen, Natascha Santacroce, Elodie Burcklen, Christiane Beckmann, Maurice Redondo, Olivier Kobel, Christoph Noppen, Niko Beerenwinkel, Tanja Stadler |
| EPI_ISL_3295949 | Wales Specialist Virology Centre Sequencing lab: Pathogen Genomics Unit | Public Health Wales Microbiology Cardiff Wales Specialist Virology Centre | Catherine Moore, Johnathan Evans, Laura Gifford, Malorie Perry, Simon Cottrell, Angela Marchbank, Alec Birchley, Alexander Adams, Amy Gaskin, Bree Gatica-Wilcox, Jason Coombes, Joel Southgate, Lauren Gilbert, Lee Graham, Nicole Pacchiarini, Sara Kumziene-Summerhayes, Sarah Taylor, Sophie Jones, Sara Rey, Matthew Bull, Joanne Watkins, Sally Corden, Tom Connor |
| EPI_ISL_3295950 | Viollier AG | Department of Biosystems Science and Engineering, ETH Zürich | Christian Beisel, Sarah Nadeau, Chaoran Chen, Ivan Topolsky, Kim Philipp Jablonski, Lara Fuhrmann, Rebecca Denes, Mirjam Feldkamp, Ina Nissen, Natascha Santacroce, Elodie Burcklen, Christiane Beckmann, Maurice Redondo, Olivier Kobel, Christoph Noppen, Niko Beerenwinkel, Tanja Stadler |
| EPI_ISL_3295951 | Wales Specialist Virology Centre Sequencing lab: Pathogen Genomics Unit | Public Health Wales Microbiology Cardiff Wales Specialist Virology Centre | Catherine Moore, Johnathan Evans, Laura Gifford, Malorie Perry, Simon Cottrell, Angela Marchbank, Alec Birchley, Alexander Adams, Amy Gaskin, Bree Gatica-Wilcox, Jason Coombes, Joel Southgate, Lauren Gilbert, Lee Graham, Nicole Pacchiarini, Sara Kumziene-Summerhayes, Sarah Taylor, Sophie Jones, Sara Rey, Matthew Bull, Joanne Watkins, Sally Corden, Tom Connor |
| EPI_ISL_3295952 | Viollier AG | Department of Biosystems Science and Engineering, ETH Zürich | Christian Beisel, Sarah Nadeau, Chaoran Chen, Ivan Topolsky, Kim Philipp Jablonski, Lara Fuhrmann, Rebecca Denes, Mirjam Feldkamp, Ina Nissen, Natascha Santacroce, Elodie Burcklen, Christiane Beckmann, Maurice Redondo, Olivier Kobel, Christoph Noppen, Niko Beerenwinkel, Tanja Stadler |
| EPI_ISL_3295953 | Wales Specialist Virology Centre Sequencing lab: Pathogen Genomics Unit | Public Health Wales Microbiology Cardiff Wales Specialist Virology Centre | Catherine Moore, Johnathan Evans, Laura Gifford, Malorie Perry, Simon Cottrell, Angela Marchbank, Alec Birchley, Alexander Adams, Amy Gaskin, Bree Gatica-Wilcox, Jason Coombes, Joel Southgate, Lauren Gilbert, Lee Graham, Nicole Pacchiarini, Sara Kumziene-Summerhayes, Sarah Taylor, Sophie Jones, Sara Rey, Matthew Bull, Joanne Watkins, Sally Corden, Tom Connor |
| EPI_ISL_3295954 | Viollier AG | Department of Biosystems Science and Engineering, ETH Zürich | Christian Beisel, Sarah Nadeau, Chaoran Chen, Ivan Topolsky, Kim Philipp Jablonski, Lara Fuhrmann, Rebecca Denes, Mirjam Feldkamp, Ina Nissen, Natascha Santacroce, Elodie Burcklen, Christiane Beckmann, Maurice Redondo, Olivier Kobel, Christoph Noppen, Niko Beerenwinkel, Tanja Stadler |
| EPI_ISL_3295955 | Wales Specialist Virology Centre Sequencing lab: Pathogen Genomics Unit | Public Health Wales Microbiology Cardiff Wales Specialist Virology Centre | Catherine Moore, Johnathan Evans, Laura Gifford, Malorie Perry, Simon Cottrell, Angela Marchbank, Alec Birchley, Alexander Adams, Amy Gaskin, Bree Gatica-Wilcox, Jason Coombes, Joel Southgate, Lauren Gilbert, Lee Graham, Nicole Pacchiarini, Sara Kumziene-Summerhayes, Sarah Taylor, Sophie Jones, Sara Rey, Matthew Bull, Joanne Watkins, Sally Corden, Tom Connor |
| EPI_ISL_3295956 | Viollier AG | Department of Biosystems Science and Engineering, ETH Zürich | Christian Beisel, Sarah Nadeau, Chaoran Chen, Ivan Topolsky, Kim Philipp Jablonski, Lara Fuhrmann, Rebecca Denes, Mirjam Feldkamp, Ina Nissen, Natascha Santacroce, Elodie Burcklen, Christiane Beckmann, Maurice Redondo, Olivier Kobel, Christoph Noppen, Niko Beerenwinkel, Tanja Stadler |
| EPI_ISL_3295957 | Wales Specialist Virology Centre Sequencing lab: Pathogen Genomics Unit | Public Health Wales Microbiology Cardiff Wales Specialist Virology Centre | Catherine Moore, Johnathan Evans, Laura Gifford, Malorie Perry, Simon Cottrell, Angela Marchbank, Alec Birchley, Alexander Adams, Amy Gaskin, Bree Gatica-Wilcox, Jason Coombes, Joel Southgate, Lauren Gilbert, Lee Graham, Nicole Pacchiarini, Sara Kumziene-Summerhayes, Sarah Taylor, Sophie Jones, Sara Rey, Matthew Bull, Joanne Watkins, Sally Corden, Tom Connor |
| EPI_ISL_3295958 | Viollier AG | Department of Biosystems Science and Engineering, ETH Zürich | Christian Beisel, Sarah Nadeau, Chaoran Chen, Ivan Topolsky, Kim Philipp Jablonski, Lara Fuhrmann, Rebecca Denes, Mirjam Feldkamp, Ina Nissen, Natascha Santacroce, Elodie Burcklen, Christiane Beckmann, Maurice Redondo, Olivier Kobel, Christoph Noppen, Niko Beerenwinkel, Tanja Stadler |
| EPI_ISL_3295959 | Wales Specialist Virology Centre Sequencing lab: Pathogen Genomics Unit | Public Health Wales Microbiology Cardiff Wales Specialist Virology Centre | Catherine Moore, Johnathan Evans, Laura Gifford, Malorie Perry, Simon Cottrell, Angela Marchbank, Alec Birchley, Alexander Adams, Amy Gaskin, Bree Gatica-Wilcox, Jason Coombes, Joel Southgate, Lauren Gilbert, Lee Graham, Nicole Pacchiarini, Sara Kumziene-Summerhayes, Sarah Taylor, Sophie Jones, Sara Rey, Matthew Bull, Joanne Watkins, Sally Corden, Tom Connor |
| EPI_ISL_3295960 | Viollier AG | Department of Biosystems Science and Engineering, ETH Zürich | Christian Beisel, Sarah Nadeau, Chaoran Chen, Ivan Topolsky, Kim Philipp Jablonski, Lara Fuhrmann, Rebecca Denes, Mirjam Feldkamp, Ina Nissen, Natascha Santacroce, Elodie Burcklen, Christiane Beckmann, Maurice Redondo, Olivier Kobel, Christoph Noppen, Niko Beerenwinkel, Tanja Stadler |
| EPI_ISL_3295961 | Wales Specialist Virology Centre Sequencing lab: Pathogen Genomics Unit | Public Health Wales Microbiology Cardiff Wales Specialist Virology Centre | Catherine Moore, Johnathan Evans, Laura Gifford, Malorie Perry, Simon Cottrell, Angela Marchbank, Alec Birchley, Alexander Adams, Amy Gaskin, Bree Gatica-Wilcox, Jason Coombes, Joel Southgate, Lauren Gilbert, Lee Graham, Nicole Pacchiarini, Sara Kumziene-Summerhayes, Sarah Taylor, Sophie Jones, Sara Rey, Matthew Bull, Joanne Watkins, Sally Corden, Tom Connor |
| EPI_ISL_3295962 | Viollier AG | Department of Biosystems Science and Engineering, ETH Zürich | Christian Beisel, Sarah Nadeau, Chaoran Chen, Ivan Topolsky, Kim Philipp Jablonski, Lara Fuhrmann, Rebecca Denes, Mirjam Feldkamp, Ina Nissen, Natascha Santacroce, Elodie Burcklen, Christiane Beckmann, Maurice Redondo, Olivier Kobel, Christoph Noppen, Niko Beerenwinkel, Tanja Stadler |
| EPI_ISL_3295963 | Wales Specialist Virology Centre Sequencing lab: Pathogen Genomics Unit | Public Health Wales Microbiology Cardiff Wales Specialist Virology Centre | Catherine Moore, Johnathan Evans, Laura Gifford, Malorie Perry, Simon Cottrell, Angela Marchbank, Alec Birchley, Alexander Adams, Amy Gaskin, Bree Gatica-Wilcox, Jason Coombes, Joel Southgate, Lauren Gilbert, Lee Graham, Nicole Pacchiarini, Sara Kumziene-Summerhayes, Sarah Taylor, Sophie Jones, Sara Rey, Matthew Bull, Joanne Watkins, Sally Corden, Tom Connor |
| EPI_ISL_3295964 | Viollier AG | Department of Biosystems Science and Engineering, ETH Zürich | Christian Beisel, Sarah Nadeau, Chaoran Chen, Ivan Topolsky, Kim Philipp Jablonski, Lara Fuhrmann, Rebecca Denes, Mirjam Feldkamp, Ina Nissen, Natascha Santacroce, Elodie Burcklen, Christiane Beckmann, Maurice Redondo, Olivier Kobel, |

| | | | |
|---|---|---|---|
| | | | Christoph Noppen, Niko Beerenwinkel, Tanja Stadler |
| EPI_ISL_3295965 | Wales Specialist Virology Centre Sequencing lab: Pathogen Genomics Unit | Public Health Wales Microbiology Cardiff Wales Specialist Virology Centre | Catherine Moore, Johnathan Evans, Laura Gifford, Malorie Perry, Simon Cottrell, Angela Marchbank, Alec Birchley, Alexander Adams, Amy Gaskin, Bree Gatica-Wilcox, Jason Coombes, Joel Southgate, Lauren Gilbert, Lee Graham, Nicole Pacchiarini, Sara Kumziene-Summerhayes, Sarah Taylor, Sophie Jones, Sara Rey, Matthew Bull, Joanne Watkins, Sally Corden, Tom Connor |
| EPI_ISL_3295966 | Viollier AG | Department of Biosystems Science and Engineering, ETH Zürich | Christian Beisel, Sarah Nadeau, Chaoran Chen, Ivan Topolsky, Kim Philipp Jablonski, Lara Fuhrmann, Rebecca Denes, Mirjam Feldkamp, Ina Nissen, Natascha Santacroce, Elodie Burcklen, Christiane Beckmann, Maurice Redondo, Olivier Kobel, Christoph Noppen, Niko Beerenwinkel, Tanja Stadler |
| EPI_ISL_3295967 | Wales Specialist Virology Centre Sequencing lab: Pathogen Genomics Unit | Public Health Wales Microbiology Cardiff Wales Specialist Virology Centre | Catherine Moore, Johnathan Evans, Laura Gifford, Malorie Perry, Simon Cottrell, Angela Marchbank, Alec Birchley, Alexander Adams, Amy Gaskin, Bree Gatica-Wilcox, Jason Coombes, Joel Southgate, Lauren Gilbert, Lee Graham, Nicole Pacchiarini, Sara Kumziene-Summerhayes, Sarah Taylor, Sophie Jones, Sara Rey, Matthew Bull, Joanne Watkins, Sally Corden, Tom Connor |
| EPI_ISL_3295968 | Viollier AG | Department of Biosystems Science and Engineering, ETH Zürich | Christian Beisel, Sarah Nadeau, Chaoran Chen, Ivan Topolsky, Kim Philipp Jablonski, Lara Fuhrmann, Rebecca Denes, Mirjam Feldkamp, Ina Nissen, Natascha Santacroce, Elodie Burcklen, Christiane Beckmann, Maurice Redondo, Olivier Kobel, Christoph Noppen, Niko Beerenwinkel, Tanja Stadler |
| EPI_ISL_3295969, EPI_ISL_3295970 | Wales Specialist Virology Centre Sequencing lab: Pathogen Genomics Unit | Public Health Wales Microbiology Cardiff Wales Specialist Virology Centre | Catherine Moore, Johnathan Evans, Laura Gifford, Malorie Perry, Simon Cottrell, Angela Marchbank, Alec Birchley, Alexander Adams, Amy Gaskin, Bree Gatica-Wilcox, Jason Coombes, Joel Southgate, Lauren Gilbert, Lee Graham, Nicole Pacchiarini, Sara Kumziene-Summerhayes, Sarah Taylor, Sophie Jones, Sara Rey, Matthew Bull, Joanne Watkins, Sally Corden, Tom Connor |
| EPI_ISL_3295971 | Viollier AG | Department of Biosystems Science and Engineering, ETH Zürich | Christian Beisel, Sarah Nadeau, Chaoran Chen, Ivan Topolsky, Kim Philipp Jablonski, Lara Fuhrmann, Rebecca Denes, Mirjam Feldkamp, Ina Nissen, Natascha Santacroce, Elodie Burcklen, Christiane Beckmann, Maurice Redondo, Olivier Kobel, Christoph Noppen, Niko Beerenwinkel, Tanja Stadler |
| EPI_ISL_3295972 | Wales Specialist Virology Centre Sequencing lab: Pathogen Genomics Unit | Public Health Wales Microbiology Cardiff Wales Specialist Virology Centre | Catherine Moore, Johnathan Evans, Laura Gifford, Malorie Perry, Simon Cottrell, Angela Marchbank, Alec Birchley, Alexander Adams, Amy Gaskin, Bree Gatica-Wilcox, Jason Coombes, Joel Southgate, Lauren Gilbert, Lee Graham, Nicole Pacchiarini, Sara Kumziene-Summerhayes, Sarah Taylor, Sophie Jones, Sara Rey, Matthew Bull, Joanne Watkins, Sally Corden, Tom Connor |
| EPI_ISL_3295973 | Viollier AG | Department of Biosystems Science and Engineering, ETH Zürich | Christian Beisel, Sarah Nadeau, Chaoran Chen, Ivan Topolsky, Kim Philipp Jablonski, Lara Fuhrmann, Rebecca Denes, Mirjam Feldkamp, Ina Nissen, Natascha Santacroce, Elodie Burcklen, Christiane Beckmann, Maurice Redondo, Olivier Kobel, Christoph Noppen, Niko Beerenwinkel, Tanja Stadler |
| EPI_ISL_3295974 | Wales Specialist Virology Centre Sequencing lab: Pathogen Genomics Unit | Public Health Wales Microbiology Cardiff Wales Specialist Virology Centre | Catherine Moore, Johnathan Evans, Laura Gifford, Malorie Perry, Simon Cottrell, Angela Marchbank, Alec Birchley, Alexander Adams, Amy Gaskin, Bree Gatica-Wilcox, Jason Coombes, Joel Southgate, Lauren Gilbert, Lee Graham, Nicole Pacchiarini, Sara Kumziene-Summerhayes, Sarah Taylor, Sophie Jones, Sara Rey, Matthew Bull, Joanne Watkins, Sally Corden, Tom Connor |
| EPI_ISL_3295975 | Viollier AG | Department of Biosystems Science and Engineering, ETH Zürich | Christian Beisel, Sarah Nadeau, Chaoran Chen, Ivan Topolsky, Kim Philipp Jablonski, Lara Fuhrmann, Rebecca Denes, Mirjam Feldkamp, Ina Nissen, Natascha Santacroce, Elodie Burcklen, Christiane Beckmann, Maurice Redondo, Olivier Kobel, Christoph Noppen, Niko Beerenwinkel, Tanja Stadler |
| EPI_ISL_3295976 | Wales Specialist Virology Centre Sequencing lab: Pathogen Genomics Unit | Public Health Wales Microbiology Cardiff Wales Specialist Virology Centre | Catherine Moore, Johnathan Evans, Laura Gifford, Malorie Perry, Simon Cottrell, Angela Marchbank, Alec Birchley, Alexander Adams, Amy Gaskin, Bree Gatica-Wilcox, Jason Coombes, Joel Southgate, Lauren Gilbert, Lee Graham, Nicole Pacchiarini, Sara Kumziene-Summerhayes, Sarah Taylor, Sophie Jones, Sara Rey, Matthew Bull, Joanne Watkins, Sally Corden, Tom Connor |
| EPI_ISL_3295977 | Viollier AG | Department of Biosystems Science and Engineering, ETH Zürich | Christian Beisel, Sarah Nadeau, Chaoran Chen, Ivan Topolsky, Kim Philipp Jablonski, Lara Fuhrmann, Rebecca Denes, Mirjam Feldkamp, Ina Nissen, Natascha Santacroce, Elodie Burcklen, Christiane Beckmann, Maurice Redondo, Olivier Kobel, Christoph Noppen, Niko Beerenwinkel, Tanja Stadler |
| EPI_ISL_3295978 | Wales Specialist Virology Centre Sequencing lab: Pathogen Genomics Unit | Public Health Wales Microbiology Cardiff Wales Specialist Virology Centre | Catherine Moore, Johnathan Evans, Laura Gifford, Malorie Perry, Simon Cottrell, Angela Marchbank, Alec Birchley, Alexander Adams, Amy Gaskin, Bree Gatica-Wilcox, Jason Coombes, Joel Southgate, Lauren Gilbert, Lee Graham, Nicole Pacchiarini, Sara Kumziene-Summerhayes, Sarah Taylor, Sophie Jones, Sara Rey, Matthew Bull, Joanne Watkins, Sally Corden, Tom Connor |
| EPI_ISL_3295979 | Viollier AG | Department of Biosystems Science and Engineering, ETH Zürich | Christian Beisel, Sarah Nadeau, Chaoran Chen, Ivan Topolsky, Kim Philipp Jablonski, Lara Fuhrmann, Rebecca Denes, Mirjam Feldkamp, Ina Nissen, Natascha Santacroce, Elodie Burcklen, Christiane Beckmann, Maurice Redondo, Olivier Kobel, Christoph Noppen, Niko Beerenwinkel, Tanja Stadler |
| EPI_ISL_3295980 | Wales Specialist Virology Centre Sequencing lab: Pathogen Genomics Unit | Public Health Wales Microbiology Cardiff Wales Specialist Virology Centre | Catherine Moore, Johnathan Evans, Laura Gifford, Malorie Perry, Simon Cottrell, Angela Marchbank, Alec Birchley, Alexander Adams, Amy Gaskin, Bree Gatica-Wilcox, Jason Coombes, Joel Southgate, Lauren Gilbert, Lee Graham, Nicole Pacchiarini, Sara Kumziene-Summerhayes, Sarah Taylor, Sophie Jones, Sara Rey, Matthew Bull, Joanne Watkins, Sally Corden, Tom Connor |
| EPI_ISL_3295981 | Viollier AG | Department of Biosystems Science and Engineering, ETH Zürich | Christian Beisel, Sarah Nadeau, Chaoran Chen, Ivan Topolsky, Kim Philipp Jablonski, Lara Fuhrmann, Rebecca Denes, Mirjam Feldkamp, Ina Nissen, Natascha Santacroce, Elodie Burcklen, Christiane Beckmann, Maurice Redondo, Olivier Kobel, Christoph Noppen, Niko Beerenwinkel, Tanja Stadler |
| EPI_ISL_3295982 | Wales Specialist Virology Centre Sequencing lab: Pathogen Genomics Unit | Public Health Wales Microbiology Cardiff Wales Specialist Virology Centre | Catherine Moore, Johnathan Evans, Laura Gifford, Malorie Perry, Simon Cottrell, Angela Marchbank, Alec Birchley, Alexander Adams, Amy Gaskin, Bree Gatica-Wilcox, Jason Coombes, Joel Southgate, Lauren Gilbert, Lee Graham, Nicole Pacchiarini, Sara Kumziene-Summerhayes, Sarah Taylor, Sophie Jones, Sara Rey, Matthew Bull, Joanne Watkins, Sally Corden, Tom Connor |
| EPI_ISL_3295983 | Viollier AG | Department of Biosystems Science and Engineering, ETH Zürich | Christian Beisel, Sarah Nadeau, Chaoran Chen, Ivan Topolsky, Kim Philipp Jablonski, Lara Fuhrmann, Rebecca Denes, Mirjam Feldkamp, Ina Nissen, Natascha Santacroce, Elodie Burcklen, Christiane Beckmann, Maurice Redondo, Olivier Kobel, Christoph Noppen, Niko Beerenwinkel, Tanja Stadler |
| EPI_ISL_3295984 | Wales Specialist Virology Centre Sequencing lab: Pathogen Genomics Unit | Public Health Wales Microbiology Cardiff Wales Specialist Virology Centre | Catherine Moore, Johnathan Evans, Laura Gifford, Malorie Perry, Simon Cottrell, Angela Marchbank, Alec Birchley, Alexander Adams, Amy Gaskin, Bree Gatica-Wilcox, Jason Coombes, Joel Southgate, Lauren Gilbert, Lee Graham, Nicole Pacchiarini, Sara Kumziene-Summerhayes, Sarah Taylor, Sophie Jones, Sara Rey, Matthew Bull, Joanne Watkins, Sally Corden, Tom Connor |
| EPI_ISL_3295985 | Viollier AG | Department of Biosystems Science and Engineering, ETH Zürich | Christian Beisel, Sarah Nadeau, Chaoran Chen, Ivan Topolsky, Kim Philipp Jablonski, Lara Fuhrmann, Rebecca Denes, Mirjam Feldkamp, Ina Nissen, Natascha Santacroce, Elodie Burcklen, Christiane Beckmann, Maurice Redondo, Olivier Kobel, Christoph Noppen, Niko Beerenwinkel, Tanja Stadler |
| EPI_ISL_3295986, EPI_ISL_3295987 | Wales Specialist Virology Centre Sequencing lab: Pathogen Genomics Unit | Public Health Wales Microbiology Cardiff Wales Specialist Virology Centre | Catherine Moore, Johnathan Evans, Laura Gifford, Malorie Perry, Simon Cottrell, Angela Marchbank, Alec Birchley, Alexander Adams, Amy Gaskin, Bree Gatica-Wilcox, Jason Coombes, Joel Southgate, Lauren Gilbert, Lee Graham, Nicole Pacchiarini, Sara Kumziene-Summerhayes, Sarah Taylor, Sophie Jones, Sara Rey, Matthew Bull, Joanne Watkins, Sally Corden, Tom Connor |
| EPI_ISL_3295988 | Viollier AG | Department of Biosystems Science and Engineering, ETH Zürich | Christian Beisel, Sarah Nadeau, Chaoran Chen, Ivan Topolsky, Kim Philipp Jablonski, Lara Fuhrmann, Rebecca Denes, Mirjam Feldkamp, Ina Nissen, Natascha Santacroce, Elodie Burcklen, Christiane Beckmann, Maurice Redondo, Olivier Kobel, Christoph Noppen, Niko Beerenwinkel, Tanja Stadler |
| EPI_ISL_3295989 | Wales Specialist Virology Centre Sequencing lab: Pathogen Genomics Unit | Public Health Wales Microbiology Cardiff Wales Specialist Virology Centre | Catherine Moore, Johnathan Evans, Laura Gifford, Malorie Perry, Simon Cottrell, Angela Marchbank, Alec Birchley, Alexander Adams, Amy Gaskin, Bree Gatica-Wilcox, Jason Coombes, Joel Southgate, Lauren Gilbert, Lee Graham, Nicole Pacchiarini, Sara Kumziene-Summerhayes, Sarah Taylor, Sophie Jones, Sara Rey, Matthew Bull, Joanne Watkins, Sally Corden, Tom Connor |
| EPI_ISL_3295990 | Viollier AG | Department of Biosystems Science and Engineering, ETH Zürich | Christian Beisel, Sarah Nadeau, Chaoran Chen, Ivan Topolsky, Kim Philipp Jablonski, Lara Fuhrmann, Rebecca Denes, Mirjam Feldkamp, Ina Nissen, Natascha Santacroce, Elodie Burcklen, Christiane Beckmann, Maurice Redondo, Olivier Kobel, Christoph Noppen, Niko Beerenwinkel, Tanja Stadler |
| EPI_ISL_3295991 | Wales Specialist Virology Centre Sequencing lab: Pathogen Genomics Unit | Public Health Wales Microbiology Cardiff Wales Specialist Virology Centre | Catherine Moore, Johnathan Evans, Laura Gifford, Malorie Perry, Simon Cottrell, Angela Marchbank, Alec Birchley, Alexander Adams, Amy Gaskin, Bree Gatica-Wilcox, Jason Coombes, Joel Southgate, Lauren Gilbert, Lee Graham, Nicole Pacchiarini, Sara Kumziene-Summerhayes, Sarah Taylor, Sophie Jones, Sara Rey, Matthew Bull, Joanne Watkins, Sally Corden, Tom Connor |
| EPI_ISL_3295992 | Viollier AG | Department of Biosystems Science and Engineering, ETH Zürich | Christian Beisel, Sarah Nadeau, Chaoran Chen, Ivan Topolsky, Kim Philipp Jablonski, Lara Fuhrmann, Rebecca Denes, Mirjam Feldkamp, Ina Nissen, Natascha Santacroce, Elodie Burcklen, Christiane Beckmann, Maurice Redondo, Olivier Kobel, Christoph Noppen, Niko Beerenwinkel, Tanja Stadler |
| EPI_ISL_3295993 | Wales Specialist Virology Centre Sequencing lab: Pathogen Genomics Unit | Public Health Wales Microbiology Cardiff Wales Specialist Virology Centre | Catherine Moore, Johnathan Evans, Laura Gifford, Malorie Perry, Simon Cottrell, Angela Marchbank, Alec Birchley, Alexander Adams, Amy Gaskin, Bree Gatica-Wilcox, Jason Coombes, Joel Southgate, Lauren Gilbert, Lee Graham, Nicole Pacchiarini, Sara Kumziene-Summerhayes, Sarah Taylor, Sophie Jones, Sara Rey, Matthew Bull, Joanne Watkins, Sally Corden, Tom Connor |
| EPI_ISL_3295994 | Viollier AG | Department of Biosystems Science and Engineering, ETH Zürich | Christian Beisel, Sarah Nadeau, Chaoran Chen, Ivan Topolsky, Kim Philipp Jablonski, Lara Fuhrmann, Rebecca Denes, Mirjam Feldkamp, Ina Nissen, Natascha Santacroce, Elodie Burcklen, Christiane Beckmann, Maurice Redondo, Olivier Kobel, Christoph Noppen, Niko Beerenwinkel, Tanja Stadler |
| EPI_ISL_3295995 | Wales Specialist Virology Centre Sequencing lab: Pathogen Genomics Unit | Public Health Wales Microbiology Cardiff Wales Specialist Virology Centre | Catherine Moore, Johnathan Evans, Laura Gifford, Malorie Perry, Simon Cottrell, Angela Marchbank, Alec Birchley, Alexander Adams, Amy Gaskin, Bree Gatica-Wilcox, Jason Coombes, Joel Southgate, Lauren Gilbert, Lee Graham, Nicole Pacchiarini, Sara Kumziene-Summerhayes, Sarah Taylor, Sophie Jones, Sara Rey, Matthew Bull, Joanne Watkins, Sally Corden, Tom Connor |
| EPI_ISL_3295996 | Viollier AG | Department of Biosystems Science and Engineering, ETH Zürich | Christian Beisel, Sarah Nadeau, Chaoran Chen, Ivan Topolsky, Kim Philipp Jablonski, Lara Fuhrmann, Rebecca Denes, Mirjam Feldkamp, Ina Nissen, Natascha Santacroce, Elodie Burcklen, Christiane Beckmann, Maurice Redondo, Olivier Kobel, Christoph Noppen, Niko Beerenwinkel, Tanja Stadler |
| EPI_ISL_3295997 | Wales Specialist Virology Centre Sequencing lab: Pathogen Genomics Unit | Public Health Wales Microbiology Cardiff Wales Specialist Virology Centre | Catherine Moore, Johnathan Evans, Laura Gifford, Malorie Perry, Simon Cottrell, Angela Marchbank, Alec Birchley, Alexander Adams, Amy Gaskin, Bree Gatica-Wilcox, Jason Coombes, Joel Southgate, Lauren Gilbert, Lee Graham, Nicole Pacchiarini, Sara Kumziene-Summerhayes, Sarah Taylor, Sophie Jones, Sara Rey, Matthew Bull, Joanne Watkins, Sally Corden, Tom Connor |
| EPI_ISL_3295998 | Viollier AG | Department of Biosystems Science and Engineering, ETH Zürich | Christian Beisel, Sarah Nadeau, Chaoran Chen, Ivan Topolsky, Kim Philipp Jablonski, Lara Fuhrmann, Rebecca Denes, Mirjam Feldkamp, Ina Nissen, Natascha Santacroce, Elodie Burcklen, Christiane Beckmann, Maurice Redondo, Olivier Kobel, Christoph Noppen, Niko Beerenwinkel, Tanja Stadler |
| EPI_ISL_3295999 | Wales Specialist Virology Centre Sequencing lab: Pathogen Genomics Unit | Public Health Wales Microbiology Cardiff Wales Specialist Virology Centre | Catherine Moore, Johnathan Evans, Laura Gifford, Malorie Perry, Simon Cottrell, Angela Marchbank, Alec Birchley, Alexander Adams, Amy Gaskin, Bree Gatica-Wilcox, Jason Coombes, Joel Southgate, Lauren Gilbert, Lee Graham, Nicole Pacchiarini, Sara Kumziene-Summerhayes, Sarah Taylor, Sophie Jones, Sara Rey, Matthew Bull, Joanne Watkins, Sally Corden, Tom Connor |
| EPI_ISL_3296000 | Viollier AG | Department of Biosystems Science and Engineering, ETH Zürich | Christian Beisel, Sarah Nadeau, Chaoran Chen, Ivan Topolsky, Kim Philipp Jablonski, Lara Fuhrmann, Rebecca Denes, Mirjam Feldkamp, Ina Nissen, Natascha Santacroce, Elodie Burcklen, Christiane Beckmann, Maurice Redondo, Olivier Kobel, Christoph Noppen, Niko Beerenwinkel, Tanja Stadler |
| EPI_ISL_3296001, EPI_ISL_3296002 | Wales Specialist Virology Centre Sequencing lab: Pathogen Genomics Unit | Public Health Wales Microbiology Cardiff Wales Specialist Virology Centre | Catherine Moore, Johnathan Evans, Laura Gifford, Malorie Perry, Simon Cottrell, Angela Marchbank, Alec Birchley, Alexander Adams, Amy Gaskin, Bree Gatica-Wilcox, Jason Coombes, Joel Southgate, Lauren Gilbert, Lee Graham, Nicole Pacchiarini, Sara Kumziene-Summerhayes, Sarah Taylor, Sophie Jones, Sara Rey, Matthew Bull, Joanne Watkins, Sally Corden, Tom Connor |
| EPI_ISL_3296003 | Viollier AG | Department of Biosystems Science and Engineering, ETH Zürich | Christian Beisel, Sarah Nadeau, Chaoran Chen, Ivan Topolsky, Kim Philipp Jablonski, Lara Fuhrmann, Rebecca Denes, Mirjam Feldkamp, Ina Nissen, Natascha Santacroce, Elodie Burcklen, Christiane Beckmann, Maurice Redondo, Olivier Kobel, Christoph Noppen, Niko Beerenwinkel, Tanja Stadler |
| EPI_ISL_3296004 | Wales Specialist Virology Centre Sequencing lab: Pathogen Genomics Unit | Public Health Wales Microbiology Cardiff Wales Specialist Virology Centre | Catherine Moore, Johnathan Evans, Laura Gifford, Malorie Perry, Simon Cottrell, Angela Marchbank, Alec Birchley, Alexander Adams, Amy Gaskin, Bree Gatica-Wilcox, Jason Coombes, Joel Southgate, Lauren Gilbert, Lee Graham, Nicole Pacchiarini, Sara Kumziene-Summerhayes, Sarah Taylor, Sophie Jones, Sara Rey, Matthew Bull, Joanne Watkins, Sally Corden, Tom Connor |
| EPI_ISL_3296005 | Viollier AG | Department of Biosystems Science and Engineering, ETH Zürich | Christian Beisel, Sarah Nadeau, Chaoran Chen, Ivan Topolsky, Kim Philipp Jablonski, Lara Fuhrmann, Rebecca Denes, Mirjam Feldkamp, Ina Nissen, Natascha Santacroce, Elodie Burcklen, Christiane Beckmann, Maurice Redondo, Olivier Kobel, Christoph Noppen, Niko Beerenwinkel, Tanja Stadler |
| EPI_ISL_3296006 | Wales Specialist Virology Centre Sequencing lab: Pathogen Genomics Unit | Public Health Wales Microbiology Cardiff Wales Specialist Virology Centre | Catherine Moore, Johnathan Evans, Laura Gifford, Malorie Perry, Simon Cottrell, Angela Marchbank, Alec Birchley, Alexander Adams, Amy Gaskin, Bree Gatica-Wilcox, Jason Coombes, Joel Southgate, Lauren Gilbert, Lee Graham, Nicole Pacchiarini, Sara Kumziene-Summerhayes, Sarah Taylor, Sophie Jones, Sara Rey, Matthew Bull, Joanne Watkins, Sally Corden, Tom Connor |
| EPI_ISL_3296007 | Viollier AG | Department of Biosystems Science and Engineering, ETH Zürich | Christian Beisel, Sarah Nadeau, Chaoran Chen, Ivan Topolsky, Kim Philipp Jablonski, Lara Fuhrmann, Rebecca Denes, Mirjam Feldkamp, Ina Nissen, Natascha Santacroce, Elodie Burcklen, Christiane Beckmann, Maurice Redondo, Olivier Kobel, Christoph Noppen, Niko Beerenwinkel, Tanja Stadler |
| EPI_ISL_3296008 | Wales Specialist Virology Centre Sequencing lab: Pathogen Genomics Unit | Public Health Wales Microbiology Cardiff Wales Specialist Virology Centre | Catherine Moore, Johnathan Evans, Laura Gifford, Malorie Perry, Simon Cottrell, Angela Marchbank, Alec Birchley, Alexander Adams, Amy Gaskin, Bree Gatica-Wilcox, Jason Coombes, Joel Southgate, Lauren Gilbert, Lee Graham, Nicole Pacchiarini, Sara Kumziene-Summerhayes, Sarah Taylor, Sophie Jones, Sara Rey, Matthew Bull, Joanne Watkins, Sally Corden, Tom Connor |
| EPI_ISL_3296009 | Viollier AG | Department of Biosystems Science and Engineering, ETH Zürich | Christian Beisel, Sarah Nadeau, Chaoran Chen, Ivan Topolsky, Kim Philipp Jablonski, Lara Fuhrmann, Rebecca Denes, Mirjam Feldkamp, Ina Nissen, Natascha Santacroce, Elodie Burcklen, Christiane Beckmann, Maurice Redondo, Olivier Kobel, Christoph Noppen, Niko Beerenwinkel, Tanja Stadler |
| EPI_ISL_3296010, EPI_ISL_3296012 | Wales Specialist Virology Centre Sequencing lab: Pathogen Genomics Unit | Public Health Wales Microbiology Cardiff Wales Specialist Virology Centre | Catherine Moore, Johnathan Evans, Laura Gifford, Malorie Perry, Simon Cottrell, Angela Marchbank, Alec Birchley, Alexander Adams, Amy Gaskin, Bree Gatica-Wilcox, Jason Coombes, Joel Southgate, Lauren Gilbert, Lee Graham, Nicole Pacchiarini, Sara Kumziene-Summerhayes, Sarah Taylor, Sophie Jones, Sara Rey, Matthew Bull, Joanne Watkins, Sally Corden, Tom Connor |
| EPI_ISL_3296013 | Viollier AG | Department of Biosystems Science and Engineering, ETH Zürich | Christian Beisel, Sarah Nadeau, Chaoran Chen, Ivan Topolsky, Kim Philipp Jablonski, Lara Fuhrmann, Rebecca Denes, Mirjam Feldkamp, Ina Nissen, Natascha Santacroce, Elodie Burcklen, Christiane Beckmann, Maurice Redondo, Olivier Kobel, Christoph Noppen, Niko Beerenwinkel, Tanja Stadler |
| EPI_ISL_3296014 | Wales Specialist Virology Centre Sequencing lab: Pathogen | Public Health Wales Microbiology Cardiff Wales Specialist Virology | Catherine Moore, Johnathan Evans, Laura Gifford, Malorie Perry, Simon Cottrell, Angela Marchbank, Alec Birchley, Alexander Adams, Amy Gaskin, Bree Gatica-Wilcox, Jason Coombes, Joel Southgate, Lauren Gilbert, Lee Graham, Nicole |

| | | | |
|---|---|---|---|
| | Genomics Unit | Centre | Pacchiarini, Sara Kumziene-Summerhayes, Sarah Taylor, Sophie Jones, Sara Rey, Matthew Bull, Joanne Watkins, Sally Corden, Tom Connor |
| EPI_ISL_3296015 | Viollier AG | Department of Biosystems Science and Engineering, ETH Zürich | Christian Beisel, Sarah Nadeau, Chaoran Chen, Ivan Topolsky, Kim Philipp Jablonski, Lara Fuhrmann, Rebecca Denes, Mirjam Feldkamp, Ina Nissen, Natascha Santacroce, Elodie Burcklen, Christiane Beckmann, Maurice Redondo, Olivier Kobel, Christoph Noppen, Niko Beerenwinkel, Tanja Stadler |
| EPI_ISL_3296016 | Wales Specialist Virology Centre Sequencing lab: Pathogen Genomics Unit | Public Health Wales Microbiology Cardiff Wales Specialist Virology Centre | Catherine Moore, Johnathan Evans, Laura Gifford, Malorie Perry, Simon Cottrell, Angela Marchbank, Alec Birchley, Alexander Adams, Amy Gaskin, Bree Gatica-Wilcox, Jason Coombes, Joel Southgate, Lauren Gilbert, Lee Graham, Nicole Pacchiarini, Sara Kumziene-Summerhayes, Sarah Taylor, Sophie Jones, Sara Rey, Matthew Bull, Joanne Watkins, Sally Corden, Tom Connor |
| EPI_ISL_3296017 | Viollier AG | Department of Biosystems Science and Engineering, ETH Zürich | Christian Beisel, Sarah Nadeau, Chaoran Chen, Ivan Topolsky, Kim Philipp Jablonski, Lara Fuhrmann, Rebecca Denes, Mirjam Feldkamp, Ina Nissen, Natascha Santacroce, Elodie Burcklen, Christiane Beckmann, Maurice Redondo, Olivier Kobel, Christoph Noppen, Niko Beerenwinkel, Tanja Stadler |
| EPI_ISL_3296018 | Wales Specialist Virology Centre Sequencing lab: Pathogen Genomics Unit | Public Health Wales Microbiology Cardiff Wales Specialist Virology Centre | Catherine Moore, Johnathan Evans, Laura Gifford, Malorie Perry, Simon Cottrell, Angela Marchbank, Alec Birchley, Alexander Adams, Amy Gaskin, Bree Gatica-Wilcox, Jason Coombes, Joel Southgate, Lauren Gilbert, Lee Graham, Nicole Pacchiarini, Sara Kumziene-Summerhayes, Sarah Taylor, Sophie Jones, Sara Rey, Matthew Bull, Joanne Watkins, Sally Corden, Tom Connor |
| EPI_ISL_3296019 | Viollier AG | Department of Biosystems Science and Engineering, ETH Zürich | Christian Beisel, Sarah Nadeau, Chaoran Chen, Ivan Topolsky, Kim Philipp Jablonski, Lara Fuhrmann, Rebecca Denes, Mirjam Feldkamp, Ina Nissen, Natascha Santacroce, Elodie Burcklen, Christiane Beckmann, Maurice Redondo, Olivier Kobel, Christoph Noppen, Niko Beerenwinkel, Tanja Stadler |
| EPI_ISL_3296020 | Wales Specialist Virology Centre Sequencing lab: Pathogen Genomics Unit | Public Health Wales Microbiology Cardiff Wales Specialist Virology Centre | Catherine Moore, Johnathan Evans, Laura Gifford, Malorie Perry, Simon Cottrell, Angela Marchbank, Alec Birchley, Alexander Adams, Amy Gaskin, Bree Gatica-Wilcox, Jason Coombes, Joel Southgate, Lauren Gilbert, Lee Graham, Nicole Pacchiarini, Sara Kumziene-Summerhayes, Sarah Taylor, Sophie Jones, Sara Rey, Matthew Bull, Joanne Watkins, Sally Corden, Tom Connor |
| EPI_ISL_3296021 | Viollier AG | Department of Biosystems Science and Engineering, ETH Zürich | Christian Beisel, Sarah Nadeau, Chaoran Chen, Ivan Topolsky, Kim Philipp Jablonski, Lara Fuhrmann, Rebecca Denes, Mirjam Feldkamp, Ina Nissen, Natascha Santacroce, Elodie Burcklen, Christiane Beckmann, Maurice Redondo, Olivier Kobel, Christoph Noppen, Niko Beerenwinkel, Tanja Stadler |
| EPI_ISL_3296022 | Wales Specialist Virology Centre Sequencing lab: Pathogen Genomics Unit | Public Health Wales Microbiology Cardiff Wales Specialist Virology Centre | Catherine Moore, Johnathan Evans, Laura Gifford, Malorie Perry, Simon Cottrell, Angela Marchbank, Alec Birchley, Alexander Adams, Amy Gaskin, Bree Gatica-Wilcox, Jason Coombes, Joel Southgate, Lauren Gilbert, Lee Graham, Nicole Pacchiarini, Sara Kumziene-Summerhayes, Sarah Taylor, Sophie Jones, Sara Rey, Matthew Bull, Joanne Watkins, Sally Corden, Tom Connor |
| EPI_ISL_3296023 | Viollier AG | Department of Biosystems Science and Engineering, ETH Zürich | Christian Beisel, Sarah Nadeau, Chaoran Chen, Ivan Topolsky, Kim Philipp Jablonski, Lara Fuhrmann, Rebecca Denes, Mirjam Feldkamp, Ina Nissen, Natascha Santacroce, Elodie Burcklen, Christiane Beckmann, Maurice Redondo, Olivier Kobel, Christoph Noppen, Niko Beerenwinkel, Tanja Stadler |
| EPI_ISL_3296024 | Wales Specialist Virology Centre Sequencing lab: Pathogen Genomics Unit | Public Health Wales Microbiology Cardiff Wales Specialist Virology Centre | Catherine Moore, Johnathan Evans, Laura Gifford, Malorie Perry, Simon Cottrell, Angela Marchbank, Alec Birchley, Alexander Adams, Amy Gaskin, Bree Gatica-Wilcox, Jason Coombes, Joel Southgate, Lauren Gilbert, Lee Graham, Nicole Pacchiarini, Sara Kumziene-Summerhayes, Sarah Taylor, Sophie Jones, Sara Rey, Matthew Bull, Joanne Watkins, Sally Corden, Tom Connor |
| EPI_ISL_3296025 | Viollier AG | Department of Biosystems Science and Engineering, ETH Zürich | Christian Beisel, Sarah Nadeau, Chaoran Chen, Ivan Topolsky, Kim Philipp Jablonski, Lara Fuhrmann, Rebecca Denes, Mirjam Feldkamp, Ina Nissen, Natascha Santacroce, Elodie Burcklen, Christiane Beckmann, Maurice Redondo, Olivier Kobel, Christoph Noppen, Niko Beerenwinkel, Tanja Stadler |
| EPI_ISL_3296026 | Wales Specialist Virology Centre Sequencing lab: Pathogen Genomics Unit | Public Health Wales Microbiology Cardiff Wales Specialist Virology Centre | Catherine Moore, Johnathan Evans, Laura Gifford, Malorie Perry, Simon Cottrell, Angela Marchbank, Alec Birchley, Alexander Adams, Amy Gaskin, Bree Gatica-Wilcox, Jason Coombes, Joel Southgate, Lauren Gilbert, Lee Graham, Nicole Pacchiarini, Sara Kumziene-Summerhayes, Sarah Taylor, Sophie Jones, Sara Rey, Matthew Bull, Joanne Watkins, Sally Corden, Tom Connor |
| EPI_ISL_3296027 | Viollier AG | Department of Biosystems Science and Engineering, ETH Zürich | Christian Beisel, Sarah Nadeau, Chaoran Chen, Ivan Topolsky, Kim Philipp Jablonski, Lara Fuhrmann, Rebecca Denes, Mirjam Feldkamp, Ina Nissen, Natascha Santacroce, Elodie Burcklen, Christiane Beckmann, Maurice Redondo, Olivier Kobel, Christoph Noppen, Niko Beerenwinkel, Tanja Stadler |
| EPI_ISL_3296028 | Wales Specialist Virology Centre Sequencing lab: Pathogen Genomics Unit | Public Health Wales Microbiology Cardiff Wales Specialist Virology Centre | Catherine Moore, Johnathan Evans, Laura Gifford, Malorie Perry, Simon Cottrell, Angela Marchbank, Alec Birchley, Alexander Adams, Amy Gaskin, Bree Gatica-Wilcox, Jason Coombes, Joel Southgate, Lauren Gilbert, Lee Graham, Nicole Pacchiarini, Sara Kumziene-Summerhayes, Sarah Taylor, Sophie Jones, Sara Rey, Matthew Bull, Joanne Watkins, Sally Corden, Tom Connor |
| EPI_ISL_3296029 | Viollier AG | Department of Biosystems Science and Engineering, ETH Zürich | Christian Beisel, Sarah Nadeau, Chaoran Chen, Ivan Topolsky, Kim Philipp Jablonski, Lara Fuhrmann, Rebecca Denes, Mirjam Feldkamp, Ina Nissen, Natascha Santacroce, Elodie Burcklen, Christiane Beckmann, Maurice Redondo, Olivier Kobel, Christoph Noppen, Niko Beerenwinkel, Tanja Stadler |
| EPI_ISL_3296030 | Wales Specialist Virology Centre Sequencing lab: Pathogen Genomics Unit | Public Health Wales Microbiology Cardiff Wales Specialist Virology Centre | Catherine Moore, Johnathan Evans, Laura Gifford, Malorie Perry, Simon Cottrell, Angela Marchbank, Alec Birchley, Alexander Adams, Amy Gaskin, Bree Gatica-Wilcox, Jason Coombes, Joel Southgate, Lauren Gilbert, Lee Graham, Nicole Pacchiarini, Sara Kumziene-Summerhayes, Sarah Taylor, Sophie Jones, Sara Rey, Matthew Bull, Joanne Watkins, Sally Corden, Tom Connor |
| EPI_ISL_3296031 | Viollier AG | Department of Biosystems Science and Engineering, ETH Zürich | Christian Beisel, Sarah Nadeau, Chaoran Chen, Ivan Topolsky, Kim Philipp Jablonski, Lara Fuhrmann, Rebecca Denes, Mirjam Feldkamp, Ina Nissen, Natascha Santacroce, Elodie Burcklen, Christiane Beckmann, Maurice Redondo, Olivier Kobel, Christoph Noppen, Niko Beerenwinkel, Tanja Stadler |
| EPI_ISL_3296032 | Wales Specialist Virology Centre Sequencing lab: Pathogen Genomics Unit | Public Health Wales Microbiology Cardiff Wales Specialist Virology Centre | Catherine Moore, Johnathan Evans, Laura Gifford, Malorie Perry, Simon Cottrell, Angela Marchbank, Alec Birchley, Alexander Adams, Amy Gaskin, Bree Gatica-Wilcox, Jason Coombes, Joel Southgate, Lauren Gilbert, Lee Graham, Nicole Pacchiarini, Sara Kumziene-Summerhayes, Sarah Taylor, Sophie Jones, Sara Rey, Matthew Bull, Joanne Watkins, Sally Corden, Tom Connor |
| EPI_ISL_3296033 | Viollier AG | Department of Biosystems Science and Engineering, ETH Zürich | Christian Beisel, Sarah Nadeau, Chaoran Chen, Ivan Topolsky, Kim Philipp Jablonski, Lara Fuhrmann, Rebecca Denes, Mirjam Feldkamp, Ina Nissen, Natascha Santacroce, Elodie Burcklen, Christiane Beckmann, Maurice Redondo, Olivier Kobel, Christoph Noppen, Niko Beerenwinkel, Tanja Stadler |
| EPI_ISL_3296034 | Wales Specialist Virology Centre Sequencing lab: Pathogen Genomics Unit | Public Health Wales Microbiology Cardiff Wales Specialist Virology Centre | Catherine Moore, Johnathan Evans, Laura Gifford, Malorie Perry, Simon Cottrell, Angela Marchbank, Alec Birchley, Alexander Adams, Amy Gaskin, Bree Gatica-Wilcox, Jason Coombes, Joel Southgate, Lauren Gilbert, Lee Graham, Nicole Pacchiarini, Sara Kumziene-Summerhayes, Sarah Taylor, Sophie Jones, Sara Rey, Matthew Bull, Joanne Watkins, Sally Corden, Tom Connor |
| EPI_ISL_3296035 | Viollier AG | Department of Biosystems Science and Engineering, ETH Zürich | Christian Beisel, Sarah Nadeau, Chaoran Chen, Ivan Topolsky, Kim Philipp Jablonski, Lara Fuhrmann, Rebecca Denes, Mirjam Feldkamp, Ina Nissen, Natascha Santacroce, Elodie Burcklen, Christiane Beckmann, Maurice Redondo, Olivier Kobel, Christoph Noppen, Niko Beerenwinkel, Tanja Stadler |
| EPI_ISL_3296036 | Wales Specialist Virology Centre Sequencing lab: Pathogen Genomics Unit | Public Health Wales Microbiology Cardiff Wales Specialist Virology Centre | Catherine Moore, Johnathan Evans, Laura Gifford, Malorie Perry, Simon Cottrell, Angela Marchbank, Alec Birchley, Alexander Adams, Amy Gaskin, Bree Gatica-Wilcox, Jason Coombes, Joel Southgate, Lauren Gilbert, Lee Graham, Nicole Pacchiarini, Sara Kumziene-Summerhayes, Sarah Taylor, Sophie Jones, Sara Rey, Matthew Bull, Joanne Watkins, Sally Corden, Tom Connor |
| EPI_ISL_3296037 | Viollier AG | Department of Biosystems Science and Engineering, ETH Zürich | Christian Beisel, Sarah Nadeau, Chaoran Chen, Ivan Topolsky, Kim Philipp Jablonski, Lara Fuhrmann, Rebecca Denes, Mirjam Feldkamp, Ina Nissen, Natascha Santacroce, Elodie Burcklen, Christiane Beckmann, Maurice Redondo, Olivier Kobel, Christoph Noppen, Niko Beerenwinkel, Tanja Stadler |
| EPI_ISL_3296038, EPI_ISL_3296039 | Wales Specialist Virology Centre Sequencing lab: Pathogen Genomics Unit | Public Health Wales Microbiology Cardiff Wales Specialist Virology Centre | Catherine Moore, Johnathan Evans, Laura Gifford, Malorie Perry, Simon Cottrell, Angela Marchbank, Alec Birchley, Alexander Adams, Amy Gaskin, Bree Gatica-Wilcox, Jason Coombes, Joel Southgate, Lauren Gilbert, Lee Graham, Nicole Pacchiarini, Sara Kumziene-Summerhayes, Sarah Taylor, Sophie Jones, Sara Rey, Matthew Bull, Joanne Watkins, Sally Corden, Tom Connor |
| EPI_ISL_3296040 | Viollier AG | Department of Biosystems Science and Engineering, ETH Zürich | Christian Beisel, Sarah Nadeau, Chaoran Chen, Ivan Topolsky, Kim Philipp Jablonski, Lara Fuhrmann, Rebecca Denes, Mirjam Feldkamp, Ina Nissen, Natascha Santacroce, Elodie Burcklen, Christiane Beckmann, Maurice Redondo, Olivier Kobel, Christoph Noppen, Niko Beerenwinkel, Tanja Stadler |
| EPI_ISL_3296041 | Wales Specialist Virology Centre Sequencing lab: Pathogen Genomics Unit | Public Health Wales Microbiology Cardiff Wales Specialist Virology Centre | Catherine Moore, Johnathan Evans, Laura Gifford, Malorie Perry, Simon Cottrell, Angela Marchbank, Alec Birchley, Alexander Adams, Amy Gaskin, Bree Gatica-Wilcox, Jason Coombes, Joel Southgate, Lauren Gilbert, Lee Graham, Nicole Pacchiarini, Sara Kumziene-Summerhayes, Sarah Taylor, Sophie Jones, Sara Rey, Matthew Bull, Joanne Watkins, Sally Corden, Tom Connor |
| EPI_ISL_3296042 | Viollier AG | Department of Biosystems Science and Engineering, ETH Zürich | Christian Beisel, Sarah Nadeau, Chaoran Chen, Ivan Topolsky, Kim Philipp Jablonski, Lara Fuhrmann, Rebecca Denes, Mirjam Feldkamp, Ina Nissen, Natascha Santacroce, Elodie Burcklen, Christiane Beckmann, Maurice Redondo, Olivier Kobel, Christoph Noppen, Niko Beerenwinkel, Tanja Stadler |
| EPI_ISL_3296043 | Wales Specialist Virology Centre Sequencing lab: Pathogen Genomics Unit | Public Health Wales Microbiology Cardiff Wales Specialist Virology Centre | Catherine Moore, Johnathan Evans, Laura Gifford, Malorie Perry, Simon Cottrell, Angela Marchbank, Alec Birchley, Alexander Adams, Amy Gaskin, Bree Gatica-Wilcox, Jason Coombes, Joel Southgate, Lauren Gilbert, Lee Graham, Nicole Pacchiarini, Sara Kumziene-Summerhayes, Sarah Taylor, Sophie Jones, Sara Rey, Matthew Bull, Joanne Watkins, Sally Corden, Tom Connor |
| EPI_ISL_3296044 | Viollier AG | Department of Biosystems Science and Engineering, ETH Zürich | Christian Beisel, Sarah Nadeau, Chaoran Chen, Ivan Topolsky, Kim Philipp Jablonski, Lara Fuhrmann, Rebecca Denes, Mirjam Feldkamp, Ina Nissen, Natascha Santacroce, Elodie Burcklen, Christiane Beckmann, Maurice Redondo, Olivier Kobel, Christoph Noppen, Niko Beerenwinkel, Tanja Stadler |
| EPI_ISL_3296045 | Wales Specialist Virology Centre Sequencing lab: Pathogen Genomics Unit | Public Health Wales Microbiology Cardiff Wales Specialist Virology Centre | Catherine Moore, Johnathan Evans, Laura Gifford, Malorie Perry, Simon Cottrell, Angela Marchbank, Alec Birchley, Alexander Adams, Amy Gaskin, Bree Gatica-Wilcox, Jason Coombes, Joel Southgate, Lauren Gilbert, Lee Graham, Nicole Pacchiarini, Sara Kumziene-Summerhayes, Sarah Taylor, Sophie Jones, Sara Rey, Matthew Bull, Joanne Watkins, Sally Corden, Tom Connor |
| EPI_ISL_3296046 | Viollier AG | Department of Biosystems Science and Engineering, ETH Zürich | Christian Beisel, Sarah Nadeau, Chaoran Chen, Ivan Topolsky, Kim Philipp Jablonski, Lara Fuhrmann, Rebecca Denes, Mirjam Feldkamp, Ina Nissen, Natascha Santacroce, Elodie Burcklen, Christiane Beckmann, Maurice Redondo, Olivier Kobel, Christoph Noppen, Niko Beerenwinkel, Tanja Stadler |
| EPI_ISL_3296047 | Wales Specialist Virology Centre Sequencing lab: Pathogen Genomics Unit | Public Health Wales Microbiology Cardiff Wales Specialist Virology Centre | Catherine Moore, Johnathan Evans, Laura Gifford, Malorie Perry, Simon Cottrell, Angela Marchbank, Alec Birchley, Alexander Adams, Amy Gaskin, Bree Gatica-Wilcox, Jason Coombes, Joel Southgate, Lauren Gilbert, Lee Graham, Nicole Pacchiarini, Sara Kumziene-Summerhayes, Sarah Taylor, Sophie Jones, Sara Rey, Matthew Bull, Joanne Watkins, Sally Corden, Tom Connor |
| EPI_ISL_3296048 | Viollier AG | Department of Biosystems Science and Engineering, ETH Zürich | Christian Beisel, Sarah Nadeau, Chaoran Chen, Ivan Topolsky, Kim Philipp Jablonski, Lara Fuhrmann, Rebecca Denes, Mirjam Feldkamp, Ina Nissen, Natascha Santacroce, Elodie Burcklen, Christiane Beckmann, Maurice Redondo, Olivier Kobel, Christoph Noppen, Niko Beerenwinkel, Tanja Stadler |
| EPI_ISL_3296049 | Wales Specialist Virology Centre Sequencing lab: Pathogen Genomics Unit | Public Health Wales Microbiology Cardiff Wales Specialist Virology Centre | Catherine Moore, Johnathan Evans, Laura Gifford, Malorie Perry, Simon Cottrell, Angela Marchbank, Alec Birchley, Alexander Adams, Amy Gaskin, Bree Gatica-Wilcox, Jason Coombes, Joel Southgate, Lauren Gilbert, Lee Graham, Nicole Pacchiarini, Sara Kumziene-Summerhayes, Sarah Taylor, Sophie Jones, Sara Rey, Matthew Bull, Joanne Watkins, Sally Corden, Tom Connor |
| EPI_ISL_3296050 | Viollier AG | Department of Biosystems Science and Engineering, ETH Zürich | Christian Beisel, Sarah Nadeau, Chaoran Chen, Ivan Topolsky, Kim Philipp Jablonski, Lara Fuhrmann, Rebecca Denes, Mirjam Feldkamp, Ina Nissen, Natascha Santacroce, Elodie Burcklen, Christiane Beckmann, Maurice Redondo, Olivier Kobel, Christoph Noppen, Niko Beerenwinkel, Tanja Stadler |
| EPI_ISL_3296051 | Wales Specialist Virology Centre Sequencing lab: Pathogen Genomics Unit | Public Health Wales Microbiology Cardiff Wales Specialist Virology Centre | Catherine Moore, Johnathan Evans, Laura Gifford, Malorie Perry, Simon Cottrell, Angela Marchbank, Alec Birchley, Alexander Adams, Amy Gaskin, Bree Gatica-Wilcox, Jason Coombes, Joel Southgate, Lauren Gilbert, Lee Graham, Nicole Pacchiarini, Sara Kumziene-Summerhayes, Sarah Taylor, Sophie Jones, Sara Rey, Matthew Bull, Joanne Watkins, Sally Corden, Tom Connor |
| EPI_ISL_3296052 | Viollier AG | Department of Biosystems Science and Engineering, ETH Zürich | Christian Beisel, Sarah Nadeau, Chaoran Chen, Ivan Topolsky, Kim Philipp Jablonski, Lara Fuhrmann, Rebecca Denes, Mirjam Feldkamp, Ina Nissen, Natascha Santacroce, Elodie Burcklen, Christiane Beckmann, Maurice Redondo, Olivier Kobel, Christoph Noppen, Niko Beerenwinkel, Tanja Stadler |
| EPI_ISL_3296053 | Wales Specialist Virology Centre Sequencing lab: Pathogen Genomics Unit | Public Health Wales Microbiology Cardiff Wales Specialist Virology Centre | Catherine Moore, Johnathan Evans, Laura Gifford, Malorie Perry, Simon Cottrell, Angela Marchbank, Alec Birchley, Alexander Adams, Amy Gaskin, Bree Gatica-Wilcox, Jason Coombes, Joel Southgate, Lauren Gilbert, Lee Graham, Nicole Pacchiarini, Sara Kumziene-Summerhayes, Sarah Taylor, Sophie Jones, Sara Rey, Matthew Bull, Joanne Watkins, Sally Corden, Tom Connor |
| EPI_ISL_3296054 | Viollier AG | Department of Biosystems Science and Engineering, ETH Zürich | Christian Beisel, Sarah Nadeau, Chaoran Chen, Ivan Topolsky, Kim Philipp Jablonski, Lara Fuhrmann, Rebecca Denes, Mirjam Feldkamp, Ina Nissen, Natascha Santacroce, Elodie Burcklen, Christiane Beckmann, Maurice Redondo, Olivier Kobel, Christoph Noppen, Niko Beerenwinkel, Tanja Stadler |
| EPI_ISL_3296055 | Wales Specialist Virology Centre Sequencing lab: Pathogen Genomics Unit | Public Health Wales Microbiology Cardiff Wales Specialist Virology Centre | Catherine Moore, Johnathan Evans, Laura Gifford, Malorie Perry, Simon Cottrell, Angela Marchbank, Alec Birchley, Alexander Adams, Amy Gaskin, Bree Gatica-Wilcox, Jason Coombes, Joel Southgate, Lauren Gilbert, Lee Graham, Nicole Pacchiarini, Sara Kumziene-Summerhayes, Sarah Taylor, Sophie Jones, Sara Rey, Matthew Bull, Joanne Watkins, Sally Corden, Tom Connor |
| EPI_ISL_3296056 | Viollier AG | Department of Biosystems Science and Engineering, ETH Zürich | Christian Beisel, Sarah Nadeau, Chaoran Chen, Ivan Topolsky, Kim Philipp Jablonski, Lara Fuhrmann, Rebecca Denes, Mirjam Feldkamp, Ina Nissen, Natascha Santacroce, Elodie Burcklen, Christiane Beckmann, Maurice Redondo, Olivier Kobel, Christoph Noppen, Niko Beerenwinkel, Tanja Stadler |
| EPI_ISL_3296057 | Wales Specialist Virology Centre Sequencing lab: Pathogen Genomics Unit | Public Health Wales Microbiology Cardiff Wales Specialist Virology Centre | Catherine Moore, Johnathan Evans, Laura Gifford, Malorie Perry, Simon Cottrell, Angela Marchbank, Alec Birchley, Alexander Adams, Amy Gaskin, Bree Gatica-Wilcox, Jason Coombes, Joel Southgate, Lauren Gilbert, Lee Graham, Nicole Pacchiarini, Sara Kumziene-Summerhayes, Sarah Taylor, Sophie Jones, Sara Rey, Matthew Bull, Joanne Watkins, Sally Corden, Tom Connor |
| EPI_ISL_3296058 | Viollier AG | Department of Biosystems Science and Engineering, ETH Zürich | Christian Beisel, Sarah Nadeau, Chaoran Chen, Ivan Topolsky, Kim Philipp Jablonski, Lara Fuhrmann, Rebecca Denes, Mirjam Feldkamp, Ina Nissen, Natascha Santacroce, Elodie Burcklen, Christiane Beckmann, Maurice Redondo, Olivier Kobel, Christoph Noppen, Niko Beerenwinkel, Tanja Stadler |
| EPI_ISL_3296059 | Wales Specialist Virology Centre Sequencing lab: Pathogen Genomics Unit | Public Health Wales Microbiology Cardiff Wales Specialist Virology Centre | Catherine Moore, Johnathan Evans, Laura Gifford, Malorie Perry, Simon Cottrell, Angela Marchbank, Alec Birchley, Alexander Adams, Amy Gaskin, Bree Gatica-Wilcox, Jason Coombes, Joel Southgate, Lauren Gilbert, Lee Graham, Nicole Pacchiarini, Sara Kumziene-Summerhayes, Sarah Taylor, Sophie Jones, Sara Rey, Matthew Bull, Joanne Watkins, Sally Corden, Tom Connor |
| EPI_ISL_3296060 | Viollier AG | Department of Biosystems Science and Engineering, ETH Zürich | Christian Beisel, Sarah Nadeau, Chaoran Chen, Ivan Topolsky, Kim Philipp Jablonski, Lara Fuhrmann, Rebecca Denes, Mirjam Feldkamp, Ina Nissen, Natascha Santacroce, Elodie Burcklen, Christiane Beckmann, Maurice Redondo, Olivier Kobel, |

| | | | |
|---|---|---|---|
| | | | Christoph Noppen, Niko Beerenwinkel, Tanja Stadler |
| EPI_ISL_3296061 | Wales Specialist Virology Centre Sequencing lab: Pathogen Genomics Unit | Public Health Wales Microbiology Cardiff Wales Specialist Virology Centre | Catherine Moore, Johnathan Evans, Laura Gifford, Malorie Perry, Simon Cottrell, Angela Marchbank, Alec Birchley, Alexander Adams, Amy Gaskin, Bree Gatica-Wilcox, Jason Coombes, Joel Southgate, Lauren Gilbert, Lee Graham, Nicole Pacchiarini, Sara Kumziene-Summerhayes, Sarah Taylor, Sophie Jones, Sara Rey, Matthew Bull, Joanne Watkins, Sally Corden, Tom Connor |
| EPI_ISL_3296062 | Viollier AG | Department of Biosystems Science and Engineering, ETH Zürich | Christian Beisel, Sarah Nadeau, Chaoran Chen, Ivan Topolsky, Kim Philipp Jablonski, Lara Fuhrmann, Rebecca Denes, Mirjam Feldkamp, Ina Nissen, Natascha Santacroce, Elodie Burcklen, Christiane Beckmann, Maurice Redondo, Olivier Kobel, Christoph Noppen, Niko Beerenwinkel, Tanja Stadler |
| EPI_ISL_3296063 | Wales Specialist Virology Centre Sequencing lab: Pathogen Genomics Unit | Public Health Wales Microbiology Cardiff Wales Specialist Virology Centre | Catherine Moore, Johnathan Evans, Laura Gifford, Malorie Perry, Simon Cottrell, Angela Marchbank, Alec Birchley, Alexander Adams, Amy Gaskin, Bree Gatica-Wilcox, Jason Coombes, Joel Southgate, Lauren Gilbert, Lee Graham, Nicole Pacchiarini, Sara Kumziene-Summerhayes, Sarah Taylor, Sophie Jones, Sara Rey, Matthew Bull, Joanne Watkins, Sally Corden, Tom Connor |
| EPI_ISL_3296064 | Viollier AG | Department of Biosystems Science and Engineering, ETH Zürich | Christian Beisel, Sarah Nadeau, Chaoran Chen, Ivan Topolsky, Kim Philipp Jablonski, Lara Fuhrmann, Rebecca Denes, Mirjam Feldkamp, Ina Nissen, Natascha Santacroce, Elodie Burcklen, Christiane Beckmann, Maurice Redondo, Olivier Kobel, Christoph Noppen, Niko Beerenwinkel, Tanja Stadler |
| EPI_ISL_3296065 | Wales Specialist Virology Centre Sequencing lab: Pathogen Genomics Unit | Public Health Wales Microbiology Cardiff Wales Specialist Virology Centre | Catherine Moore, Johnathan Evans, Laura Gifford, Malorie Perry, Simon Cottrell, Angela Marchbank, Alec Birchley, Alexander Adams, Amy Gaskin, Bree Gatica-Wilcox, Jason Coombes, Joel Southgate, Lauren Gilbert, Lee Graham, Nicole Pacchiarini, Sara Kumziene-Summerhayes, Sarah Taylor, Sophie Jones, Sara Rey, Matthew Bull, Joanne Watkins, Sally Corden, Tom Connor |
| EPI_ISL_3296066 | Viollier AG | Department of Biosystems Science and Engineering, ETH Zürich | Christian Beisel, Sarah Nadeau, Chaoran Chen, Ivan Topolsky, Kim Philipp Jablonski, Lara Fuhrmann, Rebecca Denes, Mirjam Feldkamp, Ina Nissen, Natascha Santacroce, Elodie Burcklen, Christiane Beckmann, Maurice Redondo, Olivier Kobel, Christoph Noppen, Niko Beerenwinkel, Tanja Stadler |
| EPI_ISL_3296067 | Wales Specialist Virology Centre Sequencing lab: Pathogen Genomics Unit | Public Health Wales Microbiology Cardiff Wales Specialist Virology Centre | Catherine Moore, Johnathan Evans, Laura Gifford, Malorie Perry, Simon Cottrell, Angela Marchbank, Alec Birchley, Alexander Adams, Amy Gaskin, Bree Gatica-Wilcox, Jason Coombes, Joel Southgate, Lauren Gilbert, Lee Graham, Nicole Pacchiarini, Sara Kumziene-Summerhayes, Sarah Taylor, Sophie Jones, Sara Rey, Matthew Bull, Joanne Watkins, Sally Corden, Tom Connor |
| EPI_ISL_3296068 | Viollier AG | Department of Biosystems Science and Engineering, ETH Zürich | Christian Beisel, Sarah Nadeau, Chaoran Chen, Ivan Topolsky, Kim Philipp Jablonski, Lara Fuhrmann, Rebecca Denes, Mirjam Feldkamp, Ina Nissen, Natascha Santacroce, Elodie Burcklen, Christiane Beckmann, Maurice Redondo, Olivier Kobel, Christoph Noppen, Niko Beerenwinkel, Tanja Stadler |
| EPI_ISL_3296069 | Wales Specialist Virology Centre Sequencing lab: Pathogen Genomics Unit | Public Health Wales Microbiology Cardiff Wales Specialist Virology Centre | Catherine Moore, Johnathan Evans, Laura Gifford, Malorie Perry, Simon Cottrell, Angela Marchbank, Alec Birchley, Alexander Adams, Amy Gaskin, Bree Gatica-Wilcox, Jason Coombes, Joel Southgate, Lauren Gilbert, Lee Graham, Nicole Pacchiarini, Sara Kumziene-Summerhayes, Sarah Taylor, Sophie Jones, Sara Rey, Matthew Bull, Joanne Watkins, Sally Corden, Tom Connor |
| EPI_ISL_3296070 | Viollier AG | Department of Biosystems Science and Engineering, ETH Zürich | Christian Beisel, Sarah Nadeau, Chaoran Chen, Ivan Topolsky, Kim Philipp Jablonski, Lara Fuhrmann, Rebecca Denes, Mirjam Feldkamp, Ina Nissen, Natascha Santacroce, Elodie Burcklen, Christiane Beckmann, Maurice Redondo, Olivier Kobel, Christoph Noppen, Niko Beerenwinkel, Tanja Stadler |
| EPI_ISL_3296071 | Wales Specialist Virology Centre Sequencing lab: Pathogen Genomics Unit | Public Health Wales Microbiology Cardiff Wales Specialist Virology Centre | Catherine Moore, Johnathan Evans, Laura Gifford, Malorie Perry, Simon Cottrell, Angela Marchbank, Alec Birchley, Alexander Adams, Amy Gaskin, Bree Gatica-Wilcox, Jason Coombes, Joel Southgate, Lauren Gilbert, Lee Graham, Nicole Pacchiarini, Sara Kumziene-Summerhayes, Sarah Taylor, Sophie Jones, Sara Rey, Matthew Bull, Joanne Watkins, Sally Corden, Tom Connor |
| EPI_ISL_3296072 | Viollier AG | Department of Biosystems Science and Engineering, ETH Zürich | Christian Beisel, Sarah Nadeau, Chaoran Chen, Ivan Topolsky, Kim Philipp Jablonski, Lara Fuhrmann, Rebecca Denes, Mirjam Feldkamp, Ina Nissen, Natascha Santacroce, Elodie Burcklen, Christiane Beckmann, Maurice Redondo, Olivier Kobel, Christoph Noppen, Niko Beerenwinkel, Tanja Stadler |
| EPI_ISL_3296073 | Wales Specialist Virology Centre Sequencing lab: Pathogen Genomics Unit | Public Health Wales Microbiology Cardiff Wales Specialist Virology Centre | Catherine Moore, Johnathan Evans, Laura Gifford, Malorie Perry, Simon Cottrell, Angela Marchbank, Alec Birchley, Alexander Adams, Amy Gaskin, Bree Gatica-Wilcox, Jason Coombes, Joel Southgate, Lauren Gilbert, Lee Graham, Nicole Pacchiarini, Sara Kumziene-Summerhayes, Sarah Taylor, Sophie Jones, Sara Rey, Matthew Bull, Joanne Watkins, Sally Corden, Tom Connor |
| EPI_ISL_3296074 | Viollier AG | Department of Biosystems Science and Engineering, ETH Zürich | Christian Beisel, Sarah Nadeau, Chaoran Chen, Ivan Topolsky, Kim Philipp Jablonski, Lara Fuhrmann, Rebecca Denes, Mirjam Feldkamp, Ina Nissen, Natascha Santacroce, Elodie Burcklen, Christiane Beckmann, Maurice Redondo, Olivier Kobel, Christoph Noppen, Niko Beerenwinkel, Tanja Stadler |
| EPI_ISL_3296075, EPI_ISL_3296076 | Wales Specialist Virology Centre Sequencing lab: Pathogen Genomics Unit | Public Health Wales Microbiology Cardiff Wales Specialist Virology Centre | Catherine Moore, Johnathan Evans, Laura Gifford, Malorie Perry, Simon Cottrell, Angela Marchbank, Alec Birchley, Alexander Adams, Amy Gaskin, Bree Gatica-Wilcox, Jason Coombes, Joel Southgate, Lauren Gilbert, Lee Graham, Nicole Pacchiarini, Sara Kumziene-Summerhayes, Sarah Taylor, Sophie Jones, Sara Rey, Matthew Bull, Joanne Watkins, Sally Corden, Tom Connor |
| EPI_ISL_3296077 | Viollier AG | Department of Biosystems Science and Engineering, ETH Zürich | Christian Beisel, Sarah Nadeau, Chaoran Chen, Ivan Topolsky, Kim Philipp Jablonski, Lara Fuhrmann, Rebecca Denes, Mirjam Feldkamp, Ina Nissen, Natascha Santacroce, Elodie Burcklen, Christiane Beckmann, Maurice Redondo, Olivier Kobel, Christoph Noppen, Niko Beerenwinkel, Tanja Stadler |
| EPI_ISL_3296078 | Wales Specialist Virology Centre Sequencing lab: Pathogen Genomics Unit | Public Health Wales Microbiology Cardiff Wales Specialist Virology Centre | Catherine Moore, Johnathan Evans, Laura Gifford, Malorie Perry, Simon Cottrell, Angela Marchbank, Alec Birchley, Alexander Adams, Amy Gaskin, Bree Gatica-Wilcox, Jason Coombes, Joel Southgate, Lauren Gilbert, Lee Graham, Nicole Pacchiarini, Sara Kumziene-Summerhayes, Sarah Taylor, Sophie Jones, Sara Rey, Matthew Bull, Joanne Watkins, Sally Corden, Tom Connor |
| EPI_ISL_3296079 | Viollier AG | Department of Biosystems Science and Engineering, ETH Zürich | Christian Beisel, Sarah Nadeau, Chaoran Chen, Ivan Topolsky, Kim Philipp Jablonski, Lara Fuhrmann, Rebecca Denes, Mirjam Feldkamp, Ina Nissen, Natascha Santacroce, Elodie Burcklen, Christiane Beckmann, Maurice Redondo, Olivier Kobel, Christoph Noppen, Niko Beerenwinkel, Tanja Stadler |
| EPI_ISL_3296080 | Wales Specialist Virology Centre Sequencing lab: Pathogen Genomics Unit | Public Health Wales Microbiology Cardiff Wales Specialist Virology Centre | Catherine Moore, Johnathan Evans, Laura Gifford, Malorie Perry, Simon Cottrell, Angela Marchbank, Alec Birchley, Alexander Adams, Amy Gaskin, Bree Gatica-Wilcox, Jason Coombes, Joel Southgate, Lauren Gilbert, Lee Graham, Nicole Pacchiarini, Sara Kumziene-Summerhayes, Sarah Taylor, Sophie Jones, Sara Rey, Matthew Bull, Joanne Watkins, Sally Corden, Tom Connor |
| EPI_ISL_3296081 | Viollier AG | Department of Biosystems Science and Engineering, ETH Zürich | Christian Beisel, Sarah Nadeau, Chaoran Chen, Ivan Topolsky, Kim Philipp Jablonski, Lara Fuhrmann, Rebecca Denes, Mirjam Feldkamp, Ina Nissen, Natascha Santacroce, Elodie Burcklen, Christiane Beckmann, Maurice Redondo, Olivier Kobel, Christoph Noppen, Niko Beerenwinkel, Tanja Stadler |
| EPI_ISL_3296082 | Wales Specialist Virology Centre Sequencing lab: Pathogen Genomics Unit | Public Health Wales Microbiology Cardiff Wales Specialist Virology Centre | Catherine Moore, Johnathan Evans, Laura Gifford, Malorie Perry, Simon Cottrell, Angela Marchbank, Alec Birchley, Alexander Adams, Amy Gaskin, Bree Gatica-Wilcox, Jason Coombes, Joel Southgate, Lauren Gilbert, Lee Graham, Nicole Pacchiarini, Sara Kumziene-Summerhayes, Sarah Taylor, Sophie Jones, Sara Rey, Matthew Bull, Joanne Watkins, Sally Corden, Tom Connor |
| EPI_ISL_3296083 | Viollier AG | Department of Biosystems Science and Engineering, ETH Zürich | Christian Beisel, Sarah Nadeau, Chaoran Chen, Ivan Topolsky, Kim Philipp Jablonski, Lara Fuhrmann, Rebecca Denes, Mirjam Feldkamp, Ina Nissen, Natascha Santacroce, Elodie Burcklen, Christiane Beckmann, Maurice Redondo, Olivier Kobel, Christoph Noppen, Niko Beerenwinkel, Tanja Stadler |
| EPI_ISL_3296084 | Wales Specialist Virology Centre Sequencing lab: Pathogen Genomics Unit | Public Health Wales Microbiology Cardiff Wales Specialist Virology Centre | Catherine Moore, Johnathan Evans, Laura Gifford, Malorie Perry, Simon Cottrell, Angela Marchbank, Alec Birchley, Alexander Adams, Amy Gaskin, Bree Gatica-Wilcox, Jason Coombes, Joel Southgate, Lauren Gilbert, Lee Graham, Nicole Pacchiarini, Sara Kumziene-Summerhayes, Sarah Taylor, Sophie Jones, Sara Rey, Matthew Bull, Joanne Watkins, Sally Corden, Tom Connor |
| EPI_ISL_3296085 | Viollier AG | Department of Biosystems Science and Engineering, ETH Zürich | Christian Beisel, Sarah Nadeau, Chaoran Chen, Ivan Topolsky, Kim Philipp Jablonski, Lara Fuhrmann, Rebecca Denes, Mirjam Feldkamp, Ina Nissen, Natascha Santacroce, Elodie Burcklen, Christiane Beckmann, Maurice Redondo, Olivier Kobel, Christoph Noppen, Niko Beerenwinkel, Tanja Stadler |
| EPI_ISL_3296086 | Wales Specialist Virology Centre Sequencing lab: Pathogen Genomics Unit | Public Health Wales Microbiology Cardiff Wales Specialist Virology Centre | Catherine Moore, Johnathan Evans, Laura Gifford, Malorie Perry, Simon Cottrell, Angela Marchbank, Alec Birchley, Alexander Adams, Amy Gaskin, Bree Gatica-Wilcox, Jason Coombes, Joel Southgate, Lauren Gilbert, Lee Graham, Nicole Pacchiarini, Sara Kumziene-Summerhayes, Sarah Taylor, Sophie Jones, Sara Rey, Matthew Bull, Joanne Watkins, Sally Corden, Tom Connor |
| EPI_ISL_3296087 | Viollier AG | Department of Biosystems Science and Engineering, ETH Zürich | Christian Beisel, Sarah Nadeau, Chaoran Chen, Ivan Topolsky, Kim Philipp Jablonski, Lara Fuhrmann, Rebecca Denes, Mirjam Feldkamp, Ina Nissen, Natascha Santacroce, Elodie Burcklen, Christiane Beckmann, Maurice Redondo, Olivier Kobel, Christoph Noppen, Niko Beerenwinkel, Tanja Stadler |
| EPI_ISL_3296088 | Wales Specialist Virology Centre Sequencing lab: Pathogen Genomics Unit | Public Health Wales Microbiology Cardiff Wales Specialist Virology Centre | Catherine Moore, Johnathan Evans, Laura Gifford, Malorie Perry, Simon Cottrell, Angela Marchbank, Alec Birchley, Alexander Adams, Amy Gaskin, Bree Gatica-Wilcox, Jason Coombes, Joel Southgate, Lauren Gilbert, Lee Graham, Nicole Pacchiarini, Sara Kumziene-Summerhayes, Sarah Taylor, Sophie Jones, Sara Rey, Matthew Bull, Joanne Watkins, Sally Corden, Tom Connor |
| EPI_ISL_3296089 | Viollier AG | Department of Biosystems Science and Engineering, ETH Zürich | Christian Beisel, Sarah Nadeau, Chaoran Chen, Ivan Topolsky, Kim Philipp Jablonski, Lara Fuhrmann, Rebecca Denes, Mirjam Feldkamp, Ina Nissen, Natascha Santacroce, Elodie Burcklen, Christiane Beckmann, Maurice Redondo, Olivier Kobel, Christoph Noppen, Niko Beerenwinkel, Tanja Stadler |
| EPI_ISL_3296090 | Wales Specialist Virology Centre Sequencing lab: Pathogen Genomics Unit | Public Health Wales Microbiology Cardiff Wales Specialist Virology Centre | Catherine Moore, Johnathan Evans, Laura Gifford, Malorie Perry, Simon Cottrell, Angela Marchbank, Alec Birchley, Alexander Adams, Amy Gaskin, Bree Gatica-Wilcox, Jason Coombes, Joel Southgate, Lauren Gilbert, Lee Graham, Nicole Pacchiarini, Sara Kumziene-Summerhayes, Sarah Taylor, Sophie Jones, Sara Rey, Matthew Bull, Joanne Watkins, Sally Corden, Tom Connor |
| EPI_ISL_3296091 | Viollier AG | Department of Biosystems Science and Engineering, ETH Zürich | Christian Beisel, Sarah Nadeau, Chaoran Chen, Ivan Topolsky, Kim Philipp Jablonski, Lara Fuhrmann, Rebecca Denes, Mirjam Feldkamp, Ina Nissen, Natascha Santacroce, Elodie Burcklen, Christiane Beckmann, Maurice Redondo, Olivier Kobel, Christoph Noppen, Niko Beerenwinkel, Tanja Stadler |
| EPI_ISL_3296092 | Wales Specialist Virology Centre Sequencing lab: Pathogen Genomics Unit | Public Health Wales Microbiology Cardiff Wales Specialist Virology Centre | Catherine Moore, Johnathan Evans, Laura Gifford, Malorie Perry, Simon Cottrell, Angela Marchbank, Alec Birchley, Alexander Adams, Amy Gaskin, Bree Gatica-Wilcox, Jason Coombes, Joel Southgate, Lauren Gilbert, Lee Graham, Nicole Pacchiarini, Sara Kumziene-Summerhayes, Sarah Taylor, Sophie Jones, Sara Rey, Matthew Bull, Joanne Watkins, Sally Corden, Tom Connor |
| EPI_ISL_3296093 | Viollier AG | Department of Biosystems Science and Engineering, ETH Zürich | Christian Beisel, Sarah Nadeau, Chaoran Chen, Ivan Topolsky, Kim Philipp Jablonski, Lara Fuhrmann, Rebecca Denes, Mirjam Feldkamp, Ina Nissen, Natascha Santacroce, Elodie Burcklen, Christiane Beckmann, Maurice Redondo, Olivier Kobel, Christoph Noppen, Niko Beerenwinkel, Tanja Stadler |
| EPI_ISL_3296094 | Wales Specialist Virology Centre Sequencing lab: Pathogen Genomics Unit | Public Health Wales Microbiology Cardiff Wales Specialist Virology Centre | Catherine Moore, Johnathan Evans, Laura Gifford, Malorie Perry, Simon Cottrell, Angela Marchbank, Alec Birchley, Alexander Adams, Amy Gaskin, Bree Gatica-Wilcox, Jason Coombes, Joel Southgate, Lauren Gilbert, Lee Graham, Nicole Pacchiarini, Sara Kumziene-Summerhayes, Sarah Taylor, Sophie Jones, Sara Rey, Matthew Bull, Joanne Watkins, Sally Corden, Tom Connor |
| EPI_ISL_3296095 | Viollier AG | Department of Biosystems Science and Engineering, ETH Zürich | Christian Beisel, Sarah Nadeau, Chaoran Chen, Ivan Topolsky, Kim Philipp Jablonski, Lara Fuhrmann, Rebecca Denes, Mirjam Feldkamp, Ina Nissen, Natascha Santacroce, Elodie Burcklen, Christiane Beckmann, Maurice Redondo, Olivier Kobel, Christoph Noppen, Niko Beerenwinkel, Tanja Stadler |
| EPI_ISL_3296096 | Wales Specialist Virology Centre Sequencing lab: Pathogen Genomics Unit | Public Health Wales Microbiology Cardiff Wales Specialist Virology Centre | Catherine Moore, Johnathan Evans, Laura Gifford, Malorie Perry, Simon Cottrell, Angela Marchbank, Alec Birchley, Alexander Adams, Amy Gaskin, Bree Gatica-Wilcox, Jason Coombes, Joel Southgate, Lauren Gilbert, Lee Graham, Nicole Pacchiarini, Sara Kumziene-Summerhayes, Sarah Taylor, Sophie Jones, Sara Rey, Matthew Bull, Joanne Watkins, Sally Corden, Tom Connor |
| EPI_ISL_3296097 | Viollier AG | Department of Biosystems Science and Engineering, ETH Zürich | Christian Beisel, Sarah Nadeau, Chaoran Chen, Ivan Topolsky, Kim Philipp Jablonski, Lara Fuhrmann, Rebecca Denes, Mirjam Feldkamp, Ina Nissen, Natascha Santacroce, Elodie Burcklen, Christiane Beckmann, Maurice Redondo, Olivier Kobel, Christoph Noppen, Niko Beerenwinkel, Tanja Stadler |
| EPI_ISL_3296098 | Wales Specialist Virology Centre Sequencing lab: Pathogen Genomics Unit | Public Health Wales Microbiology Cardiff Wales Specialist Virology Centre | Catherine Moore, Johnathan Evans, Laura Gifford, Malorie Perry, Simon Cottrell, Angela Marchbank, Alec Birchley, Alexander Adams, Amy Gaskin, Bree Gatica-Wilcox, Jason Coombes, Joel Southgate, Lauren Gilbert, Lee Graham, Nicole Pacchiarini, Sara Kumziene-Summerhayes, Sarah Taylor, Sophie Jones, Sara Rey, Matthew Bull, Joanne Watkins, Sally Corden, Tom Connor |
| EPI_ISL_3296099 | Viollier AG | Department of Biosystems Science and Engineering, ETH Zürich | Christian Beisel, Sarah Nadeau, Chaoran Chen, Ivan Topolsky, Kim Philipp Jablonski, Lara Fuhrmann, Rebecca Denes, Mirjam Feldkamp, Ina Nissen, Natascha Santacroce, Elodie Burcklen, Christiane Beckmann, Maurice Redondo, Olivier Kobel, Christoph Noppen, Niko Beerenwinkel, Tanja Stadler |
| EPI_ISL_3296100 | Wales Specialist Virology Centre Sequencing lab: Pathogen Genomics Unit | Public Health Wales Microbiology Cardiff Wales Specialist Virology Centre | Catherine Moore, Johnathan Evans, Laura Gifford, Malorie Perry, Simon Cottrell, Angela Marchbank, Alec Birchley, Alexander Adams, Amy Gaskin, Bree Gatica-Wilcox, Jason Coombes, Joel Southgate, Lauren Gilbert, Lee Graham, Nicole Pacchiarini, Sara Kumziene-Summerhayes, Sarah Taylor, Sophie Jones, Sara Rey, Matthew Bull, Joanne Watkins, Sally Corden, Tom Connor |
| EPI_ISL_3296101 | Viollier AG | Department of Biosystems Science and Engineering, ETH Zürich | Christian Beisel, Sarah Nadeau, Chaoran Chen, Ivan Topolsky, Kim Philipp Jablonski, Lara Fuhrmann, Rebecca Denes, Mirjam Feldkamp, Ina Nissen, Natascha Santacroce, Elodie Burcklen, Christiane Beckmann, Maurice Redondo, Olivier Kobel, Christoph Noppen, Niko Beerenwinkel, Tanja Stadler |
| EPI_ISL_3296102 | Wales Specialist Virology Centre Sequencing lab: Pathogen Genomics Unit | Public Health Wales Microbiology Cardiff Wales Specialist Virology Centre | Catherine Moore, Johnathan Evans, Laura Gifford, Malorie Perry, Simon Cottrell, Angela Marchbank, Alec Birchley, Alexander Adams, Amy Gaskin, Bree Gatica-Wilcox, Jason Coombes, Joel Southgate, Lauren Gilbert, Lee Graham, Nicole Pacchiarini, Sara Kumziene-Summerhayes, Sarah Taylor, Sophie Jones, Sara Rey, Matthew Bull, Joanne Watkins, Sally Corden, Tom Connor |
| EPI_ISL_3296103 | Viollier AG | Department of Biosystems Science and Engineering, ETH Zürich | Christian Beisel, Sarah Nadeau, Chaoran Chen, Ivan Topolsky, Kim Philipp Jablonski, Lara Fuhrmann, Rebecca Denes, Mirjam Feldkamp, Ina Nissen, Natascha Santacroce, Elodie Burcklen, Christiane Beckmann, Maurice Redondo, Olivier Kobel, Christoph Noppen, Niko Beerenwinkel, Tanja Stadler |
| EPI_ISL_3296104 | Wales Specialist Virology Centre Sequencing lab: Pathogen Genomics Unit | Public Health Wales Microbiology Cardiff Wales Specialist Virology Centre | Catherine Moore, Johnathan Evans, Laura Gifford, Malorie Perry, Simon Cottrell, Angela Marchbank, Alec Birchley, Alexander Adams, Amy Gaskin, Bree Gatica-Wilcox, Jason Coombes, Joel Southgate, Lauren Gilbert, Lee Graham, Nicole Pacchiarini, Sara Kumziene-Summerhayes, Sarah Taylor, Sophie Jones, Sara Rey, Matthew Bull, Joanne Watkins, Sally Corden, Tom Connor |
| EPI_ISL_3296105 | Viollier AG | Department of Biosystems Science and Engineering, ETH Zürich | Christian Beisel, Sarah Nadeau, Chaoran Chen, Ivan Topolsky, Kim Philipp Jablonski, Lara Fuhrmann, Rebecca Denes, Mirjam Feldkamp, Ina Nissen, Natascha Santacroce, Elodie Burcklen, Christiane Beckmann, Maurice Redondo, Olivier Kobel, Christoph Noppen, Niko Beerenwinkel, Tanja Stadler |
| EPI_ISL_3296106, | Wales Specialist Virology Centre Sequencing lab: Pathogen | Public Health Wales Microbiology Cardiff Wales Specialist Virology | Catherine Moore, Johnathan Evans, Laura Gifford, Malorie Perry, Simon Cottrell, Angela Marchbank, Alec Birchley, Alexander Adams, Amy Gaskin, Bree Gatica-Wilcox, Jason Coombes, Joel Southgate, Lauren Gilbert, Lee Graham, Nicole |

| | | | |
|---|---|---|---|
| EPI_ISL_3296107 | Genomics Unit | Centre | Pacchiarini, Sara Kumziene-Summerhayes, Sarah Taylor, Sophie Jones, Sara Rey, Matthew Bull, Joanne Watkins, Sally Corden, Tom Connor |
| EPI_ISL_3296108 | Viollier AG | Department of Biosystems Science and Engineering, ETH Zürich | Christian Beisel, Sarah Nadeau, Chaoran Chen, Ivan Topolsky, Kim Philipp Jablonski, Lara Fuhrmann, Rebecca Denes, Mirjam Feldkamp, Ina Nissen, Natascha Santacroce, Elodie Burcklen, Christiane Beckmann, Maurice Redondo, Olivier Kobel, Christoph Noppen, Niko Beerenwinkel, Tanja Stadler |
| EPI_ISL_3296109 | Wales Specialist Virology Centre Sequencing lab: Pathogen Genomics Unit | Public Health Wales Microbiology Cardiff Wales Specialist Virology Centre | Catherine Moore, Johnathan Evans, Laura Gifford, Malorie Perry, Simon Cottrell, Angela Marchbank, Alec Birchley, Alexander Adams, Amy Gaskin, Bree Gatica-Wilcox, Jason Coombes, Joel Southgate, Lauren Gilbert, Lee Graham, Nicole Pacchiarini, Sara Kumziene-Summerhayes, Sarah Taylor, Sophie Jones, Sara Rey, Matthew Bull, Joanne Watkins, Sally Corden, Tom Connor |
| EPI_ISL_3296110 | Viollier AG | Department of Biosystems Science and Engineering, ETH Zürich | Christian Beisel, Sarah Nadeau, Chaoran Chen, Ivan Topolsky, Kim Philipp Jablonski, Lara Fuhrmann, Rebecca Denes, Mirjam Feldkamp, Ina Nissen, Natascha Santacroce, Elodie Burcklen, Christiane Beckmann, Maurice Redondo, Olivier Kobel, Christoph Noppen, Niko Beerenwinkel, Tanja Stadler |
| EPI_ISL_3296111 | Wales Specialist Virology Centre Sequencing lab: Pathogen Genomics Unit | Public Health Wales Microbiology Cardiff Wales Specialist Virology Centre | Catherine Moore, Johnathan Evans, Laura Gifford, Malorie Perry, Simon Cottrell, Angela Marchbank, Alec Birchley, Alexander Adams, Amy Gaskin, Bree Gatica-Wilcox, Jason Coombes, Joel Southgate, Lauren Gilbert, Lee Graham, Nicole Pacchiarini, Sara Kumziene-Summerhayes, Sarah Taylor, Sophie Jones, Sara Rey, Matthew Bull, Joanne Watkins, Sally Corden, Tom Connor |
| EPI_ISL_3296112 | Viollier AG | Department of Biosystems Science and Engineering, ETH Zürich | Christian Beisel, Sarah Nadeau, Chaoran Chen, Ivan Topolsky, Kim Philipp Jablonski, Lara Fuhrmann, Rebecca Denes, Mirjam Feldkamp, Ina Nissen, Natascha Santacroce, Elodie Burcklen, Christiane Beckmann, Maurice Redondo, Olivier Kobel, Christoph Noppen, Niko Beerenwinkel, Tanja Stadler |
| EPI_ISL_3296113 | Wales Specialist Virology Centre Sequencing lab: Pathogen Genomics Unit | Public Health Wales Microbiology Cardiff Wales Specialist Virology Centre | Catherine Moore, Johnathan Evans, Laura Gifford, Malorie Perry, Simon Cottrell, Angela Marchbank, Alec Birchley, Alexander Adams, Amy Gaskin, Bree Gatica-Wilcox, Jason Coombes, Joel Southgate, Lauren Gilbert, Lee Graham, Nicole Pacchiarini, Sara Kumziene-Summerhayes, Sarah Taylor, Sophie Jones, Sara Rey, Matthew Bull, Joanne Watkins, Sally Corden, Tom Connor |
| EPI_ISL_3296114 | Viollier AG | Department of Biosystems Science and Engineering, ETH Zürich | Christian Beisel, Sarah Nadeau, Chaoran Chen, Ivan Topolsky, Kim Philipp Jablonski, Lara Fuhrmann, Rebecca Denes, Mirjam Feldkamp, Ina Nissen, Natascha Santacroce, Elodie Burcklen, Christiane Beckmann, Maurice Redondo, Olivier Kobel, Christoph Noppen, Niko Beerenwinkel, Tanja Stadler |
| EPI_ISL_3296115 | Wales Specialist Virology Centre Sequencing lab: Pathogen Genomics Unit | Public Health Wales Microbiology Cardiff Wales Specialist Virology Centre | Catherine Moore, Johnathan Evans, Laura Gifford, Malorie Perry, Simon Cottrell, Angela Marchbank, Alec Birchley, Alexander Adams, Amy Gaskin, Bree Gatica-Wilcox, Jason Coombes, Joel Southgate, Lauren Gilbert, Lee Graham, Nicole Pacchiarini, Sara Kumziene-Summerhayes, Sarah Taylor, Sophie Jones, Sara Rey, Matthew Bull, Joanne Watkins, Sally Corden, Tom Connor |
| EPI_ISL_3296116 | Viollier AG | Department of Biosystems Science and Engineering, ETH Zürich | Christian Beisel, Sarah Nadeau, Chaoran Chen, Ivan Topolsky, Kim Philipp Jablonski, Lara Fuhrmann, Rebecca Denes, Mirjam Feldkamp, Ina Nissen, Natascha Santacroce, Elodie Burcklen, Christiane Beckmann, Maurice Redondo, Olivier Kobel, Christoph Noppen, Niko Beerenwinkel, Tanja Stadler |
| EPI_ISL_3296117 | Wales Specialist Virology Centre Sequencing lab: Pathogen Genomics Unit | Public Health Wales Microbiology Cardiff Wales Specialist Virology Centre | Catherine Moore, Johnathan Evans, Laura Gifford, Malorie Perry, Simon Cottrell, Angela Marchbank, Alec Birchley, Alexander Adams, Amy Gaskin, Bree Gatica-Wilcox, Jason Coombes, Joel Southgate, Lauren Gilbert, Lee Graham, Nicole Pacchiarini, Sara Kumziene-Summerhayes, Sarah Taylor, Sophie Jones, Sara Rey, Matthew Bull, Joanne Watkins, Sally Corden, Tom Connor |
| EPI_ISL_3296118 | Viollier AG | Department of Biosystems Science and Engineering, ETH Zürich | Christian Beisel, Sarah Nadeau, Chaoran Chen, Ivan Topolsky, Kim Philipp Jablonski, Lara Fuhrmann, Rebecca Denes, Mirjam Feldkamp, Ina Nissen, Natascha Santacroce, Elodie Burcklen, Christiane Beckmann, Maurice Redondo, Olivier Kobel, Christoph Noppen, Niko Beerenwinkel, Tanja Stadler |
| EPI_ISL_3296119 | Wales Specialist Virology Centre Sequencing lab: Pathogen Genomics Unit | Public Health Wales Microbiology Cardiff Wales Specialist Virology Centre | Catherine Moore, Johnathan Evans, Laura Gifford, Malorie Perry, Simon Cottrell, Angela Marchbank, Alec Birchley, Alexander Adams, Amy Gaskin, Bree Gatica-Wilcox, Jason Coombes, Joel Southgate, Lauren Gilbert, Lee Graham, Nicole Pacchiarini, Sara Kumziene-Summerhayes, Sarah Taylor, Sophie Jones, Sara Rey, Matthew Bull, Joanne Watkins, Sally Corden, Tom Connor |
| EPI_ISL_3296120 | Viollier AG | Department of Biosystems Science and Engineering, ETH Zürich | Christian Beisel, Sarah Nadeau, Chaoran Chen, Ivan Topolsky, Kim Philipp Jablonski, Lara Fuhrmann, Rebecca Denes, Mirjam Feldkamp, Ina Nissen, Natascha Santacroce, Elodie Burcklen, Christiane Beckmann, Maurice Redondo, Olivier Kobel, Christoph Noppen, Niko Beerenwinkel, Tanja Stadler |
| EPI_ISL_3296121, EPI_ISL_3296122 | Wales Specialist Virology Centre Sequencing lab: Pathogen Genomics Unit | Public Health Wales Microbiology Cardiff Wales Specialist Virology Centre | Catherine Moore, Johnathan Evans, Laura Gifford, Malorie Perry, Simon Cottrell, Angela Marchbank, Alec Birchley, Alexander Adams, Amy Gaskin, Bree Gatica-Wilcox, Jason Coombes, Joel Southgate, Lauren Gilbert, Lee Graham, Nicole Pacchiarini, Sara Kumziene-Summerhayes, Sarah Taylor, Sophie Jones, Sara Rey, Matthew Bull, Joanne Watkins, Sally Corden, Tom Connor |
| EPI_ISL_3296123 | Viollier AG | Department of Biosystems Science and Engineering, ETH Zürich | Christian Beisel, Sarah Nadeau, Chaoran Chen, Ivan Topolsky, Kim Philipp Jablonski, Lara Fuhrmann, Rebecca Denes, Mirjam Feldkamp, Ina Nissen, Natascha Santacroce, Elodie Burcklen, Christiane Beckmann, Maurice Redondo, Olivier Kobel, Christoph Noppen, Niko Beerenwinkel, Tanja Stadler |
| EPI_ISL_3296124 | Wales Specialist Virology Centre Sequencing lab: Pathogen Genomics Unit | Public Health Wales Microbiology Cardiff Wales Specialist Virology Centre | Catherine Moore, Johnathan Evans, Laura Gifford, Malorie Perry, Simon Cottrell, Angela Marchbank, Alec Birchley, Alexander Adams, Amy Gaskin, Bree Gatica-Wilcox, Jason Coombes, Joel Southgate, Lauren Gilbert, Lee Graham, Nicole Pacchiarini, Sara Kumziene-Summerhayes, Sarah Taylor, Sophie Jones, Sara Rey, Matthew Bull, Joanne Watkins, Sally Corden, Tom Connor |
| EPI_ISL_3296125 | Viollier AG | Department of Biosystems Science and Engineering, ETH Zürich | Christian Beisel, Sarah Nadeau, Chaoran Chen, Ivan Topolsky, Kim Philipp Jablonski, Lara Fuhrmann, Rebecca Denes, Mirjam Feldkamp, Ina Nissen, Natascha Santacroce, Elodie Burcklen, Christiane Beckmann, Maurice Redondo, Olivier Kobel, Christoph Noppen, Niko Beerenwinkel, Tanja Stadler |
| EPI_ISL_3296126 | Wales Specialist Virology Centre Sequencing lab: Pathogen Genomics Unit | Public Health Wales Microbiology Cardiff Wales Specialist Virology Centre | Catherine Moore, Johnathan Evans, Laura Gifford, Malorie Perry, Simon Cottrell, Angela Marchbank, Alec Birchley, Alexander Adams, Amy Gaskin, Bree Gatica-Wilcox, Jason Coombes, Joel Southgate, Lauren Gilbert, Lee Graham, Nicole Pacchiarini, Sara Kumziene-Summerhayes, Sarah Taylor, Sophie Jones, Sara Rey, Matthew Bull, Joanne Watkins, Sally Corden, Tom Connor |
| EPI_ISL_3296127 | Viollier AG | Department of Biosystems Science and Engineering, ETH Zürich | Christian Beisel, Sarah Nadeau, Chaoran Chen, Ivan Topolsky, Kim Philipp Jablonski, Lara Fuhrmann, Rebecca Denes, Mirjam Feldkamp, Ina Nissen, Natascha Santacroce, Elodie Burcklen, Christiane Beckmann, Maurice Redondo, Olivier Kobel, Christoph Noppen, Niko Beerenwinkel, Tanja Stadler |
| EPI_ISL_3296128 | Wales Specialist Virology Centre Sequencing lab: Pathogen Genomics Unit | Public Health Wales Microbiology Cardiff Wales Specialist Virology Centre | Catherine Moore, Johnathan Evans, Laura Gifford, Malorie Perry, Simon Cottrell, Angela Marchbank, Alec Birchley, Alexander Adams, Amy Gaskin, Bree Gatica-Wilcox, Jason Coombes, Joel Southgate, Lauren Gilbert, Lee Graham, Nicole Pacchiarini, Sara Kumziene-Summerhayes, Sarah Taylor, Sophie Jones, Sara Rey, Matthew Bull, Joanne Watkins, Sally Corden, Tom Connor |
| EPI_ISL_3296129 | Viollier AG | Department of Biosystems Science and Engineering, ETH Zürich | Christian Beisel, Sarah Nadeau, Chaoran Chen, Ivan Topolsky, Kim Philipp Jablonski, Lara Fuhrmann, Rebecca Denes, Mirjam Feldkamp, Ina Nissen, Natascha Santacroce, Elodie Burcklen, Christiane Beckmann, Maurice Redondo, Olivier Kobel, Christoph Noppen, Niko Beerenwinkel, Tanja Stadler |
| EPI_ISL_3296130 | Wales Specialist Virology Centre Sequencing lab: Pathogen Genomics Unit | Public Health Wales Microbiology Cardiff Wales Specialist Virology Centre | Catherine Moore, Johnathan Evans, Laura Gifford, Malorie Perry, Simon Cottrell, Angela Marchbank, Alec Birchley, Alexander Adams, Amy Gaskin, Bree Gatica-Wilcox, Jason Coombes, Joel Southgate, Lauren Gilbert, Lee Graham, Nicole Pacchiarini, Sara Kumziene-Summerhayes, Sarah Taylor, Sophie Jones, Sara Rey, Matthew Bull, Joanne Watkins, Sally Corden, Tom Connor |
| EPI_ISL_3296131 | Viollier AG | Department of Biosystems Science and Engineering, ETH Zürich | Christian Beisel, Sarah Nadeau, Chaoran Chen, Ivan Topolsky, Kim Philipp Jablonski, Lara Fuhrmann, Rebecca Denes, Mirjam Feldkamp, Ina Nissen, Natascha Santacroce, Elodie Burcklen, Christiane Beckmann, Maurice Redondo, Olivier Kobel, Christoph Noppen, Niko Beerenwinkel, Tanja Stadler |
| EPI_ISL_3296132, EPI_ISL_3296133 | Wales Specialist Virology Centre Sequencing lab: Pathogen Genomics Unit | Public Health Wales Microbiology Cardiff Wales Specialist Virology Centre | Catherine Moore, Johnathan Evans, Laura Gifford, Malorie Perry, Simon Cottrell, Angela Marchbank, Alec Birchley, Alexander Adams, Amy Gaskin, Bree Gatica-Wilcox, Jason Coombes, Joel Southgate, Lauren Gilbert, Lee Graham, Nicole Pacchiarini, Sara Kumziene-Summerhayes, Sarah Taylor, Sophie Jones, Sara Rey, Matthew Bull, Joanne Watkins, Sally Corden, Tom Connor |
| EPI_ISL_3296134 | Viollier AG | Department of Biosystems Science and Engineering, ETH Zürich | Christian Beisel, Sarah Nadeau, Chaoran Chen, Ivan Topolsky, Kim Philipp Jablonski, Lara Fuhrmann, Rebecca Denes, Mirjam Feldkamp, Ina Nissen, Natascha Santacroce, Elodie Burcklen, Christiane Beckmann, Maurice Redondo, Olivier Kobel, Christoph Noppen, Niko Beerenwinkel, Tanja Stadler |
| EPI_ISL_3296135 | Wales Specialist Virology Centre Sequencing lab: Pathogen Genomics Unit | Public Health Wales Microbiology Cardiff Wales Specialist Virology Centre | Catherine Moore, Johnathan Evans, Laura Gifford, Malorie Perry, Simon Cottrell, Angela Marchbank, Alec Birchley, Alexander Adams, Amy Gaskin, Bree Gatica-Wilcox, Jason Coombes, Joel Southgate, Lauren Gilbert, Lee Graham, Nicole Pacchiarini, Sara Kumziene-Summerhayes, Sarah Taylor, Sophie Jones, Sara Rey, Matthew Bull, Joanne Watkins, Sally Corden, Tom Connor |
| EPI_ISL_3296136, EPI_ISL_3296137 | Viollier AG | Department of Biosystems Science and Engineering, ETH Zürich | Christian Beisel, Sarah Nadeau, Chaoran Chen, Ivan Topolsky, Kim Philipp Jablonski, Lara Fuhrmann, Rebecca Denes, Mirjam Feldkamp, Ina Nissen, Natascha Santacroce, Elodie Burcklen, Christiane Beckmann, Maurice Redondo, Olivier Kobel, Christoph Noppen, Niko Beerenwinkel, Tanja Stadler |
| EPI_ISL_3296138, EPI_ISL_3296139 | Wales Specialist Virology Centre Sequencing lab: Pathogen Genomics Unit | Public Health Wales Microbiology Cardiff Wales Specialist Virology Centre | Catherine Moore, Johnathan Evans, Laura Gifford, Malorie Perry, Simon Cottrell, Angela Marchbank, Alec Birchley, Alexander Adams, Amy Gaskin, Bree Gatica-Wilcox, Jason Coombes, Joel Southgate, Lauren Gilbert, Lee Graham, Nicole Pacchiarini, Sara Kumziene-Summerhayes, Sarah Taylor, Sophie Jones, Sara Rey, Matthew Bull, Joanne Watkins, Sally Corden, Tom Connor |
| EPI_ISL_3296140 | Viollier AG | Department of Biosystems Science and Engineering, ETH Zürich | Christian Beisel, Sarah Nadeau, Chaoran Chen, Ivan Topolsky, Kim Philipp Jablonski, Lara Fuhrmann, Rebecca Denes, Mirjam Feldkamp, Ina Nissen, Natascha Santacroce, Elodie Burcklen, Christiane Beckmann, Maurice Redondo, Olivier Kobel, Christoph Noppen, Niko Beerenwinkel, Tanja Stadler |
| EPI_ISL_3296141 | Wales Specialist Virology Centre Sequencing lab: Pathogen Genomics Unit | Public Health Wales Microbiology Cardiff Wales Specialist Virology Centre | Catherine Moore, Johnathan Evans, Laura Gifford, Malorie Perry, Simon Cottrell, Angela Marchbank, Alec Birchley, Alexander Adams, Amy Gaskin, Bree Gatica-Wilcox, Jason Coombes, Joel Southgate, Lauren Gilbert, Lee Graham, Nicole Pacchiarini, Sara Kumziene-Summerhayes, Sarah Taylor, Sophie Jones, Sara Rey, Matthew Bull, Joanne Watkins, Sally Corden, Tom Connor |
| EPI_ISL_3296142 | Viollier AG | Department of Biosystems Science and Engineering, ETH Zürich | Christian Beisel, Sarah Nadeau, Chaoran Chen, Ivan Topolsky, Kim Philipp Jablonski, Lara Fuhrmann, Rebecca Denes, Mirjam Feldkamp, Ina Nissen, Natascha Santacroce, Elodie Burcklen, Christiane Beckmann, Maurice Redondo, Olivier Kobel, Christoph Noppen, Niko Beerenwinkel, Tanja Stadler |
| EPI_ISL_3296143 | Wales Specialist Virology Centre Sequencing lab: Pathogen Genomics Unit | Public Health Wales Microbiology Cardiff Wales Specialist Virology Centre | Catherine Moore, Johnathan Evans, Laura Gifford, Malorie Perry, Simon Cottrell, Angela Marchbank, Alec Birchley, Alexander Adams, Amy Gaskin, Bree Gatica-Wilcox, Jason Coombes, Joel Southgate, Lauren Gilbert, Lee Graham, Nicole Pacchiarini, Sara Kumziene-Summerhayes, Sarah Taylor, Sophie Jones, Sara Rey, Matthew Bull, Joanne Watkins, Sally Corden, Tom Connor |
| EPI_ISL_3296144 | Viollier AG | Department of Biosystems Science and Engineering, ETH Zürich | Christian Beisel, Sarah Nadeau, Chaoran Chen, Ivan Topolsky, Kim Philipp Jablonski, Lara Fuhrmann, Rebecca Denes, Mirjam Feldkamp, Ina Nissen, Natascha Santacroce, Elodie Burcklen, Christiane Beckmann, Maurice Redondo, Olivier Kobel, Christoph Noppen, Niko Beerenwinkel, Tanja Stadler |
| EPI_ISL_3296145 | Wales Specialist Virology Centre Sequencing lab: Pathogen Genomics Unit | Public Health Wales Microbiology Cardiff Wales Specialist Virology Centre | Catherine Moore, Johnathan Evans, Laura Gifford, Malorie Perry, Simon Cottrell, Angela Marchbank, Alec Birchley, Alexander Adams, Amy Gaskin, Bree Gatica-Wilcox, Jason Coombes, Joel Southgate, Lauren Gilbert, Lee Graham, Nicole Pacchiarini, Sara Kumziene-Summerhayes, Sarah Taylor, Sophie Jones, Sara Rey, Matthew Bull, Joanne Watkins, Sally Corden, Tom Connor |
| EPI_ISL_3296146 | Viollier AG | Department of Biosystems Science and Engineering, ETH Zürich | Christian Beisel, Sarah Nadeau, Chaoran Chen, Ivan Topolsky, Kim Philipp Jablonski, Lara Fuhrmann, Rebecca Denes, Mirjam Feldkamp, Ina Nissen, Natascha Santacroce, Elodie Burcklen, Christiane Beckmann, Maurice Redondo, Olivier Kobel, Christoph Noppen, Niko Beerenwinkel, Tanja Stadler |
| EPI_ISL_3296147 | Wales Specialist Virology Centre Sequencing lab: Pathogen Genomics Unit | Public Health Wales Microbiology Cardiff Wales Specialist Virology Centre | Catherine Moore, Johnathan Evans, Laura Gifford, Malorie Perry, Simon Cottrell, Angela Marchbank, Alec Birchley, Alexander Adams, Amy Gaskin, Bree Gatica-Wilcox, Jason Coombes, Joel Southgate, Lauren Gilbert, Lee Graham, Nicole Pacchiarini, Sara Kumziene-Summerhayes, Sarah Taylor, Sophie Jones, Sara Rey, Matthew Bull, Joanne Watkins, Sally Corden, Tom Connor |
| EPI_ISL_3296148 | Viollier AG | Department of Biosystems Science and Engineering, ETH Zürich | Christian Beisel, Sarah Nadeau, Chaoran Chen, Ivan Topolsky, Kim Philipp Jablonski, Lara Fuhrmann, Rebecca Denes, Mirjam Feldkamp, Ina Nissen, Natascha Santacroce, Elodie Burcklen, Christiane Beckmann, Maurice Redondo, Olivier Kobel, Christoph Noppen, Niko Beerenwinkel, Tanja Stadler |
| EPI_ISL_3296149 | Wales Specialist Virology Centre Sequencing lab: Pathogen Genomics Unit | Public Health Wales Microbiology Cardiff Wales Specialist Virology Centre | Catherine Moore, Johnathan Evans, Laura Gifford, Malorie Perry, Simon Cottrell, Angela Marchbank, Alec Birchley, Alexander Adams, Amy Gaskin, Bree Gatica-Wilcox, Jason Coombes, Joel Southgate, Lauren Gilbert, Lee Graham, Nicole Pacchiarini, Sara Kumziene-Summerhayes, Sarah Taylor, Sophie Jones, Sara Rey, Matthew Bull, Joanne Watkins, Sally Corden, Tom Connor |
| EPI_ISL_3296150 | Viollier AG | Department of Biosystems Science and Engineering, ETH Zürich | Christian Beisel, Sarah Nadeau, Chaoran Chen, Ivan Topolsky, Kim Philipp Jablonski, Lara Fuhrmann, Rebecca Denes, Mirjam Feldkamp, Ina Nissen, Natascha Santacroce, Elodie Burcklen, Christiane Beckmann, Maurice Redondo, Olivier Kobel, Christoph Noppen, Niko Beerenwinkel, Tanja Stadler |
| EPI_ISL_3296151 | Wales Specialist Virology Centre Sequencing lab: Pathogen Genomics Unit | Public Health Wales Microbiology Cardiff Wales Specialist Virology Centre | Catherine Moore, Johnathan Evans, Laura Gifford, Malorie Perry, Simon Cottrell, Angela Marchbank, Alec Birchley, Alexander Adams, Amy Gaskin, Bree Gatica-Wilcox, Jason Coombes, Joel Southgate, Lauren Gilbert, Lee Graham, Nicole Pacchiarini, Sara Kumziene-Summerhayes, Sarah Taylor, Sophie Jones, Sara Rey, Matthew Bull, Joanne Watkins, Sally Corden, Tom Connor |
| EPI_ISL_3296152 | Viollier AG | Department of Biosystems Science and Engineering, ETH Zürich | Christian Beisel, Sarah Nadeau, Chaoran Chen, Ivan Topolsky, Kim Philipp Jablonski, Lara Fuhrmann, Rebecca Denes, Mirjam Feldkamp, Ina Nissen, Natascha Santacroce, Elodie Burcklen, Christiane Beckmann, Maurice Redondo, Olivier Kobel, Christoph Noppen, Niko Beerenwinkel, Tanja Stadler |
| EPI_ISL_3296153 | Wales Specialist Virology Centre Sequencing lab: Pathogen Genomics Unit | Public Health Wales Microbiology Cardiff Wales Specialist Virology Centre | Catherine Moore, Johnathan Evans, Laura Gifford, Malorie Perry, Simon Cottrell, Angela Marchbank, Alec Birchley, Alexander Adams, Amy Gaskin, Bree Gatica-Wilcox, Jason Coombes, Joel Southgate, Lauren Gilbert, Lee Graham, Nicole Pacchiarini, Sara Kumziene-Summerhayes, Sarah Taylor, Sophie Jones, Sara Rey, Matthew Bull, Joanne Watkins, Sally Corden, Tom Connor |
| EPI_ISL_3296154 | Viollier AG | Department of Biosystems Science and Engineering, ETH Zürich | Christian Beisel, Sarah Nadeau, Chaoran Chen, Ivan Topolsky, Kim Philipp Jablonski, Lara Fuhrmann, Rebecca Denes, Mirjam Feldkamp, Ina Nissen, Natascha Santacroce, Elodie Burcklen, Christiane Beckmann, Maurice Redondo, Olivier Kobel, Christoph Noppen, Niko Beerenwinkel, Tanja Stadler |
| EPI_ISL_3296155 | Wales Specialist Virology Centre Sequencing lab: Pathogen Genomics Unit | Public Health Wales Microbiology Cardiff Wales Specialist Virology Centre | Catherine Moore, Johnathan Evans, Laura Gifford, Malorie Perry, Simon Cottrell, Angela Marchbank, Alec Birchley, Alexander Adams, Amy Gaskin, Bree Gatica-Wilcox, Jason Coombes, Joel Southgate, Lauren Gilbert, Lee Graham, Nicole Pacchiarini, Sara Kumziene-Summerhayes, Sarah Taylor, Sophie Jones, Sara Rey, Matthew Bull, Joanne Watkins, Sally Corden, Tom Connor |
| EPI_ISL_3296156 | Viollier AG | Department of Biosystems Science and Engineering, ETH Zürich | Christian Beisel, Sarah Nadeau, Chaoran Chen, Ivan Topolsky, Kim Philipp Jablonski, Lara Fuhrmann, Rebecca Denes, Mirjam Feldkamp, Ina Nissen, Natascha Santacroce, Elodie Burcklen, Christiane Beckmann, Maurice Redondo, Olivier Kobel, |

| | | | |
|---|---|---|---|
| | | | Christoph Noppen, Niko Beerenwinkel, Tanja Stadler |
| EPI_ISL_3296157 | Wales Specialist Virology Centre Sequencing lab: Pathogen Genomics Unit | Public Health Wales Microbiology Cardiff Wales Specialist Virology Centre | Catherine Moore, Johnathan Evans, Laura Gifford, Malorie Perry, Simon Cottrell, Angela Marchbank, Alec Birchley, Alexander Adams, Amy Gaskin, Bree Gatica-Wilcox, Jason Coombes, Joel Southgate, Lauren Gilbert, Lee Graham, Nicole Pacchiarini, Sara Kumziene-Summerhayes, Sarah Taylor, Sophie Jones, Sara Rey, Matthew Bull, Joanne Watkins, Sally Corden, Tom Connor |
| EPI_ISL_3296158 | Viollier AG | Department of Biosystems Science and Engineering, ETH Zürich | Christian Beisel, Sarah Nadeau, Chaoran Chen, Ivan Topolsky, Kim Philipp Jablonski, Lara Fuhrmann, Rebecca Denes, Mirjam Feldkamp, Ina Nissen, Natascha Santacroce, Elodie Burcklen, Christiane Beckmann, Maurice Redondo, Olivier Kobel, Christoph Noppen, Niko Beerenwinkel, Tanja Stadler |
| EPI_ISL_3296159 | Wales Specialist Virology Centre Sequencing lab: Pathogen Genomics Unit | Public Health Wales Microbiology Cardiff Wales Specialist Virology Centre | Catherine Moore, Johnathan Evans, Laura Gifford, Malorie Perry, Simon Cottrell, Angela Marchbank, Alec Birchley, Alexander Adams, Amy Gaskin, Bree Gatica-Wilcox, Jason Coombes, Joel Southgate, Lauren Gilbert, Lee Graham, Nicole Pacchiarini, Sara Kumziene-Summerhayes, Sarah Taylor, Sophie Jones, Sara Rey, Matthew Bull, Joanne Watkins, Sally Corden, Tom Connor |
| EPI_ISL_3296160 | Viollier AG | Department of Biosystems Science and Engineering, ETH Zürich | Christian Beisel, Sarah Nadeau, Chaoran Chen, Ivan Topolsky, Kim Philipp Jablonski, Lara Fuhrmann, Rebecca Denes, Mirjam Feldkamp, Ina Nissen, Natascha Santacroce, Elodie Burcklen, Christiane Beckmann, Maurice Redondo, Olivier Kobel, Christoph Noppen, Niko Beerenwinkel, Tanja Stadler |
| EPI_ISL_3296161 | Wales Specialist Virology Centre Sequencing lab: Pathogen Genomics Unit | Public Health Wales Microbiology Cardiff Wales Specialist Virology Centre | Catherine Moore, Johnathan Evans, Laura Gifford, Malorie Perry, Simon Cottrell, Angela Marchbank, Alec Birchley, Alexander Adams, Amy Gaskin, Bree Gatica-Wilcox, Jason Coombes, Joel Southgate, Lauren Gilbert, Lee Graham, Nicole Pacchiarini, Sara Kumziene-Summerhayes, Sarah Taylor, Sophie Jones, Sara Rey, Matthew Bull, Joanne Watkins, Sally Corden, Tom Connor |
| EPI_ISL_3296162 | Viollier AG | Department of Biosystems Science and Engineering, ETH Zürich | Christian Beisel, Sarah Nadeau, Chaoran Chen, Ivan Topolsky, Kim Philipp Jablonski, Lara Fuhrmann, Rebecca Denes, Mirjam Feldkamp, Ina Nissen, Natascha Santacroce, Elodie Burcklen, Christiane Beckmann, Maurice Redondo, Olivier Kobel, Christoph Noppen, Niko Beerenwinkel, Tanja Stadler |
| EPI_ISL_3296163, EPI_ISL_3296164 | Wales Specialist Virology Centre Sequencing lab: Pathogen Genomics Unit | Public Health Wales Microbiology Cardiff Wales Specialist Virology Centre | Catherine Moore, Johnathan Evans, Laura Gifford, Malorie Perry, Simon Cottrell, Angela Marchbank, Alec Birchley, Alexander Adams, Amy Gaskin, Bree Gatica-Wilcox, Jason Coombes, Joel Southgate, Lauren Gilbert, Lee Graham, Nicole Pacchiarini, Sara Kumziene-Summerhayes, Sarah Taylor, Sophie Jones, Sara Rey, Matthew Bull, Joanne Watkins, Sally Corden, Tom Connor |
| EPI_ISL_3296165 | Viollier AG | Department of Biosystems Science and Engineering, ETH Zürich | Christian Beisel, Sarah Nadeau, Chaoran Chen, Ivan Topolsky, Kim Philipp Jablonski, Lara Fuhrmann, Rebecca Denes, Mirjam Feldkamp, Ina Nissen, Natascha Santacroce, Elodie Burcklen, Christiane Beckmann, Maurice Redondo, Olivier Kobel, Christoph Noppen, Niko Beerenwinkel, Tanja Stadler |
| EPI_ISL_3296166 | Wales Specialist Virology Centre Sequencing lab: Pathogen Genomics Unit | Public Health Wales Microbiology Cardiff Wales Specialist Virology Centre | Catherine Moore, Johnathan Evans, Laura Gifford, Malorie Perry, Simon Cottrell, Angela Marchbank, Alec Birchley, Alexander Adams, Amy Gaskin, Bree Gatica-Wilcox, Jason Coombes, Joel Southgate, Lauren Gilbert, Lee Graham, Nicole Pacchiarini, Sara Kumziene-Summerhayes, Sarah Taylor, Sophie Jones, Sara Rey, Matthew Bull, Joanne Watkins, Sally Corden, Tom Connor |
| EPI_ISL_3296167 | Viollier AG | Department of Biosystems Science and Engineering, ETH Zürich | Christian Beisel, Sarah Nadeau, Chaoran Chen, Ivan Topolsky, Kim Philipp Jablonski, Lara Fuhrmann, Rebecca Denes, Mirjam Feldkamp, Ina Nissen, Natascha Santacroce, Elodie Burcklen, Christiane Beckmann, Maurice Redondo, Olivier Kobel, Christoph Noppen, Niko Beerenwinkel, Tanja Stadler |
| EPI_ISL_3296168 | Wales Specialist Virology Centre Sequencing lab: Pathogen Genomics Unit | Public Health Wales Microbiology Cardiff Wales Specialist Virology Centre | Catherine Moore, Johnathan Evans, Laura Gifford, Malorie Perry, Simon Cottrell, Angela Marchbank, Alec Birchley, Alexander Adams, Amy Gaskin, Bree Gatica-Wilcox, Jason Coombes, Joel Southgate, Lauren Gilbert, Lee Graham, Nicole Pacchiarini, Sara Kumziene-Summerhayes, Sarah Taylor, Sophie Jones, Sara Rey, Matthew Bull, Joanne Watkins, Sally Corden, Tom Connor |
| EPI_ISL_3296169 | Viollier AG | Department of Biosystems Science and Engineering, ETH Zürich | Christian Beisel, Sarah Nadeau, Chaoran Chen, Ivan Topolsky, Kim Philipp Jablonski, Lara Fuhrmann, Rebecca Denes, Mirjam Feldkamp, Ina Nissen, Natascha Santacroce, Elodie Burcklen, Christiane Beckmann, Maurice Redondo, Olivier Kobel, Christoph Noppen, Niko Beerenwinkel, Tanja Stadler |
| EPI_ISL_3296170 | Wales Specialist Virology Centre Sequencing lab: Pathogen Genomics Unit | Public Health Wales Microbiology Cardiff Wales Specialist Virology Centre | Catherine Moore, Johnathan Evans, Laura Gifford, Malorie Perry, Simon Cottrell, Angela Marchbank, Alec Birchley, Alexander Adams, Amy Gaskin, Bree Gatica-Wilcox, Jason Coombes, Joel Southgate, Lauren Gilbert, Lee Graham, Nicole Pacchiarini, Sara Kumziene-Summerhayes, Sarah Taylor, Sophie Jones, Sara Rey, Matthew Bull, Joanne Watkins, Sally Corden, Tom Connor |
| EPI_ISL_3296171 | Viollier AG | Department of Biosystems Science and Engineering, ETH Zürich | Christian Beisel, Sarah Nadeau, Chaoran Chen, Ivan Topolsky, Kim Philipp Jablonski, Lara Fuhrmann, Rebecca Denes, Mirjam Feldkamp, Ina Nissen, Natascha Santacroce, Elodie Burcklen, Christiane Beckmann, Maurice Redondo, Olivier Kobel, Christoph Noppen, Niko Beerenwinkel, Tanja Stadler |
| EPI_ISL_3296172 | Wales Specialist Virology Centre Sequencing lab: Pathogen Genomics Unit | Public Health Wales Microbiology Cardiff Wales Specialist Virology Centre | Catherine Moore, Johnathan Evans, Laura Gifford, Malorie Perry, Simon Cottrell, Angela Marchbank, Alec Birchley, Alexander Adams, Amy Gaskin, Bree Gatica-Wilcox, Jason Coombes, Joel Southgate, Lauren Gilbert, Lee Graham, Nicole Pacchiarini, Sara Kumziene-Summerhayes, Sarah Taylor, Sophie Jones, Sara Rey, Matthew Bull, Joanne Watkins, Sally Corden, Tom Connor |
| EPI_ISL_3296173 | Viollier AG | Department of Biosystems Science and Engineering, ETH Zürich | Christian Beisel, Sarah Nadeau, Chaoran Chen, Ivan Topolsky, Kim Philipp Jablonski, Lara Fuhrmann, Rebecca Denes, Mirjam Feldkamp, Ina Nissen, Natascha Santacroce, Elodie Burcklen, Christiane Beckmann, Maurice Redondo, Olivier Kobel, Christoph Noppen, Niko Beerenwinkel, Tanja Stadler |
| EPI_ISL_3296174 | Wales Specialist Virology Centre Sequencing lab: Pathogen Genomics Unit | Public Health Wales Microbiology Cardiff Wales Specialist Virology Centre | Catherine Moore, Johnathan Evans, Laura Gifford, Malorie Perry, Simon Cottrell, Angela Marchbank, Alec Birchley, Alexander Adams, Amy Gaskin, Bree Gatica-Wilcox, Jason Coombes, Joel Southgate, Lauren Gilbert, Lee Graham, Nicole Pacchiarini, Sara Kumziene-Summerhayes, Sarah Taylor, Sophie Jones, Sara Rey, Matthew Bull, Joanne Watkins, Sally Corden, Tom Connor |
| EPI_ISL_3296175 | Viollier AG | Department of Biosystems Science and Engineering, ETH Zürich | Christian Beisel, Sarah Nadeau, Chaoran Chen, Ivan Topolsky, Kim Philipp Jablonski, Lara Fuhrmann, Rebecca Denes, Mirjam Feldkamp, Ina Nissen, Natascha Santacroce, Elodie Burcklen, Christiane Beckmann, Maurice Redondo, Olivier Kobel, Christoph Noppen, Niko Beerenwinkel, Tanja Stadler |
| EPI_ISL_3296176 | Wales Specialist Virology Centre Sequencing lab: Pathogen Genomics Unit | Public Health Wales Microbiology Cardiff Wales Specialist Virology Centre | Catherine Moore, Johnathan Evans, Laura Gifford, Malorie Perry, Simon Cottrell, Angela Marchbank, Alec Birchley, Alexander Adams, Amy Gaskin, Bree Gatica-Wilcox, Jason Coombes, Joel Southgate, Lauren Gilbert, Lee Graham, Nicole Pacchiarini, Sara Kumziene-Summerhayes, Sarah Taylor, Sophie Jones, Sara Rey, Matthew Bull, Joanne Watkins, Sally Corden, Tom Connor |
| EPI_ISL_3296177 | Viollier AG | Department of Biosystems Science and Engineering, ETH Zürich | Christian Beisel, Sarah Nadeau, Chaoran Chen, Ivan Topolsky, Kim Philipp Jablonski, Lara Fuhrmann, Rebecca Denes, Mirjam Feldkamp, Ina Nissen, Natascha Santacroce, Elodie Burcklen, Christiane Beckmann, Maurice Redondo, Olivier Kobel, Christoph Noppen, Niko Beerenwinkel, Tanja Stadler |
| EPI_ISL_3296178, EPI_ISL_3296179 | Wales Specialist Virology Centre Sequencing lab: Pathogen Genomics Unit | Public Health Wales Microbiology Cardiff Wales Specialist Virology Centre | Catherine Moore, Johnathan Evans, Laura Gifford, Malorie Perry, Simon Cottrell, Angela Marchbank, Alec Birchley, Alexander Adams, Amy Gaskin, Bree Gatica-Wilcox, Jason Coombes, Joel Southgate, Lauren Gilbert, Lee Graham, Nicole Pacchiarini, Sara Kumziene-Summerhayes, Sarah Taylor, Sophie Jones, Sara Rey, Matthew Bull, Joanne Watkins, Sally Corden, Tom Connor |
| EPI_ISL_3296180 | Viollier AG | Department of Biosystems Science and Engineering, ETH Zürich | Christian Beisel, Sarah Nadeau, Chaoran Chen, Ivan Topolsky, Kim Philipp Jablonski, Lara Fuhrmann, Rebecca Denes, Mirjam Feldkamp, Ina Nissen, Natascha Santacroce, Elodie Burcklen, Christiane Beckmann, Maurice Redondo, Olivier Kobel, Christoph Noppen, Niko Beerenwinkel, Tanja Stadler |
| EPI_ISL_3296181 | Wales Specialist Virology Centre Sequencing lab: Pathogen Genomics Unit | Public Health Wales Microbiology Cardiff Wales Specialist Virology Centre | Catherine Moore, Johnathan Evans, Laura Gifford, Malorie Perry, Simon Cottrell, Angela Marchbank, Alec Birchley, Alexander Adams, Amy Gaskin, Bree Gatica-Wilcox, Jason Coombes, Joel Southgate, Lauren Gilbert, Lee Graham, Nicole Pacchiarini, Sara Kumziene-Summerhayes, Sarah Taylor, Sophie Jones, Sara Rey, Matthew Bull, Joanne Watkins, Sally Corden, Tom Connor |
| EPI_ISL_3296182 | Viollier AG | Department of Biosystems Science and Engineering, ETH Zürich | Christian Beisel, Sarah Nadeau, Chaoran Chen, Ivan Topolsky, Kim Philipp Jablonski, Lara Fuhrmann, Rebecca Denes, Mirjam Feldkamp, Ina Nissen, Natascha Santacroce, Elodie Burcklen, Christiane Beckmann, Maurice Redondo, Olivier Kobel, Christoph Noppen, Niko Beerenwinkel, Tanja Stadler |
| EPI_ISL_3296183 | Wales Specialist Virology Centre Sequencing lab: Pathogen Genomics Unit | Public Health Wales Microbiology Cardiff Wales Specialist Virology Centre | Catherine Moore, Johnathan Evans, Laura Gifford, Malorie Perry, Simon Cottrell, Angela Marchbank, Alec Birchley, Alexander Adams, Amy Gaskin, Bree Gatica-Wilcox, Jason Coombes, Joel Southgate, Lauren Gilbert, Lee Graham, Nicole Pacchiarini, Sara Kumziene-Summerhayes, Sarah Taylor, Sophie Jones, Sara Rey, Matthew Bull, Joanne Watkins, Sally Corden, Tom Connor |
| EPI_ISL_3296184 | Viollier AG | Department of Biosystems Science and Engineering, ETH Zürich | Christian Beisel, Sarah Nadeau, Chaoran Chen, Ivan Topolsky, Kim Philipp Jablonski, Lara Fuhrmann, Rebecca Denes, Mirjam Feldkamp, Ina Nissen, Natascha Santacroce, Elodie Burcklen, Christiane Beckmann, Maurice Redondo, Olivier Kobel, Christoph Noppen, Niko Beerenwinkel, Tanja Stadler |
| EPI_ISL_3296185 | Wales Specialist Virology Centre Sequencing lab: Pathogen Genomics Unit | Public Health Wales Microbiology Cardiff Wales Specialist Virology Centre | Catherine Moore, Johnathan Evans, Laura Gifford, Malorie Perry, Simon Cottrell, Angela Marchbank, Alec Birchley, Alexander Adams, Amy Gaskin, Bree Gatica-Wilcox, Jason Coombes, Joel Southgate, Lauren Gilbert, Lee Graham, Nicole Pacchiarini, Sara Kumziene-Summerhayes, Sarah Taylor, Sophie Jones, Sara Rey, Matthew Bull, Joanne Watkins, Sally Corden, Tom Connor |
| EPI_ISL_3296186 | Viollier AG | Department of Biosystems Science and Engineering, ETH Zürich | Christian Beisel, Sarah Nadeau, Chaoran Chen, Ivan Topolsky, Kim Philipp Jablonski, Lara Fuhrmann, Rebecca Denes, Mirjam Feldkamp, Ina Nissen, Natascha Santacroce, Elodie Burcklen, Christiane Beckmann, Maurice Redondo, Olivier Kobel, Christoph Noppen, Niko Beerenwinkel, Tanja Stadler |
| EPI_ISL_3296187 | Wales Specialist Virology Centre Sequencing lab: Pathogen Genomics Unit | Public Health Wales Microbiology Cardiff Wales Specialist Virology Centre | Catherine Moore, Johnathan Evans, Laura Gifford, Malorie Perry, Simon Cottrell, Angela Marchbank, Alec Birchley, Alexander Adams, Amy Gaskin, Bree Gatica-Wilcox, Jason Coombes, Joel Southgate, Lauren Gilbert, Lee Graham, Nicole Pacchiarini, Sara Kumziene-Summerhayes, Sarah Taylor, Sophie Jones, Sara Rey, Matthew Bull, Joanne Watkins, Sally Corden, Tom Connor |
| EPI_ISL_3296188 | Viollier AG | Department of Biosystems Science and Engineering, ETH Zürich | Christian Beisel, Sarah Nadeau, Chaoran Chen, Ivan Topolsky, Kim Philipp Jablonski, Lara Fuhrmann, Rebecca Denes, Mirjam Feldkamp, Ina Nissen, Natascha Santacroce, Elodie Burcklen, Christiane Beckmann, Maurice Redondo, Olivier Kobel, Christoph Noppen, Niko Beerenwinkel, Tanja Stadler |
| EPI_ISL_3296189 | Wales Specialist Virology Centre Sequencing lab: Pathogen Genomics Unit | Public Health Wales Microbiology Cardiff Wales Specialist Virology Centre | Catherine Moore, Johnathan Evans, Laura Gifford, Malorie Perry, Simon Cottrell, Angela Marchbank, Alec Birchley, Alexander Adams, Amy Gaskin, Bree Gatica-Wilcox, Jason Coombes, Joel Southgate, Lauren Gilbert, Lee Graham, Nicole Pacchiarini, Sara Kumziene-Summerhayes, Sarah Taylor, Sophie Jones, Sara Rey, Matthew Bull, Joanne Watkins, Sally Corden, Tom Connor |
| EPI_ISL_3296190 | Viollier AG | Department of Biosystems Science and Engineering, ETH Zürich | Christian Beisel, Sarah Nadeau, Chaoran Chen, Ivan Topolsky, Kim Philipp Jablonski, Lara Fuhrmann, Rebecca Denes, Mirjam Feldkamp, Ina Nissen, Natascha Santacroce, Elodie Burcklen, Christiane Beckmann, Maurice Redondo, Olivier Kobel, Christoph Noppen, Niko Beerenwinkel, Tanja Stadler |
| EPI_ISL_3296191 | Wales Specialist Virology Centre Sequencing lab: Pathogen Genomics Unit | Public Health Wales Microbiology Cardiff Wales Specialist Virology Centre | Catherine Moore, Johnathan Evans, Laura Gifford, Malorie Perry, Simon Cottrell, Angela Marchbank, Alec Birchley, Alexander Adams, Amy Gaskin, Bree Gatica-Wilcox, Jason Coombes, Joel Southgate, Lauren Gilbert, Lee Graham, Nicole Pacchiarini, Sara Kumziene-Summerhayes, Sarah Taylor, Sophie Jones, Sara Rey, Matthew Bull, Joanne Watkins, Sally Corden, Tom Connor |
| EPI_ISL_3296192 | Viollier AG | Department of Biosystems Science and Engineering, ETH Zürich | Christian Beisel, Sarah Nadeau, Chaoran Chen, Ivan Topolsky, Kim Philipp Jablonski, Lara Fuhrmann, Rebecca Denes, Mirjam Feldkamp, Ina Nissen, Natascha Santacroce, Elodie Burcklen, Christiane Beckmann, Maurice Redondo, Olivier Kobel, Christoph Noppen, Niko Beerenwinkel, Tanja Stadler |
| EPI_ISL_3296193 | Wales Specialist Virology Centre Sequencing lab: Pathogen Genomics Unit | Public Health Wales Microbiology Cardiff Wales Specialist Virology Centre | Catherine Moore, Johnathan Evans, Laura Gifford, Malorie Perry, Simon Cottrell, Angela Marchbank, Alec Birchley, Alexander Adams, Amy Gaskin, Bree Gatica-Wilcox, Jason Coombes, Joel Southgate, Lauren Gilbert, Lee Graham, Nicole Pacchiarini, Sara Kumziene-Summerhayes, Sarah Taylor, Sophie Jones, Sara Rey, Matthew Bull, Joanne Watkins, Sally Corden, Tom Connor |
| EPI_ISL_3296194 | Viollier AG | Department of Biosystems Science and Engineering, ETH Zürich | Christian Beisel, Sarah Nadeau, Chaoran Chen, Ivan Topolsky, Kim Philipp Jablonski, Lara Fuhrmann, Rebecca Denes, Mirjam Feldkamp, Ina Nissen, Natascha Santacroce, Elodie Burcklen, Christiane Beckmann, Maurice Redondo, Olivier Kobel, Christoph Noppen, Niko Beerenwinkel, Tanja Stadler |
| EPI_ISL_3296195 | Wales Specialist Virology Centre Sequencing lab: Pathogen Genomics Unit | Public Health Wales Microbiology Cardiff Wales Specialist Virology Centre | Catherine Moore, Johnathan Evans, Laura Gifford, Malorie Perry, Simon Cottrell, Angela Marchbank, Alec Birchley, Alexander Adams, Amy Gaskin, Bree Gatica-Wilcox, Jason Coombes, Joel Southgate, Lauren Gilbert, Lee Graham, Nicole Pacchiarini, Sara Kumziene-Summerhayes, Sarah Taylor, Sophie Jones, Sara Rey, Matthew Bull, Joanne Watkins, Sally Corden, Tom Connor |
| EPI_ISL_3296196 | Viollier AG | Department of Biosystems Science and Engineering, ETH Zürich | Christian Beisel, Sarah Nadeau, Chaoran Chen, Ivan Topolsky, Kim Philipp Jablonski, Lara Fuhrmann, Rebecca Denes, Mirjam Feldkamp, Ina Nissen, Natascha Santacroce, Elodie Burcklen, Christiane Beckmann, Maurice Redondo, Olivier Kobel, Christoph Noppen, Niko Beerenwinkel, Tanja Stadler |
| EPI_ISL_3296197, EPI_ISL_3296198 | Wales Specialist Virology Centre Sequencing lab: Pathogen Genomics Unit | Public Health Wales Microbiology Cardiff Wales Specialist Virology Centre | Catherine Moore, Johnathan Evans, Laura Gifford, Malorie Perry, Simon Cottrell, Angela Marchbank, Alec Birchley, Alexander Adams, Amy Gaskin, Bree Gatica-Wilcox, Jason Coombes, Joel Southgate, Lauren Gilbert, Lee Graham, Nicole Pacchiarini, Sara Kumziene-Summerhayes, Sarah Taylor, Sophie Jones, Sara Rey, Matthew Bull, Joanne Watkins, Sally Corden, Tom Connor |
| EPI_ISL_3296199 | Viollier AG | Department of Biosystems Science and Engineering, ETH Zürich | Christian Beisel, Sarah Nadeau, Chaoran Chen, Ivan Topolsky, Kim Philipp Jablonski, Lara Fuhrmann, Rebecca Denes, Mirjam Feldkamp, Ina Nissen, Natascha Santacroce, Elodie Burcklen, Christiane Beckmann, Maurice Redondo, Olivier Kobel, Christoph Noppen, Niko Beerenwinkel, Tanja Stadler |
| EPI_ISL_3296200 | Wales Specialist Virology Centre Sequencing lab: Pathogen Genomics Unit | Public Health Wales Microbiology Cardiff Wales Specialist Virology Centre | Catherine Moore, Johnathan Evans, Laura Gifford, Malorie Perry, Simon Cottrell, Angela Marchbank, Alec Birchley, Alexander Adams, Amy Gaskin, Bree Gatica-Wilcox, Jason Coombes, Joel Southgate, Lauren Gilbert, Lee Graham, Nicole Pacchiarini, Sara Kumziene-Summerhayes, Sarah Taylor, Sophie Jones, Sara Rey, Matthew Bull, Joanne Watkins, Sally Corden, Tom Connor |
| EPI_ISL_3296201 | Viollier AG | Department of Biosystems Science and Engineering, ETH Zürich | Christian Beisel, Sarah Nadeau, Chaoran Chen, Ivan Topolsky, Kim Philipp Jablonski, Lara Fuhrmann, Rebecca Denes, Mirjam Feldkamp, Ina Nissen, Natascha Santacroce, Elodie Burcklen, Christiane Beckmann, Maurice Redondo, Olivier Kobel, Christoph Noppen, Niko Beerenwinkel, Tanja Stadler |
| EPI_ISL_3296202 | Wales Specialist Virology Centre Sequencing lab: Pathogen Genomics Unit | Public Health Wales Microbiology Cardiff Wales Specialist Virology Centre | Catherine Moore, Johnathan Evans, Laura Gifford, Malorie Perry, Simon Cottrell, Angela Marchbank, Alec Birchley, Alexander Adams, Amy Gaskin, Bree Gatica-Wilcox, Jason Coombes, Joel Southgate, Lauren Gilbert, Lee Graham, Nicole Pacchiarini, Sara Kumziene-Summerhayes, Sarah Taylor, Sophie Jones, Sara Rey, Matthew Bull, Joanne Watkins, Sally Corden, Tom Connor |
| EPI_ISL_3296203 | Viollier AG | Department of Biosystems Science and Engineering, ETH Zürich | Christian Beisel, Sarah Nadeau, Chaoran Chen, Ivan Topolsky, Kim Philipp Jablonski, Lara Fuhrmann, Rebecca Denes, Mirjam Feldkamp, Ina Nissen, Natascha Santacroce, Elodie Burcklen, Christiane Beckmann, Maurice Redondo, Olivier Kobel, Christoph Noppen, Niko Beerenwinkel, Tanja Stadler |

| | | | |
|---|---|---|---|
| EPI_ISL_3296204 | Wales Specialist Virology Centre Sequencing lab: Pathogen Genomics Unit | Public Health Wales Microbiology Cardiff Wales Specialist Virology Centre | Catherine Moore, Johnathan Evans, Laura Gifford, Malorie Perry, Simon Cottrell, Angela Marchbank, Alec Birchley, Alexander Adams, Amy Gaskin, Bree Gatica-Wilcox, Jason Coombes, Joel Southgate, Lauren Gilbert, Lee Graham, Nicole Pacchiarini, Sara Kumziene-Summerhayes, Sarah Taylor, Sophie Jones, Sara Rey, Matthew Bull, Joanne Watkins, Sally Corden, Tom Connor |
| EPI_ISL_3296205 | Viollier AG | Department of Biosystems Science and Engineering, ETH Zürich | Christian Beisel, Sarah Nadeau, Chaoran Chen, Ivan Topolsky, Kim Philipp Jablonski, Lara Fuhrmann, Rebecca Denes, Mirjam Feldkamp, Ina Nissen, Natascha Santacroce, Elodie Burcklen, Christiane Beckmann, Maurice Redondo, Olivier Kobel, Christoph Noppen, Niko Beerenwinkel, Tanja Stadler |
| EPI_ISL_3296206 | Wales Specialist Virology Centre Sequencing lab: Pathogen Genomics Unit | Public Health Wales Microbiology Cardiff Wales Specialist Virology Centre | Catherine Moore, Johnathan Evans, Laura Gifford, Malorie Perry, Simon Cottrell, Angela Marchbank, Alec Birchley, Alexander Adams, Amy Gaskin, Bree Gatica-Wilcox, Jason Coombes, Joel Southgate, Lauren Gilbert, Lee Graham, Nicole Pacchiarini, Sara Kumziene-Summerhayes, Sarah Taylor, Sophie Jones, Sara Rey, Matthew Bull, Joanne Watkins, Sally Corden, Tom Connor |
| EPI_ISL_3296207 | Viollier AG | Department of Biosystems Science and Engineering, ETH Zürich | Christian Beisel, Sarah Nadeau, Chaoran Chen, Ivan Topolsky, Kim Philipp Jablonski, Lara Fuhrmann, Rebecca Denes, Mirjam Feldkamp, Ina Nissen, Natascha Santacroce, Elodie Burcklen, Christiane Beckmann, Maurice Redondo, Olivier Kobel, Christoph Noppen, Niko Beerenwinkel, Tanja Stadler |
| EPI_ISL_3296208 | Wales Specialist Virology Centre Sequencing lab: Pathogen Genomics Unit | Public Health Wales Microbiology Cardiff Wales Specialist Virology Centre | Catherine Moore, Johnathan Evans, Laura Gifford, Malorie Perry, Simon Cottrell, Angela Marchbank, Alec Birchley, Alexander Adams, Amy Gaskin, Bree Gatica-Wilcox, Jason Coombes, Joel Southgate, Lauren Gilbert, Lee Graham, Nicole Pacchiarini, Sara Kumziene-Summerhayes, Sarah Taylor, Sophie Jones, Sara Rey, Matthew Bull, Joanne Watkins, Sally Corden, Tom Connor |
| EPI_ISL_3296209 | Viollier AG | Department of Biosystems Science and Engineering, ETH Zürich | Christian Beisel, Sarah Nadeau, Chaoran Chen, Ivan Topolsky, Kim Philipp Jablonski, Lara Fuhrmann, Rebecca Denes, Mirjam Feldkamp, Ina Nissen, Natascha Santacroce, Elodie Burcklen, Christiane Beckmann, Maurice Redondo, Olivier Kobel, Christoph Noppen, Niko Beerenwinkel, Tanja Stadler |
| EPI_ISL_3296210 | Wales Specialist Virology Centre Sequencing lab: Pathogen Genomics Unit | Public Health Wales Microbiology Cardiff Wales Specialist Virology Centre | Catherine Moore, Johnathan Evans, Laura Gifford, Malorie Perry, Simon Cottrell, Angela Marchbank, Alec Birchley, Alexander Adams, Amy Gaskin, Bree Gatica-Wilcox, Jason Coombes, Joel Southgate, Lauren Gilbert, Lee Graham, Nicole Pacchiarini, Sara Kumziene-Summerhayes, Sarah Taylor, Sophie Jones, Sara Rey, Matthew Bull, Joanne Watkins, Sally Corden, Tom Connor |
| EPI_ISL_3296211 | Viollier AG | Department of Biosystems Science and Engineering, ETH Zürich | Christian Beisel, Sarah Nadeau, Chaoran Chen, Ivan Topolsky, Kim Philipp Jablonski, Lara Fuhrmann, Rebecca Denes, Mirjam Feldkamp, Ina Nissen, Natascha Santacroce, Elodie Burcklen, Christiane Beckmann, Maurice Redondo, Olivier Kobel, Christoph Noppen, Niko Beerenwinkel, Tanja Stadler |
| EPI_ISL_3296212 | Wales Specialist Virology Centre Sequencing lab: Pathogen Genomics Unit | Public Health Wales Microbiology Cardiff Wales Specialist Virology Centre | Catherine Moore, Johnathan Evans, Laura Gifford, Malorie Perry, Simon Cottrell, Angela Marchbank, Alec Birchley, Alexander Adams, Amy Gaskin, Bree Gatica-Wilcox, Jason Coombes, Joel Southgate, Lauren Gilbert, Lee Graham, Nicole Pacchiarini, Sara Kumziene-Summerhayes, Sarah Taylor, Sophie Jones, Sara Rey, Matthew Bull, Joanne Watkins, Sally Corden, Tom Connor |
| EPI_ISL_3296213 | Viollier AG | Department of Biosystems Science and Engineering, ETH Zürich | Christian Beisel, Sarah Nadeau, Chaoran Chen, Ivan Topolsky, Kim Philipp Jablonski, Lara Fuhrmann, Rebecca Denes, Mirjam Feldkamp, Ina Nissen, Natascha Santacroce, Elodie Burcklen, Christiane Beckmann, Maurice Redondo, Olivier Kobel, Christoph Noppen, Niko Beerenwinkel, Tanja Stadler |
| EPI_ISL_3296214 | Wales Specialist Virology Centre Sequencing lab: Pathogen Genomics Unit | Public Health Wales Microbiology Cardiff Wales Specialist Virology Centre | Catherine Moore, Johnathan Evans, Laura Gifford, Malorie Perry, Simon Cottrell, Angela Marchbank, Alec Birchley, Alexander Adams, Amy Gaskin, Bree Gatica-Wilcox, Jason Coombes, Joel Southgate, Lauren Gilbert, Lee Graham, Nicole Pacchiarini, Sara Kumziene-Summerhayes, Sarah Taylor, Sophie Jones, Sara Rey, Matthew Bull, Joanne Watkins, Sally Corden, Tom Connor |
| EPI_ISL_3296215 | Viollier AG | Department of Biosystems Science and Engineering, ETH Zürich | Christian Beisel, Sarah Nadeau, Chaoran Chen, Ivan Topolsky, Kim Philipp Jablonski, Lara Fuhrmann, Rebecca Denes, Mirjam Feldkamp, Ina Nissen, Natascha Santacroce, Elodie Burcklen, Christiane Beckmann, Maurice Redondo, Olivier Kobel, Christoph Noppen, Niko Beerenwinkel, Tanja Stadler |
| EPI_ISL_3296216 | Wales Specialist Virology Centre Sequencing lab: Pathogen Genomics Unit | Public Health Wales Microbiology Cardiff Wales Specialist Virology Centre | Catherine Moore, Johnathan Evans, Laura Gifford, Malorie Perry, Simon Cottrell, Angela Marchbank, Alec Birchley, Alexander Adams, Amy Gaskin, Bree Gatica-Wilcox, Jason Coombes, Joel Southgate, Lauren Gilbert, Lee Graham, Nicole Pacchiarini, Sara Kumziene-Summerhayes, Sarah Taylor, Sophie Jones, Sara Rey, Matthew Bull, Joanne Watkins, Sally Corden, Tom Connor |
| EPI_ISL_3296217 | Viollier AG | Department of Biosystems Science and Engineering, ETH Zürich | Christian Beisel, Sarah Nadeau, Chaoran Chen, Ivan Topolsky, Kim Philipp Jablonski, Lara Fuhrmann, Rebecca Denes, Mirjam Feldkamp, Ina Nissen, Natascha Santacroce, Elodie Burcklen, Christiane Beckmann, Maurice Redondo, Olivier Kobel, Christoph Noppen, Niko Beerenwinkel, Tanja Stadler |
| EPI_ISL_3296218 | Wales Specialist Virology Centre Sequencing lab: Pathogen Genomics Unit | Public Health Wales Microbiology Cardiff Wales Specialist Virology Centre | Catherine Moore, Johnathan Evans, Laura Gifford, Malorie Perry, Simon Cottrell, Angela Marchbank, Alec Birchley, Alexander Adams, Amy Gaskin, Bree Gatica-Wilcox, Jason Coombes, Joel Southgate, Lauren Gilbert, Lee Graham, Nicole Pacchiarini, Sara Kumziene-Summerhayes, Sarah Taylor, Sophie Jones, Sara Rey, Matthew Bull, Joanne Watkins, Sally Corden, Tom Connor |
| EPI_ISL_3296219 | Viollier AG | Department of Biosystems Science and Engineering, ETH Zürich | Christian Beisel, Sarah Nadeau, Chaoran Chen, Ivan Topolsky, Kim Philipp Jablonski, Lara Fuhrmann, Rebecca Denes, Mirjam Feldkamp, Ina Nissen, Natascha Santacroce, Elodie Burcklen, Christiane Beckmann, Maurice Redondo, Olivier Kobel, Christoph Noppen, Niko Beerenwinkel, Tanja Stadler |
| EPI_ISL_3296220 | Wales Specialist Virology Centre Sequencing lab: Pathogen Genomics Unit | Public Health Wales Microbiology Cardiff Wales Specialist Virology Centre | Catherine Moore, Johnathan Evans, Laura Gifford, Malorie Perry, Simon Cottrell, Angela Marchbank, Alec Birchley, Alexander Adams, Amy Gaskin, Bree Gatica-Wilcox, Jason Coombes, Joel Southgate, Lauren Gilbert, Lee Graham, Nicole Pacchiarini, Sara Kumziene-Summerhayes, Sarah Taylor, Sophie Jones, Sara Rey, Matthew Bull, Joanne Watkins, Sally Corden, Tom Connor |
| EPI_ISL_3296221 | Viollier AG | Department of Biosystems Science and Engineering, ETH Zürich | Christian Beisel, Sarah Nadeau, Chaoran Chen, Ivan Topolsky, Kim Philipp Jablonski, Lara Fuhrmann, Rebecca Denes, Mirjam Feldkamp, Ina Nissen, Natascha Santacroce, Elodie Burcklen, Christiane Beckmann, Maurice Redondo, Olivier Kobel, Christoph Noppen, Niko Beerenwinkel, Tanja Stadler |
| EPI_ISL_3296222, EPI_ISL_3296223 | Wales Specialist Virology Centre Sequencing lab: Pathogen Genomics Unit | Public Health Wales Microbiology Cardiff Wales Specialist Virology Centre | Catherine Moore, Johnathan Evans, Laura Gifford, Malorie Perry, Simon Cottrell, Angela Marchbank, Alec Birchley, Alexander Adams, Amy Gaskin, Bree Gatica-Wilcox, Jason Coombes, Joel Southgate, Lauren Gilbert, Lee Graham, Nicole Pacchiarini, Sara Kumziene-Summerhayes, Sarah Taylor, Sophie Jones, Sara Rey, Matthew Bull, Joanne Watkins, Sally Corden, Tom Connor |
| EPI_ISL_3296224 | Viollier AG | Department of Biosystems Science and Engineering, ETH Zürich | Christian Beisel, Sarah Nadeau, Chaoran Chen, Ivan Topolsky, Kim Philipp Jablonski, Lara Fuhrmann, Rebecca Denes, Mirjam Feldkamp, Ina Nissen, Natascha Santacroce, Elodie Burcklen, Christiane Beckmann, Maurice Redondo, Olivier Kobel, Christoph Noppen, Niko Beerenwinkel, Tanja Stadler |
| EPI_ISL_3296225 | Wales Specialist Virology Centre Sequencing lab: Pathogen Genomics Unit | Public Health Wales Microbiology Cardiff Wales Specialist Virology Centre | Catherine Moore, Johnathan Evans, Laura Gifford, Malorie Perry, Simon Cottrell, Angela Marchbank, Alec Birchley, Alexander Adams, Amy Gaskin, Bree Gatica-Wilcox, Jason Coombes, Joel Southgate, Lauren Gilbert, Lee Graham, Nicole Pacchiarini, Sara Kumziene-Summerhayes, Sarah Taylor, Sophie Jones, Sara Rey, Matthew Bull, Joanne Watkins, Sally Corden, Tom Connor |
| EPI_ISL_3296226 | Viollier AG | Department of Biosystems Science and Engineering, ETH Zürich | Christian Beisel, Sarah Nadeau, Chaoran Chen, Ivan Topolsky, Kim Philipp Jablonski, Lara Fuhrmann, Rebecca Denes, Mirjam Feldkamp, Ina Nissen, Natascha Santacroce, Elodie Burcklen, Christiane Beckmann, Maurice Redondo, Olivier Kobel, Christoph Noppen, Niko Beerenwinkel, Tanja Stadler |
| EPI_ISL_3296227 | Wales Specialist Virology Centre Sequencing lab: Pathogen Genomics Unit | Public Health Wales Microbiology Cardiff Wales Specialist Virology Centre | Catherine Moore, Johnathan Evans, Laura Gifford, Malorie Perry, Simon Cottrell, Angela Marchbank, Alec Birchley, Alexander Adams, Amy Gaskin, Bree Gatica-Wilcox, Jason Coombes, Joel Southgate, Lauren Gilbert, Lee Graham, Nicole Pacchiarini, Sara Kumziene-Summerhayes, Sarah Taylor, Sophie Jones, Sara Rey, Matthew Bull, Joanne Watkins, Sally Corden, Tom Connor |
| EPI_ISL_3296228 | Viollier AG | Department of Biosystems Science and Engineering, ETH Zürich | Christian Beisel, Sarah Nadeau, Chaoran Chen, Ivan Topolsky, Kim Philipp Jablonski, Lara Fuhrmann, Rebecca Denes, Mirjam Feldkamp, Ina Nissen, Natascha Santacroce, Elodie Burcklen, Christiane Beckmann, Maurice Redondo, Olivier Kobel, Christoph Noppen, Niko Beerenwinkel, Tanja Stadler |
| EPI_ISL_3296229 | Wales Specialist Virology Centre Sequencing lab: Pathogen Genomics Unit | Public Health Wales Microbiology Cardiff Wales Specialist Virology Centre | Catherine Moore, Johnathan Evans, Laura Gifford, Malorie Perry, Simon Cottrell, Angela Marchbank, Alec Birchley, Alexander Adams, Amy Gaskin, Bree Gatica-Wilcox, Jason Coombes, Joel Southgate, Lauren Gilbert, Lee Graham, Nicole Pacchiarini, Sara Kumziene-Summerhayes, Sarah Taylor, Sophie Jones, Sara Rey, Matthew Bull, Joanne Watkins, Sally Corden, Tom Connor |
| EPI_ISL_3296230 | Viollier AG | Department of Biosystems Science and Engineering, ETH Zürich | Christian Beisel, Sarah Nadeau, Chaoran Chen, Ivan Topolsky, Kim Philipp Jablonski, Lara Fuhrmann, Rebecca Denes, Mirjam Feldkamp, Ina Nissen, Natascha Santacroce, Elodie Burcklen, Christiane Beckmann, Maurice Redondo, Olivier Kobel, Christoph Noppen, Niko Beerenwinkel, Tanja Stadler |
| EPI_ISL_3296231 | Wales Specialist Virology Centre Sequencing lab: Pathogen Genomics Unit | Public Health Wales Microbiology Cardiff Wales Specialist Virology Centre | Catherine Moore, Johnathan Evans, Laura Gifford, Malorie Perry, Simon Cottrell, Angela Marchbank, Alec Birchley, Alexander Adams, Amy Gaskin, Bree Gatica-Wilcox, Jason Coombes, Joel Southgate, Lauren Gilbert, Lee Graham, Nicole Pacchiarini, Sara Kumziene-Summerhayes, Sarah Taylor, Sophie Jones, Sara Rey, Matthew Bull, Joanne Watkins, Sally Corden, Tom Connor |
| EPI_ISL_3296232 | Viollier AG | Department of Biosystems Science and Engineering, ETH Zürich | Christian Beisel, Sarah Nadeau, Chaoran Chen, Ivan Topolsky, Kim Philipp Jablonski, Lara Fuhrmann, Rebecca Denes, Mirjam Feldkamp, Ina Nissen, Natascha Santacroce, Elodie Burcklen, Christiane Beckmann, Maurice Redondo, Olivier Kobel, Christoph Noppen, Niko Beerenwinkel, Tanja Stadler |
| EPI_ISL_3296233 | Wales Specialist Virology Centre Sequencing lab: Pathogen Genomics Unit | Public Health Wales Microbiology Cardiff Wales Specialist Virology Centre | Catherine Moore, Johnathan Evans, Laura Gifford, Malorie Perry, Simon Cottrell, Angela Marchbank, Alec Birchley, Alexander Adams, Amy Gaskin, Bree Gatica-Wilcox, Jason Coombes, Joel Southgate, Lauren Gilbert, Lee Graham, Nicole Pacchiarini, Sara Kumziene-Summerhayes, Sarah Taylor, Sophie Jones, Sara Rey, Matthew Bull, Joanne Watkins, Sally Corden, Tom Connor |
| EPI_ISL_3296234 | Viollier AG | Department of Biosystems Science and Engineering, ETH Zürich | Christian Beisel, Sarah Nadeau, Chaoran Chen, Ivan Topolsky, Kim Philipp Jablonski, Lara Fuhrmann, Rebecca Denes, Mirjam Feldkamp, Ina Nissen, Natascha Santacroce, Elodie Burcklen, Christiane Beckmann, Maurice Redondo, Olivier Kobel, Christoph Noppen, Niko Beerenwinkel, Tanja Stadler |
| EPI_ISL_3296235 | Wales Specialist Virology Centre Sequencing lab: Pathogen Genomics Unit | Public Health Wales Microbiology Cardiff Wales Specialist Virology Centre | Catherine Moore, Johnathan Evans, Laura Gifford, Malorie Perry, Simon Cottrell, Angela Marchbank, Alec Birchley, Alexander Adams, Amy Gaskin, Bree Gatica-Wilcox, Jason Coombes, Joel Southgate, Lauren Gilbert, Lee Graham, Nicole Pacchiarini, Sara Kumziene-Summerhayes, Sarah Taylor, Sophie Jones, Sara Rey, Matthew Bull, Joanne Watkins, Sally Corden, Tom Connor |
| EPI_ISL_3296236 | Viollier AG | Department of Biosystems Science and Engineering, ETH Zürich | Christian Beisel, Sarah Nadeau, Chaoran Chen, Ivan Topolsky, Kim Philipp Jablonski, Lara Fuhrmann, Rebecca Denes, Mirjam Feldkamp, Ina Nissen, Natascha Santacroce, Elodie Burcklen, Christiane Beckmann, Maurice Redondo, Olivier Kobel, Christoph Noppen, Niko Beerenwinkel, Tanja Stadler |
| EPI_ISL_3296237 | Wales Specialist Virology Centre Sequencing lab: Pathogen Genomics Unit | Public Health Wales Microbiology Cardiff Wales Specialist Virology Centre | Catherine Moore, Johnathan Evans, Laura Gifford, Malorie Perry, Simon Cottrell, Angela Marchbank, Alec Birchley, Alexander Adams, Amy Gaskin, Bree Gatica-Wilcox, Jason Coombes, Joel Southgate, Lauren Gilbert, Lee Graham, Nicole Pacchiarini, Sara Kumziene-Summerhayes, Sarah Taylor, Sophie Jones, Sara Rey, Matthew Bull, Joanne Watkins, Sally Corden, Tom Connor |
| EPI_ISL_3296238 | Viollier AG | Department of Biosystems Science and Engineering, ETH Zürich | Christian Beisel, Sarah Nadeau, Chaoran Chen, Ivan Topolsky, Kim Philipp Jablonski, Lara Fuhrmann, Rebecca Denes, Mirjam Feldkamp, Ina Nissen, Natascha Santacroce, Elodie Burcklen, Christiane Beckmann, Maurice Redondo, Olivier Kobel, Christoph Noppen, Niko Beerenwinkel, Tanja Stadler |
| EPI_ISL_3296239, EPI_ISL_3296241 | Wales Specialist Virology Centre Sequencing lab: Pathogen Genomics Unit | Public Health Wales Microbiology Cardiff Wales Specialist Virology Centre | Catherine Moore, Johnathan Evans, Laura Gifford, Malorie Perry, Simon Cottrell, Angela Marchbank, Alec Birchley, Alexander Adams, Amy Gaskin, Bree Gatica-Wilcox, Jason Coombes, Joel Southgate, Lauren Gilbert, Lee Graham, Nicole Pacchiarini, Sara Kumziene-Summerhayes, Sarah Taylor, Sophie Jones, Sara Rey, Matthew Bull, Joanne Watkins, Sally Corden, Tom Connor |
| EPI_ISL_3296242 | Viollier AG | Department of Biosystems Science and Engineering, ETH Zürich | Christian Beisel, Sarah Nadeau, Chaoran Chen, Ivan Topolsky, Kim Philipp Jablonski, Lara Fuhrmann, Rebecca Denes, Mirjam Feldkamp, Ina Nissen, Natascha Santacroce, Elodie Burcklen, Christiane Beckmann, Maurice Redondo, Olivier Kobel, Christoph Noppen, Niko Beerenwinkel, Tanja Stadler |
| EPI_ISL_3296243 | Wales Specialist Virology Centre Sequencing lab: Pathogen Genomics Unit | Public Health Wales Microbiology Cardiff Wales Specialist Virology Centre | Catherine Moore, Johnathan Evans, Laura Gifford, Malorie Perry, Simon Cottrell, Angela Marchbank, Alec Birchley, Alexander Adams, Amy Gaskin, Bree Gatica-Wilcox, Jason Coombes, Joel Southgate, Lauren Gilbert, Lee Graham, Nicole Pacchiarini, Sara Kumziene-Summerhayes, Sarah Taylor, Sophie Jones, Sara Rey, Matthew Bull, Joanne Watkins, Sally Corden, Tom Connor |
| EPI_ISL_3296244 | Viollier AG | Department of Biosystems Science and Engineering, ETH Zürich | Christian Beisel, Sarah Nadeau, Chaoran Chen, Ivan Topolsky, Kim Philipp Jablonski, Lara Fuhrmann, Rebecca Denes, Mirjam Feldkamp, Ina Nissen, Natascha Santacroce, Elodie Burcklen, Christiane Beckmann, Maurice Redondo, Olivier Kobel, Christoph Noppen, Niko Beerenwinkel, Tanja Stadler |
| EPI_ISL_3296245, EPI_ISL_3296247, EPI_ISL_3296249 | Wales Specialist Virology Centre Sequencing lab: Pathogen Genomics Unit | Public Health Wales Microbiology Cardiff Wales Specialist Virology Centre | Catherine Moore, Johnathan Evans, Laura Gifford, Malorie Perry, Simon Cottrell, Angela Marchbank, Alec Birchley, Alexander Adams, Amy Gaskin, Bree Gatica-Wilcox, Jason Coombes, Joel Southgate, Lauren Gilbert, Lee Graham, Nicole Pacchiarini, Sara Kumziene-Summerhayes, Sarah Taylor, Sophie Jones, Sara Rey, Matthew Bull, Joanne Watkins, Sally Corden, Tom Connor |
| EPI_ISL_3296250 | Viollier AG | Department of Biosystems Science and Engineering, ETH Zürich | Christian Beisel, Sarah Nadeau, Chaoran Chen, Ivan Topolsky, Kim Philipp Jablonski, Lara Fuhrmann, Rebecca Denes, Mirjam Feldkamp, Ina Nissen, Natascha Santacroce, Elodie Burcklen, Christiane Beckmann, Maurice Redondo, Olivier Kobel, Christoph Noppen, Niko Beerenwinkel, Tanja Stadler |
| EPI_ISL_3296251 | Wales Specialist Virology Centre Sequencing lab: Pathogen Genomics Unit | Public Health Wales Microbiology Cardiff Wales Specialist Virology Centre | Catherine Moore, Johnathan Evans, Laura Gifford, Malorie Perry, Simon Cottrell, Angela Marchbank, Alec Birchley, Alexander Adams, Amy Gaskin, Bree Gatica-Wilcox, Jason Coombes, Joel Southgate, Lauren Gilbert, Lee Graham, Nicole Pacchiarini, Sara Kumziene-Summerhayes, Sarah Taylor, Sophie Jones, Sara Rey, Matthew Bull, Joanne Watkins, Sally Corden, Tom Connor |
| EPI_ISL_3296252 | Viollier AG | Department of Biosystems Science and Engineering, ETH Zürich | Christian Beisel, Sarah Nadeau, Chaoran Chen, Ivan Topolsky, Kim Philipp Jablonski, Lara Fuhrmann, Rebecca Denes, Mirjam Feldkamp, Ina Nissen, Natascha Santacroce, Elodie Burcklen, Christiane Beckmann, Maurice Redondo, Olivier Kobel, Christoph Noppen, Niko Beerenwinkel, Tanja Stadler |
| EPI_ISL_3296253 | Wales Specialist Virology Centre Sequencing lab: Pathogen Genomics Unit | Public Health Wales Microbiology Cardiff Wales Specialist Virology Centre | Catherine Moore, Johnathan Evans, Laura Gifford, Malorie Perry, Simon Cottrell, Angela Marchbank, Alec Birchley, Alexander Adams, Amy Gaskin, Bree Gatica-Wilcox, Jason Coombes, Joel Southgate, Lauren Gilbert, Lee Graham, Nicole Pacchiarini, Sara Kumziene-Summerhayes, Sarah Taylor, Sophie Jones, Sara Rey, Matthew Bull, Joanne Watkins, Sally Corden, Tom Connor |
| EPI_ISL_3296254 | Viollier AG | Department of Biosystems Science and Engineering, ETH Zürich | Christian Beisel, Sarah Nadeau, Chaoran Chen, Ivan Topolsky, Kim Philipp Jablonski, Lara Fuhrmann, Rebecca Denes, Mirjam Feldkamp, Ina Nissen, Natascha Santacroce, Elodie Burcklen, Christiane Beckmann, Maurice Redondo, Olivier Kobel, Christoph Noppen, Niko Beerenwinkel, Tanja Stadler |
| EPI_ISL_3296255, | Wales Specialist Virology Centre Sequencing lab: Pathogen | Public Health Wales Microbiology Cardiff Wales Specialist Virology | Catherine Moore, Johnathan Evans, Laura Gifford, Malorie Perry, Simon Cottrell, Angela Marchbank, Alec Birchley, Alexander Adams, Amy Gaskin, Bree Gatica-Wilcox, Jason Coombes, Joel Southgate, Lauren Gilbert, Lee Graham, Nicole |

| | | | |
|---|---|---|---|
| EPI_ISL_3296256 | Genomics Unit | Centre | Pacchiarini, Sara Kumziene-Summerhayes, Sarah Taylor, Sophie Jones, Sara Rey, Matthew Bull, Joanne Watkins, Sally Corden, Tom Connor |
| EPI_ISL_3296257, EPI_ISL_3296258 | Viollier AG | Department of Biosystems Science and Engineering, ETH Zürich | Christian Beisel, Sarah Nadeau, Chaoran Chen, Ivan Topolsky, Kim Philipp Jablonski, Lara Fuhrmann, Rebecca Denes, Mirjam Feldkamp, Ina Nissen, Natascha Santacroce, Elodie Burcklen, Christiane Beckmann, Maurice Redondo, Olivier Kobel, Christoph Noppen, Niko Beerenwinkel, Tanja Stadler |
| EPI_ISL_3296259, EPI_ISL_3296261, EPI_ISL_3296262 | Wales Specialist Virology Centre Sequencing lab: Pathogen Genomics Unit | Public Health Wales Microbiology Cardiff Wales Specialist Virology Centre | Catherine Moore, Johnathan Evans, Laura Gifford, Malorie Perry, Simon Cottrell, Angela Marchbank, Alec Birchley, Alexander Adams, Amy Gaskin, Bree Gatica-Wilcox, Jason Coombes, Joel Southgate, Lauren Gilbert, Lee Graham, Nicole Pacchiarini, Sara Kumziene-Summerhayes, Sarah Taylor, Sophie Jones, Sara Rey, Matthew Bull, Joanne Watkins, Sally Corden, Tom Connor |
| EPI_ISL_3296263 | Viollier AG | Department of Biosystems Science and Engineering, ETH Zürich | Christian Beisel, Sarah Nadeau, Chaoran Chen, Ivan Topolsky, Kim Philipp Jablonski, Lara Fuhrmann, Rebecca Denes, Mirjam Feldkamp, Ina Nissen, Natascha Santacroce, Elodie Burcklen, Christiane Beckmann, Maurice Redondo, Olivier Kobel, Christoph Noppen, Niko Beerenwinkel, Tanja Stadler |
| EPI_ISL_3296264 | Wales Specialist Virology Centre Sequencing lab: Pathogen Genomics Unit | Public Health Wales Microbiology Cardiff Wales Specialist Virology Centre | Catherine Moore, Johnathan Evans, Laura Gifford, Malorie Perry, Simon Cottrell, Angela Marchbank, Alec Birchley, Alexander Adams, Amy Gaskin, Bree Gatica-Wilcox, Jason Coombes, Joel Southgate, Lauren Gilbert, Lee Graham, Nicole Pacchiarini, Sara Kumziene-Summerhayes, Sarah Taylor, Sophie Jones, Sara Rey, Matthew Bull, Joanne Watkins, Sally Corden, Tom Connor |
| EPI_ISL_3296265 | Viollier AG | Department of Biosystems Science and Engineering, ETH Zürich | Christian Beisel, Sarah Nadeau, Chaoran Chen, Ivan Topolsky, Kim Philipp Jablonski, Lara Fuhrmann, Rebecca Denes, Mirjam Feldkamp, Ina Nissen, Natascha Santacroce, Elodie Burcklen, Christiane Beckmann, Maurice Redondo, Olivier Kobel, Christoph Noppen, Niko Beerenwinkel, Tanja Stadler |
| EPI_ISL_3296266, EPI_ISL_3296268 | Wales Specialist Virology Centre Sequencing lab: Pathogen Genomics Unit | Public Health Wales Microbiology Cardiff Wales Specialist Virology Centre | Catherine Moore, Johnathan Evans, Laura Gifford, Malorie Perry, Simon Cottrell, Angela Marchbank, Alec Birchley, Alexander Adams, Amy Gaskin, Bree Gatica-Wilcox, Jason Coombes, Joel Southgate, Lauren Gilbert, Lee Graham, Nicole Pacchiarini, Sara Kumziene-Summerhayes, Sarah Taylor, Sophie Jones, Sara Rey, Matthew Bull, Joanne Watkins, Sally Corden, Tom Connor |
| EPI_ISL_3296269 | Viollier AG | Department of Biosystems Science and Engineering, ETH Zürich | Christian Beisel, Sarah Nadeau, Chaoran Chen, Ivan Topolsky, Kim Philipp Jablonski, Lara Fuhrmann, Rebecca Denes, Mirjam Feldkamp, Ina Nissen, Natascha Santacroce, Elodie Burcklen, Christiane Beckmann, Maurice Redondo, Olivier Kobel, Christoph Noppen, Niko Beerenwinkel, Tanja Stadler |
| EPI_ISL_3296270 | Wales Specialist Virology Centre Sequencing lab: Pathogen Genomics Unit | Public Health Wales Microbiology Cardiff Wales Specialist Virology Centre | Catherine Moore, Johnathan Evans, Laura Gifford, Malorie Perry, Simon Cottrell, Angela Marchbank, Alec Birchley, Alexander Adams, Amy Gaskin, Bree Gatica-Wilcox, Jason Coombes, Joel Southgate, Lauren Gilbert, Lee Graham, Nicole Pacchiarini, Sara Kumziene-Summerhayes, Sarah Taylor, Sophie Jones, Sara Rey, Matthew Bull, Joanne Watkins, Sally Corden, Tom Connor |
| EPI_ISL_3296271 | Viollier AG | Department of Biosystems Science and Engineering, ETH Zürich | Christian Beisel, Sarah Nadeau, Chaoran Chen, Ivan Topolsky, Kim Philipp Jablonski, Lara Fuhrmann, Rebecca Denes, Mirjam Feldkamp, Ina Nissen, Natascha Santacroce, Elodie Burcklen, Christiane Beckmann, Maurice Redondo, Olivier Kobel, Christoph Noppen, Niko Beerenwinkel, Tanja Stadler |
| EPI_ISL_3296272 | Wales Specialist Virology Centre Sequencing lab: Pathogen Genomics Unit | Public Health Wales Microbiology Cardiff Wales Specialist Virology Centre | Catherine Moore, Johnathan Evans, Laura Gifford, Malorie Perry, Simon Cottrell, Angela Marchbank, Alec Birchley, Alexander Adams, Amy Gaskin, Bree Gatica-Wilcox, Jason Coombes, Joel Southgate, Lauren Gilbert, Lee Graham, Nicole Pacchiarini, Sara Kumziene-Summerhayes, Sarah Taylor, Sophie Jones, Sara Rey, Matthew Bull, Joanne Watkins, Sally Corden, Tom Connor |
| EPI_ISL_3296273 | Viollier AG | Department of Biosystems Science and Engineering, ETH Zürich | Christian Beisel, Sarah Nadeau, Chaoran Chen, Ivan Topolsky, Kim Philipp Jablonski, Lara Fuhrmann, Rebecca Denes, Mirjam Feldkamp, Ina Nissen, Natascha Santacroce, Elodie Burcklen, Christiane Beckmann, Maurice Redondo, Olivier Kobel, Christoph Noppen, Niko Beerenwinkel, Tanja Stadler |
| EPI_ISL_3296274 | Wales Specialist Virology Centre Sequencing lab: Pathogen Genomics Unit | Public Health Wales Microbiology Cardiff Wales Specialist Virology Centre | Catherine Moore, Johnathan Evans, Laura Gifford, Malorie Perry, Simon Cottrell, Angela Marchbank, Alec Birchley, Alexander Adams, Amy Gaskin, Bree Gatica-Wilcox, Jason Coombes, Joel Southgate, Lauren Gilbert, Lee Graham, Nicole Pacchiarini, Sara Kumziene-Summerhayes, Sarah Taylor, Sophie Jones, Sara Rey, Matthew Bull, Joanne Watkins, Sally Corden, Tom Connor |
| EPI_ISL_3296275 | Viollier AG | Department of Biosystems Science and Engineering, ETH Zürich | Christian Beisel, Sarah Nadeau, Chaoran Chen, Ivan Topolsky, Kim Philipp Jablonski, Lara Fuhrmann, Rebecca Denes, Mirjam Feldkamp, Ina Nissen, Natascha Santacroce, Elodie Burcklen, Christiane Beckmann, Maurice Redondo, Olivier Kobel, Christoph Noppen, Niko Beerenwinkel, Tanja Stadler |
| EPI_ISL_3296276 | Wales Specialist Virology Centre Sequencing lab: Pathogen Genomics Unit | Public Health Wales Microbiology Cardiff Wales Specialist Virology Centre | Catherine Moore, Johnathan Evans, Laura Gifford, Malorie Perry, Simon Cottrell, Angela Marchbank, Alec Birchley, Alexander Adams, Amy Gaskin, Bree Gatica-Wilcox, Jason Coombes, Joel Southgate, Lauren Gilbert, Lee Graham, Nicole Pacchiarini, Sara Kumziene-Summerhayes, Sarah Taylor, Sophie Jones, Sara Rey, Matthew Bull, Joanne Watkins, Sally Corden, Tom Connor |
| EPI_ISL_3296277 | Viollier AG | Department of Biosystems Science and Engineering, ETH Zürich | Christian Beisel, Sarah Nadeau, Chaoran Chen, Ivan Topolsky, Kim Philipp Jablonski, Lara Fuhrmann, Rebecca Denes, Mirjam Feldkamp, Ina Nissen, Natascha Santacroce, Elodie Burcklen, Christiane Beckmann, Maurice Redondo, Olivier Kobel, Christoph Noppen, Niko Beerenwinkel, Tanja Stadler |
| EPI_ISL_3296278 | Wales Specialist Virology Centre Sequencing lab: Pathogen Genomics Unit | Public Health Wales Microbiology Cardiff Wales Specialist Virology Centre | Catherine Moore, Johnathan Evans, Laura Gifford, Malorie Perry, Simon Cottrell, Angela Marchbank, Alec Birchley, Alexander Adams, Amy Gaskin, Bree Gatica-Wilcox, Jason Coombes, Joel Southgate, Lauren Gilbert, Lee Graham, Nicole Pacchiarini, Sara Kumziene-Summerhayes, Sarah Taylor, Sophie Jones, Sara Rey, Matthew Bull, Joanne Watkins, Sally Corden, Tom Connor |
| EPI_ISL_3296279 | Viollier AG | Department of Biosystems Science and Engineering, ETH Zürich | Christian Beisel, Sarah Nadeau, Chaoran Chen, Ivan Topolsky, Kim Philipp Jablonski, Lara Fuhrmann, Rebecca Denes, Mirjam Feldkamp, Ina Nissen, Natascha Santacroce, Elodie Burcklen, Christiane Beckmann, Maurice Redondo, Olivier Kobel, Christoph Noppen, Niko Beerenwinkel, Tanja Stadler |
| EPI_ISL_3296280 | Wales Specialist Virology Centre Sequencing lab: Pathogen Genomics Unit | Public Health Wales Microbiology Cardiff Wales Specialist Virology Centre | Catherine Moore, Johnathan Evans, Laura Gifford, Malorie Perry, Simon Cottrell, Angela Marchbank, Alec Birchley, Alexander Adams, Amy Gaskin, Bree Gatica-Wilcox, Jason Coombes, Joel Southgate, Lauren Gilbert, Lee Graham, Nicole Pacchiarini, Sara Kumziene-Summerhayes, Sarah Taylor, Sophie Jones, Sara Rey, Matthew Bull, Joanne Watkins, Sally Corden, Tom Connor |
| EPI_ISL_3296281 | Viollier AG | Department of Biosystems Science and Engineering, ETH Zürich | Christian Beisel, Sarah Nadeau, Chaoran Chen, Ivan Topolsky, Kim Philipp Jablonski, Lara Fuhrmann, Rebecca Denes, Mirjam Feldkamp, Ina Nissen, Natascha Santacroce, Elodie Burcklen, Christiane Beckmann, Maurice Redondo, Olivier Kobel, Christoph Noppen, Niko Beerenwinkel, Tanja Stadler |
| EPI_ISL_3296282, EPI_ISL_3296283 | Wales Specialist Virology Centre Sequencing lab: Pathogen Genomics Unit | Public Health Wales Microbiology Cardiff Wales Specialist Virology Centre | Catherine Moore, Johnathan Evans, Laura Gifford, Malorie Perry, Simon Cottrell, Angela Marchbank, Alec Birchley, Alexander Adams, Amy Gaskin, Bree Gatica-Wilcox, Jason Coombes, Joel Southgate, Lauren Gilbert, Lee Graham, Nicole Pacchiarini, Sara Kumziene-Summerhayes, Sarah Taylor, Sophie Jones, Sara Rey, Matthew Bull, Joanne Watkins, Sally Corden, Tom Connor |
| EPI_ISL_3296284 | Viollier AG | Department of Biosystems Science and Engineering, ETH Zürich | Christian Beisel, Sarah Nadeau, Chaoran Chen, Ivan Topolsky, Kim Philipp Jablonski, Lara Fuhrmann, Rebecca Denes, Mirjam Feldkamp, Ina Nissen, Natascha Santacroce, Elodie Burcklen, Christiane Beckmann, Maurice Redondo, Olivier Kobel, Christoph Noppen, Niko Beerenwinkel, Tanja Stadler |
| EPI_ISL_3296285 | Wales Specialist Virology Centre Sequencing lab: Pathogen Genomics Unit | Public Health Wales Microbiology Cardiff Wales Specialist Virology Centre | Catherine Moore, Johnathan Evans, Laura Gifford, Malorie Perry, Simon Cottrell, Angela Marchbank, Alec Birchley, Alexander Adams, Amy Gaskin, Bree Gatica-Wilcox, Jason Coombes, Joel Southgate, Lauren Gilbert, Lee Graham, Nicole Pacchiarini, Sara Kumziene-Summerhayes, Sarah Taylor, Sophie Jones, Sara Rey, Matthew Bull, Joanne Watkins, Sally Corden, Tom Connor |
| EPI_ISL_3296286 | Viollier AG | Department of Biosystems Science and Engineering, ETH Zürich | Christian Beisel, Sarah Nadeau, Chaoran Chen, Ivan Topolsky, Kim Philipp Jablonski, Lara Fuhrmann, Rebecca Denes, Mirjam Feldkamp, Ina Nissen, Natascha Santacroce, Elodie Burcklen, Christiane Beckmann, Maurice Redondo, Olivier Kobel, Christoph Noppen, Niko Beerenwinkel, Tanja Stadler |
| EPI_ISL_3296287 | Wales Specialist Virology Centre Sequencing lab: Pathogen Genomics Unit | Public Health Wales Microbiology Cardiff Wales Specialist Virology Centre | Catherine Moore, Johnathan Evans, Laura Gifford, Malorie Perry, Simon Cottrell, Angela Marchbank, Alec Birchley, Alexander Adams, Amy Gaskin, Bree Gatica-Wilcox, Jason Coombes, Joel Southgate, Lauren Gilbert, Lee Graham, Nicole Pacchiarini, Sara Kumziene-Summerhayes, Sarah Taylor, Sophie Jones, Sara Rey, Matthew Bull, Joanne Watkins, Sally Corden, Tom Connor |
| EPI_ISL_3296288 | Viollier AG | Department of Biosystems Science and Engineering, ETH Zürich | Christian Beisel, Sarah Nadeau, Chaoran Chen, Ivan Topolsky, Kim Philipp Jablonski, Lara Fuhrmann, Rebecca Denes, Mirjam Feldkamp, Ina Nissen, Natascha Santacroce, Elodie Burcklen, Christiane Beckmann, Maurice Redondo, Olivier Kobel, Christoph Noppen, Niko Beerenwinkel, Tanja Stadler |
| EPI_ISL_3296289 | Wales Specialist Virology Centre Sequencing lab: Pathogen Genomics Unit | Public Health Wales Microbiology Cardiff Wales Specialist Virology Centre | Catherine Moore, Johnathan Evans, Laura Gifford, Malorie Perry, Simon Cottrell, Angela Marchbank, Alec Birchley, Alexander Adams, Amy Gaskin, Bree Gatica-Wilcox, Jason Coombes, Joel Southgate, Lauren Gilbert, Lee Graham, Nicole Pacchiarini, Sara Kumziene-Summerhayes, Sarah Taylor, Sophie Jones, Sara Rey, Matthew Bull, Joanne Watkins, Sally Corden, Tom Connor |
| EPI_ISL_3296290 | Viollier AG | Department of Biosystems Science and Engineering, ETH Zürich | Christian Beisel, Sarah Nadeau, Chaoran Chen, Ivan Topolsky, Kim Philipp Jablonski, Lara Fuhrmann, Rebecca Denes, Mirjam Feldkamp, Ina Nissen, Natascha Santacroce, Elodie Burcklen, Christiane Beckmann, Maurice Redondo, Olivier Kobel, Christoph Noppen, Niko Beerenwinkel, Tanja Stadler |
| EPI_ISL_3296291 | Wales Specialist Virology Centre Sequencing lab: Pathogen Genomics Unit | Public Health Wales Microbiology Cardiff Wales Specialist Virology Centre | Catherine Moore, Johnathan Evans, Laura Gifford, Malorie Perry, Simon Cottrell, Angela Marchbank, Alec Birchley, Alexander Adams, Amy Gaskin, Bree Gatica-Wilcox, Jason Coombes, Joel Southgate, Lauren Gilbert, Lee Graham, Nicole Pacchiarini, Sara Kumziene-Summerhayes, Sarah Taylor, Sophie Jones, Sara Rey, Matthew Bull, Joanne Watkins, Sally Corden, Tom Connor |
| EPI_ISL_3296292 | Viollier AG | Department of Biosystems Science and Engineering, ETH Zürich | Christian Beisel, Sarah Nadeau, Chaoran Chen, Ivan Topolsky, Kim Philipp Jablonski, Lara Fuhrmann, Rebecca Denes, Mirjam Feldkamp, Ina Nissen, Natascha Santacroce, Elodie Burcklen, Christiane Beckmann, Maurice Redondo, Olivier Kobel, Christoph Noppen, Niko Beerenwinkel, Tanja Stadler |
| EPI_ISL_3296293 | Wales Specialist Virology Centre Sequencing lab: Pathogen Genomics Unit | Public Health Wales Microbiology Cardiff Wales Specialist Virology Centre | Catherine Moore, Johnathan Evans, Laura Gifford, Malorie Perry, Simon Cottrell, Angela Marchbank, Alec Birchley, Alexander Adams, Amy Gaskin, Bree Gatica-Wilcox, Jason Coombes, Joel Southgate, Lauren Gilbert, Lee Graham, Nicole Pacchiarini, Sara Kumziene-Summerhayes, Sarah Taylor, Sophie Jones, Sara Rey, Matthew Bull, Joanne Watkins, Sally Corden, Tom Connor |
| EPI_ISL_3296294 | Viollier AG | Department of Biosystems Science and Engineering, ETH Zürich | Christian Beisel, Sarah Nadeau, Chaoran Chen, Ivan Topolsky, Kim Philipp Jablonski, Lara Fuhrmann, Rebecca Denes, Mirjam Feldkamp, Ina Nissen, Natascha Santacroce, Elodie Burcklen, Christiane Beckmann, Maurice Redondo, Olivier Kobel, Christoph Noppen, Niko Beerenwinkel, Tanja Stadler |
| EPI_ISL_3296295, EPI_ISL_3296297 | Wales Specialist Virology Centre Sequencing lab: Pathogen Genomics Unit | Public Health Wales Microbiology Cardiff Wales Specialist Virology Centre | Catherine Moore, Johnathan Evans, Laura Gifford, Malorie Perry, Simon Cottrell, Angela Marchbank, Alec Birchley, Alexander Adams, Amy Gaskin, Bree Gatica-Wilcox, Jason Coombes, Joel Southgate, Lauren Gilbert, Lee Graham, Nicole Pacchiarini, Sara Kumziene-Summerhayes, Sarah Taylor, Sophie Jones, Sara Rey, Matthew Bull, Joanne Watkins, Sally Corden, Tom Connor |
| EPI_ISL_3296298 | Viollier AG | Department of Biosystems Science and Engineering, ETH Zürich | Christian Beisel, Sarah Nadeau, Chaoran Chen, Ivan Topolsky, Kim Philipp Jablonski, Lara Fuhrmann, Rebecca Denes, Mirjam Feldkamp, Ina Nissen, Natascha Santacroce, Elodie Burcklen, Christiane Beckmann, Maurice Redondo, Olivier Kobel, Christoph Noppen, Niko Beerenwinkel, Tanja Stadler |
| EPI_ISL_3296299, EPI_ISL_3296301, EPI_ISL_3296303 | Wales Specialist Virology Centre Sequencing lab: Pathogen Genomics Unit | Public Health Wales Microbiology Cardiff Wales Specialist Virology Centre | Catherine Moore, Johnathan Evans, Laura Gifford, Malorie Perry, Simon Cottrell, Angela Marchbank, Alec Birchley, Alexander Adams, Amy Gaskin, Bree Gatica-Wilcox, Jason Coombes, Joel Southgate, Lauren Gilbert, Lee Graham, Nicole Pacchiarini, Sara Kumziene-Summerhayes, Sarah Taylor, Sophie Jones, Sara Rey, Matthew Bull, Joanne Watkins, Sally Corden, Tom Connor |
| EPI_ISL_3296304 | Viollier AG | Department of Biosystems Science and Engineering, ETH Zürich | Christian Beisel, Sarah Nadeau, Chaoran Chen, Ivan Topolsky, Kim Philipp Jablonski, Lara Fuhrmann, Rebecca Denes, Mirjam Feldkamp, Ina Nissen, Natascha Santacroce, Elodie Burcklen, Christiane Beckmann, Maurice Redondo, Olivier Kobel, Christoph Noppen, Niko Beerenwinkel, Tanja Stadler |
| EPI_ISL_3296305 | Wales Specialist Virology Centre Sequencing lab: Pathogen Genomics Unit | Public Health Wales Microbiology Cardiff Wales Specialist Virology Centre | Catherine Moore, Johnathan Evans, Laura Gifford, Malorie Perry, Simon Cottrell, Angela Marchbank, Alec Birchley, Alexander Adams, Amy Gaskin, Bree Gatica-Wilcox, Jason Coombes, Joel Southgate, Lauren Gilbert, Lee Graham, Nicole Pacchiarini, Sara Kumziene-Summerhayes, Sarah Taylor, Sophie Jones, Sara Rey, Matthew Bull, Joanne Watkins, Sally Corden, Tom Connor |
| EPI_ISL_3296306 | Viollier AG | Department of Biosystems Science and Engineering, ETH Zürich | Christian Beisel, Sarah Nadeau, Chaoran Chen, Ivan Topolsky, Kim Philipp Jablonski, Lara Fuhrmann, Rebecca Denes, Mirjam Feldkamp, Ina Nissen, Natascha Santacroce, Elodie Burcklen, Christiane Beckmann, Maurice Redondo, Olivier Kobel, Christoph Noppen, Niko Beerenwinkel, Tanja Stadler |
| EPI_ISL_3296307 | Wales Specialist Virology Centre Sequencing lab: Pathogen Genomics Unit | Public Health Wales Microbiology Cardiff Wales Specialist Virology Centre | Catherine Moore, Johnathan Evans, Laura Gifford, Malorie Perry, Simon Cottrell, Angela Marchbank, Alec Birchley, Alexander Adams, Amy Gaskin, Bree Gatica-Wilcox, Jason Coombes, Joel Southgate, Lauren Gilbert, Lee Graham, Nicole Pacchiarini, Sara Kumziene-Summerhayes, Sarah Taylor, Sophie Jones, Sara Rey, Matthew Bull, Joanne Watkins, Sally Corden, Tom Connor |
| EPI_ISL_3296308 | Viollier AG | Department of Biosystems Science and Engineering, ETH Zürich | Christian Beisel, Sarah Nadeau, Chaoran Chen, Ivan Topolsky, Kim Philipp Jablonski, Lara Fuhrmann, Rebecca Denes, Mirjam Feldkamp, Ina Nissen, Natascha Santacroce, Elodie Burcklen, Christiane Beckmann, Maurice Redondo, Olivier Kobel, Christoph Noppen, Niko Beerenwinkel, Tanja Stadler |
| EPI_ISL_3296309 | Wales Specialist Virology Centre Sequencing lab: Pathogen Genomics Unit | Public Health Wales Microbiology Cardiff Wales Specialist Virology Centre | Catherine Moore, Johnathan Evans, Laura Gifford, Malorie Perry, Simon Cottrell, Angela Marchbank, Alec Birchley, Alexander Adams, Amy Gaskin, Bree Gatica-Wilcox, Jason Coombes, Joel Southgate, Lauren Gilbert, Lee Graham, Nicole Pacchiarini, Sara Kumziene-Summerhayes, Sarah Taylor, Sophie Jones, Sara Rey, Matthew Bull, Joanne Watkins, Sally Corden, Tom Connor |
| EPI_ISL_3296310 | Viollier AG | Department of Biosystems Science and Engineering, ETH Zürich | Christian Beisel, Sarah Nadeau, Chaoran Chen, Ivan Topolsky, Kim Philipp Jablonski, Lara Fuhrmann, Rebecca Denes, Mirjam Feldkamp, Ina Nissen, Natascha Santacroce, Elodie Burcklen, Christiane Beckmann, Maurice Redondo, Olivier Kobel, Christoph Noppen, Niko Beerenwinkel, Tanja Stadler |
| EPI_ISL_3296311 | Wales Specialist Virology Centre Sequencing lab: Pathogen Genomics Unit | Public Health Wales Microbiology Cardiff Wales Specialist Virology Centre | Catherine Moore, Johnathan Evans, Laura Gifford, Malorie Perry, Simon Cottrell, Angela Marchbank, Alec Birchley, Alexander Adams, Amy Gaskin, Bree Gatica-Wilcox, Jason Coombes, Joel Southgate, Lauren Gilbert, Lee Graham, Nicole Pacchiarini, Sara Kumziene-Summerhayes, Sarah Taylor, Sophie Jones, Sara Rey, Matthew Bull, Joanne Watkins, Sally Corden, Tom Connor |
| EPI_ISL_3296312 | Viollier AG | Department of Biosystems Science and Engineering, ETH Zürich | Christian Beisel, Sarah Nadeau, Chaoran Chen, Ivan Topolsky, Kim Philipp Jablonski, Lara Fuhrmann, Rebecca Denes, Mirjam Feldkamp, Ina Nissen, Natascha Santacroce, Elodie Burcklen, Christiane Beckmann, Maurice Redondo, Olivier Kobel, Christoph Noppen, Niko Beerenwinkel, Tanja Stadler |
| EPI_ISL_3296313 | Wales Specialist Virology Centre Sequencing lab: Pathogen | Public Health Wales Microbiology Cardiff Wales Specialist Virology | Catherine Moore, Johnathan Evans, Laura Gifford, Malorie Perry, Simon Cottrell, Angela Marchbank, Alec Birchley, Alexander Adams, Amy Gaskin, Bree Gatica-Wilcox, Jason Coombes, Joel Southgate, Lauren Gilbert, Lee Graham, Nicole |

| | | | |
|---|---|---|---|
| | Genomics Unit | Centre | Pacchiarini, Sara Kumziene-Summerhayes, Sarah Taylor, Sophie Jones, Sara Rey, Matthew Bull, Joanne Watkins, Sally Corden, Tom Connor |
| EPI_ISL_3296314 | Viollier AG | Department of Biosystems Science and Engineering, ETH Zürich | Christian Beisel, Sarah Nadeau, Chaoran Chen, Ivan Topolsky, Kim Philipp Jablonski, Lara Fuhrmann, Rebecca Denes, Mirjam Feldkamp, Ina Nissen, Natascha Santacroce, Elodie Burcklen, Christiane Beckmann, Maurice Redondo, Olivier Kobel, Christoph Noppen, Niko Beerenwinkel, Tanja Stadler |
| EPI_ISL_3296315 | Wales Specialist Virology Centre Sequencing lab: Pathogen Genomics Unit | Public Health Wales Microbiology Cardiff Wales Specialist Virology Centre | Catherine Moore, Johnathan Evans, Laura Gifford, Malorie Perry, Simon Cottrell, Angela Marchbank, Alec Birchley, Alexander Adams, Amy Gaskin, Bree Gatica-Wilcox, Jason Coombes, Joel Southgate, Lauren Gilbert, Lee Graham, Nicole Pacchiarini, Sara Kumziene-Summerhayes, Sarah Taylor, Sophie Jones, Sara Rey, Matthew Bull, Joanne Watkins, Sally Corden, Tom Connor |
| EPI_ISL_3296316 | Viollier AG | Department of Biosystems Science and Engineering, ETH Zürich | Christian Beisel, Sarah Nadeau, Chaoran Chen, Ivan Topolsky, Kim Philipp Jablonski, Lara Fuhrmann, Rebecca Denes, Mirjam Feldkamp, Ina Nissen, Natascha Santacroce, Elodie Burcklen, Christiane Beckmann, Maurice Redondo, Olivier Kobel, Christoph Noppen, Niko Beerenwinkel, Tanja Stadler |
| EPI_ISL_3296317 | Wales Specialist Virology Centre Sequencing lab: Pathogen Genomics Unit | Public Health Wales Microbiology Cardiff Wales Specialist Virology Centre | Catherine Moore, Johnathan Evans, Laura Gifford, Malorie Perry, Simon Cottrell, Angela Marchbank, Alec Birchley, Alexander Adams, Amy Gaskin, Bree Gatica-Wilcox, Jason Coombes, Joel Southgate, Lauren Gilbert, Lee Graham, Nicole Pacchiarini, Sara Kumziene-Summerhayes, Sarah Taylor, Sophie Jones, Sara Rey, Matthew Bull, Joanne Watkins, Sally Corden, Tom Connor |
| EPI_ISL_3296318 | Viollier AG | Department of Biosystems Science and Engineering, ETH Zürich | Christian Beisel, Sarah Nadeau, Chaoran Chen, Ivan Topolsky, Kim Philipp Jablonski, Lara Fuhrmann, Rebecca Denes, Mirjam Feldkamp, Ina Nissen, Natascha Santacroce, Elodie Burcklen, Christiane Beckmann, Maurice Redondo, Olivier Kobel, Christoph Noppen, Niko Beerenwinkel, Tanja Stadler |
| EPI_ISL_3296319 | Wales Specialist Virology Centre Sequencing lab: Pathogen Genomics Unit | Public Health Wales Microbiology Cardiff Wales Specialist Virology Centre | Catherine Moore, Johnathan Evans, Laura Gifford, Malorie Perry, Simon Cottrell, Angela Marchbank, Alec Birchley, Alexander Adams, Amy Gaskin, Bree Gatica-Wilcox, Jason Coombes, Joel Southgate, Lauren Gilbert, Lee Graham, Nicole Pacchiarini, Sara Kumziene-Summerhayes, Sarah Taylor, Sophie Jones, Sara Rey, Matthew Bull, Joanne Watkins, Sally Corden, Tom Connor |
| EPI_ISL_3296320 | Viollier AG | Department of Biosystems Science and Engineering, ETH Zürich | Christian Beisel, Sarah Nadeau, Chaoran Chen, Ivan Topolsky, Kim Philipp Jablonski, Lara Fuhrmann, Rebecca Denes, Mirjam Feldkamp, Ina Nissen, Natascha Santacroce, Elodie Burcklen, Christiane Beckmann, Maurice Redondo, Olivier Kobel, Christoph Noppen, Niko Beerenwinkel, Tanja Stadler |
| EPI_ISL_3296321 | Wales Specialist Virology Centre Sequencing lab: Pathogen Genomics Unit | Public Health Wales Microbiology Cardiff Wales Specialist Virology Centre | Catherine Moore, Johnathan Evans, Laura Gifford, Malorie Perry, Simon Cottrell, Angela Marchbank, Alec Birchley, Alexander Adams, Amy Gaskin, Bree Gatica-Wilcox, Jason Coombes, Joel Southgate, Lauren Gilbert, Lee Graham, Nicole Pacchiarini, Sara Kumziene-Summerhayes, Sarah Taylor, Sophie Jones, Sara Rey, Matthew Bull, Joanne Watkins, Sally Corden, Tom Connor |
| EPI_ISL_3296322, EPI_ISL_3296323, EPI_ISL_3296324, EPI_ISL_3296325, EPI_ISL_3296326 | Viollier AG | Department of Biosystems Science and Engineering, ETH Zürich | Christian Beisel, Sarah Nadeau, Chaoran Chen, Ivan Topolsky, Kim Philipp Jablonski, Lara Fuhrmann, Rebecca Denes, Mirjam Feldkamp, Ina Nissen, Natascha Santacroce, Elodie Burcklen, Christiane Beckmann, Maurice Redondo, Olivier Kobel, Christoph Noppen, Niko Beerenwinkel, Tanja Stadler |
| EPI_ISL_3296327 | Wales Specialist Virology Centre Sequencing lab: Pathogen Genomics Unit | Public Health Wales Microbiology Cardiff Wales Specialist Virology Centre | Catherine Moore, Johnathan Evans, Laura Gifford, Malorie Perry, Simon Cottrell, Angela Marchbank, Alec Birchley, Alexander Adams, Amy Gaskin, Bree Gatica-Wilcox, Jason Coombes, Joel Southgate, Lauren Gilbert, Lee Graham, Nicole Pacchiarini, Sara Kumziene-Summerhayes, Sarah Taylor, Sophie Jones, Sara Rey, Matthew Bull, Joanne Watkins, Sally Corden, Tom Connor |
| EPI_ISL_3296328 | Viollier AG | Department of Biosystems Science and Engineering, ETH Zürich | Christian Beisel, Sarah Nadeau, Chaoran Chen, Ivan Topolsky, Kim Philipp Jablonski, Lara Fuhrmann, Rebecca Denes, Mirjam Feldkamp, Ina Nissen, Natascha Santacroce, Elodie Burcklen, Christiane Beckmann, Maurice Redondo, Olivier Kobel, Christoph Noppen, Niko Beerenwinkel, Tanja Stadler |
| EPI_ISL_3296329 | Wales Specialist Virology Centre Sequencing lab: Pathogen Genomics Unit | Public Health Wales Microbiology Cardiff Wales Specialist Virology Centre | Catherine Moore, Johnathan Evans, Laura Gifford, Malorie Perry, Simon Cottrell, Angela Marchbank, Alec Birchley, Alexander Adams, Amy Gaskin, Bree Gatica-Wilcox, Jason Coombes, Joel Southgate, Lauren Gilbert, Lee Graham, Nicole Pacchiarini, Sara Kumziene-Summerhayes, Sarah Taylor, Sophie Jones, Sara Rey, Matthew Bull, Joanne Watkins, Sally Corden, Tom Connor |
| EPI_ISL_3296330 | Viollier AG | Department of Biosystems Science and Engineering, ETH Zürich | Christian Beisel, Sarah Nadeau, Chaoran Chen, Ivan Topolsky, Kim Philipp Jablonski, Lara Fuhrmann, Rebecca Denes, Mirjam Feldkamp, Ina Nissen, Natascha Santacroce, Elodie Burcklen, Christiane Beckmann, Maurice Redondo, Olivier Kobel, Christoph Noppen, Niko Beerenwinkel, Tanja Stadler |
| EPI_ISL_3296331 | Wales Specialist Virology Centre Sequencing lab: Pathogen Genomics Unit | Public Health Wales Microbiology Cardiff Wales Specialist Virology Centre | Catherine Moore, Johnathan Evans, Laura Gifford, Malorie Perry, Simon Cottrell, Angela Marchbank, Alec Birchley, Alexander Adams, Amy Gaskin, Bree Gatica-Wilcox, Jason Coombes, Joel Southgate, Lauren Gilbert, Lee Graham, Nicole Pacchiarini, Sara Kumziene-Summerhayes, Sarah Taylor, Sophie Jones, Sara Rey, Matthew Bull, Joanne Watkins, Sally Corden, Tom Connor |
| EPI_ISL_3296332 | Viollier AG | Department of Biosystems Science and Engineering, ETH Zürich | Christian Beisel, Sarah Nadeau, Chaoran Chen, Ivan Topolsky, Kim Philipp Jablonski, Lara Fuhrmann, Rebecca Denes, Mirjam Feldkamp, Ina Nissen, Natascha Santacroce, Elodie Burcklen, Christiane Beckmann, Maurice Redondo, Olivier Kobel, Christoph Noppen, Niko Beerenwinkel, Tanja Stadler |
| EPI_ISL_3296333 | Wales Specialist Virology Centre Sequencing lab: Pathogen Genomics Unit | Public Health Wales Microbiology Cardiff Wales Specialist Virology Centre | Catherine Moore, Johnathan Evans, Laura Gifford, Malorie Perry, Simon Cottrell, Angela Marchbank, Alec Birchley, Alexander Adams, Amy Gaskin, Bree Gatica-Wilcox, Jason Coombes, Joel Southgate, Lauren Gilbert, Lee Graham, Nicole Pacchiarini, Sara Kumziene-Summerhayes, Sarah Taylor, Sophie Jones, Sara Rey, Matthew Bull, Joanne Watkins, Sally Corden, Tom Connor |
| EPI_ISL_3296334 | Viollier AG | Department of Biosystems Science and Engineering, ETH Zürich | Christian Beisel, Sarah Nadeau, Chaoran Chen, Ivan Topolsky, Kim Philipp Jablonski, Lara Fuhrmann, Rebecca Denes, Mirjam Feldkamp, Ina Nissen, Natascha Santacroce, Elodie Burcklen, Christiane Beckmann, Maurice Redondo, Olivier Kobel, Christoph Noppen, Niko Beerenwinkel, Tanja Stadler |
| EPI_ISL_3296335 | Wales Specialist Virology Centre Sequencing lab: Pathogen Genomics Unit | Public Health Wales Microbiology Cardiff Wales Specialist Virology Centre | Catherine Moore, Johnathan Evans, Laura Gifford, Malorie Perry, Simon Cottrell, Angela Marchbank, Alec Birchley, Alexander Adams, Amy Gaskin, Bree Gatica-Wilcox, Jason Coombes, Joel Southgate, Lauren Gilbert, Lee Graham, Nicole Pacchiarini, Sara Kumziene-Summerhayes, Sarah Taylor, Sophie Jones, Sara Rey, Matthew Bull, Joanne Watkins, Sally Corden, Tom Connor |
| EPI_ISL_3296336 | Viollier AG | Department of Biosystems Science and Engineering, ETH Zürich | Christian Beisel, Sarah Nadeau, Chaoran Chen, Ivan Topolsky, Kim Philipp Jablonski, Lara Fuhrmann, Rebecca Denes, Mirjam Feldkamp, Ina Nissen, Natascha Santacroce, Elodie Burcklen, Christiane Beckmann, Maurice Redondo, Olivier Kobel, Christoph Noppen, Niko Beerenwinkel, Tanja Stadler |
| EPI_ISL_3296337, EPI_ISL_3296338 | Wales Specialist Virology Centre Sequencing lab: Pathogen Genomics Unit | Public Health Wales Microbiology Cardiff Wales Specialist Virology Centre | Catherine Moore, Johnathan Evans, Laura Gifford, Malorie Perry, Simon Cottrell, Angela Marchbank, Alec Birchley, Alexander Adams, Amy Gaskin, Bree Gatica-Wilcox, Jason Coombes, Joel Southgate, Lauren Gilbert, Lee Graham, Nicole Pacchiarini, Sara Kumziene-Summerhayes, Sarah Taylor, Sophie Jones, Sara Rey, Matthew Bull, Joanne Watkins, Sally Corden, Tom Connor |
| EPI_ISL_3296339 | Viollier AG | Department of Biosystems Science and Engineering, ETH Zürich | Christian Beisel, Sarah Nadeau, Chaoran Chen, Ivan Topolsky, Kim Philipp Jablonski, Lara Fuhrmann, Rebecca Denes, Mirjam Feldkamp, Ina Nissen, Natascha Santacroce, Elodie Burcklen, Christiane Beckmann, Maurice Redondo, Olivier Kobel, Christoph Noppen, Niko Beerenwinkel, Tanja Stadler |
| EPI_ISL_3296340 | Wales Specialist Virology Centre Sequencing lab: Pathogen Genomics Unit | Public Health Wales Microbiology Cardiff Wales Specialist Virology Centre | Catherine Moore, Johnathan Evans, Laura Gifford, Malorie Perry, Simon Cottrell, Angela Marchbank, Alec Birchley, Alexander Adams, Amy Gaskin, Bree Gatica-Wilcox, Jason Coombes, Joel Southgate, Lauren Gilbert, Lee Graham, Nicole Pacchiarini, Sara Kumziene-Summerhayes, Sarah Taylor, Sophie Jones, Sara Rey, Matthew Bull, Joanne Watkins, Sally Corden, Tom Connor |
| EPI_ISL_3296341 | Viollier AG | Department of Biosystems Science and Engineering, ETH Zürich | Christian Beisel, Sarah Nadeau, Chaoran Chen, Ivan Topolsky, Kim Philipp Jablonski, Lara Fuhrmann, Rebecca Denes, Mirjam Feldkamp, Ina Nissen, Natascha Santacroce, Elodie Burcklen, Christiane Beckmann, Maurice Redondo, Olivier Kobel, Christoph Noppen, Niko Beerenwinkel, Tanja Stadler |
| EPI_ISL_3296342 | Wales Specialist Virology Centre Sequencing lab: Pathogen Genomics Unit | Public Health Wales Microbiology Cardiff Wales Specialist Virology Centre | Catherine Moore, Johnathan Evans, Laura Gifford, Malorie Perry, Simon Cottrell, Angela Marchbank, Alec Birchley, Alexander Adams, Amy Gaskin, Bree Gatica-Wilcox, Jason Coombes, Joel Southgate, Lauren Gilbert, Lee Graham, Nicole Pacchiarini, Sara Kumziene-Summerhayes, Sarah Taylor, Sophie Jones, Sara Rey, Matthew Bull, Joanne Watkins, Sally Corden, Tom Connor |
| EPI_ISL_3296343 | Viollier AG | Department of Biosystems Science and Engineering, ETH Zürich | Christian Beisel, Sarah Nadeau, Chaoran Chen, Ivan Topolsky, Kim Philipp Jablonski, Lara Fuhrmann, Rebecca Denes, Mirjam Feldkamp, Ina Nissen, Natascha Santacroce, Elodie Burcklen, Christiane Beckmann, Maurice Redondo, Olivier Kobel, Christoph Noppen, Niko Beerenwinkel, Tanja Stadler |
| EPI_ISL_3296344 | Wales Specialist Virology Centre Sequencing lab: Pathogen Genomics Unit | Public Health Wales Microbiology Cardiff Wales Specialist Virology Centre | Catherine Moore, Johnathan Evans, Laura Gifford, Malorie Perry, Simon Cottrell, Angela Marchbank, Alec Birchley, Alexander Adams, Amy Gaskin, Bree Gatica-Wilcox, Jason Coombes, Joel Southgate, Lauren Gilbert, Lee Graham, Nicole Pacchiarini, Sara Kumziene-Summerhayes, Sarah Taylor, Sophie Jones, Sara Rey, Matthew Bull, Joanne Watkins, Sally Corden, Tom Connor |
| EPI_ISL_3296345 | Viollier AG | Department of Biosystems Science and Engineering, ETH Zürich | Christian Beisel, Sarah Nadeau, Chaoran Chen, Ivan Topolsky, Kim Philipp Jablonski, Lara Fuhrmann, Rebecca Denes, Mirjam Feldkamp, Ina Nissen, Natascha Santacroce, Elodie Burcklen, Christiane Beckmann, Maurice Redondo, Olivier Kobel, Christoph Noppen, Niko Beerenwinkel, Tanja Stadler |
| EPI_ISL_3296346, EPI_ISL_3296348 | Wales Specialist Virology Centre Sequencing lab: Pathogen Genomics Unit | Public Health Wales Microbiology Cardiff Wales Specialist Virology Centre | Catherine Moore, Johnathan Evans, Laura Gifford, Malorie Perry, Simon Cottrell, Angela Marchbank, Alec Birchley, Alexander Adams, Amy Gaskin, Bree Gatica-Wilcox, Jason Coombes, Joel Southgate, Lauren Gilbert, Lee Graham, Nicole Pacchiarini, Sara Kumziene-Summerhayes, Sarah Taylor, Sophie Jones, Sara Rey, Matthew Bull, Joanne Watkins, Sally Corden, Tom Connor |
| EPI_ISL_3296349 | Viollier AG | Department of Biosystems Science and Engineering, ETH Zürich | Christian Beisel, Sarah Nadeau, Chaoran Chen, Ivan Topolsky, Kim Philipp Jablonski, Lara Fuhrmann, Rebecca Denes, Mirjam Feldkamp, Ina Nissen, Natascha Santacroce, Elodie Burcklen, Christiane Beckmann, Maurice Redondo, Olivier Kobel, Christoph Noppen, Niko Beerenwinkel, Tanja Stadler |
| EPI_ISL_3296350, EPI_ISL_3296351 | Wales Specialist Virology Centre Sequencing lab: Pathogen Genomics Unit | Public Health Wales Microbiology Cardiff Wales Specialist Virology Centre | Catherine Moore, Johnathan Evans, Laura Gifford, Malorie Perry, Simon Cottrell, Angela Marchbank, Alec Birchley, Alexander Adams, Amy Gaskin, Bree Gatica-Wilcox, Jason Coombes, Joel Southgate, Lauren Gilbert, Lee Graham, Nicole Pacchiarini, Sara Kumziene-Summerhayes, Sarah Taylor, Sophie Jones, Sara Rey, Matthew Bull, Joanne Watkins, Sally Corden, Tom Connor |
| EPI_ISL_3296352 | Viollier AG | Department of Biosystems Science and Engineering, ETH Zürich | Christian Beisel, Sarah Nadeau, Chaoran Chen, Ivan Topolsky, Kim Philipp Jablonski, Lara Fuhrmann, Rebecca Denes, Mirjam Feldkamp, Ina Nissen, Natascha Santacroce, Elodie Burcklen, Christiane Beckmann, Maurice Redondo, Olivier Kobel, Christoph Noppen, Niko Beerenwinkel, Tanja Stadler |
| EPI_ISL_3296353 | Wales Specialist Virology Centre Sequencing lab: Pathogen Genomics Unit | Public Health Wales Microbiology Cardiff Wales Specialist Virology Centre | Catherine Moore, Johnathan Evans, Laura Gifford, Malorie Perry, Simon Cottrell, Angela Marchbank, Alec Birchley, Alexander Adams, Amy Gaskin, Bree Gatica-Wilcox, Jason Coombes, Joel Southgate, Lauren Gilbert, Lee Graham, Nicole Pacchiarini, Sara Kumziene-Summerhayes, Sarah Taylor, Sophie Jones, Sara Rey, Matthew Bull, Joanne Watkins, Sally Corden, Tom Connor |
| EPI_ISL_3296354 | Viollier AG | Department of Biosystems Science and Engineering, ETH Zürich | Christian Beisel, Sarah Nadeau, Chaoran Chen, Ivan Topolsky, Kim Philipp Jablonski, Lara Fuhrmann, Rebecca Denes, Mirjam Feldkamp, Ina Nissen, Natascha Santacroce, Elodie Burcklen, Christiane Beckmann, Maurice Redondo, Olivier Kobel, Christoph Noppen, Niko Beerenwinkel, Tanja Stadler |
| EPI_ISL_3296355, EPI_ISL_3296357 | Wales Specialist Virology Centre Sequencing lab: Pathogen Genomics Unit | Public Health Wales Microbiology Cardiff Wales Specialist Virology Centre | Catherine Moore, Johnathan Evans, Laura Gifford, Malorie Perry, Simon Cottrell, Angela Marchbank, Alec Birchley, Alexander Adams, Amy Gaskin, Bree Gatica-Wilcox, Jason Coombes, Joel Southgate, Lauren Gilbert, Lee Graham, Nicole Pacchiarini, Sara Kumziene-Summerhayes, Sarah Taylor, Sophie Jones, Sara Rey, Matthew Bull, Joanne Watkins, Sally Corden, Tom Connor |
| EPI_ISL_3296358 | Viollier AG | Department of Biosystems Science and Engineering, ETH Zürich | Christian Beisel, Sarah Nadeau, Chaoran Chen, Ivan Topolsky, Kim Philipp Jablonski, Lara Fuhrmann, Rebecca Denes, Mirjam Feldkamp, Ina Nissen, Natascha Santacroce, Elodie Burcklen, Christiane Beckmann, Maurice Redondo, Olivier Kobel, Christoph Noppen, Niko Beerenwinkel, Tanja Stadler |
| EPI_ISL_3296359 | Wales Specialist Virology Centre Sequencing lab: Pathogen Genomics Unit | Public Health Wales Microbiology Cardiff Wales Specialist Virology Centre | Catherine Moore, Johnathan Evans, Laura Gifford, Malorie Perry, Simon Cottrell, Angela Marchbank, Alec Birchley, Alexander Adams, Amy Gaskin, Bree Gatica-Wilcox, Jason Coombes, Joel Southgate, Lauren Gilbert, Lee Graham, Nicole Pacchiarini, Sara Kumziene-Summerhayes, Sarah Taylor, Sophie Jones, Sara Rey, Matthew Bull, Joanne Watkins, Sally Corden, Tom Connor |
| EPI_ISL_3296360 | Viollier AG | Department of Biosystems Science and Engineering, ETH Zürich | Christian Beisel, Sarah Nadeau, Chaoran Chen, Ivan Topolsky, Kim Philipp Jablonski, Lara Fuhrmann, Rebecca Denes, Mirjam Feldkamp, Ina Nissen, Natascha Santacroce, Elodie Burcklen, Christiane Beckmann, Maurice Redondo, Olivier Kobel, Christoph Noppen, Niko Beerenwinkel, Tanja Stadler |
| EPI_ISL_3296361 | Wales Specialist Virology Centre Sequencing lab: Pathogen Genomics Unit | Public Health Wales Microbiology Cardiff Wales Specialist Virology Centre | Catherine Moore, Johnathan Evans, Laura Gifford, Malorie Perry, Simon Cottrell, Angela Marchbank, Alec Birchley, Alexander Adams, Amy Gaskin, Bree Gatica-Wilcox, Jason Coombes, Joel Southgate, Lauren Gilbert, Lee Graham, Nicole Pacchiarini, Sara Kumziene-Summerhayes, Sarah Taylor, Sophie Jones, Sara Rey, Matthew Bull, Joanne Watkins, Sally Corden, Tom Connor |
| EPI_ISL_3296362 | Viollier AG | Department of Biosystems Science and Engineering, ETH Zürich | Christian Beisel, Sarah Nadeau, Chaoran Chen, Ivan Topolsky, Kim Philipp Jablonski, Lara Fuhrmann, Rebecca Denes, Mirjam Feldkamp, Ina Nissen, Natascha Santacroce, Elodie Burcklen, Christiane Beckmann, Maurice Redondo, Olivier Kobel, Christoph Noppen, Niko Beerenwinkel, Tanja Stadler |
| EPI_ISL_3296363 | Wales Specialist Virology Centre Sequencing lab: Pathogen Genomics Unit | Public Health Wales Microbiology Cardiff Wales Specialist Virology Centre | Catherine Moore, Johnathan Evans, Laura Gifford, Malorie Perry, Simon Cottrell, Angela Marchbank, Alec Birchley, Alexander Adams, Amy Gaskin, Bree Gatica-Wilcox, Jason Coombes, Joel Southgate, Lauren Gilbert, Lee Graham, Nicole Pacchiarini, Sara Kumziene-Summerhayes, Sarah Taylor, Sophie Jones, Sara Rey, Matthew Bull, Joanne Watkins, Sally Corden, Tom Connor |
| EPI_ISL_3296364 | Viollier AG | Department of Biosystems Science and Engineering, ETH Zürich | Christian Beisel, Sarah Nadeau, Chaoran Chen, Ivan Topolsky, Kim Philipp Jablonski, Lara Fuhrmann, Rebecca Denes, Mirjam Feldkamp, Ina Nissen, Natascha Santacroce, Elodie Burcklen, Christiane Beckmann, Maurice Redondo, Olivier Kobel, Christoph Noppen, Niko Beerenwinkel, Tanja Stadler |
| EPI_ISL_3296365 | Wales Specialist Virology Centre Sequencing lab: Pathogen Genomics Unit | Public Health Wales Microbiology Cardiff Wales Specialist Virology Centre | Catherine Moore, Johnathan Evans, Laura Gifford, Malorie Perry, Simon Cottrell, Angela Marchbank, Alec Birchley, Alexander Adams, Amy Gaskin, Bree Gatica-Wilcox, Jason Coombes, Joel Southgate, Lauren Gilbert, Lee Graham, Nicole Pacchiarini, Sara Kumziene-Summerhayes, Sarah Taylor, Sophie Jones, Sara Rey, Matthew Bull, Joanne Watkins, Sally Corden, Tom Connor |
| EPI_ISL_3296366 | Viollier AG | Department of Biosystems Science and Engineering, ETH Zürich | Christian Beisel, Sarah Nadeau, Chaoran Chen, Ivan Topolsky, Kim Philipp Jablonski, Lara Fuhrmann, Rebecca Denes, Mirjam Feldkamp, Ina Nissen, Natascha Santacroce, Elodie Burcklen, Christiane Beckmann, Maurice Redondo, Olivier Kobel, Christoph Noppen, Niko Beerenwinkel, Tanja Stadler |

| | | | |
|---|---|---|---|
| EPI_ISL_3296367 | Wales Specialist Virology Centre Sequencing lab: Pathogen Genomics Unit | Public Health Wales Microbiology Cardiff Wales Specialist Virology Centre | Catherine Moore, Johnathan Evans, Laura Gifford, Malorie Perry, Simon Cottrell, Angela Marchbank, Alec Birchley, Alexander Adams, Amy Gaskin, Bree Gatica-Wilcox, Jason Coombes, Joel Southgate, Lauren Gilbert, Lee Graham, Nicole Pacchiarini, Sara Kumziene-Summerhayes, Sarah Taylor, Sophie Jones, Sara Rey, Matthew Bull, Joanne Watkins, Sally Corden, Tom Connor |
| EPI_ISL_3296368 | Viollier AG | Department of Biosystems Science and Engineering, ETH Zürich | Christian Beisel, Sarah Nadeau, Chaoran Chen, Ivan Topolsky, Kim Philipp Jablonski, Lara Fuhrmann, Rebecca Denes, Mirjam Feldkamp, Ina Nissen, Natascha Santacroce, Elodie Burcklen, Christiane Beckmann, Maurice Redondo, Olivier Kobel, Christoph Noppen, Niko Beerenwinkel, Tanja Stadler |
| EPI_ISL_3296369 | Wales Specialist Virology Centre Sequencing lab: Pathogen Genomics Unit | Public Health Wales Microbiology Cardiff Wales Specialist Virology Centre | Catherine Moore, Johnathan Evans, Laura Gifford, Malorie Perry, Simon Cottrell, Angela Marchbank, Alec Birchley, Alexander Adams, Amy Gaskin, Bree Gatica-Wilcox, Jason Coombes, Joel Southgate, Lauren Gilbert, Lee Graham, Nicole Pacchiarini, Sara Kumziene-Summerhayes, Sarah Taylor, Sophie Jones, Sara Rey, Matthew Bull, Joanne Watkins, Sally Corden, Tom Connor |
| EPI_ISL_3296370 | Viollier AG | Department of Biosystems Science and Engineering, ETH Zürich | Christian Beisel, Sarah Nadeau, Chaoran Chen, Ivan Topolsky, Kim Philipp Jablonski, Lara Fuhrmann, Rebecca Denes, Mirjam Feldkamp, Ina Nissen, Natascha Santacroce, Elodie Burcklen, Christiane Beckmann, Maurice Redondo, Olivier Kobel, Christoph Noppen, Niko Beerenwinkel, Tanja Stadler |
| EPI_ISL_3296371 | Wales Specialist Virology Centre Sequencing lab: Pathogen Genomics Unit | Public Health Wales Microbiology Cardiff Wales Specialist Virology Centre | Catherine Moore, Johnathan Evans, Laura Gifford, Malorie Perry, Simon Cottrell, Angela Marchbank, Alec Birchley, Alexander Adams, Amy Gaskin, Bree Gatica-Wilcox, Jason Coombes, Joel Southgate, Lauren Gilbert, Lee Graham, Nicole Pacchiarini, Sara Kumziene-Summerhayes, Sarah Taylor, Sophie Jones, Sara Rey, Matthew Bull, Joanne Watkins, Sally Corden, Tom Connor |
| EPI_ISL_3296372 | Viollier AG | Department of Biosystems Science and Engineering, ETH Zürich | Christian Beisel, Sarah Nadeau, Chaoran Chen, Ivan Topolsky, Kim Philipp Jablonski, Lara Fuhrmann, Rebecca Denes, Mirjam Feldkamp, Ina Nissen, Natascha Santacroce, Elodie Burcklen, Christiane Beckmann, Maurice Redondo, Olivier Kobel, Christoph Noppen, Niko Beerenwinkel, Tanja Stadler |
| EPI_ISL_3296373 | Wales Specialist Virology Centre Sequencing lab: Pathogen Genomics Unit | Public Health Wales Microbiology Cardiff Wales Specialist Virology Centre | Catherine Moore, Johnathan Evans, Laura Gifford, Malorie Perry, Simon Cottrell, Angela Marchbank, Alec Birchley, Alexander Adams, Amy Gaskin, Bree Gatica-Wilcox, Jason Coombes, Joel Southgate, Lauren Gilbert, Lee Graham, Nicole Pacchiarini, Sara Kumziene-Summerhayes, Sarah Taylor, Sophie Jones, Sara Rey, Matthew Bull, Joanne Watkins, Sally Corden, Tom Connor |
| EPI_ISL_3296374 | Viollier AG | Department of Biosystems Science and Engineering, ETH Zürich | Christian Beisel, Sarah Nadeau, Chaoran Chen, Ivan Topolsky, Kim Philipp Jablonski, Lara Fuhrmann, Rebecca Denes, Mirjam Feldkamp, Ina Nissen, Natascha Santacroce, Elodie Burcklen, Christiane Beckmann, Maurice Redondo, Olivier Kobel, Christoph Noppen, Niko Beerenwinkel, Tanja Stadler |
| EPI_ISL_3296375, EPI_ISL_3296376, EPI_ISL_3296378 | Wales Specialist Virology Centre Sequencing lab: Pathogen Genomics Unit | Public Health Wales Microbiology Cardiff Wales Specialist Virology Centre | Catherine Moore, Johnathan Evans, Laura Gifford, Malorie Perry, Simon Cottrell, Angela Marchbank, Alec Birchley, Alexander Adams, Amy Gaskin, Bree Gatica-Wilcox, Jason Coombes, Joel Southgate, Lauren Gilbert, Lee Graham, Nicole Pacchiarini, Sara Kumziene-Summerhayes, Sarah Taylor, Sophie Jones, Sara Rey, Matthew Bull, Joanne Watkins, Sally Corden, Tom Connor |
| EPI_ISL_3296379 | Viollier AG | Department of Biosystems Science and Engineering, ETH Zürich | Christian Beisel, Sarah Nadeau, Chaoran Chen, Ivan Topolsky, Kim Philipp Jablonski, Lara Fuhrmann, Rebecca Denes, Mirjam Feldkamp, Ina Nissen, Natascha Santacroce, Elodie Burcklen, Christiane Beckmann, Maurice Redondo, Olivier Kobel, Christoph Noppen, Niko Beerenwinkel, Tanja Stadler |
| EPI_ISL_3296380 | Wales Specialist Virology Centre Sequencing lab: Pathogen Genomics Unit | Public Health Wales Microbiology Cardiff Wales Specialist Virology Centre | Catherine Moore, Johnathan Evans, Laura Gifford, Malorie Perry, Simon Cottrell, Angela Marchbank, Alec Birchley, Alexander Adams, Amy Gaskin, Bree Gatica-Wilcox, Jason Coombes, Joel Southgate, Lauren Gilbert, Lee Graham, Nicole Pacchiarini, Sara Kumziene-Summerhayes, Sarah Taylor, Sophie Jones, Sara Rey, Matthew Bull, Joanne Watkins, Sally Corden, Tom Connor |
| EPI_ISL_3296381 | Viollier AG | Department of Biosystems Science and Engineering, ETH Zürich | Christian Beisel, Sarah Nadeau, Chaoran Chen, Ivan Topolsky, Kim Philipp Jablonski, Lara Fuhrmann, Rebecca Denes, Mirjam Feldkamp, Ina Nissen, Natascha Santacroce, Elodie Burcklen, Christiane Beckmann, Maurice Redondo, Olivier Kobel, Christoph Noppen, Niko Beerenwinkel, Tanja Stadler |
| EPI_ISL_3296382 | Wales Specialist Virology Centre Sequencing lab: Pathogen Genomics Unit | Public Health Wales Microbiology Cardiff Wales Specialist Virology Centre | Catherine Moore, Johnathan Evans, Laura Gifford, Malorie Perry, Simon Cottrell, Angela Marchbank, Alec Birchley, Alexander Adams, Amy Gaskin, Bree Gatica-Wilcox, Jason Coombes, Joel Southgate, Lauren Gilbert, Lee Graham, Nicole Pacchiarini, Sara Kumziene-Summerhayes, Sarah Taylor, Sophie Jones, Sara Rey, Matthew Bull, Joanne Watkins, Sally Corden, Tom Connor |
| EPI_ISL_3296383 | Viollier AG | Department of Biosystems Science and Engineering, ETH Zürich | Christian Beisel, Sarah Nadeau, Chaoran Chen, Ivan Topolsky, Kim Philipp Jablonski, Lara Fuhrmann, Rebecca Denes, Mirjam Feldkamp, Ina Nissen, Natascha Santacroce, Elodie Burcklen, Christiane Beckmann, Maurice Redondo, Olivier Kobel, Christoph Noppen, Niko Beerenwinkel, Tanja Stadler |
| EPI_ISL_3296384 | Wales Specialist Virology Centre Sequencing lab: Pathogen Genomics Unit | Public Health Wales Microbiology Cardiff Wales Specialist Virology Centre | Catherine Moore, Johnathan Evans, Laura Gifford, Malorie Perry, Simon Cottrell, Angela Marchbank, Alec Birchley, Alexander Adams, Amy Gaskin, Bree Gatica-Wilcox, Jason Coombes, Joel Southgate, Lauren Gilbert, Lee Graham, Nicole Pacchiarini, Sara Kumziene-Summerhayes, Sarah Taylor, Sophie Jones, Sara Rey, Matthew Bull, Joanne Watkins, Sally Corden, Tom Connor |
| EPI_ISL_3296385 | Viollier AG | Department of Biosystems Science and Engineering, ETH Zürich | Christian Beisel, Sarah Nadeau, Chaoran Chen, Ivan Topolsky, Kim Philipp Jablonski, Lara Fuhrmann, Rebecca Denes, Mirjam Feldkamp, Ina Nissen, Natascha Santacroce, Elodie Burcklen, Christiane Beckmann, Maurice Redondo, Olivier Kobel, Christoph Noppen, Niko Beerenwinkel, Tanja Stadler |
| EPI_ISL_3296386 | Wales Specialist Virology Centre Sequencing lab: Pathogen Genomics Unit | Public Health Wales Microbiology Cardiff Wales Specialist Virology Centre | Catherine Moore, Johnathan Evans, Laura Gifford, Malorie Perry, Simon Cottrell, Angela Marchbank, Alec Birchley, Alexander Adams, Amy Gaskin, Bree Gatica-Wilcox, Jason Coombes, Joel Southgate, Lauren Gilbert, Lee Graham, Nicole Pacchiarini, Sara Kumziene-Summerhayes, Sarah Taylor, Sophie Jones, Sara Rey, Matthew Bull, Joanne Watkins, Sally Corden, Tom Connor |
| EPI_ISL_3296387 | Viollier AG | Department of Biosystems Science and Engineering, ETH Zürich | Christian Beisel, Sarah Nadeau, Chaoran Chen, Ivan Topolsky, Kim Philipp Jablonski, Lara Fuhrmann, Rebecca Denes, Mirjam Feldkamp, Ina Nissen, Natascha Santacroce, Elodie Burcklen, Christiane Beckmann, Maurice Redondo, Olivier Kobel, Christoph Noppen, Niko Beerenwinkel, Tanja Stadler |
| EPI_ISL_3296388 | Wales Specialist Virology Centre Sequencing lab: Pathogen Genomics Unit | Public Health Wales Microbiology Cardiff Wales Specialist Virology Centre | Catherine Moore, Johnathan Evans, Laura Gifford, Malorie Perry, Simon Cottrell, Angela Marchbank, Alec Birchley, Alexander Adams, Amy Gaskin, Bree Gatica-Wilcox, Jason Coombes, Joel Southgate, Lauren Gilbert, Lee Graham, Nicole Pacchiarini, Sara Kumziene-Summerhayes, Sarah Taylor, Sophie Jones, Sara Rey, Matthew Bull, Joanne Watkins, Sally Corden, Tom Connor |
| EPI_ISL_3296389 | Viollier AG | Department of Biosystems Science and Engineering, ETH Zürich | Christian Beisel, Sarah Nadeau, Chaoran Chen, Ivan Topolsky, Kim Philipp Jablonski, Lara Fuhrmann, Rebecca Denes, Mirjam Feldkamp, Ina Nissen, Natascha Santacroce, Elodie Burcklen, Christiane Beckmann, Maurice Redondo, Olivier Kobel, Christoph Noppen, Niko Beerenwinkel, Tanja Stadler |
| EPI_ISL_3296390 | Wales Specialist Virology Centre Sequencing lab: Pathogen Genomics Unit | Public Health Wales Microbiology Cardiff Wales Specialist Virology Centre | Catherine Moore, Johnathan Evans, Laura Gifford, Malorie Perry, Simon Cottrell, Angela Marchbank, Alec Birchley, Alexander Adams, Amy Gaskin, Bree Gatica-Wilcox, Jason Coombes, Joel Southgate, Lauren Gilbert, Lee Graham, Nicole Pacchiarini, Sara Kumziene-Summerhayes, Sarah Taylor, Sophie Jones, Sara Rey, Matthew Bull, Joanne Watkins, Sally Corden, Tom Connor |
| EPI_ISL_3296391 | Viollier AG | Department of Biosystems Science and Engineering, ETH Zürich | Christian Beisel, Sarah Nadeau, Chaoran Chen, Ivan Topolsky, Kim Philipp Jablonski, Lara Fuhrmann, Rebecca Denes, Mirjam Feldkamp, Ina Nissen, Natascha Santacroce, Elodie Burcklen, Christiane Beckmann, Maurice Redondo, Olivier Kobel, Christoph Noppen, Niko Beerenwinkel, Tanja Stadler |
| EPI_ISL_3296392 | Wales Specialist Virology Centre Sequencing lab: Pathogen Genomics Unit | Public Health Wales Microbiology Cardiff Wales Specialist Virology Centre | Catherine Moore, Johnathan Evans, Laura Gifford, Malorie Perry, Simon Cottrell, Angela Marchbank, Alec Birchley, Alexander Adams, Amy Gaskin, Bree Gatica-Wilcox, Jason Coombes, Joel Southgate, Lauren Gilbert, Lee Graham, Nicole Pacchiarini, Sara Kumziene-Summerhayes, Sarah Taylor, Sophie Jones, Sara Rey, Matthew Bull, Joanne Watkins, Sally Corden, Tom Connor |
| EPI_ISL_3296393 | Viollier AG | Department of Biosystems Science and Engineering, ETH Zürich | Christian Beisel, Sarah Nadeau, Chaoran Chen, Ivan Topolsky, Kim Philipp Jablonski, Lara Fuhrmann, Rebecca Denes, Mirjam Feldkamp, Ina Nissen, Natascha Santacroce, Elodie Burcklen, Christiane Beckmann, Maurice Redondo, Olivier Kobel, Christoph Noppen, Niko Beerenwinkel, Tanja Stadler |
| EPI_ISL_3296394, EPI_ISL_3296395 | Wales Specialist Virology Centre Sequencing lab: Pathogen Genomics Unit | Public Health Wales Microbiology Cardiff Wales Specialist Virology Centre | Catherine Moore, Johnathan Evans, Laura Gifford, Malorie Perry, Simon Cottrell, Angela Marchbank, Alec Birchley, Alexander Adams, Amy Gaskin, Bree Gatica-Wilcox, Jason Coombes, Joel Southgate, Lauren Gilbert, Lee Graham, Nicole Pacchiarini, Sara Kumziene-Summerhayes, Sarah Taylor, Sophie Jones, Sara Rey, Matthew Bull, Joanne Watkins, Sally Corden, Tom Connor |
| EPI_ISL_3296396 | Viollier AG | Department of Biosystems Science and Engineering, ETH Zürich | Christian Beisel, Sarah Nadeau, Chaoran Chen, Ivan Topolsky, Kim Philipp Jablonski, Lara Fuhrmann, Rebecca Denes, Mirjam Feldkamp, Ina Nissen, Natascha Santacroce, Elodie Burcklen, Christiane Beckmann, Maurice Redondo, Olivier Kobel, Christoph Noppen, Niko Beerenwinkel, Tanja Stadler |
| EPI_ISL_3296397 | Wales Specialist Virology Centre Sequencing lab: Pathogen Genomics Unit | Public Health Wales Microbiology Cardiff Wales Specialist Virology Centre | Catherine Moore, Johnathan Evans, Laura Gifford, Malorie Perry, Simon Cottrell, Angela Marchbank, Alec Birchley, Alexander Adams, Amy Gaskin, Bree Gatica-Wilcox, Jason Coombes, Joel Southgate, Lauren Gilbert, Lee Graham, Nicole Pacchiarini, Sara Kumziene-Summerhayes, Sarah Taylor, Sophie Jones, Sara Rey, Matthew Bull, Joanne Watkins, Sally Corden, Tom Connor |
| EPI_ISL_3296398 | Viollier AG | Department of Biosystems Science and Engineering, ETH Zürich | Christian Beisel, Sarah Nadeau, Chaoran Chen, Ivan Topolsky, Kim Philipp Jablonski, Lara Fuhrmann, Rebecca Denes, Mirjam Feldkamp, Ina Nissen, Natascha Santacroce, Elodie Burcklen, Christiane Beckmann, Maurice Redondo, Olivier Kobel, Christoph Noppen, Niko Beerenwinkel, Tanja Stadler |
| EPI_ISL_3296399 | Wales Specialist Virology Centre Sequencing lab: Pathogen Genomics Unit | Public Health Wales Microbiology Cardiff Wales Specialist Virology Centre | Catherine Moore, Johnathan Evans, Laura Gifford, Malorie Perry, Simon Cottrell, Angela Marchbank, Alec Birchley, Alexander Adams, Amy Gaskin, Bree Gatica-Wilcox, Jason Coombes, Joel Southgate, Lauren Gilbert, Lee Graham, Nicole Pacchiarini, Sara Kumziene-Summerhayes, Sarah Taylor, Sophie Jones, Sara Rey, Matthew Bull, Joanne Watkins, Sally Corden, Tom Connor |
| EPI_ISL_3296400 | Viollier AG | Department of Biosystems Science and Engineering, ETH Zürich | Christian Beisel, Sarah Nadeau, Chaoran Chen, Ivan Topolsky, Kim Philipp Jablonski, Lara Fuhrmann, Rebecca Denes, Mirjam Feldkamp, Ina Nissen, Natascha Santacroce, Elodie Burcklen, Christiane Beckmann, Maurice Redondo, Olivier Kobel, Christoph Noppen, Niko Beerenwinkel, Tanja Stadler |
| EPI_ISL_3296401 | Wales Specialist Virology Centre Sequencing lab: Pathogen Genomics Unit | Public Health Wales Microbiology Cardiff Wales Specialist Virology Centre | Catherine Moore, Johnathan Evans, Laura Gifford, Malorie Perry, Simon Cottrell, Angela Marchbank, Alec Birchley, Alexander Adams, Amy Gaskin, Bree Gatica-Wilcox, Jason Coombes, Joel Southgate, Lauren Gilbert, Lee Graham, Nicole Pacchiarini, Sara Kumziene-Summerhayes, Sarah Taylor, Sophie Jones, Sara Rey, Matthew Bull, Joanne Watkins, Sally Corden, Tom Connor |
| EPI_ISL_3296402 | Viollier AG | Department of Biosystems Science and Engineering, ETH Zürich | Christian Beisel, Sarah Nadeau, Chaoran Chen, Ivan Topolsky, Kim Philipp Jablonski, Lara Fuhrmann, Rebecca Denes, Mirjam Feldkamp, Ina Nissen, Natascha Santacroce, Elodie Burcklen, Christiane Beckmann, Maurice Redondo, Olivier Kobel, Christoph Noppen, Niko Beerenwinkel, Tanja Stadler |
| EPI_ISL_3296403 | Wales Specialist Virology Centre Sequencing lab: Pathogen Genomics Unit | Public Health Wales Microbiology Cardiff Wales Specialist Virology Centre | Catherine Moore, Johnathan Evans, Laura Gifford, Malorie Perry, Simon Cottrell, Angela Marchbank, Alec Birchley, Alexander Adams, Amy Gaskin, Bree Gatica-Wilcox, Jason Coombes, Joel Southgate, Lauren Gilbert, Lee Graham, Nicole Pacchiarini, Sara Kumziene-Summerhayes, Sarah Taylor, Sophie Jones, Sara Rey, Matthew Bull, Joanne Watkins, Sally Corden, Tom Connor |
| EPI_ISL_3296404 | Viollier AG | Department of Biosystems Science and Engineering, ETH Zürich | Christian Beisel, Sarah Nadeau, Chaoran Chen, Ivan Topolsky, Kim Philipp Jablonski, Lara Fuhrmann, Rebecca Denes, Mirjam Feldkamp, Ina Nissen, Natascha Santacroce, Elodie Burcklen, Christiane Beckmann, Maurice Redondo, Olivier Kobel, Christoph Noppen, Niko Beerenwinkel, Tanja Stadler |
| EPI_ISL_3296405 | Wales Specialist Virology Centre Sequencing lab: Pathogen Genomics Unit | Public Health Wales Microbiology Cardiff Wales Specialist Virology Centre | Catherine Moore, Johnathan Evans, Laura Gifford, Malorie Perry, Simon Cottrell, Angela Marchbank, Alec Birchley, Alexander Adams, Amy Gaskin, Bree Gatica-Wilcox, Jason Coombes, Joel Southgate, Lauren Gilbert, Lee Graham, Nicole Pacchiarini, Sara Kumziene-Summerhayes, Sarah Taylor, Sophie Jones, Sara Rey, Matthew Bull, Joanne Watkins, Sally Corden, Tom Connor |
| EPI_ISL_3296406 | Viollier AG | Department of Biosystems Science and Engineering, ETH Zürich | Christian Beisel, Sarah Nadeau, Chaoran Chen, Ivan Topolsky, Kim Philipp Jablonski, Lara Fuhrmann, Rebecca Denes, Mirjam Feldkamp, Ina Nissen, Natascha Santacroce, Elodie Burcklen, Christiane Beckmann, Maurice Redondo, Olivier Kobel, Christoph Noppen, Niko Beerenwinkel, Tanja Stadler |
| EPI_ISL_3296407 | Wales Specialist Virology Centre Sequencing lab: Pathogen Genomics Unit | Public Health Wales Microbiology Cardiff Wales Specialist Virology Centre | Catherine Moore, Johnathan Evans, Laura Gifford, Malorie Perry, Simon Cottrell, Angela Marchbank, Alec Birchley, Alexander Adams, Amy Gaskin, Bree Gatica-Wilcox, Jason Coombes, Joel Southgate, Lauren Gilbert, Lee Graham, Nicole Pacchiarini, Sara Kumziene-Summerhayes, Sarah Taylor, Sophie Jones, Sara Rey, Matthew Bull, Joanne Watkins, Sally Corden, Tom Connor |
| EPI_ISL_3296408 | Viollier AG | Department of Biosystems Science and Engineering, ETH Zürich | Christian Beisel, Sarah Nadeau, Chaoran Chen, Ivan Topolsky, Kim Philipp Jablonski, Lara Fuhrmann, Rebecca Denes, Mirjam Feldkamp, Ina Nissen, Natascha Santacroce, Elodie Burcklen, Christiane Beckmann, Maurice Redondo, Olivier Kobel, Christoph Noppen, Niko Beerenwinkel, Tanja Stadler |
| EPI_ISL_3296409 | Wales Specialist Virology Centre Sequencing lab: Pathogen Genomics Unit | Public Health Wales Microbiology Cardiff Wales Specialist Virology Centre | Catherine Moore, Johnathan Evans, Laura Gifford, Malorie Perry, Simon Cottrell, Angela Marchbank, Alec Birchley, Alexander Adams, Amy Gaskin, Bree Gatica-Wilcox, Jason Coombes, Joel Southgate, Lauren Gilbert, Lee Graham, Nicole Pacchiarini, Sara Kumziene-Summerhayes, Sarah Taylor, Sophie Jones, Sara Rey, Matthew Bull, Joanne Watkins, Sally Corden, Tom Connor |
| EPI_ISL_3296410 | Viollier AG | Department of Biosystems Science and Engineering, ETH Zürich | Christian Beisel, Sarah Nadeau, Chaoran Chen, Ivan Topolsky, Kim Philipp Jablonski, Lara Fuhrmann, Rebecca Denes, Mirjam Feldkamp, Ina Nissen, Natascha Santacroce, Elodie Burcklen, Christiane Beckmann, Maurice Redondo, Olivier Kobel, Christoph Noppen, Niko Beerenwinkel, Tanja Stadler |
| EPI_ISL_3296411 | Wales Specialist Virology Centre Sequencing lab: Pathogen Genomics Unit | Public Health Wales Microbiology Cardiff Wales Specialist Virology Centre | Catherine Moore, Johnathan Evans, Laura Gifford, Malorie Perry, Simon Cottrell, Angela Marchbank, Alec Birchley, Alexander Adams, Amy Gaskin, Bree Gatica-Wilcox, Jason Coombes, Joel Southgate, Lauren Gilbert, Lee Graham, Nicole Pacchiarini, Sara Kumziene-Summerhayes, Sarah Taylor, Sophie Jones, Sara Rey, Matthew Bull, Joanne Watkins, Sally Corden, Tom Connor |
| EPI_ISL_3296412 | Viollier AG | Department of Biosystems Science and Engineering, ETH Zürich | Christian Beisel, Sarah Nadeau, Chaoran Chen, Ivan Topolsky, Kim Philipp Jablonski, Lara Fuhrmann, Rebecca Denes, Mirjam Feldkamp, Ina Nissen, Natascha Santacroce, Elodie Burcklen, Christiane Beckmann, Maurice Redondo, Olivier Kobel, Christoph Noppen, Niko Beerenwinkel, Tanja Stadler |
| EPI_ISL_3296413 | Wales Specialist Virology Centre Sequencing lab: Pathogen Genomics Unit | Public Health Wales Microbiology Cardiff Wales Specialist Virology Centre | Catherine Moore, Johnathan Evans, Laura Gifford, Malorie Perry, Simon Cottrell, Angela Marchbank, Alec Birchley, Alexander Adams, Amy Gaskin, Bree Gatica-Wilcox, Jason Coombes, Joel Southgate, Lauren Gilbert, Lee Graham, Nicole Pacchiarini, Sara Kumziene-Summerhayes, Sarah Taylor, Sophie Jones, Sara Rey, Matthew Bull, Joanne Watkins, Sally Corden, Tom Connor |
| EPI_ISL_3296414 | Viollier AG | Department of Biosystems Science and Engineering, ETH Zürich | Christian Beisel, Sarah Nadeau, Chaoran Chen, Ivan Topolsky, Kim Philipp Jablonski, Lara Fuhrmann, Rebecca Denes, Mirjam Feldkamp, Ina Nissen, Natascha Santacroce, Elodie Burcklen, Christiane Beckmann, Maurice Redondo, Olivier Kobel, Christoph Noppen, Niko Beerenwinkel, Tanja Stadler |
| EPI_ISL_3296415 | Wales Specialist Virology Centre Sequencing lab: Pathogen | Public Health Wales Microbiology Cardiff Wales Specialist Virology | Catherine Moore, Johnathan Evans, Laura Gifford, Malorie Perry, Simon Cottrell, Angela Marchbank, Alec Birchley, Alexander Adams, Amy Gaskin, Bree Gatica-Wilcox, Jason Coombes, Joel Southgate, Lauren Gilbert, Lee Graham, Nicole |

| | | | |
|---|---|---|---|
| | Genomics Unit | Centre | Pacchiarini, Sara Kumziene-Summerhayes, Sarah Taylor, Sophie Jones, Sara Rey, Matthew Bull, Joanne Watkins, Sally Corden, Tom Connor |
| EPI_ISL_3296416 | Viollier AG | Department of Biosystems Science and Engineering, ETH Zürich | Christian Beisel, Sarah Nadeau, Chaoran Chen, Ivan Topolsky, Kim Philipp Jablonski, Lara Fuhrmann, Rebecca Denes, Mirjam Feldkamp, Ina Nissen, Natascha Santacroce, Elodie Burcklen, Christiane Beckmann, Maurice Redondo, Olivier Kobel, Christoph Noppen, Niko Beerenwinkel, Tanja Stadler |
| EPI_ISL_3296417 | Wales Specialist Virology Centre Sequencing lab: Pathogen Genomics Unit | Public Health Wales Microbiology Cardiff Wales Specialist Virology Centre | Catherine Moore, Johnathan Evans, Laura Gifford, Malorie Perry, Simon Cottrell, Angela Marchbank, Alec Birchley, Alexander Adams, Amy Gaskin, Bree Gatica-Wilcox, Jason Coombes, Joel Southgate, Lauren Gilbert, Lee Graham, Nicole Pacchiarini, Sara Kumziene-Summerhayes, Sarah Taylor, Sophie Jones, Sara Rey, Matthew Bull, Joanne Watkins, Sally Corden, Tom Connor |
| EPI_ISL_3296418 | Viollier AG | Department of Biosystems Science and Engineering, ETH Zürich | Christian Beisel, Sarah Nadeau, Chaoran Chen, Ivan Topolsky, Kim Philipp Jablonski, Lara Fuhrmann, Rebecca Denes, Mirjam Feldkamp, Ina Nissen, Natascha Santacroce, Elodie Burcklen, Christiane Beckmann, Maurice Redondo, Olivier Kobel, Christoph Noppen, Niko Beerenwinkel, Tanja Stadler |
| EPI_ISL_3296419 | Wales Specialist Virology Centre Sequencing lab: Pathogen Genomics Unit | Public Health Wales Microbiology Cardiff Wales Specialist Virology Centre | Catherine Moore, Johnathan Evans, Laura Gifford, Malorie Perry, Simon Cottrell, Angela Marchbank, Alec Birchley, Alexander Adams, Amy Gaskin, Bree Gatica-Wilcox, Jason Coombes, Joel Southgate, Lauren Gilbert, Lee Graham, Nicole Pacchiarini, Sara Kumziene-Summerhayes, Sarah Taylor, Sophie Jones, Sara Rey, Matthew Bull, Joanne Watkins, Sally Corden, Tom Connor |
| EPI_ISL_3296420 | Viollier AG | Department of Biosystems Science and Engineering, ETH Zürich | Christian Beisel, Sarah Nadeau, Chaoran Chen, Ivan Topolsky, Kim Philipp Jablonski, Lara Fuhrmann, Rebecca Denes, Mirjam Feldkamp, Ina Nissen, Natascha Santacroce, Elodie Burcklen, Christiane Beckmann, Maurice Redondo, Olivier Kobel, Christoph Noppen, Niko Beerenwinkel, Tanja Stadler |
| EPI_ISL_3296421 | Wales Specialist Virology Centre Sequencing lab: Pathogen Genomics Unit | Public Health Wales Microbiology Cardiff Wales Specialist Virology Centre | Catherine Moore, Johnathan Evans, Laura Gifford, Malorie Perry, Simon Cottrell, Angela Marchbank, Alec Birchley, Alexander Adams, Amy Gaskin, Bree Gatica-Wilcox, Jason Coombes, Joel Southgate, Lauren Gilbert, Lee Graham, Nicole Pacchiarini, Sara Kumziene-Summerhayes, Sarah Taylor, Sophie Jones, Sara Rey, Matthew Bull, Joanne Watkins, Sally Corden, Tom Connor |
| EPI_ISL_3296422 | Viollier AG | Department of Biosystems Science and Engineering, ETH Zürich | Christian Beisel, Sarah Nadeau, Chaoran Chen, Ivan Topolsky, Kim Philipp Jablonski, Lara Fuhrmann, Rebecca Denes, Mirjam Feldkamp, Ina Nissen, Natascha Santacroce, Elodie Burcklen, Christiane Beckmann, Maurice Redondo, Olivier Kobel, Christoph Noppen, Niko Beerenwinkel, Tanja Stadler |
| EPI_ISL_3296423 | Wales Specialist Virology Centre Sequencing lab: Pathogen Genomics Unit | Public Health Wales Microbiology Cardiff Wales Specialist Virology Centre | Catherine Moore, Johnathan Evans, Laura Gifford, Malorie Perry, Simon Cottrell, Angela Marchbank, Alec Birchley, Alexander Adams, Amy Gaskin, Bree Gatica-Wilcox, Jason Coombes, Joel Southgate, Lauren Gilbert, Lee Graham, Nicole Pacchiarini, Sara Kumziene-Summerhayes, Sarah Taylor, Sophie Jones, Sara Rey, Matthew Bull, Joanne Watkins, Sally Corden, Tom Connor |
| EPI_ISL_3296424 | Viollier AG | Department of Biosystems Science and Engineering, ETH Zürich | Christian Beisel, Sarah Nadeau, Chaoran Chen, Ivan Topolsky, Kim Philipp Jablonski, Lara Fuhrmann, Rebecca Denes, Mirjam Feldkamp, Ina Nissen, Natascha Santacroce, Elodie Burcklen, Christiane Beckmann, Maurice Redondo, Olivier Kobel, Christoph Noppen, Niko Beerenwinkel, Tanja Stadler |
| EPI_ISL_3296425 | Wales Specialist Virology Centre Sequencing lab: Pathogen Genomics Unit | Public Health Wales Microbiology Cardiff Wales Specialist Virology Centre | Catherine Moore, Johnathan Evans, Laura Gifford, Malorie Perry, Simon Cottrell, Angela Marchbank, Alec Birchley, Alexander Adams, Amy Gaskin, Bree Gatica-Wilcox, Jason Coombes, Joel Southgate, Lauren Gilbert, Lee Graham, Nicole Pacchiarini, Sara Kumziene-Summerhayes, Sarah Taylor, Sophie Jones, Sara Rey, Matthew Bull, Joanne Watkins, Sally Corden, Tom Connor |
| EPI_ISL_3296426 | Viollier AG | Department of Biosystems Science and Engineering, ETH Zürich | Christian Beisel, Sarah Nadeau, Chaoran Chen, Ivan Topolsky, Kim Philipp Jablonski, Lara Fuhrmann, Rebecca Denes, Mirjam Feldkamp, Ina Nissen, Natascha Santacroce, Elodie Burcklen, Christiane Beckmann, Maurice Redondo, Olivier Kobel, Christoph Noppen, Niko Beerenwinkel, Tanja Stadler |
| EPI_ISL_3296427 | Wales Specialist Virology Centre Sequencing lab: Pathogen Genomics Unit | Public Health Wales Microbiology Cardiff Wales Specialist Virology Centre | Catherine Moore, Johnathan Evans, Laura Gifford, Malorie Perry, Simon Cottrell, Angela Marchbank, Alec Birchley, Alexander Adams, Amy Gaskin, Bree Gatica-Wilcox, Jason Coombes, Joel Southgate, Lauren Gilbert, Lee Graham, Nicole Pacchiarini, Sara Kumziene-Summerhayes, Sarah Taylor, Sophie Jones, Sara Rey, Matthew Bull, Joanne Watkins, Sally Corden, Tom Connor |
| EPI_ISL_3296428 | Viollier AG | Department of Biosystems Science and Engineering, ETH Zürich | Christian Beisel, Sarah Nadeau, Chaoran Chen, Ivan Topolsky, Kim Philipp Jablonski, Lara Fuhrmann, Rebecca Denes, Mirjam Feldkamp, Ina Nissen, Natascha Santacroce, Elodie Burcklen, Christiane Beckmann, Maurice Redondo, Olivier Kobel, Christoph Noppen, Niko Beerenwinkel, Tanja Stadler |
| EPI_ISL_3296429 | Wales Specialist Virology Centre Sequencing lab: Pathogen Genomics Unit | Public Health Wales Microbiology Cardiff Wales Specialist Virology Centre | Catherine Moore, Johnathan Evans, Laura Gifford, Malorie Perry, Simon Cottrell, Angela Marchbank, Alec Birchley, Alexander Adams, Amy Gaskin, Bree Gatica-Wilcox, Jason Coombes, Joel Southgate, Lauren Gilbert, Lee Graham, Nicole Pacchiarini, Sara Kumziene-Summerhayes, Sarah Taylor, Sophie Jones, Sara Rey, Matthew Bull, Joanne Watkins, Sally Corden, Tom Connor |
| EPI_ISL_3296430 | Viollier AG | Department of Biosystems Science and Engineering, ETH Zürich | Christian Beisel, Sarah Nadeau, Chaoran Chen, Ivan Topolsky, Kim Philipp Jablonski, Lara Fuhrmann, Rebecca Denes, Mirjam Feldkamp, Ina Nissen, Natascha Santacroce, Elodie Burcklen, Christiane Beckmann, Maurice Redondo, Olivier Kobel, Christoph Noppen, Niko Beerenwinkel, Tanja Stadler |
| EPI_ISL_3296431, EPI_ISL_3296432 | Wales Specialist Virology Centre Sequencing lab: Pathogen Genomics Unit | Public Health Wales Microbiology Cardiff Wales Specialist Virology Centre | Catherine Moore, Johnathan Evans, Laura Gifford, Malorie Perry, Simon Cottrell, Angela Marchbank, Alec Birchley, Alexander Adams, Amy Gaskin, Bree Gatica-Wilcox, Jason Coombes, Joel Southgate, Lauren Gilbert, Lee Graham, Nicole Pacchiarini, Sara Kumziene-Summerhayes, Sarah Taylor, Sophie Jones, Sara Rey, Matthew Bull, Joanne Watkins, Sally Corden, Tom Connor |
| EPI_ISL_3296433 | Viollier AG | Department of Biosystems Science and Engineering, ETH Zürich | Christian Beisel, Sarah Nadeau, Chaoran Chen, Ivan Topolsky, Kim Philipp Jablonski, Lara Fuhrmann, Rebecca Denes, Mirjam Feldkamp, Ina Nissen, Natascha Santacroce, Elodie Burcklen, Christiane Beckmann, Maurice Redondo, Olivier Kobel, Christoph Noppen, Niko Beerenwinkel, Tanja Stadler |
| EPI_ISL_3296434 | Wales Specialist Virology Centre Sequencing lab: Pathogen Genomics Unit | Public Health Wales Microbiology Cardiff Wales Specialist Virology Centre | Catherine Moore, Johnathan Evans, Laura Gifford, Malorie Perry, Simon Cottrell, Angela Marchbank, Alec Birchley, Alexander Adams, Amy Gaskin, Bree Gatica-Wilcox, Jason Coombes, Joel Southgate, Lauren Gilbert, Lee Graham, Nicole Pacchiarini, Sara Kumziene-Summerhayes, Sarah Taylor, Sophie Jones, Sara Rey, Matthew Bull, Joanne Watkins, Sally Corden, Tom Connor |
| EPI_ISL_3296435 | Viollier AG | Department of Biosystems Science and Engineering, ETH Zürich | Christian Beisel, Sarah Nadeau, Chaoran Chen, Ivan Topolsky, Kim Philipp Jablonski, Lara Fuhrmann, Rebecca Denes, Mirjam Feldkamp, Ina Nissen, Natascha Santacroce, Elodie Burcklen, Christiane Beckmann, Maurice Redondo, Olivier Kobel, Christoph Noppen, Niko Beerenwinkel, Tanja Stadler |
| EPI_ISL_3296436 | Wales Specialist Virology Centre Sequencing lab: Pathogen Genomics Unit | Public Health Wales Microbiology Cardiff Wales Specialist Virology Centre | Catherine Moore, Johnathan Evans, Laura Gifford, Malorie Perry, Simon Cottrell, Angela Marchbank, Alec Birchley, Alexander Adams, Amy Gaskin, Bree Gatica-Wilcox, Jason Coombes, Joel Southgate, Lauren Gilbert, Lee Graham, Nicole Pacchiarini, Sara Kumziene-Summerhayes, Sarah Taylor, Sophie Jones, Sara Rey, Matthew Bull, Joanne Watkins, Sally Corden, Tom Connor |
| EPI_ISL_3296437 | Viollier AG | Department of Biosystems Science and Engineering, ETH Zürich | Christian Beisel, Sarah Nadeau, Chaoran Chen, Ivan Topolsky, Kim Philipp Jablonski, Lara Fuhrmann, Rebecca Denes, Mirjam Feldkamp, Ina Nissen, Natascha Santacroce, Elodie Burcklen, Christiane Beckmann, Maurice Redondo, Olivier Kobel, Christoph Noppen, Niko Beerenwinkel, Tanja Stadler |
| EPI_ISL_3296438 | Wales Specialist Virology Centre Sequencing lab: Pathogen Genomics Unit | Public Health Wales Microbiology Cardiff Wales Specialist Virology Centre | Catherine Moore, Johnathan Evans, Laura Gifford, Malorie Perry, Simon Cottrell, Angela Marchbank, Alec Birchley, Alexander Adams, Amy Gaskin, Bree Gatica-Wilcox, Jason Coombes, Joel Southgate, Lauren Gilbert, Lee Graham, Nicole Pacchiarini, Sara Kumziene-Summerhayes, Sarah Taylor, Sophie Jones, Sara Rey, Matthew Bull, Joanne Watkins, Sally Corden, Tom Connor |
| EPI_ISL_3296439 | Viollier AG | Department of Biosystems Science and Engineering, ETH Zürich | Christian Beisel, Sarah Nadeau, Chaoran Chen, Ivan Topolsky, Kim Philipp Jablonski, Lara Fuhrmann, Rebecca Denes, Mirjam Feldkamp, Ina Nissen, Natascha Santacroce, Elodie Burcklen, Christiane Beckmann, Maurice Redondo, Olivier Kobel, Christoph Noppen, Niko Beerenwinkel, Tanja Stadler |
| EPI_ISL_3296440 | Wales Specialist Virology Centre Sequencing lab: Pathogen Genomics Unit | Public Health Wales Microbiology Cardiff Wales Specialist Virology Centre | Catherine Moore, Johnathan Evans, Laura Gifford, Malorie Perry, Simon Cottrell, Angela Marchbank, Alec Birchley, Alexander Adams, Amy Gaskin, Bree Gatica-Wilcox, Jason Coombes, Joel Southgate, Lauren Gilbert, Lee Graham, Nicole Pacchiarini, Sara Kumziene-Summerhayes, Sarah Taylor, Sophie Jones, Sara Rey, Matthew Bull, Joanne Watkins, Sally Corden, Tom Connor |
| EPI_ISL_3296441 | Viollier AG | Department of Biosystems Science and Engineering, ETH Zürich | Christian Beisel, Sarah Nadeau, Chaoran Chen, Ivan Topolsky, Kim Philipp Jablonski, Lara Fuhrmann, Rebecca Denes, Mirjam Feldkamp, Ina Nissen, Natascha Santacroce, Elodie Burcklen, Christiane Beckmann, Maurice Redondo, Olivier Kobel, Christoph Noppen, Niko Beerenwinkel, Tanja Stadler |
| EPI_ISL_3296442 | Wales Specialist Virology Centre Sequencing lab: Pathogen Genomics Unit | Public Health Wales Microbiology Cardiff Wales Specialist Virology Centre | Catherine Moore, Johnathan Evans, Laura Gifford, Malorie Perry, Simon Cottrell, Angela Marchbank, Alec Birchley, Alexander Adams, Amy Gaskin, Bree Gatica-Wilcox, Jason Coombes, Joel Southgate, Lauren Gilbert, Lee Graham, Nicole Pacchiarini, Sara Kumziene-Summerhayes, Sarah Taylor, Sophie Jones, Sara Rey, Matthew Bull, Joanne Watkins, Sally Corden, Tom Connor |
| EPI_ISL_3296443 | Viollier AG | Department of Biosystems Science and Engineering, ETH Zürich | Christian Beisel, Sarah Nadeau, Chaoran Chen, Ivan Topolsky, Kim Philipp Jablonski, Lara Fuhrmann, Rebecca Denes, Mirjam Feldkamp, Ina Nissen, Natascha Santacroce, Elodie Burcklen, Christiane Beckmann, Maurice Redondo, Olivier Kobel, Christoph Noppen, Niko Beerenwinkel, Tanja Stadler |
| EPI_ISL_3296444 | Wales Specialist Virology Centre Sequencing lab: Pathogen Genomics Unit | Public Health Wales Microbiology Cardiff Wales Specialist Virology Centre | Catherine Moore, Johnathan Evans, Laura Gifford, Malorie Perry, Simon Cottrell, Angela Marchbank, Alec Birchley, Alexander Adams, Amy Gaskin, Bree Gatica-Wilcox, Jason Coombes, Joel Southgate, Lauren Gilbert, Lee Graham, Nicole Pacchiarini, Sara Kumziene-Summerhayes, Sarah Taylor, Sophie Jones, Sara Rey, Matthew Bull, Joanne Watkins, Sally Corden, Tom Connor |
| EPI_ISL_3296445 | Viollier AG | Department of Biosystems Science and Engineering, ETH Zürich | Christian Beisel, Sarah Nadeau, Chaoran Chen, Ivan Topolsky, Kim Philipp Jablonski, Lara Fuhrmann, Rebecca Denes, Mirjam Feldkamp, Ina Nissen, Natascha Santacroce, Elodie Burcklen, Christiane Beckmann, Maurice Redondo, Olivier Kobel, Christoph Noppen, Niko Beerenwinkel, Tanja Stadler |
| EPI_ISL_3296446 | Wales Specialist Virology Centre Sequencing lab: Pathogen Genomics Unit | Public Health Wales Microbiology Cardiff Wales Specialist Virology Centre | Catherine Moore, Johnathan Evans, Laura Gifford, Malorie Perry, Simon Cottrell, Angela Marchbank, Alec Birchley, Alexander Adams, Amy Gaskin, Bree Gatica-Wilcox, Jason Coombes, Joel Southgate, Lauren Gilbert, Lee Graham, Nicole Pacchiarini, Sara Kumziene-Summerhayes, Sarah Taylor, Sophie Jones, Sara Rey, Matthew Bull, Joanne Watkins, Sally Corden, Tom Connor |
| EPI_ISL_3296447 | Viollier AG | Department of Biosystems Science and Engineering, ETH Zürich | Christian Beisel, Sarah Nadeau, Chaoran Chen, Ivan Topolsky, Kim Philipp Jablonski, Lara Fuhrmann, Rebecca Denes, Mirjam Feldkamp, Ina Nissen, Natascha Santacroce, Elodie Burcklen, Christiane Beckmann, Maurice Redondo, Olivier Kobel, Christoph Noppen, Niko Beerenwinkel, Tanja Stadler |
| EPI_ISL_3296448 | Wales Specialist Virology Centre Sequencing lab: Pathogen Genomics Unit | Public Health Wales Microbiology Cardiff Wales Specialist Virology Centre | Catherine Moore, Johnathan Evans, Laura Gifford, Malorie Perry, Simon Cottrell, Angela Marchbank, Alec Birchley, Alexander Adams, Amy Gaskin, Bree Gatica-Wilcox, Jason Coombes, Joel Southgate, Lauren Gilbert, Lee Graham, Nicole Pacchiarini, Sara Kumziene-Summerhayes, Sarah Taylor, Sophie Jones, Sara Rey, Matthew Bull, Joanne Watkins, Sally Corden, Tom Connor |
| EPI_ISL_3296449 | Viollier AG | Department of Biosystems Science and Engineering, ETH Zürich | Christian Beisel, Sarah Nadeau, Chaoran Chen, Ivan Topolsky, Kim Philipp Jablonski, Lara Fuhrmann, Rebecca Denes, Mirjam Feldkamp, Ina Nissen, Natascha Santacroce, Elodie Burcklen, Christiane Beckmann, Maurice Redondo, Olivier Kobel, Christoph Noppen, Niko Beerenwinkel, Tanja Stadler |
| EPI_ISL_3296450 | Wales Specialist Virology Centre Sequencing lab: Pathogen Genomics Unit | Public Health Wales Microbiology Cardiff Wales Specialist Virology Centre | Catherine Moore, Johnathan Evans, Laura Gifford, Malorie Perry, Simon Cottrell, Angela Marchbank, Alec Birchley, Alexander Adams, Amy Gaskin, Bree Gatica-Wilcox, Jason Coombes, Joel Southgate, Lauren Gilbert, Lee Graham, Nicole Pacchiarini, Sara Kumziene-Summerhayes, Sarah Taylor, Sophie Jones, Sara Rey, Matthew Bull, Joanne Watkins, Sally Corden, Tom Connor |
| EPI_ISL_3296451 | Viollier AG | Department of Biosystems Science and Engineering, ETH Zürich | Christian Beisel, Sarah Nadeau, Chaoran Chen, Ivan Topolsky, Kim Philipp Jablonski, Lara Fuhrmann, Rebecca Denes, Mirjam Feldkamp, Ina Nissen, Natascha Santacroce, Elodie Burcklen, Christiane Beckmann, Maurice Redondo, Olivier Kobel, Christoph Noppen, Niko Beerenwinkel, Tanja Stadler |
| EPI_ISL_3296452 | Wales Specialist Virology Centre Sequencing lab: Pathogen Genomics Unit | Public Health Wales Microbiology Cardiff Wales Specialist Virology Centre | Catherine Moore, Johnathan Evans, Laura Gifford, Malorie Perry, Simon Cottrell, Angela Marchbank, Alec Birchley, Alexander Adams, Amy Gaskin, Bree Gatica-Wilcox, Jason Coombes, Joel Southgate, Lauren Gilbert, Lee Graham, Nicole Pacchiarini, Sara Kumziene-Summerhayes, Sarah Taylor, Sophie Jones, Sara Rey, Matthew Bull, Joanne Watkins, Sally Corden, Tom Connor |
| EPI_ISL_3296453 | Viollier AG | Department of Biosystems Science and Engineering, ETH Zürich | Christian Beisel, Sarah Nadeau, Chaoran Chen, Ivan Topolsky, Kim Philipp Jablonski, Lara Fuhrmann, Rebecca Denes, Mirjam Feldkamp, Ina Nissen, Natascha Santacroce, Elodie Burcklen, Christiane Beckmann, Maurice Redondo, Olivier Kobel, Christoph Noppen, Niko Beerenwinkel, Tanja Stadler |
| EPI_ISL_3296454 | Wales Specialist Virology Centre Sequencing lab: Pathogen Genomics Unit | Public Health Wales Microbiology Cardiff Wales Specialist Virology Centre | Catherine Moore, Johnathan Evans, Laura Gifford, Malorie Perry, Simon Cottrell, Angela Marchbank, Alec Birchley, Alexander Adams, Amy Gaskin, Bree Gatica-Wilcox, Jason Coombes, Joel Southgate, Lauren Gilbert, Lee Graham, Nicole Pacchiarini, Sara Kumziene-Summerhayes, Sarah Taylor, Sophie Jones, Sara Rey, Matthew Bull, Joanne Watkins, Sally Corden, Tom Connor |
| EPI_ISL_3296455 | Viollier AG | Department of Biosystems Science and Engineering, ETH Zürich | Christian Beisel, Sarah Nadeau, Chaoran Chen, Ivan Topolsky, Kim Philipp Jablonski, Lara Fuhrmann, Rebecca Denes, Mirjam Feldkamp, Ina Nissen, Natascha Santacroce, Elodie Burcklen, Christiane Beckmann, Maurice Redondo, Olivier Kobel, Christoph Noppen, Niko Beerenwinkel, Tanja Stadler |
| EPI_ISL_3296456 | Wales Specialist Virology Centre Sequencing lab: Pathogen Genomics Unit | Public Health Wales Microbiology Cardiff Wales Specialist Virology Centre | Catherine Moore, Johnathan Evans, Laura Gifford, Malorie Perry, Simon Cottrell, Angela Marchbank, Alec Birchley, Alexander Adams, Amy Gaskin, Bree Gatica-Wilcox, Jason Coombes, Joel Southgate, Lauren Gilbert, Lee Graham, Nicole Pacchiarini, Sara Kumziene-Summerhayes, Sarah Taylor, Sophie Jones, Sara Rey, Matthew Bull, Joanne Watkins, Sally Corden, Tom Connor |
| EPI_ISL_3296457 | Viollier AG | Department of Biosystems Science and Engineering, ETH Zürich | Christian Beisel, Sarah Nadeau, Chaoran Chen, Ivan Topolsky, Kim Philipp Jablonski, Lara Fuhrmann, Rebecca Denes, Mirjam Feldkamp, Ina Nissen, Natascha Santacroce, Elodie Burcklen, Christiane Beckmann, Maurice Redondo, Olivier Kobel, Christoph Noppen, Niko Beerenwinkel, Tanja Stadler |
| EPI_ISL_3296458, EPI_ISL_3296459 | Wales Specialist Virology Centre Sequencing lab: Pathogen Genomics Unit | Public Health Wales Microbiology Cardiff Wales Specialist Virology Centre | Catherine Moore, Johnathan Evans, Laura Gifford, Malorie Perry, Simon Cottrell, Angela Marchbank, Alec Birchley, Alexander Adams, Amy Gaskin, Bree Gatica-Wilcox, Jason Coombes, Joel Southgate, Lauren Gilbert, Lee Graham, Nicole Pacchiarini, Sara Kumziene-Summerhayes, Sarah Taylor, Sophie Jones, Sara Rey, Matthew Bull, Joanne Watkins, Sally Corden, Tom Connor |
| EPI_ISL_3296460 | Viollier AG | Department of Biosystems Science and Engineering, ETH Zürich | Christian Beisel, Sarah Nadeau, Chaoran Chen, Ivan Topolsky, Kim Philipp Jablonski, Lara Fuhrmann, Rebecca Denes, Mirjam Feldkamp, Ina Nissen, Natascha Santacroce, Elodie Burcklen, Christiane Beckmann, Maurice Redondo, Olivier Kobel, Christoph Noppen, Niko Beerenwinkel, Tanja Stadler |
| EPI_ISL_3296461 | Wales Specialist Virology Centre Sequencing lab: Pathogen Genomics Unit | Public Health Wales Microbiology Cardiff Wales Specialist Virology Centre | Catherine Moore, Johnathan Evans, Laura Gifford, Malorie Perry, Simon Cottrell, Angela Marchbank, Alec Birchley, Alexander Adams, Amy Gaskin, Bree Gatica-Wilcox, Jason Coombes, Joel Southgate, Lauren Gilbert, Lee Graham, Nicole Pacchiarini, Sara Kumziene-Summerhayes, Sarah Taylor, Sophie Jones, Sara Rey, Matthew Bull, Joanne Watkins, Sally Corden, Tom Connor |
| EPI_ISL_3296462 | Viollier AG | Department of Biosystems Science and Engineering, ETH Zürich | Christian Beisel, Sarah Nadeau, Chaoran Chen, Ivan Topolsky, Kim Philipp Jablonski, Lara Fuhrmann, Rebecca Denes, Mirjam Feldkamp, Ina Nissen, Natascha Santacroce, Elodie Burcklen, Christiane Beckmann, Maurice Redondo, Olivier Kobel, |

| | | | |
|---|---|---|---|
| | | | Christoph Noppen, Niko Beerenwinkel, Tanja Stadler |
| EPI_ISL_3296463, EPI_ISL_3296465 | Wales Specialist Virology Centre Sequencing lab: Pathogen Genomics Unit | Public Health Wales Microbiology Cardiff Wales Specialist Virology Centre | Catherine Moore, Johnathan Evans, Laura Gifford, Malorie Perry, Simon Cottrell, Angela Marchbank, Alec Birchley, Alexander Adams, Amy Gaskin, Bree Gatica-Wilcox, Jason Coombes, Joel Southgate, Lauren Gilbert, Lee Graham, Nicole Pacchiarini, Sara Kumziene-Summerhayes, Sarah Taylor, Sophie Jones, Sara Rey, Matthew Bull, Joanne Watkins, Sally Corden, Tom Connor |
| EPI_ISL_3296466 | Viollier AG | Department of Biosystems Science and Engineering, ETH Zürich | Christian Beisel, Sarah Nadeau, Chaoran Chen, Ivan Topolsky, Kim Philipp Jablonski, Lara Fuhrmann, Rebecca Denes, Mirjam Feldkamp, Ina Nissen, Natascha Santacroce, Elodie Burcklen, Christiane Beckmann, Maurice Redondo, Olivier Kobel, Christoph Noppen, Niko Beerenwinkel, Tanja Stadler |
| EPI_ISL_3296467 | Wales Specialist Virology Centre Sequencing lab: Pathogen Genomics Unit | Public Health Wales Microbiology Cardiff Wales Specialist Virology Centre | Catherine Moore, Johnathan Evans, Laura Gifford, Malorie Perry, Simon Cottrell, Angela Marchbank, Alec Birchley, Alexander Adams, Amy Gaskin, Bree Gatica-Wilcox, Jason Coombes, Joel Southgate, Lauren Gilbert, Lee Graham, Nicole Pacchiarini, Sara Kumziene-Summerhayes, Sarah Taylor, Sophie Jones, Sara Rey, Matthew Bull, Joanne Watkins, Sally Corden, Tom Connor |
| EPI_ISL_3296468 | Viollier AG | Department of Biosystems Science and Engineering, ETH Zürich | Christian Beisel, Sarah Nadeau, Chaoran Chen, Ivan Topolsky, Kim Philipp Jablonski, Lara Fuhrmann, Rebecca Denes, Mirjam Feldkamp, Ina Nissen, Natascha Santacroce, Elodie Burcklen, Christiane Beckmann, Maurice Redondo, Olivier Kobel, Christoph Noppen, Niko Beerenwinkel, Tanja Stadler |
| EPI_ISL_3296469 | Wales Specialist Virology Centre Sequencing lab: Pathogen Genomics Unit | Public Health Wales Microbiology Cardiff Wales Specialist Virology Centre | Catherine Moore, Johnathan Evans, Laura Gifford, Malorie Perry, Simon Cottrell, Angela Marchbank, Alec Birchley, Alexander Adams, Amy Gaskin, Bree Gatica-Wilcox, Jason Coombes, Joel Southgate, Lauren Gilbert, Lee Graham, Nicole Pacchiarini, Sara Kumziene-Summerhayes, Sarah Taylor, Sophie Jones, Sara Rey, Matthew Bull, Joanne Watkins, Sally Corden, Tom Connor |
| EPI_ISL_3296470 | Viollier AG | Department of Biosystems Science and Engineering, ETH Zürich | Christian Beisel, Sarah Nadeau, Chaoran Chen, Ivan Topolsky, Kim Philipp Jablonski, Lara Fuhrmann, Rebecca Denes, Mirjam Feldkamp, Ina Nissen, Natascha Santacroce, Elodie Burcklen, Christiane Beckmann, Maurice Redondo, Olivier Kobel, Christoph Noppen, Niko Beerenwinkel, Tanja Stadler |
| EPI_ISL_3296471 | Wales Specialist Virology Centre Sequencing lab: Pathogen Genomics Unit | Public Health Wales Microbiology Cardiff Wales Specialist Virology Centre | Catherine Moore, Johnathan Evans, Laura Gifford, Malorie Perry, Simon Cottrell, Angela Marchbank, Alec Birchley, Alexander Adams, Amy Gaskin, Bree Gatica-Wilcox, Jason Coombes, Joel Southgate, Lauren Gilbert, Lee Graham, Nicole Pacchiarini, Sara Kumziene-Summerhayes, Sarah Taylor, Sophie Jones, Sara Rey, Matthew Bull, Joanne Watkins, Sally Corden, Tom Connor |
| EPI_ISL_3296472 | Viollier AG | Department of Biosystems Science and Engineering, ETH Zürich | Christian Beisel, Sarah Nadeau, Chaoran Chen, Ivan Topolsky, Kim Philipp Jablonski, Lara Fuhrmann, Rebecca Denes, Mirjam Feldkamp, Ina Nissen, Natascha Santacroce, Elodie Burcklen, Christiane Beckmann, Maurice Redondo, Olivier Kobel, Christoph Noppen, Niko Beerenwinkel, Tanja Stadler |
| EPI_ISL_3296473 | Wales Specialist Virology Centre Sequencing lab: Pathogen Genomics Unit | Public Health Wales Microbiology Cardiff Wales Specialist Virology Centre | Catherine Moore, Johnathan Evans, Laura Gifford, Malorie Perry, Simon Cottrell, Angela Marchbank, Alec Birchley, Alexander Adams, Amy Gaskin, Bree Gatica-Wilcox, Jason Coombes, Joel Southgate, Lauren Gilbert, Lee Graham, Nicole Pacchiarini, Sara Kumziene-Summerhayes, Sarah Taylor, Sophie Jones, Sara Rey, Matthew Bull, Joanne Watkins, Sally Corden, Tom Connor |
| EPI_ISL_3296474 | Viollier AG | Department of Biosystems Science and Engineering, ETH Zürich | Christian Beisel, Sarah Nadeau, Chaoran Chen, Ivan Topolsky, Kim Philipp Jablonski, Lara Fuhrmann, Rebecca Denes, Mirjam Feldkamp, Ina Nissen, Natascha Santacroce, Elodie Burcklen, Christiane Beckmann, Maurice Redondo, Olivier Kobel, Christoph Noppen, Niko Beerenwinkel, Tanja Stadler |
| EPI_ISL_3296475 | Wales Specialist Virology Centre Sequencing lab: Pathogen Genomics Unit | Public Health Wales Microbiology Cardiff Wales Specialist Virology Centre | Catherine Moore, Johnathan Evans, Laura Gifford, Malorie Perry, Simon Cottrell, Angela Marchbank, Alec Birchley, Alexander Adams, Amy Gaskin, Bree Gatica-Wilcox, Jason Coombes, Joel Southgate, Lauren Gilbert, Lee Graham, Nicole Pacchiarini, Sara Kumziene-Summerhayes, Sarah Taylor, Sophie Jones, Sara Rey, Matthew Bull, Joanne Watkins, Sally Corden, Tom Connor |
| EPI_ISL_3296476 | Viollier AG | Department of Biosystems Science and Engineering, ETH Zürich | Christian Beisel, Sarah Nadeau, Chaoran Chen, Ivan Topolsky, Kim Philipp Jablonski, Lara Fuhrmann, Rebecca Denes, Mirjam Feldkamp, Ina Nissen, Natascha Santacroce, Elodie Burcklen, Christiane Beckmann, Maurice Redondo, Olivier Kobel, Christoph Noppen, Niko Beerenwinkel, Tanja Stadler |
| EPI_ISL_3296477 | Wales Specialist Virology Centre Sequencing lab: Pathogen Genomics Unit | Public Health Wales Microbiology Cardiff Wales Specialist Virology Centre | Catherine Moore, Johnathan Evans, Laura Gifford, Malorie Perry, Simon Cottrell, Angela Marchbank, Alec Birchley, Alexander Adams, Amy Gaskin, Bree Gatica-Wilcox, Jason Coombes, Joel Southgate, Lauren Gilbert, Lee Graham, Nicole Pacchiarini, Sara Kumziene-Summerhayes, Sarah Taylor, Sophie Jones, Sara Rey, Matthew Bull, Joanne Watkins, Sally Corden, Tom Connor |
| EPI_ISL_3296478 | Viollier AG | Department of Biosystems Science and Engineering, ETH Zürich | Christian Beisel, Sarah Nadeau, Chaoran Chen, Ivan Topolsky, Kim Philipp Jablonski, Lara Fuhrmann, Rebecca Denes, Mirjam Feldkamp, Ina Nissen, Natascha Santacroce, Elodie Burcklen, Christiane Beckmann, Maurice Redondo, Olivier Kobel, Christoph Noppen, Niko Beerenwinkel, Tanja Stadler |
| EPI_ISL_3296479 | Wales Specialist Virology Centre Sequencing lab: Pathogen Genomics Unit | Public Health Wales Microbiology Cardiff Wales Specialist Virology Centre | Catherine Moore, Johnathan Evans, Laura Gifford, Malorie Perry, Simon Cottrell, Angela Marchbank, Alec Birchley, Alexander Adams, Amy Gaskin, Bree Gatica-Wilcox, Jason Coombes, Joel Southgate, Lauren Gilbert, Lee Graham, Nicole Pacchiarini, Sara Kumziene-Summerhayes, Sarah Taylor, Sophie Jones, Sara Rey, Matthew Bull, Joanne Watkins, Sally Corden, Tom Connor |
| EPI_ISL_3296480 | Viollier AG | Department of Biosystems Science and Engineering, ETH Zürich | Christian Beisel, Sarah Nadeau, Chaoran Chen, Ivan Topolsky, Kim Philipp Jablonski, Lara Fuhrmann, Rebecca Denes, Mirjam Feldkamp, Ina Nissen, Natascha Santacroce, Elodie Burcklen, Christiane Beckmann, Maurice Redondo, Olivier Kobel, Christoph Noppen, Niko Beerenwinkel, Tanja Stadler |
| EPI_ISL_3296481 | Wales Specialist Virology Centre Sequencing lab: Pathogen Genomics Unit | Public Health Wales Microbiology Cardiff Wales Specialist Virology Centre | Catherine Moore, Johnathan Evans, Laura Gifford, Malorie Perry, Simon Cottrell, Angela Marchbank, Alec Birchley, Alexander Adams, Amy Gaskin, Bree Gatica-Wilcox, Jason Coombes, Joel Southgate, Lauren Gilbert, Lee Graham, Nicole Pacchiarini, Sara Kumziene-Summerhayes, Sarah Taylor, Sophie Jones, Sara Rey, Matthew Bull, Joanne Watkins, Sally Corden, Tom Connor |
| EPI_ISL_3296482 | Viollier AG | Department of Biosystems Science and Engineering, ETH Zürich | Christian Beisel, Sarah Nadeau, Chaoran Chen, Ivan Topolsky, Kim Philipp Jablonski, Lara Fuhrmann, Rebecca Denes, Mirjam Feldkamp, Ina Nissen, Natascha Santacroce, Elodie Burcklen, Christiane Beckmann, Maurice Redondo, Olivier Kobel, Christoph Noppen, Niko Beerenwinkel, Tanja Stadler |
| EPI_ISL_3296483, EPI_ISL_3296484 | Wales Specialist Virology Centre Sequencing lab: Pathogen Genomics Unit | Public Health Wales Microbiology Cardiff Wales Specialist Virology Centre | Catherine Moore, Johnathan Evans, Laura Gifford, Malorie Perry, Simon Cottrell, Angela Marchbank, Alec Birchley, Alexander Adams, Amy Gaskin, Bree Gatica-Wilcox, Jason Coombes, Joel Southgate, Lauren Gilbert, Lee Graham, Nicole Pacchiarini, Sara Kumziene-Summerhayes, Sarah Taylor, Sophie Jones, Sara Rey, Matthew Bull, Joanne Watkins, Sally Corden, Tom Connor |
| EPI_ISL_3296485 | Viollier AG | Department of Biosystems Science and Engineering, ETH Zürich | Christian Beisel, Sarah Nadeau, Chaoran Chen, Ivan Topolsky, Kim Philipp Jablonski, Lara Fuhrmann, Rebecca Denes, Mirjam Feldkamp, Ina Nissen, Natascha Santacroce, Elodie Burcklen, Christiane Beckmann, Maurice Redondo, Olivier Kobel, Christoph Noppen, Niko Beerenwinkel, Tanja Stadler |
| EPI_ISL_3296486 | Wales Specialist Virology Centre Sequencing lab: Pathogen Genomics Unit | Public Health Wales Microbiology Cardiff Wales Specialist Virology Centre | Catherine Moore, Johnathan Evans, Laura Gifford, Malorie Perry, Simon Cottrell, Angela Marchbank, Alec Birchley, Alexander Adams, Amy Gaskin, Bree Gatica-Wilcox, Jason Coombes, Joel Southgate, Lauren Gilbert, Lee Graham, Nicole Pacchiarini, Sara Kumziene-Summerhayes, Sarah Taylor, Sophie Jones, Sara Rey, Matthew Bull, Joanne Watkins, Sally Corden, Tom Connor |
| EPI_ISL_3296487 | Viollier AG | Department of Biosystems Science and Engineering, ETH Zürich | Christian Beisel, Sarah Nadeau, Chaoran Chen, Ivan Topolsky, Kim Philipp Jablonski, Lara Fuhrmann, Rebecca Denes, Mirjam Feldkamp, Ina Nissen, Natascha Santacroce, Elodie Burcklen, Christiane Beckmann, Maurice Redondo, Olivier Kobel, Christoph Noppen, Niko Beerenwinkel, Tanja Stadler |
| EPI_ISL_3296488 | Wales Specialist Virology Centre Sequencing lab: Pathogen Genomics Unit | Public Health Wales Microbiology Cardiff Wales Specialist Virology Centre | Catherine Moore, Johnathan Evans, Laura Gifford, Malorie Perry, Simon Cottrell, Angela Marchbank, Alec Birchley, Alexander Adams, Amy Gaskin, Bree Gatica-Wilcox, Jason Coombes, Joel Southgate, Lauren Gilbert, Lee Graham, Nicole Pacchiarini, Sara Kumziene-Summerhayes, Sarah Taylor, Sophie Jones, Sara Rey, Matthew Bull, Joanne Watkins, Sally Corden, Tom Connor |
| EPI_ISL_3296489 | Viollier AG | Department of Biosystems Science and Engineering, ETH Zürich | Christian Beisel, Sarah Nadeau, Chaoran Chen, Ivan Topolsky, Kim Philipp Jablonski, Lara Fuhrmann, Rebecca Denes, Mirjam Feldkamp, Ina Nissen, Natascha Santacroce, Elodie Burcklen, Christiane Beckmann, Maurice Redondo, Olivier Kobel, Christoph Noppen, Niko Beerenwinkel, Tanja Stadler |
| EPI_ISL_3296490 | Wales Specialist Virology Centre Sequencing lab: Pathogen Genomics Unit | Public Health Wales Microbiology Cardiff Wales Specialist Virology Centre | Catherine Moore, Johnathan Evans, Laura Gifford, Malorie Perry, Simon Cottrell, Angela Marchbank, Alec Birchley, Alexander Adams, Amy Gaskin, Bree Gatica-Wilcox, Jason Coombes, Joel Southgate, Lauren Gilbert, Lee Graham, Nicole Pacchiarini, Sara Kumziene-Summerhayes, Sarah Taylor, Sophie Jones, Sara Rey, Matthew Bull, Joanne Watkins, Sally Corden, Tom Connor |
| EPI_ISL_3296491 | Viollier AG | Department of Biosystems Science and Engineering, ETH Zürich | Christian Beisel, Sarah Nadeau, Chaoran Chen, Ivan Topolsky, Kim Philipp Jablonski, Lara Fuhrmann, Rebecca Denes, Mirjam Feldkamp, Ina Nissen, Natascha Santacroce, Elodie Burcklen, Christiane Beckmann, Maurice Redondo, Olivier Kobel, Christoph Noppen, Niko Beerenwinkel, Tanja Stadler |
| EPI_ISL_3296492, EPI_ISL_3296494 | Wales Specialist Virology Centre Sequencing lab: Pathogen Genomics Unit | Public Health Wales Microbiology Cardiff Wales Specialist Virology Centre | Catherine Moore, Johnathan Evans, Laura Gifford, Malorie Perry, Simon Cottrell, Angela Marchbank, Alec Birchley, Alexander Adams, Amy Gaskin, Bree Gatica-Wilcox, Jason Coombes, Joel Southgate, Lauren Gilbert, Lee Graham, Nicole Pacchiarini, Sara Kumziene-Summerhayes, Sarah Taylor, Sophie Jones, Sara Rey, Matthew Bull, Joanne Watkins, Sally Corden, Tom Connor |
| EPI_ISL_3296495 | Viollier AG | Department of Biosystems Science and Engineering, ETH Zürich | Christian Beisel, Sarah Nadeau, Chaoran Chen, Ivan Topolsky, Kim Philipp Jablonski, Lara Fuhrmann, Rebecca Denes, Mirjam Feldkamp, Ina Nissen, Natascha Santacroce, Elodie Burcklen, Christiane Beckmann, Maurice Redondo, Olivier Kobel, Christoph Noppen, Niko Beerenwinkel, Tanja Stadler |
| EPI_ISL_3296496 | Wales Specialist Virology Centre Sequencing lab: Pathogen Genomics Unit | Public Health Wales Microbiology Cardiff Wales Specialist Virology Centre | Catherine Moore, Johnathan Evans, Laura Gifford, Malorie Perry, Simon Cottrell, Angela Marchbank, Alec Birchley, Alexander Adams, Amy Gaskin, Bree Gatica-Wilcox, Jason Coombes, Joel Southgate, Lauren Gilbert, Lee Graham, Nicole Pacchiarini, Sara Kumziene-Summerhayes, Sarah Taylor, Sophie Jones, Sara Rey, Matthew Bull, Joanne Watkins, Sally Corden, Tom Connor |
| EPI_ISL_3296497 | Viollier AG | Department of Biosystems Science and Engineering, ETH Zürich | Christian Beisel, Sarah Nadeau, Chaoran Chen, Ivan Topolsky, Kim Philipp Jablonski, Lara Fuhrmann, Rebecca Denes, Mirjam Feldkamp, Ina Nissen, Natascha Santacroce, Elodie Burcklen, Christiane Beckmann, Maurice Redondo, Olivier Kobel, Christoph Noppen, Niko Beerenwinkel, Tanja Stadler |
| EPI_ISL_3296498 | Wales Specialist Virology Centre Sequencing lab: Pathogen Genomics Unit | Public Health Wales Microbiology Cardiff Wales Specialist Virology Centre | Catherine Moore, Johnathan Evans, Laura Gifford, Malorie Perry, Simon Cottrell, Angela Marchbank, Alec Birchley, Alexander Adams, Amy Gaskin, Bree Gatica-Wilcox, Jason Coombes, Joel Southgate, Lauren Gilbert, Lee Graham, Nicole Pacchiarini, Sara Kumziene-Summerhayes, Sarah Taylor, Sophie Jones, Sara Rey, Matthew Bull, Joanne Watkins, Sally Corden, Tom Connor |
| EPI_ISL_3296499 | Viollier AG | Department of Biosystems Science and Engineering, ETH Zürich | Christian Beisel, Sarah Nadeau, Chaoran Chen, Ivan Topolsky, Kim Philipp Jablonski, Lara Fuhrmann, Rebecca Denes, Mirjam Feldkamp, Ina Nissen, Natascha Santacroce, Elodie Burcklen, Christiane Beckmann, Maurice Redondo, Olivier Kobel, Christoph Noppen, Niko Beerenwinkel, Tanja Stadler |
| EPI_ISL_3296500 | Wales Specialist Virology Centre Sequencing lab: Pathogen Genomics Unit | Public Health Wales Microbiology Cardiff Wales Specialist Virology Centre | Catherine Moore, Johnathan Evans, Laura Gifford, Malorie Perry, Simon Cottrell, Angela Marchbank, Alec Birchley, Alexander Adams, Amy Gaskin, Bree Gatica-Wilcox, Jason Coombes, Joel Southgate, Lauren Gilbert, Lee Graham, Nicole Pacchiarini, Sara Kumziene-Summerhayes, Sarah Taylor, Sophie Jones, Sara Rey, Matthew Bull, Joanne Watkins, Sally Corden, Tom Connor |
| EPI_ISL_3296501 | Viollier AG | Department of Biosystems Science and Engineering, ETH Zürich | Christian Beisel, Sarah Nadeau, Chaoran Chen, Ivan Topolsky, Kim Philipp Jablonski, Lara Fuhrmann, Rebecca Denes, Mirjam Feldkamp, Ina Nissen, Natascha Santacroce, Elodie Burcklen, Christiane Beckmann, Maurice Redondo, Olivier Kobel, Christoph Noppen, Niko Beerenwinkel, Tanja Stadler |
| EPI_ISL_3296502 | Wales Specialist Virology Centre Sequencing lab: Pathogen Genomics Unit | Public Health Wales Microbiology Cardiff Wales Specialist Virology Centre | Catherine Moore, Johnathan Evans, Laura Gifford, Malorie Perry, Simon Cottrell, Angela Marchbank, Alec Birchley, Alexander Adams, Amy Gaskin, Bree Gatica-Wilcox, Jason Coombes, Joel Southgate, Lauren Gilbert, Lee Graham, Nicole Pacchiarini, Sara Kumziene-Summerhayes, Sarah Taylor, Sophie Jones, Sara Rey, Matthew Bull, Joanne Watkins, Sally Corden, Tom Connor |
| EPI_ISL_3296503 | Viollier AG | Department of Biosystems Science and Engineering, ETH Zürich | Christian Beisel, Sarah Nadeau, Chaoran Chen, Ivan Topolsky, Kim Philipp Jablonski, Lara Fuhrmann, Rebecca Denes, Mirjam Feldkamp, Ina Nissen, Natascha Santacroce, Elodie Burcklen, Christiane Beckmann, Maurice Redondo, Olivier Kobel, Christoph Noppen, Niko Beerenwinkel, Tanja Stadler |
| EPI_ISL_3296504 | Wales Specialist Virology Centre Sequencing lab: Pathogen Genomics Unit | Public Health Wales Microbiology Cardiff Wales Specialist Virology Centre | Catherine Moore, Johnathan Evans, Laura Gifford, Malorie Perry, Simon Cottrell, Angela Marchbank, Alec Birchley, Alexander Adams, Amy Gaskin, Bree Gatica-Wilcox, Jason Coombes, Joel Southgate, Lauren Gilbert, Lee Graham, Nicole Pacchiarini, Sara Kumziene-Summerhayes, Sarah Taylor, Sophie Jones, Sara Rey, Matthew Bull, Joanne Watkins, Sally Corden, Tom Connor |
| EPI_ISL_3296505 | Viollier AG | Department of Biosystems Science and Engineering, ETH Zürich | Christian Beisel, Sarah Nadeau, Chaoran Chen, Ivan Topolsky, Kim Philipp Jablonski, Lara Fuhrmann, Rebecca Denes, Mirjam Feldkamp, Ina Nissen, Natascha Santacroce, Elodie Burcklen, Christiane Beckmann, Maurice Redondo, Olivier Kobel, Christoph Noppen, Niko Beerenwinkel, Tanja Stadler |
| EPI_ISL_3296506 | Wales Specialist Virology Centre Sequencing lab: Pathogen Genomics Unit | Public Health Wales Microbiology Cardiff Wales Specialist Virology Centre | Catherine Moore, Johnathan Evans, Laura Gifford, Malorie Perry, Simon Cottrell, Angela Marchbank, Alec Birchley, Alexander Adams, Amy Gaskin, Bree Gatica-Wilcox, Jason Coombes, Joel Southgate, Lauren Gilbert, Lee Graham, Nicole Pacchiarini, Sara Kumziene-Summerhayes, Sarah Taylor, Sophie Jones, Sara Rey, Matthew Bull, Joanne Watkins, Sally Corden, Tom Connor |
| EPI_ISL_3296507 | Viollier AG | Department of Biosystems Science and Engineering, ETH Zürich | Christian Beisel, Sarah Nadeau, Chaoran Chen, Ivan Topolsky, Kim Philipp Jablonski, Lara Fuhrmann, Rebecca Denes, Mirjam Feldkamp, Ina Nissen, Natascha Santacroce, Elodie Burcklen, Christiane Beckmann, Maurice Redondo, Olivier Kobel, Christoph Noppen, Niko Beerenwinkel, Tanja Stadler |
| EPI_ISL_3296508, EPI_ISL_3296509 | Wales Specialist Virology Centre Sequencing lab: Pathogen Genomics Unit | Public Health Wales Microbiology Cardiff Wales Specialist Virology Centre | Catherine Moore, Johnathan Evans, Laura Gifford, Malorie Perry, Simon Cottrell, Angela Marchbank, Alec Birchley, Alexander Adams, Amy Gaskin, Bree Gatica-Wilcox, Jason Coombes, Joel Southgate, Lauren Gilbert, Lee Graham, Nicole Pacchiarini, Sara Kumziene-Summerhayes, Sarah Taylor, Sophie Jones, Sara Rey, Matthew Bull, Joanne Watkins, Sally Corden, Tom Connor |
| EPI_ISL_3296510 | Viollier AG | Department of Biosystems Science and Engineering, ETH Zürich | Christian Beisel, Sarah Nadeau, Chaoran Chen, Ivan Topolsky, Kim Philipp Jablonski, Lara Fuhrmann, Rebecca Denes, Mirjam Feldkamp, Ina Nissen, Natascha Santacroce, Elodie Burcklen, Christiane Beckmann, Maurice Redondo, Olivier Kobel, Christoph Noppen, Niko Beerenwinkel, Tanja Stadler |
| EPI_ISL_3296511 | Wales Specialist Virology Centre Sequencing lab: Pathogen Genomics Unit | Public Health Wales Microbiology Cardiff Wales Specialist Virology Centre | Catherine Moore, Johnathan Evans, Laura Gifford, Malorie Perry, Simon Cottrell, Angela Marchbank, Alec Birchley, Alexander Adams, Amy Gaskin, Bree Gatica-Wilcox, Jason Coombes, Joel Southgate, Lauren Gilbert, Lee Graham, Nicole Pacchiarini, Sara Kumziene-Summerhayes, Sarah Taylor, Sophie Jones, Sara Rey, Matthew Bull, Joanne Watkins, Sally Corden, Tom Connor |
| EPI_ISL_3296512 | Viollier AG | Department of Biosystems Science and Engineering, ETH Zürich | Christian Beisel, Sarah Nadeau, Chaoran Chen, Ivan Topolsky, Kim Philipp Jablonski, Lara Fuhrmann, Rebecca Denes, Mirjam Feldkamp, Ina Nissen, Natascha Santacroce, Elodie Burcklen, Christiane Beckmann, Maurice Redondo, Olivier Kobel, Christoph Noppen, Niko Beerenwinkel, Tanja Stadler |

| | | | |
|---|---|---|---|
| EPI_ISL_3296513 | Wales Specialist Virology Centre Sequencing lab: Pathogen Genomics Unit | Public Health Wales Microbiology Cardiff Wales Specialist Virology Centre | Catherine Moore, Johnathan Evans, Laura Gifford, Malorie Perry, Simon Cottrell, Angela Marchbank, Alec Birchley, Alexander Adams, Amy Gaskin, Bree Gatica-Wilcox, Jason Coombes, Joel Southgate, Lauren Gilbert, Lee Graham, Nicole Pacchiarini, Sara Kumziene-Summerhayes, Sarah Taylor, Sophie Jones, Sara Rey, Matthew Bull, Joanne Watkins, Sally Corden, Tom Connor |
| EPI_ISL_3296514 | Viollier AG | Department of Biosystems Science and Engineering, ETH Zürich | Christian Beisel, Sarah Nadeau, Chaoran Chen, Ivan Topolsky, Kim Philipp Jablonski, Lara Fuhrmann, Rebecca Denes, Mirjam Feldkamp, Ina Nissen, Natascha Santacroce, Elodie Burcklen, Christiane Beckmann, Maurice Redondo, Olivier Kobel, Christoph Noppen, Niko Beerenwinkel, Tanja Stadler |
| EPI_ISL_3296515 | Wales Specialist Virology Centre Sequencing lab: Pathogen Genomics Unit | Public Health Wales Microbiology Cardiff Wales Specialist Virology Centre | Catherine Moore, Johnathan Evans, Laura Gifford, Malorie Perry, Simon Cottrell, Angela Marchbank, Alec Birchley, Alexander Adams, Amy Gaskin, Bree Gatica-Wilcox, Jason Coombes, Joel Southgate, Lauren Gilbert, Lee Graham, Nicole Pacchiarini, Sara Kumziene-Summerhayes, Sarah Taylor, Sophie Jones, Sara Rey, Matthew Bull, Joanne Watkins, Sally Corden, Tom Connor |
| EPI_ISL_3296516 | Viollier AG | Department of Biosystems Science and Engineering, ETH Zürich | Christian Beisel, Sarah Nadeau, Chaoran Chen, Ivan Topolsky, Kim Philipp Jablonski, Lara Fuhrmann, Rebecca Denes, Mirjam Feldkamp, Ina Nissen, Natascha Santacroce, Elodie Burcklen, Christiane Beckmann, Maurice Redondo, Olivier Kobel, Christoph Noppen, Niko Beerenwinkel, Tanja Stadler |
| EPI_ISL_3296517 | Wales Specialist Virology Centre Sequencing lab: Pathogen Genomics Unit | Public Health Wales Microbiology Cardiff Wales Specialist Virology Centre | Catherine Moore, Johnathan Evans, Laura Gifford, Malorie Perry, Simon Cottrell, Angela Marchbank, Alec Birchley, Alexander Adams, Amy Gaskin, Bree Gatica-Wilcox, Jason Coombes, Joel Southgate, Lauren Gilbert, Lee Graham, Nicole Pacchiarini, Sara Kumziene-Summerhayes, Sarah Taylor, Sophie Jones, Sara Rey, Matthew Bull, Joanne Watkins, Sally Corden, Tom Connor |
| EPI_ISL_3296518 | Viollier AG | Department of Biosystems Science and Engineering, ETH Zürich | Christian Beisel, Sarah Nadeau, Chaoran Chen, Ivan Topolsky, Kim Philipp Jablonski, Lara Fuhrmann, Rebecca Denes, Mirjam Feldkamp, Ina Nissen, Natascha Santacroce, Elodie Burcklen, Christiane Beckmann, Maurice Redondo, Olivier Kobel, Christoph Noppen, Niko Beerenwinkel, Tanja Stadler |
| EPI_ISL_3296519, EPI_ISL_3296520 | Wales Specialist Virology Centre Sequencing lab: Pathogen Genomics Unit | Public Health Wales Microbiology Cardiff Wales Specialist Virology Centre | Catherine Moore, Johnathan Evans, Laura Gifford, Malorie Perry, Simon Cottrell, Angela Marchbank, Alec Birchley, Alexander Adams, Amy Gaskin, Bree Gatica-Wilcox, Jason Coombes, Joel Southgate, Lauren Gilbert, Lee Graham, Nicole Pacchiarini, Sara Kumziene-Summerhayes, Sarah Taylor, Sophie Jones, Sara Rey, Matthew Bull, Joanne Watkins, Sally Corden, Tom Connor |
| EPI_ISL_3296521 | Viollier AG | Department of Biosystems Science and Engineering, ETH Zürich | Christian Beisel, Sarah Nadeau, Chaoran Chen, Ivan Topolsky, Kim Philipp Jablonski, Lara Fuhrmann, Rebecca Denes, Mirjam Feldkamp, Ina Nissen, Natascha Santacroce, Elodie Burcklen, Christiane Beckmann, Maurice Redondo, Olivier Kobel, Christoph Noppen, Niko Beerenwinkel, Tanja Stadler |
| EPI_ISL_3296522 | Wales Specialist Virology Centre Sequencing lab: Pathogen Genomics Unit | Public Health Wales Microbiology Cardiff Wales Specialist Virology Centre | Catherine Moore, Johnathan Evans, Laura Gifford, Malorie Perry, Simon Cottrell, Angela Marchbank, Alec Birchley, Alexander Adams, Amy Gaskin, Bree Gatica-Wilcox, Jason Coombes, Joel Southgate, Lauren Gilbert, Lee Graham, Nicole Pacchiarini, Sara Kumziene-Summerhayes, Sarah Taylor, Sophie Jones, Sara Rey, Matthew Bull, Joanne Watkins, Sally Corden, Tom Connor |
| EPI_ISL_3296523 | Viollier AG | Department of Biosystems Science and Engineering, ETH Zürich | Christian Beisel, Sarah Nadeau, Chaoran Chen, Ivan Topolsky, Kim Philipp Jablonski, Lara Fuhrmann, Rebecca Denes, Mirjam Feldkamp, Ina Nissen, Natascha Santacroce, Elodie Burcklen, Christiane Beckmann, Maurice Redondo, Olivier Kobel, Christoph Noppen, Niko Beerenwinkel, Tanja Stadler |
| EPI_ISL_3296524 | Wales Specialist Virology Centre Sequencing lab: Pathogen Genomics Unit | Public Health Wales Microbiology Cardiff Wales Specialist Virology Centre | Catherine Moore, Johnathan Evans, Laura Gifford, Malorie Perry, Simon Cottrell, Angela Marchbank, Alec Birchley, Alexander Adams, Amy Gaskin, Bree Gatica-Wilcox, Jason Coombes, Joel Southgate, Lauren Gilbert, Lee Graham, Nicole Pacchiarini, Sara Kumziene-Summerhayes, Sarah Taylor, Sophie Jones, Sara Rey, Matthew Bull, Joanne Watkins, Sally Corden, Tom Connor |
| EPI_ISL_3296525 | Viollier AG | Department of Biosystems Science and Engineering, ETH Zürich | Christian Beisel, Sarah Nadeau, Chaoran Chen, Ivan Topolsky, Kim Philipp Jablonski, Lara Fuhrmann, Rebecca Denes, Mirjam Feldkamp, Ina Nissen, Natascha Santacroce, Elodie Burcklen, Christiane Beckmann, Maurice Redondo, Olivier Kobel, Christoph Noppen, Niko Beerenwinkel, Tanja Stadler |
| EPI_ISL_3296526 | Wales Specialist Virology Centre Sequencing lab: Pathogen Genomics Unit | Public Health Wales Microbiology Cardiff Wales Specialist Virology Centre | Catherine Moore, Johnathan Evans, Laura Gifford, Malorie Perry, Simon Cottrell, Angela Marchbank, Alec Birchley, Alexander Adams, Amy Gaskin, Bree Gatica-Wilcox, Jason Coombes, Joel Southgate, Lauren Gilbert, Lee Graham, Nicole Pacchiarini, Sara Kumziene-Summerhayes, Sarah Taylor, Sophie Jones, Sara Rey, Matthew Bull, Joanne Watkins, Sally Corden, Tom Connor |
| EPI_ISL_3296527 | Viollier AG | Department of Biosystems Science and Engineering, ETH Zürich | Christian Beisel, Sarah Nadeau, Chaoran Chen, Ivan Topolsky, Kim Philipp Jablonski, Lara Fuhrmann, Rebecca Denes, Mirjam Feldkamp, Ina Nissen, Natascha Santacroce, Elodie Burcklen, Christiane Beckmann, Maurice Redondo, Olivier Kobel, Christoph Noppen, Niko Beerenwinkel, Tanja Stadler |
| EPI_ISL_3296528 | Wales Specialist Virology Centre Sequencing lab: Pathogen Genomics Unit | Public Health Wales Microbiology Cardiff Wales Specialist Virology Centre | Catherine Moore, Johnathan Evans, Laura Gifford, Malorie Perry, Simon Cottrell, Angela Marchbank, Alec Birchley, Alexander Adams, Amy Gaskin, Bree Gatica-Wilcox, Jason Coombes, Joel Southgate, Lauren Gilbert, Lee Graham, Nicole Pacchiarini, Sara Kumziene-Summerhayes, Sarah Taylor, Sophie Jones, Sara Rey, Matthew Bull, Joanne Watkins, Sally Corden, Tom Connor |
| EPI_ISL_3296529 | Viollier AG | Department of Biosystems Science and Engineering, ETH Zürich | Christian Beisel, Sarah Nadeau, Chaoran Chen, Ivan Topolsky, Kim Philipp Jablonski, Lara Fuhrmann, Rebecca Denes, Mirjam Feldkamp, Ina Nissen, Natascha Santacroce, Elodie Burcklen, Christiane Beckmann, Maurice Redondo, Olivier Kobel, Christoph Noppen, Niko Beerenwinkel, Tanja Stadler |
| EPI_ISL_3296530 | Wales Specialist Virology Centre Sequencing lab: Pathogen Genomics Unit | Public Health Wales Microbiology Cardiff Wales Specialist Virology Centre | Catherine Moore, Johnathan Evans, Laura Gifford, Malorie Perry, Simon Cottrell, Angela Marchbank, Alec Birchley, Alexander Adams, Amy Gaskin, Bree Gatica-Wilcox, Jason Coombes, Joel Southgate, Lauren Gilbert, Lee Graham, Nicole Pacchiarini, Sara Kumziene-Summerhayes, Sarah Taylor, Sophie Jones, Sara Rey, Matthew Bull, Joanne Watkins, Sally Corden, Tom Connor |
| EPI_ISL_3296531 | Viollier AG | Department of Biosystems Science and Engineering, ETH Zürich | Christian Beisel, Sarah Nadeau, Chaoran Chen, Ivan Topolsky, Kim Philipp Jablonski, Lara Fuhrmann, Rebecca Denes, Mirjam Feldkamp, Ina Nissen, Natascha Santacroce, Elodie Burcklen, Christiane Beckmann, Maurice Redondo, Olivier Kobel, Christoph Noppen, Niko Beerenwinkel, Tanja Stadler |
| EPI_ISL_3296532 | Wales Specialist Virology Centre Sequencing lab: Pathogen Genomics Unit | Public Health Wales Microbiology Cardiff Wales Specialist Virology Centre | Catherine Moore, Johnathan Evans, Laura Gifford, Malorie Perry, Simon Cottrell, Angela Marchbank, Alec Birchley, Alexander Adams, Amy Gaskin, Bree Gatica-Wilcox, Jason Coombes, Joel Southgate, Lauren Gilbert, Lee Graham, Nicole Pacchiarini, Sara Kumziene-Summerhayes, Sarah Taylor, Sophie Jones, Sara Rey, Matthew Bull, Joanne Watkins, Sally Corden, Tom Connor |
| EPI_ISL_3296533 | Viollier AG | Department of Biosystems Science and Engineering, ETH Zürich | Christian Beisel, Sarah Nadeau, Chaoran Chen, Ivan Topolsky, Kim Philipp Jablonski, Lara Fuhrmann, Rebecca Denes, Mirjam Feldkamp, Ina Nissen, Natascha Santacroce, Elodie Burcklen, Christiane Beckmann, Maurice Redondo, Olivier Kobel, Christoph Noppen, Niko Beerenwinkel, Tanja Stadler |
| EPI_ISL_3296534 | Wales Specialist Virology Centre Sequencing lab: Pathogen Genomics Unit | Public Health Wales Microbiology Cardiff Wales Specialist Virology Centre | Catherine Moore, Johnathan Evans, Laura Gifford, Malorie Perry, Simon Cottrell, Angela Marchbank, Alec Birchley, Alexander Adams, Amy Gaskin, Bree Gatica-Wilcox, Jason Coombes, Joel Southgate, Lauren Gilbert, Lee Graham, Nicole Pacchiarini, Sara Kumziene-Summerhayes, Sarah Taylor, Sophie Jones, Sara Rey, Matthew Bull, Joanne Watkins, Sally Corden, Tom Connor |
| EPI_ISL_3296535 | Viollier AG | Department of Biosystems Science and Engineering, ETH Zürich | Christian Beisel, Sarah Nadeau, Chaoran Chen, Ivan Topolsky, Kim Philipp Jablonski, Lara Fuhrmann, Rebecca Denes, Mirjam Feldkamp, Ina Nissen, Natascha Santacroce, Elodie Burcklen, Christiane Beckmann, Maurice Redondo, Olivier Kobel, Christoph Noppen, Niko Beerenwinkel, Tanja Stadler |
| EPI_ISL_3296536 | Wales Specialist Virology Centre Sequencing lab: Pathogen Genomics Unit | Public Health Wales Microbiology Cardiff Wales Specialist Virology Centre | Catherine Moore, Johnathan Evans, Laura Gifford, Malorie Perry, Simon Cottrell, Angela Marchbank, Alec Birchley, Alexander Adams, Amy Gaskin, Bree Gatica-Wilcox, Jason Coombes, Joel Southgate, Lauren Gilbert, Lee Graham, Nicole Pacchiarini, Sara Kumziene-Summerhayes, Sarah Taylor, Sophie Jones, Sara Rey, Matthew Bull, Joanne Watkins, Sally Corden, Tom Connor |
| EPI_ISL_3296537 | Viollier AG | Department of Biosystems Science and Engineering, ETH Zürich | Christian Beisel, Sarah Nadeau, Chaoran Chen, Ivan Topolsky, Kim Philipp Jablonski, Lara Fuhrmann, Rebecca Denes, Mirjam Feldkamp, Ina Nissen, Natascha Santacroce, Elodie Burcklen, Christiane Beckmann, Maurice Redondo, Olivier Kobel, Christoph Noppen, Niko Beerenwinkel, Tanja Stadler |
| EPI_ISL_3296538, EPI_ISL_3296539, EPI_ISL_3296540, EPI_ISL_3296541, EPI_ISL_3296542, EPI_ISL_3296544 | Wales Specialist Virology Centre Sequencing lab: Pathogen Genomics Unit | Public Health Wales Microbiology Cardiff Wales Specialist Virology Centre | Catherine Moore, Johnathan Evans, Laura Gifford, Malorie Perry, Simon Cottrell, Angela Marchbank, Alec Birchley, Alexander Adams, Amy Gaskin, Bree Gatica-Wilcox, Jason Coombes, Joel Southgate, Lauren Gilbert, Lee Graham, Nicole Pacchiarini, Sara Kumziene-Summerhayes, Sarah Taylor, Sophie Jones, Sara Rey, Matthew Bull, Joanne Watkins, Sally Corden, Tom Connor |
| EPI_ISL_3296545 | Viollier AG | Department of Biosystems Science and Engineering, ETH Zürich | Christian Beisel, Sarah Nadeau, Chaoran Chen, Ivan Topolsky, Kim Philipp Jablonski, Lara Fuhrmann, Rebecca Denes, Mirjam Feldkamp, Ina Nissen, Natascha Santacroce, Elodie Burcklen, Christiane Beckmann, Maurice Redondo, Olivier Kobel, Christoph Noppen, Niko Beerenwinkel, Tanja Stadler |
| EPI_ISL_3296546 | Wales Specialist Virology Centre Sequencing lab: Pathogen Genomics Unit | Public Health Wales Microbiology Cardiff Wales Specialist Virology Centre | Catherine Moore, Johnathan Evans, Laura Gifford, Malorie Perry, Simon Cottrell, Angela Marchbank, Alec Birchley, Alexander Adams, Amy Gaskin, Bree Gatica-Wilcox, Jason Coombes, Joel Southgate, Lauren Gilbert, Lee Graham, Nicole Pacchiarini, Sara Kumziene-Summerhayes, Sarah Taylor, Sophie Jones, Sara Rey, Matthew Bull, Joanne Watkins, Sally Corden, Tom Connor |
| EPI_ISL_3296547 | Viollier AG | Department of Biosystems Science and Engineering, ETH Zürich | Christian Beisel, Sarah Nadeau, Chaoran Chen, Ivan Topolsky, Kim Philipp Jablonski, Lara Fuhrmann, Rebecca Denes, Mirjam Feldkamp, Ina Nissen, Natascha Santacroce, Elodie Burcklen, Christiane Beckmann, Maurice Redondo, Olivier Kobel, Christoph Noppen, Niko Beerenwinkel, Tanja Stadler |
| EPI_ISL_3296548 | Wales Specialist Virology Centre Sequencing lab: Pathogen Genomics Unit | Public Health Wales Microbiology Cardiff Wales Specialist Virology Centre | Catherine Moore, Johnathan Evans, Laura Gifford, Malorie Perry, Simon Cottrell, Angela Marchbank, Alec Birchley, Alexander Adams, Amy Gaskin, Bree Gatica-Wilcox, Jason Coombes, Joel Southgate, Lauren Gilbert, Lee Graham, Nicole Pacchiarini, Sara Kumziene-Summerhayes, Sarah Taylor, Sophie Jones, Sara Rey, Matthew Bull, Joanne Watkins, Sally Corden, Tom Connor |
| EPI_ISL_3296549 | Viollier AG | Department of Biosystems Science and Engineering, ETH Zürich | Christian Beisel, Sarah Nadeau, Chaoran Chen, Ivan Topolsky, Kim Philipp Jablonski, Lara Fuhrmann, Rebecca Denes, Mirjam Feldkamp, Ina Nissen, Natascha Santacroce, Elodie Burcklen, Christiane Beckmann, Maurice Redondo, Olivier Kobel, Christoph Noppen, Niko Beerenwinkel, Tanja Stadler |
| EPI_ISL_3296550 | Wales Specialist Virology Centre Sequencing lab: Pathogen Genomics Unit | Public Health Wales Microbiology Cardiff Wales Specialist Virology Centre | Catherine Moore, Johnathan Evans, Laura Gifford, Malorie Perry, Simon Cottrell, Angela Marchbank, Alec Birchley, Alexander Adams, Amy Gaskin, Bree Gatica-Wilcox, Jason Coombes, Joel Southgate, Lauren Gilbert, Lee Graham, Nicole Pacchiarini, Sara Kumziene-Summerhayes, Sarah Taylor, Sophie Jones, Sara Rey, Matthew Bull, Joanne Watkins, Sally Corden, Tom Connor |
| EPI_ISL_3296551 | Viollier AG | Department of Biosystems Science and Engineering, ETH Zürich | Christian Beisel, Sarah Nadeau, Chaoran Chen, Ivan Topolsky, Kim Philipp Jablonski, Lara Fuhrmann, Rebecca Denes, Mirjam Feldkamp, Ina Nissen, Natascha Santacroce, Elodie Burcklen, Christiane Beckmann, Maurice Redondo, Olivier Kobel, Christoph Noppen, Niko Beerenwinkel, Tanja Stadler |
| EPI_ISL_3296552 | Wales Specialist Virology Centre Sequencing lab: Pathogen Genomics Unit | Public Health Wales Microbiology Cardiff Wales Specialist Virology Centre | Catherine Moore, Johnathan Evans, Laura Gifford, Malorie Perry, Simon Cottrell, Angela Marchbank, Alec Birchley, Alexander Adams, Amy Gaskin, Bree Gatica-Wilcox, Jason Coombes, Joel Southgate, Lauren Gilbert, Lee Graham, Nicole Pacchiarini, Sara Kumziene-Summerhayes, Sarah Taylor, Sophie Jones, Sara Rey, Matthew Bull, Joanne Watkins, Sally Corden, Tom Connor |
| EPI_ISL_3296553 | Viollier AG | Department of Biosystems Science and Engineering, ETH Zürich | Christian Beisel, Sarah Nadeau, Chaoran Chen, Ivan Topolsky, Kim Philipp Jablonski, Lara Fuhrmann, Rebecca Denes, Mirjam Feldkamp, Ina Nissen, Natascha Santacroce, Elodie Burcklen, Christiane Beckmann, Maurice Redondo, Olivier Kobel, Christoph Noppen, Niko Beerenwinkel, Tanja Stadler |
| EPI_ISL_3296554 | Wales Specialist Virology Centre Sequencing lab: Pathogen Genomics Unit | Public Health Wales Microbiology Cardiff Wales Specialist Virology Centre | Catherine Moore, Johnathan Evans, Laura Gifford, Malorie Perry, Simon Cottrell, Angela Marchbank, Alec Birchley, Alexander Adams, Amy Gaskin, Bree Gatica-Wilcox, Jason Coombes, Joel Southgate, Lauren Gilbert, Lee Graham, Nicole Pacchiarini, Sara Kumziene-Summerhayes, Sarah Taylor, Sophie Jones, Sara Rey, Matthew Bull, Joanne Watkins, Sally Corden, Tom Connor |
| EPI_ISL_3296555 | Viollier AG | Department of Biosystems Science and Engineering, ETH Zürich | Christian Beisel, Sarah Nadeau, Chaoran Chen, Ivan Topolsky, Kim Philipp Jablonski, Lara Fuhrmann, Rebecca Denes, Mirjam Feldkamp, Ina Nissen, Natascha Santacroce, Elodie Burcklen, Christiane Beckmann, Maurice Redondo, Olivier Kobel, Christoph Noppen, Niko Beerenwinkel, Tanja Stadler |
| EPI_ISL_3296556 | Wales Specialist Virology Centre Sequencing lab: Pathogen Genomics Unit | Public Health Wales Microbiology Cardiff Wales Specialist Virology Centre | Catherine Moore, Johnathan Evans, Laura Gifford, Malorie Perry, Simon Cottrell, Angela Marchbank, Alec Birchley, Alexander Adams, Amy Gaskin, Bree Gatica-Wilcox, Jason Coombes, Joel Southgate, Lauren Gilbert, Lee Graham, Nicole Pacchiarini, Sara Kumziene-Summerhayes, Sarah Taylor, Sophie Jones, Sara Rey, Matthew Bull, Joanne Watkins, Sally Corden, Tom Connor |
| EPI_ISL_3296557 | Viollier AG | Department of Biosystems Science and Engineering, ETH Zürich | Christian Beisel, Sarah Nadeau, Chaoran Chen, Ivan Topolsky, Kim Philipp Jablonski, Lara Fuhrmann, Rebecca Denes, Mirjam Feldkamp, Ina Nissen, Natascha Santacroce, Elodie Burcklen, Christiane Beckmann, Maurice Redondo, Olivier Kobel, Christoph Noppen, Niko Beerenwinkel, Tanja Stadler |
| EPI_ISL_3296558 | Wales Specialist Virology Centre Sequencing lab: Pathogen Genomics Unit | Public Health Wales Microbiology Cardiff Wales Specialist Virology Centre | Catherine Moore, Johnathan Evans, Laura Gifford, Malorie Perry, Simon Cottrell, Angela Marchbank, Alec Birchley, Alexander Adams, Amy Gaskin, Bree Gatica-Wilcox, Jason Coombes, Joel Southgate, Lauren Gilbert, Lee Graham, Nicole Pacchiarini, Sara Kumziene-Summerhayes, Sarah Taylor, Sophie Jones, Sara Rey, Matthew Bull, Joanne Watkins, Sally Corden, Tom Connor |
| EPI_ISL_3296559 | Viollier AG | Department of Biosystems Science and Engineering, ETH Zürich | Christian Beisel, Sarah Nadeau, Chaoran Chen, Ivan Topolsky, Kim Philipp Jablonski, Lara Fuhrmann, Rebecca Denes, Mirjam Feldkamp, Ina Nissen, Natascha Santacroce, Elodie Burcklen, Christiane Beckmann, Maurice Redondo, Olivier Kobel, Christoph Noppen, Niko Beerenwinkel, Tanja Stadler |
| EPI_ISL_3296560, EPI_ISL_3296561 | Wales Specialist Virology Centre Sequencing lab: Pathogen Genomics Unit | Public Health Wales Microbiology Cardiff Wales Specialist Virology Centre | Catherine Moore, Johnathan Evans, Laura Gifford, Malorie Perry, Simon Cottrell, Angela Marchbank, Alec Birchley, Alexander Adams, Amy Gaskin, Bree Gatica-Wilcox, Jason Coombes, Joel Southgate, Lauren Gilbert, Lee Graham, Nicole Pacchiarini, Sara Kumziene-Summerhayes, Sarah Taylor, Sophie Jones, Sara Rey, Matthew Bull, Joanne Watkins, Sally Corden, Tom Connor |
| EPI_ISL_3296562 | Viollier AG | Department of Biosystems Science and Engineering, ETH Zürich | Christian Beisel, Sarah Nadeau, Chaoran Chen, Ivan Topolsky, Kim Philipp Jablonski, Lara Fuhrmann, Rebecca Denes, Mirjam Feldkamp, Ina Nissen, Natascha Santacroce, Elodie Burcklen, Christiane Beckmann, Maurice Redondo, Olivier Kobel, Christoph Noppen, Niko Beerenwinkel, Tanja Stadler |
| EPI_ISL_3296563 | Wales Specialist Virology Centre Sequencing lab: Pathogen Genomics Unit | Public Health Wales Microbiology Cardiff Wales Specialist Virology Centre | Catherine Moore, Johnathan Evans, Laura Gifford, Malorie Perry, Simon Cottrell, Angela Marchbank, Alec Birchley, Alexander Adams, Amy Gaskin, Bree Gatica-Wilcox, Jason Coombes, Joel Southgate, Lauren Gilbert, Lee Graham, Nicole Pacchiarini, Sara Kumziene-Summerhayes, Sarah Taylor, Sophie Jones, Sara Rey, Matthew Bull, Joanne Watkins, Sally Corden, Tom Connor |
| EPI_ISL_3296564 | Viollier AG | Department of Biosystems Science and Engineering, ETH Zürich | Christian Beisel, Sarah Nadeau, Chaoran Chen, Ivan Topolsky, Kim Philipp Jablonski, Lara Fuhrmann, Rebecca Denes, Mirjam Feldkamp, Ina Nissen, Natascha Santacroce, Elodie Burcklen, Christiane Beckmann, Maurice Redondo, Olivier Kobel, |

| | | | |
|---|---|---|---|
| | | | Christoph Noppen, Niko Beerenwinkel, Tanja Stadler |
| EPI_ISL_3296565 | Wales Specialist Virology Centre Sequencing lab: Pathogen Genomics Unit | Public Health Wales Microbiology Cardiff Wales Specialist Virology Centre | Catherine Moore, Johnathan Evans, Laura Gifford, Malorie Perry, Simon Cottrell, Angela Marchbank, Alec Birchley, Alexander Adams, Amy Gaskin, Bree Gatica-Wilcox, Jason Coombes, Joel Southgate, Lauren Gilbert, Lee Graham, Nicole Pacchiarini, Sara Kumziene-Summerhayes, Sarah Taylor, Sophie Jones, Sara Rey, Matthew Bull, Joanne Watkins, Sally Corden, Tom Connor |
| EPI_ISL_3296566 | Viollier AG | Department of Biosystems Science and Engineering, ETH Zürich | Christian Beisel, Sarah Nadeau, Chaoran Chen, Ivan Topolsky, Kim Philipp Jablonski, Lara Fuhrmann, Rebecca Denes, Mirjam Feldkamp, Ina Nissen, Natascha Santacroce, Elodie Burcklen, Christiane Beckmann, Maurice Redondo, Olivier Kobel, Christoph Noppen, Niko Beerenwinkel, Tanja Stadler |
| EPI_ISL_3296567 | Wales Specialist Virology Centre Sequencing lab: Pathogen Genomics Unit | Public Health Wales Microbiology Cardiff Wales Specialist Virology Centre | Catherine Moore, Johnathan Evans, Laura Gifford, Malorie Perry, Simon Cottrell, Angela Marchbank, Alec Birchley, Alexander Adams, Amy Gaskin, Bree Gatica-Wilcox, Jason Coombes, Joel Southgate, Lauren Gilbert, Lee Graham, Nicole Pacchiarini, Sara Kumziene-Summerhayes, Sarah Taylor, Sophie Jones, Sara Rey, Matthew Bull, Joanne Watkins, Sally Corden, Tom Connor |
| EPI_ISL_3296568 | Viollier AG | Department of Biosystems Science and Engineering, ETH Zürich | Christian Beisel, Sarah Nadeau, Chaoran Chen, Ivan Topolsky, Kim Philipp Jablonski, Lara Fuhrmann, Rebecca Denes, Mirjam Feldkamp, Ina Nissen, Natascha Santacroce, Elodie Burcklen, Christiane Beckmann, Maurice Redondo, Olivier Kobel, Christoph Noppen, Niko Beerenwinkel, Tanja Stadler |
| EPI_ISL_3296569 | Wales Specialist Virology Centre Sequencing lab: Pathogen Genomics Unit | Public Health Wales Microbiology Cardiff Wales Specialist Virology Centre | Catherine Moore, Johnathan Evans, Laura Gifford, Malorie Perry, Simon Cottrell, Angela Marchbank, Alec Birchley, Alexander Adams, Amy Gaskin, Bree Gatica-Wilcox, Jason Coombes, Joel Southgate, Lauren Gilbert, Lee Graham, Nicole Pacchiarini, Sara Kumziene-Summerhayes, Sarah Taylor, Sophie Jones, Sara Rey, Matthew Bull, Joanne Watkins, Sally Corden, Tom Connor |
| EPI_ISL_3296570 | Viollier AG | Department of Biosystems Science and Engineering, ETH Zürich | Christian Beisel, Sarah Nadeau, Chaoran Chen, Ivan Topolsky, Kim Philipp Jablonski, Lara Fuhrmann, Rebecca Denes, Mirjam Feldkamp, Ina Nissen, Natascha Santacroce, Elodie Burcklen, Christiane Beckmann, Maurice Redondo, Olivier Kobel, Christoph Noppen, Niko Beerenwinkel, Tanja Stadler |
| EPI_ISL_3296571 | Wales Specialist Virology Centre Sequencing lab: Pathogen Genomics Unit | Public Health Wales Microbiology Cardiff Wales Specialist Virology Centre | Catherine Moore, Johnathan Evans, Laura Gifford, Malorie Perry, Simon Cottrell, Angela Marchbank, Alec Birchley, Alexander Adams, Amy Gaskin, Bree Gatica-Wilcox, Jason Coombes, Joel Southgate, Lauren Gilbert, Lee Graham, Nicole Pacchiarini, Sara Kumziene-Summerhayes, Sarah Taylor, Sophie Jones, Sara Rey, Matthew Bull, Joanne Watkins, Sally Corden, Tom Connor |
| EPI_ISL_3296572 | Viollier AG | Department of Biosystems Science and Engineering, ETH Zürich | Christian Beisel, Sarah Nadeau, Chaoran Chen, Ivan Topolsky, Kim Philipp Jablonski, Lara Fuhrmann, Rebecca Denes, Mirjam Feldkamp, Ina Nissen, Natascha Santacroce, Elodie Burcklen, Christiane Beckmann, Maurice Redondo, Olivier Kobel, Christoph Noppen, Niko Beerenwinkel, Tanja Stadler |
| EPI_ISL_3296573 | Wales Specialist Virology Centre Sequencing lab: Pathogen Genomics Unit | Public Health Wales Microbiology Cardiff Wales Specialist Virology Centre | Catherine Moore, Johnathan Evans, Laura Gifford, Malorie Perry, Simon Cottrell, Angela Marchbank, Alec Birchley, Alexander Adams, Amy Gaskin, Bree Gatica-Wilcox, Jason Coombes, Joel Southgate, Lauren Gilbert, Lee Graham, Nicole Pacchiarini, Sara Kumziene-Summerhayes, Sarah Taylor, Sophie Jones, Sara Rey, Matthew Bull, Joanne Watkins, Sally Corden, Tom Connor |
| EPI_ISL_3296574 | Viollier AG | Department of Biosystems Science and Engineering, ETH Zürich | Christian Beisel, Sarah Nadeau, Chaoran Chen, Ivan Topolsky, Kim Philipp Jablonski, Lara Fuhrmann, Rebecca Denes, Mirjam Feldkamp, Ina Nissen, Natascha Santacroce, Elodie Burcklen, Christiane Beckmann, Maurice Redondo, Olivier Kobel, Christoph Noppen, Niko Beerenwinkel, Tanja Stadler |
| EPI_ISL_3296575 | Wales Specialist Virology Centre Sequencing lab: Pathogen Genomics Unit | Public Health Wales Microbiology Cardiff Wales Specialist Virology Centre | Catherine Moore, Johnathan Evans, Laura Gifford, Malorie Perry, Simon Cottrell, Angela Marchbank, Alec Birchley, Alexander Adams, Amy Gaskin, Bree Gatica-Wilcox, Jason Coombes, Joel Southgate, Lauren Gilbert, Lee Graham, Nicole Pacchiarini, Sara Kumziene-Summerhayes, Sarah Taylor, Sophie Jones, Sara Rey, Matthew Bull, Joanne Watkins, Sally Corden, Tom Connor |
| EPI_ISL_3296576 | Viollier AG | Department of Biosystems Science and Engineering, ETH Zürich | Christian Beisel, Sarah Nadeau, Chaoran Chen, Ivan Topolsky, Kim Philipp Jablonski, Lara Fuhrmann, Rebecca Denes, Mirjam Feldkamp, Ina Nissen, Natascha Santacroce, Elodie Burcklen, Christiane Beckmann, Maurice Redondo, Olivier Kobel, Christoph Noppen, Niko Beerenwinkel, Tanja Stadler |
| EPI_ISL_3296577, EPI_ISL_3296579 | Wales Specialist Virology Centre Sequencing lab: Pathogen Genomics Unit | Public Health Wales Microbiology Cardiff Wales Specialist Virology Centre | Catherine Moore, Johnathan Evans, Laura Gifford, Malorie Perry, Simon Cottrell, Angela Marchbank, Alec Birchley, Alexander Adams, Amy Gaskin, Bree Gatica-Wilcox, Jason Coombes, Joel Southgate, Lauren Gilbert, Lee Graham, Nicole Pacchiarini, Sara Kumziene-Summerhayes, Sarah Taylor, Sophie Jones, Sara Rey, Matthew Bull, Joanne Watkins, Sally Corden, Tom Connor |
| EPI_ISL_3296580 | Viollier AG | Department of Biosystems Science and Engineering, ETH Zürich | Christian Beisel, Sarah Nadeau, Chaoran Chen, Ivan Topolsky, Kim Philipp Jablonski, Lara Fuhrmann, Rebecca Denes, Mirjam Feldkamp, Ina Nissen, Natascha Santacroce, Elodie Burcklen, Christiane Beckmann, Maurice Redondo, Olivier Kobel, Christoph Noppen, Niko Beerenwinkel, Tanja Stadler |
| EPI_ISL_3296581 | Wales Specialist Virology Centre Sequencing lab: Pathogen Genomics Unit | Public Health Wales Microbiology Cardiff Wales Specialist Virology Centre | Catherine Moore, Johnathan Evans, Laura Gifford, Malorie Perry, Simon Cottrell, Angela Marchbank, Alec Birchley, Alexander Adams, Amy Gaskin, Bree Gatica-Wilcox, Jason Coombes, Joel Southgate, Lauren Gilbert, Lee Graham, Nicole Pacchiarini, Sara Kumziene-Summerhayes, Sarah Taylor, Sophie Jones, Sara Rey, Matthew Bull, Joanne Watkins, Sally Corden, Tom Connor |
| EPI_ISL_3296582 | Viollier AG | Department of Biosystems Science and Engineering, ETH Zürich | Christian Beisel, Sarah Nadeau, Chaoran Chen, Ivan Topolsky, Kim Philipp Jablonski, Lara Fuhrmann, Rebecca Denes, Mirjam Feldkamp, Ina Nissen, Natascha Santacroce, Elodie Burcklen, Christiane Beckmann, Maurice Redondo, Olivier Kobel, Christoph Noppen, Niko Beerenwinkel, Tanja Stadler |
| EPI_ISL_3296583, EPI_ISL_3296585, EPI_ISL_3296586 | Wales Specialist Virology Centre Sequencing lab: Pathogen Genomics Unit | Public Health Wales Microbiology Cardiff Wales Specialist Virology Centre | Catherine Moore, Johnathan Evans, Laura Gifford, Malorie Perry, Simon Cottrell, Angela Marchbank, Alec Birchley, Alexander Adams, Amy Gaskin, Bree Gatica-Wilcox, Jason Coombes, Joel Southgate, Lauren Gilbert, Lee Graham, Nicole Pacchiarini, Sara Kumziene-Summerhayes, Sarah Taylor, Sophie Jones, Sara Rey, Matthew Bull, Joanne Watkins, Sally Corden, Tom Connor |
| EPI_ISL_3296587 | Viollier AG | Department of Biosystems Science and Engineering, ETH Zürich | Christian Beisel, Sarah Nadeau, Chaoran Chen, Ivan Topolsky, Kim Philipp Jablonski, Lara Fuhrmann, Rebecca Denes, Mirjam Feldkamp, Ina Nissen, Natascha Santacroce, Elodie Burcklen, Christiane Beckmann, Maurice Redondo, Olivier Kobel, Christoph Noppen, Niko Beerenwinkel, Tanja Stadler |
| EPI_ISL_3296588 | Wales Specialist Virology Centre Sequencing lab: Pathogen Genomics Unit | Public Health Wales Microbiology Cardiff Wales Specialist Virology Centre | Catherine Moore, Johnathan Evans, Laura Gifford, Malorie Perry, Simon Cottrell, Angela Marchbank, Alec Birchley, Alexander Adams, Amy Gaskin, Bree Gatica-Wilcox, Jason Coombes, Joel Southgate, Lauren Gilbert, Lee Graham, Nicole Pacchiarini, Sara Kumziene-Summerhayes, Sarah Taylor, Sophie Jones, Sara Rey, Matthew Bull, Joanne Watkins, Sally Corden, Tom Connor |
| EPI_ISL_3296589 | Viollier AG | Department of Biosystems Science and Engineering, ETH Zürich | Christian Beisel, Sarah Nadeau, Chaoran Chen, Ivan Topolsky, Kim Philipp Jablonski, Lara Fuhrmann, Rebecca Denes, Mirjam Feldkamp, Ina Nissen, Natascha Santacroce, Elodie Burcklen, Christiane Beckmann, Maurice Redondo, Olivier Kobel, Christoph Noppen, Niko Beerenwinkel, Tanja Stadler |
| EPI_ISL_3296590 | Wales Specialist Virology Centre Sequencing lab: Pathogen Genomics Unit | Public Health Wales Microbiology Cardiff Wales Specialist Virology Centre | Catherine Moore, Johnathan Evans, Laura Gifford, Malorie Perry, Simon Cottrell, Angela Marchbank, Alec Birchley, Alexander Adams, Amy Gaskin, Bree Gatica-Wilcox, Jason Coombes, Joel Southgate, Lauren Gilbert, Lee Graham, Nicole Pacchiarini, Sara Kumziene-Summerhayes, Sarah Taylor, Sophie Jones, Sara Rey, Matthew Bull, Joanne Watkins, Sally Corden, Tom Connor |
| EPI_ISL_3296591 | Viollier AG | Department of Biosystems Science and Engineering, ETH Zürich | Christian Beisel, Sarah Nadeau, Chaoran Chen, Ivan Topolsky, Kim Philipp Jablonski, Lara Fuhrmann, Rebecca Denes, Mirjam Feldkamp, Ina Nissen, Natascha Santacroce, Elodie Burcklen, Christiane Beckmann, Maurice Redondo, Olivier Kobel, Christoph Noppen, Niko Beerenwinkel, Tanja Stadler |
| EPI_ISL_3296592 | Wales Specialist Virology Centre Sequencing lab: Pathogen Genomics Unit | Public Health Wales Microbiology Cardiff Wales Specialist Virology Centre | Catherine Moore, Johnathan Evans, Laura Gifford, Malorie Perry, Simon Cottrell, Angela Marchbank, Alec Birchley, Alexander Adams, Amy Gaskin, Bree Gatica-Wilcox, Jason Coombes, Joel Southgate, Lauren Gilbert, Lee Graham, Nicole Pacchiarini, Sara Kumziene-Summerhayes, Sarah Taylor, Sophie Jones, Sara Rey, Matthew Bull, Joanne Watkins, Sally Corden, Tom Connor |
| EPI_ISL_3296593 | Viollier AG | Department of Biosystems Science and Engineering, ETH Zürich | Christian Beisel, Sarah Nadeau, Chaoran Chen, Ivan Topolsky, Kim Philipp Jablonski, Lara Fuhrmann, Rebecca Denes, Mirjam Feldkamp, Ina Nissen, Natascha Santacroce, Elodie Burcklen, Christiane Beckmann, Maurice Redondo, Olivier Kobel, Christoph Noppen, Niko Beerenwinkel, Tanja Stadler |
| EPI_ISL_3296594 | Wales Specialist Virology Centre Sequencing lab: Pathogen Genomics Unit | Public Health Wales Microbiology Cardiff Wales Specialist Virology Centre | Catherine Moore, Johnathan Evans, Laura Gifford, Malorie Perry, Simon Cottrell, Angela Marchbank, Alec Birchley, Alexander Adams, Amy Gaskin, Bree Gatica-Wilcox, Jason Coombes, Joel Southgate, Lauren Gilbert, Lee Graham, Nicole Pacchiarini, Sara Kumziene-Summerhayes, Sarah Taylor, Sophie Jones, Sara Rey, Matthew Bull, Joanne Watkins, Sally Corden, Tom Connor |
| EPI_ISL_3296595 | Viollier AG | Department of Biosystems Science and Engineering, ETH Zürich | Christian Beisel, Sarah Nadeau, Chaoran Chen, Ivan Topolsky, Kim Philipp Jablonski, Lara Fuhrmann, Rebecca Denes, Mirjam Feldkamp, Ina Nissen, Natascha Santacroce, Elodie Burcklen, Christiane Beckmann, Maurice Redondo, Olivier Kobel, Christoph Noppen, Niko Beerenwinkel, Tanja Stadler |
| EPI_ISL_3296596, EPI_ISL_3296598 | Wales Specialist Virology Centre Sequencing lab: Pathogen Genomics Unit | Public Health Wales Microbiology Cardiff Wales Specialist Virology Centre | Catherine Moore, Johnathan Evans, Laura Gifford, Malorie Perry, Simon Cottrell, Angela Marchbank, Alec Birchley, Alexander Adams, Amy Gaskin, Bree Gatica-Wilcox, Jason Coombes, Joel Southgate, Lauren Gilbert, Lee Graham, Nicole Pacchiarini, Sara Kumziene-Summerhayes, Sarah Taylor, Sophie Jones, Sara Rey, Matthew Bull, Joanne Watkins, Sally Corden, Tom Connor |
| EPI_ISL_3296599 | Viollier AG | Department of Biosystems Science and Engineering, ETH Zürich | Christian Beisel, Sarah Nadeau, Chaoran Chen, Ivan Topolsky, Kim Philipp Jablonski, Lara Fuhrmann, Rebecca Denes, Mirjam Feldkamp, Ina Nissen, Natascha Santacroce, Elodie Burcklen, Christiane Beckmann, Maurice Redondo, Olivier Kobel, Christoph Noppen, Niko Beerenwinkel, Tanja Stadler |
| EPI_ISL_3296600 | Wales Specialist Virology Centre Sequencing lab: Pathogen Genomics Unit | Public Health Wales Microbiology Cardiff Wales Specialist Virology Centre | Catherine Moore, Johnathan Evans, Laura Gifford, Malorie Perry, Simon Cottrell, Angela Marchbank, Alec Birchley, Alexander Adams, Amy Gaskin, Bree Gatica-Wilcox, Jason Coombes, Joel Southgate, Lauren Gilbert, Lee Graham, Nicole Pacchiarini, Sara Kumziene-Summerhayes, Sarah Taylor, Sophie Jones, Sara Rey, Matthew Bull, Joanne Watkins, Sally Corden, Tom Connor |
| EPI_ISL_3296601 | Viollier AG | Department of Biosystems Science and Engineering, ETH Zürich | Christian Beisel, Sarah Nadeau, Chaoran Chen, Ivan Topolsky, Kim Philipp Jablonski, Lara Fuhrmann, Rebecca Denes, Mirjam Feldkamp, Ina Nissen, Natascha Santacroce, Elodie Burcklen, Christiane Beckmann, Maurice Redondo, Olivier Kobel, Christoph Noppen, Niko Beerenwinkel, Tanja Stadler |
| EPI_ISL_3296602 | Wales Specialist Virology Centre Sequencing lab: Pathogen Genomics Unit | Public Health Wales Microbiology Cardiff Wales Specialist Virology Centre | Catherine Moore, Johnathan Evans, Laura Gifford, Malorie Perry, Simon Cottrell, Angela Marchbank, Alec Birchley, Alexander Adams, Amy Gaskin, Bree Gatica-Wilcox, Jason Coombes, Joel Southgate, Lauren Gilbert, Lee Graham, Nicole Pacchiarini, Sara Kumziene-Summerhayes, Sarah Taylor, Sophie Jones, Sara Rey, Matthew Bull, Joanne Watkins, Sally Corden, Tom Connor |
| EPI_ISL_3296603 | Viollier AG | Department of Biosystems Science and Engineering, ETH Zürich | Christian Beisel, Sarah Nadeau, Chaoran Chen, Ivan Topolsky, Kim Philipp Jablonski, Lara Fuhrmann, Rebecca Denes, Mirjam Feldkamp, Ina Nissen, Natascha Santacroce, Elodie Burcklen, Christiane Beckmann, Maurice Redondo, Olivier Kobel, Christoph Noppen, Niko Beerenwinkel, Tanja Stadler |
| EPI_ISL_3296604 | Wales Specialist Virology Centre Sequencing lab: Pathogen Genomics Unit | Public Health Wales Microbiology Cardiff Wales Specialist Virology Centre | Catherine Moore, Johnathan Evans, Laura Gifford, Malorie Perry, Simon Cottrell, Angela Marchbank, Alec Birchley, Alexander Adams, Amy Gaskin, Bree Gatica-Wilcox, Jason Coombes, Joel Southgate, Lauren Gilbert, Lee Graham, Nicole Pacchiarini, Sara Kumziene-Summerhayes, Sarah Taylor, Sophie Jones, Sara Rey, Matthew Bull, Joanne Watkins, Sally Corden, Tom Connor |
| EPI_ISL_3296605 | Viollier AG | Department of Biosystems Science and Engineering, ETH Zürich | Christian Beisel, Sarah Nadeau, Chaoran Chen, Ivan Topolsky, Kim Philipp Jablonski, Lara Fuhrmann, Rebecca Denes, Mirjam Feldkamp, Ina Nissen, Natascha Santacroce, Elodie Burcklen, Christiane Beckmann, Maurice Redondo, Olivier Kobel, Christoph Noppen, Niko Beerenwinkel, Tanja Stadler |
| EPI_ISL_3296606 | Wales Specialist Virology Centre Sequencing lab: Pathogen Genomics Unit | Public Health Wales Microbiology Cardiff Wales Specialist Virology Centre | Catherine Moore, Johnathan Evans, Laura Gifford, Malorie Perry, Simon Cottrell, Angela Marchbank, Alec Birchley, Alexander Adams, Amy Gaskin, Bree Gatica-Wilcox, Jason Coombes, Joel Southgate, Lauren Gilbert, Lee Graham, Nicole Pacchiarini, Sara Kumziene-Summerhayes, Sarah Taylor, Sophie Jones, Sara Rey, Matthew Bull, Joanne Watkins, Sally Corden, Tom Connor |
| EPI_ISL_3296607 | Viollier AG | Department of Biosystems Science and Engineering, ETH Zürich | Christian Beisel, Sarah Nadeau, Chaoran Chen, Ivan Topolsky, Kim Philipp Jablonski, Lara Fuhrmann, Rebecca Denes, Mirjam Feldkamp, Ina Nissen, Natascha Santacroce, Elodie Burcklen, Christiane Beckmann, Maurice Redondo, Olivier Kobel, Christoph Noppen, Niko Beerenwinkel, Tanja Stadler |
| EPI_ISL_3296608 | Wales Specialist Virology Centre Sequencing lab: Pathogen Genomics Unit | Public Health Wales Microbiology Cardiff Wales Specialist Virology Centre | Catherine Moore, Johnathan Evans, Laura Gifford, Malorie Perry, Simon Cottrell, Angela Marchbank, Alec Birchley, Alexander Adams, Amy Gaskin, Bree Gatica-Wilcox, Jason Coombes, Joel Southgate, Lauren Gilbert, Lee Graham, Nicole Pacchiarini, Sara Kumziene-Summerhayes, Sarah Taylor, Sophie Jones, Sara Rey, Matthew Bull, Joanne Watkins, Sally Corden, Tom Connor |
| EPI_ISL_3296609 | Viollier AG | Department of Biosystems Science and Engineering, ETH Zürich | Christian Beisel, Sarah Nadeau, Chaoran Chen, Ivan Topolsky, Kim Philipp Jablonski, Lara Fuhrmann, Rebecca Denes, Mirjam Feldkamp, Ina Nissen, Natascha Santacroce, Elodie Burcklen, Christiane Beckmann, Maurice Redondo, Olivier Kobel, Christoph Noppen, Niko Beerenwinkel, Tanja Stadler |
| EPI_ISL_3296610 | Wales Specialist Virology Centre Sequencing lab: Pathogen Genomics Unit | Public Health Wales Microbiology Cardiff Wales Specialist Virology Centre | Catherine Moore, Johnathan Evans, Laura Gifford, Malorie Perry, Simon Cottrell, Angela Marchbank, Alec Birchley, Alexander Adams, Amy Gaskin, Bree Gatica-Wilcox, Jason Coombes, Joel Southgate, Lauren Gilbert, Lee Graham, Nicole Pacchiarini, Sara Kumziene-Summerhayes, Sarah Taylor, Sophie Jones, Sara Rey, Matthew Bull, Joanne Watkins, Sally Corden, Tom Connor |
| EPI_ISL_3296611 | Viollier AG | Department of Biosystems Science and Engineering, ETH Zürich | Christian Beisel, Sarah Nadeau, Chaoran Chen, Ivan Topolsky, Kim Philipp Jablonski, Lara Fuhrmann, Rebecca Denes, Mirjam Feldkamp, Ina Nissen, Natascha Santacroce, Elodie Burcklen, Christiane Beckmann, Maurice Redondo, Olivier Kobel, Christoph Noppen, Niko Beerenwinkel, Tanja Stadler |
| EPI_ISL_3296612 | Wales Specialist Virology Centre Sequencing lab: Pathogen Genomics Unit | Public Health Wales Microbiology Cardiff Wales Specialist Virology Centre | Catherine Moore, Johnathan Evans, Laura Gifford, Malorie Perry, Simon Cottrell, Angela Marchbank, Alec Birchley, Alexander Adams, Amy Gaskin, Bree Gatica-Wilcox, Jason Coombes, Joel Southgate, Lauren Gilbert, Lee Graham, Nicole Pacchiarini, Sara Kumziene-Summerhayes, Sarah Taylor, Sophie Jones, Sara Rey, Matthew Bull, Joanne Watkins, Sally Corden, Tom Connor |
| EPI_ISL_3296613 | Viollier AG | Department of Biosystems Science and Engineering, ETH Zürich | Christian Beisel, Sarah Nadeau, Chaoran Chen, Ivan Topolsky, Kim Philipp Jablonski, Lara Fuhrmann, Rebecca Denes, Mirjam Feldkamp, Ina Nissen, Natascha Santacroce, Elodie Burcklen, Christiane Beckmann, Maurice Redondo, Olivier Kobel, Christoph Noppen, Niko Beerenwinkel, Tanja Stadler |
| EPI_ISL_3296614 | Wales Specialist Virology Centre Sequencing lab: Pathogen Genomics Unit | Public Health Wales Microbiology Cardiff Wales Specialist Virology Centre | Catherine Moore, Johnathan Evans, Laura Gifford, Malorie Perry, Simon Cottrell, Angela Marchbank, Alec Birchley, Alexander Adams, Amy Gaskin, Bree Gatica-Wilcox, Jason Coombes, Joel Southgate, Lauren Gilbert, Lee Graham, Nicole Pacchiarini, Sara Kumziene-Summerhayes, Sarah Taylor, Sophie Jones, Sara Rey, Matthew Bull, Joanne Watkins, Sally Corden, Tom Connor |
| EPI_ISL_3296615 | Viollier AG | Department of Biosystems Science and Engineering, ETH Zürich | Christian Beisel, Sarah Nadeau, Chaoran Chen, Ivan Topolsky, Kim Philipp Jablonski, Lara Fuhrmann, Rebecca Denes, Mirjam Feldkamp, Ina Nissen, Natascha Santacroce, Elodie Burcklen, Christiane Beckmann, Maurice Redondo, Olivier Kobel, Christoph Noppen, Niko Beerenwinkel, Tanja Stadler |

| | | | |
|---|---|---|---|
| EPI_ISL_3296616 | Wales Specialist Virology Centre Sequencing lab: Pathogen Genomics Unit | Public Health Wales Microbiology Cardiff Wales Specialist Virology Centre | Catherine Moore, Johnathan Evans, Laura Gifford, Malorie Perry, Simon Cottrell, Angela Marchbank, Alec Birchley, Alexander Adams, Amy Gaskin, Bree Gatica-Wilcox, Jason Coombes, Joel Southgate, Lauren Gilbert, Lee Graham, Nicole Pacchiarini, Sara Kumziene-Summerhayes, Sarah Taylor, Sophie Jones, Sara Rey, Matthew Bull, Joanne Watkins, Sally Corden, Tom Connor |
| EPI_ISL_3296617 | Viollier AG | Department of Biosystems Science and Engineering, ETH Zürich | Christian Beisel, Sarah Nadeau, Chaoran Chen, Ivan Topolsky, Kim Philipp Jablonski, Lara Fuhrmann, Rebecca Denes, Mirjam Feldkamp, Ina Nissen, Natascha Santacroce, Elodie Burcklen, Christiane Beckmann, Maurice Redondo, Olivier Kobel, Christoph Noppen, Niko Beerenwinkel, Tanja Stadler |
| EPI_ISL_3296618 | Wales Specialist Virology Centre Sequencing lab: Pathogen Genomics Unit | Public Health Wales Microbiology Cardiff Wales Specialist Virology Centre | Catherine Moore, Johnathan Evans, Laura Gifford, Malorie Perry, Simon Cottrell, Angela Marchbank, Alec Birchley, Alexander Adams, Amy Gaskin, Bree Gatica-Wilcox, Jason Coombes, Joel Southgate, Lauren Gilbert, Lee Graham, Nicole Pacchiarini, Sara Kumziene-Summerhayes, Sarah Taylor, Sophie Jones, Sara Rey, Matthew Bull, Joanne Watkins, Sally Corden, Tom Connor |
| EPI_ISL_3296619 | Viollier AG | Department of Biosystems Science and Engineering, ETH Zürich | Christian Beisel, Sarah Nadeau, Chaoran Chen, Ivan Topolsky, Kim Philipp Jablonski, Lara Fuhrmann, Rebecca Denes, Mirjam Feldkamp, Ina Nissen, Natascha Santacroce, Elodie Burcklen, Christiane Beckmann, Maurice Redondo, Olivier Kobel, Christoph Noppen, Niko Beerenwinkel, Tanja Stadler |
| EPI_ISL_3296620 | Wales Specialist Virology Centre Sequencing lab: Pathogen Genomics Unit | Public Health Wales Microbiology Cardiff Wales Specialist Virology Centre | Catherine Moore, Johnathan Evans, Laura Gifford, Malorie Perry, Simon Cottrell, Angela Marchbank, Alec Birchley, Alexander Adams, Amy Gaskin, Bree Gatica-Wilcox, Jason Coombes, Joel Southgate, Lauren Gilbert, Lee Graham, Nicole Pacchiarini, Sara Kumziene-Summerhayes, Sarah Taylor, Sophie Jones, Sara Rey, Matthew Bull, Joanne Watkins, Sally Corden, Tom Connor |
| EPI_ISL_3296621 | Viollier AG | Department of Biosystems Science and Engineering, ETH Zürich | Christian Beisel, Sarah Nadeau, Chaoran Chen, Ivan Topolsky, Kim Philipp Jablonski, Lara Fuhrmann, Rebecca Denes, Mirjam Feldkamp, Ina Nissen, Natascha Santacroce, Elodie Burcklen, Christiane Beckmann, Maurice Redondo, Olivier Kobel, Christoph Noppen, Niko Beerenwinkel, Tanja Stadler |
| EPI_ISL_3296622 | Wales Specialist Virology Centre Sequencing lab: Pathogen Genomics Unit | Public Health Wales Microbiology Cardiff Wales Specialist Virology Centre | Catherine Moore, Johnathan Evans, Laura Gifford, Malorie Perry, Simon Cottrell, Angela Marchbank, Alec Birchley, Alexander Adams, Amy Gaskin, Bree Gatica-Wilcox, Jason Coombes, Joel Southgate, Lauren Gilbert, Lee Graham, Nicole Pacchiarini, Sara Kumziene-Summerhayes, Sarah Taylor, Sophie Jones, Sara Rey, Matthew Bull, Joanne Watkins, Sally Corden, Tom Connor |
| EPI_ISL_3296623 | Viollier AG | Department of Biosystems Science and Engineering, ETH Zürich | Christian Beisel, Sarah Nadeau, Chaoran Chen, Ivan Topolsky, Kim Philipp Jablonski, Lara Fuhrmann, Rebecca Denes, Mirjam Feldkamp, Ina Nissen, Natascha Santacroce, Elodie Burcklen, Christiane Beckmann, Maurice Redondo, Olivier Kobel, Christoph Noppen, Niko Beerenwinkel, Tanja Stadler |
| EPI_ISL_3296624 | Wales Specialist Virology Centre Sequencing lab: Pathogen Genomics Unit | Public Health Wales Microbiology Cardiff Wales Specialist Virology Centre | Catherine Moore, Johnathan Evans, Laura Gifford, Malorie Perry, Simon Cottrell, Angela Marchbank, Alec Birchley, Alexander Adams, Amy Gaskin, Bree Gatica-Wilcox, Jason Coombes, Joel Southgate, Lauren Gilbert, Lee Graham, Nicole Pacchiarini, Sara Kumziene-Summerhayes, Sarah Taylor, Sophie Jones, Sara Rey, Matthew Bull, Joanne Watkins, Sally Corden, Tom Connor |
| EPI_ISL_3296625 | Viollier AG | Department of Biosystems Science and Engineering, ETH Zürich | Christian Beisel, Sarah Nadeau, Chaoran Chen, Ivan Topolsky, Kim Philipp Jablonski, Lara Fuhrmann, Rebecca Denes, Mirjam Feldkamp, Ina Nissen, Natascha Santacroce, Elodie Burcklen, Christiane Beckmann, Maurice Redondo, Olivier Kobel, Christoph Noppen, Niko Beerenwinkel, Tanja Stadler |
| EPI_ISL_3296626 | Wales Specialist Virology Centre Sequencing lab: Pathogen Genomics Unit | Public Health Wales Microbiology Cardiff Wales Specialist Virology Centre | Catherine Moore, Johnathan Evans, Laura Gifford, Malorie Perry, Simon Cottrell, Angela Marchbank, Alec Birchley, Alexander Adams, Amy Gaskin, Bree Gatica-Wilcox, Jason Coombes, Joel Southgate, Lauren Gilbert, Lee Graham, Nicole Pacchiarini, Sara Kumziene-Summerhayes, Sarah Taylor, Sophie Jones, Sara Rey, Matthew Bull, Joanne Watkins, Sally Corden, Tom Connor |
| EPI_ISL_3296627 | Viollier AG | Department of Biosystems Science and Engineering, ETH Zürich | Christian Beisel, Sarah Nadeau, Chaoran Chen, Ivan Topolsky, Kim Philipp Jablonski, Lara Fuhrmann, Rebecca Denes, Mirjam Feldkamp, Ina Nissen, Natascha Santacroce, Elodie Burcklen, Christiane Beckmann, Maurice Redondo, Olivier Kobel, Christoph Noppen, Niko Beerenwinkel, Tanja Stadler |
| EPI_ISL_3296628 | Wales Specialist Virology Centre Sequencing lab: Pathogen Genomics Unit | Public Health Wales Microbiology Cardiff Wales Specialist Virology Centre | Catherine Moore, Johnathan Evans, Laura Gifford, Malorie Perry, Simon Cottrell, Angela Marchbank, Alec Birchley, Alexander Adams, Amy Gaskin, Bree Gatica-Wilcox, Jason Coombes, Joel Southgate, Lauren Gilbert, Lee Graham, Nicole Pacchiarini, Sara Kumziene-Summerhayes, Sarah Taylor, Sophie Jones, Sara Rey, Matthew Bull, Joanne Watkins, Sally Corden, Tom Connor |
| EPI_ISL_3296629 | Viollier AG | Department of Biosystems Science and Engineering, ETH Zürich | Christian Beisel, Sarah Nadeau, Chaoran Chen, Ivan Topolsky, Kim Philipp Jablonski, Lara Fuhrmann, Rebecca Denes, Mirjam Feldkamp, Ina Nissen, Natascha Santacroce, Elodie Burcklen, Christiane Beckmann, Maurice Redondo, Olivier Kobel, Christoph Noppen, Niko Beerenwinkel, Tanja Stadler |
| EPI_ISL_3296630 | Wales Specialist Virology Centre Sequencing lab: Pathogen Genomics Unit | Public Health Wales Microbiology Cardiff Wales Specialist Virology Centre | Catherine Moore, Johnathan Evans, Laura Gifford, Malorie Perry, Simon Cottrell, Angela Marchbank, Alec Birchley, Alexander Adams, Amy Gaskin, Bree Gatica-Wilcox, Jason Coombes, Joel Southgate, Lauren Gilbert, Lee Graham, Nicole Pacchiarini, Sara Kumziene-Summerhayes, Sarah Taylor, Sophie Jones, Sara Rey, Matthew Bull, Joanne Watkins, Sally Corden, Tom Connor |
| EPI_ISL_3296631 | Viollier AG | Department of Biosystems Science and Engineering, ETH Zürich | Christian Beisel, Sarah Nadeau, Chaoran Chen, Ivan Topolsky, Kim Philipp Jablonski, Lara Fuhrmann, Rebecca Denes, Mirjam Feldkamp, Ina Nissen, Natascha Santacroce, Elodie Burcklen, Christiane Beckmann, Maurice Redondo, Olivier Kobel, Christoph Noppen, Niko Beerenwinkel, Tanja Stadler |
| EPI_ISL_3296632 | Wales Specialist Virology Centre Sequencing lab: Pathogen Genomics Unit | Public Health Wales Microbiology Cardiff Wales Specialist Virology Centre | Catherine Moore, Johnathan Evans, Laura Gifford, Malorie Perry, Simon Cottrell, Angela Marchbank, Alec Birchley, Alexander Adams, Amy Gaskin, Bree Gatica-Wilcox, Jason Coombes, Joel Southgate, Lauren Gilbert, Lee Graham, Nicole Pacchiarini, Sara Kumziene-Summerhayes, Sarah Taylor, Sophie Jones, Sara Rey, Matthew Bull, Joanne Watkins, Sally Corden, Tom Connor |
| EPI_ISL_3296633 | Viollier AG | Department of Biosystems Science and Engineering, ETH Zürich | Christian Beisel, Sarah Nadeau, Chaoran Chen, Ivan Topolsky, Kim Philipp Jablonski, Lara Fuhrmann, Rebecca Denes, Mirjam Feldkamp, Ina Nissen, Natascha Santacroce, Elodie Burcklen, Christiane Beckmann, Maurice Redondo, Olivier Kobel, Christoph Noppen, Niko Beerenwinkel, Tanja Stadler |
| EPI_ISL_3296634 | Wales Specialist Virology Centre Sequencing lab: Pathogen Genomics Unit | Public Health Wales Microbiology Cardiff Wales Specialist Virology Centre | Catherine Moore, Johnathan Evans, Laura Gifford, Malorie Perry, Simon Cottrell, Angela Marchbank, Alec Birchley, Alexander Adams, Amy Gaskin, Bree Gatica-Wilcox, Jason Coombes, Joel Southgate, Lauren Gilbert, Lee Graham, Nicole Pacchiarini, Sara Kumziene-Summerhayes, Sarah Taylor, Sophie Jones, Sara Rey, Matthew Bull, Joanne Watkins, Sally Corden, Tom Connor |
| EPI_ISL_3296635 | Viollier AG | Department of Biosystems Science and Engineering, ETH Zürich | Christian Beisel, Sarah Nadeau, Chaoran Chen, Ivan Topolsky, Kim Philipp Jablonski, Lara Fuhrmann, Rebecca Denes, Mirjam Feldkamp, Ina Nissen, Natascha Santacroce, Elodie Burcklen, Christiane Beckmann, Maurice Redondo, Olivier Kobel, Christoph Noppen, Niko Beerenwinkel, Tanja Stadler |
| EPI_ISL_3296636 | Wales Specialist Virology Centre Sequencing lab: Pathogen Genomics Unit | Public Health Wales Microbiology Cardiff Wales Specialist Virology Centre | Catherine Moore, Johnathan Evans, Laura Gifford, Malorie Perry, Simon Cottrell, Angela Marchbank, Alec Birchley, Alexander Adams, Amy Gaskin, Bree Gatica-Wilcox, Jason Coombes, Joel Southgate, Lauren Gilbert, Lee Graham, Nicole Pacchiarini, Sara Kumziene-Summerhayes, Sarah Taylor, Sophie Jones, Sara Rey, Matthew Bull, Joanne Watkins, Sally Corden, Tom Connor |
| EPI_ISL_3296637 | Viollier AG | Department of Biosystems Science and Engineering, ETH Zürich | Christian Beisel, Sarah Nadeau, Chaoran Chen, Ivan Topolsky, Kim Philipp Jablonski, Lara Fuhrmann, Rebecca Denes, Mirjam Feldkamp, Ina Nissen, Natascha Santacroce, Elodie Burcklen, Christiane Beckmann, Maurice Redondo, Olivier Kobel, Christoph Noppen, Niko Beerenwinkel, Tanja Stadler |
| EPI_ISL_3296638 | Wales Specialist Virology Centre Sequencing lab: Pathogen Genomics Unit | Public Health Wales Microbiology Cardiff Wales Specialist Virology Centre | Catherine Moore, Johnathan Evans, Laura Gifford, Malorie Perry, Simon Cottrell, Angela Marchbank, Alec Birchley, Alexander Adams, Amy Gaskin, Bree Gatica-Wilcox, Jason Coombes, Joel Southgate, Lauren Gilbert, Lee Graham, Nicole Pacchiarini, Sara Kumziene-Summerhayes, Sarah Taylor, Sophie Jones, Sara Rey, Matthew Bull, Joanne Watkins, Sally Corden, Tom Connor |
| EPI_ISL_3296639 | Viollier AG | Department of Biosystems Science and Engineering, ETH Zürich | Christian Beisel, Sarah Nadeau, Chaoran Chen, Ivan Topolsky, Kim Philipp Jablonski, Lara Fuhrmann, Rebecca Denes, Mirjam Feldkamp, Ina Nissen, Natascha Santacroce, Elodie Burcklen, Christiane Beckmann, Maurice Redondo, Olivier Kobel, Christoph Noppen, Niko Beerenwinkel, Tanja Stadler |
| EPI_ISL_3296640 | Wales Specialist Virology Centre Sequencing lab: Pathogen Genomics Unit | Public Health Wales Microbiology Cardiff Wales Specialist Virology Centre | Catherine Moore, Johnathan Evans, Laura Gifford, Malorie Perry, Simon Cottrell, Angela Marchbank, Alec Birchley, Alexander Adams, Amy Gaskin, Bree Gatica-Wilcox, Jason Coombes, Joel Southgate, Lauren Gilbert, Lee Graham, Nicole Pacchiarini, Sara Kumziene-Summerhayes, Sarah Taylor, Sophie Jones, Sara Rey, Matthew Bull, Joanne Watkins, Sally Corden, Tom Connor |
| EPI_ISL_3296641 | Viollier AG | Department of Biosystems Science and Engineering, ETH Zürich | Christian Beisel, Sarah Nadeau, Chaoran Chen, Ivan Topolsky, Kim Philipp Jablonski, Lara Fuhrmann, Rebecca Denes, Mirjam Feldkamp, Ina Nissen, Natascha Santacroce, Elodie Burcklen, Christiane Beckmann, Maurice Redondo, Olivier Kobel, Christoph Noppen, Niko Beerenwinkel, Tanja Stadler |
| EPI_ISL_3296642 | Wales Specialist Virology Centre Sequencing lab: Pathogen Genomics Unit | Public Health Wales Microbiology Cardiff Wales Specialist Virology Centre | Catherine Moore, Johnathan Evans, Laura Gifford, Malorie Perry, Simon Cottrell, Angela Marchbank, Alec Birchley, Alexander Adams, Amy Gaskin, Bree Gatica-Wilcox, Jason Coombes, Joel Southgate, Lauren Gilbert, Lee Graham, Nicole Pacchiarini, Sara Kumziene-Summerhayes, Sarah Taylor, Sophie Jones, Sara Rey, Matthew Bull, Joanne Watkins, Sally Corden, Tom Connor |
| EPI_ISL_3296643 | Viollier AG | Department of Biosystems Science and Engineering, ETH Zürich | Christian Beisel, Sarah Nadeau, Chaoran Chen, Ivan Topolsky, Kim Philipp Jablonski, Lara Fuhrmann, Rebecca Denes, Mirjam Feldkamp, Ina Nissen, Natascha Santacroce, Elodie Burcklen, Christiane Beckmann, Maurice Redondo, Olivier Kobel, Christoph Noppen, Niko Beerenwinkel, Tanja Stadler |
| EPI_ISL_3296644 | Wales Specialist Virology Centre Sequencing lab: Pathogen Genomics Unit | Public Health Wales Microbiology Cardiff Wales Specialist Virology Centre | Catherine Moore, Johnathan Evans, Laura Gifford, Malorie Perry, Simon Cottrell, Angela Marchbank, Alec Birchley, Alexander Adams, Amy Gaskin, Bree Gatica-Wilcox, Jason Coombes, Joel Southgate, Lauren Gilbert, Lee Graham, Nicole Pacchiarini, Sara Kumziene-Summerhayes, Sarah Taylor, Sophie Jones, Sara Rey, Matthew Bull, Joanne Watkins, Sally Corden, Tom Connor |
| EPI_ISL_3296645 | Viollier AG | Department of Biosystems Science and Engineering, ETH Zürich | Christian Beisel, Sarah Nadeau, Chaoran Chen, Ivan Topolsky, Kim Philipp Jablonski, Lara Fuhrmann, Rebecca Denes, Mirjam Feldkamp, Ina Nissen, Natascha Santacroce, Elodie Burcklen, Christiane Beckmann, Maurice Redondo, Olivier Kobel, Christoph Noppen, Niko Beerenwinkel, Tanja Stadler |
| EPI_ISL_3296646, EPI_ISL_3296648 | Wales Specialist Virology Centre Sequencing lab: Pathogen Genomics Unit | Public Health Wales Microbiology Cardiff Wales Specialist Virology Centre | Catherine Moore, Johnathan Evans, Laura Gifford, Malorie Perry, Simon Cottrell, Angela Marchbank, Alec Birchley, Alexander Adams, Amy Gaskin, Bree Gatica-Wilcox, Jason Coombes, Joel Southgate, Lauren Gilbert, Lee Graham, Nicole Pacchiarini, Sara Kumziene-Summerhayes, Sarah Taylor, Sophie Jones, Sara Rey, Matthew Bull, Joanne Watkins, Sally Corden, Tom Connor |
| EPI_ISL_3296649 | Viollier AG | Department of Biosystems Science and Engineering, ETH Zürich | Christian Beisel, Sarah Nadeau, Chaoran Chen, Ivan Topolsky, Kim Philipp Jablonski, Lara Fuhrmann, Rebecca Denes, Mirjam Feldkamp, Ina Nissen, Natascha Santacroce, Elodie Burcklen, Christiane Beckmann, Maurice Redondo, Olivier Kobel, Christoph Noppen, Niko Beerenwinkel, Tanja Stadler |
| EPI_ISL_3296650 | Wales Specialist Virology Centre Sequencing lab: Pathogen Genomics Unit | Public Health Wales Microbiology Cardiff Wales Specialist Virology Centre | Catherine Moore, Johnathan Evans, Laura Gifford, Malorie Perry, Simon Cottrell, Angela Marchbank, Alec Birchley, Alexander Adams, Amy Gaskin, Bree Gatica-Wilcox, Jason Coombes, Joel Southgate, Lauren Gilbert, Lee Graham, Nicole Pacchiarini, Sara Kumziene-Summerhayes, Sarah Taylor, Sophie Jones, Sara Rey, Matthew Bull, Joanne Watkins, Sally Corden, Tom Connor |
| EPI_ISL_3296651 | Viollier AG | Department of Biosystems Science and Engineering, ETH Zürich | Christian Beisel, Sarah Nadeau, Chaoran Chen, Ivan Topolsky, Kim Philipp Jablonski, Lara Fuhrmann, Rebecca Denes, Mirjam Feldkamp, Ina Nissen, Natascha Santacroce, Elodie Burcklen, Christiane Beckmann, Maurice Redondo, Olivier Kobel, Christoph Noppen, Niko Beerenwinkel, Tanja Stadler |
| EPI_ISL_3296652 | Wales Specialist Virology Centre Sequencing lab: Pathogen Genomics Unit | Public Health Wales Microbiology Cardiff Wales Specialist Virology Centre | Catherine Moore, Johnathan Evans, Laura Gifford, Malorie Perry, Simon Cottrell, Angela Marchbank, Alec Birchley, Alexander Adams, Amy Gaskin, Bree Gatica-Wilcox, Jason Coombes, Joel Southgate, Lauren Gilbert, Lee Graham, Nicole Pacchiarini, Sara Kumziene-Summerhayes, Sarah Taylor, Sophie Jones, Sara Rey, Matthew Bull, Joanne Watkins, Sally Corden, Tom Connor |
| EPI_ISL_3296653 | Viollier AG | Department of Biosystems Science and Engineering, ETH Zürich | Christian Beisel, Sarah Nadeau, Chaoran Chen, Ivan Topolsky, Kim Philipp Jablonski, Lara Fuhrmann, Rebecca Denes, Mirjam Feldkamp, Ina Nissen, Natascha Santacroce, Elodie Burcklen, Christiane Beckmann, Maurice Redondo, Olivier Kobel, Christoph Noppen, Niko Beerenwinkel, Tanja Stadler |
| EPI_ISL_3296654 | Wales Specialist Virology Centre Sequencing lab: Pathogen Genomics Unit | Public Health Wales Microbiology Cardiff Wales Specialist Virology Centre | Catherine Moore, Johnathan Evans, Laura Gifford, Malorie Perry, Simon Cottrell, Angela Marchbank, Alec Birchley, Alexander Adams, Amy Gaskin, Bree Gatica-Wilcox, Jason Coombes, Joel Southgate, Lauren Gilbert, Lee Graham, Nicole Pacchiarini, Sara Kumziene-Summerhayes, Sarah Taylor, Sophie Jones, Sara Rey, Matthew Bull, Joanne Watkins, Sally Corden, Tom Connor |
| EPI_ISL_3296655 | Viollier AG | Department of Biosystems Science and Engineering, ETH Zürich | Christian Beisel, Sarah Nadeau, Chaoran Chen, Ivan Topolsky, Kim Philipp Jablonski, Lara Fuhrmann, Rebecca Denes, Mirjam Feldkamp, Ina Nissen, Natascha Santacroce, Elodie Burcklen, Christiane Beckmann, Maurice Redondo, Olivier Kobel, Christoph Noppen, Niko Beerenwinkel, Tanja Stadler |
| EPI_ISL_3296656 | Wales Specialist Virology Centre Sequencing lab: Pathogen Genomics Unit | Public Health Wales Microbiology Cardiff Wales Specialist Virology Centre | Catherine Moore, Johnathan Evans, Laura Gifford, Malorie Perry, Simon Cottrell, Angela Marchbank, Alec Birchley, Alexander Adams, Amy Gaskin, Bree Gatica-Wilcox, Jason Coombes, Joel Southgate, Lauren Gilbert, Lee Graham, Nicole Pacchiarini, Sara Kumziene-Summerhayes, Sarah Taylor, Sophie Jones, Sara Rey, Matthew Bull, Joanne Watkins, Sally Corden, Tom Connor |
| EPI_ISL_3296657 | Viollier AG | Department of Biosystems Science and Engineering, ETH Zürich | Christian Beisel, Sarah Nadeau, Chaoran Chen, Ivan Topolsky, Kim Philipp Jablonski, Lara Fuhrmann, Rebecca Denes, Mirjam Feldkamp, Ina Nissen, Natascha Santacroce, Elodie Burcklen, Christiane Beckmann, Maurice Redondo, Olivier Kobel, Christoph Noppen, Niko Beerenwinkel, Tanja Stadler |
| EPI_ISL_3296658 | Wales Specialist Virology Centre Sequencing lab: Pathogen Genomics Unit | Public Health Wales Microbiology Cardiff Wales Specialist Virology Centre | Catherine Moore, Johnathan Evans, Laura Gifford, Malorie Perry, Simon Cottrell, Angela Marchbank, Alec Birchley, Alexander Adams, Amy Gaskin, Bree Gatica-Wilcox, Jason Coombes, Joel Southgate, Lauren Gilbert, Lee Graham, Nicole Pacchiarini, Sara Kumziene-Summerhayes, Sarah Taylor, Sophie Jones, Sara Rey, Matthew Bull, Joanne Watkins, Sally Corden, Tom Connor |
| EPI_ISL_3296659 | Viollier AG | Department of Biosystems Science and Engineering, ETH Zürich | Christian Beisel, Sarah Nadeau, Chaoran Chen, Ivan Topolsky, Kim Philipp Jablonski, Lara Fuhrmann, Rebecca Denes, Mirjam Feldkamp, Ina Nissen, Natascha Santacroce, Elodie Burcklen, Christiane Beckmann, Maurice Redondo, Olivier Kobel, Christoph Noppen, Niko Beerenwinkel, Tanja Stadler |
| EPI_ISL_3296660, EPI_ISL_3296662, EPI_ISL_3296663 | Wales Specialist Virology Centre Sequencing lab: Pathogen Genomics Unit | Public Health Wales Microbiology Cardiff Wales Specialist Virology Centre | Catherine Moore, Johnathan Evans, Laura Gifford, Malorie Perry, Simon Cottrell, Angela Marchbank, Alec Birchley, Alexander Adams, Amy Gaskin, Bree Gatica-Wilcox, Jason Coombes, Joel Southgate, Lauren Gilbert, Lee Graham, Nicole Pacchiarini, Sara Kumziene-Summerhayes, Sarah Taylor, Sophie Jones, Sara Rey, Matthew Bull, Joanne Watkins, Sally Corden, Tom Connor |
| EPI_ISL_3296664 | Viollier AG | Department of Biosystems Science and Engineering, ETH Zürich | Christian Beisel, Sarah Nadeau, Chaoran Chen, Ivan Topolsky, Kim Philipp Jablonski, Lara Fuhrmann, Rebecca Denes, Mirjam Feldkamp, Ina Nissen, Natascha Santacroce, Elodie Burcklen, Christiane Beckmann, Maurice Redondo, Olivier Kobel, Christoph Noppen, Niko Beerenwinkel, Tanja Stadler |

| | | | |
|---|---|---|---|
| EPI_ISL_3296665 | Wales Specialist Virology Centre Sequencing lab: Pathogen Genomics Unit | Public Health Wales Microbiology Cardiff Wales Specialist Virology Centre | Catherine Moore, Johnathan Evans, Laura Gifford, Malorie Perry, Simon Cottrell, Angela Marchbank, Alec Birchley, Alexander Adams, Amy Gaskin, Bree Gatica-Wilcox, Jason Coombes, Joel Southgate, Lauren Gilbert, Lee Graham, Nicole Pacchiarini, Sara Kumziene-Summerhayes, Sarah Taylor, Sophie Jones, Sara Rey, Matthew Bull, Joanne Watkins, Sally Corden, Tom Connor |
| EPI_ISL_3296666, EPI_ISL_3296667 | Viollier AG | Department of Biosystems Science and Engineering, ETH Zürich | Christian Beisel, Sarah Nadeau, Chaoran Chen, Ivan Topolsky, Kim Philipp Jablonski, Lara Fuhrmann, Rebecca Denes, Mirjam Feldkamp, Ina Nissen, Natascha Santacroce, Elodie Burcklen, Christiane Beckmann, Maurice Redondo, Olivier Kobel, Christoph Noppen, Niko Beerenwinkel, Tanja Stadler |
| EPI_ISL_3296668 | Wales Specialist Virology Centre Sequencing lab: Pathogen Genomics Unit | Public Health Wales Microbiology Cardiff Wales Specialist Virology Centre | Catherine Moore, Johnathan Evans, Laura Gifford, Malorie Perry, Simon Cottrell, Angela Marchbank, Alec Birchley, Alexander Adams, Amy Gaskin, Bree Gatica-Wilcox, Jason Coombes, Joel Southgate, Lauren Gilbert, Lee Graham, Nicole Pacchiarini, Sara Kumziene-Summerhayes, Sarah Taylor, Sophie Jones, Sara Rey, Matthew Bull, Joanne Watkins, Sally Corden, Tom Connor |
| EPI_ISL_3296669 | Viollier AG | Department of Biosystems Science and Engineering, ETH Zürich | Christian Beisel, Sarah Nadeau, Chaoran Chen, Ivan Topolsky, Kim Philipp Jablonski, Lara Fuhrmann, Rebecca Denes, Mirjam Feldkamp, Ina Nissen, Natascha Santacroce, Elodie Burcklen, Christiane Beckmann, Maurice Redondo, Olivier Kobel, Christoph Noppen, Niko Beerenwinkel, Tanja Stadler |
| EPI_ISL_3296670 | Wales Specialist Virology Centre Sequencing lab: Pathogen Genomics Unit | Public Health Wales Microbiology Cardiff Wales Specialist Virology Centre | Catherine Moore, Johnathan Evans, Laura Gifford, Malorie Perry, Simon Cottrell, Angela Marchbank, Alec Birchley, Alexander Adams, Amy Gaskin, Bree Gatica-Wilcox, Jason Coombes, Joel Southgate, Lauren Gilbert, Lee Graham, Nicole Pacchiarini, Sara Kumziene-Summerhayes, Sarah Taylor, Sophie Jones, Sara Rey, Matthew Bull, Joanne Watkins, Sally Corden, Tom Connor |
| EPI_ISL_3296671 | Viollier AG | Department of Biosystems Science and Engineering, ETH Zürich | Christian Beisel, Sarah Nadeau, Chaoran Chen, Ivan Topolsky, Kim Philipp Jablonski, Lara Fuhrmann, Rebecca Denes, Mirjam Feldkamp, Ina Nissen, Natascha Santacroce, Elodie Burcklen, Christiane Beckmann, Maurice Redondo, Olivier Kobel, Christoph Noppen, Niko Beerenwinkel, Tanja Stadler |
| EPI_ISL_3296672 | Wales Specialist Virology Centre Sequencing lab: Pathogen Genomics Unit | Public Health Wales Microbiology Cardiff Wales Specialist Virology Centre | Catherine Moore, Johnathan Evans, Laura Gifford, Malorie Perry, Simon Cottrell, Angela Marchbank, Alec Birchley, Alexander Adams, Amy Gaskin, Bree Gatica-Wilcox, Jason Coombes, Joel Southgate, Lauren Gilbert, Lee Graham, Nicole Pacchiarini, Sara Kumziene-Summerhayes, Sarah Taylor, Sophie Jones, Sara Rey, Matthew Bull, Joanne Watkins, Sally Corden, Tom Connor |
| EPI_ISL_3296673 | Viollier AG | Department of Biosystems Science and Engineering, ETH Zürich | Christian Beisel, Sarah Nadeau, Chaoran Chen, Ivan Topolsky, Kim Philipp Jablonski, Lara Fuhrmann, Rebecca Denes, Mirjam Feldkamp, Ina Nissen, Natascha Santacroce, Elodie Burcklen, Christiane Beckmann, Maurice Redondo, Olivier Kobel, Christoph Noppen, Niko Beerenwinkel, Tanja Stadler |
| EPI_ISL_3296674, EPI_ISL_3296676 | Wales Specialist Virology Centre Sequencing lab: Pathogen Genomics Unit | Public Health Wales Microbiology Cardiff Wales Specialist Virology Centre | Catherine Moore, Johnathan Evans, Laura Gifford, Malorie Perry, Simon Cottrell, Angela Marchbank, Alec Birchley, Alexander Adams, Amy Gaskin, Bree Gatica-Wilcox, Jason Coombes, Joel Southgate, Lauren Gilbert, Lee Graham, Nicole Pacchiarini, Sara Kumziene-Summerhayes, Sarah Taylor, Sophie Jones, Sara Rey, Matthew Bull, Joanne Watkins, Sally Corden, Tom Connor |
| EPI_ISL_3296677 | Viollier AG | Department of Biosystems Science and Engineering, ETH Zürich | Christian Beisel, Sarah Nadeau, Chaoran Chen, Ivan Topolsky, Kim Philipp Jablonski, Lara Fuhrmann, Rebecca Denes, Mirjam Feldkamp, Ina Nissen, Natascha Santacroce, Elodie Burcklen, Christiane Beckmann, Maurice Redondo, Olivier Kobel, Christoph Noppen, Niko Beerenwinkel, Tanja Stadler |
| EPI_ISL_3296678 | Wales Specialist Virology Centre Sequencing lab: Pathogen Genomics Unit | Public Health Wales Microbiology Cardiff Wales Specialist Virology Centre | Catherine Moore, Johnathan Evans, Laura Gifford, Malorie Perry, Simon Cottrell, Angela Marchbank, Alec Birchley, Alexander Adams, Amy Gaskin, Bree Gatica-Wilcox, Jason Coombes, Joel Southgate, Lauren Gilbert, Lee Graham, Nicole Pacchiarini, Sara Kumziene-Summerhayes, Sarah Taylor, Sophie Jones, Sara Rey, Matthew Bull, Joanne Watkins, Sally Corden, Tom Connor |
| EPI_ISL_3296679 | Viollier AG | Department of Biosystems Science and Engineering, ETH Zürich | Christian Beisel, Sarah Nadeau, Chaoran Chen, Ivan Topolsky, Kim Philipp Jablonski, Lara Fuhrmann, Rebecca Denes, Mirjam Feldkamp, Ina Nissen, Natascha Santacroce, Elodie Burcklen, Christiane Beckmann, Maurice Redondo, Olivier Kobel, Christoph Noppen, Niko Beerenwinkel, Tanja Stadler |
| EPI_ISL_3296680, EPI_ISL_3296681 | Wales Specialist Virology Centre Sequencing lab: Pathogen Genomics Unit | Public Health Wales Microbiology Cardiff Wales Specialist Virology Centre | Catherine Moore, Johnathan Evans, Laura Gifford, Malorie Perry, Simon Cottrell, Angela Marchbank, Alec Birchley, Alexander Adams, Amy Gaskin, Bree Gatica-Wilcox, Jason Coombes, Joel Southgate, Lauren Gilbert, Lee Graham, Nicole Pacchiarini, Sara Kumziene-Summerhayes, Sarah Taylor, Sophie Jones, Sara Rey, Matthew Bull, Joanne Watkins, Sally Corden, Tom Connor |
| EPI_ISL_3296682 | Viollier AG | Department of Biosystems Science and Engineering, ETH Zürich | Christian Beisel, Sarah Nadeau, Chaoran Chen, Ivan Topolsky, Kim Philipp Jablonski, Lara Fuhrmann, Rebecca Denes, Mirjam Feldkamp, Ina Nissen, Natascha Santacroce, Elodie Burcklen, Christiane Beckmann, Maurice Redondo, Olivier Kobel, Christoph Noppen, Niko Beerenwinkel, Tanja Stadler |
| EPI_ISL_3296683 | Wales Specialist Virology Centre Sequencing lab: Pathogen Genomics Unit | Public Health Wales Microbiology Cardiff Wales Specialist Virology Centre | Catherine Moore, Johnathan Evans, Laura Gifford, Malorie Perry, Simon Cottrell, Angela Marchbank, Alec Birchley, Alexander Adams, Amy Gaskin, Bree Gatica-Wilcox, Jason Coombes, Joel Southgate, Lauren Gilbert, Lee Graham, Nicole Pacchiarini, Sara Kumziene-Summerhayes, Sarah Taylor, Sophie Jones, Sara Rey, Matthew Bull, Joanne Watkins, Sally Corden, Tom Connor |
| EPI_ISL_3296684 | Viollier AG | Department of Biosystems Science and Engineering, ETH Zürich | Christian Beisel, Sarah Nadeau, Chaoran Chen, Ivan Topolsky, Kim Philipp Jablonski, Lara Fuhrmann, Rebecca Denes, Mirjam Feldkamp, Ina Nissen, Natascha Santacroce, Elodie Burcklen, Christiane Beckmann, Maurice Redondo, Olivier Kobel, Christoph Noppen, Niko Beerenwinkel, Tanja Stadler |
| EPI_ISL_3296685 | Wales Specialist Virology Centre Sequencing lab: Pathogen Genomics Unit | Public Health Wales Microbiology Cardiff Wales Specialist Virology Centre | Catherine Moore, Johnathan Evans, Laura Gifford, Malorie Perry, Simon Cottrell, Angela Marchbank, Alec Birchley, Alexander Adams, Amy Gaskin, Bree Gatica-Wilcox, Jason Coombes, Joel Southgate, Lauren Gilbert, Lee Graham, Nicole Pacchiarini, Sara Kumziene-Summerhayes, Sarah Taylor, Sophie Jones, Sara Rey, Matthew Bull, Joanne Watkins, Sally Corden, Tom Connor |
| EPI_ISL_3296686 | Viollier AG | Department of Biosystems Science and Engineering, ETH Zürich | Christian Beisel, Sarah Nadeau, Chaoran Chen, Ivan Topolsky, Kim Philipp Jablonski, Lara Fuhrmann, Rebecca Denes, Mirjam Feldkamp, Ina Nissen, Natascha Santacroce, Elodie Burcklen, Christiane Beckmann, Maurice Redondo, Olivier Kobel, Christoph Noppen, Niko Beerenwinkel, Tanja Stadler |
| EPI_ISL_3296687, EPI_ISL_3296688 | Wales Specialist Virology Centre Sequencing lab: Pathogen Genomics Unit | Public Health Wales Microbiology Cardiff Wales Specialist Virology Centre | Catherine Moore, Johnathan Evans, Laura Gifford, Malorie Perry, Simon Cottrell, Angela Marchbank, Alec Birchley, Alexander Adams, Amy Gaskin, Bree Gatica-Wilcox, Jason Coombes, Joel Southgate, Lauren Gilbert, Lee Graham, Nicole Pacchiarini, Sara Kumziene-Summerhayes, Sarah Taylor, Sophie Jones, Sara Rey, Matthew Bull, Joanne Watkins, Sally Corden, Tom Connor |
| EPI_ISL_3296689 | Viollier AG | Department of Biosystems Science and Engineering, ETH Zürich | Christian Beisel, Sarah Nadeau, Chaoran Chen, Ivan Topolsky, Kim Philipp Jablonski, Lara Fuhrmann, Rebecca Denes, Mirjam Feldkamp, Ina Nissen, Natascha Santacroce, Elodie Burcklen, Christiane Beckmann, Maurice Redondo, Olivier Kobel, Christoph Noppen, Niko Beerenwinkel, Tanja Stadler |
| EPI_ISL_3296690 | Wales Specialist Virology Centre Sequencing lab: Pathogen Genomics Unit | Public Health Wales Microbiology Cardiff Wales Specialist Virology Centre | Catherine Moore, Johnathan Evans, Laura Gifford, Malorie Perry, Simon Cottrell, Angela Marchbank, Alec Birchley, Alexander Adams, Amy Gaskin, Bree Gatica-Wilcox, Jason Coombes, Joel Southgate, Lauren Gilbert, Lee Graham, Nicole Pacchiarini, Sara Kumziene-Summerhayes, Sarah Taylor, Sophie Jones, Sara Rey, Matthew Bull, Joanne Watkins, Sally Corden, Tom Connor |
| EPI_ISL_3296691 | Viollier AG | Department of Biosystems Science and Engineering, ETH Zürich | Christian Beisel, Sarah Nadeau, Chaoran Chen, Ivan Topolsky, Kim Philipp Jablonski, Lara Fuhrmann, Rebecca Denes, Mirjam Feldkamp, Ina Nissen, Natascha Santacroce, Elodie Burcklen, Christiane Beckmann, Maurice Redondo, Olivier Kobel, Christoph Noppen, Niko Beerenwinkel, Tanja Stadler |
| EPI_ISL_3296692, EPI_ISL_3296694, EPI_ISL_3296695, EPI_ISL_3296697 | Wales Specialist Virology Centre Sequencing lab: Pathogen Genomics Unit | Public Health Wales Microbiology Cardiff Wales Specialist Virology Centre | Catherine Moore, Johnathan Evans, Laura Gifford, Malorie Perry, Simon Cottrell, Angela Marchbank, Alec Birchley, Alexander Adams, Amy Gaskin, Bree Gatica-Wilcox, Jason Coombes, Joel Southgate, Lauren Gilbert, Lee Graham, Nicole Pacchiarini, Sara Kumziene-Summerhayes, Sarah Taylor, Sophie Jones, Sara Rey, Matthew Bull, Joanne Watkins, Sally Corden, Tom Connor |
| EPI_ISL_3296698 | Viollier AG | Department of Biosystems Science and Engineering, ETH Zürich | Christian Beisel, Sarah Nadeau, Chaoran Chen, Ivan Topolsky, Kim Philipp Jablonski, Lara Fuhrmann, Rebecca Denes, Mirjam Feldkamp, Ina Nissen, Natascha Santacroce, Elodie Burcklen, Christiane Beckmann, Maurice Redondo, Olivier Kobel, Christoph Noppen, Niko Beerenwinkel, Tanja Stadler |
| EPI_ISL_3296699 | Wales Specialist Virology Centre Sequencing lab: Pathogen Genomics Unit | Public Health Wales Microbiology Cardiff Wales Specialist Virology Centre | Catherine Moore, Johnathan Evans, Laura Gifford, Malorie Perry, Simon Cottrell, Angela Marchbank, Alec Birchley, Alexander Adams, Amy Gaskin, Bree Gatica-Wilcox, Jason Coombes, Joel Southgate, Lauren Gilbert, Lee Graham, Nicole Pacchiarini, Sara Kumziene-Summerhayes, Sarah Taylor, Sophie Jones, Sara Rey, Matthew Bull, Joanne Watkins, Sally Corden, Tom Connor |
| EPI_ISL_3296700 | Viollier AG | Department of Biosystems Science and Engineering, ETH Zürich | Christian Beisel, Sarah Nadeau, Chaoran Chen, Ivan Topolsky, Kim Philipp Jablonski, Lara Fuhrmann, Rebecca Denes, Mirjam Feldkamp, Ina Nissen, Natascha Santacroce, Elodie Burcklen, Christiane Beckmann, Maurice Redondo, Olivier Kobel, Christoph Noppen, Niko Beerenwinkel, Tanja Stadler |
| EPI_ISL_3296701, EPI_ISL_3296703 | Wales Specialist Virology Centre Sequencing lab: Pathogen Genomics Unit | Public Health Wales Microbiology Cardiff Wales Specialist Virology Centre | Catherine Moore, Johnathan Evans, Laura Gifford, Malorie Perry, Simon Cottrell, Angela Marchbank, Alec Birchley, Alexander Adams, Amy Gaskin, Bree Gatica-Wilcox, Jason Coombes, Joel Southgate, Lauren Gilbert, Lee Graham, Nicole Pacchiarini, Sara Kumziene-Summerhayes, Sarah Taylor, Sophie Jones, Sara Rey, Matthew Bull, Joanne Watkins, Sally Corden, Tom Connor |
| EPI_ISL_3296704 | Viollier AG | Department of Biosystems Science and Engineering, ETH Zürich | Christian Beisel, Sarah Nadeau, Chaoran Chen, Ivan Topolsky, Kim Philipp Jablonski, Lara Fuhrmann, Rebecca Denes, Mirjam Feldkamp, Ina Nissen, Natascha Santacroce, Elodie Burcklen, Christiane Beckmann, Maurice Redondo, Olivier Kobel, Christoph Noppen, Niko Beerenwinkel, Tanja Stadler |
| EPI_ISL_3296705 | Wales Specialist Virology Centre Sequencing lab: Pathogen Genomics Unit | Public Health Wales Microbiology Cardiff Wales Specialist Virology Centre | Catherine Moore, Johnathan Evans, Laura Gifford, Malorie Perry, Simon Cottrell, Angela Marchbank, Alec Birchley, Alexander Adams, Amy Gaskin, Bree Gatica-Wilcox, Jason Coombes, Joel Southgate, Lauren Gilbert, Lee Graham, Nicole Pacchiarini, Sara Kumziene-Summerhayes, Sarah Taylor, Sophie Jones, Sara Rey, Matthew Bull, Joanne Watkins, Sally Corden, Tom Connor |
| EPI_ISL_3296706 | Viollier AG | Department of Biosystems Science and Engineering, ETH Zürich | Christian Beisel, Sarah Nadeau, Chaoran Chen, Ivan Topolsky, Kim Philipp Jablonski, Lara Fuhrmann, Rebecca Denes, Mirjam Feldkamp, Ina Nissen, Natascha Santacroce, Elodie Burcklen, Christiane Beckmann, Maurice Redondo, Olivier Kobel, Christoph Noppen, Niko Beerenwinkel, Tanja Stadler |
| EPI_ISL_3296707 | Wales Specialist Virology Centre Sequencing lab: Pathogen Genomics Unit | Public Health Wales Microbiology Cardiff Wales Specialist Virology Centre | Catherine Moore, Johnathan Evans, Laura Gifford, Malorie Perry, Simon Cottrell, Angela Marchbank, Alec Birchley, Alexander Adams, Amy Gaskin, Bree Gatica-Wilcox, Jason Coombes, Joel Southgate, Lauren Gilbert, Lee Graham, Nicole Pacchiarini, Sara Kumziene-Summerhayes, Sarah Taylor, Sophie Jones, Sara Rey, Matthew Bull, Joanne Watkins, Sally Corden, Tom Connor |
| EPI_ISL_3296708 | Viollier AG | Department of Biosystems Science and Engineering, ETH Zürich | Christian Beisel, Sarah Nadeau, Chaoran Chen, Ivan Topolsky, Kim Philipp Jablonski, Lara Fuhrmann, Rebecca Denes, Mirjam Feldkamp, Ina Nissen, Natascha Santacroce, Elodie Burcklen, Christiane Beckmann, Maurice Redondo, Olivier Kobel, Christoph Noppen, Niko Beerenwinkel, Tanja Stadler |
| EPI_ISL_3296709 | Wales Specialist Virology Centre Sequencing lab: Pathogen Genomics Unit | Public Health Wales Microbiology Cardiff Wales Specialist Virology Centre | Catherine Moore, Johnathan Evans, Laura Gifford, Malorie Perry, Simon Cottrell, Angela Marchbank, Alec Birchley, Alexander Adams, Amy Gaskin, Bree Gatica-Wilcox, Jason Coombes, Joel Southgate, Lauren Gilbert, Lee Graham, Nicole Pacchiarini, Sara Kumziene-Summerhayes, Sarah Taylor, Sophie Jones, Sara Rey, Matthew Bull, Joanne Watkins, Sally Corden, Tom Connor |
| EPI_ISL_3296710 | Viollier AG | Department of Biosystems Science and Engineering, ETH Zürich | Christian Beisel, Sarah Nadeau, Chaoran Chen, Ivan Topolsky, Kim Philipp Jablonski, Lara Fuhrmann, Rebecca Denes, Mirjam Feldkamp, Ina Nissen, Natascha Santacroce, Elodie Burcklen, Christiane Beckmann, Maurice Redondo, Olivier Kobel, Christoph Noppen, Niko Beerenwinkel, Tanja Stadler |
| EPI_ISL_3296711, EPI_ISL_3296712 | Wales Specialist Virology Centre Sequencing lab: Pathogen Genomics Unit | Public Health Wales Microbiology Cardiff Wales Specialist Virology Centre | Catherine Moore, Johnathan Evans, Laura Gifford, Malorie Perry, Simon Cottrell, Angela Marchbank, Alec Birchley, Alexander Adams, Amy Gaskin, Bree Gatica-Wilcox, Jason Coombes, Joel Southgate, Lauren Gilbert, Lee Graham, Nicole Pacchiarini, Sara Kumziene-Summerhayes, Sarah Taylor, Sophie Jones, Sara Rey, Matthew Bull, Joanne Watkins, Sally Corden, Tom Connor |
| EPI_ISL_3296713 | Viollier AG | Department of Biosystems Science and Engineering, ETH Zürich | Christian Beisel, Sarah Nadeau, Chaoran Chen, Ivan Topolsky, Kim Philipp Jablonski, Lara Fuhrmann, Rebecca Denes, Mirjam Feldkamp, Ina Nissen, Natascha Santacroce, Elodie Burcklen, Christiane Beckmann, Maurice Redondo, Olivier Kobel, Christoph Noppen, Niko Beerenwinkel, Tanja Stadler |
| EPI_ISL_3296714 | Wales Specialist Virology Centre Sequencing lab: Pathogen Genomics Unit | Public Health Wales Microbiology Cardiff Wales Specialist Virology Centre | Catherine Moore, Johnathan Evans, Laura Gifford, Malorie Perry, Simon Cottrell, Angela Marchbank, Alec Birchley, Alexander Adams, Amy Gaskin, Bree Gatica-Wilcox, Jason Coombes, Joel Southgate, Lauren Gilbert, Lee Graham, Nicole Pacchiarini, Sara Kumziene-Summerhayes, Sarah Taylor, Sophie Jones, Sara Rey, Matthew Bull, Joanne Watkins, Sally Corden, Tom Connor |
| EPI_ISL_3296715 | Viollier AG | Department of Biosystems Science and Engineering, ETH Zürich | Christian Beisel, Sarah Nadeau, Chaoran Chen, Ivan Topolsky, Kim Philipp Jablonski, Lara Fuhrmann, Rebecca Denes, Mirjam Feldkamp, Ina Nissen, Natascha Santacroce, Elodie Burcklen, Christiane Beckmann, Maurice Redondo, Olivier Kobel, Christoph Noppen, Niko Beerenwinkel, Tanja Stadler |
| EPI_ISL_3296716 | Wales Specialist Virology Centre Sequencing lab: Pathogen Genomics Unit | Public Health Wales Microbiology Cardiff Wales Specialist Virology Centre | Catherine Moore, Johnathan Evans, Laura Gifford, Malorie Perry, Simon Cottrell, Angela Marchbank, Alec Birchley, Alexander Adams, Amy Gaskin, Bree Gatica-Wilcox, Jason Coombes, Joel Southgate, Lauren Gilbert, Lee Graham, Nicole Pacchiarini, Sara Kumziene-Summerhayes, Sarah Taylor, Sophie Jones, Sara Rey, Matthew Bull, Joanne Watkins, Sally Corden, Tom Connor |
| EPI_ISL_3296717 | Viollier AG | Department of Biosystems Science and Engineering, ETH Zürich | Christian Beisel, Sarah Nadeau, Chaoran Chen, Ivan Topolsky, Kim Philipp Jablonski, Lara Fuhrmann, Rebecca Denes, Mirjam Feldkamp, Ina Nissen, Natascha Santacroce, Elodie Burcklen, Christiane Beckmann, Maurice Redondo, Olivier Kobel, Christoph Noppen, Niko Beerenwinkel, Tanja Stadler |
| EPI_ISL_3296718 | Wales Specialist Virology Centre Sequencing lab: Pathogen Genomics Unit | Public Health Wales Microbiology Cardiff Wales Specialist Virology Centre | Catherine Moore, Johnathan Evans, Laura Gifford, Malorie Perry, Simon Cottrell, Angela Marchbank, Alec Birchley, Alexander Adams, Amy Gaskin, Bree Gatica-Wilcox, Jason Coombes, Joel Southgate, Lauren Gilbert, Lee Graham, Nicole Pacchiarini, Sara Kumziene-Summerhayes, Sarah Taylor, Sophie Jones, Sara Rey, Matthew Bull, Joanne Watkins, Sally Corden, Tom Connor |
| EPI_ISL_3296719 | Viollier AG | Department of Biosystems Science and Engineering, ETH Zürich | Christian Beisel, Sarah Nadeau, Chaoran Chen, Ivan Topolsky, Kim Philipp Jablonski, Lara Fuhrmann, Rebecca Denes, Mirjam Feldkamp, Ina Nissen, Natascha Santacroce, Elodie Burcklen, Christiane Beckmann, Maurice Redondo, Olivier Kobel, Christoph Noppen, Niko Beerenwinkel, Tanja Stadler |
| EPI_ISL_3296720 | Wales Specialist Virology Centre Sequencing lab: Pathogen Genomics Unit | Public Health Wales Microbiology Cardiff Wales Specialist Virology Centre | Catherine Moore, Johnathan Evans, Laura Gifford, Malorie Perry, Simon Cottrell, Angela Marchbank, Alec Birchley, Alexander Adams, Amy Gaskin, Bree Gatica-Wilcox, Jason Coombes, Joel Southgate, Lauren Gilbert, Lee Graham, Nicole Pacchiarini, Sara Kumziene-Summerhayes, Sarah Taylor, Sophie Jones, Sara Rey, Matthew Bull, Joanne Watkins, Sally Corden, Tom Connor |
| EPI_ISL_3296721 | Viollier AG | Department of Biosystems Science and Engineering, ETH Zürich | Christian Beisel, Sarah Nadeau, Chaoran Chen, Ivan Topolsky, Kim Philipp Jablonski, Lara Fuhrmann, Rebecca Denes, Mirjam Feldkamp, Ina Nissen, Natascha Santacroce, Elodie Burcklen, Christiane Beckmann, Maurice Redondo, Olivier Kobel, Christoph Noppen, Niko Beerenwinkel, Tanja Stadler |
| EPI_ISL_3296722 | Wales Specialist Virology Centre Sequencing lab: Pathogen | Public Health Wales Microbiology Cardiff Wales Specialist Virology | Catherine Moore, Johnathan Evans, Laura Gifford, Malorie Perry, Simon Cottrell, Angela Marchbank, Alec Birchley, Alexander Adams, Amy Gaskin, Bree Gatica-Wilcox, Jason Coombes, Joel Southgate, Lauren Gilbert, Lee Graham, Nicole |

| | | | |
|---|---|---|---|
| | Genomics Unit | Centre | Pacchiarini, Sara Kumziene-Summerhayes, Sarah Taylor, Sophie Jones, Sara Rey, Matthew Bull, Joanne Watkins, Sally Corden, Tom Connor |
| EPI_ISL_3296723 | Viollier AG | Department of Biosystems Science and Engineering, ETH Zürich | Christian Beisel, Sarah Nadeau, Chaoran Chen, Ivan Topolsky, Kim Philipp Jablonski, Lara Fuhrmann, Rebecca Denes, Mirjam Feldkamp, Ina Nissen, Natascha Santacroce, Elodie Burcklen, Christiane Beckmann, Maurice Redondo, Olivier Kobel, Christoph Noppen, Niko Beerenwinkel, Tanja Stadler |
| EPI_ISL_3296724 | Wales Specialist Virology Centre Sequencing lab: Pathogen Genomics Unit | Public Health Wales Microbiology Cardiff Wales Specialist Virology Centre | Catherine Moore, Johnathan Evans, Laura Gifford, Malorie Perry, Simon Cottrell, Angela Marchbank, Alec Birchley, Alexander Adams, Amy Gaskin, Bree Gatica-Wilcox, Jason Coombes, Joel Southgate, Lauren Gilbert, Lee Graham, Nicole Pacchiarini, Sara Kumziene-Summerhayes, Sarah Taylor, Sophie Jones, Sara Rey, Matthew Bull, Joanne Watkins, Sally Corden, Tom Connor |
| EPI_ISL_3296725 | Viollier AG | Department of Biosystems Science and Engineering, ETH Zürich | Christian Beisel, Sarah Nadeau, Chaoran Chen, Ivan Topolsky, Kim Philipp Jablonski, Lara Fuhrmann, Rebecca Denes, Mirjam Feldkamp, Ina Nissen, Natascha Santacroce, Elodie Burcklen, Christiane Beckmann, Maurice Redondo, Olivier Kobel, Christoph Noppen, Niko Beerenwinkel, Tanja Stadler |
| EPI_ISL_3296726 | Wales Specialist Virology Centre Sequencing lab: Pathogen Genomics Unit | Public Health Wales Microbiology Cardiff Wales Specialist Virology Centre | Catherine Moore, Johnathan Evans, Laura Gifford, Malorie Perry, Simon Cottrell, Angela Marchbank, Alec Birchley, Alexander Adams, Amy Gaskin, Bree Gatica-Wilcox, Jason Coombes, Joel Southgate, Lauren Gilbert, Lee Graham, Nicole Pacchiarini, Sara Kumziene-Summerhayes, Sarah Taylor, Sophie Jones, Sara Rey, Matthew Bull, Joanne Watkins, Sally Corden, Tom Connor |
| EPI_ISL_3296727 | Viollier AG | Department of Biosystems Science and Engineering, ETH Zürich | Christian Beisel, Sarah Nadeau, Chaoran Chen, Ivan Topolsky, Kim Philipp Jablonski, Lara Fuhrmann, Rebecca Denes, Mirjam Feldkamp, Ina Nissen, Natascha Santacroce, Elodie Burcklen, Christiane Beckmann, Maurice Redondo, Olivier Kobel, Christoph Noppen, Niko Beerenwinkel, Tanja Stadler |
| EPI_ISL_3296728 | Wales Specialist Virology Centre Sequencing lab: Pathogen Genomics Unit | Public Health Wales Microbiology Cardiff Wales Specialist Virology Centre | Catherine Moore, Johnathan Evans, Laura Gifford, Malorie Perry, Simon Cottrell, Angela Marchbank, Alec Birchley, Alexander Adams, Amy Gaskin, Bree Gatica-Wilcox, Jason Coombes, Joel Southgate, Lauren Gilbert, Lee Graham, Nicole Pacchiarini, Sara Kumziene-Summerhayes, Sarah Taylor, Sophie Jones, Sara Rey, Matthew Bull, Joanne Watkins, Sally Corden, Tom Connor |
| EPI_ISL_3296729 | Viollier AG | Department of Biosystems Science and Engineering, ETH Zürich | Christian Beisel, Sarah Nadeau, Chaoran Chen, Ivan Topolsky, Kim Philipp Jablonski, Lara Fuhrmann, Rebecca Denes, Mirjam Feldkamp, Ina Nissen, Natascha Santacroce, Elodie Burcklen, Christiane Beckmann, Maurice Redondo, Olivier Kobel, Christoph Noppen, Niko Beerenwinkel, Tanja Stadler |
| EPI_ISL_3296730 | Wales Specialist Virology Centre Sequencing lab: Pathogen Genomics Unit | Public Health Wales Microbiology Cardiff Wales Specialist Virology Centre | Catherine Moore, Johnathan Evans, Laura Gifford, Malorie Perry, Simon Cottrell, Angela Marchbank, Alec Birchley, Alexander Adams, Amy Gaskin, Bree Gatica-Wilcox, Jason Coombes, Joel Southgate, Lauren Gilbert, Lee Graham, Nicole Pacchiarini, Sara Kumziene-Summerhayes, Sarah Taylor, Sophie Jones, Sara Rey, Matthew Bull, Joanne Watkins, Sally Corden, Tom Connor |
| EPI_ISL_3296731 | Viollier AG | Department of Biosystems Science and Engineering, ETH Zürich | Christian Beisel, Sarah Nadeau, Chaoran Chen, Ivan Topolsky, Kim Philipp Jablonski, Lara Fuhrmann, Rebecca Denes, Mirjam Feldkamp, Ina Nissen, Natascha Santacroce, Elodie Burcklen, Christiane Beckmann, Maurice Redondo, Olivier Kobel, Christoph Noppen, Niko Beerenwinkel, Tanja Stadler |
| EPI_ISL_3296732 | Wales Specialist Virology Centre Sequencing lab: Pathogen Genomics Unit | Public Health Wales Microbiology Cardiff Wales Specialist Virology Centre | Catherine Moore, Johnathan Evans, Laura Gifford, Malorie Perry, Simon Cottrell, Angela Marchbank, Alec Birchley, Alexander Adams, Amy Gaskin, Bree Gatica-Wilcox, Jason Coombes, Joel Southgate, Lauren Gilbert, Lee Graham, Nicole Pacchiarini, Sara Kumziene-Summerhayes, Sarah Taylor, Sophie Jones, Sara Rey, Matthew Bull, Joanne Watkins, Sally Corden, Tom Connor |
| EPI_ISL_3296733 | Viollier AG | Department of Biosystems Science and Engineering, ETH Zürich | Christian Beisel, Sarah Nadeau, Chaoran Chen, Ivan Topolsky, Kim Philipp Jablonski, Lara Fuhrmann, Rebecca Denes, Mirjam Feldkamp, Ina Nissen, Natascha Santacroce, Elodie Burcklen, Christiane Beckmann, Maurice Redondo, Olivier Kobel, Christoph Noppen, Niko Beerenwinkel, Tanja Stadler |
| EPI_ISL_3296734, EPI_ISL_3296736 | Wales Specialist Virology Centre Sequencing lab: Pathogen Genomics Unit | Public Health Wales Microbiology Cardiff Wales Specialist Virology Centre | Catherine Moore, Johnathan Evans, Laura Gifford, Malorie Perry, Simon Cottrell, Angela Marchbank, Alec Birchley, Alexander Adams, Amy Gaskin, Bree Gatica-Wilcox, Jason Coombes, Joel Southgate, Lauren Gilbert, Lee Graham, Nicole Pacchiarini, Sara Kumziene-Summerhayes, Sarah Taylor, Sophie Jones, Sara Rey, Matthew Bull, Joanne Watkins, Sally Corden, Tom Connor |
| EPI_ISL_3296737 | Viollier AG | Department of Biosystems Science and Engineering, ETH Zürich | Christian Beisel, Sarah Nadeau, Chaoran Chen, Ivan Topolsky, Kim Philipp Jablonski, Lara Fuhrmann, Rebecca Denes, Mirjam Feldkamp, Ina Nissen, Natascha Santacroce, Elodie Burcklen, Christiane Beckmann, Maurice Redondo, Olivier Kobel, Christoph Noppen, Niko Beerenwinkel, Tanja Stadler |
| EPI_ISL_3296738 | Wales Specialist Virology Centre Sequencing lab: Pathogen Genomics Unit | Public Health Wales Microbiology Cardiff Wales Specialist Virology Centre | Catherine Moore, Johnathan Evans, Laura Gifford, Malorie Perry, Simon Cottrell, Angela Marchbank, Alec Birchley, Alexander Adams, Amy Gaskin, Bree Gatica-Wilcox, Jason Coombes, Joel Southgate, Lauren Gilbert, Lee Graham, Nicole Pacchiarini, Sara Kumziene-Summerhayes, Sarah Taylor, Sophie Jones, Sara Rey, Matthew Bull, Joanne Watkins, Sally Corden, Tom Connor |
| EPI_ISL_3296739 | Viollier AG | Department of Biosystems Science and Engineering, ETH Zürich | Christian Beisel, Sarah Nadeau, Chaoran Chen, Ivan Topolsky, Kim Philipp Jablonski, Lara Fuhrmann, Rebecca Denes, Mirjam Feldkamp, Ina Nissen, Natascha Santacroce, Elodie Burcklen, Christiane Beckmann, Maurice Redondo, Olivier Kobel, Christoph Noppen, Niko Beerenwinkel, Tanja Stadler |
| EPI_ISL_3296740 | Wales Specialist Virology Centre Sequencing lab: Pathogen Genomics Unit | Public Health Wales Microbiology Cardiff Wales Specialist Virology Centre | Catherine Moore, Johnathan Evans, Laura Gifford, Malorie Perry, Simon Cottrell, Angela Marchbank, Alec Birchley, Alexander Adams, Amy Gaskin, Bree Gatica-Wilcox, Jason Coombes, Joel Southgate, Lauren Gilbert, Lee Graham, Nicole Pacchiarini, Sara Kumziene-Summerhayes, Sarah Taylor, Sophie Jones, Sara Rey, Matthew Bull, Joanne Watkins, Sally Corden, Tom Connor |
| EPI_ISL_3296741 | Viollier AG | Department of Biosystems Science and Engineering, ETH Zürich | Christian Beisel, Sarah Nadeau, Chaoran Chen, Ivan Topolsky, Kim Philipp Jablonski, Lara Fuhrmann, Rebecca Denes, Mirjam Feldkamp, Ina Nissen, Natascha Santacroce, Elodie Burcklen, Christiane Beckmann, Maurice Redondo, Olivier Kobel, Christoph Noppen, Niko Beerenwinkel, Tanja Stadler |
| EPI_ISL_3296742 | Wales Specialist Virology Centre Sequencing lab: Pathogen Genomics Unit | Public Health Wales Microbiology Cardiff Wales Specialist Virology Centre | Catherine Moore, Johnathan Evans, Laura Gifford, Malorie Perry, Simon Cottrell, Angela Marchbank, Alec Birchley, Alexander Adams, Amy Gaskin, Bree Gatica-Wilcox, Jason Coombes, Joel Southgate, Lauren Gilbert, Lee Graham, Nicole Pacchiarini, Sara Kumziene-Summerhayes, Sarah Taylor, Sophie Jones, Sara Rey, Matthew Bull, Joanne Watkins, Sally Corden, Tom Connor |
| EPI_ISL_3296743 | Viollier AG | Department of Biosystems Science and Engineering, ETH Zürich | Christian Beisel, Sarah Nadeau, Chaoran Chen, Ivan Topolsky, Kim Philipp Jablonski, Lara Fuhrmann, Rebecca Denes, Mirjam Feldkamp, Ina Nissen, Natascha Santacroce, Elodie Burcklen, Christiane Beckmann, Maurice Redondo, Olivier Kobel, Christoph Noppen, Niko Beerenwinkel, Tanja Stadler |
| EPI_ISL_3296744 | Wales Specialist Virology Centre Sequencing lab: Pathogen Genomics Unit | Public Health Wales Microbiology Cardiff Wales Specialist Virology Centre | Catherine Moore, Johnathan Evans, Laura Gifford, Malorie Perry, Simon Cottrell, Angela Marchbank, Alec Birchley, Alexander Adams, Amy Gaskin, Bree Gatica-Wilcox, Jason Coombes, Joel Southgate, Lauren Gilbert, Lee Graham, Nicole Pacchiarini, Sara Kumziene-Summerhayes, Sarah Taylor, Sophie Jones, Sara Rey, Matthew Bull, Joanne Watkins, Sally Corden, Tom Connor |
| EPI_ISL_3296745 | Viollier AG | Department of Biosystems Science and Engineering, ETH Zürich | Christian Beisel, Sarah Nadeau, Chaoran Chen, Ivan Topolsky, Kim Philipp Jablonski, Lara Fuhrmann, Rebecca Denes, Mirjam Feldkamp, Ina Nissen, Natascha Santacroce, Elodie Burcklen, Christiane Beckmann, Maurice Redondo, Olivier Kobel, Christoph Noppen, Niko Beerenwinkel, Tanja Stadler |
| EPI_ISL_3296746 | Wales Specialist Virology Centre Sequencing lab: Pathogen Genomics Unit | Public Health Wales Microbiology Cardiff Wales Specialist Virology Centre | Catherine Moore, Johnathan Evans, Laura Gifford, Malorie Perry, Simon Cottrell, Angela Marchbank, Alec Birchley, Alexander Adams, Amy Gaskin, Bree Gatica-Wilcox, Jason Coombes, Joel Southgate, Lauren Gilbert, Lee Graham, Nicole Pacchiarini, Sara Kumziene-Summerhayes, Sarah Taylor, Sophie Jones, Sara Rey, Matthew Bull, Joanne Watkins, Sally Corden, Tom Connor |
| EPI_ISL_3296747 | Viollier AG | Department of Biosystems Science and Engineering, ETH Zürich | Christian Beisel, Sarah Nadeau, Chaoran Chen, Ivan Topolsky, Kim Philipp Jablonski, Lara Fuhrmann, Rebecca Denes, Mirjam Feldkamp, Ina Nissen, Natascha Santacroce, Elodie Burcklen, Christiane Beckmann, Maurice Redondo, Olivier Kobel, Christoph Noppen, Niko Beerenwinkel, Tanja Stadler |
| EPI_ISL_3296748 | Wales Specialist Virology Centre Sequencing lab: Pathogen Genomics Unit | Public Health Wales Microbiology Cardiff Wales Specialist Virology Centre | Catherine Moore, Johnathan Evans, Laura Gifford, Malorie Perry, Simon Cottrell, Angela Marchbank, Alec Birchley, Alexander Adams, Amy Gaskin, Bree Gatica-Wilcox, Jason Coombes, Joel Southgate, Lauren Gilbert, Lee Graham, Nicole Pacchiarini, Sara Kumziene-Summerhayes, Sarah Taylor, Sophie Jones, Sara Rey, Matthew Bull, Joanne Watkins, Sally Corden, Tom Connor |
| EPI_ISL_3296749 | Viollier AG | Department of Biosystems Science and Engineering, ETH Zürich | Christian Beisel, Sarah Nadeau, Chaoran Chen, Ivan Topolsky, Kim Philipp Jablonski, Lara Fuhrmann, Rebecca Denes, Mirjam Feldkamp, Ina Nissen, Natascha Santacroce, Elodie Burcklen, Christiane Beckmann, Maurice Redondo, Olivier Kobel, Christoph Noppen, Niko Beerenwinkel, Tanja Stadler |
| EPI_ISL_3296750 | Wales Specialist Virology Centre Sequencing lab: Pathogen Genomics Unit | Public Health Wales Microbiology Cardiff Wales Specialist Virology Centre | Catherine Moore, Johnathan Evans, Laura Gifford, Malorie Perry, Simon Cottrell, Angela Marchbank, Alec Birchley, Alexander Adams, Amy Gaskin, Bree Gatica-Wilcox, Jason Coombes, Joel Southgate, Lauren Gilbert, Lee Graham, Nicole Pacchiarini, Sara Kumziene-Summerhayes, Sarah Taylor, Sophie Jones, Sara Rey, Matthew Bull, Joanne Watkins, Sally Corden, Tom Connor |
| EPI_ISL_3296751 | Viollier AG | Department of Biosystems Science and Engineering, ETH Zürich | Christian Beisel, Sarah Nadeau, Chaoran Chen, Ivan Topolsky, Kim Philipp Jablonski, Lara Fuhrmann, Rebecca Denes, Mirjam Feldkamp, Ina Nissen, Natascha Santacroce, Elodie Burcklen, Christiane Beckmann, Maurice Redondo, Olivier Kobel, Christoph Noppen, Niko Beerenwinkel, Tanja Stadler |
| EPI_ISL_3296752 | Wales Specialist Virology Centre Sequencing lab: Pathogen Genomics Unit | Public Health Wales Microbiology Cardiff Wales Specialist Virology Centre | Catherine Moore, Johnathan Evans, Laura Gifford, Malorie Perry, Simon Cottrell, Angela Marchbank, Alec Birchley, Alexander Adams, Amy Gaskin, Bree Gatica-Wilcox, Jason Coombes, Joel Southgate, Lauren Gilbert, Lee Graham, Nicole Pacchiarini, Sara Kumziene-Summerhayes, Sarah Taylor, Sophie Jones, Sara Rey, Matthew Bull, Joanne Watkins, Sally Corden, Tom Connor |
| EPI_ISL_3296753 | Viollier AG | Department of Biosystems Science and Engineering, ETH Zürich | Christian Beisel, Sarah Nadeau, Chaoran Chen, Ivan Topolsky, Kim Philipp Jablonski, Lara Fuhrmann, Rebecca Denes, Mirjam Feldkamp, Ina Nissen, Natascha Santacroce, Elodie Burcklen, Christiane Beckmann, Maurice Redondo, Olivier Kobel, Christoph Noppen, Niko Beerenwinkel, Tanja Stadler |
| EPI_ISL_3296754, EPI_ISL_3296756 | Wales Specialist Virology Centre Sequencing lab: Pathogen Genomics Unit | Public Health Wales Microbiology Cardiff Wales Specialist Virology Centre | Catherine Moore, Johnathan Evans, Laura Gifford, Malorie Perry, Simon Cottrell, Angela Marchbank, Alec Birchley, Alexander Adams, Amy Gaskin, Bree Gatica-Wilcox, Jason Coombes, Joel Southgate, Lauren Gilbert, Lee Graham, Nicole Pacchiarini, Sara Kumziene-Summerhayes, Sarah Taylor, Sophie Jones, Sara Rey, Matthew Bull, Joanne Watkins, Sally Corden, Tom Connor |
| EPI_ISL_3296757 | Viollier AG | Department of Biosystems Science and Engineering, ETH Zürich | Christian Beisel, Sarah Nadeau, Chaoran Chen, Ivan Topolsky, Kim Philipp Jablonski, Lara Fuhrmann, Rebecca Denes, Mirjam Feldkamp, Ina Nissen, Natascha Santacroce, Elodie Burcklen, Christiane Beckmann, Maurice Redondo, Olivier Kobel, Christoph Noppen, Niko Beerenwinkel, Tanja Stadler |
| EPI_ISL_3296758, EPI_ISL_3296760 | Wales Specialist Virology Centre Sequencing lab: Pathogen Genomics Unit | Public Health Wales Microbiology Cardiff Wales Specialist Virology Centre | Catherine Moore, Johnathan Evans, Laura Gifford, Malorie Perry, Simon Cottrell, Angela Marchbank, Alec Birchley, Alexander Adams, Amy Gaskin, Bree Gatica-Wilcox, Jason Coombes, Joel Southgate, Lauren Gilbert, Lee Graham, Nicole Pacchiarini, Sara Kumziene-Summerhayes, Sarah Taylor, Sophie Jones, Sara Rey, Matthew Bull, Joanne Watkins, Sally Corden, Tom Connor |
| EPI_ISL_3296761 | Viollier AG | Department of Biosystems Science and Engineering, ETH Zürich | Christian Beisel, Sarah Nadeau, Chaoran Chen, Ivan Topolsky, Kim Philipp Jablonski, Lara Fuhrmann, Rebecca Denes, Mirjam Feldkamp, Ina Nissen, Natascha Santacroce, Elodie Burcklen, Christiane Beckmann, Maurice Redondo, Olivier Kobel, Christoph Noppen, Niko Beerenwinkel, Tanja Stadler |
| EPI_ISL_3296762 | Wales Specialist Virology Centre Sequencing lab: Pathogen Genomics Unit | Public Health Wales Microbiology Cardiff Wales Specialist Virology Centre | Catherine Moore, Johnathan Evans, Laura Gifford, Malorie Perry, Simon Cottrell, Angela Marchbank, Alec Birchley, Alexander Adams, Amy Gaskin, Bree Gatica-Wilcox, Jason Coombes, Joel Southgate, Lauren Gilbert, Lee Graham, Nicole Pacchiarini, Sara Kumziene-Summerhayes, Sarah Taylor, Sophie Jones, Sara Rey, Matthew Bull, Joanne Watkins, Sally Corden, Tom Connor |
| EPI_ISL_3296763 | Viollier AG | Department of Biosystems Science and Engineering, ETH Zürich | Christian Beisel, Sarah Nadeau, Chaoran Chen, Ivan Topolsky, Kim Philipp Jablonski, Lara Fuhrmann, Rebecca Denes, Mirjam Feldkamp, Ina Nissen, Natascha Santacroce, Elodie Burcklen, Christiane Beckmann, Maurice Redondo, Olivier Kobel, Christoph Noppen, Niko Beerenwinkel, Tanja Stadler |
| EPI_ISL_3296764 | Wales Specialist Virology Centre Sequencing lab: Pathogen Genomics Unit | Public Health Wales Microbiology Cardiff Wales Specialist Virology Centre | Catherine Moore, Johnathan Evans, Laura Gifford, Malorie Perry, Simon Cottrell, Angela Marchbank, Alec Birchley, Alexander Adams, Amy Gaskin, Bree Gatica-Wilcox, Jason Coombes, Joel Southgate, Lauren Gilbert, Lee Graham, Nicole Pacchiarini, Sara Kumziene-Summerhayes, Sarah Taylor, Sophie Jones, Sara Rey, Matthew Bull, Joanne Watkins, Sally Corden, Tom Connor |
| EPI_ISL_3296765 | Viollier AG | Department of Biosystems Science and Engineering, ETH Zürich | Christian Beisel, Sarah Nadeau, Chaoran Chen, Ivan Topolsky, Kim Philipp Jablonski, Lara Fuhrmann, Rebecca Denes, Mirjam Feldkamp, Ina Nissen, Natascha Santacroce, Elodie Burcklen, Christiane Beckmann, Maurice Redondo, Olivier Kobel, Christoph Noppen, Niko Beerenwinkel, Tanja Stadler |
| EPI_ISL_3296766 | Wales Specialist Virology Centre Sequencing lab: Pathogen Genomics Unit | Public Health Wales Microbiology Cardiff Wales Specialist Virology Centre | Catherine Moore, Johnathan Evans, Laura Gifford, Malorie Perry, Simon Cottrell, Angela Marchbank, Alec Birchley, Alexander Adams, Amy Gaskin, Bree Gatica-Wilcox, Jason Coombes, Joel Southgate, Lauren Gilbert, Lee Graham, Nicole Pacchiarini, Sara Kumziene-Summerhayes, Sarah Taylor, Sophie Jones, Sara Rey, Matthew Bull, Joanne Watkins, Sally Corden, Tom Connor |
| EPI_ISL_3296767 | Viollier AG | Department of Biosystems Science and Engineering, ETH Zürich | Christian Beisel, Sarah Nadeau, Chaoran Chen, Ivan Topolsky, Kim Philipp Jablonski, Lara Fuhrmann, Rebecca Denes, Mirjam Feldkamp, Ina Nissen, Natascha Santacroce, Elodie Burcklen, Christiane Beckmann, Maurice Redondo, Olivier Kobel, Christoph Noppen, Niko Beerenwinkel, Tanja Stadler |
| EPI_ISL_3296768 | Wales Specialist Virology Centre Sequencing lab: Pathogen Genomics Unit | Public Health Wales Microbiology Cardiff Wales Specialist Virology Centre | Catherine Moore, Johnathan Evans, Laura Gifford, Malorie Perry, Simon Cottrell, Angela Marchbank, Alec Birchley, Alexander Adams, Amy Gaskin, Bree Gatica-Wilcox, Jason Coombes, Joel Southgate, Lauren Gilbert, Lee Graham, Nicole Pacchiarini, Sara Kumziene-Summerhayes, Sarah Taylor, Sophie Jones, Sara Rey, Matthew Bull, Joanne Watkins, Sally Corden, Tom Connor |
| EPI_ISL_3296769 | Viollier AG | Department of Biosystems Science and Engineering, ETH Zürich | Christian Beisel, Sarah Nadeau, Chaoran Chen, Ivan Topolsky, Kim Philipp Jablonski, Lara Fuhrmann, Rebecca Denes, Mirjam Feldkamp, Ina Nissen, Natascha Santacroce, Elodie Burcklen, Christiane Beckmann, Maurice Redondo, Olivier Kobel, Christoph Noppen, Niko Beerenwinkel, Tanja Stadler |
| EPI_ISL_3296770 | Wales Specialist Virology Centre Sequencing lab: Pathogen Genomics Unit | Public Health Wales Microbiology Cardiff Wales Specialist Virology Centre | Catherine Moore, Johnathan Evans, Laura Gifford, Malorie Perry, Simon Cottrell, Angela Marchbank, Alec Birchley, Alexander Adams, Amy Gaskin, Bree Gatica-Wilcox, Jason Coombes, Joel Southgate, Lauren Gilbert, Lee Graham, Nicole Pacchiarini, Sara Kumziene-Summerhayes, Sarah Taylor, Sophie Jones, Sara Rey, Matthew Bull, Joanne Watkins, Sally Corden, Tom Connor |
| EPI_ISL_3296771 | Viollier AG | Department of Biosystems Science and Engineering, ETH Zürich | Christian Beisel, Sarah Nadeau, Chaoran Chen, Ivan Topolsky, Kim Philipp Jablonski, Lara Fuhrmann, Rebecca Denes, Mirjam Feldkamp, Ina Nissen, Natascha Santacroce, Elodie Burcklen, Christiane Beckmann, Maurice Redondo, Olivier Kobel, Christoph Noppen, Niko Beerenwinkel, Tanja Stadler |
| EPI_ISL_3296772 | Wales Specialist Virology Centre Sequencing lab: Pathogen Genomics Unit | Public Health Wales Microbiology Cardiff Wales Specialist Virology Centre | Catherine Moore, Johnathan Evans, Laura Gifford, Malorie Perry, Simon Cottrell, Angela Marchbank, Alec Birchley, Alexander Adams, Amy Gaskin, Bree Gatica-Wilcox, Jason Coombes, Joel Southgate, Lauren Gilbert, Lee Graham, Nicole Pacchiarini, Sara Kumziene-Summerhayes, Sarah Taylor, Sophie Jones, Sara Rey, Matthew Bull, Joanne Watkins, Sally Corden, Tom Connor |
| EPI_ISL_3296773 | Viollier AG | Department of Biosystems Science and Engineering, ETH Zürich | Christian Beisel, Sarah Nadeau, Chaoran Chen, Ivan Topolsky, Kim Philipp Jablonski, Lara Fuhrmann, Rebecca Denes, Mirjam Feldkamp, Ina Nissen, Natascha Santacroce, Elodie Burcklen, Christiane Beckmann, Maurice Redondo, Olivier Kobel, |

| | | | |
|---|---|---|---|
| | | | Christoph Noppen, Niko Beerenwinkel, Tanja Stadler |
| EPI_ISL_3296774, EPI_ISL_3296775 | Wales Specialist Virology Centre Sequencing lab: Pathogen Genomics Unit | Public Health Wales Microbiology Cardiff Wales Specialist Virology Centre | Catherine Moore, Johnathan Evans, Laura Gifford, Malorie Perry, Simon Cottrell, Angela Marchbank, Alec Birchley, Alexander Adams, Amy Gaskin, Bree Gatica-Wilcox, Jason Coombes, Joel Southgate, Lauren Gilbert, Lee Graham, Nicole Pacchiarini, Sara Kumziene-Summerhayes, Sarah Taylor, Sophie Jones, Sara Rey, Matthew Bull, Joanne Watkins, Sally Corden, Tom Connor |
| EPI_ISL_3296776 | Viollier AG | Department of Biosystems Science and Engineering, ETH Zürich | Christian Beisel, Sarah Nadeau, Chaoran Chen, Ivan Topolsky, Kim Philipp Jablonski, Lara Fuhrmann, Rebecca Denes, Mirjam Feldkamp, Ina Nissen, Natascha Santacroce, Elodie Burcklen, Christiane Beckmann, Maurice Redondo, Olivier Kobel, Christoph Noppen, Niko Beerenwinkel, Tanja Stadler |
| EPI_ISL_3296777 | Wales Specialist Virology Centre Sequencing lab: Pathogen Genomics Unit | Public Health Wales Microbiology Cardiff Wales Specialist Virology Centre | Catherine Moore, Johnathan Evans, Laura Gifford, Malorie Perry, Simon Cottrell, Angela Marchbank, Alec Birchley, Alexander Adams, Amy Gaskin, Bree Gatica-Wilcox, Jason Coombes, Joel Southgate, Lauren Gilbert, Lee Graham, Nicole Pacchiarini, Sara Kumziene-Summerhayes, Sarah Taylor, Sophie Jones, Sara Rey, Matthew Bull, Joanne Watkins, Sally Corden, Tom Connor |
| EPI_ISL_3296778 | Viollier AG | Department of Biosystems Science and Engineering, ETH Zürich | Christian Beisel, Sarah Nadeau, Chaoran Chen, Ivan Topolsky, Kim Philipp Jablonski, Lara Fuhrmann, Rebecca Denes, Mirjam Feldkamp, Ina Nissen, Natascha Santacroce, Elodie Burcklen, Christiane Beckmann, Maurice Redondo, Olivier Kobel, Christoph Noppen, Niko Beerenwinkel, Tanja Stadler |
| EPI_ISL_3296779 | Wales Specialist Virology Centre Sequencing lab: Pathogen Genomics Unit | Public Health Wales Microbiology Cardiff Wales Specialist Virology Centre | Catherine Moore, Johnathan Evans, Laura Gifford, Malorie Perry, Simon Cottrell, Angela Marchbank, Alec Birchley, Alexander Adams, Amy Gaskin, Bree Gatica-Wilcox, Jason Coombes, Joel Southgate, Lauren Gilbert, Lee Graham, Nicole Pacchiarini, Sara Kumziene-Summerhayes, Sarah Taylor, Sophie Jones, Sara Rey, Matthew Bull, Joanne Watkins, Sally Corden, Tom Connor |
| EPI_ISL_3296780 | Viollier AG | Department of Biosystems Science and Engineering, ETH Zürich | Christian Beisel, Sarah Nadeau, Chaoran Chen, Ivan Topolsky, Kim Philipp Jablonski, Lara Fuhrmann, Rebecca Denes, Mirjam Feldkamp, Ina Nissen, Natascha Santacroce, Elodie Burcklen, Christiane Beckmann, Maurice Redondo, Olivier Kobel, Christoph Noppen, Niko Beerenwinkel, Tanja Stadler |
| EPI_ISL_3296781 | Wales Specialist Virology Centre Sequencing lab: Pathogen Genomics Unit | Public Health Wales Microbiology Cardiff Wales Specialist Virology Centre | Catherine Moore, Johnathan Evans, Laura Gifford, Malorie Perry, Simon Cottrell, Angela Marchbank, Alec Birchley, Alexander Adams, Amy Gaskin, Bree Gatica-Wilcox, Jason Coombes, Joel Southgate, Lauren Gilbert, Lee Graham, Nicole Pacchiarini, Sara Kumziene-Summerhayes, Sarah Taylor, Sophie Jones, Sara Rey, Matthew Bull, Joanne Watkins, Sally Corden, Tom Connor |
| EPI_ISL_3296782 | Viollier AG | Department of Biosystems Science and Engineering, ETH Zürich | Christian Beisel, Sarah Nadeau, Chaoran Chen, Ivan Topolsky, Kim Philipp Jablonski, Lara Fuhrmann, Rebecca Denes, Mirjam Feldkamp, Ina Nissen, Natascha Santacroce, Elodie Burcklen, Christiane Beckmann, Maurice Redondo, Olivier Kobel, Christoph Noppen, Niko Beerenwinkel, Tanja Stadler |
| EPI_ISL_3296783 | Wales Specialist Virology Centre Sequencing lab: Pathogen Genomics Unit | Public Health Wales Microbiology Cardiff Wales Specialist Virology Centre | Catherine Moore, Johnathan Evans, Laura Gifford, Malorie Perry, Simon Cottrell, Angela Marchbank, Alec Birchley, Alexander Adams, Amy Gaskin, Bree Gatica-Wilcox, Jason Coombes, Joel Southgate, Lauren Gilbert, Lee Graham, Nicole Pacchiarini, Sara Kumziene-Summerhayes, Sarah Taylor, Sophie Jones, Sara Rey, Matthew Bull, Joanne Watkins, Sally Corden, Tom Connor |
| EPI_ISL_3296784 | Viollier AG | Department of Biosystems Science and Engineering, ETH Zürich | Christian Beisel, Sarah Nadeau, Chaoran Chen, Ivan Topolsky, Kim Philipp Jablonski, Lara Fuhrmann, Rebecca Denes, Mirjam Feldkamp, Ina Nissen, Natascha Santacroce, Elodie Burcklen, Christiane Beckmann, Maurice Redondo, Olivier Kobel, Christoph Noppen, Niko Beerenwinkel, Tanja Stadler |
| EPI_ISL_3296785 | Wales Specialist Virology Centre Sequencing lab: Pathogen Genomics Unit | Public Health Wales Microbiology Cardiff Wales Specialist Virology Centre | Catherine Moore, Johnathan Evans, Laura Gifford, Malorie Perry, Simon Cottrell, Angela Marchbank, Alec Birchley, Alexander Adams, Amy Gaskin, Bree Gatica-Wilcox, Jason Coombes, Joel Southgate, Lauren Gilbert, Lee Graham, Nicole Pacchiarini, Sara Kumziene-Summerhayes, Sarah Taylor, Sophie Jones, Sara Rey, Matthew Bull, Joanne Watkins, Sally Corden, Tom Connor |
| EPI_ISL_3296786 | Viollier AG | Department of Biosystems Science and Engineering, ETH Zürich | Christian Beisel, Sarah Nadeau, Chaoran Chen, Ivan Topolsky, Kim Philipp Jablonski, Lara Fuhrmann, Rebecca Denes, Mirjam Feldkamp, Ina Nissen, Natascha Santacroce, Elodie Burcklen, Christiane Beckmann, Maurice Redondo, Olivier Kobel, Christoph Noppen, Niko Beerenwinkel, Tanja Stadler |
| EPI_ISL_3296787 | Wales Specialist Virology Centre Sequencing lab: Pathogen Genomics Unit | Public Health Wales Microbiology Cardiff Wales Specialist Virology Centre | Catherine Moore, Johnathan Evans, Laura Gifford, Malorie Perry, Simon Cottrell, Angela Marchbank, Alec Birchley, Alexander Adams, Amy Gaskin, Bree Gatica-Wilcox, Jason Coombes, Joel Southgate, Lauren Gilbert, Lee Graham, Nicole Pacchiarini, Sara Kumziene-Summerhayes, Sarah Taylor, Sophie Jones, Sara Rey, Matthew Bull, Joanne Watkins, Sally Corden, Tom Connor |
| EPI_ISL_3296788 | Viollier AG | Department of Biosystems Science and Engineering, ETH Zürich | Christian Beisel, Sarah Nadeau, Chaoran Chen, Ivan Topolsky, Kim Philipp Jablonski, Lara Fuhrmann, Rebecca Denes, Mirjam Feldkamp, Ina Nissen, Natascha Santacroce, Elodie Burcklen, Christiane Beckmann, Maurice Redondo, Olivier Kobel, Christoph Noppen, Niko Beerenwinkel, Tanja Stadler |
| EPI_ISL_3296789 | Wales Specialist Virology Centre Sequencing lab: Pathogen Genomics Unit | Public Health Wales Microbiology Cardiff Wales Specialist Virology Centre | Catherine Moore, Johnathan Evans, Laura Gifford, Malorie Perry, Simon Cottrell, Angela Marchbank, Alec Birchley, Alexander Adams, Amy Gaskin, Bree Gatica-Wilcox, Jason Coombes, Joel Southgate, Lauren Gilbert, Lee Graham, Nicole Pacchiarini, Sara Kumziene-Summerhayes, Sarah Taylor, Sophie Jones, Sara Rey, Matthew Bull, Joanne Watkins, Sally Corden, Tom Connor |
| EPI_ISL_3296790 | Viollier AG | Department of Biosystems Science and Engineering, ETH Zürich | Christian Beisel, Sarah Nadeau, Chaoran Chen, Ivan Topolsky, Kim Philipp Jablonski, Lara Fuhrmann, Rebecca Denes, Mirjam Feldkamp, Ina Nissen, Natascha Santacroce, Elodie Burcklen, Christiane Beckmann, Maurice Redondo, Olivier Kobel, Christoph Noppen, Niko Beerenwinkel, Tanja Stadler |
| EPI_ISL_3296791 | Wales Specialist Virology Centre Sequencing lab: Pathogen Genomics Unit | Public Health Wales Microbiology Cardiff Wales Specialist Virology Centre | Catherine Moore, Johnathan Evans, Laura Gifford, Malorie Perry, Simon Cottrell, Angela Marchbank, Alec Birchley, Alexander Adams, Amy Gaskin, Bree Gatica-Wilcox, Jason Coombes, Joel Southgate, Lauren Gilbert, Lee Graham, Nicole Pacchiarini, Sara Kumziene-Summerhayes, Sarah Taylor, Sophie Jones, Sara Rey, Matthew Bull, Joanne Watkins, Sally Corden, Tom Connor |
| EPI_ISL_3296792 | Viollier AG | Department of Biosystems Science and Engineering, ETH Zürich | Christian Beisel, Sarah Nadeau, Chaoran Chen, Ivan Topolsky, Kim Philipp Jablonski, Lara Fuhrmann, Rebecca Denes, Mirjam Feldkamp, Ina Nissen, Natascha Santacroce, Elodie Burcklen, Christiane Beckmann, Maurice Redondo, Olivier Kobel, Christoph Noppen, Niko Beerenwinkel, Tanja Stadler |
| EPI_ISL_3296793 | Wales Specialist Virology Centre Sequencing lab: Pathogen Genomics Unit | Public Health Wales Microbiology Cardiff Wales Specialist Virology Centre | Catherine Moore, Johnathan Evans, Laura Gifford, Malorie Perry, Simon Cottrell, Angela Marchbank, Alec Birchley, Alexander Adams, Amy Gaskin, Bree Gatica-Wilcox, Jason Coombes, Joel Southgate, Lauren Gilbert, Lee Graham, Nicole Pacchiarini, Sara Kumziene-Summerhayes, Sarah Taylor, Sophie Jones, Sara Rey, Matthew Bull, Joanne Watkins, Sally Corden, Tom Connor |
| EPI_ISL_3296794, EPI_ISL_3296795 | Viollier AG | Department of Biosystems Science and Engineering, ETH Zürich | Christian Beisel, Sarah Nadeau, Chaoran Chen, Ivan Topolsky, Kim Philipp Jablonski, Lara Fuhrmann, Rebecca Denes, Mirjam Feldkamp, Ina Nissen, Natascha Santacroce, Elodie Burcklen, Christiane Beckmann, Maurice Redondo, Olivier Kobel, Christoph Noppen, Niko Beerenwinkel, Tanja Stadler |
| EPI_ISL_3296796, EPI_ISL_3296797 | Wales Specialist Virology Centre Sequencing lab: Pathogen Genomics Unit | Public Health Wales Microbiology Cardiff Wales Specialist Virology Centre | Catherine Moore, Johnathan Evans, Laura Gifford, Malorie Perry, Simon Cottrell, Angela Marchbank, Alec Birchley, Alexander Adams, Amy Gaskin, Bree Gatica-Wilcox, Jason Coombes, Joel Southgate, Lauren Gilbert, Lee Graham, Nicole Pacchiarini, Sara Kumziene-Summerhayes, Sarah Taylor, Sophie Jones, Sara Rey, Matthew Bull, Joanne Watkins, Sally Corden, Tom Connor |
| EPI_ISL_3296798 | Viollier AG | Department of Biosystems Science and Engineering, ETH Zürich | Christian Beisel, Sarah Nadeau, Chaoran Chen, Ivan Topolsky, Kim Philipp Jablonski, Lara Fuhrmann, Rebecca Denes, Mirjam Feldkamp, Ina Nissen, Natascha Santacroce, Elodie Burcklen, Christiane Beckmann, Maurice Redondo, Olivier Kobel, Christoph Noppen, Niko Beerenwinkel, Tanja Stadler |
| EPI_ISL_3296799 | Wales Specialist Virology Centre Sequencing lab: Pathogen Genomics Unit | Public Health Wales Microbiology Cardiff Wales Specialist Virology Centre | Catherine Moore, Johnathan Evans, Laura Gifford, Malorie Perry, Simon Cottrell, Angela Marchbank, Alec Birchley, Alexander Adams, Amy Gaskin, Bree Gatica-Wilcox, Jason Coombes, Joel Southgate, Lauren Gilbert, Lee Graham, Nicole Pacchiarini, Sara Kumziene-Summerhayes, Sarah Taylor, Sophie Jones, Sara Rey, Matthew Bull, Joanne Watkins, Sally Corden, Tom Connor |
| EPI_ISL_3296800 | Viollier AG | Department of Biosystems Science and Engineering, ETH Zürich | Christian Beisel, Sarah Nadeau, Chaoran Chen, Ivan Topolsky, Kim Philipp Jablonski, Lara Fuhrmann, Rebecca Denes, Mirjam Feldkamp, Ina Nissen, Natascha Santacroce, Elodie Burcklen, Christiane Beckmann, Maurice Redondo, Olivier Kobel, Christoph Noppen, Niko Beerenwinkel, Tanja Stadler |
| EPI_ISL_3296801 | Wales Specialist Virology Centre Sequencing lab: Pathogen Genomics Unit | Public Health Wales Microbiology Cardiff Wales Specialist Virology Centre | Catherine Moore, Johnathan Evans, Laura Gifford, Malorie Perry, Simon Cottrell, Angela Marchbank, Alec Birchley, Alexander Adams, Amy Gaskin, Bree Gatica-Wilcox, Jason Coombes, Joel Southgate, Lauren Gilbert, Lee Graham, Nicole Pacchiarini, Sara Kumziene-Summerhayes, Sarah Taylor, Sophie Jones, Sara Rey, Matthew Bull, Joanne Watkins, Sally Corden, Tom Connor |
| EPI_ISL_3296802 | Viollier AG | Department of Biosystems Science and Engineering, ETH Zürich | Christian Beisel, Sarah Nadeau, Chaoran Chen, Ivan Topolsky, Kim Philipp Jablonski, Lara Fuhrmann, Rebecca Denes, Mirjam Feldkamp, Ina Nissen, Natascha Santacroce, Elodie Burcklen, Christiane Beckmann, Maurice Redondo, Olivier Kobel, Christoph Noppen, Niko Beerenwinkel, Tanja Stadler |
| EPI_ISL_3296803 | Wales Specialist Virology Centre Sequencing lab: Pathogen Genomics Unit | Public Health Wales Microbiology Cardiff Wales Specialist Virology Centre | Catherine Moore, Johnathan Evans, Laura Gifford, Malorie Perry, Simon Cottrell, Angela Marchbank, Alec Birchley, Alexander Adams, Amy Gaskin, Bree Gatica-Wilcox, Jason Coombes, Joel Southgate, Lauren Gilbert, Lee Graham, Nicole Pacchiarini, Sara Kumziene-Summerhayes, Sarah Taylor, Sophie Jones, Sara Rey, Matthew Bull, Joanne Watkins, Sally Corden, Tom Connor |
| EPI_ISL_3296804 | Viollier AG | Department of Biosystems Science and Engineering, ETH Zürich | Christian Beisel, Sarah Nadeau, Chaoran Chen, Ivan Topolsky, Kim Philipp Jablonski, Lara Fuhrmann, Rebecca Denes, Mirjam Feldkamp, Ina Nissen, Natascha Santacroce, Elodie Burcklen, Christiane Beckmann, Maurice Redondo, Olivier Kobel, Christoph Noppen, Niko Beerenwinkel, Tanja Stadler |
| EPI_ISL_3296805 | Wales Specialist Virology Centre Sequencing lab: Pathogen Genomics Unit | Public Health Wales Microbiology Cardiff Wales Specialist Virology Centre | Catherine Moore, Johnathan Evans, Laura Gifford, Malorie Perry, Simon Cottrell, Angela Marchbank, Alec Birchley, Alexander Adams, Amy Gaskin, Bree Gatica-Wilcox, Jason Coombes, Joel Southgate, Lauren Gilbert, Lee Graham, Nicole Pacchiarini, Sara Kumziene-Summerhayes, Sarah Taylor, Sophie Jones, Sara Rey, Matthew Bull, Joanne Watkins, Sally Corden, Tom Connor |
| EPI_ISL_3296806 | Viollier AG | Department of Biosystems Science and Engineering, ETH Zürich | Christian Beisel, Sarah Nadeau, Chaoran Chen, Ivan Topolsky, Kim Philipp Jablonski, Lara Fuhrmann, Rebecca Denes, Mirjam Feldkamp, Ina Nissen, Natascha Santacroce, Elodie Burcklen, Christiane Beckmann, Maurice Redondo, Olivier Kobel, Christoph Noppen, Niko Beerenwinkel, Tanja Stadler |
| EPI_ISL_3296807 | Wales Specialist Virology Centre Sequencing lab: Pathogen Genomics Unit | Public Health Wales Microbiology Cardiff Wales Specialist Virology Centre | Catherine Moore, Johnathan Evans, Laura Gifford, Malorie Perry, Simon Cottrell, Angela Marchbank, Alec Birchley, Alexander Adams, Amy Gaskin, Bree Gatica-Wilcox, Jason Coombes, Joel Southgate, Lauren Gilbert, Lee Graham, Nicole Pacchiarini, Sara Kumziene-Summerhayes, Sarah Taylor, Sophie Jones, Sara Rey, Matthew Bull, Joanne Watkins, Sally Corden, Tom Connor |
| EPI_ISL_3296808 | Viollier AG | Department of Biosystems Science and Engineering, ETH Zürich | Christian Beisel, Sarah Nadeau, Chaoran Chen, Ivan Topolsky, Kim Philipp Jablonski, Lara Fuhrmann, Rebecca Denes, Mirjam Feldkamp, Ina Nissen, Natascha Santacroce, Elodie Burcklen, Christiane Beckmann, Maurice Redondo, Olivier Kobel, Christoph Noppen, Niko Beerenwinkel, Tanja Stadler |
| EPI_ISL_3296809, EPI_ISL_3296810 | Wales Specialist Virology Centre Sequencing lab: Pathogen Genomics Unit | Public Health Wales Microbiology Cardiff Wales Specialist Virology Centre | Catherine Moore, Johnathan Evans, Laura Gifford, Malorie Perry, Simon Cottrell, Angela Marchbank, Alec Birchley, Alexander Adams, Amy Gaskin, Bree Gatica-Wilcox, Jason Coombes, Joel Southgate, Lauren Gilbert, Lee Graham, Nicole Pacchiarini, Sara Kumziene-Summerhayes, Sarah Taylor, Sophie Jones, Sara Rey, Matthew Bull, Joanne Watkins, Sally Corden, Tom Connor |
| EPI_ISL_3296811 | Viollier AG | Department of Biosystems Science and Engineering, ETH Zürich | Christian Beisel, Sarah Nadeau, Chaoran Chen, Ivan Topolsky, Kim Philipp Jablonski, Lara Fuhrmann, Rebecca Denes, Mirjam Feldkamp, Ina Nissen, Natascha Santacroce, Elodie Burcklen, Christiane Beckmann, Maurice Redondo, Olivier Kobel, Christoph Noppen, Niko Beerenwinkel, Tanja Stadler |
| EPI_ISL_3296812 | Wales Specialist Virology Centre Sequencing lab: Pathogen Genomics Unit | Public Health Wales Microbiology Cardiff Wales Specialist Virology Centre | Catherine Moore, Johnathan Evans, Laura Gifford, Malorie Perry, Simon Cottrell, Angela Marchbank, Alec Birchley, Alexander Adams, Amy Gaskin, Bree Gatica-Wilcox, Jason Coombes, Joel Southgate, Lauren Gilbert, Lee Graham, Nicole Pacchiarini, Sara Kumziene-Summerhayes, Sarah Taylor, Sophie Jones, Sara Rey, Matthew Bull, Joanne Watkins, Sally Corden, Tom Connor |
| EPI_ISL_3296813 | Viollier AG | Department of Biosystems Science and Engineering, ETH Zürich | Christian Beisel, Sarah Nadeau, Chaoran Chen, Ivan Topolsky, Kim Philipp Jablonski, Lara Fuhrmann, Rebecca Denes, Mirjam Feldkamp, Ina Nissen, Natascha Santacroce, Elodie Burcklen, Christiane Beckmann, Maurice Redondo, Olivier Kobel, Christoph Noppen, Niko Beerenwinkel, Tanja Stadler |
| EPI_ISL_3296814 | Wales Specialist Virology Centre Sequencing lab: Pathogen Genomics Unit | Public Health Wales Microbiology Cardiff Wales Specialist Virology Centre | Catherine Moore, Johnathan Evans, Laura Gifford, Malorie Perry, Simon Cottrell, Angela Marchbank, Alec Birchley, Alexander Adams, Amy Gaskin, Bree Gatica-Wilcox, Jason Coombes, Joel Southgate, Lauren Gilbert, Lee Graham, Nicole Pacchiarini, Sara Kumziene-Summerhayes, Sarah Taylor, Sophie Jones, Sara Rey, Matthew Bull, Joanne Watkins, Sally Corden, Tom Connor |
| EPI_ISL_3296815 | Viollier AG | Department of Biosystems Science and Engineering, ETH Zürich | Christian Beisel, Sarah Nadeau, Chaoran Chen, Ivan Topolsky, Kim Philipp Jablonski, Lara Fuhrmann, Rebecca Denes, Mirjam Feldkamp, Ina Nissen, Natascha Santacroce, Elodie Burcklen, Christiane Beckmann, Maurice Redondo, Olivier Kobel, Christoph Noppen, Niko Beerenwinkel, Tanja Stadler |
| EPI_ISL_3296816 | Wales Specialist Virology Centre Sequencing lab: Pathogen Genomics Unit | Public Health Wales Microbiology Cardiff Wales Specialist Virology Centre | Catherine Moore, Johnathan Evans, Laura Gifford, Malorie Perry, Simon Cottrell, Angela Marchbank, Alec Birchley, Alexander Adams, Amy Gaskin, Bree Gatica-Wilcox, Jason Coombes, Joel Southgate, Lauren Gilbert, Lee Graham, Nicole Pacchiarini, Sara Kumziene-Summerhayes, Sarah Taylor, Sophie Jones, Sara Rey, Matthew Bull, Joanne Watkins, Sally Corden, Tom Connor |
| EPI_ISL_3296817 | Viollier AG | Department of Biosystems Science and Engineering, ETH Zürich | Christian Beisel, Sarah Nadeau, Chaoran Chen, Ivan Topolsky, Kim Philipp Jablonski, Lara Fuhrmann, Rebecca Denes, Mirjam Feldkamp, Ina Nissen, Natascha Santacroce, Elodie Burcklen, Christiane Beckmann, Maurice Redondo, Olivier Kobel, Christoph Noppen, Niko Beerenwinkel, Tanja Stadler |
| EPI_ISL_3296818 | Wales Specialist Virology Centre Sequencing lab: Pathogen Genomics Unit | Public Health Wales Microbiology Cardiff Wales Specialist Virology Centre | Catherine Moore, Johnathan Evans, Laura Gifford, Malorie Perry, Simon Cottrell, Angela Marchbank, Alec Birchley, Alexander Adams, Amy Gaskin, Bree Gatica-Wilcox, Jason Coombes, Joel Southgate, Lauren Gilbert, Lee Graham, Nicole Pacchiarini, Sara Kumziene-Summerhayes, Sarah Taylor, Sophie Jones, Sara Rey, Matthew Bull, Joanne Watkins, Sally Corden, Tom Connor |
| EPI_ISL_3296819 | Viollier AG | Department of Biosystems Science and Engineering, ETH Zürich | Christian Beisel, Sarah Nadeau, Chaoran Chen, Ivan Topolsky, Kim Philipp Jablonski, Lara Fuhrmann, Rebecca Denes, Mirjam Feldkamp, Ina Nissen, Natascha Santacroce, Elodie Burcklen, Christiane Beckmann, Maurice Redondo, Olivier Kobel, Christoph Noppen, Niko Beerenwinkel, Tanja Stadler |
| EPI_ISL_3296820 | Wales Specialist Virology Centre Sequencing lab: Pathogen Genomics Unit | Public Health Wales Microbiology Cardiff Wales Specialist Virology Centre | Catherine Moore, Johnathan Evans, Laura Gifford, Malorie Perry, Simon Cottrell, Angela Marchbank, Alec Birchley, Alexander Adams, Amy Gaskin, Bree Gatica-Wilcox, Jason Coombes, Joel Southgate, Lauren Gilbert, Lee Graham, Nicole Pacchiarini, Sara Kumziene-Summerhayes, Sarah Taylor, Sophie Jones, Sara Rey, Matthew Bull, Joanne Watkins, Sally Corden, Tom Connor |
| EPI_ISL_3296821 | Viollier AG | Department of Biosystems Science and Engineering, ETH Zürich | Christian Beisel, Sarah Nadeau, Chaoran Chen, Ivan Topolsky, Kim Philipp Jablonski, Lara Fuhrmann, Rebecca Denes, Mirjam Feldkamp, Ina Nissen, Natascha Santacroce, Elodie Burcklen, Christiane Beckmann, Maurice Redondo, Olivier Kobel, Christoph Noppen, Niko Beerenwinkel, Tanja Stadler |
| EPI_ISL_3296822 | Wales Specialist Virology Centre Sequencing lab: Pathogen | Public Health Wales Microbiology Cardiff Wales Specialist Virology | Catherine Moore, Johnathan Evans, Laura Gifford, Malorie Perry, Simon Cottrell, Angela Marchbank, Alec Birchley, Alexander Adams, Amy Gaskin, Bree Gatica-Wilcox, Jason Coombes, Joel Southgate, Lauren Gilbert, Lee Graham, Nicole |

| | | | |
|---|---|---|---|
| | Genomics Unit | Centre | Pacchiarini, Sara Kumziene-Summerhayes, Sarah Taylor, Sophie Jones, Sara Rey, Matthew Bull, Joanne Watkins, Sally Corden, Tom Connor |
| EPI_ISL_3296823 | Viollier AG | Department of Biosystems Science and Engineering, ETH Zürich | Christian Beisel, Sarah Nadeau, Chaoran Chen, Ivan Topolsky, Kim Philipp Jablonski, Lara Fuhrmann, Rebecca Denes, Mirjam Feldkamp, Ina Nissen, Natascha Santacroce, Elodie Burcklen, Christiane Beckmann, Maurice Redondo, Olivier Kobel, Christoph Noppen, Niko Beerenwinkel, Tanja Stadler |
| EPI_ISL_3296824 | Wales Specialist Virology Centre Sequencing lab: Pathogen Genomics Unit | Public Health Wales Microbiology Cardiff Wales Specialist Virology Centre | Catherine Moore, Johnathan Evans, Laura Gifford, Malorie Perry, Simon Cottrell, Angela Marchbank, Alec Birchley, Alexander Adams, Amy Gaskin, Bree Gatica-Wilcox, Jason Coombes, Joel Southgate, Lauren Gilbert, Lee Graham, Nicole Pacchiarini, Sara Kumziene-Summerhayes, Sarah Taylor, Sophie Jones, Sara Rey, Matthew Bull, Joanne Watkins, Sally Corden, Tom Connor |
| EPI_ISL_3296825 | Viollier AG | Department of Biosystems Science and Engineering, ETH Zürich | Christian Beisel, Sarah Nadeau, Chaoran Chen, Ivan Topolsky, Kim Philipp Jablonski, Lara Fuhrmann, Rebecca Denes, Mirjam Feldkamp, Ina Nissen, Natascha Santacroce, Elodie Burcklen, Christiane Beckmann, Maurice Redondo, Olivier Kobel, Christoph Noppen, Niko Beerenwinkel, Tanja Stadler |
| EPI_ISL_3296826 | Wales Specialist Virology Centre Sequencing lab: Pathogen Genomics Unit | Public Health Wales Microbiology Cardiff Wales Specialist Virology Centre | Catherine Moore, Johnathan Evans, Laura Gifford, Malorie Perry, Simon Cottrell, Angela Marchbank, Alec Birchley, Alexander Adams, Amy Gaskin, Bree Gatica-Wilcox, Jason Coombes, Joel Southgate, Lauren Gilbert, Lee Graham, Nicole Pacchiarini, Sara Kumziene-Summerhayes, Sarah Taylor, Sophie Jones, Sara Rey, Matthew Bull, Joanne Watkins, Sally Corden, Tom Connor |
| EPI_ISL_3296827 | Viollier AG | Department of Biosystems Science and Engineering, ETH Zürich | Christian Beisel, Sarah Nadeau, Chaoran Chen, Ivan Topolsky, Kim Philipp Jablonski, Lara Fuhrmann, Rebecca Denes, Mirjam Feldkamp, Ina Nissen, Natascha Santacroce, Elodie Burcklen, Christiane Beckmann, Maurice Redondo, Olivier Kobel, Christoph Noppen, Niko Beerenwinkel, Tanja Stadler |
| EPI_ISL_3296828 | Wales Specialist Virology Centre Sequencing lab: Pathogen Genomics Unit | Public Health Wales Microbiology Cardiff Wales Specialist Virology Centre | Catherine Moore, Johnathan Evans, Laura Gifford, Malorie Perry, Simon Cottrell, Angela Marchbank, Alec Birchley, Alexander Adams, Amy Gaskin, Bree Gatica-Wilcox, Jason Coombes, Joel Southgate, Lauren Gilbert, Lee Graham, Nicole Pacchiarini, Sara Kumziene-Summerhayes, Sarah Taylor, Sophie Jones, Sara Rey, Matthew Bull, Joanne Watkins, Sally Corden, Tom Connor |
| EPI_ISL_3296829 | Viollier AG | Department of Biosystems Science and Engineering, ETH Zürich | Christian Beisel, Sarah Nadeau, Chaoran Chen, Ivan Topolsky, Kim Philipp Jablonski, Lara Fuhrmann, Rebecca Denes, Mirjam Feldkamp, Ina Nissen, Natascha Santacroce, Elodie Burcklen, Christiane Beckmann, Maurice Redondo, Olivier Kobel, Christoph Noppen, Niko Beerenwinkel, Tanja Stadler |
| EPI_ISL_3296830 | Wales Specialist Virology Centre Sequencing lab: Pathogen Genomics Unit | Public Health Wales Microbiology Cardiff Wales Specialist Virology Centre | Catherine Moore, Johnathan Evans, Laura Gifford, Malorie Perry, Simon Cottrell, Angela Marchbank, Alec Birchley, Alexander Adams, Amy Gaskin, Bree Gatica-Wilcox, Jason Coombes, Joel Southgate, Lauren Gilbert, Lee Graham, Nicole Pacchiarini, Sara Kumziene-Summerhayes, Sarah Taylor, Sophie Jones, Sara Rey, Matthew Bull, Joanne Watkins, Sally Corden, Tom Connor |
| EPI_ISL_3296831 | Viollier AG | Department of Biosystems Science and Engineering, ETH Zürich | Christian Beisel, Sarah Nadeau, Chaoran Chen, Ivan Topolsky, Kim Philipp Jablonski, Lara Fuhrmann, Rebecca Denes, Mirjam Feldkamp, Ina Nissen, Natascha Santacroce, Elodie Burcklen, Christiane Beckmann, Maurice Redondo, Olivier Kobel, Christoph Noppen, Niko Beerenwinkel, Tanja Stadler |
| EPI_ISL_3296832 | Wales Specialist Virology Centre Sequencing lab: Pathogen Genomics Unit | Public Health Wales Microbiology Cardiff Wales Specialist Virology Centre | Catherine Moore, Johnathan Evans, Laura Gifford, Malorie Perry, Simon Cottrell, Angela Marchbank, Alec Birchley, Alexander Adams, Amy Gaskin, Bree Gatica-Wilcox, Jason Coombes, Joel Southgate, Lauren Gilbert, Lee Graham, Nicole Pacchiarini, Sara Kumziene-Summerhayes, Sarah Taylor, Sophie Jones, Sara Rey, Matthew Bull, Joanne Watkins, Sally Corden, Tom Connor |
| EPI_ISL_3296833 | Viollier AG | Department of Biosystems Science and Engineering, ETH Zürich | Christian Beisel, Sarah Nadeau, Chaoran Chen, Ivan Topolsky, Kim Philipp Jablonski, Lara Fuhrmann, Rebecca Denes, Mirjam Feldkamp, Ina Nissen, Natascha Santacroce, Elodie Burcklen, Christiane Beckmann, Maurice Redondo, Olivier Kobel, Christoph Noppen, Niko Beerenwinkel, Tanja Stadler |
| EPI_ISL_3296834 | Wales Specialist Virology Centre Sequencing lab: Pathogen Genomics Unit | Public Health Wales Microbiology Cardiff Wales Specialist Virology Centre | Catherine Moore, Johnathan Evans, Laura Gifford, Malorie Perry, Simon Cottrell, Angela Marchbank, Alec Birchley, Alexander Adams, Amy Gaskin, Bree Gatica-Wilcox, Jason Coombes, Joel Southgate, Lauren Gilbert, Lee Graham, Nicole Pacchiarini, Sara Kumziene-Summerhayes, Sarah Taylor, Sophie Jones, Sara Rey, Matthew Bull, Joanne Watkins, Sally Corden, Tom Connor |
| EPI_ISL_3296835 | Viollier AG | Department of Biosystems Science and Engineering, ETH Zürich | Christian Beisel, Sarah Nadeau, Chaoran Chen, Ivan Topolsky, Kim Philipp Jablonski, Lara Fuhrmann, Rebecca Denes, Mirjam Feldkamp, Ina Nissen, Natascha Santacroce, Elodie Burcklen, Christiane Beckmann, Maurice Redondo, Olivier Kobel, Christoph Noppen, Niko Beerenwinkel, Tanja Stadler |
| EPI_ISL_3296836 | Wales Specialist Virology Centre Sequencing lab: Pathogen Genomics Unit | Public Health Wales Microbiology Cardiff Wales Specialist Virology Centre | Catherine Moore, Johnathan Evans, Laura Gifford, Malorie Perry, Simon Cottrell, Angela Marchbank, Alec Birchley, Alexander Adams, Amy Gaskin, Bree Gatica-Wilcox, Jason Coombes, Joel Southgate, Lauren Gilbert, Lee Graham, Nicole Pacchiarini, Sara Kumziene-Summerhayes, Sarah Taylor, Sophie Jones, Sara Rey, Matthew Bull, Joanne Watkins, Sally Corden, Tom Connor |
| EPI_ISL_3296837 | Viollier AG | Department of Biosystems Science and Engineering, ETH Zürich | Christian Beisel, Sarah Nadeau, Chaoran Chen, Ivan Topolsky, Kim Philipp Jablonski, Lara Fuhrmann, Rebecca Denes, Mirjam Feldkamp, Ina Nissen, Natascha Santacroce, Elodie Burcklen, Christiane Beckmann, Maurice Redondo, Olivier Kobel, Christoph Noppen, Niko Beerenwinkel, Tanja Stadler |
| EPI_ISL_3296838, EPI_ISL_3296840 | Wales Specialist Virology Centre Sequencing lab: Pathogen Genomics Unit | Public Health Wales Microbiology Cardiff Wales Specialist Virology Centre | Catherine Moore, Johnathan Evans, Laura Gifford, Malorie Perry, Simon Cottrell, Angela Marchbank, Alec Birchley, Alexander Adams, Amy Gaskin, Bree Gatica-Wilcox, Jason Coombes, Joel Southgate, Lauren Gilbert, Lee Graham, Nicole Pacchiarini, Sara Kumziene-Summerhayes, Sarah Taylor, Sophie Jones, Sara Rey, Matthew Bull, Joanne Watkins, Sally Corden, Tom Connor |
| EPI_ISL_3296841 | Viollier AG | Department of Biosystems Science and Engineering, ETH Zürich | Christian Beisel, Sarah Nadeau, Chaoran Chen, Ivan Topolsky, Kim Philipp Jablonski, Lara Fuhrmann, Rebecca Denes, Mirjam Feldkamp, Ina Nissen, Natascha Santacroce, Elodie Burcklen, Christiane Beckmann, Maurice Redondo, Olivier Kobel, Christoph Noppen, Niko Beerenwinkel, Tanja Stadler |
| EPI_ISL_3296842 | Wales Specialist Virology Centre Sequencing lab: Pathogen Genomics Unit | Public Health Wales Microbiology Cardiff Wales Specialist Virology Centre | Catherine Moore, Johnathan Evans, Laura Gifford, Malorie Perry, Simon Cottrell, Angela Marchbank, Alec Birchley, Alexander Adams, Amy Gaskin, Bree Gatica-Wilcox, Jason Coombes, Joel Southgate, Lauren Gilbert, Lee Graham, Nicole Pacchiarini, Sara Kumziene-Summerhayes, Sarah Taylor, Sophie Jones, Sara Rey, Matthew Bull, Joanne Watkins, Sally Corden, Tom Connor |
| EPI_ISL_3296843 | Viollier AG | Department of Biosystems Science and Engineering, ETH Zürich | Christian Beisel, Sarah Nadeau, Chaoran Chen, Ivan Topolsky, Kim Philipp Jablonski, Lara Fuhrmann, Rebecca Denes, Mirjam Feldkamp, Ina Nissen, Natascha Santacroce, Elodie Burcklen, Christiane Beckmann, Maurice Redondo, Olivier Kobel, Christoph Noppen, Niko Beerenwinkel, Tanja Stadler |
| EPI_ISL_3296844, EPI_ISL_3296845 | Wales Specialist Virology Centre Sequencing lab: Pathogen Genomics Unit | Public Health Wales Microbiology Cardiff Wales Specialist Virology Centre | Catherine Moore, Johnathan Evans, Laura Gifford, Malorie Perry, Simon Cottrell, Angela Marchbank, Alec Birchley, Alexander Adams, Amy Gaskin, Bree Gatica-Wilcox, Jason Coombes, Joel Southgate, Lauren Gilbert, Lee Graham, Nicole Pacchiarini, Sara Kumziene-Summerhayes, Sarah Taylor, Sophie Jones, Sara Rey, Matthew Bull, Joanne Watkins, Sally Corden, Tom Connor |
| EPI_ISL_3296846, EPI_ISL_3296847 | Viollier AG | Department of Biosystems Science and Engineering, ETH Zürich | Christian Beisel, Sarah Nadeau, Chaoran Chen, Ivan Topolsky, Kim Philipp Jablonski, Lara Fuhrmann, Rebecca Denes, Mirjam Feldkamp, Ina Nissen, Natascha Santacroce, Elodie Burcklen, Christiane Beckmann, Maurice Redondo, Olivier Kobel, Christoph Noppen, Niko Beerenwinkel, Tanja Stadler |
| EPI_ISL_3296848 | Wales Specialist Virology Centre Sequencing lab: Pathogen Genomics Unit | Public Health Wales Microbiology Cardiff Wales Specialist Virology Centre | Catherine Moore, Johnathan Evans, Laura Gifford, Malorie Perry, Simon Cottrell, Angela Marchbank, Alec Birchley, Alexander Adams, Amy Gaskin, Bree Gatica-Wilcox, Jason Coombes, Joel Southgate, Lauren Gilbert, Lee Graham, Nicole Pacchiarini, Sara Kumziene-Summerhayes, Sarah Taylor, Sophie Jones, Sara Rey, Matthew Bull, Joanne Watkins, Sally Corden, Tom Connor |
| EPI_ISL_3296849 | Viollier AG | Department of Biosystems Science and Engineering, ETH Zürich | Christian Beisel, Sarah Nadeau, Chaoran Chen, Ivan Topolsky, Kim Philipp Jablonski, Lara Fuhrmann, Rebecca Denes, Mirjam Feldkamp, Ina Nissen, Natascha Santacroce, Elodie Burcklen, Christiane Beckmann, Maurice Redondo, Olivier Kobel, Christoph Noppen, Niko Beerenwinkel, Tanja Stadler |
| EPI_ISL_3296850, EPI_ISL_3296851 | Wales Specialist Virology Centre Sequencing lab: Pathogen Genomics Unit | Public Health Wales Microbiology Cardiff Wales Specialist Virology Centre | Catherine Moore, Johnathan Evans, Laura Gifford, Malorie Perry, Simon Cottrell, Angela Marchbank, Alec Birchley, Alexander Adams, Amy Gaskin, Bree Gatica-Wilcox, Jason Coombes, Joel Southgate, Lauren Gilbert, Lee Graham, Nicole Pacchiarini, Sara Kumziene-Summerhayes, Sarah Taylor, Sophie Jones, Sara Rey, Matthew Bull, Joanne Watkins, Sally Corden, Tom Connor |
| EPI_ISL_3296852 | Viollier AG | Department of Biosystems Science and Engineering, ETH Zürich | Christian Beisel, Sarah Nadeau, Chaoran Chen, Ivan Topolsky, Kim Philipp Jablonski, Lara Fuhrmann, Rebecca Denes, Mirjam Feldkamp, Ina Nissen, Natascha Santacroce, Elodie Burcklen, Christiane Beckmann, Maurice Redondo, Olivier Kobel, Christoph Noppen, Niko Beerenwinkel, Tanja Stadler |
| EPI_ISL_3296853 | Wales Specialist Virology Centre Sequencing lab: Pathogen Genomics Unit | Public Health Wales Microbiology Cardiff Wales Specialist Virology Centre | Catherine Moore, Johnathan Evans, Laura Gifford, Malorie Perry, Simon Cottrell, Angela Marchbank, Alec Birchley, Alexander Adams, Amy Gaskin, Bree Gatica-Wilcox, Jason Coombes, Joel Southgate, Lauren Gilbert, Lee Graham, Nicole Pacchiarini, Sara Kumziene-Summerhayes, Sarah Taylor, Sophie Jones, Sara Rey, Matthew Bull, Joanne Watkins, Sally Corden, Tom Connor |
| EPI_ISL_3296854, EPI_ISL_3296855 | Viollier AG | Department of Biosystems Science and Engineering, ETH Zürich | Christian Beisel, Sarah Nadeau, Chaoran Chen, Ivan Topolsky, Kim Philipp Jablonski, Lara Fuhrmann, Rebecca Denes, Mirjam Feldkamp, Ina Nissen, Natascha Santacroce, Elodie Burcklen, Christiane Beckmann, Maurice Redondo, Olivier Kobel, Christoph Noppen, Niko Beerenwinkel, Tanja Stadler |
| EPI_ISL_3296856 | Wales Specialist Virology Centre Sequencing lab: Pathogen Genomics Unit | Public Health Wales Microbiology Cardiff Wales Specialist Virology Centre | Catherine Moore, Johnathan Evans, Laura Gifford, Malorie Perry, Simon Cottrell, Angela Marchbank, Alec Birchley, Alexander Adams, Amy Gaskin, Bree Gatica-Wilcox, Jason Coombes, Joel Southgate, Lauren Gilbert, Lee Graham, Nicole Pacchiarini, Sara Kumziene-Summerhayes, Sarah Taylor, Sophie Jones, Sara Rey, Matthew Bull, Joanne Watkins, Sally Corden, Tom Connor |
| EPI_ISL_3296857 | Viollier AG | Department of Biosystems Science and Engineering, ETH Zürich | Christian Beisel, Sarah Nadeau, Chaoran Chen, Ivan Topolsky, Kim Philipp Jablonski, Lara Fuhrmann, Rebecca Denes, Mirjam Feldkamp, Ina Nissen, Natascha Santacroce, Elodie Burcklen, Christiane Beckmann, Maurice Redondo, Olivier Kobel, Christoph Noppen, Niko Beerenwinkel, Tanja Stadler |
| EPI_ISL_3296858 | Wales Specialist Virology Centre Sequencing lab: Pathogen Genomics Unit | Public Health Wales Microbiology Cardiff Wales Specialist Virology Centre | Catherine Moore, Johnathan Evans, Laura Gifford, Malorie Perry, Simon Cottrell, Angela Marchbank, Alec Birchley, Alexander Adams, Amy Gaskin, Bree Gatica-Wilcox, Jason Coombes, Joel Southgate, Lauren Gilbert, Lee Graham, Nicole Pacchiarini, Sara Kumziene-Summerhayes, Sarah Taylor, Sophie Jones, Sara Rey, Matthew Bull, Joanne Watkins, Sally Corden, Tom Connor |
| EPI_ISL_3296859 | Viollier AG | Department of Biosystems Science and Engineering, ETH Zürich | Christian Beisel, Sarah Nadeau, Chaoran Chen, Ivan Topolsky, Kim Philipp Jablonski, Lara Fuhrmann, Rebecca Denes, Mirjam Feldkamp, Ina Nissen, Natascha Santacroce, Elodie Burcklen, Christiane Beckmann, Maurice Redondo, Olivier Kobel, Christoph Noppen, Niko Beerenwinkel, Tanja Stadler |
| EPI_ISL_3296860 | Wales Specialist Virology Centre Sequencing lab: Pathogen Genomics Unit | Public Health Wales Microbiology Cardiff Wales Specialist Virology Centre | Catherine Moore, Johnathan Evans, Laura Gifford, Malorie Perry, Simon Cottrell, Angela Marchbank, Alec Birchley, Alexander Adams, Amy Gaskin, Bree Gatica-Wilcox, Jason Coombes, Joel Southgate, Lauren Gilbert, Lee Graham, Nicole Pacchiarini, Sara Kumziene-Summerhayes, Sarah Taylor, Sophie Jones, Sara Rey, Matthew Bull, Joanne Watkins, Sally Corden, Tom Connor |
| EPI_ISL_3296861 | Viollier AG | Department of Biosystems Science and Engineering, ETH Zürich | Christian Beisel, Sarah Nadeau, Chaoran Chen, Ivan Topolsky, Kim Philipp Jablonski, Lara Fuhrmann, Rebecca Denes, Mirjam Feldkamp, Ina Nissen, Natascha Santacroce, Elodie Burcklen, Christiane Beckmann, Maurice Redondo, Olivier Kobel, Christoph Noppen, Niko Beerenwinkel, Tanja Stadler |
| EPI_ISL_3296862 | Wales Specialist Virology Centre Sequencing lab: Pathogen Genomics Unit | Public Health Wales Microbiology Cardiff Wales Specialist Virology Centre | Catherine Moore, Johnathan Evans, Laura Gifford, Malorie Perry, Simon Cottrell, Angela Marchbank, Alec Birchley, Alexander Adams, Amy Gaskin, Bree Gatica-Wilcox, Jason Coombes, Joel Southgate, Lauren Gilbert, Lee Graham, Nicole Pacchiarini, Sara Kumziene-Summerhayes, Sarah Taylor, Sophie Jones, Sara Rey, Matthew Bull, Joanne Watkins, Sally Corden, Tom Connor |
| EPI_ISL_3296863 | Viollier AG | Department of Biosystems Science and Engineering, ETH Zürich | Christian Beisel, Sarah Nadeau, Chaoran Chen, Ivan Topolsky, Kim Philipp Jablonski, Lara Fuhrmann, Rebecca Denes, Mirjam Feldkamp, Ina Nissen, Natascha Santacroce, Elodie Burcklen, Christiane Beckmann, Maurice Redondo, Olivier Kobel, Christoph Noppen, Niko Beerenwinkel, Tanja Stadler |
| EPI_ISL_3296864 | Wales Specialist Virology Centre Sequencing lab: Pathogen Genomics Unit | Public Health Wales Microbiology Cardiff Wales Specialist Virology Centre | Catherine Moore, Johnathan Evans, Laura Gifford, Malorie Perry, Simon Cottrell, Angela Marchbank, Alec Birchley, Alexander Adams, Amy Gaskin, Bree Gatica-Wilcox, Jason Coombes, Joel Southgate, Lauren Gilbert, Lee Graham, Nicole Pacchiarini, Sara Kumziene-Summerhayes, Sarah Taylor, Sophie Jones, Sara Rey, Matthew Bull, Joanne Watkins, Sally Corden, Tom Connor |
| EPI_ISL_3296865 | Viollier AG | Department of Biosystems Science and Engineering, ETH Zürich | Christian Beisel, Sarah Nadeau, Chaoran Chen, Ivan Topolsky, Kim Philipp Jablonski, Lara Fuhrmann, Rebecca Denes, Mirjam Feldkamp, Ina Nissen, Natascha Santacroce, Elodie Burcklen, Christiane Beckmann, Maurice Redondo, Olivier Kobel, Christoph Noppen, Niko Beerenwinkel, Tanja Stadler |
| EPI_ISL_3296866, EPI_ISL_3296868 | Wales Specialist Virology Centre Sequencing lab: Pathogen Genomics Unit | Public Health Wales Microbiology Cardiff Wales Specialist Virology Centre | Catherine Moore, Johnathan Evans, Laura Gifford, Malorie Perry, Simon Cottrell, Angela Marchbank, Alec Birchley, Alexander Adams, Amy Gaskin, Bree Gatica-Wilcox, Jason Coombes, Joel Southgate, Lauren Gilbert, Lee Graham, Nicole Pacchiarini, Sara Kumziene-Summerhayes, Sarah Taylor, Sophie Jones, Sara Rey, Matthew Bull, Joanne Watkins, Sally Corden, Tom Connor |
| EPI_ISL_3296869 | Viollier AG | Department of Biosystems Science and Engineering, ETH Zürich | Christian Beisel, Sarah Nadeau, Chaoran Chen, Ivan Topolsky, Kim Philipp Jablonski, Lara Fuhrmann, Rebecca Denes, Mirjam Feldkamp, Ina Nissen, Natascha Santacroce, Elodie Burcklen, Christiane Beckmann, Maurice Redondo, Olivier Kobel, Christoph Noppen, Niko Beerenwinkel, Tanja Stadler |
| EPI_ISL_3296870, EPI_ISL_3296872 | Wales Specialist Virology Centre Sequencing lab: Pathogen Genomics Unit | Public Health Wales Microbiology Cardiff Wales Specialist Virology Centre | Catherine Moore, Johnathan Evans, Laura Gifford, Malorie Perry, Simon Cottrell, Angela Marchbank, Alec Birchley, Alexander Adams, Amy Gaskin, Bree Gatica-Wilcox, Jason Coombes, Joel Southgate, Lauren Gilbert, Lee Graham, Nicole Pacchiarini, Sara Kumziene-Summerhayes, Sarah Taylor, Sophie Jones, Sara Rey, Matthew Bull, Joanne Watkins, Sally Corden, Tom Connor |
| EPI_ISL_3296873 | Viollier AG | Department of Biosystems Science and Engineering, ETH Zürich | Christian Beisel, Sarah Nadeau, Chaoran Chen, Ivan Topolsky, Kim Philipp Jablonski, Lara Fuhrmann, Rebecca Denes, Mirjam Feldkamp, Ina Nissen, Natascha Santacroce, Elodie Burcklen, Christiane Beckmann, Maurice Redondo, Olivier Kobel, Christoph Noppen, Niko Beerenwinkel, Tanja Stadler |
| EPI_ISL_3296874 | Wales Specialist Virology Centre Sequencing lab: Pathogen Genomics Unit | Public Health Wales Microbiology Cardiff Wales Specialist Virology Centre | Catherine Moore, Johnathan Evans, Laura Gifford, Malorie Perry, Simon Cottrell, Angela Marchbank, Alec Birchley, Alexander Adams, Amy Gaskin, Bree Gatica-Wilcox, Jason Coombes, Joel Southgate, Lauren Gilbert, Lee Graham, Nicole Pacchiarini, Sara Kumziene-Summerhayes, Sarah Taylor, Sophie Jones, Sara Rey, Matthew Bull, Joanne Watkins, Sally Corden, Tom Connor |
| EPI_ISL_3296875 | Viollier AG | Department of Biosystems Science and Engineering, ETH Zürich | Christian Beisel, Sarah Nadeau, Chaoran Chen, Ivan Topolsky, Kim Philipp Jablonski, Lara Fuhrmann, Rebecca Denes, Mirjam Feldkamp, Ina Nissen, Natascha Santacroce, Elodie Burcklen, Christiane Beckmann, Maurice Redondo, Olivier Kobel, Christoph Noppen, Niko Beerenwinkel, Tanja Stadler |
| EPI_ISL_3296876 | Wales Specialist Virology Centre Sequencing lab: Pathogen Genomics Unit | Public Health Wales Microbiology Cardiff Wales Specialist Virology Centre | Catherine Moore, Johnathan Evans, Laura Gifford, Malorie Perry, Simon Cottrell, Angela Marchbank, Alec Birchley, Alexander Adams, Amy Gaskin, Bree Gatica-Wilcox, Jason Coombes, Joel Southgate, Lauren Gilbert, Lee Graham, Nicole Pacchiarini, Sara Kumziene-Summerhayes, Sarah Taylor, Sophie Jones, Sara Rey, Matthew Bull, Joanne Watkins, Sally Corden, Tom Connor |
| EPI_ISL_3296877 | Viollier AG | Department of Biosystems Science and Engineering, ETH Zürich | Christian Beisel, Sarah Nadeau, Chaoran Chen, Ivan Topolsky, Kim Philipp Jablonski, Lara Fuhrmann, Rebecca Denes, Mirjam Feldkamp, Ina Nissen, Natascha Santacroce, Elodie Burcklen, Christiane Beckmann, Maurice Redondo, Olivier Kobel, |

| | | | |
|---|---|---|---|
| | | | Christoph Noppen, Niko Beerenwinkel, Tanja Stadler |
| EPI_ISL_3296878, EPI_ISL_3296879 | Wales Specialist Virology Centre Sequencing lab: Pathogen Genomics Unit | Public Health Wales Microbiology Cardiff Wales Specialist Virology Centre | Catherine Moore, Johnathan Evans, Laura Gifford, Malorie Perry, Simon Cottrell, Angela Marchbank, Alec Birchley, Alexander Adams, Amy Gaskin, Bree Gatica-Wilcox, Jason Coombes, Joel Southgate, Lauren Gilbert, Lee Graham, Nicole Pacchiarini, Sara Kumziene-Summerhayes, Sarah Taylor, Sophie Jones, Sara Rey, Matthew Bull, Joanne Watkins, Sally Corden, Tom Connor |
| EPI_ISL_3296880 | Viollier AG | Department of Biosystems Science and Engineering, ETH Zürich | Christian Beisel, Sarah Nadeau, Chaoran Chen, Ivan Topolsky, Kim Philipp Jablonski, Lara Fuhrmann, Rebecca Denes, Mirjam Feldkamp, Ina Nissen, Natascha Santacroce, Elodie Burcklen, Christiane Beckmann, Maurice Redondo, Olivier Kobel, Christoph Noppen, Niko Beerenwinkel, Tanja Stadler |
| EPI_ISL_3296881 | Wales Specialist Virology Centre Sequencing lab: Pathogen Genomics Unit | Public Health Wales Microbiology Cardiff Wales Specialist Virology Centre | Catherine Moore, Johnathan Evans, Laura Gifford, Malorie Perry, Simon Cottrell, Angela Marchbank, Alec Birchley, Alexander Adams, Amy Gaskin, Bree Gatica-Wilcox, Jason Coombes, Joel Southgate, Lauren Gilbert, Lee Graham, Nicole Pacchiarini, Sara Kumziene-Summerhayes, Sarah Taylor, Sophie Jones, Sara Rey, Matthew Bull, Joanne Watkins, Sally Corden, Tom Connor |
| EPI_ISL_3296882 | Viollier AG | Department of Biosystems Science and Engineering, ETH Zürich | Christian Beisel, Sarah Nadeau, Chaoran Chen, Ivan Topolsky, Kim Philipp Jablonski, Lara Fuhrmann, Rebecca Denes, Mirjam Feldkamp, Ina Nissen, Natascha Santacroce, Elodie Burcklen, Christiane Beckmann, Maurice Redondo, Olivier Kobel, Christoph Noppen, Niko Beerenwinkel, Tanja Stadler |
| EPI_ISL_3296883 | Wales Specialist Virology Centre Sequencing lab: Pathogen Genomics Unit | Public Health Wales Microbiology Cardiff Wales Specialist Virology Centre | Catherine Moore, Johnathan Evans, Laura Gifford, Malorie Perry, Simon Cottrell, Angela Marchbank, Alec Birchley, Alexander Adams, Amy Gaskin, Bree Gatica-Wilcox, Jason Coombes, Joel Southgate, Lauren Gilbert, Lee Graham, Nicole Pacchiarini, Sara Kumziene-Summerhayes, Sarah Taylor, Sophie Jones, Sara Rey, Matthew Bull, Joanne Watkins, Sally Corden, Tom Connor |
| EPI_ISL_3296884 | Viollier AG | Department of Biosystems Science and Engineering, ETH Zürich | Christian Beisel, Sarah Nadeau, Chaoran Chen, Ivan Topolsky, Kim Philipp Jablonski, Lara Fuhrmann, Rebecca Denes, Mirjam Feldkamp, Ina Nissen, Natascha Santacroce, Elodie Burcklen, Christiane Beckmann, Maurice Redondo, Olivier Kobel, Christoph Noppen, Niko Beerenwinkel, Tanja Stadler |
| EPI_ISL_3296885, EPI_ISL_3296887 | Wales Specialist Virology Centre Sequencing lab: Pathogen Genomics Unit | Public Health Wales Microbiology Cardiff Wales Specialist Virology Centre | Catherine Moore, Johnathan Evans, Laura Gifford, Malorie Perry, Simon Cottrell, Angela Marchbank, Alec Birchley, Alexander Adams, Amy Gaskin, Bree Gatica-Wilcox, Jason Coombes, Joel Southgate, Lauren Gilbert, Lee Graham, Nicole Pacchiarini, Sara Kumziene-Summerhayes, Sarah Taylor, Sophie Jones, Sara Rey, Matthew Bull, Joanne Watkins, Sally Corden, Tom Connor |
| EPI_ISL_3296888 | Viollier AG | Department of Biosystems Science and Engineering, ETH Zürich | Christian Beisel, Sarah Nadeau, Chaoran Chen, Ivan Topolsky, Kim Philipp Jablonski, Lara Fuhrmann, Rebecca Denes, Mirjam Feldkamp, Ina Nissen, Natascha Santacroce, Elodie Burcklen, Christiane Beckmann, Maurice Redondo, Olivier Kobel, Christoph Noppen, Niko Beerenwinkel, Tanja Stadler |
| EPI_ISL_3296889, EPI_ISL_3296891, EPI_ISL_3296893 | Wales Specialist Virology Centre Sequencing lab: Pathogen Genomics Unit | Public Health Wales Microbiology Cardiff Wales Specialist Virology Centre | Catherine Moore, Johnathan Evans, Laura Gifford, Malorie Perry, Simon Cottrell, Angela Marchbank, Alec Birchley, Alexander Adams, Amy Gaskin, Bree Gatica-Wilcox, Jason Coombes, Joel Southgate, Lauren Gilbert, Lee Graham, Nicole Pacchiarini, Sara Kumziene-Summerhayes, Sarah Taylor, Sophie Jones, Sara Rey, Matthew Bull, Joanne Watkins, Sally Corden, Tom Connor |
| EPI_ISL_3296894 | Viollier AG | Department of Biosystems Science and Engineering, ETH Zürich | Christian Beisel, Sarah Nadeau, Chaoran Chen, Ivan Topolsky, Kim Philipp Jablonski, Lara Fuhrmann, Rebecca Denes, Mirjam Feldkamp, Ina Nissen, Natascha Santacroce, Elodie Burcklen, Christiane Beckmann, Maurice Redondo, Olivier Kobel, Christoph Noppen, Niko Beerenwinkel, Tanja Stadler |
| EPI_ISL_3296895 | Wales Specialist Virology Centre Sequencing lab: Pathogen Genomics Unit | Public Health Wales Microbiology Cardiff Wales Specialist Virology Centre | Catherine Moore, Johnathan Evans, Laura Gifford, Malorie Perry, Simon Cottrell, Angela Marchbank, Alec Birchley, Alexander Adams, Amy Gaskin, Bree Gatica-Wilcox, Jason Coombes, Joel Southgate, Lauren Gilbert, Lee Graham, Nicole Pacchiarini, Sara Kumziene-Summerhayes, Sarah Taylor, Sophie Jones, Sara Rey, Matthew Bull, Joanne Watkins, Sally Corden, Tom Connor |
| EPI_ISL_3296896 | Viollier AG | Department of Biosystems Science and Engineering, ETH Zürich | Christian Beisel, Sarah Nadeau, Chaoran Chen, Ivan Topolsky, Kim Philipp Jablonski, Lara Fuhrmann, Rebecca Denes, Mirjam Feldkamp, Ina Nissen, Natascha Santacroce, Elodie Burcklen, Christiane Beckmann, Maurice Redondo, Olivier Kobel, Christoph Noppen, Niko Beerenwinkel, Tanja Stadler |
| EPI_ISL_3296897, EPI_ISL_3296899, EPI_ISL_3296900, EPI_ISL_3296902 | Wales Specialist Virology Centre Sequencing lab: Pathogen Genomics Unit | Public Health Wales Microbiology Cardiff Wales Specialist Virology Centre | Catherine Moore, Johnathan Evans, Laura Gifford, Malorie Perry, Simon Cottrell, Angela Marchbank, Alec Birchley, Alexander Adams, Amy Gaskin, Bree Gatica-Wilcox, Jason Coombes, Joel Southgate, Lauren Gilbert, Lee Graham, Nicole Pacchiarini, Sara Kumziene-Summerhayes, Sarah Taylor, Sophie Jones, Sara Rey, Matthew Bull, Joanne Watkins, Sally Corden, Tom Connor |
| EPI_ISL_3296903 | Viollier AG | Department of Biosystems Science and Engineering, ETH Zürich | Christian Beisel, Sarah Nadeau, Chaoran Chen, Ivan Topolsky, Kim Philipp Jablonski, Lara Fuhrmann, Rebecca Denes, Mirjam Feldkamp, Ina Nissen, Natascha Santacroce, Elodie Burcklen, Christiane Beckmann, Maurice Redondo, Olivier Kobel, Christoph Noppen, Niko Beerenwinkel, Tanja Stadler |
| EPI_ISL_3296904 | Wales Specialist Virology Centre Sequencing lab: Pathogen Genomics Unit | Public Health Wales Microbiology Cardiff Wales Specialist Virology Centre | Catherine Moore, Johnathan Evans, Laura Gifford, Malorie Perry, Simon Cottrell, Angela Marchbank, Alec Birchley, Alexander Adams, Amy Gaskin, Bree Gatica-Wilcox, Jason Coombes, Joel Southgate, Lauren Gilbert, Lee Graham, Nicole Pacchiarini, Sara Kumziene-Summerhayes, Sarah Taylor, Sophie Jones, Sara Rey, Matthew Bull, Joanne Watkins, Sally Corden, Tom Connor |
| EPI_ISL_3296905 | Viollier AG | Department of Biosystems Science and Engineering, ETH Zürich | Christian Beisel, Sarah Nadeau, Chaoran Chen, Ivan Topolsky, Kim Philipp Jablonski, Lara Fuhrmann, Rebecca Denes, Mirjam Feldkamp, Ina Nissen, Natascha Santacroce, Elodie Burcklen, Christiane Beckmann, Maurice Redondo, Olivier Kobel, Christoph Noppen, Niko Beerenwinkel, Tanja Stadler |
| EPI_ISL_3296906 | Wales Specialist Virology Centre Sequencing lab: Pathogen Genomics Unit | Public Health Wales Microbiology Cardiff Wales Specialist Virology Centre | Catherine Moore, Johnathan Evans, Laura Gifford, Malorie Perry, Simon Cottrell, Angela Marchbank, Alec Birchley, Alexander Adams, Amy Gaskin, Bree Gatica-Wilcox, Jason Coombes, Joel Southgate, Lauren Gilbert, Lee Graham, Nicole Pacchiarini, Sara Kumziene-Summerhayes, Sarah Taylor, Sophie Jones, Sara Rey, Matthew Bull, Joanne Watkins, Sally Corden, Tom Connor |
| EPI_ISL_3296907 | Viollier AG | Department of Biosystems Science and Engineering, ETH Zürich | Christian Beisel, Sarah Nadeau, Chaoran Chen, Ivan Topolsky, Kim Philipp Jablonski, Lara Fuhrmann, Rebecca Denes, Mirjam Feldkamp, Ina Nissen, Natascha Santacroce, Elodie Burcklen, Christiane Beckmann, Maurice Redondo, Olivier Kobel, Christoph Noppen, Niko Beerenwinkel, Tanja Stadler |
| EPI_ISL_3296908 | Wales Specialist Virology Centre Sequencing lab: Pathogen Genomics Unit | Public Health Wales Microbiology Cardiff Wales Specialist Virology Centre | Catherine Moore, Johnathan Evans, Laura Gifford, Malorie Perry, Simon Cottrell, Angela Marchbank, Alec Birchley, Alexander Adams, Amy Gaskin, Bree Gatica-Wilcox, Jason Coombes, Joel Southgate, Lauren Gilbert, Lee Graham, Nicole Pacchiarini, Sara Kumziene-Summerhayes, Sarah Taylor, Sophie Jones, Sara Rey, Matthew Bull, Joanne Watkins, Sally Corden, Tom Connor |
| EPI_ISL_3296909 | Viollier AG | Department of Biosystems Science and Engineering, ETH Zürich | Christian Beisel, Sarah Nadeau, Chaoran Chen, Ivan Topolsky, Kim Philipp Jablonski, Lara Fuhrmann, Rebecca Denes, Mirjam Feldkamp, Ina Nissen, Natascha Santacroce, Elodie Burcklen, Christiane Beckmann, Maurice Redondo, Olivier Kobel, Christoph Noppen, Niko Beerenwinkel, Tanja Stadler |
| EPI_ISL_3296910 | Wales Specialist Virology Centre Sequencing lab: Pathogen Genomics Unit | Public Health Wales Microbiology Cardiff Wales Specialist Virology Centre | Catherine Moore, Johnathan Evans, Laura Gifford, Malorie Perry, Simon Cottrell, Angela Marchbank, Alec Birchley, Alexander Adams, Amy Gaskin, Bree Gatica-Wilcox, Jason Coombes, Joel Southgate, Lauren Gilbert, Lee Graham, Nicole Pacchiarini, Sara Kumziene-Summerhayes, Sarah Taylor, Sophie Jones, Sara Rey, Matthew Bull, Joanne Watkins, Sally Corden, Tom Connor |
| EPI_ISL_3296911 | Viollier AG | Department of Biosystems Science and Engineering, ETH Zürich | Christian Beisel, Sarah Nadeau, Chaoran Chen, Ivan Topolsky, Kim Philipp Jablonski, Lara Fuhrmann, Rebecca Denes, Mirjam Feldkamp, Ina Nissen, Natascha Santacroce, Elodie Burcklen, Christiane Beckmann, Maurice Redondo, Olivier Kobel, Christoph Noppen, Niko Beerenwinkel, Tanja Stadler |
| EPI_ISL_3296912 | Wales Specialist Virology Centre Sequencing lab: Pathogen Genomics Unit | Public Health Wales Microbiology Cardiff Wales Specialist Virology Centre | Catherine Moore, Johnathan Evans, Laura Gifford, Malorie Perry, Simon Cottrell, Angela Marchbank, Alec Birchley, Alexander Adams, Amy Gaskin, Bree Gatica-Wilcox, Jason Coombes, Joel Southgate, Lauren Gilbert, Lee Graham, Nicole Pacchiarini, Sara Kumziene-Summerhayes, Sarah Taylor, Sophie Jones, Sara Rey, Matthew Bull, Joanne Watkins, Sally Corden, Tom Connor |
| EPI_ISL_3296913 | Viollier AG | Department of Biosystems Science and Engineering, ETH Zürich | Christian Beisel, Sarah Nadeau, Chaoran Chen, Ivan Topolsky, Kim Philipp Jablonski, Lara Fuhrmann, Rebecca Denes, Mirjam Feldkamp, Ina Nissen, Natascha Santacroce, Elodie Burcklen, Christiane Beckmann, Maurice Redondo, Olivier Kobel, Christoph Noppen, Niko Beerenwinkel, Tanja Stadler |
| EPI_ISL_3296914, EPI_ISL_3296916 | Wales Specialist Virology Centre Sequencing lab: Pathogen Genomics Unit | Public Health Wales Microbiology Cardiff Wales Specialist Virology Centre | Catherine Moore, Johnathan Evans, Laura Gifford, Malorie Perry, Simon Cottrell, Angela Marchbank, Alec Birchley, Alexander Adams, Amy Gaskin, Bree Gatica-Wilcox, Jason Coombes, Joel Southgate, Lauren Gilbert, Lee Graham, Nicole Pacchiarini, Sara Kumziene-Summerhayes, Sarah Taylor, Sophie Jones, Sara Rey, Matthew Bull, Joanne Watkins, Sally Corden, Tom Connor |
| EPI_ISL_3296917 | Viollier AG | Department of Biosystems Science and Engineering, ETH Zürich | Christian Beisel, Sarah Nadeau, Chaoran Chen, Ivan Topolsky, Kim Philipp Jablonski, Lara Fuhrmann, Rebecca Denes, Mirjam Feldkamp, Ina Nissen, Natascha Santacroce, Elodie Burcklen, Christiane Beckmann, Maurice Redondo, Olivier Kobel, Christoph Noppen, Niko Beerenwinkel, Tanja Stadler |
| EPI_ISL_3296918 | Wales Specialist Virology Centre Sequencing lab: Pathogen Genomics Unit | Public Health Wales Microbiology Cardiff Wales Specialist Virology Centre | Catherine Moore, Johnathan Evans, Laura Gifford, Malorie Perry, Simon Cottrell, Angela Marchbank, Alec Birchley, Alexander Adams, Amy Gaskin, Bree Gatica-Wilcox, Jason Coombes, Joel Southgate, Lauren Gilbert, Lee Graham, Nicole Pacchiarini, Sara Kumziene-Summerhayes, Sarah Taylor, Sophie Jones, Sara Rey, Matthew Bull, Joanne Watkins, Sally Corden, Tom Connor |
| EPI_ISL_3296919 | Viollier AG | Department of Biosystems Science and Engineering, ETH Zürich | Christian Beisel, Sarah Nadeau, Chaoran Chen, Ivan Topolsky, Kim Philipp Jablonski, Lara Fuhrmann, Rebecca Denes, Mirjam Feldkamp, Ina Nissen, Natascha Santacroce, Elodie Burcklen, Christiane Beckmann, Maurice Redondo, Olivier Kobel, Christoph Noppen, Niko Beerenwinkel, Tanja Stadler |
| EPI_ISL_3296920 | Wales Specialist Virology Centre Sequencing lab: Pathogen Genomics Unit | Public Health Wales Microbiology Cardiff Wales Specialist Virology Centre | Catherine Moore, Johnathan Evans, Laura Gifford, Malorie Perry, Simon Cottrell, Angela Marchbank, Alec Birchley, Alexander Adams, Amy Gaskin, Bree Gatica-Wilcox, Jason Coombes, Joel Southgate, Lauren Gilbert, Lee Graham, Nicole Pacchiarini, Sara Kumziene-Summerhayes, Sarah Taylor, Sophie Jones, Sara Rey, Matthew Bull, Joanne Watkins, Sally Corden, Tom Connor |
| EPI_ISL_3296921 | Viollier AG | Department of Biosystems Science and Engineering, ETH Zürich | Christian Beisel, Sarah Nadeau, Chaoran Chen, Ivan Topolsky, Kim Philipp Jablonski, Lara Fuhrmann, Rebecca Denes, Mirjam Feldkamp, Ina Nissen, Natascha Santacroce, Elodie Burcklen, Christiane Beckmann, Maurice Redondo, Olivier Kobel, Christoph Noppen, Niko Beerenwinkel, Tanja Stadler |
| EPI_ISL_3296922 | Wales Specialist Virology Centre Sequencing lab: Pathogen Genomics Unit | Public Health Wales Microbiology Cardiff Wales Specialist Virology Centre | Catherine Moore, Johnathan Evans, Laura Gifford, Malorie Perry, Simon Cottrell, Angela Marchbank, Alec Birchley, Alexander Adams, Amy Gaskin, Bree Gatica-Wilcox, Jason Coombes, Joel Southgate, Lauren Gilbert, Lee Graham, Nicole Pacchiarini, Sara Kumziene-Summerhayes, Sarah Taylor, Sophie Jones, Sara Rey, Matthew Bull, Joanne Watkins, Sally Corden, Tom Connor |
| EPI_ISL_3296923 | Viollier AG | Department of Biosystems Science and Engineering, ETH Zürich | Christian Beisel, Sarah Nadeau, Chaoran Chen, Ivan Topolsky, Kim Philipp Jablonski, Lara Fuhrmann, Rebecca Denes, Mirjam Feldkamp, Ina Nissen, Natascha Santacroce, Elodie Burcklen, Christiane Beckmann, Maurice Redondo, Olivier Kobel, Christoph Noppen, Niko Beerenwinkel, Tanja Stadler |
| EPI_ISL_3296924 | Wales Specialist Virology Centre Sequencing lab: Pathogen Genomics Unit | Public Health Wales Microbiology Cardiff Wales Specialist Virology Centre | Catherine Moore, Johnathan Evans, Laura Gifford, Malorie Perry, Simon Cottrell, Angela Marchbank, Alec Birchley, Alexander Adams, Amy Gaskin, Bree Gatica-Wilcox, Jason Coombes, Joel Southgate, Lauren Gilbert, Lee Graham, Nicole Pacchiarini, Sara Kumziene-Summerhayes, Sarah Taylor, Sophie Jones, Sara Rey, Matthew Bull, Joanne Watkins, Sally Corden, Tom Connor |
| EPI_ISL_3296925 | Viollier AG | Department of Biosystems Science and Engineering, ETH Zürich | Christian Beisel, Sarah Nadeau, Chaoran Chen, Ivan Topolsky, Kim Philipp Jablonski, Lara Fuhrmann, Rebecca Denes, Mirjam Feldkamp, Ina Nissen, Natascha Santacroce, Elodie Burcklen, Christiane Beckmann, Maurice Redondo, Olivier Kobel, Christoph Noppen, Niko Beerenwinkel, Tanja Stadler |
| EPI_ISL_3296926 | Wales Specialist Virology Centre Sequencing lab: Pathogen Genomics Unit | Public Health Wales Microbiology Cardiff Wales Specialist Virology Centre | Catherine Moore, Johnathan Evans, Laura Gifford, Malorie Perry, Simon Cottrell, Angela Marchbank, Alec Birchley, Alexander Adams, Amy Gaskin, Bree Gatica-Wilcox, Jason Coombes, Joel Southgate, Lauren Gilbert, Lee Graham, Nicole Pacchiarini, Sara Kumziene-Summerhayes, Sarah Taylor, Sophie Jones, Sara Rey, Matthew Bull, Joanne Watkins, Sally Corden, Tom Connor |
| EPI_ISL_3296927 | Viollier AG | Department of Biosystems Science and Engineering, ETH Zürich | Christian Beisel, Sarah Nadeau, Chaoran Chen, Ivan Topolsky, Kim Philipp Jablonski, Lara Fuhrmann, Rebecca Denes, Mirjam Feldkamp, Ina Nissen, Natascha Santacroce, Elodie Burcklen, Christiane Beckmann, Maurice Redondo, Olivier Kobel, Christoph Noppen, Niko Beerenwinkel, Tanja Stadler |
| EPI_ISL_3296928, EPI_ISL_3296929 | Wales Specialist Virology Centre Sequencing lab: Pathogen Genomics Unit | Public Health Wales Microbiology Cardiff Wales Specialist Virology Centre | Catherine Moore, Johnathan Evans, Laura Gifford, Malorie Perry, Simon Cottrell, Angela Marchbank, Alec Birchley, Alexander Adams, Amy Gaskin, Bree Gatica-Wilcox, Jason Coombes, Joel Southgate, Lauren Gilbert, Lee Graham, Nicole Pacchiarini, Sara Kumziene-Summerhayes, Sarah Taylor, Sophie Jones, Sara Rey, Matthew Bull, Joanne Watkins, Sally Corden, Tom Connor |
| EPI_ISL_3296930 | Viollier AG | Department of Biosystems Science and Engineering, ETH Zürich | Christian Beisel, Sarah Nadeau, Chaoran Chen, Ivan Topolsky, Kim Philipp Jablonski, Lara Fuhrmann, Rebecca Denes, Mirjam Feldkamp, Ina Nissen, Natascha Santacroce, Elodie Burcklen, Christiane Beckmann, Maurice Redondo, Olivier Kobel, Christoph Noppen, Niko Beerenwinkel, Tanja Stadler |
| EPI_ISL_3296931 | Wales Specialist Virology Centre Sequencing lab: Pathogen Genomics Unit | Public Health Wales Microbiology Cardiff Wales Specialist Virology Centre | Catherine Moore, Johnathan Evans, Laura Gifford, Malorie Perry, Simon Cottrell, Angela Marchbank, Alec Birchley, Alexander Adams, Amy Gaskin, Bree Gatica-Wilcox, Jason Coombes, Joel Southgate, Lauren Gilbert, Lee Graham, Nicole Pacchiarini, Sara Kumziene-Summerhayes, Sarah Taylor, Sophie Jones, Sara Rey, Matthew Bull, Joanne Watkins, Sally Corden, Tom Connor |
| EPI_ISL_3296932 | Viollier AG | Department of Biosystems Science and Engineering, ETH Zürich | Christian Beisel, Sarah Nadeau, Chaoran Chen, Ivan Topolsky, Kim Philipp Jablonski, Lara Fuhrmann, Rebecca Denes, Mirjam Feldkamp, Ina Nissen, Natascha Santacroce, Elodie Burcklen, Christiane Beckmann, Maurice Redondo, Olivier Kobel, Christoph Noppen, Niko Beerenwinkel, Tanja Stadler |
| EPI_ISL_3296933 | Wales Specialist Virology Centre Sequencing lab: Pathogen Genomics Unit | Public Health Wales Microbiology Cardiff Wales Specialist Virology Centre | Catherine Moore, Johnathan Evans, Laura Gifford, Malorie Perry, Simon Cottrell, Angela Marchbank, Alec Birchley, Alexander Adams, Amy Gaskin, Bree Gatica-Wilcox, Jason Coombes, Joel Southgate, Lauren Gilbert, Lee Graham, Nicole Pacchiarini, Sara Kumziene-Summerhayes, Sarah Taylor, Sophie Jones, Sara Rey, Matthew Bull, Joanne Watkins, Sally Corden, Tom Connor |
| EPI_ISL_3296934 | Viollier AG | Department of Biosystems Science and Engineering, ETH Zürich | Christian Beisel, Sarah Nadeau, Chaoran Chen, Ivan Topolsky, Kim Philipp Jablonski, Lara Fuhrmann, Rebecca Denes, Mirjam Feldkamp, Ina Nissen, Natascha Santacroce, Elodie Burcklen, Christiane Beckmann, Maurice Redondo, Olivier Kobel, Christoph Noppen, Niko Beerenwinkel, Tanja Stadler |
| EPI_ISL_3296935 | Wales Specialist Virology Centre Sequencing lab: Pathogen Genomics Unit | Public Health Wales Microbiology Cardiff Wales Specialist Virology Centre | Catherine Moore, Johnathan Evans, Laura Gifford, Malorie Perry, Simon Cottrell, Angela Marchbank, Alec Birchley, Alexander Adams, Amy Gaskin, Bree Gatica-Wilcox, Jason Coombes, Joel Southgate, Lauren Gilbert, Lee Graham, Nicole Pacchiarini, Sara Kumziene-Summerhayes, Sarah Taylor, Sophie Jones, Sara Rey, Matthew Bull, Joanne Watkins, Sally Corden, Tom Connor |

| | | | |
|---|---|---|---|
| EPI_ISL_3296936 | Viollier AG | Department of Biosystems Science and Engineering, ETH Zürich | Christian Beisel, Sarah Nadeau, Chaoran Chen, Ivan Topolsky, Kim Philipp Jablonski, Lara Fuhrmann, Rebecca Denes, Mirjam Feldkamp, Ina Nissen, Natascha Santacroce, Elodie Burcklen, Christiane Beckmann, Maurice Redondo, Olivier Kobel, Christoph Noppen, Niko Beerenwinkel, Tanja Stadler |
| EPI_ISL_3296937 | Wales Specialist Virology Centre Sequencing lab: Pathogen Genomics Unit | Public Health Wales Microbiology Cardiff Wales Specialist Virology Centre | Catherine Moore, Johnathan Evans, Laura Gifford, Malorie Perry, Simon Cottrell, Angela Marchbank, Alec Birchley, Alexander Adams, Amy Gaskin, Bree Gatica-Wilcox, Jason Coombes, Joel Southgate, Lauren Gilbert, Lee Graham, Nicole Pacchiarini, Sara Kumziene-Summerhayes, Sarah Taylor, Sophie Jones, Sara Rey, Matthew Bull, Joanne Watkins, Sally Corden, Tom Connor |
| EPI_ISL_3296938 | Viollier AG | Department of Biosystems Science and Engineering, ETH Zürich | Christian Beisel, Sarah Nadeau, Chaoran Chen, Ivan Topolsky, Kim Philipp Jablonski, Lara Fuhrmann, Rebecca Denes, Mirjam Feldkamp, Ina Nissen, Natascha Santacroce, Elodie Burcklen, Christiane Beckmann, Maurice Redondo, Olivier Kobel, Christoph Noppen, Niko Beerenwinkel, Tanja Stadler |
| EPI_ISL_3296939 | Wales Specialist Virology Centre Sequencing lab: Pathogen Genomics Unit | Public Health Wales Microbiology Cardiff Wales Specialist Virology Centre | Catherine Moore, Johnathan Evans, Laura Gifford, Malorie Perry, Simon Cottrell, Angela Marchbank, Alec Birchley, Alexander Adams, Amy Gaskin, Bree Gatica-Wilcox, Jason Coombes, Joel Southgate, Lauren Gilbert, Lee Graham, Nicole Pacchiarini, Sara Kumziene-Summerhayes, Sarah Taylor, Sophie Jones, Sara Rey, Matthew Bull, Joanne Watkins, Sally Corden, Tom Connor |
| EPI_ISL_3296940 | Viollier AG | Department of Biosystems Science and Engineering, ETH Zürich | Christian Beisel, Sarah Nadeau, Chaoran Chen, Ivan Topolsky, Kim Philipp Jablonski, Lara Fuhrmann, Rebecca Denes, Mirjam Feldkamp, Ina Nissen, Natascha Santacroce, Elodie Burcklen, Christiane Beckmann, Maurice Redondo, Olivier Kobel, Christoph Noppen, Niko Beerenwinkel, Tanja Stadler |
| EPI_ISL_3296941 | Wales Specialist Virology Centre Sequencing lab: Pathogen Genomics Unit | Public Health Wales Microbiology Cardiff Wales Specialist Virology Centre | Catherine Moore, Johnathan Evans, Laura Gifford, Malorie Perry, Simon Cottrell, Angela Marchbank, Alec Birchley, Alexander Adams, Amy Gaskin, Bree Gatica-Wilcox, Jason Coombes, Joel Southgate, Lauren Gilbert, Lee Graham, Nicole Pacchiarini, Sara Kumziene-Summerhayes, Sarah Taylor, Sophie Jones, Sara Rey, Matthew Bull, Joanne Watkins, Sally Corden, Tom Connor |
| EPI_ISL_3296942 | Viollier AG | Department of Biosystems Science and Engineering, ETH Zürich | Christian Beisel, Sarah Nadeau, Chaoran Chen, Ivan Topolsky, Kim Philipp Jablonski, Lara Fuhrmann, Rebecca Denes, Mirjam Feldkamp, Ina Nissen, Natascha Santacroce, Elodie Burcklen, Christiane Beckmann, Maurice Redondo, Olivier Kobel, Christoph Noppen, Niko Beerenwinkel, Tanja Stadler |
| EPI_ISL_3296943, EPI_ISL_3296944 | Wales Specialist Virology Centre Sequencing lab: Pathogen Genomics Unit | Public Health Wales Microbiology Cardiff Wales Specialist Virology Centre | Catherine Moore, Johnathan Evans, Laura Gifford, Malorie Perry, Simon Cottrell, Angela Marchbank, Alec Birchley, Alexander Adams, Amy Gaskin, Bree Gatica-Wilcox, Jason Coombes, Joel Southgate, Lauren Gilbert, Lee Graham, Nicole Pacchiarini, Sara Kumziene-Summerhayes, Sarah Taylor, Sophie Jones, Sara Rey, Matthew Bull, Joanne Watkins, Sally Corden, Tom Connor |
| EPI_ISL_3296945 | Viollier AG | Department of Biosystems Science and Engineering, ETH Zürich | Christian Beisel, Sarah Nadeau, Chaoran Chen, Ivan Topolsky, Kim Philipp Jablonski, Lara Fuhrmann, Rebecca Denes, Mirjam Feldkamp, Ina Nissen, Natascha Santacroce, Elodie Burcklen, Christiane Beckmann, Maurice Redondo, Olivier Kobel, Christoph Noppen, Niko Beerenwinkel, Tanja Stadler |
| EPI_ISL_3296946 | Wales Specialist Virology Centre Sequencing lab: Pathogen Genomics Unit | Public Health Wales Microbiology Cardiff Wales Specialist Virology Centre | Catherine Moore, Johnathan Evans, Laura Gifford, Malorie Perry, Simon Cottrell, Angela Marchbank, Alec Birchley, Alexander Adams, Amy Gaskin, Bree Gatica-Wilcox, Jason Coombes, Joel Southgate, Lauren Gilbert, Lee Graham, Nicole Pacchiarini, Sara Kumziene-Summerhayes, Sarah Taylor, Sophie Jones, Sara Rey, Matthew Bull, Joanne Watkins, Sally Corden, Tom Connor |
| EPI_ISL_3296947 | Viollier AG | Department of Biosystems Science and Engineering, ETH Zürich | Christian Beisel, Sarah Nadeau, Chaoran Chen, Ivan Topolsky, Kim Philipp Jablonski, Lara Fuhrmann, Rebecca Denes, Mirjam Feldkamp, Ina Nissen, Natascha Santacroce, Elodie Burcklen, Christiane Beckmann, Maurice Redondo, Olivier Kobel, Christoph Noppen, Niko Beerenwinkel, Tanja Stadler |
| EPI_ISL_3296948 | Wales Specialist Virology Centre Sequencing lab: Pathogen Genomics Unit | Public Health Wales Microbiology Cardiff Wales Specialist Virology Centre | Catherine Moore, Johnathan Evans, Laura Gifford, Malorie Perry, Simon Cottrell, Angela Marchbank, Alec Birchley, Alexander Adams, Amy Gaskin, Bree Gatica-Wilcox, Jason Coombes, Joel Southgate, Lauren Gilbert, Lee Graham, Nicole Pacchiarini, Sara Kumziene-Summerhayes, Sarah Taylor, Sophie Jones, Sara Rey, Matthew Bull, Joanne Watkins, Sally Corden, Tom Connor |
| EPI_ISL_3296949 | Viollier AG | Department of Biosystems Science and Engineering, ETH Zürich | Christian Beisel, Sarah Nadeau, Chaoran Chen, Ivan Topolsky, Kim Philipp Jablonski, Lara Fuhrmann, Rebecca Denes, Mirjam Feldkamp, Ina Nissen, Natascha Santacroce, Elodie Burcklen, Christiane Beckmann, Maurice Redondo, Olivier Kobel, Christoph Noppen, Niko Beerenwinkel, Tanja Stadler |
| EPI_ISL_3296950 | Wales Specialist Virology Centre Sequencing lab: Pathogen Genomics Unit | Public Health Wales Microbiology Cardiff Wales Specialist Virology Centre | Catherine Moore, Johnathan Evans, Laura Gifford, Malorie Perry, Simon Cottrell, Angela Marchbank, Alec Birchley, Alexander Adams, Amy Gaskin, Bree Gatica-Wilcox, Jason Coombes, Joel Southgate, Lauren Gilbert, Lee Graham, Nicole Pacchiarini, Sara Kumziene-Summerhayes, Sarah Taylor, Sophie Jones, Sara Rey, Matthew Bull, Joanne Watkins, Sally Corden, Tom Connor |
| EPI_ISL_3296951 | Viollier AG | Department of Biosystems Science and Engineering, ETH Zürich | Christian Beisel, Sarah Nadeau, Chaoran Chen, Ivan Topolsky, Kim Philipp Jablonski, Lara Fuhrmann, Rebecca Denes, Mirjam Feldkamp, Ina Nissen, Natascha Santacroce, Elodie Burcklen, Christiane Beckmann, Maurice Redondo, Olivier Kobel, Christoph Noppen, Niko Beerenwinkel, Tanja Stadler |
| EPI_ISL_3296952 | Wales Specialist Virology Centre Sequencing lab: Pathogen Genomics Unit | Public Health Wales Microbiology Cardiff Wales Specialist Virology Centre | Catherine Moore, Johnathan Evans, Laura Gifford, Malorie Perry, Simon Cottrell, Angela Marchbank, Alec Birchley, Alexander Adams, Amy Gaskin, Bree Gatica-Wilcox, Jason Coombes, Joel Southgate, Lauren Gilbert, Lee Graham, Nicole Pacchiarini, Sara Kumziene-Summerhayes, Sarah Taylor, Sophie Jones, Sara Rey, Matthew Bull, Joanne Watkins, Sally Corden, Tom Connor |
| EPI_ISL_3296953 | Viollier AG | Department of Biosystems Science and Engineering, ETH Zürich | Christian Beisel, Sarah Nadeau, Chaoran Chen, Ivan Topolsky, Kim Philipp Jablonski, Lara Fuhrmann, Rebecca Denes, Mirjam Feldkamp, Ina Nissen, Natascha Santacroce, Elodie Burcklen, Christiane Beckmann, Maurice Redondo, Olivier Kobel, Christoph Noppen, Niko Beerenwinkel, Tanja Stadler |
| EPI_ISL_3296954 | Wales Specialist Virology Centre Sequencing lab: Pathogen Genomics Unit | Public Health Wales Microbiology Cardiff Wales Specialist Virology Centre | Catherine Moore, Johnathan Evans, Laura Gifford, Malorie Perry, Simon Cottrell, Angela Marchbank, Alec Birchley, Alexander Adams, Amy Gaskin, Bree Gatica-Wilcox, Jason Coombes, Joel Southgate, Lauren Gilbert, Lee Graham, Nicole Pacchiarini, Sara Kumziene-Summerhayes, Sarah Taylor, Sophie Jones, Sara Rey, Matthew Bull, Joanne Watkins, Sally Corden, Tom Connor |
| EPI_ISL_3296955 | Viollier AG | Department of Biosystems Science and Engineering, ETH Zürich | Christian Beisel, Sarah Nadeau, Chaoran Chen, Ivan Topolsky, Kim Philipp Jablonski, Lara Fuhrmann, Rebecca Denes, Mirjam Feldkamp, Ina Nissen, Natascha Santacroce, Elodie Burcklen, Christiane Beckmann, Maurice Redondo, Olivier Kobel, Christoph Noppen, Niko Beerenwinkel, Tanja Stadler |
| EPI_ISL_3296956 | Wales Specialist Virology Centre Sequencing lab: Pathogen Genomics Unit | Public Health Wales Microbiology Cardiff Wales Specialist Virology Centre | Catherine Moore, Johnathan Evans, Laura Gifford, Malorie Perry, Simon Cottrell, Angela Marchbank, Alec Birchley, Alexander Adams, Amy Gaskin, Bree Gatica-Wilcox, Jason Coombes, Joel Southgate, Lauren Gilbert, Lee Graham, Nicole Pacchiarini, Sara Kumziene-Summerhayes, Sarah Taylor, Sophie Jones, Sara Rey, Matthew Bull, Joanne Watkins, Sally Corden, Tom Connor |
| EPI_ISL_3296957 | Viollier AG | Department of Biosystems Science and Engineering, ETH Zürich | Christian Beisel, Sarah Nadeau, Chaoran Chen, Ivan Topolsky, Kim Philipp Jablonski, Lara Fuhrmann, Rebecca Denes, Mirjam Feldkamp, Ina Nissen, Natascha Santacroce, Elodie Burcklen, Christiane Beckmann, Maurice Redondo, Olivier Kobel, Christoph Noppen, Niko Beerenwinkel, Tanja Stadler |
| EPI_ISL_3296958 | Wales Specialist Virology Centre Sequencing lab: Pathogen Genomics Unit | Public Health Wales Microbiology Cardiff Wales Specialist Virology Centre | Catherine Moore, Johnathan Evans, Laura Gifford, Malorie Perry, Simon Cottrell, Angela Marchbank, Alec Birchley, Alexander Adams, Amy Gaskin, Bree Gatica-Wilcox, Jason Coombes, Joel Southgate, Lauren Gilbert, Lee Graham, Nicole Pacchiarini, Sara Kumziene-Summerhayes, Sarah Taylor, Sophie Jones, Sara Rey, Matthew Bull, Joanne Watkins, Sally Corden, Tom Connor |
| EPI_ISL_3296959 | Viollier AG | Department of Biosystems Science and Engineering, ETH Zürich | Christian Beisel, Sarah Nadeau, Chaoran Chen, Ivan Topolsky, Kim Philipp Jablonski, Lara Fuhrmann, Rebecca Denes, Mirjam Feldkamp, Ina Nissen, Natascha Santacroce, Elodie Burcklen, Christiane Beckmann, Maurice Redondo, Olivier Kobel, Christoph Noppen, Niko Beerenwinkel, Tanja Stadler |
| EPI_ISL_3296960 | Wales Specialist Virology Centre Sequencing lab: Pathogen Genomics Unit | Public Health Wales Microbiology Cardiff Wales Specialist Virology Centre | Catherine Moore, Johnathan Evans, Laura Gifford, Malorie Perry, Simon Cottrell, Angela Marchbank, Alec Birchley, Alexander Adams, Amy Gaskin, Bree Gatica-Wilcox, Jason Coombes, Joel Southgate, Lauren Gilbert, Lee Graham, Nicole Pacchiarini, Sara Kumziene-Summerhayes, Sarah Taylor, Sophie Jones, Sara Rey, Matthew Bull, Joanne Watkins, Sally Corden, Tom Connor |
| EPI_ISL_3296961 | Viollier AG | Department of Biosystems Science and Engineering, ETH Zürich | Christian Beisel, Sarah Nadeau, Chaoran Chen, Ivan Topolsky, Kim Philipp Jablonski, Lara Fuhrmann, Rebecca Denes, Mirjam Feldkamp, Ina Nissen, Natascha Santacroce, Elodie Burcklen, Christiane Beckmann, Maurice Redondo, Olivier Kobel, Christoph Noppen, Niko Beerenwinkel, Tanja Stadler |
| EPI_ISL_3296962 | Wales Specialist Virology Centre Sequencing lab: Pathogen Genomics Unit | Public Health Wales Microbiology Cardiff Wales Specialist Virology Centre | Catherine Moore, Johnathan Evans, Laura Gifford, Malorie Perry, Simon Cottrell, Angela Marchbank, Alec Birchley, Alexander Adams, Amy Gaskin, Bree Gatica-Wilcox, Jason Coombes, Joel Southgate, Lauren Gilbert, Lee Graham, Nicole Pacchiarini, Sara Kumziene-Summerhayes, Sarah Taylor, Sophie Jones, Sara Rey, Matthew Bull, Joanne Watkins, Sally Corden, Tom Connor |
| EPI_ISL_3296963 | Viollier AG | Department of Biosystems Science and Engineering, ETH Zürich | Christian Beisel, Sarah Nadeau, Chaoran Chen, Ivan Topolsky, Kim Philipp Jablonski, Lara Fuhrmann, Rebecca Denes, Mirjam Feldkamp, Ina Nissen, Natascha Santacroce, Elodie Burcklen, Christiane Beckmann, Maurice Redondo, Olivier Kobel, Christoph Noppen, Niko Beerenwinkel, Tanja Stadler |
| EPI_ISL_3296964 | Wales Specialist Virology Centre Sequencing lab: Pathogen Genomics Unit | Public Health Wales Microbiology Cardiff Wales Specialist Virology Centre | Catherine Moore, Johnathan Evans, Laura Gifford, Malorie Perry, Simon Cottrell, Angela Marchbank, Alec Birchley, Alexander Adams, Amy Gaskin, Bree Gatica-Wilcox, Jason Coombes, Joel Southgate, Lauren Gilbert, Lee Graham, Nicole Pacchiarini, Sara Kumziene-Summerhayes, Sarah Taylor, Sophie Jones, Sara Rey, Matthew Bull, Joanne Watkins, Sally Corden, Tom Connor |
| EPI_ISL_3296965 | Viollier AG | Department of Biosystems Science and Engineering, ETH Zürich | Christian Beisel, Sarah Nadeau, Chaoran Chen, Ivan Topolsky, Kim Philipp Jablonski, Lara Fuhrmann, Rebecca Denes, Mirjam Feldkamp, Ina Nissen, Natascha Santacroce, Elodie Burcklen, Christiane Beckmann, Maurice Redondo, Olivier Kobel, Christoph Noppen, Niko Beerenwinkel, Tanja Stadler |
| EPI_ISL_3296966 | Wales Specialist Virology Centre Sequencing lab: Pathogen Genomics Unit | Public Health Wales Microbiology Cardiff Wales Specialist Virology Centre | Catherine Moore, Johnathan Evans, Laura Gifford, Malorie Perry, Simon Cottrell, Angela Marchbank, Alec Birchley, Alexander Adams, Amy Gaskin, Bree Gatica-Wilcox, Jason Coombes, Joel Southgate, Lauren Gilbert, Lee Graham, Nicole Pacchiarini, Sara Kumziene-Summerhayes, Sarah Taylor, Sophie Jones, Sara Rey, Matthew Bull, Joanne Watkins, Sally Corden, Tom Connor |
| EPI_ISL_3296967 | Viollier AG | Department of Biosystems Science and Engineering, ETH Zürich | Christian Beisel, Sarah Nadeau, Chaoran Chen, Ivan Topolsky, Kim Philipp Jablonski, Lara Fuhrmann, Rebecca Denes, Mirjam Feldkamp, Ina Nissen, Natascha Santacroce, Elodie Burcklen, Christiane Beckmann, Maurice Redondo, Olivier Kobel, Christoph Noppen, Niko Beerenwinkel, Tanja Stadler |
| EPI_ISL_3296968 | Wales Specialist Virology Centre Sequencing lab: Pathogen Genomics Unit | Public Health Wales Microbiology Cardiff Wales Specialist Virology Centre | Catherine Moore, Johnathan Evans, Laura Gifford, Malorie Perry, Simon Cottrell, Angela Marchbank, Alec Birchley, Alexander Adams, Amy Gaskin, Bree Gatica-Wilcox, Jason Coombes, Joel Southgate, Lauren Gilbert, Lee Graham, Nicole Pacchiarini, Sara Kumziene-Summerhayes, Sarah Taylor, Sophie Jones, Sara Rey, Matthew Bull, Joanne Watkins, Sally Corden, Tom Connor |
| EPI_ISL_3296969 | Viollier AG | Department of Biosystems Science and Engineering, ETH Zürich | Christian Beisel, Sarah Nadeau, Chaoran Chen, Ivan Topolsky, Kim Philipp Jablonski, Lara Fuhrmann, Rebecca Denes, Mirjam Feldkamp, Ina Nissen, Natascha Santacroce, Elodie Burcklen, Christiane Beckmann, Maurice Redondo, Olivier Kobel, Christoph Noppen, Niko Beerenwinkel, Tanja Stadler |
| EPI_ISL_3296970, EPI_ISL_3296971 | Wales Specialist Virology Centre Sequencing lab: Pathogen Genomics Unit | Public Health Wales Microbiology Cardiff Wales Specialist Virology Centre | Catherine Moore, Johnathan Evans, Laura Gifford, Malorie Perry, Simon Cottrell, Angela Marchbank, Alec Birchley, Alexander Adams, Amy Gaskin, Bree Gatica-Wilcox, Jason Coombes, Joel Southgate, Lauren Gilbert, Lee Graham, Nicole Pacchiarini, Sara Kumziene-Summerhayes, Sarah Taylor, Sophie Jones, Sara Rey, Matthew Bull, Joanne Watkins, Sally Corden, Tom Connor |
| EPI_ISL_3296972 | Viollier AG | Department of Biosystems Science and Engineering, ETH Zürich | Christian Beisel, Sarah Nadeau, Chaoran Chen, Ivan Topolsky, Kim Philipp Jablonski, Lara Fuhrmann, Rebecca Denes, Mirjam Feldkamp, Ina Nissen, Natascha Santacroce, Elodie Burcklen, Christiane Beckmann, Maurice Redondo, Olivier Kobel, Christoph Noppen, Niko Beerenwinkel, Tanja Stadler |
| EPI_ISL_3296973 | Wales Specialist Virology Centre Sequencing lab: Pathogen Genomics Unit | Public Health Wales Microbiology Cardiff Wales Specialist Virology Centre | Catherine Moore, Johnathan Evans, Laura Gifford, Malorie Perry, Simon Cottrell, Angela Marchbank, Alec Birchley, Alexander Adams, Amy Gaskin, Bree Gatica-Wilcox, Jason Coombes, Joel Southgate, Lauren Gilbert, Lee Graham, Nicole Pacchiarini, Sara Kumziene-Summerhayes, Sarah Taylor, Sophie Jones, Sara Rey, Matthew Bull, Joanne Watkins, Sally Corden, Tom Connor |
| EPI_ISL_3296974 | Viollier AG | Department of Biosystems Science and Engineering, ETH Zürich | Christian Beisel, Sarah Nadeau, Chaoran Chen, Ivan Topolsky, Kim Philipp Jablonski, Lara Fuhrmann, Rebecca Denes, Mirjam Feldkamp, Ina Nissen, Natascha Santacroce, Elodie Burcklen, Christiane Beckmann, Maurice Redondo, Olivier Kobel, Christoph Noppen, Niko Beerenwinkel, Tanja Stadler |
| EPI_ISL_3296975 | Wales Specialist Virology Centre Sequencing lab: Pathogen Genomics Unit | Public Health Wales Microbiology Cardiff Wales Specialist Virology Centre | Catherine Moore, Johnathan Evans, Laura Gifford, Malorie Perry, Simon Cottrell, Angela Marchbank, Alec Birchley, Alexander Adams, Amy Gaskin, Bree Gatica-Wilcox, Jason Coombes, Joel Southgate, Lauren Gilbert, Lee Graham, Nicole Pacchiarini, Sara Kumziene-Summerhayes, Sarah Taylor, Sophie Jones, Sara Rey, Matthew Bull, Joanne Watkins, Sally Corden, Tom Connor |
| EPI_ISL_3296976 | Viollier AG | Department of Biosystems Science and Engineering, ETH Zürich | Christian Beisel, Sarah Nadeau, Chaoran Chen, Ivan Topolsky, Kim Philipp Jablonski, Lara Fuhrmann, Rebecca Denes, Mirjam Feldkamp, Ina Nissen, Natascha Santacroce, Elodie Burcklen, Christiane Beckmann, Maurice Redondo, Olivier Kobel, Christoph Noppen, Niko Beerenwinkel, Tanja Stadler |
| EPI_ISL_3296977 | Wales Specialist Virology Centre Sequencing lab: Pathogen Genomics Unit | Public Health Wales Microbiology Cardiff Wales Specialist Virology Centre | Catherine Moore, Johnathan Evans, Laura Gifford, Malorie Perry, Simon Cottrell, Angela Marchbank, Alec Birchley, Alexander Adams, Amy Gaskin, Bree Gatica-Wilcox, Jason Coombes, Joel Southgate, Lauren Gilbert, Lee Graham, Nicole Pacchiarini, Sara Kumziene-Summerhayes, Sarah Taylor, Sophie Jones, Sara Rey, Matthew Bull, Joanne Watkins, Sally Corden, Tom Connor |
| EPI_ISL_3296978 | Viollier AG | Department of Biosystems Science and Engineering, ETH Zürich | Christian Beisel, Sarah Nadeau, Chaoran Chen, Ivan Topolsky, Kim Philipp Jablonski, Lara Fuhrmann, Rebecca Denes, Mirjam Feldkamp, Ina Nissen, Natascha Santacroce, Elodie Burcklen, Christiane Beckmann, Maurice Redondo, Olivier Kobel, Christoph Noppen, Niko Beerenwinkel, Tanja Stadler |
| EPI_ISL_3296979 | Wales Specialist Virology Centre Sequencing lab: Pathogen Genomics Unit | Public Health Wales Microbiology Cardiff Wales Specialist Virology Centre | Catherine Moore, Johnathan Evans, Laura Gifford, Malorie Perry, Simon Cottrell, Angela Marchbank, Alec Birchley, Alexander Adams, Amy Gaskin, Bree Gatica-Wilcox, Jason Coombes, Joel Southgate, Lauren Gilbert, Lee Graham, Nicole Pacchiarini, Sara Kumziene-Summerhayes, Sarah Taylor, Sophie Jones, Sara Rey, Matthew Bull, Joanne Watkins, Sally Corden, Tom Connor |
| EPI_ISL_3296980 | Viollier AG | Department of Biosystems Science and Engineering, ETH Zürich | Christian Beisel, Sarah Nadeau, Chaoran Chen, Ivan Topolsky, Kim Philipp Jablonski, Lara Fuhrmann, Rebecca Denes, Mirjam Feldkamp, Ina Nissen, Natascha Santacroce, Elodie Burcklen, Christiane Beckmann, Maurice Redondo, Olivier Kobel, Christoph Noppen, Niko Beerenwinkel, Tanja Stadler |
| EPI_ISL_3296981 | Wales Specialist Virology Centre Sequencing lab: Pathogen Genomics Unit | Public Health Wales Microbiology Cardiff Wales Specialist Virology Centre | Catherine Moore, Johnathan Evans, Laura Gifford, Malorie Perry, Simon Cottrell, Angela Marchbank, Alec Birchley, Alexander Adams, Amy Gaskin, Bree Gatica-Wilcox, Jason Coombes, Joel Southgate, Lauren Gilbert, Lee Graham, Nicole Pacchiarini, Sara Kumziene-Summerhayes, Sarah Taylor, Sophie Jones, Sara Rey, Matthew Bull, Joanne Watkins, Sally Corden, Tom Connor |
| EPI_ISL_3296982 | Viollier AG | Department of Biosystems Science and Engineering, ETH Zürich | Christian Beisel, Sarah Nadeau, Chaoran Chen, Ivan Topolsky, Kim Philipp Jablonski, Lara Fuhrmann, Rebecca Denes, Mirjam Feldkamp, Ina Nissen, Natascha Santacroce, Elodie Burcklen, Christiane Beckmann, Maurice Redondo, Olivier Kobel, Christoph Noppen, Niko Beerenwinkel, Tanja Stadler |

| | | | |
|---|---|---|---|
| EPI_ISL_3296983 | Wales Specialist Virology Centre Sequencing lab: Pathogen Genomics Unit | Public Health Wales Microbiology Cardiff Wales Specialist Virology Centre | Catherine Moore, Johnathan Evans, Laura Gifford, Malorie Perry, Simon Cottrell, Angela Marchbank, Alec Birchley, Alexander Adams, Amy Gaskin, Bree Gatica-Wilcox, Jason Coombes, Joel Southgate, Lauren Gilbert, Lee Graham, Nicole Pacchiarini, Sara Kumziene-Summerhayes, Sarah Taylor, Sophie Jones, Sara Rey, Matthew Bull, Joanne Watkins, Sally Corden, Tom Connor |
| EPI_ISL_3296984 | Viollier AG | Department of Biosystems Science and Engineering, ETH Zürich | Christian Beisel, Sarah Nadeau, Chaoran Chen, Ivan Topolsky, Kim Philipp Jablonski, Lara Fuhrmann, Rebecca Denes, Mirjam Feldkamp, Ina Nissen, Natascha Santacroce, Elodie Burcklen, Christiane Beckmann, Maurice Redondo, Olivier Kobel, Christoph Noppen, Niko Beerenwinkel, Tanja Stadler |
| EPI_ISL_3296985 | Wales Specialist Virology Centre Sequencing lab: Pathogen Genomics Unit | Public Health Wales Microbiology Cardiff Wales Specialist Virology Centre | Catherine Moore, Johnathan Evans, Laura Gifford, Malorie Perry, Simon Cottrell, Angela Marchbank, Alec Birchley, Alexander Adams, Amy Gaskin, Bree Gatica-Wilcox, Jason Coombes, Joel Southgate, Lauren Gilbert, Lee Graham, Nicole Pacchiarini, Sara Kumziene-Summerhayes, Sarah Taylor, Sophie Jones, Sara Rey, Matthew Bull, Joanne Watkins, Sally Corden, Tom Connor |
| EPI_ISL_3296986 | Viollier AG | Department of Biosystems Science and Engineering, ETH Zürich | Christian Beisel, Sarah Nadeau, Chaoran Chen, Ivan Topolsky, Kim Philipp Jablonski, Lara Fuhrmann, Rebecca Denes, Mirjam Feldkamp, Ina Nissen, Natascha Santacroce, Elodie Burcklen, Christiane Beckmann, Maurice Redondo, Olivier Kobel, Christoph Noppen, Niko Beerenwinkel, Tanja Stadler |
| EPI_ISL_3296987 | Wales Specialist Virology Centre Sequencing lab: Pathogen Genomics Unit | Public Health Wales Microbiology Cardiff Wales Specialist Virology Centre | Catherine Moore, Johnathan Evans, Laura Gifford, Malorie Perry, Simon Cottrell, Angela Marchbank, Alec Birchley, Alexander Adams, Amy Gaskin, Bree Gatica-Wilcox, Jason Coombes, Joel Southgate, Lauren Gilbert, Lee Graham, Nicole Pacchiarini, Sara Kumziene-Summerhayes, Sarah Taylor, Sophie Jones, Sara Rey, Matthew Bull, Joanne Watkins, Sally Corden, Tom Connor |
| EPI_ISL_3296988 | Viollier AG | Department of Biosystems Science and Engineering, ETH Zürich | Christian Beisel, Sarah Nadeau, Chaoran Chen, Ivan Topolsky, Kim Philipp Jablonski, Lara Fuhrmann, Rebecca Denes, Mirjam Feldkamp, Ina Nissen, Natascha Santacroce, Elodie Burcklen, Christiane Beckmann, Maurice Redondo, Olivier Kobel, Christoph Noppen, Niko Beerenwinkel, Tanja Stadler |
| EPI_ISL_3296989 | Wales Specialist Virology Centre Sequencing lab: Pathogen Genomics Unit | Public Health Wales Microbiology Cardiff Wales Specialist Virology Centre | Catherine Moore, Johnathan Evans, Laura Gifford, Malorie Perry, Simon Cottrell, Angela Marchbank, Alec Birchley, Alexander Adams, Amy Gaskin, Bree Gatica-Wilcox, Jason Coombes, Joel Southgate, Lauren Gilbert, Lee Graham, Nicole Pacchiarini, Sara Kumziene-Summerhayes, Sarah Taylor, Sophie Jones, Sara Rey, Matthew Bull, Joanne Watkins, Sally Corden, Tom Connor |
| EPI_ISL_3296990 | Viollier AG | Department of Biosystems Science and Engineering, ETH Zürich | Christian Beisel, Sarah Nadeau, Chaoran Chen, Ivan Topolsky, Kim Philipp Jablonski, Lara Fuhrmann, Rebecca Denes, Mirjam Feldkamp, Ina Nissen, Natascha Santacroce, Elodie Burcklen, Christiane Beckmann, Maurice Redondo, Olivier Kobel, Christoph Noppen, Niko Beerenwinkel, Tanja Stadler |
| EPI_ISL_3296991 | Wales Specialist Virology Centre Sequencing lab: Pathogen Genomics Unit | Public Health Wales Microbiology Cardiff Wales Specialist Virology Centre | Catherine Moore, Johnathan Evans, Laura Gifford, Malorie Perry, Simon Cottrell, Angela Marchbank, Alec Birchley, Alexander Adams, Amy Gaskin, Bree Gatica-Wilcox, Jason Coombes, Joel Southgate, Lauren Gilbert, Lee Graham, Nicole Pacchiarini, Sara Kumziene-Summerhayes, Sarah Taylor, Sophie Jones, Sara Rey, Matthew Bull, Joanne Watkins, Sally Corden, Tom Connor |
| EPI_ISL_3296992 | Viollier AG | Department of Biosystems Science and Engineering, ETH Zürich | Christian Beisel, Sarah Nadeau, Chaoran Chen, Ivan Topolsky, Kim Philipp Jablonski, Lara Fuhrmann, Rebecca Denes, Mirjam Feldkamp, Ina Nissen, Natascha Santacroce, Elodie Burcklen, Christiane Beckmann, Maurice Redondo, Olivier Kobel, Christoph Noppen, Niko Beerenwinkel, Tanja Stadler |
| EPI_ISL_3296993, EPI_ISL_3296995 | Wales Specialist Virology Centre Sequencing lab: Pathogen Genomics Unit | Public Health Wales Microbiology Cardiff Wales Specialist Virology Centre | Catherine Moore, Johnathan Evans, Laura Gifford, Malorie Perry, Simon Cottrell, Angela Marchbank, Alec Birchley, Alexander Adams, Amy Gaskin, Bree Gatica-Wilcox, Jason Coombes, Joel Southgate, Lauren Gilbert, Lee Graham, Nicole Pacchiarini, Sara Kumziene-Summerhayes, Sarah Taylor, Sophie Jones, Sara Rey, Matthew Bull, Joanne Watkins, Sally Corden, Tom Connor |
| EPI_ISL_3296996 | Viollier AG | Department of Biosystems Science and Engineering, ETH Zürich | Christian Beisel, Sarah Nadeau, Chaoran Chen, Ivan Topolsky, Kim Philipp Jablonski, Lara Fuhrmann, Rebecca Denes, Mirjam Feldkamp, Ina Nissen, Natascha Santacroce, Elodie Burcklen, Christiane Beckmann, Maurice Redondo, Olivier Kobel, Christoph Noppen, Niko Beerenwinkel, Tanja Stadler |
| EPI_ISL_3296997, EPI_ISL_3296998 | Wales Specialist Virology Centre Sequencing lab: Pathogen Genomics Unit | Public Health Wales Microbiology Cardiff Wales Specialist Virology Centre | Catherine Moore, Johnathan Evans, Laura Gifford, Malorie Perry, Simon Cottrell, Angela Marchbank, Alec Birchley, Alexander Adams, Amy Gaskin, Bree Gatica-Wilcox, Jason Coombes, Joel Southgate, Lauren Gilbert, Lee Graham, Nicole Pacchiarini, Sara Kumziene-Summerhayes, Sarah Taylor, Sophie Jones, Sara Rey, Matthew Bull, Joanne Watkins, Sally Corden, Tom Connor |
| EPI_ISL_3296999 | Viollier AG | Department of Biosystems Science and Engineering, ETH Zürich | Christian Beisel, Sarah Nadeau, Chaoran Chen, Ivan Topolsky, Kim Philipp Jablonski, Lara Fuhrmann, Rebecca Denes, Mirjam Feldkamp, Ina Nissen, Natascha Santacroce, Elodie Burcklen, Christiane Beckmann, Maurice Redondo, Olivier Kobel, Christoph Noppen, Niko Beerenwinkel, Tanja Stadler |
| EPI_ISL_3297000 | Wales Specialist Virology Centre Sequencing lab: Pathogen Genomics Unit | Public Health Wales Microbiology Cardiff Wales Specialist Virology Centre | Catherine Moore, Johnathan Evans, Laura Gifford, Malorie Perry, Simon Cottrell, Angela Marchbank, Alec Birchley, Alexander Adams, Amy Gaskin, Bree Gatica-Wilcox, Jason Coombes, Joel Southgate, Lauren Gilbert, Lee Graham, Nicole Pacchiarini, Sara Kumziene-Summerhayes, Sarah Taylor, Sophie Jones, Sara Rey, Matthew Bull, Joanne Watkins, Sally Corden, Tom Connor |
| EPI_ISL_3297001 | Viollier AG | Department of Biosystems Science and Engineering, ETH Zürich | Christian Beisel, Sarah Nadeau, Chaoran Chen, Ivan Topolsky, Kim Philipp Jablonski, Lara Fuhrmann, Rebecca Denes, Mirjam Feldkamp, Ina Nissen, Natascha Santacroce, Elodie Burcklen, Christiane Beckmann, Maurice Redondo, Olivier Kobel, Christoph Noppen, Niko Beerenwinkel, Tanja Stadler |
| EPI_ISL_3297002 | Wales Specialist Virology Centre Sequencing lab: Pathogen Genomics Unit | Public Health Wales Microbiology Cardiff Wales Specialist Virology Centre | Catherine Moore, Johnathan Evans, Laura Gifford, Malorie Perry, Simon Cottrell, Angela Marchbank, Alec Birchley, Alexander Adams, Amy Gaskin, Bree Gatica-Wilcox, Jason Coombes, Joel Southgate, Lauren Gilbert, Lee Graham, Nicole Pacchiarini, Sara Kumziene-Summerhayes, Sarah Taylor, Sophie Jones, Sara Rey, Matthew Bull, Joanne Watkins, Sally Corden, Tom Connor |
| EPI_ISL_3297003 | Viollier AG | Department of Biosystems Science and Engineering, ETH Zürich | Christian Beisel, Sarah Nadeau, Chaoran Chen, Ivan Topolsky, Kim Philipp Jablonski, Lara Fuhrmann, Rebecca Denes, Mirjam Feldkamp, Ina Nissen, Natascha Santacroce, Elodie Burcklen, Christiane Beckmann, Maurice Redondo, Olivier Kobel, Christoph Noppen, Niko Beerenwinkel, Tanja Stadler |
| EPI_ISL_3297004 | Wales Specialist Virology Centre Sequencing lab: Pathogen Genomics Unit | Public Health Wales Microbiology Cardiff Wales Specialist Virology Centre | Catherine Moore, Johnathan Evans, Laura Gifford, Malorie Perry, Simon Cottrell, Angela Marchbank, Alec Birchley, Alexander Adams, Amy Gaskin, Bree Gatica-Wilcox, Jason Coombes, Joel Southgate, Lauren Gilbert, Lee Graham, Nicole Pacchiarini, Sara Kumziene-Summerhayes, Sarah Taylor, Sophie Jones, Sara Rey, Matthew Bull, Joanne Watkins, Sally Corden, Tom Connor |
| EPI_ISL_3297005 | Viollier AG | Department of Biosystems Science and Engineering, ETH Zürich | Christian Beisel, Sarah Nadeau, Chaoran Chen, Ivan Topolsky, Kim Philipp Jablonski, Lara Fuhrmann, Rebecca Denes, Mirjam Feldkamp, Ina Nissen, Natascha Santacroce, Elodie Burcklen, Christiane Beckmann, Maurice Redondo, Olivier Kobel, Christoph Noppen, Niko Beerenwinkel, Tanja Stadler |
| EPI_ISL_3297006 | Wales Specialist Virology Centre Sequencing lab: Pathogen Genomics Unit | Public Health Wales Microbiology Cardiff Wales Specialist Virology Centre | Catherine Moore, Johnathan Evans, Laura Gifford, Malorie Perry, Simon Cottrell, Angela Marchbank, Alec Birchley, Alexander Adams, Amy Gaskin, Bree Gatica-Wilcox, Jason Coombes, Joel Southgate, Lauren Gilbert, Lee Graham, Nicole Pacchiarini, Sara Kumziene-Summerhayes, Sarah Taylor, Sophie Jones, Sara Rey, Matthew Bull, Joanne Watkins, Sally Corden, Tom Connor |
| EPI_ISL_3297007 | Viollier AG | Department of Biosystems Science and Engineering, ETH Zürich | Christian Beisel, Sarah Nadeau, Chaoran Chen, Ivan Topolsky, Kim Philipp Jablonski, Lara Fuhrmann, Rebecca Denes, Mirjam Feldkamp, Ina Nissen, Natascha Santacroce, Elodie Burcklen, Christiane Beckmann, Maurice Redondo, Olivier Kobel, Christoph Noppen, Niko Beerenwinkel, Tanja Stadler |
| EPI_ISL_3297008 | Wales Specialist Virology Centre Sequencing lab: Pathogen Genomics Unit | Public Health Wales Microbiology Cardiff Wales Specialist Virology Centre | Catherine Moore, Johnathan Evans, Laura Gifford, Malorie Perry, Simon Cottrell, Angela Marchbank, Alec Birchley, Alexander Adams, Amy Gaskin, Bree Gatica-Wilcox, Jason Coombes, Joel Southgate, Lauren Gilbert, Lee Graham, Nicole Pacchiarini, Sara Kumziene-Summerhayes, Sarah Taylor, Sophie Jones, Sara Rey, Matthew Bull, Joanne Watkins, Sally Corden, Tom Connor |
| EPI_ISL_3297009 | Viollier AG | Department of Biosystems Science and Engineering, ETH Zürich | Christian Beisel, Sarah Nadeau, Chaoran Chen, Ivan Topolsky, Kim Philipp Jablonski, Lara Fuhrmann, Rebecca Denes, Mirjam Feldkamp, Ina Nissen, Natascha Santacroce, Elodie Burcklen, Christiane Beckmann, Maurice Redondo, Olivier Kobel, Christoph Noppen, Niko Beerenwinkel, Tanja Stadler |
| EPI_ISL_3297010 | Wales Specialist Virology Centre Sequencing lab: Pathogen Genomics Unit | Public Health Wales Microbiology Cardiff Wales Specialist Virology Centre | Catherine Moore, Johnathan Evans, Laura Gifford, Malorie Perry, Simon Cottrell, Angela Marchbank, Alec Birchley, Alexander Adams, Amy Gaskin, Bree Gatica-Wilcox, Jason Coombes, Joel Southgate, Lauren Gilbert, Lee Graham, Nicole Pacchiarini, Sara Kumziene-Summerhayes, Sarah Taylor, Sophie Jones, Sara Rey, Matthew Bull, Joanne Watkins, Sally Corden, Tom Connor |
| EPI_ISL_3297011 | Viollier AG | Department of Biosystems Science and Engineering, ETH Zürich | Christian Beisel, Sarah Nadeau, Chaoran Chen, Ivan Topolsky, Kim Philipp Jablonski, Lara Fuhrmann, Rebecca Denes, Mirjam Feldkamp, Ina Nissen, Natascha Santacroce, Elodie Burcklen, Christiane Beckmann, Maurice Redondo, Olivier Kobel, Christoph Noppen, Niko Beerenwinkel, Tanja Stadler |
| EPI_ISL_3297012 | Wales Specialist Virology Centre Sequencing lab: Pathogen Genomics Unit | Public Health Wales Microbiology Cardiff Wales Specialist Virology Centre | Catherine Moore, Johnathan Evans, Laura Gifford, Malorie Perry, Simon Cottrell, Angela Marchbank, Alec Birchley, Alexander Adams, Amy Gaskin, Bree Gatica-Wilcox, Jason Coombes, Joel Southgate, Lauren Gilbert, Lee Graham, Nicole Pacchiarini, Sara Kumziene-Summerhayes, Sarah Taylor, Sophie Jones, Sara Rey, Matthew Bull, Joanne Watkins, Sally Corden, Tom Connor |
| EPI_ISL_3297013 | Viollier AG | Department of Biosystems Science and Engineering, ETH Zürich | Christian Beisel, Sarah Nadeau, Chaoran Chen, Ivan Topolsky, Kim Philipp Jablonski, Lara Fuhrmann, Rebecca Denes, Mirjam Feldkamp, Ina Nissen, Natascha Santacroce, Elodie Burcklen, Christiane Beckmann, Maurice Redondo, Olivier Kobel, Christoph Noppen, Niko Beerenwinkel, Tanja Stadler |
| EPI_ISL_3297014, EPI_ISL_3297015 | Wales Specialist Virology Centre Sequencing lab: Pathogen Genomics Unit | Public Health Wales Microbiology Cardiff Wales Specialist Virology Centre | Catherine Moore, Johnathan Evans, Laura Gifford, Malorie Perry, Simon Cottrell, Angela Marchbank, Alec Birchley, Alexander Adams, Amy Gaskin, Bree Gatica-Wilcox, Jason Coombes, Joel Southgate, Lauren Gilbert, Lee Graham, Nicole Pacchiarini, Sara Kumziene-Summerhayes, Sarah Taylor, Sophie Jones, Sara Rey, Matthew Bull, Joanne Watkins, Sally Corden, Tom Connor |
| EPI_ISL_3297016 | Viollier AG | Department of Biosystems Science and Engineering, ETH Zürich | Christian Beisel, Sarah Nadeau, Chaoran Chen, Ivan Topolsky, Kim Philipp Jablonski, Lara Fuhrmann, Rebecca Denes, Mirjam Feldkamp, Ina Nissen, Natascha Santacroce, Elodie Burcklen, Christiane Beckmann, Maurice Redondo, Olivier Kobel, Christoph Noppen, Niko Beerenwinkel, Tanja Stadler |
| EPI_ISL_3297017 | Wales Specialist Virology Centre Sequencing lab: Pathogen Genomics Unit | Public Health Wales Microbiology Cardiff Wales Specialist Virology Centre | Catherine Moore, Johnathan Evans, Laura Gifford, Malorie Perry, Simon Cottrell, Angela Marchbank, Alec Birchley, Alexander Adams, Amy Gaskin, Bree Gatica-Wilcox, Jason Coombes, Joel Southgate, Lauren Gilbert, Lee Graham, Nicole Pacchiarini, Sara Kumziene-Summerhayes, Sarah Taylor, Sophie Jones, Sara Rey, Matthew Bull, Joanne Watkins, Sally Corden, Tom Connor |
| EPI_ISL_3297018 | Viollier AG | Department of Biosystems Science and Engineering, ETH Zürich | Christian Beisel, Sarah Nadeau, Chaoran Chen, Ivan Topolsky, Kim Philipp Jablonski, Lara Fuhrmann, Rebecca Denes, Mirjam Feldkamp, Ina Nissen, Natascha Santacroce, Elodie Burcklen, Christiane Beckmann, Maurice Redondo, Olivier Kobel, Christoph Noppen, Niko Beerenwinkel, Tanja Stadler |
| EPI_ISL_3297019, EPI_ISL_3297021 | Wales Specialist Virology Centre Sequencing lab: Pathogen Genomics Unit | Public Health Wales Microbiology Cardiff Wales Specialist Virology Centre | Catherine Moore, Johnathan Evans, Laura Gifford, Malorie Perry, Simon Cottrell, Angela Marchbank, Alec Birchley, Alexander Adams, Amy Gaskin, Bree Gatica-Wilcox, Jason Coombes, Joel Southgate, Lauren Gilbert, Lee Graham, Nicole Pacchiarini, Sara Kumziene-Summerhayes, Sarah Taylor, Sophie Jones, Sara Rey, Matthew Bull, Joanne Watkins, Sally Corden, Tom Connor |
| EPI_ISL_3297022 | Viollier AG | Department of Biosystems Science and Engineering, ETH Zürich | Christian Beisel, Sarah Nadeau, Chaoran Chen, Ivan Topolsky, Kim Philipp Jablonski, Lara Fuhrmann, Rebecca Denes, Mirjam Feldkamp, Ina Nissen, Natascha Santacroce, Elodie Burcklen, Christiane Beckmann, Maurice Redondo, Olivier Kobel, Christoph Noppen, Niko Beerenwinkel, Tanja Stadler |
| EPI_ISL_3297023 | Wales Specialist Virology Centre Sequencing lab: Pathogen Genomics Unit | Public Health Wales Microbiology Cardiff Wales Specialist Virology Centre | Catherine Moore, Johnathan Evans, Laura Gifford, Malorie Perry, Simon Cottrell, Angela Marchbank, Alec Birchley, Alexander Adams, Amy Gaskin, Bree Gatica-Wilcox, Jason Coombes, Joel Southgate, Lauren Gilbert, Lee Graham, Nicole Pacchiarini, Sara Kumziene-Summerhayes, Sarah Taylor, Sophie Jones, Sara Rey, Matthew Bull, Joanne Watkins, Sally Corden, Tom Connor |
| EPI_ISL_3297024 | Viollier AG | Department of Biosystems Science and Engineering, ETH Zürich | Christian Beisel, Sarah Nadeau, Chaoran Chen, Ivan Topolsky, Kim Philipp Jablonski, Lara Fuhrmann, Rebecca Denes, Mirjam Feldkamp, Ina Nissen, Natascha Santacroce, Elodie Burcklen, Christiane Beckmann, Maurice Redondo, Olivier Kobel, Christoph Noppen, Niko Beerenwinkel, Tanja Stadler |
| EPI_ISL_3297025 | Wales Specialist Virology Centre Sequencing lab: Pathogen Genomics Unit | Public Health Wales Microbiology Cardiff Wales Specialist Virology Centre | Catherine Moore, Johnathan Evans, Laura Gifford, Malorie Perry, Simon Cottrell, Angela Marchbank, Alec Birchley, Alexander Adams, Amy Gaskin, Bree Gatica-Wilcox, Jason Coombes, Joel Southgate, Lauren Gilbert, Lee Graham, Nicole Pacchiarini, Sara Kumziene-Summerhayes, Sarah Taylor, Sophie Jones, Sara Rey, Matthew Bull, Joanne Watkins, Sally Corden, Tom Connor |
| EPI_ISL_3297026 | Viollier AG | Department of Biosystems Science and Engineering, ETH Zürich | Christian Beisel, Sarah Nadeau, Chaoran Chen, Ivan Topolsky, Kim Philipp Jablonski, Lara Fuhrmann, Rebecca Denes, Mirjam Feldkamp, Ina Nissen, Natascha Santacroce, Elodie Burcklen, Christiane Beckmann, Maurice Redondo, Olivier Kobel, Christoph Noppen, Niko Beerenwinkel, Tanja Stadler |
| EPI_ISL_3297027 | Wales Specialist Virology Centre Sequencing lab: Pathogen Genomics Unit | Public Health Wales Microbiology Cardiff Wales Specialist Virology Centre | Catherine Moore, Johnathan Evans, Laura Gifford, Malorie Perry, Simon Cottrell, Angela Marchbank, Alec Birchley, Alexander Adams, Amy Gaskin, Bree Gatica-Wilcox, Jason Coombes, Joel Southgate, Lauren Gilbert, Lee Graham, Nicole Pacchiarini, Sara Kumziene-Summerhayes, Sarah Taylor, Sophie Jones, Sara Rey, Matthew Bull, Joanne Watkins, Sally Corden, Tom Connor |
| EPI_ISL_3297028 | Viollier AG | Department of Biosystems Science and Engineering, ETH Zürich | Christian Beisel, Sarah Nadeau, Chaoran Chen, Ivan Topolsky, Kim Philipp Jablonski, Lara Fuhrmann, Rebecca Denes, Mirjam Feldkamp, Ina Nissen, Natascha Santacroce, Elodie Burcklen, Christiane Beckmann, Maurice Redondo, Olivier Kobel, Christoph Noppen, Niko Beerenwinkel, Tanja Stadler |
| EPI_ISL_3297029 | Wales Specialist Virology Centre Sequencing lab: Pathogen Genomics Unit | Public Health Wales Microbiology Cardiff Wales Specialist Virology Centre | Catherine Moore, Johnathan Evans, Laura Gifford, Malorie Perry, Simon Cottrell, Angela Marchbank, Alec Birchley, Alexander Adams, Amy Gaskin, Bree Gatica-Wilcox, Jason Coombes, Joel Southgate, Lauren Gilbert, Lee Graham, Nicole Pacchiarini, Sara Kumziene-Summerhayes, Sarah Taylor, Sophie Jones, Sara Rey, Matthew Bull, Joanne Watkins, Sally Corden, Tom Connor |
| EPI_ISL_3297030 | Viollier AG | Department of Biosystems Science and Engineering, ETH Zürich | Christian Beisel, Sarah Nadeau, Chaoran Chen, Ivan Topolsky, Kim Philipp Jablonski, Lara Fuhrmann, Rebecca Denes, Mirjam Feldkamp, Ina Nissen, Natascha Santacroce, Elodie Burcklen, Christiane Beckmann, Maurice Redondo, Olivier Kobel, Christoph Noppen, Niko Beerenwinkel, Tanja Stadler |
| EPI_ISL_3297031 | Wales Specialist Virology Centre Sequencing lab: Pathogen Genomics Unit | Public Health Wales Microbiology Cardiff Wales Specialist Virology Centre | Catherine Moore, Johnathan Evans, Laura Gifford, Malorie Perry, Simon Cottrell, Angela Marchbank, Alec Birchley, Alexander Adams, Amy Gaskin, Bree Gatica-Wilcox, Jason Coombes, Joel Southgate, Lauren Gilbert, Lee Graham, Nicole Pacchiarini, Sara Kumziene-Summerhayes, Sarah Taylor, Sophie Jones, Sara Rey, Matthew Bull, Joanne Watkins, Sally Corden, Tom Connor |
| EPI_ISL_3297032 | Viollier AG | Department of Biosystems Science and Engineering, ETH Zürich | Christian Beisel, Sarah Nadeau, Chaoran Chen, Ivan Topolsky, Kim Philipp Jablonski, Lara Fuhrmann, Rebecca Denes, Mirjam Feldkamp, Ina Nissen, Natascha Santacroce, Elodie Burcklen, Christiane Beckmann, Maurice Redondo, Olivier Kobel, Christoph Noppen, Niko Beerenwinkel, Tanja Stadler |
| EPI_ISL_3297033 | Wales Specialist Virology Centre Sequencing lab: Pathogen Genomics Unit | Public Health Wales Microbiology Cardiff Wales Specialist Virology Centre | Catherine Moore, Johnathan Evans, Laura Gifford, Malorie Perry, Simon Cottrell, Angela Marchbank, Alec Birchley, Alexander Adams, Amy Gaskin, Bree Gatica-Wilcox, Jason Coombes, Joel Southgate, Lauren Gilbert, Lee Graham, Nicole Pacchiarini, Sara Kumziene-Summerhayes, Sarah Taylor, Sophie Jones, Sara Rey, Matthew Bull, Joanne Watkins, Sally Corden, Tom Connor |

| | | | |
|---|---|---|---|
| EPI_ISL_3297034 | Viollier AG | Department of Biosystems Science and Engineering, ETH Zürich | Christian Beisel, Sarah Nadeau, Chaoran Chen, Ivan Topolsky, Kim Philipp Jablonski, Lara Fuhrmann, Rebecca Denes, Mirjam Feldkamp, Ina Nissen, Natascha Santacroce, Elodie Burcklen, Christiane Beckmann, Maurice Redondo, Olivier Kobel, Christoph Noppen, Niko Beerenwinkel, Tanja Stadler |
| EPI_ISL_3297035 | Wales Specialist Virology Centre Sequencing lab: Pathogen Genomics Unit | Public Health Wales Microbiology Cardiff Wales Specialist Virology Centre | Catherine Moore, Johnathan Evans, Laura Gifford, Malorie Perry, Simon Cottrell, Angela Marchbank, Alec Birchley, Alexander Adams, Amy Gaskin, Bree Gatica-Wilcox, Jason Coombes, Joel Southgate, Lauren Gilbert, Lee Graham, Nicole Pacchiarini, Sara Kumziene-Summerhayes, Sarah Taylor, Sophie Jones, Sara Rey, Matthew Bull, Joanne Watkins, Sally Corden, Tom Connor |
| EPI_ISL_3297036 | Viollier AG | Department of Biosystems Science and Engineering, ETH Zürich | Christian Beisel, Sarah Nadeau, Chaoran Chen, Ivan Topolsky, Kim Philipp Jablonski, Lara Fuhrmann, Rebecca Denes, Mirjam Feldkamp, Ina Nissen, Natascha Santacroce, Elodie Burcklen, Christiane Beckmann, Maurice Redondo, Olivier Kobel, Christoph Noppen, Niko Beerenwinkel, Tanja Stadler |
| EPI_ISL_3297037, EPI_ISL_3297038 | Wales Specialist Virology Centre Sequencing lab: Pathogen Genomics Unit | Public Health Wales Microbiology Cardiff Wales Specialist Virology Centre | Catherine Moore, Johnathan Evans, Laura Gifford, Malorie Perry, Simon Cottrell, Angela Marchbank, Alec Birchley, Alexander Adams, Amy Gaskin, Bree Gatica-Wilcox, Jason Coombes, Joel Southgate, Lauren Gilbert, Lee Graham, Nicole Pacchiarini, Sara Kumziene-Summerhayes, Sarah Taylor, Sophie Jones, Sara Rey, Matthew Bull, Joanne Watkins, Sally Corden, Tom Connor |
| EPI_ISL_3297039 | Viollier AG | Department of Biosystems Science and Engineering, ETH Zürich | Christian Beisel, Sarah Nadeau, Chaoran Chen, Ivan Topolsky, Kim Philipp Jablonski, Lara Fuhrmann, Rebecca Denes, Mirjam Feldkamp, Ina Nissen, Natascha Santacroce, Elodie Burcklen, Christiane Beckmann, Maurice Redondo, Olivier Kobel, Christoph Noppen, Niko Beerenwinkel, Tanja Stadler |
| EPI_ISL_3297040 | Wales Specialist Virology Centre Sequencing lab: Pathogen Genomics Unit | Public Health Wales Microbiology Cardiff Wales Specialist Virology Centre | Catherine Moore, Johnathan Evans, Laura Gifford, Malorie Perry, Simon Cottrell, Angela Marchbank, Alec Birchley, Alexander Adams, Amy Gaskin, Bree Gatica-Wilcox, Jason Coombes, Joel Southgate, Lauren Gilbert, Lee Graham, Nicole Pacchiarini, Sara Kumziene-Summerhayes, Sarah Taylor, Sophie Jones, Sara Rey, Matthew Bull, Joanne Watkins, Sally Corden, Tom Connor |
| EPI_ISL_3297041 | Viollier AG | Department of Biosystems Science and Engineering, ETH Zürich | Christian Beisel, Sarah Nadeau, Chaoran Chen, Ivan Topolsky, Kim Philipp Jablonski, Lara Fuhrmann, Rebecca Denes, Mirjam Feldkamp, Ina Nissen, Natascha Santacroce, Elodie Burcklen, Christiane Beckmann, Maurice Redondo, Olivier Kobel, Christoph Noppen, Niko Beerenwinkel, Tanja Stadler |
| EPI_ISL_3297042 | Wales Specialist Virology Centre Sequencing lab: Pathogen Genomics Unit | Public Health Wales Microbiology Cardiff Wales Specialist Virology Centre | Catherine Moore, Johnathan Evans, Laura Gifford, Malorie Perry, Simon Cottrell, Angela Marchbank, Alec Birchley, Alexander Adams, Amy Gaskin, Bree Gatica-Wilcox, Jason Coombes, Joel Southgate, Lauren Gilbert, Lee Graham, Nicole Pacchiarini, Sara Kumziene-Summerhayes, Sarah Taylor, Sophie Jones, Sara Rey, Matthew Bull, Joanne Watkins, Sally Corden, Tom Connor |
| EPI_ISL_3297043 | Viollier AG | Department of Biosystems Science and Engineering, ETH Zürich | Christian Beisel, Sarah Nadeau, Chaoran Chen, Ivan Topolsky, Kim Philipp Jablonski, Lara Fuhrmann, Rebecca Denes, Mirjam Feldkamp, Ina Nissen, Natascha Santacroce, Elodie Burcklen, Christiane Beckmann, Maurice Redondo, Olivier Kobel, Christoph Noppen, Niko Beerenwinkel, Tanja Stadler |
| EPI_ISL_3297044 | Wales Specialist Virology Centre Sequencing lab: Pathogen Genomics Unit | Public Health Wales Microbiology Cardiff Wales Specialist Virology Centre | Catherine Moore, Johnathan Evans, Laura Gifford, Malorie Perry, Simon Cottrell, Angela Marchbank, Alec Birchley, Alexander Adams, Amy Gaskin, Bree Gatica-Wilcox, Jason Coombes, Joel Southgate, Lauren Gilbert, Lee Graham, Nicole Pacchiarini, Sara Kumziene-Summerhayes, Sarah Taylor, Sophie Jones, Sara Rey, Matthew Bull, Joanne Watkins, Sally Corden, Tom Connor |
| EPI_ISL_3297045 | Viollier AG | Department of Biosystems Science and Engineering, ETH Zürich | Christian Beisel, Sarah Nadeau, Chaoran Chen, Ivan Topolsky, Kim Philipp Jablonski, Lara Fuhrmann, Rebecca Denes, Mirjam Feldkamp, Ina Nissen, Natascha Santacroce, Elodie Burcklen, Christiane Beckmann, Maurice Redondo, Olivier Kobel, Christoph Noppen, Niko Beerenwinkel, Tanja Stadler |
| EPI_ISL_3297046 | Wales Specialist Virology Centre Sequencing lab: Pathogen Genomics Unit | Public Health Wales Microbiology Cardiff Wales Specialist Virology Centre | Catherine Moore, Johnathan Evans, Laura Gifford, Malorie Perry, Simon Cottrell, Angela Marchbank, Alec Birchley, Alexander Adams, Amy Gaskin, Bree Gatica-Wilcox, Jason Coombes, Joel Southgate, Lauren Gilbert, Lee Graham, Nicole Pacchiarini, Sara Kumziene-Summerhayes, Sarah Taylor, Sophie Jones, Sara Rey, Matthew Bull, Joanne Watkins, Sally Corden, Tom Connor |
| EPI_ISL_3297047 | Viollier AG | Department of Biosystems Science and Engineering, ETH Zürich | Christian Beisel, Sarah Nadeau, Chaoran Chen, Ivan Topolsky, Kim Philipp Jablonski, Lara Fuhrmann, Rebecca Denes, Mirjam Feldkamp, Ina Nissen, Natascha Santacroce, Elodie Burcklen, Christiane Beckmann, Maurice Redondo, Olivier Kobel, Christoph Noppen, Niko Beerenwinkel, Tanja Stadler |
| EPI_ISL_3297049 | Wales Specialist Virology Centre Sequencing lab: Pathogen Genomics Unit | Public Health Wales Microbiology Cardiff Wales Specialist Virology Centre | Catherine Moore, Johnathan Evans, Laura Gifford, Malorie Perry, Simon Cottrell, Angela Marchbank, Alec Birchley, Alexander Adams, Amy Gaskin, Bree Gatica-Wilcox, Jason Coombes, Joel Southgate, Lauren Gilbert, Lee Graham, Nicole Pacchiarini, Sara Kumziene-Summerhayes, Sarah Taylor, Sophie Jones, Sara Rey, Matthew Bull, Joanne Watkins, Sally Corden, Tom Connor |
| EPI_ISL_3297050 | Viollier AG | Department of Biosystems Science and Engineering, ETH Zürich | Christian Beisel, Sarah Nadeau, Chaoran Chen, Ivan Topolsky, Kim Philipp Jablonski, Lara Fuhrmann, Rebecca Denes, Mirjam Feldkamp, Ina Nissen, Natascha Santacroce, Elodie Burcklen, Christiane Beckmann, Maurice Redondo, Olivier Kobel, Christoph Noppen, Niko Beerenwinkel, Tanja Stadler |
| EPI_ISL_3297051 | Wales Specialist Virology Centre Sequencing lab: Pathogen Genomics Unit | Public Health Wales Microbiology Cardiff Wales Specialist Virology Centre | Catherine Moore, Johnathan Evans, Laura Gifford, Malorie Perry, Simon Cottrell, Angela Marchbank, Alec Birchley, Alexander Adams, Amy Gaskin, Bree Gatica-Wilcox, Jason Coombes, Joel Southgate, Lauren Gilbert, Lee Graham, Nicole Pacchiarini, Sara Kumziene-Summerhayes, Sarah Taylor, Sophie Jones, Sara Rey, Matthew Bull, Joanne Watkins, Sally Corden, Tom Connor |
| EPI_ISL_3297052 | Viollier AG | Department of Biosystems Science and Engineering, ETH Zürich | Christian Beisel, Sarah Nadeau, Chaoran Chen, Ivan Topolsky, Kim Philipp Jablonski, Lara Fuhrmann, Rebecca Denes, Mirjam Feldkamp, Ina Nissen, Natascha Santacroce, Elodie Burcklen, Christiane Beckmann, Maurice Redondo, Olivier Kobel, Christoph Noppen, Niko Beerenwinkel, Tanja Stadler |
| EPI_ISL_3297053 | Wales Specialist Virology Centre Sequencing lab: Pathogen Genomics Unit | Public Health Wales Microbiology Cardiff Wales Specialist Virology Centre | Catherine Moore, Johnathan Evans, Laura Gifford, Malorie Perry, Simon Cottrell, Angela Marchbank, Alec Birchley, Alexander Adams, Amy Gaskin, Bree Gatica-Wilcox, Jason Coombes, Joel Southgate, Lauren Gilbert, Lee Graham, Nicole Pacchiarini, Sara Kumziene-Summerhayes, Sarah Taylor, Sophie Jones, Sara Rey, Matthew Bull, Joanne Watkins, Sally Corden, Tom Connor |
| EPI_ISL_3297054 | Viollier AG | Department of Biosystems Science and Engineering, ETH Zürich | Christian Beisel, Sarah Nadeau, Chaoran Chen, Ivan Topolsky, Kim Philipp Jablonski, Lara Fuhrmann, Rebecca Denes, Mirjam Feldkamp, Ina Nissen, Natascha Santacroce, Elodie Burcklen, Christiane Beckmann, Maurice Redondo, Olivier Kobel, Christoph Noppen, Niko Beerenwinkel, Tanja Stadler |
| EPI_ISL_3297055 | Wales Specialist Virology Centre Sequencing lab: Pathogen Genomics Unit | Public Health Wales Microbiology Cardiff Wales Specialist Virology Centre | Catherine Moore, Johnathan Evans, Laura Gifford, Malorie Perry, Simon Cottrell, Angela Marchbank, Alec Birchley, Alexander Adams, Amy Gaskin, Bree Gatica-Wilcox, Jason Coombes, Joel Southgate, Lauren Gilbert, Lee Graham, Nicole Pacchiarini, Sara Kumziene-Summerhayes, Sarah Taylor, Sophie Jones, Sara Rey, Matthew Bull, Joanne Watkins, Sally Corden, Tom Connor |
| EPI_ISL_3297056 | Viollier AG | Department of Biosystems Science and Engineering, ETH Zürich | Christian Beisel, Sarah Nadeau, Chaoran Chen, Ivan Topolsky, Kim Philipp Jablonski, Lara Fuhrmann, Rebecca Denes, Mirjam Feldkamp, Ina Nissen, Natascha Santacroce, Elodie Burcklen, Christiane Beckmann, Maurice Redondo, Olivier Kobel, Christoph Noppen, Niko Beerenwinkel, Tanja Stadler |
| EPI_ISL_3297057 | Wales Specialist Virology Centre Sequencing lab: Pathogen Genomics Unit | Public Health Wales Microbiology Cardiff Wales Specialist Virology Centre | Catherine Moore, Johnathan Evans, Laura Gifford, Malorie Perry, Simon Cottrell, Angela Marchbank, Alec Birchley, Alexander Adams, Amy Gaskin, Bree Gatica-Wilcox, Jason Coombes, Joel Southgate, Lauren Gilbert, Lee Graham, Nicole Pacchiarini, Sara Kumziene-Summerhayes, Sarah Taylor, Sophie Jones, Sara Rey, Matthew Bull, Joanne Watkins, Sally Corden, Tom Connor |
| EPI_ISL_3297058 | Viollier AG | Department of Biosystems Science and Engineering, ETH Zürich | Christian Beisel, Sarah Nadeau, Chaoran Chen, Ivan Topolsky, Kim Philipp Jablonski, Lara Fuhrmann, Rebecca Denes, Mirjam Feldkamp, Ina Nissen, Natascha Santacroce, Elodie Burcklen, Christiane Beckmann, Maurice Redondo, Olivier Kobel, Christoph Noppen, Niko Beerenwinkel, Tanja Stadler |
| EPI_ISL_3297059, EPI_ISL_3297060 | Wales Specialist Virology Centre Sequencing lab: Pathogen Genomics Unit | Public Health Wales Microbiology Cardiff Wales Specialist Virology Centre | Catherine Moore, Johnathan Evans, Laura Gifford, Malorie Perry, Simon Cottrell, Angela Marchbank, Alec Birchley, Alexander Adams, Amy Gaskin, Bree Gatica-Wilcox, Jason Coombes, Joel Southgate, Lauren Gilbert, Lee Graham, Nicole Pacchiarini, Sara Kumziene-Summerhayes, Sarah Taylor, Sophie Jones, Sara Rey, Matthew Bull, Joanne Watkins, Sally Corden, Tom Connor |
| EPI_ISL_3297061 | Viollier AG | Department of Biosystems Science and Engineering, ETH Zürich | Christian Beisel, Sarah Nadeau, Chaoran Chen, Ivan Topolsky, Kim Philipp Jablonski, Lara Fuhrmann, Rebecca Denes, Mirjam Feldkamp, Ina Nissen, Natascha Santacroce, Elodie Burcklen, Christiane Beckmann, Maurice Redondo, Olivier Kobel, Christoph Noppen, Niko Beerenwinkel, Tanja Stadler |
| EPI_ISL_3297062 | Wales Specialist Virology Centre Sequencing lab: Pathogen Genomics Unit | Public Health Wales Microbiology Cardiff Wales Specialist Virology Centre | Catherine Moore, Johnathan Evans, Laura Gifford, Malorie Perry, Simon Cottrell, Angela Marchbank, Alec Birchley, Alexander Adams, Amy Gaskin, Bree Gatica-Wilcox, Jason Coombes, Joel Southgate, Lauren Gilbert, Lee Graham, Nicole Pacchiarini, Sara Kumziene-Summerhayes, Sarah Taylor, Sophie Jones, Sara Rey, Matthew Bull, Joanne Watkins, Sally Corden, Tom Connor |
| EPI_ISL_3297063 | Viollier AG | Department of Biosystems Science and Engineering, ETH Zürich | Christian Beisel, Sarah Nadeau, Chaoran Chen, Ivan Topolsky, Kim Philipp Jablonski, Lara Fuhrmann, Rebecca Denes, Mirjam Feldkamp, Ina Nissen, Natascha Santacroce, Elodie Burcklen, Christiane Beckmann, Maurice Redondo, Olivier Kobel, Christoph Noppen, Niko Beerenwinkel, Tanja Stadler |
| EPI_ISL_3297064 | Wales Specialist Virology Centre Sequencing lab: Pathogen Genomics Unit | Public Health Wales Microbiology Cardiff Wales Specialist Virology Centre | Catherine Moore, Johnathan Evans, Laura Gifford, Malorie Perry, Simon Cottrell, Angela Marchbank, Alec Birchley, Alexander Adams, Amy Gaskin, Bree Gatica-Wilcox, Jason Coombes, Joel Southgate, Lauren Gilbert, Lee Graham, Nicole Pacchiarini, Sara Kumziene-Summerhayes, Sarah Taylor, Sophie Jones, Sara Rey, Matthew Bull, Joanne Watkins, Sally Corden, Tom Connor |
| EPI_ISL_3297065 | Viollier AG | Department of Biosystems Science and Engineering, ETH Zürich | Christian Beisel, Sarah Nadeau, Chaoran Chen, Ivan Topolsky, Kim Philipp Jablonski, Lara Fuhrmann, Rebecca Denes, Mirjam Feldkamp, Ina Nissen, Natascha Santacroce, Elodie Burcklen, Christiane Beckmann, Maurice Redondo, Olivier Kobel, Christoph Noppen, Niko Beerenwinkel, Tanja Stadler |
| EPI_ISL_3297066 | Wales Specialist Virology Centre Sequencing lab: Pathogen Genomics Unit | Public Health Wales Microbiology Cardiff Wales Specialist Virology Centre | Catherine Moore, Johnathan Evans, Laura Gifford, Malorie Perry, Simon Cottrell, Angela Marchbank, Alec Birchley, Alexander Adams, Amy Gaskin, Bree Gatica-Wilcox, Jason Coombes, Joel Southgate, Lauren Gilbert, Lee Graham, Nicole Pacchiarini, Sara Kumziene-Summerhayes, Sarah Taylor, Sophie Jones, Sara Rey, Matthew Bull, Joanne Watkins, Sally Corden, Tom Connor |
| EPI_ISL_3297067 | Viollier AG | Department of Biosystems Science and Engineering, ETH Zürich | Christian Beisel, Sarah Nadeau, Chaoran Chen, Ivan Topolsky, Kim Philipp Jablonski, Lara Fuhrmann, Rebecca Denes, Mirjam Feldkamp, Ina Nissen, Natascha Santacroce, Elodie Burcklen, Christiane Beckmann, Maurice Redondo, Olivier Kobel, Christoph Noppen, Niko Beerenwinkel, Tanja Stadler |
| EPI_ISL_3297068 | Wales Specialist Virology Centre Sequencing lab: Pathogen Genomics Unit | Public Health Wales Microbiology Cardiff Wales Specialist Virology Centre | Catherine Moore, Johnathan Evans, Laura Gifford, Malorie Perry, Simon Cottrell, Angela Marchbank, Alec Birchley, Alexander Adams, Amy Gaskin, Bree Gatica-Wilcox, Jason Coombes, Joel Southgate, Lauren Gilbert, Lee Graham, Nicole Pacchiarini, Sara Kumziene-Summerhayes, Sarah Taylor, Sophie Jones, Sara Rey, Matthew Bull, Joanne Watkins, Sally Corden, Tom Connor |
| EPI_ISL_3297069 | Viollier AG | Department of Biosystems Science and Engineering, ETH Zürich | Christian Beisel, Sarah Nadeau, Chaoran Chen, Ivan Topolsky, Kim Philipp Jablonski, Lara Fuhrmann, Rebecca Denes, Mirjam Feldkamp, Ina Nissen, Natascha Santacroce, Elodie Burcklen, Christiane Beckmann, Maurice Redondo, Olivier Kobel, Christoph Noppen, Niko Beerenwinkel, Tanja Stadler |
| EPI_ISL_3297070 | Wales Specialist Virology Centre Sequencing lab: Pathogen Genomics Unit | Public Health Wales Microbiology Cardiff Wales Specialist Virology Centre | Catherine Moore, Johnathan Evans, Laura Gifford, Malorie Perry, Simon Cottrell, Angela Marchbank, Alec Birchley, Alexander Adams, Amy Gaskin, Bree Gatica-Wilcox, Jason Coombes, Joel Southgate, Lauren Gilbert, Lee Graham, Nicole Pacchiarini, Sara Kumziene-Summerhayes, Sarah Taylor, Sophie Jones, Sara Rey, Matthew Bull, Joanne Watkins, Sally Corden, Tom Connor |
| EPI_ISL_3297071 | Viollier AG | Department of Biosystems Science and Engineering, ETH Zürich | Christian Beisel, Sarah Nadeau, Chaoran Chen, Ivan Topolsky, Kim Philipp Jablonski, Lara Fuhrmann, Rebecca Denes, Mirjam Feldkamp, Ina Nissen, Natascha Santacroce, Elodie Burcklen, Christiane Beckmann, Maurice Redondo, Olivier Kobel, Christoph Noppen, Niko Beerenwinkel, Tanja Stadler |
| EPI_ISL_3297072 | Wales Specialist Virology Centre Sequencing lab: Pathogen Genomics Unit | Public Health Wales Microbiology Cardiff Wales Specialist Virology Centre | Catherine Moore, Johnathan Evans, Laura Gifford, Malorie Perry, Simon Cottrell, Angela Marchbank, Alec Birchley, Alexander Adams, Amy Gaskin, Bree Gatica-Wilcox, Jason Coombes, Joel Southgate, Lauren Gilbert, Lee Graham, Nicole Pacchiarini, Sara Kumziene-Summerhayes, Sarah Taylor, Sophie Jones, Sara Rey, Matthew Bull, Joanne Watkins, Sally Corden, Tom Connor |
| EPI_ISL_3297073 | Viollier AG | Department of Biosystems Science and Engineering, ETH Zürich | Christian Beisel, Sarah Nadeau, Chaoran Chen, Ivan Topolsky, Kim Philipp Jablonski, Lara Fuhrmann, Rebecca Denes, Mirjam Feldkamp, Ina Nissen, Natascha Santacroce, Elodie Burcklen, Christiane Beckmann, Maurice Redondo, Olivier Kobel, Christoph Noppen, Niko Beerenwinkel, Tanja Stadler |
| EPI_ISL_3297074, EPI_ISL_3297075 | Wales Specialist Virology Centre Sequencing lab: Pathogen Genomics Unit | Public Health Wales Microbiology Cardiff Wales Specialist Virology Centre | Catherine Moore, Johnathan Evans, Laura Gifford, Malorie Perry, Simon Cottrell, Angela Marchbank, Alec Birchley, Alexander Adams, Amy Gaskin, Bree Gatica-Wilcox, Jason Coombes, Joel Southgate, Lauren Gilbert, Lee Graham, Nicole Pacchiarini, Sara Kumziene-Summerhayes, Sarah Taylor, Sophie Jones, Sara Rey, Matthew Bull, Joanne Watkins, Sally Corden, Tom Connor |
| EPI_ISL_3297076 | Viollier AG | Department of Biosystems Science and Engineering, ETH Zürich | Christian Beisel, Sarah Nadeau, Chaoran Chen, Ivan Topolsky, Kim Philipp Jablonski, Lara Fuhrmann, Rebecca Denes, Mirjam Feldkamp, Ina Nissen, Natascha Santacroce, Elodie Burcklen, Christiane Beckmann, Maurice Redondo, Olivier Kobel, Christoph Noppen, Niko Beerenwinkel, Tanja Stadler |
| EPI_ISL_3297077 | Wales Specialist Virology Centre Sequencing lab: Pathogen Genomics Unit | Public Health Wales Microbiology Cardiff Wales Specialist Virology Centre | Catherine Moore, Johnathan Evans, Laura Gifford, Malorie Perry, Simon Cottrell, Angela Marchbank, Alec Birchley, Alexander Adams, Amy Gaskin, Bree Gatica-Wilcox, Jason Coombes, Joel Southgate, Lauren Gilbert, Lee Graham, Nicole Pacchiarini, Sara Kumziene-Summerhayes, Sarah Taylor, Sophie Jones, Sara Rey, Matthew Bull, Joanne Watkins, Sally Corden, Tom Connor |
| EPI_ISL_3297078 | Viollier AG | Department of Biosystems Science and Engineering, ETH Zürich | Christian Beisel, Sarah Nadeau, Chaoran Chen, Ivan Topolsky, Kim Philipp Jablonski, Lara Fuhrmann, Rebecca Denes, Mirjam Feldkamp, Ina Nissen, Natascha Santacroce, Elodie Burcklen, Christiane Beckmann, Maurice Redondo, Olivier Kobel, Christoph Noppen, Niko Beerenwinkel, Tanja Stadler |
| EPI_ISL_3297079 | Wales Specialist Virology Centre Sequencing lab: Pathogen Genomics Unit | Public Health Wales Microbiology Cardiff Wales Specialist Virology Centre | Catherine Moore, Johnathan Evans, Laura Gifford, Malorie Perry, Simon Cottrell, Angela Marchbank, Alec Birchley, Alexander Adams, Amy Gaskin, Bree Gatica-Wilcox, Jason Coombes, Joel Southgate, Lauren Gilbert, Lee Graham, Nicole Pacchiarini, Sara Kumziene-Summerhayes, Sarah Taylor, Sophie Jones, Sara Rey, Matthew Bull, Joanne Watkins, Sally Corden, Tom Connor |
| EPI_ISL_3297080 | Viollier AG | Department of Biosystems Science and Engineering, ETH Zürich | Christian Beisel, Sarah Nadeau, Chaoran Chen, Ivan Topolsky, Kim Philipp Jablonski, Lara Fuhrmann, Rebecca Denes, Mirjam Feldkamp, Ina Nissen, Natascha Santacroce, Elodie Burcklen, Christiane Beckmann, Maurice Redondo, Olivier Kobel, Christoph Noppen, Niko Beerenwinkel, Tanja Stadler |
| EPI_ISL_3297081 | Wales Specialist Virology Centre Sequencing lab: Pathogen Genomics Unit | Public Health Wales Microbiology Cardiff Wales Specialist Virology Centre | Catherine Moore, Johnathan Evans, Laura Gifford, Malorie Perry, Simon Cottrell, Angela Marchbank, Alec Birchley, Alexander Adams, Amy Gaskin, Bree Gatica-Wilcox, Jason Coombes, Joel Southgate, Lauren Gilbert, Lee Graham, Nicole Pacchiarini, Sara Kumziene-Summerhayes, Sarah Taylor, Sophie Jones, Sara Rey, Matthew Bull, Joanne Watkins, Sally Corden, Tom Connor |
| EPI_ISL_3297082 | Viollier AG | Department of Biosystems Science and Engineering, ETH Zürich | Christian Beisel, Sarah Nadeau, Chaoran Chen, Ivan Topolsky, Kim Philipp Jablonski, Lara Fuhrmann, Rebecca Denes, Mirjam Feldkamp, Ina Nissen, Natascha Santacroce, Elodie Burcklen, Christiane Beckmann, Maurice Redondo, Olivier Kobel, Christoph Noppen, Niko Beerenwinkel, Tanja Stadler |

| | | | |
|---|---|---|---|
| EPI_ISL_3297083, EPI_ISL_3297084 | Wales Specialist Virology Centre Sequencing lab: Pathogen Genomics Unit | Public Health Wales Microbiology Cardiff Wales Specialist Virology Centre | Catherine Moore, Johnathan Evans, Laura Gifford, Malorie Perry, Simon Cottrell, Angela Marchbank, Alec Birchley, Alexander Adams, Amy Gaskin, Bree Gatica-Wilcox, Jason Coombes, Joel Southgate, Lauren Gilbert, Lee Graham, Nicole Pacchiarini, Sara Kumziene-Summerhayes, Sarah Taylor, Sophie Jones, Sara Rey, Matthew Bull, Joanne Watkins, Sally Corden, Tom Connor |
| EPI_ISL_3297085 | Viollier AG | Department of Biosystems Science and Engineering, ETH Zürich | Christian Beisel, Sarah Nadeau, Chaoran Chen, Ivan Topolsky, Kim Philipp Jablonski, Lara Fuhrmann, Rebecca Denes, Mirjam Feldkamp, Ina Nissen, Natascha Santacroce, Elodie Burcklen, Christiane Beckmann, Maurice Redondo, Olivier Kobel, Christoph Noppen, Niko Beerenwinkel, Tanja Stadler |
| EPI_ISL_3297086 | Wales Specialist Virology Centre Sequencing lab: Pathogen Genomics Unit | Public Health Wales Microbiology Cardiff Wales Specialist Virology Centre | Catherine Moore, Johnathan Evans, Laura Gifford, Malorie Perry, Simon Cottrell, Angela Marchbank, Alec Birchley, Alexander Adams, Amy Gaskin, Bree Gatica-Wilcox, Jason Coombes, Joel Southgate, Lauren Gilbert, Lee Graham, Nicole Pacchiarini, Sara Kumziene-Summerhayes, Sarah Taylor, Sophie Jones, Sara Rey, Matthew Bull, Joanne Watkins, Sally Corden, Tom Connor |
| EPI_ISL_3297087 | Viollier AG | Department of Biosystems Science and Engineering, ETH Zürich | Christian Beisel, Sarah Nadeau, Chaoran Chen, Ivan Topolsky, Kim Philipp Jablonski, Lara Fuhrmann, Rebecca Denes, Mirjam Feldkamp, Ina Nissen, Natascha Santacroce, Elodie Burcklen, Christiane Beckmann, Maurice Redondo, Olivier Kobel, Christoph Noppen, Niko Beerenwinkel, Tanja Stadler |
| EPI_ISL_3297088 | Wales Specialist Virology Centre Sequencing lab: Pathogen Genomics Unit | Public Health Wales Microbiology Cardiff Wales Specialist Virology Centre | Catherine Moore, Johnathan Evans, Laura Gifford, Malorie Perry, Simon Cottrell, Angela Marchbank, Alec Birchley, Alexander Adams, Amy Gaskin, Bree Gatica-Wilcox, Jason Coombes, Joel Southgate, Lauren Gilbert, Lee Graham, Nicole Pacchiarini, Sara Kumziene-Summerhayes, Sarah Taylor, Sophie Jones, Sara Rey, Matthew Bull, Joanne Watkins, Sally Corden, Tom Connor |
| EPI_ISL_3297089 | Viollier AG | Department of Biosystems Science and Engineering, ETH Zürich | Christian Beisel, Sarah Nadeau, Chaoran Chen, Ivan Topolsky, Kim Philipp Jablonski, Lara Fuhrmann, Rebecca Denes, Mirjam Feldkamp, Ina Nissen, Natascha Santacroce, Elodie Burcklen, Christiane Beckmann, Maurice Redondo, Olivier Kobel, Christoph Noppen, Niko Beerenwinkel, Tanja Stadler |
| EPI_ISL_3297090, EPI_ISL_3297091 | Wales Specialist Virology Centre Sequencing lab: Pathogen Genomics Unit | Public Health Wales Microbiology Cardiff Wales Specialist Virology Centre | Catherine Moore, Johnathan Evans, Laura Gifford, Malorie Perry, Simon Cottrell, Angela Marchbank, Alec Birchley, Alexander Adams, Amy Gaskin, Bree Gatica-Wilcox, Jason Coombes, Joel Southgate, Lauren Gilbert, Lee Graham, Nicole Pacchiarini, Sara Kumziene-Summerhayes, Sarah Taylor, Sophie Jones, Sara Rey, Matthew Bull, Joanne Watkins, Sally Corden, Tom Connor |
| EPI_ISL_3297092 | Viollier AG | Department of Biosystems Science and Engineering, ETH Zürich | Christian Beisel, Sarah Nadeau, Chaoran Chen, Ivan Topolsky, Kim Philipp Jablonski, Lara Fuhrmann, Rebecca Denes, Mirjam Feldkamp, Ina Nissen, Natascha Santacroce, Elodie Burcklen, Christiane Beckmann, Maurice Redondo, Olivier Kobel, Christoph Noppen, Niko Beerenwinkel, Tanja Stadler |
| EPI_ISL_3297093 | Wales Specialist Virology Centre Sequencing lab: Pathogen Genomics Unit | Public Health Wales Microbiology Cardiff Wales Specialist Virology Centre | Catherine Moore, Johnathan Evans, Laura Gifford, Malorie Perry, Simon Cottrell, Angela Marchbank, Alec Birchley, Alexander Adams, Amy Gaskin, Bree Gatica-Wilcox, Jason Coombes, Joel Southgate, Lauren Gilbert, Lee Graham, Nicole Pacchiarini, Sara Kumziene-Summerhayes, Sarah Taylor, Sophie Jones, Sara Rey, Matthew Bull, Joanne Watkins, Sally Corden, Tom Connor |
| EPI_ISL_3297094 | Viollier AG | Department of Biosystems Science and Engineering, ETH Zürich | Christian Beisel, Sarah Nadeau, Chaoran Chen, Ivan Topolsky, Kim Philipp Jablonski, Lara Fuhrmann, Rebecca Denes, Mirjam Feldkamp, Ina Nissen, Natascha Santacroce, Elodie Burcklen, Christiane Beckmann, Maurice Redondo, Olivier Kobel, Christoph Noppen, Niko Beerenwinkel, Tanja Stadler |
| EPI_ISL_3297095 | Wales Specialist Virology Centre Sequencing lab: Pathogen Genomics Unit | Public Health Wales Microbiology Cardiff Wales Specialist Virology Centre | Catherine Moore, Johnathan Evans, Laura Gifford, Malorie Perry, Simon Cottrell, Angela Marchbank, Alec Birchley, Alexander Adams, Amy Gaskin, Bree Gatica-Wilcox, Jason Coombes, Joel Southgate, Lauren Gilbert, Lee Graham, Nicole Pacchiarini, Sara Kumziene-Summerhayes, Sarah Taylor, Sophie Jones, Sara Rey, Matthew Bull, Joanne Watkins, Sally Corden, Tom Connor |
| EPI_ISL_3297096 | Viollier AG | Department of Biosystems Science and Engineering, ETH Zürich | Christian Beisel, Sarah Nadeau, Chaoran Chen, Ivan Topolsky, Kim Philipp Jablonski, Lara Fuhrmann, Rebecca Denes, Mirjam Feldkamp, Ina Nissen, Natascha Santacroce, Elodie Burcklen, Christiane Beckmann, Maurice Redondo, Olivier Kobel, Christoph Noppen, Niko Beerenwinkel, Tanja Stadler |
| EPI_ISL_3297097 | Wales Specialist Virology Centre Sequencing lab: Pathogen Genomics Unit | Public Health Wales Microbiology Cardiff Wales Specialist Virology Centre | Catherine Moore, Johnathan Evans, Laura Gifford, Malorie Perry, Simon Cottrell, Angela Marchbank, Alec Birchley, Alexander Adams, Amy Gaskin, Bree Gatica-Wilcox, Jason Coombes, Joel Southgate, Lauren Gilbert, Lee Graham, Nicole Pacchiarini, Sara Kumziene-Summerhayes, Sarah Taylor, Sophie Jones, Sara Rey, Matthew Bull, Joanne Watkins, Sally Corden, Tom Connor |
| EPI_ISL_3297098 | Viollier AG | Department of Biosystems Science and Engineering, ETH Zürich | Christian Beisel, Sarah Nadeau, Chaoran Chen, Ivan Topolsky, Kim Philipp Jablonski, Lara Fuhrmann, Rebecca Denes, Mirjam Feldkamp, Ina Nissen, Natascha Santacroce, Elodie Burcklen, Christiane Beckmann, Maurice Redondo, Olivier Kobel, Christoph Noppen, Niko Beerenwinkel, Tanja Stadler |
| EPI_ISL_3297099 | Wales Specialist Virology Centre Sequencing lab: Pathogen Genomics Unit | Public Health Wales Microbiology Cardiff Wales Specialist Virology Centre | Catherine Moore, Johnathan Evans, Laura Gifford, Malorie Perry, Simon Cottrell, Angela Marchbank, Alec Birchley, Alexander Adams, Amy Gaskin, Bree Gatica-Wilcox, Jason Coombes, Joel Southgate, Lauren Gilbert, Lee Graham, Nicole Pacchiarini, Sara Kumziene-Summerhayes, Sarah Taylor, Sophie Jones, Sara Rey, Matthew Bull, Joanne Watkins, Sally Corden, Tom Connor |
| EPI_ISL_3297100 | Viollier AG | Department of Biosystems Science and Engineering, ETH Zürich | Christian Beisel, Sarah Nadeau, Chaoran Chen, Ivan Topolsky, Kim Philipp Jablonski, Lara Fuhrmann, Rebecca Denes, Mirjam Feldkamp, Ina Nissen, Natascha Santacroce, Elodie Burcklen, Christiane Beckmann, Maurice Redondo, Olivier Kobel, Christoph Noppen, Niko Beerenwinkel, Tanja Stadler |
| EPI_ISL_3297101 | Wales Specialist Virology Centre Sequencing lab: Pathogen Genomics Unit | Public Health Wales Microbiology Cardiff Wales Specialist Virology Centre | Catherine Moore, Johnathan Evans, Laura Gifford, Malorie Perry, Simon Cottrell, Angela Marchbank, Alec Birchley, Alexander Adams, Amy Gaskin, Bree Gatica-Wilcox, Jason Coombes, Joel Southgate, Lauren Gilbert, Lee Graham, Nicole Pacchiarini, Sara Kumziene-Summerhayes, Sarah Taylor, Sophie Jones, Sara Rey, Matthew Bull, Joanne Watkins, Sally Corden, Tom Connor |
| EPI_ISL_3297102 | Viollier AG | Department of Biosystems Science and Engineering, ETH Zürich | Christian Beisel, Sarah Nadeau, Chaoran Chen, Ivan Topolsky, Kim Philipp Jablonski, Lara Fuhrmann, Rebecca Denes, Mirjam Feldkamp, Ina Nissen, Natascha Santacroce, Elodie Burcklen, Christiane Beckmann, Maurice Redondo, Olivier Kobel, Christoph Noppen, Niko Beerenwinkel, Tanja Stadler |
| EPI_ISL_3297103 | Wales Specialist Virology Centre Sequencing lab: Pathogen Genomics Unit | Public Health Wales Microbiology Cardiff Wales Specialist Virology Centre | Catherine Moore, Johnathan Evans, Laura Gifford, Malorie Perry, Simon Cottrell, Angela Marchbank, Alec Birchley, Alexander Adams, Amy Gaskin, Bree Gatica-Wilcox, Jason Coombes, Joel Southgate, Lauren Gilbert, Lee Graham, Nicole Pacchiarini, Sara Kumziene-Summerhayes, Sarah Taylor, Sophie Jones, Sara Rey, Matthew Bull, Joanne Watkins, Sally Corden, Tom Connor |
| EPI_ISL_3297104 | Viollier AG | Department of Biosystems Science and Engineering, ETH Zürich | Christian Beisel, Sarah Nadeau, Chaoran Chen, Ivan Topolsky, Kim Philipp Jablonski, Lara Fuhrmann, Rebecca Denes, Mirjam Feldkamp, Ina Nissen, Natascha Santacroce, Elodie Burcklen, Christiane Beckmann, Maurice Redondo, Olivier Kobel, Christoph Noppen, Niko Beerenwinkel, Tanja Stadler |
| EPI_ISL_3297105 | Wales Specialist Virology Centre Sequencing lab: Pathogen Genomics Unit | Public Health Wales Microbiology Cardiff Wales Specialist Virology Centre | Catherine Moore, Johnathan Evans, Laura Gifford, Malorie Perry, Simon Cottrell, Angela Marchbank, Alec Birchley, Alexander Adams, Amy Gaskin, Bree Gatica-Wilcox, Jason Coombes, Joel Southgate, Lauren Gilbert, Lee Graham, Nicole Pacchiarini, Sara Kumziene-Summerhayes, Sarah Taylor, Sophie Jones, Sara Rey, Matthew Bull, Joanne Watkins, Sally Corden, Tom Connor |
| EPI_ISL_3297106 | Viollier AG | Department of Biosystems Science and Engineering, ETH Zürich | Christian Beisel, Sarah Nadeau, Chaoran Chen, Ivan Topolsky, Kim Philipp Jablonski, Lara Fuhrmann, Rebecca Denes, Mirjam Feldkamp, Ina Nissen, Natascha Santacroce, Elodie Burcklen, Christiane Beckmann, Maurice Redondo, Olivier Kobel, Christoph Noppen, Niko Beerenwinkel, Tanja Stadler |
| EPI_ISL_3297107 | Wales Specialist Virology Centre Sequencing lab: Pathogen Genomics Unit | Public Health Wales Microbiology Cardiff Wales Specialist Virology Centre | Catherine Moore, Johnathan Evans, Laura Gifford, Malorie Perry, Simon Cottrell, Angela Marchbank, Alec Birchley, Alexander Adams, Amy Gaskin, Bree Gatica-Wilcox, Jason Coombes, Joel Southgate, Lauren Gilbert, Lee Graham, Nicole Pacchiarini, Sara Kumziene-Summerhayes, Sarah Taylor, Sophie Jones, Sara Rey, Matthew Bull, Joanne Watkins, Sally Corden, Tom Connor |
| EPI_ISL_3297108 | Viollier AG | Department of Biosystems Science and Engineering, ETH Zürich | Christian Beisel, Sarah Nadeau, Chaoran Chen, Ivan Topolsky, Kim Philipp Jablonski, Lara Fuhrmann, Rebecca Denes, Mirjam Feldkamp, Ina Nissen, Natascha Santacroce, Elodie Burcklen, Christiane Beckmann, Maurice Redondo, Olivier Kobel, Christoph Noppen, Niko Beerenwinkel, Tanja Stadler |
| EPI_ISL_3297109 | Wales Specialist Virology Centre Sequencing lab: Pathogen Genomics Unit | Public Health Wales Microbiology Cardiff Wales Specialist Virology Centre | Catherine Moore, Johnathan Evans, Laura Gifford, Malorie Perry, Simon Cottrell, Angela Marchbank, Alec Birchley, Alexander Adams, Amy Gaskin, Bree Gatica-Wilcox, Jason Coombes, Joel Southgate, Lauren Gilbert, Lee Graham, Nicole Pacchiarini, Sara Kumziene-Summerhayes, Sarah Taylor, Sophie Jones, Sara Rey, Matthew Bull, Joanne Watkins, Sally Corden, Tom Connor |
| EPI_ISL_3297110 | Viollier AG | Department of Biosystems Science and Engineering, ETH Zürich | Christian Beisel, Sarah Nadeau, Chaoran Chen, Ivan Topolsky, Kim Philipp Jablonski, Lara Fuhrmann, Rebecca Denes, Mirjam Feldkamp, Ina Nissen, Natascha Santacroce, Elodie Burcklen, Christiane Beckmann, Maurice Redondo, Olivier Kobel, Christoph Noppen, Niko Beerenwinkel, Tanja Stadler |
| EPI_ISL_3297111 | Wales Specialist Virology Centre Sequencing lab: Pathogen Genomics Unit | Public Health Wales Microbiology Cardiff Wales Specialist Virology Centre | Catherine Moore, Johnathan Evans, Laura Gifford, Malorie Perry, Simon Cottrell, Angela Marchbank, Alec Birchley, Alexander Adams, Amy Gaskin, Bree Gatica-Wilcox, Jason Coombes, Joel Southgate, Lauren Gilbert, Lee Graham, Nicole Pacchiarini, Sara Kumziene-Summerhayes, Sarah Taylor, Sophie Jones, Sara Rey, Matthew Bull, Joanne Watkins, Sally Corden, Tom Connor |
| EPI_ISL_3297112 | Viollier AG | Department of Biosystems Science and Engineering, ETH Zürich | Christian Beisel, Sarah Nadeau, Chaoran Chen, Ivan Topolsky, Kim Philipp Jablonski, Lara Fuhrmann, Rebecca Denes, Mirjam Feldkamp, Ina Nissen, Natascha Santacroce, Elodie Burcklen, Christiane Beckmann, Maurice Redondo, Olivier Kobel, Christoph Noppen, Niko Beerenwinkel, Tanja Stadler |
| EPI_ISL_3297113 | Wales Specialist Virology Centre Sequencing lab: Pathogen Genomics Unit | Public Health Wales Microbiology Cardiff Wales Specialist Virology Centre | Catherine Moore, Johnathan Evans, Laura Gifford, Malorie Perry, Simon Cottrell, Angela Marchbank, Alec Birchley, Alexander Adams, Amy Gaskin, Bree Gatica-Wilcox, Jason Coombes, Joel Southgate, Lauren Gilbert, Lee Graham, Nicole Pacchiarini, Sara Kumziene-Summerhayes, Sarah Taylor, Sophie Jones, Sara Rey, Matthew Bull, Joanne Watkins, Sally Corden, Tom Connor |
| EPI_ISL_3297114 | Viollier AG | Department of Biosystems Science and Engineering, ETH Zürich | Christian Beisel, Sarah Nadeau, Chaoran Chen, Ivan Topolsky, Kim Philipp Jablonski, Lara Fuhrmann, Rebecca Denes, Mirjam Feldkamp, Ina Nissen, Natascha Santacroce, Elodie Burcklen, Christiane Beckmann, Maurice Redondo, Olivier Kobel, Christoph Noppen, Niko Beerenwinkel, Tanja Stadler |
| EPI_ISL_3297115 | Wales Specialist Virology Centre Sequencing lab: Pathogen Genomics Unit | Public Health Wales Microbiology Cardiff Wales Specialist Virology Centre | Catherine Moore, Johnathan Evans, Laura Gifford, Malorie Perry, Simon Cottrell, Angela Marchbank, Alec Birchley, Alexander Adams, Amy Gaskin, Bree Gatica-Wilcox, Jason Coombes, Joel Southgate, Lauren Gilbert, Lee Graham, Nicole Pacchiarini, Sara Kumziene-Summerhayes, Sarah Taylor, Sophie Jones, Sara Rey, Matthew Bull, Joanne Watkins, Sally Corden, Tom Connor |
| EPI_ISL_3297116 | Viollier AG | Department of Biosystems Science and Engineering, ETH Zürich | Christian Beisel, Sarah Nadeau, Chaoran Chen, Ivan Topolsky, Kim Philipp Jablonski, Lara Fuhrmann, Rebecca Denes, Mirjam Feldkamp, Ina Nissen, Natascha Santacroce, Elodie Burcklen, Christiane Beckmann, Maurice Redondo, Olivier Kobel, Christoph Noppen, Niko Beerenwinkel, Tanja Stadler |
| EPI_ISL_3297117 | Wales Specialist Virology Centre Sequencing lab: Pathogen Genomics Unit | Public Health Wales Microbiology Cardiff Wales Specialist Virology Centre | Catherine Moore, Johnathan Evans, Laura Gifford, Malorie Perry, Simon Cottrell, Angela Marchbank, Alec Birchley, Alexander Adams, Amy Gaskin, Bree Gatica-Wilcox, Jason Coombes, Joel Southgate, Lauren Gilbert, Lee Graham, Nicole Pacchiarini, Sara Kumziene-Summerhayes, Sarah Taylor, Sophie Jones, Sara Rey, Matthew Bull, Joanne Watkins, Sally Corden, Tom Connor |
| EPI_ISL_3297118 | Viollier AG | Department of Biosystems Science and Engineering, ETH Zürich | Christian Beisel, Sarah Nadeau, Chaoran Chen, Ivan Topolsky, Kim Philipp Jablonski, Lara Fuhrmann, Rebecca Denes, Mirjam Feldkamp, Ina Nissen, Natascha Santacroce, Elodie Burcklen, Christiane Beckmann, Maurice Redondo, Olivier Kobel, Christoph Noppen, Niko Beerenwinkel, Tanja Stadler |
| EPI_ISL_3297119, EPI_ISL_3297121 | Wales Specialist Virology Centre Sequencing lab: Pathogen Genomics Unit | Public Health Wales Microbiology Cardiff Wales Specialist Virology Centre | Catherine Moore, Johnathan Evans, Laura Gifford, Malorie Perry, Simon Cottrell, Angela Marchbank, Alec Birchley, Alexander Adams, Amy Gaskin, Bree Gatica-Wilcox, Jason Coombes, Joel Southgate, Lauren Gilbert, Lee Graham, Nicole Pacchiarini, Sara Kumziene-Summerhayes, Sarah Taylor, Sophie Jones, Sara Rey, Matthew Bull, Joanne Watkins, Sally Corden, Tom Connor |
| EPI_ISL_3297122 | Viollier AG | Department of Biosystems Science and Engineering, ETH Zürich | Christian Beisel, Sarah Nadeau, Chaoran Chen, Ivan Topolsky, Kim Philipp Jablonski, Lara Fuhrmann, Rebecca Denes, Mirjam Feldkamp, Ina Nissen, Natascha Santacroce, Elodie Burcklen, Christiane Beckmann, Maurice Redondo, Olivier Kobel, Christoph Noppen, Niko Beerenwinkel, Tanja Stadler |
| EPI_ISL_3297123 | Wales Specialist Virology Centre Sequencing lab: Pathogen Genomics Unit | Public Health Wales Microbiology Cardiff Wales Specialist Virology Centre | Catherine Moore, Johnathan Evans, Laura Gifford, Malorie Perry, Simon Cottrell, Angela Marchbank, Alec Birchley, Alexander Adams, Amy Gaskin, Bree Gatica-Wilcox, Jason Coombes, Joel Southgate, Lauren Gilbert, Lee Graham, Nicole Pacchiarini, Sara Kumziene-Summerhayes, Sarah Taylor, Sophie Jones, Sara Rey, Matthew Bull, Joanne Watkins, Sally Corden, Tom Connor |
| EPI_ISL_3297124 | Viollier AG | Department of Biosystems Science and Engineering, ETH Zürich | Christian Beisel, Sarah Nadeau, Chaoran Chen, Ivan Topolsky, Kim Philipp Jablonski, Lara Fuhrmann, Rebecca Denes, Mirjam Feldkamp, Ina Nissen, Natascha Santacroce, Elodie Burcklen, Christiane Beckmann, Maurice Redondo, Olivier Kobel, Christoph Noppen, Niko Beerenwinkel, Tanja Stadler |
| EPI_ISL_3297125 | Wales Specialist Virology Centre Sequencing lab: Pathogen Genomics Unit | Public Health Wales Microbiology Cardiff Wales Specialist Virology Centre | Catherine Moore, Johnathan Evans, Laura Gifford, Malorie Perry, Simon Cottrell, Angela Marchbank, Alec Birchley, Alexander Adams, Amy Gaskin, Bree Gatica-Wilcox, Jason Coombes, Joel Southgate, Lauren Gilbert, Lee Graham, Nicole Pacchiarini, Sara Kumziene-Summerhayes, Sarah Taylor, Sophie Jones, Sara Rey, Matthew Bull, Joanne Watkins, Sally Corden, Tom Connor |
| EPI_ISL_3297126 | Viollier AG | Department of Biosystems Science and Engineering, ETH Zürich | Christian Beisel, Sarah Nadeau, Chaoran Chen, Ivan Topolsky, Kim Philipp Jablonski, Lara Fuhrmann, Rebecca Denes, Mirjam Feldkamp, Ina Nissen, Natascha Santacroce, Elodie Burcklen, Christiane Beckmann, Maurice Redondo, Olivier Kobel, Christoph Noppen, Niko Beerenwinkel, Tanja Stadler |
| EPI_ISL_3297127 | Wales Specialist Virology Centre Sequencing lab: Pathogen Genomics Unit | Public Health Wales Microbiology Cardiff Wales Specialist Virology Centre | Catherine Moore, Johnathan Evans, Laura Gifford, Malorie Perry, Simon Cottrell, Angela Marchbank, Alec Birchley, Alexander Adams, Amy Gaskin, Bree Gatica-Wilcox, Jason Coombes, Joel Southgate, Lauren Gilbert, Lee Graham, Nicole Pacchiarini, Sara Kumziene-Summerhayes, Sarah Taylor, Sophie Jones, Sara Rey, Matthew Bull, Joanne Watkins, Sally Corden, Tom Connor |
| EPI_ISL_3297128 | Viollier AG | Department of Biosystems Science and Engineering, ETH Zürich | Christian Beisel, Sarah Nadeau, Chaoran Chen, Ivan Topolsky, Kim Philipp Jablonski, Lara Fuhrmann, Rebecca Denes, Mirjam Feldkamp, Ina Nissen, Natascha Santacroce, Elodie Burcklen, Christiane Beckmann, Maurice Redondo, Olivier Kobel, Christoph Noppen, Niko Beerenwinkel, Tanja Stadler |
| EPI_ISL_3297129 | Wales Specialist Virology Centre Sequencing lab: Pathogen Genomics Unit | Public Health Wales Microbiology Cardiff Wales Specialist Virology Centre | Catherine Moore, Johnathan Evans, Laura Gifford, Malorie Perry, Simon Cottrell, Angela Marchbank, Alec Birchley, Alexander Adams, Amy Gaskin, Bree Gatica-Wilcox, Jason Coombes, Joel Southgate, Lauren Gilbert, Lee Graham, Nicole Pacchiarini, Sara Kumziene-Summerhayes, Sarah Taylor, Sophie Jones, Sara Rey, Matthew Bull, Joanne Watkins, Sally Corden, Tom Connor |
| EPI_ISL_3297130 | Viollier AG | Department of Biosystems Science and Engineering, ETH Zürich | Christian Beisel, Sarah Nadeau, Chaoran Chen, Ivan Topolsky, Kim Philipp Jablonski, Lara Fuhrmann, Rebecca Denes, Mirjam Feldkamp, Ina Nissen, Natascha Santacroce, Elodie Burcklen, Christiane Beckmann, Maurice Redondo, Olivier Kobel, Christoph Noppen, Niko Beerenwinkel, Tanja Stadler |
| EPI_ISL_3297131 | Wales Specialist Virology Centre Sequencing lab: Pathogen Genomics Unit | Public Health Wales Microbiology Cardiff Wales Specialist Virology Centre | Catherine Moore, Johnathan Evans, Laura Gifford, Malorie Perry, Simon Cottrell, Angela Marchbank, Alec Birchley, Alexander Adams, Amy Gaskin, Bree Gatica-Wilcox, Jason Coombes, Joel Southgate, Lauren Gilbert, Lee Graham, Nicole Pacchiarini, Sara Kumziene-Summerhayes, Sarah Taylor, Sophie Jones, Sara Rey, Matthew Bull, Joanne Watkins, Sally Corden, Tom Connor |

| | | | |
|---|---|---|---|
| EPI_ISL_3297132 | Viollier AG | Department of Biosystems Science and Engineering, ETH Zürich | Christian Beisel, Sarah Nadeau, Chaoran Chen, Ivan Topolsky, Kim Philipp Jablonski, Lara Fuhrmann, Rebecca Denes, Mirjam Feldkamp, Ina Nissen, Natascha Santacroce, Elodie Burcklen, Christiane Beckmann, Maurice Redondo, Olivier Kobel, Christoph Noppen, Niko Beerenwinkel, Tanja Stadler |
| EPI_ISL_3297133 | Wales Specialist Virology Centre Sequencing lab: Pathogen Genomics Unit | Public Health Wales Microbiology Cardiff Wales Specialist Virology Centre | Catherine Moore, Johnathan Evans, Laura Gifford, Malorie Perry, Simon Cottrell, Angela Marchbank, Alec Birchley, Alexander Adams, Amy Gaskin, Bree Gatica-Wilcox, Jason Coombes, Joel Southgate, Lauren Gilbert, Lee Graham, Nicole Pacchiarini, Sara Kumziene-Summerhayes, Sarah Taylor, Sophie Jones, Sara Rey, Matthew Bull, Joanne Watkins, Sally Corden, Tom Connor |
| EPI_ISL_3297134 | Viollier AG | Department of Biosystems Science and Engineering, ETH Zürich | Christian Beisel, Sarah Nadeau, Chaoran Chen, Ivan Topolsky, Kim Philipp Jablonski, Lara Fuhrmann, Rebecca Denes, Mirjam Feldkamp, Ina Nissen, Natascha Santacroce, Elodie Burcklen, Christiane Beckmann, Maurice Redondo, Olivier Kobel, Christoph Noppen, Niko Beerenwinkel, Tanja Stadler |
| EPI_ISL_3297135 | Wales Specialist Virology Centre Sequencing lab: Pathogen Genomics Unit | Public Health Wales Microbiology Cardiff Wales Specialist Virology Centre | Catherine Moore, Johnathan Evans, Laura Gifford, Malorie Perry, Simon Cottrell, Angela Marchbank, Alec Birchley, Alexander Adams, Amy Gaskin, Bree Gatica-Wilcox, Jason Coombes, Joel Southgate, Lauren Gilbert, Lee Graham, Nicole Pacchiarini, Sara Kumziene-Summerhayes, Sarah Taylor, Sophie Jones, Sara Rey, Matthew Bull, Joanne Watkins, Sally Corden, Tom Connor |
| EPI_ISL_3297136 | Viollier AG | Department of Biosystems Science and Engineering, ETH Zürich | Christian Beisel, Sarah Nadeau, Chaoran Chen, Ivan Topolsky, Kim Philipp Jablonski, Lara Fuhrmann, Rebecca Denes, Mirjam Feldkamp, Ina Nissen, Natascha Santacroce, Elodie Burcklen, Christiane Beckmann, Maurice Redondo, Olivier Kobel, Christoph Noppen, Niko Beerenwinkel, Tanja Stadler |
| EPI_ISL_3297137, EPI_ISL_3297138 | Wales Specialist Virology Centre Sequencing lab: Pathogen Genomics Unit | Public Health Wales Microbiology Cardiff Wales Specialist Virology Centre | Catherine Moore, Johnathan Evans, Laura Gifford, Malorie Perry, Simon Cottrell, Angela Marchbank, Alec Birchley, Alexander Adams, Amy Gaskin, Bree Gatica-Wilcox, Jason Coombes, Joel Southgate, Lauren Gilbert, Lee Graham, Nicole Pacchiarini, Sara Kumziene-Summerhayes, Sarah Taylor, Sophie Jones, Sara Rey, Matthew Bull, Joanne Watkins, Sally Corden, Tom Connor |
| EPI_ISL_3297139 | Viollier AG | Department of Biosystems Science and Engineering, ETH Zürich | Christian Beisel, Sarah Nadeau, Chaoran Chen, Ivan Topolsky, Kim Philipp Jablonski, Lara Fuhrmann, Rebecca Denes, Mirjam Feldkamp, Ina Nissen, Natascha Santacroce, Elodie Burcklen, Christiane Beckmann, Maurice Redondo, Olivier Kobel, Christoph Noppen, Niko Beerenwinkel, Tanja Stadler |
| EPI_ISL_3297140 | Wales Specialist Virology Centre Sequencing lab: Pathogen Genomics Unit | Public Health Wales Microbiology Cardiff Wales Specialist Virology Centre | Catherine Moore, Johnathan Evans, Laura Gifford, Malorie Perry, Simon Cottrell, Angela Marchbank, Alec Birchley, Alexander Adams, Amy Gaskin, Bree Gatica-Wilcox, Jason Coombes, Joel Southgate, Lauren Gilbert, Lee Graham, Nicole Pacchiarini, Sara Kumziene-Summerhayes, Sarah Taylor, Sophie Jones, Sara Rey, Matthew Bull, Joanne Watkins, Sally Corden, Tom Connor |
| EPI_ISL_3297141 | Viollier AG | Department of Biosystems Science and Engineering, ETH Zürich | Christian Beisel, Sarah Nadeau, Chaoran Chen, Ivan Topolsky, Kim Philipp Jablonski, Lara Fuhrmann, Rebecca Denes, Mirjam Feldkamp, Ina Nissen, Natascha Santacroce, Elodie Burcklen, Christiane Beckmann, Maurice Redondo, Olivier Kobel, Christoph Noppen, Niko Beerenwinkel, Tanja Stadler |
| EPI_ISL_3297142 | Wales Specialist Virology Centre Sequencing lab: Pathogen Genomics Unit | Public Health Wales Microbiology Cardiff Wales Specialist Virology Centre | Catherine Moore, Johnathan Evans, Laura Gifford, Malorie Perry, Simon Cottrell, Angela Marchbank, Alec Birchley, Alexander Adams, Amy Gaskin, Bree Gatica-Wilcox, Jason Coombes, Joel Southgate, Lauren Gilbert, Lee Graham, Nicole Pacchiarini, Sara Kumziene-Summerhayes, Sarah Taylor, Sophie Jones, Sara Rey, Matthew Bull, Joanne Watkins, Sally Corden, Tom Connor |
| EPI_ISL_3297143 | Viollier AG | Department of Biosystems Science and Engineering, ETH Zürich | Christian Beisel, Sarah Nadeau, Chaoran Chen, Ivan Topolsky, Kim Philipp Jablonski, Lara Fuhrmann, Rebecca Denes, Mirjam Feldkamp, Ina Nissen, Natascha Santacroce, Elodie Burcklen, Christiane Beckmann, Maurice Redondo, Olivier Kobel, Christoph Noppen, Niko Beerenwinkel, Tanja Stadler |
| EPI_ISL_3297144 | Wales Specialist Virology Centre Sequencing lab: Pathogen Genomics Unit | Public Health Wales Microbiology Cardiff Wales Specialist Virology Centre | Catherine Moore, Johnathan Evans, Laura Gifford, Malorie Perry, Simon Cottrell, Angela Marchbank, Alec Birchley, Alexander Adams, Amy Gaskin, Bree Gatica-Wilcox, Jason Coombes, Joel Southgate, Lauren Gilbert, Lee Graham, Nicole Pacchiarini, Sara Kumziene-Summerhayes, Sarah Taylor, Sophie Jones, Sara Rey, Matthew Bull, Joanne Watkins, Sally Corden, Tom Connor |
| EPI_ISL_3297145 | Viollier AG | Department of Biosystems Science and Engineering, ETH Zürich | Christian Beisel, Sarah Nadeau, Chaoran Chen, Ivan Topolsky, Kim Philipp Jablonski, Lara Fuhrmann, Rebecca Denes, Mirjam Feldkamp, Ina Nissen, Natascha Santacroce, Elodie Burcklen, Christiane Beckmann, Maurice Redondo, Olivier Kobel, Christoph Noppen, Niko Beerenwinkel, Tanja Stadler |
| EPI_ISL_3297146 | Wales Specialist Virology Centre Sequencing lab: Pathogen Genomics Unit | Public Health Wales Microbiology Cardiff Wales Specialist Virology Centre | Catherine Moore, Johnathan Evans, Laura Gifford, Malorie Perry, Simon Cottrell, Angela Marchbank, Alec Birchley, Alexander Adams, Amy Gaskin, Bree Gatica-Wilcox, Jason Coombes, Joel Southgate, Lauren Gilbert, Lee Graham, Nicole Pacchiarini, Sara Kumziene-Summerhayes, Sarah Taylor, Sophie Jones, Sara Rey, Matthew Bull, Joanne Watkins, Sally Corden, Tom Connor |
| EPI_ISL_3297147 | Viollier AG | Department of Biosystems Science and Engineering, ETH Zürich | Christian Beisel, Sarah Nadeau, Chaoran Chen, Ivan Topolsky, Kim Philipp Jablonski, Lara Fuhrmann, Rebecca Denes, Mirjam Feldkamp, Ina Nissen, Natascha Santacroce, Elodie Burcklen, Christiane Beckmann, Maurice Redondo, Olivier Kobel, Christoph Noppen, Niko Beerenwinkel, Tanja Stadler |
| EPI_ISL_3297148 | Wales Specialist Virology Centre Sequencing lab: Pathogen Genomics Unit | Public Health Wales Microbiology Cardiff Wales Specialist Virology Centre | Catherine Moore, Johnathan Evans, Laura Gifford, Malorie Perry, Simon Cottrell, Angela Marchbank, Alec Birchley, Alexander Adams, Amy Gaskin, Bree Gatica-Wilcox, Jason Coombes, Joel Southgate, Lauren Gilbert, Lee Graham, Nicole Pacchiarini, Sara Kumziene-Summerhayes, Sarah Taylor, Sophie Jones, Sara Rey, Matthew Bull, Joanne Watkins, Sally Corden, Tom Connor |
| EPI_ISL_3297149 | Viollier AG | Department of Biosystems Science and Engineering, ETH Zürich | Christian Beisel, Sarah Nadeau, Chaoran Chen, Ivan Topolsky, Kim Philipp Jablonski, Lara Fuhrmann, Rebecca Denes, Mirjam Feldkamp, Ina Nissen, Natascha Santacroce, Elodie Burcklen, Christiane Beckmann, Maurice Redondo, Olivier Kobel, Christoph Noppen, Niko Beerenwinkel, Tanja Stadler |
| EPI_ISL_3297150 | Wales Specialist Virology Centre Sequencing lab: Pathogen Genomics Unit | Public Health Wales Microbiology Cardiff Wales Specialist Virology Centre | Catherine Moore, Johnathan Evans, Laura Gifford, Malorie Perry, Simon Cottrell, Angela Marchbank, Alec Birchley, Alexander Adams, Amy Gaskin, Bree Gatica-Wilcox, Jason Coombes, Joel Southgate, Lauren Gilbert, Lee Graham, Nicole Pacchiarini, Sara Kumziene-Summerhayes, Sarah Taylor, Sophie Jones, Sara Rey, Matthew Bull, Joanne Watkins, Sally Corden, Tom Connor |
| EPI_ISL_3297151 | Viollier AG | Department of Biosystems Science and Engineering, ETH Zürich | Christian Beisel, Sarah Nadeau, Chaoran Chen, Ivan Topolsky, Kim Philipp Jablonski, Lara Fuhrmann, Rebecca Denes, Mirjam Feldkamp, Ina Nissen, Natascha Santacroce, Elodie Burcklen, Christiane Beckmann, Maurice Redondo, Olivier Kobel, Christoph Noppen, Niko Beerenwinkel, Tanja Stadler |
| EPI_ISL_3297152 | Wales Specialist Virology Centre Sequencing lab: Pathogen Genomics Unit | Public Health Wales Microbiology Cardiff Wales Specialist Virology Centre | Catherine Moore, Johnathan Evans, Laura Gifford, Malorie Perry, Simon Cottrell, Angela Marchbank, Alec Birchley, Alexander Adams, Amy Gaskin, Bree Gatica-Wilcox, Jason Coombes, Joel Southgate, Lauren Gilbert, Lee Graham, Nicole Pacchiarini, Sara Kumziene-Summerhayes, Sarah Taylor, Sophie Jones, Sara Rey, Matthew Bull, Joanne Watkins, Sally Corden, Tom Connor |
| EPI_ISL_3297153 | Viollier AG | Department of Biosystems Science and Engineering, ETH Zürich | Christian Beisel, Sarah Nadeau, Chaoran Chen, Ivan Topolsky, Kim Philipp Jablonski, Lara Fuhrmann, Rebecca Denes, Mirjam Feldkamp, Ina Nissen, Natascha Santacroce, Elodie Burcklen, Christiane Beckmann, Maurice Redondo, Olivier Kobel, Christoph Noppen, Niko Beerenwinkel, Tanja Stadler |
| EPI_ISL_3297154 | Wales Specialist Virology Centre Sequencing lab: Pathogen Genomics Unit | Public Health Wales Microbiology Cardiff Wales Specialist Virology Centre | Catherine Moore, Johnathan Evans, Laura Gifford, Malorie Perry, Simon Cottrell, Angela Marchbank, Alec Birchley, Alexander Adams, Amy Gaskin, Bree Gatica-Wilcox, Jason Coombes, Joel Southgate, Lauren Gilbert, Lee Graham, Nicole Pacchiarini, Sara Kumziene-Summerhayes, Sarah Taylor, Sophie Jones, Sara Rey, Matthew Bull, Joanne Watkins, Sally Corden, Tom Connor |
| EPI_ISL_3297155 | Viollier AG | Department of Biosystems Science and Engineering, ETH Zürich | Christian Beisel, Sarah Nadeau, Chaoran Chen, Ivan Topolsky, Kim Philipp Jablonski, Lara Fuhrmann, Rebecca Denes, Mirjam Feldkamp, Ina Nissen, Natascha Santacroce, Elodie Burcklen, Christiane Beckmann, Maurice Redondo, Olivier Kobel, Christoph Noppen, Niko Beerenwinkel, Tanja Stadler |
| EPI_ISL_3297156 | Wales Specialist Virology Centre Sequencing lab: Pathogen Genomics Unit | Public Health Wales Microbiology Cardiff Wales Specialist Virology Centre | Catherine Moore, Johnathan Evans, Laura Gifford, Malorie Perry, Simon Cottrell, Angela Marchbank, Alec Birchley, Alexander Adams, Amy Gaskin, Bree Gatica-Wilcox, Jason Coombes, Joel Southgate, Lauren Gilbert, Lee Graham, Nicole Pacchiarini, Sara Kumziene-Summerhayes, Sarah Taylor, Sophie Jones, Sara Rey, Matthew Bull, Joanne Watkins, Sally Corden, Tom Connor |
| EPI_ISL_3297157 | Viollier AG | Department of Biosystems Science and Engineering, ETH Zürich | Christian Beisel, Sarah Nadeau, Chaoran Chen, Ivan Topolsky, Kim Philipp Jablonski, Lara Fuhrmann, Rebecca Denes, Mirjam Feldkamp, Ina Nissen, Natascha Santacroce, Elodie Burcklen, Christiane Beckmann, Maurice Redondo, Olivier Kobel, Christoph Noppen, Niko Beerenwinkel, Tanja Stadler |
| EPI_ISL_3297158 | Wales Specialist Virology Centre Sequencing lab: Pathogen Genomics Unit | Public Health Wales Microbiology Cardiff Wales Specialist Virology Centre | Catherine Moore, Johnathan Evans, Laura Gifford, Malorie Perry, Simon Cottrell, Angela Marchbank, Alec Birchley, Alexander Adams, Amy Gaskin, Bree Gatica-Wilcox, Jason Coombes, Joel Southgate, Lauren Gilbert, Lee Graham, Nicole Pacchiarini, Sara Kumziene-Summerhayes, Sarah Taylor, Sophie Jones, Sara Rey, Matthew Bull, Joanne Watkins, Sally Corden, Tom Connor |
| EPI_ISL_3297159 | Viollier AG | Department of Biosystems Science and Engineering, ETH Zürich | Christian Beisel, Sarah Nadeau, Chaoran Chen, Ivan Topolsky, Kim Philipp Jablonski, Lara Fuhrmann, Rebecca Denes, Mirjam Feldkamp, Ina Nissen, Natascha Santacroce, Elodie Burcklen, Christiane Beckmann, Maurice Redondo, Olivier Kobel, Christoph Noppen, Niko Beerenwinkel, Tanja Stadler |
| EPI_ISL_3297160 | Wales Specialist Virology Centre Sequencing lab: Pathogen Genomics Unit | Public Health Wales Microbiology Cardiff Wales Specialist Virology Centre | Catherine Moore, Johnathan Evans, Laura Gifford, Malorie Perry, Simon Cottrell, Angela Marchbank, Alec Birchley, Alexander Adams, Amy Gaskin, Bree Gatica-Wilcox, Jason Coombes, Joel Southgate, Lauren Gilbert, Lee Graham, Nicole Pacchiarini, Sara Kumziene-Summerhayes, Sarah Taylor, Sophie Jones, Sara Rey, Matthew Bull, Joanne Watkins, Sally Corden, Tom Connor |
| EPI_ISL_3297161, EPI_ISL_3297162, EPI_ISL_3297163, EPI_ISL_3297164, EPI_ISL_3297165, EPI_ISL_3297166, EPI_ISL_3297168, EPI_ISL_3297169, EPI_ISL_3297170, EPI_ISL_3297171 | Viollier AG | Department of Biosystems Science and Engineering, ETH Zürich | Christian Beisel, Sarah Nadeau, Chaoran Chen, Ivan Topolsky, Kim Philipp Jablonski, Lara Fuhrmann, Rebecca Denes, Mirjam Feldkamp, Ina Nissen, Natascha Santacroce, Elodie Burcklen, Christiane Beckmann, Maurice Redondo, Olivier Kobel, Christoph Noppen, Niko Beerenwinkel, Tanja Stadler |
| EPI_ISL_3297172 | Wales Specialist Virology Centre Sequencing lab: Pathogen Genomics Unit | Public Health Wales Microbiology Cardiff Wales Specialist Virology Centre | Catherine Moore, Johnathan Evans, Laura Gifford, Malorie Perry, Simon Cottrell, Angela Marchbank, Alec Birchley, Alexander Adams, Amy Gaskin, Bree Gatica-Wilcox, Jason Coombes, Joel Southgate, Lauren Gilbert, Lee Graham, Nicole Pacchiarini, Sara Kumziene-Summerhayes, Sarah Taylor, Sophie Jones, Sara Rey, Matthew Bull, Joanne Watkins, Sally Corden, Tom Connor |
| EPI_ISL_3297173 | Viollier AG | Department of Biosystems Science and Engineering, ETH Zürich | Christian Beisel, Sarah Nadeau, Chaoran Chen, Ivan Topolsky, Kim Philipp Jablonski, Lara Fuhrmann, Rebecca Denes, Mirjam Feldkamp, Ina Nissen, Natascha Santacroce, Elodie Burcklen, Christiane Beckmann, Maurice Redondo, Olivier Kobel, Christoph Noppen, Niko Beerenwinkel, Tanja Stadler |
| EPI_ISL_3297174 | Wales Specialist Virology Centre Sequencing lab: Pathogen Genomics Unit | Public Health Wales Microbiology Cardiff Wales Specialist Virology Centre | Catherine Moore, Johnathan Evans, Laura Gifford, Malorie Perry, Simon Cottrell, Angela Marchbank, Alec Birchley, Alexander Adams, Amy Gaskin, Bree Gatica-Wilcox, Jason Coombes, Joel Southgate, Lauren Gilbert, Lee Graham, Nicole Pacchiarini, Sara Kumziene-Summerhayes, Sarah Taylor, Sophie Jones, Sara Rey, Matthew Bull, Joanne Watkins, Sally Corden, Tom Connor |
| EPI_ISL_3297175 | Viollier AG | Department of Biosystems Science and Engineering, ETH Zürich | Christian Beisel, Sarah Nadeau, Chaoran Chen, Ivan Topolsky, Kim Philipp Jablonski, Lara Fuhrmann, Rebecca Denes, Mirjam Feldkamp, Ina Nissen, Natascha Santacroce, Elodie Burcklen, Christiane Beckmann, Maurice Redondo, Olivier Kobel, Christoph Noppen, Niko Beerenwinkel, Tanja Stadler |
| EPI_ISL_3297176 | Wales Specialist Virology Centre Sequencing lab: Pathogen Genomics Unit | Public Health Wales Microbiology Cardiff Wales Specialist Virology Centre | Catherine Moore, Johnathan Evans, Laura Gifford, Malorie Perry, Simon Cottrell, Angela Marchbank, Alec Birchley, Alexander Adams, Amy Gaskin, Bree Gatica-Wilcox, Jason Coombes, Joel Southgate, Lauren Gilbert, Lee Graham, Nicole Pacchiarini, Sara Kumziene-Summerhayes, Sarah Taylor, Sophie Jones, Sara Rey, Matthew Bull, Joanne Watkins, Sally Corden, Tom Connor |
| EPI_ISL_3297177 | Viollier AG | Department of Biosystems Science and Engineering, ETH Zürich | Christian Beisel, Sarah Nadeau, Chaoran Chen, Ivan Topolsky, Kim Philipp Jablonski, Lara Fuhrmann, Rebecca Denes, Mirjam Feldkamp, Ina Nissen, Natascha Santacroce, Elodie Burcklen, Christiane Beckmann, Maurice Redondo, Olivier Kobel, Christoph Noppen, Niko Beerenwinkel, Tanja Stadler |
| EPI_ISL_3297178 | Wales Specialist Virology Centre Sequencing lab: Pathogen Genomics Unit | Public Health Wales Microbiology Cardiff Wales Specialist Virology Centre | Catherine Moore, Johnathan Evans, Laura Gifford, Malorie Perry, Simon Cottrell, Angela Marchbank, Alec Birchley, Alexander Adams, Amy Gaskin, Bree Gatica-Wilcox, Jason Coombes, Joel Southgate, Lauren Gilbert, Lee Graham, Nicole Pacchiarini, Sara Kumziene-Summerhayes, Sarah Taylor, Sophie Jones, Sara Rey, Matthew Bull, Joanne Watkins, Sally Corden, Tom Connor |
| EPI_ISL_3297179 | Viollier AG | Department of Biosystems Science and Engineering, ETH Zürich | Christian Beisel, Sarah Nadeau, Chaoran Chen, Ivan Topolsky, Kim Philipp Jablonski, Lara Fuhrmann, Rebecca Denes, Mirjam Feldkamp, Ina Nissen, Natascha Santacroce, Elodie Burcklen, Christiane Beckmann, Maurice Redondo, Olivier Kobel, Christoph Noppen, Niko Beerenwinkel, Tanja Stadler |
| EPI_ISL_3297180 | Wales Specialist Virology Centre Sequencing lab: Pathogen Genomics Unit | Public Health Wales Microbiology Cardiff Wales Specialist Virology Centre | Catherine Moore, Johnathan Evans, Laura Gifford, Malorie Perry, Simon Cottrell, Angela Marchbank, Alec Birchley, Alexander Adams, Amy Gaskin, Bree Gatica-Wilcox, Jason Coombes, Joel Southgate, Lauren Gilbert, Lee Graham, Nicole Pacchiarini, Sara Kumziene-Summerhayes, Sarah Taylor, Sophie Jones, Sara Rey, Matthew Bull, Joanne Watkins, Sally Corden, Tom Connor |
| EPI_ISL_3297181 | Viollier AG | Department of Biosystems Science and Engineering, ETH Zürich | Christian Beisel, Sarah Nadeau, Chaoran Chen, Ivan Topolsky, Kim Philipp Jablonski, Lara Fuhrmann, Rebecca Denes, Mirjam Feldkamp, Ina Nissen, Natascha Santacroce, Elodie Burcklen, Christiane Beckmann, Maurice Redondo, Olivier Kobel, Christoph Noppen, Niko Beerenwinkel, Tanja Stadler |
| EPI_ISL_3297182 | Wales Specialist Virology Centre Sequencing lab: Pathogen Genomics Unit | Public Health Wales Microbiology Cardiff Wales Specialist Virology Centre | Catherine Moore, Johnathan Evans, Laura Gifford, Malorie Perry, Simon Cottrell, Angela Marchbank, Alec Birchley, Alexander Adams, Amy Gaskin, Bree Gatica-Wilcox, Jason Coombes, Joel Southgate, Lauren Gilbert, Lee Graham, Nicole Pacchiarini, Sara Kumziene-Summerhayes, Sarah Taylor, Sophie Jones, Sara Rey, Matthew Bull, Joanne Watkins, Sally Corden, Tom Connor |
| EPI_ISL_3297183 | Viollier AG | Department of Biosystems Science and Engineering, ETH Zürich | Christian Beisel, Sarah Nadeau, Chaoran Chen, Ivan Topolsky, Kim Philipp Jablonski, Lara Fuhrmann, Rebecca Denes, Mirjam Feldkamp, Ina Nissen, Natascha Santacroce, Elodie Burcklen, Christiane Beckmann, Maurice Redondo, Olivier Kobel, Christoph Noppen, Niko Beerenwinkel, Tanja Stadler |
| EPI_ISL_3297184, EPI_ISL_3297186 | Wales Specialist Virology Centre Sequencing lab: Pathogen Genomics Unit | Public Health Wales Microbiology Cardiff Wales Specialist Virology Centre | Catherine Moore, Johnathan Evans, Laura Gifford, Malorie Perry, Simon Cottrell, Angela Marchbank, Alec Birchley, Alexander Adams, Amy Gaskin, Bree Gatica-Wilcox, Jason Coombes, Joel Southgate, Lauren Gilbert, Lee Graham, Nicole Pacchiarini, Sara Kumziene-Summerhayes, Sarah Taylor, Sophie Jones, Sara Rey, Matthew Bull, Joanne Watkins, Sally Corden, Tom Connor |

| | | | |
|---|---|---|---|
| EPI_ISL_3297187 | Viollier AG | Department of Biosystems Science and Engineering, ETH Zürich | Christian Beisel, Sarah Nadeau, Chaoran Chen, Ivan Topolsky, Kim Philipp Jablonski, Lara Fuhrmann, Rebecca Denes, Mirjam Feldkamp, Ina Nissen, Natascha Santacroce, Elodie Burcklen, Christiane Beckmann, Maurice Redondo, Olivier Kobel, Christoph Noppen, Niko Beerenwinkel, Tanja Stadler |
| EPI_ISL_3297188, EPI_ISL_3297190 | Wales Specialist Virology Centre Sequencing lab: Pathogen Genomics Unit | Public Health Wales Microbiology Cardiff Wales Specialist Virology Centre | Catherine Moore, Johnathan Evans, Laura Gifford, Malorie Perry, Simon Cottrell, Angela Marchbank, Alec Birchley, Alexander Adams, Amy Gaskin, Bree Gatica-Wilcox, Jason Coombes, Joel Southgate, Lauren Gilbert, Lee Graham, Nicole Pacchiarini, Sara Kumziene-Summerhayes, Sarah Taylor, Sophie Jones, Sara Rey, Matthew Bull, Joanne Watkins, Sally Corden, Tom Connor |
| EPI_ISL_3297191 | Viollier AG | Department of Biosystems Science and Engineering, ETH Zürich | Christian Beisel, Sarah Nadeau, Chaoran Chen, Ivan Topolsky, Kim Philipp Jablonski, Lara Fuhrmann, Rebecca Denes, Mirjam Feldkamp, Ina Nissen, Natascha Santacroce, Elodie Burcklen, Christiane Beckmann, Maurice Redondo, Olivier Kobel, Christoph Noppen, Niko Beerenwinkel, Tanja Stadler |
| EPI_ISL_3297192 | Wales Specialist Virology Centre Sequencing lab: Pathogen Genomics Unit | Public Health Wales Microbiology Cardiff Wales Specialist Virology Centre | Catherine Moore, Johnathan Evans, Laura Gifford, Malorie Perry, Simon Cottrell, Angela Marchbank, Alec Birchley, Alexander Adams, Amy Gaskin, Bree Gatica-Wilcox, Jason Coombes, Joel Southgate, Lauren Gilbert, Lee Graham, Nicole Pacchiarini, Sara Kumziene-Summerhayes, Sarah Taylor, Sophie Jones, Sara Rey, Matthew Bull, Joanne Watkins, Sally Corden, Tom Connor |
| EPI_ISL_3297193 | Viollier AG | Department of Biosystems Science and Engineering, ETH Zürich | Christian Beisel, Sarah Nadeau, Chaoran Chen, Ivan Topolsky, Kim Philipp Jablonski, Lara Fuhrmann, Rebecca Denes, Mirjam Feldkamp, Ina Nissen, Natascha Santacroce, Elodie Burcklen, Christiane Beckmann, Maurice Redondo, Olivier Kobel, Christoph Noppen, Niko Beerenwinkel, Tanja Stadler |
| EPI_ISL_3297194, EPI_ISL_3297196, EPI_ISL_3297198 | Wales Specialist Virology Centre Sequencing lab: Pathogen Genomics Unit | Public Health Wales Microbiology Cardiff Wales Specialist Virology Centre | Catherine Moore, Johnathan Evans, Laura Gifford, Malorie Perry, Simon Cottrell, Angela Marchbank, Alec Birchley, Alexander Adams, Amy Gaskin, Bree Gatica-Wilcox, Jason Coombes, Joel Southgate, Lauren Gilbert, Lee Graham, Nicole Pacchiarini, Sara Kumziene-Summerhayes, Sarah Taylor, Sophie Jones, Sara Rey, Matthew Bull, Joanne Watkins, Sally Corden, Tom Connor |
| EPI_ISL_3297199 | Viollier AG | Department of Biosystems Science and Engineering, ETH Zürich | Christian Beisel, Sarah Nadeau, Chaoran Chen, Ivan Topolsky, Kim Philipp Jablonski, Lara Fuhrmann, Rebecca Denes, Mirjam Feldkamp, Ina Nissen, Natascha Santacroce, Elodie Burcklen, Christiane Beckmann, Maurice Redondo, Olivier Kobel, Christoph Noppen, Niko Beerenwinkel, Tanja Stadler |
| EPI_ISL_3297200 | Wales Specialist Virology Centre Sequencing lab: Pathogen Genomics Unit | Public Health Wales Microbiology Cardiff Wales Specialist Virology Centre | Catherine Moore, Johnathan Evans, Laura Gifford, Malorie Perry, Simon Cottrell, Angela Marchbank, Alec Birchley, Alexander Adams, Amy Gaskin, Bree Gatica-Wilcox, Jason Coombes, Joel Southgate, Lauren Gilbert, Lee Graham, Nicole Pacchiarini, Sara Kumziene-Summerhayes, Sarah Taylor, Sophie Jones, Sara Rey, Matthew Bull, Joanne Watkins, Sally Corden, Tom Connor |
| EPI_ISL_3297201 | Viollier AG | Department of Biosystems Science and Engineering, ETH Zürich | Christian Beisel, Sarah Nadeau, Chaoran Chen, Ivan Topolsky, Kim Philipp Jablonski, Lara Fuhrmann, Rebecca Denes, Mirjam Feldkamp, Ina Nissen, Natascha Santacroce, Elodie Burcklen, Christiane Beckmann, Maurice Redondo, Olivier Kobel, Christoph Noppen, Niko Beerenwinkel, Tanja Stadler |
| EPI_ISL_3297202 | Wales Specialist Virology Centre Sequencing lab: Pathogen Genomics Unit | Public Health Wales Microbiology Cardiff Wales Specialist Virology Centre | Catherine Moore, Johnathan Evans, Laura Gifford, Malorie Perry, Simon Cottrell, Angela Marchbank, Alec Birchley, Alexander Adams, Amy Gaskin, Bree Gatica-Wilcox, Jason Coombes, Joel Southgate, Lauren Gilbert, Lee Graham, Nicole Pacchiarini, Sara Kumziene-Summerhayes, Sarah Taylor, Sophie Jones, Sara Rey, Matthew Bull, Joanne Watkins, Sally Corden, Tom Connor |
| EPI_ISL_3297203 | Viollier AG | Department of Biosystems Science and Engineering, ETH Zürich | Christian Beisel, Sarah Nadeau, Chaoran Chen, Ivan Topolsky, Kim Philipp Jablonski, Lara Fuhrmann, Rebecca Denes, Mirjam Feldkamp, Ina Nissen, Natascha Santacroce, Elodie Burcklen, Christiane Beckmann, Maurice Redondo, Olivier Kobel, Christoph Noppen, Niko Beerenwinkel, Tanja Stadler |
| EPI_ISL_3297204 | Wales Specialist Virology Centre Sequencing lab: Pathogen Genomics Unit | Public Health Wales Microbiology Cardiff Wales Specialist Virology Centre | Catherine Moore, Johnathan Evans, Laura Gifford, Malorie Perry, Simon Cottrell, Angela Marchbank, Alec Birchley, Alexander Adams, Amy Gaskin, Bree Gatica-Wilcox, Jason Coombes, Joel Southgate, Lauren Gilbert, Lee Graham, Nicole Pacchiarini, Sara Kumziene-Summerhayes, Sarah Taylor, Sophie Jones, Sara Rey, Matthew Bull, Joanne Watkins, Sally Corden, Tom Connor |
| EPI_ISL_3297205 | Viollier AG | Department of Biosystems Science and Engineering, ETH Zürich | Christian Beisel, Sarah Nadeau, Chaoran Chen, Ivan Topolsky, Kim Philipp Jablonski, Lara Fuhrmann, Rebecca Denes, Mirjam Feldkamp, Ina Nissen, Natascha Santacroce, Elodie Burcklen, Christiane Beckmann, Maurice Redondo, Olivier Kobel, Christoph Noppen, Niko Beerenwinkel, Tanja Stadler |
| EPI_ISL_3297206 | Wales Specialist Virology Centre Sequencing lab: Pathogen Genomics Unit | Public Health Wales Microbiology Cardiff Wales Specialist Virology Centre | Catherine Moore, Johnathan Evans, Laura Gifford, Malorie Perry, Simon Cottrell, Angela Marchbank, Alec Birchley, Alexander Adams, Amy Gaskin, Bree Gatica-Wilcox, Jason Coombes, Joel Southgate, Lauren Gilbert, Lee Graham, Nicole Pacchiarini, Sara Kumziene-Summerhayes, Sarah Taylor, Sophie Jones, Sara Rey, Matthew Bull, Joanne Watkins, Sally Corden, Tom Connor |
| EPI_ISL_3297207 | Viollier AG | Department of Biosystems Science and Engineering, ETH Zürich | Christian Beisel, Sarah Nadeau, Chaoran Chen, Ivan Topolsky, Kim Philipp Jablonski, Lara Fuhrmann, Rebecca Denes, Mirjam Feldkamp, Ina Nissen, Natascha Santacroce, Elodie Burcklen, Christiane Beckmann, Maurice Redondo, Olivier Kobel, Christoph Noppen, Niko Beerenwinkel, Tanja Stadler |
| EPI_ISL_3297208, EPI_ISL_3297209, EPI_ISL_3297212, EPI_ISL_3297213 | Wales Specialist Virology Centre Sequencing lab: Pathogen Genomics Unit | Public Health Wales Microbiology Cardiff Wales Specialist Virology Centre | Catherine Moore, Johnathan Evans, Laura Gifford, Malorie Perry, Simon Cottrell, Angela Marchbank, Alec Birchley, Alexander Adams, Amy Gaskin, Bree Gatica-Wilcox, Jason Coombes, Joel Southgate, Lauren Gilbert, Lee Graham, Nicole Pacchiarini, Sara Kumziene-Summerhayes, Sarah Taylor, Sophie Jones, Sara Rey, Matthew Bull, Joanne Watkins, Sally Corden, Tom Connor |
| EPI_ISL_3297214 | Viollier AG | Department of Biosystems Science and Engineering, ETH Zürich | Christian Beisel, Sarah Nadeau, Chaoran Chen, Ivan Topolsky, Kim Philipp Jablonski, Lara Fuhrmann, Rebecca Denes, Mirjam Feldkamp, Ina Nissen, Natascha Santacroce, Elodie Burcklen, Christiane Beckmann, Maurice Redondo, Olivier Kobel, Christoph Noppen, Niko Beerenwinkel, Tanja Stadler |
| EPI_ISL_3297215 | Wales Specialist Virology Centre Sequencing lab: Pathogen Genomics Unit | Public Health Wales Microbiology Cardiff Wales Specialist Virology Centre | Catherine Moore, Johnathan Evans, Laura Gifford, Malorie Perry, Simon Cottrell, Angela Marchbank, Alec Birchley, Alexander Adams, Amy Gaskin, Bree Gatica-Wilcox, Jason Coombes, Joel Southgate, Lauren Gilbert, Lee Graham, Nicole Pacchiarini, Sara Kumziene-Summerhayes, Sarah Taylor, Sophie Jones, Sara Rey, Matthew Bull, Joanne Watkins, Sally Corden, Tom Connor |
| EPI_ISL_3297216 | Viollier AG | Department of Biosystems Science and Engineering, ETH Zürich | Christian Beisel, Sarah Nadeau, Chaoran Chen, Ivan Topolsky, Kim Philipp Jablonski, Lara Fuhrmann, Rebecca Denes, Mirjam Feldkamp, Ina Nissen, Natascha Santacroce, Elodie Burcklen, Christiane Beckmann, Maurice Redondo, Olivier Kobel, Christoph Noppen, Niko Beerenwinkel, Tanja Stadler |
| EPI_ISL_3297217 | Wales Specialist Virology Centre Sequencing lab: Pathogen Genomics Unit | Public Health Wales Microbiology Cardiff Wales Specialist Virology Centre | Catherine Moore, Johnathan Evans, Laura Gifford, Malorie Perry, Simon Cottrell, Angela Marchbank, Alec Birchley, Alexander Adams, Amy Gaskin, Bree Gatica-Wilcox, Jason Coombes, Joel Southgate, Lauren Gilbert, Lee Graham, Nicole Pacchiarini, Sara Kumziene-Summerhayes, Sarah Taylor, Sophie Jones, Sara Rey, Matthew Bull, Joanne Watkins, Sally Corden, Tom Connor |
| EPI_ISL_3297218 | Viollier AG | Department of Biosystems Science and Engineering, ETH Zürich | Christian Beisel, Sarah Nadeau, Chaoran Chen, Ivan Topolsky, Kim Philipp Jablonski, Lara Fuhrmann, Rebecca Denes, Mirjam Feldkamp, Ina Nissen, Natascha Santacroce, Elodie Burcklen, Christiane Beckmann, Maurice Redondo, Olivier Kobel, Christoph Noppen, Niko Beerenwinkel, Tanja Stadler |
| EPI_ISL_3297219 | Wales Specialist Virology Centre Sequencing lab: Pathogen Genomics Unit | Public Health Wales Microbiology Cardiff Wales Specialist Virology Centre | Catherine Moore, Johnathan Evans, Laura Gifford, Malorie Perry, Simon Cottrell, Angela Marchbank, Alec Birchley, Alexander Adams, Amy Gaskin, Bree Gatica-Wilcox, Jason Coombes, Joel Southgate, Lauren Gilbert, Lee Graham, Nicole Pacchiarini, Sara Kumziene-Summerhayes, Sarah Taylor, Sophie Jones, Sara Rey, Matthew Bull, Joanne Watkins, Sally Corden, Tom Connor |
| EPI_ISL_3297220 | Viollier AG | Department of Biosystems Science and Engineering, ETH Zürich | Christian Beisel, Sarah Nadeau, Chaoran Chen, Ivan Topolsky, Kim Philipp Jablonski, Lara Fuhrmann, Rebecca Denes, Mirjam Feldkamp, Ina Nissen, Natascha Santacroce, Elodie Burcklen, Christiane Beckmann, Maurice Redondo, Olivier Kobel, Christoph Noppen, Niko Beerenwinkel, Tanja Stadler |
| EPI_ISL_3297221 | Wales Specialist Virology Centre Sequencing lab: Pathogen Genomics Unit | Public Health Wales Microbiology Cardiff Wales Specialist Virology Centre | Catherine Moore, Johnathan Evans, Laura Gifford, Malorie Perry, Simon Cottrell, Angela Marchbank, Alec Birchley, Alexander Adams, Amy Gaskin, Bree Gatica-Wilcox, Jason Coombes, Joel Southgate, Lauren Gilbert, Lee Graham, Nicole Pacchiarini, Sara Kumziene-Summerhayes, Sarah Taylor, Sophie Jones, Sara Rey, Matthew Bull, Joanne Watkins, Sally Corden, Tom Connor |
| EPI_ISL_3297222 | Viollier AG | Department of Biosystems Science and Engineering, ETH Zürich | Christian Beisel, Sarah Nadeau, Chaoran Chen, Ivan Topolsky, Kim Philipp Jablonski, Lara Fuhrmann, Rebecca Denes, Mirjam Feldkamp, Ina Nissen, Natascha Santacroce, Elodie Burcklen, Christiane Beckmann, Maurice Redondo, Olivier Kobel, Christoph Noppen, Niko Beerenwinkel, Tanja Stadler |
| EPI_ISL_3297223 | Wales Specialist Virology Centre Sequencing lab: Pathogen Genomics Unit | Public Health Wales Microbiology Cardiff Wales Specialist Virology Centre | Catherine Moore, Johnathan Evans, Laura Gifford, Malorie Perry, Simon Cottrell, Angela Marchbank, Alec Birchley, Alexander Adams, Amy Gaskin, Bree Gatica-Wilcox, Jason Coombes, Joel Southgate, Lauren Gilbert, Lee Graham, Nicole Pacchiarini, Sara Kumziene-Summerhayes, Sarah Taylor, Sophie Jones, Sara Rey, Matthew Bull, Joanne Watkins, Sally Corden, Tom Connor |
| EPI_ISL_3297224 | Viollier AG | Department of Biosystems Science and Engineering, ETH Zürich | Christian Beisel, Sarah Nadeau, Chaoran Chen, Ivan Topolsky, Kim Philipp Jablonski, Lara Fuhrmann, Rebecca Denes, Mirjam Feldkamp, Ina Nissen, Natascha Santacroce, Elodie Burcklen, Christiane Beckmann, Maurice Redondo, Olivier Kobel, Christoph Noppen, Niko Beerenwinkel, Tanja Stadler |
| EPI_ISL_3297225, EPI_ISL_3297226 | Wales Specialist Virology Centre Sequencing lab: Pathogen Genomics Unit | Public Health Wales Microbiology Cardiff Wales Specialist Virology Centre | Catherine Moore, Johnathan Evans, Laura Gifford, Malorie Perry, Simon Cottrell, Angela Marchbank, Alec Birchley, Alexander Adams, Amy Gaskin, Bree Gatica-Wilcox, Jason Coombes, Joel Southgate, Lauren Gilbert, Lee Graham, Nicole Pacchiarini, Sara Kumziene-Summerhayes, Sarah Taylor, Sophie Jones, Sara Rey, Matthew Bull, Joanne Watkins, Sally Corden, Tom Connor |
| EPI_ISL_3297227, EPI_ISL_3297228 | Viollier AG | Department of Biosystems Science and Engineering, ETH Zürich | Christian Beisel, Sarah Nadeau, Chaoran Chen, Ivan Topolsky, Kim Philipp Jablonski, Lara Fuhrmann, Rebecca Denes, Mirjam Feldkamp, Ina Nissen, Natascha Santacroce, Elodie Burcklen, Christiane Beckmann, Maurice Redondo, Olivier Kobel, Christoph Noppen, Niko Beerenwinkel, Tanja Stadler |
| EPI_ISL_3297229, EPI_ISL_3297230 | Wales Specialist Virology Centre Sequencing lab: Pathogen Genomics Unit | Public Health Wales Microbiology Cardiff Wales Specialist Virology Centre | Catherine Moore, Johnathan Evans, Laura Gifford, Malorie Perry, Simon Cottrell, Angela Marchbank, Alec Birchley, Alexander Adams, Amy Gaskin, Bree Gatica-Wilcox, Jason Coombes, Joel Southgate, Lauren Gilbert, Lee Graham, Nicole Pacchiarini, Sara Kumziene-Summerhayes, Sarah Taylor, Sophie Jones, Sara Rey, Matthew Bull, Joanne Watkins, Sally Corden, Tom Connor |
| EPI_ISL_3297231 | Viollier AG | Department of Biosystems Science and Engineering, ETH Zürich | Christian Beisel, Sarah Nadeau, Chaoran Chen, Ivan Topolsky, Kim Philipp Jablonski, Lara Fuhrmann, Rebecca Denes, Mirjam Feldkamp, Ina Nissen, Natascha Santacroce, Elodie Burcklen, Christiane Beckmann, Maurice Redondo, Olivier Kobel, Christoph Noppen, Niko Beerenwinkel, Tanja Stadler |
| EPI_ISL_3297232 | Wales Specialist Virology Centre Sequencing lab: Pathogen Genomics Unit | Public Health Wales Microbiology Cardiff Wales Specialist Virology Centre | Catherine Moore, Johnathan Evans, Laura Gifford, Malorie Perry, Simon Cottrell, Angela Marchbank, Alec Birchley, Alexander Adams, Amy Gaskin, Bree Gatica-Wilcox, Jason Coombes, Joel Southgate, Lauren Gilbert, Lee Graham, Nicole Pacchiarini, Sara Kumziene-Summerhayes, Sarah Taylor, Sophie Jones, Sara Rey, Matthew Bull, Joanne Watkins, Sally Corden, Tom Connor |
| EPI_ISL_3297233 | Viollier AG | Department of Biosystems Science and Engineering, ETH Zürich | Christian Beisel, Sarah Nadeau, Chaoran Chen, Ivan Topolsky, Kim Philipp Jablonski, Lara Fuhrmann, Rebecca Denes, Mirjam Feldkamp, Ina Nissen, Natascha Santacroce, Elodie Burcklen, Christiane Beckmann, Maurice Redondo, Olivier Kobel, Christoph Noppen, Niko Beerenwinkel, Tanja Stadler |
| EPI_ISL_3297234 | Wales Specialist Virology Centre Sequencing lab: Pathogen Genomics Unit | Public Health Wales Microbiology Cardiff Wales Specialist Virology Centre | Catherine Moore, Johnathan Evans, Laura Gifford, Malorie Perry, Simon Cottrell, Angela Marchbank, Alec Birchley, Alexander Adams, Amy Gaskin, Bree Gatica-Wilcox, Jason Coombes, Joel Southgate, Lauren Gilbert, Lee Graham, Nicole Pacchiarini, Sara Kumziene-Summerhayes, Sarah Taylor, Sophie Jones, Sara Rey, Matthew Bull, Joanne Watkins, Sally Corden, Tom Connor |
| EPI_ISL_3297235 | Viollier AG | Department of Biosystems Science and Engineering, ETH Zürich | Christian Beisel, Sarah Nadeau, Chaoran Chen, Ivan Topolsky, Kim Philipp Jablonski, Lara Fuhrmann, Rebecca Denes, Mirjam Feldkamp, Ina Nissen, Natascha Santacroce, Elodie Burcklen, Christiane Beckmann, Maurice Redondo, Olivier Kobel, Christoph Noppen, Niko Beerenwinkel, Tanja Stadler |
| EPI_ISL_3297236 | Wales Specialist Virology Centre Sequencing lab: Pathogen Genomics Unit | Public Health Wales Microbiology Cardiff Wales Specialist Virology Centre | Catherine Moore, Johnathan Evans, Laura Gifford, Malorie Perry, Simon Cottrell, Angela Marchbank, Alec Birchley, Alexander Adams, Amy Gaskin, Bree Gatica-Wilcox, Jason Coombes, Joel Southgate, Lauren Gilbert, Lee Graham, Nicole Pacchiarini, Sara Kumziene-Summerhayes, Sarah Taylor, Sophie Jones, Sara Rey, Matthew Bull, Joanne Watkins, Sally Corden, Tom Connor |
| EPI_ISL_3297237 | Viollier AG | Department of Biosystems Science and Engineering, ETH Zürich | Christian Beisel, Sarah Nadeau, Chaoran Chen, Ivan Topolsky, Kim Philipp Jablonski, Lara Fuhrmann, Rebecca Denes, Mirjam Feldkamp, Ina Nissen, Natascha Santacroce, Elodie Burcklen, Christiane Beckmann, Maurice Redondo, Olivier Kobel, Christoph Noppen, Niko Beerenwinkel, Tanja Stadler |
| EPI_ISL_3297238 | Wales Specialist Virology Centre Sequencing lab: Pathogen Genomics Unit | Public Health Wales Microbiology Cardiff Wales Specialist Virology Centre | Catherine Moore, Johnathan Evans, Laura Gifford, Malorie Perry, Simon Cottrell, Angela Marchbank, Alec Birchley, Alexander Adams, Amy Gaskin, Bree Gatica-Wilcox, Jason Coombes, Joel Southgate, Lauren Gilbert, Lee Graham, Nicole Pacchiarini, Sara Kumziene-Summerhayes, Sarah Taylor, Sophie Jones, Sara Rey, Matthew Bull, Joanne Watkins, Sally Corden, Tom Connor |
| EPI_ISL_3297239 | Viollier AG | Department of Biosystems Science and Engineering, ETH Zürich | Christian Beisel, Sarah Nadeau, Chaoran Chen, Ivan Topolsky, Kim Philipp Jablonski, Lara Fuhrmann, Rebecca Denes, Mirjam Feldkamp, Ina Nissen, Natascha Santacroce, Elodie Burcklen, Christiane Beckmann, Maurice Redondo, Olivier Kobel, Christoph Noppen, Niko Beerenwinkel, Tanja Stadler |
| EPI_ISL_3297240, EPI_ISL_3297241 | Wales Specialist Virology Centre Sequencing lab: Pathogen Genomics Unit | Public Health Wales Microbiology Cardiff Wales Specialist Virology Centre | Catherine Moore, Johnathan Evans, Laura Gifford, Malorie Perry, Simon Cottrell, Angela Marchbank, Alec Birchley, Alexander Adams, Amy Gaskin, Bree Gatica-Wilcox, Jason Coombes, Joel Southgate, Lauren Gilbert, Lee Graham, Nicole Pacchiarini, Sara Kumziene-Summerhayes, Sarah Taylor, Sophie Jones, Sara Rey, Matthew Bull, Joanne Watkins, Sally Corden, Tom Connor |
| EPI_ISL_3297242 | Viollier AG | Department of Biosystems Science and Engineering, ETH Zürich | Christian Beisel, Sarah Nadeau, Chaoran Chen, Ivan Topolsky, Kim Philipp Jablonski, Lara Fuhrmann, Rebecca Denes, Mirjam Feldkamp, Ina Nissen, Natascha Santacroce, Elodie Burcklen, Christiane Beckmann, Maurice Redondo, Olivier Kobel, Christoph Noppen, Niko Beerenwinkel, Tanja Stadler |
| EPI_ISL_3297243 | Wales Specialist Virology Centre Sequencing lab: Pathogen Genomics Unit | Public Health Wales Microbiology Cardiff Wales Specialist Virology Centre | Catherine Moore, Johnathan Evans, Laura Gifford, Malorie Perry, Simon Cottrell, Angela Marchbank, Alec Birchley, Alexander Adams, Amy Gaskin, Bree Gatica-Wilcox, Jason Coombes, Joel Southgate, Lauren Gilbert, Lee Graham, Nicole Pacchiarini, Sara Kumziene-Summerhayes, Sarah Taylor, Sophie Jones, Sara Rey, Matthew Bull, Joanne Watkins, Sally Corden, Tom Connor |
| EPI_ISL_3297244 | Viollier AG | Department of Biosystems Science and Engineering, ETH Zürich | Christian Beisel, Sarah Nadeau, Chaoran Chen, Ivan Topolsky, Kim Philipp Jablonski, Lara Fuhrmann, Rebecca Denes, Mirjam Feldkamp, Ina Nissen, Natascha Santacroce, Elodie Burcklen, Christiane Beckmann, Maurice Redondo, Olivier Kobel, Christoph Noppen, Niko Beerenwinkel, Tanja Stadler |
| EPI_ISL_3297245 | Wales Specialist Virology Centre Sequencing lab: Pathogen | Public Health Wales Microbiology Cardiff Wales Specialist Virology | Catherine Moore, Johnathan Evans, Laura Gifford, Malorie Perry, Simon Cottrell, Angela Marchbank, Alec Birchley, Alexander Adams, Amy Gaskin, Bree Gatica-Wilcox, Jason Coombes, Joel Southgate, Lauren Gilbert, Lee Graham, Nicole |

| | | | |
|---|---|---|---|
| | Genomics Unit | Centre | Pacchiarini, Sara Kumziene-Summerhayes, Sarah Taylor, Sophie Jones, Sara Rey, Matthew Bull, Joanne Watkins, Sally Corden, Tom Connor |
| EPI_ISL_3297246 | Viollier AG | Department of Biosystems Science and Engineering, ETH Zürich | Christian Beisel, Sarah Nadeau, Chaoran Chen, Ivan Topolsky, Kim Philipp Jablonski, Lara Fuhrmann, Rebecca Denes, Mirjam Feldkamp, Ina Nissen, Natascha Santacroce, Elodie Burcklen, Christiane Beckmann, Maurice Redondo, Olivier Kobel, Christoph Noppen, Niko Beerenwinkel, Tanja Stadler |
| EPI_ISL_3297247 | Wales Specialist Virology Centre Sequencing lab: Pathogen Genomics Unit | Public Health Wales Microbiology Cardiff Wales Specialist Virology Centre | Catherine Moore, Johnathan Evans, Laura Gifford, Malorie Perry, Simon Cottrell, Angela Marchbank, Alec Birchley, Alexander Adams, Amy Gaskin, Bree Gatica-Wilcox, Jason Coombes, Joel Southgate, Lauren Gilbert, Lee Graham, Nicole Pacchiarini, Sara Kumziene-Summerhayes, Sarah Taylor, Sophie Jones, Sara Rey, Matthew Bull, Joanne Watkins, Sally Corden, Tom Connor |
| EPI_ISL_3297248 | Viollier AG | Department of Biosystems Science and Engineering, ETH Zürich | Christian Beisel, Sarah Nadeau, Chaoran Chen, Ivan Topolsky, Kim Philipp Jablonski, Lara Fuhrmann, Rebecca Denes, Mirjam Feldkamp, Ina Nissen, Natascha Santacroce, Elodie Burcklen, Christiane Beckmann, Maurice Redondo, Olivier Kobel, Christoph Noppen, Niko Beerenwinkel, Tanja Stadler |
| EPI_ISL_3297249, EPI_ISL_3297251 | Wales Specialist Virology Centre Sequencing lab: Pathogen Genomics Unit | Public Health Wales Microbiology Cardiff Wales Specialist Virology Centre | Catherine Moore, Johnathan Evans, Laura Gifford, Malorie Perry, Simon Cottrell, Angela Marchbank, Alec Birchley, Alexander Adams, Amy Gaskin, Bree Gatica-Wilcox, Jason Coombes, Joel Southgate, Lauren Gilbert, Lee Graham, Nicole Pacchiarini, Sara Kumziene-Summerhayes, Sarah Taylor, Sophie Jones, Sara Rey, Matthew Bull, Joanne Watkins, Sally Corden, Tom Connor |
| EPI_ISL_3297252 | Viollier AG | Department of Biosystems Science and Engineering, ETH Zürich | Christian Beisel, Sarah Nadeau, Chaoran Chen, Ivan Topolsky, Kim Philipp Jablonski, Lara Fuhrmann, Rebecca Denes, Mirjam Feldkamp, Ina Nissen, Natascha Santacroce, Elodie Burcklen, Christiane Beckmann, Maurice Redondo, Olivier Kobel, Christoph Noppen, Niko Beerenwinkel, Tanja Stadler |
| EPI_ISL_3297253 | Wales Specialist Virology Centre Sequencing lab: Pathogen Genomics Unit | Public Health Wales Microbiology Cardiff Wales Specialist Virology Centre | Catherine Moore, Johnathan Evans, Laura Gifford, Malorie Perry, Simon Cottrell, Angela Marchbank, Alec Birchley, Alexander Adams, Amy Gaskin, Bree Gatica-Wilcox, Jason Coombes, Joel Southgate, Lauren Gilbert, Lee Graham, Nicole Pacchiarini, Sara Kumziene-Summerhayes, Sarah Taylor, Sophie Jones, Sara Rey, Matthew Bull, Joanne Watkins, Sally Corden, Tom Connor |
| EPI_ISL_3297254 | Viollier AG | Department of Biosystems Science and Engineering, ETH Zürich | Christian Beisel, Sarah Nadeau, Chaoran Chen, Ivan Topolsky, Kim Philipp Jablonski, Lara Fuhrmann, Rebecca Denes, Mirjam Feldkamp, Ina Nissen, Natascha Santacroce, Elodie Burcklen, Christiane Beckmann, Maurice Redondo, Olivier Kobel, Christoph Noppen, Niko Beerenwinkel, Tanja Stadler |
| EPI_ISL_3297257 | Wales Specialist Virology Centre Sequencing lab: Pathogen Genomics Unit | Public Health Wales Microbiology Cardiff Wales Specialist Virology Centre | Catherine Moore, Johnathan Evans, Laura Gifford, Malorie Perry, Simon Cottrell, Angela Marchbank, Alec Birchley, Alexander Adams, Amy Gaskin, Bree Gatica-Wilcox, Jason Coombes, Joel Southgate, Lauren Gilbert, Lee Graham, Nicole Pacchiarini, Sara Kumziene-Summerhayes, Sarah Taylor, Sophie Jones, Sara Rey, Matthew Bull, Joanne Watkins, Sally Corden, Tom Connor |
| EPI_ISL_3297258 | Viollier AG | Department of Biosystems Science and Engineering, ETH Zürich | Christian Beisel, Sarah Nadeau, Chaoran Chen, Ivan Topolsky, Kim Philipp Jablonski, Lara Fuhrmann, Rebecca Denes, Mirjam Feldkamp, Ina Nissen, Natascha Santacroce, Elodie Burcklen, Christiane Beckmann, Maurice Redondo, Olivier Kobel, Christoph Noppen, Niko Beerenwinkel, Tanja Stadler |
| EPI_ISL_3297259 | Wales Specialist Virology Centre Sequencing lab: Pathogen Genomics Unit | Public Health Wales Microbiology Cardiff Wales Specialist Virology Centre | Catherine Moore, Johnathan Evans, Laura Gifford, Malorie Perry, Simon Cottrell, Angela Marchbank, Alec Birchley, Alexander Adams, Amy Gaskin, Bree Gatica-Wilcox, Jason Coombes, Joel Southgate, Lauren Gilbert, Lee Graham, Nicole Pacchiarini, Sara Kumziene-Summerhayes, Sarah Taylor, Sophie Jones, Sara Rey, Matthew Bull, Joanne Watkins, Sally Corden, Tom Connor |
| EPI_ISL_3297260, EPI_ISL_3297261, EPI_ISL_3297262, EPI_ISL_3297263 | Viollier AG | Department of Biosystems Science and Engineering, ETH Zürich | Christian Beisel, Sarah Nadeau, Chaoran Chen, Ivan Topolsky, Kim Philipp Jablonski, Lara Fuhrmann, Rebecca Denes, Mirjam Feldkamp, Ina Nissen, Natascha Santacroce, Elodie Burcklen, Christiane Beckmann, Maurice Redondo, Olivier Kobel, Christoph Noppen, Niko Beerenwinkel, Tanja Stadler |
| EPI_ISL_3297264 | Wales Specialist Virology Centre Sequencing lab: Pathogen Genomics Unit | Public Health Wales Microbiology Cardiff Wales Specialist Virology Centre | Catherine Moore, Johnathan Evans, Laura Gifford, Malorie Perry, Simon Cottrell, Angela Marchbank, Alec Birchley, Alexander Adams, Amy Gaskin, Bree Gatica-Wilcox, Jason Coombes, Joel Southgate, Lauren Gilbert, Lee Graham, Nicole Pacchiarini, Sara Kumziene-Summerhayes, Sarah Taylor, Sophie Jones, Sara Rey, Matthew Bull, Joanne Watkins, Sally Corden, Tom Connor |
| EPI_ISL_3297265 | Viollier AG | Department of Biosystems Science and Engineering, ETH Zürich | Christian Beisel, Sarah Nadeau, Chaoran Chen, Ivan Topolsky, Kim Philipp Jablonski, Lara Fuhrmann, Rebecca Denes, Mirjam Feldkamp, Ina Nissen, Natascha Santacroce, Elodie Burcklen, Christiane Beckmann, Maurice Redondo, Olivier Kobel, Christoph Noppen, Niko Beerenwinkel, Tanja Stadler |
| EPI_ISL_3297266 | Wales Specialist Virology Centre Sequencing lab: Pathogen Genomics Unit | Public Health Wales Microbiology Cardiff Wales Specialist Virology Centre | Catherine Moore, Johnathan Evans, Laura Gifford, Malorie Perry, Simon Cottrell, Angela Marchbank, Alec Birchley, Alexander Adams, Amy Gaskin, Bree Gatica-Wilcox, Jason Coombes, Joel Southgate, Lauren Gilbert, Lee Graham, Nicole Pacchiarini, Sara Kumziene-Summerhayes, Sarah Taylor, Sophie Jones, Sara Rey, Matthew Bull, Joanne Watkins, Sally Corden, Tom Connor |
| EPI_ISL_3297267 | Viollier AG | Department of Biosystems Science and Engineering, ETH Zürich | Christian Beisel, Sarah Nadeau, Chaoran Chen, Ivan Topolsky, Kim Philipp Jablonski, Lara Fuhrmann, Rebecca Denes, Mirjam Feldkamp, Ina Nissen, Natascha Santacroce, Elodie Burcklen, Christiane Beckmann, Maurice Redondo, Olivier Kobel, Christoph Noppen, Niko Beerenwinkel, Tanja Stadler |
| EPI_ISL_3297268 | Wales Specialist Virology Centre Sequencing lab: Pathogen Genomics Unit | Public Health Wales Microbiology Cardiff Wales Specialist Virology Centre | Catherine Moore, Johnathan Evans, Laura Gifford, Malorie Perry, Simon Cottrell, Angela Marchbank, Alec Birchley, Alexander Adams, Amy Gaskin, Bree Gatica-Wilcox, Jason Coombes, Joel Southgate, Lauren Gilbert, Lee Graham, Nicole Pacchiarini, Sara Kumziene-Summerhayes, Sarah Taylor, Sophie Jones, Sara Rey, Matthew Bull, Joanne Watkins, Sally Corden, Tom Connor |
| EPI_ISL_3297269 | Viollier AG | Department of Biosystems Science and Engineering, ETH Zürich | Christian Beisel, Sarah Nadeau, Chaoran Chen, Ivan Topolsky, Kim Philipp Jablonski, Lara Fuhrmann, Rebecca Denes, Mirjam Feldkamp, Ina Nissen, Natascha Santacroce, Elodie Burcklen, Christiane Beckmann, Maurice Redondo, Olivier Kobel, Christoph Noppen, Niko Beerenwinkel, Tanja Stadler |
| EPI_ISL_3297270, EPI_ISL_3297271 | Wales Specialist Virology Centre Sequencing lab: Pathogen Genomics Unit | Public Health Wales Microbiology Cardiff Wales Specialist Virology Centre | Catherine Moore, Johnathan Evans, Laura Gifford, Malorie Perry, Simon Cottrell, Angela Marchbank, Alec Birchley, Alexander Adams, Amy Gaskin, Bree Gatica-Wilcox, Jason Coombes, Joel Southgate, Lauren Gilbert, Lee Graham, Nicole Pacchiarini, Sara Kumziene-Summerhayes, Sarah Taylor, Sophie Jones, Sara Rey, Matthew Bull, Joanne Watkins, Sally Corden, Tom Connor |
| EPI_ISL_3297272 | Viollier AG | Department of Biosystems Science and Engineering, ETH Zürich | Christian Beisel, Sarah Nadeau, Chaoran Chen, Ivan Topolsky, Kim Philipp Jablonski, Lara Fuhrmann, Rebecca Denes, Mirjam Feldkamp, Ina Nissen, Natascha Santacroce, Elodie Burcklen, Christiane Beckmann, Maurice Redondo, Olivier Kobel, Christoph Noppen, Niko Beerenwinkel, Tanja Stadler |
| EPI_ISL_3297273 | Wales Specialist Virology Centre Sequencing lab: Pathogen Genomics Unit | Public Health Wales Microbiology Cardiff Wales Specialist Virology Centre | Catherine Moore, Johnathan Evans, Laura Gifford, Malorie Perry, Simon Cottrell, Angela Marchbank, Alec Birchley, Alexander Adams, Amy Gaskin, Bree Gatica-Wilcox, Jason Coombes, Joel Southgate, Lauren Gilbert, Lee Graham, Nicole Pacchiarini, Sara Kumziene-Summerhayes, Sarah Taylor, Sophie Jones, Sara Rey, Matthew Bull, Joanne Watkins, Sally Corden, Tom Connor |
| EPI_ISL_3297274 | Viollier AG | Department of Biosystems Science and Engineering, ETH Zürich | Christian Beisel, Sarah Nadeau, Chaoran Chen, Ivan Topolsky, Kim Philipp Jablonski, Lara Fuhrmann, Rebecca Denes, Mirjam Feldkamp, Ina Nissen, Natascha Santacroce, Elodie Burcklen, Christiane Beckmann, Maurice Redondo, Olivier Kobel, Christoph Noppen, Niko Beerenwinkel, Tanja Stadler |
| EPI_ISL_3297275 | Wales Specialist Virology Centre Sequencing lab: Pathogen Genomics Unit | Public Health Wales Microbiology Cardiff Wales Specialist Virology Centre | Catherine Moore, Johnathan Evans, Laura Gifford, Malorie Perry, Simon Cottrell, Angela Marchbank, Alec Birchley, Alexander Adams, Amy Gaskin, Bree Gatica-Wilcox, Jason Coombes, Joel Southgate, Lauren Gilbert, Lee Graham, Nicole Pacchiarini, Sara Kumziene-Summerhayes, Sarah Taylor, Sophie Jones, Sara Rey, Matthew Bull, Joanne Watkins, Sally Corden, Tom Connor |
| EPI_ISL_3297276 | Viollier AG | Department of Biosystems Science and Engineering, ETH Zürich | Christian Beisel, Sarah Nadeau, Chaoran Chen, Ivan Topolsky, Kim Philipp Jablonski, Lara Fuhrmann, Rebecca Denes, Mirjam Feldkamp, Ina Nissen, Natascha Santacroce, Elodie Burcklen, Christiane Beckmann, Maurice Redondo, Olivier Kobel, Christoph Noppen, Niko Beerenwinkel, Tanja Stadler |
| EPI_ISL_3297277 | Wales Specialist Virology Centre Sequencing lab: Pathogen Genomics Unit | Public Health Wales Microbiology Cardiff Wales Specialist Virology Centre | Catherine Moore, Johnathan Evans, Laura Gifford, Malorie Perry, Simon Cottrell, Angela Marchbank, Alec Birchley, Alexander Adams, Amy Gaskin, Bree Gatica-Wilcox, Jason Coombes, Joel Southgate, Lauren Gilbert, Lee Graham, Nicole Pacchiarini, Sara Kumziene-Summerhayes, Sarah Taylor, Sophie Jones, Sara Rey, Matthew Bull, Joanne Watkins, Sally Corden, Tom Connor |
| EPI_ISL_3297278 | Viollier AG | Department of Biosystems Science and Engineering, ETH Zürich | Christian Beisel, Sarah Nadeau, Chaoran Chen, Ivan Topolsky, Kim Philipp Jablonski, Lara Fuhrmann, Rebecca Denes, Mirjam Feldkamp, Ina Nissen, Natascha Santacroce, Elodie Burcklen, Christiane Beckmann, Maurice Redondo, Olivier Kobel, Christoph Noppen, Niko Beerenwinkel, Tanja Stadler |
| EPI_ISL_3297279 | Wales Specialist Virology Centre Sequencing lab: Pathogen Genomics Unit | Public Health Wales Microbiology Cardiff Wales Specialist Virology Centre | Catherine Moore, Johnathan Evans, Laura Gifford, Malorie Perry, Simon Cottrell, Angela Marchbank, Alec Birchley, Alexander Adams, Amy Gaskin, Bree Gatica-Wilcox, Jason Coombes, Joel Southgate, Lauren Gilbert, Lee Graham, Nicole Pacchiarini, Sara Kumziene-Summerhayes, Sarah Taylor, Sophie Jones, Sara Rey, Matthew Bull, Joanne Watkins, Sally Corden, Tom Connor |
| EPI_ISL_3297280 | Viollier AG | Department of Biosystems Science and Engineering, ETH Zürich | Christian Beisel, Sarah Nadeau, Chaoran Chen, Ivan Topolsky, Kim Philipp Jablonski, Lara Fuhrmann, Rebecca Denes, Mirjam Feldkamp, Ina Nissen, Natascha Santacroce, Elodie Burcklen, Christiane Beckmann, Maurice Redondo, Olivier Kobel, Christoph Noppen, Niko Beerenwinkel, Tanja Stadler |
| EPI_ISL_3297281 | Wales Specialist Virology Centre Sequencing lab: Pathogen Genomics Unit | Public Health Wales Microbiology Cardiff Wales Specialist Virology Centre | Catherine Moore, Johnathan Evans, Laura Gifford, Malorie Perry, Simon Cottrell, Angela Marchbank, Alec Birchley, Alexander Adams, Amy Gaskin, Bree Gatica-Wilcox, Jason Coombes, Joel Southgate, Lauren Gilbert, Lee Graham, Nicole Pacchiarini, Sara Kumziene-Summerhayes, Sarah Taylor, Sophie Jones, Sara Rey, Matthew Bull, Joanne Watkins, Sally Corden, Tom Connor |
| EPI_ISL_3297282 | Viollier AG | Department of Biosystems Science and Engineering, ETH Zürich | Christian Beisel, Sarah Nadeau, Chaoran Chen, Ivan Topolsky, Kim Philipp Jablonski, Lara Fuhrmann, Rebecca Denes, Mirjam Feldkamp, Ina Nissen, Natascha Santacroce, Elodie Burcklen, Christiane Beckmann, Maurice Redondo, Olivier Kobel, Christoph Noppen, Niko Beerenwinkel, Tanja Stadler |
| EPI_ISL_3297283 | Wales Specialist Virology Centre Sequencing lab: Pathogen Genomics Unit | Public Health Wales Microbiology Cardiff Wales Specialist Virology Centre | Catherine Moore, Johnathan Evans, Laura Gifford, Malorie Perry, Simon Cottrell, Angela Marchbank, Alec Birchley, Alexander Adams, Amy Gaskin, Bree Gatica-Wilcox, Jason Coombes, Joel Southgate, Lauren Gilbert, Lee Graham, Nicole Pacchiarini, Sara Kumziene-Summerhayes, Sarah Taylor, Sophie Jones, Sara Rey, Matthew Bull, Joanne Watkins, Sally Corden, Tom Connor |
| EPI_ISL_3297284 | Viollier AG | Department of Biosystems Science and Engineering, ETH Zürich | Christian Beisel, Sarah Nadeau, Chaoran Chen, Ivan Topolsky, Kim Philipp Jablonski, Lara Fuhrmann, Rebecca Denes, Mirjam Feldkamp, Ina Nissen, Natascha Santacroce, Elodie Burcklen, Christiane Beckmann, Maurice Redondo, Olivier Kobel, Christoph Noppen, Niko Beerenwinkel, Tanja Stadler |
| EPI_ISL_3297285 | Wales Specialist Virology Centre Sequencing lab: Pathogen Genomics Unit | Public Health Wales Microbiology Cardiff Wales Specialist Virology Centre | Catherine Moore, Johnathan Evans, Laura Gifford, Malorie Perry, Simon Cottrell, Angela Marchbank, Alec Birchley, Alexander Adams, Amy Gaskin, Bree Gatica-Wilcox, Jason Coombes, Joel Southgate, Lauren Gilbert, Lee Graham, Nicole Pacchiarini, Sara Kumziene-Summerhayes, Sarah Taylor, Sophie Jones, Sara Rey, Matthew Bull, Joanne Watkins, Sally Corden, Tom Connor |
| EPI_ISL_3297286 | Viollier AG | Department of Biosystems Science and Engineering, ETH Zürich | Christian Beisel, Sarah Nadeau, Chaoran Chen, Ivan Topolsky, Kim Philipp Jablonski, Lara Fuhrmann, Rebecca Denes, Mirjam Feldkamp, Ina Nissen, Natascha Santacroce, Elodie Burcklen, Christiane Beckmann, Maurice Redondo, Olivier Kobel, Christoph Noppen, Niko Beerenwinkel, Tanja Stadler |
| EPI_ISL_3297287, EPI_ISL_3297288 | Wales Specialist Virology Centre Sequencing lab: Pathogen Genomics Unit | Public Health Wales Microbiology Cardiff Wales Specialist Virology Centre | Catherine Moore, Johnathan Evans, Laura Gifford, Malorie Perry, Simon Cottrell, Angela Marchbank, Alec Birchley, Alexander Adams, Amy Gaskin, Bree Gatica-Wilcox, Jason Coombes, Joel Southgate, Lauren Gilbert, Lee Graham, Nicole Pacchiarini, Sara Kumziene-Summerhayes, Sarah Taylor, Sophie Jones, Sara Rey, Matthew Bull, Joanne Watkins, Sally Corden, Tom Connor |
| EPI_ISL_3297289 | Viollier AG | Department of Biosystems Science and Engineering, ETH Zürich | Christian Beisel, Sarah Nadeau, Chaoran Chen, Ivan Topolsky, Kim Philipp Jablonski, Lara Fuhrmann, Rebecca Denes, Mirjam Feldkamp, Ina Nissen, Natascha Santacroce, Elodie Burcklen, Christiane Beckmann, Maurice Redondo, Olivier Kobel, Christoph Noppen, Niko Beerenwinkel, Tanja Stadler |
| EPI_ISL_3297290 | Wales Specialist Virology Centre Sequencing lab: Pathogen Genomics Unit | Public Health Wales Microbiology Cardiff Wales Specialist Virology Centre | Catherine Moore, Johnathan Evans, Laura Gifford, Malorie Perry, Simon Cottrell, Angela Marchbank, Alec Birchley, Alexander Adams, Amy Gaskin, Bree Gatica-Wilcox, Jason Coombes, Joel Southgate, Lauren Gilbert, Lee Graham, Nicole Pacchiarini, Sara Kumziene-Summerhayes, Sarah Taylor, Sophie Jones, Sara Rey, Matthew Bull, Joanne Watkins, Sally Corden, Tom Connor |
| EPI_ISL_3297291 | Viollier AG | Department of Biosystems Science and Engineering, ETH Zürich | Christian Beisel, Sarah Nadeau, Chaoran Chen, Ivan Topolsky, Kim Philipp Jablonski, Lara Fuhrmann, Rebecca Denes, Mirjam Feldkamp, Ina Nissen, Natascha Santacroce, Elodie Burcklen, Christiane Beckmann, Maurice Redondo, Olivier Kobel, Christoph Noppen, Niko Beerenwinkel, Tanja Stadler |
| EPI_ISL_3297292 | Wales Specialist Virology Centre Sequencing lab: Pathogen Genomics Unit | Public Health Wales Microbiology Cardiff Wales Specialist Virology Centre | Catherine Moore, Johnathan Evans, Laura Gifford, Malorie Perry, Simon Cottrell, Angela Marchbank, Alec Birchley, Alexander Adams, Amy Gaskin, Bree Gatica-Wilcox, Jason Coombes, Joel Southgate, Lauren Gilbert, Lee Graham, Nicole Pacchiarini, Sara Kumziene-Summerhayes, Sarah Taylor, Sophie Jones, Sara Rey, Matthew Bull, Joanne Watkins, Sally Corden, Tom Connor |
| EPI_ISL_3297293 | Viollier AG | Department of Biosystems Science and Engineering, ETH Zürich | Christian Beisel, Sarah Nadeau, Chaoran Chen, Ivan Topolsky, Kim Philipp Jablonski, Lara Fuhrmann, Rebecca Denes, Mirjam Feldkamp, Ina Nissen, Natascha Santacroce, Elodie Burcklen, Christiane Beckmann, Maurice Redondo, Olivier Kobel, Christoph Noppen, Niko Beerenwinkel, Tanja Stadler |
| EPI_ISL_3297294 | Wales Specialist Virology Centre Sequencing lab: Pathogen Genomics Unit | Public Health Wales Microbiology Cardiff Wales Specialist Virology Centre | Catherine Moore, Johnathan Evans, Laura Gifford, Malorie Perry, Simon Cottrell, Angela Marchbank, Alec Birchley, Alexander Adams, Amy Gaskin, Bree Gatica-Wilcox, Jason Coombes, Joel Southgate, Lauren Gilbert, Lee Graham, Nicole Pacchiarini, Sara Kumziene-Summerhayes, Sarah Taylor, Sophie Jones, Sara Rey, Matthew Bull, Joanne Watkins, Sally Corden, Tom Connor |
| EPI_ISL_3297295 | Viollier AG | Department of Biosystems Science and Engineering, ETH Zürich | Christian Beisel, Sarah Nadeau, Chaoran Chen, Ivan Topolsky, Kim Philipp Jablonski, Lara Fuhrmann, Rebecca Denes, Mirjam Feldkamp, Ina Nissen, Natascha Santacroce, Elodie Burcklen, Christiane Beckmann, Maurice Redondo, Olivier Kobel, Christoph Noppen, Niko Beerenwinkel, Tanja Stadler |
| EPI_ISL_3297296 | Wales Specialist Virology Centre Sequencing lab: Pathogen Genomics Unit | Public Health Wales Microbiology Cardiff Wales Specialist Virology Centre | Catherine Moore, Johnathan Evans, Laura Gifford, Malorie Perry, Simon Cottrell, Angela Marchbank, Alec Birchley, Alexander Adams, Amy Gaskin, Bree Gatica-Wilcox, Jason Coombes, Joel Southgate, Lauren Gilbert, Lee Graham, Nicole Pacchiarini, Sara Kumziene-Summerhayes, Sarah Taylor, Sophie Jones, Sara Rey, Matthew Bull, Joanne Watkins, Sally Corden, Tom Connor |
| EPI_ISL_3297297 | Viollier AG | Department of Biosystems Science and Engineering, ETH Zürich | Christian Beisel, Sarah Nadeau, Chaoran Chen, Ivan Topolsky, Kim Philipp Jablonski, Lara Fuhrmann, Rebecca Denes, Mirjam Feldkamp, Ina Nissen, Natascha Santacroce, Elodie Burcklen, Christiane Beckmann, Maurice Redondo, Olivier Kobel, Christoph Noppen, Niko Beerenwinkel, Tanja Stadler |
| EPI_ISL_3297298 | Wales Specialist Virology Centre Sequencing lab: Pathogen Genomics Unit | Public Health Wales Microbiology Cardiff Wales Specialist Virology Centre | Catherine Moore, Johnathan Evans, Laura Gifford, Malorie Perry, Simon Cottrell, Angela Marchbank, Alec Birchley, Alexander Adams, Amy Gaskin, Bree Gatica-Wilcox, Jason Coombes, Joel Southgate, Lauren Gilbert, Lee Graham, Nicole Pacchiarini, Sara Kumziene-Summerhayes, Sarah Taylor, Sophie Jones, Sara Rey, Matthew Bull, Joanne Watkins, Sally Corden, Tom Connor |

| | | | |
|---|---|---|---|
| EPI_ISL_3297299 | Viollier AG | Department of Biosystems Science and Engineering, ETH Zürich | Christian Beisel, Sarah Nadeau, Chaoran Chen, Ivan Topolsky, Kim Philipp Jablonski, Lara Fuhrmann, Rebecca Denes, Mirjam Feldkamp, Ina Nissen, Natascha Santacroce, Elodie Burcklen, Christiane Beckmann, Maurice Redondo, Olivier Kobel, Christoph Noppen, Niko Beerenwinkel, Tanja Stadler |
| EPI_ISL_3297300, EPI_ISL_3297301 | Wales Specialist Virology Centre Sequencing lab: Pathogen Genomics Unit | Public Health Wales Microbiology Cardiff Wales Specialist Virology Centre | Catherine Moore, Johnathan Evans, Laura Gifford, Malorie Perry, Simon Cottrell, Angela Marchbank, Alec Birchley, Alexander Adams, Amy Gaskin, Bree Gatica-Wilcox, Jason Coombes, Joel Southgate, Lauren Gilbert, Lee Graham, Nicole Pacchiarini, Sara Kumziene-Summerhayes, Sarah Taylor, Sophie Jones, Sara Rey, Matthew Bull, Joanne Watkins, Sally Corden, Tom Connor |
| EPI_ISL_3297302 | Viollier AG | Department of Biosystems Science and Engineering, ETH Zürich | Christian Beisel, Sarah Nadeau, Chaoran Chen, Ivan Topolsky, Kim Philipp Jablonski, Lara Fuhrmann, Rebecca Denes, Mirjam Feldkamp, Ina Nissen, Natascha Santacroce, Elodie Burcklen, Christiane Beckmann, Maurice Redondo, Olivier Kobel, Christoph Noppen, Niko Beerenwinkel, Tanja Stadler |
| EPI_ISL_3297303 | Wales Specialist Virology Centre Sequencing lab: Pathogen Genomics Unit | Public Health Wales Microbiology Cardiff Wales Specialist Virology Centre | Catherine Moore, Johnathan Evans, Laura Gifford, Malorie Perry, Simon Cottrell, Angela Marchbank, Alec Birchley, Alexander Adams, Amy Gaskin, Bree Gatica-Wilcox, Jason Coombes, Joel Southgate, Lauren Gilbert, Lee Graham, Nicole Pacchiarini, Sara Kumziene-Summerhayes, Sarah Taylor, Sophie Jones, Sara Rey, Matthew Bull, Joanne Watkins, Sally Corden, Tom Connor |
| EPI_ISL_3297304 | Viollier AG | Department of Biosystems Science and Engineering, ETH Zürich | Christian Beisel, Sarah Nadeau, Chaoran Chen, Ivan Topolsky, Kim Philipp Jablonski, Lara Fuhrmann, Rebecca Denes, Mirjam Feldkamp, Ina Nissen, Natascha Santacroce, Elodie Burcklen, Christiane Beckmann, Maurice Redondo, Olivier Kobel, Christoph Noppen, Niko Beerenwinkel, Tanja Stadler |
| EPI_ISL_3297305 | Wales Specialist Virology Centre Sequencing lab: Pathogen Genomics Unit | Public Health Wales Microbiology Cardiff Wales Specialist Virology Centre | Catherine Moore, Johnathan Evans, Laura Gifford, Malorie Perry, Simon Cottrell, Angela Marchbank, Alec Birchley, Alexander Adams, Amy Gaskin, Bree Gatica-Wilcox, Jason Coombes, Joel Southgate, Lauren Gilbert, Lee Graham, Nicole Pacchiarini, Sara Kumziene-Summerhayes, Sarah Taylor, Sophie Jones, Sara Rey, Matthew Bull, Joanne Watkins, Sally Corden, Tom Connor |
| EPI_ISL_3297306 | Viollier AG | Department of Biosystems Science and Engineering, ETH Zürich | Christian Beisel, Sarah Nadeau, Chaoran Chen, Ivan Topolsky, Kim Philipp Jablonski, Lara Fuhrmann, Rebecca Denes, Mirjam Feldkamp, Ina Nissen, Natascha Santacroce, Elodie Burcklen, Christiane Beckmann, Maurice Redondo, Olivier Kobel, Christoph Noppen, Niko Beerenwinkel, Tanja Stadler |
| EPI_ISL_3297307 | Wales Specialist Virology Centre Sequencing lab: Pathogen Genomics Unit | Public Health Wales Microbiology Cardiff Wales Specialist Virology Centre | Catherine Moore, Johnathan Evans, Laura Gifford, Malorie Perry, Simon Cottrell, Angela Marchbank, Alec Birchley, Alexander Adams, Amy Gaskin, Bree Gatica-Wilcox, Jason Coombes, Joel Southgate, Lauren Gilbert, Lee Graham, Nicole Pacchiarini, Sara Kumziene-Summerhayes, Sarah Taylor, Sophie Jones, Sara Rey, Matthew Bull, Joanne Watkins, Sally Corden, Tom Connor |
| EPI_ISL_3297308 | Viollier AG | Department of Biosystems Science and Engineering, ETH Zürich | Christian Beisel, Sarah Nadeau, Chaoran Chen, Ivan Topolsky, Kim Philipp Jablonski, Lara Fuhrmann, Rebecca Denes, Mirjam Feldkamp, Ina Nissen, Natascha Santacroce, Elodie Burcklen, Christiane Beckmann, Maurice Redondo, Olivier Kobel, Christoph Noppen, Niko Beerenwinkel, Tanja Stadler |
| EPI_ISL_3297309 | Wales Specialist Virology Centre Sequencing lab: Pathogen Genomics Unit | Public Health Wales Microbiology Cardiff Wales Specialist Virology Centre | Catherine Moore, Johnathan Evans, Laura Gifford, Malorie Perry, Simon Cottrell, Angela Marchbank, Alec Birchley, Alexander Adams, Amy Gaskin, Bree Gatica-Wilcox, Jason Coombes, Joel Southgate, Lauren Gilbert, Lee Graham, Nicole Pacchiarini, Sara Kumziene-Summerhayes, Sarah Taylor, Sophie Jones, Sara Rey, Matthew Bull, Joanne Watkins, Sally Corden, Tom Connor |
| EPI_ISL_3297310 | Viollier AG | Department of Biosystems Science and Engineering, ETH Zürich | Christian Beisel, Sarah Nadeau, Chaoran Chen, Ivan Topolsky, Kim Philipp Jablonski, Lara Fuhrmann, Rebecca Denes, Mirjam Feldkamp, Ina Nissen, Natascha Santacroce, Elodie Burcklen, Christiane Beckmann, Maurice Redondo, Olivier Kobel, Christoph Noppen, Niko Beerenwinkel, Tanja Stadler |
| EPI_ISL_3297311 | Wales Specialist Virology Centre Sequencing lab: Pathogen Genomics Unit | Public Health Wales Microbiology Cardiff Wales Specialist Virology Centre | Catherine Moore, Johnathan Evans, Laura Gifford, Malorie Perry, Simon Cottrell, Angela Marchbank, Alec Birchley, Alexander Adams, Amy Gaskin, Bree Gatica-Wilcox, Jason Coombes, Joel Southgate, Lauren Gilbert, Lee Graham, Nicole Pacchiarini, Sara Kumziene-Summerhayes, Sarah Taylor, Sophie Jones, Sara Rey, Matthew Bull, Joanne Watkins, Sally Corden, Tom Connor |
| EPI_ISL_3297312 | Viollier AG | Department of Biosystems Science and Engineering, ETH Zürich | Christian Beisel, Sarah Nadeau, Chaoran Chen, Ivan Topolsky, Kim Philipp Jablonski, Lara Fuhrmann, Rebecca Denes, Mirjam Feldkamp, Ina Nissen, Natascha Santacroce, Elodie Burcklen, Christiane Beckmann, Maurice Redondo, Olivier Kobel, Christoph Noppen, Niko Beerenwinkel, Tanja Stadler |
| EPI_ISL_3297313 | Wales Specialist Virology Centre Sequencing lab: Pathogen Genomics Unit | Public Health Wales Microbiology Cardiff Wales Specialist Virology Centre | Catherine Moore, Johnathan Evans, Laura Gifford, Malorie Perry, Simon Cottrell, Angela Marchbank, Alec Birchley, Alexander Adams, Amy Gaskin, Bree Gatica-Wilcox, Jason Coombes, Joel Southgate, Lauren Gilbert, Lee Graham, Nicole Pacchiarini, Sara Kumziene-Summerhayes, Sarah Taylor, Sophie Jones, Sara Rey, Matthew Bull, Joanne Watkins, Sally Corden, Tom Connor |
| EPI_ISL_3297314 | Viollier AG | Department of Biosystems Science and Engineering, ETH Zürich | Christian Beisel, Sarah Nadeau, Chaoran Chen, Ivan Topolsky, Kim Philipp Jablonski, Lara Fuhrmann, Rebecca Denes, Mirjam Feldkamp, Ina Nissen, Natascha Santacroce, Elodie Burcklen, Christiane Beckmann, Maurice Redondo, Olivier Kobel, Christoph Noppen, Niko Beerenwinkel, Tanja Stadler |
| EPI_ISL_3297315 | Wales Specialist Virology Centre Sequencing lab: Pathogen Genomics Unit | Public Health Wales Microbiology Cardiff Wales Specialist Virology Centre | Catherine Moore, Johnathan Evans, Laura Gifford, Malorie Perry, Simon Cottrell, Angela Marchbank, Alec Birchley, Alexander Adams, Amy Gaskin, Bree Gatica-Wilcox, Jason Coombes, Joel Southgate, Lauren Gilbert, Lee Graham, Nicole Pacchiarini, Sara Kumziene-Summerhayes, Sarah Taylor, Sophie Jones, Sara Rey, Matthew Bull, Joanne Watkins, Sally Corden, Tom Connor |
| EPI_ISL_3297316 | Viollier AG | Department of Biosystems Science and Engineering, ETH Zürich | Christian Beisel, Sarah Nadeau, Chaoran Chen, Ivan Topolsky, Kim Philipp Jablonski, Lara Fuhrmann, Rebecca Denes, Mirjam Feldkamp, Ina Nissen, Natascha Santacroce, Elodie Burcklen, Christiane Beckmann, Maurice Redondo, Olivier Kobel, Christoph Noppen, Niko Beerenwinkel, Tanja Stadler |
| EPI_ISL_3297317 | Wales Specialist Virology Centre Sequencing lab: Pathogen Genomics Unit | Public Health Wales Microbiology Cardiff Wales Specialist Virology Centre | Catherine Moore, Johnathan Evans, Laura Gifford, Malorie Perry, Simon Cottrell, Angela Marchbank, Alec Birchley, Alexander Adams, Amy Gaskin, Bree Gatica-Wilcox, Jason Coombes, Joel Southgate, Lauren Gilbert, Lee Graham, Nicole Pacchiarini, Sara Kumziene-Summerhayes, Sarah Taylor, Sophie Jones, Sara Rey, Matthew Bull, Joanne Watkins, Sally Corden, Tom Connor |
| EPI_ISL_3297318 | Viollier AG | Department of Biosystems Science and Engineering, ETH Zürich | Christian Beisel, Sarah Nadeau, Chaoran Chen, Ivan Topolsky, Kim Philipp Jablonski, Lara Fuhrmann, Rebecca Denes, Mirjam Feldkamp, Ina Nissen, Natascha Santacroce, Elodie Burcklen, Christiane Beckmann, Maurice Redondo, Olivier Kobel, Christoph Noppen, Niko Beerenwinkel, Tanja Stadler |
| EPI_ISL_3297319, EPI_ISL_3297320 | Wales Specialist Virology Centre Sequencing lab: Pathogen Genomics Unit | Public Health Wales Microbiology Cardiff Wales Specialist Virology Centre | Catherine Moore, Johnathan Evans, Laura Gifford, Malorie Perry, Simon Cottrell, Angela Marchbank, Alec Birchley, Alexander Adams, Amy Gaskin, Bree Gatica-Wilcox, Jason Coombes, Joel Southgate, Lauren Gilbert, Lee Graham, Nicole Pacchiarini, Sara Kumziene-Summerhayes, Sarah Taylor, Sophie Jones, Sara Rey, Matthew Bull, Joanne Watkins, Sally Corden, Tom Connor |
| EPI_ISL_3297321 | Viollier AG | Department of Biosystems Science and Engineering, ETH Zürich | Christian Beisel, Sarah Nadeau, Chaoran Chen, Ivan Topolsky, Kim Philipp Jablonski, Lara Fuhrmann, Rebecca Denes, Mirjam Feldkamp, Ina Nissen, Natascha Santacroce, Elodie Burcklen, Christiane Beckmann, Maurice Redondo, Olivier Kobel, Christoph Noppen, Niko Beerenwinkel, Tanja Stadler |
| EPI_ISL_3297322 | Wales Specialist Virology Centre Sequencing lab: Pathogen Genomics Unit | Public Health Wales Microbiology Cardiff Wales Specialist Virology Centre | Catherine Moore, Johnathan Evans, Laura Gifford, Malorie Perry, Simon Cottrell, Angela Marchbank, Alec Birchley, Alexander Adams, Amy Gaskin, Bree Gatica-Wilcox, Jason Coombes, Joel Southgate, Lauren Gilbert, Lee Graham, Nicole Pacchiarini, Sara Kumziene-Summerhayes, Sarah Taylor, Sophie Jones, Sara Rey, Matthew Bull, Joanne Watkins, Sally Corden, Tom Connor |
| EPI_ISL_3297323 | Viollier AG | Department of Biosystems Science and Engineering, ETH Zürich | Christian Beisel, Sarah Nadeau, Chaoran Chen, Ivan Topolsky, Kim Philipp Jablonski, Lara Fuhrmann, Rebecca Denes, Mirjam Feldkamp, Ina Nissen, Natascha Santacroce, Elodie Burcklen, Christiane Beckmann, Maurice Redondo, Olivier Kobel, Christoph Noppen, Niko Beerenwinkel, Tanja Stadler |
| EPI_ISL_3297324 | Wales Specialist Virology Centre Sequencing lab: Pathogen Genomics Unit | Public Health Wales Microbiology Cardiff Wales Specialist Virology Centre | Catherine Moore, Johnathan Evans, Laura Gifford, Malorie Perry, Simon Cottrell, Angela Marchbank, Alec Birchley, Alexander Adams, Amy Gaskin, Bree Gatica-Wilcox, Jason Coombes, Joel Southgate, Lauren Gilbert, Lee Graham, Nicole Pacchiarini, Sara Kumziene-Summerhayes, Sarah Taylor, Sophie Jones, Sara Rey, Matthew Bull, Joanne Watkins, Sally Corden, Tom Connor |
| EPI_ISL_3297325 | Viollier AG | Department of Biosystems Science and Engineering, ETH Zürich | Christian Beisel, Sarah Nadeau, Chaoran Chen, Ivan Topolsky, Kim Philipp Jablonski, Lara Fuhrmann, Rebecca Denes, Mirjam Feldkamp, Ina Nissen, Natascha Santacroce, Elodie Burcklen, Christiane Beckmann, Maurice Redondo, Olivier Kobel, Christoph Noppen, Niko Beerenwinkel, Tanja Stadler |
| EPI_ISL_3297326 | Wales Specialist Virology Centre Sequencing lab: Pathogen Genomics Unit | Public Health Wales Microbiology Cardiff Wales Specialist Virology Centre | Catherine Moore, Johnathan Evans, Laura Gifford, Malorie Perry, Simon Cottrell, Angela Marchbank, Alec Birchley, Alexander Adams, Amy Gaskin, Bree Gatica-Wilcox, Jason Coombes, Joel Southgate, Lauren Gilbert, Lee Graham, Nicole Pacchiarini, Sara Kumziene-Summerhayes, Sarah Taylor, Sophie Jones, Sara Rey, Matthew Bull, Joanne Watkins, Sally Corden, Tom Connor |
| EPI_ISL_3297327 | Viollier AG | Department of Biosystems Science and Engineering, ETH Zürich | Christian Beisel, Sarah Nadeau, Chaoran Chen, Ivan Topolsky, Kim Philipp Jablonski, Lara Fuhrmann, Rebecca Denes, Mirjam Feldkamp, Ina Nissen, Natascha Santacroce, Elodie Burcklen, Christiane Beckmann, Maurice Redondo, Olivier Kobel, Christoph Noppen, Niko Beerenwinkel, Tanja Stadler |
| EPI_ISL_3297328 | Wales Specialist Virology Centre Sequencing lab: Pathogen Genomics Unit | Public Health Wales Microbiology Cardiff Wales Specialist Virology Centre | Catherine Moore, Johnathan Evans, Laura Gifford, Malorie Perry, Simon Cottrell, Angela Marchbank, Alec Birchley, Alexander Adams, Amy Gaskin, Bree Gatica-Wilcox, Jason Coombes, Joel Southgate, Lauren Gilbert, Lee Graham, Nicole Pacchiarini, Sara Kumziene-Summerhayes, Sarah Taylor, Sophie Jones, Sara Rey, Matthew Bull, Joanne Watkins, Sally Corden, Tom Connor |
| EPI_ISL_3297329 | Viollier AG | Department of Biosystems Science and Engineering, ETH Zürich | Christian Beisel, Sarah Nadeau, Chaoran Chen, Ivan Topolsky, Kim Philipp Jablonski, Lara Fuhrmann, Rebecca Denes, Mirjam Feldkamp, Ina Nissen, Natascha Santacroce, Elodie Burcklen, Christiane Beckmann, Maurice Redondo, Olivier Kobel, Christoph Noppen, Niko Beerenwinkel, Tanja Stadler |
| EPI_ISL_3297330 | Wales Specialist Virology Centre Sequencing lab: Pathogen Genomics Unit | Public Health Wales Microbiology Cardiff Wales Specialist Virology Centre | Catherine Moore, Johnathan Evans, Laura Gifford, Malorie Perry, Simon Cottrell, Angela Marchbank, Alec Birchley, Alexander Adams, Amy Gaskin, Bree Gatica-Wilcox, Jason Coombes, Joel Southgate, Lauren Gilbert, Lee Graham, Nicole Pacchiarini, Sara Kumziene-Summerhayes, Sarah Taylor, Sophie Jones, Sara Rey, Matthew Bull, Joanne Watkins, Sally Corden, Tom Connor |
| EPI_ISL_3297331 | Viollier AG | Department of Biosystems Science and Engineering, ETH Zürich | Christian Beisel, Sarah Nadeau, Chaoran Chen, Ivan Topolsky, Kim Philipp Jablonski, Lara Fuhrmann, Rebecca Denes, Mirjam Feldkamp, Ina Nissen, Natascha Santacroce, Elodie Burcklen, Christiane Beckmann, Maurice Redondo, Olivier Kobel, Christoph Noppen, Niko Beerenwinkel, Tanja Stadler |
| EPI_ISL_3297332 | Wales Specialist Virology Centre Sequencing lab: Pathogen Genomics Unit | Public Health Wales Microbiology Cardiff Wales Specialist Virology Centre | Catherine Moore, Johnathan Evans, Laura Gifford, Malorie Perry, Simon Cottrell, Angela Marchbank, Alec Birchley, Alexander Adams, Amy Gaskin, Bree Gatica-Wilcox, Jason Coombes, Joel Southgate, Lauren Gilbert, Lee Graham, Nicole Pacchiarini, Sara Kumziene-Summerhayes, Sarah Taylor, Sophie Jones, Sara Rey, Matthew Bull, Joanne Watkins, Sally Corden, Tom Connor |
| EPI_ISL_3297333 | Viollier AG | Department of Biosystems Science and Engineering, ETH Zürich | Christian Beisel, Sarah Nadeau, Chaoran Chen, Ivan Topolsky, Kim Philipp Jablonski, Lara Fuhrmann, Rebecca Denes, Mirjam Feldkamp, Ina Nissen, Natascha Santacroce, Elodie Burcklen, Christiane Beckmann, Maurice Redondo, Olivier Kobel, Christoph Noppen, Niko Beerenwinkel, Tanja Stadler |
| EPI_ISL_3297334 | Wales Specialist Virology Centre Sequencing lab: Pathogen Genomics Unit | Public Health Wales Microbiology Cardiff Wales Specialist Virology Centre | Catherine Moore, Johnathan Evans, Laura Gifford, Malorie Perry, Simon Cottrell, Angela Marchbank, Alec Birchley, Alexander Adams, Amy Gaskin, Bree Gatica-Wilcox, Jason Coombes, Joel Southgate, Lauren Gilbert, Lee Graham, Nicole Pacchiarini, Sara Kumziene-Summerhayes, Sarah Taylor, Sophie Jones, Sara Rey, Matthew Bull, Joanne Watkins, Sally Corden, Tom Connor |
| EPI_ISL_3297335 | Viollier AG | Department of Biosystems Science and Engineering, ETH Zürich | Christian Beisel, Sarah Nadeau, Chaoran Chen, Ivan Topolsky, Kim Philipp Jablonski, Lara Fuhrmann, Rebecca Denes, Mirjam Feldkamp, Ina Nissen, Natascha Santacroce, Elodie Burcklen, Christiane Beckmann, Maurice Redondo, Olivier Kobel, Christoph Noppen, Niko Beerenwinkel, Tanja Stadler |
| EPI_ISL_3297336 | Wales Specialist Virology Centre Sequencing lab: Pathogen Genomics Unit | Public Health Wales Microbiology Cardiff Wales Specialist Virology Centre | Catherine Moore, Johnathan Evans, Laura Gifford, Malorie Perry, Simon Cottrell, Angela Marchbank, Alec Birchley, Alexander Adams, Amy Gaskin, Bree Gatica-Wilcox, Jason Coombes, Joel Southgate, Lauren Gilbert, Lee Graham, Nicole Pacchiarini, Sara Kumziene-Summerhayes, Sarah Taylor, Sophie Jones, Sara Rey, Matthew Bull, Joanne Watkins, Sally Corden, Tom Connor |
| EPI_ISL_3297337 | Viollier AG | Department of Biosystems Science and Engineering, ETH Zürich | Christian Beisel, Sarah Nadeau, Chaoran Chen, Ivan Topolsky, Kim Philipp Jablonski, Lara Fuhrmann, Rebecca Denes, Mirjam Feldkamp, Ina Nissen, Natascha Santacroce, Elodie Burcklen, Christiane Beckmann, Maurice Redondo, Olivier Kobel, Christoph Noppen, Niko Beerenwinkel, Tanja Stadler |
| EPI_ISL_3297338, EPI_ISL_3297339 | Wales Specialist Virology Centre Sequencing lab: Pathogen Genomics Unit | Public Health Wales Microbiology Cardiff Wales Specialist Virology Centre | Catherine Moore, Johnathan Evans, Laura Gifford, Malorie Perry, Simon Cottrell, Angela Marchbank, Alec Birchley, Alexander Adams, Amy Gaskin, Bree Gatica-Wilcox, Jason Coombes, Joel Southgate, Lauren Gilbert, Lee Graham, Nicole Pacchiarini, Sara Kumziene-Summerhayes, Sarah Taylor, Sophie Jones, Sara Rey, Matthew Bull, Joanne Watkins, Sally Corden, Tom Connor |
| EPI_ISL_3297340 | Viollier AG | Department of Biosystems Science and Engineering, ETH Zürich | Christian Beisel, Sarah Nadeau, Chaoran Chen, Ivan Topolsky, Kim Philipp Jablonski, Lara Fuhrmann, Rebecca Denes, Mirjam Feldkamp, Ina Nissen, Natascha Santacroce, Elodie Burcklen, Christiane Beckmann, Maurice Redondo, Olivier Kobel, Christoph Noppen, Niko Beerenwinkel, Tanja Stadler |
| EPI_ISL_3297341, EPI_ISL_3297343 | Wales Specialist Virology Centre Sequencing lab: Pathogen Genomics Unit | Public Health Wales Microbiology Cardiff Wales Specialist Virology Centre | Catherine Moore, Johnathan Evans, Laura Gifford, Malorie Perry, Simon Cottrell, Angela Marchbank, Alec Birchley, Alexander Adams, Amy Gaskin, Bree Gatica-Wilcox, Jason Coombes, Joel Southgate, Lauren Gilbert, Lee Graham, Nicole Pacchiarini, Sara Kumziene-Summerhayes, Sarah Taylor, Sophie Jones, Sara Rey, Matthew Bull, Joanne Watkins, Sally Corden, Tom Connor |
| EPI_ISL_3297344 | Viollier AG | Department of Biosystems Science and Engineering, ETH Zürich | Christian Beisel, Sarah Nadeau, Chaoran Chen, Ivan Topolsky, Kim Philipp Jablonski, Lara Fuhrmann, Rebecca Denes, Mirjam Feldkamp, Ina Nissen, Natascha Santacroce, Elodie Burcklen, Christiane Beckmann, Maurice Redondo, Olivier Kobel, Christoph Noppen, Niko Beerenwinkel, Tanja Stadler |
| EPI_ISL_3297345, EPI_ISL_3297347 | Wales Specialist Virology Centre Sequencing lab: Pathogen Genomics Unit | Public Health Wales Microbiology Cardiff Wales Specialist Virology Centre | Catherine Moore, Johnathan Evans, Laura Gifford, Malorie Perry, Simon Cottrell, Angela Marchbank, Alec Birchley, Alexander Adams, Amy Gaskin, Bree Gatica-Wilcox, Jason Coombes, Joel Southgate, Lauren Gilbert, Lee Graham, Nicole Pacchiarini, Sara Kumziene-Summerhayes, Sarah Taylor, Sophie Jones, Sara Rey, Matthew Bull, Joanne Watkins, Sally Corden, Tom Connor |
| EPI_ISL_3297348 | Viollier AG | Department of Biosystems Science and Engineering, ETH Zürich | Christian Beisel, Sarah Nadeau, Chaoran Chen, Ivan Topolsky, Kim Philipp Jablonski, Lara Fuhrmann, Rebecca Denes, Mirjam Feldkamp, Ina Nissen, Natascha Santacroce, Elodie Burcklen, Christiane Beckmann, Maurice Redondo, Olivier Kobel, Christoph Noppen, Niko Beerenwinkel, Tanja Stadler |
| EPI_ISL_3297349 | Wales Specialist Virology Centre Sequencing lab: Pathogen Genomics Unit | Public Health Wales Microbiology Cardiff Wales Specialist Virology Centre | Catherine Moore, Johnathan Evans, Laura Gifford, Malorie Perry, Simon Cottrell, Angela Marchbank, Alec Birchley, Alexander Adams, Amy Gaskin, Bree Gatica-Wilcox, Jason Coombes, Joel Southgate, Lauren Gilbert, Lee Graham, Nicole Pacchiarini, Sara Kumziene-Summerhayes, Sarah Taylor, Sophie Jones, Sara Rey, Matthew Bull, Joanne Watkins, Sally Corden, Tom Connor |
| EPI_ISL_3297350 | Viollier AG | Department of Biosystems Science and Engineering, ETH Zürich | Christian Beisel, Sarah Nadeau, Chaoran Chen, Ivan Topolsky, Kim Philipp Jablonski, Lara Fuhrmann, Rebecca Denes, Mirjam Feldkamp, Ina Nissen, Natascha Santacroce, Elodie Burcklen, Christiane Beckmann, Maurice Redondo, Olivier Kobel, Christoph Noppen, Niko Beerenwinkel, Tanja Stadler |

| | | | |
|---|---|---|---|
| EPI_ISL_3297351 | Wales Specialist Virology Centre Sequencing lab: Pathogen Genomics Unit | Public Health Wales Microbiology Cardiff Wales Specialist Virology Centre | Catherine Moore, Johnathan Evans, Laura Gifford, Malorie Perry, Simon Cottrell, Angela Marchbank, Alec Birchley, Alexander Adams, Amy Gaskin, Bree Gatica-Wilcox, Jason Coombes, Joel Southgate, Lauren Gilbert, Lee Graham, Nicole Pacchiarini, Sara Kumziene-Summerhayes, Sarah Taylor, Sophie Jones, Sara Rey, Matthew Bull, Joanne Watkins, Sally Corden, Tom Connor |
| EPI_ISL_3297352 | Viollier AG | Department of Biosystems Science and Engineering, ETH Zürich | Christian Beisel, Sarah Nadeau, Chaoran Chen, Ivan Topolsky, Kim Philipp Jablonski, Lara Fuhrmann, Rebecca Denes, Mirjam Feldkamp, Ina Nissen, Natascha Santacroce, Elodie Burcklen, Christiane Beckmann, Maurice Redondo, Olivier Kobel, Christoph Noppen, Niko Beerenwinkel, Tanja Stadler |
| EPI_ISL_3297353 | Wales Specialist Virology Centre Sequencing lab: Pathogen Genomics Unit | Public Health Wales Microbiology Cardiff Wales Specialist Virology Centre | Catherine Moore, Johnathan Evans, Laura Gifford, Malorie Perry, Simon Cottrell, Angela Marchbank, Alec Birchley, Alexander Adams, Amy Gaskin, Bree Gatica-Wilcox, Jason Coombes, Joel Southgate, Lauren Gilbert, Lee Graham, Nicole Pacchiarini, Sara Kumziene-Summerhayes, Sarah Taylor, Sophie Jones, Sara Rey, Matthew Bull, Joanne Watkins, Sally Corden, Tom Connor |
| EPI_ISL_3297354 | Viollier AG | Department of Biosystems Science and Engineering, ETH Zürich | Christian Beisel, Sarah Nadeau, Chaoran Chen, Ivan Topolsky, Kim Philipp Jablonski, Lara Fuhrmann, Rebecca Denes, Mirjam Feldkamp, Ina Nissen, Natascha Santacroce, Elodie Burcklen, Christiane Beckmann, Maurice Redondo, Olivier Kobel, Christoph Noppen, Niko Beerenwinkel, Tanja Stadler |
| EPI_ISL_3297355, EPI_ISL_3297356 | Wales Specialist Virology Centre Sequencing lab: Pathogen Genomics Unit | Public Health Wales Microbiology Cardiff Wales Specialist Virology Centre | Catherine Moore, Johnathan Evans, Laura Gifford, Malorie Perry, Simon Cottrell, Angela Marchbank, Alec Birchley, Alexander Adams, Amy Gaskin, Bree Gatica-Wilcox, Jason Coombes, Joel Southgate, Lauren Gilbert, Lee Graham, Nicole Pacchiarini, Sara Kumziene-Summerhayes, Sarah Taylor, Sophie Jones, Sara Rey, Matthew Bull, Joanne Watkins, Sally Corden, Tom Connor |
| EPI_ISL_3297357 | Viollier AG | Department of Biosystems Science and Engineering, ETH Zürich | Christian Beisel, Sarah Nadeau, Chaoran Chen, Ivan Topolsky, Kim Philipp Jablonski, Lara Fuhrmann, Rebecca Denes, Mirjam Feldkamp, Ina Nissen, Natascha Santacroce, Elodie Burcklen, Christiane Beckmann, Maurice Redondo, Olivier Kobel, Christoph Noppen, Niko Beerenwinkel, Tanja Stadler |
| EPI_ISL_3297358 | Wales Specialist Virology Centre Sequencing lab: Pathogen Genomics Unit | Public Health Wales Microbiology Cardiff Wales Specialist Virology Centre | Catherine Moore, Johnathan Evans, Laura Gifford, Malorie Perry, Simon Cottrell, Angela Marchbank, Alec Birchley, Alexander Adams, Amy Gaskin, Bree Gatica-Wilcox, Jason Coombes, Joel Southgate, Lauren Gilbert, Lee Graham, Nicole Pacchiarini, Sara Kumziene-Summerhayes, Sarah Taylor, Sophie Jones, Sara Rey, Matthew Bull, Joanne Watkins, Sally Corden, Tom Connor |
| EPI_ISL_3297359 | Viollier AG | Department of Biosystems Science and Engineering, ETH Zürich | Christian Beisel, Sarah Nadeau, Chaoran Chen, Ivan Topolsky, Kim Philipp Jablonski, Lara Fuhrmann, Rebecca Denes, Mirjam Feldkamp, Ina Nissen, Natascha Santacroce, Elodie Burcklen, Christiane Beckmann, Maurice Redondo, Olivier Kobel, Christoph Noppen, Niko Beerenwinkel, Tanja Stadler |
| EPI_ISL_3297360 | Wales Specialist Virology Centre Sequencing lab: Pathogen Genomics Unit | Public Health Wales Microbiology Cardiff Wales Specialist Virology Centre | Catherine Moore, Johnathan Evans, Laura Gifford, Malorie Perry, Simon Cottrell, Angela Marchbank, Alec Birchley, Alexander Adams, Amy Gaskin, Bree Gatica-Wilcox, Jason Coombes, Joel Southgate, Lauren Gilbert, Lee Graham, Nicole Pacchiarini, Sara Kumziene-Summerhayes, Sarah Taylor, Sophie Jones, Sara Rey, Matthew Bull, Joanne Watkins, Sally Corden, Tom Connor |
| EPI_ISL_3297361 | Viollier AG | Department of Biosystems Science and Engineering, ETH Zürich | Christian Beisel, Sarah Nadeau, Chaoran Chen, Ivan Topolsky, Kim Philipp Jablonski, Lara Fuhrmann, Rebecca Denes, Mirjam Feldkamp, Ina Nissen, Natascha Santacroce, Elodie Burcklen, Christiane Beckmann, Maurice Redondo, Olivier Kobel, Christoph Noppen, Niko Beerenwinkel, Tanja Stadler |
| EPI_ISL_3297362 | Wales Specialist Virology Centre Sequencing lab: Pathogen Genomics Unit | Public Health Wales Microbiology Cardiff Wales Specialist Virology Centre | Catherine Moore, Johnathan Evans, Laura Gifford, Malorie Perry, Simon Cottrell, Angela Marchbank, Alec Birchley, Alexander Adams, Amy Gaskin, Bree Gatica-Wilcox, Jason Coombes, Joel Southgate, Lauren Gilbert, Lee Graham, Nicole Pacchiarini, Sara Kumziene-Summerhayes, Sarah Taylor, Sophie Jones, Sara Rey, Matthew Bull, Joanne Watkins, Sally Corden, Tom Connor |
| EPI_ISL_3297363 | Viollier AG | Department of Biosystems Science and Engineering, ETH Zürich | Christian Beisel, Sarah Nadeau, Chaoran Chen, Ivan Topolsky, Kim Philipp Jablonski, Lara Fuhrmann, Rebecca Denes, Mirjam Feldkamp, Ina Nissen, Natascha Santacroce, Elodie Burcklen, Christiane Beckmann, Maurice Redondo, Olivier Kobel, Christoph Noppen, Niko Beerenwinkel, Tanja Stadler |
| EPI_ISL_3297364, EPI_ISL_3297365 | Wales Specialist Virology Centre Sequencing lab: Pathogen Genomics Unit | Public Health Wales Microbiology Cardiff Wales Specialist Virology Centre | Catherine Moore, Johnathan Evans, Laura Gifford, Malorie Perry, Simon Cottrell, Angela Marchbank, Alec Birchley, Alexander Adams, Amy Gaskin, Bree Gatica-Wilcox, Jason Coombes, Joel Southgate, Lauren Gilbert, Lee Graham, Nicole Pacchiarini, Sara Kumziene-Summerhayes, Sarah Taylor, Sophie Jones, Sara Rey, Matthew Bull, Joanne Watkins, Sally Corden, Tom Connor |
| EPI_ISL_3297366 | Viollier AG | Department of Biosystems Science and Engineering, ETH Zürich | Christian Beisel, Sarah Nadeau, Chaoran Chen, Ivan Topolsky, Kim Philipp Jablonski, Lara Fuhrmann, Rebecca Denes, Mirjam Feldkamp, Ina Nissen, Natascha Santacroce, Elodie Burcklen, Christiane Beckmann, Maurice Redondo, Olivier Kobel, Christoph Noppen, Niko Beerenwinkel, Tanja Stadler |
| EPI_ISL_3297367 | Wales Specialist Virology Centre Sequencing lab: Pathogen Genomics Unit | Public Health Wales Microbiology Cardiff Wales Specialist Virology Centre | Catherine Moore, Johnathan Evans, Laura Gifford, Malorie Perry, Simon Cottrell, Angela Marchbank, Alec Birchley, Alexander Adams, Amy Gaskin, Bree Gatica-Wilcox, Jason Coombes, Joel Southgate, Lauren Gilbert, Lee Graham, Nicole Pacchiarini, Sara Kumziene-Summerhayes, Sarah Taylor, Sophie Jones, Sara Rey, Matthew Bull, Joanne Watkins, Sally Corden, Tom Connor |
| EPI_ISL_3297368 | Viollier AG | Department of Biosystems Science and Engineering, ETH Zürich | Christian Beisel, Sarah Nadeau, Chaoran Chen, Ivan Topolsky, Kim Philipp Jablonski, Lara Fuhrmann, Rebecca Denes, Mirjam Feldkamp, Ina Nissen, Natascha Santacroce, Elodie Burcklen, Christiane Beckmann, Maurice Redondo, Olivier Kobel, Christoph Noppen, Niko Beerenwinkel, Tanja Stadler |
| EPI_ISL_3297369, EPI_ISL_3297371 | Wales Specialist Virology Centre Sequencing lab: Pathogen Genomics Unit | Public Health Wales Microbiology Cardiff Wales Specialist Virology Centre | Catherine Moore, Johnathan Evans, Laura Gifford, Malorie Perry, Simon Cottrell, Angela Marchbank, Alec Birchley, Alexander Adams, Amy Gaskin, Bree Gatica-Wilcox, Jason Coombes, Joel Southgate, Lauren Gilbert, Lee Graham, Nicole Pacchiarini, Sara Kumziene-Summerhayes, Sarah Taylor, Sophie Jones, Sara Rey, Matthew Bull, Joanne Watkins, Sally Corden, Tom Connor |
| EPI_ISL_3297372 | Viollier AG | Department of Biosystems Science and Engineering, ETH Zürich | Christian Beisel, Sarah Nadeau, Chaoran Chen, Ivan Topolsky, Kim Philipp Jablonski, Lara Fuhrmann, Rebecca Denes, Mirjam Feldkamp, Ina Nissen, Natascha Santacroce, Elodie Burcklen, Christiane Beckmann, Maurice Redondo, Olivier Kobel, Christoph Noppen, Niko Beerenwinkel, Tanja Stadler |
| EPI_ISL_3297373 | Wales Specialist Virology Centre Sequencing lab: Pathogen Genomics Unit | Public Health Wales Microbiology Cardiff Wales Specialist Virology Centre | Catherine Moore, Johnathan Evans, Laura Gifford, Malorie Perry, Simon Cottrell, Angela Marchbank, Alec Birchley, Alexander Adams, Amy Gaskin, Bree Gatica-Wilcox, Jason Coombes, Joel Southgate, Lauren Gilbert, Lee Graham, Nicole Pacchiarini, Sara Kumziene-Summerhayes, Sarah Taylor, Sophie Jones, Sara Rey, Matthew Bull, Joanne Watkins, Sally Corden, Tom Connor |
| EPI_ISL_3297374 | Viollier AG | Department of Biosystems Science and Engineering, ETH Zürich | Christian Beisel, Sarah Nadeau, Chaoran Chen, Ivan Topolsky, Kim Philipp Jablonski, Lara Fuhrmann, Rebecca Denes, Mirjam Feldkamp, Ina Nissen, Natascha Santacroce, Elodie Burcklen, Christiane Beckmann, Maurice Redondo, Olivier Kobel, Christoph Noppen, Niko Beerenwinkel, Tanja Stadler |
| EPI_ISL_3297375, EPI_ISL_3297376 | Wales Specialist Virology Centre Sequencing lab: Pathogen Genomics Unit | Public Health Wales Microbiology Cardiff Wales Specialist Virology Centre | Catherine Moore, Johnathan Evans, Laura Gifford, Malorie Perry, Simon Cottrell, Angela Marchbank, Alec Birchley, Alexander Adams, Amy Gaskin, Bree Gatica-Wilcox, Jason Coombes, Joel Southgate, Lauren Gilbert, Lee Graham, Nicole Pacchiarini, Sara Kumziene-Summerhayes, Sarah Taylor, Sophie Jones, Sara Rey, Matthew Bull, Joanne Watkins, Sally Corden, Tom Connor |
| EPI_ISL_3297377 | Viollier AG | Department of Biosystems Science and Engineering, ETH Zürich | Christian Beisel, Sarah Nadeau, Chaoran Chen, Ivan Topolsky, Kim Philipp Jablonski, Lara Fuhrmann, Rebecca Denes, Mirjam Feldkamp, Ina Nissen, Natascha Santacroce, Elodie Burcklen, Christiane Beckmann, Maurice Redondo, Olivier Kobel, Christoph Noppen, Niko Beerenwinkel, Tanja Stadler |
| EPI_ISL_3297378 | Wales Specialist Virology Centre Sequencing lab: Pathogen Genomics Unit | Public Health Wales Microbiology Cardiff Wales Specialist Virology Centre | Catherine Moore, Johnathan Evans, Laura Gifford, Malorie Perry, Simon Cottrell, Angela Marchbank, Alec Birchley, Alexander Adams, Amy Gaskin, Bree Gatica-Wilcox, Jason Coombes, Joel Southgate, Lauren Gilbert, Lee Graham, Nicole Pacchiarini, Sara Kumziene-Summerhayes, Sarah Taylor, Sophie Jones, Sara Rey, Matthew Bull, Joanne Watkins, Sally Corden, Tom Connor |
| EPI_ISL_3297379 | Viollier AG | Department of Biosystems Science and Engineering, ETH Zürich | Christian Beisel, Sarah Nadeau, Chaoran Chen, Ivan Topolsky, Kim Philipp Jablonski, Lara Fuhrmann, Rebecca Denes, Mirjam Feldkamp, Ina Nissen, Natascha Santacroce, Elodie Burcklen, Christiane Beckmann, Maurice Redondo, Olivier Kobel, Christoph Noppen, Niko Beerenwinkel, Tanja Stadler |
| EPI_ISL_3297380 | Wales Specialist Virology Centre Sequencing lab: Pathogen Genomics Unit | Public Health Wales Microbiology Cardiff Wales Specialist Virology Centre | Catherine Moore, Johnathan Evans, Laura Gifford, Malorie Perry, Simon Cottrell, Angela Marchbank, Alec Birchley, Alexander Adams, Amy Gaskin, Bree Gatica-Wilcox, Jason Coombes, Joel Southgate, Lauren Gilbert, Lee Graham, Nicole Pacchiarini, Sara Kumziene-Summerhayes, Sarah Taylor, Sophie Jones, Sara Rey, Matthew Bull, Joanne Watkins, Sally Corden, Tom Connor |
| EPI_ISL_3297381 | Viollier AG | Department of Biosystems Science and Engineering, ETH Zürich | Christian Beisel, Sarah Nadeau, Chaoran Chen, Ivan Topolsky, Kim Philipp Jablonski, Lara Fuhrmann, Rebecca Denes, Mirjam Feldkamp, Ina Nissen, Natascha Santacroce, Elodie Burcklen, Christiane Beckmann, Maurice Redondo, Olivier Kobel, Christoph Noppen, Niko Beerenwinkel, Tanja Stadler |
| EPI_ISL_3297382 | Wales Specialist Virology Centre Sequencing lab: Pathogen Genomics Unit | Public Health Wales Microbiology Cardiff Wales Specialist Virology Centre | Catherine Moore, Johnathan Evans, Laura Gifford, Malorie Perry, Simon Cottrell, Angela Marchbank, Alec Birchley, Alexander Adams, Amy Gaskin, Bree Gatica-Wilcox, Jason Coombes, Joel Southgate, Lauren Gilbert, Lee Graham, Nicole Pacchiarini, Sara Kumziene-Summerhayes, Sarah Taylor, Sophie Jones, Sara Rey, Matthew Bull, Joanne Watkins, Sally Corden, Tom Connor |
| EPI_ISL_3297383 | Viollier AG | Department of Biosystems Science and Engineering, ETH Zürich | Christian Beisel, Sarah Nadeau, Chaoran Chen, Ivan Topolsky, Kim Philipp Jablonski, Lara Fuhrmann, Rebecca Denes, Mirjam Feldkamp, Ina Nissen, Natascha Santacroce, Elodie Burcklen, Christiane Beckmann, Maurice Redondo, Olivier Kobel, Christoph Noppen, Niko Beerenwinkel, Tanja Stadler |
| EPI_ISL_3297384 | Wales Specialist Virology Centre Sequencing lab: Pathogen Genomics Unit | Public Health Wales Microbiology Cardiff Wales Specialist Virology Centre | Catherine Moore, Johnathan Evans, Laura Gifford, Malorie Perry, Simon Cottrell, Angela Marchbank, Alec Birchley, Alexander Adams, Amy Gaskin, Bree Gatica-Wilcox, Jason Coombes, Joel Southgate, Lauren Gilbert, Lee Graham, Nicole Pacchiarini, Sara Kumziene-Summerhayes, Sarah Taylor, Sophie Jones, Sara Rey, Matthew Bull, Joanne Watkins, Sally Corden, Tom Connor |
| EPI_ISL_3297385 | Viollier AG | Department of Biosystems Science and Engineering, ETH Zürich | Christian Beisel, Sarah Nadeau, Chaoran Chen, Ivan Topolsky, Kim Philipp Jablonski, Lara Fuhrmann, Rebecca Denes, Mirjam Feldkamp, Ina Nissen, Natascha Santacroce, Elodie Burcklen, Christiane Beckmann, Maurice Redondo, Olivier Kobel, Christoph Noppen, Niko Beerenwinkel, Tanja Stadler |
| EPI_ISL_3297386 | Wales Specialist Virology Centre Sequencing lab: Pathogen Genomics Unit | Public Health Wales Microbiology Cardiff Wales Specialist Virology Centre | Catherine Moore, Johnathan Evans, Laura Gifford, Malorie Perry, Simon Cottrell, Angela Marchbank, Alec Birchley, Alexander Adams, Amy Gaskin, Bree Gatica-Wilcox, Jason Coombes, Joel Southgate, Lauren Gilbert, Lee Graham, Nicole Pacchiarini, Sara Kumziene-Summerhayes, Sarah Taylor, Sophie Jones, Sara Rey, Matthew Bull, Joanne Watkins, Sally Corden, Tom Connor |
| EPI_ISL_3297387 | Viollier AG | Department of Biosystems Science and Engineering, ETH Zürich | Christian Beisel, Sarah Nadeau, Chaoran Chen, Ivan Topolsky, Kim Philipp Jablonski, Lara Fuhrmann, Rebecca Denes, Mirjam Feldkamp, Ina Nissen, Natascha Santacroce, Elodie Burcklen, Christiane Beckmann, Maurice Redondo, Olivier Kobel, Christoph Noppen, Niko Beerenwinkel, Tanja Stadler |
| EPI_ISL_3297388 | Wales Specialist Virology Centre Sequencing lab: Pathogen Genomics Unit | Public Health Wales Microbiology Cardiff Wales Specialist Virology Centre | Catherine Moore, Johnathan Evans, Laura Gifford, Malorie Perry, Simon Cottrell, Angela Marchbank, Alec Birchley, Alexander Adams, Amy Gaskin, Bree Gatica-Wilcox, Jason Coombes, Joel Southgate, Lauren Gilbert, Lee Graham, Nicole Pacchiarini, Sara Kumziene-Summerhayes, Sarah Taylor, Sophie Jones, Sara Rey, Matthew Bull, Joanne Watkins, Sally Corden, Tom Connor |
| EPI_ISL_3297389 | Viollier AG | Department of Biosystems Science and Engineering, ETH Zürich | Christian Beisel, Sarah Nadeau, Chaoran Chen, Ivan Topolsky, Kim Philipp Jablonski, Lara Fuhrmann, Rebecca Denes, Mirjam Feldkamp, Ina Nissen, Natascha Santacroce, Elodie Burcklen, Christiane Beckmann, Maurice Redondo, Olivier Kobel, Christoph Noppen, Niko Beerenwinkel, Tanja Stadler |
| EPI_ISL_3297390 | Wales Specialist Virology Centre Sequencing lab: Pathogen Genomics Unit | Public Health Wales Microbiology Cardiff Wales Specialist Virology Centre | Catherine Moore, Johnathan Evans, Laura Gifford, Malorie Perry, Simon Cottrell, Angela Marchbank, Alec Birchley, Alexander Adams, Amy Gaskin, Bree Gatica-Wilcox, Jason Coombes, Joel Southgate, Lauren Gilbert, Lee Graham, Nicole Pacchiarini, Sara Kumziene-Summerhayes, Sarah Taylor, Sophie Jones, Sara Rey, Matthew Bull, Joanne Watkins, Sally Corden, Tom Connor |
| EPI_ISL_3297391, EPI_ISL_3297392 | Viollier AG | Department of Biosystems Science and Engineering, ETH Zürich | Christian Beisel, Sarah Nadeau, Chaoran Chen, Ivan Topolsky, Kim Philipp Jablonski, Lara Fuhrmann, Rebecca Denes, Mirjam Feldkamp, Ina Nissen, Natascha Santacroce, Elodie Burcklen, Christiane Beckmann, Maurice Redondo, Olivier Kobel, Christoph Noppen, Niko Beerenwinkel, Tanja Stadler |
| EPI_ISL_3297393 | Wales Specialist Virology Centre Sequencing lab: Pathogen Genomics Unit | Public Health Wales Microbiology Cardiff Wales Specialist Virology Centre | Catherine Moore, Johnathan Evans, Laura Gifford, Malorie Perry, Simon Cottrell, Angela Marchbank, Alec Birchley, Alexander Adams, Amy Gaskin, Bree Gatica-Wilcox, Jason Coombes, Joel Southgate, Lauren Gilbert, Lee Graham, Nicole Pacchiarini, Sara Kumziene-Summerhayes, Sarah Taylor, Sophie Jones, Sara Rey, Matthew Bull, Joanne Watkins, Sally Corden, Tom Connor |
| EPI_ISL_3297394 | Viollier AG | Department of Biosystems Science and Engineering, ETH Zürich | Christian Beisel, Sarah Nadeau, Chaoran Chen, Ivan Topolsky, Kim Philipp Jablonski, Lara Fuhrmann, Rebecca Denes, Mirjam Feldkamp, Ina Nissen, Natascha Santacroce, Elodie Burcklen, Christiane Beckmann, Maurice Redondo, Olivier Kobel, Christoph Noppen, Niko Beerenwinkel, Tanja Stadler |
| EPI_ISL_3297395, EPI_ISL_3297396 | Wales Specialist Virology Centre Sequencing lab: Pathogen Genomics Unit | Public Health Wales Microbiology Cardiff Wales Specialist Virology Centre | Catherine Moore, Johnathan Evans, Laura Gifford, Malorie Perry, Simon Cottrell, Angela Marchbank, Alec Birchley, Alexander Adams, Amy Gaskin, Bree Gatica-Wilcox, Jason Coombes, Joel Southgate, Lauren Gilbert, Lee Graham, Nicole Pacchiarini, Sara Kumziene-Summerhayes, Sarah Taylor, Sophie Jones, Sara Rey, Matthew Bull, Joanne Watkins, Sally Corden, Tom Connor |
| EPI_ISL_3297397, EPI_ISL_3297398 | Viollier AG | Department of Biosystems Science and Engineering, ETH Zürich | Christian Beisel, Sarah Nadeau, Chaoran Chen, Ivan Topolsky, Kim Philipp Jablonski, Lara Fuhrmann, Rebecca Denes, Mirjam Feldkamp, Ina Nissen, Natascha Santacroce, Elodie Burcklen, Christiane Beckmann, Maurice Redondo, Olivier Kobel, Christoph Noppen, Niko Beerenwinkel, Tanja Stadler |
| EPI_ISL_3297399, EPI_ISL_3297400 | Wales Specialist Virology Centre Sequencing lab: Pathogen Genomics Unit | Public Health Wales Microbiology Cardiff Wales Specialist Virology Centre | Catherine Moore, Johnathan Evans, Laura Gifford, Malorie Perry, Simon Cottrell, Angela Marchbank, Alec Birchley, Alexander Adams, Amy Gaskin, Bree Gatica-Wilcox, Jason Coombes, Joel Southgate, Lauren Gilbert, Lee Graham, Nicole Pacchiarini, Sara Kumziene-Summerhayes, Sarah Taylor, Sophie Jones, Sara Rey, Matthew Bull, Joanne Watkins, Sally Corden, Tom Connor |
| EPI_ISL_3297401 | Viollier AG | Department of Biosystems Science and Engineering, ETH Zürich | Christian Beisel, Sarah Nadeau, Chaoran Chen, Ivan Topolsky, Kim Philipp Jablonski, Lara Fuhrmann, Rebecca Denes, Mirjam Feldkamp, Ina Nissen, Natascha Santacroce, Elodie Burcklen, Christiane Beckmann, Maurice Redondo, Olivier Kobel, Christoph Noppen, Niko Beerenwinkel, Tanja Stadler |
| EPI_ISL_3297402 | Wales Specialist Virology Centre Sequencing lab: Pathogen Genomics Unit | Public Health Wales Microbiology Cardiff Wales Specialist Virology Centre | Catherine Moore, Johnathan Evans, Laura Gifford, Malorie Perry, Simon Cottrell, Angela Marchbank, Alec Birchley, Alexander Adams, Amy Gaskin, Bree Gatica-Wilcox, Jason Coombes, Joel Southgate, Lauren Gilbert, Lee Graham, Nicole Pacchiarini, Sara Kumziene-Summerhayes, Sarah Taylor, Sophie Jones, Sara Rey, Matthew Bull, Joanne Watkins, Sally Corden, Tom Connor |
| EPI_ISL_3297403 | Viollier AG | Department of Biosystems Science and Engineering, ETH Zürich | Christian Beisel, Sarah Nadeau, Chaoran Chen, Ivan Topolsky, Kim Philipp Jablonski, Lara Fuhrmann, Rebecca Denes, Mirjam Feldkamp, Ina Nissen, Natascha Santacroce, Elodie Burcklen, Christiane Beckmann, Maurice Redondo, Olivier Kobel, Christoph Noppen, Niko Beerenwinkel, Tanja Stadler |
| EPI_ISL_3297404 | Wales Specialist Virology Centre Sequencing lab: Pathogen | Public Health Wales Microbiology Cardiff Wales Specialist Virology | Catherine Moore, Johnathan Evans, Laura Gifford, Malorie Perry, Simon Cottrell, Angela Marchbank, Alec Birchley, Alexander Adams, Amy Gaskin, Bree Gatica-Wilcox, Jason Coombes, Joel Southgate, Lauren Gilbert, Lee Graham, Nicole |

| | | | |
|---|---|---|---|
| | Genomics Unit | Centre | Pacchiarini, Sara Kumziene-Summerhayes, Sarah Taylor, Sophie Jones, Sara Rey, Matthew Bull, Joanne Watkins, Sally Corden, Tom Connor |
| EPI_ISL_3297405 | Viollier AG | Department of Biosystems Science and Engineering, ETH Zürich | Christian Beisel, Sarah Nadeau, Chaoran Chen, Ivan Topolsky, Kim Philipp Jablonski, Lara Fuhrmann, Rebecca Denes, Mirjam Feldkamp, Ina Nissen, Natascha Santacroce, Elodie Burcklen, Christiane Beckmann, Maurice Redondo, Olivier Kobel, Christoph Noppen, Niko Beerenwinkel, Tanja Stadler |
| EPI_ISL_3297406 | Wales Specialist Virology Centre Sequencing lab: Pathogen Genomics Unit | Public Health Wales Microbiology Cardiff Wales Specialist Virology Centre | Catherine Moore, Johnathan Evans, Laura Gifford, Malorie Perry, Simon Cottrell, Angela Marchbank, Alec Birchley, Alexander Adams, Amy Gaskin, Bree Gatica-Wilcox, Jason Coombes, Joel Southgate, Lauren Gilbert, Lee Graham, Nicole Pacchiarini, Sara Kumziene-Summerhayes, Sarah Taylor, Sophie Jones, Sara Rey, Matthew Bull, Joanne Watkins, Sally Corden, Tom Connor |
| EPI_ISL_3297407 | Viollier AG | Department of Biosystems Science and Engineering, ETH Zürich | Christian Beisel, Sarah Nadeau, Chaoran Chen, Ivan Topolsky, Kim Philipp Jablonski, Lara Fuhrmann, Rebecca Denes, Mirjam Feldkamp, Ina Nissen, Natascha Santacroce, Elodie Burcklen, Christiane Beckmann, Maurice Redondo, Olivier Kobel, Christoph Noppen, Niko Beerenwinkel, Tanja Stadler |
| EPI_ISL_3297408 | Wales Specialist Virology Centre Sequencing lab: Pathogen Genomics Unit | Public Health Wales Microbiology Cardiff Wales Specialist Virology Centre | Catherine Moore, Johnathan Evans, Laura Gifford, Malorie Perry, Simon Cottrell, Angela Marchbank, Alec Birchley, Alexander Adams, Amy Gaskin, Bree Gatica-Wilcox, Jason Coombes, Joel Southgate, Lauren Gilbert, Lee Graham, Nicole Pacchiarini, Sara Kumziene-Summerhayes, Sarah Taylor, Sophie Jones, Sara Rey, Matthew Bull, Joanne Watkins, Sally Corden, Tom Connor |
| EPI_ISL_3297409 | Viollier AG | Department of Biosystems Science and Engineering, ETH Zürich | Christian Beisel, Sarah Nadeau, Chaoran Chen, Ivan Topolsky, Kim Philipp Jablonski, Lara Fuhrmann, Rebecca Denes, Mirjam Feldkamp, Ina Nissen, Natascha Santacroce, Elodie Burcklen, Christiane Beckmann, Maurice Redondo, Olivier Kobel, Christoph Noppen, Niko Beerenwinkel, Tanja Stadler |
| EPI_ISL_3297410 | Wales Specialist Virology Centre Sequencing lab: Pathogen Genomics Unit | Public Health Wales Microbiology Cardiff Wales Specialist Virology Centre | Catherine Moore, Johnathan Evans, Laura Gifford, Malorie Perry, Simon Cottrell, Angela Marchbank, Alec Birchley, Alexander Adams, Amy Gaskin, Bree Gatica-Wilcox, Jason Coombes, Joel Southgate, Lauren Gilbert, Lee Graham, Nicole Pacchiarini, Sara Kumziene-Summerhayes, Sarah Taylor, Sophie Jones, Sara Rey, Matthew Bull, Joanne Watkins, Sally Corden, Tom Connor |
| EPI_ISL_3297411 | Viollier AG | Department of Biosystems Science and Engineering, ETH Zürich | Christian Beisel, Sarah Nadeau, Chaoran Chen, Ivan Topolsky, Kim Philipp Jablonski, Lara Fuhrmann, Rebecca Denes, Mirjam Feldkamp, Ina Nissen, Natascha Santacroce, Elodie Burcklen, Christiane Beckmann, Maurice Redondo, Olivier Kobel, Christoph Noppen, Niko Beerenwinkel, Tanja Stadler |
| EPI_ISL_3297412 | Wales Specialist Virology Centre Sequencing lab: Pathogen Genomics Unit | Public Health Wales Microbiology Cardiff Wales Specialist Virology Centre | Catherine Moore, Johnathan Evans, Laura Gifford, Malorie Perry, Simon Cottrell, Angela Marchbank, Alec Birchley, Alexander Adams, Amy Gaskin, Bree Gatica-Wilcox, Jason Coombes, Joel Southgate, Lauren Gilbert, Lee Graham, Nicole Pacchiarini, Sara Kumziene-Summerhayes, Sarah Taylor, Sophie Jones, Sara Rey, Matthew Bull, Joanne Watkins, Sally Corden, Tom Connor |
| EPI_ISL_3297413 | Viollier AG | Department of Biosystems Science and Engineering, ETH Zürich | Christian Beisel, Sarah Nadeau, Chaoran Chen, Ivan Topolsky, Kim Philipp Jablonski, Lara Fuhrmann, Rebecca Denes, Mirjam Feldkamp, Ina Nissen, Natascha Santacroce, Elodie Burcklen, Christiane Beckmann, Maurice Redondo, Olivier Kobel, Christoph Noppen, Niko Beerenwinkel, Tanja Stadler |
| EPI_ISL_3297414 | Wales Specialist Virology Centre Sequencing lab: Pathogen Genomics Unit | Public Health Wales Microbiology Cardiff Wales Specialist Virology Centre | Catherine Moore, Johnathan Evans, Laura Gifford, Malorie Perry, Simon Cottrell, Angela Marchbank, Alec Birchley, Alexander Adams, Amy Gaskin, Bree Gatica-Wilcox, Jason Coombes, Joel Southgate, Lauren Gilbert, Lee Graham, Nicole Pacchiarini, Sara Kumziene-Summerhayes, Sarah Taylor, Sophie Jones, Sara Rey, Matthew Bull, Joanne Watkins, Sally Corden, Tom Connor |
| EPI_ISL_3297415 | Viollier AG | Department of Biosystems Science and Engineering, ETH Zürich | Christian Beisel, Sarah Nadeau, Chaoran Chen, Ivan Topolsky, Kim Philipp Jablonski, Lara Fuhrmann, Rebecca Denes, Mirjam Feldkamp, Ina Nissen, Natascha Santacroce, Elodie Burcklen, Christiane Beckmann, Maurice Redondo, Olivier Kobel, Christoph Noppen, Niko Beerenwinkel, Tanja Stadler |
| EPI_ISL_3297416 | Wales Specialist Virology Centre Sequencing lab: Pathogen Genomics Unit | Public Health Wales Microbiology Cardiff Wales Specialist Virology Centre | Catherine Moore, Johnathan Evans, Laura Gifford, Malorie Perry, Simon Cottrell, Angela Marchbank, Alec Birchley, Alexander Adams, Amy Gaskin, Bree Gatica-Wilcox, Jason Coombes, Joel Southgate, Lauren Gilbert, Lee Graham, Nicole Pacchiarini, Sara Kumziene-Summerhayes, Sarah Taylor, Sophie Jones, Sara Rey, Matthew Bull, Joanne Watkins, Sally Corden, Tom Connor |
| EPI_ISL_3297417 | Viollier AG | Department of Biosystems Science and Engineering, ETH Zürich | Christian Beisel, Sarah Nadeau, Chaoran Chen, Ivan Topolsky, Kim Philipp Jablonski, Lara Fuhrmann, Rebecca Denes, Mirjam Feldkamp, Ina Nissen, Natascha Santacroce, Elodie Burcklen, Christiane Beckmann, Maurice Redondo, Olivier Kobel, Christoph Noppen, Niko Beerenwinkel, Tanja Stadler |
| EPI_ISL_3297418 | Wales Specialist Virology Centre Sequencing lab: Pathogen Genomics Unit | Public Health Wales Microbiology Cardiff Wales Specialist Virology Centre | Catherine Moore, Johnathan Evans, Laura Gifford, Malorie Perry, Simon Cottrell, Angela Marchbank, Alec Birchley, Alexander Adams, Amy Gaskin, Bree Gatica-Wilcox, Jason Coombes, Joel Southgate, Lauren Gilbert, Lee Graham, Nicole Pacchiarini, Sara Kumziene-Summerhayes, Sarah Taylor, Sophie Jones, Sara Rey, Matthew Bull, Joanne Watkins, Sally Corden, Tom Connor |
| EPI_ISL_3297419 | Viollier AG | Department of Biosystems Science and Engineering, ETH Zürich | Christian Beisel, Sarah Nadeau, Chaoran Chen, Ivan Topolsky, Kim Philipp Jablonski, Lara Fuhrmann, Rebecca Denes, Mirjam Feldkamp, Ina Nissen, Natascha Santacroce, Elodie Burcklen, Christiane Beckmann, Maurice Redondo, Olivier Kobel, Christoph Noppen, Niko Beerenwinkel, Tanja Stadler |
| EPI_ISL_3297420 | Wales Specialist Virology Centre Sequencing lab: Pathogen Genomics Unit | Public Health Wales Microbiology Cardiff Wales Specialist Virology Centre | Catherine Moore, Johnathan Evans, Laura Gifford, Malorie Perry, Simon Cottrell, Angela Marchbank, Alec Birchley, Alexander Adams, Amy Gaskin, Bree Gatica-Wilcox, Jason Coombes, Joel Southgate, Lauren Gilbert, Lee Graham, Nicole Pacchiarini, Sara Kumziene-Summerhayes, Sarah Taylor, Sophie Jones, Sara Rey, Matthew Bull, Joanne Watkins, Sally Corden, Tom Connor |
| EPI_ISL_3297421 | Viollier AG | Department of Biosystems Science and Engineering, ETH Zürich | Christian Beisel, Sarah Nadeau, Chaoran Chen, Ivan Topolsky, Kim Philipp Jablonski, Lara Fuhrmann, Rebecca Denes, Mirjam Feldkamp, Ina Nissen, Natascha Santacroce, Elodie Burcklen, Christiane Beckmann, Maurice Redondo, Olivier Kobel, Christoph Noppen, Niko Beerenwinkel, Tanja Stadler |
| EPI_ISL_3297422 | Wales Specialist Virology Centre Sequencing lab: Pathogen Genomics Unit | Public Health Wales Microbiology Cardiff Wales Specialist Virology Centre | Catherine Moore, Johnathan Evans, Laura Gifford, Malorie Perry, Simon Cottrell, Angela Marchbank, Alec Birchley, Alexander Adams, Amy Gaskin, Bree Gatica-Wilcox, Jason Coombes, Joel Southgate, Lauren Gilbert, Lee Graham, Nicole Pacchiarini, Sara Kumziene-Summerhayes, Sarah Taylor, Sophie Jones, Sara Rey, Matthew Bull, Joanne Watkins, Sally Corden, Tom Connor |
| EPI_ISL_3297423 | Viollier AG | Department of Biosystems Science and Engineering, ETH Zürich | Christian Beisel, Sarah Nadeau, Chaoran Chen, Ivan Topolsky, Kim Philipp Jablonski, Lara Fuhrmann, Rebecca Denes, Mirjam Feldkamp, Ina Nissen, Natascha Santacroce, Elodie Burcklen, Christiane Beckmann, Maurice Redondo, Olivier Kobel, Christoph Noppen, Niko Beerenwinkel, Tanja Stadler |
| EPI_ISL_3297424, EPI_ISL_3297425 | Wales Specialist Virology Centre Sequencing lab: Pathogen Genomics Unit | Public Health Wales Microbiology Cardiff Wales Specialist Virology Centre | Catherine Moore, Johnathan Evans, Laura Gifford, Malorie Perry, Simon Cottrell, Angela Marchbank, Alec Birchley, Alexander Adams, Amy Gaskin, Bree Gatica-Wilcox, Jason Coombes, Joel Southgate, Lauren Gilbert, Lee Graham, Nicole Pacchiarini, Sara Kumziene-Summerhayes, Sarah Taylor, Sophie Jones, Sara Rey, Matthew Bull, Joanne Watkins, Sally Corden, Tom Connor |
| EPI_ISL_3297426 | Viollier AG | Department of Biosystems Science and Engineering, ETH Zürich | Christian Beisel, Sarah Nadeau, Chaoran Chen, Ivan Topolsky, Kim Philipp Jablonski, Lara Fuhrmann, Rebecca Denes, Mirjam Feldkamp, Ina Nissen, Natascha Santacroce, Elodie Burcklen, Christiane Beckmann, Maurice Redondo, Olivier Kobel, Christoph Noppen, Niko Beerenwinkel, Tanja Stadler |
| EPI_ISL_3297427 | Wales Specialist Virology Centre Sequencing lab: Pathogen Genomics Unit | Public Health Wales Microbiology Cardiff Wales Specialist Virology Centre | Catherine Moore, Johnathan Evans, Laura Gifford, Malorie Perry, Simon Cottrell, Angela Marchbank, Alec Birchley, Alexander Adams, Amy Gaskin, Bree Gatica-Wilcox, Jason Coombes, Joel Southgate, Lauren Gilbert, Lee Graham, Nicole Pacchiarini, Sara Kumziene-Summerhayes, Sarah Taylor, Sophie Jones, Sara Rey, Matthew Bull, Joanne Watkins, Sally Corden, Tom Connor |
| EPI_ISL_3297428 | Viollier AG | Department of Biosystems Science and Engineering, ETH Zürich | Christian Beisel, Sarah Nadeau, Chaoran Chen, Ivan Topolsky, Kim Philipp Jablonski, Lara Fuhrmann, Rebecca Denes, Mirjam Feldkamp, Ina Nissen, Natascha Santacroce, Elodie Burcklen, Christiane Beckmann, Maurice Redondo, Olivier Kobel, Christoph Noppen, Niko Beerenwinkel, Tanja Stadler |
| EPI_ISL_3297429 | Wales Specialist Virology Centre Sequencing lab: Pathogen Genomics Unit | Public Health Wales Microbiology Cardiff Wales Specialist Virology Centre | Catherine Moore, Johnathan Evans, Laura Gifford, Malorie Perry, Simon Cottrell, Angela Marchbank, Alec Birchley, Alexander Adams, Amy Gaskin, Bree Gatica-Wilcox, Jason Coombes, Joel Southgate, Lauren Gilbert, Lee Graham, Nicole Pacchiarini, Sara Kumziene-Summerhayes, Sarah Taylor, Sophie Jones, Sara Rey, Matthew Bull, Joanne Watkins, Sally Corden, Tom Connor |
| EPI_ISL_3297430 | Viollier AG | Department of Biosystems Science and Engineering, ETH Zürich | Christian Beisel, Sarah Nadeau, Chaoran Chen, Ivan Topolsky, Kim Philipp Jablonski, Lara Fuhrmann, Rebecca Denes, Mirjam Feldkamp, Ina Nissen, Natascha Santacroce, Elodie Burcklen, Christiane Beckmann, Maurice Redondo, Olivier Kobel, Christoph Noppen, Niko Beerenwinkel, Tanja Stadler |
| EPI_ISL_3297431 | Wales Specialist Virology Centre Sequencing lab: Pathogen Genomics Unit | Public Health Wales Microbiology Cardiff Wales Specialist Virology Centre | Catherine Moore, Johnathan Evans, Laura Gifford, Malorie Perry, Simon Cottrell, Angela Marchbank, Alec Birchley, Alexander Adams, Amy Gaskin, Bree Gatica-Wilcox, Jason Coombes, Joel Southgate, Lauren Gilbert, Lee Graham, Nicole Pacchiarini, Sara Kumziene-Summerhayes, Sarah Taylor, Sophie Jones, Sara Rey, Matthew Bull, Joanne Watkins, Sally Corden, Tom Connor |
| EPI_ISL_3297432 | Viollier AG | Department of Biosystems Science and Engineering, ETH Zürich | Christian Beisel, Sarah Nadeau, Chaoran Chen, Ivan Topolsky, Kim Philipp Jablonski, Lara Fuhrmann, Rebecca Denes, Mirjam Feldkamp, Ina Nissen, Natascha Santacroce, Elodie Burcklen, Christiane Beckmann, Maurice Redondo, Olivier Kobel, Christoph Noppen, Niko Beerenwinkel, Tanja Stadler |
| EPI_ISL_3297433 | Wales Specialist Virology Centre Sequencing lab: Pathogen Genomics Unit | Public Health Wales Microbiology Cardiff Wales Specialist Virology Centre | Catherine Moore, Johnathan Evans, Laura Gifford, Malorie Perry, Simon Cottrell, Angela Marchbank, Alec Birchley, Alexander Adams, Amy Gaskin, Bree Gatica-Wilcox, Jason Coombes, Joel Southgate, Lauren Gilbert, Lee Graham, Nicole Pacchiarini, Sara Kumziene-Summerhayes, Sarah Taylor, Sophie Jones, Sara Rey, Matthew Bull, Joanne Watkins, Sally Corden, Tom Connor |
| EPI_ISL_3297434 | Viollier AG | Department of Biosystems Science and Engineering, ETH Zürich | Christian Beisel, Sarah Nadeau, Chaoran Chen, Ivan Topolsky, Kim Philipp Jablonski, Lara Fuhrmann, Rebecca Denes, Mirjam Feldkamp, Ina Nissen, Natascha Santacroce, Elodie Burcklen, Christiane Beckmann, Maurice Redondo, Olivier Kobel, Christoph Noppen, Niko Beerenwinkel, Tanja Stadler |
| EPI_ISL_3297435 | Wales Specialist Virology Centre Sequencing lab: Pathogen Genomics Unit | Public Health Wales Microbiology Cardiff Wales Specialist Virology Centre | Catherine Moore, Johnathan Evans, Laura Gifford, Malorie Perry, Simon Cottrell, Angela Marchbank, Alec Birchley, Alexander Adams, Amy Gaskin, Bree Gatica-Wilcox, Jason Coombes, Joel Southgate, Lauren Gilbert, Lee Graham, Nicole Pacchiarini, Sara Kumziene-Summerhayes, Sarah Taylor, Sophie Jones, Sara Rey, Matthew Bull, Joanne Watkins, Sally Corden, Tom Connor |
| EPI_ISL_3297436 | Viollier AG | Department of Biosystems Science and Engineering, ETH Zürich | Christian Beisel, Sarah Nadeau, Chaoran Chen, Ivan Topolsky, Kim Philipp Jablonski, Lara Fuhrmann, Rebecca Denes, Mirjam Feldkamp, Ina Nissen, Natascha Santacroce, Elodie Burcklen, Christiane Beckmann, Maurice Redondo, Olivier Kobel, Christoph Noppen, Niko Beerenwinkel, Tanja Stadler |
| EPI_ISL_3297437 | Wales Specialist Virology Centre Sequencing lab: Pathogen Genomics Unit | Public Health Wales Microbiology Cardiff Wales Specialist Virology Centre | Catherine Moore, Johnathan Evans, Laura Gifford, Malorie Perry, Simon Cottrell, Angela Marchbank, Alec Birchley, Alexander Adams, Amy Gaskin, Bree Gatica-Wilcox, Jason Coombes, Joel Southgate, Lauren Gilbert, Lee Graham, Nicole Pacchiarini, Sara Kumziene-Summerhayes, Sarah Taylor, Sophie Jones, Sara Rey, Matthew Bull, Joanne Watkins, Sally Corden, Tom Connor |
| EPI_ISL_3297438 | Viollier AG | Department of Biosystems Science and Engineering, ETH Zürich | Christian Beisel, Sarah Nadeau, Chaoran Chen, Ivan Topolsky, Kim Philipp Jablonski, Lara Fuhrmann, Rebecca Denes, Mirjam Feldkamp, Ina Nissen, Natascha Santacroce, Elodie Burcklen, Christiane Beckmann, Maurice Redondo, Olivier Kobel, Christoph Noppen, Niko Beerenwinkel, Tanja Stadler |
| EPI_ISL_3297439, EPI_ISL_3297440 | Wales Specialist Virology Centre Sequencing lab: Pathogen Genomics Unit | Public Health Wales Microbiology Cardiff Wales Specialist Virology Centre | Catherine Moore, Johnathan Evans, Laura Gifford, Malorie Perry, Simon Cottrell, Angela Marchbank, Alec Birchley, Alexander Adams, Amy Gaskin, Bree Gatica-Wilcox, Jason Coombes, Joel Southgate, Lauren Gilbert, Lee Graham, Nicole Pacchiarini, Sara Kumziene-Summerhayes, Sarah Taylor, Sophie Jones, Sara Rey, Matthew Bull, Joanne Watkins, Sally Corden, Tom Connor |
| EPI_ISL_3297441 | Viollier AG | Department of Biosystems Science and Engineering, ETH Zürich | Christian Beisel, Sarah Nadeau, Chaoran Chen, Ivan Topolsky, Kim Philipp Jablonski, Lara Fuhrmann, Rebecca Denes, Mirjam Feldkamp, Ina Nissen, Natascha Santacroce, Elodie Burcklen, Christiane Beckmann, Maurice Redondo, Olivier Kobel, Christoph Noppen, Niko Beerenwinkel, Tanja Stadler |
| EPI_ISL_3297442, EPI_ISL_3297444 | Wales Specialist Virology Centre Sequencing lab: Pathogen Genomics Unit | Public Health Wales Microbiology Cardiff Wales Specialist Virology Centre | Catherine Moore, Johnathan Evans, Laura Gifford, Malorie Perry, Simon Cottrell, Angela Marchbank, Alec Birchley, Alexander Adams, Amy Gaskin, Bree Gatica-Wilcox, Jason Coombes, Joel Southgate, Lauren Gilbert, Lee Graham, Nicole Pacchiarini, Sara Kumziene-Summerhayes, Sarah Taylor, Sophie Jones, Sara Rey, Matthew Bull, Joanne Watkins, Sally Corden, Tom Connor |
| EPI_ISL_3297445 | Viollier AG | Department of Biosystems Science and Engineering, ETH Zürich | Christian Beisel, Sarah Nadeau, Chaoran Chen, Ivan Topolsky, Kim Philipp Jablonski, Lara Fuhrmann, Rebecca Denes, Mirjam Feldkamp, Ina Nissen, Natascha Santacroce, Elodie Burcklen, Christiane Beckmann, Maurice Redondo, Olivier Kobel, Christoph Noppen, Niko Beerenwinkel, Tanja Stadler |
| EPI_ISL_3297446 | Wales Specialist Virology Centre Sequencing lab: Pathogen Genomics Unit | Public Health Wales Microbiology Cardiff Wales Specialist Virology Centre | Catherine Moore, Johnathan Evans, Laura Gifford, Malorie Perry, Simon Cottrell, Angela Marchbank, Alec Birchley, Alexander Adams, Amy Gaskin, Bree Gatica-Wilcox, Jason Coombes, Joel Southgate, Lauren Gilbert, Lee Graham, Nicole Pacchiarini, Sara Kumziene-Summerhayes, Sarah Taylor, Sophie Jones, Sara Rey, Matthew Bull, Joanne Watkins, Sally Corden, Tom Connor |
| EPI_ISL_3297447 | Viollier AG | Department of Biosystems Science and Engineering, ETH Zürich | Christian Beisel, Sarah Nadeau, Chaoran Chen, Ivan Topolsky, Kim Philipp Jablonski, Lara Fuhrmann, Rebecca Denes, Mirjam Feldkamp, Ina Nissen, Natascha Santacroce, Elodie Burcklen, Christiane Beckmann, Maurice Redondo, Olivier Kobel, Christoph Noppen, Niko Beerenwinkel, Tanja Stadler |
| EPI_ISL_3297448 | Wales Specialist Virology Centre Sequencing lab: Pathogen Genomics Unit | Public Health Wales Microbiology Cardiff Wales Specialist Virology Centre | Catherine Moore, Johnathan Evans, Laura Gifford, Malorie Perry, Simon Cottrell, Angela Marchbank, Alec Birchley, Alexander Adams, Amy Gaskin, Bree Gatica-Wilcox, Jason Coombes, Joel Southgate, Lauren Gilbert, Lee Graham, Nicole Pacchiarini, Sara Kumziene-Summerhayes, Sarah Taylor, Sophie Jones, Sara Rey, Matthew Bull, Joanne Watkins, Sally Corden, Tom Connor |
| EPI_ISL_3297449 | Viollier AG | Department of Biosystems Science and Engineering, ETH Zürich | Christian Beisel, Sarah Nadeau, Chaoran Chen, Ivan Topolsky, Kim Philipp Jablonski, Lara Fuhrmann, Rebecca Denes, Mirjam Feldkamp, Ina Nissen, Natascha Santacroce, Elodie Burcklen, Christiane Beckmann, Maurice Redondo, Olivier Kobel, Christoph Noppen, Niko Beerenwinkel, Tanja Stadler |
| EPI_ISL_3297450 | Wales Specialist Virology Centre Sequencing lab: Pathogen Genomics Unit | Public Health Wales Microbiology Cardiff Wales Specialist Virology Centre | Catherine Moore, Johnathan Evans, Laura Gifford, Malorie Perry, Simon Cottrell, Angela Marchbank, Alec Birchley, Alexander Adams, Amy Gaskin, Bree Gatica-Wilcox, Jason Coombes, Joel Southgate, Lauren Gilbert, Lee Graham, Nicole Pacchiarini, Sara Kumziene-Summerhayes, Sarah Taylor, Sophie Jones, Sara Rey, Matthew Bull, Joanne Watkins, Sally Corden, Tom Connor |
| EPI_ISL_3297451 | Viollier AG | Department of Biosystems Science and Engineering, ETH Zürich | Christian Beisel, Sarah Nadeau, Chaoran Chen, Ivan Topolsky, Kim Philipp Jablonski, Lara Fuhrmann, Rebecca Denes, Mirjam Feldkamp, Ina Nissen, Natascha Santacroce, Elodie Burcklen, Christiane Beckmann, Maurice Redondo, Olivier Kobel, Christoph Noppen, Niko Beerenwinkel, Tanja Stadler |
| EPI_ISL_3297452 | Wales Specialist Virology Centre Sequencing lab: Pathogen Genomics Unit | Public Health Wales Microbiology Cardiff Wales Specialist Virology Centre | Catherine Moore, Johnathan Evans, Laura Gifford, Malorie Perry, Simon Cottrell, Angela Marchbank, Alec Birchley, Alexander Adams, Amy Gaskin, Bree Gatica-Wilcox, Jason Coombes, Joel Southgate, Lauren Gilbert, Lee Graham, Nicole Pacchiarini, Sara Kumziene-Summerhayes, Sarah Taylor, Sophie Jones, Sara Rey, Matthew Bull, Joanne Watkins, Sally Corden, Tom Connor |
| EPI_ISL_3297453 | Viollier AG | Department of Biosystems Science and Engineering, ETH Zürich | Christian Beisel, Sarah Nadeau, Chaoran Chen, Ivan Topolsky, Kim Philipp Jablonski, Lara Fuhrmann, Rebecca Denes, Mirjam Feldkamp, Ina Nissen, Natascha Santacroce, Elodie Burcklen, Christiane Beckmann, Maurice Redondo, Olivier Kobel, |

| | | | |
|---|---|---|---|
| | | | Christoph Noppen, Niko Beerenwinkel, Tanja Stadler |
| EPI_ISL_3297454 | Wales Specialist Virology Centre Sequencing lab: Pathogen Genomics Unit | Public Health Wales Microbiology Cardiff Wales Specialist Virology Centre | Catherine Moore, Johnathan Evans, Laura Gifford, Malorie Perry, Simon Cottrell, Angela Marchbank, Alec Birchley, Alexander Adams, Amy Gaskin, Bree Gatica-Wilcox, Jason Coombes, Joel Southgate, Lauren Gilbert, Lee Graham, Nicole Pacchiarini, Sara Kumziene-Summerhayes, Sarah Taylor, Sophie Jones, Sara Rey, Matthew Bull, Joanne Watkins, Sally Corden, Tom Connor |
| EPI_ISL_3297455 | Viollier AG | Department of Biosystems Science and Engineering, ETH Zürich | Christian Beisel, Sarah Nadeau, Chaoran Chen, Ivan Topolsky, Kim Philipp Jablonski, Lara Fuhrmann, Rebecca Denes, Mirjam Feldkamp, Ina Nissen, Natascha Santacroce, Elodie Burcklen, Christiane Beckmann, Maurice Redondo, Olivier Kobel, Christoph Noppen, Niko Beerenwinkel, Tanja Stadler |
| EPI_ISL_3297456 | Wales Specialist Virology Centre Sequencing lab: Pathogen Genomics Unit | Public Health Wales Microbiology Cardiff Wales Specialist Virology Centre | Catherine Moore, Johnathan Evans, Laura Gifford, Malorie Perry, Simon Cottrell, Angela Marchbank, Alec Birchley, Alexander Adams, Amy Gaskin, Bree Gatica-Wilcox, Jason Coombes, Joel Southgate, Lauren Gilbert, Lee Graham, Nicole Pacchiarini, Sara Kumziene-Summerhayes, Sarah Taylor, Sophie Jones, Sara Rey, Matthew Bull, Joanne Watkins, Sally Corden, Tom Connor |
| EPI_ISL_3297457 | Viollier AG | Department of Biosystems Science and Engineering, ETH Zürich | Christian Beisel, Sarah Nadeau, Chaoran Chen, Ivan Topolsky, Kim Philipp Jablonski, Lara Fuhrmann, Rebecca Denes, Mirjam Feldkamp, Ina Nissen, Natascha Santacroce, Elodie Burcklen, Christiane Beckmann, Maurice Redondo, Olivier Kobel, Christoph Noppen, Niko Beerenwinkel, Tanja Stadler |
| EPI_ISL_3297458 | Wales Specialist Virology Centre Sequencing lab: Pathogen Genomics Unit | Public Health Wales Microbiology Cardiff Wales Specialist Virology Centre | Catherine Moore, Johnathan Evans, Laura Gifford, Malorie Perry, Simon Cottrell, Angela Marchbank, Alec Birchley, Alexander Adams, Amy Gaskin, Bree Gatica-Wilcox, Jason Coombes, Joel Southgate, Lauren Gilbert, Lee Graham, Nicole Pacchiarini, Sara Kumziene-Summerhayes, Sarah Taylor, Sophie Jones, Sara Rey, Matthew Bull, Joanne Watkins, Sally Corden, Tom Connor |
| EPI_ISL_3297459 | Viollier AG | Department of Biosystems Science and Engineering, ETH Zürich | Christian Beisel, Sarah Nadeau, Chaoran Chen, Ivan Topolsky, Kim Philipp Jablonski, Lara Fuhrmann, Rebecca Denes, Mirjam Feldkamp, Ina Nissen, Natascha Santacroce, Elodie Burcklen, Christiane Beckmann, Maurice Redondo, Olivier Kobel, Christoph Noppen, Niko Beerenwinkel, Tanja Stadler |
| EPI_ISL_3297460 | Wales Specialist Virology Centre Sequencing lab: Pathogen Genomics Unit | Public Health Wales Microbiology Cardiff Wales Specialist Virology Centre | Catherine Moore, Johnathan Evans, Laura Gifford, Malorie Perry, Simon Cottrell, Angela Marchbank, Alec Birchley, Alexander Adams, Amy Gaskin, Bree Gatica-Wilcox, Jason Coombes, Joel Southgate, Lauren Gilbert, Lee Graham, Nicole Pacchiarini, Sara Kumziene-Summerhayes, Sarah Taylor, Sophie Jones, Sara Rey, Matthew Bull, Joanne Watkins, Sally Corden, Tom Connor |
| EPI_ISL_3297461 | Viollier AG | Department of Biosystems Science and Engineering, ETH Zürich | Christian Beisel, Sarah Nadeau, Chaoran Chen, Ivan Topolsky, Kim Philipp Jablonski, Lara Fuhrmann, Rebecca Denes, Mirjam Feldkamp, Ina Nissen, Natascha Santacroce, Elodie Burcklen, Christiane Beckmann, Maurice Redondo, Olivier Kobel, Christoph Noppen, Niko Beerenwinkel, Tanja Stadler |
| EPI_ISL_3297462 | Wales Specialist Virology Centre Sequencing lab: Pathogen Genomics Unit | Public Health Wales Microbiology Cardiff Wales Specialist Virology Centre | Catherine Moore, Johnathan Evans, Laura Gifford, Malorie Perry, Simon Cottrell, Angela Marchbank, Alec Birchley, Alexander Adams, Amy Gaskin, Bree Gatica-Wilcox, Jason Coombes, Joel Southgate, Lauren Gilbert, Lee Graham, Nicole Pacchiarini, Sara Kumziene-Summerhayes, Sarah Taylor, Sophie Jones, Sara Rey, Matthew Bull, Joanne Watkins, Sally Corden, Tom Connor |
| EPI_ISL_3297463 | Viollier AG | Department of Biosystems Science and Engineering, ETH Zürich | Christian Beisel, Sarah Nadeau, Chaoran Chen, Ivan Topolsky, Kim Philipp Jablonski, Lara Fuhrmann, Rebecca Denes, Mirjam Feldkamp, Ina Nissen, Natascha Santacroce, Elodie Burcklen, Christiane Beckmann, Maurice Redondo, Olivier Kobel, Christoph Noppen, Niko Beerenwinkel, Tanja Stadler |
| EPI_ISL_3297464 | Wales Specialist Virology Centre Sequencing lab: Pathogen Genomics Unit | Public Health Wales Microbiology Cardiff Wales Specialist Virology Centre | Catherine Moore, Johnathan Evans, Laura Gifford, Malorie Perry, Simon Cottrell, Angela Marchbank, Alec Birchley, Alexander Adams, Amy Gaskin, Bree Gatica-Wilcox, Jason Coombes, Joel Southgate, Lauren Gilbert, Lee Graham, Nicole Pacchiarini, Sara Kumziene-Summerhayes, Sarah Taylor, Sophie Jones, Sara Rey, Matthew Bull, Joanne Watkins, Sally Corden, Tom Connor |
| EPI_ISL_3297465 | Viollier AG | Department of Biosystems Science and Engineering, ETH Zürich | Christian Beisel, Sarah Nadeau, Chaoran Chen, Ivan Topolsky, Kim Philipp Jablonski, Lara Fuhrmann, Rebecca Denes, Mirjam Feldkamp, Ina Nissen, Natascha Santacroce, Elodie Burcklen, Christiane Beckmann, Maurice Redondo, Olivier Kobel, Christoph Noppen, Niko Beerenwinkel, Tanja Stadler |
| EPI_ISL_3297466, EPI_ISL_3297467 | Wales Specialist Virology Centre Sequencing lab: Pathogen Genomics Unit | Public Health Wales Microbiology Cardiff Wales Specialist Virology Centre | Catherine Moore, Johnathan Evans, Laura Gifford, Malorie Perry, Simon Cottrell, Angela Marchbank, Alec Birchley, Alexander Adams, Amy Gaskin, Bree Gatica-Wilcox, Jason Coombes, Joel Southgate, Lauren Gilbert, Lee Graham, Nicole Pacchiarini, Sara Kumziene-Summerhayes, Sarah Taylor, Sophie Jones, Sara Rey, Matthew Bull, Joanne Watkins, Sally Corden, Tom Connor |
| EPI_ISL_3297468 | Viollier AG | Department of Biosystems Science and Engineering, ETH Zürich | Christian Beisel, Sarah Nadeau, Chaoran Chen, Ivan Topolsky, Kim Philipp Jablonski, Lara Fuhrmann, Rebecca Denes, Mirjam Feldkamp, Ina Nissen, Natascha Santacroce, Elodie Burcklen, Christiane Beckmann, Maurice Redondo, Olivier Kobel, Christoph Noppen, Niko Beerenwinkel, Tanja Stadler |
| EPI_ISL_3297469 | Wales Specialist Virology Centre Sequencing lab: Pathogen Genomics Unit | Public Health Wales Microbiology Cardiff Wales Specialist Virology Centre | Catherine Moore, Johnathan Evans, Laura Gifford, Malorie Perry, Simon Cottrell, Angela Marchbank, Alec Birchley, Alexander Adams, Amy Gaskin, Bree Gatica-Wilcox, Jason Coombes, Joel Southgate, Lauren Gilbert, Lee Graham, Nicole Pacchiarini, Sara Kumziene-Summerhayes, Sarah Taylor, Sophie Jones, Sara Rey, Matthew Bull, Joanne Watkins, Sally Corden, Tom Connor |
| EPI_ISL_3297470 | Viollier AG | Department of Biosystems Science and Engineering, ETH Zürich | Christian Beisel, Sarah Nadeau, Chaoran Chen, Ivan Topolsky, Kim Philipp Jablonski, Lara Fuhrmann, Rebecca Denes, Mirjam Feldkamp, Ina Nissen, Natascha Santacroce, Elodie Burcklen, Christiane Beckmann, Maurice Redondo, Olivier Kobel, Christoph Noppen, Niko Beerenwinkel, Tanja Stadler |
| EPI_ISL_3297471 | Wales Specialist Virology Centre Sequencing lab: Pathogen Genomics Unit | Public Health Wales Microbiology Cardiff Wales Specialist Virology Centre | Catherine Moore, Johnathan Evans, Laura Gifford, Malorie Perry, Simon Cottrell, Angela Marchbank, Alec Birchley, Alexander Adams, Amy Gaskin, Bree Gatica-Wilcox, Jason Coombes, Joel Southgate, Lauren Gilbert, Lee Graham, Nicole Pacchiarini, Sara Kumziene-Summerhayes, Sarah Taylor, Sophie Jones, Sara Rey, Matthew Bull, Joanne Watkins, Sally Corden, Tom Connor |
| EPI_ISL_3297472 | Viollier AG | Department of Biosystems Science and Engineering, ETH Zürich | Christian Beisel, Sarah Nadeau, Chaoran Chen, Ivan Topolsky, Kim Philipp Jablonski, Lara Fuhrmann, Rebecca Denes, Mirjam Feldkamp, Ina Nissen, Natascha Santacroce, Elodie Burcklen, Christiane Beckmann, Maurice Redondo, Olivier Kobel, Christoph Noppen, Niko Beerenwinkel, Tanja Stadler |
| EPI_ISL_3297473 | Wales Specialist Virology Centre Sequencing lab: Pathogen Genomics Unit | Public Health Wales Microbiology Cardiff Wales Specialist Virology Centre | Catherine Moore, Johnathan Evans, Laura Gifford, Malorie Perry, Simon Cottrell, Angela Marchbank, Alec Birchley, Alexander Adams, Amy Gaskin, Bree Gatica-Wilcox, Jason Coombes, Joel Southgate, Lauren Gilbert, Lee Graham, Nicole Pacchiarini, Sara Kumziene-Summerhayes, Sarah Taylor, Sophie Jones, Sara Rey, Matthew Bull, Joanne Watkins, Sally Corden, Tom Connor |
| EPI_ISL_3297474 | Viollier AG | Department of Biosystems Science and Engineering, ETH Zürich | Christian Beisel, Sarah Nadeau, Chaoran Chen, Ivan Topolsky, Kim Philipp Jablonski, Lara Fuhrmann, Rebecca Denes, Mirjam Feldkamp, Ina Nissen, Natascha Santacroce, Elodie Burcklen, Christiane Beckmann, Maurice Redondo, Olivier Kobel, Christoph Noppen, Niko Beerenwinkel, Tanja Stadler |
| EPI_ISL_3297475 | Wales Specialist Virology Centre Sequencing lab: Pathogen Genomics Unit | Public Health Wales Microbiology Cardiff Wales Specialist Virology Centre | Catherine Moore, Johnathan Evans, Laura Gifford, Malorie Perry, Simon Cottrell, Angela Marchbank, Alec Birchley, Alexander Adams, Amy Gaskin, Bree Gatica-Wilcox, Jason Coombes, Joel Southgate, Lauren Gilbert, Lee Graham, Nicole Pacchiarini, Sara Kumziene-Summerhayes, Sarah Taylor, Sophie Jones, Sara Rey, Matthew Bull, Joanne Watkins, Sally Corden, Tom Connor |
| EPI_ISL_3297476 | Viollier AG | Department of Biosystems Science and Engineering, ETH Zürich | Christian Beisel, Sarah Nadeau, Chaoran Chen, Ivan Topolsky, Kim Philipp Jablonski, Lara Fuhrmann, Rebecca Denes, Mirjam Feldkamp, Ina Nissen, Natascha Santacroce, Elodie Burcklen, Christiane Beckmann, Maurice Redondo, Olivier Kobel, Christoph Noppen, Niko Beerenwinkel, Tanja Stadler |
| EPI_ISL_3297477 | Wales Specialist Virology Centre Sequencing lab: Pathogen Genomics Unit | Public Health Wales Microbiology Cardiff Wales Specialist Virology Centre | Catherine Moore, Johnathan Evans, Laura Gifford, Malorie Perry, Simon Cottrell, Angela Marchbank, Alec Birchley, Alexander Adams, Amy Gaskin, Bree Gatica-Wilcox, Jason Coombes, Joel Southgate, Lauren Gilbert, Lee Graham, Nicole Pacchiarini, Sara Kumziene-Summerhayes, Sarah Taylor, Sophie Jones, Sara Rey, Matthew Bull, Joanne Watkins, Sally Corden, Tom Connor |
| EPI_ISL_3297478 | Viollier AG | Department of Biosystems Science and Engineering, ETH Zürich | Christian Beisel, Sarah Nadeau, Chaoran Chen, Ivan Topolsky, Kim Philipp Jablonski, Lara Fuhrmann, Rebecca Denes, Mirjam Feldkamp, Ina Nissen, Natascha Santacroce, Elodie Burcklen, Christiane Beckmann, Maurice Redondo, Olivier Kobel, Christoph Noppen, Niko Beerenwinkel, Tanja Stadler |
| EPI_ISL_3297479, EPI_ISL_3297480 | Wales Specialist Virology Centre Sequencing lab: Pathogen Genomics Unit | Public Health Wales Microbiology Cardiff Wales Specialist Virology Centre | Catherine Moore, Johnathan Evans, Laura Gifford, Malorie Perry, Simon Cottrell, Angela Marchbank, Alec Birchley, Alexander Adams, Amy Gaskin, Bree Gatica-Wilcox, Jason Coombes, Joel Southgate, Lauren Gilbert, Lee Graham, Nicole Pacchiarini, Sara Kumziene-Summerhayes, Sarah Taylor, Sophie Jones, Sara Rey, Matthew Bull, Joanne Watkins, Sally Corden, Tom Connor |
| EPI_ISL_3297481 | Viollier AG | Department of Biosystems Science and Engineering, ETH Zürich | Christian Beisel, Sarah Nadeau, Chaoran Chen, Ivan Topolsky, Kim Philipp Jablonski, Lara Fuhrmann, Rebecca Denes, Mirjam Feldkamp, Ina Nissen, Natascha Santacroce, Elodie Burcklen, Christiane Beckmann, Maurice Redondo, Olivier Kobel, Christoph Noppen, Niko Beerenwinkel, Tanja Stadler |
| EPI_ISL_3297482 | Wales Specialist Virology Centre Sequencing lab: Pathogen Genomics Unit | Public Health Wales Microbiology Cardiff Wales Specialist Virology Centre | Catherine Moore, Johnathan Evans, Laura Gifford, Malorie Perry, Simon Cottrell, Angela Marchbank, Alec Birchley, Alexander Adams, Amy Gaskin, Bree Gatica-Wilcox, Jason Coombes, Joel Southgate, Lauren Gilbert, Lee Graham, Nicole Pacchiarini, Sara Kumziene-Summerhayes, Sarah Taylor, Sophie Jones, Sara Rey, Matthew Bull, Joanne Watkins, Sally Corden, Tom Connor |
| EPI_ISL_3297483 | Viollier AG | Department of Biosystems Science and Engineering, ETH Zürich | Christian Beisel, Sarah Nadeau, Chaoran Chen, Ivan Topolsky, Kim Philipp Jablonski, Lara Fuhrmann, Rebecca Denes, Mirjam Feldkamp, Ina Nissen, Natascha Santacroce, Elodie Burcklen, Christiane Beckmann, Maurice Redondo, Olivier Kobel, Christoph Noppen, Niko Beerenwinkel, Tanja Stadler |
| EPI_ISL_3297484 | Wales Specialist Virology Centre Sequencing lab: Pathogen Genomics Unit | Public Health Wales Microbiology Cardiff Wales Specialist Virology Centre | Catherine Moore, Johnathan Evans, Laura Gifford, Malorie Perry, Simon Cottrell, Angela Marchbank, Alec Birchley, Alexander Adams, Amy Gaskin, Bree Gatica-Wilcox, Jason Coombes, Joel Southgate, Lauren Gilbert, Lee Graham, Nicole Pacchiarini, Sara Kumziene-Summerhayes, Sarah Taylor, Sophie Jones, Sara Rey, Matthew Bull, Joanne Watkins, Sally Corden, Tom Connor |
| EPI_ISL_3297485 | Viollier AG | Department of Biosystems Science and Engineering, ETH Zürich | Christian Beisel, Sarah Nadeau, Chaoran Chen, Ivan Topolsky, Kim Philipp Jablonski, Lara Fuhrmann, Rebecca Denes, Mirjam Feldkamp, Ina Nissen, Natascha Santacroce, Elodie Burcklen, Christiane Beckmann, Maurice Redondo, Olivier Kobel, Christoph Noppen, Niko Beerenwinkel, Tanja Stadler |
| EPI_ISL_3297486 | Wales Specialist Virology Centre Sequencing lab: Pathogen Genomics Unit | Public Health Wales Microbiology Cardiff Wales Specialist Virology Centre | Catherine Moore, Johnathan Evans, Laura Gifford, Malorie Perry, Simon Cottrell, Angela Marchbank, Alec Birchley, Alexander Adams, Amy Gaskin, Bree Gatica-Wilcox, Jason Coombes, Joel Southgate, Lauren Gilbert, Lee Graham, Nicole Pacchiarini, Sara Kumziene-Summerhayes, Sarah Taylor, Sophie Jones, Sara Rey, Matthew Bull, Joanne Watkins, Sally Corden, Tom Connor |
| EPI_ISL_3297487 | Viollier AG | Department of Biosystems Science and Engineering, ETH Zürich | Christian Beisel, Sarah Nadeau, Chaoran Chen, Ivan Topolsky, Kim Philipp Jablonski, Lara Fuhrmann, Rebecca Denes, Mirjam Feldkamp, Ina Nissen, Natascha Santacroce, Elodie Burcklen, Christiane Beckmann, Maurice Redondo, Olivier Kobel, Christoph Noppen, Niko Beerenwinkel, Tanja Stadler |
| EPI_ISL_3297488, EPI_ISL_3297489 | Wales Specialist Virology Centre Sequencing lab: Pathogen Genomics Unit | Public Health Wales Microbiology Cardiff Wales Specialist Virology Centre | Catherine Moore, Johnathan Evans, Laura Gifford, Malorie Perry, Simon Cottrell, Angela Marchbank, Alec Birchley, Alexander Adams, Amy Gaskin, Bree Gatica-Wilcox, Jason Coombes, Joel Southgate, Lauren Gilbert, Lee Graham, Nicole Pacchiarini, Sara Kumziene-Summerhayes, Sarah Taylor, Sophie Jones, Sara Rey, Matthew Bull, Joanne Watkins, Sally Corden, Tom Connor |
| EPI_ISL_3297490 | Viollier AG | Department of Biosystems Science and Engineering, ETH Zürich | Christian Beisel, Sarah Nadeau, Chaoran Chen, Ivan Topolsky, Kim Philipp Jablonski, Lara Fuhrmann, Rebecca Denes, Mirjam Feldkamp, Ina Nissen, Natascha Santacroce, Elodie Burcklen, Christiane Beckmann, Maurice Redondo, Olivier Kobel, Christoph Noppen, Niko Beerenwinkel, Tanja Stadler |
| EPI_ISL_3297491 | Wales Specialist Virology Centre Sequencing lab: Pathogen Genomics Unit | Public Health Wales Microbiology Cardiff Wales Specialist Virology Centre | Catherine Moore, Johnathan Evans, Laura Gifford, Malorie Perry, Simon Cottrell, Angela Marchbank, Alec Birchley, Alexander Adams, Amy Gaskin, Bree Gatica-Wilcox, Jason Coombes, Joel Southgate, Lauren Gilbert, Lee Graham, Nicole Pacchiarini, Sara Kumziene-Summerhayes, Sarah Taylor, Sophie Jones, Sara Rey, Matthew Bull, Joanne Watkins, Sally Corden, Tom Connor |
| EPI_ISL_3297492 | Viollier AG | Department of Biosystems Science and Engineering, ETH Zürich | Christian Beisel, Sarah Nadeau, Chaoran Chen, Ivan Topolsky, Kim Philipp Jablonski, Lara Fuhrmann, Rebecca Denes, Mirjam Feldkamp, Ina Nissen, Natascha Santacroce, Elodie Burcklen, Christiane Beckmann, Maurice Redondo, Olivier Kobel, Christoph Noppen, Niko Beerenwinkel, Tanja Stadler |
| EPI_ISL_3297493 | Wales Specialist Virology Centre Sequencing lab: Pathogen Genomics Unit | Public Health Wales Microbiology Cardiff Wales Specialist Virology Centre | Catherine Moore, Johnathan Evans, Laura Gifford, Malorie Perry, Simon Cottrell, Angela Marchbank, Alec Birchley, Alexander Adams, Amy Gaskin, Bree Gatica-Wilcox, Jason Coombes, Joel Southgate, Lauren Gilbert, Lee Graham, Nicole Pacchiarini, Sara Kumziene-Summerhayes, Sarah Taylor, Sophie Jones, Sara Rey, Matthew Bull, Joanne Watkins, Sally Corden, Tom Connor |
| EPI_ISL_3297494 | Viollier AG | Department of Biosystems Science and Engineering, ETH Zürich | Christian Beisel, Sarah Nadeau, Chaoran Chen, Ivan Topolsky, Kim Philipp Jablonski, Lara Fuhrmann, Rebecca Denes, Mirjam Feldkamp, Ina Nissen, Natascha Santacroce, Elodie Burcklen, Christiane Beckmann, Maurice Redondo, Olivier Kobel, Christoph Noppen, Niko Beerenwinkel, Tanja Stadler |
| EPI_ISL_3297495 | Wales Specialist Virology Centre Sequencing lab: Pathogen Genomics Unit | Public Health Wales Microbiology Cardiff Wales Specialist Virology Centre | Catherine Moore, Johnathan Evans, Laura Gifford, Malorie Perry, Simon Cottrell, Angela Marchbank, Alec Birchley, Alexander Adams, Amy Gaskin, Bree Gatica-Wilcox, Jason Coombes, Joel Southgate, Lauren Gilbert, Lee Graham, Nicole Pacchiarini, Sara Kumziene-Summerhayes, Sarah Taylor, Sophie Jones, Sara Rey, Matthew Bull, Joanne Watkins, Sally Corden, Tom Connor |
| EPI_ISL_3297496 | Viollier AG | Department of Biosystems Science and Engineering, ETH Zürich | Christian Beisel, Sarah Nadeau, Chaoran Chen, Ivan Topolsky, Kim Philipp Jablonski, Lara Fuhrmann, Rebecca Denes, Mirjam Feldkamp, Ina Nissen, Natascha Santacroce, Elodie Burcklen, Christiane Beckmann, Maurice Redondo, Olivier Kobel, Christoph Noppen, Niko Beerenwinkel, Tanja Stadler |
| EPI_ISL_3297497 | Wales Specialist Virology Centre Sequencing lab: Pathogen Genomics Unit | Public Health Wales Microbiology Cardiff Wales Specialist Virology Centre | Catherine Moore, Johnathan Evans, Laura Gifford, Malorie Perry, Simon Cottrell, Angela Marchbank, Alec Birchley, Alexander Adams, Amy Gaskin, Bree Gatica-Wilcox, Jason Coombes, Joel Southgate, Lauren Gilbert, Lee Graham, Nicole Pacchiarini, Sara Kumziene-Summerhayes, Sarah Taylor, Sophie Jones, Sara Rey, Matthew Bull, Joanne Watkins, Sally Corden, Tom Connor |
| EPI_ISL_3297498 | Viollier AG | Department of Biosystems Science and Engineering, ETH Zürich | Christian Beisel, Sarah Nadeau, Chaoran Chen, Ivan Topolsky, Kim Philipp Jablonski, Lara Fuhrmann, Rebecca Denes, Mirjam Feldkamp, Ina Nissen, Natascha Santacroce, Elodie Burcklen, Christiane Beckmann, Maurice Redondo, Olivier Kobel, Christoph Noppen, Niko Beerenwinkel, Tanja Stadler |
| EPI_ISL_3297499, EPI_ISL_3297500 | Wales Specialist Virology Centre Sequencing lab: Pathogen Genomics Unit | Public Health Wales Microbiology Cardiff Wales Specialist Virology Centre | Catherine Moore, Johnathan Evans, Laura Gifford, Malorie Perry, Simon Cottrell, Angela Marchbank, Alec Birchley, Alexander Adams, Amy Gaskin, Bree Gatica-Wilcox, Jason Coombes, Joel Southgate, Lauren Gilbert, Lee Graham, Nicole Pacchiarini, Sara Kumziene-Summerhayes, Sarah Taylor, Sophie Jones, Sara Rey, Matthew Bull, Joanne Watkins, Sally Corden, Tom Connor |
| EPI_ISL_3297501 | Viollier AG | Department of Biosystems Science and Engineering, ETH Zürich | Christian Beisel, Sarah Nadeau, Chaoran Chen, Ivan Topolsky, Kim Philipp Jablonski, Lara Fuhrmann, Rebecca Denes, Mirjam Feldkamp, Ina Nissen, Natascha Santacroce, Elodie Burcklen, Christiane Beckmann, Maurice Redondo, Olivier Kobel, Christoph Noppen, Niko Beerenwinkel, Tanja Stadler |

| | | | |
|---|---|---|---|
| EPI_ISL_3297502 | Wales Specialist Virology Centre Sequencing lab: Pathogen Genomics Unit | Public Health Wales Microbiology Cardiff Wales Specialist Virology Centre | Catherine Moore, Johnathan Evans, Laura Gifford, Malorie Perry, Simon Cottrell, Angela Marchbank, Alec Birchley, Alexander Adams, Amy Gaskin, Bree Gatica-Wilcox, Jason Coombes, Joel Southgate, Lauren Gilbert, Lee Graham, Nicole Pacchiarini, Sara Kumziene-Summerhayes, Sarah Taylor, Sophie Jones, Sara Rey, Matthew Bull, Joanne Watkins, Sally Corden, Tom Connor |
| EPI_ISL_3297503 | Viollier AG | Department of Biosystems Science and Engineering, ETH Zürich | Christian Beisel, Sarah Nadeau, Chaoran Chen, Ivan Topolsky, Kim Philipp Jablonski, Lara Fuhrmann, Rebecca Denes, Mirjam Feldkamp, Ina Nissen, Natascha Santacroce, Elodie Burcklen, Christiane Beckmann, Maurice Redondo, Olivier Kobel, Christoph Noppen, Niko Beerenwinkel, Tanja Stadler |
| EPI_ISL_3297504 | Wales Specialist Virology Centre Sequencing lab: Pathogen Genomics Unit | Public Health Wales Microbiology Cardiff Wales Specialist Virology Centre | Catherine Moore, Johnathan Evans, Laura Gifford, Malorie Perry, Simon Cottrell, Angela Marchbank, Alec Birchley, Alexander Adams, Amy Gaskin, Bree Gatica-Wilcox, Jason Coombes, Joel Southgate, Lauren Gilbert, Lee Graham, Nicole Pacchiarini, Sara Kumziene-Summerhayes, Sarah Taylor, Sophie Jones, Sara Rey, Matthew Bull, Joanne Watkins, Sally Corden, Tom Connor |
| EPI_ISL_3297505 | Viollier AG | Department of Biosystems Science and Engineering, ETH Zürich | Christian Beisel, Sarah Nadeau, Chaoran Chen, Ivan Topolsky, Kim Philipp Jablonski, Lara Fuhrmann, Rebecca Denes, Mirjam Feldkamp, Ina Nissen, Natascha Santacroce, Elodie Burcklen, Christiane Beckmann, Maurice Redondo, Olivier Kobel, Christoph Noppen, Niko Beerenwinkel, Tanja Stadler |
| EPI_ISL_3297506 | Wales Specialist Virology Centre Sequencing lab: Pathogen Genomics Unit | Public Health Wales Microbiology Cardiff Wales Specialist Virology Centre | Catherine Moore, Johnathan Evans, Laura Gifford, Malorie Perry, Simon Cottrell, Angela Marchbank, Alec Birchley, Alexander Adams, Amy Gaskin, Bree Gatica-Wilcox, Jason Coombes, Joel Southgate, Lauren Gilbert, Lee Graham, Nicole Pacchiarini, Sara Kumziene-Summerhayes, Sarah Taylor, Sophie Jones, Sara Rey, Matthew Bull, Joanne Watkins, Sally Corden, Tom Connor |
| EPI_ISL_3297507 | Viollier AG | Department of Biosystems Science and Engineering, ETH Zürich | Christian Beisel, Sarah Nadeau, Chaoran Chen, Ivan Topolsky, Kim Philipp Jablonski, Lara Fuhrmann, Rebecca Denes, Mirjam Feldkamp, Ina Nissen, Natascha Santacroce, Elodie Burcklen, Christiane Beckmann, Maurice Redondo, Olivier Kobel, Christoph Noppen, Niko Beerenwinkel, Tanja Stadler |
| EPI_ISL_3297508, EPI_ISL_3297509 | Wales Specialist Virology Centre Sequencing lab: Pathogen Genomics Unit | Public Health Wales Microbiology Cardiff Wales Specialist Virology Centre | Catherine Moore, Johnathan Evans, Laura Gifford, Malorie Perry, Simon Cottrell, Angela Marchbank, Alec Birchley, Alexander Adams, Amy Gaskin, Bree Gatica-Wilcox, Jason Coombes, Joel Southgate, Lauren Gilbert, Lee Graham, Nicole Pacchiarini, Sara Kumziene-Summerhayes, Sarah Taylor, Sophie Jones, Sara Rey, Matthew Bull, Joanne Watkins, Sally Corden, Tom Connor |
| EPI_ISL_3297510 | Viollier AG | Department of Biosystems Science and Engineering, ETH Zürich | Christian Beisel, Sarah Nadeau, Chaoran Chen, Ivan Topolsky, Kim Philipp Jablonski, Lara Fuhrmann, Rebecca Denes, Mirjam Feldkamp, Ina Nissen, Natascha Santacroce, Elodie Burcklen, Christiane Beckmann, Maurice Redondo, Olivier Kobel, Christoph Noppen, Niko Beerenwinkel, Tanja Stadler |
| EPI_ISL_3297511 | Wales Specialist Virology Centre Sequencing lab: Pathogen Genomics Unit | Public Health Wales Microbiology Cardiff Wales Specialist Virology Centre | Catherine Moore, Johnathan Evans, Laura Gifford, Malorie Perry, Simon Cottrell, Angela Marchbank, Alec Birchley, Alexander Adams, Amy Gaskin, Bree Gatica-Wilcox, Jason Coombes, Joel Southgate, Lauren Gilbert, Lee Graham, Nicole Pacchiarini, Sara Kumziene-Summerhayes, Sarah Taylor, Sophie Jones, Sara Rey, Matthew Bull, Joanne Watkins, Sally Corden, Tom Connor |
| EPI_ISL_3297512 | Viollier AG | Department of Biosystems Science and Engineering, ETH Zürich | Christian Beisel, Sarah Nadeau, Chaoran Chen, Ivan Topolsky, Kim Philipp Jablonski, Lara Fuhrmann, Rebecca Denes, Mirjam Feldkamp, Ina Nissen, Natascha Santacroce, Elodie Burcklen, Christiane Beckmann, Maurice Redondo, Olivier Kobel, Christoph Noppen, Niko Beerenwinkel, Tanja Stadler |
| EPI_ISL_3297513 | Wales Specialist Virology Centre Sequencing lab: Pathogen Genomics Unit | Public Health Wales Microbiology Cardiff Wales Specialist Virology Centre | Catherine Moore, Johnathan Evans, Laura Gifford, Malorie Perry, Simon Cottrell, Angela Marchbank, Alec Birchley, Alexander Adams, Amy Gaskin, Bree Gatica-Wilcox, Jason Coombes, Joel Southgate, Lauren Gilbert, Lee Graham, Nicole Pacchiarini, Sara Kumziene-Summerhayes, Sarah Taylor, Sophie Jones, Sara Rey, Matthew Bull, Joanne Watkins, Sally Corden, Tom Connor |
| EPI_ISL_3297514 | Viollier AG | Department of Biosystems Science and Engineering, ETH Zürich | Christian Beisel, Sarah Nadeau, Chaoran Chen, Ivan Topolsky, Kim Philipp Jablonski, Lara Fuhrmann, Rebecca Denes, Mirjam Feldkamp, Ina Nissen, Natascha Santacroce, Elodie Burcklen, Christiane Beckmann, Maurice Redondo, Olivier Kobel, Christoph Noppen, Niko Beerenwinkel, Tanja Stadler |
| EPI_ISL_3297515 | Wales Specialist Virology Centre Sequencing lab: Pathogen Genomics Unit | Public Health Wales Microbiology Cardiff Wales Specialist Virology Centre | Catherine Moore, Johnathan Evans, Laura Gifford, Malorie Perry, Simon Cottrell, Angela Marchbank, Alec Birchley, Alexander Adams, Amy Gaskin, Bree Gatica-Wilcox, Jason Coombes, Joel Southgate, Lauren Gilbert, Lee Graham, Nicole Pacchiarini, Sara Kumziene-Summerhayes, Sarah Taylor, Sophie Jones, Sara Rey, Matthew Bull, Joanne Watkins, Sally Corden, Tom Connor |
| EPI_ISL_3297516 | Viollier AG | Department of Biosystems Science and Engineering, ETH Zürich | Christian Beisel, Sarah Nadeau, Chaoran Chen, Ivan Topolsky, Kim Philipp Jablonski, Lara Fuhrmann, Rebecca Denes, Mirjam Feldkamp, Ina Nissen, Natascha Santacroce, Elodie Burcklen, Christiane Beckmann, Maurice Redondo, Olivier Kobel, Christoph Noppen, Niko Beerenwinkel, Tanja Stadler |
| EPI_ISL_3297517 | Wales Specialist Virology Centre Sequencing lab: Pathogen Genomics Unit | Public Health Wales Microbiology Cardiff Wales Specialist Virology Centre | Catherine Moore, Johnathan Evans, Laura Gifford, Malorie Perry, Simon Cottrell, Angela Marchbank, Alec Birchley, Alexander Adams, Amy Gaskin, Bree Gatica-Wilcox, Jason Coombes, Joel Southgate, Lauren Gilbert, Lee Graham, Nicole Pacchiarini, Sara Kumziene-Summerhayes, Sarah Taylor, Sophie Jones, Sara Rey, Matthew Bull, Joanne Watkins, Sally Corden, Tom Connor |
| EPI_ISL_3297518 | Viollier AG | Department of Biosystems Science and Engineering, ETH Zürich | Christian Beisel, Sarah Nadeau, Chaoran Chen, Ivan Topolsky, Kim Philipp Jablonski, Lara Fuhrmann, Rebecca Denes, Mirjam Feldkamp, Ina Nissen, Natascha Santacroce, Elodie Burcklen, Christiane Beckmann, Maurice Redondo, Olivier Kobel, Christoph Noppen, Niko Beerenwinkel, Tanja Stadler |
| EPI_ISL_3297519 | Wales Specialist Virology Centre Sequencing lab: Pathogen Genomics Unit | Public Health Wales Microbiology Cardiff Wales Specialist Virology Centre | Catherine Moore, Johnathan Evans, Laura Gifford, Malorie Perry, Simon Cottrell, Angela Marchbank, Alec Birchley, Alexander Adams, Amy Gaskin, Bree Gatica-Wilcox, Jason Coombes, Joel Southgate, Lauren Gilbert, Lee Graham, Nicole Pacchiarini, Sara Kumziene-Summerhayes, Sarah Taylor, Sophie Jones, Sara Rey, Matthew Bull, Joanne Watkins, Sally Corden, Tom Connor |
| EPI_ISL_3297520 | Viollier AG | Department of Biosystems Science and Engineering, ETH Zürich | Christian Beisel, Sarah Nadeau, Chaoran Chen, Ivan Topolsky, Kim Philipp Jablonski, Lara Fuhrmann, Rebecca Denes, Mirjam Feldkamp, Ina Nissen, Natascha Santacroce, Elodie Burcklen, Christiane Beckmann, Maurice Redondo, Olivier Kobel, Christoph Noppen, Niko Beerenwinkel, Tanja Stadler |
| EPI_ISL_3297521 | Wales Specialist Virology Centre Sequencing lab: Pathogen Genomics Unit | Public Health Wales Microbiology Cardiff Wales Specialist Virology Centre | Catherine Moore, Johnathan Evans, Laura Gifford, Malorie Perry, Simon Cottrell, Angela Marchbank, Alec Birchley, Alexander Adams, Amy Gaskin, Bree Gatica-Wilcox, Jason Coombes, Joel Southgate, Lauren Gilbert, Lee Graham, Nicole Pacchiarini, Sara Kumziene-Summerhayes, Sarah Taylor, Sophie Jones, Sara Rey, Matthew Bull, Joanne Watkins, Sally Corden, Tom Connor |
| EPI_ISL_3297522 | Viollier AG | Department of Biosystems Science and Engineering, ETH Zürich | Christian Beisel, Sarah Nadeau, Chaoran Chen, Ivan Topolsky, Kim Philipp Jablonski, Lara Fuhrmann, Rebecca Denes, Mirjam Feldkamp, Ina Nissen, Natascha Santacroce, Elodie Burcklen, Christiane Beckmann, Maurice Redondo, Olivier Kobel, Christoph Noppen, Niko Beerenwinkel, Tanja Stadler |
| EPI_ISL_3297523 | Wales Specialist Virology Centre Sequencing lab: Pathogen Genomics Unit | Public Health Wales Microbiology Cardiff Wales Specialist Virology Centre | Catherine Moore, Johnathan Evans, Laura Gifford, Malorie Perry, Simon Cottrell, Angela Marchbank, Alec Birchley, Alexander Adams, Amy Gaskin, Bree Gatica-Wilcox, Jason Coombes, Joel Southgate, Lauren Gilbert, Lee Graham, Nicole Pacchiarini, Sara Kumziene-Summerhayes, Sarah Taylor, Sophie Jones, Sara Rey, Matthew Bull, Joanne Watkins, Sally Corden, Tom Connor |
| EPI_ISL_3297524 | Viollier AG | Department of Biosystems Science and Engineering, ETH Zürich | Christian Beisel, Sarah Nadeau, Chaoran Chen, Ivan Topolsky, Kim Philipp Jablonski, Lara Fuhrmann, Rebecca Denes, Mirjam Feldkamp, Ina Nissen, Natascha Santacroce, Elodie Burcklen, Christiane Beckmann, Maurice Redondo, Olivier Kobel, Christoph Noppen, Niko Beerenwinkel, Tanja Stadler |
| EPI_ISL_3297525 | Wales Specialist Virology Centre Sequencing lab: Pathogen Genomics Unit | Public Health Wales Microbiology Cardiff Wales Specialist Virology Centre | Catherine Moore, Johnathan Evans, Laura Gifford, Malorie Perry, Simon Cottrell, Angela Marchbank, Alec Birchley, Alexander Adams, Amy Gaskin, Bree Gatica-Wilcox, Jason Coombes, Joel Southgate, Lauren Gilbert, Lee Graham, Nicole Pacchiarini, Sara Kumziene-Summerhayes, Sarah Taylor, Sophie Jones, Sara Rey, Matthew Bull, Joanne Watkins, Sally Corden, Tom Connor |
| EPI_ISL_3297526 | Viollier AG | Department of Biosystems Science and Engineering, ETH Zürich | Christian Beisel, Sarah Nadeau, Chaoran Chen, Ivan Topolsky, Kim Philipp Jablonski, Lara Fuhrmann, Rebecca Denes, Mirjam Feldkamp, Ina Nissen, Natascha Santacroce, Elodie Burcklen, Christiane Beckmann, Maurice Redondo, Olivier Kobel, Christoph Noppen, Niko Beerenwinkel, Tanja Stadler |
| EPI_ISL_3297527 | Wales Specialist Virology Centre Sequencing lab: Pathogen Genomics Unit | Public Health Wales Microbiology Cardiff Wales Specialist Virology Centre | Catherine Moore, Johnathan Evans, Laura Gifford, Malorie Perry, Simon Cottrell, Angela Marchbank, Alec Birchley, Alexander Adams, Amy Gaskin, Bree Gatica-Wilcox, Jason Coombes, Joel Southgate, Lauren Gilbert, Lee Graham, Nicole Pacchiarini, Sara Kumziene-Summerhayes, Sarah Taylor, Sophie Jones, Sara Rey, Matthew Bull, Joanne Watkins, Sally Corden, Tom Connor |
| EPI_ISL_3297528 | Viollier AG | Department of Biosystems Science and Engineering, ETH Zürich | Christian Beisel, Sarah Nadeau, Chaoran Chen, Ivan Topolsky, Kim Philipp Jablonski, Lara Fuhrmann, Rebecca Denes, Mirjam Feldkamp, Ina Nissen, Natascha Santacroce, Elodie Burcklen, Christiane Beckmann, Maurice Redondo, Olivier Kobel, Christoph Noppen, Niko Beerenwinkel, Tanja Stadler |
| EPI_ISL_3297529 | Wales Specialist Virology Centre Sequencing lab: Pathogen Genomics Unit | Public Health Wales Microbiology Cardiff Wales Specialist Virology Centre | Catherine Moore, Johnathan Evans, Laura Gifford, Malorie Perry, Simon Cottrell, Angela Marchbank, Alec Birchley, Alexander Adams, Amy Gaskin, Bree Gatica-Wilcox, Jason Coombes, Joel Southgate, Lauren Gilbert, Lee Graham, Nicole Pacchiarini, Sara Kumziene-Summerhayes, Sarah Taylor, Sophie Jones, Sara Rey, Matthew Bull, Joanne Watkins, Sally Corden, Tom Connor |
| EPI_ISL_3297530 | Viollier AG | Department of Biosystems Science and Engineering, ETH Zürich | Christian Beisel, Sarah Nadeau, Chaoran Chen, Ivan Topolsky, Kim Philipp Jablonski, Lara Fuhrmann, Rebecca Denes, Mirjam Feldkamp, Ina Nissen, Natascha Santacroce, Elodie Burcklen, Christiane Beckmann, Maurice Redondo, Olivier Kobel, Christoph Noppen, Niko Beerenwinkel, Tanja Stadler |
| EPI_ISL_3297531 | Wales Specialist Virology Centre Sequencing lab: Pathogen Genomics Unit | Public Health Wales Microbiology Cardiff Wales Specialist Virology Centre | Catherine Moore, Johnathan Evans, Laura Gifford, Malorie Perry, Simon Cottrell, Angela Marchbank, Alec Birchley, Alexander Adams, Amy Gaskin, Bree Gatica-Wilcox, Jason Coombes, Joel Southgate, Lauren Gilbert, Lee Graham, Nicole Pacchiarini, Sara Kumziene-Summerhayes, Sarah Taylor, Sophie Jones, Sara Rey, Matthew Bull, Joanne Watkins, Sally Corden, Tom Connor |
| EPI_ISL_3297532 | Viollier AG | Department of Biosystems Science and Engineering, ETH Zürich | Christian Beisel, Sarah Nadeau, Chaoran Chen, Ivan Topolsky, Kim Philipp Jablonski, Lara Fuhrmann, Rebecca Denes, Mirjam Feldkamp, Ina Nissen, Natascha Santacroce, Elodie Burcklen, Christiane Beckmann, Maurice Redondo, Olivier Kobel, Christoph Noppen, Niko Beerenwinkel, Tanja Stadler |
| EPI_ISL_3297533, EPI_ISL_3297534 | Wales Specialist Virology Centre Sequencing lab: Pathogen Genomics Unit | Public Health Wales Microbiology Cardiff Wales Specialist Virology Centre | Catherine Moore, Johnathan Evans, Laura Gifford, Malorie Perry, Simon Cottrell, Angela Marchbank, Alec Birchley, Alexander Adams, Amy Gaskin, Bree Gatica-Wilcox, Jason Coombes, Joel Southgate, Lauren Gilbert, Lee Graham, Nicole Pacchiarini, Sara Kumziene-Summerhayes, Sarah Taylor, Sophie Jones, Sara Rey, Matthew Bull, Joanne Watkins, Sally Corden, Tom Connor |
| EPI_ISL_3297535 | Viollier AG | Department of Biosystems Science and Engineering, ETH Zürich | Christian Beisel, Sarah Nadeau, Chaoran Chen, Ivan Topolsky, Kim Philipp Jablonski, Lara Fuhrmann, Rebecca Denes, Mirjam Feldkamp, Ina Nissen, Natascha Santacroce, Elodie Burcklen, Christiane Beckmann, Maurice Redondo, Olivier Kobel, Christoph Noppen, Niko Beerenwinkel, Tanja Stadler |
| EPI_ISL_3297536 | Wales Specialist Virology Centre Sequencing lab: Pathogen Genomics Unit | Public Health Wales Microbiology Cardiff Wales Specialist Virology Centre | Catherine Moore, Johnathan Evans, Laura Gifford, Malorie Perry, Simon Cottrell, Angela Marchbank, Alec Birchley, Alexander Adams, Amy Gaskin, Bree Gatica-Wilcox, Jason Coombes, Joel Southgate, Lauren Gilbert, Lee Graham, Nicole Pacchiarini, Sara Kumziene-Summerhayes, Sarah Taylor, Sophie Jones, Sara Rey, Matthew Bull, Joanne Watkins, Sally Corden, Tom Connor |
| EPI_ISL_3297537 | Viollier AG | Department of Biosystems Science and Engineering, ETH Zürich | Christian Beisel, Sarah Nadeau, Chaoran Chen, Ivan Topolsky, Kim Philipp Jablonski, Lara Fuhrmann, Rebecca Denes, Mirjam Feldkamp, Ina Nissen, Natascha Santacroce, Elodie Burcklen, Christiane Beckmann, Maurice Redondo, Olivier Kobel, Christoph Noppen, Niko Beerenwinkel, Tanja Stadler |
| EPI_ISL_3297538 | Wales Specialist Virology Centre Sequencing lab: Pathogen Genomics Unit | Public Health Wales Microbiology Cardiff Wales Specialist Virology Centre | Catherine Moore, Johnathan Evans, Laura Gifford, Malorie Perry, Simon Cottrell, Angela Marchbank, Alec Birchley, Alexander Adams, Amy Gaskin, Bree Gatica-Wilcox, Jason Coombes, Joel Southgate, Lauren Gilbert, Lee Graham, Nicole Pacchiarini, Sara Kumziene-Summerhayes, Sarah Taylor, Sophie Jones, Sara Rey, Matthew Bull, Joanne Watkins, Sally Corden, Tom Connor |
| EPI_ISL_3297539 | Viollier AG | Department of Biosystems Science and Engineering, ETH Zürich | Christian Beisel, Sarah Nadeau, Chaoran Chen, Ivan Topolsky, Kim Philipp Jablonski, Lara Fuhrmann, Rebecca Denes, Mirjam Feldkamp, Ina Nissen, Natascha Santacroce, Elodie Burcklen, Christiane Beckmann, Maurice Redondo, Olivier Kobel, Christoph Noppen, Niko Beerenwinkel, Tanja Stadler |
| EPI_ISL_3297540 | Wales Specialist Virology Centre Sequencing lab: Pathogen Genomics Unit | Public Health Wales Microbiology Cardiff Wales Specialist Virology Centre | Catherine Moore, Johnathan Evans, Laura Gifford, Malorie Perry, Simon Cottrell, Angela Marchbank, Alec Birchley, Alexander Adams, Amy Gaskin, Bree Gatica-Wilcox, Jason Coombes, Joel Southgate, Lauren Gilbert, Lee Graham, Nicole Pacchiarini, Sara Kumziene-Summerhayes, Sarah Taylor, Sophie Jones, Sara Rey, Matthew Bull, Joanne Watkins, Sally Corden, Tom Connor |
| EPI_ISL_3297541 | Viollier AG | Department of Biosystems Science and Engineering, ETH Zürich | Christian Beisel, Sarah Nadeau, Chaoran Chen, Ivan Topolsky, Kim Philipp Jablonski, Lara Fuhrmann, Rebecca Denes, Mirjam Feldkamp, Ina Nissen, Natascha Santacroce, Elodie Burcklen, Christiane Beckmann, Maurice Redondo, Olivier Kobel, Christoph Noppen, Niko Beerenwinkel, Tanja Stadler |
| EPI_ISL_3297542 | Wales Specialist Virology Centre Sequencing lab: Pathogen Genomics Unit | Public Health Wales Microbiology Cardiff Wales Specialist Virology Centre | Catherine Moore, Johnathan Evans, Laura Gifford, Malorie Perry, Simon Cottrell, Angela Marchbank, Alec Birchley, Alexander Adams, Amy Gaskin, Bree Gatica-Wilcox, Jason Coombes, Joel Southgate, Lauren Gilbert, Lee Graham, Nicole Pacchiarini, Sara Kumziene-Summerhayes, Sarah Taylor, Sophie Jones, Sara Rey, Matthew Bull, Joanne Watkins, Sally Corden, Tom Connor |
| EPI_ISL_3297543, EPI_ISL_3297544 | Viollier AG | Department of Biosystems Science and Engineering, ETH Zürich | Christian Beisel, Sarah Nadeau, Chaoran Chen, Ivan Topolsky, Kim Philipp Jablonski, Lara Fuhrmann, Rebecca Denes, Mirjam Feldkamp, Ina Nissen, Natascha Santacroce, Elodie Burcklen, Christiane Beckmann, Maurice Redondo, Olivier Kobel, Christoph Noppen, Niko Beerenwinkel, Tanja Stadler |
| EPI_ISL_3297545 | Wales Specialist Virology Centre Sequencing lab: Pathogen Genomics Unit | Public Health Wales Microbiology Cardiff Wales Specialist Virology Centre | Catherine Moore, Johnathan Evans, Laura Gifford, Malorie Perry, Simon Cottrell, Angela Marchbank, Alec Birchley, Alexander Adams, Amy Gaskin, Bree Gatica-Wilcox, Jason Coombes, Joel Southgate, Lauren Gilbert, Lee Graham, Nicole Pacchiarini, Sara Kumziene-Summerhayes, Sarah Taylor, Sophie Jones, Sara Rey, Matthew Bull, Joanne Watkins, Sally Corden, Tom Connor |
| EPI_ISL_3297546 | Viollier AG | Department of Biosystems Science and Engineering, ETH Zürich | Christian Beisel, Sarah Nadeau, Chaoran Chen, Ivan Topolsky, Kim Philipp Jablonski, Lara Fuhrmann, Rebecca Denes, Mirjam Feldkamp, Ina Nissen, Natascha Santacroce, Elodie Burcklen, Christiane Beckmann, Maurice Redondo, Olivier Kobel, Christoph Noppen, Niko Beerenwinkel, Tanja Stadler |
| EPI_ISL_3297547 | Wales Specialist Virology Centre Sequencing lab: Pathogen Genomics Unit | Public Health Wales Microbiology Cardiff Wales Specialist Virology Centre | Catherine Moore, Johnathan Evans, Laura Gifford, Malorie Perry, Simon Cottrell, Angela Marchbank, Alec Birchley, Alexander Adams, Amy Gaskin, Bree Gatica-Wilcox, Jason Coombes, Joel Southgate, Lauren Gilbert, Lee Graham, Nicole Pacchiarini, Sara Kumziene-Summerhayes, Sarah Taylor, Sophie Jones, Sara Rey, Matthew Bull, Joanne Watkins, Sally Corden, Tom Connor |
| EPI_ISL_3297548 | Viollier AG | Department of Biosystems Science and Engineering, ETH Zürich | Christian Beisel, Sarah Nadeau, Chaoran Chen, Ivan Topolsky, Kim Philipp Jablonski, Lara Fuhrmann, Rebecca Denes, Mirjam Feldkamp, Ina Nissen, Natascha Santacroce, Elodie Burcklen, Christiane Beckmann, Maurice Redondo, Olivier Kobel, Christoph Noppen, Niko Beerenwinkel, Tanja Stadler |
| EPI_ISL_3297549 | Wales Specialist Virology Centre Sequencing lab: Pathogen Genomics Unit | Public Health Wales Microbiology Cardiff Wales Specialist Virology Centre | Catherine Moore, Johnathan Evans, Laura Gifford, Malorie Perry, Simon Cottrell, Angela Marchbank, Alec Birchley, Alexander Adams, Amy Gaskin, Bree Gatica-Wilcox, Jason Coombes, Joel Southgate, Lauren Gilbert, Lee Graham, Nicole Pacchiarini, Sara Kumziene-Summerhayes, Sarah Taylor, Sophie Jones, Sara Rey, Matthew Bull, Joanne Watkins, Sally Corden, Tom Connor |

| | | | |
|---|---|---|---|
| EPI_ISL_3297550 | Viollier AG | Department of Biosystems Science and Engineering, ETH Zürich | Christian Beisel, Sarah Nadeau, Chaoran Chen, Ivan Topolsky, Kim Philipp Jablonski, Lara Fuhrmann, Rebecca Denes, Mirjam Feldkamp, Ina Nissen, Natascha Santacroce, Elodie Burcklen, Christiane Beckmann, Maurice Redondo, Olivier Kobel, Christoph Noppen, Niko Beerenwinkel, Tanja Stadler |
| EPI_ISL_3297551 | Wales Specialist Virology Centre Sequencing lab: Pathogen Genomics Unit | Public Health Wales Microbiology Cardiff Wales Specialist Virology Centre | Catherine Moore, Johnathan Evans, Laura Gifford, Malorie Perry, Simon Cottrell, Angela Marchbank, Alec Birchley, Alexander Adams, Amy Gaskin, Bree Gatica-Wilcox, Jason Coombes, Joel Southgate, Lauren Gilbert, Lee Graham, Nicole Pacchiarini, Sara Kumziene-Summerhayes, Sarah Taylor, Sophie Jones, Sara Rey, Matthew Bull, Joanne Watkins, Sally Corden, Tom Connor |
| EPI_ISL_3297552 | Viollier AG | Department of Biosystems Science and Engineering, ETH Zürich | Christian Beisel, Sarah Nadeau, Chaoran Chen, Ivan Topolsky, Kim Philipp Jablonski, Lara Fuhrmann, Rebecca Denes, Mirjam Feldkamp, Ina Nissen, Natascha Santacroce, Elodie Burcklen, Christiane Beckmann, Maurice Redondo, Olivier Kobel, Christoph Noppen, Niko Beerenwinkel, Tanja Stadler |
| EPI_ISL_3297553 | Wales Specialist Virology Centre Sequencing lab: Pathogen Genomics Unit | Public Health Wales Microbiology Cardiff Wales Specialist Virology Centre | Catherine Moore, Johnathan Evans, Laura Gifford, Malorie Perry, Simon Cottrell, Angela Marchbank, Alec Birchley, Alexander Adams, Amy Gaskin, Bree Gatica-Wilcox, Jason Coombes, Joel Southgate, Lauren Gilbert, Lee Graham, Nicole Pacchiarini, Sara Kumziene-Summerhayes, Sarah Taylor, Sophie Jones, Sara Rey, Matthew Bull, Joanne Watkins, Sally Corden, Tom Connor |
| EPI_ISL_3297554 | Viollier AG | Department of Biosystems Science and Engineering, ETH Zürich | Christian Beisel, Sarah Nadeau, Chaoran Chen, Ivan Topolsky, Kim Philipp Jablonski, Lara Fuhrmann, Rebecca Denes, Mirjam Feldkamp, Ina Nissen, Natascha Santacroce, Elodie Burcklen, Christiane Beckmann, Maurice Redondo, Olivier Kobel, Christoph Noppen, Niko Beerenwinkel, Tanja Stadler |
| EPI_ISL_3297555 | Wales Specialist Virology Centre Sequencing lab: Pathogen Genomics Unit | Public Health Wales Microbiology Cardiff Wales Specialist Virology Centre | Catherine Moore, Johnathan Evans, Laura Gifford, Malorie Perry, Simon Cottrell, Angela Marchbank, Alec Birchley, Alexander Adams, Amy Gaskin, Bree Gatica-Wilcox, Jason Coombes, Joel Southgate, Lauren Gilbert, Lee Graham, Nicole Pacchiarini, Sara Kumziene-Summerhayes, Sarah Taylor, Sophie Jones, Sara Rey, Matthew Bull, Joanne Watkins, Sally Corden, Tom Connor |
| EPI_ISL_3297556 | Viollier AG | Department of Biosystems Science and Engineering, ETH Zürich | Christian Beisel, Sarah Nadeau, Chaoran Chen, Ivan Topolsky, Kim Philipp Jablonski, Lara Fuhrmann, Rebecca Denes, Mirjam Feldkamp, Ina Nissen, Natascha Santacroce, Elodie Burcklen, Christiane Beckmann, Maurice Redondo, Olivier Kobel, Christoph Noppen, Niko Beerenwinkel, Tanja Stadler |
| EPI_ISL_3297557, EPI_ISL_3297558 | Wales Specialist Virology Centre Sequencing lab: Pathogen Genomics Unit | Public Health Wales Microbiology Cardiff Wales Specialist Virology Centre | Catherine Moore, Johnathan Evans, Laura Gifford, Malorie Perry, Simon Cottrell, Angela Marchbank, Alec Birchley, Alexander Adams, Amy Gaskin, Bree Gatica-Wilcox, Jason Coombes, Joel Southgate, Lauren Gilbert, Lee Graham, Nicole Pacchiarini, Sara Kumziene-Summerhayes, Sarah Taylor, Sophie Jones, Sara Rey, Matthew Bull, Joanne Watkins, Sally Corden, Tom Connor |
| EPI_ISL_3297559 | Viollier AG | Department of Biosystems Science and Engineering, ETH Zürich | Christian Beisel, Sarah Nadeau, Chaoran Chen, Ivan Topolsky, Kim Philipp Jablonski, Lara Fuhrmann, Rebecca Denes, Mirjam Feldkamp, Ina Nissen, Natascha Santacroce, Elodie Burcklen, Christiane Beckmann, Maurice Redondo, Olivier Kobel, Christoph Noppen, Niko Beerenwinkel, Tanja Stadler |
| EPI_ISL_3297560 | Wales Specialist Virology Centre Sequencing lab: Pathogen Genomics Unit | Public Health Wales Microbiology Cardiff Wales Specialist Virology Centre | Catherine Moore, Johnathan Evans, Laura Gifford, Malorie Perry, Simon Cottrell, Angela Marchbank, Alec Birchley, Alexander Adams, Amy Gaskin, Bree Gatica-Wilcox, Jason Coombes, Joel Southgate, Lauren Gilbert, Lee Graham, Nicole Pacchiarini, Sara Kumziene-Summerhayes, Sarah Taylor, Sophie Jones, Sara Rey, Matthew Bull, Joanne Watkins, Sally Corden, Tom Connor |
| EPI_ISL_3297561 | Viollier AG | Department of Biosystems Science and Engineering, ETH Zürich | Christian Beisel, Sarah Nadeau, Chaoran Chen, Ivan Topolsky, Kim Philipp Jablonski, Lara Fuhrmann, Rebecca Denes, Mirjam Feldkamp, Ina Nissen, Natascha Santacroce, Elodie Burcklen, Christiane Beckmann, Maurice Redondo, Olivier Kobel, Christoph Noppen, Niko Beerenwinkel, Tanja Stadler |
| EPI_ISL_3297562 | Wales Specialist Virology Centre Sequencing lab: Pathogen Genomics Unit | Public Health Wales Microbiology Cardiff Wales Specialist Virology Centre | Catherine Moore, Johnathan Evans, Laura Gifford, Malorie Perry, Simon Cottrell, Angela Marchbank, Alec Birchley, Alexander Adams, Amy Gaskin, Bree Gatica-Wilcox, Jason Coombes, Joel Southgate, Lauren Gilbert, Lee Graham, Nicole Pacchiarini, Sara Kumziene-Summerhayes, Sarah Taylor, Sophie Jones, Sara Rey, Matthew Bull, Joanne Watkins, Sally Corden, Tom Connor |
| EPI_ISL_3297563 | Viollier AG | Department of Biosystems Science and Engineering, ETH Zürich | Christian Beisel, Sarah Nadeau, Chaoran Chen, Ivan Topolsky, Kim Philipp Jablonski, Lara Fuhrmann, Rebecca Denes, Mirjam Feldkamp, Ina Nissen, Natascha Santacroce, Elodie Burcklen, Christiane Beckmann, Maurice Redondo, Olivier Kobel, Christoph Noppen, Niko Beerenwinkel, Tanja Stadler |
| EPI_ISL_3297564, EPI_ISL_3297565 | Wales Specialist Virology Centre Sequencing lab: Pathogen Genomics Unit | Public Health Wales Microbiology Cardiff Wales Specialist Virology Centre | Catherine Moore, Johnathan Evans, Laura Gifford, Malorie Perry, Simon Cottrell, Angela Marchbank, Alec Birchley, Alexander Adams, Amy Gaskin, Bree Gatica-Wilcox, Jason Coombes, Joel Southgate, Lauren Gilbert, Lee Graham, Nicole Pacchiarini, Sara Kumziene-Summerhayes, Sarah Taylor, Sophie Jones, Sara Rey, Matthew Bull, Joanne Watkins, Sally Corden, Tom Connor |
| EPI_ISL_3297566 | Viollier AG | Department of Biosystems Science and Engineering, ETH Zürich | Christian Beisel, Sarah Nadeau, Chaoran Chen, Ivan Topolsky, Kim Philipp Jablonski, Lara Fuhrmann, Rebecca Denes, Mirjam Feldkamp, Ina Nissen, Natascha Santacroce, Elodie Burcklen, Christiane Beckmann, Maurice Redondo, Olivier Kobel, Christoph Noppen, Niko Beerenwinkel, Tanja Stadler |
| EPI_ISL_3297567 | Wales Specialist Virology Centre Sequencing lab: Pathogen Genomics Unit | Public Health Wales Microbiology Cardiff Wales Specialist Virology Centre | Catherine Moore, Johnathan Evans, Laura Gifford, Malorie Perry, Simon Cottrell, Angela Marchbank, Alec Birchley, Alexander Adams, Amy Gaskin, Bree Gatica-Wilcox, Jason Coombes, Joel Southgate, Lauren Gilbert, Lee Graham, Nicole Pacchiarini, Sara Kumziene-Summerhayes, Sarah Taylor, Sophie Jones, Sara Rey, Matthew Bull, Joanne Watkins, Sally Corden, Tom Connor |
| EPI_ISL_3297568 | Viollier AG | Department of Biosystems Science and Engineering, ETH Zürich | Christian Beisel, Sarah Nadeau, Chaoran Chen, Ivan Topolsky, Kim Philipp Jablonski, Lara Fuhrmann, Rebecca Denes, Mirjam Feldkamp, Ina Nissen, Natascha Santacroce, Elodie Burcklen, Christiane Beckmann, Maurice Redondo, Olivier Kobel, Christoph Noppen, Niko Beerenwinkel, Tanja Stadler |
| EPI_ISL_3297569, EPI_ISL_3297570, EPI_ISL_3297571, EPI_ISL_3297572, EPI_ISL_3297573 | Wales Specialist Virology Centre Sequencing lab: Pathogen Genomics Unit | Public Health Wales Microbiology Cardiff Wales Specialist Virology Centre | Catherine Moore, Johnathan Evans, Laura Gifford, Malorie Perry, Simon Cottrell, Angela Marchbank, Alec Birchley, Alexander Adams, Amy Gaskin, Bree Gatica-Wilcox, Jason Coombes, Joel Southgate, Lauren Gilbert, Lee Graham, Nicole Pacchiarini, Sara Kumziene-Summerhayes, Sarah Taylor, Sophie Jones, Sara Rey, Matthew Bull, Joanne Watkins, Sally Corden, Tom Connor |
| EPI_ISL_3297574 | Viollier AG | Department of Biosystems Science and Engineering, ETH Zürich | Christian Beisel, Sarah Nadeau, Chaoran Chen, Ivan Topolsky, Kim Philipp Jablonski, Lara Fuhrmann, Rebecca Denes, Mirjam Feldkamp, Ina Nissen, Natascha Santacroce, Elodie Burcklen, Christiane Beckmann, Maurice Redondo, Olivier Kobel, Christoph Noppen, Niko Beerenwinkel, Tanja Stadler |
| EPI_ISL_3297575 | Wales Specialist Virology Centre Sequencing lab: Pathogen Genomics Unit | Public Health Wales Microbiology Cardiff Wales Specialist Virology Centre | Catherine Moore, Johnathan Evans, Laura Gifford, Malorie Perry, Simon Cottrell, Angela Marchbank, Alec Birchley, Alexander Adams, Amy Gaskin, Bree Gatica-Wilcox, Jason Coombes, Joel Southgate, Lauren Gilbert, Lee Graham, Nicole Pacchiarini, Sara Kumziene-Summerhayes, Sarah Taylor, Sophie Jones, Sara Rey, Matthew Bull, Joanne Watkins, Sally Corden, Tom Connor |
| EPI_ISL_3297576, EPI_ISL_3297577 | Viollier AG | Department of Biosystems Science and Engineering, ETH Zürich | Christian Beisel, Sarah Nadeau, Chaoran Chen, Ivan Topolsky, Kim Philipp Jablonski, Lara Fuhrmann, Rebecca Denes, Mirjam Feldkamp, Ina Nissen, Natascha Santacroce, Elodie Burcklen, Christiane Beckmann, Maurice Redondo, Olivier Kobel, Christoph Noppen, Niko Beerenwinkel, Tanja Stadler |
| EPI_ISL_3297578, EPI_ISL_3297580, EPI_ISL_3297581, EPI_ISL_3297582, EPI_ISL_3297583, EPI_ISL_3297584, EPI_ISL_3297585, EPI_ISL_3297586, EPI_ISL_3297587, EPI_ISL_3297588, EPI_ISL_3297589, EPI_ISL_3297590, EPI_ISL_3297591, EPI_ISL_3297592, EPI_ISL_3297593, EPI_ISL_3297594, EPI_ISL_3297595, EPI_ISL_3297596, EPI_ISL_3297597, EPI_ISL_3297598, EPI_ISL_3297599, EPI_ISL_3297600, EPI_ISL_3297601, EPI_ISL_3297602, EPI_ISL_3297603, EPI_ISL_3297604, EPI_ISL_3297605, EPI_ISL_3297606, EPI_ISL_3297607, EPI_ISL_3297608, EPI_ISL_3297609, EPI_ISL_3297610, EPI_ISL_3297611, EPI_ISL_3297612, EPI_ISL_3297613, EPI_ISL_3297614, EPI_ISL_3297615, EPI_ISL_3297616, EPI_ISL_3297617, EPI_ISL_3297618, EPI_ISL_3297619, EPI_ISL_3297620, EPI_ISL_3297621, EPI_ISL_3297622, EPI_ISL_3297623, EPI_ISL_3297624, EPI_ISL_3297625, EPI_ISL_3297626, EPI_ISL_3297627, EPI_ISL_3297628, EPI_ISL_3297629, EPI_ISL_3297630, EPI_ISL_3297631, EPI_ISL_3297632, EPI_ISL_3297633, EPI_ISL_3297634, EPI_ISL_3297635, EPI_ISL_3297636, EPI_ISL_3297637, EPI_ISL_3297638, EPI_ISL_3297639, EPI_ISL_3297640, EPI_ISL_3297641, EPI_ISL_3297642, EPI_ISL_3297643, EPI_ISL_3297644, EPI_ISL_3297645, EPI_ISL_3297646, EPI_ISL_3297647, EPI_ISL_3297648, EPI_ISL_3297649, EPI_ISL_3297650, EPI_ISL_3297651, EPI_ISL_3297652, EPI_ISL_3297653, EPI_ISL_3297654, EPI_ISL_3297655, EPI_ISL_3297656, EPI_ISL_3297657, EPI_ISL_3297658, EPI_ISL_3297659, EPI_ISL_3297660, EPI_ISL_3297661, EPI_ISL_3297662, EPI_ISL_3297663, EPI_ISL_3297664, EPI_ISL_3297665, EPI_ISL_3297666, EPI_ISL_3297667, EPI_ISL_3297668, EPI_ISL_3297669, EPI_ISL_3297670, EPI_ISL_3297671, EPI_ISL_3297672, EPI_ISL_3297673, EPI_ISL_3297674, EPI_ISL_3297675, EPI_ISL_3297676, EPI_ISL_3297677, EPI_ISL_3297678, EPI_ISL_3297679, EPI_ISL_3297680, EPI_ISL_3297681, EPI_ISL_3297682, EPI_ISL_3297683, EPI_ISL_3297684, EPI_ISL_3297685, EPI_ISL_3297686, EPI_ISL_3297687, EPI_ISL_3297688, EPI_ISL_3297689, EPI_ISL_3297690, EPI_ISL_3297691, EPI_ISL_3297692, EPI_ISL_3297693, EPI_ISL_3297694, EPI_ISL_3297695, EPI_ISL_3297696, EPI_ISL_3297697, EPI_ISL_3297698, EPI_ISL_3297699, EPI_ISL_3297700, EPI_ISL_3297701, EPI_ISL_3297702, EPI_ISL_3297703, EPI_ISL_3297704, EPI_ISL_3297705, EPI_ISL_3297706, EPI_ISL_3297707, EPI_ISL_3297708, EPI_ISL_3297709, EPI_ISL_3297710, EPI_ISL_3297711, EPI_ISL_3297712, EPI_ISL_3297713, EPI_ISL_3297714, EPI_ISL_3297715, EPI_ISL_3297716, EPI_ISL_3297717, EPI_ISL_3297718, EPI_ISL_3297719, EPI_ISL_3297720, EPI_ISL_3297721, EPI_ISL_3297722, EPI_ISL_3297723, EPI_ISL_3297724, EPI_ISL_3297725, EPI_ISL_3297726, EPI_ISL_3297727, EPI_ISL_3297728, EPI_ISL_3297729, EPI_ISL_3297730, EPI_ISL_3297731, EPI_ISL_3297732, EPI_ISL_3297733, EPI_ISL_3297734, EPI_ISL_3297735, EPI_ISL_3297736, EPI_ISL_3297737, EPI_ISL_3297738, EPI_ISL_3297739, EPI_ISL_3297740, EPI_ISL_3297741, EPI_ISL_3297742, EPI_ISL_3297743, EPI_ISL_3297744, EPI_ISL_3297745, EPI_ISL_3297746, EPI_ISL_3297747, EPI_ISL_3297748, EPI_ISL_3297749, EPI_ISL_3297750, EPI_ISL_3297751, EPI_ISL_3297752, EPI_ISL_3297753, EPI_ISL_3297754, EPI_ISL_3297755, EPI_ISL_3297756, EPI_ISL_3297757, EPI_ISL_3297758, EPI_ISL_3297759, EPI_ISL_3297760, EPI_ISL_3297761, EPI_ISL_3297762, EPI_ISL_3297763, EPI_ISL_3297764, EPI_ISL_3297765, EPI_ISL_3297766, EPI_ISL_3297767, EPI_ISL_3297768, EPI_ISL_3297769, EPI_ISL_3297770, EPI_ISL_3297771, EPI_ISL_3297772, EPI_ISL_3297773, EPI_ISL_3297774, EPI_ISL_3297775, EPI_ISL_3297776, EPI_ISL_3297777, EPI_ISL_3297778, EPI_ISL_3297779, EPI_ISL_3297780, EPI_ISL_3297781, EPI_ISL_3297782, EPI_ISL_3297783, EPI_ISL_3297784, EPI_ISL_3297785, EPI_ISL_3297786, EPI_ISL_3297787, EPI_ISL_3297788, EPI_ISL_3297789, EPI_ISL_3297790, EPI_ISL_3297791, EPI_ISL_3297792, EPI_ISL_3297793, EPI_ISL_3297794, EPI_ISL_3297795, EPI_ISL_3297796, EPI_ISL_3297797, EPI_ISL_3297798, EPI_ISL_3297799, EPI_ISL_3297800, EPI_ISL_3297801, EPI_ISL_3297802, EPI_ISL_3297803, EPI_ISL_3297804, EPI_ISL_3297805, EPI_ISL_3297806, EPI_ISL_3297807, EPI_ISL_3297808, EPI_ISL_3297809, EPI_ISL_3297810, EPI_ISL_3297811, EPI_ISL_3297812, EPI_ISL_3297813, EPI_ISL_3297814, EPI_ISL_3297815, EPI_ISL_3297816, EPI_ISL_3297817, EPI_ISL_3297818, EPI_ISL_3297819, EPI_ISL_3297820, EPI_ISL_3297821, EPI_ISL_3297822, EPI_ISL_3297823, EPI_ISL_3297824, EPI_ISL_3297825, EPI_ISL_3297826, EPI_ISL_3297827, EPI_ISL_3297828, EPI_ISL_3297829, EPI_ISL_3297830, EPI_ISL_3297831, EPI_ISL_3297832, EPI_ISL_3297833, EPI_ISL_3297834, EPI_ISL_3297835, EPI_ISL_3297836, EPI_ISL_3297837, EPI_ISL_3297838, EPI_ISL_3297839, EPI_ISL_3297840, EPI_ISL_3297841, EPI_ISL_3297842, EPI_ISL_3297843, EPI_ISL_3297844, EPI_ISL_3297845, EPI_ISL_3297846, EPI_ISL_3297847, EPI_ISL_3297848, EPI_ISL_3297849, EPI_ISL_3297850, EPI_ISL_3297851, EPI_ISL_3297852, EPI_ISL_3297853, EPI_ISL_3297854, EPI_ISL_3297855, EPI_ISL_3297856, EPI_ISL_3297857, EPI_ISL_3297858, EPI_ISL_3297859, EPI_ISL_3297860, EPI_ISL_3297861, EPI_ISL_3297862, EPI_ISL_3297863, EPI_ISL_3297864, EPI_ISL_3297865, EPI_ISL_3297866, EPI_ISL_3297867, EPI_ISL_3297868, EPI_ISL_3297869, EPI_ISL_3297870, EPI_ISL_3297871, EPI_ISL_3297872, EPI_ISL_3297873, EPI_ISL_3297874, EPI_ISL_3297875, EPI_ISL_3297876, EPI_ISL_3297877, EPI_ISL_3297878, EPI_ISL_3297879, EPI_ISL_3297880, EPI_ISL_3297881, EPI_ISL_3297882, EPI_ISL_3297883 | | | |
| see above | Wales Specialist Virology Centre Sequencing lab: Pathogen Genomics Unit | Public Health Wales Microbiology Cardiff Wales Specialist Virology Centre | Catherine Moore, Johnathan Evans, Laura Gifford, Malorie Perry, Simon Cottrell, Angela Marchbank, Alec Birchley, Alexander Adams, Amy Gaskin, Bree Gatica-Wilcox, Jason Coombes, Joel Southgate, Lauren Gilbert, Lee Graham, Nicole Pacchiarini, Sara Kumziene-Summerhayes, Sarah Taylor, Sophie Jones, Sara Rey, Matthew Bull, Joanne Watkins, Sally Corden, Tom Connor |
| EPI_ISL_3297884, EPI_ISL_3297885 | Originating lab: Wales Specialist Virology Centre Sequencing lab: Pathogen Genomics Unit | Public Health Wales Microbiology Cardiff Wales Specialist Virology Centre | Catherine Moore, Johnathan Evans, Laura Gifford, Malorie Perry, Simon Cottrell, Angela Marchbank, Alec Birchley, Alexander Adams, Amy Gaskin, Bree Gatica-Wilcox, Jason Coombes, Joel Southgate, Lauren Gilbert, Lee Graham, Nicole Pacchiarini, Sara Kumziene-Summerhayes, Sarah Taylor, Sophie Jones, Sara Rey, Matthew Bull, Joanne Watkins, Sally Corden, Tom Connor |
| EPI_ISL_3297886, EPI_ISL_3297887, EPI_ISL_3297888, EPI_ISL_3297889, EPI_ISL_3297890, EPI_ISL_3297891, EPI_ISL_3297892, EPI_ISL_3297893, EPI_ISL_3297894, EPI_ISL_3297895, EPI_ISL_3297896, EPI_ISL_3297897, EPI_ISL_3297898, EPI_ISL_3297899, EPI_ISL_3297900, EPI_ISL_3297901, EPI_ISL_3297902, EPI_ISL_3297903, EPI_ISL_3297904, EPI_ISL_3297905, EPI_ISL_3297906, EPI_ISL_3297907, EPI_ISL_3297908, EPI_ISL_3297909, EPI_ISL_3297910, EPI_ISL_3297911, EPI_ISL_3297912, EPI_ISL_3297913, EPI_ISL_3297914, EPI_ISL_3297915, EPI_ISL_3297916, EPI_ISL_3297917, EPI_ISL_3297918, EPI_ISL_3297919, EPI_ISL_3297920, EPI_ISL_3297921, EPI_ISL_3297922, EPI_ISL_3297923, EPI_ISL_3297924, EPI_ISL_3297925, EPI_ISL_3297926, EPI_ISL_3297927, EPI_ISL_3297928, EPI_ISL_3297929, EPI_ISL_3297930, EPI_ISL_3297931, EPI_ISL_3297932, EPI_ISL_3297933, EPI_ISL_3297934, EPI_ISL_3297935, EPI_ISL_3297936, EPI_ISL_3297937, EPI_ISL_3297938, EPI_ISL_3297939, EPI_ISL_3297940, EPI_ISL_3297941, EPI_ISL_3297942, EPI_ISL_3297943, EPI_ISL_3297944, EPI_ISL_3297945, EPI_ISL_3297946, EPI_ISL_3297947, EPI_ISL_3297948, EPI_ISL_3297949, EPI_ISL_3297950, EPI_ISL_3297951, EPI_ISL_3297952, EPI_ISL_3297953, EPI_ISL_3297954, EPI_ISL_3297955, EPI_ISL_3297956, EPI_ISL_3297957, EPI_ISL_3297958, EPI_ISL_3297959, EPI_ISL_3297960, EPI_ISL_3297961, EPI_ISL_3297962, EPI_ISL_3297963, EPI_ISL_3297964, EPI_ISL_3297965, EPI_ISL_3297966, EPI_ISL_3297967, EPI_ISL_3297968, EPI_ISL_3297969, EPI_ISL_3297970, EPI_ISL_3297971, EPI_ISL_3297972, EPI_ISL_3297973, EPI_ISL_3297974, EPI_ISL_3297975, EPI_ISL_3297976, EPI_ISL_3297977, EPI_ISL_3297978, EPI_ISL_3297979, EPI_ISL_3297980, EPI_ISL_3297981, EPI_ISL_3297982, EPI_ISL_3297983, EPI_ISL_3297984, EPI_ISL_3297985, EPI_ISL_3297986, EPI_ISL_3297987, EPI_ISL_3297988, EPI_ISL_3297989, EPI_ISL_3297990, EPI_ISL_3297991, EPI_ISL_3297992, EPI_ISL_3297993, EPI_ISL_3297994, EPI_ISL_3297995, EPI_ISL_3297996, EPI_ISL_3297997, EPI_ISL_3297998, EPI_ISL_3297999, EPI_ISL_3298000 | | | |
| see above | Centre for Enzyme Innovation, University of Portsmouth / Translational Research Laboratory, Portsmouth Hospitals NHS Trust / Hampshire Hospitals NHS Foundation Trust / Bournemouth University / University Hospitals Dorset NHS Foundation Trust / Brighton and Sussex University Hospitals NHS Trust / Dartford and Gravesham NHS Trust / Isle of Wight NHS Trust / Maidstone and Tunbridge Wells NHS Trust / University Hospital Southampton NHS Foundation Trust | COVID-19 Genomics UK (COG-UK) Consortium | Samuel Robson, Angela Beckett, Salman Goudarzi, Christopher Fearn, Kate Cook, Katie Loveson, Sharon Glaysher, Scott Elliott, Kelly Bicknell, Sarah Wyllie, Allyson Lloyd, Robert Impey, Anoop Chauhan, Stephen Kidd, Nathan Moore, Nick Cortes, Claire Thomas, Anna Mantzaouratou, Sarah Buchan, Magdalena Barrow, Andrew Butt, Liz Sheridan, Jonnie Seymour, Dorian Crudgington, Ben Macklin, Mohammed Hassan-Ibrahim, Cassandra Malone, Benjamin Cogger, Kevin Tucker, Samirakhon Raupova, Rachael Jeremiah, Anibolina Castigador, Emily Macnaughton, Karen Withell, Kordo Saeed, Jacqui Prieto, Adhyana Mahanama, Buddhini Samaraweera, Siona Silvieri, Emanuela Pelosi, Eleri Wilson-Davies, Sarah Jeremiah, Helen Wheeler, Matthew Harvey, Thea Sass, Helen Umpleby, Stephen Aplin |
| EPI_ISL_3298001, EPI_ISL_3298002, EPI_ISL_3298003, EPI_ISL_3298004, EPI_ISL_3298005, EPI_ISL_3298006, EPI_ISL_3298007, EPI_ISL_3298008, EPI_ISL_3298009, EPI_ISL_3298010, EPI_ISL_3298011, EPI_ISL_3298012, EPI_ISL_3298013, EPI_ISL_3298014, EPI_ISL_3298015, EPI_ISL_3298016, EPI_ISL_3298017, EPI_ISL_3298018, EPI_ISL_3298019, EPI_ISL_3298020, EPI_ISL_3298021, EPI_ISL_3298022, EPI_ISL_3298023, EPI_ISL_3298024, EPI_ISL_3298025, EPI_ISL_3298026, EPI_ISL_3298027, EPI_ISL_3298028, EPI_ISL_3298029, EPI_ISL_3298030, EPI_ISL_3298031, EPI_ISL_3298032, EPI_ISL_3298033, EPI_ISL_3298034, EPI_ISL_3298035, EPI_ISL_3298036, EPI_ISL_3298037, EPI_ISL_3298038, EPI_ISL_3298039, EPI_ISL_3298040, EPI_ISL_3298041, EPI_ISL_3298042, EPI_ISL_3298043, EPI_ISL_3298044, EPI_ISL_3298045, EPI_ISL_3298046, EPI_ISL_3298047, EPI_ISL_3298048, EPI_ISL_3298049, EPI_ISL_3298050, EPI_ISL_3298051, EPI_ISL_3298052, EPI_ISL_3298053, EPI_ISL_3298054, EPI_ISL_3298055, EPI_ISL_3298056, EPI_ISL_3298057, EPI_ISL_3298058, EPI_ISL_3298059, EPI_ISL_3298060, EPI_ISL_3298061, EPI_ISL_3298062, EPI_ISL_3298064, EPI_ISL_3298067, EPI_ISL_3298068, EPI_ISL_3298069, EPI_ISL_3298070, EPI_ISL_3298071, EPI_ISL_3298072, EPI_ISL_3298073, EPI_ISL_3298074, EPI_ISL_3298075, EPI_ISL_3298076, EPI_ISL_3298077, EPI_ISL_3298078, EPI_ISL_3298079, EPI_ISL_3298080, EPI_ISL_3298081, EPI_ISL_3298082, EPI_ISL_3298083, EPI_ISL_3298084, EPI_ISL_3298085, EPI_ISL_3298086, EPI_ISL_3298087, EPI_ISL_3298088, EPI_ISL_3298089, EPI_ISL_3298090, EPI_ISL_3298091, EPI_ISL_3298092, EPI_ISL_3298093, EPI_ISL_3298094, EPI_ISL_3298095, EPI_ISL_3298096, EPI_ISL_3298097, EPI_ISL_3298098, EPI_ISL_3298099, EPI_ISL_3298100, EPI_ISL_3298101, EPI_ISL_3298102, EPI_ISL_3298103, EPI_ISL_3298104, EPI_ISL_3298105, EPI_ISL_3298106, EPI_ISL_3298107, EPI_ISL_3298108, EPI_ISL_3298109, EPI_ISL_3298110, EPI_ISL_3298111, EPI_ISL_3298112, EPI_ISL_3298114, EPI_ISL_3298115, EPI_ISL_3298116, EPI_ISL_3298117, EPI_ISL_3298118, EPI_ISL_3298120, EPI_ISL_3298121, EPI_ISL_3298122, EPI_ISL_3298123, EPI_ISL_3298124, EPI_ISL_3298125, EPI_ISL_3298127, EPI_ISL_3298128, EPI_ISL_3298129, EPI_ISL_3298130, EPI_ISL_3298131, EPI_ISL_3298132, EPI_ISL_3298133, EPI_ISL_3298134, EPI_ISL_3298135, EPI_ISL_3298136, EPI_ISL_3298137, EPI_ISL_3298138, EPI_ISL_3298139, EPI_ISL_3298140, EPI_ISL_3298141, EPI_ISL_3298142, EPI_ISL_3298143, EPI_ISL_3298144, EPI_ISL_3298145, EPI_ISL_3298146, EPI_ISL_3298147, EPI_ISL_3298148, EPI_ISL_3298149, EPI_ISL_3298150, EPI_ISL_3298151, EPI_ISL_3298152, EPI_ISL_3298153, EPI_ISL_3298154, EPI_ISL_3298155, EPI_ISL_3298156, EPI_ISL_3298157, EPI_ISL_3298158, EPI_ISL_3298159, EPI_ISL_3298160, EPI_ISL_3298161, EPI_ISL_3298162, EPI_ISL_3298163, EPI_ISL_3298164, EPI_ISL_3298165, EPI_ISL_3298166, EPI_ISL_3298167, EPI_ISL_3298168, EPI_ISL_3298169, EPI_ISL_3298170, EPI_ISL_3298171, EPI_ISL_3298172, EPI_ISL_3298173, EPI_ISL_3298174, EPI_ISL_3298175, EPI_ISL_3298176, EPI_ISL_3298177, EPI_ISL_3298178, EPI_ISL_3298179, EPI_ISL_3298180, EPI_ISL_3298181, EPI_ISL_3298182, EPI_ISL_3298183, EPI_ISL_3298184, EPI_ISL_3298185, EPI_ISL_3298186, EPI_ISL_3298187, EPI_ISL_3298188, EPI_ISL_3298189, EPI_ISL_3298190, EPI_ISL_3298191, EPI_ISL_3298192, EPI_ISL_3298193, EPI_ISL_3298194 | | | |
| see above | Virology Department, Sheffield Teaching Hospitals NHS Foundation Trust/Department of Infection, Immunity and Cardiovascular Disease, The Medical School, University of Sheffield | COVID-19 Genomics UK (COG-UK) Consortium | Thushan de Silva, Matthew Parker, Nikki Smith, Adri Angyal, Rebecca Brown, Luke Green, Rachel Tucker, Paul Parsons, Danielle Groves, Katie Johnson, Laura Carrilero, Alex Keeley, Dave Partridge, Matthew Wyles, Benjamin Lindsey, Mehmet Yavuz, Mohammad Raza, Cariad Evans |
| EPI_ISL_3298196, EPI_ISL_3298197, EPI_ISL_3298198, EPI_ISL_3298199, EPI_ISL_3298200, EPI_ISL_3298201, EPI_ISL_3298202, EPI_ISL_3298203, EPI_ISL_3298204, EPI_ISL_3298205, EPI_ISL_3298206, EPI_ISL_3298207, EPI_ISL_3298208, EPI_ISL_3298214, EPI_ISL_3298215, EPI_ISL_3298216, EPI_ISL_3298217, EPI_ISL_3298218, EPI_ISL_3298219, EPI_ISL_3298220, EPI_ISL_3298221, EPI_ISL_3298222, EPI_ISL_3298223, EPI_ISL_3298224, EPI_ISL_3298225, EPI_ISL_3298226, EPI_ISL_3298228, EPI_ISL_3298229, EPI_ISL_3298230, EPI_ISL_3298231, EPI_ISL_3298232, EPI_ISL_3298233 | | | |
| see above | U.O. Microbiologia Laboratorio Unico Centro Servizi - AUSL della Romagna | U.O. Microbiologia, Laboratorio Unico Centro Servizi - AUSL della Romagna | Giorgio Dirani |
| EPI_ISL_3298235, EPI_ISL_3298236, EPI_ISL_3298237, EPI_ISL_3298238, EPI_ISL_3298239, EPI_ISL_3298240, EPI_ISL_3298241, EPI_ISL_3298242, EPI_ISL_3298244, EPI_ISL_3298245, EPI_ISL_3298246, EPI_ISL_3298247, EPI_ISL_3298248, EPI_ISL_3298249, EPI_ISL_3298251, EPI_ISL_3298252, EPI_ISL_3298253, EPI_ISL_3298254, EPI_ISL_3298255, EPI_ISL_3298294, EPI_ISL_3298295, EPI_ISL_3298296, EPI_ISL_3298297 | | | |
| see above | Wyoming Public Health Laboratory | Wyoming Public Health Laboratory | Jim Mildenberger, Taylor Fearing, Channing Weber, Ashley Norberg, Chayse Rowley, Marley Goetz, Brian Dominguez, Elliot Thomasson, Sam Britz, Cari Sloma, Robert Petit, and Rob Christensen |

| | | | |
|---|---|---|---|
| EPI_ISL_3298318 | HOSPITAL DR. FERNANDO ESCALANTE PRADILLA | Inciensa, Instituto Costarricense de Investigación y Enseñanza en Nutrición y Salud | Francisco Duarte, Hebleen Porras, Claudio Soto-Garita, Estela Cordero, Adriana Godínez, Melany Calderón, José Luis Vargas, Mariela Gutiérrez, Joselyn Prado & Lizeth Blanco |
| EPI_ISL_3298329 | HOSPITAL MEXICO | Inciensa, Instituto Costarricense de Investigación y Enseñanza en Nutrición y Salud | Francisco Duarte, Hebleen Porras, Claudio Soto-Garita, Estela Cordero, Adriana Godínez, Melany Calderón, José Luis Vargas, Mariela Gutiérrez, Joselyn Prado & Juan Carlos Villalobos |
| EPI_ISL_3298339 | AREA DE SALUD GUATUSO | Inciensa, Instituto Costarricense de Investigación y Enseñanza en Nutrición y Salud | Francisco Duarte, Hebleen Porras, Claudio Soto-Garita, Estela Cordero, Adriana Godínez, Melany Calderón, José Luis Vargas, Mariela Gutiérrez, Joselyn Prado & Francisco Chacón |
| EPI_ISL_3298343 | AREA DE SALUD MATA REDONDA-HOSPITAL - CLINICA DR. MORENO CAÑAS | Inciensa, Instituto Costarricense de Investigación y Enseñanza en Nutrición y Salud | Francisco Duarte, Hebleen Porras, Claudio Soto-Garita, Estela Cordero, Adriana Godínez, Melany Calderón, José Luis Vargas, Mariela Gutiérrez, Joselyn Prado & Ricardo González |
| EPI_ISL_3298344 | AREA DE SALUD GOICOECHEA 2 - CLINICA DR. JIMENEZ NUÑEZ | Inciensa, Instituto Costarricense de Investigación y Enseñanza en Nutrición y Salud | Francisco Duarte, Hebleen Porras, Claudio Soto-Garita, Estela Cordero, Adriana Godínez, Melany Calderón, José Luis Vargas, Mariela Gutiérrez, Joselyn Prado & Juan Carlos Cartes |
| EPI_ISL_3298345 | HOSPITAL SAN VICENTE DE PAUL | Inciensa, Instituto Costarricense de Investigación y Enseñanza en Nutrición y Salud | Francisco Duarte, Hebleen Porras, Claudio Soto-Garita, Estela Cordero, Adriana Godínez, Melany Calderón, José Luis Vargas, Mariela Gutiérrez, Joselyn Prado & Silvia Sáenz |
| EPI_ISL_3298346 | HOSPITAL DR. CARLOS LUIS VALVERDE VEGA | Inciensa, Instituto Costarricense de Investigación y Enseñanza en Nutrición y Salud | Francisco Duarte, Hebleen Porras, Claudio Soto-Garita, Estela Cordero, Adriana Godínez, Melany Calderón, José Luis Vargas, Mariela Gutiérrez, Joselyn Prado & Charbel Vargas |
| EPI_ISL_3298348 | HOSPITAL SAN VITO DE COTO BRUS | Inciensa, Instituto Costarricense de Investigación y Enseñanza en Nutrición y Salud | Francisco Duarte, Hebleen Porras, Claudio Soto-Garita, Estela Cordero, Adriana Godínez, Melany Calderón, José Luis Vargas, Mariela Gutiérrez, Joselyn Prado & Yendry Ramírez |
| EPI_ISL_3298350 | AREA DE SALUD HEREDIA-VIRILLA | Inciensa, Instituto Costarricense de Investigación y Enseñanza en Nutrición y Salud | Francisco Duarte, Hebleen Porras, Claudio Soto-Garita, Estela Cordero, Adriana Godínez, Melany Calderón, José Luis Vargas, Mariela Gutiérrez, Joselyn Prado & Angie Retana |
| EPI_ISL_3298352 | HOSPITAL METROPOLITANO | Inciensa, Instituto Costarricense de Investigación y Enseñanza en Nutrición y Salud | Francisco Duarte, Hebleen Porras, Claudio Soto-Garita, Estela Cordero, Adriana Godínez, Melany Calderón, José Luis Vargas, Mariela Gutiérrez, Joselyn Prado & Margarita Lee-Lui |
| EPI_ISL_3298354 | AREA DE SALUD FORTUNA | Inciensa, Instituto Costarricense de Investigación y Enseñanza en Nutrición y Salud | Francisco Duarte, Hebleen Porras, Claudio Soto-Garita, Estela Cordero, Adriana Godínez, Melany Calderón, José Luis Vargas, Mariela Gutiérrez, Joselyn Prado & Carolina Arrieta |
| EPI_ISL_3298357 | AREA DE SALUD ALAJUELITA | Inciensa, Instituto Costarricense de Investigación y Enseñanza en Nutrición y Salud | Francisco Duarte, Hebleen Porras, Claudio Soto-Garita, Estela Cordero, Adriana Godínez, Melany Calderón, José Luis Vargas, Mariela Gutiérrez & Joselyn Prado |
| EPI_ISL_3298362 | AREA DE SALUD SANTA CRUZ | Inciensa, Instituto Costarricense de Investigación y Enseñanza en Nutrición y Salud | Francisco Duarte, Hebleen Porras, Claudio Soto-Garita, Estela Cordero, Adriana Godínez, Melany Calderón, José Luis Vargas, Mariela Gutiérrez, Joselyn Prado & Fabiola Gutiérrez |
| EPI_ISL_3298364 | AREA DE SALUD PAQUERA | Inciensa, Instituto Costarricense de Investigación y Enseñanza en Nutrición y Salud | Francisco Duarte, Hebleen Porras, Claudio Soto-Garita, Estela Cordero, Adriana Godínez, Melany Calderón, José Luis Vargas, Mariela Gutiérrez, Joselyn Prado & María José Gómez |
| EPI_ISL_3298367 | HOSPITAL SAN RAFAEL DE ALAJUELA | Inciensa, Instituto Costarricense de Investigación y Enseñanza en Nutrición y Salud | Francisco Duarte, Hebleen Porras, Claudio Soto-Garita, Estela Cordero, Adriana Godínez, Melany Calderón, José Luis Vargas, Mariela Gutiérrez, Joselyn Prado & Christian Sánchez |
| EPI_ISL_3298368, EPI_ISL_3298369 | AREA DE SALUD GUATUSO | Inciensa, Instituto Costarricense de Investigación y Enseñanza en Nutrición y Salud | Francisco Duarte, Hebleen Porras, Claudio Soto-Garita, Estela Cordero, Adriana Godínez, Melany Calderón, José Luis Vargas, Mariela Gutiérrez, Joselyn Prado & Francisco Chacón |
| EPI_ISL_3298370 | AREA DE SALUD FORTUNA | Inciensa, Instituto Costarricense de Investigación y Enseñanza en Nutrición y Salud | Francisco Duarte, Hebleen Porras, Claudio Soto-Garita, Estela Cordero, Adriana Godínez, Melany Calderón, José Luis Vargas, Mariela Gutiérrez, Joselyn Prado & Carolina Arrieta |
| EPI_ISL_3298372, EPI_ISL_3298373 | AREA DE SALUD GUATUSO | Inciensa, Instituto Costarricense de Investigación y Enseñanza en Nutrición y Salud | Francisco Duarte, Hebleen Porras, Claudio Soto-Garita, Estela Cordero, Adriana Godínez, Melany Calderón, José Luis Vargas, Mariela Gutiérrez, Joselyn Prado & Francisco Chacón |
| EPI_ISL_3298384, EPI_ISL_3298385 | Dipartimento di Medicina di Laboratorio, Azienda sanitaria universitaria Friuli Centrale (ASU FC) | Dipartimento di Medicina di Laboratorio, Azienda sanitaria universitaria Friuli Centrale (ASU FC) | Corrado Pipan, Catia Mio, Chiara Dal Secco, Stefania Marzinotto, Francesco Curcio |
| EPI_ISL_3298387, EPI_ISL_3298388, EPI_ISL_3298389, EPI_ISL_3298390, EPI_ISL_3298391, EPI_ISL_3298392, EPI_ISL_3298393, EPI_ISL_3298394, EPI_ISL_3298395, EPI_ISL_3298396, EPI_ISL_3298397, EPI_ISL_3298398, EPI_ISL_3298399, EPI_ISL_3298400, EPI_ISL_3298401, EPI_ISL_3298402, EPI_ISL_3298403, EPI_ISL_3298404, EPI_ISL_3298405, EPI_ISL_3298406, EPI_ISL_3298407, EPI_ISL_3298408, EPI_ISL_3298409, EPI_ISL_3298410, EPI_ISL_3298411, EPI_ISL_3298412, EPI_ISL_3298413, EPI_ISL_3298414, EPI_ISL_3298415, EPI_ISL_3298416, EPI_ISL_3298417, EPI_ISL_3298418, EPI_ISL_3298419, EPI_ISL_3298420, EPI_ISL_3298421, EPI_ISL_3298422, EPI_ISL_3298423, EPI_ISL_3298424, EPI_ISL_3298425, EPI_ISL_3298426, EPI_ISL_3298427, EPI_ISL_3298428, EPI_ISL_3298429, EPI_ISL_3298430, EPI_ISL_3298431, EPI_ISL_3298432, EPI_ISL_3298433, EPI_ISL_3298434, EPI_ISL_3298435, EPI_ISL_3298436, EPI_ISL_3298437, EPI_ISL_3298438, EPI_ISL_3298439, EPI_ISL_3298440, EPI_ISL_3298441, EPI_ISL_3298442, EPI_ISL_3298443, EPI_ISL_3298444, EPI_ISL_3298445, EPI_ISL_3298446, EPI_ISL_3298447, EPI_ISL_3298448, EPI_ISL_3298449, EPI_ISL_3298450, EPI_ISL_3298451, EPI_ISL_3298452, EPI_ISL_3298453, EPI_ISL_3298454, EPI_ISL_3298455, EPI_ISL_3298456, EPI_ISL_3298457, EPI_ISL_3298458, EPI_ISL_3298459, EPI_ISL_3298460, EPI_ISL_3298461, EPI_ISL_3298462, EPI_ISL_3298463, EPI_ISL_3298464, EPI_ISL_3298465, EPI_ISL_3298466, EPI_ISL_3298467, EPI_ISL_3298468, EPI_ISL_3298469, EPI_ISL_3298470, EPI_ISL_3298471, EPI_ISL_3298472, EPI_ISL_3298473, EPI_ISL_3298474, EPI_ISL_3298475, EPI_ISL_3298476, EPI_ISL_3298477, EPI_ISL_3298478, EPI_ISL_3298479, EPI_ISL_3298480, EPI_ISL_3298481, EPI_ISL_3298482, EPI_ISL_3298483, EPI_ISL_3298484, EPI_ISL_3298485, EPI_ISL_3298486, EPI_ISL_3298487, EPI_ISL_3298488, EPI_ISL_3298489, EPI_ISL_3298490, EPI_ISL_3298491, EPI_ISL_3298492, EPI_ISL_3298493, EPI_ISL_3298494, EPI_ISL_3298495, EPI_ISL_3298496, EPI_ISL_3298497, EPI_ISL_3298498, EPI_ISL_3298499, EPI_ISL_3298500, EPI_ISL_3298501, EPI_ISL_3298502, EPI_ISL_3298503, EPI_ISL_3298504, EPI_ISL_3298505, EPI_ISL_3298506, EPI_ISL_3298507, EPI_ISL_3298508, EPI_ISL_3298509, EPI_ISL_3298510, EPI_ISL_3298511 | | | |
| see above | National Virus Reference Laboratory | National Virus Reference Laboratory | Zoe Yandle, Charlene Bennett, Gabriel Gonzalez, Michael Carr, Jonathan Dean, Cillian F De Gascun |
| EPI_ISL_3298512 | City of Milwaukee Health Department Laboratory | City of Milwaukee Health Department Laboratory | Manjeet Khubbar, Amy Bauer, Jennifer Lentz, Samantha Scott, Sanjib Bhattacharyya |
| EPI_ISL_3298513, EPI_ISL_3298514, EPI_ISL_3298515, EPI_ISL_3298516, EPI_ISL_3298517, EPI_ISL_3298518, EPI_ISL_3298519, EPI_ISL_3298520, EPI_ISL_3298521, EPI_ISL_3298522, EPI_ISL_3298523, EPI_ISL_3298524, EPI_ISL_3298525, EPI_ISL_3298526, EPI_ISL_3298527, EPI_ISL_3298528, EPI_ISL_3298529, EPI_ISL_3298530, EPI_ISL_3298531, EPI_ISL_3298532, EPI_ISL_3298533, EPI_ISL_3298534, EPI_ISL_3298535, EPI_ISL_3298536, EPI_ISL_3298537, EPI_ISL_3298538, EPI_ISL_3298539, EPI_ISL_3298540, EPI_ISL_3298541, EPI_ISL_3298542, EPI_ISL_3298543, EPI_ISL_3298544, EPI_ISL_3298545, EPI_ISL_3298546, EPI_ISL_3298547, EPI_ISL_3298548, EPI_ISL_3298549, EPI_ISL_3298550, EPI_ISL_3298551, EPI_ISL_3298552, EPI_ISL_3298553, EPI_ISL_3298554, EPI_ISL_3298555, EPI_ISL_3298556, EPI_ISL_3298557, EPI_ISL_3298558, EPI_ISL_3298559, EPI_ISL_3298560, EPI_ISL_3298561, EPI_ISL_3298562, EPI_ISL_3298563, EPI_ISL_3298564, EPI_ISL_3298565, EPI_ISL_3298566, EPI_ISL_3298567, EPI_ISL_3298568, EPI_ISL_3298569, EPI_ISL_3298570, EPI_ISL_3298571, EPI_ISL_3298572, EPI_ISL_3298573, EPI_ISL_3298574, EPI_ISL_3298575, EPI_ISL_3298576 | | | |
| see above | National Virus Reference Laboratory | National Virus Reference Laboratory | Zoe Yandle, Charlene Bennett, Gabriel Gonzalez, Michael Carr, Jonathan Dean, Cillian F De Gascun |
| EPI_ISL_3298577 | Ohio Department of Health Laboratory | Ohio Department of Health Laboratory | Holmes, Jennifer; Eric Brandt, Keoni Omura, Glen McGillivary, Caitlin McDonnell, Jade Mowery, Stephanie Mccracken, Tyler Payne, Kirtana Ramadugu, Erica Leasure, Brent Lee, Kelsey Florek, Heather Blankenship, Allison Black, Quanta Brown, and Tammy Bannerman |
| EPI_ISL_3298578, EPI_ISL_3298579, EPI_ISL_3298580, EPI_ISL_3298581, EPI_ISL_3298582, EPI_ISL_3298583, EPI_ISL_3298584, EPI_ISL_3298585, EPI_ISL_3298586, EPI_ISL_3298587, EPI_ISL_3298588, EPI_ISL_3298589, EPI_ISL_3298590, EPI_ISL_3298591, EPI_ISL_3298592, EPI_ISL_3298593, EPI_ISL_3298594, EPI_ISL_3298595, EPI_ISL_3298596, EPI_ISL_3298597, EPI_ISL_3298598, EPI_ISL_3298599, EPI_ISL_3298600, EPI_ISL_3298601, EPI_ISL_3298602, EPI_ISL_3298603, EPI_ISL_3298604, EPI_ISL_3298605, EPI_ISL_3298606, EPI_ISL_3298607, EPI_ISL_3298608, EPI_ISL_3298609, EPI_ISL_3298610, EPI_ISL_3298611, EPI_ISL_3298612, EPI_ISL_3298613, EPI_ISL_3298614, EPI_ISL_3298615, EPI_ISL_3298616, EPI_ISL_3298617, EPI_ISL_3298618, EPI_ISL_3298626, EPI_ISL_3298627, EPI_ISL_3298628, EPI_ISL_3298629, EPI_ISL_3298630, EPI_ISL_3298631, EPI_ISL_3298632, EPI_ISL_3298633, EPI_ISL_3298634, EPI_ISL_3298635, EPI_ISL_3298636 | | | |
| see above | National Virus Reference Laboratory | National Virus Reference Laboratory | Zoe Yandle, Charlene Bennett, Gabriel Gonzalez, Michael Carr, Jonathan Dean, Cillian F De Gascun |
| EPI_ISL_3298638 | Ohio Department of Health Laboratory | Ohio Department of Health Laboratory | Holmes, Jennifer; Eric Brandt, Keoni Omura, Glen McGillivary, Caitlin McDonnell, Jade Mowery, Stephanie Mccracken, Tyler Payne, Kirtana Ramadugu, Erica Leasure, Brent Lee, Kelsey Florek, Heather Blankenship, Allison Black, Quanta Brown, and Tammy Bannerman |
| EPI_ISL_3298639, EPI_ISL_3298640, EPI_ISL_3298641, EPI_ISL_3298642, EPI_ISL_3298643, EPI_ISL_3298644 | Instituto Adolfo Lutz - Regional de Santos | Instituto Adolfo Lutz, Interdiciplinary Procedures Center, Strategic Laboratory | Claudio Tavares Sacchi, Claudia Regina Gonçalves, Erica Valessa Ramos Gomes, Karoline Rodrigues Campos, Caio Vinicius Dias Lopes, Leonardo Tadeu de Araujo, Marlon Benedito Nascomento Santos |
| EPI_ISL_3298645, EPI_ISL_3298646, EPI_ISL_3298647, EPI_ISL_3298648, EPI_ISL_3298649, EPI_ISL_3298650, EPI_ISL_3298651, EPI_ISL_3298652, EPI_ISL_3298653, EPI_ISL_3298654, EPI_ISL_3298655, EPI_ISL_3298656, EPI_ISL_3298657, EPI_ISL_3298658, EPI_ISL_3298659, EPI_ISL_3298660, EPI_ISL_3298661, EPI_ISL_3298662, EPI_ISL_3298663, EPI_ISL_3298664, EPI_ISL_3298665, EPI_ISL_3298666, EPI_ISL_3298667, EPI_ISL_3298668, EPI_ISL_3298669, EPI_ISL_3298670, EPI_ISL_3298671, EPI_ISL_3298672, EPI_ISL_3298673, EPI_ISL_3298674, EPI_ISL_3298675, EPI_ISL_3298676, EPI_ISL_3298677, EPI_ISL_3298678, EPI_ISL_3298679, EPI_ISL_3298680, EPI_ISL_3298681, EPI_ISL_3298682 | | | |
| see above | U.O. Microbiologia Laboratorio Unico Centro Servizi - AUSL della Romagna | U.O. Microbiologia, Laboratorio Unico Centro Servizi - AUSL della Romagna | Giorgio Dirani |
| EPI_ISL_3298770 | IVIC | Laboratorio de Virología Molecular | Rossana C Jaspe, Carmen L Loureiro, Yoneira Sulbaran, Pierina D´Angelo, Zoila Moros, Lieska Rodríguez, José Luis Zambrano, Mariana Hidalgo, Esmeralda Vizzi, Víctor Alarcón, Domingo J Garzaro, CoViMol Group, Héctor R Rangel, Flor H Pujol |
| EPI_ISL_3298925, EPI_ISL_3298926, EPI_ISL_3298927, EPI_ISL_3298928, EPI_ISL_3298929, EPI_ISL_3298930, EPI_ISL_3298932, EPI_ISL_3298933, EPI_ISL_3298935, EPI_ISL_3298936, EPI_ISL_3298937, EPI_ISL_3298938, EPI_ISL_3298939, EPI_ISL_3298940, EPI_ISL_3298941, EPI_ISL_3298942, EPI_ISL_3298943, EPI_ISL_3298944, EPI_ISL_3298945, EPI_ISL_3298946, EPI_ISL_3298947, EPI_ISL_3298948, EPI_ISL_3298949, EPI_ISL_3298950, EPI_ISL_3298951, EPI_ISL_3298952, EPI_ISL_3298953, EPI_ISL_3298954, EPI_ISL_3298955, EPI_ISL_3298956, EPI_ISL_3298957, EPI_ISL_3298958, EPI_ISL_3298959, EPI_ISL_3298960, EPI_ISL_3298961, EPI_ISL_3298962, EPI_ISL_3298963, EPI_ISL_3298964, EPI_ISL_3298965, EPI_ISL_3298966, EPI_ISL_3298967, EPI_ISL_3298968, EPI_ISL_3298969, EPI_ISL_3298970, EPI_ISL_3298971, EPI_ISL_3298972, EPI_ISL_3298973, EPI_ISL_3298974, EPI_ISL_3298976, EPI_ISL_3298977, EPI_ISL_3298978, EPI_ISL_3298979, EPI_ISL_3298980, EPI_ISL_3298981, EPI_ISL_3298982, EPI_ISL_3298983, EPI_ISL_3298984, EPI_ISL_3298985, EPI_ISL_3298986, EPI_ISL_3298987, EPI_ISL_3298988, EPI_ISL_3298989, EPI_ISL_3298990, EPI_ISL_3298991, EPI_ISL_3298992, EPI_ISL_3298993, EPI_ISL_3298994, EPI_ISL_3298995, EPI_ISL_3298996 | | | |
| see above | Maine Health and Environmental Testing Laboratory | Tewhey Lab, The Jackson Laboratory | Matluk,N., Dewey,H., Iosue,F., Barter,M., Lynch,R., Munger,H. and Tewhey,R. |
| EPI_ISL_3299023 | Genomik Solidaritas Indonesia Laboratorium | Genomik Solidaritas Indonesia Laboratorium | Dhahlia Agustina Cahyono, Himawan Masyhuri, Vania Gavrila Wikasa, Carissa Sintca Wijaya, Gracia Felias Enos Korompis, Annisa Muthiah Sukirman, Anuraj Shankar, Louisa Markus, Meutia Ayuputeri Kumaheri, Ariel Pradipta |
| EPI_ISL_3299024 | Martin Luther King, Jr. Outpatient Center | Los Angeles County Public Health Laboratories | P. Hemarajata et al. |
| EPI_ISL_3299025 | Palmdale Regional Medical Center | Los Angeles County Public Health Laboratories | P. Hemarajata et al. |
| EPI_ISL_3299026 | DxTerity | Los Angeles County Public Health Laboratories | P. Hemarajata et al. |
| EPI_ISL_3299027, EPI_ISL_3299028 | Martin Luther King, Jr. Outpatient Center | Los Angeles County Public Health Laboratories | P. Hemarajata et al. |
| EPI_ISL_3299029 | Palmdale Regional Medical Center | Los Angeles County Public Health Laboratories | P. Hemarajata et al. |
| EPI_ISL_3299030 | USC Clinical Lab | Los Angeles County Public Health Laboratories | P. Hemarajata et al. |
| EPI_ISL_3299031 | Palmdale Regional Medical Center | Los Angeles County Public Health Laboratories | P. Hemarajata et al. |
| EPI_ISL_3299032 | Martin Luther King Jr. Community Hospital | Los Angeles County Public Health Laboratories | P. Hemarajata et al. |
| EPI_ISL_3299033, EPI_ISL_3299034, EPI_ISL_3299036 | Palmdale Regional Medical Center | Los Angeles County Public Health Laboratories | P. Hemarajata et al. |
| EPI_ISL_3299037 | USC Clinical Lab | Los Angeles County Public Health Laboratories | P. Hemarajata et al. |
| EPI_ISL_3299038, | Palmdale Regional Medical Center | Los Angeles County Public Health Laboratories | P. Hemarajata et al. |

| | | | |
|---|---|---|---|
| EPI_ISL_3299040 | | | |
| EPI_ISL_3299044 | USC Verdugo Hills Hospital | Los Angeles County Public Health Laboratories | P. Hemarajata et al. |
| EPI_ISL_3299045, EPI_ISL_3299046, EPI_ISL_3299047, EPI_ISL_3299048 | Palmdale Regional Medical Center | Los Angeles County Public Health Laboratories | P. Hemarajata et al. |
| EPI_ISL_3299210 | Genomik Solidaritas Indonesia Laboratorium | Genomik Solidaritas Indonesia Laboratorium | Himawan Masyhuri, Dhahlia Agustina Cahyono, Andani Eka Putra, Dede Rahman Agustian, Linosefa, Vania Gavrila Wikasa, Carissa Sintca Wijaya, Gracia Felias Enos Korompis, Annisa Muthiah Sukirman, Anuraj Shankar, Louisa Markus, Meutia Ayuputeri Kumaheri, Ariel Pradipta |
| EPI_ISL_3299595, EPI_ISL_3299596, EPI_ISL_3299597, EPI_ISL_3299599, EPI_ISL_3299600, EPI_ISL_3299601, EPI_ISL_3299602, EPI_ISL_3299604, EPI_ISL_3299605, EPI_ISL_3299606, EPI_ISL_3299607, EPI_ISL_3299608, EPI_ISL_3299609, EPI_ISL_3299610, EPI_ISL_3299611, EPI_ISL_3299612, EPI_ISL_3299613, EPI_ISL_3299615, EPI_ISL_3299616, EPI_ISL_3299617, EPI_ISL_3299618, EPI_ISL_3299620, EPI_ISL_3299621, EPI_ISL_3299622, EPI_ISL_3299623, EPI_ISL_3299624, EPI_ISL_3299625, EPI_ISL_3299626, EPI_ISL_3299627, EPI_ISL_3299628, EPI_ISL_3299629, EPI_ISL_3299630, EPI_ISL_3299631, EPI_ISL_3299632, EPI_ISL_3299633, EPI_ISL_3299634, EPI_ISL_3299635, EPI_ISL_3299636, EPI_ISL_3299638, EPI_ISL_3299639, EPI_ISL_3299640, EPI_ISL_3299642, EPI_ISL_3299643, EPI_ISL_3299644, EPI_ISL_3299645, EPI_ISL_3299646, EPI_ISL_3299647, EPI_ISL_3299648, EPI_ISL_3299649, EPI_ISL_3299650, EPI_ISL_3299651, EPI_ISL_3299652, EPI_ISL_3299653, EPI_ISL_3299654, EPI_ISL_3299656, EPI_ISL_3299658, EPI_ISL_3299659, EPI_ISL_3299660, EPI_ISL_3299661, EPI_ISL_3299662, EPI_ISL_3299663, EPI_ISL_3299664, EPI_ISL_3299665, EPI_ISL_3299666, EPI_ISL_3299667, EPI_ISL_3299668, EPI_ISL_3299669, EPI_ISL_3299670, EPI_ISL_3299671, EPI_ISL_3299672, EPI_ISL_3299673, EPI_ISL_3299674, EPI_ISL_3299675, EPI_ISL_3299676, EPI_ISL_3299677, EPI_ISL_3299678, EPI_ISL_3299679, EPI_ISL_3299681, EPI_ISL_3299682, EPI_ISL_3299683, EPI_ISL_3299684, EPI_ISL_3299685, EPI_ISL_3299686, EPI_ISL_3299687, EPI_ISL_3299688, EPI_ISL_3299689, EPI_ISL_3299690, EPI_ISL_3299691, EPI_ISL_3299692, EPI_ISL_3299693, EPI_ISL_3299694, EPI_ISL_3299695, EPI_ISL_3299696, EPI_ISL_3299697, EPI_ISL_3299698, EPI_ISL_3299699, EPI_ISL_3299701, EPI_ISL_3299702, EPI_ISL_3299703, EPI_ISL_3299704, EPI_ISL_3299705, EPI_ISL_3299706, EPI_ISL_3299707, EPI_ISL_3299709, EPI_ISL_3299710, EPI_ISL_3299712, EPI_ISL_3299713, EPI_ISL_3299715, EPI_ISL_3299716, EPI_ISL_3299717, EPI_ISL_3299718, EPI_ISL_3299719, EPI_ISL_3299720, EPI_ISL_3299722, EPI_ISL_3299723, EPI_ISL_3299724, EPI_ISL_3299725, EPI_ISL_3299726, EPI_ISL_3299727, EPI_ISL_3299728, EPI_ISL_3299729, EPI_ISL_3299730, EPI_ISL_3299731, EPI_ISL_3299732, EPI_ISL_3299733, EPI_ISL_3299734, EPI_ISL_3299735, EPI_ISL_3299736, EPI_ISL_3299737, EPI_ISL_3299738, EPI_ISL_3299739, EPI_ISL_3299740, EPI_ISL_3299741, EPI_ISL_3299742, EPI_ISL_3299743, EPI_ISL_3299744, EPI_ISL_3299745, EPI_ISL_3299747, EPI_ISL_3299748, EPI_ISL_3299749, EPI_ISL_3299750, EPI_ISL_3299751, EPI_ISL_3299752, EPI_ISL_3299753, EPI_ISL_3299754, EPI_ISL_3299755, EPI_ISL_3299756, EPI_ISL_3299757, EPI_ISL_3299758, EPI_ISL_3299759, EPI_ISL_3299760, EPI_ISL_3299761, EPI_ISL_3299762, EPI_ISL_3299763, EPI_ISL_3299764, EPI_ISL_3299765, EPI_ISL_3299766, EPI_ISL_3299767, EPI_ISL_3299768, EPI_ISL_3299769, EPI_ISL_3299770, EPI_ISL_3299771, EPI_ISL_3299772, EPI_ISL_3299773, EPI_ISL_3299774, EPI_ISL_3299776, EPI_ISL_3299777, EPI_ISL_3299778 | | | |
| see above | Infinity Biologix | Centers for Disease Control and Prevention Division of Viral Diseases, Pathogen Discovery | Dakota Howard, Dhwani Batra, Peter W. Cook, Kara Moser, Adrian Paskey, Jason Caravas, Benjamin Rambo-Martin, Shatavia Morrison, Christopher Gulvick, Scott Sammons, Yvette Unoarumhi, Darlene Wagner, Matthew Schmerer, Christian Bixby, Yihe Wang, Jonathan Schultz, Chirayu Goswami, Russ Hager, Robin Grimwood, Clinton R. Paden, Duncan MacCannell |
| EPI_ISL_3299779, EPI_ISL_3299784, EPI_ISL_3299785, EPI_ISL_3299786, EPI_ISL_3299789, EPI_ISL_3299791, EPI_ISL_3299792, EPI_ISL_3299794, EPI_ISL_3299795, EPI_ISL_3299796, EPI_ISL_3299797, EPI_ISL_3299798, EPI_ISL_3299800, EPI_ISL_3299801, EPI_ISL_3299802, EPI_ISL_3299803, EPI_ISL_3299804, EPI_ISL_3299806, EPI_ISL_3299807, EPI_ISL_3299808, EPI_ISL_3299809, EPI_ISL_3299810, EPI_ISL_3299811, EPI_ISL_3299813, EPI_ISL_3299815, EPI_ISL_3299816, EPI_ISL_3299817, EPI_ISL_3299818, EPI_ISL_3299819, EPI_ISL_3299820, EPI_ISL_3299821, EPI_ISL_3299822, EPI_ISL_3299823, EPI_ISL_3299824, EPI_ISL_3299825, EPI_ISL_3299826, EPI_ISL_3299827, EPI_ISL_3299828, EPI_ISL_3299829, EPI_ISL_3299830, EPI_ISL_3299831, EPI_ISL_3299832, EPI_ISL_3299833, EPI_ISL_3299834, EPI_ISL_3299835, EPI_ISL_3299836, EPI_ISL_3299837, EPI_ISL_3299839, EPI_ISL_3299840, EPI_ISL_3299842, EPI_ISL_3299843, EPI_ISL_3299844, EPI_ISL_3299845, EPI_ISL_3299846, EPI_ISL_3299848, EPI_ISL_3299849, EPI_ISL_3299850, EPI_ISL_3299851, EPI_ISL_3299852, EPI_ISL_3299853, EPI_ISL_3299854, EPI_ISL_3299855, EPI_ISL_3299856, EPI_ISL_3299860, EPI_ISL_3299861, EPI_ISL_3299862, EPI_ISL_3299863, EPI_ISL_3299865, EPI_ISL_3299868, EPI_ISL_3299869, EPI_ISL_3299870, EPI_ISL_3299872, EPI_ISL_3299873, EPI_ISL_3299874, EPI_ISL_3299875, EPI_ISL_3299876, EPI_ISL_3299878, EPI_ISL_3299879, EPI_ISL_3299880, EPI_ISL_3299881, EPI_ISL_3299883, EPI_ISL_3299884, EPI_ISL_3299885, EPI_ISL_3299887, EPI_ISL_3299888, EPI_ISL_3299891, EPI_ISL_3299892, EPI_ISL_3299895, EPI_ISL_3299896, EPI_ISL_3299897, EPI_ISL_3299898, EPI_ISL_3299900, EPI_ISL_3299901, EPI_ISL_3299903, EPI_ISL_3299904, EPI_ISL_3299905, EPI_ISL_3299906, EPI_ISL_3299907, EPI_ISL_3299908, EPI_ISL_3299909, EPI_ISL_3299910, EPI_ISL_3299911, EPI_ISL_3299912, EPI_ISL_3299914, EPI_ISL_3299915, EPI_ISL_3299916, EPI_ISL_3299917, EPI_ISL_3299918, EPI_ISL_3299919, EPI_ISL_3299920, EPI_ISL_3299921, EPI_ISL_3299922, EPI_ISL_3299923, EPI_ISL_3299925, EPI_ISL_3299927, EPI_ISL_3299928, EPI_ISL_3299929, EPI_ISL_3299930, EPI_ISL_3299931, EPI_ISL_3299932, EPI_ISL_3299933, EPI_ISL_3299934, EPI_ISL_3299935, EPI_ISL_3299936, EPI_ISL_3299937, EPI_ISL_3299938, EPI_ISL_3299940, EPI_ISL_3299941, EPI_ISL_3299942, EPI_ISL_3299943, EPI_ISL_3299944, EPI_ISL_3299945, EPI_ISL_3299946, EPI_ISL_3299947, EPI_ISL_3299948, EPI_ISL_3299949, EPI_ISL_3299950, EPI_ISL_3299951, EPI_ISL_3299952, EPI_ISL_3299953, EPI_ISL_3299954, EPI_ISL_3299955, EPI_ISL_3299956, EPI_ISL_3299957, EPI_ISL_3299958, EPI_ISL_3299959, EPI_ISL_3299960, EPI_ISL_3299961, EPI_ISL_3299962, EPI_ISL_3299963, EPI_ISL_3299964, EPI_ISL_3299966, EPI_ISL_3299969, EPI_ISL_3299970, EPI_ISL_3299972, EPI_ISL_3299973, EPI_ISL_3299975, EPI_ISL_3299977, EPI_ISL_3299978, EPI_ISL_3299979, EPI_ISL_3299980, EPI_ISL_3299981, EPI_ISL_3299982, EPI_ISL_3299985, EPI_ISL_3299986, EPI_ISL_3299987, EPI_ISL_3299988, EPI_ISL_3299989, EPI_ISL_3299992, EPI_ISL_3299993, EPI_ISL_3299994, EPI_ISL_3299995, EPI_ISL_3299996, EPI_ISL_3299997, EPI_ISL_3299998, EPI_ISL_3299999, EPI_ISL_3300000, EPI_ISL_3300001, EPI_ISL_3300003, EPI_ISL_3300004, EPI_ISL_3300006, EPI_ISL_3300007, EPI_ISL_3300008, EPI_ISL_3300010, EPI_ISL_3300011, EPI_ISL_3300012, EPI_ISL_3300013, EPI_ISL_3300014, EPI_ISL_3300015, EPI_ISL_3300016, EPI_ISL_3300017, EPI_ISL_3300018, EPI_ISL_3300019, EPI_ISL_3300020, EPI_ISL_3300022, EPI_ISL_3300023, EPI_ISL_3300024, EPI_ISL_3300025, EPI_ISL_3300026, EPI_ISL_3300027, EPI_ISL_3300028, EPI_ISL_3300029, EPI_ISL_3300030, EPI_ISL_3300031, EPI_ISL_3300032, EPI_ISL_3300033, EPI_ISL_3300035, EPI_ISL_3300036, EPI_ISL_3300038, EPI_ISL_3300039, EPI_ISL_3300040, EPI_ISL_3300041, EPI_ISL_3300042, EPI_ISL_3300043, EPI_ISL_3300044, EPI_ISL_3300046, EPI_ISL_3300047, EPI_ISL_3300048, EPI_ISL_3300049, EPI_ISL_3300050, EPI_ISL_3300051, EPI_ISL_3300052, EPI_ISL_3300053, EPI_ISL_3300054, EPI_ISL_3300055, EPI_ISL_3300056, EPI_ISL_3300057, EPI_ISL_3300058, EPI_ISL_3300059, EPI_ISL_3300060, EPI_ISL_3300061, EPI_ISL_3300062, EPI_ISL_3300063, EPI_ISL_3300064, EPI_ISL_3300065, EPI_ISL_3300066, EPI_ISL_3300067, EPI_ISL_3300068, EPI_ISL_3300069, EPI_ISL_3300070, EPI_ISL_3300072, EPI_ISL_3300073, EPI_ISL_3300074, EPI_ISL_3300075, EPI_ISL_3300076, EPI_ISL_3300077, EPI_ISL_3300078, EPI_ISL_3300079, EPI_ISL_3300080, EPI_ISL_3300081, EPI_ISL_3300082, EPI_ISL_3300083, EPI_ISL_3300084, EPI_ISL_3300085, EPI_ISL_3300086, EPI_ISL_3300087, EPI_ISL_3300088, EPI_ISL_3300089, EPI_ISL_3300090, EPI_ISL_3300091, EPI_ISL_3300092, EPI_ISL_3300093, EPI_ISL_3300094, EPI_ISL_3300095, EPI_ISL_3300096, EPI_ISL_3300097, EPI_ISL_3300098, EPI_ISL_3300099, EPI_ISL_3300100, EPI_ISL_3300101, EPI_ISL_3300102, EPI_ISL_3300103, EPI_ISL_3300104, EPI_ISL_3300105, EPI_ISL_3300106, EPI_ISL_3300107, EPI_ISL_3300108, EPI_ISL_3300109, EPI_ISL_3300110, EPI_ISL_3300111, EPI_ISL_3300112, EPI_ISL_3300113, EPI_ISL_3300114, EPI_ISL_3300115, EPI_ISL_3300116, EPI_ISL_3300118, EPI_ISL_3300119, EPI_ISL_3300120, EPI_ISL_3300121, EPI_ISL_3300122, EPI_ISL_3300123, EPI_ISL_3300124, EPI_ISL_3300125, EPI_ISL_3300126, EPI_ISL_3300127, EPI_ISL_3300128, EPI_ISL_3300129, EPI_ISL_3300130, EPI_ISL_3300131, EPI_ISL_3300132, EPI_ISL_3300133, EPI_ISL_3300134, EPI_ISL_3300135, EPI_ISL_3300136, EPI_ISL_3300137, EPI_ISL_3300138, EPI_ISL_3300139, EPI_ISL_3300140, EPI_ISL_3300141, EPI_ISL_3300142, EPI_ISL_3300143, EPI_ISL_3300144, EPI_ISL_3300145, EPI_ISL_3300146, EPI_ISL_3300147, EPI_ISL_3300148, EPI_ISL_3300149, EPI_ISL_3300150, EPI_ISL_3300151, EPI_ISL_3300152, EPI_ISL_3300153, EPI_ISL_3300154, EPI_ISL_3300155, EPI_ISL_3300156, EPI_ISL_3300157, EPI_ISL_3300158, EPI_ISL_3300159, EPI_ISL_3300160, EPI_ISL_3300161, EPI_ISL_3300162, EPI_ISL_3300163, EPI_ISL_3300164, EPI_ISL_3300165, EPI_ISL_3300166, EPI_ISL_3300167, EPI_ISL_3300168, EPI_ISL_3300169, EPI_ISL_3300170, EPI_ISL_3300171, EPI_ISL_3300172, EPI_ISL_3300173, EPI_ISL_3300174, EPI_ISL_3300175, EPI_ISL_3300176, EPI_ISL_3300177, EPI_ISL_3300178, EPI_ISL_3300179, EPI_ISL_3300180, EPI_ISL_3300181, EPI_ISL_3300182, EPI_ISL_3300183, EPI_ISL_3300184, EPI_ISL_3300185, EPI_ISL_3300186, EPI_ISL_3300187, EPI_ISL_3300188, EPI_ISL_3300189, EPI_ISL_3300190, EPI_ISL_3300191, EPI_ISL_3300192, EPI_ISL_3300193, EPI_ISL_3300194, EPI_ISL_3300195, EPI_ISL_3300196, EPI_ISL_3300197, EPI_ISL_3300198, EPI_ISL_3300199, EPI_ISL_3300200, EPI_ISL_3300201, EPI_ISL_3300202, EPI_ISL_3300203, EPI_ISL_3300204, EPI_ISL_3300205, EPI_ISL_3300206, EPI_ISL_3300207, EPI_ISL_3300208, EPI_ISL_3300209, EPI_ISL_3300210, EPI_ISL_3300211, EPI_ISL_3300212, EPI_ISL_3300213, EPI_ISL_3300214, EPI_ISL_3300215, EPI_ISL_3300216, EPI_ISL_3300217, EPI_ISL_3300218, EPI_ISL_3300220, EPI_ISL_3300221, EPI_ISL_3300222, EPI_ISL_3300223, EPI_ISL_3300224, EPI_ISL_3300225, EPI_ISL_3300226, EPI_ISL_3300227, EPI_ISL_3300228 | | | |
| see above | Fulgent Genetics | Centers for Disease Control and Prevention Division of Viral Diseases, Pathogen Discovery | Dakota Howard, Dhwani Batra, Peter W. Cook, Kara Moser, Adrian Paskey, Jason Caravas, Benjamin Rambo-Martin, Shatavia Morrison, Christopher Gulvick, Scott Sammons, Yvette Unoarumhi, Darlene Wagner, Matthew Schmerer, Harry Gao, Mickey Li, John Gao, Joseph Fierro, Benafsh Sapra, Becky Tsai, Yan Meng, Doreen Ng, James Xie, Clinton R. Paden, Duncan MacCannell |
| EPI_ISL_3300229 | Centinela Hospital Medical Center | Los Angeles County Public Health Laboratories | P. Hemarajata et al. |
| EPI_ISL_3300230, EPI_ISL_3300231, EPI_ISL_3300232, EPI_ISL_3300234 | Fulgent Genetics | Centers for Disease Control and Prevention Division of Viral Diseases, Pathogen Discovery | Dakota Howard, Dhwani Batra, Peter W. Cook, Kara Moser, Adrian Paskey, Jason Caravas, Benjamin Rambo-Martin, Shatavia Morrison, Christopher Gulvick, Scott Sammons, Yvette Unoarumhi, Darlene Wagner, Matthew Schmerer, Harry Gao, Mickey Li, John Gao, Joseph Fierro, Benafsh Sapra, Becky Tsai, Yan Meng, Doreen Ng, James Xie, Clinton R. Paden, Duncan MacCannell |
| EPI_ISL_3300236 | Centinela Hospital Medical Center | Los Angeles County Public Health Laboratories | P. Hemarajata et al. |
| EPI_ISL_3300237, EPI_ISL_3300238 | Fulgent Genetics | Centers for Disease Control and Prevention Division of Viral Diseases, Pathogen Discovery | Dakota Howard, Dhwani Batra, Peter W. Cook, Kara Moser, Adrian Paskey, Jason Caravas, Benjamin Rambo-Martin, Shatavia Morrison, Christopher Gulvick, Scott Sammons, Yvette Unoarumhi, Darlene Wagner, Matthew Schmerer, Harry Gao, Mickey Li, John Gao, Joseph Fierro, Benafsh Sapra, Becky Tsai, Yan Meng, Doreen Ng, James Xie, Clinton R. Paden, Duncan MacCannell |
| EPI_ISL_3300239 | Centinela Hospital Medical Center | Los Angeles County Public Health Laboratories | P. Hemarajata et al. |
| EPI_ISL_3300240, EPI_ISL_3300241 | Fulgent Genetics | Centers for Disease Control and Prevention Division of Viral Diseases, Pathogen Discovery | Dakota Howard, Dhwani Batra, Peter W. Cook, Kara Moser, Adrian Paskey, Jason Caravas, Benjamin Rambo-Martin, Shatavia Morrison, Christopher Gulvick, Scott Sammons, Yvette Unoarumhi, Darlene Wagner, Matthew Schmerer, Harry Gao, Mickey Li, John Gao, Joseph Fierro, Benafsh Sapra, Becky Tsai, Yan Meng, Doreen Ng, James Xie, Clinton R. Paden, Duncan MacCannell |
| EPI_ISL_3300242 | Centinela Hospital Medical Center | Los Angeles County Public Health Laboratories | P. Hemarajata et al. |
| EPI_ISL_3300243, EPI_ISL_3300244, EPI_ISL_3300245, EPI_ISL_3300246, EPI_ISL_3300247, EPI_ISL_3300248, EPI_ISL_3300250, EPI_ISL_3300251 | Fulgent Genetics | Centers for Disease Control and Prevention Division of Viral Diseases, Pathogen Discovery | Dakota Howard, Dhwani Batra, Peter W. Cook, Kara Moser, Adrian Paskey, Jason Caravas, Benjamin Rambo-Martin, Shatavia Morrison, Christopher Gulvick, Scott Sammons, Yvette Unoarumhi, Darlene Wagner, Matthew Schmerer, Harry Gao, Mickey Li, John Gao, Joseph Fierro, Benafsh Sapra, Becky Tsai, Yan Meng, Doreen Ng, James Xie, Clinton R. Paden, Duncan MacCannell |
| EPI_ISL_3300252 | Centinela Hospital Medical Center | Los Angeles County Public Health Laboratories | P. Hemarajata et al. |
| EPI_ISL_3300253, EPI_ISL_3300254, EPI_ISL_3300255, EPI_ISL_3300256, EPI_ISL_3300257, EPI_ISL_3300258, EPI_ISL_3300260, EPI_ISL_3300261, EPI_ISL_3300262, EPI_ISL_3300263 | Fulgent Genetics | Centers for Disease Control and Prevention Division of Viral Diseases, Pathogen Discovery | Dakota Howard, Dhwani Batra, Peter W. Cook, Kara Moser, Adrian Paskey, Jason Caravas, Benjamin Rambo-Martin, Shatavia Morrison, Christopher Gulvick, Scott Sammons, Yvette Unoarumhi, Darlene Wagner, Matthew Schmerer, Harry Gao, Mickey Li, John Gao, Joseph Fierro, Benafsh Sapra, Becky Tsai, Yan Meng, Doreen Ng, James Xie, Clinton R. Paden, Duncan MacCannell |
| EPI_ISL_3300264 | Sun Clinical Laboratory | Los Angeles County Public Health Laboratories | P. Hemarajata et al. |
| EPI_ISL_3300265, EPI_ISL_3300266, EPI_ISL_3300267 | Fulgent Genetics | Centers for Disease Control and Prevention Division of Viral Diseases, Pathogen Discovery | Dakota Howard, Dhwani Batra, Peter W. Cook, Kara Moser, Adrian Paskey, Jason Caravas, Benjamin Rambo-Martin, Shatavia Morrison, Christopher Gulvick, Scott Sammons, Yvette Unoarumhi, Darlene Wagner, Matthew Schmerer, Harry Gao, Mickey Li, John Gao, Joseph Fierro, Benafsh Sapra, Becky Tsai, Yan Meng, Doreen Ng, James Xie, Clinton R. Paden, Duncan MacCannell |
| EPI_ISL_3300268 | Sun Clinical Laboratory | Los Angeles County Public Health Laboratories | P. Hemarajata et al. |
| EPI_ISL_3300269, EPI_ISL_3300270, EPI_ISL_3300271, EPI_ISL_3300272, EPI_ISL_3300273 | Fulgent Genetics | Centers for Disease Control and Prevention Division of Viral Diseases, Pathogen Discovery | Dakota Howard, Dhwani Batra, Peter W. Cook, Kara Moser, Adrian Paskey, Jason Caravas, Benjamin Rambo-Martin, Shatavia Morrison, Christopher Gulvick, Scott Sammons, Yvette Unoarumhi, Darlene Wagner, Matthew Schmerer, Harry Gao, Mickey Li, John Gao, Joseph Fierro, Benafsh Sapra, Becky Tsai, Yan Meng, Doreen Ng, James Xie, Clinton R. Paden, Duncan MacCannell |
| EPI_ISL_3300274 | Sun Clinical Laboratory | Los Angeles County Public Health Laboratories | P. Hemarajata et al. |
| EPI_ISL_3300275, EPI_ISL_3300276 | Fulgent Genetics | Centers for Disease Control and Prevention Division of Viral Diseases, Pathogen Discovery | Dakota Howard, Dhwani Batra, Peter W. Cook, Kara Moser, Adrian Paskey, Jason Caravas, Benjamin Rambo-Martin, Shatavia Morrison, Christopher Gulvick, Scott Sammons, Yvette Unoarumhi, Darlene Wagner, Matthew Schmerer, Harry Gao, Mickey Li, John Gao, Joseph Fierro, Benafsh Sapra, Becky Tsai, Yan Meng, Doreen Ng, James Xie, Clinton R. Paden, Duncan MacCannell |
| EPI_ISL_3300277 | Sun Clinical Laboratory | Los Angeles County Public Health Laboratories | P. Hemarajata et al. |
| EPI_ISL_3300278, EPI_ISL_3300279, EPI_ISL_3300280, EPI_ISL_3300281, EPI_ISL_3300282 | Fulgent Genetics | Centers for Disease Control and Prevention Division of Viral Diseases, Pathogen Discovery | Dakota Howard, Dhwani Batra, Peter W. Cook, Kara Moser, Adrian Paskey, Jason Caravas, Benjamin Rambo-Martin, Shatavia Morrison, Christopher Gulvick, Scott Sammons, Yvette Unoarumhi, Darlene Wagner, Matthew Schmerer, Harry Gao, Mickey Li, John Gao, Joseph Fierro, Benafsh Sapra, Becky Tsai, Yan Meng, Doreen Ng, James Xie, Clinton R. Paden, Duncan MacCannell |
| EPI_ISL_3300283 | Sun Clinical Laboratory | Los Angeles County Public Health Laboratories | P. Hemarajata et al. |
| EPI_ISL_3300284, EPI_ISL_3300285, EPI_ISL_3300286 | Fulgent Genetics | Centers for Disease Control and Prevention Division of Viral Diseases, Pathogen Discovery | Dakota Howard, Dhwani Batra, Peter W. Cook, Kara Moser, Adrian Paskey, Jason Caravas, Benjamin Rambo-Martin, Shatavia Morrison, Christopher Gulvick, Scott Sammons, Yvette Unoarumhi, Darlene Wagner, Matthew Schmerer, Harry Gao, Mickey Li, John Gao, Joseph Fierro, Benafsh Sapra, Becky Tsai, Yan Meng, Doreen Ng, James Xie, Clinton R. Paden, Duncan MacCannell |
| EPI_ISL_3300287 | Sun Clinical Laboratory | Los Angeles County Public Health Laboratories | P. Hemarajata et al. |
| EPI_ISL_3300288, EPI_ISL_3300289, EPI_ISL_3300290, EPI_ISL_3300291, | Fulgent Genetics | Centers for Disease Control and Prevention Division of Viral Diseases, Pathogen Discovery | Dakota Howard, Dhwani Batra, Peter W. Cook, Kara Moser, Adrian Paskey, Jason Caravas, Benjamin Rambo-Martin, Shatavia Morrison, Christopher Gulvick, Scott Sammons, Yvette Unoarumhi, Darlene Wagner, Matthew Schmerer, Harry Gao, Mickey Li, John Gao, Joseph Fierro, Benafsh Sapra, Becky Tsai, Yan Meng, Doreen Ng, James Xie, Clinton R. Paden, Duncan MacCannell |

| | | | |
|---|---|---|---|
| EPI_ISL_3300292 | | | |
| EPI_ISL_3300293 | Sun Clinical Laboratory | Los Angeles County Public Health Laboratories | P. Hemarajata et al. |
| EPI_ISL_3300295 | Fulgent Genetics | Centers for Disease Control and Prevention Division of Viral Diseases, Pathogen Discovery | Dakota Howard, Dhwani Batra, Peter W. Cook, Kara Moser, Adrian Paskey, Jason Caravas, Benjamin Rambo-Martin, Shatavia Morrison, Christopher Gulvick, Scott Sammons, Yvette Unoarumhi, Darlene Wagner, Matthew Schmerer, Harry Gao, Mickey Li, John Gao, Joseph Fierro, Benafsh Sapra, Becky Tsai, Yan Meng, Doreen Ng, James Xie, Clinton R. Paden, Duncan MacCannell |
| EPI_ISL_3300296 | Providence Saint John's Health Center | Los Angeles County Public Health Laboratories | P. Hemarajata et al. |
| EPI_ISL_3300297, EPI_ISL_3300298 | Fulgent Genetics | Centers for Disease Control and Prevention Division of Viral Diseases, Pathogen Discovery | Dakota Howard, Dhwani Batra, Peter W. Cook, Kara Moser, Adrian Paskey, Jason Caravas, Benjamin Rambo-Martin, Shatavia Morrison, Christopher Gulvick, Scott Sammons, Yvette Unoarumhi, Darlene Wagner, Matthew Schmerer, Harry Gao, Mickey Li, John Gao, Joseph Fierro, Benafsh Sapra, Becky Tsai, Yan Meng, Doreen Ng, James Xie, Clinton R. Paden, Duncan MacCannell |
| EPI_ISL_3300299 | Providence Saint John's Health Center | Los Angeles County Public Health Laboratories | P. Hemarajata et al. |
| EPI_ISL_3300300, EPI_ISL_3300301, EPI_ISL_3300302 | Fulgent Genetics | Centers for Disease Control and Prevention Division of Viral Diseases, Pathogen Discovery | Dakota Howard, Dhwani Batra, Peter W. Cook, Kara Moser, Adrian Paskey, Jason Caravas, Benjamin Rambo-Martin, Shatavia Morrison, Christopher Gulvick, Scott Sammons, Yvette Unoarumhi, Darlene Wagner, Matthew Schmerer, Harry Gao, Mickey Li, John Gao, Joseph Fierro, Benafsh Sapra, Becky Tsai, Yan Meng, Doreen Ng, James Xie, Clinton R. Paden, Duncan MacCannell |
| EPI_ISL_3300303 | Providence Saint John's Health Center | Los Angeles County Public Health Laboratories | P. Hemarajata et al. |
| EPI_ISL_3300304, EPI_ISL_3300305, EPI_ISL_3300306, EPI_ISL_3300307, EPI_ISL_3300308 | Fulgent Genetics | Centers for Disease Control and Prevention Division of Viral Diseases, Pathogen Discovery | Dakota Howard, Dhwani Batra, Peter W. Cook, Kara Moser, Adrian Paskey, Jason Caravas, Benjamin Rambo-Martin, Shatavia Morrison, Christopher Gulvick, Scott Sammons, Yvette Unoarumhi, Darlene Wagner, Matthew Schmerer, Harry Gao, Mickey Li, John Gao, Joseph Fierro, Benafsh Sapra, Becky Tsai, Yan Meng, Doreen Ng, James Xie, Clinton R. Paden, Duncan MacCannell |
| EPI_ISL_3300309 | Providence Saint John's Health Center | Los Angeles County Public Health Laboratories | P. Hemarajata et al. |
| EPI_ISL_3300310, EPI_ISL_3300311, EPI_ISL_3300312, EPI_ISL_3300313, EPI_ISL_3300314 | Fulgent Genetics | Centers for Disease Control and Prevention Division of Viral Diseases, Pathogen Discovery | Dakota Howard, Dhwani Batra, Peter W. Cook, Kara Moser, Adrian Paskey, Jason Caravas, Benjamin Rambo-Martin, Shatavia Morrison, Christopher Gulvick, Scott Sammons, Yvette Unoarumhi, Darlene Wagner, Matthew Schmerer, Harry Gao, Mickey Li, John Gao, Joseph Fierro, Benafsh Sapra, Becky Tsai, Yan Meng, Doreen Ng, James Xie, Clinton R. Paden, Duncan MacCannell |
| EPI_ISL_3300315 | Providence Saint John's Health Center | Los Angeles County Public Health Laboratories | P. Hemarajata et al. |
| EPI_ISL_3300316, EPI_ISL_3300317, EPI_ISL_3300318, EPI_ISL_3300319 | Fulgent Genetics | Centers for Disease Control and Prevention Division of Viral Diseases, Pathogen Discovery | Dakota Howard, Dhwani Batra, Peter W. Cook, Kara Moser, Adrian Paskey, Jason Caravas, Benjamin Rambo-Martin, Shatavia Morrison, Christopher Gulvick, Scott Sammons, Yvette Unoarumhi, Darlene Wagner, Matthew Schmerer, Harry Gao, Mickey Li, John Gao, Joseph Fierro, Benafsh Sapra, Becky Tsai, Yan Meng, Doreen Ng, James Xie, Clinton R. Paden, Duncan MacCannell |
| EPI_ISL_3300320 | Providence Saint John's Health Center | Los Angeles County Public Health Laboratories | P. Hemarajata et al. |
| EPI_ISL_3300321, EPI_ISL_3300322, EPI_ISL_3300323 | Fulgent Genetics | Centers for Disease Control and Prevention Division of Viral Diseases, Pathogen Discovery | Dakota Howard, Dhwani Batra, Peter W. Cook, Kara Moser, Adrian Paskey, Jason Caravas, Benjamin Rambo-Martin, Shatavia Morrison, Christopher Gulvick, Scott Sammons, Yvette Unoarumhi, Darlene Wagner, Matthew Schmerer, Harry Gao, Mickey Li, John Gao, Joseph Fierro, Benafsh Sapra, Becky Tsai, Yan Meng, Doreen Ng, James Xie, Clinton R. Paden, Duncan MacCannell |
| EPI_ISL_3300324 | Providence Saint John's Health Center | Los Angeles County Public Health Laboratories | P. Hemarajata et al. |
| EPI_ISL_3300325, EPI_ISL_3300326, EPI_ISL_3300327, EPI_ISL_3300328, EPI_ISL_3300329, EPI_ISL_3300330, EPI_ISL_3300331, EPI_ISL_3300332, EPI_ISL_3300333, EPI_ISL_3300334, EPI_ISL_3300335, EPI_ISL_3300336, EPI_ISL_3300337, EPI_ISL_3300338, EPI_ISL_3300339, EPI_ISL_3300340, EPI_ISL_3300341, EPI_ISL_3300342, EPI_ISL_3300343, EPI_ISL_3300344, EPI_ISL_3300345, EPI_ISL_3300346, EPI_ISL_3300347, EPI_ISL_3300348, EPI_ISL_3300349, EPI_ISL_3300350, EPI_ISL_3300351, EPI_ISL_3300352, EPI_ISL_3300353, EPI_ISL_3300354, EPI_ISL_3300355, EPI_ISL_3300356, EPI_ISL_3300357, EPI_ISL_3300358, EPI_ISL_3300359, EPI_ISL_3300360, EPI_ISL_3300362, EPI_ISL_3300363, EPI_ISL_3300364, EPI_ISL_3300365, EPI_ISL_3300366, EPI_ISL_3300367, EPI_ISL_3300368, EPI_ISL_3300369, EPI_ISL_3300370, EPI_ISL_3300371, EPI_ISL_3300373, EPI_ISL_3300374, EPI_ISL_3300375, EPI_ISL_3300376, EPI_ISL_3300377, EPI_ISL_3300378, EPI_ISL_3300379, EPI_ISL_3300380, EPI_ISL_3300381, EPI_ISL_3300382, EPI_ISL_3300383, EPI_ISL_3300384, EPI_ISL_3300385, EPI_ISL_3300386, EPI_ISL_3300387, EPI_ISL_3300388, EPI_ISL_3300389, EPI_ISL_3300390, EPI_ISL_3300391, EPI_ISL_3300392, EPI_ISL_3300393, EPI_ISL_3300394, EPI_ISL_3300395, EPI_ISL_3300396, EPI_ISL_3300397, EPI_ISL_3300398, EPI_ISL_3300399, EPI_ISL_3300400, EPI_ISL_3300401, EPI_ISL_3300402, EPI_ISL_3300403, EPI_ISL_3300404, EPI_ISL_3300405, EPI_ISL_3300406, EPI_ISL_3300407, EPI_ISL_3300408 | | | |
| see above | Fulgent Genetics | Centers for Disease Control and Prevention Division of Viral Diseases, Pathogen Discovery | Dakota Howard, Dhwani Batra, Peter W. Cook, Kara Moser, Adrian Paskey, Jason Caravas, Benjamin Rambo-Martin, Shatavia Morrison, Christopher Gulvick, Scott Sammons, Yvette Unoarumhi, Darlene Wagner, Matthew Schmerer, Harry Gao, Mickey Li, John Gao, Joseph Fierro, Benafsh Sapra, Becky Tsai, Yan Meng, Doreen Ng, James Xie, Clinton R. Paden, Duncan MacCannell |
| EPI_ISL_3300409 | Gandhi Medical College | CDFD | Murali Bashyam, Asmita Gupta, Reelina Basu, Vinay Donipadi, Nagamani Kammili, Divya Vashisht, Ashwin Dalal |
| EPI_ISL_3300410, EPI_ISL_3300411, EPI_ISL_3300412, EPI_ISL_3300413 | Fulgent Genetics | Centers for Disease Control and Prevention Division of Viral Diseases, Pathogen Discovery | Dakota Howard, Dhwani Batra, Peter W. Cook, Kara Moser, Adrian Paskey, Jason Caravas, Benjamin Rambo-Martin, Shatavia Morrison, Christopher Gulvick, Scott Sammons, Yvette Unoarumhi, Darlene Wagner, Matthew Schmerer, Harry Gao, Mickey Li, John Gao, Joseph Fierro, Benafsh Sapra, Becky Tsai, Yan Meng, Doreen Ng, James Xie, Clinton R. Paden, Duncan MacCannell |
| EPI_ISL_3300414 | Gandhi Medical College | CDFD | Murali Bashyam, Asmita Gupta, Reelina Basu, Vinay Donipadi, Nagamani Kammili, Divya Vashisht, Ashwin Dalal |
| EPI_ISL_3300415, EPI_ISL_3300416, EPI_ISL_3300417, EPI_ISL_3300418, EPI_ISL_3300419 | Fulgent Genetics | Centers for Disease Control and Prevention Division of Viral Diseases, Pathogen Discovery | Dakota Howard, Dhwani Batra, Peter W. Cook, Kara Moser, Adrian Paskey, Jason Caravas, Benjamin Rambo-Martin, Shatavia Morrison, Christopher Gulvick, Scott Sammons, Yvette Unoarumhi, Darlene Wagner, Matthew Schmerer, Harry Gao, Mickey Li, John Gao, Joseph Fierro, Benafsh Sapra, Becky Tsai, Yan Meng, Doreen Ng, James Xie, Clinton R. Paden, Duncan MacCannell |
| EPI_ISL_3300420 | Gandhi Medical College | CDFD | Murali Bashyam, Asmita Gupta, Reelina Basu, Vinay Donipadi, Nagamani Kammili, Divya Vashisht, Ashwin Dalal |
| EPI_ISL_3300421, EPI_ISL_3300422, EPI_ISL_3300423, EPI_ISL_3300424 | Fulgent Genetics | Centers for Disease Control and Prevention Division of Viral Diseases, Pathogen Discovery | Dakota Howard, Dhwani Batra, Peter W. Cook, Kara Moser, Adrian Paskey, Jason Caravas, Benjamin Rambo-Martin, Shatavia Morrison, Christopher Gulvick, Scott Sammons, Yvette Unoarumhi, Darlene Wagner, Matthew Schmerer, Harry Gao, Mickey Li, John Gao, Joseph Fierro, Benafsh Sapra, Becky Tsai, Yan Meng, Doreen Ng, James Xie, Clinton R. Paden, Duncan MacCannell |
| EPI_ISL_3300425 | Gandhi Hospital | CDFD | Murali Bashyam, Asmita Gupta, Reelina Basu, Vinay Donipadi, Nagamani Kammili, Divya Vashisht, Ashwin Dalal |
| EPI_ISL_3300426, EPI_ISL_3300427, EPI_ISL_3300429, EPI_ISL_3300430 | Fulgent Genetics | Centers for Disease Control and Prevention Division of Viral Diseases, Pathogen Discovery | Dakota Howard, Dhwani Batra, Peter W. Cook, Kara Moser, Adrian Paskey, Jason Caravas, Benjamin Rambo-Martin, Shatavia Morrison, Christopher Gulvick, Scott Sammons, Yvette Unoarumhi, Darlene Wagner, Matthew Schmerer, Harry Gao, Mickey Li, John Gao, Joseph Fierro, Benafsh Sapra, Becky Tsai, Yan Meng, Doreen Ng, James Xie, Clinton R. Paden, Duncan MacCannell |
| EPI_ISL_3300431 | Gandhi Hospital | CDFD | Murali Bashyam, Asmita Gupta, Reelina Basu, Vinay Donipadi, Nagamani Kammili, Divya Vashisht, Ashwin Dalal |
| EPI_ISL_3300432, EPI_ISL_3300433, EPI_ISL_3300434 | Fulgent Genetics | Centers for Disease Control and Prevention Division of Viral Diseases, Pathogen Discovery | Dakota Howard, Dhwani Batra, Peter W. Cook, Kara Moser, Adrian Paskey, Jason Caravas, Benjamin Rambo-Martin, Shatavia Morrison, Christopher Gulvick, Scott Sammons, Yvette Unoarumhi, Darlene Wagner, Matthew Schmerer, Harry Gao, Mickey Li, John Gao, Joseph Fierro, Benafsh Sapra, Becky Tsai, Yan Meng, Doreen Ng, James Xie, Clinton R. Paden, Duncan MacCannell |
| EPI_ISL_3300435 | Gandhi Medical College | CDFD | Murali Bashyam, Asmita Gupta, Reelina Basu, Vinay Donipadi, Nagamani Kammili, Divya Vashisht, Ashwin Dalal |
| EPI_ISL_3300436, EPI_ISL_3300437 | Fulgent Genetics | Centers for Disease Control and Prevention Division of Viral Diseases, Pathogen Discovery | Dakota Howard, Dhwani Batra, Peter W. Cook, Kara Moser, Adrian Paskey, Jason Caravas, Benjamin Rambo-Martin, Shatavia Morrison, Christopher Gulvick, Scott Sammons, Yvette Unoarumhi, Darlene Wagner, Matthew Schmerer, Harry Gao, Mickey Li, John Gao, Joseph Fierro, Benafsh Sapra, Becky Tsai, Yan Meng, Doreen Ng, James Xie, Clinton R. Paden, Duncan MacCannell |
| EPI_ISL_3300438 | Gandhi Hospital | CDFD | Murali Bashyam, Asmita Gupta, Reelina Basu, Vinay Donipadi, Nagamani Kammili, Divya Vashisht, Ashwin Dalal |
| EPI_ISL_3300439, EPI_ISL_3300440, EPI_ISL_3300441 | Fulgent Genetics | Centers for Disease Control and Prevention Division of Viral Diseases, Pathogen Discovery | Dakota Howard, Dhwani Batra, Peter W. Cook, Kara Moser, Adrian Paskey, Jason Caravas, Benjamin Rambo-Martin, Shatavia Morrison, Christopher Gulvick, Scott Sammons, Yvette Unoarumhi, Darlene Wagner, Matthew Schmerer, Harry Gao, Mickey Li, John Gao, Joseph Fierro, Benafsh Sapra, Becky Tsai, Yan Meng, Doreen Ng, James Xie, Clinton R. Paden, Duncan MacCannell |
| EPI_ISL_3300442 | Gandhi Hospital | CDFD | Murali Bashyam, Asmita Gupta, Reelina Basu, Vinay Donipadi, Nagamani Kammili, Divya Vashisht, Ashwin Dalal |
| EPI_ISL_3300443, EPI_ISL_3300444 | Fulgent Genetics | Centers for Disease Control and Prevention Division of Viral Diseases, Pathogen Discovery | Dakota Howard, Dhwani Batra, Peter W. Cook, Kara Moser, Adrian Paskey, Jason Caravas, Benjamin Rambo-Martin, Shatavia Morrison, Christopher Gulvick, Scott Sammons, Yvette Unoarumhi, Darlene Wagner, Matthew Schmerer, Harry Gao, Mickey Li, John Gao, Joseph Fierro, Benafsh Sapra, Becky Tsai, Yan Meng, Doreen Ng, James Xie, Clinton R. Paden, Duncan MacCannell |
| EPI_ISL_3300445 | Gandhi Medical College | CDFD | Murali Bashyam, Asmita Gupta, Reelina Basu, Vinay Donipadi, Nagamani Kammili, Divya Vashisht, Ashwin Dalal |
| EPI_ISL_3300446, EPI_ISL_3300447 | Fulgent Genetics | Centers for Disease Control and Prevention Division of Viral Diseases, Pathogen Discovery | Dakota Howard, Dhwani Batra, Peter W. Cook, Kara Moser, Adrian Paskey, Jason Caravas, Benjamin Rambo-Martin, Shatavia Morrison, Christopher Gulvick, Scott Sammons, Yvette Unoarumhi, Darlene Wagner, Matthew Schmerer, Harry Gao, Mickey Li, John Gao, Joseph Fierro, Benafsh Sapra, Becky Tsai, Yan Meng, Doreen Ng, James Xie, Clinton R. Paden, Duncan MacCannell |
| EPI_ISL_3300448 | Gandhi Medical College | CDFD | Murali Bashyam, Asmita Gupta, Reelina Basu, Vinay Donipadi, Nagamani Kammili, Divya Vashisht, Ashwin Dalal |
| EPI_ISL_3300449, EPI_ISL_3300450 | Fulgent Genetics | Centers for Disease Control and Prevention Division of Viral Diseases, Pathogen Discovery | Dakota Howard, Dhwani Batra, Peter W. Cook, Kara Moser, Adrian Paskey, Jason Caravas, Benjamin Rambo-Martin, Shatavia Morrison, Christopher Gulvick, Scott Sammons, Yvette Unoarumhi, Darlene Wagner, Matthew Schmerer, Harry Gao, Mickey Li, John Gao, Joseph Fierro, Benafsh Sapra, Becky Tsai, Yan Meng, Doreen Ng, James Xie, Clinton R. Paden, Duncan MacCannell |
| EPI_ISL_3300451 | Gandhi Medical College | CDFD | Murali Bashyam, Asmita Gupta, Reelina Basu, Vinay Donipadi, Nagamani Kammili, Divya Vashisht, Ashwin Dalal |
| EPI_ISL_3300452, EPI_ISL_3300453, EPI_ISL_3300454 | Fulgent Genetics | Centers for Disease Control and Prevention Division of Viral Diseases, Pathogen Discovery | Dakota Howard, Dhwani Batra, Peter W. Cook, Kara Moser, Adrian Paskey, Jason Caravas, Benjamin Rambo-Martin, Shatavia Morrison, Christopher Gulvick, Scott Sammons, Yvette Unoarumhi, Darlene Wagner, Matthew Schmerer, Harry Gao, Mickey Li, John Gao, Joseph Fierro, Benafsh Sapra, Becky Tsai, Yan Meng, Doreen Ng, James Xie, Clinton R. Paden, Duncan MacCannell |
| EPI_ISL_3300455 | Gandhi Medical College | CDFD | Murali Bashyam, Asmita Gupta, Reelina Basu, Vinay Donipadi, Nagamani Kammili, Divya Vashisht, Ashwin Dalal |
| EPI_ISL_3300456, EPI_ISL_3300457 | Fulgent Genetics | Centers for Disease Control and Prevention Division of Viral Diseases, Pathogen Discovery | Dakota Howard, Dhwani Batra, Peter W. Cook, Kara Moser, Adrian Paskey, Jason Caravas, Benjamin Rambo-Martin, Shatavia Morrison, Christopher Gulvick, Scott Sammons, Yvette Unoarumhi, Darlene Wagner, Matthew Schmerer, Harry Gao, Mickey Li, John Gao, Joseph Fierro, Benafsh Sapra, Becky Tsai, Yan Meng, Doreen Ng, James Xie, Clinton R. Paden, Duncan MacCannell |
| EPI_ISL_3300458 | Gandhi Medical College | CDFD | Murali Bashyam, Asmita Gupta, Reelina Basu, Vinay Donipadi, Nagamani Kammili, Divya Vashisht, Ashwin Dalal |
| EPI_ISL_3300459, EPI_ISL_3300460, EPI_ISL_3300461 | Fulgent Genetics | Centers for Disease Control and Prevention Division of Viral Diseases, Pathogen Discovery | Dakota Howard, Dhwani Batra, Peter W. Cook, Kara Moser, Adrian Paskey, Jason Caravas, Benjamin Rambo-Martin, Shatavia Morrison, Christopher Gulvick, Scott Sammons, Yvette Unoarumhi, Darlene Wagner, Matthew Schmerer, Harry Gao, Mickey Li, John Gao, Joseph Fierro, Benafsh Sapra, Becky Tsai, Yan Meng, Doreen Ng, James Xie, Clinton R. Paden, Duncan MacCannell |

| | | | |
|---|---|---|---|
| EPI_ISL_3300462 | Gandhi Medical College | CDFD | Murali Bashyam, Asmita Gupta, Reelina Basu, Vinay Donipadi, Nagamani Kammili, Divya Vashisht, Ashwin Dalal |
| EPI_ISL_3300463, EPI_ISL_3300464 | Fulgent Genetics | Centers for Disease Control and Prevention Division of Viral Diseases, Pathogen Discovery | Dakota Howard, Dhwani Batra, Peter W. Cook, Kara Moser, Adrian Paskey, Jason Caravas, Benjamin Rambo-Martin, Shatavia Morrison, Christopher Gulvick, Scott Sammons, Yvette Unoarumhi, Darlene Wagner, Matthew Schmerer, Harry Gao, Mickey Li, John Gao, Joseph Fierro, Benafsh Sapra, Becky Tsai, Yan Meng, Doreen Ng, James Xie, Clinton R. Paden, Duncan MacCannell |
| EPI_ISL_3300465 | Gandhi Medical College | CDFD | Murali Bashyam, Asmita Gupta, Reelina Basu, Vinay Donipadi, Nagamani Kammili, Divya Vashisht, Ashwin Dalal |
| EPI_ISL_3300466, EPI_ISL_3300467 | Fulgent Genetics | Centers for Disease Control and Prevention Division of Viral Diseases, Pathogen Discovery | Dakota Howard, Dhwani Batra, Peter W. Cook, Kara Moser, Adrian Paskey, Jason Caravas, Benjamin Rambo-Martin, Shatavia Morrison, Christopher Gulvick, Scott Sammons, Yvette Unoarumhi, Darlene Wagner, Matthew Schmerer, Harry Gao, Mickey Li, John Gao, Joseph Fierro, Benafsh Sapra, Becky Tsai, Yan Meng, Doreen Ng, James Xie, Clinton R. Paden, Duncan MacCannell |
| EPI_ISL_3300468 | Gandhi Medical College | CDFD | Murali Bashyam, Asmita Gupta, Reelina Basu, Vinay Donipadi, Nagamani Kammili, Divya Vashisht, Ashwin Dalal |
| EPI_ISL_3300469, EPI_ISL_3300471 | Fulgent Genetics | Centers for Disease Control and Prevention Division of Viral Diseases, Pathogen Discovery | Dakota Howard, Dhwani Batra, Peter W. Cook, Kara Moser, Adrian Paskey, Jason Caravas, Benjamin Rambo-Martin, Shatavia Morrison, Christopher Gulvick, Scott Sammons, Yvette Unoarumhi, Darlene Wagner, Matthew Schmerer, Harry Gao, Mickey Li, John Gao, Joseph Fierro, Benafsh Sapra, Becky Tsai, Yan Meng, Doreen Ng, James Xie, Clinton R. Paden, Duncan MacCannell |
| EPI_ISL_3300472 | Gandhi Medical College | CDFD | Murali Bashyam, Asmita Gupta, Reelina Basu, Vinay Donipadi, Nagamani Kammili, Divya Vashisht, Ashwin Dalal |
| EPI_ISL_3300473, EPI_ISL_3300474 | Fulgent Genetics | Centers for Disease Control and Prevention Division of Viral Diseases, Pathogen Discovery | Dakota Howard, Dhwani Batra, Peter W. Cook, Kara Moser, Adrian Paskey, Jason Caravas, Benjamin Rambo-Martin, Shatavia Morrison, Christopher Gulvick, Scott Sammons, Yvette Unoarumhi, Darlene Wagner, Matthew Schmerer, Harry Gao, Mickey Li, John Gao, Joseph Fierro, Benafsh Sapra, Becky Tsai, Yan Meng, Doreen Ng, James Xie, Clinton R. Paden, Duncan MacCannell |
| EPI_ISL_3300475 | Gandhi Medical College | CDFD | Murali Bashyam, Asmita Gupta, Reelina Basu, Vinay Donipadi, Nagamani Kammili, Divya Vashisht, Ashwin Dalal |
| EPI_ISL_3300476, EPI_ISL_3300477 | Fulgent Genetics | Centers for Disease Control and Prevention Division of Viral Diseases, Pathogen Discovery | Dakota Howard, Dhwani Batra, Peter W. Cook, Kara Moser, Adrian Paskey, Jason Caravas, Benjamin Rambo-Martin, Shatavia Morrison, Christopher Gulvick, Scott Sammons, Yvette Unoarumhi, Darlene Wagner, Matthew Schmerer, Harry Gao, Mickey Li, John Gao, Joseph Fierro, Benafsh Sapra, Becky Tsai, Yan Meng, Doreen Ng, James Xie, Clinton R. Paden, Duncan MacCannell |
| EPI_ISL_3300478 | Gandhi Medical College | CDFD | Murali Bashyam, Asmita Gupta, Reelina Basu, Vinay Donipadi, Nagamani Kammili, Divya Vashisht, Ashwin Dalal |
| EPI_ISL_3300479, EPI_ISL_3300480, EPI_ISL_3300481 | Fulgent Genetics | Centers for Disease Control and Prevention Division of Viral Diseases, Pathogen Discovery | Dakota Howard, Dhwani Batra, Peter W. Cook, Kara Moser, Adrian Paskey, Jason Caravas, Benjamin Rambo-Martin, Shatavia Morrison, Christopher Gulvick, Scott Sammons, Yvette Unoarumhi, Darlene Wagner, Matthew Schmerer, Harry Gao, Mickey Li, John Gao, Joseph Fierro, Benafsh Sapra, Becky Tsai, Yan Meng, Doreen Ng, James Xie, Clinton R. Paden, Duncan MacCannell |
| EPI_ISL_3300482 | Gandhi Medical College | CDFD | Murali Bashyam, Asmita Gupta, Reelina Basu, Vinay Donipadi, Nagamani Kammili, Divya Vashisht, Ashwin Dalal |
| EPI_ISL_3300483, EPI_ISL_3300484 | Fulgent Genetics | Centers for Disease Control and Prevention Division of Viral Diseases, Pathogen Discovery | Dakota Howard, Dhwani Batra, Peter W. Cook, Kara Moser, Adrian Paskey, Jason Caravas, Benjamin Rambo-Martin, Shatavia Morrison, Christopher Gulvick, Scott Sammons, Yvette Unoarumhi, Darlene Wagner, Matthew Schmerer, Harry Gao, Mickey Li, John Gao, Joseph Fierro, Benafsh Sapra, Becky Tsai, Yan Meng, Doreen Ng, James Xie, Clinton R. Paden, Duncan MacCannell |
| EPI_ISL_3300485 | Gandhi Medical College | CDFD | Murali Bashyam, Asmita Gupta, Reelina Basu, Vinay Donipadi, Nagamani Kammili, Divya Vashisht, Ashwin Dalal |
| EPI_ISL_3300486, EPI_ISL_3300487 | Fulgent Genetics | Centers for Disease Control and Prevention Division of Viral Diseases, Pathogen Discovery | Dakota Howard, Dhwani Batra, Peter W. Cook, Kara Moser, Adrian Paskey, Jason Caravas, Benjamin Rambo-Martin, Shatavia Morrison, Christopher Gulvick, Scott Sammons, Yvette Unoarumhi, Darlene Wagner, Matthew Schmerer, Harry Gao, Mickey Li, John Gao, Joseph Fierro, Benafsh Sapra, Becky Tsai, Yan Meng, Doreen Ng, James Xie, Clinton R. Paden, Duncan MacCannell |
| EPI_ISL_3300488 | Gandhi Medical College | CDFD | Murali Bashyam, Asmita Gupta, Reelina Basu, Vinay Donipadi, Nagamani Kammili, Divya Vashisht, Ashwin Dalal |
| EPI_ISL_3300489, EPI_ISL_3300490, EPI_ISL_3300491 | Fulgent Genetics | Centers for Disease Control and Prevention Division of Viral Diseases, Pathogen Discovery | Dakota Howard, Dhwani Batra, Peter W. Cook, Kara Moser, Adrian Paskey, Jason Caravas, Benjamin Rambo-Martin, Shatavia Morrison, Christopher Gulvick, Scott Sammons, Yvette Unoarumhi, Darlene Wagner, Matthew Schmerer, Harry Gao, Mickey Li, John Gao, Joseph Fierro, Benafsh Sapra, Becky Tsai, Yan Meng, Doreen Ng, James Xie, Clinton R. Paden, Duncan MacCannell |
| EPI_ISL_3300492 | Gandhi Medical College | CDFD | Murali Bashyam, Asmita Gupta, Reelina Basu, Vinay Donipadi, Nagamani Kammili, Divya Vashisht, Ashwin Dalal |
| EPI_ISL_3300493, EPI_ISL_3300494 | Fulgent Genetics | Centers for Disease Control and Prevention Division of Viral Diseases, Pathogen Discovery | Dakota Howard, Dhwani Batra, Peter W. Cook, Kara Moser, Adrian Paskey, Jason Caravas, Benjamin Rambo-Martin, Shatavia Morrison, Christopher Gulvick, Scott Sammons, Yvette Unoarumhi, Darlene Wagner, Matthew Schmerer, Harry Gao, Mickey Li, John Gao, Joseph Fierro, Benafsh Sapra, Becky Tsai, Yan Meng, Doreen Ng, James Xie, Clinton R. Paden, Duncan MacCannell |
| EPI_ISL_3300495 | Gandhi Medical College | CDFD | Murali Bashyam, Asmita Gupta, Reelina Basu, Vinay Donipadi, Nagamani Kammili, Divya Vashisht, Ashwin Dalal |
| EPI_ISL_3300496, EPI_ISL_3300497 | Fulgent Genetics | Centers for Disease Control and Prevention Division of Viral Diseases, Pathogen Discovery | Dakota Howard, Dhwani Batra, Peter W. Cook, Kara Moser, Adrian Paskey, Jason Caravas, Benjamin Rambo-Martin, Shatavia Morrison, Christopher Gulvick, Scott Sammons, Yvette Unoarumhi, Darlene Wagner, Matthew Schmerer, Harry Gao, Mickey Li, John Gao, Joseph Fierro, Benafsh Sapra, Becky Tsai, Yan Meng, Doreen Ng, James Xie, Clinton R. Paden, Duncan MacCannell |
| EPI_ISL_3300498 | Gandhi Hospital | CDFD | Murali Bashyam, Asmita Gupta, Reelina Basu, Vinay Donipadi, Nagamani Kammili, Divya Vashisht, Ashwin Dalal |
| EPI_ISL_3300499, EPI_ISL_3300500, EPI_ISL_3300502, EPI_ISL_3300503 | Fulgent Genetics | Centers for Disease Control and Prevention Division of Viral Diseases, Pathogen Discovery | Dakota Howard, Dhwani Batra, Peter W. Cook, Kara Moser, Adrian Paskey, Jason Caravas, Benjamin Rambo-Martin, Shatavia Morrison, Christopher Gulvick, Scott Sammons, Yvette Unoarumhi, Darlene Wagner, Matthew Schmerer, Harry Gao, Mickey Li, John Gao, Joseph Fierro, Benafsh Sapra, Becky Tsai, Yan Meng, Doreen Ng, James Xie, Clinton R. Paden, Duncan MacCannell |
| EPI_ISL_3300504 | Gandhi Medical College | CDFD | Murali Bashyam, Asmita Gupta, Reelina Basu, Vinay Donipadi, Nagamani Kammili, Divya Vashisht, Ashwin Dalal |
| EPI_ISL_3300506 | Fulgent Genetics | Centers for Disease Control and Prevention Division of Viral Diseases, Pathogen Discovery | Dakota Howard, Dhwani Batra, Peter W. Cook, Kara Moser, Adrian Paskey, Jason Caravas, Benjamin Rambo-Martin, Shatavia Morrison, Christopher Gulvick, Scott Sammons, Yvette Unoarumhi, Darlene Wagner, Matthew Schmerer, Harry Gao, Mickey Li, John Gao, Joseph Fierro, Benafsh Sapra, Becky Tsai, Yan Meng, Doreen Ng, James Xie, Clinton R. Paden, Duncan MacCannell |
| EPI_ISL_3300507 | Gandhi Medical College | CDFD | Murali Bashyam, Asmita Gupta, Reelina Basu, Vinay Donipadi, Nagamani Kammili, Divya Vashisht, Ashwin Dalal |
| EPI_ISL_3300508, EPI_ISL_3300509 | Fulgent Genetics | Centers for Disease Control and Prevention Division of Viral Diseases, Pathogen Discovery | Dakota Howard, Dhwani Batra, Peter W. Cook, Kara Moser, Adrian Paskey, Jason Caravas, Benjamin Rambo-Martin, Shatavia Morrison, Christopher Gulvick, Scott Sammons, Yvette Unoarumhi, Darlene Wagner, Matthew Schmerer, Harry Gao, Mickey Li, John Gao, Joseph Fierro, Benafsh Sapra, Becky Tsai, Yan Meng, Doreen Ng, James Xie, Clinton R. Paden, Duncan MacCannell |
| EPI_ISL_3300510 | Gandhi Medical College | CDFD | Murali Bashyam, Asmita Gupta, Reelina Basu, Vinay Donipadi, Nagamani Kammili, Divya Vashisht, Ashwin Dalal |
| EPI_ISL_3300511, EPI_ISL_3300512 | Fulgent Genetics | Centers for Disease Control and Prevention Division of Viral Diseases, Pathogen Discovery | Dakota Howard, Dhwani Batra, Peter W. Cook, Kara Moser, Adrian Paskey, Jason Caravas, Benjamin Rambo-Martin, Shatavia Morrison, Christopher Gulvick, Scott Sammons, Yvette Unoarumhi, Darlene Wagner, Matthew Schmerer, Harry Gao, Mickey Li, John Gao, Joseph Fierro, Benafsh Sapra, Becky Tsai, Yan Meng, Doreen Ng, James Xie, Clinton R. Paden, Duncan MacCannell |
| EPI_ISL_3300513 | Gandhi Medical College | CDFD | Murali Bashyam, Asmita Gupta, Reelina Basu, Vinay Donipadi, Nagamani Kammili, Divya Vashisht, Ashwin Dalal |
| EPI_ISL_3300514, EPI_ISL_3300515 | Fulgent Genetics | Centers for Disease Control and Prevention Division of Viral Diseases, Pathogen Discovery | Dakota Howard, Dhwani Batra, Peter W. Cook, Kara Moser, Adrian Paskey, Jason Caravas, Benjamin Rambo-Martin, Shatavia Morrison, Christopher Gulvick, Scott Sammons, Yvette Unoarumhi, Darlene Wagner, Matthew Schmerer, Harry Gao, Mickey Li, John Gao, Joseph Fierro, Benafsh Sapra, Becky Tsai, Yan Meng, Doreen Ng, James Xie, Clinton R. Paden, Duncan MacCannell |
| EPI_ISL_3300516 | Gandhi Medical College | CDFD | Murali Bashyam, Asmita Gupta, Reelina Basu, Vinay Donipadi, Nagamani Kammili, Divya Vashisht, Ashwin Dalal |
| EPI_ISL_3300517, EPI_ISL_3300518 | Fulgent Genetics | Centers for Disease Control and Prevention Division of Viral Diseases, Pathogen Discovery | Dakota Howard, Dhwani Batra, Peter W. Cook, Kara Moser, Adrian Paskey, Jason Caravas, Benjamin Rambo-Martin, Shatavia Morrison, Christopher Gulvick, Scott Sammons, Yvette Unoarumhi, Darlene Wagner, Matthew Schmerer, Harry Gao, Mickey Li, John Gao, Joseph Fierro, Benafsh Sapra, Becky Tsai, Yan Meng, Doreen Ng, James Xie, Clinton R. Paden, Duncan MacCannell |
| EPI_ISL_3300519 | Gandhi Medical College | CDFD | Murali Bashyam, Asmita Gupta, Reelina Basu, Vinay Donipadi, Nagamani Kammili, Divya Vashisht, Ashwin Dalal |
| EPI_ISL_3300520, EPI_ISL_3300521 | Fulgent Genetics | Centers for Disease Control and Prevention Division of Viral Diseases, Pathogen Discovery | Dakota Howard, Dhwani Batra, Peter W. Cook, Kara Moser, Adrian Paskey, Jason Caravas, Benjamin Rambo-Martin, Shatavia Morrison, Christopher Gulvick, Scott Sammons, Yvette Unoarumhi, Darlene Wagner, Matthew Schmerer, Harry Gao, Mickey Li, John Gao, Joseph Fierro, Benafsh Sapra, Becky Tsai, Yan Meng, Doreen Ng, James Xie, Clinton R. Paden, Duncan MacCannell |
| EPI_ISL_3300522 | Gandhi Medical College | CDFD | Murali Bashyam, Asmita Gupta, Reelina Basu, Vinay Donipadi, Nagamani Kammili, Divya Vashisht, Ashwin Dalal |
| EPI_ISL_3300523, EPI_ISL_3300524, EPI_ISL_3300525, EPI_ISL_3300527, EPI_ISL_3300528 | Fulgent Genetics | Centers for Disease Control and Prevention Division of Viral Diseases, Pathogen Discovery | Dakota Howard, Dhwani Batra, Peter W. Cook, Kara Moser, Adrian Paskey, Jason Caravas, Benjamin Rambo-Martin, Shatavia Morrison, Christopher Gulvick, Scott Sammons, Yvette Unoarumhi, Darlene Wagner, Matthew Schmerer, Harry Gao, Mickey Li, John Gao, Joseph Fierro, Benafsh Sapra, Becky Tsai, Yan Meng, Doreen Ng, James Xie, Clinton R. Paden, Duncan MacCannell |
| EPI_ISL_3300529 | Gandhi Medical College | CDFD | Murali Bashyam, Asmita Gupta, Reelina Basu, Vinay Donipadi, Nagamani Kammili, Divya Vashisht, Ashwin Dalal |
| EPI_ISL_3300530 | Fulgent Genetics | Centers for Disease Control and Prevention Division of Viral Diseases, Pathogen Discovery | Dakota Howard, Dhwani Batra, Peter W. Cook, Kara Moser, Adrian Paskey, Jason Caravas, Benjamin Rambo-Martin, Shatavia Morrison, Christopher Gulvick, Scott Sammons, Yvette Unoarumhi, Darlene Wagner, Matthew Schmerer, Harry Gao, Mickey Li, John Gao, Joseph Fierro, Benafsh Sapra, Becky Tsai, Yan Meng, Doreen Ng, James Xie, Clinton R. Paden, Duncan MacCannell |
| EPI_ISL_3300531 | Gandhi Hospital | CDFD | Murali Bashyam, Asmita Gupta, Reelina Basu, Vinay Donipadi, Nagamani Kammili, Divya Vashisht, Ashwin Dalal |
| EPI_ISL_3300532, EPI_ISL_3300533 | Fulgent Genetics | Centers for Disease Control and Prevention Division of Viral Diseases, Pathogen Discovery | Dakota Howard, Dhwani Batra, Peter W. Cook, Kara Moser, Adrian Paskey, Jason Caravas, Benjamin Rambo-Martin, Shatavia Morrison, Christopher Gulvick, Scott Sammons, Yvette Unoarumhi, Darlene Wagner, Matthew Schmerer, Harry Gao, Mickey Li, John Gao, Joseph Fierro, Benafsh Sapra, Becky Tsai, Yan Meng, Doreen Ng, James Xie, Clinton R. Paden, Duncan MacCannell |
| EPI_ISL_3300534 | Gandhi Medical College | CDFD | Murali Bashyam, Asmita Gupta, Reelina Basu, Vinay Donipadi, Nagamani Kammili, Divya Vashisht, Ashwin Dalal |
| EPI_ISL_3300535, EPI_ISL_3300536, EPI_ISL_3300537 | Fulgent Genetics | Centers for Disease Control and Prevention Division of Viral Diseases, Pathogen Discovery | Dakota Howard, Dhwani Batra, Peter W. Cook, Kara Moser, Adrian Paskey, Jason Caravas, Benjamin Rambo-Martin, Shatavia Morrison, Christopher Gulvick, Scott Sammons, Yvette Unoarumhi, Darlene Wagner, Matthew Schmerer, Harry Gao, Mickey Li, John Gao, Joseph Fierro, Benafsh Sapra, Becky Tsai, Yan Meng, Doreen Ng, James Xie, Clinton R. Paden, Duncan MacCannell |
| EPI_ISL_3300538 | Gandhi Medical College | CDFD | Murali Bashyam, Asmita Gupta, Reelina Basu, Vinay Donipadi, Nagamani Kammili, Divya Vashisht, Ashwin Dalal |
| EPI_ISL_3300539, EPI_ISL_3300540 | Fulgent Genetics | Centers for Disease Control and Prevention Division of Viral Diseases, Pathogen Discovery | Dakota Howard, Dhwani Batra, Peter W. Cook, Kara Moser, Adrian Paskey, Jason Caravas, Benjamin Rambo-Martin, Shatavia Morrison, Christopher Gulvick, Scott Sammons, Yvette Unoarumhi, Darlene Wagner, Matthew Schmerer, Harry Gao, Mickey Li, John Gao, Joseph Fierro, Benafsh Sapra, Becky Tsai, Yan Meng, Doreen Ng, James Xie, Clinton R. Paden, Duncan MacCannell |
| EPI_ISL_3300541 | Gandhi Medical College | CDFD | Murali Bashyam, Asmita Gupta, Reelina Basu, Vinay Donipadi, Nagamani Kammili, Divya Vashisht, Ashwin Dalal |
| EPI_ISL_3300542 | Fulgent Genetics | Centers for Disease Control and Prevention Division of Viral Diseases, Pathogen Discovery | Dakota Howard, Dhwani Batra, Peter W. Cook, Kara Moser, Adrian Paskey, Jason Caravas, Benjamin Rambo-Martin, Shatavia Morrison, Christopher Gulvick, Scott Sammons, Yvette Unoarumhi, Darlene Wagner, Matthew Schmerer, Harry Gao, Mickey Li, John Gao, Joseph Fierro, Benafsh Sapra, Becky Tsai, Yan Meng, Doreen Ng, James Xie, Clinton R. Paden, Duncan MacCannell |
| EPI_ISL_3300543 | Gandhi Hospital | CDFD | Murali Bashyam, Asmita Gupta, Reelina Basu, Vinay Donipadi, Nagamani Kammili, Divya Vashisht, Ashwin Dalal |
| EPI_ISL_3300544, EPI_ISL_3300545 | Fulgent Genetics | Centers for Disease Control and Prevention Division of Viral Diseases, Pathogen Discovery | Dakota Howard, Dhwani Batra, Peter W. Cook, Kara Moser, Adrian Paskey, Jason Caravas, Benjamin Rambo-Martin, Shatavia Morrison, Christopher Gulvick, Scott Sammons, Yvette Unoarumhi, Darlene Wagner, Matthew Schmerer, Harry Gao, Mickey Li, John Gao, Joseph Fierro, Benafsh Sapra, Becky Tsai, Yan Meng, Doreen Ng, James Xie, Clinton R. Paden, Duncan MacCannell |
| EPI_ISL_3300546 | Gandhi Hospital | CDFD | Murali Bashyam, Asmita Gupta, Reelina Basu, Vinay Donipadi, Nagamani Kammili, Divya Vashisht, Ashwin Dalal |
| EPI_ISL_3300547, EPI_ISL_3300548, EPI_ISL_3300550, | Fulgent Genetics | Centers for Disease Control and Prevention Division of Viral Diseases, Pathogen Discovery | Dakota Howard, Dhwani Batra, Peter W. Cook, Kara Moser, Adrian Paskey, Jason Caravas, Benjamin Rambo-Martin, Shatavia Morrison, Christopher Gulvick, Scott Sammons, Yvette Unoarumhi, Darlene Wagner, Matthew Schmerer, Harry Gao, Mickey Li, John Gao, Joseph Fierro, Benafsh Sapra, Becky Tsai, Yan Meng, Doreen Ng, James Xie, Clinton R. Paden, Duncan MacCannell |

| | | | |
|---|---|---|---|
| EPI_ISL_3300551, EPI_ISL_3300553, EPI_ISL_3300554 | | | |
| EPI_ISL_3300555 | Gandhi Medical College | CDFD | Murali Bashyam, Asmita Gupta, Reelina Basu, Vinay Donipadi, Nagamani Kammili, Divya Vashisht, Ashwin Dalal |
| EPI_ISL_3300556, EPI_ISL_3300557 | Fulgent Genetics | Centers for Disease Control and Prevention Division of Viral Diseases, Pathogen Discovery | Dakota Howard, Dhwani Batra, Peter W. Cook, Kara Moser, Adrian Paskey, Jason Caravas, Benjamin Rambo-Martin, Shatavia Morrison, Christopher Gulvick, Scott Sammons, Yvette Unoarumhi, Darlene Wagner, Matthew Schmerer, Harry Gao, Mickey Li, John Gao, Joseph Fierro, Benafsh Sapra, Becky Tsai, Yan Meng, Doreen Ng, James Xie, Clinton R. Paden, Duncan MacCannell |
| EPI_ISL_3300558 | Gandhi Medical College | CDFD | Murali Bashyam, Asmita Gupta, Reelina Basu, Vinay Donipadi, Nagamani Kammili, Divya Vashisht, Ashwin Dalal |
| EPI_ISL_3300559, EPI_ISL_3300560 | Fulgent Genetics | Centers for Disease Control and Prevention Division of Viral Diseases, Pathogen Discovery | Dakota Howard, Dhwani Batra, Peter W. Cook, Kara Moser, Adrian Paskey, Jason Caravas, Benjamin Rambo-Martin, Shatavia Morrison, Christopher Gulvick, Scott Sammons, Yvette Unoarumhi, Darlene Wagner, Matthew Schmerer, Harry Gao, Mickey Li, John Gao, Joseph Fierro, Benafsh Sapra, Becky Tsai, Yan Meng, Doreen Ng, James Xie, Clinton R. Paden, Duncan MacCannell |
| EPI_ISL_3300561 | Gandhi Medical College | CDFD | Murali Bashyam, Asmita Gupta, Reelina Basu, Vinay Donipadi, Nagamani Kammili, Divya Vashisht, Ashwin Dalal |
| EPI_ISL_3300562, EPI_ISL_3300563 | Fulgent Genetics | Centers for Disease Control and Prevention Division of Viral Diseases, Pathogen Discovery | Dakota Howard, Dhwani Batra, Peter W. Cook, Kara Moser, Adrian Paskey, Jason Caravas, Benjamin Rambo-Martin, Shatavia Morrison, Christopher Gulvick, Scott Sammons, Yvette Unoarumhi, Darlene Wagner, Matthew Schmerer, Harry Gao, Mickey Li, John Gao, Joseph Fierro, Benafsh Sapra, Becky Tsai, Yan Meng, Doreen Ng, James Xie, Clinton R. Paden, Duncan MacCannell |
| EPI_ISL_3300564 | Gandhi Medical College | CDFD | Murali Bashyam, Asmita Gupta, Reelina Basu, Vinay Donipadi, Nagamani Kammili, Divya Vashisht, Ashwin Dalal |
| EPI_ISL_3300565, EPI_ISL_3300566 | Fulgent Genetics | Centers for Disease Control and Prevention Division of Viral Diseases, Pathogen Discovery | Dakota Howard, Dhwani Batra, Peter W. Cook, Kara Moser, Adrian Paskey, Jason Caravas, Benjamin Rambo-Martin, Shatavia Morrison, Christopher Gulvick, Scott Sammons, Yvette Unoarumhi, Darlene Wagner, Matthew Schmerer, Harry Gao, Mickey Li, John Gao, Joseph Fierro, Benafsh Sapra, Becky Tsai, Yan Meng, Doreen Ng, James Xie, Clinton R. Paden, Duncan MacCannell |
| EPI_ISL_3300567 | Gandhi Medical College | CDFD | Murali Bashyam, Asmita Gupta, Reelina Basu, Vinay Donipadi, Nagamani Kammili, Divya Vashisht, Ashwin Dalal |
| EPI_ISL_3300568, EPI_ISL_3300569 | Fulgent Genetics | Centers for Disease Control and Prevention Division of Viral Diseases, Pathogen Discovery | Dakota Howard, Dhwani Batra, Peter W. Cook, Kara Moser, Adrian Paskey, Jason Caravas, Benjamin Rambo-Martin, Shatavia Morrison, Christopher Gulvick, Scott Sammons, Yvette Unoarumhi, Darlene Wagner, Matthew Schmerer, Harry Gao, Mickey Li, John Gao, Joseph Fierro, Benafsh Sapra, Becky Tsai, Yan Meng, Doreen Ng, James Xie, Clinton R. Paden, Duncan MacCannell |
| EPI_ISL_3300570 | Gandhi Hospital | CDFD | Murali Bashyam, Asmita Gupta, Reelina Basu, Vinay Donipadi, Nagamani Kammili, Divya Vashisht, Ashwin Dalal |
| EPI_ISL_3300571, EPI_ISL_3300572 | Fulgent Genetics | Centers for Disease Control and Prevention Division of Viral Diseases, Pathogen Discovery | Dakota Howard, Dhwani Batra, Peter W. Cook, Kara Moser, Adrian Paskey, Jason Caravas, Benjamin Rambo-Martin, Shatavia Morrison, Christopher Gulvick, Scott Sammons, Yvette Unoarumhi, Darlene Wagner, Matthew Schmerer, Harry Gao, Mickey Li, John Gao, Joseph Fierro, Benafsh Sapra, Becky Tsai, Yan Meng, Doreen Ng, James Xie, Clinton R. Paden, Duncan MacCannell |
| EPI_ISL_3300573 | Gandhi Hospital | CDFD | Murali Bashyam, Asmita Gupta, Reelina Basu, Vinay Donipadi, Nagamani Kammili, Divya Vashisht, Ashwin Dalal |
| EPI_ISL_3300574, EPI_ISL_3300575 | Fulgent Genetics | Centers for Disease Control and Prevention Division of Viral Diseases, Pathogen Discovery | Dakota Howard, Dhwani Batra, Peter W. Cook, Kara Moser, Adrian Paskey, Jason Caravas, Benjamin Rambo-Martin, Shatavia Morrison, Christopher Gulvick, Scott Sammons, Yvette Unoarumhi, Darlene Wagner, Matthew Schmerer, Harry Gao, Mickey Li, John Gao, Joseph Fierro, Benafsh Sapra, Becky Tsai, Yan Meng, Doreen Ng, James Xie, Clinton R. Paden, Duncan MacCannell |
| EPI_ISL_3300576 | Gandhi Hospital | CDFD | Murali Bashyam, Asmita Gupta, Reelina Basu, Vinay Donipadi, Nagamani Kammili, Divya Vashisht, Ashwin Dalal |
| EPI_ISL_3300577, EPI_ISL_3300578 | Fulgent Genetics | Centers for Disease Control and Prevention Division of Viral Diseases, Pathogen Discovery | Dakota Howard, Dhwani Batra, Peter W. Cook, Kara Moser, Adrian Paskey, Jason Caravas, Benjamin Rambo-Martin, Shatavia Morrison, Christopher Gulvick, Scott Sammons, Yvette Unoarumhi, Darlene Wagner, Matthew Schmerer, Harry Gao, Mickey Li, John Gao, Joseph Fierro, Benafsh Sapra, Becky Tsai, Yan Meng, Doreen Ng, James Xie, Clinton R. Paden, Duncan MacCannell |
| EPI_ISL_3300579 | Gandhi Hospital | CDFD | Murali Bashyam, Asmita Gupta, Reelina Basu, Vinay Donipadi, Nagamani Kammili, Divya Vashisht, Ashwin Dalal |
| EPI_ISL_3300581 | Fulgent Genetics | Centers for Disease Control and Prevention Division of Viral Diseases, Pathogen Discovery | Dakota Howard, Dhwani Batra, Peter W. Cook, Kara Moser, Adrian Paskey, Jason Caravas, Benjamin Rambo-Martin, Shatavia Morrison, Christopher Gulvick, Scott Sammons, Yvette Unoarumhi, Darlene Wagner, Matthew Schmerer, Harry Gao, Mickey Li, John Gao, Joseph Fierro, Benafsh Sapra, Becky Tsai, Yan Meng, Doreen Ng, James Xie, Clinton R. Paden, Duncan MacCannell |
| EPI_ISL_3300582 | Gandhi Hospital | CDFD | Murali Bashyam, Asmita Gupta, Reelina Basu, Vinay Donipadi, Nagamani Kammili, Divya Vashisht, Ashwin Dalal |
| EPI_ISL_3300583, EPI_ISL_3300584 | Fulgent Genetics | Centers for Disease Control and Prevention Division of Viral Diseases, Pathogen Discovery | Dakota Howard, Dhwani Batra, Peter W. Cook, Kara Moser, Adrian Paskey, Jason Caravas, Benjamin Rambo-Martin, Shatavia Morrison, Christopher Gulvick, Scott Sammons, Yvette Unoarumhi, Darlene Wagner, Matthew Schmerer, Harry Gao, Mickey Li, John Gao, Joseph Fierro, Benafsh Sapra, Becky Tsai, Yan Meng, Doreen Ng, James Xie, Clinton R. Paden, Duncan MacCannell |
| EPI_ISL_3300585 | Gandhi Hospital | CDFD | Murali Bashyam, Asmita Gupta, Reelina Basu, Vinay Donipadi, Nagamani Kammili, Divya Vashisht, Ashwin Dalal |
| EPI_ISL_3300586, EPI_ISL_3300587 | Fulgent Genetics | Centers for Disease Control and Prevention Division of Viral Diseases, Pathogen Discovery | Dakota Howard, Dhwani Batra, Peter W. Cook, Kara Moser, Adrian Paskey, Jason Caravas, Benjamin Rambo-Martin, Shatavia Morrison, Christopher Gulvick, Scott Sammons, Yvette Unoarumhi, Darlene Wagner, Matthew Schmerer, Harry Gao, Mickey Li, John Gao, Joseph Fierro, Benafsh Sapra, Becky Tsai, Yan Meng, Doreen Ng, James Xie, Clinton R. Paden, Duncan MacCannell |
| EPI_ISL_3300588 | Gandhi Medical College | CDFD | Murali Bashyam, Asmita Gupta, Reelina Basu, Vinay Donipadi, Nagamani Kammili, Divya Vashisht, Ashwin Dalal |
| EPI_ISL_3300589 | Fulgent Genetics | Centers for Disease Control and Prevention Division of Viral Diseases, Pathogen Discovery | Dakota Howard, Dhwani Batra, Peter W. Cook, Kara Moser, Adrian Paskey, Jason Caravas, Benjamin Rambo-Martin, Shatavia Morrison, Christopher Gulvick, Scott Sammons, Yvette Unoarumhi, Darlene Wagner, Matthew Schmerer, Harry Gao, Mickey Li, John Gao, Joseph Fierro, Benafsh Sapra, Becky Tsai, Yan Meng, Doreen Ng, James Xie, Clinton R. Paden, Duncan MacCannell |
| EPI_ISL_3300590 | Gandhi Medical College | CDFD | Murali Bashyam, Asmita Gupta, Reelina Basu, Vinay Donipadi, Nagamani Kammili, Divya Vashisht, Ashwin Dalal |
| EPI_ISL_3300591, EPI_ISL_3300592, EPI_ISL_3300593 | Fulgent Genetics | Centers for Disease Control and Prevention Division of Viral Diseases, Pathogen Discovery | Dakota Howard, Dhwani Batra, Peter W. Cook, Kara Moser, Adrian Paskey, Jason Caravas, Benjamin Rambo-Martin, Shatavia Morrison, Christopher Gulvick, Scott Sammons, Yvette Unoarumhi, Darlene Wagner, Matthew Schmerer, Harry Gao, Mickey Li, John Gao, Joseph Fierro, Benafsh Sapra, Becky Tsai, Yan Meng, Doreen Ng, James Xie, Clinton R. Paden, Duncan MacCannell |
| EPI_ISL_3300594 | Gandhi Medical College | CDFD | Murali Bashyam, Asmita Gupta, Reelina Basu, Vinay Donipadi, Nagamani Kammili, Divya Vashisht, Ashwin Dalal |
| EPI_ISL_3300595, EPI_ISL_3300596 | Fulgent Genetics | Centers for Disease Control and Prevention Division of Viral Diseases, Pathogen Discovery | Dakota Howard, Dhwani Batra, Peter W. Cook, Kara Moser, Adrian Paskey, Jason Caravas, Benjamin Rambo-Martin, Shatavia Morrison, Christopher Gulvick, Scott Sammons, Yvette Unoarumhi, Darlene Wagner, Matthew Schmerer, Harry Gao, Mickey Li, John Gao, Joseph Fierro, Benafsh Sapra, Becky Tsai, Yan Meng, Doreen Ng, James Xie, Clinton R. Paden, Duncan MacCannell |
| EPI_ISL_3300597 | Gandhi Medical College | CDFD | Murali Bashyam, Asmita Gupta, Reelina Basu, Vinay Donipadi, Nagamani Kammili, Divya Vashisht, Ashwin Dalal |
| EPI_ISL_3300598, EPI_ISL_3300599 | Fulgent Genetics | Centers for Disease Control and Prevention Division of Viral Diseases, Pathogen Discovery | Dakota Howard, Dhwani Batra, Peter W. Cook, Kara Moser, Adrian Paskey, Jason Caravas, Benjamin Rambo-Martin, Shatavia Morrison, Christopher Gulvick, Scott Sammons, Yvette Unoarumhi, Darlene Wagner, Matthew Schmerer, Harry Gao, Mickey Li, John Gao, Joseph Fierro, Benafsh Sapra, Becky Tsai, Yan Meng, Doreen Ng, James Xie, Clinton R. Paden, Duncan MacCannell |
| EPI_ISL_3300600 | Gandhi Medical College | CDFD | Murali Bashyam, Asmita Gupta, Reelina Basu, Vinay Donipadi, Nagamani Kammili, Divya Vashisht, Ashwin Dalal |
| EPI_ISL_3300601, EPI_ISL_3300602, EPI_ISL_3300603, EPI_ISL_3300604 | Fulgent Genetics | Centers for Disease Control and Prevention Division of Viral Diseases, Pathogen Discovery | Dakota Howard, Dhwani Batra, Peter W. Cook, Kara Moser, Adrian Paskey, Jason Caravas, Benjamin Rambo-Martin, Shatavia Morrison, Christopher Gulvick, Scott Sammons, Yvette Unoarumhi, Darlene Wagner, Matthew Schmerer, Harry Gao, Mickey Li, John Gao, Joseph Fierro, Benafsh Sapra, Becky Tsai, Yan Meng, Doreen Ng, James Xie, Clinton R. Paden, Duncan MacCannell |
| EPI_ISL_3300605 | Gandhi Medical College | CDFD | Murali Bashyam, Asmita Gupta, Reelina Basu, Vinay Donipadi, Nagamani Kammili, Divya Vashisht, Ashwin Dalal |
| EPI_ISL_3300606 | Fulgent Genetics | Centers for Disease Control and Prevention Division of Viral Diseases, Pathogen Discovery | Dakota Howard, Dhwani Batra, Peter W. Cook, Kara Moser, Adrian Paskey, Jason Caravas, Benjamin Rambo-Martin, Shatavia Morrison, Christopher Gulvick, Scott Sammons, Yvette Unoarumhi, Darlene Wagner, Matthew Schmerer, Harry Gao, Mickey Li, John Gao, Joseph Fierro, Benafsh Sapra, Becky Tsai, Yan Meng, Doreen Ng, James Xie, Clinton R. Paden, Duncan MacCannell |
| EPI_ISL_3300607 | Gandhi Medical College | CDFD | Murali Bashyam, Asmita Gupta, Reelina Basu, Vinay Donipadi, Nagamani Kammili, Divya Vashisht, Ashwin Dalal |
| EPI_ISL_3300608, EPI_ISL_3300609 | Fulgent Genetics | Centers for Disease Control and Prevention Division of Viral Diseases, Pathogen Discovery | Dakota Howard, Dhwani Batra, Peter W. Cook, Kara Moser, Adrian Paskey, Jason Caravas, Benjamin Rambo-Martin, Shatavia Morrison, Christopher Gulvick, Scott Sammons, Yvette Unoarumhi, Darlene Wagner, Matthew Schmerer, Harry Gao, Mickey Li, John Gao, Joseph Fierro, Benafsh Sapra, Becky Tsai, Yan Meng, Doreen Ng, James Xie, Clinton R. Paden, Duncan MacCannell |
| EPI_ISL_3300610 | Gandhi Hospital | CDFD | Murali Bashyam, Asmita Gupta, Reelina Basu, Vinay Donipadi, Nagamani Kammili, Divya Vashisht, Ashwin Dalal |
| EPI_ISL_3300611, EPI_ISL_3300612, EPI_ISL_3300613, EPI_ISL_3300614, EPI_ISL_3300616, EPI_ISL_3300617, EPI_ISL_3300618 | Fulgent Genetics | Centers for Disease Control and Prevention Division of Viral Diseases, Pathogen Discovery | Dakota Howard, Dhwani Batra, Peter W. Cook, Kara Moser, Adrian Paskey, Jason Caravas, Benjamin Rambo-Martin, Shatavia Morrison, Christopher Gulvick, Scott Sammons, Yvette Unoarumhi, Darlene Wagner, Matthew Schmerer, Harry Gao, Mickey Li, John Gao, Joseph Fierro, Benafsh Sapra, Becky Tsai, Yan Meng, Doreen Ng, James Xie, Clinton R. Paden, Duncan MacCannell |
| EPI_ISL_3300619 | Gandhi Hospital | CDFD | Murali Bashyam, Asmita Gupta, Reelina Basu, Vinay Donipadi, Nagamani Kammili, Divya Vashisht, Ashwin Dalal |
| EPI_ISL_3300620, EPI_ISL_3300621, EPI_ISL_3300622, EPI_ISL_3300623 | Fulgent Genetics | Centers for Disease Control and Prevention Division of Viral Diseases, Pathogen Discovery | Dakota Howard, Dhwani Batra, Peter W. Cook, Kara Moser, Adrian Paskey, Jason Caravas, Benjamin Rambo-Martin, Shatavia Morrison, Christopher Gulvick, Scott Sammons, Yvette Unoarumhi, Darlene Wagner, Matthew Schmerer, Harry Gao, Mickey Li, John Gao, Joseph Fierro, Benafsh Sapra, Becky Tsai, Yan Meng, Doreen Ng, James Xie, Clinton R. Paden, Duncan MacCannell |
| EPI_ISL_3300624 | Gandhi Hospital | CDFD | Murali Bashyam, Asmita Gupta, Reelina Basu, Vinay Donipadi, Nagamani Kammili, Divya Vashisht, Ashwin Dalal |
| EPI_ISL_3300625, EPI_ISL_3300626 | Fulgent Genetics | Centers for Disease Control and Prevention Division of Viral Diseases, Pathogen Discovery | Dakota Howard, Dhwani Batra, Peter W. Cook, Kara Moser, Adrian Paskey, Jason Caravas, Benjamin Rambo-Martin, Shatavia Morrison, Christopher Gulvick, Scott Sammons, Yvette Unoarumhi, Darlene Wagner, Matthew Schmerer, Harry Gao, Mickey Li, John Gao, Joseph Fierro, Benafsh Sapra, Becky Tsai, Yan Meng, Doreen Ng, James Xie, Clinton R. Paden, Duncan MacCannell |
| EPI_ISL_3300627 | Gandhi Hospital | CDFD | Murali Bashyam, Asmita Gupta, Reelina Basu, Vinay Donipadi, Nagamani Kammili, Divya Vashisht, Ashwin Dalal |
| EPI_ISL_3300628, EPI_ISL_3300629, EPI_ISL_3300630, EPI_ISL_3300631 | Fulgent Genetics | Centers for Disease Control and Prevention Division of Viral Diseases, Pathogen Discovery | Dakota Howard, Dhwani Batra, Peter W. Cook, Kara Moser, Adrian Paskey, Jason Caravas, Benjamin Rambo-Martin, Shatavia Morrison, Christopher Gulvick, Scott Sammons, Yvette Unoarumhi, Darlene Wagner, Matthew Schmerer, Harry Gao, Mickey Li, John Gao, Joseph Fierro, Benafsh Sapra, Becky Tsai, Yan Meng, Doreen Ng, James Xie, Clinton R. Paden, Duncan MacCannell |
| EPI_ISL_3300632 | Gandhi Medical College | CDFD | Murali Bashyam, Asmita Gupta, Reelina Basu, Vinay Donipadi, Nagamani Kammili, Divya Vashisht, Ashwin Dalal |
| EPI_ISL_3300633, EPI_ISL_3300634 | Fulgent Genetics | Centers for Disease Control and Prevention Division of Viral Diseases, Pathogen Discovery | Dakota Howard, Dhwani Batra, Peter W. Cook, Kara Moser, Adrian Paskey, Jason Caravas, Benjamin Rambo-Martin, Shatavia Morrison, Christopher Gulvick, Scott Sammons, Yvette Unoarumhi, Darlene Wagner, Matthew Schmerer, Harry Gao, Mickey Li, John Gao, Joseph Fierro, Benafsh Sapra, Becky Tsai, Yan Meng, Doreen Ng, James Xie, Clinton R. Paden, Duncan MacCannell |
| EPI_ISL_3300635 | Gandhi Medical College | CDFD | Murali Bashyam, Asmita Gupta, Reelina Basu, Vinay Donipadi, Nagamani Kammili, Divya Vashisht, Ashwin Dalal |
| EPI_ISL_3300636, EPI_ISL_3300637, EPI_ISL_3300638, | Fulgent Genetics | Centers for Disease Control and Prevention Division of Viral Diseases, Pathogen Discovery | Dakota Howard, Dhwani Batra, Peter W. Cook, Kara Moser, Adrian Paskey, Jason Caravas, Benjamin Rambo-Martin, Shatavia Morrison, Christopher Gulvick, Scott Sammons, Yvette Unoarumhi, Darlene Wagner, Matthew Schmerer, Harry Gao, Mickey Li, John Gao, Joseph Fierro, Benafsh Sapra, Becky Tsai, Yan Meng, Doreen Ng, James Xie, Clinton R. Paden, Duncan MacCannell |

| | | | |
|---|---|---|---|
| EPI_ISL_3300639 | | | |
| EPI_ISL_3300640 | Gandhi Hospital | CDFD | Murali Bashyam, Asmita Gupta, Reelina Basu, Vinay Donipadi, Nagamani Kammili, Divya Vashisht, Ashwin Dalal |
| EPI_ISL_3300641, EPI_ISL_3300642, EPI_ISL_3300643, EPI_ISL_3300644, EPI_ISL_3300645 | Fulgent Genetics | Centers for Disease Control and Prevention Division of Viral Diseases, Pathogen Discovery | Dakota Howard, Dhwani Batra, Peter W. Cook, Kara Moser, Adrian Paskey, Jason Caravas, Benjamin Rambo-Martin, Shatavia Morrison, Christopher Gulvick, Scott Sammons, Yvette Unoarumhi, Darlene Wagner, Matthew Schmerer, Harry Gao, Mickey Li, John Gao, Joseph Fierro, Benafsh Sapra, Becky Tsai, Yan Meng, Doreen Ng, James Xie, Clinton R. Paden, Duncan MacCannell |
| EPI_ISL_3300646 | Government General Hospital | CDFD | Murali Bashyam, Asmita Gupta, Reelina Basu, Vinay Donipadi, Nagamani Kammili, Divya Vashisht, Ashwin Dalal |
| EPI_ISL_3300648, EPI_ISL_3300649, EPI_ISL_3300650 | Fulgent Genetics | Centers for Disease Control and Prevention Division of Viral Diseases, Pathogen Discovery | Dakota Howard, Dhwani Batra, Peter W. Cook, Kara Moser, Adrian Paskey, Jason Caravas, Benjamin Rambo-Martin, Shatavia Morrison, Christopher Gulvick, Scott Sammons, Yvette Unoarumhi, Darlene Wagner, Matthew Schmerer, Harry Gao, Mickey Li, John Gao, Joseph Fierro, Benafsh Sapra, Becky Tsai, Yan Meng, Doreen Ng, James Xie, Clinton R. Paden, Duncan MacCannell |
| EPI_ISL_3300651 | Government General Hospital | CDFD | Murali Bashyam, Asmita Gupta, Reelina Basu, Vinay Donipadi, Nagamani Kammili, Divya Vashisht, Ashwin Dalal |
| EPI_ISL_3300652, EPI_ISL_3300653, EPI_ISL_3300655, EPI_ISL_3300656 | Fulgent Genetics | Centers for Disease Control and Prevention Division of Viral Diseases, Pathogen Discovery | Dakota Howard, Dhwani Batra, Peter W. Cook, Kara Moser, Adrian Paskey, Jason Caravas, Benjamin Rambo-Martin, Shatavia Morrison, Christopher Gulvick, Scott Sammons, Yvette Unoarumhi, Darlene Wagner, Matthew Schmerer, Harry Gao, Mickey Li, John Gao, Joseph Fierro, Benafsh Sapra, Becky Tsai, Yan Meng, Doreen Ng, James Xie, Clinton R. Paden, Duncan MacCannell |
| EPI_ISL_3300657 | Government General Hospital | CDFD | Murali Bashyam, Asmita Gupta, Reelina Basu, Vinay Donipadi, Nagamani Kammili, Divya Vashisht, Ashwin Dalal |
| EPI_ISL_3300658, EPI_ISL_3300659, EPI_ISL_3300660, EPI_ISL_3300661 | Fulgent Genetics | Centers for Disease Control and Prevention Division of Viral Diseases, Pathogen Discovery | Dakota Howard, Dhwani Batra, Peter W. Cook, Kara Moser, Adrian Paskey, Jason Caravas, Benjamin Rambo-Martin, Shatavia Morrison, Christopher Gulvick, Scott Sammons, Yvette Unoarumhi, Darlene Wagner, Matthew Schmerer, Harry Gao, Mickey Li, John Gao, Joseph Fierro, Benafsh Sapra, Becky Tsai, Yan Meng, Doreen Ng, James Xie, Clinton R. Paden, Duncan MacCannell |
| EPI_ISL_3300662 | Government General Hospital | CDFD | Murali Bashyam, Asmita Gupta, Reelina Basu, Vinay Donipadi, Nagamani Kammili, Divya Vashisht, Ashwin Dalal |
| EPI_ISL_3300663, EPI_ISL_3300664, EPI_ISL_3300665, EPI_ISL_3300666 | Fulgent Genetics | Centers for Disease Control and Prevention Division of Viral Diseases, Pathogen Discovery | Dakota Howard, Dhwani Batra, Peter W. Cook, Kara Moser, Adrian Paskey, Jason Caravas, Benjamin Rambo-Martin, Shatavia Morrison, Christopher Gulvick, Scott Sammons, Yvette Unoarumhi, Darlene Wagner, Matthew Schmerer, Harry Gao, Mickey Li, John Gao, Joseph Fierro, Benafsh Sapra, Becky Tsai, Yan Meng, Doreen Ng, James Xie, Clinton R. Paden, Duncan MacCannell |
| EPI_ISL_3300667 | Government General Hospital | CDFD | Murali Bashyam, Asmita Gupta, Reelina Basu, Vinay Donipadi, Nagamani Kammili, Divya Vashisht, Ashwin Dalal |
| EPI_ISL_3300668, EPI_ISL_3300669, EPI_ISL_3300670 | Fulgent Genetics | Centers for Disease Control and Prevention Division of Viral Diseases, Pathogen Discovery | Dakota Howard, Dhwani Batra, Peter W. Cook, Kara Moser, Adrian Paskey, Jason Caravas, Benjamin Rambo-Martin, Shatavia Morrison, Christopher Gulvick, Scott Sammons, Yvette Unoarumhi, Darlene Wagner, Matthew Schmerer, Harry Gao, Mickey Li, John Gao, Joseph Fierro, Benafsh Sapra, Becky Tsai, Yan Meng, Doreen Ng, James Xie, Clinton R. Paden, Duncan MacCannell |
| EPI_ISL_3300671 | Government General Hospital | CDFD | Murali Bashyam, Asmita Gupta, Reelina Basu, Vinay Donipadi, Nagamani Kammili, Divya Vashisht, Ashwin Dalal |
| EPI_ISL_3300672, EPI_ISL_3300673 | Fulgent Genetics | Centers for Disease Control and Prevention Division of Viral Diseases, Pathogen Discovery | Dakota Howard, Dhwani Batra, Peter W. Cook, Kara Moser, Adrian Paskey, Jason Caravas, Benjamin Rambo-Martin, Shatavia Morrison, Christopher Gulvick, Scott Sammons, Yvette Unoarumhi, Darlene Wagner, Matthew Schmerer, Harry Gao, Mickey Li, John Gao, Joseph Fierro, Benafsh Sapra, Becky Tsai, Yan Meng, Doreen Ng, James Xie, Clinton R. Paden, Duncan MacCannell |
| EPI_ISL_3300674 | Government General Hospital | CDFD | Murali Bashyam, Asmita Gupta, Reelina Basu, Vinay Donipadi, Nagamani Kammili, Divya Vashisht, Ashwin Dalal |
| EPI_ISL_3300675, EPI_ISL_3300676, EPI_ISL_3300677, EPI_ISL_3300678, EPI_ISL_3300679 | Fulgent Genetics | Centers for Disease Control and Prevention Division of Viral Diseases, Pathogen Discovery | Dakota Howard, Dhwani Batra, Peter W. Cook, Kara Moser, Adrian Paskey, Jason Caravas, Benjamin Rambo-Martin, Shatavia Morrison, Christopher Gulvick, Scott Sammons, Yvette Unoarumhi, Darlene Wagner, Matthew Schmerer, Harry Gao, Mickey Li, John Gao, Joseph Fierro, Benafsh Sapra, Becky Tsai, Yan Meng, Doreen Ng, James Xie, Clinton R. Paden, Duncan MacCannell |
| EPI_ISL_3300680 | Government General Hospital | CDFD | Murali Bashyam, Asmita Gupta, Reelina Basu, Vinay Donipadi, Nagamani Kammili, Divya Vashisht, Ashwin Dalal |
| EPI_ISL_3300681, EPI_ISL_3300682, EPI_ISL_3300683 | Fulgent Genetics | Centers for Disease Control and Prevention Division of Viral Diseases, Pathogen Discovery | Dakota Howard, Dhwani Batra, Peter W. Cook, Kara Moser, Adrian Paskey, Jason Caravas, Benjamin Rambo-Martin, Shatavia Morrison, Christopher Gulvick, Scott Sammons, Yvette Unoarumhi, Darlene Wagner, Matthew Schmerer, Harry Gao, Mickey Li, John Gao, Joseph Fierro, Benafsh Sapra, Becky Tsai, Yan Meng, Doreen Ng, James Xie, Clinton R. Paden, Duncan MacCannell |
| EPI_ISL_3300684 | Government General Hospital | CDFD | Murali Bashyam, Asmita Gupta, Reelina Basu, Vinay Donipadi, Nagamani Kammili, Divya Vashisht, Ashwin Dalal |
| EPI_ISL_3300685, EPI_ISL_3300686, EPI_ISL_3300687, EPI_ISL_3300688 | Fulgent Genetics | Centers for Disease Control and Prevention Division of Viral Diseases, Pathogen Discovery | Dakota Howard, Dhwani Batra, Peter W. Cook, Kara Moser, Adrian Paskey, Jason Caravas, Benjamin Rambo-Martin, Shatavia Morrison, Christopher Gulvick, Scott Sammons, Yvette Unoarumhi, Darlene Wagner, Matthew Schmerer, Harry Gao, Mickey Li, John Gao, Joseph Fierro, Benafsh Sapra, Becky Tsai, Yan Meng, Doreen Ng, James Xie, Clinton R. Paden, Duncan MacCannell |
| EPI_ISL_3300690 | Government General Hospital | CDFD | Murali Bashyam, Asmita Gupta, Reelina Basu, Vinay Donipadi, Nagamani Kammili, Divya Vashisht, Ashwin Dalal |
| EPI_ISL_3300691, EPI_ISL_3300692 | Fulgent Genetics | Centers for Disease Control and Prevention Division of Viral Diseases, Pathogen Discovery | Dakota Howard, Dhwani Batra, Peter W. Cook, Kara Moser, Adrian Paskey, Jason Caravas, Benjamin Rambo-Martin, Shatavia Morrison, Christopher Gulvick, Scott Sammons, Yvette Unoarumhi, Darlene Wagner, Matthew Schmerer, Harry Gao, Mickey Li, John Gao, Joseph Fierro, Benafsh Sapra, Becky Tsai, Yan Meng, Doreen Ng, James Xie, Clinton R. Paden, Duncan MacCannell |
| EPI_ISL_3300693 | Government General Hospital | CDFD | Murali Bashyam, Asmita Gupta, Reelina Basu, Vinay Donipadi, Nagamani Kammili, Divya Vashisht, Ashwin Dalal |
| EPI_ISL_3300694, EPI_ISL_3300695, EPI_ISL_3300696, EPI_ISL_3300697, EPI_ISL_3300699, EPI_ISL_3300700, EPI_ISL_3300702, EPI_ISL_3300703, EPI_ISL_3300704, EPI_ISL_3300705 | Fulgent Genetics | Centers for Disease Control and Prevention Division of Viral Diseases, Pathogen Discovery | Dakota Howard, Dhwani Batra, Peter W. Cook, Kara Moser, Adrian Paskey, Jason Caravas, Benjamin Rambo-Martin, Shatavia Morrison, Christopher Gulvick, Scott Sammons, Yvette Unoarumhi, Darlene Wagner, Matthew Schmerer, Harry Gao, Mickey Li, John Gao, Joseph Fierro, Benafsh Sapra, Becky Tsai, Yan Meng, Doreen Ng, James Xie, Clinton R. Paden, Duncan MacCannell |
| EPI_ISL_3300706 | CDFD | CDFD | Murali Bashyam, Asmita Gupta, Reelina Basu, Vinay Donipadi, Nagamani Kammili, Divya Vashisht, Ashwin Dalal |
| EPI_ISL_3300707, EPI_ISL_3300708, EPI_ISL_3300709, EPI_ISL_3300710 | Fulgent Genetics | Centers for Disease Control and Prevention Division of Viral Diseases, Pathogen Discovery | Dakota Howard, Dhwani Batra, Peter W. Cook, Kara Moser, Adrian Paskey, Jason Caravas, Benjamin Rambo-Martin, Shatavia Morrison, Christopher Gulvick, Scott Sammons, Yvette Unoarumhi, Darlene Wagner, Matthew Schmerer, Harry Gao, Mickey Li, John Gao, Joseph Fierro, Benafsh Sapra, Becky Tsai, Yan Meng, Doreen Ng, James Xie, Clinton R. Paden, Duncan MacCannell |
| EPI_ISL_3300711 | CDFD | CDFD | Murali Bashyam, Asmita Gupta, Reelina Basu, Vinay Donipadi, Nagamani Kammili, Divya Vashisht, Ashwin Dalal |
| EPI_ISL_3300712, EPI_ISL_3300713, EPI_ISL_3300714 | Fulgent Genetics | Centers for Disease Control and Prevention Division of Viral Diseases, Pathogen Discovery | Dakota Howard, Dhwani Batra, Peter W. Cook, Kara Moser, Adrian Paskey, Jason Caravas, Benjamin Rambo-Martin, Shatavia Morrison, Christopher Gulvick, Scott Sammons, Yvette Unoarumhi, Darlene Wagner, Matthew Schmerer, Harry Gao, Mickey Li, John Gao, Joseph Fierro, Benafsh Sapra, Becky Tsai, Yan Meng, Doreen Ng, James Xie, Clinton R. Paden, Duncan MacCannell |
| EPI_ISL_3300715 | CDFD | CDFD | Murali Bashyam, Asmita Gupta, Reelina Basu, Vinay Donipadi, Nagamani Kammili, Divya Vashisht, Ashwin Dalal |
| EPI_ISL_3300716, EPI_ISL_3300717 | Fulgent Genetics | Centers for Disease Control and Prevention Division of Viral Diseases, Pathogen Discovery | Dakota Howard, Dhwani Batra, Peter W. Cook, Kara Moser, Adrian Paskey, Jason Caravas, Benjamin Rambo-Martin, Shatavia Morrison, Christopher Gulvick, Scott Sammons, Yvette Unoarumhi, Darlene Wagner, Matthew Schmerer, Harry Gao, Mickey Li, John Gao, Joseph Fierro, Benafsh Sapra, Becky Tsai, Yan Meng, Doreen Ng, James Xie, Clinton R. Paden, Duncan MacCannell |
| EPI_ISL_3300718 | CDFD | CDFD | Murali Bashyam, Asmita Gupta, Reelina Basu, Vinay Donipadi, Nagamani Kammili, Divya Vashisht, Ashwin Dalal |
| EPI_ISL_3300719, EPI_ISL_3300720, EPI_ISL_3300721, EPI_ISL_3300723, EPI_ISL_3300724 | Fulgent Genetics | Centers for Disease Control and Prevention Division of Viral Diseases, Pathogen Discovery | Dakota Howard, Dhwani Batra, Peter W. Cook, Kara Moser, Adrian Paskey, Jason Caravas, Benjamin Rambo-Martin, Shatavia Morrison, Christopher Gulvick, Scott Sammons, Yvette Unoarumhi, Darlene Wagner, Matthew Schmerer, Harry Gao, Mickey Li, John Gao, Joseph Fierro, Benafsh Sapra, Becky Tsai, Yan Meng, Doreen Ng, James Xie, Clinton R. Paden, Duncan MacCannell |
| EPI_ISL_3300725 | CDFD | CDFD | Murali Bashyam, Asmita Gupta, Reelina Basu, Vinay Donipadi, Nagamani Kammili, Divya Vashisht, Ashwin Dalal |
| EPI_ISL_3300726, EPI_ISL_3300727, EPI_ISL_3300729, EPI_ISL_3300730 | Fulgent Genetics | Centers for Disease Control and Prevention Division of Viral Diseases, Pathogen Discovery | Dakota Howard, Dhwani Batra, Peter W. Cook, Kara Moser, Adrian Paskey, Jason Caravas, Benjamin Rambo-Martin, Shatavia Morrison, Christopher Gulvick, Scott Sammons, Yvette Unoarumhi, Darlene Wagner, Matthew Schmerer, Harry Gao, Mickey Li, John Gao, Joseph Fierro, Benafsh Sapra, Becky Tsai, Yan Meng, Doreen Ng, James Xie, Clinton R. Paden, Duncan MacCannell |
| EPI_ISL_3300731 | CDFD | CDFD | Murali Bashyam, Asmita Gupta, Reelina Basu, Vinay Donipadi, Nagamani Kammili, Divya Vashisht, Ashwin Dalal |
| EPI_ISL_3300732, EPI_ISL_3300733 | Fulgent Genetics | Centers for Disease Control and Prevention Division of Viral Diseases, Pathogen Discovery | Dakota Howard, Dhwani Batra, Peter W. Cook, Kara Moser, Adrian Paskey, Jason Caravas, Benjamin Rambo-Martin, Shatavia Morrison, Christopher Gulvick, Scott Sammons, Yvette Unoarumhi, Darlene Wagner, Matthew Schmerer, Harry Gao, Mickey Li, John Gao, Joseph Fierro, Benafsh Sapra, Becky Tsai, Yan Meng, Doreen Ng, James Xie, Clinton R. Paden, Duncan MacCannell |
| EPI_ISL_3300734 | CDFD | CDFD | Murali Bashyam, Asmita Gupta, Reelina Basu, Vinay Donipadi, Nagamani Kammili, Divya Vashisht, Ashwin Dalal |
| EPI_ISL_3300735 | Fulgent Genetics | Centers for Disease Control and Prevention Division of Viral Diseases, Pathogen Discovery | Dakota Howard, Dhwani Batra, Peter W. Cook, Kara Moser, Adrian Paskey, Jason Caravas, Benjamin Rambo-Martin, Shatavia Morrison, Christopher Gulvick, Scott Sammons, Yvette Unoarumhi, Darlene Wagner, Matthew Schmerer, Harry Gao, Mickey Li, John Gao, Joseph Fierro, Benafsh Sapra, Becky Tsai, Yan Meng, Doreen Ng, James Xie, Clinton R. Paden, Duncan MacCannell |
| EPI_ISL_3300736 | CDFD | CDFD | Murali Bashyam, Asmita Gupta, Reelina Basu, Vinay Donipadi, Nagamani Kammili, Divya Vashisht, Ashwin Dalal |

| | | | |
|---|---|---|---|
| EPI_ISL_3300737 | Fulgent Genetics | Centers for Disease Control and Prevention Division of Viral Diseases, Pathogen Discovery | Dakota Howard, Dhwani Batra, Peter W. Cook, Kara Moser, Adrian Paskey, Jason Caravas, Benjamin Rambo-Martin, Shatavia Morrison, Christopher Gulvick, Scott Sammons, Yvette Unoarumhi, Darlene Wagner, Matthew Schmerer, Harry Gao, Mickey Li, John Gao, Joseph Fierro, Benafsh Sapra, Becky Tsai, Yan Meng, Doreen Ng, James Xie, Clinton R. Paden, Duncan MacCannell |
| EPI_ISL_3300738 | CDFD | CDFD | Murali Bashyam, Asmita Gupta, Reelina Basu, Vinay Donipadi, Nagamani Kammili, Divya Vashisht, Ashwin Dalal |
| EPI_ISL_3300739 | Fulgent Genetics | Centers for Disease Control and Prevention Division of Viral Diseases, Pathogen Discovery | Dakota Howard, Dhwani Batra, Peter W. Cook, Kara Moser, Adrian Paskey, Jason Caravas, Benjamin Rambo-Martin, Shatavia Morrison, Christopher Gulvick, Scott Sammons, Yvette Unoarumhi, Darlene Wagner, Matthew Schmerer, Harry Gao, Mickey Li, John Gao, Joseph Fierro, Benafsh Sapra, Becky Tsai, Yan Meng, Doreen Ng, James Xie, Clinton R. Paden, Duncan MacCannell |
| EPI_ISL_3300740 | CDFD | CDFD | Murali Bashyam, Asmita Gupta, Reelina Basu, Vinay Donipadi, Nagamani Kammili, Divya Vashisht, Ashwin Dalal |
| EPI_ISL_3300741 | Fulgent Genetics | Centers for Disease Control and Prevention Division of Viral Diseases, Pathogen Discovery | Dakota Howard, Dhwani Batra, Peter W. Cook, Kara Moser, Adrian Paskey, Jason Caravas, Benjamin Rambo-Martin, Shatavia Morrison, Christopher Gulvick, Scott Sammons, Yvette Unoarumhi, Darlene Wagner, Matthew Schmerer, Harry Gao, Mickey Li, John Gao, Joseph Fierro, Benafsh Sapra, Becky Tsai, Yan Meng, Doreen Ng, James Xie, Clinton R. Paden, Duncan MacCannell |
| EPI_ISL_3300742, EPI_ISL_3300743 | CDFD | CDFD | Murali Bashyam, Asmita Gupta, Reelina Basu, Vinay Donipadi, Nagamani Kammili, Divya Vashisht, Ashwin Dalal |
| EPI_ISL_3300744 | Fulgent Genetics | Centers for Disease Control and Prevention Division of Viral Diseases, Pathogen Discovery | Dakota Howard, Dhwani Batra, Peter W. Cook, Kara Moser, Adrian Paskey, Jason Caravas, Benjamin Rambo-Martin, Shatavia Morrison, Christopher Gulvick, Scott Sammons, Yvette Unoarumhi, Darlene Wagner, Matthew Schmerer, Harry Gao, Mickey Li, John Gao, Joseph Fierro, Benafsh Sapra, Becky Tsai, Yan Meng, Doreen Ng, James Xie, Clinton R. Paden, Duncan MacCannell |
| EPI_ISL_3300745 | CDFD | CDFD | Murali Bashyam, Asmita Gupta, Reelina Basu, Vinay Donipadi, Nagamani Kammili, Divya Vashisht, Ashwin Dalal |
| EPI_ISL_3300746, EPI_ISL_3300747 | Fulgent Genetics | Centers for Disease Control and Prevention Division of Viral Diseases, Pathogen Discovery | Dakota Howard, Dhwani Batra, Peter W. Cook, Kara Moser, Adrian Paskey, Jason Caravas, Benjamin Rambo-Martin, Shatavia Morrison, Christopher Gulvick, Scott Sammons, Yvette Unoarumhi, Darlene Wagner, Matthew Schmerer, Harry Gao, Mickey Li, John Gao, Joseph Fierro, Benafsh Sapra, Becky Tsai, Yan Meng, Doreen Ng, James Xie, Clinton R. Paden, Duncan MacCannell |
| EPI_ISL_3300748 | CDFD | CDFD | Murali Bashyam, Asmita Gupta, Reelina Basu, Vinay Donipadi, Nagamani Kammili, Divya Vashisht, Ashwin Dalal |
| EPI_ISL_3300749 | Fulgent Genetics | Centers for Disease Control and Prevention Division of Viral Diseases, Pathogen Discovery | Dakota Howard, Dhwani Batra, Peter W. Cook, Kara Moser, Adrian Paskey, Jason Caravas, Benjamin Rambo-Martin, Shatavia Morrison, Christopher Gulvick, Scott Sammons, Yvette Unoarumhi, Darlene Wagner, Matthew Schmerer, Harry Gao, Mickey Li, John Gao, Joseph Fierro, Benafsh Sapra, Becky Tsai, Yan Meng, Doreen Ng, James Xie, Clinton R. Paden, Duncan MacCannell |
| EPI_ISL_3300750 | CDFD | CDFD | Murali Bashyam, Asmita Gupta, Reelina Basu, Vinay Donipadi, Nagamani Kammili, Divya Vashisht, Ashwin Dalal |
| EPI_ISL_3300751 | Fulgent Genetics | Centers for Disease Control and Prevention Division of Viral Diseases, Pathogen Discovery | Dakota Howard, Dhwani Batra, Peter W. Cook, Kara Moser, Adrian Paskey, Jason Caravas, Benjamin Rambo-Martin, Shatavia Morrison, Christopher Gulvick, Scott Sammons, Yvette Unoarumhi, Darlene Wagner, Matthew Schmerer, Harry Gao, Mickey Li, John Gao, Joseph Fierro, Benafsh Sapra, Becky Tsai, Yan Meng, Doreen Ng, James Xie, Clinton R. Paden, Duncan MacCannell |
| EPI_ISL_3300752 | CDFD | CDFD | Murali Bashyam, Asmita Gupta, Reelina Basu, Vinay Donipadi, Nagamani Kammili, Divya Vashisht, Ashwin Dalal |
| EPI_ISL_3300753, EPI_ISL_3300754, EPI_ISL_3300755 | Fulgent Genetics | Centers for Disease Control and Prevention Division of Viral Diseases, Pathogen Discovery | Dakota Howard, Dhwani Batra, Peter W. Cook, Kara Moser, Adrian Paskey, Jason Caravas, Benjamin Rambo-Martin, Shatavia Morrison, Christopher Gulvick, Scott Sammons, Yvette Unoarumhi, Darlene Wagner, Matthew Schmerer, Harry Gao, Mickey Li, John Gao, Joseph Fierro, Benafsh Sapra, Becky Tsai, Yan Meng, Doreen Ng, James Xie, Clinton R. Paden, Duncan MacCannell |
| EPI_ISL_3300756 | CDFD | CDFD | Murali Bashyam, Asmita Gupta, Reelina Basu, Vinay Donipadi, Nagamani Kammili, Divya Vashisht, Ashwin Dalal |
| EPI_ISL_3300757 | Fulgent Genetics | Centers for Disease Control and Prevention Division of Viral Diseases, Pathogen Discovery | Dakota Howard, Dhwani Batra, Peter W. Cook, Kara Moser, Adrian Paskey, Jason Caravas, Benjamin Rambo-Martin, Shatavia Morrison, Christopher Gulvick, Scott Sammons, Yvette Unoarumhi, Darlene Wagner, Matthew Schmerer, Harry Gao, Mickey Li, John Gao, Joseph Fierro, Benafsh Sapra, Becky Tsai, Yan Meng, Doreen Ng, James Xie, Clinton R. Paden, Duncan MacCannell |
| EPI_ISL_3300758 | CDFD | CDFD | Murali Bashyam, Asmita Gupta, Reelina Basu, Vinay Donipadi, Nagamani Kammili, Divya Vashisht, Ashwin Dalal |
| EPI_ISL_3300759, EPI_ISL_3300761 | Fulgent Genetics | Centers for Disease Control and Prevention Division of Viral Diseases, Pathogen Discovery | Dakota Howard, Dhwani Batra, Peter W. Cook, Kara Moser, Adrian Paskey, Jason Caravas, Benjamin Rambo-Martin, Shatavia Morrison, Christopher Gulvick, Scott Sammons, Yvette Unoarumhi, Darlene Wagner, Matthew Schmerer, Harry Gao, Mickey Li, John Gao, Joseph Fierro, Benafsh Sapra, Becky Tsai, Yan Meng, Doreen Ng, James Xie, Clinton R. Paden, Duncan MacCannell |
| EPI_ISL_3300762 | CDFD | CDFD | Murali Bashyam, Asmita Gupta, Reelina Basu, Vinay Donipadi, Nagamani Kammili, Divya Vashisht, Ashwin Dalal |
| EPI_ISL_3300763 | Fulgent Genetics | Centers for Disease Control and Prevention Division of Viral Diseases, Pathogen Discovery | Dakota Howard, Dhwani Batra, Peter W. Cook, Kara Moser, Adrian Paskey, Jason Caravas, Benjamin Rambo-Martin, Shatavia Morrison, Christopher Gulvick, Scott Sammons, Yvette Unoarumhi, Darlene Wagner, Matthew Schmerer, Harry Gao, Mickey Li, John Gao, Joseph Fierro, Benafsh Sapra, Becky Tsai, Yan Meng, Doreen Ng, James Xie, Clinton R. Paden, Duncan MacCannell |
| EPI_ISL_3300764 | CDFD | CDFD | Murali Bashyam, Asmita Gupta, Reelina Basu, Vinay Donipadi, Nagamani Kammili, Divya Vashisht, Ashwin Dalal |
| EPI_ISL_3300765 | Fulgent Genetics | Centers for Disease Control and Prevention Division of Viral Diseases, Pathogen Discovery | Dakota Howard, Dhwani Batra, Peter W. Cook, Kara Moser, Adrian Paskey, Jason Caravas, Benjamin Rambo-Martin, Shatavia Morrison, Christopher Gulvick, Scott Sammons, Yvette Unoarumhi, Darlene Wagner, Matthew Schmerer, Harry Gao, Mickey Li, John Gao, Joseph Fierro, Benafsh Sapra, Becky Tsai, Yan Meng, Doreen Ng, James Xie, Clinton R. Paden, Duncan MacCannell |
| EPI_ISL_3300766 | CDFD | CDFD | Murali Bashyam, Asmita Gupta, Reelina Basu, Vinay Donipadi, Nagamani Kammili, Divya Vashisht, Ashwin Dalal |
| EPI_ISL_3300767, EPI_ISL_3300768 | Fulgent Genetics | Centers for Disease Control and Prevention Division of Viral Diseases, Pathogen Discovery | Dakota Howard, Dhwani Batra, Peter W. Cook, Kara Moser, Adrian Paskey, Jason Caravas, Benjamin Rambo-Martin, Shatavia Morrison, Christopher Gulvick, Scott Sammons, Yvette Unoarumhi, Darlene Wagner, Matthew Schmerer, Harry Gao, Mickey Li, John Gao, Joseph Fierro, Benafsh Sapra, Becky Tsai, Yan Meng, Doreen Ng, James Xie, Clinton R. Paden, Duncan MacCannell |
| EPI_ISL_3300769 | CDFD | CDFD | Murali Bashyam, Asmita Gupta, Reelina Basu, Vinay Donipadi, Nagamani Kammili, Divya Vashisht, Ashwin Dalal |
| EPI_ISL_3300770, EPI_ISL_3300771, EPI_ISL_3300772 | Fulgent Genetics | Centers for Disease Control and Prevention Division of Viral Diseases, Pathogen Discovery | Dakota Howard, Dhwani Batra, Peter W. Cook, Kara Moser, Adrian Paskey, Jason Caravas, Benjamin Rambo-Martin, Shatavia Morrison, Christopher Gulvick, Scott Sammons, Yvette Unoarumhi, Darlene Wagner, Matthew Schmerer, Harry Gao, Mickey Li, John Gao, Joseph Fierro, Benafsh Sapra, Becky Tsai, Yan Meng, Doreen Ng, James Xie, Clinton R. Paden, Duncan MacCannell |
| EPI_ISL_3300773 | CDFD | CDFD | Murali Bashyam, Asmita Gupta, Reelina Basu, Vinay Donipadi, Nagamani Kammili, Divya Vashisht, Ashwin Dalal |
| EPI_ISL_3300774 | Fulgent Genetics | Centers for Disease Control and Prevention Division of Viral Diseases, Pathogen Discovery | Dakota Howard, Dhwani Batra, Peter W. Cook, Kara Moser, Adrian Paskey, Jason Caravas, Benjamin Rambo-Martin, Shatavia Morrison, Christopher Gulvick, Scott Sammons, Yvette Unoarumhi, Darlene Wagner, Matthew Schmerer, Harry Gao, Mickey Li, John Gao, Joseph Fierro, Benafsh Sapra, Becky Tsai, Yan Meng, Doreen Ng, James Xie, Clinton R. Paden, Duncan MacCannell |
| EPI_ISL_3300775, EPI_ISL_3300776 | CDFD | CDFD | Murali Bashyam, Asmita Gupta, Reelina Basu, Vinay Donipadi, Nagamani Kammili, Divya Vashisht, Ashwin Dalal |
| EPI_ISL_3300777, EPI_ISL_3300778, EPI_ISL_3300779 | Fulgent Genetics | Centers for Disease Control and Prevention Division of Viral Diseases, Pathogen Discovery | Dakota Howard, Dhwani Batra, Peter W. Cook, Kara Moser, Adrian Paskey, Jason Caravas, Benjamin Rambo-Martin, Shatavia Morrison, Christopher Gulvick, Scott Sammons, Yvette Unoarumhi, Darlene Wagner, Matthew Schmerer, Harry Gao, Mickey Li, John Gao, Joseph Fierro, Benafsh Sapra, Becky Tsai, Yan Meng, Doreen Ng, James Xie, Clinton R. Paden, Duncan MacCannell |
| EPI_ISL_3300780 | CDFD | CDFD | Murali Bashyam, Asmita Gupta, Reelina Basu, Vinay Donipadi, Nagamani Kammili, Divya Vashisht, Ashwin Dalal |
| EPI_ISL_3300781 | Fulgent Genetics | Centers for Disease Control and Prevention Division of Viral Diseases, Pathogen Discovery | Dakota Howard, Dhwani Batra, Peter W. Cook, Kara Moser, Adrian Paskey, Jason Caravas, Benjamin Rambo-Martin, Shatavia Morrison, Christopher Gulvick, Scott Sammons, Yvette Unoarumhi, Darlene Wagner, Matthew Schmerer, Harry Gao, Mickey Li, John Gao, Joseph Fierro, Benafsh Sapra, Becky Tsai, Yan Meng, Doreen Ng, James Xie, Clinton R. Paden, Duncan MacCannell |
| EPI_ISL_3300782 | CDFD | CDFD | Murali Bashyam, Asmita Gupta, Reelina Basu, Vinay Donipadi, Nagamani Kammili, Divya Vashisht, Ashwin Dalal |
| EPI_ISL_3300783 | Fulgent Genetics | Centers for Disease Control and Prevention Division of Viral Diseases, Pathogen Discovery | Dakota Howard, Dhwani Batra, Peter W. Cook, Kara Moser, Adrian Paskey, Jason Caravas, Benjamin Rambo-Martin, Shatavia Morrison, Christopher Gulvick, Scott Sammons, Yvette Unoarumhi, Darlene Wagner, Matthew Schmerer, Harry Gao, Mickey Li, John Gao, Joseph Fierro, Benafsh Sapra, Becky Tsai, Yan Meng, Doreen Ng, James Xie, Clinton R. Paden, Duncan MacCannell |
| EPI_ISL_3300784 | CDFD | CDFD | Murali Bashyam, Asmita Gupta, Reelina Basu, Vinay Donipadi, Nagamani Kammili, Divya Vashisht, Ashwin Dalal |
| EPI_ISL_3300785 | Fulgent Genetics | Centers for Disease Control and Prevention Division of Viral Diseases, Pathogen Discovery | Dakota Howard, Dhwani Batra, Peter W. Cook, Kara Moser, Adrian Paskey, Jason Caravas, Benjamin Rambo-Martin, Shatavia Morrison, Christopher Gulvick, Scott Sammons, Yvette Unoarumhi, Darlene Wagner, Matthew Schmerer, Harry Gao, Mickey Li, John Gao, Joseph Fierro, Benafsh Sapra, Becky Tsai, Yan Meng, Doreen Ng, James Xie, Clinton R. Paden, Duncan MacCannell |
| EPI_ISL_3300786 | CDFD | CDFD | Murali Bashyam, Asmita Gupta, Reelina Basu, Vinay Donipadi, Nagamani Kammili, Divya Vashisht, Ashwin Dalal |
| EPI_ISL_3300787 | Fulgent Genetics | Centers for Disease Control and Prevention Division of Viral Diseases, Pathogen Discovery | Dakota Howard, Dhwani Batra, Peter W. Cook, Kara Moser, Adrian Paskey, Jason Caravas, Benjamin Rambo-Martin, Shatavia Morrison, Christopher Gulvick, Scott Sammons, Yvette Unoarumhi, Darlene Wagner, Matthew Schmerer, Harry Gao, Mickey Li, John Gao, Joseph Fierro, Benafsh Sapra, Becky Tsai, Yan Meng, Doreen Ng, James Xie, Clinton R. Paden, Duncan MacCannell |
| EPI_ISL_3300788, EPI_ISL_3300789 | CDFD | CDFD | Murali Bashyam, Asmita Gupta, Reelina Basu, Vinay Donipadi, Nagamani Kammili, Divya Vashisht, Ashwin Dalal |
| EPI_ISL_3300790 | Fulgent Genetics | Centers for Disease Control and Prevention Division of Viral Diseases, Pathogen Discovery | Dakota Howard, Dhwani Batra, Peter W. Cook, Kara Moser, Adrian Paskey, Jason Caravas, Benjamin Rambo-Martin, Shatavia Morrison, Christopher Gulvick, Scott Sammons, Yvette Unoarumhi, Darlene Wagner, Matthew Schmerer, Harry Gao, Mickey Li, John Gao, Joseph Fierro, Benafsh Sapra, Becky Tsai, Yan Meng, Doreen Ng, James Xie, Clinton R. Paden, Duncan MacCannell |
| EPI_ISL_3300791 | CDFD | CDFD | Murali Bashyam, Asmita Gupta, Reelina Basu, Vinay Donipadi, Nagamani Kammili, Divya Vashisht, Ashwin Dalal |
| EPI_ISL_3300792 | Fulgent Genetics | Centers for Disease Control and Prevention Division of Viral Diseases, Pathogen Discovery | Dakota Howard, Dhwani Batra, Peter W. Cook, Kara Moser, Adrian Paskey, Jason Caravas, Benjamin Rambo-Martin, Shatavia Morrison, Christopher Gulvick, Scott Sammons, Yvette Unoarumhi, Darlene Wagner, Matthew Schmerer, Harry Gao, Mickey Li, John Gao, Joseph Fierro, Benafsh Sapra, Becky Tsai, Yan Meng, Doreen Ng, James Xie, Clinton R. Paden, Duncan MacCannell |
| EPI_ISL_3300793 | CDFD | CDFD | Murali Bashyam, Asmita Gupta, Reelina Basu, Vinay Donipadi, Nagamani Kammili, Divya Vashisht, Ashwin Dalal |
| EPI_ISL_3300794 | Fulgent Genetics | Centers for Disease Control and Prevention Division of Viral Diseases, Pathogen Discovery | Dakota Howard, Dhwani Batra, Peter W. Cook, Kara Moser, Adrian Paskey, Jason Caravas, Benjamin Rambo-Martin, Shatavia Morrison, Christopher Gulvick, Scott Sammons, Yvette Unoarumhi, Darlene Wagner, Matthew Schmerer, Harry Gao, Mickey Li, John Gao, Joseph Fierro, Benafsh Sapra, Becky Tsai, Yan Meng, Doreen Ng, James Xie, Clinton R. Paden, Duncan MacCannell |
| EPI_ISL_3300795 | CDFD | CDFD | Murali Bashyam, Asmita Gupta, Reelina Basu, Vinay Donipadi, Nagamani Kammili, Divya Vashisht, Ashwin Dalal |
| EPI_ISL_3300796 | Fulgent Genetics | Centers for Disease Control and Prevention Division of Viral Diseases, Pathogen Discovery | Dakota Howard, Dhwani Batra, Peter W. Cook, Kara Moser, Adrian Paskey, Jason Caravas, Benjamin Rambo-Martin, Shatavia Morrison, Christopher Gulvick, Scott Sammons, Yvette Unoarumhi, Darlene Wagner, Matthew Schmerer, Harry Gao, Mickey Li, John Gao, Joseph Fierro, Benafsh Sapra, Becky Tsai, Yan Meng, Doreen Ng, James Xie, Clinton R. Paden, Duncan MacCannell |
| EPI_ISL_3300797 | CDFD | CDFD | Murali Bashyam, Asmita Gupta, Reelina Basu, Vinay Donipadi, Nagamani Kammili, Divya Vashisht, Ashwin Dalal |
| EPI_ISL_3300798 | Fulgent Genetics | Centers for Disease Control and Prevention Division of Viral Diseases, Pathogen Discovery | Dakota Howard, Dhwani Batra, Peter W. Cook, Kara Moser, Adrian Paskey, Jason Caravas, Benjamin Rambo-Martin, Shatavia Morrison, Christopher Gulvick, Scott Sammons, Yvette Unoarumhi, Darlene Wagner, Matthew Schmerer, Harry Gao, Mickey Li, John Gao, Joseph Fierro, Benafsh Sapra, Becky Tsai, Yan Meng, Doreen Ng, James Xie, Clinton R. Paden, Duncan MacCannell |
| EPI_ISL_3300799, EPI_ISL_3300800 | CDFD | CDFD | Murali Bashyam, Asmita Gupta, Reelina Basu, Vinay Donipadi, Nagamani Kammili, Divya Vashisht, Ashwin Dalal |
| EPI_ISL_3300801, EPI_ISL_3300803 | Fulgent Genetics | Centers for Disease Control and Prevention Division of Viral Diseases, Pathogen Discovery | Dakota Howard, Dhwani Batra, Peter W. Cook, Kara Moser, Adrian Paskey, Jason Caravas, Benjamin Rambo-Martin, Shatavia Morrison, Christopher Gulvick, Scott Sammons, Yvette Unoarumhi, Darlene Wagner, Matthew Schmerer, Harry Gao, Mickey Li, John Gao, Joseph Fierro, Benafsh Sapra, Becky Tsai, Yan Meng, Doreen Ng, James Xie, Clinton R. Paden, Duncan MacCannell |
| EPI_ISL_3300804 | CDFD | CDFD | Murali Bashyam, Asmita Gupta, Reelina Basu, Vinay Donipadi, Nagamani Kammili, Divya Vashisht, Ashwin Dalal |
| EPI_ISL_3300805 | Fulgent Genetics | Centers for Disease Control and Prevention Division of Viral | Dakota Howard, Dhwani Batra, Peter W. Cook, Kara Moser, Adrian Paskey, Jason Caravas, Benjamin Rambo-Martin, Shatavia Morrison, Christopher Gulvick, Scott Sammons, Yvette Unoarumhi, Darlene Wagner, Matthew Schmerer, Harry Gao, |

| | | | |
|---|---|---|---|
| | | Diseases, Pathogen Discovery | Mickey Li, John Gao, Joseph Fierro, Benafsh Sapra, Becky Tsai, Yan Meng, Doreen Ng, James Xie, Clinton R. Paden, Duncan MacCannell |
| EPI_ISL_3300806 | CDFD | CDFD | Murali Bashyam, Asmita Gupta, Reelina Basu, Vinay Donipadi, Nagamani Kammili, Divya Vashisht, Ashwin Dalal |
| EPI_ISL_3300807 | Fulgent Genetics | Centers for Disease Control and Prevention Division of Viral Diseases, Pathogen Discovery | Dakota Howard, Dhwani Batra, Peter W. Cook, Kara Moser, Adrian Paskey, Jason Caravas, Benjamin Rambo-Martin, Shatavia Morrison, Christopher Gulvick, Scott Sammons, Yvette Unoarumhi, Darlene Wagner, Matthew Schmerer, Harry Gao, Mickey Li, John Gao, Joseph Fierro, Benafsh Sapra, Becky Tsai, Yan Meng, Doreen Ng, James Xie, Clinton R. Paden, Duncan MacCannell |
| EPI_ISL_3300808 | CDFD | CDFD | Murali Bashyam, Asmita Gupta, Reelina Basu, Vinay Donipadi, Nagamani Kammili, Divya Vashisht, Ashwin Dalal |
| EPI_ISL_3300809, EPI_ISL_3300810 | Fulgent Genetics | Centers for Disease Control and Prevention Division of Viral Diseases, Pathogen Discovery | Dakota Howard, Dhwani Batra, Peter W. Cook, Kara Moser, Adrian Paskey, Jason Caravas, Benjamin Rambo-Martin, Shatavia Morrison, Christopher Gulvick, Scott Sammons, Yvette Unoarumhi, Darlene Wagner, Matthew Schmerer, Harry Gao, Mickey Li, John Gao, Joseph Fierro, Benafsh Sapra, Becky Tsai, Yan Meng, Doreen Ng, James Xie, Clinton R. Paden, Duncan MacCannell |
| EPI_ISL_3300811 | CDFD | CDFD | Murali Bashyam, Asmita Gupta, Reelina Basu, Vinay Donipadi, Nagamani Kammili, Divya Vashisht, Ashwin Dalal |
| EPI_ISL_3300812 | Fulgent Genetics | Centers for Disease Control and Prevention Division of Viral Diseases, Pathogen Discovery | Dakota Howard, Dhwani Batra, Peter W. Cook, Kara Moser, Adrian Paskey, Jason Caravas, Benjamin Rambo-Martin, Shatavia Morrison, Christopher Gulvick, Scott Sammons, Yvette Unoarumhi, Darlene Wagner, Matthew Schmerer, Harry Gao, Mickey Li, John Gao, Joseph Fierro, Benafsh Sapra, Becky Tsai, Yan Meng, Doreen Ng, James Xie, Clinton R. Paden, Duncan MacCannell |
| EPI_ISL_3300813 | CDFD | CDFD | Murali Bashyam, Asmita Gupta, Reelina Basu, Vinay Donipadi, Nagamani Kammili, Divya Vashisht, Ashwin Dalal |
| EPI_ISL_3300814, EPI_ISL_3300816 | Fulgent Genetics | Centers for Disease Control and Prevention Division of Viral Diseases, Pathogen Discovery | Dakota Howard, Dhwani Batra, Peter W. Cook, Kara Moser, Adrian Paskey, Jason Caravas, Benjamin Rambo-Martin, Shatavia Morrison, Christopher Gulvick, Scott Sammons, Yvette Unoarumhi, Darlene Wagner, Matthew Schmerer, Harry Gao, Mickey Li, John Gao, Joseph Fierro, Benafsh Sapra, Becky Tsai, Yan Meng, Doreen Ng, James Xie, Clinton R. Paden, Duncan MacCannell |
| EPI_ISL_3300817 | CDFD | CDFD | Murali Bashyam, Asmita Gupta, Reelina Basu, Vinay Donipadi, Nagamani Kammili, Divya Vashisht, Ashwin Dalal |
| EPI_ISL_3300818 | Fulgent Genetics | Centers for Disease Control and Prevention Division of Viral Diseases, Pathogen Discovery | Dakota Howard, Dhwani Batra, Peter W. Cook, Kara Moser, Adrian Paskey, Jason Caravas, Benjamin Rambo-Martin, Shatavia Morrison, Christopher Gulvick, Scott Sammons, Yvette Unoarumhi, Darlene Wagner, Matthew Schmerer, Harry Gao, Mickey Li, John Gao, Joseph Fierro, Benafsh Sapra, Becky Tsai, Yan Meng, Doreen Ng, James Xie, Clinton R. Paden, Duncan MacCannell |
| EPI_ISL_3300819 | CDFD | CDFD | Murali Bashyam, Asmita Gupta, Reelina Basu, Vinay Donipadi, Nagamani Kammili, Divya Vashisht, Ashwin Dalal |
| EPI_ISL_3300820 | Fulgent Genetics | Centers for Disease Control and Prevention Division of Viral Diseases, Pathogen Discovery | Dakota Howard, Dhwani Batra, Peter W. Cook, Kara Moser, Adrian Paskey, Jason Caravas, Benjamin Rambo-Martin, Shatavia Morrison, Christopher Gulvick, Scott Sammons, Yvette Unoarumhi, Darlene Wagner, Matthew Schmerer, Harry Gao, Mickey Li, John Gao, Joseph Fierro, Benafsh Sapra, Becky Tsai, Yan Meng, Doreen Ng, James Xie, Clinton R. Paden, Duncan MacCannell |
| EPI_ISL_3300821, EPI_ISL_3300822 | CDFD | CDFD | Murali Bashyam, Asmita Gupta, Reelina Basu, Vinay Donipadi, Nagamani Kammili, Divya Vashisht, Ashwin Dalal |
| EPI_ISL_3300823 | Fulgent Genetics | Centers for Disease Control and Prevention Division of Viral Diseases, Pathogen Discovery | Dakota Howard, Dhwani Batra, Peter W. Cook, Kara Moser, Adrian Paskey, Jason Caravas, Benjamin Rambo-Martin, Shatavia Morrison, Christopher Gulvick, Scott Sammons, Yvette Unoarumhi, Darlene Wagner, Matthew Schmerer, Harry Gao, Mickey Li, John Gao, Joseph Fierro, Benafsh Sapra, Becky Tsai, Yan Meng, Doreen Ng, James Xie, Clinton R. Paden, Duncan MacCannell |
| EPI_ISL_3300824 | CDFD | CDFD | Murali Bashyam, Asmita Gupta, Reelina Basu, Vinay Donipadi, Nagamani Kammili, Divya Vashisht, Ashwin Dalal |
| EPI_ISL_3300825 | Fulgent Genetics | Centers for Disease Control and Prevention Division of Viral Diseases, Pathogen Discovery | Dakota Howard, Dhwani Batra, Peter W. Cook, Kara Moser, Adrian Paskey, Jason Caravas, Benjamin Rambo-Martin, Shatavia Morrison, Christopher Gulvick, Scott Sammons, Yvette Unoarumhi, Darlene Wagner, Matthew Schmerer, Harry Gao, Mickey Li, John Gao, Joseph Fierro, Benafsh Sapra, Becky Tsai, Yan Meng, Doreen Ng, James Xie, Clinton R. Paden, Duncan MacCannell |
| EPI_ISL_3300826 | CDFD | CDFD | Murali Bashyam, Asmita Gupta, Reelina Basu, Vinay Donipadi, Nagamani Kammili, Divya Vashisht, Ashwin Dalal |
| EPI_ISL_3300827, EPI_ISL_3300828 | Fulgent Genetics | Centers for Disease Control and Prevention Division of Viral Diseases, Pathogen Discovery | Dakota Howard, Dhwani Batra, Peter W. Cook, Kara Moser, Adrian Paskey, Jason Caravas, Benjamin Rambo-Martin, Shatavia Morrison, Christopher Gulvick, Scott Sammons, Yvette Unoarumhi, Darlene Wagner, Matthew Schmerer, Harry Gao, Mickey Li, John Gao, Joseph Fierro, Benafsh Sapra, Becky Tsai, Yan Meng, Doreen Ng, James Xie, Clinton R. Paden, Duncan MacCannell |
| EPI_ISL_3300829 | CDFD | CDFD | Murali Bashyam, Asmita Gupta, Reelina Basu, Vinay Donipadi, Nagamani Kammili, Divya Vashisht, Ashwin Dalal |
| EPI_ISL_3300830, EPI_ISL_3300831, EPI_ISL_3300832 | Fulgent Genetics | Centers for Disease Control and Prevention Division of Viral Diseases, Pathogen Discovery | Dakota Howard, Dhwani Batra, Peter W. Cook, Kara Moser, Adrian Paskey, Jason Caravas, Benjamin Rambo-Martin, Shatavia Morrison, Christopher Gulvick, Scott Sammons, Yvette Unoarumhi, Darlene Wagner, Matthew Schmerer, Harry Gao, Mickey Li, John Gao, Joseph Fierro, Benafsh Sapra, Becky Tsai, Yan Meng, Doreen Ng, James Xie, Clinton R. Paden, Duncan MacCannell |
| EPI_ISL_3300833 | CDFD | CDFD | Murali Bashyam, Asmita Gupta, Reelina Basu, Vinay Donipadi, Nagamani Kammili, Divya Vashisht, Ashwin Dalal |
| EPI_ISL_3300834, EPI_ISL_3300835 | Fulgent Genetics | Centers for Disease Control and Prevention Division of Viral Diseases, Pathogen Discovery | Dakota Howard, Dhwani Batra, Peter W. Cook, Kara Moser, Adrian Paskey, Jason Caravas, Benjamin Rambo-Martin, Shatavia Morrison, Christopher Gulvick, Scott Sammons, Yvette Unoarumhi, Darlene Wagner, Matthew Schmerer, Harry Gao, Mickey Li, John Gao, Joseph Fierro, Benafsh Sapra, Becky Tsai, Yan Meng, Doreen Ng, James Xie, Clinton R. Paden, Duncan MacCannell |
| EPI_ISL_3300836 | CDFD | CDFD | Murali Bashyam, Asmita Gupta, Reelina Basu, Vinay Donipadi, Nagamani Kammili, Divya Vashisht, Ashwin Dalal |
| EPI_ISL_3300837, EPI_ISL_3300838 | Fulgent Genetics | Centers for Disease Control and Prevention Division of Viral Diseases, Pathogen Discovery | Dakota Howard, Dhwani Batra, Peter W. Cook, Kara Moser, Adrian Paskey, Jason Caravas, Benjamin Rambo-Martin, Shatavia Morrison, Christopher Gulvick, Scott Sammons, Yvette Unoarumhi, Darlene Wagner, Matthew Schmerer, Harry Gao, Mickey Li, John Gao, Joseph Fierro, Benafsh Sapra, Becky Tsai, Yan Meng, Doreen Ng, James Xie, Clinton R. Paden, Duncan MacCannell |
| EPI_ISL_3300839 | CDFD | CDFD | Murali Bashyam, Asmita Gupta, Reelina Basu, Vinay Donipadi, Nagamani Kammili, Divya Vashisht, Ashwin Dalal |
| EPI_ISL_3300840 | Fulgent Genetics | Centers for Disease Control and Prevention Division of Viral Diseases, Pathogen Discovery | Dakota Howard, Dhwani Batra, Peter W. Cook, Kara Moser, Adrian Paskey, Jason Caravas, Benjamin Rambo-Martin, Shatavia Morrison, Christopher Gulvick, Scott Sammons, Yvette Unoarumhi, Darlene Wagner, Matthew Schmerer, Harry Gao, Mickey Li, John Gao, Joseph Fierro, Benafsh Sapra, Becky Tsai, Yan Meng, Doreen Ng, James Xie, Clinton R. Paden, Duncan MacCannell |
| EPI_ISL_3300841 | CDFD | CDFD | Murali Bashyam, Asmita Gupta, Reelina Basu, Vinay Donipadi, Nagamani Kammili, Divya Vashisht, Ashwin Dalal |
| EPI_ISL_3300842 | Fulgent Genetics | Centers for Disease Control and Prevention Division of Viral Diseases, Pathogen Discovery | Dakota Howard, Dhwani Batra, Peter W. Cook, Kara Moser, Adrian Paskey, Jason Caravas, Benjamin Rambo-Martin, Shatavia Morrison, Christopher Gulvick, Scott Sammons, Yvette Unoarumhi, Darlene Wagner, Matthew Schmerer, Harry Gao, Mickey Li, John Gao, Joseph Fierro, Benafsh Sapra, Becky Tsai, Yan Meng, Doreen Ng, James Xie, Clinton R. Paden, Duncan MacCannell |
| EPI_ISL_3300843 | CDFD | CDFD | Murali Bashyam, Asmita Gupta, Reelina Basu, Vinay Donipadi, Nagamani Kammili, Divya Vashisht, Ashwin Dalal |
| EPI_ISL_3300844 | Fulgent Genetics | Centers for Disease Control and Prevention Division of Viral Diseases, Pathogen Discovery | Dakota Howard, Dhwani Batra, Peter W. Cook, Kara Moser, Adrian Paskey, Jason Caravas, Benjamin Rambo-Martin, Shatavia Morrison, Christopher Gulvick, Scott Sammons, Yvette Unoarumhi, Darlene Wagner, Matthew Schmerer, Harry Gao, Mickey Li, John Gao, Joseph Fierro, Benafsh Sapra, Becky Tsai, Yan Meng, Doreen Ng, James Xie, Clinton R. Paden, Duncan MacCannell |
| EPI_ISL_3300845 | CDFD | CDFD | Murali Bashyam, Asmita Gupta, Reelina Basu, Vinay Donipadi, Nagamani Kammili, Divya Vashisht, Ashwin Dalal |

EPI_ISL_3300846, EPI_ISL_3300847, EPI_ISL_3300848, EPI_ISL_3300849, EPI_ISL_3300850, EPI_ISL_3300851, EPI_ISL_3300852, EPI_ISL_3300853, EPI_ISL_3300854, EPI_ISL_3300855, EPI_ISL_3300856, EPI_ISL_3300857, EPI_ISL_3300858, EPI_ISL_3300859, EPI_ISL_3300860, EPI_ISL_3300861, EPI_ISL_3300863, EPI_ISL_3300864, EPI_ISL_3300865, EPI_ISL_3300866, EPI_ISL_3300867, EPI_ISL_3300868,
EPI_ISL_3300869, EPI_ISL_3300870, EPI_ISL_3300871, EPI_ISL_3300872, EPI_ISL_3300873, EPI_ISL_3300874, EPI_ISL_3300875, EPI_ISL_3300876, EPI_ISL_3300877, EPI_ISL_3300878, EPI_ISL_3300879, EPI_ISL_3300880, EPI_ISL_3300881, EPI_ISL_3300882, EPI_ISL_3300883, EPI_ISL_3300884, EPI_ISL_3300885, EPI_ISL_3300886, EPI_ISL_3300887, EPI_ISL_3300888, EPI_ISL_3300889, EPI_ISL_3300890,
EPI_ISL_3300891, EPI_ISL_3300892, EPI_ISL_3300893, EPI_ISL_3300894, EPI_ISL_3300895, EPI_ISL_3300896, EPI_ISL_3300897, EPI_ISL_3300898, EPI_ISL_3300899, EPI_ISL_3300900, EPI_ISL_3300901, EPI_ISL_3300902, EPI_ISL_3300903, EPI_ISL_3300904, EPI_ISL_3300905, EPI_ISL_3300906, EPI_ISL_3300907, EPI_ISL_3300908, EPI_ISL_3300909, EPI_ISL_3300910, EPI_ISL_3300911, EPI_ISL_3300912,
EPI_ISL_3300913, EPI_ISL_3300914, EPI_ISL_3300915, EPI_ISL_3300916, EPI_ISL_3300917, EPI_ISL_3300918, EPI_ISL_3300919, EPI_ISL_3300920, EPI_ISL_3300921, EPI_ISL_3300922, EPI_ISL_3300923, EPI_ISL_3300924, EPI_ISL_3300925, EPI_ISL_3300926, EPI_ISL_3300927, EPI_ISL_3300928, EPI_ISL_3300929, EPI_ISL_3300930, EPI_ISL_3300931, EPI_ISL_3300932, EPI_ISL_3300933, EPI_ISL_3300934,
EPI_ISL_3300935, EPI_ISL_3300936, EPI_ISL_3300937, EPI_ISL_3300938, EPI_ISL_3300939, EPI_ISL_3300940, EPI_ISL_3300941, EPI_ISL_3300942, EPI_ISL_3300943, EPI_ISL_3300944, EPI_ISL_3300945, EPI_ISL_3300946, EPI_ISL_3300947, EPI_ISL_3300948, EPI_ISL_3300949, EPI_ISL_3300950, EPI_ISL_3300951, EPI_ISL_3300952, EPI_ISL_3300953, EPI_ISL_3300954, EPI_ISL_3300955, EPI_ISL_3300956,
EPI_ISL_3300957, EPI_ISL_3300958, EPI_ISL_3300959, EPI_ISL_3300960, EPI_ISL_3300961, EPI_ISL_3300962, EPI_ISL_3300963, EPI_ISL_3300964, EPI_ISL_3300965, EPI_ISL_3300966, EPI_ISL_3300967, EPI_ISL_3300968, EPI_ISL_3300969, EPI_ISL_3300970, EPI_ISL_3300971, EPI_ISL_3300972, EPI_ISL_3300973, EPI_ISL_3300974, EPI_ISL_3300975, EPI_ISL_3300976, EPI_ISL_3300977, EPI_ISL_3300978,
EPI_ISL_3300979, EPI_ISL_3300980, EPI_ISL_3300981, EPI_ISL_3300982, EPI_ISL_3300983, EPI_ISL_3300984, EPI_ISL_3300985, EPI_ISL_3300986, EPI_ISL_3300987, EPI_ISL_3300988, EPI_ISL_3300989, EPI_ISL_3300990, EPI_ISL_3300991, EPI_ISL_3300992, EPI_ISL_3300993, EPI_ISL_3300994, EPI_ISL_3300995, EPI_ISL_3300996, EPI_ISL_3300997, EPI_ISL_3300998, EPI_ISL_3300999, EPI_ISL_3301000,
EPI_ISL_3301001, EPI_ISL_3301002, EPI_ISL_3301003, EPI_ISL_3301004, EPI_ISL_3301005, EPI_ISL_3301007, EPI_ISL_3301009, EPI_ISL_3301010, EPI_ISL_3301011, EPI_ISL_3301012, EPI_ISL_3301013, EPI_ISL_3301014, EPI_ISL_3301015, EPI_ISL_3301016, EPI_ISL_3301017, EPI_ISL_3301018, EPI_ISL_3301019, EPI_ISL_3301021, EPI_ISL_3301022, EPI_ISL_3301023, EPI_ISL_3301024, EPI_ISL_3301025,
EPI_ISL_3301026, EPI_ISL_3301027, EPI_ISL_3301028, EPI_ISL_3301029, EPI_ISL_3301030, EPI_ISL_3301031, EPI_ISL_3301032, EPI_ISL_3301033, EPI_ISL_3301034, EPI_ISL_3301035, EPI_ISL_3301036, EPI_ISL_3301037, EPI_ISL_3301038, EPI_ISL_3301039, EPI_ISL_3301040, EPI_ISL_3301041, EPI_ISL_3301043, EPI_ISL_3301044, EPI_ISL_3301045, EPI_ISL_3301046, EPI_ISL_3301047, EPI_ISL_3301048,
EPI_ISL_3301049, EPI_ISL_3301050, EPI_ISL_3301051, EPI_ISL_3301052, EPI_ISL_3301053, EPI_ISL_3301054, EPI_ISL_3301055, EPI_ISL_3301056, EPI_ISL_3301057, EPI_ISL_3301058, EPI_ISL_3301059, EPI_ISL_3301060, EPI_ISL_3301063, EPI_ISL_3301064, EPI_ISL_3301065, EPI_ISL_3301066, EPI_ISL_3301067, EPI_ISL_3301068, EPI_ISL_3301069, EPI_ISL_3301070, EPI_ISL_3301071, EPI_ISL_3301072,
EPI_ISL_3301073, EPI_ISL_3301074, EPI_ISL_3301075, EPI_ISL_3301076, EPI_ISL_3301077, EPI_ISL_3301078, EPI_ISL_3301079, EPI_ISL_3301080, EPI_ISL_3301081, EPI_ISL_3301082, EPI_ISL_3301083, EPI_ISL_3301084, EPI_ISL_3301085, EPI_ISL_3301086, EPI_ISL_3301087, EPI_ISL_3301088, EPI_ISL_3301089, EPI_ISL_3301090, EPI_ISL_3301091, EPI_ISL_3301092, EPI_ISL_3301094, EPI_ISL_3301095,
EPI_ISL_3301096, EPI_ISL_3301097, EPI_ISL_3301098, EPI_ISL_3301099, EPI_ISL_3301100, EPI_ISL_3301101, EPI_ISL_3301102, EPI_ISL_3301103, EPI_ISL_3301104, EPI_ISL_3301105, EPI_ISL_3301106, EPI_ISL_3301107, EPI_ISL_3301108, EPI_ISL_3301109, EPI_ISL_3301110, EPI_ISL_3301111, EPI_ISL_3301112, EPI_ISL_3301113, EPI_ISL_3301115, EPI_ISL_3301116, EPI_ISL_3301117, EPI_ISL_3301118,
EPI_ISL_3301119, EPI_ISL_3301120, EPI_ISL_3301121, EPI_ISL_3301122, EPI_ISL_3301123, EPI_ISL_3301124, EPI_ISL_3301125, EPI_ISL_3301126, EPI_ISL_3301127, EPI_ISL_3301128, EPI_ISL_3301129, EPI_ISL_3301130, EPI_ISL_3301131, EPI_ISL_3301133, EPI_ISL_3301134, EPI_ISL_3301135, EPI_ISL_3301136, EPI_ISL_3301137, EPI_ISL_3301138, EPI_ISL_3301139, EPI_ISL_3301140, EPI_ISL_3301141,
EPI_ISL_3301142, EPI_ISL_3301143, EPI_ISL_3301144, EPI_ISL_3301145, EPI_ISL_3301146, EPI_ISL_3301147, EPI_ISL_3301148, EPI_ISL_3301149, EPI_ISL_3301150, EPI_ISL_3301151, EPI_ISL_3301152, EPI_ISL_3301153, EPI_ISL_3301154, EPI_ISL_3301155, EPI_ISL_3301156, EPI_ISL_3301157, EPI_ISL_3301158, EPI_ISL_3301159, EPI_ISL_3301160, EPI_ISL_3301161, EPI_ISL_3301162, EPI_ISL_3301163,
EPI_ISL_3301164, EPI_ISL_3301165, EPI_ISL_3301166, EPI_ISL_3301167, EPI_ISL_3301168, EPI_ISL_3301169, EPI_ISL_3301170, EPI_ISL_3301171, EPI_ISL_3301172, EPI_ISL_3301173, EPI_ISL_3301174, EPI_ISL_3301175, EPI_ISL_3301176, EPI_ISL_3301177, EPI_ISL_3301178, EPI_ISL_3301179, EPI_ISL_3301180, EPI_ISL_3301181, EPI_ISL_3301182, EPI_ISL_3301183, EPI_ISL_3301184, EPI_ISL_3301185,
EPI_ISL_3301186, EPI_ISL_3301187, EPI_ISL_3301188, EPI_ISL_3301189, EPI_ISL_3301192, EPI_ISL_3301193, EPI_ISL_3301194, EPI_ISL_3301195, EPI_ISL_3301196, EPI_ISL_3301197, EPI_ISL_3301198, EPI_ISL_3301199, EPI_ISL_3301200, EPI_ISL_3301201, EPI_ISL_3301202, EPI_ISL_3301203, EPI_ISL_3301204, EPI_ISL_3301205, EPI_ISL_3301206, EPI_ISL_3301207, EPI_ISL_3301208, EPI_ISL_3301209,
EPI_ISL_3301210, EPI_ISL_3301211, EPI_ISL_3301212, EPI_ISL_3301214, EPI_ISL_3301215, EPI_ISL_3301216, EPI_ISL_3301217, EPI_ISL_3301218, EPI_ISL_3301219, EPI_ISL_3301220, EPI_ISL_3301221, EPI_ISL_3301222, EPI_ISL_3301223, EPI_ISL_3301224, EPI_ISL_3301225, EPI_ISL_3301226, EPI_ISL_3301227, EPI_ISL_3301228, EPI_ISL_3301229, EPI_ISL_3301230, EPI_ISL_3301231, EPI_ISL_3301232,
EPI_ISL_3301233, EPI_ISL_3301234, EPI_ISL_3301235, EPI_ISL_3301236, EPI_ISL_3301237, EPI_ISL_3301238, EPI_ISL_3301239, EPI_ISL_3301240, EPI_ISL_3301242, EPI_ISL_3301243, EPI_ISL_3301244, EPI_ISL_3301245, EPI_ISL_3301246, EPI_ISL_3301247, EPI_ISL_3301248, EPI_ISL_3301249, EPI_ISL_3301250, EPI_ISL_3301251, EPI_ISL_3301252, EPI_ISL_3301253, EPI_ISL_3301254, EPI_ISL_3301255,
EPI_ISL_3301256, EPI_ISL_3301257, EPI_ISL_3301258, EPI_ISL_3301259, EPI_ISL_3301260, EPI_ISL_3301261, EPI_ISL_3301262, EPI_ISL_3301263, EPI_ISL_3301264, EPI_ISL_3301265, EPI_ISL_3301266, EPI_ISL_3301267, EPI_ISL_3301268, EPI_ISL_3301269, EPI_ISL_3301270, EPI_ISL_3301271, EPI_ISL_3301272, EPI_ISL_3301273, EPI_ISL_3301274, EPI_ISL_3301275, EPI_ISL_3301276, EPI_ISL_3301279,
EPI_ISL_3301280, EPI_ISL_3301281, EPI_ISL_3301282, EPI_ISL_3301283, EPI_ISL_3301284, EPI_ISL_3301285, EPI_ISL_3301286, EPI_ISL_3301287, EPI_ISL_3301288, EPI_ISL_3301289, EPI_ISL_3301290, EPI_ISL_3301291, EPI_ISL_3301292, EPI_ISL_3301293, EPI_ISL_3301294, EPI_ISL_3301295, EPI_ISL_3301296, EPI_ISL_3301297, EPI_ISL_3301298, EPI_ISL_3301299, EPI_ISL_3301300, EPI_ISL_3301301,
EPI_ISL_3301302, EPI_ISL_3301303, EPI_ISL_3301304, EPI_ISL_3301305, EPI_ISL_3301306, EPI_ISL_3301307, EPI_ISL_3301308, EPI_ISL_3301309, EPI_ISL_3301310, EPI_ISL_3301311, EPI_ISL_3301312, EPI_ISL_3301313, EPI_ISL_3301314, EPI_ISL_3301315, EPI_ISL_3301316, EPI_ISL_3301317, EPI_ISL_3301318, EPI_ISL_3301319, EPI_ISL_3301320, EPI_ISL_3301321, EPI_ISL_3301322, EPI_ISL_3301323,
EPI_ISL_3301324, EPI_ISL_3301325, EPI_ISL_3301326, EPI_ISL_3301327, EPI_ISL_3301328, EPI_ISL_3301329, EPI_ISL_3301330, EPI_ISL_3301332, EPI_ISL_3301333, EPI_ISL_3301334, EPI_ISL_3301335, EPI_ISL_3301336, EPI_ISL_3301337, EPI_ISL_3301338, EPI_ISL_3301339, EPI_ISL_3301340, EPI_ISL_3301341, EPI_ISL_3301342, EPI_ISL_3301343, EPI_ISL_3301344, EPI_ISL_3301345, EPI_ISL_3301346,
EPI_ISL_3301347, EPI_ISL_3301348, EPI_ISL_3301349, EPI_ISL_3301350, EPI_ISL_3301351, EPI_ISL_3301352, EPI_ISL_3301353, EPI_ISL_3301354, EPI_ISL_3301355, EPI_ISL_3301356, EPI_ISL_3301357, EPI_ISL_3301358, EPI_ISL_3301359, EPI_ISL_3301360, EPI_ISL_3301361, EPI_ISL_3301362, EPI_ISL_3301363, EPI_ISL_3301364, EPI_ISL_3301365, EPI_ISL_3301366, EPI_ISL_3301367, EPI_ISL_3301368,
EPI_ISL_3301369, EPI_ISL_3301370, EPI_ISL_3301371, EPI_ISL_3301372, EPI_ISL_3301373, EPI_ISL_3301374, EPI_ISL_3301375, EPI_ISL_3301376, EPI_ISL_3301377, EPI_ISL_3301378, EPI_ISL_3301379, EPI_ISL_3301380, EPI_ISL_3301381, EPI_ISL_3301382, EPI_ISL_3301383, EPI_ISL_3301384, EPI_ISL_3301385, EPI_ISL_3301386, EPI_ISL_3301387, EPI_ISL_3301388, EPI_ISL_3301389, EPI_ISL_3301390,
EPI_ISL_3301391, EPI_ISL_3301392, EPI_ISL_3301393, EPI_ISL_3301394, EPI_ISL_3301395, EPI_ISL_3301396, EPI_ISL_3301397, EPI_ISL_3301398, EPI_ISL_3301399, EPI_ISL_3301400, EPI_ISL_3301401, EPI_ISL_3301402, EPI_ISL_3301403, EPI_ISL_3301404, EPI_ISL_3301405, EPI_ISL_3301406, EPI_ISL_3301407, EPI_ISL_3301408, EPI_ISL_3301409, EPI_ISL_3301410, EPI_ISL_3301411, EPI_ISL_3301412,
EPI_ISL_3301413, EPI_ISL_3301414, EPI_ISL_3301415, EPI_ISL_3301416, EPI_ISL_3301417, EPI_ISL_3301418, EPI_ISL_3301419, EPI_ISL_3301420, EPI_ISL_3301421, EPI_ISL_3301422, EPI_ISL_3301423, EPI_ISL_3301424, EPI_ISL_3301425, EPI_ISL_3301426, EPI_ISL_3301427, EPI_ISL_3301428, EPI_ISL_3301429, EPI_ISL_3301430, EPI_ISL_3301431, EPI_ISL_3301432, EPI_ISL_3301433, EPI_ISL_3301434,
EPI_ISL_3301435, EPI_ISL_3301436, EPI_ISL_3301437, EPI_ISL_3301438, EPI_ISL_3301439, EPI_ISL_3301440, EPI_ISL_3301441, EPI_ISL_3301442, EPI_ISL_3301443, EPI_ISL_3301444, EPI_ISL_3301445, EPI_ISL_3301446, EPI_ISL_3301447, EPI_ISL_3301448, EPI_ISL_3301449, EPI_ISL_3301450, EPI_ISL_3301452, EPI_ISL_3301453, EPI_ISL_3301454, EPI_ISL_3301455, EPI_ISL_3301456, EPI_ISL_3301457,
EPI_ISL_3301458, EPI_ISL_3301459, EPI_ISL_3301460, EPI_ISL_3301461, EPI_ISL_3301462, EPI_ISL_3301463, EPI_ISL_3301464, EPI_ISL_3301465, EPI_ISL_3301466, EPI_ISL_3301467, EPI_ISL_3301468, EPI_ISL_3301469, EPI_ISL_3301470, EPI_ISL_3301471, EPI_ISL_3301472, EPI_ISL_3301473, EPI_ISL_3301474, EPI_ISL_3301475, EPI_ISL_3301476, EPI_ISL_3301477, EPI_ISL_3301478, EPI_ISL_3301479,
EPI_ISL_3301480, EPI_ISL_3301481, EPI_ISL_3301482, EPI_ISL_3301483, EPI_ISL_3301484, EPI_ISL_3301485, EPI_ISL_3301486, EPI_ISL_3301487, EPI_ISL_3301489, EPI_ISL_3301490, EPI_ISL_3301491, EPI_ISL_3301492, EPI_ISL_3301493, EPI_ISL_3301494, EPI_ISL_3301495, EPI_ISL_3301496, EPI_ISL_3301497, EPI_ISL_3301498, EPI_ISL_3301499, EPI_ISL_3301500, EPI_ISL_3301501, EPI_ISL_3301502,
EPI_ISL_3301503, EPI_ISL_3301504, EPI_ISL_3301505, EPI_ISL_3301506, EPI_ISL_3301507, EPI_ISL_3301508, EPI_ISL_3301509, EPI_ISL_3301510, EPI_ISL_3301511, EPI_ISL_3301512, EPI_ISL_3301514, EPI_ISL_3301515, EPI_ISL_3301516, EPI_ISL_3301517, EPI_ISL_3301518, EPI_ISL_3301519, EPI_ISL_3301520, EPI_ISL_3301521, EPI_ISL_3301522, EPI_ISL_3301523, EPI_ISL_3301524, EPI_ISL_3301525,
EPI_ISL_3301526, EPI_ISL_3301527, EPI_ISL_3301528, EPI_ISL_3301530, EPI_ISL_3301531, EPI_ISL_3301532, EPI_ISL_3301533, EPI_ISL_3301534, EPI_ISL_3301535, EPI_ISL_3301536, EPI_ISL_3301537, EPI_ISL_3301538, EPI_ISL_3301539, EPI_ISL_3301540, EPI_ISL_3301541, EPI_ISL_3301542, EPI_ISL_3301543, EPI_ISL_3301544, EPI_ISL_3301545, EPI_ISL_3301546, EPI_ISL_3301547, EPI_ISL_3301548,
EPI_ISL_3301549, EPI_ISL_3301550, EPI_ISL_3301552, EPI_ISL_3301553, EPI_ISL_3301554, EPI_ISL_3301555, EPI_ISL_3301556, EPI_ISL_3301557, EPI_ISL_3301558, EPI_ISL_3301559, EPI_ISL_3301560, EPI_ISL_3301561, EPI_ISL_3301562, EPI_ISL_3301563, EPI_ISL_3301564, EPI_ISL_3301565, EPI_ISL_3301566, EPI_ISL_3301567, EPI_ISL_3301568, EPI_ISL_3301569, EPI_ISL_3301570, EPI_ISL_3301571,
EPI_ISL_3301572, EPI_ISL_3301574, EPI_ISL_3301575, EPI_ISL_3301577, EPI_ISL_3301578, EPI_ISL_3301579, EPI_ISL_3301580, EPI_ISL_3301581, EPI_ISL_3301582, EPI_ISL_3301583, EPI_ISL_3301584, EPI_ISL_3301585, EPI_ISL_3301586, EPI_ISL_3301587, EPI_ISL_3301588, EPI_ISL_3301589, EPI_ISL_3301590, EPI_ISL_3301591, EPI_ISL_3301593, EPI_ISL_3301594, EPI_ISL_3301595, EPI_ISL_3301596,
EPI_ISL_3301597, EPI_ISL_3301598, EPI_ISL_3301599, EPI_ISL_3301600, EPI_ISL_3301601, EPI_ISL_3301602, EPI_ISL_3301603, EPI_ISL_3301604, EPI_ISL_3301605, EPI_ISL_3301606, EPI_ISL_3301607, EPI_ISL_3301608, EPI_ISL_3301609, EPI_ISL_3301610, EPI_ISL_3301611, EPI_ISL_3301612, EPI_ISL_3301613, EPI_ISL_3301614, EPI_ISL_3301615, EPI_ISL_3301616, EPI_ISL_3301617, EPI_ISL_3301618,
EPI_ISL_3301619, EPI_ISL_3301620, EPI_ISL_3301621, EPI_ISL_3301622, EPI_ISL_3301623, EPI_ISL_3301624, EPI_ISL_3301625, EPI_ISL_3301626, EPI_ISL_3301627, EPI_ISL_3301628, EPI_ISL_3301629, EPI_ISL_3301630, EPI_ISL_3301631, EPI_ISL_3301632, EPI_ISL_3301633, EPI_ISL_3301634, EPI_ISL_3301635, EPI_ISL_3301636, EPI_ISL_3301637, EPI_ISL_3301638, EPI_ISL_3301639, EPI_ISL_3301640,
EPI_ISL_3301641, EPI_ISL_3301642, EPI_ISL_3301644, EPI_ISL_3301645, EPI_ISL_3301646, EPI_ISL_3301647, EPI_ISL_3301648, EPI_ISL_3301649, EPI_ISL_3301650, EPI_ISL_3301651, EPI_ISL_3301652, EPI_ISL_3301653, EPI_ISL_3301654, EPI_ISL_3301655, EPI_ISL_3301656, EPI_ISL_3301657, EPI_ISL_3301658, EPI_ISL_3301659, EPI_ISL_3301660, EPI_ISL_3301661, EPI_ISL_3301662, EPI_ISL_3301663,
EPI_ISL_3301664, EPI_ISL_3301665, EPI_ISL_3301666, EPI_ISL_3301667, EPI_ISL_3301668, EPI_ISL_3301669, EPI_ISL_3301670, EPI_ISL_3301671, EPI_ISL_3301672, EPI_ISL_3301673, EPI_ISL_3301674, EPI_ISL_3301675, EPI_ISL_3301676, EPI_ISL_3301677, EPI_ISL_3301678, EPI_ISL_3301679, EPI_ISL_3301680, EPI_ISL_3301681, EPI_ISL_3301682, EPI_ISL_3301683, EPI_ISL_3301684, EPI_ISL_3301685,
EPI_ISL_3301686, EPI_ISL_3301687, EPI_ISL_3301688, EPI_ISL_3301689, EPI_ISL_3301690, EPI_ISL_3301691, EPI_ISL_3301692, EPI_ISL_3301693, EPI_ISL_3301694, EPI_ISL_3301696, EPI_ISL_3301697, EPI_ISL_3301698, EPI_ISL_3301699, EPI_ISL_3301700, EPI_ISL_3301701, EPI_ISL_3301702, EPI_ISL_3301703, EPI_ISL_3301704, EPI_ISL_3301705, EPI_ISL_3301706, EPI_ISL_3301707, EPI_ISL_3301708,
EPI_ISL_3301709, EPI_ISL_3301710, EPI_ISL_3301711, EPI_ISL_3301712, EPI_ISL_3301713, EPI_ISL_3301714, EPI_ISL_3301715, EPI_ISL_3301716, EPI_ISL_3301717, EPI_ISL_3301718, EPI_ISL_3301719, EPI_ISL_3301720, EPI_ISL_3301721, EPI_ISL_3301722, EPI_ISL_3301723, EPI_ISL_3301724, EPI_ISL_3301726, EPI_ISL_3301727, EPI_ISL_3301728, EPI_ISL_3301729, EPI_ISL_3301730, EPI_ISL_3301731,
EPI_ISL_3301732, EPI_ISL_3301733, EPI_ISL_3301734, EPI_ISL_3301735, EPI_ISL_3301736, EPI_ISL_3301737, EPI_ISL_3301738, EPI_ISL_3301739, EPI_ISL_3301740, EPI_ISL_3301741, EPI_ISL_3301742, EPI_ISL_3301743, EPI_ISL_3301744, EPI_ISL_3301745, EPI_ISL_3301746, EPI_ISL_3301747, EPI_ISL_3301748, EPI_ISL_3301749, EPI_ISL_3301750, EPI_ISL_3301751, EPI_ISL_3301752, EPI_ISL_3301753,
EPI_ISL_3301754, EPI_ISL_3301755, EPI_ISL_3301756, EPI_ISL_3301757, EPI_ISL_3301758, EPI_ISL_3301759, EPI_ISL_3301760, EPI_ISL_3301761, EPI_ISL_3301762, EPI_ISL_3301763, EPI_ISL_3301764, EPI_ISL_3301765, EPI_ISL_3301766, EPI_ISL_3301767, EPI_ISL_3301768, EPI_ISL_3301769, EPI_ISL_3301770, EPI_ISL_3301771, EPI_ISL_3301772, EPI_ISL_3301773, EPI_ISL_3301774, EPI_ISL_3301775,
EPI_ISL_3301776, EPI_ISL_3301777, EPI_ISL_3301778, EPI_ISL_3301779, EPI_ISL_3301780, EPI_ISL_3301781, EPI_ISL_3301782, EPI_ISL_3301783, EPI_ISL_3301784, EPI_ISL_3301785, EPI_ISL_3301786, EPI_ISL_3301787, EPI_ISL_3301788, EPI_ISL_3301789, EPI_ISL_3301790, EPI_ISL_3301791, EPI_ISL_3301792, EPI_ISL_3301793, EPI_ISL_3301794, EPI_ISL_3301795, EPI_ISL_3301796, EPI_ISL_3301797,
EPI_ISL_3301798, EPI_ISL_3301799, EPI_ISL_3301800, EPI_ISL_3301801, EPI_ISL_3301802, EPI_ISL_3301803, EPI_ISL_3301804, EPI_ISL_3301805, EPI_ISL_3301806, EPI_ISL_3301807, EPI_ISL_3301808, EPI_ISL_3301809, EPI_ISL_3301810, EPI_ISL_3301811, EPI_ISL_3301812, EPI_ISL_3301813, EPI_ISL_3301814, EPI_ISL_3301815, EPI_ISL_3301816, EPI_ISL_3301817, EPI_ISL_3301818, EPI_ISL_3301819,
EPI_ISL_3301820, EPI_ISL_3301821, EPI_ISL_3301822, EPI_ISL_3301823, EPI_ISL_3301824, EPI_ISL_3301825, EPI_ISL_3301826, EPI_ISL_3301827, EPI_ISL_3301828, EPI_ISL_3301829, EPI_ISL_3301830, EPI_ISL_3301831, EPI_ISL_3301832, EPI_ISL_3301833, EPI_ISL_3301834, EPI_ISL_3301835, EPI_ISL_3301836, EPI_ISL_3301837, EPI_ISL_3301838, EPI_ISL_3301840, EPI_ISL_3301841, EPI_ISL_3301843,
EPI_ISL_3301844, EPI_ISL_3301845, EPI_ISL_3301847, EPI_ISL_3301848, EPI_ISL_3301849, EPI_ISL_3301850, EPI_ISL_3301851, EPI_ISL_3301852, EPI_ISL_3301853, EPI_ISL_3301854, EPI_ISL_3301856, EPI_ISL_3301860, EPI_ISL_3301861, EPI_ISL_3301862, EPI_ISL_3301863, EPI_ISL_3301864, EPI_ISL_3301865, EPI_ISL_3301866, EPI_ISL_3301867, EPI_ISL_3301868, EPI_ISL_3301869, EPI_ISL_3301870,
EPI_ISL_3301871, EPI_ISL_3301874, EPI_ISL_3301875, EPI_ISL_3301876, EPI_ISL_3301877, EPI_ISL_3301878, EPI_ISL_3301879, EPI_ISL_3301880, EPI_ISL_3301881, EPI_ISL_3301882, EPI_ISL_3301883, EPI_ISL_3301884, EPI_ISL_3301885, EPI_ISL_3301886, EPI_ISL_3301887, EPI_ISL_3301888, EPI_ISL_3301889, EPI_ISL_3301890, EPI_ISL_3301891, EPI_ISL_3301892, EPI_ISL_3301893, EPI_ISL_3301894,

| | | | |
|---|---|---|---|
| EPI_ISL_3301895, EPI_ISL_3301896, EPI_ISL_3301897, EPI_ISL_3301898, EPI_ISL_3301899, EPI_ISL_3301900, EPI_ISL_3301901, EPI_ISL_3301902, EPI_ISL_3301903, EPI_ISL_3301904, EPI_ISL_3301905, EPI_ISL_3301906, EPI_ISL_3301907, EPI_ISL_3301908, EPI_ISL_3301909, EPI_ISL_3301910, EPI_ISL_3301911, EPI_ISL_3301912, EPI_ISL_3301913, EPI_ISL_3301914, EPI_ISL_3301915, EPI_ISL_3301916, EPI_ISL_3301917, EPI_ISL_3301918, EPI_ISL_3301919, EPI_ISL_3301920, EPI_ISL_3301921, EPI_ISL_3301922, EPI_ISL_3301923, EPI_ISL_3301924, EPI_ISL_3301925, EPI_ISL_3301926, EPI_ISL_3301927, EPI_ISL_3301928, EPI_ISL_3301929, EPI_ISL_3301930, EPI_ISL_3301931, EPI_ISL_3301932, EPI_ISL_3301933, EPI_ISL_3301934, EPI_ISL_3301935, EPI_ISL_3301936, EPI_ISL_3301937, EPI_ISL_3301938, EPI_ISL_3301939, EPI_ISL_3301940, EPI_ISL_3301941, EPI_ISL_3301942, EPI_ISL_3301943, EPI_ISL_3301944, EPI_ISL_3301945, EPI_ISL_3301946, EPI_ISL_3301947, EPI_ISL_3301948, EPI_ISL_3301949, EPI_ISL_3301951, EPI_ISL_3301952, EPI_ISL_3301953, EPI_ISL_3301954, EPI_ISL_3301955, EPI_ISL_3301956, EPI_ISL_3301957, EPI_ISL_3301958, EPI_ISL_3301959, EPI_ISL_3301960, EPI_ISL_3301961, EPI_ISL_3301962, EPI_ISL_3301963, EPI_ISL_3301964, EPI_ISL_3301965, EPI_ISL_3301966, EPI_ISL_3301967, EPI_ISL_3301968, EPI_ISL_3301969, EPI_ISL_3301970, EPI_ISL_3301971, EPI_ISL_3301972, EPI_ISL_3301973, EPI_ISL_3301974, EPI_ISL_3301975, EPI_ISL_3301976, EPI_ISL_3301978, EPI_ISL_3301980, EPI_ISL_3301981, EPI_ISL_3301982, EPI_ISL_3301983, EPI_ISL_3301984, EPI_ISL_3301985, EPI_ISL_3301986, EPI_ISL_3301987, EPI_ISL_3301988, EPI_ISL_3301989, EPI_ISL_3301990, EPI_ISL_3301991, EPI_ISL_3301992, EPI_ISL_3301993, EPI_ISL_3301994, EPI_ISL_3301995, EPI_ISL_3301996, EPI_ISL_3301997, EPI_ISL_3301998, EPI_ISL_3301999, EPI_ISL_3302000, EPI_ISL_3302001, EPI_ISL_3302002, EPI_ISL_3302003, EPI_ISL_3302004, EPI_ISL_3302005, EPI_ISL_3302006, EPI_ISL_3302007, EPI_ISL_3302008, EPI_ISL_3302009, EPI_ISL_3302010, EPI_ISL_3302012, EPI_ISL_3302013, EPI_ISL_3302014, EPI_ISL_3302015, EPI_ISL_3302016, EPI_ISL_3302017, EPI_ISL_3302018, EPI_ISL_3302019, EPI_ISL_3302020, EPI_ISL_3302021, EPI_ISL_3302022, EPI_ISL_3302023, EPI_ISL_3302024, EPI_ISL_3302025, EPI_ISL_3302026, EPI_ISL_3302027, EPI_ISL_3302028, EPI_ISL_3302029, EPI_ISL_3302030, EPI_ISL_3302031, EPI_ISL_3302032, EPI_ISL_3302033, EPI_ISL_3302034, EPI_ISL_3302035, EPI_ISL_3302036, EPI_ISL_3302037, EPI_ISL_3302038, EPI_ISL_3302039, EPI_ISL_3302040, EPI_ISL_3302041, EPI_ISL_3302042, EPI_ISL_3302043, EPI_ISL_3302044, EPI_ISL_3302045, EPI_ISL_3302046, EPI_ISL_3302047, EPI_ISL_3302048, EPI_ISL_3302049, EPI_ISL_3302050, EPI_ISL_3302051, EPI_ISL_3302052, EPI_ISL_3302053, EPI_ISL_3302054, EPI_ISL_3302055, EPI_ISL_3302056, EPI_ISL_3302057, EPI_ISL_3302058, EPI_ISL_3302059, EPI_ISL_3302060, EPI_ISL_3302061, EPI_ISL_3302062, EPI_ISL_3302063, EPI_ISL_3302064, EPI_ISL_3302065, EPI_ISL_3302066, EPI_ISL_3302067, EPI_ISL_3302068, EPI_ISL_3302069, EPI_ISL_3302070, EPI_ISL_3302071, EPI_ISL_3302072, EPI_ISL_3302073, EPI_ISL_3302074, EPI_ISL_3302075, EPI_ISL_3302076, EPI_ISL_3302077, EPI_ISL_3302078, EPI_ISL_3302079, EPI_ISL_3302080, EPI_ISL_3302081, EPI_ISL_3302082, EPI_ISL_3302083, EPI_ISL_3302084, EPI_ISL_3302085, EPI_ISL_3302086, EPI_ISL_3302087, EPI_ISL_3302088, EPI_ISL_3302089, EPI_ISL_3302090, EPI_ISL_3302091, EPI_ISL_3302092, EPI_ISL_3302093, EPI_ISL_3302094, EPI_ISL_3302095, EPI_ISL_3302096, EPI_ISL_3302097, EPI_ISL_3302098, EPI_ISL_3302099, EPI_ISL_3302100, EPI_ISL_3302101, EPI_ISL_3302102, EPI_ISL_3302103, EPI_ISL_3302104, EPI_ISL_3302105, EPI_ISL_3302106, EPI_ISL_3302107, EPI_ISL_3302108, EPI_ISL_3302109, EPI_ISL_3302110, EPI_ISL_3302111, EPI_ISL_3302112, EPI_ISL_3302113, EPI_ISL_3302114, EPI_ISL_3302115, EPI_ISL_3302116, EPI_ISL_3302117, EPI_ISL_3302118, EPI_ISL_3302119, EPI_ISL_3302120, EPI_ISL_3302121, EPI_ISL_3302122, EPI_ISL_3302123, EPI_ISL_3302124, EPI_ISL_3302125, EPI_ISL_3302126, EPI_ISL_3302127, EPI_ISL_3302128, EPI_ISL_3302129, EPI_ISL_3302130, EPI_ISL_3302131, EPI_ISL_3302132, EPI_ISL_3302133, EPI_ISL_3302134, EPI_ISL_3302135, EPI_ISL_3302136, EPI_ISL_3302137, EPI_ISL_3302138, EPI_ISL_3302139, EPI_ISL_3302140, EPI_ISL_3302141, EPI_ISL_3302142, EPI_ISL_3302143, EPI_ISL_3302144, EPI_ISL_3302145, EPI_ISL_3302146, EPI_ISL_3302147, EPI_ISL_3302148, EPI_ISL_3302149, EPI_ISL_3302150, EPI_ISL_3302151, EPI_ISL_3302152, EPI_ISL_3302153, EPI_ISL_3302154, EPI_ISL_3302155, EPI_ISL_3302156, EPI_ISL_3302157, EPI_ISL_3302158, EPI_ISL_3302159, EPI_ISL_3302160, EPI_ISL_3302161, EPI_ISL_3302162, EPI_ISL_3302163, EPI_ISL_3302164, EPI_ISL_3302165, EPI_ISL_3302166, EPI_ISL_3302167, EPI_ISL_3302168, EPI_ISL_3302169, EPI_ISL_3302170, EPI_ISL_3302171, EPI_ISL_3302172, EPI_ISL_3302173, EPI_ISL_3302174, EPI_ISL_3302175, EPI_ISL_3302176, EPI_ISL_3302177, EPI_ISL_3302178, EPI_ISL_3302179, EPI_ISL_3302180, EPI_ISL_3302181, EPI_ISL_3302182, EPI_ISL_3302183, EPI_ISL_3302184, EPI_ISL_3302185, EPI_ISL_3302186, EPI_ISL_3302187, EPI_ISL_3302188, EPI_ISL_3302189, EPI_ISL_3302190, EPI_ISL_3302191, EPI_ISL_3302192, EPI_ISL_3302193, EPI_ISL_3302194, EPI_ISL_3302195, EPI_ISL_3302196, EPI_ISL_3302197, EPI_ISL_3302198, EPI_ISL_3302199, EPI_ISL_3302200, EPI_ISL_3302201, EPI_ISL_3302202, EPI_ISL_3302203, EPI_ISL_3302204, EPI_ISL_3302205, EPI_ISL_3302206, EPI_ISL_3302207, EPI_ISL_3302208, EPI_ISL_3302209, EPI_ISL_3302210, EPI_ISL_3302211, EPI_ISL_3302212, EPI_ISL_3302213, EPI_ISL_3302214, EPI_ISL_3302215, EPI_ISL_3302216, EPI_ISL_3302217, EPI_ISL_3302218, EPI_ISL_3302219, EPI_ISL_3302220, EPI_ISL_3302221, EPI_ISL_3302222, EPI_ISL_3302223, EPI_ISL_3302224, EPI_ISL_3302225, EPI_ISL_3302226, EPI_ISL_3302227, EPI_ISL_3302228, EPI_ISL_3302229, EPI_ISL_3302230, EPI_ISL_3302231, EPI_ISL_3302232, EPI_ISL_3302233, EPI_ISL_3302234, EPI_ISL_3302235, EPI_ISL_3302236, EPI_ISL_3302237, EPI_ISL_3302238, EPI_ISL_3302239, EPI_ISL_3302240, EPI_ISL_3302241, EPI_ISL_3302242, EPI_ISL_3302243, EPI_ISL_3302244, EPI_ISL_3302246, EPI_ISL_3302247, EPI_ISL_3302248, EPI_ISL_3302249, EPI_ISL_3302250, EPI_ISL_3302251, EPI_ISL_3302252, EPI_ISL_3302253, EPI_ISL_3302254, EPI_ISL_3302255, EPI_ISL_3302256, EPI_ISL_3302257, EPI_ISL_3302258, EPI_ISL_3302259, EPI_ISL_3302260, EPI_ISL_3302261, EPI_ISL_3302262, EPI_ISL_3302263, EPI_ISL_3302264, EPI_ISL_3302265, EPI_ISL_3302266, EPI_ISL_3302267, EPI_ISL_3302268, EPI_ISL_3302269, EPI_ISL_3302270, EPI_ISL_3302271, EPI_ISL_3302272, EPI_ISL_3302273, EPI_ISL_3302274, EPI_ISL_3302275, EPI_ISL_3302276, EPI_ISL_3302277, EPI_ISL_3302278, EPI_ISL_3302279, EPI_ISL_3302280, EPI_ISL_3302281, EPI_ISL_3302282, EPI_ISL_3302283, EPI_ISL_3302284, EPI_ISL_3302285, EPI_ISL_3302286, EPI_ISL_3302287, EPI_ISL_3302288, EPI_ISL_3302289, EPI_ISL_3302290, EPI_ISL_3302291, EPI_ISL_3302292, EPI_ISL_3302293, EPI_ISL_3302294, EPI_ISL_3302295, EPI_ISL_3302296, EPI_ISL_3302297, EPI_ISL_3302298, EPI_ISL_3302299, EPI_ISL_3302300, EPI_ISL_3302301, EPI_ISL_3302302, EPI_ISL_3302303, EPI_ISL_3302304, EPI_ISL_3302305, EPI_ISL_3302306, EPI_ISL_3302307, EPI_ISL_3302308, EPI_ISL_3302309, EPI_ISL_3302310, EPI_ISL_3302311, EPI_ISL_3302313, EPI_ISL_3302314, EPI_ISL_3302315, EPI_ISL_3302316, EPI_ISL_3302317, EPI_ISL_3302318, EPI_ISL_3302319, EPI_ISL_3302320, EPI_ISL_3302321, EPI_ISL_3302322, EPI_ISL_3302323, EPI_ISL_3302324, EPI_ISL_3302325, EPI_ISL_3302326, EPI_ISL_3302327, EPI_ISL_3302328, EPI_ISL_3302329, EPI_ISL_3302331, EPI_ISL_3302332, EPI_ISL_3302333, EPI_ISL_3302334, EPI_ISL_3302336, EPI_ISL_3302337, EPI_ISL_3302338, EPI_ISL_3302339, EPI_ISL_3302340, EPI_ISL_3302341, EPI_ISL_3302342, EPI_ISL_3302343, EPI_ISL_3302344, EPI_ISL_3302345, EPI_ISL_3302346, EPI_ISL_3302347, EPI_ISL_3302348, EPI_ISL_3302349, EPI_ISL_3302350, EPI_ISL_3302351, EPI_ISL_3302352, EPI_ISL_3302353, EPI_ISL_3302354, EPI_ISL_3302355, EPI_ISL_3302356, EPI_ISL_3302357, EPI_ISL_3302358, EPI_ISL_3302359, EPI_ISL_3302360, EPI_ISL_3302361, EPI_ISL_3302362, EPI_ISL_3302363, EPI_ISL_3302364, EPI_ISL_3302365, EPI_ISL_3302366, EPI_ISL_3302368, EPI_ISL_3302369, EPI_ISL_3302370, EPI_ISL_3302371, EPI_ISL_3302372, EPI_ISL_3302373, EPI_ISL_3302374, EPI_ISL_3302375, EPI_ISL_3302376, EPI_ISL_3302377, EPI_ISL_3302378, EPI_ISL_3302379, EPI_ISL_3302380, EPI_ISL_3302381, EPI_ISL_3302382, EPI_ISL_3302383, EPI_ISL_3302384, EPI_ISL_3302385, EPI_ISL_3302386, EPI_ISL_3302387, EPI_ISL_3302388, EPI_ISL_3302389, EPI_ISL_3302390, EPI_ISL_3302391, EPI_ISL_3302392, EPI_ISL_3302393, EPI_ISL_3302394, EPI_ISL_3302395, EPI_ISL_3302396, EPI_ISL_3302397, EPI_ISL_3302398, EPI_ISL_3302399, EPI_ISL_3302400, EPI_ISL_3302401, EPI_ISL_3302403, EPI_ISL_3302404, EPI_ISL_3302405, EPI_ISL_3302406, EPI_ISL_3302407, EPI_ISL_3302409, EPI_ISL_3302410, EPI_ISL_3302411, EPI_ISL_3302412, EPI_ISL_3302413, EPI_ISL_3302414, EPI_ISL_3302415, EPI_ISL_3302416, EPI_ISL_3302417, EPI_ISL_3302418, EPI_ISL_3302419, EPI_ISL_3302420, EPI_ISL_3302421, EPI_ISL_3302422, EPI_ISL_3302423, EPI_ISL_3302424, EPI_ISL_3302425, EPI_ISL_3302426, EPI_ISL_3302427, EPI_ISL_3302429, EPI_ISL_3302430, EPI_ISL_3302431, EPI_ISL_3302432, EPI_ISL_3302433, EPI_ISL_3302434, EPI_ISL_3302435, EPI_ISL_3302436, EPI_ISL_3302437, EPI_ISL_3302438, EPI_ISL_3302439, EPI_ISL_3302440, EPI_ISL_3302441, EPI_ISL_3302442, EPI_ISL_3302443, EPI_ISL_3302444, EPI_ISL_3302445, EPI_ISL_3302446, EPI_ISL_3302447, EPI_ISL_3302448, EPI_ISL_3302449, EPI_ISL_3302450, EPI_ISL_3302451, EPI_ISL_3302452, EPI_ISL_3302453, EPI_ISL_3302454, EPI_ISL_3302455, EPI_ISL_3302456, EPI_ISL_3302457, EPI_ISL_3302458, EPI_ISL_3302459, EPI_ISL_3302460, EPI_ISL_3302461, EPI_ISL_3302462, EPI_ISL_3302463, EPI_ISL_3302464, EPI_ISL_3302465, EPI_ISL_3302466, EPI_ISL_3302467, EPI_ISL_3302468, EPI_ISL_3302469, EPI_ISL_3302470, EPI_ISL_3302471, EPI_ISL_3302472, EPI_ISL_3302473, EPI_ISL_3302474, EPI_ISL_3302475, EPI_ISL_3302476, EPI_ISL_3302477, EPI_ISL_3302478, EPI_ISL_3302479, EPI_ISL_3302480, EPI_ISL_3302481, EPI_ISL_3302482, EPI_ISL_3302483, EPI_ISL_3302484, EPI_ISL_3302486, EPI_ISL_3302487, EPI_ISL_3302488, EPI_ISL_3302490, EPI_ISL_3302491, EPI_ISL_3302492, EPI_ISL_3302493, EPI_ISL_3302494, EPI_ISL_3302495, EPI_ISL_3302497, EPI_ISL_3302498, EPI_ISL_3302499, EPI_ISL_3302500, EPI_ISL_3302501, EPI_ISL_3302502, EPI_ISL_3302503, EPI_ISL_3302504, EPI_ISL_3302505, EPI_ISL_3302506, EPI_ISL_3302507, EPI_ISL_3302508, EPI_ISL_3302509, EPI_ISL_3302510, EPI_ISL_3302511, EPI_ISL_3302512, EPI_ISL_3302513, EPI_ISL_3302514, EPI_ISL_3302515, EPI_ISL_3302516, EPI_ISL_3302517, EPI_ISL_3302518, EPI_ISL_3302519, EPI_ISL_3302520, EPI_ISL_3302521, EPI_ISL_3302522, EPI_ISL_3302523, EPI_ISL_3302524, EPI_ISL_3302525, EPI_ISL_3302526, EPI_ISL_3302527, EPI_ISL_3302528, EPI_ISL_3302529, EPI_ISL_3302530, EPI_ISL_3302531, EPI_ISL_3302532, EPI_ISL_3302533, EPI_ISL_3302534, EPI_ISL_3302535, EPI_ISL_3302536, EPI_ISL_3302537, EPI_ISL_3302538, EPI_ISL_3302539, EPI_ISL_3302540, EPI_ISL_3302541, EPI_ISL_3302542, EPI_ISL_3302543, EPI_ISL_3302544, EPI_ISL_3302545, EPI_ISL_3302546, EPI_ISL_3302547, EPI_ISL_3302548, EPI_ISL_3302549, EPI_ISL_3302550, EPI_ISL_3302551, EPI_ISL_3302552, EPI_ISL_3302553, EPI_ISL_3302554, EPI_ISL_3302555, EPI_ISL_3302556, EPI_ISL_3302557, EPI_ISL_3302558, EPI_ISL_3302559, EPI_ISL_3302560, EPI_ISL_3302561, EPI_ISL_3302562, EPI_ISL_3302563, EPI_ISL_3302564, EPI_ISL_3302565, EPI_ISL_3302566, EPI_ISL_3302567, EPI_ISL_3302568, EPI_ISL_3302569, EPI_ISL_3302570, EPI_ISL_3302571, EPI_ISL_3302572, EPI_ISL_3302574, EPI_ISL_3302575, EPI_ISL_3302576, EPI_ISL_3302577, EPI_ISL_3302578, EPI_ISL_3302579, EPI_ISL_3302580, EPI_ISL_3302581, EPI_ISL_3302582, EPI_ISL_3302583, EPI_ISL_3302584, EPI_ISL_3302585, EPI_ISL_3302586, EPI_ISL_3302587, EPI_ISL_3302588, EPI_ISL_3302589, EPI_ISL_3302590, EPI_ISL_3302591, EPI_ISL_3302592, EPI_ISL_3302593, EPI_ISL_3302594, EPI_ISL_3302595, EPI_ISL_3302596, EPI_ISL_3302597, EPI_ISL_3302598, EPI_ISL_3302599, EPI_ISL_3302600, EPI_ISL_3302601, EPI_ISL_3302602, EPI_ISL_3302603, EPI_ISL_3302604, EPI_ISL_3302605, EPI_ISL_3302606, EPI_ISL_3302607, EPI_ISL_3302608, EPI_ISL_3302609, EPI_ISL_3302610, EPI_ISL_3302611, EPI_ISL_3302612, EPI_ISL_3302613, EPI_ISL_3302614, EPI_ISL_3302615, EPI_ISL_3302616, EPI_ISL_3302617, EPI_ISL_3302618, EPI_ISL_3302619, EPI_ISL_3302620, EPI_ISL_3302621, EPI_ISL_3302622, EPI_ISL_3302623, EPI_ISL_3302624, EPI_ISL_3302625, EPI_ISL_3302626, EPI_ISL_3302628, EPI_ISL_3302629, EPI_ISL_3302630, EPI_ISL_3302631, EPI_ISL_3302632, EPI_ISL_3302633, EPI_ISL_3302634, EPI_ISL_3302635, EPI_ISL_3302636, EPI_ISL_3302637, EPI_ISL_3302638, EPI_ISL_3302639, EPI_ISL_3302640, EPI_ISL_3302641, EPI_ISL_3302642, EPI_ISL_3302643, EPI_ISL_3302644, EPI_ISL_3302645, EPI_ISL_3302646, EPI_ISL_3302647, EPI_ISL_3302648, EPI_ISL_3302649, EPI_ISL_3302650, EPI_ISL_3302651, EPI_ISL_3302652, EPI_ISL_3302653, EPI_ISL_3302654, EPI_ISL_3302655, EPI_ISL_3302656, EPI_ISL_3302657, EPI_ISL_3302658, EPI_ISL_3302659, EPI_ISL_3302660, EPI_ISL_3302661, EPI_ISL_3302662, EPI_ISL_3302663, EPI_ISL_3302664, EPI_ISL_3302665, EPI_ISL_3302666, EPI_ISL_3302667, EPI_ISL_3302668, EPI_ISL_3302669, EPI_ISL_3302670, EPI_ISL_3302671, EPI_ISL_3302674, EPI_ISL_3302675, EPI_ISL_3302676, EPI_ISL_3302677, EPI_ISL_3302678, EPI_ISL_3302679, EPI_ISL_3302680, EPI_ISL_3302681, EPI_ISL_3302682, EPI_ISL_3302683, EPI_ISL_3302684, EPI_ISL_3302685, EPI_ISL_3302686, EPI_ISL_3302687, EPI_ISL_3302688, EPI_ISL_3302689, EPI_ISL_3302690, EPI_ISL_3302691, EPI_ISL_3302692, EPI_ISL_3302693, EPI_ISL_3302694, EPI_ISL_3302695, EPI_ISL_3302696, EPI_ISL_3302697, EPI_ISL_3302698, EPI_ISL_3302699, EPI_ISL_3302701, EPI_ISL_3302702, EPI_ISL_3302703, EPI_ISL_3302704, EPI_ISL_3302705, EPI_ISL_3302706, EPI_ISL_3302707, EPI_ISL_3302708, EPI_ISL_3302709, EPI_ISL_3302710, EPI_ISL_3302711, EPI_ISL_3302712, EPI_ISL_3302713, EPI_ISL_3302714, EPI_ISL_3302715, EPI_ISL_3302716, EPI_ISL_3302717, EPI_ISL_3302718, EPI_ISL_3302719, EPI_ISL_3302720, EPI_ISL_3302721, EPI_ISL_3302722, EPI_ISL_3302723, EPI_ISL_3302724, EPI_ISL_3302725, EPI_ISL_3302726, EPI_ISL_3302727, EPI_ISL_3302728, EPI_ISL_3302729, EPI_ISL_3302730, EPI_ISL_3302731, EPI_ISL_3302732, EPI_ISL_3302733, EPI_ISL_3302734, EPI_ISL_3302735, EPI_ISL_3302736, EPI_ISL_3302737, EPI_ISL_3302739, EPI_ISL_3302740, EPI_ISL_3302741, EPI_ISL_3302742, EPI_ISL_3302743, EPI_ISL_3302745, EPI_ISL_3302746, EPI_ISL_3302747, EPI_ISL_3302748, EPI_ISL_3302749, EPI_ISL_3302750, EPI_ISL_3302751, EPI_ISL_3302752, EPI_ISL_3302753, EPI_ISL_3302754, EPI_ISL_3302755, EPI_ISL_3302756, EPI_ISL_3302757, EPI_ISL_3302758, EPI_ISL_3302759, EPI_ISL_3302760, EPI_ISL_3302761, EPI_ISL_3302762, EPI_ISL_3302763, EPI_ISL_3302764, EPI_ISL_3302765, EPI_ISL_3302766, EPI_ISL_3302767, EPI_ISL_3302768, EPI_ISL_3302769, EPI_ISL_3302770, EPI_ISL_3302771, EPI_ISL_3302772, EPI_ISL_3302773, EPI_ISL_3302774, EPI_ISL_3302775, EPI_ISL_3302776, EPI_ISL_3302777, EPI_ISL_3302778, EPI_ISL_3302779, EPI_ISL_3302780, EPI_ISL_3302781, EPI_ISL_3302782, EPI_ISL_3302783, EPI_ISL_3302784, EPI_ISL_3302785, EPI_ISL_3302786, EPI_ISL_3302787, EPI_ISL_3302788, EPI_ISL_3302789, EPI_ISL_3302791, EPI_ISL_3302792, EPI_ISL_3302794, EPI_ISL_3302795, EPI_ISL_3302796, EPI_ISL_3302797, EPI_ISL_3302798, EPI_ISL_3302799, EPI_ISL_3302800, EPI_ISL_3302801, EPI_ISL_3302802, EPI_ISL_3302803, EPI_ISL_3302804, EPI_ISL_3302805, EPI_ISL_3302806, EPI_ISL_3302807, EPI_ISL_3302808, EPI_ISL_3302809, EPI_ISL_3302810, EPI_ISL_3302811, EPI_ISL_3302812, EPI_ISL_3302813, EPI_ISL_3302814, EPI_ISL_3302815, EPI_ISL_3302816, EPI_ISL_3302817, EPI_ISL_3302818, EPI_ISL_3302819, EPI_ISL_3302820, EPI_ISL_3302821, EPI_ISL_3302822, EPI_ISL_3302823, EPI_ISL_3302824, EPI_ISL_3302825, EPI_ISL_3302826, EPI_ISL_3302827, EPI_ISL_3302828, EPI_ISL_3302829, EPI_ISL_3302830, EPI_ISL_3302831, EPI_ISL_3302832, EPI_ISL_3302833, EPI_ISL_3302834, EPI_ISL_3302835, EPI_ISL_3302836, EPI_ISL_3302837, EPI_ISL_3302838, EPI_ISL_3302839, EPI_ISL_3302840, EPI_ISL_3302841, EPI_ISL_3302842, EPI_ISL_3302843, EPI_ISL_3302844, EPI_ISL_3302845, EPI_ISL_3302846, EPI_ISL_3302847, EPI_ISL_3302848, EPI_ISL_3302849, EPI_ISL_3302850, EPI_ISL_3302851, EPI_ISL_3302852, EPI_ISL_3302853, EPI_ISL_3302854, EPI_ISL_3302855, EPI_ISL_3302856, EPI_ISL_3302857, EPI_ISL_3302858, EPI_ISL_3302859, EPI_ISL_3302860, EPI_ISL_3302861, EPI_ISL_3302862, EPI_ISL_3302863, EPI_ISL_3302864, EPI_ISL_3302865, EPI_ISL_3302866, EPI_ISL_3302867, EPI_ISL_3302868, EPI_ISL_3302869, EPI_ISL_3302870, EPI_ISL_3302871, EPI_ISL_3302872, EPI_ISL_3302873, EPI_ISL_3302874, EPI_ISL_3302875, EPI_ISL_3302876, EPI_ISL_3302877, EPI_ISL_3302878, EPI_ISL_3302879, EPI_ISL_3302880, EPI_ISL_3302881, EPI_ISL_3302882, EPI_ISL_3302883, EPI_ISL_3302884, EPI_ISL_3302886, EPI_ISL_3302887, EPI_ISL_3302888, EPI_ISL_3302889, EPI_ISL_3302890, EPI_ISL_3302891, EPI_ISL_3302892, EPI_ISL_3302893, EPI_ISL_3302894, EPI_ISL_3302895, EPI_ISL_3302896, EPI_ISL_3302897, EPI_ISL_3302898, EPI_ISL_3302899, EPI_ISL_3302900, EPI_ISL_3302902, EPI_ISL_3302903, EPI_ISL_3302904, EPI_ISL_3302905, EPI_ISL_3302906, EPI_ISL_3302907, EPI_ISL_3302908, EPI_ISL_3302909, EPI_ISL_3302910, EPI_ISL_3302911, EPI_ISL_3302912, EPI_ISL_3302913, EPI_ISL_3302914, EPI_ISL_3302915, EPI_ISL_3302916, EPI_ISL_3302917, EPI_ISL_3302918, EPI_ISL_3302919, EPI_ISL_3302920, EPI_ISL_3302921, EPI_ISL_3302922, EPI_ISL_3302923, EPI_ISL_3302924, EPI_ISL_3302925, EPI_ISL_3302926, EPI_ISL_3302928, EPI_ISL_3302929, EPI_ISL_3302931, EPI_ISL_3302932, EPI_ISL_3302933, EPI_ISL_3302934, EPI_ISL_3302936, EPI_ISL_3302937, EPI_ISL_3302938, EPI_ISL_3302939, EPI_ISL_3302940, EPI_ISL_3302941, EPI_ISL_3302942, EPI_ISL_3302943, EPI_ISL_3302944, EPI_ISL_3302945, EPI_ISL_3302946, EPI_ISL_3302947, EPI_ISL_3302948, EPI_ISL_3302949, EPI_ISL_3302950, EPI_ISL_3302951, EPI_ISL_3302952, EPI_ISL_3302953, EPI_ISL_3302954, EPI_ISL_3302955, EPI_ISL_3302956, EPI_ISL_3302957, EPI_ISL_3302958, EPI_ISL_3302959, EPI_ISL_3302960, EPI_ISL_3302961, EPI_ISL_3302962, EPI_ISL_3302963, EPI_ISL_3302964, EPI_ISL_3302965, EPI_ISL_3302966, EPI_ISL_3302967, EPI_ISL_3302968, EPI_ISL_3302969, EPI_ISL_3302970, EPI_ISL_3302971, EPI_ISL_3302972, EPI_ISL_3302973, EPI_ISL_3302974, EPI_ISL_3302975, EPI_ISL_3302976, EPI_ISL_3302977, EPI_ISL_3302978, EPI_ISL_3302979, EPI_ISL_3302980, EPI_ISL_3302981, EPI_ISL_3302982, EPI_ISL_3302983, EPI_ISL_3302984, EPI_ISL_3302985, EPI_ISL_3302986, EPI_ISL_3302987, EPI_ISL_3302988, EPI_ISL_3302989, EPI_ISL_3302990, EPI_ISL_3302991, EPI_ISL_3302992, EPI_ISL_3302993, EPI_ISL_3302994, EPI_ISL_3302995, EPI_ISL_3302996, EPI_ISL_3302997, EPI_ISL_3302998, EPI_ISL_3302999, EPI_ISL_3303000, EPI_ISL_3303001, EPI_ISL_3303002, EPI_ISL_3303003, EPI_ISL_3303004, EPI_ISL_3303005, EPI_ISL_3303006, EPI_ISL_3303007, EPI_ISL_3303008, EPI_ISL_3303009, EPI_ISL_3303010, EPI_ISL_3303011, EPI_ISL_3303012, EPI_ISL_3303013, EPI_ISL_3303014, EPI_ISL_3303015, EPI_ISL_3303017, EPI_ISL_3303018, EPI_ISL_3303019, EPI_ISL_3303020, EPI_ISL_3303021, EPI_ISL_3303022, EPI_ISL_3303023, EPI_ISL_3303024, EPI_ISL_3303025, EPI_ISL_3303026, EPI_ISL_3303027, EPI_ISL_3303028, EPI_ISL_3303029, EPI_ISL_3303030, EPI_ISL_3303031, EPI_ISL_3303033, EPI_ISL_3303034, EPI_ISL_3303036, EPI_ISL_3303037, EPI_ISL_3303038, EPI_ISL_3303039, EPI_ISL_3303040, EPI_ISL_3303041, EPI_ISL_3303042, EPI_ISL_3303043, EPI_ISL_3303044, EPI_ISL_3303045, EPI_ISL_3303046, EPI_ISL_3303047, EPI_ISL_3303048, EPI_ISL_3303049, EPI_ISL_3303050, EPI_ISL_3303051, EPI_ISL_3303052, EPI_ISL_3303053, EPI_ISL_3303054, EPI_ISL_3303055, EPI_ISL_3303056, EPI_ISL_3303057, EPI_ISL_3303058, EPI_ISL_3303059, EPI_ISL_3303060, EPI_ISL_3303061, EPI_ISL_3303062, EPI_ISL_3303063, EPI_ISL_3303064, EPI_ISL_3303065, EPI_ISL_3303066, EPI_ISL_3303067, EPI_ISL_3303068, EPI_ISL_3303069, EPI_ISL_3303070, EPI_ISL_3303071, EPI_ISL_3303072, EPI_ISL_3303073, EPI_ISL_3303074, EPI_ISL_3303075, EPI_ISL_3303076, EPI_ISL_3303077, EPI_ISL_3303078, EPI_ISL_3303079, EPI_ISL_3303080, EPI_ISL_3303081, EPI_ISL_3303082, EPI_ISL_3303083, EPI_ISL_3303084, EPI_ISL_3303085, EPI_ISL_3303086, EPI_ISL_3303087, EPI_ISL_3303089, EPI_ISL_3303090, EPI_ISL_3303091, EPI_ISL_3303092, EPI_ISL_3303093, EPI_ISL_3303094, EPI_ISL_3303095, EPI_ISL_3303096, EPI_ISL_3303097, EPI_ISL_3303098, EPI_ISL_3303099, EPI_ISL_3303100, EPI_ISL_3303101, EPI_ISL_3303102, EPI_ISL_3303103, EPI_ISL_3303104, EPI_ISL_3303105, EPI_ISL_3303106, EPI_ISL_3303107, EPI_ISL_3303108, EPI_ISL_3303109, EPI_ISL_3303110, EPI_ISL_3303111, EPI_ISL_3303112, EPI_ISL_3303113, EPI_ISL_3303114, EPI_ISL_3303115, EPI_ISL_3303116, EPI_ISL_3303117, EPI_ISL_3303118, EPI_ISL_3303119, EPI_ISL_3303120, EPI_ISL_3303121, EPI_ISL_3303122, EPI_ISL_3303123, EPI_ISL_3303124, EPI_ISL_3303125, EPI_ISL_3303126, EPI_ISL_3303127, EPI_ISL_3303128, EPI_ISL_3303130, EPI_ISL_3303132, EPI_ISL_3303133, EPI_ISL_3303134, EPI_ISL_3303135, EPI_ISL_3303136, EPI_ISL_3303137, EPI_ISL_3303138, EPI_ISL_3303139, EPI_ISL_3303140, EPI_ISL_3303141, EPI_ISL_3303142, EPI_ISL_3303143, EPI_ISL_3303144, EPI_ISL_3303145, EPI_ISL_3303146, EPI_ISL_3303147, EPI_ISL_3303148, EPI_ISL_3303149, EPI_ISL_3303150, EPI_ISL_3303151, EPI_ISL_3303152, EPI_ISL_3303153, EPI_ISL_3303154, EPI_ISL_3303155, EPI_ISL_3303156, EPI_ISL_3303158, EPI_ISL_3303159, EPI_ISL_3303160, EPI_ISL_3303161, EPI_ISL_3303162, EPI_ISL_3303163, EPI_ISL_3303164, EPI_ISL_3303165, EPI_ISL_3303166, EPI_ISL_3303167, EPI_ISL_3303168, EPI_ISL_3303169, EPI_ISL_3303170, EPI_ISL_3303171, EPI_ISL_3303172, EPI_ISL_3303173, EPI_ISL_3303174, EPI_ISL_3303175, EPI_ISL_3303176, EPI_ISL_3303177, EPI_ISL_3303178, EPI_ISL_3303179, EPI_ISL_3303180, EPI_ISL_3303181, EPI_ISL_3303182, EPI_ISL_3303183, EPI_ISL_3303184, EPI_ISL_3303185, EPI_ISL_3303186, EPI_ISL_3303187, EPI_ISL_3303188, EPI_ISL_3303189, EPI_ISL_3303190, EPI_ISL_3303191, EPI_ISL_3303192, EPI_ISL_3303193, EPI_ISL_3303194, EPI_ISL_3303195, EPI_ISL_3303196, EPI_ISL_3303197, EPI_ISL_3303198, EPI_ISL_3303199, EPI_ISL_3303200, EPI_ISL_3303201, EPI_ISL_3303202, EPI_ISL_3303203, EPI_ISL_3303204, EPI_ISL_3303205, EPI_ISL_3303206, EPI_ISL_3303207, EPI_ISL_3303208, EPI_ISL_3303209, EPI_ISL_3303210, EPI_ISL_3303211, EPI_ISL_3303212, EPI_ISL_3303213, EPI_ISL_3303214, EPI_ISL_3303215, EPI_ISL_3303216, EPI_ISL_3303217, EPI_ISL_3303218, EPI_ISL_3303219, EPI_ISL_3303220, EPI_ISL_3303221, EPI_ISL_3303222, EPI_ISL_3303223, EPI_ISL_3303224, EPI_ISL_3303225, EPI_ISL_3303226, EPI_ISL_3303227, EPI_ISL_3303228, EPI_ISL_3303229, EPI_ISL_3303230, EPI_ISL_3303231, EPI_ISL_3303232, EPI_ISL_3303233, EPI_ISL_3303234, EPI_ISL_3303235, EPI_ISL_3303236, EPI_ISL_3303237, EPI_ISL_3303238, EPI_ISL_3303239, EPI_ISL_3303240, EPI_ISL_3303241, EPI_ISL_3303242, EPI_ISL_3303243, EPI_ISL_3303244, EPI_ISL_3303245, EPI_ISL_3303246, EPI_ISL_3303247, EPI_ISL_3303248, EPI_ISL_3303249, EPI_ISL_3303250, EPI_ISL_3303251, EPI_ISL_3303252, EPI_ISL_3303253, EPI_ISL_3303254, EPI_ISL_3303255, EPI_ISL_3303256, EPI_ISL_3303257, EPI_ISL_3303258, EPI_ISL_3303259, EPI_ISL_3303260, EPI_ISL_3303261, EPI_ISL_3303262, EPI_ISL_3303263, EPI_ISL_3303264, EPI_ISL_3303265, EPI_ISL_3303266, EPI_ISL_3303267, EPI_ISL_3303268, EPI_ISL_3303269, EPI_ISL_3303270, EPI_ISL_3303271, EPI_ISL_3303272, EPI_ISL_3303273, EPI_ISL_3303274, EPI_ISL_3303275, EPI_ISL_3303276, EPI_ISL_3303277, EPI_ISL_3303278, EPI_ISL_3303279, EPI_ISL_3303280, EPI_ISL_3303281, EPI_ISL_3303282, EPI_ISL_3303283, EPI_ISL_3303284, EPI_ISL_3303287, EPI_ISL_3303288, EPI_ISL_3303289, EPI_ISL_3303290, EPI_ISL_3303291, EPI_ISL_3303292, EPI_ISL_3303293, EPI_ISL_3303294, EPI_ISL_3303295, EPI_ISL_3303296, EPI_ISL_3303297, EPI_ISL_3303298, EPI_ISL_3303299, EPI_ISL_3303300, EPI_ISL_3303301, EPI_ISL_3303302, EPI_ISL_3303303, EPI_ISL_3303304, EPI_ISL_3303305, EPI_ISL_3303306, EPI_ISL_3303307, EPI_ISL_3303308, EPI_ISL_3303309, EPI_ISL_3303310, EPI_ISL_3303311, EPI_ISL_3303312, EPI_ISL_3303313, EPI_ISL_3303314, EPI_ISL_3303315, EPI_ISL_3303316, EPI_ISL_3303317, EPI_ISL_3303318, EPI_ISL_3303319, EPI_ISL_3303320, EPI_ISL_3303321, EPI_ISL_3303322, EPI_ISL_3303323, EPI_ISL_3303324, EPI_ISL_3303325, EPI_ISL_3303326, EPI_ISL_3303327, EPI_ISL_3303328, EPI_ISL_3303329, EPI_ISL_3303330, EPI_ISL_3303331, EPI_ISL_3303332, EPI_ISL_3303333, EPI_ISL_3303334, EPI_ISL_3303335, EPI_ISL_3303336, EPI_ISL_3303337, EPI_ISL_3303338, EPI_ISL_3303339, EPI_ISL_3303340, EPI_ISL_3303341, EPI_ISL_3303342, EPI_ISL_3303343, EPI_ISL_3303344, EPI_ISL_3303345, EPI_ISL_3303346, EPI_ISL_3303347, EPI_ISL_3303348, EPI_ISL_3303349, EPI_ISL_3303350, EPI_ISL_3303351, EPI_ISL_3303352, EPI_ISL_3303353, EPI_ISL_3303354, EPI_ISL_3303355, EPI_ISL_3303356, EPI_ISL_3303357, EPI_ISL_3303358, EPI_ISL_3303359, EPI_ISL_3303360, EPI_ISL_3303361, EPI_ISL_3303362, EPI_ISL_3303363, EPI_ISL_3303364, EPI_ISL_3303365, EPI_ISL_3303366, EPI_ISL_3303367, EPI_ISL_3303368, EPI_ISL_3303369, EPI_ISL_3303370, EPI_ISL_3303371, EPI_ISL_3303372, EPI_ISL_3303373, EPI_ISL_3303374, EPI_ISL_3303375, EPI_ISL_3303376, EPI_ISL_3303377, EPI_ISL_3303378, EPI_ISL_3303379, EPI_ISL_3303381, EPI_ISL_3303382, EPI_ISL_3303383, EPI_ISL_3303384, EPI_ISL_3303385, EPI_ISL_3303386, EPI_ISL_3303387, EPI_ISL_3303388, EPI_ISL_3303389, EPI_ISL_3303390, EPI_ISL_3303391, EPI_ISL_3303392, EPI_ISL_3303393, EPI_ISL_3303394, EPI_ISL_3303395, EPI_ISL_3303396, EPI_ISL_3303397, EPI_ISL_3303398, EPI_ISL_3303399, EPI_ISL_3303400, EPI_ISL_3303401, EPI_ISL_3303402, EPI_ISL_3303403, EPI_ISL_3303404, EPI_ISL_3303405, EPI_ISL_3303406, EPI_ISL_3303407, EPI_ISL_3303408, EPI_ISL_3303409, EPI_ISL_3303410, EPI_ISL_3303411, EPI_ISL_3303412, EPI_ISL_3303413, EPI_ISL_3303414, EPI_ISL_3303415, EPI_ISL_3303416, EPI_ISL_3303417, EPI_ISL_3303418, EPI_ISL_3303419, EPI_ISL_3303420, EPI_ISL_3303421, EPI_ISL_3303422, EPI_ISL_3303423, EPI_ISL_3303425, EPI_ISL_3303426, EPI_ISL_3303427, EPI_ISL_3303428, EPI_ISL_3303429, EPI_ISL_3303430, EPI_ISL_3303431, EPI_ISL_3303432, EPI_ISL_3303433, EPI_ISL_3303434, EPI_ISL_3303435, EPI_ISL_3303436, EPI_ISL_3303437, EPI_ISL_3303438, EPI_ISL_3303439, EPI_ISL_3303440, EPI_ISL_3303441, EPI_ISL_3303442, EPI_ISL_3303443, EPI_ISL_3303444, EPI_ISL_3303445, EPI_ISL_3303446, EPI_ISL_3303447, EPI_ISL_3303448, EPI_ISL_3303449, EPI_ISL_3303450, EPI_ISL_3303451, EPI_ISL_3303452, EPI_ISL_3303453, EPI_ISL_3303454, EPI_ISL_3303455, EPI_ISL_3303456, EPI_ISL_3303457, EPI_ISL_3303458, EPI_ISL_3303459, EPI_ISL_3303460, EPI_ISL_3303461, EPI_ISL_3303462, EPI_ISL_3303463, EPI_ISL_3303464, EPI_ISL_3303465, EPI_ISL_3303466, EPI_ISL_3303467, EPI_ISL_3303468, EPI_ISL_3303469, EPI_ISL_3303470, EPI_ISL_3303471, EPI_ISL_3303472, EPI_ISL_3303473, EPI_ISL_3303474, EPI_ISL_3303475, EPI_ISL_3303476, EPI_ISL_3303477, EPI_ISL_3303478, EPI_ISL_3303479, EPI_ISL_3303480, EPI_ISL_3303481, EPI_ISL_3303482, EPI_ISL_3303483, EPI_ISL_3303484, EPI_ISL_3303485, EPI_ISL_3303486, EPI_ISL_3303487, EPI_ISL_3303488, EPI_ISL_3303489, EPI_ISL_3303490, EPI_ISL_3303491, EPI_ISL_3303492, EPI_ISL_3303493, EPI_ISL_3303494, EPI_ISL_3303495, EPI_ISL_3303496, EPI_ISL_3303497, EPI_ISL_3303498, EPI_ISL_3303499, EPI_ISL_3303500, EPI_ISL_3303501, EPI_ISL_3303502, EPI_ISL_3303503, EPI_ISL_3303505, EPI_ISL_3303506, EPI_ISL_3303507, EPI_ISL_3303508, EPI_ISL_3303509, EPI_ISL_3303510, EPI_ISL_3303511, EPI_ISL_3303512, EPI_ISL_3303513, EPI_ISL_3303514, EPI_ISL_3303515, EPI_ISL_3303517, EPI_ISL_3303518, EPI_ISL_3303519, EPI_ISL_3303520, EPI_ISL_3303521, EPI_ISL_3303523, EPI_ISL_3303524, EPI_ISL_3303525, EPI_ISL_3303526, EPI_ISL_3303527, EPI_ISL_3303528, EPI_ISL_3303529, EPI_ISL_3303530, EPI_ISL_3303531, EPI_ISL_3303532, EPI_ISL_3303533, EPI_ISL_3303534, EPI_ISL_3303536, EPI_ISL_3303538, EPI_ISL_3303539, EPI_ISL_3303540, EPI_ISL_3303541, EPI_ISL_3303542, EPI_ISL_3303543, EPI_ISL_3303544, EPI_ISL_3303545, EPI_ISL_3303546, EPI_ISL_3303547, EPI_ISL_3303548, EPI_ISL_3303549, EPI_ISL_3303550, EPI_ISL_3303551, EPI_ISL_3303552, EPI_ISL_3303553, EPI_ISL_3303555, EPI_ISL_3303556, EPI_ISL_3303557, EPI_ISL_3303558, EPI_ISL_3303559, EPI_ISL_3303560, EPI_ISL_3303561, EPI_ISL_3303562, EPI_ISL_3303563, EPI_ISL_3303564, EPI_ISL_3303565, EPI_ISL_3303566, EPI_ISL_3303567, EPI_ISL_3303568, EPI_ISL_3303569, EPI_ISL_3303570, EPI_ISL_3303571, EPI_ISL_3303572, EPI_ISL_3303573, EPI_ISL_3303574, EPI_ISL_3303575, EPI_ISL_3303576, EPI_ISL_3303577, EPI_ISL_3303578, EPI_ISL_3303579, EPI_ISL_3303580, EPI_ISL_3303581, EPI_ISL_3303582, EPI_ISL_3303583, EPI_ISL_3303584, EPI_ISL_3303585, EPI_ISL_3303586, EPI_ISL_3303587, EPI_ISL_3303589, EPI_ISL_3303590, EPI_ISL_3303592, EPI_ISL_3303593, EPI_ISL_3303594, EPI_ISL_3303595, EPI_ISL_3303596, EPI_ISL_3303597, EPI_ISL_3303598, EPI_ISL_3303599, EPI_ISL_3303600, EPI_ISL_3303601, EPI_ISL_3303602, EPI_ISL_3303603, EPI_ISL_3303604, EPI_ISL_3303606, EPI_ISL_3303607, EPI_ISL_3303608, EPI_ISL_3303609, EPI_ISL_3303611, EPI_ISL_3303612, EPI_ISL_3303613, EPI_ISL_3303614, EPI_ISL_3303615, EPI_ISL_3303616, EPI_ISL_3303617, EPI_ISL_3303618, EPI_ISL_3303619, EPI_ISL_3303620, EPI_ISL_3303621, EPI_ISL_3303622, EPI_ISL_3303623, EPI_ISL_3303624, EPI_ISL_3303625, EPI_ISL_3303626, EPI_ISL_3303627, EPI_ISL_3303628, EPI_ISL_3303629, EPI_ISL_3303630, EPI_ISL_3303632, EPI_ISL_3303633, EPI_ISL_3303634, EPI_ISL_3303635, EPI_ISL_3303636, EPI_ISL_3303637, EPI_ISL_3303638, EPI_ISL_3303639, EPI_ISL_3303640, EPI_ISL_3303641, EPI_ISL_3303642, EPI_ISL_3303643, EPI_ISL_3303644, EPI_ISL_3303645, EPI_ISL_3303646, EPI_ISL_3303647, EPI_ISL_3303649, EPI_ISL_3303650, EPI_ISL_3303651, EPI_ISL_3303652, EPI_ISL_3303653, EPI_ISL_3303654, EPI_ISL_3303655, EPI_ISL_3303656, EPI_ISL_3303657, EPI_ISL_3303658, EPI_ISL_3303659, EPI_ISL_3303660, EPI_ISL_3303661, EPI_ISL_3303662, EPI_ISL_3303663, EPI_ISL_3303664, EPI_ISL_3303665, EPI_ISL_3303666, EPI_ISL_3303667, EPI_ISL_3303668, EPI_ISL_3303669, EPI_ISL_3303670, EPI_ISL_3303671, EPI_ISL_3303672, EPI_ISL_3303673, EPI_ISL_3303674, EPI_ISL_3303675, EPI_ISL_3303676, EPI_ISL_3303677, EPI_ISL_3303678, EPI_ISL_3303679, EPI_ISL_3303680, EPI_ISL_3303681, EPI_ISL_3303682, EPI_ISL_3303683, EPI_ISL_3303685, EPI_ISL_3303686, EPI_ISL_3303687, EPI_ISL_3303688, EPI_ISL_3303689, EPI_ISL_3303690, EPI_ISL_3303691, EPI_ISL_3303692, EPI_ISL_3303693, EPI_ISL_3303694, EPI_ISL_3303695, EPI_ISL_3303696, EPI_ISL_3303697, EPI_ISL_3303698, EPI_ISL_3303699, EPI_ISL_3303701, EPI_ISL_3303702, EPI_ISL_3303703, EPI_ISL_3303704, EPI_ISL_3303705, EPI_ISL_3303706, EPI_ISL_3303707, EPI_ISL_3303708, EPI_ISL_3303709, EPI_ISL_3303710, EPI_ISL_3303711, EPI_ISL_3303712, EPI_ISL_3303713, EPI_ISL_3303714, EPI_ISL_3303715, EPI_ISL_3303716, EPI_ISL_3303717, EPI_ISL_3303718, EPI_ISL_3303719, EPI_ISL_3303720, EPI_ISL_3303721, EPI_ISL_3303722, EPI_ISL_3303723, EPI_ISL_3303724, EPI_ISL_3303725 | | | |
| see above | Fulgent Genetics | Centers for Disease Control and Prevention Division of Viral Diseases, Pathogen Discovery | Dakota Howard, Dhwani Batra, Peter W. Cook, Kara Moser, Adrian Paskey, Jason Caravas, Benjamin Rambo-Martin, Shatavia Morrison, Christopher Gulvick, Scott Sammons, Yvette Unoarumhi, Darlene Wagner, Matthew Schmerer, Harry Gao, Mickey Li, John Gao, Joseph Fierro, Benafsh Sapra, Becky Tsai, Yan Meng, Doreen Ng, James Xie, Clinton R. Paden, Duncan MacCannell |
| EPI_ISL_3303728 | Ohio Department of Health Laboratory | Ohio Department of Health Laboratory | Holmes, Jennifer; Eric Brandt, Keoni Omura, Glen McGillivary, Caitlin McDonnell, Jade Mowery, Stephanie Mccracken, Tyler Payne, Kirtana Ramadugu, Erica Leasure, Brent Lee, Kelsey Florek, Heather Blankenship, Allison Black, Quanta Brown, and Tammy Bannerman |
| EPI_ISL_3303731, EPI_ISL_3303732, EPI_ISL_3303733, EPI_ISL_3303734 | Clinical Virology | Clinical Bacteriology | Tim Roloff, Fanny Wegner, Helena MB Seth-Smith, Alfredo Mari, Karoline Leuzinger, Julia Bielicki, Manuel Battegay, Hans Hirsch, Adrian Egli |
| EPI_ISL_3303735 | Kantonsarztamt Solothurn | Clinical Bacteriology | Tim Roloff, Fanny Wegner, Helena MB Seth-Smith, Alfredo Mari, Karoline Leuzinger, Julia Bielicki, Manuel Battegay, Lukas, Fenner, Hans Hirsch, Adrian Egli |
| EPI_ISL_3303736, EPI_ISL_3303737, EPI_ISL_3303738, EPI_ISL_3303739, EPI_ISL_3303740, EPI_ISL_3303741, EPI_ISL_3303742, EPI_ISL_3303743, EPI_ISL_3303744, EPI_ISL_3303745, EPI_ISL_3303746, EPI_ISL_3303747, EPI_ISL_3303748, EPI_ISL_3303749, EPI_ISL_3303750, EPI_ISL_3303751, EPI_ISL_3303752, EPI_ISL_3303753, EPI_ISL_3303754, EPI_ISL_3303755, EPI_ISL_3303756, EPI_ISL_3303757, EPI_ISL_3303758, EPI_ISL_3303759, EPI_ISL_3303760, EPI_ISL_3303761, EPI_ISL_3303762, EPI_ISL_3303763, EPI_ISL_3303764, EPI_ISL_3303765, EPI_ISL_3303766 | | | |
| see above | Clinical Virology | Clinical Bacteriology | Tim Roloff, Fanny Wegner, Helena MB Seth-Smith, Alfredo Mari, Karoline Leuzinger, Julia Bielicki, Manuel Battegay, Hans Hirsch, Adrian Egli |
| EPI_ISL_3303785, EPI_ISL_3303786, EPI_ISL_3303787 | Biolytix AG | Clinical Bacteriology | Tim Roloff, Fanny Wegner, Helena MB Seth-Smith, Alfredo Mari, Karoline Leuzinger, Julia Bielicki, Manuel Battegay, Hans Hirsch, Adrian Egli |
| EPI_ISL_3303795 | Clinical Virology | Clinical Bacteriology | Tim Roloff, Fanny Wegner, Helena MB Seth-Smith, Alfredo Mari, Karoline Leuzinger, Julia Bielicki, Manuel Battegay, Hans Hirsch, Adrian Egli |
| EPI_ISL_3303796 | Ohio Department of Health Laboratory | Ohio Department of Health Laboratory | Holmes, Jennifer; Eric Brandt, Keoni Omura, Glen McGillivary, Caitlin McDonnell, Jade Mowery, Stephanie Mccracken, Tyler Payne, Kirtana Ramadugu, Erica Leasure, Brent Lee, Kelsey Florek, Heather Blankenship, Allison Black, Quanta Brown, and Tammy Bannerman |
| EPI_ISL_3303797, EPI_ISL_3303798, EPI_ISL_3303799, EPI_ISL_3303800, EPI_ISL_3303801, EPI_ISL_3303802, EPI_ISL_3303803, EPI_ISL_3303806, EPI_ISL_3303807, EPI_ISL_3303808, EPI_ISL_3303809, EPI_ISL_3303810, EPI_ISL_3303811, EPI_ISL_3303812, EPI_ISL_3303813, EPI_ISL_3303814, EPI_ISL_3303815, EPI_ISL_3303816, EPI_ISL_3303817, EPI_ISL_3303818, EPI_ISL_3303821, EPI_ISL_3303822, EPI_ISL_3303823, EPI_ISL_3303824, EPI_ISL_3303825, EPI_ISL_3303827 | | | |
| see above | IN State Department of Health Laboratory Services | IN State Department of Health Laboratory Services | Cassandra Campion, Jamie Yeadon, Brian Pope, Lixia Liu, Kyle Brownlee, Melissa Hindenlang, Mark Glazier |
| EPI_ISL_3303828, EPI_ISL_3303829 | Infinity Biologix | Centers for Disease Control and Prevention Division of Viral Diseases, Pathogen Discovery | Dakota Howard, Dhwani Batra, Peter W. Cook, Kara Moser, Adrian Paskey, Jason Caravas, Benjamin Rambo-Martin, Shatavia Morrison, Christopher Gulvick, Scott Sammons, Yvette Unoarumhi, Darlene Wagner, Matthew Schmerer, Christian Bixby, Yihe Wang, Jonathan Schultz, Chirayu Goswami, Russ Hager, Robin Grimwood, Clinton R. Paden, Duncan MacCannell |
| EPI_ISL_3303833, | Laboratory Corporation of America | Centers for Disease Control and Prevention Division of Viral | Dakota Howard, Dhwani Batra, Peter W. Cook, Kara Moser, Adrian Paskey, Jason Caravas, Benjamin Rambo-Martin, Shatavia Morrison, Christopher Gulvick, Scott Sammons, Yvette Unoarumhi, Darlene Wagner, Matthew Schmerer, Minoo |

| | | | |
|---|---|---|---|
| EPI_ISL_3303834, EPI_ISL_3303835, EPI_ISL_3303836 | | Diseases, Pathogen Discovery | Agarwal, Eyad Almasri, Debbie Boles, Ayla Burns, Nuthawin Charoensri, Oren Cohen, Susan Countryman, Mary Ann Cristobal, Bobbi Croy, Suzanne Dale, Hrushikesh Deshmukh, Amanda Douglas, Vincent Drouillon, Marcia Eisenberg, Howard Engler, Rama Ghatti, Prashant Gupta, Susan Hicks, Jake Humphrey, Lax Iyer, Lisa Pfefferle, Manoj Jain, Matthew Robinson, Mohan Kolli, Brian Krueger, Tim Kuphal, Stanley Letovsky, Michael Levandoski, Craig Lukasik, Jonathan Meltzer, Brian Norvell, Mindy Nye, Scott Parker, Christos Petropoulos, John Pruitt, Steven Ragan, Scott Ryan, Mike Sapeta, Jana Schroth, Suresh Babu Selvaraju, Goran Stevovic, Amanda Suchanek, Andrea Throop, Lyndon Tilson, Thomas Urban, Joe Voshell, Kimberly Wagner, Jonathan Williams, Mary Williamson, Qian Zeng, Tricia Zwiefelhofer, Clinton R. Paden, Duncan MacCannell |
| EPI_ISL_3303837 | Fulgent Genetics | Centers for Disease Control and Prevention Division of Viral Diseases, Pathogen Discovery | Dakota Howard, Dhwani Batra, Peter W. Cook, Kara Moser, Adrian Paskey, Jason Caravas, Benjamin Rambo-Martin, Shatavia Morrison, Christopher Gulvick, Scott Sammons, Yvette Unoarumhi, Darlene Wagner, Matthew Schmerer, Harry Gao, Mickey Li, John Gao, Joseph Fierro, Benafsh Sapra, Becky Tsai, Yan Meng, Doreen Ng, James Xie, Clinton R. Paden, Duncan MacCannell |
| EPI_ISL_3303926 | Commonwealth Healthcare Center | Centers for Disease Control and Prevention Division of Viral Diseases, Pathogen Discovery | Mili Sheth, Sarah Nobles, Jasmine Padilla, Alex Burgin, Meghan Bentz, Mark Burroughs, Peter Cook, Clinton R. Paden, Dhwani Batra, Krista Queen, Kristen Knipe, Dakota Howard, Yvette Unoarumhi, Matthew Schmerer, Ben L. Rambo-Martin, Kristine Lacek, Sam Shepard, Dave Wentworth, Vivien Dugan, Suxiang Tong, Justin Lee |
| EPI_ISL_3303927 | "CO Dept. of Public Health and Environment, Lab Services Division" | Centers for Disease Control and Prevention Division of Viral Diseases, Pathogen Discovery | Mili Sheth, Sarah Nobles, Jasmine Padilla, Alex Burgin, Meghan Bentz, Mark Burroughs, Peter Cook, Clinton R. Paden, Dhwani Batra, Krista Queen, Kristen Knipe, Dakota Howard, Yvette Unoarumhi, Matthew Schmerer, Ben L. Rambo-Martin, Kristine Lacek, Sam Shepard, Dave Wentworth, Vivien Dugan, Suxiang Tong, Justin Lee |
| EPI_ISL_3303928 | WA State Department of Health | Centers for Disease Control and Prevention Division of Viral Diseases, Pathogen Discovery | Mili Sheth, Sarah Nobles, Jasmine Padilla, Alex Burgin, Meghan Bentz, Mark Burroughs, Peter Cook, Clinton R. Paden, Dhwani Batra, Krista Queen, Kristen Knipe, Dakota Howard, Yvette Unoarumhi, Matthew Schmerer, Ben L. Rambo-Martin, Kristine Lacek, Sam Shepard, Dave Wentworth, Vivien Dugan, Suxiang Tong, Justin Lee |
| EPI_ISL_3303931 | Fulgent Genetics | Centers for Disease Control and Prevention Division of Viral Diseases, Pathogen Discovery | Dakota Howard, Dhwani Batra, Peter W. Cook, Kara Moser, Adrian Paskey, Jason Caravas, Benjamin Rambo-Martin, Shatavia Morrison, Christopher Gulvick, Scott Sammons, Yvette Unoarumhi, Darlene Wagner, Matthew Schmerer, Harry Gao, Mickey Li, John Gao, Joseph Fierro, Benafsh Sapra, Becky Tsai, Yan Meng, Doreen Ng, James Xie, Clinton R. Paden, Duncan MacCannell |
| EPI_ISL_3303936, EPI_ISL_3303937 | Ohio Department of Health Laboratory | Ohio Department of Health Laboratory | Holmes, Jennifer; Eric Brandt, Keoni Omura, Glen McGillivary, Caitlin McDonnell, Jade Mowery, Stephanie Mccracken, Tyler Payne, Kirtana Ramadugu, Erica Leasure, Brent Lee, Kelsey Florek, Heather Blankenship, Allison Black, Quanta Brown, and Tammy Bannerman |
| EPI_ISL_3303938, EPI_ISL_3303939, EPI_ISL_3303940, EPI_ISL_3303941, EPI_ISL_3303942, EPI_ISL_3303943, EPI_ISL_3303944, EPI_ISL_3303945, EPI_ISL_3303946, EPI_ISL_3303947, EPI_ISL_3303948, EPI_ISL_3303949, EPI_ISL_3303950 | | | |
| see above | California Hospital Medical Center | Los Angeles County Public Health Laboratories | P. Hemarajata et al. |
| EPI_ISL_3303951 | Clarity Mobile Venture | Los Angeles County Public Health Laboratories | P. Hemarajata et al. |
| EPI_ISL_3303952 | Exer with Providence | Los Angeles County Public Health Laboratories | P. Hemarajata et al. |
| EPI_ISL_3303953 | USC Clinical Lab | Los Angeles County Public Health Laboratories | P. Hemarajata et al. |
| EPI_ISL_3303954, EPI_ISL_3303955 | Martin Luther King, Jr. Outpatient Center | Los Angeles County Public Health Laboratories | P. Hemarajata et al. |
| EPI_ISL_3303956, EPI_ISL_3303957, EPI_ISL_3303958, EPI_ISL_3303959, EPI_ISL_3303960, EPI_ISL_3303961, EPI_ISL_3303962, EPI_ISL_3303963, EPI_ISL_3303964 | Torrance Memorial Medical Center | Los Angeles County Public Health Laboratories | P. Hemarajata et al. |
| EPI_ISL_3303965, EPI_ISL_3303966 | Ohio Department of Health Laboratory | Ohio Department of Health Laboratory | Holmes, Jennifer; Eric Brandt, Keoni Omura, Glen McGillivary, Caitlin McDonnell, Jade Mowery, Stephanie Mccracken, Tyler Payne, Kirtana Ramadugu, Erica Leasure, Brent Lee, Kelsey Florek, Heather Blankenship, Allison Black, Quanta Brown, and Tammy Bannerman |
| EPI_ISL_3304295, EPI_ISL_3304296, EPI_ISL_3304297, EPI_ISL_3304298, EPI_ISL_3304300, EPI_ISL_3304301, EPI_ISL_3304302, EPI_ISL_3304303, EPI_ISL_3304304, EPI_ISL_3304305, EPI_ISL_3304306, EPI_ISL_3304307, EPI_ISL_3304308, EPI_ISL_3304309, EPI_ISL_3304310 | | | |
| see above | MD PHL | Maryland Department of Health Laboratories Administration | Eric N. Keller, Jillian Loomis, Kwang Low, Terence L. Moore, Ami Patel, and Robert Myers |
| EPI_ISL_3304311, EPI_ISL_3304313, EPI_ISL_3304314, EPI_ISL_3304315, EPI_ISL_3304316, EPI_ISL_3304317, EPI_ISL_3304318, EPI_ISL_3304319, EPI_ISL_3304320, EPI_ISL_3304321, EPI_ISL_3304322, EPI_ISL_3304323, EPI_ISL_3304324, EPI_ISL_3304325, EPI_ISL_3304326, EPI_ISL_3304328, EPI_ISL_3304329, EPI_ISL_3304330, EPI_ISL_3304331, EPI_ISL_3304332, EPI_ISL_3304333, EPI_ISL_3304334, EPI_ISL_3304335, EPI_ISL_3304336, EPI_ISL_3304337, EPI_ISL_3304338, EPI_ISL_3304339, EPI_ISL_3304340, EPI_ISL_3304341, EPI_ISL_3304342, EPI_ISL_3304343, EPI_ISL_3304344 | | | |
| see above | Lurie Children's Hospital of Chicago | Northwestern University - Ozer Lab | Ramon Lorenzo-Redondo, Lacy M. Simons, Taylor J. Dean, Michael G. Ison, Xiaotian, Zheng, William J. Muller, Larry K. Kociolek, Judd F. Hultquist, Egon A. Ozer |
| EPI_ISL_3304345, EPI_ISL_3304346, EPI_ISL_3304347, EPI_ISL_3304348, EPI_ISL_3304349, EPI_ISL_3304350, EPI_ISL_3304351, EPI_ISL_3304352, EPI_ISL_3304353, EPI_ISL_3304354, EPI_ISL_3304355, EPI_ISL_3304356, EPI_ISL_3304357, EPI_ISL_3304358, EPI_ISL_3304359, EPI_ISL_3304360, EPI_ISL_3304362, EPI_ISL_3304363, EPI_ISL_3304365, EPI_ISL_3304366, EPI_ISL_3304367, EPI_ISL_3304368, EPI_ISL_3304369, EPI_ISL_3304370, EPI_ISL_3304371, EPI_ISL_3304372, EPI_ISL_3304373, EPI_ISL_3304374, EPI_ISL_3304375, EPI_ISL_3304376, EPI_ISL_3304377, EPI_ISL_3304378, EPI_ISL_3304379, EPI_ISL_3304380, EPI_ISL_3304381, EPI_ISL_3304382, EPI_ISL_3304383, EPI_ISL_3304384, EPI_ISL_3304385, EPI_ISL_3304386, EPI_ISL_3304387, EPI_ISL_3304388, EPI_ISL_3304389, EPI_ISL_3304390, EPI_ISL_3304391, EPI_ISL_3304392, EPI_ISL_3304393, EPI_ISL_3304394, EPI_ISL_3304395, EPI_ISL_3304396, EPI_ISL_3304397, EPI_ISL_3304398, EPI_ISL_3304400, EPI_ISL_3304401, EPI_ISL_3304402, EPI_ISL_3304403, EPI_ISL_3304404, EPI_ISL_3304405, EPI_ISL_3304406, EPI_ISL_3304407, EPI_ISL_3304408, EPI_ISL_3304409, EPI_ISL_3304411, EPI_ISL_3304412, EPI_ISL_3304413, EPI_ISL_3304414, EPI_ISL_3304416, EPI_ISL_3304417, EPI_ISL_3304418, EPI_ISL_3304419, EPI_ISL_3304420, EPI_ISL_3304421 | | | |
| see above | MD PHL | Maryland Department of Health Laboratories Administration | Eric N. Keller, Jillian Loomis, Kwang Low, Terence L. Moore, Ami Patel, and Robert Myers |
| EPI_ISL_3304438, EPI_ISL_3304439, EPI_ISL_3304440, EPI_ISL_3304441, EPI_ISL_3304442, EPI_ISL_3304443, EPI_ISL_3304444, EPI_ISL_3304445, EPI_ISL_3304446, EPI_ISL_3304447, EPI_ISL_3304448, EPI_ISL_3304449, EPI_ISL_3304450, EPI_ISL_3304451, EPI_ISL_3304452, EPI_ISL_3304453, EPI_ISL_3304454, EPI_ISL_3304455, EPI_ISL_3304456 | | | |
| see above | OSPEDALE CIVILE TERAMO - CENTRO TRASFUSIONALE | Istituto Zooprofilattico Sperimentale dell'Abruzzo e Molise "G. Caporale" | Lorusso A, Marcacci M, Di Domenico M, Ancora M, Curini V, Di Lollo Valeria, Mangone I, Rinaldi A, Delli Compagni E, Scialabba S, Caporale M, Di Pasquale A, Cammà C, Puglia I, Calistri P, Savini G |
| EPI_ISL_3304457, EPI_ISL_3304458, EPI_ISL_3304460, EPI_ISL_3304461, EPI_ISL_3304462, EPI_ISL_3304463, EPI_ISL_3304464, EPI_ISL_3304465, EPI_ISL_3304466, EPI_ISL_3304467 | OSPEDALE SAN SALVATORE - MEDICINA DI LABORATORIO | Istituto Zooprofilattico Sperimentale dell'Abruzzo e Molise "G. Caporale" | Lorusso A, Marcacci M, Di Domenico M, Ancora M, Curini V, Di Lollo Valeria, Mangone I, Rinaldi A, Delli Compagni E, Scialabba S, Caporale M, Di Pasquale A, Cammà C, Puglia I, Calistri P, Savini G |
| EPI_ISL_3304468, EPI_ISL_3304469, EPI_ISL_3304470, EPI_ISL_3304471, EPI_ISL_3304472, EPI_ISL_3304473, EPI_ISL_3304474, EPI_ISL_3304475, EPI_ISL_3304476, EPI_ISL_3304477, EPI_ISL_3304478, EPI_ISL_3304479, EPI_ISL_3304480, EPI_ISL_3304481, EPI_ISL_3304482, EPI_ISL_3304485, EPI_ISL_3304486, EPI_ISL_3304487 | | | |
| see above | PO AVEZZANO - U.O.C. Laboratorio Analisi | Istituto Zooprofilattico Sperimentale dell'Abruzzo e Molise "G. Caporale" | Lorusso A, Marcacci M, Di Domenico M, Ancora M, Curini V, Di Lollo Valeria, Mangone I, Rinaldi A, Delli Compagni E, Scialabba S, Caporale M, Di Pasquale A, Cammà C, Puglia I, Calistri P, Savini G |
| EPI_ISL_3304488 | SIESP DIPARTIMENTO DI PREVENZIONE TERAMO | Istituto Zooprofilattico Sperimentale dell'Abruzzo e Molise "G. Caporale" | Lorusso A, Marcacci M, Di Domenico M, Ancora M, Curini V, Di Lollo Valeria, Mangone I, Rinaldi A, Delli Compagni E, Scialabba S, Caporale M, Di Pasquale A, Cammà C, Puglia I, Calistri P, Savini G |
| EPI_ISL_3304489, EPI_ISL_3304490, EPI_ISL_3304491, EPI_ISL_3304492, EPI_ISL_3304493, EPI_ISL_3304494, EPI_ISL_3304495, EPI_ISL_3304496, EPI_ISL_3304497, EPI_ISL_3304498, EPI_ISL_3304499, EPI_ISL_3304500, EPI_ISL_3304501, EPI_ISL_3304502, EPI_ISL_3304503, EPI_ISL_3304504, EPI_ISL_3304506 | | | |
| see above | OSPEDALE CIVILE TERAMO - CENTRO TRASFUSIONALE | Istituto Zooprofilattico Sperimentale dell'Abruzzo e Molise "G. Caporale" | Lorusso A, Marcacci M, Di Domenico M, Ancora M, Curini V, Di Lollo Valeria, Mangone I, Rinaldi A, Delli Compagni E, Scialabba S, Caporale M, Di Pasquale A, Cammà C, Puglia I, Calistri P, Savini G |
| EPI_ISL_3304507, EPI_ISL_3304508, EPI_ISL_3304509, EPI_ISL_3304510, EPI_ISL_3304511, EPI_ISL_3304512 | OSPEDALE SAN SALVATORE - MEDICINA DI LABORATORIO | Istituto Zooprofilattico Sperimentale dell'Abruzzo e Molise "G. Caporale" | Lorusso A, Marcacci M, Di Domenico M, Ancora M, Curini V, Di Lollo Valeria, Mangone I, Rinaldi A, Delli Compagni E, Scialabba S, Caporale M, Di Pasquale A, Cammà C, Puglia I, Calistri P, Savini G |
| EPI_ISL_3304513 | SIESP DIPARTIMENTO DI PREVENZIONE TERAMO | Istituto Zooprofilattico Sperimentale dell'Abruzzo e Molise "G. Caporale" | Lorusso A, Marcacci M, Di Domenico M, Ancora M, Curini V, Di Lollo Valeria, Mangone I, Rinaldi A, Delli Compagni E, Scialabba S, Caporale M, Di Pasquale A, Cammà C, Puglia I, Calistri P, Savini G |
| EPI_ISL_3304514, EPI_ISL_3304515, EPI_ISL_3304517, EPI_ISL_3304518, EPI_ISL_3304519, EPI_ISL_3304520, EPI_ISL_3304521, EPI_ISL_3304522, EPI_ISL_3304523 | PO AVEZZANO - U.O.C. Laboratorio Analisi | Istituto Zooprofilattico Sperimentale dell'Abruzzo e Molise "G. Caporale" | Lorusso A, Marcacci M, Di Domenico M, Ancora M, Curini V, Di Lollo Valeria, Mangone I, Rinaldi A, Delli Compagni E, Scialabba S, Caporale M, Di Pasquale A, Cammà C, Puglia I, Calistri P, Savini G |
| EPI_ISL_3304524, EPI_ISL_3304525 | OSPEDALE SAN SALVATORE - MEDICINA DI LABORATORIO | Istituto Zooprofilattico Sperimentale dell'Abruzzo e Molise "G. Caporale" | Lorusso A, Marcacci M, Di Domenico M, Ancora M, Curini V, Di Lollo Valeria, Mangone I, Rinaldi A, Delli Compagni E, Scialabba S, Caporale M, Di Pasquale A, Cammà C, Puglia I, Calistri P, Savini G |
| EPI_ISL_3304526, EPI_ISL_3304527, EPI_ISL_3304528, EPI_ISL_3304529, EPI_ISL_3304530, EPI_ISL_3304531, EPI_ISL_3304532, EPI_ISL_3304533, EPI_ISL_3304534, EPI_ISL_3304535, EPI_ISL_3304536, EPI_ISL_3304537, EPI_ISL_3304538, EPI_ISL_3304539, EPI_ISL_3304540, EPI_ISL_3304541, EPI_ISL_3304542, EPI_ISL_3304543, EPI_ISL_3304544, EPI_ISL_3304545, EPI_ISL_3304546, EPI_ISL_3304547, EPI_ISL_3304548, EPI_ISL_3304549, EPI_ISL_3304550, EPI_ISL_3304551, EPI_ISL_3304552, EPI_ISL_3304553, EPI_ISL_3304554, EPI_ISL_3304555, EPI_ISL_3304556, EPI_ISL_3304557, EPI_ISL_3304558, EPI_ISL_3304559, EPI_ISL_3304560, EPI_ISL_3304561, EPI_ISL_3304562, EPI_ISL_3304563, EPI_ISL_3304564, EPI_ISL_3304565 | | | |
| see above | OSPEDALE CIVILE TERAMO - CENTRO TRASFUSIONALE | Istituto Zooprofilattico Sperimentale dell'Abruzzo e Molise "G. Caporale" | Lorusso A, Marcacci M, Di Domenico M, Ancora M, Curini V, Di Lollo Valeria, Mangone I, Rinaldi A, Delli Compagni E, Scialabba S, Caporale M, Di Pasquale A, Cammà C, Puglia I, Calistri P, Savini G |
| EPI_ISL_3304568, EPI_ISL_3304570, EPI_ISL_3304572, EPI_ISL_3304573, EPI_ISL_3304574, EPI_ISL_3304575, EPI_ISL_3304576, EPI_ISL_3304577, EPI_ISL_3304578, EPI_ISL_3304579, EPI_ISL_3304580, EPI_ISL_3304581, EPI_ISL_3304582, EPI_ISL_3304583, EPI_ISL_3304584, EPI_ISL_3304585, EPI_ISL_3304586, EPI_ISL_3304587, EPI_ISL_3304588, EPI_ISL_3304589, EPI_ISL_3304590, EPI_ISL_3304591, EPI_ISL_3304592, EPI_ISL_3304593, EPI_ISL_3304594, EPI_ISL_3304595, EPI_ISL_3304596, EPI_ISL_3304597, EPI_ISL_3304598, EPI_ISL_3304599, EPI_ISL_3304603 | | | |
| see above | Università degli Studi di Perugia | Istituto Zooprofilattico Sperimentale dell'Abruzzo e Molise "G. Caporale" | Mencacci A, Camilloni B, Biagetti M, Giammarioli M, Lorusso A, Marcacci M, Di Domenico M, Ancora M, Curini V, Mangone I, Rinaldi A, Delli Compagni E, Scialabba S, Di Pasquale A, Cammà C, Puglia I, Calistri P, Savini G |
| EPI_ISL_3304604, EPI_ISL_3304605, EPI_ISL_3304606, EPI_ISL_3304607, EPI_ISL_3304608, EPI_ISL_3304609, EPI_ISL_3304610, EPI_ISL_3304611, EPI_ISL_3304612, EPI_ISL_3304613, EPI_ISL_3304614 | | | |

| | | | |
|---|---|---|---|
| see above | Azienda Ospedaliera Terni | Istituto Zooprofilattico Sperimentale dell'Abruzzo e Molise "G. Caporale" | Palumbo M, Scaccetti A, Lorusso A, Marcacci M, Di Domenico M, Ancora M, Curini V, Mangone I, Rinaldi A, Delli Compagni E, Scialabba S, Di Pasquale A, Cammà C, Puglia I, Calistri P, Savini G |
| EPI_ISL_3304615, EPI_ISL_3304616, EPI_ISL_3304617, EPI_ISL_3304618, EPI_ISL_3304619, EPI_ISL_3304620, EPI_ISL_3304621 | Azienda USL Umbria 2 | Istituto Zooprofilattico Sperimentale dell'Abruzzo e Molise "G. Caporale" | Proietti A, Pistoni E, Lorusso A, Marcacci M, Di Domenico M, Ancora M, Curini V, Mangone I, Rinaldi A, Delli Compagni E, Scialabba S, Di Pasquale A, Cammà C, Puglia I, Calistri P, Savini G |
| EPI_ISL_3304622 | Laboratorio Analisi Osp. Città di Castello - Azienda USL Umbria1 | Istituto Zooprofilattico Sperimentale dell'Abruzzo e Molise "G. Caporale" | Malagigi V, Tacconi P,Lorusso A, Marcacci M, Di Domenico M, Ancora M, Curini V, Mangone I, Rinaldi A, Delli Compagni E, Scialabba S, Di Pasquale A, Cammà C, Puglia I, Calistri P, Savini G |
| EPI_ISL_3304623 | Azienda USL Umbria 2 | Istituto Zooprofilattico Sperimentale dell'Abruzzo e Molise "G. Caporale" | Proietti A, Pistoni E, Lorusso A, Marcacci M, Di Domenico M, Ancora M, Curini V, Mangone I, Rinaldi A, Delli Compagni E, Scialabba S, Di Pasquale A, Cammà C, Puglia I, Calistri P, Savini G |
| EPI_ISL_3304936, EPI_ISL_3304937, EPI_ISL_3304938, EPI_ISL_3304963, EPI_ISL_3304966, EPI_ISL_3304970, EPI_ISL_3304981, EPI_ISL_3304984, EPI_ISL_3304986 | Fulgent Genetics | Centers for Disease Control and Prevention Division of Viral Diseases, Pathogen Discovery | Dakota Howard, Dhwani Batra, Peter W. Cook, Kara Moser, Adrian Paskey, Jason Caravas, Benjamin Rambo-Martin, Shatavia Morrison, Christopher Gulvick, Scott Sammons, Yvette Unoarumhi, Darlene Wagner, Matthew Schmerer, Harry Gao, Mickey Li, John Gao, Joseph Fierro, Benafsh Sapra, Becky Tsai, Yan Meng, Doreen Ng, James Xie, Clinton R. Paden, Duncan MacCannell |
| EPI_ISL_3305016, EPI_ISL_3305031, EPI_ISL_3305042, EPI_ISL_3305053, EPI_ISL_3305060, EPI_ISL_3305061, EPI_ISL_3305066, EPI_ISL_3305069, EPI_ISL_3305086, EPI_ISL_3305088, EPI_ISL_3305091, EPI_ISL_3305092, EPI_ISL_3305104, EPI_ISL_3305107, EPI_ISL_3305116, EPI_ISL_3305120, EPI_ISL_3305132, EPI_ISL_3305146, EPI_ISL_3305149, EPI_ISL_3305153, EPI_ISL_3305164, EPI_ISL_3305185, EPI_ISL_3305187, EPI_ISL_3305188 | | | |
| see above | Quest Diagnostics Incorporated | Centers for Disease Control and Prevention Division of Viral Diseases, Pathogen Discovery | Dakota Howard, Dhwani Batra, Peter W. Cook, Kara Moser, Adrian Paskey, Jason Caravas, Benjamin Rambo-Martin, Shatavia Morrison, Christopher Gulvick, Scott Sammons, Yvette Unoarumhi, Darlene Wagner, Matthew Schmerer, S. H. Rosenthal, A. Gerasimova, R. M. Kagan, B. Anderson, M. Hua, Y. Liu, L.E. Bernstein, K.E. Livingston, A. Perez, I. A. Shlyakhter, R. V. Rolando, R. Owen, P. Tanpaiboon, F. Lacbawan, Clinton R. Paden, Duncan MacCannell |
| EPI_ISL_3305288, EPI_ISL_3305289, EPI_ISL_3305290, EPI_ISL_3305291, EPI_ISL_3305292, EPI_ISL_3305293, EPI_ISL_3305294, EPI_ISL_3305295, EPI_ISL_3305296, EPI_ISL_3305297, EPI_ISL_3305298, EPI_ISL_3305300, EPI_ISL_3305301, EPI_ISL_3305304, EPI_ISL_3305306, EPI_ISL_3305307, EPI_ISL_3305309, EPI_ISL_3305310, EPI_ISL_3305311, EPI_ISL_3305317, EPI_ISL_3305318, EPI_ISL_3305319, EPI_ISL_3305320, EPI_ISL_3305323, EPI_ISL_3305324, EPI_ISL_3305325, EPI_ISL_3305326, EPI_ISL_3305329, EPI_ISL_3305330, EPI_ISL_3305331, EPI_ISL_3305332, EPI_ISL_3305334, EPI_ISL_3305335, EPI_ISL_3305336, EPI_ISL_3305337, EPI_ISL_3305338, EPI_ISL_3305339, EPI_ISL_3305341, EPI_ISL_3305343, EPI_ISL_3305345, EPI_ISL_3305346, EPI_ISL_3305347, EPI_ISL_3305348, EPI_ISL_3305349, EPI_ISL_3305352, EPI_ISL_3305353, EPI_ISL_3305355, EPI_ISL_3305356, EPI_ISL_3305357, EPI_ISL_3305358, EPI_ISL_3305359, EPI_ISL_3305360, EPI_ISL_3305361, EPI_ISL_3305362, EPI_ISL_3305363, EPI_ISL_3305364, EPI_ISL_3305365, EPI_ISL_3305366, EPI_ISL_3305367, EPI_ISL_3305368, EPI_ISL_3305369, EPI_ISL_3305371, EPI_ISL_3305373, EPI_ISL_3305374, EPI_ISL_3305375, EPI_ISL_3305376, EPI_ISL_3305377, EPI_ISL_3305378, EPI_ISL_3305379, EPI_ISL_3305380, EPI_ISL_3305381, EPI_ISL_3305382, EPI_ISL_3305385, EPI_ISL_3305388, EPI_ISL_3305389, EPI_ISL_3305391, EPI_ISL_3305392, EPI_ISL_3305393, EPI_ISL_3305394, EPI_ISL_3305395, EPI_ISL_3305396, EPI_ISL_3305397, EPI_ISL_3305398, EPI_ISL_3305399, EPI_ISL_3305402, EPI_ISL_3305403, EPI_ISL_3305404 | | | |
| see above | Aegis Sciences Corporation | Centers for Disease Control and Prevention Division of Viral Diseases, Pathogen Discovery | Dakota Howard, Dhwani Batra, Peter W. Cook, Kara Moser, Adrian Paskey, Jason Caravas, Benjamin Rambo-Martin, Shatavia Morrison, Christopher Gulvick, Scott Sammons, Yvette Unoarumhi, Darlene Wagner, Matthew Schmerer, Cyndi Clark, Patrick Campbell, Rob Case, Vikramsinha Ghorpade, Holly Houdeshell, Ola Kvalvaag, Dillon Nall, Ethan Sanders, Alec Vest, Shaun Westlund, Matthew Hardison, Clinton R. Paden, Duncan MacCannell |
| EPI_ISL_3305405 | Australian Clinical Labs | NSW Health Pathology - Institute of Clinical Pathology and Medical Research; Westmead Hospital; University of Sydney | Arnott A., Draper J., Gall M., Martinez E., Rockett R., Sintchenko V., on behalf of ICPMR |
| EPI_ISL_3305406 | Aegis Sciences Corporation | Centers for Disease Control and Prevention Division of Viral Diseases, Pathogen Discovery | Dakota Howard, Dhwani Batra, Peter W. Cook, Kara Moser, Adrian Paskey, Jason Caravas, Benjamin Rambo-Martin, Shatavia Morrison, Christopher Gulvick, Scott Sammons, Yvette Unoarumhi, Darlene Wagner, Matthew Schmerer, Cyndi Clark, Patrick Campbell, Rob Case, Vikramsinha Ghorpade, Holly Houdeshell, Ola Kvalvaag, Dillon Nall, Ethan Sanders, Alec Vest, Shaun Westlund, Matthew Hardison, Clinton R. Paden, Duncan MacCannell |
| EPI_ISL_3305407 | Pathology West - NSW Health Pathology | NSW Health Pathology - Institute of Clinical Pathology and Medical Research; Westmead Hospital; University of Sydney | Arnott A., Draper J., Gall M., Martinez E., Rockett R., Sintchenko V., on behalf of ICPMR |
| EPI_ISL_3305409 | South Eastern Area Laboratory Services (SEALS) | NSW Health Pathology - Institute of Clinical Pathology and Medical Research; Westmead Hospital; University of Sydney | Arnott A., Draper J., Gall M., Martinez E., Rockett R., Sintchenko V., on behalf of ICPMR |
| EPI_ISL_3305410 | Aegis Sciences Corporation | Centers for Disease Control and Prevention Division of Viral Diseases, Pathogen Discovery | Dakota Howard, Dhwani Batra, Peter W. Cook, Kara Moser, Adrian Paskey, Jason Caravas, Benjamin Rambo-Martin, Shatavia Morrison, Christopher Gulvick, Scott Sammons, Yvette Unoarumhi, Darlene Wagner, Matthew Schmerer, Cyndi Clark, Patrick Campbell, Rob Case, Vikramsinha Ghorpade, Holly Houdeshell, Ola Kvalvaag, Dillon Nall, Ethan Sanders, Alec Vest, Shaun Westlund, Matthew Hardison, Clinton R. Paden, Duncan MacCannell |
| EPI_ISL_3305411 | Sydney South West Pathology Service (SSWPS) - Royal Prince Alfred Hospital - NSW Health Pathology | NSW Health Pathology - Institute of Clinical Pathology and Medical Research; Westmead Hospital; University of Sydney | Arnott A., Draper J., Gall M., Martinez E., Rockett R., Sintchenko V., on behalf of ICPMR |
| EPI_ISL_3305412 | Aegis Sciences Corporation | Centers for Disease Control and Prevention Division of Viral Diseases, Pathogen Discovery | Dakota Howard, Dhwani Batra, Peter W. Cook, Kara Moser, Adrian Paskey, Jason Caravas, Benjamin Rambo-Martin, Shatavia Morrison, Christopher Gulvick, Scott Sammons, Yvette Unoarumhi, Darlene Wagner, Matthew Schmerer, Cyndi Clark, Patrick Campbell, Rob Case, Vikramsinha Ghorpade, Holly Houdeshell, Ola Kvalvaag, Dillon Nall, Ethan Sanders, Alec Vest, Shaun Westlund, Matthew Hardison, Clinton R. Paden, Duncan MacCannell |
| EPI_ISL_3305413 | Sydney South West Pathology Service (SSWPS) - Liverpool Hospital - NSW Health Pathology | NSW Health Pathology - Institute of Clinical Pathology and Medical Research; Westmead Hospital; University of Sydney | Arnott A., Draper J., Gall M., Martinez E., Rockett R., Sintchenko V., on behalf of ICPMR |
| EPI_ISL_3305414, EPI_ISL_3305415 | Aegis Sciences Corporation | Centers for Disease Control and Prevention Division of Viral Diseases, Pathogen Discovery | Dakota Howard, Dhwani Batra, Peter W. Cook, Kara Moser, Adrian Paskey, Jason Caravas, Benjamin Rambo-Martin, Shatavia Morrison, Christopher Gulvick, Scott Sammons, Yvette Unoarumhi, Darlene Wagner, Matthew Schmerer, Cyndi Clark, Patrick Campbell, Rob Case, Vikramsinha Ghorpade, Holly Houdeshell, Ola Kvalvaag, Dillon Nall, Ethan Sanders, Alec Vest, Shaun Westlund, Matthew Hardison, Clinton R. Paden, Duncan MacCannell |
| EPI_ISL_3305416 | Sydney South West Pathology Service (SSWPS) - Liverpool Hospital - NSW Health Pathology | NSW Health Pathology - Institute of Clinical Pathology and Medical Research; Westmead Hospital; University of Sydney | Arnott A., Draper J., Gall M., Martinez E., Rockett R., Sintchenko V., on behalf of ICPMR |
| EPI_ISL_3305417 | Histopath | NSW Health Pathology - Institute of Clinical Pathology and Medical Research; Westmead Hospital; University of Sydney | Arnott A., Draper J., Gall M., Martinez E., Rockett R., Sintchenko V., on behalf of ICPMR |
| EPI_ISL_3305418 | Aegis Sciences Corporation | Centers for Disease Control and Prevention Division of Viral Diseases, Pathogen Discovery | Dakota Howard, Dhwani Batra, Peter W. Cook, Kara Moser, Adrian Paskey, Jason Caravas, Benjamin Rambo-Martin, Shatavia Morrison, Christopher Gulvick, Scott Sammons, Yvette Unoarumhi, Darlene Wagner, Matthew Schmerer, Cyndi Clark, Patrick Campbell, Rob Case, Vikramsinha Ghorpade, Holly Houdeshell, Ola Kvalvaag, Dillon Nall, Ethan Sanders, Alec Vest, Shaun Westlund, Matthew Hardison, Clinton R. Paden, Duncan MacCannell |
| EPI_ISL_3305419 | 4Cyte Pathology | NSW Health Pathology - Institute of Clinical Pathology and Medical Research; Westmead Hospital; University of Sydney | Arnott A., Draper J., Gall M., Martinez E., Rockett R., Sintchenko V., on behalf of ICPMR |
| EPI_ISL_3305420 | Aegis Sciences Corporation | Centers for Disease Control and Prevention Division of Viral Diseases, Pathogen Discovery | Dakota Howard, Dhwani Batra, Peter W. Cook, Kara Moser, Adrian Paskey, Jason Caravas, Benjamin Rambo-Martin, Shatavia Morrison, Christopher Gulvick, Scott Sammons, Yvette Unoarumhi, Darlene Wagner, Matthew Schmerer, Cyndi Clark, Patrick Campbell, Rob Case, Vikramsinha Ghorpade, Holly Houdeshell, Ola Kvalvaag, Dillon Nall, Ethan Sanders, Alec Vest, Shaun Westlund, Matthew Hardison, Clinton R. Paden, Duncan MacCannell |
| EPI_ISL_3305421 | Sydney South West Pathology Service (SSWPS) - Liverpool Hospital - NSW Health Pathology | NSW Health Pathology - Institute of Clinical Pathology and Medical Research; Westmead Hospital; University of Sydney | Arnott A., Draper J., Gall M., Martinez E., Rockett R., Sintchenko V., on behalf of ICPMR |
| EPI_ISL_3305422 | Aegis Sciences Corporation | Centers for Disease Control and Prevention Division of Viral Diseases, Pathogen Discovery | Dakota Howard, Dhwani Batra, Peter W. Cook, Kara Moser, Adrian Paskey, Jason Caravas, Benjamin Rambo-Martin, Shatavia Morrison, Christopher Gulvick, Scott Sammons, Yvette Unoarumhi, Darlene Wagner, Matthew Schmerer, Cyndi Clark, Patrick Campbell, Rob Case, Vikramsinha Ghorpade, Holly Houdeshell, Ola Kvalvaag, Dillon Nall, Ethan Sanders, Alec Vest, Shaun Westlund, Matthew Hardison, Clinton R. Paden, Duncan MacCannell |
| EPI_ISL_3305424, EPI_ISL_3305426 | Douglass Hanly Moir Pathology | NSW Health Pathology - Institute of Clinical Pathology and Medical Research; Westmead Hospital; University of Sydney | Arnott A., Draper J., Gall M., Martinez E., Rockett R., Sintchenko V., on behalf of ICPMR |
| EPI_ISL_3305427 | Aegis Sciences Corporation | Centers for Disease Control and Prevention Division of Viral Diseases, Pathogen Discovery | Dakota Howard, Dhwani Batra, Peter W. Cook, Kara Moser, Adrian Paskey, Jason Caravas, Benjamin Rambo-Martin, Shatavia Morrison, Christopher Gulvick, Scott Sammons, Yvette Unoarumhi, Darlene Wagner, Matthew Schmerer, Cyndi Clark, Patrick Campbell, Rob Case, Vikramsinha Ghorpade, Holly Houdeshell, Ola Kvalvaag, Dillon Nall, Ethan Sanders, Alec Vest, Shaun Westlund, Matthew Hardison, Clinton R. Paden, Duncan MacCannell |
| EPI_ISL_3305428, EPI_ISL_3305429 | Sydney South West Pathology Service (SSWPS) - Liverpool Hospital - NSW Health Pathology | NSW Health Pathology - Institute of Clinical Pathology and Medical Research; Westmead Hospital; University of Sydney | Arnott A., Draper J., Gall M., Martinez E., Rockett R., Sintchenko V., on behalf of ICPMR |
| EPI_ISL_3305430 | Aegis Sciences Corporation | Centers for Disease Control and Prevention Division of Viral Diseases, Pathogen Discovery | Dakota Howard, Dhwani Batra, Peter W. Cook, Kara Moser, Adrian Paskey, Jason Caravas, Benjamin Rambo-Martin, Shatavia Morrison, Christopher Gulvick, Scott Sammons, Yvette Unoarumhi, Darlene Wagner, Matthew Schmerer, Cyndi Clark, Patrick Campbell, Rob Case, Vikramsinha Ghorpade, Holly Houdeshell, Ola Kvalvaag, Dillon Nall, Ethan Sanders, Alec Vest, Shaun Westlund, Matthew Hardison, Clinton R. Paden, Duncan MacCannell |
| EPI_ISL_3305431 | Sydney South West Pathology Service (SSWPS) - Liverpool Hospital - NSW Health Pathology | NSW Health Pathology - Institute of Clinical Pathology and Medical Research; Westmead Hospital; University of Sydney | Arnott A., Draper J., Gall M., Martinez E., Rockett R., Sintchenko V., on behalf of ICPMR |
| EPI_ISL_3305432 | Aegis Sciences Corporation | Centers for Disease Control and Prevention Division of Viral Diseases, Pathogen Discovery | Dakota Howard, Dhwani Batra, Peter W. Cook, Kara Moser, Adrian Paskey, Jason Caravas, Benjamin Rambo-Martin, Shatavia Morrison, Christopher Gulvick, Scott Sammons, Yvette Unoarumhi, Darlene Wagner, Matthew Schmerer, Cyndi Clark, Patrick Campbell, Rob Case, Vikramsinha Ghorpade, Holly Houdeshell, Ola Kvalvaag, Dillon Nall, Ethan Sanders, Alec Vest, Shaun Westlund, Matthew Hardison, Clinton R. Paden, Duncan MacCannell |
| EPI_ISL_3305434, EPI_ISL_3305435 | Laverty Pathology | NSW Health Pathology - Institute of Clinical Pathology and Medical Research; Westmead Hospital; University of Sydney | Arnott A., Draper J., Gall M., Martinez E., Rockett R., Sintchenko V., on behalf of ICPMR |
| EPI_ISL_3305436 | Aegis Sciences Corporation | Centers for Disease Control and Prevention Division of Viral Diseases, Pathogen Discovery | Dakota Howard, Dhwani Batra, Peter W. Cook, Kara Moser, Adrian Paskey, Jason Caravas, Benjamin Rambo-Martin, Shatavia Morrison, Christopher Gulvick, Scott Sammons, Yvette Unoarumhi, Darlene Wagner, Matthew Schmerer, Cyndi Clark, Patrick Campbell, Rob Case, Vikramsinha Ghorpade, Holly Houdeshell, Ola Kvalvaag, Dillon Nall, Ethan Sanders, Alec Vest, Shaun Westlund, Matthew Hardison, Clinton R. Paden, Duncan MacCannell |
| EPI_ISL_3305437 | Sydney South West Pathology Service (SSWPS) - Liverpool Hospital - NSW Health Pathology | NSW Health Pathology - Institute of Clinical Pathology and Medical Research; Westmead Hospital; University of Sydney | Arnott A., Draper J., Gall M., Martinez E., Rockett R., Sintchenko V., on behalf of ICPMR |
| EPI_ISL_3305438 | Aegis Sciences Corporation | Centers for Disease Control and Prevention Division of Viral Diseases, Pathogen Discovery | Dakota Howard, Dhwani Batra, Peter W. Cook, Kara Moser, Adrian Paskey, Jason Caravas, Benjamin Rambo-Martin, Shatavia Morrison, Christopher Gulvick, Scott Sammons, Yvette Unoarumhi, Darlene Wagner, Matthew Schmerer, Cyndi Clark, Patrick Campbell, Rob Case, Vikramsinha Ghorpade, Holly Houdeshell, Ola Kvalvaag, Dillon Nall, Ethan Sanders, Alec Vest, Shaun Westlund, Matthew Hardison, Clinton R. Paden, Duncan MacCannell |
| EPI_ISL_3305439 | Sydney South West Pathology Service (SSWPS) - Liverpool Hospital - NSW Health Pathology | NSW Health Pathology - Institute of Clinical Pathology and Medical Research; Westmead Hospital; University of Sydney | Arnott A., Draper J., Gall M., Martinez E., Rockett R., Sintchenko V., on behalf of ICPMR |
| EPI_ISL_3305440 | Aegis Sciences Corporation | Centers for Disease Control and Prevention Division of Viral Diseases, Pathogen Discovery | Dakota Howard, Dhwani Batra, Peter W. Cook, Kara Moser, Adrian Paskey, Jason Caravas, Benjamin Rambo-Martin, Shatavia Morrison, Christopher Gulvick, Scott Sammons, Yvette Unoarumhi, Darlene Wagner, Matthew Schmerer, Cyndi Clark, Patrick Campbell, Rob Case, Vikramsinha Ghorpade, Holly Houdeshell, Ola Kvalvaag, Dillon Nall, Ethan Sanders, Alec Vest, Shaun Westlund, Matthew Hardison, Clinton R. Paden, Duncan MacCannell |
| EPI_ISL_3305441 | Histopath | NSW Health Pathology - Institute of Clinical Pathology and Medical Research; Westmead Hospital; University of Sydney | Arnott A., Draper J., Gall M., Martinez E., Rockett R., Sintchenko V., on behalf of ICPMR |
| EPI_ISL_3305442 | Aegis Sciences Corporation | Centers for Disease Control and Prevention Division of Viral Diseases, Pathogen Discovery | Dakota Howard, Dhwani Batra, Peter W. Cook, Kara Moser, Adrian Paskey, Jason Caravas, Benjamin Rambo-Martin, Shatavia Morrison, Christopher Gulvick, Scott Sammons, Yvette Unoarumhi, Darlene Wagner, Matthew Schmerer, Cyndi Clark, Patrick Campbell, Rob Case, Vikramsinha Ghorpade, Holly Houdeshell, Ola Kvalvaag, Dillon Nall, Ethan Sanders, Alec Vest, Shaun Westlund, Matthew Hardison, Clinton R. Paden, Duncan MacCannell |

| | | | |
|---|---|---|---|
| EPI_ISL_3305443 | Sydney South West Pathology Service (SSWPS) - Liverpool Hospital - NSW Health Pathology | NSW Health Pathology - Institute of Clinical Pathology and Medical Research; Westmead Hospital; University of Sydney | Arnott A., Draper J., Gall M., Martinez E., Rockett R., Sintchenko V., on behalf of ICPMR |
| EPI_ISL_3305445 | Medlab Pathology | NSW Health Pathology - Institute of Clinical Pathology and Medical Research; Westmead Hospital; University of Sydney | Arnott A., Draper J., Gall M., Martinez E., Rockett R., Sintchenko V., on behalf of ICPMR |
| EPI_ISL_3305446 | Aegis Sciences Corporation | Centers for Disease Control and Prevention Division of Viral Diseases, Pathogen Discovery | Dakota Howard, Dhwani Batra, Peter W. Cook, Kara Moser, Adrian Paskey, Jason Caravas, Benjamin Rambo-Martin, Shatavia Morrison, Christopher Gulvick, Scott Sammons, Yvette Unoarumhi, Darlene Wagner, Matthew Schmerer, Cyndi Clark, Patrick Campbell, Rob Case, Vikramsinha Ghorpade, Holly Houdeshell, Ola Kvalvaag, Dillon Nall, Ethan Sanders, Alec Vest, Shaun Westlund, Matthew Hardison, Clinton R. Paden, Duncan MacCannell |
| EPI_ISL_3305447 | Laverty Pathology | NSW Health Pathology - Institute of Clinical Pathology and Medical Research; Westmead Hospital; University of Sydney | Arnott A., Draper J., Gall M., Martinez E., Rockett R., Sintchenko V., on behalf of ICPMR |
| EPI_ISL_3305448 | Aegis Sciences Corporation | Centers for Disease Control and Prevention Division of Viral Diseases, Pathogen Discovery | Dakota Howard, Dhwani Batra, Peter W. Cook, Kara Moser, Adrian Paskey, Jason Caravas, Benjamin Rambo-Martin, Shatavia Morrison, Christopher Gulvick, Scott Sammons, Yvette Unoarumhi, Darlene Wagner, Matthew Schmerer, Cyndi Clark, Patrick Campbell, Rob Case, Vikramsinha Ghorpade, Holly Houdeshell, Ola Kvalvaag, Dillon Nall, Ethan Sanders, Alec Vest, Shaun Westlund, Matthew Hardison, Clinton R. Paden, Duncan MacCannell |
| EPI_ISL_3305449 | Laverty Pathology | NSW Health Pathology - Institute of Clinical Pathology and Medical Research; Westmead Hospital; University of Sydney | Arnott A., Draper J., Gall M., Martinez E., Rockett R., Sintchenko V., on behalf of ICPMR |
| EPI_ISL_3305450 | Aegis Sciences Corporation | Centers for Disease Control and Prevention Division of Viral Diseases, Pathogen Discovery | Dakota Howard, Dhwani Batra, Peter W. Cook, Kara Moser, Adrian Paskey, Jason Caravas, Benjamin Rambo-Martin, Shatavia Morrison, Christopher Gulvick, Scott Sammons, Yvette Unoarumhi, Darlene Wagner, Matthew Schmerer, Cyndi Clark, Patrick Campbell, Rob Case, Vikramsinha Ghorpade, Holly Houdeshell, Ola Kvalvaag, Dillon Nall, Ethan Sanders, Alec Vest, Shaun Westlund, Matthew Hardison, Clinton R. Paden, Duncan MacCannell |
| EPI_ISL_3305451, EPI_ISL_3305453, EPI_ISL_3305454 | Histopath | NSW Health Pathology - Institute of Clinical Pathology and Medical Research; Westmead Hospital; University of Sydney | Arnott A., Draper J., Gall M., Martinez E., Rockett R., Sintchenko V., on behalf of ICPMR |
| EPI_ISL_3305455 | Aegis Sciences Corporation | Centers for Disease Control and Prevention Division of Viral Diseases, Pathogen Discovery | Dakota Howard, Dhwani Batra, Peter W. Cook, Kara Moser, Adrian Paskey, Jason Caravas, Benjamin Rambo-Martin, Shatavia Morrison, Christopher Gulvick, Scott Sammons, Yvette Unoarumhi, Darlene Wagner, Matthew Schmerer, Cyndi Clark, Patrick Campbell, Rob Case, Vikramsinha Ghorpade, Holly Houdeshell, Ola Kvalvaag, Dillon Nall, Ethan Sanders, Alec Vest, Shaun Westlund, Matthew Hardison, Clinton R. Paden, Duncan MacCannell |
| EPI_ISL_3305457, EPI_ISL_3305459, EPI_ISL_3305460 | Sydney South West Pathology Service (SSWPS) - Liverpool Hospital - NSW Health Pathology | NSW Health Pathology - Institute of Clinical Pathology and Medical Research; Westmead Hospital; University of Sydney | Arnott A., Draper J., Gall M., Martinez E., Rockett R., Sintchenko V., on behalf of ICPMR |
| EPI_ISL_3305462 | St Vincent's Pathology (SydPath) | NSW Health Pathology - Institute of Clinical Pathology and Medical Research; Westmead Hospital; University of Sydney | Arnott A., Draper J., Gall M., Martinez E., Rockett R., Sintchenko V., on behalf of ICPMR |
| EPI_ISL_3305463 | Aegis Sciences Corporation | Centers for Disease Control and Prevention Division of Viral Diseases, Pathogen Discovery | Dakota Howard, Dhwani Batra, Peter W. Cook, Kara Moser, Adrian Paskey, Jason Caravas, Benjamin Rambo-Martin, Shatavia Morrison, Christopher Gulvick, Scott Sammons, Yvette Unoarumhi, Darlene Wagner, Matthew Schmerer, Cyndi Clark, Patrick Campbell, Rob Case, Vikramsinha Ghorpade, Holly Houdeshell, Ola Kvalvaag, Dillon Nall, Ethan Sanders, Alec Vest, Shaun Westlund, Matthew Hardison, Clinton R. Paden, Duncan MacCannell |
| EPI_ISL_3305464 | Histopath | NSW Health Pathology - Institute of Clinical Pathology and Medical Research; Westmead Hospital; University of Sydney | Arnott A., Draper J., Gall M., Martinez E., Rockett R., Sintchenko V., on behalf of ICPMR |
| EPI_ISL_3305465 | Aegis Sciences Corporation | Centers for Disease Control and Prevention Division of Viral Diseases, Pathogen Discovery | Dakota Howard, Dhwani Batra, Peter W. Cook, Kara Moser, Adrian Paskey, Jason Caravas, Benjamin Rambo-Martin, Shatavia Morrison, Christopher Gulvick, Scott Sammons, Yvette Unoarumhi, Darlene Wagner, Matthew Schmerer, Cyndi Clark, Patrick Campbell, Rob Case, Vikramsinha Ghorpade, Holly Houdeshell, Ola Kvalvaag, Dillon Nall, Ethan Sanders, Alec Vest, Shaun Westlund, Matthew Hardison, Clinton R. Paden, Duncan MacCannell |
| EPI_ISL_3305466 | Douglass Hanly Moir Pathology | NSW Health Pathology - Institute of Clinical Pathology and Medical Research; Westmead Hospital; University of Sydney | Arnott A., Draper J., Gall M., Martinez E., Rockett R., Sintchenko V., on behalf of ICPMR |
| EPI_ISL_3305467 | Sydney South West Pathology Service (SSWPS) - Liverpool Hospital - NSW Health Pathology | NSW Health Pathology - Institute of Clinical Pathology and Medical Research; Westmead Hospital; University of Sydney | Arnott A., Draper J., Gall M., Martinez E., Rockett R., Sintchenko V., on behalf of ICPMR |
| EPI_ISL_3305468, EPI_ISL_3305469, EPI_ISL_3305470 | Histopath | NSW Health Pathology - Institute of Clinical Pathology and Medical Research; Westmead Hospital; University of Sydney | Arnott A., Draper J., Gall M., Martinez E., Rockett R., Sintchenko V., on behalf of ICPMR |
| EPI_ISL_3305471 | Sydney South West Pathology Service (SSWPS) - Liverpool Hospital - NSW Health Pathology | NSW Health Pathology - Institute of Clinical Pathology and Medical Research; Westmead Hospital; University of Sydney | Arnott A., Draper J., Gall M., Martinez E., Rockett R., Sintchenko V., on behalf of ICPMR |
| EPI_ISL_3305472 | Medlab Pathology | NSW Health Pathology - Institute of Clinical Pathology and Medical Research; Westmead Hospital; University of Sydney | Arnott A., Draper J., Gall M., Martinez E., Rockett R., Sintchenko V., on behalf of ICPMR |
| EPI_ISL_3305473 | Aegis Sciences Corporation | Centers for Disease Control and Prevention Division of Viral Diseases, Pathogen Discovery | Dakota Howard, Dhwani Batra, Peter W. Cook, Kara Moser, Adrian Paskey, Jason Caravas, Benjamin Rambo-Martin, Shatavia Morrison, Christopher Gulvick, Scott Sammons, Yvette Unoarumhi, Darlene Wagner, Matthew Schmerer, Cyndi Clark, Patrick Campbell, Rob Case, Vikramsinha Ghorpade, Holly Houdeshell, Ola Kvalvaag, Dillon Nall, Ethan Sanders, Alec Vest, Shaun Westlund, Matthew Hardison, Clinton R. Paden, Duncan MacCannell |
| EPI_ISL_3305474 | St Vincent's Pathology (SydPath) | NSW Health Pathology - Institute of Clinical Pathology and Medical Research; Westmead Hospital; University of Sydney | Arnott A., Draper J., Gall M., Martinez E., Rockett R., Sintchenko V., on behalf of ICPMR |
| EPI_ISL_3305476 | Medlab Pathology | NSW Health Pathology - Institute of Clinical Pathology and Medical Research; Westmead Hospital; University of Sydney | Arnott A., Draper J., Gall M., Martinez E., Rockett R., Sintchenko V., on behalf of ICPMR |
| EPI_ISL_3305477 | Aegis Sciences Corporation | Centers for Disease Control and Prevention Division of Viral Diseases, Pathogen Discovery | Dakota Howard, Dhwani Batra, Peter W. Cook, Kara Moser, Adrian Paskey, Jason Caravas, Benjamin Rambo-Martin, Shatavia Morrison, Christopher Gulvick, Scott Sammons, Yvette Unoarumhi, Darlene Wagner, Matthew Schmerer, Cyndi Clark, Patrick Campbell, Rob Case, Vikramsinha Ghorpade, Holly Houdeshell, Ola Kvalvaag, Dillon Nall, Ethan Sanders, Alec Vest, Shaun Westlund, Matthew Hardison, Clinton R. Paden, Duncan MacCannell |
| EPI_ISL_3305478 | Medlab Pathology | NSW Health Pathology - Institute of Clinical Pathology and Medical Research; Westmead Hospital; University of Sydney | Arnott A., Draper J., Gall M., Martinez E., Rockett R., Sintchenko V., on behalf of ICPMR |
| EPI_ISL_3305479 | Aegis Sciences Corporation | Centers for Disease Control and Prevention Division of Viral Diseases, Pathogen Discovery | Dakota Howard, Dhwani Batra, Peter W. Cook, Kara Moser, Adrian Paskey, Jason Caravas, Benjamin Rambo-Martin, Shatavia Morrison, Christopher Gulvick, Scott Sammons, Yvette Unoarumhi, Darlene Wagner, Matthew Schmerer, Cyndi Clark, Patrick Campbell, Rob Case, Vikramsinha Ghorpade, Holly Houdeshell, Ola Kvalvaag, Dillon Nall, Ethan Sanders, Alec Vest, Shaun Westlund, Matthew Hardison, Clinton R. Paden, Duncan MacCannell |
| EPI_ISL_3305480 | Medlab Pathology | NSW Health Pathology - Institute of Clinical Pathology and Medical Research; Westmead Hospital; University of Sydney | Arnott A., Draper J., Gall M., Martinez E., Rockett R., Sintchenko V., on behalf of ICPMR |
| EPI_ISL_3305481 | Aegis Sciences Corporation | Centers for Disease Control and Prevention Division of Viral Diseases, Pathogen Discovery | Dakota Howard, Dhwani Batra, Peter W. Cook, Kara Moser, Adrian Paskey, Jason Caravas, Benjamin Rambo-Martin, Shatavia Morrison, Christopher Gulvick, Scott Sammons, Yvette Unoarumhi, Darlene Wagner, Matthew Schmerer, Cyndi Clark, Patrick Campbell, Rob Case, Vikramsinha Ghorpade, Holly Houdeshell, Ola Kvalvaag, Dillon Nall, Ethan Sanders, Alec Vest, Shaun Westlund, Matthew Hardison, Clinton R. Paden, Duncan MacCannell |
| EPI_ISL_3305482 | Australian Clinical Labs (formerly Healthscope Pathology) | NSW Health Pathology - Institute of Clinical Pathology and Medical Research; Westmead Hospital; University of Sydney | Arnott A., Draper J., Gall M., Martinez E., Rockett R., Sintchenko V., on behalf of ICPMR |
| EPI_ISL_3305483 | Aegis Sciences Corporation | Centers for Disease Control and Prevention Division of Viral Diseases, Pathogen Discovery | Dakota Howard, Dhwani Batra, Peter W. Cook, Kara Moser, Adrian Paskey, Jason Caravas, Benjamin Rambo-Martin, Shatavia Morrison, Christopher Gulvick, Scott Sammons, Yvette Unoarumhi, Darlene Wagner, Matthew Schmerer, Cyndi Clark, Patrick Campbell, Rob Case, Vikramsinha Ghorpade, Holly Houdeshell, Ola Kvalvaag, Dillon Nall, Ethan Sanders, Alec Vest, Shaun Westlund, Matthew Hardison, Clinton R. Paden, Duncan MacCannell |
| EPI_ISL_3305485 | Sydney South West Pathology Service (SSWPS) - Liverpool Hospital - NSW Health Pathology | NSW Health Pathology - Institute of Clinical Pathology and Medical Research; Westmead Hospital; University of Sydney | Arnott A., Draper J., Gall M., Martinez E., Rockett R., Sintchenko V., on behalf of ICPMR |
| EPI_ISL_3305486 | Aegis Sciences Corporation | Centers for Disease Control and Prevention Division of Viral Diseases, Pathogen Discovery | Dakota Howard, Dhwani Batra, Peter W. Cook, Kara Moser, Adrian Paskey, Jason Caravas, Benjamin Rambo-Martin, Shatavia Morrison, Christopher Gulvick, Scott Sammons, Yvette Unoarumhi, Darlene Wagner, Matthew Schmerer, Cyndi Clark, Patrick Campbell, Rob Case, Vikramsinha Ghorpade, Holly Houdeshell, Ola Kvalvaag, Dillon Nall, Ethan Sanders, Alec Vest, Shaun Westlund, Matthew Hardison, Clinton R. Paden, Duncan MacCannell |
| EPI_ISL_3305487 | Sydney South West Pathology Service (SSWPS) - Liverpool Hospital - NSW Health Pathology | NSW Health Pathology - Institute of Clinical Pathology and Medical Research; Westmead Hospital; University of Sydney | Arnott A., Draper J., Gall M., Martinez E., Rockett R., Sintchenko V., on behalf of ICPMR |
| EPI_ISL_3305488 | Aegis Sciences Corporation | Centers for Disease Control and Prevention Division of Viral Diseases, Pathogen Discovery | Dakota Howard, Dhwani Batra, Peter W. Cook, Kara Moser, Adrian Paskey, Jason Caravas, Benjamin Rambo-Martin, Shatavia Morrison, Christopher Gulvick, Scott Sammons, Yvette Unoarumhi, Darlene Wagner, Matthew Schmerer, Cyndi Clark, Patrick Campbell, Rob Case, Vikramsinha Ghorpade, Holly Houdeshell, Ola Kvalvaag, Dillon Nall, Ethan Sanders, Alec Vest, Shaun Westlund, Matthew Hardison, Clinton R. Paden, Duncan MacCannell |
| EPI_ISL_3305489 | Sydney South West Pathology Service (SSWPS) - Liverpool Hospital - NSW Health Pathology | NSW Health Pathology - Institute of Clinical Pathology and Medical Research; Westmead Hospital; University of Sydney | Arnott A., Draper J., Gall M., Martinez E., Rockett R., Sintchenko V., on behalf of ICPMR |
| EPI_ISL_3305490 | Aegis Sciences Corporation | Centers for Disease Control and Prevention Division of Viral Diseases, Pathogen Discovery | Dakota Howard, Dhwani Batra, Peter W. Cook, Kara Moser, Adrian Paskey, Jason Caravas, Benjamin Rambo-Martin, Shatavia Morrison, Christopher Gulvick, Scott Sammons, Yvette Unoarumhi, Darlene Wagner, Matthew Schmerer, Cyndi Clark, Patrick Campbell, Rob Case, Vikramsinha Ghorpade, Holly Houdeshell, Ola Kvalvaag, Dillon Nall, Ethan Sanders, Alec Vest, Shaun Westlund, Matthew Hardison, Clinton R. Paden, Duncan MacCannell |
| EPI_ISL_3305491 | Laverty Pathology | NSW Health Pathology - Institute of Clinical Pathology and Medical Research; Westmead Hospital; University of Sydney | Arnott A., Draper J., Gall M., Martinez E., Rockett R., Sintchenko V., on behalf of ICPMR |
| EPI_ISL_3305492, EPI_ISL_3305493 | Aegis Sciences Corporation | Centers for Disease Control and Prevention Division of Viral Diseases, Pathogen Discovery | Dakota Howard, Dhwani Batra, Peter W. Cook, Kara Moser, Adrian Paskey, Jason Caravas, Benjamin Rambo-Martin, Shatavia Morrison, Christopher Gulvick, Scott Sammons, Yvette Unoarumhi, Darlene Wagner, Matthew Schmerer, Cyndi Clark, Patrick Campbell, Rob Case, Vikramsinha Ghorpade, Holly Houdeshell, Ola Kvalvaag, Dillon Nall, Ethan Sanders, Alec Vest, Shaun Westlund, Matthew Hardison, Clinton R. Paden, Duncan MacCannell |
| EPI_ISL_3305494 | Douglass Hanly Moir Pathology | NSW Health Pathology - Institute of Clinical Pathology and Medical Research; Westmead Hospital; University of Sydney | Arnott A., Draper J., Gall M., Martinez E., Rockett R., Sintchenko V., on behalf of ICPMR |
| EPI_ISL_3305495 | Aegis Sciences Corporation | Centers for Disease Control and Prevention Division of Viral Diseases, Pathogen Discovery | Dakota Howard, Dhwani Batra, Peter W. Cook, Kara Moser, Adrian Paskey, Jason Caravas, Benjamin Rambo-Martin, Shatavia Morrison, Christopher Gulvick, Scott Sammons, Yvette Unoarumhi, Darlene Wagner, Matthew Schmerer, Cyndi Clark, Patrick Campbell, Rob Case, Vikramsinha Ghorpade, Holly Houdeshell, Ola Kvalvaag, Dillon Nall, Ethan Sanders, Alec Vest, Shaun Westlund, Matthew Hardison, Clinton R. Paden, Duncan MacCannell |
| EPI_ISL_3305496 | Laverty Pathology | NSW Health Pathology - Institute of Clinical Pathology and Medical Research; Westmead Hospital; University of Sydney | Arnott A., Draper J., Gall M., Martinez E., Rockett R., Sintchenko V., on behalf of ICPMR |
| EPI_ISL_3305497 | Aegis Sciences Corporation | Centers for Disease Control and Prevention Division of Viral Diseases, Pathogen Discovery | Dakota Howard, Dhwani Batra, Peter W. Cook, Kara Moser, Adrian Paskey, Jason Caravas, Benjamin Rambo-Martin, Shatavia Morrison, Christopher Gulvick, Scott Sammons, Yvette Unoarumhi, Darlene Wagner, Matthew Schmerer, Cyndi Clark, Patrick Campbell, Rob Case, Vikramsinha Ghorpade, Holly Houdeshell, Ola Kvalvaag, Dillon Nall, Ethan Sanders, Alec Vest, Shaun Westlund, Matthew Hardison, Clinton R. Paden, Duncan MacCannell |
| EPI_ISL_3305498 | Laverty Pathology | NSW Health Pathology - Institute of Clinical Pathology and Medical Research; Westmead Hospital; University of Sydney | Arnott A., Draper J., Gall M., Martinez E., Rockett R., Sintchenko V., on behalf of ICPMR |
| EPI_ISL_3305500 | Australian Clinical Labs (formerly Healthscope Pathology) | NSW Health Pathology - Institute of Clinical Pathology and Medical Research; Westmead Hospital; University of Sydney | Arnott A., Draper J., Gall M., Martinez E., Rockett R., Sintchenko V., on behalf of ICPMR |
| EPI_ISL_3305501 | Aegis Sciences Corporation | Centers for Disease Control and Prevention Division of Viral | Dakota Howard, Dhwani Batra, Peter W. Cook, Kara Moser, Adrian Paskey, Jason Caravas, Benjamin Rambo-Martin, Shatavia Morrison, Christopher Gulvick, Scott Sammons, Yvette Unoarumhi, Darlene Wagner, Matthew Schmerer, Cyndi Clark, |

| | | | |
|---|---|---|---|
| | | Diseases, Pathogen Discovery | Patrick Campbell, Rob Case, Vikramsinha Ghorpade, Holly Houdeshell, Ola Kvalvaag, Dillon Nall, Ethan Sanders, Alec Vest, Shaun Westlund, Matthew Hardison, Clinton R. Paden, Duncan MacCannell |
| EPI_ISL_3305502 | Australian Clinical Labs (formerly Healthscope Pathology) | NSW Health Pathology - Institute of Clinical Pathology and Medical Research; Westmead Hospital; University of Sydney | Arnott A., Draper J., Gall M., Martinez E., Rockett R., Sintchenko V., on behalf of ICPMR |
| EPI_ISL_3305503 | Aegis Sciences Corporation | Centers for Disease Control and Prevention Division of Viral Diseases, Pathogen Discovery | Dakota Howard, Dhwani Batra, Peter W. Cook, Kara Moser, Adrian Paskey, Jason Caravas, Benjamin Rambo-Martin, Shatavia Morrison, Christopher Gulvick, Scott Sammons, Yvette Unoarumhi, Darlene Wagner, Matthew Schmerer, Cyndi Clark, Patrick Campbell, Rob Case, Vikramsinha Ghorpade, Holly Houdeshell, Ola Kvalvaag, Dillon Nall, Ethan Sanders, Alec Vest, Shaun Westlund, Matthew Hardison, Clinton R. Paden, Duncan MacCannell |
| EPI_ISL_3305504, EPI_ISL_3305505 | Medlab Pathology | NSW Health Pathology - Institute of Clinical Pathology and Medical Research; Westmead Hospital; University of Sydney | Arnott A., Draper J., Gall M., Martinez E., Rockett R., Sintchenko V., on behalf of ICPMR |
| EPI_ISL_3305506 | Aegis Sciences Corporation | Centers for Disease Control and Prevention Division of Viral Diseases, Pathogen Discovery | Dakota Howard, Dhwani Batra, Peter W. Cook, Kara Moser, Adrian Paskey, Jason Caravas, Benjamin Rambo-Martin, Shatavia Morrison, Christopher Gulvick, Scott Sammons, Yvette Unoarumhi, Darlene Wagner, Matthew Schmerer, Cyndi Clark, Patrick Campbell, Rob Case, Vikramsinha Ghorpade, Holly Houdeshell, Ola Kvalvaag, Dillon Nall, Ethan Sanders, Alec Vest, Shaun Westlund, Matthew Hardison, Clinton R. Paden, Duncan MacCannell |
| EPI_ISL_3305507 | St Vincent's Pathology (SydPath) | NSW Health Pathology - Institute of Clinical Pathology and Medical Research; Westmead Hospital; University of Sydney | Arnott A., Draper J., Gall M., Martinez E., Rockett R., Sintchenko V., on behalf of ICPMR |
| EPI_ISL_3305508 | Aegis Sciences Corporation | Centers for Disease Control and Prevention Division of Viral Diseases, Pathogen Discovery | Dakota Howard, Dhwani Batra, Peter W. Cook, Kara Moser, Adrian Paskey, Jason Caravas, Benjamin Rambo-Martin, Shatavia Morrison, Christopher Gulvick, Scott Sammons, Yvette Unoarumhi, Darlene Wagner, Matthew Schmerer, Cyndi Clark, Patrick Campbell, Rob Case, Vikramsinha Ghorpade, Holly Houdeshell, Ola Kvalvaag, Dillon Nall, Ethan Sanders, Alec Vest, Shaun Westlund, Matthew Hardison, Clinton R. Paden, Duncan MacCannell |
| EPI_ISL_3305509 | St Vincent's Pathology (SydPath) | NSW Health Pathology - Institute of Clinical Pathology and Medical Research; Westmead Hospital; University of Sydney | Arnott A., Draper J., Gall M., Martinez E., Rockett R., Sintchenko V., on behalf of ICPMR |
| EPI_ISL_3305510 | Aegis Sciences Corporation | Centers for Disease Control and Prevention Division of Viral Diseases, Pathogen Discovery | Dakota Howard, Dhwani Batra, Peter W. Cook, Kara Moser, Adrian Paskey, Jason Caravas, Benjamin Rambo-Martin, Shatavia Morrison, Christopher Gulvick, Scott Sammons, Yvette Unoarumhi, Darlene Wagner, Matthew Schmerer, Cyndi Clark, Patrick Campbell, Rob Case, Vikramsinha Ghorpade, Holly Houdeshell, Ola Kvalvaag, Dillon Nall, Ethan Sanders, Alec Vest, Shaun Westlund, Matthew Hardison, Clinton R. Paden, Duncan MacCannell |
| EPI_ISL_3305511, EPI_ISL_3305513 | St Vincent's Pathology (SydPath) | NSW Health Pathology - Institute of Clinical Pathology and Medical Research; Westmead Hospital; University of Sydney | Arnott A., Draper J., Gall M., Martinez E., Rockett R., Sintchenko V., on behalf of ICPMR |
| EPI_ISL_3305514 | Aegis Sciences Corporation | Centers for Disease Control and Prevention Division of Viral Diseases, Pathogen Discovery | Dakota Howard, Dhwani Batra, Peter W. Cook, Kara Moser, Adrian Paskey, Jason Caravas, Benjamin Rambo-Martin, Shatavia Morrison, Christopher Gulvick, Scott Sammons, Yvette Unoarumhi, Darlene Wagner, Matthew Schmerer, Cyndi Clark, Patrick Campbell, Rob Case, Vikramsinha Ghorpade, Holly Houdeshell, Ola Kvalvaag, Dillon Nall, Ethan Sanders, Alec Vest, Shaun Westlund, Matthew Hardison, Clinton R. Paden, Duncan MacCannell |
| EPI_ISL_3305515 | Histopath | NSW Health Pathology - Institute of Clinical Pathology and Medical Research; Westmead Hospital; University of Sydney | Arnott A., Draper J., Gall M., Martinez E., Rockett R., Sintchenko V., on behalf of ICPMR |
| EPI_ISL_3305516 | Aegis Sciences Corporation | Centers for Disease Control and Prevention Division of Viral Diseases, Pathogen Discovery | Dakota Howard, Dhwani Batra, Peter W. Cook, Kara Moser, Adrian Paskey, Jason Caravas, Benjamin Rambo-Martin, Shatavia Morrison, Christopher Gulvick, Scott Sammons, Yvette Unoarumhi, Darlene Wagner, Matthew Schmerer, Cyndi Clark, Patrick Campbell, Rob Case, Vikramsinha Ghorpade, Holly Houdeshell, Ola Kvalvaag, Dillon Nall, Ethan Sanders, Alec Vest, Shaun Westlund, Matthew Hardison, Clinton R. Paden, Duncan MacCannell |
| EPI_ISL_3305517, EPI_ISL_3305518 | Histopath | NSW Health Pathology - Institute of Clinical Pathology and Medical Research; Westmead Hospital; University of Sydney | Arnott A., Draper J., Gall M., Martinez E., Rockett R., Sintchenko V., on behalf of ICPMR |
| EPI_ISL_3305519 | Aegis Sciences Corporation | Centers for Disease Control and Prevention Division of Viral Diseases, Pathogen Discovery | Dakota Howard, Dhwani Batra, Peter W. Cook, Kara Moser, Adrian Paskey, Jason Caravas, Benjamin Rambo-Martin, Shatavia Morrison, Christopher Gulvick, Scott Sammons, Yvette Unoarumhi, Darlene Wagner, Matthew Schmerer, Cyndi Clark, Patrick Campbell, Rob Case, Vikramsinha Ghorpade, Holly Houdeshell, Ola Kvalvaag, Dillon Nall, Ethan Sanders, Alec Vest, Shaun Westlund, Matthew Hardison, Clinton R. Paden, Duncan MacCannell |
| EPI_ISL_3305520 | Histopath | NSW Health Pathology - Institute of Clinical Pathology and Medical Research; Westmead Hospital; University of Sydney | Arnott A., Draper J., Gall M., Martinez E., Rockett R., Sintchenko V., on behalf of ICPMR |
| EPI_ISL_3305521 | Aegis Sciences Corporation | Centers for Disease Control and Prevention Division of Viral Diseases, Pathogen Discovery | Dakota Howard, Dhwani Batra, Peter W. Cook, Kara Moser, Adrian Paskey, Jason Caravas, Benjamin Rambo-Martin, Shatavia Morrison, Christopher Gulvick, Scott Sammons, Yvette Unoarumhi, Darlene Wagner, Matthew Schmerer, Cyndi Clark, Patrick Campbell, Rob Case, Vikramsinha Ghorpade, Holly Houdeshell, Ola Kvalvaag, Dillon Nall, Ethan Sanders, Alec Vest, Shaun Westlund, Matthew Hardison, Clinton R. Paden, Duncan MacCannell |
| EPI_ISL_3305522 | Histopath | NSW Health Pathology - Institute of Clinical Pathology and Medical Research; Westmead Hospital; University of Sydney | Arnott A., Draper J., Gall M., Martinez E., Rockett R., Sintchenko V., on behalf of ICPMR |
| EPI_ISL_3305523 | Aegis Sciences Corporation | Centers for Disease Control and Prevention Division of Viral Diseases, Pathogen Discovery | Dakota Howard, Dhwani Batra, Peter W. Cook, Kara Moser, Adrian Paskey, Jason Caravas, Benjamin Rambo-Martin, Shatavia Morrison, Christopher Gulvick, Scott Sammons, Yvette Unoarumhi, Darlene Wagner, Matthew Schmerer, Cyndi Clark, Patrick Campbell, Rob Case, Vikramsinha Ghorpade, Holly Houdeshell, Ola Kvalvaag, Dillon Nall, Ethan Sanders, Alec Vest, Shaun Westlund, Matthew Hardison, Clinton R. Paden, Duncan MacCannell |
| EPI_ISL_3305524 | Histopath | NSW Health Pathology - Institute of Clinical Pathology and Medical Research; Westmead Hospital; University of Sydney | Arnott A., Draper J., Gall M., Martinez E., Rockett R., Sintchenko V., on behalf of ICPMR |
| EPI_ISL_3305525 | Aegis Sciences Corporation | Centers for Disease Control and Prevention Division of Viral Diseases, Pathogen Discovery | Dakota Howard, Dhwani Batra, Peter W. Cook, Kara Moser, Adrian Paskey, Jason Caravas, Benjamin Rambo-Martin, Shatavia Morrison, Christopher Gulvick, Scott Sammons, Yvette Unoarumhi, Darlene Wagner, Matthew Schmerer, Cyndi Clark, Patrick Campbell, Rob Case, Vikramsinha Ghorpade, Holly Houdeshell, Ola Kvalvaag, Dillon Nall, Ethan Sanders, Alec Vest, Shaun Westlund, Matthew Hardison, Clinton R. Paden, Duncan MacCannell |
| EPI_ISL_3305526 | Histopath | NSW Health Pathology - Institute of Clinical Pathology and Medical Research; Westmead Hospital; University of Sydney | Arnott A., Draper J., Gall M., Martinez E., Rockett R., Sintchenko V., on behalf of ICPMR |
| EPI_ISL_3305527 | Aegis Sciences Corporation | Centers for Disease Control and Prevention Division of Viral Diseases, Pathogen Discovery | Dakota Howard, Dhwani Batra, Peter W. Cook, Kara Moser, Adrian Paskey, Jason Caravas, Benjamin Rambo-Martin, Shatavia Morrison, Christopher Gulvick, Scott Sammons, Yvette Unoarumhi, Darlene Wagner, Matthew Schmerer, Cyndi Clark, Patrick Campbell, Rob Case, Vikramsinha Ghorpade, Holly Houdeshell, Ola Kvalvaag, Dillon Nall, Ethan Sanders, Alec Vest, Shaun Westlund, Matthew Hardison, Clinton R. Paden, Duncan MacCannell |
| EPI_ISL_3305528, EPI_ISL_3305529 | Histopath | NSW Health Pathology - Institute of Clinical Pathology and Medical Research; Westmead Hospital; University of Sydney | Arnott A., Draper J., Gall M., Martinez E., Rockett R., Sintchenko V., on behalf of ICPMR |
| EPI_ISL_3305530 | Aegis Sciences Corporation | Centers for Disease Control and Prevention Division of Viral Diseases, Pathogen Discovery | Dakota Howard, Dhwani Batra, Peter W. Cook, Kara Moser, Adrian Paskey, Jason Caravas, Benjamin Rambo-Martin, Shatavia Morrison, Christopher Gulvick, Scott Sammons, Yvette Unoarumhi, Darlene Wagner, Matthew Schmerer, Cyndi Clark, Patrick Campbell, Rob Case, Vikramsinha Ghorpade, Holly Houdeshell, Ola Kvalvaag, Dillon Nall, Ethan Sanders, Alec Vest, Shaun Westlund, Matthew Hardison, Clinton R. Paden, Duncan MacCannell |
| EPI_ISL_3305531, EPI_ISL_3305533, EPI_ISL_3305535 | Histopath | NSW Health Pathology - Institute of Clinical Pathology and Medical Research; Westmead Hospital; University of Sydney | Arnott A., Draper J., Gall M., Martinez E., Rockett R., Sintchenko V., on behalf of ICPMR |
| EPI_ISL_3305536 | Aegis Sciences Corporation | Centers for Disease Control and Prevention Division of Viral Diseases, Pathogen Discovery | Dakota Howard, Dhwani Batra, Peter W. Cook, Kara Moser, Adrian Paskey, Jason Caravas, Benjamin Rambo-Martin, Shatavia Morrison, Christopher Gulvick, Scott Sammons, Yvette Unoarumhi, Darlene Wagner, Matthew Schmerer, Cyndi Clark, Patrick Campbell, Rob Case, Vikramsinha Ghorpade, Holly Houdeshell, Ola Kvalvaag, Dillon Nall, Ethan Sanders, Alec Vest, Shaun Westlund, Matthew Hardison, Clinton R. Paden, Duncan MacCannell |
| EPI_ISL_3305537 | Histopath | NSW Health Pathology - Institute of Clinical Pathology and Medical Research; Westmead Hospital; University of Sydney | Arnott A., Draper J., Gall M., Martinez E., Rockett R., Sintchenko V., on behalf of ICPMR |
| EPI_ISL_3305538 | Aegis Sciences Corporation | Centers for Disease Control and Prevention Division of Viral Diseases, Pathogen Discovery | Dakota Howard, Dhwani Batra, Peter W. Cook, Kara Moser, Adrian Paskey, Jason Caravas, Benjamin Rambo-Martin, Shatavia Morrison, Christopher Gulvick, Scott Sammons, Yvette Unoarumhi, Darlene Wagner, Matthew Schmerer, Cyndi Clark, Patrick Campbell, Rob Case, Vikramsinha Ghorpade, Holly Houdeshell, Ola Kvalvaag, Dillon Nall, Ethan Sanders, Alec Vest, Shaun Westlund, Matthew Hardison, Clinton R. Paden, Duncan MacCannell |
| EPI_ISL_3305539 | Histopath | NSW Health Pathology - Institute of Clinical Pathology and Medical Research; Westmead Hospital; University of Sydney | Arnott A., Draper J., Gall M., Martinez E., Rockett R., Sintchenko V., on behalf of ICPMR |
| EPI_ISL_3305540 | Aegis Sciences Corporation | Centers for Disease Control and Prevention Division of Viral Diseases, Pathogen Discovery | Dakota Howard, Dhwani Batra, Peter W. Cook, Kara Moser, Adrian Paskey, Jason Caravas, Benjamin Rambo-Martin, Shatavia Morrison, Christopher Gulvick, Scott Sammons, Yvette Unoarumhi, Darlene Wagner, Matthew Schmerer, Cyndi Clark, Patrick Campbell, Rob Case, Vikramsinha Ghorpade, Holly Houdeshell, Ola Kvalvaag, Dillon Nall, Ethan Sanders, Alec Vest, Shaun Westlund, Matthew Hardison, Clinton R. Paden, Duncan MacCannell |
| EPI_ISL_3305541, EPI_ISL_3305542 | Histopath | NSW Health Pathology - Institute of Clinical Pathology and Medical Research; Westmead Hospital; University of Sydney | Arnott A., Draper J., Gall M., Martinez E., Rockett R., Sintchenko V., on behalf of ICPMR |
| EPI_ISL_3305543, EPI_ISL_3305544, EPI_ISL_3305545, EPI_ISL_3305546, EPI_ISL_3305547, EPI_ISL_3305548, EPI_ISL_3305549, EPI_ISL_3305550, EPI_ISL_3305551, EPI_ISL_3305552, EPI_ISL_3305555, EPI_ISL_3305556, EPI_ISL_3305557, EPI_ISL_3305558, EPI_ISL_3305559, EPI_ISL_3305560, EPI_ISL_3305561, EPI_ISL_3305562, EPI_ISL_3305563, EPI_ISL_3305564, EPI_ISL_3305565, EPI_ISL_3305566, EPI_ISL_3305568, EPI_ISL_3305569, EPI_ISL_3305571, EPI_ISL_3305572, EPI_ISL_3305573, EPI_ISL_3305574, EPI_ISL_3305575, EPI_ISL_3305576, EPI_ISL_3305577, EPI_ISL_3305580, EPI_ISL_3305581, EPI_ISL_3305582, EPI_ISL_3305583, EPI_ISL_3305584, EPI_ISL_3305585 | | | |
| see above | Aegis Sciences Corporation | Centers for Disease Control and Prevention Division of Viral Diseases, Pathogen Discovery | Dakota Howard, Dhwani Batra, Peter W. Cook, Kara Moser, Adrian Paskey, Jason Caravas, Benjamin Rambo-Martin, Shatavia Morrison, Christopher Gulvick, Scott Sammons, Yvette Unoarumhi, Darlene Wagner, Matthew Schmerer, Cyndi Clark, Patrick Campbell, Rob Case, Vikramsinha Ghorpade, Holly Houdeshell, Ola Kvalvaag, Dillon Nall, Ethan Sanders, Alec Vest, Shaun Westlund, Matthew Hardison, Clinton R. Paden, Duncan MacCannell |
| EPI_ISL_3305599, EPI_ISL_3305606, EPI_ISL_3305610, EPI_ISL_3305634, EPI_ISL_3305636, EPI_ISL_3305637, EPI_ISL_3305638, EPI_ISL_3305639, EPI_ISL_3305640, EPI_ISL_3305643, EPI_ISL_3305645, EPI_ISL_3305646, EPI_ISL_3305649, EPI_ISL_3305651, EPI_ISL_3305654, EPI_ISL_3305655, EPI_ISL_3305658, EPI_ISL_3305659, EPI_ISL_3305661, EPI_ISL_3305665, EPI_ISL_3305668, EPI_ISL_3305669, EPI_ISL_3305670, EPI_ISL_3305671, EPI_ISL_3305672, EPI_ISL_3305673, EPI_ISL_3305677, EPI_ISL_3305679, EPI_ISL_3305680, EPI_ISL_3305683, EPI_ISL_3305685, EPI_ISL_3305686, EPI_ISL_3305689, EPI_ISL_3305693, EPI_ISL_3305698, EPI_ISL_3305699, EPI_ISL_3305700, EPI_ISL_3305702, EPI_ISL_3305705, EPI_ISL_3305706, EPI_ISL_3305708, EPI_ISL_3305711, EPI_ISL_3305712, EPI_ISL_3305715, EPI_ISL_3305717, EPI_ISL_3305718, EPI_ISL_3305720, EPI_ISL_3305721, EPI_ISL_3305722, EPI_ISL_3305723, EPI_ISL_3305725, EPI_ISL_3305726, EPI_ISL_3305728, EPI_ISL_3305729, EPI_ISL_3305730, EPI_ISL_3305731, EPI_ISL_3305733, EPI_ISL_3305734, EPI_ISL_3305735, EPI_ISL_3305736, EPI_ISL_3305737, EPI_ISL_3305739, EPI_ISL_3305741 | | | |
| see above | Fulgent Genetics | Centers for Disease Control and Prevention Division of Viral Diseases, Pathogen Discovery | Dakota Howard, Dhwani Batra, Peter W. Cook, Kara Moser, Adrian Paskey, Jason Caravas, Benjamin Rambo-Martin, Shatavia Morrison, Christopher Gulvick, Scott Sammons, Yvette Unoarumhi, Darlene Wagner, Matthew Schmerer, Harry Gao, Mickey Li, John Gao, Joseph Fierro, Benafsh Sapra, Becky Tsai, Yan Meng, Doreen Ng, James Xie, Clinton R. Paden, Duncan MacCannell |
| EPI_ISL_3305745, EPI_ISL_3305746, EPI_ISL_3305749, EPI_ISL_3305750, EPI_ISL_3305751, EPI_ISL_3305752, EPI_ISL_3305755, EPI_ISL_3305758 | Infinity Biologix | Centers for Disease Control and Prevention Division of Viral Diseases, Pathogen Discovery | Dakota Howard, Dhwani Batra, Peter W. Cook, Kara Moser, Adrian Paskey, Jason Caravas, Benjamin Rambo-Martin, Shatavia Morrison, Christopher Gulvick, Scott Sammons, Yvette Unoarumhi, Darlene Wagner, Matthew Schmerer, Christian Bixby, Yihe Wang, Jonathan Schultz, Chirayu Goswami, Russ Hager, Robin Grimwood, Clinton R. Paden, Duncan MacCannell |
| EPI_ISL_3305767, EPI_ISL_3305768 | IL Dept. of Public Health Springfield Laboratory | Centers for Disease Control and Prevention Division of Viral Diseases, Pathogen Discovery | Mili Sheth, Sarah Nobles, Jasmine Padilla, Alex Burgin, Meghan Bentz, Mark Burroughs, Peter Cook, Clinton R. Paden, Dhwani Batra, Krista Queen, Kristen Knipe, Dakota Howard, Yvette Unoarumhi, Matthew Schmerer, Ben L. Rambo-Martin, Kristine Lacek, Sam Shepard, Dave Wentworth, Vivien Dugan, Suxiang Tong, Justin Lee |
| EPI_ISL_3305770 | MI - Michigan Department of Health and Human Services - Bureau of Laboratories | Centers for Disease Control and Prevention Division of Viral Diseases, Pathogen Discovery | Mili Sheth, Sarah Nobles, Jasmine Padilla, Alex Burgin, Meghan Bentz, Mark Burroughs, Peter Cook, Clinton R. Paden, Dhwani Batra, Krista Queen, Kristen Knipe, Dakota Howard, Yvette Unoarumhi, Matthew Schmerer, Ben L. Rambo-Martin, Kristine Lacek, Sam Shepard, Dave Wentworth, Vivien Dugan, Suxiang Tong, Justin Lee |
| EPI_ISL_3305774, EPI_ISL_3305775, EPI_ISL_3305776 | SD Public Health Laboratory | Centers for Disease Control and Prevention Division of Viral Diseases, Pathogen Discovery | Mili Sheth, Sarah Nobles, Jasmine Padilla, Alex Burgin, Meghan Bentz, Mark Burroughs, Peter Cook, Clinton R. Paden, Dhwani Batra, Krista Queen, Kristen Knipe, Dakota Howard, Yvette Unoarumhi, Matthew Schmerer, Ben L. Rambo-Martin, Kristine Lacek, Sam Shepard, Dave Wentworth, Vivien Dugan, Suxiang Tong, Justin Lee |
| EPI_ISL_3305779 | ND Dept. of Health Laboratory Services-Microbiology | Centers for Disease Control and Prevention Division of Viral Diseases, Pathogen Discovery | Mili Sheth, Sarah Nobles, Jasmine Padilla, Alex Burgin, Meghan Bentz, Mark Burroughs, Peter Cook, Clinton R. Paden, Dhwani Batra, Krista Queen, Kristen Knipe, Dakota Howard, Yvette Unoarumhi, Matthew Schmerer, Ben L. Rambo-Martin, Kristine Lacek, Sam Shepard, Dave Wentworth, Vivien Dugan, Suxiang Tong, Justin Lee |
| EPI_ISL_3305783 | RI State Health Laboratories | Centers for Disease Control and Prevention Division of Viral Diseases, Pathogen Discovery | Mili Sheth, Sarah Nobles, Jasmine Padilla, Alex Burgin, Meghan Bentz, Mark Burroughs, Peter Cook, Clinton R. Paden, Dhwani Batra, Krista Queen, Kristen Knipe, Dakota Howard, Yvette Unoarumhi, Matthew Schmerer, Ben L. Rambo-Martin, Kristine Lacek, Sam Shepard, Dave Wentworth, Vivien Dugan, Suxiang Tong, Justin Lee |

| | | | |
|---|---|---|---|
| EPI_ISL_3305842 | Department of Acute Infectious Diseases Control and Prevention, Yunnan Provincial Center for Disease Control and Prevention | Department of Acute Infectious Diseases Control and Prevention, Yunnan Provincial Center for Disease Control and Prevention | Meiling Zhang, Duo Li, Wenpeng Gu, Xiaonan Zhao, Senquan Jia, Jienan Zhou, Zhaosheng Liu, Xiaoqing Fu |
| EPI_ISL_3305843, EPI_ISL_3305844, EPI_ISL_3305845, EPI_ISL_3305846, EPI_ISL_3305847, EPI_ISL_3305848, EPI_ISL_3305849, EPI_ISL_3305850, EPI_ISL_3305851, EPI_ISL_3305852 | Department of Acute Infectious Diseases Control and Prevention, Yunnan Provincial Center for Disease Control and Prevention | Department of Acute Infectious Diseases Control and Prevention, Yunnan Provincial Center for Disease Control and Prevention | Meiling Zhang, Jienan Zhou, Zhaosheng Liu, Xiaoqing Fu |
| EPI_ISL_3305854 | Labo Analyses Med | National Reference Center for Viruses of Respiratory Infections, Institut Pasteur, Paris | Marion Barbet, Sylvie Behillil, Méline Bizard, Angela Brisebarre, Camille Capel, Vincent Enouf, Louise Lefrançois, Frédéric Lemoine, Christophe Malabat, Corinne Maufrais, Hub de Bioinformatique et Biostatistique, Etienne Simon-Lorière, Maud Vanpeene, Sylvie Van der Werf ,GréGoire Potiron |
| EPI_ISL_3305855, EPI_ISL_3305856, EPI_ISL_3305858, EPI_ISL_3305860, EPI_ISL_3305861, EPI_ISL_3305862, EPI_ISL_3305863, EPI_ISL_3305864, EPI_ISL_3305865, EPI_ISL_3305866, EPI_ISL_3305867, EPI_ISL_3305868, EPI_ISL_3305869, EPI_ISL_3305870, EPI_ISL_3305871, EPI_ISL_3305872, EPI_ISL_3305873, EPI_ISL_3305874, EPI_ISL_3305875, EPI_ISL_3305876, EPI_ISL_3305877, EPI_ISL_3305878, EPI_ISL_3305879, EPI_ISL_3305880, EPI_ISL_3305881, EPI_ISL_3305882, EPI_ISL_3305883, EPI_ISL_3305884, EPI_ISL_3305885, EPI_ISL_3305886, EPI_ISL_3305887, EPI_ISL_3305888, EPI_ISL_3305889, EPI_ISL_3305890, EPI_ISL_3305891, EPI_ISL_3305892, EPI_ISL_3305893, EPI_ISL_3305894, EPI_ISL_3305895, EPI_ISL_3305896 | | | |
| see above | Labo Analyses Med | National Reference Center for Viruses of Respiratory Infections, Institut Pasteur, Paris | Marion Barbet, Sylvie Behillil, Méline Bizard, Angela Brisebarre, Camille Capel, Vincent Enouf, Louise Lefrançois, Frédéric Lemoine, Christophe Malabat, Corinne Maufrais, Hub de Bioinformatique et Biostatistique, Etienne Simon-Lorière, Maud Vanpeene, Sylvie Van der Werf ,S. Cocco |
| EPI_ISL_3305897, EPI_ISL_3305898, EPI_ISL_3305899, EPI_ISL_3305900, EPI_ISL_3305901, EPI_ISL_3305902, EPI_ISL_3305903, EPI_ISL_3305905, EPI_ISL_3305906, EPI_ISL_3305907, EPI_ISL_3305908, EPI_ISL_3305909, EPI_ISL_3305910, EPI_ISL_3305911, EPI_ISL_3305912, EPI_ISL_3305913, EPI_ISL_3305914, EPI_ISL_3305915, EPI_ISL_3305916, EPI_ISL_3305917, EPI_ISL_3305918, EPI_ISL_3305919, EPI_ISL_3305921, EPI_ISL_3305922, EPI_ISL_3305923, EPI_ISL_3305924, EPI_ISL_3305925, EPI_ISL_3305926, EPI_ISL_3305927, EPI_ISL_3305928, EPI_ISL_3305929, EPI_ISL_3305930, EPI_ISL_3305931, EPI_ISL_3305932, EPI_ISL_3305933, EPI_ISL_3305934, EPI_ISL_3305935, EPI_ISL_3305936, EPI_ISL_3305937 | | | |
| see above | Labo Analyses Med | National Reference Center for Viruses of Respiratory Infections, Institut Pasteur, Paris | Marion Barbet, Sylvie Behillil, Méline Bizard, Angela Brisebarre, Camille Capel, Vincent Enouf, Louise Lefrançois, Frédéric Lemoine, Christophe Malabat, Corinne Maufrais, Hub de Bioinformatique et Biostatistique, Etienne Simon-Lorière, Maud Vanpeene, Sylvie Van der Werf ,Anne Holstein |
| EPI_ISL_3305938 | Labo Analyses Med | National Reference Center for Viruses of Respiratory Infections, Institut Pasteur, Paris | Marion Barbet, Sylvie Behillil, Méline Bizard, Angela Brisebarre, Camille Capel, Vincent Enouf, Louise Lefrançois, Frédéric Lemoine, Christophe Malabat, Corinne Maufrais, Hub de Bioinformatique et Biostatistique, Etienne Simon-Lorière, Maud Vanpeene, Sylvie Van der Werf ,ValéRie Serazin |
| EPI_ISL_3305939, EPI_ISL_3305940, EPI_ISL_3305941, EPI_ISL_3305942 | Labo Analyses Med | National Reference Center for Viruses of Respiratory Infections, Institut Pasteur, Paris | Marion Barbet, Sylvie Behillil, Méline Bizard, Angela Brisebarre, Camille Capel, Vincent Enouf, Louise Lefrançois, Frédéric Lemoine, Christophe Malabat, Corinne Maufrais, Hub de Bioinformatique et Biostatistique, Etienne Simon-Lorière, Maud Vanpeene, Sylvie Van der Werf ,ValéRie Weber |
| EPI_ISL_3305943, EPI_ISL_3305944, EPI_ISL_3305945, EPI_ISL_3305946, EPI_ISL_3305947, EPI_ISL_3305948, EPI_ISL_3305949, EPI_ISL_3305950, EPI_ISL_3305951, EPI_ISL_3305952, EPI_ISL_3305953 | | | |
| see above | Labo Analyses Med | National Reference Center for Viruses of Respiratory Infections, Institut Pasteur, Paris | Marion Barbet, Sylvie Behillil, Méline Bizard, Angela Brisebarre, Camille Capel, Vincent Enouf, Louise Lefrançois, Frédéric Lemoine, Christophe Malabat, Corinne Maufrais, Hub de Bioinformatique et Biostatistique, Etienne Simon-Lorière, Maud Vanpeene, Sylvie Van der Werf ,Emmanuel Pasquier |
| EPI_ISL_3305954 | Hospital | National Reference Center for Viruses of Respiratory Infections, Institut Pasteur, Paris | Marion Barbet, Sylvie Behillil, Méline Bizard, Angela Brisebarre, Camille Capel, Vincent Enouf, Louise Lefrançois, Frédéric Lemoine, Christophe Malabat, Corinne Maufrais, Hub de Bioinformatique et Biostatistique, Etienne Simon-Lorière, Maud Vanpeene, Sylvie Van der Werf ,Anne Vachee |
| EPI_ISL_3305955, EPI_ISL_3305956, EPI_ISL_3305957, EPI_ISL_3305958, EPI_ISL_3305959, EPI_ISL_3305960, EPI_ISL_3305961, EPI_ISL_3305962, EPI_ISL_3305964, EPI_ISL_3305965, EPI_ISL_3305966 | | | |
| see above | Labo Analyses Med | National Reference Center for Viruses of Respiratory Infections, Institut Pasteur, Paris | Marion Barbet, Sylvie Behillil, Méline Bizard, Angela Brisebarre, Camille Capel, Vincent Enouf, Louise Lefrançois, Frédéric Lemoine, Christophe Malabat, Corinne Maufrais, Hub de Bioinformatique et Biostatistique, Etienne Simon-Lorière, Maud Vanpeene, Sylvie Van der Werf ,Guillaume Aubin |
| EPI_ISL_3305967, EPI_ISL_3305968, EPI_ISL_3305969, EPI_ISL_3305971, EPI_ISL_3305972, EPI_ISL_3305973, EPI_ISL_3305974, EPI_ISL_3305975, EPI_ISL_3305976, EPI_ISL_3305978, EPI_ISL_3305979, EPI_ISL_3305980, EPI_ISL_3305981, EPI_ISL_3305982, EPI_ISL_3305983, EPI_ISL_3305984, EPI_ISL_3305985, EPI_ISL_3305986, EPI_ISL_3305987, EPI_ISL_3305988, EPI_ISL_3305989, EPI_ISL_3305991 | | | |
| see above | Labo Analyses Med | National Reference Center for Viruses of Respiratory Infections, Institut Pasteur, Paris | Marion Barbet, Sylvie Behillil, Méline Bizard, Angela Brisebarre, Camille Capel, Vincent Enouf, Louise Lefrançois, Frédéric Lemoine, Christophe Malabat, Corinne Maufrais, Hub de Bioinformatique et Biostatistique, Etienne Simon-Lorière, Maud Vanpeene, Sylvie Van der Werf ,Denis Le Minous |
| EPI_ISL_3305993, EPI_ISL_3305994, EPI_ISL_3305995 | Hospital | National Reference Center for Viruses of Respiratory Infections, Institut Pasteur, Paris | Marion Barbet, Sylvie Behillil, Méline Bizard, Angela Brisebarre, Camille Capel, Vincent Enouf, Louise Lefrançois, Frédéric Lemoine, Christophe Malabat, Corinne Maufrais, Hub de Bioinformatique et Biostatistique, Etienne Simon-Lorière, Maud Vanpeene, Sylvie Van der Werf ,Pascale Martres |
| EPI_ISL_3305996, EPI_ISL_3305997, EPI_ISL_3305998 | Hospital | National Reference Center for Viruses of Respiratory Infections, Institut Pasteur, Paris | Marion Barbet, Sylvie Behillil, Méline Bizard, Angela Brisebarre, Camille Capel, Vincent Enouf, Louise Lefrançois, Frédéric Lemoine, Christophe Malabat, Corinne Maufrais, Hub de Bioinformatique et Biostatistique, Etienne Simon-Lorière, Maud Vanpeene, Sylvie Van der Werf ,Axelle Paquin |
| EPI_ISL_3305999, EPI_ISL_3306000 | Labo Analyses Med | National Reference Center for Viruses of Respiratory Infections, Institut Pasteur, Paris | Marion Barbet, Sylvie Behillil, Méline Bizard, Angela Brisebarre, Camille Capel, Vincent Enouf, Louise Lefrançois, Frédéric Lemoine, Christophe Malabat, Corinne Maufrais, Hub de Bioinformatique et Biostatistique, Etienne Simon-Lorière, Maud Vanpeene, Sylvie Van der Werf ,Caroline Malderet |
| EPI_ISL_3306001, EPI_ISL_3306002, EPI_ISL_3306003, EPI_ISL_3306005, EPI_ISL_3306006, EPI_ISL_3306007 | Hospital | National Reference Center for Viruses of Respiratory Infections, Institut Pasteur, Paris | Marion Barbet, Sylvie Behillil, Méline Bizard, Angela Brisebarre, Camille Capel, Vincent Enouf, Louise Lefrançois, Frédéric Lemoine, Christophe Malabat, Corinne Maufrais, Hub de Bioinformatique et Biostatistique, Etienne Simon-Lorière, Maud Vanpeene, Sylvie Van der Werf ,L. Courdavault |
| EPI_ISL_3306008, EPI_ISL_3306009, EPI_ISL_3306010, EPI_ISL_3306011, EPI_ISL_3306012, EPI_ISL_3306013 | Labo Analyses Med | National Reference Center for Viruses of Respiratory Infections, Institut Pasteur, Paris | Marion Barbet, Sylvie Behillil, Méline Bizard, Angela Brisebarre, Camille Capel, Vincent Enouf, Louise Lefrançois, Frédéric Lemoine, Christophe Malabat, Corinne Maufrais, Hub de Bioinformatique et Biostatistique, Etienne Simon-Lorière, Maud Vanpeene, Sylvie Van der Werf ,Arnaud Jean |
| EPI_ISL_3306014 | Hospital | National Reference Center for Viruses of Respiratory Infections, Institut Pasteur, Paris | Marion Barbet, Sylvie Behillil, Méline Bizard, Angela Brisebarre, Camille Capel, Vincent Enouf, Louise Lefrançois, Frédéric Lemoine, Christophe Malabat, Corinne Maufrais, Hub de Bioinformatique et Biostatistique, Etienne Simon-Lorière, Maud Vanpeene, Sylvie Van der Werf ,Marianne Burgard |
| EPI_ISL_3306016, EPI_ISL_3306017 | Labo Analyses Med | National Reference Center for Viruses of Respiratory Infections, Institut Pasteur, Paris | Marion Barbet, Sylvie Behillil, Méline Bizard, Angela Brisebarre, Camille Capel, Vincent Enouf, Louise Lefrançois, Frédéric Lemoine, Christophe Malabat, Corinne Maufrais, Hub de Bioinformatique et Biostatistique, Etienne Simon-Lorière, Maud Vanpeene, Sylvie Van der Werf |
| EPI_ISL_3306018, EPI_ISL_3306019, EPI_ISL_3306020, EPI_ISL_3306021 | Labo Analyses Med | National Reference Center for Viruses of Respiratory Infections, Institut Pasteur, Paris | Marion Barbet, Sylvie Behillil, Méline Bizard, Angela Brisebarre, Camille Capel, Vincent Enouf, Louise Lefrançois, Frédéric Lemoine, Christophe Malabat, Corinne Maufrais, Hub de Bioinformatique et Biostatistique, Etienne Simon-Lorière, Maud Vanpeene, Sylvie Van der Werf ,Beate Heym |
| EPI_ISL_3306022, EPI_ISL_3306023, EPI_ISL_3306024, EPI_ISL_3306025, EPI_ISL_3306026, EPI_ISL_3306027 | Labo Analyses Med | National Reference Center for Viruses of Respiratory Infections, Institut Pasteur, Paris | Marion Barbet, Sylvie Behillil, Méline Bizard, Angela Brisebarre, Camille Capel, Vincent Enouf, Louise Lefrançois, Frédéric Lemoine, Christophe Malabat, Corinne Maufrais, Hub de Bioinformatique et Biostatistique, Etienne Simon-Lorière, Maud Vanpeene, Sylvie Van der Werf ,Sylvie Robert |
| EPI_ISL_3306028, EPI_ISL_3306029, EPI_ISL_3306030, EPI_ISL_3306031, EPI_ISL_3306032, EPI_ISL_3306033, EPI_ISL_3306034, EPI_ISL_3306035, EPI_ISL_3306036, EPI_ISL_3306037, EPI_ISL_3306038, EPI_ISL_3306039, EPI_ISL_3306040, EPI_ISL_3306041, EPI_ISL_3306042, EPI_ISL_3306043, EPI_ISL_3306044, EPI_ISL_3306045 | | | |
| see above | Labo Analyses Med | National Reference Center for Viruses of Respiratory Infections, Institut Pasteur, Paris | Marion Barbet, Sylvie Behillil, Méline Bizard, Angela Brisebarre, Camille Capel, Vincent Enouf, Louise Lefrançois, Frédéric Lemoine, Christophe Malabat, Corinne Maufrais, Hub de Bioinformatique et Biostatistique, Etienne Simon-Lorière, Maud Vanpeene, Sylvie Van der Werf ,Claire Felloni |
| EPI_ISL_3306046, EPI_ISL_3306047, EPI_ISL_3306048 | Hospital | National Reference Center for Viruses of Respiratory Infections, Institut Pasteur, Paris | Marion Barbet, Sylvie Behillil, Méline Bizard, Angela Brisebarre, Camille Capel, Vincent Enouf, Louise Lefrançois, Frédéric Lemoine, Christophe Malabat, Corinne Maufrais, Hub de Bioinformatique et Biostatistique, Etienne Simon-Lorière, Maud Vanpeene, Sylvie Van der Werf ,Esther Gyde |
| EPI_ISL_3306049, EPI_ISL_3306050, EPI_ISL_3306051, EPI_ISL_3306052 | Hospital | National Reference Center for Viruses of Respiratory Infections, Institut Pasteur, Paris | Marion Barbet, Sylvie Behillil, Méline Bizard, Angela Brisebarre, Camille Capel, Vincent Enouf, Louise Lefrançois, Frédéric Lemoine, Christophe Malabat, Corinne Maufrais, Hub de Bioinformatique et Biostatistique, Etienne Simon-Lorière, Maud Vanpeene, Sylvie Van der Werf ,CéCile Farrugia |
| EPI_ISL_3306053 | Hospital | National Reference Center for Viruses of Respiratory Infections, Institut Pasteur, Paris | Marion Barbet, Sylvie Behillil, Méline Bizard, Angela Brisebarre, Camille Capel, Vincent Enouf, Louise Lefrançois, Frédéric Lemoine, Christophe Malabat, Corinne Maufrais, Hub de Bioinformatique et Biostatistique, Etienne Simon-Lorière, Maud Vanpeene, Sylvie Van der Werf ,StéPhanie Marque Juillet |
| EPI_ISL_3306054, EPI_ISL_3306055, EPI_ISL_3306056, EPI_ISL_3306057, EPI_ISL_3306058, EPI_ISL_3306059, EPI_ISL_3306060, EPI_ISL_3306061, EPI_ISL_3306062, EPI_ISL_3306063, EPI_ISL_3306065, EPI_ISL_3306066, EPI_ISL_3306067, EPI_ISL_3306068, EPI_ISL_3306070, EPI_ISL_3306071, EPI_ISL_3306072, EPI_ISL_3306074, EPI_ISL_3306075, EPI_ISL_3306076, EPI_ISL_3306077, EPI_ISL_3306078, EPI_ISL_3306083 | | | |
| see above | Labo Analyses Med | National Reference Center for Viruses of Respiratory Infections, Institut Pasteur, Paris | Marion Barbet, Sylvie Behillil, Méline Bizard, Angela Brisebarre, Camille Capel, Vincent Enouf, Louise Lefrançois, Frédéric Lemoine, Christophe Malabat, Corinne Maufrais, Hub de Bioinformatique et Biostatistique, Etienne Simon-Lorière, Maud Vanpeene, Sylvie Van der Werf ,Karine Michez |
| EPI_ISL_3306084, EPI_ISL_3306085, EPI_ISL_3306086, EPI_ISL_3306087, EPI_ISL_3306089, EPI_ISL_3306091, | Labo Analyses Med | National Reference Center for Viruses of Respiratory Infections, Institut Pasteur, Paris | Marion Barbet, Sylvie Behillil, Méline Bizard, Angela Brisebarre, Camille Capel, Vincent Enouf, Louise Lefrançois, Frédéric Lemoine, Christophe Malabat, Corinne Maufrais, Hub de Bioinformatique et Biostatistique, Etienne Simon-Lorière, Maud Vanpeene, Sylvie Van der Werf ,Nabila Belhouachi |

| | | | |
|---|---|---|---|
| EPI_ISL_3306092, EPI_ISL_3306093 | | | |
| EPI_ISL_3306104 | Labo Analyses Med | National Reference Center for Viruses of Respiratory Infections, Institut Pasteur, Paris | Marion Barbet, Sylvie Behillil, Méline Bizard, Angela Brisebarre, Camille Capel, Vincent Enouf, Louise Lefrançois, Frédéric Lemoine, Christophe Malabat, Corinne Maufrais, Hub de Bioinformatique et Biostatistique, Etienne Simon-Lorière, Maud Vanpeene, Sylvie Van der Werf ,FrançOis Nestour |
| EPI_ISL_3306118 | Hospital | National Reference Center for Viruses of Respiratory Infections, Institut Pasteur, Paris | Marion Barbet, Sylvie Behillil, Méline Bizard, Angela Brisebarre, Camille Capel, Vincent Enouf, Louise Lefrançois, Frédéric Lemoine, Christophe Malabat, Corinne Maufrais, Hub de Bioinformatique et Biostatistique, Etienne Simon-Lorière, Maud Vanpeene, Sylvie Van der Werf ,Dominique Descamps |
| EPI_ISL_3306119 | Hospital | National Reference Center for Viruses of Respiratory Infections, Institut Pasteur, Paris | Marion Barbet, Sylvie Behillil, Méline Bizard, Angela Brisebarre, Camille Capel, Vincent Enouf, Louise Lefrançois, Frédéric Lemoine, Christophe Malabat, Corinne Maufrais, Hub de Bioinformatique et Biostatistique, Etienne Simon-Lorière, Maud Vanpeene, Sylvie Van der Werf ,StéPhanie Van Agt |
| EPI_ISL_3306120, EPI_ISL_3306121, EPI_ISL_3306122, EPI_ISL_3306123, EPI_ISL_3306124, EPI_ISL_3306125, EPI_ISL_3306126, EPI_ISL_3306127, EPI_ISL_3306128, EPI_ISL_3306129, EPI_ISL_3306130, EPI_ISL_3306132, EPI_ISL_3306133, EPI_ISL_3306134, EPI_ISL_3306135, EPI_ISL_3306136, EPI_ISL_3306137, EPI_ISL_3306138, EPI_ISL_3306139, EPI_ISL_3306140, EPI_ISL_3306141, EPI_ISL_3306142, EPI_ISL_3306143, EPI_ISL_3306144, EPI_ISL_3306146, EPI_ISL_3306147, EPI_ISL_3306148, EPI_ISL_3306149, EPI_ISL_3306150, EPI_ISL_3306151, EPI_ISL_3306152, EPI_ISL_3306153, EPI_ISL_3306154, EPI_ISL_3306155, EPI_ISL_3306156, EPI_ISL_3306157, EPI_ISL_3306158, EPI_ISL_3306159, EPI_ISL_3306160, EPI_ISL_3306161, EPI_ISL_3306162, EPI_ISL_3306163, EPI_ISL_3306164, EPI_ISL_3306165, EPI_ISL_3306166, EPI_ISL_3306167, EPI_ISL_3306168, EPI_ISL_3306169, EPI_ISL_3306170, EPI_ISL_3306171, EPI_ISL_3306172, EPI_ISL_3306173, EPI_ISL_3306174, EPI_ISL_3306175, EPI_ISL_3306176, EPI_ISL_3306177, EPI_ISL_3306178, EPI_ISL_3306179, EPI_ISL_3306180, EPI_ISL_3306181 | | | |
| see above | Labo Analyses Med | National Reference Center for Viruses of Respiratory Infections, Institut Pasteur, Paris | Marion Barbet, Sylvie Behillil, Méline Bizard, Angela Brisebarre, Camille Capel, Vincent Enouf, Louise Lefrançois, Frédéric Lemoine, Christophe Malabat, Corinne Maufrais, Hub de Bioinformatique et Biostatistique, Etienne Simon-Lorière, Maud Vanpeene, Sylvie Van der Werf ,Jean-FrançOis Comes |
| EPI_ISL_3306183, EPI_ISL_3306184, EPI_ISL_3306185, EPI_ISL_3306186, EPI_ISL_3306187, EPI_ISL_3306188, EPI_ISL_3306189, EPI_ISL_3306190, EPI_ISL_3306191, EPI_ISL_3306192, EPI_ISL_3306193, EPI_ISL_3306194, EPI_ISL_3306195, EPI_ISL_3306196, EPI_ISL_3306197, EPI_ISL_3306198, EPI_ISL_3306199, EPI_ISL_3306200, EPI_ISL_3306201, EPI_ISL_3306202, EPI_ISL_3306203, EPI_ISL_3306204, EPI_ISL_3306205, EPI_ISL_3306206, EPI_ISL_3306207, EPI_ISL_3306208, EPI_ISL_3306209, EPI_ISL_3306210, EPI_ISL_3306211, EPI_ISL_3306212, EPI_ISL_3306213, EPI_ISL_3306214, EPI_ISL_3306215, EPI_ISL_3306216, EPI_ISL_3306217, EPI_ISL_3306218, EPI_ISL_3306219, EPI_ISL_3306220, EPI_ISL_3306221, EPI_ISL_3306222, EPI_ISL_3306223, EPI_ISL_3306224, EPI_ISL_3306225, EPI_ISL_3306226, EPI_ISL_3306228, EPI_ISL_3306229, EPI_ISL_3306230, EPI_ISL_3306231, EPI_ISL_3306232, EPI_ISL_3306233, EPI_ISL_3306234, EPI_ISL_3306235, EPI_ISL_3306236, EPI_ISL_3306237, EPI_ISL_3306238, EPI_ISL_3306239, EPI_ISL_3306240, EPI_ISL_3306241, EPI_ISL_3306242, EPI_ISL_3306243, EPI_ISL_3306244, EPI_ISL_3306245, EPI_ISL_3306246, EPI_ISL_3306247, EPI_ISL_3306248, EPI_ISL_3306249, EPI_ISL_3306250, EPI_ISL_3306251, EPI_ISL_3306252, EPI_ISL_3306253, EPI_ISL_3306254, EPI_ISL_3306255, EPI_ISL_3306256, EPI_ISL_3306257, EPI_ISL_3306258, EPI_ISL_3306260, EPI_ISL_3306261, EPI_ISL_3306262, EPI_ISL_3306263, EPI_ISL_3306264, EPI_ISL_3306265, EPI_ISL_3306266, EPI_ISL_3306268, EPI_ISL_3306269, EPI_ISL_3306270, EPI_ISL_3306271, EPI_ISL_3306272, EPI_ISL_3306273, EPI_ISL_3306274, EPI_ISL_3306275, EPI_ISL_3306276, EPI_ISL_3306277, EPI_ISL_3306278, EPI_ISL_3306279, EPI_ISL_3306280, EPI_ISL_3306281, EPI_ISL_3306282, EPI_ISL_3306283, EPI_ISL_3306284, EPI_ISL_3306285, EPI_ISL_3306286, EPI_ISL_3306287, EPI_ISL_3306288, EPI_ISL_3306289, EPI_ISL_3306290, EPI_ISL_3306291, EPI_ISL_3306292, EPI_ISL_3306293, EPI_ISL_3306294, EPI_ISL_3306295, EPI_ISL_3306297, EPI_ISL_3306298, EPI_ISL_3306299, EPI_ISL_3306300, EPI_ISL_3306301, EPI_ISL_3306302, EPI_ISL_3306303, EPI_ISL_3306304, EPI_ISL_3306305, EPI_ISL_3306306, EPI_ISL_3306307, EPI_ISL_3306308, EPI_ISL_3306309, EPI_ISL_3306310, EPI_ISL_3306311, EPI_ISL_3306312, EPI_ISL_3306313, EPI_ISL_3306314, EPI_ISL_3306315 | | | |
| see above | Labo Analyses Med | National Reference Center for Viruses of Respiratory Infections, Institut Pasteur, Paris | Marion Barbet, Sylvie Behillil, Méline Bizard, Angela Brisebarre, Camille Capel, Vincent Enouf, Louise Lefrançois, Frédéric Lemoine, Christophe Malabat, Corinne Maufrais, Hub de Bioinformatique et Biostatistique, Etienne Simon-Lorière, Maud Vanpeene, Sylvie Van der Werf ,OphéLie Said-Delattre |
| EPI_ISL_3306316 | Hospital | National Reference Center for Viruses of Respiratory Infections, Institut Pasteur, Paris | Marion Barbet, Sylvie Behillil, Méline Bizard, Angela Brisebarre, Camille Capel, Vincent Enouf, Louise Lefrançois, Frédéric Lemoine, Christophe Malabat, Corinne Maufrais, Hub de Bioinformatique et Biostatistique, Etienne Simon-Lorière, Maud Vanpeene, Sylvie Van der Werf ,Catherine Gaudry Graffin |
| EPI_ISL_3306317, EPI_ISL_3306318, EPI_ISL_3306319 | Hospital | National Reference Center for Viruses of Respiratory Infections, Institut Pasteur, Paris | Marion Barbet, Sylvie Behillil, Méline Bizard, Angela Brisebarre, Camille Capel, Vincent Enouf, Louise Lefrançois, Frédéric Lemoine, Christophe Malabat, Corinne Maufrais, Hub de Bioinformatique et Biostatistique, Etienne Simon-Lorière, Maud Vanpeene, Sylvie Van der Werf ,Anne Cady |
| EPI_ISL_3306320, EPI_ISL_3306321, EPI_ISL_3306322 | Labo Analyses Med | National Reference Center for Viruses of Respiratory Infections, Institut Pasteur, Paris | Marion Barbet, Sylvie Behillil, Méline Bizard, Angela Brisebarre, Camille Capel, Vincent Enouf, Louise Lefrançois, Frédéric Lemoine, Christophe Malabat, Corinne Maufrais, Hub de Bioinformatique et Biostatistique, Etienne Simon-Lorière, Maud Vanpeene, Sylvie Van der Werf ,Caroline Malderet |
| EPI_ISL_3306323, EPI_ISL_3306324, EPI_ISL_3306325, EPI_ISL_3306326, EPI_ISL_3306327, EPI_ISL_3306328, EPI_ISL_3306329, EPI_ISL_3306331 | Labo Analyses Med | National Reference Center for Viruses of Respiratory Infections, Institut Pasteur, Paris | Marion Barbet, Sylvie Behillil, Méline Bizard, Angela Brisebarre, Camille Capel, Vincent Enouf, Louise Lefrançois, Frédéric Lemoine, Christophe Malabat, Corinne Maufrais, Hub de Bioinformatique et Biostatistique, Etienne Simon-Lorière, Maud Vanpeene, Sylvie Van der Werf ,Baptiste Moreira |
| EPI_ISL_3306332, EPI_ISL_3306333 | Emergency County Hospital Targu Mures | National Institute of Infectious Diseases-Prof. Dr. Matei Bals Molecular Diagnostics Laboratory | Corina Casangiu, Leontina Banica, Marius Surleac, Ovidiu Vlaicu, Petre Milu, Robert Hohan, Simona Paraschiv, Dan Otelea |
| EPI_ISL_3306334 | Department of Public Health Gorj | National Institute of Infectious Diseases-Prof. Dr. Matei Bals Molecular Diagnostics Laboratory | Corina Casangiu, Leontina Banica, Marius Surleac, Ovidiu Vlaicu, Petre Milu, Robert Hohan, Simona Paraschiv, Dan Otelea |
| EPI_ISL_3306335, EPI_ISL_3306336, EPI_ISL_3306337, EPI_ISL_3306338, EPI_ISL_3306339, EPI_ISL_3306340, EPI_ISL_3306341, EPI_ISL_3306342, EPI_ISL_3306343, EPI_ISL_3306344, EPI_ISL_3306345, EPI_ISL_3306346, EPI_ISL_3306348, EPI_ISL_3306349, EPI_ISL_3306350, EPI_ISL_3306351, EPI_ISL_3306352, EPI_ISL_3306353, EPI_ISL_3306354, EPI_ISL_3306355, EPI_ISL_3306357, EPI_ISL_3306358, EPI_ISL_3306359, EPI_ISL_3306360, EPI_ISL_3306361, EPI_ISL_3306362, EPI_ISL_3306363, EPI_ISL_3306364, EPI_ISL_3306365, EPI_ISL_3306366, EPI_ISL_3306367, EPI_ISL_3306368, EPI_ISL_3306369, EPI_ISL_3306370, EPI_ISL_3306371, EPI_ISL_3306372, EPI_ISL_3306373, EPI_ISL_3306374, EPI_ISL_3306375, EPI_ISL_3306376, EPI_ISL_3306377, EPI_ISL_3306378, EPI_ISL_3306379, EPI_ISL_3306380, EPI_ISL_3306381, EPI_ISL_3306382, EPI_ISL_3306383, EPI_ISL_3306384, EPI_ISL_3306385, EPI_ISL_3306386, EPI_ISL_3306387, EPI_ISL_3306388, EPI_ISL_3306389, EPI_ISL_3306390 | | | |
| see above | Hospital General Universitario Gregorio Marañón | Hospital General Universitario Gregorio Marañón | Sergio Buenestado Serrano, Pedro Sola Campoy, Laura Pérez-Lago, Cristina Rodriguez-Grande, Marta Herranz Martin, Victor Manuel de la Cueva, Julia Suárez, Pilar Catalán, Patricia Muñoz, Darío García de Viedma |
| EPI_ISL_3306392 | Public Health Authority of the Slovak Republic | Laboratory of Genomics and Bioinformatics, Comenius University Science Park | Tatiana Sedláčková, Diana Rusňáková, Miroslav Böhmer, Anna Gičová, Jaroslav Budiš, Tomáš Szemes, Jakub Styk |
| EPI_ISL_3306394, EPI_ISL_3306395, EPI_ISL_3306396 | CHU Pontchaillou | CHU Pontchaillou | GROLHIER Claire, DENOUAL Florent, ETCHEVERRY Amandine, SASSI Mohamed, JAGLINE Steven, FEBREAU Christine, PRONIER Charlotte, GALIBERT Marie Dominique, DE TAYRAC Marie, THIBAULT Vincent |
| EPI_ISL_3306397, EPI_ISL_3306398, EPI_ISL_3306399, EPI_ISL_3306400, EPI_ISL_3306401, EPI_ISL_3306402, EPI_ISL_3306403, EPI_ISL_3306404, EPI_ISL_3306405, EPI_ISL_3306406, EPI_ISL_3306407, EPI_ISL_3306408, EPI_ISL_3306409, EPI_ISL_3306410, EPI_ISL_3306411, EPI_ISL_3306412, EPI_ISL_3306413, EPI_ISL_3306414, EPI_ISL_3306415, EPI_ISL_3306416, EPI_ISL_3306417, EPI_ISL_3306418, EPI_ISL_3306419, EPI_ISL_3306420, EPI_ISL_3306421, EPI_ISL_3306422, EPI_ISL_3306423, EPI_ISL_3306424, EPI_ISL_3306426, EPI_ISL_3306427, EPI_ISL_3306428, EPI_ISL_3306429, EPI_ISL_3306430, EPI_ISL_3306431, EPI_ISL_3306432, EPI_ISL_3306433, EPI_ISL_3306434, EPI_ISL_3306435, EPI_ISL_3306436, EPI_ISL_3306437, EPI_ISL_3306438, EPI_ISL_3306439, EPI_ISL_3306440, EPI_ISL_3306441, EPI_ISL_3306442, EPI_ISL_3306443, EPI_ISL_3306444, EPI_ISL_3306445, EPI_ISL_3306446, EPI_ISL_3306447, EPI_ISL_3306448, EPI_ISL_3306449, EPI_ISL_3306450, EPI_ISL_3306451, EPI_ISL_3306452, EPI_ISL_3306453, EPI_ISL_3306454, EPI_ISL_3306455, EPI_ISL_3306456, EPI_ISL_3306457, EPI_ISL_3306458, EPI_ISL_3306459, EPI_ISL_3306460, EPI_ISL_3306461, EPI_ISL_3306462, EPI_ISL_3306463, EPI_ISL_3306464, EPI_ISL_3306465, EPI_ISL_3306466, EPI_ISL_3306467 | | | |
| see above | Hospital Universitari Vall d'Hebron - Vall d'Hebron Institut de Recerca | Hospital Universitari Vall d'Hebron - Vall d'Hebron Institut de Recerca | Cristina Andrés, Maria Piñana, Alejandra González-Sánchez, Damir Garcia-Cehic, Ariadna Rando, Juliana Esperalba, Maria Gema Codina, Carla Castillo, Karen García, Rodrigo Vásquez, Maria Carmen Martin, Tomàs Pumarola, Josep Quer, Andrés Antón |
| EPI_ISL_3306468, EPI_ISL_3306469, EPI_ISL_3306470, EPI_ISL_3306471, EPI_ISL_3306472, EPI_ISL_3306473 | Hospital Universitari Arnau de Vilanova | Hospital Universitari Vall d'Hebron - Vall d'Hebron Institut de Recerca | Cristina Andrés, Maria Piñana, Alejandra González-Sánchez, Damir Garcia-Cehic, Ariadna Rando, Juliana Esperalba, Maria Gema Codina, Carla Castillo, Karen García, Rodrigo Vásquez, Maria Carmen Martin, Tomàs Pumarola, Josep Quer, Andrés Antón |
| EPI_ISL_3306474, EPI_ISL_3306475, EPI_ISL_3306476, EPI_ISL_3306477, EPI_ISL_3306478, EPI_ISL_3306479, EPI_ISL_3306480, EPI_ISL_3306481, EPI_ISL_3306482, EPI_ISL_3306483, EPI_ISL_3306484, EPI_ISL_3306485, EPI_ISL_3306487, EPI_ISL_3306488, EPI_ISL_3306489, EPI_ISL_3306490, EPI_ISL_3306491, EPI_ISL_3306492, EPI_ISL_3306493, EPI_ISL_3306494, EPI_ISL_3306495, EPI_ISL_3306496, EPI_ISL_3306497, EPI_ISL_3306498, EPI_ISL_3306499, EPI_ISL_3306500, EPI_ISL_3306501, EPI_ISL_3306502, EPI_ISL_3306503, EPI_ISL_3306504, EPI_ISL_3306505, EPI_ISL_3306507, EPI_ISL_3306508, EPI_ISL_3306509, EPI_ISL_3306510, EPI_ISL_3306511, EPI_ISL_3306512, EPI_ISL_3306513, EPI_ISL_3306514, EPI_ISL_3306516, EPI_ISL_3306517, EPI_ISL_3306518, EPI_ISL_3306519, EPI_ISL_3306520, EPI_ISL_3306521, EPI_ISL_3306522, EPI_ISL_3306523, EPI_ISL_3306524, EPI_ISL_3306525, EPI_ISL_3306526, EPI_ISL_3306527, EPI_ISL_3306528, EPI_ISL_3306529, EPI_ISL_3306530, EPI_ISL_3306531, EPI_ISL_3306533, EPI_ISL_3306534, EPI_ISL_3306535, EPI_ISL_3306536, EPI_ISL_3306537, EPI_ISL_3306538, EPI_ISL_3306539, EPI_ISL_3306540, EPI_ISL_3306541, EPI_ISL_3306542, EPI_ISL_3306543, EPI_ISL_3306544, EPI_ISL_3306545, EPI_ISL_3306546, EPI_ISL_3306547, EPI_ISL_3306548, EPI_ISL_3306549, EPI_ISL_3306550, EPI_ISL_3306551, EPI_ISL_3306552, EPI_ISL_3306553, EPI_ISL_3306554, EPI_ISL_3306555, EPI_ISL_3306556, EPI_ISL_3306557, EPI_ISL_3306558, EPI_ISL_3306559, EPI_ISL_3306560, EPI_ISL_3306561, EPI_ISL_3306562, EPI_ISL_3306563, EPI_ISL_3306564, EPI_ISL_3306565, EPI_ISL_3306566, EPI_ISL_3306567, EPI_ISL_3306568, EPI_ISL_3306569, EPI_ISL_3306570, EPI_ISL_3306571, EPI_ISL_3306572, EPI_ISL_3306573, EPI_ISL_3306574, EPI_ISL_3306575, EPI_ISL_3306576, EPI_ISL_3306577, EPI_ISL_3306578, EPI_ISL_3306579, EPI_ISL_3306580, EPI_ISL_3306581, EPI_ISL_3306582, EPI_ISL_3306583, EPI_ISL_3306584, EPI_ISL_3306585, EPI_ISL_3306586, EPI_ISL_3306587, EPI_ISL_3306588, EPI_ISL_3306589, EPI_ISL_3306590, EPI_ISL_3306591, EPI_ISL_3306592, EPI_ISL_3306593, EPI_ISL_3306594, EPI_ISL_3306595, EPI_ISL_3306596, EPI_ISL_3306597, EPI_ISL_3306598, EPI_ISL_3306599, EPI_ISL_3306600, EPI_ISL_3306601, EPI_ISL_3306602, EPI_ISL_3306603, EPI_ISL_3306604, EPI_ISL_3306605, EPI_ISL_3306606, EPI_ISL_3306607, EPI_ISL_3306608, EPI_ISL_3306609, EPI_ISL_3306610, EPI_ISL_3306611, EPI_ISL_3306612, EPI_ISL_3306613, EPI_ISL_3306614, EPI_ISL_3306615, EPI_ISL_3306616, EPI_ISL_3306617, EPI_ISL_3306618, EPI_ISL_3306619, EPI_ISL_3306620, EPI_ISL_3306621, EPI_ISL_3306622, EPI_ISL_3306623, EPI_ISL_3306624, EPI_ISL_3306625, EPI_ISL_3306626, EPI_ISL_3306627, EPI_ISL_3306628, EPI_ISL_3306629, EPI_ISL_3306630, EPI_ISL_3306631, EPI_ISL_3306632, EPI_ISL_3306633, EPI_ISL_3306634, EPI_ISL_3306635, EPI_ISL_3306636, EPI_ISL_3306637, EPI_ISL_3306638, EPI_ISL_3306639, EPI_ISL_3306640, EPI_ISL_3306641, EPI_ISL_3306642, EPI_ISL_3306643, EPI_ISL_3306644, EPI_ISL_3306645, EPI_ISL_3306646, EPI_ISL_3306647, EPI_ISL_3306648, EPI_ISL_3306649, EPI_ISL_3306650, EPI_ISL_3306651, EPI_ISL_3306652, EPI_ISL_3306653, EPI_ISL_3306654, EPI_ISL_3306655, EPI_ISL_3306656, EPI_ISL_3306657, EPI_ISL_3306658, EPI_ISL_3306659, EPI_ISL_3306660, EPI_ISL_3306661, EPI_ISL_3306662, EPI_ISL_3306663, EPI_ISL_3306664, EPI_ISL_3306665, EPI_ISL_3306666, EPI_ISL_3306667, EPI_ISL_3306668, EPI_ISL_3306669, EPI_ISL_3306670, EPI_ISL_3306671, EPI_ISL_3306672, EPI_ISL_3306673, EPI_ISL_3306674, EPI_ISL_3306675, EPI_ISL_3306676, EPI_ISL_3306677, EPI_ISL_3306678, EPI_ISL_3306680, EPI_ISL_3306681, EPI_ISL_3306682, EPI_ISL_3306683, EPI_ISL_3306684, EPI_ISL_3306685, EPI_ISL_3306686, EPI_ISL_3306687, EPI_ISL_3306688, EPI_ISL_3306689, EPI_ISL_3306690, EPI_ISL_3306691, EPI_ISL_3306692, EPI_ISL_3306693, EPI_ISL_3306694, EPI_ISL_3306695, EPI_ISL_3306696, EPI_ISL_3306697, EPI_ISL_3306698, EPI_ISL_3306699, EPI_ISL_3306701, EPI_ISL_3306702, EPI_ISL_3306703, EPI_ISL_3306704, EPI_ISL_3306705, EPI_ISL_3306706, EPI_ISL_3306707, EPI_ISL_3306708, EPI_ISL_3306709, EPI_ISL_3306710, EPI_ISL_3306711, EPI_ISL_3306712, EPI_ISL_3306713, EPI_ISL_3306714, EPI_ISL_3306715, EPI_ISL_3306716, EPI_ISL_3306717, EPI_ISL_3306718, EPI_ISL_3306719, EPI_ISL_3306720, EPI_ISL_3306721, EPI_ISL_3306722, EPI_ISL_3306723, EPI_ISL_3306724, EPI_ISL_3306726, EPI_ISL_3306727, EPI_ISL_3306728, EPI_ISL_3306729, EPI_ISL_3306731, EPI_ISL_3306732, EPI_ISL_3306733, EPI_ISL_3306734, EPI_ISL_3306735, EPI_ISL_3306736, EPI_ISL_3306737, EPI_ISL_3306738, EPI_ISL_3306739, EPI_ISL_3306740, EPI_ISL_3306741, EPI_ISL_3306742, EPI_ISL_3306743, EPI_ISL_3306744, EPI_ISL_3306745, EPI_ISL_3306746, EPI_ISL_3306747, EPI_ISL_3306748, EPI_ISL_3306749, EPI_ISL_3306750, EPI_ISL_3306751, EPI_ISL_3306752, EPI_ISL_3306753, EPI_ISL_3306754, EPI_ISL_3306755, EPI_ISL_3306756, EPI_ISL_3306757, EPI_ISL_3306758, EPI_ISL_3306759, EPI_ISL_3306760, EPI_ISL_3306761, EPI_ISL_3306762, EPI_ISL_3306763, EPI_ISL_3306764, EPI_ISL_3306765, EPI_ISL_3306766, EPI_ISL_3306767, EPI_ISL_3306768, EPI_ISL_3306769, EPI_ISL_3306770, EPI_ISL_3306771, EPI_ISL_3306772, EPI_ISL_3306773, EPI_ISL_3306774, EPI_ISL_3306775, EPI_ISL_3306776, EPI_ISL_3306777, EPI_ISL_3306778, EPI_ISL_3306779, EPI_ISL_3306780, EPI_ISL_3306781, EPI_ISL_3306782, EPI_ISL_3306783, EPI_ISL_3306784, EPI_ISL_3306785, EPI_ISL_3306786, EPI_ISL_3306787, EPI_ISL_3306788, EPI_ISL_3306789, EPI_ISL_3306790, EPI_ISL_3306791, EPI_ISL_3306792, EPI_ISL_3306793, EPI_ISL_3306794, EPI_ISL_3306795, EPI_ISL_3306796, EPI_ISL_3306797, EPI_ISL_3306798, EPI_ISL_3306800, EPI_ISL_3306801, EPI_ISL_3306802, EPI_ISL_3306803, EPI_ISL_3306804, EPI_ISL_3306805, EPI_ISL_3306806, EPI_ISL_3306807, EPI_ISL_3306808, EPI_ISL_3306809, EPI_ISL_3306810, EPI_ISL_3306811, EPI_ISL_3306812, EPI_ISL_3306813, EPI_ISL_3306814, EPI_ISL_3306815, EPI_ISL_3306816, EPI_ISL_3306817, EPI_ISL_3306818, EPI_ISL_3306819, EPI_ISL_3306820, EPI_ISL_3306822, EPI_ISL_3306823, EPI_ISL_3306824, EPI_ISL_3306825, EPI_ISL_3306826, EPI_ISL_3306827, EPI_ISL_3306828, EPI_ISL_3306829, EPI_ISL_3306830 | | | |
| see above | Ministry of Health Turkey | Ministry of Health Turkey | Fatma Bayrakdar, Yasemin Coşgun, Süleyman Yalcin, Gülay Korukluoğlu |
| EPI_ISL_3306832, EPI_ISL_3306833, EPI_ISL_3306834 | Public Health Authority of the Slovak Republic | Laboratory of Genomics and Bioinformatics, Comenius University Science Park | Tatiana Sedláčková, Diana Rusňáková, Miroslav Böhmer, Anna Gičová, Jaroslav Budiš, Tomáš Szemes, Jakub Styk |
| EPI_ISL_3306836, EPI_ISL_3306837, EPI_ISL_3306838, EPI_ISL_3306839, EPI_ISL_3306841, EPI_ISL_3306842, EPI_ISL_3306843, EPI_ISL_3306844, EPI_ISL_3306845, EPI_ISL_3306846, EPI_ISL_3306847, EPI_ISL_3306848, EPI_ISL_3306849 | | | |
| see above | Tulare County Public Health Laboratory | Tulare County Public Health Laboratory | Denise L Lopez, Joshua Kootstra |
| EPI_ISL_3306862, EPI_ISL_3306863, EPI_ISL_3306864, EPI_ISL_3306865, EPI_ISL_3306867 | Department of Virology, Istituto Zooprofilattico Sperimentale del Lazio e della Toscana (IZSLT) | Department of General Diagnostics; Department of Virology; Istituto Zooprofilattico Sperimentale del Lazio e della Toscana (IZSLT) | Patricia Alba, Giuseppe Manna, Elena L. Diaconu, Fabiola Feltrin, Raffaela Conti, Teresa Scicluna, Virginia Carfora, Antonella Cersini, Alessia Franco, Antonio Battisti. |
| EPI_ISL_3306868, EPI_ISL_3306869, EPI_ISL_3306870 | University of Rome Tor Vergata: Departm Experim Medicine Chair of Virology | Department of General Diagnostics; Department of Virology; Istituto Zooprofilattico Sperimentale del Lazio e della Toscana (IZSLT) | Patricia Alba, Giuseppe Manna, Elena L. Diaconu, Fabiola Feltrin, Raffaela Conti, Teresa Scicluna, Virginia Carfora, Antonella Cersini, Alessia Franco, Antonio Battisti. |
| EPI_ISL_3306873 | AOU Policlinico Umberto I; Sapienza Università di Roma | Department of General Diagnostics; Department of Virology; Istituto Zooprofilattico Sperimentale del Lazio e della Toscana (IZSLT) | Patricia Alba, Giuseppe Manna, Elena L. Diaconu, Fabiola Feltrin, Raffaela Conti, Teresa Scicluna, Virginia Carfora, Antonella Cersini, Alessia Franco, Antonio Battisti. |
| EPI_ISL_3306874, EPI_ISL_3306877 | UOC PATOLOGIA CLINICA - OSPEDALE F. SPAZIANI | Department of General Diagnostics; Department of Virology; Istituto Zooprofilattico Sperimentale del Lazio e della Toscana (IZSLT) | Patricia Alba, Giuseppe Manna, Elena L. Diaconu, Fabiola Feltrin, Raffaela Conti, Teresa Scicluna, Virginia Carfora, Antonella Cersini, Alessia Franco, Antonio Battisti. |
| EPI_ISL_3306879 | OSPEDALE SAN CAMILLO DE LELLIS | Department of General Diagnostics; Department of Virology; Istituto | Patricia Alba, Giuseppe Manna, Elena L. Diaconu, Fabiola Feltrin, Raffaela Conti, Teresa Scicluna, Virginia Carfora, Antonella Cersini, Alessia Franco, Antonio Battisti. |

| | | | |
|---|---|---|---|
| | | Zooprofilattico Sperimentale del Lazio e della Toscana (IZSLT) | |
| EPI_ISL_3306880, EPI_ISL_3306881 | Department of Virology, Istituto Zooprofilattico Sperimentale del Lazio e della Toscana (IZSLT) | Department of General Diagnostics; Department of Virology; Istituto Zooprofilattico Sperimentale del Lazio e della Toscana (IZSLT) | Patricia Alba, Giuseppe Manna, Elena L. Diaconu, Fabiola Feltrin, Raffaela Conti, Teresa Scicluna, Virginia Carfora, Antonella Cersini, Alessia Franco, Antonio Battisti. |
| EPI_ISL_3306882, EPI_ISL_3306883 | University of Rome Tor Vergata: Departm Experim Medicine Chair of Virology | Department of General Diagnostics; Department of Virology; Istituto Zooprofilattico Sperimentale del Lazio e della Toscana (IZSLT) | Patricia Alba, Giuseppe Manna, Elena L. Diaconu, Fabiola Feltrin, Raffaela Conti, Teresa Scicluna, Virginia Carfora, Antonella Cersini, Alessia Franco, Antonio Battisti. |
| EPI_ISL_3306884, EPI_ISL_3306885, EPI_ISL_3306886, EPI_ISL_3306888, EPI_ISL_3306889, EPI_ISL_3306890 | Department of Virology, Istituto Zooprofilattico Sperimentale del Lazio e della Toscana (IZSLT) | Department of General Diagnostics; Department of Virology; Istituto Zooprofilattico Sperimentale del Lazio e della Toscana (IZSLT) | Patricia Alba, Giuseppe Manna, Elena L. Diaconu, Fabiola Feltrin, Raffaela Conti, Teresa Scicluna, Virginia Carfora, Antonella Cersini, Alessia Franco, Antonio Battisti. |
| EPI_ISL_3306891, EPI_ISL_3306892, EPI_ISL_3306893, EPI_ISL_3306894, EPI_ISL_3306895, EPI_ISL_3306896, EPI_ISL_3306897, EPI_ISL_3306898, EPI_ISL_3306899, EPI_ISL_3306900, EPI_ISL_3306901, EPI_ISL_3306902, EPI_ISL_3306903, EPI_ISL_3306904, EPI_ISL_3306905 | | | |
| see above | CHULN - H Santa Maria | Instituto Nacional de Saude (INSA) | Borges et al |
| EPI_ISL_3306906, EPI_ISL_3306907, EPI_ISL_3306908, EPI_ISL_3306909, EPI_ISL_3306910 | H Santarem | Instituto Nacional de Saude (INSA) | Borges et al |
| EPI_ISL_3306911, EPI_ISL_3306912, EPI_ISL_3306913, EPI_ISL_3306914, EPI_ISL_3306915, EPI_ISL_3306916, EPI_ISL_3306917, EPI_ISL_3306918, EPI_ISL_3306919, EPI_ISL_3306920, EPI_ISL_3306921, EPI_ISL_3306922, EPI_ISL_3306923, EPI_ISL_3306924 | | | |
| see above | ULS Litoral Alentejano | Instituto Nacional de Saude (INSA) | Borges et al |
| EPI_ISL_3306926, EPI_ISL_3306927, EPI_ISL_3306928, EPI_ISL_3306929, EPI_ISL_3306930, EPI_ISL_3306931, EPI_ISL_3306932, EPI_ISL_3306933, EPI_ISL_3306934, EPI_ISL_3306935, EPI_ISL_3306936, EPI_ISL_3306937, EPI_ISL_3306938, EPI_ISL_3306939, EPI_ISL_3306940, EPI_ISL_3306941, EPI_ISL_3306942, EPI_ISL_3306943, EPI_ISL_3306944, EPI_ISL_3306945, EPI_ISL_3306946, EPI_ISL_3306947 | | | |
| see above | ULS Norte Alentejano | Instituto Nacional de Saude (INSA) | Borges et al |
| EPI_ISL_3306948, EPI_ISL_3306949, EPI_ISL_3306950, EPI_ISL_3306951, EPI_ISL_3306952, EPI_ISL_3306953, EPI_ISL_3306954, EPI_ISL_3306955, EPI_ISL_3306956, EPI_ISL_3306957, EPI_ISL_3306958, EPI_ISL_3306959, EPI_ISL_3306960, EPI_ISL_3306961, EPI_ISL_3306962, EPI_ISL_3306963, EPI_ISL_3306964 | | | |
| see above | CHLC | Instituto Nacional de Saude (INSA) | Borges et al |
| EPI_ISL_3306965, EPI_ISL_3306966, EPI_ISL_3306967, EPI_ISL_3306968, EPI_ISL_3306969, EPI_ISL_3306970, EPI_ISL_3306971, EPI_ISL_3306972, EPI_ISL_3306973, EPI_ISL_3306974, EPI_ISL_3306975, EPI_ISL_3306976, EPI_ISL_3306977, EPI_ISL_3306978, EPI_ISL_3306979 | | | |
| see above | ULS Castelo Branco | Instituto Nacional de Saude (INSA) | Borges et al |
| EPI_ISL_3306980, EPI_ISL_3306981, EPI_ISL_3306982, EPI_ISL_3306983, EPI_ISL_3306984, EPI_ISL_3306985, EPI_ISL_3306986, EPI_ISL_3306987, EPI_ISL_3306988, EPI_ISL_3306989, EPI_ISL_3306990, EPI_ISL_3306991, EPI_ISL_3306992, EPI_ISL_3306993 | | | |
| see above | LAC UCoimbra | Instituto Nacional de Saude (INSA) | Borges et al |
| EPI_ISL_3306994, EPI_ISL_3306995, EPI_ISL_3306996, EPI_ISL_3306997, EPI_ISL_3306998, EPI_ISL_3306999, EPI_ISL_3307000, EPI_ISL_3307001, EPI_ISL_3307002, EPI_ISL_3307003, EPI_ISL_3307004, EPI_ISL_3307005, EPI_ISL_3307006, EPI_ISL_3307007, EPI_ISL_3307008, EPI_ISL_3307009, EPI_ISL_3307010, EPI_ISL_3307011, EPI_ISL_3307012, EPI_ISL_3307013, EPI_ISL_3307014 | | | |
| see above | CHMT | Instituto Nacional de Saude (INSA) | Borges et al |
| EPI_ISL_3307015, EPI_ISL_3307016, EPI_ISL_3307017, EPI_ISL_3307018, EPI_ISL_3307019, EPI_ISL_3307020, EPI_ISL_3307021, EPI_ISL_3307022, EPI_ISL_3307023, EPI_ISL_3307024, EPI_ISL_3307025, EPI_ISL_3307026, EPI_ISL_3307027, EPI_ISL_3307028, EPI_ISL_3307029, EPI_ISL_3307030, EPI_ISL_3307031, EPI_ISL_3307032, EPI_ISL_3307033, EPI_ISL_3307034, EPI_ISL_3307035, EPI_ISL_3307036, EPI_ISL_3307037, EPI_ISL_3307038, EPI_ISL_3307039, EPI_ISL_3307040, EPI_ISL_3307041, EPI_ISL_3307042, EPI_ISL_3307043, EPI_ISL_3307044, EPI_ISL_3307045, EPI_ISL_3307046, EPI_ISL_3307047, EPI_ISL_3307048, EPI_ISL_3307049, EPI_ISL_3307050, EPI_ISL_3307051, EPI_ISL_3307052, EPI_ISL_3307053, EPI_ISL_3307054, EPI_ISL_3307055, EPI_ISL_3307056, EPI_ISL_3307057, EPI_ISL_3307058, EPI_ISL_3307059, EPI_ISL_3307060, EPI_ISL_3307061, EPI_ISL_3307062 | | | |
| see above | ARS Algarve - Laboratorio Laura Ayres | Instituto Nacional de Saude (INSA) | Borges et al |
| EPI_ISL_3307063, EPI_ISL_3307064, EPI_ISL_3307065, EPI_ISL_3307066, EPI_ISL_3307067, EPI_ISL_3307068, EPI_ISL_3307069, EPI_ISL_3307070, EPI_ISL_3307071, EPI_ISL_3307072, EPI_ISL_3307073, EPI_ISL_3307074, EPI_ISL_3307075 | | | |
| see above | H Fernando Fonseca | Instituto Nacional de Saude (INSA) | Borges et al |
| EPI_ISL_3307076, EPI_ISL_3307077, EPI_ISL_3307078, EPI_ISL_3307079, EPI_ISL_3307080, EPI_ISL_3307081, EPI_ISL_3307082, EPI_ISL_3307083, EPI_ISL_3307084, EPI_ISL_3307085, EPI_ISL_3307086, EPI_ISL_3307087, EPI_ISL_3307089, EPI_ISL_3307090 | | | |
| see above | CHU Sao Joao, Porto | Instituto Nacional de Saude (INSA) | Borges et al |
| EPI_ISL_3307091, EPI_ISL_3307092, EPI_ISL_3307093, EPI_ISL_3307094, EPI_ISL_3307095, EPI_ISL_3307096, EPI_ISL_3307097, EPI_ISL_3307098, EPI_ISL_3307099, EPI_ISL_3307100, EPI_ISL_3307101, EPI_ISL_3307102, EPI_ISL_3307103, EPI_ISL_3307104, EPI_ISL_3307105, EPI_ISL_3307106, EPI_ISL_3307107, EPI_ISL_3307108, EPI_ISL_3307109, EPI_ISL_3307110, EPI_ISL_3307111, EPI_ISL_3307112, EPI_ISL_3307113, EPI_ISL_3307114, EPI_ISL_3307115, EPI_ISL_3307116, EPI_ISL_3307117, EPI_ISL_3307118, EPI_ISL_3307119 | | | |
| see above | CHTMAD | Instituto Nacional de Saude (INSA) | Borges et al |
| EPI_ISL_3307120, EPI_ISL_3307121, EPI_ISL_3307122, EPI_ISL_3307123, EPI_ISL_3307124, EPI_ISL_3307125, EPI_ISL_3307126, EPI_ISL_3307127, EPI_ISL_3307128, EPI_ISL_3307129, EPI_ISL_3307130, EPI_ISL_3307131, EPI_ISL_3307132 | | | |
| see above | CHEDV | Instituto Nacional de Saude (INSA) | Borges et al |
| EPI_ISL_3307133, EPI_ISL_3307134, EPI_ISL_3307135, EPI_ISL_3307136, EPI_ISL_3307137, EPI_ISL_3307138, EPI_ISL_3307139, EPI_ISL_3307140, EPI_ISL_3307141, EPI_ISL_3307142, EPI_ISL_3307143 | | | |
| see above | CH Tondela Viseu | Instituto Nacional de Saude (INSA) | Borges et al |
| EPI_ISL_3307144, EPI_ISL_3307145, EPI_ISL_3307146, EPI_ISL_3307147, EPI_ISL_3307148, EPI_ISL_3307149, EPI_ISL_3307150, EPI_ISL_3307151, EPI_ISL_3307152, EPI_ISL_3307153, EPI_ISL_3307154, EPI_ISL_3307155, EPI_ISL_3307156, EPI_ISL_3307157, EPI_ISL_3307158 | | | |
| see above | H Braga | Instituto Nacional de Saude (INSA) | Borges et al |
| EPI_ISL_3307159, EPI_ISL_3307160, EPI_ISL_3307161, EPI_ISL_3307163, EPI_ISL_3307164, EPI_ISL_3307165, EPI_ISL_3307166, EPI_ISL_3307167, EPI_ISL_3307168 | H Guimaraes | Instituto Nacional de Saude (INSA) | Borges et al |
| EPI_ISL_3307170, EPI_ISL_3307171, EPI_ISL_3307172 | INSA | Instituto Nacional de Saude (INSA) | Borges et al |
| EPI_ISL_3307173, EPI_ISL_3307174 | H Guimaraes | Instituto Nacional de Saude (INSA) | Borges et al |
| EPI_ISL_3307175, EPI_ISL_3307176, EPI_ISL_3307177, EPI_ISL_3307178, EPI_ISL_3307179, EPI_ISL_3307180, EPI_ISL_3307181, EPI_ISL_3307182, EPI_ISL_3307183, EPI_ISL_3307184, EPI_ISL_3307185, EPI_ISL_3307186, EPI_ISL_3307187, EPI_ISL_3307188, EPI_ISL_3307189, EPI_ISL_3307190, EPI_ISL_3307191, EPI_ISL_3307192, EPI_ISL_3307193, EPI_ISL_3307194, EPI_ISL_3307195, EPI_ISL_3307196, EPI_ISL_3307197, EPI_ISL_3307198, EPI_ISL_3307199, EPI_ISL_3307200, EPI_ISL_3307201, EPI_ISL_3307202, EPI_ISL_3307203, EPI_ISL_3307204, EPI_ISL_3307205, EPI_ISL_3307206, EPI_ISL_3307207, EPI_ISL_3307208 | | | |
| see above | SYNLAB | Instituto Nacional de Saude (INSA) | Borges et al |
| EPI_ISL_3307209, EPI_ISL_3307210, EPI_ISL_3307211, EPI_ISL_3307212, EPI_ISL_3307213, EPI_ISL_3307214, EPI_ISL_3307215, EPI_ISL_3307216, EPI_ISL_3307217, EPI_ISL_3307218, EPI_ISL_3307219, EPI_ISL_3307220, EPI_ISL_3307221, EPI_ISL_3307222, EPI_ISL_3307223 | | | |
| see above | Labeto - CAB - Leiria | Instituto Nacional de Saude (INSA) | Borges et al |
| EPI_ISL_3307224, EPI_ISL_3307225, EPI_ISL_3307226, EPI_ISL_3307227, EPI_ISL_3307228, EPI_ISL_3307229, EPI_ISL_3307230, EPI_ISL_3307231, EPI_ISL_3307232, EPI_ISL_3307233, EPI_ISL_3307234, EPI_ISL_3307235, EPI_ISL_3307236, EPI_ISL_3307237, EPI_ISL_3307238 | | | |
| see above | CH Porto - H Sto Antonio | Instituto Nacional de Saude (INSA) | Borges et al |
| EPI_ISL_3307239, EPI_ISL_3307240, EPI_ISL_3307241, EPI_ISL_3307242, EPI_ISL_3307243, EPI_ISL_3307244, EPI_ISL_3307245, EPI_ISL_3307246, EPI_ISL_3307247, EPI_ISL_3307248, EPI_ISL_3307249, EPI_ISL_3307250, EPI_ISL_3307251, EPI_ISL_3307252, EPI_ISL_3307253 | | | |
| see above | H Evora | Instituto Nacional de Saude (INSA) | Borges et al |
| EPI_ISL_3307254, EPI_ISL_3307255, EPI_ISL_3307256, EPI_ISL_3307257, EPI_ISL_3307258, EPI_ISL_3307259, EPI_ISL_3307260, EPI_ISL_3307261, EPI_ISL_3307262, EPI_ISL_3307263, EPI_ISL_3307264, EPI_ISL_3307265, EPI_ISL_3307266, EPI_ISL_3307267, EPI_ISL_3307268, EPI_ISL_3307269, EPI_ISL_3307270, EPI_ISL_3307271, EPI_ISL_3307272, EPI_ISL_3307273, EPI_ISL_3307274, EPI_ISL_3307275, EPI_ISL_3307276, EPI_ISL_3307277, EPI_ISL_3307278, EPI_ISL_3307279, EPI_ISL_3307280, EPI_ISL_3307281, EPI_ISL_3307282, EPI_ISL_3307283, EPI_ISL_3307284, EPI_ISL_3307285 | | | |
| see above | SEEBMO | Instituto Nacional de Saude (INSA) | Borges et al |
| EPI_ISL_3307286, EPI_ISL_3307287, EPI_ISL_3307288 | CH Setubal | Instituto Nacional de Saude (INSA) | Borges et al |
| EPI_ISL_3307289, EPI_ISL_3307290, EPI_ISL_3307291, EPI_ISL_3307292, EPI_ISL_3307293, EPI_ISL_3307294, EPI_ISL_3307295, EPI_ISL_3307296, EPI_ISL_3307297, EPI_ISL_3307298, EPI_ISL_3307299, EPI_ISL_3307300, EPI_ISL_3307301, EPI_ISL_3307302, EPI_ISL_3307303, EPI_ISL_3307304, EPI_ISL_3307305, EPI_ISL_3307306, EPI_ISL_3307307, EPI_ISL_3307308, EPI_ISL_3307309, EPI_ISL_3307310, EPI_ISL_3307311, EPI_ISL_3307312, EPI_ISL_3307313 | | | |
| see above | H Divino Espirito Santo - Ponta Delgada | Instituto Nacional de Saude (INSA) | Borges et al |
| EPI_ISL_3307314, EPI_ISL_3307315, EPI_ISL_3307316, EPI_ISL_3307317, EPI_ISL_3307318, EPI_ISL_3307319, EPI_ISL_3307320, EPI_ISL_3307321, EPI_ISL_3307322, EPI_ISL_3307323, EPI_ISL_3307324, EPI_ISL_3307325, EPI_ISL_3307326, EPI_ISL_3307327, EPI_ISL_3307328, EPI_ISL_3307329, EPI_ISL_3307330, EPI_ISL_3307331, EPI_ISL_3307332, EPI_ISL_3307333, EPI_ISL_3307334, EPI_ISL_3307335, EPI_ISL_3307336, EPI_ISL_3307337, EPI_ISL_3307338, EPI_ISL_3307339, EPI_ISL_3307340, EPI_ISL_3307341, EPI_ISL_3307342, EPI_ISL_3307343, EPI_ISL_3307344, EPI_ISL_3307345, EPI_ISL_3307346, EPI_ISL_3307347, EPI_ISL_3307348, EPI_ISL_3307349, EPI_ISL_3307350, EPI_ISL_3307351, EPI_ISL_3307352, EPI_ISL_3307353, EPI_ISL_3307354, EPI_ISL_3307355, EPI_ISL_3307356, EPI_ISL_3307357 | | | |
| see above | SESARAM | Instituto Nacional de Saude (INSA) | Borges et al |
| EPI_ISL_3307358, EPI_ISL_3307359, EPI_ISL_3307360, EPI_ISL_3307361, EPI_ISL_3307362, EPI_ISL_3307363, EPI_ISL_3307364, EPI_ISL_3307365, EPI_ISL_3307366, EPI_ISL_3307367, EPI_ISL_3307368, EPI_ISL_3307369, EPI_ISL_3307370, EPI_ISL_3307371 | | | |
| see above | CHULN - H Santa Maria | Instituto Nacional de Saude (INSA) | Borges et al |
| EPI_ISL_3307372, EPI_ISL_3307373, EPI_ISL_3307374, EPI_ISL_3307375, EPI_ISL_3307376, EPI_ISL_3307377, EPI_ISL_3307378, EPI_ISL_3307379, EPI_ISL_3307380, EPI_ISL_3307381, EPI_ISL_3307382, EPI_ISL_3307383 | | | |

| | | | |
|---|---|---|---|
| see above | Avelab | Instituto Nacional de Saude (INSA) | Borges et al |
| EPI_ISL_3307384, EPI_ISL_3307385, EPI_ISL_3307387, EPI_ISL_3307388, EPI_ISL_3307389, EPI_ISL_3307391, EPI_ISL_3307392, EPI_ISL_3307393, EPI_ISL_3307394 | ULS Guarda | Instituto Nacional de Saude (INSA) | Borges et al |
| EPI_ISL_3307395, EPI_ISL_3307396, EPI_ISL_3307397, EPI_ISL_3307398, EPI_ISL_3307399, EPI_ISL_3307400, EPI_ISL_3307401, EPI_ISL_3307402, EPI_ISL_3307403, EPI_ISL_3307404, EPI_ISL_3307405, EPI_ISL_3307406, EPI_ISL_3307407, EPI_ISL_3307408, EPI_ISL_3307409 | | | |
| see above | CH Tamega e Sousa | Instituto Nacional de Saude (INSA) | Borges et al |
| EPI_ISL_3307410, EPI_ISL_3307411, EPI_ISL_3307412, EPI_ISL_3307413, EPI_ISL_3307414, EPI_ISL_3307415, EPI_ISL_3307416, EPI_ISL_3307417, EPI_ISL_3307418, EPI_ISL_3307419, EPI_ISL_3307420, EPI_ISL_3307421, EPI_ISL_3307422, EPI_ISL_3307423, EPI_ISL_3307424, EPI_ISL_3307425, EPI_ISL_3307426, EPI_ISL_3307427, EPI_ISL_3307428, EPI_ISL_3307429, EPI_ISL_3307430, EPI_ISL_3307431, EPI_ISL_3307432, EPI_ISL_3307433, EPI_ISL_3307434, EPI_ISL_3307435, EPI_ISL_3307436, EPI_ISL_3307437, EPI_ISL_3307438, EPI_ISL_3307439, EPI_ISL_3307440, EPI_ISL_3307441 | | | |
| see above | UNILABS | Instituto Nacional de Saude (INSA) | Borges et al |
| EPI_ISL_3307459 | Tripler Army Medical Center Microbiology Department | Department of Clinical Investigation | Rachael Downham, Kendra Carter, Jonathan D'Ambrozio, Catherine Uyehara |
| EPI_ISL_3307460, EPI_ISL_3307461, EPI_ISL_3307462 | Tripler Army Medical Center Microbiology Department | Department of Clinical Investigation | Rachael Downham, Kendra Carter, Jonathan D'Ambrozio, Catherine Uyehara |
| EPI_ISL_3307463, EPI_ISL_3307464, EPI_ISL_3307465, EPI_ISL_3307466 | Austrian Agency for Health and Food Safety (AGES) | Bergthaler laboratory, CeMM Research Center for Molecular Medicine of the Austrian Academy of Sciences | Lukas Endler, Anna Schedl, Fabian Amman, Petr Triska, Matthew Thornton, Thomas Penz, Benedikt Agerer, Maelle Le Moing, Michael Schuster, Bekir Erguner, Jan Laine, Martin Senekowitsch, Christoph Bock, Andreas Bergthaler |
| EPI_ISL_3307467 | Tripler Army Medical Center Microbiology Department | Department of Clinical Investigation | Rachael Downham, Kendra Carter, Jonathan D'Ambrozio, Catherine Uyehara |
| EPI_ISL_3307468, EPI_ISL_3307469, EPI_ISL_3307470, EPI_ISL_3307471, EPI_ISL_3307472, EPI_ISL_3307473, EPI_ISL_3307474, EPI_ISL_3307475, EPI_ISL_3307477, EPI_ISL_3307478, EPI_ISL_3307479, EPI_ISL_3307480, EPI_ISL_3307481, EPI_ISL_3307482, EPI_ISL_3307483, EPI_ISL_3307484, EPI_ISL_3307485, EPI_ISL_3307486, EPI_ISL_3307487, EPI_ISL_3307488, EPI_ISL_3307489, EPI_ISL_3307490, EPI_ISL_3307491, EPI_ISL_3307492, EPI_ISL_3307493, EPI_ISL_3307494, EPI_ISL_3307495, EPI_ISL_3307496, EPI_ISL_3307497, EPI_ISL_3307498, EPI_ISL_3307499, EPI_ISL_3307500, EPI_ISL_3307501, EPI_ISL_3307502, EPI_ISL_3307503, EPI_ISL_3307504, EPI_ISL_3307505, EPI_ISL_3307506, EPI_ISL_3307507, EPI_ISL_3307508, EPI_ISL_3307509, EPI_ISL_3307510, EPI_ISL_3307511, EPI_ISL_3307512, EPI_ISL_3307513, EPI_ISL_3307514, EPI_ISL_3307515, EPI_ISL_3307516, EPI_ISL_3307517, EPI_ISL_3307518, EPI_ISL_3307519, EPI_ISL_3307520, EPI_ISL_3307521, EPI_ISL_3307522, EPI_ISL_3307523, EPI_ISL_3307524, EPI_ISL_3307525, EPI_ISL_3307526, EPI_ISL_3307527, EPI_ISL_3307528, EPI_ISL_3307529, EPI_ISL_3307530, EPI_ISL_3307531, EPI_ISL_3307532, EPI_ISL_3307533, EPI_ISL_3307534, EPI_ISL_3307535, EPI_ISL_3307536, EPI_ISL_3307537, EPI_ISL_3307538, EPI_ISL_3307539, EPI_ISL_3307540, EPI_ISL_3307541, EPI_ISL_3307542, EPI_ISL_3307543, EPI_ISL_3307544, EPI_ISL_3307545, EPI_ISL_3307546, EPI_ISL_3307548, EPI_ISL_3307549, EPI_ISL_3307550, EPI_ISL_3307551, EPI_ISL_3307552, EPI_ISL_3307553, EPI_ISL_3307554, EPI_ISL_3307555, EPI_ISL_3307556, EPI_ISL_3307557, EPI_ISL_3307558, EPI_ISL_3307559, EPI_ISL_3307560, EPI_ISL_3307561, EPI_ISL_3307562, EPI_ISL_3307563, EPI_ISL_3307564, EPI_ISL_3307565, EPI_ISL_3307566, EPI_ISL_3307567, EPI_ISL_3307568, EPI_ISL_3307569, EPI_ISL_3307570, EPI_ISL_3307571, EPI_ISL_3307572, EPI_ISL_3307573, EPI_ISL_3307574, EPI_ISL_3307575, EPI_ISL_3307576, EPI_ISL_3307577, EPI_ISL_3307578, EPI_ISL_3307579, EPI_ISL_3307580, EPI_ISL_3307581, EPI_ISL_3307582, EPI_ISL_3307583, EPI_ISL_3307584, EPI_ISL_3307585, EPI_ISL_3307586, EPI_ISL_3307587, EPI_ISL_3307588, EPI_ISL_3307589, EPI_ISL_3307590, EPI_ISL_3307591, EPI_ISL_3307592, EPI_ISL_3307593, EPI_ISL_3307594, EPI_ISL_3307595, EPI_ISL_3307596, EPI_ISL_3307597, EPI_ISL_3307598, EPI_ISL_3307599, EPI_ISL_3307600, EPI_ISL_3307601, EPI_ISL_3307602, EPI_ISL_3307603, EPI_ISL_3307604, EPI_ISL_3307605, EPI_ISL_3307606, EPI_ISL_3307607, EPI_ISL_3307608, EPI_ISL_3307609, EPI_ISL_3307610, EPI_ISL_3307612, EPI_ISL_3307613, EPI_ISL_3307614, EPI_ISL_3307615, EPI_ISL_3307616, EPI_ISL_3307617, EPI_ISL_3307618, EPI_ISL_3307619, EPI_ISL_3307621, EPI_ISL_3307622, EPI_ISL_3307623, EPI_ISL_3307624, EPI_ISL_3307625, EPI_ISL_3307626, EPI_ISL_3307627, EPI_ISL_3307628, EPI_ISL_3307629, EPI_ISL_3307630, EPI_ISL_3307632, EPI_ISL_3307633, EPI_ISL_3307634, EPI_ISL_3307635, EPI_ISL_3307636, EPI_ISL_3307637, EPI_ISL_3307638, EPI_ISL_3307639, EPI_ISL_3307640, EPI_ISL_3307641, EPI_ISL_3307642, EPI_ISL_3307643, EPI_ISL_3307644, EPI_ISL_3307645, EPI_ISL_3307646, EPI_ISL_3307647, EPI_ISL_3307648, EPI_ISL_3307649, EPI_ISL_3307650, EPI_ISL_3307651, EPI_ISL_3307652, EPI_ISL_3307653, EPI_ISL_3307654, EPI_ISL_3307655, EPI_ISL_3307656, EPI_ISL_3307657, EPI_ISL_3307658, EPI_ISL_3307659, EPI_ISL_3307660, EPI_ISL_3307661, EPI_ISL_3307662, EPI_ISL_3307663, EPI_ISL_3307664, EPI_ISL_3307665, EPI_ISL_3307666, EPI_ISL_3307667, EPI_ISL_3307668, EPI_ISL_3307669, EPI_ISL_3307670, EPI_ISL_3307671, EPI_ISL_3307672, EPI_ISL_3307673, EPI_ISL_3307674, EPI_ISL_3307675, EPI_ISL_3307676, EPI_ISL_3307677, EPI_ISL_3307678, EPI_ISL_3307679, EPI_ISL_3307680, EPI_ISL_3307681, EPI_ISL_3307683, EPI_ISL_3307685, EPI_ISL_3307686, EPI_ISL_3307687, EPI_ISL_3307688, EPI_ISL_3307689, EPI_ISL_3307690, EPI_ISL_3307691, EPI_ISL_3307692, EPI_ISL_3307694, EPI_ISL_3307695, EPI_ISL_3307696, EPI_ISL_3307697, EPI_ISL_3307698, EPI_ISL_3307699, EPI_ISL_3307700, EPI_ISL_3307701, EPI_ISL_3307702, EPI_ISL_3307703, EPI_ISL_3307704, EPI_ISL_3307705, EPI_ISL_3307706, EPI_ISL_3307707, EPI_ISL_3307708, EPI_ISL_3307709, EPI_ISL_3307710, EPI_ISL_3307711, EPI_ISL_3307712, EPI_ISL_3307713, EPI_ISL_3307714, EPI_ISL_3307715, EPI_ISL_3307716, EPI_ISL_3307717, EPI_ISL_3307718, EPI_ISL_3307719, EPI_ISL_3307720, EPI_ISL_3307721, EPI_ISL_3307722, EPI_ISL_3307723, EPI_ISL_3307724, EPI_ISL_3307725, EPI_ISL_3307726, EPI_ISL_3307727, EPI_ISL_3307728, EPI_ISL_3307729, EPI_ISL_3307730, EPI_ISL_3307731, EPI_ISL_3307733, EPI_ISL_3307734, EPI_ISL_3307735, EPI_ISL_3307736, EPI_ISL_3307737, EPI_ISL_3307738 | | | |
| see above | Austrian Agency for Health and Food Safety (AGES) | Bergthaler laboratory, CeMM Research Center for Molecular Medicine of the Austrian Academy of Sciences | Lukas Endler, Anna Schedl, Fabian Amman, Petr Triska, Matthew Thornton, Thomas Penz, Benedikt Agerer, Maelle Le Moing, Michael Schuster, Bekir Erguner, Jan Laine, Martin Senekowitsch, Christoph Bock, Andreas Bergthaler |
| EPI_ISL_3307739 | Tripler Army Medical Center Microbiology Department | Department of Clinical Investigation | Rachael Downham, Kendra Carter, Jonathan D'Ambrozio, Catherine Uyehara |
| EPI_ISL_3307740, EPI_ISL_3307741, EPI_ISL_3307742 | Austrian Agency for Health and Food Safety (AGES) | Bergthaler laboratory, CeMM Research Center for Molecular Medicine of the Austrian Academy of Sciences | Lukas Endler, Anna Schedl, Fabian Amman, Petr Triska, Matthew Thornton, Thomas Penz, Benedikt Agerer, Maelle Le Moing, Michael Schuster, Bekir Erguner, Jan Laine, Martin Senekowitsch, Christoph Bock, Andreas Bergthaler |
| EPI_ISL_3307743 | ILV Kärnten | Bergthaler laboratory, CeMM Research Center for Molecular Medicine of the Austrian Academy of Sciences | Lukas Endler, Anna Schedl, Fabian Amman, Petr Triska, Matthew Thornton, Thomas Penz, Benedikt Agerer, Maelle Le Moing, Michael Schuster, Bekir Erguner, Jan Laine, Martin Senekowitsch, Christoph Bock, Andreas Bergthaler |
| EPI_ISL_3307744, EPI_ISL_3307745, EPI_ISL_3307746, EPI_ISL_3307747, EPI_ISL_3307748, EPI_ISL_3307749, EPI_ISL_3307750, EPI_ISL_3307751, EPI_ISL_3307752, EPI_ISL_3307753, EPI_ISL_3307755, EPI_ISL_3307756, EPI_ISL_3307757, EPI_ISL_3307758, EPI_ISL_3307759 | | | |
| see above | Universität Innsbruck, Institut für Mikrobiologie | Bergthaler laboratory, CeMM Research Center for Molecular Medicine of the Austrian Academy of Sciences | Lukas Endler, Anna Schedl, Fabian Amman, Petr Triska, Matthew Thornton, Thomas Penz, Benedikt Agerer, Maelle Le Moing, Michael Schuster, Bekir Erguner, Jan Laine, Martin Senekowitsch, Christoph Bock, Andreas Bergthaler |
| EPI_ISL_3307760, EPI_ISL_3307761, EPI_ISL_3307762, EPI_ISL_3307763, EPI_ISL_3307764, EPI_ISL_3307765, EPI_ISL_3307766, EPI_ISL_3307767, EPI_ISL_3307768, EPI_ISL_3307769, EPI_ISL_3307770, EPI_ISL_3307771, EPI_ISL_3307772, EPI_ISL_3307773, EPI_ISL_3307774, EPI_ISL_3307775, EPI_ISL_3307776, EPI_ISL_3307777, EPI_ISL_3307778, EPI_ISL_3307779, EPI_ISL_3307780 | | | |
| see above | Klinikum Wels-Grieskirchen | Bergthaler laboratory, CeMM Research Center for Molecular Medicine of the Austrian Academy of Sciences | Lukas Endler, Anna Schedl, Fabian Amman, Petr Triska, Matthew Thornton, Thomas Penz, Benedikt Agerer, Maelle Le Moing, Michael Schuster, Bekir Erguner, Jan Laine, Martin Senekowitsch, Christoph Bock, Andreas Bergthaler |
| EPI_ISL_3307781 | Institute of Legal Medicine, Medical University of Innsbruck | Bergthaler laboratory, CeMM Research Center for Molecular Medicine of the Austrian Academy of Sciences | Lukas Endler, Anna Schedl, Fabian Amman, Petr Triska, Matthew Thornton, Thomas Penz, Benedikt Agerer, Maelle Le Moing, Michael Schuster, Bekir Erguner, Jan Laine, Martin Senekowitsch, Christoph Bock, Andreas Bergthaler |
| EPI_ISL_3307783, EPI_ISL_3307784, EPI_ISL_3307785 | Center for Virology, Medical University of Vienna | Bergthaler laboratory, CeMM Research Center for Molecular Medicine of the Austrian Academy of Sciences | Lukas Endler, Anna Schedl, Fabian Amman, Petr Triska, Matthew Thornton, Thomas Penz, Benedikt Agerer, Maelle Le Moing, Michael Schuster, Bekir Erguner, Jan Laine, Martin Senekowitsch, Christoph Bock, Andreas Bergthaler |
| EPI_ISL_3307788, EPI_ISL_3307789, EPI_ISL_3307791, EPI_ISL_3307792, EPI_ISL_3307793 | Institute for Water Quality and Resource Management, Technical University Vienna | Bergthaler laboratory, CeMM Research Center for Molecular Medicine of the Austrian Academy of Sciences | Lukas Endler, Anna Schedl, Fabian Amman, Petr Triska, Matthew Thornton, Thomas Penz, Benedikt Agerer, Maelle Le Moing, Michael Schuster, Bekir Erguner, Jan Laine, Martin Senekowitsch, Christoph Bock, Andreas Bergthaler |
| EPI_ISL_3307809, EPI_ISL_3307810, EPI_ISL_3307811, EPI_ISL_3307814, EPI_ISL_3307815, EPI_ISL_3307816, EPI_ISL_3307818, EPI_ISL_3307819, EPI_ISL_3307820, EPI_ISL_3307821, EPI_ISL_3307822, EPI_ISL_3307823, EPI_ISL_3307824, EPI_ISL_3307826, EPI_ISL_3307827, EPI_ISL_3307828, EPI_ISL_3307829, EPI_ISL_3307830, EPI_ISL_3307831, EPI_ISL_3307832, EPI_ISL_3307833, EPI_ISL_3307834, EPI_ISL_3307835, EPI_ISL_3307836, EPI_ISL_3307837, EPI_ISL_3307839, EPI_ISL_3307840, EPI_ISL_3307842, EPI_ISL_3307843, EPI_ISL_3307844, EPI_ISL_3307845, EPI_ISL_3307846, EPI_ISL_3307847, EPI_ISL_3307848, EPI_ISL_3307849, EPI_ISL_3307850, EPI_ISL_3307851, EPI_ISL_3307852, EPI_ISL_3307853 | | | |
| see above | State Laboratories Division, Hawaii State Department of Health | State Laboratories Division, Hawaii State Department of Health | Pamela O'Brien, Drew Kuwazaki, Ayana Garnet, Daniel Strange, Razvan Sultana, Edward Desmond |
| EPI_ISL_3307855 | Tripler Army Medical Center Microbiology Department | Department of Clinical Investigation | Rachael Downham, Kendra Carter, Jonathan D'Ambrozio, Catherine Uyehara |
| EPI_ISL_3307856, EPI_ISL_3307857, EPI_ISL_3307858, EPI_ISL_3307859, EPI_ISL_3307860, EPI_ISL_3307861, EPI_ISL_3307862, EPI_ISL_3307863, EPI_ISL_3307865, EPI_ISL_3307866, EPI_ISL_3307867, EPI_ISL_3307868, EPI_ISL_3307869, EPI_ISL_3307870, EPI_ISL_3307871, EPI_ISL_3307872, EPI_ISL_3307874, EPI_ISL_3307875, EPI_ISL_3307876, EPI_ISL_3307877, EPI_ISL_3307878, EPI_ISL_3307880, EPI_ISL_3307881, EPI_ISL_3307882, EPI_ISL_3307883, EPI_ISL_3307884, EPI_ISL_3307885, EPI_ISL_3307886, EPI_ISL_3307887, EPI_ISL_3307888, EPI_ISL_3307889, EPI_ISL_3307891, EPI_ISL_3307892, EPI_ISL_3307893, EPI_ISL_3307894, EPI_ISL_3307895, EPI_ISL_3307896, EPI_ISL_3307897, EPI_ISL_3307899, EPI_ISL_3307901, EPI_ISL_3307902, EPI_ISL_3307903, EPI_ISL_3307905, EPI_ISL_3307907, EPI_ISL_3307908, EPI_ISL_3307909, EPI_ISL_3307910, EPI_ISL_3307913, EPI_ISL_3307915, EPI_ISL_3307916, EPI_ISL_3307917, EPI_ISL_3307918, EPI_ISL_3307919, EPI_ISL_3307920, EPI_ISL_3307923, EPI_ISL_3307924, EPI_ISL_3307925, EPI_ISL_3307928, EPI_ISL_3307929, EPI_ISL_3307930, EPI_ISL_3307931, EPI_ISL_3307932, EPI_ISL_3307933, EPI_ISL_3307934, EPI_ISL_3307935, EPI_ISL_3307936, EPI_ISL_3307937, EPI_ISL_3307938, EPI_ISL_3307940, EPI_ISL_3307941, EPI_ISL_3307942, EPI_ISL_3307944, EPI_ISL_3307945, EPI_ISL_3307946, EPI_ISL_3307947, EPI_ISL_3307948, EPI_ISL_3307949, EPI_ISL_3307950, EPI_ISL_3307951, EPI_ISL_3307952, EPI_ISL_3307954, EPI_ISL_3307955, EPI_ISL_3307956, EPI_ISL_3307958, EPI_ISL_3307959, EPI_ISL_3307960, EPI_ISL_3307961 | | | |
| see above | State Laboratories Division, Hawaii State Department of Health | State Laboratories Division, Hawaii State Department of Health | Pamela O'Brien, Drew Kuwazaki, Ayana Garnet, Daniel Strange, Razvan Sultana, Edward Desmond |
| EPI_ISL_3307963 | Tripler Army Medical Center Microbiology Department | Department of Clinical Investigation | Rachael Downham, Kendra Carter, Jonathan D'Ambrozio, Catherine Uyehara |
| EPI_ISL_3307964, EPI_ISL_3307965, EPI_ISL_3307966, EPI_ISL_3307967, EPI_ISL_3307968, EPI_ISL_3307969, EPI_ISL_3307970, EPI_ISL_3307971, EPI_ISL_3307972, EPI_ISL_3307973, EPI_ISL_3307974, EPI_ISL_3307975, EPI_ISL_3307976, EPI_ISL_3307977, EPI_ISL_3307978, EPI_ISL_3307979, EPI_ISL_3307980, EPI_ISL_3307981, EPI_ISL_3307982, EPI_ISL_3307983, EPI_ISL_3307984, EPI_ISL_3307985, EPI_ISL_3307986, EPI_ISL_3307987, EPI_ISL_3307988, EPI_ISL_3307989, EPI_ISL_3307990, EPI_ISL_3307991, EPI_ISL_3307992, EPI_ISL_3307993, EPI_ISL_3307994, EPI_ISL_3307995, EPI_ISL_3307996, EPI_ISL_3307997, EPI_ISL_3307998, EPI_ISL_3307999, EPI_ISL_3308000, EPI_ISL_3308001, EPI_ISL_3308002, EPI_ISL_3308003, EPI_ISL_3308004, EPI_ISL_3308005, EPI_ISL_3308006, EPI_ISL_3308007, EPI_ISL_3308008, EPI_ISL_3308009, EPI_ISL_3308010, EPI_ISL_3308011, EPI_ISL_3308012, EPI_ISL_3308013, EPI_ISL_3308014, EPI_ISL_3308015, EPI_ISL_3308016, EPI_ISL_3308017, EPI_ISL_3308018, EPI_ISL_3308019, EPI_ISL_3308020, EPI_ISL_3308021, EPI_ISL_3308022, EPI_ISL_3308023, EPI_ISL_3308024, EPI_ISL_3308025, EPI_ISL_3308026, EPI_ISL_3308027, EPI_ISL_3308028, EPI_ISL_3308029, EPI_ISL_3308030, EPI_ISL_3308031, EPI_ISL_3308032, EPI_ISL_3308033, EPI_ISL_3308034, EPI_ISL_3308035, EPI_ISL_3308036, EPI_ISL_3308037, EPI_ISL_3308038, EPI_ISL_3308039, EPI_ISL_3308040, EPI_ISL_3308041, EPI_ISL_3308042, EPI_ISL_3308043, EPI_ISL_3308044, EPI_ISL_3308045, EPI_ISL_3308046, EPI_ISL_3308047, EPI_ISL_3308048, EPI_ISL_3308049, EPI_ISL_3308050, EPI_ISL_3308051, EPI_ISL_3308052, EPI_ISL_3308053, EPI_ISL_3308054, EPI_ISL_3308055, EPI_ISL_3308056, EPI_ISL_3308057, EPI_ISL_3308058, EPI_ISL_3308059, EPI_ISL_3308060, EPI_ISL_3308061, EPI_ISL_3308062, EPI_ISL_3308063, EPI_ISL_3308064, EPI_ISL_3308065, EPI_ISL_3308066, EPI_ISL_3308067, EPI_ISL_3308068, EPI_ISL_3308069, EPI_ISL_3308070, EPI_ISL_3308071, EPI_ISL_3308072, EPI_ISL_3308073, EPI_ISL_3308074, EPI_ISL_3308075, EPI_ISL_3308076, EPI_ISL_3308077, EPI_ISL_3308078, EPI_ISL_3308079, EPI_ISL_3308080, EPI_ISL_3308081, EPI_ISL_3308082, EPI_ISL_3308083, EPI_ISL_3308084, EPI_ISL_3308085, EPI_ISL_3308086, EPI_ISL_3308087, EPI_ISL_3308088, EPI_ISL_3308089, EPI_ISL_3308090, EPI_ISL_3308091, EPI_ISL_3308092, EPI_ISL_3308093, EPI_ISL_3308094, EPI_ISL_3308095, EPI_ISL_3308096, EPI_ISL_3308097, EPI_ISL_3308098, EPI_ISL_3308099, EPI_ISL_3308100, EPI_ISL_3308101, EPI_ISL_3308102, EPI_ISL_3308103, EPI_ISL_3308104, EPI_ISL_3308105, EPI_ISL_3308106, EPI_ISL_3308107, EPI_ISL_3308108, EPI_ISL_3308109, EPI_ISL_3308110, EPI_ISL_3308111, EPI_ISL_3308112, EPI_ISL_3308113, EPI_ISL_3308114, EPI_ISL_3308115, EPI_ISL_3308116, EPI_ISL_3308117, EPI_ISL_3308118, EPI_ISL_3308119, EPI_ISL_3308120, EPI_ISL_3308121, EPI_ISL_3308122, EPI_ISL_3308123, EPI_ISL_3308124, EPI_ISL_3308125, EPI_ISL_3308126, EPI_ISL_3308127, EPI_ISL_3308128, EPI_ISL_3308129, EPI_ISL_3308130, EPI_ISL_3308131, EPI_ISL_3308132, EPI_ISL_3308133, EPI_ISL_3308134, EPI_ISL_3308135, EPI_ISL_3308136, EPI_ISL_3308137, EPI_ISL_3308138, EPI_ISL_3308139, EPI_ISL_3308140, EPI_ISL_3308141, EPI_ISL_3308142, EPI_ISL_3308143, EPI_ISL_3308144, EPI_ISL_3308145, EPI_ISL_3308146, EPI_ISL_3308147, EPI_ISL_3308148 | | | |
| see above | National Public Health Laboratory, National Centre for Infectious Diseases | National Public Health Laboratory, National Centre for Infectious Diseases | Tze Minn Mak, Katherine Ching, Zhenyang Zhou, Royce Ang, Lin Cui, Raymond Tzer Pin Lin |
| EPI_ISL_3308149 | Tripler Army Medical Center Microbiology Department | Department of Clinical Investigation | Rachael Downham, Kendra Carter, Jonathan D'Ambrozio, Catherine Uyehara |
| EPI_ISL_3308150, EPI_ISL_3308151, EPI_ISL_3308152, EPI_ISL_3308153, EPI_ISL_3308154, EPI_ISL_3308155, EPI_ISL_3308156, EPI_ISL_3308157, EPI_ISL_3308158, EPI_ISL_3308159, EPI_ISL_3308160, EPI_ISL_3308161, EPI_ISL_3308162, EPI_ISL_3308163, EPI_ISL_3308164, EPI_ISL_3308165, EPI_ISL_3308166, EPI_ISL_3308167 | | | |
| see above | National Public Health Laboratory, National Centre for Infectious Diseases | National Public Health Laboratory, National Centre for Infectious Diseases | Tze Minn Mak, Katherine Ching, Zhenyang Zhou, Royce Ang, Lin Cui, Raymond Tzer Pin Lin |
| EPI_ISL_3308168, EPI_ISL_3308169 | HOSPITAL MANACOR | HOSPITAL UNIVERSITARIO SON ESPASES | Dr. Antonio Oliver, Dr. Carla López-Causapé, Dr. Gabriel Cabot, on behalf of Servicio de Microbiología, Hospital Universitario Son Espases |
| EPI_ISL_3308170 | HOSPITAL UNIVERSITARIO SON ESPASES | HOSPITAL UNIVERSITARIO SON ESPASES | Dr. Antonio Oliver, Dr. Carla López-Causapé, Dr. Gabriel Cabot, on behalf of Servicio de Microbiología, Hospital Universitario Son Espases |
| EPI_ISL_3308171, EPI_ISL_3308172 | HOSPITAL MATEU ORFILA | HOSPITAL UNIVERSITARIO SON ESPASES | Dr. Antonio Oliver, Dr. Carla López-Causapé, Dr. Gabriel Cabot, on behalf of Servicio de Microbiología, Hospital Universitario Son Espases |
| EPI_ISL_3308173 | HOSPITAL UNIVERSITARIO SON ESPASES | HOSPITAL UNIVERSITARIO SON ESPASES | Dr. Antonio Oliver, Dr. Carla López-Causapé, Dr. Gabriel Cabot, on behalf of Servicio de Microbiología, Hospital Universitario Son Espases |
| EPI_ISL_3308174, EPI_ISL_3308175, EPI_ISL_3308176 | HOSPITAL CAN MISSES | HOSPITAL UNIVERSITARIO SON ESPASES | Dr. Antonio Oliver, Dr. Carla López-Causapé, Dr. Gabriel Cabot, on behalf of Servicio de Microbiología, Hospital Universitario Son Espases |

| | | | |
|---|---|---|---|
| EPI_ISL_3308177 | HOSPITAL UNIVERSITARIO SON ESPASES | HOSPITAL UNIVERSITARIO SON ESPASES | Dr. Antonio Oliver, Dr. Carla López-Causapé, Dr. Gabriel Cabot, on behalf of Servicio de Microbiología, Hospital Universitario Son Espases |
| EPI_ISL_3308178, EPI_ISL_3308179 | HOSPITAL MANACOR | HOSPITAL UNIVERSITARIO SON ESPASES | Dr. Antonio Oliver, Dr. Carla López-Causapé, Dr. Gabriel Cabot, on behalf of Servicio de Microbiología, Hospital Universitario Son Espases |
| EPI_ISL_3308180 | HOSPITAL UNIVERSITARIO SON ESPASES | HOSPITAL UNIVERSITARIO SON ESPASES | Dr. Antonio Oliver, Dr. Carla López-Causapé, Dr. Gabriel Cabot, on behalf of Servicio de Microbiología, Hospital Universitario Son Espases |
| EPI_ISL_3308181 | HOSPITAL MATEU ORFILA | HOSPITAL UNIVERSITARIO SON ESPASES | Dr. Antonio Oliver, Dr. Carla López-Causapé, Dr. Gabriel Cabot, on behalf of Servicio de Microbiología, Hospital Universitario Son Espases |
| EPI_ISL_3308182, EPI_ISL_3308183, EPI_ISL_3308184, EPI_ISL_3308186, EPI_ISL_3308187 | HOSPITAL UNIVERSITARIO SON ESPASES | HOSPITAL UNIVERSITARIO SON ESPASES | Dr. Antonio Oliver, Dr. Carla López-Causapé, Dr. Gabriel Cabot, on behalf of Servicio de Microbiología, Hospital Universitario Son Espases |
| EPI_ISL_3308188 | HOSPITAL MANACOR | HOSPITAL UNIVERSITARIO SON ESPASES | Dr. Antonio Oliver, Dr. Carla López-Causapé, Dr. Gabriel Cabot, on behalf of Servicio de Microbiología, Hospital Universitario Son Espases |
| EPI_ISL_3308189, EPI_ISL_3308190 | HOSPITAL UNIVERSITARIO SON ESPASES | HOSPITAL UNIVERSITARIO SON ESPASES | Dr. Antonio Oliver, Dr. Carla López-Causapé, Dr. Gabriel Cabot, on behalf of Servicio de Microbiología, Hospital Universitario Son Espases |
| EPI_ISL_3308191 | HOSPITAL MATEU ORFILA | HOSPITAL UNIVERSITARIO SON ESPASES | Dr. Antonio Oliver, Dr. Carla López-Causapé, Dr. Gabriel Cabot, on behalf of Servicio de Microbiología, Hospital Universitario Son Espases |
| EPI_ISL_3308192, EPI_ISL_3308194, EPI_ISL_3308195 | HOSPITAL UNIVERSITARIO SON ESPASES | HOSPITAL UNIVERSITARIO SON ESPASES | Dr. Antonio Oliver, Dr. Carla López-Causapé, Dr. Gabriel Cabot, on behalf of Servicio de Microbiología, Hospital Universitario Son Espases |
| EPI_ISL_3308196 | HOSPITAL UNIVERSITARIO SON LLATZER | HOSPITAL UNIVERSITARIO SON ESPASES | Dr. Antonio Oliver, Dr. Carla López-Causapé, Dr. Gabriel Cabot, on behalf of Servicio de Microbiología, Hospital Universitario Son Espases |
| EPI_ISL_3308197, EPI_ISL_3308198, EPI_ISL_3308199 | HOSPITAL CAN MISSES | HOSPITAL UNIVERSITARIO SON ESPASES | Dr. Antonio Oliver, Dr. Carla López-Causapé, Dr. Gabriel Cabot, on behalf of Servicio de Microbiología, Hospital Universitario Son Espases |
| EPI_ISL_3308200 | HOSPITAL UNIVERSITARIO SON ESPASES | HOSPITAL UNIVERSITARIO SON ESPASES | Dr. Antonio Oliver, Dr. Carla López-Causapé, Dr. Gabriel Cabot, on behalf of Servicio de Microbiología, Hospital Universitario Son Espases |
| EPI_ISL_3308201 | HOSPITAL MANACOR | HOSPITAL UNIVERSITARIO SON ESPASES | Dr. Antonio Oliver, Dr. Carla López-Causapé, Dr. Gabriel Cabot, on behalf of Servicio de Microbiología, Hospital Universitario Son Espases |
| EPI_ISL_3308202, EPI_ISL_3308203 | HOSPITAL UNIVERSITARIO SON ESPASES | HOSPITAL UNIVERSITARIO SON ESPASES | Dr. Antonio Oliver, Dr. Carla López-Causapé, Dr. Gabriel Cabot, on behalf of Servicio de Microbiología, Hospital Universitario Son Espases |
| EPI_ISL_3308204 | HOSPITAL MATEU ORFILA | HOSPITAL UNIVERSITARIO SON ESPASES | Dr. Antonio Oliver, Dr. Carla López-Causapé, Dr. Gabriel Cabot, on behalf of Servicio de Microbiología, Hospital Universitario Son Espases |
| EPI_ISL_3308205, EPI_ISL_3308206, EPI_ISL_3308207 | HOSPITAL UNIVERSITARIO SON ESPASES | HOSPITAL UNIVERSITARIO SON ESPASES | Dr. Antonio Oliver, Dr. Carla López-Causapé, Dr. Gabriel Cabot, on behalf of Servicio de Microbiología, Hospital Universitario Son Espases |
| EPI_ISL_3308208 | HOSPITAL UNIVERSITARIO SON LLATZER | HOSPITAL UNIVERSITARIO SON ESPASES | Dr. Antonio Oliver, Dr. Carla López-Causapé, Dr. Gabriel Cabot, on behalf of Servicio de Microbiología, Hospital Universitario Son Espases |
| EPI_ISL_3308209 | HOSPITAL CAN MISSES | HOSPITAL UNIVERSITARIO SON ESPASES | Dr. Antonio Oliver, Dr. Carla López-Causapé, Dr. Gabriel Cabot, on behalf of Servicio de Microbiología, Hospital Universitario Son Espases |
| EPI_ISL_3308210 | HOSPITAL UNIVERSITARIO SON ESPASES | HOSPITAL UNIVERSITARIO SON ESPASES | Dr. Antonio Oliver, Dr. Carla López-Causapé, Dr. Gabriel Cabot, on behalf of Servicio de Microbiología, Hospital Universitario Son Espases |
| EPI_ISL_3308211 | HOSPITAL MANACOR | HOSPITAL UNIVERSITARIO SON ESPASES | Dr. Antonio Oliver, Dr. Carla López-Causapé, Dr. Gabriel Cabot, on behalf of Servicio de Microbiología, Hospital Universitario Son Espases |
| EPI_ISL_3308212, EPI_ISL_3308213, EPI_ISL_3308214, EPI_ISL_3308215, EPI_ISL_3308216 | HOSPITAL UNIVERSITARIO SON ESPASES | HOSPITAL UNIVERSITARIO SON ESPASES | Dr. Antonio Oliver, Dr. Carla López-Causapé, Dr. Gabriel Cabot, on behalf of Servicio de Microbiología, Hospital Universitario Son Espases |
| EPI_ISL_3308217 | HOSPITAL CAN MISSES | HOSPITAL UNIVERSITARIO SON ESPASES | Dr. Antonio Oliver, Dr. Carla López-Causapé, Dr. Gabriel Cabot, on behalf of Servicio de Microbiología, Hospital Universitario Son Espases |
| EPI_ISL_3308218 | HOSPITAL UNIVERSITARIO SON ESPASES | HOSPITAL UNIVERSITARIO SON ESPASES | Dr. Antonio Oliver, Dr. Carla López-Causapé, Dr. Gabriel Cabot, on behalf of Servicio de Microbiología, Hospital Universitario Son Espases |
| EPI_ISL_3308219 | HOSPITAL MANACOR | HOSPITAL UNIVERSITARIO SON ESPASES | Dr. Antonio Oliver, Dr. Carla López-Causapé, Dr. Gabriel Cabot, on behalf of Servicio de Microbiología, Hospital Universitario Son Espases |
| EPI_ISL_3308220, EPI_ISL_3308221, EPI_ISL_3308222, EPI_ISL_3308223, EPI_ISL_3308224, EPI_ISL_3308225, EPI_ISL_3308226 | HOSPITAL UNIVERSITARIO SON ESPASES | HOSPITAL UNIVERSITARIO SON ESPASES | Dr. Antonio Oliver, Dr. Carla López-Causapé, Dr. Gabriel Cabot, on behalf of Servicio de Microbiología, Hospital Universitario Son Espases |
| EPI_ISL_3308227 | HOSPITAL CAN MISSES | HOSPITAL UNIVERSITARIO SON ESPASES | Dr. Antonio Oliver, Dr. Carla López-Causapé, Dr. Gabriel Cabot, on behalf of Servicio de Microbiología, Hospital Universitario Son Espases |
| EPI_ISL_3308228 | HOSPITAL UNIVERSITARIO SON ESPASES | HOSPITAL UNIVERSITARIO SON ESPASES | Dr. Antonio Oliver, Dr. Carla López-Causapé, Dr. Gabriel Cabot, on behalf of Servicio de Microbiología, Hospital Universitario Son Espases |
| EPI_ISL_3308229 | HOSPITAL MANACOR | HOSPITAL UNIVERSITARIO SON ESPASES | Dr. Antonio Oliver, Dr. Carla López-Causapé, Dr. Gabriel Cabot, on behalf of Servicio de Microbiología, Hospital Universitario Son Espases |
| EPI_ISL_3308230, EPI_ISL_3308231 | HOSPITAL UNIVERSITARIO SON ESPASES | HOSPITAL UNIVERSITARIO SON ESPASES | Dr. Antonio Oliver, Dr. Carla López-Causapé, Dr. Gabriel Cabot, on behalf of Servicio de Microbiología, Hospital Universitario Son Espases |
| EPI_ISL_3308232 | HOSPITAL MATEU ORFILA | HOSPITAL UNIVERSITARIO SON ESPASES | Dr. Antonio Oliver, Dr. Carla López-Causapé, Dr. Gabriel Cabot, on behalf of Servicio de Microbiología, Hospital Universitario Son Espases |
| EPI_ISL_3308233, EPI_ISL_3308234 | HOSPITAL UNIVERSITARIO SON ESPASES | HOSPITAL UNIVERSITARIO SON ESPASES | Dr. Antonio Oliver, Dr. Carla López-Causapé, Dr. Gabriel Cabot, on behalf of Servicio de Microbiología, Hospital Universitario Son Espases |
| EPI_ISL_3308236, EPI_ISL_3308237 | HOSPITAL CAN MISSES | HOSPITAL UNIVERSITARIO SON ESPASES | Dr. Antonio Oliver, Dr. Carla López-Causapé, Dr. Gabriel Cabot, on behalf of Servicio de Microbiología, Hospital Universitario Son Espases |
| EPI_ISL_3308238, EPI_ISL_3308239 | HOSPITAL UNIVERSITARIO SON ESPASES | HOSPITAL UNIVERSITARIO SON ESPASES | Dr. Antonio Oliver, Dr. Carla López-Causapé, Dr. Gabriel Cabot, on behalf of Servicio de Microbiología, Hospital Universitario Son Espases |
| EPI_ISL_3308240 | HOSPITAL MANACOR | HOSPITAL UNIVERSITARIO SON ESPASES | Dr. Antonio Oliver, Dr. Carla López-Causapé, Dr. Gabriel Cabot, on behalf of Servicio de Microbiología, Hospital Universitario Son Espases |
| EPI_ISL_3308241, EPI_ISL_3308242, EPI_ISL_3308243, EPI_ISL_3308244, EPI_ISL_3308245, EPI_ISL_3308246, EPI_ISL_3308247 | HOSPITAL UNIVERSITARIO SON ESPASES | HOSPITAL UNIVERSITARIO SON ESPASES | Dr. Antonio Oliver, Dr. Carla López-Causapé, Dr. Gabriel Cabot, on behalf of Servicio de Microbiología, Hospital Universitario Son Espases |
| EPI_ISL_3308248 | HOSPITAL CAN MISSES | HOSPITAL UNIVERSITARIO SON ESPASES | Dr. Antonio Oliver, Dr. Carla López-Causapé, Dr. Gabriel Cabot, on behalf of Servicio de Microbiología, Hospital Universitario Son Espases |
| EPI_ISL_3308249 | HOSPITAL UNIVERSITARIO SON ESPASES | HOSPITAL UNIVERSITARIO SON ESPASES | Dr. Antonio Oliver, Dr. Carla López-Causapé, Dr. Gabriel Cabot, on behalf of Servicio de Microbiología, Hospital Universitario Son Espases |
| EPI_ISL_3308251 | Tripler Army Medical Center Microbiology Department | Department of Clinical Investigation | Rachael Downham, Kendra Carter, Jonathan D'Ambrozio, Catherine Uyehara |
| EPI_ISL_3308252, EPI_ISL_3308253, EPI_ISL_3308254, EPI_ISL_3308255, EPI_ISL_3308256, EPI_ISL_3308257, EPI_ISL_3308258, EPI_ISL_3308259, EPI_ISL_3308260, EPI_ISL_3308261, EPI_ISL_3308262, EPI_ISL_3308263, EPI_ISL_3308264, EPI_ISL_3308265, EPI_ISL_3308266, EPI_ISL_3308267, EPI_ISL_3308268, EPI_ISL_3308269, EPI_ISL_3308270, EPI_ISL_3308271, EPI_ISL_3308272, EPI_ISL_3308273, EPI_ISL_3308274, EPI_ISL_3308275, EPI_ISL_3308276, EPI_ISL_3308277, EPI_ISL_3308278, EPI_ISL_3308279, EPI_ISL_3308280, EPI_ISL_3308281, EPI_ISL_3308282, EPI_ISL_3308283, EPI_ISL_3308284, EPI_ISL_3308285, EPI_ISL_3308286, EPI_ISL_3308287, EPI_ISL_3308288, EPI_ISL_3308289, EPI_ISL_3308290, EPI_ISL_3308291, EPI_ISL_3308292, EPI_ISL_3308293, EPI_ISL_3308294, EPI_ISL_3308295, EPI_ISL_3308296, EPI_ISL_3308297, EPI_ISL_3308298, EPI_ISL_3308299, EPI_ISL_3308300, EPI_ISL_3308301, EPI_ISL_3308302, EPI_ISL_3308303, EPI_ISL_3308304, EPI_ISL_3308305, EPI_ISL_3308306, EPI_ISL_3308307, EPI_ISL_3308308, EPI_ISL_3308309, EPI_ISL_3308310, EPI_ISL_3308311, EPI_ISL_3308312, EPI_ISL_3308313, EPI_ISL_3308314, EPI_ISL_3308315, EPI_ISL_3308317, EPI_ISL_3308318, EPI_ISL_3308319, EPI_ISL_3308320, EPI_ISL_3308321, EPI_ISL_3308322, EPI_ISL_3308323, EPI_ISL_3308324, EPI_ISL_3308325, EPI_ISL_3308326, EPI_ISL_3308327, EPI_ISL_3308328, EPI_ISL_3308329, EPI_ISL_3308330 | | | |
| see above | Dipartimento di Medicina di Laboratorio, Azienda sanitaria universitaria Friuli Centrale (ASU FC) | Dipartimento di Medicina di Laboratorio, Azienda sanitaria universitaria Friuli Centrale (ASU FC) | Corrado Pipan, Catia Mio, Chiara Dal Secco, Stefania Marzinotto, Francesco Curcio |
| EPI_ISL_3308410, EPI_ISL_3308411 | Tripler Army Medical Center Microbiology Department | Department of Clinical Investigation | Rachael Downham, Kendra Carter, Jonathan D'Ambrozio, Catherine Uyehara |
| EPI_ISL_3308412, EPI_ISL_3308413, EPI_ISL_3308414, EPI_ISL_3308415, EPI_ISL_3308416, EPI_ISL_3308417, EPI_ISL_3308418, EPI_ISL_3308419, EPI_ISL_3308420, EPI_ISL_3308421, EPI_ISL_3308422, EPI_ISL_3308423, EPI_ISL_3308424, EPI_ISL_3308425, EPI_ISL_3308426 | | | |
| see above | Centracare Laboratory Services | Minnesota Department of Health, Public Health Laboratory | Alexandra Lorentz, Jacob Garfin, Matt Plumb, and Xiong Wang |
| EPI_ISL_3308427 | Minnesota Department of Health, Public Health Laboratory | Minnesota Department of Health, Public Health Laboratory | Alexandra Lorentz, Jacob Garfin, Matt Plumb, and Xiong Wang |
| EPI_ISL_3308428, EPI_ISL_3308429, EPI_ISL_3308430, EPI_ISL_3308432, EPI_ISL_3308433, EPI_ISL_3308434, EPI_ISL_3308435, EPI_ISL_3308436, EPI_ISL_3308437, EPI_ISL_3308438, EPI_ISL_3308439, EPI_ISL_3308440, EPI_ISL_3308441, EPI_ISL_3308442, EPI_ISL_3308443, EPI_ISL_3308444, EPI_ISL_3308445, EPI_ISL_3308446, EPI_ISL_3308447, EPI_ISL_3308448, EPI_ISL_3308449, EPI_ISL_3308450, EPI_ISL_3308451, EPI_ISL_3308452, EPI_ISL_3308453, EPI_ISL_3308454 | | | |
| see above | Centracare Laboratory Services | Minnesota Department of Health, Public Health Laboratory | Alexandra Lorentz, Jacob Garfin, Matt Plumb, and Xiong Wang |
| EPI_ISL_3308455, EPI_ISL_3308456, EPI_ISL_3308457, EPI_ISL_3308458, EPI_ISL_3308459, EPI_ISL_3308460, EPI_ISL_3308461, EPI_ISL_3308462, EPI_ISL_3308463, EPI_ISL_3308464, EPI_ISL_3308465, EPI_ISL_3308466 | | | |
| see above | Minnesota Department of Health, Public Health Laboratory | Minnesota Department of Health, Public Health Laboratory | Alexandra Lorentz, Jacob Garfin, Matt Plumb, and Xiong Wang |
| EPI_ISL_3308467 | M Health Fairview | Minnesota Department of Health, Public Health Laboratory | Alexandra Lorentz, Jacob Garfin, Matt Plumb, and Xiong Wang |

| | | | |
|---|---|---|---|
| EPI_ISL_3308468, EPI_ISL_3308469, EPI_ISL_3308470, EPI_ISL_3308471, EPI_ISL_3308472, EPI_ISL_3308473 | Minnesota Department of Health, Public Health Laboratory | Minnesota Department of Health, Public Health Laboratory | Alexandra Lorentz, Jacob Garfin, Matt Plumb, and Xiong Wang |
| EPI_ISL_3308474 | Tripler Army Medical Center Microbiology Department | Department of Clinical Investigation | Rachael Downham, Kendra Carter, Jonathan D'Ambrozio, Catherine Uyehara |
| EPI_ISL_3308475, EPI_ISL_3308477, EPI_ISL_3308478, EPI_ISL_3308479, EPI_ISL_3308480, EPI_ISL_3308481, EPI_ISL_3308482, EPI_ISL_3308483, EPI_ISL_3308484, EPI_ISL_3308485, EPI_ISL_3308486, EPI_ISL_3308487, EPI_ISL_3308488 | | | |
| see above | Minnesota Department of Health, Public Health Laboratory | Minnesota Department of Health, Public Health Laboratory | Alexandra Lorentz, Jacob Garfin, Matt Plumb, and Xiong Wang |
| EPI_ISL_3308489 | M Health Fairview | Minnesota Department of Health, Public Health Laboratory | Alexandra Lorentz, Jacob Garfin, Matt Plumb, and Xiong Wang |
| EPI_ISL_3308490, EPI_ISL_3308491, EPI_ISL_3308492, EPI_ISL_3308493, EPI_ISL_3308494, EPI_ISL_3308495, EPI_ISL_3308496, EPI_ISL_3308497, EPI_ISL_3308498, EPI_ISL_3308499, EPI_ISL_3308500, EPI_ISL_3308501, EPI_ISL_3308502, EPI_ISL_3308503, EPI_ISL_3308504, EPI_ISL_3308505 | | | |
| see above | Minnesota Department of Health, Public Health Laboratory | Minnesota Department of Health, Public Health Laboratory | Alexandra Lorentz, Jacob Garfin, Matt Plumb, and Xiong Wang |
| EPI_ISL_3308506, EPI_ISL_3308586 | Tripler Army Medical Center Microbiology Department | Department of Clinical Investigation | Rachael Downham, Kendra Carter, Jonathan D'Ambrozio, Catherine Uyehara |
| EPI_ISL_3308587 | CHU Pontchaillou | CHU Pontchaillou | GROLHIER Claire, DENOUAL Florent, ETCHEVERRY Amandine, SASSI Mohamed, JAGLINE Steven, FEBREAU Christine, PRONIER Charlotte, GALIBERT Marie Dominique, DE TAYRAC Marie, THIBAULT Vincent |
| EPI_ISL_3308588, EPI_ISL_3308589 | Tripler Army Medical Center Microbiology Department | Department of Clinical Investigation | Rachael Downham, Kendra Carter, Jonathan D'Ambrozio, Catherine Uyehara |
| EPI_ISL_3308590, EPI_ISL_3308591, EPI_ISL_3308592, EPI_ISL_3308593, EPI_ISL_3308594, EPI_ISL_3308595 | Department of Clinical Microbiology | GIGA Medical Genomics | Keith Durkin, Maria Artesi, Bouchra Boujemla, Nathalie Renotte, Nadine Cambisano, Sébastien Bontems, Cécile Meex, Claire Gourzonès, Olivier Ek, Laurent Gillet, Marie-Pierre Hayette, Vincent Bours |
| EPI_ISL_3308597, EPI_ISL_3308598, EPI_ISL_3308599, EPI_ISL_3308600, EPI_ISL_3308601, EPI_ISL_3308602, EPI_ISL_3308603, EPI_ISL_3308604, EPI_ISL_3308605, EPI_ISL_3308606, EPI_ISL_3308607, EPI_ISL_3308608, EPI_ISL_3308609, EPI_ISL_3308610 | | | |
| see above | SYNLAB | GIGA Medical Genomics | Keith Durkin, Maria Artesi, Bouchra Boujemla, Nathalie Renotte, Nadine Cambisano, Sébastien Bontems, Cécile Meex, Claire Gourzonès, Olivier Ek, Laurent Gillet, Marie-Pierre Hayette, Vincent Bours |
| EPI_ISL_3308611, EPI_ISL_3308612, EPI_ISL_3308613, EPI_ISL_3308614 | Department of Clinical Microbiology | GIGA Medical Genomics | Keith Durkin, Maria Artesi, Bouchra Boujemla, Nathalie Renotte, Nadine Cambisano, Sébastien Bontems, Cécile Meex, Claire Gourzonès, Olivier Ek, Laurent Gillet, Marie-Pierre Hayette, Vincent Bours |
| EPI_ISL_3308615, EPI_ISL_3308616, EPI_ISL_3308617, EPI_ISL_3308618, EPI_ISL_3308619, EPI_ISL_3308620 | SYNLAB | GIGA Medical Genomics | Keith Durkin, Maria Artesi, Bouchra Boujemla, Nathalie Renotte, Nadine Cambisano, Sébastien Bontems, Cécile Meex, Claire Gourzonès, Olivier Ek, Laurent Gillet, Marie-Pierre Hayette, Vincent Bours |
| EPI_ISL_3308621, EPI_ISL_3308622, EPI_ISL_3308623, EPI_ISL_3308624, EPI_ISL_3308625, EPI_ISL_3308626, EPI_ISL_3308627, EPI_ISL_3308628, EPI_ISL_3308629, EPI_ISL_3308630, EPI_ISL_3308632, EPI_ISL_3308633, EPI_ISL_3308634, EPI_ISL_3308635, EPI_ISL_3308636, EPI_ISL_3308637, EPI_ISL_3308638, EPI_ISL_3308639, EPI_ISL_3308640, EPI_ISL_3308641, EPI_ISL_3308642, EPI_ISL_3308644, EPI_ISL_3308645, EPI_ISL_3308646, EPI_ISL_3308647, EPI_ISL_3308652, EPI_ISL_3308653, EPI_ISL_3308654, EPI_ISL_3308655, EPI_ISL_3308656, EPI_ISL_3308657, EPI_ISL_3308658, EPI_ISL_3308659 | | | |
| see above | University of Liège COVID-19 testing center | GIGA Medical Genomics | Keith Durkin, Maria Artesi, Bouchra Boujemla, Nathalie Renotte, Nadine Cambisano, Sébastien Bontems, Cécile Meex, Claire Gourzonès, Olivier Ek, Laurent Gillet, Marie-Pierre Hayette, Vincent Bours |
| EPI_ISL_3308660, EPI_ISL_3308664 | Tripler Army Medical Center Microbiology Department | Department of Clinical Investigation | Rachael Downham, Kendra Carter, Jonathan D'Ambrozio, Catherine Uyehara |
| EPI_ISL_3308665, EPI_ISL_3308666, EPI_ISL_3308667, EPI_ISL_3308668, EPI_ISL_3308669, EPI_ISL_3308670, EPI_ISL_3308671, EPI_ISL_3308672, EPI_ISL_3308673, EPI_ISL_3308674 | Hospital Universitari Joan XXIII | Hospital Universitari Joan XXIII | Gemma Recio, Cristina Gutiérrez, Clara Benavent, Carla Martín, Margarida Terrón, Ester Picó, Natalia Bastón |
| EPI_ISL_3308675, EPI_ISL_3308676, EPI_ISL_3308687, EPI_ISL_3308688 | Tripler Army Medical Center Microbiology Department | Department of Clinical Investigation | Rachael Downham, Kendra Carter, Jonathan D'Ambrozio, Catherine Uyehara |
| EPI_ISL_3308689, EPI_ISL_3308690, EPI_ISL_3308691, EPI_ISL_3308693, EPI_ISL_3308694, EPI_ISL_3308695, EPI_ISL_3308696, EPI_ISL_3308697, EPI_ISL_3308699, EPI_ISL_3308701, EPI_ISL_3308702, EPI_ISL_3308703, EPI_ISL_3308704, EPI_ISL_3308705, EPI_ISL_3308707, EPI_ISL_3308709, EPI_ISL_3308710, EPI_ISL_3308711, EPI_ISL_3308712, EPI_ISL_3308713, EPI_ISL_3308714, EPI_ISL_3308715, EPI_ISL_3308717, EPI_ISL_3308718, EPI_ISL_3308720, EPI_ISL_3308721, EPI_ISL_3308722, EPI_ISL_3308724, EPI_ISL_3308726, EPI_ISL_3308727, EPI_ISL_3308728, EPI_ISL_3308730, EPI_ISL_3308731, EPI_ISL_3308732, EPI_ISL_3308733, EPI_ISL_3308734, EPI_ISL_3308735, EPI_ISL_3308736, EPI_ISL_3308737, EPI_ISL_3308738, EPI_ISL_3308740, EPI_ISL_3308741, EPI_ISL_3308742, EPI_ISL_3308743, EPI_ISL_3308744, EPI_ISL_3308745, EPI_ISL_3308746, EPI_ISL_3308747, EPI_ISL_3308748, EPI_ISL_3308749, EPI_ISL_3308750 | | | |
| see above | Servicio Microbiología Hospital La Paz | Servicio Microbiología Hospital La Paz | Fernando Lázaro, Rubén Cáceres, Iván Bloise, Jesús Mingorance Cruz, Elie Dahdouh |
| EPI_ISL_3308753, EPI_ISL_3308756, EPI_ISL_3308757, EPI_ISL_3308759, EPI_ISL_3308760, EPI_ISL_3308761, EPI_ISL_3308762, EPI_ISL_3308763, EPI_ISL_3308764, EPI_ISL_3308765, EPI_ISL_3308766, EPI_ISL_3308767, EPI_ISL_3308768, EPI_ISL_3308769, EPI_ISL_3308771 | | | |
| see above | Fakultni nemocnice Ostrava | Fakultni nemocnice Ostrava | Chorzempa |
| EPI_ISL_3308776, EPI_ISL_3308777, EPI_ISL_3308778, EPI_ISL_3308779, EPI_ISL_3308780, EPI_ISL_3308782 | Tripler Army Medical Center Microbiology Department | Department of Clinical Investigation | Rachael Downham, Kendra Carter, Jonathan D'Ambrozio, Catherine Uyehara |
| EPI_ISL_3308783, EPI_ISL_3308784, EPI_ISL_3308785, EPI_ISL_3308786, EPI_ISL_3308787, EPI_ISL_3308788, EPI_ISL_3308789, EPI_ISL_3308790 | AULSS 6 Euganea | Istituto Zooprofilattico Sperimentale delle Venezie | Adelaide Milani, Alessia Schivo, Annalisa Salviato, Elisa Palumbo, Erika Giorgia Quaranta, Luca Tassoni, Ambra Pastori, Edoardo Giussani, Alice Fusaro, Isabella Monne, Calogero Terregino, Antonia Ricci |
| EPI_ISL_3308791 | AULSS 9 Scaligera - San Bonifacio | Istituto Zooprofilattico Sperimentale delle Venezie | Adelaide Milani, Alessia Schivo, Annalisa Salviato, Elisa Palumbo, Erika Giorgia Quaranta, Luca Tassoni, Ambra Pastori, Edoardo Giussani, Alice Fusaro, Isabella Monne, Calogero Terregino, Antonia Ricci |
| EPI_ISL_3308792, EPI_ISL_3308793 | AULSS 6 Euganea | Istituto Zooprofilattico Sperimentale delle Venezie | Adelaide Milani, Alessia Schivo, Annalisa Salviato, Elisa Palumbo, Erika Giorgia Quaranta, Luca Tassoni, Ambra Pastori, Edoardo Giussani, Alice Fusaro, Isabella Monne, Calogero Terregino, Antonia Ricci |
| EPI_ISL_3308794, EPI_ISL_3308795, EPI_ISL_3308796, EPI_ISL_3308797, EPI_ISL_3308798, EPI_ISL_3308799, EPI_ISL_3308800, EPI_ISL_3308801, EPI_ISL_3308802, EPI_ISL_3308803 | AULSS 9 Scaligera - San Bonifacio | Istituto Zooprofilattico Sperimentale delle Venezie | Adelaide Milani, Alessia Schivo, Annalisa Salviato, Elisa Palumbo, Erika Giorgia Quaranta, Luca Tassoni, Ambra Pastori, Edoardo Giussani, Alice Fusaro, Isabella Monne, Calogero Terregino, Antonia Ricci |
| EPI_ISL_3308804 | AULSS 9 Scaligera - Villafranca | Istituto Zooprofilattico Sperimentale delle Venezie | Adelaide Milani, Alessia Schivo, Annalisa Salviato, Elisa Palumbo, Erika Giorgia Quaranta, Luca Tassoni, Ambra Pastori, Edoardo Giussani, Alice Fusaro, Isabella Monne, Calogero Terregino, Antonia Ricci |
| EPI_ISL_3308805 | AULSS 9 Scaligera - San Bonifacio | Istituto Zooprofilattico Sperimentale delle Venezie | Adelaide Milani, Alessia Schivo, Annalisa Salviato, Elisa Palumbo, Erika Giorgia Quaranta, Luca Tassoni, Ambra Pastori, Edoardo Giussani, Alice Fusaro, Isabella Monne, Calogero Terregino, Antonia Ricci |
| EPI_ISL_3308806, EPI_ISL_3308807, EPI_ISL_3308808 | AULSS 9 Scaligera - Villafranca | Istituto Zooprofilattico Sperimentale delle Venezie | Adelaide Milani, Alessia Schivo, Annalisa Salviato, Elisa Palumbo, Erika Giorgia Quaranta, Luca Tassoni, Ambra Pastori, Edoardo Giussani, Alice Fusaro, Isabella Monne, Calogero Terregino, Antonia Ricci |
| EPI_ISL_3308809 | AULSS 7 Pedemontana - Santorso | Istituto Zooprofilattico Sperimentale delle Venezie | Adelaide Milani, Alessia Schivo, Annalisa Salviato, Elisa Palumbo, Erika Giorgia Quaranta, Luca Tassoni, Ambra Pastori, Edoardo Giussani, Alice Fusaro, Isabella Monne, Calogero Terregino, Antonia Ricci |
| EPI_ISL_3308810 | AULSS 9 Scaligera - Villafranca | Istituto Zooprofilattico Sperimentale delle Venezie | Adelaide Milani, Alessia Schivo, Annalisa Salviato, Elisa Palumbo, Erika Giorgia Quaranta, Luca Tassoni, Ambra Pastori, Edoardo Giussani, Alice Fusaro, Isabella Monne, Calogero Terregino, Antonia Ricci |
| EPI_ISL_3308811, | AULSS 7 Pedemontana - Santorso | Istituto Zooprofilattico Sperimentale delle Venezie | Adelaide Milani, Alessia Schivo, Annalisa Salviato, Elisa Palumbo, Erika Giorgia Quaranta, Luca Tassoni, Ambra Pastori, Edoardo Giussani, Alice Fusaro, Isabella Monne, Calogero Terregino, Antonia Ricci |

| | | | |
|---|---|---|---|
| EPI_ISL_3308812, EPI_ISL_3308813, EPI_ISL_3308814 | | | |
| EPI_ISL_3308815 | AULSS 1 Dolomiti | Istituto Zooprofilattico Sperimentale delle Venezie | Adelaide Milani, Alessia Schivo, Annalisa Salviato, Elisa Palumbo, Erika Giorgia Quaranta, Luca Tassoni, Ambra Pastori, Edoardo Giussani, Alice Fusaro, Isabella Monne, Calogero Terregino, Antonia Ricci |
| EPI_ISL_3308816 | AULSS 5 Polesana | Istituto Zooprofilattico Sperimentale delle Venezie | Adelaide Milani, Alessia Schivo, Annalisa Salviato, Elisa Palumbo, Erika Giorgia Quaranta, Luca Tassoni, Ambra Pastori, Edoardo Giussani, Alice Fusaro, Isabella Monne, Calogero Terregino, Antonia Ricci |
| EPI_ISL_3308817, EPI_ISL_3308818 | AULSS 1 Dolomiti | Istituto Zooprofilattico Sperimentale delle Venezie | Adelaide Milani, Alessia Schivo, Annalisa Salviato, Elisa Palumbo, Erika Giorgia Quaranta, Luca Tassoni, Ambra Pastori, Edoardo Giussani, Alice Fusaro, Isabella Monne, Calogero Terregino, Antonia Ricci |
| EPI_ISL_3308819, EPI_ISL_3308820, EPI_ISL_3308821, EPI_ISL_3308822, EPI_ISL_3308823, EPI_ISL_3308825 | AULSS 5 Polesana | Istituto Zooprofilattico Sperimentale delle Venezie | Adelaide Milani, Alessia Schivo, Annalisa Salviato, Elisa Palumbo, Erika Giorgia Quaranta, Luca Tassoni, Ambra Pastori, Edoardo Giussani, Alice Fusaro, Isabella Monne, Calogero Terregino, Antonia Ricci |
| EPI_ISL_3308826, EPI_ISL_3308827, EPI_ISL_3308829, EPI_ISL_3308830, EPI_ISL_3308831, EPI_ISL_3308832, EPI_ISL_3308833, EPI_ISL_3308834, EPI_ISL_3308835 | AULSS 2 Marca Trevigiana | Istituto Zooprofilattico Sperimentale delle Venezie | Adelaide Milani, Alessia Schivo, Annalisa Salviato, Elisa Palumbo, Erika Giorgia Quaranta, Luca Tassoni, Ambra Pastori, Edoardo Giussani, Alice Fusaro, Isabella Monne, Calogero Terregino, Antonia Ricci |
| EPI_ISL_3308836 | AOPD | Istituto Zooprofilattico Sperimentale delle Venezie | Adelaide Milani, Alessia Schivo, Annalisa Salviato, Elisa Palumbo, Erika Giorgia Quaranta, Luca Tassoni, Ambra Pastori, Edoardo Giussani, Alice Fusaro, Isabella Monne, Calogero Terregino, Antonia Ricci |
| EPI_ISL_3308837, EPI_ISL_3308838, EPI_ISL_3308839, EPI_ISL_3308840, EPI_ISL_3308841 | AOUIVR | Istituto Zooprofilattico Sperimentale delle Venezie | Adelaide Milani, Alessia Schivo, Annalisa Salviato, Elisa Palumbo, Erika Giorgia Quaranta, Luca Tassoni, Ambra Pastori, Edoardo Giussani, Alice Fusaro, Isabella Monne, Calogero Terregino, Antonia Ricci |
| EPI_ISL_3308842, EPI_ISL_3308843, EPI_ISL_3308844, EPI_ISL_3308845, EPI_ISL_3308846, EPI_ISL_3308847, EPI_ISL_3308848, EPI_ISL_3308849, EPI_ISL_3308850, EPI_ISL_3308851 | AULSS 9 Scaligera - San Bonifacio | Istituto Zooprofilattico Sperimentale delle Venezie | Adelaide Milani, Alessia Schivo, Annalisa Salviato, Elisa Palumbo, Erika Giorgia Quaranta, Luca Tassoni, Ambra Pastori, Edoardo Giussani, Alice Fusaro, Isabella Monne, Calogero Terregino, Antonia Ricci |
| EPI_ISL_3308852, EPI_ISL_3308853, EPI_ISL_3308855, EPI_ISL_3308856, EPI_ISL_3308857, EPI_ISL_3308858 | AULSS 7 Pedemontana - Santorso | Istituto Zooprofilattico Sperimentale delle Venezie | Adelaide Milani, Alessia Schivo, Annalisa Salviato, Elisa Palumbo, Erika Giorgia Quaranta, Luca Tassoni, Ambra Pastori, Edoardo Giussani, Alice Fusaro, Isabella Monne, Calogero Terregino, Antonia Ricci |
| EPI_ISL_3308859, EPI_ISL_3308860, EPI_ISL_3308861, EPI_ISL_3308862, EPI_ISL_3308863, EPI_ISL_3308864, EPI_ISL_3308865, EPI_ISL_3308866, EPI_ISL_3308867, EPI_ISL_3308868 | AULSS 7 Pedemontana - Bassano | Istituto Zooprofilattico Sperimentale delle Venezie | Adelaide Milani, Alessia Schivo, Annalisa Salviato, Elisa Palumbo, Erika Giorgia Quaranta, Luca Tassoni, Ambra Pastori, Edoardo Giussani, Alice Fusaro, Isabella Monne, Calogero Terregino, Antonia Ricci |
| EPI_ISL_3308869 | AULSS 8 Berica | Istituto Zooprofilattico Sperimentale delle Venezie | Adelaide Milani, Alessia Schivo, Annalisa Salviato, Elisa Palumbo, Erika Giorgia Quaranta, Luca Tassoni, Ambra Pastori, Edoardo Giussani, Alice Fusaro, Isabella Monne, Calogero Terregino, Antonia Ricci |
| EPI_ISL_3308871, EPI_ISL_3308872, EPI_ISL_3308873, EPI_ISL_3308874, EPI_ISL_3308875, EPI_ISL_3308876, EPI_ISL_3308877, EPI_ISL_3308878, EPI_ISL_3308879, EPI_ISL_3308880, EPI_ISL_3308881, EPI_ISL_3308882 | | | |
| see above | AULSS 1 Dolomiti | Istituto Zooprofilattico Sperimentale delle Venezie | Adelaide Milani, Alessia Schivo, Annalisa Salviato, Elisa Palumbo, Erika Giorgia Quaranta, Luca Tassoni, Ambra Pastori, Edoardo Giussani, Alice Fusaro, Isabella Monne, Calogero Terregino, Antonia Ricci |
| EPI_ISL_3308883 | AOPD | Istituto Zooprofilattico Sperimentale delle Venezie | Adelaide Milani, Alessia Schivo, Annalisa Salviato, Elisa Palumbo, Erika Giorgia Quaranta, Luca Tassoni, Ambra Pastori, Edoardo Giussani, Alice Fusaro, Isabella Monne, Calogero Terregino, Antonia Ricci |
| EPI_ISL_3308884 | AULSS 2 Marca Trevigiana | Istituto Zooprofilattico Sperimentale delle Venezie | Adelaide Milani, Alessia Schivo, Annalisa Salviato, Elisa Palumbo, Erika Giorgia Quaranta, Luca Tassoni, Ambra Pastori, Edoardo Giussani, Alice Fusaro, Isabella Monne, Calogero Terregino, Antonia Ricci |
| EPI_ISL_3308885 | AOPD | Istituto Zooprofilattico Sperimentale delle Venezie | Adelaide Milani, Alessia Schivo, Annalisa Salviato, Elisa Palumbo, Erika Giorgia Quaranta, Luca Tassoni, Ambra Pastori, Edoardo Giussani, Alice Fusaro, Isabella Monne, Calogero Terregino, Antonia Ricci |
| EPI_ISL_3308887 | AULSS 2 Marca Trevigiana | Istituto Zooprofilattico Sperimentale delle Venezie | Adelaide Milani, Alessia Schivo, Annalisa Salviato, Elisa Palumbo, Erika Giorgia Quaranta, Luca Tassoni, Ambra Pastori, Edoardo Giussani, Alice Fusaro, Isabella Monne, Calogero Terregino, Antonia Ricci |
| EPI_ISL_3308888 | AOPD | Istituto Zooprofilattico Sperimentale delle Venezie | Adelaide Milani, Alessia Schivo, Annalisa Salviato, Elisa Palumbo, Erika Giorgia Quaranta, Luca Tassoni, Ambra Pastori, Edoardo Giussani, Alice Fusaro, Isabella Monne, Calogero Terregino, Antonia Ricci |
| EPI_ISL_3308889, EPI_ISL_3308890, EPI_ISL_3308891, EPI_ISL_3308892, EPI_ISL_3308893, EPI_ISL_3308894, EPI_ISL_3308895 | AULSS 2 Marca Trevigiana | Istituto Zooprofilattico Sperimentale delle Venezie | Adelaide Milani, Alessia Schivo, Annalisa Salviato, Elisa Palumbo, Erika Giorgia Quaranta, Luca Tassoni, Ambra Pastori, Edoardo Giussani, Alice Fusaro, Isabella Monne, Calogero Terregino, Antonia Ricci |
| EPI_ISL_3308896, EPI_ISL_3308897 | AULSS 9 Scaligera - Legnago | Istituto Zooprofilattico Sperimentale delle Venezie | Adelaide Milani, Alessia Schivo, Annalisa Salviato, Elisa Palumbo, Erika Giorgia Quaranta, Luca Tassoni, Ambra Pastori, Edoardo Giussani, Alice Fusaro, Isabella Monne, Calogero Terregino, Antonia Ricci |
| EPI_ISL_3308898 | AULSS 2 Marca Trevigiana | Istituto Zooprofilattico Sperimentale delle Venezie | Adelaide Milani, Alessia Schivo, Annalisa Salviato, Elisa Palumbo, Erika Giorgia Quaranta, Luca Tassoni, Ambra Pastori, Edoardo Giussani, Alice Fusaro, Isabella Monne, Calogero Terregino, Antonia Ricci |
| EPI_ISL_3308899, EPI_ISL_3308900, EPI_ISL_3308901 | AULSS 9 Scaligera - Legnago | Istituto Zooprofilattico Sperimentale delle Venezie | Adelaide Milani, Alessia Schivo, Annalisa Salviato, Elisa Palumbo, Erika Giorgia Quaranta, Luca Tassoni, Ambra Pastori, Edoardo Giussani, Alice Fusaro, Isabella Monne, Calogero Terregino, Antonia Ricci |
| EPI_ISL_3308902 | AULSS 9 Scaligera - San Bonifacio | Istituto Zooprofilattico Sperimentale delle Venezie | Adelaide Milani, Alessia Schivo, Annalisa Salviato, Elisa Palumbo, Erika Giorgia Quaranta, Luca Tassoni, Ambra Pastori, Edoardo Giussani, Alice Fusaro, Isabella Monne, Calogero Terregino, Antonia Ricci |
| EPI_ISL_3308903 | AULSS 9 Scaligera - Legnago | Istituto Zooprofilattico Sperimentale delle Venezie | Adelaide Milani, Alessia Schivo, Annalisa Salviato, Elisa Palumbo, Erika Giorgia Quaranta, Luca Tassoni, Ambra Pastori, Edoardo Giussani, Alice Fusaro, Isabella Monne, Calogero Terregino, Antonia Ricci |
| EPI_ISL_3308904, EPI_ISL_3308905, EPI_ISL_3308906, EPI_ISL_3308907, EPI_ISL_3308908, EPI_ISL_3308909, EPI_ISL_3308910, EPI_ISL_3308911, EPI_ISL_3308912, EPI_ISL_3308913 | AULSS 9 Scaligera - San Bonifacio | Istituto Zooprofilattico Sperimentale delle Venezie | Adelaide Milani, Alessia Schivo, Annalisa Salviato, Elisa Palumbo, Erika Giorgia Quaranta, Luca Tassoni, Ambra Pastori, Edoardo Giussani, Alice Fusaro, Isabella Monne, Calogero Terregino, Antonia Ricci |
| EPI_ISL_3308914 | AULSS 6 Euganea | Istituto Zooprofilattico Sperimentale delle Venezie | Adelaide Milani, Alessia Schivo, Annalisa Salviato, Elisa Palumbo, Erika Giorgia Quaranta, Luca Tassoni, Ambra Pastori, Edoardo Giussani, Alice Fusaro, Isabella Monne, Calogero Terregino, Antonia Ricci |
| EPI_ISL_3308915 | AULSS 9 Scaligera - San Bonifacio | Istituto Zooprofilattico Sperimentale delle Venezie | Adelaide Milani, Alessia Schivo, Annalisa Salviato, Elisa Palumbo, Erika Giorgia Quaranta, Luca Tassoni, Ambra Pastori, Edoardo Giussani, Alice Fusaro, Isabella Monne, Calogero Terregino, Antonia Ricci |
| EPI_ISL_3308916, EPI_ISL_3308917, EPI_ISL_3308918, EPI_ISL_3308919, EPI_ISL_3308920, EPI_ISL_3308921, EPI_ISL_3308922, EPI_ISL_3308923 | AULSS 6 Euganea | Istituto Zooprofilattico Sperimentale delle Venezie | Adelaide Milani, Alessia Schivo, Annalisa Salviato, Elisa Palumbo, Erika Giorgia Quaranta, Luca Tassoni, Ambra Pastori, Edoardo Giussani, Alice Fusaro, Isabella Monne, Calogero Terregino, Antonia Ricci |

| | | | |
|---|---|---|---|
| EPI_ISL_3308924, EPI_ISL_3308925 | AULSS 7 Pedemontana - Santorso | Istituto Zooprofilattico Sperimentale delle Venezie | Adelaide Milani, Alessia Schivo, Annalisa Salviato, Elisa Palumbo, Erika Giorgia Quaranta, Luca Tassoni, Ambra Pastori, Edoardo Giussani, Alice Fusaro, Isabella Monne, Calogero Terregino, Antonia Ricci |
| EPI_ISL_3308926 | AULSS 6 Euganea | Istituto Zooprofilattico Sperimentale delle Venezie | Adelaide Milani, Alessia Schivo, Annalisa Salviato, Elisa Palumbo, Erika Giorgia Quaranta, Luca Tassoni, Ambra Pastori, Edoardo Giussani, Alice Fusaro, Isabella Monne, Calogero Terregino, Antonia Ricci |
| EPI_ISL_3308927, EPI_ISL_3308928, EPI_ISL_3308929, EPI_ISL_3308930, EPI_ISL_3308931 | AULSS 7 Pedemontana - Santorso | Istituto Zooprofilattico Sperimentale delle Venezie | Adelaide Milani, Alessia Schivo, Annalisa Salviato, Elisa Palumbo, Erika Giorgia Quaranta, Luca Tassoni, Ambra Pastori, Edoardo Giussani, Alice Fusaro, Isabella Monne, Calogero Terregino, Antonia Ricci |
| EPI_ISL_3308932, EPI_ISL_3308933, EPI_ISL_3308934, EPI_ISL_3308935, EPI_ISL_3308936, EPI_ISL_3308937, EPI_ISL_3308938, EPI_ISL_3308939, EPI_ISL_3308940 | AULSS 7 Pedemontana - Bassano | Istituto Zooprofilattico Sperimentale delle Venezie | Adelaide Milani, Alessia Schivo, Annalisa Salviato, Elisa Palumbo, Erika Giorgia Quaranta, Luca Tassoni, Ambra Pastori, Edoardo Giussani, Alice Fusaro, Isabella Monne, Calogero Terregino, Antonia Ricci |
| EPI_ISL_3308941 | AULSS 8 Berica | Istituto Zooprofilattico Sperimentale delle Venezie | Adelaide Milani, Alessia Schivo, Annalisa Salviato, Elisa Palumbo, Erika Giorgia Quaranta, Luca Tassoni, Ambra Pastori, Edoardo Giussani, Alice Fusaro, Isabella Monne, Calogero Terregino, Antonia Ricci |
| EPI_ISL_3308942 | AULSS 7 Pedemontana - Bassano | Istituto Zooprofilattico Sperimentale delle Venezie | Adelaide Milani, Alessia Schivo, Annalisa Salviato, Elisa Palumbo, Erika Giorgia Quaranta, Luca Tassoni, Ambra Pastori, Edoardo Giussani, Alice Fusaro, Isabella Monne, Calogero Terregino, Antonia Ricci |
| EPI_ISL_3308943, EPI_ISL_3308944, EPI_ISL_3308945, EPI_ISL_3308946, EPI_ISL_3308947, EPI_ISL_3308948, EPI_ISL_3308949, EPI_ISL_3308950, EPI_ISL_3308951, EPI_ISL_3308952, EPI_ISL_3308953, EPI_ISL_3308954, EPI_ISL_3308955 | | | |
| see above | AULSS 8 Berica | Istituto Zooprofilattico Sperimentale delle Venezie | Adelaide Milani, Alessia Schivo, Annalisa Salviato, Elisa Palumbo, Erika Giorgia Quaranta, Luca Tassoni, Ambra Pastori, Edoardo Giussani, Alice Fusaro, Isabella Monne, Calogero Terregino, Antonia Ricci |
| EPI_ISL_3308956 | AULSS 5 Polesana | Istituto Zooprofilattico Sperimentale delle Venezie | Adelaide Milani, Alessia Schivo, Annalisa Salviato, Elisa Palumbo, Erika Giorgia Quaranta, Luca Tassoni, Ambra Pastori, Edoardo Giussani, Alice Fusaro, Isabella Monne, Calogero Terregino, Antonia Ricci |
| EPI_ISL_3308957, EPI_ISL_3308958, EPI_ISL_3308959 | AULSS 8 Berica | Istituto Zooprofilattico Sperimentale delle Venezie | Adelaide Milani, Alessia Schivo, Annalisa Salviato, Elisa Palumbo, Erika Giorgia Quaranta, Luca Tassoni, Ambra Pastori, Edoardo Giussani, Alice Fusaro, Isabella Monne, Calogero Terregino, Antonia Ricci |
| EPI_ISL_3308960, EPI_ISL_3308961, EPI_ISL_3308962 | AULSS 5 Polesana | Istituto Zooprofilattico Sperimentale delle Venezie | Adelaide Milani, Alessia Schivo, Annalisa Salviato, Elisa Palumbo, Erika Giorgia Quaranta, Luca Tassoni, Ambra Pastori, Edoardo Giussani, Alice Fusaro, Isabella Monne, Calogero Terregino, Antonia Ricci |
| EPI_ISL_3308969 | Lighthouse Laboratory Plymouth | Wellcome Sanger Institute for the COVID-19 Genomics UK (COG-UK) Consortium | Lighthouse Laboratory Plymouth and Alex Alderton, Roberto Amato, Jeffrey Barrett, Sonia Goncalves, Ewan Harrison, David K. Jackson, Ian Johnston, Dominic Kwiatkowski, Cordelia Langford, John Sillitoe on behalf of the Wellcome Sanger Institute COVID-19 Surveillance Team |
| EPI_ISL_3308970, EPI_ISL_3308971, EPI_ISL_3308972, EPI_ISL_3308973, EPI_ISL_3308974 | Lighthouse Lab in Milton Keynes | Wellcome Sanger Institute for the COVID-19 Genomics UK (COG-UK) Consortium | The Lighthouse Lab in Milton Keynes and Alex Alderton, Roberto Amato, Jeffrey Barrett, Sonia Goncalves, Ewan Harrison, David K. Jackson, Ian Johnston, Dominic Kwiatkowski, Cordelia Langford, John Sillitoe on behalf of the Wellcome Sanger Institute COVID-19 Surveillance Team |
| EPI_ISL_3308975 | Lighthouse Laboratory Plymouth | Wellcome Sanger Institute for the COVID-19 Genomics UK (COG-UK) Consortium | Lighthouse Laboratory Plymouth and Alex Alderton, Roberto Amato, Jeffrey Barrett, Sonia Goncalves, Ewan Harrison, David K. Jackson, Ian Johnston, Dominic Kwiatkowski, Cordelia Langford, John Sillitoe on behalf of the Wellcome Sanger Institute COVID-19 Surveillance Team |
| EPI_ISL_3308976, EPI_ISL_3308977, EPI_ISL_3308978, EPI_ISL_3308979, EPI_ISL_3308980, EPI_ISL_3308981, EPI_ISL_3308982 | Lighthouse Lab in Milton Keynes | Wellcome Sanger Institute for the COVID-19 Genomics UK (COG-UK) Consortium | The Lighthouse Lab in Milton Keynes and Alex Alderton, Roberto Amato, Jeffrey Barrett, Sonia Goncalves, Ewan Harrison, David K. Jackson, Ian Johnston, Dominic Kwiatkowski, Cordelia Langford, John Sillitoe on behalf of the Wellcome Sanger Institute COVID-19 Surveillance Team |
| EPI_ISL_3308983 | Lighthouse Laboratory Plymouth | Wellcome Sanger Institute for the COVID-19 Genomics UK (COG-UK) Consortium | Lighthouse Laboratory Plymouth and Alex Alderton, Roberto Amato, Jeffrey Barrett, Sonia Goncalves, Ewan Harrison, David K. Jackson, Ian Johnston, Dominic Kwiatkowski, Cordelia Langford, John Sillitoe on behalf of the Wellcome Sanger Institute COVID-19 Surveillance Team |
| EPI_ISL_3308984 | Lighthouse Lab in Milton Keynes | Wellcome Sanger Institute for the COVID-19 Genomics UK (COG-UK) Consortium | The Lighthouse Lab in Milton Keynes and Alex Alderton, Roberto Amato, Jeffrey Barrett, Sonia Goncalves, Ewan Harrison, David K. Jackson, Ian Johnston, Dominic Kwiatkowski, Cordelia Langford, John Sillitoe on behalf of the Wellcome Sanger Institute COVID-19 Surveillance Team |
| EPI_ISL_3308985 | Lighthouse Laboratory Plymouth | Wellcome Sanger Institute for the COVID-19 Genomics UK (COG-UK) Consortium | Lighthouse Laboratory Plymouth and Alex Alderton, Roberto Amato, Jeffrey Barrett, Sonia Goncalves, Ewan Harrison, David K. Jackson, Ian Johnston, Dominic Kwiatkowski, Cordelia Langford, John Sillitoe on behalf of the Wellcome Sanger Institute COVID-19 Surveillance Team |
| EPI_ISL_3308986, EPI_ISL_3308987 | Lighthouse Lab in Milton Keynes | Wellcome Sanger Institute for the COVID-19 Genomics UK (COG-UK) Consortium | The Lighthouse Lab in Milton Keynes and Alex Alderton, Roberto Amato, Jeffrey Barrett, Sonia Goncalves, Ewan Harrison, David K. Jackson, Ian Johnston, Dominic Kwiatkowski, Cordelia Langford, John Sillitoe on behalf of the Wellcome Sanger Institute COVID-19 Surveillance Team |
| EPI_ISL_3308988 | Lighthouse Laboratory Plymouth | Wellcome Sanger Institute for the COVID-19 Genomics UK (COG-UK) Consortium | Lighthouse Laboratory Plymouth and Alex Alderton, Roberto Amato, Jeffrey Barrett, Sonia Goncalves, Ewan Harrison, David K. Jackson, Ian Johnston, Dominic Kwiatkowski, Cordelia Langford, John Sillitoe on behalf of the Wellcome Sanger Institute COVID-19 Surveillance Team |
| EPI_ISL_3308989, EPI_ISL_3308990, EPI_ISL_3308991 | Lighthouse Lab in Milton Keynes | Wellcome Sanger Institute for the COVID-19 Genomics UK (COG-UK) Consortium | The Lighthouse Lab in Milton Keynes and Alex Alderton, Roberto Amato, Jeffrey Barrett, Sonia Goncalves, Ewan Harrison, David K. Jackson, Ian Johnston, Dominic Kwiatkowski, Cordelia Langford, John Sillitoe on behalf of the Wellcome Sanger Institute COVID-19 Surveillance Team |
| EPI_ISL_3308992 | Lighthouse Laboratory Plymouth | Wellcome Sanger Institute for the COVID-19 Genomics UK (COG-UK) Consortium | Lighthouse Laboratory Plymouth and Alex Alderton, Roberto Amato, Jeffrey Barrett, Sonia Goncalves, Ewan Harrison, David K. Jackson, Ian Johnston, Dominic Kwiatkowski, Cordelia Langford, John Sillitoe on behalf of the Wellcome Sanger Institute COVID-19 Surveillance Team |
| EPI_ISL_3308993, EPI_ISL_3308994, EPI_ISL_3308995, EPI_ISL_3308996, EPI_ISL_3308997, EPI_ISL_3308998, EPI_ISL_3308999, EPI_ISL_3309000, EPI_ISL_3309001 | Lighthouse Lab in Milton Keynes | Wellcome Sanger Institute for the COVID-19 Genomics UK (COG-UK) Consortium | The Lighthouse Lab in Milton Keynes and Alex Alderton, Roberto Amato, Jeffrey Barrett, Sonia Goncalves, Ewan Harrison, David K. Jackson, Ian Johnston, Dominic Kwiatkowski, Cordelia Langford, John Sillitoe on behalf of the Wellcome Sanger Institute COVID-19 Surveillance Team |
| EPI_ISL_3309002 | Lighthouse Laboratory Plymouth | Wellcome Sanger Institute for the COVID-19 Genomics UK (COG-UK) Consortium | Lighthouse Laboratory Plymouth and Alex Alderton, Roberto Amato, Jeffrey Barrett, Sonia Goncalves, Ewan Harrison, David K. Jackson, Ian Johnston, Dominic Kwiatkowski, Cordelia Langford, John Sillitoe on behalf of the Wellcome Sanger Institute COVID-19 Surveillance Team |
| EPI_ISL_3309003 | Lighthouse Lab in Milton Keynes | Wellcome Sanger Institute for the COVID-19 Genomics UK (COG-UK) Consortium | The Lighthouse Lab in Milton Keynes and Alex Alderton, Roberto Amato, Jeffrey Barrett, Sonia Goncalves, Ewan Harrison, David K. Jackson, Ian Johnston, Dominic Kwiatkowski, Cordelia Langford, John Sillitoe on behalf of the Wellcome Sanger Institute COVID-19 Surveillance Team |
| EPI_ISL_3309004 | Lighthouse Laboratory Plymouth | Wellcome Sanger Institute for the COVID-19 Genomics UK (COG-UK) Consortium | Lighthouse Laboratory Plymouth and Alex Alderton, Roberto Amato, Jeffrey Barrett, Sonia Goncalves, Ewan Harrison, David K. Jackson, Ian Johnston, Dominic Kwiatkowski, Cordelia Langford, John Sillitoe on behalf of the Wellcome Sanger Institute COVID-19 Surveillance Team |
| EPI_ISL_3309005, EPI_ISL_3309006, EPI_ISL_3309007 | Lighthouse Lab in Milton Keynes | Wellcome Sanger Institute for the COVID-19 Genomics UK (COG-UK) Consortium | The Lighthouse Lab in Milton Keynes and Alex Alderton, Roberto Amato, Jeffrey Barrett, Sonia Goncalves, Ewan Harrison, David K. Jackson, Ian Johnston, Dominic Kwiatkowski, Cordelia Langford, John Sillitoe on behalf of the Wellcome Sanger Institute COVID-19 Surveillance Team |
| EPI_ISL_3309008 | Lighthouse Laboratory Plymouth | Wellcome Sanger Institute for the COVID-19 Genomics UK (COG-UK) Consortium | Lighthouse Laboratory Plymouth and Alex Alderton, Roberto Amato, Jeffrey Barrett, Sonia Goncalves, Ewan Harrison, David K. Jackson, Ian Johnston, Dominic Kwiatkowski, Cordelia Langford, John Sillitoe on behalf of the Wellcome Sanger Institute COVID-19 Surveillance Team |
| EPI_ISL_3309009, EPI_ISL_3309010, EPI_ISL_3309011 | Lighthouse Lab in Milton Keynes | Wellcome Sanger Institute for the COVID-19 Genomics UK (COG-UK) Consortium | The Lighthouse Lab in Milton Keynes and Alex Alderton, Roberto Amato, Jeffrey Barrett, Sonia Goncalves, Ewan Harrison, David K. Jackson, Ian Johnston, Dominic Kwiatkowski, Cordelia Langford, John Sillitoe on behalf of the Wellcome Sanger Institute COVID-19 Surveillance Team |
| EPI_ISL_3309012 | Lighthouse Laboratory Plymouth | Wellcome Sanger Institute for the COVID-19 Genomics UK (COG-UK) Consortium | Lighthouse Laboratory Plymouth and Alex Alderton, Roberto Amato, Jeffrey Barrett, Sonia Goncalves, Ewan Harrison, David K. Jackson, Ian Johnston, Dominic Kwiatkowski, Cordelia Langford, John Sillitoe on behalf of the Wellcome Sanger Institute COVID-19 Surveillance Team |
| EPI_ISL_3309013, EPI_ISL_3309014 | Lighthouse Lab in Milton Keynes | Wellcome Sanger Institute for the COVID-19 Genomics UK (COG-UK) Consortium | The Lighthouse Lab in Milton Keynes and Alex Alderton, Roberto Amato, Jeffrey Barrett, Sonia Goncalves, Ewan Harrison, David K. Jackson, Ian Johnston, Dominic Kwiatkowski, Cordelia Langford, John Sillitoe on behalf of the Wellcome Sanger Institute COVID-19 Surveillance Team |
| EPI_ISL_3309015, EPI_ISL_3309016 | Lighthouse Laboratory Plymouth | Wellcome Sanger Institute for the COVID-19 Genomics UK (COG-UK) Consortium | Lighthouse Laboratory Plymouth and Alex Alderton, Roberto Amato, Jeffrey Barrett, Sonia Goncalves, Ewan Harrison, David K. Jackson, Ian Johnston, Dominic Kwiatkowski, Cordelia Langford, John Sillitoe on behalf of the Wellcome Sanger Institute COVID-19 Surveillance Team |
| EPI_ISL_3309017, EPI_ISL_3309018, EPI_ISL_3309019 | Lighthouse Lab in Milton Keynes | Wellcome Sanger Institute for the COVID-19 Genomics UK (COG-UK) Consortium | The Lighthouse Lab in Milton Keynes and Alex Alderton, Roberto Amato, Jeffrey Barrett, Sonia Goncalves, Ewan Harrison, David K. Jackson, Ian Johnston, Dominic Kwiatkowski, Cordelia Langford, John Sillitoe on behalf of the Wellcome Sanger Institute COVID-19 Surveillance Team |
| EPI_ISL_3309020 | Lighthouse Laboratory Plymouth | Wellcome Sanger Institute for the COVID-19 Genomics UK (COG-UK) Consortium | Lighthouse Laboratory Plymouth and Alex Alderton, Roberto Amato, Jeffrey Barrett, Sonia Goncalves, Ewan Harrison, David K. Jackson, Ian Johnston, Dominic Kwiatkowski, Cordelia Langford, John Sillitoe on behalf of the Wellcome Sanger Institute COVID-19 Surveillance Team |
| EPI_ISL_3309021, EPI_ISL_3309022, | Lighthouse Lab in Milton Keynes | Wellcome Sanger Institute for the COVID-19 Genomics UK (COG-UK) Consortium | The Lighthouse Lab in Milton Keynes and Alex Alderton, Roberto Amato, Jeffrey Barrett, Sonia Goncalves, Ewan Harrison, David K. Jackson, Ian Johnston, Dominic Kwiatkowski, Cordelia Langford, John Sillitoe on behalf of the Wellcome Sanger Institute COVID-19 Surveillance Team |

| | | | |
|---|---|---|---|
| EPI_ISL_3309023, EPI_ISL_3309024, EPI_ISL_3309025, EPI_ISL_3309026, EPI_ISL_3309027, EPI_ISL_3309028, EPI_ISL_3309029 | | | |
| EPI_ISL_3309030, EPI_ISL_3309031 | Lighthouse Laboratory Plymouth | Wellcome Sanger Institute for the COVID-19 Genomics UK (COG-UK) Consortium | Lighthouse Laboratory Plymouth and Alex Alderton, Roberto Amato, Jeffrey Barrett, Sonia Goncalves, Ewan Harrison, David K. Jackson, Ian Johnston, Dominic Kwiatkowski, Cordelia Langford, John Sillitoe on behalf of the Wellcome Sanger Institute COVID-19 Surveillance Team |
| EPI_ISL_3309032 | Lighthouse Lab in Milton Keynes | Wellcome Sanger Institute for the COVID-19 Genomics UK (COG-UK) Consortium | The Lighthouse Lab in Milton Keynes and Alex Alderton, Roberto Amato, Jeffrey Barrett, Sonia Goncalves, Ewan Harrison, David K. Jackson, Ian Johnston, Dominic Kwiatkowski, Cordelia Langford, John Sillitoe on behalf of the Wellcome Sanger Institute COVID-19 Surveillance Team |
| EPI_ISL_3309033 | Lighthouse Laboratory Plymouth | Wellcome Sanger Institute for the COVID-19 Genomics UK (COG-UK) Consortium | Lighthouse Laboratory Plymouth and Alex Alderton, Roberto Amato, Jeffrey Barrett, Sonia Goncalves, Ewan Harrison, David K. Jackson, Ian Johnston, Dominic Kwiatkowski, Cordelia Langford, John Sillitoe on behalf of the Wellcome Sanger Institute COVID-19 Surveillance Team |
| EPI_ISL_3309034, EPI_ISL_3309035, EPI_ISL_3309036, EPI_ISL_3309037, EPI_ISL_3309038, EPI_ISL_3309039, EPI_ISL_3309040, EPI_ISL_3309041 | Lighthouse Lab in Milton Keynes | Wellcome Sanger Institute for the COVID-19 Genomics UK (COG-UK) Consortium | The Lighthouse Lab in Milton Keynes and Alex Alderton, Roberto Amato, Jeffrey Barrett, Sonia Goncalves, Ewan Harrison, David K. Jackson, Ian Johnston, Dominic Kwiatkowski, Cordelia Langford, John Sillitoe on behalf of the Wellcome Sanger Institute COVID-19 Surveillance Team |
| EPI_ISL_3309042 | Lighthouse Laboratory Plymouth | Wellcome Sanger Institute for the COVID-19 Genomics UK (COG-UK) Consortium | Lighthouse Laboratory Plymouth and Alex Alderton, Roberto Amato, Jeffrey Barrett, Sonia Goncalves, Ewan Harrison, David K. Jackson, Ian Johnston, Dominic Kwiatkowski, Cordelia Langford, John Sillitoe on behalf of the Wellcome Sanger Institute COVID-19 Surveillance Team |
| EPI_ISL_3309043 | Berkshire and Surrey Pathology Services Lighthouse Laboratory | Wellcome Sanger Institute for the COVID-19 Genomics UK (COG-UK) Consortium | Berkshire and Surrey Pathology Services Lighthouse Laboratory and Alex Alderton, Roberto Amato, Jeffrey Barrett, Sonia Goncalves, Ewan Harrison, David K. Jackson, Ian Johnston, Dominic Kwiatkowski, Cordelia Langford, John Sillitoe on behalf of the Wellcome Sanger Institute COVID-19 Surveillance Team |
| EPI_ISL_3309044 | Lighthouse Lab in Milton Keynes | Wellcome Sanger Institute for the COVID-19 Genomics UK (COG-UK) Consortium | The Lighthouse Lab in Milton Keynes and Alex Alderton, Roberto Amato, Jeffrey Barrett, Sonia Goncalves, Ewan Harrison, David K. Jackson, Ian Johnston, Dominic Kwiatkowski, Cordelia Langford, John Sillitoe on behalf of the Wellcome Sanger Institute COVID-19 Surveillance Team |
| EPI_ISL_3309045, EPI_ISL_3309046 | Lighthouse Laboratory Plymouth | Wellcome Sanger Institute for the COVID-19 Genomics UK (COG-UK) Consortium | Lighthouse Laboratory Plymouth and Alex Alderton, Roberto Amato, Jeffrey Barrett, Sonia Goncalves, Ewan Harrison, David K. Jackson, Ian Johnston, Dominic Kwiatkowski, Cordelia Langford, John Sillitoe on behalf of the Wellcome Sanger Institute COVID-19 Surveillance Team |
| EPI_ISL_3309047, EPI_ISL_3309048 | Lighthouse Lab in Milton Keynes | Wellcome Sanger Institute for the COVID-19 Genomics UK (COG-UK) Consortium | The Lighthouse Lab in Milton Keynes and Alex Alderton, Roberto Amato, Jeffrey Barrett, Sonia Goncalves, Ewan Harrison, David K. Jackson, Ian Johnston, Dominic Kwiatkowski, Cordelia Langford, John Sillitoe on behalf of the Wellcome Sanger Institute COVID-19 Surveillance Team |
| EPI_ISL_3309049 | Lighthouse Laboratory Plymouth | Wellcome Sanger Institute for the COVID-19 Genomics UK (COG-UK) Consortium | Lighthouse Laboratory Plymouth and Alex Alderton, Roberto Amato, Jeffrey Barrett, Sonia Goncalves, Ewan Harrison, David K. Jackson, Ian Johnston, Dominic Kwiatkowski, Cordelia Langford, John Sillitoe on behalf of the Wellcome Sanger Institute COVID-19 Surveillance Team |
| EPI_ISL_3309050 | Lighthouse Lab in Milton Keynes | Wellcome Sanger Institute for the COVID-19 Genomics UK (COG-UK) Consortium | The Lighthouse Lab in Milton Keynes and Alex Alderton, Roberto Amato, Jeffrey Barrett, Sonia Goncalves, Ewan Harrison, David K. Jackson, Ian Johnston, Dominic Kwiatkowski, Cordelia Langford, John Sillitoe on behalf of the Wellcome Sanger Institute COVID-19 Surveillance Team |
| EPI_ISL_3309051 | Lighthouse Laboratory Plymouth | Wellcome Sanger Institute for the COVID-19 Genomics UK (COG-UK) Consortium | Lighthouse Laboratory Plymouth and Alex Alderton, Roberto Amato, Jeffrey Barrett, Sonia Goncalves, Ewan Harrison, David K. Jackson, Ian Johnston, Dominic Kwiatkowski, Cordelia Langford, John Sillitoe on behalf of the Wellcome Sanger Institute COVID-19 Surveillance Team |
| EPI_ISL_3309052, EPI_ISL_3309053 | Lighthouse Lab in Milton Keynes | Wellcome Sanger Institute for the COVID-19 Genomics UK (COG-UK) Consortium | The Lighthouse Lab in Milton Keynes and Alex Alderton, Roberto Amato, Jeffrey Barrett, Sonia Goncalves, Ewan Harrison, David K. Jackson, Ian Johnston, Dominic Kwiatkowski, Cordelia Langford, John Sillitoe on behalf of the Wellcome Sanger Institute COVID-19 Surveillance Team |
| EPI_ISL_3309054, EPI_ISL_3309055 | Lighthouse Laboratory Plymouth | Wellcome Sanger Institute for the COVID-19 Genomics UK (COG-UK) Consortium | Lighthouse Laboratory Plymouth and Alex Alderton, Roberto Amato, Jeffrey Barrett, Sonia Goncalves, Ewan Harrison, David K. Jackson, Ian Johnston, Dominic Kwiatkowski, Cordelia Langford, John Sillitoe on behalf of the Wellcome Sanger Institute COVID-19 Surveillance Team |
| EPI_ISL_3309056, EPI_ISL_3309057, EPI_ISL_3309058, EPI_ISL_3309059, EPI_ISL_3309060, EPI_ISL_3309061, EPI_ISL_3309062, EPI_ISL_3309063, EPI_ISL_3309064 | Lighthouse Lab in Milton Keynes | Wellcome Sanger Institute for the COVID-19 Genomics UK (COG-UK) Consortium | The Lighthouse Lab in Milton Keynes and Alex Alderton, Roberto Amato, Jeffrey Barrett, Sonia Goncalves, Ewan Harrison, David K. Jackson, Ian Johnston, Dominic Kwiatkowski, Cordelia Langford, John Sillitoe on behalf of the Wellcome Sanger Institute COVID-19 Surveillance Team |
| EPI_ISL_3309065 | Lighthouse Laboratory Plymouth | Wellcome Sanger Institute for the COVID-19 Genomics UK (COG-UK) Consortium | Lighthouse Laboratory Plymouth and Alex Alderton, Roberto Amato, Jeffrey Barrett, Sonia Goncalves, Ewan Harrison, David K. Jackson, Ian Johnston, Dominic Kwiatkowski, Cordelia Langford, John Sillitoe on behalf of the Wellcome Sanger Institute COVID-19 Surveillance Team |
| EPI_ISL_3309066, EPI_ISL_3309067 | Lighthouse Lab in Milton Keynes | Wellcome Sanger Institute for the COVID-19 Genomics UK (COG-UK) Consortium | The Lighthouse Lab in Milton Keynes and Alex Alderton, Roberto Amato, Jeffrey Barrett, Sonia Goncalves, Ewan Harrison, David K. Jackson, Ian Johnston, Dominic Kwiatkowski, Cordelia Langford, John Sillitoe on behalf of the Wellcome Sanger Institute COVID-19 Surveillance Team |
| EPI_ISL_3309068 | Lighthouse Laboratory Plymouth | Wellcome Sanger Institute for the COVID-19 Genomics UK (COG-UK) Consortium | Lighthouse Laboratory Plymouth and Alex Alderton, Roberto Amato, Jeffrey Barrett, Sonia Goncalves, Ewan Harrison, David K. Jackson, Ian Johnston, Dominic Kwiatkowski, Cordelia Langford, John Sillitoe on behalf of the Wellcome Sanger Institute COVID-19 Surveillance Team |
| EPI_ISL_3309069, EPI_ISL_3309070, EPI_ISL_3309071, EPI_ISL_3309072, EPI_ISL_3309073, EPI_ISL_3309074, EPI_ISL_3309075 | Lighthouse Lab in Milton Keynes | Wellcome Sanger Institute for the COVID-19 Genomics UK (COG-UK) Consortium | The Lighthouse Lab in Milton Keynes and Alex Alderton, Roberto Amato, Jeffrey Barrett, Sonia Goncalves, Ewan Harrison, David K. Jackson, Ian Johnston, Dominic Kwiatkowski, Cordelia Langford, John Sillitoe on behalf of the Wellcome Sanger Institute COVID-19 Surveillance Team |
| EPI_ISL_3309076 | Lighthouse Laboratory Plymouth | Wellcome Sanger Institute for the COVID-19 Genomics UK (COG-UK) Consortium | Lighthouse Laboratory Plymouth and Alex Alderton, Roberto Amato, Jeffrey Barrett, Sonia Goncalves, Ewan Harrison, David K. Jackson, Ian Johnston, Dominic Kwiatkowski, Cordelia Langford, John Sillitoe on behalf of the Wellcome Sanger Institute COVID-19 Surveillance Team |
| EPI_ISL_3309077, EPI_ISL_3309078 | Lighthouse Lab in Milton Keynes | Wellcome Sanger Institute for the COVID-19 Genomics UK (COG-UK) Consortium | The Lighthouse Lab in Milton Keynes and Alex Alderton, Roberto Amato, Jeffrey Barrett, Sonia Goncalves, Ewan Harrison, David K. Jackson, Ian Johnston, Dominic Kwiatkowski, Cordelia Langford, John Sillitoe on behalf of the Wellcome Sanger Institute COVID-19 Surveillance Team |
| EPI_ISL_3309079 | Lighthouse Laboratory Plymouth | Wellcome Sanger Institute for the COVID-19 Genomics UK (COG-UK) Consortium | Lighthouse Laboratory Plymouth and Alex Alderton, Roberto Amato, Jeffrey Barrett, Sonia Goncalves, Ewan Harrison, David K. Jackson, Ian Johnston, Dominic Kwiatkowski, Cordelia Langford, John Sillitoe on behalf of the Wellcome Sanger Institute COVID-19 Surveillance Team |
| EPI_ISL_3309080, EPI_ISL_3309081, EPI_ISL_3309082, EPI_ISL_3309083, EPI_ISL_3309084, EPI_ISL_3309085, EPI_ISL_3309086, EPI_ISL_3309087 | Lighthouse Lab in Milton Keynes | Wellcome Sanger Institute for the COVID-19 Genomics UK (COG-UK) Consortium | The Lighthouse Lab in Milton Keynes and Alex Alderton, Roberto Amato, Jeffrey Barrett, Sonia Goncalves, Ewan Harrison, David K. Jackson, Ian Johnston, Dominic Kwiatkowski, Cordelia Langford, John Sillitoe on behalf of the Wellcome Sanger Institute COVID-19 Surveillance Team |
| EPI_ISL_3309088 | Lighthouse Laboratory Plymouth | Wellcome Sanger Institute for the COVID-19 Genomics UK (COG-UK) Consortium | Lighthouse Laboratory Plymouth and Alex Alderton, Roberto Amato, Jeffrey Barrett, Sonia Goncalves, Ewan Harrison, David K. Jackson, Ian Johnston, Dominic Kwiatkowski, Cordelia Langford, John Sillitoe on behalf of the Wellcome Sanger Institute COVID-19 Surveillance Team |
| EPI_ISL_3309089, EPI_ISL_3309090, EPI_ISL_3309091, EPI_ISL_3309092, EPI_ISL_3309093, EPI_ISL_3309094, EPI_ISL_3309095, EPI_ISL_3309096, EPI_ISL_3309097, EPI_ISL_3309099, EPI_ISL_3309100, EPI_ISL_3309101 | | | |
| see above | Lighthouse Lab in Milton Keynes | Wellcome Sanger Institute for the COVID-19 Genomics UK (COG-UK) Consortium | The Lighthouse Lab in Milton Keynes and Alex Alderton, Roberto Amato, Jeffrey Barrett, Sonia Goncalves, Ewan Harrison, David K. Jackson, Ian Johnston, Dominic Kwiatkowski, Cordelia Langford, John Sillitoe on behalf of the Wellcome Sanger Institute COVID-19 Surveillance Team |
| EPI_ISL_3309102 | Lighthouse Laboratory Plymouth | Wellcome Sanger Institute for the COVID-19 Genomics UK (COG-UK) Consortium | Lighthouse Laboratory Plymouth and Alex Alderton, Roberto Amato, Jeffrey Barrett, Sonia Goncalves, Ewan Harrison, David K. Jackson, Ian Johnston, Dominic Kwiatkowski, Cordelia Langford, John Sillitoe on behalf of the Wellcome Sanger Institute COVID-19 Surveillance Team |
| EPI_ISL_3309103 | Lighthouse Lab in Milton Keynes | Wellcome Sanger Institute for the COVID-19 Genomics UK (COG-UK) Consortium | The Lighthouse Lab in Milton Keynes and Alex Alderton, Roberto Amato, Jeffrey Barrett, Sonia Goncalves, Ewan Harrison, David K. Jackson, Ian Johnston, Dominic Kwiatkowski, Cordelia Langford, John Sillitoe on behalf of the Wellcome Sanger Institute COVID-19 Surveillance Team |
| EPI_ISL_3309104 | Lighthouse Laboratory Plymouth | Wellcome Sanger Institute for the COVID-19 Genomics UK (COG-UK) Consortium | Lighthouse Laboratory Plymouth and Alex Alderton, Roberto Amato, Jeffrey Barrett, Sonia Goncalves, Ewan Harrison, David K. Jackson, Ian Johnston, Dominic Kwiatkowski, Cordelia Langford, John Sillitoe on behalf of the Wellcome Sanger Institute COVID-19 Surveillance Team |
| EPI_ISL_3309105, EPI_ISL_3309106, EPI_ISL_3309107, EPI_ISL_3309108, EPI_ISL_3309109, EPI_ISL_3309110 | Lighthouse Lab in Milton Keynes | Wellcome Sanger Institute for the COVID-19 Genomics UK (COG-UK) Consortium | The Lighthouse Lab in Milton Keynes and Alex Alderton, Roberto Amato, Jeffrey Barrett, Sonia Goncalves, Ewan Harrison, David K. Jackson, Ian Johnston, Dominic Kwiatkowski, Cordelia Langford, John Sillitoe on behalf of the Wellcome Sanger Institute COVID-19 Surveillance Team |
| EPI_ISL_3309111 | Lighthouse Laboratory Plymouth | Wellcome Sanger Institute for the COVID-19 Genomics UK (COG-UK) Consortium | Lighthouse Laboratory Plymouth and Alex Alderton, Roberto Amato, Jeffrey Barrett, Sonia Goncalves, Ewan Harrison, David K. Jackson, Ian Johnston, Dominic Kwiatkowski, Cordelia Langford, John Sillitoe on behalf of the Wellcome Sanger Institute COVID-19 Surveillance Team |

| | | | |
|---|---|---|---|
| EPI_ISL_3309112, EPI_ISL_3309113 | Lighthouse Lab in Milton Keynes | Wellcome Sanger Institute for the COVID-19 Genomics UK (COG-UK) Consortium | The Lighthouse Lab in Milton Keynes and Alex Alderton, Roberto Amato, Jeffrey Barrett, Sonia Goncalves, Ewan Harrison, David K. Jackson, Ian Johnston, Dominic Kwiatkowski, Cordelia Langford, John Sillitoe on behalf of the Wellcome Sanger Institute COVID-19 Surveillance Team |
| EPI_ISL_3309114, EPI_ISL_3309115 | Lighthouse Laboratory Plymouth | Wellcome Sanger Institute for the COVID-19 Genomics UK (COG-UK) Consortium | Lighthouse Laboratory Plymouth and Alex Alderton, Roberto Amato, Jeffrey Barrett, Sonia Goncalves, Ewan Harrison, David K. Jackson, Ian Johnston, Dominic Kwiatkowski, Cordelia Langford, John Sillitoe on behalf of the Wellcome Sanger Institute COVID-19 Surveillance Team |
| EPI_ISL_3309116, EPI_ISL_3309117, EPI_ISL_3309118 | Lighthouse Lab in Milton Keynes | Wellcome Sanger Institute for the COVID-19 Genomics UK (COG-UK) Consortium | The Lighthouse Lab in Milton Keynes and Alex Alderton, Roberto Amato, Jeffrey Barrett, Sonia Goncalves, Ewan Harrison, David K. Jackson, Ian Johnston, Dominic Kwiatkowski, Cordelia Langford, John Sillitoe on behalf of the Wellcome Sanger Institute COVID-19 Surveillance Team |
| EPI_ISL_3309119, EPI_ISL_3309120 | Lighthouse Laboratory Plymouth | Wellcome Sanger Institute for the COVID-19 Genomics UK (COG-UK) Consortium | Lighthouse Laboratory Plymouth and Alex Alderton, Roberto Amato, Jeffrey Barrett, Sonia Goncalves, Ewan Harrison, David K. Jackson, Ian Johnston, Dominic Kwiatkowski, Cordelia Langford, John Sillitoe on behalf of the Wellcome Sanger Institute COVID-19 Surveillance Team |
| EPI_ISL_3309121, EPI_ISL_3309122, EPI_ISL_3309123 | Lighthouse Lab in Milton Keynes | Wellcome Sanger Institute for the COVID-19 Genomics UK (COG-UK) Consortium | The Lighthouse Lab in Milton Keynes and Alex Alderton, Roberto Amato, Jeffrey Barrett, Sonia Goncalves, Ewan Harrison, David K. Jackson, Ian Johnston, Dominic Kwiatkowski, Cordelia Langford, John Sillitoe on behalf of the Wellcome Sanger Institute COVID-19 Surveillance Team |
| EPI_ISL_3309124, EPI_ISL_3309125, EPI_ISL_3309126 | Lighthouse Laboratory Plymouth | Wellcome Sanger Institute for the COVID-19 Genomics UK (COG-UK) Consortium | Lighthouse Laboratory Plymouth and Alex Alderton, Roberto Amato, Jeffrey Barrett, Sonia Goncalves, Ewan Harrison, David K. Jackson, Ian Johnston, Dominic Kwiatkowski, Cordelia Langford, John Sillitoe on behalf of the Wellcome Sanger Institute COVID-19 Surveillance Team |
| EPI_ISL_3309127 | Lighthouse Lab in Milton Keynes | Wellcome Sanger Institute for the COVID-19 Genomics UK (COG-UK) Consortium | The Lighthouse Lab in Milton Keynes and Alex Alderton, Roberto Amato, Jeffrey Barrett, Sonia Goncalves, Ewan Harrison, David K. Jackson, Ian Johnston, Dominic Kwiatkowski, Cordelia Langford, John Sillitoe on behalf of the Wellcome Sanger Institute COVID-19 Surveillance Team |
| EPI_ISL_3309128 | Lighthouse Laboratory Plymouth | Wellcome Sanger Institute for the COVID-19 Genomics UK (COG-UK) Consortium | Lighthouse Laboratory Plymouth and Alex Alderton, Roberto Amato, Jeffrey Barrett, Sonia Goncalves, Ewan Harrison, David K. Jackson, Ian Johnston, Dominic Kwiatkowski, Cordelia Langford, John Sillitoe on behalf of the Wellcome Sanger Institute COVID-19 Surveillance Team |
| EPI_ISL_3309129 | Lighthouse Lab in Milton Keynes | Wellcome Sanger Institute for the COVID-19 Genomics UK (COG-UK) Consortium | The Lighthouse Lab in Milton Keynes and Alex Alderton, Roberto Amato, Jeffrey Barrett, Sonia Goncalves, Ewan Harrison, David K. Jackson, Ian Johnston, Dominic Kwiatkowski, Cordelia Langford, John Sillitoe on behalf of the Wellcome Sanger Institute COVID-19 Surveillance Team |
| EPI_ISL_3309130 | Lighthouse Laboratory Plymouth | Wellcome Sanger Institute for the COVID-19 Genomics UK (COG-UK) Consortium | Lighthouse Laboratory Plymouth and Alex Alderton, Roberto Amato, Jeffrey Barrett, Sonia Goncalves, Ewan Harrison, David K. Jackson, Ian Johnston, Dominic Kwiatkowski, Cordelia Langford, John Sillitoe on behalf of the Wellcome Sanger Institute COVID-19 Surveillance Team |
| EPI_ISL_3309131, EPI_ISL_3309132, EPI_ISL_3309133, EPI_ISL_3309134, EPI_ISL_3309135 | Lighthouse Lab in Milton Keynes | Wellcome Sanger Institute for the COVID-19 Genomics UK (COG-UK) Consortium | The Lighthouse Lab in Milton Keynes and Alex Alderton, Roberto Amato, Jeffrey Barrett, Sonia Goncalves, Ewan Harrison, David K. Jackson, Ian Johnston, Dominic Kwiatkowski, Cordelia Langford, John Sillitoe on behalf of the Wellcome Sanger Institute COVID-19 Surveillance Team |
| EPI_ISL_3309136 | Lighthouse Laboratory Plymouth | Wellcome Sanger Institute for the COVID-19 Genomics UK (COG-UK) Consortium | Lighthouse Laboratory Plymouth and Alex Alderton, Roberto Amato, Jeffrey Barrett, Sonia Goncalves, Ewan Harrison, David K. Jackson, Ian Johnston, Dominic Kwiatkowski, Cordelia Langford, John Sillitoe on behalf of the Wellcome Sanger Institute COVID-19 Surveillance Team |
| EPI_ISL_3309137 | Lighthouse Lab in Milton Keynes | Wellcome Sanger Institute for the COVID-19 Genomics UK (COG-UK) Consortium | The Lighthouse Lab in Milton Keynes and Alex Alderton, Roberto Amato, Jeffrey Barrett, Sonia Goncalves, Ewan Harrison, David K. Jackson, Ian Johnston, Dominic Kwiatkowski, Cordelia Langford, John Sillitoe on behalf of the Wellcome Sanger Institute COVID-19 Surveillance Team |
| EPI_ISL_3309138, EPI_ISL_3309139 | Lighthouse Laboratory Plymouth | Wellcome Sanger Institute for the COVID-19 Genomics UK (COG-UK) Consortium | Lighthouse Laboratory Plymouth and Alex Alderton, Roberto Amato, Jeffrey Barrett, Sonia Goncalves, Ewan Harrison, David K. Jackson, Ian Johnston, Dominic Kwiatkowski, Cordelia Langford, John Sillitoe on behalf of the Wellcome Sanger Institute COVID-19 Surveillance Team |
| EPI_ISL_3309140, EPI_ISL_3309141, EPI_ISL_3309142, EPI_ISL_3309143, EPI_ISL_3309144 | Lighthouse Lab in Milton Keynes | Wellcome Sanger Institute for the COVID-19 Genomics UK (COG-UK) Consortium | The Lighthouse Lab in Milton Keynes and Alex Alderton, Roberto Amato, Jeffrey Barrett, Sonia Goncalves, Ewan Harrison, David K. Jackson, Ian Johnston, Dominic Kwiatkowski, Cordelia Langford, John Sillitoe on behalf of the Wellcome Sanger Institute COVID-19 Surveillance Team |
| EPI_ISL_3309145 | Lighthouse Laboratory Plymouth | Wellcome Sanger Institute for the COVID-19 Genomics UK (COG-UK) Consortium | Lighthouse Laboratory Plymouth and Alex Alderton, Roberto Amato, Jeffrey Barrett, Sonia Goncalves, Ewan Harrison, David K. Jackson, Ian Johnston, Dominic Kwiatkowski, Cordelia Langford, John Sillitoe on behalf of the Wellcome Sanger Institute COVID-19 Surveillance Team |
| EPI_ISL_3309146 | Lighthouse Lab in Milton Keynes | Wellcome Sanger Institute for the COVID-19 Genomics UK (COG-UK) Consortium | The Lighthouse Lab in Milton Keynes and Alex Alderton, Roberto Amato, Jeffrey Barrett, Sonia Goncalves, Ewan Harrison, David K. Jackson, Ian Johnston, Dominic Kwiatkowski, Cordelia Langford, John Sillitoe on behalf of the Wellcome Sanger Institute COVID-19 Surveillance Team |
| EPI_ISL_3309147, EPI_ISL_3309148 | Lighthouse Laboratory Plymouth | Wellcome Sanger Institute for the COVID-19 Genomics UK (COG-UK) Consortium | Lighthouse Laboratory Plymouth and Alex Alderton, Roberto Amato, Jeffrey Barrett, Sonia Goncalves, Ewan Harrison, David K. Jackson, Ian Johnston, Dominic Kwiatkowski, Cordelia Langford, John Sillitoe on behalf of the Wellcome Sanger Institute COVID-19 Surveillance Team |
| EPI_ISL_3309149, EPI_ISL_3309150, EPI_ISL_3309151, EPI_ISL_3309152, EPI_ISL_3309153, EPI_ISL_3309154, EPI_ISL_3309155, EPI_ISL_3309156, EPI_ISL_3309157, EPI_ISL_3309158, EPI_ISL_3309159, EPI_ISL_3309160 | | | |
| see above | Lighthouse Lab in Milton Keynes | Wellcome Sanger Institute for the COVID-19 Genomics UK (COG-UK) Consortium | The Lighthouse Lab in Milton Keynes and Alex Alderton, Roberto Amato, Jeffrey Barrett, Sonia Goncalves, Ewan Harrison, David K. Jackson, Ian Johnston, Dominic Kwiatkowski, Cordelia Langford, John Sillitoe on behalf of the Wellcome Sanger Institute COVID-19 Surveillance Team |
| EPI_ISL_3309161 | Lighthouse Laboratory Plymouth | Wellcome Sanger Institute for the COVID-19 Genomics UK (COG-UK) Consortium | Lighthouse Laboratory Plymouth and Alex Alderton, Roberto Amato, Jeffrey Barrett, Sonia Goncalves, Ewan Harrison, David K. Jackson, Ian Johnston, Dominic Kwiatkowski, Cordelia Langford, John Sillitoe on behalf of the Wellcome Sanger Institute COVID-19 Surveillance Team |
| EPI_ISL_3309162, EPI_ISL_3309163, EPI_ISL_3309164 | Lighthouse Lab in Milton Keynes | Wellcome Sanger Institute for the COVID-19 Genomics UK (COG-UK) Consortium | The Lighthouse Lab in Milton Keynes and Alex Alderton, Roberto Amato, Jeffrey Barrett, Sonia Goncalves, Ewan Harrison, David K. Jackson, Ian Johnston, Dominic Kwiatkowski, Cordelia Langford, John Sillitoe on behalf of the Wellcome Sanger Institute COVID-19 Surveillance Team |
| EPI_ISL_3309165 | Lighthouse Laboratory Plymouth | Wellcome Sanger Institute for the COVID-19 Genomics UK (COG-UK) Consortium | Lighthouse Laboratory Plymouth and Alex Alderton, Roberto Amato, Jeffrey Barrett, Sonia Goncalves, Ewan Harrison, David K. Jackson, Ian Johnston, Dominic Kwiatkowski, Cordelia Langford, John Sillitoe on behalf of the Wellcome Sanger Institute COVID-19 Surveillance Team |
| EPI_ISL_3309166, EPI_ISL_3309167, EPI_ISL_3309168 | Lighthouse Lab in Milton Keynes | Wellcome Sanger Institute for the COVID-19 Genomics UK (COG-UK) Consortium | The Lighthouse Lab in Milton Keynes and Alex Alderton, Roberto Amato, Jeffrey Barrett, Sonia Goncalves, Ewan Harrison, David K. Jackson, Ian Johnston, Dominic Kwiatkowski, Cordelia Langford, John Sillitoe on behalf of the Wellcome Sanger Institute COVID-19 Surveillance Team |
| EPI_ISL_3309169 | Lighthouse Laboratory Plymouth | Wellcome Sanger Institute for the COVID-19 Genomics UK (COG-UK) Consortium | Lighthouse Laboratory Plymouth and Alex Alderton, Roberto Amato, Jeffrey Barrett, Sonia Goncalves, Ewan Harrison, David K. Jackson, Ian Johnston, Dominic Kwiatkowski, Cordelia Langford, John Sillitoe on behalf of the Wellcome Sanger Institute COVID-19 Surveillance Team |
| EPI_ISL_3309170, EPI_ISL_3309171, EPI_ISL_3309172 | Lighthouse Lab in Milton Keynes | Wellcome Sanger Institute for the COVID-19 Genomics UK (COG-UK) Consortium | The Lighthouse Lab in Milton Keynes and Alex Alderton, Roberto Amato, Jeffrey Barrett, Sonia Goncalves, Ewan Harrison, David K. Jackson, Ian Johnston, Dominic Kwiatkowski, Cordelia Langford, John Sillitoe on behalf of the Wellcome Sanger Institute COVID-19 Surveillance Team |
| EPI_ISL_3309173 | Lighthouse Laboratory Plymouth | Wellcome Sanger Institute for the COVID-19 Genomics UK (COG-UK) Consortium | Lighthouse Laboratory Plymouth and Alex Alderton, Roberto Amato, Jeffrey Barrett, Sonia Goncalves, Ewan Harrison, David K. Jackson, Ian Johnston, Dominic Kwiatkowski, Cordelia Langford, John Sillitoe on behalf of the Wellcome Sanger Institute COVID-19 Surveillance Team |
| EPI_ISL_3309174, EPI_ISL_3309175, EPI_ISL_3309176 | Lighthouse Lab in Milton Keynes | Wellcome Sanger Institute for the COVID-19 Genomics UK (COG-UK) Consortium | The Lighthouse Lab in Milton Keynes and Alex Alderton, Roberto Amato, Jeffrey Barrett, Sonia Goncalves, Ewan Harrison, David K. Jackson, Ian Johnston, Dominic Kwiatkowski, Cordelia Langford, John Sillitoe on behalf of the Wellcome Sanger Institute COVID-19 Surveillance Team |
| EPI_ISL_3309177 | Lighthouse Laboratory Plymouth | Wellcome Sanger Institute for the COVID-19 Genomics UK (COG-UK) Consortium | Lighthouse Laboratory Plymouth and Alex Alderton, Roberto Amato, Jeffrey Barrett, Sonia Goncalves, Ewan Harrison, David K. Jackson, Ian Johnston, Dominic Kwiatkowski, Cordelia Langford, John Sillitoe on behalf of the Wellcome Sanger Institute COVID-19 Surveillance Team |
| EPI_ISL_3309178 | Lighthouse Lab in Milton Keynes | Wellcome Sanger Institute for the COVID-19 Genomics UK (COG-UK) Consortium | The Lighthouse Lab in Milton Keynes and Alex Alderton, Roberto Amato, Jeffrey Barrett, Sonia Goncalves, Ewan Harrison, David K. Jackson, Ian Johnston, Dominic Kwiatkowski, Cordelia Langford, John Sillitoe on behalf of the Wellcome Sanger Institute COVID-19 Surveillance Team |
| EPI_ISL_3309179, EPI_ISL_3309180 | Lighthouse Laboratory Plymouth | Wellcome Sanger Institute for the COVID-19 Genomics UK (COG-UK) Consortium | Lighthouse Laboratory Plymouth and Alex Alderton, Roberto Amato, Jeffrey Barrett, Sonia Goncalves, Ewan Harrison, David K. Jackson, Ian Johnston, Dominic Kwiatkowski, Cordelia Langford, John Sillitoe on behalf of the Wellcome Sanger Institute COVID-19 Surveillance Team |
| EPI_ISL_3309181, EPI_ISL_3309182, EPI_ISL_3309183 | Lighthouse Lab in Milton Keynes | Wellcome Sanger Institute for the COVID-19 Genomics UK (COG-UK) Consortium | The Lighthouse Lab in Milton Keynes and Alex Alderton, Roberto Amato, Jeffrey Barrett, Sonia Goncalves, Ewan Harrison, David K. Jackson, Ian Johnston, Dominic Kwiatkowski, Cordelia Langford, John Sillitoe on behalf of the Wellcome Sanger Institute COVID-19 Surveillance Team |
| EPI_ISL_3309184 | Lighthouse Laboratory Plymouth | Wellcome Sanger Institute for the COVID-19 Genomics UK (COG-UK) Consortium | Lighthouse Laboratory Plymouth and Alex Alderton, Roberto Amato, Jeffrey Barrett, Sonia Goncalves, Ewan Harrison, David K. Jackson, Ian Johnston, Dominic Kwiatkowski, Cordelia Langford, John Sillitoe on behalf of the Wellcome Sanger Institute COVID-19 Surveillance Team |
| EPI_ISL_3309185 | Lighthouse Lab in Milton Keynes | Wellcome Sanger Institute for the COVID-19 Genomics UK (COG-UK) Consortium | The Lighthouse Lab in Milton Keynes and Alex Alderton, Roberto Amato, Jeffrey Barrett, Sonia Goncalves, Ewan Harrison, David K. Jackson, Ian Johnston, Dominic Kwiatkowski, Cordelia Langford, John Sillitoe on behalf of the Wellcome Sanger Institute COVID-19 Surveillance Team |
| EPI_ISL_3309186, EPI_ISL_3309187 | Lighthouse Laboratory Plymouth | Wellcome Sanger Institute for the COVID-19 Genomics UK (COG-UK) Consortium | Lighthouse Laboratory Plymouth and Alex Alderton, Roberto Amato, Jeffrey Barrett, Sonia Goncalves, Ewan Harrison, David K. Jackson, Ian Johnston, Dominic Kwiatkowski, Cordelia Langford, John Sillitoe on behalf of the Wellcome Sanger Institute COVID-19 Surveillance Team |
| EPI_ISL_3309188 | Lighthouse Lab in Milton Keynes | Wellcome Sanger Institute for the COVID-19 Genomics UK (COG-UK) Consortium | The Lighthouse Lab in Milton Keynes and Alex Alderton, Roberto Amato, Jeffrey Barrett, Sonia Goncalves, Ewan Harrison, David K. Jackson, Ian Johnston, Dominic Kwiatkowski, Cordelia Langford, John Sillitoe on behalf of the Wellcome Sanger Institute COVID-19 Surveillance Team |
| EPI_ISL_3309189 | Lighthouse Laboratory Plymouth | Wellcome Sanger Institute for the COVID-19 Genomics UK (COG-UK) Consortium | Lighthouse Laboratory Plymouth and Alex Alderton, Roberto Amato, Jeffrey Barrett, Sonia Goncalves, Ewan Harrison, David K. Jackson, Ian Johnston, Dominic Kwiatkowski, Cordelia Langford, John Sillitoe on behalf of the Wellcome Sanger Institute COVID-19 Surveillance Team |
| EPI_ISL_3309190, EPI_ISL_3309191, EPI_ISL_3309192, EPI_ISL_3309193 | Lighthouse Lab in Milton Keynes | Wellcome Sanger Institute for the COVID-19 Genomics UK (COG-UK) Consortium | The Lighthouse Lab in Milton Keynes and Alex Alderton, Roberto Amato, Jeffrey Barrett, Sonia Goncalves, Ewan Harrison, David K. Jackson, Ian Johnston, Dominic Kwiatkowski, Cordelia Langford, John Sillitoe on behalf of the Wellcome Sanger Institute COVID-19 Surveillance Team |
| EPI_ISL_3309194 | Lighthouse Laboratory Plymouth | Wellcome Sanger Institute for the COVID-19 Genomics UK (COG-UK) Consortium | Lighthouse Laboratory Plymouth and Alex Alderton, Roberto Amato, Jeffrey Barrett, Sonia Goncalves, Ewan Harrison, David K. Jackson, Ian Johnston, Dominic Kwiatkowski, Cordelia Langford, John Sillitoe on behalf of the Wellcome Sanger Institute COVID-19 Surveillance Team |

| | | | |
|---|---|---|---|
| EPI_ISL_3309195, EPI_ISL_3309196, EPI_ISL_3309197 | Lighthouse Lab in Milton Keynes | Wellcome Sanger Institute for the COVID-19 Genomics UK (COG-UK) Consortium | The Lighthouse Lab in Milton Keynes and Alex Alderton, Roberto Amato, Jeffrey Barrett, Sonia Goncalves, Ewan Harrison, David K. Jackson, Ian Johnston, Dominic Kwiatkowski, Cordelia Langford, John Sillitoe on behalf of the Wellcome Sanger Institute COVID-19 Surveillance Team |
| EPI_ISL_3309198 | Lighthouse Laboratory Plymouth | Wellcome Sanger Institute for the COVID-19 Genomics UK (COG-UK) Consortium | Lighthouse Laboratory Plymouth and Alex Alderton, Roberto Amato, Jeffrey Barrett, Sonia Goncalves, Ewan Harrison, David K. Jackson, Ian Johnston, Dominic Kwiatkowski, Cordelia Langford, John Sillitoe on behalf of the Wellcome Sanger Institute COVID-19 Surveillance Team |
| EPI_ISL_3309199, EPI_ISL_3309200, EPI_ISL_3309201, EPI_ISL_3309202 | Lighthouse Lab in Milton Keynes | Wellcome Sanger Institute for the COVID-19 Genomics UK (COG-UK) Consortium | The Lighthouse Lab in Milton Keynes and Alex Alderton, Roberto Amato, Jeffrey Barrett, Sonia Goncalves, Ewan Harrison, David K. Jackson, Ian Johnston, Dominic Kwiatkowski, Cordelia Langford, John Sillitoe on behalf of the Wellcome Sanger Institute COVID-19 Surveillance Team |
| EPI_ISL_3309203 | Lighthouse Laboratory Plymouth | Wellcome Sanger Institute for the COVID-19 Genomics UK (COG-UK) Consortium | Lighthouse Laboratory Plymouth and Alex Alderton, Roberto Amato, Jeffrey Barrett, Sonia Goncalves, Ewan Harrison, David K. Jackson, Ian Johnston, Dominic Kwiatkowski, Cordelia Langford, John Sillitoe on behalf of the Wellcome Sanger Institute COVID-19 Surveillance Team |
| EPI_ISL_3309204 | Lighthouse Lab in Milton Keynes | Wellcome Sanger Institute for the COVID-19 Genomics UK (COG-UK) Consortium | The Lighthouse Lab in Milton Keynes and Alex Alderton, Roberto Amato, Jeffrey Barrett, Sonia Goncalves, Ewan Harrison, David K. Jackson, Ian Johnston, Dominic Kwiatkowski, Cordelia Langford, John Sillitoe on behalf of the Wellcome Sanger Institute COVID-19 Surveillance Team |
| EPI_ISL_3309205 | Lighthouse Laboratory Plymouth | Wellcome Sanger Institute for the COVID-19 Genomics UK (COG-UK) Consortium | Lighthouse Laboratory Plymouth and Alex Alderton, Roberto Amato, Jeffrey Barrett, Sonia Goncalves, Ewan Harrison, David K. Jackson, Ian Johnston, Dominic Kwiatkowski, Cordelia Langford, John Sillitoe on behalf of the Wellcome Sanger Institute COVID-19 Surveillance Team |
| EPI_ISL_3309206, EPI_ISL_3309207, EPI_ISL_3309208, EPI_ISL_3309209, EPI_ISL_3309210, EPI_ISL_3309211, EPI_ISL_3309212, EPI_ISL_3309213 | Lighthouse Lab in Milton Keynes | Wellcome Sanger Institute for the COVID-19 Genomics UK (COG-UK) Consortium | The Lighthouse Lab in Milton Keynes and Alex Alderton, Roberto Amato, Jeffrey Barrett, Sonia Goncalves, Ewan Harrison, David K. Jackson, Ian Johnston, Dominic Kwiatkowski, Cordelia Langford, John Sillitoe on behalf of the Wellcome Sanger Institute COVID-19 Surveillance Team |
| EPI_ISL_3309214 | Lighthouse Laboratory Plymouth | Wellcome Sanger Institute for the COVID-19 Genomics UK (COG-UK) Consortium | Lighthouse Laboratory Plymouth and Alex Alderton, Roberto Amato, Jeffrey Barrett, Sonia Goncalves, Ewan Harrison, David K. Jackson, Ian Johnston, Dominic Kwiatkowski, Cordelia Langford, John Sillitoe on behalf of the Wellcome Sanger Institute COVID-19 Surveillance Team |
| EPI_ISL_3309215 | Lighthouse Lab in Milton Keynes | Wellcome Sanger Institute for the COVID-19 Genomics UK (COG-UK) Consortium | The Lighthouse Lab in Milton Keynes and Alex Alderton, Roberto Amato, Jeffrey Barrett, Sonia Goncalves, Ewan Harrison, David K. Jackson, Ian Johnston, Dominic Kwiatkowski, Cordelia Langford, John Sillitoe on behalf of the Wellcome Sanger Institute COVID-19 Surveillance Team |
| EPI_ISL_3309216 | Lighthouse Laboratory Plymouth | Wellcome Sanger Institute for the COVID-19 Genomics UK (COG-UK) Consortium | Lighthouse Laboratory Plymouth and Alex Alderton, Roberto Amato, Jeffrey Barrett, Sonia Goncalves, Ewan Harrison, David K. Jackson, Ian Johnston, Dominic Kwiatkowski, Cordelia Langford, John Sillitoe on behalf of the Wellcome Sanger Institute COVID-19 Surveillance Team |
| EPI_ISL_3309217, EPI_ISL_3309218 | Lighthouse Lab in Milton Keynes | Wellcome Sanger Institute for the COVID-19 Genomics UK (COG-UK) Consortium | The Lighthouse Lab in Milton Keynes and Alex Alderton, Roberto Amato, Jeffrey Barrett, Sonia Goncalves, Ewan Harrison, David K. Jackson, Ian Johnston, Dominic Kwiatkowski, Cordelia Langford, John Sillitoe on behalf of the Wellcome Sanger Institute COVID-19 Surveillance Team |
| EPI_ISL_3309219 | Lighthouse Laboratory Plymouth | Wellcome Sanger Institute for the COVID-19 Genomics UK (COG-UK) Consortium | Lighthouse Laboratory Plymouth and Alex Alderton, Roberto Amato, Jeffrey Barrett, Sonia Goncalves, Ewan Harrison, David K. Jackson, Ian Johnston, Dominic Kwiatkowski, Cordelia Langford, John Sillitoe on behalf of the Wellcome Sanger Institute COVID-19 Surveillance Team |
| EPI_ISL_3309220, EPI_ISL_3309221, EPI_ISL_3309222, EPI_ISL_3309223, EPI_ISL_3309224, EPI_ISL_3309225, EPI_ISL_3309226, EPI_ISL_3309227 | Lighthouse Lab in Milton Keynes | Wellcome Sanger Institute for the COVID-19 Genomics UK (COG-UK) Consortium | The Lighthouse Lab in Milton Keynes and Alex Alderton, Roberto Amato, Jeffrey Barrett, Sonia Goncalves, Ewan Harrison, David K. Jackson, Ian Johnston, Dominic Kwiatkowski, Cordelia Langford, John Sillitoe on behalf of the Wellcome Sanger Institute COVID-19 Surveillance Team |
| EPI_ISL_3309228 | Lighthouse Laboratory Plymouth | Wellcome Sanger Institute for the COVID-19 Genomics UK (COG-UK) Consortium | Lighthouse Laboratory Plymouth and Alex Alderton, Roberto Amato, Jeffrey Barrett, Sonia Goncalves, Ewan Harrison, David K. Jackson, Ian Johnston, Dominic Kwiatkowski, Cordelia Langford, John Sillitoe on behalf of the Wellcome Sanger Institute COVID-19 Surveillance Team |
| EPI_ISL_3309229, EPI_ISL_3309230, EPI_ISL_3309231, EPI_ISL_3309232, EPI_ISL_3309233, EPI_ISL_3309234, EPI_ISL_3309235, EPI_ISL_3309236, EPI_ISL_3309237, EPI_ISL_3309238, EPI_ISL_3309239, EPI_ISL_3309240, EPI_ISL_3309241 | | | |
| see above | Lighthouse Lab in Milton Keynes | Wellcome Sanger Institute for the COVID-19 Genomics UK (COG-UK) Consortium | The Lighthouse Lab in Milton Keynes and Alex Alderton, Roberto Amato, Jeffrey Barrett, Sonia Goncalves, Ewan Harrison, David K. Jackson, Ian Johnston, Dominic Kwiatkowski, Cordelia Langford, John Sillitoe on behalf of the Wellcome Sanger Institute COVID-19 Surveillance Team |
| EPI_ISL_3309242 | Lighthouse Laboratory Plymouth | Wellcome Sanger Institute for the COVID-19 Genomics UK (COG-UK) Consortium | Lighthouse Laboratory Plymouth and Alex Alderton, Roberto Amato, Jeffrey Barrett, Sonia Goncalves, Ewan Harrison, David K. Jackson, Ian Johnston, Dominic Kwiatkowski, Cordelia Langford, John Sillitoe on behalf of the Wellcome Sanger Institute COVID-19 Surveillance Team |
| EPI_ISL_3309243 | Lighthouse Lab in Milton Keynes | Wellcome Sanger Institute for the COVID-19 Genomics UK (COG-UK) Consortium | The Lighthouse Lab in Milton Keynes and Alex Alderton, Roberto Amato, Jeffrey Barrett, Sonia Goncalves, Ewan Harrison, David K. Jackson, Ian Johnston, Dominic Kwiatkowski, Cordelia Langford, John Sillitoe on behalf of the Wellcome Sanger Institute COVID-19 Surveillance Team |
| EPI_ISL_3309244 | Lighthouse Laboratory Plymouth | Wellcome Sanger Institute for the COVID-19 Genomics UK (COG-UK) Consortium | Lighthouse Laboratory Plymouth and Alex Alderton, Roberto Amato, Jeffrey Barrett, Sonia Goncalves, Ewan Harrison, David K. Jackson, Ian Johnston, Dominic Kwiatkowski, Cordelia Langford, John Sillitoe on behalf of the Wellcome Sanger Institute COVID-19 Surveillance Team |
| EPI_ISL_3309245 | Lighthouse Lab in Milton Keynes | Wellcome Sanger Institute for the COVID-19 Genomics UK (COG-UK) Consortium | The Lighthouse Lab in Milton Keynes and Alex Alderton, Roberto Amato, Jeffrey Barrett, Sonia Goncalves, Ewan Harrison, David K. Jackson, Ian Johnston, Dominic Kwiatkowski, Cordelia Langford, John Sillitoe on behalf of the Wellcome Sanger Institute COVID-19 Surveillance Team |
| EPI_ISL_3309246 | Lighthouse Laboratory Plymouth | Wellcome Sanger Institute for the COVID-19 Genomics UK (COG-UK) Consortium | Lighthouse Laboratory Plymouth and Alex Alderton, Roberto Amato, Jeffrey Barrett, Sonia Goncalves, Ewan Harrison, David K. Jackson, Ian Johnston, Dominic Kwiatkowski, Cordelia Langford, John Sillitoe on behalf of the Wellcome Sanger Institute COVID-19 Surveillance Team |
| EPI_ISL_3309247, EPI_ISL_3309248, EPI_ISL_3309249, EPI_ISL_3309250, EPI_ISL_3309251, EPI_ISL_3309252, EPI_ISL_3309253, EPI_ISL_3309254 | Lighthouse Lab in Milton Keynes | Wellcome Sanger Institute for the COVID-19 Genomics UK (COG-UK) Consortium | The Lighthouse Lab in Milton Keynes and Alex Alderton, Roberto Amato, Jeffrey Barrett, Sonia Goncalves, Ewan Harrison, David K. Jackson, Ian Johnston, Dominic Kwiatkowski, Cordelia Langford, John Sillitoe on behalf of the Wellcome Sanger Institute COVID-19 Surveillance Team |
| EPI_ISL_3309255 | Lighthouse Laboratory Plymouth | Wellcome Sanger Institute for the COVID-19 Genomics UK (COG-UK) Consortium | Lighthouse Laboratory Plymouth and Alex Alderton, Roberto Amato, Jeffrey Barrett, Sonia Goncalves, Ewan Harrison, David K. Jackson, Ian Johnston, Dominic Kwiatkowski, Cordelia Langford, John Sillitoe on behalf of the Wellcome Sanger Institute COVID-19 Surveillance Team |
| EPI_ISL_3309256, EPI_ISL_3309257, EPI_ISL_3309258, EPI_ISL_3309259, EPI_ISL_3309260, EPI_ISL_3309261, EPI_ISL_3309262 | Lighthouse Lab in Milton Keynes | Wellcome Sanger Institute for the COVID-19 Genomics UK (COG-UK) Consortium | The Lighthouse Lab in Milton Keynes and Alex Alderton, Roberto Amato, Jeffrey Barrett, Sonia Goncalves, Ewan Harrison, David K. Jackson, Ian Johnston, Dominic Kwiatkowski, Cordelia Langford, John Sillitoe on behalf of the Wellcome Sanger Institute COVID-19 Surveillance Team |
| EPI_ISL_3309263 | Lighthouse Laboratory Plymouth | Wellcome Sanger Institute for the COVID-19 Genomics UK (COG-UK) Consortium | Lighthouse Laboratory Plymouth and Alex Alderton, Roberto Amato, Jeffrey Barrett, Sonia Goncalves, Ewan Harrison, David K. Jackson, Ian Johnston, Dominic Kwiatkowski, Cordelia Langford, John Sillitoe on behalf of the Wellcome Sanger Institute COVID-19 Surveillance Team |
| EPI_ISL_3309264, EPI_ISL_3309265 | Lighthouse Lab in Milton Keynes | Wellcome Sanger Institute for the COVID-19 Genomics UK (COG-UK) Consortium | The Lighthouse Lab in Milton Keynes and Alex Alderton, Roberto Amato, Jeffrey Barrett, Sonia Goncalves, Ewan Harrison, David K. Jackson, Ian Johnston, Dominic Kwiatkowski, Cordelia Langford, John Sillitoe on behalf of the Wellcome Sanger Institute COVID-19 Surveillance Team |
| EPI_ISL_3309266 | Lighthouse Laboratory Plymouth | Wellcome Sanger Institute for the COVID-19 Genomics UK (COG-UK) Consortium | Lighthouse Laboratory Plymouth and Alex Alderton, Roberto Amato, Jeffrey Barrett, Sonia Goncalves, Ewan Harrison, David K. Jackson, Ian Johnston, Dominic Kwiatkowski, Cordelia Langford, John Sillitoe on behalf of the Wellcome Sanger Institute COVID-19 Surveillance Team |
| EPI_ISL_3309267, EPI_ISL_3309268 | Lighthouse Lab in Milton Keynes | Wellcome Sanger Institute for the COVID-19 Genomics UK (COG-UK) Consortium | The Lighthouse Lab in Milton Keynes and Alex Alderton, Roberto Amato, Jeffrey Barrett, Sonia Goncalves, Ewan Harrison, David K. Jackson, Ian Johnston, Dominic Kwiatkowski, Cordelia Langford, John Sillitoe on behalf of the Wellcome Sanger Institute COVID-19 Surveillance Team |
| EPI_ISL_3309269 | Lighthouse Laboratory Plymouth | Wellcome Sanger Institute for the COVID-19 Genomics UK (COG-UK) Consortium | Lighthouse Laboratory Plymouth and Alex Alderton, Roberto Amato, Jeffrey Barrett, Sonia Goncalves, Ewan Harrison, David K. Jackson, Ian Johnston, Dominic Kwiatkowski, Cordelia Langford, John Sillitoe on behalf of the Wellcome Sanger Institute COVID-19 Surveillance Team |
| EPI_ISL_3309270, EPI_ISL_3309271, EPI_ISL_3309272, EPI_ISL_3309273, EPI_ISL_3309274, EPI_ISL_3309275 | Lighthouse Lab in Milton Keynes | Wellcome Sanger Institute for the COVID-19 Genomics UK (COG-UK) Consortium | The Lighthouse Lab in Milton Keynes and Alex Alderton, Roberto Amato, Jeffrey Barrett, Sonia Goncalves, Ewan Harrison, David K. Jackson, Ian Johnston, Dominic Kwiatkowski, Cordelia Langford, John Sillitoe on behalf of the Wellcome Sanger Institute COVID-19 Surveillance Team |

EPI_ISL_3309276, EPI_ISL_3309277, EPI_ISL_3309278, EPI_ISL_3309279, EPI_ISL_3309280, EPI_ISL_3309281, EPI_ISL_3309282, EPI_ISL_3309283, EPI_ISL_3309284, EPI_ISL_3309285, EPI_ISL_3309286, EPI_ISL_3309287, EPI_ISL_3309288, EPI_ISL_3309289, EPI_ISL_3309290, EPI_ISL_3309291, EPI_ISL_3309292, EPI_ISL_3309293, EPI_ISL_3309294, EPI_ISL_3309295, EPI_ISL_3309296, EPI_ISL_3309297, EPI_ISL_3309298, EPI_ISL_3309299, EPI_ISL_3309300, EPI_ISL_3309301, EPI_ISL_3309302, EPI_ISL_3309303, EPI_ISL_3309304, EPI_ISL_3309305, EPI_ISL_3309306, EPI_ISL_3309307, EPI_ISL_3309308, EPI_ISL_3309309, EPI_ISL_3309310, EPI_ISL_3309311, EPI_ISL_3309312, EPI_ISL_3309313, EPI_ISL_3309314, EPI_ISL_3309315, EPI_ISL_3309316, EPI_ISL_3309317, EPI_ISL_3309318, EPI_ISL_3309319, EPI_ISL_3309320, EPI_ISL_3309321, EPI_ISL_3309322, EPI_ISL_3309323, EPI_ISL_3309324, EPI_ISL_3309325, EPI_ISL_3309326, EPI_ISL_3309327, EPI_ISL_3309328, EPI_ISL_3309329, EPI_ISL_3309330, EPI_ISL_3309331, EPI_ISL_3309332, EPI_ISL_3309333, EPI_ISL_3309334, EPI_ISL_3309335, EPI_ISL_3309336, EPI_ISL_3309337, EPI_ISL_3309338, EPI_ISL_3309339, EPI_ISL_3309340, EPI_ISL_3309341, EPI_ISL_3309342, EPI_ISL_3309343, EPI_ISL_3309344, EPI_ISL_3309345, EPI_ISL_3309346, EPI_ISL_3309347, EPI_ISL_3309348, EPI_ISL_3309349, EPI_ISL_3309350, EPI_ISL_3309351, EPI_ISL_3309352, EPI_ISL_3309353, EPI_ISL_3309354, EPI_ISL_3309355, EPI_ISL_3309356, EPI_ISL_3309357, EPI_ISL_3309358, EPI_ISL_3309359, EPI_ISL_3309360, EPI_ISL_3309361, EPI_ISL_3309362, EPI_ISL_3309363, EPI_ISL_3309364, EPI_ISL_3309365, EPI_ISL_3309366, EPI_ISL_3309367, EPI_ISL_3309368, EPI_ISL_3309369, EPI_ISL_3309370, EPI_ISL_3309371, EPI_ISL_3309372, EPI_ISL_3309373, EPI_ISL_3309374, EPI_ISL_3309375, EPI_ISL_3309376, EPI_ISL_3309377, EPI_ISL_3309378, EPI_ISL_3309379, EPI_ISL_3309380, EPI_ISL_3309381, EPI_ISL_3309382, EPI_ISL_3309383, EPI_ISL_3309384, EPI_ISL_3309385, EPI_ISL_3309386, EPI_ISL_3309387, EPI_ISL_3309388, EPI_ISL_3309389, EPI_ISL_3309390, EPI_ISL_3309391, EPI_ISL_3309392, EPI_ISL_3309393, EPI_ISL_3309394, EPI_ISL_3309395, EPI_ISL_3309396, EPI_ISL_3309397, EPI_ISL_3309398, EPI_ISL_3309399, EPI_ISL_3309400, EPI_ISL_3309401, EPI_ISL_3309402, EPI_ISL_3309403, EPI_ISL_3309404, EPI_ISL_3309405, EPI_ISL_3309406, EPI_ISL_3309407, EPI_ISL_3309408, EPI_ISL_3309409, EPI_ISL_3309410, EPI_ISL_3309411, EPI_ISL_3309412, EPI_ISL_3309413, EPI_ISL_3309414, EPI_ISL_3309415, EPI_ISL_3309416, EPI_ISL_3309417, EPI_ISL_3309418, EPI_ISL_3309419, EPI_ISL_3309420, EPI_ISL_3309421, EPI_ISL_3309422, EPI_ISL_3309423, EPI_ISL_3309424, EPI_ISL_3309425, EPI_ISL_3309426, EPI_ISL_3309427, EPI_ISL_3309428, EPI_ISL_3309429,

EPI_ISL_3309430, EPI_ISL_3309431, EPI_ISL_3309432, EPI_ISL_3309433, EPI_ISL_3309434, EPI_ISL_3309435, EPI_ISL_3309436, EPI_ISL_3309437, EPI_ISL_3309438, EPI_ISL_3309439, EPI_ISL_3309440, EPI_ISL_3309441, EPI_ISL_3309442, EPI_ISL_3309443, EPI_ISL_3309444, EPI_ISL_3309445, EPI_ISL_3309446, EPI_ISL_3309447, EPI_ISL_3309448, EPI_ISL_3309449, EPI_ISL_3309450, EPI_ISL_3309451, EPI_ISL_3309452, EPI_ISL_3309453, EPI_ISL_3309454, EPI_ISL_3309455, EPI_ISL_3309456, EPI_ISL_3309457, EPI_ISL_3309458, EPI_ISL_3309459, EPI_ISL_3309460, EPI_ISL_3309461, EPI_ISL_3309462, EPI_ISL_3309463, EPI_ISL_3309464, EPI_ISL_3309465, EPI_ISL_3309466, EPI_ISL_3309467, EPI_ISL_3309468, EPI_ISL_3309469, EPI_ISL_3309470, EPI_ISL_3309471, EPI_ISL_3309472, EPI_ISL_3309473, EPI_ISL_3309474, EPI_ISL_3309475, EPI_ISL_3309476, EPI_ISL_3309477, EPI_ISL_3309478, EPI_ISL_3309479, EPI_ISL_3309480, EPI_ISL_3309481, EPI_ISL_3309482, EPI_ISL_3309483, EPI_ISL_3309484, EPI_ISL_3309485, EPI_ISL_3309486, EPI_ISL_3309487, EPI_ISL_3309488, EPI_ISL_3309489, EPI_ISL_3309490, EPI_ISL_3309491, EPI_ISL_3309492, EPI_ISL_3309493, EPI_ISL_3309494, EPI_ISL_3309495, EPI_ISL_3309496, EPI_ISL_3309497, EPI_ISL_3309498, EPI_ISL_3309499, EPI_ISL_3309500, EPI_ISL_3309501, EPI_ISL_3309502, EPI_ISL_3309503, EPI_ISL_3309504, EPI_ISL_3309505, EPI_ISL_3309506, EPI_ISL_3309507, EPI_ISL_3309508, EPI_ISL_3309509, EPI_ISL_3309510, EPI_ISL_3309511, EPI_ISL_3309512, EPI_ISL_3309513, EPI_ISL_3309514, EPI_ISL_3309515, EPI_ISL_3309516, EPI_ISL_3309517, EPI_ISL_3309518, EPI_ISL_3309519, EPI_ISL_3309520, EPI_ISL_3309521, EPI_ISL_3309522, EPI_ISL_3309523, EPI_ISL_3309524, EPI_ISL_3309525, EPI_ISL_3309526, EPI_ISL_3309527, EPI_ISL_3309528, EPI_ISL_3309529, EPI_ISL_3309530, EPI_ISL_3309531, EPI_ISL_3309532, EPI_ISL_3309533, EPI_ISL_3309534, EPI_ISL_3309535, EPI_ISL_3309536, EPI_ISL_3309537, EPI_ISL_3309538, EPI_ISL_3309539, EPI_ISL_3309540, EPI_ISL_3309541, EPI_ISL_3309542, EPI_ISL_3309543, EPI_ISL_3309544, EPI_ISL_3309545, EPI_ISL_3309546, EPI_ISL_3309547, EPI_ISL_3309548, EPI_ISL_3309549, EPI_ISL_3309550, EPI_ISL_3309551, EPI_ISL_3309552, EPI_ISL_3309553, EPI_ISL_3309554, EPI_ISL_3309555, EPI_ISL_3309556, EPI_ISL_3309557, EPI_ISL_3309558, EPI_ISL_3309559, EPI_ISL_3309560, EPI_ISL_3309561, EPI_ISL_3309562, EPI_ISL_3309563, EPI_ISL_3309564, EPI_ISL_3309565, EPI_ISL_3309566, EPI_ISL_3309567, EPI_ISL_3309568, EPI_ISL_3309569, EPI_ISL_3309570, EPI_ISL_3309571, EPI_ISL_3309572, EPI_ISL_3309573, EPI_ISL_3309574, EPI_ISL_3309575, EPI_ISL_3309576, EPI_ISL_3309577, EPI_ISL_3309578, EPI_ISL_3309579, EPI_ISL_3309580, EPI_ISL_3309581, EPI_ISL_3309582, EPI_ISL_3309583, EPI_ISL_3309584, EPI_ISL_3309585, EPI_ISL_3309586, EPI_ISL_3309587, EPI_ISL_3309588, EPI_ISL_3309589, EPI_ISL_3309590, EPI_ISL_3309591, EPI_ISL_3309592, EPI_ISL_3309593, EPI_ISL_3309594, EPI_ISL_3309595, EPI_ISL_3309596, EPI_ISL_3309597, EPI_ISL_3309598, EPI_ISL_3309599, EPI_ISL_3309600, EPI_ISL_3309601, EPI_ISL_3309602, EPI_ISL_3309603, EPI_ISL_3309604, EPI_ISL_3309605, EPI_ISL_3309606, EPI_ISL_3309607, EPI_ISL_3309608, EPI_ISL_3309609, EPI_ISL_3309610, EPI_ISL_3309611, EPI_ISL_3309612, EPI_ISL_3309613, EPI_ISL_3309614, EPI_ISL_3309615, EPI_ISL_3309616, EPI_ISL_3309617, EPI_ISL_3309618

| | | | |
|---|---|---|---|
| see above | Lighthouse Laboratory Plymouth | Wellcome Sanger Institute for the COVID-19 Genomics UK (COG-UK) Consortium | Lighthouse Laboratory Plymouth and Alex Alderton, Roberto Amato, Jeffrey Barrett, Sonia Goncalves, Ewan Harrison, David K. Jackson, Ian Johnston, Dominic Kwiatkowski, Cordelia Langford, John Sillitoe on behalf of the Wellcome Sanger Institute COVID-19 Surveillance Team |
| EPI_ISL_3309619 | Lighthouse Lab in Glasgow | Wellcome Sanger Institute for the COVID-19 Genomics UK (COG-UK) Consortium | Harper VanSteenhouse, Yumi Kasai, David Gray, Carol Clugston, Anna Dominiczak and Alex Alderton, Roberto Amato, Jeffrey Barrett, Sonia Goncalves, Ewan Harrison, David K. Jackson, Ian Johnston, Dominic Kwiatkowski, Cordelia Langford, John Sillitoe on behalf of the Wellcome Sanger Institute COVID-19 Surveillance Team |
| EPI_ISL_3309620, EPI_ISL_3309621, EPI_ISL_3309622, EPI_ISL_3309623 | Lighthouse Laboratory Plymouth | Wellcome Sanger Institute for the COVID-19 Genomics UK (COG-UK) Consortium | Lighthouse Laboratory Plymouth and Alex Alderton, Roberto Amato, Jeffrey Barrett, Sonia Goncalves, Ewan Harrison, David K. Jackson, Ian Johnston, Dominic Kwiatkowski, Cordelia Langford, John Sillitoe on behalf of the Wellcome Sanger Institute COVID-19 Surveillance Team |
| EPI_ISL_3309624 | Health Services Laboratories | Wellcome Sanger Institute for the COVID-19 Genomics UK (COG-UK) Consortium | Health Services Laboratories and Alex Alderton, Roberto Amato, Jeffrey Barrett, Sonia Goncalves, Ewan Harrison, David K. Jackson, Ian Johnston, Dominic Kwiatkowski, Cordelia Langford, John Sillitoe on behalf of the Wellcome Sanger Institute COVID-19 Surveillance Team |
| EPI_ISL_3309625, EPI_ISL_3309626, EPI_ISL_3309627 | Lighthouse Laboratory Plymouth | Wellcome Sanger Institute for the COVID-19 Genomics UK (COG-UK) Consortium | Lighthouse Laboratory Plymouth and Alex Alderton, Roberto Amato, Jeffrey Barrett, Sonia Goncalves, Ewan Harrison, David K. Jackson, Ian Johnston, Dominic Kwiatkowski, Cordelia Langford, John Sillitoe on behalf of the Wellcome Sanger Institute COVID-19 Surveillance Team |
| EPI_ISL_3309628, EPI_ISL_3309629, EPI_ISL_3309630 | Health Services Laboratories | Wellcome Sanger Institute for the COVID-19 Genomics UK (COG-UK) Consortium | Health Services Laboratories and Alex Alderton, Roberto Amato, Jeffrey Barrett, Sonia Goncalves, Ewan Harrison, David K. Jackson, Ian Johnston, Dominic Kwiatkowski, Cordelia Langford, John Sillitoe on behalf of the Wellcome Sanger Institute COVID-19 Surveillance Team |
| EPI_ISL_3309631, EPI_ISL_3309632, EPI_ISL_3309633 | Lighthouse Laboratory Plymouth | Wellcome Sanger Institute for the COVID-19 Genomics UK (COG-UK) Consortium | Lighthouse Laboratory Plymouth and Alex Alderton, Roberto Amato, Jeffrey Barrett, Sonia Goncalves, Ewan Harrison, David K. Jackson, Ian Johnston, Dominic Kwiatkowski, Cordelia Langford, John Sillitoe on behalf of the Wellcome Sanger Institute COVID-19 Surveillance Team |
| EPI_ISL_3309634, EPI_ISL_3309635, EPI_ISL_3309636 | Health Services Laboratories | Wellcome Sanger Institute for the COVID-19 Genomics UK (COG-UK) Consortium | Health Services Laboratories and Alex Alderton, Roberto Amato, Jeffrey Barrett, Sonia Goncalves, Ewan Harrison, David K. Jackson, Ian Johnston, Dominic Kwiatkowski, Cordelia Langford, John Sillitoe on behalf of the Wellcome Sanger Institute COVID-19 Surveillance Team |
| EPI_ISL_3309637, EPI_ISL_3309638, EPI_ISL_3309639, EPI_ISL_3309640 | Lighthouse Laboratory Plymouth | Wellcome Sanger Institute for the COVID-19 Genomics UK (COG-UK) Consortium | Lighthouse Laboratory Plymouth and Alex Alderton, Roberto Amato, Jeffrey Barrett, Sonia Goncalves, Ewan Harrison, David K. Jackson, Ian Johnston, Dominic Kwiatkowski, Cordelia Langford, John Sillitoe on behalf of the Wellcome Sanger Institute COVID-19 Surveillance Team |
| EPI_ISL_3309641 | Health Services Laboratories | Wellcome Sanger Institute for the COVID-19 Genomics UK (COG-UK) Consortium | Health Services Laboratories and Alex Alderton, Roberto Amato, Jeffrey Barrett, Sonia Goncalves, Ewan Harrison, David K. Jackson, Ian Johnston, Dominic Kwiatkowski, Cordelia Langford, John Sillitoe on behalf of the Wellcome Sanger Institute COVID-19 Surveillance Team |
| EPI_ISL_3309642, EPI_ISL_3309643, EPI_ISL_3309644 | Lighthouse Laboratory Plymouth | Wellcome Sanger Institute for the COVID-19 Genomics UK (COG-UK) Consortium | Lighthouse Laboratory Plymouth and Alex Alderton, Roberto Amato, Jeffrey Barrett, Sonia Goncalves, Ewan Harrison, David K. Jackson, Ian Johnston, Dominic Kwiatkowski, Cordelia Langford, John Sillitoe on behalf of the Wellcome Sanger Institute COVID-19 Surveillance Team |
| EPI_ISL_3309645 | Health Services Laboratories | Wellcome Sanger Institute for the COVID-19 Genomics UK (COG-UK) Consortium | Health Services Laboratories and Alex Alderton, Roberto Amato, Jeffrey Barrett, Sonia Goncalves, Ewan Harrison, David K. Jackson, Ian Johnston, Dominic Kwiatkowski, Cordelia Langford, John Sillitoe on behalf of the Wellcome Sanger Institute COVID-19 Surveillance Team |
| EPI_ISL_3309646 | Lighthouse Laboratory Plymouth | Wellcome Sanger Institute for the COVID-19 Genomics UK (COG-UK) Consortium | Lighthouse Laboratory Plymouth and Alex Alderton, Roberto Amato, Jeffrey Barrett, Sonia Goncalves, Ewan Harrison, David K. Jackson, Ian Johnston, Dominic Kwiatkowski, Cordelia Langford, John Sillitoe on behalf of the Wellcome Sanger Institute COVID-19 Surveillance Team |
| EPI_ISL_3309647 | Health Services Laboratories | Wellcome Sanger Institute for the COVID-19 Genomics UK (COG-UK) Consortium | Health Services Laboratories and Alex Alderton, Roberto Amato, Jeffrey Barrett, Sonia Goncalves, Ewan Harrison, David K. Jackson, Ian Johnston, Dominic Kwiatkowski, Cordelia Langford, John Sillitoe on behalf of the Wellcome Sanger Institute COVID-19 Surveillance Team |
| EPI_ISL_3309648 | Lighthouse Laboratory Plymouth | Wellcome Sanger Institute for the COVID-19 Genomics UK (COG-UK) Consortium | Lighthouse Laboratory Plymouth and Alex Alderton, Roberto Amato, Jeffrey Barrett, Sonia Goncalves, Ewan Harrison, David K. Jackson, Ian Johnston, Dominic Kwiatkowski, Cordelia Langford, John Sillitoe on behalf of the Wellcome Sanger Institute COVID-19 Surveillance Team |
| EPI_ISL_3309649 | Health Services Laboratories | Wellcome Sanger Institute for the COVID-19 Genomics UK (COG-UK) Consortium | Health Services Laboratories and Alex Alderton, Roberto Amato, Jeffrey Barrett, Sonia Goncalves, Ewan Harrison, David K. Jackson, Ian Johnston, Dominic Kwiatkowski, Cordelia Langford, John Sillitoe on behalf of the Wellcome Sanger Institute COVID-19 Surveillance Team |
| EPI_ISL_3309650, EPI_ISL_3309651 | Lighthouse Laboratory Plymouth | Wellcome Sanger Institute for the COVID-19 Genomics UK (COG-UK) Consortium | Lighthouse Laboratory Plymouth and Alex Alderton, Roberto Amato, Jeffrey Barrett, Sonia Goncalves, Ewan Harrison, David K. Jackson, Ian Johnston, Dominic Kwiatkowski, Cordelia Langford, John Sillitoe on behalf of the Wellcome Sanger Institute COVID-19 Surveillance Team |
| EPI_ISL_3309652, EPI_ISL_3309653, EPI_ISL_3309654 | Health Services Laboratories | Wellcome Sanger Institute for the COVID-19 Genomics UK (COG-UK) Consortium | Health Services Laboratories and Alex Alderton, Roberto Amato, Jeffrey Barrett, Sonia Goncalves, Ewan Harrison, David K. Jackson, Ian Johnston, Dominic Kwiatkowski, Cordelia Langford, John Sillitoe on behalf of the Wellcome Sanger Institute COVID-19 Surveillance Team |
| EPI_ISL_3309655 | Lighthouse Laboratory Plymouth | Wellcome Sanger Institute for the COVID-19 Genomics UK (COG-UK) Consortium | Lighthouse Laboratory Plymouth and Alex Alderton, Roberto Amato, Jeffrey Barrett, Sonia Goncalves, Ewan Harrison, David K. Jackson, Ian Johnston, Dominic Kwiatkowski, Cordelia Langford, John Sillitoe on behalf of the Wellcome Sanger Institute COVID-19 Surveillance Team |
| EPI_ISL_3309656 | Health Services Laboratories | Wellcome Sanger Institute for the COVID-19 Genomics UK (COG-UK) Consortium | Health Services Laboratories and Alex Alderton, Roberto Amato, Jeffrey Barrett, Sonia Goncalves, Ewan Harrison, David K. Jackson, Ian Johnston, Dominic Kwiatkowski, Cordelia Langford, John Sillitoe on behalf of the Wellcome Sanger Institute COVID-19 Surveillance Team |
| EPI_ISL_3309657, EPI_ISL_3309658, EPI_ISL_3309659 | Lighthouse Laboratory Plymouth | Wellcome Sanger Institute for the COVID-19 Genomics UK (COG-UK) Consortium | Lighthouse Laboratory Plymouth and Alex Alderton, Roberto Amato, Jeffrey Barrett, Sonia Goncalves, Ewan Harrison, David K. Jackson, Ian Johnston, Dominic Kwiatkowski, Cordelia Langford, John Sillitoe on behalf of the Wellcome Sanger Institute COVID-19 Surveillance Team |
| EPI_ISL_3309660, EPI_ISL_3309661 | Health Services Laboratories | Wellcome Sanger Institute for the COVID-19 Genomics UK (COG-UK) Consortium | Health Services Laboratories and Alex Alderton, Roberto Amato, Jeffrey Barrett, Sonia Goncalves, Ewan Harrison, David K. Jackson, Ian Johnston, Dominic Kwiatkowski, Cordelia Langford, John Sillitoe on behalf of the Wellcome Sanger Institute COVID-19 Surveillance Team |
| EPI_ISL_3309662, EPI_ISL_3309663 | Lighthouse Laboratory Plymouth | Wellcome Sanger Institute for the COVID-19 Genomics UK (COG-UK) Consortium | Lighthouse Laboratory Plymouth and Alex Alderton, Roberto Amato, Jeffrey Barrett, Sonia Goncalves, Ewan Harrison, David K. Jackson, Ian Johnston, Dominic Kwiatkowski, Cordelia Langford, John Sillitoe on behalf of the Wellcome Sanger Institute COVID-19 Surveillance Team |
| EPI_ISL_3309664, EPI_ISL_3309665 | Health Services Laboratories | Wellcome Sanger Institute for the COVID-19 Genomics UK (COG-UK) Consortium | Health Services Laboratories and Alex Alderton, Roberto Amato, Jeffrey Barrett, Sonia Goncalves, Ewan Harrison, David K. Jackson, Ian Johnston, Dominic Kwiatkowski, Cordelia Langford, John Sillitoe on behalf of the Wellcome Sanger Institute COVID-19 Surveillance Team |
| EPI_ISL_3309666, EPI_ISL_3309667, EPI_ISL_3309668 | Lighthouse Laboratory Plymouth | Wellcome Sanger Institute for the COVID-19 Genomics UK (COG-UK) Consortium | Lighthouse Laboratory Plymouth and Alex Alderton, Roberto Amato, Jeffrey Barrett, Sonia Goncalves, Ewan Harrison, David K. Jackson, Ian Johnston, Dominic Kwiatkowski, Cordelia Langford, John Sillitoe on behalf of the Wellcome Sanger Institute COVID-19 Surveillance Team |
| EPI_ISL_3309669, EPI_ISL_3309670, EPI_ISL_3309671 | Health Services Laboratories | Wellcome Sanger Institute for the COVID-19 Genomics UK (COG-UK) Consortium | Health Services Laboratories and Alex Alderton, Roberto Amato, Jeffrey Barrett, Sonia Goncalves, Ewan Harrison, David K. Jackson, Ian Johnston, Dominic Kwiatkowski, Cordelia Langford, John Sillitoe on behalf of the Wellcome Sanger Institute COVID-19 Surveillance Team |
| EPI_ISL_3309672 | Lighthouse Laboratory Plymouth | Wellcome Sanger Institute for the COVID-19 Genomics UK (COG-UK) Consortium | Lighthouse Laboratory Plymouth and Alex Alderton, Roberto Amato, Jeffrey Barrett, Sonia Goncalves, Ewan Harrison, David K. Jackson, Ian Johnston, Dominic Kwiatkowski, Cordelia Langford, John Sillitoe on behalf of the Wellcome Sanger Institute COVID-19 Surveillance Team |
| EPI_ISL_3309673 | Health Services Laboratories | Wellcome Sanger Institute for the COVID-19 Genomics UK (COG-UK) Consortium | Health Services Laboratories and Alex Alderton, Roberto Amato, Jeffrey Barrett, Sonia Goncalves, Ewan Harrison, David K. Jackson, Ian Johnston, Dominic Kwiatkowski, Cordelia Langford, John Sillitoe on behalf of the Wellcome Sanger Institute COVID-19 Surveillance Team |
| EPI_ISL_3309674 | Lighthouse Laboratory Plymouth | Wellcome Sanger Institute for the COVID-19 Genomics UK (COG-UK) Consortium | Lighthouse Laboratory Plymouth and Alex Alderton, Roberto Amato, Jeffrey Barrett, Sonia Goncalves, Ewan Harrison, David K. Jackson, Ian Johnston, Dominic Kwiatkowski, Cordelia Langford, John Sillitoe on behalf of the Wellcome Sanger Institute COVID-19 Surveillance Team |
| EPI_ISL_3309675, EPI_ISL_3309676, EPI_ISL_3309677 | Health Services Laboratories | Wellcome Sanger Institute for the COVID-19 Genomics UK (COG-UK) Consortium | Health Services Laboratories and Alex Alderton, Roberto Amato, Jeffrey Barrett, Sonia Goncalves, Ewan Harrison, David K. Jackson, Ian Johnston, Dominic Kwiatkowski, Cordelia Langford, John Sillitoe on behalf of the Wellcome Sanger Institute COVID-19 Surveillance Team |
| EPI_ISL_3309678 | Lighthouse Laboratory Plymouth | Wellcome Sanger Institute for the COVID-19 Genomics UK (COG-UK) Consortium | Lighthouse Laboratory Plymouth and Alex Alderton, Roberto Amato, Jeffrey Barrett, Sonia Goncalves, Ewan Harrison, David K. Jackson, Ian Johnston, Dominic Kwiatkowski, Cordelia Langford, John Sillitoe on behalf of the Wellcome Sanger Institute COVID-19 Surveillance Team |
| EPI_ISL_3309679 | Lighthouse Lab in Glasgow | Wellcome Sanger Institute for the COVID-19 Genomics UK (COG-UK) Consortium | Harper VanSteenhouse, Yumi Kasai, David Gray, Carol Clugston, Anna Dominiczak and Alex Alderton, Roberto Amato, Jeffrey Barrett, Sonia Goncalves, Ewan Harrison, David K. Jackson, Ian Johnston, Dominic Kwiatkowski, Cordelia Langford, John Sillitoe on behalf of the Wellcome Sanger Institute COVID-19 Surveillance Team |
| EPI_ISL_3309680, EPI_ISL_3309681, EPI_ISL_3309682, EPI_ISL_3309683 | Lighthouse Laboratory Plymouth | Wellcome Sanger Institute for the COVID-19 Genomics UK (COG-UK) Consortium | Lighthouse Laboratory Plymouth and Alex Alderton, Roberto Amato, Jeffrey Barrett, Sonia Goncalves, Ewan Harrison, David K. Jackson, Ian Johnston, Dominic Kwiatkowski, Cordelia Langford, John Sillitoe on behalf of the Wellcome Sanger Institute COVID-19 Surveillance Team |
| EPI_ISL_3309684, EPI_ISL_3309685, EPI_ISL_3309686 | Health Services Laboratories | Wellcome Sanger Institute for the COVID-19 Genomics UK (COG-UK) Consortium | Health Services Laboratories and Alex Alderton, Roberto Amato, Jeffrey Barrett, Sonia Goncalves, Ewan Harrison, David K. Jackson, Ian Johnston, Dominic Kwiatkowski, Cordelia Langford, John Sillitoe on behalf of the Wellcome Sanger Institute COVID-19 Surveillance Team |
| EPI_ISL_3309687, | Lighthouse Laboratory Plymouth | Wellcome Sanger Institute for the COVID-19 Genomics UK (COG-UK) | Lighthouse Laboratory Plymouth and Alex Alderton, Roberto Amato, Jeffrey Barrett, Sonia Goncalves, Ewan Harrison, David K. Jackson, Ian Johnston, Dominic Kwiatkowski, Cordelia Langford, John Sillitoe on behalf of the Wellcome Sanger |

| | | | |
|---|---|---|---|
| EPI_ISL_3309688, EPI_ISL_3309689, EPI_ISL_3309690, EPI_ISL_3309691, EPI_ISL_3309692 | | Consortium | Institute COVID-19 Surveillance Team |
| EPI_ISL_3309693 | Health Services Laboratories | Wellcome Sanger Institute for the COVID-19 Genomics UK (COG-UK) Consortium | Health Services Laboratories and Alex Alderton, Roberto Amato, Jeffrey Barrett, Sonia Goncalves, Ewan Harrison, David K. Jackson, Ian Johnston, Dominic Kwiatkowski, Cordelia Langford, John Sillitoe on behalf of the Wellcome Sanger Institute COVID-19 Surveillance Team |
| EPI_ISL_3309694, EPI_ISL_3309695, EPI_ISL_3309696, EPI_ISL_3309697, EPI_ISL_3309698, EPI_ISL_3309699, EPI_ISL_3309700, EPI_ISL_3309701 | Lighthouse Laboratory Plymouth | Wellcome Sanger Institute for the COVID-19 Genomics UK (COG-UK) Consortium | Lighthouse Laboratory Plymouth and Alex Alderton, Roberto Amato, Jeffrey Barrett, Sonia Goncalves, Ewan Harrison, David K. Jackson, Ian Johnston, Dominic Kwiatkowski, Cordelia Langford, John Sillitoe on behalf of the Wellcome Sanger Institute COVID-19 Surveillance Team |
| EPI_ISL_3309702, EPI_ISL_3309703 | Health Services Laboratories | Wellcome Sanger Institute for the COVID-19 Genomics UK (COG-UK) Consortium | Health Services Laboratories and Alex Alderton, Roberto Amato, Jeffrey Barrett, Sonia Goncalves, Ewan Harrison, David K. Jackson, Ian Johnston, Dominic Kwiatkowski, Cordelia Langford, John Sillitoe on behalf of the Wellcome Sanger Institute COVID-19 Surveillance Team |
| EPI_ISL_3309704, EPI_ISL_3309705, EPI_ISL_3309706, EPI_ISL_3309707, EPI_ISL_3309708, EPI_ISL_3309709, EPI_ISL_3309710, EPI_ISL_3309711 | Lighthouse Laboratory Plymouth | Wellcome Sanger Institute for the COVID-19 Genomics UK (COG-UK) Consortium | Lighthouse Laboratory Plymouth and Alex Alderton, Roberto Amato, Jeffrey Barrett, Sonia Goncalves, Ewan Harrison, David K. Jackson, Ian Johnston, Dominic Kwiatkowski, Cordelia Langford, John Sillitoe on behalf of the Wellcome Sanger Institute COVID-19 Surveillance Team |
| EPI_ISL_3309712, EPI_ISL_3309713, EPI_ISL_3309714 | Health Services Laboratories | Wellcome Sanger Institute for the COVID-19 Genomics UK (COG-UK) Consortium | Health Services Laboratories and Alex Alderton, Roberto Amato, Jeffrey Barrett, Sonia Goncalves, Ewan Harrison, David K. Jackson, Ian Johnston, Dominic Kwiatkowski, Cordelia Langford, John Sillitoe on behalf of the Wellcome Sanger Institute COVID-19 Surveillance Team |
| EPI_ISL_3309715, EPI_ISL_3309716 | Lighthouse Laboratory Plymouth | Wellcome Sanger Institute for the COVID-19 Genomics UK (COG-UK) Consortium | Lighthouse Laboratory Plymouth and Alex Alderton, Roberto Amato, Jeffrey Barrett, Sonia Goncalves, Ewan Harrison, David K. Jackson, Ian Johnston, Dominic Kwiatkowski, Cordelia Langford, John Sillitoe on behalf of the Wellcome Sanger Institute COVID-19 Surveillance Team |
| EPI_ISL_3309717, EPI_ISL_3309718 | Health Services Laboratories | Wellcome Sanger Institute for the COVID-19 Genomics UK (COG-UK) Consortium | Health Services Laboratories and Alex Alderton, Roberto Amato, Jeffrey Barrett, Sonia Goncalves, Ewan Harrison, David K. Jackson, Ian Johnston, Dominic Kwiatkowski, Cordelia Langford, John Sillitoe on behalf of the Wellcome Sanger Institute COVID-19 Surveillance Team |
| EPI_ISL_3309719, EPI_ISL_3309720 | Lighthouse Laboratory Plymouth | Wellcome Sanger Institute for the COVID-19 Genomics UK (COG-UK) Consortium | Lighthouse Laboratory Plymouth and Alex Alderton, Roberto Amato, Jeffrey Barrett, Sonia Goncalves, Ewan Harrison, David K. Jackson, Ian Johnston, Dominic Kwiatkowski, Cordelia Langford, John Sillitoe on behalf of the Wellcome Sanger Institute COVID-19 Surveillance Team |
| EPI_ISL_3309721 | Health Services Laboratories | Wellcome Sanger Institute for the COVID-19 Genomics UK (COG-UK) Consortium | Health Services Laboratories and Alex Alderton, Roberto Amato, Jeffrey Barrett, Sonia Goncalves, Ewan Harrison, David K. Jackson, Ian Johnston, Dominic Kwiatkowski, Cordelia Langford, John Sillitoe on behalf of the Wellcome Sanger Institute COVID-19 Surveillance Team |
| EPI_ISL_3309722 | Lighthouse Laboratory Plymouth | Wellcome Sanger Institute for the COVID-19 Genomics UK (COG-UK) Consortium | Lighthouse Laboratory Plymouth and Alex Alderton, Roberto Amato, Jeffrey Barrett, Sonia Goncalves, Ewan Harrison, David K. Jackson, Ian Johnston, Dominic Kwiatkowski, Cordelia Langford, John Sillitoe on behalf of the Wellcome Sanger Institute COVID-19 Surveillance Team |
| EPI_ISL_3309723 | Health Services Laboratories | Wellcome Sanger Institute for the COVID-19 Genomics UK (COG-UK) Consortium | Health Services Laboratories and Alex Alderton, Roberto Amato, Jeffrey Barrett, Sonia Goncalves, Ewan Harrison, David K. Jackson, Ian Johnston, Dominic Kwiatkowski, Cordelia Langford, John Sillitoe on behalf of the Wellcome Sanger Institute COVID-19 Surveillance Team |
| EPI_ISL_3309724 | Lighthouse Laboratory Plymouth | Wellcome Sanger Institute for the COVID-19 Genomics UK (COG-UK) Consortium | Lighthouse Laboratory Plymouth and Alex Alderton, Roberto Amato, Jeffrey Barrett, Sonia Goncalves, Ewan Harrison, David K. Jackson, Ian Johnston, Dominic Kwiatkowski, Cordelia Langford, John Sillitoe on behalf of the Wellcome Sanger Institute COVID-19 Surveillance Team |
| EPI_ISL_3309725 | Health Services Laboratories | Wellcome Sanger Institute for the COVID-19 Genomics UK (COG-UK) Consortium | Health Services Laboratories and Alex Alderton, Roberto Amato, Jeffrey Barrett, Sonia Goncalves, Ewan Harrison, David K. Jackson, Ian Johnston, Dominic Kwiatkowski, Cordelia Langford, John Sillitoe on behalf of the Wellcome Sanger Institute COVID-19 Surveillance Team |
| EPI_ISL_3309726, EPI_ISL_3309727, EPI_ISL_3309728 | Lighthouse Laboratory Plymouth | Wellcome Sanger Institute for the COVID-19 Genomics UK (COG-UK) Consortium | Lighthouse Laboratory Plymouth and Alex Alderton, Roberto Amato, Jeffrey Barrett, Sonia Goncalves, Ewan Harrison, David K. Jackson, Ian Johnston, Dominic Kwiatkowski, Cordelia Langford, John Sillitoe on behalf of the Wellcome Sanger Institute COVID-19 Surveillance Team |
| EPI_ISL_3309729 | Health Services Laboratories | Wellcome Sanger Institute for the COVID-19 Genomics UK (COG-UK) Consortium | Health Services Laboratories and Alex Alderton, Roberto Amato, Jeffrey Barrett, Sonia Goncalves, Ewan Harrison, David K. Jackson, Ian Johnston, Dominic Kwiatkowski, Cordelia Langford, John Sillitoe on behalf of the Wellcome Sanger Institute COVID-19 Surveillance Team |
| EPI_ISL_3309730, EPI_ISL_3309731 | Lighthouse Laboratory Plymouth | Wellcome Sanger Institute for the COVID-19 Genomics UK (COG-UK) Consortium | Lighthouse Laboratory Plymouth and Alex Alderton, Roberto Amato, Jeffrey Barrett, Sonia Goncalves, Ewan Harrison, David K. Jackson, Ian Johnston, Dominic Kwiatkowski, Cordelia Langford, John Sillitoe on behalf of the Wellcome Sanger Institute COVID-19 Surveillance Team |
| EPI_ISL_3309732, EPI_ISL_3309733, EPI_ISL_3309734 | Health Services Laboratories | Wellcome Sanger Institute for the COVID-19 Genomics UK (COG-UK) Consortium | Health Services Laboratories and Alex Alderton, Roberto Amato, Jeffrey Barrett, Sonia Goncalves, Ewan Harrison, David K. Jackson, Ian Johnston, Dominic Kwiatkowski, Cordelia Langford, John Sillitoe on behalf of the Wellcome Sanger Institute COVID-19 Surveillance Team |
| EPI_ISL_3309735 | Lighthouse Laboratory Plymouth | Wellcome Sanger Institute for the COVID-19 Genomics UK (COG-UK) Consortium | Lighthouse Laboratory Plymouth and Alex Alderton, Roberto Amato, Jeffrey Barrett, Sonia Goncalves, Ewan Harrison, David K. Jackson, Ian Johnston, Dominic Kwiatkowski, Cordelia Langford, John Sillitoe on behalf of the Wellcome Sanger Institute COVID-19 Surveillance Team |
| EPI_ISL_3309736, EPI_ISL_3309737 | Health Services Laboratories | Wellcome Sanger Institute for the COVID-19 Genomics UK (COG-UK) Consortium | Health Services Laboratories and Alex Alderton, Roberto Amato, Jeffrey Barrett, Sonia Goncalves, Ewan Harrison, David K. Jackson, Ian Johnston, Dominic Kwiatkowski, Cordelia Langford, John Sillitoe on behalf of the Wellcome Sanger Institute COVID-19 Surveillance Team |
| EPI_ISL_3309738 | Lighthouse Laboratory Plymouth | Wellcome Sanger Institute for the COVID-19 Genomics UK (COG-UK) Consortium | Lighthouse Laboratory Plymouth and Alex Alderton, Roberto Amato, Jeffrey Barrett, Sonia Goncalves, Ewan Harrison, David K. Jackson, Ian Johnston, Dominic Kwiatkowski, Cordelia Langford, John Sillitoe on behalf of the Wellcome Sanger Institute COVID-19 Surveillance Team |
| EPI_ISL_3309739, EPI_ISL_3309740 | Health Services Laboratories | Wellcome Sanger Institute for the COVID-19 Genomics UK (COG-UK) Consortium | Health Services Laboratories and Alex Alderton, Roberto Amato, Jeffrey Barrett, Sonia Goncalves, Ewan Harrison, David K. Jackson, Ian Johnston, Dominic Kwiatkowski, Cordelia Langford, John Sillitoe on behalf of the Wellcome Sanger Institute COVID-19 Surveillance Team |
| EPI_ISL_3309741 | Lighthouse Laboratory Plymouth | Wellcome Sanger Institute for the COVID-19 Genomics UK (COG-UK) Consortium | Lighthouse Laboratory Plymouth and Alex Alderton, Roberto Amato, Jeffrey Barrett, Sonia Goncalves, Ewan Harrison, David K. Jackson, Ian Johnston, Dominic Kwiatkowski, Cordelia Langford, John Sillitoe on behalf of the Wellcome Sanger Institute COVID-19 Surveillance Team |
| EPI_ISL_3309742, EPI_ISL_3309743 | Health Services Laboratories | Wellcome Sanger Institute for the COVID-19 Genomics UK (COG-UK) Consortium | Health Services Laboratories and Alex Alderton, Roberto Amato, Jeffrey Barrett, Sonia Goncalves, Ewan Harrison, David K. Jackson, Ian Johnston, Dominic Kwiatkowski, Cordelia Langford, John Sillitoe on behalf of the Wellcome Sanger Institute COVID-19 Surveillance Team |
| EPI_ISL_3309744 | Lighthouse Laboratory Plymouth | Wellcome Sanger Institute for the COVID-19 Genomics UK (COG-UK) Consortium | Lighthouse Laboratory Plymouth and Alex Alderton, Roberto Amato, Jeffrey Barrett, Sonia Goncalves, Ewan Harrison, David K. Jackson, Ian Johnston, Dominic Kwiatkowski, Cordelia Langford, John Sillitoe on behalf of the Wellcome Sanger Institute COVID-19 Surveillance Team |
| EPI_ISL_3309745 | Lighthouse Lab in Glasgow | Wellcome Sanger Institute for the COVID-19 Genomics UK (COG-UK) Consortium | Harper VanSteenhouse, Yumi Kasai, David Gray, Carol Clugston, Anna Dominiczak and Alex Alderton, Roberto Amato, Jeffrey Barrett, Sonia Goncalves, Ewan Harrison, David K. Jackson, Ian Johnston, Dominic Kwiatkowski, Cordelia Langford, John Sillitoe on behalf of the Wellcome Sanger Institute COVID-19 Surveillance Team |
| EPI_ISL_3309746, EPI_ISL_3309747, EPI_ISL_3309748, EPI_ISL_3309749 | Lighthouse Laboratory Plymouth | Wellcome Sanger Institute for the COVID-19 Genomics UK (COG-UK) Consortium | Lighthouse Laboratory Plymouth and Alex Alderton, Roberto Amato, Jeffrey Barrett, Sonia Goncalves, Ewan Harrison, David K. Jackson, Ian Johnston, Dominic Kwiatkowski, Cordelia Langford, John Sillitoe on behalf of the Wellcome Sanger Institute COVID-19 Surveillance Team |
| EPI_ISL_3309750 | Health Services Laboratories | Wellcome Sanger Institute for the COVID-19 Genomics UK (COG-UK) Consortium | Health Services Laboratories and Alex Alderton, Roberto Amato, Jeffrey Barrett, Sonia Goncalves, Ewan Harrison, David K. Jackson, Ian Johnston, Dominic Kwiatkowski, Cordelia Langford, John Sillitoe on behalf of the Wellcome Sanger Institute COVID-19 Surveillance Team |
| EPI_ISL_3309751, EPI_ISL_3309752 | Lighthouse Laboratory Plymouth | Wellcome Sanger Institute for the COVID-19 Genomics UK (COG-UK) Consortium | Lighthouse Laboratory Plymouth and Alex Alderton, Roberto Amato, Jeffrey Barrett, Sonia Goncalves, Ewan Harrison, David K. Jackson, Ian Johnston, Dominic Kwiatkowski, Cordelia Langford, John Sillitoe on behalf of the Wellcome Sanger Institute COVID-19 Surveillance Team |
| EPI_ISL_3309753 | Health Services Laboratories | Wellcome Sanger Institute for the COVID-19 Genomics UK (COG-UK) Consortium | Health Services Laboratories and Alex Alderton, Roberto Amato, Jeffrey Barrett, Sonia Goncalves, Ewan Harrison, David K. Jackson, Ian Johnston, Dominic Kwiatkowski, Cordelia Langford, John Sillitoe on behalf of the Wellcome Sanger Institute COVID-19 Surveillance Team |
| EPI_ISL_3309754 | Lighthouse Laboratory Plymouth | Wellcome Sanger Institute for the COVID-19 Genomics UK (COG-UK) Consortium | Lighthouse Laboratory Plymouth and Alex Alderton, Roberto Amato, Jeffrey Barrett, Sonia Goncalves, Ewan Harrison, David K. Jackson, Ian Johnston, Dominic Kwiatkowski, Cordelia Langford, John Sillitoe on behalf of the Wellcome Sanger Institute COVID-19 Surveillance Team |
| EPI_ISL_3309755 | Health Services Laboratories | Wellcome Sanger Institute for the COVID-19 Genomics UK (COG-UK) Consortium | Health Services Laboratories and Alex Alderton, Roberto Amato, Jeffrey Barrett, Sonia Goncalves, Ewan Harrison, David K. Jackson, Ian Johnston, Dominic Kwiatkowski, Cordelia Langford, John Sillitoe on behalf of the Wellcome Sanger Institute COVID-19 Surveillance Team |
| EPI_ISL_3309756 | Lighthouse Laboratory Plymouth | Wellcome Sanger Institute for the COVID-19 Genomics UK (COG-UK) Consortium | Lighthouse Laboratory Plymouth and Alex Alderton, Roberto Amato, Jeffrey Barrett, Sonia Goncalves, Ewan Harrison, David K. Jackson, Ian Johnston, Dominic Kwiatkowski, Cordelia Langford, John Sillitoe on behalf of the Wellcome Sanger Institute COVID-19 Surveillance Team |
| EPI_ISL_3309757, EPI_ISL_3309758 | Health Services Laboratories | Wellcome Sanger Institute for the COVID-19 Genomics UK (COG-UK) Consortium | Health Services Laboratories and Alex Alderton, Roberto Amato, Jeffrey Barrett, Sonia Goncalves, Ewan Harrison, David K. Jackson, Ian Johnston, Dominic Kwiatkowski, Cordelia Langford, John Sillitoe on behalf of the Wellcome Sanger Institute COVID-19 Surveillance Team |
| EPI_ISL_3309759, EPI_ISL_3309760, EPI_ISL_3309761, EPI_ISL_3309762 | Lighthouse Laboratory Plymouth | Wellcome Sanger Institute for the COVID-19 Genomics UK (COG-UK) Consortium | Lighthouse Laboratory Plymouth and Alex Alderton, Roberto Amato, Jeffrey Barrett, Sonia Goncalves, Ewan Harrison, David K. Jackson, Ian Johnston, Dominic Kwiatkowski, Cordelia Langford, John Sillitoe on behalf of the Wellcome Sanger Institute COVID-19 Surveillance Team |
| EPI_ISL_3309763 | Lighthouse Lab in Glasgow | Wellcome Sanger Institute for the COVID-19 Genomics UK (COG-UK) Consortium | Harper VanSteenhouse, Yumi Kasai, David Gray, Carol Clugston, Anna Dominiczak and Alex Alderton, Roberto Amato, Jeffrey Barrett, Sonia Goncalves, Ewan Harrison, David K. Jackson, Ian Johnston, Dominic Kwiatkowski, Cordelia Langford, John Sillitoe on behalf of the Wellcome Sanger Institute COVID-19 Surveillance Team |

| | | | |
|---|---|---|---|
| EPI_ISL_3309764, EPI_ISL_3309765, EPI_ISL_3309766 | Health Services Laboratories | Wellcome Sanger Institute for the COVID-19 Genomics UK (COG-UK) Consortium | Health Services Laboratories and Alex Alderton, Roberto Amato, Jeffrey Barrett, Sonia Goncalves, Ewan Harrison, David K. Jackson, Ian Johnston, Dominic Kwiatkowski, Cordelia Langford, John Sillitoe on behalf of the Wellcome Sanger Institute COVID-19 Surveillance Team |
| EPI_ISL_3309767 | Lighthouse Laboratory Plymouth | Wellcome Sanger Institute for the COVID-19 Genomics UK (COG-UK) Consortium | Lighthouse Laboratory Plymouth and Alex Alderton, Roberto Amato, Jeffrey Barrett, Sonia Goncalves, Ewan Harrison, David K. Jackson, Ian Johnston, Dominic Kwiatkowski, Cordelia Langford, John Sillitoe on behalf of the Wellcome Sanger Institute COVID-19 Surveillance Team |
| EPI_ISL_3309768, EPI_ISL_3309769 | Health Services Laboratories | Wellcome Sanger Institute for the COVID-19 Genomics UK (COG-UK) Consortium | Health Services Laboratories and Alex Alderton, Roberto Amato, Jeffrey Barrett, Sonia Goncalves, Ewan Harrison, David K. Jackson, Ian Johnston, Dominic Kwiatkowski, Cordelia Langford, John Sillitoe on behalf of the Wellcome Sanger Institute COVID-19 Surveillance Team |
| EPI_ISL_3309770, EPI_ISL_3309771, EPI_ISL_3309772 | Lighthouse Laboratory Plymouth | Wellcome Sanger Institute for the COVID-19 Genomics UK (COG-UK) Consortium | Lighthouse Laboratory Plymouth and Alex Alderton, Roberto Amato, Jeffrey Barrett, Sonia Goncalves, Ewan Harrison, David K. Jackson, Ian Johnston, Dominic Kwiatkowski, Cordelia Langford, John Sillitoe on behalf of the Wellcome Sanger Institute COVID-19 Surveillance Team |
| EPI_ISL_3309773 | Health Services Laboratories | Wellcome Sanger Institute for the COVID-19 Genomics UK (COG-UK) Consortium | Health Services Laboratories and Alex Alderton, Roberto Amato, Jeffrey Barrett, Sonia Goncalves, Ewan Harrison, David K. Jackson, Ian Johnston, Dominic Kwiatkowski, Cordelia Langford, John Sillitoe on behalf of the Wellcome Sanger Institute COVID-19 Surveillance Team |
| EPI_ISL_3309774 | Lighthouse Laboratory Plymouth | Wellcome Sanger Institute for the COVID-19 Genomics UK (COG-UK) Consortium | Lighthouse Laboratory Plymouth and Alex Alderton, Roberto Amato, Jeffrey Barrett, Sonia Goncalves, Ewan Harrison, David K. Jackson, Ian Johnston, Dominic Kwiatkowski, Cordelia Langford, John Sillitoe on behalf of the Wellcome Sanger Institute COVID-19 Surveillance Team |
| EPI_ISL_3309775 | Health Services Laboratories | Wellcome Sanger Institute for the COVID-19 Genomics UK (COG-UK) Consortium | Health Services Laboratories and Alex Alderton, Roberto Amato, Jeffrey Barrett, Sonia Goncalves, Ewan Harrison, David K. Jackson, Ian Johnston, Dominic Kwiatkowski, Cordelia Langford, John Sillitoe on behalf of the Wellcome Sanger Institute COVID-19 Surveillance Team |
| EPI_ISL_3309776 | Lighthouse Laboratory Plymouth | Wellcome Sanger Institute for the COVID-19 Genomics UK (COG-UK) Consortium | Lighthouse Laboratory Plymouth and Alex Alderton, Roberto Amato, Jeffrey Barrett, Sonia Goncalves, Ewan Harrison, David K. Jackson, Ian Johnston, Dominic Kwiatkowski, Cordelia Langford, John Sillitoe on behalf of the Wellcome Sanger Institute COVID-19 Surveillance Team |
| EPI_ISL_3309777, EPI_ISL_3309778 | Health Services Laboratories | Wellcome Sanger Institute for the COVID-19 Genomics UK (COG-UK) Consortium | Health Services Laboratories and Alex Alderton, Roberto Amato, Jeffrey Barrett, Sonia Goncalves, Ewan Harrison, David K. Jackson, Ian Johnston, Dominic Kwiatkowski, Cordelia Langford, John Sillitoe on behalf of the Wellcome Sanger Institute COVID-19 Surveillance Team |
| EPI_ISL_3309779 | Lighthouse Laboratory Plymouth | Wellcome Sanger Institute for the COVID-19 Genomics UK (COG-UK) Consortium | Lighthouse Laboratory Plymouth and Alex Alderton, Roberto Amato, Jeffrey Barrett, Sonia Goncalves, Ewan Harrison, David K. Jackson, Ian Johnston, Dominic Kwiatkowski, Cordelia Langford, John Sillitoe on behalf of the Wellcome Sanger Institute COVID-19 Surveillance Team |
| EPI_ISL_3309780, EPI_ISL_3309781, EPI_ISL_3309782, EPI_ISL_3309783 | Health Services Laboratories | Wellcome Sanger Institute for the COVID-19 Genomics UK (COG-UK) Consortium | Health Services Laboratories and Alex Alderton, Roberto Amato, Jeffrey Barrett, Sonia Goncalves, Ewan Harrison, David K. Jackson, Ian Johnston, Dominic Kwiatkowski, Cordelia Langford, John Sillitoe on behalf of the Wellcome Sanger Institute COVID-19 Surveillance Team |
| EPI_ISL_3309784, EPI_ISL_3309785, EPI_ISL_3309786 | Lighthouse Laboratory Plymouth | Wellcome Sanger Institute for the COVID-19 Genomics UK (COG-UK) Consortium | Lighthouse Laboratory Plymouth and Alex Alderton, Roberto Amato, Jeffrey Barrett, Sonia Goncalves, Ewan Harrison, David K. Jackson, Ian Johnston, Dominic Kwiatkowski, Cordelia Langford, John Sillitoe on behalf of the Wellcome Sanger Institute COVID-19 Surveillance Team |
| EPI_ISL_3309787 | Health Services Laboratories | Wellcome Sanger Institute for the COVID-19 Genomics UK (COG-UK) Consortium | Health Services Laboratories and Alex Alderton, Roberto Amato, Jeffrey Barrett, Sonia Goncalves, Ewan Harrison, David K. Jackson, Ian Johnston, Dominic Kwiatkowski, Cordelia Langford, John Sillitoe on behalf of the Wellcome Sanger Institute COVID-19 Surveillance Team |
| EPI_ISL_3309788 | Lighthouse Laboratory Plymouth | Wellcome Sanger Institute for the COVID-19 Genomics UK (COG-UK) Consortium | Lighthouse Laboratory Plymouth and Alex Alderton, Roberto Amato, Jeffrey Barrett, Sonia Goncalves, Ewan Harrison, David K. Jackson, Ian Johnston, Dominic Kwiatkowski, Cordelia Langford, John Sillitoe on behalf of the Wellcome Sanger Institute COVID-19 Surveillance Team |
| EPI_ISL_3309789 | Lighthouse Lab in Glasgow | Wellcome Sanger Institute for the COVID-19 Genomics UK (COG-UK) Consortium | Harper VanSteenhouse, Yumi Kasai, David Gray, Carol Clugston, Anna Dominiczak and Alex Alderton, Roberto Amato, Jeffrey Barrett, Sonia Goncalves, Ewan Harrison, David K. Jackson, Ian Johnston, Dominic Kwiatkowski, Cordelia Langford, John Sillitoe on behalf of the Wellcome Sanger Institute COVID-19 Surveillance Team |
| EPI_ISL_3309790 | Health Services Laboratories | Wellcome Sanger Institute for the COVID-19 Genomics UK (COG-UK) Consortium | Health Services Laboratories and Alex Alderton, Roberto Amato, Jeffrey Barrett, Sonia Goncalves, Ewan Harrison, David K. Jackson, Ian Johnston, Dominic Kwiatkowski, Cordelia Langford, John Sillitoe on behalf of the Wellcome Sanger Institute COVID-19 Surveillance Team |
| EPI_ISL_3309791 | Lighthouse Laboratory Plymouth | Wellcome Sanger Institute for the COVID-19 Genomics UK (COG-UK) Consortium | Lighthouse Laboratory Plymouth and Alex Alderton, Roberto Amato, Jeffrey Barrett, Sonia Goncalves, Ewan Harrison, David K. Jackson, Ian Johnston, Dominic Kwiatkowski, Cordelia Langford, John Sillitoe on behalf of the Wellcome Sanger Institute COVID-19 Surveillance Team |
| EPI_ISL_3309792, EPI_ISL_3309793, EPI_ISL_3309794, EPI_ISL_3309795, EPI_ISL_3309796 | Health Services Laboratories | Wellcome Sanger Institute for the COVID-19 Genomics UK (COG-UK) Consortium | Health Services Laboratories and Alex Alderton, Roberto Amato, Jeffrey Barrett, Sonia Goncalves, Ewan Harrison, David K. Jackson, Ian Johnston, Dominic Kwiatkowski, Cordelia Langford, John Sillitoe on behalf of the Wellcome Sanger Institute COVID-19 Surveillance Team |
| EPI_ISL_3309797 | Lighthouse Laboratory Plymouth | Wellcome Sanger Institute for the COVID-19 Genomics UK (COG-UK) Consortium | Lighthouse Laboratory Plymouth and Alex Alderton, Roberto Amato, Jeffrey Barrett, Sonia Goncalves, Ewan Harrison, David K. Jackson, Ian Johnston, Dominic Kwiatkowski, Cordelia Langford, John Sillitoe on behalf of the Wellcome Sanger Institute COVID-19 Surveillance Team |
| EPI_ISL_3309798, EPI_ISL_3309799, EPI_ISL_3309800 | Health Services Laboratories | Wellcome Sanger Institute for the COVID-19 Genomics UK (COG-UK) Consortium | Health Services Laboratories and Alex Alderton, Roberto Amato, Jeffrey Barrett, Sonia Goncalves, Ewan Harrison, David K. Jackson, Ian Johnston, Dominic Kwiatkowski, Cordelia Langford, John Sillitoe on behalf of the Wellcome Sanger Institute COVID-19 Surveillance Team |
| EPI_ISL_3309801 | Lighthouse Laboratory Plymouth | Wellcome Sanger Institute for the COVID-19 Genomics UK (COG-UK) Consortium | Lighthouse Laboratory Plymouth and Alex Alderton, Roberto Amato, Jeffrey Barrett, Sonia Goncalves, Ewan Harrison, David K. Jackson, Ian Johnston, Dominic Kwiatkowski, Cordelia Langford, John Sillitoe on behalf of the Wellcome Sanger Institute COVID-19 Surveillance Team |
| EPI_ISL_3309802 | Lighthouse Lab in Glasgow | Wellcome Sanger Institute for the COVID-19 Genomics UK (COG-UK) Consortium | Harper VanSteenhouse, Yumi Kasai, David Gray, Carol Clugston, Anna Dominiczak and Alex Alderton, Roberto Amato, Jeffrey Barrett, Sonia Goncalves, Ewan Harrison, David K. Jackson, Ian Johnston, Dominic Kwiatkowski, Cordelia Langford, John Sillitoe on behalf of the Wellcome Sanger Institute COVID-19 Surveillance Team |
| EPI_ISL_3309803, EPI_ISL_3309804 | Health Services Laboratories | Wellcome Sanger Institute for the COVID-19 Genomics UK (COG-UK) Consortium | Health Services Laboratories and Alex Alderton, Roberto Amato, Jeffrey Barrett, Sonia Goncalves, Ewan Harrison, David K. Jackson, Ian Johnston, Dominic Kwiatkowski, Cordelia Langford, John Sillitoe on behalf of the Wellcome Sanger Institute COVID-19 Surveillance Team |
| EPI_ISL_3309805 | Lighthouse Lab in Glasgow | Wellcome Sanger Institute for the COVID-19 Genomics UK (COG-UK) Consortium | Harper VanSteenhouse, Yumi Kasai, David Gray, Carol Clugston, Anna Dominiczak and Alex Alderton, Roberto Amato, Jeffrey Barrett, Sonia Goncalves, Ewan Harrison, David K. Jackson, Ian Johnston, Dominic Kwiatkowski, Cordelia Langford, John Sillitoe on behalf of the Wellcome Sanger Institute COVID-19 Surveillance Team |
| EPI_ISL_3309806, EPI_ISL_3309807, EPI_ISL_3309808, EPI_ISL_3309809, EPI_ISL_3309810, EPI_ISL_3309811, EPI_ISL_3309812, EPI_ISL_3309813, EPI_ISL_3309814, EPI_ISL_3309815, EPI_ISL_3309816, EPI_ISL_3309817, EPI_ISL_3309818 | | | |
| see above | Lighthouse Laboratory Plymouth | Wellcome Sanger Institute for the COVID-19 Genomics UK (COG-UK) Consortium | Lighthouse Laboratory Plymouth and Alex Alderton, Roberto Amato, Jeffrey Barrett, Sonia Goncalves, Ewan Harrison, David K. Jackson, Ian Johnston, Dominic Kwiatkowski, Cordelia Langford, John Sillitoe on behalf of the Wellcome Sanger Institute COVID-19 Surveillance Team |
| EPI_ISL_3309819 | Health Services Laboratories | Wellcome Sanger Institute for the COVID-19 Genomics UK (COG-UK) Consortium | Health Services Laboratories and Alex Alderton, Roberto Amato, Jeffrey Barrett, Sonia Goncalves, Ewan Harrison, David K. Jackson, Ian Johnston, Dominic Kwiatkowski, Cordelia Langford, John Sillitoe on behalf of the Wellcome Sanger Institute COVID-19 Surveillance Team |
| EPI_ISL_3309820, EPI_ISL_3309821, EPI_ISL_3309822, EPI_ISL_3309823, EPI_ISL_3309824, EPI_ISL_3309825, EPI_ISL_3309826, EPI_ISL_3309827 | Lighthouse Laboratory Plymouth | Wellcome Sanger Institute for the COVID-19 Genomics UK (COG-UK) Consortium | Lighthouse Laboratory Plymouth and Alex Alderton, Roberto Amato, Jeffrey Barrett, Sonia Goncalves, Ewan Harrison, David K. Jackson, Ian Johnston, Dominic Kwiatkowski, Cordelia Langford, John Sillitoe on behalf of the Wellcome Sanger Institute COVID-19 Surveillance Team |
| EPI_ISL_3309828, EPI_ISL_3309829, EPI_ISL_3309830 | Health Services Laboratories | Wellcome Sanger Institute for the COVID-19 Genomics UK (COG-UK) Consortium | Health Services Laboratories and Alex Alderton, Roberto Amato, Jeffrey Barrett, Sonia Goncalves, Ewan Harrison, David K. Jackson, Ian Johnston, Dominic Kwiatkowski, Cordelia Langford, John Sillitoe on behalf of the Wellcome Sanger Institute COVID-19 Surveillance Team |
| EPI_ISL_3309831, EPI_ISL_3309832 | Lighthouse Laboratory Plymouth | Wellcome Sanger Institute for the COVID-19 Genomics UK (COG-UK) Consortium | Lighthouse Laboratory Plymouth and Alex Alderton, Roberto Amato, Jeffrey Barrett, Sonia Goncalves, Ewan Harrison, David K. Jackson, Ian Johnston, Dominic Kwiatkowski, Cordelia Langford, John Sillitoe on behalf of the Wellcome Sanger Institute COVID-19 Surveillance Team |
| EPI_ISL_3309833, EPI_ISL_3309834, EPI_ISL_3309835 | Health Services Laboratories | Wellcome Sanger Institute for the COVID-19 Genomics UK (COG-UK) Consortium | Health Services Laboratories and Alex Alderton, Roberto Amato, Jeffrey Barrett, Sonia Goncalves, Ewan Harrison, David K. Jackson, Ian Johnston, Dominic Kwiatkowski, Cordelia Langford, John Sillitoe on behalf of the Wellcome Sanger Institute COVID-19 Surveillance Team |
| EPI_ISL_3309836 | Lighthouse Lab in Glasgow | Wellcome Sanger Institute for the COVID-19 Genomics UK (COG-UK) Consortium | Harper VanSteenhouse, Yumi Kasai, David Gray, Carol Clugston, Anna Dominiczak and Alex Alderton, Roberto Amato, Jeffrey Barrett, Sonia Goncalves, Ewan Harrison, David K. Jackson, Ian Johnston, Dominic Kwiatkowski, Cordelia Langford, John Sillitoe on behalf of the Wellcome Sanger Institute COVID-19 Surveillance Team |
| EPI_ISL_3309837, EPI_ISL_3309838, EPI_ISL_3309839, EPI_ISL_3309840, EPI_ISL_3309841, EPI_ISL_3309842 | Lighthouse Laboratory Plymouth | Wellcome Sanger Institute for the COVID-19 Genomics UK (COG-UK) Consortium | Lighthouse Laboratory Plymouth and Alex Alderton, Roberto Amato, Jeffrey Barrett, Sonia Goncalves, Ewan Harrison, David K. Jackson, Ian Johnston, Dominic Kwiatkowski, Cordelia Langford, John Sillitoe on behalf of the Wellcome Sanger Institute COVID-19 Surveillance Team |
| EPI_ISL_3309843, EPI_ISL_3309844 | Health Services Laboratories | Wellcome Sanger Institute for the COVID-19 Genomics UK (COG-UK) Consortium | Health Services Laboratories and Alex Alderton, Roberto Amato, Jeffrey Barrett, Sonia Goncalves, Ewan Harrison, David K. Jackson, Ian Johnston, Dominic Kwiatkowski, Cordelia Langford, John Sillitoe on behalf of the Wellcome Sanger Institute COVID-19 Surveillance Team |
| EPI_ISL_3309845 | Lighthouse Laboratory Plymouth | Wellcome Sanger Institute for the COVID-19 Genomics UK (COG-UK) Consortium | Lighthouse Laboratory Plymouth and Alex Alderton, Roberto Amato, Jeffrey Barrett, Sonia Goncalves, Ewan Harrison, David K. Jackson, Ian Johnston, Dominic Kwiatkowski, Cordelia Langford, John Sillitoe on behalf of the Wellcome Sanger Institute COVID-19 Surveillance Team |
| EPI_ISL_3309846 | Health Services Laboratories | Wellcome Sanger Institute for the COVID-19 Genomics UK (COG-UK) Consortium | Health Services Laboratories and Alex Alderton, Roberto Amato, Jeffrey Barrett, Sonia Goncalves, Ewan Harrison, David K. Jackson, Ian Johnston, Dominic Kwiatkowski, Cordelia Langford, John Sillitoe on behalf of the Wellcome Sanger Institute COVID-19 Surveillance Team |
| EPI_ISL_3309847, EPI_ISL_3309848 | Lighthouse Laboratory Plymouth | Wellcome Sanger Institute for the COVID-19 Genomics UK (COG-UK) Consortium | Lighthouse Laboratory Plymouth and Alex Alderton, Roberto Amato, Jeffrey Barrett, Sonia Goncalves, Ewan Harrison, David K. Jackson, Ian Johnston, Dominic Kwiatkowski, Cordelia Langford, John Sillitoe on behalf of the Wellcome Sanger Institute COVID-19 Surveillance Team |

| | | | |
|---|---|---|---|
| EPI_ISL_3309849 | Health Services Laboratories | Wellcome Sanger Institute for the COVID-19 Genomics UK (COG-UK) Consortium | Health Services Laboratories and Alex Alderton, Roberto Amato, Jeffrey Barrett, Sonia Goncalves, Ewan Harrison, David K. Jackson, Ian Johnston, Dominic Kwiatkowski, Cordelia Langford, John Sillitoe on behalf of the Wellcome Sanger Institute COVID-19 Surveillance Team |
| EPI_ISL_3309850, EPI_ISL_3309851 | Lighthouse Laboratory Plymouth | Wellcome Sanger Institute for the COVID-19 Genomics UK (COG-UK) Consortium | Lighthouse Laboratory Plymouth and Alex Alderton, Roberto Amato, Jeffrey Barrett, Sonia Goncalves, Ewan Harrison, David K. Jackson, Ian Johnston, Dominic Kwiatkowski, Cordelia Langford, John Sillitoe on behalf of the Wellcome Sanger Institute COVID-19 Surveillance Team |
| EPI_ISL_3309852 | Health Services Laboratories | Wellcome Sanger Institute for the COVID-19 Genomics UK (COG-UK) Consortium | Health Services Laboratories and Alex Alderton, Roberto Amato, Jeffrey Barrett, Sonia Goncalves, Ewan Harrison, David K. Jackson, Ian Johnston, Dominic Kwiatkowski, Cordelia Langford, John Sillitoe on behalf of the Wellcome Sanger Institute COVID-19 Surveillance Team |
| EPI_ISL_3309853 | Lighthouse Lab in Glasgow | Wellcome Sanger Institute for the COVID-19 Genomics UK (COG-UK) Consortium | Harper VanSteenhouse, Yumi Kasai, David Gray, Carol Clugston, Anna Dominiczak and Alex Alderton, Roberto Amato, Jeffrey Barrett, Sonia Goncalves, Ewan Harrison, David K. Jackson, Ian Johnston, Dominic Kwiatkowski, Cordelia Langford, John Sillitoe on behalf of the Wellcome Sanger Institute COVID-19 Surveillance Team |
| EPI_ISL_3309854, EPI_ISL_3309855 | Lighthouse Laboratory Plymouth | Wellcome Sanger Institute for the COVID-19 Genomics UK (COG-UK) Consortium | Lighthouse Laboratory Plymouth and Alex Alderton, Roberto Amato, Jeffrey Barrett, Sonia Goncalves, Ewan Harrison, David K. Jackson, Ian Johnston, Dominic Kwiatkowski, Cordelia Langford, John Sillitoe on behalf of the Wellcome Sanger Institute COVID-19 Surveillance Team |
| EPI_ISL_3309856 | Health Services Laboratories | Wellcome Sanger Institute for the COVID-19 Genomics UK (COG-UK) Consortium | Health Services Laboratories and Alex Alderton, Roberto Amato, Jeffrey Barrett, Sonia Goncalves, Ewan Harrison, David K. Jackson, Ian Johnston, Dominic Kwiatkowski, Cordelia Langford, John Sillitoe on behalf of the Wellcome Sanger Institute COVID-19 Surveillance Team |
| EPI_ISL_3309857 | Lighthouse Laboratory Plymouth | Wellcome Sanger Institute for the COVID-19 Genomics UK (COG-UK) Consortium | Lighthouse Laboratory Plymouth and Alex Alderton, Roberto Amato, Jeffrey Barrett, Sonia Goncalves, Ewan Harrison, David K. Jackson, Ian Johnston, Dominic Kwiatkowski, Cordelia Langford, John Sillitoe on behalf of the Wellcome Sanger Institute COVID-19 Surveillance Team |
| EPI_ISL_3309858 | Health Services Laboratories | Wellcome Sanger Institute for the COVID-19 Genomics UK (COG-UK) Consortium | Health Services Laboratories and Alex Alderton, Roberto Amato, Jeffrey Barrett, Sonia Goncalves, Ewan Harrison, David K. Jackson, Ian Johnston, Dominic Kwiatkowski, Cordelia Langford, John Sillitoe on behalf of the Wellcome Sanger Institute COVID-19 Surveillance Team |
| EPI_ISL_3309859 | Lighthouse Laboratory Plymouth | Wellcome Sanger Institute for the COVID-19 Genomics UK (COG-UK) Consortium | Lighthouse Laboratory Plymouth and Alex Alderton, Roberto Amato, Jeffrey Barrett, Sonia Goncalves, Ewan Harrison, David K. Jackson, Ian Johnston, Dominic Kwiatkowski, Cordelia Langford, John Sillitoe on behalf of the Wellcome Sanger Institute COVID-19 Surveillance Team |
| EPI_ISL_3309860 | Health Services Laboratories | Wellcome Sanger Institute for the COVID-19 Genomics UK (COG-UK) Consortium | Health Services Laboratories and Alex Alderton, Roberto Amato, Jeffrey Barrett, Sonia Goncalves, Ewan Harrison, David K. Jackson, Ian Johnston, Dominic Kwiatkowski, Cordelia Langford, John Sillitoe on behalf of the Wellcome Sanger Institute COVID-19 Surveillance Team |
| EPI_ISL_3309861 | Lighthouse Laboratory Plymouth | Wellcome Sanger Institute for the COVID-19 Genomics UK (COG-UK) Consortium | Lighthouse Laboratory Plymouth and Alex Alderton, Roberto Amato, Jeffrey Barrett, Sonia Goncalves, Ewan Harrison, David K. Jackson, Ian Johnston, Dominic Kwiatkowski, Cordelia Langford, John Sillitoe on behalf of the Wellcome Sanger Institute COVID-19 Surveillance Team |
| EPI_ISL_3309862 | Health Services Laboratories | Wellcome Sanger Institute for the COVID-19 Genomics UK (COG-UK) Consortium | Health Services Laboratories and Alex Alderton, Roberto Amato, Jeffrey Barrett, Sonia Goncalves, Ewan Harrison, David K. Jackson, Ian Johnston, Dominic Kwiatkowski, Cordelia Langford, John Sillitoe on behalf of the Wellcome Sanger Institute COVID-19 Surveillance Team |
| EPI_ISL_3309863, EPI_ISL_3309864 | Lighthouse Laboratory Plymouth | Wellcome Sanger Institute for the COVID-19 Genomics UK (COG-UK) Consortium | Lighthouse Laboratory Plymouth and Alex Alderton, Roberto Amato, Jeffrey Barrett, Sonia Goncalves, Ewan Harrison, David K. Jackson, Ian Johnston, Dominic Kwiatkowski, Cordelia Langford, John Sillitoe on behalf of the Wellcome Sanger Institute COVID-19 Surveillance Team |
| EPI_ISL_3309865, EPI_ISL_3309866 | Health Services Laboratories | Wellcome Sanger Institute for the COVID-19 Genomics UK (COG-UK) Consortium | Health Services Laboratories and Alex Alderton, Roberto Amato, Jeffrey Barrett, Sonia Goncalves, Ewan Harrison, David K. Jackson, Ian Johnston, Dominic Kwiatkowski, Cordelia Langford, John Sillitoe on behalf of the Wellcome Sanger Institute COVID-19 Surveillance Team |
| EPI_ISL_3309867 | Lighthouse Lab in Glasgow | Wellcome Sanger Institute for the COVID-19 Genomics UK (COG-UK) Consortium | Harper VanSteenhouse, Yumi Kasai, David Gray, Carol Clugston, Anna Dominiczak and Alex Alderton, Roberto Amato, Jeffrey Barrett, Sonia Goncalves, Ewan Harrison, David K. Jackson, Ian Johnston, Dominic Kwiatkowski, Cordelia Langford, John Sillitoe on behalf of the Wellcome Sanger Institute COVID-19 Surveillance Team |
| EPI_ISL_3309868, EPI_ISL_3309869, EPI_ISL_3309870, EPI_ISL_3309871, EPI_ISL_3309872 | Lighthouse Laboratory Plymouth | Wellcome Sanger Institute for the COVID-19 Genomics UK (COG-UK) Consortium | Lighthouse Laboratory Plymouth and Alex Alderton, Roberto Amato, Jeffrey Barrett, Sonia Goncalves, Ewan Harrison, David K. Jackson, Ian Johnston, Dominic Kwiatkowski, Cordelia Langford, John Sillitoe on behalf of the Wellcome Sanger Institute COVID-19 Surveillance Team |
| EPI_ISL_3309873 | Health Services Laboratories | Wellcome Sanger Institute for the COVID-19 Genomics UK (COG-UK) Consortium | Health Services Laboratories and Alex Alderton, Roberto Amato, Jeffrey Barrett, Sonia Goncalves, Ewan Harrison, David K. Jackson, Ian Johnston, Dominic Kwiatkowski, Cordelia Langford, John Sillitoe on behalf of the Wellcome Sanger Institute COVID-19 Surveillance Team |
| EPI_ISL_3309874, EPI_ISL_3309875 | Lighthouse Laboratory Plymouth | Wellcome Sanger Institute for the COVID-19 Genomics UK (COG-UK) Consortium | Lighthouse Laboratory Plymouth and Alex Alderton, Roberto Amato, Jeffrey Barrett, Sonia Goncalves, Ewan Harrison, David K. Jackson, Ian Johnston, Dominic Kwiatkowski, Cordelia Langford, John Sillitoe on behalf of the Wellcome Sanger Institute COVID-19 Surveillance Team |
| EPI_ISL_3309876 | Health Services Laboratories | Wellcome Sanger Institute for the COVID-19 Genomics UK (COG-UK) Consortium | Health Services Laboratories and Alex Alderton, Roberto Amato, Jeffrey Barrett, Sonia Goncalves, Ewan Harrison, David K. Jackson, Ian Johnston, Dominic Kwiatkowski, Cordelia Langford, John Sillitoe on behalf of the Wellcome Sanger Institute COVID-19 Surveillance Team |
| EPI_ISL_3309877, EPI_ISL_3309878 | Lighthouse Laboratory Plymouth | Wellcome Sanger Institute for the COVID-19 Genomics UK (COG-UK) Consortium | Lighthouse Laboratory Plymouth and Alex Alderton, Roberto Amato, Jeffrey Barrett, Sonia Goncalves, Ewan Harrison, David K. Jackson, Ian Johnston, Dominic Kwiatkowski, Cordelia Langford, John Sillitoe on behalf of the Wellcome Sanger Institute COVID-19 Surveillance Team |
| EPI_ISL_3309879 | Lighthouse Lab in Alderley Park | Wellcome Sanger Institute for the COVID-19 Genomics UK (COG-UK) Consortium | Jacquelyn Wynn, Mairead Hyland, The Lighthouse Lab in Alderley Park and Alex Alderton, Roberto Amato, Jeffrey Barrett, Sonia Goncalves, Ewan Harrison, David K. Jackson, Ian Johnston, Dominic Kwiatkowski, Cordelia Langford, John Sillitoe on behalf of the Wellcome Sanger Institute COVID-19 Surveillance Team |
| EPI_ISL_3309880 | Health Services Laboratories | Wellcome Sanger Institute for the COVID-19 Genomics UK (COG-UK) Consortium | Health Services Laboratories and Alex Alderton, Roberto Amato, Jeffrey Barrett, Sonia Goncalves, Ewan Harrison, David K. Jackson, Ian Johnston, Dominic Kwiatkowski, Cordelia Langford, John Sillitoe on behalf of the Wellcome Sanger Institute COVID-19 Surveillance Team |
| EPI_ISL_3309881 | Lighthouse Laboratory Plymouth | Wellcome Sanger Institute for the COVID-19 Genomics UK (COG-UK) Consortium | Lighthouse Laboratory Plymouth and Alex Alderton, Roberto Amato, Jeffrey Barrett, Sonia Goncalves, Ewan Harrison, David K. Jackson, Ian Johnston, Dominic Kwiatkowski, Cordelia Langford, John Sillitoe on behalf of the Wellcome Sanger Institute COVID-19 Surveillance Team |
| EPI_ISL_3309882 | Lighthouse Lab in Milton Keynes | Wellcome Sanger Institute for the COVID-19 Genomics UK (COG-UK) Consortium | The Lighthouse Lab in Milton Keynes and Alex Alderton, Roberto Amato, Jeffrey Barrett, Sonia Goncalves, Ewan Harrison, David K. Jackson, Ian Johnston, Dominic Kwiatkowski, Cordelia Langford, John Sillitoe on behalf of the Wellcome Sanger Institute COVID-19 Surveillance Team |
| EPI_ISL_3309883, EPI_ISL_3309884, EPI_ISL_3309885 | Lighthouse Lab in Alderley Park | Wellcome Sanger Institute for the COVID-19 Genomics UK (COG-UK) Consortium | Jacquelyn Wynn, Mairead Hyland, The Lighthouse Lab in Alderley Park and Alex Alderton, Roberto Amato, Jeffrey Barrett, Sonia Goncalves, Ewan Harrison, David K. Jackson, Ian Johnston, Dominic Kwiatkowski, Cordelia Langford, John Sillitoe on behalf of the Wellcome Sanger Institute COVID-19 Surveillance Team |
| EPI_ISL_3309886, EPI_ISL_3309887 | Lighthouse Lab in Milton Keynes | Wellcome Sanger Institute for the COVID-19 Genomics UK (COG-UK) Consortium | The Lighthouse Lab in Milton Keynes and Alex Alderton, Roberto Amato, Jeffrey Barrett, Sonia Goncalves, Ewan Harrison, David K. Jackson, Ian Johnston, Dominic Kwiatkowski, Cordelia Langford, John Sillitoe on behalf of the Wellcome Sanger Institute COVID-19 Surveillance Team |
| EPI_ISL_3309888 | Lighthouse Laboratory Plymouth | Wellcome Sanger Institute for the COVID-19 Genomics UK (COG-UK) Consortium | Lighthouse Laboratory Plymouth and Alex Alderton, Roberto Amato, Jeffrey Barrett, Sonia Goncalves, Ewan Harrison, David K. Jackson, Ian Johnston, Dominic Kwiatkowski, Cordelia Langford, John Sillitoe on behalf of the Wellcome Sanger Institute COVID-19 Surveillance Team |
| EPI_ISL_3309889 | Health Services Laboratories | Wellcome Sanger Institute for the COVID-19 Genomics UK (COG-UK) Consortium | Health Services Laboratories and Alex Alderton, Roberto Amato, Jeffrey Barrett, Sonia Goncalves, Ewan Harrison, David K. Jackson, Ian Johnston, Dominic Kwiatkowski, Cordelia Langford, John Sillitoe on behalf of the Wellcome Sanger Institute COVID-19 Surveillance Team |
| EPI_ISL_3309890, EPI_ISL_3309891 | Lighthouse Lab in Milton Keynes | Wellcome Sanger Institute for the COVID-19 Genomics UK (COG-UK) Consortium | The Lighthouse Lab in Milton Keynes and Alex Alderton, Roberto Amato, Jeffrey Barrett, Sonia Goncalves, Ewan Harrison, David K. Jackson, Ian Johnston, Dominic Kwiatkowski, Cordelia Langford, John Sillitoe on behalf of the Wellcome Sanger Institute COVID-19 Surveillance Team |
| EPI_ISL_3309892 | Lighthouse Lab in Alderley Park | Wellcome Sanger Institute for the COVID-19 Genomics UK (COG-UK) Consortium | Jacquelyn Wynn, Mairead Hyland, The Lighthouse Lab in Alderley Park and Alex Alderton, Roberto Amato, Jeffrey Barrett, Sonia Goncalves, Ewan Harrison, David K. Jackson, Ian Johnston, Dominic Kwiatkowski, Cordelia Langford, John Sillitoe on behalf of the Wellcome Sanger Institute COVID-19 Surveillance Team |
| EPI_ISL_3309893, EPI_ISL_3309894 | Health Services Laboratories | Wellcome Sanger Institute for the COVID-19 Genomics UK (COG-UK) Consortium | Health Services Laboratories and Alex Alderton, Roberto Amato, Jeffrey Barrett, Sonia Goncalves, Ewan Harrison, David K. Jackson, Ian Johnston, Dominic Kwiatkowski, Cordelia Langford, John Sillitoe on behalf of the Wellcome Sanger Institute COVID-19 Surveillance Team |
| EPI_ISL_3309895 | Lighthouse Lab in Alderley Park | Wellcome Sanger Institute for the COVID-19 Genomics UK (COG-UK) Consortium | Jacquelyn Wynn, Mairead Hyland, The Lighthouse Lab in Alderley Park and Alex Alderton, Roberto Amato, Jeffrey Barrett, Sonia Goncalves, Ewan Harrison, David K. Jackson, Ian Johnston, Dominic Kwiatkowski, Cordelia Langford, John Sillitoe on behalf of the Wellcome Sanger Institute COVID-19 Surveillance Team |
| EPI_ISL_3309896 | Lighthouse Lab in Milton Keynes | Wellcome Sanger Institute for the COVID-19 Genomics UK (COG-UK) Consortium | The Lighthouse Lab in Milton Keynes and Alex Alderton, Roberto Amato, Jeffrey Barrett, Sonia Goncalves, Ewan Harrison, David K. Jackson, Ian Johnston, Dominic Kwiatkowski, Cordelia Langford, John Sillitoe on behalf of the Wellcome Sanger Institute COVID-19 Surveillance Team |
| EPI_ISL_3309897 | Lighthouse Lab in Alderley Park | Wellcome Sanger Institute for the COVID-19 Genomics UK (COG-UK) Consortium | Jacquelyn Wynn, Mairead Hyland, The Lighthouse Lab in Alderley Park and Alex Alderton, Roberto Amato, Jeffrey Barrett, Sonia Goncalves, Ewan Harrison, David K. Jackson, Ian Johnston, Dominic Kwiatkowski, Cordelia Langford, John Sillitoe on behalf of the Wellcome Sanger Institute COVID-19 Surveillance Team |
| EPI_ISL_3309898 | Lighthouse Lab in Milton Keynes | Wellcome Sanger Institute for the COVID-19 Genomics UK (COG-UK) Consortium | The Lighthouse Lab in Milton Keynes and Alex Alderton, Roberto Amato, Jeffrey Barrett, Sonia Goncalves, Ewan Harrison, David K. Jackson, Ian Johnston, Dominic Kwiatkowski, Cordelia Langford, John Sillitoe on behalf of the Wellcome Sanger Institute COVID-19 Surveillance Team |
| EPI_ISL_3309899 | Lighthouse Lab in Alderley Park | Wellcome Sanger Institute for the COVID-19 Genomics UK (COG-UK) Consortium | Jacquelyn Wynn, Mairead Hyland, The Lighthouse Lab in Alderley Park and Alex Alderton, Roberto Amato, Jeffrey Barrett, Sonia Goncalves, Ewan Harrison, David K. Jackson, Ian Johnston, Dominic Kwiatkowski, Cordelia Langford, John Sillitoe on behalf of the Wellcome Sanger Institute COVID-19 Surveillance Team |
| EPI_ISL_3309900 | Lighthouse Lab in Milton Keynes | Wellcome Sanger Institute for the COVID-19 Genomics UK (COG-UK) Consortium | The Lighthouse Lab in Milton Keynes and Alex Alderton, Roberto Amato, Jeffrey Barrett, Sonia Goncalves, Ewan Harrison, David K. Jackson, Ian Johnston, Dominic Kwiatkowski, Cordelia Langford, John Sillitoe on behalf of the Wellcome Sanger Institute COVID-19 Surveillance Team |
| EPI_ISL_3309901 | Lighthouse Laboratory Plymouth | Wellcome Sanger Institute for the COVID-19 Genomics UK (COG-UK) Consortium | Lighthouse Laboratory Plymouth and Alex Alderton, Roberto Amato, Jeffrey Barrett, Sonia Goncalves, Ewan Harrison, David K. Jackson, Ian Johnston, Dominic Kwiatkowski, Cordelia Langford, John Sillitoe on behalf of the Wellcome Sanger Institute COVID-19 Surveillance Team |
| EPI_ISL_3309902 | Lighthouse Lab in Alderley Park | Wellcome Sanger Institute for the COVID-19 Genomics UK (COG-UK) Consortium | Jacquelyn Wynn, Mairead Hyland, The Lighthouse Lab in Alderley Park and Alex Alderton, Roberto Amato, Jeffrey Barrett, Sonia Goncalves, Ewan Harrison, David K. Jackson, Ian Johnston, Dominic Kwiatkowski, Cordelia Langford, John Sillitoe on behalf of the Wellcome Sanger Institute COVID-19 Surveillance Team |
| EPI_ISL_3309903 | Lighthouse Lab in Milton Keynes | Wellcome Sanger Institute for the COVID-19 Genomics UK (COG-UK) Consortium | The Lighthouse Lab in Milton Keynes and Alex Alderton, Roberto Amato, Jeffrey Barrett, Sonia Goncalves, Ewan Harrison, David K. Jackson, Ian Johnston, Dominic Kwiatkowski, Cordelia Langford, John Sillitoe on behalf of the Wellcome Sanger Institute COVID-19 Surveillance Team |
| EPI_ISL_3309904 | Lighthouse Lab in Alderley Park | Wellcome Sanger Institute for the COVID-19 Genomics UK (COG-UK) Consortium | Jacquelyn Wynn, Mairead Hyland, The Lighthouse Lab in Alderley Park and Alex Alderton, Roberto Amato, Jeffrey Barrett, Sonia Goncalves, Ewan Harrison, David K. Jackson, Ian Johnston, Dominic Kwiatkowski, Cordelia Langford, John Sillitoe on behalf of the Wellcome Sanger Institute COVID-19 Surveillance Team |
| EPI_ISL_3309905 | Lighthouse Lab in Milton Keynes | Wellcome Sanger Institute for the COVID-19 Genomics UK (COG-UK) Consortium | The Lighthouse Lab in Milton Keynes and Alex Alderton, Roberto Amato, Jeffrey Barrett, Sonia Goncalves, Ewan Harrison, David K. Jackson, Ian Johnston, Dominic Kwiatkowski, Cordelia Langford, John Sillitoe on behalf of the Wellcome Sanger Institute COVID-19 Surveillance Team |
| EPI_ISL_3309906 | Health Services Laboratories | Wellcome Sanger Institute for the COVID-19 Genomics UK (COG-UK) Consortium | Health Services Laboratories and Alex Alderton, Roberto Amato, Jeffrey Barrett, Sonia Goncalves, Ewan Harrison, David K. Jackson, Ian Johnston, Dominic Kwiatkowski, Cordelia Langford, John Sillitoe on behalf of the Wellcome Sanger Institute COVID-19 Surveillance Team |

| | | | |
|---|---|---|---|
| EPI_ISL_3309907, EPI_ISL_3309908 | Lighthouse Laboratory Plymouth | Wellcome Sanger Institute for the COVID-19 Genomics UK (COG-UK) Consortium | Lighthouse Laboratory Plymouth and Alex Alderton, Roberto Amato, Jeffrey Barrett, Sonia Goncalves, Ewan Harrison, David K. Jackson, Ian Johnston, Dominic Kwiatkowski, Cordelia Langford, John Sillitoe on behalf of the Wellcome Sanger Institute COVID-19 Surveillance Team |
| EPI_ISL_3309909, EPI_ISL_3309910 | Lighthouse Lab in Milton Keynes | Wellcome Sanger Institute for the COVID-19 Genomics UK (COG-UK) Consortium | The Lighthouse Lab in Milton Keynes and Alex Alderton, Roberto Amato, Jeffrey Barrett, Sonia Goncalves, Ewan Harrison, David K. Jackson, Ian Johnston, Dominic Kwiatkowski, Cordelia Langford, John Sillitoe on behalf of the Wellcome Sanger Institute COVID-19 Surveillance Team |
| EPI_ISL_3309911 | Lighthouse Lab in Alderley Park | Wellcome Sanger Institute for the COVID-19 Genomics UK (COG-UK) Consortium | Jacquelyn Wynn, Mairead Hyland, The Lighthouse Lab in Alderley Park and Alex Alderton, Roberto Amato, Jeffrey Barrett, Sonia Goncalves, Ewan Harrison, David K. Jackson, Ian Johnston, Dominic Kwiatkowski, Cordelia Langford, John Sillitoe on behalf of the Wellcome Sanger Institute COVID-19 Surveillance Team |
| EPI_ISL_3309912, EPI_ISL_3309913, EPI_ISL_3309914 | Lighthouse Laboratory Plymouth | Wellcome Sanger Institute for the COVID-19 Genomics UK (COG-UK) Consortium | Lighthouse Laboratory Plymouth and Alex Alderton, Roberto Amato, Jeffrey Barrett, Sonia Goncalves, Ewan Harrison, David K. Jackson, Ian Johnston, Dominic Kwiatkowski, Cordelia Langford, John Sillitoe on behalf of the Wellcome Sanger Institute COVID-19 Surveillance Team |
| EPI_ISL_3309915 | Health Services Laboratories | Wellcome Sanger Institute for the COVID-19 Genomics UK (COG-UK) Consortium | Health Services Laboratories and Alex Alderton, Roberto Amato, Jeffrey Barrett, Sonia Goncalves, Ewan Harrison, David K. Jackson, Ian Johnston, Dominic Kwiatkowski, Cordelia Langford, John Sillitoe on behalf of the Wellcome Sanger Institute COVID-19 Surveillance Team |
| EPI_ISL_3309916 | Lighthouse Lab in Alderley Park | Wellcome Sanger Institute for the COVID-19 Genomics UK (COG-UK) Consortium | Jacquelyn Wynn, Mairead Hyland, The Lighthouse Lab in Alderley Park and Alex Alderton, Roberto Amato, Jeffrey Barrett, Sonia Goncalves, Ewan Harrison, David K. Jackson, Ian Johnston, Dominic Kwiatkowski, Cordelia Langford, John Sillitoe on behalf of the Wellcome Sanger Institute COVID-19 Surveillance Team |
| EPI_ISL_3309917 | Health Services Laboratories | Wellcome Sanger Institute for the COVID-19 Genomics UK (COG-UK) Consortium | Health Services Laboratories and Alex Alderton, Roberto Amato, Jeffrey Barrett, Sonia Goncalves, Ewan Harrison, David K. Jackson, Ian Johnston, Dominic Kwiatkowski, Cordelia Langford, John Sillitoe on behalf of the Wellcome Sanger Institute COVID-19 Surveillance Team |
| EPI_ISL_3309918 | Lighthouse Laboratory Plymouth | Wellcome Sanger Institute for the COVID-19 Genomics UK (COG-UK) Consortium | Lighthouse Laboratory Plymouth and Alex Alderton, Roberto Amato, Jeffrey Barrett, Sonia Goncalves, Ewan Harrison, David K. Jackson, Ian Johnston, Dominic Kwiatkowski, Cordelia Langford, John Sillitoe on behalf of the Wellcome Sanger Institute COVID-19 Surveillance Team |
| EPI_ISL_3309919, EPI_ISL_3309920, EPI_ISL_3309921, EPI_ISL_3309922 | Health Services Laboratories | Wellcome Sanger Institute for the COVID-19 Genomics UK (COG-UK) Consortium | Health Services Laboratories and Alex Alderton, Roberto Amato, Jeffrey Barrett, Sonia Goncalves, Ewan Harrison, David K. Jackson, Ian Johnston, Dominic Kwiatkowski, Cordelia Langford, John Sillitoe on behalf of the Wellcome Sanger Institute COVID-19 Surveillance Team |
| EPI_ISL_3309923 | Lighthouse Laboratory Plymouth | Wellcome Sanger Institute for the COVID-19 Genomics UK (COG-UK) Consortium | Lighthouse Laboratory Plymouth and Alex Alderton, Roberto Amato, Jeffrey Barrett, Sonia Goncalves, Ewan Harrison, David K. Jackson, Ian Johnston, Dominic Kwiatkowski, Cordelia Langford, John Sillitoe on behalf of the Wellcome Sanger Institute COVID-19 Surveillance Team |
| EPI_ISL_3309924 | Lighthouse Lab in Alderley Park | Wellcome Sanger Institute for the COVID-19 Genomics UK (COG-UK) Consortium | Jacquelyn Wynn, Mairead Hyland, The Lighthouse Lab in Alderley Park and Alex Alderton, Roberto Amato, Jeffrey Barrett, Sonia Goncalves, Ewan Harrison, David K. Jackson, Ian Johnston, Dominic Kwiatkowski, Cordelia Langford, John Sillitoe on behalf of the Wellcome Sanger Institute COVID-19 Surveillance Team |
| EPI_ISL_3309925 | Lighthouse Laboratory Plymouth | Wellcome Sanger Institute for the COVID-19 Genomics UK (COG-UK) Consortium | Lighthouse Laboratory Plymouth and Alex Alderton, Roberto Amato, Jeffrey Barrett, Sonia Goncalves, Ewan Harrison, David K. Jackson, Ian Johnston, Dominic Kwiatkowski, Cordelia Langford, John Sillitoe on behalf of the Wellcome Sanger Institute COVID-19 Surveillance Team |
| EPI_ISL_3309926, EPI_ISL_3309927 | Lighthouse Lab in Alderley Park | Wellcome Sanger Institute for the COVID-19 Genomics UK (COG-UK) Consortium | Jacquelyn Wynn, Mairead Hyland, The Lighthouse Lab in Alderley Park and Alex Alderton, Roberto Amato, Jeffrey Barrett, Sonia Goncalves, Ewan Harrison, David K. Jackson, Ian Johnston, Dominic Kwiatkowski, Cordelia Langford, John Sillitoe on behalf of the Wellcome Sanger Institute COVID-19 Surveillance Team |
| EPI_ISL_3309928, EPI_ISL_3309929, EPI_ISL_3309930, EPI_ISL_3309931, EPI_ISL_3309932 | Health Services Laboratories | Wellcome Sanger Institute for the COVID-19 Genomics UK (COG-UK) Consortium | Health Services Laboratories and Alex Alderton, Roberto Amato, Jeffrey Barrett, Sonia Goncalves, Ewan Harrison, David K. Jackson, Ian Johnston, Dominic Kwiatkowski, Cordelia Langford, John Sillitoe on behalf of the Wellcome Sanger Institute COVID-19 Surveillance Team |
| EPI_ISL_3309933, EPI_ISL_3309934 | Lighthouse Laboratory Plymouth | Wellcome Sanger Institute for the COVID-19 Genomics UK (COG-UK) Consortium | Lighthouse Laboratory Plymouth and Alex Alderton, Roberto Amato, Jeffrey Barrett, Sonia Goncalves, Ewan Harrison, David K. Jackson, Ian Johnston, Dominic Kwiatkowski, Cordelia Langford, John Sillitoe on behalf of the Wellcome Sanger Institute COVID-19 Surveillance Team |
| EPI_ISL_3309935 | Health Services Laboratories | Wellcome Sanger Institute for the COVID-19 Genomics UK (COG-UK) Consortium | Health Services Laboratories and Alex Alderton, Roberto Amato, Jeffrey Barrett, Sonia Goncalves, Ewan Harrison, David K. Jackson, Ian Johnston, Dominic Kwiatkowski, Cordelia Langford, John Sillitoe on behalf of the Wellcome Sanger Institute COVID-19 Surveillance Team |
| EPI_ISL_3309936 | Lighthouse Laboratory Plymouth | Wellcome Sanger Institute for the COVID-19 Genomics UK (COG-UK) Consortium | Lighthouse Laboratory Plymouth and Alex Alderton, Roberto Amato, Jeffrey Barrett, Sonia Goncalves, Ewan Harrison, David K. Jackson, Ian Johnston, Dominic Kwiatkowski, Cordelia Langford, John Sillitoe on behalf of the Wellcome Sanger Institute COVID-19 Surveillance Team |
| EPI_ISL_3309937 | Lighthouse Lab in Alderley Park | Wellcome Sanger Institute for the COVID-19 Genomics UK (COG-UK) Consortium | Jacquelyn Wynn, Mairead Hyland, The Lighthouse Lab in Alderley Park and Alex Alderton, Roberto Amato, Jeffrey Barrett, Sonia Goncalves, Ewan Harrison, David K. Jackson, Ian Johnston, Dominic Kwiatkowski, Cordelia Langford, John Sillitoe on behalf of the Wellcome Sanger Institute COVID-19 Surveillance Team |
| EPI_ISL_3309938 | Health Services Laboratories | Wellcome Sanger Institute for the COVID-19 Genomics UK (COG-UK) Consortium | Health Services Laboratories and Alex Alderton, Roberto Amato, Jeffrey Barrett, Sonia Goncalves, Ewan Harrison, David K. Jackson, Ian Johnston, Dominic Kwiatkowski, Cordelia Langford, John Sillitoe on behalf of the Wellcome Sanger Institute COVID-19 Surveillance Team |
| EPI_ISL_3309939 | Lighthouse Laboratory Plymouth | Wellcome Sanger Institute for the COVID-19 Genomics UK (COG-UK) Consortium | Lighthouse Laboratory Plymouth and Alex Alderton, Roberto Amato, Jeffrey Barrett, Sonia Goncalves, Ewan Harrison, David K. Jackson, Ian Johnston, Dominic Kwiatkowski, Cordelia Langford, John Sillitoe on behalf of the Wellcome Sanger Institute COVID-19 Surveillance Team |
| EPI_ISL_3309940, EPI_ISL_3309941, EPI_ISL_3309942 | Lighthouse Lab in Alderley Park | Wellcome Sanger Institute for the COVID-19 Genomics UK (COG-UK) Consortium | Jacquelyn Wynn, Mairead Hyland, The Lighthouse Lab in Alderley Park and Alex Alderton, Roberto Amato, Jeffrey Barrett, Sonia Goncalves, Ewan Harrison, David K. Jackson, Ian Johnston, Dominic Kwiatkowski, Cordelia Langford, John Sillitoe on behalf of the Wellcome Sanger Institute COVID-19 Surveillance Team |
| EPI_ISL_3309943, EPI_ISL_3309944 | Lighthouse Lab in Milton Keynes | Wellcome Sanger Institute for the COVID-19 Genomics UK (COG-UK) Consortium | The Lighthouse Lab in Milton Keynes and Alex Alderton, Roberto Amato, Jeffrey Barrett, Sonia Goncalves, Ewan Harrison, David K. Jackson, Ian Johnston, Dominic Kwiatkowski, Cordelia Langford, John Sillitoe on behalf of the Wellcome Sanger Institute COVID-19 Surveillance Team |
| EPI_ISL_3309945 | Health Services Laboratories | Wellcome Sanger Institute for the COVID-19 Genomics UK (COG-UK) Consortium | Health Services Laboratories and Alex Alderton, Roberto Amato, Jeffrey Barrett, Sonia Goncalves, Ewan Harrison, David K. Jackson, Ian Johnston, Dominic Kwiatkowski, Cordelia Langford, John Sillitoe on behalf of the Wellcome Sanger Institute COVID-19 Surveillance Team |
| EPI_ISL_3309946 | Lighthouse Lab in Alderley Park | Wellcome Sanger Institute for the COVID-19 Genomics UK (COG-UK) Consortium | Jacquelyn Wynn, Mairead Hyland, The Lighthouse Lab in Alderley Park and Alex Alderton, Roberto Amato, Jeffrey Barrett, Sonia Goncalves, Ewan Harrison, David K. Jackson, Ian Johnston, Dominic Kwiatkowski, Cordelia Langford, John Sillitoe on behalf of the Wellcome Sanger Institute COVID-19 Surveillance Team |
| EPI_ISL_3309947 | Lighthouse Laboratory Plymouth | Wellcome Sanger Institute for the COVID-19 Genomics UK (COG-UK) Consortium | Lighthouse Laboratory Plymouth and Alex Alderton, Roberto Amato, Jeffrey Barrett, Sonia Goncalves, Ewan Harrison, David K. Jackson, Ian Johnston, Dominic Kwiatkowski, Cordelia Langford, John Sillitoe on behalf of the Wellcome Sanger Institute COVID-19 Surveillance Team |
| EPI_ISL_3309948 | Lighthouse Lab in Milton Keynes | Wellcome Sanger Institute for the COVID-19 Genomics UK (COG-UK) Consortium | The Lighthouse Lab in Milton Keynes and Alex Alderton, Roberto Amato, Jeffrey Barrett, Sonia Goncalves, Ewan Harrison, David K. Jackson, Ian Johnston, Dominic Kwiatkowski, Cordelia Langford, John Sillitoe on behalf of the Wellcome Sanger Institute COVID-19 Surveillance Team |
| EPI_ISL_3309949 | Lighthouse Laboratory Plymouth | Wellcome Sanger Institute for the COVID-19 Genomics UK (COG-UK) Consortium | Lighthouse Laboratory Plymouth and Alex Alderton, Roberto Amato, Jeffrey Barrett, Sonia Goncalves, Ewan Harrison, David K. Jackson, Ian Johnston, Dominic Kwiatkowski, Cordelia Langford, John Sillitoe on behalf of the Wellcome Sanger Institute COVID-19 Surveillance Team |
| EPI_ISL_3309950 | Lighthouse Lab in Alderley Park | Wellcome Sanger Institute for the COVID-19 Genomics UK (COG-UK) Consortium | Jacquelyn Wynn, Mairead Hyland, The Lighthouse Lab in Alderley Park and Alex Alderton, Roberto Amato, Jeffrey Barrett, Sonia Goncalves, Ewan Harrison, David K. Jackson, Ian Johnston, Dominic Kwiatkowski, Cordelia Langford, John Sillitoe on behalf of the Wellcome Sanger Institute COVID-19 Surveillance Team |
| EPI_ISL_3309951 | Lighthouse Lab in Milton Keynes | Wellcome Sanger Institute for the COVID-19 Genomics UK (COG-UK) Consortium | The Lighthouse Lab in Milton Keynes and Alex Alderton, Roberto Amato, Jeffrey Barrett, Sonia Goncalves, Ewan Harrison, David K. Jackson, Ian Johnston, Dominic Kwiatkowski, Cordelia Langford, John Sillitoe on behalf of the Wellcome Sanger Institute COVID-19 Surveillance Team |
| EPI_ISL_3309952, EPI_ISL_3309953 | Lighthouse Laboratory Plymouth | Wellcome Sanger Institute for the COVID-19 Genomics UK (COG-UK) Consortium | Lighthouse Laboratory Plymouth and Alex Alderton, Roberto Amato, Jeffrey Barrett, Sonia Goncalves, Ewan Harrison, David K. Jackson, Ian Johnston, Dominic Kwiatkowski, Cordelia Langford, John Sillitoe on behalf of the Wellcome Sanger Institute COVID-19 Surveillance Team |
| EPI_ISL_3309954 | Health Services Laboratories | Wellcome Sanger Institute for the COVID-19 Genomics UK (COG-UK) Consortium | Health Services Laboratories and Alex Alderton, Roberto Amato, Jeffrey Barrett, Sonia Goncalves, Ewan Harrison, David K. Jackson, Ian Johnston, Dominic Kwiatkowski, Cordelia Langford, John Sillitoe on behalf of the Wellcome Sanger Institute COVID-19 Surveillance Team |
| EPI_ISL_3309955 | Lighthouse Laboratory Plymouth | Wellcome Sanger Institute for the COVID-19 Genomics UK (COG-UK) Consortium | Lighthouse Laboratory Plymouth and Alex Alderton, Roberto Amato, Jeffrey Barrett, Sonia Goncalves, Ewan Harrison, David K. Jackson, Ian Johnston, Dominic Kwiatkowski, Cordelia Langford, John Sillitoe on behalf of the Wellcome Sanger Institute COVID-19 Surveillance Team |
| EPI_ISL_3309956 | Health Services Laboratories | Wellcome Sanger Institute for the COVID-19 Genomics UK (COG-UK) Consortium | Health Services Laboratories and Alex Alderton, Roberto Amato, Jeffrey Barrett, Sonia Goncalves, Ewan Harrison, David K. Jackson, Ian Johnston, Dominic Kwiatkowski, Cordelia Langford, John Sillitoe on behalf of the Wellcome Sanger Institute COVID-19 Surveillance Team |
| EPI_ISL_3309957 | Lighthouse Lab in Milton Keynes | Wellcome Sanger Institute for the COVID-19 Genomics UK (COG-UK) Consortium | The Lighthouse Lab in Milton Keynes and Alex Alderton, Roberto Amato, Jeffrey Barrett, Sonia Goncalves, Ewan Harrison, David K. Jackson, Ian Johnston, Dominic Kwiatkowski, Cordelia Langford, John Sillitoe on behalf of the Wellcome Sanger Institute COVID-19 Surveillance Team |
| EPI_ISL_3309958 | Health Services Laboratories | Wellcome Sanger Institute for the COVID-19 Genomics UK (COG-UK) Consortium | Health Services Laboratories and Alex Alderton, Roberto Amato, Jeffrey Barrett, Sonia Goncalves, Ewan Harrison, David K. Jackson, Ian Johnston, Dominic Kwiatkowski, Cordelia Langford, John Sillitoe on behalf of the Wellcome Sanger Institute COVID-19 Surveillance Team |
| EPI_ISL_3309959 | Lighthouse Lab in Milton Keynes | Wellcome Sanger Institute for the COVID-19 Genomics UK (COG-UK) Consortium | The Lighthouse Lab in Milton Keynes and Alex Alderton, Roberto Amato, Jeffrey Barrett, Sonia Goncalves, Ewan Harrison, David K. Jackson, Ian Johnston, Dominic Kwiatkowski, Cordelia Langford, John Sillitoe on behalf of the Wellcome Sanger Institute COVID-19 Surveillance Team |
| EPI_ISL_3309960 | Lighthouse Laboratory Plymouth | Wellcome Sanger Institute for the COVID-19 Genomics UK (COG-UK) Consortium | Lighthouse Laboratory Plymouth and Alex Alderton, Roberto Amato, Jeffrey Barrett, Sonia Goncalves, Ewan Harrison, David K. Jackson, Ian Johnston, Dominic Kwiatkowski, Cordelia Langford, John Sillitoe on behalf of the Wellcome Sanger Institute COVID-19 Surveillance Team |
| EPI_ISL_3309961 | Lighthouse Lab in Alderley Park | Wellcome Sanger Institute for the COVID-19 Genomics UK (COG-UK) Consortium | Jacquelyn Wynn, Mairead Hyland, The Lighthouse Lab in Alderley Park and Alex Alderton, Roberto Amato, Jeffrey Barrett, Sonia Goncalves, Ewan Harrison, David K. Jackson, Ian Johnston, Dominic Kwiatkowski, Cordelia Langford, John Sillitoe on behalf of the Wellcome Sanger Institute COVID-19 Surveillance Team |
| EPI_ISL_3309962 | Lighthouse Lab in Milton Keynes | Wellcome Sanger Institute for the COVID-19 Genomics UK (COG-UK) Consortium | The Lighthouse Lab in Milton Keynes and Alex Alderton, Roberto Amato, Jeffrey Barrett, Sonia Goncalves, Ewan Harrison, David K. Jackson, Ian Johnston, Dominic Kwiatkowski, Cordelia Langford, John Sillitoe on behalf of the Wellcome Sanger Institute COVID-19 Surveillance Team |
| EPI_ISL_3309963 | Lighthouse Laboratory Plymouth | Wellcome Sanger Institute for the COVID-19 Genomics UK (COG-UK) Consortium | Lighthouse Laboratory Plymouth and Alex Alderton, Roberto Amato, Jeffrey Barrett, Sonia Goncalves, Ewan Harrison, David K. Jackson, Ian Johnston, Dominic Kwiatkowski, Cordelia Langford, John Sillitoe on behalf of the Wellcome Sanger Institute COVID-19 Surveillance Team |
| EPI_ISL_3309964 | Health Services Laboratories | Wellcome Sanger Institute for the COVID-19 Genomics UK (COG-UK) Consortium | Health Services Laboratories and Alex Alderton, Roberto Amato, Jeffrey Barrett, Sonia Goncalves, Ewan Harrison, David K. Jackson, Ian Johnston, Dominic Kwiatkowski, Cordelia Langford, John Sillitoe on behalf of the Wellcome Sanger Institute COVID-19 Surveillance Team |
| EPI_ISL_3309965, EPI_ISL_3309966 | Lighthouse Lab in Milton Keynes | Wellcome Sanger Institute for the COVID-19 Genomics UK (COG-UK) Consortium | The Lighthouse Lab in Milton Keynes and Alex Alderton, Roberto Amato, Jeffrey Barrett, Sonia Goncalves, Ewan Harrison, David K. Jackson, Ian Johnston, Dominic Kwiatkowski, Cordelia Langford, John Sillitoe on behalf of the Wellcome Sanger Institute COVID-19 Surveillance Team |
| EPI_ISL_3309967 | Health Services Laboratories | Wellcome Sanger Institute for the COVID-19 Genomics UK (COG-UK) | Health Services Laboratories and Alex Alderton, Roberto Amato, Jeffrey Barrett, Sonia Goncalves, Ewan Harrison, David K. Jackson, Ian Johnston, Dominic Kwiatkowski, Cordelia Langford, John Sillitoe on behalf of the Wellcome Sanger Institute |

| | | | |
|---|---|---|---|
| | | Consortium | COVID-19 Surveillance Team |
| EPI_ISL_3309968 | Lighthouse Laboratory Plymouth | Wellcome Sanger Institute for the COVID-19 Genomics UK (COG-UK) Consortium | Lighthouse Laboratory Plymouth and Alex Alderton, Roberto Amato, Jeffrey Barrett, Sonia Goncalves, Ewan Harrison, David K. Jackson, Ian Johnston, Dominic Kwiatkowski, Cordelia Langford, John Sillitoe on behalf of the Wellcome Sanger Institute COVID-19 Surveillance Team |
| EPI_ISL_3309969, EPI_ISL_3309970 | Lighthouse Lab in Alderley Park | Wellcome Sanger Institute for the COVID-19 Genomics UK (COG-UK) Consortium | Jacquelyn Wynn, Mairead Hyland, The Lighthouse Lab in Alderley Park and Alex Alderton, Roberto Amato, Jeffrey Barrett, Sonia Goncalves, Ewan Harrison, David K. Jackson, Ian Johnston, Dominic Kwiatkowski, Cordelia Langford, John Sillitoe on behalf of the Wellcome Sanger Institute COVID-19 Surveillance Team |
| EPI_ISL_3309971 | Lighthouse Laboratory Plymouth | Wellcome Sanger Institute for the COVID-19 Genomics UK (COG-UK) Consortium | Lighthouse Laboratory Plymouth and Alex Alderton, Roberto Amato, Jeffrey Barrett, Sonia Goncalves, Ewan Harrison, David K. Jackson, Ian Johnston, Dominic Kwiatkowski, Cordelia Langford, John Sillitoe on behalf of the Wellcome Sanger Institute COVID-19 Surveillance Team |
| EPI_ISL_3309972, EPI_ISL_3309973 | Health Services Laboratories | Wellcome Sanger Institute for the COVID-19 Genomics UK (COG-UK) Consortium | Health Services Laboratories and Alex Alderton, Roberto Amato, Jeffrey Barrett, Sonia Goncalves, Ewan Harrison, David K. Jackson, Ian Johnston, Dominic Kwiatkowski, Cordelia Langford, John Sillitoe on behalf of the Wellcome Sanger Institute COVID-19 Surveillance Team |
| EPI_ISL_3309974 | Lighthouse Laboratory Plymouth | Wellcome Sanger Institute for the COVID-19 Genomics UK (COG-UK) Consortium | Lighthouse Laboratory Plymouth and Alex Alderton, Roberto Amato, Jeffrey Barrett, Sonia Goncalves, Ewan Harrison, David K. Jackson, Ian Johnston, Dominic Kwiatkowski, Cordelia Langford, John Sillitoe on behalf of the Wellcome Sanger Institute COVID-19 Surveillance Team |
| EPI_ISL_3309975, EPI_ISL_3309976 | Lighthouse Lab in Milton Keynes | Wellcome Sanger Institute for the COVID-19 Genomics UK (COG-UK) Consortium | The Lighthouse Lab in Milton Keynes and Alex Alderton, Roberto Amato, Jeffrey Barrett, Sonia Goncalves, Ewan Harrison, David K. Jackson, Ian Johnston, Dominic Kwiatkowski, Cordelia Langford, John Sillitoe on behalf of the Wellcome Sanger Institute COVID-19 Surveillance Team |
| EPI_ISL_3309977 | Health Services Laboratories | Wellcome Sanger Institute for the COVID-19 Genomics UK (COG-UK) Consortium | Health Services Laboratories and Alex Alderton, Roberto Amato, Jeffrey Barrett, Sonia Goncalves, Ewan Harrison, David K. Jackson, Ian Johnston, Dominic Kwiatkowski, Cordelia Langford, John Sillitoe on behalf of the Wellcome Sanger Institute COVID-19 Surveillance Team |
| EPI_ISL_3309978 | Lighthouse Lab in Milton Keynes | Wellcome Sanger Institute for the COVID-19 Genomics UK (COG-UK) Consortium | The Lighthouse Lab in Milton Keynes and Alex Alderton, Roberto Amato, Jeffrey Barrett, Sonia Goncalves, Ewan Harrison, David K. Jackson, Ian Johnston, Dominic Kwiatkowski, Cordelia Langford, John Sillitoe on behalf of the Wellcome Sanger Institute COVID-19 Surveillance Team |
| EPI_ISL_3309979 | Lighthouse Lab in Alderley Park | Wellcome Sanger Institute for the COVID-19 Genomics UK (COG-UK) Consortium | Jacquelyn Wynn, Mairead Hyland, The Lighthouse Lab in Alderley Park and Alex Alderton, Roberto Amato, Jeffrey Barrett, Sonia Goncalves, Ewan Harrison, David K. Jackson, Ian Johnston, Dominic Kwiatkowski, Cordelia Langford, John Sillitoe on behalf of the Wellcome Sanger Institute COVID-19 Surveillance Team |
| EPI_ISL_3309980 | Lighthouse Laboratory Plymouth | Wellcome Sanger Institute for the COVID-19 Genomics UK (COG-UK) Consortium | Lighthouse Laboratory Plymouth and Alex Alderton, Roberto Amato, Jeffrey Barrett, Sonia Goncalves, Ewan Harrison, David K. Jackson, Ian Johnston, Dominic Kwiatkowski, Cordelia Langford, John Sillitoe on behalf of the Wellcome Sanger Institute COVID-19 Surveillance Team |
| EPI_ISL_3309981 | Health Services Laboratories | Wellcome Sanger Institute for the COVID-19 Genomics UK (COG-UK) Consortium | Health Services Laboratories and Alex Alderton, Roberto Amato, Jeffrey Barrett, Sonia Goncalves, Ewan Harrison, David K. Jackson, Ian Johnston, Dominic Kwiatkowski, Cordelia Langford, John Sillitoe on behalf of the Wellcome Sanger Institute COVID-19 Surveillance Team |
| EPI_ISL_3309982 | Lighthouse Laboratory Plymouth | Wellcome Sanger Institute for the COVID-19 Genomics UK (COG-UK) Consortium | Lighthouse Laboratory Plymouth and Alex Alderton, Roberto Amato, Jeffrey Barrett, Sonia Goncalves, Ewan Harrison, David K. Jackson, Ian Johnston, Dominic Kwiatkowski, Cordelia Langford, John Sillitoe on behalf of the Wellcome Sanger Institute COVID-19 Surveillance Team |
| EPI_ISL_3309983, EPI_ISL_3309984, EPI_ISL_3309985, EPI_ISL_3309986, EPI_ISL_3309987, EPI_ISL_3309988 | Lighthouse Lab in Milton Keynes | Wellcome Sanger Institute for the COVID-19 Genomics UK (COG-UK) Consortium | The Lighthouse Lab in Milton Keynes and Alex Alderton, Roberto Amato, Jeffrey Barrett, Sonia Goncalves, Ewan Harrison, David K. Jackson, Ian Johnston, Dominic Kwiatkowski, Cordelia Langford, John Sillitoe on behalf of the Wellcome Sanger Institute COVID-19 Surveillance Team |
| EPI_ISL_3309989 | Lighthouse Lab in Alderley Park | Wellcome Sanger Institute for the COVID-19 Genomics UK (COG-UK) Consortium | Jacquelyn Wynn, Mairead Hyland, The Lighthouse Lab in Alderley Park and Alex Alderton, Roberto Amato, Jeffrey Barrett, Sonia Goncalves, Ewan Harrison, David K. Jackson, Ian Johnston, Dominic Kwiatkowski, Cordelia Langford, John Sillitoe on behalf of the Wellcome Sanger Institute COVID-19 Surveillance Team |
| EPI_ISL_3309990 | Lighthouse Laboratory Plymouth | Wellcome Sanger Institute for the COVID-19 Genomics UK (COG-UK) Consortium | Lighthouse Laboratory Plymouth and Alex Alderton, Roberto Amato, Jeffrey Barrett, Sonia Goncalves, Ewan Harrison, David K. Jackson, Ian Johnston, Dominic Kwiatkowski, Cordelia Langford, John Sillitoe on behalf of the Wellcome Sanger Institute COVID-19 Surveillance Team |
| EPI_ISL_3309991, EPI_ISL_3309992 | Health Services Laboratories | Wellcome Sanger Institute for the COVID-19 Genomics UK (COG-UK) Consortium | Health Services Laboratories and Alex Alderton, Roberto Amato, Jeffrey Barrett, Sonia Goncalves, Ewan Harrison, David K. Jackson, Ian Johnston, Dominic Kwiatkowski, Cordelia Langford, John Sillitoe on behalf of the Wellcome Sanger Institute COVID-19 Surveillance Team |
| EPI_ISL_3309993 | Lighthouse Lab in Milton Keynes | Wellcome Sanger Institute for the COVID-19 Genomics UK (COG-UK) Consortium | The Lighthouse Lab in Milton Keynes and Alex Alderton, Roberto Amato, Jeffrey Barrett, Sonia Goncalves, Ewan Harrison, David K. Jackson, Ian Johnston, Dominic Kwiatkowski, Cordelia Langford, John Sillitoe on behalf of the Wellcome Sanger Institute COVID-19 Surveillance Team |
| EPI_ISL_3309994 | Lighthouse Lab in Alderley Park | Wellcome Sanger Institute for the COVID-19 Genomics UK (COG-UK) Consortium | Jacquelyn Wynn, Mairead Hyland, The Lighthouse Lab in Alderley Park and Alex Alderton, Roberto Amato, Jeffrey Barrett, Sonia Goncalves, Ewan Harrison, David K. Jackson, Ian Johnston, Dominic Kwiatkowski, Cordelia Langford, John Sillitoe on behalf of the Wellcome Sanger Institute COVID-19 Surveillance Team |
| EPI_ISL_3309995 | Lighthouse Laboratory Plymouth | Wellcome Sanger Institute for the COVID-19 Genomics UK (COG-UK) Consortium | Lighthouse Laboratory Plymouth and Alex Alderton, Roberto Amato, Jeffrey Barrett, Sonia Goncalves, Ewan Harrison, David K. Jackson, Ian Johnston, Dominic Kwiatkowski, Cordelia Langford, John Sillitoe on behalf of the Wellcome Sanger Institute COVID-19 Surveillance Team |
| EPI_ISL_3309996 | Lighthouse Lab in Milton Keynes | Wellcome Sanger Institute for the COVID-19 Genomics UK (COG-UK) Consortium | The Lighthouse Lab in Milton Keynes and Alex Alderton, Roberto Amato, Jeffrey Barrett, Sonia Goncalves, Ewan Harrison, David K. Jackson, Ian Johnston, Dominic Kwiatkowski, Cordelia Langford, John Sillitoe on behalf of the Wellcome Sanger Institute COVID-19 Surveillance Team |
| EPI_ISL_3309997 | Health Services Laboratories | Wellcome Sanger Institute for the COVID-19 Genomics UK (COG-UK) Consortium | Health Services Laboratories and Alex Alderton, Roberto Amato, Jeffrey Barrett, Sonia Goncalves, Ewan Harrison, David K. Jackson, Ian Johnston, Dominic Kwiatkowski, Cordelia Langford, John Sillitoe on behalf of the Wellcome Sanger Institute COVID-19 Surveillance Team |
| EPI_ISL_3309998 | Lighthouse Lab in Milton Keynes | Wellcome Sanger Institute for the COVID-19 Genomics UK (COG-UK) Consortium | The Lighthouse Lab in Milton Keynes and Alex Alderton, Roberto Amato, Jeffrey Barrett, Sonia Goncalves, Ewan Harrison, David K. Jackson, Ian Johnston, Dominic Kwiatkowski, Cordelia Langford, John Sillitoe on behalf of the Wellcome Sanger Institute COVID-19 Surveillance Team |
| EPI_ISL_3309999 | Health Services Laboratories | Wellcome Sanger Institute for the COVID-19 Genomics UK (COG-UK) Consortium | Health Services Laboratories and Alex Alderton, Roberto Amato, Jeffrey Barrett, Sonia Goncalves, Ewan Harrison, David K. Jackson, Ian Johnston, Dominic Kwiatkowski, Cordelia Langford, John Sillitoe on behalf of the Wellcome Sanger Institute COVID-19 Surveillance Team |
| EPI_ISL_3310000 | Lighthouse Laboratory Plymouth | Wellcome Sanger Institute for the COVID-19 Genomics UK (COG-UK) Consortium | Lighthouse Laboratory Plymouth and Alex Alderton, Roberto Amato, Jeffrey Barrett, Sonia Goncalves, Ewan Harrison, David K. Jackson, Ian Johnston, Dominic Kwiatkowski, Cordelia Langford, John Sillitoe on behalf of the Wellcome Sanger Institute COVID-19 Surveillance Team |
| EPI_ISL_3310001 | Health Services Laboratories | Wellcome Sanger Institute for the COVID-19 Genomics UK (COG-UK) Consortium | Health Services Laboratories and Alex Alderton, Roberto Amato, Jeffrey Barrett, Sonia Goncalves, Ewan Harrison, David K. Jackson, Ian Johnston, Dominic Kwiatkowski, Cordelia Langford, John Sillitoe on behalf of the Wellcome Sanger Institute COVID-19 Surveillance Team |
| EPI_ISL_3310002 | Lighthouse Lab in Alderley Park | Wellcome Sanger Institute for the COVID-19 Genomics UK (COG-UK) Consortium | Jacquelyn Wynn, Mairead Hyland, The Lighthouse Lab in Alderley Park and Alex Alderton, Roberto Amato, Jeffrey Barrett, Sonia Goncalves, Ewan Harrison, David K. Jackson, Ian Johnston, Dominic Kwiatkowski, Cordelia Langford, John Sillitoe on behalf of the Wellcome Sanger Institute COVID-19 Surveillance Team |
| EPI_ISL_3310003, EPI_ISL_3310004, EPI_ISL_3310005 | Lighthouse Lab in Milton Keynes | Wellcome Sanger Institute for the COVID-19 Genomics UK (COG-UK) Consortium | The Lighthouse Lab in Milton Keynes and Alex Alderton, Roberto Amato, Jeffrey Barrett, Sonia Goncalves, Ewan Harrison, David K. Jackson, Ian Johnston, Dominic Kwiatkowski, Cordelia Langford, John Sillitoe on behalf of the Wellcome Sanger Institute COVID-19 Surveillance Team |
| EPI_ISL_3310006 | Lighthouse Laboratory Plymouth | Wellcome Sanger Institute for the COVID-19 Genomics UK (COG-UK) Consortium | Lighthouse Laboratory Plymouth and Alex Alderton, Roberto Amato, Jeffrey Barrett, Sonia Goncalves, Ewan Harrison, David K. Jackson, Ian Johnston, Dominic Kwiatkowski, Cordelia Langford, John Sillitoe on behalf of the Wellcome Sanger Institute COVID-19 Surveillance Team |
| EPI_ISL_3310007 | Lighthouse Lab in Alderley Park | Wellcome Sanger Institute for the COVID-19 Genomics UK (COG-UK) Consortium | Jacquelyn Wynn, Mairead Hyland, The Lighthouse Lab in Alderley Park and Alex Alderton, Roberto Amato, Jeffrey Barrett, Sonia Goncalves, Ewan Harrison, David K. Jackson, Ian Johnston, Dominic Kwiatkowski, Cordelia Langford, John Sillitoe on behalf of the Wellcome Sanger Institute COVID-19 Surveillance Team |
| EPI_ISL_3310008, EPI_ISL_3310009 | Lighthouse Lab in Milton Keynes | Wellcome Sanger Institute for the COVID-19 Genomics UK (COG-UK) Consortium | The Lighthouse Lab in Milton Keynes and Alex Alderton, Roberto Amato, Jeffrey Barrett, Sonia Goncalves, Ewan Harrison, David K. Jackson, Ian Johnston, Dominic Kwiatkowski, Cordelia Langford, John Sillitoe on behalf of the Wellcome Sanger Institute COVID-19 Surveillance Team |
| EPI_ISL_3310010 | Lighthouse Laboratory Plymouth | Wellcome Sanger Institute for the COVID-19 Genomics UK (COG-UK) Consortium | Lighthouse Laboratory Plymouth and Alex Alderton, Roberto Amato, Jeffrey Barrett, Sonia Goncalves, Ewan Harrison, David K. Jackson, Ian Johnston, Dominic Kwiatkowski, Cordelia Langford, John Sillitoe on behalf of the Wellcome Sanger Institute COVID-19 Surveillance Team |
| EPI_ISL_3310011 | Lighthouse Lab in Alderley Park | Wellcome Sanger Institute for the COVID-19 Genomics UK (COG-UK) Consortium | Jacquelyn Wynn, Mairead Hyland, The Lighthouse Lab in Alderley Park and Alex Alderton, Roberto Amato, Jeffrey Barrett, Sonia Goncalves, Ewan Harrison, David K. Jackson, Ian Johnston, Dominic Kwiatkowski, Cordelia Langford, John Sillitoe on behalf of the Wellcome Sanger Institute COVID-19 Surveillance Team |
| EPI_ISL_3310012, EPI_ISL_3310013, EPI_ISL_3310014, EPI_ISL_3310015, EPI_ISL_3310016 | Lighthouse Lab in Milton Keynes | Wellcome Sanger Institute for the COVID-19 Genomics UK (COG-UK) Consortium | The Lighthouse Lab in Milton Keynes and Alex Alderton, Roberto Amato, Jeffrey Barrett, Sonia Goncalves, Ewan Harrison, David K. Jackson, Ian Johnston, Dominic Kwiatkowski, Cordelia Langford, John Sillitoe on behalf of the Wellcome Sanger Institute COVID-19 Surveillance Team |
| EPI_ISL_3310017 | Lighthouse Lab in Alderley Park | Wellcome Sanger Institute for the COVID-19 Genomics UK (COG-UK) Consortium | Jacquelyn Wynn, Mairead Hyland, The Lighthouse Lab in Alderley Park and Alex Alderton, Roberto Amato, Jeffrey Barrett, Sonia Goncalves, Ewan Harrison, David K. Jackson, Ian Johnston, Dominic Kwiatkowski, Cordelia Langford, John Sillitoe on behalf of the Wellcome Sanger Institute COVID-19 Surveillance Team |
| EPI_ISL_3310018, EPI_ISL_3310019, EPI_ISL_3310020, EPI_ISL_3310021 | Lighthouse Lab in Milton Keynes | Wellcome Sanger Institute for the COVID-19 Genomics UK (COG-UK) Consortium | The Lighthouse Lab in Milton Keynes and Alex Alderton, Roberto Amato, Jeffrey Barrett, Sonia Goncalves, Ewan Harrison, David K. Jackson, Ian Johnston, Dominic Kwiatkowski, Cordelia Langford, John Sillitoe on behalf of the Wellcome Sanger Institute COVID-19 Surveillance Team |
| EPI_ISL_3310022 | Lighthouse Lab in Alderley Park | Wellcome Sanger Institute for the COVID-19 Genomics UK (COG-UK) Consortium | Jacquelyn Wynn, Mairead Hyland, The Lighthouse Lab in Alderley Park and Alex Alderton, Roberto Amato, Jeffrey Barrett, Sonia Goncalves, Ewan Harrison, David K. Jackson, Ian Johnston, Dominic Kwiatkowski, Cordelia Langford, John Sillitoe on behalf of the Wellcome Sanger Institute COVID-19 Surveillance Team |
| EPI_ISL_3310023 | Health Services Laboratories | Wellcome Sanger Institute for the COVID-19 Genomics UK (COG-UK) Consortium | Health Services Laboratories and Alex Alderton, Roberto Amato, Jeffrey Barrett, Sonia Goncalves, Ewan Harrison, David K. Jackson, Ian Johnston, Dominic Kwiatkowski, Cordelia Langford, John Sillitoe on behalf of the Wellcome Sanger Institute COVID-19 Surveillance Team |
| EPI_ISL_3310024 | Lighthouse Lab in Alderley Park | Wellcome Sanger Institute for the COVID-19 Genomics UK (COG-UK) Consortium | Jacquelyn Wynn, Mairead Hyland, The Lighthouse Lab in Alderley Park and Alex Alderton, Roberto Amato, Jeffrey Barrett, Sonia Goncalves, Ewan Harrison, David K. Jackson, Ian Johnston, Dominic Kwiatkowski, Cordelia Langford, John Sillitoe on behalf of the Wellcome Sanger Institute COVID-19 Surveillance Team |
| EPI_ISL_3310025 | Health Services Laboratories | Wellcome Sanger Institute for the COVID-19 Genomics UK (COG-UK) Consortium | Health Services Laboratories and Alex Alderton, Roberto Amato, Jeffrey Barrett, Sonia Goncalves, Ewan Harrison, David K. Jackson, Ian Johnston, Dominic Kwiatkowski, Cordelia Langford, John Sillitoe on behalf of the Wellcome Sanger Institute COVID-19 Surveillance Team |
| EPI_ISL_3310026 | Lighthouse Laboratory Plymouth | Wellcome Sanger Institute for the COVID-19 Genomics UK (COG-UK) Consortium | Lighthouse Laboratory Plymouth and Alex Alderton, Roberto Amato, Jeffrey Barrett, Sonia Goncalves, Ewan Harrison, David K. Jackson, Ian Johnston, Dominic Kwiatkowski, Cordelia Langford, John Sillitoe on behalf of the Wellcome Sanger Institute COVID-19 Surveillance Team |
| EPI_ISL_3310027 | Lighthouse Lab in Milton Keynes | Wellcome Sanger Institute for the COVID-19 Genomics UK (COG-UK) | The Lighthouse Lab in Milton Keynes and Alex Alderton, Roberto Amato, Jeffrey Barrett, Sonia Goncalves, Ewan Harrison, David K. Jackson, Ian Johnston, Dominic Kwiatkowski, Cordelia Langford, John Sillitoe on behalf of the Wellcome Sanger |

| | | | |
|---|---|---|---|
| | | Consortium | Institute COVID-19 Surveillance Team |
| EPI_ISL_3310028, EPI_ISL_3310029, EPI_ISL_3310030 | Health Services Laboratories | Wellcome Sanger Institute for the COVID-19 Genomics UK (COG-UK) Consortium | Health Services Laboratories and Alex Alderton, Roberto Amato, Jeffrey Barrett, Sonia Goncalves, Ewan Harrison, David K. Jackson, Ian Johnston, Dominic Kwiatkowski, Cordelia Langford, John Sillitoe on behalf of the Wellcome Sanger Institute COVID-19 Surveillance Team |
| EPI_ISL_3310031, EPI_ISL_3310032 | Lighthouse Laboratory Plymouth | Wellcome Sanger Institute for the COVID-19 Genomics UK (COG-UK) Consortium | Lighthouse Laboratory Plymouth and Alex Alderton, Roberto Amato, Jeffrey Barrett, Sonia Goncalves, Ewan Harrison, David K. Jackson, Ian Johnston, Dominic Kwiatkowski, Cordelia Langford, John Sillitoe on behalf of the Wellcome Sanger Institute COVID-19 Surveillance Team |
| EPI_ISL_3310033 | Lighthouse Lab in Milton Keynes | Wellcome Sanger Institute for the COVID-19 Genomics UK (COG-UK) Consortium | The Lighthouse Lab in Milton Keynes and Alex Alderton, Roberto Amato, Jeffrey Barrett, Sonia Goncalves, Ewan Harrison, David K. Jackson, Ian Johnston, Dominic Kwiatkowski, Cordelia Langford, John Sillitoe on behalf of the Wellcome Sanger Institute COVID-19 Surveillance Team |
| EPI_ISL_3310034 | Lighthouse Lab in Alderley Park | Wellcome Sanger Institute for the COVID-19 Genomics UK (COG-UK) Consortium | Jacquelyn Wynn, Mairead Hyland, The Lighthouse Lab in Alderley Park and Alex Alderton, Roberto Amato, Jeffrey Barrett, Sonia Goncalves, Ewan Harrison, David K. Jackson, Ian Johnston, Dominic Kwiatkowski, Cordelia Langford, John Sillitoe on behalf of the Wellcome Sanger Institute COVID-19 Surveillance Team |
| EPI_ISL_3310035, EPI_ISL_3310036 | Lighthouse Laboratory Plymouth | Wellcome Sanger Institute for the COVID-19 Genomics UK (COG-UK) Consortium | Lighthouse Laboratory Plymouth and Alex Alderton, Roberto Amato, Jeffrey Barrett, Sonia Goncalves, Ewan Harrison, David K. Jackson, Ian Johnston, Dominic Kwiatkowski, Cordelia Langford, John Sillitoe on behalf of the Wellcome Sanger Institute COVID-19 Surveillance Team |
| EPI_ISL_3310037 | Lighthouse Lab in Alderley Park | Wellcome Sanger Institute for the COVID-19 Genomics UK (COG-UK) Consortium | Jacquelyn Wynn, Mairead Hyland, The Lighthouse Lab in Alderley Park and Alex Alderton, Roberto Amato, Jeffrey Barrett, Sonia Goncalves, Ewan Harrison, David K. Jackson, Ian Johnston, Dominic Kwiatkowski, Cordelia Langford, John Sillitoe on behalf of the Wellcome Sanger Institute COVID-19 Surveillance Team |
| EPI_ISL_3310038 | Lighthouse Laboratory Plymouth | Wellcome Sanger Institute for the COVID-19 Genomics UK (COG-UK) Consortium | Lighthouse Laboratory Plymouth and Alex Alderton, Roberto Amato, Jeffrey Barrett, Sonia Goncalves, Ewan Harrison, David K. Jackson, Ian Johnston, Dominic Kwiatkowski, Cordelia Langford, John Sillitoe on behalf of the Wellcome Sanger Institute COVID-19 Surveillance Team |
| EPI_ISL_3310039 | Lighthouse Lab in Milton Keynes | Wellcome Sanger Institute for the COVID-19 Genomics UK (COG-UK) Consortium | The Lighthouse Lab in Milton Keynes and Alex Alderton, Roberto Amato, Jeffrey Barrett, Sonia Goncalves, Ewan Harrison, David K. Jackson, Ian Johnston, Dominic Kwiatkowski, Cordelia Langford, John Sillitoe on behalf of the Wellcome Sanger Institute COVID-19 Surveillance Team |
| EPI_ISL_3310040, EPI_ISL_3310041, EPI_ISL_3310042, EPI_ISL_3310043, EPI_ISL_3310044 | Lighthouse Laboratory Plymouth | Wellcome Sanger Institute for the COVID-19 Genomics UK (COG-UK) Consortium | Lighthouse Laboratory Plymouth and Alex Alderton, Roberto Amato, Jeffrey Barrett, Sonia Goncalves, Ewan Harrison, David K. Jackson, Ian Johnston, Dominic Kwiatkowski, Cordelia Langford, John Sillitoe on behalf of the Wellcome Sanger Institute COVID-19 Surveillance Team |
| EPI_ISL_3310045 | Health Services Laboratories | Wellcome Sanger Institute for the COVID-19 Genomics UK (COG-UK) Consortium | Health Services Laboratories and Alex Alderton, Roberto Amato, Jeffrey Barrett, Sonia Goncalves, Ewan Harrison, David K. Jackson, Ian Johnston, Dominic Kwiatkowski, Cordelia Langford, John Sillitoe on behalf of the Wellcome Sanger Institute COVID-19 Surveillance Team |
| EPI_ISL_3310046 | Lighthouse Lab in Alderley Park | Wellcome Sanger Institute for the COVID-19 Genomics UK (COG-UK) Consortium | Jacquelyn Wynn, Mairead Hyland, The Lighthouse Lab in Alderley Park and Alex Alderton, Roberto Amato, Jeffrey Barrett, Sonia Goncalves, Ewan Harrison, David K. Jackson, Ian Johnston, Dominic Kwiatkowski, Cordelia Langford, John Sillitoe on behalf of the Wellcome Sanger Institute COVID-19 Surveillance Team |
| EPI_ISL_3310047 | Lighthouse Laboratory Plymouth | Wellcome Sanger Institute for the COVID-19 Genomics UK (COG-UK) Consortium | Lighthouse Laboratory Plymouth and Alex Alderton, Roberto Amato, Jeffrey Barrett, Sonia Goncalves, Ewan Harrison, David K. Jackson, Ian Johnston, Dominic Kwiatkowski, Cordelia Langford, John Sillitoe on behalf of the Wellcome Sanger Institute COVID-19 Surveillance Team |
| EPI_ISL_3310048, EPI_ISL_3310049, EPI_ISL_3310050 | Lighthouse Lab in Alderley Park | Wellcome Sanger Institute for the COVID-19 Genomics UK (COG-UK) Consortium | Jacquelyn Wynn, Mairead Hyland, The Lighthouse Lab in Alderley Park and Alex Alderton, Roberto Amato, Jeffrey Barrett, Sonia Goncalves, Ewan Harrison, David K. Jackson, Ian Johnston, Dominic Kwiatkowski, Cordelia Langford, John Sillitoe on behalf of the Wellcome Sanger Institute COVID-19 Surveillance Team |
| EPI_ISL_3310051 | Lighthouse Laboratory Plymouth | Wellcome Sanger Institute for the COVID-19 Genomics UK (COG-UK) Consortium | Lighthouse Laboratory Plymouth and Alex Alderton, Roberto Amato, Jeffrey Barrett, Sonia Goncalves, Ewan Harrison, David K. Jackson, Ian Johnston, Dominic Kwiatkowski, Cordelia Langford, John Sillitoe on behalf of the Wellcome Sanger Institute COVID-19 Surveillance Team |
| EPI_ISL_3310052 | Lighthouse Lab in Milton Keynes | Wellcome Sanger Institute for the COVID-19 Genomics UK (COG-UK) Consortium | The Lighthouse Lab in Milton Keynes and Alex Alderton, Roberto Amato, Jeffrey Barrett, Sonia Goncalves, Ewan Harrison, David K. Jackson, Ian Johnston, Dominic Kwiatkowski, Cordelia Langford, John Sillitoe on behalf of the Wellcome Sanger Institute COVID-19 Surveillance Team |
| EPI_ISL_3310053, EPI_ISL_3310054 | Lighthouse Lab in Alderley Park | Wellcome Sanger Institute for the COVID-19 Genomics UK (COG-UK) Consortium | Jacquelyn Wynn, Mairead Hyland, The Lighthouse Lab in Alderley Park and Alex Alderton, Roberto Amato, Jeffrey Barrett, Sonia Goncalves, Ewan Harrison, David K. Jackson, Ian Johnston, Dominic Kwiatkowski, Cordelia Langford, John Sillitoe on behalf of the Wellcome Sanger Institute COVID-19 Surveillance Team |
| EPI_ISL_3310055 | Lighthouse Lab in Milton Keynes | Wellcome Sanger Institute for the COVID-19 Genomics UK (COG-UK) Consortium | The Lighthouse Lab in Milton Keynes and Alex Alderton, Roberto Amato, Jeffrey Barrett, Sonia Goncalves, Ewan Harrison, David K. Jackson, Ian Johnston, Dominic Kwiatkowski, Cordelia Langford, John Sillitoe on behalf of the Wellcome Sanger Institute COVID-19 Surveillance Team |
| EPI_ISL_3310056 | Health Services Laboratories | Wellcome Sanger Institute for the COVID-19 Genomics UK (COG-UK) Consortium | Health Services Laboratories and Alex Alderton, Roberto Amato, Jeffrey Barrett, Sonia Goncalves, Ewan Harrison, David K. Jackson, Ian Johnston, Dominic Kwiatkowski, Cordelia Langford, John Sillitoe on behalf of the Wellcome Sanger Institute COVID-19 Surveillance Team |
| EPI_ISL_3310057 | Lighthouse Laboratory Plymouth | Wellcome Sanger Institute for the COVID-19 Genomics UK (COG-UK) Consortium | Lighthouse Laboratory Plymouth and Alex Alderton, Roberto Amato, Jeffrey Barrett, Sonia Goncalves, Ewan Harrison, David K. Jackson, Ian Johnston, Dominic Kwiatkowski, Cordelia Langford, John Sillitoe on behalf of the Wellcome Sanger Institute COVID-19 Surveillance Team |
| EPI_ISL_3310058 | Health Services Laboratories | Wellcome Sanger Institute for the COVID-19 Genomics UK (COG-UK) Consortium | Health Services Laboratories and Alex Alderton, Roberto Amato, Jeffrey Barrett, Sonia Goncalves, Ewan Harrison, David K. Jackson, Ian Johnston, Dominic Kwiatkowski, Cordelia Langford, John Sillitoe on behalf of the Wellcome Sanger Institute COVID-19 Surveillance Team |
| EPI_ISL_3310059 | Lighthouse Lab in Milton Keynes | Wellcome Sanger Institute for the COVID-19 Genomics UK (COG-UK) Consortium | The Lighthouse Lab in Milton Keynes and Alex Alderton, Roberto Amato, Jeffrey Barrett, Sonia Goncalves, Ewan Harrison, David K. Jackson, Ian Johnston, Dominic Kwiatkowski, Cordelia Langford, John Sillitoe on behalf of the Wellcome Sanger Institute COVID-19 Surveillance Team |
| EPI_ISL_3310060 | Lighthouse Lab in Alderley Park | Wellcome Sanger Institute for the COVID-19 Genomics UK (COG-UK) Consortium | Jacquelyn Wynn, Mairead Hyland, The Lighthouse Lab in Alderley Park and Alex Alderton, Roberto Amato, Jeffrey Barrett, Sonia Goncalves, Ewan Harrison, David K. Jackson, Ian Johnston, Dominic Kwiatkowski, Cordelia Langford, John Sillitoe on behalf of the Wellcome Sanger Institute COVID-19 Surveillance Team |
| EPI_ISL_3310061 | Lighthouse Lab in Milton Keynes | Wellcome Sanger Institute for the COVID-19 Genomics UK (COG-UK) Consortium | The Lighthouse Lab in Milton Keynes and Alex Alderton, Roberto Amato, Jeffrey Barrett, Sonia Goncalves, Ewan Harrison, David K. Jackson, Ian Johnston, Dominic Kwiatkowski, Cordelia Langford, John Sillitoe on behalf of the Wellcome Sanger Institute COVID-19 Surveillance Team |
| EPI_ISL_3310062, EPI_ISL_3310063, EPI_ISL_3310064 | Health Services Laboratories | Wellcome Sanger Institute for the COVID-19 Genomics UK (COG-UK) Consortium | Health Services Laboratories and Alex Alderton, Roberto Amato, Jeffrey Barrett, Sonia Goncalves, Ewan Harrison, David K. Jackson, Ian Johnston, Dominic Kwiatkowski, Cordelia Langford, John Sillitoe on behalf of the Wellcome Sanger Institute COVID-19 Surveillance Team |
| EPI_ISL_3310065, EPI_ISL_3310066 | Lighthouse Laboratory Plymouth | Wellcome Sanger Institute for the COVID-19 Genomics UK (COG-UK) Consortium | Lighthouse Laboratory Plymouth and Alex Alderton, Roberto Amato, Jeffrey Barrett, Sonia Goncalves, Ewan Harrison, David K. Jackson, Ian Johnston, Dominic Kwiatkowski, Cordelia Langford, John Sillitoe on behalf of the Wellcome Sanger Institute COVID-19 Surveillance Team |
| EPI_ISL_3310067, EPI_ISL_3310068 | Health Services Laboratories | Wellcome Sanger Institute for the COVID-19 Genomics UK (COG-UK) Consortium | Health Services Laboratories and Alex Alderton, Roberto Amato, Jeffrey Barrett, Sonia Goncalves, Ewan Harrison, David K. Jackson, Ian Johnston, Dominic Kwiatkowski, Cordelia Langford, John Sillitoe on behalf of the Wellcome Sanger Institute COVID-19 Surveillance Team |
| EPI_ISL_3310069 | Lighthouse Laboratory Plymouth | Wellcome Sanger Institute for the COVID-19 Genomics UK (COG-UK) Consortium | Lighthouse Laboratory Plymouth and Alex Alderton, Roberto Amato, Jeffrey Barrett, Sonia Goncalves, Ewan Harrison, David K. Jackson, Ian Johnston, Dominic Kwiatkowski, Cordelia Langford, John Sillitoe on behalf of the Wellcome Sanger Institute COVID-19 Surveillance Team |
| EPI_ISL_3310070, EPI_ISL_3310071, EPI_ISL_3310072, EPI_ISL_3310073, EPI_ISL_3310074, EPI_ISL_3310075 | Lighthouse Lab in Alderley Park | Wellcome Sanger Institute for the COVID-19 Genomics UK (COG-UK) Consortium | Jacquelyn Wynn, Mairead Hyland, The Lighthouse Lab in Alderley Park and Alex Alderton, Roberto Amato, Jeffrey Barrett, Sonia Goncalves, Ewan Harrison, David K. Jackson, Ian Johnston, Dominic Kwiatkowski, Cordelia Langford, John Sillitoe on behalf of the Wellcome Sanger Institute COVID-19 Surveillance Team |
| EPI_ISL_3310076 | Lighthouse Lab in Milton Keynes | Wellcome Sanger Institute for the COVID-19 Genomics UK (COG-UK) Consortium | The Lighthouse Lab in Milton Keynes and Alex Alderton, Roberto Amato, Jeffrey Barrett, Sonia Goncalves, Ewan Harrison, David K. Jackson, Ian Johnston, Dominic Kwiatkowski, Cordelia Langford, John Sillitoe on behalf of the Wellcome Sanger Institute COVID-19 Surveillance Team |
| EPI_ISL_3310077 | Lighthouse Laboratory Plymouth | Wellcome Sanger Institute for the COVID-19 Genomics UK (COG-UK) Consortium | Lighthouse Laboratory Plymouth and Alex Alderton, Roberto Amato, Jeffrey Barrett, Sonia Goncalves, Ewan Harrison, David K. Jackson, Ian Johnston, Dominic Kwiatkowski, Cordelia Langford, John Sillitoe on behalf of the Wellcome Sanger Institute COVID-19 Surveillance Team |
| EPI_ISL_3310078, EPI_ISL_3310079 | Lighthouse Lab in Alderley Park | Wellcome Sanger Institute for the COVID-19 Genomics UK (COG-UK) Consortium | Jacquelyn Wynn, Mairead Hyland, The Lighthouse Lab in Alderley Park and Alex Alderton, Roberto Amato, Jeffrey Barrett, Sonia Goncalves, Ewan Harrison, David K. Jackson, Ian Johnston, Dominic Kwiatkowski, Cordelia Langford, John Sillitoe on behalf of the Wellcome Sanger Institute COVID-19 Surveillance Team |
| EPI_ISL_3310080 | Lighthouse Lab in Milton Keynes | Wellcome Sanger Institute for the COVID-19 Genomics UK (COG-UK) Consortium | The Lighthouse Lab in Milton Keynes and Alex Alderton, Roberto Amato, Jeffrey Barrett, Sonia Goncalves, Ewan Harrison, David K. Jackson, Ian Johnston, Dominic Kwiatkowski, Cordelia Langford, John Sillitoe on behalf of the Wellcome Sanger Institute COVID-19 Surveillance Team |
| EPI_ISL_3310081 | Health Services Laboratories | Wellcome Sanger Institute for the COVID-19 Genomics UK (COG-UK) Consortium | Health Services Laboratories and Alex Alderton, Roberto Amato, Jeffrey Barrett, Sonia Goncalves, Ewan Harrison, David K. Jackson, Ian Johnston, Dominic Kwiatkowski, Cordelia Langford, John Sillitoe on behalf of the Wellcome Sanger Institute COVID-19 Surveillance Team |
| EPI_ISL_3310082, EPI_ISL_3310083 | Lighthouse Laboratory Plymouth | Wellcome Sanger Institute for the COVID-19 Genomics UK (COG-UK) Consortium | Lighthouse Laboratory Plymouth and Alex Alderton, Roberto Amato, Jeffrey Barrett, Sonia Goncalves, Ewan Harrison, David K. Jackson, Ian Johnston, Dominic Kwiatkowski, Cordelia Langford, John Sillitoe on behalf of the Wellcome Sanger Institute COVID-19 Surveillance Team |
| EPI_ISL_3310084 | Health Services Laboratories | Wellcome Sanger Institute for the COVID-19 Genomics UK (COG-UK) Consortium | Health Services Laboratories and Alex Alderton, Roberto Amato, Jeffrey Barrett, Sonia Goncalves, Ewan Harrison, David K. Jackson, Ian Johnston, Dominic Kwiatkowski, Cordelia Langford, John Sillitoe on behalf of the Wellcome Sanger Institute COVID-19 Surveillance Team |
| EPI_ISL_3310085, EPI_ISL_3310086 | Lighthouse Laboratory Plymouth | Wellcome Sanger Institute for the COVID-19 Genomics UK (COG-UK) Consortium | Lighthouse Laboratory Plymouth and Alex Alderton, Roberto Amato, Jeffrey Barrett, Sonia Goncalves, Ewan Harrison, David K. Jackson, Ian Johnston, Dominic Kwiatkowski, Cordelia Langford, John Sillitoe on behalf of the Wellcome Sanger Institute COVID-19 Surveillance Team |
| EPI_ISL_3310087 | Health Services Laboratories | Wellcome Sanger Institute for the COVID-19 Genomics UK (COG-UK) Consortium | Health Services Laboratories and Alex Alderton, Roberto Amato, Jeffrey Barrett, Sonia Goncalves, Ewan Harrison, David K. Jackson, Ian Johnston, Dominic Kwiatkowski, Cordelia Langford, John Sillitoe on behalf of the Wellcome Sanger Institute COVID-19 Surveillance Team |
| EPI_ISL_3310088, EPI_ISL_3310089, EPI_ISL_3310090 | Lighthouse Lab in Milton Keynes | Wellcome Sanger Institute for the COVID-19 Genomics UK (COG-UK) Consortium | The Lighthouse Lab in Milton Keynes and Alex Alderton, Roberto Amato, Jeffrey Barrett, Sonia Goncalves, Ewan Harrison, David K. Jackson, Ian Johnston, Dominic Kwiatkowski, Cordelia Langford, John Sillitoe on behalf of the Wellcome Sanger Institute COVID-19 Surveillance Team |
| EPI_ISL_3310091 | Health Services Laboratories | Wellcome Sanger Institute for the COVID-19 Genomics UK (COG-UK) Consortium | Health Services Laboratories and Alex Alderton, Roberto Amato, Jeffrey Barrett, Sonia Goncalves, Ewan Harrison, David K. Jackson, Ian Johnston, Dominic Kwiatkowski, Cordelia Langford, John Sillitoe on behalf of the Wellcome Sanger Institute COVID-19 Surveillance Team |
| EPI_ISL_3310092 | Lighthouse Lab in Alderley Park | Wellcome Sanger Institute for the COVID-19 Genomics UK (COG-UK) Consortium | Jacquelyn Wynn, Mairead Hyland, The Lighthouse Lab in Alderley Park and Alex Alderton, Roberto Amato, Jeffrey Barrett, Sonia Goncalves, Ewan Harrison, David K. Jackson, Ian Johnston, Dominic Kwiatkowski, Cordelia Langford, John Sillitoe on behalf of the Wellcome Sanger Institute COVID-19 Surveillance Team |

| | | | |
|---|---|---|---|
| EPI_ISL_3310093 | Lighthouse Lab in Milton Keynes | Wellcome Sanger Institute for the COVID-19 Genomics UK (COG-UK) Consortium | The Lighthouse Lab in Milton Keynes and Alex Alderton, Roberto Amato, Jeffrey Barrett, Sonia Goncalves, Ewan Harrison, David K. Jackson, Ian Johnston, Dominic Kwiatkowski, Cordelia Langford, John Sillitoe on behalf of the Wellcome Sanger Institute COVID-19 Surveillance Team |
| EPI_ISL_3310094 | Lighthouse Laboratory Plymouth | Wellcome Sanger Institute for the COVID-19 Genomics UK (COG-UK) Consortium | Lighthouse Laboratory Plymouth and Alex Alderton, Roberto Amato, Jeffrey Barrett, Sonia Goncalves, Ewan Harrison, David K. Jackson, Ian Johnston, Dominic Kwiatkowski, Cordelia Langford, John Sillitoe on behalf of the Wellcome Sanger Institute COVID-19 Surveillance Team |
| EPI_ISL_3310095 | Health Services Laboratories | Wellcome Sanger Institute for the COVID-19 Genomics UK (COG-UK) Consortium | Health Services Laboratories and Alex Alderton, Roberto Amato, Jeffrey Barrett, Sonia Goncalves, Ewan Harrison, David K. Jackson, Ian Johnston, Dominic Kwiatkowski, Cordelia Langford, John Sillitoe on behalf of the Wellcome Sanger Institute COVID-19 Surveillance Team |
| EPI_ISL_3310096, EPI_ISL_3310097 | Lighthouse Laboratory Plymouth | Wellcome Sanger Institute for the COVID-19 Genomics UK (COG-UK) Consortium | Lighthouse Laboratory Plymouth and Alex Alderton, Roberto Amato, Jeffrey Barrett, Sonia Goncalves, Ewan Harrison, David K. Jackson, Ian Johnston, Dominic Kwiatkowski, Cordelia Langford, John Sillitoe on behalf of the Wellcome Sanger Institute COVID-19 Surveillance Team |
| EPI_ISL_3310098 | Lighthouse Lab in Alderley Park | Wellcome Sanger Institute for the COVID-19 Genomics UK (COG-UK) Consortium | Jacquelyn Wynn, Mairead Hyland, The Lighthouse Lab in Alderley Park and Alex Alderton, Roberto Amato, Jeffrey Barrett, Sonia Goncalves, Ewan Harrison, David K. Jackson, Ian Johnston, Dominic Kwiatkowski, Cordelia Langford, John Sillitoe on behalf of the Wellcome Sanger Institute COVID-19 Surveillance Team |
| EPI_ISL_3310099 | Lighthouse Laboratory Plymouth | Wellcome Sanger Institute for the COVID-19 Genomics UK (COG-UK) Consortium | Lighthouse Laboratory Plymouth and Alex Alderton, Roberto Amato, Jeffrey Barrett, Sonia Goncalves, Ewan Harrison, David K. Jackson, Ian Johnston, Dominic Kwiatkowski, Cordelia Langford, John Sillitoe on behalf of the Wellcome Sanger Institute COVID-19 Surveillance Team |
| EPI_ISL_3310100, EPI_ISL_3310101, EPI_ISL_3310102 | Health Services Laboratories | Wellcome Sanger Institute for the COVID-19 Genomics UK (COG-UK) Consortium | Health Services Laboratories and Alex Alderton, Roberto Amato, Jeffrey Barrett, Sonia Goncalves, Ewan Harrison, David K. Jackson, Ian Johnston, Dominic Kwiatkowski, Cordelia Langford, John Sillitoe on behalf of the Wellcome Sanger Institute COVID-19 Surveillance Team |
| EPI_ISL_3310103 | Lighthouse Lab in Alderley Park | Wellcome Sanger Institute for the COVID-19 Genomics UK (COG-UK) Consortium | Jacquelyn Wynn, Mairead Hyland, The Lighthouse Lab in Alderley Park and Alex Alderton, Roberto Amato, Jeffrey Barrett, Sonia Goncalves, Ewan Harrison, David K. Jackson, Ian Johnston, Dominic Kwiatkowski, Cordelia Langford, John Sillitoe on behalf of the Wellcome Sanger Institute COVID-19 Surveillance Team |
| EPI_ISL_3310104 | Lighthouse Lab in Milton Keynes | Wellcome Sanger Institute for the COVID-19 Genomics UK (COG-UK) Consortium | The Lighthouse Lab in Milton Keynes and Alex Alderton, Roberto Amato, Jeffrey Barrett, Sonia Goncalves, Ewan Harrison, David K. Jackson, Ian Johnston, Dominic Kwiatkowski, Cordelia Langford, John Sillitoe on behalf of the Wellcome Sanger Institute COVID-19 Surveillance Team |
| EPI_ISL_3310105 | Lighthouse Lab in Alderley Park | Wellcome Sanger Institute for the COVID-19 Genomics UK (COG-UK) Consortium | Jacquelyn Wynn, Mairead Hyland, The Lighthouse Lab in Alderley Park and Alex Alderton, Roberto Amato, Jeffrey Barrett, Sonia Goncalves, Ewan Harrison, David K. Jackson, Ian Johnston, Dominic Kwiatkowski, Cordelia Langford, John Sillitoe on behalf of the Wellcome Sanger Institute COVID-19 Surveillance Team |
| EPI_ISL_3310106 | Health Services Laboratories | Wellcome Sanger Institute for the COVID-19 Genomics UK (COG-UK) Consortium | Health Services Laboratories and Alex Alderton, Roberto Amato, Jeffrey Barrett, Sonia Goncalves, Ewan Harrison, David K. Jackson, Ian Johnston, Dominic Kwiatkowski, Cordelia Langford, John Sillitoe on behalf of the Wellcome Sanger Institute COVID-19 Surveillance Team |
| EPI_ISL_3310107 | Lighthouse Lab in Milton Keynes | Wellcome Sanger Institute for the COVID-19 Genomics UK (COG-UK) Consortium | The Lighthouse Lab in Milton Keynes and Alex Alderton, Roberto Amato, Jeffrey Barrett, Sonia Goncalves, Ewan Harrison, David K. Jackson, Ian Johnston, Dominic Kwiatkowski, Cordelia Langford, John Sillitoe on behalf of the Wellcome Sanger Institute COVID-19 Surveillance Team |
| EPI_ISL_3310108 | Lighthouse Laboratory Plymouth | Wellcome Sanger Institute for the COVID-19 Genomics UK (COG-UK) Consortium | Lighthouse Laboratory Plymouth and Alex Alderton, Roberto Amato, Jeffrey Barrett, Sonia Goncalves, Ewan Harrison, David K. Jackson, Ian Johnston, Dominic Kwiatkowski, Cordelia Langford, John Sillitoe on behalf of the Wellcome Sanger Institute COVID-19 Surveillance Team |
| EPI_ISL_3310109, EPI_ISL_3310110, EPI_ISL_3310111, EPI_ISL_3310112 | Lighthouse Lab in Milton Keynes | Wellcome Sanger Institute for the COVID-19 Genomics UK (COG-UK) Consortium | The Lighthouse Lab in Milton Keynes and Alex Alderton, Roberto Amato, Jeffrey Barrett, Sonia Goncalves, Ewan Harrison, David K. Jackson, Ian Johnston, Dominic Kwiatkowski, Cordelia Langford, John Sillitoe on behalf of the Wellcome Sanger Institute COVID-19 Surveillance Team |
| EPI_ISL_3310113, EPI_ISL_3310114, EPI_ISL_3310115 | Lighthouse Lab in Alderley Park | Wellcome Sanger Institute for the COVID-19 Genomics UK (COG-UK) Consortium | Jacquelyn Wynn, Mairead Hyland, The Lighthouse Lab in Alderley Park and Alex Alderton, Roberto Amato, Jeffrey Barrett, Sonia Goncalves, Ewan Harrison, David K. Jackson, Ian Johnston, Dominic Kwiatkowski, Cordelia Langford, John Sillitoe on behalf of the Wellcome Sanger Institute COVID-19 Surveillance Team |
| EPI_ISL_3310116 | Lighthouse Laboratory Plymouth | Wellcome Sanger Institute for the COVID-19 Genomics UK (COG-UK) Consortium | Lighthouse Laboratory Plymouth and Alex Alderton, Roberto Amato, Jeffrey Barrett, Sonia Goncalves, Ewan Harrison, David K. Jackson, Ian Johnston, Dominic Kwiatkowski, Cordelia Langford, John Sillitoe on behalf of the Wellcome Sanger Institute COVID-19 Surveillance Team |
| EPI_ISL_3310117, EPI_ISL_3310118 | Lighthouse Lab in Alderley Park | Wellcome Sanger Institute for the COVID-19 Genomics UK (COG-UK) Consortium | Jacquelyn Wynn, Mairead Hyland, The Lighthouse Lab in Alderley Park and Alex Alderton, Roberto Amato, Jeffrey Barrett, Sonia Goncalves, Ewan Harrison, David K. Jackson, Ian Johnston, Dominic Kwiatkowski, Cordelia Langford, John Sillitoe on behalf of the Wellcome Sanger Institute COVID-19 Surveillance Team |
| EPI_ISL_3310119 | Health Services Laboratories | Wellcome Sanger Institute for the COVID-19 Genomics UK (COG-UK) Consortium | Health Services Laboratories and Alex Alderton, Roberto Amato, Jeffrey Barrett, Sonia Goncalves, Ewan Harrison, David K. Jackson, Ian Johnston, Dominic Kwiatkowski, Cordelia Langford, John Sillitoe on behalf of the Wellcome Sanger Institute COVID-19 Surveillance Team |
| EPI_ISL_3310120, EPI_ISL_3310121, EPI_ISL_3310122 | Lighthouse Lab in Alderley Park | Wellcome Sanger Institute for the COVID-19 Genomics UK (COG-UK) Consortium | Jacquelyn Wynn, Mairead Hyland, The Lighthouse Lab in Alderley Park and Alex Alderton, Roberto Amato, Jeffrey Barrett, Sonia Goncalves, Ewan Harrison, David K. Jackson, Ian Johnston, Dominic Kwiatkowski, Cordelia Langford, John Sillitoe on behalf of the Wellcome Sanger Institute COVID-19 Surveillance Team |
| EPI_ISL_3310123 | Health Services Laboratories | Wellcome Sanger Institute for the COVID-19 Genomics UK (COG-UK) Consortium | Health Services Laboratories and Alex Alderton, Roberto Amato, Jeffrey Barrett, Sonia Goncalves, Ewan Harrison, David K. Jackson, Ian Johnston, Dominic Kwiatkowski, Cordelia Langford, John Sillitoe on behalf of the Wellcome Sanger Institute COVID-19 Surveillance Team |
| EPI_ISL_3310124 | Lighthouse Lab in Milton Keynes | Wellcome Sanger Institute for the COVID-19 Genomics UK (COG-UK) Consortium | The Lighthouse Lab in Milton Keynes and Alex Alderton, Roberto Amato, Jeffrey Barrett, Sonia Goncalves, Ewan Harrison, David K. Jackson, Ian Johnston, Dominic Kwiatkowski, Cordelia Langford, John Sillitoe on behalf of the Wellcome Sanger Institute COVID-19 Surveillance Team |
| EPI_ISL_3310125 | Health Services Laboratories | Wellcome Sanger Institute for the COVID-19 Genomics UK (COG-UK) Consortium | Health Services Laboratories and Alex Alderton, Roberto Amato, Jeffrey Barrett, Sonia Goncalves, Ewan Harrison, David K. Jackson, Ian Johnston, Dominic Kwiatkowski, Cordelia Langford, John Sillitoe on behalf of the Wellcome Sanger Institute COVID-19 Surveillance Team |
| EPI_ISL_3310126 | Lighthouse Lab in Milton Keynes | Wellcome Sanger Institute for the COVID-19 Genomics UK (COG-UK) Consortium | The Lighthouse Lab in Milton Keynes and Alex Alderton, Roberto Amato, Jeffrey Barrett, Sonia Goncalves, Ewan Harrison, David K. Jackson, Ian Johnston, Dominic Kwiatkowski, Cordelia Langford, John Sillitoe on behalf of the Wellcome Sanger Institute COVID-19 Surveillance Team |
| EPI_ISL_3310127 | Lighthouse Laboratory Plymouth | Wellcome Sanger Institute for the COVID-19 Genomics UK (COG-UK) Consortium | Lighthouse Laboratory Plymouth and Alex Alderton, Roberto Amato, Jeffrey Barrett, Sonia Goncalves, Ewan Harrison, David K. Jackson, Ian Johnston, Dominic Kwiatkowski, Cordelia Langford, John Sillitoe on behalf of the Wellcome Sanger Institute COVID-19 Surveillance Team |
| EPI_ISL_3310128 | Lighthouse Lab in Alderley Park | Wellcome Sanger Institute for the COVID-19 Genomics UK (COG-UK) Consortium | Jacquelyn Wynn, Mairead Hyland, The Lighthouse Lab in Alderley Park and Alex Alderton, Roberto Amato, Jeffrey Barrett, Sonia Goncalves, Ewan Harrison, David K. Jackson, Ian Johnston, Dominic Kwiatkowski, Cordelia Langford, John Sillitoe on behalf of the Wellcome Sanger Institute COVID-19 Surveillance Team |
| EPI_ISL_3310129 | Lighthouse Lab in Milton Keynes | Wellcome Sanger Institute for the COVID-19 Genomics UK (COG-UK) Consortium | The Lighthouse Lab in Milton Keynes and Alex Alderton, Roberto Amato, Jeffrey Barrett, Sonia Goncalves, Ewan Harrison, David K. Jackson, Ian Johnston, Dominic Kwiatkowski, Cordelia Langford, John Sillitoe on behalf of the Wellcome Sanger Institute COVID-19 Surveillance Team |
| EPI_ISL_3310130 | Lighthouse Lab in Alderley Park | Wellcome Sanger Institute for the COVID-19 Genomics UK (COG-UK) Consortium | Jacquelyn Wynn, Mairead Hyland, The Lighthouse Lab in Alderley Park and Alex Alderton, Roberto Amato, Jeffrey Barrett, Sonia Goncalves, Ewan Harrison, David K. Jackson, Ian Johnston, Dominic Kwiatkowski, Cordelia Langford, John Sillitoe on behalf of the Wellcome Sanger Institute COVID-19 Surveillance Team |
| EPI_ISL_3310131 | Lighthouse Laboratory Plymouth | Wellcome Sanger Institute for the COVID-19 Genomics UK (COG-UK) Consortium | Lighthouse Laboratory Plymouth and Alex Alderton, Roberto Amato, Jeffrey Barrett, Sonia Goncalves, Ewan Harrison, David K. Jackson, Ian Johnston, Dominic Kwiatkowski, Cordelia Langford, John Sillitoe on behalf of the Wellcome Sanger Institute COVID-19 Surveillance Team |
| EPI_ISL_3310132 | Lighthouse Lab in Milton Keynes | Wellcome Sanger Institute for the COVID-19 Genomics UK (COG-UK) Consortium | The Lighthouse Lab in Milton Keynes and Alex Alderton, Roberto Amato, Jeffrey Barrett, Sonia Goncalves, Ewan Harrison, David K. Jackson, Ian Johnston, Dominic Kwiatkowski, Cordelia Langford, John Sillitoe on behalf of the Wellcome Sanger Institute COVID-19 Surveillance Team |
| EPI_ISL_3310133, EPI_ISL_3310134 | Lighthouse Lab in Alderley Park | Wellcome Sanger Institute for the COVID-19 Genomics UK (COG-UK) Consortium | Jacquelyn Wynn, Mairead Hyland, The Lighthouse Lab in Alderley Park and Alex Alderton, Roberto Amato, Jeffrey Barrett, Sonia Goncalves, Ewan Harrison, David K. Jackson, Ian Johnston, Dominic Kwiatkowski, Cordelia Langford, John Sillitoe on behalf of the Wellcome Sanger Institute COVID-19 Surveillance Team |
| EPI_ISL_3310135 | Lighthouse Lab in Milton Keynes | Wellcome Sanger Institute for the COVID-19 Genomics UK (COG-UK) Consortium | The Lighthouse Lab in Milton Keynes and Alex Alderton, Roberto Amato, Jeffrey Barrett, Sonia Goncalves, Ewan Harrison, David K. Jackson, Ian Johnston, Dominic Kwiatkowski, Cordelia Langford, John Sillitoe on behalf of the Wellcome Sanger Institute COVID-19 Surveillance Team |
| EPI_ISL_3310136 | Lighthouse Lab in Alderley Park | Wellcome Sanger Institute for the COVID-19 Genomics UK (COG-UK) Consortium | Jacquelyn Wynn, Mairead Hyland, The Lighthouse Lab in Alderley Park and Alex Alderton, Roberto Amato, Jeffrey Barrett, Sonia Goncalves, Ewan Harrison, David K. Jackson, Ian Johnston, Dominic Kwiatkowski, Cordelia Langford, John Sillitoe on behalf of the Wellcome Sanger Institute COVID-19 Surveillance Team |
| EPI_ISL_3310137 | Lighthouse Laboratory Plymouth | Wellcome Sanger Institute for the COVID-19 Genomics UK (COG-UK) Consortium | Lighthouse Laboratory Plymouth and Alex Alderton, Roberto Amato, Jeffrey Barrett, Sonia Goncalves, Ewan Harrison, David K. Jackson, Ian Johnston, Dominic Kwiatkowski, Cordelia Langford, John Sillitoe on behalf of the Wellcome Sanger Institute COVID-19 Surveillance Team |
| EPI_ISL_3310138, EPI_ISL_3310139 | Lighthouse Lab in Milton Keynes | Wellcome Sanger Institute for the COVID-19 Genomics UK (COG-UK) Consortium | The Lighthouse Lab in Milton Keynes and Alex Alderton, Roberto Amato, Jeffrey Barrett, Sonia Goncalves, Ewan Harrison, David K. Jackson, Ian Johnston, Dominic Kwiatkowski, Cordelia Langford, John Sillitoe on behalf of the Wellcome Sanger Institute COVID-19 Surveillance Team |
| EPI_ISL_3310140 | Lighthouse Lab in Alderley Park | Wellcome Sanger Institute for the COVID-19 Genomics UK (COG-UK) Consortium | Jacquelyn Wynn, Mairead Hyland, The Lighthouse Lab in Alderley Park and Alex Alderton, Roberto Amato, Jeffrey Barrett, Sonia Goncalves, Ewan Harrison, David K. Jackson, Ian Johnston, Dominic Kwiatkowski, Cordelia Langford, John Sillitoe on behalf of the Wellcome Sanger Institute COVID-19 Surveillance Team |
| EPI_ISL_3310141 | Lighthouse Lab in Milton Keynes | Wellcome Sanger Institute for the COVID-19 Genomics UK (COG-UK) Consortium | The Lighthouse Lab in Milton Keynes and Alex Alderton, Roberto Amato, Jeffrey Barrett, Sonia Goncalves, Ewan Harrison, David K. Jackson, Ian Johnston, Dominic Kwiatkowski, Cordelia Langford, John Sillitoe on behalf of the Wellcome Sanger Institute COVID-19 Surveillance Team |
| EPI_ISL_3310142 | Lighthouse Laboratory Plymouth | Wellcome Sanger Institute for the COVID-19 Genomics UK (COG-UK) Consortium | Lighthouse Laboratory Plymouth and Alex Alderton, Roberto Amato, Jeffrey Barrett, Sonia Goncalves, Ewan Harrison, David K. Jackson, Ian Johnston, Dominic Kwiatkowski, Cordelia Langford, John Sillitoe on behalf of the Wellcome Sanger Institute COVID-19 Surveillance Team |
| EPI_ISL_3310143 | Lighthouse Lab in Milton Keynes | Wellcome Sanger Institute for the COVID-19 Genomics UK (COG-UK) Consortium | The Lighthouse Lab in Milton Keynes and Alex Alderton, Roberto Amato, Jeffrey Barrett, Sonia Goncalves, Ewan Harrison, David K. Jackson, Ian Johnston, Dominic Kwiatkowski, Cordelia Langford, John Sillitoe on behalf of the Wellcome Sanger Institute COVID-19 Surveillance Team |
| EPI_ISL_3310144, EPI_ISL_3310145 | Health Services Laboratories | Wellcome Sanger Institute for the COVID-19 Genomics UK (COG-UK) Consortium | Health Services Laboratories and Alex Alderton, Roberto Amato, Jeffrey Barrett, Sonia Goncalves, Ewan Harrison, David K. Jackson, Ian Johnston, Dominic Kwiatkowski, Cordelia Langford, John Sillitoe on behalf of the Wellcome Sanger Institute COVID-19 Surveillance Team |
| EPI_ISL_3310146, EPI_ISL_3310147 | Lighthouse Laboratory Plymouth | Wellcome Sanger Institute for the COVID-19 Genomics UK (COG-UK) Consortium | Lighthouse Laboratory Plymouth and Alex Alderton, Roberto Amato, Jeffrey Barrett, Sonia Goncalves, Ewan Harrison, David K. Jackson, Ian Johnston, Dominic Kwiatkowski, Cordelia Langford, John Sillitoe on behalf of the Wellcome Sanger Institute COVID-19 Surveillance Team |
| EPI_ISL_3310148 | Health Services Laboratories | Wellcome Sanger Institute for the COVID-19 Genomics UK (COG-UK) Consortium | Health Services Laboratories and Alex Alderton, Roberto Amato, Jeffrey Barrett, Sonia Goncalves, Ewan Harrison, David K. Jackson, Ian Johnston, Dominic Kwiatkowski, Cordelia Langford, John Sillitoe on behalf of the Wellcome Sanger Institute COVID-19 Surveillance Team |
| EPI_ISL_3310149, EPI_ISL_3310150 | Lighthouse Lab in Milton Keynes | Wellcome Sanger Institute for the COVID-19 Genomics UK (COG-UK) Consortium | The Lighthouse Lab in Milton Keynes and Alex Alderton, Roberto Amato, Jeffrey Barrett, Sonia Goncalves, Ewan Harrison, David K. Jackson, Ian Johnston, Dominic Kwiatkowski, Cordelia Langford, John Sillitoe on behalf of the Wellcome Sanger Institute COVID-19 Surveillance Team |
| EPI_ISL_3310151 | Lighthouse Lab in Alderley Park | Wellcome Sanger Institute for the COVID-19 Genomics UK (COG-UK) Consortium | Jacquelyn Wynn, Mairead Hyland, The Lighthouse Lab in Alderley Park and Alex Alderton, Roberto Amato, Jeffrey Barrett, Sonia Goncalves, Ewan Harrison, David K. Jackson, Ian Johnston, Dominic Kwiatkowski, Cordelia Langford, John Sillitoe on behalf of the Wellcome Sanger Institute COVID-19 Surveillance Team |

| | | | |
|---|---|---|---|
| EPI_ISL_3310152 | Health Services Laboratories | Wellcome Sanger Institute for the COVID-19 Genomics UK (COG-UK) Consortium | Health Services Laboratories and Alex Alderton, Roberto Amato, Jeffrey Barrett, Sonia Goncalves, Ewan Harrison, David K. Jackson, Ian Johnston, Dominic Kwiatkowski, Cordelia Langford, John Sillitoe on behalf of the Wellcome Sanger Institute COVID-19 Surveillance Team |
| EPI_ISL_3310153 | Lighthouse Lab in Alderley Park | Wellcome Sanger Institute for the COVID-19 Genomics UK (COG-UK) Consortium | Jacquelyn Wynn, Mairead Hyland, The Lighthouse Lab in Alderley Park and Alex Alderton, Roberto Amato, Jeffrey Barrett, Sonia Goncalves, Ewan Harrison, David K. Jackson, Ian Johnston, Dominic Kwiatkowski, Cordelia Langford, John Sillitoe on behalf of the Wellcome Sanger Institute COVID-19 Surveillance Team |
| EPI_ISL_3310154 | Lighthouse Laboratory Plymouth | Wellcome Sanger Institute for the COVID-19 Genomics UK (COG-UK) Consortium | Lighthouse Laboratory Plymouth and Alex Alderton, Roberto Amato, Jeffrey Barrett, Sonia Goncalves, Ewan Harrison, David K. Jackson, Ian Johnston, Dominic Kwiatkowski, Cordelia Langford, John Sillitoe on behalf of the Wellcome Sanger Institute COVID-19 Surveillance Team |
| EPI_ISL_3310155 | Health Services Laboratories | Wellcome Sanger Institute for the COVID-19 Genomics UK (COG-UK) Consortium | Health Services Laboratories and Alex Alderton, Roberto Amato, Jeffrey Barrett, Sonia Goncalves, Ewan Harrison, David K. Jackson, Ian Johnston, Dominic Kwiatkowski, Cordelia Langford, John Sillitoe on behalf of the Wellcome Sanger Institute COVID-19 Surveillance Team |
| EPI_ISL_3310156 | Lighthouse Laboratory Plymouth | Wellcome Sanger Institute for the COVID-19 Genomics UK (COG-UK) Consortium | Lighthouse Laboratory Plymouth and Alex Alderton, Roberto Amato, Jeffrey Barrett, Sonia Goncalves, Ewan Harrison, David K. Jackson, Ian Johnston, Dominic Kwiatkowski, Cordelia Langford, John Sillitoe on behalf of the Wellcome Sanger Institute COVID-19 Surveillance Team |
| EPI_ISL_3310157, EPI_ISL_3310158 | Lighthouse Lab in Alderley Park | Wellcome Sanger Institute for the COVID-19 Genomics UK (COG-UK) Consortium | Jacquelyn Wynn, Mairead Hyland, The Lighthouse Lab in Alderley Park and Alex Alderton, Roberto Amato, Jeffrey Barrett, Sonia Goncalves, Ewan Harrison, David K. Jackson, Ian Johnston, Dominic Kwiatkowski, Cordelia Langford, John Sillitoe on behalf of the Wellcome Sanger Institute COVID-19 Surveillance Team |
| EPI_ISL_3310159 | Health Services Laboratories | Wellcome Sanger Institute for the COVID-19 Genomics UK (COG-UK) Consortium | Health Services Laboratories and Alex Alderton, Roberto Amato, Jeffrey Barrett, Sonia Goncalves, Ewan Harrison, David K. Jackson, Ian Johnston, Dominic Kwiatkowski, Cordelia Langford, John Sillitoe on behalf of the Wellcome Sanger Institute COVID-19 Surveillance Team |
| EPI_ISL_3310160 | Lighthouse Laboratory Plymouth | Wellcome Sanger Institute for the COVID-19 Genomics UK (COG-UK) Consortium | Lighthouse Laboratory Plymouth and Alex Alderton, Roberto Amato, Jeffrey Barrett, Sonia Goncalves, Ewan Harrison, David K. Jackson, Ian Johnston, Dominic Kwiatkowski, Cordelia Langford, John Sillitoe on behalf of the Wellcome Sanger Institute COVID-19 Surveillance Team |
| EPI_ISL_3310161 | Health Services Laboratories | Wellcome Sanger Institute for the COVID-19 Genomics UK (COG-UK) Consortium | Health Services Laboratories and Alex Alderton, Roberto Amato, Jeffrey Barrett, Sonia Goncalves, Ewan Harrison, David K. Jackson, Ian Johnston, Dominic Kwiatkowski, Cordelia Langford, John Sillitoe on behalf of the Wellcome Sanger Institute COVID-19 Surveillance Team |
| EPI_ISL_3310162 | Lighthouse Lab in Alderley Park | Wellcome Sanger Institute for the COVID-19 Genomics UK (COG-UK) Consortium | Jacquelyn Wynn, Mairead Hyland, The Lighthouse Lab in Alderley Park and Alex Alderton, Roberto Amato, Jeffrey Barrett, Sonia Goncalves, Ewan Harrison, David K. Jackson, Ian Johnston, Dominic Kwiatkowski, Cordelia Langford, John Sillitoe on behalf of the Wellcome Sanger Institute COVID-19 Surveillance Team |
| EPI_ISL_3310163, EPI_ISL_3310164 | Lighthouse Laboratory Plymouth | Wellcome Sanger Institute for the COVID-19 Genomics UK (COG-UK) Consortium | Lighthouse Laboratory Plymouth and Alex Alderton, Roberto Amato, Jeffrey Barrett, Sonia Goncalves, Ewan Harrison, David K. Jackson, Ian Johnston, Dominic Kwiatkowski, Cordelia Langford, John Sillitoe on behalf of the Wellcome Sanger Institute COVID-19 Surveillance Team |
| EPI_ISL_3310165 | Lighthouse Lab in Milton Keynes | Wellcome Sanger Institute for the COVID-19 Genomics UK (COG-UK) Consortium | The Lighthouse Lab in Milton Keynes and Alex Alderton, Roberto Amato, Jeffrey Barrett, Sonia Goncalves, Ewan Harrison, David K. Jackson, Ian Johnston, Dominic Kwiatkowski, Cordelia Langford, John Sillitoe on behalf of the Wellcome Sanger Institute COVID-19 Surveillance Team |
| EPI_ISL_3310166 | Lighthouse Lab in Glasgow | Wellcome Sanger Institute for the COVID-19 Genomics UK (COG-UK) Consortium | Harper VanSteenhouse, Yumi Kasai, David Gray, Carol Clugston, Anna Dominiczak and Alex Alderton, Roberto Amato, Jeffrey Barrett, Sonia Goncalves, Ewan Harrison, David K. Jackson, Ian Johnston, Dominic Kwiatkowski, Cordelia Langford, John Sillitoe on behalf of the Wellcome Sanger Institute COVID-19 Surveillance Team |
| EPI_ISL_3310167, EPI_ISL_3310168 | Lighthouse Lab in Milton Keynes | Wellcome Sanger Institute for the COVID-19 Genomics UK (COG-UK) Consortium | The Lighthouse Lab in Milton Keynes and Alex Alderton, Roberto Amato, Jeffrey Barrett, Sonia Goncalves, Ewan Harrison, David K. Jackson, Ian Johnston, Dominic Kwiatkowski, Cordelia Langford, John Sillitoe on behalf of the Wellcome Sanger Institute COVID-19 Surveillance Team |
| EPI_ISL_3310169 | Lighthouse Lab in Glasgow | Wellcome Sanger Institute for the COVID-19 Genomics UK (COG-UK) Consortium | Harper VanSteenhouse, Yumi Kasai, David Gray, Carol Clugston, Anna Dominiczak and Alex Alderton, Roberto Amato, Jeffrey Barrett, Sonia Goncalves, Ewan Harrison, David K. Jackson, Ian Johnston, Dominic Kwiatkowski, Cordelia Langford, John Sillitoe on behalf of the Wellcome Sanger Institute COVID-19 Surveillance Team |
| EPI_ISL_3310170 | Lighthouse Lab in Alderley Park | Wellcome Sanger Institute for the COVID-19 Genomics UK (COG-UK) Consortium | Jacquelyn Wynn, Mairead Hyland, The Lighthouse Lab in Alderley Park and Alex Alderton, Roberto Amato, Jeffrey Barrett, Sonia Goncalves, Ewan Harrison, David K. Jackson, Ian Johnston, Dominic Kwiatkowski, Cordelia Langford, John Sillitoe on behalf of the Wellcome Sanger Institute COVID-19 Surveillance Team |
| EPI_ISL_3310171, EPI_ISL_3310172 | Lighthouse Lab in Glasgow | Wellcome Sanger Institute for the COVID-19 Genomics UK (COG-UK) Consortium | Harper VanSteenhouse, Yumi Kasai, David Gray, Carol Clugston, Anna Dominiczak and Alex Alderton, Roberto Amato, Jeffrey Barrett, Sonia Goncalves, Ewan Harrison, David K. Jackson, Ian Johnston, Dominic Kwiatkowski, Cordelia Langford, John Sillitoe on behalf of the Wellcome Sanger Institute COVID-19 Surveillance Team |
| EPI_ISL_3310173, EPI_ISL_3310174 | Lighthouse Lab in Milton Keynes | Wellcome Sanger Institute for the COVID-19 Genomics UK (COG-UK) Consortium | The Lighthouse Lab in Milton Keynes and Alex Alderton, Roberto Amato, Jeffrey Barrett, Sonia Goncalves, Ewan Harrison, David K. Jackson, Ian Johnston, Dominic Kwiatkowski, Cordelia Langford, John Sillitoe on behalf of the Wellcome Sanger Institute COVID-19 Surveillance Team |
| EPI_ISL_3310175 | Lighthouse Lab in Glasgow | Wellcome Sanger Institute for the COVID-19 Genomics UK (COG-UK) Consortium | Harper VanSteenhouse, Yumi Kasai, David Gray, Carol Clugston, Anna Dominiczak and Alex Alderton, Roberto Amato, Jeffrey Barrett, Sonia Goncalves, Ewan Harrison, David K. Jackson, Ian Johnston, Dominic Kwiatkowski, Cordelia Langford, John Sillitoe on behalf of the Wellcome Sanger Institute COVID-19 Surveillance Team |
| EPI_ISL_3310176 | Lighthouse Lab in Alderley Park | Wellcome Sanger Institute for the COVID-19 Genomics UK (COG-UK) Consortium | Jacquelyn Wynn, Mairead Hyland, The Lighthouse Lab in Alderley Park and Alex Alderton, Roberto Amato, Jeffrey Barrett, Sonia Goncalves, Ewan Harrison, David K. Jackson, Ian Johnston, Dominic Kwiatkowski, Cordelia Langford, John Sillitoe on behalf of the Wellcome Sanger Institute COVID-19 Surveillance Team |
| EPI_ISL_3310177 | Lighthouse Lab in Milton Keynes | Wellcome Sanger Institute for the COVID-19 Genomics UK (COG-UK) Consortium | The Lighthouse Lab in Milton Keynes and Alex Alderton, Roberto Amato, Jeffrey Barrett, Sonia Goncalves, Ewan Harrison, David K. Jackson, Ian Johnston, Dominic Kwiatkowski, Cordelia Langford, John Sillitoe on behalf of the Wellcome Sanger Institute COVID-19 Surveillance Team |
| EPI_ISL_3310178, EPI_ISL_3310179 | Lighthouse Lab in Alderley Park | Wellcome Sanger Institute for the COVID-19 Genomics UK (COG-UK) Consortium | Jacquelyn Wynn, Mairead Hyland, The Lighthouse Lab in Alderley Park and Alex Alderton, Roberto Amato, Jeffrey Barrett, Sonia Goncalves, Ewan Harrison, David K. Jackson, Ian Johnston, Dominic Kwiatkowski, Cordelia Langford, John Sillitoe on behalf of the Wellcome Sanger Institute COVID-19 Surveillance Team |
| EPI_ISL_3310180, EPI_ISL_3310181 | Lighthouse Lab in Milton Keynes | Wellcome Sanger Institute for the COVID-19 Genomics UK (COG-UK) Consortium | The Lighthouse Lab in Milton Keynes and Alex Alderton, Roberto Amato, Jeffrey Barrett, Sonia Goncalves, Ewan Harrison, David K. Jackson, Ian Johnston, Dominic Kwiatkowski, Cordelia Langford, John Sillitoe on behalf of the Wellcome Sanger Institute COVID-19 Surveillance Team |
| EPI_ISL_3310182 | Lighthouse Lab in Alderley Park | Wellcome Sanger Institute for the COVID-19 Genomics UK (COG-UK) Consortium | Jacquelyn Wynn, Mairead Hyland, The Lighthouse Lab in Alderley Park and Alex Alderton, Roberto Amato, Jeffrey Barrett, Sonia Goncalves, Ewan Harrison, David K. Jackson, Ian Johnston, Dominic Kwiatkowski, Cordelia Langford, John Sillitoe on behalf of the Wellcome Sanger Institute COVID-19 Surveillance Team |
| EPI_ISL_3310183 | Lighthouse Lab in Milton Keynes | Wellcome Sanger Institute for the COVID-19 Genomics UK (COG-UK) Consortium | The Lighthouse Lab in Milton Keynes and Alex Alderton, Roberto Amato, Jeffrey Barrett, Sonia Goncalves, Ewan Harrison, David K. Jackson, Ian Johnston, Dominic Kwiatkowski, Cordelia Langford, John Sillitoe on behalf of the Wellcome Sanger Institute COVID-19 Surveillance Team |
| EPI_ISL_3310184 | Lighthouse Lab in Glasgow | Wellcome Sanger Institute for the COVID-19 Genomics UK (COG-UK) Consortium | Harper VanSteenhouse, Yumi Kasai, David Gray, Carol Clugston, Anna Dominiczak and Alex Alderton, Roberto Amato, Jeffrey Barrett, Sonia Goncalves, Ewan Harrison, David K. Jackson, Ian Johnston, Dominic Kwiatkowski, Cordelia Langford, John Sillitoe on behalf of the Wellcome Sanger Institute COVID-19 Surveillance Team |
| EPI_ISL_3310185 | Lighthouse Lab in Milton Keynes | Wellcome Sanger Institute for the COVID-19 Genomics UK (COG-UK) Consortium | The Lighthouse Lab in Milton Keynes and Alex Alderton, Roberto Amato, Jeffrey Barrett, Sonia Goncalves, Ewan Harrison, David K. Jackson, Ian Johnston, Dominic Kwiatkowski, Cordelia Langford, John Sillitoe on behalf of the Wellcome Sanger Institute COVID-19 Surveillance Team |
| EPI_ISL_3310186, EPI_ISL_3310187 | Lighthouse Lab in Alderley Park | Wellcome Sanger Institute for the COVID-19 Genomics UK (COG-UK) Consortium | Jacquelyn Wynn, Mairead Hyland, The Lighthouse Lab in Alderley Park and Alex Alderton, Roberto Amato, Jeffrey Barrett, Sonia Goncalves, Ewan Harrison, David K. Jackson, Ian Johnston, Dominic Kwiatkowski, Cordelia Langford, John Sillitoe on behalf of the Wellcome Sanger Institute COVID-19 Surveillance Team |
| EPI_ISL_3310188 | Lighthouse Lab in Glasgow | Wellcome Sanger Institute for the COVID-19 Genomics UK (COG-UK) Consortium | Harper VanSteenhouse, Yumi Kasai, David Gray, Carol Clugston, Anna Dominiczak and Alex Alderton, Roberto Amato, Jeffrey Barrett, Sonia Goncalves, Ewan Harrison, David K. Jackson, Ian Johnston, Dominic Kwiatkowski, Cordelia Langford, John Sillitoe on behalf of the Wellcome Sanger Institute COVID-19 Surveillance Team |
| EPI_ISL_3310189 | Lighthouse Lab in Milton Keynes | Wellcome Sanger Institute for the COVID-19 Genomics UK (COG-UK) Consortium | The Lighthouse Lab in Milton Keynes and Alex Alderton, Roberto Amato, Jeffrey Barrett, Sonia Goncalves, Ewan Harrison, David K. Jackson, Ian Johnston, Dominic Kwiatkowski, Cordelia Langford, John Sillitoe on behalf of the Wellcome Sanger Institute COVID-19 Surveillance Team |
| EPI_ISL_3310190, EPI_ISL_3310191 | Lighthouse Lab in Glasgow | Wellcome Sanger Institute for the COVID-19 Genomics UK (COG-UK) Consortium | Harper VanSteenhouse, Yumi Kasai, David Gray, Carol Clugston, Anna Dominiczak and Alex Alderton, Roberto Amato, Jeffrey Barrett, Sonia Goncalves, Ewan Harrison, David K. Jackson, Ian Johnston, Dominic Kwiatkowski, Cordelia Langford, John Sillitoe on behalf of the Wellcome Sanger Institute COVID-19 Surveillance Team |
| EPI_ISL_3310192, EPI_ISL_3310193, EPI_ISL_3310194 | Lighthouse Lab in Alderley Park | Wellcome Sanger Institute for the COVID-19 Genomics UK (COG-UK) Consortium | Jacquelyn Wynn, Mairead Hyland, The Lighthouse Lab in Alderley Park and Alex Alderton, Roberto Amato, Jeffrey Barrett, Sonia Goncalves, Ewan Harrison, David K. Jackson, Ian Johnston, Dominic Kwiatkowski, Cordelia Langford, John Sillitoe on behalf of the Wellcome Sanger Institute COVID-19 Surveillance Team |
| EPI_ISL_3310195 | Lighthouse Lab in Glasgow | Wellcome Sanger Institute for the COVID-19 Genomics UK (COG-UK) Consortium | Harper VanSteenhouse, Yumi Kasai, David Gray, Carol Clugston, Anna Dominiczak and Alex Alderton, Roberto Amato, Jeffrey Barrett, Sonia Goncalves, Ewan Harrison, David K. Jackson, Ian Johnston, Dominic Kwiatkowski, Cordelia Langford, John Sillitoe on behalf of the Wellcome Sanger Institute COVID-19 Surveillance Team |
| EPI_ISL_3310196 | Lighthouse Lab in Milton Keynes | Wellcome Sanger Institute for the COVID-19 Genomics UK (COG-UK) Consortium | The Lighthouse Lab in Milton Keynes and Alex Alderton, Roberto Amato, Jeffrey Barrett, Sonia Goncalves, Ewan Harrison, David K. Jackson, Ian Johnston, Dominic Kwiatkowski, Cordelia Langford, John Sillitoe on behalf of the Wellcome Sanger Institute COVID-19 Surveillance Team |
| EPI_ISL_3310197 | Lighthouse Lab in Alderley Park | Wellcome Sanger Institute for the COVID-19 Genomics UK (COG-UK) Consortium | Jacquelyn Wynn, Mairead Hyland, The Lighthouse Lab in Alderley Park and Alex Alderton, Roberto Amato, Jeffrey Barrett, Sonia Goncalves, Ewan Harrison, David K. Jackson, Ian Johnston, Dominic Kwiatkowski, Cordelia Langford, John Sillitoe on behalf of the Wellcome Sanger Institute COVID-19 Surveillance Team |
| EPI_ISL_3310198, EPI_ISL_3310199, EPI_ISL_3310200 | Lighthouse Lab in Glasgow | Wellcome Sanger Institute for the COVID-19 Genomics UK (COG-UK) Consortium | Harper VanSteenhouse, Yumi Kasai, David Gray, Carol Clugston, Anna Dominiczak and Alex Alderton, Roberto Amato, Jeffrey Barrett, Sonia Goncalves, Ewan Harrison, David K. Jackson, Ian Johnston, Dominic Kwiatkowski, Cordelia Langford, John Sillitoe on behalf of the Wellcome Sanger Institute COVID-19 Surveillance Team |
| EPI_ISL_3310201 | Lighthouse Lab in Milton Keynes | Wellcome Sanger Institute for the COVID-19 Genomics UK (COG-UK) Consortium | The Lighthouse Lab in Milton Keynes and Alex Alderton, Roberto Amato, Jeffrey Barrett, Sonia Goncalves, Ewan Harrison, David K. Jackson, Ian Johnston, Dominic Kwiatkowski, Cordelia Langford, John Sillitoe on behalf of the Wellcome Sanger Institute COVID-19 Surveillance Team |
| EPI_ISL_3310202 | Lighthouse Lab in Alderley Park | Wellcome Sanger Institute for the COVID-19 Genomics UK (COG-UK) Consortium | Jacquelyn Wynn, Mairead Hyland, The Lighthouse Lab in Alderley Park and Alex Alderton, Roberto Amato, Jeffrey Barrett, Sonia Goncalves, Ewan Harrison, David K. Jackson, Ian Johnston, Dominic Kwiatkowski, Cordelia Langford, John Sillitoe on behalf of the Wellcome Sanger Institute COVID-19 Surveillance Team |
| EPI_ISL_3310203, EPI_ISL_3310204 | Lighthouse Lab in Milton Keynes | Wellcome Sanger Institute for the COVID-19 Genomics UK (COG-UK) Consortium | The Lighthouse Lab in Milton Keynes and Alex Alderton, Roberto Amato, Jeffrey Barrett, Sonia Goncalves, Ewan Harrison, David K. Jackson, Ian Johnston, Dominic Kwiatkowski, Cordelia Langford, John Sillitoe on behalf of the Wellcome Sanger Institute COVID-19 Surveillance Team |
| EPI_ISL_3310205 | Lighthouse Lab in Alderley Park | Wellcome Sanger Institute for the COVID-19 Genomics UK (COG-UK) Consortium | Jacquelyn Wynn, Mairead Hyland, The Lighthouse Lab in Alderley Park and Alex Alderton, Roberto Amato, Jeffrey Barrett, Sonia Goncalves, Ewan Harrison, David K. Jackson, Ian Johnston, Dominic Kwiatkowski, Cordelia Langford, John Sillitoe on behalf of the Wellcome Sanger Institute COVID-19 Surveillance Team |
| EPI_ISL_3310206 | Lighthouse Lab in Milton Keynes | Wellcome Sanger Institute for the COVID-19 Genomics UK (COG-UK) Consortium | The Lighthouse Lab in Milton Keynes and Alex Alderton, Roberto Amato, Jeffrey Barrett, Sonia Goncalves, Ewan Harrison, David K. Jackson, Ian Johnston, Dominic Kwiatkowski, Cordelia Langford, John Sillitoe on behalf of the Wellcome Sanger Institute COVID-19 Surveillance Team |
| EPI_ISL_3310207, EPI_ISL_3310208 | Lighthouse Lab in Glasgow | Wellcome Sanger Institute for the COVID-19 Genomics UK (COG-UK) Consortium | Harper VanSteenhouse, Yumi Kasai, David Gray, Carol Clugston, Anna Dominiczak and Alex Alderton, Roberto Amato, Jeffrey Barrett, Sonia Goncalves, Ewan Harrison, David K. Jackson, Ian Johnston, Dominic Kwiatkowski, Cordelia Langford, John Sillitoe on behalf of the Wellcome Sanger Institute COVID-19 Surveillance Team |
| EPI_ISL_3310209, EPI_ISL_3310210 | Lighthouse Lab in Alderley Park | Wellcome Sanger Institute for the COVID-19 Genomics UK (COG-UK) Consortium | Jacquelyn Wynn, Mairead Hyland, The Lighthouse Lab in Alderley Park and Alex Alderton, Roberto Amato, Jeffrey Barrett, Sonia Goncalves, Ewan Harrison, David K. Jackson, Ian Johnston, Dominic Kwiatkowski, Cordelia Langford, John Sillitoe on behalf of the Wellcome Sanger Institute COVID-19 Surveillance Team |
| EPI_ISL_3310211, EPI_ISL_3310212, | Lighthouse Lab in Glasgow | Wellcome Sanger Institute for the COVID-19 Genomics UK (COG-UK) Consortium | Harper VanSteenhouse, Yumi Kasai, David Gray, Carol Clugston, Anna Dominiczak and Alex Alderton, Roberto Amato, Jeffrey Barrett, Sonia Goncalves, Ewan Harrison, David K. Jackson, Ian Johnston, Dominic Kwiatkowski, Cordelia Langford, John Sillitoe on behalf of the Wellcome Sanger Institute COVID-19 Surveillance Team |

| | | | |
|---|---|---|---|
| EPI_ISL_3310213 | | | |
| EPI_ISL_3310214 | Lighthouse Lab in Milton Keynes | Wellcome Sanger Institute for the COVID-19 Genomics UK (COG-UK) Consortium | The Lighthouse Lab in Milton Keynes and Alex Alderton, Roberto Amato, Jeffrey Barrett, Sonia Goncalves, Ewan Harrison, David K. Jackson, Ian Johnston, Dominic Kwiatkowski, Cordelia Langford, John Sillitoe on behalf of the Wellcome Sanger Institute COVID-19 Surveillance Team |
| EPI_ISL_3310215, EPI_ISL_3310216 | Lighthouse Lab in Glasgow | Wellcome Sanger Institute for the COVID-19 Genomics UK (COG-UK) Consortium | Harper VanSteenhouse, Yumi Kasai, David Gray, Carol Clugston, Anna Dominiczak and Alex Alderton, Roberto Amato, Jeffrey Barrett, Sonia Goncalves, Ewan Harrison, David K. Jackson, Ian Johnston, Dominic Kwiatkowski, Cordelia Langford, John Sillitoe on behalf of the Wellcome Sanger Institute COVID-19 Surveillance Team |
| EPI_ISL_3310217, EPI_ISL_3310218 | Lighthouse Lab in Milton Keynes | Wellcome Sanger Institute for the COVID-19 Genomics UK (COG-UK) Consortium | The Lighthouse Lab in Milton Keynes and Alex Alderton, Roberto Amato, Jeffrey Barrett, Sonia Goncalves, Ewan Harrison, David K. Jackson, Ian Johnston, Dominic Kwiatkowski, Cordelia Langford, John Sillitoe on behalf of the Wellcome Sanger Institute COVID-19 Surveillance Team |
| EPI_ISL_3310219 | Lighthouse Lab in Alderley Park | Wellcome Sanger Institute for the COVID-19 Genomics UK (COG-UK) Consortium | Jacquelyn Wynn, Mairead Hyland, The Lighthouse Lab in Alderley Park and Alex Alderton, Roberto Amato, Jeffrey Barrett, Sonia Goncalves, Ewan Harrison, David K. Jackson, Ian Johnston, Dominic Kwiatkowski, Cordelia Langford, John Sillitoe on behalf of the Wellcome Sanger Institute COVID-19 Surveillance Team |
| EPI_ISL_3310220 | Lighthouse Lab in Glasgow | Wellcome Sanger Institute for the COVID-19 Genomics UK (COG-UK) Consortium | Harper VanSteenhouse, Yumi Kasai, David Gray, Carol Clugston, Anna Dominiczak and Alex Alderton, Roberto Amato, Jeffrey Barrett, Sonia Goncalves, Ewan Harrison, David K. Jackson, Ian Johnston, Dominic Kwiatkowski, Cordelia Langford, John Sillitoe on behalf of the Wellcome Sanger Institute COVID-19 Surveillance Team |
| EPI_ISL_3310221, EPI_ISL_3310222, EPI_ISL_3310223 | Lighthouse Lab in Milton Keynes | Wellcome Sanger Institute for the COVID-19 Genomics UK (COG-UK) Consortium | The Lighthouse Lab in Milton Keynes and Alex Alderton, Roberto Amato, Jeffrey Barrett, Sonia Goncalves, Ewan Harrison, David K. Jackson, Ian Johnston, Dominic Kwiatkowski, Cordelia Langford, John Sillitoe on behalf of the Wellcome Sanger Institute COVID-19 Surveillance Team |
| EPI_ISL_3310224, EPI_ISL_3310225, EPI_ISL_3310226 | Lighthouse Lab in Alderley Park | Wellcome Sanger Institute for the COVID-19 Genomics UK (COG-UK) Consortium | Jacquelyn Wynn, Mairead Hyland, The Lighthouse Lab in Alderley Park and Alex Alderton, Roberto Amato, Jeffrey Barrett, Sonia Goncalves, Ewan Harrison, David K. Jackson, Ian Johnston, Dominic Kwiatkowski, Cordelia Langford, John Sillitoe on behalf of the Wellcome Sanger Institute COVID-19 Surveillance Team |
| EPI_ISL_3310227 | Lighthouse Lab in Milton Keynes | Wellcome Sanger Institute for the COVID-19 Genomics UK (COG-UK) Consortium | The Lighthouse Lab in Milton Keynes and Alex Alderton, Roberto Amato, Jeffrey Barrett, Sonia Goncalves, Ewan Harrison, David K. Jackson, Ian Johnston, Dominic Kwiatkowski, Cordelia Langford, John Sillitoe on behalf of the Wellcome Sanger Institute COVID-19 Surveillance Team |
| EPI_ISL_3310228, EPI_ISL_3310229, EPI_ISL_3310230, EPI_ISL_3310231, EPI_ISL_3310232 | Lighthouse Lab in Glasgow | Wellcome Sanger Institute for the COVID-19 Genomics UK (COG-UK) Consortium | Harper VanSteenhouse, Yumi Kasai, David Gray, Carol Clugston, Anna Dominiczak and Alex Alderton, Roberto Amato, Jeffrey Barrett, Sonia Goncalves, Ewan Harrison, David K. Jackson, Ian Johnston, Dominic Kwiatkowski, Cordelia Langford, John Sillitoe on behalf of the Wellcome Sanger Institute COVID-19 Surveillance Team |
| EPI_ISL_3310233, EPI_ISL_3310234 | Lighthouse Lab in Alderley Park | Wellcome Sanger Institute for the COVID-19 Genomics UK (COG-UK) Consortium | Jacquelyn Wynn, Mairead Hyland, The Lighthouse Lab in Alderley Park and Alex Alderton, Roberto Amato, Jeffrey Barrett, Sonia Goncalves, Ewan Harrison, David K. Jackson, Ian Johnston, Dominic Kwiatkowski, Cordelia Langford, John Sillitoe on behalf of the Wellcome Sanger Institute COVID-19 Surveillance Team |
| EPI_ISL_3310235 | Lighthouse Lab in Milton Keynes | Wellcome Sanger Institute for the COVID-19 Genomics UK (COG-UK) Consortium | The Lighthouse Lab in Milton Keynes and Alex Alderton, Roberto Amato, Jeffrey Barrett, Sonia Goncalves, Ewan Harrison, David K. Jackson, Ian Johnston, Dominic Kwiatkowski, Cordelia Langford, John Sillitoe on behalf of the Wellcome Sanger Institute COVID-19 Surveillance Team |
| EPI_ISL_3310236 | Lighthouse Lab in Glasgow | Wellcome Sanger Institute for the COVID-19 Genomics UK (COG-UK) Consortium | Harper VanSteenhouse, Yumi Kasai, David Gray, Carol Clugston, Anna Dominiczak and Alex Alderton, Roberto Amato, Jeffrey Barrett, Sonia Goncalves, Ewan Harrison, David K. Jackson, Ian Johnston, Dominic Kwiatkowski, Cordelia Langford, John Sillitoe on behalf of the Wellcome Sanger Institute COVID-19 Surveillance Team |
| EPI_ISL_3310237 | Lighthouse Lab in Milton Keynes | Wellcome Sanger Institute for the COVID-19 Genomics UK (COG-UK) Consortium | The Lighthouse Lab in Milton Keynes and Alex Alderton, Roberto Amato, Jeffrey Barrett, Sonia Goncalves, Ewan Harrison, David K. Jackson, Ian Johnston, Dominic Kwiatkowski, Cordelia Langford, John Sillitoe on behalf of the Wellcome Sanger Institute COVID-19 Surveillance Team |
| EPI_ISL_3310238 | Lighthouse Lab in Glasgow | Wellcome Sanger Institute for the COVID-19 Genomics UK (COG-UK) Consortium | Harper VanSteenhouse, Yumi Kasai, David Gray, Carol Clugston, Anna Dominiczak and Alex Alderton, Roberto Amato, Jeffrey Barrett, Sonia Goncalves, Ewan Harrison, David K. Jackson, Ian Johnston, Dominic Kwiatkowski, Cordelia Langford, John Sillitoe on behalf of the Wellcome Sanger Institute COVID-19 Surveillance Team |
| EPI_ISL_3310239 | Lighthouse Lab in Alderley Park | Wellcome Sanger Institute for the COVID-19 Genomics UK (COG-UK) Consortium | Jacquelyn Wynn, Mairead Hyland, The Lighthouse Lab in Alderley Park and Alex Alderton, Roberto Amato, Jeffrey Barrett, Sonia Goncalves, Ewan Harrison, David K. Jackson, Ian Johnston, Dominic Kwiatkowski, Cordelia Langford, John Sillitoe on behalf of the Wellcome Sanger Institute COVID-19 Surveillance Team |
| EPI_ISL_3310240, EPI_ISL_3310241, EPI_ISL_3310242 | Lighthouse Lab in Glasgow | Wellcome Sanger Institute for the COVID-19 Genomics UK (COG-UK) Consortium | Harper VanSteenhouse, Yumi Kasai, David Gray, Carol Clugston, Anna Dominiczak and Alex Alderton, Roberto Amato, Jeffrey Barrett, Sonia Goncalves, Ewan Harrison, David K. Jackson, Ian Johnston, Dominic Kwiatkowski, Cordelia Langford, John Sillitoe on behalf of the Wellcome Sanger Institute COVID-19 Surveillance Team |
| EPI_ISL_3310243 | Lighthouse Lab in Milton Keynes | Wellcome Sanger Institute for the COVID-19 Genomics UK (COG-UK) Consortium | The Lighthouse Lab in Milton Keynes and Alex Alderton, Roberto Amato, Jeffrey Barrett, Sonia Goncalves, Ewan Harrison, David K. Jackson, Ian Johnston, Dominic Kwiatkowski, Cordelia Langford, John Sillitoe on behalf of the Wellcome Sanger Institute COVID-19 Surveillance Team |
| EPI_ISL_3310244, EPI_ISL_3310245 | Lighthouse Lab in Glasgow | Wellcome Sanger Institute for the COVID-19 Genomics UK (COG-UK) Consortium | Harper VanSteenhouse, Yumi Kasai, David Gray, Carol Clugston, Anna Dominiczak and Alex Alderton, Roberto Amato, Jeffrey Barrett, Sonia Goncalves, Ewan Harrison, David K. Jackson, Ian Johnston, Dominic Kwiatkowski, Cordelia Langford, John Sillitoe on behalf of the Wellcome Sanger Institute COVID-19 Surveillance Team |
| EPI_ISL_3310246 | Lighthouse Lab in Alderley Park | Wellcome Sanger Institute for the COVID-19 Genomics UK (COG-UK) Consortium | Jacquelyn Wynn, Mairead Hyland, The Lighthouse Lab in Alderley Park and Alex Alderton, Roberto Amato, Jeffrey Barrett, Sonia Goncalves, Ewan Harrison, David K. Jackson, Ian Johnston, Dominic Kwiatkowski, Cordelia Langford, John Sillitoe on behalf of the Wellcome Sanger Institute COVID-19 Surveillance Team |
| EPI_ISL_3310247 | Lighthouse Lab in Milton Keynes | Wellcome Sanger Institute for the COVID-19 Genomics UK (COG-UK) Consortium | The Lighthouse Lab in Milton Keynes and Alex Alderton, Roberto Amato, Jeffrey Barrett, Sonia Goncalves, Ewan Harrison, David K. Jackson, Ian Johnston, Dominic Kwiatkowski, Cordelia Langford, John Sillitoe on behalf of the Wellcome Sanger Institute COVID-19 Surveillance Team |
| EPI_ISL_3310248 | Lighthouse Lab in Glasgow | Wellcome Sanger Institute for the COVID-19 Genomics UK (COG-UK) Consortium | Harper VanSteenhouse, Yumi Kasai, David Gray, Carol Clugston, Anna Dominiczak and Alex Alderton, Roberto Amato, Jeffrey Barrett, Sonia Goncalves, Ewan Harrison, David K. Jackson, Ian Johnston, Dominic Kwiatkowski, Cordelia Langford, John Sillitoe on behalf of the Wellcome Sanger Institute COVID-19 Surveillance Team |
| EPI_ISL_3310249 | Lighthouse Lab in Alderley Park | Wellcome Sanger Institute for the COVID-19 Genomics UK (COG-UK) Consortium | Jacquelyn Wynn, Mairead Hyland, The Lighthouse Lab in Alderley Park and Alex Alderton, Roberto Amato, Jeffrey Barrett, Sonia Goncalves, Ewan Harrison, David K. Jackson, Ian Johnston, Dominic Kwiatkowski, Cordelia Langford, John Sillitoe on behalf of the Wellcome Sanger Institute COVID-19 Surveillance Team |
| EPI_ISL_3310250 | Lighthouse Lab in Glasgow | Wellcome Sanger Institute for the COVID-19 Genomics UK (COG-UK) Consortium | Harper VanSteenhouse, Yumi Kasai, David Gray, Carol Clugston, Anna Dominiczak and Alex Alderton, Roberto Amato, Jeffrey Barrett, Sonia Goncalves, Ewan Harrison, David K. Jackson, Ian Johnston, Dominic Kwiatkowski, Cordelia Langford, John Sillitoe on behalf of the Wellcome Sanger Institute COVID-19 Surveillance Team |
| EPI_ISL_3310251 | Lighthouse Lab in Alderley Park | Wellcome Sanger Institute for the COVID-19 Genomics UK (COG-UK) Consortium | Jacquelyn Wynn, Mairead Hyland, The Lighthouse Lab in Alderley Park and Alex Alderton, Roberto Amato, Jeffrey Barrett, Sonia Goncalves, Ewan Harrison, David K. Jackson, Ian Johnston, Dominic Kwiatkowski, Cordelia Langford, John Sillitoe on behalf of the Wellcome Sanger Institute COVID-19 Surveillance Team |
| EPI_ISL_3310252, EPI_ISL_3310253 | Lighthouse Lab in Glasgow | Wellcome Sanger Institute for the COVID-19 Genomics UK (COG-UK) Consortium | Harper VanSteenhouse, Yumi Kasai, David Gray, Carol Clugston, Anna Dominiczak and Alex Alderton, Roberto Amato, Jeffrey Barrett, Sonia Goncalves, Ewan Harrison, David K. Jackson, Ian Johnston, Dominic Kwiatkowski, Cordelia Langford, John Sillitoe on behalf of the Wellcome Sanger Institute COVID-19 Surveillance Team |
| EPI_ISL_3310254 | Lighthouse Lab in Milton Keynes | Wellcome Sanger Institute for the COVID-19 Genomics UK (COG-UK) Consortium | The Lighthouse Lab in Milton Keynes and Alex Alderton, Roberto Amato, Jeffrey Barrett, Sonia Goncalves, Ewan Harrison, David K. Jackson, Ian Johnston, Dominic Kwiatkowski, Cordelia Langford, John Sillitoe on behalf of the Wellcome Sanger Institute COVID-19 Surveillance Team |
| EPI_ISL_3310255, EPI_ISL_3310256, EPI_ISL_3310257, EPI_ISL_3310258 | Lighthouse Lab in Glasgow | Wellcome Sanger Institute for the COVID-19 Genomics UK (COG-UK) Consortium | Harper VanSteenhouse, Yumi Kasai, David Gray, Carol Clugston, Anna Dominiczak and Alex Alderton, Roberto Amato, Jeffrey Barrett, Sonia Goncalves, Ewan Harrison, David K. Jackson, Ian Johnston, Dominic Kwiatkowski, Cordelia Langford, John Sillitoe on behalf of the Wellcome Sanger Institute COVID-19 Surveillance Team |
| EPI_ISL_3310259, EPI_ISL_3310260 | Lighthouse Lab in Milton Keynes | Wellcome Sanger Institute for the COVID-19 Genomics UK (COG-UK) Consortium | The Lighthouse Lab in Milton Keynes and Alex Alderton, Roberto Amato, Jeffrey Barrett, Sonia Goncalves, Ewan Harrison, David K. Jackson, Ian Johnston, Dominic Kwiatkowski, Cordelia Langford, John Sillitoe on behalf of the Wellcome Sanger Institute COVID-19 Surveillance Team |
| EPI_ISL_3310261 | Lighthouse Lab in Glasgow | Wellcome Sanger Institute for the COVID-19 Genomics UK (COG-UK) Consortium | Harper VanSteenhouse, Yumi Kasai, David Gray, Carol Clugston, Anna Dominiczak and Alex Alderton, Roberto Amato, Jeffrey Barrett, Sonia Goncalves, Ewan Harrison, David K. Jackson, Ian Johnston, Dominic Kwiatkowski, Cordelia Langford, John Sillitoe on behalf of the Wellcome Sanger Institute COVID-19 Surveillance Team |
| EPI_ISL_3310262 | Lighthouse Lab in Milton Keynes | Wellcome Sanger Institute for the COVID-19 Genomics UK (COG-UK) Consortium | The Lighthouse Lab in Milton Keynes and Alex Alderton, Roberto Amato, Jeffrey Barrett, Sonia Goncalves, Ewan Harrison, David K. Jackson, Ian Johnston, Dominic Kwiatkowski, Cordelia Langford, John Sillitoe on behalf of the Wellcome Sanger Institute COVID-19 Surveillance Team |
| EPI_ISL_3310263, EPI_ISL_3310264, EPI_ISL_3310265 | Lighthouse Lab in Glasgow | Wellcome Sanger Institute for the COVID-19 Genomics UK (COG-UK) Consortium | Harper VanSteenhouse, Yumi Kasai, David Gray, Carol Clugston, Anna Dominiczak and Alex Alderton, Roberto Amato, Jeffrey Barrett, Sonia Goncalves, Ewan Harrison, David K. Jackson, Ian Johnston, Dominic Kwiatkowski, Cordelia Langford, John Sillitoe on behalf of the Wellcome Sanger Institute COVID-19 Surveillance Team |
| EPI_ISL_3310266 | Lighthouse Lab in Milton Keynes | Wellcome Sanger Institute for the COVID-19 Genomics UK (COG-UK) Consortium | The Lighthouse Lab in Milton Keynes and Alex Alderton, Roberto Amato, Jeffrey Barrett, Sonia Goncalves, Ewan Harrison, David K. Jackson, Ian Johnston, Dominic Kwiatkowski, Cordelia Langford, John Sillitoe on behalf of the Wellcome Sanger Institute COVID-19 Surveillance Team |
| EPI_ISL_3310267 | Lighthouse Lab in Glasgow | Wellcome Sanger Institute for the COVID-19 Genomics UK (COG-UK) Consortium | Harper VanSteenhouse, Yumi Kasai, David Gray, Carol Clugston, Anna Dominiczak and Alex Alderton, Roberto Amato, Jeffrey Barrett, Sonia Goncalves, Ewan Harrison, David K. Jackson, Ian Johnston, Dominic Kwiatkowski, Cordelia Langford, John Sillitoe on behalf of the Wellcome Sanger Institute COVID-19 Surveillance Team |
| EPI_ISL_3310268 | Lighthouse Lab in Alderley Park | Wellcome Sanger Institute for the COVID-19 Genomics UK (COG-UK) Consortium | Jacquelyn Wynn, Mairead Hyland, The Lighthouse Lab in Alderley Park and Alex Alderton, Roberto Amato, Jeffrey Barrett, Sonia Goncalves, Ewan Harrison, David K. Jackson, Ian Johnston, Dominic Kwiatkowski, Cordelia Langford, John Sillitoe on behalf of the Wellcome Sanger Institute COVID-19 Surveillance Team |
| EPI_ISL_3310269 | Lighthouse Lab in Milton Keynes | Wellcome Sanger Institute for the COVID-19 Genomics UK (COG-UK) Consortium | The Lighthouse Lab in Milton Keynes and Alex Alderton, Roberto Amato, Jeffrey Barrett, Sonia Goncalves, Ewan Harrison, David K. Jackson, Ian Johnston, Dominic Kwiatkowski, Cordelia Langford, John Sillitoe on behalf of the Wellcome Sanger Institute COVID-19 Surveillance Team |
| EPI_ISL_3310270 | Lighthouse Lab in Alderley Park | Wellcome Sanger Institute for the COVID-19 Genomics UK (COG-UK) Consortium | Jacquelyn Wynn, Mairead Hyland, The Lighthouse Lab in Alderley Park and Alex Alderton, Roberto Amato, Jeffrey Barrett, Sonia Goncalves, Ewan Harrison, David K. Jackson, Ian Johnston, Dominic Kwiatkowski, Cordelia Langford, John Sillitoe on behalf of the Wellcome Sanger Institute COVID-19 Surveillance Team |
| EPI_ISL_3310271 | Lighthouse Lab in Milton Keynes | Wellcome Sanger Institute for the COVID-19 Genomics UK (COG-UK) Consortium | The Lighthouse Lab in Milton Keynes and Alex Alderton, Roberto Amato, Jeffrey Barrett, Sonia Goncalves, Ewan Harrison, David K. Jackson, Ian Johnston, Dominic Kwiatkowski, Cordelia Langford, John Sillitoe on behalf of the Wellcome Sanger Institute COVID-19 Surveillance Team |
| EPI_ISL_3310272, EPI_ISL_3310273 | Lighthouse Lab in Glasgow | Wellcome Sanger Institute for the COVID-19 Genomics UK (COG-UK) Consortium | Harper VanSteenhouse, Yumi Kasai, David Gray, Carol Clugston, Anna Dominiczak and Alex Alderton, Roberto Amato, Jeffrey Barrett, Sonia Goncalves, Ewan Harrison, David K. Jackson, Ian Johnston, Dominic Kwiatkowski, Cordelia Langford, John Sillitoe on behalf of the Wellcome Sanger Institute COVID-19 Surveillance Team |
| EPI_ISL_3310274 | Lighthouse Lab in Alderley Park | Wellcome Sanger Institute for the COVID-19 Genomics UK (COG-UK) Consortium | Jacquelyn Wynn, Mairead Hyland, The Lighthouse Lab in Alderley Park and Alex Alderton, Roberto Amato, Jeffrey Barrett, Sonia Goncalves, Ewan Harrison, David K. Jackson, Ian Johnston, Dominic Kwiatkowski, Cordelia Langford, John Sillitoe on behalf of the Wellcome Sanger Institute COVID-19 Surveillance Team |
| EPI_ISL_3310275 | Lighthouse Lab in Glasgow | Wellcome Sanger Institute for the COVID-19 Genomics UK (COG-UK) Consortium | Harper VanSteenhouse, Yumi Kasai, David Gray, Carol Clugston, Anna Dominiczak and Alex Alderton, Roberto Amato, Jeffrey Barrett, Sonia Goncalves, Ewan Harrison, David K. Jackson, Ian Johnston, Dominic Kwiatkowski, Cordelia Langford, John Sillitoe on behalf of the Wellcome Sanger Institute COVID-19 Surveillance Team |
| EPI_ISL_3310276 | Lighthouse Lab in Alderley Park | Wellcome Sanger Institute for the COVID-19 Genomics UK (COG-UK) | Jacquelyn Wynn, Mairead Hyland, The Lighthouse Lab in Alderley Park and Alex Alderton, Roberto Amato, Jeffrey Barrett, Sonia Goncalves, Ewan Harrison, David K. Jackson, Ian Johnston, Dominic Kwiatkowski, Cordelia Langford, John Sillitoe on |

| | | | |
|---|---|---|---|
| | | Consortium | behalf of the Wellcome Sanger Institute COVID-19 Surveillance Team |
| EPI_ISL_3310277 | Lighthouse Lab in Glasgow | Wellcome Sanger Institute for the COVID-19 Genomics UK (COG-UK) Consortium | Harper VanSteenhouse, Yumi Kasai, David Gray, Carol Clugston, Anna Dominiczak and Alex Alderton, Roberto Amato, Jeffrey Barrett, Sonia Goncalves, Ewan Harrison, David K. Jackson, Ian Johnston, Dominic Kwiatkowski, Cordelia Langford, John Sillitoe on behalf of the Wellcome Sanger Institute COVID-19 Surveillance Team |
| EPI_ISL_3310278 | Lighthouse Lab in Alderley Park | Wellcome Sanger Institute for the COVID-19 Genomics UK (COG-UK) Consortium | Jacquelyn Wynn, Mairead Hyland, The Lighthouse Lab in Alderley Park and Alex Alderton, Roberto Amato, Jeffrey Barrett, Sonia Goncalves, Ewan Harrison, David K. Jackson, Ian Johnston, Dominic Kwiatkowski, Cordelia Langford, John Sillitoe on behalf of the Wellcome Sanger Institute COVID-19 Surveillance Team |
| EPI_ISL_3310279 | Lighthouse Lab in Milton Keynes | Wellcome Sanger Institute for the COVID-19 Genomics UK (COG-UK) Consortium | The Lighthouse Lab in Milton Keynes and Alex Alderton, Roberto Amato, Jeffrey Barrett, Sonia Goncalves, Ewan Harrison, David K. Jackson, Ian Johnston, Dominic Kwiatkowski, Cordelia Langford, John Sillitoe on behalf of the Wellcome Sanger Institute COVID-19 Surveillance Team |
| EPI_ISL_3310280 | Lighthouse Lab in Glasgow | Wellcome Sanger Institute for the COVID-19 Genomics UK (COG-UK) Consortium | Harper VanSteenhouse, Yumi Kasai, David Gray, Carol Clugston, Anna Dominiczak and Alex Alderton, Roberto Amato, Jeffrey Barrett, Sonia Goncalves, Ewan Harrison, David K. Jackson, Ian Johnston, Dominic Kwiatkowski, Cordelia Langford, John Sillitoe on behalf of the Wellcome Sanger Institute COVID-19 Surveillance Team |
| EPI_ISL_3310281 | Lighthouse Lab in Alderley Park | Wellcome Sanger Institute for the COVID-19 Genomics UK (COG-UK) Consortium | Jacquelyn Wynn, Mairead Hyland, The Lighthouse Lab in Alderley Park and Alex Alderton, Roberto Amato, Jeffrey Barrett, Sonia Goncalves, Ewan Harrison, David K. Jackson, Ian Johnston, Dominic Kwiatkowski, Cordelia Langford, John Sillitoe on behalf of the Wellcome Sanger Institute COVID-19 Surveillance Team |
| EPI_ISL_3310282 | Lighthouse Lab in Glasgow | Wellcome Sanger Institute for the COVID-19 Genomics UK (COG-UK) Consortium | Harper VanSteenhouse, Yumi Kasai, David Gray, Carol Clugston, Anna Dominiczak and Alex Alderton, Roberto Amato, Jeffrey Barrett, Sonia Goncalves, Ewan Harrison, David K. Jackson, Ian Johnston, Dominic Kwiatkowski, Cordelia Langford, John Sillitoe on behalf of the Wellcome Sanger Institute COVID-19 Surveillance Team |
| EPI_ISL_3310283 | Lighthouse Lab in Milton Keynes | Wellcome Sanger Institute for the COVID-19 Genomics UK (COG-UK) Consortium | The Lighthouse Lab in Milton Keynes and Alex Alderton, Roberto Amato, Jeffrey Barrett, Sonia Goncalves, Ewan Harrison, David K. Jackson, Ian Johnston, Dominic Kwiatkowski, Cordelia Langford, John Sillitoe on behalf of the Wellcome Sanger Institute COVID-19 Surveillance Team |
| EPI_ISL_3310284, EPI_ISL_3310285 | Lighthouse Lab in Glasgow | Wellcome Sanger Institute for the COVID-19 Genomics UK (COG-UK) Consortium | Harper VanSteenhouse, Yumi Kasai, David Gray, Carol Clugston, Anna Dominiczak and Alex Alderton, Roberto Amato, Jeffrey Barrett, Sonia Goncalves, Ewan Harrison, David K. Jackson, Ian Johnston, Dominic Kwiatkowski, Cordelia Langford, John Sillitoe on behalf of the Wellcome Sanger Institute COVID-19 Surveillance Team |
| EPI_ISL_3310286 | Lighthouse Lab in Milton Keynes | Wellcome Sanger Institute for the COVID-19 Genomics UK (COG-UK) Consortium | The Lighthouse Lab in Milton Keynes and Alex Alderton, Roberto Amato, Jeffrey Barrett, Sonia Goncalves, Ewan Harrison, David K. Jackson, Ian Johnston, Dominic Kwiatkowski, Cordelia Langford, John Sillitoe on behalf of the Wellcome Sanger Institute COVID-19 Surveillance Team |
| EPI_ISL_3310287 | Lighthouse Lab in Alderley Park | Wellcome Sanger Institute for the COVID-19 Genomics UK (COG-UK) Consortium | Jacquelyn Wynn, Mairead Hyland, The Lighthouse Lab in Alderley Park and Alex Alderton, Roberto Amato, Jeffrey Barrett, Sonia Goncalves, Ewan Harrison, David K. Jackson, Ian Johnston, Dominic Kwiatkowski, Cordelia Langford, John Sillitoe on behalf of the Wellcome Sanger Institute COVID-19 Surveillance Team |
| EPI_ISL_3310288 | Lighthouse Lab in Milton Keynes | Wellcome Sanger Institute for the COVID-19 Genomics UK (COG-UK) Consortium | The Lighthouse Lab in Milton Keynes and Alex Alderton, Roberto Amato, Jeffrey Barrett, Sonia Goncalves, Ewan Harrison, David K. Jackson, Ian Johnston, Dominic Kwiatkowski, Cordelia Langford, John Sillitoe on behalf of the Wellcome Sanger Institute COVID-19 Surveillance Team |
| EPI_ISL_3310289 | Lighthouse Lab in Glasgow | Wellcome Sanger Institute for the COVID-19 Genomics UK (COG-UK) Consortium | Harper VanSteenhouse, Yumi Kasai, David Gray, Carol Clugston, Anna Dominiczak and Alex Alderton, Roberto Amato, Jeffrey Barrett, Sonia Goncalves, Ewan Harrison, David K. Jackson, Ian Johnston, Dominic Kwiatkowski, Cordelia Langford, John Sillitoe on behalf of the Wellcome Sanger Institute COVID-19 Surveillance Team |
| EPI_ISL_3310290 | Lighthouse Lab in Alderley Park | Wellcome Sanger Institute for the COVID-19 Genomics UK (COG-UK) Consortium | Jacquelyn Wynn, Mairead Hyland, The Lighthouse Lab in Alderley Park and Alex Alderton, Roberto Amato, Jeffrey Barrett, Sonia Goncalves, Ewan Harrison, David K. Jackson, Ian Johnston, Dominic Kwiatkowski, Cordelia Langford, John Sillitoe on behalf of the Wellcome Sanger Institute COVID-19 Surveillance Team |
| EPI_ISL_3310291 | Lighthouse Lab in Glasgow | Wellcome Sanger Institute for the COVID-19 Genomics UK (COG-UK) Consortium | Harper VanSteenhouse, Yumi Kasai, David Gray, Carol Clugston, Anna Dominiczak and Alex Alderton, Roberto Amato, Jeffrey Barrett, Sonia Goncalves, Ewan Harrison, David K. Jackson, Ian Johnston, Dominic Kwiatkowski, Cordelia Langford, John Sillitoe on behalf of the Wellcome Sanger Institute COVID-19 Surveillance Team |
| EPI_ISL_3310292 | Lighthouse Lab in Milton Keynes | Wellcome Sanger Institute for the COVID-19 Genomics UK (COG-UK) Consortium | The Lighthouse Lab in Milton Keynes and Alex Alderton, Roberto Amato, Jeffrey Barrett, Sonia Goncalves, Ewan Harrison, David K. Jackson, Ian Johnston, Dominic Kwiatkowski, Cordelia Langford, John Sillitoe on behalf of the Wellcome Sanger Institute COVID-19 Surveillance Team |
| EPI_ISL_3310293 | Lighthouse Lab in Alderley Park | Wellcome Sanger Institute for the COVID-19 Genomics UK (COG-UK) Consortium | Jacquelyn Wynn, Mairead Hyland, The Lighthouse Lab in Alderley Park and Alex Alderton, Roberto Amato, Jeffrey Barrett, Sonia Goncalves, Ewan Harrison, David K. Jackson, Ian Johnston, Dominic Kwiatkowski, Cordelia Langford, John Sillitoe on behalf of the Wellcome Sanger Institute COVID-19 Surveillance Team |
| EPI_ISL_3310294 | Lighthouse Lab in Glasgow | Wellcome Sanger Institute for the COVID-19 Genomics UK (COG-UK) Consortium | Harper VanSteenhouse, Yumi Kasai, David Gray, Carol Clugston, Anna Dominiczak and Alex Alderton, Roberto Amato, Jeffrey Barrett, Sonia Goncalves, Ewan Harrison, David K. Jackson, Ian Johnston, Dominic Kwiatkowski, Cordelia Langford, John Sillitoe on behalf of the Wellcome Sanger Institute COVID-19 Surveillance Team |
| EPI_ISL_3310295 | Lighthouse Lab in Alderley Park | Wellcome Sanger Institute for the COVID-19 Genomics UK (COG-UK) Consortium | Jacquelyn Wynn, Mairead Hyland, The Lighthouse Lab in Alderley Park and Alex Alderton, Roberto Amato, Jeffrey Barrett, Sonia Goncalves, Ewan Harrison, David K. Jackson, Ian Johnston, Dominic Kwiatkowski, Cordelia Langford, John Sillitoe on behalf of the Wellcome Sanger Institute COVID-19 Surveillance Team |
| EPI_ISL_3310296 | Lighthouse Lab in Milton Keynes | Wellcome Sanger Institute for the COVID-19 Genomics UK (COG-UK) Consortium | The Lighthouse Lab in Milton Keynes and Alex Alderton, Roberto Amato, Jeffrey Barrett, Sonia Goncalves, Ewan Harrison, David K. Jackson, Ian Johnston, Dominic Kwiatkowski, Cordelia Langford, John Sillitoe on behalf of the Wellcome Sanger Institute COVID-19 Surveillance Team |
| EPI_ISL_3310297 | Lighthouse Lab in Alderley Park | Wellcome Sanger Institute for the COVID-19 Genomics UK (COG-UK) Consortium | Jacquelyn Wynn, Mairead Hyland, The Lighthouse Lab in Alderley Park and Alex Alderton, Roberto Amato, Jeffrey Barrett, Sonia Goncalves, Ewan Harrison, David K. Jackson, Ian Johnston, Dominic Kwiatkowski, Cordelia Langford, John Sillitoe on behalf of the Wellcome Sanger Institute COVID-19 Surveillance Team |
| EPI_ISL_3310298 | Lighthouse Lab in Milton Keynes | Wellcome Sanger Institute for the COVID-19 Genomics UK (COG-UK) Consortium | The Lighthouse Lab in Milton Keynes and Alex Alderton, Roberto Amato, Jeffrey Barrett, Sonia Goncalves, Ewan Harrison, David K. Jackson, Ian Johnston, Dominic Kwiatkowski, Cordelia Langford, John Sillitoe on behalf of the Wellcome Sanger Institute COVID-19 Surveillance Team |
| EPI_ISL_3310299 | Lighthouse Lab in Alderley Park | Wellcome Sanger Institute for the COVID-19 Genomics UK (COG-UK) Consortium | Jacquelyn Wynn, Mairead Hyland, The Lighthouse Lab in Alderley Park and Alex Alderton, Roberto Amato, Jeffrey Barrett, Sonia Goncalves, Ewan Harrison, David K. Jackson, Ian Johnston, Dominic Kwiatkowski, Cordelia Langford, John Sillitoe on behalf of the Wellcome Sanger Institute COVID-19 Surveillance Team |
| EPI_ISL_3310300, EPI_ISL_3310301 | Lighthouse Lab in Glasgow | Wellcome Sanger Institute for the COVID-19 Genomics UK (COG-UK) Consortium | Harper VanSteenhouse, Yumi Kasai, David Gray, Carol Clugston, Anna Dominiczak and Alex Alderton, Roberto Amato, Jeffrey Barrett, Sonia Goncalves, Ewan Harrison, David K. Jackson, Ian Johnston, Dominic Kwiatkowski, Cordelia Langford, John Sillitoe on behalf of the Wellcome Sanger Institute COVID-19 Surveillance Team |
| EPI_ISL_3310302 | Lighthouse Lab in Alderley Park | Wellcome Sanger Institute for the COVID-19 Genomics UK (COG-UK) Consortium | Jacquelyn Wynn, Mairead Hyland, The Lighthouse Lab in Alderley Park and Alex Alderton, Roberto Amato, Jeffrey Barrett, Sonia Goncalves, Ewan Harrison, David K. Jackson, Ian Johnston, Dominic Kwiatkowski, Cordelia Langford, John Sillitoe on behalf of the Wellcome Sanger Institute COVID-19 Surveillance Team |
| EPI_ISL_3310303, EPI_ISL_3310304, EPI_ISL_3310305, EPI_ISL_3310306, EPI_ISL_3310307 | Lighthouse Lab in Glasgow | Wellcome Sanger Institute for the COVID-19 Genomics UK (COG-UK) Consortium | Harper VanSteenhouse, Yumi Kasai, David Gray, Carol Clugston, Anna Dominiczak and Alex Alderton, Roberto Amato, Jeffrey Barrett, Sonia Goncalves, Ewan Harrison, David K. Jackson, Ian Johnston, Dominic Kwiatkowski, Cordelia Langford, John Sillitoe on behalf of the Wellcome Sanger Institute COVID-19 Surveillance Team |
| EPI_ISL_3310308 | Lighthouse Lab in Milton Keynes | Wellcome Sanger Institute for the COVID-19 Genomics UK (COG-UK) Consortium | The Lighthouse Lab in Milton Keynes and Alex Alderton, Roberto Amato, Jeffrey Barrett, Sonia Goncalves, Ewan Harrison, David K. Jackson, Ian Johnston, Dominic Kwiatkowski, Cordelia Langford, John Sillitoe on behalf of the Wellcome Sanger Institute COVID-19 Surveillance Team |
| EPI_ISL_3310309 | Lighthouse Lab in Glasgow | Wellcome Sanger Institute for the COVID-19 Genomics UK (COG-UK) Consortium | Harper VanSteenhouse, Yumi Kasai, David Gray, Carol Clugston, Anna Dominiczak and Alex Alderton, Roberto Amato, Jeffrey Barrett, Sonia Goncalves, Ewan Harrison, David K. Jackson, Ian Johnston, Dominic Kwiatkowski, Cordelia Langford, John Sillitoe on behalf of the Wellcome Sanger Institute COVID-19 Surveillance Team |
| EPI_ISL_3310310 | Lighthouse Lab in Alderley Park | Wellcome Sanger Institute for the COVID-19 Genomics UK (COG-UK) Consortium | Jacquelyn Wynn, Mairead Hyland, The Lighthouse Lab in Alderley Park and Alex Alderton, Roberto Amato, Jeffrey Barrett, Sonia Goncalves, Ewan Harrison, David K. Jackson, Ian Johnston, Dominic Kwiatkowski, Cordelia Langford, John Sillitoe on behalf of the Wellcome Sanger Institute COVID-19 Surveillance Team |
| EPI_ISL_3310311 | Lighthouse Lab in Milton Keynes | Wellcome Sanger Institute for the COVID-19 Genomics UK (COG-UK) Consortium | The Lighthouse Lab in Milton Keynes and Alex Alderton, Roberto Amato, Jeffrey Barrett, Sonia Goncalves, Ewan Harrison, David K. Jackson, Ian Johnston, Dominic Kwiatkowski, Cordelia Langford, John Sillitoe on behalf of the Wellcome Sanger Institute COVID-19 Surveillance Team |
| EPI_ISL_3310312 | Lighthouse Lab in Alderley Park | Wellcome Sanger Institute for the COVID-19 Genomics UK (COG-UK) Consortium | Jacquelyn Wynn, Mairead Hyland, The Lighthouse Lab in Alderley Park and Alex Alderton, Roberto Amato, Jeffrey Barrett, Sonia Goncalves, Ewan Harrison, David K. Jackson, Ian Johnston, Dominic Kwiatkowski, Cordelia Langford, John Sillitoe on behalf of the Wellcome Sanger Institute COVID-19 Surveillance Team |
| EPI_ISL_3310313 | Lighthouse Lab in Glasgow | Wellcome Sanger Institute for the COVID-19 Genomics UK (COG-UK) Consortium | Harper VanSteenhouse, Yumi Kasai, David Gray, Carol Clugston, Anna Dominiczak and Alex Alderton, Roberto Amato, Jeffrey Barrett, Sonia Goncalves, Ewan Harrison, David K. Jackson, Ian Johnston, Dominic Kwiatkowski, Cordelia Langford, John Sillitoe on behalf of the Wellcome Sanger Institute COVID-19 Surveillance Team |
| EPI_ISL_3310314 | Lighthouse Lab in Milton Keynes | Wellcome Sanger Institute for the COVID-19 Genomics UK (COG-UK) Consortium | The Lighthouse Lab in Milton Keynes and Alex Alderton, Roberto Amato, Jeffrey Barrett, Sonia Goncalves, Ewan Harrison, David K. Jackson, Ian Johnston, Dominic Kwiatkowski, Cordelia Langford, John Sillitoe on behalf of the Wellcome Sanger Institute COVID-19 Surveillance Team |
| EPI_ISL_3310315 | Lighthouse Lab in Glasgow | Wellcome Sanger Institute for the COVID-19 Genomics UK (COG-UK) Consortium | Harper VanSteenhouse, Yumi Kasai, David Gray, Carol Clugston, Anna Dominiczak and Alex Alderton, Roberto Amato, Jeffrey Barrett, Sonia Goncalves, Ewan Harrison, David K. Jackson, Ian Johnston, Dominic Kwiatkowski, Cordelia Langford, John Sillitoe on behalf of the Wellcome Sanger Institute COVID-19 Surveillance Team |
| EPI_ISL_3310316 | Lighthouse Lab in Milton Keynes | Wellcome Sanger Institute for the COVID-19 Genomics UK (COG-UK) Consortium | The Lighthouse Lab in Milton Keynes and Alex Alderton, Roberto Amato, Jeffrey Barrett, Sonia Goncalves, Ewan Harrison, David K. Jackson, Ian Johnston, Dominic Kwiatkowski, Cordelia Langford, John Sillitoe on behalf of the Wellcome Sanger Institute COVID-19 Surveillance Team |
| EPI_ISL_3310317, EPI_ISL_3310318 | Lighthouse Lab in Glasgow | Wellcome Sanger Institute for the COVID-19 Genomics UK (COG-UK) Consortium | Harper VanSteenhouse, Yumi Kasai, David Gray, Carol Clugston, Anna Dominiczak and Alex Alderton, Roberto Amato, Jeffrey Barrett, Sonia Goncalves, Ewan Harrison, David K. Jackson, Ian Johnston, Dominic Kwiatkowski, Cordelia Langford, John Sillitoe on behalf of the Wellcome Sanger Institute COVID-19 Surveillance Team |
| EPI_ISL_3310319 | Lighthouse Lab in Milton Keynes | Wellcome Sanger Institute for the COVID-19 Genomics UK (COG-UK) Consortium | The Lighthouse Lab in Milton Keynes and Alex Alderton, Roberto Amato, Jeffrey Barrett, Sonia Goncalves, Ewan Harrison, David K. Jackson, Ian Johnston, Dominic Kwiatkowski, Cordelia Langford, John Sillitoe on behalf of the Wellcome Sanger Institute COVID-19 Surveillance Team |
| EPI_ISL_3310320 | Lighthouse Lab in Alderley Park | Wellcome Sanger Institute for the COVID-19 Genomics UK (COG-UK) Consortium | Jacquelyn Wynn, Mairead Hyland, The Lighthouse Lab in Alderley Park and Alex Alderton, Roberto Amato, Jeffrey Barrett, Sonia Goncalves, Ewan Harrison, David K. Jackson, Ian Johnston, Dominic Kwiatkowski, Cordelia Langford, John Sillitoe on behalf of the Wellcome Sanger Institute COVID-19 Surveillance Team |
| EPI_ISL_3310321 | Lighthouse Lab in Milton Keynes | Wellcome Sanger Institute for the COVID-19 Genomics UK (COG-UK) Consortium | The Lighthouse Lab in Milton Keynes and Alex Alderton, Roberto Amato, Jeffrey Barrett, Sonia Goncalves, Ewan Harrison, David K. Jackson, Ian Johnston, Dominic Kwiatkowski, Cordelia Langford, John Sillitoe on behalf of the Wellcome Sanger Institute COVID-19 Surveillance Team |
| EPI_ISL_3310322 | Lighthouse Lab in Alderley Park | Wellcome Sanger Institute for the COVID-19 Genomics UK (COG-UK) Consortium | Jacquelyn Wynn, Mairead Hyland, The Lighthouse Lab in Alderley Park and Alex Alderton, Roberto Amato, Jeffrey Barrett, Sonia Goncalves, Ewan Harrison, David K. Jackson, Ian Johnston, Dominic Kwiatkowski, Cordelia Langford, John Sillitoe on behalf of the Wellcome Sanger Institute COVID-19 Surveillance Team |
| EPI_ISL_3310323, EPI_ISL_3310324 | Lighthouse Lab in Milton Keynes | Wellcome Sanger Institute for the COVID-19 Genomics UK (COG-UK) Consortium | The Lighthouse Lab in Milton Keynes and Alex Alderton, Roberto Amato, Jeffrey Barrett, Sonia Goncalves, Ewan Harrison, David K. Jackson, Ian Johnston, Dominic Kwiatkowski, Cordelia Langford, John Sillitoe on behalf of the Wellcome Sanger Institute COVID-19 Surveillance Team |
| EPI_ISL_3310325, EPI_ISL_3310326, EPI_ISL_3310327, EPI_ISL_3310328, EPI_ISL_3310329, EPI_ISL_3310330, EPI_ISL_3310331 | Lighthouse Lab in Glasgow | Wellcome Sanger Institute for the COVID-19 Genomics UK (COG-UK) Consortium | Harper VanSteenhouse, Yumi Kasai, David Gray, Carol Clugston, Anna Dominiczak and Alex Alderton, Roberto Amato, Jeffrey Barrett, Sonia Goncalves, Ewan Harrison, David K. Jackson, Ian Johnston, Dominic Kwiatkowski, Cordelia Langford, John Sillitoe on behalf of the Wellcome Sanger Institute COVID-19 Surveillance Team |

| | | | |
|---|---|---|---|
| EPI_ISL_3310332 | Lighthouse Lab in Alderley Park | Wellcome Sanger Institute for the COVID-19 Genomics UK (COG-UK) Consortium | Jacquelyn Wynn, Mairead Hyland, The Lighthouse Lab in Alderley Park and Alex Alderton, Roberto Amato, Jeffrey Barrett, Sonia Goncalves, Ewan Harrison, David K. Jackson, Ian Johnston, Dominic Kwiatkowski, Cordelia Langford, John Sillitoe on behalf of the Wellcome Sanger Institute COVID-19 Surveillance Team |
| EPI_ISL_3310333 | Lighthouse Lab in Milton Keynes | Wellcome Sanger Institute for the COVID-19 Genomics UK (COG-UK) Consortium | The Lighthouse Lab in Milton Keynes and Alex Alderton, Roberto Amato, Jeffrey Barrett, Sonia Goncalves, Ewan Harrison, David K. Jackson, Ian Johnston, Dominic Kwiatkowski, Cordelia Langford, John Sillitoe on behalf of the Wellcome Sanger Institute COVID-19 Surveillance Team |
| EPI_ISL_3310334 | Lighthouse Lab in Glasgow | Wellcome Sanger Institute for the COVID-19 Genomics UK (COG-UK) Consortium | Harper VanSteenhouse, Yumi Kasai, David Gray, Carol Clugston, Anna Dominiczak and Alex Alderton, Roberto Amato, Jeffrey Barrett, Sonia Goncalves, Ewan Harrison, David K. Jackson, Ian Johnston, Dominic Kwiatkowski, Cordelia Langford, John Sillitoe on behalf of the Wellcome Sanger Institute COVID-19 Surveillance Team |
| EPI_ISL_3310335 | Lighthouse Lab in Alderley Park | Wellcome Sanger Institute for the COVID-19 Genomics UK (COG-UK) Consortium | Jacquelyn Wynn, Mairead Hyland, The Lighthouse Lab in Alderley Park and Alex Alderton, Roberto Amato, Jeffrey Barrett, Sonia Goncalves, Ewan Harrison, David K. Jackson, Ian Johnston, Dominic Kwiatkowski, Cordelia Langford, John Sillitoe on behalf of the Wellcome Sanger Institute COVID-19 Surveillance Team |
| EPI_ISL_3310336 | Lighthouse Lab in Milton Keynes | Wellcome Sanger Institute for the COVID-19 Genomics UK (COG-UK) Consortium | The Lighthouse Lab in Milton Keynes and Alex Alderton, Roberto Amato, Jeffrey Barrett, Sonia Goncalves, Ewan Harrison, David K. Jackson, Ian Johnston, Dominic Kwiatkowski, Cordelia Langford, John Sillitoe on behalf of the Wellcome Sanger Institute COVID-19 Surveillance Team |
| EPI_ISL_3310337 | Lighthouse Lab in Alderley Park | Wellcome Sanger Institute for the COVID-19 Genomics UK (COG-UK) Consortium | Jacquelyn Wynn, Mairead Hyland, The Lighthouse Lab in Alderley Park and Alex Alderton, Roberto Amato, Jeffrey Barrett, Sonia Goncalves, Ewan Harrison, David K. Jackson, Ian Johnston, Dominic Kwiatkowski, Cordelia Langford, John Sillitoe on behalf of the Wellcome Sanger Institute COVID-19 Surveillance Team |
| EPI_ISL_3310338, EPI_ISL_3310339, EPI_ISL_3310340, EPI_ISL_3310341 | Lighthouse Lab in Glasgow | Wellcome Sanger Institute for the COVID-19 Genomics UK (COG-UK) Consortium | Harper VanSteenhouse, Yumi Kasai, David Gray, Carol Clugston, Anna Dominiczak and Alex Alderton, Roberto Amato, Jeffrey Barrett, Sonia Goncalves, Ewan Harrison, David K. Jackson, Ian Johnston, Dominic Kwiatkowski, Cordelia Langford, John Sillitoe on behalf of the Wellcome Sanger Institute COVID-19 Surveillance Team |
| EPI_ISL_3310342, EPI_ISL_3310343 | Lighthouse Lab in Alderley Park | Wellcome Sanger Institute for the COVID-19 Genomics UK (COG-UK) Consortium | Jacquelyn Wynn, Mairead Hyland, The Lighthouse Lab in Alderley Park and Alex Alderton, Roberto Amato, Jeffrey Barrett, Sonia Goncalves, Ewan Harrison, David K. Jackson, Ian Johnston, Dominic Kwiatkowski, Cordelia Langford, John Sillitoe on behalf of the Wellcome Sanger Institute COVID-19 Surveillance Team |
| EPI_ISL_3310344 | Lighthouse Lab in Glasgow | Wellcome Sanger Institute for the COVID-19 Genomics UK (COG-UK) Consortium | Harper VanSteenhouse, Yumi Kasai, David Gray, Carol Clugston, Anna Dominiczak and Alex Alderton, Roberto Amato, Jeffrey Barrett, Sonia Goncalves, Ewan Harrison, David K. Jackson, Ian Johnston, Dominic Kwiatkowski, Cordelia Langford, John Sillitoe on behalf of the Wellcome Sanger Institute COVID-19 Surveillance Team |
| EPI_ISL_3310345 | Lighthouse Lab in Alderley Park | Wellcome Sanger Institute for the COVID-19 Genomics UK (COG-UK) Consortium | Jacquelyn Wynn, Mairead Hyland, The Lighthouse Lab in Alderley Park and Alex Alderton, Roberto Amato, Jeffrey Barrett, Sonia Goncalves, Ewan Harrison, David K. Jackson, Ian Johnston, Dominic Kwiatkowski, Cordelia Langford, John Sillitoe on behalf of the Wellcome Sanger Institute COVID-19 Surveillance Team |
| EPI_ISL_3310346, EPI_ISL_3310347 | Lighthouse Lab in Glasgow | Wellcome Sanger Institute for the COVID-19 Genomics UK (COG-UK) Consortium | Harper VanSteenhouse, Yumi Kasai, David Gray, Carol Clugston, Anna Dominiczak and Alex Alderton, Roberto Amato, Jeffrey Barrett, Sonia Goncalves, Ewan Harrison, David K. Jackson, Ian Johnston, Dominic Kwiatkowski, Cordelia Langford, John Sillitoe on behalf of the Wellcome Sanger Institute COVID-19 Surveillance Team |
| EPI_ISL_3310348, EPI_ISL_3310349 | Lighthouse Lab in Alderley Park | Wellcome Sanger Institute for the COVID-19 Genomics UK (COG-UK) Consortium | Jacquelyn Wynn, Mairead Hyland, The Lighthouse Lab in Alderley Park and Alex Alderton, Roberto Amato, Jeffrey Barrett, Sonia Goncalves, Ewan Harrison, David K. Jackson, Ian Johnston, Dominic Kwiatkowski, Cordelia Langford, John Sillitoe on behalf of the Wellcome Sanger Institute COVID-19 Surveillance Team |
| EPI_ISL_3310350 | Lighthouse Lab in Milton Keynes | Wellcome Sanger Institute for the COVID-19 Genomics UK (COG-UK) Consortium | The Lighthouse Lab in Milton Keynes and Alex Alderton, Roberto Amato, Jeffrey Barrett, Sonia Goncalves, Ewan Harrison, David K. Jackson, Ian Johnston, Dominic Kwiatkowski, Cordelia Langford, John Sillitoe on behalf of the Wellcome Sanger Institute COVID-19 Surveillance Team |
| EPI_ISL_3310351, EPI_ISL_3310352 | Lighthouse Lab in Glasgow | Wellcome Sanger Institute for the COVID-19 Genomics UK (COG-UK) Consortium | Harper VanSteenhouse, Yumi Kasai, David Gray, Carol Clugston, Anna Dominiczak and Alex Alderton, Roberto Amato, Jeffrey Barrett, Sonia Goncalves, Ewan Harrison, David K. Jackson, Ian Johnston, Dominic Kwiatkowski, Cordelia Langford, John Sillitoe on behalf of the Wellcome Sanger Institute COVID-19 Surveillance Team |
| EPI_ISL_3310353 | Lighthouse Lab in Milton Keynes | Wellcome Sanger Institute for the COVID-19 Genomics UK (COG-UK) Consortium | The Lighthouse Lab in Milton Keynes and Alex Alderton, Roberto Amato, Jeffrey Barrett, Sonia Goncalves, Ewan Harrison, David K. Jackson, Ian Johnston, Dominic Kwiatkowski, Cordelia Langford, John Sillitoe on behalf of the Wellcome Sanger Institute COVID-19 Surveillance Team |
| EPI_ISL_3310354 | Lighthouse Lab in Alderley Park | Wellcome Sanger Institute for the COVID-19 Genomics UK (COG-UK) Consortium | Jacquelyn Wynn, Mairead Hyland, The Lighthouse Lab in Alderley Park and Alex Alderton, Roberto Amato, Jeffrey Barrett, Sonia Goncalves, Ewan Harrison, David K. Jackson, Ian Johnston, Dominic Kwiatkowski, Cordelia Langford, John Sillitoe on behalf of the Wellcome Sanger Institute COVID-19 Surveillance Team |
| EPI_ISL_3310355 | Lighthouse Lab in Glasgow | Wellcome Sanger Institute for the COVID-19 Genomics UK (COG-UK) Consortium | Harper VanSteenhouse, Yumi Kasai, David Gray, Carol Clugston, Anna Dominiczak and Alex Alderton, Roberto Amato, Jeffrey Barrett, Sonia Goncalves, Ewan Harrison, David K. Jackson, Ian Johnston, Dominic Kwiatkowski, Cordelia Langford, John Sillitoe on behalf of the Wellcome Sanger Institute COVID-19 Surveillance Team |
| EPI_ISL_3310356 | Lighthouse Lab in Alderley Park | Wellcome Sanger Institute for the COVID-19 Genomics UK (COG-UK) Consortium | Jacquelyn Wynn, Mairead Hyland, The Lighthouse Lab in Alderley Park and Alex Alderton, Roberto Amato, Jeffrey Barrett, Sonia Goncalves, Ewan Harrison, David K. Jackson, Ian Johnston, Dominic Kwiatkowski, Cordelia Langford, John Sillitoe on behalf of the Wellcome Sanger Institute COVID-19 Surveillance Team |
| EPI_ISL_3310357, EPI_ISL_3310358, EPI_ISL_3310359 | Lighthouse Lab in Glasgow | Wellcome Sanger Institute for the COVID-19 Genomics UK (COG-UK) Consortium | Harper VanSteenhouse, Yumi Kasai, David Gray, Carol Clugston, Anna Dominiczak and Alex Alderton, Roberto Amato, Jeffrey Barrett, Sonia Goncalves, Ewan Harrison, David K. Jackson, Ian Johnston, Dominic Kwiatkowski, Cordelia Langford, John Sillitoe on behalf of the Wellcome Sanger Institute COVID-19 Surveillance Team |
| EPI_ISL_3310360 | Lighthouse Lab in Alderley Park | Wellcome Sanger Institute for the COVID-19 Genomics UK (COG-UK) Consortium | Jacquelyn Wynn, Mairead Hyland, The Lighthouse Lab in Alderley Park and Alex Alderton, Roberto Amato, Jeffrey Barrett, Sonia Goncalves, Ewan Harrison, David K. Jackson, Ian Johnston, Dominic Kwiatkowski, Cordelia Langford, John Sillitoe on behalf of the Wellcome Sanger Institute COVID-19 Surveillance Team |
| EPI_ISL_3310361 | Lighthouse Lab in Glasgow | Wellcome Sanger Institute for the COVID-19 Genomics UK (COG-UK) Consortium | Harper VanSteenhouse, Yumi Kasai, David Gray, Carol Clugston, Anna Dominiczak and Alex Alderton, Roberto Amato, Jeffrey Barrett, Sonia Goncalves, Ewan Harrison, David K. Jackson, Ian Johnston, Dominic Kwiatkowski, Cordelia Langford, John Sillitoe on behalf of the Wellcome Sanger Institute COVID-19 Surveillance Team |
| EPI_ISL_3310362, EPI_ISL_3310363, EPI_ISL_3310364 | Lighthouse Lab in Alderley Park | Wellcome Sanger Institute for the COVID-19 Genomics UK (COG-UK) Consortium | Jacquelyn Wynn, Mairead Hyland, The Lighthouse Lab in Alderley Park and Alex Alderton, Roberto Amato, Jeffrey Barrett, Sonia Goncalves, Ewan Harrison, David K. Jackson, Ian Johnston, Dominic Kwiatkowski, Cordelia Langford, John Sillitoe on behalf of the Wellcome Sanger Institute COVID-19 Surveillance Team |
| EPI_ISL_3310365 | Lighthouse Lab in Glasgow | Wellcome Sanger Institute for the COVID-19 Genomics UK (COG-UK) Consortium | Harper VanSteenhouse, Yumi Kasai, David Gray, Carol Clugston, Anna Dominiczak and Alex Alderton, Roberto Amato, Jeffrey Barrett, Sonia Goncalves, Ewan Harrison, David K. Jackson, Ian Johnston, Dominic Kwiatkowski, Cordelia Langford, John Sillitoe on behalf of the Wellcome Sanger Institute COVID-19 Surveillance Team |
| EPI_ISL_3310366 | Lighthouse Lab in Alderley Park | Wellcome Sanger Institute for the COVID-19 Genomics UK (COG-UK) Consortium | Jacquelyn Wynn, Mairead Hyland, The Lighthouse Lab in Alderley Park and Alex Alderton, Roberto Amato, Jeffrey Barrett, Sonia Goncalves, Ewan Harrison, David K. Jackson, Ian Johnston, Dominic Kwiatkowski, Cordelia Langford, John Sillitoe on behalf of the Wellcome Sanger Institute COVID-19 Surveillance Team |
| EPI_ISL_3310367, EPI_ISL_3310368, EPI_ISL_3310369 | Lighthouse Lab in Glasgow | Wellcome Sanger Institute for the COVID-19 Genomics UK (COG-UK) Consortium | Harper VanSteenhouse, Yumi Kasai, David Gray, Carol Clugston, Anna Dominiczak and Alex Alderton, Roberto Amato, Jeffrey Barrett, Sonia Goncalves, Ewan Harrison, David K. Jackson, Ian Johnston, Dominic Kwiatkowski, Cordelia Langford, John Sillitoe on behalf of the Wellcome Sanger Institute COVID-19 Surveillance Team |
| EPI_ISL_3310370 | Lighthouse Lab in Alderley Park | Wellcome Sanger Institute for the COVID-19 Genomics UK (COG-UK) Consortium | Jacquelyn Wynn, Mairead Hyland, The Lighthouse Lab in Alderley Park and Alex Alderton, Roberto Amato, Jeffrey Barrett, Sonia Goncalves, Ewan Harrison, David K. Jackson, Ian Johnston, Dominic Kwiatkowski, Cordelia Langford, John Sillitoe on behalf of the Wellcome Sanger Institute COVID-19 Surveillance Team |
| EPI_ISL_3310371 | Lighthouse Lab in Milton Keynes | Wellcome Sanger Institute for the COVID-19 Genomics UK (COG-UK) Consortium | The Lighthouse Lab in Milton Keynes and Alex Alderton, Roberto Amato, Jeffrey Barrett, Sonia Goncalves, Ewan Harrison, David K. Jackson, Ian Johnston, Dominic Kwiatkowski, Cordelia Langford, John Sillitoe on behalf of the Wellcome Sanger Institute COVID-19 Surveillance Team |
| EPI_ISL_3310372, EPI_ISL_3310373, EPI_ISL_3310374, EPI_ISL_3310375 | Lighthouse Lab in Glasgow | Wellcome Sanger Institute for the COVID-19 Genomics UK (COG-UK) Consortium | Harper VanSteenhouse, Yumi Kasai, David Gray, Carol Clugston, Anna Dominiczak and Alex Alderton, Roberto Amato, Jeffrey Barrett, Sonia Goncalves, Ewan Harrison, David K. Jackson, Ian Johnston, Dominic Kwiatkowski, Cordelia Langford, John Sillitoe on behalf of the Wellcome Sanger Institute COVID-19 Surveillance Team |
| EPI_ISL_3310376 | Lighthouse Lab in Milton Keynes | Wellcome Sanger Institute for the COVID-19 Genomics UK (COG-UK) Consortium | The Lighthouse Lab in Milton Keynes and Alex Alderton, Roberto Amato, Jeffrey Barrett, Sonia Goncalves, Ewan Harrison, David K. Jackson, Ian Johnston, Dominic Kwiatkowski, Cordelia Langford, John Sillitoe on behalf of the Wellcome Sanger Institute COVID-19 Surveillance Team |
| EPI_ISL_3310377 | Lighthouse Lab in Alderley Park | Wellcome Sanger Institute for the COVID-19 Genomics UK (COG-UK) Consortium | Jacquelyn Wynn, Mairead Hyland, The Lighthouse Lab in Alderley Park and Alex Alderton, Roberto Amato, Jeffrey Barrett, Sonia Goncalves, Ewan Harrison, David K. Jackson, Ian Johnston, Dominic Kwiatkowski, Cordelia Langford, John Sillitoe on behalf of the Wellcome Sanger Institute COVID-19 Surveillance Team |
| EPI_ISL_3310378 | Lighthouse Lab in Glasgow | Wellcome Sanger Institute for the COVID-19 Genomics UK (COG-UK) Consortium | Harper VanSteenhouse, Yumi Kasai, David Gray, Carol Clugston, Anna Dominiczak and Alex Alderton, Roberto Amato, Jeffrey Barrett, Sonia Goncalves, Ewan Harrison, David K. Jackson, Ian Johnston, Dominic Kwiatkowski, Cordelia Langford, John Sillitoe on behalf of the Wellcome Sanger Institute COVID-19 Surveillance Team |
| EPI_ISL_3310379, EPI_ISL_3310380 | Lighthouse Lab in Alderley Park | Wellcome Sanger Institute for the COVID-19 Genomics UK (COG-UK) Consortium | Jacquelyn Wynn, Mairead Hyland, The Lighthouse Lab in Alderley Park and Alex Alderton, Roberto Amato, Jeffrey Barrett, Sonia Goncalves, Ewan Harrison, David K. Jackson, Ian Johnston, Dominic Kwiatkowski, Cordelia Langford, John Sillitoe on behalf of the Wellcome Sanger Institute COVID-19 Surveillance Team |
| EPI_ISL_3310381 | Lighthouse Lab in Glasgow | Wellcome Sanger Institute for the COVID-19 Genomics UK (COG-UK) Consortium | Harper VanSteenhouse, Yumi Kasai, David Gray, Carol Clugston, Anna Dominiczak and Alex Alderton, Roberto Amato, Jeffrey Barrett, Sonia Goncalves, Ewan Harrison, David K. Jackson, Ian Johnston, Dominic Kwiatkowski, Cordelia Langford, John Sillitoe on behalf of the Wellcome Sanger Institute COVID-19 Surveillance Team |
| EPI_ISL_3310382, EPI_ISL_3310383, EPI_ISL_3310384 | Lighthouse Lab in Alderley Park | Wellcome Sanger Institute for the COVID-19 Genomics UK (COG-UK) Consortium | Jacquelyn Wynn, Mairead Hyland, The Lighthouse Lab in Alderley Park and Alex Alderton, Roberto Amato, Jeffrey Barrett, Sonia Goncalves, Ewan Harrison, David K. Jackson, Ian Johnston, Dominic Kwiatkowski, Cordelia Langford, John Sillitoe on behalf of the Wellcome Sanger Institute COVID-19 Surveillance Team |
| EPI_ISL_3310385, EPI_ISL_3310386 | Lighthouse Lab in Glasgow | Wellcome Sanger Institute for the COVID-19 Genomics UK (COG-UK) Consortium | Harper VanSteenhouse, Yumi Kasai, David Gray, Carol Clugston, Anna Dominiczak and Alex Alderton, Roberto Amato, Jeffrey Barrett, Sonia Goncalves, Ewan Harrison, David K. Jackson, Ian Johnston, Dominic Kwiatkowski, Cordelia Langford, John Sillitoe on behalf of the Wellcome Sanger Institute COVID-19 Surveillance Team |
| EPI_ISL_3310387 | Lighthouse Lab in Milton Keynes | Wellcome Sanger Institute for the COVID-19 Genomics UK (COG-UK) Consortium | The Lighthouse Lab in Milton Keynes and Alex Alderton, Roberto Amato, Jeffrey Barrett, Sonia Goncalves, Ewan Harrison, David K. Jackson, Ian Johnston, Dominic Kwiatkowski, Cordelia Langford, John Sillitoe on behalf of the Wellcome Sanger Institute COVID-19 Surveillance Team |
| EPI_ISL_3310388 | Lighthouse Lab in Glasgow | Wellcome Sanger Institute for the COVID-19 Genomics UK (COG-UK) Consortium | Harper VanSteenhouse, Yumi Kasai, David Gray, Carol Clugston, Anna Dominiczak and Alex Alderton, Roberto Amato, Jeffrey Barrett, Sonia Goncalves, Ewan Harrison, David K. Jackson, Ian Johnston, Dominic Kwiatkowski, Cordelia Langford, John Sillitoe on behalf of the Wellcome Sanger Institute COVID-19 Surveillance Team |
| EPI_ISL_3310389 | Lighthouse Lab in Milton Keynes | Wellcome Sanger Institute for the COVID-19 Genomics UK (COG-UK) Consortium | The Lighthouse Lab in Milton Keynes and Alex Alderton, Roberto Amato, Jeffrey Barrett, Sonia Goncalves, Ewan Harrison, David K. Jackson, Ian Johnston, Dominic Kwiatkowski, Cordelia Langford, John Sillitoe on behalf of the Wellcome Sanger Institute COVID-19 Surveillance Team |
| EPI_ISL_3310390 | Lighthouse Lab in Alderley Park | Wellcome Sanger Institute for the COVID-19 Genomics UK (COG-UK) Consortium | Jacquelyn Wynn, Mairead Hyland, The Lighthouse Lab in Alderley Park and Alex Alderton, Roberto Amato, Jeffrey Barrett, Sonia Goncalves, Ewan Harrison, David K. Jackson, Ian Johnston, Dominic Kwiatkowski, Cordelia Langford, John Sillitoe on behalf of the Wellcome Sanger Institute COVID-19 Surveillance Team |
| EPI_ISL_3310391, EPI_ISL_3310392, EPI_ISL_3310393 | Lighthouse Lab in Glasgow | Wellcome Sanger Institute for the COVID-19 Genomics UK (COG-UK) Consortium | Harper VanSteenhouse, Yumi Kasai, David Gray, Carol Clugston, Anna Dominiczak and Alex Alderton, Roberto Amato, Jeffrey Barrett, Sonia Goncalves, Ewan Harrison, David K. Jackson, Ian Johnston, Dominic Kwiatkowski, Cordelia Langford, John Sillitoe on behalf of the Wellcome Sanger Institute COVID-19 Surveillance Team |
| EPI_ISL_3310394 | Lighthouse Lab in Milton Keynes | Wellcome Sanger Institute for the COVID-19 Genomics UK (COG-UK) Consortium | The Lighthouse Lab in Milton Keynes and Alex Alderton, Roberto Amato, Jeffrey Barrett, Sonia Goncalves, Ewan Harrison, David K. Jackson, Ian Johnston, Dominic Kwiatkowski, Cordelia Langford, John Sillitoe on behalf of the Wellcome Sanger Institute COVID-19 Surveillance Team |
| EPI_ISL_3310395 | Lighthouse Lab in Alderley Park | Wellcome Sanger Institute for the COVID-19 Genomics UK (COG-UK) | Jacquelyn Wynn, Mairead Hyland, The Lighthouse Lab in Alderley Park and Alex Alderton, Roberto Amato, Jeffrey Barrett, Sonia Goncalves, Ewan Harrison, David K. Jackson, Ian Johnston, Dominic Kwiatkowski, Cordelia Langford, John Sillitoe on |

| | | | |
|---|---|---|---|
| | | Consortium | behalf of the Wellcome Sanger Institute COVID-19 Surveillance Team |
| EPI_ISL_3310396 | Lighthouse Lab in Glasgow | Wellcome Sanger Institute for the COVID-19 Genomics UK (COG-UK) Consortium | Harper VanSteenhouse, Yumi Kasai, David Gray, Carol Clugston, Anna Dominiczak and Alex Alderton, Roberto Amato, Jeffrey Barrett, Sonia Goncalves, Ewan Harrison, David K. Jackson, Ian Johnston, Dominic Kwiatkowski, Cordelia Langford, John Sillitoe on behalf of the Wellcome Sanger Institute COVID-19 Surveillance Team |
| EPI_ISL_3310397 | Lighthouse Lab in Alderley Park | Wellcome Sanger Institute for the COVID-19 Genomics UK (COG-UK) Consortium | Jacquelyn Wynn, Mairead Hyland, The Lighthouse Lab in Alderley Park and Alex Alderton, Roberto Amato, Jeffrey Barrett, Sonia Goncalves, Ewan Harrison, David K. Jackson, Ian Johnston, Dominic Kwiatkowski, Cordelia Langford, John Sillitoe on behalf of the Wellcome Sanger Institute COVID-19 Surveillance Team |
| EPI_ISL_3310398 | Lighthouse Lab in Glasgow | Wellcome Sanger Institute for the COVID-19 Genomics UK (COG-UK) Consortium | Harper VanSteenhouse, Yumi Kasai, David Gray, Carol Clugston, Anna Dominiczak and Alex Alderton, Roberto Amato, Jeffrey Barrett, Sonia Goncalves, Ewan Harrison, David K. Jackson, Ian Johnston, Dominic Kwiatkowski, Cordelia Langford, John Sillitoe on behalf of the Wellcome Sanger Institute COVID-19 Surveillance Team |
| EPI_ISL_3310399 | Lighthouse Lab in Milton Keynes | Wellcome Sanger Institute for the COVID-19 Genomics UK (COG-UK) Consortium | The Lighthouse Lab in Milton Keynes and Alex Alderton, Roberto Amato, Jeffrey Barrett, Sonia Goncalves, Ewan Harrison, David K. Jackson, Ian Johnston, Dominic Kwiatkowski, Cordelia Langford, John Sillitoe on behalf of the Wellcome Sanger Institute COVID-19 Surveillance Team |
| EPI_ISL_3310400 | Lighthouse Lab in Alderley Park | Wellcome Sanger Institute for the COVID-19 Genomics UK (COG-UK) Consortium | Jacquelyn Wynn, Mairead Hyland, The Lighthouse Lab in Alderley Park and Alex Alderton, Roberto Amato, Jeffrey Barrett, Sonia Goncalves, Ewan Harrison, David K. Jackson, Ian Johnston, Dominic Kwiatkowski, Cordelia Langford, John Sillitoe on behalf of the Wellcome Sanger Institute COVID-19 Surveillance Team |
| EPI_ISL_3310401 | Lighthouse Lab in Milton Keynes | Wellcome Sanger Institute for the COVID-19 Genomics UK (COG-UK) Consortium | The Lighthouse Lab in Milton Keynes and Alex Alderton, Roberto Amato, Jeffrey Barrett, Sonia Goncalves, Ewan Harrison, David K. Jackson, Ian Johnston, Dominic Kwiatkowski, Cordelia Langford, John Sillitoe on behalf of the Wellcome Sanger Institute COVID-19 Surveillance Team |
| EPI_ISL_3310402 | Lighthouse Lab in Alderley Park | Wellcome Sanger Institute for the COVID-19 Genomics UK (COG-UK) Consortium | Jacquelyn Wynn, Mairead Hyland, The Lighthouse Lab in Alderley Park and Alex Alderton, Roberto Amato, Jeffrey Barrett, Sonia Goncalves, Ewan Harrison, David K. Jackson, Ian Johnston, Dominic Kwiatkowski, Cordelia Langford, John Sillitoe on behalf of the Wellcome Sanger Institute COVID-19 Surveillance Team |
| EPI_ISL_3310403 | Lighthouse Lab in Glasgow | Wellcome Sanger Institute for the COVID-19 Genomics UK (COG-UK) Consortium | Harper VanSteenhouse, Yumi Kasai, David Gray, Carol Clugston, Anna Dominiczak and Alex Alderton, Roberto Amato, Jeffrey Barrett, Sonia Goncalves, Ewan Harrison, David K. Jackson, Ian Johnston, Dominic Kwiatkowski, Cordelia Langford, John Sillitoe on behalf of the Wellcome Sanger Institute COVID-19 Surveillance Team |
| EPI_ISL_3310404 | Lighthouse Lab in Milton Keynes | Wellcome Sanger Institute for the COVID-19 Genomics UK (COG-UK) Consortium | The Lighthouse Lab in Milton Keynes and Alex Alderton, Roberto Amato, Jeffrey Barrett, Sonia Goncalves, Ewan Harrison, David K. Jackson, Ian Johnston, Dominic Kwiatkowski, Cordelia Langford, John Sillitoe on behalf of the Wellcome Sanger Institute COVID-19 Surveillance Team |
| EPI_ISL_3310405, EPI_ISL_3310406, EPI_ISL_3310407 | Lighthouse Lab in Glasgow | Wellcome Sanger Institute for the COVID-19 Genomics UK (COG-UK) Consortium | Harper VanSteenhouse, Yumi Kasai, David Gray, Carol Clugston, Anna Dominiczak and Alex Alderton, Roberto Amato, Jeffrey Barrett, Sonia Goncalves, Ewan Harrison, David K. Jackson, Ian Johnston, Dominic Kwiatkowski, Cordelia Langford, John Sillitoe on behalf of the Wellcome Sanger Institute COVID-19 Surveillance Team |
| EPI_ISL_3310408, EPI_ISL_3310409 | Lighthouse Lab in Milton Keynes | Wellcome Sanger Institute for the COVID-19 Genomics UK (COG-UK) Consortium | The Lighthouse Lab in Milton Keynes and Alex Alderton, Roberto Amato, Jeffrey Barrett, Sonia Goncalves, Ewan Harrison, David K. Jackson, Ian Johnston, Dominic Kwiatkowski, Cordelia Langford, John Sillitoe on behalf of the Wellcome Sanger Institute COVID-19 Surveillance Team |
| EPI_ISL_3310410 | Lighthouse Lab in Glasgow | Wellcome Sanger Institute for the COVID-19 Genomics UK (COG-UK) Consortium | Harper VanSteenhouse, Yumi Kasai, David Gray, Carol Clugston, Anna Dominiczak and Alex Alderton, Roberto Amato, Jeffrey Barrett, Sonia Goncalves, Ewan Harrison, David K. Jackson, Ian Johnston, Dominic Kwiatkowski, Cordelia Langford, John Sillitoe on behalf of the Wellcome Sanger Institute COVID-19 Surveillance Team |
| EPI_ISL_3310411 | Lighthouse Lab in Alderley Park | Wellcome Sanger Institute for the COVID-19 Genomics UK (COG-UK) Consortium | Jacquelyn Wynn, Mairead Hyland, The Lighthouse Lab in Alderley Park and Alex Alderton, Roberto Amato, Jeffrey Barrett, Sonia Goncalves, Ewan Harrison, David K. Jackson, Ian Johnston, Dominic Kwiatkowski, Cordelia Langford, John Sillitoe on behalf of the Wellcome Sanger Institute COVID-19 Surveillance Team |
| EPI_ISL_3310412 | Lighthouse Lab in Milton Keynes | Wellcome Sanger Institute for the COVID-19 Genomics UK (COG-UK) Consortium | The Lighthouse Lab in Milton Keynes and Alex Alderton, Roberto Amato, Jeffrey Barrett, Sonia Goncalves, Ewan Harrison, David K. Jackson, Ian Johnston, Dominic Kwiatkowski, Cordelia Langford, John Sillitoe on behalf of the Wellcome Sanger Institute COVID-19 Surveillance Team |
| EPI_ISL_3310413 | Lighthouse Lab in Alderley Park | Wellcome Sanger Institute for the COVID-19 Genomics UK (COG-UK) Consortium | Jacquelyn Wynn, Mairead Hyland, The Lighthouse Lab in Alderley Park and Alex Alderton, Roberto Amato, Jeffrey Barrett, Sonia Goncalves, Ewan Harrison, David K. Jackson, Ian Johnston, Dominic Kwiatkowski, Cordelia Langford, John Sillitoe on behalf of the Wellcome Sanger Institute COVID-19 Surveillance Team |
| EPI_ISL_3310414 | Lighthouse Lab in Milton Keynes | Wellcome Sanger Institute for the COVID-19 Genomics UK (COG-UK) Consortium | The Lighthouse Lab in Milton Keynes and Alex Alderton, Roberto Amato, Jeffrey Barrett, Sonia Goncalves, Ewan Harrison, David K. Jackson, Ian Johnston, Dominic Kwiatkowski, Cordelia Langford, John Sillitoe on behalf of the Wellcome Sanger Institute COVID-19 Surveillance Team |
| EPI_ISL_3310415, EPI_ISL_3310416 | Lighthouse Lab in Alderley Park | Wellcome Sanger Institute for the COVID-19 Genomics UK (COG-UK) Consortium | Jacquelyn Wynn, Mairead Hyland, The Lighthouse Lab in Alderley Park and Alex Alderton, Roberto Amato, Jeffrey Barrett, Sonia Goncalves, Ewan Harrison, David K. Jackson, Ian Johnston, Dominic Kwiatkowski, Cordelia Langford, John Sillitoe on behalf of the Wellcome Sanger Institute COVID-19 Surveillance Team |
| EPI_ISL_3310417 | Lighthouse Lab in Milton Keynes | Wellcome Sanger Institute for the COVID-19 Genomics UK (COG-UK) Consortium | The Lighthouse Lab in Milton Keynes and Alex Alderton, Roberto Amato, Jeffrey Barrett, Sonia Goncalves, Ewan Harrison, David K. Jackson, Ian Johnston, Dominic Kwiatkowski, Cordelia Langford, John Sillitoe on behalf of the Wellcome Sanger Institute COVID-19 Surveillance Team |
| EPI_ISL_3310418, EPI_ISL_3310419 | Lighthouse Lab in Glasgow | Wellcome Sanger Institute for the COVID-19 Genomics UK (COG-UK) Consortium | Harper VanSteenhouse, Yumi Kasai, David Gray, Carol Clugston, Anna Dominiczak and Alex Alderton, Roberto Amato, Jeffrey Barrett, Sonia Goncalves, Ewan Harrison, David K. Jackson, Ian Johnston, Dominic Kwiatkowski, Cordelia Langford, John Sillitoe on behalf of the Wellcome Sanger Institute COVID-19 Surveillance Team |
| EPI_ISL_3310420 | Lighthouse Lab in Alderley Park | Wellcome Sanger Institute for the COVID-19 Genomics UK (COG-UK) Consortium | Jacquelyn Wynn, Mairead Hyland, The Lighthouse Lab in Alderley Park and Alex Alderton, Roberto Amato, Jeffrey Barrett, Sonia Goncalves, Ewan Harrison, David K. Jackson, Ian Johnston, Dominic Kwiatkowski, Cordelia Langford, John Sillitoe on behalf of the Wellcome Sanger Institute COVID-19 Surveillance Team |
| EPI_ISL_3310421, EPI_ISL_3310422 | Lighthouse Lab in Glasgow | Wellcome Sanger Institute for the COVID-19 Genomics UK (COG-UK) Consortium | Harper VanSteenhouse, Yumi Kasai, David Gray, Carol Clugston, Anna Dominiczak and Alex Alderton, Roberto Amato, Jeffrey Barrett, Sonia Goncalves, Ewan Harrison, David K. Jackson, Ian Johnston, Dominic Kwiatkowski, Cordelia Langford, John Sillitoe on behalf of the Wellcome Sanger Institute COVID-19 Surveillance Team |
| EPI_ISL_3310423 | Lighthouse Lab in Alderley Park | Wellcome Sanger Institute for the COVID-19 Genomics UK (COG-UK) Consortium | Jacquelyn Wynn, Mairead Hyland, The Lighthouse Lab in Alderley Park and Alex Alderton, Roberto Amato, Jeffrey Barrett, Sonia Goncalves, Ewan Harrison, David K. Jackson, Ian Johnston, Dominic Kwiatkowski, Cordelia Langford, John Sillitoe on behalf of the Wellcome Sanger Institute COVID-19 Surveillance Team |
| EPI_ISL_3310424 | Lighthouse Lab in Glasgow | Wellcome Sanger Institute for the COVID-19 Genomics UK (COG-UK) Consortium | Harper VanSteenhouse, Yumi Kasai, David Gray, Carol Clugston, Anna Dominiczak and Alex Alderton, Roberto Amato, Jeffrey Barrett, Sonia Goncalves, Ewan Harrison, David K. Jackson, Ian Johnston, Dominic Kwiatkowski, Cordelia Langford, John Sillitoe on behalf of the Wellcome Sanger Institute COVID-19 Surveillance Team |
| EPI_ISL_3310425, EPI_ISL_3310426, EPI_ISL_3310427 | Lighthouse Lab in Milton Keynes | Wellcome Sanger Institute for the COVID-19 Genomics UK (COG-UK) Consortium | The Lighthouse Lab in Milton Keynes and Alex Alderton, Roberto Amato, Jeffrey Barrett, Sonia Goncalves, Ewan Harrison, David K. Jackson, Ian Johnston, Dominic Kwiatkowski, Cordelia Langford, John Sillitoe on behalf of the Wellcome Sanger Institute COVID-19 Surveillance Team |
| EPI_ISL_3310428, EPI_ISL_3310429 | Lighthouse Lab in Alderley Park | Wellcome Sanger Institute for the COVID-19 Genomics UK (COG-UK) Consortium | Jacquelyn Wynn, Mairead Hyland, The Lighthouse Lab in Alderley Park and Alex Alderton, Roberto Amato, Jeffrey Barrett, Sonia Goncalves, Ewan Harrison, David K. Jackson, Ian Johnston, Dominic Kwiatkowski, Cordelia Langford, John Sillitoe on behalf of the Wellcome Sanger Institute COVID-19 Surveillance Team |
| EPI_ISL_3310430 | Lighthouse Lab in Glasgow | Wellcome Sanger Institute for the COVID-19 Genomics UK (COG-UK) Consortium | Harper VanSteenhouse, Yumi Kasai, David Gray, Carol Clugston, Anna Dominiczak and Alex Alderton, Roberto Amato, Jeffrey Barrett, Sonia Goncalves, Ewan Harrison, David K. Jackson, Ian Johnston, Dominic Kwiatkowski, Cordelia Langford, John Sillitoe on behalf of the Wellcome Sanger Institute COVID-19 Surveillance Team |
| EPI_ISL_3310431 | Lighthouse Lab in Alderley Park | Wellcome Sanger Institute for the COVID-19 Genomics UK (COG-UK) Consortium | Jacquelyn Wynn, Mairead Hyland, The Lighthouse Lab in Alderley Park and Alex Alderton, Roberto Amato, Jeffrey Barrett, Sonia Goncalves, Ewan Harrison, David K. Jackson, Ian Johnston, Dominic Kwiatkowski, Cordelia Langford, John Sillitoe on behalf of the Wellcome Sanger Institute COVID-19 Surveillance Team |
| EPI_ISL_3310432 | Lighthouse Lab in Milton Keynes | Wellcome Sanger Institute for the COVID-19 Genomics UK (COG-UK) Consortium | The Lighthouse Lab in Milton Keynes and Alex Alderton, Roberto Amato, Jeffrey Barrett, Sonia Goncalves, Ewan Harrison, David K. Jackson, Ian Johnston, Dominic Kwiatkowski, Cordelia Langford, John Sillitoe on behalf of the Wellcome Sanger Institute COVID-19 Surveillance Team |
| EPI_ISL_3310433, EPI_ISL_3310434 | Lighthouse Lab in Glasgow | Wellcome Sanger Institute for the COVID-19 Genomics UK (COG-UK) Consortium | Harper VanSteenhouse, Yumi Kasai, David Gray, Carol Clugston, Anna Dominiczak and Alex Alderton, Roberto Amato, Jeffrey Barrett, Sonia Goncalves, Ewan Harrison, David K. Jackson, Ian Johnston, Dominic Kwiatkowski, Cordelia Langford, John Sillitoe on behalf of the Wellcome Sanger Institute COVID-19 Surveillance Team |
| EPI_ISL_3310435, EPI_ISL_3310436 | Lighthouse Lab in Milton Keynes | Wellcome Sanger Institute for the COVID-19 Genomics UK (COG-UK) Consortium | The Lighthouse Lab in Milton Keynes and Alex Alderton, Roberto Amato, Jeffrey Barrett, Sonia Goncalves, Ewan Harrison, David K. Jackson, Ian Johnston, Dominic Kwiatkowski, Cordelia Langford, John Sillitoe on behalf of the Wellcome Sanger Institute COVID-19 Surveillance Team |
| EPI_ISL_3310437 | Lighthouse Lab in Glasgow | Wellcome Sanger Institute for the COVID-19 Genomics UK (COG-UK) Consortium | Harper VanSteenhouse, Yumi Kasai, David Gray, Carol Clugston, Anna Dominiczak and Alex Alderton, Roberto Amato, Jeffrey Barrett, Sonia Goncalves, Ewan Harrison, David K. Jackson, Ian Johnston, Dominic Kwiatkowski, Cordelia Langford, John Sillitoe on behalf of the Wellcome Sanger Institute COVID-19 Surveillance Team |
| EPI_ISL_3310438 | Lighthouse Lab in Milton Keynes | Wellcome Sanger Institute for the COVID-19 Genomics UK (COG-UK) Consortium | The Lighthouse Lab in Milton Keynes and Alex Alderton, Roberto Amato, Jeffrey Barrett, Sonia Goncalves, Ewan Harrison, David K. Jackson, Ian Johnston, Dominic Kwiatkowski, Cordelia Langford, John Sillitoe on behalf of the Wellcome Sanger Institute COVID-19 Surveillance Team |
| EPI_ISL_3310439 | Lighthouse Lab in Glasgow | Wellcome Sanger Institute for the COVID-19 Genomics UK (COG-UK) Consortium | Harper VanSteenhouse, Yumi Kasai, David Gray, Carol Clugston, Anna Dominiczak and Alex Alderton, Roberto Amato, Jeffrey Barrett, Sonia Goncalves, Ewan Harrison, David K. Jackson, Ian Johnston, Dominic Kwiatkowski, Cordelia Langford, John Sillitoe on behalf of the Wellcome Sanger Institute COVID-19 Surveillance Team |
| EPI_ISL_3310440, EPI_ISL_3310441 | Lighthouse Lab in Milton Keynes | Wellcome Sanger Institute for the COVID-19 Genomics UK (COG-UK) Consortium | The Lighthouse Lab in Milton Keynes and Alex Alderton, Roberto Amato, Jeffrey Barrett, Sonia Goncalves, Ewan Harrison, David K. Jackson, Ian Johnston, Dominic Kwiatkowski, Cordelia Langford, John Sillitoe on behalf of the Wellcome Sanger Institute COVID-19 Surveillance Team |
| EPI_ISL_3310442 | Lighthouse Lab in Glasgow | Wellcome Sanger Institute for the COVID-19 Genomics UK (COG-UK) Consortium | Harper VanSteenhouse, Yumi Kasai, David Gray, Carol Clugston, Anna Dominiczak and Alex Alderton, Roberto Amato, Jeffrey Barrett, Sonia Goncalves, Ewan Harrison, David K. Jackson, Ian Johnston, Dominic Kwiatkowski, Cordelia Langford, John Sillitoe on behalf of the Wellcome Sanger Institute COVID-19 Surveillance Team |
| EPI_ISL_3310443 | Lighthouse Lab in Alderley Park | Wellcome Sanger Institute for the COVID-19 Genomics UK (COG-UK) Consortium | Jacquelyn Wynn, Mairead Hyland, The Lighthouse Lab in Alderley Park and Alex Alderton, Roberto Amato, Jeffrey Barrett, Sonia Goncalves, Ewan Harrison, David K. Jackson, Ian Johnston, Dominic Kwiatkowski, Cordelia Langford, John Sillitoe on behalf of the Wellcome Sanger Institute COVID-19 Surveillance Team |
| EPI_ISL_3310444 | Lighthouse Lab in Milton Keynes | Wellcome Sanger Institute for the COVID-19 Genomics UK (COG-UK) Consortium | The Lighthouse Lab in Milton Keynes and Alex Alderton, Roberto Amato, Jeffrey Barrett, Sonia Goncalves, Ewan Harrison, David K. Jackson, Ian Johnston, Dominic Kwiatkowski, Cordelia Langford, John Sillitoe on behalf of the Wellcome Sanger Institute COVID-19 Surveillance Team |
| EPI_ISL_3310445, EPI_ISL_3310446 | Lighthouse Lab in Alderley Park | Wellcome Sanger Institute for the COVID-19 Genomics UK (COG-UK) Consortium | Jacquelyn Wynn, Mairead Hyland, The Lighthouse Lab in Alderley Park and Alex Alderton, Roberto Amato, Jeffrey Barrett, Sonia Goncalves, Ewan Harrison, David K. Jackson, Ian Johnston, Dominic Kwiatkowski, Cordelia Langford, John Sillitoe on behalf of the Wellcome Sanger Institute COVID-19 Surveillance Team |
| EPI_ISL_3310447 | Berkshire and Surrey Pathology Services Lighthouse Laboratory | Wellcome Sanger Institute for the COVID-19 Genomics UK (COG-UK) Consortium | Berkshire and Surrey Pathology Services Lighthouse Laboratory and Alex Alderton, Roberto Amato, Jeffrey Barrett, Sonia Goncalves, Ewan Harrison, David K. Jackson, Ian Johnston, Dominic Kwiatkowski, Cordelia Langford, John Sillitoe on behalf of the Wellcome Sanger Institute COVID-19 Surveillance Team |
| EPI_ISL_3310448 | Lighthouse Lab in Alderley Park | Wellcome Sanger Institute for the COVID-19 Genomics UK (COG-UK) Consortium | Jacquelyn Wynn, Mairead Hyland, The Lighthouse Lab in Alderley Park and Alex Alderton, Roberto Amato, Jeffrey Barrett, Sonia Goncalves, Ewan Harrison, David K. Jackson, Ian Johnston, Dominic Kwiatkowski, Cordelia Langford, John Sillitoe on behalf of the Wellcome Sanger Institute COVID-19 Surveillance Team |
| EPI_ISL_3310449 | Berkshire and Surrey Pathology Services Lighthouse Laboratory | Wellcome Sanger Institute for the COVID-19 Genomics UK (COG-UK) Consortium | Berkshire and Surrey Pathology Services Lighthouse Laboratory and Alex Alderton, Roberto Amato, Jeffrey Barrett, Sonia Goncalves, Ewan Harrison, David K. Jackson, Ian Johnston, Dominic Kwiatkowski, Cordelia Langford, John Sillitoe on behalf of the Wellcome Sanger Institute COVID-19 Surveillance Team |
| EPI_ISL_3310450, EPI_ISL_3310451 | Lighthouse Lab in Alderley Park | Wellcome Sanger Institute for the COVID-19 Genomics UK (COG-UK) Consortium | Jacquelyn Wynn, Mairead Hyland, The Lighthouse Lab in Alderley Park and Alex Alderton, Roberto Amato, Jeffrey Barrett, Sonia Goncalves, Ewan Harrison, David K. Jackson, Ian Johnston, Dominic Kwiatkowski, Cordelia Langford, John Sillitoe on behalf of the Wellcome Sanger Institute COVID-19 Surveillance Team |
| EPI_ISL_3310452, EPI_ISL_3310453 | Berkshire and Surrey Pathology Services Lighthouse Laboratory | Wellcome Sanger Institute for the COVID-19 Genomics UK (COG-UK) Consortium | Berkshire and Surrey Pathology Services Lighthouse Laboratory and Alex Alderton, Roberto Amato, Jeffrey Barrett, Sonia Goncalves, Ewan Harrison, David K. Jackson, Ian Johnston, Dominic Kwiatkowski, Cordelia Langford, John Sillitoe on behalf of the Wellcome Sanger Institute COVID-19 Surveillance Team |
| EPI_ISL_3310454, EPI_ISL_3310455, | Lighthouse Lab in Alderley Park | Wellcome Sanger Institute for the COVID-19 Genomics UK (COG-UK) Consortium | Jacquelyn Wynn, Mairead Hyland, The Lighthouse Lab in Alderley Park and Alex Alderton, Roberto Amato, Jeffrey Barrett, Sonia Goncalves, Ewan Harrison, David K. Jackson, Ian Johnston, Dominic Kwiatkowski, Cordelia Langford, John Sillitoe on behalf of the Wellcome Sanger Institute COVID-19 Surveillance Team |

| | | | |
|---|---|---|---|
| EPI_ISL_3310456, EPI_ISL_3310457 | | | |
| EPI_ISL_3310458 | Berkshire and Surrey Pathology Services Lighthouse Laboratory | Wellcome Sanger Institute for the COVID-19 Genomics UK (COG-UK) Consortium | Berkshire and Surrey Pathology Services Lighthouse Laboratory and Alex Alderton, Roberto Amato, Jeffrey Barrett, Sonia Goncalves, Ewan Harrison, David K. Jackson, Ian Johnston, Dominic Kwiatkowski, Cordelia Langford, John Sillitoe on behalf of the Wellcome Sanger Institute COVID-19 Surveillance Team |
| EPI_ISL_3310459, EPI_ISL_3310460, EPI_ISL_3310461 | Lighthouse Lab in Alderley Park | Wellcome Sanger Institute for the COVID-19 Genomics UK (COG-UK) Consortium | Jacquelyn Wynn, Mairead Hyland, The Lighthouse Lab in Alderley Park and Alex Alderton, Roberto Amato, Jeffrey Barrett, Sonia Goncalves, Ewan Harrison, David K. Jackson, Ian Johnston, Dominic Kwiatkowski, Cordelia Langford, John Sillitoe on behalf of the Wellcome Sanger Institute COVID-19 Surveillance Team |
| EPI_ISL_3310462, EPI_ISL_3310463 | Berkshire and Surrey Pathology Services Lighthouse Laboratory | Wellcome Sanger Institute for the COVID-19 Genomics UK (COG-UK) Consortium | Berkshire and Surrey Pathology Services Lighthouse Laboratory and Alex Alderton, Roberto Amato, Jeffrey Barrett, Sonia Goncalves, Ewan Harrison, David K. Jackson, Ian Johnston, Dominic Kwiatkowski, Cordelia Langford, John Sillitoe on behalf of the Wellcome Sanger Institute COVID-19 Surveillance Team |
| EPI_ISL_3310464, EPI_ISL_3310465, EPI_ISL_3310466, EPI_ISL_3310467, EPI_ISL_3310468 | Lighthouse Lab in Alderley Park | Wellcome Sanger Institute for the COVID-19 Genomics UK (COG-UK) Consortium | Jacquelyn Wynn, Mairead Hyland, The Lighthouse Lab in Alderley Park and Alex Alderton, Roberto Amato, Jeffrey Barrett, Sonia Goncalves, Ewan Harrison, David K. Jackson, Ian Johnston, Dominic Kwiatkowski, Cordelia Langford, John Sillitoe on behalf of the Wellcome Sanger Institute COVID-19 Surveillance Team |
| EPI_ISL_3310469 | Berkshire and Surrey Pathology Services Lighthouse Laboratory | Wellcome Sanger Institute for the COVID-19 Genomics UK (COG-UK) Consortium | Berkshire and Surrey Pathology Services Lighthouse Laboratory and Alex Alderton, Roberto Amato, Jeffrey Barrett, Sonia Goncalves, Ewan Harrison, David K. Jackson, Ian Johnston, Dominic Kwiatkowski, Cordelia Langford, John Sillitoe on behalf of the Wellcome Sanger Institute COVID-19 Surveillance Team |
| EPI_ISL_3310470, EPI_ISL_3310471, EPI_ISL_3310472, EPI_ISL_3310473, EPI_ISL_3310474, EPI_ISL_3310475 | Lighthouse Lab in Alderley Park | Wellcome Sanger Institute for the COVID-19 Genomics UK (COG-UK) Consortium | Jacquelyn Wynn, Mairead Hyland, The Lighthouse Lab in Alderley Park and Alex Alderton, Roberto Amato, Jeffrey Barrett, Sonia Goncalves, Ewan Harrison, David K. Jackson, Ian Johnston, Dominic Kwiatkowski, Cordelia Langford, John Sillitoe on behalf of the Wellcome Sanger Institute COVID-19 Surveillance Team |
| EPI_ISL_3310476 | Berkshire and Surrey Pathology Services Lighthouse Laboratory | Wellcome Sanger Institute for the COVID-19 Genomics UK (COG-UK) Consortium | Berkshire and Surrey Pathology Services Lighthouse Laboratory and Alex Alderton, Roberto Amato, Jeffrey Barrett, Sonia Goncalves, Ewan Harrison, David K. Jackson, Ian Johnston, Dominic Kwiatkowski, Cordelia Langford, John Sillitoe on behalf of the Wellcome Sanger Institute COVID-19 Surveillance Team |
| EPI_ISL_3310477 | Lighthouse Lab in Alderley Park | Wellcome Sanger Institute for the COVID-19 Genomics UK (COG-UK) Consortium | Jacquelyn Wynn, Mairead Hyland, The Lighthouse Lab in Alderley Park and Alex Alderton, Roberto Amato, Jeffrey Barrett, Sonia Goncalves, Ewan Harrison, David K. Jackson, Ian Johnston, Dominic Kwiatkowski, Cordelia Langford, John Sillitoe on behalf of the Wellcome Sanger Institute COVID-19 Surveillance Team |
| EPI_ISL_3310478 | Berkshire and Surrey Pathology Services Lighthouse Laboratory | Wellcome Sanger Institute for the COVID-19 Genomics UK (COG-UK) Consortium | Berkshire and Surrey Pathology Services Lighthouse Laboratory and Alex Alderton, Roberto Amato, Jeffrey Barrett, Sonia Goncalves, Ewan Harrison, David K. Jackson, Ian Johnston, Dominic Kwiatkowski, Cordelia Langford, John Sillitoe on behalf of the Wellcome Sanger Institute COVID-19 Surveillance Team |
| EPI_ISL_3310479, EPI_ISL_3310480 | Lighthouse Lab in Alderley Park | Wellcome Sanger Institute for the COVID-19 Genomics UK (COG-UK) Consortium | Jacquelyn Wynn, Mairead Hyland, The Lighthouse Lab in Alderley Park and Alex Alderton, Roberto Amato, Jeffrey Barrett, Sonia Goncalves, Ewan Harrison, David K. Jackson, Ian Johnston, Dominic Kwiatkowski, Cordelia Langford, John Sillitoe on behalf of the Wellcome Sanger Institute COVID-19 Surveillance Team |
| EPI_ISL_3310481 | Berkshire and Surrey Pathology Services Lighthouse Laboratory | Wellcome Sanger Institute for the COVID-19 Genomics UK (COG-UK) Consortium | Berkshire and Surrey Pathology Services Lighthouse Laboratory and Alex Alderton, Roberto Amato, Jeffrey Barrett, Sonia Goncalves, Ewan Harrison, David K. Jackson, Ian Johnston, Dominic Kwiatkowski, Cordelia Langford, John Sillitoe on behalf of the Wellcome Sanger Institute COVID-19 Surveillance Team |
| EPI_ISL_3310482, EPI_ISL_3310483, EPI_ISL_3310484, EPI_ISL_3310485, EPI_ISL_3310486, EPI_ISL_3310487 | Lighthouse Lab in Alderley Park | Wellcome Sanger Institute for the COVID-19 Genomics UK (COG-UK) Consortium | Jacquelyn Wynn, Mairead Hyland, The Lighthouse Lab in Alderley Park and Alex Alderton, Roberto Amato, Jeffrey Barrett, Sonia Goncalves, Ewan Harrison, David K. Jackson, Ian Johnston, Dominic Kwiatkowski, Cordelia Langford, John Sillitoe on behalf of the Wellcome Sanger Institute COVID-19 Surveillance Team |
| EPI_ISL_3310488, EPI_ISL_3310489 | Berkshire and Surrey Pathology Services Lighthouse Laboratory | Wellcome Sanger Institute for the COVID-19 Genomics UK (COG-UK) Consortium | Berkshire and Surrey Pathology Services Lighthouse Laboratory and Alex Alderton, Roberto Amato, Jeffrey Barrett, Sonia Goncalves, Ewan Harrison, David K. Jackson, Ian Johnston, Dominic Kwiatkowski, Cordelia Langford, John Sillitoe on behalf of the Wellcome Sanger Institute COVID-19 Surveillance Team |
| EPI_ISL_3310490 | Lighthouse Lab in Alderley Park | Wellcome Sanger Institute for the COVID-19 Genomics UK (COG-UK) Consortium | Jacquelyn Wynn, Mairead Hyland, The Lighthouse Lab in Alderley Park and Alex Alderton, Roberto Amato, Jeffrey Barrett, Sonia Goncalves, Ewan Harrison, David K. Jackson, Ian Johnston, Dominic Kwiatkowski, Cordelia Langford, John Sillitoe on behalf of the Wellcome Sanger Institute COVID-19 Surveillance Team |
| EPI_ISL_3310491, EPI_ISL_3310492 | Berkshire and Surrey Pathology Services Lighthouse Laboratory | Wellcome Sanger Institute for the COVID-19 Genomics UK (COG-UK) Consortium | Berkshire and Surrey Pathology Services Lighthouse Laboratory and Alex Alderton, Roberto Amato, Jeffrey Barrett, Sonia Goncalves, Ewan Harrison, David K. Jackson, Ian Johnston, Dominic Kwiatkowski, Cordelia Langford, John Sillitoe on behalf of the Wellcome Sanger Institute COVID-19 Surveillance Team |
| EPI_ISL_3310493, EPI_ISL_3310494 | Lighthouse Lab in Alderley Park | Wellcome Sanger Institute for the COVID-19 Genomics UK (COG-UK) Consortium | Jacquelyn Wynn, Mairead Hyland, The Lighthouse Lab in Alderley Park and Alex Alderton, Roberto Amato, Jeffrey Barrett, Sonia Goncalves, Ewan Harrison, David K. Jackson, Ian Johnston, Dominic Kwiatkowski, Cordelia Langford, John Sillitoe on behalf of the Wellcome Sanger Institute COVID-19 Surveillance Team |
| EPI_ISL_3310495, EPI_ISL_3310496 | Berkshire and Surrey Pathology Services Lighthouse Laboratory | Wellcome Sanger Institute for the COVID-19 Genomics UK (COG-UK) Consortium | Berkshire and Surrey Pathology Services Lighthouse Laboratory and Alex Alderton, Roberto Amato, Jeffrey Barrett, Sonia Goncalves, Ewan Harrison, David K. Jackson, Ian Johnston, Dominic Kwiatkowski, Cordelia Langford, John Sillitoe on behalf of the Wellcome Sanger Institute COVID-19 Surveillance Team |
| EPI_ISL_3310497, EPI_ISL_3310498 | Lighthouse Lab in Alderley Park | Wellcome Sanger Institute for the COVID-19 Genomics UK (COG-UK) Consortium | Jacquelyn Wynn, Mairead Hyland, The Lighthouse Lab in Alderley Park and Alex Alderton, Roberto Amato, Jeffrey Barrett, Sonia Goncalves, Ewan Harrison, David K. Jackson, Ian Johnston, Dominic Kwiatkowski, Cordelia Langford, John Sillitoe on behalf of the Wellcome Sanger Institute COVID-19 Surveillance Team |
| EPI_ISL_3310499 | Berkshire and Surrey Pathology Services Lighthouse Laboratory | Wellcome Sanger Institute for the COVID-19 Genomics UK (COG-UK) Consortium | Berkshire and Surrey Pathology Services Lighthouse Laboratory and Alex Alderton, Roberto Amato, Jeffrey Barrett, Sonia Goncalves, Ewan Harrison, David K. Jackson, Ian Johnston, Dominic Kwiatkowski, Cordelia Langford, John Sillitoe on behalf of the Wellcome Sanger Institute COVID-19 Surveillance Team |
| EPI_ISL_3310500, EPI_ISL_3310501, EPI_ISL_3310502, EPI_ISL_3310503, EPI_ISL_3310504, EPI_ISL_3310505, EPI_ISL_3310506, EPI_ISL_3310507, EPI_ISL_3310508, EPI_ISL_3310509, EPI_ISL_3310510, EPI_ISL_3310511, EPI_ISL_3310512, EPI_ISL_3310513, EPI_ISL_3310514 | | | |
| see above | Lighthouse Lab in Alderley Park | Wellcome Sanger Institute for the COVID-19 Genomics UK (COG-UK) Consortium | Jacquelyn Wynn, Mairead Hyland, The Lighthouse Lab in Alderley Park and Alex Alderton, Roberto Amato, Jeffrey Barrett, Sonia Goncalves, Ewan Harrison, David K. Jackson, Ian Johnston, Dominic Kwiatkowski, Cordelia Langford, John Sillitoe on behalf of the Wellcome Sanger Institute COVID-19 Surveillance Team |
| EPI_ISL_3310515 | Berkshire and Surrey Pathology Services Lighthouse Laboratory | Wellcome Sanger Institute for the COVID-19 Genomics UK (COG-UK) Consortium | Berkshire and Surrey Pathology Services Lighthouse Laboratory and Alex Alderton, Roberto Amato, Jeffrey Barrett, Sonia Goncalves, Ewan Harrison, David K. Jackson, Ian Johnston, Dominic Kwiatkowski, Cordelia Langford, John Sillitoe on behalf of the Wellcome Sanger Institute COVID-19 Surveillance Team |
| EPI_ISL_3310516, EPI_ISL_3310517, EPI_ISL_3310518, EPI_ISL_3310519, EPI_ISL_3310520, EPI_ISL_3310521 | Lighthouse Lab in Alderley Park | Wellcome Sanger Institute for the COVID-19 Genomics UK (COG-UK) Consortium | Jacquelyn Wynn, Mairead Hyland, The Lighthouse Lab in Alderley Park and Alex Alderton, Roberto Amato, Jeffrey Barrett, Sonia Goncalves, Ewan Harrison, David K. Jackson, Ian Johnston, Dominic Kwiatkowski, Cordelia Langford, John Sillitoe on behalf of the Wellcome Sanger Institute COVID-19 Surveillance Team |
| EPI_ISL_3310522 | Berkshire and Surrey Pathology Services Lighthouse Laboratory | Wellcome Sanger Institute for the COVID-19 Genomics UK (COG-UK) Consortium | Berkshire and Surrey Pathology Services Lighthouse Laboratory and Alex Alderton, Roberto Amato, Jeffrey Barrett, Sonia Goncalves, Ewan Harrison, David K. Jackson, Ian Johnston, Dominic Kwiatkowski, Cordelia Langford, John Sillitoe on behalf of the Wellcome Sanger Institute COVID-19 Surveillance Team |
| EPI_ISL_3310523, EPI_ISL_3310524, EPI_ISL_3310525 | Lighthouse Lab in Alderley Park | Wellcome Sanger Institute for the COVID-19 Genomics UK (COG-UK) Consortium | Jacquelyn Wynn, Mairead Hyland, The Lighthouse Lab in Alderley Park and Alex Alderton, Roberto Amato, Jeffrey Barrett, Sonia Goncalves, Ewan Harrison, David K. Jackson, Ian Johnston, Dominic Kwiatkowski, Cordelia Langford, John Sillitoe on behalf of the Wellcome Sanger Institute COVID-19 Surveillance Team |
| EPI_ISL_3310526 | Berkshire and Surrey Pathology Services Lighthouse Laboratory | Wellcome Sanger Institute for the COVID-19 Genomics UK (COG-UK) Consortium | Berkshire and Surrey Pathology Services Lighthouse Laboratory and Alex Alderton, Roberto Amato, Jeffrey Barrett, Sonia Goncalves, Ewan Harrison, David K. Jackson, Ian Johnston, Dominic Kwiatkowski, Cordelia Langford, John Sillitoe on behalf of the Wellcome Sanger Institute COVID-19 Surveillance Team |
| EPI_ISL_3310527, EPI_ISL_3310528, EPI_ISL_3310529, EPI_ISL_3310530 | Lighthouse Lab in Alderley Park | Wellcome Sanger Institute for the COVID-19 Genomics UK (COG-UK) Consortium | Jacquelyn Wynn, Mairead Hyland, The Lighthouse Lab in Alderley Park and Alex Alderton, Roberto Amato, Jeffrey Barrett, Sonia Goncalves, Ewan Harrison, David K. Jackson, Ian Johnston, Dominic Kwiatkowski, Cordelia Langford, John Sillitoe on behalf of the Wellcome Sanger Institute COVID-19 Surveillance Team |
| EPI_ISL_3310531 | Berkshire and Surrey Pathology Services Lighthouse Laboratory | Wellcome Sanger Institute for the COVID-19 Genomics UK (COG-UK) Consortium | Berkshire and Surrey Pathology Services Lighthouse Laboratory and Alex Alderton, Roberto Amato, Jeffrey Barrett, Sonia Goncalves, Ewan Harrison, David K. Jackson, Ian Johnston, Dominic Kwiatkowski, Cordelia Langford, John Sillitoe on behalf of the Wellcome Sanger Institute COVID-19 Surveillance Team |
| EPI_ISL_3310532, EPI_ISL_3310533, EPI_ISL_3310534 | Lighthouse Lab in Alderley Park | Wellcome Sanger Institute for the COVID-19 Genomics UK (COG-UK) Consortium | Jacquelyn Wynn, Mairead Hyland, The Lighthouse Lab in Alderley Park and Alex Alderton, Roberto Amato, Jeffrey Barrett, Sonia Goncalves, Ewan Harrison, David K. Jackson, Ian Johnston, Dominic Kwiatkowski, Cordelia Langford, John Sillitoe on behalf of the Wellcome Sanger Institute COVID-19 Surveillance Team |
| EPI_ISL_3310535 | Berkshire and Surrey Pathology Services Lighthouse Laboratory | Wellcome Sanger Institute for the COVID-19 Genomics UK (COG-UK) Consortium | Berkshire and Surrey Pathology Services Lighthouse Laboratory and Alex Alderton, Roberto Amato, Jeffrey Barrett, Sonia Goncalves, Ewan Harrison, David K. Jackson, Ian Johnston, Dominic Kwiatkowski, Cordelia Langford, John Sillitoe on behalf of the Wellcome Sanger Institute COVID-19 Surveillance Team |
| EPI_ISL_3310536 | Lighthouse Lab in Alderley Park | Wellcome Sanger Institute for the COVID-19 Genomics UK (COG-UK) Consortium | Jacquelyn Wynn, Mairead Hyland, The Lighthouse Lab in Alderley Park and Alex Alderton, Roberto Amato, Jeffrey Barrett, Sonia Goncalves, Ewan Harrison, David K. Jackson, Ian Johnston, Dominic Kwiatkowski, Cordelia Langford, John Sillitoe on behalf of the Wellcome Sanger Institute COVID-19 Surveillance Team |
| EPI_ISL_3310537 | Berkshire and Surrey Pathology Services Lighthouse Laboratory | Wellcome Sanger Institute for the COVID-19 Genomics UK (COG-UK) Consortium | Berkshire and Surrey Pathology Services Lighthouse Laboratory and Alex Alderton, Roberto Amato, Jeffrey Barrett, Sonia Goncalves, Ewan Harrison, David K. Jackson, Ian Johnston, Dominic Kwiatkowski, Cordelia Langford, John Sillitoe on behalf of the Wellcome Sanger Institute COVID-19 Surveillance Team |
| EPI_ISL_3310538, EPI_ISL_3310539, EPI_ISL_3310540, EPI_ISL_3310541, EPI_ISL_3310542, EPI_ISL_3310543, EPI_ISL_3310544, EPI_ISL_3310545 | Lighthouse Lab in Alderley Park | Wellcome Sanger Institute for the COVID-19 Genomics UK (COG-UK) Consortium | Jacquelyn Wynn, Mairead Hyland, The Lighthouse Lab in Alderley Park and Alex Alderton, Roberto Amato, Jeffrey Barrett, Sonia Goncalves, Ewan Harrison, David K. Jackson, Ian Johnston, Dominic Kwiatkowski, Cordelia Langford, John Sillitoe on behalf of the Wellcome Sanger Institute COVID-19 Surveillance Team |
| EPI_ISL_3310546 | Berkshire and Surrey Pathology Services Lighthouse Laboratory | Wellcome Sanger Institute for the COVID-19 Genomics UK (COG-UK) Consortium | Berkshire and Surrey Pathology Services Lighthouse Laboratory and Alex Alderton, Roberto Amato, Jeffrey Barrett, Sonia Goncalves, Ewan Harrison, David K. Jackson, Ian Johnston, Dominic Kwiatkowski, Cordelia Langford, John Sillitoe on behalf of the Wellcome Sanger Institute COVID-19 Surveillance Team |

| | | | |
|---|---|---|---|
| EPI_ISL_3310547, EPI_ISL_3310548, EPI_ISL_3310549 | Lighthouse Lab in Alderley Park | Wellcome Sanger Institute for the COVID-19 Genomics UK (COG-UK) Consortium | Jacquelyn Wynn, Mairead Hyland, The Lighthouse Lab in Alderley Park and Alex Alderton, Roberto Amato, Jeffrey Barrett, Sonia Goncalves, Ewan Harrison, David K. Jackson, Ian Johnston, Dominic Kwiatkowski, Cordelia Langford, John Sillitoe on behalf of the Wellcome Sanger Institute COVID-19 Surveillance Team |
| EPI_ISL_3310550 | Berkshire and Surrey Pathology Services Lighthouse Laboratory | Wellcome Sanger Institute for the COVID-19 Genomics UK (COG-UK) Consortium | Berkshire and Surrey Pathology Services Lighthouse Laboratory and Alex Alderton, Roberto Amato, Jeffrey Barrett, Sonia Goncalves, Ewan Harrison, David K. Jackson, Ian Johnston, Dominic Kwiatkowski, Cordelia Langford, John Sillitoe on behalf of the Wellcome Sanger Institute COVID-19 Surveillance Team |
| EPI_ISL_3310551, EPI_ISL_3310552, EPI_ISL_3310553, EPI_ISL_3310554, EPI_ISL_3310555, EPI_ISL_3310556, EPI_ISL_3310557, EPI_ISL_3310558, EPI_ISL_3310559, EPI_ISL_3310560, EPI_ISL_3310561, EPI_ISL_3310562, EPI_ISL_3310563 | | | |
| see above | Lighthouse Lab in Alderley Park | Wellcome Sanger Institute for the COVID-19 Genomics UK (COG-UK) Consortium | Jacquelyn Wynn, Mairead Hyland, The Lighthouse Lab in Alderley Park and Alex Alderton, Roberto Amato, Jeffrey Barrett, Sonia Goncalves, Ewan Harrison, David K. Jackson, Ian Johnston, Dominic Kwiatkowski, Cordelia Langford, John Sillitoe on behalf of the Wellcome Sanger Institute COVID-19 Surveillance Team |
| EPI_ISL_3310564 | Berkshire and Surrey Pathology Services Lighthouse Laboratory | Wellcome Sanger Institute for the COVID-19 Genomics UK (COG-UK) Consortium | Berkshire and Surrey Pathology Services Lighthouse Laboratory and Alex Alderton, Roberto Amato, Jeffrey Barrett, Sonia Goncalves, Ewan Harrison, David K. Jackson, Ian Johnston, Dominic Kwiatkowski, Cordelia Langford, John Sillitoe on behalf of the Wellcome Sanger Institute COVID-19 Surveillance Team |
| EPI_ISL_3310565, EPI_ISL_3310566, EPI_ISL_3310567, EPI_ISL_3310568, EPI_ISL_3310569, EPI_ISL_3310570, EPI_ISL_3310571, EPI_ISL_3310572, EPI_ISL_3310573, EPI_ISL_3310574, EPI_ISL_3310575, EPI_ISL_3310576, EPI_ISL_3310577, EPI_ISL_3310578, EPI_ISL_3310579, EPI_ISL_3310580, EPI_ISL_3310581, EPI_ISL_3310582, EPI_ISL_3310583 | | | |
| see above | Lighthouse Lab in Alderley Park | Wellcome Sanger Institute for the COVID-19 Genomics UK (COG-UK) Consortium | Jacquelyn Wynn, Mairead Hyland, The Lighthouse Lab in Alderley Park and Alex Alderton, Roberto Amato, Jeffrey Barrett, Sonia Goncalves, Ewan Harrison, David K. Jackson, Ian Johnston, Dominic Kwiatkowski, Cordelia Langford, John Sillitoe on behalf of the Wellcome Sanger Institute COVID-19 Surveillance Team |
| EPI_ISL_3310584 | Berkshire and Surrey Pathology Services Lighthouse Laboratory | Wellcome Sanger Institute for the COVID-19 Genomics UK (COG-UK) Consortium | Berkshire and Surrey Pathology Services Lighthouse Laboratory and Alex Alderton, Roberto Amato, Jeffrey Barrett, Sonia Goncalves, Ewan Harrison, David K. Jackson, Ian Johnston, Dominic Kwiatkowski, Cordelia Langford, John Sillitoe on behalf of the Wellcome Sanger Institute COVID-19 Surveillance Team |
| EPI_ISL_3310585, EPI_ISL_3310586 | Lighthouse Lab in Alderley Park | Wellcome Sanger Institute for the COVID-19 Genomics UK (COG-UK) Consortium | Jacquelyn Wynn, Mairead Hyland, The Lighthouse Lab in Alderley Park and Alex Alderton, Roberto Amato, Jeffrey Barrett, Sonia Goncalves, Ewan Harrison, David K. Jackson, Ian Johnston, Dominic Kwiatkowski, Cordelia Langford, John Sillitoe on behalf of the Wellcome Sanger Institute COVID-19 Surveillance Team |
| EPI_ISL_3310587 | Berkshire and Surrey Pathology Services Lighthouse Laboratory | Wellcome Sanger Institute for the COVID-19 Genomics UK (COG-UK) Consortium | Berkshire and Surrey Pathology Services Lighthouse Laboratory and Alex Alderton, Roberto Amato, Jeffrey Barrett, Sonia Goncalves, Ewan Harrison, David K. Jackson, Ian Johnston, Dominic Kwiatkowski, Cordelia Langford, John Sillitoe on behalf of the Wellcome Sanger Institute COVID-19 Surveillance Team |
| EPI_ISL_3310588, EPI_ISL_3310589 | Lighthouse Lab in Alderley Park | Wellcome Sanger Institute for the COVID-19 Genomics UK (COG-UK) Consortium | Jacquelyn Wynn, Mairead Hyland, The Lighthouse Lab in Alderley Park and Alex Alderton, Roberto Amato, Jeffrey Barrett, Sonia Goncalves, Ewan Harrison, David K. Jackson, Ian Johnston, Dominic Kwiatkowski, Cordelia Langford, John Sillitoe on behalf of the Wellcome Sanger Institute COVID-19 Surveillance Team |
| EPI_ISL_3310590 | Berkshire and Surrey Pathology Services Lighthouse Laboratory | Wellcome Sanger Institute for the COVID-19 Genomics UK (COG-UK) Consortium | Berkshire and Surrey Pathology Services Lighthouse Laboratory and Alex Alderton, Roberto Amato, Jeffrey Barrett, Sonia Goncalves, Ewan Harrison, David K. Jackson, Ian Johnston, Dominic Kwiatkowski, Cordelia Langford, John Sillitoe on behalf of the Wellcome Sanger Institute COVID-19 Surveillance Team |
| EPI_ISL_3310591, EPI_ISL_3310592, EPI_ISL_3310593, EPI_ISL_3310594, EPI_ISL_3310595, EPI_ISL_3310596, EPI_ISL_3310597 | Lighthouse Lab in Alderley Park | Wellcome Sanger Institute for the COVID-19 Genomics UK (COG-UK) Consortium | Jacquelyn Wynn, Mairead Hyland, The Lighthouse Lab in Alderley Park and Alex Alderton, Roberto Amato, Jeffrey Barrett, Sonia Goncalves, Ewan Harrison, David K. Jackson, Ian Johnston, Dominic Kwiatkowski, Cordelia Langford, John Sillitoe on behalf of the Wellcome Sanger Institute COVID-19 Surveillance Team |
| EPI_ISL_3310598 | Berkshire and Surrey Pathology Services Lighthouse Laboratory | Wellcome Sanger Institute for the COVID-19 Genomics UK (COG-UK) Consortium | Berkshire and Surrey Pathology Services Lighthouse Laboratory and Alex Alderton, Roberto Amato, Jeffrey Barrett, Sonia Goncalves, Ewan Harrison, David K. Jackson, Ian Johnston, Dominic Kwiatkowski, Cordelia Langford, John Sillitoe on behalf of the Wellcome Sanger Institute COVID-19 Surveillance Team |
| EPI_ISL_3310599, EPI_ISL_3310600, EPI_ISL_3310601, EPI_ISL_3310602, EPI_ISL_3310603, EPI_ISL_3310604, EPI_ISL_3310605, EPI_ISL_3310606, EPI_ISL_3310607, EPI_ISL_3310608, EPI_ISL_3310609 | | | |
| see above | Lighthouse Lab in Alderley Park | Wellcome Sanger Institute for the COVID-19 Genomics UK (COG-UK) Consortium | Jacquelyn Wynn, Mairead Hyland, The Lighthouse Lab in Alderley Park and Alex Alderton, Roberto Amato, Jeffrey Barrett, Sonia Goncalves, Ewan Harrison, David K. Jackson, Ian Johnston, Dominic Kwiatkowski, Cordelia Langford, John Sillitoe on behalf of the Wellcome Sanger Institute COVID-19 Surveillance Team |
| EPI_ISL_3310610, EPI_ISL_3310611, EPI_ISL_3310612 | Berkshire and Surrey Pathology Services Lighthouse Laboratory | Wellcome Sanger Institute for the COVID-19 Genomics UK (COG-UK) Consortium | Berkshire and Surrey Pathology Services Lighthouse Laboratory and Alex Alderton, Roberto Amato, Jeffrey Barrett, Sonia Goncalves, Ewan Harrison, David K. Jackson, Ian Johnston, Dominic Kwiatkowski, Cordelia Langford, John Sillitoe on behalf of the Wellcome Sanger Institute COVID-19 Surveillance Team |
| EPI_ISL_3310613 | Lighthouse Lab in Alderley Park | Wellcome Sanger Institute for the COVID-19 Genomics UK (COG-UK) Consortium | Jacquelyn Wynn, Mairead Hyland, The Lighthouse Lab in Alderley Park and Alex Alderton, Roberto Amato, Jeffrey Barrett, Sonia Goncalves, Ewan Harrison, David K. Jackson, Ian Johnston, Dominic Kwiatkowski, Cordelia Langford, John Sillitoe on behalf of the Wellcome Sanger Institute COVID-19 Surveillance Team |
| EPI_ISL_3310614 | Berkshire and Surrey Pathology Services Lighthouse Laboratory | Wellcome Sanger Institute for the COVID-19 Genomics UK (COG-UK) Consortium | Berkshire and Surrey Pathology Services Lighthouse Laboratory and Alex Alderton, Roberto Amato, Jeffrey Barrett, Sonia Goncalves, Ewan Harrison, David K. Jackson, Ian Johnston, Dominic Kwiatkowski, Cordelia Langford, John Sillitoe on behalf of the Wellcome Sanger Institute COVID-19 Surveillance Team |
| EPI_ISL_3310615, EPI_ISL_3310616, EPI_ISL_3310617, EPI_ISL_3310618, EPI_ISL_3310619, EPI_ISL_3310620 | Lighthouse Lab in Alderley Park | Wellcome Sanger Institute for the COVID-19 Genomics UK (COG-UK) Consortium | Jacquelyn Wynn, Mairead Hyland, The Lighthouse Lab in Alderley Park and Alex Alderton, Roberto Amato, Jeffrey Barrett, Sonia Goncalves, Ewan Harrison, David K. Jackson, Ian Johnston, Dominic Kwiatkowski, Cordelia Langford, John Sillitoe on behalf of the Wellcome Sanger Institute COVID-19 Surveillance Team |
| EPI_ISL_3310621, EPI_ISL_3310622 | Berkshire and Surrey Pathology Services Lighthouse Laboratory | Wellcome Sanger Institute for the COVID-19 Genomics UK (COG-UK) Consortium | Berkshire and Surrey Pathology Services Lighthouse Laboratory and Alex Alderton, Roberto Amato, Jeffrey Barrett, Sonia Goncalves, Ewan Harrison, David K. Jackson, Ian Johnston, Dominic Kwiatkowski, Cordelia Langford, John Sillitoe on behalf of the Wellcome Sanger Institute COVID-19 Surveillance Team |
| EPI_ISL_3310623, EPI_ISL_3310624 | Lighthouse Lab in Alderley Park | Wellcome Sanger Institute for the COVID-19 Genomics UK (COG-UK) Consortium | Jacquelyn Wynn, Mairead Hyland, The Lighthouse Lab in Alderley Park and Alex Alderton, Roberto Amato, Jeffrey Barrett, Sonia Goncalves, Ewan Harrison, David K. Jackson, Ian Johnston, Dominic Kwiatkowski, Cordelia Langford, John Sillitoe on behalf of the Wellcome Sanger Institute COVID-19 Surveillance Team |
| EPI_ISL_3310625, EPI_ISL_3310626 | Berkshire and Surrey Pathology Services Lighthouse Laboratory | Wellcome Sanger Institute for the COVID-19 Genomics UK (COG-UK) Consortium | Berkshire and Surrey Pathology Services Lighthouse Laboratory and Alex Alderton, Roberto Amato, Jeffrey Barrett, Sonia Goncalves, Ewan Harrison, David K. Jackson, Ian Johnston, Dominic Kwiatkowski, Cordelia Langford, John Sillitoe on behalf of the Wellcome Sanger Institute COVID-19 Surveillance Team |
| EPI_ISL_3310627 | Lighthouse Lab in Alderley Park | Wellcome Sanger Institute for the COVID-19 Genomics UK (COG-UK) Consortium | Jacquelyn Wynn, Mairead Hyland, The Lighthouse Lab in Alderley Park and Alex Alderton, Roberto Amato, Jeffrey Barrett, Sonia Goncalves, Ewan Harrison, David K. Jackson, Ian Johnston, Dominic Kwiatkowski, Cordelia Langford, John Sillitoe on behalf of the Wellcome Sanger Institute COVID-19 Surveillance Team |
| EPI_ISL_3310628 | Berkshire and Surrey Pathology Services Lighthouse Laboratory | Wellcome Sanger Institute for the COVID-19 Genomics UK (COG-UK) Consortium | Berkshire and Surrey Pathology Services Lighthouse Laboratory and Alex Alderton, Roberto Amato, Jeffrey Barrett, Sonia Goncalves, Ewan Harrison, David K. Jackson, Ian Johnston, Dominic Kwiatkowski, Cordelia Langford, John Sillitoe on behalf of the Wellcome Sanger Institute COVID-19 Surveillance Team |
| EPI_ISL_3310629, EPI_ISL_3310630, EPI_ISL_3310631, EPI_ISL_3310632 | Lighthouse Lab in Alderley Park | Wellcome Sanger Institute for the COVID-19 Genomics UK (COG-UK) Consortium | Jacquelyn Wynn, Mairead Hyland, The Lighthouse Lab in Alderley Park and Alex Alderton, Roberto Amato, Jeffrey Barrett, Sonia Goncalves, Ewan Harrison, David K. Jackson, Ian Johnston, Dominic Kwiatkowski, Cordelia Langford, John Sillitoe on behalf of the Wellcome Sanger Institute COVID-19 Surveillance Team |
| EPI_ISL_3310633 | Berkshire and Surrey Pathology Services Lighthouse Laboratory | Wellcome Sanger Institute for the COVID-19 Genomics UK (COG-UK) Consortium | Berkshire and Surrey Pathology Services Lighthouse Laboratory and Alex Alderton, Roberto Amato, Jeffrey Barrett, Sonia Goncalves, Ewan Harrison, David K. Jackson, Ian Johnston, Dominic Kwiatkowski, Cordelia Langford, John Sillitoe on behalf of the Wellcome Sanger Institute COVID-19 Surveillance Team |
| EPI_ISL_3310634, EPI_ISL_3310635, EPI_ISL_3310636, EPI_ISL_3310637, EPI_ISL_3310638, EPI_ISL_3310639, EPI_ISL_3310640, EPI_ISL_3310641, EPI_ISL_3310642, EPI_ISL_3310643, EPI_ISL_3310644 | | | |
| see above | Lighthouse Lab in Alderley Park | Wellcome Sanger Institute for the COVID-19 Genomics UK (COG-UK) Consortium | Jacquelyn Wynn, Mairead Hyland, The Lighthouse Lab in Alderley Park and Alex Alderton, Roberto Amato, Jeffrey Barrett, Sonia Goncalves, Ewan Harrison, David K. Jackson, Ian Johnston, Dominic Kwiatkowski, Cordelia Langford, John Sillitoe on behalf of the Wellcome Sanger Institute COVID-19 Surveillance Team |
| EPI_ISL_3310645, EPI_ISL_3310646 | Berkshire and Surrey Pathology Services Lighthouse Laboratory | Wellcome Sanger Institute for the COVID-19 Genomics UK (COG-UK) Consortium | Berkshire and Surrey Pathology Services Lighthouse Laboratory and Alex Alderton, Roberto Amato, Jeffrey Barrett, Sonia Goncalves, Ewan Harrison, David K. Jackson, Ian Johnston, Dominic Kwiatkowski, Cordelia Langford, John Sillitoe on behalf of the Wellcome Sanger Institute COVID-19 Surveillance Team |
| EPI_ISL_3310647, EPI_ISL_3310648 | Lighthouse Lab in Alderley Park | Wellcome Sanger Institute for the COVID-19 Genomics UK (COG-UK) Consortium | Jacquelyn Wynn, Mairead Hyland, The Lighthouse Lab in Alderley Park and Alex Alderton, Roberto Amato, Jeffrey Barrett, Sonia Goncalves, Ewan Harrison, David K. Jackson, Ian Johnston, Dominic Kwiatkowski, Cordelia Langford, John Sillitoe on behalf of the Wellcome Sanger Institute COVID-19 Surveillance Team |
| EPI_ISL_3310649, EPI_ISL_3310650 | Berkshire and Surrey Pathology Services Lighthouse Laboratory | Wellcome Sanger Institute for the COVID-19 Genomics UK (COG-UK) Consortium | Berkshire and Surrey Pathology Services Lighthouse Laboratory and Alex Alderton, Roberto Amato, Jeffrey Barrett, Sonia Goncalves, Ewan Harrison, David K. Jackson, Ian Johnston, Dominic Kwiatkowski, Cordelia Langford, John Sillitoe on behalf of the Wellcome Sanger Institute COVID-19 Surveillance Team |
| EPI_ISL_3310651, EPI_ISL_3310652, EPI_ISL_3310653, EPI_ISL_3310654, EPI_ISL_3310655, EPI_ISL_3310656, EPI_ISL_3310657 | Lighthouse Lab in Alderley Park | Wellcome Sanger Institute for the COVID-19 Genomics UK (COG-UK) Consortium | Jacquelyn Wynn, Mairead Hyland, The Lighthouse Lab in Alderley Park and Alex Alderton, Roberto Amato, Jeffrey Barrett, Sonia Goncalves, Ewan Harrison, David K. Jackson, Ian Johnston, Dominic Kwiatkowski, Cordelia Langford, John Sillitoe on behalf of the Wellcome Sanger Institute COVID-19 Surveillance Team |
| EPI_ISL_3310658 | Berkshire and Surrey Pathology Services Lighthouse Laboratory | Wellcome Sanger Institute for the COVID-19 Genomics UK (COG-UK) Consortium | Berkshire and Surrey Pathology Services Lighthouse Laboratory and Alex Alderton, Roberto Amato, Jeffrey Barrett, Sonia Goncalves, Ewan Harrison, David K. Jackson, Ian Johnston, Dominic Kwiatkowski, Cordelia Langford, John Sillitoe on behalf of the Wellcome Sanger Institute COVID-19 Surveillance Team |
| EPI_ISL_3310659, EPI_ISL_3310660, EPI_ISL_3310661, EPI_ISL_3310662, EPI_ISL_3310663, EPI_ISL_3310664, EPI_ISL_3310665 | Lighthouse Lab in Alderley Park | Wellcome Sanger Institute for the COVID-19 Genomics UK (COG-UK) Consortium | Jacquelyn Wynn, Mairead Hyland, The Lighthouse Lab in Alderley Park and Alex Alderton, Roberto Amato, Jeffrey Barrett, Sonia Goncalves, Ewan Harrison, David K. Jackson, Ian Johnston, Dominic Kwiatkowski, Cordelia Langford, John Sillitoe on behalf of the Wellcome Sanger Institute COVID-19 Surveillance Team |
| EPI_ISL_3310666 | Berkshire and Surrey Pathology Services Lighthouse Laboratory | Wellcome Sanger Institute for the COVID-19 Genomics UK (COG-UK) Consortium | Berkshire and Surrey Pathology Services Lighthouse Laboratory and Alex Alderton, Roberto Amato, Jeffrey Barrett, Sonia Goncalves, Ewan Harrison, David K. Jackson, Ian Johnston, Dominic Kwiatkowski, Cordelia Langford, John Sillitoe on behalf of the Wellcome Sanger Institute COVID-19 Surveillance Team |
| EPI_ISL_3310667, EPI_ISL_3310668, | Lighthouse Lab in Alderley Park | Wellcome Sanger Institute for the COVID-19 Genomics UK (COG-UK) Consortium | Jacquelyn Wynn, Mairead Hyland, The Lighthouse Lab in Alderley Park and Alex Alderton, Roberto Amato, Jeffrey Barrett, Sonia Goncalves, Ewan Harrison, David K. Jackson, Ian Johnston, Dominic Kwiatkowski, Cordelia Langford, John Sillitoe on behalf of the Wellcome Sanger Institute COVID-19 Surveillance Team |

| | | | |
|---|---|---|---|
| EPI_ISL_3310669, EPI_ISL_3310670, EPI_ISL_3310671 | | | |
| EPI_ISL_3310672 | Berkshire and Surrey Pathology Services Lighthouse Laboratory | Wellcome Sanger Institute for the COVID-19 Genomics UK (COG-UK) Consortium | Berkshire and Surrey Pathology Services Lighthouse Laboratory and Alex Alderton, Roberto Amato, Jeffrey Barrett, Sonia Goncalves, Ewan Harrison, David K. Jackson, Ian Johnston, Dominic Kwiatkowski, Cordelia Langford, John Sillitoe on behalf of the Wellcome Sanger Institute COVID-19 Surveillance Team |
| EPI_ISL_3310673 | Lighthouse Lab in Alderley Park | Wellcome Sanger Institute for the COVID-19 Genomics UK (COG-UK) Consortium | Jacquelyn Wynn, Mairead Hyland, The Lighthouse Lab in Alderley Park and Alex Alderton, Roberto Amato, Jeffrey Barrett, Sonia Goncalves, Ewan Harrison, David K. Jackson, Ian Johnston, Dominic Kwiatkowski, Cordelia Langford, John Sillitoe on behalf of the Wellcome Sanger Institute COVID-19 Surveillance Team |
| EPI_ISL_3310674, EPI_ISL_3310675 | Berkshire and Surrey Pathology Services Lighthouse Laboratory | Wellcome Sanger Institute for the COVID-19 Genomics UK (COG-UK) Consortium | Berkshire and Surrey Pathology Services Lighthouse Laboratory and Alex Alderton, Roberto Amato, Jeffrey Barrett, Sonia Goncalves, Ewan Harrison, David K. Jackson, Ian Johnston, Dominic Kwiatkowski, Cordelia Langford, John Sillitoe on behalf of the Wellcome Sanger Institute COVID-19 Surveillance Team |
| EPI_ISL_3310676, EPI_ISL_3310677, EPI_ISL_3310678, EPI_ISL_3310679, EPI_ISL_3310680, EPI_ISL_3310681, EPI_ISL_3310682, EPI_ISL_3310683, EPI_ISL_3310684 | Lighthouse Lab in Alderley Park | Wellcome Sanger Institute for the COVID-19 Genomics UK (COG-UK) Consortium | Jacquelyn Wynn, Mairead Hyland, The Lighthouse Lab in Alderley Park and Alex Alderton, Roberto Amato, Jeffrey Barrett, Sonia Goncalves, Ewan Harrison, David K. Jackson, Ian Johnston, Dominic Kwiatkowski, Cordelia Langford, John Sillitoe on behalf of the Wellcome Sanger Institute COVID-19 Surveillance Team |
| EPI_ISL_3310685, EPI_ISL_3310686, EPI_ISL_3310687 | Berkshire and Surrey Pathology Services Lighthouse Laboratory | Wellcome Sanger Institute for the COVID-19 Genomics UK (COG-UK) Consortium | Berkshire and Surrey Pathology Services Lighthouse Laboratory and Alex Alderton, Roberto Amato, Jeffrey Barrett, Sonia Goncalves, Ewan Harrison, David K. Jackson, Ian Johnston, Dominic Kwiatkowski, Cordelia Langford, John Sillitoe on behalf of the Wellcome Sanger Institute COVID-19 Surveillance Team |
| EPI_ISL_3310688, EPI_ISL_3310689, EPI_ISL_3310690 | Lighthouse Lab in Alderley Park | Wellcome Sanger Institute for the COVID-19 Genomics UK (COG-UK) Consortium | Jacquelyn Wynn, Mairead Hyland, The Lighthouse Lab in Alderley Park and Alex Alderton, Roberto Amato, Jeffrey Barrett, Sonia Goncalves, Ewan Harrison, David K. Jackson, Ian Johnston, Dominic Kwiatkowski, Cordelia Langford, John Sillitoe on behalf of the Wellcome Sanger Institute COVID-19 Surveillance Team |
| EPI_ISL_3310691, EPI_ISL_3310692, EPI_ISL_3310693 | Berkshire and Surrey Pathology Services Lighthouse Laboratory | Wellcome Sanger Institute for the COVID-19 Genomics UK (COG-UK) Consortium | Berkshire and Surrey Pathology Services Lighthouse Laboratory and Alex Alderton, Roberto Amato, Jeffrey Barrett, Sonia Goncalves, Ewan Harrison, David K. Jackson, Ian Johnston, Dominic Kwiatkowski, Cordelia Langford, John Sillitoe on behalf of the Wellcome Sanger Institute COVID-19 Surveillance Team |
| EPI_ISL_3310694, EPI_ISL_3310695, EPI_ISL_3310696, EPI_ISL_3310697, EPI_ISL_3310698, EPI_ISL_3310699 | Lighthouse Lab in Alderley Park | Wellcome Sanger Institute for the COVID-19 Genomics UK (COG-UK) Consortium | Jacquelyn Wynn, Mairead Hyland, The Lighthouse Lab in Alderley Park and Alex Alderton, Roberto Amato, Jeffrey Barrett, Sonia Goncalves, Ewan Harrison, David K. Jackson, Ian Johnston, Dominic Kwiatkowski, Cordelia Langford, John Sillitoe on behalf of the Wellcome Sanger Institute COVID-19 Surveillance Team |
| EPI_ISL_3310700 | Berkshire and Surrey Pathology Services Lighthouse Laboratory | Wellcome Sanger Institute for the COVID-19 Genomics UK (COG-UK) Consortium | Berkshire and Surrey Pathology Services Lighthouse Laboratory and Alex Alderton, Roberto Amato, Jeffrey Barrett, Sonia Goncalves, Ewan Harrison, David K. Jackson, Ian Johnston, Dominic Kwiatkowski, Cordelia Langford, John Sillitoe on behalf of the Wellcome Sanger Institute COVID-19 Surveillance Team |
| EPI_ISL_3310701 | Lighthouse Lab in Alderley Park | Wellcome Sanger Institute for the COVID-19 Genomics UK (COG-UK) Consortium | Jacquelyn Wynn, Mairead Hyland, The Lighthouse Lab in Alderley Park and Alex Alderton, Roberto Amato, Jeffrey Barrett, Sonia Goncalves, Ewan Harrison, David K. Jackson, Ian Johnston, Dominic Kwiatkowski, Cordelia Langford, John Sillitoe on behalf of the Wellcome Sanger Institute COVID-19 Surveillance Team |
| EPI_ISL_3310702 | Berkshire and Surrey Pathology Services Lighthouse Laboratory | Wellcome Sanger Institute for the COVID-19 Genomics UK (COG-UK) Consortium | Berkshire and Surrey Pathology Services Lighthouse Laboratory and Alex Alderton, Roberto Amato, Jeffrey Barrett, Sonia Goncalves, Ewan Harrison, David K. Jackson, Ian Johnston, Dominic Kwiatkowski, Cordelia Langford, John Sillitoe on behalf of the Wellcome Sanger Institute COVID-19 Surveillance Team |
| EPI_ISL_3310703 | Lighthouse Lab in Alderley Park | Wellcome Sanger Institute for the COVID-19 Genomics UK (COG-UK) Consortium | Jacquelyn Wynn, Mairead Hyland, The Lighthouse Lab in Alderley Park and Alex Alderton, Roberto Amato, Jeffrey Barrett, Sonia Goncalves, Ewan Harrison, David K. Jackson, Ian Johnston, Dominic Kwiatkowski, Cordelia Langford, John Sillitoe on behalf of the Wellcome Sanger Institute COVID-19 Surveillance Team |
| EPI_ISL_3310704 | Berkshire and Surrey Pathology Services Lighthouse Laboratory | Wellcome Sanger Institute for the COVID-19 Genomics UK (COG-UK) Consortium | Berkshire and Surrey Pathology Services Lighthouse Laboratory and Alex Alderton, Roberto Amato, Jeffrey Barrett, Sonia Goncalves, Ewan Harrison, David K. Jackson, Ian Johnston, Dominic Kwiatkowski, Cordelia Langford, John Sillitoe on behalf of the Wellcome Sanger Institute COVID-19 Surveillance Team |
| EPI_ISL_3310705, EPI_ISL_3310706, EPI_ISL_3310707 | Lighthouse Lab in Alderley Park | Wellcome Sanger Institute for the COVID-19 Genomics UK (COG-UK) Consortium | Jacquelyn Wynn, Mairead Hyland, The Lighthouse Lab in Alderley Park and Alex Alderton, Roberto Amato, Jeffrey Barrett, Sonia Goncalves, Ewan Harrison, David K. Jackson, Ian Johnston, Dominic Kwiatkowski, Cordelia Langford, John Sillitoe on behalf of the Wellcome Sanger Institute COVID-19 Surveillance Team |
| EPI_ISL_3310708 | Berkshire and Surrey Pathology Services Lighthouse Laboratory | Wellcome Sanger Institute for the COVID-19 Genomics UK (COG-UK) Consortium | Berkshire and Surrey Pathology Services Lighthouse Laboratory and Alex Alderton, Roberto Amato, Jeffrey Barrett, Sonia Goncalves, Ewan Harrison, David K. Jackson, Ian Johnston, Dominic Kwiatkowski, Cordelia Langford, John Sillitoe on behalf of the Wellcome Sanger Institute COVID-19 Surveillance Team |
| EPI_ISL_3310709, EPI_ISL_3310710, EPI_ISL_3310711 | Lighthouse Lab in Alderley Park | Wellcome Sanger Institute for the COVID-19 Genomics UK (COG-UK) Consortium | Jacquelyn Wynn, Mairead Hyland, The Lighthouse Lab in Alderley Park and Alex Alderton, Roberto Amato, Jeffrey Barrett, Sonia Goncalves, Ewan Harrison, David K. Jackson, Ian Johnston, Dominic Kwiatkowski, Cordelia Langford, John Sillitoe on behalf of the Wellcome Sanger Institute COVID-19 Surveillance Team |
| EPI_ISL_3310712 | Berkshire and Surrey Pathology Services Lighthouse Laboratory | Wellcome Sanger Institute for the COVID-19 Genomics UK (COG-UK) Consortium | Berkshire and Surrey Pathology Services Lighthouse Laboratory and Alex Alderton, Roberto Amato, Jeffrey Barrett, Sonia Goncalves, Ewan Harrison, David K. Jackson, Ian Johnston, Dominic Kwiatkowski, Cordelia Langford, John Sillitoe on behalf of the Wellcome Sanger Institute COVID-19 Surveillance Team |
| EPI_ISL_3310713 | Lighthouse Lab in Alderley Park | Wellcome Sanger Institute for the COVID-19 Genomics UK (COG-UK) Consortium | Jacquelyn Wynn, Mairead Hyland, The Lighthouse Lab in Alderley Park and Alex Alderton, Roberto Amato, Jeffrey Barrett, Sonia Goncalves, Ewan Harrison, David K. Jackson, Ian Johnston, Dominic Kwiatkowski, Cordelia Langford, John Sillitoe on behalf of the Wellcome Sanger Institute COVID-19 Surveillance Team |
| EPI_ISL_3310714 | Berkshire and Surrey Pathology Services Lighthouse Laboratory | Wellcome Sanger Institute for the COVID-19 Genomics UK (COG-UK) Consortium | Berkshire and Surrey Pathology Services Lighthouse Laboratory and Alex Alderton, Roberto Amato, Jeffrey Barrett, Sonia Goncalves, Ewan Harrison, David K. Jackson, Ian Johnston, Dominic Kwiatkowski, Cordelia Langford, John Sillitoe on behalf of the Wellcome Sanger Institute COVID-19 Surveillance Team |
| EPI_ISL_3310715 | Lighthouse Lab in Alderley Park | Wellcome Sanger Institute for the COVID-19 Genomics UK (COG-UK) Consortium | Jacquelyn Wynn, Mairead Hyland, The Lighthouse Lab in Alderley Park and Alex Alderton, Roberto Amato, Jeffrey Barrett, Sonia Goncalves, Ewan Harrison, David K. Jackson, Ian Johnston, Dominic Kwiatkowski, Cordelia Langford, John Sillitoe on behalf of the Wellcome Sanger Institute COVID-19 Surveillance Team |
| EPI_ISL_3310716 | Berkshire and Surrey Pathology Services Lighthouse Laboratory | Wellcome Sanger Institute for the COVID-19 Genomics UK (COG-UK) Consortium | Berkshire and Surrey Pathology Services Lighthouse Laboratory and Alex Alderton, Roberto Amato, Jeffrey Barrett, Sonia Goncalves, Ewan Harrison, David K. Jackson, Ian Johnston, Dominic Kwiatkowski, Cordelia Langford, John Sillitoe on behalf of the Wellcome Sanger Institute COVID-19 Surveillance Team |
| EPI_ISL_3310717 | Lighthouse Lab in Alderley Park | Wellcome Sanger Institute for the COVID-19 Genomics UK (COG-UK) Consortium | Jacquelyn Wynn, Mairead Hyland, The Lighthouse Lab in Alderley Park and Alex Alderton, Roberto Amato, Jeffrey Barrett, Sonia Goncalves, Ewan Harrison, David K. Jackson, Ian Johnston, Dominic Kwiatkowski, Cordelia Langford, John Sillitoe on behalf of the Wellcome Sanger Institute COVID-19 Surveillance Team |
| EPI_ISL_3310718 | Berkshire and Surrey Pathology Services Lighthouse Laboratory | Wellcome Sanger Institute for the COVID-19 Genomics UK (COG-UK) Consortium | Berkshire and Surrey Pathology Services Lighthouse Laboratory and Alex Alderton, Roberto Amato, Jeffrey Barrett, Sonia Goncalves, Ewan Harrison, David K. Jackson, Ian Johnston, Dominic Kwiatkowski, Cordelia Langford, John Sillitoe on behalf of the Wellcome Sanger Institute COVID-19 Surveillance Team |
| EPI_ISL_3310719, EPI_ISL_3310720, EPI_ISL_3310721, EPI_ISL_3310722, EPI_ISL_3310723, EPI_ISL_3310724 | Lighthouse Lab in Alderley Park | Wellcome Sanger Institute for the COVID-19 Genomics UK (COG-UK) Consortium | Jacquelyn Wynn, Mairead Hyland, The Lighthouse Lab in Alderley Park and Alex Alderton, Roberto Amato, Jeffrey Barrett, Sonia Goncalves, Ewan Harrison, David K. Jackson, Ian Johnston, Dominic Kwiatkowski, Cordelia Langford, John Sillitoe on behalf of the Wellcome Sanger Institute COVID-19 Surveillance Team |
| EPI_ISL_3310725 | Berkshire and Surrey Pathology Services Lighthouse Laboratory | Wellcome Sanger Institute for the COVID-19 Genomics UK (COG-UK) Consortium | Berkshire and Surrey Pathology Services Lighthouse Laboratory and Alex Alderton, Roberto Amato, Jeffrey Barrett, Sonia Goncalves, Ewan Harrison, David K. Jackson, Ian Johnston, Dominic Kwiatkowski, Cordelia Langford, John Sillitoe on behalf of the Wellcome Sanger Institute COVID-19 Surveillance Team |
| EPI_ISL_3310726, EPI_ISL_3310727, EPI_ISL_3310728, EPI_ISL_3310729 | Lighthouse Lab in Alderley Park | Wellcome Sanger Institute for the COVID-19 Genomics UK (COG-UK) Consortium | Jacquelyn Wynn, Mairead Hyland, The Lighthouse Lab in Alderley Park and Alex Alderton, Roberto Amato, Jeffrey Barrett, Sonia Goncalves, Ewan Harrison, David K. Jackson, Ian Johnston, Dominic Kwiatkowski, Cordelia Langford, John Sillitoe on behalf of the Wellcome Sanger Institute COVID-19 Surveillance Team |
| EPI_ISL_3310730 | Berkshire and Surrey Pathology Services Lighthouse Laboratory | Wellcome Sanger Institute for the COVID-19 Genomics UK (COG-UK) Consortium | Berkshire and Surrey Pathology Services Lighthouse Laboratory and Alex Alderton, Roberto Amato, Jeffrey Barrett, Sonia Goncalves, Ewan Harrison, David K. Jackson, Ian Johnston, Dominic Kwiatkowski, Cordelia Langford, John Sillitoe on behalf of the Wellcome Sanger Institute COVID-19 Surveillance Team |
| EPI_ISL_3310731, EPI_ISL_3310732, EPI_ISL_3310733, EPI_ISL_3310734 | Lighthouse Lab in Alderley Park | Wellcome Sanger Institute for the COVID-19 Genomics UK (COG-UK) Consortium | Jacquelyn Wynn, Mairead Hyland, The Lighthouse Lab in Alderley Park and Alex Alderton, Roberto Amato, Jeffrey Barrett, Sonia Goncalves, Ewan Harrison, David K. Jackson, Ian Johnston, Dominic Kwiatkowski, Cordelia Langford, John Sillitoe on behalf of the Wellcome Sanger Institute COVID-19 Surveillance Team |
| EPI_ISL_3310735 | Berkshire and Surrey Pathology Services Lighthouse Laboratory | Wellcome Sanger Institute for the COVID-19 Genomics UK (COG-UK) Consortium | Berkshire and Surrey Pathology Services Lighthouse Laboratory and Alex Alderton, Roberto Amato, Jeffrey Barrett, Sonia Goncalves, Ewan Harrison, David K. Jackson, Ian Johnston, Dominic Kwiatkowski, Cordelia Langford, John Sillitoe on behalf of the Wellcome Sanger Institute COVID-19 Surveillance Team |
| EPI_ISL_3310736, EPI_ISL_3310737 | Lighthouse Lab in Alderley Park | Wellcome Sanger Institute for the COVID-19 Genomics UK (COG-UK) Consortium | Jacquelyn Wynn, Mairead Hyland, The Lighthouse Lab in Alderley Park and Alex Alderton, Roberto Amato, Jeffrey Barrett, Sonia Goncalves, Ewan Harrison, David K. Jackson, Ian Johnston, Dominic Kwiatkowski, Cordelia Langford, John Sillitoe on behalf of the Wellcome Sanger Institute COVID-19 Surveillance Team |
| EPI_ISL_3310738 | Berkshire and Surrey Pathology Services Lighthouse Laboratory | Wellcome Sanger Institute for the COVID-19 Genomics UK (COG-UK) Consortium | Berkshire and Surrey Pathology Services Lighthouse Laboratory and Alex Alderton, Roberto Amato, Jeffrey Barrett, Sonia Goncalves, Ewan Harrison, David K. Jackson, Ian Johnston, Dominic Kwiatkowski, Cordelia Langford, John Sillitoe on behalf of the Wellcome Sanger Institute COVID-19 Surveillance Team |
| EPI_ISL_3310739, EPI_ISL_3310740 | Lighthouse Lab in Alderley Park | Wellcome Sanger Institute for the COVID-19 Genomics UK (COG-UK) Consortium | Jacquelyn Wynn, Mairead Hyland, The Lighthouse Lab in Alderley Park and Alex Alderton, Roberto Amato, Jeffrey Barrett, Sonia Goncalves, Ewan Harrison, David K. Jackson, Ian Johnston, Dominic Kwiatkowski, Cordelia Langford, John Sillitoe on behalf of the Wellcome Sanger Institute COVID-19 Surveillance Team |
| EPI_ISL_3310741 | Berkshire and Surrey Pathology Services Lighthouse Laboratory | Wellcome Sanger Institute for the COVID-19 Genomics UK (COG-UK) Consortium | Berkshire and Surrey Pathology Services Lighthouse Laboratory and Alex Alderton, Roberto Amato, Jeffrey Barrett, Sonia Goncalves, Ewan Harrison, David K. Jackson, Ian Johnston, Dominic Kwiatkowski, Cordelia Langford, John Sillitoe on behalf of the Wellcome Sanger Institute COVID-19 Surveillance Team |
| EPI_ISL_3310742, | Lighthouse Lab in Alderley Park | Wellcome Sanger Institute for the COVID-19 Genomics UK (COG-UK) | Jacquelyn Wynn, Mairead Hyland, The Lighthouse Lab in Alderley Park and Alex Alderton, Roberto Amato, Jeffrey Barrett, Sonia Goncalves, Ewan Harrison, David K. Jackson, Ian Johnston, Dominic Kwiatkowski, Cordelia Langford, John Sillitoe on |

| | | | |
|---|---|---|---|
| EPI_ISL_3310743 | | Consortium | behalf of the Wellcome Sanger Institute COVID-19 Surveillance Team |
| EPI_ISL_3310744 | Berkshire and Surrey Pathology Services Lighthouse Laboratory | Wellcome Sanger Institute for the COVID-19 Genomics UK (COG-UK) Consortium | Berkshire and Surrey Pathology Services Lighthouse Laboratory and Alex Alderton, Roberto Amato, Jeffrey Barrett, Sonia Goncalves, Ewan Harrison, David K. Jackson, Ian Johnston, Dominic Kwiatkowski, Cordelia Langford, John Sillitoe on behalf of the Wellcome Sanger Institute COVID-19 Surveillance Team |
| EPI_ISL_3310745, EPI_ISL_3310746, EPI_ISL_3310747, EPI_ISL_3310748, EPI_ISL_3310749, EPI_ISL_3310750 | Lighthouse Lab in Alderley Park | Wellcome Sanger Institute for the COVID-19 Genomics UK (COG-UK) Consortium | Jacquelyn Wynn, Mairead Hyland, The Lighthouse Lab in Alderley Park and Alex Alderton, Roberto Amato, Jeffrey Barrett, Sonia Goncalves, Ewan Harrison, David K. Jackson, Ian Johnston, Dominic Kwiatkowski, Cordelia Langford, John Sillitoe on behalf of the Wellcome Sanger Institute COVID-19 Surveillance Team |
| EPI_ISL_3310751, EPI_ISL_3310752, EPI_ISL_3310753 | Berkshire and Surrey Pathology Services Lighthouse Laboratory | Wellcome Sanger Institute for the COVID-19 Genomics UK (COG-UK) Consortium | Berkshire and Surrey Pathology Services Lighthouse Laboratory and Alex Alderton, Roberto Amato, Jeffrey Barrett, Sonia Goncalves, Ewan Harrison, David K. Jackson, Ian Johnston, Dominic Kwiatkowski, Cordelia Langford, John Sillitoe on behalf of the Wellcome Sanger Institute COVID-19 Surveillance Team |
| EPI_ISL_3310754, EPI_ISL_3310755 | Lighthouse Lab in Alderley Park | Wellcome Sanger Institute for the COVID-19 Genomics UK (COG-UK) Consortium | Jacquelyn Wynn, Mairead Hyland, The Lighthouse Lab in Alderley Park and Alex Alderton, Roberto Amato, Jeffrey Barrett, Sonia Goncalves, Ewan Harrison, David K. Jackson, Ian Johnston, Dominic Kwiatkowski, Cordelia Langford, John Sillitoe on behalf of the Wellcome Sanger Institute COVID-19 Surveillance Team |
| EPI_ISL_3310756 | Berkshire and Surrey Pathology Services Lighthouse Laboratory | Wellcome Sanger Institute for the COVID-19 Genomics UK (COG-UK) Consortium | Berkshire and Surrey Pathology Services Lighthouse Laboratory and Alex Alderton, Roberto Amato, Jeffrey Barrett, Sonia Goncalves, Ewan Harrison, David K. Jackson, Ian Johnston, Dominic Kwiatkowski, Cordelia Langford, John Sillitoe on behalf of the Wellcome Sanger Institute COVID-19 Surveillance Team |
| EPI_ISL_3310757, EPI_ISL_3310758, EPI_ISL_3310759 | Lighthouse Lab in Alderley Park | Wellcome Sanger Institute for the COVID-19 Genomics UK (COG-UK) Consortium | Jacquelyn Wynn, Mairead Hyland, The Lighthouse Lab in Alderley Park and Alex Alderton, Roberto Amato, Jeffrey Barrett, Sonia Goncalves, Ewan Harrison, David K. Jackson, Ian Johnston, Dominic Kwiatkowski, Cordelia Langford, John Sillitoe on behalf of the Wellcome Sanger Institute COVID-19 Surveillance Team |
| EPI_ISL_3310760 | Berkshire and Surrey Pathology Services Lighthouse Laboratory | Wellcome Sanger Institute for the COVID-19 Genomics UK (COG-UK) Consortium | Berkshire and Surrey Pathology Services Lighthouse Laboratory and Alex Alderton, Roberto Amato, Jeffrey Barrett, Sonia Goncalves, Ewan Harrison, David K. Jackson, Ian Johnston, Dominic Kwiatkowski, Cordelia Langford, John Sillitoe on behalf of the Wellcome Sanger Institute COVID-19 Surveillance Team |
| EPI_ISL_3310761 | Lighthouse Lab in Alderley Park | Wellcome Sanger Institute for the COVID-19 Genomics UK (COG-UK) Consortium | Jacquelyn Wynn, Mairead Hyland, The Lighthouse Lab in Alderley Park and Alex Alderton, Roberto Amato, Jeffrey Barrett, Sonia Goncalves, Ewan Harrison, David K. Jackson, Ian Johnston, Dominic Kwiatkowski, Cordelia Langford, John Sillitoe on behalf of the Wellcome Sanger Institute COVID-19 Surveillance Team |
| EPI_ISL_3310762 | Berkshire and Surrey Pathology Services Lighthouse Laboratory | Wellcome Sanger Institute for the COVID-19 Genomics UK (COG-UK) Consortium | Berkshire and Surrey Pathology Services Lighthouse Laboratory and Alex Alderton, Roberto Amato, Jeffrey Barrett, Sonia Goncalves, Ewan Harrison, David K. Jackson, Ian Johnston, Dominic Kwiatkowski, Cordelia Langford, John Sillitoe on behalf of the Wellcome Sanger Institute COVID-19 Surveillance Team |
| EPI_ISL_3310763, EPI_ISL_3310764, EPI_ISL_3310765 | Lighthouse Lab in Alderley Park | Wellcome Sanger Institute for the COVID-19 Genomics UK (COG-UK) Consortium | Jacquelyn Wynn, Mairead Hyland, The Lighthouse Lab in Alderley Park and Alex Alderton, Roberto Amato, Jeffrey Barrett, Sonia Goncalves, Ewan Harrison, David K. Jackson, Ian Johnston, Dominic Kwiatkowski, Cordelia Langford, John Sillitoe on behalf of the Wellcome Sanger Institute COVID-19 Surveillance Team |
| EPI_ISL_3310766 | Berkshire and Surrey Pathology Services Lighthouse Laboratory | Wellcome Sanger Institute for the COVID-19 Genomics UK (COG-UK) Consortium | Berkshire and Surrey Pathology Services Lighthouse Laboratory and Alex Alderton, Roberto Amato, Jeffrey Barrett, Sonia Goncalves, Ewan Harrison, David K. Jackson, Ian Johnston, Dominic Kwiatkowski, Cordelia Langford, John Sillitoe on behalf of the Wellcome Sanger Institute COVID-19 Surveillance Team |
| EPI_ISL_3310767, EPI_ISL_3310768, EPI_ISL_3310769 | Lighthouse Lab in Alderley Park | Wellcome Sanger Institute for the COVID-19 Genomics UK (COG-UK) Consortium | Jacquelyn Wynn, Mairead Hyland, The Lighthouse Lab in Alderley Park and Alex Alderton, Roberto Amato, Jeffrey Barrett, Sonia Goncalves, Ewan Harrison, David K. Jackson, Ian Johnston, Dominic Kwiatkowski, Cordelia Langford, John Sillitoe on behalf of the Wellcome Sanger Institute COVID-19 Surveillance Team |
| EPI_ISL_3310770, EPI_ISL_3310771 | Lighthouse Lab in Glasgow | Wellcome Sanger Institute for the COVID-19 Genomics UK (COG-UK) Consortium | Harper VanSteenhouse, Yumi Kasai, David Gray, Carol Clugston, Anna Dominiczak and Alex Alderton, Roberto Amato, Jeffrey Barrett, Sonia Goncalves, Ewan Harrison, David K. Jackson, Ian Johnston, Dominic Kwiatkowski, Cordelia Langford, John Sillitoe on behalf of the Wellcome Sanger Institute COVID-19 Surveillance Team |
| EPI_ISL_3310772, EPI_ISL_3310773, EPI_ISL_3310774 | Lighthouse Lab in Alderley Park | Wellcome Sanger Institute for the COVID-19 Genomics UK (COG-UK) Consortium | Jacquelyn Wynn, Mairead Hyland, The Lighthouse Lab in Alderley Park and Alex Alderton, Roberto Amato, Jeffrey Barrett, Sonia Goncalves, Ewan Harrison, David K. Jackson, Ian Johnston, Dominic Kwiatkowski, Cordelia Langford, John Sillitoe on behalf of the Wellcome Sanger Institute COVID-19 Surveillance Team |
| EPI_ISL_3310775 | Lighthouse Lab in Glasgow | Wellcome Sanger Institute for the COVID-19 Genomics UK (COG-UK) Consortium | Harper VanSteenhouse, Yumi Kasai, David Gray, Carol Clugston, Anna Dominiczak and Alex Alderton, Roberto Amato, Jeffrey Barrett, Sonia Goncalves, Ewan Harrison, David K. Jackson, Ian Johnston, Dominic Kwiatkowski, Cordelia Langford, John Sillitoe on behalf of the Wellcome Sanger Institute COVID-19 Surveillance Team |
| EPI_ISL_3310776, EPI_ISL_3310777 | Lighthouse Lab in Alderley Park | Wellcome Sanger Institute for the COVID-19 Genomics UK (COG-UK) Consortium | Jacquelyn Wynn, Mairead Hyland, The Lighthouse Lab in Alderley Park and Alex Alderton, Roberto Amato, Jeffrey Barrett, Sonia Goncalves, Ewan Harrison, David K. Jackson, Ian Johnston, Dominic Kwiatkowski, Cordelia Langford, John Sillitoe on behalf of the Wellcome Sanger Institute COVID-19 Surveillance Team |
| EPI_ISL_3310778 | Lighthouse Lab in Glasgow | Wellcome Sanger Institute for the COVID-19 Genomics UK (COG-UK) Consortium | Harper VanSteenhouse, Yumi Kasai, David Gray, Carol Clugston, Anna Dominiczak and Alex Alderton, Roberto Amato, Jeffrey Barrett, Sonia Goncalves, Ewan Harrison, David K. Jackson, Ian Johnston, Dominic Kwiatkowski, Cordelia Langford, John Sillitoe on behalf of the Wellcome Sanger Institute COVID-19 Surveillance Team |
| EPI_ISL_3310779, EPI_ISL_3310780, EPI_ISL_3310781 | Lighthouse Lab in Alderley Park | Wellcome Sanger Institute for the COVID-19 Genomics UK (COG-UK) Consortium | Jacquelyn Wynn, Mairead Hyland, The Lighthouse Lab in Alderley Park and Alex Alderton, Roberto Amato, Jeffrey Barrett, Sonia Goncalves, Ewan Harrison, David K. Jackson, Ian Johnston, Dominic Kwiatkowski, Cordelia Langford, John Sillitoe on behalf of the Wellcome Sanger Institute COVID-19 Surveillance Team |
| EPI_ISL_3310782 | Lighthouse Lab in Glasgow | Wellcome Sanger Institute for the COVID-19 Genomics UK (COG-UK) Consortium | Harper VanSteenhouse, Yumi Kasai, David Gray, Carol Clugston, Anna Dominiczak and Alex Alderton, Roberto Amato, Jeffrey Barrett, Sonia Goncalves, Ewan Harrison, David K. Jackson, Ian Johnston, Dominic Kwiatkowski, Cordelia Langford, John Sillitoe on behalf of the Wellcome Sanger Institute COVID-19 Surveillance Team |
| EPI_ISL_3310783, EPI_ISL_3310784, EPI_ISL_3310785, EPI_ISL_3310786 | Lighthouse Lab in Alderley Park | Wellcome Sanger Institute for the COVID-19 Genomics UK (COG-UK) Consortium | Jacquelyn Wynn, Mairead Hyland, The Lighthouse Lab in Alderley Park and Alex Alderton, Roberto Amato, Jeffrey Barrett, Sonia Goncalves, Ewan Harrison, David K. Jackson, Ian Johnston, Dominic Kwiatkowski, Cordelia Langford, John Sillitoe on behalf of the Wellcome Sanger Institute COVID-19 Surveillance Team |
| EPI_ISL_3310787 | Lighthouse Lab in Glasgow | Wellcome Sanger Institute for the COVID-19 Genomics UK (COG-UK) Consortium | Harper VanSteenhouse, Yumi Kasai, David Gray, Carol Clugston, Anna Dominiczak and Alex Alderton, Roberto Amato, Jeffrey Barrett, Sonia Goncalves, Ewan Harrison, David K. Jackson, Ian Johnston, Dominic Kwiatkowski, Cordelia Langford, John Sillitoe on behalf of the Wellcome Sanger Institute COVID-19 Surveillance Team |
| EPI_ISL_3310788, EPI_ISL_3310789 | Lighthouse Lab in Alderley Park | Wellcome Sanger Institute for the COVID-19 Genomics UK (COG-UK) Consortium | Jacquelyn Wynn, Mairead Hyland, The Lighthouse Lab in Alderley Park and Alex Alderton, Roberto Amato, Jeffrey Barrett, Sonia Goncalves, Ewan Harrison, David K. Jackson, Ian Johnston, Dominic Kwiatkowski, Cordelia Langford, John Sillitoe on behalf of the Wellcome Sanger Institute COVID-19 Surveillance Team |
| EPI_ISL_3310790 | Lighthouse Lab in Glasgow | Wellcome Sanger Institute for the COVID-19 Genomics UK (COG-UK) Consortium | Harper VanSteenhouse, Yumi Kasai, David Gray, Carol Clugston, Anna Dominiczak and Alex Alderton, Roberto Amato, Jeffrey Barrett, Sonia Goncalves, Ewan Harrison, David K. Jackson, Ian Johnston, Dominic Kwiatkowski, Cordelia Langford, John Sillitoe on behalf of the Wellcome Sanger Institute COVID-19 Surveillance Team |
| EPI_ISL_3310791, EPI_ISL_3310792 | Lighthouse Lab in Alderley Park | Wellcome Sanger Institute for the COVID-19 Genomics UK (COG-UK) Consortium | Jacquelyn Wynn, Mairead Hyland, The Lighthouse Lab in Alderley Park and Alex Alderton, Roberto Amato, Jeffrey Barrett, Sonia Goncalves, Ewan Harrison, David K. Jackson, Ian Johnston, Dominic Kwiatkowski, Cordelia Langford, John Sillitoe on behalf of the Wellcome Sanger Institute COVID-19 Surveillance Team |
| EPI_ISL_3310793 | Lighthouse Lab in Glasgow | Wellcome Sanger Institute for the COVID-19 Genomics UK (COG-UK) Consortium | Harper VanSteenhouse, Yumi Kasai, David Gray, Carol Clugston, Anna Dominiczak and Alex Alderton, Roberto Amato, Jeffrey Barrett, Sonia Goncalves, Ewan Harrison, David K. Jackson, Ian Johnston, Dominic Kwiatkowski, Cordelia Langford, John Sillitoe on behalf of the Wellcome Sanger Institute COVID-19 Surveillance Team |
| EPI_ISL_3310794, EPI_ISL_3310795, EPI_ISL_3310796, EPI_ISL_3310797, EPI_ISL_3310798, EPI_ISL_3310799, EPI_ISL_3310800, EPI_ISL_3310801, EPI_ISL_3310802 | Lighthouse Lab in Alderley Park | Wellcome Sanger Institute for the COVID-19 Genomics UK (COG-UK) Consortium | Jacquelyn Wynn, Mairead Hyland, The Lighthouse Lab in Alderley Park and Alex Alderton, Roberto Amato, Jeffrey Barrett, Sonia Goncalves, Ewan Harrison, David K. Jackson, Ian Johnston, Dominic Kwiatkowski, Cordelia Langford, John Sillitoe on behalf of the Wellcome Sanger Institute COVID-19 Surveillance Team |
| EPI_ISL_3310803, EPI_ISL_3310804 | Lighthouse Lab in Glasgow | Wellcome Sanger Institute for the COVID-19 Genomics UK (COG-UK) Consortium | Harper VanSteenhouse, Yumi Kasai, David Gray, Carol Clugston, Anna Dominiczak and Alex Alderton, Roberto Amato, Jeffrey Barrett, Sonia Goncalves, Ewan Harrison, David K. Jackson, Ian Johnston, Dominic Kwiatkowski, Cordelia Langford, John Sillitoe on behalf of the Wellcome Sanger Institute COVID-19 Surveillance Team |
| EPI_ISL_3310805, EPI_ISL_3310806 | Lighthouse Lab in Alderley Park | Wellcome Sanger Institute for the COVID-19 Genomics UK (COG-UK) Consortium | Jacquelyn Wynn, Mairead Hyland, The Lighthouse Lab in Alderley Park and Alex Alderton, Roberto Amato, Jeffrey Barrett, Sonia Goncalves, Ewan Harrison, David K. Jackson, Ian Johnston, Dominic Kwiatkowski, Cordelia Langford, John Sillitoe on behalf of the Wellcome Sanger Institute COVID-19 Surveillance Team |
| EPI_ISL_3310807 | Lighthouse Lab in Glasgow | Wellcome Sanger Institute for the COVID-19 Genomics UK (COG-UK) Consortium | Harper VanSteenhouse, Yumi Kasai, David Gray, Carol Clugston, Anna Dominiczak and Alex Alderton, Roberto Amato, Jeffrey Barrett, Sonia Goncalves, Ewan Harrison, David K. Jackson, Ian Johnston, Dominic Kwiatkowski, Cordelia Langford, John Sillitoe on behalf of the Wellcome Sanger Institute COVID-19 Surveillance Team |
| EPI_ISL_3310808 | Lighthouse Lab in Alderley Park | Wellcome Sanger Institute for the COVID-19 Genomics UK (COG-UK) Consortium | Jacquelyn Wynn, Mairead Hyland, The Lighthouse Lab in Alderley Park and Alex Alderton, Roberto Amato, Jeffrey Barrett, Sonia Goncalves, Ewan Harrison, David K. Jackson, Ian Johnston, Dominic Kwiatkowski, Cordelia Langford, John Sillitoe on behalf of the Wellcome Sanger Institute COVID-19 Surveillance Team |
| EPI_ISL_3310809 | Lighthouse Lab in Glasgow | Wellcome Sanger Institute for the COVID-19 Genomics UK (COG-UK) Consortium | Harper VanSteenhouse, Yumi Kasai, David Gray, Carol Clugston, Anna Dominiczak and Alex Alderton, Roberto Amato, Jeffrey Barrett, Sonia Goncalves, Ewan Harrison, David K. Jackson, Ian Johnston, Dominic Kwiatkowski, Cordelia Langford, John Sillitoe on behalf of the Wellcome Sanger Institute COVID-19 Surveillance Team |
| EPI_ISL_3310810, EPI_ISL_3310811, EPI_ISL_3310812, EPI_ISL_3310813, EPI_ISL_3310814, EPI_ISL_3310815, EPI_ISL_3310816 | Lighthouse Lab in Alderley Park | Wellcome Sanger Institute for the COVID-19 Genomics UK (COG-UK) Consortium | Jacquelyn Wynn, Mairead Hyland, The Lighthouse Lab in Alderley Park and Alex Alderton, Roberto Amato, Jeffrey Barrett, Sonia Goncalves, Ewan Harrison, David K. Jackson, Ian Johnston, Dominic Kwiatkowski, Cordelia Langford, John Sillitoe on behalf of the Wellcome Sanger Institute COVID-19 Surveillance Team |
| EPI_ISL_3310817 | Lighthouse Lab in Glasgow | Wellcome Sanger Institute for the COVID-19 Genomics UK (COG-UK) Consortium | Harper VanSteenhouse, Yumi Kasai, David Gray, Carol Clugston, Anna Dominiczak and Alex Alderton, Roberto Amato, Jeffrey Barrett, Sonia Goncalves, Ewan Harrison, David K. Jackson, Ian Johnston, Dominic Kwiatkowski, Cordelia Langford, John Sillitoe on behalf of the Wellcome Sanger Institute COVID-19 Surveillance Team |
| EPI_ISL_3310818, EPI_ISL_3310819, EPI_ISL_3310820, EPI_ISL_3310821, | Lighthouse Lab in Alderley Park | Wellcome Sanger Institute for the COVID-19 Genomics UK (COG-UK) Consortium | Jacquelyn Wynn, Mairead Hyland, The Lighthouse Lab in Alderley Park and Alex Alderton, Roberto Amato, Jeffrey Barrett, Sonia Goncalves, Ewan Harrison, David K. Jackson, Ian Johnston, Dominic Kwiatkowski, Cordelia Langford, John Sillitoe on behalf of the Wellcome Sanger Institute COVID-19 Surveillance Team |

| | | | |
|---|---|---|---|
| EPI_ISL_3310822 | | | |
| EPI_ISL_3310823 | Lighthouse Lab in Glasgow | Wellcome Sanger Institute for the COVID-19 Genomics UK (COG-UK) Consortium | Harper VanSteenhouse, Yumi Kasai, David Gray, Carol Clugston, Anna Dominiczak and Alex Alderton, Roberto Amato, Jeffrey Barrett, Sonia Goncalves, Ewan Harrison, David K. Jackson, Ian Johnston, Dominic Kwiatkowski, Cordelia Langford, John Sillitoe on behalf of the Wellcome Sanger Institute COVID-19 Surveillance Team |
| EPI_ISL_3310824, EPI_ISL_3310825, EPI_ISL_3310826, EPI_ISL_3310827, EPI_ISL_3310828 | Lighthouse Lab in Alderley Park | Wellcome Sanger Institute for the COVID-19 Genomics UK (COG-UK) Consortium | Jacquelyn Wynn, Mairead Hyland, The Lighthouse Lab in Alderley Park and Alex Alderton, Roberto Amato, Jeffrey Barrett, Sonia Goncalves, Ewan Harrison, David K. Jackson, Ian Johnston, Dominic Kwiatkowski, Cordelia Langford, John Sillitoe on behalf of the Wellcome Sanger Institute COVID-19 Surveillance Team |
| EPI_ISL_3310829 | Lighthouse Lab in Glasgow | Wellcome Sanger Institute for the COVID-19 Genomics UK (COG-UK) Consortium | Harper VanSteenhouse, Yumi Kasai, David Gray, Carol Clugston, Anna Dominiczak and Alex Alderton, Roberto Amato, Jeffrey Barrett, Sonia Goncalves, Ewan Harrison, David K. Jackson, Ian Johnston, Dominic Kwiatkowski, Cordelia Langford, John Sillitoe on behalf of the Wellcome Sanger Institute COVID-19 Surveillance Team |
| EPI_ISL_3310830, EPI_ISL_3310831 | Lighthouse Lab in Alderley Park | Wellcome Sanger Institute for the COVID-19 Genomics UK (COG-UK) Consortium | Jacquelyn Wynn, Mairead Hyland, The Lighthouse Lab in Alderley Park and Alex Alderton, Roberto Amato, Jeffrey Barrett, Sonia Goncalves, Ewan Harrison, David K. Jackson, Ian Johnston, Dominic Kwiatkowski, Cordelia Langford, John Sillitoe on behalf of the Wellcome Sanger Institute COVID-19 Surveillance Team |
| EPI_ISL_3310832 | Lighthouse Lab in Glasgow | Wellcome Sanger Institute for the COVID-19 Genomics UK (COG-UK) Consortium | Harper VanSteenhouse, Yumi Kasai, David Gray, Carol Clugston, Anna Dominiczak and Alex Alderton, Roberto Amato, Jeffrey Barrett, Sonia Goncalves, Ewan Harrison, David K. Jackson, Ian Johnston, Dominic Kwiatkowski, Cordelia Langford, John Sillitoe on behalf of the Wellcome Sanger Institute COVID-19 Surveillance Team |
| EPI_ISL_3310833 | Lighthouse Lab in Alderley Park | Wellcome Sanger Institute for the COVID-19 Genomics UK (COG-UK) Consortium | Jacquelyn Wynn, Mairead Hyland, The Lighthouse Lab in Alderley Park and Alex Alderton, Roberto Amato, Jeffrey Barrett, Sonia Goncalves, Ewan Harrison, David K. Jackson, Ian Johnston, Dominic Kwiatkowski, Cordelia Langford, John Sillitoe on behalf of the Wellcome Sanger Institute COVID-19 Surveillance Team |
| EPI_ISL_3310834 | Lighthouse Lab in Glasgow | Wellcome Sanger Institute for the COVID-19 Genomics UK (COG-UK) Consortium | Harper VanSteenhouse, Yumi Kasai, David Gray, Carol Clugston, Anna Dominiczak and Alex Alderton, Roberto Amato, Jeffrey Barrett, Sonia Goncalves, Ewan Harrison, David K. Jackson, Ian Johnston, Dominic Kwiatkowski, Cordelia Langford, John Sillitoe on behalf of the Wellcome Sanger Institute COVID-19 Surveillance Team |
| EPI_ISL_3310835, EPI_ISL_3310836, EPI_ISL_3310837 | Lighthouse Lab in Alderley Park | Wellcome Sanger Institute for the COVID-19 Genomics UK (COG-UK) Consortium | Jacquelyn Wynn, Mairead Hyland, The Lighthouse Lab in Alderley Park and Alex Alderton, Roberto Amato, Jeffrey Barrett, Sonia Goncalves, Ewan Harrison, David K. Jackson, Ian Johnston, Dominic Kwiatkowski, Cordelia Langford, John Sillitoe on behalf of the Wellcome Sanger Institute COVID-19 Surveillance Team |
| EPI_ISL_3310839 | Lighthouse Lab in Glasgow | Wellcome Sanger Institute for the COVID-19 Genomics UK (COG-UK) Consortium | Harper VanSteenhouse, Yumi Kasai, David Gray, Carol Clugston, Anna Dominiczak and Alex Alderton, Roberto Amato, Jeffrey Barrett, Sonia Goncalves, Ewan Harrison, David K. Jackson, Ian Johnston, Dominic Kwiatkowski, Cordelia Langford, John Sillitoe on behalf of the Wellcome Sanger Institute COVID-19 Surveillance Team |
| EPI_ISL_3310840, EPI_ISL_3310841, EPI_ISL_3310842, EPI_ISL_3310843, EPI_ISL_3310844 | Lighthouse Lab in Alderley Park | Wellcome Sanger Institute for the COVID-19 Genomics UK (COG-UK) Consortium | Jacquelyn Wynn, Mairead Hyland, The Lighthouse Lab in Alderley Park and Alex Alderton, Roberto Amato, Jeffrey Barrett, Sonia Goncalves, Ewan Harrison, David K. Jackson, Ian Johnston, Dominic Kwiatkowski, Cordelia Langford, John Sillitoe on behalf of the Wellcome Sanger Institute COVID-19 Surveillance Team |
| EPI_ISL_3310845 | Lighthouse Lab in Glasgow | Wellcome Sanger Institute for the COVID-19 Genomics UK (COG-UK) Consortium | Harper VanSteenhouse, Yumi Kasai, David Gray, Carol Clugston, Anna Dominiczak and Alex Alderton, Roberto Amato, Jeffrey Barrett, Sonia Goncalves, Ewan Harrison, David K. Jackson, Ian Johnston, Dominic Kwiatkowski, Cordelia Langford, John Sillitoe on behalf of the Wellcome Sanger Institute COVID-19 Surveillance Team |
| EPI_ISL_3310846 | Lighthouse Lab in Alderley Park | Wellcome Sanger Institute for the COVID-19 Genomics UK (COG-UK) Consortium | Jacquelyn Wynn, Mairead Hyland, The Lighthouse Lab in Alderley Park and Alex Alderton, Roberto Amato, Jeffrey Barrett, Sonia Goncalves, Ewan Harrison, David K. Jackson, Ian Johnston, Dominic Kwiatkowski, Cordelia Langford, John Sillitoe on behalf of the Wellcome Sanger Institute COVID-19 Surveillance Team |
| EPI_ISL_3310847 | Lighthouse Lab in Glasgow | Wellcome Sanger Institute for the COVID-19 Genomics UK (COG-UK) Consortium | Harper VanSteenhouse, Yumi Kasai, David Gray, Carol Clugston, Anna Dominiczak and Alex Alderton, Roberto Amato, Jeffrey Barrett, Sonia Goncalves, Ewan Harrison, David K. Jackson, Ian Johnston, Dominic Kwiatkowski, Cordelia Langford, John Sillitoe on behalf of the Wellcome Sanger Institute COVID-19 Surveillance Team |
| EPI_ISL_3310848 | Lighthouse Lab in Alderley Park | Wellcome Sanger Institute for the COVID-19 Genomics UK (COG-UK) Consortium | Jacquelyn Wynn, Mairead Hyland, The Lighthouse Lab in Alderley Park and Alex Alderton, Roberto Amato, Jeffrey Barrett, Sonia Goncalves, Ewan Harrison, David K. Jackson, Ian Johnston, Dominic Kwiatkowski, Cordelia Langford, John Sillitoe on behalf of the Wellcome Sanger Institute COVID-19 Surveillance Team |
| EPI_ISL_3310849 | Lighthouse Lab in Glasgow | Wellcome Sanger Institute for the COVID-19 Genomics UK (COG-UK) Consortium | Harper VanSteenhouse, Yumi Kasai, David Gray, Carol Clugston, Anna Dominiczak and Alex Alderton, Roberto Amato, Jeffrey Barrett, Sonia Goncalves, Ewan Harrison, David K. Jackson, Ian Johnston, Dominic Kwiatkowski, Cordelia Langford, John Sillitoe on behalf of the Wellcome Sanger Institute COVID-19 Surveillance Team |
| EPI_ISL_3310850, EPI_ISL_3310851, EPI_ISL_3310852, EPI_ISL_3310853, EPI_ISL_3310854 | Lighthouse Lab in Alderley Park | Wellcome Sanger Institute for the COVID-19 Genomics UK (COG-UK) Consortium | Jacquelyn Wynn, Mairead Hyland, The Lighthouse Lab in Alderley Park and Alex Alderton, Roberto Amato, Jeffrey Barrett, Sonia Goncalves, Ewan Harrison, David K. Jackson, Ian Johnston, Dominic Kwiatkowski, Cordelia Langford, John Sillitoe on behalf of the Wellcome Sanger Institute COVID-19 Surveillance Team |
| EPI_ISL_3310855, EPI_ISL_3310856 | Lighthouse Lab in Glasgow | Wellcome Sanger Institute for the COVID-19 Genomics UK (COG-UK) Consortium | Harper VanSteenhouse, Yumi Kasai, David Gray, Carol Clugston, Anna Dominiczak and Alex Alderton, Roberto Amato, Jeffrey Barrett, Sonia Goncalves, Ewan Harrison, David K. Jackson, Ian Johnston, Dominic Kwiatkowski, Cordelia Langford, John Sillitoe on behalf of the Wellcome Sanger Institute COVID-19 Surveillance Team |
| EPI_ISL_3310857 | Lighthouse Lab in Alderley Park | Wellcome Sanger Institute for the COVID-19 Genomics UK (COG-UK) Consortium | Jacquelyn Wynn, Mairead Hyland, The Lighthouse Lab in Alderley Park and Alex Alderton, Roberto Amato, Jeffrey Barrett, Sonia Goncalves, Ewan Harrison, David K. Jackson, Ian Johnston, Dominic Kwiatkowski, Cordelia Langford, John Sillitoe on behalf of the Wellcome Sanger Institute COVID-19 Surveillance Team |
| EPI_ISL_3310858 | Lighthouse Lab in Glasgow | Wellcome Sanger Institute for the COVID-19 Genomics UK (COG-UK) Consortium | Harper VanSteenhouse, Yumi Kasai, David Gray, Carol Clugston, Anna Dominiczak and Alex Alderton, Roberto Amato, Jeffrey Barrett, Sonia Goncalves, Ewan Harrison, David K. Jackson, Ian Johnston, Dominic Kwiatkowski, Cordelia Langford, John Sillitoe on behalf of the Wellcome Sanger Institute COVID-19 Surveillance Team |
| EPI_ISL_3310859, EPI_ISL_3310860, EPI_ISL_3310861 | Lighthouse Lab in Alderley Park | Wellcome Sanger Institute for the COVID-19 Genomics UK (COG-UK) Consortium | Jacquelyn Wynn, Mairead Hyland, The Lighthouse Lab in Alderley Park and Alex Alderton, Roberto Amato, Jeffrey Barrett, Sonia Goncalves, Ewan Harrison, David K. Jackson, Ian Johnston, Dominic Kwiatkowski, Cordelia Langford, John Sillitoe on behalf of the Wellcome Sanger Institute COVID-19 Surveillance Team |
| EPI_ISL_3310862, EPI_ISL_3310863 | Lighthouse Lab in Glasgow | Wellcome Sanger Institute for the COVID-19 Genomics UK (COG-UK) Consortium | Harper VanSteenhouse, Yumi Kasai, David Gray, Carol Clugston, Anna Dominiczak and Alex Alderton, Roberto Amato, Jeffrey Barrett, Sonia Goncalves, Ewan Harrison, David K. Jackson, Ian Johnston, Dominic Kwiatkowski, Cordelia Langford, John Sillitoe on behalf of the Wellcome Sanger Institute COVID-19 Surveillance Team |
| EPI_ISL_3310864, EPI_ISL_3310865, EPI_ISL_3310866 | Lighthouse Lab in Alderley Park | Wellcome Sanger Institute for the COVID-19 Genomics UK (COG-UK) Consortium | Jacquelyn Wynn, Mairead Hyland, The Lighthouse Lab in Alderley Park and Alex Alderton, Roberto Amato, Jeffrey Barrett, Sonia Goncalves, Ewan Harrison, David K. Jackson, Ian Johnston, Dominic Kwiatkowski, Cordelia Langford, John Sillitoe on behalf of the Wellcome Sanger Institute COVID-19 Surveillance Team |
| EPI_ISL_3310867 | Lighthouse Lab in Glasgow | Wellcome Sanger Institute for the COVID-19 Genomics UK (COG-UK) Consortium | Harper VanSteenhouse, Yumi Kasai, David Gray, Carol Clugston, Anna Dominiczak and Alex Alderton, Roberto Amato, Jeffrey Barrett, Sonia Goncalves, Ewan Harrison, David K. Jackson, Ian Johnston, Dominic Kwiatkowski, Cordelia Langford, John Sillitoe on behalf of the Wellcome Sanger Institute COVID-19 Surveillance Team |
| EPI_ISL_3310868 | Lighthouse Lab in Alderley Park | Wellcome Sanger Institute for the COVID-19 Genomics UK (COG-UK) Consortium | Jacquelyn Wynn, Mairead Hyland, The Lighthouse Lab in Alderley Park and Alex Alderton, Roberto Amato, Jeffrey Barrett, Sonia Goncalves, Ewan Harrison, David K. Jackson, Ian Johnston, Dominic Kwiatkowski, Cordelia Langford, John Sillitoe on behalf of the Wellcome Sanger Institute COVID-19 Surveillance Team |
| EPI_ISL_3310869 | Lighthouse Lab in Glasgow | Wellcome Sanger Institute for the COVID-19 Genomics UK (COG-UK) Consortium | Harper VanSteenhouse, Yumi Kasai, David Gray, Carol Clugston, Anna Dominiczak and Alex Alderton, Roberto Amato, Jeffrey Barrett, Sonia Goncalves, Ewan Harrison, David K. Jackson, Ian Johnston, Dominic Kwiatkowski, Cordelia Langford, John Sillitoe on behalf of the Wellcome Sanger Institute COVID-19 Surveillance Team |
| EPI_ISL_3310870, EPI_ISL_3310871, EPI_ISL_3310872 | Lighthouse Lab in Alderley Park | Wellcome Sanger Institute for the COVID-19 Genomics UK (COG-UK) Consortium | Jacquelyn Wynn, Mairead Hyland, The Lighthouse Lab in Alderley Park and Alex Alderton, Roberto Amato, Jeffrey Barrett, Sonia Goncalves, Ewan Harrison, David K. Jackson, Ian Johnston, Dominic Kwiatkowski, Cordelia Langford, John Sillitoe on behalf of the Wellcome Sanger Institute COVID-19 Surveillance Team |
| EPI_ISL_3310873 | Lighthouse Lab in Glasgow | Wellcome Sanger Institute for the COVID-19 Genomics UK (COG-UK) Consortium | Harper VanSteenhouse, Yumi Kasai, David Gray, Carol Clugston, Anna Dominiczak and Alex Alderton, Roberto Amato, Jeffrey Barrett, Sonia Goncalves, Ewan Harrison, David K. Jackson, Ian Johnston, Dominic Kwiatkowski, Cordelia Langford, John Sillitoe on behalf of the Wellcome Sanger Institute COVID-19 Surveillance Team |
| EPI_ISL_3310874, EPI_ISL_3310875, EPI_ISL_3310876, EPI_ISL_3310877, EPI_ISL_3310878, EPI_ISL_3310879, EPI_ISL_3310880 | Lighthouse Lab in Alderley Park | Wellcome Sanger Institute for the COVID-19 Genomics UK (COG-UK) Consortium | Jacquelyn Wynn, Mairead Hyland, The Lighthouse Lab in Alderley Park and Alex Alderton, Roberto Amato, Jeffrey Barrett, Sonia Goncalves, Ewan Harrison, David K. Jackson, Ian Johnston, Dominic Kwiatkowski, Cordelia Langford, John Sillitoe on behalf of the Wellcome Sanger Institute COVID-19 Surveillance Team |
| EPI_ISL_3310881, EPI_ISL_3310882 | Lighthouse Lab in Glasgow | Wellcome Sanger Institute for the COVID-19 Genomics UK (COG-UK) Consortium | Harper VanSteenhouse, Yumi Kasai, David Gray, Carol Clugston, Anna Dominiczak and Alex Alderton, Roberto Amato, Jeffrey Barrett, Sonia Goncalves, Ewan Harrison, David K. Jackson, Ian Johnston, Dominic Kwiatkowski, Cordelia Langford, John Sillitoe on behalf of the Wellcome Sanger Institute COVID-19 Surveillance Team |
| EPI_ISL_3310883, EPI_ISL_3310884, EPI_ISL_3310885 | Lighthouse Lab in Alderley Park | Wellcome Sanger Institute for the COVID-19 Genomics UK (COG-UK) Consortium | Jacquelyn Wynn, Mairead Hyland, The Lighthouse Lab in Alderley Park and Alex Alderton, Roberto Amato, Jeffrey Barrett, Sonia Goncalves, Ewan Harrison, David K. Jackson, Ian Johnston, Dominic Kwiatkowski, Cordelia Langford, John Sillitoe on behalf of the Wellcome Sanger Institute COVID-19 Surveillance Team |
| EPI_ISL_3310886, EPI_ISL_3310887 | Lighthouse Lab in Glasgow | Wellcome Sanger Institute for the COVID-19 Genomics UK (COG-UK) Consortium | Harper VanSteenhouse, Yumi Kasai, David Gray, Carol Clugston, Anna Dominiczak and Alex Alderton, Roberto Amato, Jeffrey Barrett, Sonia Goncalves, Ewan Harrison, David K. Jackson, Ian Johnston, Dominic Kwiatkowski, Cordelia Langford, John Sillitoe on behalf of the Wellcome Sanger Institute COVID-19 Surveillance Team |
| EPI_ISL_3310888, EPI_ISL_3310889, EPI_ISL_3310890, EPI_ISL_3310891, EPI_ISL_3310892, EPI_ISL_3310893, EPI_ISL_3310894, EPI_ISL_3310895, EPI_ISL_3310896, EPI_ISL_3310897, EPI_ISL_3310898, EPI_ISL_3310899, EPI_ISL_3310900, EPI_ISL_3310901, EPI_ISL_3310902, EPI_ISL_3310903, EPI_ISL_3310904, EPI_ISL_3310905, EPI_ISL_3310906, EPI_ISL_3310907, EPI_ISL_3310908, EPI_ISL_3310909, EPI_ISL_3310910 | | | |
| see above | Lighthouse Lab in Alderley Park | Wellcome Sanger Institute for the COVID-19 Genomics UK (COG-UK) Consortium | Jacquelyn Wynn, Mairead Hyland, The Lighthouse Lab in Alderley Park and Alex Alderton, Roberto Amato, Jeffrey Barrett, Sonia Goncalves, Ewan Harrison, David K. Jackson, Ian Johnston, Dominic Kwiatkowski, Cordelia Langford, John Sillitoe on behalf of the Wellcome Sanger Institute COVID-19 Surveillance Team |
| EPI_ISL_3310911 | Lighthouse Lab in Glasgow | Wellcome Sanger Institute for the COVID-19 Genomics UK (COG-UK) Consortium | Harper VanSteenhouse, Yumi Kasai, David Gray, Carol Clugston, Anna Dominiczak and Alex Alderton, Roberto Amato, Jeffrey Barrett, Sonia Goncalves, Ewan Harrison, David K. Jackson, Ian Johnston, Dominic Kwiatkowski, Cordelia Langford, John Sillitoe on behalf of the Wellcome Sanger Institute COVID-19 Surveillance Team |
| EPI_ISL_3310912 | Lighthouse Lab in Alderley Park | Wellcome Sanger Institute for the COVID-19 Genomics UK (COG-UK) Consortium | Jacquelyn Wynn, Mairead Hyland, The Lighthouse Lab in Alderley Park and Alex Alderton, Roberto Amato, Jeffrey Barrett, Sonia Goncalves, Ewan Harrison, David K. Jackson, Ian Johnston, Dominic Kwiatkowski, Cordelia Langford, John Sillitoe on behalf of the Wellcome Sanger Institute COVID-19 Surveillance Team |
| EPI_ISL_3310913 | Lighthouse Lab in Glasgow | Wellcome Sanger Institute for the COVID-19 Genomics UK (COG-UK) Consortium | Harper VanSteenhouse, Yumi Kasai, David Gray, Carol Clugston, Anna Dominiczak and Alex Alderton, Roberto Amato, Jeffrey Barrett, Sonia Goncalves, Ewan Harrison, David K. Jackson, Ian Johnston, Dominic Kwiatkowski, Cordelia Langford, John Sillitoe on behalf of the Wellcome Sanger Institute COVID-19 Surveillance Team |
| EPI_ISL_3310914 | Lighthouse Lab in Alderley Park | Wellcome Sanger Institute for the COVID-19 Genomics UK (COG-UK) Consortium | Jacquelyn Wynn, Mairead Hyland, The Lighthouse Lab in Alderley Park and Alex Alderton, Roberto Amato, Jeffrey Barrett, Sonia Goncalves, Ewan Harrison, David K. Jackson, Ian Johnston, Dominic Kwiatkowski, Cordelia Langford, John Sillitoe on behalf of the Wellcome Sanger Institute COVID-19 Surveillance Team |

| | | | |
|---|---|---|---|
| EPI_ISL_3310915 | Lighthouse Lab in Glasgow | Wellcome Sanger Institute for the COVID-19 Genomics UK (COG-UK) Consortium | Harper VanSteenhouse, Yumi Kasai, David Gray, Carol Clugston, Anna Dominiczak and Alex Alderton, Roberto Amato, Jeffrey Barrett, Sonia Goncalves, Ewan Harrison, David K. Jackson, Ian Johnston, Dominic Kwiatkowski, Cordelia Langford, John Sillitoe on behalf of the Wellcome Sanger Institute COVID-19 Surveillance Team |
| EPI_ISL_3310916, EPI_ISL_3310917 | Lighthouse Lab in Alderley Park | Wellcome Sanger Institute for the COVID-19 Genomics UK (COG-UK) Consortium | Jacquelyn Wynn, Mairead Hyland, The Lighthouse Lab in Alderley Park and Alex Alderton, Roberto Amato, Jeffrey Barrett, Sonia Goncalves, Ewan Harrison, David K. Jackson, Ian Johnston, Dominic Kwiatkowski, Cordelia Langford, John Sillitoe on behalf of the Wellcome Sanger Institute COVID-19 Surveillance Team |
| EPI_ISL_3310918, EPI_ISL_3310919, EPI_ISL_3310920 | Lighthouse Lab in Glasgow | Wellcome Sanger Institute for the COVID-19 Genomics UK (COG-UK) Consortium | Harper VanSteenhouse, Yumi Kasai, David Gray, Carol Clugston, Anna Dominiczak and Alex Alderton, Roberto Amato, Jeffrey Barrett, Sonia Goncalves, Ewan Harrison, David K. Jackson, Ian Johnston, Dominic Kwiatkowski, Cordelia Langford, John Sillitoe on behalf of the Wellcome Sanger Institute COVID-19 Surveillance Team |
| EPI_ISL_3310921, EPI_ISL_3310922, EPI_ISL_3310923, EPI_ISL_3310924, EPI_ISL_3310925, EPI_ISL_3310926 | Lighthouse Lab in Alderley Park | Wellcome Sanger Institute for the COVID-19 Genomics UK (COG-UK) Consortium | Jacquelyn Wynn, Mairead Hyland, The Lighthouse Lab in Alderley Park and Alex Alderton, Roberto Amato, Jeffrey Barrett, Sonia Goncalves, Ewan Harrison, David K. Jackson, Ian Johnston, Dominic Kwiatkowski, Cordelia Langford, John Sillitoe on behalf of the Wellcome Sanger Institute COVID-19 Surveillance Team |
| EPI_ISL_3310927 | Lighthouse Lab in Glasgow | Wellcome Sanger Institute for the COVID-19 Genomics UK (COG-UK) Consortium | Harper VanSteenhouse, Yumi Kasai, David Gray, Carol Clugston, Anna Dominiczak and Alex Alderton, Roberto Amato, Jeffrey Barrett, Sonia Goncalves, Ewan Harrison, David K. Jackson, Ian Johnston, Dominic Kwiatkowski, Cordelia Langford, John Sillitoe on behalf of the Wellcome Sanger Institute COVID-19 Surveillance Team |
| EPI_ISL_3310928 | Lighthouse Lab in Alderley Park | Wellcome Sanger Institute for the COVID-19 Genomics UK (COG-UK) Consortium | Jacquelyn Wynn, Mairead Hyland, The Lighthouse Lab in Alderley Park and Alex Alderton, Roberto Amato, Jeffrey Barrett, Sonia Goncalves, Ewan Harrison, David K. Jackson, Ian Johnston, Dominic Kwiatkowski, Cordelia Langford, John Sillitoe on behalf of the Wellcome Sanger Institute COVID-19 Surveillance Team |
| EPI_ISL_3310929 | Lighthouse Lab in Glasgow | Wellcome Sanger Institute for the COVID-19 Genomics UK (COG-UK) Consortium | Harper VanSteenhouse, Yumi Kasai, David Gray, Carol Clugston, Anna Dominiczak and Alex Alderton, Roberto Amato, Jeffrey Barrett, Sonia Goncalves, Ewan Harrison, David K. Jackson, Ian Johnston, Dominic Kwiatkowski, Cordelia Langford, John Sillitoe on behalf of the Wellcome Sanger Institute COVID-19 Surveillance Team |
| EPI_ISL_3310930, EPI_ISL_3310931, EPI_ISL_3310932, EPI_ISL_3310933, EPI_ISL_3310934, EPI_ISL_3310935, EPI_ISL_3310936, EPI_ISL_3310937, EPI_ISL_3310938, EPI_ISL_3310939, EPI_ISL_3310940, EPI_ISL_3310941, EPI_ISL_3310942, EPI_ISL_3310943, EPI_ISL_3310944 | | | |
| see above | Lighthouse Lab in Alderley Park | Wellcome Sanger Institute for the COVID-19 Genomics UK (COG-UK) Consortium | Jacquelyn Wynn, Mairead Hyland, The Lighthouse Lab in Alderley Park and Alex Alderton, Roberto Amato, Jeffrey Barrett, Sonia Goncalves, Ewan Harrison, David K. Jackson, Ian Johnston, Dominic Kwiatkowski, Cordelia Langford, John Sillitoe on behalf of the Wellcome Sanger Institute COVID-19 Surveillance Team |
| EPI_ISL_3310945 | Lighthouse Lab in Glasgow | Wellcome Sanger Institute for the COVID-19 Genomics UK (COG-UK) Consortium | Harper VanSteenhouse, Yumi Kasai, David Gray, Carol Clugston, Anna Dominiczak and Alex Alderton, Roberto Amato, Jeffrey Barrett, Sonia Goncalves, Ewan Harrison, David K. Jackson, Ian Johnston, Dominic Kwiatkowski, Cordelia Langford, John Sillitoe on behalf of the Wellcome Sanger Institute COVID-19 Surveillance Team |
| EPI_ISL_3310946, EPI_ISL_3310947, EPI_ISL_3310948, EPI_ISL_3310949, EPI_ISL_3310950, EPI_ISL_3310951, EPI_ISL_3310952, EPI_ISL_3310953, EPI_ISL_3310954, EPI_ISL_3310955 | Lighthouse Lab in Alderley Park | Wellcome Sanger Institute for the COVID-19 Genomics UK (COG-UK) Consortium | Jacquelyn Wynn, Mairead Hyland, The Lighthouse Lab in Alderley Park and Alex Alderton, Roberto Amato, Jeffrey Barrett, Sonia Goncalves, Ewan Harrison, David K. Jackson, Ian Johnston, Dominic Kwiatkowski, Cordelia Langford, John Sillitoe on behalf of the Wellcome Sanger Institute COVID-19 Surveillance Team |
| EPI_ISL_3310956 | Lighthouse Lab in Glasgow | Wellcome Sanger Institute for the COVID-19 Genomics UK (COG-UK) Consortium | Harper VanSteenhouse, Yumi Kasai, David Gray, Carol Clugston, Anna Dominiczak and Alex Alderton, Roberto Amato, Jeffrey Barrett, Sonia Goncalves, Ewan Harrison, David K. Jackson, Ian Johnston, Dominic Kwiatkowski, Cordelia Langford, John Sillitoe on behalf of the Wellcome Sanger Institute COVID-19 Surveillance Team |
| EPI_ISL_3310957, EPI_ISL_3310958, EPI_ISL_3310959, EPI_ISL_3310960, EPI_ISL_3310961, EPI_ISL_3310962 | Lighthouse Lab in Alderley Park | Wellcome Sanger Institute for the COVID-19 Genomics UK (COG-UK) Consortium | Jacquelyn Wynn, Mairead Hyland, The Lighthouse Lab in Alderley Park and Alex Alderton, Roberto Amato, Jeffrey Barrett, Sonia Goncalves, Ewan Harrison, David K. Jackson, Ian Johnston, Dominic Kwiatkowski, Cordelia Langford, John Sillitoe on behalf of the Wellcome Sanger Institute COVID-19 Surveillance Team |
| EPI_ISL_3310963 | Lighthouse Lab in Glasgow | Wellcome Sanger Institute for the COVID-19 Genomics UK (COG-UK) Consortium | Harper VanSteenhouse, Yumi Kasai, David Gray, Carol Clugston, Anna Dominiczak and Alex Alderton, Roberto Amato, Jeffrey Barrett, Sonia Goncalves, Ewan Harrison, David K. Jackson, Ian Johnston, Dominic Kwiatkowski, Cordelia Langford, John Sillitoe on behalf of the Wellcome Sanger Institute COVID-19 Surveillance Team |
| EPI_ISL_3310964, EPI_ISL_3310965, EPI_ISL_3310966, EPI_ISL_3310967, EPI_ISL_3310968, EPI_ISL_3310969, EPI_ISL_3310970, EPI_ISL_3310971, EPI_ISL_3310972, EPI_ISL_3310973 | Lighthouse Lab in Alderley Park | Wellcome Sanger Institute for the COVID-19 Genomics UK (COG-UK) Consortium | Jacquelyn Wynn, Mairead Hyland, The Lighthouse Lab in Alderley Park and Alex Alderton, Roberto Amato, Jeffrey Barrett, Sonia Goncalves, Ewan Harrison, David K. Jackson, Ian Johnston, Dominic Kwiatkowski, Cordelia Langford, John Sillitoe on behalf of the Wellcome Sanger Institute COVID-19 Surveillance Team |
| EPI_ISL_3310974 | Lighthouse Lab in Glasgow | Wellcome Sanger Institute for the COVID-19 Genomics UK (COG-UK) Consortium | Harper VanSteenhouse, Yumi Kasai, David Gray, Carol Clugston, Anna Dominiczak and Alex Alderton, Roberto Amato, Jeffrey Barrett, Sonia Goncalves, Ewan Harrison, David K. Jackson, Ian Johnston, Dominic Kwiatkowski, Cordelia Langford, John Sillitoe on behalf of the Wellcome Sanger Institute COVID-19 Surveillance Team |
| EPI_ISL_3310975, EPI_ISL_3310976, EPI_ISL_3310977, EPI_ISL_3310978, EPI_ISL_3310979, EPI_ISL_3310980, EPI_ISL_3310981, EPI_ISL_3310982, EPI_ISL_3310983 | Lighthouse Lab in Alderley Park | Wellcome Sanger Institute for the COVID-19 Genomics UK (COG-UK) Consortium | Jacquelyn Wynn, Mairead Hyland, The Lighthouse Lab in Alderley Park and Alex Alderton, Roberto Amato, Jeffrey Barrett, Sonia Goncalves, Ewan Harrison, David K. Jackson, Ian Johnston, Dominic Kwiatkowski, Cordelia Langford, John Sillitoe on behalf of the Wellcome Sanger Institute COVID-19 Surveillance Team |
| EPI_ISL_3310984 | Lighthouse Lab in Glasgow | Wellcome Sanger Institute for the COVID-19 Genomics UK (COG-UK) Consortium | Harper VanSteenhouse, Yumi Kasai, David Gray, Carol Clugston, Anna Dominiczak and Alex Alderton, Roberto Amato, Jeffrey Barrett, Sonia Goncalves, Ewan Harrison, David K. Jackson, Ian Johnston, Dominic Kwiatkowski, Cordelia Langford, John Sillitoe on behalf of the Wellcome Sanger Institute COVID-19 Surveillance Team |
| EPI_ISL_3310985, EPI_ISL_3310986 | Lighthouse Lab in Alderley Park | Wellcome Sanger Institute for the COVID-19 Genomics UK (COG-UK) Consortium | Jacquelyn Wynn, Mairead Hyland, The Lighthouse Lab in Alderley Park and Alex Alderton, Roberto Amato, Jeffrey Barrett, Sonia Goncalves, Ewan Harrison, David K. Jackson, Ian Johnston, Dominic Kwiatkowski, Cordelia Langford, John Sillitoe on behalf of the Wellcome Sanger Institute COVID-19 Surveillance Team |
| EPI_ISL_3310987 | Lighthouse Lab in Glasgow | Wellcome Sanger Institute for the COVID-19 Genomics UK (COG-UK) Consortium | Harper VanSteenhouse, Yumi Kasai, David Gray, Carol Clugston, Anna Dominiczak and Alex Alderton, Roberto Amato, Jeffrey Barrett, Sonia Goncalves, Ewan Harrison, David K. Jackson, Ian Johnston, Dominic Kwiatkowski, Cordelia Langford, John Sillitoe on behalf of the Wellcome Sanger Institute COVID-19 Surveillance Team |
| EPI_ISL_3310988, EPI_ISL_3310989, EPI_ISL_3310990, EPI_ISL_3310991, EPI_ISL_3310992, EPI_ISL_3310993, EPI_ISL_3310994, EPI_ISL_3310995, EPI_ISL_3310996, EPI_ISL_3310997, EPI_ISL_3310998 | | | |
| see above | Lighthouse Lab in Alderley Park | Wellcome Sanger Institute for the COVID-19 Genomics UK (COG-UK) Consortium | Jacquelyn Wynn, Mairead Hyland, The Lighthouse Lab in Alderley Park and Alex Alderton, Roberto Amato, Jeffrey Barrett, Sonia Goncalves, Ewan Harrison, David K. Jackson, Ian Johnston, Dominic Kwiatkowski, Cordelia Langford, John Sillitoe on behalf of the Wellcome Sanger Institute COVID-19 Surveillance Team |
| EPI_ISL_3310999, EPI_ISL_3311000 | Lighthouse Lab in Glasgow | Wellcome Sanger Institute for the COVID-19 Genomics UK (COG-UK) Consortium | Harper VanSteenhouse, Yumi Kasai, David Gray, Carol Clugston, Anna Dominiczak and Alex Alderton, Roberto Amato, Jeffrey Barrett, Sonia Goncalves, Ewan Harrison, David K. Jackson, Ian Johnston, Dominic Kwiatkowski, Cordelia Langford, John Sillitoe on behalf of the Wellcome Sanger Institute COVID-19 Surveillance Team |
| EPI_ISL_3311001, EPI_ISL_3311002, EPI_ISL_3311003 | Lighthouse Lab in Alderley Park | Wellcome Sanger Institute for the COVID-19 Genomics UK (COG-UK) Consortium | Jacquelyn Wynn, Mairead Hyland, The Lighthouse Lab in Alderley Park and Alex Alderton, Roberto Amato, Jeffrey Barrett, Sonia Goncalves, Ewan Harrison, David K. Jackson, Ian Johnston, Dominic Kwiatkowski, Cordelia Langford, John Sillitoe on behalf of the Wellcome Sanger Institute COVID-19 Surveillance Team |
| EPI_ISL_3311004 | Lighthouse Lab in Glasgow | Wellcome Sanger Institute for the COVID-19 Genomics UK (COG-UK) Consortium | Harper VanSteenhouse, Yumi Kasai, David Gray, Carol Clugston, Anna Dominiczak and Alex Alderton, Roberto Amato, Jeffrey Barrett, Sonia Goncalves, Ewan Harrison, David K. Jackson, Ian Johnston, Dominic Kwiatkowski, Cordelia Langford, John Sillitoe on behalf of the Wellcome Sanger Institute COVID-19 Surveillance Team |
| EPI_ISL_3311005, EPI_ISL_3311006, EPI_ISL_3311007 | Lighthouse Lab in Alderley Park | Wellcome Sanger Institute for the COVID-19 Genomics UK (COG-UK) Consortium | Jacquelyn Wynn, Mairead Hyland, The Lighthouse Lab in Alderley Park and Alex Alderton, Roberto Amato, Jeffrey Barrett, Sonia Goncalves, Ewan Harrison, David K. Jackson, Ian Johnston, Dominic Kwiatkowski, Cordelia Langford, John Sillitoe on behalf of the Wellcome Sanger Institute COVID-19 Surveillance Team |
| EPI_ISL_3311008, EPI_ISL_3311009 | Lighthouse Lab in Glasgow | Wellcome Sanger Institute for the COVID-19 Genomics UK (COG-UK) Consortium | Harper VanSteenhouse, Yumi Kasai, David Gray, Carol Clugston, Anna Dominiczak and Alex Alderton, Roberto Amato, Jeffrey Barrett, Sonia Goncalves, Ewan Harrison, David K. Jackson, Ian Johnston, Dominic Kwiatkowski, Cordelia Langford, John Sillitoe on behalf of the Wellcome Sanger Institute COVID-19 Surveillance Team |
| EPI_ISL_3311010, EPI_ISL_3311011, EPI_ISL_3311012, EPI_ISL_3311013, EPI_ISL_3311014 | Lighthouse Lab in Alderley Park | Wellcome Sanger Institute for the COVID-19 Genomics UK (COG-UK) Consortium | Jacquelyn Wynn, Mairead Hyland, The Lighthouse Lab in Alderley Park and Alex Alderton, Roberto Amato, Jeffrey Barrett, Sonia Goncalves, Ewan Harrison, David K. Jackson, Ian Johnston, Dominic Kwiatkowski, Cordelia Langford, John Sillitoe on behalf of the Wellcome Sanger Institute COVID-19 Surveillance Team |
| EPI_ISL_3311015 | Lighthouse Lab in Glasgow | Wellcome Sanger Institute for the COVID-19 Genomics UK (COG-UK) Consortium | Harper VanSteenhouse, Yumi Kasai, David Gray, Carol Clugston, Anna Dominiczak and Alex Alderton, Roberto Amato, Jeffrey Barrett, Sonia Goncalves, Ewan Harrison, David K. Jackson, Ian Johnston, Dominic Kwiatkowski, Cordelia Langford, John Sillitoe on behalf of the Wellcome Sanger Institute COVID-19 Surveillance Team |
| EPI_ISL_3311016, EPI_ISL_3311017, EPI_ISL_3311018, EPI_ISL_3311019, EPI_ISL_3311020, EPI_ISL_3311021, EPI_ISL_3311022, EPI_ISL_3311023, EPI_ISL_3311024, EPI_ISL_3311025, EPI_ISL_3311026, EPI_ISL_3311027, EPI_ISL_3311028, EPI_ISL_3311029 | | | |
| see above | Lighthouse Lab in Alderley Park | Wellcome Sanger Institute for the COVID-19 Genomics UK (COG-UK) Consortium | Jacquelyn Wynn, Mairead Hyland, The Lighthouse Lab in Alderley Park and Alex Alderton, Roberto Amato, Jeffrey Barrett, Sonia Goncalves, Ewan Harrison, David K. Jackson, Ian Johnston, Dominic Kwiatkowski, Cordelia Langford, John Sillitoe on behalf of the Wellcome Sanger Institute COVID-19 Surveillance Team |

| | | | |
|---|---|---|---|
| EPI_ISL_3311030, EPI_ISL_3311031 | Lighthouse Lab in Glasgow | Wellcome Sanger Institute for the COVID-19 Genomics UK (COG-UK) Consortium | Harper VanSteenhouse, Yumi Kasai, David Gray, Carol Clugston, Anna Dominiczak and Alex Alderton, Roberto Amato, Jeffrey Barrett, Sonia Goncalves, Ewan Harrison, David K. Jackson, Ian Johnston, Dominic Kwiatkowski, Cordelia Langford, John Sillitoe on behalf of the Wellcome Sanger Institute COVID-19 Surveillance Team |
| EPI_ISL_3311032, EPI_ISL_3311033 | Lighthouse Lab in Alderley Park | Wellcome Sanger Institute for the COVID-19 Genomics UK (COG-UK) Consortium | Jacquelyn Wynn, Mairead Hyland, The Lighthouse Lab in Alderley Park and Alex Alderton, Roberto Amato, Jeffrey Barrett, Sonia Goncalves, Ewan Harrison, David K. Jackson, Ian Johnston, Dominic Kwiatkowski, Cordelia Langford, John Sillitoe on behalf of the Wellcome Sanger Institute COVID-19 Surveillance Team |
| EPI_ISL_3311034 | Lighthouse Lab in Glasgow | Wellcome Sanger Institute for the COVID-19 Genomics UK (COG-UK) Consortium | Harper VanSteenhouse, Yumi Kasai, David Gray, Carol Clugston, Anna Dominiczak and Alex Alderton, Roberto Amato, Jeffrey Barrett, Sonia Goncalves, Ewan Harrison, David K. Jackson, Ian Johnston, Dominic Kwiatkowski, Cordelia Langford, John Sillitoe on behalf of the Wellcome Sanger Institute COVID-19 Surveillance Team |
| EPI_ISL_3311035 | Lighthouse Lab in Alderley Park | Wellcome Sanger Institute for the COVID-19 Genomics UK (COG-UK) Consortium | Jacquelyn Wynn, Mairead Hyland, The Lighthouse Lab in Alderley Park and Alex Alderton, Roberto Amato, Jeffrey Barrett, Sonia Goncalves, Ewan Harrison, David K. Jackson, Ian Johnston, Dominic Kwiatkowski, Cordelia Langford, John Sillitoe on behalf of the Wellcome Sanger Institute COVID-19 Surveillance Team |
| EPI_ISL_3311036 | Lighthouse Lab in Glasgow | Wellcome Sanger Institute for the COVID-19 Genomics UK (COG-UK) Consortium | Harper VanSteenhouse, Yumi Kasai, David Gray, Carol Clugston, Anna Dominiczak and Alex Alderton, Roberto Amato, Jeffrey Barrett, Sonia Goncalves, Ewan Harrison, David K. Jackson, Ian Johnston, Dominic Kwiatkowski, Cordelia Langford, John Sillitoe on behalf of the Wellcome Sanger Institute COVID-19 Surveillance Team |
| EPI_ISL_3311037, EPI_ISL_3311038, EPI_ISL_3311039, EPI_ISL_3311040, EPI_ISL_3311041, EPI_ISL_3311042 | Lighthouse Lab in Alderley Park | Wellcome Sanger Institute for the COVID-19 Genomics UK (COG-UK) Consortium | Jacquelyn Wynn, Mairead Hyland, The Lighthouse Lab in Alderley Park and Alex Alderton, Roberto Amato, Jeffrey Barrett, Sonia Goncalves, Ewan Harrison, David K. Jackson, Ian Johnston, Dominic Kwiatkowski, Cordelia Langford, John Sillitoe on behalf of the Wellcome Sanger Institute COVID-19 Surveillance Team |
| EPI_ISL_3311043 | Lighthouse Lab in Glasgow | Wellcome Sanger Institute for the COVID-19 Genomics UK (COG-UK) Consortium | Harper VanSteenhouse, Yumi Kasai, David Gray, Carol Clugston, Anna Dominiczak and Alex Alderton, Roberto Amato, Jeffrey Barrett, Sonia Goncalves, Ewan Harrison, David K. Jackson, Ian Johnston, Dominic Kwiatkowski, Cordelia Langford, John Sillitoe on behalf of the Wellcome Sanger Institute COVID-19 Surveillance Team |
| EPI_ISL_3311044, EPI_ISL_3311045 | Lighthouse Lab in Alderley Park | Wellcome Sanger Institute for the COVID-19 Genomics UK (COG-UK) Consortium | Jacquelyn Wynn, Mairead Hyland, The Lighthouse Lab in Alderley Park and Alex Alderton, Roberto Amato, Jeffrey Barrett, Sonia Goncalves, Ewan Harrison, David K. Jackson, Ian Johnston, Dominic Kwiatkowski, Cordelia Langford, John Sillitoe on behalf of the Wellcome Sanger Institute COVID-19 Surveillance Team |
| EPI_ISL_3311046, EPI_ISL_3311047, EPI_ISL_3311048 | Lighthouse Lab in Milton Keynes | Wellcome Sanger Institute for the COVID-19 Genomics UK (COG-UK) Consortium | The Lighthouse Lab in Milton Keynes and Alex Alderton, Roberto Amato, Jeffrey Barrett, Sonia Goncalves, Ewan Harrison, David K. Jackson, Ian Johnston, Dominic Kwiatkowski, Cordelia Langford, John Sillitoe on behalf of the Wellcome Sanger Institute COVID-19 Surveillance Team |
| EPI_ISL_3311049 | Lighthouse Lab in Alderley Park | Wellcome Sanger Institute for the COVID-19 Genomics UK (COG-UK) Consortium | Jacquelyn Wynn, Mairead Hyland, The Lighthouse Lab in Alderley Park and Alex Alderton, Roberto Amato, Jeffrey Barrett, Sonia Goncalves, Ewan Harrison, David K. Jackson, Ian Johnston, Dominic Kwiatkowski, Cordelia Langford, John Sillitoe on behalf of the Wellcome Sanger Institute COVID-19 Surveillance Team |
| EPI_ISL_3311050, EPI_ISL_3311051, EPI_ISL_3311052 | Lighthouse Lab in Milton Keynes | Wellcome Sanger Institute for the COVID-19 Genomics UK (COG-UK) Consortium | The Lighthouse Lab in Milton Keynes and Alex Alderton, Roberto Amato, Jeffrey Barrett, Sonia Goncalves, Ewan Harrison, David K. Jackson, Ian Johnston, Dominic Kwiatkowski, Cordelia Langford, John Sillitoe on behalf of the Wellcome Sanger Institute COVID-19 Surveillance Team |
| EPI_ISL_3311053 | Lighthouse Lab in Alderley Park | Wellcome Sanger Institute for the COVID-19 Genomics UK (COG-UK) Consortium | Jacquelyn Wynn, Mairead Hyland, The Lighthouse Lab in Alderley Park and Alex Alderton, Roberto Amato, Jeffrey Barrett, Sonia Goncalves, Ewan Harrison, David K. Jackson, Ian Johnston, Dominic Kwiatkowski, Cordelia Langford, John Sillitoe on behalf of the Wellcome Sanger Institute COVID-19 Surveillance Team |
| EPI_ISL_3311054, EPI_ISL_3311055, EPI_ISL_3311056, EPI_ISL_3311057, EPI_ISL_3311058, EPI_ISL_3311059 | Lighthouse Lab in Milton Keynes | Wellcome Sanger Institute for the COVID-19 Genomics UK (COG-UK) Consortium | The Lighthouse Lab in Milton Keynes and Alex Alderton, Roberto Amato, Jeffrey Barrett, Sonia Goncalves, Ewan Harrison, David K. Jackson, Ian Johnston, Dominic Kwiatkowski, Cordelia Langford, John Sillitoe on behalf of the Wellcome Sanger Institute COVID-19 Surveillance Team |
| EPI_ISL_3311060 | Lighthouse Lab in Alderley Park | Wellcome Sanger Institute for the COVID-19 Genomics UK (COG-UK) Consortium | Jacquelyn Wynn, Mairead Hyland, The Lighthouse Lab in Alderley Park and Alex Alderton, Roberto Amato, Jeffrey Barrett, Sonia Goncalves, Ewan Harrison, David K. Jackson, Ian Johnston, Dominic Kwiatkowski, Cordelia Langford, John Sillitoe on behalf of the Wellcome Sanger Institute COVID-19 Surveillance Team |
| EPI_ISL_3311061, EPI_ISL_3311062, EPI_ISL_3311063, EPI_ISL_3311064, EPI_ISL_3311065 | Lighthouse Lab in Milton Keynes | Wellcome Sanger Institute for the COVID-19 Genomics UK (COG-UK) Consortium | The Lighthouse Lab in Milton Keynes and Alex Alderton, Roberto Amato, Jeffrey Barrett, Sonia Goncalves, Ewan Harrison, David K. Jackson, Ian Johnston, Dominic Kwiatkowski, Cordelia Langford, John Sillitoe on behalf of the Wellcome Sanger Institute COVID-19 Surveillance Team |
| EPI_ISL_3311066 | Lighthouse Lab in Alderley Park | Wellcome Sanger Institute for the COVID-19 Genomics UK (COG-UK) Consortium | Jacquelyn Wynn, Mairead Hyland, The Lighthouse Lab in Alderley Park and Alex Alderton, Roberto Amato, Jeffrey Barrett, Sonia Goncalves, Ewan Harrison, David K. Jackson, Ian Johnston, Dominic Kwiatkowski, Cordelia Langford, John Sillitoe on behalf of the Wellcome Sanger Institute COVID-19 Surveillance Team |
| EPI_ISL_3311067, EPI_ISL_3311068, EPI_ISL_3311069, EPI_ISL_3311070, EPI_ISL_3311071, EPI_ISL_3311072, EPI_ISL_3311073 | Lighthouse Lab in Milton Keynes | Wellcome Sanger Institute for the COVID-19 Genomics UK (COG-UK) Consortium | The Lighthouse Lab in Milton Keynes and Alex Alderton, Roberto Amato, Jeffrey Barrett, Sonia Goncalves, Ewan Harrison, David K. Jackson, Ian Johnston, Dominic Kwiatkowski, Cordelia Langford, John Sillitoe on behalf of the Wellcome Sanger Institute COVID-19 Surveillance Team |
| EPI_ISL_3311074 | Lighthouse Lab in Alderley Park | Wellcome Sanger Institute for the COVID-19 Genomics UK (COG-UK) Consortium | Jacquelyn Wynn, Mairead Hyland, The Lighthouse Lab in Alderley Park and Alex Alderton, Roberto Amato, Jeffrey Barrett, Sonia Goncalves, Ewan Harrison, David K. Jackson, Ian Johnston, Dominic Kwiatkowski, Cordelia Langford, John Sillitoe on behalf of the Wellcome Sanger Institute COVID-19 Surveillance Team |
| EPI_ISL_3311075, EPI_ISL_3311076, EPI_ISL_3311077 | Lighthouse Lab in Milton Keynes | Wellcome Sanger Institute for the COVID-19 Genomics UK (COG-UK) Consortium | The Lighthouse Lab in Milton Keynes and Alex Alderton, Roberto Amato, Jeffrey Barrett, Sonia Goncalves, Ewan Harrison, David K. Jackson, Ian Johnston, Dominic Kwiatkowski, Cordelia Langford, John Sillitoe on behalf of the Wellcome Sanger Institute COVID-19 Surveillance Team |
| EPI_ISL_3311078 | Lighthouse Lab in Alderley Park | Wellcome Sanger Institute for the COVID-19 Genomics UK (COG-UK) Consortium | Jacquelyn Wynn, Mairead Hyland, The Lighthouse Lab in Alderley Park and Alex Alderton, Roberto Amato, Jeffrey Barrett, Sonia Goncalves, Ewan Harrison, David K. Jackson, Ian Johnston, Dominic Kwiatkowski, Cordelia Langford, John Sillitoe on behalf of the Wellcome Sanger Institute COVID-19 Surveillance Team |
| EPI_ISL_3311079, EPI_ISL_3311080, EPI_ISL_3311081, EPI_ISL_3311082, EPI_ISL_3311083, EPI_ISL_3311084, EPI_ISL_3311085 | Lighthouse Lab in Milton Keynes | Wellcome Sanger Institute for the COVID-19 Genomics UK (COG-UK) Consortium | The Lighthouse Lab in Milton Keynes and Alex Alderton, Roberto Amato, Jeffrey Barrett, Sonia Goncalves, Ewan Harrison, David K. Jackson, Ian Johnston, Dominic Kwiatkowski, Cordelia Langford, John Sillitoe on behalf of the Wellcome Sanger Institute COVID-19 Surveillance Team |
| EPI_ISL_3311086 | Lighthouse Lab in Alderley Park | Wellcome Sanger Institute for the COVID-19 Genomics UK (COG-UK) Consortium | Jacquelyn Wynn, Mairead Hyland, The Lighthouse Lab in Alderley Park and Alex Alderton, Roberto Amato, Jeffrey Barrett, Sonia Goncalves, Ewan Harrison, David K. Jackson, Ian Johnston, Dominic Kwiatkowski, Cordelia Langford, John Sillitoe on behalf of the Wellcome Sanger Institute COVID-19 Surveillance Team |
| EPI_ISL_3311087, EPI_ISL_3311088 | Lighthouse Lab in Milton Keynes | Wellcome Sanger Institute for the COVID-19 Genomics UK (COG-UK) Consortium | The Lighthouse Lab in Milton Keynes and Alex Alderton, Roberto Amato, Jeffrey Barrett, Sonia Goncalves, Ewan Harrison, David K. Jackson, Ian Johnston, Dominic Kwiatkowski, Cordelia Langford, John Sillitoe on behalf of the Wellcome Sanger Institute COVID-19 Surveillance Team |
| EPI_ISL_3311089 | Lighthouse Lab in Alderley Park | Wellcome Sanger Institute for the COVID-19 Genomics UK (COG-UK) Consortium | Jacquelyn Wynn, Mairead Hyland, The Lighthouse Lab in Alderley Park and Alex Alderton, Roberto Amato, Jeffrey Barrett, Sonia Goncalves, Ewan Harrison, David K. Jackson, Ian Johnston, Dominic Kwiatkowski, Cordelia Langford, John Sillitoe on behalf of the Wellcome Sanger Institute COVID-19 Surveillance Team |
| EPI_ISL_3311090, EPI_ISL_3311091, EPI_ISL_3311092, EPI_ISL_3311093 | Lighthouse Lab in Milton Keynes | Wellcome Sanger Institute for the COVID-19 Genomics UK (COG-UK) Consortium | The Lighthouse Lab in Milton Keynes and Alex Alderton, Roberto Amato, Jeffrey Barrett, Sonia Goncalves, Ewan Harrison, David K. Jackson, Ian Johnston, Dominic Kwiatkowski, Cordelia Langford, John Sillitoe on behalf of the Wellcome Sanger Institute COVID-19 Surveillance Team |
| EPI_ISL_3311094 | Lighthouse Lab in Alderley Park | Wellcome Sanger Institute for the COVID-19 Genomics UK (COG-UK) Consortium | Jacquelyn Wynn, Mairead Hyland, The Lighthouse Lab in Alderley Park and Alex Alderton, Roberto Amato, Jeffrey Barrett, Sonia Goncalves, Ewan Harrison, David K. Jackson, Ian Johnston, Dominic Kwiatkowski, Cordelia Langford, John Sillitoe on behalf of the Wellcome Sanger Institute COVID-19 Surveillance Team |
| EPI_ISL_3311095, EPI_ISL_3311096, EPI_ISL_3311097, EPI_ISL_3311098, EPI_ISL_3311099, EPI_ISL_3311100, EPI_ISL_3311101, EPI_ISL_3311102, EPI_ISL_3311103 | Lighthouse Lab in Milton Keynes | Wellcome Sanger Institute for the COVID-19 Genomics UK (COG-UK) Consortium | The Lighthouse Lab in Milton Keynes and Alex Alderton, Roberto Amato, Jeffrey Barrett, Sonia Goncalves, Ewan Harrison, David K. Jackson, Ian Johnston, Dominic Kwiatkowski, Cordelia Langford, John Sillitoe on behalf of the Wellcome Sanger Institute COVID-19 Surveillance Team |
| EPI_ISL_3311104 | Lighthouse Lab in Alderley Park | Wellcome Sanger Institute for the COVID-19 Genomics UK (COG-UK) Consortium | Jacquelyn Wynn, Mairead Hyland, The Lighthouse Lab in Alderley Park and Alex Alderton, Roberto Amato, Jeffrey Barrett, Sonia Goncalves, Ewan Harrison, David K. Jackson, Ian Johnston, Dominic Kwiatkowski, Cordelia Langford, John Sillitoe on behalf of the Wellcome Sanger Institute COVID-19 Surveillance Team |
| EPI_ISL_3311105, EPI_ISL_3311106, EPI_ISL_3311107, EPI_ISL_3311108 | Lighthouse Lab in Milton Keynes | Wellcome Sanger Institute for the COVID-19 Genomics UK (COG-UK) Consortium | The Lighthouse Lab in Milton Keynes and Alex Alderton, Roberto Amato, Jeffrey Barrett, Sonia Goncalves, Ewan Harrison, David K. Jackson, Ian Johnston, Dominic Kwiatkowski, Cordelia Langford, John Sillitoe on behalf of the Wellcome Sanger Institute COVID-19 Surveillance Team |
| EPI_ISL_3311109 | Lighthouse Lab in Alderley Park | Wellcome Sanger Institute for the COVID-19 Genomics UK (COG-UK) Consortium | Jacquelyn Wynn, Mairead Hyland, The Lighthouse Lab in Alderley Park and Alex Alderton, Roberto Amato, Jeffrey Barrett, Sonia Goncalves, Ewan Harrison, David K. Jackson, Ian Johnston, Dominic Kwiatkowski, Cordelia Langford, John Sillitoe on behalf of the Wellcome Sanger Institute COVID-19 Surveillance Team |
| EPI_ISL_3311110, EPI_ISL_3311111, | Lighthouse Lab in Milton Keynes | Wellcome Sanger Institute for the COVID-19 Genomics UK (COG-UK) Consortium | The Lighthouse Lab in Milton Keynes and Alex Alderton, Roberto Amato, Jeffrey Barrett, Sonia Goncalves, Ewan Harrison, David K. Jackson, Ian Johnston, Dominic Kwiatkowski, Cordelia Langford, John Sillitoe on behalf of the Wellcome Sanger Institute COVID-19 Surveillance Team |

| | | | |
|---|---|---|---|
| EPI_ISL_3311112, EPI_ISL_3311113, EPI_ISL_3311114, EPI_ISL_3311115, EPI_ISL_3311116 | | | |
| EPI_ISL_3311117 | Lighthouse Lab in Alderley Park | Wellcome Sanger Institute for the COVID-19 Genomics UK (COG-UK) Consortium | Jacquelyn Wynn, Mairead Hyland, The Lighthouse Lab in Alderley Park and Alex Alderton, Roberto Amato, Jeffrey Barrett, Sonia Goncalves, Ewan Harrison, David K. Jackson, Ian Johnston, Dominic Kwiatkowski, Cordelia Langford, John Sillitoe on behalf of the Wellcome Sanger Institute COVID-19 Surveillance Team |
| EPI_ISL_3311118 | Lighthouse Lab in Milton Keynes | Wellcome Sanger Institute for the COVID-19 Genomics UK (COG-UK) Consortium | The Lighthouse Lab in Milton Keynes and Alex Alderton, Roberto Amato, Jeffrey Barrett, Sonia Goncalves, Ewan Harrison, David K. Jackson, Ian Johnston, Dominic Kwiatkowski, Cordelia Langford, John Sillitoe on behalf of the Wellcome Sanger Institute COVID-19 Surveillance Team |
| EPI_ISL_3311119 | Lighthouse Lab in Alderley Park | Wellcome Sanger Institute for the COVID-19 Genomics UK (COG-UK) Consortium | Jacquelyn Wynn, Mairead Hyland, The Lighthouse Lab in Alderley Park and Alex Alderton, Roberto Amato, Jeffrey Barrett, Sonia Goncalves, Ewan Harrison, David K. Jackson, Ian Johnston, Dominic Kwiatkowski, Cordelia Langford, John Sillitoe on behalf of the Wellcome Sanger Institute COVID-19 Surveillance Team |
| EPI_ISL_3311120, EPI_ISL_3311121 | Lighthouse Lab in Milton Keynes | Wellcome Sanger Institute for the COVID-19 Genomics UK (COG-UK) Consortium | The Lighthouse Lab in Milton Keynes and Alex Alderton, Roberto Amato, Jeffrey Barrett, Sonia Goncalves, Ewan Harrison, David K. Jackson, Ian Johnston, Dominic Kwiatkowski, Cordelia Langford, John Sillitoe on behalf of the Wellcome Sanger Institute COVID-19 Surveillance Team |
| EPI_ISL_3311122 | Lighthouse Lab in Alderley Park | Wellcome Sanger Institute for the COVID-19 Genomics UK (COG-UK) Consortium | Jacquelyn Wynn, Mairead Hyland, The Lighthouse Lab in Alderley Park and Alex Alderton, Roberto Amato, Jeffrey Barrett, Sonia Goncalves, Ewan Harrison, David K. Jackson, Ian Johnston, Dominic Kwiatkowski, Cordelia Langford, John Sillitoe on behalf of the Wellcome Sanger Institute COVID-19 Surveillance Team |
| EPI_ISL_3311123, EPI_ISL_3311124, EPI_ISL_3311125, EPI_ISL_3311126, EPI_ISL_3311127, EPI_ISL_3311128, EPI_ISL_3311129 | Lighthouse Lab in Milton Keynes | Wellcome Sanger Institute for the COVID-19 Genomics UK (COG-UK) Consortium | The Lighthouse Lab in Milton Keynes and Alex Alderton, Roberto Amato, Jeffrey Barrett, Sonia Goncalves, Ewan Harrison, David K. Jackson, Ian Johnston, Dominic Kwiatkowski, Cordelia Langford, John Sillitoe on behalf of the Wellcome Sanger Institute COVID-19 Surveillance Team |
| EPI_ISL_3311130 | Lighthouse Lab in Alderley Park | Wellcome Sanger Institute for the COVID-19 Genomics UK (COG-UK) Consortium | Jacquelyn Wynn, Mairead Hyland, The Lighthouse Lab in Alderley Park and Alex Alderton, Roberto Amato, Jeffrey Barrett, Sonia Goncalves, Ewan Harrison, David K. Jackson, Ian Johnston, Dominic Kwiatkowski, Cordelia Langford, John Sillitoe on behalf of the Wellcome Sanger Institute COVID-19 Surveillance Team |
| EPI_ISL_3311131, EPI_ISL_3311132, EPI_ISL_3311133, EPI_ISL_3311134, EPI_ISL_3311135, EPI_ISL_3311136, EPI_ISL_3311137 | Lighthouse Lab in Milton Keynes | Wellcome Sanger Institute for the COVID-19 Genomics UK (COG-UK) Consortium | The Lighthouse Lab in Milton Keynes and Alex Alderton, Roberto Amato, Jeffrey Barrett, Sonia Goncalves, Ewan Harrison, David K. Jackson, Ian Johnston, Dominic Kwiatkowski, Cordelia Langford, John Sillitoe on behalf of the Wellcome Sanger Institute COVID-19 Surveillance Team |
| EPI_ISL_3311138 | Lighthouse Lab in Alderley Park | Wellcome Sanger Institute for the COVID-19 Genomics UK (COG-UK) Consortium | Jacquelyn Wynn, Mairead Hyland, The Lighthouse Lab in Alderley Park and Alex Alderton, Roberto Amato, Jeffrey Barrett, Sonia Goncalves, Ewan Harrison, David K. Jackson, Ian Johnston, Dominic Kwiatkowski, Cordelia Langford, John Sillitoe on behalf of the Wellcome Sanger Institute COVID-19 Surveillance Team |
| EPI_ISL_3311139 | Lighthouse Lab in Milton Keynes | Wellcome Sanger Institute for the COVID-19 Genomics UK (COG-UK) Consortium | The Lighthouse Lab in Milton Keynes and Alex Alderton, Roberto Amato, Jeffrey Barrett, Sonia Goncalves, Ewan Harrison, David K. Jackson, Ian Johnston, Dominic Kwiatkowski, Cordelia Langford, John Sillitoe on behalf of the Wellcome Sanger Institute COVID-19 Surveillance Team |
| EPI_ISL_3311140 | Lighthouse Lab in Alderley Park | Wellcome Sanger Institute for the COVID-19 Genomics UK (COG-UK) Consortium | Jacquelyn Wynn, Mairead Hyland, The Lighthouse Lab in Alderley Park and Alex Alderton, Roberto Amato, Jeffrey Barrett, Sonia Goncalves, Ewan Harrison, David K. Jackson, Ian Johnston, Dominic Kwiatkowski, Cordelia Langford, John Sillitoe on behalf of the Wellcome Sanger Institute COVID-19 Surveillance Team |
| EPI_ISL_3311141, EPI_ISL_3311142, EPI_ISL_3311143, EPI_ISL_3311144, EPI_ISL_3311145 | Lighthouse Lab in Milton Keynes | Wellcome Sanger Institute for the COVID-19 Genomics UK (COG-UK) Consortium | The Lighthouse Lab in Milton Keynes and Alex Alderton, Roberto Amato, Jeffrey Barrett, Sonia Goncalves, Ewan Harrison, David K. Jackson, Ian Johnston, Dominic Kwiatkowski, Cordelia Langford, John Sillitoe on behalf of the Wellcome Sanger Institute COVID-19 Surveillance Team |
| EPI_ISL_3311146, EPI_ISL_3311147 | Lighthouse Lab in Alderley Park | Wellcome Sanger Institute for the COVID-19 Genomics UK (COG-UK) Consortium | Jacquelyn Wynn, Mairead Hyland, The Lighthouse Lab in Alderley Park and Alex Alderton, Roberto Amato, Jeffrey Barrett, Sonia Goncalves, Ewan Harrison, David K. Jackson, Ian Johnston, Dominic Kwiatkowski, Cordelia Langford, John Sillitoe on behalf of the Wellcome Sanger Institute COVID-19 Surveillance Team |
| EPI_ISL_3311148, EPI_ISL_3311149, EPI_ISL_3311150 | Lighthouse Lab in Milton Keynes | Wellcome Sanger Institute for the COVID-19 Genomics UK (COG-UK) Consortium | The Lighthouse Lab in Milton Keynes and Alex Alderton, Roberto Amato, Jeffrey Barrett, Sonia Goncalves, Ewan Harrison, David K. Jackson, Ian Johnston, Dominic Kwiatkowski, Cordelia Langford, John Sillitoe on behalf of the Wellcome Sanger Institute COVID-19 Surveillance Team |
| EPI_ISL_3311151, EPI_ISL_3311152 | Lighthouse Lab in Alderley Park | Wellcome Sanger Institute for the COVID-19 Genomics UK (COG-UK) Consortium | Jacquelyn Wynn, Mairead Hyland, The Lighthouse Lab in Alderley Park and Alex Alderton, Roberto Amato, Jeffrey Barrett, Sonia Goncalves, Ewan Harrison, David K. Jackson, Ian Johnston, Dominic Kwiatkowski, Cordelia Langford, John Sillitoe on behalf of the Wellcome Sanger Institute COVID-19 Surveillance Team |
| EPI_ISL_3311153 | Lighthouse Lab in Milton Keynes | Wellcome Sanger Institute for the COVID-19 Genomics UK (COG-UK) Consortium | The Lighthouse Lab in Milton Keynes and Alex Alderton, Roberto Amato, Jeffrey Barrett, Sonia Goncalves, Ewan Harrison, David K. Jackson, Ian Johnston, Dominic Kwiatkowski, Cordelia Langford, John Sillitoe on behalf of the Wellcome Sanger Institute COVID-19 Surveillance Team |
| EPI_ISL_3311154 | Lighthouse Lab in Alderley Park | Wellcome Sanger Institute for the COVID-19 Genomics UK (COG-UK) Consortium | Jacquelyn Wynn, Mairead Hyland, The Lighthouse Lab in Alderley Park and Alex Alderton, Roberto Amato, Jeffrey Barrett, Sonia Goncalves, Ewan Harrison, David K. Jackson, Ian Johnston, Dominic Kwiatkowski, Cordelia Langford, John Sillitoe on behalf of the Wellcome Sanger Institute COVID-19 Surveillance Team |
| EPI_ISL_3311155, EPI_ISL_3311156, EPI_ISL_3311157 | Lighthouse Lab in Milton Keynes | Wellcome Sanger Institute for the COVID-19 Genomics UK (COG-UK) Consortium | The Lighthouse Lab in Milton Keynes and Alex Alderton, Roberto Amato, Jeffrey Barrett, Sonia Goncalves, Ewan Harrison, David K. Jackson, Ian Johnston, Dominic Kwiatkowski, Cordelia Langford, John Sillitoe on behalf of the Wellcome Sanger Institute COVID-19 Surveillance Team |
| EPI_ISL_3311158 | Lighthouse Lab in Alderley Park | Wellcome Sanger Institute for the COVID-19 Genomics UK (COG-UK) Consortium | Jacquelyn Wynn, Mairead Hyland, The Lighthouse Lab in Alderley Park and Alex Alderton, Roberto Amato, Jeffrey Barrett, Sonia Goncalves, Ewan Harrison, David K. Jackson, Ian Johnston, Dominic Kwiatkowski, Cordelia Langford, John Sillitoe on behalf of the Wellcome Sanger Institute COVID-19 Surveillance Team |
| EPI_ISL_3311159 | Lighthouse Lab in Milton Keynes | Wellcome Sanger Institute for the COVID-19 Genomics UK (COG-UK) Consortium | The Lighthouse Lab in Milton Keynes and Alex Alderton, Roberto Amato, Jeffrey Barrett, Sonia Goncalves, Ewan Harrison, David K. Jackson, Ian Johnston, Dominic Kwiatkowski, Cordelia Langford, John Sillitoe on behalf of the Wellcome Sanger Institute COVID-19 Surveillance Team |
| EPI_ISL_3311160 | Lighthouse Lab in Alderley Park | Wellcome Sanger Institute for the COVID-19 Genomics UK (COG-UK) Consortium | Jacquelyn Wynn, Mairead Hyland, The Lighthouse Lab in Alderley Park and Alex Alderton, Roberto Amato, Jeffrey Barrett, Sonia Goncalves, Ewan Harrison, David K. Jackson, Ian Johnston, Dominic Kwiatkowski, Cordelia Langford, John Sillitoe on behalf of the Wellcome Sanger Institute COVID-19 Surveillance Team |
| EPI_ISL_3311161, EPI_ISL_3311162, EPI_ISL_3311163 | Lighthouse Lab in Milton Keynes | Wellcome Sanger Institute for the COVID-19 Genomics UK (COG-UK) Consortium | The Lighthouse Lab in Milton Keynes and Alex Alderton, Roberto Amato, Jeffrey Barrett, Sonia Goncalves, Ewan Harrison, David K. Jackson, Ian Johnston, Dominic Kwiatkowski, Cordelia Langford, John Sillitoe on behalf of the Wellcome Sanger Institute COVID-19 Surveillance Team |
| EPI_ISL_3311164, EPI_ISL_3311165, EPI_ISL_3311166 | Lighthouse Lab in Alderley Park | Wellcome Sanger Institute for the COVID-19 Genomics UK (COG-UK) Consortium | Jacquelyn Wynn, Mairead Hyland, The Lighthouse Lab in Alderley Park and Alex Alderton, Roberto Amato, Jeffrey Barrett, Sonia Goncalves, Ewan Harrison, David K. Jackson, Ian Johnston, Dominic Kwiatkowski, Cordelia Langford, John Sillitoe on behalf of the Wellcome Sanger Institute COVID-19 Surveillance Team |
| EPI_ISL_3311167, EPI_ISL_3311168 | Lighthouse Lab in Milton Keynes | Wellcome Sanger Institute for the COVID-19 Genomics UK (COG-UK) Consortium | The Lighthouse Lab in Milton Keynes and Alex Alderton, Roberto Amato, Jeffrey Barrett, Sonia Goncalves, Ewan Harrison, David K. Jackson, Ian Johnston, Dominic Kwiatkowski, Cordelia Langford, John Sillitoe on behalf of the Wellcome Sanger Institute COVID-19 Surveillance Team |
| EPI_ISL_3311169 | Lighthouse Lab in Alderley Park | Wellcome Sanger Institute for the COVID-19 Genomics UK (COG-UK) Consortium | Jacquelyn Wynn, Mairead Hyland, The Lighthouse Lab in Alderley Park and Alex Alderton, Roberto Amato, Jeffrey Barrett, Sonia Goncalves, Ewan Harrison, David K. Jackson, Ian Johnston, Dominic Kwiatkowski, Cordelia Langford, John Sillitoe on behalf of the Wellcome Sanger Institute COVID-19 Surveillance Team |
| EPI_ISL_3311170, EPI_ISL_3311171, EPI_ISL_3311172, EPI_ISL_3311173, EPI_ISL_3311174, EPI_ISL_3311175, EPI_ISL_3311176, EPI_ISL_3311177, EPI_ISL_3311178, EPI_ISL_3311179, EPI_ISL_3311180 | | | |
| see above | Lighthouse Lab in Milton Keynes | Wellcome Sanger Institute for the COVID-19 Genomics UK (COG-UK) Consortium | The Lighthouse Lab in Milton Keynes and Alex Alderton, Roberto Amato, Jeffrey Barrett, Sonia Goncalves, Ewan Harrison, David K. Jackson, Ian Johnston, Dominic Kwiatkowski, Cordelia Langford, John Sillitoe on behalf of the Wellcome Sanger Institute COVID-19 Surveillance Team |
| EPI_ISL_3311181 | Lighthouse Lab in Alderley Park | Wellcome Sanger Institute for the COVID-19 Genomics UK (COG-UK) Consortium | Jacquelyn Wynn, Mairead Hyland, The Lighthouse Lab in Alderley Park and Alex Alderton, Roberto Amato, Jeffrey Barrett, Sonia Goncalves, Ewan Harrison, David K. Jackson, Ian Johnston, Dominic Kwiatkowski, Cordelia Langford, John Sillitoe on behalf of the Wellcome Sanger Institute COVID-19 Surveillance Team |
| EPI_ISL_3311182, EPI_ISL_3311183, EPI_ISL_3311184, EPI_ISL_3311185, EPI_ISL_3311186, EPI_ISL_3311187 | Lighthouse Lab in Milton Keynes | Wellcome Sanger Institute for the COVID-19 Genomics UK (COG-UK) Consortium | The Lighthouse Lab in Milton Keynes and Alex Alderton, Roberto Amato, Jeffrey Barrett, Sonia Goncalves, Ewan Harrison, David K. Jackson, Ian Johnston, Dominic Kwiatkowski, Cordelia Langford, John Sillitoe on behalf of the Wellcome Sanger Institute COVID-19 Surveillance Team |
| EPI_ISL_3311188, EPI_ISL_3311189 | Lighthouse Lab in Alderley Park | Wellcome Sanger Institute for the COVID-19 Genomics UK (COG-UK) Consortium | Jacquelyn Wynn, Mairead Hyland, The Lighthouse Lab in Alderley Park and Alex Alderton, Roberto Amato, Jeffrey Barrett, Sonia Goncalves, Ewan Harrison, David K. Jackson, Ian Johnston, Dominic Kwiatkowski, Cordelia Langford, John Sillitoe on behalf of the Wellcome Sanger Institute COVID-19 Surveillance Team |
| EPI_ISL_3311190 | Lighthouse Lab in Milton Keynes | Wellcome Sanger Institute for the COVID-19 Genomics UK (COG-UK) Consortium | The Lighthouse Lab in Milton Keynes and Alex Alderton, Roberto Amato, Jeffrey Barrett, Sonia Goncalves, Ewan Harrison, David K. Jackson, Ian Johnston, Dominic Kwiatkowski, Cordelia Langford, John Sillitoe on behalf of the Wellcome Sanger Institute COVID-19 Surveillance Team |
| EPI_ISL_3311191 | Lighthouse Lab in Alderley Park | Wellcome Sanger Institute for the COVID-19 Genomics UK (COG-UK) Consortium | Jacquelyn Wynn, Mairead Hyland, The Lighthouse Lab in Alderley Park and Alex Alderton, Roberto Amato, Jeffrey Barrett, Sonia Goncalves, Ewan Harrison, David K. Jackson, Ian Johnston, Dominic Kwiatkowski, Cordelia Langford, John Sillitoe on behalf of the Wellcome Sanger Institute COVID-19 Surveillance Team |
| EPI_ISL_3311192 | Lighthouse Lab in Milton Keynes | Wellcome Sanger Institute for the COVID-19 Genomics UK (COG-UK) Consortium | The Lighthouse Lab in Milton Keynes and Alex Alderton, Roberto Amato, Jeffrey Barrett, Sonia Goncalves, Ewan Harrison, David K. Jackson, Ian Johnston, Dominic Kwiatkowski, Cordelia Langford, John Sillitoe on behalf of the Wellcome Sanger Institute COVID-19 Surveillance Team |
| EPI_ISL_3311193 | Lighthouse Lab in Alderley Park | Wellcome Sanger Institute for the COVID-19 Genomics UK (COG-UK) Consortium | Jacquelyn Wynn, Mairead Hyland, The Lighthouse Lab in Alderley Park and Alex Alderton, Roberto Amato, Jeffrey Barrett, Sonia Goncalves, Ewan Harrison, David K. Jackson, Ian Johnston, Dominic Kwiatkowski, Cordelia Langford, John Sillitoe on behalf of the Wellcome Sanger Institute COVID-19 Surveillance Team |
| EPI_ISL_3311194, EPI_ISL_3311195, | Lighthouse Lab in Milton Keynes | Wellcome Sanger Institute for the COVID-19 Genomics UK (COG-UK) Consortium | The Lighthouse Lab in Milton Keynes and Alex Alderton, Roberto Amato, Jeffrey Barrett, Sonia Goncalves, Ewan Harrison, David K. Jackson, Ian Johnston, Dominic Kwiatkowski, Cordelia Langford, John Sillitoe on behalf of the Wellcome Sanger Institute COVID-19 Surveillance Team |

| | | | |
|---|---|---|---|
| EPI_ISL_3311196, EPI_ISL_3311197, EPI_ISL_3311198, EPI_ISL_3311199, EPI_ISL_3311200 | | | |
| EPI_ISL_3311201 | Lighthouse Lab in Alderley Park | Wellcome Sanger Institute for the COVID-19 Genomics UK (COG-UK) Consortium | Jacquelyn Wynn, Mairead Hyland, The Lighthouse Lab in Alderley Park and Alex Alderton, Roberto Amato, Jeffrey Barrett, Sonia Goncalves, Ewan Harrison, David K. Jackson, Ian Johnston, Dominic Kwiatkowski, Cordelia Langford, John Sillitoe on behalf of the Wellcome Sanger Institute COVID-19 Surveillance Team |
| EPI_ISL_3311202, EPI_ISL_3311203, EPI_ISL_3311204, EPI_ISL_3311205 | Lighthouse Lab in Milton Keynes | Wellcome Sanger Institute for the COVID-19 Genomics UK (COG-UK) Consortium | The Lighthouse Lab in Milton Keynes and Alex Alderton, Roberto Amato, Jeffrey Barrett, Sonia Goncalves, Ewan Harrison, David K. Jackson, Ian Johnston, Dominic Kwiatkowski, Cordelia Langford, John Sillitoe on behalf of the Wellcome Sanger Institute COVID-19 Surveillance Team |
| EPI_ISL_3311206 | Lighthouse Lab in Alderley Park | Wellcome Sanger Institute for the COVID-19 Genomics UK (COG-UK) Consortium | Jacquelyn Wynn, Mairead Hyland, The Lighthouse Lab in Alderley Park and Alex Alderton, Roberto Amato, Jeffrey Barrett, Sonia Goncalves, Ewan Harrison, David K. Jackson, Ian Johnston, Dominic Kwiatkowski, Cordelia Langford, John Sillitoe on behalf of the Wellcome Sanger Institute COVID-19 Surveillance Team |
| EPI_ISL_3311207, EPI_ISL_3311208, EPI_ISL_3311209, EPI_ISL_3311210, EPI_ISL_3311211 | Lighthouse Lab in Milton Keynes | Wellcome Sanger Institute for the COVID-19 Genomics UK (COG-UK) Consortium | The Lighthouse Lab in Milton Keynes and Alex Alderton, Roberto Amato, Jeffrey Barrett, Sonia Goncalves, Ewan Harrison, David K. Jackson, Ian Johnston, Dominic Kwiatkowski, Cordelia Langford, John Sillitoe on behalf of the Wellcome Sanger Institute COVID-19 Surveillance Team |
| EPI_ISL_3311212 | Lighthouse Lab in Alderley Park | Wellcome Sanger Institute for the COVID-19 Genomics UK (COG-UK) Consortium | Jacquelyn Wynn, Mairead Hyland, The Lighthouse Lab in Alderley Park and Alex Alderton, Roberto Amato, Jeffrey Barrett, Sonia Goncalves, Ewan Harrison, David K. Jackson, Ian Johnston, Dominic Kwiatkowski, Cordelia Langford, John Sillitoe on behalf of the Wellcome Sanger Institute COVID-19 Surveillance Team |
| EPI_ISL_3311213, EPI_ISL_3311214, EPI_ISL_3311215, EPI_ISL_3311216 | Lighthouse Lab in Milton Keynes | Wellcome Sanger Institute for the COVID-19 Genomics UK (COG-UK) Consortium | The Lighthouse Lab in Milton Keynes and Alex Alderton, Roberto Amato, Jeffrey Barrett, Sonia Goncalves, Ewan Harrison, David K. Jackson, Ian Johnston, Dominic Kwiatkowski, Cordelia Langford, John Sillitoe on behalf of the Wellcome Sanger Institute COVID-19 Surveillance Team |
| EPI_ISL_3311217 | Lighthouse Lab in Alderley Park | Wellcome Sanger Institute for the COVID-19 Genomics UK (COG-UK) Consortium | Jacquelyn Wynn, Mairead Hyland, The Lighthouse Lab in Alderley Park and Alex Alderton, Roberto Amato, Jeffrey Barrett, Sonia Goncalves, Ewan Harrison, David K. Jackson, Ian Johnston, Dominic Kwiatkowski, Cordelia Langford, John Sillitoe on behalf of the Wellcome Sanger Institute COVID-19 Surveillance Team |
| EPI_ISL_3311218, EPI_ISL_3311219, EPI_ISL_3311220, EPI_ISL_3311221, EPI_ISL_3311222 | Lighthouse Lab in Milton Keynes | Wellcome Sanger Institute for the COVID-19 Genomics UK (COG-UK) Consortium | The Lighthouse Lab in Milton Keynes and Alex Alderton, Roberto Amato, Jeffrey Barrett, Sonia Goncalves, Ewan Harrison, David K. Jackson, Ian Johnston, Dominic Kwiatkowski, Cordelia Langford, John Sillitoe on behalf of the Wellcome Sanger Institute COVID-19 Surveillance Team |
| EPI_ISL_3311223, EPI_ISL_3311224, EPI_ISL_3311225 | Lighthouse Lab in Alderley Park | Wellcome Sanger Institute for the COVID-19 Genomics UK (COG-UK) Consortium | Jacquelyn Wynn, Mairead Hyland, The Lighthouse Lab in Alderley Park and Alex Alderton, Roberto Amato, Jeffrey Barrett, Sonia Goncalves, Ewan Harrison, David K. Jackson, Ian Johnston, Dominic Kwiatkowski, Cordelia Langford, John Sillitoe on behalf of the Wellcome Sanger Institute COVID-19 Surveillance Team |
| EPI_ISL_3311226 | Lighthouse Lab in Milton Keynes | Wellcome Sanger Institute for the COVID-19 Genomics UK (COG-UK) Consortium | The Lighthouse Lab in Milton Keynes and Alex Alderton, Roberto Amato, Jeffrey Barrett, Sonia Goncalves, Ewan Harrison, David K. Jackson, Ian Johnston, Dominic Kwiatkowski, Cordelia Langford, John Sillitoe on behalf of the Wellcome Sanger Institute COVID-19 Surveillance Team |
| EPI_ISL_3311227 | Lighthouse Lab in Alderley Park | Wellcome Sanger Institute for the COVID-19 Genomics UK (COG-UK) Consortium | Jacquelyn Wynn, Mairead Hyland, The Lighthouse Lab in Alderley Park and Alex Alderton, Roberto Amato, Jeffrey Barrett, Sonia Goncalves, Ewan Harrison, David K. Jackson, Ian Johnston, Dominic Kwiatkowski, Cordelia Langford, John Sillitoe on behalf of the Wellcome Sanger Institute COVID-19 Surveillance Team |
| EPI_ISL_3311228, EPI_ISL_3311229, EPI_ISL_3311230, EPI_ISL_3311231, EPI_ISL_3311232, EPI_ISL_3311233, EPI_ISL_3311234 | Lighthouse Lab in Milton Keynes | Wellcome Sanger Institute for the COVID-19 Genomics UK (COG-UK) Consortium | The Lighthouse Lab in Milton Keynes and Alex Alderton, Roberto Amato, Jeffrey Barrett, Sonia Goncalves, Ewan Harrison, David K. Jackson, Ian Johnston, Dominic Kwiatkowski, Cordelia Langford, John Sillitoe on behalf of the Wellcome Sanger Institute COVID-19 Surveillance Team |
| EPI_ISL_3311235, EPI_ISL_3311236 | Lighthouse Lab in Alderley Park | Wellcome Sanger Institute for the COVID-19 Genomics UK (COG-UK) Consortium | Jacquelyn Wynn, Mairead Hyland, The Lighthouse Lab in Alderley Park and Alex Alderton, Roberto Amato, Jeffrey Barrett, Sonia Goncalves, Ewan Harrison, David K. Jackson, Ian Johnston, Dominic Kwiatkowski, Cordelia Langford, John Sillitoe on behalf of the Wellcome Sanger Institute COVID-19 Surveillance Team |
| EPI_ISL_3311237, EPI_ISL_3311238, EPI_ISL_3311239, EPI_ISL_3311240, EPI_ISL_3311241, EPI_ISL_3311242, EPI_ISL_3311243, EPI_ISL_3311244, EPI_ISL_3311245, EPI_ISL_3311246, EPI_ISL_3311247 | | | |
| see above | Lighthouse Lab in Milton Keynes | Wellcome Sanger Institute for the COVID-19 Genomics UK (COG-UK) Consortium | The Lighthouse Lab in Milton Keynes and Alex Alderton, Roberto Amato, Jeffrey Barrett, Sonia Goncalves, Ewan Harrison, David K. Jackson, Ian Johnston, Dominic Kwiatkowski, Cordelia Langford, John Sillitoe on behalf of the Wellcome Sanger Institute COVID-19 Surveillance Team |
| EPI_ISL_3311248 | Lighthouse Lab in Alderley Park | Wellcome Sanger Institute for the COVID-19 Genomics UK (COG-UK) Consortium | Jacquelyn Wynn, Mairead Hyland, The Lighthouse Lab in Alderley Park and Alex Alderton, Roberto Amato, Jeffrey Barrett, Sonia Goncalves, Ewan Harrison, David K. Jackson, Ian Johnston, Dominic Kwiatkowski, Cordelia Langford, John Sillitoe on behalf of the Wellcome Sanger Institute COVID-19 Surveillance Team |
| EPI_ISL_3311249, EPI_ISL_3311250 | Lighthouse Lab in Milton Keynes | Wellcome Sanger Institute for the COVID-19 Genomics UK (COG-UK) Consortium | The Lighthouse Lab in Milton Keynes and Alex Alderton, Roberto Amato, Jeffrey Barrett, Sonia Goncalves, Ewan Harrison, David K. Jackson, Ian Johnston, Dominic Kwiatkowski, Cordelia Langford, John Sillitoe on behalf of the Wellcome Sanger Institute COVID-19 Surveillance Team |
| EPI_ISL_3311251 | Lighthouse Lab in Alderley Park | Wellcome Sanger Institute for the COVID-19 Genomics UK (COG-UK) Consortium | Jacquelyn Wynn, Mairead Hyland, The Lighthouse Lab in Alderley Park and Alex Alderton, Roberto Amato, Jeffrey Barrett, Sonia Goncalves, Ewan Harrison, David K. Jackson, Ian Johnston, Dominic Kwiatkowski, Cordelia Langford, John Sillitoe on behalf of the Wellcome Sanger Institute COVID-19 Surveillance Team |
| EPI_ISL_3311252, EPI_ISL_3311253 | Lighthouse Lab in Milton Keynes | Wellcome Sanger Institute for the COVID-19 Genomics UK (COG-UK) Consortium | The Lighthouse Lab in Milton Keynes and Alex Alderton, Roberto Amato, Jeffrey Barrett, Sonia Goncalves, Ewan Harrison, David K. Jackson, Ian Johnston, Dominic Kwiatkowski, Cordelia Langford, John Sillitoe on behalf of the Wellcome Sanger Institute COVID-19 Surveillance Team |
| EPI_ISL_3311254, EPI_ISL_3311255 | Lighthouse Lab in Alderley Park | Wellcome Sanger Institute for the COVID-19 Genomics UK (COG-UK) Consortium | Jacquelyn Wynn, Mairead Hyland, The Lighthouse Lab in Alderley Park and Alex Alderton, Roberto Amato, Jeffrey Barrett, Sonia Goncalves, Ewan Harrison, David K. Jackson, Ian Johnston, Dominic Kwiatkowski, Cordelia Langford, John Sillitoe on behalf of the Wellcome Sanger Institute COVID-19 Surveillance Team |
| EPI_ISL_3311256, EPI_ISL_3311257, EPI_ISL_3311258, EPI_ISL_3311259 | Lighthouse Lab in Milton Keynes | Wellcome Sanger Institute for the COVID-19 Genomics UK (COG-UK) Consortium | The Lighthouse Lab in Milton Keynes and Alex Alderton, Roberto Amato, Jeffrey Barrett, Sonia Goncalves, Ewan Harrison, David K. Jackson, Ian Johnston, Dominic Kwiatkowski, Cordelia Langford, John Sillitoe on behalf of the Wellcome Sanger Institute COVID-19 Surveillance Team |
| EPI_ISL_3311260 | Lighthouse Lab in Alderley Park | Wellcome Sanger Institute for the COVID-19 Genomics UK (COG-UK) Consortium | Jacquelyn Wynn, Mairead Hyland, The Lighthouse Lab in Alderley Park and Alex Alderton, Roberto Amato, Jeffrey Barrett, Sonia Goncalves, Ewan Harrison, David K. Jackson, Ian Johnston, Dominic Kwiatkowski, Cordelia Langford, John Sillitoe on behalf of the Wellcome Sanger Institute COVID-19 Surveillance Team |
| EPI_ISL_3311261, EPI_ISL_3311262, EPI_ISL_3311263, EPI_ISL_3311264, EPI_ISL_3311265 | Lighthouse Lab in Milton Keynes | Wellcome Sanger Institute for the COVID-19 Genomics UK (COG-UK) Consortium | The Lighthouse Lab in Milton Keynes and Alex Alderton, Roberto Amato, Jeffrey Barrett, Sonia Goncalves, Ewan Harrison, David K. Jackson, Ian Johnston, Dominic Kwiatkowski, Cordelia Langford, John Sillitoe on behalf of the Wellcome Sanger Institute COVID-19 Surveillance Team |
| EPI_ISL_3311266 | Lighthouse Lab in Alderley Park | Wellcome Sanger Institute for the COVID-19 Genomics UK (COG-UK) Consortium | Jacquelyn Wynn, Mairead Hyland, The Lighthouse Lab in Alderley Park and Alex Alderton, Roberto Amato, Jeffrey Barrett, Sonia Goncalves, Ewan Harrison, David K. Jackson, Ian Johnston, Dominic Kwiatkowski, Cordelia Langford, John Sillitoe on behalf of the Wellcome Sanger Institute COVID-19 Surveillance Team |
| EPI_ISL_3311267, EPI_ISL_3311268, EPI_ISL_3311269 | Lighthouse Lab in Milton Keynes | Wellcome Sanger Institute for the COVID-19 Genomics UK (COG-UK) Consortium | The Lighthouse Lab in Milton Keynes and Alex Alderton, Roberto Amato, Jeffrey Barrett, Sonia Goncalves, Ewan Harrison, David K. Jackson, Ian Johnston, Dominic Kwiatkowski, Cordelia Langford, John Sillitoe on behalf of the Wellcome Sanger Institute COVID-19 Surveillance Team |
| EPI_ISL_3311270 | Lighthouse Lab in Alderley Park | Wellcome Sanger Institute for the COVID-19 Genomics UK (COG-UK) Consortium | Jacquelyn Wynn, Mairead Hyland, The Lighthouse Lab in Alderley Park and Alex Alderton, Roberto Amato, Jeffrey Barrett, Sonia Goncalves, Ewan Harrison, David K. Jackson, Ian Johnston, Dominic Kwiatkowski, Cordelia Langford, John Sillitoe on behalf of the Wellcome Sanger Institute COVID-19 Surveillance Team |
| EPI_ISL_3311271, EPI_ISL_3311272, EPI_ISL_3311273 | Lighthouse Lab in Milton Keynes | Wellcome Sanger Institute for the COVID-19 Genomics UK (COG-UK) Consortium | The Lighthouse Lab in Milton Keynes and Alex Alderton, Roberto Amato, Jeffrey Barrett, Sonia Goncalves, Ewan Harrison, David K. Jackson, Ian Johnston, Dominic Kwiatkowski, Cordelia Langford, John Sillitoe on behalf of the Wellcome Sanger Institute COVID-19 Surveillance Team |
| EPI_ISL_3311274, EPI_ISL_3311275 | Lighthouse Lab in Alderley Park | Wellcome Sanger Institute for the COVID-19 Genomics UK (COG-UK) Consortium | Jacquelyn Wynn, Mairead Hyland, The Lighthouse Lab in Alderley Park and Alex Alderton, Roberto Amato, Jeffrey Barrett, Sonia Goncalves, Ewan Harrison, David K. Jackson, Ian Johnston, Dominic Kwiatkowski, Cordelia Langford, John Sillitoe on behalf of the Wellcome Sanger Institute COVID-19 Surveillance Team |
| EPI_ISL_3311276, EPI_ISL_3311277, EPI_ISL_3311278, EPI_ISL_3311279, EPI_ISL_3311280 | Lighthouse Lab in Milton Keynes | Wellcome Sanger Institute for the COVID-19 Genomics UK (COG-UK) Consortium | The Lighthouse Lab in Milton Keynes and Alex Alderton, Roberto Amato, Jeffrey Barrett, Sonia Goncalves, Ewan Harrison, David K. Jackson, Ian Johnston, Dominic Kwiatkowski, Cordelia Langford, John Sillitoe on behalf of the Wellcome Sanger Institute COVID-19 Surveillance Team |
| EPI_ISL_3311281 | Lighthouse Lab in Alderley Park | Wellcome Sanger Institute for the COVID-19 Genomics UK (COG-UK) Consortium | Jacquelyn Wynn, Mairead Hyland, The Lighthouse Lab in Alderley Park and Alex Alderton, Roberto Amato, Jeffrey Barrett, Sonia Goncalves, Ewan Harrison, David K. Jackson, Ian Johnston, Dominic Kwiatkowski, Cordelia Langford, John Sillitoe on behalf of the Wellcome Sanger Institute COVID-19 Surveillance Team |
| EPI_ISL_3311282, EPI_ISL_3311283, EPI_ISL_3311284, EPI_ISL_3311285 | Lighthouse Lab in Milton Keynes | Wellcome Sanger Institute for the COVID-19 Genomics UK (COG-UK) Consortium | The Lighthouse Lab in Milton Keynes and Alex Alderton, Roberto Amato, Jeffrey Barrett, Sonia Goncalves, Ewan Harrison, David K. Jackson, Ian Johnston, Dominic Kwiatkowski, Cordelia Langford, John Sillitoe on behalf of the Wellcome Sanger Institute COVID-19 Surveillance Team |
| EPI_ISL_3311286 | Lighthouse Lab in Alderley Park | Wellcome Sanger Institute for the COVID-19 Genomics UK (COG-UK) Consortium | Jacquelyn Wynn, Mairead Hyland, The Lighthouse Lab in Alderley Park and Alex Alderton, Roberto Amato, Jeffrey Barrett, Sonia Goncalves, Ewan Harrison, David K. Jackson, Ian Johnston, Dominic Kwiatkowski, Cordelia Langford, John Sillitoe on behalf of the Wellcome Sanger Institute COVID-19 Surveillance Team |

| | | | |
|---|---|---|---|
| EPI_ISL_3311287, EPI_ISL_3311288, EPI_ISL_3311289, EPI_ISL_3311290 | Lighthouse Lab in Milton Keynes | Wellcome Sanger Institute for the COVID-19 Genomics UK (COG-UK) Consortium | The Lighthouse Lab in Milton Keynes and Alex Alderton, Roberto Amato, Jeffrey Barrett, Sonia Goncalves, Ewan Harrison, David K. Jackson, Ian Johnston, Dominic Kwiatkowski, Cordelia Langford, John Sillitoe on behalf of the Wellcome Sanger Institute COVID-19 Surveillance Team |
| EPI_ISL_3311291 | Lighthouse Lab in Alderley Park | Wellcome Sanger Institute for the COVID-19 Genomics UK (COG-UK) Consortium | Jacquelyn Wynn, Mairead Hyland, The Lighthouse Lab in Alderley Park and Alex Alderton, Roberto Amato, Jeffrey Barrett, Sonia Goncalves, Ewan Harrison, David K. Jackson, Ian Johnston, Dominic Kwiatkowski, Cordelia Langford, John Sillitoe on behalf of the Wellcome Sanger Institute COVID-19 Surveillance Team |
| EPI_ISL_3311292, EPI_ISL_3311293, EPI_ISL_3311294, EPI_ISL_3311295, EPI_ISL_3311296, EPI_ISL_3311297, EPI_ISL_3311298, EPI_ISL_3311299, EPI_ISL_3311300 | Lighthouse Lab in Milton Keynes | Wellcome Sanger Institute for the COVID-19 Genomics UK (COG-UK) Consortium | The Lighthouse Lab in Milton Keynes and Alex Alderton, Roberto Amato, Jeffrey Barrett, Sonia Goncalves, Ewan Harrison, David K. Jackson, Ian Johnston, Dominic Kwiatkowski, Cordelia Langford, John Sillitoe on behalf of the Wellcome Sanger Institute COVID-19 Surveillance Team |
| EPI_ISL_3311301, EPI_ISL_3311302, EPI_ISL_3311303 | Lighthouse Lab in Alderley Park | Wellcome Sanger Institute for the COVID-19 Genomics UK (COG-UK) Consortium | Jacquelyn Wynn, Mairead Hyland, The Lighthouse Lab in Alderley Park and Alex Alderton, Roberto Amato, Jeffrey Barrett, Sonia Goncalves, Ewan Harrison, David K. Jackson, Ian Johnston, Dominic Kwiatkowski, Cordelia Langford, John Sillitoe on behalf of the Wellcome Sanger Institute COVID-19 Surveillance Team |
| EPI_ISL_3311304, EPI_ISL_3311305, EPI_ISL_3311306, EPI_ISL_3311307, EPI_ISL_3311308, EPI_ISL_3311309, EPI_ISL_3311310, EPI_ISL_3311311 | Lighthouse Lab in Milton Keynes | Wellcome Sanger Institute for the COVID-19 Genomics UK (COG-UK) Consortium | The Lighthouse Lab in Milton Keynes and Alex Alderton, Roberto Amato, Jeffrey Barrett, Sonia Goncalves, Ewan Harrison, David K. Jackson, Ian Johnston, Dominic Kwiatkowski, Cordelia Langford, John Sillitoe on behalf of the Wellcome Sanger Institute COVID-19 Surveillance Team |
| EPI_ISL_3311312 | Lighthouse Lab in Alderley Park | Wellcome Sanger Institute for the COVID-19 Genomics UK (COG-UK) Consortium | Jacquelyn Wynn, Mairead Hyland, The Lighthouse Lab in Alderley Park and Alex Alderton, Roberto Amato, Jeffrey Barrett, Sonia Goncalves, Ewan Harrison, David K. Jackson, Ian Johnston, Dominic Kwiatkowski, Cordelia Langford, John Sillitoe on behalf of the Wellcome Sanger Institute COVID-19 Surveillance Team |
| EPI_ISL_3311313, EPI_ISL_3311314, EPI_ISL_3311315 | Lighthouse Lab in Milton Keynes | Wellcome Sanger Institute for the COVID-19 Genomics UK (COG-UK) Consortium | The Lighthouse Lab in Milton Keynes and Alex Alderton, Roberto Amato, Jeffrey Barrett, Sonia Goncalves, Ewan Harrison, David K. Jackson, Ian Johnston, Dominic Kwiatkowski, Cordelia Langford, John Sillitoe on behalf of the Wellcome Sanger Institute COVID-19 Surveillance Team |
| EPI_ISL_3311316 | Lighthouse Lab in Alderley Park | Wellcome Sanger Institute for the COVID-19 Genomics UK (COG-UK) Consortium | Jacquelyn Wynn, Mairead Hyland, The Lighthouse Lab in Alderley Park and Alex Alderton, Roberto Amato, Jeffrey Barrett, Sonia Goncalves, Ewan Harrison, David K. Jackson, Ian Johnston, Dominic Kwiatkowski, Cordelia Langford, John Sillitoe on behalf of the Wellcome Sanger Institute COVID-19 Surveillance Team |
| EPI_ISL_3311317, EPI_ISL_3311318 | Lighthouse Lab in Milton Keynes | Wellcome Sanger Institute for the COVID-19 Genomics UK (COG-UK) Consortium | The Lighthouse Lab in Milton Keynes and Alex Alderton, Roberto Amato, Jeffrey Barrett, Sonia Goncalves, Ewan Harrison, David K. Jackson, Ian Johnston, Dominic Kwiatkowski, Cordelia Langford, John Sillitoe on behalf of the Wellcome Sanger Institute COVID-19 Surveillance Team |
| EPI_ISL_3311319, EPI_ISL_3311320 | Lighthouse Lab in Alderley Park | Wellcome Sanger Institute for the COVID-19 Genomics UK (COG-UK) Consortium | Jacquelyn Wynn, Mairead Hyland, The Lighthouse Lab in Alderley Park and Alex Alderton, Roberto Amato, Jeffrey Barrett, Sonia Goncalves, Ewan Harrison, David K. Jackson, Ian Johnston, Dominic Kwiatkowski, Cordelia Langford, John Sillitoe on behalf of the Wellcome Sanger Institute COVID-19 Surveillance Team |
| EPI_ISL_3311321, EPI_ISL_3311322, EPI_ISL_3311323, EPI_ISL_3311324, EPI_ISL_3311325, EPI_ISL_3311326 | Lighthouse Lab in Milton Keynes | Wellcome Sanger Institute for the COVID-19 Genomics UK (COG-UK) Consortium | The Lighthouse Lab in Milton Keynes and Alex Alderton, Roberto Amato, Jeffrey Barrett, Sonia Goncalves, Ewan Harrison, David K. Jackson, Ian Johnston, Dominic Kwiatkowski, Cordelia Langford, John Sillitoe on behalf of the Wellcome Sanger Institute COVID-19 Surveillance Team |
| EPI_ISL_3311327 | Lighthouse Lab in Alderley Park | Wellcome Sanger Institute for the COVID-19 Genomics UK (COG-UK) Consortium | Jacquelyn Wynn, Mairead Hyland, The Lighthouse Lab in Alderley Park and Alex Alderton, Roberto Amato, Jeffrey Barrett, Sonia Goncalves, Ewan Harrison, David K. Jackson, Ian Johnston, Dominic Kwiatkowski, Cordelia Langford, John Sillitoe on behalf of the Wellcome Sanger Institute COVID-19 Surveillance Team |
| EPI_ISL_3311328, EPI_ISL_3311329, EPI_ISL_3311330 | Lighthouse Lab in Milton Keynes | Wellcome Sanger Institute for the COVID-19 Genomics UK (COG-UK) Consortium | The Lighthouse Lab in Milton Keynes and Alex Alderton, Roberto Amato, Jeffrey Barrett, Sonia Goncalves, Ewan Harrison, David K. Jackson, Ian Johnston, Dominic Kwiatkowski, Cordelia Langford, John Sillitoe on behalf of the Wellcome Sanger Institute COVID-19 Surveillance Team |
| EPI_ISL_3311331 | Lighthouse Lab in Alderley Park | Wellcome Sanger Institute for the COVID-19 Genomics UK (COG-UK) Consortium | Jacquelyn Wynn, Mairead Hyland, The Lighthouse Lab in Alderley Park and Alex Alderton, Roberto Amato, Jeffrey Barrett, Sonia Goncalves, Ewan Harrison, David K. Jackson, Ian Johnston, Dominic Kwiatkowski, Cordelia Langford, John Sillitoe on behalf of the Wellcome Sanger Institute COVID-19 Surveillance Team |
| EPI_ISL_3311332, EPI_ISL_3311333, EPI_ISL_3311334 | Lighthouse Lab in Milton Keynes | Wellcome Sanger Institute for the COVID-19 Genomics UK (COG-UK) Consortium | The Lighthouse Lab in Milton Keynes and Alex Alderton, Roberto Amato, Jeffrey Barrett, Sonia Goncalves, Ewan Harrison, David K. Jackson, Ian Johnston, Dominic Kwiatkowski, Cordelia Langford, John Sillitoe on behalf of the Wellcome Sanger Institute COVID-19 Surveillance Team |
| EPI_ISL_3311335 | Lighthouse Lab in Alderley Park | Wellcome Sanger Institute for the COVID-19 Genomics UK (COG-UK) Consortium | Jacquelyn Wynn, Mairead Hyland, The Lighthouse Lab in Alderley Park and Alex Alderton, Roberto Amato, Jeffrey Barrett, Sonia Goncalves, Ewan Harrison, David K. Jackson, Ian Johnston, Dominic Kwiatkowski, Cordelia Langford, John Sillitoe on behalf of the Wellcome Sanger Institute COVID-19 Surveillance Team |
| EPI_ISL_3311336 | Lighthouse Lab in Milton Keynes | Wellcome Sanger Institute for the COVID-19 Genomics UK (COG-UK) Consortium | The Lighthouse Lab in Milton Keynes and Alex Alderton, Roberto Amato, Jeffrey Barrett, Sonia Goncalves, Ewan Harrison, David K. Jackson, Ian Johnston, Dominic Kwiatkowski, Cordelia Langford, John Sillitoe on behalf of the Wellcome Sanger Institute COVID-19 Surveillance Team |
| EPI_ISL_3311337 | Lighthouse Lab in Alderley Park | Wellcome Sanger Institute for the COVID-19 Genomics UK (COG-UK) Consortium | Jacquelyn Wynn, Mairead Hyland, The Lighthouse Lab in Alderley Park and Alex Alderton, Roberto Amato, Jeffrey Barrett, Sonia Goncalves, Ewan Harrison, David K. Jackson, Ian Johnston, Dominic Kwiatkowski, Cordelia Langford, John Sillitoe on behalf of the Wellcome Sanger Institute COVID-19 Surveillance Team |
| EPI_ISL_3311338, EPI_ISL_3311339, EPI_ISL_3311340, EPI_ISL_3311341 | Lighthouse Lab in Milton Keynes | Wellcome Sanger Institute for the COVID-19 Genomics UK (COG-UK) Consortium | The Lighthouse Lab in Milton Keynes and Alex Alderton, Roberto Amato, Jeffrey Barrett, Sonia Goncalves, Ewan Harrison, David K. Jackson, Ian Johnston, Dominic Kwiatkowski, Cordelia Langford, John Sillitoe on behalf of the Wellcome Sanger Institute COVID-19 Surveillance Team |
| EPI_ISL_3311342 | Lighthouse Lab in Alderley Park | Wellcome Sanger Institute for the COVID-19 Genomics UK (COG-UK) Consortium | Jacquelyn Wynn, Mairead Hyland, The Lighthouse Lab in Alderley Park and Alex Alderton, Roberto Amato, Jeffrey Barrett, Sonia Goncalves, Ewan Harrison, David K. Jackson, Ian Johnston, Dominic Kwiatkowski, Cordelia Langford, John Sillitoe on behalf of the Wellcome Sanger Institute COVID-19 Surveillance Team |
| EPI_ISL_3311343 | Lighthouse Lab in Milton Keynes | Wellcome Sanger Institute for the COVID-19 Genomics UK (COG-UK) Consortium | The Lighthouse Lab in Milton Keynes and Alex Alderton, Roberto Amato, Jeffrey Barrett, Sonia Goncalves, Ewan Harrison, David K. Jackson, Ian Johnston, Dominic Kwiatkowski, Cordelia Langford, John Sillitoe on behalf of the Wellcome Sanger Institute COVID-19 Surveillance Team |
| EPI_ISL_3311344, EPI_ISL_3311345 | Lighthouse Lab in Alderley Park | Wellcome Sanger Institute for the COVID-19 Genomics UK (COG-UK) Consortium | Jacquelyn Wynn, Mairead Hyland, The Lighthouse Lab in Alderley Park and Alex Alderton, Roberto Amato, Jeffrey Barrett, Sonia Goncalves, Ewan Harrison, David K. Jackson, Ian Johnston, Dominic Kwiatkowski, Cordelia Langford, John Sillitoe on behalf of the Wellcome Sanger Institute COVID-19 Surveillance Team |
| EPI_ISL_3311346 | Lighthouse Lab in Milton Keynes | Wellcome Sanger Institute for the COVID-19 Genomics UK (COG-UK) Consortium | The Lighthouse Lab in Milton Keynes and Alex Alderton, Roberto Amato, Jeffrey Barrett, Sonia Goncalves, Ewan Harrison, David K. Jackson, Ian Johnston, Dominic Kwiatkowski, Cordelia Langford, John Sillitoe on behalf of the Wellcome Sanger Institute COVID-19 Surveillance Team |
| EPI_ISL_3311347 | Lighthouse Lab in Alderley Park | Wellcome Sanger Institute for the COVID-19 Genomics UK (COG-UK) Consortium | Jacquelyn Wynn, Mairead Hyland, The Lighthouse Lab in Alderley Park and Alex Alderton, Roberto Amato, Jeffrey Barrett, Sonia Goncalves, Ewan Harrison, David K. Jackson, Ian Johnston, Dominic Kwiatkowski, Cordelia Langford, John Sillitoe on behalf of the Wellcome Sanger Institute COVID-19 Surveillance Team |
| EPI_ISL_3311348, EPI_ISL_3311349 | Lighthouse Lab in Milton Keynes | Wellcome Sanger Institute for the COVID-19 Genomics UK (COG-UK) Consortium | The Lighthouse Lab in Milton Keynes and Alex Alderton, Roberto Amato, Jeffrey Barrett, Sonia Goncalves, Ewan Harrison, David K. Jackson, Ian Johnston, Dominic Kwiatkowski, Cordelia Langford, John Sillitoe on behalf of the Wellcome Sanger Institute COVID-19 Surveillance Team |
| EPI_ISL_3311350 | Lighthouse Lab in Alderley Park | Wellcome Sanger Institute for the COVID-19 Genomics UK (COG-UK) Consortium | Jacquelyn Wynn, Mairead Hyland, The Lighthouse Lab in Alderley Park and Alex Alderton, Roberto Amato, Jeffrey Barrett, Sonia Goncalves, Ewan Harrison, David K. Jackson, Ian Johnston, Dominic Kwiatkowski, Cordelia Langford, John Sillitoe on behalf of the Wellcome Sanger Institute COVID-19 Surveillance Team |
| EPI_ISL_3311351 | Lighthouse Lab in Milton Keynes | Wellcome Sanger Institute for the COVID-19 Genomics UK (COG-UK) Consortium | The Lighthouse Lab in Milton Keynes and Alex Alderton, Roberto Amato, Jeffrey Barrett, Sonia Goncalves, Ewan Harrison, David K. Jackson, Ian Johnston, Dominic Kwiatkowski, Cordelia Langford, John Sillitoe on behalf of the Wellcome Sanger Institute COVID-19 Surveillance Team |
| EPI_ISL_3311352 | Lighthouse Lab in Alderley Park | Wellcome Sanger Institute for the COVID-19 Genomics UK (COG-UK) Consortium | Jacquelyn Wynn, Mairead Hyland, The Lighthouse Lab in Alderley Park and Alex Alderton, Roberto Amato, Jeffrey Barrett, Sonia Goncalves, Ewan Harrison, David K. Jackson, Ian Johnston, Dominic Kwiatkowski, Cordelia Langford, John Sillitoe on behalf of the Wellcome Sanger Institute COVID-19 Surveillance Team |
| EPI_ISL_3311353, EPI_ISL_3311354 | Lighthouse Lab in Milton Keynes | Wellcome Sanger Institute for the COVID-19 Genomics UK (COG-UK) Consortium | The Lighthouse Lab in Milton Keynes and Alex Alderton, Roberto Amato, Jeffrey Barrett, Sonia Goncalves, Ewan Harrison, David K. Jackson, Ian Johnston, Dominic Kwiatkowski, Cordelia Langford, John Sillitoe on behalf of the Wellcome Sanger Institute COVID-19 Surveillance Team |
| EPI_ISL_3311355, EPI_ISL_3311356 | Lighthouse Lab in Alderley Park | Wellcome Sanger Institute for the COVID-19 Genomics UK (COG-UK) Consortium | Jacquelyn Wynn, Mairead Hyland, The Lighthouse Lab in Alderley Park and Alex Alderton, Roberto Amato, Jeffrey Barrett, Sonia Goncalves, Ewan Harrison, David K. Jackson, Ian Johnston, Dominic Kwiatkowski, Cordelia Langford, John Sillitoe on behalf of the Wellcome Sanger Institute COVID-19 Surveillance Team |
| EPI_ISL_3311357, EPI_ISL_3311358, EPI_ISL_3311359, EPI_ISL_3311360 | Lighthouse Lab in Milton Keynes | Wellcome Sanger Institute for the COVID-19 Genomics UK (COG-UK) Consortium | The Lighthouse Lab in Milton Keynes and Alex Alderton, Roberto Amato, Jeffrey Barrett, Sonia Goncalves, Ewan Harrison, David K. Jackson, Ian Johnston, Dominic Kwiatkowski, Cordelia Langford, John Sillitoe on behalf of the Wellcome Sanger Institute COVID-19 Surveillance Team |
| EPI_ISL_3311361 | Lighthouse Lab in Alderley Park | Wellcome Sanger Institute for the COVID-19 Genomics UK (COG-UK) Consortium | Jacquelyn Wynn, Mairead Hyland, The Lighthouse Lab in Alderley Park and Alex Alderton, Roberto Amato, Jeffrey Barrett, Sonia Goncalves, Ewan Harrison, David K. Jackson, Ian Johnston, Dominic Kwiatkowski, Cordelia Langford, John Sillitoe on behalf of the Wellcome Sanger Institute COVID-19 Surveillance Team |
| EPI_ISL_3311362 | Lighthouse Lab in Milton Keynes | Wellcome Sanger Institute for the COVID-19 Genomics UK (COG-UK) Consortium | The Lighthouse Lab in Milton Keynes and Alex Alderton, Roberto Amato, Jeffrey Barrett, Sonia Goncalves, Ewan Harrison, David K. Jackson, Ian Johnston, Dominic Kwiatkowski, Cordelia Langford, John Sillitoe on behalf of the Wellcome Sanger Institute COVID-19 Surveillance Team |
| EPI_ISL_3311363 | Lighthouse Lab in Alderley Park | Wellcome Sanger Institute for the COVID-19 Genomics UK (COG-UK) Consortium | Jacquelyn Wynn, Mairead Hyland, The Lighthouse Lab in Alderley Park and Alex Alderton, Roberto Amato, Jeffrey Barrett, Sonia Goncalves, Ewan Harrison, David K. Jackson, Ian Johnston, Dominic Kwiatkowski, Cordelia Langford, John Sillitoe on behalf of the Wellcome Sanger Institute COVID-19 Surveillance Team |

| | | | |
|---|---|---|---|
| EPI_ISL_3311364 | Lighthouse Lab in Milton Keynes | Wellcome Sanger Institute for the COVID-19 Genomics UK (COG-UK) Consortium | The Lighthouse Lab in Milton Keynes and Alex Alderton, Roberto Amato, Jeffrey Barrett, Sonia Goncalves, Ewan Harrison, David K. Jackson, Ian Johnston, Dominic Kwiatkowski, Cordelia Langford, John Sillitoe on behalf of the Wellcome Sanger Institute COVID-19 Surveillance Team |
| EPI_ISL_3311365 | Lighthouse Lab in Alderley Park | Wellcome Sanger Institute for the COVID-19 Genomics UK (COG-UK) Consortium | Jacquelyn Wynn, Mairead Hyland, The Lighthouse Lab in Alderley Park and Alex Alderton, Roberto Amato, Jeffrey Barrett, Sonia Goncalves, Ewan Harrison, David K. Jackson, Ian Johnston, Dominic Kwiatkowski, Cordelia Langford, John Sillitoe on behalf of the Wellcome Sanger Institute COVID-19 Surveillance Team |
| EPI_ISL_3311366, EPI_ISL_3311367 | Lighthouse Lab in Milton Keynes | Wellcome Sanger Institute for the COVID-19 Genomics UK (COG-UK) Consortium | The Lighthouse Lab in Milton Keynes and Alex Alderton, Roberto Amato, Jeffrey Barrett, Sonia Goncalves, Ewan Harrison, David K. Jackson, Ian Johnston, Dominic Kwiatkowski, Cordelia Langford, John Sillitoe on behalf of the Wellcome Sanger Institute COVID-19 Surveillance Team |
| EPI_ISL_3311368 | Lighthouse Lab in Alderley Park | Wellcome Sanger Institute for the COVID-19 Genomics UK (COG-UK) Consortium | Jacquelyn Wynn, Mairead Hyland, The Lighthouse Lab in Alderley Park and Alex Alderton, Roberto Amato, Jeffrey Barrett, Sonia Goncalves, Ewan Harrison, David K. Jackson, Ian Johnston, Dominic Kwiatkowski, Cordelia Langford, John Sillitoe on behalf of the Wellcome Sanger Institute COVID-19 Surveillance Team |
| EPI_ISL_3311369 | Lighthouse Lab in Milton Keynes | Wellcome Sanger Institute for the COVID-19 Genomics UK (COG-UK) Consortium | The Lighthouse Lab in Milton Keynes and Alex Alderton, Roberto Amato, Jeffrey Barrett, Sonia Goncalves, Ewan Harrison, David K. Jackson, Ian Johnston, Dominic Kwiatkowski, Cordelia Langford, John Sillitoe on behalf of the Wellcome Sanger Institute COVID-19 Surveillance Team |
| EPI_ISL_3311370, EPI_ISL_3311371 | Lighthouse Lab in Glasgow | Wellcome Sanger Institute for the COVID-19 Genomics UK (COG-UK) Consortium | Harper VanSteenhouse, Yumi Kasai, David Gray, Carol Clugston, Anna Dominiczak and Alex Alderton, Roberto Amato, Jeffrey Barrett, Sonia Goncalves, Ewan Harrison, David K. Jackson, Ian Johnston, Dominic Kwiatkowski, Cordelia Langford, John Sillitoe on behalf of the Wellcome Sanger Institute COVID-19 Surveillance Team |
| EPI_ISL_3311372 | Lighthouse Lab in Milton Keynes | Wellcome Sanger Institute for the COVID-19 Genomics UK (COG-UK) Consortium | The Lighthouse Lab in Milton Keynes and Alex Alderton, Roberto Amato, Jeffrey Barrett, Sonia Goncalves, Ewan Harrison, David K. Jackson, Ian Johnston, Dominic Kwiatkowski, Cordelia Langford, John Sillitoe on behalf of the Wellcome Sanger Institute COVID-19 Surveillance Team |
| EPI_ISL_3311373, EPI_ISL_3311374, EPI_ISL_3311375 | Lighthouse Lab in Alderley Park | Wellcome Sanger Institute for the COVID-19 Genomics UK (COG-UK) Consortium | Jacquelyn Wynn, Mairead Hyland, The Lighthouse Lab in Alderley Park and Alex Alderton, Roberto Amato, Jeffrey Barrett, Sonia Goncalves, Ewan Harrison, David K. Jackson, Ian Johnston, Dominic Kwiatkowski, Cordelia Langford, John Sillitoe on behalf of the Wellcome Sanger Institute COVID-19 Surveillance Team |
| EPI_ISL_3311376 | Lighthouse Lab in Milton Keynes | Wellcome Sanger Institute for the COVID-19 Genomics UK (COG-UK) Consortium | The Lighthouse Lab in Milton Keynes and Alex Alderton, Roberto Amato, Jeffrey Barrett, Sonia Goncalves, Ewan Harrison, David K. Jackson, Ian Johnston, Dominic Kwiatkowski, Cordelia Langford, John Sillitoe on behalf of the Wellcome Sanger Institute COVID-19 Surveillance Team |
| EPI_ISL_3311377 | Health Services Laboratories | Wellcome Sanger Institute for the COVID-19 Genomics UK (COG-UK) Consortium | Health Services Laboratories and Alex Alderton, Roberto Amato, Jeffrey Barrett, Sonia Goncalves, Ewan Harrison, David K. Jackson, Ian Johnston, Dominic Kwiatkowski, Cordelia Langford, John Sillitoe on behalf of the Wellcome Sanger Institute COVID-19 Surveillance Team |
| EPI_ISL_3311378 | Lighthouse Lab in Alderley Park | Wellcome Sanger Institute for the COVID-19 Genomics UK (COG-UK) Consortium | Jacquelyn Wynn, Mairead Hyland, The Lighthouse Lab in Alderley Park and Alex Alderton, Roberto Amato, Jeffrey Barrett, Sonia Goncalves, Ewan Harrison, David K. Jackson, Ian Johnston, Dominic Kwiatkowski, Cordelia Langford, John Sillitoe on behalf of the Wellcome Sanger Institute COVID-19 Surveillance Team |
| EPI_ISL_3311379 | Lighthouse Lab in Glasgow | Wellcome Sanger Institute for the COVID-19 Genomics UK (COG-UK) Consortium | Harper VanSteenhouse, Yumi Kasai, David Gray, Carol Clugston, Anna Dominiczak and Alex Alderton, Roberto Amato, Jeffrey Barrett, Sonia Goncalves, Ewan Harrison, David K. Jackson, Ian Johnston, Dominic Kwiatkowski, Cordelia Langford, John Sillitoe on behalf of the Wellcome Sanger Institute COVID-19 Surveillance Team |
| EPI_ISL_3311380, EPI_ISL_3311381, EPI_ISL_3311382 | Lighthouse Lab in Alderley Park | Wellcome Sanger Institute for the COVID-19 Genomics UK (COG-UK) Consortium | Jacquelyn Wynn, Mairead Hyland, The Lighthouse Lab in Alderley Park and Alex Alderton, Roberto Amato, Jeffrey Barrett, Sonia Goncalves, Ewan Harrison, David K. Jackson, Ian Johnston, Dominic Kwiatkowski, Cordelia Langford, John Sillitoe on behalf of the Wellcome Sanger Institute COVID-19 Surveillance Team |
| EPI_ISL_3311383 | Lighthouse Lab in Glasgow | Wellcome Sanger Institute for the COVID-19 Genomics UK (COG-UK) Consortium | Harper VanSteenhouse, Yumi Kasai, David Gray, Carol Clugston, Anna Dominiczak and Alex Alderton, Roberto Amato, Jeffrey Barrett, Sonia Goncalves, Ewan Harrison, David K. Jackson, Ian Johnston, Dominic Kwiatkowski, Cordelia Langford, John Sillitoe on behalf of the Wellcome Sanger Institute COVID-19 Surveillance Team |
| EPI_ISL_3311384, EPI_ISL_3311385 | Lighthouse Lab in Alderley Park | Wellcome Sanger Institute for the COVID-19 Genomics UK (COG-UK) Consortium | Jacquelyn Wynn, Mairead Hyland, The Lighthouse Lab in Alderley Park and Alex Alderton, Roberto Amato, Jeffrey Barrett, Sonia Goncalves, Ewan Harrison, David K. Jackson, Ian Johnston, Dominic Kwiatkowski, Cordelia Langford, John Sillitoe on behalf of the Wellcome Sanger Institute COVID-19 Surveillance Team |
| EPI_ISL_3311386 | Lighthouse Lab in Glasgow | Wellcome Sanger Institute for the COVID-19 Genomics UK (COG-UK) Consortium | Harper VanSteenhouse, Yumi Kasai, David Gray, Carol Clugston, Anna Dominiczak and Alex Alderton, Roberto Amato, Jeffrey Barrett, Sonia Goncalves, Ewan Harrison, David K. Jackson, Ian Johnston, Dominic Kwiatkowski, Cordelia Langford, John Sillitoe on behalf of the Wellcome Sanger Institute COVID-19 Surveillance Team |
| EPI_ISL_3311387, EPI_ISL_3311388, EPI_ISL_3311389, EPI_ISL_3311390 | Lighthouse Lab in Milton Keynes | Wellcome Sanger Institute for the COVID-19 Genomics UK (COG-UK) Consortium | The Lighthouse Lab in Milton Keynes and Alex Alderton, Roberto Amato, Jeffrey Barrett, Sonia Goncalves, Ewan Harrison, David K. Jackson, Ian Johnston, Dominic Kwiatkowski, Cordelia Langford, John Sillitoe on behalf of the Wellcome Sanger Institute COVID-19 Surveillance Team |
| EPI_ISL_3311391, EPI_ISL_3311392, EPI_ISL_3311393 | Lighthouse Lab in Alderley Park | Wellcome Sanger Institute for the COVID-19 Genomics UK (COG-UK) Consortium | Jacquelyn Wynn, Mairead Hyland, The Lighthouse Lab in Alderley Park and Alex Alderton, Roberto Amato, Jeffrey Barrett, Sonia Goncalves, Ewan Harrison, David K. Jackson, Ian Johnston, Dominic Kwiatkowski, Cordelia Langford, John Sillitoe on behalf of the Wellcome Sanger Institute COVID-19 Surveillance Team |
| EPI_ISL_3311394, EPI_ISL_3311395, EPI_ISL_3311396 | Lighthouse Lab in Glasgow | Wellcome Sanger Institute for the COVID-19 Genomics UK (COG-UK) Consortium | Harper VanSteenhouse, Yumi Kasai, David Gray, Carol Clugston, Anna Dominiczak and Alex Alderton, Roberto Amato, Jeffrey Barrett, Sonia Goncalves, Ewan Harrison, David K. Jackson, Ian Johnston, Dominic Kwiatkowski, Cordelia Langford, John Sillitoe on behalf of the Wellcome Sanger Institute COVID-19 Surveillance Team |
| EPI_ISL_3311397 | Lighthouse Lab in Alderley Park | Wellcome Sanger Institute for the COVID-19 Genomics UK (COG-UK) Consortium | Jacquelyn Wynn, Mairead Hyland, The Lighthouse Lab in Alderley Park and Alex Alderton, Roberto Amato, Jeffrey Barrett, Sonia Goncalves, Ewan Harrison, David K. Jackson, Ian Johnston, Dominic Kwiatkowski, Cordelia Langford, John Sillitoe on behalf of the Wellcome Sanger Institute COVID-19 Surveillance Team |
| EPI_ISL_3311398 | Lighthouse Lab in Milton Keynes | Wellcome Sanger Institute for the COVID-19 Genomics UK (COG-UK) Consortium | The Lighthouse Lab in Milton Keynes and Alex Alderton, Roberto Amato, Jeffrey Barrett, Sonia Goncalves, Ewan Harrison, David K. Jackson, Ian Johnston, Dominic Kwiatkowski, Cordelia Langford, John Sillitoe on behalf of the Wellcome Sanger Institute COVID-19 Surveillance Team |
| EPI_ISL_3311399, EPI_ISL_3311400 | Lighthouse Lab in Alderley Park | Wellcome Sanger Institute for the COVID-19 Genomics UK (COG-UK) Consortium | Jacquelyn Wynn, Mairead Hyland, The Lighthouse Lab in Alderley Park and Alex Alderton, Roberto Amato, Jeffrey Barrett, Sonia Goncalves, Ewan Harrison, David K. Jackson, Ian Johnston, Dominic Kwiatkowski, Cordelia Langford, John Sillitoe on behalf of the Wellcome Sanger Institute COVID-19 Surveillance Team |
| EPI_ISL_3311401 | Lighthouse Lab in Glasgow | Wellcome Sanger Institute for the COVID-19 Genomics UK (COG-UK) Consortium | Harper VanSteenhouse, Yumi Kasai, David Gray, Carol Clugston, Anna Dominiczak and Alex Alderton, Roberto Amato, Jeffrey Barrett, Sonia Goncalves, Ewan Harrison, David K. Jackson, Ian Johnston, Dominic Kwiatkowski, Cordelia Langford, John Sillitoe on behalf of the Wellcome Sanger Institute COVID-19 Surveillance Team |
| EPI_ISL_3311402 | Lighthouse Lab in Alderley Park | Wellcome Sanger Institute for the COVID-19 Genomics UK (COG-UK) Consortium | Jacquelyn Wynn, Mairead Hyland, The Lighthouse Lab in Alderley Park and Alex Alderton, Roberto Amato, Jeffrey Barrett, Sonia Goncalves, Ewan Harrison, David K. Jackson, Ian Johnston, Dominic Kwiatkowski, Cordelia Langford, John Sillitoe on behalf of the Wellcome Sanger Institute COVID-19 Surveillance Team |
| EPI_ISL_3311403 | Lighthouse Lab in Glasgow | Wellcome Sanger Institute for the COVID-19 Genomics UK (COG-UK) Consortium | Harper VanSteenhouse, Yumi Kasai, David Gray, Carol Clugston, Anna Dominiczak and Alex Alderton, Roberto Amato, Jeffrey Barrett, Sonia Goncalves, Ewan Harrison, David K. Jackson, Ian Johnston, Dominic Kwiatkowski, Cordelia Langford, John Sillitoe on behalf of the Wellcome Sanger Institute COVID-19 Surveillance Team |
| EPI_ISL_3311404, EPI_ISL_3311405 | Lighthouse Lab in Alderley Park | Wellcome Sanger Institute for the COVID-19 Genomics UK (COG-UK) Consortium | Jacquelyn Wynn, Mairead Hyland, The Lighthouse Lab in Alderley Park and Alex Alderton, Roberto Amato, Jeffrey Barrett, Sonia Goncalves, Ewan Harrison, David K. Jackson, Ian Johnston, Dominic Kwiatkowski, Cordelia Langford, John Sillitoe on behalf of the Wellcome Sanger Institute COVID-19 Surveillance Team |
| EPI_ISL_3311406 | Lighthouse Lab in Milton Keynes | Wellcome Sanger Institute for the COVID-19 Genomics UK (COG-UK) Consortium | The Lighthouse Lab in Milton Keynes and Alex Alderton, Roberto Amato, Jeffrey Barrett, Sonia Goncalves, Ewan Harrison, David K. Jackson, Ian Johnston, Dominic Kwiatkowski, Cordelia Langford, John Sillitoe on behalf of the Wellcome Sanger Institute COVID-19 Surveillance Team |
| EPI_ISL_3311407 | Lighthouse Lab in Glasgow | Wellcome Sanger Institute for the COVID-19 Genomics UK (COG-UK) Consortium | Harper VanSteenhouse, Yumi Kasai, David Gray, Carol Clugston, Anna Dominiczak and Alex Alderton, Roberto Amato, Jeffrey Barrett, Sonia Goncalves, Ewan Harrison, David K. Jackson, Ian Johnston, Dominic Kwiatkowski, Cordelia Langford, John Sillitoe on behalf of the Wellcome Sanger Institute COVID-19 Surveillance Team |
| EPI_ISL_3311408 | Lighthouse Lab in Milton Keynes | Wellcome Sanger Institute for the COVID-19 Genomics UK (COG-UK) Consortium | The Lighthouse Lab in Milton Keynes and Alex Alderton, Roberto Amato, Jeffrey Barrett, Sonia Goncalves, Ewan Harrison, David K. Jackson, Ian Johnston, Dominic Kwiatkowski, Cordelia Langford, John Sillitoe on behalf of the Wellcome Sanger Institute COVID-19 Surveillance Team |
| EPI_ISL_3311409, EPI_ISL_3311410 | Lighthouse Lab in Glasgow | Wellcome Sanger Institute for the COVID-19 Genomics UK (COG-UK) Consortium | Harper VanSteenhouse, Yumi Kasai, David Gray, Carol Clugston, Anna Dominiczak and Alex Alderton, Roberto Amato, Jeffrey Barrett, Sonia Goncalves, Ewan Harrison, David K. Jackson, Ian Johnston, Dominic Kwiatkowski, Cordelia Langford, John Sillitoe on behalf of the Wellcome Sanger Institute COVID-19 Surveillance Team |
| EPI_ISL_3311411, EPI_ISL_3311412 | Lighthouse Lab in Milton Keynes | Wellcome Sanger Institute for the COVID-19 Genomics UK (COG-UK) Consortium | The Lighthouse Lab in Milton Keynes and Alex Alderton, Roberto Amato, Jeffrey Barrett, Sonia Goncalves, Ewan Harrison, David K. Jackson, Ian Johnston, Dominic Kwiatkowski, Cordelia Langford, John Sillitoe on behalf of the Wellcome Sanger Institute COVID-19 Surveillance Team |
| EPI_ISL_3311413 | Lighthouse Lab in Alderley Park | Wellcome Sanger Institute for the COVID-19 Genomics UK (COG-UK) Consortium | Jacquelyn Wynn, Mairead Hyland, The Lighthouse Lab in Alderley Park and Alex Alderton, Roberto Amato, Jeffrey Barrett, Sonia Goncalves, Ewan Harrison, David K. Jackson, Ian Johnston, Dominic Kwiatkowski, Cordelia Langford, John Sillitoe on behalf of the Wellcome Sanger Institute COVID-19 Surveillance Team |
| EPI_ISL_3311414, EPI_ISL_3311415 | Lighthouse Lab in Milton Keynes | Wellcome Sanger Institute for the COVID-19 Genomics UK (COG-UK) Consortium | The Lighthouse Lab in Milton Keynes and Alex Alderton, Roberto Amato, Jeffrey Barrett, Sonia Goncalves, Ewan Harrison, David K. Jackson, Ian Johnston, Dominic Kwiatkowski, Cordelia Langford, John Sillitoe on behalf of the Wellcome Sanger Institute COVID-19 Surveillance Team |
| EPI_ISL_3311416 | Lighthouse Lab in Glasgow | Wellcome Sanger Institute for the COVID-19 Genomics UK (COG-UK) Consortium | Harper VanSteenhouse, Yumi Kasai, David Gray, Carol Clugston, Anna Dominiczak and Alex Alderton, Roberto Amato, Jeffrey Barrett, Sonia Goncalves, Ewan Harrison, David K. Jackson, Ian Johnston, Dominic Kwiatkowski, Cordelia Langford, John Sillitoe on behalf of the Wellcome Sanger Institute COVID-19 Surveillance Team |
| EPI_ISL_3311417 | Lighthouse Lab in Alderley Park | Wellcome Sanger Institute for the COVID-19 Genomics UK (COG-UK) Consortium | Jacquelyn Wynn, Mairead Hyland, The Lighthouse Lab in Alderley Park and Alex Alderton, Roberto Amato, Jeffrey Barrett, Sonia Goncalves, Ewan Harrison, David K. Jackson, Ian Johnston, Dominic Kwiatkowski, Cordelia Langford, John Sillitoe on behalf of the Wellcome Sanger Institute COVID-19 Surveillance Team |
| EPI_ISL_3311418 | Lighthouse Lab in Milton Keynes | Wellcome Sanger Institute for the COVID-19 Genomics UK (COG-UK) Consortium | The Lighthouse Lab in Milton Keynes and Alex Alderton, Roberto Amato, Jeffrey Barrett, Sonia Goncalves, Ewan Harrison, David K. Jackson, Ian Johnston, Dominic Kwiatkowski, Cordelia Langford, John Sillitoe on behalf of the Wellcome Sanger Institute COVID-19 Surveillance Team |
| EPI_ISL_3311419 | Lighthouse Lab in Alderley Park | Wellcome Sanger Institute for the COVID-19 Genomics UK (COG-UK) Consortium | Jacquelyn Wynn, Mairead Hyland, The Lighthouse Lab in Alderley Park and Alex Alderton, Roberto Amato, Jeffrey Barrett, Sonia Goncalves, Ewan Harrison, David K. Jackson, Ian Johnston, Dominic Kwiatkowski, Cordelia Langford, John Sillitoe on behalf of the Wellcome Sanger Institute COVID-19 Surveillance Team |
| EPI_ISL_3311420 | Lighthouse Lab in Milton Keynes | Wellcome Sanger Institute for the COVID-19 Genomics UK (COG-UK) Consortium | The Lighthouse Lab in Milton Keynes and Alex Alderton, Roberto Amato, Jeffrey Barrett, Sonia Goncalves, Ewan Harrison, David K. Jackson, Ian Johnston, Dominic Kwiatkowski, Cordelia Langford, John Sillitoe on behalf of the Wellcome Sanger Institute COVID-19 Surveillance Team |
| EPI_ISL_3311421 | Lighthouse Lab in Alderley Park | Wellcome Sanger Institute for the COVID-19 Genomics UK (COG-UK) Consortium | Jacquelyn Wynn, Mairead Hyland, The Lighthouse Lab in Alderley Park and Alex Alderton, Roberto Amato, Jeffrey Barrett, Sonia Goncalves, Ewan Harrison, David K. Jackson, Ian Johnston, Dominic Kwiatkowski, Cordelia Langford, John Sillitoe on behalf of the Wellcome Sanger Institute COVID-19 Surveillance Team |
| EPI_ISL_3311422 | Lighthouse Lab in Milton Keynes | Wellcome Sanger Institute for the COVID-19 Genomics UK (COG-UK) Consortium | The Lighthouse Lab in Milton Keynes and Alex Alderton, Roberto Amato, Jeffrey Barrett, Sonia Goncalves, Ewan Harrison, David K. Jackson, Ian Johnston, Dominic Kwiatkowski, Cordelia Langford, John Sillitoe on behalf of the Wellcome Sanger Institute COVID-19 Surveillance Team |
| EPI_ISL_3311423 | Lighthouse Lab in Glasgow | Wellcome Sanger Institute for the COVID-19 Genomics UK (COG-UK) Consortium | Harper VanSteenhouse, Yumi Kasai, David Gray, Carol Clugston, Anna Dominiczak and Alex Alderton, Roberto Amato, Jeffrey Barrett, Sonia Goncalves, Ewan Harrison, David K. Jackson, Ian Johnston, Dominic Kwiatkowski, Cordelia Langford, John Sillitoe on behalf of the Wellcome Sanger Institute COVID-19 Surveillance Team |
| EPI_ISL_3311424 | Lighthouse Lab in Alderley Park | Wellcome Sanger Institute for the COVID-19 Genomics UK (COG-UK) Consortium | Jacquelyn Wynn, Mairead Hyland, The Lighthouse Lab in Alderley Park and Alex Alderton, Roberto Amato, Jeffrey Barrett, Sonia Goncalves, Ewan Harrison, David K. Jackson, Ian Johnston, Dominic Kwiatkowski, Cordelia Langford, John Sillitoe on behalf of the Wellcome Sanger Institute COVID-19 Surveillance Team |
| EPI_ISL_3311425 | Lighthouse Lab in Milton Keynes | Wellcome Sanger Institute for the COVID-19 Genomics UK (COG-UK) | The Lighthouse Lab in Milton Keynes and Alex Alderton, Roberto Amato, Jeffrey Barrett, Sonia Goncalves, Ewan Harrison, David K. Jackson, Ian Johnston, Dominic Kwiatkowski, Cordelia Langford, John Sillitoe on behalf of the Wellcome Sanger |

| | | | |
|---|---|---|---|
| | | Consortium | Institute COVID-19 Surveillance Team |
| EPI_ISL_3311426 | Lighthouse Lab in Glasgow | Wellcome Sanger Institute for the COVID-19 Genomics UK (COG-UK) Consortium | Harper VanSteenhouse, Yumi Kasai, David Gray, Carol Clugston, Anna Dominiczak and Alex Alderton, Roberto Amato, Jeffrey Barrett, Sonia Goncalves, Ewan Harrison, David K. Jackson, Ian Johnston, Dominic Kwiatkowski, Cordelia Langford, John Sillitoe on behalf of the Wellcome Sanger Institute COVID-19 Surveillance Team |
| EPI_ISL_3311427 | Lighthouse Lab in Alderley Park | Wellcome Sanger Institute for the COVID-19 Genomics UK (COG-UK) Consortium | Jacquelyn Wynn, Mairead Hyland, The Lighthouse Lab in Alderley Park and Alex Alderton, Roberto Amato, Jeffrey Barrett, Sonia Goncalves, Ewan Harrison, David K. Jackson, Ian Johnston, Dominic Kwiatkowski, Cordelia Langford, John Sillitoe on behalf of the Wellcome Sanger Institute COVID-19 Surveillance Team |
| EPI_ISL_3311428, EPI_ISL_3311429 | Lighthouse Lab in Glasgow | Wellcome Sanger Institute for the COVID-19 Genomics UK (COG-UK) Consortium | Harper VanSteenhouse, Yumi Kasai, David Gray, Carol Clugston, Anna Dominiczak and Alex Alderton, Roberto Amato, Jeffrey Barrett, Sonia Goncalves, Ewan Harrison, David K. Jackson, Ian Johnston, Dominic Kwiatkowski, Cordelia Langford, John Sillitoe on behalf of the Wellcome Sanger Institute COVID-19 Surveillance Team |
| EPI_ISL_3311430 | Lighthouse Lab in Alderley Park | Wellcome Sanger Institute for the COVID-19 Genomics UK (COG-UK) Consortium | Jacquelyn Wynn, Mairead Hyland, The Lighthouse Lab in Alderley Park and Alex Alderton, Roberto Amato, Jeffrey Barrett, Sonia Goncalves, Ewan Harrison, David K. Jackson, Ian Johnston, Dominic Kwiatkowski, Cordelia Langford, John Sillitoe on behalf of the Wellcome Sanger Institute COVID-19 Surveillance Team |
| EPI_ISL_3311431 | Lighthouse Lab in Glasgow | Wellcome Sanger Institute for the COVID-19 Genomics UK (COG-UK) Consortium | Harper VanSteenhouse, Yumi Kasai, David Gray, Carol Clugston, Anna Dominiczak and Alex Alderton, Roberto Amato, Jeffrey Barrett, Sonia Goncalves, Ewan Harrison, David K. Jackson, Ian Johnston, Dominic Kwiatkowski, Cordelia Langford, John Sillitoe on behalf of the Wellcome Sanger Institute COVID-19 Surveillance Team |
| EPI_ISL_3311432, EPI_ISL_3311433, EPI_ISL_3311434, EPI_ISL_3311435 | Lighthouse Lab in Milton Keynes | Wellcome Sanger Institute for the COVID-19 Genomics UK (COG-UK) Consortium | The Lighthouse Lab in Milton Keynes and Alex Alderton, Roberto Amato, Jeffrey Barrett, Sonia Goncalves, Ewan Harrison, David K. Jackson, Ian Johnston, Dominic Kwiatkowski, Cordelia Langford, John Sillitoe on behalf of the Wellcome Sanger Institute COVID-19 Surveillance Team |
| EPI_ISL_3311436 | Lighthouse Lab in Glasgow | Wellcome Sanger Institute for the COVID-19 Genomics UK (COG-UK) Consortium | Harper VanSteenhouse, Yumi Kasai, David Gray, Carol Clugston, Anna Dominiczak and Alex Alderton, Roberto Amato, Jeffrey Barrett, Sonia Goncalves, Ewan Harrison, David K. Jackson, Ian Johnston, Dominic Kwiatkowski, Cordelia Langford, John Sillitoe on behalf of the Wellcome Sanger Institute COVID-19 Surveillance Team |
| EPI_ISL_3311437 | Lighthouse Lab in Milton Keynes | Wellcome Sanger Institute for the COVID-19 Genomics UK (COG-UK) Consortium | The Lighthouse Lab in Milton Keynes and Alex Alderton, Roberto Amato, Jeffrey Barrett, Sonia Goncalves, Ewan Harrison, David K. Jackson, Ian Johnston, Dominic Kwiatkowski, Cordelia Langford, John Sillitoe on behalf of the Wellcome Sanger Institute COVID-19 Surveillance Team |
| EPI_ISL_3311438 | Lighthouse Lab in Glasgow | Wellcome Sanger Institute for the COVID-19 Genomics UK (COG-UK) Consortium | Harper VanSteenhouse, Yumi Kasai, David Gray, Carol Clugston, Anna Dominiczak and Alex Alderton, Roberto Amato, Jeffrey Barrett, Sonia Goncalves, Ewan Harrison, David K. Jackson, Ian Johnston, Dominic Kwiatkowski, Cordelia Langford, John Sillitoe on behalf of the Wellcome Sanger Institute COVID-19 Surveillance Team |
| EPI_ISL_3311439 | Lighthouse Lab in Alderley Park | Wellcome Sanger Institute for the COVID-19 Genomics UK (COG-UK) Consortium | Jacquelyn Wynn, Mairead Hyland, The Lighthouse Lab in Alderley Park and Alex Alderton, Roberto Amato, Jeffrey Barrett, Sonia Goncalves, Ewan Harrison, David K. Jackson, Ian Johnston, Dominic Kwiatkowski, Cordelia Langford, John Sillitoe on behalf of the Wellcome Sanger Institute COVID-19 Surveillance Team |
| EPI_ISL_3311440 | Health Services Laboratories | Wellcome Sanger Institute for the COVID-19 Genomics UK (COG-UK) Consortium | Health Services Laboratories and Alex Alderton, Roberto Amato, Jeffrey Barrett, Sonia Goncalves, Ewan Harrison, David K. Jackson, Ian Johnston, Dominic Kwiatkowski, Cordelia Langford, John Sillitoe on behalf of the Wellcome Sanger Institute COVID-19 Surveillance Team |
| EPI_ISL_3311441 | Lighthouse Lab in Glasgow | Wellcome Sanger Institute for the COVID-19 Genomics UK (COG-UK) Consortium | Harper VanSteenhouse, Yumi Kasai, David Gray, Carol Clugston, Anna Dominiczak and Alex Alderton, Roberto Amato, Jeffrey Barrett, Sonia Goncalves, Ewan Harrison, David K. Jackson, Ian Johnston, Dominic Kwiatkowski, Cordelia Langford, John Sillitoe on behalf of the Wellcome Sanger Institute COVID-19 Surveillance Team |
| EPI_ISL_3311442 | Lighthouse Lab in Milton Keynes | Wellcome Sanger Institute for the COVID-19 Genomics UK (COG-UK) Consortium | The Lighthouse Lab in Milton Keynes and Alex Alderton, Roberto Amato, Jeffrey Barrett, Sonia Goncalves, Ewan Harrison, David K. Jackson, Ian Johnston, Dominic Kwiatkowski, Cordelia Langford, John Sillitoe on behalf of the Wellcome Sanger Institute COVID-19 Surveillance Team |
| EPI_ISL_3311443 | Lighthouse Lab in Glasgow | Wellcome Sanger Institute for the COVID-19 Genomics UK (COG-UK) Consortium | Harper VanSteenhouse, Yumi Kasai, David Gray, Carol Clugston, Anna Dominiczak and Alex Alderton, Roberto Amato, Jeffrey Barrett, Sonia Goncalves, Ewan Harrison, David K. Jackson, Ian Johnston, Dominic Kwiatkowski, Cordelia Langford, John Sillitoe on behalf of the Wellcome Sanger Institute COVID-19 Surveillance Team |
| EPI_ISL_3311444 | Lighthouse Lab in Alderley Park | Wellcome Sanger Institute for the COVID-19 Genomics UK (COG-UK) Consortium | Jacquelyn Wynn, Mairead Hyland, The Lighthouse Lab in Alderley Park and Alex Alderton, Roberto Amato, Jeffrey Barrett, Sonia Goncalves, Ewan Harrison, David K. Jackson, Ian Johnston, Dominic Kwiatkowski, Cordelia Langford, John Sillitoe on behalf of the Wellcome Sanger Institute COVID-19 Surveillance Team |
| EPI_ISL_3311445, EPI_ISL_3311446 | Lighthouse Lab in Milton Keynes | Wellcome Sanger Institute for the COVID-19 Genomics UK (COG-UK) Consortium | The Lighthouse Lab in Milton Keynes and Alex Alderton, Roberto Amato, Jeffrey Barrett, Sonia Goncalves, Ewan Harrison, David K. Jackson, Ian Johnston, Dominic Kwiatkowski, Cordelia Langford, John Sillitoe on behalf of the Wellcome Sanger Institute COVID-19 Surveillance Team |
| EPI_ISL_3311447 | Lighthouse Lab in Glasgow | Wellcome Sanger Institute for the COVID-19 Genomics UK (COG-UK) Consortium | Harper VanSteenhouse, Yumi Kasai, David Gray, Carol Clugston, Anna Dominiczak and Alex Alderton, Roberto Amato, Jeffrey Barrett, Sonia Goncalves, Ewan Harrison, David K. Jackson, Ian Johnston, Dominic Kwiatkowski, Cordelia Langford, John Sillitoe on behalf of the Wellcome Sanger Institute COVID-19 Surveillance Team |
| EPI_ISL_3311448 | Lighthouse Lab in Alderley Park | Wellcome Sanger Institute for the COVID-19 Genomics UK (COG-UK) Consortium | Jacquelyn Wynn, Mairead Hyland, The Lighthouse Lab in Alderley Park and Alex Alderton, Roberto Amato, Jeffrey Barrett, Sonia Goncalves, Ewan Harrison, David K. Jackson, Ian Johnston, Dominic Kwiatkowski, Cordelia Langford, John Sillitoe on behalf of the Wellcome Sanger Institute COVID-19 Surveillance Team |
| EPI_ISL_3311449, EPI_ISL_3311450 | Lighthouse Lab in Milton Keynes | Wellcome Sanger Institute for the COVID-19 Genomics UK (COG-UK) Consortium | The Lighthouse Lab in Milton Keynes and Alex Alderton, Roberto Amato, Jeffrey Barrett, Sonia Goncalves, Ewan Harrison, David K. Jackson, Ian Johnston, Dominic Kwiatkowski, Cordelia Langford, John Sillitoe on behalf of the Wellcome Sanger Institute COVID-19 Surveillance Team |
| EPI_ISL_3311451 | Lighthouse Lab in Glasgow | Wellcome Sanger Institute for the COVID-19 Genomics UK (COG-UK) Consortium | Harper VanSteenhouse, Yumi Kasai, David Gray, Carol Clugston, Anna Dominiczak and Alex Alderton, Roberto Amato, Jeffrey Barrett, Sonia Goncalves, Ewan Harrison, David K. Jackson, Ian Johnston, Dominic Kwiatkowski, Cordelia Langford, John Sillitoe on behalf of the Wellcome Sanger Institute COVID-19 Surveillance Team |
| EPI_ISL_3311452 | Lighthouse Lab in Milton Keynes | Wellcome Sanger Institute for the COVID-19 Genomics UK (COG-UK) Consortium | The Lighthouse Lab in Milton Keynes and Alex Alderton, Roberto Amato, Jeffrey Barrett, Sonia Goncalves, Ewan Harrison, David K. Jackson, Ian Johnston, Dominic Kwiatkowski, Cordelia Langford, John Sillitoe on behalf of the Wellcome Sanger Institute COVID-19 Surveillance Team |
| EPI_ISL_3311453 | Lighthouse Lab in Alderley Park | Wellcome Sanger Institute for the COVID-19 Genomics UK (COG-UK) Consortium | Jacquelyn Wynn, Mairead Hyland, The Lighthouse Lab in Alderley Park and Alex Alderton, Roberto Amato, Jeffrey Barrett, Sonia Goncalves, Ewan Harrison, David K. Jackson, Ian Johnston, Dominic Kwiatkowski, Cordelia Langford, John Sillitoe on behalf of the Wellcome Sanger Institute COVID-19 Surveillance Team |
| EPI_ISL_3311454 | Lighthouse Lab in Milton Keynes | Wellcome Sanger Institute for the COVID-19 Genomics UK (COG-UK) Consortium | The Lighthouse Lab in Milton Keynes and Alex Alderton, Roberto Amato, Jeffrey Barrett, Sonia Goncalves, Ewan Harrison, David K. Jackson, Ian Johnston, Dominic Kwiatkowski, Cordelia Langford, John Sillitoe on behalf of the Wellcome Sanger Institute COVID-19 Surveillance Team |
| EPI_ISL_3311455 | Lighthouse Lab in Alderley Park | Wellcome Sanger Institute for the COVID-19 Genomics UK (COG-UK) Consortium | Jacquelyn Wynn, Mairead Hyland, The Lighthouse Lab in Alderley Park and Alex Alderton, Roberto Amato, Jeffrey Barrett, Sonia Goncalves, Ewan Harrison, David K. Jackson, Ian Johnston, Dominic Kwiatkowski, Cordelia Langford, John Sillitoe on behalf of the Wellcome Sanger Institute COVID-19 Surveillance Team |
| EPI_ISL_3311456 | Lighthouse Lab in Milton Keynes | Wellcome Sanger Institute for the COVID-19 Genomics UK (COG-UK) Consortium | The Lighthouse Lab in Milton Keynes and Alex Alderton, Roberto Amato, Jeffrey Barrett, Sonia Goncalves, Ewan Harrison, David K. Jackson, Ian Johnston, Dominic Kwiatkowski, Cordelia Langford, John Sillitoe on behalf of the Wellcome Sanger Institute COVID-19 Surveillance Team |
| EPI_ISL_3311457 | Lighthouse Lab in Glasgow | Wellcome Sanger Institute for the COVID-19 Genomics UK (COG-UK) Consortium | Harper VanSteenhouse, Yumi Kasai, David Gray, Carol Clugston, Anna Dominiczak and Alex Alderton, Roberto Amato, Jeffrey Barrett, Sonia Goncalves, Ewan Harrison, David K. Jackson, Ian Johnston, Dominic Kwiatkowski, Cordelia Langford, John Sillitoe on behalf of the Wellcome Sanger Institute COVID-19 Surveillance Team |
| EPI_ISL_3311458 | Lighthouse Lab in Alderley Park | Wellcome Sanger Institute for the COVID-19 Genomics UK (COG-UK) Consortium | Jacquelyn Wynn, Mairead Hyland, The Lighthouse Lab in Alderley Park and Alex Alderton, Roberto Amato, Jeffrey Barrett, Sonia Goncalves, Ewan Harrison, David K. Jackson, Ian Johnston, Dominic Kwiatkowski, Cordelia Langford, John Sillitoe on behalf of the Wellcome Sanger Institute COVID-19 Surveillance Team |
| EPI_ISL_3311459, EPI_ISL_3311460, EPI_ISL_3311461 | Lighthouse Lab in Glasgow | Wellcome Sanger Institute for the COVID-19 Genomics UK (COG-UK) Consortium | Harper VanSteenhouse, Yumi Kasai, David Gray, Carol Clugston, Anna Dominiczak and Alex Alderton, Roberto Amato, Jeffrey Barrett, Sonia Goncalves, Ewan Harrison, David K. Jackson, Ian Johnston, Dominic Kwiatkowski, Cordelia Langford, John Sillitoe on behalf of the Wellcome Sanger Institute COVID-19 Surveillance Team |
| EPI_ISL_3311462, EPI_ISL_3311463, EPI_ISL_3311464 | Lighthouse Lab in Milton Keynes | Wellcome Sanger Institute for the COVID-19 Genomics UK (COG-UK) Consortium | The Lighthouse Lab in Milton Keynes and Alex Alderton, Roberto Amato, Jeffrey Barrett, Sonia Goncalves, Ewan Harrison, David K. Jackson, Ian Johnston, Dominic Kwiatkowski, Cordelia Langford, John Sillitoe on behalf of the Wellcome Sanger Institute COVID-19 Surveillance Team |
| EPI_ISL_3311465 | Lighthouse Lab in Alderley Park | Wellcome Sanger Institute for the COVID-19 Genomics UK (COG-UK) Consortium | Jacquelyn Wynn, Mairead Hyland, The Lighthouse Lab in Alderley Park and Alex Alderton, Roberto Amato, Jeffrey Barrett, Sonia Goncalves, Ewan Harrison, David K. Jackson, Ian Johnston, Dominic Kwiatkowski, Cordelia Langford, John Sillitoe on behalf of the Wellcome Sanger Institute COVID-19 Surveillance Team |
| EPI_ISL_3311466 | Lighthouse Lab in Glasgow | Wellcome Sanger Institute for the COVID-19 Genomics UK (COG-UK) Consortium | Harper VanSteenhouse, Yumi Kasai, David Gray, Carol Clugston, Anna Dominiczak and Alex Alderton, Roberto Amato, Jeffrey Barrett, Sonia Goncalves, Ewan Harrison, David K. Jackson, Ian Johnston, Dominic Kwiatkowski, Cordelia Langford, John Sillitoe on behalf of the Wellcome Sanger Institute COVID-19 Surveillance Team |
| EPI_ISL_3311467, EPI_ISL_3311468, EPI_ISL_3311469 | Lighthouse Lab in Milton Keynes | Wellcome Sanger Institute for the COVID-19 Genomics UK (COG-UK) Consortium | The Lighthouse Lab in Milton Keynes and Alex Alderton, Roberto Amato, Jeffrey Barrett, Sonia Goncalves, Ewan Harrison, David K. Jackson, Ian Johnston, Dominic Kwiatkowski, Cordelia Langford, John Sillitoe on behalf of the Wellcome Sanger Institute COVID-19 Surveillance Team |
| EPI_ISL_3311470 | Lighthouse Lab in Alderley Park | Wellcome Sanger Institute for the COVID-19 Genomics UK (COG-UK) Consortium | Jacquelyn Wynn, Mairead Hyland, The Lighthouse Lab in Alderley Park and Alex Alderton, Roberto Amato, Jeffrey Barrett, Sonia Goncalves, Ewan Harrison, David K. Jackson, Ian Johnston, Dominic Kwiatkowski, Cordelia Langford, John Sillitoe on behalf of the Wellcome Sanger Institute COVID-19 Surveillance Team |
| EPI_ISL_3311471, EPI_ISL_3311472 | Lighthouse Lab in Milton Keynes | Wellcome Sanger Institute for the COVID-19 Genomics UK (COG-UK) Consortium | The Lighthouse Lab in Milton Keynes and Alex Alderton, Roberto Amato, Jeffrey Barrett, Sonia Goncalves, Ewan Harrison, David K. Jackson, Ian Johnston, Dominic Kwiatkowski, Cordelia Langford, John Sillitoe on behalf of the Wellcome Sanger Institute COVID-19 Surveillance Team |
| EPI_ISL_3311473 | Lighthouse Lab in Alderley Park | Wellcome Sanger Institute for the COVID-19 Genomics UK (COG-UK) Consortium | Jacquelyn Wynn, Mairead Hyland, The Lighthouse Lab in Alderley Park and Alex Alderton, Roberto Amato, Jeffrey Barrett, Sonia Goncalves, Ewan Harrison, David K. Jackson, Ian Johnston, Dominic Kwiatkowski, Cordelia Langford, John Sillitoe on behalf of the Wellcome Sanger Institute COVID-19 Surveillance Team |
| EPI_ISL_3311474 | Lighthouse Lab in Glasgow | Wellcome Sanger Institute for the COVID-19 Genomics UK (COG-UK) Consortium | Harper VanSteenhouse, Yumi Kasai, David Gray, Carol Clugston, Anna Dominiczak and Alex Alderton, Roberto Amato, Jeffrey Barrett, Sonia Goncalves, Ewan Harrison, David K. Jackson, Ian Johnston, Dominic Kwiatkowski, Cordelia Langford, John Sillitoe on behalf of the Wellcome Sanger Institute COVID-19 Surveillance Team |
| EPI_ISL_3311475 | Lighthouse Lab in Milton Keynes | Wellcome Sanger Institute for the COVID-19 Genomics UK (COG-UK) Consortium | The Lighthouse Lab in Milton Keynes and Alex Alderton, Roberto Amato, Jeffrey Barrett, Sonia Goncalves, Ewan Harrison, David K. Jackson, Ian Johnston, Dominic Kwiatkowski, Cordelia Langford, John Sillitoe on behalf of the Wellcome Sanger Institute COVID-19 Surveillance Team |
| EPI_ISL_3311476 | Lighthouse Lab in Glasgow | Wellcome Sanger Institute for the COVID-19 Genomics UK (COG-UK) Consortium | Harper VanSteenhouse, Yumi Kasai, David Gray, Carol Clugston, Anna Dominiczak and Alex Alderton, Roberto Amato, Jeffrey Barrett, Sonia Goncalves, Ewan Harrison, David K. Jackson, Ian Johnston, Dominic Kwiatkowski, Cordelia Langford, John Sillitoe on behalf of the Wellcome Sanger Institute COVID-19 Surveillance Team |
| EPI_ISL_3311477, EPI_ISL_3311478 | Lighthouse Lab in Milton Keynes | Wellcome Sanger Institute for the COVID-19 Genomics UK (COG-UK) Consortium | The Lighthouse Lab in Milton Keynes and Alex Alderton, Roberto Amato, Jeffrey Barrett, Sonia Goncalves, Ewan Harrison, David K. Jackson, Ian Johnston, Dominic Kwiatkowski, Cordelia Langford, John Sillitoe on behalf of the Wellcome Sanger Institute COVID-19 Surveillance Team |
| EPI_ISL_3311479 | Lighthouse Lab in Alderley Park | Wellcome Sanger Institute for the COVID-19 Genomics UK (COG-UK) Consortium | Jacquelyn Wynn, Mairead Hyland, The Lighthouse Lab in Alderley Park and Alex Alderton, Roberto Amato, Jeffrey Barrett, Sonia Goncalves, Ewan Harrison, David K. Jackson, Ian Johnston, Dominic Kwiatkowski, Cordelia Langford, John Sillitoe on behalf of the Wellcome Sanger Institute COVID-19 Surveillance Team |
| EPI_ISL_3311480 | Lighthouse Lab in Glasgow | Wellcome Sanger Institute for the COVID-19 Genomics UK (COG-UK) Consortium | Harper VanSteenhouse, Yumi Kasai, David Gray, Carol Clugston, Anna Dominiczak and Alex Alderton, Roberto Amato, Jeffrey Barrett, Sonia Goncalves, Ewan Harrison, David K. Jackson, Ian Johnston, Dominic Kwiatkowski, Cordelia Langford, John Sillitoe on behalf of the Wellcome Sanger Institute COVID-19 Surveillance Team |
| EPI_ISL_3311481, EPI_ISL_3311482 | Lighthouse Lab in Milton Keynes | Wellcome Sanger Institute for the COVID-19 Genomics UK (COG-UK) Consortium | The Lighthouse Lab in Milton Keynes and Alex Alderton, Roberto Amato, Jeffrey Barrett, Sonia Goncalves, Ewan Harrison, David K. Jackson, Ian Johnston, Dominic Kwiatkowski, Cordelia Langford, John Sillitoe on behalf of the Wellcome Sanger Institute COVID-19 Surveillance Team |
| EPI_ISL_3311483, | Lighthouse Lab in Alderley Park | Wellcome Sanger Institute for the COVID-19 Genomics UK (COG-UK) | Jacquelyn Wynn, Mairead Hyland, The Lighthouse Lab in Alderley Park and Alex Alderton, Roberto Amato, Jeffrey Barrett, Sonia Goncalves, Ewan Harrison, David K. Jackson, Ian Johnston, Dominic Kwiatkowski, Cordelia Langford, John Sillitoe on |

| | | | |
|---|---|---|---|
| EPI_ISL_3311484 | | Consortium | behalf of the Wellcome Sanger Institute COVID-19 Surveillance Team |
| EPI_ISL_3311485, EPI_ISL_3311486 | Lighthouse Lab in Glasgow | Wellcome Sanger Institute for the COVID-19 Genomics UK (COG-UK) Consortium | Harper VanSteenhouse, Yumi Kasai, David Gray, Carol Clugston, Anna Dominiczak and Alex Alderton, Roberto Amato, Jeffrey Barrett, Sonia Goncalves, Ewan Harrison, David K. Jackson, Ian Johnston, Dominic Kwiatkowski, Cordelia Langford, John Sillitoe on behalf of the Wellcome Sanger Institute COVID-19 Surveillance Team |
| EPI_ISL_3311487 | Lighthouse Lab in Milton Keynes | Wellcome Sanger Institute for the COVID-19 Genomics UK (COG-UK) Consortium | The Lighthouse Lab in Milton Keynes and Alex Alderton, Roberto Amato, Jeffrey Barrett, Sonia Goncalves, Ewan Harrison, David K. Jackson, Ian Johnston, Dominic Kwiatkowski, Cordelia Langford, John Sillitoe on behalf of the Wellcome Sanger Institute COVID-19 Surveillance Team |
| EPI_ISL_3311488, EPI_ISL_3311489 | Lighthouse Lab in Alderley Park | Wellcome Sanger Institute for the COVID-19 Genomics UK (COG-UK) Consortium | Jacquelyn Wynn, Mairead Hyland, The Lighthouse Lab in Alderley Park and Alex Alderton, Roberto Amato, Jeffrey Barrett, Sonia Goncalves, Ewan Harrison, David K. Jackson, Ian Johnston, Dominic Kwiatkowski, Cordelia Langford, John Sillitoe on behalf of the Wellcome Sanger Institute COVID-19 Surveillance Team |
| EPI_ISL_3311490, EPI_ISL_3311491, EPI_ISL_3311492 | Lighthouse Lab in Milton Keynes | Wellcome Sanger Institute for the COVID-19 Genomics UK (COG-UK) Consortium | The Lighthouse Lab in Milton Keynes and Alex Alderton, Roberto Amato, Jeffrey Barrett, Sonia Goncalves, Ewan Harrison, David K. Jackson, Ian Johnston, Dominic Kwiatkowski, Cordelia Langford, John Sillitoe on behalf of the Wellcome Sanger Institute COVID-19 Surveillance Team |
| EPI_ISL_3311493, EPI_ISL_3311494 | Lighthouse Lab in Alderley Park | Wellcome Sanger Institute for the COVID-19 Genomics UK (COG-UK) Consortium | Jacquelyn Wynn, Mairead Hyland, The Lighthouse Lab in Alderley Park and Alex Alderton, Roberto Amato, Jeffrey Barrett, Sonia Goncalves, Ewan Harrison, David K. Jackson, Ian Johnston, Dominic Kwiatkowski, Cordelia Langford, John Sillitoe on behalf of the Wellcome Sanger Institute COVID-19 Surveillance Team |
| EPI_ISL_3311495, EPI_ISL_3311496 | Lighthouse Lab in Milton Keynes | Wellcome Sanger Institute for the COVID-19 Genomics UK (COG-UK) Consortium | The Lighthouse Lab in Milton Keynes and Alex Alderton, Roberto Amato, Jeffrey Barrett, Sonia Goncalves, Ewan Harrison, David K. Jackson, Ian Johnston, Dominic Kwiatkowski, Cordelia Langford, John Sillitoe on behalf of the Wellcome Sanger Institute COVID-19 Surveillance Team |
| EPI_ISL_3311497 | Lighthouse Lab in Alderley Park | Wellcome Sanger Institute for the COVID-19 Genomics UK (COG-UK) Consortium | Jacquelyn Wynn, Mairead Hyland, The Lighthouse Lab in Alderley Park and Alex Alderton, Roberto Amato, Jeffrey Barrett, Sonia Goncalves, Ewan Harrison, David K. Jackson, Ian Johnston, Dominic Kwiatkowski, Cordelia Langford, John Sillitoe on behalf of the Wellcome Sanger Institute COVID-19 Surveillance Team |
| EPI_ISL_3311498, EPI_ISL_3311499 | Lighthouse Lab in Glasgow | Wellcome Sanger Institute for the COVID-19 Genomics UK (COG-UK) Consortium | Harper VanSteenhouse, Yumi Kasai, David Gray, Carol Clugston, Anna Dominiczak and Alex Alderton, Roberto Amato, Jeffrey Barrett, Sonia Goncalves, Ewan Harrison, David K. Jackson, Ian Johnston, Dominic Kwiatkowski, Cordelia Langford, John Sillitoe on behalf of the Wellcome Sanger Institute COVID-19 Surveillance Team |
| EPI_ISL_3311500 | Lighthouse Lab in Alderley Park | Wellcome Sanger Institute for the COVID-19 Genomics UK (COG-UK) Consortium | Jacquelyn Wynn, Mairead Hyland, The Lighthouse Lab in Alderley Park and Alex Alderton, Roberto Amato, Jeffrey Barrett, Sonia Goncalves, Ewan Harrison, David K. Jackson, Ian Johnston, Dominic Kwiatkowski, Cordelia Langford, John Sillitoe on behalf of the Wellcome Sanger Institute COVID-19 Surveillance Team |
| EPI_ISL_3311501, EPI_ISL_3311502 | Lighthouse Lab in Milton Keynes | Wellcome Sanger Institute for the COVID-19 Genomics UK (COG-UK) Consortium | The Lighthouse Lab in Milton Keynes and Alex Alderton, Roberto Amato, Jeffrey Barrett, Sonia Goncalves, Ewan Harrison, David K. Jackson, Ian Johnston, Dominic Kwiatkowski, Cordelia Langford, John Sillitoe on behalf of the Wellcome Sanger Institute COVID-19 Surveillance Team |
| EPI_ISL_3311503 | Lighthouse Lab in Glasgow | Wellcome Sanger Institute for the COVID-19 Genomics UK (COG-UK) Consortium | Harper VanSteenhouse, Yumi Kasai, David Gray, Carol Clugston, Anna Dominiczak and Alex Alderton, Roberto Amato, Jeffrey Barrett, Sonia Goncalves, Ewan Harrison, David K. Jackson, Ian Johnston, Dominic Kwiatkowski, Cordelia Langford, John Sillitoe on behalf of the Wellcome Sanger Institute COVID-19 Surveillance Team |
| EPI_ISL_3311504 | Health Services Laboratories | Wellcome Sanger Institute for the COVID-19 Genomics UK (COG-UK) Consortium | Health Services Laboratories and Alex Alderton, Roberto Amato, Jeffrey Barrett, Sonia Goncalves, Ewan Harrison, David K. Jackson, Ian Johnston, Dominic Kwiatkowski, Cordelia Langford, John Sillitoe on behalf of the Wellcome Sanger Institute COVID-19 Surveillance Team |
| EPI_ISL_3311505 | Lighthouse Lab in Glasgow | Wellcome Sanger Institute for the COVID-19 Genomics UK (COG-UK) Consortium | Harper VanSteenhouse, Yumi Kasai, David Gray, Carol Clugston, Anna Dominiczak and Alex Alderton, Roberto Amato, Jeffrey Barrett, Sonia Goncalves, Ewan Harrison, David K. Jackson, Ian Johnston, Dominic Kwiatkowski, Cordelia Langford, John Sillitoe on behalf of the Wellcome Sanger Institute COVID-19 Surveillance Team |
| EPI_ISL_3311506 | Lighthouse Lab in Milton Keynes | Wellcome Sanger Institute for the COVID-19 Genomics UK (COG-UK) Consortium | The Lighthouse Lab in Milton Keynes and Alex Alderton, Roberto Amato, Jeffrey Barrett, Sonia Goncalves, Ewan Harrison, David K. Jackson, Ian Johnston, Dominic Kwiatkowski, Cordelia Langford, John Sillitoe on behalf of the Wellcome Sanger Institute COVID-19 Surveillance Team |
| EPI_ISL_3311507, EPI_ISL_3311508 | Lighthouse Lab in Glasgow | Wellcome Sanger Institute for the COVID-19 Genomics UK (COG-UK) Consortium | Harper VanSteenhouse, Yumi Kasai, David Gray, Carol Clugston, Anna Dominiczak and Alex Alderton, Roberto Amato, Jeffrey Barrett, Sonia Goncalves, Ewan Harrison, David K. Jackson, Ian Johnston, Dominic Kwiatkowski, Cordelia Langford, John Sillitoe on behalf of the Wellcome Sanger Institute COVID-19 Surveillance Team |
| EPI_ISL_3311509, EPI_ISL_3311510, EPI_ISL_3311511 | Lighthouse Lab in Alderley Park | Wellcome Sanger Institute for the COVID-19 Genomics UK (COG-UK) Consortium | Jacquelyn Wynn, Mairead Hyland, The Lighthouse Lab in Alderley Park and Alex Alderton, Roberto Amato, Jeffrey Barrett, Sonia Goncalves, Ewan Harrison, David K. Jackson, Ian Johnston, Dominic Kwiatkowski, Cordelia Langford, John Sillitoe on behalf of the Wellcome Sanger Institute COVID-19 Surveillance Team |
| EPI_ISL_3311512, EPI_ISL_3311513 | Lighthouse Lab in Glasgow | Wellcome Sanger Institute for the COVID-19 Genomics UK (COG-UK) Consortium | Harper VanSteenhouse, Yumi Kasai, David Gray, Carol Clugston, Anna Dominiczak and Alex Alderton, Roberto Amato, Jeffrey Barrett, Sonia Goncalves, Ewan Harrison, David K. Jackson, Ian Johnston, Dominic Kwiatkowski, Cordelia Langford, John Sillitoe on behalf of the Wellcome Sanger Institute COVID-19 Surveillance Team |
| EPI_ISL_3311514 | Lighthouse Lab in Milton Keynes | Wellcome Sanger Institute for the COVID-19 Genomics UK (COG-UK) Consortium | The Lighthouse Lab in Milton Keynes and Alex Alderton, Roberto Amato, Jeffrey Barrett, Sonia Goncalves, Ewan Harrison, David K. Jackson, Ian Johnston, Dominic Kwiatkowski, Cordelia Langford, John Sillitoe on behalf of the Wellcome Sanger Institute COVID-19 Surveillance Team |
| EPI_ISL_3311515 | Health Services Laboratories | Wellcome Sanger Institute for the COVID-19 Genomics UK (COG-UK) Consortium | Health Services Laboratories and Alex Alderton, Roberto Amato, Jeffrey Barrett, Sonia Goncalves, Ewan Harrison, David K. Jackson, Ian Johnston, Dominic Kwiatkowski, Cordelia Langford, John Sillitoe on behalf of the Wellcome Sanger Institute COVID-19 Surveillance Team |
| EPI_ISL_3311516 | Lighthouse Lab in Glasgow | Wellcome Sanger Institute for the COVID-19 Genomics UK (COG-UK) Consortium | Harper VanSteenhouse, Yumi Kasai, David Gray, Carol Clugston, Anna Dominiczak and Alex Alderton, Roberto Amato, Jeffrey Barrett, Sonia Goncalves, Ewan Harrison, David K. Jackson, Ian Johnston, Dominic Kwiatkowski, Cordelia Langford, John Sillitoe on behalf of the Wellcome Sanger Institute COVID-19 Surveillance Team |
| EPI_ISL_3311517 | Lighthouse Lab in Milton Keynes | Wellcome Sanger Institute for the COVID-19 Genomics UK (COG-UK) Consortium | The Lighthouse Lab in Milton Keynes and Alex Alderton, Roberto Amato, Jeffrey Barrett, Sonia Goncalves, Ewan Harrison, David K. Jackson, Ian Johnston, Dominic Kwiatkowski, Cordelia Langford, John Sillitoe on behalf of the Wellcome Sanger Institute COVID-19 Surveillance Team |
| EPI_ISL_3311518 | Lighthouse Lab in Glasgow | Wellcome Sanger Institute for the COVID-19 Genomics UK (COG-UK) Consortium | Harper VanSteenhouse, Yumi Kasai, David Gray, Carol Clugston, Anna Dominiczak and Alex Alderton, Roberto Amato, Jeffrey Barrett, Sonia Goncalves, Ewan Harrison, David K. Jackson, Ian Johnston, Dominic Kwiatkowski, Cordelia Langford, John Sillitoe on behalf of the Wellcome Sanger Institute COVID-19 Surveillance Team |
| EPI_ISL_3311519 | Lighthouse Lab in Alderley Park | Wellcome Sanger Institute for the COVID-19 Genomics UK (COG-UK) Consortium | Jacquelyn Wynn, Mairead Hyland, The Lighthouse Lab in Alderley Park and Alex Alderton, Roberto Amato, Jeffrey Barrett, Sonia Goncalves, Ewan Harrison, David K. Jackson, Ian Johnston, Dominic Kwiatkowski, Cordelia Langford, John Sillitoe on behalf of the Wellcome Sanger Institute COVID-19 Surveillance Team |
| EPI_ISL_3311520 | Lighthouse Lab in Milton Keynes | Wellcome Sanger Institute for the COVID-19 Genomics UK (COG-UK) Consortium | The Lighthouse Lab in Milton Keynes and Alex Alderton, Roberto Amato, Jeffrey Barrett, Sonia Goncalves, Ewan Harrison, David K. Jackson, Ian Johnston, Dominic Kwiatkowski, Cordelia Langford, John Sillitoe on behalf of the Wellcome Sanger Institute COVID-19 Surveillance Team |
| EPI_ISL_3311521 | Lighthouse Lab in Alderley Park | Wellcome Sanger Institute for the COVID-19 Genomics UK (COG-UK) Consortium | Jacquelyn Wynn, Mairead Hyland, The Lighthouse Lab in Alderley Park and Alex Alderton, Roberto Amato, Jeffrey Barrett, Sonia Goncalves, Ewan Harrison, David K. Jackson, Ian Johnston, Dominic Kwiatkowski, Cordelia Langford, John Sillitoe on behalf of the Wellcome Sanger Institute COVID-19 Surveillance Team |
| EPI_ISL_3311522 | Lighthouse Lab in Milton Keynes | Wellcome Sanger Institute for the COVID-19 Genomics UK (COG-UK) Consortium | The Lighthouse Lab in Milton Keynes and Alex Alderton, Roberto Amato, Jeffrey Barrett, Sonia Goncalves, Ewan Harrison, David K. Jackson, Ian Johnston, Dominic Kwiatkowski, Cordelia Langford, John Sillitoe on behalf of the Wellcome Sanger Institute COVID-19 Surveillance Team |
| EPI_ISL_3311523 | Health Services Laboratories | Wellcome Sanger Institute for the COVID-19 Genomics UK (COG-UK) Consortium | Health Services Laboratories and Alex Alderton, Roberto Amato, Jeffrey Barrett, Sonia Goncalves, Ewan Harrison, David K. Jackson, Ian Johnston, Dominic Kwiatkowski, Cordelia Langford, John Sillitoe on behalf of the Wellcome Sanger Institute COVID-19 Surveillance Team |
| EPI_ISL_3311524, EPI_ISL_3311525, EPI_ISL_3311526 | Lighthouse Lab in Alderley Park | Wellcome Sanger Institute for the COVID-19 Genomics UK (COG-UK) Consortium | Jacquelyn Wynn, Mairead Hyland, The Lighthouse Lab in Alderley Park and Alex Alderton, Roberto Amato, Jeffrey Barrett, Sonia Goncalves, Ewan Harrison, David K. Jackson, Ian Johnston, Dominic Kwiatkowski, Cordelia Langford, John Sillitoe on behalf of the Wellcome Sanger Institute COVID-19 Surveillance Team |
| EPI_ISL_3311527 | Lighthouse Lab in Glasgow | Wellcome Sanger Institute for the COVID-19 Genomics UK (COG-UK) Consortium | Harper VanSteenhouse, Yumi Kasai, David Gray, Carol Clugston, Anna Dominiczak and Alex Alderton, Roberto Amato, Jeffrey Barrett, Sonia Goncalves, Ewan Harrison, David K. Jackson, Ian Johnston, Dominic Kwiatkowski, Cordelia Langford, John Sillitoe on behalf of the Wellcome Sanger Institute COVID-19 Surveillance Team |
| EPI_ISL_3311528, EPI_ISL_3311529 | Lighthouse Lab in Alderley Park | Wellcome Sanger Institute for the COVID-19 Genomics UK (COG-UK) Consortium | Jacquelyn Wynn, Mairead Hyland, The Lighthouse Lab in Alderley Park and Alex Alderton, Roberto Amato, Jeffrey Barrett, Sonia Goncalves, Ewan Harrison, David K. Jackson, Ian Johnston, Dominic Kwiatkowski, Cordelia Langford, John Sillitoe on behalf of the Wellcome Sanger Institute COVID-19 Surveillance Team |
| EPI_ISL_3311530 | Lighthouse Lab in Milton Keynes | Wellcome Sanger Institute for the COVID-19 Genomics UK (COG-UK) Consortium | The Lighthouse Lab in Milton Keynes and Alex Alderton, Roberto Amato, Jeffrey Barrett, Sonia Goncalves, Ewan Harrison, David K. Jackson, Ian Johnston, Dominic Kwiatkowski, Cordelia Langford, John Sillitoe on behalf of the Wellcome Sanger Institute COVID-19 Surveillance Team |
| EPI_ISL_3311531 | Lighthouse Lab in Glasgow | Wellcome Sanger Institute for the COVID-19 Genomics UK (COG-UK) Consortium | Harper VanSteenhouse, Yumi Kasai, David Gray, Carol Clugston, Anna Dominiczak and Alex Alderton, Roberto Amato, Jeffrey Barrett, Sonia Goncalves, Ewan Harrison, David K. Jackson, Ian Johnston, Dominic Kwiatkowski, Cordelia Langford, John Sillitoe on behalf of the Wellcome Sanger Institute COVID-19 Surveillance Team |
| EPI_ISL_3311532, EPI_ISL_3311533 | Lighthouse Lab in Milton Keynes | Wellcome Sanger Institute for the COVID-19 Genomics UK (COG-UK) Consortium | The Lighthouse Lab in Milton Keynes and Alex Alderton, Roberto Amato, Jeffrey Barrett, Sonia Goncalves, Ewan Harrison, David K. Jackson, Ian Johnston, Dominic Kwiatkowski, Cordelia Langford, John Sillitoe on behalf of the Wellcome Sanger Institute COVID-19 Surveillance Team |
| EPI_ISL_3311534 | Lighthouse Lab in Alderley Park | Wellcome Sanger Institute for the COVID-19 Genomics UK (COG-UK) Consortium | Jacquelyn Wynn, Mairead Hyland, The Lighthouse Lab in Alderley Park and Alex Alderton, Roberto Amato, Jeffrey Barrett, Sonia Goncalves, Ewan Harrison, David K. Jackson, Ian Johnston, Dominic Kwiatkowski, Cordelia Langford, John Sillitoe on behalf of the Wellcome Sanger Institute COVID-19 Surveillance Team |
| EPI_ISL_3311535 | Lighthouse Lab in Milton Keynes | Wellcome Sanger Institute for the COVID-19 Genomics UK (COG-UK) Consortium | The Lighthouse Lab in Milton Keynes and Alex Alderton, Roberto Amato, Jeffrey Barrett, Sonia Goncalves, Ewan Harrison, David K. Jackson, Ian Johnston, Dominic Kwiatkowski, Cordelia Langford, John Sillitoe on behalf of the Wellcome Sanger Institute COVID-19 Surveillance Team |
| EPI_ISL_3311536, EPI_ISL_3311537 | Lighthouse Lab in Alderley Park | Wellcome Sanger Institute for the COVID-19 Genomics UK (COG-UK) Consortium | Jacquelyn Wynn, Mairead Hyland, The Lighthouse Lab in Alderley Park and Alex Alderton, Roberto Amato, Jeffrey Barrett, Sonia Goncalves, Ewan Harrison, David K. Jackson, Ian Johnston, Dominic Kwiatkowski, Cordelia Langford, John Sillitoe on behalf of the Wellcome Sanger Institute COVID-19 Surveillance Team |
| EPI_ISL_3311538, EPI_ISL_3311539, EPI_ISL_3311540 | Lighthouse Lab in Milton Keynes | Wellcome Sanger Institute for the COVID-19 Genomics UK (COG-UK) Consortium | The Lighthouse Lab in Milton Keynes and Alex Alderton, Roberto Amato, Jeffrey Barrett, Sonia Goncalves, Ewan Harrison, David K. Jackson, Ian Johnston, Dominic Kwiatkowski, Cordelia Langford, John Sillitoe on behalf of the Wellcome Sanger Institute COVID-19 Surveillance Team |
| EPI_ISL_3311541 | Lighthouse Lab in Glasgow | Wellcome Sanger Institute for the COVID-19 Genomics UK (COG-UK) Consortium | Harper VanSteenhouse, Yumi Kasai, David Gray, Carol Clugston, Anna Dominiczak and Alex Alderton, Roberto Amato, Jeffrey Barrett, Sonia Goncalves, Ewan Harrison, David K. Jackson, Ian Johnston, Dominic Kwiatkowski, Cordelia Langford, John Sillitoe on behalf of the Wellcome Sanger Institute COVID-19 Surveillance Team |
| EPI_ISL_3311542 | Lighthouse Lab in Milton Keynes | Wellcome Sanger Institute for the COVID-19 Genomics UK (COG-UK) Consortium | The Lighthouse Lab in Milton Keynes and Alex Alderton, Roberto Amato, Jeffrey Barrett, Sonia Goncalves, Ewan Harrison, David K. Jackson, Ian Johnston, Dominic Kwiatkowski, Cordelia Langford, John Sillitoe on behalf of the Wellcome Sanger Institute COVID-19 Surveillance Team |
| EPI_ISL_3311543 | Lighthouse Lab in Alderley Park | Wellcome Sanger Institute for the COVID-19 Genomics UK (COG-UK) Consortium | Jacquelyn Wynn, Mairead Hyland, The Lighthouse Lab in Alderley Park and Alex Alderton, Roberto Amato, Jeffrey Barrett, Sonia Goncalves, Ewan Harrison, David K. Jackson, Ian Johnston, Dominic Kwiatkowski, Cordelia Langford, John Sillitoe on behalf of the Wellcome Sanger Institute COVID-19 Surveillance Team |
| EPI_ISL_3311544 | Lighthouse Lab in Glasgow | Wellcome Sanger Institute for the COVID-19 Genomics UK (COG-UK) Consortium | Harper VanSteenhouse, Yumi Kasai, David Gray, Carol Clugston, Anna Dominiczak and Alex Alderton, Roberto Amato, Jeffrey Barrett, Sonia Goncalves, Ewan Harrison, David K. Jackson, Ian Johnston, Dominic Kwiatkowski, Cordelia Langford, John Sillitoe on behalf of the Wellcome Sanger Institute COVID-19 Surveillance Team |
| EPI_ISL_3311545 | Lighthouse Lab in Milton Keynes | Wellcome Sanger Institute for the COVID-19 Genomics UK (COG-UK) Consortium | The Lighthouse Lab in Milton Keynes and Alex Alderton, Roberto Amato, Jeffrey Barrett, Sonia Goncalves, Ewan Harrison, David K. Jackson, Ian Johnston, Dominic Kwiatkowski, Cordelia Langford, John Sillitoe on behalf of the Wellcome Sanger Institute COVID-19 Surveillance Team |
| EPI_ISL_3311546 | Lighthouse Lab in Glasgow | Wellcome Sanger Institute for the COVID-19 Genomics UK (COG-UK) Consortium | Harper VanSteenhouse, Yumi Kasai, David Gray, Carol Clugston, Anna Dominiczak and Alex Alderton, Roberto Amato, Jeffrey Barrett, Sonia Goncalves, Ewan Harrison, David K. Jackson, Ian Johnston, Dominic Kwiatkowski, Cordelia Langford, John Sillitoe on behalf of the Wellcome Sanger Institute COVID-19 Surveillance Team |

| | | | |
|---|---|---|---|
| EPI_ISL_3311547, EPI_ISL_3311548 | Lighthouse Lab in Alderley Park | Wellcome Sanger Institute for the COVID-19 Genomics UK (COG-UK) Consortium | Jacquelyn Wynn, Mairead Hyland, The Lighthouse Lab in Alderley Park and Alex Alderton, Roberto Amato, Jeffrey Barrett, Sonia Goncalves, Ewan Harrison, David K. Jackson, Ian Johnston, Dominic Kwiatkowski, Cordelia Langford, John Sillitoe on behalf of the Wellcome Sanger Institute COVID-19 Surveillance Team |
| EPI_ISL_3311549 | Lighthouse Lab in Milton Keynes | Wellcome Sanger Institute for the COVID-19 Genomics UK (COG-UK) Consortium | The Lighthouse Lab in Milton Keynes and Alex Alderton, Roberto Amato, Jeffrey Barrett, Sonia Goncalves, Ewan Harrison, David K. Jackson, Ian Johnston, Dominic Kwiatkowski, Cordelia Langford, John Sillitoe on behalf of the Wellcome Sanger Institute COVID-19 Surveillance Team |
| EPI_ISL_3311550 | Lighthouse Lab in Glasgow | Wellcome Sanger Institute for the COVID-19 Genomics UK (COG-UK) Consortium | Harper VanSteenhouse, Yumi Kasai, David Gray, Carol Clugston, Anna Dominiczak and Alex Alderton, Roberto Amato, Jeffrey Barrett, Sonia Goncalves, Ewan Harrison, David K. Jackson, Ian Johnston, Dominic Kwiatkowski, Cordelia Langford, John Sillitoe on behalf of the Wellcome Sanger Institute COVID-19 Surveillance Team |
| EPI_ISL_3311551, EPI_ISL_3311552 | Lighthouse Lab in Alderley Park | Wellcome Sanger Institute for the COVID-19 Genomics UK (COG-UK) Consortium | Jacquelyn Wynn, Mairead Hyland, The Lighthouse Lab in Alderley Park and Alex Alderton, Roberto Amato, Jeffrey Barrett, Sonia Goncalves, Ewan Harrison, David K. Jackson, Ian Johnston, Dominic Kwiatkowski, Cordelia Langford, John Sillitoe on behalf of the Wellcome Sanger Institute COVID-19 Surveillance Team |
| EPI_ISL_3311553, EPI_ISL_3311554 | Lighthouse Lab in Milton Keynes | Wellcome Sanger Institute for the COVID-19 Genomics UK (COG-UK) Consortium | The Lighthouse Lab in Milton Keynes and Alex Alderton, Roberto Amato, Jeffrey Barrett, Sonia Goncalves, Ewan Harrison, David K. Jackson, Ian Johnston, Dominic Kwiatkowski, Cordelia Langford, John Sillitoe on behalf of the Wellcome Sanger Institute COVID-19 Surveillance Team |
| EPI_ISL_3311555 | Lighthouse Lab in Glasgow | Wellcome Sanger Institute for the COVID-19 Genomics UK (COG-UK) Consortium | Harper VanSteenhouse, Yumi Kasai, David Gray, Carol Clugston, Anna Dominiczak and Alex Alderton, Roberto Amato, Jeffrey Barrett, Sonia Goncalves, Ewan Harrison, David K. Jackson, Ian Johnston, Dominic Kwiatkowski, Cordelia Langford, John Sillitoe on behalf of the Wellcome Sanger Institute COVID-19 Surveillance Team |
| EPI_ISL_3311556 | Lighthouse Lab in Milton Keynes | Wellcome Sanger Institute for the COVID-19 Genomics UK (COG-UK) Consortium | The Lighthouse Lab in Milton Keynes and Alex Alderton, Roberto Amato, Jeffrey Barrett, Sonia Goncalves, Ewan Harrison, David K. Jackson, Ian Johnston, Dominic Kwiatkowski, Cordelia Langford, John Sillitoe on behalf of the Wellcome Sanger Institute COVID-19 Surveillance Team |
| EPI_ISL_3311557 | Lighthouse Lab in Glasgow | Wellcome Sanger Institute for the COVID-19 Genomics UK (COG-UK) Consortium | Harper VanSteenhouse, Yumi Kasai, David Gray, Carol Clugston, Anna Dominiczak and Alex Alderton, Roberto Amato, Jeffrey Barrett, Sonia Goncalves, Ewan Harrison, David K. Jackson, Ian Johnston, Dominic Kwiatkowski, Cordelia Langford, John Sillitoe on behalf of the Wellcome Sanger Institute COVID-19 Surveillance Team |
| EPI_ISL_3311558 | Lighthouse Lab in Alderley Park | Wellcome Sanger Institute for the COVID-19 Genomics UK (COG-UK) Consortium | Jacquelyn Wynn, Mairead Hyland, The Lighthouse Lab in Alderley Park and Alex Alderton, Roberto Amato, Jeffrey Barrett, Sonia Goncalves, Ewan Harrison, David K. Jackson, Ian Johnston, Dominic Kwiatkowski, Cordelia Langford, John Sillitoe on behalf of the Wellcome Sanger Institute COVID-19 Surveillance Team |
| EPI_ISL_3311559 | Lighthouse Lab in Glasgow | Wellcome Sanger Institute for the COVID-19 Genomics UK (COG-UK) Consortium | Harper VanSteenhouse, Yumi Kasai, David Gray, Carol Clugston, Anna Dominiczak and Alex Alderton, Roberto Amato, Jeffrey Barrett, Sonia Goncalves, Ewan Harrison, David K. Jackson, Ian Johnston, Dominic Kwiatkowski, Cordelia Langford, John Sillitoe on behalf of the Wellcome Sanger Institute COVID-19 Surveillance Team |
| EPI_ISL_3311560, EPI_ISL_3311561, EPI_ISL_3311562, EPI_ISL_3311563, EPI_ISL_3311564 | Lighthouse Lab in Milton Keynes | Wellcome Sanger Institute for the COVID-19 Genomics UK (COG-UK) Consortium | The Lighthouse Lab in Milton Keynes and Alex Alderton, Roberto Amato, Jeffrey Barrett, Sonia Goncalves, Ewan Harrison, David K. Jackson, Ian Johnston, Dominic Kwiatkowski, Cordelia Langford, John Sillitoe on behalf of the Wellcome Sanger Institute COVID-19 Surveillance Team |
| EPI_ISL_3311565, EPI_ISL_3311566 | Lighthouse Lab in Glasgow | Wellcome Sanger Institute for the COVID-19 Genomics UK (COG-UK) Consortium | Harper VanSteenhouse, Yumi Kasai, David Gray, Carol Clugston, Anna Dominiczak and Alex Alderton, Roberto Amato, Jeffrey Barrett, Sonia Goncalves, Ewan Harrison, David K. Jackson, Ian Johnston, Dominic Kwiatkowski, Cordelia Langford, John Sillitoe on behalf of the Wellcome Sanger Institute COVID-19 Surveillance Team |
| EPI_ISL_3311567 | Lighthouse Lab in Milton Keynes | Wellcome Sanger Institute for the COVID-19 Genomics UK (COG-UK) Consortium | The Lighthouse Lab in Milton Keynes and Alex Alderton, Roberto Amato, Jeffrey Barrett, Sonia Goncalves, Ewan Harrison, David K. Jackson, Ian Johnston, Dominic Kwiatkowski, Cordelia Langford, John Sillitoe on behalf of the Wellcome Sanger Institute COVID-19 Surveillance Team |
| EPI_ISL_3311568 | Lighthouse Lab in Alderley Park | Wellcome Sanger Institute for the COVID-19 Genomics UK (COG-UK) Consortium | Jacquelyn Wynn, Mairead Hyland, The Lighthouse Lab in Alderley Park and Alex Alderton, Roberto Amato, Jeffrey Barrett, Sonia Goncalves, Ewan Harrison, David K. Jackson, Ian Johnston, Dominic Kwiatkowski, Cordelia Langford, John Sillitoe on behalf of the Wellcome Sanger Institute COVID-19 Surveillance Team |
| EPI_ISL_3311569 | Lighthouse Lab in Glasgow | Wellcome Sanger Institute for the COVID-19 Genomics UK (COG-UK) Consortium | Harper VanSteenhouse, Yumi Kasai, David Gray, Carol Clugston, Anna Dominiczak and Alex Alderton, Roberto Amato, Jeffrey Barrett, Sonia Goncalves, Ewan Harrison, David K. Jackson, Ian Johnston, Dominic Kwiatkowski, Cordelia Langford, John Sillitoe on behalf of the Wellcome Sanger Institute COVID-19 Surveillance Team |
| EPI_ISL_3311570 | Health Services Laboratories | Wellcome Sanger Institute for the COVID-19 Genomics UK (COG-UK) Consortium | Health Services Laboratories and Alex Alderton, Roberto Amato, Jeffrey Barrett, Sonia Goncalves, Ewan Harrison, David K. Jackson, Ian Johnston, Dominic Kwiatkowski, Cordelia Langford, John Sillitoe on behalf of the Wellcome Sanger Institute COVID-19 Surveillance Team |
| EPI_ISL_3311571, EPI_ISL_3311572 | Lighthouse Lab in Glasgow | Wellcome Sanger Institute for the COVID-19 Genomics UK (COG-UK) Consortium | Harper VanSteenhouse, Yumi Kasai, David Gray, Carol Clugston, Anna Dominiczak and Alex Alderton, Roberto Amato, Jeffrey Barrett, Sonia Goncalves, Ewan Harrison, David K. Jackson, Ian Johnston, Dominic Kwiatkowski, Cordelia Langford, John Sillitoe on behalf of the Wellcome Sanger Institute COVID-19 Surveillance Team |
| EPI_ISL_3311573, EPI_ISL_3311574 | Lighthouse Lab in Milton Keynes | Wellcome Sanger Institute for the COVID-19 Genomics UK (COG-UK) Consortium | The Lighthouse Lab in Milton Keynes and Alex Alderton, Roberto Amato, Jeffrey Barrett, Sonia Goncalves, Ewan Harrison, David K. Jackson, Ian Johnston, Dominic Kwiatkowski, Cordelia Langford, John Sillitoe on behalf of the Wellcome Sanger Institute COVID-19 Surveillance Team |
| EPI_ISL_3311575 | Lighthouse Lab in Alderley Park | Wellcome Sanger Institute for the COVID-19 Genomics UK (COG-UK) Consortium | Jacquelyn Wynn, Mairead Hyland, The Lighthouse Lab in Alderley Park and Alex Alderton, Roberto Amato, Jeffrey Barrett, Sonia Goncalves, Ewan Harrison, David K. Jackson, Ian Johnston, Dominic Kwiatkowski, Cordelia Langford, John Sillitoe on behalf of the Wellcome Sanger Institute COVID-19 Surveillance Team |
| EPI_ISL_3311576 | Lighthouse Lab in Glasgow | Wellcome Sanger Institute for the COVID-19 Genomics UK (COG-UK) Consortium | Harper VanSteenhouse, Yumi Kasai, David Gray, Carol Clugston, Anna Dominiczak and Alex Alderton, Roberto Amato, Jeffrey Barrett, Sonia Goncalves, Ewan Harrison, David K. Jackson, Ian Johnston, Dominic Kwiatkowski, Cordelia Langford, John Sillitoe on behalf of the Wellcome Sanger Institute COVID-19 Surveillance Team |
| EPI_ISL_3311577, EPI_ISL_3311578 | Lighthouse Lab in Milton Keynes | Wellcome Sanger Institute for the COVID-19 Genomics UK (COG-UK) Consortium | The Lighthouse Lab in Milton Keynes and Alex Alderton, Roberto Amato, Jeffrey Barrett, Sonia Goncalves, Ewan Harrison, David K. Jackson, Ian Johnston, Dominic Kwiatkowski, Cordelia Langford, John Sillitoe on behalf of the Wellcome Sanger Institute COVID-19 Surveillance Team |
| EPI_ISL_3311579 | Lighthouse Lab in Alderley Park | Wellcome Sanger Institute for the COVID-19 Genomics UK (COG-UK) Consortium | Jacquelyn Wynn, Mairead Hyland, The Lighthouse Lab in Alderley Park and Alex Alderton, Roberto Amato, Jeffrey Barrett, Sonia Goncalves, Ewan Harrison, David K. Jackson, Ian Johnston, Dominic Kwiatkowski, Cordelia Langford, John Sillitoe on behalf of the Wellcome Sanger Institute COVID-19 Surveillance Team |
| EPI_ISL_3311580 | Lighthouse Lab in Milton Keynes | Wellcome Sanger Institute for the COVID-19 Genomics UK (COG-UK) Consortium | The Lighthouse Lab in Milton Keynes and Alex Alderton, Roberto Amato, Jeffrey Barrett, Sonia Goncalves, Ewan Harrison, David K. Jackson, Ian Johnston, Dominic Kwiatkowski, Cordelia Langford, John Sillitoe on behalf of the Wellcome Sanger Institute COVID-19 Surveillance Team |
| EPI_ISL_3311581 | Lighthouse Lab in Alderley Park | Wellcome Sanger Institute for the COVID-19 Genomics UK (COG-UK) Consortium | Jacquelyn Wynn, Mairead Hyland, The Lighthouse Lab in Alderley Park and Alex Alderton, Roberto Amato, Jeffrey Barrett, Sonia Goncalves, Ewan Harrison, David K. Jackson, Ian Johnston, Dominic Kwiatkowski, Cordelia Langford, John Sillitoe on behalf of the Wellcome Sanger Institute COVID-19 Surveillance Team |
| EPI_ISL_3311582 | Health Services Laboratories | Wellcome Sanger Institute for the COVID-19 Genomics UK (COG-UK) Consortium | Health Services Laboratories and Alex Alderton, Roberto Amato, Jeffrey Barrett, Sonia Goncalves, Ewan Harrison, David K. Jackson, Ian Johnston, Dominic Kwiatkowski, Cordelia Langford, John Sillitoe on behalf of the Wellcome Sanger Institute COVID-19 Surveillance Team |
| EPI_ISL_3311583 | Lighthouse Lab in Milton Keynes | Wellcome Sanger Institute for the COVID-19 Genomics UK (COG-UK) Consortium | The Lighthouse Lab in Milton Keynes and Alex Alderton, Roberto Amato, Jeffrey Barrett, Sonia Goncalves, Ewan Harrison, David K. Jackson, Ian Johnston, Dominic Kwiatkowski, Cordelia Langford, John Sillitoe on behalf of the Wellcome Sanger Institute COVID-19 Surveillance Team |
| EPI_ISL_3311584, EPI_ISL_3311585, EPI_ISL_3311586 | Lighthouse Lab in Alderley Park | Wellcome Sanger Institute for the COVID-19 Genomics UK (COG-UK) Consortium | Jacquelyn Wynn, Mairead Hyland, The Lighthouse Lab in Alderley Park and Alex Alderton, Roberto Amato, Jeffrey Barrett, Sonia Goncalves, Ewan Harrison, David K. Jackson, Ian Johnston, Dominic Kwiatkowski, Cordelia Langford, John Sillitoe on behalf of the Wellcome Sanger Institute COVID-19 Surveillance Team |
| EPI_ISL_3311587 | Lighthouse Lab in Milton Keynes | Wellcome Sanger Institute for the COVID-19 Genomics UK (COG-UK) Consortium | The Lighthouse Lab in Milton Keynes and Alex Alderton, Roberto Amato, Jeffrey Barrett, Sonia Goncalves, Ewan Harrison, David K. Jackson, Ian Johnston, Dominic Kwiatkowski, Cordelia Langford, John Sillitoe on behalf of the Wellcome Sanger Institute COVID-19 Surveillance Team |
| EPI_ISL_3311588 | Lighthouse Lab in Glasgow | Wellcome Sanger Institute for the COVID-19 Genomics UK (COG-UK) Consortium | Harper VanSteenhouse, Yumi Kasai, David Gray, Carol Clugston, Anna Dominiczak and Alex Alderton, Roberto Amato, Jeffrey Barrett, Sonia Goncalves, Ewan Harrison, David K. Jackson, Ian Johnston, Dominic Kwiatkowski, Cordelia Langford, John Sillitoe on behalf of the Wellcome Sanger Institute COVID-19 Surveillance Team |
| EPI_ISL_3311589, EPI_ISL_3311590, EPI_ISL_3311591, EPI_ISL_3311592, EPI_ISL_3311593 | Lighthouse Lab in Alderley Park | Wellcome Sanger Institute for the COVID-19 Genomics UK (COG-UK) Consortium | Jacquelyn Wynn, Mairead Hyland, The Lighthouse Lab in Alderley Park and Alex Alderton, Roberto Amato, Jeffrey Barrett, Sonia Goncalves, Ewan Harrison, David K. Jackson, Ian Johnston, Dominic Kwiatkowski, Cordelia Langford, John Sillitoe on behalf of the Wellcome Sanger Institute COVID-19 Surveillance Team |
| EPI_ISL_3311594 | Lighthouse Lab in Milton Keynes | Wellcome Sanger Institute for the COVID-19 Genomics UK (COG-UK) Consortium | The Lighthouse Lab in Milton Keynes and Alex Alderton, Roberto Amato, Jeffrey Barrett, Sonia Goncalves, Ewan Harrison, David K. Jackson, Ian Johnston, Dominic Kwiatkowski, Cordelia Langford, John Sillitoe on behalf of the Wellcome Sanger Institute COVID-19 Surveillance Team |
| EPI_ISL_3311595, EPI_ISL_3311596 | Lighthouse Lab in Alderley Park | Wellcome Sanger Institute for the COVID-19 Genomics UK (COG-UK) Consortium | Jacquelyn Wynn, Mairead Hyland, The Lighthouse Lab in Alderley Park and Alex Alderton, Roberto Amato, Jeffrey Barrett, Sonia Goncalves, Ewan Harrison, David K. Jackson, Ian Johnston, Dominic Kwiatkowski, Cordelia Langford, John Sillitoe on behalf of the Wellcome Sanger Institute COVID-19 Surveillance Team |
| EPI_ISL_3311597 | Lighthouse Lab in Milton Keynes | Wellcome Sanger Institute for the COVID-19 Genomics UK (COG-UK) Consortium | The Lighthouse Lab in Milton Keynes and Alex Alderton, Roberto Amato, Jeffrey Barrett, Sonia Goncalves, Ewan Harrison, David K. Jackson, Ian Johnston, Dominic Kwiatkowski, Cordelia Langford, John Sillitoe on behalf of the Wellcome Sanger Institute COVID-19 Surveillance Team |
| EPI_ISL_3311598, EPI_ISL_3311599 | Lighthouse Lab in Glasgow | Wellcome Sanger Institute for the COVID-19 Genomics UK (COG-UK) Consortium | Harper VanSteenhouse, Yumi Kasai, David Gray, Carol Clugston, Anna Dominiczak and Alex Alderton, Roberto Amato, Jeffrey Barrett, Sonia Goncalves, Ewan Harrison, David K. Jackson, Ian Johnston, Dominic Kwiatkowski, Cordelia Langford, John Sillitoe on behalf of the Wellcome Sanger Institute COVID-19 Surveillance Team |
| EPI_ISL_3311600, EPI_ISL_3311601 | Lighthouse Lab in Milton Keynes | Wellcome Sanger Institute for the COVID-19 Genomics UK (COG-UK) Consortium | The Lighthouse Lab in Milton Keynes and Alex Alderton, Roberto Amato, Jeffrey Barrett, Sonia Goncalves, Ewan Harrison, David K. Jackson, Ian Johnston, Dominic Kwiatkowski, Cordelia Langford, John Sillitoe on behalf of the Wellcome Sanger Institute COVID-19 Surveillance Team |
| EPI_ISL_3311602 | Lighthouse Laboratory Plymouth | Wellcome Sanger Institute for the COVID-19 Genomics UK (COG-UK) Consortium | Lighthouse Laboratory Plymouth and Alex Alderton, Roberto Amato, Jeffrey Barrett, Sonia Goncalves, Ewan Harrison, David K. Jackson, Ian Johnston, Dominic Kwiatkowski, Cordelia Langford, John Sillitoe on behalf of the Wellcome Sanger Institute COVID-19 Surveillance Team |
| EPI_ISL_3311603 | Lighthouse Lab in Glasgow | Wellcome Sanger Institute for the COVID-19 Genomics UK (COG-UK) Consortium | Harper VanSteenhouse, Yumi Kasai, David Gray, Carol Clugston, Anna Dominiczak and Alex Alderton, Roberto Amato, Jeffrey Barrett, Sonia Goncalves, Ewan Harrison, David K. Jackson, Ian Johnston, Dominic Kwiatkowski, Cordelia Langford, John Sillitoe on behalf of the Wellcome Sanger Institute COVID-19 Surveillance Team |
| EPI_ISL_3311604, EPI_ISL_3311605, EPI_ISL_3311606, EPI_ISL_3311607, EPI_ISL_3311608 | Lighthouse Laboratory Plymouth | Wellcome Sanger Institute for the COVID-19 Genomics UK (COG-UK) Consortium | Lighthouse Laboratory Plymouth and Alex Alderton, Roberto Amato, Jeffrey Barrett, Sonia Goncalves, Ewan Harrison, David K. Jackson, Ian Johnston, Dominic Kwiatkowski, Cordelia Langford, John Sillitoe on behalf of the Wellcome Sanger Institute COVID-19 Surveillance Team |
| EPI_ISL_3311609, EPI_ISL_3311610 | Lighthouse Lab in Glasgow | Wellcome Sanger Institute for the COVID-19 Genomics UK (COG-UK) Consortium | Harper VanSteenhouse, Yumi Kasai, David Gray, Carol Clugston, Anna Dominiczak and Alex Alderton, Roberto Amato, Jeffrey Barrett, Sonia Goncalves, Ewan Harrison, David K. Jackson, Ian Johnston, Dominic Kwiatkowski, Cordelia Langford, John Sillitoe on behalf of the Wellcome Sanger Institute COVID-19 Surveillance Team |
| EPI_ISL_3311611, EPI_ISL_3311612, EPI_ISL_3311613, EPI_ISL_3311614 | Lighthouse Lab in Milton Keynes | Wellcome Sanger Institute for the COVID-19 Genomics UK (COG-UK) Consortium | The Lighthouse Lab in Milton Keynes and Alex Alderton, Roberto Amato, Jeffrey Barrett, Sonia Goncalves, Ewan Harrison, David K. Jackson, Ian Johnston, Dominic Kwiatkowski, Cordelia Langford, John Sillitoe on behalf of the Wellcome Sanger Institute COVID-19 Surveillance Team |

| | | | |
|---|---|---|---|
| EPI_ISL_3311615 | Lighthouse Laboratory Plymouth | Wellcome Sanger Institute for the COVID-19 Genomics UK (COG-UK) Consortium | Lighthouse Laboratory Plymouth and Alex Alderton, Roberto Amato, Jeffrey Barrett, Sonia Goncalves, Ewan Harrison, David K. Jackson, Ian Johnston, Dominic Kwiatkowski, Cordelia Langford, John Sillitoe on behalf of the Wellcome Sanger Institute COVID-19 Surveillance Team |
| EPI_ISL_3311616 | Lighthouse Lab in Glasgow | Wellcome Sanger Institute for the COVID-19 Genomics UK (COG-UK) Consortium | Harper VanSteenhouse, Yumi Kasai, David Gray, Carol Clugston, Anna Dominiczak and Alex Alderton, Roberto Amato, Jeffrey Barrett, Sonia Goncalves, Ewan Harrison, David K. Jackson, Ian Johnston, Dominic Kwiatkowski, Cordelia Langford, John Sillitoe on behalf of the Wellcome Sanger Institute COVID-19 Surveillance Team |
| EPI_ISL_3311617 | Lighthouse Laboratory Plymouth | Wellcome Sanger Institute for the COVID-19 Genomics UK (COG-UK) Consortium | Lighthouse Laboratory Plymouth and Alex Alderton, Roberto Amato, Jeffrey Barrett, Sonia Goncalves, Ewan Harrison, David K. Jackson, Ian Johnston, Dominic Kwiatkowski, Cordelia Langford, John Sillitoe on behalf of the Wellcome Sanger Institute COVID-19 Surveillance Team |
| EPI_ISL_3311618 | Lighthouse Lab in Milton Keynes | Wellcome Sanger Institute for the COVID-19 Genomics UK (COG-UK) Consortium | The Lighthouse Lab in Milton Keynes and Alex Alderton, Roberto Amato, Jeffrey Barrett, Sonia Goncalves, Ewan Harrison, David K. Jackson, Ian Johnston, Dominic Kwiatkowski, Cordelia Langford, John Sillitoe on behalf of the Wellcome Sanger Institute COVID-19 Surveillance Team |
| EPI_ISL_3311619 | Lighthouse Laboratory Plymouth | Wellcome Sanger Institute for the COVID-19 Genomics UK (COG-UK) Consortium | Lighthouse Laboratory Plymouth and Alex Alderton, Roberto Amato, Jeffrey Barrett, Sonia Goncalves, Ewan Harrison, David K. Jackson, Ian Johnston, Dominic Kwiatkowski, Cordelia Langford, John Sillitoe on behalf of the Wellcome Sanger Institute COVID-19 Surveillance Team |
| EPI_ISL_3311620 | Lighthouse Lab in Glasgow | Wellcome Sanger Institute for the COVID-19 Genomics UK (COG-UK) Consortium | Harper VanSteenhouse, Yumi Kasai, David Gray, Carol Clugston, Anna Dominiczak and Alex Alderton, Roberto Amato, Jeffrey Barrett, Sonia Goncalves, Ewan Harrison, David K. Jackson, Ian Johnston, Dominic Kwiatkowski, Cordelia Langford, John Sillitoe on behalf of the Wellcome Sanger Institute COVID-19 Surveillance Team |
| EPI_ISL_3311621 | Lighthouse Lab in Milton Keynes | Wellcome Sanger Institute for the COVID-19 Genomics UK (COG-UK) Consortium | The Lighthouse Lab in Milton Keynes and Alex Alderton, Roberto Amato, Jeffrey Barrett, Sonia Goncalves, Ewan Harrison, David K. Jackson, Ian Johnston, Dominic Kwiatkowski, Cordelia Langford, John Sillitoe on behalf of the Wellcome Sanger Institute COVID-19 Surveillance Team |
| EPI_ISL_3311622 | Lighthouse Lab in Glasgow | Wellcome Sanger Institute for the COVID-19 Genomics UK (COG-UK) Consortium | Harper VanSteenhouse, Yumi Kasai, David Gray, Carol Clugston, Anna Dominiczak and Alex Alderton, Roberto Amato, Jeffrey Barrett, Sonia Goncalves, Ewan Harrison, David K. Jackson, Ian Johnston, Dominic Kwiatkowski, Cordelia Langford, John Sillitoe on behalf of the Wellcome Sanger Institute COVID-19 Surveillance Team |
| EPI_ISL_3311623 | Lighthouse Laboratory Plymouth | Wellcome Sanger Institute for the COVID-19 Genomics UK (COG-UK) Consortium | Lighthouse Laboratory Plymouth and Alex Alderton, Roberto Amato, Jeffrey Barrett, Sonia Goncalves, Ewan Harrison, David K. Jackson, Ian Johnston, Dominic Kwiatkowski, Cordelia Langford, John Sillitoe on behalf of the Wellcome Sanger Institute COVID-19 Surveillance Team |
| EPI_ISL_3311624 | Lighthouse Lab in Glasgow | Wellcome Sanger Institute for the COVID-19 Genomics UK (COG-UK) Consortium | Harper VanSteenhouse, Yumi Kasai, David Gray, Carol Clugston, Anna Dominiczak and Alex Alderton, Roberto Amato, Jeffrey Barrett, Sonia Goncalves, Ewan Harrison, David K. Jackson, Ian Johnston, Dominic Kwiatkowski, Cordelia Langford, John Sillitoe on behalf of the Wellcome Sanger Institute COVID-19 Surveillance Team |
| EPI_ISL_3311625, EPI_ISL_3311626, EPI_ISL_3311627 | Lighthouse Lab in Milton Keynes | Wellcome Sanger Institute for the COVID-19 Genomics UK (COG-UK) Consortium | The Lighthouse Lab in Milton Keynes and Alex Alderton, Roberto Amato, Jeffrey Barrett, Sonia Goncalves, Ewan Harrison, David K. Jackson, Ian Johnston, Dominic Kwiatkowski, Cordelia Langford, John Sillitoe on behalf of the Wellcome Sanger Institute COVID-19 Surveillance Team |
| EPI_ISL_3311628, EPI_ISL_3311629, EPI_ISL_3311630 | Lighthouse Laboratory Plymouth | Wellcome Sanger Institute for the COVID-19 Genomics UK (COG-UK) Consortium | Lighthouse Laboratory Plymouth and Alex Alderton, Roberto Amato, Jeffrey Barrett, Sonia Goncalves, Ewan Harrison, David K. Jackson, Ian Johnston, Dominic Kwiatkowski, Cordelia Langford, John Sillitoe on behalf of the Wellcome Sanger Institute COVID-19 Surveillance Team |
| EPI_ISL_3311631 | Lighthouse Lab in Glasgow | Wellcome Sanger Institute for the COVID-19 Genomics UK (COG-UK) Consortium | Harper VanSteenhouse, Yumi Kasai, David Gray, Carol Clugston, Anna Dominiczak and Alex Alderton, Roberto Amato, Jeffrey Barrett, Sonia Goncalves, Ewan Harrison, David K. Jackson, Ian Johnston, Dominic Kwiatkowski, Cordelia Langford, John Sillitoe on behalf of the Wellcome Sanger Institute COVID-19 Surveillance Team |
| EPI_ISL_3311632, EPI_ISL_3311633, EPI_ISL_3311634 | Lighthouse Laboratory Plymouth | Wellcome Sanger Institute for the COVID-19 Genomics UK (COG-UK) Consortium | Lighthouse Laboratory Plymouth and Alex Alderton, Roberto Amato, Jeffrey Barrett, Sonia Goncalves, Ewan Harrison, David K. Jackson, Ian Johnston, Dominic Kwiatkowski, Cordelia Langford, John Sillitoe on behalf of the Wellcome Sanger Institute COVID-19 Surveillance Team |
| EPI_ISL_3311635 | Lighthouse Lab in Milton Keynes | Wellcome Sanger Institute for the COVID-19 Genomics UK (COG-UK) Consortium | The Lighthouse Lab in Milton Keynes and Alex Alderton, Roberto Amato, Jeffrey Barrett, Sonia Goncalves, Ewan Harrison, David K. Jackson, Ian Johnston, Dominic Kwiatkowski, Cordelia Langford, John Sillitoe on behalf of the Wellcome Sanger Institute COVID-19 Surveillance Team |
| EPI_ISL_3311636 | Lighthouse Laboratory Plymouth | Wellcome Sanger Institute for the COVID-19 Genomics UK (COG-UK) Consortium | Lighthouse Laboratory Plymouth and Alex Alderton, Roberto Amato, Jeffrey Barrett, Sonia Goncalves, Ewan Harrison, David K. Jackson, Ian Johnston, Dominic Kwiatkowski, Cordelia Langford, John Sillitoe on behalf of the Wellcome Sanger Institute COVID-19 Surveillance Team |
| EPI_ISL_3311637, EPI_ISL_3311638 | Lighthouse Lab in Milton Keynes | Wellcome Sanger Institute for the COVID-19 Genomics UK (COG-UK) Consortium | The Lighthouse Lab in Milton Keynes and Alex Alderton, Roberto Amato, Jeffrey Barrett, Sonia Goncalves, Ewan Harrison, David K. Jackson, Ian Johnston, Dominic Kwiatkowski, Cordelia Langford, John Sillitoe on behalf of the Wellcome Sanger Institute COVID-19 Surveillance Team |
| EPI_ISL_3311639, EPI_ISL_3311640, EPI_ISL_3311641 | Lighthouse Lab in Glasgow | Wellcome Sanger Institute for the COVID-19 Genomics UK (COG-UK) Consortium | Harper VanSteenhouse, Yumi Kasai, David Gray, Carol Clugston, Anna Dominiczak and Alex Alderton, Roberto Amato, Jeffrey Barrett, Sonia Goncalves, Ewan Harrison, David K. Jackson, Ian Johnston, Dominic Kwiatkowski, Cordelia Langford, John Sillitoe on behalf of the Wellcome Sanger Institute COVID-19 Surveillance Team |
| EPI_ISL_3311642 | Lighthouse Laboratory Plymouth | Wellcome Sanger Institute for the COVID-19 Genomics UK (COG-UK) Consortium | Lighthouse Laboratory Plymouth and Alex Alderton, Roberto Amato, Jeffrey Barrett, Sonia Goncalves, Ewan Harrison, David K. Jackson, Ian Johnston, Dominic Kwiatkowski, Cordelia Langford, John Sillitoe on behalf of the Wellcome Sanger Institute COVID-19 Surveillance Team |
| EPI_ISL_3311643 | Lighthouse Lab in Milton Keynes | Wellcome Sanger Institute for the COVID-19 Genomics UK (COG-UK) Consortium | The Lighthouse Lab in Milton Keynes and Alex Alderton, Roberto Amato, Jeffrey Barrett, Sonia Goncalves, Ewan Harrison, David K. Jackson, Ian Johnston, Dominic Kwiatkowski, Cordelia Langford, John Sillitoe on behalf of the Wellcome Sanger Institute COVID-19 Surveillance Team |
| EPI_ISL_3311644 | Lighthouse Lab in Glasgow | Wellcome Sanger Institute for the COVID-19 Genomics UK (COG-UK) Consortium | Harper VanSteenhouse, Yumi Kasai, David Gray, Carol Clugston, Anna Dominiczak and Alex Alderton, Roberto Amato, Jeffrey Barrett, Sonia Goncalves, Ewan Harrison, David K. Jackson, Ian Johnston, Dominic Kwiatkowski, Cordelia Langford, John Sillitoe on behalf of the Wellcome Sanger Institute COVID-19 Surveillance Team |
| EPI_ISL_3311645 | Lighthouse Laboratory Plymouth | Wellcome Sanger Institute for the COVID-19 Genomics UK (COG-UK) Consortium | Lighthouse Laboratory Plymouth and Alex Alderton, Roberto Amato, Jeffrey Barrett, Sonia Goncalves, Ewan Harrison, David K. Jackson, Ian Johnston, Dominic Kwiatkowski, Cordelia Langford, John Sillitoe on behalf of the Wellcome Sanger Institute COVID-19 Surveillance Team |
| EPI_ISL_3311646, EPI_ISL_3311647 | Lighthouse Lab in Glasgow | Wellcome Sanger Institute for the COVID-19 Genomics UK (COG-UK) Consortium | Harper VanSteenhouse, Yumi Kasai, David Gray, Carol Clugston, Anna Dominiczak and Alex Alderton, Roberto Amato, Jeffrey Barrett, Sonia Goncalves, Ewan Harrison, David K. Jackson, Ian Johnston, Dominic Kwiatkowski, Cordelia Langford, John Sillitoe on behalf of the Wellcome Sanger Institute COVID-19 Surveillance Team |
| EPI_ISL_3311648 | Lighthouse Lab in Milton Keynes | Wellcome Sanger Institute for the COVID-19 Genomics UK (COG-UK) Consortium | The Lighthouse Lab in Milton Keynes and Alex Alderton, Roberto Amato, Jeffrey Barrett, Sonia Goncalves, Ewan Harrison, David K. Jackson, Ian Johnston, Dominic Kwiatkowski, Cordelia Langford, John Sillitoe on behalf of the Wellcome Sanger Institute COVID-19 Surveillance Team |
| EPI_ISL_3311649, EPI_ISL_3311650 | Lighthouse Lab in Glasgow | Wellcome Sanger Institute for the COVID-19 Genomics UK (COG-UK) Consortium | Harper VanSteenhouse, Yumi Kasai, David Gray, Carol Clugston, Anna Dominiczak and Alex Alderton, Roberto Amato, Jeffrey Barrett, Sonia Goncalves, Ewan Harrison, David K. Jackson, Ian Johnston, Dominic Kwiatkowski, Cordelia Langford, John Sillitoe on behalf of the Wellcome Sanger Institute COVID-19 Surveillance Team |
| EPI_ISL_3311651 | Lighthouse Laboratory Plymouth | Wellcome Sanger Institute for the COVID-19 Genomics UK (COG-UK) Consortium | Lighthouse Laboratory Plymouth and Alex Alderton, Roberto Amato, Jeffrey Barrett, Sonia Goncalves, Ewan Harrison, David K. Jackson, Ian Johnston, Dominic Kwiatkowski, Cordelia Langford, John Sillitoe on behalf of the Wellcome Sanger Institute COVID-19 Surveillance Team |
| EPI_ISL_3311652 | Lighthouse Lab in Glasgow | Wellcome Sanger Institute for the COVID-19 Genomics UK (COG-UK) Consortium | Harper VanSteenhouse, Yumi Kasai, David Gray, Carol Clugston, Anna Dominiczak and Alex Alderton, Roberto Amato, Jeffrey Barrett, Sonia Goncalves, Ewan Harrison, David K. Jackson, Ian Johnston, Dominic Kwiatkowski, Cordelia Langford, John Sillitoe on behalf of the Wellcome Sanger Institute COVID-19 Surveillance Team |
| EPI_ISL_3311653 | Lighthouse Lab in Milton Keynes | Wellcome Sanger Institute for the COVID-19 Genomics UK (COG-UK) Consortium | The Lighthouse Lab in Milton Keynes and Alex Alderton, Roberto Amato, Jeffrey Barrett, Sonia Goncalves, Ewan Harrison, David K. Jackson, Ian Johnston, Dominic Kwiatkowski, Cordelia Langford, John Sillitoe on behalf of the Wellcome Sanger Institute COVID-19 Surveillance Team |
| EPI_ISL_3311654, EPI_ISL_3311655 | Lighthouse Lab in Glasgow | Wellcome Sanger Institute for the COVID-19 Genomics UK (COG-UK) Consortium | Harper VanSteenhouse, Yumi Kasai, David Gray, Carol Clugston, Anna Dominiczak and Alex Alderton, Roberto Amato, Jeffrey Barrett, Sonia Goncalves, Ewan Harrison, David K. Jackson, Ian Johnston, Dominic Kwiatkowski, Cordelia Langford, John Sillitoe on behalf of the Wellcome Sanger Institute COVID-19 Surveillance Team |
| EPI_ISL_3311656 | Lighthouse Laboratory Plymouth | Wellcome Sanger Institute for the COVID-19 Genomics UK (COG-UK) Consortium | Lighthouse Laboratory Plymouth and Alex Alderton, Roberto Amato, Jeffrey Barrett, Sonia Goncalves, Ewan Harrison, David K. Jackson, Ian Johnston, Dominic Kwiatkowski, Cordelia Langford, John Sillitoe on behalf of the Wellcome Sanger Institute COVID-19 Surveillance Team |
| EPI_ISL_3311657, EPI_ISL_3311658 | Lighthouse Lab in Milton Keynes | Wellcome Sanger Institute for the COVID-19 Genomics UK (COG-UK) Consortium | The Lighthouse Lab in Milton Keynes and Alex Alderton, Roberto Amato, Jeffrey Barrett, Sonia Goncalves, Ewan Harrison, David K. Jackson, Ian Johnston, Dominic Kwiatkowski, Cordelia Langford, John Sillitoe on behalf of the Wellcome Sanger Institute COVID-19 Surveillance Team |
| EPI_ISL_3311659 | Lighthouse Lab in Glasgow | Wellcome Sanger Institute for the COVID-19 Genomics UK (COG-UK) Consortium | Harper VanSteenhouse, Yumi Kasai, David Gray, Carol Clugston, Anna Dominiczak and Alex Alderton, Roberto Amato, Jeffrey Barrett, Sonia Goncalves, Ewan Harrison, David K. Jackson, Ian Johnston, Dominic Kwiatkowski, Cordelia Langford, John Sillitoe on behalf of the Wellcome Sanger Institute COVID-19 Surveillance Team |
| EPI_ISL_3311660, EPI_ISL_3311661 | Lighthouse Lab in Milton Keynes | Wellcome Sanger Institute for the COVID-19 Genomics UK (COG-UK) Consortium | The Lighthouse Lab in Milton Keynes and Alex Alderton, Roberto Amato, Jeffrey Barrett, Sonia Goncalves, Ewan Harrison, David K. Jackson, Ian Johnston, Dominic Kwiatkowski, Cordelia Langford, John Sillitoe on behalf of the Wellcome Sanger Institute COVID-19 Surveillance Team |
| EPI_ISL_3311662 | Lighthouse Lab in Glasgow | Wellcome Sanger Institute for the COVID-19 Genomics UK (COG-UK) Consortium | Harper VanSteenhouse, Yumi Kasai, David Gray, Carol Clugston, Anna Dominiczak and Alex Alderton, Roberto Amato, Jeffrey Barrett, Sonia Goncalves, Ewan Harrison, David K. Jackson, Ian Johnston, Dominic Kwiatkowski, Cordelia Langford, John Sillitoe on behalf of the Wellcome Sanger Institute COVID-19 Surveillance Team |
| EPI_ISL_3311663, EPI_ISL_3311664 | Lighthouse Laboratory Plymouth | Wellcome Sanger Institute for the COVID-19 Genomics UK (COG-UK) Consortium | Lighthouse Laboratory Plymouth and Alex Alderton, Roberto Amato, Jeffrey Barrett, Sonia Goncalves, Ewan Harrison, David K. Jackson, Ian Johnston, Dominic Kwiatkowski, Cordelia Langford, John Sillitoe on behalf of the Wellcome Sanger Institute COVID-19 Surveillance Team |
| EPI_ISL_3311665 | Lighthouse Lab in Milton Keynes | Wellcome Sanger Institute for the COVID-19 Genomics UK (COG-UK) Consortium | The Lighthouse Lab in Milton Keynes and Alex Alderton, Roberto Amato, Jeffrey Barrett, Sonia Goncalves, Ewan Harrison, David K. Jackson, Ian Johnston, Dominic Kwiatkowski, Cordelia Langford, John Sillitoe on behalf of the Wellcome Sanger Institute COVID-19 Surveillance Team |
| EPI_ISL_3311666 | Lighthouse Laboratory Plymouth | Wellcome Sanger Institute for the COVID-19 Genomics UK (COG-UK) Consortium | Lighthouse Laboratory Plymouth and Alex Alderton, Roberto Amato, Jeffrey Barrett, Sonia Goncalves, Ewan Harrison, David K. Jackson, Ian Johnston, Dominic Kwiatkowski, Cordelia Langford, John Sillitoe on behalf of the Wellcome Sanger Institute COVID-19 Surveillance Team |
| EPI_ISL_3311667, EPI_ISL_3311668, EPI_ISL_3311669, EPI_ISL_3311670 | Lighthouse Lab in Milton Keynes | Wellcome Sanger Institute for the COVID-19 Genomics UK (COG-UK) Consortium | The Lighthouse Lab in Milton Keynes and Alex Alderton, Roberto Amato, Jeffrey Barrett, Sonia Goncalves, Ewan Harrison, David K. Jackson, Ian Johnston, Dominic Kwiatkowski, Cordelia Langford, John Sillitoe on behalf of the Wellcome Sanger Institute COVID-19 Surveillance Team |
| EPI_ISL_3311671, EPI_ISL_3311672 | Lighthouse Laboratory Plymouth | Wellcome Sanger Institute for the COVID-19 Genomics UK (COG-UK) Consortium | Lighthouse Laboratory Plymouth and Alex Alderton, Roberto Amato, Jeffrey Barrett, Sonia Goncalves, Ewan Harrison, David K. Jackson, Ian Johnston, Dominic Kwiatkowski, Cordelia Langford, John Sillitoe on behalf of the Wellcome Sanger Institute COVID-19 Surveillance Team |
| EPI_ISL_3311673 | Lighthouse Lab in Milton Keynes | Wellcome Sanger Institute for the COVID-19 Genomics UK (COG-UK) Consortium | The Lighthouse Lab in Milton Keynes and Alex Alderton, Roberto Amato, Jeffrey Barrett, Sonia Goncalves, Ewan Harrison, David K. Jackson, Ian Johnston, Dominic Kwiatkowski, Cordelia Langford, John Sillitoe on behalf of the Wellcome Sanger Institute COVID-19 Surveillance Team |
| EPI_ISL_3311674, EPI_ISL_3311675, EPI_ISL_3311676, EPI_ISL_3311677 | Lighthouse Laboratory Plymouth | Wellcome Sanger Institute for the COVID-19 Genomics UK (COG-UK) Consortium | Lighthouse Laboratory Plymouth and Alex Alderton, Roberto Amato, Jeffrey Barrett, Sonia Goncalves, Ewan Harrison, David K. Jackson, Ian Johnston, Dominic Kwiatkowski, Cordelia Langford, John Sillitoe on behalf of the Wellcome Sanger Institute COVID-19 Surveillance Team |
| EPI_ISL_3311678 | Lighthouse Lab in Milton Keynes | Wellcome Sanger Institute for the COVID-19 Genomics UK (COG-UK) Consortium | The Lighthouse Lab in Milton Keynes and Alex Alderton, Roberto Amato, Jeffrey Barrett, Sonia Goncalves, Ewan Harrison, David K. Jackson, Ian Johnston, Dominic Kwiatkowski, Cordelia Langford, John Sillitoe on behalf of the Wellcome Sanger Institute COVID-19 Surveillance Team |

| | | | |
|---|---|---|---|
| EPI_ISL_3311679 | Lighthouse Lab in Glasgow | Wellcome Sanger Institute for the COVID-19 Genomics UK (COG-UK) Consortium | Harper VanSteenhouse, Yumi Kasai, David Gray, Carol Clugston, Anna Dominiczak and Alex Alderton, Roberto Amato, Jeffrey Barrett, Sonia Goncalves, Ewan Harrison, David K. Jackson, Ian Johnston, Dominic Kwiatkowski, Cordelia Langford, John Sillitoe on behalf of the Wellcome Sanger Institute COVID-19 Surveillance Team |
| EPI_ISL_3311680 | Lighthouse Laboratory Plymouth | Wellcome Sanger Institute for the COVID-19 Genomics UK (COG-UK) Consortium | Lighthouse Laboratory Plymouth and Alex Alderton, Roberto Amato, Jeffrey Barrett, Sonia Goncalves, Ewan Harrison, David K. Jackson, Ian Johnston, Dominic Kwiatkowski, Cordelia Langford, John Sillitoe on behalf of the Wellcome Sanger Institute COVID-19 Surveillance Team |
| EPI_ISL_3311681, EPI_ISL_3311682 | Lighthouse Lab in Glasgow | Wellcome Sanger Institute for the COVID-19 Genomics UK (COG-UK) Consortium | Harper VanSteenhouse, Yumi Kasai, David Gray, Carol Clugston, Anna Dominiczak and Alex Alderton, Roberto Amato, Jeffrey Barrett, Sonia Goncalves, Ewan Harrison, David K. Jackson, Ian Johnston, Dominic Kwiatkowski, Cordelia Langford, John Sillitoe on behalf of the Wellcome Sanger Institute COVID-19 Surveillance Team |
| EPI_ISL_3311683 | Lighthouse Lab in Milton Keynes | Wellcome Sanger Institute for the COVID-19 Genomics UK (COG-UK) Consortium | The Lighthouse Lab in Milton Keynes and Alex Alderton, Roberto Amato, Jeffrey Barrett, Sonia Goncalves, Ewan Harrison, David K. Jackson, Ian Johnston, Dominic Kwiatkowski, Cordelia Langford, John Sillitoe on behalf of the Wellcome Sanger Institute COVID-19 Surveillance Team |
| EPI_ISL_3311684, EPI_ISL_3311685 | Lighthouse Laboratory Plymouth | Wellcome Sanger Institute for the COVID-19 Genomics UK (COG-UK) Consortium | Lighthouse Laboratory Plymouth and Alex Alderton, Roberto Amato, Jeffrey Barrett, Sonia Goncalves, Ewan Harrison, David K. Jackson, Ian Johnston, Dominic Kwiatkowski, Cordelia Langford, John Sillitoe on behalf of the Wellcome Sanger Institute COVID-19 Surveillance Team |
| EPI_ISL_3311686, EPI_ISL_3311687, EPI_ISL_3311688 | Lighthouse Lab in Milton Keynes | Wellcome Sanger Institute for the COVID-19 Genomics UK (COG-UK) Consortium | The Lighthouse Lab in Milton Keynes and Alex Alderton, Roberto Amato, Jeffrey Barrett, Sonia Goncalves, Ewan Harrison, David K. Jackson, Ian Johnston, Dominic Kwiatkowski, Cordelia Langford, John Sillitoe on behalf of the Wellcome Sanger Institute COVID-19 Surveillance Team |
| EPI_ISL_3311689, EPI_ISL_3311690 | Lighthouse Lab in Glasgow | Wellcome Sanger Institute for the COVID-19 Genomics UK (COG-UK) Consortium | Harper VanSteenhouse, Yumi Kasai, David Gray, Carol Clugston, Anna Dominiczak and Alex Alderton, Roberto Amato, Jeffrey Barrett, Sonia Goncalves, Ewan Harrison, David K. Jackson, Ian Johnston, Dominic Kwiatkowski, Cordelia Langford, John Sillitoe on behalf of the Wellcome Sanger Institute COVID-19 Surveillance Team |
| EPI_ISL_3311691, EPI_ISL_3311692 | Lighthouse Laboratory Plymouth | Wellcome Sanger Institute for the COVID-19 Genomics UK (COG-UK) Consortium | Lighthouse Laboratory Plymouth and Alex Alderton, Roberto Amato, Jeffrey Barrett, Sonia Goncalves, Ewan Harrison, David K. Jackson, Ian Johnston, Dominic Kwiatkowski, Cordelia Langford, John Sillitoe on behalf of the Wellcome Sanger Institute COVID-19 Surveillance Team |
| EPI_ISL_3311693 | Lighthouse Lab in Glasgow | Wellcome Sanger Institute for the COVID-19 Genomics UK (COG-UK) Consortium | Harper VanSteenhouse, Yumi Kasai, David Gray, Carol Clugston, Anna Dominiczak and Alex Alderton, Roberto Amato, Jeffrey Barrett, Sonia Goncalves, Ewan Harrison, David K. Jackson, Ian Johnston, Dominic Kwiatkowski, Cordelia Langford, John Sillitoe on behalf of the Wellcome Sanger Institute COVID-19 Surveillance Team |
| EPI_ISL_3311694 | Lighthouse Lab in Milton Keynes | Wellcome Sanger Institute for the COVID-19 Genomics UK (COG-UK) Consortium | The Lighthouse Lab in Milton Keynes and Alex Alderton, Roberto Amato, Jeffrey Barrett, Sonia Goncalves, Ewan Harrison, David K. Jackson, Ian Johnston, Dominic Kwiatkowski, Cordelia Langford, John Sillitoe on behalf of the Wellcome Sanger Institute COVID-19 Surveillance Team |
| EPI_ISL_3311695, EPI_ISL_3311696 | Lighthouse Laboratory Plymouth | Wellcome Sanger Institute for the COVID-19 Genomics UK (COG-UK) Consortium | Lighthouse Laboratory Plymouth and Alex Alderton, Roberto Amato, Jeffrey Barrett, Sonia Goncalves, Ewan Harrison, David K. Jackson, Ian Johnston, Dominic Kwiatkowski, Cordelia Langford, John Sillitoe on behalf of the Wellcome Sanger Institute COVID-19 Surveillance Team |
| EPI_ISL_3311697 | Lighthouse Lab in Glasgow | Wellcome Sanger Institute for the COVID-19 Genomics UK (COG-UK) Consortium | Harper VanSteenhouse, Yumi Kasai, David Gray, Carol Clugston, Anna Dominiczak and Alex Alderton, Roberto Amato, Jeffrey Barrett, Sonia Goncalves, Ewan Harrison, David K. Jackson, Ian Johnston, Dominic Kwiatkowski, Cordelia Langford, John Sillitoe on behalf of the Wellcome Sanger Institute COVID-19 Surveillance Team |
| EPI_ISL_3311698 | Lighthouse Lab in Milton Keynes | Wellcome Sanger Institute for the COVID-19 Genomics UK (COG-UK) Consortium | The Lighthouse Lab in Milton Keynes and Alex Alderton, Roberto Amato, Jeffrey Barrett, Sonia Goncalves, Ewan Harrison, David K. Jackson, Ian Johnston, Dominic Kwiatkowski, Cordelia Langford, John Sillitoe on behalf of the Wellcome Sanger Institute COVID-19 Surveillance Team |
| EPI_ISL_3311699 | Lighthouse Lab in Glasgow | Wellcome Sanger Institute for the COVID-19 Genomics UK (COG-UK) Consortium | Harper VanSteenhouse, Yumi Kasai, David Gray, Carol Clugston, Anna Dominiczak and Alex Alderton, Roberto Amato, Jeffrey Barrett, Sonia Goncalves, Ewan Harrison, David K. Jackson, Ian Johnston, Dominic Kwiatkowski, Cordelia Langford, John Sillitoe on behalf of the Wellcome Sanger Institute COVID-19 Surveillance Team |
| EPI_ISL_3311700 | Lighthouse Laboratory Plymouth | Wellcome Sanger Institute for the COVID-19 Genomics UK (COG-UK) Consortium | Lighthouse Laboratory Plymouth and Alex Alderton, Roberto Amato, Jeffrey Barrett, Sonia Goncalves, Ewan Harrison, David K. Jackson, Ian Johnston, Dominic Kwiatkowski, Cordelia Langford, John Sillitoe on behalf of the Wellcome Sanger Institute COVID-19 Surveillance Team |
| EPI_ISL_3311701 | Lighthouse Lab in Milton Keynes | Wellcome Sanger Institute for the COVID-19 Genomics UK (COG-UK) Consortium | The Lighthouse Lab in Milton Keynes and Alex Alderton, Roberto Amato, Jeffrey Barrett, Sonia Goncalves, Ewan Harrison, David K. Jackson, Ian Johnston, Dominic Kwiatkowski, Cordelia Langford, John Sillitoe on behalf of the Wellcome Sanger Institute COVID-19 Surveillance Team |
| EPI_ISL_3311702, EPI_ISL_3311703, EPI_ISL_3311704 | Lighthouse Lab in Glasgow | Wellcome Sanger Institute for the COVID-19 Genomics UK (COG-UK) Consortium | Harper VanSteenhouse, Yumi Kasai, David Gray, Carol Clugston, Anna Dominiczak and Alex Alderton, Roberto Amato, Jeffrey Barrett, Sonia Goncalves, Ewan Harrison, David K. Jackson, Ian Johnston, Dominic Kwiatkowski, Cordelia Langford, John Sillitoe on behalf of the Wellcome Sanger Institute COVID-19 Surveillance Team |
| EPI_ISL_3311705 | Lighthouse Laboratory Plymouth | Wellcome Sanger Institute for the COVID-19 Genomics UK (COG-UK) Consortium | Lighthouse Laboratory Plymouth and Alex Alderton, Roberto Amato, Jeffrey Barrett, Sonia Goncalves, Ewan Harrison, David K. Jackson, Ian Johnston, Dominic Kwiatkowski, Cordelia Langford, John Sillitoe on behalf of the Wellcome Sanger Institute COVID-19 Surveillance Team |
| EPI_ISL_3311706, EPI_ISL_3311707 | Lighthouse Lab in Milton Keynes | Wellcome Sanger Institute for the COVID-19 Genomics UK (COG-UK) Consortium | The Lighthouse Lab in Milton Keynes and Alex Alderton, Roberto Amato, Jeffrey Barrett, Sonia Goncalves, Ewan Harrison, David K. Jackson, Ian Johnston, Dominic Kwiatkowski, Cordelia Langford, John Sillitoe on behalf of the Wellcome Sanger Institute COVID-19 Surveillance Team |
| EPI_ISL_3311708, EPI_ISL_3311709 | Lighthouse Laboratory Plymouth | Wellcome Sanger Institute for the COVID-19 Genomics UK (COG-UK) Consortium | Lighthouse Laboratory Plymouth and Alex Alderton, Roberto Amato, Jeffrey Barrett, Sonia Goncalves, Ewan Harrison, David K. Jackson, Ian Johnston, Dominic Kwiatkowski, Cordelia Langford, John Sillitoe on behalf of the Wellcome Sanger Institute COVID-19 Surveillance Team |
| EPI_ISL_3311710 | Lighthouse Lab in Milton Keynes | Wellcome Sanger Institute for the COVID-19 Genomics UK (COG-UK) Consortium | The Lighthouse Lab in Milton Keynes and Alex Alderton, Roberto Amato, Jeffrey Barrett, Sonia Goncalves, Ewan Harrison, David K. Jackson, Ian Johnston, Dominic Kwiatkowski, Cordelia Langford, John Sillitoe on behalf of the Wellcome Sanger Institute COVID-19 Surveillance Team |
| EPI_ISL_3311711 | Lighthouse Laboratory Plymouth | Wellcome Sanger Institute for the COVID-19 Genomics UK (COG-UK) Consortium | Lighthouse Laboratory Plymouth and Alex Alderton, Roberto Amato, Jeffrey Barrett, Sonia Goncalves, Ewan Harrison, David K. Jackson, Ian Johnston, Dominic Kwiatkowski, Cordelia Langford, John Sillitoe on behalf of the Wellcome Sanger Institute COVID-19 Surveillance Team |
| EPI_ISL_3311712 | Lighthouse Lab in Milton Keynes | Wellcome Sanger Institute for the COVID-19 Genomics UK (COG-UK) Consortium | The Lighthouse Lab in Milton Keynes and Alex Alderton, Roberto Amato, Jeffrey Barrett, Sonia Goncalves, Ewan Harrison, David K. Jackson, Ian Johnston, Dominic Kwiatkowski, Cordelia Langford, John Sillitoe on behalf of the Wellcome Sanger Institute COVID-19 Surveillance Team |
| EPI_ISL_3311713 | Lighthouse Lab in Glasgow | Wellcome Sanger Institute for the COVID-19 Genomics UK (COG-UK) Consortium | Harper VanSteenhouse, Yumi Kasai, David Gray, Carol Clugston, Anna Dominiczak and Alex Alderton, Roberto Amato, Jeffrey Barrett, Sonia Goncalves, Ewan Harrison, David K. Jackson, Ian Johnston, Dominic Kwiatkowski, Cordelia Langford, John Sillitoe on behalf of the Wellcome Sanger Institute COVID-19 Surveillance Team |
| EPI_ISL_3311714 | Lighthouse Laboratory Plymouth | Wellcome Sanger Institute for the COVID-19 Genomics UK (COG-UK) Consortium | Lighthouse Laboratory Plymouth and Alex Alderton, Roberto Amato, Jeffrey Barrett, Sonia Goncalves, Ewan Harrison, David K. Jackson, Ian Johnston, Dominic Kwiatkowski, Cordelia Langford, John Sillitoe on behalf of the Wellcome Sanger Institute COVID-19 Surveillance Team |
| EPI_ISL_3311715, EPI_ISL_3311716, EPI_ISL_3311717 | Lighthouse Lab in Milton Keynes | Wellcome Sanger Institute for the COVID-19 Genomics UK (COG-UK) Consortium | The Lighthouse Lab in Milton Keynes and Alex Alderton, Roberto Amato, Jeffrey Barrett, Sonia Goncalves, Ewan Harrison, David K. Jackson, Ian Johnston, Dominic Kwiatkowski, Cordelia Langford, John Sillitoe on behalf of the Wellcome Sanger Institute COVID-19 Surveillance Team |
| EPI_ISL_3311718 | Lighthouse Laboratory Plymouth | Wellcome Sanger Institute for the COVID-19 Genomics UK (COG-UK) Consortium | Lighthouse Laboratory Plymouth and Alex Alderton, Roberto Amato, Jeffrey Barrett, Sonia Goncalves, Ewan Harrison, David K. Jackson, Ian Johnston, Dominic Kwiatkowski, Cordelia Langford, John Sillitoe on behalf of the Wellcome Sanger Institute COVID-19 Surveillance Team |
| EPI_ISL_3311719 | Lighthouse Lab in Milton Keynes | Wellcome Sanger Institute for the COVID-19 Genomics UK (COG-UK) Consortium | The Lighthouse Lab in Milton Keynes and Alex Alderton, Roberto Amato, Jeffrey Barrett, Sonia Goncalves, Ewan Harrison, David K. Jackson, Ian Johnston, Dominic Kwiatkowski, Cordelia Langford, John Sillitoe on behalf of the Wellcome Sanger Institute COVID-19 Surveillance Team |
| EPI_ISL_3311720, EPI_ISL_3311721 | Lighthouse Laboratory Plymouth | Wellcome Sanger Institute for the COVID-19 Genomics UK (COG-UK) Consortium | Lighthouse Laboratory Plymouth and Alex Alderton, Roberto Amato, Jeffrey Barrett, Sonia Goncalves, Ewan Harrison, David K. Jackson, Ian Johnston, Dominic Kwiatkowski, Cordelia Langford, John Sillitoe on behalf of the Wellcome Sanger Institute COVID-19 Surveillance Team |
| EPI_ISL_3311722 | Lighthouse Lab in Milton Keynes | Wellcome Sanger Institute for the COVID-19 Genomics UK (COG-UK) Consortium | The Lighthouse Lab in Milton Keynes and Alex Alderton, Roberto Amato, Jeffrey Barrett, Sonia Goncalves, Ewan Harrison, David K. Jackson, Ian Johnston, Dominic Kwiatkowski, Cordelia Langford, John Sillitoe on behalf of the Wellcome Sanger Institute COVID-19 Surveillance Team |
| EPI_ISL_3311723 | Lighthouse Lab in Glasgow | Wellcome Sanger Institute for the COVID-19 Genomics UK (COG-UK) Consortium | Harper VanSteenhouse, Yumi Kasai, David Gray, Carol Clugston, Anna Dominiczak and Alex Alderton, Roberto Amato, Jeffrey Barrett, Sonia Goncalves, Ewan Harrison, David K. Jackson, Ian Johnston, Dominic Kwiatkowski, Cordelia Langford, John Sillitoe on behalf of the Wellcome Sanger Institute COVID-19 Surveillance Team |
| EPI_ISL_3311724, EPI_ISL_3311725, EPI_ISL_3311726, EPI_ISL_3311727 | Lighthouse Laboratory Plymouth | Wellcome Sanger Institute for the COVID-19 Genomics UK (COG-UK) Consortium | Lighthouse Laboratory Plymouth and Alex Alderton, Roberto Amato, Jeffrey Barrett, Sonia Goncalves, Ewan Harrison, David K. Jackson, Ian Johnston, Dominic Kwiatkowski, Cordelia Langford, John Sillitoe on behalf of the Wellcome Sanger Institute COVID-19 Surveillance Team |
| EPI_ISL_3311728, EPI_ISL_3311729, EPI_ISL_3311730, EPI_ISL_3311731 | Lighthouse Lab in Milton Keynes | Wellcome Sanger Institute for the COVID-19 Genomics UK (COG-UK) Consortium | The Lighthouse Lab in Milton Keynes and Alex Alderton, Roberto Amato, Jeffrey Barrett, Sonia Goncalves, Ewan Harrison, David K. Jackson, Ian Johnston, Dominic Kwiatkowski, Cordelia Langford, John Sillitoe on behalf of the Wellcome Sanger Institute COVID-19 Surveillance Team |
| EPI_ISL_3311732 | Lighthouse Laboratory Plymouth | Wellcome Sanger Institute for the COVID-19 Genomics UK (COG-UK) Consortium | Lighthouse Laboratory Plymouth and Alex Alderton, Roberto Amato, Jeffrey Barrett, Sonia Goncalves, Ewan Harrison, David K. Jackson, Ian Johnston, Dominic Kwiatkowski, Cordelia Langford, John Sillitoe on behalf of the Wellcome Sanger Institute COVID-19 Surveillance Team |
| EPI_ISL_3311733 | Lighthouse Lab in Glasgow | Wellcome Sanger Institute for the COVID-19 Genomics UK (COG-UK) Consortium | Harper VanSteenhouse, Yumi Kasai, David Gray, Carol Clugston, Anna Dominiczak and Alex Alderton, Roberto Amato, Jeffrey Barrett, Sonia Goncalves, Ewan Harrison, David K. Jackson, Ian Johnston, Dominic Kwiatkowski, Cordelia Langford, John Sillitoe on behalf of the Wellcome Sanger Institute COVID-19 Surveillance Team |
| EPI_ISL_3311734, EPI_ISL_3311735 | Lighthouse Lab in Milton Keynes | Wellcome Sanger Institute for the COVID-19 Genomics UK (COG-UK) Consortium | The Lighthouse Lab in Milton Keynes and Alex Alderton, Roberto Amato, Jeffrey Barrett, Sonia Goncalves, Ewan Harrison, David K. Jackson, Ian Johnston, Dominic Kwiatkowski, Cordelia Langford, John Sillitoe on behalf of the Wellcome Sanger Institute COVID-19 Surveillance Team |
| EPI_ISL_3311736 | Lighthouse Laboratory Plymouth | Wellcome Sanger Institute for the COVID-19 Genomics UK (COG-UK) Consortium | Lighthouse Laboratory Plymouth and Alex Alderton, Roberto Amato, Jeffrey Barrett, Sonia Goncalves, Ewan Harrison, David K. Jackson, Ian Johnston, Dominic Kwiatkowski, Cordelia Langford, John Sillitoe on behalf of the Wellcome Sanger Institute COVID-19 Surveillance Team |
| EPI_ISL_3311737 | Lighthouse Lab in Milton Keynes | Wellcome Sanger Institute for the COVID-19 Genomics UK (COG-UK) Consortium | The Lighthouse Lab in Milton Keynes and Alex Alderton, Roberto Amato, Jeffrey Barrett, Sonia Goncalves, Ewan Harrison, David K. Jackson, Ian Johnston, Dominic Kwiatkowski, Cordelia Langford, John Sillitoe on behalf of the Wellcome Sanger Institute COVID-19 Surveillance Team |
| EPI_ISL_3311738 | Lighthouse Laboratory Plymouth | Wellcome Sanger Institute for the COVID-19 Genomics UK (COG-UK) Consortium | Lighthouse Laboratory Plymouth and Alex Alderton, Roberto Amato, Jeffrey Barrett, Sonia Goncalves, Ewan Harrison, David K. Jackson, Ian Johnston, Dominic Kwiatkowski, Cordelia Langford, John Sillitoe on behalf of the Wellcome Sanger Institute COVID-19 Surveillance Team |
| EPI_ISL_3311739, EPI_ISL_3311740 | Lighthouse Lab in Milton Keynes | Wellcome Sanger Institute for the COVID-19 Genomics UK (COG-UK) Consortium | The Lighthouse Lab in Milton Keynes and Alex Alderton, Roberto Amato, Jeffrey Barrett, Sonia Goncalves, Ewan Harrison, David K. Jackson, Ian Johnston, Dominic Kwiatkowski, Cordelia Langford, John Sillitoe on behalf of the Wellcome Sanger Institute COVID-19 Surveillance Team |
| EPI_ISL_3311741, EPI_ISL_3311742, EPI_ISL_3311743 | Lighthouse Laboratory Plymouth | Wellcome Sanger Institute for the COVID-19 Genomics UK (COG-UK) Consortium | Lighthouse Laboratory Plymouth and Alex Alderton, Roberto Amato, Jeffrey Barrett, Sonia Goncalves, Ewan Harrison, David K. Jackson, Ian Johnston, Dominic Kwiatkowski, Cordelia Langford, John Sillitoe on behalf of the Wellcome Sanger Institute COVID-19 Surveillance Team |
| EPI_ISL_3311744 | Lighthouse Lab in Glasgow | Wellcome Sanger Institute for the COVID-19 Genomics UK (COG-UK) Consortium | Harper VanSteenhouse, Yumi Kasai, David Gray, Carol Clugston, Anna Dominiczak and Alex Alderton, Roberto Amato, Jeffrey Barrett, Sonia Goncalves, Ewan Harrison, David K. Jackson, Ian Johnston, Dominic Kwiatkowski, Cordelia Langford, John Sillitoe on behalf of the Wellcome Sanger Institute COVID-19 Surveillance Team |

| | | | |
|---|---|---|---|
| EPI_ISL_3311745 | Lighthouse Lab in Milton Keynes | Wellcome Sanger Institute for the COVID-19 Genomics UK (COG-UK) Consortium | The Lighthouse Lab in Milton Keynes and Alex Alderton, Roberto Amato, Jeffrey Barrett, Sonia Goncalves, Ewan Harrison, David K. Jackson, Ian Johnston, Dominic Kwiatkowski, Cordelia Langford, John Sillitoe on behalf of the Wellcome Sanger Institute COVID-19 Surveillance Team |
| EPI_ISL_3311746 | Lighthouse Laboratory Plymouth | Wellcome Sanger Institute for the COVID-19 Genomics UK (COG-UK) Consortium | Lighthouse Laboratory Plymouth and Alex Alderton, Roberto Amato, Jeffrey Barrett, Sonia Goncalves, Ewan Harrison, David K. Jackson, Ian Johnston, Dominic Kwiatkowski, Cordelia Langford, John Sillitoe on behalf of the Wellcome Sanger Institute COVID-19 Surveillance Team |
| EPI_ISL_3311747 | Lighthouse Lab in Milton Keynes | Wellcome Sanger Institute for the COVID-19 Genomics UK (COG-UK) Consortium | The Lighthouse Lab in Milton Keynes and Alex Alderton, Roberto Amato, Jeffrey Barrett, Sonia Goncalves, Ewan Harrison, David K. Jackson, Ian Johnston, Dominic Kwiatkowski, Cordelia Langford, John Sillitoe on behalf of the Wellcome Sanger Institute COVID-19 Surveillance Team |
| EPI_ISL_3311748, EPI_ISL_3311749, EPI_ISL_3311750, EPI_ISL_3311751, EPI_ISL_3311752, EPI_ISL_3311753, EPI_ISL_3311754 | Lighthouse Laboratory Plymouth | Wellcome Sanger Institute for the COVID-19 Genomics UK (COG-UK) Consortium | Lighthouse Laboratory Plymouth and Alex Alderton, Roberto Amato, Jeffrey Barrett, Sonia Goncalves, Ewan Harrison, David K. Jackson, Ian Johnston, Dominic Kwiatkowski, Cordelia Langford, John Sillitoe on behalf of the Wellcome Sanger Institute COVID-19 Surveillance Team |
| EPI_ISL_3311755, EPI_ISL_3311756, EPI_ISL_3311757, EPI_ISL_3311758 | Lighthouse Lab in Milton Keynes | Wellcome Sanger Institute for the COVID-19 Genomics UK (COG-UK) Consortium | The Lighthouse Lab in Milton Keynes and Alex Alderton, Roberto Amato, Jeffrey Barrett, Sonia Goncalves, Ewan Harrison, David K. Jackson, Ian Johnston, Dominic Kwiatkowski, Cordelia Langford, John Sillitoe on behalf of the Wellcome Sanger Institute COVID-19 Surveillance Team |
| EPI_ISL_3311759 | Lighthouse Laboratory Plymouth | Wellcome Sanger Institute for the COVID-19 Genomics UK (COG-UK) Consortium | Lighthouse Laboratory Plymouth and Alex Alderton, Roberto Amato, Jeffrey Barrett, Sonia Goncalves, Ewan Harrison, David K. Jackson, Ian Johnston, Dominic Kwiatkowski, Cordelia Langford, John Sillitoe on behalf of the Wellcome Sanger Institute COVID-19 Surveillance Team |
| EPI_ISL_3311760, EPI_ISL_3311761 | Lighthouse Lab in Milton Keynes | Wellcome Sanger Institute for the COVID-19 Genomics UK (COG-UK) Consortium | The Lighthouse Lab in Milton Keynes and Alex Alderton, Roberto Amato, Jeffrey Barrett, Sonia Goncalves, Ewan Harrison, David K. Jackson, Ian Johnston, Dominic Kwiatkowski, Cordelia Langford, John Sillitoe on behalf of the Wellcome Sanger Institute COVID-19 Surveillance Team |
| EPI_ISL_3311762 | Lighthouse Laboratory Plymouth | Wellcome Sanger Institute for the COVID-19 Genomics UK (COG-UK) Consortium | Lighthouse Laboratory Plymouth and Alex Alderton, Roberto Amato, Jeffrey Barrett, Sonia Goncalves, Ewan Harrison, David K. Jackson, Ian Johnston, Dominic Kwiatkowski, Cordelia Langford, John Sillitoe on behalf of the Wellcome Sanger Institute COVID-19 Surveillance Team |
| EPI_ISL_3311763, EPI_ISL_3311764 | Lighthouse Lab in Milton Keynes | Wellcome Sanger Institute for the COVID-19 Genomics UK (COG-UK) Consortium | The Lighthouse Lab in Milton Keynes and Alex Alderton, Roberto Amato, Jeffrey Barrett, Sonia Goncalves, Ewan Harrison, David K. Jackson, Ian Johnston, Dominic Kwiatkowski, Cordelia Langford, John Sillitoe on behalf of the Wellcome Sanger Institute COVID-19 Surveillance Team |
| EPI_ISL_3311765 | Lighthouse Laboratory Plymouth | Wellcome Sanger Institute for the COVID-19 Genomics UK (COG-UK) Consortium | Lighthouse Laboratory Plymouth and Alex Alderton, Roberto Amato, Jeffrey Barrett, Sonia Goncalves, Ewan Harrison, David K. Jackson, Ian Johnston, Dominic Kwiatkowski, Cordelia Langford, John Sillitoe on behalf of the Wellcome Sanger Institute COVID-19 Surveillance Team |
| EPI_ISL_3311766, EPI_ISL_3311767 | Lighthouse Lab in Milton Keynes | Wellcome Sanger Institute for the COVID-19 Genomics UK (COG-UK) Consortium | The Lighthouse Lab in Milton Keynes and Alex Alderton, Roberto Amato, Jeffrey Barrett, Sonia Goncalves, Ewan Harrison, David K. Jackson, Ian Johnston, Dominic Kwiatkowski, Cordelia Langford, John Sillitoe on behalf of the Wellcome Sanger Institute COVID-19 Surveillance Team |
| EPI_ISL_3311768 | Lighthouse Lab in Glasgow | Wellcome Sanger Institute for the COVID-19 Genomics UK (COG-UK) Consortium | Harper VanSteenhouse, Yumi Kasai, David Gray, Carol Clugston, Anna Dominiczak and Alex Alderton, Roberto Amato, Jeffrey Barrett, Sonia Goncalves, Ewan Harrison, David K. Jackson, Ian Johnston, Dominic Kwiatkowski, Cordelia Langford, John Sillitoe on behalf of the Wellcome Sanger Institute COVID-19 Surveillance Team |
| EPI_ISL_3311769 | Lighthouse Lab in Milton Keynes | Wellcome Sanger Institute for the COVID-19 Genomics UK (COG-UK) Consortium | The Lighthouse Lab in Milton Keynes and Alex Alderton, Roberto Amato, Jeffrey Barrett, Sonia Goncalves, Ewan Harrison, David K. Jackson, Ian Johnston, Dominic Kwiatkowski, Cordelia Langford, John Sillitoe on behalf of the Wellcome Sanger Institute COVID-19 Surveillance Team |
| EPI_ISL_3311770 | Lighthouse Laboratory Plymouth | Wellcome Sanger Institute for the COVID-19 Genomics UK (COG-UK) Consortium | Lighthouse Laboratory Plymouth and Alex Alderton, Roberto Amato, Jeffrey Barrett, Sonia Goncalves, Ewan Harrison, David K. Jackson, Ian Johnston, Dominic Kwiatkowski, Cordelia Langford, John Sillitoe on behalf of the Wellcome Sanger Institute COVID-19 Surveillance Team |
| EPI_ISL_3311771 | Lighthouse Lab in Milton Keynes | Wellcome Sanger Institute for the COVID-19 Genomics UK (COG-UK) Consortium | The Lighthouse Lab in Milton Keynes and Alex Alderton, Roberto Amato, Jeffrey Barrett, Sonia Goncalves, Ewan Harrison, David K. Jackson, Ian Johnston, Dominic Kwiatkowski, Cordelia Langford, John Sillitoe on behalf of the Wellcome Sanger Institute COVID-19 Surveillance Team |
| EPI_ISL_3311772 | Lighthouse Laboratory Plymouth | Wellcome Sanger Institute for the COVID-19 Genomics UK (COG-UK) Consortium | Lighthouse Laboratory Plymouth and Alex Alderton, Roberto Amato, Jeffrey Barrett, Sonia Goncalves, Ewan Harrison, David K. Jackson, Ian Johnston, Dominic Kwiatkowski, Cordelia Langford, John Sillitoe on behalf of the Wellcome Sanger Institute COVID-19 Surveillance Team |
| EPI_ISL_3311773 | Lighthouse Lab in Glasgow | Wellcome Sanger Institute for the COVID-19 Genomics UK (COG-UK) Consortium | Harper VanSteenhouse, Yumi Kasai, David Gray, Carol Clugston, Anna Dominiczak and Alex Alderton, Roberto Amato, Jeffrey Barrett, Sonia Goncalves, Ewan Harrison, David K. Jackson, Ian Johnston, Dominic Kwiatkowski, Cordelia Langford, John Sillitoe on behalf of the Wellcome Sanger Institute COVID-19 Surveillance Team |
| EPI_ISL_3311774, EPI_ISL_3311775, EPI_ISL_3311776 | Lighthouse Lab in Milton Keynes | Wellcome Sanger Institute for the COVID-19 Genomics UK (COG-UK) Consortium | The Lighthouse Lab in Milton Keynes and Alex Alderton, Roberto Amato, Jeffrey Barrett, Sonia Goncalves, Ewan Harrison, David K. Jackson, Ian Johnston, Dominic Kwiatkowski, Cordelia Langford, John Sillitoe on behalf of the Wellcome Sanger Institute COVID-19 Surveillance Team |
| EPI_ISL_3311777 | Lighthouse Laboratory Plymouth | Wellcome Sanger Institute for the COVID-19 Genomics UK (COG-UK) Consortium | Lighthouse Laboratory Plymouth and Alex Alderton, Roberto Amato, Jeffrey Barrett, Sonia Goncalves, Ewan Harrison, David K. Jackson, Ian Johnston, Dominic Kwiatkowski, Cordelia Langford, John Sillitoe on behalf of the Wellcome Sanger Institute COVID-19 Surveillance Team |
| EPI_ISL_3311778, EPI_ISL_3311779 | Lighthouse Lab in Glasgow | Wellcome Sanger Institute for the COVID-19 Genomics UK (COG-UK) Consortium | Harper VanSteenhouse, Yumi Kasai, David Gray, Carol Clugston, Anna Dominiczak and Alex Alderton, Roberto Amato, Jeffrey Barrett, Sonia Goncalves, Ewan Harrison, David K. Jackson, Ian Johnston, Dominic Kwiatkowski, Cordelia Langford, John Sillitoe on behalf of the Wellcome Sanger Institute COVID-19 Surveillance Team |
| EPI_ISL_3311780 | Lighthouse Laboratory Plymouth | Wellcome Sanger Institute for the COVID-19 Genomics UK (COG-UK) Consortium | Lighthouse Laboratory Plymouth and Alex Alderton, Roberto Amato, Jeffrey Barrett, Sonia Goncalves, Ewan Harrison, David K. Jackson, Ian Johnston, Dominic Kwiatkowski, Cordelia Langford, John Sillitoe on behalf of the Wellcome Sanger Institute COVID-19 Surveillance Team |
| EPI_ISL_3311781 | Lighthouse Lab in Milton Keynes | Wellcome Sanger Institute for the COVID-19 Genomics UK (COG-UK) Consortium | The Lighthouse Lab in Milton Keynes and Alex Alderton, Roberto Amato, Jeffrey Barrett, Sonia Goncalves, Ewan Harrison, David K. Jackson, Ian Johnston, Dominic Kwiatkowski, Cordelia Langford, John Sillitoe on behalf of the Wellcome Sanger Institute COVID-19 Surveillance Team |
| EPI_ISL_3311782 | Lighthouse Lab in Glasgow | Wellcome Sanger Institute for the COVID-19 Genomics UK (COG-UK) Consortium | Harper VanSteenhouse, Yumi Kasai, David Gray, Carol Clugston, Anna Dominiczak and Alex Alderton, Roberto Amato, Jeffrey Barrett, Sonia Goncalves, Ewan Harrison, David K. Jackson, Ian Johnston, Dominic Kwiatkowski, Cordelia Langford, John Sillitoe on behalf of the Wellcome Sanger Institute COVID-19 Surveillance Team |
| EPI_ISL_3311783, EPI_ISL_3311784, EPI_ISL_3311785 | Lighthouse Lab in Milton Keynes | Wellcome Sanger Institute for the COVID-19 Genomics UK (COG-UK) Consortium | The Lighthouse Lab in Milton Keynes and Alex Alderton, Roberto Amato, Jeffrey Barrett, Sonia Goncalves, Ewan Harrison, David K. Jackson, Ian Johnston, Dominic Kwiatkowski, Cordelia Langford, John Sillitoe on behalf of the Wellcome Sanger Institute COVID-19 Surveillance Team |
| EPI_ISL_3311786 | Lighthouse Lab in Glasgow | Wellcome Sanger Institute for the COVID-19 Genomics UK (COG-UK) Consortium | Harper VanSteenhouse, Yumi Kasai, David Gray, Carol Clugston, Anna Dominiczak and Alex Alderton, Roberto Amato, Jeffrey Barrett, Sonia Goncalves, Ewan Harrison, David K. Jackson, Ian Johnston, Dominic Kwiatkowski, Cordelia Langford, John Sillitoe on behalf of the Wellcome Sanger Institute COVID-19 Surveillance Team |
| EPI_ISL_3311787, EPI_ISL_3311788, EPI_ISL_3311789 | Lighthouse Laboratory Plymouth | Wellcome Sanger Institute for the COVID-19 Genomics UK (COG-UK) Consortium | Lighthouse Laboratory Plymouth and Alex Alderton, Roberto Amato, Jeffrey Barrett, Sonia Goncalves, Ewan Harrison, David K. Jackson, Ian Johnston, Dominic Kwiatkowski, Cordelia Langford, John Sillitoe on behalf of the Wellcome Sanger Institute COVID-19 Surveillance Team |
| EPI_ISL_3311790 | Lighthouse Lab in Milton Keynes | Wellcome Sanger Institute for the COVID-19 Genomics UK (COG-UK) Consortium | The Lighthouse Lab in Milton Keynes and Alex Alderton, Roberto Amato, Jeffrey Barrett, Sonia Goncalves, Ewan Harrison, David K. Jackson, Ian Johnston, Dominic Kwiatkowski, Cordelia Langford, John Sillitoe on behalf of the Wellcome Sanger Institute COVID-19 Surveillance Team |
| EPI_ISL_3311791, EPI_ISL_3311792 | Lighthouse Laboratory Plymouth | Wellcome Sanger Institute for the COVID-19 Genomics UK (COG-UK) Consortium | Lighthouse Laboratory Plymouth and Alex Alderton, Roberto Amato, Jeffrey Barrett, Sonia Goncalves, Ewan Harrison, David K. Jackson, Ian Johnston, Dominic Kwiatkowski, Cordelia Langford, John Sillitoe on behalf of the Wellcome Sanger Institute COVID-19 Surveillance Team |
| EPI_ISL_3311793 | Lighthouse Lab in Milton Keynes | Wellcome Sanger Institute for the COVID-19 Genomics UK (COG-UK) Consortium | The Lighthouse Lab in Milton Keynes and Alex Alderton, Roberto Amato, Jeffrey Barrett, Sonia Goncalves, Ewan Harrison, David K. Jackson, Ian Johnston, Dominic Kwiatkowski, Cordelia Langford, John Sillitoe on behalf of the Wellcome Sanger Institute COVID-19 Surveillance Team |
| EPI_ISL_3311794 | Lighthouse Lab in Glasgow | Wellcome Sanger Institute for the COVID-19 Genomics UK (COG-UK) Consortium | Harper VanSteenhouse, Yumi Kasai, David Gray, Carol Clugston, Anna Dominiczak and Alex Alderton, Roberto Amato, Jeffrey Barrett, Sonia Goncalves, Ewan Harrison, David K. Jackson, Ian Johnston, Dominic Kwiatkowski, Cordelia Langford, John Sillitoe on behalf of the Wellcome Sanger Institute COVID-19 Surveillance Team |
| EPI_ISL_3311795 | Lighthouse Laboratory Plymouth | Wellcome Sanger Institute for the COVID-19 Genomics UK (COG-UK) Consortium | Lighthouse Laboratory Plymouth and Alex Alderton, Roberto Amato, Jeffrey Barrett, Sonia Goncalves, Ewan Harrison, David K. Jackson, Ian Johnston, Dominic Kwiatkowski, Cordelia Langford, John Sillitoe on behalf of the Wellcome Sanger Institute COVID-19 Surveillance Team |
| EPI_ISL_3311796, EPI_ISL_3311797 | Lighthouse Lab in Milton Keynes | Wellcome Sanger Institute for the COVID-19 Genomics UK (COG-UK) Consortium | The Lighthouse Lab in Milton Keynes and Alex Alderton, Roberto Amato, Jeffrey Barrett, Sonia Goncalves, Ewan Harrison, David K. Jackson, Ian Johnston, Dominic Kwiatkowski, Cordelia Langford, John Sillitoe on behalf of the Wellcome Sanger Institute COVID-19 Surveillance Team |
| EPI_ISL_3311798 | Lighthouse Lab in Alderley Park | Wellcome Sanger Institute for the COVID-19 Genomics UK (COG-UK) Consortium | Jacquelyn Wynn, Mairead Hyland, The Lighthouse Lab in Alderley Park and Alex Alderton, Roberto Amato, Jeffrey Barrett, Sonia Goncalves, Ewan Harrison, David K. Jackson, Ian Johnston, Dominic Kwiatkowski, Cordelia Langford, John Sillitoe on behalf of the Wellcome Sanger Institute COVID-19 Surveillance Team |
| EPI_ISL_3311799, EPI_ISL_3311800, EPI_ISL_3311801, EPI_ISL_3311802 | Lighthouse Lab in Milton Keynes | Wellcome Sanger Institute for the COVID-19 Genomics UK (COG-UK) Consortium | The Lighthouse Lab in Milton Keynes and Alex Alderton, Roberto Amato, Jeffrey Barrett, Sonia Goncalves, Ewan Harrison, David K. Jackson, Ian Johnston, Dominic Kwiatkowski, Cordelia Langford, John Sillitoe on behalf of the Wellcome Sanger Institute COVID-19 Surveillance Team |
| EPI_ISL_3311803 | Lighthouse Lab in Alderley Park | Wellcome Sanger Institute for the COVID-19 Genomics UK (COG-UK) Consortium | Jacquelyn Wynn, Mairead Hyland, The Lighthouse Lab in Alderley Park and Alex Alderton, Roberto Amato, Jeffrey Barrett, Sonia Goncalves, Ewan Harrison, David K. Jackson, Ian Johnston, Dominic Kwiatkowski, Cordelia Langford, John Sillitoe on behalf of the Wellcome Sanger Institute COVID-19 Surveillance Team |
| EPI_ISL_3311804, EPI_ISL_3311805, EPI_ISL_3311806, EPI_ISL_3311807 | Lighthouse Lab in Milton Keynes | Wellcome Sanger Institute for the COVID-19 Genomics UK (COG-UK) Consortium | The Lighthouse Lab in Milton Keynes and Alex Alderton, Roberto Amato, Jeffrey Barrett, Sonia Goncalves, Ewan Harrison, David K. Jackson, Ian Johnston, Dominic Kwiatkowski, Cordelia Langford, John Sillitoe on behalf of the Wellcome Sanger Institute COVID-19 Surveillance Team |
| EPI_ISL_3311808 | Lighthouse Lab in Alderley Park | Wellcome Sanger Institute for the COVID-19 Genomics UK (COG-UK) Consortium | Jacquelyn Wynn, Mairead Hyland, The Lighthouse Lab in Alderley Park and Alex Alderton, Roberto Amato, Jeffrey Barrett, Sonia Goncalves, Ewan Harrison, David K. Jackson, Ian Johnston, Dominic Kwiatkowski, Cordelia Langford, John Sillitoe on behalf of the Wellcome Sanger Institute COVID-19 Surveillance Team |
| EPI_ISL_3311809, EPI_ISL_3311810, EPI_ISL_3311811, EPI_ISL_3311812 | Lighthouse Lab in Milton Keynes | Wellcome Sanger Institute for the COVID-19 Genomics UK (COG-UK) Consortium | The Lighthouse Lab in Milton Keynes and Alex Alderton, Roberto Amato, Jeffrey Barrett, Sonia Goncalves, Ewan Harrison, David K. Jackson, Ian Johnston, Dominic Kwiatkowski, Cordelia Langford, John Sillitoe on behalf of the Wellcome Sanger Institute COVID-19 Surveillance Team |
| EPI_ISL_3311813 | Lighthouse Lab in Alderley Park | Wellcome Sanger Institute for the COVID-19 Genomics UK (COG-UK) | Jacquelyn Wynn, Mairead Hyland, The Lighthouse Lab in Alderley Park and Alex Alderton, Roberto Amato, Jeffrey Barrett, Sonia Goncalves, Ewan Harrison, David K. Jackson, Ian Johnston, Dominic Kwiatkowski, Cordelia Langford, John Sillitoe on |

| | | | |
|---|---|---|---|
| | | Consortium | behalf of the Wellcome Sanger Institute COVID-19 Surveillance Team |
| EPI_ISL_3311814, EPI_ISL_3311815, EPI_ISL_3311816, EPI_ISL_3311817, EPI_ISL_3311818, EPI_ISL_3311819, EPI_ISL_3311820 | Lighthouse Lab in Milton Keynes | Wellcome Sanger Institute for the COVID-19 Genomics UK (COG-UK) Consortium | The Lighthouse Lab in Milton Keynes and Alex Alderton, Roberto Amato, Jeffrey Barrett, Sonia Goncalves, Ewan Harrison, David K. Jackson, Ian Johnston, Dominic Kwiatkowski, Cordelia Langford, John Sillitoe on behalf of the Wellcome Sanger Institute COVID-19 Surveillance Team |
| EPI_ISL_3311821 | Lighthouse Lab in Alderley Park | Wellcome Sanger Institute for the COVID-19 Genomics UK (COG-UK) Consortium | Jacquelyn Wynn, Mairead Hyland, The Lighthouse Lab in Alderley Park and Alex Alderton, Roberto Amato, Jeffrey Barrett, Sonia Goncalves, Ewan Harrison, David K. Jackson, Ian Johnston, Dominic Kwiatkowski, Cordelia Langford, John Sillitoe on behalf of the Wellcome Sanger Institute COVID-19 Surveillance Team |
| EPI_ISL_3311822 | Lighthouse Lab in Milton Keynes | Wellcome Sanger Institute for the COVID-19 Genomics UK (COG-UK) Consortium | The Lighthouse Lab in Milton Keynes and Alex Alderton, Roberto Amato, Jeffrey Barrett, Sonia Goncalves, Ewan Harrison, David K. Jackson, Ian Johnston, Dominic Kwiatkowski, Cordelia Langford, John Sillitoe on behalf of the Wellcome Sanger Institute COVID-19 Surveillance Team |
| EPI_ISL_3311823 | Lighthouse Lab in Alderley Park | Wellcome Sanger Institute for the COVID-19 Genomics UK (COG-UK) Consortium | Jacquelyn Wynn, Mairead Hyland, The Lighthouse Lab in Alderley Park and Alex Alderton, Roberto Amato, Jeffrey Barrett, Sonia Goncalves, Ewan Harrison, David K. Jackson, Ian Johnston, Dominic Kwiatkowski, Cordelia Langford, John Sillitoe on behalf of the Wellcome Sanger Institute COVID-19 Surveillance Team |
| EPI_ISL_3311824, EPI_ISL_3311825 | Lighthouse Lab in Milton Keynes | Wellcome Sanger Institute for the COVID-19 Genomics UK (COG-UK) Consortium | The Lighthouse Lab in Milton Keynes and Alex Alderton, Roberto Amato, Jeffrey Barrett, Sonia Goncalves, Ewan Harrison, David K. Jackson, Ian Johnston, Dominic Kwiatkowski, Cordelia Langford, John Sillitoe on behalf of the Wellcome Sanger Institute COVID-19 Surveillance Team |
| EPI_ISL_3311826 | Lighthouse Lab in Alderley Park | Wellcome Sanger Institute for the COVID-19 Genomics UK (COG-UK) Consortium | Jacquelyn Wynn, Mairead Hyland, The Lighthouse Lab in Alderley Park and Alex Alderton, Roberto Amato, Jeffrey Barrett, Sonia Goncalves, Ewan Harrison, David K. Jackson, Ian Johnston, Dominic Kwiatkowski, Cordelia Langford, John Sillitoe on behalf of the Wellcome Sanger Institute COVID-19 Surveillance Team |
| EPI_ISL_3311827, EPI_ISL_3311828, EPI_ISL_3311829 | Lighthouse Lab in Milton Keynes | Wellcome Sanger Institute for the COVID-19 Genomics UK (COG-UK) Consortium | The Lighthouse Lab in Milton Keynes and Alex Alderton, Roberto Amato, Jeffrey Barrett, Sonia Goncalves, Ewan Harrison, David K. Jackson, Ian Johnston, Dominic Kwiatkowski, Cordelia Langford, John Sillitoe on behalf of the Wellcome Sanger Institute COVID-19 Surveillance Team |
| EPI_ISL_3311830 | Lighthouse Lab in Alderley Park | Wellcome Sanger Institute for the COVID-19 Genomics UK (COG-UK) Consortium | Jacquelyn Wynn, Mairead Hyland, The Lighthouse Lab in Alderley Park and Alex Alderton, Roberto Amato, Jeffrey Barrett, Sonia Goncalves, Ewan Harrison, David K. Jackson, Ian Johnston, Dominic Kwiatkowski, Cordelia Langford, John Sillitoe on behalf of the Wellcome Sanger Institute COVID-19 Surveillance Team |
| EPI_ISL_3311831, EPI_ISL_3311832, EPI_ISL_3311833 | Lighthouse Lab in Milton Keynes | Wellcome Sanger Institute for the COVID-19 Genomics UK (COG-UK) Consortium | The Lighthouse Lab in Milton Keynes and Alex Alderton, Roberto Amato, Jeffrey Barrett, Sonia Goncalves, Ewan Harrison, David K. Jackson, Ian Johnston, Dominic Kwiatkowski, Cordelia Langford, John Sillitoe on behalf of the Wellcome Sanger Institute COVID-19 Surveillance Team |
| EPI_ISL_3311834 | Lighthouse Lab in Alderley Park | Wellcome Sanger Institute for the COVID-19 Genomics UK (COG-UK) Consortium | Jacquelyn Wynn, Mairead Hyland, The Lighthouse Lab in Alderley Park and Alex Alderton, Roberto Amato, Jeffrey Barrett, Sonia Goncalves, Ewan Harrison, David K. Jackson, Ian Johnston, Dominic Kwiatkowski, Cordelia Langford, John Sillitoe on behalf of the Wellcome Sanger Institute COVID-19 Surveillance Team |
| EPI_ISL_3311835 | Lighthouse Lab in Milton Keynes | Wellcome Sanger Institute for the COVID-19 Genomics UK (COG-UK) Consortium | The Lighthouse Lab in Milton Keynes and Alex Alderton, Roberto Amato, Jeffrey Barrett, Sonia Goncalves, Ewan Harrison, David K. Jackson, Ian Johnston, Dominic Kwiatkowski, Cordelia Langford, John Sillitoe on behalf of the Wellcome Sanger Institute COVID-19 Surveillance Team |
| EPI_ISL_3311836 | Lighthouse Lab in Alderley Park | Wellcome Sanger Institute for the COVID-19 Genomics UK (COG-UK) Consortium | Jacquelyn Wynn, Mairead Hyland, The Lighthouse Lab in Alderley Park and Alex Alderton, Roberto Amato, Jeffrey Barrett, Sonia Goncalves, Ewan Harrison, David K. Jackson, Ian Johnston, Dominic Kwiatkowski, Cordelia Langford, John Sillitoe on behalf of the Wellcome Sanger Institute COVID-19 Surveillance Team |
| EPI_ISL_3311837, EPI_ISL_3311838, EPI_ISL_3311839, EPI_ISL_3311840, EPI_ISL_3311841 | Lighthouse Lab in Milton Keynes | Wellcome Sanger Institute for the COVID-19 Genomics UK (COG-UK) Consortium | The Lighthouse Lab in Milton Keynes and Alex Alderton, Roberto Amato, Jeffrey Barrett, Sonia Goncalves, Ewan Harrison, David K. Jackson, Ian Johnston, Dominic Kwiatkowski, Cordelia Langford, John Sillitoe on behalf of the Wellcome Sanger Institute COVID-19 Surveillance Team |
| EPI_ISL_3311842, EPI_ISL_3311843 | Lighthouse Lab in Alderley Park | Wellcome Sanger Institute for the COVID-19 Genomics UK (COG-UK) Consortium | Jacquelyn Wynn, Mairead Hyland, The Lighthouse Lab in Alderley Park and Alex Alderton, Roberto Amato, Jeffrey Barrett, Sonia Goncalves, Ewan Harrison, David K. Jackson, Ian Johnston, Dominic Kwiatkowski, Cordelia Langford, John Sillitoe on behalf of the Wellcome Sanger Institute COVID-19 Surveillance Team |
| EPI_ISL_3311844, EPI_ISL_3311845, EPI_ISL_3311846, EPI_ISL_3311847 | Lighthouse Lab in Milton Keynes | Wellcome Sanger Institute for the COVID-19 Genomics UK (COG-UK) Consortium | The Lighthouse Lab in Milton Keynes and Alex Alderton, Roberto Amato, Jeffrey Barrett, Sonia Goncalves, Ewan Harrison, David K. Jackson, Ian Johnston, Dominic Kwiatkowski, Cordelia Langford, John Sillitoe on behalf of the Wellcome Sanger Institute COVID-19 Surveillance Team |
| EPI_ISL_3311848 | Lighthouse Lab in Alderley Park | Wellcome Sanger Institute for the COVID-19 Genomics UK (COG-UK) Consortium | Jacquelyn Wynn, Mairead Hyland, The Lighthouse Lab in Alderley Park and Alex Alderton, Roberto Amato, Jeffrey Barrett, Sonia Goncalves, Ewan Harrison, David K. Jackson, Ian Johnston, Dominic Kwiatkowski, Cordelia Langford, John Sillitoe on behalf of the Wellcome Sanger Institute COVID-19 Surveillance Team |
| EPI_ISL_3311849, EPI_ISL_3311850, EPI_ISL_3311851 | Lighthouse Lab in Milton Keynes | Wellcome Sanger Institute for the COVID-19 Genomics UK (COG-UK) Consortium | The Lighthouse Lab in Milton Keynes and Alex Alderton, Roberto Amato, Jeffrey Barrett, Sonia Goncalves, Ewan Harrison, David K. Jackson, Ian Johnston, Dominic Kwiatkowski, Cordelia Langford, John Sillitoe on behalf of the Wellcome Sanger Institute COVID-19 Surveillance Team |
| EPI_ISL_3311852 | Lighthouse Lab in Alderley Park | Wellcome Sanger Institute for the COVID-19 Genomics UK (COG-UK) Consortium | Jacquelyn Wynn, Mairead Hyland, The Lighthouse Lab in Alderley Park and Alex Alderton, Roberto Amato, Jeffrey Barrett, Sonia Goncalves, Ewan Harrison, David K. Jackson, Ian Johnston, Dominic Kwiatkowski, Cordelia Langford, John Sillitoe on behalf of the Wellcome Sanger Institute COVID-19 Surveillance Team |
| EPI_ISL_3311853 | Lighthouse Lab in Milton Keynes | Wellcome Sanger Institute for the COVID-19 Genomics UK (COG-UK) Consortium | The Lighthouse Lab in Milton Keynes and Alex Alderton, Roberto Amato, Jeffrey Barrett, Sonia Goncalves, Ewan Harrison, David K. Jackson, Ian Johnston, Dominic Kwiatkowski, Cordelia Langford, John Sillitoe on behalf of the Wellcome Sanger Institute COVID-19 Surveillance Team |
| EPI_ISL_3311854 | Lighthouse Lab in Alderley Park | Wellcome Sanger Institute for the COVID-19 Genomics UK (COG-UK) Consortium | Jacquelyn Wynn, Mairead Hyland, The Lighthouse Lab in Alderley Park and Alex Alderton, Roberto Amato, Jeffrey Barrett, Sonia Goncalves, Ewan Harrison, David K. Jackson, Ian Johnston, Dominic Kwiatkowski, Cordelia Langford, John Sillitoe on behalf of the Wellcome Sanger Institute COVID-19 Surveillance Team |
| EPI_ISL_3311855, EPI_ISL_3311856, EPI_ISL_3311857, EPI_ISL_3311858, EPI_ISL_3311859, EPI_ISL_3311860, EPI_ISL_3311862, EPI_ISL_3311863, EPI_ISL_3311864 | Lighthouse Lab in Milton Keynes | Wellcome Sanger Institute for the COVID-19 Genomics UK (COG-UK) Consortium | The Lighthouse Lab in Milton Keynes and Alex Alderton, Roberto Amato, Jeffrey Barrett, Sonia Goncalves, Ewan Harrison, David K. Jackson, Ian Johnston, Dominic Kwiatkowski, Cordelia Langford, John Sillitoe on behalf of the Wellcome Sanger Institute COVID-19 Surveillance Team |
| EPI_ISL_3311865, EPI_ISL_3311866 | Lighthouse Lab in Alderley Park | Wellcome Sanger Institute for the COVID-19 Genomics UK (COG-UK) Consortium | Jacquelyn Wynn, Mairead Hyland, The Lighthouse Lab in Alderley Park and Alex Alderton, Roberto Amato, Jeffrey Barrett, Sonia Goncalves, Ewan Harrison, David K. Jackson, Ian Johnston, Dominic Kwiatkowski, Cordelia Langford, John Sillitoe on behalf of the Wellcome Sanger Institute COVID-19 Surveillance Team |
| EPI_ISL_3311867 | Lighthouse Lab in Milton Keynes | Wellcome Sanger Institute for the COVID-19 Genomics UK (COG-UK) Consortium | The Lighthouse Lab in Milton Keynes and Alex Alderton, Roberto Amato, Jeffrey Barrett, Sonia Goncalves, Ewan Harrison, David K. Jackson, Ian Johnston, Dominic Kwiatkowski, Cordelia Langford, John Sillitoe on behalf of the Wellcome Sanger Institute COVID-19 Surveillance Team |
| EPI_ISL_3311868 | Lighthouse Lab in Alderley Park | Wellcome Sanger Institute for the COVID-19 Genomics UK (COG-UK) Consortium | Jacquelyn Wynn, Mairead Hyland, The Lighthouse Lab in Alderley Park and Alex Alderton, Roberto Amato, Jeffrey Barrett, Sonia Goncalves, Ewan Harrison, David K. Jackson, Ian Johnston, Dominic Kwiatkowski, Cordelia Langford, John Sillitoe on behalf of the Wellcome Sanger Institute COVID-19 Surveillance Team |
| EPI_ISL_3311869 | Lighthouse Lab in Milton Keynes | Wellcome Sanger Institute for the COVID-19 Genomics UK (COG-UK) Consortium | The Lighthouse Lab in Milton Keynes and Alex Alderton, Roberto Amato, Jeffrey Barrett, Sonia Goncalves, Ewan Harrison, David K. Jackson, Ian Johnston, Dominic Kwiatkowski, Cordelia Langford, John Sillitoe on behalf of the Wellcome Sanger Institute COVID-19 Surveillance Team |
| EPI_ISL_3311870, EPI_ISL_3311871, EPI_ISL_3311872, EPI_ISL_3311873, EPI_ISL_3311874 | Lighthouse Lab in Alderley Park | Wellcome Sanger Institute for the COVID-19 Genomics UK (COG-UK) Consortium | Jacquelyn Wynn, Mairead Hyland, The Lighthouse Lab in Alderley Park and Alex Alderton, Roberto Amato, Jeffrey Barrett, Sonia Goncalves, Ewan Harrison, David K. Jackson, Ian Johnston, Dominic Kwiatkowski, Cordelia Langford, John Sillitoe on behalf of the Wellcome Sanger Institute COVID-19 Surveillance Team |
| EPI_ISL_3311875, EPI_ISL_3311876 | Lighthouse Lab in Milton Keynes | Wellcome Sanger Institute for the COVID-19 Genomics UK (COG-UK) Consortium | The Lighthouse Lab in Milton Keynes and Alex Alderton, Roberto Amato, Jeffrey Barrett, Sonia Goncalves, Ewan Harrison, David K. Jackson, Ian Johnston, Dominic Kwiatkowski, Cordelia Langford, John Sillitoe on behalf of the Wellcome Sanger Institute COVID-19 Surveillance Team |
| EPI_ISL_3311877 | Lighthouse Lab in Alderley Park | Wellcome Sanger Institute for the COVID-19 Genomics UK (COG-UK) Consortium | Jacquelyn Wynn, Mairead Hyland, The Lighthouse Lab in Alderley Park and Alex Alderton, Roberto Amato, Jeffrey Barrett, Sonia Goncalves, Ewan Harrison, David K. Jackson, Ian Johnston, Dominic Kwiatkowski, Cordelia Langford, John Sillitoe on behalf of the Wellcome Sanger Institute COVID-19 Surveillance Team |
| EPI_ISL_3311878, EPI_ISL_3311879, EPI_ISL_3311880, EPI_ISL_3311881, EPI_ISL_3311882, EPI_ISL_3311883, EPI_ISL_3311884, EPI_ISL_3311885 | Lighthouse Lab in Milton Keynes | Wellcome Sanger Institute for the COVID-19 Genomics UK (COG-UK) Consortium | The Lighthouse Lab in Milton Keynes and Alex Alderton, Roberto Amato, Jeffrey Barrett, Sonia Goncalves, Ewan Harrison, David K. Jackson, Ian Johnston, Dominic Kwiatkowski, Cordelia Langford, John Sillitoe on behalf of the Wellcome Sanger Institute COVID-19 Surveillance Team |
| EPI_ISL_3311886 | Lighthouse Lab in Alderley Park | Wellcome Sanger Institute for the COVID-19 Genomics UK (COG-UK) Consortium | Jacquelyn Wynn, Mairead Hyland, The Lighthouse Lab in Alderley Park and Alex Alderton, Roberto Amato, Jeffrey Barrett, Sonia Goncalves, Ewan Harrison, David K. Jackson, Ian Johnston, Dominic Kwiatkowski, Cordelia Langford, John Sillitoe on behalf of the Wellcome Sanger Institute COVID-19 Surveillance Team |
| EPI_ISL_3311887, EPI_ISL_3311888 | Lighthouse Lab in Milton Keynes | Wellcome Sanger Institute for the COVID-19 Genomics UK (COG-UK) Consortium | The Lighthouse Lab in Milton Keynes and Alex Alderton, Roberto Amato, Jeffrey Barrett, Sonia Goncalves, Ewan Harrison, David K. Jackson, Ian Johnston, Dominic Kwiatkowski, Cordelia Langford, John Sillitoe on behalf of the Wellcome Sanger Institute COVID-19 Surveillance Team |
| EPI_ISL_3311889 | Lighthouse Lab in Alderley Park | Wellcome Sanger Institute for the COVID-19 Genomics UK (COG-UK) Consortium | Jacquelyn Wynn, Mairead Hyland, The Lighthouse Lab in Alderley Park and Alex Alderton, Roberto Amato, Jeffrey Barrett, Sonia Goncalves, Ewan Harrison, David K. Jackson, Ian Johnston, Dominic Kwiatkowski, Cordelia Langford, John Sillitoe on behalf of the Wellcome Sanger Institute COVID-19 Surveillance Team |
| EPI_ISL_3311890, | Lighthouse Lab in Milton Keynes | Wellcome Sanger Institute for the COVID-19 Genomics UK (COG-UK) | The Lighthouse Lab in Milton Keynes and Alex Alderton, Roberto Amato, Jeffrey Barrett, Sonia Goncalves, Ewan Harrison, David K. Jackson, Ian Johnston, Dominic Kwiatkowski, Cordelia Langford, John Sillitoe on behalf of the Wellcome Sanger |

| | | | |
|---|---|---|---|
| EPI_ISL_3311891, EPI_ISL_3311892 | | Consortium | Institute COVID-19 Surveillance Team |
| EPI_ISL_3311893 | Lighthouse Lab in Alderley Park | Wellcome Sanger Institute for the COVID-19 Genomics UK (COG-UK) Consortium | Jacquelyn Wynn, Mairead Hyland, The Lighthouse Lab in Alderley Park and Alex Alderton, Roberto Amato, Jeffrey Barrett, Sonia Goncalves, Ewan Harrison, David K. Jackson, Ian Johnston, Dominic Kwiatkowski, Cordelia Langford, John Sillitoe on behalf of the Wellcome Sanger Institute COVID-19 Surveillance Team |
| EPI_ISL_3311894 | Lighthouse Lab in Milton Keynes | Wellcome Sanger Institute for the COVID-19 Genomics UK (COG-UK) Consortium | The Lighthouse Lab in Milton Keynes and Alex Alderton, Roberto Amato, Jeffrey Barrett, Sonia Goncalves, Ewan Harrison, David K. Jackson, Ian Johnston, Dominic Kwiatkowski, Cordelia Langford, John Sillitoe on behalf of the Wellcome Sanger Institute COVID-19 Surveillance Team |
| EPI_ISL_3311895 | Lighthouse Lab in Alderley Park | Wellcome Sanger Institute for the COVID-19 Genomics UK (COG-UK) Consortium | Jacquelyn Wynn, Mairead Hyland, The Lighthouse Lab in Alderley Park and Alex Alderton, Roberto Amato, Jeffrey Barrett, Sonia Goncalves, Ewan Harrison, David K. Jackson, Ian Johnston, Dominic Kwiatkowski, Cordelia Langford, John Sillitoe on behalf of the Wellcome Sanger Institute COVID-19 Surveillance Team |
| EPI_ISL_3311896 | Lighthouse Lab in Milton Keynes | Wellcome Sanger Institute for the COVID-19 Genomics UK (COG-UK) Consortium | The Lighthouse Lab in Milton Keynes and Alex Alderton, Roberto Amato, Jeffrey Barrett, Sonia Goncalves, Ewan Harrison, David K. Jackson, Ian Johnston, Dominic Kwiatkowski, Cordelia Langford, John Sillitoe on behalf of the Wellcome Sanger Institute COVID-19 Surveillance Team |
| EPI_ISL_3311897, EPI_ISL_3311898 | Lighthouse Lab in Alderley Park | Wellcome Sanger Institute for the COVID-19 Genomics UK (COG-UK) Consortium | Jacquelyn Wynn, Mairead Hyland, The Lighthouse Lab in Alderley Park and Alex Alderton, Roberto Amato, Jeffrey Barrett, Sonia Goncalves, Ewan Harrison, David K. Jackson, Ian Johnston, Dominic Kwiatkowski, Cordelia Langford, John Sillitoe on behalf of the Wellcome Sanger Institute COVID-19 Surveillance Team |
| EPI_ISL_3311899, EPI_ISL_3311900, EPI_ISL_3311901, EPI_ISL_3311902, EPI_ISL_3311903, EPI_ISL_3311904 | Lighthouse Lab in Milton Keynes | Wellcome Sanger Institute for the COVID-19 Genomics UK (COG-UK) Consortium | The Lighthouse Lab in Milton Keynes and Alex Alderton, Roberto Amato, Jeffrey Barrett, Sonia Goncalves, Ewan Harrison, David K. Jackson, Ian Johnston, Dominic Kwiatkowski, Cordelia Langford, John Sillitoe on behalf of the Wellcome Sanger Institute COVID-19 Surveillance Team |
| EPI_ISL_3311905, EPI_ISL_3311906 | Lighthouse Lab in Alderley Park | Wellcome Sanger Institute for the COVID-19 Genomics UK (COG-UK) Consortium | Jacquelyn Wynn, Mairead Hyland, The Lighthouse Lab in Alderley Park and Alex Alderton, Roberto Amato, Jeffrey Barrett, Sonia Goncalves, Ewan Harrison, David K. Jackson, Ian Johnston, Dominic Kwiatkowski, Cordelia Langford, John Sillitoe on behalf of the Wellcome Sanger Institute COVID-19 Surveillance Team |
| EPI_ISL_3311907 | Lighthouse Lab in Milton Keynes | Wellcome Sanger Institute for the COVID-19 Genomics UK (COG-UK) Consortium | The Lighthouse Lab in Milton Keynes and Alex Alderton, Roberto Amato, Jeffrey Barrett, Sonia Goncalves, Ewan Harrison, David K. Jackson, Ian Johnston, Dominic Kwiatkowski, Cordelia Langford, John Sillitoe on behalf of the Wellcome Sanger Institute COVID-19 Surveillance Team |
| EPI_ISL_3311908 | Lighthouse Lab in Alderley Park | Wellcome Sanger Institute for the COVID-19 Genomics UK (COG-UK) Consortium | Jacquelyn Wynn, Mairead Hyland, The Lighthouse Lab in Alderley Park and Alex Alderton, Roberto Amato, Jeffrey Barrett, Sonia Goncalves, Ewan Harrison, David K. Jackson, Ian Johnston, Dominic Kwiatkowski, Cordelia Langford, John Sillitoe on behalf of the Wellcome Sanger Institute COVID-19 Surveillance Team |
| EPI_ISL_3311909, EPI_ISL_3311910 | Lighthouse Lab in Milton Keynes | Wellcome Sanger Institute for the COVID-19 Genomics UK (COG-UK) Consortium | The Lighthouse Lab in Milton Keynes and Alex Alderton, Roberto Amato, Jeffrey Barrett, Sonia Goncalves, Ewan Harrison, David K. Jackson, Ian Johnston, Dominic Kwiatkowski, Cordelia Langford, John Sillitoe on behalf of the Wellcome Sanger Institute COVID-19 Surveillance Team |
| EPI_ISL_3311911, EPI_ISL_3311912, EPI_ISL_3311913 | Lighthouse Lab in Alderley Park | Wellcome Sanger Institute for the COVID-19 Genomics UK (COG-UK) Consortium | Jacquelyn Wynn, Mairead Hyland, The Lighthouse Lab in Alderley Park and Alex Alderton, Roberto Amato, Jeffrey Barrett, Sonia Goncalves, Ewan Harrison, David K. Jackson, Ian Johnston, Dominic Kwiatkowski, Cordelia Langford, John Sillitoe on behalf of the Wellcome Sanger Institute COVID-19 Surveillance Team |
| EPI_ISL_3311914 | Lighthouse Lab in Milton Keynes | Wellcome Sanger Institute for the COVID-19 Genomics UK (COG-UK) Consortium | The Lighthouse Lab in Milton Keynes and Alex Alderton, Roberto Amato, Jeffrey Barrett, Sonia Goncalves, Ewan Harrison, David K. Jackson, Ian Johnston, Dominic Kwiatkowski, Cordelia Langford, John Sillitoe on behalf of the Wellcome Sanger Institute COVID-19 Surveillance Team |
| EPI_ISL_3311915 | Lighthouse Lab in Alderley Park | Wellcome Sanger Institute for the COVID-19 Genomics UK (COG-UK) Consortium | Jacquelyn Wynn, Mairead Hyland, The Lighthouse Lab in Alderley Park and Alex Alderton, Roberto Amato, Jeffrey Barrett, Sonia Goncalves, Ewan Harrison, David K. Jackson, Ian Johnston, Dominic Kwiatkowski, Cordelia Langford, John Sillitoe on behalf of the Wellcome Sanger Institute COVID-19 Surveillance Team |
| EPI_ISL_3311916 | Lighthouse Lab in Milton Keynes | Wellcome Sanger Institute for the COVID-19 Genomics UK (COG-UK) Consortium | The Lighthouse Lab in Milton Keynes and Alex Alderton, Roberto Amato, Jeffrey Barrett, Sonia Goncalves, Ewan Harrison, David K. Jackson, Ian Johnston, Dominic Kwiatkowski, Cordelia Langford, John Sillitoe on behalf of the Wellcome Sanger Institute COVID-19 Surveillance Team |
| EPI_ISL_3311917 | Lighthouse Lab in Alderley Park | Wellcome Sanger Institute for the COVID-19 Genomics UK (COG-UK) Consortium | Jacquelyn Wynn, Mairead Hyland, The Lighthouse Lab in Alderley Park and Alex Alderton, Roberto Amato, Jeffrey Barrett, Sonia Goncalves, Ewan Harrison, David K. Jackson, Ian Johnston, Dominic Kwiatkowski, Cordelia Langford, John Sillitoe on behalf of the Wellcome Sanger Institute COVID-19 Surveillance Team |
| EPI_ISL_3311918, EPI_ISL_3311919 | Lighthouse Lab in Milton Keynes | Wellcome Sanger Institute for the COVID-19 Genomics UK (COG-UK) Consortium | The Lighthouse Lab in Milton Keynes and Alex Alderton, Roberto Amato, Jeffrey Barrett, Sonia Goncalves, Ewan Harrison, David K. Jackson, Ian Johnston, Dominic Kwiatkowski, Cordelia Langford, John Sillitoe on behalf of the Wellcome Sanger Institute COVID-19 Surveillance Team |
| EPI_ISL_3311920 | Lighthouse Lab in Alderley Park | Wellcome Sanger Institute for the COVID-19 Genomics UK (COG-UK) Consortium | Jacquelyn Wynn, Mairead Hyland, The Lighthouse Lab in Alderley Park and Alex Alderton, Roberto Amato, Jeffrey Barrett, Sonia Goncalves, Ewan Harrison, David K. Jackson, Ian Johnston, Dominic Kwiatkowski, Cordelia Langford, John Sillitoe on behalf of the Wellcome Sanger Institute COVID-19 Surveillance Team |
| EPI_ISL_3311921, EPI_ISL_3311922, EPI_ISL_3311923 | Lighthouse Lab in Milton Keynes | Wellcome Sanger Institute for the COVID-19 Genomics UK (COG-UK) Consortium | The Lighthouse Lab in Milton Keynes and Alex Alderton, Roberto Amato, Jeffrey Barrett, Sonia Goncalves, Ewan Harrison, David K. Jackson, Ian Johnston, Dominic Kwiatkowski, Cordelia Langford, John Sillitoe on behalf of the Wellcome Sanger Institute COVID-19 Surveillance Team |
| EPI_ISL_3311924 | Lighthouse Lab in Alderley Park | Wellcome Sanger Institute for the COVID-19 Genomics UK (COG-UK) Consortium | Jacquelyn Wynn, Mairead Hyland, The Lighthouse Lab in Alderley Park and Alex Alderton, Roberto Amato, Jeffrey Barrett, Sonia Goncalves, Ewan Harrison, David K. Jackson, Ian Johnston, Dominic Kwiatkowski, Cordelia Langford, John Sillitoe on behalf of the Wellcome Sanger Institute COVID-19 Surveillance Team |
| EPI_ISL_3311925, EPI_ISL_3311926, EPI_ISL_3311927, EPI_ISL_3311928, EPI_ISL_3311929, EPI_ISL_3311930 | Lighthouse Lab in Milton Keynes | Wellcome Sanger Institute for the COVID-19 Genomics UK (COG-UK) Consortium | The Lighthouse Lab in Milton Keynes and Alex Alderton, Roberto Amato, Jeffrey Barrett, Sonia Goncalves, Ewan Harrison, David K. Jackson, Ian Johnston, Dominic Kwiatkowski, Cordelia Langford, John Sillitoe on behalf of the Wellcome Sanger Institute COVID-19 Surveillance Team |
| EPI_ISL_3311931 | Lighthouse Lab in Alderley Park | Wellcome Sanger Institute for the COVID-19 Genomics UK (COG-UK) Consortium | Jacquelyn Wynn, Mairead Hyland, The Lighthouse Lab in Alderley Park and Alex Alderton, Roberto Amato, Jeffrey Barrett, Sonia Goncalves, Ewan Harrison, David K. Jackson, Ian Johnston, Dominic Kwiatkowski, Cordelia Langford, John Sillitoe on behalf of the Wellcome Sanger Institute COVID-19 Surveillance Team |
| EPI_ISL_3311932, EPI_ISL_3311933, EPI_ISL_3311934, EPI_ISL_3311935, EPI_ISL_3311936, EPI_ISL_3311937, EPI_ISL_3311938, EPI_ISL_3311939 | Lighthouse Lab in Milton Keynes | Wellcome Sanger Institute for the COVID-19 Genomics UK (COG-UK) Consortium | The Lighthouse Lab in Milton Keynes and Alex Alderton, Roberto Amato, Jeffrey Barrett, Sonia Goncalves, Ewan Harrison, David K. Jackson, Ian Johnston, Dominic Kwiatkowski, Cordelia Langford, John Sillitoe on behalf of the Wellcome Sanger Institute COVID-19 Surveillance Team |
| EPI_ISL_3311940 | Lighthouse Lab in Alderley Park | Wellcome Sanger Institute for the COVID-19 Genomics UK (COG-UK) Consortium | Jacquelyn Wynn, Mairead Hyland, The Lighthouse Lab in Alderley Park and Alex Alderton, Roberto Amato, Jeffrey Barrett, Sonia Goncalves, Ewan Harrison, David K. Jackson, Ian Johnston, Dominic Kwiatkowski, Cordelia Langford, John Sillitoe on behalf of the Wellcome Sanger Institute COVID-19 Surveillance Team |
| EPI_ISL_3311941 | Lighthouse Lab in Milton Keynes | Wellcome Sanger Institute for the COVID-19 Genomics UK (COG-UK) Consortium | The Lighthouse Lab in Milton Keynes and Alex Alderton, Roberto Amato, Jeffrey Barrett, Sonia Goncalves, Ewan Harrison, David K. Jackson, Ian Johnston, Dominic Kwiatkowski, Cordelia Langford, John Sillitoe on behalf of the Wellcome Sanger Institute COVID-19 Surveillance Team |
| EPI_ISL_3311942, EPI_ISL_3311943 | Lighthouse Lab in Alderley Park | Wellcome Sanger Institute for the COVID-19 Genomics UK (COG-UK) Consortium | Jacquelyn Wynn, Mairead Hyland, The Lighthouse Lab in Alderley Park and Alex Alderton, Roberto Amato, Jeffrey Barrett, Sonia Goncalves, Ewan Harrison, David K. Jackson, Ian Johnston, Dominic Kwiatkowski, Cordelia Langford, John Sillitoe on behalf of the Wellcome Sanger Institute COVID-19 Surveillance Team |
| EPI_ISL_3311944 | Lighthouse Lab in Milton Keynes | Wellcome Sanger Institute for the COVID-19 Genomics UK (COG-UK) Consortium | The Lighthouse Lab in Milton Keynes and Alex Alderton, Roberto Amato, Jeffrey Barrett, Sonia Goncalves, Ewan Harrison, David K. Jackson, Ian Johnston, Dominic Kwiatkowski, Cordelia Langford, John Sillitoe on behalf of the Wellcome Sanger Institute COVID-19 Surveillance Team |
| EPI_ISL_3311945 | Lighthouse Lab in Alderley Park | Wellcome Sanger Institute for the COVID-19 Genomics UK (COG-UK) Consortium | Jacquelyn Wynn, Mairead Hyland, The Lighthouse Lab in Alderley Park and Alex Alderton, Roberto Amato, Jeffrey Barrett, Sonia Goncalves, Ewan Harrison, David K. Jackson, Ian Johnston, Dominic Kwiatkowski, Cordelia Langford, John Sillitoe on behalf of the Wellcome Sanger Institute COVID-19 Surveillance Team |
| EPI_ISL_3311946 | Lighthouse Lab in Milton Keynes | Wellcome Sanger Institute for the COVID-19 Genomics UK (COG-UK) Consortium | The Lighthouse Lab in Milton Keynes and Alex Alderton, Roberto Amato, Jeffrey Barrett, Sonia Goncalves, Ewan Harrison, David K. Jackson, Ian Johnston, Dominic Kwiatkowski, Cordelia Langford, John Sillitoe on behalf of the Wellcome Sanger Institute COVID-19 Surveillance Team |
| EPI_ISL_3311947 | Lighthouse Lab in Alderley Park | Wellcome Sanger Institute for the COVID-19 Genomics UK (COG-UK) Consortium | Jacquelyn Wynn, Mairead Hyland, The Lighthouse Lab in Alderley Park and Alex Alderton, Roberto Amato, Jeffrey Barrett, Sonia Goncalves, Ewan Harrison, David K. Jackson, Ian Johnston, Dominic Kwiatkowski, Cordelia Langford, John Sillitoe on behalf of the Wellcome Sanger Institute COVID-19 Surveillance Team |
| EPI_ISL_3311948, EPI_ISL_3311949, EPI_ISL_3311950, EPI_ISL_3311951, EPI_ISL_3311952, EPI_ISL_3311953, EPI_ISL_3311954 | Lighthouse Lab in Milton Keynes | Wellcome Sanger Institute for the COVID-19 Genomics UK (COG-UK) Consortium | The Lighthouse Lab in Milton Keynes and Alex Alderton, Roberto Amato, Jeffrey Barrett, Sonia Goncalves, Ewan Harrison, David K. Jackson, Ian Johnston, Dominic Kwiatkowski, Cordelia Langford, John Sillitoe on behalf of the Wellcome Sanger Institute COVID-19 Surveillance Team |
| EPI_ISL_3311955, EPI_ISL_3311956, EPI_ISL_3311957 | Lighthouse Lab in Alderley Park | Wellcome Sanger Institute for the COVID-19 Genomics UK (COG-UK) Consortium | Jacquelyn Wynn, Mairead Hyland, The Lighthouse Lab in Alderley Park and Alex Alderton, Roberto Amato, Jeffrey Barrett, Sonia Goncalves, Ewan Harrison, David K. Jackson, Ian Johnston, Dominic Kwiatkowski, Cordelia Langford, John Sillitoe on behalf of the Wellcome Sanger Institute COVID-19 Surveillance Team |
| EPI_ISL_3311958, EPI_ISL_3311959, EPI_ISL_3311960, EPI_ISL_3311961, EPI_ISL_3311962 | Lighthouse Lab in Milton Keynes | Wellcome Sanger Institute for the COVID-19 Genomics UK (COG-UK) Consortium | The Lighthouse Lab in Milton Keynes and Alex Alderton, Roberto Amato, Jeffrey Barrett, Sonia Goncalves, Ewan Harrison, David K. Jackson, Ian Johnston, Dominic Kwiatkowski, Cordelia Langford, John Sillitoe on behalf of the Wellcome Sanger Institute COVID-19 Surveillance Team |
| EPI_ISL_3311963 | Lighthouse Lab in Alderley Park | Wellcome Sanger Institute for the COVID-19 Genomics UK (COG-UK) Consortium | Jacquelyn Wynn, Mairead Hyland, The Lighthouse Lab in Alderley Park and Alex Alderton, Roberto Amato, Jeffrey Barrett, Sonia Goncalves, Ewan Harrison, David K. Jackson, Ian Johnston, Dominic Kwiatkowski, Cordelia Langford, John Sillitoe on behalf of the Wellcome Sanger Institute COVID-19 Surveillance Team |
| EPI_ISL_3311964, | Lighthouse Lab in Milton Keynes | Wellcome Sanger Institute for the COVID-19 Genomics UK (COG-UK) | The Lighthouse Lab in Milton Keynes and Alex Alderton, Roberto Amato, Jeffrey Barrett, Sonia Goncalves, Ewan Harrison, David K. Jackson, Ian Johnston, Dominic Kwiatkowski, Cordelia Langford, John Sillitoe on behalf of the Wellcome Sanger |

| | | | |
|---|---|---|---|
| EPI_ISL_3311965 | | Consortium | Institute COVID-19 Surveillance Team |
| EPI_ISL_3311966 | Lighthouse Lab in Alderley Park | Wellcome Sanger Institute for the COVID-19 Genomics UK (COG-UK) Consortium | Jacquelyn Wynn, Mairead Hyland, The Lighthouse Lab in Alderley Park and Alex Alderton, Roberto Amato, Jeffrey Barrett, Sonia Goncalves, Ewan Harrison, David K. Jackson, Ian Johnston, Dominic Kwiatkowski, Cordelia Langford, John Sillitoe on behalf of the Wellcome Sanger Institute COVID-19 Surveillance Team |
| EPI_ISL_3311967, EPI_ISL_3311968, EPI_ISL_3311969 | Lighthouse Lab in Milton Keynes | Wellcome Sanger Institute for the COVID-19 Genomics UK (COG-UK) Consortium | The Lighthouse Lab in Milton Keynes and Alex Alderton, Roberto Amato, Jeffrey Barrett, Sonia Goncalves, Ewan Harrison, David K. Jackson, Ian Johnston, Dominic Kwiatkowski, Cordelia Langford, John Sillitoe on behalf of the Wellcome Sanger Institute COVID-19 Surveillance Team |
| EPI_ISL_3311970, EPI_ISL_3311971 | Lighthouse Lab in Alderley Park | Wellcome Sanger Institute for the COVID-19 Genomics UK (COG-UK) Consortium | Jacquelyn Wynn, Mairead Hyland, The Lighthouse Lab in Alderley Park and Alex Alderton, Roberto Amato, Jeffrey Barrett, Sonia Goncalves, Ewan Harrison, David K. Jackson, Ian Johnston, Dominic Kwiatkowski, Cordelia Langford, John Sillitoe on behalf of the Wellcome Sanger Institute COVID-19 Surveillance Team |
| EPI_ISL_3311972, EPI_ISL_3311973, EPI_ISL_3311974, EPI_ISL_3311975 | Lighthouse Lab in Milton Keynes | Wellcome Sanger Institute for the COVID-19 Genomics UK (COG-UK) Consortium | The Lighthouse Lab in Milton Keynes and Alex Alderton, Roberto Amato, Jeffrey Barrett, Sonia Goncalves, Ewan Harrison, David K. Jackson, Ian Johnston, Dominic Kwiatkowski, Cordelia Langford, John Sillitoe on behalf of the Wellcome Sanger Institute COVID-19 Surveillance Team |
| EPI_ISL_3311976 | Lighthouse Lab in Alderley Park | Wellcome Sanger Institute for the COVID-19 Genomics UK (COG-UK) Consortium | Jacquelyn Wynn, Mairead Hyland, The Lighthouse Lab in Alderley Park and Alex Alderton, Roberto Amato, Jeffrey Barrett, Sonia Goncalves, Ewan Harrison, David K. Jackson, Ian Johnston, Dominic Kwiatkowski, Cordelia Langford, John Sillitoe on behalf of the Wellcome Sanger Institute COVID-19 Surveillance Team |
| EPI_ISL_3311977, EPI_ISL_3311978, EPI_ISL_3311979 | Lighthouse Lab in Milton Keynes | Wellcome Sanger Institute for the COVID-19 Genomics UK (COG-UK) Consortium | The Lighthouse Lab in Milton Keynes and Alex Alderton, Roberto Amato, Jeffrey Barrett, Sonia Goncalves, Ewan Harrison, David K. Jackson, Ian Johnston, Dominic Kwiatkowski, Cordelia Langford, John Sillitoe on behalf of the Wellcome Sanger Institute COVID-19 Surveillance Team |
| EPI_ISL_3311980, EPI_ISL_3311981 | Lighthouse Lab in Alderley Park | Wellcome Sanger Institute for the COVID-19 Genomics UK (COG-UK) Consortium | Jacquelyn Wynn, Mairead Hyland, The Lighthouse Lab in Alderley Park and Alex Alderton, Roberto Amato, Jeffrey Barrett, Sonia Goncalves, Ewan Harrison, David K. Jackson, Ian Johnston, Dominic Kwiatkowski, Cordelia Langford, John Sillitoe on behalf of the Wellcome Sanger Institute COVID-19 Surveillance Team |
| EPI_ISL_3311982, EPI_ISL_3311983 | Lighthouse Lab in Milton Keynes | Wellcome Sanger Institute for the COVID-19 Genomics UK (COG-UK) Consortium | The Lighthouse Lab in Milton Keynes and Alex Alderton, Roberto Amato, Jeffrey Barrett, Sonia Goncalves, Ewan Harrison, David K. Jackson, Ian Johnston, Dominic Kwiatkowski, Cordelia Langford, John Sillitoe on behalf of the Wellcome Sanger Institute COVID-19 Surveillance Team |
| EPI_ISL_3311984 | Lighthouse Lab in Alderley Park | Wellcome Sanger Institute for the COVID-19 Genomics UK (COG-UK) Consortium | Jacquelyn Wynn, Mairead Hyland, The Lighthouse Lab in Alderley Park and Alex Alderton, Roberto Amato, Jeffrey Barrett, Sonia Goncalves, Ewan Harrison, David K. Jackson, Ian Johnston, Dominic Kwiatkowski, Cordelia Langford, John Sillitoe on behalf of the Wellcome Sanger Institute COVID-19 Surveillance Team |
| EPI_ISL_3311985, EPI_ISL_3311986, EPI_ISL_3311987, EPI_ISL_3311988, EPI_ISL_3311989, EPI_ISL_3311990 | Lighthouse Lab in Milton Keynes | Wellcome Sanger Institute for the COVID-19 Genomics UK (COG-UK) Consortium | The Lighthouse Lab in Milton Keynes and Alex Alderton, Roberto Amato, Jeffrey Barrett, Sonia Goncalves, Ewan Harrison, David K. Jackson, Ian Johnston, Dominic Kwiatkowski, Cordelia Langford, John Sillitoe on behalf of the Wellcome Sanger Institute COVID-19 Surveillance Team |
| EPI_ISL_3311991 | Lighthouse Lab in Alderley Park | Wellcome Sanger Institute for the COVID-19 Genomics UK (COG-UK) Consortium | Jacquelyn Wynn, Mairead Hyland, The Lighthouse Lab in Alderley Park and Alex Alderton, Roberto Amato, Jeffrey Barrett, Sonia Goncalves, Ewan Harrison, David K. Jackson, Ian Johnston, Dominic Kwiatkowski, Cordelia Langford, John Sillitoe on behalf of the Wellcome Sanger Institute COVID-19 Surveillance Team |
| EPI_ISL_3311992 | Lighthouse Lab in Milton Keynes | Wellcome Sanger Institute for the COVID-19 Genomics UK (COG-UK) Consortium | The Lighthouse Lab in Milton Keynes and Alex Alderton, Roberto Amato, Jeffrey Barrett, Sonia Goncalves, Ewan Harrison, David K. Jackson, Ian Johnston, Dominic Kwiatkowski, Cordelia Langford, John Sillitoe on behalf of the Wellcome Sanger Institute COVID-19 Surveillance Team |
| EPI_ISL_3311993 | Lighthouse Lab in Alderley Park | Wellcome Sanger Institute for the COVID-19 Genomics UK (COG-UK) Consortium | Jacquelyn Wynn, Mairead Hyland, The Lighthouse Lab in Alderley Park and Alex Alderton, Roberto Amato, Jeffrey Barrett, Sonia Goncalves, Ewan Harrison, David K. Jackson, Ian Johnston, Dominic Kwiatkowski, Cordelia Langford, John Sillitoe on behalf of the Wellcome Sanger Institute COVID-19 Surveillance Team |
| EPI_ISL_3311994, EPI_ISL_3311995 | Lighthouse Lab in Milton Keynes | Wellcome Sanger Institute for the COVID-19 Genomics UK (COG-UK) Consortium | The Lighthouse Lab in Milton Keynes and Alex Alderton, Roberto Amato, Jeffrey Barrett, Sonia Goncalves, Ewan Harrison, David K. Jackson, Ian Johnston, Dominic Kwiatkowski, Cordelia Langford, John Sillitoe on behalf of the Wellcome Sanger Institute COVID-19 Surveillance Team |
| EPI_ISL_3311996, EPI_ISL_3311997, EPI_ISL_3311998, EPI_ISL_3311999 | Lighthouse Lab in Alderley Park | Wellcome Sanger Institute for the COVID-19 Genomics UK (COG-UK) Consortium | Jacquelyn Wynn, Mairead Hyland, The Lighthouse Lab in Alderley Park and Alex Alderton, Roberto Amato, Jeffrey Barrett, Sonia Goncalves, Ewan Harrison, David K. Jackson, Ian Johnston, Dominic Kwiatkowski, Cordelia Langford, John Sillitoe on behalf of the Wellcome Sanger Institute COVID-19 Surveillance Team |
| EPI_ISL_3312000, EPI_ISL_3312001, EPI_ISL_3312002, EPI_ISL_3312003, EPI_ISL_3312004 | Lighthouse Lab in Milton Keynes | Wellcome Sanger Institute for the COVID-19 Genomics UK (COG-UK) Consortium | The Lighthouse Lab in Milton Keynes and Alex Alderton, Roberto Amato, Jeffrey Barrett, Sonia Goncalves, Ewan Harrison, David K. Jackson, Ian Johnston, Dominic Kwiatkowski, Cordelia Langford, John Sillitoe on behalf of the Wellcome Sanger Institute COVID-19 Surveillance Team |
| EPI_ISL_3312005 | Lighthouse Lab in Alderley Park | Wellcome Sanger Institute for the COVID-19 Genomics UK (COG-UK) Consortium | Jacquelyn Wynn, Mairead Hyland, The Lighthouse Lab in Alderley Park and Alex Alderton, Roberto Amato, Jeffrey Barrett, Sonia Goncalves, Ewan Harrison, David K. Jackson, Ian Johnston, Dominic Kwiatkowski, Cordelia Langford, John Sillitoe on behalf of the Wellcome Sanger Institute COVID-19 Surveillance Team |
| EPI_ISL_3312006, EPI_ISL_3312007, EPI_ISL_3312008, EPI_ISL_3312009, EPI_ISL_3312010, EPI_ISL_3312011, EPI_ISL_3312012, EPI_ISL_3312013, EPI_ISL_3312014 | Lighthouse Lab in Milton Keynes | Wellcome Sanger Institute for the COVID-19 Genomics UK (COG-UK) Consortium | The Lighthouse Lab in Milton Keynes and Alex Alderton, Roberto Amato, Jeffrey Barrett, Sonia Goncalves, Ewan Harrison, David K. Jackson, Ian Johnston, Dominic Kwiatkowski, Cordelia Langford, John Sillitoe on behalf of the Wellcome Sanger Institute COVID-19 Surveillance Team |
| EPI_ISL_3312015 | Lighthouse Lab in Alderley Park | Wellcome Sanger Institute for the COVID-19 Genomics UK (COG-UK) Consortium | Jacquelyn Wynn, Mairead Hyland, The Lighthouse Lab in Alderley Park and Alex Alderton, Roberto Amato, Jeffrey Barrett, Sonia Goncalves, Ewan Harrison, David K. Jackson, Ian Johnston, Dominic Kwiatkowski, Cordelia Langford, John Sillitoe on behalf of the Wellcome Sanger Institute COVID-19 Surveillance Team |
| EPI_ISL_3312016 | Lighthouse Lab in Milton Keynes | Wellcome Sanger Institute for the COVID-19 Genomics UK (COG-UK) Consortium | The Lighthouse Lab in Milton Keynes and Alex Alderton, Roberto Amato, Jeffrey Barrett, Sonia Goncalves, Ewan Harrison, David K. Jackson, Ian Johnston, Dominic Kwiatkowski, Cordelia Langford, John Sillitoe on behalf of the Wellcome Sanger Institute COVID-19 Surveillance Team |
| EPI_ISL_3312017 | Lighthouse Lab in Alderley Park | Wellcome Sanger Institute for the COVID-19 Genomics UK (COG-UK) Consortium | Jacquelyn Wynn, Mairead Hyland, The Lighthouse Lab in Alderley Park and Alex Alderton, Roberto Amato, Jeffrey Barrett, Sonia Goncalves, Ewan Harrison, David K. Jackson, Ian Johnston, Dominic Kwiatkowski, Cordelia Langford, John Sillitoe on behalf of the Wellcome Sanger Institute COVID-19 Surveillance Team |
| EPI_ISL_3312018, EPI_ISL_3312019 | Lighthouse Lab in Milton Keynes | Wellcome Sanger Institute for the COVID-19 Genomics UK (COG-UK) Consortium | The Lighthouse Lab in Milton Keynes and Alex Alderton, Roberto Amato, Jeffrey Barrett, Sonia Goncalves, Ewan Harrison, David K. Jackson, Ian Johnston, Dominic Kwiatkowski, Cordelia Langford, John Sillitoe on behalf of the Wellcome Sanger Institute COVID-19 Surveillance Team |
| EPI_ISL_3312020 | Lighthouse Lab in Alderley Park | Wellcome Sanger Institute for the COVID-19 Genomics UK (COG-UK) Consortium | Jacquelyn Wynn, Mairead Hyland, The Lighthouse Lab in Alderley Park and Alex Alderton, Roberto Amato, Jeffrey Barrett, Sonia Goncalves, Ewan Harrison, David K. Jackson, Ian Johnston, Dominic Kwiatkowski, Cordelia Langford, John Sillitoe on behalf of the Wellcome Sanger Institute COVID-19 Surveillance Team |
| EPI_ISL_3312021, EPI_ISL_3312022, EPI_ISL_3312023 | Lighthouse Lab in Milton Keynes | Wellcome Sanger Institute for the COVID-19 Genomics UK (COG-UK) Consortium | The Lighthouse Lab in Milton Keynes and Alex Alderton, Roberto Amato, Jeffrey Barrett, Sonia Goncalves, Ewan Harrison, David K. Jackson, Ian Johnston, Dominic Kwiatkowski, Cordelia Langford, John Sillitoe on behalf of the Wellcome Sanger Institute COVID-19 Surveillance Team |
| EPI_ISL_3312024 | Lighthouse Lab in Alderley Park | Wellcome Sanger Institute for the COVID-19 Genomics UK (COG-UK) Consortium | Jacquelyn Wynn, Mairead Hyland, The Lighthouse Lab in Alderley Park and Alex Alderton, Roberto Amato, Jeffrey Barrett, Sonia Goncalves, Ewan Harrison, David K. Jackson, Ian Johnston, Dominic Kwiatkowski, Cordelia Langford, John Sillitoe on behalf of the Wellcome Sanger Institute COVID-19 Surveillance Team |
| EPI_ISL_3312025, EPI_ISL_3312026 | Lighthouse Lab in Milton Keynes | Wellcome Sanger Institute for the COVID-19 Genomics UK (COG-UK) Consortium | The Lighthouse Lab in Milton Keynes and Alex Alderton, Roberto Amato, Jeffrey Barrett, Sonia Goncalves, Ewan Harrison, David K. Jackson, Ian Johnston, Dominic Kwiatkowski, Cordelia Langford, John Sillitoe on behalf of the Wellcome Sanger Institute COVID-19 Surveillance Team |
| EPI_ISL_3312027 | Lighthouse Lab in Alderley Park | Wellcome Sanger Institute for the COVID-19 Genomics UK (COG-UK) Consortium | Jacquelyn Wynn, Mairead Hyland, The Lighthouse Lab in Alderley Park and Alex Alderton, Roberto Amato, Jeffrey Barrett, Sonia Goncalves, Ewan Harrison, David K. Jackson, Ian Johnston, Dominic Kwiatkowski, Cordelia Langford, John Sillitoe on behalf of the Wellcome Sanger Institute COVID-19 Surveillance Team |
| EPI_ISL_3312028, EPI_ISL_3312029, EPI_ISL_3312030, EPI_ISL_3312031 | Lighthouse Lab in Milton Keynes | Wellcome Sanger Institute for the COVID-19 Genomics UK (COG-UK) Consortium | The Lighthouse Lab in Milton Keynes and Alex Alderton, Roberto Amato, Jeffrey Barrett, Sonia Goncalves, Ewan Harrison, David K. Jackson, Ian Johnston, Dominic Kwiatkowski, Cordelia Langford, John Sillitoe on behalf of the Wellcome Sanger Institute COVID-19 Surveillance Team |
| EPI_ISL_3312032, EPI_ISL_3312033 | Lighthouse Lab in Alderley Park | Wellcome Sanger Institute for the COVID-19 Genomics UK (COG-UK) Consortium | Jacquelyn Wynn, Mairead Hyland, The Lighthouse Lab in Alderley Park and Alex Alderton, Roberto Amato, Jeffrey Barrett, Sonia Goncalves, Ewan Harrison, David K. Jackson, Ian Johnston, Dominic Kwiatkowski, Cordelia Langford, John Sillitoe on behalf of the Wellcome Sanger Institute COVID-19 Surveillance Team |
| EPI_ISL_3312034, EPI_ISL_3312035, EPI_ISL_3312036, EPI_ISL_3312037, EPI_ISL_3312038, EPI_ISL_3312039, EPI_ISL_3312040, EPI_ISL_3312041 | Lighthouse Lab in Milton Keynes | Wellcome Sanger Institute for the COVID-19 Genomics UK (COG-UK) Consortium | The Lighthouse Lab in Milton Keynes and Alex Alderton, Roberto Amato, Jeffrey Barrett, Sonia Goncalves, Ewan Harrison, David K. Jackson, Ian Johnston, Dominic Kwiatkowski, Cordelia Langford, John Sillitoe on behalf of the Wellcome Sanger Institute COVID-19 Surveillance Team |
| EPI_ISL_3312042 | Lighthouse Lab in Alderley Park | Wellcome Sanger Institute for the COVID-19 Genomics UK (COG-UK) Consortium | Jacquelyn Wynn, Mairead Hyland, The Lighthouse Lab in Alderley Park and Alex Alderton, Roberto Amato, Jeffrey Barrett, Sonia Goncalves, Ewan Harrison, David K. Jackson, Ian Johnston, Dominic Kwiatkowski, Cordelia Langford, John Sillitoe on behalf of the Wellcome Sanger Institute COVID-19 Surveillance Team |
| EPI_ISL_3312043, EPI_ISL_3312044, EPI_ISL_3312045 | Lighthouse Lab in Milton Keynes | Wellcome Sanger Institute for the COVID-19 Genomics UK (COG-UK) Consortium | The Lighthouse Lab in Milton Keynes and Alex Alderton, Roberto Amato, Jeffrey Barrett, Sonia Goncalves, Ewan Harrison, David K. Jackson, Ian Johnston, Dominic Kwiatkowski, Cordelia Langford, John Sillitoe on behalf of the Wellcome Sanger Institute COVID-19 Surveillance Team |
| EPI_ISL_3312046 | Lighthouse Lab in Alderley Park | Wellcome Sanger Institute for the COVID-19 Genomics UK (COG-UK) | Jacquelyn Wynn, Mairead Hyland, The Lighthouse Lab in Alderley Park and Alex Alderton, Roberto Amato, Jeffrey Barrett, Sonia Goncalves, Ewan Harrison, David K. Jackson, Ian Johnston, Dominic Kwiatkowski, Cordelia Langford, John Sillitoe on |

| | | | |
|---|---|---|---|
| | | Consortium | behalf of the Wellcome Sanger Institute COVID-19 Surveillance Team |
| EPI_ISL_3312047, EPI_ISL_3312048 | Lighthouse Lab in Milton Keynes | Wellcome Sanger Institute for the COVID-19 Genomics UK (COG-UK) Consortium | The Lighthouse Lab in Milton Keynes and Alex Alderton, Roberto Amato, Jeffrey Barrett, Sonia Goncalves, Ewan Harrison, David K. Jackson, Ian Johnston, Dominic Kwiatkowski, Cordelia Langford, John Sillitoe on behalf of the Wellcome Sanger Institute COVID-19 Surveillance Team |
| EPI_ISL_3312049 | Lighthouse Lab in Alderley Park | Wellcome Sanger Institute for the COVID-19 Genomics UK (COG-UK) Consortium | Jacquelyn Wynn, Mairead Hyland, The Lighthouse Lab in Alderley Park and Alex Alderton, Roberto Amato, Jeffrey Barrett, Sonia Goncalves, Ewan Harrison, David K. Jackson, Ian Johnston, Dominic Kwiatkowski, Cordelia Langford, John Sillitoe on behalf of the Wellcome Sanger Institute COVID-19 Surveillance Team |
| EPI_ISL_3312050, EPI_ISL_3312051 | Lighthouse Lab in Milton Keynes | Wellcome Sanger Institute for the COVID-19 Genomics UK (COG-UK) Consortium | The Lighthouse Lab in Milton Keynes and Alex Alderton, Roberto Amato, Jeffrey Barrett, Sonia Goncalves, Ewan Harrison, David K. Jackson, Ian Johnston, Dominic Kwiatkowski, Cordelia Langford, John Sillitoe on behalf of the Wellcome Sanger Institute COVID-19 Surveillance Team |
| EPI_ISL_3312052, EPI_ISL_3312053 | Lighthouse Lab in Alderley Park | Wellcome Sanger Institute for the COVID-19 Genomics UK (COG-UK) Consortium | Jacquelyn Wynn, Mairead Hyland, The Lighthouse Lab in Alderley Park and Alex Alderton, Roberto Amato, Jeffrey Barrett, Sonia Goncalves, Ewan Harrison, David K. Jackson, Ian Johnston, Dominic Kwiatkowski, Cordelia Langford, John Sillitoe on behalf of the Wellcome Sanger Institute COVID-19 Surveillance Team |
| EPI_ISL_3312054, EPI_ISL_3312055, EPI_ISL_3312056 | Lighthouse Lab in Milton Keynes | Wellcome Sanger Institute for the COVID-19 Genomics UK (COG-UK) Consortium | The Lighthouse Lab in Milton Keynes and Alex Alderton, Roberto Amato, Jeffrey Barrett, Sonia Goncalves, Ewan Harrison, David K. Jackson, Ian Johnston, Dominic Kwiatkowski, Cordelia Langford, John Sillitoe on behalf of the Wellcome Sanger Institute COVID-19 Surveillance Team |
| EPI_ISL_3312057, EPI_ISL_3312058 | Lighthouse Lab in Alderley Park | Wellcome Sanger Institute for the COVID-19 Genomics UK (COG-UK) Consortium | Jacquelyn Wynn, Mairead Hyland, The Lighthouse Lab in Alderley Park and Alex Alderton, Roberto Amato, Jeffrey Barrett, Sonia Goncalves, Ewan Harrison, David K. Jackson, Ian Johnston, Dominic Kwiatkowski, Cordelia Langford, John Sillitoe on behalf of the Wellcome Sanger Institute COVID-19 Surveillance Team |
| EPI_ISL_3312059, EPI_ISL_3312060, EPI_ISL_3312061, EPI_ISL_3312062, EPI_ISL_3312063, EPI_ISL_3312064, EPI_ISL_3312065, EPI_ISL_3312066, EPI_ISL_3312067 | Lighthouse Lab in Glasgow | Wellcome Sanger Institute for the COVID-19 Genomics UK (COG-UK) Consortium | Harper VanSteenhouse, Yumi Kasai, David Gray, Carol Clugston, Anna Dominiczak and Alex Alderton, Roberto Amato, Jeffrey Barrett, Sonia Goncalves, Ewan Harrison, David K. Jackson, Ian Johnston, Dominic Kwiatkowski, Cordelia Langford, John Sillitoe on behalf of the Wellcome Sanger Institute COVID-19 Surveillance Team |
| EPI_ISL_3312068 | Berkshire and Surrey Pathology Services Lighthouse Laboratory | Wellcome Sanger Institute for the COVID-19 Genomics UK (COG-UK) Consortium | Berkshire and Surrey Pathology Services Lighthouse Laboratory and Alex Alderton, Roberto Amato, Jeffrey Barrett, Sonia Goncalves, Ewan Harrison, David K. Jackson, Ian Johnston, Dominic Kwiatkowski, Cordelia Langford, John Sillitoe on behalf of the Wellcome Sanger Institute COVID-19 Surveillance Team |
| EPI_ISL_3312069 | Lighthouse Lab in Glasgow | Wellcome Sanger Institute for the COVID-19 Genomics UK (COG-UK) Consortium | Harper VanSteenhouse, Yumi Kasai, David Gray, Carol Clugston, Anna Dominiczak and Alex Alderton, Roberto Amato, Jeffrey Barrett, Sonia Goncalves, Ewan Harrison, David K. Jackson, Ian Johnston, Dominic Kwiatkowski, Cordelia Langford, John Sillitoe on behalf of the Wellcome Sanger Institute COVID-19 Surveillance Team |
| EPI_ISL_3312070 | Berkshire and Surrey Pathology Services Lighthouse Laboratory | Wellcome Sanger Institute for the COVID-19 Genomics UK (COG-UK) Consortium | Berkshire and Surrey Pathology Services Lighthouse Laboratory and Alex Alderton, Roberto Amato, Jeffrey Barrett, Sonia Goncalves, Ewan Harrison, David K. Jackson, Ian Johnston, Dominic Kwiatkowski, Cordelia Langford, John Sillitoe on behalf of the Wellcome Sanger Institute COVID-19 Surveillance Team |
| EPI_ISL_3312071, EPI_ISL_3312072 | Lighthouse Lab in Glasgow | Wellcome Sanger Institute for the COVID-19 Genomics UK (COG-UK) Consortium | Harper VanSteenhouse, Yumi Kasai, David Gray, Carol Clugston, Anna Dominiczak and Alex Alderton, Roberto Amato, Jeffrey Barrett, Sonia Goncalves, Ewan Harrison, David K. Jackson, Ian Johnston, Dominic Kwiatkowski, Cordelia Langford, John Sillitoe on behalf of the Wellcome Sanger Institute COVID-19 Surveillance Team |
| EPI_ISL_3312073 | Berkshire and Surrey Pathology Services Lighthouse Laboratory | Wellcome Sanger Institute for the COVID-19 Genomics UK (COG-UK) Consortium | Berkshire and Surrey Pathology Services Lighthouse Laboratory and Alex Alderton, Roberto Amato, Jeffrey Barrett, Sonia Goncalves, Ewan Harrison, David K. Jackson, Ian Johnston, Dominic Kwiatkowski, Cordelia Langford, John Sillitoe on behalf of the Wellcome Sanger Institute COVID-19 Surveillance Team |
| EPI_ISL_3312074, EPI_ISL_3312075, EPI_ISL_3312076 | Lighthouse Lab in Glasgow | Wellcome Sanger Institute for the COVID-19 Genomics UK (COG-UK) Consortium | Harper VanSteenhouse, Yumi Kasai, David Gray, Carol Clugston, Anna Dominiczak and Alex Alderton, Roberto Amato, Jeffrey Barrett, Sonia Goncalves, Ewan Harrison, David K. Jackson, Ian Johnston, Dominic Kwiatkowski, Cordelia Langford, John Sillitoe on behalf of the Wellcome Sanger Institute COVID-19 Surveillance Team |
| EPI_ISL_3312077 | Berkshire and Surrey Pathology Services Lighthouse Laboratory | Wellcome Sanger Institute for the COVID-19 Genomics UK (COG-UK) Consortium | Berkshire and Surrey Pathology Services Lighthouse Laboratory and Alex Alderton, Roberto Amato, Jeffrey Barrett, Sonia Goncalves, Ewan Harrison, David K. Jackson, Ian Johnston, Dominic Kwiatkowski, Cordelia Langford, John Sillitoe on behalf of the Wellcome Sanger Institute COVID-19 Surveillance Team |
| EPI_ISL_3312078, EPI_ISL_3312079 | Lighthouse Lab in Glasgow | Wellcome Sanger Institute for the COVID-19 Genomics UK (COG-UK) Consortium | Harper VanSteenhouse, Yumi Kasai, David Gray, Carol Clugston, Anna Dominiczak and Alex Alderton, Roberto Amato, Jeffrey Barrett, Sonia Goncalves, Ewan Harrison, David K. Jackson, Ian Johnston, Dominic Kwiatkowski, Cordelia Langford, John Sillitoe on behalf of the Wellcome Sanger Institute COVID-19 Surveillance Team |
| EPI_ISL_3312080 | Berkshire and Surrey Pathology Services Lighthouse Laboratory | Wellcome Sanger Institute for the COVID-19 Genomics UK (COG-UK) Consortium | Berkshire and Surrey Pathology Services Lighthouse Laboratory and Alex Alderton, Roberto Amato, Jeffrey Barrett, Sonia Goncalves, Ewan Harrison, David K. Jackson, Ian Johnston, Dominic Kwiatkowski, Cordelia Langford, John Sillitoe on behalf of the Wellcome Sanger Institute COVID-19 Surveillance Team |
| EPI_ISL_3312081, EPI_ISL_3312082, EPI_ISL_3312083, EPI_ISL_3312084, EPI_ISL_3312085, EPI_ISL_3312086, EPI_ISL_3312087, EPI_ISL_3312088, EPI_ISL_3312089, EPI_ISL_3312090, EPI_ISL_3312091, EPI_ISL_3312092, EPI_ISL_3312093, EPI_ISL_3312094, EPI_ISL_3312095, EPI_ISL_3312096, EPI_ISL_3312097, EPI_ISL_3312098, EPI_ISL_3312099, EPI_ISL_3312100, EPI_ISL_3312101, EPI_ISL_3312102, EPI_ISL_3312103 | | | |
| see above | Lighthouse Lab in Glasgow | Wellcome Sanger Institute for the COVID-19 Genomics UK (COG-UK) Consortium | Harper VanSteenhouse, Yumi Kasai, David Gray, Carol Clugston, Anna Dominiczak and Alex Alderton, Roberto Amato, Jeffrey Barrett, Sonia Goncalves, Ewan Harrison, David K. Jackson, Ian Johnston, Dominic Kwiatkowski, Cordelia Langford, John Sillitoe on behalf of the Wellcome Sanger Institute COVID-19 Surveillance Team |
| EPI_ISL_3312104 | Berkshire and Surrey Pathology Services Lighthouse Laboratory | Wellcome Sanger Institute for the COVID-19 Genomics UK (COG-UK) Consortium | Berkshire and Surrey Pathology Services Lighthouse Laboratory and Alex Alderton, Roberto Amato, Jeffrey Barrett, Sonia Goncalves, Ewan Harrison, David K. Jackson, Ian Johnston, Dominic Kwiatkowski, Cordelia Langford, John Sillitoe on behalf of the Wellcome Sanger Institute COVID-19 Surveillance Team |
| EPI_ISL_3312105, EPI_ISL_3312106, EPI_ISL_3312107, EPI_ISL_3312108, EPI_ISL_3312109, EPI_ISL_3312110, EPI_ISL_3312111, EPI_ISL_3312112, EPI_ISL_3312113, EPI_ISL_3312114, EPI_ISL_3312115 | | | |
| see above | Lighthouse Lab in Glasgow | Wellcome Sanger Institute for the COVID-19 Genomics UK (COG-UK) Consortium | Harper VanSteenhouse, Yumi Kasai, David Gray, Carol Clugston, Anna Dominiczak and Alex Alderton, Roberto Amato, Jeffrey Barrett, Sonia Goncalves, Ewan Harrison, David K. Jackson, Ian Johnston, Dominic Kwiatkowski, Cordelia Langford, John Sillitoe on behalf of the Wellcome Sanger Institute COVID-19 Surveillance Team |
| EPI_ISL_3312116 | Berkshire and Surrey Pathology Services Lighthouse Laboratory | Wellcome Sanger Institute for the COVID-19 Genomics UK (COG-UK) Consortium | Berkshire and Surrey Pathology Services Lighthouse Laboratory and Alex Alderton, Roberto Amato, Jeffrey Barrett, Sonia Goncalves, Ewan Harrison, David K. Jackson, Ian Johnston, Dominic Kwiatkowski, Cordelia Langford, John Sillitoe on behalf of the Wellcome Sanger Institute COVID-19 Surveillance Team |
| EPI_ISL_3312117 | Lighthouse Lab in Glasgow | Wellcome Sanger Institute for the COVID-19 Genomics UK (COG-UK) Consortium | Harper VanSteenhouse, Yumi Kasai, David Gray, Carol Clugston, Anna Dominiczak and Alex Alderton, Roberto Amato, Jeffrey Barrett, Sonia Goncalves, Ewan Harrison, David K. Jackson, Ian Johnston, Dominic Kwiatkowski, Cordelia Langford, John Sillitoe on behalf of the Wellcome Sanger Institute COVID-19 Surveillance Team |
| EPI_ISL_3312118 | Berkshire and Surrey Pathology Services Lighthouse Laboratory | Wellcome Sanger Institute for the COVID-19 Genomics UK (COG-UK) Consortium | Berkshire and Surrey Pathology Services Lighthouse Laboratory and Alex Alderton, Roberto Amato, Jeffrey Barrett, Sonia Goncalves, Ewan Harrison, David K. Jackson, Ian Johnston, Dominic Kwiatkowski, Cordelia Langford, John Sillitoe on behalf of the Wellcome Sanger Institute COVID-19 Surveillance Team |
| EPI_ISL_3312119 | Lighthouse Lab in Glasgow | Wellcome Sanger Institute for the COVID-19 Genomics UK (COG-UK) Consortium | Harper VanSteenhouse, Yumi Kasai, David Gray, Carol Clugston, Anna Dominiczak and Alex Alderton, Roberto Amato, Jeffrey Barrett, Sonia Goncalves, Ewan Harrison, David K. Jackson, Ian Johnston, Dominic Kwiatkowski, Cordelia Langford, John Sillitoe on behalf of the Wellcome Sanger Institute COVID-19 Surveillance Team |
| EPI_ISL_3312120 | Berkshire and Surrey Pathology Services Lighthouse Laboratory | Wellcome Sanger Institute for the COVID-19 Genomics UK (COG-UK) Consortium | Berkshire and Surrey Pathology Services Lighthouse Laboratory and Alex Alderton, Roberto Amato, Jeffrey Barrett, Sonia Goncalves, Ewan Harrison, David K. Jackson, Ian Johnston, Dominic Kwiatkowski, Cordelia Langford, John Sillitoe on behalf of the Wellcome Sanger Institute COVID-19 Surveillance Team |
| EPI_ISL_3312121, EPI_ISL_3312122 | Lighthouse Lab in Glasgow | Wellcome Sanger Institute for the COVID-19 Genomics UK (COG-UK) Consortium | Harper VanSteenhouse, Yumi Kasai, David Gray, Carol Clugston, Anna Dominiczak and Alex Alderton, Roberto Amato, Jeffrey Barrett, Sonia Goncalves, Ewan Harrison, David K. Jackson, Ian Johnston, Dominic Kwiatkowski, Cordelia Langford, John Sillitoe on behalf of the Wellcome Sanger Institute COVID-19 Surveillance Team |
| EPI_ISL_3312123 | Berkshire and Surrey Pathology Services Lighthouse Laboratory | Wellcome Sanger Institute for the COVID-19 Genomics UK (COG-UK) Consortium | Berkshire and Surrey Pathology Services Lighthouse Laboratory and Alex Alderton, Roberto Amato, Jeffrey Barrett, Sonia Goncalves, Ewan Harrison, David K. Jackson, Ian Johnston, Dominic Kwiatkowski, Cordelia Langford, John Sillitoe on behalf of the Wellcome Sanger Institute COVID-19 Surveillance Team |
| EPI_ISL_3312124, EPI_ISL_3312125, EPI_ISL_3312126, EPI_ISL_3312127, EPI_ISL_3312128, EPI_ISL_3312129, EPI_ISL_3312130, EPI_ISL_3312131, EPI_ISL_3312132, EPI_ISL_3312133 | Lighthouse Lab in Glasgow | Wellcome Sanger Institute for the COVID-19 Genomics UK (COG-UK) Consortium | Harper VanSteenhouse, Yumi Kasai, David Gray, Carol Clugston, Anna Dominiczak and Alex Alderton, Roberto Amato, Jeffrey Barrett, Sonia Goncalves, Ewan Harrison, David K. Jackson, Ian Johnston, Dominic Kwiatkowski, Cordelia Langford, John Sillitoe on behalf of the Wellcome Sanger Institute COVID-19 Surveillance Team |
| EPI_ISL_3312134 | Berkshire and Surrey Pathology Services Lighthouse Laboratory | Wellcome Sanger Institute for the COVID-19 Genomics UK (COG-UK) Consortium | Berkshire and Surrey Pathology Services Lighthouse Laboratory and Alex Alderton, Roberto Amato, Jeffrey Barrett, Sonia Goncalves, Ewan Harrison, David K. Jackson, Ian Johnston, Dominic Kwiatkowski, Cordelia Langford, John Sillitoe on behalf of the Wellcome Sanger Institute COVID-19 Surveillance Team |
| EPI_ISL_3312135, EPI_ISL_3312136 | Lighthouse Lab in Glasgow | Wellcome Sanger Institute for the COVID-19 Genomics UK (COG-UK) Consortium | Harper VanSteenhouse, Yumi Kasai, David Gray, Carol Clugston, Anna Dominiczak and Alex Alderton, Roberto Amato, Jeffrey Barrett, Sonia Goncalves, Ewan Harrison, David K. Jackson, Ian Johnston, Dominic Kwiatkowski, Cordelia Langford, John Sillitoe on behalf of the Wellcome Sanger Institute COVID-19 Surveillance Team |
| EPI_ISL_3312137 | Berkshire and Surrey Pathology Services Lighthouse Laboratory | Wellcome Sanger Institute for the COVID-19 Genomics UK (COG-UK) Consortium | Berkshire and Surrey Pathology Services Lighthouse Laboratory and Alex Alderton, Roberto Amato, Jeffrey Barrett, Sonia Goncalves, Ewan Harrison, David K. Jackson, Ian Johnston, Dominic Kwiatkowski, Cordelia Langford, John Sillitoe on behalf of the Wellcome Sanger Institute COVID-19 Surveillance Team |
| EPI_ISL_3312138, EPI_ISL_3312139, EPI_ISL_3312140 | Lighthouse Lab in Glasgow | Wellcome Sanger Institute for the COVID-19 Genomics UK (COG-UK) Consortium | Harper VanSteenhouse, Yumi Kasai, David Gray, Carol Clugston, Anna Dominiczak and Alex Alderton, Roberto Amato, Jeffrey Barrett, Sonia Goncalves, Ewan Harrison, David K. Jackson, Ian Johnston, Dominic Kwiatkowski, Cordelia Langford, John Sillitoe on behalf of the Wellcome Sanger Institute COVID-19 Surveillance Team |
| EPI_ISL_3312141 | Berkshire and Surrey Pathology Services Lighthouse Laboratory | Wellcome Sanger Institute for the COVID-19 Genomics UK (COG-UK) Consortium | Berkshire and Surrey Pathology Services Lighthouse Laboratory and Alex Alderton, Roberto Amato, Jeffrey Barrett, Sonia Goncalves, Ewan Harrison, David K. Jackson, Ian Johnston, Dominic Kwiatkowski, Cordelia Langford, John Sillitoe on behalf of the Wellcome Sanger Institute COVID-19 Surveillance Team |
| EPI_ISL_3312142, EPI_ISL_3312143 | Lighthouse Lab in Glasgow | Wellcome Sanger Institute for the COVID-19 Genomics UK (COG-UK) Consortium | Harper VanSteenhouse, Yumi Kasai, David Gray, Carol Clugston, Anna Dominiczak and Alex Alderton, Roberto Amato, Jeffrey Barrett, Sonia Goncalves, Ewan Harrison, David K. Jackson, Ian Johnston, Dominic Kwiatkowski, Cordelia Langford, John Sillitoe on behalf of the Wellcome Sanger Institute COVID-19 Surveillance Team |
| EPI_ISL_3312144, EPI_ISL_3312145 | Berkshire and Surrey Pathology Services Lighthouse Laboratory | Wellcome Sanger Institute for the COVID-19 Genomics UK (COG-UK) Consortium | Berkshire and Surrey Pathology Services Lighthouse Laboratory and Alex Alderton, Roberto Amato, Jeffrey Barrett, Sonia Goncalves, Ewan Harrison, David K. Jackson, Ian Johnston, Dominic Kwiatkowski, Cordelia Langford, John Sillitoe on behalf of the Wellcome Sanger Institute COVID-19 Surveillance Team |
| EPI_ISL_3312146 | Lighthouse Lab in Glasgow | Wellcome Sanger Institute for the COVID-19 Genomics UK (COG-UK) Consortium | Harper VanSteenhouse, Yumi Kasai, David Gray, Carol Clugston, Anna Dominiczak and Alex Alderton, Roberto Amato, Jeffrey Barrett, Sonia Goncalves, Ewan Harrison, David K. Jackson, Ian Johnston, Dominic Kwiatkowski, Cordelia Langford, John Sillitoe on behalf of the Wellcome Sanger Institute COVID-19 Surveillance Team |
| EPI_ISL_3312147, | Berkshire and Surrey Pathology Services Lighthouse Laboratory | Wellcome Sanger Institute for the COVID-19 Genomics UK (COG-UK) | Berkshire and Surrey Pathology Services Lighthouse Laboratory and Alex Alderton, Roberto Amato, Jeffrey Barrett, Sonia Goncalves, Ewan Harrison, David K. Jackson, Ian Johnston, Dominic Kwiatkowski, Cordelia Langford, John Sillitoe on behalf |

| | | | |
|---|---|---|---|
| EPI_ISL_3312148 | | Consortium | of the Wellcome Sanger Institute COVID-19 Surveillance Team |
| EPI_ISL_3312149, EPI_ISL_3312150 | Lighthouse Lab in Glasgow | Wellcome Sanger Institute for the COVID-19 Genomics UK (COG-UK) Consortium | Harper VanSteenhouse, Yumi Kasai, David Gray, Carol Clugston, Anna Dominiczak and Alex Alderton, Roberto Amato, Jeffrey Barrett, Sonia Goncalves, Ewan Harrison, David K. Jackson, Ian Johnston, Dominic Kwiatkowski, Cordelia Langford, John Sillitoe on behalf of the Wellcome Sanger Institute COVID-19 Surveillance Team |
| EPI_ISL_3312151 | Berkshire and Surrey Pathology Services Lighthouse Laboratory | Wellcome Sanger Institute for the COVID-19 Genomics UK (COG-UK) Consortium | Berkshire and Surrey Pathology Services Lighthouse Laboratory and Alex Alderton, Roberto Amato, Jeffrey Barrett, Sonia Goncalves, Ewan Harrison, David K. Jackson, Ian Johnston, Dominic Kwiatkowski, Cordelia Langford, John Sillitoe on behalf of the Wellcome Sanger Institute COVID-19 Surveillance Team |
| EPI_ISL_3312152, EPI_ISL_3312153, EPI_ISL_3312154, EPI_ISL_3312155 | Lighthouse Lab in Glasgow | Wellcome Sanger Institute for the COVID-19 Genomics UK (COG-UK) Consortium | Harper VanSteenhouse, Yumi Kasai, David Gray, Carol Clugston, Anna Dominiczak and Alex Alderton, Roberto Amato, Jeffrey Barrett, Sonia Goncalves, Ewan Harrison, David K. Jackson, Ian Johnston, Dominic Kwiatkowski, Cordelia Langford, John Sillitoe on behalf of the Wellcome Sanger Institute COVID-19 Surveillance Team |
| EPI_ISL_3312156 | Berkshire and Surrey Pathology Services Lighthouse Laboratory | Wellcome Sanger Institute for the COVID-19 Genomics UK (COG-UK) Consortium | Berkshire and Surrey Pathology Services Lighthouse Laboratory and Alex Alderton, Roberto Amato, Jeffrey Barrett, Sonia Goncalves, Ewan Harrison, David K. Jackson, Ian Johnston, Dominic Kwiatkowski, Cordelia Langford, John Sillitoe on behalf of the Wellcome Sanger Institute COVID-19 Surveillance Team |
| EPI_ISL_3312157, EPI_ISL_3312158 | Lighthouse Lab in Glasgow | Wellcome Sanger Institute for the COVID-19 Genomics UK (COG-UK) Consortium | Harper VanSteenhouse, Yumi Kasai, David Gray, Carol Clugston, Anna Dominiczak and Alex Alderton, Roberto Amato, Jeffrey Barrett, Sonia Goncalves, Ewan Harrison, David K. Jackson, Ian Johnston, Dominic Kwiatkowski, Cordelia Langford, John Sillitoe on behalf of the Wellcome Sanger Institute COVID-19 Surveillance Team |
| EPI_ISL_3312159 | Berkshire and Surrey Pathology Services Lighthouse Laboratory | Wellcome Sanger Institute for the COVID-19 Genomics UK (COG-UK) Consortium | Berkshire and Surrey Pathology Services Lighthouse Laboratory and Alex Alderton, Roberto Amato, Jeffrey Barrett, Sonia Goncalves, Ewan Harrison, David K. Jackson, Ian Johnston, Dominic Kwiatkowski, Cordelia Langford, John Sillitoe on behalf of the Wellcome Sanger Institute COVID-19 Surveillance Team |
| EPI_ISL_3312160 | Lighthouse Lab in Glasgow | Wellcome Sanger Institute for the COVID-19 Genomics UK (COG-UK) Consortium | Harper VanSteenhouse, Yumi Kasai, David Gray, Carol Clugston, Anna Dominiczak and Alex Alderton, Roberto Amato, Jeffrey Barrett, Sonia Goncalves, Ewan Harrison, David K. Jackson, Ian Johnston, Dominic Kwiatkowski, Cordelia Langford, John Sillitoe on behalf of the Wellcome Sanger Institute COVID-19 Surveillance Team |
| EPI_ISL_3312161 | Berkshire and Surrey Pathology Services Lighthouse Laboratory | Wellcome Sanger Institute for the COVID-19 Genomics UK (COG-UK) Consortium | Berkshire and Surrey Pathology Services Lighthouse Laboratory and Alex Alderton, Roberto Amato, Jeffrey Barrett, Sonia Goncalves, Ewan Harrison, David K. Jackson, Ian Johnston, Dominic Kwiatkowski, Cordelia Langford, John Sillitoe on behalf of the Wellcome Sanger Institute COVID-19 Surveillance Team |
| EPI_ISL_3312162, EPI_ISL_3312163, EPI_ISL_3312164, EPI_ISL_3312165, EPI_ISL_3312166, EPI_ISL_3312167, EPI_ISL_3312168, EPI_ISL_3312169, EPI_ISL_3312170, EPI_ISL_3312171, EPI_ISL_3312172, EPI_ISL_3312173, EPI_ISL_3312174, EPI_ISL_3312175, EPI_ISL_3312176, EPI_ISL_3312177, EPI_ISL_3312178 | | | |
| see above | Lighthouse Lab in Glasgow | Wellcome Sanger Institute for the COVID-19 Genomics UK (COG-UK) Consortium | Harper VanSteenhouse, Yumi Kasai, David Gray, Carol Clugston, Anna Dominiczak and Alex Alderton, Roberto Amato, Jeffrey Barrett, Sonia Goncalves, Ewan Harrison, David K. Jackson, Ian Johnston, Dominic Kwiatkowski, Cordelia Langford, John Sillitoe on behalf of the Wellcome Sanger Institute COVID-19 Surveillance Team |
| EPI_ISL_3312179 | Berkshire and Surrey Pathology Services Lighthouse Laboratory | Wellcome Sanger Institute for the COVID-19 Genomics UK (COG-UK) Consortium | Berkshire and Surrey Pathology Services Lighthouse Laboratory and Alex Alderton, Roberto Amato, Jeffrey Barrett, Sonia Goncalves, Ewan Harrison, David K. Jackson, Ian Johnston, Dominic Kwiatkowski, Cordelia Langford, John Sillitoe on behalf of the Wellcome Sanger Institute COVID-19 Surveillance Team |
| EPI_ISL_3312180, EPI_ISL_3312181, EPI_ISL_3312182, EPI_ISL_3312183, EPI_ISL_3312184, EPI_ISL_3312185, EPI_ISL_3312186, EPI_ISL_3312187, EPI_ISL_3312188, EPI_ISL_3312189, EPI_ISL_3312190, EPI_ISL_3312191, EPI_ISL_3312192, EPI_ISL_3312193, EPI_ISL_3312194, EPI_ISL_3312195 | | | |
| see above | Lighthouse Lab in Glasgow | Wellcome Sanger Institute for the COVID-19 Genomics UK (COG-UK) Consortium | Harper VanSteenhouse, Yumi Kasai, David Gray, Carol Clugston, Anna Dominiczak and Alex Alderton, Roberto Amato, Jeffrey Barrett, Sonia Goncalves, Ewan Harrison, David K. Jackson, Ian Johnston, Dominic Kwiatkowski, Cordelia Langford, John Sillitoe on behalf of the Wellcome Sanger Institute COVID-19 Surveillance Team |
| EPI_ISL_3312196 | Berkshire and Surrey Pathology Services Lighthouse Laboratory | Wellcome Sanger Institute for the COVID-19 Genomics UK (COG-UK) Consortium | Berkshire and Surrey Pathology Services Lighthouse Laboratory and Alex Alderton, Roberto Amato, Jeffrey Barrett, Sonia Goncalves, Ewan Harrison, David K. Jackson, Ian Johnston, Dominic Kwiatkowski, Cordelia Langford, John Sillitoe on behalf of the Wellcome Sanger Institute COVID-19 Surveillance Team |
| EPI_ISL_3312197, EPI_ISL_3312198, EPI_ISL_3312199 | Lighthouse Lab in Glasgow | Wellcome Sanger Institute for the COVID-19 Genomics UK (COG-UK) Consortium | Harper VanSteenhouse, Yumi Kasai, David Gray, Carol Clugston, Anna Dominiczak and Alex Alderton, Roberto Amato, Jeffrey Barrett, Sonia Goncalves, Ewan Harrison, David K. Jackson, Ian Johnston, Dominic Kwiatkowski, Cordelia Langford, John Sillitoe on behalf of the Wellcome Sanger Institute COVID-19 Surveillance Team |
| EPI_ISL_3312200 | Berkshire and Surrey Pathology Services Lighthouse Laboratory | Wellcome Sanger Institute for the COVID-19 Genomics UK (COG-UK) Consortium | Berkshire and Surrey Pathology Services Lighthouse Laboratory and Alex Alderton, Roberto Amato, Jeffrey Barrett, Sonia Goncalves, Ewan Harrison, David K. Jackson, Ian Johnston, Dominic Kwiatkowski, Cordelia Langford, John Sillitoe on behalf of the Wellcome Sanger Institute COVID-19 Surveillance Team |
| EPI_ISL_3312201, EPI_ISL_3312202, EPI_ISL_3312203, EPI_ISL_3312204, EPI_ISL_3312205, EPI_ISL_3312206, EPI_ISL_3312207 | Lighthouse Lab in Glasgow | Wellcome Sanger Institute for the COVID-19 Genomics UK (COG-UK) Consortium | Harper VanSteenhouse, Yumi Kasai, David Gray, Carol Clugston, Anna Dominiczak and Alex Alderton, Roberto Amato, Jeffrey Barrett, Sonia Goncalves, Ewan Harrison, David K. Jackson, Ian Johnston, Dominic Kwiatkowski, Cordelia Langford, John Sillitoe on behalf of the Wellcome Sanger Institute COVID-19 Surveillance Team |
| EPI_ISL_3312208 | Berkshire and Surrey Pathology Services Lighthouse Laboratory | Wellcome Sanger Institute for the COVID-19 Genomics UK (COG-UK) Consortium | Berkshire and Surrey Pathology Services Lighthouse Laboratory and Alex Alderton, Roberto Amato, Jeffrey Barrett, Sonia Goncalves, Ewan Harrison, David K. Jackson, Ian Johnston, Dominic Kwiatkowski, Cordelia Langford, John Sillitoe on behalf of the Wellcome Sanger Institute COVID-19 Surveillance Team |
| EPI_ISL_3312209, EPI_ISL_3312210, EPI_ISL_3312211 | Lighthouse Lab in Glasgow | Wellcome Sanger Institute for the COVID-19 Genomics UK (COG-UK) Consortium | Harper VanSteenhouse, Yumi Kasai, David Gray, Carol Clugston, Anna Dominiczak and Alex Alderton, Roberto Amato, Jeffrey Barrett, Sonia Goncalves, Ewan Harrison, David K. Jackson, Ian Johnston, Dominic Kwiatkowski, Cordelia Langford, John Sillitoe on behalf of the Wellcome Sanger Institute COVID-19 Surveillance Team |
| EPI_ISL_3312212 | Berkshire and Surrey Pathology Services Lighthouse Laboratory | Wellcome Sanger Institute for the COVID-19 Genomics UK (COG-UK) Consortium | Berkshire and Surrey Pathology Services Lighthouse Laboratory and Alex Alderton, Roberto Amato, Jeffrey Barrett, Sonia Goncalves, Ewan Harrison, David K. Jackson, Ian Johnston, Dominic Kwiatkowski, Cordelia Langford, John Sillitoe on behalf of the Wellcome Sanger Institute COVID-19 Surveillance Team |
| EPI_ISL_3312213, EPI_ISL_3312214, EPI_ISL_3312215 | Lighthouse Lab in Glasgow | Wellcome Sanger Institute for the COVID-19 Genomics UK (COG-UK) Consortium | Harper VanSteenhouse, Yumi Kasai, David Gray, Carol Clugston, Anna Dominiczak and Alex Alderton, Roberto Amato, Jeffrey Barrett, Sonia Goncalves, Ewan Harrison, David K. Jackson, Ian Johnston, Dominic Kwiatkowski, Cordelia Langford, John Sillitoe on behalf of the Wellcome Sanger Institute COVID-19 Surveillance Team |
| EPI_ISL_3312216 | Berkshire and Surrey Pathology Services Lighthouse Laboratory | Wellcome Sanger Institute for the COVID-19 Genomics UK (COG-UK) Consortium | Berkshire and Surrey Pathology Services Lighthouse Laboratory and Alex Alderton, Roberto Amato, Jeffrey Barrett, Sonia Goncalves, Ewan Harrison, David K. Jackson, Ian Johnston, Dominic Kwiatkowski, Cordelia Langford, John Sillitoe on behalf of the Wellcome Sanger Institute COVID-19 Surveillance Team |
| EPI_ISL_3312217, EPI_ISL_3312218, EPI_ISL_3312219, EPI_ISL_3312220, EPI_ISL_3312221 | Lighthouse Lab in Glasgow | Wellcome Sanger Institute for the COVID-19 Genomics UK (COG-UK) Consortium | Harper VanSteenhouse, Yumi Kasai, David Gray, Carol Clugston, Anna Dominiczak and Alex Alderton, Roberto Amato, Jeffrey Barrett, Sonia Goncalves, Ewan Harrison, David K. Jackson, Ian Johnston, Dominic Kwiatkowski, Cordelia Langford, John Sillitoe on behalf of the Wellcome Sanger Institute COVID-19 Surveillance Team |
| EPI_ISL_3312222 | Berkshire and Surrey Pathology Services Lighthouse Laboratory | Wellcome Sanger Institute for the COVID-19 Genomics UK (COG-UK) Consortium | Berkshire and Surrey Pathology Services Lighthouse Laboratory and Alex Alderton, Roberto Amato, Jeffrey Barrett, Sonia Goncalves, Ewan Harrison, David K. Jackson, Ian Johnston, Dominic Kwiatkowski, Cordelia Langford, John Sillitoe on behalf of the Wellcome Sanger Institute COVID-19 Surveillance Team |
| EPI_ISL_3312223, EPI_ISL_3312224, EPI_ISL_3312225, EPI_ISL_3312226, EPI_ISL_3312227, EPI_ISL_3312228, EPI_ISL_3312229, EPI_ISL_3312230, EPI_ISL_3312231, EPI_ISL_3312232 | Lighthouse Lab in Glasgow | Wellcome Sanger Institute for the COVID-19 Genomics UK (COG-UK) Consortium | Harper VanSteenhouse, Yumi Kasai, David Gray, Carol Clugston, Anna Dominiczak and Alex Alderton, Roberto Amato, Jeffrey Barrett, Sonia Goncalves, Ewan Harrison, David K. Jackson, Ian Johnston, Dominic Kwiatkowski, Cordelia Langford, John Sillitoe on behalf of the Wellcome Sanger Institute COVID-19 Surveillance Team |
| EPI_ISL_3312233 | Berkshire and Surrey Pathology Services Lighthouse Laboratory | Wellcome Sanger Institute for the COVID-19 Genomics UK (COG-UK) Consortium | Berkshire and Surrey Pathology Services Lighthouse Laboratory and Alex Alderton, Roberto Amato, Jeffrey Barrett, Sonia Goncalves, Ewan Harrison, David K. Jackson, Ian Johnston, Dominic Kwiatkowski, Cordelia Langford, John Sillitoe on behalf of the Wellcome Sanger Institute COVID-19 Surveillance Team |
| EPI_ISL_3312234, EPI_ISL_3312235, EPI_ISL_3312236, EPI_ISL_3312237, EPI_ISL_3312238, EPI_ISL_3312239, EPI_ISL_3312240, EPI_ISL_3312241, EPI_ISL_3312242, EPI_ISL_3312243, EPI_ISL_3312244, EPI_ISL_3312245, EPI_ISL_3312246 | | | |
| see above | Lighthouse Lab in Glasgow | Wellcome Sanger Institute for the COVID-19 Genomics UK (COG-UK) Consortium | Harper VanSteenhouse, Yumi Kasai, David Gray, Carol Clugston, Anna Dominiczak and Alex Alderton, Roberto Amato, Jeffrey Barrett, Sonia Goncalves, Ewan Harrison, David K. Jackson, Ian Johnston, Dominic Kwiatkowski, Cordelia Langford, John Sillitoe on behalf of the Wellcome Sanger Institute COVID-19 Surveillance Team |
| EPI_ISL_3312247 | Berkshire and Surrey Pathology Services Lighthouse Laboratory | Wellcome Sanger Institute for the COVID-19 Genomics UK (COG-UK) Consortium | Berkshire and Surrey Pathology Services Lighthouse Laboratory and Alex Alderton, Roberto Amato, Jeffrey Barrett, Sonia Goncalves, Ewan Harrison, David K. Jackson, Ian Johnston, Dominic Kwiatkowski, Cordelia Langford, John Sillitoe on behalf of the Wellcome Sanger Institute COVID-19 Surveillance Team |
| EPI_ISL_3312248, EPI_ISL_3312249, EPI_ISL_3312250, EPI_ISL_3312251, EPI_ISL_3312252, EPI_ISL_3312253 | Lighthouse Lab in Glasgow | Wellcome Sanger Institute for the COVID-19 Genomics UK (COG-UK) Consortium | Harper VanSteenhouse, Yumi Kasai, David Gray, Carol Clugston, Anna Dominiczak and Alex Alderton, Roberto Amato, Jeffrey Barrett, Sonia Goncalves, Ewan Harrison, David K. Jackson, Ian Johnston, Dominic Kwiatkowski, Cordelia Langford, John Sillitoe on behalf of the Wellcome Sanger Institute COVID-19 Surveillance Team |
| EPI_ISL_3312254, EPI_ISL_3312255, EPI_ISL_3312256, EPI_ISL_3312257, EPI_ISL_3312258, EPI_ISL_3312259, EPI_ISL_3312260, EPI_ISL_3312261 | Berkshire and Surrey Pathology Services Lighthouse Laboratory | Wellcome Sanger Institute for the COVID-19 Genomics UK (COG-UK) Consortium | Berkshire and Surrey Pathology Services Lighthouse Laboratory and Alex Alderton, Roberto Amato, Jeffrey Barrett, Sonia Goncalves, Ewan Harrison, David K. Jackson, Ian Johnston, Dominic Kwiatkowski, Cordelia Langford, John Sillitoe on behalf of the Wellcome Sanger Institute COVID-19 Surveillance Team |
| EPI_ISL_3312262, EPI_ISL_3312263, EPI_ISL_3312264 | Lighthouse Lab in Glasgow | Wellcome Sanger Institute for the COVID-19 Genomics UK (COG-UK) Consortium | Harper VanSteenhouse, Yumi Kasai, David Gray, Carol Clugston, Anna Dominiczak and Alex Alderton, Roberto Amato, Jeffrey Barrett, Sonia Goncalves, Ewan Harrison, David K. Jackson, Ian Johnston, Dominic Kwiatkowski, Cordelia Langford, John Sillitoe on behalf of the Wellcome Sanger Institute COVID-19 Surveillance Team |
| EPI_ISL_3312265 | Berkshire and Surrey Pathology Services Lighthouse Laboratory | Wellcome Sanger Institute for the COVID-19 Genomics UK (COG-UK) | Berkshire and Surrey Pathology Services Lighthouse Laboratory and Alex Alderton, Roberto Amato, Jeffrey Barrett, Sonia Goncalves, Ewan Harrison, David K. Jackson, Ian Johnston, Dominic Kwiatkowski, Cordelia Langford, John Sillitoe on behalf |

| | | | |
|---|---|---|---|
| | | Consortium | of the Wellcome Sanger Institute COVID-19 Surveillance Team |
| EPI_ISL_3312266, EPI_ISL_3312267 | Lighthouse Lab in Glasgow | Wellcome Sanger Institute for the COVID-19 Genomics UK (COG-UK) Consortium | Harper VanSteenhouse, Yumi Kasai, David Gray, Carol Clugston, Anna Dominiczak and Alex Alderton, Roberto Amato, Jeffrey Barrett, Sonia Goncalves, Ewan Harrison, David K. Jackson, Ian Johnston, Dominic Kwiatkowski, Cordelia Langford, John Sillitoe on behalf of the Wellcome Sanger Institute COVID-19 Surveillance Team |
| EPI_ISL_3312268 | Berkshire and Surrey Pathology Services Lighthouse Laboratory | Wellcome Sanger Institute for the COVID-19 Genomics UK (COG-UK) Consortium | Berkshire and Surrey Pathology Services Lighthouse Laboratory and Alex Alderton, Roberto Amato, Jeffrey Barrett, Sonia Goncalves, Ewan Harrison, David K. Jackson, Ian Johnston, Dominic Kwiatkowski, Cordelia Langford, John Sillitoe on behalf of the Wellcome Sanger Institute COVID-19 Surveillance Team |
| EPI_ISL_3312269, EPI_ISL_3312270, EPI_ISL_3312271 | Lighthouse Lab in Glasgow | Wellcome Sanger Institute for the COVID-19 Genomics UK (COG-UK) Consortium | Harper VanSteenhouse, Yumi Kasai, David Gray, Carol Clugston, Anna Dominiczak and Alex Alderton, Roberto Amato, Jeffrey Barrett, Sonia Goncalves, Ewan Harrison, David K. Jackson, Ian Johnston, Dominic Kwiatkowski, Cordelia Langford, John Sillitoe on behalf of the Wellcome Sanger Institute COVID-19 Surveillance Team |
| EPI_ISL_3312272 | Berkshire and Surrey Pathology Services Lighthouse Laboratory | Wellcome Sanger Institute for the COVID-19 Genomics UK (COG-UK) Consortium | Berkshire and Surrey Pathology Services Lighthouse Laboratory and Alex Alderton, Roberto Amato, Jeffrey Barrett, Sonia Goncalves, Ewan Harrison, David K. Jackson, Ian Johnston, Dominic Kwiatkowski, Cordelia Langford, John Sillitoe on behalf of the Wellcome Sanger Institute COVID-19 Surveillance Team |
| EPI_ISL_3312273, EPI_ISL_3312274, EPI_ISL_3312275, EPI_ISL_3312276 | Lighthouse Lab in Glasgow | Wellcome Sanger Institute for the COVID-19 Genomics UK (COG-UK) Consortium | Harper VanSteenhouse, Yumi Kasai, David Gray, Carol Clugston, Anna Dominiczak and Alex Alderton, Roberto Amato, Jeffrey Barrett, Sonia Goncalves, Ewan Harrison, David K. Jackson, Ian Johnston, Dominic Kwiatkowski, Cordelia Langford, John Sillitoe on behalf of the Wellcome Sanger Institute COVID-19 Surveillance Team |
| EPI_ISL_3312277, EPI_ISL_3312278 | Berkshire and Surrey Pathology Services Lighthouse Laboratory | Wellcome Sanger Institute for the COVID-19 Genomics UK (COG-UK) Consortium | Berkshire and Surrey Pathology Services Lighthouse Laboratory and Alex Alderton, Roberto Amato, Jeffrey Barrett, Sonia Goncalves, Ewan Harrison, David K. Jackson, Ian Johnston, Dominic Kwiatkowski, Cordelia Langford, John Sillitoe on behalf of the Wellcome Sanger Institute COVID-19 Surveillance Team |
| EPI_ISL_3312279, EPI_ISL_3312280, EPI_ISL_3312281 | Lighthouse Lab in Glasgow | Wellcome Sanger Institute for the COVID-19 Genomics UK (COG-UK) Consortium | Harper VanSteenhouse, Yumi Kasai, David Gray, Carol Clugston, Anna Dominiczak and Alex Alderton, Roberto Amato, Jeffrey Barrett, Sonia Goncalves, Ewan Harrison, David K. Jackson, Ian Johnston, Dominic Kwiatkowski, Cordelia Langford, John Sillitoe on behalf of the Wellcome Sanger Institute COVID-19 Surveillance Team |
| EPI_ISL_3312282 | Berkshire and Surrey Pathology Services Lighthouse Laboratory | Wellcome Sanger Institute for the COVID-19 Genomics UK (COG-UK) Consortium | Berkshire and Surrey Pathology Services Lighthouse Laboratory and Alex Alderton, Roberto Amato, Jeffrey Barrett, Sonia Goncalves, Ewan Harrison, David K. Jackson, Ian Johnston, Dominic Kwiatkowski, Cordelia Langford, John Sillitoe on behalf of the Wellcome Sanger Institute COVID-19 Surveillance Team |
| EPI_ISL_3312283, EPI_ISL_3312284 | Lighthouse Lab in Glasgow | Wellcome Sanger Institute for the COVID-19 Genomics UK (COG-UK) Consortium | Harper VanSteenhouse, Yumi Kasai, David Gray, Carol Clugston, Anna Dominiczak and Alex Alderton, Roberto Amato, Jeffrey Barrett, Sonia Goncalves, Ewan Harrison, David K. Jackson, Ian Johnston, Dominic Kwiatkowski, Cordelia Langford, John Sillitoe on behalf of the Wellcome Sanger Institute COVID-19 Surveillance Team |
| EPI_ISL_3312285 | Berkshire and Surrey Pathology Services Lighthouse Laboratory | Wellcome Sanger Institute for the COVID-19 Genomics UK (COG-UK) Consortium | Berkshire and Surrey Pathology Services Lighthouse Laboratory and Alex Alderton, Roberto Amato, Jeffrey Barrett, Sonia Goncalves, Ewan Harrison, David K. Jackson, Ian Johnston, Dominic Kwiatkowski, Cordelia Langford, John Sillitoe on behalf of the Wellcome Sanger Institute COVID-19 Surveillance Team |
| EPI_ISL_3312286 | Lighthouse Lab in Glasgow | Wellcome Sanger Institute for the COVID-19 Genomics UK (COG-UK) Consortium | Harper VanSteenhouse, Yumi Kasai, David Gray, Carol Clugston, Anna Dominiczak and Alex Alderton, Roberto Amato, Jeffrey Barrett, Sonia Goncalves, Ewan Harrison, David K. Jackson, Ian Johnston, Dominic Kwiatkowski, Cordelia Langford, John Sillitoe on behalf of the Wellcome Sanger Institute COVID-19 Surveillance Team |
| EPI_ISL_3312287 | Berkshire and Surrey Pathology Services Lighthouse Laboratory | Wellcome Sanger Institute for the COVID-19 Genomics UK (COG-UK) Consortium | Berkshire and Surrey Pathology Services Lighthouse Laboratory and Alex Alderton, Roberto Amato, Jeffrey Barrett, Sonia Goncalves, Ewan Harrison, David K. Jackson, Ian Johnston, Dominic Kwiatkowski, Cordelia Langford, John Sillitoe on behalf of the Wellcome Sanger Institute COVID-19 Surveillance Team |
| EPI_ISL_3312288 | Lighthouse Lab in Glasgow | Wellcome Sanger Institute for the COVID-19 Genomics UK (COG-UK) Consortium | Harper VanSteenhouse, Yumi Kasai, David Gray, Carol Clugston, Anna Dominiczak and Alex Alderton, Roberto Amato, Jeffrey Barrett, Sonia Goncalves, Ewan Harrison, David K. Jackson, Ian Johnston, Dominic Kwiatkowski, Cordelia Langford, John Sillitoe on behalf of the Wellcome Sanger Institute COVID-19 Surveillance Team |
| EPI_ISL_3312289 | Lighthouse Lab in Alderley Park | Wellcome Sanger Institute for the COVID-19 Genomics UK (COG-UK) Consortium | Jacquelyn Wynn, Mairead Hyland, The Lighthouse Lab in Alderley Park and Alex Alderton, Roberto Amato, Jeffrey Barrett, Sonia Goncalves, Ewan Harrison, David K. Jackson, Ian Johnston, Dominic Kwiatkowski, Cordelia Langford, John Sillitoe on behalf of the Wellcome Sanger Institute COVID-19 Surveillance Team |
| EPI_ISL_3312290, EPI_ISL_3312291 | Lighthouse Laboratory Plymouth | Wellcome Sanger Institute for the COVID-19 Genomics UK (COG-UK) Consortium | Lighthouse Laboratory Plymouth and Alex Alderton, Roberto Amato, Jeffrey Barrett, Sonia Goncalves, Ewan Harrison, David K. Jackson, Ian Johnston, Dominic Kwiatkowski, Cordelia Langford, John Sillitoe on behalf of the Wellcome Sanger Institute COVID-19 Surveillance Team |
| EPI_ISL_3312292 | Lighthouse Lab in Glasgow | Wellcome Sanger Institute for the COVID-19 Genomics UK (COG-UK) Consortium | Harper VanSteenhouse, Yumi Kasai, David Gray, Carol Clugston, Anna Dominiczak and Alex Alderton, Roberto Amato, Jeffrey Barrett, Sonia Goncalves, Ewan Harrison, David K. Jackson, Ian Johnston, Dominic Kwiatkowski, Cordelia Langford, John Sillitoe on behalf of the Wellcome Sanger Institute COVID-19 Surveillance Team |
| EPI_ISL_3312293 | Lighthouse Lab in Alderley Park | Wellcome Sanger Institute for the COVID-19 Genomics UK (COG-UK) Consortium | Jacquelyn Wynn, Mairead Hyland, The Lighthouse Lab in Alderley Park and Alex Alderton, Roberto Amato, Jeffrey Barrett, Sonia Goncalves, Ewan Harrison, David K. Jackson, Ian Johnston, Dominic Kwiatkowski, Cordelia Langford, John Sillitoe on behalf of the Wellcome Sanger Institute COVID-19 Surveillance Team |
| EPI_ISL_3312294 | Lighthouse Lab in Glasgow | Wellcome Sanger Institute for the COVID-19 Genomics UK (COG-UK) Consortium | Harper VanSteenhouse, Yumi Kasai, David Gray, Carol Clugston, Anna Dominiczak and Alex Alderton, Roberto Amato, Jeffrey Barrett, Sonia Goncalves, Ewan Harrison, David K. Jackson, Ian Johnston, Dominic Kwiatkowski, Cordelia Langford, John Sillitoe on behalf of the Wellcome Sanger Institute COVID-19 Surveillance Team |
| EPI_ISL_3312295 | Lighthouse Lab in Alderley Park | Wellcome Sanger Institute for the COVID-19 Genomics UK (COG-UK) Consortium | Jacquelyn Wynn, Mairead Hyland, The Lighthouse Lab in Alderley Park and Alex Alderton, Roberto Amato, Jeffrey Barrett, Sonia Goncalves, Ewan Harrison, David K. Jackson, Ian Johnston, Dominic Kwiatkowski, Cordelia Langford, John Sillitoe on behalf of the Wellcome Sanger Institute COVID-19 Surveillance Team |
| EPI_ISL_3312296, EPI_ISL_3312297, EPI_ISL_3312298, EPI_ISL_3312299, EPI_ISL_3312300 | Lighthouse Lab in Glasgow | Wellcome Sanger Institute for the COVID-19 Genomics UK (COG-UK) Consortium | Harper VanSteenhouse, Yumi Kasai, David Gray, Carol Clugston, Anna Dominiczak and Alex Alderton, Roberto Amato, Jeffrey Barrett, Sonia Goncalves, Ewan Harrison, David K. Jackson, Ian Johnston, Dominic Kwiatkowski, Cordelia Langford, John Sillitoe on behalf of the Wellcome Sanger Institute COVID-19 Surveillance Team |
| EPI_ISL_3312301 | Lighthouse Laboratory Plymouth | Wellcome Sanger Institute for the COVID-19 Genomics UK (COG-UK) Consortium | Lighthouse Laboratory Plymouth and Alex Alderton, Roberto Amato, Jeffrey Barrett, Sonia Goncalves, Ewan Harrison, David K. Jackson, Ian Johnston, Dominic Kwiatkowski, Cordelia Langford, John Sillitoe on behalf of the Wellcome Sanger Institute COVID-19 Surveillance Team |
| EPI_ISL_3312302 | Lighthouse Lab in Alderley Park | Wellcome Sanger Institute for the COVID-19 Genomics UK (COG-UK) Consortium | Jacquelyn Wynn, Mairead Hyland, The Lighthouse Lab in Alderley Park and Alex Alderton, Roberto Amato, Jeffrey Barrett, Sonia Goncalves, Ewan Harrison, David K. Jackson, Ian Johnston, Dominic Kwiatkowski, Cordelia Langford, John Sillitoe on behalf of the Wellcome Sanger Institute COVID-19 Surveillance Team |
| EPI_ISL_3312303 | Lighthouse Laboratory Plymouth | Wellcome Sanger Institute for the COVID-19 Genomics UK (COG-UK) Consortium | Lighthouse Laboratory Plymouth and Alex Alderton, Roberto Amato, Jeffrey Barrett, Sonia Goncalves, Ewan Harrison, David K. Jackson, Ian Johnston, Dominic Kwiatkowski, Cordelia Langford, John Sillitoe on behalf of the Wellcome Sanger Institute COVID-19 Surveillance Team |
| EPI_ISL_3312304 | Lighthouse Lab in Alderley Park | Wellcome Sanger Institute for the COVID-19 Genomics UK (COG-UK) Consortium | Jacquelyn Wynn, Mairead Hyland, The Lighthouse Lab in Alderley Park and Alex Alderton, Roberto Amato, Jeffrey Barrett, Sonia Goncalves, Ewan Harrison, David K. Jackson, Ian Johnston, Dominic Kwiatkowski, Cordelia Langford, John Sillitoe on behalf of the Wellcome Sanger Institute COVID-19 Surveillance Team |
| EPI_ISL_3312305 | Lighthouse Lab in Glasgow | Wellcome Sanger Institute for the COVID-19 Genomics UK (COG-UK) Consortium | Harper VanSteenhouse, Yumi Kasai, David Gray, Carol Clugston, Anna Dominiczak and Alex Alderton, Roberto Amato, Jeffrey Barrett, Sonia Goncalves, Ewan Harrison, David K. Jackson, Ian Johnston, Dominic Kwiatkowski, Cordelia Langford, John Sillitoe on behalf of the Wellcome Sanger Institute COVID-19 Surveillance Team |
| EPI_ISL_3312306 | Lighthouse Laboratory Plymouth | Wellcome Sanger Institute for the COVID-19 Genomics UK (COG-UK) Consortium | Lighthouse Laboratory Plymouth and Alex Alderton, Roberto Amato, Jeffrey Barrett, Sonia Goncalves, Ewan Harrison, David K. Jackson, Ian Johnston, Dominic Kwiatkowski, Cordelia Langford, John Sillitoe on behalf of the Wellcome Sanger Institute COVID-19 Surveillance Team |
| EPI_ISL_3312307, EPI_ISL_3312308, EPI_ISL_3312309 | Lighthouse Lab in Glasgow | Wellcome Sanger Institute for the COVID-19 Genomics UK (COG-UK) Consortium | Harper VanSteenhouse, Yumi Kasai, David Gray, Carol Clugston, Anna Dominiczak and Alex Alderton, Roberto Amato, Jeffrey Barrett, Sonia Goncalves, Ewan Harrison, David K. Jackson, Ian Johnston, Dominic Kwiatkowski, Cordelia Langford, John Sillitoe on behalf of the Wellcome Sanger Institute COVID-19 Surveillance Team |
| EPI_ISL_3312310 | Lighthouse Lab in Alderley Park | Wellcome Sanger Institute for the COVID-19 Genomics UK (COG-UK) Consortium | Jacquelyn Wynn, Mairead Hyland, The Lighthouse Lab in Alderley Park and Alex Alderton, Roberto Amato, Jeffrey Barrett, Sonia Goncalves, Ewan Harrison, David K. Jackson, Ian Johnston, Dominic Kwiatkowski, Cordelia Langford, John Sillitoe on behalf of the Wellcome Sanger Institute COVID-19 Surveillance Team |
| EPI_ISL_3312311, EPI_ISL_3312312 | Lighthouse Laboratory Plymouth | Wellcome Sanger Institute for the COVID-19 Genomics UK (COG-UK) Consortium | Lighthouse Laboratory Plymouth and Alex Alderton, Roberto Amato, Jeffrey Barrett, Sonia Goncalves, Ewan Harrison, David K. Jackson, Ian Johnston, Dominic Kwiatkowski, Cordelia Langford, John Sillitoe on behalf of the Wellcome Sanger Institute COVID-19 Surveillance Team |
| EPI_ISL_3312313 | Lighthouse Lab in Glasgow | Wellcome Sanger Institute for the COVID-19 Genomics UK (COG-UK) Consortium | Harper VanSteenhouse, Yumi Kasai, David Gray, Carol Clugston, Anna Dominiczak and Alex Alderton, Roberto Amato, Jeffrey Barrett, Sonia Goncalves, Ewan Harrison, David K. Jackson, Ian Johnston, Dominic Kwiatkowski, Cordelia Langford, John Sillitoe on behalf of the Wellcome Sanger Institute COVID-19 Surveillance Team |
| EPI_ISL_3312314 | Lighthouse Lab in Alderley Park | Wellcome Sanger Institute for the COVID-19 Genomics UK (COG-UK) Consortium | Jacquelyn Wynn, Mairead Hyland, The Lighthouse Lab in Alderley Park and Alex Alderton, Roberto Amato, Jeffrey Barrett, Sonia Goncalves, Ewan Harrison, David K. Jackson, Ian Johnston, Dominic Kwiatkowski, Cordelia Langford, John Sillitoe on behalf of the Wellcome Sanger Institute COVID-19 Surveillance Team |
| EPI_ISL_3312315, EPI_ISL_3312316 | Lighthouse Laboratory Plymouth | Wellcome Sanger Institute for the COVID-19 Genomics UK (COG-UK) Consortium | Lighthouse Laboratory Plymouth and Alex Alderton, Roberto Amato, Jeffrey Barrett, Sonia Goncalves, Ewan Harrison, David K. Jackson, Ian Johnston, Dominic Kwiatkowski, Cordelia Langford, John Sillitoe on behalf of the Wellcome Sanger Institute COVID-19 Surveillance Team |
| EPI_ISL_3312317, EPI_ISL_3312318 | Lighthouse Lab in Alderley Park | Wellcome Sanger Institute for the COVID-19 Genomics UK (COG-UK) Consortium | Jacquelyn Wynn, Mairead Hyland, The Lighthouse Lab in Alderley Park and Alex Alderton, Roberto Amato, Jeffrey Barrett, Sonia Goncalves, Ewan Harrison, David K. Jackson, Ian Johnston, Dominic Kwiatkowski, Cordelia Langford, John Sillitoe on behalf of the Wellcome Sanger Institute COVID-19 Surveillance Team |
| EPI_ISL_3312319 | Lighthouse Laboratory Plymouth | Wellcome Sanger Institute for the COVID-19 Genomics UK (COG-UK) Consortium | Lighthouse Laboratory Plymouth and Alex Alderton, Roberto Amato, Jeffrey Barrett, Sonia Goncalves, Ewan Harrison, David K. Jackson, Ian Johnston, Dominic Kwiatkowski, Cordelia Langford, John Sillitoe on behalf of the Wellcome Sanger Institute COVID-19 Surveillance Team |
| EPI_ISL_3312320, EPI_ISL_3312321, EPI_ISL_3312322 | Lighthouse Lab in Glasgow | Wellcome Sanger Institute for the COVID-19 Genomics UK (COG-UK) Consortium | Harper VanSteenhouse, Yumi Kasai, David Gray, Carol Clugston, Anna Dominiczak and Alex Alderton, Roberto Amato, Jeffrey Barrett, Sonia Goncalves, Ewan Harrison, David K. Jackson, Ian Johnston, Dominic Kwiatkowski, Cordelia Langford, John Sillitoe on behalf of the Wellcome Sanger Institute COVID-19 Surveillance Team |
| EPI_ISL_3312323 | Lighthouse Laboratory Plymouth | Wellcome Sanger Institute for the COVID-19 Genomics UK (COG-UK) Consortium | Lighthouse Laboratory Plymouth and Alex Alderton, Roberto Amato, Jeffrey Barrett, Sonia Goncalves, Ewan Harrison, David K. Jackson, Ian Johnston, Dominic Kwiatkowski, Cordelia Langford, John Sillitoe on behalf of the Wellcome Sanger Institute COVID-19 Surveillance Team |
| EPI_ISL_3312324, EPI_ISL_3312325 | Lighthouse Lab in Glasgow | Wellcome Sanger Institute for the COVID-19 Genomics UK (COG-UK) Consortium | Harper VanSteenhouse, Yumi Kasai, David Gray, Carol Clugston, Anna Dominiczak and Alex Alderton, Roberto Amato, Jeffrey Barrett, Sonia Goncalves, Ewan Harrison, David K. Jackson, Ian Johnston, Dominic Kwiatkowski, Cordelia Langford, John Sillitoe on behalf of the Wellcome Sanger Institute COVID-19 Surveillance Team |
| EPI_ISL_3312326 | Lighthouse Laboratory Plymouth | Wellcome Sanger Institute for the COVID-19 Genomics UK (COG-UK) Consortium | Lighthouse Laboratory Plymouth and Alex Alderton, Roberto Amato, Jeffrey Barrett, Sonia Goncalves, Ewan Harrison, David K. Jackson, Ian Johnston, Dominic Kwiatkowski, Cordelia Langford, John Sillitoe on behalf of the Wellcome Sanger Institute COVID-19 Surveillance Team |
| EPI_ISL_3312327, EPI_ISL_3312328 | Lighthouse Lab in Glasgow | Wellcome Sanger Institute for the COVID-19 Genomics UK (COG-UK) Consortium | Harper VanSteenhouse, Yumi Kasai, David Gray, Carol Clugston, Anna Dominiczak and Alex Alderton, Roberto Amato, Jeffrey Barrett, Sonia Goncalves, Ewan Harrison, David K. Jackson, Ian Johnston, Dominic Kwiatkowski, Cordelia Langford, John Sillitoe on behalf of the Wellcome Sanger Institute COVID-19 Surveillance Team |
| EPI_ISL_3312329 | Lighthouse Laboratory Plymouth | Wellcome Sanger Institute for the COVID-19 Genomics UK (COG-UK) Consortium | Lighthouse Laboratory Plymouth and Alex Alderton, Roberto Amato, Jeffrey Barrett, Sonia Goncalves, Ewan Harrison, David K. Jackson, Ian Johnston, Dominic Kwiatkowski, Cordelia Langford, John Sillitoe on behalf of the Wellcome Sanger Institute COVID-19 Surveillance Team |
| EPI_ISL_3312330, EPI_ISL_3312331 | Lighthouse Lab in Alderley Park | Wellcome Sanger Institute for the COVID-19 Genomics UK (COG-UK) Consortium | Jacquelyn Wynn, Mairead Hyland, The Lighthouse Lab in Alderley Park and Alex Alderton, Roberto Amato, Jeffrey Barrett, Sonia Goncalves, Ewan Harrison, David K. Jackson, Ian Johnston, Dominic Kwiatkowski, Cordelia Langford, John Sillitoe on behalf of the Wellcome Sanger Institute COVID-19 Surveillance Team |

| | | | |
|---|---|---|---|
| EPI_ISL_3312332 | Lighthouse Lab in Glasgow | Wellcome Sanger Institute for the COVID-19 Genomics UK (COG-UK) Consortium | Harper VanSteenhouse, Yumi Kasai, David Gray, Carol Clugston, Anna Dominiczak and Alex Alderton, Roberto Amato, Jeffrey Barrett, Sonia Goncalves, Ewan Harrison, David K. Jackson, Ian Johnston, Dominic Kwiatkowski, Cordelia Langford, John Sillitoe on behalf of the Wellcome Sanger Institute COVID-19 Surveillance Team |
| EPI_ISL_3312333 | Lighthouse Lab in Alderley Park | Wellcome Sanger Institute for the COVID-19 Genomics UK (COG-UK) Consortium | Jacquelyn Wynn, Mairead Hyland, The Lighthouse Lab in Alderley Park and Alex Alderton, Roberto Amato, Jeffrey Barrett, Sonia Goncalves, Ewan Harrison, David K. Jackson, Ian Johnston, Dominic Kwiatkowski, Cordelia Langford, John Sillitoe on behalf of the Wellcome Sanger Institute COVID-19 Surveillance Team |
| EPI_ISL_3312334 | Lighthouse Laboratory Plymouth | Wellcome Sanger Institute for the COVID-19 Genomics UK (COG-UK) Consortium | Lighthouse Laboratory Plymouth and Alex Alderton, Roberto Amato, Jeffrey Barrett, Sonia Goncalves, Ewan Harrison, David K. Jackson, Ian Johnston, Dominic Kwiatkowski, Cordelia Langford, John Sillitoe on behalf of the Wellcome Sanger Institute COVID-19 Surveillance Team |
| EPI_ISL_3312335, EPI_ISL_3312336, EPI_ISL_3312337, EPI_ISL_3312338, EPI_ISL_3312339 | Lighthouse Lab in Glasgow | Wellcome Sanger Institute for the COVID-19 Genomics UK (COG-UK) Consortium | Harper VanSteenhouse, Yumi Kasai, David Gray, Carol Clugston, Anna Dominiczak and Alex Alderton, Roberto Amato, Jeffrey Barrett, Sonia Goncalves, Ewan Harrison, David K. Jackson, Ian Johnston, Dominic Kwiatkowski, Cordelia Langford, John Sillitoe on behalf of the Wellcome Sanger Institute COVID-19 Surveillance Team |
| EPI_ISL_3312340 | Lighthouse Lab in Alderley Park | Wellcome Sanger Institute for the COVID-19 Genomics UK (COG-UK) Consortium | Jacquelyn Wynn, Mairead Hyland, The Lighthouse Lab in Alderley Park and Alex Alderton, Roberto Amato, Jeffrey Barrett, Sonia Goncalves, Ewan Harrison, David K. Jackson, Ian Johnston, Dominic Kwiatkowski, Cordelia Langford, John Sillitoe on behalf of the Wellcome Sanger Institute COVID-19 Surveillance Team |
| EPI_ISL_3312341 | Lighthouse Laboratory Plymouth | Wellcome Sanger Institute for the COVID-19 Genomics UK (COG-UK) Consortium | Lighthouse Laboratory Plymouth and Alex Alderton, Roberto Amato, Jeffrey Barrett, Sonia Goncalves, Ewan Harrison, David K. Jackson, Ian Johnston, Dominic Kwiatkowski, Cordelia Langford, John Sillitoe on behalf of the Wellcome Sanger Institute COVID-19 Surveillance Team |
| EPI_ISL_3312342, EPI_ISL_3312343, EPI_ISL_3312344, EPI_ISL_3312345 | Lighthouse Lab in Glasgow | Wellcome Sanger Institute for the COVID-19 Genomics UK (COG-UK) Consortium | Harper VanSteenhouse, Yumi Kasai, David Gray, Carol Clugston, Anna Dominiczak and Alex Alderton, Roberto Amato, Jeffrey Barrett, Sonia Goncalves, Ewan Harrison, David K. Jackson, Ian Johnston, Dominic Kwiatkowski, Cordelia Langford, John Sillitoe on behalf of the Wellcome Sanger Institute COVID-19 Surveillance Team |
| EPI_ISL_3312346, EPI_ISL_3312347 | Lighthouse Laboratory Plymouth | Wellcome Sanger Institute for the COVID-19 Genomics UK (COG-UK) Consortium | Lighthouse Laboratory Plymouth and Alex Alderton, Roberto Amato, Jeffrey Barrett, Sonia Goncalves, Ewan Harrison, David K. Jackson, Ian Johnston, Dominic Kwiatkowski, Cordelia Langford, John Sillitoe on behalf of the Wellcome Sanger Institute COVID-19 Surveillance Team |
| EPI_ISL_3312348 | Lighthouse Lab in Alderley Park | Wellcome Sanger Institute for the COVID-19 Genomics UK (COG-UK) Consortium | Jacquelyn Wynn, Mairead Hyland, The Lighthouse Lab in Alderley Park and Alex Alderton, Roberto Amato, Jeffrey Barrett, Sonia Goncalves, Ewan Harrison, David K. Jackson, Ian Johnston, Dominic Kwiatkowski, Cordelia Langford, John Sillitoe on behalf of the Wellcome Sanger Institute COVID-19 Surveillance Team |
| EPI_ISL_3312349 | Lighthouse Lab in Glasgow | Wellcome Sanger Institute for the COVID-19 Genomics UK (COG-UK) Consortium | Harper VanSteenhouse, Yumi Kasai, David Gray, Carol Clugston, Anna Dominiczak and Alex Alderton, Roberto Amato, Jeffrey Barrett, Sonia Goncalves, Ewan Harrison, David K. Jackson, Ian Johnston, Dominic Kwiatkowski, Cordelia Langford, John Sillitoe on behalf of the Wellcome Sanger Institute COVID-19 Surveillance Team |
| EPI_ISL_3312350 | Lighthouse Lab in Alderley Park | Wellcome Sanger Institute for the COVID-19 Genomics UK (COG-UK) Consortium | Jacquelyn Wynn, Mairead Hyland, The Lighthouse Lab in Alderley Park and Alex Alderton, Roberto Amato, Jeffrey Barrett, Sonia Goncalves, Ewan Harrison, David K. Jackson, Ian Johnston, Dominic Kwiatkowski, Cordelia Langford, John Sillitoe on behalf of the Wellcome Sanger Institute COVID-19 Surveillance Team |
| EPI_ISL_3312351 | Lighthouse Laboratory Plymouth | Wellcome Sanger Institute for the COVID-19 Genomics UK (COG-UK) Consortium | Lighthouse Laboratory Plymouth and Alex Alderton, Roberto Amato, Jeffrey Barrett, Sonia Goncalves, Ewan Harrison, David K. Jackson, Ian Johnston, Dominic Kwiatkowski, Cordelia Langford, John Sillitoe on behalf of the Wellcome Sanger Institute COVID-19 Surveillance Team |
| EPI_ISL_3312352 | Lighthouse Lab in Glasgow | Wellcome Sanger Institute for the COVID-19 Genomics UK (COG-UK) Consortium | Harper VanSteenhouse, Yumi Kasai, David Gray, Carol Clugston, Anna Dominiczak and Alex Alderton, Roberto Amato, Jeffrey Barrett, Sonia Goncalves, Ewan Harrison, David K. Jackson, Ian Johnston, Dominic Kwiatkowski, Cordelia Langford, John Sillitoe on behalf of the Wellcome Sanger Institute COVID-19 Surveillance Team |
| EPI_ISL_3312353, EPI_ISL_3312354 | Lighthouse Laboratory Plymouth | Wellcome Sanger Institute for the COVID-19 Genomics UK (COG-UK) Consortium | Lighthouse Laboratory Plymouth and Alex Alderton, Roberto Amato, Jeffrey Barrett, Sonia Goncalves, Ewan Harrison, David K. Jackson, Ian Johnston, Dominic Kwiatkowski, Cordelia Langford, John Sillitoe on behalf of the Wellcome Sanger Institute COVID-19 Surveillance Team |
| EPI_ISL_3312355 | Lighthouse Lab in Alderley Park | Wellcome Sanger Institute for the COVID-19 Genomics UK (COG-UK) Consortium | Jacquelyn Wynn, Mairead Hyland, The Lighthouse Lab in Alderley Park and Alex Alderton, Roberto Amato, Jeffrey Barrett, Sonia Goncalves, Ewan Harrison, David K. Jackson, Ian Johnston, Dominic Kwiatkowski, Cordelia Langford, John Sillitoe on behalf of the Wellcome Sanger Institute COVID-19 Surveillance Team |
| EPI_ISL_3312356, EPI_ISL_3312357, EPI_ISL_3312358 | Lighthouse Laboratory Plymouth | Wellcome Sanger Institute for the COVID-19 Genomics UK (COG-UK) Consortium | Lighthouse Laboratory Plymouth and Alex Alderton, Roberto Amato, Jeffrey Barrett, Sonia Goncalves, Ewan Harrison, David K. Jackson, Ian Johnston, Dominic Kwiatkowski, Cordelia Langford, John Sillitoe on behalf of the Wellcome Sanger Institute COVID-19 Surveillance Team |
| EPI_ISL_3312359, EPI_ISL_3312360 | Lighthouse Lab in Glasgow | Wellcome Sanger Institute for the COVID-19 Genomics UK (COG-UK) Consortium | Harper VanSteenhouse, Yumi Kasai, David Gray, Carol Clugston, Anna Dominiczak and Alex Alderton, Roberto Amato, Jeffrey Barrett, Sonia Goncalves, Ewan Harrison, David K. Jackson, Ian Johnston, Dominic Kwiatkowski, Cordelia Langford, John Sillitoe on behalf of the Wellcome Sanger Institute COVID-19 Surveillance Team |
| EPI_ISL_3312361 | Lighthouse Lab in Alderley Park | Wellcome Sanger Institute for the COVID-19 Genomics UK (COG-UK) Consortium | Jacquelyn Wynn, Mairead Hyland, The Lighthouse Lab in Alderley Park and Alex Alderton, Roberto Amato, Jeffrey Barrett, Sonia Goncalves, Ewan Harrison, David K. Jackson, Ian Johnston, Dominic Kwiatkowski, Cordelia Langford, John Sillitoe on behalf of the Wellcome Sanger Institute COVID-19 Surveillance Team |
| EPI_ISL_3312362 | Lighthouse Laboratory Plymouth | Wellcome Sanger Institute for the COVID-19 Genomics UK (COG-UK) Consortium | Lighthouse Laboratory Plymouth and Alex Alderton, Roberto Amato, Jeffrey Barrett, Sonia Goncalves, Ewan Harrison, David K. Jackson, Ian Johnston, Dominic Kwiatkowski, Cordelia Langford, John Sillitoe on behalf of the Wellcome Sanger Institute COVID-19 Surveillance Team |
| EPI_ISL_3312363 | Lighthouse Lab in Glasgow | Wellcome Sanger Institute for the COVID-19 Genomics UK (COG-UK) Consortium | Harper VanSteenhouse, Yumi Kasai, David Gray, Carol Clugston, Anna Dominiczak and Alex Alderton, Roberto Amato, Jeffrey Barrett, Sonia Goncalves, Ewan Harrison, David K. Jackson, Ian Johnston, Dominic Kwiatkowski, Cordelia Langford, John Sillitoe on behalf of the Wellcome Sanger Institute COVID-19 Surveillance Team |
| EPI_ISL_3312364 | Lighthouse Lab in Alderley Park | Wellcome Sanger Institute for the COVID-19 Genomics UK (COG-UK) Consortium | Jacquelyn Wynn, Mairead Hyland, The Lighthouse Lab in Alderley Park and Alex Alderton, Roberto Amato, Jeffrey Barrett, Sonia Goncalves, Ewan Harrison, David K. Jackson, Ian Johnston, Dominic Kwiatkowski, Cordelia Langford, John Sillitoe on behalf of the Wellcome Sanger Institute COVID-19 Surveillance Team |
| EPI_ISL_3312365 | Lighthouse Lab in Glasgow | Wellcome Sanger Institute for the COVID-19 Genomics UK (COG-UK) Consortium | Harper VanSteenhouse, Yumi Kasai, David Gray, Carol Clugston, Anna Dominiczak and Alex Alderton, Roberto Amato, Jeffrey Barrett, Sonia Goncalves, Ewan Harrison, David K. Jackson, Ian Johnston, Dominic Kwiatkowski, Cordelia Langford, John Sillitoe on behalf of the Wellcome Sanger Institute COVID-19 Surveillance Team |
| EPI_ISL_3312366 | Lighthouse Laboratory Plymouth | Wellcome Sanger Institute for the COVID-19 Genomics UK (COG-UK) Consortium | Lighthouse Laboratory Plymouth and Alex Alderton, Roberto Amato, Jeffrey Barrett, Sonia Goncalves, Ewan Harrison, David K. Jackson, Ian Johnston, Dominic Kwiatkowski, Cordelia Langford, John Sillitoe on behalf of the Wellcome Sanger Institute COVID-19 Surveillance Team |
| EPI_ISL_3312367 | Lighthouse Lab in Glasgow | Wellcome Sanger Institute for the COVID-19 Genomics UK (COG-UK) Consortium | Harper VanSteenhouse, Yumi Kasai, David Gray, Carol Clugston, Anna Dominiczak and Alex Alderton, Roberto Amato, Jeffrey Barrett, Sonia Goncalves, Ewan Harrison, David K. Jackson, Ian Johnston, Dominic Kwiatkowski, Cordelia Langford, John Sillitoe on behalf of the Wellcome Sanger Institute COVID-19 Surveillance Team |
| EPI_ISL_3312368 | Lighthouse Lab in Alderley Park | Wellcome Sanger Institute for the COVID-19 Genomics UK (COG-UK) Consortium | Jacquelyn Wynn, Mairead Hyland, The Lighthouse Lab in Alderley Park and Alex Alderton, Roberto Amato, Jeffrey Barrett, Sonia Goncalves, Ewan Harrison, David K. Jackson, Ian Johnston, Dominic Kwiatkowski, Cordelia Langford, John Sillitoe on behalf of the Wellcome Sanger Institute COVID-19 Surveillance Team |
| EPI_ISL_3312369 | Lighthouse Lab in Glasgow | Wellcome Sanger Institute for the COVID-19 Genomics UK (COG-UK) Consortium | Harper VanSteenhouse, Yumi Kasai, David Gray, Carol Clugston, Anna Dominiczak and Alex Alderton, Roberto Amato, Jeffrey Barrett, Sonia Goncalves, Ewan Harrison, David K. Jackson, Ian Johnston, Dominic Kwiatkowski, Cordelia Langford, John Sillitoe on behalf of the Wellcome Sanger Institute COVID-19 Surveillance Team |
| EPI_ISL_3312370 | Lighthouse Lab in Alderley Park | Wellcome Sanger Institute for the COVID-19 Genomics UK (COG-UK) Consortium | Jacquelyn Wynn, Mairead Hyland, The Lighthouse Lab in Alderley Park and Alex Alderton, Roberto Amato, Jeffrey Barrett, Sonia Goncalves, Ewan Harrison, David K. Jackson, Ian Johnston, Dominic Kwiatkowski, Cordelia Langford, John Sillitoe on behalf of the Wellcome Sanger Institute COVID-19 Surveillance Team |
| EPI_ISL_3312371, EPI_ISL_3312372 | Lighthouse Lab in Glasgow | Wellcome Sanger Institute for the COVID-19 Genomics UK (COG-UK) Consortium | Harper VanSteenhouse, Yumi Kasai, David Gray, Carol Clugston, Anna Dominiczak and Alex Alderton, Roberto Amato, Jeffrey Barrett, Sonia Goncalves, Ewan Harrison, David K. Jackson, Ian Johnston, Dominic Kwiatkowski, Cordelia Langford, John Sillitoe on behalf of the Wellcome Sanger Institute COVID-19 Surveillance Team |
| EPI_ISL_3312373 | Lighthouse Laboratory Plymouth | Wellcome Sanger Institute for the COVID-19 Genomics UK (COG-UK) Consortium | Lighthouse Laboratory Plymouth and Alex Alderton, Roberto Amato, Jeffrey Barrett, Sonia Goncalves, Ewan Harrison, David K. Jackson, Ian Johnston, Dominic Kwiatkowski, Cordelia Langford, John Sillitoe on behalf of the Wellcome Sanger Institute COVID-19 Surveillance Team |
| EPI_ISL_3312374, EPI_ISL_3312375, EPI_ISL_3312376 | Lighthouse Lab in Glasgow | Wellcome Sanger Institute for the COVID-19 Genomics UK (COG-UK) Consortium | Harper VanSteenhouse, Yumi Kasai, David Gray, Carol Clugston, Anna Dominiczak and Alex Alderton, Roberto Amato, Jeffrey Barrett, Sonia Goncalves, Ewan Harrison, David K. Jackson, Ian Johnston, Dominic Kwiatkowski, Cordelia Langford, John Sillitoe on behalf of the Wellcome Sanger Institute COVID-19 Surveillance Team |
| EPI_ISL_3312377 | Lighthouse Laboratory Plymouth | Wellcome Sanger Institute for the COVID-19 Genomics UK (COG-UK) Consortium | Lighthouse Laboratory Plymouth and Alex Alderton, Roberto Amato, Jeffrey Barrett, Sonia Goncalves, Ewan Harrison, David K. Jackson, Ian Johnston, Dominic Kwiatkowski, Cordelia Langford, John Sillitoe on behalf of the Wellcome Sanger Institute COVID-19 Surveillance Team |
| EPI_ISL_3312378 | Lighthouse Lab in Alderley Park | Wellcome Sanger Institute for the COVID-19 Genomics UK (COG-UK) Consortium | Jacquelyn Wynn, Mairead Hyland, The Lighthouse Lab in Alderley Park and Alex Alderton, Roberto Amato, Jeffrey Barrett, Sonia Goncalves, Ewan Harrison, David K. Jackson, Ian Johnston, Dominic Kwiatkowski, Cordelia Langford, John Sillitoe on behalf of the Wellcome Sanger Institute COVID-19 Surveillance Team |
| EPI_ISL_3312379, EPI_ISL_3312380, EPI_ISL_3312381, EPI_ISL_3312382, EPI_ISL_3312383, EPI_ISL_3312384, EPI_ISL_3312385 | Lighthouse Lab in Glasgow | Wellcome Sanger Institute for the COVID-19 Genomics UK (COG-UK) Consortium | Harper VanSteenhouse, Yumi Kasai, David Gray, Carol Clugston, Anna Dominiczak and Alex Alderton, Roberto Amato, Jeffrey Barrett, Sonia Goncalves, Ewan Harrison, David K. Jackson, Ian Johnston, Dominic Kwiatkowski, Cordelia Langford, John Sillitoe on behalf of the Wellcome Sanger Institute COVID-19 Surveillance Team |
| EPI_ISL_3312386 | Lighthouse Laboratory Plymouth | Wellcome Sanger Institute for the COVID-19 Genomics UK (COG-UK) Consortium | Lighthouse Laboratory Plymouth and Alex Alderton, Roberto Amato, Jeffrey Barrett, Sonia Goncalves, Ewan Harrison, David K. Jackson, Ian Johnston, Dominic Kwiatkowski, Cordelia Langford, John Sillitoe on behalf of the Wellcome Sanger Institute COVID-19 Surveillance Team |
| EPI_ISL_3312387 | Lighthouse Lab in Glasgow | Wellcome Sanger Institute for the COVID-19 Genomics UK (COG-UK) Consortium | Harper VanSteenhouse, Yumi Kasai, David Gray, Carol Clugston, Anna Dominiczak and Alex Alderton, Roberto Amato, Jeffrey Barrett, Sonia Goncalves, Ewan Harrison, David K. Jackson, Ian Johnston, Dominic Kwiatkowski, Cordelia Langford, John Sillitoe on behalf of the Wellcome Sanger Institute COVID-19 Surveillance Team |
| EPI_ISL_3312388 | Lighthouse Lab in Alderley Park | Wellcome Sanger Institute for the COVID-19 Genomics UK (COG-UK) Consortium | Jacquelyn Wynn, Mairead Hyland, The Lighthouse Lab in Alderley Park and Alex Alderton, Roberto Amato, Jeffrey Barrett, Sonia Goncalves, Ewan Harrison, David K. Jackson, Ian Johnston, Dominic Kwiatkowski, Cordelia Langford, John Sillitoe on behalf of the Wellcome Sanger Institute COVID-19 Surveillance Team |
| EPI_ISL_3312389 | Lighthouse Lab in Glasgow | Wellcome Sanger Institute for the COVID-19 Genomics UK (COG-UK) Consortium | Harper VanSteenhouse, Yumi Kasai, David Gray, Carol Clugston, Anna Dominiczak and Alex Alderton, Roberto Amato, Jeffrey Barrett, Sonia Goncalves, Ewan Harrison, David K. Jackson, Ian Johnston, Dominic Kwiatkowski, Cordelia Langford, John Sillitoe on behalf of the Wellcome Sanger Institute COVID-19 Surveillance Team |
| EPI_ISL_3312390 | Lighthouse Lab in Alderley Park | Wellcome Sanger Institute for the COVID-19 Genomics UK (COG-UK) Consortium | Jacquelyn Wynn, Mairead Hyland, The Lighthouse Lab in Alderley Park and Alex Alderton, Roberto Amato, Jeffrey Barrett, Sonia Goncalves, Ewan Harrison, David K. Jackson, Ian Johnston, Dominic Kwiatkowski, Cordelia Langford, John Sillitoe on behalf of the Wellcome Sanger Institute COVID-19 Surveillance Team |
| EPI_ISL_3312391 | Lighthouse Lab in Glasgow | Wellcome Sanger Institute for the COVID-19 Genomics UK (COG-UK) Consortium | Harper VanSteenhouse, Yumi Kasai, David Gray, Carol Clugston, Anna Dominiczak and Alex Alderton, Roberto Amato, Jeffrey Barrett, Sonia Goncalves, Ewan Harrison, David K. Jackson, Ian Johnston, Dominic Kwiatkowski, Cordelia Langford, John Sillitoe on behalf of the Wellcome Sanger Institute COVID-19 Surveillance Team |
| EPI_ISL_3312392, EPI_ISL_3312393 | Lighthouse Laboratory Plymouth | Wellcome Sanger Institute for the COVID-19 Genomics UK (COG-UK) Consortium | Lighthouse Laboratory Plymouth and Alex Alderton, Roberto Amato, Jeffrey Barrett, Sonia Goncalves, Ewan Harrison, David K. Jackson, Ian Johnston, Dominic Kwiatkowski, Cordelia Langford, John Sillitoe on behalf of the Wellcome Sanger Institute COVID-19 Surveillance Team |

| | | | |
|---|---|---|---|
| EPI_ISL_3312394 | Lighthouse Lab in Alderley Park | Wellcome Sanger Institute for the COVID-19 Genomics UK (COG-UK) Consortium | Jacquelyn Wynn, Mairead Hyland, The Lighthouse Lab in Alderley Park and Alex Alderton, Roberto Amato, Jeffrey Barrett, Sonia Goncalves, Ewan Harrison, David K. Jackson, Ian Johnston, Dominic Kwiatkowski, Cordelia Langford, John Sillitoe on behalf of the Wellcome Sanger Institute COVID-19 Surveillance Team |
| EPI_ISL_3312395 | Lighthouse Lab in Glasgow | Wellcome Sanger Institute for the COVID-19 Genomics UK (COG-UK) Consortium | Harper VanSteenhouse, Yumi Kasai, David Gray, Carol Clugston, Anna Dominiczak and Alex Alderton, Roberto Amato, Jeffrey Barrett, Sonia Goncalves, Ewan Harrison, David K. Jackson, Ian Johnston, Dominic Kwiatkowski, Cordelia Langford, John Sillitoe on behalf of the Wellcome Sanger Institute COVID-19 Surveillance Team |
| EPI_ISL_3312396 | Lighthouse Laboratory Plymouth | Wellcome Sanger Institute for the COVID-19 Genomics UK (COG-UK) Consortium | Lighthouse Laboratory Plymouth and Alex Alderton, Roberto Amato, Jeffrey Barrett, Sonia Goncalves, Ewan Harrison, David K. Jackson, Ian Johnston, Dominic Kwiatkowski, Cordelia Langford, John Sillitoe on behalf of the Wellcome Sanger Institute COVID-19 Surveillance Team |
| EPI_ISL_3312397, EPI_ISL_3312398 | Lighthouse Lab in Glasgow | Wellcome Sanger Institute for the COVID-19 Genomics UK (COG-UK) Consortium | Harper VanSteenhouse, Yumi Kasai, David Gray, Carol Clugston, Anna Dominiczak and Alex Alderton, Roberto Amato, Jeffrey Barrett, Sonia Goncalves, Ewan Harrison, David K. Jackson, Ian Johnston, Dominic Kwiatkowski, Cordelia Langford, John Sillitoe on behalf of the Wellcome Sanger Institute COVID-19 Surveillance Team |
| EPI_ISL_3312399 | Lighthouse Lab in Alderley Park | Wellcome Sanger Institute for the COVID-19 Genomics UK (COG-UK) Consortium | Jacquelyn Wynn, Mairead Hyland, The Lighthouse Lab in Alderley Park and Alex Alderton, Roberto Amato, Jeffrey Barrett, Sonia Goncalves, Ewan Harrison, David K. Jackson, Ian Johnston, Dominic Kwiatkowski, Cordelia Langford, John Sillitoe on behalf of the Wellcome Sanger Institute COVID-19 Surveillance Team |
| EPI_ISL_3312400 | Lighthouse Lab in Glasgow | Wellcome Sanger Institute for the COVID-19 Genomics UK (COG-UK) Consortium | Harper VanSteenhouse, Yumi Kasai, David Gray, Carol Clugston, Anna Dominiczak and Alex Alderton, Roberto Amato, Jeffrey Barrett, Sonia Goncalves, Ewan Harrison, David K. Jackson, Ian Johnston, Dominic Kwiatkowski, Cordelia Langford, John Sillitoe on behalf of the Wellcome Sanger Institute COVID-19 Surveillance Team |
| EPI_ISL_3312401, EPI_ISL_3312402 | Lighthouse Laboratory Plymouth | Wellcome Sanger Institute for the COVID-19 Genomics UK (COG-UK) Consortium | Lighthouse Laboratory Plymouth and Alex Alderton, Roberto Amato, Jeffrey Barrett, Sonia Goncalves, Ewan Harrison, David K. Jackson, Ian Johnston, Dominic Kwiatkowski, Cordelia Langford, John Sillitoe on behalf of the Wellcome Sanger Institute COVID-19 Surveillance Team |
| EPI_ISL_3312403, EPI_ISL_3312404 | Lighthouse Lab in Alderley Park | Wellcome Sanger Institute for the COVID-19 Genomics UK (COG-UK) Consortium | Jacquelyn Wynn, Mairead Hyland, The Lighthouse Lab in Alderley Park and Alex Alderton, Roberto Amato, Jeffrey Barrett, Sonia Goncalves, Ewan Harrison, David K. Jackson, Ian Johnston, Dominic Kwiatkowski, Cordelia Langford, John Sillitoe on behalf of the Wellcome Sanger Institute COVID-19 Surveillance Team |
| EPI_ISL_3312405, EPI_ISL_3312406, EPI_ISL_3312407, EPI_ISL_3312408, EPI_ISL_3312409, EPI_ISL_3312410 | Lighthouse Lab in Glasgow | Wellcome Sanger Institute for the COVID-19 Genomics UK (COG-UK) Consortium | Harper VanSteenhouse, Yumi Kasai, David Gray, Carol Clugston, Anna Dominiczak and Alex Alderton, Roberto Amato, Jeffrey Barrett, Sonia Goncalves, Ewan Harrison, David K. Jackson, Ian Johnston, Dominic Kwiatkowski, Cordelia Langford, John Sillitoe on behalf of the Wellcome Sanger Institute COVID-19 Surveillance Team |
| EPI_ISL_3312411 | Lighthouse Lab in Alderley Park | Wellcome Sanger Institute for the COVID-19 Genomics UK (COG-UK) Consortium | Jacquelyn Wynn, Mairead Hyland, The Lighthouse Lab in Alderley Park and Alex Alderton, Roberto Amato, Jeffrey Barrett, Sonia Goncalves, Ewan Harrison, David K. Jackson, Ian Johnston, Dominic Kwiatkowski, Cordelia Langford, John Sillitoe on behalf of the Wellcome Sanger Institute COVID-19 Surveillance Team |
| EPI_ISL_3312412 | Lighthouse Lab in Glasgow | Wellcome Sanger Institute for the COVID-19 Genomics UK (COG-UK) Consortium | Harper VanSteenhouse, Yumi Kasai, David Gray, Carol Clugston, Anna Dominiczak and Alex Alderton, Roberto Amato, Jeffrey Barrett, Sonia Goncalves, Ewan Harrison, David K. Jackson, Ian Johnston, Dominic Kwiatkowski, Cordelia Langford, John Sillitoe on behalf of the Wellcome Sanger Institute COVID-19 Surveillance Team |
| EPI_ISL_3312413 | Lighthouse Laboratory Plymouth | Wellcome Sanger Institute for the COVID-19 Genomics UK (COG-UK) Consortium | Lighthouse Laboratory Plymouth and Alex Alderton, Roberto Amato, Jeffrey Barrett, Sonia Goncalves, Ewan Harrison, David K. Jackson, Ian Johnston, Dominic Kwiatkowski, Cordelia Langford, John Sillitoe on behalf of the Wellcome Sanger Institute COVID-19 Surveillance Team |
| EPI_ISL_3312414, EPI_ISL_3312415, EPI_ISL_3312416, EPI_ISL_3312417, EPI_ISL_3312418 | Lighthouse Lab in Glasgow | Wellcome Sanger Institute for the COVID-19 Genomics UK (COG-UK) Consortium | Harper VanSteenhouse, Yumi Kasai, David Gray, Carol Clugston, Anna Dominiczak and Alex Alderton, Roberto Amato, Jeffrey Barrett, Sonia Goncalves, Ewan Harrison, David K. Jackson, Ian Johnston, Dominic Kwiatkowski, Cordelia Langford, John Sillitoe on behalf of the Wellcome Sanger Institute COVID-19 Surveillance Team |
| EPI_ISL_3312419, EPI_ISL_3312420 | Lighthouse Lab in Alderley Park | Wellcome Sanger Institute for the COVID-19 Genomics UK (COG-UK) Consortium | Jacquelyn Wynn, Mairead Hyland, The Lighthouse Lab in Alderley Park and Alex Alderton, Roberto Amato, Jeffrey Barrett, Sonia Goncalves, Ewan Harrison, David K. Jackson, Ian Johnston, Dominic Kwiatkowski, Cordelia Langford, John Sillitoe on behalf of the Wellcome Sanger Institute COVID-19 Surveillance Team |
| EPI_ISL_3312421, EPI_ISL_3312422 | Lighthouse Lab in Glasgow | Wellcome Sanger Institute for the COVID-19 Genomics UK (COG-UK) Consortium | Harper VanSteenhouse, Yumi Kasai, David Gray, Carol Clugston, Anna Dominiczak and Alex Alderton, Roberto Amato, Jeffrey Barrett, Sonia Goncalves, Ewan Harrison, David K. Jackson, Ian Johnston, Dominic Kwiatkowski, Cordelia Langford, John Sillitoe on behalf of the Wellcome Sanger Institute COVID-19 Surveillance Team |
| EPI_ISL_3312423 | Lighthouse Lab in Alderley Park | Wellcome Sanger Institute for the COVID-19 Genomics UK (COG-UK) Consortium | Jacquelyn Wynn, Mairead Hyland, The Lighthouse Lab in Alderley Park and Alex Alderton, Roberto Amato, Jeffrey Barrett, Sonia Goncalves, Ewan Harrison, David K. Jackson, Ian Johnston, Dominic Kwiatkowski, Cordelia Langford, John Sillitoe on behalf of the Wellcome Sanger Institute COVID-19 Surveillance Team |
| EPI_ISL_3312424, EPI_ISL_3312425 | Lighthouse Laboratory Plymouth | Wellcome Sanger Institute for the COVID-19 Genomics UK (COG-UK) Consortium | Lighthouse Laboratory Plymouth and Alex Alderton, Roberto Amato, Jeffrey Barrett, Sonia Goncalves, Ewan Harrison, David K. Jackson, Ian Johnston, Dominic Kwiatkowski, Cordelia Langford, John Sillitoe on behalf of the Wellcome Sanger Institute COVID-19 Surveillance Team |
| EPI_ISL_3312426 | Lighthouse Lab in Alderley Park | Wellcome Sanger Institute for the COVID-19 Genomics UK (COG-UK) Consortium | Jacquelyn Wynn, Mairead Hyland, The Lighthouse Lab in Alderley Park and Alex Alderton, Roberto Amato, Jeffrey Barrett, Sonia Goncalves, Ewan Harrison, David K. Jackson, Ian Johnston, Dominic Kwiatkowski, Cordelia Langford, John Sillitoe on behalf of the Wellcome Sanger Institute COVID-19 Surveillance Team |
| EPI_ISL_3312427 | Lighthouse Laboratory Plymouth | Wellcome Sanger Institute for the COVID-19 Genomics UK (COG-UK) Consortium | Lighthouse Laboratory Plymouth and Alex Alderton, Roberto Amato, Jeffrey Barrett, Sonia Goncalves, Ewan Harrison, David K. Jackson, Ian Johnston, Dominic Kwiatkowski, Cordelia Langford, John Sillitoe on behalf of the Wellcome Sanger Institute COVID-19 Surveillance Team |
| EPI_ISL_3312428 | Lighthouse Lab in Alderley Park | Wellcome Sanger Institute for the COVID-19 Genomics UK (COG-UK) Consortium | Jacquelyn Wynn, Mairead Hyland, The Lighthouse Lab in Alderley Park and Alex Alderton, Roberto Amato, Jeffrey Barrett, Sonia Goncalves, Ewan Harrison, David K. Jackson, Ian Johnston, Dominic Kwiatkowski, Cordelia Langford, John Sillitoe on behalf of the Wellcome Sanger Institute COVID-19 Surveillance Team |
| EPI_ISL_3312429 | Lighthouse Lab in Glasgow | Wellcome Sanger Institute for the COVID-19 Genomics UK (COG-UK) Consortium | Harper VanSteenhouse, Yumi Kasai, David Gray, Carol Clugston, Anna Dominiczak and Alex Alderton, Roberto Amato, Jeffrey Barrett, Sonia Goncalves, Ewan Harrison, David K. Jackson, Ian Johnston, Dominic Kwiatkowski, Cordelia Langford, John Sillitoe on behalf of the Wellcome Sanger Institute COVID-19 Surveillance Team |
| EPI_ISL_3312430 | Lighthouse Lab in Alderley Park | Wellcome Sanger Institute for the COVID-19 Genomics UK (COG-UK) Consortium | Jacquelyn Wynn, Mairead Hyland, The Lighthouse Lab in Alderley Park and Alex Alderton, Roberto Amato, Jeffrey Barrett, Sonia Goncalves, Ewan Harrison, David K. Jackson, Ian Johnston, Dominic Kwiatkowski, Cordelia Langford, John Sillitoe on behalf of the Wellcome Sanger Institute COVID-19 Surveillance Team |
| EPI_ISL_3312431 | Lighthouse Lab in Glasgow | Wellcome Sanger Institute for the COVID-19 Genomics UK (COG-UK) Consortium | Harper VanSteenhouse, Yumi Kasai, David Gray, Carol Clugston, Anna Dominiczak and Alex Alderton, Roberto Amato, Jeffrey Barrett, Sonia Goncalves, Ewan Harrison, David K. Jackson, Ian Johnston, Dominic Kwiatkowski, Cordelia Langford, John Sillitoe on behalf of the Wellcome Sanger Institute COVID-19 Surveillance Team |
| EPI_ISL_3312432 | Lighthouse Laboratory Plymouth | Wellcome Sanger Institute for the COVID-19 Genomics UK (COG-UK) Consortium | Lighthouse Laboratory Plymouth and Alex Alderton, Roberto Amato, Jeffrey Barrett, Sonia Goncalves, Ewan Harrison, David K. Jackson, Ian Johnston, Dominic Kwiatkowski, Cordelia Langford, John Sillitoe on behalf of the Wellcome Sanger Institute COVID-19 Surveillance Team |
| EPI_ISL_3312433, EPI_ISL_3312434 | Lighthouse Lab in Glasgow | Wellcome Sanger Institute for the COVID-19 Genomics UK (COG-UK) Consortium | Harper VanSteenhouse, Yumi Kasai, David Gray, Carol Clugston, Anna Dominiczak and Alex Alderton, Roberto Amato, Jeffrey Barrett, Sonia Goncalves, Ewan Harrison, David K. Jackson, Ian Johnston, Dominic Kwiatkowski, Cordelia Langford, John Sillitoe on behalf of the Wellcome Sanger Institute COVID-19 Surveillance Team |
| EPI_ISL_3312435 | Lighthouse Laboratory Plymouth | Wellcome Sanger Institute for the COVID-19 Genomics UK (COG-UK) Consortium | Lighthouse Laboratory Plymouth and Alex Alderton, Roberto Amato, Jeffrey Barrett, Sonia Goncalves, Ewan Harrison, David K. Jackson, Ian Johnston, Dominic Kwiatkowski, Cordelia Langford, John Sillitoe on behalf of the Wellcome Sanger Institute COVID-19 Surveillance Team |
| EPI_ISL_3312436 | Lighthouse Lab in Alderley Park | Wellcome Sanger Institute for the COVID-19 Genomics UK (COG-UK) Consortium | Jacquelyn Wynn, Mairead Hyland, The Lighthouse Lab in Alderley Park and Alex Alderton, Roberto Amato, Jeffrey Barrett, Sonia Goncalves, Ewan Harrison, David K. Jackson, Ian Johnston, Dominic Kwiatkowski, Cordelia Langford, John Sillitoe on behalf of the Wellcome Sanger Institute COVID-19 Surveillance Team |
| EPI_ISL_3312437 | Lighthouse Lab in Glasgow | Wellcome Sanger Institute for the COVID-19 Genomics UK (COG-UK) Consortium | Harper VanSteenhouse, Yumi Kasai, David Gray, Carol Clugston, Anna Dominiczak and Alex Alderton, Roberto Amato, Jeffrey Barrett, Sonia Goncalves, Ewan Harrison, David K. Jackson, Ian Johnston, Dominic Kwiatkowski, Cordelia Langford, John Sillitoe on behalf of the Wellcome Sanger Institute COVID-19 Surveillance Team |
| EPI_ISL_3312438, EPI_ISL_3312439 | Lighthouse Lab in Alderley Park | Wellcome Sanger Institute for the COVID-19 Genomics UK (COG-UK) Consortium | Jacquelyn Wynn, Mairead Hyland, The Lighthouse Lab in Alderley Park and Alex Alderton, Roberto Amato, Jeffrey Barrett, Sonia Goncalves, Ewan Harrison, David K. Jackson, Ian Johnston, Dominic Kwiatkowski, Cordelia Langford, John Sillitoe on behalf of the Wellcome Sanger Institute COVID-19 Surveillance Team |
| EPI_ISL_3312440 | Lighthouse Lab in Glasgow | Wellcome Sanger Institute for the COVID-19 Genomics UK (COG-UK) Consortium | Harper VanSteenhouse, Yumi Kasai, David Gray, Carol Clugston, Anna Dominiczak and Alex Alderton, Roberto Amato, Jeffrey Barrett, Sonia Goncalves, Ewan Harrison, David K. Jackson, Ian Johnston, Dominic Kwiatkowski, Cordelia Langford, John Sillitoe on behalf of the Wellcome Sanger Institute COVID-19 Surveillance Team |
| EPI_ISL_3312441, EPI_ISL_3312442 | Lighthouse Lab in Alderley Park | Wellcome Sanger Institute for the COVID-19 Genomics UK (COG-UK) Consortium | Jacquelyn Wynn, Mairead Hyland, The Lighthouse Lab in Alderley Park and Alex Alderton, Roberto Amato, Jeffrey Barrett, Sonia Goncalves, Ewan Harrison, David K. Jackson, Ian Johnston, Dominic Kwiatkowski, Cordelia Langford, John Sillitoe on behalf of the Wellcome Sanger Institute COVID-19 Surveillance Team |
| EPI_ISL_3312443 | Lighthouse Lab in Glasgow | Wellcome Sanger Institute for the COVID-19 Genomics UK (COG-UK) Consortium | Harper VanSteenhouse, Yumi Kasai, David Gray, Carol Clugston, Anna Dominiczak and Alex Alderton, Roberto Amato, Jeffrey Barrett, Sonia Goncalves, Ewan Harrison, David K. Jackson, Ian Johnston, Dominic Kwiatkowski, Cordelia Langford, John Sillitoe on behalf of the Wellcome Sanger Institute COVID-19 Surveillance Team |
| EPI_ISL_3312444 | Lighthouse Laboratory Plymouth | Wellcome Sanger Institute for the COVID-19 Genomics UK (COG-UK) Consortium | Lighthouse Laboratory Plymouth and Alex Alderton, Roberto Amato, Jeffrey Barrett, Sonia Goncalves, Ewan Harrison, David K. Jackson, Ian Johnston, Dominic Kwiatkowski, Cordelia Langford, John Sillitoe on behalf of the Wellcome Sanger Institute COVID-19 Surveillance Team |
| EPI_ISL_3312445, EPI_ISL_3312446 | Lighthouse Lab in Alderley Park | Wellcome Sanger Institute for the COVID-19 Genomics UK (COG-UK) Consortium | Jacquelyn Wynn, Mairead Hyland, The Lighthouse Lab in Alderley Park and Alex Alderton, Roberto Amato, Jeffrey Barrett, Sonia Goncalves, Ewan Harrison, David K. Jackson, Ian Johnston, Dominic Kwiatkowski, Cordelia Langford, John Sillitoe on behalf of the Wellcome Sanger Institute COVID-19 Surveillance Team |
| EPI_ISL_3312447 | Lighthouse Laboratory Plymouth | Wellcome Sanger Institute for the COVID-19 Genomics UK (COG-UK) Consortium | Lighthouse Laboratory Plymouth and Alex Alderton, Roberto Amato, Jeffrey Barrett, Sonia Goncalves, Ewan Harrison, David K. Jackson, Ian Johnston, Dominic Kwiatkowski, Cordelia Langford, John Sillitoe on behalf of the Wellcome Sanger Institute COVID-19 Surveillance Team |
| EPI_ISL_3312448, EPI_ISL_3312449 | Lighthouse Lab in Glasgow | Wellcome Sanger Institute for the COVID-19 Genomics UK (COG-UK) Consortium | Harper VanSteenhouse, Yumi Kasai, David Gray, Carol Clugston, Anna Dominiczak and Alex Alderton, Roberto Amato, Jeffrey Barrett, Sonia Goncalves, Ewan Harrison, David K. Jackson, Ian Johnston, Dominic Kwiatkowski, Cordelia Langford, John Sillitoe on behalf of the Wellcome Sanger Institute COVID-19 Surveillance Team |
| EPI_ISL_3312450 | Lighthouse Lab in Alderley Park | Wellcome Sanger Institute for the COVID-19 Genomics UK (COG-UK) Consortium | Jacquelyn Wynn, Mairead Hyland, The Lighthouse Lab in Alderley Park and Alex Alderton, Roberto Amato, Jeffrey Barrett, Sonia Goncalves, Ewan Harrison, David K. Jackson, Ian Johnston, Dominic Kwiatkowski, Cordelia Langford, John Sillitoe on behalf of the Wellcome Sanger Institute COVID-19 Surveillance Team |
| EPI_ISL_3312451, EPI_ISL_3312452 | Lighthouse Laboratory Plymouth | Wellcome Sanger Institute for the COVID-19 Genomics UK (COG-UK) Consortium | Lighthouse Laboratory Plymouth and Alex Alderton, Roberto Amato, Jeffrey Barrett, Sonia Goncalves, Ewan Harrison, David K. Jackson, Ian Johnston, Dominic Kwiatkowski, Cordelia Langford, John Sillitoe on behalf of the Wellcome Sanger Institute COVID-19 Surveillance Team |
| EPI_ISL_3312453, EPI_ISL_3312454 | Lighthouse Lab in Glasgow | Wellcome Sanger Institute for the COVID-19 Genomics UK (COG-UK) Consortium | Harper VanSteenhouse, Yumi Kasai, David Gray, Carol Clugston, Anna Dominiczak and Alex Alderton, Roberto Amato, Jeffrey Barrett, Sonia Goncalves, Ewan Harrison, David K. Jackson, Ian Johnston, Dominic Kwiatkowski, Cordelia Langford, John Sillitoe on behalf of the Wellcome Sanger Institute COVID-19 Surveillance Team |
| EPI_ISL_3312455 | Lighthouse Lab in Alderley Park | Wellcome Sanger Institute for the COVID-19 Genomics UK (COG-UK) Consortium | Jacquelyn Wynn, Mairead Hyland, The Lighthouse Lab in Alderley Park and Alex Alderton, Roberto Amato, Jeffrey Barrett, Sonia Goncalves, Ewan Harrison, David K. Jackson, Ian Johnston, Dominic Kwiatkowski, Cordelia Langford, John Sillitoe on behalf of the Wellcome Sanger Institute COVID-19 Surveillance Team |
| EPI_ISL_3312456 | Lighthouse Lab in Glasgow | Wellcome Sanger Institute for the COVID-19 Genomics UK (COG-UK) Consortium | Harper VanSteenhouse, Yumi Kasai, David Gray, Carol Clugston, Anna Dominiczak and Alex Alderton, Roberto Amato, Jeffrey Barrett, Sonia Goncalves, Ewan Harrison, David K. Jackson, Ian Johnston, Dominic Kwiatkowski, Cordelia Langford, John Sillitoe on behalf of the Wellcome Sanger Institute COVID-19 Surveillance Team |
| EPI_ISL_3312457 | Lighthouse Lab in Alderley Park | Wellcome Sanger Institute for the COVID-19 Genomics UK (COG-UK) Consortium | Jacquelyn Wynn, Mairead Hyland, The Lighthouse Lab in Alderley Park and Alex Alderton, Roberto Amato, Jeffrey Barrett, Sonia Goncalves, Ewan Harrison, David K. Jackson, Ian Johnston, Dominic Kwiatkowski, Cordelia Langford, John Sillitoe on behalf of the Wellcome Sanger Institute COVID-19 Surveillance Team |

| | | | |
|---|---|---|---|
| EPI_ISL_3312458 | Lighthouse Lab in Glasgow | Wellcome Sanger Institute for the COVID-19 Genomics UK (COG-UK) Consortium | Harper VanSteenhouse, Yumi Kasai, David Gray, Carol Clugston, Anna Dominiczak and Alex Alderton, Roberto Amato, Jeffrey Barrett, Sonia Goncalves, Ewan Harrison, David K. Jackson, Ian Johnston, Dominic Kwiatkowski, Cordelia Langford, John Sillitoe on behalf of the Wellcome Sanger Institute COVID-19 Surveillance Team |
| EPI_ISL_3312459 | Lighthouse Laboratory Plymouth | Wellcome Sanger Institute for the COVID-19 Genomics UK (COG-UK) Consortium | Lighthouse Laboratory Plymouth and Alex Alderton, Roberto Amato, Jeffrey Barrett, Sonia Goncalves, Ewan Harrison, David K. Jackson, Ian Johnston, Dominic Kwiatkowski, Cordelia Langford, John Sillitoe on behalf of the Wellcome Sanger Institute COVID-19 Surveillance Team |
| EPI_ISL_3312460, EPI_ISL_3312461 | Lighthouse Lab in Glasgow | Wellcome Sanger Institute for the COVID-19 Genomics UK (COG-UK) Consortium | Harper VanSteenhouse, Yumi Kasai, David Gray, Carol Clugston, Anna Dominiczak and Alex Alderton, Roberto Amato, Jeffrey Barrett, Sonia Goncalves, Ewan Harrison, David K. Jackson, Ian Johnston, Dominic Kwiatkowski, Cordelia Langford, John Sillitoe on behalf of the Wellcome Sanger Institute COVID-19 Surveillance Team |
| EPI_ISL_3312462 | Lighthouse Laboratory Plymouth | Wellcome Sanger Institute for the COVID-19 Genomics UK (COG-UK) Consortium | Lighthouse Laboratory Plymouth and Alex Alderton, Roberto Amato, Jeffrey Barrett, Sonia Goncalves, Ewan Harrison, David K. Jackson, Ian Johnston, Dominic Kwiatkowski, Cordelia Langford, John Sillitoe on behalf of the Wellcome Sanger Institute COVID-19 Surveillance Team |
| EPI_ISL_3312463 | Lighthouse Lab in Alderley Park | Wellcome Sanger Institute for the COVID-19 Genomics UK (COG-UK) Consortium | Jacquelyn Wynn, Mairead Hyland, The Lighthouse Lab in Alderley Park and Alex Alderton, Roberto Amato, Jeffrey Barrett, Sonia Goncalves, Ewan Harrison, David K. Jackson, Ian Johnston, Dominic Kwiatkowski, Cordelia Langford, John Sillitoe on behalf of the Wellcome Sanger Institute COVID-19 Surveillance Team |
| EPI_ISL_3312464, EPI_ISL_3312465 | Lighthouse Lab in Glasgow | Wellcome Sanger Institute for the COVID-19 Genomics UK (COG-UK) Consortium | Harper VanSteenhouse, Yumi Kasai, David Gray, Carol Clugston, Anna Dominiczak and Alex Alderton, Roberto Amato, Jeffrey Barrett, Sonia Goncalves, Ewan Harrison, David K. Jackson, Ian Johnston, Dominic Kwiatkowski, Cordelia Langford, John Sillitoe on behalf of the Wellcome Sanger Institute COVID-19 Surveillance Team |
| EPI_ISL_3312466 | Lighthouse Lab in Alderley Park | Wellcome Sanger Institute for the COVID-19 Genomics UK (COG-UK) Consortium | Jacquelyn Wynn, Mairead Hyland, The Lighthouse Lab in Alderley Park and Alex Alderton, Roberto Amato, Jeffrey Barrett, Sonia Goncalves, Ewan Harrison, David K. Jackson, Ian Johnston, Dominic Kwiatkowski, Cordelia Langford, John Sillitoe on behalf of the Wellcome Sanger Institute COVID-19 Surveillance Team |
| EPI_ISL_3312467, EPI_ISL_3312468 | Lighthouse Lab in Glasgow | Wellcome Sanger Institute for the COVID-19 Genomics UK (COG-UK) Consortium | Harper VanSteenhouse, Yumi Kasai, David Gray, Carol Clugston, Anna Dominiczak and Alex Alderton, Roberto Amato, Jeffrey Barrett, Sonia Goncalves, Ewan Harrison, David K. Jackson, Ian Johnston, Dominic Kwiatkowski, Cordelia Langford, John Sillitoe on behalf of the Wellcome Sanger Institute COVID-19 Surveillance Team |
| EPI_ISL_3312469, EPI_ISL_3312470 | Lighthouse Laboratory Plymouth | Wellcome Sanger Institute for the COVID-19 Genomics UK (COG-UK) Consortium | Lighthouse Laboratory Plymouth and Alex Alderton, Roberto Amato, Jeffrey Barrett, Sonia Goncalves, Ewan Harrison, David K. Jackson, Ian Johnston, Dominic Kwiatkowski, Cordelia Langford, John Sillitoe on behalf of the Wellcome Sanger Institute COVID-19 Surveillance Team |
| EPI_ISL_3312471 | Lighthouse Lab in Glasgow | Wellcome Sanger Institute for the COVID-19 Genomics UK (COG-UK) Consortium | Harper VanSteenhouse, Yumi Kasai, David Gray, Carol Clugston, Anna Dominiczak and Alex Alderton, Roberto Amato, Jeffrey Barrett, Sonia Goncalves, Ewan Harrison, David K. Jackson, Ian Johnston, Dominic Kwiatkowski, Cordelia Langford, John Sillitoe on behalf of the Wellcome Sanger Institute COVID-19 Surveillance Team |
| EPI_ISL_3312472, EPI_ISL_3312473 | Lighthouse Lab in Alderley Park | Wellcome Sanger Institute for the COVID-19 Genomics UK (COG-UK) Consortium | Jacquelyn Wynn, Mairead Hyland, The Lighthouse Lab in Alderley Park and Alex Alderton, Roberto Amato, Jeffrey Barrett, Sonia Goncalves, Ewan Harrison, David K. Jackson, Ian Johnston, Dominic Kwiatkowski, Cordelia Langford, John Sillitoe on behalf of the Wellcome Sanger Institute COVID-19 Surveillance Team |
| EPI_ISL_3312474 | Lighthouse Laboratory Plymouth | Wellcome Sanger Institute for the COVID-19 Genomics UK (COG-UK) Consortium | Lighthouse Laboratory Plymouth and Alex Alderton, Roberto Amato, Jeffrey Barrett, Sonia Goncalves, Ewan Harrison, David K. Jackson, Ian Johnston, Dominic Kwiatkowski, Cordelia Langford, John Sillitoe on behalf of the Wellcome Sanger Institute COVID-19 Surveillance Team |
| EPI_ISL_3312475 | Lighthouse Lab in Alderley Park | Wellcome Sanger Institute for the COVID-19 Genomics UK (COG-UK) Consortium | Jacquelyn Wynn, Mairead Hyland, The Lighthouse Lab in Alderley Park and Alex Alderton, Roberto Amato, Jeffrey Barrett, Sonia Goncalves, Ewan Harrison, David K. Jackson, Ian Johnston, Dominic Kwiatkowski, Cordelia Langford, John Sillitoe on behalf of the Wellcome Sanger Institute COVID-19 Surveillance Team |
| EPI_ISL_3312476, EPI_ISL_3312477, EPI_ISL_3312478 | Lighthouse Lab in Glasgow | Wellcome Sanger Institute for the COVID-19 Genomics UK (COG-UK) Consortium | Harper VanSteenhouse, Yumi Kasai, David Gray, Carol Clugston, Anna Dominiczak and Alex Alderton, Roberto Amato, Jeffrey Barrett, Sonia Goncalves, Ewan Harrison, David K. Jackson, Ian Johnston, Dominic Kwiatkowski, Cordelia Langford, John Sillitoe on behalf of the Wellcome Sanger Institute COVID-19 Surveillance Team |
| EPI_ISL_3312479 | Lighthouse Laboratory Plymouth | Wellcome Sanger Institute for the COVID-19 Genomics UK (COG-UK) Consortium | Lighthouse Laboratory Plymouth and Alex Alderton, Roberto Amato, Jeffrey Barrett, Sonia Goncalves, Ewan Harrison, David K. Jackson, Ian Johnston, Dominic Kwiatkowski, Cordelia Langford, John Sillitoe on behalf of the Wellcome Sanger Institute COVID-19 Surveillance Team |
| EPI_ISL_3312480, EPI_ISL_3312481, EPI_ISL_3312482, EPI_ISL_3312483, EPI_ISL_3312484 | Lighthouse Lab in Alderley Park | Wellcome Sanger Institute for the COVID-19 Genomics UK (COG-UK) Consortium | Jacquelyn Wynn, Mairead Hyland, The Lighthouse Lab in Alderley Park and Alex Alderton, Roberto Amato, Jeffrey Barrett, Sonia Goncalves, Ewan Harrison, David K. Jackson, Ian Johnston, Dominic Kwiatkowski, Cordelia Langford, John Sillitoe on behalf of the Wellcome Sanger Institute COVID-19 Surveillance Team |
| EPI_ISL_3312485, EPI_ISL_3312486 | Lighthouse Lab in Glasgow | Wellcome Sanger Institute for the COVID-19 Genomics UK (COG-UK) Consortium | Harper VanSteenhouse, Yumi Kasai, David Gray, Carol Clugston, Anna Dominiczak and Alex Alderton, Roberto Amato, Jeffrey Barrett, Sonia Goncalves, Ewan Harrison, David K. Jackson, Ian Johnston, Dominic Kwiatkowski, Cordelia Langford, John Sillitoe on behalf of the Wellcome Sanger Institute COVID-19 Surveillance Team |
| EPI_ISL_3312487, EPI_ISL_3312488, EPI_ISL_3312489, EPI_ISL_3312490 | Lighthouse Laboratory Plymouth | Wellcome Sanger Institute for the COVID-19 Genomics UK (COG-UK) Consortium | Lighthouse Laboratory Plymouth and Alex Alderton, Roberto Amato, Jeffrey Barrett, Sonia Goncalves, Ewan Harrison, David K. Jackson, Ian Johnston, Dominic Kwiatkowski, Cordelia Langford, John Sillitoe on behalf of the Wellcome Sanger Institute COVID-19 Surveillance Team |
| EPI_ISL_3312491, EPI_ISL_3312492 | Lighthouse Lab in Glasgow | Wellcome Sanger Institute for the COVID-19 Genomics UK (COG-UK) Consortium | Harper VanSteenhouse, Yumi Kasai, David Gray, Carol Clugston, Anna Dominiczak and Alex Alderton, Roberto Amato, Jeffrey Barrett, Sonia Goncalves, Ewan Harrison, David K. Jackson, Ian Johnston, Dominic Kwiatkowski, Cordelia Langford, John Sillitoe on behalf of the Wellcome Sanger Institute COVID-19 Surveillance Team |
| EPI_ISL_3312493 | Lighthouse Lab in Alderley Park | Wellcome Sanger Institute for the COVID-19 Genomics UK (COG-UK) Consortium | Jacquelyn Wynn, Mairead Hyland, The Lighthouse Lab in Alderley Park and Alex Alderton, Roberto Amato, Jeffrey Barrett, Sonia Goncalves, Ewan Harrison, David K. Jackson, Ian Johnston, Dominic Kwiatkowski, Cordelia Langford, John Sillitoe on behalf of the Wellcome Sanger Institute COVID-19 Surveillance Team |
| EPI_ISL_3312494 | Lighthouse Laboratory Plymouth | Wellcome Sanger Institute for the COVID-19 Genomics UK (COG-UK) Consortium | Lighthouse Laboratory Plymouth and Alex Alderton, Roberto Amato, Jeffrey Barrett, Sonia Goncalves, Ewan Harrison, David K. Jackson, Ian Johnston, Dominic Kwiatkowski, Cordelia Langford, John Sillitoe on behalf of the Wellcome Sanger Institute COVID-19 Surveillance Team |
| EPI_ISL_3312495 | Lighthouse Lab in Glasgow | Wellcome Sanger Institute for the COVID-19 Genomics UK (COG-UK) Consortium | Harper VanSteenhouse, Yumi Kasai, David Gray, Carol Clugston, Anna Dominiczak and Alex Alderton, Roberto Amato, Jeffrey Barrett, Sonia Goncalves, Ewan Harrison, David K. Jackson, Ian Johnston, Dominic Kwiatkowski, Cordelia Langford, John Sillitoe on behalf of the Wellcome Sanger Institute COVID-19 Surveillance Team |
| EPI_ISL_3312496 | Lighthouse Lab in Alderley Park | Wellcome Sanger Institute for the COVID-19 Genomics UK (COG-UK) Consortium | Jacquelyn Wynn, Mairead Hyland, The Lighthouse Lab in Alderley Park and Alex Alderton, Roberto Amato, Jeffrey Barrett, Sonia Goncalves, Ewan Harrison, David K. Jackson, Ian Johnston, Dominic Kwiatkowski, Cordelia Langford, John Sillitoe on behalf of the Wellcome Sanger Institute COVID-19 Surveillance Team |
| EPI_ISL_3312497 | Lighthouse Lab in Glasgow | Wellcome Sanger Institute for the COVID-19 Genomics UK (COG-UK) Consortium | Harper VanSteenhouse, Yumi Kasai, David Gray, Carol Clugston, Anna Dominiczak and Alex Alderton, Roberto Amato, Jeffrey Barrett, Sonia Goncalves, Ewan Harrison, David K. Jackson, Ian Johnston, Dominic Kwiatkowski, Cordelia Langford, John Sillitoe on behalf of the Wellcome Sanger Institute COVID-19 Surveillance Team |
| EPI_ISL_3312498, EPI_ISL_3312499 | Lighthouse Laboratory Plymouth | Wellcome Sanger Institute for the COVID-19 Genomics UK (COG-UK) Consortium | Lighthouse Laboratory Plymouth and Alex Alderton, Roberto Amato, Jeffrey Barrett, Sonia Goncalves, Ewan Harrison, David K. Jackson, Ian Johnston, Dominic Kwiatkowski, Cordelia Langford, John Sillitoe on behalf of the Wellcome Sanger Institute COVID-19 Surveillance Team |
| EPI_ISL_3312500, EPI_ISL_3312501, EPI_ISL_3312502 | Lighthouse Lab in Glasgow | Wellcome Sanger Institute for the COVID-19 Genomics UK (COG-UK) Consortium | Harper VanSteenhouse, Yumi Kasai, David Gray, Carol Clugston, Anna Dominiczak and Alex Alderton, Roberto Amato, Jeffrey Barrett, Sonia Goncalves, Ewan Harrison, David K. Jackson, Ian Johnston, Dominic Kwiatkowski, Cordelia Langford, John Sillitoe on behalf of the Wellcome Sanger Institute COVID-19 Surveillance Team |
| EPI_ISL_3312503, EPI_ISL_3312504 | Lighthouse Lab in Alderley Park | Wellcome Sanger Institute for the COVID-19 Genomics UK (COG-UK) Consortium | Jacquelyn Wynn, Mairead Hyland, The Lighthouse Lab in Alderley Park and Alex Alderton, Roberto Amato, Jeffrey Barrett, Sonia Goncalves, Ewan Harrison, David K. Jackson, Ian Johnston, Dominic Kwiatkowski, Cordelia Langford, John Sillitoe on behalf of the Wellcome Sanger Institute COVID-19 Surveillance Team |
| EPI_ISL_3312505, EPI_ISL_3312506, EPI_ISL_3312507, EPI_ISL_3312508, EPI_ISL_3312509 | Lighthouse Lab in Glasgow | Wellcome Sanger Institute for the COVID-19 Genomics UK (COG-UK) Consortium | Harper VanSteenhouse, Yumi Kasai, David Gray, Carol Clugston, Anna Dominiczak and Alex Alderton, Roberto Amato, Jeffrey Barrett, Sonia Goncalves, Ewan Harrison, David K. Jackson, Ian Johnston, Dominic Kwiatkowski, Cordelia Langford, John Sillitoe on behalf of the Wellcome Sanger Institute COVID-19 Surveillance Team |
| EPI_ISL_3312510 | Lighthouse Laboratory Plymouth | Wellcome Sanger Institute for the COVID-19 Genomics UK (COG-UK) Consortium | Lighthouse Laboratory Plymouth and Alex Alderton, Roberto Amato, Jeffrey Barrett, Sonia Goncalves, Ewan Harrison, David K. Jackson, Ian Johnston, Dominic Kwiatkowski, Cordelia Langford, John Sillitoe on behalf of the Wellcome Sanger Institute COVID-19 Surveillance Team |
| EPI_ISL_3312511, EPI_ISL_3312512, EPI_ISL_3312513 | Lighthouse Lab in Alderley Park | Wellcome Sanger Institute for the COVID-19 Genomics UK (COG-UK) Consortium | Jacquelyn Wynn, Mairead Hyland, The Lighthouse Lab in Alderley Park and Alex Alderton, Roberto Amato, Jeffrey Barrett, Sonia Goncalves, Ewan Harrison, David K. Jackson, Ian Johnston, Dominic Kwiatkowski, Cordelia Langford, John Sillitoe on behalf of the Wellcome Sanger Institute COVID-19 Surveillance Team |
| EPI_ISL_3312514 | Lighthouse Laboratory Plymouth | Wellcome Sanger Institute for the COVID-19 Genomics UK (COG-UK) Consortium | Lighthouse Laboratory Plymouth and Alex Alderton, Roberto Amato, Jeffrey Barrett, Sonia Goncalves, Ewan Harrison, David K. Jackson, Ian Johnston, Dominic Kwiatkowski, Cordelia Langford, John Sillitoe on behalf of the Wellcome Sanger Institute COVID-19 Surveillance Team |
| EPI_ISL_3312515, EPI_ISL_3312516 | Lighthouse Lab in Glasgow | Wellcome Sanger Institute for the COVID-19 Genomics UK (COG-UK) Consortium | Harper VanSteenhouse, Yumi Kasai, David Gray, Carol Clugston, Anna Dominiczak and Alex Alderton, Roberto Amato, Jeffrey Barrett, Sonia Goncalves, Ewan Harrison, David K. Jackson, Ian Johnston, Dominic Kwiatkowski, Cordelia Langford, John Sillitoe on behalf of the Wellcome Sanger Institute COVID-19 Surveillance Team |
| EPI_ISL_3312517 | Lighthouse Lab in Alderley Park | Wellcome Sanger Institute for the COVID-19 Genomics UK (COG-UK) Consortium | Jacquelyn Wynn, Mairead Hyland, The Lighthouse Lab in Alderley Park and Alex Alderton, Roberto Amato, Jeffrey Barrett, Sonia Goncalves, Ewan Harrison, David K. Jackson, Ian Johnston, Dominic Kwiatkowski, Cordelia Langford, John Sillitoe on behalf of the Wellcome Sanger Institute COVID-19 Surveillance Team |
| EPI_ISL_3312518 | Lighthouse Lab in Glasgow | Wellcome Sanger Institute for the COVID-19 Genomics UK (COG-UK) Consortium | Harper VanSteenhouse, Yumi Kasai, David Gray, Carol Clugston, Anna Dominiczak and Alex Alderton, Roberto Amato, Jeffrey Barrett, Sonia Goncalves, Ewan Harrison, David K. Jackson, Ian Johnston, Dominic Kwiatkowski, Cordelia Langford, John Sillitoe on behalf of the Wellcome Sanger Institute COVID-19 Surveillance Team |
| EPI_ISL_3312519 | Lighthouse Laboratory Plymouth | Wellcome Sanger Institute for the COVID-19 Genomics UK (COG-UK) Consortium | Lighthouse Laboratory Plymouth and Alex Alderton, Roberto Amato, Jeffrey Barrett, Sonia Goncalves, Ewan Harrison, David K. Jackson, Ian Johnston, Dominic Kwiatkowski, Cordelia Langford, John Sillitoe on behalf of the Wellcome Sanger Institute COVID-19 Surveillance Team |
| EPI_ISL_3312520 | Lighthouse Lab in Alderley Park | Wellcome Sanger Institute for the COVID-19 Genomics UK (COG-UK) Consortium | Jacquelyn Wynn, Mairead Hyland, The Lighthouse Lab in Alderley Park and Alex Alderton, Roberto Amato, Jeffrey Barrett, Sonia Goncalves, Ewan Harrison, David K. Jackson, Ian Johnston, Dominic Kwiatkowski, Cordelia Langford, John Sillitoe on behalf of the Wellcome Sanger Institute COVID-19 Surveillance Team |
| EPI_ISL_3312521 | Lighthouse Laboratory Plymouth | Wellcome Sanger Institute for the COVID-19 Genomics UK (COG-UK) Consortium | Lighthouse Laboratory Plymouth and Alex Alderton, Roberto Amato, Jeffrey Barrett, Sonia Goncalves, Ewan Harrison, David K. Jackson, Ian Johnston, Dominic Kwiatkowski, Cordelia Langford, John Sillitoe on behalf of the Wellcome Sanger Institute COVID-19 Surveillance Team |
| EPI_ISL_3312522, EPI_ISL_3312523 | Lighthouse Lab in Glasgow | Wellcome Sanger Institute for the COVID-19 Genomics UK (COG-UK) Consortium | Harper VanSteenhouse, Yumi Kasai, David Gray, Carol Clugston, Anna Dominiczak and Alex Alderton, Roberto Amato, Jeffrey Barrett, Sonia Goncalves, Ewan Harrison, David K. Jackson, Ian Johnston, Dominic Kwiatkowski, Cordelia Langford, John Sillitoe on behalf of the Wellcome Sanger Institute COVID-19 Surveillance Team |
| EPI_ISL_3312524 | Lighthouse Lab in Alderley Park | Wellcome Sanger Institute for the COVID-19 Genomics UK (COG-UK) Consortium | Jacquelyn Wynn, Mairead Hyland, The Lighthouse Lab in Alderley Park and Alex Alderton, Roberto Amato, Jeffrey Barrett, Sonia Goncalves, Ewan Harrison, David K. Jackson, Ian Johnston, Dominic Kwiatkowski, Cordelia Langford, John Sillitoe on behalf of the Wellcome Sanger Institute COVID-19 Surveillance Team |
| EPI_ISL_3312525 | Lighthouse Lab in Glasgow | Wellcome Sanger Institute for the COVID-19 Genomics UK (COG-UK) Consortium | Harper VanSteenhouse, Yumi Kasai, David Gray, Carol Clugston, Anna Dominiczak and Alex Alderton, Roberto Amato, Jeffrey Barrett, Sonia Goncalves, Ewan Harrison, David K. Jackson, Ian Johnston, Dominic Kwiatkowski, Cordelia Langford, John Sillitoe on behalf of the Wellcome Sanger Institute COVID-19 Surveillance Team |
| EPI_ISL_3312526 | Lighthouse Laboratory Plymouth | Wellcome Sanger Institute for the COVID-19 Genomics UK (COG-UK) | Lighthouse Laboratory Plymouth and Alex Alderton, Roberto Amato, Jeffrey Barrett, Sonia Goncalves, Ewan Harrison, David K. Jackson, Ian Johnston, Dominic Kwiatkowski, Cordelia Langford, John Sillitoe on behalf of the Wellcome Sanger |

| | | | |
|---|---|---|---|
| | | Consortium | Institute COVID-19 Surveillance Team |
| EPI_ISL_3312527 | Lighthouse Lab in Alderley Park | Wellcome Sanger Institute for the COVID-19 Genomics UK (COG-UK) Consortium | Jacquelyn Wynn, Mairead Hyland, The Lighthouse Lab in Alderley Park and Alex Alderton, Roberto Amato, Jeffrey Barrett, Sonia Goncalves, Ewan Harrison, David K. Jackson, Ian Johnston, Dominic Kwiatkowski, Cordelia Langford, John Sillitoe on behalf of the Wellcome Sanger Institute COVID-19 Surveillance Team |
| EPI_ISL_3312528 | Lighthouse Lab in Glasgow | Wellcome Sanger Institute for the COVID-19 Genomics UK (COG-UK) Consortium | Harper VanSteenhouse, Yumi Kasai, David Gray, Carol Clugston, Anna Dominiczak and Alex Alderton, Roberto Amato, Jeffrey Barrett, Sonia Goncalves, Ewan Harrison, David K. Jackson, Ian Johnston, Dominic Kwiatkowski, Cordelia Langford, John Sillitoe on behalf of the Wellcome Sanger Institute COVID-19 Surveillance Team |
| EPI_ISL_3312529 | Lighthouse Laboratory Plymouth | Wellcome Sanger Institute for the COVID-19 Genomics UK (COG-UK) Consortium | Lighthouse Laboratory Plymouth and Alex Alderton, Roberto Amato, Jeffrey Barrett, Sonia Goncalves, Ewan Harrison, David K. Jackson, Ian Johnston, Dominic Kwiatkowski, Cordelia Langford, John Sillitoe on behalf of the Wellcome Sanger Institute COVID-19 Surveillance Team |
| EPI_ISL_3312530, EPI_ISL_3312531 | Lighthouse Lab in Glasgow | Wellcome Sanger Institute for the COVID-19 Genomics UK (COG-UK) Consortium | Harper VanSteenhouse, Yumi Kasai, David Gray, Carol Clugston, Anna Dominiczak and Alex Alderton, Roberto Amato, Jeffrey Barrett, Sonia Goncalves, Ewan Harrison, David K. Jackson, Ian Johnston, Dominic Kwiatkowski, Cordelia Langford, John Sillitoe on behalf of the Wellcome Sanger Institute COVID-19 Surveillance Team |
| EPI_ISL_3312532, EPI_ISL_3312533 | Lighthouse Laboratory Plymouth | Wellcome Sanger Institute for the COVID-19 Genomics UK (COG-UK) Consortium | Lighthouse Laboratory Plymouth and Alex Alderton, Roberto Amato, Jeffrey Barrett, Sonia Goncalves, Ewan Harrison, David K. Jackson, Ian Johnston, Dominic Kwiatkowski, Cordelia Langford, John Sillitoe on behalf of the Wellcome Sanger Institute COVID-19 Surveillance Team |
| EPI_ISL_3312534 | Lighthouse Lab in Glasgow | Wellcome Sanger Institute for the COVID-19 Genomics UK (COG-UK) Consortium | Harper VanSteenhouse, Yumi Kasai, David Gray, Carol Clugston, Anna Dominiczak and Alex Alderton, Roberto Amato, Jeffrey Barrett, Sonia Goncalves, Ewan Harrison, David K. Jackson, Ian Johnston, Dominic Kwiatkowski, Cordelia Langford, John Sillitoe on behalf of the Wellcome Sanger Institute COVID-19 Surveillance Team |
| EPI_ISL_3312535 | Lighthouse Lab in Milton Keynes | Wellcome Sanger Institute for the COVID-19 Genomics UK (COG-UK) Consortium | The Lighthouse Lab in Milton Keynes and Alex Alderton, Roberto Amato, Jeffrey Barrett, Sonia Goncalves, Ewan Harrison, David K. Jackson, Ian Johnston, Dominic Kwiatkowski, Cordelia Langford, John Sillitoe on behalf of the Wellcome Sanger Institute COVID-19 Surveillance Team |
| EPI_ISL_3312536 | Lighthouse Lab in Alderley Park | Wellcome Sanger Institute for the COVID-19 Genomics UK (COG-UK) Consortium | Jacquelyn Wynn, Mairead Hyland, The Lighthouse Lab in Alderley Park and Alex Alderton, Roberto Amato, Jeffrey Barrett, Sonia Goncalves, Ewan Harrison, David K. Jackson, Ian Johnston, Dominic Kwiatkowski, Cordelia Langford, John Sillitoe on behalf of the Wellcome Sanger Institute COVID-19 Surveillance Team |
| EPI_ISL_3312537, EPI_ISL_3312538 | Lighthouse Lab in Milton Keynes | Wellcome Sanger Institute for the COVID-19 Genomics UK (COG-UK) Consortium | The Lighthouse Lab in Milton Keynes and Alex Alderton, Roberto Amato, Jeffrey Barrett, Sonia Goncalves, Ewan Harrison, David K. Jackson, Ian Johnston, Dominic Kwiatkowski, Cordelia Langford, John Sillitoe on behalf of the Wellcome Sanger Institute COVID-19 Surveillance Team |
| EPI_ISL_3312539, EPI_ISL_3312540, EPI_ISL_3312541, EPI_ISL_3312542 | Lighthouse Lab in Alderley Park | Wellcome Sanger Institute for the COVID-19 Genomics UK (COG-UK) Consortium | Jacquelyn Wynn, Mairead Hyland, The Lighthouse Lab in Alderley Park and Alex Alderton, Roberto Amato, Jeffrey Barrett, Sonia Goncalves, Ewan Harrison, David K. Jackson, Ian Johnston, Dominic Kwiatkowski, Cordelia Langford, John Sillitoe on behalf of the Wellcome Sanger Institute COVID-19 Surveillance Team |
| EPI_ISL_3312543, EPI_ISL_3312544, EPI_ISL_3312545 | Lighthouse Lab in Milton Keynes | Wellcome Sanger Institute for the COVID-19 Genomics UK (COG-UK) Consortium | The Lighthouse Lab in Milton Keynes and Alex Alderton, Roberto Amato, Jeffrey Barrett, Sonia Goncalves, Ewan Harrison, David K. Jackson, Ian Johnston, Dominic Kwiatkowski, Cordelia Langford, John Sillitoe on behalf of the Wellcome Sanger Institute COVID-19 Surveillance Team |
| EPI_ISL_3312546, EPI_ISL_3312547, EPI_ISL_3312548, EPI_ISL_3312549 | Lighthouse Lab in Alderley Park | Wellcome Sanger Institute for the COVID-19 Genomics UK (COG-UK) Consortium | Jacquelyn Wynn, Mairead Hyland, The Lighthouse Lab in Alderley Park and Alex Alderton, Roberto Amato, Jeffrey Barrett, Sonia Goncalves, Ewan Harrison, David K. Jackson, Ian Johnston, Dominic Kwiatkowski, Cordelia Langford, John Sillitoe on behalf of the Wellcome Sanger Institute COVID-19 Surveillance Team |
| EPI_ISL_3312550 | Lighthouse Lab in Milton Keynes | Wellcome Sanger Institute for the COVID-19 Genomics UK (COG-UK) Consortium | The Lighthouse Lab in Milton Keynes and Alex Alderton, Roberto Amato, Jeffrey Barrett, Sonia Goncalves, Ewan Harrison, David K. Jackson, Ian Johnston, Dominic Kwiatkowski, Cordelia Langford, John Sillitoe on behalf of the Wellcome Sanger Institute COVID-19 Surveillance Team |
| EPI_ISL_3312551, EPI_ISL_3312552 | Lighthouse Lab in Alderley Park | Wellcome Sanger Institute for the COVID-19 Genomics UK (COG-UK) Consortium | Jacquelyn Wynn, Mairead Hyland, The Lighthouse Lab in Alderley Park and Alex Alderton, Roberto Amato, Jeffrey Barrett, Sonia Goncalves, Ewan Harrison, David K. Jackson, Ian Johnston, Dominic Kwiatkowski, Cordelia Langford, John Sillitoe on behalf of the Wellcome Sanger Institute COVID-19 Surveillance Team |
| EPI_ISL_3312553, EPI_ISL_3312554, EPI_ISL_3312555 | Lighthouse Lab in Milton Keynes | Wellcome Sanger Institute for the COVID-19 Genomics UK (COG-UK) Consortium | The Lighthouse Lab in Milton Keynes and Alex Alderton, Roberto Amato, Jeffrey Barrett, Sonia Goncalves, Ewan Harrison, David K. Jackson, Ian Johnston, Dominic Kwiatkowski, Cordelia Langford, John Sillitoe on behalf of the Wellcome Sanger Institute COVID-19 Surveillance Team |
| EPI_ISL_3312556 | Lighthouse Lab in Alderley Park | Wellcome Sanger Institute for the COVID-19 Genomics UK (COG-UK) Consortium | Jacquelyn Wynn, Mairead Hyland, The Lighthouse Lab in Alderley Park and Alex Alderton, Roberto Amato, Jeffrey Barrett, Sonia Goncalves, Ewan Harrison, David K. Jackson, Ian Johnston, Dominic Kwiatkowski, Cordelia Langford, John Sillitoe on behalf of the Wellcome Sanger Institute COVID-19 Surveillance Team |
| EPI_ISL_3312557, EPI_ISL_3312558, EPI_ISL_3312559, EPI_ISL_3312560 | Lighthouse Lab in Milton Keynes | Wellcome Sanger Institute for the COVID-19 Genomics UK (COG-UK) Consortium | The Lighthouse Lab in Milton Keynes and Alex Alderton, Roberto Amato, Jeffrey Barrett, Sonia Goncalves, Ewan Harrison, David K. Jackson, Ian Johnston, Dominic Kwiatkowski, Cordelia Langford, John Sillitoe on behalf of the Wellcome Sanger Institute COVID-19 Surveillance Team |
| EPI_ISL_3312561 | Lighthouse Lab in Alderley Park | Wellcome Sanger Institute for the COVID-19 Genomics UK (COG-UK) Consortium | Jacquelyn Wynn, Mairead Hyland, The Lighthouse Lab in Alderley Park and Alex Alderton, Roberto Amato, Jeffrey Barrett, Sonia Goncalves, Ewan Harrison, David K. Jackson, Ian Johnston, Dominic Kwiatkowski, Cordelia Langford, John Sillitoe on behalf of the Wellcome Sanger Institute COVID-19 Surveillance Team |
| EPI_ISL_3312562 | Lighthouse Lab in Milton Keynes | Wellcome Sanger Institute for the COVID-19 Genomics UK (COG-UK) Consortium | The Lighthouse Lab in Milton Keynes and Alex Alderton, Roberto Amato, Jeffrey Barrett, Sonia Goncalves, Ewan Harrison, David K. Jackson, Ian Johnston, Dominic Kwiatkowski, Cordelia Langford, John Sillitoe on behalf of the Wellcome Sanger Institute COVID-19 Surveillance Team |
| EPI_ISL_3312563, EPI_ISL_3312564, EPI_ISL_3312565 | Lighthouse Lab in Alderley Park | Wellcome Sanger Institute for the COVID-19 Genomics UK (COG-UK) Consortium | Jacquelyn Wynn, Mairead Hyland, The Lighthouse Lab in Alderley Park and Alex Alderton, Roberto Amato, Jeffrey Barrett, Sonia Goncalves, Ewan Harrison, David K. Jackson, Ian Johnston, Dominic Kwiatkowski, Cordelia Langford, John Sillitoe on behalf of the Wellcome Sanger Institute COVID-19 Surveillance Team |
| EPI_ISL_3312566 | Lighthouse Lab in Milton Keynes | Wellcome Sanger Institute for the COVID-19 Genomics UK (COG-UK) Consortium | The Lighthouse Lab in Milton Keynes and Alex Alderton, Roberto Amato, Jeffrey Barrett, Sonia Goncalves, Ewan Harrison, David K. Jackson, Ian Johnston, Dominic Kwiatkowski, Cordelia Langford, John Sillitoe on behalf of the Wellcome Sanger Institute COVID-19 Surveillance Team |
| EPI_ISL_3312567 | Lighthouse Lab in Alderley Park | Wellcome Sanger Institute for the COVID-19 Genomics UK (COG-UK) Consortium | Jacquelyn Wynn, Mairead Hyland, The Lighthouse Lab in Alderley Park and Alex Alderton, Roberto Amato, Jeffrey Barrett, Sonia Goncalves, Ewan Harrison, David K. Jackson, Ian Johnston, Dominic Kwiatkowski, Cordelia Langford, John Sillitoe on behalf of the Wellcome Sanger Institute COVID-19 Surveillance Team |
| EPI_ISL_3312568, EPI_ISL_3312569, EPI_ISL_3312570, EPI_ISL_3312571 | Lighthouse Lab in Milton Keynes | Wellcome Sanger Institute for the COVID-19 Genomics UK (COG-UK) Consortium | The Lighthouse Lab in Milton Keynes and Alex Alderton, Roberto Amato, Jeffrey Barrett, Sonia Goncalves, Ewan Harrison, David K. Jackson, Ian Johnston, Dominic Kwiatkowski, Cordelia Langford, John Sillitoe on behalf of the Wellcome Sanger Institute COVID-19 Surveillance Team |
| EPI_ISL_3312572, EPI_ISL_3312573, EPI_ISL_3312574 | Lighthouse Lab in Alderley Park | Wellcome Sanger Institute for the COVID-19 Genomics UK (COG-UK) Consortium | Jacquelyn Wynn, Mairead Hyland, The Lighthouse Lab in Alderley Park and Alex Alderton, Roberto Amato, Jeffrey Barrett, Sonia Goncalves, Ewan Harrison, David K. Jackson, Ian Johnston, Dominic Kwiatkowski, Cordelia Langford, John Sillitoe on behalf of the Wellcome Sanger Institute COVID-19 Surveillance Team |
| EPI_ISL_3312575, EPI_ISL_3312576, EPI_ISL_3312577, EPI_ISL_3312578, EPI_ISL_3312579, EPI_ISL_3312580 | Lighthouse Lab in Milton Keynes | Wellcome Sanger Institute for the COVID-19 Genomics UK (COG-UK) Consortium | The Lighthouse Lab in Milton Keynes and Alex Alderton, Roberto Amato, Jeffrey Barrett, Sonia Goncalves, Ewan Harrison, David K. Jackson, Ian Johnston, Dominic Kwiatkowski, Cordelia Langford, John Sillitoe on behalf of the Wellcome Sanger Institute COVID-19 Surveillance Team |
| EPI_ISL_3312581, EPI_ISL_3312582 | Lighthouse Lab in Alderley Park | Wellcome Sanger Institute for the COVID-19 Genomics UK (COG-UK) Consortium | Jacquelyn Wynn, Mairead Hyland, The Lighthouse Lab in Alderley Park and Alex Alderton, Roberto Amato, Jeffrey Barrett, Sonia Goncalves, Ewan Harrison, David K. Jackson, Ian Johnston, Dominic Kwiatkowski, Cordelia Langford, John Sillitoe on behalf of the Wellcome Sanger Institute COVID-19 Surveillance Team |
| EPI_ISL_3312583 | Lighthouse Lab in Milton Keynes | Wellcome Sanger Institute for the COVID-19 Genomics UK (COG-UK) Consortium | The Lighthouse Lab in Milton Keynes and Alex Alderton, Roberto Amato, Jeffrey Barrett, Sonia Goncalves, Ewan Harrison, David K. Jackson, Ian Johnston, Dominic Kwiatkowski, Cordelia Langford, John Sillitoe on behalf of the Wellcome Sanger Institute COVID-19 Surveillance Team |
| EPI_ISL_3312584, EPI_ISL_3312585, EPI_ISL_3312586, EPI_ISL_3312587, EPI_ISL_3312588, EPI_ISL_3312589, EPI_ISL_3312590 | Lighthouse Lab in Alderley Park | Wellcome Sanger Institute for the COVID-19 Genomics UK (COG-UK) Consortium | Jacquelyn Wynn, Mairead Hyland, The Lighthouse Lab in Alderley Park and Alex Alderton, Roberto Amato, Jeffrey Barrett, Sonia Goncalves, Ewan Harrison, David K. Jackson, Ian Johnston, Dominic Kwiatkowski, Cordelia Langford, John Sillitoe on behalf of the Wellcome Sanger Institute COVID-19 Surveillance Team |
| EPI_ISL_3312591, EPI_ISL_3312592, EPI_ISL_3312593, EPI_ISL_3312594, EPI_ISL_3312595, EPI_ISL_3312596, EPI_ISL_3312597, EPI_ISL_3312598 | Lighthouse Lab in Milton Keynes | Wellcome Sanger Institute for the COVID-19 Genomics UK (COG-UK) Consortium | The Lighthouse Lab in Milton Keynes and Alex Alderton, Roberto Amato, Jeffrey Barrett, Sonia Goncalves, Ewan Harrison, David K. Jackson, Ian Johnston, Dominic Kwiatkowski, Cordelia Langford, John Sillitoe on behalf of the Wellcome Sanger Institute COVID-19 Surveillance Team |
| EPI_ISL_3312599, EPI_ISL_3312600 | Lighthouse Lab in Alderley Park | Wellcome Sanger Institute for the COVID-19 Genomics UK (COG-UK) Consortium | Jacquelyn Wynn, Mairead Hyland, The Lighthouse Lab in Alderley Park and Alex Alderton, Roberto Amato, Jeffrey Barrett, Sonia Goncalves, Ewan Harrison, David K. Jackson, Ian Johnston, Dominic Kwiatkowski, Cordelia Langford, John Sillitoe on behalf of the Wellcome Sanger Institute COVID-19 Surveillance Team |
| EPI_ISL_3312601 | Lighthouse Lab in Milton Keynes | Wellcome Sanger Institute for the COVID-19 Genomics UK (COG-UK) Consortium | The Lighthouse Lab in Milton Keynes and Alex Alderton, Roberto Amato, Jeffrey Barrett, Sonia Goncalves, Ewan Harrison, David K. Jackson, Ian Johnston, Dominic Kwiatkowski, Cordelia Langford, John Sillitoe on behalf of the Wellcome Sanger Institute COVID-19 Surveillance Team |
| EPI_ISL_3312602, EPI_ISL_3312603 | Lighthouse Lab in Alderley Park | Wellcome Sanger Institute for the COVID-19 Genomics UK (COG-UK) Consortium | Jacquelyn Wynn, Mairead Hyland, The Lighthouse Lab in Alderley Park and Alex Alderton, Roberto Amato, Jeffrey Barrett, Sonia Goncalves, Ewan Harrison, David K. Jackson, Ian Johnston, Dominic Kwiatkowski, Cordelia Langford, John Sillitoe on behalf of the Wellcome Sanger Institute COVID-19 Surveillance Team |
| EPI_ISL_3312604, EPI_ISL_3312605, EPI_ISL_3312606, | Lighthouse Lab in Milton Keynes | Wellcome Sanger Institute for the COVID-19 Genomics UK (COG-UK) Consortium | The Lighthouse Lab in Milton Keynes and Alex Alderton, Roberto Amato, Jeffrey Barrett, Sonia Goncalves, Ewan Harrison, David K. Jackson, Ian Johnston, Dominic Kwiatkowski, Cordelia Langford, John Sillitoe on behalf of the Wellcome Sanger Institute COVID-19 Surveillance Team |

| | | | |
|---|---|---|---|
| EPI_ISL_3312607, EPI_ISL_3312608 | | | |
| EPI_ISL_3312609 | Lighthouse Lab in Alderley Park | Wellcome Sanger Institute for the COVID-19 Genomics UK (COG-UK) Consortium | Jacquelyn Wynn, Mairead Hyland, The Lighthouse Lab in Alderley Park and Alex Alderton, Roberto Amato, Jeffrey Barrett, Sonia Goncalves, Ewan Harrison, David K. Jackson, Ian Johnston, Dominic Kwiatkowski, Cordelia Langford, John Sillitoe on behalf of the Wellcome Sanger Institute COVID-19 Surveillance Team |
| EPI_ISL_3312610 | Lighthouse Lab in Milton Keynes | Wellcome Sanger Institute for the COVID-19 Genomics UK (COG-UK) Consortium | The Lighthouse Lab in Milton Keynes and Alex Alderton, Roberto Amato, Jeffrey Barrett, Sonia Goncalves, Ewan Harrison, David K. Jackson, Ian Johnston, Dominic Kwiatkowski, Cordelia Langford, John Sillitoe on behalf of the Wellcome Sanger Institute COVID-19 Surveillance Team |
| EPI_ISL_3312611, EPI_ISL_3312612 | Lighthouse Lab in Alderley Park | Wellcome Sanger Institute for the COVID-19 Genomics UK (COG-UK) Consortium | Jacquelyn Wynn, Mairead Hyland, The Lighthouse Lab in Alderley Park and Alex Alderton, Roberto Amato, Jeffrey Barrett, Sonia Goncalves, Ewan Harrison, David K. Jackson, Ian Johnston, Dominic Kwiatkowski, Cordelia Langford, John Sillitoe on behalf of the Wellcome Sanger Institute COVID-19 Surveillance Team |
| EPI_ISL_3312613 | Lighthouse Lab in Milton Keynes | Wellcome Sanger Institute for the COVID-19 Genomics UK (COG-UK) Consortium | The Lighthouse Lab in Milton Keynes and Alex Alderton, Roberto Amato, Jeffrey Barrett, Sonia Goncalves, Ewan Harrison, David K. Jackson, Ian Johnston, Dominic Kwiatkowski, Cordelia Langford, John Sillitoe on behalf of the Wellcome Sanger Institute COVID-19 Surveillance Team |
| EPI_ISL_3312614 | Lighthouse Lab in Alderley Park | Wellcome Sanger Institute for the COVID-19 Genomics UK (COG-UK) Consortium | Jacquelyn Wynn, Mairead Hyland, The Lighthouse Lab in Alderley Park and Alex Alderton, Roberto Amato, Jeffrey Barrett, Sonia Goncalves, Ewan Harrison, David K. Jackson, Ian Johnston, Dominic Kwiatkowski, Cordelia Langford, John Sillitoe on behalf of the Wellcome Sanger Institute COVID-19 Surveillance Team |
| EPI_ISL_3312615 | Lighthouse Lab in Milton Keynes | Wellcome Sanger Institute for the COVID-19 Genomics UK (COG-UK) Consortium | The Lighthouse Lab in Milton Keynes and Alex Alderton, Roberto Amato, Jeffrey Barrett, Sonia Goncalves, Ewan Harrison, David K. Jackson, Ian Johnston, Dominic Kwiatkowski, Cordelia Langford, John Sillitoe on behalf of the Wellcome Sanger Institute COVID-19 Surveillance Team |
| EPI_ISL_3312616 | Lighthouse Lab in Alderley Park | Wellcome Sanger Institute for the COVID-19 Genomics UK (COG-UK) Consortium | Jacquelyn Wynn, Mairead Hyland, The Lighthouse Lab in Alderley Park and Alex Alderton, Roberto Amato, Jeffrey Barrett, Sonia Goncalves, Ewan Harrison, David K. Jackson, Ian Johnston, Dominic Kwiatkowski, Cordelia Langford, John Sillitoe on behalf of the Wellcome Sanger Institute COVID-19 Surveillance Team |
| EPI_ISL_3312617, EPI_ISL_3312618, EPI_ISL_3312619 | Lighthouse Lab in Milton Keynes | Wellcome Sanger Institute for the COVID-19 Genomics UK (COG-UK) Consortium | The Lighthouse Lab in Milton Keynes and Alex Alderton, Roberto Amato, Jeffrey Barrett, Sonia Goncalves, Ewan Harrison, David K. Jackson, Ian Johnston, Dominic Kwiatkowski, Cordelia Langford, John Sillitoe on behalf of the Wellcome Sanger Institute COVID-19 Surveillance Team |
| EPI_ISL_3312620, EPI_ISL_3312621 | Lighthouse Lab in Alderley Park | Wellcome Sanger Institute for the COVID-19 Genomics UK (COG-UK) Consortium | Jacquelyn Wynn, Mairead Hyland, The Lighthouse Lab in Alderley Park and Alex Alderton, Roberto Amato, Jeffrey Barrett, Sonia Goncalves, Ewan Harrison, David K. Jackson, Ian Johnston, Dominic Kwiatkowski, Cordelia Langford, John Sillitoe on behalf of the Wellcome Sanger Institute COVID-19 Surveillance Team |
| EPI_ISL_3312622 | Lighthouse Lab in Milton Keynes | Wellcome Sanger Institute for the COVID-19 Genomics UK (COG-UK) Consortium | The Lighthouse Lab in Milton Keynes and Alex Alderton, Roberto Amato, Jeffrey Barrett, Sonia Goncalves, Ewan Harrison, David K. Jackson, Ian Johnston, Dominic Kwiatkowski, Cordelia Langford, John Sillitoe on behalf of the Wellcome Sanger Institute COVID-19 Surveillance Team |
| EPI_ISL_3312623, EPI_ISL_3312624, EPI_ISL_3312625 | Lighthouse Lab in Alderley Park | Wellcome Sanger Institute for the COVID-19 Genomics UK (COG-UK) Consortium | Jacquelyn Wynn, Mairead Hyland, The Lighthouse Lab in Alderley Park and Alex Alderton, Roberto Amato, Jeffrey Barrett, Sonia Goncalves, Ewan Harrison, David K. Jackson, Ian Johnston, Dominic Kwiatkowski, Cordelia Langford, John Sillitoe on behalf of the Wellcome Sanger Institute COVID-19 Surveillance Team |
| EPI_ISL_3312626 | Lighthouse Lab in Milton Keynes | Wellcome Sanger Institute for the COVID-19 Genomics UK (COG-UK) Consortium | The Lighthouse Lab in Milton Keynes and Alex Alderton, Roberto Amato, Jeffrey Barrett, Sonia Goncalves, Ewan Harrison, David K. Jackson, Ian Johnston, Dominic Kwiatkowski, Cordelia Langford, John Sillitoe on behalf of the Wellcome Sanger Institute COVID-19 Surveillance Team |
| EPI_ISL_3312627 | Lighthouse Lab in Alderley Park | Wellcome Sanger Institute for the COVID-19 Genomics UK (COG-UK) Consortium | Jacquelyn Wynn, Mairead Hyland, The Lighthouse Lab in Alderley Park and Alex Alderton, Roberto Amato, Jeffrey Barrett, Sonia Goncalves, Ewan Harrison, David K. Jackson, Ian Johnston, Dominic Kwiatkowski, Cordelia Langford, John Sillitoe on behalf of the Wellcome Sanger Institute COVID-19 Surveillance Team |
| EPI_ISL_3312628 | Lighthouse Lab in Milton Keynes | Wellcome Sanger Institute for the COVID-19 Genomics UK (COG-UK) Consortium | The Lighthouse Lab in Milton Keynes and Alex Alderton, Roberto Amato, Jeffrey Barrett, Sonia Goncalves, Ewan Harrison, David K. Jackson, Ian Johnston, Dominic Kwiatkowski, Cordelia Langford, John Sillitoe on behalf of the Wellcome Sanger Institute COVID-19 Surveillance Team |
| EPI_ISL_3312629 | Lighthouse Lab in Alderley Park | Wellcome Sanger Institute for the COVID-19 Genomics UK (COG-UK) Consortium | Jacquelyn Wynn, Mairead Hyland, The Lighthouse Lab in Alderley Park and Alex Alderton, Roberto Amato, Jeffrey Barrett, Sonia Goncalves, Ewan Harrison, David K. Jackson, Ian Johnston, Dominic Kwiatkowski, Cordelia Langford, John Sillitoe on behalf of the Wellcome Sanger Institute COVID-19 Surveillance Team |
| EPI_ISL_3312630 | Lighthouse Lab in Milton Keynes | Wellcome Sanger Institute for the COVID-19 Genomics UK (COG-UK) Consortium | The Lighthouse Lab in Milton Keynes and Alex Alderton, Roberto Amato, Jeffrey Barrett, Sonia Goncalves, Ewan Harrison, David K. Jackson, Ian Johnston, Dominic Kwiatkowski, Cordelia Langford, John Sillitoe on behalf of the Wellcome Sanger Institute COVID-19 Surveillance Team |
| EPI_ISL_3312631, EPI_ISL_3312632, EPI_ISL_3312633 | Lighthouse Lab in Alderley Park | Wellcome Sanger Institute for the COVID-19 Genomics UK (COG-UK) Consortium | Jacquelyn Wynn, Mairead Hyland, The Lighthouse Lab in Alderley Park and Alex Alderton, Roberto Amato, Jeffrey Barrett, Sonia Goncalves, Ewan Harrison, David K. Jackson, Ian Johnston, Dominic Kwiatkowski, Cordelia Langford, John Sillitoe on behalf of the Wellcome Sanger Institute COVID-19 Surveillance Team |
| EPI_ISL_3312634, EPI_ISL_3312635, EPI_ISL_3312636 | Lighthouse Lab in Milton Keynes | Wellcome Sanger Institute for the COVID-19 Genomics UK (COG-UK) Consortium | The Lighthouse Lab in Milton Keynes and Alex Alderton, Roberto Amato, Jeffrey Barrett, Sonia Goncalves, Ewan Harrison, David K. Jackson, Ian Johnston, Dominic Kwiatkowski, Cordelia Langford, John Sillitoe on behalf of the Wellcome Sanger Institute COVID-19 Surveillance Team |
| EPI_ISL_3312637, EPI_ISL_3312638, EPI_ISL_3312639, EPI_ISL_3312640, EPI_ISL_3312641 | Lighthouse Lab in Alderley Park | Wellcome Sanger Institute for the COVID-19 Genomics UK (COG-UK) Consortium | Jacquelyn Wynn, Mairead Hyland, The Lighthouse Lab in Alderley Park and Alex Alderton, Roberto Amato, Jeffrey Barrett, Sonia Goncalves, Ewan Harrison, David K. Jackson, Ian Johnston, Dominic Kwiatkowski, Cordelia Langford, John Sillitoe on behalf of the Wellcome Sanger Institute COVID-19 Surveillance Team |
| EPI_ISL_3312642 | Lighthouse Lab in Milton Keynes | Wellcome Sanger Institute for the COVID-19 Genomics UK (COG-UK) Consortium | The Lighthouse Lab in Milton Keynes and Alex Alderton, Roberto Amato, Jeffrey Barrett, Sonia Goncalves, Ewan Harrison, David K. Jackson, Ian Johnston, Dominic Kwiatkowski, Cordelia Langford, John Sillitoe on behalf of the Wellcome Sanger Institute COVID-19 Surveillance Team |
| EPI_ISL_3312643, EPI_ISL_3312644, EPI_ISL_3312645 | Lighthouse Lab in Alderley Park | Wellcome Sanger Institute for the COVID-19 Genomics UK (COG-UK) Consortium | Jacquelyn Wynn, Mairead Hyland, The Lighthouse Lab in Alderley Park and Alex Alderton, Roberto Amato, Jeffrey Barrett, Sonia Goncalves, Ewan Harrison, David K. Jackson, Ian Johnston, Dominic Kwiatkowski, Cordelia Langford, John Sillitoe on behalf of the Wellcome Sanger Institute COVID-19 Surveillance Team |
| EPI_ISL_3312646, EPI_ISL_3312647 | Lighthouse Lab in Milton Keynes | Wellcome Sanger Institute for the COVID-19 Genomics UK (COG-UK) Consortium | The Lighthouse Lab in Milton Keynes and Alex Alderton, Roberto Amato, Jeffrey Barrett, Sonia Goncalves, Ewan Harrison, David K. Jackson, Ian Johnston, Dominic Kwiatkowski, Cordelia Langford, John Sillitoe on behalf of the Wellcome Sanger Institute COVID-19 Surveillance Team |
| EPI_ISL_3312648 | Lighthouse Lab in Alderley Park | Wellcome Sanger Institute for the COVID-19 Genomics UK (COG-UK) Consortium | Jacquelyn Wynn, Mairead Hyland, The Lighthouse Lab in Alderley Park and Alex Alderton, Roberto Amato, Jeffrey Barrett, Sonia Goncalves, Ewan Harrison, David K. Jackson, Ian Johnston, Dominic Kwiatkowski, Cordelia Langford, John Sillitoe on behalf of the Wellcome Sanger Institute COVID-19 Surveillance Team |
| EPI_ISL_3312649, EPI_ISL_3312650 | Lighthouse Lab in Milton Keynes | Wellcome Sanger Institute for the COVID-19 Genomics UK (COG-UK) Consortium | The Lighthouse Lab in Milton Keynes and Alex Alderton, Roberto Amato, Jeffrey Barrett, Sonia Goncalves, Ewan Harrison, David K. Jackson, Ian Johnston, Dominic Kwiatkowski, Cordelia Langford, John Sillitoe on behalf of the Wellcome Sanger Institute COVID-19 Surveillance Team |
| EPI_ISL_3312651 | Lighthouse Lab in Alderley Park | Wellcome Sanger Institute for the COVID-19 Genomics UK (COG-UK) Consortium | Jacquelyn Wynn, Mairead Hyland, The Lighthouse Lab in Alderley Park and Alex Alderton, Roberto Amato, Jeffrey Barrett, Sonia Goncalves, Ewan Harrison, David K. Jackson, Ian Johnston, Dominic Kwiatkowski, Cordelia Langford, John Sillitoe on behalf of the Wellcome Sanger Institute COVID-19 Surveillance Team |
| EPI_ISL_3312652, EPI_ISL_3312653, EPI_ISL_3312654, EPI_ISL_3312655 | Lighthouse Lab in Milton Keynes | Wellcome Sanger Institute for the COVID-19 Genomics UK (COG-UK) Consortium | The Lighthouse Lab in Milton Keynes and Alex Alderton, Roberto Amato, Jeffrey Barrett, Sonia Goncalves, Ewan Harrison, David K. Jackson, Ian Johnston, Dominic Kwiatkowski, Cordelia Langford, John Sillitoe on behalf of the Wellcome Sanger Institute COVID-19 Surveillance Team |
| EPI_ISL_3312656 | Lighthouse Lab in Alderley Park | Wellcome Sanger Institute for the COVID-19 Genomics UK (COG-UK) Consortium | Jacquelyn Wynn, Mairead Hyland, The Lighthouse Lab in Alderley Park and Alex Alderton, Roberto Amato, Jeffrey Barrett, Sonia Goncalves, Ewan Harrison, David K. Jackson, Ian Johnston, Dominic Kwiatkowski, Cordelia Langford, John Sillitoe on behalf of the Wellcome Sanger Institute COVID-19 Surveillance Team |
| EPI_ISL_3312657, EPI_ISL_3312658 | Lighthouse Lab in Milton Keynes | Wellcome Sanger Institute for the COVID-19 Genomics UK (COG-UK) Consortium | The Lighthouse Lab in Milton Keynes and Alex Alderton, Roberto Amato, Jeffrey Barrett, Sonia Goncalves, Ewan Harrison, David K. Jackson, Ian Johnston, Dominic Kwiatkowski, Cordelia Langford, John Sillitoe on behalf of the Wellcome Sanger Institute COVID-19 Surveillance Team |
| EPI_ISL_3312659, EPI_ISL_3312660 | Lighthouse Lab in Alderley Park | Wellcome Sanger Institute for the COVID-19 Genomics UK (COG-UK) Consortium | Jacquelyn Wynn, Mairead Hyland, The Lighthouse Lab in Alderley Park and Alex Alderton, Roberto Amato, Jeffrey Barrett, Sonia Goncalves, Ewan Harrison, David K. Jackson, Ian Johnston, Dominic Kwiatkowski, Cordelia Langford, John Sillitoe on behalf of the Wellcome Sanger Institute COVID-19 Surveillance Team |
| EPI_ISL_3312661 | Lighthouse Lab in Milton Keynes | Wellcome Sanger Institute for the COVID-19 Genomics UK (COG-UK) Consortium | The Lighthouse Lab in Milton Keynes and Alex Alderton, Roberto Amato, Jeffrey Barrett, Sonia Goncalves, Ewan Harrison, David K. Jackson, Ian Johnston, Dominic Kwiatkowski, Cordelia Langford, John Sillitoe on behalf of the Wellcome Sanger Institute COVID-19 Surveillance Team |
| EPI_ISL_3312662 | Lighthouse Lab in Alderley Park | Wellcome Sanger Institute for the COVID-19 Genomics UK (COG-UK) Consortium | Jacquelyn Wynn, Mairead Hyland, The Lighthouse Lab in Alderley Park and Alex Alderton, Roberto Amato, Jeffrey Barrett, Sonia Goncalves, Ewan Harrison, David K. Jackson, Ian Johnston, Dominic Kwiatkowski, Cordelia Langford, John Sillitoe on behalf of the Wellcome Sanger Institute COVID-19 Surveillance Team |
| EPI_ISL_3312663 | Lighthouse Lab in Milton Keynes | Wellcome Sanger Institute for the COVID-19 Genomics UK (COG-UK) Consortium | The Lighthouse Lab in Milton Keynes and Alex Alderton, Roberto Amato, Jeffrey Barrett, Sonia Goncalves, Ewan Harrison, David K. Jackson, Ian Johnston, Dominic Kwiatkowski, Cordelia Langford, John Sillitoe on behalf of the Wellcome Sanger Institute COVID-19 Surveillance Team |
| EPI_ISL_3312664 | Lighthouse Lab in Alderley Park | Wellcome Sanger Institute for the COVID-19 Genomics UK (COG-UK) Consortium | Jacquelyn Wynn, Mairead Hyland, The Lighthouse Lab in Alderley Park and Alex Alderton, Roberto Amato, Jeffrey Barrett, Sonia Goncalves, Ewan Harrison, David K. Jackson, Ian Johnston, Dominic Kwiatkowski, Cordelia Langford, John Sillitoe on behalf of the Wellcome Sanger Institute COVID-19 Surveillance Team |
| EPI_ISL_3312665 | Lighthouse Lab in Milton Keynes | Wellcome Sanger Institute for the COVID-19 Genomics UK (COG-UK) Consortium | The Lighthouse Lab in Milton Keynes and Alex Alderton, Roberto Amato, Jeffrey Barrett, Sonia Goncalves, Ewan Harrison, David K. Jackson, Ian Johnston, Dominic Kwiatkowski, Cordelia Langford, John Sillitoe on behalf of the Wellcome Sanger Institute COVID-19 Surveillance Team |
| EPI_ISL_3312666, EPI_ISL_3312667, EPI_ISL_3312668 | Lighthouse Lab in Alderley Park | Wellcome Sanger Institute for the COVID-19 Genomics UK (COG-UK) Consortium | Jacquelyn Wynn, Mairead Hyland, The Lighthouse Lab in Alderley Park and Alex Alderton, Roberto Amato, Jeffrey Barrett, Sonia Goncalves, Ewan Harrison, David K. Jackson, Ian Johnston, Dominic Kwiatkowski, Cordelia Langford, John Sillitoe on behalf of the Wellcome Sanger Institute COVID-19 Surveillance Team |
| EPI_ISL_3312669, EPI_ISL_3312670 | Lighthouse Lab in Milton Keynes | Wellcome Sanger Institute for the COVID-19 Genomics UK (COG-UK) Consortium | The Lighthouse Lab in Milton Keynes and Alex Alderton, Roberto Amato, Jeffrey Barrett, Sonia Goncalves, Ewan Harrison, David K. Jackson, Ian Johnston, Dominic Kwiatkowski, Cordelia Langford, John Sillitoe on behalf of the Wellcome Sanger Institute COVID-19 Surveillance Team |
| EPI_ISL_3312671 | Lighthouse Lab in Alderley Park | Wellcome Sanger Institute for the COVID-19 Genomics UK (COG-UK) Consortium | Jacquelyn Wynn, Mairead Hyland, The Lighthouse Lab in Alderley Park and Alex Alderton, Roberto Amato, Jeffrey Barrett, Sonia Goncalves, Ewan Harrison, David K. Jackson, Ian Johnston, Dominic Kwiatkowski, Cordelia Langford, John Sillitoe on behalf of the Wellcome Sanger Institute COVID-19 Surveillance Team |
| EPI_ISL_3312672 | Lighthouse Lab in Milton Keynes | Wellcome Sanger Institute for the COVID-19 Genomics UK (COG-UK) Consortium | The Lighthouse Lab in Milton Keynes and Alex Alderton, Roberto Amato, Jeffrey Barrett, Sonia Goncalves, Ewan Harrison, David K. Jackson, Ian Johnston, Dominic Kwiatkowski, Cordelia Langford, John Sillitoe on behalf of the Wellcome Sanger Institute COVID-19 Surveillance Team |
| EPI_ISL_3312673 | Lighthouse Lab in Alderley Park | Wellcome Sanger Institute for the COVID-19 Genomics UK (COG-UK) Consortium | Jacquelyn Wynn, Mairead Hyland, The Lighthouse Lab in Alderley Park and Alex Alderton, Roberto Amato, Jeffrey Barrett, Sonia Goncalves, Ewan Harrison, David K. Jackson, Ian Johnston, Dominic Kwiatkowski, Cordelia Langford, John Sillitoe on behalf of the Wellcome Sanger Institute COVID-19 Surveillance Team |
| EPI_ISL_3312674, | Lighthouse Lab in Milton Keynes | Wellcome Sanger Institute for the COVID-19 Genomics UK (COG-UK) | The Lighthouse Lab in Milton Keynes and Alex Alderton, Roberto Amato, Jeffrey Barrett, Sonia Goncalves, Ewan Harrison, David K. Jackson, Ian Johnston, Dominic Kwiatkowski, Cordelia Langford, John Sillitoe on behalf of the Wellcome Sanger |

| | | | |
|---|---|---|---|
| EPI_ISL_3312675, EPI_ISL_3312676, EPI_ISL_3312677, EPI_ISL_3312678 | | Consortium | Institute COVID-19 Surveillance Team |
| EPI_ISL_3312679, EPI_ISL_3312680, EPI_ISL_3312681, EPI_ISL_3312682 | Lighthouse Lab in Alderley Park | Wellcome Sanger Institute for the COVID-19 Genomics UK (COG-UK) Consortium | Jacquelyn Wynn, Mairead Hyland, The Lighthouse Lab in Alderley Park and Alex Alderton, Roberto Amato, Jeffrey Barrett, Sonia Goncalves, Ewan Harrison, David K. Jackson, Ian Johnston, Dominic Kwiatkowski, Cordelia Langford, John Sillitoe on behalf of the Wellcome Sanger Institute COVID-19 Surveillance Team |
| EPI_ISL_3312683, EPI_ISL_3312684, EPI_ISL_3312685 | Lighthouse Lab in Milton Keynes | Wellcome Sanger Institute for the COVID-19 Genomics UK (COG-UK) Consortium | The Lighthouse Lab in Milton Keynes and Alex Alderton, Roberto Amato, Jeffrey Barrett, Sonia Goncalves, Ewan Harrison, David K. Jackson, Ian Johnston, Dominic Kwiatkowski, Cordelia Langford, John Sillitoe on behalf of the Wellcome Sanger Institute COVID-19 Surveillance Team |
| EPI_ISL_3312686, EPI_ISL_3312687 | Lighthouse Lab in Alderley Park | Wellcome Sanger Institute for the COVID-19 Genomics UK (COG-UK) Consortium | Jacquelyn Wynn, Mairead Hyland, The Lighthouse Lab in Alderley Park and Alex Alderton, Roberto Amato, Jeffrey Barrett, Sonia Goncalves, Ewan Harrison, David K. Jackson, Ian Johnston, Dominic Kwiatkowski, Cordelia Langford, John Sillitoe on behalf of the Wellcome Sanger Institute COVID-19 Surveillance Team |
| EPI_ISL_3312688, EPI_ISL_3312689 | Lighthouse Lab in Milton Keynes | Wellcome Sanger Institute for the COVID-19 Genomics UK (COG-UK) Consortium | The Lighthouse Lab in Milton Keynes and Alex Alderton, Roberto Amato, Jeffrey Barrett, Sonia Goncalves, Ewan Harrison, David K. Jackson, Ian Johnston, Dominic Kwiatkowski, Cordelia Langford, John Sillitoe on behalf of the Wellcome Sanger Institute COVID-19 Surveillance Team |
| EPI_ISL_3312690, EPI_ISL_3312691 | Lighthouse Lab in Alderley Park | Wellcome Sanger Institute for the COVID-19 Genomics UK (COG-UK) Consortium | Jacquelyn Wynn, Mairead Hyland, The Lighthouse Lab in Alderley Park and Alex Alderton, Roberto Amato, Jeffrey Barrett, Sonia Goncalves, Ewan Harrison, David K. Jackson, Ian Johnston, Dominic Kwiatkowski, Cordelia Langford, John Sillitoe on behalf of the Wellcome Sanger Institute COVID-19 Surveillance Team |
| EPI_ISL_3312692, EPI_ISL_3312693, EPI_ISL_3312694, EPI_ISL_3312695, EPI_ISL_3312696, EPI_ISL_3312697, EPI_ISL_3312698, EPI_ISL_3312699, EPI_ISL_3312700 | Lighthouse Lab in Milton Keynes | Wellcome Sanger Institute for the COVID-19 Genomics UK (COG-UK) Consortium | The Lighthouse Lab in Milton Keynes and Alex Alderton, Roberto Amato, Jeffrey Barrett, Sonia Goncalves, Ewan Harrison, David K. Jackson, Ian Johnston, Dominic Kwiatkowski, Cordelia Langford, John Sillitoe on behalf of the Wellcome Sanger Institute COVID-19 Surveillance Team |
| EPI_ISL_3312701, EPI_ISL_3312702, EPI_ISL_3312703 | Lighthouse Lab in Alderley Park | Wellcome Sanger Institute for the COVID-19 Genomics UK (COG-UK) Consortium | Jacquelyn Wynn, Mairead Hyland, The Lighthouse Lab in Alderley Park and Alex Alderton, Roberto Amato, Jeffrey Barrett, Sonia Goncalves, Ewan Harrison, David K. Jackson, Ian Johnston, Dominic Kwiatkowski, Cordelia Langford, John Sillitoe on behalf of the Wellcome Sanger Institute COVID-19 Surveillance Team |
| EPI_ISL_3312704, EPI_ISL_3312705, EPI_ISL_3312706 | Lighthouse Lab in Milton Keynes | Wellcome Sanger Institute for the COVID-19 Genomics UK (COG-UK) Consortium | The Lighthouse Lab in Milton Keynes and Alex Alderton, Roberto Amato, Jeffrey Barrett, Sonia Goncalves, Ewan Harrison, David K. Jackson, Ian Johnston, Dominic Kwiatkowski, Cordelia Langford, John Sillitoe on behalf of the Wellcome Sanger Institute COVID-19 Surveillance Team |
| EPI_ISL_3312707, EPI_ISL_3312708, EPI_ISL_3312709 | Lighthouse Lab in Alderley Park | Wellcome Sanger Institute for the COVID-19 Genomics UK (COG-UK) Consortium | Jacquelyn Wynn, Mairead Hyland, The Lighthouse Lab in Alderley Park and Alex Alderton, Roberto Amato, Jeffrey Barrett, Sonia Goncalves, Ewan Harrison, David K. Jackson, Ian Johnston, Dominic Kwiatkowski, Cordelia Langford, John Sillitoe on behalf of the Wellcome Sanger Institute COVID-19 Surveillance Team |
| EPI_ISL_3312710 | Lighthouse Lab in Milton Keynes | Wellcome Sanger Institute for the COVID-19 Genomics UK (COG-UK) Consortium | The Lighthouse Lab in Milton Keynes and Alex Alderton, Roberto Amato, Jeffrey Barrett, Sonia Goncalves, Ewan Harrison, David K. Jackson, Ian Johnston, Dominic Kwiatkowski, Cordelia Langford, John Sillitoe on behalf of the Wellcome Sanger Institute COVID-19 Surveillance Team |
| EPI_ISL_3312711, EPI_ISL_3312712 | Lighthouse Lab in Alderley Park | Wellcome Sanger Institute for the COVID-19 Genomics UK (COG-UK) Consortium | Jacquelyn Wynn, Mairead Hyland, The Lighthouse Lab in Alderley Park and Alex Alderton, Roberto Amato, Jeffrey Barrett, Sonia Goncalves, Ewan Harrison, David K. Jackson, Ian Johnston, Dominic Kwiatkowski, Cordelia Langford, John Sillitoe on behalf of the Wellcome Sanger Institute COVID-19 Surveillance Team |
| EPI_ISL_3312713, EPI_ISL_3312714 | Lighthouse Lab in Milton Keynes | Wellcome Sanger Institute for the COVID-19 Genomics UK (COG-UK) Consortium | The Lighthouse Lab in Milton Keynes and Alex Alderton, Roberto Amato, Jeffrey Barrett, Sonia Goncalves, Ewan Harrison, David K. Jackson, Ian Johnston, Dominic Kwiatkowski, Cordelia Langford, John Sillitoe on behalf of the Wellcome Sanger Institute COVID-19 Surveillance Team |
| EPI_ISL_3312715, EPI_ISL_3312716, EPI_ISL_3312717, EPI_ISL_3312718, EPI_ISL_3312719, EPI_ISL_3312720 | Lighthouse Lab in Alderley Park | Wellcome Sanger Institute for the COVID-19 Genomics UK (COG-UK) Consortium | Jacquelyn Wynn, Mairead Hyland, The Lighthouse Lab in Alderley Park and Alex Alderton, Roberto Amato, Jeffrey Barrett, Sonia Goncalves, Ewan Harrison, David K. Jackson, Ian Johnston, Dominic Kwiatkowski, Cordelia Langford, John Sillitoe on behalf of the Wellcome Sanger Institute COVID-19 Surveillance Team |
| EPI_ISL_3312721, EPI_ISL_3312722, EPI_ISL_3312723, EPI_ISL_3312724 | Lighthouse Lab in Milton Keynes | Wellcome Sanger Institute for the COVID-19 Genomics UK (COG-UK) Consortium | The Lighthouse Lab in Milton Keynes and Alex Alderton, Roberto Amato, Jeffrey Barrett, Sonia Goncalves, Ewan Harrison, David K. Jackson, Ian Johnston, Dominic Kwiatkowski, Cordelia Langford, John Sillitoe on behalf of the Wellcome Sanger Institute COVID-19 Surveillance Team |
| EPI_ISL_3312725 | Lighthouse Lab in Alderley Park | Wellcome Sanger Institute for the COVID-19 Genomics UK (COG-UK) Consortium | Jacquelyn Wynn, Mairead Hyland, The Lighthouse Lab in Alderley Park and Alex Alderton, Roberto Amato, Jeffrey Barrett, Sonia Goncalves, Ewan Harrison, David K. Jackson, Ian Johnston, Dominic Kwiatkowski, Cordelia Langford, John Sillitoe on behalf of the Wellcome Sanger Institute COVID-19 Surveillance Team |
| EPI_ISL_3312726, EPI_ISL_3312727, EPI_ISL_3312728, EPI_ISL_3312729 | Lighthouse Lab in Milton Keynes | Wellcome Sanger Institute for the COVID-19 Genomics UK (COG-UK) Consortium | The Lighthouse Lab in Milton Keynes and Alex Alderton, Roberto Amato, Jeffrey Barrett, Sonia Goncalves, Ewan Harrison, David K. Jackson, Ian Johnston, Dominic Kwiatkowski, Cordelia Langford, John Sillitoe on behalf of the Wellcome Sanger Institute COVID-19 Surveillance Team |
| EPI_ISL_3312730, EPI_ISL_3312731, EPI_ISL_3312732, EPI_ISL_3312733, EPI_ISL_3312734, EPI_ISL_3312735, EPI_ISL_3312736 | Lighthouse Lab in Alderley Park | Wellcome Sanger Institute for the COVID-19 Genomics UK (COG-UK) Consortium | Jacquelyn Wynn, Mairead Hyland, The Lighthouse Lab in Alderley Park and Alex Alderton, Roberto Amato, Jeffrey Barrett, Sonia Goncalves, Ewan Harrison, David K. Jackson, Ian Johnston, Dominic Kwiatkowski, Cordelia Langford, John Sillitoe on behalf of the Wellcome Sanger Institute COVID-19 Surveillance Team |
| EPI_ISL_3312737, EPI_ISL_3312738 | Lighthouse Lab in Milton Keynes | Wellcome Sanger Institute for the COVID-19 Genomics UK (COG-UK) Consortium | The Lighthouse Lab in Milton Keynes and Alex Alderton, Roberto Amato, Jeffrey Barrett, Sonia Goncalves, Ewan Harrison, David K. Jackson, Ian Johnston, Dominic Kwiatkowski, Cordelia Langford, John Sillitoe on behalf of the Wellcome Sanger Institute COVID-19 Surveillance Team |
| EPI_ISL_3312739 | Lighthouse Lab in Alderley Park | Wellcome Sanger Institute for the COVID-19 Genomics UK (COG-UK) Consortium | Jacquelyn Wynn, Mairead Hyland, The Lighthouse Lab in Alderley Park and Alex Alderton, Roberto Amato, Jeffrey Barrett, Sonia Goncalves, Ewan Harrison, David K. Jackson, Ian Johnston, Dominic Kwiatkowski, Cordelia Langford, John Sillitoe on behalf of the Wellcome Sanger Institute COVID-19 Surveillance Team |
| EPI_ISL_3312740 | Lighthouse Lab in Milton Keynes | Wellcome Sanger Institute for the COVID-19 Genomics UK (COG-UK) Consortium | The Lighthouse Lab in Milton Keynes and Alex Alderton, Roberto Amato, Jeffrey Barrett, Sonia Goncalves, Ewan Harrison, David K. Jackson, Ian Johnston, Dominic Kwiatkowski, Cordelia Langford, John Sillitoe on behalf of the Wellcome Sanger Institute COVID-19 Surveillance Team |
| EPI_ISL_3312741 | Lighthouse Lab in Alderley Park | Wellcome Sanger Institute for the COVID-19 Genomics UK (COG-UK) Consortium | Jacquelyn Wynn, Mairead Hyland, The Lighthouse Lab in Alderley Park and Alex Alderton, Roberto Amato, Jeffrey Barrett, Sonia Goncalves, Ewan Harrison, David K. Jackson, Ian Johnston, Dominic Kwiatkowski, Cordelia Langford, John Sillitoe on behalf of the Wellcome Sanger Institute COVID-19 Surveillance Team |
| EPI_ISL_3312742 | Lighthouse Lab in Milton Keynes | Wellcome Sanger Institute for the COVID-19 Genomics UK (COG-UK) Consortium | The Lighthouse Lab in Milton Keynes and Alex Alderton, Roberto Amato, Jeffrey Barrett, Sonia Goncalves, Ewan Harrison, David K. Jackson, Ian Johnston, Dominic Kwiatkowski, Cordelia Langford, John Sillitoe on behalf of the Wellcome Sanger Institute COVID-19 Surveillance Team |
| EPI_ISL_3312743 | Lighthouse Lab in Alderley Park | Wellcome Sanger Institute for the COVID-19 Genomics UK (COG-UK) Consortium | Jacquelyn Wynn, Mairead Hyland, The Lighthouse Lab in Alderley Park and Alex Alderton, Roberto Amato, Jeffrey Barrett, Sonia Goncalves, Ewan Harrison, David K. Jackson, Ian Johnston, Dominic Kwiatkowski, Cordelia Langford, John Sillitoe on behalf of the Wellcome Sanger Institute COVID-19 Surveillance Team |
| EPI_ISL_3312744 | Lighthouse Lab in Milton Keynes | Wellcome Sanger Institute for the COVID-19 Genomics UK (COG-UK) Consortium | The Lighthouse Lab in Milton Keynes and Alex Alderton, Roberto Amato, Jeffrey Barrett, Sonia Goncalves, Ewan Harrison, David K. Jackson, Ian Johnston, Dominic Kwiatkowski, Cordelia Langford, John Sillitoe on behalf of the Wellcome Sanger Institute COVID-19 Surveillance Team |
| EPI_ISL_3312745, EPI_ISL_3312746 | Lighthouse Lab in Alderley Park | Wellcome Sanger Institute for the COVID-19 Genomics UK (COG-UK) Consortium | Jacquelyn Wynn, Mairead Hyland, The Lighthouse Lab in Alderley Park and Alex Alderton, Roberto Amato, Jeffrey Barrett, Sonia Goncalves, Ewan Harrison, David K. Jackson, Ian Johnston, Dominic Kwiatkowski, Cordelia Langford, John Sillitoe on behalf of the Wellcome Sanger Institute COVID-19 Surveillance Team |
| EPI_ISL_3312747 | Lighthouse Lab in Milton Keynes | Wellcome Sanger Institute for the COVID-19 Genomics UK (COG-UK) Consortium | The Lighthouse Lab in Milton Keynes and Alex Alderton, Roberto Amato, Jeffrey Barrett, Sonia Goncalves, Ewan Harrison, David K. Jackson, Ian Johnston, Dominic Kwiatkowski, Cordelia Langford, John Sillitoe on behalf of the Wellcome Sanger Institute COVID-19 Surveillance Team |
| EPI_ISL_3312748, EPI_ISL_3312749 | Lighthouse Lab in Alderley Park | Wellcome Sanger Institute for the COVID-19 Genomics UK (COG-UK) Consortium | Jacquelyn Wynn, Mairead Hyland, The Lighthouse Lab in Alderley Park and Alex Alderton, Roberto Amato, Jeffrey Barrett, Sonia Goncalves, Ewan Harrison, David K. Jackson, Ian Johnston, Dominic Kwiatkowski, Cordelia Langford, John Sillitoe on behalf of the Wellcome Sanger Institute COVID-19 Surveillance Team |
| EPI_ISL_3312750 | Lighthouse Lab in Milton Keynes | Wellcome Sanger Institute for the COVID-19 Genomics UK (COG-UK) Consortium | The Lighthouse Lab in Milton Keynes and Alex Alderton, Roberto Amato, Jeffrey Barrett, Sonia Goncalves, Ewan Harrison, David K. Jackson, Ian Johnston, Dominic Kwiatkowski, Cordelia Langford, John Sillitoe on behalf of the Wellcome Sanger Institute COVID-19 Surveillance Team |
| EPI_ISL_3312751, EPI_ISL_3312752 | Lighthouse Lab in Alderley Park | Wellcome Sanger Institute for the COVID-19 Genomics UK (COG-UK) Consortium | Jacquelyn Wynn, Mairead Hyland, The Lighthouse Lab in Alderley Park and Alex Alderton, Roberto Amato, Jeffrey Barrett, Sonia Goncalves, Ewan Harrison, David K. Jackson, Ian Johnston, Dominic Kwiatkowski, Cordelia Langford, John Sillitoe on behalf of the Wellcome Sanger Institute COVID-19 Surveillance Team |
| EPI_ISL_3312753 | Lighthouse Lab in Milton Keynes | Wellcome Sanger Institute for the COVID-19 Genomics UK (COG-UK) Consortium | The Lighthouse Lab in Milton Keynes and Alex Alderton, Roberto Amato, Jeffrey Barrett, Sonia Goncalves, Ewan Harrison, David K. Jackson, Ian Johnston, Dominic Kwiatkowski, Cordelia Langford, John Sillitoe on behalf of the Wellcome Sanger Institute COVID-19 Surveillance Team |
| EPI_ISL_3312754 | Lighthouse Lab in Alderley Park | Wellcome Sanger Institute for the COVID-19 Genomics UK (COG-UK) Consortium | Jacquelyn Wynn, Mairead Hyland, The Lighthouse Lab in Alderley Park and Alex Alderton, Roberto Amato, Jeffrey Barrett, Sonia Goncalves, Ewan Harrison, David K. Jackson, Ian Johnston, Dominic Kwiatkowski, Cordelia Langford, John Sillitoe on behalf of the Wellcome Sanger Institute COVID-19 Surveillance Team |
| EPI_ISL_3312755 | Lighthouse Lab in Milton Keynes | Wellcome Sanger Institute for the COVID-19 Genomics UK (COG-UK) Consortium | The Lighthouse Lab in Milton Keynes and Alex Alderton, Roberto Amato, Jeffrey Barrett, Sonia Goncalves, Ewan Harrison, David K. Jackson, Ian Johnston, Dominic Kwiatkowski, Cordelia Langford, John Sillitoe on behalf of the Wellcome Sanger Institute COVID-19 Surveillance Team |
| EPI_ISL_3312756, | Lighthouse Lab in Alderley Park | Wellcome Sanger Institute for the COVID-19 Genomics UK (COG-UK) | Jacquelyn Wynn, Mairead Hyland, The Lighthouse Lab in Alderley Park and Alex Alderton, Roberto Amato, Jeffrey Barrett, Sonia Goncalves, Ewan Harrison, David K. Jackson, Ian Johnston, Dominic Kwiatkowski, Cordelia Langford, John Sillitoe on |

| | | | |
|---|---|---|---|
| EPI_ISL_3312757, EPI_ISL_3312758, EPI_ISL_3312759, EPI_ISL_3312760 | | Consortium | behalf of the Wellcome Sanger Institute COVID-19 Surveillance Team |
| EPI_ISL_3312761, EPI_ISL_3312762, EPI_ISL_3312763 | Lighthouse Lab in Milton Keynes | Wellcome Sanger Institute for the COVID-19 Genomics UK (COG-UK) Consortium | The Lighthouse Lab in Milton Keynes and Alex Alderton, Roberto Amato, Jeffrey Barrett, Sonia Goncalves, Ewan Harrison, David K. Jackson, Ian Johnston, Dominic Kwiatkowski, Cordelia Langford, John Sillitoe on behalf of the Wellcome Sanger Institute COVID-19 Surveillance Team |
| EPI_ISL_3312764, EPI_ISL_3312765, EPI_ISL_3312766 | Lighthouse Lab in Alderley Park | Wellcome Sanger Institute for the COVID-19 Genomics UK (COG-UK) Consortium | Jacquelyn Wynn, Mairead Hyland, The Lighthouse Lab in Alderley Park and Alex Alderton, Roberto Amato, Jeffrey Barrett, Sonia Goncalves, Ewan Harrison, David K. Jackson, Ian Johnston, Dominic Kwiatkowski, Cordelia Langford, John Sillitoe on behalf of the Wellcome Sanger Institute COVID-19 Surveillance Team |
| EPI_ISL_3312767 | Lighthouse Lab in Milton Keynes | Wellcome Sanger Institute for the COVID-19 Genomics UK (COG-UK) Consortium | The Lighthouse Lab in Milton Keynes and Alex Alderton, Roberto Amato, Jeffrey Barrett, Sonia Goncalves, Ewan Harrison, David K. Jackson, Ian Johnston, Dominic Kwiatkowski, Cordelia Langford, John Sillitoe on behalf of the Wellcome Sanger Institute COVID-19 Surveillance Team |
| EPI_ISL_3312768 | Lighthouse Lab in Alderley Park | Wellcome Sanger Institute for the COVID-19 Genomics UK (COG-UK) Consortium | Jacquelyn Wynn, Mairead Hyland, The Lighthouse Lab in Alderley Park and Alex Alderton, Roberto Amato, Jeffrey Barrett, Sonia Goncalves, Ewan Harrison, David K. Jackson, Ian Johnston, Dominic Kwiatkowski, Cordelia Langford, John Sillitoe on behalf of the Wellcome Sanger Institute COVID-19 Surveillance Team |
| EPI_ISL_3312769, EPI_ISL_3312770 | Lighthouse Lab in Milton Keynes | Wellcome Sanger Institute for the COVID-19 Genomics UK (COG-UK) Consortium | The Lighthouse Lab in Milton Keynes and Alex Alderton, Roberto Amato, Jeffrey Barrett, Sonia Goncalves, Ewan Harrison, David K. Jackson, Ian Johnston, Dominic Kwiatkowski, Cordelia Langford, John Sillitoe on behalf of the Wellcome Sanger Institute COVID-19 Surveillance Team |
| EPI_ISL_3312771, EPI_ISL_3312772 | Lighthouse Lab in Alderley Park | Wellcome Sanger Institute for the COVID-19 Genomics UK (COG-UK) Consortium | Jacquelyn Wynn, Mairead Hyland, The Lighthouse Lab in Alderley Park and Alex Alderton, Roberto Amato, Jeffrey Barrett, Sonia Goncalves, Ewan Harrison, David K. Jackson, Ian Johnston, Dominic Kwiatkowski, Cordelia Langford, John Sillitoe on behalf of the Wellcome Sanger Institute COVID-19 Surveillance Team |
| EPI_ISL_3312773 | Lighthouse Lab in Milton Keynes | Wellcome Sanger Institute for the COVID-19 Genomics UK (COG-UK) Consortium | The Lighthouse Lab in Milton Keynes and Alex Alderton, Roberto Amato, Jeffrey Barrett, Sonia Goncalves, Ewan Harrison, David K. Jackson, Ian Johnston, Dominic Kwiatkowski, Cordelia Langford, John Sillitoe on behalf of the Wellcome Sanger Institute COVID-19 Surveillance Team |
| EPI_ISL_3312774 | Lighthouse Lab in Alderley Park | Wellcome Sanger Institute for the COVID-19 Genomics UK (COG-UK) Consortium | Jacquelyn Wynn, Mairead Hyland, The Lighthouse Lab in Alderley Park and Alex Alderton, Roberto Amato, Jeffrey Barrett, Sonia Goncalves, Ewan Harrison, David K. Jackson, Ian Johnston, Dominic Kwiatkowski, Cordelia Langford, John Sillitoe on behalf of the Wellcome Sanger Institute COVID-19 Surveillance Team |
| EPI_ISL_3312775 | Lighthouse Lab in Milton Keynes | Wellcome Sanger Institute for the COVID-19 Genomics UK (COG-UK) Consortium | The Lighthouse Lab in Milton Keynes and Alex Alderton, Roberto Amato, Jeffrey Barrett, Sonia Goncalves, Ewan Harrison, David K. Jackson, Ian Johnston, Dominic Kwiatkowski, Cordelia Langford, John Sillitoe on behalf of the Wellcome Sanger Institute COVID-19 Surveillance Team |
| EPI_ISL_3312776 | Lighthouse Lab in Alderley Park | Wellcome Sanger Institute for the COVID-19 Genomics UK (COG-UK) Consortium | Jacquelyn Wynn, Mairead Hyland, The Lighthouse Lab in Alderley Park and Alex Alderton, Roberto Amato, Jeffrey Barrett, Sonia Goncalves, Ewan Harrison, David K. Jackson, Ian Johnston, Dominic Kwiatkowski, Cordelia Langford, John Sillitoe on behalf of the Wellcome Sanger Institute COVID-19 Surveillance Team |
| EPI_ISL_3312777 | Lighthouse Lab in Milton Keynes | Wellcome Sanger Institute for the COVID-19 Genomics UK (COG-UK) Consortium | The Lighthouse Lab in Milton Keynes and Alex Alderton, Roberto Amato, Jeffrey Barrett, Sonia Goncalves, Ewan Harrison, David K. Jackson, Ian Johnston, Dominic Kwiatkowski, Cordelia Langford, John Sillitoe on behalf of the Wellcome Sanger Institute COVID-19 Surveillance Team |
| EPI_ISL_3312778 | Lighthouse Lab in Alderley Park | Wellcome Sanger Institute for the COVID-19 Genomics UK (COG-UK) Consortium | Jacquelyn Wynn, Mairead Hyland, The Lighthouse Lab in Alderley Park and Alex Alderton, Roberto Amato, Jeffrey Barrett, Sonia Goncalves, Ewan Harrison, David K. Jackson, Ian Johnston, Dominic Kwiatkowski, Cordelia Langford, John Sillitoe on behalf of the Wellcome Sanger Institute COVID-19 Surveillance Team |
| EPI_ISL_3312779 | Lighthouse Lab in Milton Keynes | Wellcome Sanger Institute for the COVID-19 Genomics UK (COG-UK) Consortium | The Lighthouse Lab in Milton Keynes and Alex Alderton, Roberto Amato, Jeffrey Barrett, Sonia Goncalves, Ewan Harrison, David K. Jackson, Ian Johnston, Dominic Kwiatkowski, Cordelia Langford, John Sillitoe on behalf of the Wellcome Sanger Institute COVID-19 Surveillance Team |
| EPI_ISL_3312780 | Lighthouse Lab in Alderley Park | Wellcome Sanger Institute for the COVID-19 Genomics UK (COG-UK) Consortium | Jacquelyn Wynn, Mairead Hyland, The Lighthouse Lab in Alderley Park and Alex Alderton, Roberto Amato, Jeffrey Barrett, Sonia Goncalves, Ewan Harrison, David K. Jackson, Ian Johnston, Dominic Kwiatkowski, Cordelia Langford, John Sillitoe on behalf of the Wellcome Sanger Institute COVID-19 Surveillance Team |
| EPI_ISL_3312781 | Lighthouse Lab in Milton Keynes | Wellcome Sanger Institute for the COVID-19 Genomics UK (COG-UK) Consortium | The Lighthouse Lab in Milton Keynes and Alex Alderton, Roberto Amato, Jeffrey Barrett, Sonia Goncalves, Ewan Harrison, David K. Jackson, Ian Johnston, Dominic Kwiatkowski, Cordelia Langford, John Sillitoe on behalf of the Wellcome Sanger Institute COVID-19 Surveillance Team |
| EPI_ISL_3312782 | Lighthouse Lab in Alderley Park | Wellcome Sanger Institute for the COVID-19 Genomics UK (COG-UK) Consortium | Jacquelyn Wynn, Mairead Hyland, The Lighthouse Lab in Alderley Park and Alex Alderton, Roberto Amato, Jeffrey Barrett, Sonia Goncalves, Ewan Harrison, David K. Jackson, Ian Johnston, Dominic Kwiatkowski, Cordelia Langford, John Sillitoe on behalf of the Wellcome Sanger Institute COVID-19 Surveillance Team |
| EPI_ISL_3312783 | Lighthouse Lab in Milton Keynes | Wellcome Sanger Institute for the COVID-19 Genomics UK (COG-UK) Consortium | The Lighthouse Lab in Milton Keynes and Alex Alderton, Roberto Amato, Jeffrey Barrett, Sonia Goncalves, Ewan Harrison, David K. Jackson, Ian Johnston, Dominic Kwiatkowski, Cordelia Langford, John Sillitoe on behalf of the Wellcome Sanger Institute COVID-19 Surveillance Team |
| EPI_ISL_3312784, EPI_ISL_3312785, EPI_ISL_3312786, EPI_ISL_3312787 | Lighthouse Lab in Alderley Park | Wellcome Sanger Institute for the COVID-19 Genomics UK (COG-UK) Consortium | Jacquelyn Wynn, Mairead Hyland, The Lighthouse Lab in Alderley Park and Alex Alderton, Roberto Amato, Jeffrey Barrett, Sonia Goncalves, Ewan Harrison, David K. Jackson, Ian Johnston, Dominic Kwiatkowski, Cordelia Langford, John Sillitoe on behalf of the Wellcome Sanger Institute COVID-19 Surveillance Team |
| EPI_ISL_3312788 | Lighthouse Lab in Milton Keynes | Wellcome Sanger Institute for the COVID-19 Genomics UK (COG-UK) Consortium | The Lighthouse Lab in Milton Keynes and Alex Alderton, Roberto Amato, Jeffrey Barrett, Sonia Goncalves, Ewan Harrison, David K. Jackson, Ian Johnston, Dominic Kwiatkowski, Cordelia Langford, John Sillitoe on behalf of the Wellcome Sanger Institute COVID-19 Surveillance Team |
| EPI_ISL_3312789, EPI_ISL_3312790, EPI_ISL_3312791 | Lighthouse Lab in Alderley Park | Wellcome Sanger Institute for the COVID-19 Genomics UK (COG-UK) Consortium | Jacquelyn Wynn, Mairead Hyland, The Lighthouse Lab in Alderley Park and Alex Alderton, Roberto Amato, Jeffrey Barrett, Sonia Goncalves, Ewan Harrison, David K. Jackson, Ian Johnston, Dominic Kwiatkowski, Cordelia Langford, John Sillitoe on behalf of the Wellcome Sanger Institute COVID-19 Surveillance Team |
| EPI_ISL_3312792 | Lighthouse Lab in Milton Keynes | Wellcome Sanger Institute for the COVID-19 Genomics UK (COG-UK) Consortium | The Lighthouse Lab in Milton Keynes and Alex Alderton, Roberto Amato, Jeffrey Barrett, Sonia Goncalves, Ewan Harrison, David K. Jackson, Ian Johnston, Dominic Kwiatkowski, Cordelia Langford, John Sillitoe on behalf of the Wellcome Sanger Institute COVID-19 Surveillance Team |
| EPI_ISL_3312793, EPI_ISL_3312794, EPI_ISL_3312795 | Lighthouse Lab in Alderley Park | Wellcome Sanger Institute for the COVID-19 Genomics UK (COG-UK) Consortium | Jacquelyn Wynn, Mairead Hyland, The Lighthouse Lab in Alderley Park and Alex Alderton, Roberto Amato, Jeffrey Barrett, Sonia Goncalves, Ewan Harrison, David K. Jackson, Ian Johnston, Dominic Kwiatkowski, Cordelia Langford, John Sillitoe on behalf of the Wellcome Sanger Institute COVID-19 Surveillance Team |
| EPI_ISL_3312796, EPI_ISL_3312797, EPI_ISL_3312798, EPI_ISL_3312799 | Lighthouse Lab in Milton Keynes | Wellcome Sanger Institute for the COVID-19 Genomics UK (COG-UK) Consortium | The Lighthouse Lab in Milton Keynes and Alex Alderton, Roberto Amato, Jeffrey Barrett, Sonia Goncalves, Ewan Harrison, David K. Jackson, Ian Johnston, Dominic Kwiatkowski, Cordelia Langford, John Sillitoe on behalf of the Wellcome Sanger Institute COVID-19 Surveillance Team |
| EPI_ISL_3312800 | Lighthouse Lab in Alderley Park | Wellcome Sanger Institute for the COVID-19 Genomics UK (COG-UK) Consortium | Jacquelyn Wynn, Mairead Hyland, The Lighthouse Lab in Alderley Park and Alex Alderton, Roberto Amato, Jeffrey Barrett, Sonia Goncalves, Ewan Harrison, David K. Jackson, Ian Johnston, Dominic Kwiatkowski, Cordelia Langford, John Sillitoe on behalf of the Wellcome Sanger Institute COVID-19 Surveillance Team |
| EPI_ISL_3312801, EPI_ISL_3312802, EPI_ISL_3312803 | Lighthouse Lab in Milton Keynes | Wellcome Sanger Institute for the COVID-19 Genomics UK (COG-UK) Consortium | The Lighthouse Lab in Milton Keynes and Alex Alderton, Roberto Amato, Jeffrey Barrett, Sonia Goncalves, Ewan Harrison, David K. Jackson, Ian Johnston, Dominic Kwiatkowski, Cordelia Langford, John Sillitoe on behalf of the Wellcome Sanger Institute COVID-19 Surveillance Team |
| EPI_ISL_3312804, EPI_ISL_3312805 | Lighthouse Lab in Alderley Park | Wellcome Sanger Institute for the COVID-19 Genomics UK (COG-UK) Consortium | Jacquelyn Wynn, Mairead Hyland, The Lighthouse Lab in Alderley Park and Alex Alderton, Roberto Amato, Jeffrey Barrett, Sonia Goncalves, Ewan Harrison, David K. Jackson, Ian Johnston, Dominic Kwiatkowski, Cordelia Langford, John Sillitoe on behalf of the Wellcome Sanger Institute COVID-19 Surveillance Team |
| EPI_ISL_3312806 | Lighthouse Lab in Milton Keynes | Wellcome Sanger Institute for the COVID-19 Genomics UK (COG-UK) Consortium | The Lighthouse Lab in Milton Keynes and Alex Alderton, Roberto Amato, Jeffrey Barrett, Sonia Goncalves, Ewan Harrison, David K. Jackson, Ian Johnston, Dominic Kwiatkowski, Cordelia Langford, John Sillitoe on behalf of the Wellcome Sanger Institute COVID-19 Surveillance Team |
| EPI_ISL_3312807 | Lighthouse Lab in Alderley Park | Wellcome Sanger Institute for the COVID-19 Genomics UK (COG-UK) Consortium | Jacquelyn Wynn, Mairead Hyland, The Lighthouse Lab in Alderley Park and Alex Alderton, Roberto Amato, Jeffrey Barrett, Sonia Goncalves, Ewan Harrison, David K. Jackson, Ian Johnston, Dominic Kwiatkowski, Cordelia Langford, John Sillitoe on behalf of the Wellcome Sanger Institute COVID-19 Surveillance Team |
| EPI_ISL_3312808, EPI_ISL_3312809 | Lighthouse Lab in Milton Keynes | Wellcome Sanger Institute for the COVID-19 Genomics UK (COG-UK) Consortium | The Lighthouse Lab in Milton Keynes and Alex Alderton, Roberto Amato, Jeffrey Barrett, Sonia Goncalves, Ewan Harrison, David K. Jackson, Ian Johnston, Dominic Kwiatkowski, Cordelia Langford, John Sillitoe on behalf of the Wellcome Sanger Institute COVID-19 Surveillance Team |
| EPI_ISL_3312810 | Lighthouse Lab in Alderley Park | Wellcome Sanger Institute for the COVID-19 Genomics UK (COG-UK) Consortium | Jacquelyn Wynn, Mairead Hyland, The Lighthouse Lab in Alderley Park and Alex Alderton, Roberto Amato, Jeffrey Barrett, Sonia Goncalves, Ewan Harrison, David K. Jackson, Ian Johnston, Dominic Kwiatkowski, Cordelia Langford, John Sillitoe on behalf of the Wellcome Sanger Institute COVID-19 Surveillance Team |
| EPI_ISL_3312811 | Lighthouse Lab in Milton Keynes | Wellcome Sanger Institute for the COVID-19 Genomics UK (COG-UK) Consortium | The Lighthouse Lab in Milton Keynes and Alex Alderton, Roberto Amato, Jeffrey Barrett, Sonia Goncalves, Ewan Harrison, David K. Jackson, Ian Johnston, Dominic Kwiatkowski, Cordelia Langford, John Sillitoe on behalf of the Wellcome Sanger Institute COVID-19 Surveillance Team |
| EPI_ISL_3312812 | Lighthouse Lab in Alderley Park | Wellcome Sanger Institute for the COVID-19 Genomics UK (COG-UK) Consortium | Jacquelyn Wynn, Mairead Hyland, The Lighthouse Lab in Alderley Park and Alex Alderton, Roberto Amato, Jeffrey Barrett, Sonia Goncalves, Ewan Harrison, David K. Jackson, Ian Johnston, Dominic Kwiatkowski, Cordelia Langford, John Sillitoe on behalf of the Wellcome Sanger Institute COVID-19 Surveillance Team |
| EPI_ISL_3312813 | Lighthouse Lab in Milton Keynes | Wellcome Sanger Institute for the COVID-19 Genomics UK (COG-UK) Consortium | The Lighthouse Lab in Milton Keynes and Alex Alderton, Roberto Amato, Jeffrey Barrett, Sonia Goncalves, Ewan Harrison, David K. Jackson, Ian Johnston, Dominic Kwiatkowski, Cordelia Langford, John Sillitoe on behalf of the Wellcome Sanger Institute COVID-19 Surveillance Team |
| EPI_ISL_3312814, EPI_ISL_3312815 | Lighthouse Lab in Alderley Park | Wellcome Sanger Institute for the COVID-19 Genomics UK (COG-UK) Consortium | Jacquelyn Wynn, Mairead Hyland, The Lighthouse Lab in Alderley Park and Alex Alderton, Roberto Amato, Jeffrey Barrett, Sonia Goncalves, Ewan Harrison, David K. Jackson, Ian Johnston, Dominic Kwiatkowski, Cordelia Langford, John Sillitoe on behalf of the Wellcome Sanger Institute COVID-19 Surveillance Team |
| EPI_ISL_3312816 | Lighthouse Lab in Milton Keynes | Wellcome Sanger Institute for the COVID-19 Genomics UK (COG-UK) Consortium | The Lighthouse Lab in Milton Keynes and Alex Alderton, Roberto Amato, Jeffrey Barrett, Sonia Goncalves, Ewan Harrison, David K. Jackson, Ian Johnston, Dominic Kwiatkowski, Cordelia Langford, John Sillitoe on behalf of the Wellcome Sanger Institute COVID-19 Surveillance Team |
| EPI_ISL_3312817, EPI_ISL_3312818, EPI_ISL_3312819 | Lighthouse Lab in Alderley Park | Wellcome Sanger Institute for the COVID-19 Genomics UK (COG-UK) Consortium | Jacquelyn Wynn, Mairead Hyland, The Lighthouse Lab in Alderley Park and Alex Alderton, Roberto Amato, Jeffrey Barrett, Sonia Goncalves, Ewan Harrison, David K. Jackson, Ian Johnston, Dominic Kwiatkowski, Cordelia Langford, John Sillitoe on behalf of the Wellcome Sanger Institute COVID-19 Surveillance Team |
| EPI_ISL_3312820, EPI_ISL_3312821 | Lighthouse Lab in Milton Keynes | Wellcome Sanger Institute for the COVID-19 Genomics UK (COG-UK) Consortium | The Lighthouse Lab in Milton Keynes and Alex Alderton, Roberto Amato, Jeffrey Barrett, Sonia Goncalves, Ewan Harrison, David K. Jackson, Ian Johnston, Dominic Kwiatkowski, Cordelia Langford, John Sillitoe on behalf of the Wellcome Sanger Institute COVID-19 Surveillance Team |
| EPI_ISL_3312822, EPI_ISL_3312823 | Lighthouse Lab in Alderley Park | Wellcome Sanger Institute for the COVID-19 Genomics UK (COG-UK) Consortium | Jacquelyn Wynn, Mairead Hyland, The Lighthouse Lab in Alderley Park and Alex Alderton, Roberto Amato, Jeffrey Barrett, Sonia Goncalves, Ewan Harrison, David K. Jackson, Ian Johnston, Dominic Kwiatkowski, Cordelia Langford, John Sillitoe on behalf of the Wellcome Sanger Institute COVID-19 Surveillance Team |
| EPI_ISL_3312824, EPI_ISL_3312825 | Lighthouse Lab in Milton Keynes | Wellcome Sanger Institute for the COVID-19 Genomics UK (COG-UK) Consortium | The Lighthouse Lab in Milton Keynes and Alex Alderton, Roberto Amato, Jeffrey Barrett, Sonia Goncalves, Ewan Harrison, David K. Jackson, Ian Johnston, Dominic Kwiatkowski, Cordelia Langford, John Sillitoe on behalf of the Wellcome Sanger Institute COVID-19 Surveillance Team |

| | | | |
|---|---|---|---|
| EPI_ISL_3312826 | Lighthouse Lab in Alderley Park | Wellcome Sanger Institute for the COVID-19 Genomics UK (COG-UK) Consortium | Jacquelyn Wynn, Mairead Hyland, The Lighthouse Lab in Alderley Park and Alex Alderton, Roberto Amato, Jeffrey Barrett, Sonia Goncalves, Ewan Harrison, David K. Jackson, Ian Johnston, Dominic Kwiatkowski, Cordelia Langford, John Sillitoe on behalf of the Wellcome Sanger Institute COVID-19 Surveillance Team |
| EPI_ISL_3312827 | Lighthouse Lab in Milton Keynes | Wellcome Sanger Institute for the COVID-19 Genomics UK (COG-UK) Consortium | The Lighthouse Lab in Milton Keynes and Alex Alderton, Roberto Amato, Jeffrey Barrett, Sonia Goncalves, Ewan Harrison, David K. Jackson, Ian Johnston, Dominic Kwiatkowski, Cordelia Langford, John Sillitoe on behalf of the Wellcome Sanger Institute COVID-19 Surveillance Team |
| EPI_ISL_3312828, EPI_ISL_3312829, EPI_ISL_3312830 | Lighthouse Lab in Alderley Park | Wellcome Sanger Institute for the COVID-19 Genomics UK (COG-UK) Consortium | Jacquelyn Wynn, Mairead Hyland, The Lighthouse Lab in Alderley Park and Alex Alderton, Roberto Amato, Jeffrey Barrett, Sonia Goncalves, Ewan Harrison, David K. Jackson, Ian Johnston, Dominic Kwiatkowski, Cordelia Langford, John Sillitoe on behalf of the Wellcome Sanger Institute COVID-19 Surveillance Team |
| EPI_ISL_3312831 | Lighthouse Lab in Milton Keynes | Wellcome Sanger Institute for the COVID-19 Genomics UK (COG-UK) Consortium | The Lighthouse Lab in Milton Keynes and Alex Alderton, Roberto Amato, Jeffrey Barrett, Sonia Goncalves, Ewan Harrison, David K. Jackson, Ian Johnston, Dominic Kwiatkowski, Cordelia Langford, John Sillitoe on behalf of the Wellcome Sanger Institute COVID-19 Surveillance Team |
| EPI_ISL_3312832, EPI_ISL_3312833, EPI_ISL_3312834 | Lighthouse Lab in Alderley Park | Wellcome Sanger Institute for the COVID-19 Genomics UK (COG-UK) Consortium | Jacquelyn Wynn, Mairead Hyland, The Lighthouse Lab in Alderley Park and Alex Alderton, Roberto Amato, Jeffrey Barrett, Sonia Goncalves, Ewan Harrison, David K. Jackson, Ian Johnston, Dominic Kwiatkowski, Cordelia Langford, John Sillitoe on behalf of the Wellcome Sanger Institute COVID-19 Surveillance Team |
| EPI_ISL_3312835, EPI_ISL_3312836, EPI_ISL_3312837 | Lighthouse Lab in Milton Keynes | Wellcome Sanger Institute for the COVID-19 Genomics UK (COG-UK) Consortium | The Lighthouse Lab in Milton Keynes and Alex Alderton, Roberto Amato, Jeffrey Barrett, Sonia Goncalves, Ewan Harrison, David K. Jackson, Ian Johnston, Dominic Kwiatkowski, Cordelia Langford, John Sillitoe on behalf of the Wellcome Sanger Institute COVID-19 Surveillance Team |
| EPI_ISL_3312838 | Lighthouse Lab in Alderley Park | Wellcome Sanger Institute for the COVID-19 Genomics UK (COG-UK) Consortium | Jacquelyn Wynn, Mairead Hyland, The Lighthouse Lab in Alderley Park and Alex Alderton, Roberto Amato, Jeffrey Barrett, Sonia Goncalves, Ewan Harrison, David K. Jackson, Ian Johnston, Dominic Kwiatkowski, Cordelia Langford, John Sillitoe on behalf of the Wellcome Sanger Institute COVID-19 Surveillance Team |
| EPI_ISL_3312839, EPI_ISL_3312840 | Lighthouse Lab in Milton Keynes | Wellcome Sanger Institute for the COVID-19 Genomics UK (COG-UK) Consortium | The Lighthouse Lab in Milton Keynes and Alex Alderton, Roberto Amato, Jeffrey Barrett, Sonia Goncalves, Ewan Harrison, David K. Jackson, Ian Johnston, Dominic Kwiatkowski, Cordelia Langford, John Sillitoe on behalf of the Wellcome Sanger Institute COVID-19 Surveillance Team |
| EPI_ISL_3312841, EPI_ISL_3312842 | Lighthouse Lab in Alderley Park | Wellcome Sanger Institute for the COVID-19 Genomics UK (COG-UK) Consortium | Jacquelyn Wynn, Mairead Hyland, The Lighthouse Lab in Alderley Park and Alex Alderton, Roberto Amato, Jeffrey Barrett, Sonia Goncalves, Ewan Harrison, David K. Jackson, Ian Johnston, Dominic Kwiatkowski, Cordelia Langford, John Sillitoe on behalf of the Wellcome Sanger Institute COVID-19 Surveillance Team |
| EPI_ISL_3312843 | Lighthouse Lab in Milton Keynes | Wellcome Sanger Institute for the COVID-19 Genomics UK (COG-UK) Consortium | The Lighthouse Lab in Milton Keynes and Alex Alderton, Roberto Amato, Jeffrey Barrett, Sonia Goncalves, Ewan Harrison, David K. Jackson, Ian Johnston, Dominic Kwiatkowski, Cordelia Langford, John Sillitoe on behalf of the Wellcome Sanger Institute COVID-19 Surveillance Team |
| EPI_ISL_3312844 | Lighthouse Lab in Alderley Park | Wellcome Sanger Institute for the COVID-19 Genomics UK (COG-UK) Consortium | Jacquelyn Wynn, Mairead Hyland, The Lighthouse Lab in Alderley Park and Alex Alderton, Roberto Amato, Jeffrey Barrett, Sonia Goncalves, Ewan Harrison, David K. Jackson, Ian Johnston, Dominic Kwiatkowski, Cordelia Langford, John Sillitoe on behalf of the Wellcome Sanger Institute COVID-19 Surveillance Team |
| EPI_ISL_3312845 | Lighthouse Lab in Milton Keynes | Wellcome Sanger Institute for the COVID-19 Genomics UK (COG-UK) Consortium | The Lighthouse Lab in Milton Keynes and Alex Alderton, Roberto Amato, Jeffrey Barrett, Sonia Goncalves, Ewan Harrison, David K. Jackson, Ian Johnston, Dominic Kwiatkowski, Cordelia Langford, John Sillitoe on behalf of the Wellcome Sanger Institute COVID-19 Surveillance Team |
| EPI_ISL_3312846, EPI_ISL_3312847 | Lighthouse Lab in Alderley Park | Wellcome Sanger Institute for the COVID-19 Genomics UK (COG-UK) Consortium | Jacquelyn Wynn, Mairead Hyland, The Lighthouse Lab in Alderley Park and Alex Alderton, Roberto Amato, Jeffrey Barrett, Sonia Goncalves, Ewan Harrison, David K. Jackson, Ian Johnston, Dominic Kwiatkowski, Cordelia Langford, John Sillitoe on behalf of the Wellcome Sanger Institute COVID-19 Surveillance Team |
| EPI_ISL_3312848 | Lighthouse Lab in Milton Keynes | Wellcome Sanger Institute for the COVID-19 Genomics UK (COG-UK) Consortium | The Lighthouse Lab in Milton Keynes and Alex Alderton, Roberto Amato, Jeffrey Barrett, Sonia Goncalves, Ewan Harrison, David K. Jackson, Ian Johnston, Dominic Kwiatkowski, Cordelia Langford, John Sillitoe on behalf of the Wellcome Sanger Institute COVID-19 Surveillance Team |
| EPI_ISL_3312849, EPI_ISL_3312850, EPI_ISL_3312851, EPI_ISL_3312852, EPI_ISL_3312853, EPI_ISL_3312854 | Lighthouse Lab in Alderley Park | Wellcome Sanger Institute for the COVID-19 Genomics UK (COG-UK) Consortium | Jacquelyn Wynn, Mairead Hyland, The Lighthouse Lab in Alderley Park and Alex Alderton, Roberto Amato, Jeffrey Barrett, Sonia Goncalves, Ewan Harrison, David K. Jackson, Ian Johnston, Dominic Kwiatkowski, Cordelia Langford, John Sillitoe on behalf of the Wellcome Sanger Institute COVID-19 Surveillance Team |
| EPI_ISL_3312855, EPI_ISL_3312856, EPI_ISL_3312857 | Lighthouse Lab in Milton Keynes | Wellcome Sanger Institute for the COVID-19 Genomics UK (COG-UK) Consortium | The Lighthouse Lab in Milton Keynes and Alex Alderton, Roberto Amato, Jeffrey Barrett, Sonia Goncalves, Ewan Harrison, David K. Jackson, Ian Johnston, Dominic Kwiatkowski, Cordelia Langford, John Sillitoe on behalf of the Wellcome Sanger Institute COVID-19 Surveillance Team |
| EPI_ISL_3312858 | Lighthouse Lab in Alderley Park | Wellcome Sanger Institute for the COVID-19 Genomics UK (COG-UK) Consortium | Jacquelyn Wynn, Mairead Hyland, The Lighthouse Lab in Alderley Park and Alex Alderton, Roberto Amato, Jeffrey Barrett, Sonia Goncalves, Ewan Harrison, David K. Jackson, Ian Johnston, Dominic Kwiatkowski, Cordelia Langford, John Sillitoe on behalf of the Wellcome Sanger Institute COVID-19 Surveillance Team |
| EPI_ISL_3312859 | Lighthouse Lab in Milton Keynes | Wellcome Sanger Institute for the COVID-19 Genomics UK (COG-UK) Consortium | The Lighthouse Lab in Milton Keynes and Alex Alderton, Roberto Amato, Jeffrey Barrett, Sonia Goncalves, Ewan Harrison, David K. Jackson, Ian Johnston, Dominic Kwiatkowski, Cordelia Langford, John Sillitoe on behalf of the Wellcome Sanger Institute COVID-19 Surveillance Team |
| EPI_ISL_3312860, EPI_ISL_3312861 | Lighthouse Lab in Alderley Park | Wellcome Sanger Institute for the COVID-19 Genomics UK (COG-UK) Consortium | Jacquelyn Wynn, Mairead Hyland, The Lighthouse Lab in Alderley Park and Alex Alderton, Roberto Amato, Jeffrey Barrett, Sonia Goncalves, Ewan Harrison, David K. Jackson, Ian Johnston, Dominic Kwiatkowski, Cordelia Langford, John Sillitoe on behalf of the Wellcome Sanger Institute COVID-19 Surveillance Team |
| EPI_ISL_3312862, EPI_ISL_3312863, EPI_ISL_3312864, EPI_ISL_3312865, EPI_ISL_3312866, EPI_ISL_3312867, EPI_ISL_3312868, EPI_ISL_3312869, EPI_ISL_3312870, EPI_ISL_3312871, EPI_ISL_3312872, EPI_ISL_3312873, EPI_ISL_3312874, EPI_ISL_3312875, EPI_ISL_3312876, EPI_ISL_3312877, EPI_ISL_3312878, EPI_ISL_3312879, EPI_ISL_3312880, EPI_ISL_3312881, EPI_ISL_3312882, EPI_ISL_3312883, EPI_ISL_3312884, EPI_ISL_3312885, EPI_ISL_3312886, EPI_ISL_3312887, EPI_ISL_3312888, EPI_ISL_3312889, EPI_ISL_3312890, EPI_ISL_3312891, EPI_ISL_3312892, EPI_ISL_3312893, EPI_ISL_3312894, EPI_ISL_3312895, EPI_ISL_3312896, EPI_ISL_3312897, EPI_ISL_3312898, EPI_ISL_3312899, EPI_ISL_3312900, EPI_ISL_3312901, EPI_ISL_3312902, EPI_ISL_3312903, EPI_ISL_3312904, EPI_ISL_3312905, EPI_ISL_3312906, EPI_ISL_3312907, EPI_ISL_3312908, EPI_ISL_3312909, EPI_ISL_3312910, EPI_ISL_3312911, EPI_ISL_3312912, EPI_ISL_3312913, EPI_ISL_3312914, EPI_ISL_3312915, EPI_ISL_3312916, EPI_ISL_3312917, EPI_ISL_3312918, EPI_ISL_3312919, EPI_ISL_3312920, EPI_ISL_3312921, EPI_ISL_3312922, EPI_ISL_3312923, EPI_ISL_3312924, EPI_ISL_3312925, EPI_ISL_3312926, EPI_ISL_3312927, EPI_ISL_3312928, EPI_ISL_3312929, EPI_ISL_3312930, EPI_ISL_3312931, EPI_ISL_3312932, EPI_ISL_3312933, EPI_ISL_3312934, EPI_ISL_3312935, EPI_ISL_3312936, EPI_ISL_3312937, EPI_ISL_3312938, EPI_ISL_3312939, EPI_ISL_3312940, EPI_ISL_3312941, EPI_ISL_3312942, EPI_ISL_3312943, EPI_ISL_3312944, EPI_ISL_3312945, EPI_ISL_3312946, EPI_ISL_3312947, EPI_ISL_3312948, EPI_ISL_3312949, EPI_ISL_3312950, EPI_ISL_3312951, EPI_ISL_3312952, EPI_ISL_3312953, EPI_ISL_3312954, EPI_ISL_3312955, EPI_ISL_3312956, EPI_ISL_3312957, EPI_ISL_3312958, EPI_ISL_3312959, EPI_ISL_3312960, EPI_ISL_3312961, EPI_ISL_3312962, EPI_ISL_3312963, EPI_ISL_3312964, EPI_ISL_3312965, EPI_ISL_3312966, EPI_ISL_3312967, EPI_ISL_3312968, EPI_ISL_3312969, EPI_ISL_3312970, EPI_ISL_3312971, EPI_ISL_3312972, EPI_ISL_3312973, EPI_ISL_3312974, EPI_ISL_3312975, EPI_ISL_3312976, EPI_ISL_3312977, EPI_ISL_3312978, EPI_ISL_3312979, EPI_ISL_3312980, EPI_ISL_3312981, EPI_ISL_3312982, EPI_ISL_3312983, EPI_ISL_3312984, EPI_ISL_3312985, EPI_ISL_3312986, EPI_ISL_3312987, EPI_ISL_3312988, EPI_ISL_3312989, EPI_ISL_3312990, EPI_ISL_3312991, EPI_ISL_3312992, EPI_ISL_3312993, EPI_ISL_3312994, EPI_ISL_3312995, EPI_ISL_3312996, EPI_ISL_3312997, EPI_ISL_3312998, EPI_ISL_3312999, EPI_ISL_3313000, EPI_ISL_3313001, EPI_ISL_3313002, EPI_ISL_3313003, EPI_ISL_3313004, EPI_ISL_3313005, EPI_ISL_3313006, EPI_ISL_3313007, EPI_ISL_3313008, EPI_ISL_3313009, EPI_ISL_3313010, EPI_ISL_3313011, EPI_ISL_3313012, EPI_ISL_3313013, EPI_ISL_3313014, EPI_ISL_3313015, EPI_ISL_3313016, EPI_ISL_3313017, EPI_ISL_3313018, EPI_ISL_3313019, EPI_ISL_3313020, EPI_ISL_3313021, EPI_ISL_3313022, EPI_ISL_3313023, EPI_ISL_3313024, EPI_ISL_3313025, EPI_ISL_3313026, EPI_ISL_3313027, EPI_ISL_3313028, EPI_ISL_3313029, EPI_ISL_3313030, EPI_ISL_3313031, EPI_ISL_3313032, EPI_ISL_3313033, EPI_ISL_3313034, EPI_ISL_3313035, EPI_ISL_3313036, EPI_ISL_3313037, EPI_ISL_3313038, EPI_ISL_3313039, EPI_ISL_3313040, EPI_ISL_3313041, EPI_ISL_3313042, EPI_ISL_3313043, EPI_ISL_3313044, EPI_ISL_3313045, EPI_ISL_3313046, EPI_ISL_3313047, EPI_ISL_3313048, EPI_ISL_3313049, EPI_ISL_3313050, EPI_ISL_3313051, EPI_ISL_3313052, EPI_ISL_3313053, EPI_ISL_3313054, EPI_ISL_3313055, EPI_ISL_3313056, EPI_ISL_3313057, EPI_ISL_3313058, EPI_ISL_3313059, EPI_ISL_3313060, EPI_ISL_3313061, EPI_ISL_3313062, EPI_ISL_3313063, EPI_ISL_3313064, EPI_ISL_3313065, EPI_ISL_3313066, EPI_ISL_3313067, EPI_ISL_3313068, EPI_ISL_3313069, EPI_ISL_3313070, EPI_ISL_3313071, EPI_ISL_3313072, EPI_ISL_3313073, EPI_ISL_3313074, EPI_ISL_3313075, EPI_ISL_3313076, EPI_ISL_3313077, EPI_ISL_3313078, EPI_ISL_3313079, EPI_ISL_3313080, EPI_ISL_3313081, EPI_ISL_3313082, EPI_ISL_3313083, EPI_ISL_3313084, EPI_ISL_3313085, EPI_ISL_3313086, EPI_ISL_3313087, EPI_ISL_3313088, EPI_ISL_3313089, EPI_ISL_3313090, EPI_ISL_3313091, EPI_ISL_3313092, EPI_ISL_3313093, EPI_ISL_3313094, EPI_ISL_3313095, EPI_ISL_3313096, EPI_ISL_3313097, EPI_ISL_3313098, EPI_ISL_3313099, EPI_ISL_3313100, EPI_ISL_3313101, EPI_ISL_3313102, EPI_ISL_3313103, EPI_ISL_3313104, EPI_ISL_3313105, EPI_ISL_3313106, EPI_ISL_3313107, EPI_ISL_3313108, EPI_ISL_3313109, EPI_ISL_3313110, EPI_ISL_3313111, EPI_ISL_3313112, EPI_ISL_3313113, EPI_ISL_3313114, EPI_ISL_3313115, EPI_ISL_3313116, EPI_ISL_3313117, EPI_ISL_3313118, EPI_ISL_3313119, EPI_ISL_3313120, EPI_ISL_3313121, EPI_ISL_3313122, EPI_ISL_3313123, EPI_ISL_3313124, EPI_ISL_3313125, EPI_ISL_3313126, EPI_ISL_3313127, EPI_ISL_3313128, EPI_ISL_3313129, EPI_ISL_3313130, EPI_ISL_3313131, EPI_ISL_3313132, EPI_ISL_3313133, EPI_ISL_3313134, EPI_ISL_3313135, EPI_ISL_3313136, EPI_ISL_3313137, EPI_ISL_3313138, EPI_ISL_3313139, EPI_ISL_3313140, EPI_ISL_3313141, EPI_ISL_3313142, EPI_ISL_3313143, EPI_ISL_3313144, EPI_ISL_3313145, EPI_ISL_3313146, EPI_ISL_3313147, EPI_ISL_3313148, EPI_ISL_3313149, EPI_ISL_3313150, EPI_ISL_3313151, EPI_ISL_3313152, EPI_ISL_3313153, EPI_ISL_3313154, EPI_ISL_3313155, EPI_ISL_3313156, EPI_ISL_3313157, EPI_ISL_3313158, EPI_ISL_3313159, EPI_ISL_3313160, EPI_ISL_3313161, EPI_ISL_3313162, EPI_ISL_3313163, EPI_ISL_3313164, EPI_ISL_3313165, EPI_ISL_3313166, EPI_ISL_3313167, EPI_ISL_3313168, EPI_ISL_3313169, EPI_ISL_3313170, EPI_ISL_3313171, EPI_ISL_3313172, EPI_ISL_3313173, EPI_ISL_3313174, EPI_ISL_3313175, EPI_ISL_3313176, EPI_ISL_3313177, EPI_ISL_3313178, EPI_ISL_3313179, EPI_ISL_3313180, EPI_ISL_3313181, EPI_ISL_3313182, EPI_ISL_3313183, EPI_ISL_3313184, EPI_ISL_3313185, EPI_ISL_3313186, EPI_ISL_3313187, EPI_ISL_3313188, EPI_ISL_3313189, EPI_ISL_3313190, EPI_ISL_3313191, EPI_ISL_3313192, EPI_ISL_3313193, EPI_ISL_3313194, EPI_ISL_3313195, EPI_ISL_3313196, EPI_ISL_3313197, EPI_ISL_3313198, EPI_ISL_3313199, EPI_ISL_3313200, EPI_ISL_3313201, EPI_ISL_3313202, EPI_ISL_3313203, EPI_ISL_3313204, EPI_ISL_3313205, EPI_ISL_3313206, EPI_ISL_3313207, EPI_ISL_3313208, EPI_ISL_3313209, EPI_ISL_3313210, EPI_ISL_3313211, EPI_ISL_3313212, EPI_ISL_3313213, EPI_ISL_3313214, EPI_ISL_3313215, EPI_ISL_3313216, EPI_ISL_3313217, EPI_ISL_3313218, EPI_ISL_3313219, EPI_ISL_3313220, EPI_ISL_3313221, EPI_ISL_3313222, EPI_ISL_3313223, EPI_ISL_3313224, EPI_ISL_3313225, EPI_ISL_3313226, EPI_ISL_3313227, EPI_ISL_3313228, EPI_ISL_3313229, EPI_ISL_3313230, EPI_ISL_3313231, EPI_ISL_3313232, EPI_ISL_3313233, EPI_ISL_3313234, EPI_ISL_3313235, EPI_ISL_3313236, EPI_ISL_3313237, EPI_ISL_3313238, EPI_ISL_3313239, EPI_ISL_3313240, EPI_ISL_3313241, EPI_ISL_3313242, EPI_ISL_3313243, EPI_ISL_3313244, EPI_ISL_3313245, EPI_ISL_3313246, EPI_ISL_3313247, EPI_ISL_3313248, EPI_ISL_3313249, EPI_ISL_3313250, EPI_ISL_3313251, EPI_ISL_3313252, EPI_ISL_3313253, EPI_ISL_3313254, EPI_ISL_3313255, EPI_ISL_3313256, EPI_ISL_3313257, EPI_ISL_3313258, EPI_ISL_3313259, EPI_ISL_3313260, EPI_ISL_3313261, EPI_ISL_3313262, EPI_ISL_3313263, EPI_ISL_3313264, EPI_ISL_3313265, EPI_ISL_3313266, EPI_ISL_3313267, EPI_ISL_3313268, EPI_ISL_3313269, EPI_ISL_3313270, EPI_ISL_3313271, EPI_ISL_3313272, EPI_ISL_3313273, EPI_ISL_3313274, EPI_ISL_3313275, EPI_ISL_3313276, EPI_ISL_3313277, EPI_ISL_3313278, EPI_ISL_3313279, EPI_ISL_3313280, EPI_ISL_3313281, EPI_ISL_3313282, EPI_ISL_3313283, EPI_ISL_3313284, EPI_ISL_3313285, EPI_ISL_3313286, EPI_ISL_3313287, EPI_ISL_3313288, EPI_ISL_3313289, EPI_ISL_3313290, EPI_ISL_3313291, EPI_ISL_3313292, EPI_ISL_3313293, EPI_ISL_3313294, EPI_ISL_3313295, EPI_ISL_3313296, EPI_ISL_3313297, EPI_ISL_3313298, EPI_ISL_3313299, EPI_ISL_3313300, EPI_ISL_3313301, EPI_ISL_3313302, EPI_ISL_3313303, EPI_ISL_3313304, EPI_ISL_3313305, EPI_ISL_3313306, EPI_ISL_3313307, EPI_ISL_3313308, EPI_ISL_3313309, EPI_ISL_3313310, EPI_ISL_3313311, EPI_ISL_3313312, EPI_ISL_3313313, EPI_ISL_3313314, EPI_ISL_3313315, EPI_ISL_3313316, EPI_ISL_3313317, EPI_ISL_3313318, EPI_ISL_3313319, EPI_ISL_3313320, EPI_ISL_3313321, EPI_ISL_3313322, EPI_ISL_3313323, EPI_ISL_3313324, EPI_ISL_3313325, EPI_ISL_3313326, EPI_ISL_3313327, EPI_ISL_3313328, EPI_ISL_3313329, EPI_ISL_3313330, EPI_ISL_3313331, EPI_ISL_3313332, EPI_ISL_3313333, EPI_ISL_3313334, EPI_ISL_3313335, EPI_ISL_3313336, EPI_ISL_3313337, EPI_ISL_3313338, EPI_ISL_3313339, EPI_ISL_3313340, EPI_ISL_3313341, EPI_ISL_3313342, EPI_ISL_3313343, EPI_ISL_3313344, EPI_ISL_3313345, EPI_ISL_3313346, EPI_ISL_3313347, EPI_ISL_3313348, EPI_ISL_3313349, EPI_ISL_3313350, EPI_ISL_3313351, EPI_ISL_3313352, EPI_ISL_3313353, EPI_ISL_3313354, EPI_ISL_3313355, EPI_ISL_3313356, EPI_ISL_3313357, EPI_ISL_3313358, EPI_ISL_3313359, EPI_ISL_3313360, EPI_ISL_3313361, EPI_ISL_3313362, EPI_ISL_3313363, EPI_ISL_3313364, EPI_ISL_3313365, EPI_ISL_3313366, EPI_ISL_3313367, EPI_ISL_3313368, EPI_ISL_3313369, EPI_ISL_3313370, EPI_ISL_3313371, EPI_ISL_3313372, EPI_ISL_3313373, EPI_ISL_3313374, EPI_ISL_3313375, EPI_ISL_3313376, EPI_ISL_3313377, EPI_ISL_3313378, EPI_ISL_3313379, EPI_ISL_3313380, EPI_ISL_3313381, EPI_ISL_3313382, EPI_ISL_3313383, EPI_ISL_3313384, EPI_ISL_3313385, EPI_ISL_3313386, EPI_ISL_3313387, EPI_ISL_3313388, EPI_ISL_3313389, EPI_ISL_3313390, EPI_ISL_3313391, EPI_ISL_3313392, EPI_ISL_3313393, EPI_ISL_3313394, EPI_ISL_3313395, EPI_ISL_3313396, EPI_ISL_3313397, EPI_ISL_3313398, EPI_ISL_3313399, EPI_ISL_3313400, EPI_ISL_3313401, EPI_ISL_3313402, EPI_ISL_3313403, EPI_ISL_3313404, EPI_ISL_3313405, EPI_ISL_3313406, EPI_ISL_3313407, EPI_ISL_3313408, EPI_ISL_3313409, EPI_ISL_3313410, EPI_ISL_3313411, EPI_ISL_3313412, EPI_ISL_3313413, EPI_ISL_3313414, EPI_ISL_3313415, EPI_ISL_3313416, EPI_ISL_3313417, EPI_ISL_3313418, EPI_ISL_3313419, EPI_ISL_3313420, EPI_ISL_3313421, EPI_ISL_3313422, EPI_ISL_3313423, EPI_ISL_3313424, EPI_ISL_3313425, EPI_ISL_3313426, EPI_ISL_3313427, EPI_ISL_3313428, EPI_ISL_3313429, EPI_ISL_3313430, EPI_ISL_3313431, EPI_ISL_3313432, EPI_ISL_3313433, EPI_ISL_3313434, EPI_ISL_3313435, EPI_ISL_3313436, EPI_ISL_3313437, EPI_ISL_3313438, EPI_ISL_3313439, EPI_ISL_3313440, EPI_ISL_3313441, EPI_ISL_3313442, EPI_ISL_3313443, EPI_ISL_3313444, EPI_ISL_3313445, EPI_ISL_3313446, EPI_ISL_3313447, EPI_ISL_3313448, EPI_ISL_3313449, EPI_ISL_3313450, EPI_ISL_3313451, EPI_ISL_3313452, EPI_ISL_3313453, EPI_ISL_3313454, EPI_ISL_3313455, EPI_ISL_3313456, EPI_ISL_3313457, EPI_ISL_3313458, EPI_ISL_3313459, EPI_ISL_3313460, EPI_ISL_3313461, EPI_ISL_3313462, EPI_ISL_3313463, EPI_ISL_3313464, EPI_ISL_3313465, EPI_ISL_3313466, EPI_ISL_3313467, EPI_ISL_3313468, EPI_ISL_3313469, EPI_ISL_3313470, EPI_ISL_3313471, EPI_ISL_3313472, EPI_ISL_3313473, EPI_ISL_3313474, EPI_ISL_3313475, EPI_ISL_3313476, EPI_ISL_3313477, EPI_ISL_3313478, EPI_ISL_3313479, EPI_ISL_3313480, EPI_ISL_3313481, EPI_ISL_3313482, EPI_ISL_3313483, EPI_ISL_3313484, EPI_ISL_3313485, EPI_ISL_3313486, EPI_ISL_3313487, EPI_ISL_3313488, EPI_ISL_3313489, EPI_ISL_3313490, EPI_ISL_3313491, EPI_ISL_3313492, EPI_ISL_3313493, EPI_ISL_3313494, EPI_ISL_3313495, EPI_ISL_3313496, EPI_ISL_3313497 | | | |
| see above | Lighthouse Lab in Milton Keynes | Wellcome Sanger Institute for the COVID-19 Genomics UK (COG-UK) Consortium | The Lighthouse Lab in Milton Keynes and Alex Alderton, Roberto Amato, Jeffrey Barrett, Sonia Goncalves, Ewan Harrison, David K. Jackson, Ian Johnston, Dominic Kwiatkowski, Cordelia Langford, John Sillitoe on behalf of the Wellcome Sanger Institute COVID-19 Surveillance Team |
| EPI_ISL_3313498 | Berkshire and Surrey Pathology Services Lighthouse Laboratory | Wellcome Sanger Institute for the COVID-19 Genomics UK (COG-UK) Consortium | Berkshire and Surrey Pathology Services Lighthouse Laboratory and Alex Alderton, Roberto Amato, Jeffrey Barrett, Sonia Goncalves, Ewan Harrison, David K. Jackson, Ian Johnston, Dominic Kwiatkowski, Cordelia Langford, John Sillitoe on behalf of the Wellcome Sanger Institute COVID-19 Surveillance Team |
| EPI_ISL_3313499 | Lighthouse Lab in Milton Keynes | Wellcome Sanger Institute for the COVID-19 Genomics UK (COG-UK) Consortium | The Lighthouse Lab in Milton Keynes and Alex Alderton, Roberto Amato, Jeffrey Barrett, Sonia Goncalves, Ewan Harrison, David K. Jackson, Ian Johnston, Dominic Kwiatkowski, Cordelia Langford, John Sillitoe on behalf of the Wellcome Sanger Institute COVID-19 Surveillance Team |
| EPI_ISL_3313500 | Berkshire and Surrey Pathology Services Lighthouse Laboratory | Wellcome Sanger Institute for the COVID-19 Genomics UK (COG-UK) Consortium | Berkshire and Surrey Pathology Services Lighthouse Laboratory and Alex Alderton, Roberto Amato, Jeffrey Barrett, Sonia Goncalves, Ewan Harrison, David K. Jackson, Ian Johnston, Dominic Kwiatkowski, Cordelia Langford, John Sillitoe on behalf of the Wellcome Sanger Institute COVID-19 Surveillance Team |
| EPI_ISL_3313501 | Lighthouse Lab in Milton Keynes | Wellcome Sanger Institute for the COVID-19 Genomics UK (COG-UK) Consortium | The Lighthouse Lab in Milton Keynes and Alex Alderton, Roberto Amato, Jeffrey Barrett, Sonia Goncalves, Ewan Harrison, David K. Jackson, Ian Johnston, Dominic Kwiatkowski, Cordelia Langford, John Sillitoe on behalf of the Wellcome Sanger Institute COVID-19 Surveillance Team |
| EPI_ISL_3313502 | Berkshire and Surrey Pathology Services Lighthouse Laboratory | Wellcome Sanger Institute for the COVID-19 Genomics UK (COG-UK) Consortium | Berkshire and Surrey Pathology Services Lighthouse Laboratory and Alex Alderton, Roberto Amato, Jeffrey Barrett, Sonia Goncalves, Ewan Harrison, David K. Jackson, Ian Johnston, Dominic Kwiatkowski, Cordelia Langford, John Sillitoe on behalf of the Wellcome Sanger Institute COVID-19 Surveillance Team |
| EPI_ISL_3313503 | Lighthouse Lab in Glasgow | Wellcome Sanger Institute for the COVID-19 Genomics UK (COG-UK) Consortium | Harper VanSteenhouse, Yumi Kasai, David Gray, Carol Clugston, Anna Dominiczak and Alex Alderton, Roberto Amato, Jeffrey Barrett, Sonia Goncalves, Ewan Harrison, David K. Jackson, Ian Johnston, Dominic Kwiatkowski, Cordelia Langford, John Sillitoe on behalf of the Wellcome Sanger Institute COVID-19 Surveillance Team |
| EPI_ISL_3313504 | Berkshire and Surrey Pathology Services Lighthouse Laboratory | Wellcome Sanger Institute for the COVID-19 Genomics UK (COG-UK) Consortium | Berkshire and Surrey Pathology Services Lighthouse Laboratory and Alex Alderton, Roberto Amato, Jeffrey Barrett, Sonia Goncalves, Ewan Harrison, David K. Jackson, Ian Johnston, Dominic Kwiatkowski, Cordelia Langford, John Sillitoe on behalf of the Wellcome Sanger Institute COVID-19 Surveillance Team |
| EPI_ISL_3313505 | Lighthouse Lab in Glasgow | Wellcome Sanger Institute for the COVID-19 Genomics UK (COG-UK) Consortium | Harper VanSteenhouse, Yumi Kasai, David Gray, Carol Clugston, Anna Dominiczak and Alex Alderton, Roberto Amato, Jeffrey Barrett, Sonia Goncalves, Ewan Harrison, David K. Jackson, Ian Johnston, Dominic Kwiatkowski, Cordelia Langford, John Sillitoe on behalf of the Wellcome Sanger Institute COVID-19 Surveillance Team |
| EPI_ISL_3313506 | Berkshire and Surrey Pathology Services Lighthouse Laboratory | Wellcome Sanger Institute for the COVID-19 Genomics UK (COG-UK) Consortium | Berkshire and Surrey Pathology Services Lighthouse Laboratory and Alex Alderton, Roberto Amato, Jeffrey Barrett, Sonia Goncalves, Ewan Harrison, David K. Jackson, Ian Johnston, Dominic Kwiatkowski, Cordelia Langford, John Sillitoe on behalf of the Wellcome Sanger Institute COVID-19 Surveillance Team |
| EPI_ISL_3313507 | Lighthouse Lab in Glasgow | Wellcome Sanger Institute for the COVID-19 Genomics UK (COG-UK) | Harper VanSteenhouse, Yumi Kasai, David Gray, Carol Clugston, Anna Dominiczak and Alex Alderton, Roberto Amato, Jeffrey Barrett, Sonia Goncalves, Ewan Harrison, David K. Jackson, Ian Johnston, Dominic Kwiatkowski, Cordelia Langford, |

| | | | |
|---|---|---|---|
| | | Consortium | John Sillitoe on behalf of the Wellcome Sanger Institute COVID-19 Surveillance Team |
| EPI_ISL_3313508, EPI_ISL_3313509, EPI_ISL_3313510 | Berkshire and Surrey Pathology Services Lighthouse Laboratory | Wellcome Sanger Institute for the COVID-19 Genomics UK (COG-UK) Consortium | Berkshire and Surrey Pathology Services Lighthouse Laboratory and Alex Alderton, Roberto Amato, Jeffrey Barrett, Sonia Goncalves, Ewan Harrison, David K. Jackson, Ian Johnston, Dominic Kwiatkowski, Cordelia Langford, John Sillitoe on behalf of the Wellcome Sanger Institute COVID-19 Surveillance Team |
| EPI_ISL_3313511 | Lighthouse Lab in Glasgow | Wellcome Sanger Institute for the COVID-19 Genomics UK (COG-UK) Consortium | Harper VanSteenhouse, Yumi Kasai, David Gray, Carol Clugston, Anna Dominiczak and Alex Alderton, Roberto Amato, Jeffrey Barrett, Sonia Goncalves, Ewan Harrison, David K. Jackson, Ian Johnston, Dominic Kwiatkowski, Cordelia Langford, John Sillitoe on behalf of the Wellcome Sanger Institute COVID-19 Surveillance Team |
| EPI_ISL_3313512 | Berkshire and Surrey Pathology Services Lighthouse Laboratory | Wellcome Sanger Institute for the COVID-19 Genomics UK (COG-UK) Consortium | Berkshire and Surrey Pathology Services Lighthouse Laboratory and Alex Alderton, Roberto Amato, Jeffrey Barrett, Sonia Goncalves, Ewan Harrison, David K. Jackson, Ian Johnston, Dominic Kwiatkowski, Cordelia Langford, John Sillitoe on behalf of the Wellcome Sanger Institute COVID-19 Surveillance Team |
| EPI_ISL_3313513, EPI_ISL_3313514, EPI_ISL_3313515 | Lighthouse Lab in Glasgow | Wellcome Sanger Institute for the COVID-19 Genomics UK (COG-UK) Consortium | Harper VanSteenhouse, Yumi Kasai, David Gray, Carol Clugston, Anna Dominiczak and Alex Alderton, Roberto Amato, Jeffrey Barrett, Sonia Goncalves, Ewan Harrison, David K. Jackson, Ian Johnston, Dominic Kwiatkowski, Cordelia Langford, John Sillitoe on behalf of the Wellcome Sanger Institute COVID-19 Surveillance Team |
| EPI_ISL_3313516 | Lighthouse Lab in Milton Keynes | Wellcome Sanger Institute for the COVID-19 Genomics UK (COG-UK) Consortium | The Lighthouse Lab in Milton Keynes and Alex Alderton, Roberto Amato, Jeffrey Barrett, Sonia Goncalves, Ewan Harrison, David K. Jackson, Ian Johnston, Dominic Kwiatkowski, Cordelia Langford, John Sillitoe on behalf of the Wellcome Sanger Institute COVID-19 Surveillance Team |
| EPI_ISL_3313517 | Berkshire and Surrey Pathology Services Lighthouse Laboratory | Wellcome Sanger Institute for the COVID-19 Genomics UK (COG-UK) Consortium | Berkshire and Surrey Pathology Services Lighthouse Laboratory and Alex Alderton, Roberto Amato, Jeffrey Barrett, Sonia Goncalves, Ewan Harrison, David K. Jackson, Ian Johnston, Dominic Kwiatkowski, Cordelia Langford, John Sillitoe on behalf of the Wellcome Sanger Institute COVID-19 Surveillance Team |
| EPI_ISL_3313518 | Lighthouse Lab in Glasgow | Wellcome Sanger Institute for the COVID-19 Genomics UK (COG-UK) Consortium | Harper VanSteenhouse, Yumi Kasai, David Gray, Carol Clugston, Anna Dominiczak and Alex Alderton, Roberto Amato, Jeffrey Barrett, Sonia Goncalves, Ewan Harrison, David K. Jackson, Ian Johnston, Dominic Kwiatkowski, Cordelia Langford, John Sillitoe on behalf of the Wellcome Sanger Institute COVID-19 Surveillance Team |
| EPI_ISL_3313519 | Berkshire and Surrey Pathology Services Lighthouse Laboratory | Wellcome Sanger Institute for the COVID-19 Genomics UK (COG-UK) Consortium | Berkshire and Surrey Pathology Services Lighthouse Laboratory and Alex Alderton, Roberto Amato, Jeffrey Barrett, Sonia Goncalves, Ewan Harrison, David K. Jackson, Ian Johnston, Dominic Kwiatkowski, Cordelia Langford, John Sillitoe on behalf of the Wellcome Sanger Institute COVID-19 Surveillance Team |
| EPI_ISL_3313520 | Lighthouse Lab in Milton Keynes | Wellcome Sanger Institute for the COVID-19 Genomics UK (COG-UK) Consortium | The Lighthouse Lab in Milton Keynes and Alex Alderton, Roberto Amato, Jeffrey Barrett, Sonia Goncalves, Ewan Harrison, David K. Jackson, Ian Johnston, Dominic Kwiatkowski, Cordelia Langford, John Sillitoe on behalf of the Wellcome Sanger Institute COVID-19 Surveillance Team |
| EPI_ISL_3313521, EPI_ISL_3313522, EPI_ISL_3313523 | Berkshire and Surrey Pathology Services Lighthouse Laboratory | Wellcome Sanger Institute for the COVID-19 Genomics UK (COG-UK) Consortium | Berkshire and Surrey Pathology Services Lighthouse Laboratory and Alex Alderton, Roberto Amato, Jeffrey Barrett, Sonia Goncalves, Ewan Harrison, David K. Jackson, Ian Johnston, Dominic Kwiatkowski, Cordelia Langford, John Sillitoe on behalf of the Wellcome Sanger Institute COVID-19 Surveillance Team |
| EPI_ISL_3313524 | Lighthouse Lab in Milton Keynes | Wellcome Sanger Institute for the COVID-19 Genomics UK (COG-UK) Consortium | The Lighthouse Lab in Milton Keynes and Alex Alderton, Roberto Amato, Jeffrey Barrett, Sonia Goncalves, Ewan Harrison, David K. Jackson, Ian Johnston, Dominic Kwiatkowski, Cordelia Langford, John Sillitoe on behalf of the Wellcome Sanger Institute COVID-19 Surveillance Team |
| EPI_ISL_3313525 | Lighthouse Lab in Glasgow | Wellcome Sanger Institute for the COVID-19 Genomics UK (COG-UK) Consortium | Harper VanSteenhouse, Yumi Kasai, David Gray, Carol Clugston, Anna Dominiczak and Alex Alderton, Roberto Amato, Jeffrey Barrett, Sonia Goncalves, Ewan Harrison, David K. Jackson, Ian Johnston, Dominic Kwiatkowski, Cordelia Langford, John Sillitoe on behalf of the Wellcome Sanger Institute COVID-19 Surveillance Team |
| EPI_ISL_3313526, EPI_ISL_3313527 | Berkshire and Surrey Pathology Services Lighthouse Laboratory | Wellcome Sanger Institute for the COVID-19 Genomics UK (COG-UK) Consortium | Berkshire and Surrey Pathology Services Lighthouse Laboratory and Alex Alderton, Roberto Amato, Jeffrey Barrett, Sonia Goncalves, Ewan Harrison, David K. Jackson, Ian Johnston, Dominic Kwiatkowski, Cordelia Langford, John Sillitoe on behalf of the Wellcome Sanger Institute COVID-19 Surveillance Team |
| EPI_ISL_3313528, EPI_ISL_3313529, EPI_ISL_3313530 | Lighthouse Lab in Glasgow | Wellcome Sanger Institute for the COVID-19 Genomics UK (COG-UK) Consortium | Harper VanSteenhouse, Yumi Kasai, David Gray, Carol Clugston, Anna Dominiczak and Alex Alderton, Roberto Amato, Jeffrey Barrett, Sonia Goncalves, Ewan Harrison, David K. Jackson, Ian Johnston, Dominic Kwiatkowski, Cordelia Langford, John Sillitoe on behalf of the Wellcome Sanger Institute COVID-19 Surveillance Team |
| EPI_ISL_3313531, EPI_ISL_3313532 | Berkshire and Surrey Pathology Services Lighthouse Laboratory | Wellcome Sanger Institute for the COVID-19 Genomics UK (COG-UK) Consortium | Berkshire and Surrey Pathology Services Lighthouse Laboratory and Alex Alderton, Roberto Amato, Jeffrey Barrett, Sonia Goncalves, Ewan Harrison, David K. Jackson, Ian Johnston, Dominic Kwiatkowski, Cordelia Langford, John Sillitoe on behalf of the Wellcome Sanger Institute COVID-19 Surveillance Team |
| EPI_ISL_3313533 | Lighthouse Lab in Milton Keynes | Wellcome Sanger Institute for the COVID-19 Genomics UK (COG-UK) Consortium | The Lighthouse Lab in Milton Keynes and Alex Alderton, Roberto Amato, Jeffrey Barrett, Sonia Goncalves, Ewan Harrison, David K. Jackson, Ian Johnston, Dominic Kwiatkowski, Cordelia Langford, John Sillitoe on behalf of the Wellcome Sanger Institute COVID-19 Surveillance Team |
| EPI_ISL_3313534 | Berkshire and Surrey Pathology Services Lighthouse Laboratory | Wellcome Sanger Institute for the COVID-19 Genomics UK (COG-UK) Consortium | Berkshire and Surrey Pathology Services Lighthouse Laboratory and Alex Alderton, Roberto Amato, Jeffrey Barrett, Sonia Goncalves, Ewan Harrison, David K. Jackson, Ian Johnston, Dominic Kwiatkowski, Cordelia Langford, John Sillitoe on behalf of the Wellcome Sanger Institute COVID-19 Surveillance Team |
| EPI_ISL_3313535 | Lighthouse Lab in Glasgow | Wellcome Sanger Institute for the COVID-19 Genomics UK (COG-UK) Consortium | Harper VanSteenhouse, Yumi Kasai, David Gray, Carol Clugston, Anna Dominiczak and Alex Alderton, Roberto Amato, Jeffrey Barrett, Sonia Goncalves, Ewan Harrison, David K. Jackson, Ian Johnston, Dominic Kwiatkowski, Cordelia Langford, John Sillitoe on behalf of the Wellcome Sanger Institute COVID-19 Surveillance Team |
| EPI_ISL_3313536 | Lighthouse Lab in Milton Keynes | Wellcome Sanger Institute for the COVID-19 Genomics UK (COG-UK) Consortium | The Lighthouse Lab in Milton Keynes and Alex Alderton, Roberto Amato, Jeffrey Barrett, Sonia Goncalves, Ewan Harrison, David K. Jackson, Ian Johnston, Dominic Kwiatkowski, Cordelia Langford, John Sillitoe on behalf of the Wellcome Sanger Institute COVID-19 Surveillance Team |
| EPI_ISL_3313537, EPI_ISL_3313538, EPI_ISL_3313539, EPI_ISL_3313540, EPI_ISL_3313541 | Berkshire and Surrey Pathology Services Lighthouse Laboratory | Wellcome Sanger Institute for the COVID-19 Genomics UK (COG-UK) Consortium | Berkshire and Surrey Pathology Services Lighthouse Laboratory and Alex Alderton, Roberto Amato, Jeffrey Barrett, Sonia Goncalves, Ewan Harrison, David K. Jackson, Ian Johnston, Dominic Kwiatkowski, Cordelia Langford, John Sillitoe on behalf of the Wellcome Sanger Institute COVID-19 Surveillance Team |
| EPI_ISL_3313542 | Lighthouse Lab in Milton Keynes | Wellcome Sanger Institute for the COVID-19 Genomics UK (COG-UK) Consortium | The Lighthouse Lab in Milton Keynes and Alex Alderton, Roberto Amato, Jeffrey Barrett, Sonia Goncalves, Ewan Harrison, David K. Jackson, Ian Johnston, Dominic Kwiatkowski, Cordelia Langford, John Sillitoe on behalf of the Wellcome Sanger Institute COVID-19 Surveillance Team |
| EPI_ISL_3313543 | Berkshire and Surrey Pathology Services Lighthouse Laboratory | Wellcome Sanger Institute for the COVID-19 Genomics UK (COG-UK) Consortium | Berkshire and Surrey Pathology Services Lighthouse Laboratory and Alex Alderton, Roberto Amato, Jeffrey Barrett, Sonia Goncalves, Ewan Harrison, David K. Jackson, Ian Johnston, Dominic Kwiatkowski, Cordelia Langford, John Sillitoe on behalf of the Wellcome Sanger Institute COVID-19 Surveillance Team |
| EPI_ISL_3313544 | Lighthouse Lab in Milton Keynes | Wellcome Sanger Institute for the COVID-19 Genomics UK (COG-UK) Consortium | The Lighthouse Lab in Milton Keynes and Alex Alderton, Roberto Amato, Jeffrey Barrett, Sonia Goncalves, Ewan Harrison, David K. Jackson, Ian Johnston, Dominic Kwiatkowski, Cordelia Langford, John Sillitoe on behalf of the Wellcome Sanger Institute COVID-19 Surveillance Team |
| EPI_ISL_3313545 | Berkshire and Surrey Pathology Services Lighthouse Laboratory | Wellcome Sanger Institute for the COVID-19 Genomics UK (COG-UK) Consortium | Berkshire and Surrey Pathology Services Lighthouse Laboratory and Alex Alderton, Roberto Amato, Jeffrey Barrett, Sonia Goncalves, Ewan Harrison, David K. Jackson, Ian Johnston, Dominic Kwiatkowski, Cordelia Langford, John Sillitoe on behalf of the Wellcome Sanger Institute COVID-19 Surveillance Team |
| EPI_ISL_3313546 | Lighthouse Lab in Glasgow | Wellcome Sanger Institute for the COVID-19 Genomics UK (COG-UK) Consortium | Harper VanSteenhouse, Yumi Kasai, David Gray, Carol Clugston, Anna Dominiczak and Alex Alderton, Roberto Amato, Jeffrey Barrett, Sonia Goncalves, Ewan Harrison, David K. Jackson, Ian Johnston, Dominic Kwiatkowski, Cordelia Langford, John Sillitoe on behalf of the Wellcome Sanger Institute COVID-19 Surveillance Team |
| EPI_ISL_3313547, EPI_ISL_3313548, EPI_ISL_3313549, EPI_ISL_3313550 | Berkshire and Surrey Pathology Services Lighthouse Laboratory | Wellcome Sanger Institute for the COVID-19 Genomics UK (COG-UK) Consortium | Berkshire and Surrey Pathology Services Lighthouse Laboratory and Alex Alderton, Roberto Amato, Jeffrey Barrett, Sonia Goncalves, Ewan Harrison, David K. Jackson, Ian Johnston, Dominic Kwiatkowski, Cordelia Langford, John Sillitoe on behalf of the Wellcome Sanger Institute COVID-19 Surveillance Team |
| EPI_ISL_3313551 | Lighthouse Lab in Glasgow | Wellcome Sanger Institute for the COVID-19 Genomics UK (COG-UK) Consortium | Harper VanSteenhouse, Yumi Kasai, David Gray, Carol Clugston, Anna Dominiczak and Alex Alderton, Roberto Amato, Jeffrey Barrett, Sonia Goncalves, Ewan Harrison, David K. Jackson, Ian Johnston, Dominic Kwiatkowski, Cordelia Langford, John Sillitoe on behalf of the Wellcome Sanger Institute COVID-19 Surveillance Team |
| EPI_ISL_3313552 | Berkshire and Surrey Pathology Services Lighthouse Laboratory | Wellcome Sanger Institute for the COVID-19 Genomics UK (COG-UK) Consortium | Berkshire and Surrey Pathology Services Lighthouse Laboratory and Alex Alderton, Roberto Amato, Jeffrey Barrett, Sonia Goncalves, Ewan Harrison, David K. Jackson, Ian Johnston, Dominic Kwiatkowski, Cordelia Langford, John Sillitoe on behalf of the Wellcome Sanger Institute COVID-19 Surveillance Team |
| EPI_ISL_3313553 | Lighthouse Lab in Glasgow | Wellcome Sanger Institute for the COVID-19 Genomics UK (COG-UK) Consortium | Harper VanSteenhouse, Yumi Kasai, David Gray, Carol Clugston, Anna Dominiczak and Alex Alderton, Roberto Amato, Jeffrey Barrett, Sonia Goncalves, Ewan Harrison, David K. Jackson, Ian Johnston, Dominic Kwiatkowski, Cordelia Langford, John Sillitoe on behalf of the Wellcome Sanger Institute COVID-19 Surveillance Team |
| EPI_ISL_3313554, EPI_ISL_3313555 | Lighthouse Lab in Milton Keynes | Wellcome Sanger Institute for the COVID-19 Genomics UK (COG-UK) Consortium | The Lighthouse Lab in Milton Keynes and Alex Alderton, Roberto Amato, Jeffrey Barrett, Sonia Goncalves, Ewan Harrison, David K. Jackson, Ian Johnston, Dominic Kwiatkowski, Cordelia Langford, John Sillitoe on behalf of the Wellcome Sanger Institute COVID-19 Surveillance Team |
| EPI_ISL_3313556 | Berkshire and Surrey Pathology Services Lighthouse Laboratory | Wellcome Sanger Institute for the COVID-19 Genomics UK (COG-UK) Consortium | Berkshire and Surrey Pathology Services Lighthouse Laboratory and Alex Alderton, Roberto Amato, Jeffrey Barrett, Sonia Goncalves, Ewan Harrison, David K. Jackson, Ian Johnston, Dominic Kwiatkowski, Cordelia Langford, John Sillitoe on behalf of the Wellcome Sanger Institute COVID-19 Surveillance Team |
| EPI_ISL_3313557, EPI_ISL_3313558, EPI_ISL_3313559 | Lighthouse Lab in Milton Keynes | Wellcome Sanger Institute for the COVID-19 Genomics UK (COG-UK) Consortium | The Lighthouse Lab in Milton Keynes and Alex Alderton, Roberto Amato, Jeffrey Barrett, Sonia Goncalves, Ewan Harrison, David K. Jackson, Ian Johnston, Dominic Kwiatkowski, Cordelia Langford, John Sillitoe on behalf of the Wellcome Sanger Institute COVID-19 Surveillance Team |
| EPI_ISL_3313560 | Lighthouse Lab in Glasgow | Wellcome Sanger Institute for the COVID-19 Genomics UK (COG-UK) Consortium | Harper VanSteenhouse, Yumi Kasai, David Gray, Carol Clugston, Anna Dominiczak and Alex Alderton, Roberto Amato, Jeffrey Barrett, Sonia Goncalves, Ewan Harrison, David K. Jackson, Ian Johnston, Dominic Kwiatkowski, Cordelia Langford, John Sillitoe on behalf of the Wellcome Sanger Institute COVID-19 Surveillance Team |
| EPI_ISL_3313561 | Berkshire and Surrey Pathology Services Lighthouse Laboratory | Wellcome Sanger Institute for the COVID-19 Genomics UK (COG-UK) Consortium | Berkshire and Surrey Pathology Services Lighthouse Laboratory and Alex Alderton, Roberto Amato, Jeffrey Barrett, Sonia Goncalves, Ewan Harrison, David K. Jackson, Ian Johnston, Dominic Kwiatkowski, Cordelia Langford, John Sillitoe on behalf of the Wellcome Sanger Institute COVID-19 Surveillance Team |
| EPI_ISL_3313562 | Lighthouse Lab in Glasgow | Wellcome Sanger Institute for the COVID-19 Genomics UK (COG-UK) Consortium | Harper VanSteenhouse, Yumi Kasai, David Gray, Carol Clugston, Anna Dominiczak and Alex Alderton, Roberto Amato, Jeffrey Barrett, Sonia Goncalves, Ewan Harrison, David K. Jackson, Ian Johnston, Dominic Kwiatkowski, Cordelia Langford, John Sillitoe on behalf of the Wellcome Sanger Institute COVID-19 Surveillance Team |
| EPI_ISL_3313563 | Lighthouse Lab in Milton Keynes | Wellcome Sanger Institute for the COVID-19 Genomics UK (COG-UK) Consortium | The Lighthouse Lab in Milton Keynes and Alex Alderton, Roberto Amato, Jeffrey Barrett, Sonia Goncalves, Ewan Harrison, David K. Jackson, Ian Johnston, Dominic Kwiatkowski, Cordelia Langford, John Sillitoe on behalf of the Wellcome Sanger Institute COVID-19 Surveillance Team |
| EPI_ISL_3313564 | Berkshire and Surrey Pathology Services Lighthouse Laboratory | Wellcome Sanger Institute for the COVID-19 Genomics UK (COG-UK) Consortium | Berkshire and Surrey Pathology Services Lighthouse Laboratory and Alex Alderton, Roberto Amato, Jeffrey Barrett, Sonia Goncalves, Ewan Harrison, David K. Jackson, Ian Johnston, Dominic Kwiatkowski, Cordelia Langford, John Sillitoe on behalf of the Wellcome Sanger Institute COVID-19 Surveillance Team |
| EPI_ISL_3313565 | Lighthouse Lab in Milton Keynes | Wellcome Sanger Institute for the COVID-19 Genomics UK (COG-UK) Consortium | The Lighthouse Lab in Milton Keynes and Alex Alderton, Roberto Amato, Jeffrey Barrett, Sonia Goncalves, Ewan Harrison, David K. Jackson, Ian Johnston, Dominic Kwiatkowski, Cordelia Langford, John Sillitoe on behalf of the Wellcome Sanger Institute COVID-19 Surveillance Team |
| EPI_ISL_3313566 | Berkshire and Surrey Pathology Services Lighthouse Laboratory | Wellcome Sanger Institute for the COVID-19 Genomics UK (COG-UK) Consortium | Berkshire and Surrey Pathology Services Lighthouse Laboratory and Alex Alderton, Roberto Amato, Jeffrey Barrett, Sonia Goncalves, Ewan Harrison, David K. Jackson, Ian Johnston, Dominic Kwiatkowski, Cordelia Langford, John Sillitoe on behalf of the Wellcome Sanger Institute COVID-19 Surveillance Team |
| EPI_ISL_3313567 | Lighthouse Lab in Milton Keynes | Wellcome Sanger Institute for the COVID-19 Genomics UK (COG-UK) Consortium | The Lighthouse Lab in Milton Keynes and Alex Alderton, Roberto Amato, Jeffrey Barrett, Sonia Goncalves, Ewan Harrison, David K. Jackson, Ian Johnston, Dominic Kwiatkowski, Cordelia Langford, John Sillitoe on behalf of the Wellcome Sanger Institute COVID-19 Surveillance Team |
| EPI_ISL_3313568 | Lighthouse Lab in Glasgow | Wellcome Sanger Institute for the COVID-19 Genomics UK (COG-UK) | Harper VanSteenhouse, Yumi Kasai, David Gray, Carol Clugston, Anna Dominiczak and Alex Alderton, Roberto Amato, Jeffrey Barrett, Sonia Goncalves, Ewan Harrison, David K. Jackson, Ian Johnston, Dominic Kwiatkowski, Cordelia Langford, |

| | | | |
|---|---|---|---|
| | | Consortium | John Sillitoe on behalf of the Wellcome Sanger Institute COVID-19 Surveillance Team |
| EPI_ISL_3313569, EPI_ISL_3313570, EPI_ISL_3313571 | Berkshire and Surrey Pathology Services Lighthouse Laboratory | Wellcome Sanger Institute for the COVID-19 Genomics UK (COG-UK) Consortium | Berkshire and Surrey Pathology Services Lighthouse Laboratory and Alex Alderton, Roberto Amato, Jeffrey Barrett, Sonia Goncalves, Ewan Harrison, David K. Jackson, Ian Johnston, Dominic Kwiatkowski, Cordelia Langford, John Sillitoe on behalf of the Wellcome Sanger Institute COVID-19 Surveillance Team |
| EPI_ISL_3313572 | Lighthouse Lab in Milton Keynes | Wellcome Sanger Institute for the COVID-19 Genomics UK (COG-UK) Consortium | The Lighthouse Lab in Milton Keynes and Alex Alderton, Roberto Amato, Jeffrey Barrett, Sonia Goncalves, Ewan Harrison, David K. Jackson, Ian Johnston, Dominic Kwiatkowski, Cordelia Langford, John Sillitoe on behalf of the Wellcome Sanger Institute COVID-19 Surveillance Team |
| EPI_ISL_3313573, EPI_ISL_3313574, EPI_ISL_3313575, EPI_ISL_3313576 | Berkshire and Surrey Pathology Services Lighthouse Laboratory | Wellcome Sanger Institute for the COVID-19 Genomics UK (COG-UK) Consortium | Berkshire and Surrey Pathology Services Lighthouse Laboratory and Alex Alderton, Roberto Amato, Jeffrey Barrett, Sonia Goncalves, Ewan Harrison, David K. Jackson, Ian Johnston, Dominic Kwiatkowski, Cordelia Langford, John Sillitoe on behalf of the Wellcome Sanger Institute COVID-19 Surveillance Team |
| EPI_ISL_3313577, EPI_ISL_3313578 | Lighthouse Lab in Glasgow | Wellcome Sanger Institute for the COVID-19 Genomics UK (COG-UK) Consortium | Harper VanSteenhouse, Yumi Kasai, David Gray, Carol Clugston, Anna Dominiczak and Alex Alderton, Roberto Amato, Jeffrey Barrett, Sonia Goncalves, Ewan Harrison, David K. Jackson, Ian Johnston, Dominic Kwiatkowski, Cordelia Langford, John Sillitoe on behalf of the Wellcome Sanger Institute COVID-19 Surveillance Team |
| EPI_ISL_3313579, EPI_ISL_3313580, EPI_ISL_3313581 | Berkshire and Surrey Pathology Services Lighthouse Laboratory | Wellcome Sanger Institute for the COVID-19 Genomics UK (COG-UK) Consortium | Berkshire and Surrey Pathology Services Lighthouse Laboratory and Alex Alderton, Roberto Amato, Jeffrey Barrett, Sonia Goncalves, Ewan Harrison, David K. Jackson, Ian Johnston, Dominic Kwiatkowski, Cordelia Langford, John Sillitoe on behalf of the Wellcome Sanger Institute COVID-19 Surveillance Team |
| EPI_ISL_3313582 | Lighthouse Lab in Milton Keynes | Wellcome Sanger Institute for the COVID-19 Genomics UK (COG-UK) Consortium | The Lighthouse Lab in Milton Keynes and Alex Alderton, Roberto Amato, Jeffrey Barrett, Sonia Goncalves, Ewan Harrison, David K. Jackson, Ian Johnston, Dominic Kwiatkowski, Cordelia Langford, John Sillitoe on behalf of the Wellcome Sanger Institute COVID-19 Surveillance Team |
| EPI_ISL_3313583 | Berkshire and Surrey Pathology Services Lighthouse Laboratory | Wellcome Sanger Institute for the COVID-19 Genomics UK (COG-UK) Consortium | Berkshire and Surrey Pathology Services Lighthouse Laboratory and Alex Alderton, Roberto Amato, Jeffrey Barrett, Sonia Goncalves, Ewan Harrison, David K. Jackson, Ian Johnston, Dominic Kwiatkowski, Cordelia Langford, John Sillitoe on behalf of the Wellcome Sanger Institute COVID-19 Surveillance Team |
| EPI_ISL_3313584, EPI_ISL_3313585 | Lighthouse Lab in Milton Keynes | Wellcome Sanger Institute for the COVID-19 Genomics UK (COG-UK) Consortium | The Lighthouse Lab in Milton Keynes and Alex Alderton, Roberto Amato, Jeffrey Barrett, Sonia Goncalves, Ewan Harrison, David K. Jackson, Ian Johnston, Dominic Kwiatkowski, Cordelia Langford, John Sillitoe on behalf of the Wellcome Sanger Institute COVID-19 Surveillance Team |
| EPI_ISL_3313586 | Berkshire and Surrey Pathology Services Lighthouse Laboratory | Wellcome Sanger Institute for the COVID-19 Genomics UK (COG-UK) Consortium | Berkshire and Surrey Pathology Services Lighthouse Laboratory and Alex Alderton, Roberto Amato, Jeffrey Barrett, Sonia Goncalves, Ewan Harrison, David K. Jackson, Ian Johnston, Dominic Kwiatkowski, Cordelia Langford, John Sillitoe on behalf of the Wellcome Sanger Institute COVID-19 Surveillance Team |
| EPI_ISL_3313587 | Lighthouse Lab in Milton Keynes | Wellcome Sanger Institute for the COVID-19 Genomics UK (COG-UK) Consortium | The Lighthouse Lab in Milton Keynes and Alex Alderton, Roberto Amato, Jeffrey Barrett, Sonia Goncalves, Ewan Harrison, David K. Jackson, Ian Johnston, Dominic Kwiatkowski, Cordelia Langford, John Sillitoe on behalf of the Wellcome Sanger Institute COVID-19 Surveillance Team |
| EPI_ISL_3313588, EPI_ISL_3313589, EPI_ISL_3313590, EPI_ISL_3313591 | Berkshire and Surrey Pathology Services Lighthouse Laboratory | Wellcome Sanger Institute for the COVID-19 Genomics UK (COG-UK) Consortium | Berkshire and Surrey Pathology Services Lighthouse Laboratory and Alex Alderton, Roberto Amato, Jeffrey Barrett, Sonia Goncalves, Ewan Harrison, David K. Jackson, Ian Johnston, Dominic Kwiatkowski, Cordelia Langford, John Sillitoe on behalf of the Wellcome Sanger Institute COVID-19 Surveillance Team |
| EPI_ISL_3313592, EPI_ISL_3313593 | Lighthouse Lab in Milton Keynes | Wellcome Sanger Institute for the COVID-19 Genomics UK (COG-UK) Consortium | The Lighthouse Lab in Milton Keynes and Alex Alderton, Roberto Amato, Jeffrey Barrett, Sonia Goncalves, Ewan Harrison, David K. Jackson, Ian Johnston, Dominic Kwiatkowski, Cordelia Langford, John Sillitoe on behalf of the Wellcome Sanger Institute COVID-19 Surveillance Team |
| EPI_ISL_3313594 | Berkshire and Surrey Pathology Services Lighthouse Laboratory | Wellcome Sanger Institute for the COVID-19 Genomics UK (COG-UK) Consortium | Berkshire and Surrey Pathology Services Lighthouse Laboratory and Alex Alderton, Roberto Amato, Jeffrey Barrett, Sonia Goncalves, Ewan Harrison, David K. Jackson, Ian Johnston, Dominic Kwiatkowski, Cordelia Langford, John Sillitoe on behalf of the Wellcome Sanger Institute COVID-19 Surveillance Team |
| EPI_ISL_3313595, EPI_ISL_3313596, EPI_ISL_3313597 | Lighthouse Lab in Milton Keynes | Wellcome Sanger Institute for the COVID-19 Genomics UK (COG-UK) Consortium | The Lighthouse Lab in Milton Keynes and Alex Alderton, Roberto Amato, Jeffrey Barrett, Sonia Goncalves, Ewan Harrison, David K. Jackson, Ian Johnston, Dominic Kwiatkowski, Cordelia Langford, John Sillitoe on behalf of the Wellcome Sanger Institute COVID-19 Surveillance Team |
| EPI_ISL_3313598, EPI_ISL_3313599, EPI_ISL_3313600, EPI_ISL_3313601 | Berkshire and Surrey Pathology Services Lighthouse Laboratory | Wellcome Sanger Institute for the COVID-19 Genomics UK (COG-UK) Consortium | Berkshire and Surrey Pathology Services Lighthouse Laboratory and Alex Alderton, Roberto Amato, Jeffrey Barrett, Sonia Goncalves, Ewan Harrison, David K. Jackson, Ian Johnston, Dominic Kwiatkowski, Cordelia Langford, John Sillitoe on behalf of the Wellcome Sanger Institute COVID-19 Surveillance Team |
| EPI_ISL_3313602 | Lighthouse Lab in Milton Keynes | Wellcome Sanger Institute for the COVID-19 Genomics UK (COG-UK) Consortium | The Lighthouse Lab in Milton Keynes and Alex Alderton, Roberto Amato, Jeffrey Barrett, Sonia Goncalves, Ewan Harrison, David K. Jackson, Ian Johnston, Dominic Kwiatkowski, Cordelia Langford, John Sillitoe on behalf of the Wellcome Sanger Institute COVID-19 Surveillance Team |
| EPI_ISL_3313603, EPI_ISL_3313604, EPI_ISL_3313605 | Berkshire and Surrey Pathology Services Lighthouse Laboratory | Wellcome Sanger Institute for the COVID-19 Genomics UK (COG-UK) Consortium | Berkshire and Surrey Pathology Services Lighthouse Laboratory and Alex Alderton, Roberto Amato, Jeffrey Barrett, Sonia Goncalves, Ewan Harrison, David K. Jackson, Ian Johnston, Dominic Kwiatkowski, Cordelia Langford, John Sillitoe on behalf of the Wellcome Sanger Institute COVID-19 Surveillance Team |
| EPI_ISL_3313606, EPI_ISL_3313607, EPI_ISL_3313608 | Lighthouse Lab in Milton Keynes | Wellcome Sanger Institute for the COVID-19 Genomics UK (COG-UK) Consortium | The Lighthouse Lab in Milton Keynes and Alex Alderton, Roberto Amato, Jeffrey Barrett, Sonia Goncalves, Ewan Harrison, David K. Jackson, Ian Johnston, Dominic Kwiatkowski, Cordelia Langford, John Sillitoe on behalf of the Wellcome Sanger Institute COVID-19 Surveillance Team |
| EPI_ISL_3313609 | Berkshire and Surrey Pathology Services Lighthouse Laboratory | Wellcome Sanger Institute for the COVID-19 Genomics UK (COG-UK) Consortium | Berkshire and Surrey Pathology Services Lighthouse Laboratory and Alex Alderton, Roberto Amato, Jeffrey Barrett, Sonia Goncalves, Ewan Harrison, David K. Jackson, Ian Johnston, Dominic Kwiatkowski, Cordelia Langford, John Sillitoe on behalf of the Wellcome Sanger Institute COVID-19 Surveillance Team |
| EPI_ISL_3313610 | Lighthouse Lab in Glasgow | Wellcome Sanger Institute for the COVID-19 Genomics UK (COG-UK) Consortium | Harper VanSteenhouse, Yumi Kasai, David Gray, Carol Clugston, Anna Dominiczak and Alex Alderton, Roberto Amato, Jeffrey Barrett, Sonia Goncalves, Ewan Harrison, David K. Jackson, Ian Johnston, Dominic Kwiatkowski, Cordelia Langford, John Sillitoe on behalf of the Wellcome Sanger Institute COVID-19 Surveillance Team |
| EPI_ISL_3313611 | Berkshire and Surrey Pathology Services Lighthouse Laboratory | Wellcome Sanger Institute for the COVID-19 Genomics UK (COG-UK) Consortium | Berkshire and Surrey Pathology Services Lighthouse Laboratory and Alex Alderton, Roberto Amato, Jeffrey Barrett, Sonia Goncalves, Ewan Harrison, David K. Jackson, Ian Johnston, Dominic Kwiatkowski, Cordelia Langford, John Sillitoe on behalf of the Wellcome Sanger Institute COVID-19 Surveillance Team |
| EPI_ISL_3313612 | Lighthouse Lab in Glasgow | Wellcome Sanger Institute for the COVID-19 Genomics UK (COG-UK) Consortium | Harper VanSteenhouse, Yumi Kasai, David Gray, Carol Clugston, Anna Dominiczak and Alex Alderton, Roberto Amato, Jeffrey Barrett, Sonia Goncalves, Ewan Harrison, David K. Jackson, Ian Johnston, Dominic Kwiatkowski, Cordelia Langford, John Sillitoe on behalf of the Wellcome Sanger Institute COVID-19 Surveillance Team |
| EPI_ISL_3313613 | Berkshire and Surrey Pathology Services Lighthouse Laboratory | Wellcome Sanger Institute for the COVID-19 Genomics UK (COG-UK) Consortium | Berkshire and Surrey Pathology Services Lighthouse Laboratory and Alex Alderton, Roberto Amato, Jeffrey Barrett, Sonia Goncalves, Ewan Harrison, David K. Jackson, Ian Johnston, Dominic Kwiatkowski, Cordelia Langford, John Sillitoe on behalf of the Wellcome Sanger Institute COVID-19 Surveillance Team |
| EPI_ISL_3313614 | Lighthouse Lab in Milton Keynes | Wellcome Sanger Institute for the COVID-19 Genomics UK (COG-UK) Consortium | The Lighthouse Lab in Milton Keynes and Alex Alderton, Roberto Amato, Jeffrey Barrett, Sonia Goncalves, Ewan Harrison, David K. Jackson, Ian Johnston, Dominic Kwiatkowski, Cordelia Langford, John Sillitoe on behalf of the Wellcome Sanger Institute COVID-19 Surveillance Team |
| EPI_ISL_3313615 | Lighthouse Lab in Glasgow | Wellcome Sanger Institute for the COVID-19 Genomics UK (COG-UK) Consortium | Harper VanSteenhouse, Yumi Kasai, David Gray, Carol Clugston, Anna Dominiczak and Alex Alderton, Roberto Amato, Jeffrey Barrett, Sonia Goncalves, Ewan Harrison, David K. Jackson, Ian Johnston, Dominic Kwiatkowski, Cordelia Langford, John Sillitoe on behalf of the Wellcome Sanger Institute COVID-19 Surveillance Team |
| EPI_ISL_3313616, EPI_ISL_3313617, EPI_ISL_3313618 | Berkshire and Surrey Pathology Services Lighthouse Laboratory | Wellcome Sanger Institute for the COVID-19 Genomics UK (COG-UK) Consortium | Berkshire and Surrey Pathology Services Lighthouse Laboratory and Alex Alderton, Roberto Amato, Jeffrey Barrett, Sonia Goncalves, Ewan Harrison, David K. Jackson, Ian Johnston, Dominic Kwiatkowski, Cordelia Langford, John Sillitoe on behalf of the Wellcome Sanger Institute COVID-19 Surveillance Team |
| EPI_ISL_3313619 | Lighthouse Lab in Glasgow | Wellcome Sanger Institute for the COVID-19 Genomics UK (COG-UK) Consortium | Harper VanSteenhouse, Yumi Kasai, David Gray, Carol Clugston, Anna Dominiczak and Alex Alderton, Roberto Amato, Jeffrey Barrett, Sonia Goncalves, Ewan Harrison, David K. Jackson, Ian Johnston, Dominic Kwiatkowski, Cordelia Langford, John Sillitoe on behalf of the Wellcome Sanger Institute COVID-19 Surveillance Team |
| EPI_ISL_3313620 | Lighthouse Lab in Milton Keynes | Wellcome Sanger Institute for the COVID-19 Genomics UK (COG-UK) Consortium | The Lighthouse Lab in Milton Keynes and Alex Alderton, Roberto Amato, Jeffrey Barrett, Sonia Goncalves, Ewan Harrison, David K. Jackson, Ian Johnston, Dominic Kwiatkowski, Cordelia Langford, John Sillitoe on behalf of the Wellcome Sanger Institute COVID-19 Surveillance Team |
| EPI_ISL_3313621, EPI_ISL_3313622, EPI_ISL_3313623, EPI_ISL_3313624 | Berkshire and Surrey Pathology Services Lighthouse Laboratory | Wellcome Sanger Institute for the COVID-19 Genomics UK (COG-UK) Consortium | Berkshire and Surrey Pathology Services Lighthouse Laboratory and Alex Alderton, Roberto Amato, Jeffrey Barrett, Sonia Goncalves, Ewan Harrison, David K. Jackson, Ian Johnston, Dominic Kwiatkowski, Cordelia Langford, John Sillitoe on behalf of the Wellcome Sanger Institute COVID-19 Surveillance Team |
| EPI_ISL_3313625 | Lighthouse Lab in Glasgow | Wellcome Sanger Institute for the COVID-19 Genomics UK (COG-UK) Consortium | Harper VanSteenhouse, Yumi Kasai, David Gray, Carol Clugston, Anna Dominiczak and Alex Alderton, Roberto Amato, Jeffrey Barrett, Sonia Goncalves, Ewan Harrison, David K. Jackson, Ian Johnston, Dominic Kwiatkowski, Cordelia Langford, John Sillitoe on behalf of the Wellcome Sanger Institute COVID-19 Surveillance Team |
| EPI_ISL_3313626 | Lighthouse Lab in Milton Keynes | Wellcome Sanger Institute for the COVID-19 Genomics UK (COG-UK) Consortium | The Lighthouse Lab in Milton Keynes and Alex Alderton, Roberto Amato, Jeffrey Barrett, Sonia Goncalves, Ewan Harrison, David K. Jackson, Ian Johnston, Dominic Kwiatkowski, Cordelia Langford, John Sillitoe on behalf of the Wellcome Sanger Institute COVID-19 Surveillance Team |
| EPI_ISL_3313627 | Berkshire and Surrey Pathology Services Lighthouse Laboratory | Wellcome Sanger Institute for the COVID-19 Genomics UK (COG-UK) Consortium | Berkshire and Surrey Pathology Services Lighthouse Laboratory and Alex Alderton, Roberto Amato, Jeffrey Barrett, Sonia Goncalves, Ewan Harrison, David K. Jackson, Ian Johnston, Dominic Kwiatkowski, Cordelia Langford, John Sillitoe on behalf of the Wellcome Sanger Institute COVID-19 Surveillance Team |
| EPI_ISL_3313628 | Lighthouse Lab in Glasgow | Wellcome Sanger Institute for the COVID-19 Genomics UK (COG-UK) Consortium | Harper VanSteenhouse, Yumi Kasai, David Gray, Carol Clugston, Anna Dominiczak and Alex Alderton, Roberto Amato, Jeffrey Barrett, Sonia Goncalves, Ewan Harrison, David K. Jackson, Ian Johnston, Dominic Kwiatkowski, Cordelia Langford, John Sillitoe on behalf of the Wellcome Sanger Institute COVID-19 Surveillance Team |
| EPI_ISL_3313629, EPI_ISL_3313630 | Berkshire and Surrey Pathology Services Lighthouse Laboratory | Wellcome Sanger Institute for the COVID-19 Genomics UK (COG-UK) Consortium | Berkshire and Surrey Pathology Services Lighthouse Laboratory and Alex Alderton, Roberto Amato, Jeffrey Barrett, Sonia Goncalves, Ewan Harrison, David K. Jackson, Ian Johnston, Dominic Kwiatkowski, Cordelia Langford, John Sillitoe on behalf of the Wellcome Sanger Institute COVID-19 Surveillance Team |
| EPI_ISL_3313631 | Lighthouse Lab in Milton Keynes | Wellcome Sanger Institute for the COVID-19 Genomics UK (COG-UK) Consortium | The Lighthouse Lab in Milton Keynes and Alex Alderton, Roberto Amato, Jeffrey Barrett, Sonia Goncalves, Ewan Harrison, David K. Jackson, Ian Johnston, Dominic Kwiatkowski, Cordelia Langford, John Sillitoe on behalf of the Wellcome Sanger Institute COVID-19 Surveillance Team |
| EPI_ISL_3313632, EPI_ISL_3313633 | Lighthouse Lab in Glasgow | Wellcome Sanger Institute for the COVID-19 Genomics UK (COG-UK) Consortium | Harper VanSteenhouse, Yumi Kasai, David Gray, Carol Clugston, Anna Dominiczak and Alex Alderton, Roberto Amato, Jeffrey Barrett, Sonia Goncalves, Ewan Harrison, David K. Jackson, Ian Johnston, Dominic Kwiatkowski, Cordelia Langford, John Sillitoe on behalf of the Wellcome Sanger Institute COVID-19 Surveillance Team |
| EPI_ISL_3313634 | Berkshire and Surrey Pathology Services Lighthouse Laboratory | Wellcome Sanger Institute for the COVID-19 Genomics UK (COG-UK) Consortium | Berkshire and Surrey Pathology Services Lighthouse Laboratory and Alex Alderton, Roberto Amato, Jeffrey Barrett, Sonia Goncalves, Ewan Harrison, David K. Jackson, Ian Johnston, Dominic Kwiatkowski, Cordelia Langford, John Sillitoe on behalf of the Wellcome Sanger Institute COVID-19 Surveillance Team |
| EPI_ISL_3313635, EPI_ISL_3313636, EPI_ISL_3313637, EPI_ISL_3313638 | Lighthouse Lab in Milton Keynes | Wellcome Sanger Institute for the COVID-19 Genomics UK (COG-UK) Consortium | The Lighthouse Lab in Milton Keynes and Alex Alderton, Roberto Amato, Jeffrey Barrett, Sonia Goncalves, Ewan Harrison, David K. Jackson, Ian Johnston, Dominic Kwiatkowski, Cordelia Langford, John Sillitoe on behalf of the Wellcome Sanger Institute COVID-19 Surveillance Team |

| | | | |
|---|---|---|---|
| EPI_ISL_3313639 | Lighthouse Lab in Glasgow | Wellcome Sanger Institute for the COVID-19 Genomics UK (COG-UK) Consortium | Harper VanSteenhouse, Yumi Kasai, David Gray, Carol Clugston, Anna Dominiczak and Alex Alderton, Roberto Amato, Jeffrey Barrett, Sonia Goncalves, Ewan Harrison, David K. Jackson, Ian Johnston, Dominic Kwiatkowski, Cordelia Langford, John Sillitoe on behalf of the Wellcome Sanger Institute COVID-19 Surveillance Team |
| EPI_ISL_3313640 | Berkshire and Surrey Pathology Services Lighthouse Laboratory | Wellcome Sanger Institute for the COVID-19 Genomics UK (COG-UK) Consortium | Berkshire and Surrey Pathology Services Lighthouse Laboratory and Alex Alderton, Roberto Amato, Jeffrey Barrett, Sonia Goncalves, Ewan Harrison, David K. Jackson, Ian Johnston, Dominic Kwiatkowski, Cordelia Langford, John Sillitoe on behalf of the Wellcome Sanger Institute COVID-19 Surveillance Team |
| EPI_ISL_3313641 | Lighthouse Lab in Milton Keynes | Wellcome Sanger Institute for the COVID-19 Genomics UK (COG-UK) Consortium | The Lighthouse Lab in Milton Keynes and Alex Alderton, Roberto Amato, Jeffrey Barrett, Sonia Goncalves, Ewan Harrison, David K. Jackson, Ian Johnston, Dominic Kwiatkowski, Cordelia Langford, John Sillitoe on behalf of the Wellcome Sanger Institute COVID-19 Surveillance Team |
| EPI_ISL_3313642 | Lighthouse Lab in Glasgow | Wellcome Sanger Institute for the COVID-19 Genomics UK (COG-UK) Consortium | Harper VanSteenhouse, Yumi Kasai, David Gray, Carol Clugston, Anna Dominiczak and Alex Alderton, Roberto Amato, Jeffrey Barrett, Sonia Goncalves, Ewan Harrison, David K. Jackson, Ian Johnston, Dominic Kwiatkowski, Cordelia Langford, John Sillitoe on behalf of the Wellcome Sanger Institute COVID-19 Surveillance Team |
| EPI_ISL_3313643 | Berkshire and Surrey Pathology Services Lighthouse Laboratory | Wellcome Sanger Institute for the COVID-19 Genomics UK (COG-UK) Consortium | Berkshire and Surrey Pathology Services Lighthouse Laboratory and Alex Alderton, Roberto Amato, Jeffrey Barrett, Sonia Goncalves, Ewan Harrison, David K. Jackson, Ian Johnston, Dominic Kwiatkowski, Cordelia Langford, John Sillitoe on behalf of the Wellcome Sanger Institute COVID-19 Surveillance Team |
| EPI_ISL_3313644, EPI_ISL_3313645 | Lighthouse Lab in Glasgow | Wellcome Sanger Institute for the COVID-19 Genomics UK (COG-UK) Consortium | Harper VanSteenhouse, Yumi Kasai, David Gray, Carol Clugston, Anna Dominiczak and Alex Alderton, Roberto Amato, Jeffrey Barrett, Sonia Goncalves, Ewan Harrison, David K. Jackson, Ian Johnston, Dominic Kwiatkowski, Cordelia Langford, John Sillitoe on behalf of the Wellcome Sanger Institute COVID-19 Surveillance Team |
| EPI_ISL_3313646 | Berkshire and Surrey Pathology Services Lighthouse Laboratory | Wellcome Sanger Institute for the COVID-19 Genomics UK (COG-UK) Consortium | Berkshire and Surrey Pathology Services Lighthouse Laboratory and Alex Alderton, Roberto Amato, Jeffrey Barrett, Sonia Goncalves, Ewan Harrison, David K. Jackson, Ian Johnston, Dominic Kwiatkowski, Cordelia Langford, John Sillitoe on behalf of the Wellcome Sanger Institute COVID-19 Surveillance Team |
| EPI_ISL_3313647 | Lighthouse Lab in Milton Keynes | Wellcome Sanger Institute for the COVID-19 Genomics UK (COG-UK) Consortium | The Lighthouse Lab in Milton Keynes and Alex Alderton, Roberto Amato, Jeffrey Barrett, Sonia Goncalves, Ewan Harrison, David K. Jackson, Ian Johnston, Dominic Kwiatkowski, Cordelia Langford, John Sillitoe on behalf of the Wellcome Sanger Institute COVID-19 Surveillance Team |
| EPI_ISL_3313648 | Berkshire and Surrey Pathology Services Lighthouse Laboratory | Wellcome Sanger Institute for the COVID-19 Genomics UK (COG-UK) Consortium | Berkshire and Surrey Pathology Services Lighthouse Laboratory and Alex Alderton, Roberto Amato, Jeffrey Barrett, Sonia Goncalves, Ewan Harrison, David K. Jackson, Ian Johnston, Dominic Kwiatkowski, Cordelia Langford, John Sillitoe on behalf of the Wellcome Sanger Institute COVID-19 Surveillance Team |
| EPI_ISL_3313649 | Lighthouse Lab in Milton Keynes | Wellcome Sanger Institute for the COVID-19 Genomics UK (COG-UK) Consortium | The Lighthouse Lab in Milton Keynes and Alex Alderton, Roberto Amato, Jeffrey Barrett, Sonia Goncalves, Ewan Harrison, David K. Jackson, Ian Johnston, Dominic Kwiatkowski, Cordelia Langford, John Sillitoe on behalf of the Wellcome Sanger Institute COVID-19 Surveillance Team |
| EPI_ISL_3313650, EPI_ISL_3313651, EPI_ISL_3313652, EPI_ISL_3313653, EPI_ISL_3313654 | Berkshire and Surrey Pathology Services Lighthouse Laboratory | Wellcome Sanger Institute for the COVID-19 Genomics UK (COG-UK) Consortium | Berkshire and Surrey Pathology Services Lighthouse Laboratory and Alex Alderton, Roberto Amato, Jeffrey Barrett, Sonia Goncalves, Ewan Harrison, David K. Jackson, Ian Johnston, Dominic Kwiatkowski, Cordelia Langford, John Sillitoe on behalf of the Wellcome Sanger Institute COVID-19 Surveillance Team |
| EPI_ISL_3313655 | Lighthouse Lab in Milton Keynes | Wellcome Sanger Institute for the COVID-19 Genomics UK (COG-UK) Consortium | The Lighthouse Lab in Milton Keynes and Alex Alderton, Roberto Amato, Jeffrey Barrett, Sonia Goncalves, Ewan Harrison, David K. Jackson, Ian Johnston, Dominic Kwiatkowski, Cordelia Langford, John Sillitoe on behalf of the Wellcome Sanger Institute COVID-19 Surveillance Team |
| EPI_ISL_3313656 | Berkshire and Surrey Pathology Services Lighthouse Laboratory | Wellcome Sanger Institute for the COVID-19 Genomics UK (COG-UK) Consortium | Berkshire and Surrey Pathology Services Lighthouse Laboratory and Alex Alderton, Roberto Amato, Jeffrey Barrett, Sonia Goncalves, Ewan Harrison, David K. Jackson, Ian Johnston, Dominic Kwiatkowski, Cordelia Langford, John Sillitoe on behalf of the Wellcome Sanger Institute COVID-19 Surveillance Team |
| EPI_ISL_3313657 | Lighthouse Lab in Glasgow | Wellcome Sanger Institute for the COVID-19 Genomics UK (COG-UK) Consortium | Harper VanSteenhouse, Yumi Kasai, David Gray, Carol Clugston, Anna Dominiczak and Alex Alderton, Roberto Amato, Jeffrey Barrett, Sonia Goncalves, Ewan Harrison, David K. Jackson, Ian Johnston, Dominic Kwiatkowski, Cordelia Langford, John Sillitoe on behalf of the Wellcome Sanger Institute COVID-19 Surveillance Team |
| EPI_ISL_3313658, EPI_ISL_3313659, EPI_ISL_3313660 | Berkshire and Surrey Pathology Services Lighthouse Laboratory | Wellcome Sanger Institute for the COVID-19 Genomics UK (COG-UK) Consortium | Berkshire and Surrey Pathology Services Lighthouse Laboratory and Alex Alderton, Roberto Amato, Jeffrey Barrett, Sonia Goncalves, Ewan Harrison, David K. Jackson, Ian Johnston, Dominic Kwiatkowski, Cordelia Langford, John Sillitoe on behalf of the Wellcome Sanger Institute COVID-19 Surveillance Team |
| EPI_ISL_3313661 | Lighthouse Lab in Glasgow | Wellcome Sanger Institute for the COVID-19 Genomics UK (COG-UK) Consortium | Harper VanSteenhouse, Yumi Kasai, David Gray, Carol Clugston, Anna Dominiczak and Alex Alderton, Roberto Amato, Jeffrey Barrett, Sonia Goncalves, Ewan Harrison, David K. Jackson, Ian Johnston, Dominic Kwiatkowski, Cordelia Langford, John Sillitoe on behalf of the Wellcome Sanger Institute COVID-19 Surveillance Team |
| EPI_ISL_3313662, EPI_ISL_3313663 | Berkshire and Surrey Pathology Services Lighthouse Laboratory | Wellcome Sanger Institute for the COVID-19 Genomics UK (COG-UK) Consortium | Berkshire and Surrey Pathology Services Lighthouse Laboratory and Alex Alderton, Roberto Amato, Jeffrey Barrett, Sonia Goncalves, Ewan Harrison, David K. Jackson, Ian Johnston, Dominic Kwiatkowski, Cordelia Langford, John Sillitoe on behalf of the Wellcome Sanger Institute COVID-19 Surveillance Team |
| EPI_ISL_3313664 | Lighthouse Lab in Glasgow | Wellcome Sanger Institute for the COVID-19 Genomics UK (COG-UK) Consortium | Harper VanSteenhouse, Yumi Kasai, David Gray, Carol Clugston, Anna Dominiczak and Alex Alderton, Roberto Amato, Jeffrey Barrett, Sonia Goncalves, Ewan Harrison, David K. Jackson, Ian Johnston, Dominic Kwiatkowski, Cordelia Langford, John Sillitoe on behalf of the Wellcome Sanger Institute COVID-19 Surveillance Team |
| EPI_ISL_3313665, EPI_ISL_3313666, EPI_ISL_3313667, EPI_ISL_3313668 | Berkshire and Surrey Pathology Services Lighthouse Laboratory | Wellcome Sanger Institute for the COVID-19 Genomics UK (COG-UK) Consortium | Berkshire and Surrey Pathology Services Lighthouse Laboratory and Alex Alderton, Roberto Amato, Jeffrey Barrett, Sonia Goncalves, Ewan Harrison, David K. Jackson, Ian Johnston, Dominic Kwiatkowski, Cordelia Langford, John Sillitoe on behalf of the Wellcome Sanger Institute COVID-19 Surveillance Team |
| EPI_ISL_3313669 | Lighthouse Lab in Glasgow | Wellcome Sanger Institute for the COVID-19 Genomics UK (COG-UK) Consortium | Harper VanSteenhouse, Yumi Kasai, David Gray, Carol Clugston, Anna Dominiczak and Alex Alderton, Roberto Amato, Jeffrey Barrett, Sonia Goncalves, Ewan Harrison, David K. Jackson, Ian Johnston, Dominic Kwiatkowski, Cordelia Langford, John Sillitoe on behalf of the Wellcome Sanger Institute COVID-19 Surveillance Team |
| EPI_ISL_3313670 | Lighthouse Lab in Milton Keynes | Wellcome Sanger Institute for the COVID-19 Genomics UK (COG-UK) Consortium | The Lighthouse Lab in Milton Keynes and Alex Alderton, Roberto Amato, Jeffrey Barrett, Sonia Goncalves, Ewan Harrison, David K. Jackson, Ian Johnston, Dominic Kwiatkowski, Cordelia Langford, John Sillitoe on behalf of the Wellcome Sanger Institute COVID-19 Surveillance Team |
| EPI_ISL_3313671 | Berkshire and Surrey Pathology Services Lighthouse Laboratory | Wellcome Sanger Institute for the COVID-19 Genomics UK (COG-UK) Consortium | Berkshire and Surrey Pathology Services Lighthouse Laboratory and Alex Alderton, Roberto Amato, Jeffrey Barrett, Sonia Goncalves, Ewan Harrison, David K. Jackson, Ian Johnston, Dominic Kwiatkowski, Cordelia Langford, John Sillitoe on behalf of the Wellcome Sanger Institute COVID-19 Surveillance Team |
| EPI_ISL_3313672 | Lighthouse Lab in Milton Keynes | Wellcome Sanger Institute for the COVID-19 Genomics UK (COG-UK) Consortium | The Lighthouse Lab in Milton Keynes and Alex Alderton, Roberto Amato, Jeffrey Barrett, Sonia Goncalves, Ewan Harrison, David K. Jackson, Ian Johnston, Dominic Kwiatkowski, Cordelia Langford, John Sillitoe on behalf of the Wellcome Sanger Institute COVID-19 Surveillance Team |
| EPI_ISL_3313673, EPI_ISL_3313674 | Berkshire and Surrey Pathology Services Lighthouse Laboratory | Wellcome Sanger Institute for the COVID-19 Genomics UK (COG-UK) Consortium | Berkshire and Surrey Pathology Services Lighthouse Laboratory and Alex Alderton, Roberto Amato, Jeffrey Barrett, Sonia Goncalves, Ewan Harrison, David K. Jackson, Ian Johnston, Dominic Kwiatkowski, Cordelia Langford, John Sillitoe on behalf of the Wellcome Sanger Institute COVID-19 Surveillance Team |
| EPI_ISL_3313675 | Lighthouse Lab in Glasgow | Wellcome Sanger Institute for the COVID-19 Genomics UK (COG-UK) Consortium | Harper VanSteenhouse, Yumi Kasai, David Gray, Carol Clugston, Anna Dominiczak and Alex Alderton, Roberto Amato, Jeffrey Barrett, Sonia Goncalves, Ewan Harrison, David K. Jackson, Ian Johnston, Dominic Kwiatkowski, Cordelia Langford, John Sillitoe on behalf of the Wellcome Sanger Institute COVID-19 Surveillance Team |
| EPI_ISL_3313676 | Berkshire and Surrey Pathology Services Lighthouse Laboratory | Wellcome Sanger Institute for the COVID-19 Genomics UK (COG-UK) Consortium | Berkshire and Surrey Pathology Services Lighthouse Laboratory and Alex Alderton, Roberto Amato, Jeffrey Barrett, Sonia Goncalves, Ewan Harrison, David K. Jackson, Ian Johnston, Dominic Kwiatkowski, Cordelia Langford, John Sillitoe on behalf of the Wellcome Sanger Institute COVID-19 Surveillance Team |
| EPI_ISL_3313677, EPI_ISL_3313678 | Lighthouse Lab in Milton Keynes | Wellcome Sanger Institute for the COVID-19 Genomics UK (COG-UK) Consortium | The Lighthouse Lab in Milton Keynes and Alex Alderton, Roberto Amato, Jeffrey Barrett, Sonia Goncalves, Ewan Harrison, David K. Jackson, Ian Johnston, Dominic Kwiatkowski, Cordelia Langford, John Sillitoe on behalf of the Wellcome Sanger Institute COVID-19 Surveillance Team |
| EPI_ISL_3313679, EPI_ISL_3313680, EPI_ISL_3313681, EPI_ISL_3313682 | Berkshire and Surrey Pathology Services Lighthouse Laboratory | Wellcome Sanger Institute for the COVID-19 Genomics UK (COG-UK) Consortium | Berkshire and Surrey Pathology Services Lighthouse Laboratory and Alex Alderton, Roberto Amato, Jeffrey Barrett, Sonia Goncalves, Ewan Harrison, David K. Jackson, Ian Johnston, Dominic Kwiatkowski, Cordelia Langford, John Sillitoe on behalf of the Wellcome Sanger Institute COVID-19 Surveillance Team |
| EPI_ISL_3313683 | Lighthouse Lab in Milton Keynes | Wellcome Sanger Institute for the COVID-19 Genomics UK (COG-UK) Consortium | The Lighthouse Lab in Milton Keynes and Alex Alderton, Roberto Amato, Jeffrey Barrett, Sonia Goncalves, Ewan Harrison, David K. Jackson, Ian Johnston, Dominic Kwiatkowski, Cordelia Langford, John Sillitoe on behalf of the Wellcome Sanger Institute COVID-19 Surveillance Team |
| EPI_ISL_3313684, EPI_ISL_3313685 | Berkshire and Surrey Pathology Services Lighthouse Laboratory | Wellcome Sanger Institute for the COVID-19 Genomics UK (COG-UK) Consortium | Berkshire and Surrey Pathology Services Lighthouse Laboratory and Alex Alderton, Roberto Amato, Jeffrey Barrett, Sonia Goncalves, Ewan Harrison, David K. Jackson, Ian Johnston, Dominic Kwiatkowski, Cordelia Langford, John Sillitoe on behalf of the Wellcome Sanger Institute COVID-19 Surveillance Team |
| EPI_ISL_3313686 | Lighthouse Lab in Milton Keynes | Wellcome Sanger Institute for the COVID-19 Genomics UK (COG-UK) Consortium | The Lighthouse Lab in Milton Keynes and Alex Alderton, Roberto Amato, Jeffrey Barrett, Sonia Goncalves, Ewan Harrison, David K. Jackson, Ian Johnston, Dominic Kwiatkowski, Cordelia Langford, John Sillitoe on behalf of the Wellcome Sanger Institute COVID-19 Surveillance Team |
| EPI_ISL_3313687 | Lighthouse Lab in Glasgow | Wellcome Sanger Institute for the COVID-19 Genomics UK (COG-UK) Consortium | Harper VanSteenhouse, Yumi Kasai, David Gray, Carol Clugston, Anna Dominiczak and Alex Alderton, Roberto Amato, Jeffrey Barrett, Sonia Goncalves, Ewan Harrison, David K. Jackson, Ian Johnston, Dominic Kwiatkowski, Cordelia Langford, John Sillitoe on behalf of the Wellcome Sanger Institute COVID-19 Surveillance Team |
| EPI_ISL_3313688 | Lighthouse Lab in Milton Keynes | Wellcome Sanger Institute for the COVID-19 Genomics UK (COG-UK) Consortium | The Lighthouse Lab in Milton Keynes and Alex Alderton, Roberto Amato, Jeffrey Barrett, Sonia Goncalves, Ewan Harrison, David K. Jackson, Ian Johnston, Dominic Kwiatkowski, Cordelia Langford, John Sillitoe on behalf of the Wellcome Sanger Institute COVID-19 Surveillance Team |
| EPI_ISL_3313689 | Lighthouse Lab in Glasgow | Wellcome Sanger Institute for the COVID-19 Genomics UK (COG-UK) Consortium | Harper VanSteenhouse, Yumi Kasai, David Gray, Carol Clugston, Anna Dominiczak and Alex Alderton, Roberto Amato, Jeffrey Barrett, Sonia Goncalves, Ewan Harrison, David K. Jackson, Ian Johnston, Dominic Kwiatkowski, Cordelia Langford, John Sillitoe on behalf of the Wellcome Sanger Institute COVID-19 Surveillance Team |
| EPI_ISL_3313690, EPI_ISL_3313691 | Lighthouse Lab in Milton Keynes | Wellcome Sanger Institute for the COVID-19 Genomics UK (COG-UK) Consortium | The Lighthouse Lab in Milton Keynes and Alex Alderton, Roberto Amato, Jeffrey Barrett, Sonia Goncalves, Ewan Harrison, David K. Jackson, Ian Johnston, Dominic Kwiatkowski, Cordelia Langford, John Sillitoe on behalf of the Wellcome Sanger Institute COVID-19 Surveillance Team |
| EPI_ISL_3313692 | Lighthouse Lab in Glasgow | Wellcome Sanger Institute for the COVID-19 Genomics UK (COG-UK) Consortium | Harper VanSteenhouse, Yumi Kasai, David Gray, Carol Clugston, Anna Dominiczak and Alex Alderton, Roberto Amato, Jeffrey Barrett, Sonia Goncalves, Ewan Harrison, David K. Jackson, Ian Johnston, Dominic Kwiatkowski, Cordelia Langford, John Sillitoe on behalf of the Wellcome Sanger Institute COVID-19 Surveillance Team |
| EPI_ISL_3313693, EPI_ISL_3313694, EPI_ISL_3313695, EPI_ISL_3313696 | Berkshire and Surrey Pathology Services Lighthouse Laboratory | Wellcome Sanger Institute for the COVID-19 Genomics UK (COG-UK) Consortium | Berkshire and Surrey Pathology Services Lighthouse Laboratory and Alex Alderton, Roberto Amato, Jeffrey Barrett, Sonia Goncalves, Ewan Harrison, David K. Jackson, Ian Johnston, Dominic Kwiatkowski, Cordelia Langford, John Sillitoe on behalf of the Wellcome Sanger Institute COVID-19 Surveillance Team |
| EPI_ISL_3313697 | Lighthouse Lab in Milton Keynes | Wellcome Sanger Institute for the COVID-19 Genomics UK (COG-UK) Consortium | The Lighthouse Lab in Milton Keynes and Alex Alderton, Roberto Amato, Jeffrey Barrett, Sonia Goncalves, Ewan Harrison, David K. Jackson, Ian Johnston, Dominic Kwiatkowski, Cordelia Langford, John Sillitoe on behalf of the Wellcome Sanger Institute COVID-19 Surveillance Team |
| EPI_ISL_3313698 | Lighthouse Lab in Glasgow | Wellcome Sanger Institute for the COVID-19 Genomics UK (COG-UK) Consortium | Harper VanSteenhouse, Yumi Kasai, David Gray, Carol Clugston, Anna Dominiczak and Alex Alderton, Roberto Amato, Jeffrey Barrett, Sonia Goncalves, Ewan Harrison, David K. Jackson, Ian Johnston, Dominic Kwiatkowski, Cordelia Langford, John Sillitoe on behalf of the Wellcome Sanger Institute COVID-19 Surveillance Team |
| EPI_ISL_3313699 | Lighthouse Lab in Milton Keynes | Wellcome Sanger Institute for the COVID-19 Genomics UK (COG-UK) Consortium | The Lighthouse Lab in Milton Keynes and Alex Alderton, Roberto Amato, Jeffrey Barrett, Sonia Goncalves, Ewan Harrison, David K. Jackson, Ian Johnston, Dominic Kwiatkowski, Cordelia Langford, John Sillitoe on behalf of the Wellcome Sanger Institute COVID-19 Surveillance Team |
| EPI_ISL_3313700 | Berkshire and Surrey Pathology Services Lighthouse Laboratory | Wellcome Sanger Institute for the COVID-19 Genomics UK (COG-UK) Consortium | Berkshire and Surrey Pathology Services Lighthouse Laboratory and Alex Alderton, Roberto Amato, Jeffrey Barrett, Sonia Goncalves, Ewan Harrison, David K. Jackson, Ian Johnston, Dominic Kwiatkowski, Cordelia Langford, John Sillitoe on behalf of the Wellcome Sanger Institute COVID-19 Surveillance Team |

| | | | |
|---|---|---|---|
| EPI_ISL_3313701 | Lighthouse Lab in Glasgow | Wellcome Sanger Institute for the COVID-19 Genomics UK (COG-UK) Consortium | Harper VanSteenhouse, Yumi Kasai, David Gray, Carol Clugston, Anna Dominiczak and Alex Alderton, Roberto Amato, Jeffrey Barrett, Sonia Goncalves, Ewan Harrison, David K. Jackson, Ian Johnston, Dominic Kwiatkowski, Cordelia Langford, John Sillitoe on behalf of the Wellcome Sanger Institute COVID-19 Surveillance Team |
| EPI_ISL_3313702, EPI_ISL_3313703, EPI_ISL_3313704, EPI_ISL_3313705, EPI_ISL_3313706 | Berkshire and Surrey Pathology Services Lighthouse Laboratory | Wellcome Sanger Institute for the COVID-19 Genomics UK (COG-UK) Consortium | Berkshire and Surrey Pathology Services Lighthouse Laboratory and Alex Alderton, Roberto Amato, Jeffrey Barrett, Sonia Goncalves, Ewan Harrison, David K. Jackson, Ian Johnston, Dominic Kwiatkowski, Cordelia Langford, John Sillitoe on behalf of the Wellcome Sanger Institute COVID-19 Surveillance Team |
| EPI_ISL_3313707 | Lighthouse Lab in Milton Keynes | Wellcome Sanger Institute for the COVID-19 Genomics UK (COG-UK) Consortium | The Lighthouse Lab in Milton Keynes and Alex Alderton, Roberto Amato, Jeffrey Barrett, Sonia Goncalves, Ewan Harrison, David K. Jackson, Ian Johnston, Dominic Kwiatkowski, Cordelia Langford, John Sillitoe on behalf of the Wellcome Sanger Institute COVID-19 Surveillance Team |
| EPI_ISL_3313708 | Berkshire and Surrey Pathology Services Lighthouse Laboratory | Wellcome Sanger Institute for the COVID-19 Genomics UK (COG-UK) Consortium | Berkshire and Surrey Pathology Services Lighthouse Laboratory and Alex Alderton, Roberto Amato, Jeffrey Barrett, Sonia Goncalves, Ewan Harrison, David K. Jackson, Ian Johnston, Dominic Kwiatkowski, Cordelia Langford, John Sillitoe on behalf of the Wellcome Sanger Institute COVID-19 Surveillance Team |
| EPI_ISL_3313709, EPI_ISL_3313710 | Lighthouse Lab in Milton Keynes | Wellcome Sanger Institute for the COVID-19 Genomics UK (COG-UK) Consortium | The Lighthouse Lab in Milton Keynes and Alex Alderton, Roberto Amato, Jeffrey Barrett, Sonia Goncalves, Ewan Harrison, David K. Jackson, Ian Johnston, Dominic Kwiatkowski, Cordelia Langford, John Sillitoe on behalf of the Wellcome Sanger Institute COVID-19 Surveillance Team |
| EPI_ISL_3313711, EPI_ISL_3313712 | Lighthouse Lab in Glasgow | Wellcome Sanger Institute for the COVID-19 Genomics UK (COG-UK) Consortium | Harper VanSteenhouse, Yumi Kasai, David Gray, Carol Clugston, Anna Dominiczak and Alex Alderton, Roberto Amato, Jeffrey Barrett, Sonia Goncalves, Ewan Harrison, David K. Jackson, Ian Johnston, Dominic Kwiatkowski, Cordelia Langford, John Sillitoe on behalf of the Wellcome Sanger Institute COVID-19 Surveillance Team |
| EPI_ISL_3313713 | Lighthouse Lab in Milton Keynes | Wellcome Sanger Institute for the COVID-19 Genomics UK (COG-UK) Consortium | The Lighthouse Lab in Milton Keynes and Alex Alderton, Roberto Amato, Jeffrey Barrett, Sonia Goncalves, Ewan Harrison, David K. Jackson, Ian Johnston, Dominic Kwiatkowski, Cordelia Langford, John Sillitoe on behalf of the Wellcome Sanger Institute COVID-19 Surveillance Team |
| EPI_ISL_3313714, EPI_ISL_3313715 | Berkshire and Surrey Pathology Services Lighthouse Laboratory | Wellcome Sanger Institute for the COVID-19 Genomics UK (COG-UK) Consortium | Berkshire and Surrey Pathology Services Lighthouse Laboratory and Alex Alderton, Roberto Amato, Jeffrey Barrett, Sonia Goncalves, Ewan Harrison, David K. Jackson, Ian Johnston, Dominic Kwiatkowski, Cordelia Langford, John Sillitoe on behalf of the Wellcome Sanger Institute COVID-19 Surveillance Team |
| EPI_ISL_3313716 | Lighthouse Lab in Glasgow | Wellcome Sanger Institute for the COVID-19 Genomics UK (COG-UK) Consortium | Harper VanSteenhouse, Yumi Kasai, David Gray, Carol Clugston, Anna Dominiczak and Alex Alderton, Roberto Amato, Jeffrey Barrett, Sonia Goncalves, Ewan Harrison, David K. Jackson, Ian Johnston, Dominic Kwiatkowski, Cordelia Langford, John Sillitoe on behalf of the Wellcome Sanger Institute COVID-19 Surveillance Team |
| EPI_ISL_3313717 | Lighthouse Lab in Milton Keynes | Wellcome Sanger Institute for the COVID-19 Genomics UK (COG-UK) Consortium | The Lighthouse Lab in Milton Keynes and Alex Alderton, Roberto Amato, Jeffrey Barrett, Sonia Goncalves, Ewan Harrison, David K. Jackson, Ian Johnston, Dominic Kwiatkowski, Cordelia Langford, John Sillitoe on behalf of the Wellcome Sanger Institute COVID-19 Surveillance Team |
| EPI_ISL_3313718, EPI_ISL_3313719, EPI_ISL_3313720 | Berkshire and Surrey Pathology Services Lighthouse Laboratory | Wellcome Sanger Institute for the COVID-19 Genomics UK (COG-UK) Consortium | Berkshire and Surrey Pathology Services Lighthouse Laboratory and Alex Alderton, Roberto Amato, Jeffrey Barrett, Sonia Goncalves, Ewan Harrison, David K. Jackson, Ian Johnston, Dominic Kwiatkowski, Cordelia Langford, John Sillitoe on behalf of the Wellcome Sanger Institute COVID-19 Surveillance Team |
| EPI_ISL_3313721 | Lighthouse Lab in Milton Keynes | Wellcome Sanger Institute for the COVID-19 Genomics UK (COG-UK) Consortium | The Lighthouse Lab in Milton Keynes and Alex Alderton, Roberto Amato, Jeffrey Barrett, Sonia Goncalves, Ewan Harrison, David K. Jackson, Ian Johnston, Dominic Kwiatkowski, Cordelia Langford, John Sillitoe on behalf of the Wellcome Sanger Institute COVID-19 Surveillance Team |
| EPI_ISL_3313722 | Berkshire and Surrey Pathology Services Lighthouse Laboratory | Wellcome Sanger Institute for the COVID-19 Genomics UK (COG-UK) Consortium | Berkshire and Surrey Pathology Services Lighthouse Laboratory and Alex Alderton, Roberto Amato, Jeffrey Barrett, Sonia Goncalves, Ewan Harrison, David K. Jackson, Ian Johnston, Dominic Kwiatkowski, Cordelia Langford, John Sillitoe on behalf of the Wellcome Sanger Institute COVID-19 Surveillance Team |
| EPI_ISL_3313723 | Lighthouse Lab in Glasgow | Wellcome Sanger Institute for the COVID-19 Genomics UK (COG-UK) Consortium | Harper VanSteenhouse, Yumi Kasai, David Gray, Carol Clugston, Anna Dominiczak and Alex Alderton, Roberto Amato, Jeffrey Barrett, Sonia Goncalves, Ewan Harrison, David K. Jackson, Ian Johnston, Dominic Kwiatkowski, Cordelia Langford, John Sillitoe on behalf of the Wellcome Sanger Institute COVID-19 Surveillance Team |
| EPI_ISL_3313724 | Lighthouse Lab in Milton Keynes | Wellcome Sanger Institute for the COVID-19 Genomics UK (COG-UK) Consortium | The Lighthouse Lab in Milton Keynes and Alex Alderton, Roberto Amato, Jeffrey Barrett, Sonia Goncalves, Ewan Harrison, David K. Jackson, Ian Johnston, Dominic Kwiatkowski, Cordelia Langford, John Sillitoe on behalf of the Wellcome Sanger Institute COVID-19 Surveillance Team |
| EPI_ISL_3313725, EPI_ISL_3313726 | Berkshire and Surrey Pathology Services Lighthouse Laboratory | Wellcome Sanger Institute for the COVID-19 Genomics UK (COG-UK) Consortium | Berkshire and Surrey Pathology Services Lighthouse Laboratory and Alex Alderton, Roberto Amato, Jeffrey Barrett, Sonia Goncalves, Ewan Harrison, David K. Jackson, Ian Johnston, Dominic Kwiatkowski, Cordelia Langford, John Sillitoe on behalf of the Wellcome Sanger Institute COVID-19 Surveillance Team |
| EPI_ISL_3313727, EPI_ISL_3313728 | Lighthouse Lab in Glasgow | Wellcome Sanger Institute for the COVID-19 Genomics UK (COG-UK) Consortium | Harper VanSteenhouse, Yumi Kasai, David Gray, Carol Clugston, Anna Dominiczak and Alex Alderton, Roberto Amato, Jeffrey Barrett, Sonia Goncalves, Ewan Harrison, David K. Jackson, Ian Johnston, Dominic Kwiatkowski, Cordelia Langford, John Sillitoe on behalf of the Wellcome Sanger Institute COVID-19 Surveillance Team |
| EPI_ISL_3313729 | Lighthouse Lab in Milton Keynes | Wellcome Sanger Institute for the COVID-19 Genomics UK (COG-UK) Consortium | The Lighthouse Lab in Milton Keynes and Alex Alderton, Roberto Amato, Jeffrey Barrett, Sonia Goncalves, Ewan Harrison, David K. Jackson, Ian Johnston, Dominic Kwiatkowski, Cordelia Langford, John Sillitoe on behalf of the Wellcome Sanger Institute COVID-19 Surveillance Team |
| EPI_ISL_3313730 | Lighthouse Lab in Glasgow | Wellcome Sanger Institute for the COVID-19 Genomics UK (COG-UK) Consortium | Harper VanSteenhouse, Yumi Kasai, David Gray, Carol Clugston, Anna Dominiczak and Alex Alderton, Roberto Amato, Jeffrey Barrett, Sonia Goncalves, Ewan Harrison, David K. Jackson, Ian Johnston, Dominic Kwiatkowski, Cordelia Langford, John Sillitoe on behalf of the Wellcome Sanger Institute COVID-19 Surveillance Team |
| EPI_ISL_3313731, EPI_ISL_3313732 | Berkshire and Surrey Pathology Services Lighthouse Laboratory | Wellcome Sanger Institute for the COVID-19 Genomics UK (COG-UK) Consortium | Berkshire and Surrey Pathology Services Lighthouse Laboratory and Alex Alderton, Roberto Amato, Jeffrey Barrett, Sonia Goncalves, Ewan Harrison, David K. Jackson, Ian Johnston, Dominic Kwiatkowski, Cordelia Langford, John Sillitoe on behalf of the Wellcome Sanger Institute COVID-19 Surveillance Team |
| EPI_ISL_3313733, EPI_ISL_3313734, EPI_ISL_3313735, EPI_ISL_3313736 | Lighthouse Lab in Milton Keynes | Wellcome Sanger Institute for the COVID-19 Genomics UK (COG-UK) Consortium | The Lighthouse Lab in Milton Keynes and Alex Alderton, Roberto Amato, Jeffrey Barrett, Sonia Goncalves, Ewan Harrison, David K. Jackson, Ian Johnston, Dominic Kwiatkowski, Cordelia Langford, John Sillitoe on behalf of the Wellcome Sanger Institute COVID-19 Surveillance Team |
| EPI_ISL_3313737, EPI_ISL_3313738 | Berkshire and Surrey Pathology Services Lighthouse Laboratory | Wellcome Sanger Institute for the COVID-19 Genomics UK (COG-UK) Consortium | Berkshire and Surrey Pathology Services Lighthouse Laboratory and Alex Alderton, Roberto Amato, Jeffrey Barrett, Sonia Goncalves, Ewan Harrison, David K. Jackson, Ian Johnston, Dominic Kwiatkowski, Cordelia Langford, John Sillitoe on behalf of the Wellcome Sanger Institute COVID-19 Surveillance Team |
| EPI_ISL_3313739, EPI_ISL_3313740, EPI_ISL_3313741 | Lighthouse Lab in Milton Keynes | Wellcome Sanger Institute for the COVID-19 Genomics UK (COG-UK) Consortium | The Lighthouse Lab in Milton Keynes and Alex Alderton, Roberto Amato, Jeffrey Barrett, Sonia Goncalves, Ewan Harrison, David K. Jackson, Ian Johnston, Dominic Kwiatkowski, Cordelia Langford, John Sillitoe on behalf of the Wellcome Sanger Institute COVID-19 Surveillance Team |
| EPI_ISL_3313742, EPI_ISL_3313743 | Berkshire and Surrey Pathology Services Lighthouse Laboratory | Wellcome Sanger Institute for the COVID-19 Genomics UK (COG-UK) Consortium | Berkshire and Surrey Pathology Services Lighthouse Laboratory and Alex Alderton, Roberto Amato, Jeffrey Barrett, Sonia Goncalves, Ewan Harrison, David K. Jackson, Ian Johnston, Dominic Kwiatkowski, Cordelia Langford, John Sillitoe on behalf of the Wellcome Sanger Institute COVID-19 Surveillance Team |
| EPI_ISL_3313744 | Lighthouse Lab in Milton Keynes | Wellcome Sanger Institute for the COVID-19 Genomics UK (COG-UK) Consortium | The Lighthouse Lab in Milton Keynes and Alex Alderton, Roberto Amato, Jeffrey Barrett, Sonia Goncalves, Ewan Harrison, David K. Jackson, Ian Johnston, Dominic Kwiatkowski, Cordelia Langford, John Sillitoe on behalf of the Wellcome Sanger Institute COVID-19 Surveillance Team |
| EPI_ISL_3313745 | Berkshire and Surrey Pathology Services Lighthouse Laboratory | Wellcome Sanger Institute for the COVID-19 Genomics UK (COG-UK) Consortium | Berkshire and Surrey Pathology Services Lighthouse Laboratory and Alex Alderton, Roberto Amato, Jeffrey Barrett, Sonia Goncalves, Ewan Harrison, David K. Jackson, Ian Johnston, Dominic Kwiatkowski, Cordelia Langford, John Sillitoe on behalf of the Wellcome Sanger Institute COVID-19 Surveillance Team |
| EPI_ISL_3313746 | Lighthouse Lab in Glasgow | Wellcome Sanger Institute for the COVID-19 Genomics UK (COG-UK) Consortium | Harper VanSteenhouse, Yumi Kasai, David Gray, Carol Clugston, Anna Dominiczak and Alex Alderton, Roberto Amato, Jeffrey Barrett, Sonia Goncalves, Ewan Harrison, David K. Jackson, Ian Johnston, Dominic Kwiatkowski, Cordelia Langford, John Sillitoe on behalf of the Wellcome Sanger Institute COVID-19 Surveillance Team |
| EPI_ISL_3313747, EPI_ISL_3313748 | Berkshire and Surrey Pathology Services Lighthouse Laboratory | Wellcome Sanger Institute for the COVID-19 Genomics UK (COG-UK) Consortium | Berkshire and Surrey Pathology Services Lighthouse Laboratory and Alex Alderton, Roberto Amato, Jeffrey Barrett, Sonia Goncalves, Ewan Harrison, David K. Jackson, Ian Johnston, Dominic Kwiatkowski, Cordelia Langford, John Sillitoe on behalf of the Wellcome Sanger Institute COVID-19 Surveillance Team |
| EPI_ISL_3313749 | Lighthouse Lab in Milton Keynes | Wellcome Sanger Institute for the COVID-19 Genomics UK (COG-UK) Consortium | The Lighthouse Lab in Milton Keynes and Alex Alderton, Roberto Amato, Jeffrey Barrett, Sonia Goncalves, Ewan Harrison, David K. Jackson, Ian Johnston, Dominic Kwiatkowski, Cordelia Langford, John Sillitoe on behalf of the Wellcome Sanger Institute COVID-19 Surveillance Team |
| EPI_ISL_3313750 | Berkshire and Surrey Pathology Services Lighthouse Laboratory | Wellcome Sanger Institute for the COVID-19 Genomics UK (COG-UK) Consortium | Berkshire and Surrey Pathology Services Lighthouse Laboratory and Alex Alderton, Roberto Amato, Jeffrey Barrett, Sonia Goncalves, Ewan Harrison, David K. Jackson, Ian Johnston, Dominic Kwiatkowski, Cordelia Langford, John Sillitoe on behalf of the Wellcome Sanger Institute COVID-19 Surveillance Team |
| EPI_ISL_3313751, EPI_ISL_3313752 | Lighthouse Lab in Milton Keynes | Wellcome Sanger Institute for the COVID-19 Genomics UK (COG-UK) Consortium | The Lighthouse Lab in Milton Keynes and Alex Alderton, Roberto Amato, Jeffrey Barrett, Sonia Goncalves, Ewan Harrison, David K. Jackson, Ian Johnston, Dominic Kwiatkowski, Cordelia Langford, John Sillitoe on behalf of the Wellcome Sanger Institute COVID-19 Surveillance Team |
| EPI_ISL_3313753, EPI_ISL_3313754 | Berkshire and Surrey Pathology Services Lighthouse Laboratory | Wellcome Sanger Institute for the COVID-19 Genomics UK (COG-UK) Consortium | Berkshire and Surrey Pathology Services Lighthouse Laboratory and Alex Alderton, Roberto Amato, Jeffrey Barrett, Sonia Goncalves, Ewan Harrison, David K. Jackson, Ian Johnston, Dominic Kwiatkowski, Cordelia Langford, John Sillitoe on behalf of the Wellcome Sanger Institute COVID-19 Surveillance Team |
| EPI_ISL_3313755 | Lighthouse Lab in Milton Keynes | Wellcome Sanger Institute for the COVID-19 Genomics UK (COG-UK) Consortium | The Lighthouse Lab in Milton Keynes and Alex Alderton, Roberto Amato, Jeffrey Barrett, Sonia Goncalves, Ewan Harrison, David K. Jackson, Ian Johnston, Dominic Kwiatkowski, Cordelia Langford, John Sillitoe on behalf of the Wellcome Sanger Institute COVID-19 Surveillance Team |
| EPI_ISL_3313756 | Berkshire and Surrey Pathology Services Lighthouse Laboratory | Wellcome Sanger Institute for the COVID-19 Genomics UK (COG-UK) Consortium | Berkshire and Surrey Pathology Services Lighthouse Laboratory and Alex Alderton, Roberto Amato, Jeffrey Barrett, Sonia Goncalves, Ewan Harrison, David K. Jackson, Ian Johnston, Dominic Kwiatkowski, Cordelia Langford, John Sillitoe on behalf of the Wellcome Sanger Institute COVID-19 Surveillance Team |
| EPI_ISL_3313757, EPI_ISL_3313758 | Lighthouse Lab in Glasgow | Wellcome Sanger Institute for the COVID-19 Genomics UK (COG-UK) Consortium | Harper VanSteenhouse, Yumi Kasai, David Gray, Carol Clugston, Anna Dominiczak and Alex Alderton, Roberto Amato, Jeffrey Barrett, Sonia Goncalves, Ewan Harrison, David K. Jackson, Ian Johnston, Dominic Kwiatkowski, Cordelia Langford, John Sillitoe on behalf of the Wellcome Sanger Institute COVID-19 Surveillance Team |
| EPI_ISL_3313759 | Lighthouse Lab in Milton Keynes | Wellcome Sanger Institute for the COVID-19 Genomics UK (COG-UK) Consortium | The Lighthouse Lab in Milton Keynes and Alex Alderton, Roberto Amato, Jeffrey Barrett, Sonia Goncalves, Ewan Harrison, David K. Jackson, Ian Johnston, Dominic Kwiatkowski, Cordelia Langford, John Sillitoe on behalf of the Wellcome Sanger Institute COVID-19 Surveillance Team |
| EPI_ISL_3313760, EPI_ISL_3313761 | Berkshire and Surrey Pathology Services Lighthouse Laboratory | Wellcome Sanger Institute for the COVID-19 Genomics UK (COG-UK) Consortium | Berkshire and Surrey Pathology Services Lighthouse Laboratory and Alex Alderton, Roberto Amato, Jeffrey Barrett, Sonia Goncalves, Ewan Harrison, David K. Jackson, Ian Johnston, Dominic Kwiatkowski, Cordelia Langford, John Sillitoe on behalf of the Wellcome Sanger Institute COVID-19 Surveillance Team |
| EPI_ISL_3313762, EPI_ISL_3313763 | Lighthouse Lab in Milton Keynes | Wellcome Sanger Institute for the COVID-19 Genomics UK (COG-UK) Consortium | The Lighthouse Lab in Milton Keynes and Alex Alderton, Roberto Amato, Jeffrey Barrett, Sonia Goncalves, Ewan Harrison, David K. Jackson, Ian Johnston, Dominic Kwiatkowski, Cordelia Langford, John Sillitoe on behalf of the Wellcome Sanger Institute COVID-19 Surveillance Team |
| EPI_ISL_3313764 | Lighthouse Lab in Glasgow | Wellcome Sanger Institute for the COVID-19 Genomics UK (COG-UK) Consortium | Harper VanSteenhouse, Yumi Kasai, David Gray, Carol Clugston, Anna Dominiczak and Alex Alderton, Roberto Amato, Jeffrey Barrett, Sonia Goncalves, Ewan Harrison, David K. Jackson, Ian Johnston, Dominic Kwiatkowski, Cordelia Langford, John Sillitoe on behalf of the Wellcome Sanger Institute COVID-19 Surveillance Team |
| EPI_ISL_3313765, EPI_ISL_3313766 | Berkshire and Surrey Pathology Services Lighthouse Laboratory | Wellcome Sanger Institute for the COVID-19 Genomics UK (COG-UK) Consortium | Berkshire and Surrey Pathology Services Lighthouse Laboratory and Alex Alderton, Roberto Amato, Jeffrey Barrett, Sonia Goncalves, Ewan Harrison, David K. Jackson, Ian Johnston, Dominic Kwiatkowski, Cordelia Langford, John Sillitoe on behalf of the Wellcome Sanger Institute COVID-19 Surveillance Team |
| EPI_ISL_3313767 | Lighthouse Lab in Glasgow | Wellcome Sanger Institute for the COVID-19 Genomics UK (COG-UK) Consortium | Harper VanSteenhouse, Yumi Kasai, David Gray, Carol Clugston, Anna Dominiczak and Alex Alderton, Roberto Amato, Jeffrey Barrett, Sonia Goncalves, Ewan Harrison, David K. Jackson, Ian Johnston, Dominic Kwiatkowski, Cordelia Langford, John Sillitoe on behalf of the Wellcome Sanger Institute COVID-19 Surveillance Team |
| EPI_ISL_3313768 | Lighthouse Lab in Milton Keynes | Wellcome Sanger Institute for the COVID-19 Genomics UK (COG-UK) Consortium | The Lighthouse Lab in Milton Keynes and Alex Alderton, Roberto Amato, Jeffrey Barrett, Sonia Goncalves, Ewan Harrison, David K. Jackson, Ian Johnston, Dominic Kwiatkowski, Cordelia Langford, John Sillitoe on behalf of the Wellcome Sanger Institute COVID-19 Surveillance Team |
| EPI_ISL_3313769, | Berkshire and Surrey Pathology Services Lighthouse Laboratory | Wellcome Sanger Institute for the COVID-19 Genomics UK (COG-UK) | Berkshire and Surrey Pathology Services Lighthouse Laboratory and Alex Alderton, Roberto Amato, Jeffrey Barrett, Sonia Goncalves, Ewan Harrison, David K. Jackson, Ian Johnston, Dominic Kwiatkowski, Cordelia Langford, John Sillitoe on behalf |

| | | | |
|---|---|---|---|
| EPI_ISL_3313770, EPI_ISL_3313771, EPI_ISL_3313772, EPI_ISL_3313773 | | Consortium | of the Wellcome Sanger Institute COVID-19 Surveillance Team |
| EPI_ISL_3313774 | Lighthouse Lab in Glasgow | Wellcome Sanger Institute for the COVID-19 Genomics UK (COG-UK) Consortium | Harper VanSteenhouse, Yumi Kasai, David Gray, Carol Clugston, Anna Dominiczak and Alex Alderton, Roberto Amato, Jeffrey Barrett, Sonia Goncalves, Ewan Harrison, David K. Jackson, Ian Johnston, Dominic Kwiatkowski, Cordelia Langford, John Sillitoe on behalf of the Wellcome Sanger Institute COVID-19 Surveillance Team |
| EPI_ISL_3313775 | Berkshire and Surrey Pathology Services Lighthouse Laboratory | Wellcome Sanger Institute for the COVID-19 Genomics UK (COG-UK) Consortium | Berkshire and Surrey Pathology Services Lighthouse Laboratory and Alex Alderton, Roberto Amato, Jeffrey Barrett, Sonia Goncalves, Ewan Harrison, David K. Jackson, Ian Johnston, Dominic Kwiatkowski, Cordelia Langford, John Sillitoe on behalf of the Wellcome Sanger Institute COVID-19 Surveillance Team |
| EPI_ISL_3313776 | Lighthouse Lab in Glasgow | Wellcome Sanger Institute for the COVID-19 Genomics UK (COG-UK) Consortium | Harper VanSteenhouse, Yumi Kasai, David Gray, Carol Clugston, Anna Dominiczak and Alex Alderton, Roberto Amato, Jeffrey Barrett, Sonia Goncalves, Ewan Harrison, David K. Jackson, Ian Johnston, Dominic Kwiatkowski, Cordelia Langford, John Sillitoe on behalf of the Wellcome Sanger Institute COVID-19 Surveillance Team |
| EPI_ISL_3313777, EPI_ISL_3313778 | Berkshire and Surrey Pathology Services Lighthouse Laboratory | Wellcome Sanger Institute for the COVID-19 Genomics UK (COG-UK) Consortium | Berkshire and Surrey Pathology Services Lighthouse Laboratory and Alex Alderton, Roberto Amato, Jeffrey Barrett, Sonia Goncalves, Ewan Harrison, David K. Jackson, Ian Johnston, Dominic Kwiatkowski, Cordelia Langford, John Sillitoe on behalf of the Wellcome Sanger Institute COVID-19 Surveillance Team |
| EPI_ISL_3313779 | Lighthouse Lab in Milton Keynes | Wellcome Sanger Institute for the COVID-19 Genomics UK (COG-UK) Consortium | The Lighthouse Lab in Milton Keynes and Alex Alderton, Roberto Amato, Jeffrey Barrett, Sonia Goncalves, Ewan Harrison, David K. Jackson, Ian Johnston, Dominic Kwiatkowski, Cordelia Langford, John Sillitoe on behalf of the Wellcome Sanger Institute COVID-19 Surveillance Team |
| EPI_ISL_3313780, EPI_ISL_3313781, EPI_ISL_3313782, EPI_ISL_3313783 | Berkshire and Surrey Pathology Services Lighthouse Laboratory | Wellcome Sanger Institute for the COVID-19 Genomics UK (COG-UK) Consortium | Berkshire and Surrey Pathology Services Lighthouse Laboratory and Alex Alderton, Roberto Amato, Jeffrey Barrett, Sonia Goncalves, Ewan Harrison, David K. Jackson, Ian Johnston, Dominic Kwiatkowski, Cordelia Langford, John Sillitoe on behalf of the Wellcome Sanger Institute COVID-19 Surveillance Team |
| EPI_ISL_3313784, EPI_ISL_3313785, EPI_ISL_3313786 | Lighthouse Lab in Milton Keynes | Wellcome Sanger Institute for the COVID-19 Genomics UK (COG-UK) Consortium | The Lighthouse Lab in Milton Keynes and Alex Alderton, Roberto Amato, Jeffrey Barrett, Sonia Goncalves, Ewan Harrison, David K. Jackson, Ian Johnston, Dominic Kwiatkowski, Cordelia Langford, John Sillitoe on behalf of the Wellcome Sanger Institute COVID-19 Surveillance Team |
| EPI_ISL_3313787, EPI_ISL_3313788 | Berkshire and Surrey Pathology Services Lighthouse Laboratory | Wellcome Sanger Institute for the COVID-19 Genomics UK (COG-UK) Consortium | Berkshire and Surrey Pathology Services Lighthouse Laboratory and Alex Alderton, Roberto Amato, Jeffrey Barrett, Sonia Goncalves, Ewan Harrison, David K. Jackson, Ian Johnston, Dominic Kwiatkowski, Cordelia Langford, John Sillitoe on behalf of the Wellcome Sanger Institute COVID-19 Surveillance Team |
| EPI_ISL_3313789 | Lighthouse Lab in Glasgow | Wellcome Sanger Institute for the COVID-19 Genomics UK (COG-UK) Consortium | Harper VanSteenhouse, Yumi Kasai, David Gray, Carol Clugston, Anna Dominiczak and Alex Alderton, Roberto Amato, Jeffrey Barrett, Sonia Goncalves, Ewan Harrison, David K. Jackson, Ian Johnston, Dominic Kwiatkowski, Cordelia Langford, John Sillitoe on behalf of the Wellcome Sanger Institute COVID-19 Surveillance Team |
| EPI_ISL_3313790, EPI_ISL_3313791, EPI_ISL_3313792 | Berkshire and Surrey Pathology Services Lighthouse Laboratory | Wellcome Sanger Institute for the COVID-19 Genomics UK (COG-UK) Consortium | Berkshire and Surrey Pathology Services Lighthouse Laboratory and Alex Alderton, Roberto Amato, Jeffrey Barrett, Sonia Goncalves, Ewan Harrison, David K. Jackson, Ian Johnston, Dominic Kwiatkowski, Cordelia Langford, John Sillitoe on behalf of the Wellcome Sanger Institute COVID-19 Surveillance Team |
| EPI_ISL_3313793 | Lighthouse Lab in Glasgow | Wellcome Sanger Institute for the COVID-19 Genomics UK (COG-UK) Consortium | Harper VanSteenhouse, Yumi Kasai, David Gray, Carol Clugston, Anna Dominiczak and Alex Alderton, Roberto Amato, Jeffrey Barrett, Sonia Goncalves, Ewan Harrison, David K. Jackson, Ian Johnston, Dominic Kwiatkowski, Cordelia Langford, John Sillitoe on behalf of the Wellcome Sanger Institute COVID-19 Surveillance Team |
| EPI_ISL_3313794 | Lighthouse Lab in Milton Keynes | Wellcome Sanger Institute for the COVID-19 Genomics UK (COG-UK) Consortium | The Lighthouse Lab in Milton Keynes and Alex Alderton, Roberto Amato, Jeffrey Barrett, Sonia Goncalves, Ewan Harrison, David K. Jackson, Ian Johnston, Dominic Kwiatkowski, Cordelia Langford, John Sillitoe on behalf of the Wellcome Sanger Institute COVID-19 Surveillance Team |
| EPI_ISL_3313795 | Lighthouse Lab in Glasgow | Wellcome Sanger Institute for the COVID-19 Genomics UK (COG-UK) Consortium | Harper VanSteenhouse, Yumi Kasai, David Gray, Carol Clugston, Anna Dominiczak and Alex Alderton, Roberto Amato, Jeffrey Barrett, Sonia Goncalves, Ewan Harrison, David K. Jackson, Ian Johnston, Dominic Kwiatkowski, Cordelia Langford, John Sillitoe on behalf of the Wellcome Sanger Institute COVID-19 Surveillance Team |
| EPI_ISL_3313796, EPI_ISL_3313797 | Berkshire and Surrey Pathology Services Lighthouse Laboratory | Wellcome Sanger Institute for the COVID-19 Genomics UK (COG-UK) Consortium | Berkshire and Surrey Pathology Services Lighthouse Laboratory and Alex Alderton, Roberto Amato, Jeffrey Barrett, Sonia Goncalves, Ewan Harrison, David K. Jackson, Ian Johnston, Dominic Kwiatkowski, Cordelia Langford, John Sillitoe on behalf of the Wellcome Sanger Institute COVID-19 Surveillance Team |
| EPI_ISL_3313798 | Lighthouse Lab in Milton Keynes | Wellcome Sanger Institute for the COVID-19 Genomics UK (COG-UK) Consortium | The Lighthouse Lab in Milton Keynes and Alex Alderton, Roberto Amato, Jeffrey Barrett, Sonia Goncalves, Ewan Harrison, David K. Jackson, Ian Johnston, Dominic Kwiatkowski, Cordelia Langford, John Sillitoe on behalf of the Wellcome Sanger Institute COVID-19 Surveillance Team |
| EPI_ISL_3313799 | Lighthouse Lab in Glasgow | Wellcome Sanger Institute for the COVID-19 Genomics UK (COG-UK) Consortium | Harper VanSteenhouse, Yumi Kasai, David Gray, Carol Clugston, Anna Dominiczak and Alex Alderton, Roberto Amato, Jeffrey Barrett, Sonia Goncalves, Ewan Harrison, David K. Jackson, Ian Johnston, Dominic Kwiatkowski, Cordelia Langford, John Sillitoe on behalf of the Wellcome Sanger Institute COVID-19 Surveillance Team |
| EPI_ISL_3313800, EPI_ISL_3313801 | Lighthouse Lab in Milton Keynes | Wellcome Sanger Institute for the COVID-19 Genomics UK (COG-UK) Consortium | The Lighthouse Lab in Milton Keynes and Alex Alderton, Roberto Amato, Jeffrey Barrett, Sonia Goncalves, Ewan Harrison, David K. Jackson, Ian Johnston, Dominic Kwiatkowski, Cordelia Langford, John Sillitoe on behalf of the Wellcome Sanger Institute COVID-19 Surveillance Team |
| EPI_ISL_3313802 | Berkshire and Surrey Pathology Services Lighthouse Laboratory | Wellcome Sanger Institute for the COVID-19 Genomics UK (COG-UK) Consortium | Berkshire and Surrey Pathology Services Lighthouse Laboratory and Alex Alderton, Roberto Amato, Jeffrey Barrett, Sonia Goncalves, Ewan Harrison, David K. Jackson, Ian Johnston, Dominic Kwiatkowski, Cordelia Langford, John Sillitoe on behalf of the Wellcome Sanger Institute COVID-19 Surveillance Team |
| EPI_ISL_3313803 | Lighthouse Lab in Milton Keynes | Wellcome Sanger Institute for the COVID-19 Genomics UK (COG-UK) Consortium | The Lighthouse Lab in Milton Keynes and Alex Alderton, Roberto Amato, Jeffrey Barrett, Sonia Goncalves, Ewan Harrison, David K. Jackson, Ian Johnston, Dominic Kwiatkowski, Cordelia Langford, John Sillitoe on behalf of the Wellcome Sanger Institute COVID-19 Surveillance Team |
| EPI_ISL_3313804, EPI_ISL_3313805, EPI_ISL_3313806, EPI_ISL_3313807 | Berkshire and Surrey Pathology Services Lighthouse Laboratory | Wellcome Sanger Institute for the COVID-19 Genomics UK (COG-UK) Consortium | Berkshire and Surrey Pathology Services Lighthouse Laboratory and Alex Alderton, Roberto Amato, Jeffrey Barrett, Sonia Goncalves, Ewan Harrison, David K. Jackson, Ian Johnston, Dominic Kwiatkowski, Cordelia Langford, John Sillitoe on behalf of the Wellcome Sanger Institute COVID-19 Surveillance Team |
| EPI_ISL_3313808 | Lighthouse Lab in Milton Keynes | Wellcome Sanger Institute for the COVID-19 Genomics UK (COG-UK) Consortium | The Lighthouse Lab in Milton Keynes and Alex Alderton, Roberto Amato, Jeffrey Barrett, Sonia Goncalves, Ewan Harrison, David K. Jackson, Ian Johnston, Dominic Kwiatkowski, Cordelia Langford, John Sillitoe on behalf of the Wellcome Sanger Institute COVID-19 Surveillance Team |
| EPI_ISL_3313809 | Berkshire and Surrey Pathology Services Lighthouse Laboratory | Wellcome Sanger Institute for the COVID-19 Genomics UK (COG-UK) Consortium | Berkshire and Surrey Pathology Services Lighthouse Laboratory and Alex Alderton, Roberto Amato, Jeffrey Barrett, Sonia Goncalves, Ewan Harrison, David K. Jackson, Ian Johnston, Dominic Kwiatkowski, Cordelia Langford, John Sillitoe on behalf of the Wellcome Sanger Institute COVID-19 Surveillance Team |
| EPI_ISL_3313810 | Lighthouse Lab in Milton Keynes | Wellcome Sanger Institute for the COVID-19 Genomics UK (COG-UK) Consortium | The Lighthouse Lab in Milton Keynes and Alex Alderton, Roberto Amato, Jeffrey Barrett, Sonia Goncalves, Ewan Harrison, David K. Jackson, Ian Johnston, Dominic Kwiatkowski, Cordelia Langford, John Sillitoe on behalf of the Wellcome Sanger Institute COVID-19 Surveillance Team |
| EPI_ISL_3313811 | Berkshire and Surrey Pathology Services Lighthouse Laboratory | Wellcome Sanger Institute for the COVID-19 Genomics UK (COG-UK) Consortium | Berkshire and Surrey Pathology Services Lighthouse Laboratory and Alex Alderton, Roberto Amato, Jeffrey Barrett, Sonia Goncalves, Ewan Harrison, David K. Jackson, Ian Johnston, Dominic Kwiatkowski, Cordelia Langford, John Sillitoe on behalf of the Wellcome Sanger Institute COVID-19 Surveillance Team |
| EPI_ISL_3313812 | Lighthouse Lab in Milton Keynes | Wellcome Sanger Institute for the COVID-19 Genomics UK (COG-UK) Consortium | The Lighthouse Lab in Milton Keynes and Alex Alderton, Roberto Amato, Jeffrey Barrett, Sonia Goncalves, Ewan Harrison, David K. Jackson, Ian Johnston, Dominic Kwiatkowski, Cordelia Langford, John Sillitoe on behalf of the Wellcome Sanger Institute COVID-19 Surveillance Team |
| EPI_ISL_3313813 | Berkshire and Surrey Pathology Services Lighthouse Laboratory | Wellcome Sanger Institute for the COVID-19 Genomics UK (COG-UK) Consortium | Berkshire and Surrey Pathology Services Lighthouse Laboratory and Alex Alderton, Roberto Amato, Jeffrey Barrett, Sonia Goncalves, Ewan Harrison, David K. Jackson, Ian Johnston, Dominic Kwiatkowski, Cordelia Langford, John Sillitoe on behalf of the Wellcome Sanger Institute COVID-19 Surveillance Team |
| EPI_ISL_3313814, EPI_ISL_3313815 | Lighthouse Lab in Glasgow | Wellcome Sanger Institute for the COVID-19 Genomics UK (COG-UK) Consortium | Harper VanSteenhouse, Yumi Kasai, David Gray, Carol Clugston, Anna Dominiczak and Alex Alderton, Roberto Amato, Jeffrey Barrett, Sonia Goncalves, Ewan Harrison, David K. Jackson, Ian Johnston, Dominic Kwiatkowski, Cordelia Langford, John Sillitoe on behalf of the Wellcome Sanger Institute COVID-19 Surveillance Team |
| EPI_ISL_3313816 | Berkshire and Surrey Pathology Services Lighthouse Laboratory | Wellcome Sanger Institute for the COVID-19 Genomics UK (COG-UK) Consortium | Berkshire and Surrey Pathology Services Lighthouse Laboratory and Alex Alderton, Roberto Amato, Jeffrey Barrett, Sonia Goncalves, Ewan Harrison, David K. Jackson, Ian Johnston, Dominic Kwiatkowski, Cordelia Langford, John Sillitoe on behalf of the Wellcome Sanger Institute COVID-19 Surveillance Team |
| EPI_ISL_3313817, EPI_ISL_3313818 | Lighthouse Lab in Milton Keynes | Wellcome Sanger Institute for the COVID-19 Genomics UK (COG-UK) Consortium | The Lighthouse Lab in Milton Keynes and Alex Alderton, Roberto Amato, Jeffrey Barrett, Sonia Goncalves, Ewan Harrison, David K. Jackson, Ian Johnston, Dominic Kwiatkowski, Cordelia Langford, John Sillitoe on behalf of the Wellcome Sanger Institute COVID-19 Surveillance Team |
| EPI_ISL_3313819 | Berkshire and Surrey Pathology Services Lighthouse Laboratory | Wellcome Sanger Institute for the COVID-19 Genomics UK (COG-UK) Consortium | Berkshire and Surrey Pathology Services Lighthouse Laboratory and Alex Alderton, Roberto Amato, Jeffrey Barrett, Sonia Goncalves, Ewan Harrison, David K. Jackson, Ian Johnston, Dominic Kwiatkowski, Cordelia Langford, John Sillitoe on behalf of the Wellcome Sanger Institute COVID-19 Surveillance Team |
| EPI_ISL_3313820 | Lighthouse Lab in Glasgow | Wellcome Sanger Institute for the COVID-19 Genomics UK (COG-UK) Consortium | Harper VanSteenhouse, Yumi Kasai, David Gray, Carol Clugston, Anna Dominiczak and Alex Alderton, Roberto Amato, Jeffrey Barrett, Sonia Goncalves, Ewan Harrison, David K. Jackson, Ian Johnston, Dominic Kwiatkowski, Cordelia Langford, John Sillitoe on behalf of the Wellcome Sanger Institute COVID-19 Surveillance Team |
| EPI_ISL_3313821 | Lighthouse Lab in Milton Keynes | Wellcome Sanger Institute for the COVID-19 Genomics UK (COG-UK) Consortium | The Lighthouse Lab in Milton Keynes and Alex Alderton, Roberto Amato, Jeffrey Barrett, Sonia Goncalves, Ewan Harrison, David K. Jackson, Ian Johnston, Dominic Kwiatkowski, Cordelia Langford, John Sillitoe on behalf of the Wellcome Sanger Institute COVID-19 Surveillance Team |
| EPI_ISL_3313822 | Berkshire and Surrey Pathology Services Lighthouse Laboratory | Wellcome Sanger Institute for the COVID-19 Genomics UK (COG-UK) Consortium | Berkshire and Surrey Pathology Services Lighthouse Laboratory and Alex Alderton, Roberto Amato, Jeffrey Barrett, Sonia Goncalves, Ewan Harrison, David K. Jackson, Ian Johnston, Dominic Kwiatkowski, Cordelia Langford, John Sillitoe on behalf of the Wellcome Sanger Institute COVID-19 Surveillance Team |
| EPI_ISL_3313823, EPI_ISL_3313824, EPI_ISL_3313825, EPI_ISL_3313826, EPI_ISL_3313827, EPI_ISL_3313828 | Lighthouse Lab in Milton Keynes | Wellcome Sanger Institute for the COVID-19 Genomics UK (COG-UK) Consortium | The Lighthouse Lab in Milton Keynes and Alex Alderton, Roberto Amato, Jeffrey Barrett, Sonia Goncalves, Ewan Harrison, David K. Jackson, Ian Johnston, Dominic Kwiatkowski, Cordelia Langford, John Sillitoe on behalf of the Wellcome Sanger Institute COVID-19 Surveillance Team |
| EPI_ISL_3313829 | Lighthouse Lab in Glasgow | Wellcome Sanger Institute for the COVID-19 Genomics UK (COG-UK) Consortium | Harper VanSteenhouse, Yumi Kasai, David Gray, Carol Clugston, Anna Dominiczak and Alex Alderton, Roberto Amato, Jeffrey Barrett, Sonia Goncalves, Ewan Harrison, David K. Jackson, Ian Johnston, Dominic Kwiatkowski, Cordelia Langford, John Sillitoe on behalf of the Wellcome Sanger Institute COVID-19 Surveillance Team |
| EPI_ISL_3313830 | Lighthouse Lab in Milton Keynes | Wellcome Sanger Institute for the COVID-19 Genomics UK (COG-UK) Consortium | The Lighthouse Lab in Milton Keynes and Alex Alderton, Roberto Amato, Jeffrey Barrett, Sonia Goncalves, Ewan Harrison, David K. Jackson, Ian Johnston, Dominic Kwiatkowski, Cordelia Langford, John Sillitoe on behalf of the Wellcome Sanger Institute COVID-19 Surveillance Team |
| EPI_ISL_3313831 | Lighthouse Lab in Glasgow | Wellcome Sanger Institute for the COVID-19 Genomics UK (COG-UK) Consortium | Harper VanSteenhouse, Yumi Kasai, David Gray, Carol Clugston, Anna Dominiczak and Alex Alderton, Roberto Amato, Jeffrey Barrett, Sonia Goncalves, Ewan Harrison, David K. Jackson, Ian Johnston, Dominic Kwiatkowski, Cordelia Langford, John Sillitoe on behalf of the Wellcome Sanger Institute COVID-19 Surveillance Team |
| EPI_ISL_3313832 | Berkshire and Surrey Pathology Services Lighthouse Laboratory | Wellcome Sanger Institute for the COVID-19 Genomics UK (COG-UK) Consortium | Berkshire and Surrey Pathology Services Lighthouse Laboratory and Alex Alderton, Roberto Amato, Jeffrey Barrett, Sonia Goncalves, Ewan Harrison, David K. Jackson, Ian Johnston, Dominic Kwiatkowski, Cordelia Langford, John Sillitoe on behalf of the Wellcome Sanger Institute COVID-19 Surveillance Team |
| EPI_ISL_3313833 | Lighthouse Lab in Glasgow | Wellcome Sanger Institute for the COVID-19 Genomics UK (COG-UK) Consortium | Harper VanSteenhouse, Yumi Kasai, David Gray, Carol Clugston, Anna Dominiczak and Alex Alderton, Roberto Amato, Jeffrey Barrett, Sonia Goncalves, Ewan Harrison, David K. Jackson, Ian Johnston, Dominic Kwiatkowski, Cordelia Langford, John Sillitoe on behalf of the Wellcome Sanger Institute COVID-19 Surveillance Team |

| | | | |
|---|---|---|---|
| EPI_ISL_3313834 | Berkshire and Surrey Pathology Services Lighthouse Laboratory | Wellcome Sanger Institute for the COVID-19 Genomics UK (COG-UK) Consortium | Berkshire and Surrey Pathology Services Lighthouse Laboratory and Alex Alderton, Roberto Amato, Jeffrey Barrett, Sonia Goncalves, Ewan Harrison, David K. Jackson, Ian Johnston, Dominic Kwiatkowski, Cordelia Langford, John Sillitoe on behalf of the Wellcome Sanger Institute COVID-19 Surveillance Team |
| EPI_ISL_3313835 | Lighthouse Lab in Milton Keynes | Wellcome Sanger Institute for the COVID-19 Genomics UK (COG-UK) Consortium | The Lighthouse Lab in Milton Keynes and Alex Alderton, Roberto Amato, Jeffrey Barrett, Sonia Goncalves, Ewan Harrison, David K. Jackson, Ian Johnston, Dominic Kwiatkowski, Cordelia Langford, John Sillitoe on behalf of the Wellcome Sanger Institute COVID-19 Surveillance Team |
| EPI_ISL_3313836 | Lighthouse Lab in Glasgow | Wellcome Sanger Institute for the COVID-19 Genomics UK (COG-UK) Consortium | Harper VanSteenhouse, Yumi Kasai, David Gray, Carol Clugston, Anna Dominiczak and Alex Alderton, Roberto Amato, Jeffrey Barrett, Sonia Goncalves, Ewan Harrison, David K. Jackson, Ian Johnston, Dominic Kwiatkowski, Cordelia Langford, John Sillitoe on behalf of the Wellcome Sanger Institute COVID-19 Surveillance Team |
| EPI_ISL_3313837 | Berkshire and Surrey Pathology Services Lighthouse Laboratory | Wellcome Sanger Institute for the COVID-19 Genomics UK (COG-UK) Consortium | Berkshire and Surrey Pathology Services Lighthouse Laboratory and Alex Alderton, Roberto Amato, Jeffrey Barrett, Sonia Goncalves, Ewan Harrison, David K. Jackson, Ian Johnston, Dominic Kwiatkowski, Cordelia Langford, John Sillitoe on behalf of the Wellcome Sanger Institute COVID-19 Surveillance Team |
| EPI_ISL_3313838, EPI_ISL_3313839, EPI_ISL_3313840 | Lighthouse Lab in Milton Keynes | Wellcome Sanger Institute for the COVID-19 Genomics UK (COG-UK) Consortium | The Lighthouse Lab in Milton Keynes and Alex Alderton, Roberto Amato, Jeffrey Barrett, Sonia Goncalves, Ewan Harrison, David K. Jackson, Ian Johnston, Dominic Kwiatkowski, Cordelia Langford, John Sillitoe on behalf of the Wellcome Sanger Institute COVID-19 Surveillance Team |
| EPI_ISL_3313841 | Berkshire and Surrey Pathology Services Lighthouse Laboratory | Wellcome Sanger Institute for the COVID-19 Genomics UK (COG-UK) Consortium | Berkshire and Surrey Pathology Services Lighthouse Laboratory and Alex Alderton, Roberto Amato, Jeffrey Barrett, Sonia Goncalves, Ewan Harrison, David K. Jackson, Ian Johnston, Dominic Kwiatkowski, Cordelia Langford, John Sillitoe on behalf of the Wellcome Sanger Institute COVID-19 Surveillance Team |
| EPI_ISL_3313842 | Lighthouse Lab in Glasgow | Wellcome Sanger Institute for the COVID-19 Genomics UK (COG-UK) Consortium | Harper VanSteenhouse, Yumi Kasai, David Gray, Carol Clugston, Anna Dominiczak and Alex Alderton, Roberto Amato, Jeffrey Barrett, Sonia Goncalves, Ewan Harrison, David K. Jackson, Ian Johnston, Dominic Kwiatkowski, Cordelia Langford, John Sillitoe on behalf of the Wellcome Sanger Institute COVID-19 Surveillance Team |
| EPI_ISL_3313843, EPI_ISL_3313844, EPI_ISL_3313845 | Lighthouse Lab in Milton Keynes | Wellcome Sanger Institute for the COVID-19 Genomics UK (COG-UK) Consortium | The Lighthouse Lab in Milton Keynes and Alex Alderton, Roberto Amato, Jeffrey Barrett, Sonia Goncalves, Ewan Harrison, David K. Jackson, Ian Johnston, Dominic Kwiatkowski, Cordelia Langford, John Sillitoe on behalf of the Wellcome Sanger Institute COVID-19 Surveillance Team |
| EPI_ISL_3313846 | Lighthouse Lab in Glasgow | Wellcome Sanger Institute for the COVID-19 Genomics UK (COG-UK) Consortium | Harper VanSteenhouse, Yumi Kasai, David Gray, Carol Clugston, Anna Dominiczak and Alex Alderton, Roberto Amato, Jeffrey Barrett, Sonia Goncalves, Ewan Harrison, David K. Jackson, Ian Johnston, Dominic Kwiatkowski, Cordelia Langford, John Sillitoe on behalf of the Wellcome Sanger Institute COVID-19 Surveillance Team |
| EPI_ISL_3313847, EPI_ISL_3313848 | Lighthouse Lab in Milton Keynes | Wellcome Sanger Institute for the COVID-19 Genomics UK (COG-UK) Consortium | The Lighthouse Lab in Milton Keynes and Alex Alderton, Roberto Amato, Jeffrey Barrett, Sonia Goncalves, Ewan Harrison, David K. Jackson, Ian Johnston, Dominic Kwiatkowski, Cordelia Langford, John Sillitoe on behalf of the Wellcome Sanger Institute COVID-19 Surveillance Team |
| EPI_ISL_3313849 | Lighthouse Lab in Glasgow | Wellcome Sanger Institute for the COVID-19 Genomics UK (COG-UK) Consortium | Harper VanSteenhouse, Yumi Kasai, David Gray, Carol Clugston, Anna Dominiczak and Alex Alderton, Roberto Amato, Jeffrey Barrett, Sonia Goncalves, Ewan Harrison, David K. Jackson, Ian Johnston, Dominic Kwiatkowski, Cordelia Langford, John Sillitoe on behalf of the Wellcome Sanger Institute COVID-19 Surveillance Team |
| EPI_ISL_3313850, EPI_ISL_3313851, EPI_ISL_3313852 | Lighthouse Lab in Milton Keynes | Wellcome Sanger Institute for the COVID-19 Genomics UK (COG-UK) Consortium | The Lighthouse Lab in Milton Keynes and Alex Alderton, Roberto Amato, Jeffrey Barrett, Sonia Goncalves, Ewan Harrison, David K. Jackson, Ian Johnston, Dominic Kwiatkowski, Cordelia Langford, John Sillitoe on behalf of the Wellcome Sanger Institute COVID-19 Surveillance Team |
| EPI_ISL_3313853 | Berkshire and Surrey Pathology Services Lighthouse Laboratory | Wellcome Sanger Institute for the COVID-19 Genomics UK (COG-UK) Consortium | Berkshire and Surrey Pathology Services Lighthouse Laboratory and Alex Alderton, Roberto Amato, Jeffrey Barrett, Sonia Goncalves, Ewan Harrison, David K. Jackson, Ian Johnston, Dominic Kwiatkowski, Cordelia Langford, John Sillitoe on behalf of the Wellcome Sanger Institute COVID-19 Surveillance Team |
| EPI_ISL_3313854 | Lighthouse Lab in Milton Keynes | Wellcome Sanger Institute for the COVID-19 Genomics UK (COG-UK) Consortium | The Lighthouse Lab in Milton Keynes and Alex Alderton, Roberto Amato, Jeffrey Barrett, Sonia Goncalves, Ewan Harrison, David K. Jackson, Ian Johnston, Dominic Kwiatkowski, Cordelia Langford, John Sillitoe on behalf of the Wellcome Sanger Institute COVID-19 Surveillance Team |
| EPI_ISL_3313855 | Lighthouse Lab in Glasgow | Wellcome Sanger Institute for the COVID-19 Genomics UK (COG-UK) Consortium | Harper VanSteenhouse, Yumi Kasai, David Gray, Carol Clugston, Anna Dominiczak and Alex Alderton, Roberto Amato, Jeffrey Barrett, Sonia Goncalves, Ewan Harrison, David K. Jackson, Ian Johnston, Dominic Kwiatkowski, Cordelia Langford, John Sillitoe on behalf of the Wellcome Sanger Institute COVID-19 Surveillance Team |
| EPI_ISL_3313856 | Berkshire and Surrey Pathology Services Lighthouse Laboratory | Wellcome Sanger Institute for the COVID-19 Genomics UK (COG-UK) Consortium | Berkshire and Surrey Pathology Services Lighthouse Laboratory and Alex Alderton, Roberto Amato, Jeffrey Barrett, Sonia Goncalves, Ewan Harrison, David K. Jackson, Ian Johnston, Dominic Kwiatkowski, Cordelia Langford, John Sillitoe on behalf of the Wellcome Sanger Institute COVID-19 Surveillance Team |
| EPI_ISL_3313857 | Lighthouse Lab in Glasgow | Wellcome Sanger Institute for the COVID-19 Genomics UK (COG-UK) Consortium | Harper VanSteenhouse, Yumi Kasai, David Gray, Carol Clugston, Anna Dominiczak and Alex Alderton, Roberto Amato, Jeffrey Barrett, Sonia Goncalves, Ewan Harrison, David K. Jackson, Ian Johnston, Dominic Kwiatkowski, Cordelia Langford, John Sillitoe on behalf of the Wellcome Sanger Institute COVID-19 Surveillance Team |
| EPI_ISL_3313858 | Berkshire and Surrey Pathology Services Lighthouse Laboratory | Wellcome Sanger Institute for the COVID-19 Genomics UK (COG-UK) Consortium | Berkshire and Surrey Pathology Services Lighthouse Laboratory and Alex Alderton, Roberto Amato, Jeffrey Barrett, Sonia Goncalves, Ewan Harrison, David K. Jackson, Ian Johnston, Dominic Kwiatkowski, Cordelia Langford, John Sillitoe on behalf of the Wellcome Sanger Institute COVID-19 Surveillance Team |
| EPI_ISL_3313859 | Lighthouse Lab in Milton Keynes | Wellcome Sanger Institute for the COVID-19 Genomics UK (COG-UK) Consortium | The Lighthouse Lab in Milton Keynes and Alex Alderton, Roberto Amato, Jeffrey Barrett, Sonia Goncalves, Ewan Harrison, David K. Jackson, Ian Johnston, Dominic Kwiatkowski, Cordelia Langford, John Sillitoe on behalf of the Wellcome Sanger Institute COVID-19 Surveillance Team |
| EPI_ISL_3313860, EPI_ISL_3313861, EPI_ISL_3313862 | Lighthouse Lab in Glasgow | Wellcome Sanger Institute for the COVID-19 Genomics UK (COG-UK) Consortium | Harper VanSteenhouse, Yumi Kasai, David Gray, Carol Clugston, Anna Dominiczak and Alex Alderton, Roberto Amato, Jeffrey Barrett, Sonia Goncalves, Ewan Harrison, David K. Jackson, Ian Johnston, Dominic Kwiatkowski, Cordelia Langford, John Sillitoe on behalf of the Wellcome Sanger Institute COVID-19 Surveillance Team |
| EPI_ISL_3313863, EPI_ISL_3313864, EPI_ISL_3313865 | Lighthouse Lab in Milton Keynes | Wellcome Sanger Institute for the COVID-19 Genomics UK (COG-UK) Consortium | The Lighthouse Lab in Milton Keynes and Alex Alderton, Roberto Amato, Jeffrey Barrett, Sonia Goncalves, Ewan Harrison, David K. Jackson, Ian Johnston, Dominic Kwiatkowski, Cordelia Langford, John Sillitoe on behalf of the Wellcome Sanger Institute COVID-19 Surveillance Team |
| EPI_ISL_3313866 | Lighthouse Lab in Glasgow | Wellcome Sanger Institute for the COVID-19 Genomics UK (COG-UK) Consortium | Harper VanSteenhouse, Yumi Kasai, David Gray, Carol Clugston, Anna Dominiczak and Alex Alderton, Roberto Amato, Jeffrey Barrett, Sonia Goncalves, Ewan Harrison, David K. Jackson, Ian Johnston, Dominic Kwiatkowski, Cordelia Langford, John Sillitoe on behalf of the Wellcome Sanger Institute COVID-19 Surveillance Team |
| EPI_ISL_3313867, EPI_ISL_3313868 | Lighthouse Lab in Milton Keynes | Wellcome Sanger Institute for the COVID-19 Genomics UK (COG-UK) Consortium | The Lighthouse Lab in Milton Keynes and Alex Alderton, Roberto Amato, Jeffrey Barrett, Sonia Goncalves, Ewan Harrison, David K. Jackson, Ian Johnston, Dominic Kwiatkowski, Cordelia Langford, John Sillitoe on behalf of the Wellcome Sanger Institute COVID-19 Surveillance Team |
| EPI_ISL_3313869 | Berkshire and Surrey Pathology Services Lighthouse Laboratory | Wellcome Sanger Institute for the COVID-19 Genomics UK (COG-UK) Consortium | Berkshire and Surrey Pathology Services Lighthouse Laboratory and Alex Alderton, Roberto Amato, Jeffrey Barrett, Sonia Goncalves, Ewan Harrison, David K. Jackson, Ian Johnston, Dominic Kwiatkowski, Cordelia Langford, John Sillitoe on behalf of the Wellcome Sanger Institute COVID-19 Surveillance Team |
| EPI_ISL_3313870 | Lighthouse Lab in Milton Keynes | Wellcome Sanger Institute for the COVID-19 Genomics UK (COG-UK) Consortium | The Lighthouse Lab in Milton Keynes and Alex Alderton, Roberto Amato, Jeffrey Barrett, Sonia Goncalves, Ewan Harrison, David K. Jackson, Ian Johnston, Dominic Kwiatkowski, Cordelia Langford, John Sillitoe on behalf of the Wellcome Sanger Institute COVID-19 Surveillance Team |
| EPI_ISL_3313871 | Berkshire and Surrey Pathology Services Lighthouse Laboratory | Wellcome Sanger Institute for the COVID-19 Genomics UK (COG-UK) Consortium | Berkshire and Surrey Pathology Services Lighthouse Laboratory and Alex Alderton, Roberto Amato, Jeffrey Barrett, Sonia Goncalves, Ewan Harrison, David K. Jackson, Ian Johnston, Dominic Kwiatkowski, Cordelia Langford, John Sillitoe on behalf of the Wellcome Sanger Institute COVID-19 Surveillance Team |
| EPI_ISL_3313872, EPI_ISL_3313873 | Lighthouse Lab in Milton Keynes | Wellcome Sanger Institute for the COVID-19 Genomics UK (COG-UK) Consortium | The Lighthouse Lab in Milton Keynes and Alex Alderton, Roberto Amato, Jeffrey Barrett, Sonia Goncalves, Ewan Harrison, David K. Jackson, Ian Johnston, Dominic Kwiatkowski, Cordelia Langford, John Sillitoe on behalf of the Wellcome Sanger Institute COVID-19 Surveillance Team |
| EPI_ISL_3313874 | Berkshire and Surrey Pathology Services Lighthouse Laboratory | Wellcome Sanger Institute for the COVID-19 Genomics UK (COG-UK) Consortium | Berkshire and Surrey Pathology Services Lighthouse Laboratory and Alex Alderton, Roberto Amato, Jeffrey Barrett, Sonia Goncalves, Ewan Harrison, David K. Jackson, Ian Johnston, Dominic Kwiatkowski, Cordelia Langford, John Sillitoe on behalf of the Wellcome Sanger Institute COVID-19 Surveillance Team |
| EPI_ISL_3313875 | Lighthouse Lab in Glasgow | Wellcome Sanger Institute for the COVID-19 Genomics UK (COG-UK) Consortium | Harper VanSteenhouse, Yumi Kasai, David Gray, Carol Clugston, Anna Dominiczak and Alex Alderton, Roberto Amato, Jeffrey Barrett, Sonia Goncalves, Ewan Harrison, David K. Jackson, Ian Johnston, Dominic Kwiatkowski, Cordelia Langford, John Sillitoe on behalf of the Wellcome Sanger Institute COVID-19 Surveillance Team |
| EPI_ISL_3313876, EPI_ISL_3313877 | Lighthouse Lab in Milton Keynes | Wellcome Sanger Institute for the COVID-19 Genomics UK (COG-UK) Consortium | The Lighthouse Lab in Milton Keynes and Alex Alderton, Roberto Amato, Jeffrey Barrett, Sonia Goncalves, Ewan Harrison, David K. Jackson, Ian Johnston, Dominic Kwiatkowski, Cordelia Langford, John Sillitoe on behalf of the Wellcome Sanger Institute COVID-19 Surveillance Team |
| EPI_ISL_3313878 | Lighthouse Lab in Glasgow | Wellcome Sanger Institute for the COVID-19 Genomics UK (COG-UK) Consortium | Harper VanSteenhouse, Yumi Kasai, David Gray, Carol Clugston, Anna Dominiczak and Alex Alderton, Roberto Amato, Jeffrey Barrett, Sonia Goncalves, Ewan Harrison, David K. Jackson, Ian Johnston, Dominic Kwiatkowski, Cordelia Langford, John Sillitoe on behalf of the Wellcome Sanger Institute COVID-19 Surveillance Team |
| EPI_ISL_3313879 | Lighthouse Lab in Milton Keynes | Wellcome Sanger Institute for the COVID-19 Genomics UK (COG-UK) Consortium | The Lighthouse Lab in Milton Keynes and Alex Alderton, Roberto Amato, Jeffrey Barrett, Sonia Goncalves, Ewan Harrison, David K. Jackson, Ian Johnston, Dominic Kwiatkowski, Cordelia Langford, John Sillitoe on behalf of the Wellcome Sanger Institute COVID-19 Surveillance Team |
| EPI_ISL_3313880 | Berkshire and Surrey Pathology Services Lighthouse Laboratory | Wellcome Sanger Institute for the COVID-19 Genomics UK (COG-UK) Consortium | Berkshire and Surrey Pathology Services Lighthouse Laboratory and Alex Alderton, Roberto Amato, Jeffrey Barrett, Sonia Goncalves, Ewan Harrison, David K. Jackson, Ian Johnston, Dominic Kwiatkowski, Cordelia Langford, John Sillitoe on behalf of the Wellcome Sanger Institute COVID-19 Surveillance Team |
| EPI_ISL_3313881, EPI_ISL_3313882, EPI_ISL_3313883 | Lighthouse Lab in Milton Keynes | Wellcome Sanger Institute for the COVID-19 Genomics UK (COG-UK) Consortium | The Lighthouse Lab in Milton Keynes and Alex Alderton, Roberto Amato, Jeffrey Barrett, Sonia Goncalves, Ewan Harrison, David K. Jackson, Ian Johnston, Dominic Kwiatkowski, Cordelia Langford, John Sillitoe on behalf of the Wellcome Sanger Institute COVID-19 Surveillance Team |
| EPI_ISL_3313884 | Berkshire and Surrey Pathology Services Lighthouse Laboratory | Wellcome Sanger Institute for the COVID-19 Genomics UK (COG-UK) Consortium | Berkshire and Surrey Pathology Services Lighthouse Laboratory and Alex Alderton, Roberto Amato, Jeffrey Barrett, Sonia Goncalves, Ewan Harrison, David K. Jackson, Ian Johnston, Dominic Kwiatkowski, Cordelia Langford, John Sillitoe on behalf of the Wellcome Sanger Institute COVID-19 Surveillance Team |
| EPI_ISL_3313885 | Lighthouse Lab in Milton Keynes | Wellcome Sanger Institute for the COVID-19 Genomics UK (COG-UK) Consortium | The Lighthouse Lab in Milton Keynes and Alex Alderton, Roberto Amato, Jeffrey Barrett, Sonia Goncalves, Ewan Harrison, David K. Jackson, Ian Johnston, Dominic Kwiatkowski, Cordelia Langford, John Sillitoe on behalf of the Wellcome Sanger Institute COVID-19 Surveillance Team |
| EPI_ISL_3313886 | Lighthouse Lab in Glasgow | Wellcome Sanger Institute for the COVID-19 Genomics UK (COG-UK) Consortium | Harper VanSteenhouse, Yumi Kasai, David Gray, Carol Clugston, Anna Dominiczak and Alex Alderton, Roberto Amato, Jeffrey Barrett, Sonia Goncalves, Ewan Harrison, David K. Jackson, Ian Johnston, Dominic Kwiatkowski, Cordelia Langford, John Sillitoe on behalf of the Wellcome Sanger Institute COVID-19 Surveillance Team |
| EPI_ISL_3313887 | Lighthouse Lab in Milton Keynes | Wellcome Sanger Institute for the COVID-19 Genomics UK (COG-UK) Consortium | The Lighthouse Lab in Milton Keynes and Alex Alderton, Roberto Amato, Jeffrey Barrett, Sonia Goncalves, Ewan Harrison, David K. Jackson, Ian Johnston, Dominic Kwiatkowski, Cordelia Langford, John Sillitoe on behalf of the Wellcome Sanger Institute COVID-19 Surveillance Team |
| EPI_ISL_3313888 | Lighthouse Lab in Glasgow | Wellcome Sanger Institute for the COVID-19 Genomics UK (COG-UK) Consortium | Harper VanSteenhouse, Yumi Kasai, David Gray, Carol Clugston, Anna Dominiczak and Alex Alderton, Roberto Amato, Jeffrey Barrett, Sonia Goncalves, Ewan Harrison, David K. Jackson, Ian Johnston, Dominic Kwiatkowski, Cordelia Langford, John Sillitoe on behalf of the Wellcome Sanger Institute COVID-19 Surveillance Team |
| EPI_ISL_3313889, EPI_ISL_3313890 | Lighthouse Lab in Milton Keynes | Wellcome Sanger Institute for the COVID-19 Genomics UK (COG-UK) Consortium | The Lighthouse Lab in Milton Keynes and Alex Alderton, Roberto Amato, Jeffrey Barrett, Sonia Goncalves, Ewan Harrison, David K. Jackson, Ian Johnston, Dominic Kwiatkowski, Cordelia Langford, John Sillitoe on behalf of the Wellcome Sanger Institute COVID-19 Surveillance Team |
| EPI_ISL_3313891 | Berkshire and Surrey Pathology Services Lighthouse Laboratory | Wellcome Sanger Institute for the COVID-19 Genomics UK (COG-UK) Consortium | Berkshire and Surrey Pathology Services Lighthouse Laboratory and Alex Alderton, Roberto Amato, Jeffrey Barrett, Sonia Goncalves, Ewan Harrison, David K. Jackson, Ian Johnston, Dominic Kwiatkowski, Cordelia Langford, John Sillitoe on behalf of the Wellcome Sanger Institute COVID-19 Surveillance Team |
| EPI_ISL_3313892 | Lighthouse Lab in Milton Keynes | Wellcome Sanger Institute for the COVID-19 Genomics UK (COG-UK) Consortium | The Lighthouse Lab in Milton Keynes and Alex Alderton, Roberto Amato, Jeffrey Barrett, Sonia Goncalves, Ewan Harrison, David K. Jackson, Ian Johnston, Dominic Kwiatkowski, Cordelia Langford, John Sillitoe on behalf of the Wellcome Sanger Institute COVID-19 Surveillance Team |
| EPI_ISL_3313893, | Berkshire and Surrey Pathology Services Lighthouse Laboratory | Wellcome Sanger Institute for the COVID-19 Genomics UK (COG-UK) | Berkshire and Surrey Pathology Services Lighthouse Laboratory and Alex Alderton, Roberto Amato, Jeffrey Barrett, Sonia Goncalves, Ewan Harrison, David K. Jackson, Ian Johnston, Dominic Kwiatkowski, Cordelia Langford, John Sillitoe on behalf |

| | | | |
|---|---|---|---|
| EPI_ISL_3313894, EPI_ISL_3313895 | | Consortium | of the Wellcome Sanger Institute COVID-19 Surveillance Team |
| EPI_ISL_3313896, EPI_ISL_3313897 | Lighthouse Lab in Milton Keynes | Wellcome Sanger Institute for the COVID-19 Genomics UK (COG-UK) Consortium | The Lighthouse Lab in Milton Keynes and Alex Alderton, Roberto Amato, Jeffrey Barrett, Sonia Goncalves, Ewan Harrison, David K. Jackson, Ian Johnston, Dominic Kwiatkowski, Cordelia Langford, John Sillitoe on behalf of the Wellcome Sanger Institute COVID-19 Surveillance Team |
| EPI_ISL_3313898 | Berkshire and Surrey Pathology Services Lighthouse Laboratory | Wellcome Sanger Institute for the COVID-19 Genomics UK (COG-UK) Consortium | Berkshire and Surrey Pathology Services Lighthouse Laboratory and Alex Alderton, Roberto Amato, Jeffrey Barrett, Sonia Goncalves, Ewan Harrison, David K. Jackson, Ian Johnston, Dominic Kwiatkowski, Cordelia Langford, John Sillitoe on behalf of the Wellcome Sanger Institute COVID-19 Surveillance Team |
| EPI_ISL_3313899 | Lighthouse Lab in Glasgow | Wellcome Sanger Institute for the COVID-19 Genomics UK (COG-UK) Consortium | Harper VanSteenhouse, Yumi Kasai, David Gray, Carol Clugston, Anna Dominiczak and Alex Alderton, Roberto Amato, Jeffrey Barrett, Sonia Goncalves, Ewan Harrison, David K. Jackson, Ian Johnston, Dominic Kwiatkowski, Cordelia Langford, John Sillitoe on behalf of the Wellcome Sanger Institute COVID-19 Surveillance Team |
| EPI_ISL_3313900 | Berkshire and Surrey Pathology Services Lighthouse Laboratory | Wellcome Sanger Institute for the COVID-19 Genomics UK (COG-UK) Consortium | Berkshire and Surrey Pathology Services Lighthouse Laboratory and Alex Alderton, Roberto Amato, Jeffrey Barrett, Sonia Goncalves, Ewan Harrison, David K. Jackson, Ian Johnston, Dominic Kwiatkowski, Cordelia Langford, John Sillitoe on behalf of the Wellcome Sanger Institute COVID-19 Surveillance Team |
| EPI_ISL_3313901, EPI_ISL_3313902 | Lighthouse Lab in Milton Keynes | Wellcome Sanger Institute for the COVID-19 Genomics UK (COG-UK) Consortium | The Lighthouse Lab in Milton Keynes and Alex Alderton, Roberto Amato, Jeffrey Barrett, Sonia Goncalves, Ewan Harrison, David K. Jackson, Ian Johnston, Dominic Kwiatkowski, Cordelia Langford, John Sillitoe on behalf of the Wellcome Sanger Institute COVID-19 Surveillance Team |
| EPI_ISL_3313903, EPI_ISL_3313904 | Berkshire and Surrey Pathology Services Lighthouse Laboratory | Wellcome Sanger Institute for the COVID-19 Genomics UK (COG-UK) Consortium | Berkshire and Surrey Pathology Services Lighthouse Laboratory and Alex Alderton, Roberto Amato, Jeffrey Barrett, Sonia Goncalves, Ewan Harrison, David K. Jackson, Ian Johnston, Dominic Kwiatkowski, Cordelia Langford, John Sillitoe on behalf of the Wellcome Sanger Institute COVID-19 Surveillance Team |
| EPI_ISL_3313905, EPI_ISL_3313906 | Lighthouse Lab in Milton Keynes | Wellcome Sanger Institute for the COVID-19 Genomics UK (COG-UK) Consortium | The Lighthouse Lab in Milton Keynes and Alex Alderton, Roberto Amato, Jeffrey Barrett, Sonia Goncalves, Ewan Harrison, David K. Jackson, Ian Johnston, Dominic Kwiatkowski, Cordelia Langford, John Sillitoe on behalf of the Wellcome Sanger Institute COVID-19 Surveillance Team |
| EPI_ISL_3313907 | Berkshire and Surrey Pathology Services Lighthouse Laboratory | Wellcome Sanger Institute for the COVID-19 Genomics UK (COG-UK) Consortium | Berkshire and Surrey Pathology Services Lighthouse Laboratory and Alex Alderton, Roberto Amato, Jeffrey Barrett, Sonia Goncalves, Ewan Harrison, David K. Jackson, Ian Johnston, Dominic Kwiatkowski, Cordelia Langford, John Sillitoe on behalf of the Wellcome Sanger Institute COVID-19 Surveillance Team |
| EPI_ISL_3313908 | Lighthouse Lab in Glasgow | Wellcome Sanger Institute for the COVID-19 Genomics UK (COG-UK) Consortium | Harper VanSteenhouse, Yumi Kasai, David Gray, Carol Clugston, Anna Dominiczak and Alex Alderton, Roberto Amato, Jeffrey Barrett, Sonia Goncalves, Ewan Harrison, David K. Jackson, Ian Johnston, Dominic Kwiatkowski, Cordelia Langford, John Sillitoe on behalf of the Wellcome Sanger Institute COVID-19 Surveillance Team |
| EPI_ISL_3313909, EPI_ISL_3313910, EPI_ISL_3313911 | Lighthouse Lab in Milton Keynes | Wellcome Sanger Institute for the COVID-19 Genomics UK (COG-UK) Consortium | The Lighthouse Lab in Milton Keynes and Alex Alderton, Roberto Amato, Jeffrey Barrett, Sonia Goncalves, Ewan Harrison, David K. Jackson, Ian Johnston, Dominic Kwiatkowski, Cordelia Langford, John Sillitoe on behalf of the Wellcome Sanger Institute COVID-19 Surveillance Team |
| EPI_ISL_3313912 | Berkshire and Surrey Pathology Services Lighthouse Laboratory | Wellcome Sanger Institute for the COVID-19 Genomics UK (COG-UK) Consortium | Berkshire and Surrey Pathology Services Lighthouse Laboratory and Alex Alderton, Roberto Amato, Jeffrey Barrett, Sonia Goncalves, Ewan Harrison, David K. Jackson, Ian Johnston, Dominic Kwiatkowski, Cordelia Langford, John Sillitoe on behalf of the Wellcome Sanger Institute COVID-19 Surveillance Team |
| EPI_ISL_3313913, EPI_ISL_3313914 | Lighthouse Lab in Glasgow | Wellcome Sanger Institute for the COVID-19 Genomics UK (COG-UK) Consortium | Harper VanSteenhouse, Yumi Kasai, David Gray, Carol Clugston, Anna Dominiczak and Alex Alderton, Roberto Amato, Jeffrey Barrett, Sonia Goncalves, Ewan Harrison, David K. Jackson, Ian Johnston, Dominic Kwiatkowski, Cordelia Langford, John Sillitoe on behalf of the Wellcome Sanger Institute COVID-19 Surveillance Team |
| EPI_ISL_3313915, EPI_ISL_3313916, EPI_ISL_3313917, EPI_ISL_3313918 | Lighthouse Lab in Milton Keynes | Wellcome Sanger Institute for the COVID-19 Genomics UK (COG-UK) Consortium | The Lighthouse Lab in Milton Keynes and Alex Alderton, Roberto Amato, Jeffrey Barrett, Sonia Goncalves, Ewan Harrison, David K. Jackson, Ian Johnston, Dominic Kwiatkowski, Cordelia Langford, John Sillitoe on behalf of the Wellcome Sanger Institute COVID-19 Surveillance Team |
| EPI_ISL_3313919 | Berkshire and Surrey Pathology Services Lighthouse Laboratory | Wellcome Sanger Institute for the COVID-19 Genomics UK (COG-UK) Consortium | Berkshire and Surrey Pathology Services Lighthouse Laboratory and Alex Alderton, Roberto Amato, Jeffrey Barrett, Sonia Goncalves, Ewan Harrison, David K. Jackson, Ian Johnston, Dominic Kwiatkowski, Cordelia Langford, John Sillitoe on behalf of the Wellcome Sanger Institute COVID-19 Surveillance Team |
| EPI_ISL_3313920, EPI_ISL_3313921, EPI_ISL_3313922 | Lighthouse Lab in Milton Keynes | Wellcome Sanger Institute for the COVID-19 Genomics UK (COG-UK) Consortium | The Lighthouse Lab in Milton Keynes and Alex Alderton, Roberto Amato, Jeffrey Barrett, Sonia Goncalves, Ewan Harrison, David K. Jackson, Ian Johnston, Dominic Kwiatkowski, Cordelia Langford, John Sillitoe on behalf of the Wellcome Sanger Institute COVID-19 Surveillance Team |
| EPI_ISL_3313923 | Lighthouse Lab in Glasgow | Wellcome Sanger Institute for the COVID-19 Genomics UK (COG-UK) Consortium | Harper VanSteenhouse, Yumi Kasai, David Gray, Carol Clugston, Anna Dominiczak and Alex Alderton, Roberto Amato, Jeffrey Barrett, Sonia Goncalves, Ewan Harrison, David K. Jackson, Ian Johnston, Dominic Kwiatkowski, Cordelia Langford, John Sillitoe on behalf of the Wellcome Sanger Institute COVID-19 Surveillance Team |
| EPI_ISL_3313924, EPI_ISL_3313925, EPI_ISL_3313926, EPI_ISL_3313927 | Lighthouse Lab in Milton Keynes | Wellcome Sanger Institute for the COVID-19 Genomics UK (COG-UK) Consortium | The Lighthouse Lab in Milton Keynes and Alex Alderton, Roberto Amato, Jeffrey Barrett, Sonia Goncalves, Ewan Harrison, David K. Jackson, Ian Johnston, Dominic Kwiatkowski, Cordelia Langford, John Sillitoe on behalf of the Wellcome Sanger Institute COVID-19 Surveillance Team |
| EPI_ISL_3313928 | Berkshire and Surrey Pathology Services Lighthouse Laboratory | Wellcome Sanger Institute for the COVID-19 Genomics UK (COG-UK) Consortium | Berkshire and Surrey Pathology Services Lighthouse Laboratory and Alex Alderton, Roberto Amato, Jeffrey Barrett, Sonia Goncalves, Ewan Harrison, David K. Jackson, Ian Johnston, Dominic Kwiatkowski, Cordelia Langford, John Sillitoe on behalf of the Wellcome Sanger Institute COVID-19 Surveillance Team |
| EPI_ISL_3313929, EPI_ISL_3313930, EPI_ISL_3313931 | Lighthouse Lab in Milton Keynes | Wellcome Sanger Institute for the COVID-19 Genomics UK (COG-UK) Consortium | The Lighthouse Lab in Milton Keynes and Alex Alderton, Roberto Amato, Jeffrey Barrett, Sonia Goncalves, Ewan Harrison, David K. Jackson, Ian Johnston, Dominic Kwiatkowski, Cordelia Langford, John Sillitoe on behalf of the Wellcome Sanger Institute COVID-19 Surveillance Team |
| EPI_ISL_3313932 | Lighthouse Lab in Glasgow | Wellcome Sanger Institute for the COVID-19 Genomics UK (COG-UK) Consortium | Harper VanSteenhouse, Yumi Kasai, David Gray, Carol Clugston, Anna Dominiczak and Alex Alderton, Roberto Amato, Jeffrey Barrett, Sonia Goncalves, Ewan Harrison, David K. Jackson, Ian Johnston, Dominic Kwiatkowski, Cordelia Langford, John Sillitoe on behalf of the Wellcome Sanger Institute COVID-19 Surveillance Team |
| EPI_ISL_3313933 | Lighthouse Lab in Milton Keynes | Wellcome Sanger Institute for the COVID-19 Genomics UK (COG-UK) Consortium | The Lighthouse Lab in Milton Keynes and Alex Alderton, Roberto Amato, Jeffrey Barrett, Sonia Goncalves, Ewan Harrison, David K. Jackson, Ian Johnston, Dominic Kwiatkowski, Cordelia Langford, John Sillitoe on behalf of the Wellcome Sanger Institute COVID-19 Surveillance Team |
| EPI_ISL_3313934, EPI_ISL_3313935 | Lighthouse Lab in Glasgow | Wellcome Sanger Institute for the COVID-19 Genomics UK (COG-UK) Consortium | Harper VanSteenhouse, Yumi Kasai, David Gray, Carol Clugston, Anna Dominiczak and Alex Alderton, Roberto Amato, Jeffrey Barrett, Sonia Goncalves, Ewan Harrison, David K. Jackson, Ian Johnston, Dominic Kwiatkowski, Cordelia Langford, John Sillitoe on behalf of the Wellcome Sanger Institute COVID-19 Surveillance Team |
| EPI_ISL_3313936 | Lighthouse Lab in Milton Keynes | Wellcome Sanger Institute for the COVID-19 Genomics UK (COG-UK) Consortium | The Lighthouse Lab in Milton Keynes and Alex Alderton, Roberto Amato, Jeffrey Barrett, Sonia Goncalves, Ewan Harrison, David K. Jackson, Ian Johnston, Dominic Kwiatkowski, Cordelia Langford, John Sillitoe on behalf of the Wellcome Sanger Institute COVID-19 Surveillance Team |
| EPI_ISL_3313937, EPI_ISL_3313938 | Lighthouse Lab in Glasgow | Wellcome Sanger Institute for the COVID-19 Genomics UK (COG-UK) Consortium | Harper VanSteenhouse, Yumi Kasai, David Gray, Carol Clugston, Anna Dominiczak and Alex Alderton, Roberto Amato, Jeffrey Barrett, Sonia Goncalves, Ewan Harrison, David K. Jackson, Ian Johnston, Dominic Kwiatkowski, Cordelia Langford, John Sillitoe on behalf of the Wellcome Sanger Institute COVID-19 Surveillance Team |
| EPI_ISL_3313939 | Lighthouse Lab in Milton Keynes | Wellcome Sanger Institute for the COVID-19 Genomics UK (COG-UK) Consortium | The Lighthouse Lab in Milton Keynes and Alex Alderton, Roberto Amato, Jeffrey Barrett, Sonia Goncalves, Ewan Harrison, David K. Jackson, Ian Johnston, Dominic Kwiatkowski, Cordelia Langford, John Sillitoe on behalf of the Wellcome Sanger Institute COVID-19 Surveillance Team |
| EPI_ISL_3313940, EPI_ISL_3313941 | Berkshire and Surrey Pathology Services Lighthouse Laboratory | Wellcome Sanger Institute for the COVID-19 Genomics UK (COG-UK) Consortium | Berkshire and Surrey Pathology Services Lighthouse Laboratory and Alex Alderton, Roberto Amato, Jeffrey Barrett, Sonia Goncalves, Ewan Harrison, David K. Jackson, Ian Johnston, Dominic Kwiatkowski, Cordelia Langford, John Sillitoe on behalf of the Wellcome Sanger Institute COVID-19 Surveillance Team |
| EPI_ISL_3313942 | Lighthouse Lab in Milton Keynes | Wellcome Sanger Institute for the COVID-19 Genomics UK (COG-UK) Consortium | The Lighthouse Lab in Milton Keynes and Alex Alderton, Roberto Amato, Jeffrey Barrett, Sonia Goncalves, Ewan Harrison, David K. Jackson, Ian Johnston, Dominic Kwiatkowski, Cordelia Langford, John Sillitoe on behalf of the Wellcome Sanger Institute COVID-19 Surveillance Team |
| EPI_ISL_3313943 | Berkshire and Surrey Pathology Services Lighthouse Laboratory | Wellcome Sanger Institute for the COVID-19 Genomics UK (COG-UK) Consortium | Berkshire and Surrey Pathology Services Lighthouse Laboratory and Alex Alderton, Roberto Amato, Jeffrey Barrett, Sonia Goncalves, Ewan Harrison, David K. Jackson, Ian Johnston, Dominic Kwiatkowski, Cordelia Langford, John Sillitoe on behalf of the Wellcome Sanger Institute COVID-19 Surveillance Team |
| EPI_ISL_3313944 | Lighthouse Lab in Glasgow | Wellcome Sanger Institute for the COVID-19 Genomics UK (COG-UK) Consortium | Harper VanSteenhouse, Yumi Kasai, David Gray, Carol Clugston, Anna Dominiczak and Alex Alderton, Roberto Amato, Jeffrey Barrett, Sonia Goncalves, Ewan Harrison, David K. Jackson, Ian Johnston, Dominic Kwiatkowski, Cordelia Langford, John Sillitoe on behalf of the Wellcome Sanger Institute COVID-19 Surveillance Team |
| EPI_ISL_3313945, EPI_ISL_3313946, EPI_ISL_3313947 | Lighthouse Lab in Milton Keynes | Wellcome Sanger Institute for the COVID-19 Genomics UK (COG-UK) Consortium | The Lighthouse Lab in Milton Keynes and Alex Alderton, Roberto Amato, Jeffrey Barrett, Sonia Goncalves, Ewan Harrison, David K. Jackson, Ian Johnston, Dominic Kwiatkowski, Cordelia Langford, John Sillitoe on behalf of the Wellcome Sanger Institute COVID-19 Surveillance Team |
| EPI_ISL_3313948 | Lighthouse Lab in Glasgow | Wellcome Sanger Institute for the COVID-19 Genomics UK (COG-UK) Consortium | Harper VanSteenhouse, Yumi Kasai, David Gray, Carol Clugston, Anna Dominiczak and Alex Alderton, Roberto Amato, Jeffrey Barrett, Sonia Goncalves, Ewan Harrison, David K. Jackson, Ian Johnston, Dominic Kwiatkowski, Cordelia Langford, John Sillitoe on behalf of the Wellcome Sanger Institute COVID-19 Surveillance Team |
| EPI_ISL_3313949, EPI_ISL_3313950, EPI_ISL_3313951 | Berkshire and Surrey Pathology Services Lighthouse Laboratory | Wellcome Sanger Institute for the COVID-19 Genomics UK (COG-UK) Consortium | Berkshire and Surrey Pathology Services Lighthouse Laboratory and Alex Alderton, Roberto Amato, Jeffrey Barrett, Sonia Goncalves, Ewan Harrison, David K. Jackson, Ian Johnston, Dominic Kwiatkowski, Cordelia Langford, John Sillitoe on behalf of the Wellcome Sanger Institute COVID-19 Surveillance Team |
| EPI_ISL_3313952 | Lighthouse Lab in Milton Keynes | Wellcome Sanger Institute for the COVID-19 Genomics UK (COG-UK) Consortium | The Lighthouse Lab in Milton Keynes and Alex Alderton, Roberto Amato, Jeffrey Barrett, Sonia Goncalves, Ewan Harrison, David K. Jackson, Ian Johnston, Dominic Kwiatkowski, Cordelia Langford, John Sillitoe on behalf of the Wellcome Sanger Institute COVID-19 Surveillance Team |
| EPI_ISL_3313953, EPI_ISL_3313954, EPI_ISL_3313955 | Berkshire and Surrey Pathology Services Lighthouse Laboratory | Wellcome Sanger Institute for the COVID-19 Genomics UK (COG-UK) Consortium | Berkshire and Surrey Pathology Services Lighthouse Laboratory and Alex Alderton, Roberto Amato, Jeffrey Barrett, Sonia Goncalves, Ewan Harrison, David K. Jackson, Ian Johnston, Dominic Kwiatkowski, Cordelia Langford, John Sillitoe on behalf of the Wellcome Sanger Institute COVID-19 Surveillance Team |
| EPI_ISL_3313956, EPI_ISL_3313957 | Lighthouse Lab in Milton Keynes | Wellcome Sanger Institute for the COVID-19 Genomics UK (COG-UK) Consortium | The Lighthouse Lab in Milton Keynes and Alex Alderton, Roberto Amato, Jeffrey Barrett, Sonia Goncalves, Ewan Harrison, David K. Jackson, Ian Johnston, Dominic Kwiatkowski, Cordelia Langford, John Sillitoe on behalf of the Wellcome Sanger Institute COVID-19 Surveillance Team |
| EPI_ISL_3313958 | Berkshire and Surrey Pathology Services Lighthouse Laboratory | Wellcome Sanger Institute for the COVID-19 Genomics UK (COG-UK) Consortium | Berkshire and Surrey Pathology Services Lighthouse Laboratory and Alex Alderton, Roberto Amato, Jeffrey Barrett, Sonia Goncalves, Ewan Harrison, David K. Jackson, Ian Johnston, Dominic Kwiatkowski, Cordelia Langford, John Sillitoe on behalf of the Wellcome Sanger Institute COVID-19 Surveillance Team |
| EPI_ISL_3313959, EPI_ISL_3313960, EPI_ISL_3313961, EPI_ISL_3313962 | Lighthouse Lab in Milton Keynes | Wellcome Sanger Institute for the COVID-19 Genomics UK (COG-UK) Consortium | The Lighthouse Lab in Milton Keynes and Alex Alderton, Roberto Amato, Jeffrey Barrett, Sonia Goncalves, Ewan Harrison, David K. Jackson, Ian Johnston, Dominic Kwiatkowski, Cordelia Langford, John Sillitoe on behalf of the Wellcome Sanger Institute COVID-19 Surveillance Team |
| EPI_ISL_3313963, EPI_ISL_3313964 | Berkshire and Surrey Pathology Services Lighthouse Laboratory | Wellcome Sanger Institute for the COVID-19 Genomics UK (COG-UK) Consortium | Berkshire and Surrey Pathology Services Lighthouse Laboratory and Alex Alderton, Roberto Amato, Jeffrey Barrett, Sonia Goncalves, Ewan Harrison, David K. Jackson, Ian Johnston, Dominic Kwiatkowski, Cordelia Langford, John Sillitoe on behalf of the Wellcome Sanger Institute COVID-19 Surveillance Team |
| EPI_ISL_3313965, EPI_ISL_3313966 | Lighthouse Lab in Milton Keynes | Wellcome Sanger Institute for the COVID-19 Genomics UK (COG-UK) Consortium | The Lighthouse Lab in Milton Keynes and Alex Alderton, Roberto Amato, Jeffrey Barrett, Sonia Goncalves, Ewan Harrison, David K. Jackson, Ian Johnston, Dominic Kwiatkowski, Cordelia Langford, John Sillitoe on behalf of the Wellcome Sanger Institute COVID-19 Surveillance Team |

| | | | |
|---|---|---|---|
| EPI_ISL_3313967 | Berkshire and Surrey Pathology Services Lighthouse Laboratory | Wellcome Sanger Institute for the COVID-19 Genomics UK (COG-UK) Consortium | Berkshire and Surrey Pathology Services Lighthouse Laboratory and Alex Alderton, Roberto Amato, Jeffrey Barrett, Sonia Goncalves, Ewan Harrison, David K. Jackson, Ian Johnston, Dominic Kwiatkowski, Cordelia Langford, John Sillitoe on behalf of the Wellcome Sanger Institute COVID-19 Surveillance Team |
| EPI_ISL_3313968 | Lighthouse Lab in Milton Keynes | Wellcome Sanger Institute for the COVID-19 Genomics UK (COG-UK) Consortium | The Lighthouse Lab in Milton Keynes and Alex Alderton, Roberto Amato, Jeffrey Barrett, Sonia Goncalves, Ewan Harrison, David K. Jackson, Ian Johnston, Dominic Kwiatkowski, Cordelia Langford, John Sillitoe on behalf of the Wellcome Sanger Institute COVID-19 Surveillance Team |
| EPI_ISL_3313969, EPI_ISL_3313970 | Lighthouse Lab in Glasgow | Wellcome Sanger Institute for the COVID-19 Genomics UK (COG-UK) Consortium | Harper VanSteenhouse, Yumi Kasai, David Gray, Carol Clugston, Anna Dominiczak and Alex Alderton, Roberto Amato, Jeffrey Barrett, Sonia Goncalves, Ewan Harrison, David K. Jackson, Ian Johnston, Dominic Kwiatkowski, Cordelia Langford, John Sillitoe on behalf of the Wellcome Sanger Institute COVID-19 Surveillance Team |
| EPI_ISL_3313971, EPI_ISL_3313972, EPI_ISL_3313973, EPI_ISL_3313974 | Lighthouse Lab in Milton Keynes | Wellcome Sanger Institute for the COVID-19 Genomics UK (COG-UK) Consortium | The Lighthouse Lab in Milton Keynes and Alex Alderton, Roberto Amato, Jeffrey Barrett, Sonia Goncalves, Ewan Harrison, David K. Jackson, Ian Johnston, Dominic Kwiatkowski, Cordelia Langford, John Sillitoe on behalf of the Wellcome Sanger Institute COVID-19 Surveillance Team |
| EPI_ISL_3313975, EPI_ISL_3313976 | Lighthouse Lab in Glasgow | Wellcome Sanger Institute for the COVID-19 Genomics UK (COG-UK) Consortium | Harper VanSteenhouse, Yumi Kasai, David Gray, Carol Clugston, Anna Dominiczak and Alex Alderton, Roberto Amato, Jeffrey Barrett, Sonia Goncalves, Ewan Harrison, David K. Jackson, Ian Johnston, Dominic Kwiatkowski, Cordelia Langford, John Sillitoe on behalf of the Wellcome Sanger Institute COVID-19 Surveillance Team |
| EPI_ISL_3313977 | Berkshire and Surrey Pathology Services Lighthouse Laboratory | Wellcome Sanger Institute for the COVID-19 Genomics UK (COG-UK) Consortium | Berkshire and Surrey Pathology Services Lighthouse Laboratory and Alex Alderton, Roberto Amato, Jeffrey Barrett, Sonia Goncalves, Ewan Harrison, David K. Jackson, Ian Johnston, Dominic Kwiatkowski, Cordelia Langford, John Sillitoe on behalf of the Wellcome Sanger Institute COVID-19 Surveillance Team |
| EPI_ISL_3313978, EPI_ISL_3313979, EPI_ISL_3313980 | Lighthouse Lab in Milton Keynes | Wellcome Sanger Institute for the COVID-19 Genomics UK (COG-UK) Consortium | The Lighthouse Lab in Milton Keynes and Alex Alderton, Roberto Amato, Jeffrey Barrett, Sonia Goncalves, Ewan Harrison, David K. Jackson, Ian Johnston, Dominic Kwiatkowski, Cordelia Langford, John Sillitoe on behalf of the Wellcome Sanger Institute COVID-19 Surveillance Team |
| EPI_ISL_3313981 | Lighthouse Lab in Glasgow | Wellcome Sanger Institute for the COVID-19 Genomics UK (COG-UK) Consortium | Harper VanSteenhouse, Yumi Kasai, David Gray, Carol Clugston, Anna Dominiczak and Alex Alderton, Roberto Amato, Jeffrey Barrett, Sonia Goncalves, Ewan Harrison, David K. Jackson, Ian Johnston, Dominic Kwiatkowski, Cordelia Langford, John Sillitoe on behalf of the Wellcome Sanger Institute COVID-19 Surveillance Team |
| EPI_ISL_3313982 | Berkshire and Surrey Pathology Services Lighthouse Laboratory | Wellcome Sanger Institute for the COVID-19 Genomics UK (COG-UK) Consortium | Berkshire and Surrey Pathology Services Lighthouse Laboratory and Alex Alderton, Roberto Amato, Jeffrey Barrett, Sonia Goncalves, Ewan Harrison, David K. Jackson, Ian Johnston, Dominic Kwiatkowski, Cordelia Langford, John Sillitoe on behalf of the Wellcome Sanger Institute COVID-19 Surveillance Team |
| EPI_ISL_3313983 | Lighthouse Lab in Milton Keynes | Wellcome Sanger Institute for the COVID-19 Genomics UK (COG-UK) Consortium | The Lighthouse Lab in Milton Keynes and Alex Alderton, Roberto Amato, Jeffrey Barrett, Sonia Goncalves, Ewan Harrison, David K. Jackson, Ian Johnston, Dominic Kwiatkowski, Cordelia Langford, John Sillitoe on behalf of the Wellcome Sanger Institute COVID-19 Surveillance Team |
| EPI_ISL_3313984 | Lighthouse Lab in Glasgow | Wellcome Sanger Institute for the COVID-19 Genomics UK (COG-UK) Consortium | Harper VanSteenhouse, Yumi Kasai, David Gray, Carol Clugston, Anna Dominiczak and Alex Alderton, Roberto Amato, Jeffrey Barrett, Sonia Goncalves, Ewan Harrison, David K. Jackson, Ian Johnston, Dominic Kwiatkowski, Cordelia Langford, John Sillitoe on behalf of the Wellcome Sanger Institute COVID-19 Surveillance Team |
| EPI_ISL_3313985 | Lighthouse Lab in Milton Keynes | Wellcome Sanger Institute for the COVID-19 Genomics UK (COG-UK) Consortium | The Lighthouse Lab in Milton Keynes and Alex Alderton, Roberto Amato, Jeffrey Barrett, Sonia Goncalves, Ewan Harrison, David K. Jackson, Ian Johnston, Dominic Kwiatkowski, Cordelia Langford, John Sillitoe on behalf of the Wellcome Sanger Institute COVID-19 Surveillance Team |
| EPI_ISL_3313986, EPI_ISL_3313987 | Lighthouse Lab in Glasgow | Wellcome Sanger Institute for the COVID-19 Genomics UK (COG-UK) Consortium | Harper VanSteenhouse, Yumi Kasai, David Gray, Carol Clugston, Anna Dominiczak and Alex Alderton, Roberto Amato, Jeffrey Barrett, Sonia Goncalves, Ewan Harrison, David K. Jackson, Ian Johnston, Dominic Kwiatkowski, Cordelia Langford, John Sillitoe on behalf of the Wellcome Sanger Institute COVID-19 Surveillance Team |
| EPI_ISL_3313988, EPI_ISL_3313989 | Lighthouse Lab in Milton Keynes | Wellcome Sanger Institute for the COVID-19 Genomics UK (COG-UK) Consortium | The Lighthouse Lab in Milton Keynes and Alex Alderton, Roberto Amato, Jeffrey Barrett, Sonia Goncalves, Ewan Harrison, David K. Jackson, Ian Johnston, Dominic Kwiatkowski, Cordelia Langford, John Sillitoe on behalf of the Wellcome Sanger Institute COVID-19 Surveillance Team |
| EPI_ISL_3313990 | Berkshire and Surrey Pathology Services Lighthouse Laboratory | Wellcome Sanger Institute for the COVID-19 Genomics UK (COG-UK) Consortium | Berkshire and Surrey Pathology Services Lighthouse Laboratory and Alex Alderton, Roberto Amato, Jeffrey Barrett, Sonia Goncalves, Ewan Harrison, David K. Jackson, Ian Johnston, Dominic Kwiatkowski, Cordelia Langford, John Sillitoe on behalf of the Wellcome Sanger Institute COVID-19 Surveillance Team |
| EPI_ISL_3313991, EPI_ISL_3313992, EPI_ISL_3313993 | Lighthouse Lab in Milton Keynes | Wellcome Sanger Institute for the COVID-19 Genomics UK (COG-UK) Consortium | The Lighthouse Lab in Milton Keynes and Alex Alderton, Roberto Amato, Jeffrey Barrett, Sonia Goncalves, Ewan Harrison, David K. Jackson, Ian Johnston, Dominic Kwiatkowski, Cordelia Langford, John Sillitoe on behalf of the Wellcome Sanger Institute COVID-19 Surveillance Team |
| EPI_ISL_3313994 | Berkshire and Surrey Pathology Services Lighthouse Laboratory | Wellcome Sanger Institute for the COVID-19 Genomics UK (COG-UK) Consortium | Berkshire and Surrey Pathology Services Lighthouse Laboratory and Alex Alderton, Roberto Amato, Jeffrey Barrett, Sonia Goncalves, Ewan Harrison, David K. Jackson, Ian Johnston, Dominic Kwiatkowski, Cordelia Langford, John Sillitoe on behalf of the Wellcome Sanger Institute COVID-19 Surveillance Team |
| EPI_ISL_3313995, EPI_ISL_3313996 | Lighthouse Lab in Glasgow | Wellcome Sanger Institute for the COVID-19 Genomics UK (COG-UK) Consortium | Harper VanSteenhouse, Yumi Kasai, David Gray, Carol Clugston, Anna Dominiczak and Alex Alderton, Roberto Amato, Jeffrey Barrett, Sonia Goncalves, Ewan Harrison, David K. Jackson, Ian Johnston, Dominic Kwiatkowski, Cordelia Langford, John Sillitoe on behalf of the Wellcome Sanger Institute COVID-19 Surveillance Team |
| EPI_ISL_3313997, EPI_ISL_3313998, EPI_ISL_3313999, EPI_ISL_3314000, EPI_ISL_3314001, EPI_ISL_3314002 | Lighthouse Lab in Milton Keynes | Wellcome Sanger Institute for the COVID-19 Genomics UK (COG-UK) Consortium | The Lighthouse Lab in Milton Keynes and Alex Alderton, Roberto Amato, Jeffrey Barrett, Sonia Goncalves, Ewan Harrison, David K. Jackson, Ian Johnston, Dominic Kwiatkowski, Cordelia Langford, John Sillitoe on behalf of the Wellcome Sanger Institute COVID-19 Surveillance Team |
| EPI_ISL_3314003 | Berkshire and Surrey Pathology Services Lighthouse Laboratory | Wellcome Sanger Institute for the COVID-19 Genomics UK (COG-UK) Consortium | Berkshire and Surrey Pathology Services Lighthouse Laboratory and Alex Alderton, Roberto Amato, Jeffrey Barrett, Sonia Goncalves, Ewan Harrison, David K. Jackson, Ian Johnston, Dominic Kwiatkowski, Cordelia Langford, John Sillitoe on behalf of the Wellcome Sanger Institute COVID-19 Surveillance Team |
| EPI_ISL_3314004, EPI_ISL_3314005 | Lighthouse Lab in Milton Keynes | Wellcome Sanger Institute for the COVID-19 Genomics UK (COG-UK) Consortium | The Lighthouse Lab in Milton Keynes and Alex Alderton, Roberto Amato, Jeffrey Barrett, Sonia Goncalves, Ewan Harrison, David K. Jackson, Ian Johnston, Dominic Kwiatkowski, Cordelia Langford, John Sillitoe on behalf of the Wellcome Sanger Institute COVID-19 Surveillance Team |
| EPI_ISL_3314006, EPI_ISL_3314007 | Berkshire and Surrey Pathology Services Lighthouse Laboratory | Wellcome Sanger Institute for the COVID-19 Genomics UK (COG-UK) Consortium | Berkshire and Surrey Pathology Services Lighthouse Laboratory and Alex Alderton, Roberto Amato, Jeffrey Barrett, Sonia Goncalves, Ewan Harrison, David K. Jackson, Ian Johnston, Dominic Kwiatkowski, Cordelia Langford, John Sillitoe on behalf of the Wellcome Sanger Institute COVID-19 Surveillance Team |
| EPI_ISL_3314008 | Lighthouse Lab in Milton Keynes | Wellcome Sanger Institute for the COVID-19 Genomics UK (COG-UK) Consortium | The Lighthouse Lab in Milton Keynes and Alex Alderton, Roberto Amato, Jeffrey Barrett, Sonia Goncalves, Ewan Harrison, David K. Jackson, Ian Johnston, Dominic Kwiatkowski, Cordelia Langford, John Sillitoe on behalf of the Wellcome Sanger Institute COVID-19 Surveillance Team |
| EPI_ISL_3314009 | Lighthouse Lab in Glasgow | Wellcome Sanger Institute for the COVID-19 Genomics UK (COG-UK) Consortium | Harper VanSteenhouse, Yumi Kasai, David Gray, Carol Clugston, Anna Dominiczak and Alex Alderton, Roberto Amato, Jeffrey Barrett, Sonia Goncalves, Ewan Harrison, David K. Jackson, Ian Johnston, Dominic Kwiatkowski, Cordelia Langford, John Sillitoe on behalf of the Wellcome Sanger Institute COVID-19 Surveillance Team |
| EPI_ISL_3314010, EPI_ISL_3314011, EPI_ISL_3314012 | Lighthouse Lab in Milton Keynes | Wellcome Sanger Institute for the COVID-19 Genomics UK (COG-UK) Consortium | The Lighthouse Lab in Milton Keynes and Alex Alderton, Roberto Amato, Jeffrey Barrett, Sonia Goncalves, Ewan Harrison, David K. Jackson, Ian Johnston, Dominic Kwiatkowski, Cordelia Langford, John Sillitoe on behalf of the Wellcome Sanger Institute COVID-19 Surveillance Team |
| EPI_ISL_3314013, EPI_ISL_3314014 | Lighthouse Lab in Glasgow | Wellcome Sanger Institute for the COVID-19 Genomics UK (COG-UK) Consortium | Harper VanSteenhouse, Yumi Kasai, David Gray, Carol Clugston, Anna Dominiczak and Alex Alderton, Roberto Amato, Jeffrey Barrett, Sonia Goncalves, Ewan Harrison, David K. Jackson, Ian Johnston, Dominic Kwiatkowski, Cordelia Langford, John Sillitoe on behalf of the Wellcome Sanger Institute COVID-19 Surveillance Team |
| EPI_ISL_3314015 | Berkshire and Surrey Pathology Services Lighthouse Laboratory | Wellcome Sanger Institute for the COVID-19 Genomics UK (COG-UK) Consortium | Berkshire and Surrey Pathology Services Lighthouse Laboratory and Alex Alderton, Roberto Amato, Jeffrey Barrett, Sonia Goncalves, Ewan Harrison, David K. Jackson, Ian Johnston, Dominic Kwiatkowski, Cordelia Langford, John Sillitoe on behalf of the Wellcome Sanger Institute COVID-19 Surveillance Team |
| EPI_ISL_3314016, EPI_ISL_3314017, EPI_ISL_3314018, EPI_ISL_3314019, EPI_ISL_3314020, EPI_ISL_3314021, EPI_ISL_3314022 | Lighthouse Lab in Milton Keynes | Wellcome Sanger Institute for the COVID-19 Genomics UK (COG-UK) Consortium | The Lighthouse Lab in Milton Keynes and Alex Alderton, Roberto Amato, Jeffrey Barrett, Sonia Goncalves, Ewan Harrison, David K. Jackson, Ian Johnston, Dominic Kwiatkowski, Cordelia Langford, John Sillitoe on behalf of the Wellcome Sanger Institute COVID-19 Surveillance Team |
| EPI_ISL_3314023 | Berkshire and Surrey Pathology Services Lighthouse Laboratory | Wellcome Sanger Institute for the COVID-19 Genomics UK (COG-UK) Consortium | Berkshire and Surrey Pathology Services Lighthouse Laboratory and Alex Alderton, Roberto Amato, Jeffrey Barrett, Sonia Goncalves, Ewan Harrison, David K. Jackson, Ian Johnston, Dominic Kwiatkowski, Cordelia Langford, John Sillitoe on behalf of the Wellcome Sanger Institute COVID-19 Surveillance Team |
| EPI_ISL_3314024, EPI_ISL_3314025, EPI_ISL_3314026 | Lighthouse Lab in Milton Keynes | Wellcome Sanger Institute for the COVID-19 Genomics UK (COG-UK) Consortium | The Lighthouse Lab in Milton Keynes and Alex Alderton, Roberto Amato, Jeffrey Barrett, Sonia Goncalves, Ewan Harrison, David K. Jackson, Ian Johnston, Dominic Kwiatkowski, Cordelia Langford, John Sillitoe on behalf of the Wellcome Sanger Institute COVID-19 Surveillance Team |
| EPI_ISL_3314027 | Berkshire and Surrey Pathology Services Lighthouse Laboratory | Wellcome Sanger Institute for the COVID-19 Genomics UK (COG-UK) Consortium | Berkshire and Surrey Pathology Services Lighthouse Laboratory and Alex Alderton, Roberto Amato, Jeffrey Barrett, Sonia Goncalves, Ewan Harrison, David K. Jackson, Ian Johnston, Dominic Kwiatkowski, Cordelia Langford, John Sillitoe on behalf of the Wellcome Sanger Institute COVID-19 Surveillance Team |
| EPI_ISL_3314028 | Lighthouse Lab in Milton Keynes | Wellcome Sanger Institute for the COVID-19 Genomics UK (COG-UK) Consortium | The Lighthouse Lab in Milton Keynes and Alex Alderton, Roberto Amato, Jeffrey Barrett, Sonia Goncalves, Ewan Harrison, David K. Jackson, Ian Johnston, Dominic Kwiatkowski, Cordelia Langford, John Sillitoe on behalf of the Wellcome Sanger Institute COVID-19 Surveillance Team |
| EPI_ISL_3314029 | Lighthouse Lab in Glasgow | Wellcome Sanger Institute for the COVID-19 Genomics UK (COG-UK) Consortium | Harper VanSteenhouse, Yumi Kasai, David Gray, Carol Clugston, Anna Dominiczak and Alex Alderton, Roberto Amato, Jeffrey Barrett, Sonia Goncalves, Ewan Harrison, David K. Jackson, Ian Johnston, Dominic Kwiatkowski, Cordelia Langford, John Sillitoe on behalf of the Wellcome Sanger Institute COVID-19 Surveillance Team |
| EPI_ISL_3314030 | Lighthouse Lab in Milton Keynes | Wellcome Sanger Institute for the COVID-19 Genomics UK (COG-UK) Consortium | The Lighthouse Lab in Milton Keynes and Alex Alderton, Roberto Amato, Jeffrey Barrett, Sonia Goncalves, Ewan Harrison, David K. Jackson, Ian Johnston, Dominic Kwiatkowski, Cordelia Langford, John Sillitoe on behalf of the Wellcome Sanger Institute COVID-19 Surveillance Team |
| EPI_ISL_3314031 | Berkshire and Surrey Pathology Services Lighthouse Laboratory | Wellcome Sanger Institute for the COVID-19 Genomics UK (COG-UK) Consortium | Berkshire and Surrey Pathology Services Lighthouse Laboratory and Alex Alderton, Roberto Amato, Jeffrey Barrett, Sonia Goncalves, Ewan Harrison, David K. Jackson, Ian Johnston, Dominic Kwiatkowski, Cordelia Langford, John Sillitoe on behalf of the Wellcome Sanger Institute COVID-19 Surveillance Team |
| EPI_ISL_3314032 | Lighthouse Lab in Milton Keynes | Wellcome Sanger Institute for the COVID-19 Genomics UK (COG-UK) Consortium | The Lighthouse Lab in Milton Keynes and Alex Alderton, Roberto Amato, Jeffrey Barrett, Sonia Goncalves, Ewan Harrison, David K. Jackson, Ian Johnston, Dominic Kwiatkowski, Cordelia Langford, John Sillitoe on behalf of the Wellcome Sanger Institute COVID-19 Surveillance Team |
| EPI_ISL_3314033 | Lighthouse Lab in Glasgow | Wellcome Sanger Institute for the COVID-19 Genomics UK (COG-UK) Consortium | Harper VanSteenhouse, Yumi Kasai, David Gray, Carol Clugston, Anna Dominiczak and Alex Alderton, Roberto Amato, Jeffrey Barrett, Sonia Goncalves, Ewan Harrison, David K. Jackson, Ian Johnston, Dominic Kwiatkowski, Cordelia Langford, John Sillitoe on behalf of the Wellcome Sanger Institute COVID-19 Surveillance Team |
| EPI_ISL_3314034, EPI_ISL_3314035, EPI_ISL_3314036, EPI_ISL_3314037, | Lighthouse Lab in Milton Keynes | Wellcome Sanger Institute for the COVID-19 Genomics UK (COG-UK) Consortium | The Lighthouse Lab in Milton Keynes and Alex Alderton, Roberto Amato, Jeffrey Barrett, Sonia Goncalves, Ewan Harrison, David K. Jackson, Ian Johnston, Dominic Kwiatkowski, Cordelia Langford, John Sillitoe on behalf of the Wellcome Sanger Institute COVID-19 Surveillance Team |

| | | | |
|---|---|---|---|
| EPI_ISL_3314038 | | | |
| EPI_ISL_3314039, EPI_ISL_3314040 | Lighthouse Lab in Glasgow | Wellcome Sanger Institute for the COVID-19 Genomics UK (COG-UK) Consortium | Harper VanSteenhouse, Yumi Kasai, David Gray, Carol Clugston, Anna Dominiczak and Alex Alderton, Roberto Amato, Jeffrey Barrett, Sonia Goncalves, Ewan Harrison, David K. Jackson, Ian Johnston, Dominic Kwiatkowski, Cordelia Langford, John Sillitoe on behalf of the Wellcome Sanger Institute COVID-19 Surveillance Team |
| EPI_ISL_3314041 | Berkshire and Surrey Pathology Services Lighthouse Laboratory | Wellcome Sanger Institute for the COVID-19 Genomics UK (COG-UK) Consortium | Berkshire and Surrey Pathology Services Lighthouse Laboratory and Alex Alderton, Roberto Amato, Jeffrey Barrett, Sonia Goncalves, Ewan Harrison, David K. Jackson, Ian Johnston, Dominic Kwiatkowski, Cordelia Langford, John Sillitoe on behalf of the Wellcome Sanger Institute COVID-19 Surveillance Team |
| EPI_ISL_3314042, EPI_ISL_3314043 | Lighthouse Lab in Glasgow | Wellcome Sanger Institute for the COVID-19 Genomics UK (COG-UK) Consortium | Harper VanSteenhouse, Yumi Kasai, David Gray, Carol Clugston, Anna Dominiczak and Alex Alderton, Roberto Amato, Jeffrey Barrett, Sonia Goncalves, Ewan Harrison, David K. Jackson, Ian Johnston, Dominic Kwiatkowski, Cordelia Langford, John Sillitoe on behalf of the Wellcome Sanger Institute COVID-19 Surveillance Team |
| EPI_ISL_3314044, EPI_ISL_3314045 | Lighthouse Lab in Milton Keynes | Wellcome Sanger Institute for the COVID-19 Genomics UK (COG-UK) Consortium | The Lighthouse Lab in Milton Keynes and Alex Alderton, Roberto Amato, Jeffrey Barrett, Sonia Goncalves, Ewan Harrison, David K. Jackson, Ian Johnston, Dominic Kwiatkowski, Cordelia Langford, John Sillitoe on behalf of the Wellcome Sanger Institute COVID-19 Surveillance Team |
| EPI_ISL_3314046 | Berkshire and Surrey Pathology Services Lighthouse Laboratory | Wellcome Sanger Institute for the COVID-19 Genomics UK (COG-UK) Consortium | Berkshire and Surrey Pathology Services Lighthouse Laboratory and Alex Alderton, Roberto Amato, Jeffrey Barrett, Sonia Goncalves, Ewan Harrison, David K. Jackson, Ian Johnston, Dominic Kwiatkowski, Cordelia Langford, John Sillitoe on behalf of the Wellcome Sanger Institute COVID-19 Surveillance Team |
| EPI_ISL_3314047, EPI_ISL_3314048 | Lighthouse Lab in Milton Keynes | Wellcome Sanger Institute for the COVID-19 Genomics UK (COG-UK) Consortium | The Lighthouse Lab in Milton Keynes and Alex Alderton, Roberto Amato, Jeffrey Barrett, Sonia Goncalves, Ewan Harrison, David K. Jackson, Ian Johnston, Dominic Kwiatkowski, Cordelia Langford, John Sillitoe on behalf of the Wellcome Sanger Institute COVID-19 Surveillance Team |
| EPI_ISL_3314049 | Lighthouse Lab in Glasgow | Wellcome Sanger Institute for the COVID-19 Genomics UK (COG-UK) Consortium | Harper VanSteenhouse, Yumi Kasai, David Gray, Carol Clugston, Anna Dominiczak and Alex Alderton, Roberto Amato, Jeffrey Barrett, Sonia Goncalves, Ewan Harrison, David K. Jackson, Ian Johnston, Dominic Kwiatkowski, Cordelia Langford, John Sillitoe on behalf of the Wellcome Sanger Institute COVID-19 Surveillance Team |
| EPI_ISL_3314050, EPI_ISL_3314051, EPI_ISL_3314052, EPI_ISL_3314053, EPI_ISL_3314054 | Lighthouse Lab in Milton Keynes | Wellcome Sanger Institute for the COVID-19 Genomics UK (COG-UK) Consortium | The Lighthouse Lab in Milton Keynes and Alex Alderton, Roberto Amato, Jeffrey Barrett, Sonia Goncalves, Ewan Harrison, David K. Jackson, Ian Johnston, Dominic Kwiatkowski, Cordelia Langford, John Sillitoe on behalf of the Wellcome Sanger Institute COVID-19 Surveillance Team |
| EPI_ISL_3314055 | Berkshire and Surrey Pathology Services Lighthouse Laboratory | Wellcome Sanger Institute for the COVID-19 Genomics UK (COG-UK) Consortium | Berkshire and Surrey Pathology Services Lighthouse Laboratory and Alex Alderton, Roberto Amato, Jeffrey Barrett, Sonia Goncalves, Ewan Harrison, David K. Jackson, Ian Johnston, Dominic Kwiatkowski, Cordelia Langford, John Sillitoe on behalf of the Wellcome Sanger Institute COVID-19 Surveillance Team |
| EPI_ISL_3314056, EPI_ISL_3314057, EPI_ISL_3314058 | Lighthouse Lab in Milton Keynes | Wellcome Sanger Institute for the COVID-19 Genomics UK (COG-UK) Consortium | The Lighthouse Lab in Milton Keynes and Alex Alderton, Roberto Amato, Jeffrey Barrett, Sonia Goncalves, Ewan Harrison, David K. Jackson, Ian Johnston, Dominic Kwiatkowski, Cordelia Langford, John Sillitoe on behalf of the Wellcome Sanger Institute COVID-19 Surveillance Team |
| EPI_ISL_3314059 | Berkshire and Surrey Pathology Services Lighthouse Laboratory | Wellcome Sanger Institute for the COVID-19 Genomics UK (COG-UK) Consortium | Berkshire and Surrey Pathology Services Lighthouse Laboratory and Alex Alderton, Roberto Amato, Jeffrey Barrett, Sonia Goncalves, Ewan Harrison, David K. Jackson, Ian Johnston, Dominic Kwiatkowski, Cordelia Langford, John Sillitoe on behalf of the Wellcome Sanger Institute COVID-19 Surveillance Team |
| EPI_ISL_3314060, EPI_ISL_3314061, EPI_ISL_3314062, EPI_ISL_3314063, EPI_ISL_3314064, EPI_ISL_3314065 | Lighthouse Lab in Milton Keynes | Wellcome Sanger Institute for the COVID-19 Genomics UK (COG-UK) Consortium | The Lighthouse Lab in Milton Keynes and Alex Alderton, Roberto Amato, Jeffrey Barrett, Sonia Goncalves, Ewan Harrison, David K. Jackson, Ian Johnston, Dominic Kwiatkowski, Cordelia Langford, John Sillitoe on behalf of the Wellcome Sanger Institute COVID-19 Surveillance Team |
| EPI_ISL_3314066 | Berkshire and Surrey Pathology Services Lighthouse Laboratory | Wellcome Sanger Institute for the COVID-19 Genomics UK (COG-UK) Consortium | Berkshire and Surrey Pathology Services Lighthouse Laboratory and Alex Alderton, Roberto Amato, Jeffrey Barrett, Sonia Goncalves, Ewan Harrison, David K. Jackson, Ian Johnston, Dominic Kwiatkowski, Cordelia Langford, John Sillitoe on behalf of the Wellcome Sanger Institute COVID-19 Surveillance Team |
| EPI_ISL_3314067 | Lighthouse Lab in Glasgow | Wellcome Sanger Institute for the COVID-19 Genomics UK (COG-UK) Consortium | Harper VanSteenhouse, Yumi Kasai, David Gray, Carol Clugston, Anna Dominiczak and Alex Alderton, Roberto Amato, Jeffrey Barrett, Sonia Goncalves, Ewan Harrison, David K. Jackson, Ian Johnston, Dominic Kwiatkowski, Cordelia Langford, John Sillitoe on behalf of the Wellcome Sanger Institute COVID-19 Surveillance Team |
| EPI_ISL_3314068, EPI_ISL_3314069 | Berkshire and Surrey Pathology Services Lighthouse Laboratory | Wellcome Sanger Institute for the COVID-19 Genomics UK (COG-UK) Consortium | Berkshire and Surrey Pathology Services Lighthouse Laboratory and Alex Alderton, Roberto Amato, Jeffrey Barrett, Sonia Goncalves, Ewan Harrison, David K. Jackson, Ian Johnston, Dominic Kwiatkowski, Cordelia Langford, John Sillitoe on behalf of the Wellcome Sanger Institute COVID-19 Surveillance Team |
| EPI_ISL_3314070 | Lighthouse Lab in Milton Keynes | Wellcome Sanger Institute for the COVID-19 Genomics UK (COG-UK) Consortium | The Lighthouse Lab in Milton Keynes and Alex Alderton, Roberto Amato, Jeffrey Barrett, Sonia Goncalves, Ewan Harrison, David K. Jackson, Ian Johnston, Dominic Kwiatkowski, Cordelia Langford, John Sillitoe on behalf of the Wellcome Sanger Institute COVID-19 Surveillance Team |
| EPI_ISL_3314071 | Berkshire and Surrey Pathology Services Lighthouse Laboratory | Wellcome Sanger Institute for the COVID-19 Genomics UK (COG-UK) Consortium | Berkshire and Surrey Pathology Services Lighthouse Laboratory and Alex Alderton, Roberto Amato, Jeffrey Barrett, Sonia Goncalves, Ewan Harrison, David K. Jackson, Ian Johnston, Dominic Kwiatkowski, Cordelia Langford, John Sillitoe on behalf of the Wellcome Sanger Institute COVID-19 Surveillance Team |
| EPI_ISL_3314072, EPI_ISL_3314073 | Lighthouse Lab in Milton Keynes | Wellcome Sanger Institute for the COVID-19 Genomics UK (COG-UK) Consortium | The Lighthouse Lab in Milton Keynes and Alex Alderton, Roberto Amato, Jeffrey Barrett, Sonia Goncalves, Ewan Harrison, David K. Jackson, Ian Johnston, Dominic Kwiatkowski, Cordelia Langford, John Sillitoe on behalf of the Wellcome Sanger Institute COVID-19 Surveillance Team |
| EPI_ISL_3314074 | Lighthouse Lab in Glasgow | Wellcome Sanger Institute for the COVID-19 Genomics UK (COG-UK) Consortium | Harper VanSteenhouse, Yumi Kasai, David Gray, Carol Clugston, Anna Dominiczak and Alex Alderton, Roberto Amato, Jeffrey Barrett, Sonia Goncalves, Ewan Harrison, David K. Jackson, Ian Johnston, Dominic Kwiatkowski, Cordelia Langford, John Sillitoe on behalf of the Wellcome Sanger Institute COVID-19 Surveillance Team |
| EPI_ISL_3314075 | Berkshire and Surrey Pathology Services Lighthouse Laboratory | Wellcome Sanger Institute for the COVID-19 Genomics UK (COG-UK) Consortium | Berkshire and Surrey Pathology Services Lighthouse Laboratory and Alex Alderton, Roberto Amato, Jeffrey Barrett, Sonia Goncalves, Ewan Harrison, David K. Jackson, Ian Johnston, Dominic Kwiatkowski, Cordelia Langford, John Sillitoe on behalf of the Wellcome Sanger Institute COVID-19 Surveillance Team |
| EPI_ISL_3314076, EPI_ISL_3314077 | Lighthouse Lab in Milton Keynes | Wellcome Sanger Institute for the COVID-19 Genomics UK (COG-UK) Consortium | The Lighthouse Lab in Milton Keynes and Alex Alderton, Roberto Amato, Jeffrey Barrett, Sonia Goncalves, Ewan Harrison, David K. Jackson, Ian Johnston, Dominic Kwiatkowski, Cordelia Langford, John Sillitoe on behalf of the Wellcome Sanger Institute COVID-19 Surveillance Team |
| EPI_ISL_3314078 | Lighthouse Lab in Glasgow | Wellcome Sanger Institute for the COVID-19 Genomics UK (COG-UK) Consortium | Harper VanSteenhouse, Yumi Kasai, David Gray, Carol Clugston, Anna Dominiczak and Alex Alderton, Roberto Amato, Jeffrey Barrett, Sonia Goncalves, Ewan Harrison, David K. Jackson, Ian Johnston, Dominic Kwiatkowski, Cordelia Langford, John Sillitoe on behalf of the Wellcome Sanger Institute COVID-19 Surveillance Team |
| EPI_ISL_3314079, EPI_ISL_3314080 | Lighthouse Lab in Milton Keynes | Wellcome Sanger Institute for the COVID-19 Genomics UK (COG-UK) Consortium | The Lighthouse Lab in Milton Keynes and Alex Alderton, Roberto Amato, Jeffrey Barrett, Sonia Goncalves, Ewan Harrison, David K. Jackson, Ian Johnston, Dominic Kwiatkowski, Cordelia Langford, John Sillitoe on behalf of the Wellcome Sanger Institute COVID-19 Surveillance Team |
| EPI_ISL_3314081 | Lighthouse Lab in Glasgow | Wellcome Sanger Institute for the COVID-19 Genomics UK (COG-UK) Consortium | Harper VanSteenhouse, Yumi Kasai, David Gray, Carol Clugston, Anna Dominiczak and Alex Alderton, Roberto Amato, Jeffrey Barrett, Sonia Goncalves, Ewan Harrison, David K. Jackson, Ian Johnston, Dominic Kwiatkowski, Cordelia Langford, John Sillitoe on behalf of the Wellcome Sanger Institute COVID-19 Surveillance Team |
| EPI_ISL_3314082, EPI_ISL_3314083 | Lighthouse Lab in Milton Keynes | Wellcome Sanger Institute for the COVID-19 Genomics UK (COG-UK) Consortium | The Lighthouse Lab in Milton Keynes and Alex Alderton, Roberto Amato, Jeffrey Barrett, Sonia Goncalves, Ewan Harrison, David K. Jackson, Ian Johnston, Dominic Kwiatkowski, Cordelia Langford, John Sillitoe on behalf of the Wellcome Sanger Institute COVID-19 Surveillance Team |
| EPI_ISL_3314084 | Lighthouse Lab in Glasgow | Wellcome Sanger Institute for the COVID-19 Genomics UK (COG-UK) Consortium | Harper VanSteenhouse, Yumi Kasai, David Gray, Carol Clugston, Anna Dominiczak and Alex Alderton, Roberto Amato, Jeffrey Barrett, Sonia Goncalves, Ewan Harrison, David K. Jackson, Ian Johnston, Dominic Kwiatkowski, Cordelia Langford, John Sillitoe on behalf of the Wellcome Sanger Institute COVID-19 Surveillance Team |
| EPI_ISL_3314085, EPI_ISL_3314086 | Lighthouse Lab in Milton Keynes | Wellcome Sanger Institute for the COVID-19 Genomics UK (COG-UK) Consortium | The Lighthouse Lab in Milton Keynes and Alex Alderton, Roberto Amato, Jeffrey Barrett, Sonia Goncalves, Ewan Harrison, David K. Jackson, Ian Johnston, Dominic Kwiatkowski, Cordelia Langford, John Sillitoe on behalf of the Wellcome Sanger Institute COVID-19 Surveillance Team |
| EPI_ISL_3314087, EPI_ISL_3314088 | Berkshire and Surrey Pathology Services Lighthouse Laboratory | Wellcome Sanger Institute for the COVID-19 Genomics UK (COG-UK) Consortium | Berkshire and Surrey Pathology Services Lighthouse Laboratory and Alex Alderton, Roberto Amato, Jeffrey Barrett, Sonia Goncalves, Ewan Harrison, David K. Jackson, Ian Johnston, Dominic Kwiatkowski, Cordelia Langford, John Sillitoe on behalf of the Wellcome Sanger Institute COVID-19 Surveillance Team |
| EPI_ISL_3314089, EPI_ISL_3314090, EPI_ISL_3314091 | Lighthouse Lab in Milton Keynes | Wellcome Sanger Institute for the COVID-19 Genomics UK (COG-UK) Consortium | The Lighthouse Lab in Milton Keynes and Alex Alderton, Roberto Amato, Jeffrey Barrett, Sonia Goncalves, Ewan Harrison, David K. Jackson, Ian Johnston, Dominic Kwiatkowski, Cordelia Langford, John Sillitoe on behalf of the Wellcome Sanger Institute COVID-19 Surveillance Team |
| EPI_ISL_3314092 | Berkshire and Surrey Pathology Services Lighthouse Laboratory | Wellcome Sanger Institute for the COVID-19 Genomics UK (COG-UK) Consortium | Berkshire and Surrey Pathology Services Lighthouse Laboratory and Alex Alderton, Roberto Amato, Jeffrey Barrett, Sonia Goncalves, Ewan Harrison, David K. Jackson, Ian Johnston, Dominic Kwiatkowski, Cordelia Langford, John Sillitoe on behalf of the Wellcome Sanger Institute COVID-19 Surveillance Team |
| EPI_ISL_3314093 | Lighthouse Lab in Glasgow | Wellcome Sanger Institute for the COVID-19 Genomics UK (COG-UK) Consortium | Harper VanSteenhouse, Yumi Kasai, David Gray, Carol Clugston, Anna Dominiczak and Alex Alderton, Roberto Amato, Jeffrey Barrett, Sonia Goncalves, Ewan Harrison, David K. Jackson, Ian Johnston, Dominic Kwiatkowski, Cordelia Langford, John Sillitoe on behalf of the Wellcome Sanger Institute COVID-19 Surveillance Team |
| EPI_ISL_3314094, EPI_ISL_3314095 | Berkshire and Surrey Pathology Services Lighthouse Laboratory | Wellcome Sanger Institute for the COVID-19 Genomics UK (COG-UK) Consortium | Berkshire and Surrey Pathology Services Lighthouse Laboratory and Alex Alderton, Roberto Amato, Jeffrey Barrett, Sonia Goncalves, Ewan Harrison, David K. Jackson, Ian Johnston, Dominic Kwiatkowski, Cordelia Langford, John Sillitoe on behalf of the Wellcome Sanger Institute COVID-19 Surveillance Team |
| EPI_ISL_3314096, EPI_ISL_3314097 | Lighthouse Lab in Milton Keynes | Wellcome Sanger Institute for the COVID-19 Genomics UK (COG-UK) Consortium | The Lighthouse Lab in Milton Keynes and Alex Alderton, Roberto Amato, Jeffrey Barrett, Sonia Goncalves, Ewan Harrison, David K. Jackson, Ian Johnston, Dominic Kwiatkowski, Cordelia Langford, John Sillitoe on behalf of the Wellcome Sanger Institute COVID-19 Surveillance Team |
| EPI_ISL_3314098, EPI_ISL_3314099 | Berkshire and Surrey Pathology Services Lighthouse Laboratory | Wellcome Sanger Institute for the COVID-19 Genomics UK (COG-UK) Consortium | Berkshire and Surrey Pathology Services Lighthouse Laboratory and Alex Alderton, Roberto Amato, Jeffrey Barrett, Sonia Goncalves, Ewan Harrison, David K. Jackson, Ian Johnston, Dominic Kwiatkowski, Cordelia Langford, John Sillitoe on behalf of the Wellcome Sanger Institute COVID-19 Surveillance Team |
| EPI_ISL_3314100, EPI_ISL_3314101, EPI_ISL_3314102, EPI_ISL_3314103 | Lighthouse Lab in Glasgow | Wellcome Sanger Institute for the COVID-19 Genomics UK (COG-UK) Consortium | Harper VanSteenhouse, Yumi Kasai, David Gray, Carol Clugston, Anna Dominiczak and Alex Alderton, Roberto Amato, Jeffrey Barrett, Sonia Goncalves, Ewan Harrison, David K. Jackson, Ian Johnston, Dominic Kwiatkowski, Cordelia Langford, John Sillitoe on behalf of the Wellcome Sanger Institute COVID-19 Surveillance Team |
| EPI_ISL_3314104, EPI_ISL_3314105, EPI_ISL_3314106, EPI_ISL_3314107, EPI_ISL_3314108, EPI_ISL_3314109 | Lighthouse Lab in Milton Keynes | Wellcome Sanger Institute for the COVID-19 Genomics UK (COG-UK) Consortium | The Lighthouse Lab in Milton Keynes and Alex Alderton, Roberto Amato, Jeffrey Barrett, Sonia Goncalves, Ewan Harrison, David K. Jackson, Ian Johnston, Dominic Kwiatkowski, Cordelia Langford, John Sillitoe on behalf of the Wellcome Sanger Institute COVID-19 Surveillance Team |
| EPI_ISL_3314110 | Lighthouse Lab in Glasgow | Wellcome Sanger Institute for the COVID-19 Genomics UK (COG-UK) Consortium | Harper VanSteenhouse, Yumi Kasai, David Gray, Carol Clugston, Anna Dominiczak and Alex Alderton, Roberto Amato, Jeffrey Barrett, Sonia Goncalves, Ewan Harrison, David K. Jackson, Ian Johnston, Dominic Kwiatkowski, Cordelia Langford, John Sillitoe on behalf of the Wellcome Sanger Institute COVID-19 Surveillance Team |
| EPI_ISL_3314111, EPI_ISL_3314112 | Lighthouse Lab in Milton Keynes | Wellcome Sanger Institute for the COVID-19 Genomics UK (COG-UK) Consortium | The Lighthouse Lab in Milton Keynes and Alex Alderton, Roberto Amato, Jeffrey Barrett, Sonia Goncalves, Ewan Harrison, David K. Jackson, Ian Johnston, Dominic Kwiatkowski, Cordelia Langford, John Sillitoe on behalf of the Wellcome Sanger Institute COVID-19 Surveillance Team |
| EPI_ISL_3314113, | Berkshire and Surrey Pathology Services Lighthouse Laboratory | Wellcome Sanger Institute for the COVID-19 Genomics UK (COG-UK) | Berkshire and Surrey Pathology Services Lighthouse Laboratory and Alex Alderton, Roberto Amato, Jeffrey Barrett, Sonia Goncalves, Ewan Harrison, David K. Jackson, Ian Johnston, Dominic Kwiatkowski, Cordelia Langford, John Sillitoe on behalf |

| | | | |
|---|---|---|---|
| EPI_ISL_3314114 | | Consortium | of the Wellcome Sanger Institute COVID-19 Surveillance Team |
| EPI_ISL_3314115 | Lighthouse Lab in Glasgow | Wellcome Sanger Institute for the COVID-19 Genomics UK (COG-UK) Consortium | Harper VanSteenhouse, Yumi Kasai, David Gray, Carol Clugston, Anna Dominiczak and Alex Alderton, Roberto Amato, Jeffrey Barrett, Sonia Goncalves, Ewan Harrison, David K. Jackson, Ian Johnston, Dominic Kwiatkowski, Cordelia Langford, John Sillitoe on behalf of the Wellcome Sanger Institute COVID-19 Surveillance Team |
| EPI_ISL_3314116, EPI_ISL_3314117 | Lighthouse Lab in Milton Keynes | Wellcome Sanger Institute for the COVID-19 Genomics UK (COG-UK) Consortium | The Lighthouse Lab in Milton Keynes and Alex Alderton, Roberto Amato, Jeffrey Barrett, Sonia Goncalves, Ewan Harrison, David K. Jackson, Ian Johnston, Dominic Kwiatkowski, Cordelia Langford, John Sillitoe on behalf of the Wellcome Sanger Institute COVID-19 Surveillance Team |
| EPI_ISL_3314118 | Lighthouse Lab in Glasgow | Wellcome Sanger Institute for the COVID-19 Genomics UK (COG-UK) Consortium | Harper VanSteenhouse, Yumi Kasai, David Gray, Carol Clugston, Anna Dominiczak and Alex Alderton, Roberto Amato, Jeffrey Barrett, Sonia Goncalves, Ewan Harrison, David K. Jackson, Ian Johnston, Dominic Kwiatkowski, Cordelia Langford, John Sillitoe on behalf of the Wellcome Sanger Institute COVID-19 Surveillance Team |
| EPI_ISL_3314119 | Berkshire and Surrey Pathology Services Lighthouse Laboratory | Wellcome Sanger Institute for the COVID-19 Genomics UK (COG-UK) Consortium | Berkshire and Surrey Pathology Services Lighthouse Laboratory and Alex Alderton, Roberto Amato, Jeffrey Barrett, Sonia Goncalves, Ewan Harrison, David K. Jackson, Ian Johnston, Dominic Kwiatkowski, Cordelia Langford, John Sillitoe on behalf of the Wellcome Sanger Institute COVID-19 Surveillance Team |
| EPI_ISL_3314120, EPI_ISL_3314121 | Lighthouse Lab in Glasgow | Wellcome Sanger Institute for the COVID-19 Genomics UK (COG-UK) Consortium | Harper VanSteenhouse, Yumi Kasai, David Gray, Carol Clugston, Anna Dominiczak and Alex Alderton, Roberto Amato, Jeffrey Barrett, Sonia Goncalves, Ewan Harrison, David K. Jackson, Ian Johnston, Dominic Kwiatkowski, Cordelia Langford, John Sillitoe on behalf of the Wellcome Sanger Institute COVID-19 Surveillance Team |
| EPI_ISL_3314122, EPI_ISL_3314123, EPI_ISL_3314124 | Lighthouse Lab in Milton Keynes | Wellcome Sanger Institute for the COVID-19 Genomics UK (COG-UK) Consortium | The Lighthouse Lab in Milton Keynes and Alex Alderton, Roberto Amato, Jeffrey Barrett, Sonia Goncalves, Ewan Harrison, David K. Jackson, Ian Johnston, Dominic Kwiatkowski, Cordelia Langford, John Sillitoe on behalf of the Wellcome Sanger Institute COVID-19 Surveillance Team |
| EPI_ISL_3314125 | Berkshire and Surrey Pathology Services Lighthouse Laboratory | Wellcome Sanger Institute for the COVID-19 Genomics UK (COG-UK) Consortium | Berkshire and Surrey Pathology Services Lighthouse Laboratory and Alex Alderton, Roberto Amato, Jeffrey Barrett, Sonia Goncalves, Ewan Harrison, David K. Jackson, Ian Johnston, Dominic Kwiatkowski, Cordelia Langford, John Sillitoe on behalf of the Wellcome Sanger Institute COVID-19 Surveillance Team |
| EPI_ISL_3314126, EPI_ISL_3314127 | Lighthouse Lab in Glasgow | Wellcome Sanger Institute for the COVID-19 Genomics UK (COG-UK) Consortium | Harper VanSteenhouse, Yumi Kasai, David Gray, Carol Clugston, Anna Dominiczak and Alex Alderton, Roberto Amato, Jeffrey Barrett, Sonia Goncalves, Ewan Harrison, David K. Jackson, Ian Johnston, Dominic Kwiatkowski, Cordelia Langford, John Sillitoe on behalf of the Wellcome Sanger Institute COVID-19 Surveillance Team |
| EPI_ISL_3314128 | Lighthouse Lab in Alderley Park | Wellcome Sanger Institute for the COVID-19 Genomics UK (COG-UK) Consortium | Jacquelyn Wynn, Mairead Hyland, The Lighthouse Lab in Alderley Park and Alex Alderton, Roberto Amato, Jeffrey Barrett, Sonia Goncalves, Ewan Harrison, David K. Jackson, Ian Johnston, Dominic Kwiatkowski, Cordelia Langford, John Sillitoe on behalf of the Wellcome Sanger Institute COVID-19 Surveillance Team |
| EPI_ISL_3314129 | Lighthouse Lab in Glasgow | Wellcome Sanger Institute for the COVID-19 Genomics UK (COG-UK) Consortium | Harper VanSteenhouse, Yumi Kasai, David Gray, Carol Clugston, Anna Dominiczak and Alex Alderton, Roberto Amato, Jeffrey Barrett, Sonia Goncalves, Ewan Harrison, David K. Jackson, Ian Johnston, Dominic Kwiatkowski, Cordelia Langford, John Sillitoe on behalf of the Wellcome Sanger Institute COVID-19 Surveillance Team |
| EPI_ISL_3314130, EPI_ISL_3314131 | Lighthouse Lab in Alderley Park | Wellcome Sanger Institute for the COVID-19 Genomics UK (COG-UK) Consortium | Jacquelyn Wynn, Mairead Hyland, The Lighthouse Lab in Alderley Park and Alex Alderton, Roberto Amato, Jeffrey Barrett, Sonia Goncalves, Ewan Harrison, David K. Jackson, Ian Johnston, Dominic Kwiatkowski, Cordelia Langford, John Sillitoe on behalf of the Wellcome Sanger Institute COVID-19 Surveillance Team |
| EPI_ISL_3314132 | Lighthouse Lab in Glasgow | Wellcome Sanger Institute for the COVID-19 Genomics UK (COG-UK) Consortium | Harper VanSteenhouse, Yumi Kasai, David Gray, Carol Clugston, Anna Dominiczak and Alex Alderton, Roberto Amato, Jeffrey Barrett, Sonia Goncalves, Ewan Harrison, David K. Jackson, Ian Johnston, Dominic Kwiatkowski, Cordelia Langford, John Sillitoe on behalf of the Wellcome Sanger Institute COVID-19 Surveillance Team |
| EPI_ISL_3314133, EPI_ISL_3314134 | Berkshire and Surrey Pathology Services Lighthouse Laboratory | Wellcome Sanger Institute for the COVID-19 Genomics UK (COG-UK) Consortium | Berkshire and Surrey Pathology Services Lighthouse Laboratory and Alex Alderton, Roberto Amato, Jeffrey Barrett, Sonia Goncalves, Ewan Harrison, David K. Jackson, Ian Johnston, Dominic Kwiatkowski, Cordelia Langford, John Sillitoe on behalf of the Wellcome Sanger Institute COVID-19 Surveillance Team |
| EPI_ISL_3314135, EPI_ISL_3314136 | Lighthouse Lab in Glasgow | Wellcome Sanger Institute for the COVID-19 Genomics UK (COG-UK) Consortium | Harper VanSteenhouse, Yumi Kasai, David Gray, Carol Clugston, Anna Dominiczak and Alex Alderton, Roberto Amato, Jeffrey Barrett, Sonia Goncalves, Ewan Harrison, David K. Jackson, Ian Johnston, Dominic Kwiatkowski, Cordelia Langford, John Sillitoe on behalf of the Wellcome Sanger Institute COVID-19 Surveillance Team |
| EPI_ISL_3314137 | Lighthouse Lab in Alderley Park | Wellcome Sanger Institute for the COVID-19 Genomics UK (COG-UK) Consortium | Jacquelyn Wynn, Mairead Hyland, The Lighthouse Lab in Alderley Park and Alex Alderton, Roberto Amato, Jeffrey Barrett, Sonia Goncalves, Ewan Harrison, David K. Jackson, Ian Johnston, Dominic Kwiatkowski, Cordelia Langford, John Sillitoe on behalf of the Wellcome Sanger Institute COVID-19 Surveillance Team |
| EPI_ISL_3314138 | Berkshire and Surrey Pathology Services Lighthouse Laboratory | Wellcome Sanger Institute for the COVID-19 Genomics UK (COG-UK) Consortium | Berkshire and Surrey Pathology Services Lighthouse Laboratory and Alex Alderton, Roberto Amato, Jeffrey Barrett, Sonia Goncalves, Ewan Harrison, David K. Jackson, Ian Johnston, Dominic Kwiatkowski, Cordelia Langford, John Sillitoe on behalf of the Wellcome Sanger Institute COVID-19 Surveillance Team |
| EPI_ISL_3314139 | Lighthouse Lab in Glasgow | Wellcome Sanger Institute for the COVID-19 Genomics UK (COG-UK) Consortium | Harper VanSteenhouse, Yumi Kasai, David Gray, Carol Clugston, Anna Dominiczak and Alex Alderton, Roberto Amato, Jeffrey Barrett, Sonia Goncalves, Ewan Harrison, David K. Jackson, Ian Johnston, Dominic Kwiatkowski, Cordelia Langford, John Sillitoe on behalf of the Wellcome Sanger Institute COVID-19 Surveillance Team |
| EPI_ISL_3314140, EPI_ISL_3314141 | Berkshire and Surrey Pathology Services Lighthouse Laboratory | Wellcome Sanger Institute for the COVID-19 Genomics UK (COG-UK) Consortium | Berkshire and Surrey Pathology Services Lighthouse Laboratory and Alex Alderton, Roberto Amato, Jeffrey Barrett, Sonia Goncalves, Ewan Harrison, David K. Jackson, Ian Johnston, Dominic Kwiatkowski, Cordelia Langford, John Sillitoe on behalf of the Wellcome Sanger Institute COVID-19 Surveillance Team |
| EPI_ISL_3314142 | Lighthouse Lab in Alderley Park | Wellcome Sanger Institute for the COVID-19 Genomics UK (COG-UK) Consortium | Jacquelyn Wynn, Mairead Hyland, The Lighthouse Lab in Alderley Park and Alex Alderton, Roberto Amato, Jeffrey Barrett, Sonia Goncalves, Ewan Harrison, David K. Jackson, Ian Johnston, Dominic Kwiatkowski, Cordelia Langford, John Sillitoe on behalf of the Wellcome Sanger Institute COVID-19 Surveillance Team |
| EPI_ISL_3314143 | Lighthouse Lab in Glasgow | Wellcome Sanger Institute for the COVID-19 Genomics UK (COG-UK) Consortium | Harper VanSteenhouse, Yumi Kasai, David Gray, Carol Clugston, Anna Dominiczak and Alex Alderton, Roberto Amato, Jeffrey Barrett, Sonia Goncalves, Ewan Harrison, David K. Jackson, Ian Johnston, Dominic Kwiatkowski, Cordelia Langford, John Sillitoe on behalf of the Wellcome Sanger Institute COVID-19 Surveillance Team |
| EPI_ISL_3314144, EPI_ISL_3314145, EPI_ISL_3314146 | Lighthouse Lab in Alderley Park | Wellcome Sanger Institute for the COVID-19 Genomics UK (COG-UK) Consortium | Jacquelyn Wynn, Mairead Hyland, The Lighthouse Lab in Alderley Park and Alex Alderton, Roberto Amato, Jeffrey Barrett, Sonia Goncalves, Ewan Harrison, David K. Jackson, Ian Johnston, Dominic Kwiatkowski, Cordelia Langford, John Sillitoe on behalf of the Wellcome Sanger Institute COVID-19 Surveillance Team |
| EPI_ISL_3314147, EPI_ISL_3314148, EPI_ISL_3314149 | Lighthouse Lab in Glasgow | Wellcome Sanger Institute for the COVID-19 Genomics UK (COG-UK) Consortium | Harper VanSteenhouse, Yumi Kasai, David Gray, Carol Clugston, Anna Dominiczak and Alex Alderton, Roberto Amato, Jeffrey Barrett, Sonia Goncalves, Ewan Harrison, David K. Jackson, Ian Johnston, Dominic Kwiatkowski, Cordelia Langford, John Sillitoe on behalf of the Wellcome Sanger Institute COVID-19 Surveillance Team |
| EPI_ISL_3314150 | Lighthouse Lab in Alderley Park | Wellcome Sanger Institute for the COVID-19 Genomics UK (COG-UK) Consortium | Jacquelyn Wynn, Mairead Hyland, The Lighthouse Lab in Alderley Park and Alex Alderton, Roberto Amato, Jeffrey Barrett, Sonia Goncalves, Ewan Harrison, David K. Jackson, Ian Johnston, Dominic Kwiatkowski, Cordelia Langford, John Sillitoe on behalf of the Wellcome Sanger Institute COVID-19 Surveillance Team |
| EPI_ISL_3314151, EPI_ISL_3314152, EPI_ISL_3314153 | Lighthouse Lab in Glasgow | Wellcome Sanger Institute for the COVID-19 Genomics UK (COG-UK) Consortium | Harper VanSteenhouse, Yumi Kasai, David Gray, Carol Clugston, Anna Dominiczak and Alex Alderton, Roberto Amato, Jeffrey Barrett, Sonia Goncalves, Ewan Harrison, David K. Jackson, Ian Johnston, Dominic Kwiatkowski, Cordelia Langford, John Sillitoe on behalf of the Wellcome Sanger Institute COVID-19 Surveillance Team |
| EPI_ISL_3314154 | Berkshire and Surrey Pathology Services Lighthouse Laboratory | Wellcome Sanger Institute for the COVID-19 Genomics UK (COG-UK) Consortium | Berkshire and Surrey Pathology Services Lighthouse Laboratory and Alex Alderton, Roberto Amato, Jeffrey Barrett, Sonia Goncalves, Ewan Harrison, David K. Jackson, Ian Johnston, Dominic Kwiatkowski, Cordelia Langford, John Sillitoe on behalf of the Wellcome Sanger Institute COVID-19 Surveillance Team |
| EPI_ISL_3314155 | Lighthouse Lab in Glasgow | Wellcome Sanger Institute for the COVID-19 Genomics UK (COG-UK) Consortium | Harper VanSteenhouse, Yumi Kasai, David Gray, Carol Clugston, Anna Dominiczak and Alex Alderton, Roberto Amato, Jeffrey Barrett, Sonia Goncalves, Ewan Harrison, David K. Jackson, Ian Johnston, Dominic Kwiatkowski, Cordelia Langford, John Sillitoe on behalf of the Wellcome Sanger Institute COVID-19 Surveillance Team |
| EPI_ISL_3314156 | Berkshire and Surrey Pathology Services Lighthouse Laboratory | Wellcome Sanger Institute for the COVID-19 Genomics UK (COG-UK) Consortium | Berkshire and Surrey Pathology Services Lighthouse Laboratory and Alex Alderton, Roberto Amato, Jeffrey Barrett, Sonia Goncalves, Ewan Harrison, David K. Jackson, Ian Johnston, Dominic Kwiatkowski, Cordelia Langford, John Sillitoe on behalf of the Wellcome Sanger Institute COVID-19 Surveillance Team |
| EPI_ISL_3314157 | Lighthouse Lab in Alderley Park | Wellcome Sanger Institute for the COVID-19 Genomics UK (COG-UK) Consortium | Jacquelyn Wynn, Mairead Hyland, The Lighthouse Lab in Alderley Park and Alex Alderton, Roberto Amato, Jeffrey Barrett, Sonia Goncalves, Ewan Harrison, David K. Jackson, Ian Johnston, Dominic Kwiatkowski, Cordelia Langford, John Sillitoe on behalf of the Wellcome Sanger Institute COVID-19 Surveillance Team |
| EPI_ISL_3314158, EPI_ISL_3314159 | Lighthouse Lab in Glasgow | Wellcome Sanger Institute for the COVID-19 Genomics UK (COG-UK) Consortium | Harper VanSteenhouse, Yumi Kasai, David Gray, Carol Clugston, Anna Dominiczak and Alex Alderton, Roberto Amato, Jeffrey Barrett, Sonia Goncalves, Ewan Harrison, David K. Jackson, Ian Johnston, Dominic Kwiatkowski, Cordelia Langford, John Sillitoe on behalf of the Wellcome Sanger Institute COVID-19 Surveillance Team |
| EPI_ISL_3314160 | Berkshire and Surrey Pathology Services Lighthouse Laboratory | Wellcome Sanger Institute for the COVID-19 Genomics UK (COG-UK) Consortium | Berkshire and Surrey Pathology Services Lighthouse Laboratory and Alex Alderton, Roberto Amato, Jeffrey Barrett, Sonia Goncalves, Ewan Harrison, David K. Jackson, Ian Johnston, Dominic Kwiatkowski, Cordelia Langford, John Sillitoe on behalf of the Wellcome Sanger Institute COVID-19 Surveillance Team |
| EPI_ISL_3314161 | Lighthouse Lab in Alderley Park | Wellcome Sanger Institute for the COVID-19 Genomics UK (COG-UK) Consortium | Jacquelyn Wynn, Mairead Hyland, The Lighthouse Lab in Alderley Park and Alex Alderton, Roberto Amato, Jeffrey Barrett, Sonia Goncalves, Ewan Harrison, David K. Jackson, Ian Johnston, Dominic Kwiatkowski, Cordelia Langford, John Sillitoe on behalf of the Wellcome Sanger Institute COVID-19 Surveillance Team |
| EPI_ISL_3314162 | Lighthouse Lab in Glasgow | Wellcome Sanger Institute for the COVID-19 Genomics UK (COG-UK) Consortium | Harper VanSteenhouse, Yumi Kasai, David Gray, Carol Clugston, Anna Dominiczak and Alex Alderton, Roberto Amato, Jeffrey Barrett, Sonia Goncalves, Ewan Harrison, David K. Jackson, Ian Johnston, Dominic Kwiatkowski, Cordelia Langford, John Sillitoe on behalf of the Wellcome Sanger Institute COVID-19 Surveillance Team |
| EPI_ISL_3314163 | Lighthouse Lab in Alderley Park | Wellcome Sanger Institute for the COVID-19 Genomics UK (COG-UK) Consortium | Jacquelyn Wynn, Mairead Hyland, The Lighthouse Lab in Alderley Park and Alex Alderton, Roberto Amato, Jeffrey Barrett, Sonia Goncalves, Ewan Harrison, David K. Jackson, Ian Johnston, Dominic Kwiatkowski, Cordelia Langford, John Sillitoe on behalf of the Wellcome Sanger Institute COVID-19 Surveillance Team |
| EPI_ISL_3314164 | Lighthouse Lab in Glasgow | Wellcome Sanger Institute for the COVID-19 Genomics UK (COG-UK) Consortium | Harper VanSteenhouse, Yumi Kasai, David Gray, Carol Clugston, Anna Dominiczak and Alex Alderton, Roberto Amato, Jeffrey Barrett, Sonia Goncalves, Ewan Harrison, David K. Jackson, Ian Johnston, Dominic Kwiatkowski, Cordelia Langford, John Sillitoe on behalf of the Wellcome Sanger Institute COVID-19 Surveillance Team |
| EPI_ISL_3314165 | Berkshire and Surrey Pathology Services Lighthouse Laboratory | Wellcome Sanger Institute for the COVID-19 Genomics UK (COG-UK) Consortium | Berkshire and Surrey Pathology Services Lighthouse Laboratory and Alex Alderton, Roberto Amato, Jeffrey Barrett, Sonia Goncalves, Ewan Harrison, David K. Jackson, Ian Johnston, Dominic Kwiatkowski, Cordelia Langford, John Sillitoe on behalf of the Wellcome Sanger Institute COVID-19 Surveillance Team |
| EPI_ISL_3314166 | Lighthouse Lab in Glasgow | Wellcome Sanger Institute for the COVID-19 Genomics UK (COG-UK) Consortium | Harper VanSteenhouse, Yumi Kasai, David Gray, Carol Clugston, Anna Dominiczak and Alex Alderton, Roberto Amato, Jeffrey Barrett, Sonia Goncalves, Ewan Harrison, David K. Jackson, Ian Johnston, Dominic Kwiatkowski, Cordelia Langford, John Sillitoe on behalf of the Wellcome Sanger Institute COVID-19 Surveillance Team |
| EPI_ISL_3314167 | Lighthouse Lab in Alderley Park | Wellcome Sanger Institute for the COVID-19 Genomics UK (COG-UK) Consortium | Jacquelyn Wynn, Mairead Hyland, The Lighthouse Lab in Alderley Park and Alex Alderton, Roberto Amato, Jeffrey Barrett, Sonia Goncalves, Ewan Harrison, David K. Jackson, Ian Johnston, Dominic Kwiatkowski, Cordelia Langford, John Sillitoe on behalf of the Wellcome Sanger Institute COVID-19 Surveillance Team |
| EPI_ISL_3314168 | Berkshire and Surrey Pathology Services Lighthouse Laboratory | Wellcome Sanger Institute for the COVID-19 Genomics UK (COG-UK) Consortium | Berkshire and Surrey Pathology Services Lighthouse Laboratory and Alex Alderton, Roberto Amato, Jeffrey Barrett, Sonia Goncalves, Ewan Harrison, David K. Jackson, Ian Johnston, Dominic Kwiatkowski, Cordelia Langford, John Sillitoe on behalf of the Wellcome Sanger Institute COVID-19 Surveillance Team |
| EPI_ISL_3314169 | Lighthouse Lab in Alderley Park | Wellcome Sanger Institute for the COVID-19 Genomics UK (COG-UK) Consortium | Jacquelyn Wynn, Mairead Hyland, The Lighthouse Lab in Alderley Park and Alex Alderton, Roberto Amato, Jeffrey Barrett, Sonia Goncalves, Ewan Harrison, David K. Jackson, Ian Johnston, Dominic Kwiatkowski, Cordelia Langford, John Sillitoe on behalf of the Wellcome Sanger Institute COVID-19 Surveillance Team |
| EPI_ISL_3314170, EPI_ISL_3314171 | Lighthouse Lab in Glasgow | Wellcome Sanger Institute for the COVID-19 Genomics UK (COG-UK) Consortium | Harper VanSteenhouse, Yumi Kasai, David Gray, Carol Clugston, Anna Dominiczak and Alex Alderton, Roberto Amato, Jeffrey Barrett, Sonia Goncalves, Ewan Harrison, David K. Jackson, Ian Johnston, Dominic Kwiatkowski, Cordelia Langford, John Sillitoe on behalf of the Wellcome Sanger Institute COVID-19 Surveillance Team |
| EPI_ISL_3314172 | Lighthouse Lab in Alderley Park | Wellcome Sanger Institute for the COVID-19 Genomics UK (COG-UK) Consortium | Jacquelyn Wynn, Mairead Hyland, The Lighthouse Lab in Alderley Park and Alex Alderton, Roberto Amato, Jeffrey Barrett, Sonia Goncalves, Ewan Harrison, David K. Jackson, Ian Johnston, Dominic Kwiatkowski, Cordelia Langford, John Sillitoe on behalf of the Wellcome Sanger Institute COVID-19 Surveillance Team |
| EPI_ISL_3314173, EPI_ISL_3314174, EPI_ISL_3314175 | Lighthouse Lab in Glasgow | Wellcome Sanger Institute for the COVID-19 Genomics UK (COG-UK) Consortium | Harper VanSteenhouse, Yumi Kasai, David Gray, Carol Clugston, Anna Dominiczak and Alex Alderton, Roberto Amato, Jeffrey Barrett, Sonia Goncalves, Ewan Harrison, David K. Jackson, Ian Johnston, Dominic Kwiatkowski, Cordelia Langford, John Sillitoe on behalf of the Wellcome Sanger Institute COVID-19 Surveillance Team |

| | | | |
|---|---|---|---|
| EPI_ISL_3314176, EPI_ISL_3314177 | Lighthouse Lab in Alderley Park | Wellcome Sanger Institute for the COVID-19 Genomics UK (COG-UK) Consortium | Jacquelyn Wynn, Mairead Hyland, The Lighthouse Lab in Alderley Park and Alex Alderton, Roberto Amato, Jeffrey Barrett, Sonia Goncalves, Ewan Harrison, David K. Jackson, Ian Johnston, Dominic Kwiatkowski, Cordelia Langford, John Sillitoe on behalf of the Wellcome Sanger Institute COVID-19 Surveillance Team |
| EPI_ISL_3314178, EPI_ISL_3314179 | Berkshire and Surrey Pathology Services Lighthouse Laboratory | Wellcome Sanger Institute for the COVID-19 Genomics UK (COG-UK) Consortium | Berkshire and Surrey Pathology Services Lighthouse Laboratory and Alex Alderton, Roberto Amato, Jeffrey Barrett, Sonia Goncalves, Ewan Harrison, David K. Jackson, Ian Johnston, Dominic Kwiatkowski, Cordelia Langford, John Sillitoe on behalf of the Wellcome Sanger Institute COVID-19 Surveillance Team |
| EPI_ISL_3314180, EPI_ISL_3314181, EPI_ISL_3314182 | Lighthouse Lab in Glasgow | Wellcome Sanger Institute for the COVID-19 Genomics UK (COG-UK) Consortium | Harper VanSteenhouse, Yumi Kasai, David Gray, Carol Clugston, Anna Dominiczak and Alex Alderton, Roberto Amato, Jeffrey Barrett, Sonia Goncalves, Ewan Harrison, David K. Jackson, Ian Johnston, Dominic Kwiatkowski, Cordelia Langford, John Sillitoe on behalf of the Wellcome Sanger Institute COVID-19 Surveillance Team |
| EPI_ISL_3314183, EPI_ISL_3314184 | Berkshire and Surrey Pathology Services Lighthouse Laboratory | Wellcome Sanger Institute for the COVID-19 Genomics UK (COG-UK) Consortium | Berkshire and Surrey Pathology Services Lighthouse Laboratory and Alex Alderton, Roberto Amato, Jeffrey Barrett, Sonia Goncalves, Ewan Harrison, David K. Jackson, Ian Johnston, Dominic Kwiatkowski, Cordelia Langford, John Sillitoe on behalf of the Wellcome Sanger Institute COVID-19 Surveillance Team |
| EPI_ISL_3314185 | Lighthouse Lab in Glasgow | Wellcome Sanger Institute for the COVID-19 Genomics UK (COG-UK) Consortium | Harper VanSteenhouse, Yumi Kasai, David Gray, Carol Clugston, Anna Dominiczak and Alex Alderton, Roberto Amato, Jeffrey Barrett, Sonia Goncalves, Ewan Harrison, David K. Jackson, Ian Johnston, Dominic Kwiatkowski, Cordelia Langford, John Sillitoe on behalf of the Wellcome Sanger Institute COVID-19 Surveillance Team |
| EPI_ISL_3314186 | Berkshire and Surrey Pathology Services Lighthouse Laboratory | Wellcome Sanger Institute for the COVID-19 Genomics UK (COG-UK) Consortium | Berkshire and Surrey Pathology Services Lighthouse Laboratory and Alex Alderton, Roberto Amato, Jeffrey Barrett, Sonia Goncalves, Ewan Harrison, David K. Jackson, Ian Johnston, Dominic Kwiatkowski, Cordelia Langford, John Sillitoe on behalf of the Wellcome Sanger Institute COVID-19 Surveillance Team |
| EPI_ISL_3314187 | Lighthouse Lab in Glasgow | Wellcome Sanger Institute for the COVID-19 Genomics UK (COG-UK) Consortium | Harper VanSteenhouse, Yumi Kasai, David Gray, Carol Clugston, Anna Dominiczak and Alex Alderton, Roberto Amato, Jeffrey Barrett, Sonia Goncalves, Ewan Harrison, David K. Jackson, Ian Johnston, Dominic Kwiatkowski, Cordelia Langford, John Sillitoe on behalf of the Wellcome Sanger Institute COVID-19 Surveillance Team |
| EPI_ISL_3314188 | Berkshire and Surrey Pathology Services Lighthouse Laboratory | Wellcome Sanger Institute for the COVID-19 Genomics UK (COG-UK) Consortium | Berkshire and Surrey Pathology Services Lighthouse Laboratory and Alex Alderton, Roberto Amato, Jeffrey Barrett, Sonia Goncalves, Ewan Harrison, David K. Jackson, Ian Johnston, Dominic Kwiatkowski, Cordelia Langford, John Sillitoe on behalf of the Wellcome Sanger Institute COVID-19 Surveillance Team |
| EPI_ISL_3314189 | Lighthouse Lab in Glasgow | Wellcome Sanger Institute for the COVID-19 Genomics UK (COG-UK) Consortium | Harper VanSteenhouse, Yumi Kasai, David Gray, Carol Clugston, Anna Dominiczak and Alex Alderton, Roberto Amato, Jeffrey Barrett, Sonia Goncalves, Ewan Harrison, David K. Jackson, Ian Johnston, Dominic Kwiatkowski, Cordelia Langford, John Sillitoe on behalf of the Wellcome Sanger Institute COVID-19 Surveillance Team |
| EPI_ISL_3314190 | Lighthouse Lab in Alderley Park | Wellcome Sanger Institute for the COVID-19 Genomics UK (COG-UK) Consortium | Jacquelyn Wynn, Mairead Hyland, The Lighthouse Lab in Alderley Park and Alex Alderton, Roberto Amato, Jeffrey Barrett, Sonia Goncalves, Ewan Harrison, David K. Jackson, Ian Johnston, Dominic Kwiatkowski, Cordelia Langford, John Sillitoe on behalf of the Wellcome Sanger Institute COVID-19 Surveillance Team |
| EPI_ISL_3314191 | Lighthouse Lab in Glasgow | Wellcome Sanger Institute for the COVID-19 Genomics UK (COG-UK) Consortium | Harper VanSteenhouse, Yumi Kasai, David Gray, Carol Clugston, Anna Dominiczak and Alex Alderton, Roberto Amato, Jeffrey Barrett, Sonia Goncalves, Ewan Harrison, David K. Jackson, Ian Johnston, Dominic Kwiatkowski, Cordelia Langford, John Sillitoe on behalf of the Wellcome Sanger Institute COVID-19 Surveillance Team |
| EPI_ISL_3314192 | Lighthouse Lab in Alderley Park | Wellcome Sanger Institute for the COVID-19 Genomics UK (COG-UK) Consortium | Jacquelyn Wynn, Mairead Hyland, The Lighthouse Lab in Alderley Park and Alex Alderton, Roberto Amato, Jeffrey Barrett, Sonia Goncalves, Ewan Harrison, David K. Jackson, Ian Johnston, Dominic Kwiatkowski, Cordelia Langford, John Sillitoe on behalf of the Wellcome Sanger Institute COVID-19 Surveillance Team |
| EPI_ISL_3314193, EPI_ISL_3314194 | Berkshire and Surrey Pathology Services Lighthouse Laboratory | Wellcome Sanger Institute for the COVID-19 Genomics UK (COG-UK) Consortium | Berkshire and Surrey Pathology Services Lighthouse Laboratory and Alex Alderton, Roberto Amato, Jeffrey Barrett, Sonia Goncalves, Ewan Harrison, David K. Jackson, Ian Johnston, Dominic Kwiatkowski, Cordelia Langford, John Sillitoe on behalf of the Wellcome Sanger Institute COVID-19 Surveillance Team |
| EPI_ISL_3314195 | Lighthouse Lab in Glasgow | Wellcome Sanger Institute for the COVID-19 Genomics UK (COG-UK) Consortium | Harper VanSteenhouse, Yumi Kasai, David Gray, Carol Clugston, Anna Dominiczak and Alex Alderton, Roberto Amato, Jeffrey Barrett, Sonia Goncalves, Ewan Harrison, David K. Jackson, Ian Johnston, Dominic Kwiatkowski, Cordelia Langford, John Sillitoe on behalf of the Wellcome Sanger Institute COVID-19 Surveillance Team |
| EPI_ISL_3314196 | Lighthouse Lab in Alderley Park | Wellcome Sanger Institute for the COVID-19 Genomics UK (COG-UK) Consortium | Jacquelyn Wynn, Mairead Hyland, The Lighthouse Lab in Alderley Park and Alex Alderton, Roberto Amato, Jeffrey Barrett, Sonia Goncalves, Ewan Harrison, David K. Jackson, Ian Johnston, Dominic Kwiatkowski, Cordelia Langford, John Sillitoe on behalf of the Wellcome Sanger Institute COVID-19 Surveillance Team |
| EPI_ISL_3314197 | Berkshire and Surrey Pathology Services Lighthouse Laboratory | Wellcome Sanger Institute for the COVID-19 Genomics UK (COG-UK) Consortium | Berkshire and Surrey Pathology Services Lighthouse Laboratory and Alex Alderton, Roberto Amato, Jeffrey Barrett, Sonia Goncalves, Ewan Harrison, David K. Jackson, Ian Johnston, Dominic Kwiatkowski, Cordelia Langford, John Sillitoe on behalf of the Wellcome Sanger Institute COVID-19 Surveillance Team |
| EPI_ISL_3314198, EPI_ISL_3314199 | Lighthouse Lab in Alderley Park | Wellcome Sanger Institute for the COVID-19 Genomics UK (COG-UK) Consortium | Jacquelyn Wynn, Mairead Hyland, The Lighthouse Lab in Alderley Park and Alex Alderton, Roberto Amato, Jeffrey Barrett, Sonia Goncalves, Ewan Harrison, David K. Jackson, Ian Johnston, Dominic Kwiatkowski, Cordelia Langford, John Sillitoe on behalf of the Wellcome Sanger Institute COVID-19 Surveillance Team |
| EPI_ISL_3314200, EPI_ISL_3314201 | Lighthouse Lab in Glasgow | Wellcome Sanger Institute for the COVID-19 Genomics UK (COG-UK) Consortium | Harper VanSteenhouse, Yumi Kasai, David Gray, Carol Clugston, Anna Dominiczak and Alex Alderton, Roberto Amato, Jeffrey Barrett, Sonia Goncalves, Ewan Harrison, David K. Jackson, Ian Johnston, Dominic Kwiatkowski, Cordelia Langford, John Sillitoe on behalf of the Wellcome Sanger Institute COVID-19 Surveillance Team |
| EPI_ISL_3314202 | Lighthouse Lab in Alderley Park | Wellcome Sanger Institute for the COVID-19 Genomics UK (COG-UK) Consortium | Jacquelyn Wynn, Mairead Hyland, The Lighthouse Lab in Alderley Park and Alex Alderton, Roberto Amato, Jeffrey Barrett, Sonia Goncalves, Ewan Harrison, David K. Jackson, Ian Johnston, Dominic Kwiatkowski, Cordelia Langford, John Sillitoe on behalf of the Wellcome Sanger Institute COVID-19 Surveillance Team |
| EPI_ISL_3314203, EPI_ISL_3314204, EPI_ISL_3314205 | Lighthouse Lab in Glasgow | Wellcome Sanger Institute for the COVID-19 Genomics UK (COG-UK) Consortium | Harper VanSteenhouse, Yumi Kasai, David Gray, Carol Clugston, Anna Dominiczak and Alex Alderton, Roberto Amato, Jeffrey Barrett, Sonia Goncalves, Ewan Harrison, David K. Jackson, Ian Johnston, Dominic Kwiatkowski, Cordelia Langford, John Sillitoe on behalf of the Wellcome Sanger Institute COVID-19 Surveillance Team |
| EPI_ISL_3314206 | Lighthouse Lab in Alderley Park | Wellcome Sanger Institute for the COVID-19 Genomics UK (COG-UK) Consortium | Jacquelyn Wynn, Mairead Hyland, The Lighthouse Lab in Alderley Park and Alex Alderton, Roberto Amato, Jeffrey Barrett, Sonia Goncalves, Ewan Harrison, David K. Jackson, Ian Johnston, Dominic Kwiatkowski, Cordelia Langford, John Sillitoe on behalf of the Wellcome Sanger Institute COVID-19 Surveillance Team |
| EPI_ISL_3314207, EPI_ISL_3314208, EPI_ISL_3314209, EPI_ISL_3314210 | Berkshire and Surrey Pathology Services Lighthouse Laboratory | Wellcome Sanger Institute for the COVID-19 Genomics UK (COG-UK) Consortium | Berkshire and Surrey Pathology Services Lighthouse Laboratory and Alex Alderton, Roberto Amato, Jeffrey Barrett, Sonia Goncalves, Ewan Harrison, David K. Jackson, Ian Johnston, Dominic Kwiatkowski, Cordelia Langford, John Sillitoe on behalf of the Wellcome Sanger Institute COVID-19 Surveillance Team |
| EPI_ISL_3314211 | Lighthouse Lab in Alderley Park | Wellcome Sanger Institute for the COVID-19 Genomics UK (COG-UK) Consortium | Jacquelyn Wynn, Mairead Hyland, The Lighthouse Lab in Alderley Park and Alex Alderton, Roberto Amato, Jeffrey Barrett, Sonia Goncalves, Ewan Harrison, David K. Jackson, Ian Johnston, Dominic Kwiatkowski, Cordelia Langford, John Sillitoe on behalf of the Wellcome Sanger Institute COVID-19 Surveillance Team |
| EPI_ISL_3314212 | Lighthouse Lab in Glasgow | Wellcome Sanger Institute for the COVID-19 Genomics UK (COG-UK) Consortium | Harper VanSteenhouse, Yumi Kasai, David Gray, Carol Clugston, Anna Dominiczak and Alex Alderton, Roberto Amato, Jeffrey Barrett, Sonia Goncalves, Ewan Harrison, David K. Jackson, Ian Johnston, Dominic Kwiatkowski, Cordelia Langford, John Sillitoe on behalf of the Wellcome Sanger Institute COVID-19 Surveillance Team |
| EPI_ISL_3314213, EPI_ISL_3314214 | Lighthouse Lab in Alderley Park | Wellcome Sanger Institute for the COVID-19 Genomics UK (COG-UK) Consortium | Jacquelyn Wynn, Mairead Hyland, The Lighthouse Lab in Alderley Park and Alex Alderton, Roberto Amato, Jeffrey Barrett, Sonia Goncalves, Ewan Harrison, David K. Jackson, Ian Johnston, Dominic Kwiatkowski, Cordelia Langford, John Sillitoe on behalf of the Wellcome Sanger Institute COVID-19 Surveillance Team |
| EPI_ISL_3314215, EPI_ISL_3314216 | Lighthouse Lab in Glasgow | Wellcome Sanger Institute for the COVID-19 Genomics UK (COG-UK) Consortium | Harper VanSteenhouse, Yumi Kasai, David Gray, Carol Clugston, Anna Dominiczak and Alex Alderton, Roberto Amato, Jeffrey Barrett, Sonia Goncalves, Ewan Harrison, David K. Jackson, Ian Johnston, Dominic Kwiatkowski, Cordelia Langford, John Sillitoe on behalf of the Wellcome Sanger Institute COVID-19 Surveillance Team |
| EPI_ISL_3314217 | Berkshire and Surrey Pathology Services Lighthouse Laboratory | Wellcome Sanger Institute for the COVID-19 Genomics UK (COG-UK) Consortium | Berkshire and Surrey Pathology Services Lighthouse Laboratory and Alex Alderton, Roberto Amato, Jeffrey Barrett, Sonia Goncalves, Ewan Harrison, David K. Jackson, Ian Johnston, Dominic Kwiatkowski, Cordelia Langford, John Sillitoe on behalf of the Wellcome Sanger Institute COVID-19 Surveillance Team |
| EPI_ISL_3314218, EPI_ISL_3314219, EPI_ISL_3314220, EPI_ISL_3314221 | Lighthouse Lab in Glasgow | Wellcome Sanger Institute for the COVID-19 Genomics UK (COG-UK) Consortium | Harper VanSteenhouse, Yumi Kasai, David Gray, Carol Clugston, Anna Dominiczak and Alex Alderton, Roberto Amato, Jeffrey Barrett, Sonia Goncalves, Ewan Harrison, David K. Jackson, Ian Johnston, Dominic Kwiatkowski, Cordelia Langford, John Sillitoe on behalf of the Wellcome Sanger Institute COVID-19 Surveillance Team |
| EPI_ISL_3314222 | Lighthouse Lab in Alderley Park | Wellcome Sanger Institute for the COVID-19 Genomics UK (COG-UK) Consortium | Jacquelyn Wynn, Mairead Hyland, The Lighthouse Lab in Alderley Park and Alex Alderton, Roberto Amato, Jeffrey Barrett, Sonia Goncalves, Ewan Harrison, David K. Jackson, Ian Johnston, Dominic Kwiatkowski, Cordelia Langford, John Sillitoe on behalf of the Wellcome Sanger Institute COVID-19 Surveillance Team |
| EPI_ISL_3314223, EPI_ISL_3314224, EPI_ISL_3314225, EPI_ISL_3314226, EPI_ISL_3314227, EPI_ISL_3314228 | Berkshire and Surrey Pathology Services Lighthouse Laboratory | Wellcome Sanger Institute for the COVID-19 Genomics UK (COG-UK) Consortium | Berkshire and Surrey Pathology Services Lighthouse Laboratory and Alex Alderton, Roberto Amato, Jeffrey Barrett, Sonia Goncalves, Ewan Harrison, David K. Jackson, Ian Johnston, Dominic Kwiatkowski, Cordelia Langford, John Sillitoe on behalf of the Wellcome Sanger Institute COVID-19 Surveillance Team |
| EPI_ISL_3314229, EPI_ISL_3314230 | Lighthouse Lab in Glasgow | Wellcome Sanger Institute for the COVID-19 Genomics UK (COG-UK) Consortium | Harper VanSteenhouse, Yumi Kasai, David Gray, Carol Clugston, Anna Dominiczak and Alex Alderton, Roberto Amato, Jeffrey Barrett, Sonia Goncalves, Ewan Harrison, David K. Jackson, Ian Johnston, Dominic Kwiatkowski, Cordelia Langford, John Sillitoe on behalf of the Wellcome Sanger Institute COVID-19 Surveillance Team |
| EPI_ISL_3314231, EPI_ISL_3314232 | Berkshire and Surrey Pathology Services Lighthouse Laboratory | Wellcome Sanger Institute for the COVID-19 Genomics UK (COG-UK) Consortium | Berkshire and Surrey Pathology Services Lighthouse Laboratory and Alex Alderton, Roberto Amato, Jeffrey Barrett, Sonia Goncalves, Ewan Harrison, David K. Jackson, Ian Johnston, Dominic Kwiatkowski, Cordelia Langford, John Sillitoe on behalf of the Wellcome Sanger Institute COVID-19 Surveillance Team |
| EPI_ISL_3314233 | Lighthouse Lab in Glasgow | Wellcome Sanger Institute for the COVID-19 Genomics UK (COG-UK) Consortium | Harper VanSteenhouse, Yumi Kasai, David Gray, Carol Clugston, Anna Dominiczak and Alex Alderton, Roberto Amato, Jeffrey Barrett, Sonia Goncalves, Ewan Harrison, David K. Jackson, Ian Johnston, Dominic Kwiatkowski, Cordelia Langford, John Sillitoe on behalf of the Wellcome Sanger Institute COVID-19 Surveillance Team |
| EPI_ISL_3314234 | Berkshire and Surrey Pathology Services Lighthouse Laboratory | Wellcome Sanger Institute for the COVID-19 Genomics UK (COG-UK) Consortium | Berkshire and Surrey Pathology Services Lighthouse Laboratory and Alex Alderton, Roberto Amato, Jeffrey Barrett, Sonia Goncalves, Ewan Harrison, David K. Jackson, Ian Johnston, Dominic Kwiatkowski, Cordelia Langford, John Sillitoe on behalf of the Wellcome Sanger Institute COVID-19 Surveillance Team |
| EPI_ISL_3314235 | Lighthouse Lab in Glasgow | Wellcome Sanger Institute for the COVID-19 Genomics UK (COG-UK) Consortium | Harper VanSteenhouse, Yumi Kasai, David Gray, Carol Clugston, Anna Dominiczak and Alex Alderton, Roberto Amato, Jeffrey Barrett, Sonia Goncalves, Ewan Harrison, David K. Jackson, Ian Johnston, Dominic Kwiatkowski, Cordelia Langford, John Sillitoe on behalf of the Wellcome Sanger Institute COVID-19 Surveillance Team |
| EPI_ISL_3314236, EPI_ISL_3314237 | Lighthouse Lab in Alderley Park | Wellcome Sanger Institute for the COVID-19 Genomics UK (COG-UK) Consortium | Jacquelyn Wynn, Mairead Hyland, The Lighthouse Lab in Alderley Park and Alex Alderton, Roberto Amato, Jeffrey Barrett, Sonia Goncalves, Ewan Harrison, David K. Jackson, Ian Johnston, Dominic Kwiatkowski, Cordelia Langford, John Sillitoe on behalf of the Wellcome Sanger Institute COVID-19 Surveillance Team |
| EPI_ISL_3314238 | Lighthouse Lab in Glasgow | Wellcome Sanger Institute for the COVID-19 Genomics UK (COG-UK) Consortium | Harper VanSteenhouse, Yumi Kasai, David Gray, Carol Clugston, Anna Dominiczak and Alex Alderton, Roberto Amato, Jeffrey Barrett, Sonia Goncalves, Ewan Harrison, David K. Jackson, Ian Johnston, Dominic Kwiatkowski, Cordelia Langford, John Sillitoe on behalf of the Wellcome Sanger Institute COVID-19 Surveillance Team |
| EPI_ISL_3314239 | Lighthouse Lab in Alderley Park | Wellcome Sanger Institute for the COVID-19 Genomics UK (COG-UK) Consortium | Jacquelyn Wynn, Mairead Hyland, The Lighthouse Lab in Alderley Park and Alex Alderton, Roberto Amato, Jeffrey Barrett, Sonia Goncalves, Ewan Harrison, David K. Jackson, Ian Johnston, Dominic Kwiatkowski, Cordelia Langford, John Sillitoe on behalf of the Wellcome Sanger Institute COVID-19 Surveillance Team |
| EPI_ISL_3314240, EPI_ISL_3314241 | Lighthouse Lab in Glasgow | Wellcome Sanger Institute for the COVID-19 Genomics UK (COG-UK) Consortium | Harper VanSteenhouse, Yumi Kasai, David Gray, Carol Clugston, Anna Dominiczak and Alex Alderton, Roberto Amato, Jeffrey Barrett, Sonia Goncalves, Ewan Harrison, David K. Jackson, Ian Johnston, Dominic Kwiatkowski, Cordelia Langford, John Sillitoe on behalf of the Wellcome Sanger Institute COVID-19 Surveillance Team |
| EPI_ISL_3314242 | Lighthouse Lab in Alderley Park | Wellcome Sanger Institute for the COVID-19 Genomics UK (COG-UK) Consortium | Jacquelyn Wynn, Mairead Hyland, The Lighthouse Lab in Alderley Park and Alex Alderton, Roberto Amato, Jeffrey Barrett, Sonia Goncalves, Ewan Harrison, David K. Jackson, Ian Johnston, Dominic Kwiatkowski, Cordelia Langford, John Sillitoe on behalf of the Wellcome Sanger Institute COVID-19 Surveillance Team |
| EPI_ISL_3314243 | Lighthouse Lab in Glasgow | Wellcome Sanger Institute for the COVID-19 Genomics UK (COG-UK) Consortium | Harper VanSteenhouse, Yumi Kasai, David Gray, Carol Clugston, Anna Dominiczak and Alex Alderton, Roberto Amato, Jeffrey Barrett, Sonia Goncalves, Ewan Harrison, David K. Jackson, Ian Johnston, Dominic Kwiatkowski, Cordelia Langford, John Sillitoe on behalf of the Wellcome Sanger Institute COVID-19 Surveillance Team |

| | | | |
|---|---|---|---|
| EPI_ISL_3314244 | Berkshire and Surrey Pathology Services Lighthouse Laboratory | Wellcome Sanger Institute for the COVID-19 Genomics UK (COG-UK) Consortium | Berkshire and Surrey Pathology Services Lighthouse Laboratory and Alex Alderton, Roberto Amato, Jeffrey Barrett, Sonia Goncalves, Ewan Harrison, David K. Jackson, Ian Johnston, Dominic Kwiatkowski, Cordelia Langford, John Sillitoe on behalf of the Wellcome Sanger Institute COVID-19 Surveillance Team |
| EPI_ISL_3314245 | Lighthouse Lab in Alderley Park | Wellcome Sanger Institute for the COVID-19 Genomics UK (COG-UK) Consortium | Jacquelyn Wynn, Mairead Hyland, The Lighthouse Lab in Alderley Park and Alex Alderton, Roberto Amato, Jeffrey Barrett, Sonia Goncalves, Ewan Harrison, David K. Jackson, Ian Johnston, Dominic Kwiatkowski, Cordelia Langford, John Sillitoe on behalf of the Wellcome Sanger Institute COVID-19 Surveillance Team |
| EPI_ISL_3314246 | Lighthouse Lab in Glasgow | Wellcome Sanger Institute for the COVID-19 Genomics UK (COG-UK) Consortium | Harper VanSteenhouse, Yumi Kasai, David Gray, Carol Clugston, Anna Dominiczak and Alex Alderton, Roberto Amato, Jeffrey Barrett, Sonia Goncalves, Ewan Harrison, David K. Jackson, Ian Johnston, Dominic Kwiatkowski, Cordelia Langford, John Sillitoe on behalf of the Wellcome Sanger Institute COVID-19 Surveillance Team |
| EPI_ISL_3314247 | Lighthouse Lab in Alderley Park | Wellcome Sanger Institute for the COVID-19 Genomics UK (COG-UK) Consortium | Jacquelyn Wynn, Mairead Hyland, The Lighthouse Lab in Alderley Park and Alex Alderton, Roberto Amato, Jeffrey Barrett, Sonia Goncalves, Ewan Harrison, David K. Jackson, Ian Johnston, Dominic Kwiatkowski, Cordelia Langford, John Sillitoe on behalf of the Wellcome Sanger Institute COVID-19 Surveillance Team |
| EPI_ISL_3314248, EPI_ISL_3314249 | Lighthouse Lab in Glasgow | Wellcome Sanger Institute for the COVID-19 Genomics UK (COG-UK) Consortium | Harper VanSteenhouse, Yumi Kasai, David Gray, Carol Clugston, Anna Dominiczak and Alex Alderton, Roberto Amato, Jeffrey Barrett, Sonia Goncalves, Ewan Harrison, David K. Jackson, Ian Johnston, Dominic Kwiatkowski, Cordelia Langford, John Sillitoe on behalf of the Wellcome Sanger Institute COVID-19 Surveillance Team |
| EPI_ISL_3314250 | Berkshire and Surrey Pathology Services Lighthouse Laboratory | Wellcome Sanger Institute for the COVID-19 Genomics UK (COG-UK) Consortium | Berkshire and Surrey Pathology Services Lighthouse Laboratory and Alex Alderton, Roberto Amato, Jeffrey Barrett, Sonia Goncalves, Ewan Harrison, David K. Jackson, Ian Johnston, Dominic Kwiatkowski, Cordelia Langford, John Sillitoe on behalf of the Wellcome Sanger Institute COVID-19 Surveillance Team |
| EPI_ISL_3314251 | Lighthouse Lab in Glasgow | Wellcome Sanger Institute for the COVID-19 Genomics UK (COG-UK) Consortium | Harper VanSteenhouse, Yumi Kasai, David Gray, Carol Clugston, Anna Dominiczak and Alex Alderton, Roberto Amato, Jeffrey Barrett, Sonia Goncalves, Ewan Harrison, David K. Jackson, Ian Johnston, Dominic Kwiatkowski, Cordelia Langford, John Sillitoe on behalf of the Wellcome Sanger Institute COVID-19 Surveillance Team |
| EPI_ISL_3314252 | Berkshire and Surrey Pathology Services Lighthouse Laboratory | Wellcome Sanger Institute for the COVID-19 Genomics UK (COG-UK) Consortium | Berkshire and Surrey Pathology Services Lighthouse Laboratory and Alex Alderton, Roberto Amato, Jeffrey Barrett, Sonia Goncalves, Ewan Harrison, David K. Jackson, Ian Johnston, Dominic Kwiatkowski, Cordelia Langford, John Sillitoe on behalf of the Wellcome Sanger Institute COVID-19 Surveillance Team |
| EPI_ISL_3314253, EPI_ISL_3314254, EPI_ISL_3314255, EPI_ISL_3314256, EPI_ISL_3314257 | Lighthouse Lab in Glasgow | Wellcome Sanger Institute for the COVID-19 Genomics UK (COG-UK) Consortium | Harper VanSteenhouse, Yumi Kasai, David Gray, Carol Clugston, Anna Dominiczak and Alex Alderton, Roberto Amato, Jeffrey Barrett, Sonia Goncalves, Ewan Harrison, David K. Jackson, Ian Johnston, Dominic Kwiatkowski, Cordelia Langford, John Sillitoe on behalf of the Wellcome Sanger Institute COVID-19 Surveillance Team |
| EPI_ISL_3314258 | Lighthouse Lab in Alderley Park | Wellcome Sanger Institute for the COVID-19 Genomics UK (COG-UK) Consortium | Jacquelyn Wynn, Mairead Hyland, The Lighthouse Lab in Alderley Park and Alex Alderton, Roberto Amato, Jeffrey Barrett, Sonia Goncalves, Ewan Harrison, David K. Jackson, Ian Johnston, Dominic Kwiatkowski, Cordelia Langford, John Sillitoe on behalf of the Wellcome Sanger Institute COVID-19 Surveillance Team |
| EPI_ISL_3314259 | Berkshire and Surrey Pathology Services Lighthouse Laboratory | Wellcome Sanger Institute for the COVID-19 Genomics UK (COG-UK) Consortium | Berkshire and Surrey Pathology Services Lighthouse Laboratory and Alex Alderton, Roberto Amato, Jeffrey Barrett, Sonia Goncalves, Ewan Harrison, David K. Jackson, Ian Johnston, Dominic Kwiatkowski, Cordelia Langford, John Sillitoe on behalf of the Wellcome Sanger Institute COVID-19 Surveillance Team |
| EPI_ISL_3314260, EPI_ISL_3314261, EPI_ISL_3314262, EPI_ISL_3314263, EPI_ISL_3314264 | Lighthouse Lab in Alderley Park | Wellcome Sanger Institute for the COVID-19 Genomics UK (COG-UK) Consortium | Jacquelyn Wynn, Mairead Hyland, The Lighthouse Lab in Alderley Park and Alex Alderton, Roberto Amato, Jeffrey Barrett, Sonia Goncalves, Ewan Harrison, David K. Jackson, Ian Johnston, Dominic Kwiatkowski, Cordelia Langford, John Sillitoe on behalf of the Wellcome Sanger Institute COVID-19 Surveillance Team |
| EPI_ISL_3314265, EPI_ISL_3314266 | Lighthouse Lab in Glasgow | Wellcome Sanger Institute for the COVID-19 Genomics UK (COG-UK) Consortium | Harper VanSteenhouse, Yumi Kasai, David Gray, Carol Clugston, Anna Dominiczak and Alex Alderton, Roberto Amato, Jeffrey Barrett, Sonia Goncalves, Ewan Harrison, David K. Jackson, Ian Johnston, Dominic Kwiatkowski, Cordelia Langford, John Sillitoe on behalf of the Wellcome Sanger Institute COVID-19 Surveillance Team |
| EPI_ISL_3314267 | Lighthouse Lab in Alderley Park | Wellcome Sanger Institute for the COVID-19 Genomics UK (COG-UK) Consortium | Jacquelyn Wynn, Mairead Hyland, The Lighthouse Lab in Alderley Park and Alex Alderton, Roberto Amato, Jeffrey Barrett, Sonia Goncalves, Ewan Harrison, David K. Jackson, Ian Johnston, Dominic Kwiatkowski, Cordelia Langford, John Sillitoe on behalf of the Wellcome Sanger Institute COVID-19 Surveillance Team |
| EPI_ISL_3314268, EPI_ISL_3314269, EPI_ISL_3314270, EPI_ISL_3314271 | Lighthouse Lab in Glasgow | Wellcome Sanger Institute for the COVID-19 Genomics UK (COG-UK) Consortium | Harper VanSteenhouse, Yumi Kasai, David Gray, Carol Clugston, Anna Dominiczak and Alex Alderton, Roberto Amato, Jeffrey Barrett, Sonia Goncalves, Ewan Harrison, David K. Jackson, Ian Johnston, Dominic Kwiatkowski, Cordelia Langford, John Sillitoe on behalf of the Wellcome Sanger Institute COVID-19 Surveillance Team |
| EPI_ISL_3314272 | Lighthouse Lab in Alderley Park | Wellcome Sanger Institute for the COVID-19 Genomics UK (COG-UK) Consortium | Jacquelyn Wynn, Mairead Hyland, The Lighthouse Lab in Alderley Park and Alex Alderton, Roberto Amato, Jeffrey Barrett, Sonia Goncalves, Ewan Harrison, David K. Jackson, Ian Johnston, Dominic Kwiatkowski, Cordelia Langford, John Sillitoe on behalf of the Wellcome Sanger Institute COVID-19 Surveillance Team |
| EPI_ISL_3314273 | Lighthouse Lab in Glasgow | Wellcome Sanger Institute for the COVID-19 Genomics UK (COG-UK) Consortium | Harper VanSteenhouse, Yumi Kasai, David Gray, Carol Clugston, Anna Dominiczak and Alex Alderton, Roberto Amato, Jeffrey Barrett, Sonia Goncalves, Ewan Harrison, David K. Jackson, Ian Johnston, Dominic Kwiatkowski, Cordelia Langford, John Sillitoe on behalf of the Wellcome Sanger Institute COVID-19 Surveillance Team |
| EPI_ISL_3314274 | Berkshire and Surrey Pathology Services Lighthouse Laboratory | Wellcome Sanger Institute for the COVID-19 Genomics UK (COG-UK) Consortium | Berkshire and Surrey Pathology Services Lighthouse Laboratory and Alex Alderton, Roberto Amato, Jeffrey Barrett, Sonia Goncalves, Ewan Harrison, David K. Jackson, Ian Johnston, Dominic Kwiatkowski, Cordelia Langford, John Sillitoe on behalf of the Wellcome Sanger Institute COVID-19 Surveillance Team |
| EPI_ISL_3314275 | Lighthouse Lab in Alderley Park | Wellcome Sanger Institute for the COVID-19 Genomics UK (COG-UK) Consortium | Jacquelyn Wynn, Mairead Hyland, The Lighthouse Lab in Alderley Park and Alex Alderton, Roberto Amato, Jeffrey Barrett, Sonia Goncalves, Ewan Harrison, David K. Jackson, Ian Johnston, Dominic Kwiatkowski, Cordelia Langford, John Sillitoe on behalf of the Wellcome Sanger Institute COVID-19 Surveillance Team |
| EPI_ISL_3314276 | Lighthouse Lab in Glasgow | Wellcome Sanger Institute for the COVID-19 Genomics UK (COG-UK) Consortium | Harper VanSteenhouse, Yumi Kasai, David Gray, Carol Clugston, Anna Dominiczak and Alex Alderton, Roberto Amato, Jeffrey Barrett, Sonia Goncalves, Ewan Harrison, David K. Jackson, Ian Johnston, Dominic Kwiatkowski, Cordelia Langford, John Sillitoe on behalf of the Wellcome Sanger Institute COVID-19 Surveillance Team |
| EPI_ISL_3314277 | Lighthouse Lab in Alderley Park | Wellcome Sanger Institute for the COVID-19 Genomics UK (COG-UK) Consortium | Jacquelyn Wynn, Mairead Hyland, The Lighthouse Lab in Alderley Park and Alex Alderton, Roberto Amato, Jeffrey Barrett, Sonia Goncalves, Ewan Harrison, David K. Jackson, Ian Johnston, Dominic Kwiatkowski, Cordelia Langford, John Sillitoe on behalf of the Wellcome Sanger Institute COVID-19 Surveillance Team |
| EPI_ISL_3314278, EPI_ISL_3314279, EPI_ISL_3314280, EPI_ISL_3314281, EPI_ISL_3314282, EPI_ISL_3314283, EPI_ISL_3314284 | Lighthouse Lab in Glasgow | Wellcome Sanger Institute for the COVID-19 Genomics UK (COG-UK) Consortium | Harper VanSteenhouse, Yumi Kasai, David Gray, Carol Clugston, Anna Dominiczak and Alex Alderton, Roberto Amato, Jeffrey Barrett, Sonia Goncalves, Ewan Harrison, David K. Jackson, Ian Johnston, Dominic Kwiatkowski, Cordelia Langford, John Sillitoe on behalf of the Wellcome Sanger Institute COVID-19 Surveillance Team |
| EPI_ISL_3314285 | Lighthouse Lab in Alderley Park | Wellcome Sanger Institute for the COVID-19 Genomics UK (COG-UK) Consortium | Jacquelyn Wynn, Mairead Hyland, The Lighthouse Lab in Alderley Park and Alex Alderton, Roberto Amato, Jeffrey Barrett, Sonia Goncalves, Ewan Harrison, David K. Jackson, Ian Johnston, Dominic Kwiatkowski, Cordelia Langford, John Sillitoe on behalf of the Wellcome Sanger Institute COVID-19 Surveillance Team |
| EPI_ISL_3314286 | Lighthouse Lab in Glasgow | Wellcome Sanger Institute for the COVID-19 Genomics UK (COG-UK) Consortium | Harper VanSteenhouse, Yumi Kasai, David Gray, Carol Clugston, Anna Dominiczak and Alex Alderton, Roberto Amato, Jeffrey Barrett, Sonia Goncalves, Ewan Harrison, David K. Jackson, Ian Johnston, Dominic Kwiatkowski, Cordelia Langford, John Sillitoe on behalf of the Wellcome Sanger Institute COVID-19 Surveillance Team |
| EPI_ISL_3314287, EPI_ISL_3314288, EPI_ISL_3314289, EPI_ISL_3314290 | Berkshire and Surrey Pathology Services Lighthouse Laboratory | Wellcome Sanger Institute for the COVID-19 Genomics UK (COG-UK) Consortium | Berkshire and Surrey Pathology Services Lighthouse Laboratory and Alex Alderton, Roberto Amato, Jeffrey Barrett, Sonia Goncalves, Ewan Harrison, David K. Jackson, Ian Johnston, Dominic Kwiatkowski, Cordelia Langford, John Sillitoe on behalf of the Wellcome Sanger Institute COVID-19 Surveillance Team |
| EPI_ISL_3314291 | Lighthouse Lab in Glasgow | Wellcome Sanger Institute for the COVID-19 Genomics UK (COG-UK) Consortium | Harper VanSteenhouse, Yumi Kasai, David Gray, Carol Clugston, Anna Dominiczak and Alex Alderton, Roberto Amato, Jeffrey Barrett, Sonia Goncalves, Ewan Harrison, David K. Jackson, Ian Johnston, Dominic Kwiatkowski, Cordelia Langford, John Sillitoe on behalf of the Wellcome Sanger Institute COVID-19 Surveillance Team |
| EPI_ISL_3314292, EPI_ISL_3314293, EPI_ISL_3314294 | Lighthouse Lab in Alderley Park | Wellcome Sanger Institute for the COVID-19 Genomics UK (COG-UK) Consortium | Jacquelyn Wynn, Mairead Hyland, The Lighthouse Lab in Alderley Park and Alex Alderton, Roberto Amato, Jeffrey Barrett, Sonia Goncalves, Ewan Harrison, David K. Jackson, Ian Johnston, Dominic Kwiatkowski, Cordelia Langford, John Sillitoe on behalf of the Wellcome Sanger Institute COVID-19 Surveillance Team |
| EPI_ISL_3314295 | Berkshire and Surrey Pathology Services Lighthouse Laboratory | Wellcome Sanger Institute for the COVID-19 Genomics UK (COG-UK) Consortium | Berkshire and Surrey Pathology Services Lighthouse Laboratory and Alex Alderton, Roberto Amato, Jeffrey Barrett, Sonia Goncalves, Ewan Harrison, David K. Jackson, Ian Johnston, Dominic Kwiatkowski, Cordelia Langford, John Sillitoe on behalf of the Wellcome Sanger Institute COVID-19 Surveillance Team |
| EPI_ISL_3314296 | Lighthouse Lab in Glasgow | Wellcome Sanger Institute for the COVID-19 Genomics UK (COG-UK) Consortium | Harper VanSteenhouse, Yumi Kasai, David Gray, Carol Clugston, Anna Dominiczak and Alex Alderton, Roberto Amato, Jeffrey Barrett, Sonia Goncalves, Ewan Harrison, David K. Jackson, Ian Johnston, Dominic Kwiatkowski, Cordelia Langford, John Sillitoe on behalf of the Wellcome Sanger Institute COVID-19 Surveillance Team |
| EPI_ISL_3314297 | Lighthouse Lab in Alderley Park | Wellcome Sanger Institute for the COVID-19 Genomics UK (COG-UK) Consortium | Jacquelyn Wynn, Mairead Hyland, The Lighthouse Lab in Alderley Park and Alex Alderton, Roberto Amato, Jeffrey Barrett, Sonia Goncalves, Ewan Harrison, David K. Jackson, Ian Johnston, Dominic Kwiatkowski, Cordelia Langford, John Sillitoe on behalf of the Wellcome Sanger Institute COVID-19 Surveillance Team |
| EPI_ISL_3314298, EPI_ISL_3314299 | Lighthouse Lab in Glasgow | Wellcome Sanger Institute for the COVID-19 Genomics UK (COG-UK) Consortium | Harper VanSteenhouse, Yumi Kasai, David Gray, Carol Clugston, Anna Dominiczak and Alex Alderton, Roberto Amato, Jeffrey Barrett, Sonia Goncalves, Ewan Harrison, David K. Jackson, Ian Johnston, Dominic Kwiatkowski, Cordelia Langford, John Sillitoe on behalf of the Wellcome Sanger Institute COVID-19 Surveillance Team |
| EPI_ISL_3314300 | Lighthouse Lab in Alderley Park | Wellcome Sanger Institute for the COVID-19 Genomics UK (COG-UK) Consortium | Jacquelyn Wynn, Mairead Hyland, The Lighthouse Lab in Alderley Park and Alex Alderton, Roberto Amato, Jeffrey Barrett, Sonia Goncalves, Ewan Harrison, David K. Jackson, Ian Johnston, Dominic Kwiatkowski, Cordelia Langford, John Sillitoe on behalf of the Wellcome Sanger Institute COVID-19 Surveillance Team |
| EPI_ISL_3314301 | Lighthouse Lab in Glasgow | Wellcome Sanger Institute for the COVID-19 Genomics UK (COG-UK) Consortium | Harper VanSteenhouse, Yumi Kasai, David Gray, Carol Clugston, Anna Dominiczak and Alex Alderton, Roberto Amato, Jeffrey Barrett, Sonia Goncalves, Ewan Harrison, David K. Jackson, Ian Johnston, Dominic Kwiatkowski, Cordelia Langford, John Sillitoe on behalf of the Wellcome Sanger Institute COVID-19 Surveillance Team |
| EPI_ISL_3314302 | Berkshire and Surrey Pathology Services Lighthouse Laboratory | Wellcome Sanger Institute for the COVID-19 Genomics UK (COG-UK) Consortium | Berkshire and Surrey Pathology Services Lighthouse Laboratory and Alex Alderton, Roberto Amato, Jeffrey Barrett, Sonia Goncalves, Ewan Harrison, David K. Jackson, Ian Johnston, Dominic Kwiatkowski, Cordelia Langford, John Sillitoe on behalf of the Wellcome Sanger Institute COVID-19 Surveillance Team |
| EPI_ISL_3314303 | Lighthouse Lab in Alderley Park | Wellcome Sanger Institute for the COVID-19 Genomics UK (COG-UK) Consortium | Jacquelyn Wynn, Mairead Hyland, The Lighthouse Lab in Alderley Park and Alex Alderton, Roberto Amato, Jeffrey Barrett, Sonia Goncalves, Ewan Harrison, David K. Jackson, Ian Johnston, Dominic Kwiatkowski, Cordelia Langford, John Sillitoe on behalf of the Wellcome Sanger Institute COVID-19 Surveillance Team |
| EPI_ISL_3314304 | Lighthouse Lab in Glasgow | Wellcome Sanger Institute for the COVID-19 Genomics UK (COG-UK) Consortium | Harper VanSteenhouse, Yumi Kasai, David Gray, Carol Clugston, Anna Dominiczak and Alex Alderton, Roberto Amato, Jeffrey Barrett, Sonia Goncalves, Ewan Harrison, David K. Jackson, Ian Johnston, Dominic Kwiatkowski, Cordelia Langford, John Sillitoe on behalf of the Wellcome Sanger Institute COVID-19 Surveillance Team |
| EPI_ISL_3314305 | Lighthouse Lab in Alderley Park | Wellcome Sanger Institute for the COVID-19 Genomics UK (COG-UK) Consortium | Jacquelyn Wynn, Mairead Hyland, The Lighthouse Lab in Alderley Park and Alex Alderton, Roberto Amato, Jeffrey Barrett, Sonia Goncalves, Ewan Harrison, David K. Jackson, Ian Johnston, Dominic Kwiatkowski, Cordelia Langford, John Sillitoe on behalf of the Wellcome Sanger Institute COVID-19 Surveillance Team |
| EPI_ISL_3314306 | Berkshire and Surrey Pathology Services Lighthouse Laboratory | Wellcome Sanger Institute for the COVID-19 Genomics UK (COG-UK) Consortium | Berkshire and Surrey Pathology Services Lighthouse Laboratory and Alex Alderton, Roberto Amato, Jeffrey Barrett, Sonia Goncalves, Ewan Harrison, David K. Jackson, Ian Johnston, Dominic Kwiatkowski, Cordelia Langford, John Sillitoe on behalf of the Wellcome Sanger Institute COVID-19 Surveillance Team |
| EPI_ISL_3314307 | Lighthouse Lab in Glasgow | Wellcome Sanger Institute for the COVID-19 Genomics UK (COG-UK) | Harper VanSteenhouse, Yumi Kasai, David Gray, Carol Clugston, Anna Dominiczak and Alex Alderton, Roberto Amato, Jeffrey Barrett, Sonia Goncalves, Ewan Harrison, David K. Jackson, Ian Johnston, Dominic Kwiatkowski, Cordelia Langford, |

| | | | |
|---|---|---|---|
| | | Consortium | John Sillitoe on behalf of the Wellcome Sanger Institute COVID-19 Surveillance Team |
| EPI_ISL_3314308, EPI_ISL_3314309, EPI_ISL_3314310 | Berkshire and Surrey Pathology Services Lighthouse Laboratory | Wellcome Sanger Institute for the COVID-19 Genomics UK (COG-UK) Consortium | Berkshire and Surrey Pathology Services Lighthouse Laboratory and Alex Alderton, Roberto Amato, Jeffrey Barrett, Sonia Goncalves, Ewan Harrison, David K. Jackson, Ian Johnston, Dominic Kwiatkowski, Cordelia Langford, John Sillitoe on behalf of the Wellcome Sanger Institute COVID-19 Surveillance Team |
| EPI_ISL_3314311, EPI_ISL_3314312, EPI_ISL_3314313 | Lighthouse Lab in Alderley Park | Wellcome Sanger Institute for the COVID-19 Genomics UK (COG-UK) Consortium | Jacquelyn Wynn, Mairead Hyland, The Lighthouse Lab in Alderley Park and Alex Alderton, Roberto Amato, Jeffrey Barrett, Sonia Goncalves, Ewan Harrison, David K. Jackson, Ian Johnston, Dominic Kwiatkowski, Cordelia Langford, John Sillitoe on behalf of the Wellcome Sanger Institute COVID-19 Surveillance Team |
| EPI_ISL_3314314, EPI_ISL_3314315 | Lighthouse Lab in Glasgow | Wellcome Sanger Institute for the COVID-19 Genomics UK (COG-UK) Consortium | Harper VanSteenhouse, Yumi Kasai, David Gray, Carol Clugston, Anna Dominiczak and Alex Alderton, Roberto Amato, Jeffrey Barrett, Sonia Goncalves, Ewan Harrison, David K. Jackson, Ian Johnston, Dominic Kwiatkowski, Cordelia Langford, John Sillitoe on behalf of the Wellcome Sanger Institute COVID-19 Surveillance Team |
| EPI_ISL_3314316 | Lighthouse Lab in Alderley Park | Wellcome Sanger Institute for the COVID-19 Genomics UK (COG-UK) Consortium | Jacquelyn Wynn, Mairead Hyland, The Lighthouse Lab in Alderley Park and Alex Alderton, Roberto Amato, Jeffrey Barrett, Sonia Goncalves, Ewan Harrison, David K. Jackson, Ian Johnston, Dominic Kwiatkowski, Cordelia Langford, John Sillitoe on behalf of the Wellcome Sanger Institute COVID-19 Surveillance Team |
| EPI_ISL_3314317 | Lighthouse Lab in Glasgow | Wellcome Sanger Institute for the COVID-19 Genomics UK (COG-UK) Consortium | Harper VanSteenhouse, Yumi Kasai, David Gray, Carol Clugston, Anna Dominiczak and Alex Alderton, Roberto Amato, Jeffrey Barrett, Sonia Goncalves, Ewan Harrison, David K. Jackson, Ian Johnston, Dominic Kwiatkowski, Cordelia Langford, John Sillitoe on behalf of the Wellcome Sanger Institute COVID-19 Surveillance Team |
| EPI_ISL_3314318 | Lighthouse Lab in Alderley Park | Wellcome Sanger Institute for the COVID-19 Genomics UK (COG-UK) Consortium | Jacquelyn Wynn, Mairead Hyland, The Lighthouse Lab in Alderley Park and Alex Alderton, Roberto Amato, Jeffrey Barrett, Sonia Goncalves, Ewan Harrison, David K. Jackson, Ian Johnston, Dominic Kwiatkowski, Cordelia Langford, John Sillitoe on behalf of the Wellcome Sanger Institute COVID-19 Surveillance Team |
| EPI_ISL_3314319, EPI_ISL_3314320, EPI_ISL_3314321 | Lighthouse Lab in Glasgow | Wellcome Sanger Institute for the COVID-19 Genomics UK (COG-UK) Consortium | Harper VanSteenhouse, Yumi Kasai, David Gray, Carol Clugston, Anna Dominiczak and Alex Alderton, Roberto Amato, Jeffrey Barrett, Sonia Goncalves, Ewan Harrison, David K. Jackson, Ian Johnston, Dominic Kwiatkowski, Cordelia Langford, John Sillitoe on behalf of the Wellcome Sanger Institute COVID-19 Surveillance Team |
| EPI_ISL_3314322 | Lighthouse Lab in Alderley Park | Wellcome Sanger Institute for the COVID-19 Genomics UK (COG-UK) Consortium | Jacquelyn Wynn, Mairead Hyland, The Lighthouse Lab in Alderley Park and Alex Alderton, Roberto Amato, Jeffrey Barrett, Sonia Goncalves, Ewan Harrison, David K. Jackson, Ian Johnston, Dominic Kwiatkowski, Cordelia Langford, John Sillitoe on behalf of the Wellcome Sanger Institute COVID-19 Surveillance Team |
| EPI_ISL_3314323 | Lighthouse Lab in Glasgow | Wellcome Sanger Institute for the COVID-19 Genomics UK (COG-UK) Consortium | Harper VanSteenhouse, Yumi Kasai, David Gray, Carol Clugston, Anna Dominiczak and Alex Alderton, Roberto Amato, Jeffrey Barrett, Sonia Goncalves, Ewan Harrison, David K. Jackson, Ian Johnston, Dominic Kwiatkowski, Cordelia Langford, John Sillitoe on behalf of the Wellcome Sanger Institute COVID-19 Surveillance Team |
| EPI_ISL_3314324 | Berkshire and Surrey Pathology Services Lighthouse Laboratory | Wellcome Sanger Institute for the COVID-19 Genomics UK (COG-UK) Consortium | Berkshire and Surrey Pathology Services Lighthouse Laboratory and Alex Alderton, Roberto Amato, Jeffrey Barrett, Sonia Goncalves, Ewan Harrison, David K. Jackson, Ian Johnston, Dominic Kwiatkowski, Cordelia Langford, John Sillitoe on behalf of the Wellcome Sanger Institute COVID-19 Surveillance Team |
| EPI_ISL_3314325 | Lighthouse Lab in Glasgow | Wellcome Sanger Institute for the COVID-19 Genomics UK (COG-UK) Consortium | Harper VanSteenhouse, Yumi Kasai, David Gray, Carol Clugston, Anna Dominiczak and Alex Alderton, Roberto Amato, Jeffrey Barrett, Sonia Goncalves, Ewan Harrison, David K. Jackson, Ian Johnston, Dominic Kwiatkowski, Cordelia Langford, John Sillitoe on behalf of the Wellcome Sanger Institute COVID-19 Surveillance Team |
| EPI_ISL_3314326 | Berkshire and Surrey Pathology Services Lighthouse Laboratory | Wellcome Sanger Institute for the COVID-19 Genomics UK (COG-UK) Consortium | Berkshire and Surrey Pathology Services Lighthouse Laboratory and Alex Alderton, Roberto Amato, Jeffrey Barrett, Sonia Goncalves, Ewan Harrison, David K. Jackson, Ian Johnston, Dominic Kwiatkowski, Cordelia Langford, John Sillitoe on behalf of the Wellcome Sanger Institute COVID-19 Surveillance Team |
| EPI_ISL_3314327, EPI_ISL_3314328 | Lighthouse Lab in Glasgow | Wellcome Sanger Institute for the COVID-19 Genomics UK (COG-UK) Consortium | Harper VanSteenhouse, Yumi Kasai, David Gray, Carol Clugston, Anna Dominiczak and Alex Alderton, Roberto Amato, Jeffrey Barrett, Sonia Goncalves, Ewan Harrison, David K. Jackson, Ian Johnston, Dominic Kwiatkowski, Cordelia Langford, John Sillitoe on behalf of the Wellcome Sanger Institute COVID-19 Surveillance Team |
| EPI_ISL_3314329, EPI_ISL_3314330, EPI_ISL_3314331 | Lighthouse Lab in Alderley Park | Wellcome Sanger Institute for the COVID-19 Genomics UK (COG-UK) Consortium | Jacquelyn Wynn, Mairead Hyland, The Lighthouse Lab in Alderley Park and Alex Alderton, Roberto Amato, Jeffrey Barrett, Sonia Goncalves, Ewan Harrison, David K. Jackson, Ian Johnston, Dominic Kwiatkowski, Cordelia Langford, John Sillitoe on behalf of the Wellcome Sanger Institute COVID-19 Surveillance Team |
| EPI_ISL_3314332, EPI_ISL_3314333 | Berkshire and Surrey Pathology Services Lighthouse Laboratory | Wellcome Sanger Institute for the COVID-19 Genomics UK (COG-UK) Consortium | Berkshire and Surrey Pathology Services Lighthouse Laboratory and Alex Alderton, Roberto Amato, Jeffrey Barrett, Sonia Goncalves, Ewan Harrison, David K. Jackson, Ian Johnston, Dominic Kwiatkowski, Cordelia Langford, John Sillitoe on behalf of the Wellcome Sanger Institute COVID-19 Surveillance Team |
| EPI_ISL_3314334, EPI_ISL_3314335 | Lighthouse Lab in Alderley Park | Wellcome Sanger Institute for the COVID-19 Genomics UK (COG-UK) Consortium | Jacquelyn Wynn, Mairead Hyland, The Lighthouse Lab in Alderley Park and Alex Alderton, Roberto Amato, Jeffrey Barrett, Sonia Goncalves, Ewan Harrison, David K. Jackson, Ian Johnston, Dominic Kwiatkowski, Cordelia Langford, John Sillitoe on behalf of the Wellcome Sanger Institute COVID-19 Surveillance Team |
| EPI_ISL_3314336, EPI_ISL_3314337 | Lighthouse Lab in Glasgow | Wellcome Sanger Institute for the COVID-19 Genomics UK (COG-UK) Consortium | Harper VanSteenhouse, Yumi Kasai, David Gray, Carol Clugston, Anna Dominiczak and Alex Alderton, Roberto Amato, Jeffrey Barrett, Sonia Goncalves, Ewan Harrison, David K. Jackson, Ian Johnston, Dominic Kwiatkowski, Cordelia Langford, John Sillitoe on behalf of the Wellcome Sanger Institute COVID-19 Surveillance Team |
| EPI_ISL_3314338 | Lighthouse Lab in Alderley Park | Wellcome Sanger Institute for the COVID-19 Genomics UK (COG-UK) Consortium | Jacquelyn Wynn, Mairead Hyland, The Lighthouse Lab in Alderley Park and Alex Alderton, Roberto Amato, Jeffrey Barrett, Sonia Goncalves, Ewan Harrison, David K. Jackson, Ian Johnston, Dominic Kwiatkowski, Cordelia Langford, John Sillitoe on behalf of the Wellcome Sanger Institute COVID-19 Surveillance Team |
| EPI_ISL_3314339, EPI_ISL_3314340 | Berkshire and Surrey Pathology Services Lighthouse Laboratory | Wellcome Sanger Institute for the COVID-19 Genomics UK (COG-UK) Consortium | Berkshire and Surrey Pathology Services Lighthouse Laboratory and Alex Alderton, Roberto Amato, Jeffrey Barrett, Sonia Goncalves, Ewan Harrison, David K. Jackson, Ian Johnston, Dominic Kwiatkowski, Cordelia Langford, John Sillitoe on behalf of the Wellcome Sanger Institute COVID-19 Surveillance Team |
| EPI_ISL_3314341 | Lighthouse Lab in Glasgow | Wellcome Sanger Institute for the COVID-19 Genomics UK (COG-UK) Consortium | Harper VanSteenhouse, Yumi Kasai, David Gray, Carol Clugston, Anna Dominiczak and Alex Alderton, Roberto Amato, Jeffrey Barrett, Sonia Goncalves, Ewan Harrison, David K. Jackson, Ian Johnston, Dominic Kwiatkowski, Cordelia Langford, John Sillitoe on behalf of the Wellcome Sanger Institute COVID-19 Surveillance Team |
| EPI_ISL_3314342 | Lighthouse Lab in Alderley Park | Wellcome Sanger Institute for the COVID-19 Genomics UK (COG-UK) Consortium | Jacquelyn Wynn, Mairead Hyland, The Lighthouse Lab in Alderley Park and Alex Alderton, Roberto Amato, Jeffrey Barrett, Sonia Goncalves, Ewan Harrison, David K. Jackson, Ian Johnston, Dominic Kwiatkowski, Cordelia Langford, John Sillitoe on behalf of the Wellcome Sanger Institute COVID-19 Surveillance Team |
| EPI_ISL_3314343, EPI_ISL_3314344, EPI_ISL_3314345 | Berkshire and Surrey Pathology Services Lighthouse Laboratory | Wellcome Sanger Institute for the COVID-19 Genomics UK (COG-UK) Consortium | Berkshire and Surrey Pathology Services Lighthouse Laboratory and Alex Alderton, Roberto Amato, Jeffrey Barrett, Sonia Goncalves, Ewan Harrison, David K. Jackson, Ian Johnston, Dominic Kwiatkowski, Cordelia Langford, John Sillitoe on behalf of the Wellcome Sanger Institute COVID-19 Surveillance Team |
| EPI_ISL_3314346 | Lighthouse Lab in Alderley Park | Wellcome Sanger Institute for the COVID-19 Genomics UK (COG-UK) Consortium | Jacquelyn Wynn, Mairead Hyland, The Lighthouse Lab in Alderley Park and Alex Alderton, Roberto Amato, Jeffrey Barrett, Sonia Goncalves, Ewan Harrison, David K. Jackson, Ian Johnston, Dominic Kwiatkowski, Cordelia Langford, John Sillitoe on behalf of the Wellcome Sanger Institute COVID-19 Surveillance Team |
| EPI_ISL_3314347 | Lighthouse Lab in Glasgow | Wellcome Sanger Institute for the COVID-19 Genomics UK (COG-UK) Consortium | Harper VanSteenhouse, Yumi Kasai, David Gray, Carol Clugston, Anna Dominiczak and Alex Alderton, Roberto Amato, Jeffrey Barrett, Sonia Goncalves, Ewan Harrison, David K. Jackson, Ian Johnston, Dominic Kwiatkowski, Cordelia Langford, John Sillitoe on behalf of the Wellcome Sanger Institute COVID-19 Surveillance Team |
| EPI_ISL_3314348 | Lighthouse Lab in Alderley Park | Wellcome Sanger Institute for the COVID-19 Genomics UK (COG-UK) Consortium | Jacquelyn Wynn, Mairead Hyland, The Lighthouse Lab in Alderley Park and Alex Alderton, Roberto Amato, Jeffrey Barrett, Sonia Goncalves, Ewan Harrison, David K. Jackson, Ian Johnston, Dominic Kwiatkowski, Cordelia Langford, John Sillitoe on behalf of the Wellcome Sanger Institute COVID-19 Surveillance Team |
| EPI_ISL_3314349, EPI_ISL_3314350 | Lighthouse Lab in Glasgow | Wellcome Sanger Institute for the COVID-19 Genomics UK (COG-UK) Consortium | Harper VanSteenhouse, Yumi Kasai, David Gray, Carol Clugston, Anna Dominiczak and Alex Alderton, Roberto Amato, Jeffrey Barrett, Sonia Goncalves, Ewan Harrison, David K. Jackson, Ian Johnston, Dominic Kwiatkowski, Cordelia Langford, John Sillitoe on behalf of the Wellcome Sanger Institute COVID-19 Surveillance Team |
| EPI_ISL_3314351 | Berkshire and Surrey Pathology Services Lighthouse Laboratory | Wellcome Sanger Institute for the COVID-19 Genomics UK (COG-UK) Consortium | Berkshire and Surrey Pathology Services Lighthouse Laboratory and Alex Alderton, Roberto Amato, Jeffrey Barrett, Sonia Goncalves, Ewan Harrison, David K. Jackson, Ian Johnston, Dominic Kwiatkowski, Cordelia Langford, John Sillitoe on behalf of the Wellcome Sanger Institute COVID-19 Surveillance Team |
| EPI_ISL_3314352 | Lighthouse Lab in Glasgow | Wellcome Sanger Institute for the COVID-19 Genomics UK (COG-UK) Consortium | Harper VanSteenhouse, Yumi Kasai, David Gray, Carol Clugston, Anna Dominiczak and Alex Alderton, Roberto Amato, Jeffrey Barrett, Sonia Goncalves, Ewan Harrison, David K. Jackson, Ian Johnston, Dominic Kwiatkowski, Cordelia Langford, John Sillitoe on behalf of the Wellcome Sanger Institute COVID-19 Surveillance Team |
| EPI_ISL_3314353 | Berkshire and Surrey Pathology Services Lighthouse Laboratory | Wellcome Sanger Institute for the COVID-19 Genomics UK (COG-UK) Consortium | Berkshire and Surrey Pathology Services Lighthouse Laboratory and Alex Alderton, Roberto Amato, Jeffrey Barrett, Sonia Goncalves, Ewan Harrison, David K. Jackson, Ian Johnston, Dominic Kwiatkowski, Cordelia Langford, John Sillitoe on behalf of the Wellcome Sanger Institute COVID-19 Surveillance Team |
| EPI_ISL_3314354 | Lighthouse Lab in Glasgow | Wellcome Sanger Institute for the COVID-19 Genomics UK (COG-UK) Consortium | Harper VanSteenhouse, Yumi Kasai, David Gray, Carol Clugston, Anna Dominiczak and Alex Alderton, Roberto Amato, Jeffrey Barrett, Sonia Goncalves, Ewan Harrison, David K. Jackson, Ian Johnston, Dominic Kwiatkowski, Cordelia Langford, John Sillitoe on behalf of the Wellcome Sanger Institute COVID-19 Surveillance Team |
| EPI_ISL_3314355, EPI_ISL_3314356 | Berkshire and Surrey Pathology Services Lighthouse Laboratory | Wellcome Sanger Institute for the COVID-19 Genomics UK (COG-UK) Consortium | Berkshire and Surrey Pathology Services Lighthouse Laboratory and Alex Alderton, Roberto Amato, Jeffrey Barrett, Sonia Goncalves, Ewan Harrison, David K. Jackson, Ian Johnston, Dominic Kwiatkowski, Cordelia Langford, John Sillitoe on behalf of the Wellcome Sanger Institute COVID-19 Surveillance Team |
| EPI_ISL_3314357 | Lighthouse Lab in Alderley Park | Wellcome Sanger Institute for the COVID-19 Genomics UK (COG-UK) Consortium | Jacquelyn Wynn, Mairead Hyland, The Lighthouse Lab in Alderley Park and Alex Alderton, Roberto Amato, Jeffrey Barrett, Sonia Goncalves, Ewan Harrison, David K. Jackson, Ian Johnston, Dominic Kwiatkowski, Cordelia Langford, John Sillitoe on behalf of the Wellcome Sanger Institute COVID-19 Surveillance Team |
| EPI_ISL_3314358 | Berkshire and Surrey Pathology Services Lighthouse Laboratory | Wellcome Sanger Institute for the COVID-19 Genomics UK (COG-UK) Consortium | Berkshire and Surrey Pathology Services Lighthouse Laboratory and Alex Alderton, Roberto Amato, Jeffrey Barrett, Sonia Goncalves, Ewan Harrison, David K. Jackson, Ian Johnston, Dominic Kwiatkowski, Cordelia Langford, John Sillitoe on behalf of the Wellcome Sanger Institute COVID-19 Surveillance Team |
| EPI_ISL_3314359 | Lighthouse Lab in Glasgow | Wellcome Sanger Institute for the COVID-19 Genomics UK (COG-UK) Consortium | Harper VanSteenhouse, Yumi Kasai, David Gray, Carol Clugston, Anna Dominiczak and Alex Alderton, Roberto Amato, Jeffrey Barrett, Sonia Goncalves, Ewan Harrison, David K. Jackson, Ian Johnston, Dominic Kwiatkowski, Cordelia Langford, John Sillitoe on behalf of the Wellcome Sanger Institute COVID-19 Surveillance Team |
| EPI_ISL_3314360, EPI_ISL_3314361 | Berkshire and Surrey Pathology Services Lighthouse Laboratory | Wellcome Sanger Institute for the COVID-19 Genomics UK (COG-UK) Consortium | Berkshire and Surrey Pathology Services Lighthouse Laboratory and Alex Alderton, Roberto Amato, Jeffrey Barrett, Sonia Goncalves, Ewan Harrison, David K. Jackson, Ian Johnston, Dominic Kwiatkowski, Cordelia Langford, John Sillitoe on behalf of the Wellcome Sanger Institute COVID-19 Surveillance Team |
| EPI_ISL_3314362 | Lighthouse Lab in Alderley Park | Wellcome Sanger Institute for the COVID-19 Genomics UK (COG-UK) Consortium | Jacquelyn Wynn, Mairead Hyland, The Lighthouse Lab in Alderley Park and Alex Alderton, Roberto Amato, Jeffrey Barrett, Sonia Goncalves, Ewan Harrison, David K. Jackson, Ian Johnston, Dominic Kwiatkowski, Cordelia Langford, John Sillitoe on behalf of the Wellcome Sanger Institute COVID-19 Surveillance Team |
| EPI_ISL_3314363, EPI_ISL_3314364, EPI_ISL_3314365 | Lighthouse Lab in Glasgow | Wellcome Sanger Institute for the COVID-19 Genomics UK (COG-UK) Consortium | Harper VanSteenhouse, Yumi Kasai, David Gray, Carol Clugston, Anna Dominiczak and Alex Alderton, Roberto Amato, Jeffrey Barrett, Sonia Goncalves, Ewan Harrison, David K. Jackson, Ian Johnston, Dominic Kwiatkowski, Cordelia Langford, John Sillitoe on behalf of the Wellcome Sanger Institute COVID-19 Surveillance Team |
| EPI_ISL_3314366 | Lighthouse Lab in Alderley Park | Wellcome Sanger Institute for the COVID-19 Genomics UK (COG-UK) Consortium | Jacquelyn Wynn, Mairead Hyland, The Lighthouse Lab in Alderley Park and Alex Alderton, Roberto Amato, Jeffrey Barrett, Sonia Goncalves, Ewan Harrison, David K. Jackson, Ian Johnston, Dominic Kwiatkowski, Cordelia Langford, John Sillitoe on behalf of the Wellcome Sanger Institute COVID-19 Surveillance Team |
| EPI_ISL_3314367 | Berkshire and Surrey Pathology Services Lighthouse Laboratory | Wellcome Sanger Institute for the COVID-19 Genomics UK (COG-UK) Consortium | Berkshire and Surrey Pathology Services Lighthouse Laboratory and Alex Alderton, Roberto Amato, Jeffrey Barrett, Sonia Goncalves, Ewan Harrison, David K. Jackson, Ian Johnston, Dominic Kwiatkowski, Cordelia Langford, John Sillitoe on behalf of the Wellcome Sanger Institute COVID-19 Surveillance Team |
| EPI_ISL_3314368, EPI_ISL_3314369, EPI_ISL_3314370 | Lighthouse Lab in Glasgow | Wellcome Sanger Institute for the COVID-19 Genomics UK (COG-UK) Consortium | Harper VanSteenhouse, Yumi Kasai, David Gray, Carol Clugston, Anna Dominiczak and Alex Alderton, Roberto Amato, Jeffrey Barrett, Sonia Goncalves, Ewan Harrison, David K. Jackson, Ian Johnston, Dominic Kwiatkowski, Cordelia Langford, John Sillitoe on behalf of the Wellcome Sanger Institute COVID-19 Surveillance Team |
| EPI_ISL_3314371, EPI_ISL_3314372 | Berkshire and Surrey Pathology Services Lighthouse Laboratory | Wellcome Sanger Institute for the COVID-19 Genomics UK (COG-UK) Consortium | Berkshire and Surrey Pathology Services Lighthouse Laboratory and Alex Alderton, Roberto Amato, Jeffrey Barrett, Sonia Goncalves, Ewan Harrison, David K. Jackson, Ian Johnston, Dominic Kwiatkowski, Cordelia Langford, John Sillitoe on behalf of the Wellcome Sanger Institute COVID-19 Surveillance Team |
| EPI_ISL_3314373 | Lighthouse Lab in Glasgow | Wellcome Sanger Institute for the COVID-19 Genomics UK (COG-UK) Consortium | Harper VanSteenhouse, Yumi Kasai, David Gray, Carol Clugston, Anna Dominiczak and Alex Alderton, Roberto Amato, Jeffrey Barrett, Sonia Goncalves, Ewan Harrison, David K. Jackson, Ian Johnston, Dominic Kwiatkowski, Cordelia Langford, John Sillitoe on behalf of the Wellcome Sanger Institute COVID-19 Surveillance Team |

| | | | |
|---|---|---|---|
| EPI_ISL_3314374 | Lighthouse Lab in Alderley Park | Wellcome Sanger Institute for the COVID-19 Genomics UK (COG-UK) Consortium | Jacquelyn Wynn, Mairead Hyland, The Lighthouse Lab in Alderley Park and Alex Alderton, Roberto Amato, Jeffrey Barrett, Sonia Goncalves, Ewan Harrison, David K. Jackson, Ian Johnston, Dominic Kwiatkowski, Cordelia Langford, John Sillitoe on behalf of the Wellcome Sanger Institute COVID-19 Surveillance Team |
| EPI_ISL_3314375 | Berkshire and Surrey Pathology Services Lighthouse Laboratory | Wellcome Sanger Institute for the COVID-19 Genomics UK (COG-UK) Consortium | Berkshire and Surrey Pathology Services Lighthouse Laboratory and Alex Alderton, Roberto Amato, Jeffrey Barrett, Sonia Goncalves, Ewan Harrison, David K. Jackson, Ian Johnston, Dominic Kwiatkowski, Cordelia Langford, John Sillitoe on behalf of the Wellcome Sanger Institute COVID-19 Surveillance Team |
| EPI_ISL_3314376 | Lighthouse Lab in Alderley Park | Wellcome Sanger Institute for the COVID-19 Genomics UK (COG-UK) Consortium | Jacquelyn Wynn, Mairead Hyland, The Lighthouse Lab in Alderley Park and Alex Alderton, Roberto Amato, Jeffrey Barrett, Sonia Goncalves, Ewan Harrison, David K. Jackson, Ian Johnston, Dominic Kwiatkowski, Cordelia Langford, John Sillitoe on behalf of the Wellcome Sanger Institute COVID-19 Surveillance Team |
| EPI_ISL_3314377 | Lighthouse Lab in Glasgow | Wellcome Sanger Institute for the COVID-19 Genomics UK (COG-UK) Consortium | Harper VanSteenhouse, Yumi Kasai, David Gray, Carol Clugston, Anna Dominiczak and Alex Alderton, Roberto Amato, Jeffrey Barrett, Sonia Goncalves, Ewan Harrison, David K. Jackson, Ian Johnston, Dominic Kwiatkowski, Cordelia Langford, John Sillitoe on behalf of the Wellcome Sanger Institute COVID-19 Surveillance Team |
| EPI_ISL_3314378 | Berkshire and Surrey Pathology Services Lighthouse Laboratory | Wellcome Sanger Institute for the COVID-19 Genomics UK (COG-UK) Consortium | Berkshire and Surrey Pathology Services Lighthouse Laboratory and Alex Alderton, Roberto Amato, Jeffrey Barrett, Sonia Goncalves, Ewan Harrison, David K. Jackson, Ian Johnston, Dominic Kwiatkowski, Cordelia Langford, John Sillitoe on behalf of the Wellcome Sanger Institute COVID-19 Surveillance Team |
| EPI_ISL_3314379 | Lighthouse Lab in Glasgow | Wellcome Sanger Institute for the COVID-19 Genomics UK (COG-UK) Consortium | Harper VanSteenhouse, Yumi Kasai, David Gray, Carol Clugston, Anna Dominiczak and Alex Alderton, Roberto Amato, Jeffrey Barrett, Sonia Goncalves, Ewan Harrison, David K. Jackson, Ian Johnston, Dominic Kwiatkowski, Cordelia Langford, John Sillitoe on behalf of the Wellcome Sanger Institute COVID-19 Surveillance Team |
| EPI_ISL_3314380, EPI_ISL_3314381 | Lighthouse Lab in Alderley Park | Wellcome Sanger Institute for the COVID-19 Genomics UK (COG-UK) Consortium | Jacquelyn Wynn, Mairead Hyland, The Lighthouse Lab in Alderley Park and Alex Alderton, Roberto Amato, Jeffrey Barrett, Sonia Goncalves, Ewan Harrison, David K. Jackson, Ian Johnston, Dominic Kwiatkowski, Cordelia Langford, John Sillitoe on behalf of the Wellcome Sanger Institute COVID-19 Surveillance Team |
| EPI_ISL_3314382 | Lighthouse Lab in Glasgow | Wellcome Sanger Institute for the COVID-19 Genomics UK (COG-UK) Consortium | Harper VanSteenhouse, Yumi Kasai, David Gray, Carol Clugston, Anna Dominiczak and Alex Alderton, Roberto Amato, Jeffrey Barrett, Sonia Goncalves, Ewan Harrison, David K. Jackson, Ian Johnston, Dominic Kwiatkowski, Cordelia Langford, John Sillitoe on behalf of the Wellcome Sanger Institute COVID-19 Surveillance Team |
| EPI_ISL_3314383 | Lighthouse Lab in Alderley Park | Wellcome Sanger Institute for the COVID-19 Genomics UK (COG-UK) Consortium | Jacquelyn Wynn, Mairead Hyland, The Lighthouse Lab in Alderley Park and Alex Alderton, Roberto Amato, Jeffrey Barrett, Sonia Goncalves, Ewan Harrison, David K. Jackson, Ian Johnston, Dominic Kwiatkowski, Cordelia Langford, John Sillitoe on behalf of the Wellcome Sanger Institute COVID-19 Surveillance Team |
| EPI_ISL_3314384, EPI_ISL_3314385, EPI_ISL_3314386 | Lighthouse Lab in Glasgow | Wellcome Sanger Institute for the COVID-19 Genomics UK (COG-UK) Consortium | Harper VanSteenhouse, Yumi Kasai, David Gray, Carol Clugston, Anna Dominiczak and Alex Alderton, Roberto Amato, Jeffrey Barrett, Sonia Goncalves, Ewan Harrison, David K. Jackson, Ian Johnston, Dominic Kwiatkowski, Cordelia Langford, John Sillitoe on behalf of the Wellcome Sanger Institute COVID-19 Surveillance Team |
| EPI_ISL_3314387 | Lighthouse Lab in Alderley Park | Wellcome Sanger Institute for the COVID-19 Genomics UK (COG-UK) Consortium | Jacquelyn Wynn, Mairead Hyland, The Lighthouse Lab in Alderley Park and Alex Alderton, Roberto Amato, Jeffrey Barrett, Sonia Goncalves, Ewan Harrison, David K. Jackson, Ian Johnston, Dominic Kwiatkowski, Cordelia Langford, John Sillitoe on behalf of the Wellcome Sanger Institute COVID-19 Surveillance Team |
| EPI_ISL_3314388 | Lighthouse Lab in Glasgow | Wellcome Sanger Institute for the COVID-19 Genomics UK (COG-UK) Consortium | Harper VanSteenhouse, Yumi Kasai, David Gray, Carol Clugston, Anna Dominiczak and Alex Alderton, Roberto Amato, Jeffrey Barrett, Sonia Goncalves, Ewan Harrison, David K. Jackson, Ian Johnston, Dominic Kwiatkowski, Cordelia Langford, John Sillitoe on behalf of the Wellcome Sanger Institute COVID-19 Surveillance Team |
| EPI_ISL_3314389, EPI_ISL_3314390, EPI_ISL_3314391 | Lighthouse Lab in Alderley Park | Wellcome Sanger Institute for the COVID-19 Genomics UK (COG-UK) Consortium | Jacquelyn Wynn, Mairead Hyland, The Lighthouse Lab in Alderley Park and Alex Alderton, Roberto Amato, Jeffrey Barrett, Sonia Goncalves, Ewan Harrison, David K. Jackson, Ian Johnston, Dominic Kwiatkowski, Cordelia Langford, John Sillitoe on behalf of the Wellcome Sanger Institute COVID-19 Surveillance Team |
| EPI_ISL_3314392, EPI_ISL_3314393, EPI_ISL_3314394 | Berkshire and Surrey Pathology Services Lighthouse Laboratory | Wellcome Sanger Institute for the COVID-19 Genomics UK (COG-UK) Consortium | Berkshire and Surrey Pathology Services Lighthouse Laboratory and Alex Alderton, Roberto Amato, Jeffrey Barrett, Sonia Goncalves, Ewan Harrison, David K. Jackson, Ian Johnston, Dominic Kwiatkowski, Cordelia Langford, John Sillitoe on behalf of the Wellcome Sanger Institute COVID-19 Surveillance Team |
| EPI_ISL_3314395 | Lighthouse Lab in Glasgow | Wellcome Sanger Institute for the COVID-19 Genomics UK (COG-UK) Consortium | Harper VanSteenhouse, Yumi Kasai, David Gray, Carol Clugston, Anna Dominiczak and Alex Alderton, Roberto Amato, Jeffrey Barrett, Sonia Goncalves, Ewan Harrison, David K. Jackson, Ian Johnston, Dominic Kwiatkowski, Cordelia Langford, John Sillitoe on behalf of the Wellcome Sanger Institute COVID-19 Surveillance Team |
| EPI_ISL_3314396, EPI_ISL_3314397 | Berkshire and Surrey Pathology Services Lighthouse Laboratory | Wellcome Sanger Institute for the COVID-19 Genomics UK (COG-UK) Consortium | Berkshire and Surrey Pathology Services Lighthouse Laboratory and Alex Alderton, Roberto Amato, Jeffrey Barrett, Sonia Goncalves, Ewan Harrison, David K. Jackson, Ian Johnston, Dominic Kwiatkowski, Cordelia Langford, John Sillitoe on behalf of the Wellcome Sanger Institute COVID-19 Surveillance Team |
| EPI_ISL_3314398, EPI_ISL_3314399 | Lighthouse Lab in Glasgow | Wellcome Sanger Institute for the COVID-19 Genomics UK (COG-UK) Consortium | Harper VanSteenhouse, Yumi Kasai, David Gray, Carol Clugston, Anna Dominiczak and Alex Alderton, Roberto Amato, Jeffrey Barrett, Sonia Goncalves, Ewan Harrison, David K. Jackson, Ian Johnston, Dominic Kwiatkowski, Cordelia Langford, John Sillitoe on behalf of the Wellcome Sanger Institute COVID-19 Surveillance Team |
| EPI_ISL_3314400, EPI_ISL_3314401 | Berkshire and Surrey Pathology Services Lighthouse Laboratory | Wellcome Sanger Institute for the COVID-19 Genomics UK (COG-UK) Consortium | Berkshire and Surrey Pathology Services Lighthouse Laboratory and Alex Alderton, Roberto Amato, Jeffrey Barrett, Sonia Goncalves, Ewan Harrison, David K. Jackson, Ian Johnston, Dominic Kwiatkowski, Cordelia Langford, John Sillitoe on behalf of the Wellcome Sanger Institute COVID-19 Surveillance Team |
| EPI_ISL_3314402 | Lighthouse Lab in Alderley Park | Wellcome Sanger Institute for the COVID-19 Genomics UK (COG-UK) Consortium | Jacquelyn Wynn, Mairead Hyland, The Lighthouse Lab in Alderley Park and Alex Alderton, Roberto Amato, Jeffrey Barrett, Sonia Goncalves, Ewan Harrison, David K. Jackson, Ian Johnston, Dominic Kwiatkowski, Cordelia Langford, John Sillitoe on behalf of the Wellcome Sanger Institute COVID-19 Surveillance Team |
| EPI_ISL_3314403, EPI_ISL_3314404, EPI_ISL_3314405, EPI_ISL_3314406, EPI_ISL_3314407, EPI_ISL_3314408, EPI_ISL_3314409 | Lighthouse Lab in Glasgow | Wellcome Sanger Institute for the COVID-19 Genomics UK (COG-UK) Consortium | Harper VanSteenhouse, Yumi Kasai, David Gray, Carol Clugston, Anna Dominiczak and Alex Alderton, Roberto Amato, Jeffrey Barrett, Sonia Goncalves, Ewan Harrison, David K. Jackson, Ian Johnston, Dominic Kwiatkowski, Cordelia Langford, John Sillitoe on behalf of the Wellcome Sanger Institute COVID-19 Surveillance Team |
| EPI_ISL_3314410 | Berkshire and Surrey Pathology Services Lighthouse Laboratory | Wellcome Sanger Institute for the COVID-19 Genomics UK (COG-UK) Consortium | Berkshire and Surrey Pathology Services Lighthouse Laboratory and Alex Alderton, Roberto Amato, Jeffrey Barrett, Sonia Goncalves, Ewan Harrison, David K. Jackson, Ian Johnston, Dominic Kwiatkowski, Cordelia Langford, John Sillitoe on behalf of the Wellcome Sanger Institute COVID-19 Surveillance Team |
| EPI_ISL_3314411 | Lighthouse Lab in Alderley Park | Wellcome Sanger Institute for the COVID-19 Genomics UK (COG-UK) Consortium | Jacquelyn Wynn, Mairead Hyland, The Lighthouse Lab in Alderley Park and Alex Alderton, Roberto Amato, Jeffrey Barrett, Sonia Goncalves, Ewan Harrison, David K. Jackson, Ian Johnston, Dominic Kwiatkowski, Cordelia Langford, John Sillitoe on behalf of the Wellcome Sanger Institute COVID-19 Surveillance Team |
| EPI_ISL_3314412, EPI_ISL_3314413 | Berkshire and Surrey Pathology Services Lighthouse Laboratory | Wellcome Sanger Institute for the COVID-19 Genomics UK (COG-UK) Consortium | Berkshire and Surrey Pathology Services Lighthouse Laboratory and Alex Alderton, Roberto Amato, Jeffrey Barrett, Sonia Goncalves, Ewan Harrison, David K. Jackson, Ian Johnston, Dominic Kwiatkowski, Cordelia Langford, John Sillitoe on behalf of the Wellcome Sanger Institute COVID-19 Surveillance Team |
| EPI_ISL_3314414 | Lighthouse Lab in Glasgow | Wellcome Sanger Institute for the COVID-19 Genomics UK (COG-UK) Consortium | Harper VanSteenhouse, Yumi Kasai, David Gray, Carol Clugston, Anna Dominiczak and Alex Alderton, Roberto Amato, Jeffrey Barrett, Sonia Goncalves, Ewan Harrison, David K. Jackson, Ian Johnston, Dominic Kwiatkowski, Cordelia Langford, John Sillitoe on behalf of the Wellcome Sanger Institute COVID-19 Surveillance Team |
| EPI_ISL_3314415 | Lighthouse Lab in Alderley Park | Wellcome Sanger Institute for the COVID-19 Genomics UK (COG-UK) Consortium | Jacquelyn Wynn, Mairead Hyland, The Lighthouse Lab in Alderley Park and Alex Alderton, Roberto Amato, Jeffrey Barrett, Sonia Goncalves, Ewan Harrison, David K. Jackson, Ian Johnston, Dominic Kwiatkowski, Cordelia Langford, John Sillitoe on behalf of the Wellcome Sanger Institute COVID-19 Surveillance Team |
| EPI_ISL_3314416 | Lighthouse Lab in Glasgow | Wellcome Sanger Institute for the COVID-19 Genomics UK (COG-UK) Consortium | Harper VanSteenhouse, Yumi Kasai, David Gray, Carol Clugston, Anna Dominiczak and Alex Alderton, Roberto Amato, Jeffrey Barrett, Sonia Goncalves, Ewan Harrison, David K. Jackson, Ian Johnston, Dominic Kwiatkowski, Cordelia Langford, John Sillitoe on behalf of the Wellcome Sanger Institute COVID-19 Surveillance Team |
| EPI_ISL_3314417, EPI_ISL_3314418 | Lighthouse Lab in Alderley Park | Wellcome Sanger Institute for the COVID-19 Genomics UK (COG-UK) Consortium | Jacquelyn Wynn, Mairead Hyland, The Lighthouse Lab in Alderley Park and Alex Alderton, Roberto Amato, Jeffrey Barrett, Sonia Goncalves, Ewan Harrison, David K. Jackson, Ian Johnston, Dominic Kwiatkowski, Cordelia Langford, John Sillitoe on behalf of the Wellcome Sanger Institute COVID-19 Surveillance Team |
| EPI_ISL_3314419, EPI_ISL_3314420, EPI_ISL_3314421 | Lighthouse Lab in Glasgow | Wellcome Sanger Institute for the COVID-19 Genomics UK (COG-UK) Consortium | Harper VanSteenhouse, Yumi Kasai, David Gray, Carol Clugston, Anna Dominiczak and Alex Alderton, Roberto Amato, Jeffrey Barrett, Sonia Goncalves, Ewan Harrison, David K. Jackson, Ian Johnston, Dominic Kwiatkowski, Cordelia Langford, John Sillitoe on behalf of the Wellcome Sanger Institute COVID-19 Surveillance Team |
| EPI_ISL_3314422, EPI_ISL_3314423 | Berkshire and Surrey Pathology Services Lighthouse Laboratory | Wellcome Sanger Institute for the COVID-19 Genomics UK (COG-UK) Consortium | Berkshire and Surrey Pathology Services Lighthouse Laboratory and Alex Alderton, Roberto Amato, Jeffrey Barrett, Sonia Goncalves, Ewan Harrison, David K. Jackson, Ian Johnston, Dominic Kwiatkowski, Cordelia Langford, John Sillitoe on behalf of the Wellcome Sanger Institute COVID-19 Surveillance Team |
| EPI_ISL_3314424 | Lighthouse Lab in Glasgow | Wellcome Sanger Institute for the COVID-19 Genomics UK (COG-UK) Consortium | Harper VanSteenhouse, Yumi Kasai, David Gray, Carol Clugston, Anna Dominiczak and Alex Alderton, Roberto Amato, Jeffrey Barrett, Sonia Goncalves, Ewan Harrison, David K. Jackson, Ian Johnston, Dominic Kwiatkowski, Cordelia Langford, John Sillitoe on behalf of the Wellcome Sanger Institute COVID-19 Surveillance Team |
| EPI_ISL_3314425 | Berkshire and Surrey Pathology Services Lighthouse Laboratory | Wellcome Sanger Institute for the COVID-19 Genomics UK (COG-UK) Consortium | Berkshire and Surrey Pathology Services Lighthouse Laboratory and Alex Alderton, Roberto Amato, Jeffrey Barrett, Sonia Goncalves, Ewan Harrison, David K. Jackson, Ian Johnston, Dominic Kwiatkowski, Cordelia Langford, John Sillitoe on behalf of the Wellcome Sanger Institute COVID-19 Surveillance Team |
| EPI_ISL_3314426, EPI_ISL_3314427, EPI_ISL_3314428, EPI_ISL_3314429, EPI_ISL_3314430, EPI_ISL_3314431 | Lighthouse Laboratory Plymouth | Wellcome Sanger Institute for the COVID-19 Genomics UK (COG-UK) Consortium | Lighthouse Laboratory Plymouth and Alex Alderton, Roberto Amato, Jeffrey Barrett, Sonia Goncalves, Ewan Harrison, David K. Jackson, Ian Johnston, Dominic Kwiatkowski, Cordelia Langford, John Sillitoe on behalf of the Wellcome Sanger Institute COVID-19 Surveillance Team |
| EPI_ISL_3314432 | Berkshire and Surrey Pathology Services Lighthouse Laboratory | Wellcome Sanger Institute for the COVID-19 Genomics UK (COG-UK) Consortium | Berkshire and Surrey Pathology Services Lighthouse Laboratory and Alex Alderton, Roberto Amato, Jeffrey Barrett, Sonia Goncalves, Ewan Harrison, David K. Jackson, Ian Johnston, Dominic Kwiatkowski, Cordelia Langford, John Sillitoe on behalf of the Wellcome Sanger Institute COVID-19 Surveillance Team |
| EPI_ISL_3314433 | Lighthouse Laboratory Plymouth | Wellcome Sanger Institute for the COVID-19 Genomics UK (COG-UK) Consortium | Lighthouse Laboratory Plymouth and Alex Alderton, Roberto Amato, Jeffrey Barrett, Sonia Goncalves, Ewan Harrison, David K. Jackson, Ian Johnston, Dominic Kwiatkowski, Cordelia Langford, John Sillitoe on behalf of the Wellcome Sanger Institute COVID-19 Surveillance Team |
| EPI_ISL_3314434, EPI_ISL_3314435, EPI_ISL_3314436, EPI_ISL_3314437, EPI_ISL_3314438 | Berkshire and Surrey Pathology Services Lighthouse Laboratory | Wellcome Sanger Institute for the COVID-19 Genomics UK (COG-UK) Consortium | Berkshire and Surrey Pathology Services Lighthouse Laboratory and Alex Alderton, Roberto Amato, Jeffrey Barrett, Sonia Goncalves, Ewan Harrison, David K. Jackson, Ian Johnston, Dominic Kwiatkowski, Cordelia Langford, John Sillitoe on behalf of the Wellcome Sanger Institute COVID-19 Surveillance Team |

EPI_ISL_3314439, EPI_ISL_3314440, EPI_ISL_3314441, EPI_ISL_3314442, EPI_ISL_3314443, EPI_ISL_3314444, EPI_ISL_3314445, EPI_ISL_3314446, EPI_ISL_3314447, EPI_ISL_3314448, EPI_ISL_3314449, EPI_ISL_3314450, EPI_ISL_3314451, EPI_ISL_3314452, EPI_ISL_3314453, EPI_ISL_3314454, EPI_ISL_3314455, EPI_ISL_3314456, EPI_ISL_3314457, EPI_ISL_3314458, EPI_ISL_3314459, EPI_ISL_3314460, EPI_ISL_3314461, EPI_ISL_3314462, EPI_ISL_3314463, EPI_ISL_3314464, EPI_ISL_3314465, EPI_ISL_3314466, EPI_ISL_3314467, EPI_ISL_3314468, EPI_ISL_3314469, EPI_ISL_3314470, EPI_ISL_3314471, EPI_ISL_3314472, EPI_ISL_3314473, EPI_ISL_3314474, EPI_ISL_3314475, EPI_ISL_3314476, EPI_ISL_3314477, EPI_ISL_3314478, EPI_ISL_3314479, EPI_ISL_3314480, EPI_ISL_3314481, EPI_ISL_3314482, EPI_ISL_3314483, EPI_ISL_3314484, EPI_ISL_3314485, EPI_ISL_3314486, EPI_ISL_3314487, EPI_ISL_3314488, EPI_ISL_3314489, EPI_ISL_3314490, EPI_ISL_3314491, EPI_ISL_3314492, EPI_ISL_3314493, EPI_ISL_3314494, EPI_ISL_3314495, EPI_ISL_3314496, EPI_ISL_3314497, EPI_ISL_3314498, EPI_ISL_3314499, EPI_ISL_3314500, EPI_ISL_3314501, EPI_ISL_3314502, EPI_ISL_3314503, EPI_ISL_3314504, EPI_ISL_3314505, EPI_ISL_3314506, EPI_ISL_3314507, EPI_ISL_3314508, EPI_ISL_3314509, EPI_ISL_3314510, EPI_ISL_3314511, EPI_ISL_3314512, EPI_ISL_3314513, EPI_ISL_3314514, EPI_ISL_3314515, EPI_ISL_3314516, EPI_ISL_3314517, EPI_ISL_3314518, EPI_ISL_3314519, EPI_ISL_3314520, EPI_ISL_3314521, EPI_ISL_3314522, EPI_ISL_3314523, EPI_ISL_3314524, EPI_ISL_3314525, EPI_ISL_3314526, EPI_ISL_3314527, EPI_ISL_3314528, EPI_ISL_3314529, EPI_ISL_3314530, EPI_ISL_3314531, EPI_ISL_3314532, EPI_ISL_3314533, EPI_ISL_3314534, EPI_ISL_3314535, EPI_ISL_3314536, EPI_ISL_3314537, EPI_ISL_3314538, EPI_ISL_3314539, EPI_ISL_3314540, EPI_ISL_3314541, EPI_ISL_3314542, EPI_ISL_3314543, EPI_ISL_3314544, EPI_ISL_3314545, EPI_ISL_3314546, EPI_ISL_3314547, EPI_ISL_3314548, EPI_ISL_3314549, EPI_ISL_3314550, EPI_ISL_3314551, EPI_ISL_3314552, EPI_ISL_3314553, EPI_ISL_3314554, EPI_ISL_3314555, EPI_ISL_3314556, EPI_ISL_3314557, EPI_ISL_3314558, EPI_ISL_3314559, EPI_ISL_3314560, EPI_ISL_3314561, EPI_ISL_3314562, EPI_ISL_3314563, EPI_ISL_3314564, EPI_ISL_3314565, EPI_ISL_3314566, EPI_ISL_3314567, EPI_ISL_3314568, EPI_ISL_3314569, EPI_ISL_3314570, EPI_ISL_3314571, EPI_ISL_3314572, EPI_ISL_3314573, EPI_ISL_3314574, EPI_ISL_3314575, EPI_ISL_3314576, EPI_ISL_3314577, EPI_ISL_3314578, EPI_ISL_3314579, EPI_ISL_3314580, EPI_ISL_3314581, EPI_ISL_3314582, EPI_ISL_3314583, EPI_ISL_3314584, EPI_ISL_3314585, EPI_ISL_3314586, EPI_ISL_3314587, EPI_ISL_3314588, EPI_ISL_3314589, EPI_ISL_3314590, EPI_ISL_3314591, EPI_ISL_3314592, EPI_ISL_3314593, EPI_ISL_3314594, EPI_ISL_3314595, EPI_ISL_3314596, EPI_ISL_3314597, EPI_ISL_3314598, EPI_ISL_3314599, EPI_ISL_3314600, EPI_ISL_3314601, EPI_ISL_3314602, EPI_ISL_3314603, EPI_ISL_3314604, EPI_ISL_3314605, EPI_ISL_3314606, EPI_ISL_3314607, EPI_ISL_3314608, EPI_ISL_3314609, EPI_ISL_3314610, EPI_ISL_3314611, EPI_ISL_3314612, EPI_ISL_3314613, EPI_ISL_3314614,

EPI_ISL_3314615, EPI_ISL_3314616, EPI_ISL_3314617, EPI_ISL_3314618, EPI_ISL_3314619, EPI_ISL_3314620, EPI_ISL_3314621, EPI_ISL_3314622, EPI_ISL_3314623, EPI_ISL_3314624, EPI_ISL_3314625, EPI_ISL_3314626, EPI_ISL_3314627, EPI_ISL_3314628, EPI_ISL_3314629, EPI_ISL_3314630, EPI_ISL_3314631, EPI_ISL_3314632, EPI_ISL_3314633, EPI_ISL_3314634, EPI_ISL_3314635, EPI_ISL_3314636, EPI_ISL_3314637, EPI_ISL_3314638, EPI_ISL_3314639, EPI_ISL_3314640, EPI_ISL_3314641, EPI_ISL_3314642, EPI_ISL_3314643, EPI_ISL_3314644, EPI_ISL_3314645, EPI_ISL_3314646, EPI_ISL_3314647, EPI_ISL_3314648, EPI_ISL_3314649, EPI_ISL_3314650, EPI_ISL_3314651, EPI_ISL_3314652, EPI_ISL_3314653, EPI_ISL_3314654, EPI_ISL_3314655, EPI_ISL_3314656, EPI_ISL_3314657, EPI_ISL_3314658, EPI_ISL_3314659, EPI_ISL_3314660, EPI_ISL_3314661, EPI_ISL_3314662, EPI_ISL_3314663, EPI_ISL_3314664, EPI_ISL_3314665, EPI_ISL_3314666

| | | | |
|---|---|---|---|
| see above | Lighthouse Lab in Milton Keynes | Wellcome Sanger Institute for the COVID-19 Genomics UK (COG-UK) Consortium | The Lighthouse Lab in Milton Keynes and Alex Alderton, Roberto Amato, Jeffrey Barrett, Sonia Goncalves, Ewan Harrison, David K. Jackson, Ian Johnston, Dominic Kwiatkowski, Cordelia Langford, John Sillitoe on behalf of the Wellcome Sanger Institute COVID-19 Surveillance Team |
| EPI_ISL_3314667 | Lighthouse Laboratory Plymouth | Wellcome Sanger Institute for the COVID-19 Genomics UK (COG-UK) Consortium | Lighthouse Laboratory Plymouth and Alex Alderton, Roberto Amato, Jeffrey Barrett, Sonia Goncalves, Ewan Harrison, David K. Jackson, Ian Johnston, Dominic Kwiatkowski, Cordelia Langford, John Sillitoe on behalf of the Wellcome Sanger Institute COVID-19 Surveillance Team |
| EPI_ISL_3314668 | Lighthouse Lab in Glasgow | Wellcome Sanger Institute for the COVID-19 Genomics UK (COG-UK) Consortium | Harper VanSteenhouse, Yumi Kasai, David Gray, Carol Clugston, Anna Dominiczak and Alex Alderton, Roberto Amato, Jeffrey Barrett, Sonia Goncalves, Ewan Harrison, David K. Jackson, Ian Johnston, Dominic Kwiatkowski, Cordelia Langford, John Sillitoe on behalf of the Wellcome Sanger Institute COVID-19 Surveillance Team |
| EPI_ISL_3314669, EPI_ISL_3314670 | Lighthouse Laboratory Plymouth | Wellcome Sanger Institute for the COVID-19 Genomics UK (COG-UK) Consortium | Lighthouse Laboratory Plymouth and Alex Alderton, Roberto Amato, Jeffrey Barrett, Sonia Goncalves, Ewan Harrison, David K. Jackson, Ian Johnston, Dominic Kwiatkowski, Cordelia Langford, John Sillitoe on behalf of the Wellcome Sanger Institute COVID-19 Surveillance Team |
| EPI_ISL_3314671, EPI_ISL_3314672, EPI_ISL_3314673, EPI_ISL_3314674 | Berkshire and Surrey Pathology Services Lighthouse Laboratory | Wellcome Sanger Institute for the COVID-19 Genomics UK (COG-UK) Consortium | Berkshire and Surrey Pathology Services Lighthouse Laboratory and Alex Alderton, Roberto Amato, Jeffrey Barrett, Sonia Goncalves, Ewan Harrison, David K. Jackson, Ian Johnston, Dominic Kwiatkowski, Cordelia Langford, John Sillitoe on behalf of the Wellcome Sanger Institute COVID-19 Surveillance Team |
| EPI_ISL_3314675, EPI_ISL_3314676 | Lighthouse Laboratory Plymouth | Wellcome Sanger Institute for the COVID-19 Genomics UK (COG-UK) Consortium | Lighthouse Laboratory Plymouth and Alex Alderton, Roberto Amato, Jeffrey Barrett, Sonia Goncalves, Ewan Harrison, David K. Jackson, Ian Johnston, Dominic Kwiatkowski, Cordelia Langford, John Sillitoe on behalf of the Wellcome Sanger Institute COVID-19 Surveillance Team |
| EPI_ISL_3314677 | Lighthouse Lab in Glasgow | Wellcome Sanger Institute for the COVID-19 Genomics UK (COG-UK) Consortium | Harper VanSteenhouse, Yumi Kasai, David Gray, Carol Clugston, Anna Dominiczak and Alex Alderton, Roberto Amato, Jeffrey Barrett, Sonia Goncalves, Ewan Harrison, David K. Jackson, Ian Johnston, Dominic Kwiatkowski, Cordelia Langford, John Sillitoe on behalf of the Wellcome Sanger Institute COVID-19 Surveillance Team |
| EPI_ISL_3314678, EPI_ISL_3314679, EPI_ISL_3314680, EPI_ISL_3314681, EPI_ISL_3314682, EPI_ISL_3314683, EPI_ISL_3314684 | Lighthouse Laboratory Plymouth | Wellcome Sanger Institute for the COVID-19 Genomics UK (COG-UK) Consortium | Lighthouse Laboratory Plymouth and Alex Alderton, Roberto Amato, Jeffrey Barrett, Sonia Goncalves, Ewan Harrison, David K. Jackson, Ian Johnston, Dominic Kwiatkowski, Cordelia Langford, John Sillitoe on behalf of the Wellcome Sanger Institute COVID-19 Surveillance Team |
| EPI_ISL_3314685 | Berkshire and Surrey Pathology Services Lighthouse Laboratory | Wellcome Sanger Institute for the COVID-19 Genomics UK (COG-UK) Consortium | Berkshire and Surrey Pathology Services Lighthouse Laboratory and Alex Alderton, Roberto Amato, Jeffrey Barrett, Sonia Goncalves, Ewan Harrison, David K. Jackson, Ian Johnston, Dominic Kwiatkowski, Cordelia Langford, John Sillitoe on behalf of the Wellcome Sanger Institute COVID-19 Surveillance Team |
| EPI_ISL_3314686, EPI_ISL_3314687, EPI_ISL_3314688, EPI_ISL_3314689, EPI_ISL_3314690 | Lighthouse Laboratory Plymouth | Wellcome Sanger Institute for the COVID-19 Genomics UK (COG-UK) Consortium | Lighthouse Laboratory Plymouth and Alex Alderton, Roberto Amato, Jeffrey Barrett, Sonia Goncalves, Ewan Harrison, David K. Jackson, Ian Johnston, Dominic Kwiatkowski, Cordelia Langford, John Sillitoe on behalf of the Wellcome Sanger Institute COVID-19 Surveillance Team |
| EPI_ISL_3314691, EPI_ISL_3314692 | Berkshire and Surrey Pathology Services Lighthouse Laboratory | Wellcome Sanger Institute for the COVID-19 Genomics UK (COG-UK) Consortium | Berkshire and Surrey Pathology Services Lighthouse Laboratory and Alex Alderton, Roberto Amato, Jeffrey Barrett, Sonia Goncalves, Ewan Harrison, David K. Jackson, Ian Johnston, Dominic Kwiatkowski, Cordelia Langford, John Sillitoe on behalf of the Wellcome Sanger Institute COVID-19 Surveillance Team |
| EPI_ISL_3314693, EPI_ISL_3314694, EPI_ISL_3314695, EPI_ISL_3314696, EPI_ISL_3314697, EPI_ISL_3314698 | Lighthouse Laboratory Plymouth | Wellcome Sanger Institute for the COVID-19 Genomics UK (COG-UK) Consortium | Lighthouse Laboratory Plymouth and Alex Alderton, Roberto Amato, Jeffrey Barrett, Sonia Goncalves, Ewan Harrison, David K. Jackson, Ian Johnston, Dominic Kwiatkowski, Cordelia Langford, John Sillitoe on behalf of the Wellcome Sanger Institute COVID-19 Surveillance Team |
| EPI_ISL_3314699 | Berkshire and Surrey Pathology Services Lighthouse Laboratory | Wellcome Sanger Institute for the COVID-19 Genomics UK (COG-UK) Consortium | Berkshire and Surrey Pathology Services Lighthouse Laboratory and Alex Alderton, Roberto Amato, Jeffrey Barrett, Sonia Goncalves, Ewan Harrison, David K. Jackson, Ian Johnston, Dominic Kwiatkowski, Cordelia Langford, John Sillitoe on behalf of the Wellcome Sanger Institute COVID-19 Surveillance Team |
| EPI_ISL_3314700, EPI_ISL_3314701, EPI_ISL_3314702 | Lighthouse Laboratory Plymouth | Wellcome Sanger Institute for the COVID-19 Genomics UK (COG-UK) Consortium | Lighthouse Laboratory Plymouth and Alex Alderton, Roberto Amato, Jeffrey Barrett, Sonia Goncalves, Ewan Harrison, David K. Jackson, Ian Johnston, Dominic Kwiatkowski, Cordelia Langford, John Sillitoe on behalf of the Wellcome Sanger Institute COVID-19 Surveillance Team |
| EPI_ISL_3314703 | Berkshire and Surrey Pathology Services Lighthouse Laboratory | Wellcome Sanger Institute for the COVID-19 Genomics UK (COG-UK) Consortium | Berkshire and Surrey Pathology Services Lighthouse Laboratory and Alex Alderton, Roberto Amato, Jeffrey Barrett, Sonia Goncalves, Ewan Harrison, David K. Jackson, Ian Johnston, Dominic Kwiatkowski, Cordelia Langford, John Sillitoe on behalf of the Wellcome Sanger Institute COVID-19 Surveillance Team |
| EPI_ISL_3314704 | Lighthouse Laboratory Plymouth | Wellcome Sanger Institute for the COVID-19 Genomics UK (COG-UK) Consortium | Lighthouse Laboratory Plymouth and Alex Alderton, Roberto Amato, Jeffrey Barrett, Sonia Goncalves, Ewan Harrison, David K. Jackson, Ian Johnston, Dominic Kwiatkowski, Cordelia Langford, John Sillitoe on behalf of the Wellcome Sanger Institute COVID-19 Surveillance Team |
| EPI_ISL_3314705, EPI_ISL_3314706 | Berkshire and Surrey Pathology Services Lighthouse Laboratory | Wellcome Sanger Institute for the COVID-19 Genomics UK (COG-UK) Consortium | Berkshire and Surrey Pathology Services Lighthouse Laboratory and Alex Alderton, Roberto Amato, Jeffrey Barrett, Sonia Goncalves, Ewan Harrison, David K. Jackson, Ian Johnston, Dominic Kwiatkowski, Cordelia Langford, John Sillitoe on behalf of the Wellcome Sanger Institute COVID-19 Surveillance Team |
| EPI_ISL_3314707, EPI_ISL_3314708 | Lighthouse Laboratory Plymouth | Wellcome Sanger Institute for the COVID-19 Genomics UK (COG-UK) Consortium | Lighthouse Laboratory Plymouth and Alex Alderton, Roberto Amato, Jeffrey Barrett, Sonia Goncalves, Ewan Harrison, David K. Jackson, Ian Johnston, Dominic Kwiatkowski, Cordelia Langford, John Sillitoe on behalf of the Wellcome Sanger Institute COVID-19 Surveillance Team |
| EPI_ISL_3314709 | Berkshire and Surrey Pathology Services Lighthouse Laboratory | Wellcome Sanger Institute for the COVID-19 Genomics UK (COG-UK) Consortium | Berkshire and Surrey Pathology Services Lighthouse Laboratory and Alex Alderton, Roberto Amato, Jeffrey Barrett, Sonia Goncalves, Ewan Harrison, David K. Jackson, Ian Johnston, Dominic Kwiatkowski, Cordelia Langford, John Sillitoe on behalf of the Wellcome Sanger Institute COVID-19 Surveillance Team |
| EPI_ISL_3314710, EPI_ISL_3314711 | Lighthouse Laboratory Plymouth | Wellcome Sanger Institute for the COVID-19 Genomics UK (COG-UK) Consortium | Lighthouse Laboratory Plymouth and Alex Alderton, Roberto Amato, Jeffrey Barrett, Sonia Goncalves, Ewan Harrison, David K. Jackson, Ian Johnston, Dominic Kwiatkowski, Cordelia Langford, John Sillitoe on behalf of the Wellcome Sanger Institute COVID-19 Surveillance Team |
| EPI_ISL_3314712 | Berkshire and Surrey Pathology Services Lighthouse Laboratory | Wellcome Sanger Institute for the COVID-19 Genomics UK (COG-UK) Consortium | Berkshire and Surrey Pathology Services Lighthouse Laboratory and Alex Alderton, Roberto Amato, Jeffrey Barrett, Sonia Goncalves, Ewan Harrison, David K. Jackson, Ian Johnston, Dominic Kwiatkowski, Cordelia Langford, John Sillitoe on behalf of the Wellcome Sanger Institute COVID-19 Surveillance Team |
| EPI_ISL_3314713 | Lighthouse Laboratory Plymouth | Wellcome Sanger Institute for the COVID-19 Genomics UK (COG-UK) Consortium | Lighthouse Laboratory Plymouth and Alex Alderton, Roberto Amato, Jeffrey Barrett, Sonia Goncalves, Ewan Harrison, David K. Jackson, Ian Johnston, Dominic Kwiatkowski, Cordelia Langford, John Sillitoe on behalf of the Wellcome Sanger Institute COVID-19 Surveillance Team |
| EPI_ISL_3314714 | Berkshire and Surrey Pathology Services Lighthouse Laboratory | Wellcome Sanger Institute for the COVID-19 Genomics UK (COG-UK) Consortium | Berkshire and Surrey Pathology Services Lighthouse Laboratory and Alex Alderton, Roberto Amato, Jeffrey Barrett, Sonia Goncalves, Ewan Harrison, David K. Jackson, Ian Johnston, Dominic Kwiatkowski, Cordelia Langford, John Sillitoe on behalf of the Wellcome Sanger Institute COVID-19 Surveillance Team |
| EPI_ISL_3314715 | Lighthouse Laboratory Plymouth | Wellcome Sanger Institute for the COVID-19 Genomics UK (COG-UK) Consortium | Lighthouse Laboratory Plymouth and Alex Alderton, Roberto Amato, Jeffrey Barrett, Sonia Goncalves, Ewan Harrison, David K. Jackson, Ian Johnston, Dominic Kwiatkowski, Cordelia Langford, John Sillitoe on behalf of the Wellcome Sanger Institute COVID-19 Surveillance Team |
| EPI_ISL_3314716, EPI_ISL_3314717, EPI_ISL_3314718 | Berkshire and Surrey Pathology Services Lighthouse Laboratory | Wellcome Sanger Institute for the COVID-19 Genomics UK (COG-UK) Consortium | Berkshire and Surrey Pathology Services Lighthouse Laboratory and Alex Alderton, Roberto Amato, Jeffrey Barrett, Sonia Goncalves, Ewan Harrison, David K. Jackson, Ian Johnston, Dominic Kwiatkowski, Cordelia Langford, John Sillitoe on behalf of the Wellcome Sanger Institute COVID-19 Surveillance Team |
| EPI_ISL_3314719 | Lighthouse Laboratory Plymouth | Wellcome Sanger Institute for the COVID-19 Genomics UK (COG-UK) Consortium | Lighthouse Laboratory Plymouth and Alex Alderton, Roberto Amato, Jeffrey Barrett, Sonia Goncalves, Ewan Harrison, David K. Jackson, Ian Johnston, Dominic Kwiatkowski, Cordelia Langford, John Sillitoe on behalf of the Wellcome Sanger Institute COVID-19 Surveillance Team |
| EPI_ISL_3314720 | Berkshire and Surrey Pathology Services Lighthouse Laboratory | Wellcome Sanger Institute for the COVID-19 Genomics UK (COG-UK) Consortium | Berkshire and Surrey Pathology Services Lighthouse Laboratory and Alex Alderton, Roberto Amato, Jeffrey Barrett, Sonia Goncalves, Ewan Harrison, David K. Jackson, Ian Johnston, Dominic Kwiatkowski, Cordelia Langford, John Sillitoe on behalf of the Wellcome Sanger Institute COVID-19 Surveillance Team |
| EPI_ISL_3314721 | Lighthouse Laboratory Plymouth | Wellcome Sanger Institute for the COVID-19 Genomics UK (COG-UK) Consortium | Lighthouse Laboratory Plymouth and Alex Alderton, Roberto Amato, Jeffrey Barrett, Sonia Goncalves, Ewan Harrison, David K. Jackson, Ian Johnston, Dominic Kwiatkowski, Cordelia Langford, John Sillitoe on behalf of the Wellcome Sanger Institute COVID-19 Surveillance Team |
| EPI_ISL_3314722 | Berkshire and Surrey Pathology Services Lighthouse Laboratory | Wellcome Sanger Institute for the COVID-19 Genomics UK (COG-UK) Consortium | Berkshire and Surrey Pathology Services Lighthouse Laboratory and Alex Alderton, Roberto Amato, Jeffrey Barrett, Sonia Goncalves, Ewan Harrison, David K. Jackson, Ian Johnston, Dominic Kwiatkowski, Cordelia Langford, John Sillitoe on behalf of the Wellcome Sanger Institute COVID-19 Surveillance Team |
| EPI_ISL_3314723 | Lighthouse Lab in Glasgow | Wellcome Sanger Institute for the COVID-19 Genomics UK (COG-UK) Consortium | Harper VanSteenhouse, Yumi Kasai, David Gray, Carol Clugston, Anna Dominiczak and Alex Alderton, Roberto Amato, Jeffrey Barrett, Sonia Goncalves, Ewan Harrison, David K. Jackson, Ian Johnston, Dominic Kwiatkowski, Cordelia Langford, John Sillitoe on behalf of the Wellcome Sanger Institute COVID-19 Surveillance Team |
| EPI_ISL_3314724 | Lighthouse Laboratory Plymouth | Wellcome Sanger Institute for the COVID-19 Genomics UK (COG-UK) Consortium | Lighthouse Laboratory Plymouth and Alex Alderton, Roberto Amato, Jeffrey Barrett, Sonia Goncalves, Ewan Harrison, David K. Jackson, Ian Johnston, Dominic Kwiatkowski, Cordelia Langford, John Sillitoe on behalf of the Wellcome Sanger Institute COVID-19 Surveillance Team |
| EPI_ISL_3314725 | Lighthouse Lab in Glasgow | Wellcome Sanger Institute for the COVID-19 Genomics UK (COG-UK) Consortium | Harper VanSteenhouse, Yumi Kasai, David Gray, Carol Clugston, Anna Dominiczak and Alex Alderton, Roberto Amato, Jeffrey Barrett, Sonia Goncalves, Ewan Harrison, David K. Jackson, Ian Johnston, Dominic Kwiatkowski, Cordelia Langford, John Sillitoe on behalf of the Wellcome Sanger Institute COVID-19 Surveillance Team |
| EPI_ISL_3314726, EPI_ISL_3314727, EPI_ISL_3314728 | Lighthouse Laboratory Plymouth | Wellcome Sanger Institute for the COVID-19 Genomics UK (COG-UK) Consortium | Lighthouse Laboratory Plymouth and Alex Alderton, Roberto Amato, Jeffrey Barrett, Sonia Goncalves, Ewan Harrison, David K. Jackson, Ian Johnston, Dominic Kwiatkowski, Cordelia Langford, John Sillitoe on behalf of the Wellcome Sanger Institute COVID-19 Surveillance Team |
| EPI_ISL_3314729 | Lighthouse Lab in Glasgow | Wellcome Sanger Institute for the COVID-19 Genomics UK (COG-UK) Consortium | Harper VanSteenhouse, Yumi Kasai, David Gray, Carol Clugston, Anna Dominiczak and Alex Alderton, Roberto Amato, Jeffrey Barrett, Sonia Goncalves, Ewan Harrison, David K. Jackson, Ian Johnston, Dominic Kwiatkowski, Cordelia Langford, John Sillitoe on behalf of the Wellcome Sanger Institute COVID-19 Surveillance Team |
| EPI_ISL_3314730, EPI_ISL_3314731, EPI_ISL_3314732, EPI_ISL_3314733, EPI_ISL_3314734 | Lighthouse Laboratory Plymouth | Wellcome Sanger Institute for the COVID-19 Genomics UK (COG-UK) Consortium | Lighthouse Laboratory Plymouth and Alex Alderton, Roberto Amato, Jeffrey Barrett, Sonia Goncalves, Ewan Harrison, David K. Jackson, Ian Johnston, Dominic Kwiatkowski, Cordelia Langford, John Sillitoe on behalf of the Wellcome Sanger Institute COVID-19 Surveillance Team |
| EPI_ISL_3314735 | Berkshire and Surrey Pathology Services Lighthouse Laboratory | Wellcome Sanger Institute for the COVID-19 Genomics UK (COG-UK) | Berkshire and Surrey Pathology Services Lighthouse Laboratory and Alex Alderton, Roberto Amato, Jeffrey Barrett, Sonia Goncalves, Ewan Harrison, David K. Jackson, Ian Johnston, Dominic Kwiatkowski, Cordelia Langford, John Sillitoe on behalf |

| | | | |
|---|---|---|---|
| | | Consortium | of the Wellcome Sanger Institute COVID-19 Surveillance Team |
| EPI_ISL_3314736, EPI_ISL_3314737 | Lighthouse Laboratory Plymouth | Wellcome Sanger Institute for the COVID-19 Genomics UK (COG-UK) Consortium | Lighthouse Laboratory Plymouth and Alex Alderton, Roberto Amato, Jeffrey Barrett, Sonia Goncalves, Ewan Harrison, David K. Jackson, Ian Johnston, Dominic Kwiatkowski, Cordelia Langford, John Sillitoe on behalf of the Wellcome Sanger Institute COVID-19 Surveillance Team |
| EPI_ISL_3314738 | Lighthouse Lab in Glasgow | Wellcome Sanger Institute for the COVID-19 Genomics UK (COG-UK) Consortium | Harper VanSteenhouse, Yumi Kasai, David Gray, Carol Clugston, Anna Dominiczak and Alex Alderton, Roberto Amato, Jeffrey Barrett, Sonia Goncalves, Ewan Harrison, David K. Jackson, Ian Johnston, Dominic Kwiatkowski, Cordelia Langford, John Sillitoe on behalf of the Wellcome Sanger Institute COVID-19 Surveillance Team |
| EPI_ISL_3314739 | Lighthouse Laboratory Plymouth | Wellcome Sanger Institute for the COVID-19 Genomics UK (COG-UK) Consortium | Lighthouse Laboratory Plymouth and Alex Alderton, Roberto Amato, Jeffrey Barrett, Sonia Goncalves, Ewan Harrison, David K. Jackson, Ian Johnston, Dominic Kwiatkowski, Cordelia Langford, John Sillitoe on behalf of the Wellcome Sanger Institute COVID-19 Surveillance Team |
| EPI_ISL_3314740 | Berkshire and Surrey Pathology Services Lighthouse Laboratory | Wellcome Sanger Institute for the COVID-19 Genomics UK (COG-UK) Consortium | Berkshire and Surrey Pathology Services Lighthouse Laboratory and Alex Alderton, Roberto Amato, Jeffrey Barrett, Sonia Goncalves, Ewan Harrison, David K. Jackson, Ian Johnston, Dominic Kwiatkowski, Cordelia Langford, John Sillitoe on behalf of the Wellcome Sanger Institute COVID-19 Surveillance Team |
| EPI_ISL_3314741, EPI_ISL_3314742 | Lighthouse Laboratory Plymouth | Wellcome Sanger Institute for the COVID-19 Genomics UK (COG-UK) Consortium | Lighthouse Laboratory Plymouth and Alex Alderton, Roberto Amato, Jeffrey Barrett, Sonia Goncalves, Ewan Harrison, David K. Jackson, Ian Johnston, Dominic Kwiatkowski, Cordelia Langford, John Sillitoe on behalf of the Wellcome Sanger Institute COVID-19 Surveillance Team |
| EPI_ISL_3314743 | Berkshire and Surrey Pathology Services Lighthouse Laboratory | Wellcome Sanger Institute for the COVID-19 Genomics UK (COG-UK) Consortium | Berkshire and Surrey Pathology Services Lighthouse Laboratory and Alex Alderton, Roberto Amato, Jeffrey Barrett, Sonia Goncalves, Ewan Harrison, David K. Jackson, Ian Johnston, Dominic Kwiatkowski, Cordelia Langford, John Sillitoe on behalf of the Wellcome Sanger Institute COVID-19 Surveillance Team |
| EPI_ISL_3314744, EPI_ISL_3314745, EPI_ISL_3314746 | Lighthouse Laboratory Plymouth | Wellcome Sanger Institute for the COVID-19 Genomics UK (COG-UK) Consortium | Lighthouse Laboratory Plymouth and Alex Alderton, Roberto Amato, Jeffrey Barrett, Sonia Goncalves, Ewan Harrison, David K. Jackson, Ian Johnston, Dominic Kwiatkowski, Cordelia Langford, John Sillitoe on behalf of the Wellcome Sanger Institute COVID-19 Surveillance Team |
| EPI_ISL_3314747 | Lighthouse Lab in Glasgow | Wellcome Sanger Institute for the COVID-19 Genomics UK (COG-UK) Consortium | Harper VanSteenhouse, Yumi Kasai, David Gray, Carol Clugston, Anna Dominiczak and Alex Alderton, Roberto Amato, Jeffrey Barrett, Sonia Goncalves, Ewan Harrison, David K. Jackson, Ian Johnston, Dominic Kwiatkowski, Cordelia Langford, John Sillitoe on behalf of the Wellcome Sanger Institute COVID-19 Surveillance Team |
| EPI_ISL_3314748 | Berkshire and Surrey Pathology Services Lighthouse Laboratory | Wellcome Sanger Institute for the COVID-19 Genomics UK (COG-UK) Consortium | Berkshire and Surrey Pathology Services Lighthouse Laboratory and Alex Alderton, Roberto Amato, Jeffrey Barrett, Sonia Goncalves, Ewan Harrison, David K. Jackson, Ian Johnston, Dominic Kwiatkowski, Cordelia Langford, John Sillitoe on behalf of the Wellcome Sanger Institute COVID-19 Surveillance Team |
| EPI_ISL_3314749 | Lighthouse Laboratory Plymouth | Wellcome Sanger Institute for the COVID-19 Genomics UK (COG-UK) Consortium | Lighthouse Laboratory Plymouth and Alex Alderton, Roberto Amato, Jeffrey Barrett, Sonia Goncalves, Ewan Harrison, David K. Jackson, Ian Johnston, Dominic Kwiatkowski, Cordelia Langford, John Sillitoe on behalf of the Wellcome Sanger Institute COVID-19 Surveillance Team |
| EPI_ISL_3314750, EPI_ISL_3314751, EPI_ISL_3314752 | Berkshire and Surrey Pathology Services Lighthouse Laboratory | Wellcome Sanger Institute for the COVID-19 Genomics UK (COG-UK) Consortium | Berkshire and Surrey Pathology Services Lighthouse Laboratory and Alex Alderton, Roberto Amato, Jeffrey Barrett, Sonia Goncalves, Ewan Harrison, David K. Jackson, Ian Johnston, Dominic Kwiatkowski, Cordelia Langford, John Sillitoe on behalf of the Wellcome Sanger Institute COVID-19 Surveillance Team |
| EPI_ISL_3314753, EPI_ISL_3314754, EPI_ISL_3314755, EPI_ISL_3314756 | Lighthouse Laboratory Plymouth | Wellcome Sanger Institute for the COVID-19 Genomics UK (COG-UK) Consortium | Lighthouse Laboratory Plymouth and Alex Alderton, Roberto Amato, Jeffrey Barrett, Sonia Goncalves, Ewan Harrison, David K. Jackson, Ian Johnston, Dominic Kwiatkowski, Cordelia Langford, John Sillitoe on behalf of the Wellcome Sanger Institute COVID-19 Surveillance Team |
| EPI_ISL_3314757 | Lighthouse Lab in Glasgow | Wellcome Sanger Institute for the COVID-19 Genomics UK (COG-UK) Consortium | Harper VanSteenhouse, Yumi Kasai, David Gray, Carol Clugston, Anna Dominiczak and Alex Alderton, Roberto Amato, Jeffrey Barrett, Sonia Goncalves, Ewan Harrison, David K. Jackson, Ian Johnston, Dominic Kwiatkowski, Cordelia Langford, John Sillitoe on behalf of the Wellcome Sanger Institute COVID-19 Surveillance Team |
| EPI_ISL_3314758 | Berkshire and Surrey Pathology Services Lighthouse Laboratory | Wellcome Sanger Institute for the COVID-19 Genomics UK (COG-UK) Consortium | Berkshire and Surrey Pathology Services Lighthouse Laboratory and Alex Alderton, Roberto Amato, Jeffrey Barrett, Sonia Goncalves, Ewan Harrison, David K. Jackson, Ian Johnston, Dominic Kwiatkowski, Cordelia Langford, John Sillitoe on behalf of the Wellcome Sanger Institute COVID-19 Surveillance Team |
| EPI_ISL_3314759 | Lighthouse Laboratory Plymouth | Wellcome Sanger Institute for the COVID-19 Genomics UK (COG-UK) Consortium | Lighthouse Laboratory Plymouth and Alex Alderton, Roberto Amato, Jeffrey Barrett, Sonia Goncalves, Ewan Harrison, David K. Jackson, Ian Johnston, Dominic Kwiatkowski, Cordelia Langford, John Sillitoe on behalf of the Wellcome Sanger Institute COVID-19 Surveillance Team |
| EPI_ISL_3314760, EPI_ISL_3314761, EPI_ISL_3314762 | Berkshire and Surrey Pathology Services Lighthouse Laboratory | Wellcome Sanger Institute for the COVID-19 Genomics UK (COG-UK) Consortium | Berkshire and Surrey Pathology Services Lighthouse Laboratory and Alex Alderton, Roberto Amato, Jeffrey Barrett, Sonia Goncalves, Ewan Harrison, David K. Jackson, Ian Johnston, Dominic Kwiatkowski, Cordelia Langford, John Sillitoe on behalf of the Wellcome Sanger Institute COVID-19 Surveillance Team |
| EPI_ISL_3314763 | Lighthouse Lab in Glasgow | Wellcome Sanger Institute for the COVID-19 Genomics UK (COG-UK) Consortium | Harper VanSteenhouse, Yumi Kasai, David Gray, Carol Clugston, Anna Dominiczak and Alex Alderton, Roberto Amato, Jeffrey Barrett, Sonia Goncalves, Ewan Harrison, David K. Jackson, Ian Johnston, Dominic Kwiatkowski, Cordelia Langford, John Sillitoe on behalf of the Wellcome Sanger Institute COVID-19 Surveillance Team |
| EPI_ISL_3314764 | Lighthouse Laboratory Plymouth | Wellcome Sanger Institute for the COVID-19 Genomics UK (COG-UK) Consortium | Lighthouse Laboratory Plymouth and Alex Alderton, Roberto Amato, Jeffrey Barrett, Sonia Goncalves, Ewan Harrison, David K. Jackson, Ian Johnston, Dominic Kwiatkowski, Cordelia Langford, John Sillitoe on behalf of the Wellcome Sanger Institute COVID-19 Surveillance Team |
| EPI_ISL_3314765 | Berkshire and Surrey Pathology Services Lighthouse Laboratory | Wellcome Sanger Institute for the COVID-19 Genomics UK (COG-UK) Consortium | Berkshire and Surrey Pathology Services Lighthouse Laboratory and Alex Alderton, Roberto Amato, Jeffrey Barrett, Sonia Goncalves, Ewan Harrison, David K. Jackson, Ian Johnston, Dominic Kwiatkowski, Cordelia Langford, John Sillitoe on behalf of the Wellcome Sanger Institute COVID-19 Surveillance Team |
| EPI_ISL_3314766 | Lighthouse Laboratory Plymouth | Wellcome Sanger Institute for the COVID-19 Genomics UK (COG-UK) Consortium | Lighthouse Laboratory Plymouth and Alex Alderton, Roberto Amato, Jeffrey Barrett, Sonia Goncalves, Ewan Harrison, David K. Jackson, Ian Johnston, Dominic Kwiatkowski, Cordelia Langford, John Sillitoe on behalf of the Wellcome Sanger Institute COVID-19 Surveillance Team |
| EPI_ISL_3314767, EPI_ISL_3314768 | Berkshire and Surrey Pathology Services Lighthouse Laboratory | Wellcome Sanger Institute for the COVID-19 Genomics UK (COG-UK) Consortium | Berkshire and Surrey Pathology Services Lighthouse Laboratory and Alex Alderton, Roberto Amato, Jeffrey Barrett, Sonia Goncalves, Ewan Harrison, David K. Jackson, Ian Johnston, Dominic Kwiatkowski, Cordelia Langford, John Sillitoe on behalf of the Wellcome Sanger Institute COVID-19 Surveillance Team |
| EPI_ISL_3314769, EPI_ISL_3314770, EPI_ISL_3314771, EPI_ISL_3314772, EPI_ISL_3314773 | Lighthouse Laboratory Plymouth | Wellcome Sanger Institute for the COVID-19 Genomics UK (COG-UK) Consortium | Lighthouse Laboratory Plymouth and Alex Alderton, Roberto Amato, Jeffrey Barrett, Sonia Goncalves, Ewan Harrison, David K. Jackson, Ian Johnston, Dominic Kwiatkowski, Cordelia Langford, John Sillitoe on behalf of the Wellcome Sanger Institute COVID-19 Surveillance Team |
| EPI_ISL_3314774 | Berkshire and Surrey Pathology Services Lighthouse Laboratory | Wellcome Sanger Institute for the COVID-19 Genomics UK (COG-UK) Consortium | Berkshire and Surrey Pathology Services Lighthouse Laboratory and Alex Alderton, Roberto Amato, Jeffrey Barrett, Sonia Goncalves, Ewan Harrison, David K. Jackson, Ian Johnston, Dominic Kwiatkowski, Cordelia Langford, John Sillitoe on behalf of the Wellcome Sanger Institute COVID-19 Surveillance Team |
| EPI_ISL_3314775 | Lighthouse Laboratory Plymouth | Wellcome Sanger Institute for the COVID-19 Genomics UK (COG-UK) Consortium | Lighthouse Laboratory Plymouth and Alex Alderton, Roberto Amato, Jeffrey Barrett, Sonia Goncalves, Ewan Harrison, David K. Jackson, Ian Johnston, Dominic Kwiatkowski, Cordelia Langford, John Sillitoe on behalf of the Wellcome Sanger Institute COVID-19 Surveillance Team |
| EPI_ISL_3314776, EPI_ISL_3314777 | Berkshire and Surrey Pathology Services Lighthouse Laboratory | Wellcome Sanger Institute for the COVID-19 Genomics UK (COG-UK) Consortium | Berkshire and Surrey Pathology Services Lighthouse Laboratory and Alex Alderton, Roberto Amato, Jeffrey Barrett, Sonia Goncalves, Ewan Harrison, David K. Jackson, Ian Johnston, Dominic Kwiatkowski, Cordelia Langford, John Sillitoe on behalf of the Wellcome Sanger Institute COVID-19 Surveillance Team |
| EPI_ISL_3314778, EPI_ISL_3314779 | Lighthouse Laboratory Plymouth | Wellcome Sanger Institute for the COVID-19 Genomics UK (COG-UK) Consortium | Lighthouse Laboratory Plymouth and Alex Alderton, Roberto Amato, Jeffrey Barrett, Sonia Goncalves, Ewan Harrison, David K. Jackson, Ian Johnston, Dominic Kwiatkowski, Cordelia Langford, John Sillitoe on behalf of the Wellcome Sanger Institute COVID-19 Surveillance Team |
| EPI_ISL_3314780 | Lighthouse Lab in Glasgow | Wellcome Sanger Institute for the COVID-19 Genomics UK (COG-UK) Consortium | Harper VanSteenhouse, Yumi Kasai, David Gray, Carol Clugston, Anna Dominiczak and Alex Alderton, Roberto Amato, Jeffrey Barrett, Sonia Goncalves, Ewan Harrison, David K. Jackson, Ian Johnston, Dominic Kwiatkowski, Cordelia Langford, John Sillitoe on behalf of the Wellcome Sanger Institute COVID-19 Surveillance Team |
| EPI_ISL_3314781, EPI_ISL_3314782, EPI_ISL_3314783, EPI_ISL_3314784 | Lighthouse Laboratory Plymouth | Wellcome Sanger Institute for the COVID-19 Genomics UK (COG-UK) Consortium | Lighthouse Laboratory Plymouth and Alex Alderton, Roberto Amato, Jeffrey Barrett, Sonia Goncalves, Ewan Harrison, David K. Jackson, Ian Johnston, Dominic Kwiatkowski, Cordelia Langford, John Sillitoe on behalf of the Wellcome Sanger Institute COVID-19 Surveillance Team |
| EPI_ISL_3314785 | Lighthouse Lab in Glasgow | Wellcome Sanger Institute for the COVID-19 Genomics UK (COG-UK) Consortium | Harper VanSteenhouse, Yumi Kasai, David Gray, Carol Clugston, Anna Dominiczak and Alex Alderton, Roberto Amato, Jeffrey Barrett, Sonia Goncalves, Ewan Harrison, David K. Jackson, Ian Johnston, Dominic Kwiatkowski, Cordelia Langford, John Sillitoe on behalf of the Wellcome Sanger Institute COVID-19 Surveillance Team |
| EPI_ISL_3314786 | Berkshire and Surrey Pathology Services Lighthouse Laboratory | Wellcome Sanger Institute for the COVID-19 Genomics UK (COG-UK) Consortium | Berkshire and Surrey Pathology Services Lighthouse Laboratory and Alex Alderton, Roberto Amato, Jeffrey Barrett, Sonia Goncalves, Ewan Harrison, David K. Jackson, Ian Johnston, Dominic Kwiatkowski, Cordelia Langford, John Sillitoe on behalf of the Wellcome Sanger Institute COVID-19 Surveillance Team |
| EPI_ISL_3314787, EPI_ISL_3314788 | Lighthouse Laboratory Plymouth | Wellcome Sanger Institute for the COVID-19 Genomics UK (COG-UK) Consortium | Lighthouse Laboratory Plymouth and Alex Alderton, Roberto Amato, Jeffrey Barrett, Sonia Goncalves, Ewan Harrison, David K. Jackson, Ian Johnston, Dominic Kwiatkowski, Cordelia Langford, John Sillitoe on behalf of the Wellcome Sanger Institute COVID-19 Surveillance Team |
| EPI_ISL_3314789 | Berkshire and Surrey Pathology Services Lighthouse Laboratory | Wellcome Sanger Institute for the COVID-19 Genomics UK (COG-UK) Consortium | Berkshire and Surrey Pathology Services Lighthouse Laboratory and Alex Alderton, Roberto Amato, Jeffrey Barrett, Sonia Goncalves, Ewan Harrison, David K. Jackson, Ian Johnston, Dominic Kwiatkowski, Cordelia Langford, John Sillitoe on behalf of the Wellcome Sanger Institute COVID-19 Surveillance Team |
| EPI_ISL_3314790 | Lighthouse Lab in Glasgow | Wellcome Sanger Institute for the COVID-19 Genomics UK (COG-UK) Consortium | Harper VanSteenhouse, Yumi Kasai, David Gray, Carol Clugston, Anna Dominiczak and Alex Alderton, Roberto Amato, Jeffrey Barrett, Sonia Goncalves, Ewan Harrison, David K. Jackson, Ian Johnston, Dominic Kwiatkowski, Cordelia Langford, John Sillitoe on behalf of the Wellcome Sanger Institute COVID-19 Surveillance Team |
| EPI_ISL_3314791 | Lighthouse Laboratory Plymouth | Wellcome Sanger Institute for the COVID-19 Genomics UK (COG-UK) Consortium | Lighthouse Laboratory Plymouth and Alex Alderton, Roberto Amato, Jeffrey Barrett, Sonia Goncalves, Ewan Harrison, David K. Jackson, Ian Johnston, Dominic Kwiatkowski, Cordelia Langford, John Sillitoe on behalf of the Wellcome Sanger Institute COVID-19 Surveillance Team |
| EPI_ISL_3314792 | Lighthouse Lab in Glasgow | Wellcome Sanger Institute for the COVID-19 Genomics UK (COG-UK) Consortium | Harper VanSteenhouse, Yumi Kasai, David Gray, Carol Clugston, Anna Dominiczak and Alex Alderton, Roberto Amato, Jeffrey Barrett, Sonia Goncalves, Ewan Harrison, David K. Jackson, Ian Johnston, Dominic Kwiatkowski, Cordelia Langford, John Sillitoe on behalf of the Wellcome Sanger Institute COVID-19 Surveillance Team |
| EPI_ISL_3314793, EPI_ISL_3314794, EPI_ISL_3314795, EPI_ISL_3314796, EPI_ISL_3314797, EPI_ISL_3314798, EPI_ISL_3314799, EPI_ISL_3314800, EPI_ISL_3314801, EPI_ISL_3314802, EPI_ISL_3314803 | | | |
| see above | Lighthouse Laboratory Plymouth | Wellcome Sanger Institute for the COVID-19 Genomics UK (COG-UK) Consortium | Lighthouse Laboratory Plymouth and Alex Alderton, Roberto Amato, Jeffrey Barrett, Sonia Goncalves, Ewan Harrison, David K. Jackson, Ian Johnston, Dominic Kwiatkowski, Cordelia Langford, John Sillitoe on behalf of the Wellcome Sanger Institute COVID-19 Surveillance Team |
| EPI_ISL_3314804, EPI_ISL_3314805 | Berkshire and Surrey Pathology Services Lighthouse Laboratory | Wellcome Sanger Institute for the COVID-19 Genomics UK (COG-UK) Consortium | Berkshire and Surrey Pathology Services Lighthouse Laboratory and Alex Alderton, Roberto Amato, Jeffrey Barrett, Sonia Goncalves, Ewan Harrison, David K. Jackson, Ian Johnston, Dominic Kwiatkowski, Cordelia Langford, John Sillitoe on behalf of the Wellcome Sanger Institute COVID-19 Surveillance Team |
| EPI_ISL_3314806 | Lighthouse Lab in Glasgow | Wellcome Sanger Institute for the COVID-19 Genomics UK (COG-UK) Consortium | Harper VanSteenhouse, Yumi Kasai, David Gray, Carol Clugston, Anna Dominiczak and Alex Alderton, Roberto Amato, Jeffrey Barrett, Sonia Goncalves, Ewan Harrison, David K. Jackson, Ian Johnston, Dominic Kwiatkowski, Cordelia Langford, John Sillitoe on behalf of the Wellcome Sanger Institute COVID-19 Surveillance Team |
| EPI_ISL_3314807, EPI_ISL_3314808 | Berkshire and Surrey Pathology Services Lighthouse Laboratory | Wellcome Sanger Institute for the COVID-19 Genomics UK (COG-UK) Consortium | Berkshire and Surrey Pathology Services Lighthouse Laboratory and Alex Alderton, Roberto Amato, Jeffrey Barrett, Sonia Goncalves, Ewan Harrison, David K. Jackson, Ian Johnston, Dominic Kwiatkowski, Cordelia Langford, John Sillitoe on behalf of the Wellcome Sanger Institute COVID-19 Surveillance Team |
| EPI_ISL_3314809, | Lighthouse Laboratory Plymouth | Wellcome Sanger Institute for the COVID-19 Genomics UK (COG-UK) | Lighthouse Laboratory Plymouth and Alex Alderton, Roberto Amato, Jeffrey Barrett, Sonia Goncalves, Ewan Harrison, David K. Jackson, Ian Johnston, Dominic Kwiatkowski, Cordelia Langford, John Sillitoe on behalf of the Wellcome Sanger |

| | | | |
|---|---|---|---|
| EPI_ISL_3314810, EPI_ISL_3314811, EPI_ISL_3314812, EPI_ISL_3314813 | | Consortium | Institute COVID-19 Surveillance Team |
| EPI_ISL_3314814 | Lighthouse Lab in Glasgow | Wellcome Sanger Institute for the COVID-19 Genomics UK (COG-UK) Consortium | Harper VanSteenhouse, Yumi Kasai, David Gray, Carol Clugston, Anna Dominiczak and Alex Alderton, Roberto Amato, Jeffrey Barrett, Sonia Goncalves, Ewan Harrison, David K. Jackson, Ian Johnston, Dominic Kwiatkowski, Cordelia Langford, John Sillitoe on behalf of the Wellcome Sanger Institute COVID-19 Surveillance Team |
| EPI_ISL_3314815 | Lighthouse Laboratory Plymouth | Wellcome Sanger Institute for the COVID-19 Genomics UK (COG-UK) Consortium | Lighthouse Laboratory Plymouth and Alex Alderton, Roberto Amato, Jeffrey Barrett, Sonia Goncalves, Ewan Harrison, David K. Jackson, Ian Johnston, Dominic Kwiatkowski, Cordelia Langford, John Sillitoe on behalf of the Wellcome Sanger Institute COVID-19 Surveillance Team |
| EPI_ISL_3314816, EPI_ISL_3314817 | Berkshire and Surrey Pathology Services Lighthouse Laboratory | Wellcome Sanger Institute for the COVID-19 Genomics UK (COG-UK) Consortium | Berkshire and Surrey Pathology Services Lighthouse Laboratory and Alex Alderton, Roberto Amato, Jeffrey Barrett, Sonia Goncalves, Ewan Harrison, David K. Jackson, Ian Johnston, Dominic Kwiatkowski, Cordelia Langford, John Sillitoe on behalf of the Wellcome Sanger Institute COVID-19 Surveillance Team |
| EPI_ISL_3314818 | Lighthouse Lab in Glasgow | Wellcome Sanger Institute for the COVID-19 Genomics UK (COG-UK) Consortium | Harper VanSteenhouse, Yumi Kasai, David Gray, Carol Clugston, Anna Dominiczak and Alex Alderton, Roberto Amato, Jeffrey Barrett, Sonia Goncalves, Ewan Harrison, David K. Jackson, Ian Johnston, Dominic Kwiatkowski, Cordelia Langford, John Sillitoe on behalf of the Wellcome Sanger Institute COVID-19 Surveillance Team |
| EPI_ISL_3314819, EPI_ISL_3314820 | Lighthouse Laboratory Plymouth | Wellcome Sanger Institute for the COVID-19 Genomics UK (COG-UK) Consortium | Lighthouse Laboratory Plymouth and Alex Alderton, Roberto Amato, Jeffrey Barrett, Sonia Goncalves, Ewan Harrison, David K. Jackson, Ian Johnston, Dominic Kwiatkowski, Cordelia Langford, John Sillitoe on behalf of the Wellcome Sanger Institute COVID-19 Surveillance Team |
| EPI_ISL_3314821 | Berkshire and Surrey Pathology Services Lighthouse Laboratory | Wellcome Sanger Institute for the COVID-19 Genomics UK (COG-UK) Consortium | Berkshire and Surrey Pathology Services Lighthouse Laboratory and Alex Alderton, Roberto Amato, Jeffrey Barrett, Sonia Goncalves, Ewan Harrison, David K. Jackson, Ian Johnston, Dominic Kwiatkowski, Cordelia Langford, John Sillitoe on behalf of the Wellcome Sanger Institute COVID-19 Surveillance Team |
| EPI_ISL_3314822, EPI_ISL_3314823 | Lighthouse Laboratory Plymouth | Wellcome Sanger Institute for the COVID-19 Genomics UK (COG-UK) Consortium | Lighthouse Laboratory Plymouth and Alex Alderton, Roberto Amato, Jeffrey Barrett, Sonia Goncalves, Ewan Harrison, David K. Jackson, Ian Johnston, Dominic Kwiatkowski, Cordelia Langford, John Sillitoe on behalf of the Wellcome Sanger Institute COVID-19 Surveillance Team |
| EPI_ISL_3314824 | Lighthouse Lab in Glasgow | Wellcome Sanger Institute for the COVID-19 Genomics UK (COG-UK) Consortium | Harper VanSteenhouse, Yumi Kasai, David Gray, Carol Clugston, Anna Dominiczak and Alex Alderton, Roberto Amato, Jeffrey Barrett, Sonia Goncalves, Ewan Harrison, David K. Jackson, Ian Johnston, Dominic Kwiatkowski, Cordelia Langford, John Sillitoe on behalf of the Wellcome Sanger Institute COVID-19 Surveillance Team |
| EPI_ISL_3314825, EPI_ISL_3314826, EPI_ISL_3314827 | Lighthouse Laboratory Plymouth | Wellcome Sanger Institute for the COVID-19 Genomics UK (COG-UK) Consortium | Lighthouse Laboratory Plymouth and Alex Alderton, Roberto Amato, Jeffrey Barrett, Sonia Goncalves, Ewan Harrison, David K. Jackson, Ian Johnston, Dominic Kwiatkowski, Cordelia Langford, John Sillitoe on behalf of the Wellcome Sanger Institute COVID-19 Surveillance Team |
| EPI_ISL_3314828, EPI_ISL_3314829, EPI_ISL_3314830 | Berkshire and Surrey Pathology Services Lighthouse Laboratory | Wellcome Sanger Institute for the COVID-19 Genomics UK (COG-UK) Consortium | Berkshire and Surrey Pathology Services Lighthouse Laboratory and Alex Alderton, Roberto Amato, Jeffrey Barrett, Sonia Goncalves, Ewan Harrison, David K. Jackson, Ian Johnston, Dominic Kwiatkowski, Cordelia Langford, John Sillitoe on behalf of the Wellcome Sanger Institute COVID-19 Surveillance Team |
| EPI_ISL_3314831, EPI_ISL_3314832, EPI_ISL_3314833, EPI_ISL_3314834 | Lighthouse Laboratory Plymouth | Wellcome Sanger Institute for the COVID-19 Genomics UK (COG-UK) Consortium | Lighthouse Laboratory Plymouth and Alex Alderton, Roberto Amato, Jeffrey Barrett, Sonia Goncalves, Ewan Harrison, David K. Jackson, Ian Johnston, Dominic Kwiatkowski, Cordelia Langford, John Sillitoe on behalf of the Wellcome Sanger Institute COVID-19 Surveillance Team |
| EPI_ISL_3314835 | Lighthouse Lab in Glasgow | Wellcome Sanger Institute for the COVID-19 Genomics UK (COG-UK) Consortium | Harper VanSteenhouse, Yumi Kasai, David Gray, Carol Clugston, Anna Dominiczak and Alex Alderton, Roberto Amato, Jeffrey Barrett, Sonia Goncalves, Ewan Harrison, David K. Jackson, Ian Johnston, Dominic Kwiatkowski, Cordelia Langford, John Sillitoe on behalf of the Wellcome Sanger Institute COVID-19 Surveillance Team |
| EPI_ISL_3314836 | Lighthouse Laboratory Plymouth | Wellcome Sanger Institute for the COVID-19 Genomics UK (COG-UK) Consortium | Lighthouse Laboratory Plymouth and Alex Alderton, Roberto Amato, Jeffrey Barrett, Sonia Goncalves, Ewan Harrison, David K. Jackson, Ian Johnston, Dominic Kwiatkowski, Cordelia Langford, John Sillitoe on behalf of the Wellcome Sanger Institute COVID-19 Surveillance Team |
| EPI_ISL_3314837 | Berkshire and Surrey Pathology Services Lighthouse Laboratory | Wellcome Sanger Institute for the COVID-19 Genomics UK (COG-UK) Consortium | Berkshire and Surrey Pathology Services Lighthouse Laboratory and Alex Alderton, Roberto Amato, Jeffrey Barrett, Sonia Goncalves, Ewan Harrison, David K. Jackson, Ian Johnston, Dominic Kwiatkowski, Cordelia Langford, John Sillitoe on behalf of the Wellcome Sanger Institute COVID-19 Surveillance Team |
| EPI_ISL_3314838, EPI_ISL_3314839 | Lighthouse Laboratory Plymouth | Wellcome Sanger Institute for the COVID-19 Genomics UK (COG-UK) Consortium | Lighthouse Laboratory Plymouth and Alex Alderton, Roberto Amato, Jeffrey Barrett, Sonia Goncalves, Ewan Harrison, David K. Jackson, Ian Johnston, Dominic Kwiatkowski, Cordelia Langford, John Sillitoe on behalf of the Wellcome Sanger Institute COVID-19 Surveillance Team |
| EPI_ISL_3314840, EPI_ISL_3314841 | Berkshire and Surrey Pathology Services Lighthouse Laboratory | Wellcome Sanger Institute for the COVID-19 Genomics UK (COG-UK) Consortium | Berkshire and Surrey Pathology Services Lighthouse Laboratory and Alex Alderton, Roberto Amato, Jeffrey Barrett, Sonia Goncalves, Ewan Harrison, David K. Jackson, Ian Johnston, Dominic Kwiatkowski, Cordelia Langford, John Sillitoe on behalf of the Wellcome Sanger Institute COVID-19 Surveillance Team |
| EPI_ISL_3314842, EPI_ISL_3314843 | Lighthouse Laboratory Plymouth | Wellcome Sanger Institute for the COVID-19 Genomics UK (COG-UK) Consortium | Lighthouse Laboratory Plymouth and Alex Alderton, Roberto Amato, Jeffrey Barrett, Sonia Goncalves, Ewan Harrison, David K. Jackson, Ian Johnston, Dominic Kwiatkowski, Cordelia Langford, John Sillitoe on behalf of the Wellcome Sanger Institute COVID-19 Surveillance Team |
| EPI_ISL_3314844 | Berkshire and Surrey Pathology Services Lighthouse Laboratory | Wellcome Sanger Institute for the COVID-19 Genomics UK (COG-UK) Consortium | Berkshire and Surrey Pathology Services Lighthouse Laboratory and Alex Alderton, Roberto Amato, Jeffrey Barrett, Sonia Goncalves, Ewan Harrison, David K. Jackson, Ian Johnston, Dominic Kwiatkowski, Cordelia Langford, John Sillitoe on behalf of the Wellcome Sanger Institute COVID-19 Surveillance Team |
| EPI_ISL_3314845, EPI_ISL_3314846 | Lighthouse Laboratory Plymouth | Wellcome Sanger Institute for the COVID-19 Genomics UK (COG-UK) Consortium | Lighthouse Laboratory Plymouth and Alex Alderton, Roberto Amato, Jeffrey Barrett, Sonia Goncalves, Ewan Harrison, David K. Jackson, Ian Johnston, Dominic Kwiatkowski, Cordelia Langford, John Sillitoe on behalf of the Wellcome Sanger Institute COVID-19 Surveillance Team |
| EPI_ISL_3314847 | Berkshire and Surrey Pathology Services Lighthouse Laboratory | Wellcome Sanger Institute for the COVID-19 Genomics UK (COG-UK) Consortium | Berkshire and Surrey Pathology Services Lighthouse Laboratory and Alex Alderton, Roberto Amato, Jeffrey Barrett, Sonia Goncalves, Ewan Harrison, David K. Jackson, Ian Johnston, Dominic Kwiatkowski, Cordelia Langford, John Sillitoe on behalf of the Wellcome Sanger Institute COVID-19 Surveillance Team |
| EPI_ISL_3314848, EPI_ISL_3314849, EPI_ISL_3314850 | Lighthouse Lab in Glasgow | Wellcome Sanger Institute for the COVID-19 Genomics UK (COG-UK) Consortium | Harper VanSteenhouse, Yumi Kasai, David Gray, Carol Clugston, Anna Dominiczak and Alex Alderton, Roberto Amato, Jeffrey Barrett, Sonia Goncalves, Ewan Harrison, David K. Jackson, Ian Johnston, Dominic Kwiatkowski, Cordelia Langford, John Sillitoe on behalf of the Wellcome Sanger Institute COVID-19 Surveillance Team |
| EPI_ISL_3314851 | Berkshire and Surrey Pathology Services Lighthouse Laboratory | Wellcome Sanger Institute for the COVID-19 Genomics UK (COG-UK) Consortium | Berkshire and Surrey Pathology Services Lighthouse Laboratory and Alex Alderton, Roberto Amato, Jeffrey Barrett, Sonia Goncalves, Ewan Harrison, David K. Jackson, Ian Johnston, Dominic Kwiatkowski, Cordelia Langford, John Sillitoe on behalf of the Wellcome Sanger Institute COVID-19 Surveillance Team |
| EPI_ISL_3314852 | Lighthouse Laboratory Plymouth | Wellcome Sanger Institute for the COVID-19 Genomics UK (COG-UK) Consortium | Lighthouse Laboratory Plymouth and Alex Alderton, Roberto Amato, Jeffrey Barrett, Sonia Goncalves, Ewan Harrison, David K. Jackson, Ian Johnston, Dominic Kwiatkowski, Cordelia Langford, John Sillitoe on behalf of the Wellcome Sanger Institute COVID-19 Surveillance Team |
| EPI_ISL_3314853, EPI_ISL_3314854 | Berkshire and Surrey Pathology Services Lighthouse Laboratory | Wellcome Sanger Institute for the COVID-19 Genomics UK (COG-UK) Consortium | Berkshire and Surrey Pathology Services Lighthouse Laboratory and Alex Alderton, Roberto Amato, Jeffrey Barrett, Sonia Goncalves, Ewan Harrison, David K. Jackson, Ian Johnston, Dominic Kwiatkowski, Cordelia Langford, John Sillitoe on behalf of the Wellcome Sanger Institute COVID-19 Surveillance Team |
| EPI_ISL_3314855, EPI_ISL_3314856, EPI_ISL_3314857 | Lighthouse Laboratory Plymouth | Wellcome Sanger Institute for the COVID-19 Genomics UK (COG-UK) Consortium | Lighthouse Laboratory Plymouth and Alex Alderton, Roberto Amato, Jeffrey Barrett, Sonia Goncalves, Ewan Harrison, David K. Jackson, Ian Johnston, Dominic Kwiatkowski, Cordelia Langford, John Sillitoe on behalf of the Wellcome Sanger Institute COVID-19 Surveillance Team |
| EPI_ISL_3314858 | Berkshire and Surrey Pathology Services Lighthouse Laboratory | Wellcome Sanger Institute for the COVID-19 Genomics UK (COG-UK) Consortium | Berkshire and Surrey Pathology Services Lighthouse Laboratory and Alex Alderton, Roberto Amato, Jeffrey Barrett, Sonia Goncalves, Ewan Harrison, David K. Jackson, Ian Johnston, Dominic Kwiatkowski, Cordelia Langford, John Sillitoe on behalf of the Wellcome Sanger Institute COVID-19 Surveillance Team |
| EPI_ISL_3314859, EPI_ISL_3314860, EPI_ISL_3314861, EPI_ISL_3314862 | Lighthouse Laboratory Plymouth | Wellcome Sanger Institute for the COVID-19 Genomics UK (COG-UK) Consortium | Lighthouse Laboratory Plymouth and Alex Alderton, Roberto Amato, Jeffrey Barrett, Sonia Goncalves, Ewan Harrison, David K. Jackson, Ian Johnston, Dominic Kwiatkowski, Cordelia Langford, John Sillitoe on behalf of the Wellcome Sanger Institute COVID-19 Surveillance Team |
| EPI_ISL_3314863 | Berkshire and Surrey Pathology Services Lighthouse Laboratory | Wellcome Sanger Institute for the COVID-19 Genomics UK (COG-UK) Consortium | Berkshire and Surrey Pathology Services Lighthouse Laboratory and Alex Alderton, Roberto Amato, Jeffrey Barrett, Sonia Goncalves, Ewan Harrison, David K. Jackson, Ian Johnston, Dominic Kwiatkowski, Cordelia Langford, John Sillitoe on behalf of the Wellcome Sanger Institute COVID-19 Surveillance Team |
| EPI_ISL_3314864 | Lighthouse Laboratory Plymouth | Wellcome Sanger Institute for the COVID-19 Genomics UK (COG-UK) Consortium | Lighthouse Laboratory Plymouth and Alex Alderton, Roberto Amato, Jeffrey Barrett, Sonia Goncalves, Ewan Harrison, David K. Jackson, Ian Johnston, Dominic Kwiatkowski, Cordelia Langford, John Sillitoe on behalf of the Wellcome Sanger Institute COVID-19 Surveillance Team |
| EPI_ISL_3314865 | Berkshire and Surrey Pathology Services Lighthouse Laboratory | Wellcome Sanger Institute for the COVID-19 Genomics UK (COG-UK) Consortium | Berkshire and Surrey Pathology Services Lighthouse Laboratory and Alex Alderton, Roberto Amato, Jeffrey Barrett, Sonia Goncalves, Ewan Harrison, David K. Jackson, Ian Johnston, Dominic Kwiatkowski, Cordelia Langford, John Sillitoe on behalf of the Wellcome Sanger Institute COVID-19 Surveillance Team |
| EPI_ISL_3314866 | Lighthouse Laboratory Plymouth | Wellcome Sanger Institute for the COVID-19 Genomics UK (COG-UK) Consortium | Lighthouse Laboratory Plymouth and Alex Alderton, Roberto Amato, Jeffrey Barrett, Sonia Goncalves, Ewan Harrison, David K. Jackson, Ian Johnston, Dominic Kwiatkowski, Cordelia Langford, John Sillitoe on behalf of the Wellcome Sanger Institute COVID-19 Surveillance Team |
| EPI_ISL_3314867, EPI_ISL_3314868 | Berkshire and Surrey Pathology Services Lighthouse Laboratory | Wellcome Sanger Institute for the COVID-19 Genomics UK (COG-UK) Consortium | Berkshire and Surrey Pathology Services Lighthouse Laboratory and Alex Alderton, Roberto Amato, Jeffrey Barrett, Sonia Goncalves, Ewan Harrison, David K. Jackson, Ian Johnston, Dominic Kwiatkowski, Cordelia Langford, John Sillitoe on behalf of the Wellcome Sanger Institute COVID-19 Surveillance Team |
| EPI_ISL_3314869 | Lighthouse Laboratory Plymouth | Wellcome Sanger Institute for the COVID-19 Genomics UK (COG-UK) Consortium | Lighthouse Laboratory Plymouth and Alex Alderton, Roberto Amato, Jeffrey Barrett, Sonia Goncalves, Ewan Harrison, David K. Jackson, Ian Johnston, Dominic Kwiatkowski, Cordelia Langford, John Sillitoe on behalf of the Wellcome Sanger Institute COVID-19 Surveillance Team |
| EPI_ISL_3314870 | Berkshire and Surrey Pathology Services Lighthouse Laboratory | Wellcome Sanger Institute for the COVID-19 Genomics UK (COG-UK) Consortium | Berkshire and Surrey Pathology Services Lighthouse Laboratory and Alex Alderton, Roberto Amato, Jeffrey Barrett, Sonia Goncalves, Ewan Harrison, David K. Jackson, Ian Johnston, Dominic Kwiatkowski, Cordelia Langford, John Sillitoe on behalf of the Wellcome Sanger Institute COVID-19 Surveillance Team |
| EPI_ISL_3314871, EPI_ISL_3314872, EPI_ISL_3314873 | Lighthouse Laboratory Plymouth | Wellcome Sanger Institute for the COVID-19 Genomics UK (COG-UK) Consortium | Lighthouse Laboratory Plymouth and Alex Alderton, Roberto Amato, Jeffrey Barrett, Sonia Goncalves, Ewan Harrison, David K. Jackson, Ian Johnston, Dominic Kwiatkowski, Cordelia Langford, John Sillitoe on behalf of the Wellcome Sanger Institute COVID-19 Surveillance Team |
| EPI_ISL_3314874, EPI_ISL_3314875 | Berkshire and Surrey Pathology Services Lighthouse Laboratory | Wellcome Sanger Institute for the COVID-19 Genomics UK (COG-UK) Consortium | Berkshire and Surrey Pathology Services Lighthouse Laboratory and Alex Alderton, Roberto Amato, Jeffrey Barrett, Sonia Goncalves, Ewan Harrison, David K. Jackson, Ian Johnston, Dominic Kwiatkowski, Cordelia Langford, John Sillitoe on behalf of the Wellcome Sanger Institute COVID-19 Surveillance Team |
| EPI_ISL_3314876, EPI_ISL_3314877, EPI_ISL_3314878, EPI_ISL_3314879, EPI_ISL_3314880, EPI_ISL_3314881, EPI_ISL_3314882 | Lighthouse Laboratory Plymouth | Wellcome Sanger Institute for the COVID-19 Genomics UK (COG-UK) Consortium | Lighthouse Laboratory Plymouth and Alex Alderton, Roberto Amato, Jeffrey Barrett, Sonia Goncalves, Ewan Harrison, David K. Jackson, Ian Johnston, Dominic Kwiatkowski, Cordelia Langford, John Sillitoe on behalf of the Wellcome Sanger Institute COVID-19 Surveillance Team |
| EPI_ISL_3314883 | Berkshire and Surrey Pathology Services Lighthouse Laboratory | Wellcome Sanger Institute for the COVID-19 Genomics UK (COG-UK) | Berkshire and Surrey Pathology Services Lighthouse Laboratory and Alex Alderton, Roberto Amato, Jeffrey Barrett, Sonia Goncalves, Ewan Harrison, David K. Jackson, Ian Johnston, Dominic Kwiatkowski, Cordelia Langford, John Sillitoe on behalf |

| | | | |
|---|---|---|---|
| | | Consortium | of the Wellcome Sanger Institute COVID-19 Surveillance Team |
| EPI_ISL_3314884, EPI_ISL_3314885, EPI_ISL_3314886, EPI_ISL_3314887, EPI_ISL_3314888 | Lighthouse Laboratory Plymouth | Wellcome Sanger Institute for the COVID-19 Genomics UK (COG-UK) Consortium | Lighthouse Laboratory Plymouth and Alex Alderton, Roberto Amato, Jeffrey Barrett, Sonia Goncalves, Ewan Harrison, David K. Jackson, Ian Johnston, Dominic Kwiatkowski, Cordelia Langford, John Sillitoe on behalf of the Wellcome Sanger Institute COVID-19 Surveillance Team |
| EPI_ISL_3314889 | Lighthouse Lab in Glasgow | Wellcome Sanger Institute for the COVID-19 Genomics UK (COG-UK) Consortium | Harper VanSteenhouse, Yumi Kasai, David Gray, Carol Clugston, Anna Dominiczak and Alex Alderton, Roberto Amato, Jeffrey Barrett, Sonia Goncalves, Ewan Harrison, David K. Jackson, Ian Johnston, Dominic Kwiatkowski, Cordelia Langford, John Sillitoe on behalf of the Wellcome Sanger Institute COVID-19 Surveillance Team |
| EPI_ISL_3314890 | Berkshire and Surrey Pathology Services Lighthouse Laboratory | Wellcome Sanger Institute for the COVID-19 Genomics UK (COG-UK) Consortium | Berkshire and Surrey Pathology Services Lighthouse Laboratory and Alex Alderton, Roberto Amato, Jeffrey Barrett, Sonia Goncalves, Ewan Harrison, David K. Jackson, Ian Johnston, Dominic Kwiatkowski, Cordelia Langford, John Sillitoe on behalf of the Wellcome Sanger Institute COVID-19 Surveillance Team |
| EPI_ISL_3314891, EPI_ISL_3314892, EPI_ISL_3314893 | Lighthouse Laboratory Plymouth | Wellcome Sanger Institute for the COVID-19 Genomics UK (COG-UK) Consortium | Lighthouse Laboratory Plymouth and Alex Alderton, Roberto Amato, Jeffrey Barrett, Sonia Goncalves, Ewan Harrison, David K. Jackson, Ian Johnston, Dominic Kwiatkowski, Cordelia Langford, John Sillitoe on behalf of the Wellcome Sanger Institute COVID-19 Surveillance Team |
| EPI_ISL_3314894 | Berkshire and Surrey Pathology Services Lighthouse Laboratory | Wellcome Sanger Institute for the COVID-19 Genomics UK (COG-UK) Consortium | Berkshire and Surrey Pathology Services Lighthouse Laboratory and Alex Alderton, Roberto Amato, Jeffrey Barrett, Sonia Goncalves, Ewan Harrison, David K. Jackson, Ian Johnston, Dominic Kwiatkowski, Cordelia Langford, John Sillitoe on behalf of the Wellcome Sanger Institute COVID-19 Surveillance Team |
| EPI_ISL_3314895, EPI_ISL_3314896 | Lighthouse Laboratory Plymouth | Wellcome Sanger Institute for the COVID-19 Genomics UK (COG-UK) Consortium | Lighthouse Laboratory Plymouth and Alex Alderton, Roberto Amato, Jeffrey Barrett, Sonia Goncalves, Ewan Harrison, David K. Jackson, Ian Johnston, Dominic Kwiatkowski, Cordelia Langford, John Sillitoe on behalf of the Wellcome Sanger Institute COVID-19 Surveillance Team |
| EPI_ISL_3314897, EPI_ISL_3314898 | Berkshire and Surrey Pathology Services Lighthouse Laboratory | Wellcome Sanger Institute for the COVID-19 Genomics UK (COG-UK) Consortium | Berkshire and Surrey Pathology Services Lighthouse Laboratory and Alex Alderton, Roberto Amato, Jeffrey Barrett, Sonia Goncalves, Ewan Harrison, David K. Jackson, Ian Johnston, Dominic Kwiatkowski, Cordelia Langford, John Sillitoe on behalf of the Wellcome Sanger Institute COVID-19 Surveillance Team |
| EPI_ISL_3314899, EPI_ISL_3314900, EPI_ISL_3314901, EPI_ISL_3314902, EPI_ISL_3314903, EPI_ISL_3314904, EPI_ISL_3314905 | Lighthouse Laboratory Plymouth | Wellcome Sanger Institute for the COVID-19 Genomics UK (COG-UK) Consortium | Lighthouse Laboratory Plymouth and Alex Alderton, Roberto Amato, Jeffrey Barrett, Sonia Goncalves, Ewan Harrison, David K. Jackson, Ian Johnston, Dominic Kwiatkowski, Cordelia Langford, John Sillitoe on behalf of the Wellcome Sanger Institute COVID-19 Surveillance Team |
| EPI_ISL_3314906, EPI_ISL_3314907, EPI_ISL_3314908 | Berkshire and Surrey Pathology Services Lighthouse Laboratory | Wellcome Sanger Institute for the COVID-19 Genomics UK (COG-UK) Consortium | Berkshire and Surrey Pathology Services Lighthouse Laboratory and Alex Alderton, Roberto Amato, Jeffrey Barrett, Sonia Goncalves, Ewan Harrison, David K. Jackson, Ian Johnston, Dominic Kwiatkowski, Cordelia Langford, John Sillitoe on behalf of the Wellcome Sanger Institute COVID-19 Surveillance Team |
| EPI_ISL_3314909, EPI_ISL_3314910, EPI_ISL_3314911, EPI_ISL_3314912, EPI_ISL_3314913 | Lighthouse Laboratory Plymouth | Wellcome Sanger Institute for the COVID-19 Genomics UK (COG-UK) Consortium | Lighthouse Laboratory Plymouth and Alex Alderton, Roberto Amato, Jeffrey Barrett, Sonia Goncalves, Ewan Harrison, David K. Jackson, Ian Johnston, Dominic Kwiatkowski, Cordelia Langford, John Sillitoe on behalf of the Wellcome Sanger Institute COVID-19 Surveillance Team |
| EPI_ISL_3314914 | Lighthouse Lab in Glasgow | Wellcome Sanger Institute for the COVID-19 Genomics UK (COG-UK) Consortium | Harper VanSteenhouse, Yumi Kasai, David Gray, Carol Clugston, Anna Dominiczak and Alex Alderton, Roberto Amato, Jeffrey Barrett, Sonia Goncalves, Ewan Harrison, David K. Jackson, Ian Johnston, Dominic Kwiatkowski, Cordelia Langford, John Sillitoe on behalf of the Wellcome Sanger Institute COVID-19 Surveillance Team |
| EPI_ISL_3314915 | Berkshire and Surrey Pathology Services Lighthouse Laboratory | Wellcome Sanger Institute for the COVID-19 Genomics UK (COG-UK) Consortium | Berkshire and Surrey Pathology Services Lighthouse Laboratory and Alex Alderton, Roberto Amato, Jeffrey Barrett, Sonia Goncalves, Ewan Harrison, David K. Jackson, Ian Johnston, Dominic Kwiatkowski, Cordelia Langford, John Sillitoe on behalf of the Wellcome Sanger Institute COVID-19 Surveillance Team |
| EPI_ISL_3314916, EPI_ISL_3314917, EPI_ISL_3314918 | Lighthouse Laboratory Plymouth | Wellcome Sanger Institute for the COVID-19 Genomics UK (COG-UK) Consortium | Lighthouse Laboratory Plymouth and Alex Alderton, Roberto Amato, Jeffrey Barrett, Sonia Goncalves, Ewan Harrison, David K. Jackson, Ian Johnston, Dominic Kwiatkowski, Cordelia Langford, John Sillitoe on behalf of the Wellcome Sanger Institute COVID-19 Surveillance Team |
| EPI_ISL_3314919, EPI_ISL_3314920, EPI_ISL_3314921 | Berkshire and Surrey Pathology Services Lighthouse Laboratory | Wellcome Sanger Institute for the COVID-19 Genomics UK (COG-UK) Consortium | Berkshire and Surrey Pathology Services Lighthouse Laboratory and Alex Alderton, Roberto Amato, Jeffrey Barrett, Sonia Goncalves, Ewan Harrison, David K. Jackson, Ian Johnston, Dominic Kwiatkowski, Cordelia Langford, John Sillitoe on behalf of the Wellcome Sanger Institute COVID-19 Surveillance Team |
| EPI_ISL_3314922 | Lighthouse Laboratory Plymouth | Wellcome Sanger Institute for the COVID-19 Genomics UK (COG-UK) Consortium | Lighthouse Laboratory Plymouth and Alex Alderton, Roberto Amato, Jeffrey Barrett, Sonia Goncalves, Ewan Harrison, David K. Jackson, Ian Johnston, Dominic Kwiatkowski, Cordelia Langford, John Sillitoe on behalf of the Wellcome Sanger Institute COVID-19 Surveillance Team |
| EPI_ISL_3314923, EPI_ISL_3314924 | Berkshire and Surrey Pathology Services Lighthouse Laboratory | Wellcome Sanger Institute for the COVID-19 Genomics UK (COG-UK) Consortium | Berkshire and Surrey Pathology Services Lighthouse Laboratory and Alex Alderton, Roberto Amato, Jeffrey Barrett, Sonia Goncalves, Ewan Harrison, David K. Jackson, Ian Johnston, Dominic Kwiatkowski, Cordelia Langford, John Sillitoe on behalf of the Wellcome Sanger Institute COVID-19 Surveillance Team |
| EPI_ISL_3314925, EPI_ISL_3314926, EPI_ISL_3314927, EPI_ISL_3314928 | Lighthouse Laboratory Plymouth | Wellcome Sanger Institute for the COVID-19 Genomics UK (COG-UK) Consortium | Lighthouse Laboratory Plymouth and Alex Alderton, Roberto Amato, Jeffrey Barrett, Sonia Goncalves, Ewan Harrison, David K. Jackson, Ian Johnston, Dominic Kwiatkowski, Cordelia Langford, John Sillitoe on behalf of the Wellcome Sanger Institute COVID-19 Surveillance Team |
| EPI_ISL_3314929 | Lighthouse Lab in Alderley Park | Wellcome Sanger Institute for the COVID-19 Genomics UK (COG-UK) Consortium | Jacquelyn Wynn, Mairead Hyland, The Lighthouse Lab in Alderley Park and Alex Alderton, Roberto Amato, Jeffrey Barrett, Sonia Goncalves, Ewan Harrison, David K. Jackson, Ian Johnston, Dominic Kwiatkowski, Cordelia Langford, John Sillitoe on behalf of the Wellcome Sanger Institute COVID-19 Surveillance Team |
| EPI_ISL_3314930 | Lighthouse Lab in Glasgow | Wellcome Sanger Institute for the COVID-19 Genomics UK (COG-UK) Consortium | Harper VanSteenhouse, Yumi Kasai, David Gray, Carol Clugston, Anna Dominiczak and Alex Alderton, Roberto Amato, Jeffrey Barrett, Sonia Goncalves, Ewan Harrison, David K. Jackson, Ian Johnston, Dominic Kwiatkowski, Cordelia Langford, John Sillitoe on behalf of the Wellcome Sanger Institute COVID-19 Surveillance Team |
| EPI_ISL_3314931 | Lighthouse Lab in Alderley Park | Wellcome Sanger Institute for the COVID-19 Genomics UK (COG-UK) Consortium | Jacquelyn Wynn, Mairead Hyland, The Lighthouse Lab in Alderley Park and Alex Alderton, Roberto Amato, Jeffrey Barrett, Sonia Goncalves, Ewan Harrison, David K. Jackson, Ian Johnston, Dominic Kwiatkowski, Cordelia Langford, John Sillitoe on behalf of the Wellcome Sanger Institute COVID-19 Surveillance Team |
| EPI_ISL_3314932 | Lighthouse Lab in Glasgow | Wellcome Sanger Institute for the COVID-19 Genomics UK (COG-UK) Consortium | Harper VanSteenhouse, Yumi Kasai, David Gray, Carol Clugston, Anna Dominiczak and Alex Alderton, Roberto Amato, Jeffrey Barrett, Sonia Goncalves, Ewan Harrison, David K. Jackson, Ian Johnston, Dominic Kwiatkowski, Cordelia Langford, John Sillitoe on behalf of the Wellcome Sanger Institute COVID-19 Surveillance Team |
| EPI_ISL_3314933 | Lighthouse Laboratory Plymouth | Wellcome Sanger Institute for the COVID-19 Genomics UK (COG-UK) Consortium | Lighthouse Laboratory Plymouth and Alex Alderton, Roberto Amato, Jeffrey Barrett, Sonia Goncalves, Ewan Harrison, David K. Jackson, Ian Johnston, Dominic Kwiatkowski, Cordelia Langford, John Sillitoe on behalf of the Wellcome Sanger Institute COVID-19 Surveillance Team |
| EPI_ISL_3314934 | Lighthouse Lab in Glasgow | Wellcome Sanger Institute for the COVID-19 Genomics UK (COG-UK) Consortium | Harper VanSteenhouse, Yumi Kasai, David Gray, Carol Clugston, Anna Dominiczak and Alex Alderton, Roberto Amato, Jeffrey Barrett, Sonia Goncalves, Ewan Harrison, David K. Jackson, Ian Johnston, Dominic Kwiatkowski, Cordelia Langford, John Sillitoe on behalf of the Wellcome Sanger Institute COVID-19 Surveillance Team |
| EPI_ISL_3314935 | Lighthouse Laboratory Plymouth | Wellcome Sanger Institute for the COVID-19 Genomics UK (COG-UK) Consortium | Lighthouse Laboratory Plymouth and Alex Alderton, Roberto Amato, Jeffrey Barrett, Sonia Goncalves, Ewan Harrison, David K. Jackson, Ian Johnston, Dominic Kwiatkowski, Cordelia Langford, John Sillitoe on behalf of the Wellcome Sanger Institute COVID-19 Surveillance Team |
| EPI_ISL_3314936 | Lighthouse Lab in Glasgow | Wellcome Sanger Institute for the COVID-19 Genomics UK (COG-UK) Consortium | Harper VanSteenhouse, Yumi Kasai, David Gray, Carol Clugston, Anna Dominiczak and Alex Alderton, Roberto Amato, Jeffrey Barrett, Sonia Goncalves, Ewan Harrison, David K. Jackson, Ian Johnston, Dominic Kwiatkowski, Cordelia Langford, John Sillitoe on behalf of the Wellcome Sanger Institute COVID-19 Surveillance Team |
| EPI_ISL_3314937 | Lighthouse Lab in Alderley Park | Wellcome Sanger Institute for the COVID-19 Genomics UK (COG-UK) Consortium | Jacquelyn Wynn, Mairead Hyland, The Lighthouse Lab in Alderley Park and Alex Alderton, Roberto Amato, Jeffrey Barrett, Sonia Goncalves, Ewan Harrison, David K. Jackson, Ian Johnston, Dominic Kwiatkowski, Cordelia Langford, John Sillitoe on behalf of the Wellcome Sanger Institute COVID-19 Surveillance Team |
| EPI_ISL_3314938 | Lighthouse Lab in Milton Keynes | Wellcome Sanger Institute for the COVID-19 Genomics UK (COG-UK) Consortium | The Lighthouse Lab in Milton Keynes and Alex Alderton, Roberto Amato, Jeffrey Barrett, Sonia Goncalves, Ewan Harrison, David K. Jackson, Ian Johnston, Dominic Kwiatkowski, Cordelia Langford, John Sillitoe on behalf of the Wellcome Sanger Institute COVID-19 Surveillance Team |
| EPI_ISL_3314939, EPI_ISL_3314940 | Lighthouse Lab in Alderley Park | Wellcome Sanger Institute for the COVID-19 Genomics UK (COG-UK) Consortium | Jacquelyn Wynn, Mairead Hyland, The Lighthouse Lab in Alderley Park and Alex Alderton, Roberto Amato, Jeffrey Barrett, Sonia Goncalves, Ewan Harrison, David K. Jackson, Ian Johnston, Dominic Kwiatkowski, Cordelia Langford, John Sillitoe on behalf of the Wellcome Sanger Institute COVID-19 Surveillance Team |
| EPI_ISL_3314941 | Lighthouse Lab in Milton Keynes | Wellcome Sanger Institute for the COVID-19 Genomics UK (COG-UK) Consortium | The Lighthouse Lab in Milton Keynes and Alex Alderton, Roberto Amato, Jeffrey Barrett, Sonia Goncalves, Ewan Harrison, David K. Jackson, Ian Johnston, Dominic Kwiatkowski, Cordelia Langford, John Sillitoe on behalf of the Wellcome Sanger Institute COVID-19 Surveillance Team |
| EPI_ISL_3314942, EPI_ISL_3314943, EPI_ISL_3314944 | Lighthouse Lab in Alderley Park | Wellcome Sanger Institute for the COVID-19 Genomics UK (COG-UK) Consortium | Jacquelyn Wynn, Mairead Hyland, The Lighthouse Lab in Alderley Park and Alex Alderton, Roberto Amato, Jeffrey Barrett, Sonia Goncalves, Ewan Harrison, David K. Jackson, Ian Johnston, Dominic Kwiatkowski, Cordelia Langford, John Sillitoe on behalf of the Wellcome Sanger Institute COVID-19 Surveillance Team |
| EPI_ISL_3314945 | Lighthouse Lab in Milton Keynes | Wellcome Sanger Institute for the COVID-19 Genomics UK (COG-UK) Consortium | The Lighthouse Lab in Milton Keynes and Alex Alderton, Roberto Amato, Jeffrey Barrett, Sonia Goncalves, Ewan Harrison, David K. Jackson, Ian Johnston, Dominic Kwiatkowski, Cordelia Langford, John Sillitoe on behalf of the Wellcome Sanger Institute COVID-19 Surveillance Team |
| EPI_ISL_3314951, EPI_ISL_3314952, EPI_ISL_3314954, EPI_ISL_3314955, EPI_ISL_3314956, EPI_ISL_3314957, EPI_ISL_3314958, EPI_ISL_3314959, EPI_ISL_3314960, EPI_ISL_3314961, EPI_ISL_3314963, EPI_ISL_3314964, EPI_ISL_3314966, EPI_ISL_3314967, EPI_ISL_3314968, EPI_ISL_3314969, EPI_ISL_3314970, EPI_ISL_3314971, EPI_ISL_3314972 | | | |
| see above | Molecular Diagnostics Laboratory | National Reference Laboratory, Nigeria Centre for Disease Control | Dr Ndodo Nnaemeka, Olusola Anuoluwapo Akanbi, Chimaobi Chukwu, Dr Omoare Adesuyi, Celestina Obiekea, Anthony Ahumibe, Catherine Okoi, Nwando Mba, Dr Chikwe Ihekweazu |
| EPI_ISL_3314977 | Diagnostic and Research Center of Infectious Diseases, Medical Faculty, Andalas University | Genomik Solidaritas Indonesia Laboratorium / Diagnostic and Research Center of Infectious Diseases, Medical Faculty, Andalas University | Vania Gavrila Wikasa, Dhahlia Agustina Cahyono, Andani Eka Putra, Dede Rahman Agustian, Linosefa, Reinhart Gabriel, Carissa Sintca Wijaya, Annisa Muthiah Sukirman, Gracia Felias Enos Korompis, Anuraj Shankar, Louisa Markus, Meutia Ayuputeri Kumaheri, Ariel Pradipta |
| EPI_ISL_3314978 | Diagnostic and Research Center of Infectious Diseases, Medical Faculty, Andalas University | Genomik Solidaritas Indonesia Laboratorium / Diagnostic and Research Center of Infectious Diseases, Medical Faculty, Andalas University | Reinhart Gabriel, Carissa Sintca Wijaya, Andani Eka Putra, Dede Rahman Agustian, Linosefa, Vania Gavrila Wikasa, Dhahlia Agustina Cahyono, Gracia Felias Enos Korompis, Annisa Muthiah Sukirman, Anuraj Shankar, Louisa Markus, Meutia Ayuputeri Kumaheri, Ariel Pradipta |
| EPI_ISL_3314979 | Diagnostic and Research Center of Infectious Diseases, Medical Faculty, Andalas University | Genomik Solidaritas Indonesia Laboratorium / Diagnostic and Research Center of Infectious Diseases, Medical Faculty, Andalas University | Carissa Sintca Wijaya, Vania Gavrila Wikasa, Andani Eka Putra, Dede Rahman Agustian, Linosefa, Dhahlia Agustina Cahyono, Reinhart Gabriel, Gracia Felias Enos Korompis, Annisa Muthiah Sukirman, Anuraj Shankar, Louisa Markus, Meutia Ayuputeri Kumaheri, Ariel Pradipta |
| EPI_ISL_3314983 | Diagnostic and Research Center of Infectious Diseases, Medical | Genomik Solidaritas Indonesia Laboratorium / Diagnostic and | Dhahlia Agustina Cahyono, Reinhart Gabriel, Andani Eka Putra, Dede Rahman Agustian, Linosefa, Carissa Sintca Wijaya, Vania Gavrila Wikasa, Annisa Muthiah Sukirman, Gracia Felias Enos Korompis, Anuraj Shankar, Louisa Markus, Meutia |

| | | | |
|---|---|---|---|
| | Faculty, Andalas University | Research Center of Infectious Diseases, Medical Faculty, Andalas University | Ayuputeri Kumaheri, Ariel Pradipta |
| EPI_ISL_3314984 | Diagnostic and Research Center of Infectious Diseases, Medical Faculty, Andalas University | Genomik Solidaritas Indonesia Laboratorium / Diagnostic and Research Center of Infectious Diseases, Medical Faculty, Andalas University | Reinhart Gabriel, Carissa Sintca Wijaya, Andani Eka Putra, Dede Rahman Agustian, Linosefa, Vania Gavrila Wikasa, Dhahlia Agustina Cahyono, Gracia Felias Enos Korompis, Annisa Muthiah Sukirman, Anuraj Shankar, Louisa Markus, Meutia Ayuputeri Kumaheri, Ariel Pradipta |
| EPI_ISL_3314985 | Diagnostic and Research Center of Infectious Diseases, Medical Faculty, Andalas University | Genomik Solidaritas Indonesia Laboratorium / Diagnostic and Research Center of Infectious Diseases, Medical Faculty, Andalas University | Vania Gavrila Wikasa, Dhahlia Agustina Cahyono, Andani Eka Putra, Dede Rahman Agustian, Linosefa, Reinhart Gabriel, Carissa Sintca Wijaya, Annisa Muthiah Sukirman, Gracia Felias Enos Korompis, Anuraj Shankar, Louisa Markus, Meutia Ayuputeri Kumaheri, Ariel Pradipta |
| EPI_ISL_3314986, EPI_ISL_3314987 | Diagnostic and Research Center of Infectious Diseases, Medical Faculty, Andalas University | Genomik Solidaritas Indonesia Laboratorium / Diagnostic and Research Center of Infectious Diseases, Medical Faculty, Andalas University | Dhahlia Agustina Cahyono, Reinhart Gabriel, Andani Eka Putra, Dede Rahman Agustian, Linosefa, Carissa Sintca Wijaya, Vania Gavrila Wikasa, Annisa Muthiah Sukirman, Gracia Felias Enos Korompis, Anuraj Shankar, Louisa Markus, Meutia Ayuputeri Kumaheri, Ariel Pradipta |
| EPI_ISL_3314988 | Diagnostic and Research Center of Infectious Diseases, Medical Faculty, Andalas University | Genomik Solidaritas Indonesia Laboratorium / Diagnostic and Research Center of Infectious Diseases, Medical Faculty, Andalas University | Reinhart Gabriel, Carissa Sintca Wijaya, Andani Eka Putra, Dede Rahman Agustian, Linosefa, Vania Gavrila Wikasa, Dhahlia Agustina Cahyono, Gracia Felias Enos Korompis, Annisa Muthiah Sukirman, Anuraj Shankar, Louisa Markus, Meutia Ayuputeri Kumaheri, Ariel Pradipta |
| EPI_ISL_3314989, EPI_ISL_3314990, EPI_ISL_3314991, EPI_ISL_3314992 | Diagnostic and Research Center of Infectious Diseases, Medical Faculty, Andalas University | Genomik Solidaritas Indonesia Laboratorium / Diagnostic and Research Center of Infectious Diseases, Medical Faculty, Andalas University | Dhahlia Agustina Cahyono, Reinhart Gabriel, Andani Eka Putra, Dede Rahman Agustian, Linosefa, Carissa Sintca Wijaya, Vania Gavrila Wikasa, Annisa Muthiah Sukirman, Gracia Felias Enos Korompis, Anuraj Shankar, Louisa Markus, Meutia Ayuputeri Kumaheri, Ariel Pradipta |
| EPI_ISL_3314993, EPI_ISL_3314994 | Diagnostic and Research Center of Infectious Diseases, Medical Faculty, Andalas University | Genomik Solidaritas Indonesia Laboratorium / Diagnostic and Research Center of Infectious Diseases, Medical Faculty, Andalas University | Vania Gavrila Wikasa, Dhahlia Agustina Cahyono, Andani Eka Putra, Dede Rahman Agustian, Linosefa, Reinhart Gabriel, Carissa Sintca Wijaya, Annisa Muthiah Sukirman, Gracia Felias Enos Korompis, Anuraj Shankar, Louisa Markus, Meutia Ayuputeri Kumaheri, Ariel Pradipta |
| EPI_ISL_3314995 | Diagnostic and Research Center of Infectious Diseases, Medical Faculty, Andalas University | Genomik Solidaritas Indonesia Laboratorium / Diagnostic and Research Center of Infectious Diseases, Medical Faculty, Andalas University | Carissa Sintca Wijaya, Vania Gavrila Wikasa, Andani Eka Putra, Dede Rahman Agustian, Linosefa, Dhahlia Agustina Cahyono, Reinhart Gabriel, Gracia Felias Enos Korompis, Annisa Muthiah Sukirman, Anuraj Shankar, Louisa Markus, Meutia Ayuputeri Kumaheri, Ariel Pradipta |
| EPI_ISL_3315000, EPI_ISL_3315001, EPI_ISL_3315002 | Diagnostic and Research Center of Infectious Diseases, Medical Faculty, Andalas University | Genomik Solidaritas Indonesia Laboratorium / Diagnostic and Research Center of Infectious Diseases, Medical Faculty, Andalas University | Reinhart Gabriel, Carissa Sintca Wijaya, Andani Eka Putra, Dede Rahman Agustian, Linosefa, Vania Gavrila Wikasa, Dhahlia Agustina Cahyono, Gracia Felias Enos Korompis, Annisa Muthiah Sukirman, Anuraj Shankar, Louisa Markus, Meutia Ayuputeri Kumaheri, Ariel Pradipta |
| EPI_ISL_3315003 | Diagnostic and Research Center of Infectious Diseases, Medical Faculty, Andalas University | Genomik Solidaritas Indonesia Laboratorium / Diagnostic and Research Center of Infectious Diseases, Medical Faculty, Andalas University | Vania Gavrila Wikasa, Dhahlia Agustina Cahyono, Andani Eka Putra, Dede Rahman Agustian, Linosefa, Reinhart Gabriel, Carissa Sintca Wijaya, Annisa Muthiah Sukirman, Gracia Felias Enos Korompis, Anuraj Shankar, Louisa Markus, Meutia Ayuputeri Kumaheri, Ariel Pradipta |
| EPI_ISL_3315005, EPI_ISL_3315008, EPI_ISL_3315010, EPI_ISL_3315011, EPI_ISL_3315012 | Diagnostic and Research Center of Infectious Diseases, Medical Faculty, Andalas University | Genomik Solidaritas Indonesia Laboratorium / Diagnostic and Research Center of Infectious Diseases, Medical Faculty, Andalas University | Reinhart Gabriel, Carissa Sintca Wijaya, Andani Eka Putra, Dede Rahman Agustian, Linosefa, Vania Gavrila Wikasa, Dhahlia Agustina Cahyono, Gracia Felias Enos Korompis, Annisa Muthiah Sukirman, Anuraj Shankar, Louisa Markus, Meutia Ayuputeri Kumaheri, Ariel Pradipta |
| EPI_ISL_3315013, EPI_ISL_3315014, EPI_ISL_3315015 | Diagnostic and Research Center of Infectious Diseases, Medical Faculty, Andalas University | Genomik Solidaritas Indonesia Laboratorium / Diagnostic and Research Center of Infectious Diseases, Medical Faculty, Andalas University | Carissa Sintca Wijaya, Vania Gavrila Wikasa, Andani Eka Putra, Dede Rahman Agustian, Linosefa, Dhahlia Agustina Cahyono, Reinhart Gabriel, Gracia Felias Enos Korompis, Annisa Muthiah Sukirman, Anuraj Shankar, Louisa Markus, Meutia Ayuputeri Kumaheri, Ariel Pradipta |
| EPI_ISL_3315016, EPI_ISL_3315017 | Diagnostic and Research Center of Infectious Diseases, Medical Faculty, Andalas University | Genomik Solidaritas Indonesia Laboratorium / Diagnostic and Research Center of Infectious Diseases, Medical Faculty, Andalas University | Vania Gavrila Wikasa, Dhahlia Agustina Cahyono, Andani Eka Putra, Dede Rahman Agustian, Linosefa, Reinhart Gabriel, Carissa Sintca Wijaya, Annisa Muthiah Sukirman, Gracia Felias Enos Korompis, Anuraj Shankar, Louisa Markus, Meutia Ayuputeri Kumaheri, Ariel Pradipta |
| EPI_ISL_3315018, EPI_ISL_3315019 | Diagnostic and Research Center of Infectious Diseases, Medical Faculty, Andalas University | Genomik Solidaritas Indonesia Laboratorium / Diagnostic and Research Center of Infectious Diseases, Medical Faculty, Andalas University | Dhahlia Agustina Cahyono, Reinhart Gabriel, Andani Eka Putra, Dede Rahman Agustian, Linosefa, Carissa Sintca Wijaya, Vania Gavrila Wikasa, Annisa Muthiah Sukirman, Gracia Felias Enos Korompis, Anuraj Shankar, Louisa Markus, Meutia Ayuputeri Kumaheri, Ariel Pradipta |
| EPI_ISL_3315020, EPI_ISL_3315024 | Diagnostic and Research Center of Infectious Diseases, Medical Faculty, Andalas University | Genomik Solidaritas Indonesia Laboratorium / Diagnostic and Research Center of Infectious Diseases, Medical Faculty, Andalas University | Carissa Sintca Wijaya, Vania Gavrila Wikasa, Andani Eka Putra, Dede Rahman Agustian, Linosefa, Dhahlia Agustina Cahyono, Reinhart Gabriel, Gracia Felias Enos Korompis, Annisa Muthiah Sukirman, Anuraj Shankar, Louisa Markus, Meutia Ayuputeri Kumaheri, Ariel Pradipta |
| EPI_ISL_3315050 | National Centre For Cell Science | National Centre For Cell Science – INSACOG | Dhiraj Paul, Shivang P. Bhanushali, Mitali Inamdar, Mohak P Gujare, Manoj Kumar Bhat, Ajay Pillai, INSACOG Consortium team, Yogesh Shouche |
| EPI_ISL_3315051 | IFCOR Brno | University Hospital Brno, CMBG | Matej Bezdicek, Jan Svaton, Martina Lengerova, Kristyna Dufkova |
| EPI_ISL_3315052 | Cytogeneticka laborator Brno | University Hospital Brno, CMBG | Matej Bezdicek, Jan Svaton, Martina Lengerova, Kristyna Dufkova |
| EPI_ISL_3315053 | IFCOR Brno | University Hospital Brno, CMBG | Matej Bezdicek, Jan Svaton, Martina Lengerova, Kristyna Dufkova |
| EPI_ISL_3315054 | Nemocnice Valtice | University Hospital Brno, CMBG | Matej Bezdicek, Jan Svaton, Martina Lengerova, Kristyna Dufkova |
| EPI_ISL_3315055, EPI_ISL_3315056, EPI_ISL_3315057 | Nemocnice TGM Hodonin | University Hospital Brno, CMBG | Matej Bezdicek, Jan Svaton, Martina Lengerova, Kristyna Dufkova |
| EPI_ISL_3315058, EPI_ISL_3315059, EPI_ISL_3315060, EPI_ISL_3315061, EPI_ISL_3315063, EPI_ISL_3315064, EPI_ISL_3315065, EPI_ISL_3315066 | University Hospital Brno, OKMI | University Hospital Brno, CMBG | Matej Bezdicek, Jan Svaton, Martina Lengerova, Kristyna Dufkova |
| EPI_ISL_3315067, EPI_ISL_3315068 | Hola Biolabs | University Hospital Brno, CMBG | Matej Bezdicek, Jan Svaton, Martina Lengerova, Kristyna Dufkova |
| EPI_ISL_3315069, EPI_ISL_3315070, EPI_ISL_3315071, EPI_ISL_3315072, EPI_ISL_3315073, EPI_ISL_3315074, EPI_ISL_3315075, EPI_ISL_3315076 | Nemocnice Jihlava | University Hospital Brno, CMBG | Matej Bezdicek, Jan Svaton, Martina Lengerova, Kristyna Dufkova |
| EPI_ISL_3315077, EPI_ISL_3315078, EPI_ISL_3315079, EPI_ISL_3315080 | Nemocnice Havlickuv Brod | University Hospital Brno, CMBG | Matej Bezdicek, Jan Svaton, Martina Lengerova, Kristyna Dufkova |
| EPI_ISL_3315081 | Cytogeneticka laborator Brno | University Hospital Brno, CMBG | Matej Bezdicek, Jan Svaton, Martina Lengerova, Kristyna Dufkova |
| EPI_ISL_3315082, EPI_ISL_3315083, EPI_ISL_3315084, EPI_ISL_3315085, EPI_ISL_3315086, EPI_ISL_3315087, EPI_ISL_3315088, EPI_ISL_3315089 | St. Anne´s University Hospital Brno | University Hospital Brno, CMBG | Matej Bezdicek, Jan Svaton, Martina Lengerova, Kristyna Dufkova |
| EPI_ISL_3315091 | LT/Osp. S.Maria Goretti | National Institute for Infectious Diseases (INMI) L. Spallanzani I.R.C.C.S | E Giombini, B Bartolini, M Rueca, G Bonfiglio, O Butera, CEM Gruber, F Santini, F Messina, A Di Caro, MR Capobianchi |
| EPI_ISL_3315092 | RM3/A.O. S.Camillo | National Institute for Infectious Diseases (INMI) L. Spallanzani I.R.C.C.S | E Giombini, B Bartolini, M Rueca, G Bonfiglio, O Butera, CEM Gruber, F Santini, F Messina, A Di Caro, MR Capobianchi |
| EPI_ISL_3315093 | RM2/Osp. S.Pertini | National Institute for Infectious Diseases (INMI) L. Spallanzani I.R.C.C.S | E Giombini, B Bartolini, M Rueca, G Bonfiglio, O Butera, CEM Gruber, F Santini, F Messina, A Di Caro, MR Capobianchi |
| EPI_ISL_3315094, EPI_ISL_3315095, EPI_ISL_3315096 | RM3/A.O. S.Camillo | National Institute for Infectious Diseases (INMI) L. Spallanzani I.R.C.C.S | E Giombini, B Bartolini, M Rueca, G Bonfiglio, O Butera, CEM Gruber, F Santini, F Messina, A Di Caro, MR Capobianchi |
| EPI_ISL_3315097, EPI_ISL_3315098 | RM2/Osp. S.Pertini | National Institute for Infectious Diseases (INMI) L. Spallanzani I.R.C.C.S | E Giombini, B Bartolini, M Rueca, G Bonfiglio, O Butera, CEM Gruber, F Santini, F Messina, A Di Caro, MR Capobianchi |

| | | | |
|---|---|---|---|
| EPI_ISL_3315099 | LT/Osp. S.Maria Goretti | National Institute for Infectious Diseases (INMI) L. Spallanzani I.R.C.C.S | E Giombini, B Bartolini, M Rueca, G Bonfiglio, O Butera, CEM Gruber, F Santini, F Messina, A Di Caro, MR Capobianchi |
| EPI_ISL_3315100 | RM2/Osp. S.Pertini | National Institute for Infectious Diseases (INMI) L. Spallanzani I.R.C.C.S | E Giombini, B Bartolini, M Rueca, G Bonfiglio, O Butera, CEM Gruber, F Santini, F Messina, A Di Caro, MR Capobianchi |
| EPI_ISL_3315102 | LT/Osp. S.Maria Goretti | National Institute for Infectious Diseases (INMI) L. Spallanzani I.R.C.C.S | E Giombini, B Bartolini, M Rueca, G Bonfiglio, O Butera, CEM Gruber, F Santini, F Messina, A Di Caro, MR Capobianchi |
| EPI_ISL_3315104 | RM2/Osp. S.Pertini | National Institute for Infectious Diseases (INMI) L. Spallanzani I.R.C.C.S | E Giombini, B Bartolini, M Rueca, G Bonfiglio, O Butera, CEM Gruber, F Santini, F Messina, A Di Caro, MR Capobianchi |
| EPI_ISL_3315105 | LT/Osp. S.Maria Goretti | National Institute for Infectious Diseases (INMI) L. Spallanzani I.R.C.C.S | E Giombini, B Bartolini, M Rueca, G Bonfiglio, O Butera, CEM Gruber, F Santini, F Messina, A Di Caro, MR Capobianchi |
| EPI_ISL_3315106 | RM2/Osp. S.Pertini | National Institute for Infectious Diseases (INMI) L. Spallanzani I.R.C.C.S | E Giombini, B Bartolini, M Rueca, G Bonfiglio, O Butera, CEM Gruber, F Santini, F Messina, A Di Caro, MR Capobianchi |
| EPI_ISL_3315107 | LT/Osp. S.Maria Goretti | National Institute for Infectious Diseases (INMI) L. Spallanzani I.R.C.C.S | E Giombini, B Bartolini, M Rueca, G Bonfiglio, O Butera, CEM Gruber, F Santini, F Messina, A Di Caro, MR Capobianchi |
| EPI_ISL_3315109, EPI_ISL_3315110 | RM2/Osp. S.Pertini | National Institute for Infectious Diseases (INMI) L. Spallanzani I.R.C.C.S | E Giombini, B Bartolini, M Rueca, G Bonfiglio, O Butera, CEM Gruber, F Santini, F Messina, A Di Caro, MR Capobianchi |
| EPI_ISL_3315111 | RM1/A.U. Umberto I | National Institute for Infectious Diseases (INMI) L. Spallanzani I.R.C.C.S | E Giombini, B Bartolini, M Rueca, G Bonfiglio, O Butera, CEM Gruber, F Santini, F Messina, A Di Caro, MR Capobianchi |
| EPI_ISL_3315112 | LT/Osp. S.Maria Goretti | National Institute for Infectious Diseases (INMI) L. Spallanzani I.R.C.C.S | E Giombini, B Bartolini, M Rueca, G Bonfiglio, O Butera, CEM Gruber, F Santini, F Messina, A Di Caro, MR Capobianchi |
| EPI_ISL_3315113 | RM2/Osp. S.Pertini | National Institute for Infectious Diseases (INMI) L. Spallanzani I.R.C.C.S | E Giombini, B Bartolini, M Rueca, G Bonfiglio, O Butera, CEM Gruber, F Santini, F Messina, A Di Caro, MR Capobianchi |
| EPI_ISL_3315115 | RM1/A.U. Umberto I | National Institute for Infectious Diseases (INMI) L. Spallanzani I.R.C.C.S | E Giombini, B Bartolini, M Rueca, G Bonfiglio, O Butera, CEM Gruber, F Santini, F Messina, A Di Caro, MR Capobianchi |
| EPI_ISL_3315116 | RM2/Osp. S.Pertini | National Institute for Infectious Diseases (INMI) L. Spallanzani I.R.C.C.S | E Giombini, B Bartolini, M Rueca, G Bonfiglio, O Butera, CEM Gruber, F Santini, F Messina, A Di Caro, MR Capobianchi |
| EPI_ISL_3315117 | RM1/A.U. Umberto I | National Institute for Infectious Diseases (INMI) L. Spallanzani I.R.C.C.S | E Giombini, B Bartolini, M Rueca, G Bonfiglio, O Butera, CEM Gruber, F Santini, F Messina, A Di Caro, MR Capobianchi |
| EPI_ISL_3315118, EPI_ISL_3315119, EPI_ISL_3315120, EPI_ISL_3315121, EPI_ISL_3315122 | RM2/Osp. S.Pertini | National Institute for Infectious Diseases (INMI) L. Spallanzani I.R.C.C.S | E Giombini, B Bartolini, M Rueca, G Bonfiglio, O Butera, CEM Gruber, F Santini, F Messina, A Di Caro, MR Capobianchi |
| EPI_ISL_3315123, EPI_ISL_3315124 | RM6/Genzano | National Institute for Infectious Diseases (INMI) L. Spallanzani I.R.C.C.S | E Giombini, B Bartolini, M Rueca, G Bonfiglio, O Butera, CEM Gruber, F Santini, F Messina, A Di Caro, MR Capobianchi |
| EPI_ISL_3315125, EPI_ISL_3315126, EPI_ISL_3315128 | RM2/Osp. S.Pertini | National Institute for Infectious Diseases (INMI) L. Spallanzani I.R.C.C.S | E Giombini, B Bartolini, M Rueca, G Bonfiglio, O Butera, CEM Gruber, F Santini, F Messina, A Di Caro, MR Capobianchi |
| EPI_ISL_3315129 | RM1/A.U. Umberto I | National Institute for Infectious Diseases (INMI) L. Spallanzani I.R.C.C.S | E Giombini, B Bartolini, M Rueca, G Bonfiglio, O Butera, CEM Gruber, F Santini, F Messina, A Di Caro, MR Capobianchi |
| EPI_ISL_3315130 | RM3/A.O. S.Camillo | National Institute for Infectious Diseases (INMI) L. Spallanzani I.R.C.C.S | E Giombini, B Bartolini, M Rueca, G Bonfiglio, O Butera, CEM Gruber, F Santini, F Messina, A Di Caro, MR Capobianchi |
| EPI_ISL_3315131, EPI_ISL_3315132 | RM2/Osp. S.Pertini | National Institute for Infectious Diseases (INMI) L. Spallanzani I.R.C.C.S | E Giombini, B Bartolini, M Rueca, G Bonfiglio, O Butera, CEM Gruber, F Santini, F Messina, A Di Caro, MR Capobianchi |
| EPI_ISL_3315133 | RM6/Genzano | National Institute for Infectious Diseases (INMI) L. Spallanzani I.R.C.C.S | E Giombini, B Bartolini, M Rueca, G Bonfiglio, O Butera, CEM Gruber, F Santini, F Messina, A Di Caro, MR Capobianchi |
| EPI_ISL_3315134, EPI_ISL_3315135 | RM2/Osp. S.Pertini | National Institute for Infectious Diseases (INMI) L. Spallanzani I.R.C.C.S | E Giombini, B Bartolini, M Rueca, G Bonfiglio, O Butera, CEM Gruber, F Santini, F Messina, A Di Caro, MR Capobianchi |
| EPI_ISL_3315136, EPI_ISL_3315137 | RM6/Genzano | National Institute for Infectious Diseases (INMI) L. Spallanzani I.R.C.C.S | E Giombini, B Bartolini, M Rueca, G Bonfiglio, O Butera, CEM Gruber, F Santini, F Messina, A Di Caro, MR Capobianchi |
| EPI_ISL_3315138 | RM3/A.O. S.Camillo | National Institute for Infectious Diseases (INMI) L. Spallanzani I.R.C.C.S | E Giombini, B Bartolini, M Rueca, G Bonfiglio, O Butera, CEM Gruber, F Santini, F Messina, A Di Caro, MR Capobianchi |
| EPI_ISL_3315140, EPI_ISL_3315141 | RM1/A.U. Umberto I | National Institute for Infectious Diseases (INMI) L. Spallanzani I.R.C.C.S | E Giombini, B Bartolini, M Rueca, G Bonfiglio, O Butera, CEM Gruber, F Santini, F Messina, A Di Caro, MR Capobianchi |
| EPI_ISL_3315142, EPI_ISL_3315143, EPI_ISL_3315144, EPI_ISL_3315145, EPI_ISL_3315146 | RM3/A.O. S.Camillo | National Institute for Infectious Diseases (INMI) L. Spallanzani I.R.C.C.S | E Giombini, B Bartolini, M Rueca, G Bonfiglio, O Butera, CEM Gruber, F Santini, F Messina, A Di Caro, MR Capobianchi |
| EPI_ISL_3315147 | RM6/Genzano | National Institute for Infectious Diseases (INMI) L. Spallanzani I.R.C.C.S | E Giombini, B Bartolini, M Rueca, G Bonfiglio, O Butera, CEM Gruber, F Santini, F Messina, A Di Caro, MR Capobianchi |
| EPI_ISL_3315148, EPI_ISL_3315149 | RM3/A.O. S.Camillo | National Institute for Infectious Diseases (INMI) L. Spallanzani I.R.C.C.S | E Giombini, B Bartolini, M Rueca, G Bonfiglio, O Butera, CEM Gruber, F Santini, F Messina, A Di Caro, MR Capobianchi |
| EPI_ISL_3315150 | RM6/Genzano | National Institute for Infectious Diseases (INMI) L. Spallanzani I.R.C.C.S | E Giombini, B Bartolini, M Rueca, G Bonfiglio, O Butera, CEM Gruber, F Santini, F Messina, A Di Caro, MR Capobianchi |
| EPI_ISL_3315151, EPI_ISL_3315152, EPI_ISL_3315153, EPI_ISL_3315154, EPI_ISL_3315155, EPI_ISL_3315156, EPI_ISL_3315157, EPI_ISL_3315158, EPI_ISL_3315159 | RM3/A.O. S.Camillo | National Institute for Infectious Diseases (INMI) L. Spallanzani I.R.C.C.S | E Giombini, B Bartolini, M Rueca, G Bonfiglio, O Butera, CEM Gruber, F Santini, F Messina, A Di Caro, MR Capobianchi |
| EPI_ISL_3315165, EPI_ISL_3315166, EPI_ISL_3315167, EPI_ISL_3315186, EPI_ISL_3315187, EPI_ISL_3315188, EPI_ISL_3315190, EPI_ISL_3315191, EPI_ISL_3315192, EPI_ISL_3315193, EPI_ISL_3315194, EPI_ISL_3315195, EPI_ISL_3315196, EPI_ISL_3315197, EPI_ISL_3315199, EPI_ISL_3315202, EPI_ISL_3315203, EPI_ISL_3315209, EPI_ISL_3315211, EPI_ISL_3315215, EPI_ISL_3315216, EPI_ISL_3315217, EPI_ISL_3315221, EPI_ISL_3315222, EPI_ISL_3315223, EPI_ISL_3315224, EPI_ISL_3315225, EPI_ISL_3315226, EPI_ISL_3315227, EPI_ISL_3315228, EPI_ISL_3315256, EPI_ISL_3315273, EPI_ISL_3315277, EPI_ISL_3315295, EPI_ISL_3315337, EPI_ISL_3315338, EPI_ISL_3315339, EPI_ISL_3315341, EPI_ISL_3315342, EPI_ISL_3315343, EPI_ISL_3315344, EPI_ISL_3315345, EPI_ISL_3315346, EPI_ISL_3315347, EPI_ISL_3315348, EPI_ISL_3315349, EPI_ISL_3315350, EPI_ISL_3315351, EPI_ISL_3315352, EPI_ISL_3315353, EPI_ISL_3315354, EPI_ISL_3315355, EPI_ISL_3315356, EPI_ISL_3315357, EPI_ISL_3315358, EPI_ISL_3315359, EPI_ISL_3315360, EPI_ISL_3315361, EPI_ISL_3315362, EPI_ISL_3315363, EPI_ISL_3315364, EPI_ISL_3315365, EPI_ISL_3315366, EPI_ISL_3315367, EPI_ISL_3315368, EPI_ISL_3315369, EPI_ISL_3315370, EPI_ISL_3315371, EPI_ISL_3315372, EPI_ISL_3315373, EPI_ISL_3315374, EPI_ISL_3315375, EPI_ISL_3315376, EPI_ISL_3315377, EPI_ISL_3315378, EPI_ISL_3315379, EPI_ISL_3315380, EPI_ISL_3315381, EPI_ISL_3315382, EPI_ISL_3315383, EPI_ISL_3315384, EPI_ISL_3315385, EPI_ISL_3315386, EPI_ISL_3315387, EPI_ISL_3315388, EPI_ISL_3315389, EPI_ISL_3315390, EPI_ISL_3315391, EPI_ISL_3315392, EPI_ISL_3315393, EPI_ISL_3315394, EPI_ISL_3315395, EPI_ISL_3315396, EPI_ISL_3315397, EPI_ISL_3315398, EPI_ISL_3315399, EPI_ISL_3315400, EPI_ISL_3315401, EPI_ISL_3315402, EPI_ISL_3315403, EPI_ISL_3315404, EPI_ISL_3315405, EPI_ISL_3315406, EPI_ISL_3315407, EPI_ISL_3315408, EPI_ISL_3315409, EPI_ISL_3315410, EPI_ISL_3315411, EPI_ISL_3315412, EPI_ISL_3315413, EPI_ISL_3315414, EPI_ISL_3315415, EPI_ISL_3315416, EPI_ISL_3315417, EPI_ISL_3315418, EPI_ISL_3315419, EPI_ISL_3315420, EPI_ISL_3315421, EPI_ISL_3315422, EPI_ISL_3315423, EPI_ISL_3315424, EPI_ISL_3315425, EPI_ISL_3315426, EPI_ISL_3315427, EPI_ISL_3315428, EPI_ISL_3315429, EPI_ISL_3315430, EPI_ISL_3315431, EPI_ISL_3315432, EPI_ISL_3315433, EPI_ISL_3315434, EPI_ISL_3315435, EPI_ISL_3315436, EPI_ISL_3315437, EPI_ISL_3315438, EPI_ISL_3315439, EPI_ISL_3315440, EPI_ISL_3315441, EPI_ISL_3315456, EPI_ISL_3315457, EPI_ISL_3315458, EPI_ISL_3315459, EPI_ISL_3315460, EPI_ISL_3315462, EPI_ISL_3315463, EPI_ISL_3315464, EPI_ISL_3315465, EPI_ISL_3315466, EPI_ISL_3315467, EPI_ISL_3315468, EPI_ISL_3315469, EPI_ISL_3315470, EPI_ISL_3315471, EPI_ISL_3315472, EPI_ISL_3315473, EPI_ISL_3315474, EPI_ISL_3315475, EPI_ISL_3315476, EPI_ISL_3315477, EPI_ISL_3315478, EPI_ISL_3315479, EPI_ISL_3315480, EPI_ISL_3315481, EPI_ISL_3315483, EPI_ISL_3315484, EPI_ISL_3315485, EPI_ISL_3315486, EPI_ISL_3315487, EPI_ISL_3315488, EPI_ISL_3315489, EPI_ISL_3315490, EPI_ISL_3315491, EPI_ISL_3315492, EPI_ISL_3315493, EPI_ISL_3315494, EPI_ISL_3315495, EPI_ISL_3315496, EPI_ISL_3315497, EPI_ISL_3315498, EPI_ISL_3315499, EPI_ISL_3315500, EPI_ISL_3315501, EPI_ISL_3315502, EPI_ISL_3315503, EPI_ISL_3315504, EPI_ISL_3315505, EPI_ISL_3315506, EPI_ISL_3315508, EPI_ISL_3315509, EPI_ISL_3315510, EPI_ISL_3315511, EPI_ISL_3315514, EPI_ISL_3315515, EPI_ISL_3315516, EPI_ISL_3315517, EPI_ISL_3315518, EPI_ISL_3315519, EPI_ISL_3315520, EPI_ISL_3315521, EPI_ISL_3315522, EPI_ISL_3315523, EPI_ISL_3315524, EPI_ISL_3315525, EPI_ISL_3315526, EPI_ISL_3315527, EPI_ISL_3315528, EPI_ISL_3315529, EPI_ISL_3315530, EPI_ISL_3315531, EPI_ISL_3315532, EPI_ISL_3315533, EPI_ISL_3315534, EPI_ISL_3315535, EPI_ISL_3315536, EPI_ISL_3315537, EPI_ISL_3315538, EPI_ISL_3315539, EPI_ISL_3315540, EPI_ISL_3315541, EPI_ISL_3315542, EPI_ISL_3315543, EPI_ISL_3315544, EPI_ISL_3315545, EPI_ISL_3315546, EPI_ISL_3315547, EPI_ISL_3315548, EPI_ISL_3315549, EPI_ISL_3315550, EPI_ISL_3315551, EPI_ISL_3315552, EPI_ISL_3315553, EPI_ISL_3315554, EPI_ISL_3315555, EPI_ISL_3315556, EPI_ISL_3315557, EPI_ISL_3315558, EPI_ISL_3315559, EPI_ISL_3315560, EPI_ISL_3315561, EPI_ISL_3315562, EPI_ISL_3315563, EPI_ISL_3315564, EPI_ISL_3315565, EPI_ISL_3315566, EPI_ISL_3315567, EPI_ISL_3315568, EPI_ISL_3315569, EPI_ISL_3315570, EPI_ISL_3315571, EPI_ISL_3315572, EPI_ISL_3315573, EPI_ISL_3315574, EPI_ISL_3315575, EPI_ISL_3315576, EPI_ISL_3315577, EPI_ISL_3315578, EPI_ISL_3315579, EPI_ISL_3315580, EPI_ISL_3315581, EPI_ISL_3315583, EPI_ISL_3315584, EPI_ISL_3315585, EPI_ISL_3315586, EPI_ISL_3315587, EPI_ISL_3315588, EPI_ISL_3315589, EPI_ISL_3315590, EPI_ISL_3315591, EPI_ISL_3315592, EPI_ISL_3315593, EPI_ISL_3315594, EPI_ISL_3315595, EPI_ISL_3315596, EPI_ISL_3315597, EPI_ISL_3315598, EPI_ISL_3315599, EPI_ISL_3315600, EPI_ISL_3315601, EPI_ISL_3315603, EPI_ISL_3315604, EPI_ISL_3315605, EPI_ISL_3315607, EPI_ISL_3315608, EPI_ISL_3315609, EPI_ISL_3315610, EPI_ISL_3315611, EPI_ISL_3315612, EPI_ISL_3315613, EPI_ISL_3315619, EPI_ISL_3315648, EPI_ISL_3315723, EPI_ISL_3315725, EPI_ISL_3315729, EPI_ISL_3315730, EPI_ISL_3315731, EPI_ISL_3315735, EPI_ISL_3315736, EPI_ISL_3315737, EPI_ISL_3315738, EPI_ISL_3315739, EPI_ISL_3315740, EPI_ISL_3315741, EPI_ISL_3315742, EPI_ISL_3315770, EPI_ISL_3315787 | | | |
| see above | Ministry of Health Turkey | Ministry of Health Turkey | Fatma Bayrakdar, Yasemin Coşgun, Süleyman Yalcin, Gülay Korukluoğlu |
| EPI_ISL_3315791, EPI_ISL_3315792, EPI_ISL_3315793, EPI_ISL_3315794, EPI_ISL_3315795, EPI_ISL_3315796, EPI_ISL_3315797, EPI_ISL_3315798, EPI_ISL_3315799, EPI_ISL_3315800, EPI_ISL_3315801, EPI_ISL_3315802, EPI_ISL_3315803, EPI_ISL_3315804, EPI_ISL_3315805, EPI_ISL_3315806, EPI_ISL_3315807, EPI_ISL_3315808, EPI_ISL_3315809, EPI_ISL_3315810, EPI_ISL_3315811, EPI_ISL_3315812, EPI_ISL_3315813, EPI_ISL_3315816, EPI_ISL_3315818, EPI_ISL_3315819, EPI_ISL_3315820, EPI_ISL_3315821, EPI_ISL_3315822, EPI_ISL_3315823, EPI_ISL_3315824, EPI_ISL_3315825, EPI_ISL_3315826, EPI_ISL_3315827, EPI_ISL_3315828, EPI_ISL_3315829, EPI_ISL_3315830, EPI_ISL_3315831, EPI_ISL_3315832, EPI_ISL_3315833, EPI_ISL_3315834, EPI_ISL_3315835, EPI_ISL_3315836, EPI_ISL_3315837 | | | |
| see above | Servicio de Microbiologia Hospital Ramon y Cajal | Servicio de Microbiologia Hospital Ramon y Cajal | Ponce M, Galan JC, Martinez L. Abreu M, y Gonzalez-Alba JM |
| EPI_ISL_3315860, EPI_ISL_3315861, EPI_ISL_3315862, EPI_ISL_3315863, EPI_ISL_3315864, EPI_ISL_3315865, EPI_ISL_3315866, EPI_ISL_3315867, EPI_ISL_3315868, EPI_ISL_3315869, EPI_ISL_3315871, EPI_ISL_3315872, EPI_ISL_3315873, EPI_ISL_3315874, EPI_ISL_3315875, EPI_ISL_3315876, EPI_ISL_3315877, EPI_ISL_3315878, EPI_ISL_3315879, EPI_ISL_3315880, EPI_ISL_3315881 | | | |
| see above | Indian Council of Medical Research-National Institute of | Indian Council of Medical Research-National Institute of | Pragya D. Yadav |

| | | | |
|---|---|---|---|
| | Virology, Microbial Containment Complex | Virology, Microbial Containment Complex | |
| EPI_ISL_3316317 | Public Health Authority of the Slovak Republic | Laboratory of Genomics and Bioinformatics, Comenius University Science Park | Tatiana Sedláčková, Diana Rusňáková, Miroslav Böhmer, Anna Gičová, Jaroslav Budiš, Tomáš Szemes, Jakub Styk |
| EPI_ISL_3316318, EPI_ISL_3316319, EPI_ISL_3316320, EPI_ISL_3316321, EPI_ISL_3316322, EPI_ISL_3316323, EPI_ISL_3316324, EPI_ISL_3316325, EPI_ISL_3316326, EPI_ISL_3316327, EPI_ISL_3316328, EPI_ISL_3316329, EPI_ISL_3316330, EPI_ISL_3316331, EPI_ISL_3316332, EPI_ISL_3316333, EPI_ISL_3316334, EPI_ISL_3316335, EPI_ISL_3316336, EPI_ISL_3316337, EPI_ISL_3316338, EPI_ISL_3316339, EPI_ISL_3316340, EPI_ISL_3316341, EPI_ISL_3316342, EPI_ISL_3316343, EPI_ISL_3316344, EPI_ISL_3316345, EPI_ISL_3316346, EPI_ISL_3316347, EPI_ISL_3316348, EPI_ISL_3316349, EPI_ISL_3316350, EPI_ISL_3316351, EPI_ISL_3316352, EPI_ISL_3316353, EPI_ISL_3316354, EPI_ISL_3316355, EPI_ISL_3316356, EPI_ISL_3316357, EPI_ISL_3316358, EPI_ISL_3316359, EPI_ISL_3316360, EPI_ISL_3316361, EPI_ISL_3316362, EPI_ISL_3316363, EPI_ISL_3316364, EPI_ISL_3316365, EPI_ISL_3316366, EPI_ISL_3316367, EPI_ISL_3316368, EPI_ISL_3316369, EPI_ISL_3316370, EPI_ISL_3316371, EPI_ISL_3316372, EPI_ISL_3316373, EPI_ISL_3316374, EPI_ISL_3316375, EPI_ISL_3316376, EPI_ISL_3316377, EPI_ISL_3316378, EPI_ISL_3316379, EPI_ISL_3316380, EPI_ISL_3316381, EPI_ISL_3316382, EPI_ISL_3316383, EPI_ISL_3316384, EPI_ISL_3316385, EPI_ISL_3316386, EPI_ISL_3316387, EPI_ISL_3316388, EPI_ISL_3316389, EPI_ISL_3316390, EPI_ISL_3316391, EPI_ISL_3316392 | | | |
| see above | Ministry of Health Turkey | Ministry of Health Turkey | Fatma Bayrakdar, Yasemin Coşgun, Süleyman Yalcin, Gülay Korukluoğlu |
| EPI_ISL_3316399, EPI_ISL_3316400, EPI_ISL_3316401, EPI_ISL_3316402, EPI_ISL_3316403, EPI_ISL_3316404, EPI_ISL_3316405, EPI_ISL_3316406, EPI_ISL_3316407, EPI_ISL_3316408, EPI_ISL_3316409, EPI_ISL_3316410, EPI_ISL_3316411, EPI_ISL_3316412, EPI_ISL_3316413, EPI_ISL_3316414, EPI_ISL_3316415, EPI_ISL_3316416, EPI_ISL_3316417, EPI_ISL_3316418, EPI_ISL_3316419, EPI_ISL_3316420, EPI_ISL_3316421, EPI_ISL_3316423, EPI_ISL_3316424, EPI_ISL_3316425, EPI_ISL_3316426, EPI_ISL_3316427, EPI_ISL_3316428, EPI_ISL_3316429, EPI_ISL_3316430, EPI_ISL_3316431, EPI_ISL_3316432 | | | |
| see above | AZDelta | AZ Delta Medical Laboratories in Roeselare, Belgium | Geert Martens, Dieter De Smet, Frederik Van Hoecke, on behalf of AZ Delta COVID-19 Genomics core (member of Genomic surveillance of SARS-CoV-2 in Belgium network) |
| EPI_ISL_3316544, EPI_ISL_3316545, EPI_ISL_3316547, EPI_ISL_3316548, EPI_ISL_3316550, EPI_ISL_3316551, EPI_ISL_3316553, EPI_ISL_3316554, EPI_ISL_3316555, EPI_ISL_3316556, EPI_ISL_3316557, EPI_ISL_3316558, EPI_ISL_3316559, EPI_ISL_3316560, EPI_ISL_3316561, EPI_ISL_3316562, EPI_ISL_3316563, EPI_ISL_3316564, EPI_ISL_3316565, EPI_ISL_3316566, EPI_ISL_3316567, EPI_ISL_3316568, EPI_ISL_3316569, EPI_ISL_3316570, EPI_ISL_3316571, EPI_ISL_3316572, EPI_ISL_3316574, EPI_ISL_3316575, EPI_ISL_3316576, EPI_ISL_3316577, EPI_ISL_3316578, EPI_ISL_3316580, EPI_ISL_3316581, EPI_ISL_3316582, EPI_ISL_3316583, EPI_ISL_3316584, EPI_ISL_3316585, EPI_ISL_3316586, EPI_ISL_3316587, EPI_ISL_3316588, EPI_ISL_3316589, EPI_ISL_3316590, EPI_ISL_3316594, EPI_ISL_3316595, EPI_ISL_3316596, EPI_ISL_3316597, EPI_ISL_3316598, EPI_ISL_3316599, EPI_ISL_3316600, EPI_ISL_3316601, EPI_ISL_3316602, EPI_ISL_3316603, EPI_ISL_3316604, EPI_ISL_3316605, EPI_ISL_3316606, EPI_ISL_3316607, EPI_ISL_3316608, EPI_ISL_3316609, EPI_ISL_3316610, EPI_ISL_3316611, EPI_ISL_3316612, EPI_ISL_3316613, EPI_ISL_3316614, EPI_ISL_3316615, EPI_ISL_3316616, EPI_ISL_3316617, EPI_ISL_3316618, EPI_ISL_3316619, EPI_ISL_3316620, EPI_ISL_3316621, EPI_ISL_3316622, EPI_ISL_3316623, EPI_ISL_3316624, EPI_ISL_3316625, EPI_ISL_3316626, EPI_ISL_3316627, EPI_ISL_3316628, EPI_ISL_3316629, EPI_ISL_3316630, EPI_ISL_3316631, EPI_ISL_3316632, EPI_ISL_3316633, EPI_ISL_3316634, EPI_ISL_3316635, EPI_ISL_3316636, EPI_ISL_3316637, EPI_ISL_3316638, EPI_ISL_3316639, EPI_ISL_3316640, EPI_ISL_3316641, EPI_ISL_3316642, EPI_ISL_3316643, EPI_ISL_3316644, EPI_ISL_3316645, EPI_ISL_3316646, EPI_ISL_3316647, EPI_ISL_3316648, EPI_ISL_3316649, EPI_ISL_3316650, EPI_ISL_3316651, EPI_ISL_3316652, EPI_ISL_3316653, EPI_ISL_3316654, EPI_ISL_3316655, EPI_ISL_3316656, EPI_ISL_3316657, EPI_ISL_3316658, EPI_ISL_3316659, EPI_ISL_3316660, EPI_ISL_3316661, EPI_ISL_3316662, EPI_ISL_3316663, EPI_ISL_3316664, EPI_ISL_3316665, EPI_ISL_3316666, EPI_ISL_3316667, EPI_ISL_3316668, EPI_ISL_3316669, EPI_ISL_3316670, EPI_ISL_3316671, EPI_ISL_3316672, EPI_ISL_3316673, EPI_ISL_3316674, EPI_ISL_3316675, EPI_ISL_3316676, EPI_ISL_3316677, EPI_ISL_3316678, EPI_ISL_3316679, EPI_ISL_3316680, EPI_ISL_3316681, EPI_ISL_3316682, EPI_ISL_3316683, EPI_ISL_3316684, EPI_ISL_3316685, EPI_ISL_3316686, EPI_ISL_3316687, EPI_ISL_3316688, EPI_ISL_3316689, EPI_ISL_3316691, EPI_ISL_3316692, EPI_ISL_3316693, EPI_ISL_3316694, EPI_ISL_3316695, EPI_ISL_3316696, EPI_ISL_3316697, EPI_ISL_3316698, EPI_ISL_3316699, EPI_ISL_3316700, EPI_ISL_3316701, EPI_ISL_3316702, EPI_ISL_3316703, EPI_ISL_3316704, EPI_ISL_3316705, EPI_ISL_3316706, EPI_ISL_3316707, EPI_ISL_3316708, EPI_ISL_3316709, EPI_ISL_3316710, EPI_ISL_3316711, EPI_ISL_3316712, EPI_ISL_3316713, EPI_ISL_3316714, EPI_ISL_3316715, EPI_ISL_3316716, EPI_ISL_3316717, EPI_ISL_3316718, EPI_ISL_3316719 | | | |
| see above | National Laboratory for Health, Environment and Food, OMM, Maribor | NLZOH (National Laboratory for Health, Environment and Food) / CISLD (Clinical Institute of Special Laboratory Diagnostics), University Children's Hospital, University Medical Center Ljubljana | Sandra Janezic, Aleksander Mahnic, Maja Rupnik, Tjaša Žohar Čretnik, Alenka Štorman, Nika Gobec, Aleksander Kocuvan, Kaja Tominc, Maša Jarčič, Mojca Cimerman, Nika Volmajer, Andrej Golle / Jernej Kovač, Barbara Jenko Bizjan, Tine Tesovnik, Robert Šket, Katarina Kozmos, Ana Grom, Maruša Debeljak, Marko Pokorn, Tadej Battelino |
| EPI_ISL_3316720, EPI_ISL_3316721, EPI_ISL_3316722, EPI_ISL_3316723, EPI_ISL_3316724, EPI_ISL_3316725, EPI_ISL_3316726, EPI_ISL_3316727, EPI_ISL_3316728, EPI_ISL_3316729, EPI_ISL_3316730, EPI_ISL_3316731, EPI_ISL_3316732, EPI_ISL_3316733, EPI_ISL_3316734, EPI_ISL_3316735, EPI_ISL_3316736, EPI_ISL_3316737, EPI_ISL_3316738, EPI_ISL_3316739, EPI_ISL_3316740, EPI_ISL_3316742, EPI_ISL_3316743, EPI_ISL_3316744, EPI_ISL_3316745, EPI_ISL_3316746, EPI_ISL_3316747, EPI_ISL_3316748, EPI_ISL_3316749, EPI_ISL_3316750, EPI_ISL_3316751, EPI_ISL_3316752, EPI_ISL_3316753, EPI_ISL_3316754, EPI_ISL_3316755, EPI_ISL_3316756, EPI_ISL_3316757, EPI_ISL_3316758, EPI_ISL_3316759, EPI_ISL_3316760, EPI_ISL_3316761, EPI_ISL_3316762, EPI_ISL_3316763, EPI_ISL_3316764, EPI_ISL_3316765, EPI_ISL_3316766, EPI_ISL_3316767, EPI_ISL_3316769, EPI_ISL_3316770, EPI_ISL_3316771, EPI_ISL_3316772, EPI_ISL_3316773, EPI_ISL_3316774, EPI_ISL_3316775, EPI_ISL_3316776, EPI_ISL_3316777, EPI_ISL_3316778, EPI_ISL_3316779, EPI_ISL_3316780, EPI_ISL_3316781, EPI_ISL_3316782, EPI_ISL_3316783, EPI_ISL_3316784, EPI_ISL_3316785, EPI_ISL_3316786, EPI_ISL_3316787, EPI_ISL_3316788, EPI_ISL_3316789, EPI_ISL_3316790, EPI_ISL_3316791, EPI_ISL_3316792, EPI_ISL_3316793, EPI_ISL_3316794, EPI_ISL_3316795, EPI_ISL_3316796, EPI_ISL_3316797, EPI_ISL_3316798, EPI_ISL_3316799, EPI_ISL_3316800, EPI_ISL_3316801, EPI_ISL_3316802, EPI_ISL_3316803, EPI_ISL_3316804, EPI_ISL_3316805, EPI_ISL_3316806, EPI_ISL_3316807, EPI_ISL_3316808, EPI_ISL_3316809, EPI_ISL_3316810, EPI_ISL_3316811, EPI_ISL_3316812, EPI_ISL_3316813, EPI_ISL_3316814, EPI_ISL_3316815, EPI_ISL_3316816, EPI_ISL_3316817, EPI_ISL_3316818, EPI_ISL_3316819, EPI_ISL_3316820, EPI_ISL_3316821, EPI_ISL_3316822, EPI_ISL_3316823, EPI_ISL_3316824, EPI_ISL_3316825, EPI_ISL_3316826, EPI_ISL_3316827, EPI_ISL_3316828, EPI_ISL_3316829, EPI_ISL_3316830, EPI_ISL_3316831, EPI_ISL_3316832, EPI_ISL_3316833, EPI_ISL_3316834, EPI_ISL_3316835, EPI_ISL_3316836, EPI_ISL_3316837, EPI_ISL_3316838, EPI_ISL_3316839, EPI_ISL_3316840, EPI_ISL_3316841, EPI_ISL_3316842, EPI_ISL_3316843, EPI_ISL_3316844, EPI_ISL_3316845, EPI_ISL_3316846, EPI_ISL_3316847, EPI_ISL_3316848, EPI_ISL_3316849, EPI_ISL_3316850, EPI_ISL_3316851, EPI_ISL_3316852, EPI_ISL_3316853, EPI_ISL_3316854, EPI_ISL_3316855, EPI_ISL_3316856, EPI_ISL_3316857, EPI_ISL_3316858, EPI_ISL_3316859, EPI_ISL_3316860, EPI_ISL_3316861, EPI_ISL_3316862, EPI_ISL_3316863, EPI_ISL_3316864, EPI_ISL_3316865, EPI_ISL_3316866, EPI_ISL_3316867, EPI_ISL_3316868, EPI_ISL_3316869, EPI_ISL_3316870, EPI_ISL_3316871, EPI_ISL_3316872, EPI_ISL_3316873, EPI_ISL_3316874, EPI_ISL_3316875, EPI_ISL_3316876, EPI_ISL_3316877, EPI_ISL_3316878, EPI_ISL_3316879, EPI_ISL_3316880, EPI_ISL_3316881, EPI_ISL_3316882, EPI_ISL_3316883, EPI_ISL_3316884, EPI_ISL_3316885, EPI_ISL_3316886, EPI_ISL_3316887, EPI_ISL_3316888, EPI_ISL_3316889, EPI_ISL_3316890, EPI_ISL_3316891, EPI_ISL_3316892, EPI_ISL_3316893, EPI_ISL_3316894, EPI_ISL_3316895, EPI_ISL_3316896, EPI_ISL_3316897, EPI_ISL_3316898, EPI_ISL_3316899, EPI_ISL_3316900, EPI_ISL_3316901, EPI_ISL_3316902, EPI_ISL_3316903, EPI_ISL_3316904, EPI_ISL_3316905, EPI_ISL_3316906, EPI_ISL_3316907, EPI_ISL_3316908, EPI_ISL_3316909, EPI_ISL_3316910, EPI_ISL_3316911, EPI_ISL_3316912, EPI_ISL_3316913, EPI_ISL_3316914, EPI_ISL_3316915, EPI_ISL_3316916 | | | |
| see above | National Laboratory for Health, Environment and Food, OMM, Kranj | NLZOH (National Laboratory for Health, Environment and Food) / CISLD (Clinical Institute of Special Laboratory Diagnostics), University Children's Hospital, University Medical Center Ljubljana | Sandra Janezic, Aleksander Mahnic, Maja Rupnik, Tjaša Žohar Čretnik, Alenka Štorman, Nika Gobec, Aleksander Kocuvan, Kaja Tominc, Maša Jarčič, Mateja Ravnik, Monika Korošec, Marjana Petrevčič / Jernej Kovač, Barbara Jenko Bizjan, Tine Tesovnik, Robert Šket, Katarina Kozmos, Ana Grom, Maruša Debeljak, Marko Pokorn, Tadej Battelino |
| EPI_ISL_3316918, EPI_ISL_3316919, EPI_ISL_3316920, EPI_ISL_3316921, EPI_ISL_3316922, EPI_ISL_3316923, EPI_ISL_3316924, EPI_ISL_3316925, EPI_ISL_3316926, EPI_ISL_3316929, EPI_ISL_3316930, EPI_ISL_3316935, EPI_ISL_3316937, EPI_ISL_3316938, EPI_ISL_3316939, EPI_ISL_3316940, EPI_ISL_3316941, EPI_ISL_3316942, EPI_ISL_3316943, EPI_ISL_3316944, EPI_ISL_3316945, EPI_ISL_3316946, EPI_ISL_3316947, EPI_ISL_3316948, EPI_ISL_3316949, EPI_ISL_3316950, EPI_ISL_3316951, EPI_ISL_3316952, EPI_ISL_3316953, EPI_ISL_3316954, EPI_ISL_3316955, EPI_ISL_3316956, EPI_ISL_3316957, EPI_ISL_3316958, EPI_ISL_3316959, EPI_ISL_3316960 | | | |
| see above | National Laboratory for Health, Environment and Food, OMM, Celje | NLZOH (National Laboratory for Health, Environment and Food) / CISLD (Clinical Institute of Special Laboratory Diagnostics), University Children's Hospital, University Medical Center Ljubljana | Sandra Janezic, Aleksander Mahnic, Maja Rupnik, Tjaša Žohar Čretnik, Alenka Štorman, Nika Gobec, Aleksander Kocuvan, Kaja Tominc, Maša Jarčič, Mateja Borinc, Daša Kavka / Jernej Kovač, Barbara Jenko Bizjan, Tine Tesovnik, Robert Šket, Katarina Kozmos, Ana Grom, Maruša Debeljak, Marko Pokorn, Tadej Battelino |
| EPI_ISL_3316961, EPI_ISL_3316962, EPI_ISL_3316963, EPI_ISL_3316964, EPI_ISL_3316965, EPI_ISL_3316966, EPI_ISL_3316967, EPI_ISL_3316968, EPI_ISL_3316969, EPI_ISL_3316970 | National Laboratory for Health, Environment and Food, OMM, Maribor | NLZOH (National Laboratory for Health, Environment and Food) / CISLD (Clinical Institute of Special Laboratory Diagnostics), University Children's Hospital, University Medical Center Ljubljana | Sandra Janezic, Aleksander Mahnic, Maja Rupnik, Tjaša Žohar Čretnik, Alenka Štorman, Nika Gobec, Aleksander Kocuvan, Kaja Tominc, Maša Jarčič, Mojca Cimerman, Nika Volmajer, Andrej Golle / Jernej Kovač, Barbara Jenko Bizjan, Tine Tesovnik, Robert Šket, Katarina Kozmos, Ana Grom, Maruša Debeljak, Marko Pokorn, Tadej Battelino |
| EPI_ISL_3316972, EPI_ISL_3316973, EPI_ISL_3316975, EPI_ISL_3316976, EPI_ISL_3316978, EPI_ISL_3316979, EPI_ISL_3316980, EPI_ISL_3316981, EPI_ISL_3316982, EPI_ISL_3316983, EPI_ISL_3316985, EPI_ISL_3316986, EPI_ISL_3316987, EPI_ISL_3316988, EPI_ISL_3316990, EPI_ISL_3316991, EPI_ISL_3316992, EPI_ISL_3316993, EPI_ISL_3316994, EPI_ISL_3316995, EPI_ISL_3316996, EPI_ISL_3316997, EPI_ISL_3316998, EPI_ISL_3316999, EPI_ISL_3317000, EPI_ISL_3317001, EPI_ISL_3317002, EPI_ISL_3317003, EPI_ISL_3317004, EPI_ISL_3317005, EPI_ISL_3317006, EPI_ISL_3317007, EPI_ISL_3317008, EPI_ISL_3317010, EPI_ISL_3317011, EPI_ISL_3317013, EPI_ISL_3317014, EPI_ISL_3317015, EPI_ISL_3317016, EPI_ISL_3317017, EPI_ISL_3317018, EPI_ISL_3317019, EPI_ISL_3317020, EPI_ISL_3317021, EPI_ISL_3317022, EPI_ISL_3317023, EPI_ISL_3317024, EPI_ISL_3317025, EPI_ISL_3317026, EPI_ISL_3317027, EPI_ISL_3317028, EPI_ISL_3317029, EPI_ISL_3317030, EPI_ISL_3317031 | | | |
| see above | National Laboratory for Health, Environment and Food, OMM, Koper | NLZOH (National Laboratory for Health, Environment and Food) / CISLD (Clinical Institute of Special Laboratory Diagnostics), University Children's Hospital, University Medical Center Ljubljana | Sandra Janezic, Aleksander Mahnic, Maja Rupnik, Tjaša Žohar Čretnik, Alenka Štorman, Nika Gobec, Aleksander Kocuvan, Kaja Tominc, Maša Jarčič, Gašper Strugar, Tina Cvetković, Mitja Rak / Jernej Kovač, Barbara Jenko Bizjan, Tine Tesovnik, Robert Šket, Katarina Kozmos, Ana Grom, Maruša Debeljak, Marko Pokorn, Tadej Battelino |
| EPI_ISL_3317033, EPI_ISL_3317034, EPI_ISL_3317035, EPI_ISL_3317036, EPI_ISL_3317037, EPI_ISL_3317038, EPI_ISL_3317039, EPI_ISL_3317040, EPI_ISL_3317041, EPI_ISL_3317042, EPI_ISL_3317043, EPI_ISL_3317044, EPI_ISL_3317045, EPI_ISL_3317046, EPI_ISL_3317047, EPI_ISL_3317048, EPI_ISL_3317049, EPI_ISL_3317050, EPI_ISL_3317051, EPI_ISL_3317052, EPI_ISL_3317053, EPI_ISL_3317054, EPI_ISL_3317055, EPI_ISL_3317056, EPI_ISL_3317057, EPI_ISL_3317058, EPI_ISL_3317060, EPI_ISL_3317061, EPI_ISL_3317062, EPI_ISL_3317063, EPI_ISL_3317064, EPI_ISL_3317065, EPI_ISL_3317066, EPI_ISL_3317067, EPI_ISL_3317068, EPI_ISL_3317069, EPI_ISL_3317070, EPI_ISL_3317071, EPI_ISL_3317072, EPI_ISL_3317073, EPI_ISL_3317074, EPI_ISL_3317075, EPI_ISL_3317076, EPI_ISL_3317077, EPI_ISL_3317078, EPI_ISL_3317079, EPI_ISL_3317080, EPI_ISL_3317081, EPI_ISL_3317082, EPI_ISL_3317083, EPI_ISL_3317084, EPI_ISL_3317085, EPI_ISL_3317086, EPI_ISL_3317087, EPI_ISL_3317088, EPI_ISL_3317089, EPI_ISL_3317090, EPI_ISL_3317091, EPI_ISL_3317092, EPI_ISL_3317093, EPI_ISL_3317094, EPI_ISL_3317095, EPI_ISL_3317096, EPI_ISL_3317097, EPI_ISL_3317098, EPI_ISL_3317099, EPI_ISL_3317100, EPI_ISL_3317101, EPI_ISL_3317102, EPI_ISL_3317103, EPI_ISL_3317104, EPI_ISL_3317105, EPI_ISL_3317106, EPI_ISL_3317107, EPI_ISL_3317108, EPI_ISL_3317109, EPI_ISL_3317110, EPI_ISL_3317111, EPI_ISL_3317112, EPI_ISL_3317113, EPI_ISL_3317114, EPI_ISL_3317115, EPI_ISL_3317116, EPI_ISL_3317118, EPI_ISL_3317119, EPI_ISL_3317120, EPI_ISL_3317121, EPI_ISL_3317122, EPI_ISL_3317123, EPI_ISL_3317124, EPI_ISL_3317125, EPI_ISL_3317126, EPI_ISL_3317127, EPI_ISL_3317128, EPI_ISL_3317129, EPI_ISL_3317130, EPI_ISL_3317131, EPI_ISL_3317133, EPI_ISL_3317134, EPI_ISL_3317135, EPI_ISL_3317136, EPI_ISL_3317138, EPI_ISL_3317140, EPI_ISL_3317141, EPI_ISL_3317146, EPI_ISL_3317147, EPI_ISL_3317148, EPI_ISL_3317150, EPI_ISL_3317151, EPI_ISL_3317152, EPI_ISL_3317153, EPI_ISL_3317154, EPI_ISL_3317155, EPI_ISL_3317156, EPI_ISL_3317157, EPI_ISL_3317159, EPI_ISL_3317160, EPI_ISL_3317161, EPI_ISL_3317162, EPI_ISL_3317163, EPI_ISL_3317164, EPI_ISL_3317165, EPI_ISL_3317166, EPI_ISL_3317167, EPI_ISL_3317168, EPI_ISL_3317169, EPI_ISL_3317170, EPI_ISL_3317171, EPI_ISL_3317172, EPI_ISL_3317173, EPI_ISL_3317174, EPI_ISL_3317175, EPI_ISL_3317176, EPI_ISL_3317177, EPI_ISL_3317178, EPI_ISL_3317179, EPI_ISL_3317180, EPI_ISL_3317181, EPI_ISL_3317182, EPI_ISL_3317183, EPI_ISL_3317184, EPI_ISL_3317185, EPI_ISL_3317186, EPI_ISL_3317187, EPI_ISL_3317188, EPI_ISL_3317189, EPI_ISL_3317190, EPI_ISL_3317191, EPI_ISL_3317192, EPI_ISL_3317193, EPI_ISL_3317194, EPI_ISL_3317195, EPI_ISL_3317196, EPI_ISL_3317197, EPI_ISL_3317198, EPI_ISL_3317199, EPI_ISL_3317200, EPI_ISL_3317201, EPI_ISL_3317202, EPI_ISL_3317203, EPI_ISL_3317204, EPI_ISL_3317205, EPI_ISL_3317206, EPI_ISL_3317207, EPI_ISL_3317208, EPI_ISL_3317209, EPI_ISL_3317210, EPI_ISL_3317211, EPI_ISL_3317212, EPI_ISL_3317213, EPI_ISL_3317214, EPI_ISL_3317215, EPI_ISL_3317216, EPI_ISL_3317217, EPI_ISL_3317218, EPI_ISL_3317219 | | | |
| see above | National Laboratory for Health, Environment and Food, OMM, Novo mesto | NLZOH (National Laboratory for Health, Environment and Food) / CISLD (Clinical Institute of Special Laboratory Diagnostics), University Children's Hospital, University Medical Center Ljubljana | Sandra Janezic, Aleksander Mahnic, Maja Rupnik, Tjaša Žohar Čretnik, Alenka Štorman, Nika Gobec, Aleksander Kocuvan, Kaja Tominc, Maša Jarčič, David Cvetko, Tatjana Harlander, Matjaž Retelj / Jernej Kovač, Barbara Jenko Bizjan, Tine Tesovnik, Robert Šket, Katarina Kozmos, Ana Grom, Maruša Debeljak, Marko Pokorn, Tadej Battelino |
| EPI_ISL_3317220, EPI_ISL_3317221 | ADPH | ADPH | Adrianna Walker |
| EPI_ISL_3317222 | RURAL ALABAMA PREVENTION CENTER | ADPH | Adrianna Walker |
| EPI_ISL_3317223, EPI_ISL_3317224 | DALE MEDICAL CENTER | ADPH | Adrianna Walker |
| EPI_ISL_3317225, EPI_ISL_3317226 | ADPH | ADPH | Adrianna Walker |
| EPI_ISL_3317227, EPI_ISL_3317228 | SOUTHEAST HEALTH MED CTR DOTHAN | ADPH | Adrianna Walker |
| EPI_ISL_3317229, EPI_ISL_3317230, EPI_ISL_3317231, EPI_ISL_3317232, EPI_ISL_3317233 | ADPH | ADPH | Adrianna Walker |
| EPI_ISL_3317234 | RURAL ALABAMA PREVENTION CENTER | ADPH | Adrianna Walker |
| EPI_ISL_3317235 | EAST ALABAMA MEDICAL CENTER | ADPH | Adrianna Walker |
| EPI_ISL_3317236, EPI_ISL_3317237, EPI_ISL_3317238 | ADPH | ADPH | Adrianna Walker |
| EPI_ISL_3317239 | RURAL HEALTH CENTER DALLAS COUNTY | ADPH | Adrianna Walker |
| EPI_ISL_3317240, EPI_ISL_3317241, EPI_ISL_3317242, EPI_ISL_3317243, EPI_ISL_3317244, EPI_ISL_3317245, EPI_ISL_3317246, EPI_ISL_3317247, EPI_ISL_3317248 | ADPH | ADPH | Adrianna Walker |
| EPI_ISL_3317249 | UFFORD MEDICAL ASSET GROUP CROSSVILLE | ADPH | Adrianna Walker |
| EPI_ISL_3317250, | ADPH | ADPH | Adrianna Walker |

| | | | |
|---|---|---|---|
| EPI_ISL_3317251, EPI_ISL_3317252, EPI_ISL_3317253, EPI_ISL_3317254 | | | |
| EPI_ISL_3317255 | SOUTHEAST HEALTH MED CTR DOTHAN | ADPH | Adrianna Walker |
| EPI_ISL_3317256 | UNIVERSITY MEDICAL CENTER (TUSCALOOSA) | ADPH | Adrianna Walker |
| EPI_ISL_3317257, EPI_ISL_3317258, EPI_ISL_3317259, EPI_ISL_3317260, EPI_ISL_3317261, EPI_ISL_3317262, EPI_ISL_3317263, EPI_ISL_3317264, EPI_ISL_3317265, EPI_ISL_3317266, EPI_ISL_3317267, EPI_ISL_3317268, EPI_ISL_3317269, EPI_ISL_3317270, EPI_ISL_3317271, EPI_ISL_3317272, EPI_ISL_3317273, EPI_ISL_3317274 | | | |
| see above | ADPH | ADPH | Adrianna Walker |
| EPI_ISL_3317275, EPI_ISL_3317276 | SHOALS COMMUNITY CLINIC | ADPH | Adrianna Walker |
| EPI_ISL_3317277, EPI_ISL_3317278 | EAST ALABAMA MEDICAL CENTER | ADPH | Adrianna Walker |
| EPI_ISL_3317280, EPI_ISL_3317281 | BAPTIST MED CTR SOUTH (MONTGOMERY) | ADPH | Adrianna Walker |
| EPI_ISL_3317282, EPI_ISL_3317283, EPI_ISL_3317284, EPI_ISL_3317285, EPI_ISL_3317286, EPI_ISL_3317287, EPI_ISL_3317288, EPI_ISL_3317289, EPI_ISL_3317291, EPI_ISL_3317292, EPI_ISL_3317293, EPI_ISL_3317294, EPI_ISL_3317295 | | | |
| see above | DCH REGIONAL MEDICAL CENTER | ADPH | Adrianna Walker |
| EPI_ISL_3317298 | UNIVERSITY MEDICAL CENTER (TUSCALOOSA) | ADPH | Adrianna Walker |
| EPI_ISL_3317299 | BAPTIST MED CTR (PRINCETON) | ADPH | Adrianna Walker |
| EPI_ISL_3317300 | SOUTHEAST HEALTH MED CTR DOTHAN | ADPH | Adrianna Walker |
| EPI_ISL_3317301 | ADPH | ADPH | Adrianna Walker |
| EPI_ISL_3317302, EPI_ISL_3317303, EPI_ISL_3317304, EPI_ISL_3317305, EPI_ISL_3317306, EPI_ISL_3317307, EPI_ISL_3317308, EPI_ISL_3317309, EPI_ISL_3317310, EPI_ISL_3317311, EPI_ISL_3317312, EPI_ISL_3317313, EPI_ISL_3317314, EPI_ISL_3317315, EPI_ISL_3317316, EPI_ISL_3317317, EPI_ISL_3317318, EPI_ISL_3317319, EPI_ISL_3317320, EPI_ISL_3317321, EPI_ISL_3317322, EPI_ISL_3317323, EPI_ISL_3317324, EPI_ISL_3317325, EPI_ISL_3317326, EPI_ISL_3317327, EPI_ISL_3317328, EPI_ISL_3317329, EPI_ISL_3317330, EPI_ISL_3317331, EPI_ISL_3317332, EPI_ISL_3317333, EPI_ISL_3317334, EPI_ISL_3317335, EPI_ISL_3317336, EPI_ISL_3317337, EPI_ISL_3317338, EPI_ISL_3317339, EPI_ISL_3317340, EPI_ISL_3317341, EPI_ISL_3317342, EPI_ISL_3317343, EPI_ISL_3317344, EPI_ISL_3317345, EPI_ISL_3317346, EPI_ISL_3317347, EPI_ISL_3317348, EPI_ISL_3317349, EPI_ISL_3317350, EPI_ISL_3317351, EPI_ISL_3317352, EPI_ISL_3317353, EPI_ISL_3317354, EPI_ISL_3317355, EPI_ISL_3317356, EPI_ISL_3317357, EPI_ISL_3317358, EPI_ISL_3317359, EPI_ISL_3317360, EPI_ISL_3317361, EPI_ISL_3317362, EPI_ISL_3317363, EPI_ISL_3317364, EPI_ISL_3317365, EPI_ISL_3317366, EPI_ISL_3317367, EPI_ISL_3317368, EPI_ISL_3317369, EPI_ISL_3317370, EPI_ISL_3317371, EPI_ISL_3317372, EPI_ISL_3317373, EPI_ISL_3317374, EPI_ISL_3317375, EPI_ISL_3317376, EPI_ISL_3317377, EPI_ISL_3317378, EPI_ISL_3317379, EPI_ISL_3317380, EPI_ISL_3317381, EPI_ISL_3317382, EPI_ISL_3317383, EPI_ISL_3317384, EPI_ISL_3317385, EPI_ISL_3317386, EPI_ISL_3317387, EPI_ISL_3317388, EPI_ISL_3317389, EPI_ISL_3317390, EPI_ISL_3317391, EPI_ISL_3317392, EPI_ISL_3317393, EPI_ISL_3317394, EPI_ISL_3317395, EPI_ISL_3317396, EPI_ISL_3317397, EPI_ISL_3317398, EPI_ISL_3317399, EPI_ISL_3317400, EPI_ISL_3317401, EPI_ISL_3317402, EPI_ISL_3317403, EPI_ISL_3317404, EPI_ISL_3317405, EPI_ISL_3317406, EPI_ISL_3317407, EPI_ISL_3317408, EPI_ISL_3317409, EPI_ISL_3317410, EPI_ISL_3317411, EPI_ISL_3317412, EPI_ISL_3317413, EPI_ISL_3317414, EPI_ISL_3317415, EPI_ISL_3317416, EPI_ISL_3317417, EPI_ISL_3317418, EPI_ISL_3317419, EPI_ISL_3317420, EPI_ISL_3317421, EPI_ISL_3317422, EPI_ISL_3317423, EPI_ISL_3317424, EPI_ISL_3317425, EPI_ISL_3317426, EPI_ISL_3317427, EPI_ISL_3317428, EPI_ISL_3317429, EPI_ISL_3317430, EPI_ISL_3317431, EPI_ISL_3317432, EPI_ISL_3317433, EPI_ISL_3317434, EPI_ISL_3317435, EPI_ISL_3317436, EPI_ISL_3317437, EPI_ISL_3317438, EPI_ISL_3317439, EPI_ISL_3317440, EPI_ISL_3317441, EPI_ISL_3317442, EPI_ISL_3317443, EPI_ISL_3317444, EPI_ISL_3317445, EPI_ISL_3317446, EPI_ISL_3317447, EPI_ISL_3317448, EPI_ISL_3317449, EPI_ISL_3317450, EPI_ISL_3317451, EPI_ISL_3317452, EPI_ISL_3317453, EPI_ISL_3317454, EPI_ISL_3317455, EPI_ISL_3317456, EPI_ISL_3317457, EPI_ISL_3317458, EPI_ISL_3317459, EPI_ISL_3317460, EPI_ISL_3317461, EPI_ISL_3317462, EPI_ISL_3317463, EPI_ISL_3317464, EPI_ISL_3317465, EPI_ISL_3317466, EPI_ISL_3317467, EPI_ISL_3317468, EPI_ISL_3317469, EPI_ISL_3317470, EPI_ISL_3317471, EPI_ISL_3317472, EPI_ISL_3317473, EPI_ISL_3317474, EPI_ISL_3317475, EPI_ISL_3317476, EPI_ISL_3317477, EPI_ISL_3317478, EPI_ISL_3317479, EPI_ISL_3317480, EPI_ISL_3317481, EPI_ISL_3317482, EPI_ISL_3317483, EPI_ISL_3317484, EPI_ISL_3317485, EPI_ISL_3317486, EPI_ISL_3317487, EPI_ISL_3317488, EPI_ISL_3317489, EPI_ISL_3317490, EPI_ISL_3317491, EPI_ISL_3317492, EPI_ISL_3317493, EPI_ISL_3317494, EPI_ISL_3317495, EPI_ISL_3317496, EPI_ISL_3317497, EPI_ISL_3317498, EPI_ISL_3317499, EPI_ISL_3317500, EPI_ISL_3317501, EPI_ISL_3317502, EPI_ISL_3317503, EPI_ISL_3317504, EPI_ISL_3317505, EPI_ISL_3317506, EPI_ISL_3317507, EPI_ISL_3317508, EPI_ISL_3317509, EPI_ISL_3317510, EPI_ISL_3317511, EPI_ISL_3317512, EPI_ISL_3317513, EPI_ISL_3317514, EPI_ISL_3317515, EPI_ISL_3317516, EPI_ISL_3317517, EPI_ISL_3317518, EPI_ISL_3317519, EPI_ISL_3317520, EPI_ISL_3317521, EPI_ISL_3317522, EPI_ISL_3317523, EPI_ISL_3317524, EPI_ISL_3317525, EPI_ISL_3317526, EPI_ISL_3317527, EPI_ISL_3317528, EPI_ISL_3317529, EPI_ISL_3317530, EPI_ISL_3317531, EPI_ISL_3317532, EPI_ISL_3317533, EPI_ISL_3317534, EPI_ISL_3317535, EPI_ISL_3317536, EPI_ISL_3317537, EPI_ISL_3317538, EPI_ISL_3317539, EPI_ISL_3317540, EPI_ISL_3317541, EPI_ISL_3317542, EPI_ISL_3317543, EPI_ISL_3317544, EPI_ISL_3317545, EPI_ISL_3317546, EPI_ISL_3317547, EPI_ISL_3317548, EPI_ISL_3317549, EPI_ISL_3317550, EPI_ISL_3317551, EPI_ISL_3317552, EPI_ISL_3317553, EPI_ISL_3317554, EPI_ISL_3317555, EPI_ISL_3317556, EPI_ISL_3317557, EPI_ISL_3317558, EPI_ISL_3317559, EPI_ISL_3317560, EPI_ISL_3317561, EPI_ISL_3317562, EPI_ISL_3317563, EPI_ISL_3317564, EPI_ISL_3317565, EPI_ISL_3317566, EPI_ISL_3317567, EPI_ISL_3317568, EPI_ISL_3317569, EPI_ISL_3317570, EPI_ISL_3317571, EPI_ISL_3317572, EPI_ISL_3317573, EPI_ISL_3317574, EPI_ISL_3317575, EPI_ISL_3317576, EPI_ISL_3317577, EPI_ISL_3317578, EPI_ISL_3317579, EPI_ISL_3317580, EPI_ISL_3317581, EPI_ISL_3317582, EPI_ISL_3317583, EPI_ISL_3317584, EPI_ISL_3317585, EPI_ISL_3317586, EPI_ISL_3317587, EPI_ISL_3317588, EPI_ISL_3317589, EPI_ISL_3317590, EPI_ISL_3317591, EPI_ISL_3317592, EPI_ISL_3317593, EPI_ISL_3317594, EPI_ISL_3317595, EPI_ISL_3317596, EPI_ISL_3317597, EPI_ISL_3317598, EPI_ISL_3317599, EPI_ISL_3317600, EPI_ISL_3317601, EPI_ISL_3317602, EPI_ISL_3317603, EPI_ISL_3317604, EPI_ISL_3317605, EPI_ISL_3317606, EPI_ISL_3317607, EPI_ISL_3317608, EPI_ISL_3317609, EPI_ISL_3317610, EPI_ISL_3317611, EPI_ISL_3317612, EPI_ISL_3317613, EPI_ISL_3317614, EPI_ISL_3317615, EPI_ISL_3317616, EPI_ISL_3317617, EPI_ISL_3317618, EPI_ISL_3317619, EPI_ISL_3317620, EPI_ISL_3317621, EPI_ISL_3317622, EPI_ISL_3317623, EPI_ISL_3317624, EPI_ISL_3317625, EPI_ISL_3317626, EPI_ISL_3317627, EPI_ISL_3317628, EPI_ISL_3317629, EPI_ISL_3317630, EPI_ISL_3317631, EPI_ISL_3317632, EPI_ISL_3317633, EPI_ISL_3317634, EPI_ISL_3317635, EPI_ISL_3317636, EPI_ISL_3317637, EPI_ISL_3317638, EPI_ISL_3317639, EPI_ISL_3317640, EPI_ISL_3317641, EPI_ISL_3317642, EPI_ISL_3317643, EPI_ISL_3317644, EPI_ISL_3317645, EPI_ISL_3317646, EPI_ISL_3317647, EPI_ISL_3317648, EPI_ISL_3317649, EPI_ISL_3317650, EPI_ISL_3317651, EPI_ISL_3317652, EPI_ISL_3317653, EPI_ISL_3317654, EPI_ISL_3317655, EPI_ISL_3317656, EPI_ISL_3317657, EPI_ISL_3317658, EPI_ISL_3317659, EPI_ISL_3317660, EPI_ISL_3317661, EPI_ISL_3317662, EPI_ISL_3317663, EPI_ISL_3317664, EPI_ISL_3317665, EPI_ISL_3317666, EPI_ISL_3317667, EPI_ISL_3317668, EPI_ISL_3317669, EPI_ISL_3317670, EPI_ISL_3317671, EPI_ISL_3317672, EPI_ISL_3317673, EPI_ISL_3317674, EPI_ISL_3317675, EPI_ISL_3317676, EPI_ISL_3317677, EPI_ISL_3317678, EPI_ISL_3317679, EPI_ISL_3317680, EPI_ISL_3317681, EPI_ISL_3317682, EPI_ISL_3317683, EPI_ISL_3317684, EPI_ISL_3317685, EPI_ISL_3317686, EPI_ISL_3317687, EPI_ISL_3317688, EPI_ISL_3317689, EPI_ISL_3317690, EPI_ISL_3317691, EPI_ISL_3317692, EPI_ISL_3317693, EPI_ISL_3317694, EPI_ISL_3317695, EPI_ISL_3317696, EPI_ISL_3317697, EPI_ISL_3317698, EPI_ISL_3317699, EPI_ISL_3317700, EPI_ISL_3317701, EPI_ISL_3317702, EPI_ISL_3317703, EPI_ISL_3317704, EPI_ISL_3317705, EPI_ISL_3317706, EPI_ISL_3317707, EPI_ISL_3317708, EPI_ISL_3317709, EPI_ISL_3317710, EPI_ISL_3317711, EPI_ISL_3317712, EPI_ISL_3317713, EPI_ISL_3317714, EPI_ISL_3317715, EPI_ISL_3317716, EPI_ISL_3317717, EPI_ISL_3317718, EPI_ISL_3317719, EPI_ISL_3317720, EPI_ISL_3317721, EPI_ISL_3317722, EPI_ISL_3317723, EPI_ISL_3317724, EPI_ISL_3317725, EPI_ISL_3317726, EPI_ISL_3317727, EPI_ISL_3317728, EPI_ISL_3317729, EPI_ISL_3317730, EPI_ISL_3317731, EPI_ISL_3317732, EPI_ISL_3317733, EPI_ISL_3317734, EPI_ISL_3317735, EPI_ISL_3317736, EPI_ISL_3317737, EPI_ISL_3317738, EPI_ISL_3317739, EPI_ISL_3317740, EPI_ISL_3317741, EPI_ISL_3317742, EPI_ISL_3317743, EPI_ISL_3317744, EPI_ISL_3317745, EPI_ISL_3317746, EPI_ISL_3317747, EPI_ISL_3317748, EPI_ISL_3317749, EPI_ISL_3317750, EPI_ISL_3317751, EPI_ISL_3317752, EPI_ISL_3317753, EPI_ISL_3317754, EPI_ISL_3317755, EPI_ISL_3317756, EPI_ISL_3317757, EPI_ISL_3317758, EPI_ISL_3317759, EPI_ISL_3317760, EPI_ISL_3317761, EPI_ISL_3317762, EPI_ISL_3317763, EPI_ISL_3317764, EPI_ISL_3317765, EPI_ISL_3317766, EPI_ISL_3317767, EPI_ISL_3317768, EPI_ISL_3317769, EPI_ISL_3317770, EPI_ISL_3317771, EPI_ISL_3317772, EPI_ISL_3317773, EPI_ISL_3317774, EPI_ISL_3317775, EPI_ISL_3317776, EPI_ISL_3317777, EPI_ISL_3317778, EPI_ISL_3317779, EPI_ISL_3317780, EPI_ISL_3317781, EPI_ISL_3317782, EPI_ISL_3317783, EPI_ISL_3317784, EPI_ISL_3317785, EPI_ISL_3317786, EPI_ISL_3317787, EPI_ISL_3317788, EPI_ISL_3317789, EPI_ISL_3317790, EPI_ISL_3317791, EPI_ISL_3317792, EPI_ISL_3317793, EPI_ISL_3317794, EPI_ISL_3317795, EPI_ISL_3317796, EPI_ISL_3317797, EPI_ISL_3317798, EPI_ISL_3317799, EPI_ISL_3317800, EPI_ISL_3317801, EPI_ISL_3317802, EPI_ISL_3317803, EPI_ISL_3317804, EPI_ISL_3317805, EPI_ISL_3317806, EPI_ISL_3317807, EPI_ISL_3317808, EPI_ISL_3317809, EPI_ISL_3317810, EPI_ISL_3317811, EPI_ISL_3317812, EPI_ISL_3317813, EPI_ISL_3317814, EPI_ISL_3317815, EPI_ISL_3317816, EPI_ISL_3317817, EPI_ISL_3317818, EPI_ISL_3317819, EPI_ISL_3317820, EPI_ISL_3317821, EPI_ISL_3317822, EPI_ISL_3317823, EPI_ISL_3317824, EPI_ISL_3317825, EPI_ISL_3317826, EPI_ISL_3317827, EPI_ISL_3317828, EPI_ISL_3317829, EPI_ISL_3317830, EPI_ISL_3317831, EPI_ISL_3317832, EPI_ISL_3317833, EPI_ISL_3317834, EPI_ISL_3317835, EPI_ISL_3317836, EPI_ISL_3317837, EPI_ISL_3317838, EPI_ISL_3317839, EPI_ISL_3317840, EPI_ISL_3317841 | | | |
| see above | Ministry of Health Turkey | Ministry of Health Turkey | Fatma Bayrakdar, Yasemin Coşgun, Süleyman Yalcin, Gülay Korukluoğlu |
| EPI_ISL_3317842, EPI_ISL_3317843, EPI_ISL_3317844, EPI_ISL_3317845, EPI_ISL_3317846, EPI_ISL_3317847, EPI_ISL_3317848, EPI_ISL_3317849, EPI_ISL_3317850, EPI_ISL_3317851, EPI_ISL_3317852, EPI_ISL_3317853, EPI_ISL_3317854, EPI_ISL_3317855, EPI_ISL_3317856, EPI_ISL_3317857, EPI_ISL_3317858, EPI_ISL_3317859, EPI_ISL_3317860, EPI_ISL_3317861, EPI_ISL_3317863, EPI_ISL_3317865, EPI_ISL_3317866, EPI_ISL_3317867, EPI_ISL_3317868, EPI_ISL_3317869, EPI_ISL_3317870, EPI_ISL_3317871, EPI_ISL_3317872, EPI_ISL_3317873, EPI_ISL_3317874, EPI_ISL_3317875, EPI_ISL_3317876, EPI_ISL_3317878, EPI_ISL_3317879, EPI_ISL_3317880, EPI_ISL_3317881, EPI_ISL_3317882, EPI_ISL_3317883, EPI_ISL_3317884, EPI_ISL_3317885, EPI_ISL_3317886, EPI_ISL_3317887, EPI_ISL_3317888, EPI_ISL_3317889, EPI_ISL_3317890, EPI_ISL_3317894, EPI_ISL_3317895, EPI_ISL_3317896, EPI_ISL_3317897, EPI_ISL_3317898, EPI_ISL_3317899, EPI_ISL_3317900, EPI_ISL_3317901, EPI_ISL_3317902, EPI_ISL_3317903, EPI_ISL_3317904, EPI_ISL_3317905, EPI_ISL_3317906, EPI_ISL_3317907, EPI_ISL_3317908, EPI_ISL_3317909, EPI_ISL_3317910, EPI_ISL_3317911, EPI_ISL_3317913, EPI_ISL_3317914, EPI_ISL_3317915, EPI_ISL_3317916, EPI_ISL_3317917, EPI_ISL_3317918, EPI_ISL_3317919, EPI_ISL_3317921, EPI_ISL_3317922, EPI_ISL_3317923, EPI_ISL_3317924, EPI_ISL_3317925, EPI_ISL_3317927, EPI_ISL_3317928, EPI_ISL_3317929, EPI_ISL_3317930, EPI_ISL_3317931, EPI_ISL_3317932, EPI_ISL_3317933, EPI_ISL_3317934, EPI_ISL_3317935, EPI_ISL_3317936, EPI_ISL_3317937 | | | |
| see above | NJDOH, Public Health and Environmental Laboratories | NJ_PHEL | Lindsey Bodnar, Shiv K. Verma, Ryan Pachucki, Chelsea San Filippo, Jacquelyn Deverell, Dana Woell, Allison Roder, Byeong Jeong |
| EPI_ISL_3318725, EPI_ISL_3318726, EPI_ISL_3318728, EPI_ISL_3318729, EPI_ISL_3318730, EPI_ISL_3318731, EPI_ISL_3318732, EPI_ISL_3318733, EPI_ISL_3318734, EPI_ISL_3318735, EPI_ISL_3318736, EPI_ISL_3318737, EPI_ISL_3318738, EPI_ISL_3318739, EPI_ISL_3318740, EPI_ISL_3318742, EPI_ISL_3318743, EPI_ISL_3318744, EPI_ISL_3318745, EPI_ISL_3318746, EPI_ISL_3318747, EPI_ISL_3318748, EPI_ISL_3318749, EPI_ISL_3318751, EPI_ISL_3318752, EPI_ISL_3318754, EPI_ISL_3318755, EPI_ISL_3318756, EPI_ISL_3318757, EPI_ISL_3318758, EPI_ISL_3318759, EPI_ISL_3318760, EPI_ISL_3318761, EPI_ISL_3318762, EPI_ISL_3318763, EPI_ISL_3318765, EPI_ISL_3318766, EPI_ISL_3318767, EPI_ISL_3318768, EPI_ISL_3318769, EPI_ISL_3318770, EPI_ISL_3318771, EPI_ISL_3318772, EPI_ISL_3318773, EPI_ISL_3318774, EPI_ISL_3318775, EPI_ISL_3318776, EPI_ISL_3318777, EPI_ISL_3318778, EPI_ISL_3318779, EPI_ISL_3318780, EPI_ISL_3318781, EPI_ISL_3318782, EPI_ISL_3318783, EPI_ISL_3318784, EPI_ISL_3318785, EPI_ISL_3318786, EPI_ISL_3318789, EPI_ISL_3318790, EPI_ISL_3318791, EPI_ISL_3318792, EPI_ISL_3318793, EPI_ISL_3318794, EPI_ISL_3318795, EPI_ISL_3318796, EPI_ISL_3318797, EPI_ISL_3318798, EPI_ISL_3318799, EPI_ISL_3318800, EPI_ISL_3318801, EPI_ISL_3318802, EPI_ISL_3318803, EPI_ISL_3318804, EPI_ISL_3318805, EPI_ISL_3318806, EPI_ISL_3318807, EPI_ISL_3318808, EPI_ISL_3318809, EPI_ISL_3318810, EPI_ISL_3318811, EPI_ISL_3318812, EPI_ISL_3318813, EPI_ISL_3318814, EPI_ISL_3318815, EPI_ISL_3318816, EPI_ISL_3318817, EPI_ISL_3318818, EPI_ISL_3318819, EPI_ISL_3318820, EPI_ISL_3318821, EPI_ISL_3318822, EPI_ISL_3318823, EPI_ISL_3318824, EPI_ISL_3318825, EPI_ISL_3318826, EPI_ISL_3318827, EPI_ISL_3318828, EPI_ISL_3318829, EPI_ISL_3318830, EPI_ISL_3318831, EPI_ISL_3318832, EPI_ISL_3318833, EPI_ISL_3318835, EPI_ISL_3318836, EPI_ISL_3318837, EPI_ISL_3318838 | | | |
| see above | Virginia Division of Consolidated Laboratory Services | Virginia Division of Consolidated Laboratory Services | Virginia Division of Consolidated Laboratory Services |
| EPI_ISL_3318839, EPI_ISL_3318840, EPI_ISL_3318842, EPI_ISL_3318843, EPI_ISL_3318844 | University of Michigan Clinical Microbiology Laboratory | Lauring Lab, University of Michigan, Department of Microbiology and Immunology | Valesano |
| EPI_ISL_3318846 | IVY3 Central Lab, Vanderbilt University Medical Center | Lauring Lab, University of Michigan, Department of Microbiology and Immunology | Valesano |
| EPI_ISL_3318852, EPI_ISL_3318853, EPI_ISL_3318854, EPI_ISL_3318855, EPI_ISL_3318856, EPI_ISL_3318857, EPI_ISL_3318859, EPI_ISL_3318860, EPI_ISL_3318862, EPI_ISL_3318863, EPI_ISL_3318864, EPI_ISL_3318865, EPI_ISL_3318866, EPI_ISL_3318867, EPI_ISL_3318868, EPI_ISL_3318869, EPI_ISL_3318870, EPI_ISL_3318871, EPI_ISL_3318872, EPI_ISL_3318873, EPI_ISL_3318874, EPI_ISL_3318875, EPI_ISL_3318876, EPI_ISL_3318877, EPI_ISL_3318878, EPI_ISL_3318879, EPI_ISL_3318880, EPI_ISL_3318881, EPI_ISL_3318882, EPI_ISL_3318883, EPI_ISL_3318884, EPI_ISL_3318885, EPI_ISL_3318886, EPI_ISL_3318887, EPI_ISL_3318888, EPI_ISL_3318889, EPI_ISL_3318890, EPI_ISL_3318891, EPI_ISL_3318892, EPI_ISL_3318893, EPI_ISL_3318894, EPI_ISL_3318895, EPI_ISL_3318896, EPI_ISL_3318897, EPI_ISL_3318898, EPI_ISL_3318899, EPI_ISL_3318900, EPI_ISL_3318901, EPI_ISL_3318902, EPI_ISL_3318903, EPI_ISL_3318904, EPI_ISL_3318905, EPI_ISL_3318906, EPI_ISL_3318907, EPI_ISL_3318908, EPI_ISL_3318909, EPI_ISL_3318910, EPI_ISL_3318911, EPI_ISL_3318912, EPI_ISL_3318913, EPI_ISL_3318914, EPI_ISL_3318915, EPI_ISL_3318916, EPI_ISL_3318917, EPI_ISL_3318918, EPI_ISL_3318919, EPI_ISL_3318920, EPI_ISL_3318921, EPI_ISL_3318922, EPI_ISL_3318923, EPI_ISL_3318924, EPI_ISL_3318925, EPI_ISL_3318926, EPI_ISL_3318927, EPI_ISL_3318928, EPI_ISL_3318929, EPI_ISL_3318930, EPI_ISL_3318931, EPI_ISL_3318932, EPI_ISL_3318933, EPI_ISL_3318934, EPI_ISL_3318935, EPI_ISL_3318936, EPI_ISL_3318937, EPI_ISL_3318938, EPI_ISL_3318939, EPI_ISL_3318940, EPI_ISL_3318941, EPI_ISL_3318942, EPI_ISL_3318943, EPI_ISL_3318944, EPI_ISL_3318945, EPI_ISL_3318946, EPI_ISL_3318947, EPI_ISL_3318948, EPI_ISL_3318949, EPI_ISL_3318950, EPI_ISL_3318951, EPI_ISL_3318952, EPI_ISL_3318953, EPI_ISL_3318954, EPI_ISL_3318955, EPI_ISL_3318956, EPI_ISL_3318957, EPI_ISL_3318958, EPI_ISL_3318959, EPI_ISL_3318960, EPI_ISL_3318961, EPI_ISL_3318962, EPI_ISL_3318963, EPI_ISL_3318964, EPI_ISL_3318965, EPI_ISL_3318966, EPI_ISL_3318967, EPI_ISL_3318968, EPI_ISL_3318969, EPI_ISL_3318970, EPI_ISL_3318971, EPI_ISL_3318972, EPI_ISL_3318973, EPI_ISL_3318974, EPI_ISL_3318975, EPI_ISL_3318976, EPI_ISL_3318977, EPI_ISL_3318978, EPI_ISL_3318979, EPI_ISL_3318980, EPI_ISL_3318981, EPI_ISL_3318983, EPI_ISL_3318984, EPI_ISL_3318985, EPI_ISL_3318986, EPI_ISL_3318987, EPI_ISL_3318988, EPI_ISL_3318989, EPI_ISL_3318990, EPI_ISL_3318991, EPI_ISL_3318992, EPI_ISL_3318993, EPI_ISL_3318994, EPI_ISL_3318995, EPI_ISL_3318996, EPI_ISL_3318997, EPI_ISL_3318998, EPI_ISL_3318999, EPI_ISL_3319000, EPI_ISL_3319001, EPI_ISL_3319002, EPI_ISL_3319003, EPI_ISL_3319004, EPI_ISL_3319005, EPI_ISL_3319006, EPI_ISL_3319007, EPI_ISL_3319008, EPI_ISL_3319009, EPI_ISL_3319010, EPI_ISL_3319011, EPI_ISL_3319012, EPI_ISL_3319013, EPI_ISL_3319014, EPI_ISL_3319015, EPI_ISL_3319016, EPI_ISL_3319017, EPI_ISL_3319018, EPI_ISL_3319019, EPI_ISL_3319020, EPI_ISL_3319021, EPI_ISL_3319022, EPI_ISL_3319023, EPI_ISL_3319024, EPI_ISL_3319025, EPI_ISL_3319026, EPI_ISL_3319027, EPI_ISL_3319028, EPI_ISL_3319029, EPI_ISL_3319030 | | | |
| see above | National Institute of Public Health | National Institute of Public Health | Helena Jirincova, Jaromira Vecerova, Timotej Suri, Dusan Trnka, Alexander Nagy |
| EPI_ISL_3319059, EPI_ISL_3319061, EPI_ISL_3319063, EPI_ISL_3319064, EPI_ISL_3319066 | Biolytix AG | Clinical Bacteriology | Tim Roloff, Fanny Wegner, Helena MB Seth-Smith, Alfredo Mari, Karoline Leuzinger, Julia Bielicki, Manuel Battegay, Hans Hirsch, Adrian Egli |
| EPI_ISL_3319070, EPI_ISL_3319071, EPI_ISL_3319072, EPI_ISL_3319073, EPI_ISL_3319074, EPI_ISL_3319075, EPI_ISL_3319076, EPI_ISL_3319077, EPI_ISL_3319078, EPI_ISL_3319079, EPI_ISL_3319080, EPI_ISL_3319081, EPI_ISL_3319082, EPI_ISL_3319083, EPI_ISL_3319084, EPI_ISL_3319085, EPI_ISL_3319086, EPI_ISL_3319087, EPI_ISL_3319088, EPI_ISL_3319089, EPI_ISL_3319090, EPI_ISL_3319091, EPI_ISL_3319092, EPI_ISL_3319093, EPI_ISL_3319094, EPI_ISL_3319095, EPI_ISL_3319096, EPI_ISL_3319097, EPI_ISL_3319098 | | | |
| see above | Montana Public Health Laboratory | Montana Public Health Laboratory | Joy Ritter, Michelle Mozer, Carrie Biskupiak, Deborah Gibson, Michael Dills |
| EPI_ISL_3319117 | URMC LABS | Wadsworth Center, New York State Department of Health | Kirsten St. George, Daryl M. Lamson, Alexis Russell, Matthew Shudt, Melissa A Leisner, Jonathan Plitnick, Catharine Prussing, Navjot Singh, John Kelly, Erasmus Schneider, Erica Lasek-Nesselquist |
| EPI_ISL_3319120, EPI_ISL_3319121, EPI_ISL_3319122, EPI_ISL_3319123, EPI_ISL_3319124, EPI_ISL_3319125, EPI_ISL_3319126, EPI_ISL_3319127, EPI_ISL_3319128, EPI_ISL_3319129, EPI_ISL_3319130, EPI_ISL_3319131, EPI_ISL_3319132, EPI_ISL_3319133, EPI_ISL_3319134, EPI_ISL_3319135, EPI_ISL_3319136, EPI_ISL_3319137, EPI_ISL_3319138, EPI_ISL_3319139, EPI_ISL_3319140, EPI_ISL_3319141, EPI_ISL_3319142, EPI_ISL_3319143, EPI_ISL_3319145 | | | |
| see above | The Jackson Laboratory | The Jackson Laboratory | Uvalic J, Long J, Renzette N, Adams M, Omerza G, Kelly K, Li L |
| EPI_ISL_3319146, EPI_ISL_3319147, EPI_ISL_3319148, EPI_ISL_3319149, EPI_ISL_3319150, EPI_ISL_3319151, EPI_ISL_3319152 | AULSS 3 Venezia | UOSD Genetica e Citogenetica - Azienda ULSS 3 Serenissima; Istituto Zooprofilattico Sperimentale delle Venezie | Mosé Favarato, Elisa Squarcina, Laura Squarzon, Claudia Perini, Laura Bevilacqua, Adelaide Milani, Alessia Schivo, Annalisa Salviato, Elisa Palumbo, Erika Giorgia Quaranta, Luca Tassoni, Ambra Pastori, Edoardo Giussani, Alice Fusaro, Isabella Monne, Calogero Terregino, Antonia Ricci |
| EPI_ISL_3319153, EPI_ISL_3319154, EPI_ISL_3319155, EPI_ISL_3319156, EPI_ISL_3319157, EPI_ISL_3319158, EPI_ISL_3319159, EPI_ISL_3319160, EPI_ISL_3319161, EPI_ISL_3319162, EPI_ISL_3319163, EPI_ISL_3319164, EPI_ISL_3319165, EPI_ISL_3319166, EPI_ISL_3319167, EPI_ISL_3319168, EPI_ISL_3319169, EPI_ISL_3319170, EPI_ISL_3319171, EPI_ISL_3319172, EPI_ISL_3319173, EPI_ISL_3319174, EPI_ISL_3319175, EPI_ISL_3319176, EPI_ISL_3319177 | | | |
| see above | Arizona State Public Health Laboratory | Arizona State Public Health Laboratory | Trung Huynh, Jessica Escobar, Katherine Fullerton, Nobuko Fukushima, Matthew Contursi, Stacy White, Linda Getsinger, Victor Waddell |
| EPI_ISL_3319359, EPI_ISL_3319362, EPI_ISL_3319363, EPI_ISL_3319366, EPI_ISL_3319368, EPI_ISL_3319369, EPI_ISL_3319371, EPI_ISL_3319372, EPI_ISL_3319373, EPI_ISL_3319417, EPI_ISL_3319429, EPI_ISL_3319488, EPI_ISL_3319704, EPI_ISL_3319710, EPI_ISL_3319719, EPI_ISL_3319720, EPI_ISL_3319726, EPI_ISL_3319727, EPI_ISL_3319732, EPI_ISL_3319736, EPI_ISL_3319739, EPI_ISL_3319740, | | | |

| | | | |
|---|---|---|---|
| EPI_ISL_3319741, EPI_ISL_3319742, EPI_ISL_3319744, EPI_ISL_3319746, EPI_ISL_3319748, EPI_ISL_3319751, EPI_ISL_3319753, EPI_ISL_3319755, EPI_ISL_3319786, EPI_ISL_3319788, EPI_ISL_3319789, EPI_ISL_3319791, EPI_ISL_3319792, EPI_ISL_3319793, EPI_ISL_3319795, EPI_ISL_3319799, EPI_ISL_3319800, EPI_ISL_3319801, EPI_ISL_3319802, EPI_ISL_3319803, EPI_ISL_3319804, EPI_ISL_3319806, EPI_ISL_3319807, EPI_ISL_3319808, EPI_ISL_3319810, EPI_ISL_3319811, EPI_ISL_3319812, EPI_ISL_3319813, EPI_ISL_3319814, EPI_ISL_3319815, EPI_ISL_3319816, EPI_ISL_3319817, EPI_ISL_3319818, EPI_ISL_3319819, EPI_ISL_3319820, EPI_ISL_3319821, EPI_ISL_3319822, EPI_ISL_3319823, EPI_ISL_3319824, EPI_ISL_3319825, EPI_ISL_3319826, EPI_ISL_3319827, EPI_ISL_3319828, EPI_ISL_3319830, EPI_ISL_3319831, EPI_ISL_3319832 | | | |
| see above | OHSU-MM Lab | OHSU-MM Lab | Jeannine Lama |
| EPI_ISL_3319833, EPI_ISL_3319834, EPI_ISL_3319835, EPI_ISL_3319836, EPI_ISL_3319837, EPI_ISL_3319838, EPI_ISL_3319839, EPI_ISL_3319840, EPI_ISL_3319841, EPI_ISL_3319842, EPI_ISL_3319843, EPI_ISL_3319844, EPI_ISL_3319845, EPI_ISL_3319846, EPI_ISL_3319847, EPI_ISL_3319848, EPI_ISL_3319849, EPI_ISL_3319850, EPI_ISL_3319851, EPI_ISL_3319852, EPI_ISL_3319853, EPI_ISL_3319854, EPI_ISL_3319855, EPI_ISL_3319856 | | | |
| see above | AULSS 3 Venezia | UOSD Genetica e Citogenetica - Azienda ULSS 3 Serenissima; Istituto Zooprofilattico Sperimentale delle Venezie | Mosé Favarato, Elisa Squarcina, Laura Squarzon, Claudia Perini, Laura Bevilacqua, Adelaide Milani, Alessia Schivo, Annalisa Salviato, Elisa Palumbo, Erika Giorgia Quaranta, Luca Tassoni, Ambra Pastori, Edoardo Giussani, Alice Fusaro, Isabella Monne, Calogero Terregino, Antonia Ricci |
| EPI_ISL_3319860, EPI_ISL_3319861, EPI_ISL_3319862, EPI_ISL_3319863, EPI_ISL_3319864, EPI_ISL_3319865, EPI_ISL_3319866, EPI_ISL_3319867, EPI_ISL_3319868, EPI_ISL_3319869, EPI_ISL_3319870, EPI_ISL_3319871, EPI_ISL_3319872, EPI_ISL_3319873, EPI_ISL_3319876, EPI_ISL_3319877, EPI_ISL_3319878, EPI_ISL_3319881, EPI_ISL_3319882 | | | |
| see above | Arizona State Public Health Laboratory | Arizona State Public Health Laboratory | Trung Huynh, Jessica Escobar, Katherine Fullerton, Nobuko Fukushima, Matthew Contursi, Stacy White, Linda Getsinger, Victor Waddell |
| EPI_ISL_3320128, EPI_ISL_3320129, EPI_ISL_3320130, EPI_ISL_3320131, EPI_ISL_3320132, EPI_ISL_3320133, EPI_ISL_3320134, EPI_ISL_3320135, EPI_ISL_3320136, EPI_ISL_3320137, EPI_ISL_3320138, EPI_ISL_3320139, EPI_ISL_3320140, EPI_ISL_3320141, EPI_ISL_3320142, EPI_ISL_3320143, EPI_ISL_3320144, EPI_ISL_3320145, EPI_ISL_3320146, EPI_ISL_3320147, EPI_ISL_3320148, EPI_ISL_3320149, EPI_ISL_3320150, EPI_ISL_3320151, EPI_ISL_3320154, EPI_ISL_3320155, EPI_ISL_3320156, EPI_ISL_3320157, EPI_ISL_3320158, EPI_ISL_3320159, EPI_ISL_3320160, EPI_ISL_3320161, EPI_ISL_3320162, EPI_ISL_3320163, EPI_ISL_3320164, EPI_ISL_3320165, EPI_ISL_3320166, EPI_ISL_3320167, EPI_ISL_3320168, EPI_ISL_3320169, EPI_ISL_3320170, EPI_ISL_3320171, EPI_ISL_3320172, EPI_ISL_3320173 | | | |
| see above | Tampa General Hospital Esoteric Lab | Tampa General Hospital Esoteric Research & Development Lab | Jon Faughn, Deanna Becker, Dominic Uy, Jorge Mecias-Frias, Katie Hurless, Vicki Healer, Suzane Silbert |
| EPI_ISL_3320174, EPI_ISL_3320175, EPI_ISL_3320177, EPI_ISL_3320178, EPI_ISL_3320179, EPI_ISL_3320180, EPI_ISL_3320181, EPI_ISL_3320182, EPI_ISL_3320183, EPI_ISL_3320185, EPI_ISL_3320186, EPI_ISL_3320187, EPI_ISL_3320188, EPI_ISL_3320190, EPI_ISL_3320191, EPI_ISL_3320192 | | | |
| see above | Microbiology Department, Laboratori Clínic Metropolitana Nord. Hospital Universitari Germans Trias i Pujol | Can Ruti SARS-CoV-2 Sequencing Hub (HUGTiP/IrsiCaixa/IGTP) | Marc Noguera-Julian, Pilar Armengol, Ignacio Blanco, Antoni E Bordoy, Francesc Catala-Moll, Pere-Joan Cardona, Maria Casadellà, Cristina Casañ, Gemma Clara, Bonaventura Clotet, Montserrat Giménez, Mercedes Guerrero, Anna Not, David Panisello, Roger Paredes, Mariona Parera, Alexia París, Verónica Saludes, Alba Sánchez, Laia Soler, Lauro Sumoy, and Elisa Martró on behalf of the Can Ruti SARS-CoV-2 Sequencing Hub |
| EPI_ISL_3320193 | Microbiologia CATLAB | Can Ruti SARS-CoV-2 Sequencing Hub (HUGTiP/IrsiCaixa/IGTP) | Marc Noguera-Julian, Pilar Armengol, Ignacio Blanco, Antoni E Bordoy, Francesc Catala-Moll, Pere-Joan Cardona, Maria Casadellà, Cristina Casañ, Gemma Clara, Bonaventura Clotet, Montserrat Giménez, Mercedes Guerrero, Anna Not, David Panisello, Roger Paredes, Mariona Parera, Alexia París, Verónica Saludes, Alba Sánchez, Laia Soler, Lauro Sumoy, and Elisa Martró on behalf of the Can Ruti SARS-CoV-2 Sequencing Hub |
| EPI_ISL_3320194, EPI_ISL_3320195, EPI_ISL_3320196, EPI_ISL_3320197 | Microbiology Department, Laboratori Clínic Metropolitana Nord. Hospital Universitari Germans Trias i Pujol | Can Ruti SARS-CoV-2 Sequencing Hub (HUGTiP/IrsiCaixa/IGTP) | Marc Noguera-Julian, Pilar Armengol, Ignacio Blanco, Antoni E Bordoy, Francesc Catala-Moll, Pere-Joan Cardona, Maria Casadellà, Cristina Casañ, Gemma Clara, Bonaventura Clotet, Montserrat Giménez, Mercedes Guerrero, Anna Not, David Panisello, Roger Paredes, Mariona Parera, Alexia París, Verónica Saludes, Alba Sánchez, Laia Soler, Lauro Sumoy, and Elisa Martró on behalf of the Can Ruti SARS-CoV-2 Sequencing Hub |
| EPI_ISL_3320198 | Microbiologia CATLAB | Can Ruti SARS-CoV-2 Sequencing Hub (HUGTiP/IrsiCaixa/IGTP) | Marc Noguera-Julian, Pilar Armengol, Ignacio Blanco, Antoni E Bordoy, Francesc Catala-Moll, Pere-Joan Cardona, Maria Casadellà, Cristina Casañ, Gemma Clara, Bonaventura Clotet, Montserrat Giménez, Mercedes Guerrero, Anna Not, David Panisello, Roger Paredes, Mariona Parera, Alexia París, Verónica Saludes, Alba Sánchez, Laia Soler, Lauro Sumoy, and Elisa Martró on behalf of the Can Ruti SARS-CoV-2 Sequencing Hub |
| EPI_ISL_3320199, EPI_ISL_3320200, EPI_ISL_3320203, EPI_ISL_3320204, EPI_ISL_3320205, EPI_ISL_3320206, EPI_ISL_3320207, EPI_ISL_3320208, EPI_ISL_3320209, EPI_ISL_3320210, EPI_ISL_3320211, EPI_ISL_3320212, EPI_ISL_3320213, EPI_ISL_3320214, EPI_ISL_3320215, EPI_ISL_3320216, EPI_ISL_3320218, EPI_ISL_3320219, EPI_ISL_3320220, EPI_ISL_3320221, EPI_ISL_3320222, EPI_ISL_3320223, EPI_ISL_3320224, EPI_ISL_3320225, EPI_ISL_3320226, EPI_ISL_3320227, EPI_ISL_3320228, EPI_ISL_3320229, EPI_ISL_3320230, EPI_ISL_3320231, EPI_ISL_3320232, EPI_ISL_3320233, EPI_ISL_3320234, EPI_ISL_3320235, EPI_ISL_3320236, EPI_ISL_3320237, EPI_ISL_3320238, EPI_ISL_3320239, EPI_ISL_3320240, EPI_ISL_3320243, EPI_ISL_3320244, EPI_ISL_3320245, EPI_ISL_3320246, EPI_ISL_3320247, EPI_ISL_3320248, EPI_ISL_3320249, EPI_ISL_3320250, EPI_ISL_3320251, EPI_ISL_3320252, EPI_ISL_3320253, EPI_ISL_3320254, EPI_ISL_3320255, EPI_ISL_3320256, EPI_ISL_3320257, EPI_ISL_3320258, EPI_ISL_3320259, EPI_ISL_3320260, EPI_ISL_3320261, EPI_ISL_3320262, EPI_ISL_3320264, EPI_ISL_3320265, EPI_ISL_3320266, EPI_ISL_3320267, EPI_ISL_3320268, EPI_ISL_3320269, EPI_ISL_3320270, EPI_ISL_3320271, EPI_ISL_3320272, EPI_ISL_3320273, EPI_ISL_3320274, EPI_ISL_3320275, EPI_ISL_3320276, EPI_ISL_3320277, EPI_ISL_3320278, EPI_ISL_3320280, EPI_ISL_3320281, EPI_ISL_3320282, EPI_ISL_3320283, EPI_ISL_3320284, EPI_ISL_3320285, EPI_ISL_3320287, EPI_ISL_3320288, EPI_ISL_3320289, EPI_ISL_3320290, EPI_ISL_3320291, EPI_ISL_3320292, EPI_ISL_3320294, EPI_ISL_3320295, EPI_ISL_3320296, EPI_ISL_3320297, EPI_ISL_3320299 | | | |
| see above | Microbiology Department, Laboratori Clínic Metropolitana Nord. Hospital Universitari Germans Trias i Pujol | Can Ruti SARS-CoV-2 Sequencing Hub (HUGTiP/IrsiCaixa/IGTP) | Marc Noguera-Julian, Pilar Armengol, Ignacio Blanco, Antoni E Bordoy, Francesc Catala-Moll, Pere-Joan Cardona, Maria Casadellà, Cristina Casañ, Gemma Clara, Bonaventura Clotet, Montserrat Giménez, Mercedes Guerrero, Anna Not, David Panisello, Roger Paredes, Mariona Parera, Alexia París, Verónica Saludes, Alba Sánchez, Laia Soler, Lauro Sumoy, and Elisa Martró on behalf of the Can Ruti SARS-CoV-2 Sequencing Hub |
| EPI_ISL_3320300 | AZDelta | AZ Delta Medical Laboratories in Roeselare, Belgium | Geert Martens, Dieter De Smet, Frederik Van Hoecke, on behalf of AZ Delta COVID-19 Genomics core (member of Genomic surveillance of SARS-CoV-2 in Belgium network) |
| EPI_ISL_3320301, EPI_ISL_3320302 | Microbiology Department, Laboratori Clínic Metropolitana Nord. Hospital Universitari Germans Trias i Pujol | Can Ruti SARS-CoV-2 Sequencing Hub (HUGTiP/IrsiCaixa/IGTP) | Marc Noguera-Julian, Pilar Armengol, Ignacio Blanco, Antoni E Bordoy, Francesc Catala-Moll, Pere-Joan Cardona, Maria Casadellà, Cristina Casañ, Gemma Clara, Bonaventura Clotet, Montserrat Giménez, Mercedes Guerrero, Anna Not, David Panisello, Roger Paredes, Mariona Parera, Alexia París, Verónica Saludes, Alba Sánchez, Laia Soler, Lauro Sumoy, and Elisa Martró on behalf of the Can Ruti SARS-CoV-2 Sequencing Hub |
| EPI_ISL_3320303 | AZDelta | AZ Delta Medical Laboratories in Roeselare, Belgium | Geert Martens, Dieter De Smet, Frederik Van Hoecke, on behalf of AZ Delta COVID-19 Genomics core (member of Genomic surveillance of SARS-CoV-2 in Belgium network) |
| EPI_ISL_3320304 | Microbiology Department, Laboratori Clínic Metropolitana Nord. Hospital Universitari Germans Trias i Pujol | Can Ruti SARS-CoV-2 Sequencing Hub (HUGTiP/IrsiCaixa/IGTP) | Marc Noguera-Julian, Pilar Armengol, Ignacio Blanco, Antoni E Bordoy, Francesc Catala-Moll, Pere-Joan Cardona, Maria Casadellà, Cristina Casañ, Gemma Clara, Bonaventura Clotet, Montserrat Giménez, Mercedes Guerrero, Anna Not, David Panisello, Roger Paredes, Mariona Parera, Alexia París, Verónica Saludes, Alba Sánchez, Laia Soler, Lauro Sumoy, and Elisa Martró on behalf of the Can Ruti SARS-CoV-2 Sequencing Hub |
| EPI_ISL_3320305 | AZDelta | AZ Delta Medical Laboratories in Roeselare, Belgium | Geert Martens, Dieter De Smet, Frederik Van Hoecke, on behalf of AZ Delta COVID-19 Genomics core (member of Genomic surveillance of SARS-CoV-2 in Belgium network) |
| EPI_ISL_3320306 | Microbiology Department, Laboratori Clínic Metropolitana Nord. Hospital Universitari Germans Trias i Pujol | Can Ruti SARS-CoV-2 Sequencing Hub (HUGTiP/IrsiCaixa/IGTP) | Marc Noguera-Julian, Pilar Armengol, Ignacio Blanco, Antoni E Bordoy, Francesc Catala-Moll, Pere-Joan Cardona, Maria Casadellà, Cristina Casañ, Gemma Clara, Bonaventura Clotet, Montserrat Giménez, Mercedes Guerrero, Anna Not, David Panisello, Roger Paredes, Mariona Parera, Alexia París, Verónica Saludes, Alba Sánchez, Laia Soler, Lauro Sumoy, and Elisa Martró on behalf of the Can Ruti SARS-CoV-2 Sequencing Hub |
| EPI_ISL_3320307 | AZDelta | AZ Delta Medical Laboratories in Roeselare, Belgium | Geert Martens, Dieter De Smet, Frederik Van Hoecke, on behalf of AZ Delta COVID-19 Genomics core (member of Genomic surveillance of SARS-CoV-2 in Belgium network) |
| EPI_ISL_3320308 | Microbiology Department, Laboratori Clínic Metropolitana Nord. Hospital Universitari Germans Trias i Pujol | Can Ruti SARS-CoV-2 Sequencing Hub (HUGTiP/IrsiCaixa/IGTP) | Marc Noguera-Julian, Pilar Armengol, Ignacio Blanco, Antoni E Bordoy, Francesc Catala-Moll, Pere-Joan Cardona, Maria Casadellà, Cristina Casañ, Gemma Clara, Bonaventura Clotet, Montserrat Giménez, Mercedes Guerrero, Anna Not, David Panisello, Roger Paredes, Mariona Parera, Alexia París, Verónica Saludes, Alba Sánchez, Laia Soler, Lauro Sumoy, and Elisa Martró on behalf of the Can Ruti SARS-CoV-2 Sequencing Hub |
| EPI_ISL_3320309 | AZDelta | AZ Delta Medical Laboratories in Roeselare, Belgium | Geert Martens, Dieter De Smet, Frederik Van Hoecke, on behalf of AZ Delta COVID-19 Genomics core (member of Genomic surveillance of SARS-CoV-2 in Belgium network) |
| EPI_ISL_3320310, EPI_ISL_3320311 | Microbiology Department, Laboratori Clínic Metropolitana Nord. Hospital Universitari Germans Trias i Pujol | Can Ruti SARS-CoV-2 Sequencing Hub (HUGTiP/IrsiCaixa/IGTP) | Marc Noguera-Julian, Pilar Armengol, Ignacio Blanco, Antoni E Bordoy, Francesc Catala-Moll, Pere-Joan Cardona, Maria Casadellà, Cristina Casañ, Gemma Clara, Bonaventura Clotet, Montserrat Giménez, Mercedes Guerrero, Anna Not, David Panisello, Roger Paredes, Mariona Parera, Alexia París, Verónica Saludes, Alba Sánchez, Laia Soler, Lauro Sumoy, and Elisa Martró on behalf of the Can Ruti SARS-CoV-2 Sequencing Hub |
| EPI_ISL_3320312 | AZDelta | AZ Delta Medical Laboratories in Roeselare, Belgium | Geert Martens, Dieter De Smet, Frederik Van Hoecke, on behalf of AZ Delta COVID-19 Genomics core (member of Genomic surveillance of SARS-CoV-2 in Belgium network) |
| EPI_ISL_3320313 | Microbiology Department, Laboratori Clínic Metropolitana Nord. Hospital Universitari Germans Trias i Pujol | Can Ruti SARS-CoV-2 Sequencing Hub (HUGTiP/IrsiCaixa/IGTP) | Marc Noguera-Julian, Pilar Armengol, Ignacio Blanco, Antoni E Bordoy, Francesc Catala-Moll, Pere-Joan Cardona, Maria Casadellà, Cristina Casañ, Gemma Clara, Bonaventura Clotet, Montserrat Giménez, Mercedes Guerrero, Anna Not, David Panisello, Roger Paredes, Mariona Parera, Alexia París, Verónica Saludes, Alba Sánchez, Laia Soler, Lauro Sumoy, and Elisa Martró on behalf of the Can Ruti SARS-CoV-2 Sequencing Hub |
| EPI_ISL_3320314 | AZDelta | AZ Delta Medical Laboratories in Roeselare, Belgium | Geert Martens, Dieter De Smet, Frederik Van Hoecke, on behalf of AZ Delta COVID-19 Genomics core (member of Genomic surveillance of SARS-CoV-2 in Belgium network) |
| EPI_ISL_3320315 | Microbiology Department, Laboratori Clínic Metropolitana Nord. Hospital Universitari Germans Trias i Pujol | Can Ruti SARS-CoV-2 Sequencing Hub (HUGTiP/IrsiCaixa/IGTP) | Marc Noguera-Julian, Pilar Armengol, Ignacio Blanco, Antoni E Bordoy, Francesc Catala-Moll, Pere-Joan Cardona, Maria Casadellà, Cristina Casañ, Gemma Clara, Bonaventura Clotet, Montserrat Giménez, Mercedes Guerrero, Anna Not, David Panisello, Roger Paredes, Mariona Parera, Alexia París, Verónica Saludes, Alba Sánchez, Laia Soler, Lauro Sumoy, and Elisa Martró on behalf of the Can Ruti SARS-CoV-2 Sequencing Hub |
| EPI_ISL_3320316, EPI_ISL_3320317, EPI_ISL_3320318, EPI_ISL_3320319, EPI_ISL_3320320, EPI_ISL_3320321, EPI_ISL_3320322, EPI_ISL_3320323, EPI_ISL_3320324, EPI_ISL_3320325, EPI_ISL_3320326, EPI_ISL_3320327, EPI_ISL_3320328, EPI_ISL_3320329, EPI_ISL_3320330, EPI_ISL_3320331, EPI_ISL_3320332, EPI_ISL_3320333, EPI_ISL_3320334, EPI_ISL_3320335, EPI_ISL_3320336, EPI_ISL_3320337, EPI_ISL_3320338, EPI_ISL_3320339, EPI_ISL_3320340, EPI_ISL_3320341, EPI_ISL_3320342, EPI_ISL_3320343, EPI_ISL_3320344, EPI_ISL_3320345, EPI_ISL_3320346 | | | |
| see above | AZDelta | AZ Delta Medical Laboratories in Roeselare, Belgium | Geert Martens, Dieter De Smet, Frederik Van Hoecke, on behalf of AZ Delta COVID-19 Genomics core (member of Genomic surveillance of SARS-CoV-2 in Belgium network) |
| EPI_ISL_3320349, EPI_ISL_3320350, EPI_ISL_3320351, EPI_ISL_3320352, EPI_ISL_3320353, EPI_ISL_3320354, EPI_ISL_3320355, EPI_ISL_3320356, EPI_ISL_3320358, EPI_ISL_3320359, EPI_ISL_3320360, EPI_ISL_3320361, EPI_ISL_3320362, EPI_ISL_3320363, EPI_ISL_3320364, EPI_ISL_3320365, EPI_ISL_3320366, EPI_ISL_3320367, EPI_ISL_3320368, EPI_ISL_3320369, EPI_ISL_3320370 | | | |
| see above | NJDOH, Public Health and Environmental Laboratories | NJ_PHEL | Lindsey Bodnar, Shiv K. Verma, Ryan Pachucki, Chelsea San Filippo, Jacquelyn Deverell, Dana Woell, Allison Roder, Byeong Jeong |
| EPI_ISL_3320371 | Fundacio Althaia-Manresa | Banc de Sang i Teixits | Francisco Vidal, Irene Corrales, Nina Borràs, Carlos Hobeich, Natália Comes, Lorena Ramírez, Noemí Gonzalez, Silvia Sauleda, Maria Glòria Soria |
| EPI_ISL_3320373 | CAP Folgueroles | Banc de Sang i Teixits | Francisco Vidal, Irene Corrales, Nina Borràs, Carlos Hobeich, Natália Comes, Lorena Ramírez, Noemí Gonzalez, Silvia Sauleda, Maria Glòria Soria |
| EPI_ISL_3320375 | Fundacio Althaia-Manresa | Banc de Sang i Teixits | Francisco Vidal, Irene Corrales, Nina Borràs, Carlos Hobeich, Natália Comes, Lorena Ramírez, Noemí Gonzalez, Silvia Sauleda, Maria Glòria Soria |
| EPI_ISL_3320376 | CAP Sant Quirze de Besora | Banc de Sang i Teixits | Francisco Vidal, Irene Corrales, Nina Borràs, Carlos Hobeich, Natália Comes, Lorena Ramírez, Noemí Gonzalez, Silvia Sauleda, Maria Glòria Soria |
| EPI_ISL_3320377 | CAP Vic Nord | Banc de Sang i Teixits | Francisco Vidal, Irene Corrales, Nina Borràs, Carlos Hobeich, Natália Comes, Lorena Ramírez, Noemí Gonzalez, Silvia Sauleda, Maria Glòria Soria |
| EPI_ISL_3320378 | CAP Les Hortes | Banc de Sang i Teixits | Francisco Vidal, Irene Corrales, Nina Borràs, Carlos Hobeich, Natália Comes, Lorena Ramírez, Noemí Gonzalez, Silvia Sauleda, Maria Glòria Soria |
| EPI_ISL_3320379 | CAP Tona | Banc de Sang i Teixits | Francisco Vidal, Irene Corrales, Nina Borràs, Carlos Hobeich, Natália Comes, Lorena Ramírez, Noemí Gonzalez, Silvia Sauleda, Maria Glòria Soria |
| EPI_ISL_3320380 | CAP Les Hortes | Banc de Sang i Teixits | Francisco Vidal, Irene Corrales, Nina Borràs, Carlos Hobeich, Natália Comes, Lorena Ramírez, Noemí Gonzalez, Silvia Sauleda, Maria Glòria Soria |
| EPI_ISL_3320381 | Fundacio Althaia-Manresa | Banc de Sang i Teixits | Francisco Vidal, Irene Corrales, Nina Borràs, Carlos Hobeich, Natália Comes, Lorena Ramírez, Noemí Gonzalez, Silvia Sauleda, Maria Glòria Soria |
| EPI_ISL_3320382 | CAP Les Hortes | Banc de Sang i Teixits | Francisco Vidal, Irene Corrales, Nina Borràs, Carlos Hobeich, Natália Comes, Lorena Ramírez, Noemí Gonzalez, Silvia Sauleda, Maria Glòria Soria |
| EPI_ISL_3320383 | Fundacio Althaia-Manresa | Banc de Sang i Teixits | Francisco Vidal, Irene Corrales, Nina Borràs, Carlos Hobeich, Natália Comes, Lorena Ramírez, Noemí Gonzalez, Silvia Sauleda, Maria Glòria Soria |
| EPI_ISL_3320385, EPI_ISL_3320386 | CAP Manlleu | Banc de Sang i Teixits | Francisco Vidal, Irene Corrales, Nina Borràs, Carlos Hobeich, Natália Comes, Lorena Ramírez, Noemí Gonzalez, Silvia Sauleda, Maria Glòria Soria |
| EPI_ISL_3320387 | CAP El Remei | Banc de Sang i Teixits | Francisco Vidal, Irene Corrales, Nina Borràs, Carlos Hobeich, Natália Comes, Lorena Ramírez, Noemí Gonzalez, Silvia Sauleda, Maria Glòria Soria |
| EPI_ISL_3320388 | Fundacio Althaia-Manresa | Banc de Sang i Teixits | Francisco Vidal, Irene Corrales, Nina Borràs, Carlos Hobeich, Natália Comes, Lorena Ramírez, Noemí Gonzalez, Silvia Sauleda, Maria Glòria Soria |
| EPI_ISL_3320389 | CAP El Remei | Banc de Sang i Teixits | Francisco Vidal, Irene Corrales, Nina Borràs, Carlos Hobeich, Natália Comes, Lorena Ramírez, Noemí Gonzalez, Silvia Sauleda, Maria Glòria Soria |
| EPI_ISL_3320390 | CAP Les Hortes | Banc de Sang i Teixits | Francisco Vidal, Irene Corrales, Nina Borràs, Carlos Hobeich, Natália Comes, Lorena Ramírez, Noemí Gonzalez, Silvia Sauleda, Maria Glòria Soria |
| EPI_ISL_3320391, EPI_ISL_3320392 | Fundacio Althaia-Manresa | Banc de Sang i Teixits | Francisco Vidal, Irene Corrales, Nina Borràs, Carlos Hobeich, Natália Comes, Lorena Ramírez, Noemí Gonzalez, Silvia Sauleda, Maria Glòria Soria |
| EPI_ISL_3320393 | CAP El Remei | Banc de Sang i Teixits | Francisco Vidal, Irene Corrales, Nina Borràs, Carlos Hobeich, Natália Comes, Lorena Ramírez, Noemí Gonzalez, Silvia Sauleda, Maria Glòria Soria |
| EPI_ISL_3320394, EPI_ISL_3320395 | Fundacio Althaia-Manresa | Banc de Sang i Teixits | Francisco Vidal, Irene Corrales, Nina Borràs, Carlos Hobeich, Natália Comes, Lorena Ramírez, Noemí Gonzalez, Silvia Sauleda, Maria Glòria Soria |
| EPI_ISL_3320396 | CAP El Remei | Banc de Sang i Teixits | Francisco Vidal, Irene Corrales, Nina Borràs, Carlos Hobeich, Natália Comes, Lorena Ramírez, Noemí Gonzalez, Silvia Sauleda, Maria Glòria Soria |
| EPI_ISL_3320397 | CAP Les Hortes | Banc de Sang i Teixits | Francisco Vidal, Irene Corrales, Nina Borràs, Carlos Hobeich, Natália Comes, Lorena Ramírez, Noemí Gonzalez, Silvia Sauleda, Maria Glòria Soria |
| EPI_ISL_3320398 | CAP Roda de Ter | Banc de Sang i Teixits | Francisco Vidal, Irene Corrales, Nina Borràs, Carlos Hobeich, Natália Comes, Lorena Ramírez, Noemí Gonzalez, Silvia Sauleda, Maria Glòria Soria |
| EPI_ISL_3320399 | CAP Vic Nord | Banc de Sang i Teixits | Francisco Vidal, Irene Corrales, Nina Borràs, Carlos Hobeich, Natália Comes, Lorena Ramírez, Noemí Gonzalez, Silvia Sauleda, Maria Glòria Soria |
| EPI_ISL_3320400 | CAP Roger de Flor | Banc de Sang i Teixits | Francisco Vidal, Irene Corrales, Nina Borràs, Carlos Hobeich, Natália Comes, Lorena Ramírez, Noemí Gonzalez, Silvia Sauleda, Maria Glòria Soria |
| EPI_ISL_3320401, EPI_ISL_3320402, EPI_ISL_3320403, EPI_ISL_3320404, EPI_ISL_3320405, EPI_ISL_3320406, EPI_ISL_3320407, EPI_ISL_3320408, EPI_ISL_3320409, EPI_ISL_3320410, EPI_ISL_3320411, EPI_ISL_3320412, EPI_ISL_3320413, EPI_ISL_3320414, EPI_ISL_3320415, EPI_ISL_3320417, EPI_ISL_3320418, EPI_ISL_3320419, EPI_ISL_3320420, EPI_ISL_3320421, EPI_ISL_3320422, EPI_ISL_3320423, | | | |

| | | | |
|---|---|---|---|
| EPI_ISL_3320424, EPI_ISL_3320425, EPI_ISL_3320426, EPI_ISL_3320427, EPI_ISL_3320428, EPI_ISL_3320429, EPI_ISL_3320430, EPI_ISL_3320431, EPI_ISL_3320432, EPI_ISL_3320433, EPI_ISL_3320434, EPI_ISL_3320435, EPI_ISL_3320436, EPI_ISL_3320437, EPI_ISL_3320438, EPI_ISL_3320439, EPI_ISL_3320440, EPI_ISL_3320441, EPI_ISL_3320442, EPI_ISL_3320443, EPI_ISL_3320444, EPI_ISL_3320445, EPI_ISL_3320446, EPI_ISL_3320447, EPI_ISL_3320448, EPI_ISL_3320449, EPI_ISL_3320450, EPI_ISL_3320451, EPI_ISL_3320452, EPI_ISL_3320453, EPI_ISL_3320454, EPI_ISL_3320455, EPI_ISL_3320456, EPI_ISL_3320457, EPI_ISL_3320458 | | | |
| see above | Kansas Health and Environmental Lab | Kansas Health and Environmental Lab | Mike Grose, Jonathan Barnell, Amanda Bradley, and Phil Adam |
| EPI_ISL_3320459, EPI_ISL_3320460, EPI_ISL_3320461 | Marian Medical Center Laboratory | County of San Luis Obispo Public Health Laboratory | Katrina Erwin, Frances Sidhu |
| EPI_ISL_3320462, EPI_ISL_3320463, EPI_ISL_3320464, EPI_ISL_3320465 | Sierra Vista Regional Medical Center Laboratory | County of San Luis Obispo Public Health Laboratory | Katrina Erwin, Frances Sidhu |
| EPI_ISL_3320466 | Twin Cities Community Hospital Laboratory | County of San Luis Obispo Public Health Laboratory | Katrina Erwin, Frances Sidhu |
| EPI_ISL_3320467, EPI_ISL_3320468, EPI_ISL_3320469, EPI_ISL_3320470, EPI_ISL_3320471, EPI_ISL_3320472, EPI_ISL_3320473, EPI_ISL_3320474, EPI_ISL_3320475 | Sierra Vista Regional Medical Center Laboratory | County of San Luis Obispo Public Health Laboratory | Katrina Erwin, Frances Sidhu |
| EPI_ISL_3320476 | French Hospital Laboratory | County of San Luis Obispo Public Health Laboratory | Katrina Erwin, Frances Sidhu |
| EPI_ISL_3320477 | San Luis Obispo Public Health Department | County of San Luis Obispo Public Health Laboratory | Katrina Erwin, Frances Sidhu |
| EPI_ISL_3320478, EPI_ISL_3320479 | Cal Poly University Health Center | County of San Luis Obispo Public Health Laboratory | Katrina Erwin, Frances Sidhu |
| EPI_ISL_3320480, EPI_ISL_3320481, EPI_ISL_3320482 | Marian Medical Center Laboratory | County of San Luis Obispo Public Health Laboratory | Katrina Erwin, Frances Sidhu |
| EPI_ISL_3320483, EPI_ISL_3320484, EPI_ISL_3320485, EPI_ISL_3320486, EPI_ISL_3320487, EPI_ISL_3320488 | Platform BIS UZA/UAntwerpen | Labo Klinische Biologie, UZA | Marie Le Mercier, Jasmine Coppens, Kathleen Holemans, Ines Verbesselt, Basil Britto Xavier, Christine Lammens, Veerle Matheeussen, Herman Goossens |
| EPI_ISL_3320489 | Algemeen Klinisch Labo | Labo Klinische Biologie, UZA | Marie Le Mercier, Jasmine Coppens, Kathleen Holemans, Ines Verbesselt, Basil Britto Xavier, Christine Lammens, Veerle Matheeussen, Herman Goossens |
| EPI_ISL_3320491, EPI_ISL_3320492 | Platform BIS UZA/UAntwerpen | Labo Klinische Biologie, UZA | Marie Le Mercier, Jasmine Coppens, Kathleen Holemans, Ines Verbesselt, Basil Britto Xavier, Christine Lammens, Veerle Matheeussen, Herman Goossens |
| EPI_ISL_3320493 | Centrum voor Medische Analyse | Labo Klinische Biologie, UZA | Marie Le Mercier, Jasmine Coppens, Kathleen Holemans, Ines Verbesselt, Basil Britto Xavier, Christine Lammens, Veerle Matheeussen, Herman Goossens |
| EPI_ISL_3320494, EPI_ISL_3320495, EPI_ISL_3320496, EPI_ISL_3320497, EPI_ISL_3320498 | Platform BIS UZA/UAntwerpen | Labo Klinische Biologie, UZA | Marie Le Mercier, Jasmine Coppens, Kathleen Holemans, Ines Verbesselt, Basil Britto Xavier, Christine Lammens, Veerle Matheeussen, Herman Goossens |
| EPI_ISL_3320499 | Centrum voor Medische Analyse | Labo Klinische Biologie, UZA | Marie Le Mercier, Jasmine Coppens, Kathleen Holemans, Ines Verbesselt, Basil Britto Xavier, Christine Lammens, Veerle Matheeussen, Herman Goossens |
| EPI_ISL_3320500 | Platform BIS UZA/UAntwerpen | Labo Klinische Biologie, UZA | Marie Le Mercier, Jasmine Coppens, Kathleen Holemans, Ines Verbesselt, Basil Britto Xavier, Christine Lammens, Veerle Matheeussen, Herman Goossens |
| EPI_ISL_3320501 | Centrum voor Medische Analyse | Labo Klinische Biologie, UZA | Marie Le Mercier, Jasmine Coppens, Kathleen Holemans, Ines Verbesselt, Basil Britto Xavier, Christine Lammens, Veerle Matheeussen, Herman Goossens |
| EPI_ISL_3320502 | Platform BIS UZA/UAntwerpen | Labo Klinische Biologie, UZA | Marie Le Mercier, Jasmine Coppens, Kathleen Holemans, Ines Verbesselt, Basil Britto Xavier, Christine Lammens, Veerle Matheeussen, Herman Goossens |
| EPI_ISL_3320503 | Centrum voor Medische Analyse | Labo Klinische Biologie, UZA | Marie Le Mercier, Jasmine Coppens, Kathleen Holemans, Ines Verbesselt, Basil Britto Xavier, Christine Lammens, Veerle Matheeussen, Herman Goossens |
| EPI_ISL_3320504, EPI_ISL_3320505 | Platform BIS UZA/UAntwerpen | Labo Klinische Biologie, UZA | Marie Le Mercier, Jasmine Coppens, Kathleen Holemans, Ines Verbesselt, Basil Britto Xavier, Christine Lammens, Veerle Matheeussen, Herman Goossens |
| EPI_ISL_3320506, EPI_ISL_3320507, EPI_ISL_3320508, EPI_ISL_3320509, EPI_ISL_3320510, EPI_ISL_3320511, EPI_ISL_3320512, EPI_ISL_3320513, EPI_ISL_3320514, EPI_ISL_3320515, EPI_ISL_3320516, EPI_ISL_3320517, EPI_ISL_3320518 | | | |
| see above | Centrum voor Medische Analyse | Labo Klinische Biologie, UZA | Marie Le Mercier, Jasmine Coppens, Kathleen Holemans, Ines Verbesselt, Basil Britto Xavier, Christine Lammens, Veerle Matheeussen, Herman Goossens |
| EPI_ISL_3320519 | Labo Klinische Biologie, UZA | Labo Klinische Biologie, UZA | Marie Le Mercier, Jasmine Coppens, Kathleen Holemans, Ines Verbesselt, Basil Britto Xavier, Christine Lammens, Veerle Matheeussen, Herman Goossens |
| EPI_ISL_3320520 | LABO LKO - LMC | Labo Klinische Biologie, UZA | Marie Le Mercier, Jasmine Coppens, Kathleen Holemans, Ines Verbesselt, Basil Britto Xavier, Christine Lammens, Veerle Matheeussen, Herman Goossens |
| EPI_ISL_3320521, EPI_ISL_3320522, EPI_ISL_3320523, EPI_ISL_3320524, EPI_ISL_3320525, EPI_ISL_3320526, EPI_ISL_3320527, EPI_ISL_3320528, EPI_ISL_3320529, EPI_ISL_3320530, EPI_ISL_3320531, EPI_ISL_3320532, EPI_ISL_3320533, EPI_ISL_3320534, EPI_ISL_3320535, EPI_ISL_3320536, EPI_ISL_3320537, EPI_ISL_3320538, EPI_ISL_3320539, EPI_ISL_3320540, EPI_ISL_3320541, EPI_ISL_3320542, EPI_ISL_3320543, EPI_ISL_3320544, EPI_ISL_3320545, EPI_ISL_3320546, EPI_ISL_3320547, EPI_ISL_3320548, EPI_ISL_3320549, EPI_ISL_3320550, EPI_ISL_3320551, EPI_ISL_3320552, EPI_ISL_3320553, EPI_ISL_3320554, EPI_ISL_3320555, EPI_ISL_3320556, EPI_ISL_3320557, EPI_ISL_3320558, EPI_ISL_3320559, EPI_ISL_3320560, EPI_ISL_3320561, EPI_ISL_3320562, EPI_ISL_3320563, EPI_ISL_3320564, EPI_ISL_3320565, EPI_ISL_3320566, EPI_ISL_3320567, EPI_ISL_3320568, EPI_ISL_3320569, EPI_ISL_3320570, EPI_ISL_3320571, EPI_ISL_3320572, EPI_ISL_3320573, EPI_ISL_3320574, EPI_ISL_3320575, EPI_ISL_3320576, EPI_ISL_3320577, EPI_ISL_3320578, EPI_ISL_3320579, EPI_ISL_3320580, EPI_ISL_3320581, EPI_ISL_3320582, EPI_ISL_3320583, EPI_ISL_3320584, EPI_ISL_3320585, EPI_ISL_3320586, EPI_ISL_3320587, EPI_ISL_3320588, EPI_ISL_3320589, EPI_ISL_3320590, EPI_ISL_3320591, EPI_ISL_3320592, EPI_ISL_3320593, EPI_ISL_3320594, EPI_ISL_3320595, EPI_ISL_3320596, EPI_ISL_3320597, EPI_ISL_3320598, EPI_ISL_3320599 | | | |
| see above | Platform BIS UZA/UAntwerpen | Labo Klinische Biologie, UZA | Marie Le Mercier, Jasmine Coppens, Kathleen Holemans, Ines Verbesselt, Basil Britto Xavier, Christine Lammens, Veerle Matheeussen, Herman Goossens |
| EPI_ISL_3320600, EPI_ISL_3320601, EPI_ISL_3320602, EPI_ISL_3320603, EPI_ISL_3320604, EPI_ISL_3320605, EPI_ISL_3320606, EPI_ISL_3320607, EPI_ISL_3320608, EPI_ISL_3320609, EPI_ISL_3320610, EPI_ISL_3320611, EPI_ISL_3320612, EPI_ISL_3320613, EPI_ISL_3320614, EPI_ISL_3320615, EPI_ISL_3320616, EPI_ISL_3320617, EPI_ISL_3320618, EPI_ISL_3320619, EPI_ISL_3320620, EPI_ISL_3320621, EPI_ISL_3320622, EPI_ISL_3320623, EPI_ISL_3320624, EPI_ISL_3320625, EPI_ISL_3320626, EPI_ISL_3320627, EPI_ISL_3320628, EPI_ISL_3320629, EPI_ISL_3320630, EPI_ISL_3320631, EPI_ISL_3320632, EPI_ISL_3320633, EPI_ISL_3320634, EPI_ISL_3320635, EPI_ISL_3320636 | | | |
| see above | Oregon State Public Health Laboratory | Oregon State Public Health Laboratory | Rafia Razzaque, Eugene Yeboah, Vanda Makris, Laura Tsaknaridis, John Fontana and Shane Sevey |
| EPI_ISL_3320638, EPI_ISL_3320639, EPI_ISL_3320640, EPI_ISL_3320641, EPI_ISL_3320642, EPI_ISL_3320643, EPI_ISL_3320644, EPI_ISL_3320645, EPI_ISL_3320646, EPI_ISL_3320647, EPI_ISL_3320648, EPI_ISL_3320649 | | | |
| see above | UCSC Genomics Institute | UCSC Genomics Institute | Mark Akeson, Ikenna Anigbogu, Eric Beraut, Isabel Bjork, Molly Cassatt-Johnstone, Terren Chang, Russell Corbett-Detig, Namrita Dhillon, Maximilian Haeussler, David Haussler, Angie Hinrichs, Miten Jain, Joshua Kapp, A. Marm Kilpatrick, Jakob McBroome, Hugh Olsen, Jeremy Sanford, Beth Shapiro, Michael Stone, Bryan Thornlow, Yatish Turakhia, Ciara Wanket |
| EPI_ISL_3320651, EPI_ISL_3320652, EPI_ISL_3320653, EPI_ISL_3320654, EPI_ISL_3320655, EPI_ISL_3320656, EPI_ISL_3320657, EPI_ISL_3320658, EPI_ISL_3320659, EPI_ISL_3320660, EPI_ISL_3320661, EPI_ISL_3320662, EPI_ISL_3320663, EPI_ISL_3320664, EPI_ISL_3320665, EPI_ISL_3320666, EPI_ISL_3320667, EPI_ISL_3320668, EPI_ISL_3320669, EPI_ISL_3320670, EPI_ISL_3320671, EPI_ISL_3320672, EPI_ISL_3320673, EPI_ISL_3320674, EPI_ISL_3320675, EPI_ISL_3320676, EPI_ISL_3320677, EPI_ISL_3320679, EPI_ISL_3320680, EPI_ISL_3320681, EPI_ISL_3320682, EPI_ISL_3320683, EPI_ISL_3320684, EPI_ISL_3320686, EPI_ISL_3320687, EPI_ISL_3320688, EPI_ISL_3320689, EPI_ISL_3320690, EPI_ISL_3320692, EPI_ISL_3320693, EPI_ISL_3320694, EPI_ISL_3320695, EPI_ISL_3320696, EPI_ISL_3320697, EPI_ISL_3320698, EPI_ISL_3320699, EPI_ISL_3320700, EPI_ISL_3320701, EPI_ISL_3320702, EPI_ISL_3320703, EPI_ISL_3320704, EPI_ISL_3320705, EPI_ISL_3320706, EPI_ISL_3320707, EPI_ISL_3320708, EPI_ISL_3320709, EPI_ISL_3320710, EPI_ISL_3320711, EPI_ISL_3320712, EPI_ISL_3320713, EPI_ISL_3320714, EPI_ISL_3320715, EPI_ISL_3320716 | | | |
| see above | CT Department of Public Health | CT Department of Public Health | Claire_Pearson,Tu_N_Nguyen |
| EPI_ISL_3320754 | Furst Medical Laboratory | Norwegian Institute of Public Health, Department of Virology | Kathrine Stene-Johansen, Kamilla Heddeland Instefjord, Hilde Elshaug, Garcia Llorente Ignacio, Jon Bråte, Engebretsen Serina Beate,Pedersen Benedikte Nevjen, Line Victoria Moen, Debech Nadia, Atiya R Ali,Marie Paulsen Madsen, Rasmus Riis Kopperud, Hilde Vollan, Karoline Bragstad, Olav Hungnes |
| EPI_ISL_3320756 | Vestfold Hospital, Toensberg Department of Microbiology | Norwegian Institute of Public Health, Department of Virology | Kathrine Stene-Johansen, Kamilla Heddeland Instefjord, Hilde Elshaug, Garcia Llorente Ignacio, Jon Bråte, Engebretsen Serina Beate,Pedersen Benedikte Nevjen, Line Victoria Moen, Debech Nadia, Atiya R Ali,Marie Paulsen Madsen, Rasmus Riis Kopperud, Hilde Vollan, Karoline Bragstad, Olav Hungnes |
| EPI_ISL_3320757 | Furst Medical Laboratory | Norwegian Institute of Public Health, Department of Virology | Kathrine Stene-Johansen, Kamilla Heddeland Instefjord, Hilde Elshaug, Garcia Llorente Ignacio, Jon Bråte, Engebretsen Serina Beate,Pedersen Benedikte Nevjen, Line Victoria Moen, Debech Nadia, Atiya R Ali,Marie Paulsen Madsen, Rasmus Riis Kopperud, Hilde Vollan, Karoline Bragstad, Olav Hungnes |
| EPI_ISL_3320758 | Department of Medical Microbiology - section Molde, Molde Hospital | Norwegian Institute of Public Health, Department of Virology | Kathrine Stene-Johansen, Kamilla Heddeland Instefjord, Hilde Elshaug, Garcia Llorente Ignacio, Jon Bråte, Engebretsen Serina Beate,Pedersen Benedikte Nevjen, Line Victoria Moen, Debech Nadia, Atiya R Ali,Marie Paulsen Madsen, Rasmus Riis Kopperud, Hilde Vollan, Karoline Bragstad, Olav Hungnes |
| EPI_ISL_3320759 | Vestfold Hospital, Toensberg Department of Microbiology | Norwegian Institute of Public Health, Department of Virology | Kathrine Stene-Johansen, Kamilla Heddeland Instefjord, Hilde Elshaug, Garcia Llorente Ignacio, Jon Bråte, Engebretsen Serina Beate,Pedersen Benedikte Nevjen, Line Victoria Moen, Debech Nadia, Atiya R Ali,Marie Paulsen Madsen, Rasmus Riis Kopperud, Hilde Vollan, Karoline Bragstad, Olav Hungnes |
| EPI_ISL_3320760 | Furst Medical Laboratory | Norwegian Institute of Public Health, Department of Virology | Kathrine Stene-Johansen, Kamilla Heddeland Instefjord, Hilde Elshaug, Garcia Llorente Ignacio, Jon Bråte, Engebretsen Serina Beate,Pedersen Benedikte Nevjen, Line Victoria Moen, Debech Nadia, Atiya R Ali,Marie Paulsen Madsen, Rasmus Riis Kopperud, Hilde Vollan, Karoline Bragstad, Olav Hungnes |
| EPI_ISL_3320761, EPI_ISL_3320765 | Department of Medical Microbiology - section Molde, Molde Hospital | Norwegian Institute of Public Health, Department of Virology | Kathrine Stene-Johansen, Kamilla Heddeland Instefjord, Hilde Elshaug, Garcia Llorente Ignacio, Jon Bråte, Engebretsen Serina Beate,Pedersen Benedikte Nevjen, Line Victoria Moen, Debech Nadia, Atiya R Ali,Marie Paulsen Madsen, Rasmus Riis Kopperud, Hilde Vollan, Karoline Bragstad, Olav Hungnes |
| EPI_ISL_3320766 | Vestfold Hospital, Toensberg Department of Microbiology | Norwegian Institute of Public Health, Department of Virology | Kathrine Stene-Johansen, Kamilla Heddeland Instefjord, Hilde Elshaug, Garcia Llorente Ignacio, Jon Bråte, Engebretsen Serina Beate,Pedersen Benedikte Nevjen, Line Victoria Moen, Debech Nadia, Atiya R Ali,Marie Paulsen Madsen, Rasmus Riis Kopperud, Hilde Vollan, Karoline Bragstad, Olav Hungnes |
| EPI_ISL_3320767, EPI_ISL_3320768, EPI_ISL_3320769 | Furst Medical Laboratory | Norwegian Institute of Public Health, Department of Virology | Kathrine Stene-Johansen, Kamilla Heddeland Instefjord, Hilde Elshaug, Garcia Llorente Ignacio, Jon Bråte, Engebretsen Serina Beate,Pedersen Benedikte Nevjen, Line Victoria Moen, Debech Nadia, Atiya R Ali,Marie Paulsen Madsen, Rasmus Riis Kopperud, Hilde Vollan, Karoline Bragstad, Olav Hungnes |
| EPI_ISL_3320771 | Vestfold Hospital, Toensberg Department of Microbiology | Norwegian Institute of Public Health, Department of Virology | Kathrine Stene-Johansen, Kamilla Heddeland Instefjord, Hilde Elshaug, Garcia Llorente Ignacio, Jon Bråte, Engebretsen Serina Beate,Pedersen Benedikte Nevjen, Line Victoria Moen, Debech Nadia, Atiya R Ali,Marie Paulsen Madsen, Rasmus Riis Kopperud, Hilde Vollan, Karoline Bragstad, Olav Hungnes |
| EPI_ISL_3320772 | Furst Medical Laboratory | Norwegian Institute of Public Health, Department of Virology | Kathrine Stene-Johansen, Kamilla Heddeland Instefjord, Hilde Elshaug, Garcia Llorente Ignacio, Jon Bråte, Engebretsen Serina Beate,Pedersen Benedikte Nevjen, Line Victoria Moen, Debech Nadia, Atiya R Ali,Marie Paulsen Madsen, Rasmus Riis Kopperud, Hilde Vollan, Karoline Bragstad, Olav Hungnes |

| | | | |
|---|---|---|---|
| EPI_ISL_3320773 | Vestfold Hospital, Toensberg Department of Microbiology | Norwegian Institute of Public Health, Department of Virology | Kathrine Stene-Johansen, Kamilla Heddeland Instefjord, Hilde Elshaug, Garcia Llorente Ignacio, Jon Bråte, Engebretsen Serina Beate,Pedersen Benedikte Nevjen, Line Victoria Moen, Debech Nadia, Atiya R Ali,Marie Paulsen Madsen, Rasmus Riis Kopperud, Hilde Vollan, Karoline Bragstad, Olav Hungnes |
| EPI_ISL_3320779 | Medical Microbiology Unit, Department for Laboratory Medicine, Drammen Hospital, Vestre Viken Health Trust, | Norwegian Institute of Public Health, Department of Virology | Kathrine Stene-Johansen, Kamilla Heddeland Instefjord, Hilde Elshaug, Garcia Llorente Ignacio, Jon Bråte, Engebretsen Serina Beate,Pedersen Benedikte Nevjen, Line Victoria Moen, Debech Nadia, Atiya R Ali,Marie Paulsen Madsen, Rasmus Riis Kopperud, Hilde Vollan, Karoline Bragstad, Olav Hungnes |
| EPI_ISL_3320781, EPI_ISL_3320782, EPI_ISL_3320783 | Furst Medical Laboratory | Norwegian Institute of Public Health, Department of Virology | Kathrine Stene-Johansen, Kamilla Heddeland Instefjord, Hilde Elshaug, Garcia Llorente Ignacio, Jon Bråte, Engebretsen Serina Beate,Pedersen Benedikte Nevjen, Line Victoria Moen, Debech Nadia, Atiya R Ali,Marie Paulsen Madsen, Rasmus Riis Kopperud, Hilde Vollan, Karoline Bragstad, Olav Hungnes |
| EPI_ISL_3320785 | Vestfold Hospital, Toensberg Department of Microbiology | Norwegian Institute of Public Health, Department of Virology | Kathrine Stene-Johansen, Kamilla Heddeland Instefjord, Hilde Elshaug, Garcia Llorente Ignacio, Jon Bråte, Engebretsen Serina Beate,Pedersen Benedikte Nevjen, Line Victoria Moen, Debech Nadia, Atiya R Ali,Marie Paulsen Madsen, Rasmus Riis Kopperud, Hilde Vollan, Karoline Bragstad, Olav Hungnes |
| EPI_ISL_3320787 | Department of Medical Microbiology - section Molde, Molde Hospital | Norwegian Institute of Public Health, Department of Virology | Kathrine Stene-Johansen, Kamilla Heddeland Instefjord, Hilde Elshaug, Garcia Llorente Ignacio, Jon Bråte, Engebretsen Serina Beate,Pedersen Benedikte Nevjen, Line Victoria Moen, Debech Nadia, Atiya R Ali,Marie Paulsen Madsen, Rasmus Riis Kopperud, Hilde Vollan, Karoline Bragstad, Olav Hungnes |
| EPI_ISL_3320788 | Medical Microbiology Unit, Department for Laboratory Medicine, Drammen Hospital, Vestre Viken Health Trust, | Norwegian Institute of Public Health, Department of Virology | Kathrine Stene-Johansen, Kamilla Heddeland Instefjord, Hilde Elshaug, Garcia Llorente Ignacio, Jon Bråte, Engebretsen Serina Beate,Pedersen Benedikte Nevjen, Line Victoria Moen, Debech Nadia, Atiya R Ali,Marie Paulsen Madsen, Rasmus Riis Kopperud, Hilde Vollan, Karoline Bragstad, Olav Hungnes |
| EPI_ISL_3320789 | Vestfold Hospital, Toensberg Department of Microbiology | Norwegian Institute of Public Health, Department of Virology | Kathrine Stene-Johansen, Kamilla Heddeland Instefjord, Hilde Elshaug, Garcia Llorente Ignacio, Jon Bråte, Engebretsen Serina Beate,Pedersen Benedikte Nevjen, Line Victoria Moen, Debech Nadia, Atiya R Ali,Marie Paulsen Madsen, Rasmus Riis Kopperud, Hilde Vollan, Karoline Bragstad, Olav Hungnes |
| EPI_ISL_3320790 | Medical Microbiology Unit, Department for Laboratory Medicine, Drammen Hospital, Vestre Viken Health Trust, | Norwegian Institute of Public Health, Department of Virology | Kathrine Stene-Johansen, Kamilla Heddeland Instefjord, Hilde Elshaug, Garcia Llorente Ignacio, Jon Bråte, Engebretsen Serina Beate,Pedersen Benedikte Nevjen, Line Victoria Moen, Debech Nadia, Atiya R Ali,Marie Paulsen Madsen, Rasmus Riis Kopperud, Hilde Vollan, Karoline Bragstad, Olav Hungnes |
| EPI_ISL_3320791 | Vestfold Hospital, Toensberg Department of Microbiology | Norwegian Institute of Public Health, Department of Virology | Kathrine Stene-Johansen, Kamilla Heddeland Instefjord, Hilde Elshaug, Garcia Llorente Ignacio, Jon Bråte, Engebretsen Serina Beate,Pedersen Benedikte Nevjen, Line Victoria Moen, Debech Nadia, Atiya R Ali,Marie Paulsen Madsen, Rasmus Riis Kopperud, Hilde Vollan, Karoline Bragstad, Olav Hungnes |
| EPI_ISL_3320793 | Unilabs Laboratory Medicine | Norwegian Institute of Public Health, Department of Virology | Kathrine Stene-Johansen, Kamilla Heddeland Instefjord, Hilde Elshaug, Garcia Llorente Ignacio, Jon Bråte, Engebretsen Serina Beate,Pedersen Benedikte Nevjen, Line Victoria Moen, Debech Nadia, Atiya R Ali,Marie Paulsen Madsen, Rasmus Riis Kopperud, Hilde Vollan, Karoline Bragstad, Olav Hungnes |
| EPI_ISL_3320794, EPI_ISL_3320795 | Vestfold Hospital, Toensberg Department of Microbiology | Norwegian Institute of Public Health, Department of Virology | Kathrine Stene-Johansen, Kamilla Heddeland Instefjord, Hilde Elshaug, Garcia Llorente Ignacio, Jon Bråte, Engebretsen Serina Beate,Pedersen Benedikte Nevjen, Line Victoria Moen, Debech Nadia, Atiya R Ali,Marie Paulsen Madsen, Rasmus Riis Kopperud, Hilde Vollan, Karoline Bragstad, Olav Hungnes |
| EPI_ISL_3320796 | Unilabs Laboratory Medicine | Norwegian Institute of Public Health, Department of Virology | Kathrine Stene-Johansen, Kamilla Heddeland Instefjord, Hilde Elshaug, Garcia Llorente Ignacio, Jon Bråte, Engebretsen Serina Beate,Pedersen Benedikte Nevjen, Line Victoria Moen, Debech Nadia, Atiya R Ali,Marie Paulsen Madsen, Rasmus Riis Kopperud, Hilde Vollan, Karoline Bragstad, Olav Hungnes |
| EPI_ISL_3320814 | Department of Medical Microbiology - section Molde, Molde Hospital | Norwegian Institute of Public Health, Department of Virology | Kathrine Stene-Johansen, Kamilla Heddeland Instefjord, Hilde Elshaug, Garcia Llorente Ignacio, Jon Bråte, Engebretsen Serina Beate,Pedersen Benedikte Nevjen, Line Victoria Moen, Debech Nadia, Atiya R Ali,Marie Paulsen Madsen, Rasmus Riis Kopperud, Hilde Vollan, Karoline Bragstad, Olav Hungnes |
| EPI_ISL_3320816 | Nordland Hospital - Bodo, Laboratory Department, Molecular Biology Unit | Norwegian Institute of Public Health, Department of Virology | Kathrine Stene-Johansen, Kamilla Heddeland Instefjord, Hilde Elshaug, Garcia Llorente Ignacio, Jon Bråte, Engebretsen Serina Beate,Pedersen Benedikte Nevjen, Line Victoria Moen, Debech Nadia, Atiya R Ali,Marie Paulsen Madsen, Rasmus Riis Kopperud, Hilde Vollan, Karoline Bragstad, Olav Hungnes |
| EPI_ISL_3320817, EPI_ISL_3320818 | Haukeland University Hospital, Dept. of Microbiology | Norwegian Institute of Public Health, Department of Virology | Kathrine Stene-Johansen, Kamilla Heddeland Instefjord, Hilde Elshaug, Garcia Llorente Ignacio, Jon Bråte, Engebretsen Serina Beate,Pedersen Benedikte Nevjen, Line Victoria Moen, Debech Nadia, Atiya R Ali,Marie Paulsen Madsen, Rasmus Riis Kopperud, Hilde Vollan, Karoline Bragstad, Olav Hungnes |
| EPI_ISL_3320819 | Medical Microbiology Unit, Department for Laboratory Medicine, Drammen Hospital, Vestre Viken Health Trust, | Norwegian Institute of Public Health, Department of Virology | Kathrine Stene-Johansen, Kamilla Heddeland Instefjord, Hilde Elshaug, Garcia Llorente Ignacio, Jon Bråte, Engebretsen Serina Beate,Pedersen Benedikte Nevjen, Line Victoria Moen, Debech Nadia, Atiya R Ali,Marie Paulsen Madsen, Rasmus Riis Kopperud, Hilde Vollan, Karoline Bragstad, Olav Hungnes |
| EPI_ISL_3320820 | Haukeland University Hospital, Dept. of Microbiology | Norwegian Institute of Public Health, Department of Virology | Kathrine Stene-Johansen, Kamilla Heddeland Instefjord, Hilde Elshaug, Garcia Llorente Ignacio, Jon Bråte, Engebretsen Serina Beate,Pedersen Benedikte Nevjen, Line Victoria Moen, Debech Nadia, Atiya R Ali,Marie Paulsen Madsen, Rasmus Riis Kopperud, Hilde Vollan, Karoline Bragstad, Olav Hungnes |
| EPI_ISL_3320821 | Medical Microbiology Unit, Department for Laboratory Medicine, Drammen Hospital, Vestre Viken Health Trust, | Norwegian Institute of Public Health, Department of Virology | Kathrine Stene-Johansen, Kamilla Heddeland Instefjord, Hilde Elshaug, Garcia Llorente Ignacio, Jon Bråte, Engebretsen Serina Beate,Pedersen Benedikte Nevjen, Line Victoria Moen, Debech Nadia, Atiya R Ali,Marie Paulsen Madsen, Rasmus Riis Kopperud, Hilde Vollan, Karoline Bragstad, Olav Hungnes |
| EPI_ISL_3320822 | Ostfold Hospital Trust - Kalnes, Centre for Laboratory Medicine, Section for gene technology and infection serology | Norwegian Institute of Public Health, Department of Virology | Kathrine Stene-Johansen, Kamilla Heddeland Instefjord, Hilde Elshaug, Garcia Llorente Ignacio, Jon Bråte, Engebretsen Serina Beate,Pedersen Benedikte Nevjen, Line Victoria Moen, Debech Nadia, Atiya R Ali,Marie Paulsen Madsen, Rasmus Riis Kopperud, Hilde Vollan, Karoline Bragstad, Olav Hungnes |
| EPI_ISL_3320823, EPI_ISL_3320824 | Medical Microbiology Unit, Department for Laboratory Medicine, Drammen Hospital, Vestre Viken Health Trust, | Norwegian Institute of Public Health, Department of Virology | Kathrine Stene-Johansen, Kamilla Heddeland Instefjord, Hilde Elshaug, Garcia Llorente Ignacio, Jon Bråte, Engebretsen Serina Beate,Pedersen Benedikte Nevjen, Line Victoria Moen, Debech Nadia, Atiya R Ali,Marie Paulsen Madsen, Rasmus Riis Kopperud, Hilde Vollan, Karoline Bragstad, Olav Hungnes |
| EPI_ISL_3320825, EPI_ISL_3320826, EPI_ISL_3320827, EPI_ISL_3320828, EPI_ISL_3320829 | Haukeland University Hospital, Dept. of Microbiology | Norwegian Institute of Public Health, Department of Virology | Kathrine Stene-Johansen, Kamilla Heddeland Instefjord, Hilde Elshaug, Garcia Llorente Ignacio, Jon Bråte, Engebretsen Serina Beate,Pedersen Benedikte Nevjen, Line Victoria Moen, Debech Nadia, Atiya R Ali,Marie Paulsen Madsen, Rasmus Riis Kopperud, Hilde Vollan, Karoline Bragstad, Olav Hungnes |
| EPI_ISL_3320830 | Medical Microbiology Unit, Department for Laboratory Medicine, Drammen Hospital, Vestre Viken Health Trust, | Norwegian Institute of Public Health, Department of Virology | Kathrine Stene-Johansen, Kamilla Heddeland Instefjord, Hilde Elshaug, Garcia Llorente Ignacio, Jon Bråte, Engebretsen Serina Beate,Pedersen Benedikte Nevjen, Line Victoria Moen, Debech Nadia, Atiya R Ali,Marie Paulsen Madsen, Rasmus Riis Kopperud, Hilde Vollan, Karoline Bragstad, Olav Hungnes |
| EPI_ISL_3320831 | Haukeland University Hospital, Dept. of Microbiology | Norwegian Institute of Public Health, Department of Virology | Kathrine Stene-Johansen, Kamilla Heddeland Instefjord, Hilde Elshaug, Garcia Llorente Ignacio, Jon Bråte, Engebretsen Serina Beate,Pedersen Benedikte Nevjen, Line Victoria Moen, Debech Nadia, Atiya R Ali,Marie Paulsen Madsen, Rasmus Riis Kopperud, Hilde Vollan, Karoline Bragstad, Olav Hungnes |
| EPI_ISL_3320832 | Ostfold Hospital Trust - Kalnes, Centre for Laboratory Medicine, Section for gene technology and infection serology | Norwegian Institute of Public Health, Department of Virology | Kathrine Stene-Johansen, Kamilla Heddeland Instefjord, Hilde Elshaug, Garcia Llorente Ignacio, Jon Bråte, Engebretsen Serina Beate,Pedersen Benedikte Nevjen, Line Victoria Moen, Debech Nadia, Atiya R Ali,Marie Paulsen Madsen, Rasmus Riis Kopperud, Hilde Vollan, Karoline Bragstad, Olav Hungnes |
| EPI_ISL_3320833, EPI_ISL_3320834 | Medical Microbiology Unit, Department for Laboratory Medicine, Drammen Hospital, Vestre Viken Health Trust, | Norwegian Institute of Public Health, Department of Virology | Kathrine Stene-Johansen, Kamilla Heddeland Instefjord, Hilde Elshaug, Garcia Llorente Ignacio, Jon Bråte, Engebretsen Serina Beate,Pedersen Benedikte Nevjen, Line Victoria Moen, Debech Nadia, Atiya R Ali,Marie Paulsen Madsen, Rasmus Riis Kopperud, Hilde Vollan, Karoline Bragstad, Olav Hungnes |
| EPI_ISL_3320835 | Haukeland University Hospital, Dept. of Microbiology | Norwegian Institute of Public Health, Department of Virology | Kathrine Stene-Johansen, Kamilla Heddeland Instefjord, Hilde Elshaug, Garcia Llorente Ignacio, Jon Bråte, Engebretsen Serina Beate,Pedersen Benedikte Nevjen, Line Victoria Moen, Debech Nadia, Atiya R Ali,Marie Paulsen Madsen, Rasmus Riis Kopperud, Hilde Vollan, Karoline Bragstad, Olav Hungnes |
| EPI_ISL_3320836 | Furst Medical Laboratory | Norwegian Institute of Public Health, Department of Virology | Kathrine Stene-Johansen, Kamilla Heddeland Instefjord, Hilde Elshaug, Garcia Llorente Ignacio, Jon Bråte, Engebretsen Serina Beate,Pedersen Benedikte Nevjen, Line Victoria Moen, Debech Nadia, Atiya R Ali,Marie Paulsen Madsen, Rasmus Riis Kopperud, Hilde Vollan, Karoline Bragstad, Olav Hungnes |
| EPI_ISL_3320837 | Haukeland University Hospital, Dept. of Microbiology | Norwegian Institute of Public Health, Department of Virology | Kathrine Stene-Johansen, Kamilla Heddeland Instefjord, Hilde Elshaug, Garcia Llorente Ignacio, Jon Bråte, Engebretsen Serina Beate,Pedersen Benedikte Nevjen, Line Victoria Moen, Debech Nadia, Atiya R Ali,Marie Paulsen Madsen, Rasmus Riis Kopperud, Hilde Vollan, Karoline Bragstad, Olav Hungnes |
| EPI_ISL_3320838 | Nordland Hospital - Bodo, Laboratory Department, Molecular Biology Unit | Norwegian Institute of Public Health, Department of Virology | Kathrine Stene-Johansen, Kamilla Heddeland Instefjord, Hilde Elshaug, Garcia Llorente Ignacio, Jon Bråte, Engebretsen Serina Beate,Pedersen Benedikte Nevjen, Line Victoria Moen, Debech Nadia, Atiya R Ali,Marie Paulsen Madsen, Rasmus Riis Kopperud, Hilde Vollan, Karoline Bragstad, Olav Hungnes |
| EPI_ISL_3320839 | Medical Microbiology Unit, Department for Laboratory Medicine, Drammen Hospital, Vestre Viken Health Trust, | Norwegian Institute of Public Health, Department of Virology | Kathrine Stene-Johansen, Kamilla Heddeland Instefjord, Hilde Elshaug, Garcia Llorente Ignacio, Jon Bråte, Engebretsen Serina Beate,Pedersen Benedikte Nevjen, Line Victoria Moen, Debech Nadia, Atiya R Ali,Marie Paulsen Madsen, Rasmus Riis Kopperud, Hilde Vollan, Karoline Bragstad, Olav Hungnes |
| EPI_ISL_3320840, EPI_ISL_3320841, EPI_ISL_3320842 | Nordland Hospital - Bodo, Laboratory Department, Molecular Biology Unit | Norwegian Institute of Public Health, Department of Virology | Kathrine Stene-Johansen, Kamilla Heddeland Instefjord, Hilde Elshaug, Garcia Llorente Ignacio, Jon Bråte, Engebretsen Serina Beate,Pedersen Benedikte Nevjen, Line Victoria Moen, Debech Nadia, Atiya R Ali,Marie Paulsen Madsen, Rasmus Riis Kopperud, Hilde Vollan, Karoline Bragstad, Olav Hungnes |
| EPI_ISL_3320843 | Oslo University Hospital, Department of Medical Microbiology | Norwegian Institute of Public Health, Department of Virology | Kathrine Stene-Johansen, Kamilla Heddeland Instefjord, Hilde Elshaug, Garcia Llorente Ignacio, Jon Bråte, Engebretsen Serina Beate,Pedersen Benedikte Nevjen, Line Victoria Moen, Debech Nadia, Atiya R Ali,Marie Paulsen Madsen, Rasmus Riis Kopperud, Hilde Vollan, Karoline Bragstad, Olav Hungnes |
| EPI_ISL_3320844 | Nordland Hospital - Bodo, Laboratory Department, Molecular Biology Unit | Norwegian Institute of Public Health, Department of Virology | Kathrine Stene-Johansen, Kamilla Heddeland Instefjord, Hilde Elshaug, Garcia Llorente Ignacio, Jon Bråte, Engebretsen Serina Beate,Pedersen Benedikte Nevjen, Line Victoria Moen, Debech Nadia, Atiya R Ali,Marie Paulsen Madsen, Rasmus Riis Kopperud, Hilde Vollan, Karoline Bragstad, Olav Hungnes |
| EPI_ISL_3320845, EPI_ISL_3320846 | Oslo University Hospital, Department of Medical Microbiology | Norwegian Institute of Public Health, Department of Virology | Kathrine Stene-Johansen, Kamilla Heddeland Instefjord, Hilde Elshaug, Garcia Llorente Ignacio, Jon Bråte, Engebretsen Serina Beate,Pedersen Benedikte Nevjen, Line Victoria Moen, Debech Nadia, Atiya R Ali,Marie Paulsen Madsen, Rasmus Riis Kopperud, Hilde Vollan, Karoline Bragstad, Olav Hungnes |
| EPI_ISL_3320847, EPI_ISL_3320848 | Nordland Hospital - Bodo, Laboratory Department, Molecular Biology Unit | Norwegian Institute of Public Health, Department of Virology | Kathrine Stene-Johansen, Kamilla Heddeland Instefjord, Hilde Elshaug, Garcia Llorente Ignacio, Jon Bråte, Engebretsen Serina Beate,Pedersen Benedikte Nevjen, Line Victoria Moen, Debech Nadia, Atiya R Ali,Marie Paulsen Madsen, Rasmus Riis Kopperud, Hilde Vollan, Karoline Bragstad, Olav Hungnes |
| EPI_ISL_3320849 | Furst Medical Laboratory | Norwegian Institute of Public Health, Department of Virology | Kathrine Stene-Johansen, Kamilla Heddeland Instefjord, Hilde Elshaug, Garcia Llorente Ignacio, Jon Bråte, Engebretsen Serina Beate,Pedersen Benedikte Nevjen, Line Victoria Moen, Debech Nadia, Atiya R Ali,Marie Paulsen Madsen, Rasmus Riis Kopperud, Hilde Vollan, Karoline Bragstad, Olav Hungnes |
| EPI_ISL_3320851, EPI_ISL_3320852 | Nordland Hospital - Bodo, Laboratory Department, Molecular Biology Unit | Norwegian Institute of Public Health, Department of Virology | Kathrine Stene-Johansen, Kamilla Heddeland Instefjord, Hilde Elshaug, Garcia Llorente Ignacio, Jon Bråte, Engebretsen Serina Beate,Pedersen Benedikte Nevjen, Line Victoria Moen, Debech Nadia, Atiya R Ali,Marie Paulsen Madsen, Rasmus Riis Kopperud, Hilde Vollan, Karoline Bragstad, Olav Hungnes |
| EPI_ISL_3320853, EPI_ISL_3320854, EPI_ISL_3320855, EPI_ISL_3320856, EPI_ISL_3320857 | University Hospital of Northern Norway, Department for Microbiology and Infectious Disease Control | Norwegian Institute of Public Health, Department of Virology | Kathrine Stene-Johansen, Kamilla Heddeland Instefjord, Hilde Elshaug, Garcia Llorente Ignacio, Jon Bråte, Engebretsen Serina Beate,Pedersen Benedikte Nevjen, Line Victoria Moen, Debech Nadia, Atiya R Ali,Marie Paulsen Madsen, Rasmus Riis Kopperud, Hilde Vollan, Karoline Bragstad, Olav Hungnes |
| EPI_ISL_3320859 | Innlandet Hospital Trust, Division Lillehammer, Department for Medical Microbiology | Norwegian Institute of Public Health, Department of Virology | Kathrine Stene-Johansen, Kamilla Heddeland Instefjord, Hilde Elshaug, Garcia Llorente Ignacio, Jon Bråte, Engebretsen Serina Beate,Pedersen Benedikte Nevjen, Line Victoria Moen, Debech Nadia, Atiya R Ali,Marie Paulsen Madsen, Rasmus Riis Kopperud, Hilde Vollan, Karoline Bragstad, Olav Hungnes |
| EPI_ISL_3320860 | Vestfold Hospital, Toensberg Department of Microbiology | Norwegian Institute of Public Health, Department of Virology | Kathrine Stene-Johansen, Kamilla Heddeland Instefjord, Hilde Elshaug, Garcia Llorente Ignacio, Jon Bråte, Engebretsen Serina Beate,Pedersen Benedikte Nevjen, Line Victoria Moen, Debech Nadia, Atiya R Ali,Marie Paulsen Madsen, Rasmus Riis Kopperud, Hilde Vollan, Karoline Bragstad, Olav Hungnes |
| EPI_ISL_3320861 | Innlandet Hospital Trust, Division Lillehammer, Department for Medical Microbiology | Norwegian Institute of Public Health, Department of Virology | Kathrine Stene-Johansen, Kamilla Heddeland Instefjord, Hilde Elshaug, Garcia Llorente Ignacio, Jon Bråte, Engebretsen Serina Beate,Pedersen Benedikte Nevjen, Line Victoria Moen, Debech Nadia, Atiya R Ali,Marie Paulsen Madsen, Rasmus Riis Kopperud, Hilde Vollan, Karoline Bragstad, Olav Hungnes |
| EPI_ISL_3320863 | Department of Medical Microbiology, St. Olavs hospital | Norwegian Institute of Public Health, Department of Virology | Kathrine Stene-Johansen, Kamilla Heddeland Instefjord, Hilde Elshaug, Garcia Llorente Ignacio, Jon Bråte, Engebretsen Serina Beate,Pedersen Benedikte Nevjen, Line Victoria Moen, Debech Nadia, Atiya R Ali,Marie Paulsen Madsen, Rasmus Riis Kopperud, Hilde Vollan, Karoline Bragstad, Olav Hungnes |

| | | | |
|---|---|---|---|
| EPI_ISL_3320864 | Department of Medical Microbiology - section Molde, Molde Hospital | Norwegian Institute of Public Health, Department of Virology | Kathrine Stene-Johansen, Kamilla Heddeland Instefjord, Hilde Elshaug, Garcia Llorente Ignacio, Jon Bråte, Engebretsen Serina Beate,Pedersen Benedikte Nevjen, Line Victoria Moen, Debech Nadia, Atiya R Ali,Marie Paulsen Madsen, Rasmus Riis Kopperud, Hilde Vollan, Karoline Bragstad, Olav Hungnes |
| EPI_ISL_3320865 | Department of Medical Microbiology, St. Olavs hospital | Norwegian Institute of Public Health, Department of Virology | Kathrine Stene-Johansen, Kamilla Heddeland Instefjord, Hilde Elshaug, Garcia Llorente Ignacio, Jon Bråte, Engebretsen Serina Beate,Pedersen Benedikte Nevjen, Line Victoria Moen, Debech Nadia, Atiya R Ali,Marie Paulsen Madsen, Rasmus Riis Kopperud, Hilde Vollan, Karoline Bragstad, Olav Hungnes |
| EPI_ISL_3320866 | Department of Medical Microbiology - section Molde, Molde Hospital | Norwegian Institute of Public Health, Department of Virology | Kathrine Stene-Johansen, Kamilla Heddeland Instefjord, Hilde Elshaug, Garcia Llorente Ignacio, Jon Bråte, Engebretsen Serina Beate,Pedersen Benedikte Nevjen, Line Victoria Moen, Debech Nadia, Atiya R Ali,Marie Paulsen Madsen, Rasmus Riis Kopperud, Hilde Vollan, Karoline Bragstad, Olav Hungnes |
| EPI_ISL_3320867 | Innlandet Hospital Trust, Division Lillehammer, Department for Medical Microbiology | Norwegian Institute of Public Health, Department of Virology | Kathrine Stene-Johansen, Kamilla Heddeland Instefjord, Hilde Elshaug, Garcia Llorente Ignacio, Jon Bråte, Engebretsen Serina Beate,Pedersen Benedikte Nevjen, Line Victoria Moen, Debech Nadia, Atiya R Ali,Marie Paulsen Madsen, Rasmus Riis Kopperud, Hilde Vollan, Karoline Bragstad, Olav Hungnes |
| EPI_ISL_3320868, EPI_ISL_3320869 | Department of Medical Microbiology - section Molde, Molde Hospital | Norwegian Institute of Public Health, Department of Virology | Kathrine Stene-Johansen, Kamilla Heddeland Instefjord, Hilde Elshaug, Garcia Llorente Ignacio, Jon Bråte, Engebretsen Serina Beate,Pedersen Benedikte Nevjen, Line Victoria Moen, Debech Nadia, Atiya R Ali,Marie Paulsen Madsen, Rasmus Riis Kopperud, Hilde Vollan, Karoline Bragstad, Olav Hungnes |
| EPI_ISL_3320870 | Hospital of Southern Norway - Kristiansand, Department of Medical Microbiology | Norwegian Institute of Public Health, Department of Virology | Kathrine Stene-Johansen, Kamilla Heddeland Instefjord, Hilde Elshaug, Garcia Llorente Ignacio, Jon Bråte, Engebretsen Serina Beate,Pedersen Benedikte Nevjen, Line Victoria Moen, Debech Nadia, Atiya R Ali,Marie Paulsen Madsen, Rasmus Riis Kopperud, Hilde Vollan, Karoline Bragstad, Olav Hungnes |
| EPI_ISL_3320871, EPI_ISL_3320872 | Nordland Hospital - Bodo, Laboratory Department, Molecular Biology Unit | Norwegian Institute of Public Health, Department of Virology | Kathrine Stene-Johansen, Kamilla Heddeland Instefjord, Hilde Elshaug, Garcia Llorente Ignacio, Jon Bråte, Engebretsen Serina Beate,Pedersen Benedikte Nevjen, Line Victoria Moen, Debech Nadia, Atiya R Ali,Marie Paulsen Madsen, Rasmus Riis Kopperud, Hilde Vollan, Karoline Bragstad, Olav Hungnes |
| EPI_ISL_3320874 | Department of Medical Microbiology, St. Olavs hospital | Norwegian Institute of Public Health, Department of Virology | Kathrine Stene-Johansen, Kamilla Heddeland Instefjord, Hilde Elshaug, Garcia Llorente Ignacio, Jon Bråte, Engebretsen Serina Beate,Pedersen Benedikte Nevjen, Line Victoria Moen, Debech Nadia, Atiya R Ali,Marie Paulsen Madsen, Rasmus Riis Kopperud, Hilde Vollan, Karoline Bragstad, Olav Hungnes |
| EPI_ISL_3320877 | Department of Medical Microbiology - section Molde, Molde Hospital | Norwegian Institute of Public Health, Department of Virology | Kathrine Stene-Johansen, Kamilla Heddeland Instefjord, Hilde Elshaug, Garcia Llorente Ignacio, Jon Bråte, Engebretsen Serina Beate,Pedersen Benedikte Nevjen, Line Victoria Moen, Debech Nadia, Atiya R Ali,Marie Paulsen Madsen, Rasmus Riis Kopperud, Hilde Vollan, Karoline Bragstad, Olav Hungnes |
| EPI_ISL_3320878 | Hospital of Southern Norway - Kristiansand, Department of Medical Microbiology | Norwegian Institute of Public Health, Department of Virology | Kathrine Stene-Johansen, Kamilla Heddeland Instefjord, Hilde Elshaug, Garcia Llorente Ignacio, Jon Bråte, Engebretsen Serina Beate,Pedersen Benedikte Nevjen, Line Victoria Moen, Debech Nadia, Atiya R Ali,Marie Paulsen Madsen, Rasmus Riis Kopperud, Hilde Vollan, Karoline Bragstad, Olav Hungnes |
| EPI_ISL_3320879 | Department of Medical Microbiology, St. Olavs hospital | Norwegian Institute of Public Health, Department of Virology | Kathrine Stene-Johansen, Kamilla Heddeland Instefjord, Hilde Elshaug, Garcia Llorente Ignacio, Jon Bråte, Engebretsen Serina Beate,Pedersen Benedikte Nevjen, Line Victoria Moen, Debech Nadia, Atiya R Ali,Marie Paulsen Madsen, Rasmus Riis Kopperud, Hilde Vollan, Karoline Bragstad, Olav Hungnes |
| EPI_ISL_3320880, EPI_ISL_3320881 | Hospital of Southern Norway - Kristiansand, Department of Medical Microbiology | Norwegian Institute of Public Health, Department of Virology | Kathrine Stene-Johansen, Kamilla Heddeland Instefjord, Hilde Elshaug, Garcia Llorente Ignacio, Jon Bråte, Engebretsen Serina Beate,Pedersen Benedikte Nevjen, Line Victoria Moen, Debech Nadia, Atiya R Ali,Marie Paulsen Madsen, Rasmus Riis Kopperud, Hilde Vollan, Karoline Bragstad, Olav Hungnes |
| EPI_ISL_3320882 | Nordland Hospital - Bodo, Laboratory Department, Molecular Biology Unit | Norwegian Institute of Public Health, Department of Virology | Kathrine Stene-Johansen, Kamilla Heddeland Instefjord, Hilde Elshaug, Garcia Llorente Ignacio, Jon Bråte, Engebretsen Serina Beate,Pedersen Benedikte Nevjen, Line Victoria Moen, Debech Nadia, Atiya R Ali,Marie Paulsen Madsen, Rasmus Riis Kopperud, Hilde Vollan, Karoline Bragstad, Olav Hungnes |
| EPI_ISL_3320884 | Hospital of Southern Norway - Kristiansand, Department of Medical Microbiology | Norwegian Institute of Public Health, Department of Virology | Kathrine Stene-Johansen, Kamilla Heddeland Instefjord, Hilde Elshaug, Garcia Llorente Ignacio, Jon Bråte, Engebretsen Serina Beate,Pedersen Benedikte Nevjen, Line Victoria Moen, Debech Nadia, Atiya R Ali,Marie Paulsen Madsen, Rasmus Riis Kopperud, Hilde Vollan, Karoline Bragstad, Olav Hungnes |
| EPI_ISL_3320885 | Nordland Hospital - Bodo, Laboratory Department, Molecular Biology Unit | Norwegian Institute of Public Health, Department of Virology | Kathrine Stene-Johansen, Kamilla Heddeland Instefjord, Hilde Elshaug, Garcia Llorente Ignacio, Jon Bråte, Engebretsen Serina Beate,Pedersen Benedikte Nevjen, Line Victoria Moen, Debech Nadia, Atiya R Ali,Marie Paulsen Madsen, Rasmus Riis Kopperud, Hilde Vollan, Karoline Bragstad, Olav Hungnes |
| EPI_ISL_3320886 | Haukeland University Hospital, Dept. of Microbiology | Norwegian Institute of Public Health, Department of Virology | Kathrine Stene-Johansen, Kamilla Heddeland Instefjord, Hilde Elshaug, Garcia Llorente Ignacio, Jon Bråte, Engebretsen Serina Beate,Pedersen Benedikte Nevjen, Line Victoria Moen, Debech Nadia, Atiya R Ali,Marie Paulsen Madsen, Rasmus Riis Kopperud, Hilde Vollan, Karoline Bragstad, Olav Hungnes |
| EPI_ISL_3320887 | Medical Microbiology Unit, Department for Laboratory Medicine, Drammen Hospital, Vestre Viken Health Trust, | Norwegian Institute of Public Health, Department of Virology | Kathrine Stene-Johansen, Kamilla Heddeland Instefjord, Hilde Elshaug, Garcia Llorente Ignacio, Jon Bråte, Engebretsen Serina Beate,Pedersen Benedikte Nevjen, Line Victoria Moen, Debech Nadia, Atiya R Ali,Marie Paulsen Madsen, Rasmus Riis Kopperud, Hilde Vollan, Karoline Bragstad, Olav Hungnes |
| EPI_ISL_3320888 | Ostfold Hospital Trust - Kalnes, Centre for Laboratory Medicine, Section for gene technology and infection serology | Norwegian Institute of Public Health, Department of Virology | Kathrine Stene-Johansen, Kamilla Heddeland Instefjord, Hilde Elshaug, Garcia Llorente Ignacio, Jon Bråte, Engebretsen Serina Beate,Pedersen Benedikte Nevjen, Line Victoria Moen, Debech Nadia, Atiya R Ali,Marie Paulsen Madsen, Rasmus Riis Kopperud, Hilde Vollan, Karoline Bragstad, Olav Hungnes |
| EPI_ISL_3320889, EPI_ISL_3320890 | Haukeland University Hospital, Dept. of Microbiology | Norwegian Institute of Public Health, Department of Virology | Kathrine Stene-Johansen, Kamilla Heddeland Instefjord, Hilde Elshaug, Garcia Llorente Ignacio, Jon Bråte, Engebretsen Serina Beate,Pedersen Benedikte Nevjen, Line Victoria Moen, Debech Nadia, Atiya R Ali,Marie Paulsen Madsen, Rasmus Riis Kopperud, Hilde Vollan, Karoline Bragstad, Olav Hungnes |
| EPI_ISL_3320891 | Ostfold Hospital Trust - Kalnes, Centre for Laboratory Medicine, Section for gene technology and infection serology | Norwegian Institute of Public Health, Department of Virology | Kathrine Stene-Johansen, Kamilla Heddeland Instefjord, Hilde Elshaug, Garcia Llorente Ignacio, Jon Bråte, Engebretsen Serina Beate,Pedersen Benedikte Nevjen, Line Victoria Moen, Debech Nadia, Atiya R Ali,Marie Paulsen Madsen, Rasmus Riis Kopperud, Hilde Vollan, Karoline Bragstad, Olav Hungnes |
| EPI_ISL_3320892 | Haukeland University Hospital, Dept. of Microbiology | Norwegian Institute of Public Health, Department of Virology | Kathrine Stene-Johansen, Kamilla Heddeland Instefjord, Hilde Elshaug, Garcia Llorente Ignacio, Jon Bråte, Engebretsen Serina Beate,Pedersen Benedikte Nevjen, Line Victoria Moen, Debech Nadia, Atiya R Ali,Marie Paulsen Madsen, Rasmus Riis Kopperud, Hilde Vollan, Karoline Bragstad, Olav Hungnes |
| EPI_ISL_3320893 | Medical Microbiology Unit, Department for Laboratory Medicine, Drammen Hospital, Vestre Viken Health Trust, | Norwegian Institute of Public Health, Department of Virology | Kathrine Stene-Johansen, Kamilla Heddeland Instefjord, Hilde Elshaug, Garcia Llorente Ignacio, Jon Bråte, Engebretsen Serina Beate,Pedersen Benedikte Nevjen, Line Victoria Moen, Debech Nadia, Atiya R Ali,Marie Paulsen Madsen, Rasmus Riis Kopperud, Hilde Vollan, Karoline Bragstad, Olav Hungnes |
| EPI_ISL_3320894 | Haukeland University Hospital, Dept. of Microbiology | Norwegian Institute of Public Health, Department of Virology | Kathrine Stene-Johansen, Kamilla Heddeland Instefjord, Hilde Elshaug, Garcia Llorente Ignacio, Jon Bråte, Engebretsen Serina Beate,Pedersen Benedikte Nevjen, Line Victoria Moen, Debech Nadia, Atiya R Ali,Marie Paulsen Madsen, Rasmus Riis Kopperud, Hilde Vollan, Karoline Bragstad, Olav Hungnes |
| EPI_ISL_3320895 | Unilabs Laboratory Medicine | Norwegian Institute of Public Health, Department of Virology | Kathrine Stene-Johansen, Kamilla Heddeland Instefjord, Hilde Elshaug, Garcia Llorente Ignacio, Jon Bråte, Engebretsen Serina Beate,Pedersen Benedikte Nevjen, Line Victoria Moen, Debech Nadia, Atiya R Ali,Marie Paulsen Madsen, Rasmus Riis Kopperud, Hilde Vollan, Karoline Bragstad, Olav Hungnes |
| EPI_ISL_3320896, EPI_ISL_3320897, EPI_ISL_3320898 | Haukeland University Hospital, Dept. of Microbiology | Norwegian Institute of Public Health, Department of Virology | Kathrine Stene-Johansen, Kamilla Heddeland Instefjord, Hilde Elshaug, Garcia Llorente Ignacio, Jon Bråte, Engebretsen Serina Beate,Pedersen Benedikte Nevjen, Line Victoria Moen, Debech Nadia, Atiya R Ali,Marie Paulsen Madsen, Rasmus Riis Kopperud, Hilde Vollan, Karoline Bragstad, Olav Hungnes |
| EPI_ISL_3320899 | Ostfold Hospital Trust - Kalnes, Centre for Laboratory Medicine, Section for gene technology and infection serology | Norwegian Institute of Public Health, Department of Virology | Kathrine Stene-Johansen, Kamilla Heddeland Instefjord, Hilde Elshaug, Garcia Llorente Ignacio, Jon Bråte, Engebretsen Serina Beate,Pedersen Benedikte Nevjen, Line Victoria Moen, Debech Nadia, Atiya R Ali,Marie Paulsen Madsen, Rasmus Riis Kopperud, Hilde Vollan, Karoline Bragstad, Olav Hungnes |
| EPI_ISL_3320900 | Haukeland University Hospital, Dept. of Microbiology | Norwegian Institute of Public Health, Department of Virology | Kathrine Stene-Johansen, Kamilla Heddeland Instefjord, Hilde Elshaug, Garcia Llorente Ignacio, Jon Bråte, Engebretsen Serina Beate,Pedersen Benedikte Nevjen, Line Victoria Moen, Debech Nadia, Atiya R Ali,Marie Paulsen Madsen, Rasmus Riis Kopperud, Hilde Vollan, Karoline Bragstad, Olav Hungnes |

EPI_ISL_3320923, EPI_ISL_3320926, EPI_ISL_3320929, EPI_ISL_3320933, EPI_ISL_3320935, EPI_ISL_3320939, EPI_ISL_3320942, EPI_ISL_3320945, EPI_ISL_3320947, EPI_ISL_3320950, EPI_ISL_3320954, EPI_ISL_3320957, EPI_ISL_3320961

| | | | |
|---|---|---|---|
| see above | Orebro University Hospital, Dept Laboratory Medicine, Clinical Microbiology | Orebro University Hospital | Sundqvist M et al |
| EPI_ISL_3321119, EPI_ISL_3321120 | Labo Analyses Med | National Reference Center for Viruses of Respiratory Infections, Institut Pasteur, Paris | Marion Barbet, Sylvie Behillil, Méline Bizard, Angela Brisebarre, Camille Capel, Vincent Enouf, Louise Lefrançois, Frédéric Lemoine, Christophe Malabat, Corinne Maufrais, Hub de Bioinformatique et Biostatistique, Etienne Simon-Lorière, Maud Vanpeene, Sylvie Van der Werf ,S. Cocco |
| EPI_ISL_3321121 | Labo Analyses Med | National Reference Center for Viruses of Respiratory Infections, Institut Pasteur, Paris | Marion Barbet, Sylvie Behillil, Méline Bizard, Angela Brisebarre, Camille Capel, Vincent Enouf, Louise Lefrançois, Frédéric Lemoine, Christophe Malabat, Corinne Maufrais, Hub de Bioinformatique et Biostatistique, Etienne Simon-Lorière, Maud Vanpeene, Sylvie Van der Werf ,Anne Holstein |
| EPI_ISL_3321123 | Hospital | National Reference Center for Viruses of Respiratory Infections, Institut Pasteur, Paris | Marion Barbet, Sylvie Behillil, Méline Bizard, Angela Brisebarre, Camille Capel, Vincent Enouf, Louise Lefrançois, Frédéric Lemoine, Christophe Malabat, Corinne Maufrais, Hub de Bioinformatique et Biostatistique, Etienne Simon-Lorière, Maud Vanpeene, Sylvie Van der Werf ,Pascale Martres |
| EPI_ISL_3321124 | Labo Analyses Med | National Reference Center for Viruses of Respiratory Infections, Institut Pasteur, Paris | Marion Barbet, Sylvie Behillil, Méline Bizard, Angela Brisebarre, Camille Capel, Vincent Enouf, Louise Lefrançois, Frédéric Lemoine, Christophe Malabat, Corinne Maufrais, Hub de Bioinformatique et Biostatistique, Etienne Simon-Lorière, Maud Vanpeene, Sylvie Van der Werf ,Nabila Belhouachi |
| EPI_ISL_3321126, EPI_ISL_3321127, EPI_ISL_3321128 | Labo Analyses Med | National Reference Center for Viruses of Respiratory Infections, Institut Pasteur, Paris | Marion Barbet, Sylvie Behillil, Méline Bizard, Angela Brisebarre, Camille Capel, Vincent Enouf, Louise Lefrançois, Frédéric Lemoine, Christophe Malabat, Corinne Maufrais, Hub de Bioinformatique et Biostatistique, Etienne Simon-Lorière, Maud Vanpeene, Sylvie Van der Werf ,OphéLie Said-Delattre |
| EPI_ISL_3321129, EPI_ISL_3321130, EPI_ISL_3321131, EPI_ISL_3321132, EPI_ISL_3321133, EPI_ISL_3321134, EPI_ISL_3321135, EPI_ISL_3321136 | Ochsner LSU Health | LSUHS Emerging Viral Threat Laboratory | Jeremy P. Kamil, Jennifer L. Carroll, Maarten Van Diest, Rona S. Scott, Gregory L. Ware, Alexander Mijalis, Andrew D. Yurochko, Christopher G. Kevil, John A. Vanchiere |

EPI_ISL_3321138, EPI_ISL_3321140, EPI_ISL_3321141, EPI_ISL_3321142, EPI_ISL_3321143, EPI_ISL_3321144, EPI_ISL_3321145, EPI_ISL_3321146, EPI_ISL_3321147, EPI_ISL_3321148, EPI_ISL_3321149, EPI_ISL_3321150, EPI_ISL_3321151, EPI_ISL_3321152, EPI_ISL_3321153, EPI_ISL_3321154, EPI_ISL_3321155, EPI_ISL_3321156, EPI_ISL_3321157, EPI_ISL_3321158, EPI_ISL_3321159, EPI_ISL_3321161, EPI_ISL_3321162, EPI_ISL_3321163, EPI_ISL_3321164, EPI_ISL_3321165, EPI_ISL_3321166, EPI_ISL_3321167, EPI_ISL_3321168, EPI_ISL_3321169, EPI_ISL_3321170, EPI_ISL_3321171, EPI_ISL_3321172, EPI_ISL_3321173, EPI_ISL_3321174, EPI_ISL_3321175, EPI_ISL_3321176, EPI_ISL_3321177, EPI_ISL_3321178, EPI_ISL_3321179, EPI_ISL_3321180, EPI_ISL_3321181, EPI_ISL_3321182, EPI_ISL_3321184, EPI_ISL_3321185, EPI_ISL_3321186, EPI_ISL_3321187, EPI_ISL_3321188, EPI_ISL_3321189, EPI_ISL_3321190, EPI_ISL_3321196, EPI_ISL_3321197, EPI_ISL_3321198, EPI_ISL_3321199, EPI_ISL_3321200, EPI_ISL_3321201, EPI_ISL_3321202, EPI_ISL_3321203, EPI_ISL_3321204, EPI_ISL_3321205, EPI_ISL_3321206, EPI_ISL_3321207, EPI_ISL_3321208, EPI_ISL_3321209, EPI_ISL_3321210, EPI_ISL_3321211, EPI_ISL_3321212, EPI_ISL_3321213, EPI_ISL_3321214, EPI_ISL_3321215, EPI_ISL_3321216, EPI_ISL_3321217, EPI_ISL_3321218, EPI_ISL_3321219, EPI_ISL_3321220, EPI_ISL_3321221, EPI_ISL_3321222, EPI_ISL_3321223, EPI_ISL_3321224, EPI_ISL_3321225, EPI_ISL_3321226, EPI_ISL_3321227, EPI_ISL_3321228, EPI_ISL_3321229, EPI_ISL_3321230, EPI_ISL_3321231, EPI_ISL_3321232, EPI_ISL_3321233, EPI_ISL_3321234, EPI_ISL_3321235, EPI_ISL_3321236, EPI_ISL_3321237, EPI_ISL_3321238, EPI_ISL_3321239, EPI_ISL_3321240, EPI_ISL_3321241, EPI_ISL_3321242, EPI_ISL_3321243, EPI_ISL_3321244, EPI_ISL_3321245, EPI_ISL_3321246, EPI_ISL_3321247, EPI_ISL_3321248, EPI_ISL_3321249, EPI_ISL_3321250, EPI_ISL_3321251, EPI_ISL_3321252, EPI_ISL_3321253, EPI_ISL_3321254, EPI_ISL_3321255, EPI_ISL_3321256, EPI_ISL_3321257, EPI_ISL_3321258, EPI_ISL_3321259, EPI_ISL_3321260, EPI_ISL_3321262, EPI_ISL_3321263

| | | | |
|---|---|---|---|
| see above | Willis-Knighton Medical Center Hospital Laboratory | LSUHS Emerging Viral Threat Laboratory | Lorie M. Atkins, April N. Johnson, Gregory L. Ware, Alexander Mijalis, Jeremy P. Kamil, Jennifer L. Carroll, Maarten Van Diest, Rona S. Scott, Andrew D. Yurochko, Christopher G. Kevil, John A. Vanchiere, Joseph A. Bocchini |
| EPI_ISL_3321315 | CAP Folgueroles | Banc de Sang i Teixits | Francisco Vidal, Irene Corrales, Nina Borràs, Carlos Hobeich, Natália Comes, Lorena Ramírez, Noemí Gonzalez, Silvia Sauleda, Maria Glòria Soria |

EPI_ISL_3321322, EPI_ISL_3321323, EPI_ISL_3321325, EPI_ISL_3321342, EPI_ISL_3321343, EPI_ISL_3321344, EPI_ISL_3321348, EPI_ISL_3321352, EPI_ISL_3321353, EPI_ISL_3321359, EPI_ISL_3321360, EPI_ISL_3321366, EPI_ISL_3321367, EPI_ISL_3321368, EPI_ISL_3321369, EPI_ISL_3321370, EPI_ISL_3321371, EPI_ISL_3321372, EPI_ISL_3321374, EPI_ISL_3321375, EPI_ISL_3321376, EPI_ISL_3321377, EPI_ISL_3321378, EPI_ISL_3321379, EPI_ISL_3321380, EPI_ISL_3321381, EPI_ISL_3321382, EPI_ISL_3321383, EPI_ISL_3321386, EPI_ISL_3321388, EPI_ISL_3321389, EPI_ISL_3321391, EPI_ISL_3321392, EPI_ISL_3321393, EPI_ISL_3321395, EPI_ISL_3321396, EPI_ISL_3321397, EPI_ISL_3321398, EPI_ISL_3321400, EPI_ISL_3321401, EPI_ISL_3321402, EPI_ISL_3321403, EPI_ISL_3321404, EPI_ISL_3321405, EPI_ISL_3321406, EPI_ISL_3321407, EPI_ISL_3321408, EPI_ISL_3321409, EPI_ISL_3321410, EPI_ISL_3321411, EPI_ISL_3321412, EPI_ISL_3321413, EPI_ISL_3321414, EPI_ISL_3321415, EPI_ISL_3321417, EPI_ISL_3321418, EPI_ISL_3321421, EPI_ISL_3321422, EPI_ISL_3321423, EPI_ISL_3321424, EPI_ISL_3321425, EPI_ISL_3321426, EPI_ISL_3321427, EPI_ISL_3321428, EPI_ISL_3321429, EPI_ISL_3321431, EPI_ISL_3321432, EPI_ISL_3321434, EPI_ISL_3321435, EPI_ISL_3321436, EPI_ISL_3321437, EPI_ISL_3321438, EPI_ISL_3321439, EPI_ISL_3321440, EPI_ISL_3321442, EPI_ISL_3321443, EPI_ISL_3321444, EPI_ISL_3321447, EPI_ISL_3321448, EPI_ISL_3321449, EPI_ISL_3321451, EPI_ISL_3321452, EPI_ISL_3321453, EPI_ISL_3321454, EPI_ISL_3321455, EPI_ISL_3321456, EPI_ISL_3321457, EPI_ISL_3321458, EPI_ISL_3321459, EPI_ISL_3321460, EPI_ISL_3321461, EPI_ISL_3321463, EPI_ISL_3321464, EPI_ISL_3321465, EPI_ISL_3321466, EPI_ISL_3321468, EPI_ISL_3321469, EPI_ISL_3321470, EPI_ISL_3321471, EPI_ISL_3321472, EPI_ISL_3321473, EPI_ISL_3321474, EPI_ISL_3321476, EPI_ISL_3321477, EPI_ISL_3321479, EPI_ISL_3321482, EPI_ISL_3321483, EPI_ISL_3321485, EPI_ISL_3321486, EPI_ISL_3321487, EPI_ISL_3321488, EPI_ISL_3321489, EPI_ISL_3321490, EPI_ISL_3321491, EPI_ISL_3321492, EPI_ISL_3321493, EPI_ISL_3321495, EPI_ISL_3321496, EPI_ISL_3321498, EPI_ISL_3321499, EPI_ISL_3321500, EPI_ISL_3321501, EPI_ISL_3321502, EPI_ISL_3321503, EPI_ISL_3321504, EPI_ISL_3321505, EPI_ISL_3321506, EPI_ISL_3321507, EPI_ISL_3321508, EPI_ISL_3321509, EPI_ISL_3321510, EPI_ISL_3321511, EPI_ISL_3321512, EPI_ISL_3321514, EPI_ISL_3321515, EPI_ISL_3321516, EPI_ISL_3321517, EPI_ISL_3321520, EPI_ISL_3321521, EPI_ISL_3321522, EPI_ISL_3321523, EPI_ISL_3321524, EPI_ISL_3321525, EPI_ISL_3321526, EPI_ISL_3321527, EPI_ISL_3321528, EPI_ISL_3321529, EPI_ISL_3321530, EPI_ISL_3321531, EPI_ISL_3321532, EPI_ISL_3321533, EPI_ISL_3321534, EPI_ISL_3321535, EPI_ISL_3321536, EPI_ISL_3321537, EPI_ISL_3321539, EPI_ISL_3321540, EPI_ISL_3321541, EPI_ISL_3321542, EPI_ISL_3321543, EPI_ISL_3321544, EPI_ISL_3321545, EPI_ISL_3321546, EPI_ISL_3321547, EPI_ISL_3321548, EPI_ISL_3321549, EPI_ISL_3321550, EPI_ISL_3321551, EPI_ISL_3321552, EPI_ISL_3321553, EPI_ISL_3321554, EPI_ISL_3321555, EPI_ISL_3321556, EPI_ISL_3321557, EPI_ISL_3321558, EPI_ISL_3321559,

EPI_ISL_3321560, EPI_ISL_3321561, EPI_ISL_3321562, EPI_ISL_3321563, EPI_ISL_3321564, EPI_ISL_3321565, EPI_ISL_3321566, EPI_ISL_3321568, EPI_ISL_3321570, EPI_ISL_3321572, EPI_ISL_3321573, EPI_ISL_3321574, EPI_ISL_3321575, EPI_ISL_3321576, EPI_ISL_3321577, EPI_ISL_3321578, EPI_ISL_3321579, EPI_ISL_3321580, EPI_ISL_3321581, EPI_ISL_3321582, EPI_ISL_3321583, EPI_ISL_3321584, EPI_ISL_3321586, EPI_ISL_3321587, EPI_ISL_3321588, EPI_ISL_3321589, EPI_ISL_3321590, EPI_ISL_3321591, EPI_ISL_3321592, EPI_ISL_3321594, EPI_ISL_3321595, EPI_ISL_3321596, EPI_ISL_3321597, EPI_ISL_3321598, EPI_ISL_3321599, EPI_ISL_3321602, EPI_ISL_3321603, EPI_ISL_3321604, EPI_ISL_3321605, EPI_ISL_3321606, EPI_ISL_3321607, EPI_ISL_3321608, EPI_ISL_3321609, EPI_ISL_3321610, EPI_ISL_3321611, EPI_ISL_3321612, EPI_ISL_3321613, EPI_ISL_3321614, EPI_ISL_3321616, EPI_ISL_3321617, EPI_ISL_3321619, EPI_ISL_3321621, EPI_ISL_3321622, EPI_ISL_3321623, EPI_ISL_3321624, EPI_ISL_3321625, EPI_ISL_3321626, EPI_ISL_3321627, EPI_ISL_3321628, EPI_ISL_3321629, EPI_ISL_3321630, EPI_ISL_3321631, EPI_ISL_3321632, EPI_ISL_3321633, EPI_ISL_3321634, EPI_ISL_3321638, EPI_ISL_3321639, EPI_ISL_3321640, EPI_ISL_3321641, EPI_ISL_3321642, EPI_ISL_3321643, EPI_ISL_3321644, EPI_ISL_3321645, EPI_ISL_3321646, EPI_ISL_3321647, EPI_ISL_3321648, EPI_ISL_3321650, EPI_ISL_3321651, EPI_ISL_3321652, EPI_ISL_3321653, EPI_ISL_3321654, EPI_ISL_3321655, EPI_ISL_3321656, EPI_ISL_3321657, EPI_ISL_3321658, EPI_ISL_3321659, EPI_ISL_3321660, EPI_ISL_3321661, EPI_ISL_3321663, EPI_ISL_3321664, EPI_ISL_3321666, EPI_ISL_3321667, EPI_ISL_3321668, EPI_ISL_3321669, EPI_ISL_3321670, EPI_ISL_3321671, EPI_ISL_3321672, EPI_ISL_3321673, EPI_ISL_3321674, EPI_ISL_3321675, EPI_ISL_3321677, EPI_ISL_3321678, EPI_ISL_3321679, EPI_ISL_3321680, EPI_ISL_3321681, EPI_ISL_3321683, EPI_ISL_3321684, EPI_ISL_3321685, EPI_ISL_3321686, EPI_ISL_3321687, EPI_ISL_3321689, EPI_ISL_3321690, EPI_ISL_3321691, EPI_ISL_3321692, EPI_ISL_3321693, EPI_ISL_3321695, EPI_ISL_3321696, EPI_ISL_3321697, EPI_ISL_3321699, EPI_ISL_3321701, EPI_ISL_3321702, EPI_ISL_3321703, EPI_ISL_3321704, EPI_ISL_3321705, EPI_ISL_3321706, EPI_ISL_3321707, EPI_ISL_3321708, EPI_ISL_3321709, EPI_ISL_3321710, EPI_ISL_3321711, EPI_ISL_3321712, EPI_ISL_3321713, EPI_ISL_3321714, EPI_ISL_3321715, EPI_ISL_3321716, EPI_ISL_3321717, EPI_ISL_3321719, EPI_ISL_3321720, EPI_ISL_3321721, EPI_ISL_3321722, EPI_ISL_3321723, EPI_ISL_3321724, EPI_ISL_3321726, EPI_ISL_3321727, EPI_ISL_3321728, EPI_ISL_3321729, EPI_ISL_3321730, EPI_ISL_3321731, EPI_ISL_3321732, EPI_ISL_3321733, EPI_ISL_3321735, EPI_ISL_3321736, EPI_ISL_3321737, EPI_ISL_3321738, EPI_ISL_3321739, EPI_ISL_3321740, EPI_ISL_3321742, EPI_ISL_3321743, EPI_ISL_3321744, EPI_ISL_3321745, EPI_ISL_3321746, EPI_ISL_3321747, EPI_ISL_3321748, EPI_ISL_3321749, EPI_ISL_3321750, EPI_ISL_3321751, EPI_ISL_3321752, EPI_ISL_3321753, EPI_ISL_3321754, EPI_ISL_3321755, EPI_ISL_3321756, EPI_ISL_3321757, EPI_ISL_3321758, EPI_ISL_3321759, EPI_ISL_3321760, EPI_ISL_3321761, EPI_ISL_3321762, EPI_ISL_3321764, EPI_ISL_3321765, EPI_ISL_3321766, EPI_ISL_3321767, EPI_ISL_3321768, EPI_ISL_3321769, EPI_ISL_3321770, EPI_ISL_3321771, EPI_ISL_3321772, EPI_ISL_3321773, EPI_ISL_3321774, EPI_ISL_3321775, EPI_ISL_3321776, EPI_ISL_3321777, EPI_ISL_3321778, EPI_ISL_3321779, EPI_ISL_3321780, EPI_ISL_3321781, EPI_ISL_3321782, EPI_ISL_3321783, EPI_ISL_3321784, EPI_ISL_3321785, EPI_ISL_3321786, EPI_ISL_3321787, EPI_ISL_3321788, EPI_ISL_3321789, EPI_ISL_3321790, EPI_ISL_3321791, EPI_ISL_3321792, EPI_ISL_3321793, EPI_ISL_3321795, EPI_ISL_3321796, EPI_ISL_3321797, EPI_ISL_3321798, EPI_ISL_3321799, EPI_ISL_3321801, EPI_ISL_3321802, EPI_ISL_3321803, EPI_ISL_3321804, EPI_ISL_3321805, EPI_ISL_3321806, EPI_ISL_3321808, EPI_ISL_3321809, EPI_ISL_3321810, EPI_ISL_3321811, EPI_ISL_3321812, EPI_ISL_3321813, EPI_ISL_3321814, EPI_ISL_3321815, EPI_ISL_3321816, EPI_ISL_3321817, EPI_ISL_3321818, EPI_ISL_3321819, EPI_ISL_3321820, EPI_ISL_3321821, EPI_ISL_3321822, EPI_ISL_3321823, EPI_ISL_3321826, EPI_ISL_3321827, EPI_ISL_3321828, EPI_ISL_3321829, EPI_ISL_3321830, EPI_ISL_3321831, EPI_ISL_3321832, EPI_ISL_3321833, EPI_ISL_3321834, EPI_ISL_3321835, EPI_ISL_3321836, EPI_ISL_3321837, EPI_ISL_3321838, EPI_ISL_3321839, EPI_ISL_3321840, EPI_ISL_3321841, EPI_ISL_3321842, EPI_ISL_3321843, EPI_ISL_3321844, EPI_ISL_3321845, EPI_ISL_3321846, EPI_ISL_3321847, EPI_ISL_3321848, EPI_ISL_3321850, EPI_ISL_3321851, EPI_ISL_3321852, EPI_ISL_3321853, EPI_ISL_3321854, EPI_ISL_3321855, EPI_ISL_3321856, EPI_ISL_3321858, EPI_ISL_3321859, EPI_ISL_3321860, EPI_ISL_3321861, EPI_ISL_3321862, EPI_ISL_3321863, EPI_ISL_3321864, EPI_ISL_3321866, EPI_ISL_3321867, EPI_ISL_3321869, EPI_ISL_3321870, EPI_ISL_3321871, EPI_ISL_3321872, EPI_ISL_3321873, EPI_ISL_3321874, EPI_ISL_3321875, EPI_ISL_3321876, EPI_ISL_3321878, EPI_ISL_3321879, EPI_ISL_3321880, EPI_ISL_3321881, EPI_ISL_3321883, EPI_ISL_3321885, EPI_ISL_3321886, EPI_ISL_3321887, EPI_ISL_3321888, EPI_ISL_3321890, EPI_ISL_3321891, EPI_ISL_3321895, EPI_ISL_3321897, EPI_ISL_3321900, EPI_ISL_3321902, EPI_ISL_3321904, EPI_ISL_3321908, EPI_ISL_3321911, EPI_ISL_3321913, EPI_ISL_3321915, EPI_ISL_3321917, EPI_ISL_3321919, EPI_ISL_3321920, EPI_ISL_3321921, EPI_ISL_3321925, EPI_ISL_3321926, EPI_ISL_3321929, EPI_ISL_3321930, EPI_ISL_3321931, EPI_ISL_3321932, EPI_ISL_3321933, EPI_ISL_3321935, EPI_ISL_3321936, EPI_ISL_3321937, EPI_ISL_3321938, EPI_ISL_3321939, EPI_ISL_3321940, EPI_ISL_3321941, EPI_ISL_3321942, EPI_ISL_3321943, EPI_ISL_3321944, EPI_ISL_3321945, EPI_ISL_3321946, EPI_ISL_3321947, EPI_ISL_3321948, EPI_ISL_3321949, EPI_ISL_3321950, EPI_ISL_3321951, EPI_ISL_3321952, EPI_ISL_3321953, EPI_ISL_3321954, EPI_ISL_3321955, EPI_ISL_3321956, EPI_ISL_3321958, EPI_ISL_3321959, EPI_ISL_3321961, EPI_ISL_3321964, EPI_ISL_3321965, EPI_ISL_3321966, EPI_ISL_3321967, EPI_ISL_3321968, EPI_ISL_3321969, EPI_ISL_3321970, EPI_ISL_3321971, EPI_ISL_3321972, EPI_ISL_3321973, EPI_ISL_3321974, EPI_ISL_3321975, EPI_ISL_3321976, EPI_ISL_3321978, EPI_ISL_3321979, EPI_ISL_3321980, EPI_ISL_3321981, EPI_ISL_3321982, EPI_ISL_3321983, EPI_ISL_3321984, EPI_ISL_3321985, EPI_ISL_3321986, EPI_ISL_3321987, EPI_ISL_3321988, EPI_ISL_3321989, EPI_ISL_3321990, EPI_ISL_3321991, EPI_ISL_3321992, EPI_ISL_3321993, EPI_ISL_3321994, EPI_ISL_3321995, EPI_ISL_3321996, EPI_ISL_3321997, EPI_ISL_3321998, EPI_ISL_3321999, EPI_ISL_3322000, EPI_ISL_3322001, EPI_ISL_3322002, EPI_ISL_3322003, EPI_ISL_3322005, EPI_ISL_3322006, EPI_ISL_3322007, EPI_ISL_3322009, EPI_ISL_3322011, EPI_ISL_3322012, EPI_ISL_3322013, EPI_ISL_3322014, EPI_ISL_3322016, EPI_ISL_3322017, EPI_ISL_3322018, EPI_ISL_3322019, EPI_ISL_3322020, EPI_ISL_3322021, EPI_ISL_3322022, EPI_ISL_3322023, EPI_ISL_3322024, EPI_ISL_3322025, EPI_ISL_3322026, EPI_ISL_3322028, EPI_ISL_3322029, EPI_ISL_3322030, EPI_ISL_3322031, EPI_ISL_3322032, EPI_ISL_3322033, EPI_ISL_3322034, EPI_ISL_3322036, EPI_ISL_3322037, EPI_ISL_3322038, EPI_ISL_3322039, EPI_ISL_3322041, EPI_ISL_3322042, EPI_ISL_3322043, EPI_ISL_3322044, EPI_ISL_3322045, EPI_ISL_3322046, EPI_ISL_3322047, EPI_ISL_3322048, EPI_ISL_3322050, EPI_ISL_3322051, EPI_ISL_3322052, EPI_ISL_3322053, EPI_ISL_3322054, EPI_ISL_3322058, EPI_ISL_3322060, EPI_ISL_3322062, EPI_ISL_3322063, EPI_ISL_3322064, EPI_ISL_3322065, EPI_ISL_3322066, EPI_ISL_3322067, EPI_ISL_3322068, EPI_ISL_3322069, EPI_ISL_3322070, EPI_ISL_3322071, EPI_ISL_3322073, EPI_ISL_3322075, EPI_ISL_3322076, EPI_ISL_3322077, EPI_ISL_3322078, EPI_ISL_3322079, EPI_ISL_3322080, EPI_ISL_3322081, EPI_ISL_3322082, EPI_ISL_3322083, EPI_ISL_3322084, EPI_ISL_3322085, EPI_ISL_3322086, EPI_ISL_3322087, EPI_ISL_3322088, EPI_ISL_3322089, EPI_ISL_3322090, EPI_ISL_3322091, EPI_ISL_3322092, EPI_ISL_3322093, EPI_ISL_3322094, EPI_ISL_3322095, EPI_ISL_3322097, EPI_ISL_3322098, EPI_ISL_3322099, EPI_ISL_3322100, EPI_ISL_3322101, EPI_ISL_3322102, EPI_ISL_3322103, EPI_ISL_3322104, EPI_ISL_3322105, EPI_ISL_3322106, EPI_ISL_3322107, EPI_ISL_3322108, EPI_ISL_3322109, EPI_ISL_3322110, EPI_ISL_3322111, EPI_ISL_3322112, EPI_ISL_3322113, EPI_ISL_3322114, EPI_ISL_3322115, EPI_ISL_3322116, EPI_ISL_3322117, EPI_ISL_3322118, EPI_ISL_3322120, EPI_ISL_3322121, EPI_ISL_3322122, EPI_ISL_3322124, EPI_ISL_3322125, EPI_ISL_3322126, EPI_ISL_3322127, EPI_ISL_3322128, EPI_ISL_3322129, EPI_ISL_3322130, EPI_ISL_3322131, EPI_ISL_3322132, EPI_ISL_3322133, EPI_ISL_3322134, EPI_ISL_3322135, EPI_ISL_3322136, EPI_ISL_3322137, EPI_ISL_3322138, EPI_ISL_3322139, EPI_ISL_3322140, EPI_ISL_3322143, EPI_ISL_3322144, EPI_ISL_3322145, EPI_ISL_3322146, EPI_ISL_3322147, EPI_ISL_3322149, EPI_ISL_3322150, EPI_ISL_3322151, EPI_ISL_3322152, EPI_ISL_3322153, EPI_ISL_3322154, EPI_ISL_3322155, EPI_ISL_3322156, EPI_ISL_3322157, EPI_ISL_3322158, EPI_ISL_3322159, EPI_ISL_3322160, EPI_ISL_3322161, EPI_ISL_3322162, EPI_ISL_3322164, EPI_ISL_3322165, EPI_ISL_3322166, EPI_ISL_3322168, EPI_ISL_3322169, EPI_ISL_3322171, EPI_ISL_3322172, EPI_ISL_3322173, EPI_ISL_3322175, EPI_ISL_3322176, EPI_ISL_3322177, EPI_ISL_3322178, EPI_ISL_3322179, EPI_ISL_3322180, EPI_ISL_3322181, EPI_ISL_3322182, EPI_ISL_3322183, EPI_ISL_3322184, EPI_ISL_3322185, EPI_ISL_3322186, EPI_ISL_3322189, EPI_ISL_3322190, EPI_ISL_3322191, EPI_ISL_3322192, EPI_ISL_3322196, EPI_ISL_3322197, EPI_ISL_3322198, EPI_ISL_3322199, EPI_ISL_3322200, EPI_ISL_3322201, EPI_ISL_3322203, EPI_ISL_3322204, EPI_ISL_3322205, EPI_ISL_3322206, EPI_ISL_3322207, EPI_ISL_3322208, EPI_ISL_3322209, EPI_ISL_3322210, EPI_ISL_3322211, EPI_ISL_3322213, EPI_ISL_3322215, EPI_ISL_3322216, EPI_ISL_3322217, EPI_ISL_3322218, EPI_ISL_3322219, EPI_ISL_3322220, EPI_ISL_3322221, EPI_ISL_3322222, EPI_ISL_3322223, EPI_ISL_3322224, EPI_ISL_3322226, EPI_ISL_3322227, EPI_ISL_3322228, EPI_ISL_3322229, EPI_ISL_3322230, EPI_ISL_3322231, EPI_ISL_3322232, EPI_ISL_3322233, EPI_ISL_3322234, EPI_ISL_3322235, EPI_ISL_3322236, EPI_ISL_3322237, EPI_ISL_3322238, EPI_ISL_3322239, EPI_ISL_3322240, EPI_ISL_3322241, EPI_ISL_3322242, EPI_ISL_3322243, EPI_ISL_3322244, EPI_ISL_3322245, EPI_ISL_3322246, EPI_ISL_3322247, EPI_ISL_3322248, EPI_ISL_3322249, EPI_ISL_3322250, EPI_ISL_3322251, EPI_ISL_3322253, EPI_ISL_3322254, EPI_ISL_3322255, EPI_ISL_3322257, EPI_ISL_3322258, EPI_ISL_3322260, EPI_ISL_3322262, EPI_ISL_3322263, EPI_ISL_3322264, EPI_ISL_3322265, EPI_ISL_3322266, EPI_ISL_3322267, EPI_ISL_3322268, EPI_ISL_3322269, EPI_ISL_3322270, EPI_ISL_3322271, EPI_ISL_3322272, EPI_ISL_3322273, EPI_ISL_3322274, EPI_ISL_3322275, EPI_ISL_3322276, EPI_ISL_3322277, EPI_ISL_3322278, EPI_ISL_3322279, EPI_ISL_3322280, EPI_ISL_3322281, EPI_ISL_3322282, EPI_ISL_3322283, EPI_ISL_3322284, EPI_ISL_3322285, EPI_ISL_3322286, EPI_ISL_3322287, EPI_ISL_3322288, EPI_ISL_3322290, EPI_ISL_3322291, EPI_ISL_3322293, EPI_ISL_3322294, EPI_ISL_3322295, EPI_ISL_3322296, EPI_ISL_3322297, EPI_ISL_3322299, EPI_ISL_3322300, EPI_ISL_3322301, EPI_ISL_3322302, EPI_ISL_3322303, EPI_ISL_3322304, EPI_ISL_3322305, EPI_ISL_3322306, EPI_ISL_3322307, EPI_ISL_3322308, EPI_ISL_3322309, EPI_ISL_3322310, EPI_ISL_3322311, EPI_ISL_3322312, EPI_ISL_3322313, EPI_ISL_3322314, EPI_ISL_3322316, EPI_ISL_3322317, EPI_ISL_3322318, EPI_ISL_3322321, EPI_ISL_3322322, EPI_ISL_3322324, EPI_ISL_3322325, EPI_ISL_3322326, EPI_ISL_3322327, EPI_ISL_3322329, EPI_ISL_3322331, EPI_ISL_3322332, EPI_ISL_3322333, EPI_ISL_3322334, EPI_ISL_3322335, EPI_ISL_3322336, EPI_ISL_3322337, EPI_ISL_3322338, EPI_ISL_3322339, EPI_ISL_3322340, EPI_ISL_3322342, EPI_ISL_3322343, EPI_ISL_3322344, EPI_ISL_3322345, EPI_ISL_3322346, EPI_ISL_3322347, EPI_ISL_3322348, EPI_ISL_3322349, EPI_ISL_3322350, EPI_ISL_3322351, EPI_ISL_3322352, EPI_ISL_3322353, EPI_ISL_3322354, EPI_ISL_3322355, EPI_ISL_3322356, EPI_ISL_3322357, EPI_ISL_3322361, EPI_ISL_3322362, EPI_ISL_3322363, EPI_ISL_3322366, EPI_ISL_3322367, EPI_ISL_3322368, EPI_ISL_3322369, EPI_ISL_3322370, EPI_ISL_3322371, EPI_ISL_3322372, EPI_ISL_3322373, EPI_ISL_3322374, EPI_ISL_3322375, EPI_ISL_3322376, EPI_ISL_3322377, EPI_ISL_3322378, EPI_ISL_3322379, EPI_ISL_3322380, EPI_ISL_3322381, EPI_ISL_3322382, EPI_ISL_3322383, EPI_ISL_3322384, EPI_ISL_3322385, EPI_ISL_3322386, EPI_ISL_3322387, EPI_ISL_3322388, EPI_ISL_3322389, EPI_ISL_3322390, EPI_ISL_3322391, EPI_ISL_3322392, EPI_ISL_3322393, EPI_ISL_3322394, EPI_ISL_3322395, EPI_ISL_3322396, EPI_ISL_3322397, EPI_ISL_3322398, EPI_ISL_3322400, EPI_ISL_3322401, EPI_ISL_3322402, EPI_ISL_3322403, EPI_ISL_3322404, EPI_ISL_3322405, EPI_ISL_3322406, EPI_ISL_3322407, EPI_ISL_3322408, EPI_ISL_3322409, EPI_ISL_3322411, EPI_ISL_3322412, EPI_ISL_3322413, EPI_ISL_3322415, EPI_ISL_3322416, EPI_ISL_3322417, EPI_ISL_3322418, EPI_ISL_3322420, EPI_ISL_3322421, EPI_ISL_3322422, EPI_ISL_3322423, EPI_ISL_3322424, EPI_ISL_3322425, EPI_ISL_3322426, EPI_ISL_3322427, EPI_ISL_3322428, EPI_ISL_3322429, EPI_ISL_3322430, EPI_ISL_3322431, EPI_ISL_3322432, EPI_ISL_3322433, EPI_ISL_3322436, EPI_ISL_3322437, EPI_ISL_3322438, EPI_ISL_3322439, EPI_ISL_3322440, EPI_ISL_3322442, EPI_ISL_3322443, EPI_ISL_3322444, EPI_ISL_3322446, EPI_ISL_3322448, EPI_ISL_3322449, EPI_ISL_3322450, EPI_ISL_3322451, EPI_ISL_3322452, EPI_ISL_3322453, EPI_ISL_3322454, EPI_ISL_3322455, EPI_ISL_3322456, EPI_ISL_3322457, EPI_ISL_3322459, EPI_ISL_3322461, EPI_ISL_3322462, EPI_ISL_3322464, EPI_ISL_3322465, EPI_ISL_3322466, EPI_ISL_3322467, EPI_ISL_3322468, EPI_ISL_3322469, EPI_ISL_3322470, EPI_ISL_3322471, EPI_ISL_3322472, EPI_ISL_3322474, EPI_ISL_3322475, EPI_ISL_3322476, EPI_ISL_3322478, EPI_ISL_3322479, EPI_ISL_3322480, EPI_ISL_3322481, EPI_ISL_3322482, EPI_ISL_3322483, EPI_ISL_3322484, EPI_ISL_3322485, EPI_ISL_3322486, EPI_ISL_3322487, EPI_ISL_3322488, EPI_ISL_3322489, EPI_ISL_3322490, EPI_ISL_3322491, EPI_ISL_3322492, EPI_ISL_3322493, EPI_ISL_3322496, EPI_ISL_3322497, EPI_ISL_3322498, EPI_ISL_3322499, EPI_ISL_3322501, EPI_ISL_3322502, EPI_ISL_3322503, EPI_ISL_3322504, EPI_ISL_3322505, EPI_ISL_3322506, EPI_ISL_3322507, EPI_ISL_3322508, EPI_ISL_3322509, EPI_ISL_3322510, EPI_ISL_3322512, EPI_ISL_3322513, EPI_ISL_3322514, EPI_ISL_3322515, EPI_ISL_3322516, EPI_ISL_3322517, EPI_ISL_3322518, EPI_ISL_3322519, EPI_ISL_3322520, EPI_ISL_3322521, EPI_ISL_3322522, EPI_ISL_3322523, EPI_ISL_3322524, EPI_ISL_3322525, EPI_ISL_3322527, EPI_ISL_3322528, EPI_ISL_3322529, EPI_ISL_3322530, EPI_ISL_3322531, EPI_ISL_3322532, EPI_ISL_3322533, EPI_ISL_3322534, EPI_ISL_3322535, EPI_ISL_3322536, EPI_ISL_3322537, EPI_ISL_3322538, EPI_ISL_3322539, EPI_ISL_3322540, EPI_ISL_3322541, EPI_ISL_3322543, EPI_ISL_3322544, EPI_ISL_3322545, EPI_ISL_3322546, EPI_ISL_3322547, EPI_ISL_3322548, EPI_ISL_3322549, EPI_ISL_3322550, EPI_ISL_3322551, EPI_ISL_3322552, EPI_ISL_3322553, EPI_ISL_3322554, EPI_ISL_3322555, EPI_ISL_3322556, EPI_ISL_3322557, EPI_ISL_3322558, EPI_ISL_3322559, EPI_ISL_3322560, EPI_ISL_3322561, EPI_ISL_3322562, EPI_ISL_3322564, EPI_ISL_3322565, EPI_ISL_3322566, EPI_ISL_3322567, EPI_ISL_3322568, EPI_ISL_3322569, EPI_ISL_3322570, EPI_ISL_3322572, EPI_ISL_3322573, EPI_ISL_3322574, EPI_ISL_3322575, EPI_ISL_3322576, EPI_ISL_3322577, EPI_ISL_3322578, EPI_ISL_3322579, EPI_ISL_3322580, EPI_ISL_3322581, EPI_ISL_3322582, EPI_ISL_3322583, EPI_ISL_3322584, EPI_ISL_3322585, EPI_ISL_3322586, EPI_ISL_3322587, EPI_ISL_3322588, EPI_ISL_3322589, EPI_ISL_3322590, EPI_ISL_3322591, EPI_ISL_3322592, EPI_ISL_3322593, EPI_ISL_3322594, EPI_ISL_3322595, EPI_ISL_3322597, EPI_ISL_3322598, EPI_ISL_3322599, EPI_ISL_3322601, EPI_ISL_3322602, EPI_ISL_3322603, EPI_ISL_3322604, EPI_ISL_3322605, EPI_ISL_3322606, EPI_ISL_3322607, EPI_ISL_3322609, EPI_ISL_3322610, EPI_ISL_3322611, EPI_ISL_3322612, EPI_ISL_3322614, EPI_ISL_3322615, EPI_ISL_3322617, EPI_ISL_3322618, EPI_ISL_3322619, EPI_ISL_3322620, EPI_ISL_3322621, EPI_ISL_3322622, EPI_ISL_3322623, EPI_ISL_3322624, EPI_ISL_3322625, EPI_ISL_3322626, EPI_ISL_3322627, EPI_ISL_3322628, EPI_ISL_3322629, EPI_ISL_3322630, EPI_ISL_3322631, EPI_ISL_3322632, EPI_ISL_3322633, EPI_ISL_3322634, EPI_ISL_3322635, EPI_ISL_3322636, EPI_ISL_3322637, EPI_ISL_3322638, EPI_ISL_3322639, EPI_ISL_3322640, EPI_ISL_3322641, EPI_ISL_3322642, EPI_ISL_3322643, EPI_ISL_3322644, EPI_ISL_3322645, EPI_ISL_3322646, EPI_ISL_3322647, EPI_ISL_3322648, EPI_ISL_3322649, EPI_ISL_3322650, EPI_ISL_3322651, EPI_ISL_3322652, EPI_ISL_3322653, EPI_ISL_3322654, EPI_ISL_3322655, EPI_ISL_3322656, EPI_ISL_3322658, EPI_ISL_3322659, EPI_ISL_3322660, EPI_ISL_3322662, EPI_ISL_3322664, EPI_ISL_3322665, EPI_ISL_3322666, EPI_ISL_3322667, EPI_ISL_3322668, EPI_ISL_3322670, EPI_ISL_3322671, EPI_ISL_3322672, EPI_ISL_3322673, EPI_ISL_3322674, EPI_ISL_3322675, EPI_ISL_3322676, EPI_ISL_3322677, EPI_ISL_3322678, EPI_ISL_3322679, EPI_ISL_3322680, EPI_ISL_3322681, EPI_ISL_3322682, EPI_ISL_3322685, EPI_ISL_3322687, EPI_ISL_3322689, EPI_ISL_3322691, EPI_ISL_3322694, EPI_ISL_3322696, EPI_ISL_3322700, EPI_ISL_3322705, EPI_ISL_3322707, EPI_ISL_3322709, EPI_ISL_3322712, EPI_ISL_3322716, EPI_ISL_3322719, EPI_ISL_3322721, EPI_ISL_3322723, EPI_ISL_3322725, EPI_ISL_3322727, EPI_ISL_3322730, EPI_ISL_3322732, EPI_ISL_3322733, EPI_ISL_3322736, EPI_ISL_3322738, EPI_ISL_3322740, EPI_ISL_3322742, EPI_ISL_3322744, EPI_ISL_3322747, EPI_ISL_3322749, EPI_ISL_3322753, EPI_ISL_3322755, EPI_ISL_3322762, EPI_ISL_3322765, EPI_ISL_3322767, EPI_ISL_3322769, EPI_ISL_3322772, EPI_ISL_3322774, EPI_ISL_3322776, EPI_ISL_3322778, EPI_ISL_3322781, EPI_ISL_3322783, EPI_ISL_3322785, EPI_ISL_3322787, EPI_ISL_3322789, EPI_ISL_3322791, EPI_ISL_3322794, EPI_ISL_3322796, EPI_ISL_3322798, EPI_ISL_3322802

| | | | |
|---|---|---|---|
| see above | Aegis Sciences Corporation | Centers for Disease Control and Prevention Division of Viral Diseases, Pathogen Discovery | Dakota Howard, Dhwani Batra, Peter W. Cook, Kara Moser, Adrian Paskey, Jason Caravas, Benjamin Rambo-Martin, Shatavia Morrison, Christopher Gulvick, Scott Sammons, Yvette Unoarumhi, Darlene Wagner, Matthew Schmerer, Cyndi Clark, Patrick Campbell, Rob Case, Vikramsinha Ghorpade, Holly Houdeshell, Ola Kvalvaag, Dillon Nall, Ethan Sanders, Alec Vest, Shaun Westlund, Matthew Hardison, Clinton R. Paden, Duncan MacCannell |
| EPI_ISL_3322804, EPI_ISL_3322806 | UPMC Clinical Microbiology Laboratory | Microbial Genome Sequencing Center; Microbial Genomic Epidemiology Laboratory, University of Pittsburgh | Lee H. Harrison, Jane W. Marsh, Marissa P. Griffith, Vatsala R. Srinivasa, Kady D. Waggle, Daniel J. Snyder, Vaughn S. Cooper |
| EPI_ISL_3322807 | Aegis Sciences Corporation | Centers for Disease Control and Prevention Division of Viral Diseases, Pathogen Discovery | Dakota Howard, Dhwani Batra, Peter W. Cook, Kara Moser, Adrian Paskey, Jason Caravas, Benjamin Rambo-Martin, Shatavia Morrison, Christopher Gulvick, Scott Sammons, Yvette Unoarumhi, Darlene Wagner, Matthew Schmerer, Cyndi Clark, Patrick Campbell, Rob Case, Vikramsinha Ghorpade, Holly Houdeshell, Ola Kvalvaag, Dillon Nall, Ethan Sanders, Alec Vest, Shaun Westlund, Matthew Hardison, Clinton R. Paden, Duncan MacCannell |
| EPI_ISL_3322808 | UPMC Clinical Microbiology Laboratory | Microbial Genome Sequencing Center; Microbial Genomic Epidemiology Laboratory, University of Pittsburgh | Lee H. Harrison, Jane W. Marsh, Marissa P. Griffith, Vatsala R. Srinivasa, Kady D. Waggle, Daniel J. Snyder, Vaughn S. Cooper |

EPI_ISL_3322809, EPI_ISL_3322811, EPI_ISL_3322813, EPI_ISL_3322816, EPI_ISL_3322818, EPI_ISL_3322820, EPI_ISL_3322825, EPI_ISL_3322829, EPI_ISL_3322832, EPI_ISL_3322834, EPI_ISL_3322836, EPI_ISL_3322839, EPI_ISL_3322842, EPI_ISL_3322844, EPI_ISL_3322846, EPI_ISL_3322849, EPI_ISL_3322853, EPI_ISL_3322856, EPI_ISL_3322858, EPI_ISL_3322861, EPI_ISL_3322862, EPI_ISL_3322863, EPI_ISL_3322864, EPI_ISL_3322865, EPI_ISL_3322866, EPI_ISL_3322867, EPI_ISL_3322868, EPI_ISL_3322869, EPI_ISL_3322870, EPI_ISL_3322871, EPI_ISL_3322872, EPI_ISL_3322873, EPI_ISL_3322875, EPI_ISL_3322876, EPI_ISL_3322877, EPI_ISL_3322878, EPI_ISL_3322880, EPI_ISL_3322882, EPI_ISL_3322883, EPI_ISL_3322884, EPI_ISL_3322885, EPI_ISL_3322886, EPI_ISL_3322887, EPI_ISL_3322888, EPI_ISL_3322889, EPI_ISL_3322890, EPI_ISL_3322891, EPI_ISL_3322892, EPI_ISL_3322893, EPI_ISL_3322894, EPI_ISL_3322895, EPI_ISL_3322896, EPI_ISL_3322897, EPI_ISL_3322898, EPI_ISL_3322899, EPI_ISL_3322900, EPI_ISL_3322901, EPI_ISL_3322902, EPI_ISL_3322903, EPI_ISL_3322904, EPI_ISL_3322905, EPI_ISL_3322906, EPI_ISL_3322907, EPI_ISL_3322912, EPI_ISL_3322914, EPI_ISL_3322915, EPI_ISL_3322916, EPI_ISL_3322917, EPI_ISL_3322918, EPI_ISL_3322919, EPI_ISL_3322920, EPI_ISL_3322921, EPI_ISL_3322922, EPI_ISL_3322923, EPI_ISL_3322924, EPI_ISL_3322925, EPI_ISL_3322926, EPI_ISL_3322927, EPI_ISL_3322928, EPI_ISL_3322929, EPI_ISL_3322930, EPI_ISL_3322931, EPI_ISL_3322932, EPI_ISL_3322933, EPI_ISL_3322934, EPI_ISL_3322935, EPI_ISL_3322936, EPI_ISL_3322938, EPI_ISL_3322939, EPI_ISL_3322940, EPI_ISL_3322942, EPI_ISL_3322943, EPI_ISL_3322944, EPI_ISL_3322945, EPI_ISL_3322946, EPI_ISL_3322947, EPI_ISL_3322948, EPI_ISL_3322951, EPI_ISL_3322952, EPI_ISL_3322953, EPI_ISL_3322954, EPI_ISL_3322955, EPI_ISL_3322956, EPI_ISL_3322958, EPI_ISL_3322960, EPI_ISL_3322961, EPI_ISL_3322963, EPI_ISL_3322964, EPI_ISL_3322965, EPI_ISL_3322966, EPI_ISL_3322967, EPI_ISL_3322968, EPI_ISL_3322970, EPI_ISL_3322971, EPI_ISL_3322972, EPI_ISL_3322973, EPI_ISL_3322975, EPI_ISL_3322976, EPI_ISL_3322977, EPI_ISL_3322978, EPI_ISL_3322979, EPI_ISL_3322980, EPI_ISL_3322981, EPI_ISL_3322982, EPI_ISL_3322983, EPI_ISL_3322984, EPI_ISL_3322986, EPI_ISL_3322987, EPI_ISL_3322988, EPI_ISL_3322989, EPI_ISL_3322990, EPI_ISL_3322991, EPI_ISL_3322992, EPI_ISL_3322993, EPI_ISL_3322995, EPI_ISL_3322996, EPI_ISL_3322997, EPI_ISL_3322998, EPI_ISL_3322999, EPI_ISL_3323001, EPI_ISL_3323002, EPI_ISL_3323003, EPI_ISL_3323004, EPI_ISL_3323005, EPI_ISL_3323006, EPI_ISL_3323007, EPI_ISL_3323008, EPI_ISL_3323009, EPI_ISL_3323010, EPI_ISL_3323011, EPI_ISL_3323012, EPI_ISL_3323013, EPI_ISL_3323014, EPI_ISL_3323015, EPI_ISL_3323017, EPI_ISL_3323018, EPI_ISL_3323019, EPI_ISL_3323020, EPI_ISL_3323021, EPI_ISL_3323022, EPI_ISL_3323023, EPI_ISL_3323024, EPI_ISL_3323025, EPI_ISL_3323026, EPI_ISL_3323027, EPI_ISL_3323028, EPI_ISL_3323029, EPI_ISL_3323030, EPI_ISL_3323031, EPI_ISL_3323032, EPI_ISL_3323033, EPI_ISL_3323034, EPI_ISL_3323036, EPI_ISL_3323037, EPI_ISL_3323040, EPI_ISL_3323041, EPI_ISL_3323042, EPI_ISL_3323043, EPI_ISL_3323044, EPI_ISL_3323045, EPI_ISL_3323046, EPI_ISL_3323047, EPI_ISL_3323048, EPI_ISL_3323049, EPI_ISL_3323050, EPI_ISL_3323051, EPI_ISL_3323052, EPI_ISL_3323053, EPI_ISL_3323054, EPI_ISL_3323055, EPI_ISL_3323056, EPI_ISL_3323057, EPI_ISL_3323059, EPI_ISL_3323060, EPI_ISL_3323062, EPI_ISL_3323064, EPI_ISL_3323065, EPI_ISL_3323067, EPI_ISL_3323068, EPI_ISL_3323069, EPI_ISL_3323070, EPI_ISL_3323071, EPI_ISL_3323072, EPI_ISL_3323073, EPI_ISL_3323074, EPI_ISL_3323075, EPI_ISL_3323076, EPI_ISL_3323077, EPI_ISL_3323079, EPI_ISL_3323081, EPI_ISL_3323083, EPI_ISL_3323084, EPI_ISL_3323087, EPI_ISL_3323088, EPI_ISL_3323089, EPI_ISL_3323090, EPI_ISL_3323091, EPI_ISL_3323092, EPI_ISL_3323093, EPI_ISL_3323094, EPI_ISL_3323095, EPI_ISL_3323096, EPI_ISL_3323098, EPI_ISL_3323099, EPI_ISL_3323100, EPI_ISL_3323101, EPI_ISL_3323103, EPI_ISL_3323104, EPI_ISL_3323105, EPI_ISL_3323106, EPI_ISL_3323108, EPI_ISL_3323109, EPI_ISL_3323110, EPI_ISL_3323111, EPI_ISL_3323112, EPI_ISL_3323113, EPI_ISL_3323114, EPI_ISL_3323115, EPI_ISL_3323116, EPI_ISL_3323117, EPI_ISL_3323118, EPI_ISL_3323119, EPI_ISL_3323120, EPI_ISL_3323121, EPI_ISL_3323122, EPI_ISL_3323123, EPI_ISL_3323124, EPI_ISL_3323125, EPI_ISL_3323126, EPI_ISL_3323128, EPI_ISL_3323129, EPI_ISL_3323130, EPI_ISL_3323133, EPI_ISL_3323135, EPI_ISL_3323136, EPI_ISL_3323139, EPI_ISL_3323140, EPI_ISL_3323141, EPI_ISL_3323142, EPI_ISL_3323143, EPI_ISL_3323144, EPI_ISL_3323145, EPI_ISL_3323146, EPI_ISL_3323147, EPI_ISL_3323148, EPI_ISL_3323149, EPI_ISL_3323150, EPI_ISL_3323151, EPI_ISL_3323152, EPI_ISL_3323153, EPI_ISL_3323154, EPI_ISL_3323155, EPI_ISL_3323156, EPI_ISL_3323157, EPI_ISL_3323158, EPI_ISL_3323159, EPI_ISL_3323160, EPI_ISL_3323161, EPI_ISL_3323162, EPI_ISL_3323163, EPI_ISL_3323164, EPI_ISL_3323165, EPI_ISL_3323166, EPI_ISL_3323167, EPI_ISL_3323168, EPI_ISL_3323169, EPI_ISL_3323171, EPI_ISL_3323172, EPI_ISL_3323176, EPI_ISL_3323177, EPI_ISL_3323179

| | | | |
|---|---|---|---|
| see above | Aegis Sciences Corporation | Centers for Disease Control and Prevention Division of Viral Diseases, Pathogen Discovery | Dakota Howard, Dhwani Batra, Peter W. Cook, Kara Moser, Adrian Paskey, Jason Caravas, Benjamin Rambo-Martin, Shatavia Morrison, Christopher Gulvick, Scott Sammons, Yvette Unoarumhi, Darlene Wagner, Matthew Schmerer, Cyndi Clark, Patrick Campbell, Rob Case, Vikramsinha Ghorpade, Holly Houdeshell, Ola Kvalvaag, Dillon Nall, Ethan Sanders, Alec Vest, Shaun Westlund, Matthew Hardison, Clinton R. Paden, Duncan MacCannell |
| EPI_ISL_3323181, EPI_ISL_3323182 | Azienda Sanitaria dell'Alto Adige - Laboratorio Aziendale di Microbiologia e Virologia | Azienda Sanitaria dell'Alto Adige | Irene Bianconi |
| EPI_ISL_3323183 | Aegis Sciences Corporation | Centers for Disease Control and Prevention Division of Viral Diseases, Pathogen Discovery | Dakota Howard, Dhwani Batra, Peter W. Cook, Kara Moser, Adrian Paskey, Jason Caravas, Benjamin Rambo-Martin, Shatavia Morrison, Christopher Gulvick, Scott Sammons, Yvette Unoarumhi, Darlene Wagner, Matthew Schmerer, Cyndi Clark, Patrick Campbell, Rob Case, Vikramsinha Ghorpade, Holly Houdeshell, Ola Kvalvaag, Dillon Nall, Ethan Sanders, Alec Vest, Shaun Westlund, Matthew Hardison, Clinton R. Paden, Duncan MacCannell |
| EPI_ISL_3323184, EPI_ISL_3323185 | Azienda Sanitaria dell'Alto Adige - Laboratorio Aziendale di Microbiologia e Virologia | Azienda Sanitaria dell'Alto Adige | Irene Bianconi |
| EPI_ISL_3323186 | Aegis Sciences Corporation | Centers for Disease Control and Prevention Division of Viral Diseases, Pathogen Discovery | Dakota Howard, Dhwani Batra, Peter W. Cook, Kara Moser, Adrian Paskey, Jason Caravas, Benjamin Rambo-Martin, Shatavia Morrison, Christopher Gulvick, Scott Sammons, Yvette Unoarumhi, Darlene Wagner, Matthew Schmerer, Cyndi Clark, Patrick Campbell, Rob Case, Vikramsinha Ghorpade, Holly Houdeshell, Ola Kvalvaag, Dillon Nall, Ethan Sanders, Alec Vest, Shaun Westlund, Matthew Hardison, Clinton R. Paden, Duncan MacCannell |
| EPI_ISL_3323187, EPI_ISL_3323188 | Azienda Sanitaria dell'Alto Adige - Laboratorio Aziendale di Microbiologia e Virologia | Azienda Sanitaria dell'Alto Adige | Irene Bianconi |
| EPI_ISL_3323189 | Aegis Sciences Corporation | Centers for Disease Control and Prevention Division of Viral Diseases, Pathogen Discovery | Dakota Howard, Dhwani Batra, Peter W. Cook, Kara Moser, Adrian Paskey, Jason Caravas, Benjamin Rambo-Martin, Shatavia Morrison, Christopher Gulvick, Scott Sammons, Yvette Unoarumhi, Darlene Wagner, Matthew Schmerer, Cyndi Clark, Patrick Campbell, Rob Case, Vikramsinha Ghorpade, Holly Houdeshell, Ola Kvalvaag, Dillon Nall, Ethan Sanders, Alec Vest, Shaun Westlund, Matthew Hardison, Clinton R. Paden, Duncan MacCannell |
| EPI_ISL_3323190 | Azienda Sanitaria dell'Alto Adige - Laboratorio Aziendale di Microbiologia e Virologia | Azienda Sanitaria dell'Alto Adige | Irene Bianconi |
| EPI_ISL_3323191 | Aegis Sciences Corporation | Centers for Disease Control and Prevention Division of Viral Diseases, Pathogen Discovery | Dakota Howard, Dhwani Batra, Peter W. Cook, Kara Moser, Adrian Paskey, Jason Caravas, Benjamin Rambo-Martin, Shatavia Morrison, Christopher Gulvick, Scott Sammons, Yvette Unoarumhi, Darlene Wagner, Matthew Schmerer, Cyndi Clark, Patrick Campbell, Rob Case, Vikramsinha Ghorpade, Holly Houdeshell, Ola Kvalvaag, Dillon Nall, Ethan Sanders, Alec Vest, Shaun Westlund, Matthew Hardison, Clinton R. Paden, Duncan MacCannell |
| EPI_ISL_3323192, EPI_ISL_3323193 | Azienda Sanitaria dell'Alto Adige - Laboratorio Aziendale di Microbiologia e Virologia | Azienda Sanitaria dell'Alto Adige | Irene Bianconi |
| EPI_ISL_3323194 | Aegis Sciences Corporation | Centers for Disease Control and Prevention Division of Viral Diseases, Pathogen Discovery | Dakota Howard, Dhwani Batra, Peter W. Cook, Kara Moser, Adrian Paskey, Jason Caravas, Benjamin Rambo-Martin, Shatavia Morrison, Christopher Gulvick, Scott Sammons, Yvette Unoarumhi, Darlene Wagner, Matthew Schmerer, Cyndi Clark, Patrick Campbell, Rob Case, Vikramsinha Ghorpade, Holly Houdeshell, Ola Kvalvaag, Dillon Nall, Ethan Sanders, Alec Vest, Shaun Westlund, Matthew Hardison, Clinton R. Paden, Duncan MacCannell |
| EPI_ISL_3323195 | Azienda Sanitaria dell'Alto Adige - Laboratorio Aziendale di Microbiologia e Virologia | Azienda Sanitaria dell'Alto Adige | Irene Bianconi |
| EPI_ISL_3323196 | Aegis Sciences Corporation | Centers for Disease Control and Prevention Division of Viral Diseases, Pathogen Discovery | Dakota Howard, Dhwani Batra, Peter W. Cook, Kara Moser, Adrian Paskey, Jason Caravas, Benjamin Rambo-Martin, Shatavia Morrison, Christopher Gulvick, Scott Sammons, Yvette Unoarumhi, Darlene Wagner, Matthew Schmerer, Cyndi Clark, Patrick Campbell, Rob Case, Vikramsinha Ghorpade, Holly Houdeshell, Ola Kvalvaag, Dillon Nall, Ethan Sanders, Alec Vest, Shaun Westlund, Matthew Hardison, Clinton R. Paden, Duncan MacCannell |
| EPI_ISL_3323197, EPI_ISL_3323198, EPI_ISL_3323200, EPI_ISL_3323201 | Azienda Sanitaria dell'Alto Adige - Laboratorio Aziendale di Microbiologia e Virologia | Azienda Sanitaria dell'Alto Adige | Irene Bianconi |
| EPI_ISL_3323202 | Aegis Sciences Corporation | Centers for Disease Control and Prevention Division of Viral Diseases, Pathogen Discovery | Dakota Howard, Dhwani Batra, Peter W. Cook, Kara Moser, Adrian Paskey, Jason Caravas, Benjamin Rambo-Martin, Shatavia Morrison, Christopher Gulvick, Scott Sammons, Yvette Unoarumhi, Darlene Wagner, Matthew Schmerer, Cyndi Clark, Patrick Campbell, Rob Case, Vikramsinha Ghorpade, Holly Houdeshell, Ola Kvalvaag, Dillon Nall, Ethan Sanders, Alec Vest, Shaun Westlund, Matthew Hardison, Clinton R. Paden, Duncan MacCannell |

| | | | |
|---|---|---|---|
| EPI_ISL_3323203 | Azienda Sanitaria dell'Alto Adige - Laboratorio Aziendale di Microbiologia e Virologia | Azienda Sanitaria dell'Alto Adige | Irene Bianconi |
| EPI_ISL_3323204 | Aegis Sciences Corporation | Centers for Disease Control and Prevention Division of Viral Diseases, Pathogen Discovery | Dakota Howard, Dhwani Batra, Peter W. Cook, Kara Moser, Adrian Paskey, Jason Caravas, Benjamin Rambo-Martin, Shatavia Morrison, Christopher Gulvick, Scott Sammons, Yvette Unoarumhi, Darlene Wagner, Matthew Schmerer, Cyndi Clark, Patrick Campbell, Rob Case, Vikramsinha Ghorpade, Holly Houdeshell, Ola Kvalvaag, Dillon Nall, Ethan Sanders, Alec Vest, Shaun Westlund, Matthew Hardison, Clinton R. Paden, Duncan MacCannell |
| EPI_ISL_3323205, EPI_ISL_3323206 | Azienda Sanitaria dell'Alto Adige - Laboratorio Aziendale di Microbiologia e Virologia | Azienda Sanitaria dell'Alto Adige | Irene Bianconi |
| EPI_ISL_3323207 | Aegis Sciences Corporation | Centers for Disease Control and Prevention Division of Viral Diseases, Pathogen Discovery | Dakota Howard, Dhwani Batra, Peter W. Cook, Kara Moser, Adrian Paskey, Jason Caravas, Benjamin Rambo-Martin, Shatavia Morrison, Christopher Gulvick, Scott Sammons, Yvette Unoarumhi, Darlene Wagner, Matthew Schmerer, Cyndi Clark, Patrick Campbell, Rob Case, Vikramsinha Ghorpade, Holly Houdeshell, Ola Kvalvaag, Dillon Nall, Ethan Sanders, Alec Vest, Shaun Westlund, Matthew Hardison, Clinton R. Paden, Duncan MacCannell |
| EPI_ISL_3323208, EPI_ISL_3323209 | Azienda Sanitaria dell'Alto Adige - Laboratorio Aziendale di Microbiologia e Virologia | Azienda Sanitaria dell'Alto Adige | Irene Bianconi |
| EPI_ISL_3323210 | Aegis Sciences Corporation | Centers for Disease Control and Prevention Division of Viral Diseases, Pathogen Discovery | Dakota Howard, Dhwani Batra, Peter W. Cook, Kara Moser, Adrian Paskey, Jason Caravas, Benjamin Rambo-Martin, Shatavia Morrison, Christopher Gulvick, Scott Sammons, Yvette Unoarumhi, Darlene Wagner, Matthew Schmerer, Cyndi Clark, Patrick Campbell, Rob Case, Vikramsinha Ghorpade, Holly Houdeshell, Ola Kvalvaag, Dillon Nall, Ethan Sanders, Alec Vest, Shaun Westlund, Matthew Hardison, Clinton R. Paden, Duncan MacCannell |
| EPI_ISL_3323211 | Azienda Sanitaria dell'Alto Adige - Laboratorio Aziendale di Microbiologia e Virologia | Azienda Sanitaria dell'Alto Adige | Irene Bianconi |
| EPI_ISL_3323212 | Aegis Sciences Corporation | Centers for Disease Control and Prevention Division of Viral Diseases, Pathogen Discovery | Dakota Howard, Dhwani Batra, Peter W. Cook, Kara Moser, Adrian Paskey, Jason Caravas, Benjamin Rambo-Martin, Shatavia Morrison, Christopher Gulvick, Scott Sammons, Yvette Unoarumhi, Darlene Wagner, Matthew Schmerer, Cyndi Clark, Patrick Campbell, Rob Case, Vikramsinha Ghorpade, Holly Houdeshell, Ola Kvalvaag, Dillon Nall, Ethan Sanders, Alec Vest, Shaun Westlund, Matthew Hardison, Clinton R. Paden, Duncan MacCannell |
| EPI_ISL_3323213, EPI_ISL_3323214 | Azienda Sanitaria dell'Alto Adige - Laboratorio Aziendale di Microbiologia e Virologia | Azienda Sanitaria dell'Alto Adige | Irene Bianconi |
| EPI_ISL_3323215 | Aegis Sciences Corporation | Centers for Disease Control and Prevention Division of Viral Diseases, Pathogen Discovery | Dakota Howard, Dhwani Batra, Peter W. Cook, Kara Moser, Adrian Paskey, Jason Caravas, Benjamin Rambo-Martin, Shatavia Morrison, Christopher Gulvick, Scott Sammons, Yvette Unoarumhi, Darlene Wagner, Matthew Schmerer, Cyndi Clark, Patrick Campbell, Rob Case, Vikramsinha Ghorpade, Holly Houdeshell, Ola Kvalvaag, Dillon Nall, Ethan Sanders, Alec Vest, Shaun Westlund, Matthew Hardison, Clinton R. Paden, Duncan MacCannell |
| EPI_ISL_3323216 | Azienda Sanitaria dell'Alto Adige - Laboratorio Aziendale di Microbiologia e Virologia | Azienda Sanitaria dell'Alto Adige | Irene Bianconi |
| EPI_ISL_3323217 | Aegis Sciences Corporation | Centers for Disease Control and Prevention Division of Viral Diseases, Pathogen Discovery | Dakota Howard, Dhwani Batra, Peter W. Cook, Kara Moser, Adrian Paskey, Jason Caravas, Benjamin Rambo-Martin, Shatavia Morrison, Christopher Gulvick, Scott Sammons, Yvette Unoarumhi, Darlene Wagner, Matthew Schmerer, Cyndi Clark, Patrick Campbell, Rob Case, Vikramsinha Ghorpade, Holly Houdeshell, Ola Kvalvaag, Dillon Nall, Ethan Sanders, Alec Vest, Shaun Westlund, Matthew Hardison, Clinton R. Paden, Duncan MacCannell |
| EPI_ISL_3323218, EPI_ISL_3323219 | Azienda Sanitaria dell'Alto Adige - Laboratorio Aziendale di Microbiologia e Virologia | Azienda Sanitaria dell'Alto Adige | Irene Bianconi |
| EPI_ISL_3323220 | Aegis Sciences Corporation | Centers for Disease Control and Prevention Division of Viral Diseases, Pathogen Discovery | Dakota Howard, Dhwani Batra, Peter W. Cook, Kara Moser, Adrian Paskey, Jason Caravas, Benjamin Rambo-Martin, Shatavia Morrison, Christopher Gulvick, Scott Sammons, Yvette Unoarumhi, Darlene Wagner, Matthew Schmerer, Cyndi Clark, Patrick Campbell, Rob Case, Vikramsinha Ghorpade, Holly Houdeshell, Ola Kvalvaag, Dillon Nall, Ethan Sanders, Alec Vest, Shaun Westlund, Matthew Hardison, Clinton R. Paden, Duncan MacCannell |
| EPI_ISL_3323221, EPI_ISL_3323223, EPI_ISL_3323224 | Azienda Sanitaria dell'Alto Adige - Laboratorio Aziendale di Microbiologia e Virologia | Azienda Sanitaria dell'Alto Adige | Irene Bianconi |
| EPI_ISL_3323225 | Aegis Sciences Corporation | Centers for Disease Control and Prevention Division of Viral Diseases, Pathogen Discovery | Dakota Howard, Dhwani Batra, Peter W. Cook, Kara Moser, Adrian Paskey, Jason Caravas, Benjamin Rambo-Martin, Shatavia Morrison, Christopher Gulvick, Scott Sammons, Yvette Unoarumhi, Darlene Wagner, Matthew Schmerer, Cyndi Clark, Patrick Campbell, Rob Case, Vikramsinha Ghorpade, Holly Houdeshell, Ola Kvalvaag, Dillon Nall, Ethan Sanders, Alec Vest, Shaun Westlund, Matthew Hardison, Clinton R. Paden, Duncan MacCannell |
| EPI_ISL_3323226 | Azienda Sanitaria dell'Alto Adige - Laboratorio Aziendale di Microbiologia e Virologia | Azienda Sanitaria dell'Alto Adige | Irene Bianconi |
| EPI_ISL_3323227 | Aegis Sciences Corporation | Centers for Disease Control and Prevention Division of Viral Diseases, Pathogen Discovery | Dakota Howard, Dhwani Batra, Peter W. Cook, Kara Moser, Adrian Paskey, Jason Caravas, Benjamin Rambo-Martin, Shatavia Morrison, Christopher Gulvick, Scott Sammons, Yvette Unoarumhi, Darlene Wagner, Matthew Schmerer, Cyndi Clark, Patrick Campbell, Rob Case, Vikramsinha Ghorpade, Holly Houdeshell, Ola Kvalvaag, Dillon Nall, Ethan Sanders, Alec Vest, Shaun Westlund, Matthew Hardison, Clinton R. Paden, Duncan MacCannell |
| EPI_ISL_3323228, EPI_ISL_3323229 | Azienda Sanitaria dell'Alto Adige - Laboratorio Aziendale di Microbiologia e Virologia | Azienda Sanitaria dell'Alto Adige | Irene Bianconi |
| EPI_ISL_3323230 | Aegis Sciences Corporation | Centers for Disease Control and Prevention Division of Viral Diseases, Pathogen Discovery | Dakota Howard, Dhwani Batra, Peter W. Cook, Kara Moser, Adrian Paskey, Jason Caravas, Benjamin Rambo-Martin, Shatavia Morrison, Christopher Gulvick, Scott Sammons, Yvette Unoarumhi, Darlene Wagner, Matthew Schmerer, Cyndi Clark, Patrick Campbell, Rob Case, Vikramsinha Ghorpade, Holly Houdeshell, Ola Kvalvaag, Dillon Nall, Ethan Sanders, Alec Vest, Shaun Westlund, Matthew Hardison, Clinton R. Paden, Duncan MacCannell |
| EPI_ISL_3323231, EPI_ISL_3323232 | Azienda Sanitaria dell'Alto Adige - Laboratorio Aziendale di Microbiologia e Virologia | Azienda Sanitaria dell'Alto Adige | Irene Bianconi |
| EPI_ISL_3323233 | Aegis Sciences Corporation | Centers for Disease Control and Prevention Division of Viral Diseases, Pathogen Discovery | Dakota Howard, Dhwani Batra, Peter W. Cook, Kara Moser, Adrian Paskey, Jason Caravas, Benjamin Rambo-Martin, Shatavia Morrison, Christopher Gulvick, Scott Sammons, Yvette Unoarumhi, Darlene Wagner, Matthew Schmerer, Cyndi Clark, Patrick Campbell, Rob Case, Vikramsinha Ghorpade, Holly Houdeshell, Ola Kvalvaag, Dillon Nall, Ethan Sanders, Alec Vest, Shaun Westlund, Matthew Hardison, Clinton R. Paden, Duncan MacCannell |
| EPI_ISL_3323234 | Azienda Sanitaria dell'Alto Adige - Laboratorio Aziendale di Microbiologia e Virologia | Azienda Sanitaria dell'Alto Adige | Irene Bianconi |
| EPI_ISL_3323235 | Aegis Sciences Corporation | Centers for Disease Control and Prevention Division of Viral Diseases, Pathogen Discovery | Dakota Howard, Dhwani Batra, Peter W. Cook, Kara Moser, Adrian Paskey, Jason Caravas, Benjamin Rambo-Martin, Shatavia Morrison, Christopher Gulvick, Scott Sammons, Yvette Unoarumhi, Darlene Wagner, Matthew Schmerer, Cyndi Clark, Patrick Campbell, Rob Case, Vikramsinha Ghorpade, Holly Houdeshell, Ola Kvalvaag, Dillon Nall, Ethan Sanders, Alec Vest, Shaun Westlund, Matthew Hardison, Clinton R. Paden, Duncan MacCannell |
| EPI_ISL_3323236, EPI_ISL_3323237 | Azienda Sanitaria dell'Alto Adige - Laboratorio Aziendale di Microbiologia e Virologia | Azienda Sanitaria dell'Alto Adige | Irene Bianconi |
| EPI_ISL_3323238 | Aegis Sciences Corporation | Centers for Disease Control and Prevention Division of Viral Diseases, Pathogen Discovery | Dakota Howard, Dhwani Batra, Peter W. Cook, Kara Moser, Adrian Paskey, Jason Caravas, Benjamin Rambo-Martin, Shatavia Morrison, Christopher Gulvick, Scott Sammons, Yvette Unoarumhi, Darlene Wagner, Matthew Schmerer, Cyndi Clark, Patrick Campbell, Rob Case, Vikramsinha Ghorpade, Holly Houdeshell, Ola Kvalvaag, Dillon Nall, Ethan Sanders, Alec Vest, Shaun Westlund, Matthew Hardison, Clinton R. Paden, Duncan MacCannell |
| EPI_ISL_3323239, EPI_ISL_3323240 | Azienda Sanitaria dell'Alto Adige - Laboratorio Aziendale di Microbiologia e Virologia | Azienda Sanitaria dell'Alto Adige | Irene Bianconi |
| EPI_ISL_3323241 | Aegis Sciences Corporation | Centers for Disease Control and Prevention Division of Viral Diseases, Pathogen Discovery | Dakota Howard, Dhwani Batra, Peter W. Cook, Kara Moser, Adrian Paskey, Jason Caravas, Benjamin Rambo-Martin, Shatavia Morrison, Christopher Gulvick, Scott Sammons, Yvette Unoarumhi, Darlene Wagner, Matthew Schmerer, Cyndi Clark, Patrick Campbell, Rob Case, Vikramsinha Ghorpade, Holly Houdeshell, Ola Kvalvaag, Dillon Nall, Ethan Sanders, Alec Vest, Shaun Westlund, Matthew Hardison, Clinton R. Paden, Duncan MacCannell |
| EPI_ISL_3323242, EPI_ISL_3323244, EPI_ISL_3323245 | Azienda Sanitaria dell'Alto Adige - Laboratorio Aziendale di Microbiologia e Virologia | Azienda Sanitaria dell'Alto Adige | Irene Bianconi |
| EPI_ISL_3323246 | Aegis Sciences Corporation | Centers for Disease Control and Prevention Division of Viral Diseases, Pathogen Discovery | Dakota Howard, Dhwani Batra, Peter W. Cook, Kara Moser, Adrian Paskey, Jason Caravas, Benjamin Rambo-Martin, Shatavia Morrison, Christopher Gulvick, Scott Sammons, Yvette Unoarumhi, Darlene Wagner, Matthew Schmerer, Cyndi Clark, Patrick Campbell, Rob Case, Vikramsinha Ghorpade, Holly Houdeshell, Ola Kvalvaag, Dillon Nall, Ethan Sanders, Alec Vest, Shaun Westlund, Matthew Hardison, Clinton R. Paden, Duncan MacCannell |
| EPI_ISL_3323247, EPI_ISL_3323248 | Azienda Sanitaria dell'Alto Adige - Laboratorio Aziendale di Microbiologia e Virologia | Azienda Sanitaria dell'Alto Adige | Irene Bianconi |
| EPI_ISL_3323249 | Aegis Sciences Corporation | Centers for Disease Control and Prevention Division of Viral Diseases, Pathogen Discovery | Dakota Howard, Dhwani Batra, Peter W. Cook, Kara Moser, Adrian Paskey, Jason Caravas, Benjamin Rambo-Martin, Shatavia Morrison, Christopher Gulvick, Scott Sammons, Yvette Unoarumhi, Darlene Wagner, Matthew Schmerer, Cyndi Clark, Patrick Campbell, Rob Case, Vikramsinha Ghorpade, Holly Houdeshell, Ola Kvalvaag, Dillon Nall, Ethan Sanders, Alec Vest, Shaun Westlund, Matthew Hardison, Clinton R. Paden, Duncan MacCannell |
| EPI_ISL_3323250 | Azienda Sanitaria dell'Alto Adige - Laboratorio Aziendale di Microbiologia e Virologia | Azienda Sanitaria dell'Alto Adige | Irene Bianconi |
| EPI_ISL_3323251 | Aegis Sciences Corporation | Centers for Disease Control and Prevention Division of Viral Diseases, Pathogen Discovery | Dakota Howard, Dhwani Batra, Peter W. Cook, Kara Moser, Adrian Paskey, Jason Caravas, Benjamin Rambo-Martin, Shatavia Morrison, Christopher Gulvick, Scott Sammons, Yvette Unoarumhi, Darlene Wagner, Matthew Schmerer, Cyndi Clark, Patrick Campbell, Rob Case, Vikramsinha Ghorpade, Holly Houdeshell, Ola Kvalvaag, Dillon Nall, Ethan Sanders, Alec Vest, Shaun Westlund, Matthew Hardison, Clinton R. Paden, Duncan MacCannell |
| EPI_ISL_3323252, EPI_ISL_3323253 | Azienda Sanitaria dell'Alto Adige - Laboratorio Aziendale di Microbiologia e Virologia | Azienda Sanitaria dell'Alto Adige | Irene Bianconi |
| EPI_ISL_3323254 | Aegis Sciences Corporation | Centers for Disease Control and Prevention Division of Viral Diseases, Pathogen Discovery | Dakota Howard, Dhwani Batra, Peter W. Cook, Kara Moser, Adrian Paskey, Jason Caravas, Benjamin Rambo-Martin, Shatavia Morrison, Christopher Gulvick, Scott Sammons, Yvette Unoarumhi, Darlene Wagner, Matthew Schmerer, Cyndi Clark, Patrick Campbell, Rob Case, Vikramsinha Ghorpade, Holly Houdeshell, Ola Kvalvaag, Dillon Nall, Ethan Sanders, Alec Vest, Shaun Westlund, Matthew Hardison, Clinton R. Paden, Duncan MacCannell |
| EPI_ISL_3323255, EPI_ISL_3323256 | Azienda Sanitaria dell'Alto Adige - Laboratorio Aziendale di Microbiologia e Virologia | Azienda Sanitaria dell'Alto Adige | Irene Bianconi |
| EPI_ISL_3323257 | Aegis Sciences Corporation | Centers for Disease Control and Prevention Division of Viral Diseases, Pathogen Discovery | Dakota Howard, Dhwani Batra, Peter W. Cook, Kara Moser, Adrian Paskey, Jason Caravas, Benjamin Rambo-Martin, Shatavia Morrison, Christopher Gulvick, Scott Sammons, Yvette Unoarumhi, Darlene Wagner, Matthew Schmerer, Cyndi Clark, Patrick Campbell, Rob Case, Vikramsinha Ghorpade, Holly Houdeshell, Ola Kvalvaag, Dillon Nall, Ethan Sanders, Alec Vest, Shaun Westlund, Matthew Hardison, Clinton R. Paden, Duncan MacCannell |
| EPI_ISL_3323258, EPI_ISL_3323260, EPI_ISL_3323261, EPI_ISL_3323263, EPI_ISL_3323264 | Azienda Sanitaria dell'Alto Adige - Laboratorio Aziendale di Microbiologia e Virologia | Azienda Sanitaria dell'Alto Adige | Irene Bianconi |
| EPI_ISL_3323265 | Aegis Sciences Corporation | Centers for Disease Control and Prevention Division of Viral Diseases, Pathogen Discovery | Dakota Howard, Dhwani Batra, Peter W. Cook, Kara Moser, Adrian Paskey, Jason Caravas, Benjamin Rambo-Martin, Shatavia Morrison, Christopher Gulvick, Scott Sammons, Yvette Unoarumhi, Darlene Wagner, Matthew Schmerer, Cyndi Clark, Patrick Campbell, Rob Case, Vikramsinha Ghorpade, Holly Houdeshell, Ola Kvalvaag, Dillon Nall, Ethan Sanders, Alec Vest, Shaun Westlund, Matthew Hardison, Clinton R. Paden, Duncan MacCannell |
| EPI_ISL_3323266 | Azienda Sanitaria dell'Alto Adige - Laboratorio Aziendale di Microbiologia e Virologia | Azienda Sanitaria dell'Alto Adige | Irene Bianconi |
| EPI_ISL_3323267 | Aegis Sciences Corporation | Centers for Disease Control and Prevention Division of Viral Diseases, Pathogen Discovery | Dakota Howard, Dhwani Batra, Peter W. Cook, Kara Moser, Adrian Paskey, Jason Caravas, Benjamin Rambo-Martin, Shatavia Morrison, Christopher Gulvick, Scott Sammons, Yvette Unoarumhi, Darlene Wagner, Matthew Schmerer, Cyndi Clark, Patrick Campbell, Rob Case, Vikramsinha Ghorpade, Holly Houdeshell, Ola Kvalvaag, Dillon Nall, Ethan Sanders, Alec Vest, Shaun Westlund, Matthew Hardison, Clinton R. Paden, Duncan MacCannell |
| EPI_ISL_3323268, EPI_ISL_3323269 | Azienda Sanitaria dell'Alto Adige - Laboratorio Aziendale di Microbiologia e Virologia | Azienda Sanitaria dell'Alto Adige | Irene Bianconi |

| | | | |
|---|---|---|---|
| EPI_ISL_3323270 | Aegis Sciences Corporation | Centers for Disease Control and Prevention Division of Viral Diseases, Pathogen Discovery | Dakota Howard, Dhwani Batra, Peter W. Cook, Kara Moser, Adrian Paskey, Jason Caravas, Benjamin Rambo-Martin, Shatavia Morrison, Christopher Gulvick, Scott Sammons, Yvette Unoarumhi, Darlene Wagner, Matthew Schmerer, Cyndi Clark, Patrick Campbell, Rob Case, Vikramsinha Ghorpade, Holly Houdeshell, Ola Kvalvaag, Dillon Nall, Ethan Sanders, Alec Vest, Shaun Westlund, Matthew Hardison, Clinton R. Paden, Duncan MacCannell |
| EPI_ISL_3323271, EPI_ISL_3323272 | Azienda Sanitaria dell'Alto Adige - Laboratorio Aziendale di Microbiologia e Virologia | Azienda Sanitaria dell'Alto Adige | Irene Bianconi |
| EPI_ISL_3323273 | Aegis Sciences Corporation | Centers for Disease Control and Prevention Division of Viral Diseases, Pathogen Discovery | Dakota Howard, Dhwani Batra, Peter W. Cook, Kara Moser, Adrian Paskey, Jason Caravas, Benjamin Rambo-Martin, Shatavia Morrison, Christopher Gulvick, Scott Sammons, Yvette Unoarumhi, Darlene Wagner, Matthew Schmerer, Cyndi Clark, Patrick Campbell, Rob Case, Vikramsinha Ghorpade, Holly Houdeshell, Ola Kvalvaag, Dillon Nall, Ethan Sanders, Alec Vest, Shaun Westlund, Matthew Hardison, Clinton R. Paden, Duncan MacCannell |
| EPI_ISL_3323274, EPI_ISL_3323276, EPI_ISL_3323277 | Azienda Sanitaria dell'Alto Adige - Laboratorio Aziendale di Microbiologia e Virologia | Azienda Sanitaria dell'Alto Adige | Irene Bianconi |
| EPI_ISL_3323278 | Aegis Sciences Corporation | Centers for Disease Control and Prevention Division of Viral Diseases, Pathogen Discovery | Dakota Howard, Dhwani Batra, Peter W. Cook, Kara Moser, Adrian Paskey, Jason Caravas, Benjamin Rambo-Martin, Shatavia Morrison, Christopher Gulvick, Scott Sammons, Yvette Unoarumhi, Darlene Wagner, Matthew Schmerer, Cyndi Clark, Patrick Campbell, Rob Case, Vikramsinha Ghorpade, Holly Houdeshell, Ola Kvalvaag, Dillon Nall, Ethan Sanders, Alec Vest, Shaun Westlund, Matthew Hardison, Clinton R. Paden, Duncan MacCannell |
| EPI_ISL_3323279, EPI_ISL_3323280 | Azienda Sanitaria dell'Alto Adige - Laboratorio Aziendale di Microbiologia e Virologia | Azienda Sanitaria dell'Alto Adige | Irene Bianconi |
| EPI_ISL_3323281 | Aegis Sciences Corporation | Centers for Disease Control and Prevention Division of Viral Diseases, Pathogen Discovery | Dakota Howard, Dhwani Batra, Peter W. Cook, Kara Moser, Adrian Paskey, Jason Caravas, Benjamin Rambo-Martin, Shatavia Morrison, Christopher Gulvick, Scott Sammons, Yvette Unoarumhi, Darlene Wagner, Matthew Schmerer, Cyndi Clark, Patrick Campbell, Rob Case, Vikramsinha Ghorpade, Holly Houdeshell, Ola Kvalvaag, Dillon Nall, Ethan Sanders, Alec Vest, Shaun Westlund, Matthew Hardison, Clinton R. Paden, Duncan MacCannell |
| EPI_ISL_3323282, EPI_ISL_3323284, EPI_ISL_3323285 | Azienda Sanitaria dell'Alto Adige - Laboratorio Aziendale di Microbiologia e Virologia | Azienda Sanitaria dell'Alto Adige | Irene Bianconi |
| EPI_ISL_3323286 | Aegis Sciences Corporation | Centers for Disease Control and Prevention Division of Viral Diseases, Pathogen Discovery | Dakota Howard, Dhwani Batra, Peter W. Cook, Kara Moser, Adrian Paskey, Jason Caravas, Benjamin Rambo-Martin, Shatavia Morrison, Christopher Gulvick, Scott Sammons, Yvette Unoarumhi, Darlene Wagner, Matthew Schmerer, Cyndi Clark, Patrick Campbell, Rob Case, Vikramsinha Ghorpade, Holly Houdeshell, Ola Kvalvaag, Dillon Nall, Ethan Sanders, Alec Vest, Shaun Westlund, Matthew Hardison, Clinton R. Paden, Duncan MacCannell |
| EPI_ISL_3323287, EPI_ISL_3323288 | Azienda Sanitaria dell'Alto Adige - Laboratorio Aziendale di Microbiologia e Virologia | Azienda Sanitaria dell'Alto Adige | Irene Bianconi |
| EPI_ISL_3323289 | Aegis Sciences Corporation | Centers for Disease Control and Prevention Division of Viral Diseases, Pathogen Discovery | Dakota Howard, Dhwani Batra, Peter W. Cook, Kara Moser, Adrian Paskey, Jason Caravas, Benjamin Rambo-Martin, Shatavia Morrison, Christopher Gulvick, Scott Sammons, Yvette Unoarumhi, Darlene Wagner, Matthew Schmerer, Cyndi Clark, Patrick Campbell, Rob Case, Vikramsinha Ghorpade, Holly Houdeshell, Ola Kvalvaag, Dillon Nall, Ethan Sanders, Alec Vest, Shaun Westlund, Matthew Hardison, Clinton R. Paden, Duncan MacCannell |
| EPI_ISL_3323290 | Azienda Sanitaria dell'Alto Adige - Laboratorio Aziendale di Microbiologia e Virologia | Azienda Sanitaria dell'Alto Adige | Irene Bianconi |
| EPI_ISL_3323291 | Aegis Sciences Corporation | Centers for Disease Control and Prevention Division of Viral Diseases, Pathogen Discovery | Dakota Howard, Dhwani Batra, Peter W. Cook, Kara Moser, Adrian Paskey, Jason Caravas, Benjamin Rambo-Martin, Shatavia Morrison, Christopher Gulvick, Scott Sammons, Yvette Unoarumhi, Darlene Wagner, Matthew Schmerer, Cyndi Clark, Patrick Campbell, Rob Case, Vikramsinha Ghorpade, Holly Houdeshell, Ola Kvalvaag, Dillon Nall, Ethan Sanders, Alec Vest, Shaun Westlund, Matthew Hardison, Clinton R. Paden, Duncan MacCannell |
| EPI_ISL_3323292, EPI_ISL_3323293 | Azienda Sanitaria dell'Alto Adige - Laboratorio Aziendale di Microbiologia e Virologia | Azienda Sanitaria dell'Alto Adige | Irene Bianconi |
| EPI_ISL_3323294 | Aegis Sciences Corporation | Centers for Disease Control and Prevention Division of Viral Diseases, Pathogen Discovery | Dakota Howard, Dhwani Batra, Peter W. Cook, Kara Moser, Adrian Paskey, Jason Caravas, Benjamin Rambo-Martin, Shatavia Morrison, Christopher Gulvick, Scott Sammons, Yvette Unoarumhi, Darlene Wagner, Matthew Schmerer, Cyndi Clark, Patrick Campbell, Rob Case, Vikramsinha Ghorpade, Holly Houdeshell, Ola Kvalvaag, Dillon Nall, Ethan Sanders, Alec Vest, Shaun Westlund, Matthew Hardison, Clinton R. Paden, Duncan MacCannell |
| EPI_ISL_3323295, EPI_ISL_3323296 | Azienda Sanitaria dell'Alto Adige - Laboratorio Aziendale di Microbiologia e Virologia | Azienda Sanitaria dell'Alto Adige | Irene Bianconi |
| EPI_ISL_3323297 | Aegis Sciences Corporation | Centers for Disease Control and Prevention Division of Viral Diseases, Pathogen Discovery | Dakota Howard, Dhwani Batra, Peter W. Cook, Kara Moser, Adrian Paskey, Jason Caravas, Benjamin Rambo-Martin, Shatavia Morrison, Christopher Gulvick, Scott Sammons, Yvette Unoarumhi, Darlene Wagner, Matthew Schmerer, Cyndi Clark, Patrick Campbell, Rob Case, Vikramsinha Ghorpade, Holly Houdeshell, Ola Kvalvaag, Dillon Nall, Ethan Sanders, Alec Vest, Shaun Westlund, Matthew Hardison, Clinton R. Paden, Duncan MacCannell |
| EPI_ISL_3323298, EPI_ISL_3323299 | Azienda Sanitaria dell'Alto Adige - Laboratorio Aziendale di Microbiologia e Virologia | Azienda Sanitaria dell'Alto Adige | Irene Bianconi |
| EPI_ISL_3323300 | Aegis Sciences Corporation | Centers for Disease Control and Prevention Division of Viral Diseases, Pathogen Discovery | Dakota Howard, Dhwani Batra, Peter W. Cook, Kara Moser, Adrian Paskey, Jason Caravas, Benjamin Rambo-Martin, Shatavia Morrison, Christopher Gulvick, Scott Sammons, Yvette Unoarumhi, Darlene Wagner, Matthew Schmerer, Cyndi Clark, Patrick Campbell, Rob Case, Vikramsinha Ghorpade, Holly Houdeshell, Ola Kvalvaag, Dillon Nall, Ethan Sanders, Alec Vest, Shaun Westlund, Matthew Hardison, Clinton R. Paden, Duncan MacCannell |
| EPI_ISL_3323301 | Azienda Sanitaria dell'Alto Adige - Laboratorio Aziendale di Microbiologia e Virologia | Azienda Sanitaria dell'Alto Adige | Irene Bianconi |
| EPI_ISL_3323302 | Aegis Sciences Corporation | Centers for Disease Control and Prevention Division of Viral Diseases, Pathogen Discovery | Dakota Howard, Dhwani Batra, Peter W. Cook, Kara Moser, Adrian Paskey, Jason Caravas, Benjamin Rambo-Martin, Shatavia Morrison, Christopher Gulvick, Scott Sammons, Yvette Unoarumhi, Darlene Wagner, Matthew Schmerer, Cyndi Clark, Patrick Campbell, Rob Case, Vikramsinha Ghorpade, Holly Houdeshell, Ola Kvalvaag, Dillon Nall, Ethan Sanders, Alec Vest, Shaun Westlund, Matthew Hardison, Clinton R. Paden, Duncan MacCannell |
| EPI_ISL_3323303, EPI_ISL_3323304 | Azienda Sanitaria dell'Alto Adige - Laboratorio Aziendale di Microbiologia e Virologia | Azienda Sanitaria dell'Alto Adige | Irene Bianconi |
| EPI_ISL_3323305 | Aegis Sciences Corporation | Centers for Disease Control and Prevention Division of Viral Diseases, Pathogen Discovery | Dakota Howard, Dhwani Batra, Peter W. Cook, Kara Moser, Adrian Paskey, Jason Caravas, Benjamin Rambo-Martin, Shatavia Morrison, Christopher Gulvick, Scott Sammons, Yvette Unoarumhi, Darlene Wagner, Matthew Schmerer, Cyndi Clark, Patrick Campbell, Rob Case, Vikramsinha Ghorpade, Holly Houdeshell, Ola Kvalvaag, Dillon Nall, Ethan Sanders, Alec Vest, Shaun Westlund, Matthew Hardison, Clinton R. Paden, Duncan MacCannell |
| EPI_ISL_3323306, EPI_ISL_3323307, EPI_ISL_3323309, EPI_ISL_3323310 | Azienda Sanitaria dell'Alto Adige - Laboratorio Aziendale di Microbiologia e Virologia | Azienda Sanitaria dell'Alto Adige | Irene Bianconi |
| EPI_ISL_3323311 | Aegis Sciences Corporation | Centers for Disease Control and Prevention Division of Viral Diseases, Pathogen Discovery | Dakota Howard, Dhwani Batra, Peter W. Cook, Kara Moser, Adrian Paskey, Jason Caravas, Benjamin Rambo-Martin, Shatavia Morrison, Christopher Gulvick, Scott Sammons, Yvette Unoarumhi, Darlene Wagner, Matthew Schmerer, Cyndi Clark, Patrick Campbell, Rob Case, Vikramsinha Ghorpade, Holly Houdeshell, Ola Kvalvaag, Dillon Nall, Ethan Sanders, Alec Vest, Shaun Westlund, Matthew Hardison, Clinton R. Paden, Duncan MacCannell |
| EPI_ISL_3323312 | Azienda Sanitaria dell'Alto Adige - Laboratorio Aziendale di Microbiologia e Virologia | Azienda Sanitaria dell'Alto Adige | Irene Bianconi |
| EPI_ISL_3323313 | Aegis Sciences Corporation | Centers for Disease Control and Prevention Division of Viral Diseases, Pathogen Discovery | Dakota Howard, Dhwani Batra, Peter W. Cook, Kara Moser, Adrian Paskey, Jason Caravas, Benjamin Rambo-Martin, Shatavia Morrison, Christopher Gulvick, Scott Sammons, Yvette Unoarumhi, Darlene Wagner, Matthew Schmerer, Cyndi Clark, Patrick Campbell, Rob Case, Vikramsinha Ghorpade, Holly Houdeshell, Ola Kvalvaag, Dillon Nall, Ethan Sanders, Alec Vest, Shaun Westlund, Matthew Hardison, Clinton R. Paden, Duncan MacCannell |
| EPI_ISL_3323314 | Azienda Sanitaria dell'Alto Adige - Laboratorio Aziendale di Microbiologia e Virologia | Azienda Sanitaria dell'Alto Adige | Irene Bianconi |
| EPI_ISL_3323315 | Aegis Sciences Corporation | Centers for Disease Control and Prevention Division of Viral Diseases, Pathogen Discovery | Dakota Howard, Dhwani Batra, Peter W. Cook, Kara Moser, Adrian Paskey, Jason Caravas, Benjamin Rambo-Martin, Shatavia Morrison, Christopher Gulvick, Scott Sammons, Yvette Unoarumhi, Darlene Wagner, Matthew Schmerer, Cyndi Clark, Patrick Campbell, Rob Case, Vikramsinha Ghorpade, Holly Houdeshell, Ola Kvalvaag, Dillon Nall, Ethan Sanders, Alec Vest, Shaun Westlund, Matthew Hardison, Clinton R. Paden, Duncan MacCannell |
| EPI_ISL_3323316, EPI_ISL_3323317 | Azienda Sanitaria dell'Alto Adige - Laboratorio Aziendale di Microbiologia e Virologia | Azienda Sanitaria dell'Alto Adige | Irene Bianconi |
| EPI_ISL_3323318 | Aegis Sciences Corporation | Centers for Disease Control and Prevention Division of Viral Diseases, Pathogen Discovery | Dakota Howard, Dhwani Batra, Peter W. Cook, Kara Moser, Adrian Paskey, Jason Caravas, Benjamin Rambo-Martin, Shatavia Morrison, Christopher Gulvick, Scott Sammons, Yvette Unoarumhi, Darlene Wagner, Matthew Schmerer, Cyndi Clark, Patrick Campbell, Rob Case, Vikramsinha Ghorpade, Holly Houdeshell, Ola Kvalvaag, Dillon Nall, Ethan Sanders, Alec Vest, Shaun Westlund, Matthew Hardison, Clinton R. Paden, Duncan MacCannell |
| EPI_ISL_3323319, EPI_ISL_3323320 | Azienda Sanitaria dell'Alto Adige - Laboratorio Aziendale di Microbiologia e Virologia | Azienda Sanitaria dell'Alto Adige | Irene Bianconi |
| EPI_ISL_3323321 | Aegis Sciences Corporation | Centers for Disease Control and Prevention Division of Viral Diseases, Pathogen Discovery | Dakota Howard, Dhwani Batra, Peter W. Cook, Kara Moser, Adrian Paskey, Jason Caravas, Benjamin Rambo-Martin, Shatavia Morrison, Christopher Gulvick, Scott Sammons, Yvette Unoarumhi, Darlene Wagner, Matthew Schmerer, Cyndi Clark, Patrick Campbell, Rob Case, Vikramsinha Ghorpade, Holly Houdeshell, Ola Kvalvaag, Dillon Nall, Ethan Sanders, Alec Vest, Shaun Westlund, Matthew Hardison, Clinton R. Paden, Duncan MacCannell |
| EPI_ISL_3323322, EPI_ISL_3323323 | Azienda Sanitaria dell'Alto Adige - Laboratorio Aziendale di Microbiologia e Virologia | Azienda Sanitaria dell'Alto Adige | Irene Bianconi |
| EPI_ISL_3323324 | Aegis Sciences Corporation | Centers for Disease Control and Prevention Division of Viral Diseases, Pathogen Discovery | Dakota Howard, Dhwani Batra, Peter W. Cook, Kara Moser, Adrian Paskey, Jason Caravas, Benjamin Rambo-Martin, Shatavia Morrison, Christopher Gulvick, Scott Sammons, Yvette Unoarumhi, Darlene Wagner, Matthew Schmerer, Cyndi Clark, Patrick Campbell, Rob Case, Vikramsinha Ghorpade, Holly Houdeshell, Ola Kvalvaag, Dillon Nall, Ethan Sanders, Alec Vest, Shaun Westlund, Matthew Hardison, Clinton R. Paden, Duncan MacCannell |
| EPI_ISL_3323325 | Azienda Sanitaria dell'Alto Adige - Laboratorio Aziendale di Microbiologia e Virologia | Azienda Sanitaria dell'Alto Adige | Irene Bianconi |
| EPI_ISL_3323326 | Aegis Sciences Corporation | Centers for Disease Control and Prevention Division of Viral Diseases, Pathogen Discovery | Dakota Howard, Dhwani Batra, Peter W. Cook, Kara Moser, Adrian Paskey, Jason Caravas, Benjamin Rambo-Martin, Shatavia Morrison, Christopher Gulvick, Scott Sammons, Yvette Unoarumhi, Darlene Wagner, Matthew Schmerer, Cyndi Clark, Patrick Campbell, Rob Case, Vikramsinha Ghorpade, Holly Houdeshell, Ola Kvalvaag, Dillon Nall, Ethan Sanders, Alec Vest, Shaun Westlund, Matthew Hardison, Clinton R. Paden, Duncan MacCannell |
| EPI_ISL_3323327, EPI_ISL_3323328 | Azienda Sanitaria dell'Alto Adige - Laboratorio Aziendale di Microbiologia e Virologia | Azienda Sanitaria dell'Alto Adige | Irene Bianconi |
| EPI_ISL_3323329 | Aegis Sciences Corporation | Centers for Disease Control and Prevention Division of Viral Diseases, Pathogen Discovery | Dakota Howard, Dhwani Batra, Peter W. Cook, Kara Moser, Adrian Paskey, Jason Caravas, Benjamin Rambo-Martin, Shatavia Morrison, Christopher Gulvick, Scott Sammons, Yvette Unoarumhi, Darlene Wagner, Matthew Schmerer, Cyndi Clark, Patrick Campbell, Rob Case, Vikramsinha Ghorpade, Holly Houdeshell, Ola Kvalvaag, Dillon Nall, Ethan Sanders, Alec Vest, Shaun Westlund, Matthew Hardison, Clinton R. Paden, Duncan MacCannell |
| EPI_ISL_3323330 | Azienda Sanitaria dell'Alto Adige - Laboratorio Aziendale di Microbiologia e Virologia | Azienda Sanitaria dell'Alto Adige | Irene Bianconi |

EPI_ISL_3323331, EPI_ISL_3323333, EPI_ISL_3323334, EPI_ISL_3323335, EPI_ISL_3323336, EPI_ISL_3323337, EPI_ISL_3323338, EPI_ISL_3323339, EPI_ISL_3323340, EPI_ISL_3323341, EPI_ISL_3323342, EPI_ISL_3323343, EPI_ISL_3323344, EPI_ISL_3323345, EPI_ISL_3323346, EPI_ISL_3323347, EPI_ISL_3323348, EPI_ISL_3323349, EPI_ISL_3323350, EPI_ISL_3323352, EPI_ISL_3323353, EPI_ISL_3323354, EPI_ISL_3323355, EPI_ISL_3323356, EPI_ISL_3323357, EPI_ISL_3323358, EPI_ISL_3323359, EPI_ISL_3323360, EPI_ISL_3323361, EPI_ISL_3323362, EPI_ISL_3323363, EPI_ISL_3323364, EPI_ISL_3323365, EPI_ISL_3323366, EPI_ISL_3323367, EPI_ISL_3323368, EPI_ISL_3323369, EPI_ISL_3323370, EPI_ISL_3323371, EPI_ISL_3323372, EPI_ISL_3323373, EPI_ISL_3323374, EPI_ISL_3323375, EPI_ISL_3323376, EPI_ISL_3323377, EPI_ISL_3323378, EPI_ISL_3323379, EPI_ISL_3323382, EPI_ISL_3323383, EPI_ISL_3323384, EPI_ISL_3323385, EPI_ISL_3323386, EPI_ISL_3323387, EPI_ISL_3323388, EPI_ISL_3323390, EPI_ISL_3323391, EPI_ISL_3323392, EPI_ISL_3323393, EPI_ISL_3323394, EPI_ISL_3323396, EPI_ISL_3323397, EPI_ISL_3323398, EPI_ISL_3323399, EPI_ISL_3323400, EPI_ISL_3323401, EPI_ISL_3323402, EPI_ISL_3323403, EPI_ISL_3323404, EPI_ISL_3323405, EPI_ISL_3323406, EPI_ISL_3323409, EPI_ISL_3323410, EPI_ISL_3323411, EPI_ISL_3323412, EPI_ISL_3323413, EPI_ISL_3323414, EPI_ISL_3323415, EPI_ISL_3323416, EPI_ISL_3323417, EPI_ISL_3323418, EPI_ISL_3323419, EPI_ISL_3323420, EPI_ISL_3323421, EPI_ISL_3323422, EPI_ISL_3323423, EPI_ISL_3323425, EPI_ISL_3323426, EPI_ISL_3323427, EPI_ISL_3323428, EPI_ISL_3323429, EPI_ISL_3323430, EPI_ISL_3323431, EPI_ISL_3323432, EPI_ISL_3323433, EPI_ISL_3323434, EPI_ISL_3323436, EPI_ISL_3323437, EPI_ISL_3323438, EPI_ISL_3323439, EPI_ISL_3323440, EPI_ISL_3323442, EPI_ISL_3323443, EPI_ISL_3323446, EPI_ISL_3323447, EPI_ISL_3323448, EPI_ISL_3323449, EPI_ISL_3323450, EPI_ISL_3323451, EPI_ISL_3323452, EPI_ISL_3323453, EPI_ISL_3323454, EPI_ISL_3323455, EPI_ISL_3323457, EPI_ISL_3323458, EPI_ISL_3323460, EPI_ISL_3323461, EPI_ISL_3323462, EPI_ISL_3323463, EPI_ISL_3323464, EPI_ISL_3323466, EPI_ISL_3323467, EPI_ISL_3323468, EPI_ISL_3323469, EPI_ISL_3323470, EPI_ISL_3323471, EPI_ISL_3323472, EPI_ISL_3323473, EPI_ISL_3323474, EPI_ISL_3323475, EPI_ISL_3323476, EPI_ISL_3323477, EPI_ISL_3323478, EPI_ISL_3323479, EPI_ISL_3323480, EPI_ISL_3323481, EPI_ISL_3323482, EPI_ISL_3323483, EPI_ISL_3323484, EPI_ISL_3323485, EPI_ISL_3323486, EPI_ISL_3323487, EPI_ISL_3323488, EPI_ISL_3323489, EPI_ISL_3323490, EPI_ISL_3323491, EPI_ISL_3323493, EPI_ISL_3323494, EPI_ISL_3323496, EPI_ISL_3323497, EPI_ISL_3323498, EPI_ISL_3323499, EPI_ISL_3323500, EPI_ISL_3323501, EPI_ISL_3323502, EPI_ISL_3323503, EPI_ISL_3323504, EPI_ISL_3323505, EPI_ISL_3323506, EPI_ISL_3323507, EPI_ISL_3323508, EPI_ISL_3323509, EPI_ISL_3323510, EPI_ISL_3323511, EPI_ISL_3323513, EPI_ISL_3323514, EPI_ISL_3323515, EPI_ISL_3323518, EPI_ISL_3323519, EPI_ISL_3323520, EPI_ISL_3323521, EPI_ISL_3323522, EPI_ISL_3323523, EPI_ISL_3323524, EPI_ISL_3323526, EPI_ISL_3323527, EPI_ISL_3323528,

| | | | |
|---|---|---|---|
| EPI_ISL_3323529, EPI_ISL_3323530, EPI_ISL_3323531, EPI_ISL_3323532, EPI_ISL_3323533, EPI_ISL_3323534, EPI_ISL_3323535, EPI_ISL_3323536, EPI_ISL_3323537, EPI_ISL_3323538, EPI_ISL_3323539, EPI_ISL_3323540, EPI_ISL_3323541, EPI_ISL_3323542, EPI_ISL_3323543, EPI_ISL_3323544, EPI_ISL_3323545, EPI_ISL_3323546, EPI_ISL_3323547, EPI_ISL_3323548, EPI_ISL_3323549, EPI_ISL_3323550, EPI_ISL_3323551, EPI_ISL_3323552, EPI_ISL_3323553, EPI_ISL_3323554, EPI_ISL_3323555, EPI_ISL_3323556, EPI_ISL_3323557, EPI_ISL_3323558, EPI_ISL_3323559, EPI_ISL_3323560, EPI_ISL_3323561, EPI_ISL_3323562, EPI_ISL_3323563, EPI_ISL_3323564, EPI_ISL_3323565, EPI_ISL_3323566, EPI_ISL_3323567, EPI_ISL_3323568, EPI_ISL_3323569, EPI_ISL_3323570, EPI_ISL_3323571, EPI_ISL_3323572, EPI_ISL_3323573, EPI_ISL_3323574, EPI_ISL_3323575, EPI_ISL_3323576, EPI_ISL_3323577, EPI_ISL_3323578, EPI_ISL_3323580, EPI_ISL_3323581, EPI_ISL_3323582, EPI_ISL_3323583, EPI_ISL_3323584, EPI_ISL_3323585, EPI_ISL_3323586, EPI_ISL_3323587, EPI_ISL_3323589, EPI_ISL_3323590, EPI_ISL_3323591, EPI_ISL_3323592, EPI_ISL_3323593, EPI_ISL_3323594, EPI_ISL_3323595, EPI_ISL_3323596, EPI_ISL_3323597, EPI_ISL_3323598, EPI_ISL_3323599, EPI_ISL_3323600, EPI_ISL_3323601, EPI_ISL_3323602, EPI_ISL_3323603, EPI_ISL_3323604, EPI_ISL_3323605, EPI_ISL_3323606, EPI_ISL_3323607, EPI_ISL_3323609, EPI_ISL_3323610, EPI_ISL_3323611, EPI_ISL_3323612, EPI_ISL_3323613, EPI_ISL_3323614, EPI_ISL_3323615, EPI_ISL_3323616, EPI_ISL_3323617, EPI_ISL_3323618, EPI_ISL_3323619, EPI_ISL_3323621, EPI_ISL_3323622, EPI_ISL_3323623, EPI_ISL_3323624, EPI_ISL_3323625, EPI_ISL_3323626, EPI_ISL_3323627, EPI_ISL_3323628, EPI_ISL_3323629, EPI_ISL_3323630, EPI_ISL_3323631, EPI_ISL_3323632, EPI_ISL_3323633, EPI_ISL_3323635, EPI_ISL_3323636, EPI_ISL_3323637, EPI_ISL_3323638, EPI_ISL_3323639, EPI_ISL_3323640, EPI_ISL_3323641, EPI_ISL_3323642, EPI_ISL_3323643, EPI_ISL_3323644, EPI_ISL_3323645, EPI_ISL_3323646, EPI_ISL_3323647, EPI_ISL_3323648, EPI_ISL_3323649, EPI_ISL_3323650, EPI_ISL_3323651, EPI_ISL_3323652, EPI_ISL_3323654, EPI_ISL_3323655, EPI_ISL_3323656, EPI_ISL_3323657, EPI_ISL_3323658, EPI_ISL_3323659, EPI_ISL_3323660, EPI_ISL_3323661, EPI_ISL_3323662, EPI_ISL_3323663, EPI_ISL_3323665, EPI_ISL_3323666, EPI_ISL_3323667, EPI_ISL_3323668, EPI_ISL_3323669, EPI_ISL_3323670, EPI_ISL_3323671, EPI_ISL_3323672, EPI_ISL_3323673, EPI_ISL_3323674, EPI_ISL_3323675, EPI_ISL_3323676, EPI_ISL_3323677, EPI_ISL_3323678, EPI_ISL_3323679, EPI_ISL_3323682, EPI_ISL_3323683, EPI_ISL_3323684, EPI_ISL_3323685, EPI_ISL_3323686, EPI_ISL_3323687, EPI_ISL_3323688, EPI_ISL_3323689, EPI_ISL_3323691, EPI_ISL_3323692, EPI_ISL_3323693, EPI_ISL_3323694, EPI_ISL_3323695, EPI_ISL_3323696, EPI_ISL_3323697, EPI_ISL_3323698, EPI_ISL_3323699, EPI_ISL_3323700, EPI_ISL_3323701, EPI_ISL_3323702, EPI_ISL_3323703, EPI_ISL_3323704, EPI_ISL_3323705, EPI_ISL_3323706, EPI_ISL_3323707, EPI_ISL_3323708, EPI_ISL_3323709, EPI_ISL_3323710, EPI_ISL_3323711, EPI_ISL_3323712, EPI_ISL_3323713, EPI_ISL_3323714, EPI_ISL_3323717, EPI_ISL_3323718, EPI_ISL_3323719, EPI_ISL_3323720, EPI_ISL_3323721, EPI_ISL_3323722, EPI_ISL_3323723, EPI_ISL_3323726, EPI_ISL_3323727, EPI_ISL_3323728, EPI_ISL_3323729, EPI_ISL_3323730, EPI_ISL_3323732, EPI_ISL_3323733, EPI_ISL_3323734, EPI_ISL_3323735, EPI_ISL_3323736, EPI_ISL_3323737, EPI_ISL_3323738, EPI_ISL_3323739, EPI_ISL_3323740, EPI_ISL_3323741, EPI_ISL_3323742, EPI_ISL_3323744, EPI_ISL_3323745, EPI_ISL_3323746, EPI_ISL_3323747, EPI_ISL_3323748, EPI_ISL_3323749, EPI_ISL_3323750, EPI_ISL_3323751, EPI_ISL_3323752, EPI_ISL_3323753, EPI_ISL_3323754, EPI_ISL_3323755, EPI_ISL_3323756, EPI_ISL_3323757, EPI_ISL_3323758, EPI_ISL_3323759, EPI_ISL_3323760, EPI_ISL_3323762, EPI_ISL_3323763, EPI_ISL_3323764, EPI_ISL_3323765, EPI_ISL_3323766, EPI_ISL_3323767, EPI_ISL_3323768, EPI_ISL_3323769, EPI_ISL_3323770, EPI_ISL_3323771, EPI_ISL_3323772, EPI_ISL_3323773, EPI_ISL_3323774, EPI_ISL_3323775, EPI_ISL_3323776, EPI_ISL_3323777, EPI_ISL_3323778, EPI_ISL_3323779, EPI_ISL_3323780, EPI_ISL_3323781, EPI_ISL_3323782, EPI_ISL_3323783, EPI_ISL_3323784, EPI_ISL_3323785, EPI_ISL_3323786, EPI_ISL_3323787, EPI_ISL_3323788, EPI_ISL_3323789, EPI_ISL_3323790, EPI_ISL_3323791, EPI_ISL_3323793, EPI_ISL_3323794, EPI_ISL_3323795, EPI_ISL_3323796, EPI_ISL_3323797, EPI_ISL_3323798, EPI_ISL_3323799, EPI_ISL_3323801, EPI_ISL_3323802, EPI_ISL_3323803, EPI_ISL_3323804, EPI_ISL_3323805, EPI_ISL_3323807, EPI_ISL_3323808, EPI_ISL_3323809, EPI_ISL_3323810, EPI_ISL_3323811, EPI_ISL_3323812, EPI_ISL_3323814, EPI_ISL_3323816, EPI_ISL_3323818, EPI_ISL_3323819, EPI_ISL_3323820, EPI_ISL_3323821, EPI_ISL_3323823, EPI_ISL_3323824, EPI_ISL_3323826, EPI_ISL_3323827, EPI_ISL_3323828, EPI_ISL_3323829, EPI_ISL_3323830, EPI_ISL_3323831, EPI_ISL_3323832, EPI_ISL_3323833, EPI_ISL_3323834, EPI_ISL_3323835, EPI_ISL_3323836, EPI_ISL_3323838, EPI_ISL_3323839, EPI_ISL_3323840, EPI_ISL_3323841, EPI_ISL_3323842, EPI_ISL_3323844, EPI_ISL_3323845, EPI_ISL_3323846, EPI_ISL_3323848, EPI_ISL_3323849, EPI_ISL_3323850, EPI_ISL_3323851, EPI_ISL_3323852, EPI_ISL_3323853, EPI_ISL_3323855, EPI_ISL_3323857, EPI_ISL_3323858, EPI_ISL_3323859, EPI_ISL_3323861, EPI_ISL_3323862, EPI_ISL_3323863, EPI_ISL_3323865, EPI_ISL_3323866, EPI_ISL_3323867, EPI_ISL_3323868, EPI_ISL_3323869, EPI_ISL_3323870, EPI_ISL_3323871, EPI_ISL_3323873, EPI_ISL_3323874, EPI_ISL_3323875, EPI_ISL_3323876, EPI_ISL_3323877, EPI_ISL_3323878, EPI_ISL_3323880, EPI_ISL_3323881, EPI_ISL_3323883, EPI_ISL_3323884, EPI_ISL_3323885, EPI_ISL_3323886, EPI_ISL_3323887, EPI_ISL_3323889, EPI_ISL_3323890, EPI_ISL_3323891, EPI_ISL_3323892, EPI_ISL_3323893, EPI_ISL_3323894, EPI_ISL_3323895, EPI_ISL_3323896, EPI_ISL_3323898, EPI_ISL_3323899, EPI_ISL_3323900, EPI_ISL_3323901, EPI_ISL_3323902, EPI_ISL_3323905, EPI_ISL_3323908, EPI_ISL_3323909, EPI_ISL_3323910, EPI_ISL_3323911, EPI_ISL_3323912, EPI_ISL_3323913, EPI_ISL_3323915, EPI_ISL_3323916, EPI_ISL_3323917, EPI_ISL_3323918, EPI_ISL_3323919, EPI_ISL_3323920, EPI_ISL_3323921, EPI_ISL_3323922, EPI_ISL_3323923, EPI_ISL_3323924, EPI_ISL_3323925, EPI_ISL_3323926, EPI_ISL_3323928, EPI_ISL_3323929, EPI_ISL_3323930, EPI_ISL_3323931, EPI_ISL_3323932, EPI_ISL_3323934, EPI_ISL_3323935, EPI_ISL_3323936, EPI_ISL_3323937, EPI_ISL_3323939, EPI_ISL_3323940, EPI_ISL_3323941, EPI_ISL_3323942, EPI_ISL_3323943, EPI_ISL_3323944, EPI_ISL_3323945, EPI_ISL_3323946, EPI_ISL_3323947, EPI_ISL_3323949, EPI_ISL_3323950, EPI_ISL_3323951, EPI_ISL_3323952, EPI_ISL_3323953, EPI_ISL_3323955, EPI_ISL_3323956, EPI_ISL_3323957, EPI_ISL_3323958, EPI_ISL_3323959, EPI_ISL_3323961, EPI_ISL_3323962, EPI_ISL_3323963, EPI_ISL_3323964, EPI_ISL_3323967, EPI_ISL_3323968, EPI_ISL_3323969, EPI_ISL_3323970, EPI_ISL_3323971, EPI_ISL_3323972, EPI_ISL_3323973, EPI_ISL_3323974, EPI_ISL_3323975, EPI_ISL_3323976, EPI_ISL_3323977, EPI_ISL_3323978, EPI_ISL_3323979, EPI_ISL_3323981, EPI_ISL_3323982, EPI_ISL_3323983, EPI_ISL_3323984, EPI_ISL_3323985, EPI_ISL_3323986, EPI_ISL_3323987, EPI_ISL_3323988, EPI_ISL_3323989, EPI_ISL_3323990, EPI_ISL_3323991, EPI_ISL_3323992, EPI_ISL_3323993, EPI_ISL_3323994, EPI_ISL_3323995, EPI_ISL_3323996, EPI_ISL_3323997, EPI_ISL_3323998, EPI_ISL_3323999, EPI_ISL_3324000, EPI_ISL_3324001, EPI_ISL_3324003, EPI_ISL_3324004, EPI_ISL_3324005, EPI_ISL_3324007, EPI_ISL_3324008, EPI_ISL_3324009, EPI_ISL_3324010, EPI_ISL_3324011, EPI_ISL_3324012, EPI_ISL_3324013, EPI_ISL_3324014, EPI_ISL_3324015, EPI_ISL_3324016, EPI_ISL_3324017, EPI_ISL_3324018, EPI_ISL_3324019, EPI_ISL_3324021, EPI_ISL_3324022, EPI_ISL_3324026, EPI_ISL_3324028, EPI_ISL_3324029, EPI_ISL_3324030, EPI_ISL_3324033, EPI_ISL_3324034, EPI_ISL_3324035, EPI_ISL_3324036, EPI_ISL_3324037, EPI_ISL_3324038, EPI_ISL_3324039, EPI_ISL_3324041, EPI_ISL_3324042, EPI_ISL_3324043, EPI_ISL_3324044, EPI_ISL_3324045, EPI_ISL_3324047, EPI_ISL_3324048, EPI_ISL_3324049, EPI_ISL_3324050, EPI_ISL_3324051, EPI_ISL_3324053, EPI_ISL_3324054, EPI_ISL_3324056, EPI_ISL_3324057, EPI_ISL_3324058, EPI_ISL_3324059, EPI_ISL_3324060, EPI_ISL_3324061, EPI_ISL_3324062, EPI_ISL_3324063, EPI_ISL_3324065, EPI_ISL_3324066, EPI_ISL_3324067, EPI_ISL_3324068, EPI_ISL_3324069, EPI_ISL_3324070, EPI_ISL_3324071, EPI_ISL_3324072, EPI_ISL_3324073, EPI_ISL_3324074, EPI_ISL_3324075, EPI_ISL_3324076, EPI_ISL_3324077, EPI_ISL_3324079, EPI_ISL_3324084, EPI_ISL_3324085, EPI_ISL_3324086, EPI_ISL_3324087, EPI_ISL_3324088, EPI_ISL_3324089, EPI_ISL_3324090, EPI_ISL_3324091, EPI_ISL_3324092, EPI_ISL_3324093, EPI_ISL_3324094, EPI_ISL_3324095, EPI_ISL_3324096, EPI_ISL_3324097, EPI_ISL_3324098, EPI_ISL_3324100, EPI_ISL_3324101, EPI_ISL_3324102, EPI_ISL_3324103, EPI_ISL_3324104, EPI_ISL_3324105, EPI_ISL_3324106, EPI_ISL_3324107, EPI_ISL_3324108, EPI_ISL_3324109, EPI_ISL_3324110, EPI_ISL_3324112, EPI_ISL_3324115, EPI_ISL_3324116, EPI_ISL_3324117, EPI_ISL_3324118, EPI_ISL_3324119, EPI_ISL_3324120, EPI_ISL_3324121, EPI_ISL_3324122, EPI_ISL_3324123, EPI_ISL_3324124, EPI_ISL_3324125, EPI_ISL_3324126, EPI_ISL_3324128, EPI_ISL_3324129, EPI_ISL_3324131, EPI_ISL_3324132, EPI_ISL_3324133, EPI_ISL_3324134, EPI_ISL_3324135, EPI_ISL_3324136, EPI_ISL_3324137, EPI_ISL_3324138, EPI_ISL_3324139, EPI_ISL_3324140, EPI_ISL_3324141, EPI_ISL_3324142, EPI_ISL_3324143, EPI_ISL_3324144, EPI_ISL_3324145, EPI_ISL_3324146, EPI_ISL_3324147, EPI_ISL_3324148, EPI_ISL_3324151, EPI_ISL_3324153, EPI_ISL_3324154, EPI_ISL_3324155, EPI_ISL_3324158, EPI_ISL_3324159, EPI_ISL_3324160, EPI_ISL_3324161, EPI_ISL_3324162, EPI_ISL_3324163, EPI_ISL_3324165, EPI_ISL_3324166, EPI_ISL_3324167, EPI_ISL_3324168, EPI_ISL_3324169, EPI_ISL_3324170, EPI_ISL_3324171, EPI_ISL_3324172, EPI_ISL_3324173, EPI_ISL_3324174, EPI_ISL_3324175, EPI_ISL_3324176, EPI_ISL_3324177, EPI_ISL_3324178, EPI_ISL_3324179, EPI_ISL_3324180, EPI_ISL_3324181, EPI_ISL_3324182, EPI_ISL_3324183, EPI_ISL_3324184, EPI_ISL_3324185, EPI_ISL_3324186, EPI_ISL_3324187, EPI_ISL_3324188, EPI_ISL_3324189, EPI_ISL_3324190, EPI_ISL_3324191, EPI_ISL_3324192, EPI_ISL_3324193, EPI_ISL_3324195, EPI_ISL_3324196, EPI_ISL_3324197, EPI_ISL_3324198, EPI_ISL_3324199, EPI_ISL_3324200, EPI_ISL_3324201, EPI_ISL_3324202, EPI_ISL_3324203, EPI_ISL_3324205, EPI_ISL_3324206, EPI_ISL_3324207, EPI_ISL_3324208, EPI_ISL_3324209, EPI_ISL_3324210, EPI_ISL_3324211, EPI_ISL_3324213, EPI_ISL_3324214, EPI_ISL_3324216, EPI_ISL_3324217, EPI_ISL_3324218, EPI_ISL_3324220, EPI_ISL_3324221, EPI_ISL_3324222, EPI_ISL_3324223, EPI_ISL_3324225, EPI_ISL_3324226, EPI_ISL_3324227, EPI_ISL_3324228, EPI_ISL_3324229, EPI_ISL_3324230, EPI_ISL_3324231, EPI_ISL_3324232, EPI_ISL_3324233, EPI_ISL_3324234, EPI_ISL_3324235, EPI_ISL_3324236, EPI_ISL_3324237, EPI_ISL_3324238, EPI_ISL_3324239, EPI_ISL_3324240, EPI_ISL_3324241, EPI_ISL_3324242, EPI_ISL_3324243, EPI_ISL_3324244, EPI_ISL_3324245, EPI_ISL_3324246, EPI_ISL_3324247, EPI_ISL_3324248, EPI_ISL_3324249, EPI_ISL_3324251, EPI_ISL_3324252, EPI_ISL_3324253, EPI_ISL_3324254, EPI_ISL_3324255, EPI_ISL_3324256, EPI_ISL_3324257, EPI_ISL_3324258, EPI_ISL_3324259, EPI_ISL_3324260, EPI_ISL_3324261, EPI_ISL_3324262, EPI_ISL_3324263, EPI_ISL_3324264, EPI_ISL_3324266, EPI_ISL_3324267, EPI_ISL_3324268, EPI_ISL_3324269, EPI_ISL_3324270, EPI_ISL_3324271, EPI_ISL_3324272, EPI_ISL_3324273, EPI_ISL_3324274, EPI_ISL_3324275, EPI_ISL_3324276, EPI_ISL_3324277, EPI_ISL_3324278, EPI_ISL_3324279, EPI_ISL_3324280, EPI_ISL_3324281, EPI_ISL_3324282, EPI_ISL_3324283, EPI_ISL_3324284, EPI_ISL_3324285, EPI_ISL_3324286, EPI_ISL_3324287, EPI_ISL_3324288, EPI_ISL_3324289, EPI_ISL_3324290, EPI_ISL_3324291, EPI_ISL_3324293, EPI_ISL_3324294, EPI_ISL_3324295, EPI_ISL_3324296, EPI_ISL_3324297, EPI_ISL_3324298, EPI_ISL_3324299, EPI_ISL_3324300, EPI_ISL_3324301, EPI_ISL_3324302, EPI_ISL_3324303, EPI_ISL_3324304, EPI_ISL_3324305, EPI_ISL_3324306, EPI_ISL_3324307, EPI_ISL_3324308, EPI_ISL_3324309, EPI_ISL_3324310, EPI_ISL_3324311, EPI_ISL_3324312, EPI_ISL_3324313, EPI_ISL_3324314, EPI_ISL_3324315, EPI_ISL_3324316, EPI_ISL_3324318, EPI_ISL_3324320, EPI_ISL_3324321, EPI_ISL_3324322, EPI_ISL_3324323, EPI_ISL_3324324, EPI_ISL_3324325, EPI_ISL_3324326, EPI_ISL_3324327, EPI_ISL_3324328, EPI_ISL_3324329, EPI_ISL_3324330, EPI_ISL_3324331, EPI_ISL_3324332, EPI_ISL_3324333, EPI_ISL_3324335, EPI_ISL_3324336, EPI_ISL_3324337, EPI_ISL_3324338, EPI_ISL_3324339, EPI_ISL_3324340, EPI_ISL_3324341, EPI_ISL_3324342, EPI_ISL_3324343, EPI_ISL_3324344, EPI_ISL_3324345, EPI_ISL_3324346, EPI_ISL_3324348, EPI_ISL_3324349, EPI_ISL_3324350, EPI_ISL_3324351, EPI_ISL_3324352, EPI_ISL_3324353, EPI_ISL_3324354, EPI_ISL_3324355, EPI_ISL_3324356, EPI_ISL_3324357, EPI_ISL_3324358, EPI_ISL_3324359, EPI_ISL_3324360, EPI_ISL_3324361, EPI_ISL_3324362, EPI_ISL_3324363, EPI_ISL_3324364, EPI_ISL_3324365, EPI_ISL_3324366, EPI_ISL_3324367, EPI_ISL_3324368, EPI_ISL_3324369, EPI_ISL_3324370, EPI_ISL_3324371, EPI_ISL_3324372, EPI_ISL_3324373, EPI_ISL_3324374, EPI_ISL_3324375, EPI_ISL_3324376, EPI_ISL_3324377, EPI_ISL_3324378, EPI_ISL_3324379, EPI_ISL_3324380, EPI_ISL_3324381, EPI_ISL_3324382, EPI_ISL_3324383, EPI_ISL_3324384, EPI_ISL_3324385, EPI_ISL_3324386, EPI_ISL_3324387, EPI_ISL_3324388, EPI_ISL_3324389, EPI_ISL_3324390, EPI_ISL_3324391, EPI_ISL_3324392, EPI_ISL_3324393, EPI_ISL_3324394, EPI_ISL_3324395, EPI_ISL_3324396, EPI_ISL_3324397, EPI_ISL_3324398, EPI_ISL_3324399, EPI_ISL_3324400, EPI_ISL_3324401, EPI_ISL_3324402, EPI_ISL_3324403, EPI_ISL_3324404, EPI_ISL_3324405, EPI_ISL_3324406, EPI_ISL_3324407, EPI_ISL_3324408, EPI_ISL_3324409, EPI_ISL_3324411, EPI_ISL_3324412, EPI_ISL_3324413, EPI_ISL_3324414, EPI_ISL_3324415, EPI_ISL_3324416, EPI_ISL_3324417, EPI_ISL_3324418, EPI_ISL_3324419, EPI_ISL_3324420, EPI_ISL_3324421, EPI_ISL_3324423, EPI_ISL_3324424, EPI_ISL_3324426, EPI_ISL_3324427, EPI_ISL_3324428, EPI_ISL_3324429, EPI_ISL_3324430, EPI_ISL_3324431, EPI_ISL_3324432, EPI_ISL_3324433, EPI_ISL_3324434, EPI_ISL_3324435, EPI_ISL_3324436, EPI_ISL_3324437, EPI_ISL_3324438, EPI_ISL_3324439, EPI_ISL_3324440, EPI_ISL_3324441, EPI_ISL_3324442, EPI_ISL_3324443, EPI_ISL_3324444, EPI_ISL_3324445, EPI_ISL_3324446, EPI_ISL_3324447, EPI_ISL_3324448, EPI_ISL_3324449, EPI_ISL_3324450, EPI_ISL_3324451, EPI_ISL_3324452, EPI_ISL_3324453, EPI_ISL_3324454, EPI_ISL_3324455, EPI_ISL_3324456, EPI_ISL_3324457, EPI_ISL_3324458, EPI_ISL_3324459, EPI_ISL_3324460, EPI_ISL_3324461, EPI_ISL_3324462, EPI_ISL_3324463, EPI_ISL_3324464, EPI_ISL_3324465, EPI_ISL_3324466, EPI_ISL_3324467, EPI_ISL_3324468, EPI_ISL_3324469, EPI_ISL_3324470, EPI_ISL_3324471, EPI_ISL_3324472, EPI_ISL_3324473, EPI_ISL_3324474, EPI_ISL_3324475, EPI_ISL_3324476, EPI_ISL_3324477, EPI_ISL_3324478, EPI_ISL_3324479, EPI_ISL_3324480, EPI_ISL_3324481, EPI_ISL_3324482, EPI_ISL_3324483, EPI_ISL_3324484, EPI_ISL_3324485, EPI_ISL_3324486, EPI_ISL_3324487, EPI_ISL_3324488, EPI_ISL_3324489, EPI_ISL_3324490, EPI_ISL_3324491, EPI_ISL_3324492, EPI_ISL_3324493, EPI_ISL_3324494, EPI_ISL_3324495, EPI_ISL_3324496, EPI_ISL_3324497, EPI_ISL_3324498, EPI_ISL_3324499, EPI_ISL_3324500, EPI_ISL_3324501, EPI_ISL_3324502, EPI_ISL_3324503, EPI_ISL_3324504, EPI_ISL_3324505, EPI_ISL_3324506, EPI_ISL_3324507, EPI_ISL_3324508, EPI_ISL_3324509, EPI_ISL_3324510, EPI_ISL_3324511, EPI_ISL_3324512, EPI_ISL_3324513, EPI_ISL_3324514, EPI_ISL_3324515, EPI_ISL_3324516, EPI_ISL_3324517, EPI_ISL_3324518, EPI_ISL_3324519, EPI_ISL_3324520, EPI_ISL_3324521, EPI_ISL_3324522, EPI_ISL_3324523, EPI_ISL_3324524, EPI_ISL_3324526, EPI_ISL_3324527, EPI_ISL_3324528, EPI_ISL_3324529, EPI_ISL_3324530, EPI_ISL_3324531, EPI_ISL_3324532, EPI_ISL_3324533, EPI_ISL_3324534, EPI_ISL_3324535, EPI_ISL_3324537, EPI_ISL_3324538, EPI_ISL_3324539, EPI_ISL_3324540, EPI_ISL_3324541, EPI_ISL_3324542, EPI_ISL_3324543, EPI_ISL_3324545, EPI_ISL_3324546, EPI_ISL_3324547, EPI_ISL_3324549, EPI_ISL_3324550, EPI_ISL_3324551, EPI_ISL_3324552, EPI_ISL_3324553, EPI_ISL_3324555, EPI_ISL_3324556, EPI_ISL_3324557, EPI_ISL_3324558, EPI_ISL_3324561, EPI_ISL_3324562, EPI_ISL_3324563, EPI_ISL_3324564, EPI_ISL_3324565, EPI_ISL_3324566, EPI_ISL_3324567, EPI_ISL_3324568, EPI_ISL_3324569, EPI_ISL_3324570, EPI_ISL_3324571, EPI_ISL_3324572, EPI_ISL_3324573, EPI_ISL_3324574, EPI_ISL_3324575, EPI_ISL_3324576, EPI_ISL_3324577, EPI_ISL_3324578, EPI_ISL_3324579, EPI_ISL_3324581, EPI_ISL_3324582, EPI_ISL_3324583, EPI_ISL_3324584, EPI_ISL_3324585, EPI_ISL_3324586, EPI_ISL_3324587, EPI_ISL_3324588, EPI_ISL_3324589, EPI_ISL_3324590, EPI_ISL_3324591, EPI_ISL_3324592, EPI_ISL_3324593, EPI_ISL_3324594, EPI_ISL_3324595, EPI_ISL_3324596, EPI_ISL_3324597, EPI_ISL_3324598, EPI_ISL_3324599, EPI_ISL_3324600, EPI_ISL_3324601, EPI_ISL_3324602, EPI_ISL_3324603, EPI_ISL_3324604, EPI_ISL_3324605, EPI_ISL_3324606, EPI_ISL_3324607, EPI_ISL_3324608, EPI_ISL_3324609, EPI_ISL_3324610, EPI_ISL_3324611, EPI_ISL_3324612, EPI_ISL_3324613, EPI_ISL_3324614, EPI_ISL_3324615, EPI_ISL_3324616, EPI_ISL_3324617, EPI_ISL_3324618, EPI_ISL_3324619, EPI_ISL_3324620, EPI_ISL_3324621, EPI_ISL_3324622, EPI_ISL_3324623, EPI_ISL_3324624, EPI_ISL_3324625, EPI_ISL_3324626, EPI_ISL_3324627, EPI_ISL_3324628, EPI_ISL_3324629, EPI_ISL_3324630, EPI_ISL_3324631, EPI_ISL_3324632, EPI_ISL_3324633, EPI_ISL_3324634, EPI_ISL_3324635, EPI_ISL_3324636, EPI_ISL_3324637, EPI_ISL_3324638, EPI_ISL_3324639, EPI_ISL_3324640, EPI_ISL_3324641, EPI_ISL_3324642, EPI_ISL_3324643, EPI_ISL_3324644, EPI_ISL_3324645, EPI_ISL_3324646, EPI_ISL_3324647, EPI_ISL_3324648, EPI_ISL_3324649, EPI_ISL_3324650, EPI_ISL_3324651, EPI_ISL_3324652, EPI_ISL_3324653, EPI_ISL_3324654, EPI_ISL_3324655, EPI_ISL_3324656, EPI_ISL_3324657, EPI_ISL_3324658, EPI_ISL_3324659, EPI_ISL_3324660, EPI_ISL_3324661, EPI_ISL_3324662, EPI_ISL_3324663, EPI_ISL_3324664, EPI_ISL_3324665, EPI_ISL_3324666, EPI_ISL_3324667, EPI_ISL_3324668, EPI_ISL_3324670, EPI_ISL_3324671, EPI_ISL_3324672, EPI_ISL_3324673, EPI_ISL_3324674, EPI_ISL_3324675, EPI_ISL_3324676, EPI_ISL_3324677, EPI_ISL_3324678, EPI_ISL_3324679, EPI_ISL_3324680, EPI_ISL_3324681, EPI_ISL_3324682, EPI_ISL_3324683, EPI_ISL_3324684, EPI_ISL_3324685, EPI_ISL_3324686, EPI_ISL_3324687, EPI_ISL_3324688, EPI_ISL_3324689, EPI_ISL_3324690, EPI_ISL_3324691, EPI_ISL_3324692, EPI_ISL_3324694, EPI_ISL_3324695, EPI_ISL_3324696, EPI_ISL_3324697, EPI_ISL_3324698, EPI_ISL_3324699, EPI_ISL_3324700, EPI_ISL_3324701, EPI_ISL_3324702, EPI_ISL_3324703, EPI_ISL_3324704, EPI_ISL_3324705, EPI_ISL_3324706, EPI_ISL_3324707, EPI_ISL_3324708, EPI_ISL_3324709, EPI_ISL_3324710, EPI_ISL_3324711, EPI_ISL_3324712, EPI_ISL_3324713, EPI_ISL_3324714, EPI_ISL_3324715, EPI_ISL_3324717, EPI_ISL_3324718, EPI_ISL_3324719, EPI_ISL_3324721, EPI_ISL_3324722, EPI_ISL_3324723, EPI_ISL_3324724, EPI_ISL_3324725, EPI_ISL_3324727, EPI_ISL_3324728, EPI_ISL_3324729, EPI_ISL_3324731, EPI_ISL_3324732, EPI_ISL_3324733, EPI_ISL_3324734, EPI_ISL_3324736, EPI_ISL_3324737, EPI_ISL_3324738, EPI_ISL_3324739, EPI_ISL_3324740, EPI_ISL_3324741, EPI_ISL_3324742, EPI_ISL_3324743, EPI_ISL_3324744, EPI_ISL_3324745, EPI_ISL_3324746, EPI_ISL_3324747, EPI_ISL_3324748, EPI_ISL_3324749, EPI_ISL_3324750, EPI_ISL_3324751, EPI_ISL_3324752, EPI_ISL_3324753, EPI_ISL_3324755, EPI_ISL_3324756, EPI_ISL_3324757, EPI_ISL_3324759, EPI_ISL_3324760, EPI_ISL_3324762, EPI_ISL_3324763, EPI_ISL_3324764, EPI_ISL_3324765, EPI_ISL_3324766, EPI_ISL_3324768, EPI_ISL_3324769, EPI_ISL_3324770, EPI_ISL_3324771, EPI_ISL_3324772, EPI_ISL_3324773, EPI_ISL_3324774, EPI_ISL_3324775, EPI_ISL_3324776, EPI_ISL_3324777, EPI_ISL_3324778, EPI_ISL_3324779, EPI_ISL_3324780, EPI_ISL_3324781, EPI_ISL_3324782, EPI_ISL_3324783, EPI_ISL_3324784, EPI_ISL_3324785, EPI_ISL_3324786, EPI_ISL_3324787, EPI_ISL_3324788, EPI_ISL_3324789, EPI_ISL_3324790, EPI_ISL_3324792, EPI_ISL_3324793, EPI_ISL_3324795, EPI_ISL_3324796, EPI_ISL_3324797, EPI_ISL_3324798, EPI_ISL_3324799, EPI_ISL_3324800, EPI_ISL_3324801, EPI_ISL_3324802, EPI_ISL_3324803, EPI_ISL_3324804, EPI_ISL_3324805, EPI_ISL_3324806, EPI_ISL_3324807, EPI_ISL_3324808, EPI_ISL_3324809, EPI_ISL_3324810, EPI_ISL_3324811, EPI_ISL_3324812, EPI_ISL_3324813, EPI_ISL_3324814, EPI_ISL_3324815, EPI_ISL_3324816, EPI_ISL_3324817, EPI_ISL_3324818, EPI_ISL_3324820, EPI_ISL_3324822, EPI_ISL_3324823, EPI_ISL_3324825, EPI_ISL_3324827, EPI_ISL_3324828, EPI_ISL_3324829, EPI_ISL_3324830, EPI_ISL_3324831, EPI_ISL_3324832, EPI_ISL_3324833, EPI_ISL_3324834, EPI_ISL_3324835, EPI_ISL_3324836, EPI_ISL_3324837, EPI_ISL_3324838, EPI_ISL_3324839, EPI_ISL_3324840, EPI_ISL_3324841, EPI_ISL_3324842, EPI_ISL_3324843, EPI_ISL_3324844, EPI_ISL_3324845, EPI_ISL_3324847, EPI_ISL_3324848, EPI_ISL_3324849, EPI_ISL_3324850, EPI_ISL_3324851, EPI_ISL_3324852, EPI_ISL_3324853, EPI_ISL_3324854, EPI_ISL_3324855, EPI_ISL_3324856, EPI_ISL_3324857, EPI_ISL_3324858, EPI_ISL_3324859, EPI_ISL_3324860, EPI_ISL_3324861, EPI_ISL_3324862, EPI_ISL_3324863, EPI_ISL_3324864, EPI_ISL_3324865, EPI_ISL_3324866, EPI_ISL_3324867, EPI_ISL_3324868, EPI_ISL_3324869, EPI_ISL_3324870, EPI_ISL_3324871, EPI_ISL_3324872, EPI_ISL_3324873, EPI_ISL_3324875, EPI_ISL_3324876, EPI_ISL_3324877, EPI_ISL_3324878, EPI_ISL_3324879, EPI_ISL_3324880, EPI_ISL_3324881, EPI_ISL_3324882, EPI_ISL_3324883, EPI_ISL_3324884, EPI_ISL_3324886, EPI_ISL_3324887, EPI_ISL_3324888, EPI_ISL_3324890, EPI_ISL_3324891, EPI_ISL_3324892, EPI_ISL_3324893, EPI_ISL_3324894, EPI_ISL_3324895, EPI_ISL_3324896, EPI_ISL_3324897, EPI_ISL_3324898, EPI_ISL_3324899, EPI_ISL_3324900, EPI_ISL_3324901, EPI_ISL_3324902, EPI_ISL_3324903, EPI_ISL_3324904, EPI_ISL_3324905, EPI_ISL_3324906, EPI_ISL_3324907, EPI_ISL_3324908, EPI_ISL_3324909, EPI_ISL_3324910, EPI_ISL_3324912, EPI_ISL_3324913, EPI_ISL_3324915, EPI_ISL_3324916, EPI_ISL_3324917, EPI_ISL_3324918, EPI_ISL_3324919, EPI_ISL_3324920, EPI_ISL_3324921, EPI_ISL_3324922, EPI_ISL_3324923, EPI_ISL_3324924, EPI_ISL_3324925, EPI_ISL_3324926, EPI_ISL_3324927, EPI_ISL_3324928, EPI_ISL_3324929, EPI_ISL_3324930, EPI_ISL_3324931, EPI_ISL_3324932, EPI_ISL_3324933, EPI_ISL_3324934, EPI_ISL_3324935, EPI_ISL_3324936, EPI_ISL_3324937, EPI_ISL_3324938, EPI_ISL_3324939, EPI_ISL_3324940, EPI_ISL_3324941, EPI_ISL_3324942, EPI_ISL_3324943, EPI_ISL_3324944, EPI_ISL_3324945, EPI_ISL_3324946, EPI_ISL_3324947, EPI_ISL_3324948, EPI_ISL_3324949, EPI_ISL_3324950, EPI_ISL_3324951, EPI_ISL_3324952, EPI_ISL_3324953, EPI_ISL_3324954, EPI_ISL_3324955, EPI_ISL_3324956, EPI_ISL_3324957, EPI_ISL_3324958, EPI_ISL_3324959, EPI_ISL_3324960, EPI_ISL_3324961, EPI_ISL_3324962, EPI_ISL_3324964, EPI_ISL_3324965, EPI_ISL_3324966, EPI_ISL_3324967, EPI_ISL_3324968, EPI_ISL_3324969, EPI_ISL_3324970, EPI_ISL_3324971, EPI_ISL_3324972, EPI_ISL_3324973, EPI_ISL_3324974, EPI_ISL_3324976, EPI_ISL_3324977, EPI_ISL_3324978, EPI_ISL_3324979, EPI_ISL_3324980, EPI_ISL_3324981, EPI_ISL_3324982, EPI_ISL_3324983, EPI_ISL_3324984, EPI_ISL_3324985, EPI_ISL_3324986, EPI_ISL_3324987, EPI_ISL_3324988, EPI_ISL_3324989, EPI_ISL_3324990, EPI_ISL_3324991, EPI_ISL_3324992, EPI_ISL_3324993, EPI_ISL_3324995, EPI_ISL_3324996, EPI_ISL_3324997, EPI_ISL_3324998, EPI_ISL_3324999, EPI_ISL_3325000, EPI_ISL_3325002, EPI_ISL_3325003, EPI_ISL_3325004, EPI_ISL_3325005, EPI_ISL_3325006, EPI_ISL_3325007, EPI_ISL_3325009, EPI_ISL_3325010, EPI_ISL_3325012, EPI_ISL_3325013, EPI_ISL_3325014, EPI_ISL_3325015, EPI_ISL_3325016, EPI_ISL_3325017, EPI_ISL_3325018, EPI_ISL_3325019, EPI_ISL_3325020, EPI_ISL_3325021, EPI_ISL_3325023, EPI_ISL_3325024, EPI_ISL_3325025, EPI_ISL_3325026, EPI_ISL_3325027, EPI_ISL_3325029, EPI_ISL_3325030, EPI_ISL_3325031, EPI_ISL_3325033, EPI_ISL_3325034, EPI_ISL_3325035, EPI_ISL_3325036, EPI_ISL_3325037, EPI_ISL_3325038, EPI_ISL_3325039, EPI_ISL_3325040, EPI_ISL_3325041, EPI_ISL_3325042, EPI_ISL_3325043, EPI_ISL_3325044, EPI_ISL_3325045, EPI_ISL_3325046, EPI_ISL_3325047, EPI_ISL_3325048, EPI_ISL_3325049, EPI_ISL_3325050, EPI_ISL_3325051, EPI_ISL_3325052, EPI_ISL_3325053, EPI_ISL_3325054, EPI_ISL_3325055, EPI_ISL_3325056, EPI_ISL_3325058, EPI_ISL_3325059, EPI_ISL_3325060, EPI_ISL_3325062, EPI_ISL_3325063, EPI_ISL_3325065, EPI_ISL_3325066, EPI_ISL_3325067, EPI_ISL_3325068, EPI_ISL_3325069, EPI_ISL_3325070, EPI_ISL_3325071, EPI_ISL_3325073, EPI_ISL_3325074, EPI_ISL_3325075, EPI_ISL_3325076, EPI_ISL_3325077, EPI_ISL_3325078, EPI_ISL_3325081, EPI_ISL_3325082, EPI_ISL_3325083, EPI_ISL_3325084, EPI_ISL_3325085, EPI_ISL_3325086, EPI_ISL_3325087, EPI_ISL_3325088, EPI_ISL_3325089, EPI_ISL_3325090, EPI_ISL_3325091, EPI_ISL_3325092, EPI_ISL_3325093, EPI_ISL_3325094, EPI_ISL_3325096, EPI_ISL_3325097, EPI_ISL_3325098, EPI_ISL_3325099, EPI_ISL_3325100, EPI_ISL_3325101, EPI_ISL_3325102, EPI_ISL_3325103, EPI_ISL_3325104, EPI_ISL_3325105, EPI_ISL_3325106, EPI_ISL_3325107, EPI_ISL_3325108, EPI_ISL_3325110, EPI_ISL_3325111, EPI_ISL_3325112, EPI_ISL_3325113, EPI_ISL_3325114, EPI_ISL_3325115, EPI_ISL_3325116, EPI_ISL_3325117, EPI_ISL_3325118, EPI_ISL_3325119, EPI_ISL_3325120, EPI_ISL_3325121, EPI_ISL_3325122, EPI_ISL_3325123, EPI_ISL_3325124, EPI_ISL_3325125, EPI_ISL_3325126, EPI_ISL_3325127, EPI_ISL_3325130, EPI_ISL_3325132, EPI_ISL_3325133, EPI_ISL_3325134, EPI_ISL_3325135, EPI_ISL_3325136, EPI_ISL_3325137, EPI_ISL_3325138, EPI_ISL_3325139, EPI_ISL_3325140, EPI_ISL_3325141, EPI_ISL_3325142, EPI_ISL_3325143, EPI_ISL_3325144, EPI_ISL_3325145, EPI_ISL_3325146, EPI_ISL_3325147, EPI_ISL_3325148, EPI_ISL_3325149, EPI_ISL_3325150, EPI_ISL_3325151, EPI_ISL_3325152, EPI_ISL_3325153, EPI_ISL_3325154, EPI_ISL_3325155, EPI_ISL_3325156, EPI_ISL_3325157, EPI_ISL_3325158, EPI_ISL_3325159, EPI_ISL_3325160, EPI_ISL_3325162, EPI_ISL_3325163, EPI_ISL_3325164, EPI_ISL_3325165, EPI_ISL_3325166, EPI_ISL_3325167, EPI_ISL_3325171, EPI_ISL_3325172, EPI_ISL_3325173, EPI_ISL_3325174, EPI_ISL_3325175, EPI_ISL_3325176, EPI_ISL_3325177, EPI_ISL_3325178, EPI_ISL_3325179, EPI_ISL_3325180, EPI_ISL_3325181, EPI_ISL_3325182, EPI_ISL_3325183, EPI_ISL_3325184, EPI_ISL_3325185, EPI_ISL_3325186, EPI_ISL_3325187, EPI_ISL_3325188, EPI_ISL_3325189, EPI_ISL_3325190, EPI_ISL_3325191, EPI_ISL_3325192, EPI_ISL_3325193, EPI_ISL_3325195, EPI_ISL_3325196, EPI_ISL_3325197, EPI_ISL_3325198, EPI_ISL_3325199, EPI_ISL_3325200, EPI_ISL_3325201, EPI_ISL_3325202, EPI_ISL_3325203, EPI_ISL_3325204, EPI_ISL_3325205, EPI_ISL_3325206, EPI_ISL_3325207, EPI_ISL_3325208, EPI_ISL_3325209, EPI_ISL_3325210, EPI_ISL_3325211, EPI_ISL_3325213, EPI_ISL_3325214, EPI_ISL_3325215, EPI_ISL_3325218, EPI_ISL_3325219, EPI_ISL_3325220, EPI_ISL_3325221, EPI_ISL_3325222, EPI_ISL_3325223, EPI_ISL_3325224, EPI_ISL_3325225, EPI_ISL_3325226, EPI_ISL_3325228, EPI_ISL_3325229, EPI_ISL_3325230, EPI_ISL_3325231, EPI_ISL_3325233, EPI_ISL_3325234, EPI_ISL_3325235, EPI_ISL_3325236, EPI_ISL_3325237, EPI_ISL_3325238, EPI_ISL_3325239, EPI_ISL_3325240, EPI_ISL_3325241, EPI_ISL_3325242, EPI_ISL_3325243, EPI_ISL_3325244, EPI_ISL_3325245, EPI_ISL_3325246, EPI_ISL_3325247, EPI_ISL_3325248, EPI_ISL_3325249, EPI_ISL_3325250, EPI_ISL_3325251, EPI_ISL_3325252, EPI_ISL_3325253, EPI_ISL_3325254, EPI_ISL_3325255, EPI_ISL_3325256, EPI_ISL_3325257, EPI_ISL_3325258, EPI_ISL_3325259, EPI_ISL_3325260, EPI_ISL_3325261, EPI_ISL_3325262, EPI_ISL_3325263, EPI_ISL_3325264, EPI_ISL_3325265, EPI_ISL_3325266, EPI_ISL_3325267, EPI_ISL_3325268, EPI_ISL_3325269, EPI_ISL_3325270, EPI_ISL_3325271, EPI_ISL_3325272, EPI_ISL_3325273, EPI_ISL_3325274, EPI_ISL_3325275, EPI_ISL_3325276, EPI_ISL_3325277, EPI_ISL_3325278, EPI_ISL_3325279, EPI_ISL_3325280, EPI_ISL_3325281, EPI_ISL_3325282, EPI_ISL_3325283, EPI_ISL_3325284, EPI_ISL_3325285, EPI_ISL_3325286, EPI_ISL_3325287, EPI_ISL_3325288, EPI_ISL_3325289, EPI_ISL_3325290, EPI_ISL_3325291, EPI_ISL_3325292, EPI_ISL_3325293, EPI_ISL_3325294, EPI_ISL_3325296, EPI_ISL_3325299, EPI_ISL_3325300, EPI_ISL_3325301, EPI_ISL_3325302, EPI_ISL_3325303, EPI_ISL_3325304, EPI_ISL_3325305, EPI_ISL_3325306, EPI_ISL_3325307, EPI_ISL_3325308, EPI_ISL_3325310, EPI_ISL_3325311, EPI_ISL_3325312, EPI_ISL_3325313, EPI_ISL_3325314, EPI_ISL_3325315, EPI_ISL_3325316, EPI_ISL_3325317, EPI_ISL_3325319, EPI_ISL_3325320, EPI_ISL_3325321, EPI_ISL_3325323, EPI_ISL_3325324, EPI_ISL_3325325, EPI_ISL_3325326, EPI_ISL_3325327, EPI_ISL_3325328, EPI_ISL_3325329, EPI_ISL_3325330, EPI_ISL_3325332, EPI_ISL_3325333, EPI_ISL_3325334, EPI_ISL_3325335, EPI_ISL_3325337, EPI_ISL_3325338, EPI_ISL_3325339, EPI_ISL_3325340, EPI_ISL_3325341, EPI_ISL_3325342, EPI_ISL_3325343, EPI_ISL_3325344, EPI_ISL_3325345, EPI_ISL_3325346, EPI_ISL_3325347, EPI_ISL_3325348, EPI_ISL_3325349, EPI_ISL_3325350, EPI_ISL_3325351, EPI_ISL_3325352, EPI_ISL_3325353, EPI_ISL_3325354, EPI_ISL_3325355, EPI_ISL_3325356, EPI_ISL_3325357, EPI_ISL_3325359, EPI_ISL_3325360, EPI_ISL_3325361, EPI_ISL_3325362, EPI_ISL_3325363, EPI_ISL_3325364, EPI_ISL_3325365, EPI_ISL_3325366, EPI_ISL_3325367, EPI_ISL_3325368, EPI_ISL_3325369, EPI_ISL_3325370, EPI_ISL_3325371, EPI_ISL_3325373, EPI_ISL_3325374, EPI_ISL_3325375, EPI_ISL_3325376, EPI_ISL_3325377, EPI_ISL_3325379, EPI_ISL_3325381, EPI_ISL_3325382, EPI_ISL_3325383, EPI_ISL_3325384, EPI_ISL_3325385, EPI_ISL_3325386, EPI_ISL_3325387, EPI_ISL_3325388, EPI_ISL_3325389, EPI_ISL_3325390, EPI_ISL_3325391 | | | |
| see above | Aegis Sciences Corporation | Centers for Disease Control and Prevention Division of Viral Diseases, Pathogen Discovery | Dakota Howard, Dhwani Batra, Peter W. Cook, Kara Moser, Adrian Paskey, Jason Caravas, Benjamin Rambo-Martin, Shatavia Morrison, Christopher Gulvick, Scott Sammons, Yvette Unoarumhi, Darlene Wagner, Matthew Schmerer, Cyndi Clark, Patrick Campbell, Rob Case, Vikramsinha Ghorpade, Holly Houdeshell, Ola Kvalvaag, Dillon Nall, Ethan Sanders, Alec Vest, Shaun Westlund, Matthew Hardison, Clinton R. Paden, Duncan MacCannell |
| EPI_ISL_3325392, EPI_ISL_3325393, EPI_ISL_3325394, EPI_ISL_3325395, EPI_ISL_3325396, EPI_ISL_3325397, EPI_ISL_3325398, EPI_ISL_3325399, EPI_ISL_3325400, EPI_ISL_3325401, EPI_ISL_3325402, EPI_ISL_3325403, EPI_ISL_3325404, EPI_ISL_3325405, EPI_ISL_3325406, EPI_ISL_3325407, EPI_ISL_3325408, EPI_ISL_3325409, EPI_ISL_3325410, EPI_ISL_3325411, EPI_ISL_3325412, EPI_ISL_3325413, EPI_ISL_3325414, EPI_ISL_3325415, EPI_ISL_3325416, EPI_ISL_3325417, EPI_ISL_3325418, EPI_ISL_3325419, EPI_ISL_3325420, EPI_ISL_3325421, EPI_ISL_3325422, EPI_ISL_3325423, EPI_ISL_3325424, EPI_ISL_3325425, EPI_ISL_3325426, EPI_ISL_3325427, EPI_ISL_3325428, EPI_ISL_3325429, EPI_ISL_3325430, EPI_ISL_3325431, EPI_ISL_3325432, EPI_ISL_3325433, EPI_ISL_3325434, EPI_ISL_3325435, EPI_ISL_3325436, EPI_ISL_3325437, EPI_ISL_3325438, EPI_ISL_3325439, EPI_ISL_3325440, EPI_ISL_3325441, EPI_ISL_3325442, EPI_ISL_3325443, EPI_ISL_3325444, EPI_ISL_3325445, EPI_ISL_3325446, EPI_ISL_3325447, EPI_ISL_3325448, EPI_ISL_3325449, EPI_ISL_3325450, EPI_ISL_3325451, EPI_ISL_3325452, EPI_ISL_3325453, EPI_ISL_3325454, EPI_ISL_3325455, EPI_ISL_3325456, EPI_ISL_3325457, EPI_ISL_3325458, EPI_ISL_3325459, EPI_ISL_3325460, EPI_ISL_3325461, EPI_ISL_3325462, EPI_ISL_3325463, EPI_ISL_3325464, EPI_ISL_3325465, EPI_ISL_3325466, EPI_ISL_3325467, EPI_ISL_3325468, EPI_ISL_3325469, EPI_ISL_3325470, EPI_ISL_3325471, EPI_ISL_3325472, EPI_ISL_3325473, EPI_ISL_3325474, EPI_ISL_3325475, EPI_ISL_3325476, EPI_ISL_3325477, EPI_ISL_3325478, EPI_ISL_3325479, EPI_ISL_3325480, EPI_ISL_3325481, EPI_ISL_3325482, EPI_ISL_3325483, EPI_ISL_3325484, EPI_ISL_3325485, EPI_ISL_3325486, EPI_ISL_3325487, EPI_ISL_3325488, EPI_ISL_3325489, EPI_ISL_3325490, EPI_ISL_3325491, EPI_ISL_3325492, EPI_ISL_3325493, EPI_ISL_3325494, EPI_ISL_3325495, EPI_ISL_3325496, EPI_ISL_3325497, EPI_ISL_3325498, EPI_ISL_3325499, EPI_ISL_3325500, EPI_ISL_3325501, EPI_ISL_3325502, EPI_ISL_3325503, EPI_ISL_3325504, EPI_ISL_3325505, EPI_ISL_3325506, EPI_ISL_3325507, EPI_ISL_3325508, EPI_ISL_3325509, EPI_ISL_3325510, EPI_ISL_3325511, EPI_ISL_3325512, EPI_ISL_3325513, EPI_ISL_3325514, EPI_ISL_3325515, EPI_ISL_3325516, EPI_ISL_3325517, EPI_ISL_3325518, EPI_ISL_3325519, EPI_ISL_3325520, EPI_ISL_3325521, EPI_ISL_3325522, EPI_ISL_3325523, EPI_ISL_3325524, EPI_ISL_3325525, EPI_ISL_3325526, EPI_ISL_3325527, EPI_ISL_3325528, EPI_ISL_3325529, EPI_ISL_3325530, EPI_ISL_3325531, EPI_ISL_3325533, EPI_ISL_3325534, EPI_ISL_3325535, EPI_ISL_3325536, EPI_ISL_3325537, EPI_ISL_3325538, EPI_ISL_3325539, EPI_ISL_3325540, EPI_ISL_3325541, EPI_ISL_3325542, EPI_ISL_3325543, EPI_ISL_3325545, EPI_ISL_3325546, EPI_ISL_3325547, EPI_ISL_3325549, EPI_ISL_3325550, EPI_ISL_3325551, EPI_ISL_3325552, EPI_ISL_3325553, EPI_ISL_3325554, EPI_ISL_3325555, EPI_ISL_3325558, EPI_ISL_3325560, EPI_ISL_3325561, EPI_ISL_3325562, EPI_ISL_3325563, EPI_ISL_3325564, EPI_ISL_3325565, EPI_ISL_3325566, EPI_ISL_3325567, EPI_ISL_3325568, EPI_ISL_3325569, EPI_ISL_3325570, EPI_ISL_3325571, EPI_ISL_3325572, EPI_ISL_3325573, EPI_ISL_3325574, EPI_ISL_3325575, EPI_ISL_3325576, EPI_ISL_3325578, EPI_ISL_3325579, EPI_ISL_3325580, EPI_ISL_3325581, EPI_ISL_3325582, EPI_ISL_3325583, EPI_ISL_3325585, EPI_ISL_3325586, EPI_ISL_3325587, EPI_ISL_3325588, EPI_ISL_3325589, EPI_ISL_3325590, EPI_ISL_3325591, EPI_ISL_3325592, EPI_ISL_3325593, EPI_ISL_3325594, EPI_ISL_3325595, EPI_ISL_3325596, EPI_ISL_3325597, EPI_ISL_3325598, EPI_ISL_3325599, EPI_ISL_3325600, EPI_ISL_3325601, EPI_ISL_3325602, EPI_ISL_3325603, EPI_ISL_3325604, EPI_ISL_3325605, EPI_ISL_3325606, EPI_ISL_3325607, EPI_ISL_3325608, EPI_ISL_3325609, EPI_ISL_3325610, EPI_ISL_3325612, EPI_ISL_3325613, EPI_ISL_3325614, EPI_ISL_3325616, EPI_ISL_3325617, EPI_ISL_3325618, EPI_ISL_3325619, EPI_ISL_3325620, EPI_ISL_3325621, EPI_ISL_3325622, EPI_ISL_3325624, EPI_ISL_3325625, EPI_ISL_3325626, EPI_ISL_3325627, EPI_ISL_3325628, EPI_ISL_3325629, EPI_ISL_3325630, EPI_ISL_3325631, EPI_ISL_3325632, EPI_ISL_3325633, EPI_ISL_3325634, EPI_ISL_3325635, EPI_ISL_3325636, EPI_ISL_3325637, EPI_ISL_3325639, EPI_ISL_3325640, EPI_ISL_3325641, EPI_ISL_3325642, EPI_ISL_3325643, EPI_ISL_3325644, EPI_ISL_3325646, EPI_ISL_3325647, EPI_ISL_3325648, EPI_ISL_3325650, EPI_ISL_3325651, EPI_ISL_3325652, EPI_ISL_3325653, EPI_ISL_3325654, EPI_ISL_3325655, EPI_ISL_3325656, EPI_ISL_3325657, EPI_ISL_3325658, EPI_ISL_3325659, EPI_ISL_3325660, EPI_ISL_3325661, EPI_ISL_3325662, EPI_ISL_3325663, EPI_ISL_3325664, EPI_ISL_3325665, EPI_ISL_3325666, EPI_ISL_3325667, EPI_ISL_3325668, EPI_ISL_3325669, EPI_ISL_3325670, EPI_ISL_3325671, EPI_ISL_3325672, EPI_ISL_3325673, EPI_ISL_3325674, EPI_ISL_3325675, EPI_ISL_3325676, EPI_ISL_3325677, EPI_ISL_3325678, EPI_ISL_3325679, EPI_ISL_3325680, EPI_ISL_3325681, EPI_ISL_3325682, EPI_ISL_3325683, EPI_ISL_3325684, EPI_ISL_3325685, EPI_ISL_3325686, EPI_ISL_3325687, EPI_ISL_3325688, EPI_ISL_3325689, EPI_ISL_3325690, EPI_ISL_3325692, EPI_ISL_3325693, EPI_ISL_3325694, EPI_ISL_3325695, EPI_ISL_3325696, EPI_ISL_3325697, EPI_ISL_3325698, EPI_ISL_3325699, EPI_ISL_3325700, EPI_ISL_3325701, EPI_ISL_3325702, EPI_ISL_3325703, EPI_ISL_3325705, EPI_ISL_3325706, EPI_ISL_3325707, EPI_ISL_3325708, EPI_ISL_3325710, EPI_ISL_3325711, EPI_ISL_3325712 | | | |
| see above | Laboratory Corporation of America | Centers for Disease Control and Prevention Division of Viral Diseases, Pathogen Discovery | Dakota Howard, Dhwani Batra, Peter W. Cook, Kara Moser, Adrian Paskey, Jason Caravas, Benjamin Rambo-Martin, Shatavia Morrison, Christopher Gulvick, Scott Sammons, Yvette Unoarumhi, Darlene Wagner, Matthew Schmerer, Minoo Agarwal, Eyad Almasri, Debbie Boles, Ayla Burns, Nuthawin Charoensri, Oren Cohen, Susan Countryman, Mary Ann Cristobal, Bobbi Croy, Suzanne Dale, Hrushikesh Deshmukh, Amanda Douglas, Vincent Drouillon, Marcia Eisenberg, Howard Engler, Rama Ghatti, Prashant Gupta, Susan Hicks, Jake Humphrey, Lax Iyer, Lisa Pfefferle, Manoj Jain, Matthew Robinson, Mohan Kolli, Brian Krueger, Tim Kuphal, Stanley Letovsky, Michael Levandoski, Craig Lukasik, Jonathan Meltzer, Brian Norvell, Mindy Nye, Scott Parker, Christos Petropoulos, John Pruitt, Steven Ragan, Scott Ryan, Mike Sapeta, Jana Schroth, Suresh Babu Selvaraju, Goran Stevovic, Amanda Suchanek, Andrea Throop, Lyndon Tilson, Thomas Urban, Joe Voshell, Kimberly Wagner, Jonathan Williams, Mary Williamson, Qian Zeng, Tricia Zwiefelhofer, Clinton R. Paden, Duncan MacCannell |
| EPI_ISL_3325713 | TGen North | TGen North | Jolene Bowers, Heather Centner, Brett Van Tassel, Chris French, Hayley Yaglom, Darrin Lemmer, Dave Engelthaler |
| EPI_ISL_3325714, EPI_ISL_3325715 | Laboratory Corporation of America | Centers for Disease Control and Prevention Division of Viral Diseases, Pathogen Discovery | Dakota Howard, Dhwani Batra, Peter W. Cook, Kara Moser, Adrian Paskey, Jason Caravas, Benjamin Rambo-Martin, Shatavia Morrison, Christopher Gulvick, Scott Sammons, Yvette Unoarumhi, Darlene Wagner, Matthew Schmerer, Minoo Agarwal, Eyad Almasri, Debbie Boles, Ayla Burns, Nuthawin Charoensri, Oren Cohen, Susan Countryman, Mary Ann Cristobal, Bobbi Croy, Suzanne Dale, Hrushikesh Deshmukh, Amanda Douglas, Vincent Drouillon, Marcia Eisenberg, Howard Engler, Rama Ghatti, Prashant Gupta, Susan Hicks, Jake Humphrey, Lax Iyer, Lisa Pfefferle, Manoj Jain, Matthew Robinson, Mohan Kolli, Brian Krueger, Tim Kuphal, Stanley Letovsky, Michael Levandoski, Craig Lukasik, Jonathan Meltzer, Brian Norvell, Mindy Nye, Scott Parker, Christos Petropoulos, John Pruitt, Steven Ragan, Scott Ryan, Mike Sapeta, Jana Schroth, Suresh Babu Selvaraju, Goran Stevovic, Amanda Suchanek, Andrea Throop, Lyndon Tilson, Thomas Urban, Joe Voshell, Kimberly Wagner, Jonathan Williams, Mary Williamson, Qian Zeng, Tricia Zwiefelhofer, Clinton R. Paden, Duncan MacCannell |
| EPI_ISL_3325716 | TGen North | TGen North | Jolene Bowers, Heather Centner, Brett Van Tassel, Chris French, Hayley Yaglom, Darrin Lemmer, Dave Engelthaler |
| EPI_ISL_3325717 | Laboratory Corporation of America | Centers for Disease Control and Prevention Division of Viral | Dakota Howard, Dhwani Batra, Peter W. Cook, Kara Moser, Adrian Paskey, Jason Caravas, Benjamin Rambo-Martin, Shatavia Morrison, Christopher Gulvick, Scott Sammons, Yvette Unoarumhi, Darlene Wagner, Matthew Schmerer, Minoo |

| | | | |
|---|---|---|---|
| | | Diseases, Pathogen Discovery | Agarwal, Eyad Almasri, Debbie Boles, Ayla Burns, Nuthawin Charoensri, Oren Cohen, Susan Countryman, Mary Ann Cristobal, Bobbi Croy, Suzanne Dale, Hrushikesh Deshmukh, Amanda Douglas, Vincent Drouillon, Marcia Eisenberg, Howard Engler, Rama Ghatti, Prashant Gupta, Susan Hicks, Jake Humphrey, Lax Iyer, Lisa Pfefferle, Manoj Jain, Matthew Robinson, Mohan Kolli, Brian Krueger, Tim Kuphal, Stanley Letovsky, Michael Levandoski, Craig Lukasik, Jonathan Meltzer, Brian Norvell, Mindy Nye, Scott Parker, Christos Petropoulos, John Pruitt, Steven Ragan, Scott Ryan, Mike Sapeta, Jana Schroth, Suresh Babu Selvaraju, Goran Stevovic, Amanda Suchanek, Andrea Throop, Lyndon Tilson, Thomas Urban, Joe Voshell, Kimberly Wagner, Jonathan Williams, Mary Williamson, Qian Zeng, Tricia Zwiefelhofer, Clinton R. Paden, Duncan MacCannell |
| EPI_ISL_3325718 | TGen North | TGen North | Jolene Bowers, Heather Centner, Brett Van Tassel, Chris French, Hayley Yaglom, Darrin Lemmer, Dave Engelthaler |
| EPI_ISL_3325719, EPI_ISL_3325721, EPI_ISL_3325723, EPI_ISL_3325724 | Laboratory Corporation of America | Centers for Disease Control and Prevention Division of Viral Diseases, Pathogen Discovery | Dakota Howard, Dhwani Batra, Peter W. Cook, Kara Moser, Adrian Paskey, Jason Caravas, Benjamin Rambo-Martin, Shatavia Morrison, Christopher Gulvick, Scott Sammons, Yvette Unoarumhi, Darlene Wagner, Matthew Schmerer, Minoo Agarwal, Eyad Almasri, Debbie Boles, Ayla Burns, Nuthawin Charoensri, Oren Cohen, Susan Countryman, Mary Ann Cristobal, Bobbi Croy, Suzanne Dale, Hrushikesh Deshmukh, Amanda Douglas, Vincent Drouillon, Marcia Eisenberg, Howard Engler, Rama Ghatti, Prashant Gupta, Susan Hicks, Jake Humphrey, Lax Iyer, Lisa Pfefferle, Manoj Jain, Matthew Robinson, Mohan Kolli, Brian Krueger, Tim Kuphal, Stanley Letovsky, Michael Levandoski, Craig Lukasik, Jonathan Meltzer, Brian Norvell, Mindy Nye, Scott Parker, Christos Petropoulos, John Pruitt, Steven Ragan, Scott Ryan, Mike Sapeta, Jana Schroth, Suresh Babu Selvaraju, Goran Stevovic, Amanda Suchanek, Andrea Throop, Lyndon Tilson, Thomas Urban, Joe Voshell, Kimberly Wagner, Jonathan Williams, Mary Williamson, Qian Zeng, Tricia Zwiefelhofer, Clinton R. Paden, Duncan MacCannell |
| EPI_ISL_3325725 | TGen North | TGen North | Jolene Bowers, Heather Centner, Brett Van Tassel, Chris French, Hayley Yaglom, Darrin Lemmer, Dave Engelthaler |
| EPI_ISL_3325726 | Laboratory Corporation of America | Centers for Disease Control and Prevention Division of Viral Diseases, Pathogen Discovery | Dakota Howard, Dhwani Batra, Peter W. Cook, Kara Moser, Adrian Paskey, Jason Caravas, Benjamin Rambo-Martin, Shatavia Morrison, Christopher Gulvick, Scott Sammons, Yvette Unoarumhi, Darlene Wagner, Matthew Schmerer, Minoo Agarwal, Eyad Almasri, Debbie Boles, Ayla Burns, Nuthawin Charoensri, Oren Cohen, Susan Countryman, Mary Ann Cristobal, Bobbi Croy, Suzanne Dale, Hrushikesh Deshmukh, Amanda Douglas, Vincent Drouillon, Marcia Eisenberg, Howard Engler, Rama Ghatti, Prashant Gupta, Susan Hicks, Jake Humphrey, Lax Iyer, Lisa Pfefferle, Manoj Jain, Matthew Robinson, Mohan Kolli, Brian Krueger, Tim Kuphal, Stanley Letovsky, Michael Levandoski, Craig Lukasik, Jonathan Meltzer, Brian Norvell, Mindy Nye, Scott Parker, Christos Petropoulos, John Pruitt, Steven Ragan, Scott Ryan, Mike Sapeta, Jana Schroth, Suresh Babu Selvaraju, Goran Stevovic, Amanda Suchanek, Andrea Throop, Lyndon Tilson, Thomas Urban, Joe Voshell, Kimberly Wagner, Jonathan Williams, Mary Williamson, Qian Zeng, Tricia Zwiefelhofer, Clinton R. Paden, Duncan MacCannell |
| EPI_ISL_3325727 | TGen North | TGen North | Jolene Bowers, Heather Centner, Brett Van Tassel, Chris French, Hayley Yaglom, Darrin Lemmer, Dave Engelthaler |
| EPI_ISL_3325730 | Laboratory Corporation of America | Centers for Disease Control and Prevention Division of Viral Diseases, Pathogen Discovery | Dakota Howard, Dhwani Batra, Peter W. Cook, Kara Moser, Adrian Paskey, Jason Caravas, Benjamin Rambo-Martin, Shatavia Morrison, Christopher Gulvick, Scott Sammons, Yvette Unoarumhi, Darlene Wagner, Matthew Schmerer, Minoo Agarwal, Eyad Almasri, Debbie Boles, Ayla Burns, Nuthawin Charoensri, Oren Cohen, Susan Countryman, Mary Ann Cristobal, Bobbi Croy, Suzanne Dale, Hrushikesh Deshmukh, Amanda Douglas, Vincent Drouillon, Marcia Eisenberg, Howard Engler, Rama Ghatti, Prashant Gupta, Susan Hicks, Jake Humphrey, Lax Iyer, Lisa Pfefferle, Manoj Jain, Matthew Robinson, Mohan Kolli, Brian Krueger, Tim Kuphal, Stanley Letovsky, Michael Levandoski, Craig Lukasik, Jonathan Meltzer, Brian Norvell, Mindy Nye, Scott Parker, Christos Petropoulos, John Pruitt, Steven Ragan, Scott Ryan, Mike Sapeta, Jana Schroth, Suresh Babu Selvaraju, Goran Stevovic, Amanda Suchanek, Andrea Throop, Lyndon Tilson, Thomas Urban, Joe Voshell, Kimberly Wagner, Jonathan Williams, Mary Williamson, Qian Zeng, Tricia Zwiefelhofer, Clinton R. Paden, Duncan MacCannell |
| EPI_ISL_3325731 | TGen North | TGen North | Jolene Bowers, Heather Centner, Brett Van Tassel, Chris French, Hayley Yaglom, Darrin Lemmer, Dave Engelthaler |
| EPI_ISL_3325736, EPI_ISL_3325737, EPI_ISL_3325739 | Laboratory Corporation of America | Centers for Disease Control and Prevention Division of Viral Diseases, Pathogen Discovery | Dakota Howard, Dhwani Batra, Peter W. Cook, Kara Moser, Adrian Paskey, Jason Caravas, Benjamin Rambo-Martin, Shatavia Morrison, Christopher Gulvick, Scott Sammons, Yvette Unoarumhi, Darlene Wagner, Matthew Schmerer, Minoo Agarwal, Eyad Almasri, Debbie Boles, Ayla Burns, Nuthawin Charoensri, Oren Cohen, Susan Countryman, Mary Ann Cristobal, Bobbi Croy, Suzanne Dale, Hrushikesh Deshmukh, Amanda Douglas, Vincent Drouillon, Marcia Eisenberg, Howard Engler, Rama Ghatti, Prashant Gupta, Susan Hicks, Jake Humphrey, Lax Iyer, Lisa Pfefferle, Manoj Jain, Matthew Robinson, Mohan Kolli, Brian Krueger, Tim Kuphal, Stanley Letovsky, Michael Levandoski, Craig Lukasik, Jonathan Meltzer, Brian Norvell, Mindy Nye, Scott Parker, Christos Petropoulos, John Pruitt, Steven Ragan, Scott Ryan, Mike Sapeta, Jana Schroth, Suresh Babu Selvaraju, Goran Stevovic, Amanda Suchanek, Andrea Throop, Lyndon Tilson, Thomas Urban, Joe Voshell, Kimberly Wagner, Jonathan Williams, Mary Williamson, Qian Zeng, Tricia Zwiefelhofer, Clinton R. Paden, Duncan MacCannell |
| EPI_ISL_3325740 | TGen North | TGen North | Jolene Bowers, Heather Centner, Brett Van Tassel, Chris French, Hayley Yaglom, Darrin Lemmer, Dave Engelthaler |
| EPI_ISL_3325741, EPI_ISL_3325743 | Laboratory Corporation of America | Centers for Disease Control and Prevention Division of Viral Diseases, Pathogen Discovery | Dakota Howard, Dhwani Batra, Peter W. Cook, Kara Moser, Adrian Paskey, Jason Caravas, Benjamin Rambo-Martin, Shatavia Morrison, Christopher Gulvick, Scott Sammons, Yvette Unoarumhi, Darlene Wagner, Matthew Schmerer, Minoo Agarwal, Eyad Almasri, Debbie Boles, Ayla Burns, Nuthawin Charoensri, Oren Cohen, Susan Countryman, Mary Ann Cristobal, Bobbi Croy, Suzanne Dale, Hrushikesh Deshmukh, Amanda Douglas, Vincent Drouillon, Marcia Eisenberg, Howard Engler, Rama Ghatti, Prashant Gupta, Susan Hicks, Jake Humphrey, Lax Iyer, Lisa Pfefferle, Manoj Jain, Matthew Robinson, Mohan Kolli, Brian Krueger, Tim Kuphal, Stanley Letovsky, Michael Levandoski, Craig Lukasik, Jonathan Meltzer, Brian Norvell, Mindy Nye, Scott Parker, Christos Petropoulos, John Pruitt, Steven Ragan, Scott Ryan, Mike Sapeta, Jana Schroth, Suresh Babu Selvaraju, Goran Stevovic, Amanda Suchanek, Andrea Throop, Lyndon Tilson, Thomas Urban, Joe Voshell, Kimberly Wagner, Jonathan Williams, Mary Williamson, Qian Zeng, Tricia Zwiefelhofer, Clinton R. Paden, Duncan MacCannell |
| EPI_ISL_3325744 | TGen North | TGen North | Jolene Bowers, Heather Centner, Brett Van Tassel, Chris French, Hayley Yaglom, Darrin Lemmer, Dave Engelthaler |
| EPI_ISL_3325745, EPI_ISL_3325747 | Laboratory Corporation of America | Centers for Disease Control and Prevention Division of Viral Diseases, Pathogen Discovery | Dakota Howard, Dhwani Batra, Peter W. Cook, Kara Moser, Adrian Paskey, Jason Caravas, Benjamin Rambo-Martin, Shatavia Morrison, Christopher Gulvick, Scott Sammons, Yvette Unoarumhi, Darlene Wagner, Matthew Schmerer, Minoo Agarwal, Eyad Almasri, Debbie Boles, Ayla Burns, Nuthawin Charoensri, Oren Cohen, Susan Countryman, Mary Ann Cristobal, Bobbi Croy, Suzanne Dale, Hrushikesh Deshmukh, Amanda Douglas, Vincent Drouillon, Marcia Eisenberg, Howard Engler, Rama Ghatti, Prashant Gupta, Susan Hicks, Jake Humphrey, Lax Iyer, Lisa Pfefferle, Manoj Jain, Matthew Robinson, Mohan Kolli, Brian Krueger, Tim Kuphal, Stanley Letovsky, Michael Levandoski, Craig Lukasik, Jonathan Meltzer, Brian Norvell, Mindy Nye, Scott Parker, Christos Petropoulos, John Pruitt, Steven Ragan, Scott Ryan, Mike Sapeta, Jana Schroth, Suresh Babu Selvaraju, Goran Stevovic, Amanda Suchanek, Andrea Throop, Lyndon Tilson, Thomas Urban, Joe Voshell, Kimberly Wagner, Jonathan Williams, Mary Williamson, Qian Zeng, Tricia Zwiefelhofer, Clinton R. Paden, Duncan MacCannell |
| EPI_ISL_3325748, EPI_ISL_3325750 | TGen North | TGen North | Jolene Bowers, Heather Centner, Brett Van Tassel, Chris French, Hayley Yaglom, Darrin Lemmer, Dave Engelthaler |
| EPI_ISL_3325751, EPI_ISL_3325753, EPI_ISL_3325754, EPI_ISL_3325756 | Laboratory Corporation of America | Centers for Disease Control and Prevention Division of Viral Diseases, Pathogen Discovery | Dakota Howard, Dhwani Batra, Peter W. Cook, Kara Moser, Adrian Paskey, Jason Caravas, Benjamin Rambo-Martin, Shatavia Morrison, Christopher Gulvick, Scott Sammons, Yvette Unoarumhi, Darlene Wagner, Matthew Schmerer, Minoo Agarwal, Eyad Almasri, Debbie Boles, Ayla Burns, Nuthawin Charoensri, Oren Cohen, Susan Countryman, Mary Ann Cristobal, Bobbi Croy, Suzanne Dale, Hrushikesh Deshmukh, Amanda Douglas, Vincent Drouillon, Marcia Eisenberg, Howard Engler, Rama Ghatti, Prashant Gupta, Susan Hicks, Jake Humphrey, Lax Iyer, Lisa Pfefferle, Manoj Jain, Matthew Robinson, Mohan Kolli, Brian Krueger, Tim Kuphal, Stanley Letovsky, Michael Levandoski, Craig Lukasik, Jonathan Meltzer, Brian Norvell, Mindy Nye, Scott Parker, Christos Petropoulos, John Pruitt, Steven Ragan, Scott Ryan, Mike Sapeta, Jana Schroth, Suresh Babu Selvaraju, Goran Stevovic, Amanda Suchanek, Andrea Throop, Lyndon Tilson, Thomas Urban, Joe Voshell, Kimberly Wagner, Jonathan Williams, Mary Williamson, Qian Zeng, Tricia Zwiefelhofer, Clinton R. Paden, Duncan MacCannell |
| EPI_ISL_3325757 | TGen North | TGen North | Jolene Bowers, Heather Centner, Brett Van Tassel, Chris French, Hayley Yaglom, Darrin Lemmer, Dave Engelthaler |
| EPI_ISL_3325758 | Laboratory Corporation of America | Centers for Disease Control and Prevention Division of Viral Diseases, Pathogen Discovery | Dakota Howard, Dhwani Batra, Peter W. Cook, Kara Moser, Adrian Paskey, Jason Caravas, Benjamin Rambo-Martin, Shatavia Morrison, Christopher Gulvick, Scott Sammons, Yvette Unoarumhi, Darlene Wagner, Matthew Schmerer, Minoo Agarwal, Eyad Almasri, Debbie Boles, Ayla Burns, Nuthawin Charoensri, Oren Cohen, Susan Countryman, Mary Ann Cristobal, Bobbi Croy, Suzanne Dale, Hrushikesh Deshmukh, Amanda Douglas, Vincent Drouillon, Marcia Eisenberg, Howard Engler, Rama Ghatti, Prashant Gupta, Susan Hicks, Jake Humphrey, Lax Iyer, Lisa Pfefferle, Manoj Jain, Matthew Robinson, Mohan Kolli, Brian Krueger, Tim Kuphal, Stanley Letovsky, Michael Levandoski, Craig Lukasik, Jonathan Meltzer, Brian Norvell, Mindy Nye, Scott Parker, Christos Petropoulos, John Pruitt, Steven Ragan, Scott Ryan, Mike Sapeta, Jana Schroth, Suresh Babu Selvaraju, Goran Stevovic, Amanda Suchanek, Andrea Throop, Lyndon Tilson, Thomas Urban, Joe Voshell, Kimberly Wagner, Jonathan Williams, Mary Williamson, Qian Zeng, Tricia Zwiefelhofer, Clinton R. Paden, Duncan MacCannell |
| EPI_ISL_3325759 | TGen North | TGen North | Jolene Bowers, Heather Centner, Brett Van Tassel, Chris French, Hayley Yaglom, Darrin Lemmer, Dave Engelthaler |
| EPI_ISL_3325760, EPI_ISL_3325762, EPI_ISL_3325763 | Laboratory Corporation of America | Centers for Disease Control and Prevention Division of Viral Diseases, Pathogen Discovery | Dakota Howard, Dhwani Batra, Peter W. Cook, Kara Moser, Adrian Paskey, Jason Caravas, Benjamin Rambo-Martin, Shatavia Morrison, Christopher Gulvick, Scott Sammons, Yvette Unoarumhi, Darlene Wagner, Matthew Schmerer, Minoo Agarwal, Eyad Almasri, Debbie Boles, Ayla Burns, Nuthawin Charoensri, Oren Cohen, Susan Countryman, Mary Ann Cristobal, Bobbi Croy, Suzanne Dale, Hrushikesh Deshmukh, Amanda Douglas, Vincent Drouillon, Marcia Eisenberg, Howard Engler, Rama Ghatti, Prashant Gupta, Susan Hicks, Jake Humphrey, Lax Iyer, Lisa Pfefferle, Manoj Jain, Matthew Robinson, Mohan Kolli, Brian Krueger, Tim Kuphal, Stanley Letovsky, Michael Levandoski, Craig Lukasik, Jonathan Meltzer, Brian Norvell, Mindy Nye, Scott Parker, Christos Petropoulos, John Pruitt, Steven Ragan, Scott Ryan, Mike Sapeta, Jana Schroth, Suresh Babu Selvaraju, Goran Stevovic, Amanda Suchanek, Andrea Throop, Lyndon Tilson, Thomas Urban, Joe Voshell, Kimberly Wagner, Jonathan Williams, Mary Williamson, Qian Zeng, Tricia Zwiefelhofer, Clinton R. Paden, Duncan MacCannell |
| EPI_ISL_3325764 | TGen North | TGen North | Jolene Bowers, Heather Centner, Brett Van Tassel, Chris French, Hayley Yaglom, Darrin Lemmer, Dave Engelthaler |
| EPI_ISL_3325765 | Laboratory Corporation of America | Centers for Disease Control and Prevention Division of Viral Diseases, Pathogen Discovery | Dakota Howard, Dhwani Batra, Peter W. Cook, Kara Moser, Adrian Paskey, Jason Caravas, Benjamin Rambo-Martin, Shatavia Morrison, Christopher Gulvick, Scott Sammons, Yvette Unoarumhi, Darlene Wagner, Matthew Schmerer, Minoo Agarwal, Eyad Almasri, Debbie Boles, Ayla Burns, Nuthawin Charoensri, Oren Cohen, Susan Countryman, Mary Ann Cristobal, Bobbi Croy, Suzanne Dale, Hrushikesh Deshmukh, Amanda Douglas, Vincent Drouillon, Marcia Eisenberg, Howard Engler, Rama Ghatti, Prashant Gupta, Susan Hicks, Jake Humphrey, Lax Iyer, Lisa Pfefferle, Manoj Jain, Matthew Robinson, Mohan Kolli, Brian Krueger, Tim Kuphal, Stanley Letovsky, Michael Levandoski, Craig Lukasik, Jonathan Meltzer, Brian Norvell, Mindy Nye, Scott Parker, Christos Petropoulos, John Pruitt, Steven Ragan, Scott Ryan, Mike Sapeta, Jana Schroth, Suresh Babu Selvaraju, Goran Stevovic, Amanda Suchanek, Andrea Throop, Lyndon Tilson, Thomas Urban, Joe Voshell, Kimberly Wagner, Jonathan Williams, Mary Williamson, Qian Zeng, Tricia Zwiefelhofer, Clinton R. Paden, Duncan MacCannell |
| EPI_ISL_3325766, EPI_ISL_3325768 | TGen North | TGen North | Jolene Bowers, Heather Centner, Brett Van Tassel, Chris French, Hayley Yaglom, Darrin Lemmer, Dave Engelthaler |
| EPI_ISL_3325769 | Laboratory Corporation of America | Centers for Disease Control and Prevention Division of Viral Diseases, Pathogen Discovery | Dakota Howard, Dhwani Batra, Peter W. Cook, Kara Moser, Adrian Paskey, Jason Caravas, Benjamin Rambo-Martin, Shatavia Morrison, Christopher Gulvick, Scott Sammons, Yvette Unoarumhi, Darlene Wagner, Matthew Schmerer, Minoo Agarwal, Eyad Almasri, Debbie Boles, Ayla Burns, Nuthawin Charoensri, Oren Cohen, Susan Countryman, Mary Ann Cristobal, Bobbi Croy, Suzanne Dale, Hrushikesh Deshmukh, Amanda Douglas, Vincent Drouillon, Marcia Eisenberg, Howard Engler, Rama Ghatti, Prashant Gupta, Susan Hicks, Jake Humphrey, Lax Iyer, Lisa Pfefferle, Manoj Jain, Matthew Robinson, Mohan Kolli, Brian Krueger, Tim Kuphal, Stanley Letovsky, Michael Levandoski, Craig Lukasik, Jonathan Meltzer, Brian Norvell, Mindy Nye, Scott Parker, Christos Petropoulos, John Pruitt, Steven Ragan, Scott Ryan, Mike Sapeta, Jana Schroth, Suresh Babu Selvaraju, Goran Stevovic, Amanda Suchanek, Andrea Throop, Lyndon Tilson, Thomas Urban, Joe Voshell, Kimberly Wagner, Jonathan Williams, Mary Williamson, Qian Zeng, Tricia Zwiefelhofer, Clinton R. Paden, Duncan MacCannell |
| EPI_ISL_3325770 | TGen North | TGen North | Jolene Bowers, Heather Centner, Brett Van Tassel, Chris French, Hayley Yaglom, Darrin Lemmer, Dave Engelthaler |
| EPI_ISL_3325771 | Laboratory Corporation of America | Centers for Disease Control and Prevention Division of Viral Diseases, Pathogen Discovery | Dakota Howard, Dhwani Batra, Peter W. Cook, Kara Moser, Adrian Paskey, Jason Caravas, Benjamin Rambo-Martin, Shatavia Morrison, Christopher Gulvick, Scott Sammons, Yvette Unoarumhi, Darlene Wagner, Matthew Schmerer, Minoo Agarwal, Eyad Almasri, Debbie Boles, Ayla Burns, Nuthawin Charoensri, Oren Cohen, Susan Countryman, Mary Ann Cristobal, Bobbi Croy, Suzanne Dale, Hrushikesh Deshmukh, Amanda Douglas, Vincent Drouillon, Marcia Eisenberg, Howard Engler, Rama Ghatti, Prashant Gupta, Susan Hicks, Jake Humphrey, Lax Iyer, Lisa Pfefferle, Manoj Jain, Matthew Robinson, Mohan Kolli, Brian Krueger, Tim Kuphal, Stanley Letovsky, Michael Levandoski, Craig Lukasik, Jonathan Meltzer, Brian Norvell, Mindy Nye, Scott Parker, Christos Petropoulos, John Pruitt, Steven Ragan, Scott Ryan, Mike Sapeta, Jana Schroth, Suresh Babu Selvaraju, Goran Stevovic, Amanda Suchanek, Andrea Throop, Lyndon Tilson, Thomas Urban, Joe Voshell, Kimberly Wagner, Jonathan Williams, Mary Williamson, Qian Zeng, Tricia Zwiefelhofer, Clinton R. Paden, Duncan MacCannell |
| EPI_ISL_3325772 | TGen North | TGen North | Jolene Bowers, Heather Centner, Brett Van Tassel, Chris French, Hayley Yaglom, Darrin Lemmer, Dave Engelthaler |
| EPI_ISL_3325773, EPI_ISL_3325774, EPI_ISL_3325776 | Laboratory Corporation of America | Centers for Disease Control and Prevention Division of Viral Diseases, Pathogen Discovery | Dakota Howard, Dhwani Batra, Peter W. Cook, Kara Moser, Adrian Paskey, Jason Caravas, Benjamin Rambo-Martin, Shatavia Morrison, Christopher Gulvick, Scott Sammons, Yvette Unoarumhi, Darlene Wagner, Matthew Schmerer, Minoo Agarwal, Eyad Almasri, Debbie Boles, Ayla Burns, Nuthawin Charoensri, Oren Cohen, Susan Countryman, Mary Ann Cristobal, Bobbi Croy, Suzanne Dale, Hrushikesh Deshmukh, Amanda Douglas, Vincent Drouillon, Marcia Eisenberg, Howard Engler, Rama Ghatti, Prashant Gupta, Susan Hicks, Jake Humphrey, Lax Iyer, Lisa Pfefferle, Manoj Jain, Matthew Robinson, Mohan Kolli, Brian Krueger, Tim Kuphal, Stanley Letovsky, Michael Levandoski, Craig Lukasik, Jonathan Meltzer, Brian Norvell, Mindy Nye, Scott Parker, Christos Petropoulos, John Pruitt, Steven Ragan, Scott Ryan, Mike Sapeta, Jana Schroth, Suresh Babu Selvaraju, Goran Stevovic, Amanda Suchanek, Andrea Throop, Lyndon Tilson, Thomas Urban, Joe Voshell, Kimberly Wagner, Jonathan Williams, Mary Williamson, Qian Zeng, Tricia Zwiefelhofer, Clinton R. Paden, Duncan MacCannell |
| EPI_ISL_3325777 | TGen North | TGen North | Jolene Bowers, Heather Centner, Brett Van Tassel, Chris French, Hayley Yaglom, Darrin Lemmer, Dave Engelthaler |
| EPI_ISL_3325778 | Laboratory Corporation of America | Centers for Disease Control and Prevention Division of Viral Diseases, Pathogen Discovery | Dakota Howard, Dhwani Batra, Peter W. Cook, Kara Moser, Adrian Paskey, Jason Caravas, Benjamin Rambo-Martin, Shatavia Morrison, Christopher Gulvick, Scott Sammons, Yvette Unoarumhi, Darlene Wagner, Matthew Schmerer, Minoo Agarwal, Eyad Almasri, Debbie Boles, Ayla Burns, Nuthawin Charoensri, Oren Cohen, Susan Countryman, Mary Ann Cristobal, Bobbi Croy, Suzanne Dale, Hrushikesh Deshmukh, Amanda Douglas, Vincent Drouillon, Marcia Eisenberg, Howard Engler, Rama Ghatti, Prashant Gupta, Susan Hicks, Jake Humphrey, Lax Iyer, Lisa Pfefferle, Manoj Jain, Matthew Robinson, Mohan Kolli, Brian Krueger, Tim Kuphal, Stanley Letovsky, Michael Levandoski, Craig Lukasik, Jonathan Meltzer, Brian Norvell, Mindy Nye, Scott Parker, Christos Petropoulos, John Pruitt, Steven Ragan, Scott Ryan, Mike Sapeta, Jana Schroth, Suresh Babu Selvaraju, Goran Stevovic, Amanda Suchanek, Andrea Throop, Lyndon Tilson, Thomas Urban, Joe Voshell, Kimberly Wagner, Jonathan Williams, Mary Williamson, Qian Zeng, Tricia Zwiefelhofer, Clinton R. Paden, Duncan MacCannell |
| EPI_ISL_3325779 | TGen North | TGen North | Jolene Bowers, Heather Centner, Brett Van Tassel, Chris French, Hayley Yaglom, Darrin Lemmer, Dave Engelthaler |
| EPI_ISL_3325780, EPI_ISL_3325783 | Laboratory Corporation of America | Centers for Disease Control and Prevention Division of Viral Diseases, Pathogen Discovery | Dakota Howard, Dhwani Batra, Peter W. Cook, Kara Moser, Adrian Paskey, Jason Caravas, Benjamin Rambo-Martin, Shatavia Morrison, Christopher Gulvick, Scott Sammons, Yvette Unoarumhi, Darlene Wagner, Matthew Schmerer, Minoo Agarwal, Eyad Almasri, Debbie Boles, Ayla Burns, Nuthawin Charoensri, Oren Cohen, Susan Countryman, Mary Ann Cristobal, Bobbi Croy, Suzanne Dale, Hrushikesh Deshmukh, Amanda Douglas, Vincent Drouillon, Marcia Eisenberg, Howard Engler, Rama Ghatti, Prashant Gupta, Susan Hicks, Jake Humphrey, Lax Iyer, Lisa Pfefferle, Manoj Jain, Matthew Robinson, Mohan Kolli, Brian Krueger, Tim Kuphal, Stanley Letovsky, Michael Levandoski, Craig Lukasik, Jonathan Meltzer, Brian Norvell, Mindy Nye, Scott Parker, Christos Petropoulos, John Pruitt, Steven Ragan, Scott Ryan, Mike Sapeta, Jana Schroth, Suresh Babu Selvaraju, Goran Stevovic, Amanda Suchanek, Andrea Throop, Lyndon Tilson, Thomas Urban, Joe Voshell, Kimberly Wagner, Jonathan Williams, Mary Williamson, Qian Zeng, Tricia Zwiefelhofer, Clinton R. Paden, Duncan MacCannell |

| | | | |
|---|---|---|---|
| EPI_ISL_3325784 | TGen North | TGen North | Jolene Bowers, Heather Centner, Brett Van Tassel, Chris French, Hayley Yaglom, Darrin Lemmer, Dave Engelthaler |
| EPI_ISL_3325785, EPI_ISL_3325787, EPI_ISL_3325789, EPI_ISL_3325790 | Laboratory Corporation of America | Centers for Disease Control and Prevention Division of Viral Diseases, Pathogen Discovery | Dakota Howard, Dhwani Batra, Peter W. Cook, Kara Moser, Adrian Paskey, Jason Caravas, Benjamin Rambo-Martin, Shatavia Morrison, Christopher Gulvick, Scott Sammons, Yvette Unoarumhi, Darlene Wagner, Matthew Schmerer, Minoo Agarwal, Eyad Almasri, Debbie Boles, Ayla Burns, Nuthawin Charoensri, Oren Cohen, Susan Countryman, Mary Ann Cristobal, Bobbi Croy, Suzanne Dale, Hrushikesh Deshmukh, Amanda Douglas, Vincent Drouillon, Marcia Eisenberg, Howard Engler, Rama Ghatti, Prashant Gupta, Susan Hicks, Jake Humphrey, Lax Iyer, Lisa Pfefferle, Manoj Jain, Matthew Robinson, Mohan Kolli, Brian Krueger, Tim Kuphal, Stanley Letovsky, Michael Levandoski, Craig Lukasik, Jonathan Meltzer, Brian Norvell, Mindy Nye, Scott Parker, Christos Petropoulos, John Pruitt, Steven Ragan, Scott Ryan, Mike Sapeta, Jana Schroth, Suresh Babu Selvaraju, Goran Stevovic, Amanda Suchanek, Andrea Throop, Lyndon Tilson, Thomas Urban, Joe Voshell, Kimberly Wagner, Jonathan Williams, Mary Williamson, Qian Zeng, Tricia Zwiefelhofer, Clinton R. Paden, Duncan MacCannell |
| EPI_ISL_3325791 | TGen North | TGen North | Jolene Bowers, Heather Centner, Brett Van Tassel, Chris French, Hayley Yaglom, Darrin Lemmer, Dave Engelthaler |
| EPI_ISL_3325792 | Laboratory Corporation of America | Centers for Disease Control and Prevention Division of Viral Diseases, Pathogen Discovery | Dakota Howard, Dhwani Batra, Peter W. Cook, Kara Moser, Adrian Paskey, Jason Caravas, Benjamin Rambo-Martin, Shatavia Morrison, Christopher Gulvick, Scott Sammons, Yvette Unoarumhi, Darlene Wagner, Matthew Schmerer, Minoo Agarwal, Eyad Almasri, Debbie Boles, Ayla Burns, Nuthawin Charoensri, Oren Cohen, Susan Countryman, Mary Ann Cristobal, Bobbi Croy, Suzanne Dale, Hrushikesh Deshmukh, Amanda Douglas, Vincent Drouillon, Marcia Eisenberg, Howard Engler, Rama Ghatti, Prashant Gupta, Susan Hicks, Jake Humphrey, Lax Iyer, Lisa Pfefferle, Manoj Jain, Matthew Robinson, Mohan Kolli, Brian Krueger, Tim Kuphal, Stanley Letovsky, Michael Levandoski, Craig Lukasik, Jonathan Meltzer, Brian Norvell, Mindy Nye, Scott Parker, Christos Petropoulos, John Pruitt, Steven Ragan, Scott Ryan, Mike Sapeta, Jana Schroth, Suresh Babu Selvaraju, Goran Stevovic, Amanda Suchanek, Andrea Throop, Lyndon Tilson, Thomas Urban, Joe Voshell, Kimberly Wagner, Jonathan Williams, Mary Williamson, Qian Zeng, Tricia Zwiefelhofer, Clinton R. Paden, Duncan MacCannell |
| EPI_ISL_3325793 | TGen North | TGen North | Jolene Bowers, Heather Centner, Brett Van Tassel, Chris French, Hayley Yaglom, Darrin Lemmer, Dave Engelthaler |
| EPI_ISL_3325794 | Laboratory Corporation of America | Centers for Disease Control and Prevention Division of Viral Diseases, Pathogen Discovery | Dakota Howard, Dhwani Batra, Peter W. Cook, Kara Moser, Adrian Paskey, Jason Caravas, Benjamin Rambo-Martin, Shatavia Morrison, Christopher Gulvick, Scott Sammons, Yvette Unoarumhi, Darlene Wagner, Matthew Schmerer, Minoo Agarwal, Eyad Almasri, Debbie Boles, Ayla Burns, Nuthawin Charoensri, Oren Cohen, Susan Countryman, Mary Ann Cristobal, Bobbi Croy, Suzanne Dale, Hrushikesh Deshmukh, Amanda Douglas, Vincent Drouillon, Marcia Eisenberg, Howard Engler, Rama Ghatti, Prashant Gupta, Susan Hicks, Jake Humphrey, Lax Iyer, Lisa Pfefferle, Manoj Jain, Matthew Robinson, Mohan Kolli, Brian Krueger, Tim Kuphal, Stanley Letovsky, Michael Levandoski, Craig Lukasik, Jonathan Meltzer, Brian Norvell, Mindy Nye, Scott Parker, Christos Petropoulos, John Pruitt, Steven Ragan, Scott Ryan, Mike Sapeta, Jana Schroth, Suresh Babu Selvaraju, Goran Stevovic, Amanda Suchanek, Andrea Throop, Lyndon Tilson, Thomas Urban, Joe Voshell, Kimberly Wagner, Jonathan Williams, Mary Williamson, Qian Zeng, Tricia Zwiefelhofer, Clinton R. Paden, Duncan MacCannell |
| EPI_ISL_3325795 | TGen North | TGen North | Jolene Bowers, Heather Centner, Brett Van Tassel, Chris French, Hayley Yaglom, Darrin Lemmer, Dave Engelthaler |
| EPI_ISL_3325796, EPI_ISL_3325798 | Laboratory Corporation of America | Centers for Disease Control and Prevention Division of Viral Diseases, Pathogen Discovery | Dakota Howard, Dhwani Batra, Peter W. Cook, Kara Moser, Adrian Paskey, Jason Caravas, Benjamin Rambo-Martin, Shatavia Morrison, Christopher Gulvick, Scott Sammons, Yvette Unoarumhi, Darlene Wagner, Matthew Schmerer, Minoo Agarwal, Eyad Almasri, Debbie Boles, Ayla Burns, Nuthawin Charoensri, Oren Cohen, Susan Countryman, Mary Ann Cristobal, Bobbi Croy, Suzanne Dale, Hrushikesh Deshmukh, Amanda Douglas, Vincent Drouillon, Marcia Eisenberg, Howard Engler, Rama Ghatti, Prashant Gupta, Susan Hicks, Jake Humphrey, Lax Iyer, Lisa Pfefferle, Manoj Jain, Matthew Robinson, Mohan Kolli, Brian Krueger, Tim Kuphal, Stanley Letovsky, Michael Levandoski, Craig Lukasik, Jonathan Meltzer, Brian Norvell, Mindy Nye, Scott Parker, Christos Petropoulos, John Pruitt, Steven Ragan, Scott Ryan, Mike Sapeta, Jana Schroth, Suresh Babu Selvaraju, Goran Stevovic, Amanda Suchanek, Andrea Throop, Lyndon Tilson, Thomas Urban, Joe Voshell, Kimberly Wagner, Jonathan Williams, Mary Williamson, Qian Zeng, Tricia Zwiefelhofer, Clinton R. Paden, Duncan MacCannell |
| EPI_ISL_3325799 | TGen North | TGen North | Jolene Bowers, Heather Centner, Brett Van Tassel, Chris French, Hayley Yaglom, Darrin Lemmer, Dave Engelthaler |
| EPI_ISL_3325800, EPI_ISL_3325802 | Laboratory Corporation of America | Centers for Disease Control and Prevention Division of Viral Diseases, Pathogen Discovery | Dakota Howard, Dhwani Batra, Peter W. Cook, Kara Moser, Adrian Paskey, Jason Caravas, Benjamin Rambo-Martin, Shatavia Morrison, Christopher Gulvick, Scott Sammons, Yvette Unoarumhi, Darlene Wagner, Matthew Schmerer, Minoo Agarwal, Eyad Almasri, Debbie Boles, Ayla Burns, Nuthawin Charoensri, Oren Cohen, Susan Countryman, Mary Ann Cristobal, Bobbi Croy, Suzanne Dale, Hrushikesh Deshmukh, Amanda Douglas, Vincent Drouillon, Marcia Eisenberg, Howard Engler, Rama Ghatti, Prashant Gupta, Susan Hicks, Jake Humphrey, Lax Iyer, Lisa Pfefferle, Manoj Jain, Matthew Robinson, Mohan Kolli, Brian Krueger, Tim Kuphal, Stanley Letovsky, Michael Levandoski, Craig Lukasik, Jonathan Meltzer, Brian Norvell, Mindy Nye, Scott Parker, Christos Petropoulos, John Pruitt, Steven Ragan, Scott Ryan, Mike Sapeta, Jana Schroth, Suresh Babu Selvaraju, Goran Stevovic, Amanda Suchanek, Andrea Throop, Lyndon Tilson, Thomas Urban, Joe Voshell, Kimberly Wagner, Jonathan Williams, Mary Williamson, Qian Zeng, Tricia Zwiefelhofer, Clinton R. Paden, Duncan MacCannell |
| EPI_ISL_3325803 | TGen North | TGen North | Jolene Bowers, Heather Centner, Brett Van Tassel, Chris French, Hayley Yaglom, Darrin Lemmer, Dave Engelthaler |
| EPI_ISL_3325804, EPI_ISL_3325805 | Laboratory Corporation of America | Centers for Disease Control and Prevention Division of Viral Diseases, Pathogen Discovery | Dakota Howard, Dhwani Batra, Peter W. Cook, Kara Moser, Adrian Paskey, Jason Caravas, Benjamin Rambo-Martin, Shatavia Morrison, Christopher Gulvick, Scott Sammons, Yvette Unoarumhi, Darlene Wagner, Matthew Schmerer, Minoo Agarwal, Eyad Almasri, Debbie Boles, Ayla Burns, Nuthawin Charoensri, Oren Cohen, Susan Countryman, Mary Ann Cristobal, Bobbi Croy, Suzanne Dale, Hrushikesh Deshmukh, Amanda Douglas, Vincent Drouillon, Marcia Eisenberg, Howard Engler, Rama Ghatti, Prashant Gupta, Susan Hicks, Jake Humphrey, Lax Iyer, Lisa Pfefferle, Manoj Jain, Matthew Robinson, Mohan Kolli, Brian Krueger, Tim Kuphal, Stanley Letovsky, Michael Levandoski, Craig Lukasik, Jonathan Meltzer, Brian Norvell, Mindy Nye, Scott Parker, Christos Petropoulos, John Pruitt, Steven Ragan, Scott Ryan, Mike Sapeta, Jana Schroth, Suresh Babu Selvaraju, Goran Stevovic, Amanda Suchanek, Andrea Throop, Lyndon Tilson, Thomas Urban, Joe Voshell, Kimberly Wagner, Jonathan Williams, Mary Williamson, Qian Zeng, Tricia Zwiefelhofer, Clinton R. Paden, Duncan MacCannell |
| EPI_ISL_3325806 | TGen North | TGen North | Jolene Bowers, Heather Centner, Brett Van Tassel, Chris French, Hayley Yaglom, Darrin Lemmer, Dave Engelthaler |
| EPI_ISL_3325807, EPI_ISL_3325809 | Laboratory Corporation of America | Centers for Disease Control and Prevention Division of Viral Diseases, Pathogen Discovery | Dakota Howard, Dhwani Batra, Peter W. Cook, Kara Moser, Adrian Paskey, Jason Caravas, Benjamin Rambo-Martin, Shatavia Morrison, Christopher Gulvick, Scott Sammons, Yvette Unoarumhi, Darlene Wagner, Matthew Schmerer, Minoo Agarwal, Eyad Almasri, Debbie Boles, Ayla Burns, Nuthawin Charoensri, Oren Cohen, Susan Countryman, Mary Ann Cristobal, Bobbi Croy, Suzanne Dale, Hrushikesh Deshmukh, Amanda Douglas, Vincent Drouillon, Marcia Eisenberg, Howard Engler, Rama Ghatti, Prashant Gupta, Susan Hicks, Jake Humphrey, Lax Iyer, Lisa Pfefferle, Manoj Jain, Matthew Robinson, Mohan Kolli, Brian Krueger, Tim Kuphal, Stanley Letovsky, Michael Levandoski, Craig Lukasik, Jonathan Meltzer, Brian Norvell, Mindy Nye, Scott Parker, Christos Petropoulos, John Pruitt, Steven Ragan, Scott Ryan, Mike Sapeta, Jana Schroth, Suresh Babu Selvaraju, Goran Stevovic, Amanda Suchanek, Andrea Throop, Lyndon Tilson, Thomas Urban, Joe Voshell, Kimberly Wagner, Jonathan Williams, Mary Williamson, Qian Zeng, Tricia Zwiefelhofer, Clinton R. Paden, Duncan MacCannell |
| EPI_ISL_3325810 | TGen North | TGen North | Jolene Bowers, Heather Centner, Brett Van Tassel, Chris French, Hayley Yaglom, Darrin Lemmer, Dave Engelthaler |
| EPI_ISL_3325811 | Laboratory Corporation of America | Centers for Disease Control and Prevention Division of Viral Diseases, Pathogen Discovery | Dakota Howard, Dhwani Batra, Peter W. Cook, Kara Moser, Adrian Paskey, Jason Caravas, Benjamin Rambo-Martin, Shatavia Morrison, Christopher Gulvick, Scott Sammons, Yvette Unoarumhi, Darlene Wagner, Matthew Schmerer, Minoo Agarwal, Eyad Almasri, Debbie Boles, Ayla Burns, Nuthawin Charoensri, Oren Cohen, Susan Countryman, Mary Ann Cristobal, Bobbi Croy, Suzanne Dale, Hrushikesh Deshmukh, Amanda Douglas, Vincent Drouillon, Marcia Eisenberg, Howard Engler, Rama Ghatti, Prashant Gupta, Susan Hicks, Jake Humphrey, Lax Iyer, Lisa Pfefferle, Manoj Jain, Matthew Robinson, Mohan Kolli, Brian Krueger, Tim Kuphal, Stanley Letovsky, Michael Levandoski, Craig Lukasik, Jonathan Meltzer, Brian Norvell, Mindy Nye, Scott Parker, Christos Petropoulos, John Pruitt, Steven Ragan, Scott Ryan, Mike Sapeta, Jana Schroth, Suresh Babu Selvaraju, Goran Stevovic, Amanda Suchanek, Andrea Throop, Lyndon Tilson, Thomas Urban, Joe Voshell, Kimberly Wagner, Jonathan Williams, Mary Williamson, Qian Zeng, Tricia Zwiefelhofer, Clinton R. Paden, Duncan MacCannell |
| EPI_ISL_3325812 | TGen North | TGen North | Jolene Bowers, Heather Centner, Brett Van Tassel, Chris French, Hayley Yaglom, Darrin Lemmer, Dave Engelthaler |
| EPI_ISL_3325813, EPI_ISL_3325814, EPI_ISL_3325816 | Laboratory Corporation of America | Centers for Disease Control and Prevention Division of Viral Diseases, Pathogen Discovery | Dakota Howard, Dhwani Batra, Peter W. Cook, Kara Moser, Adrian Paskey, Jason Caravas, Benjamin Rambo-Martin, Shatavia Morrison, Christopher Gulvick, Scott Sammons, Yvette Unoarumhi, Darlene Wagner, Matthew Schmerer, Minoo Agarwal, Eyad Almasri, Debbie Boles, Ayla Burns, Nuthawin Charoensri, Oren Cohen, Susan Countryman, Mary Ann Cristobal, Bobbi Croy, Suzanne Dale, Hrushikesh Deshmukh, Amanda Douglas, Vincent Drouillon, Marcia Eisenberg, Howard Engler, Rama Ghatti, Prashant Gupta, Susan Hicks, Jake Humphrey, Lax Iyer, Lisa Pfefferle, Manoj Jain, Matthew Robinson, Mohan Kolli, Brian Krueger, Tim Kuphal, Stanley Letovsky, Michael Levandoski, Craig Lukasik, Jonathan Meltzer, Brian Norvell, Mindy Nye, Scott Parker, Christos Petropoulos, John Pruitt, Steven Ragan, Scott Ryan, Mike Sapeta, Jana Schroth, Suresh Babu Selvaraju, Goran Stevovic, Amanda Suchanek, Andrea Throop, Lyndon Tilson, Thomas Urban, Joe Voshell, Kimberly Wagner, Jonathan Williams, Mary Williamson, Qian Zeng, Tricia Zwiefelhofer, Clinton R. Paden, Duncan MacCannell |
| EPI_ISL_3325817 | TGen North | TGen North | Jolene Bowers, Heather Centner, Brett Van Tassel, Chris French, Hayley Yaglom, Darrin Lemmer, Dave Engelthaler |
| EPI_ISL_3325818, EPI_ISL_3325820 | Laboratory Corporation of America | Centers for Disease Control and Prevention Division of Viral Diseases, Pathogen Discovery | Dakota Howard, Dhwani Batra, Peter W. Cook, Kara Moser, Adrian Paskey, Jason Caravas, Benjamin Rambo-Martin, Shatavia Morrison, Christopher Gulvick, Scott Sammons, Yvette Unoarumhi, Darlene Wagner, Matthew Schmerer, Minoo Agarwal, Eyad Almasri, Debbie Boles, Ayla Burns, Nuthawin Charoensri, Oren Cohen, Susan Countryman, Mary Ann Cristobal, Bobbi Croy, Suzanne Dale, Hrushikesh Deshmukh, Amanda Douglas, Vincent Drouillon, Marcia Eisenberg, Howard Engler, Rama Ghatti, Prashant Gupta, Susan Hicks, Jake Humphrey, Lax Iyer, Lisa Pfefferle, Manoj Jain, Matthew Robinson, Mohan Kolli, Brian Krueger, Tim Kuphal, Stanley Letovsky, Michael Levandoski, Craig Lukasik, Jonathan Meltzer, Brian Norvell, Mindy Nye, Scott Parker, Christos Petropoulos, John Pruitt, Steven Ragan, Scott Ryan, Mike Sapeta, Jana Schroth, Suresh Babu Selvaraju, Goran Stevovic, Amanda Suchanek, Andrea Throop, Lyndon Tilson, Thomas Urban, Joe Voshell, Kimberly Wagner, Jonathan Williams, Mary Williamson, Qian Zeng, Tricia Zwiefelhofer, Clinton R. Paden, Duncan MacCannell |
| EPI_ISL_3325821 | TGen North | TGen North | Jolene Bowers, Heather Centner, Brett Van Tassel, Chris French, Hayley Yaglom, Darrin Lemmer, Dave Engelthaler |
| EPI_ISL_3325822, EPI_ISL_3325823 | Laboratory Corporation of America | Centers for Disease Control and Prevention Division of Viral Diseases, Pathogen Discovery | Dakota Howard, Dhwani Batra, Peter W. Cook, Kara Moser, Adrian Paskey, Jason Caravas, Benjamin Rambo-Martin, Shatavia Morrison, Christopher Gulvick, Scott Sammons, Yvette Unoarumhi, Darlene Wagner, Matthew Schmerer, Minoo Agarwal, Eyad Almasri, Debbie Boles, Ayla Burns, Nuthawin Charoensri, Oren Cohen, Susan Countryman, Mary Ann Cristobal, Bobbi Croy, Suzanne Dale, Hrushikesh Deshmukh, Amanda Douglas, Vincent Drouillon, Marcia Eisenberg, Howard Engler, Rama Ghatti, Prashant Gupta, Susan Hicks, Jake Humphrey, Lax Iyer, Lisa Pfefferle, Manoj Jain, Matthew Robinson, Mohan Kolli, Brian Krueger, Tim Kuphal, Stanley Letovsky, Michael Levandoski, Craig Lukasik, Jonathan Meltzer, Brian Norvell, Mindy Nye, Scott Parker, Christos Petropoulos, John Pruitt, Steven Ragan, Scott Ryan, Mike Sapeta, Jana Schroth, Suresh Babu Selvaraju, Goran Stevovic, Amanda Suchanek, Andrea Throop, Lyndon Tilson, Thomas Urban, Joe Voshell, Kimberly Wagner, Jonathan Williams, Mary Williamson, Qian Zeng, Tricia Zwiefelhofer, Clinton R. Paden, Duncan MacCannell |
| EPI_ISL_3325824 | TGen North | TGen North | Jolene Bowers, Heather Centner, Brett Van Tassel, Chris French, Hayley Yaglom, Darrin Lemmer, Dave Engelthaler |
| EPI_ISL_3325825 | Laboratory Corporation of America | Centers for Disease Control and Prevention Division of Viral Diseases, Pathogen Discovery | Dakota Howard, Dhwani Batra, Peter W. Cook, Kara Moser, Adrian Paskey, Jason Caravas, Benjamin Rambo-Martin, Shatavia Morrison, Christopher Gulvick, Scott Sammons, Yvette Unoarumhi, Darlene Wagner, Matthew Schmerer, Minoo Agarwal, Eyad Almasri, Debbie Boles, Ayla Burns, Nuthawin Charoensri, Oren Cohen, Susan Countryman, Mary Ann Cristobal, Bobbi Croy, Suzanne Dale, Hrushikesh Deshmukh, Amanda Douglas, Vincent Drouillon, Marcia Eisenberg, Howard Engler, Rama Ghatti, Prashant Gupta, Susan Hicks, Jake Humphrey, Lax Iyer, Lisa Pfefferle, Manoj Jain, Matthew Robinson, Mohan Kolli, Brian Krueger, Tim Kuphal, Stanley Letovsky, Michael Levandoski, Craig Lukasik, Jonathan Meltzer, Brian Norvell, Mindy Nye, Scott Parker, Christos Petropoulos, John Pruitt, Steven Ragan, Scott Ryan, Mike Sapeta, Jana Schroth, Suresh Babu Selvaraju, Goran Stevovic, Amanda Suchanek, Andrea Throop, Lyndon Tilson, Thomas Urban, Joe Voshell, Kimberly Wagner, Jonathan Williams, Mary Williamson, Qian Zeng, Tricia Zwiefelhofer, Clinton R. Paden, Duncan MacCannell |
| EPI_ISL_3325826 | TGen North | TGen North | Jolene Bowers, Heather Centner, Brett Van Tassel, Chris French, Hayley Yaglom, Darrin Lemmer, Dave Engelthaler |
| EPI_ISL_3325827, EPI_ISL_3325829 | Laboratory Corporation of America | Centers for Disease Control and Prevention Division of Viral Diseases, Pathogen Discovery | Dakota Howard, Dhwani Batra, Peter W. Cook, Kara Moser, Adrian Paskey, Jason Caravas, Benjamin Rambo-Martin, Shatavia Morrison, Christopher Gulvick, Scott Sammons, Yvette Unoarumhi, Darlene Wagner, Matthew Schmerer, Minoo Agarwal, Eyad Almasri, Debbie Boles, Ayla Burns, Nuthawin Charoensri, Oren Cohen, Susan Countryman, Mary Ann Cristobal, Bobbi Croy, Suzanne Dale, Hrushikesh Deshmukh, Amanda Douglas, Vincent Drouillon, Marcia Eisenberg, Howard Engler, Rama Ghatti, Prashant Gupta, Susan Hicks, Jake Humphrey, Lax Iyer, Lisa Pfefferle, Manoj Jain, Matthew Robinson, Mohan Kolli, Brian Krueger, Tim Kuphal, Stanley Letovsky, Michael Levandoski, Craig Lukasik, Jonathan Meltzer, Brian Norvell, Mindy Nye, Scott Parker, Christos Petropoulos, John Pruitt, Steven Ragan, Scott Ryan, Mike Sapeta, Jana Schroth, Suresh Babu Selvaraju, Goran Stevovic, Amanda Suchanek, Andrea Throop, Lyndon Tilson, Thomas Urban, Joe Voshell, Kimberly Wagner, Jonathan Williams, Mary Williamson, Qian Zeng, Tricia Zwiefelhofer, Clinton R. Paden, Duncan MacCannell |
| EPI_ISL_3325830 | TGen North | TGen North | Jolene Bowers, Heather Centner, Brett Van Tassel, Chris French, Hayley Yaglom, Darrin Lemmer, Dave Engelthaler |
| EPI_ISL_3325831, EPI_ISL_3325832 | Laboratory Corporation of America | Centers for Disease Control and Prevention Division of Viral Diseases, Pathogen Discovery | Dakota Howard, Dhwani Batra, Peter W. Cook, Kara Moser, Adrian Paskey, Jason Caravas, Benjamin Rambo-Martin, Shatavia Morrison, Christopher Gulvick, Scott Sammons, Yvette Unoarumhi, Darlene Wagner, Matthew Schmerer, Minoo Agarwal, Eyad Almasri, Debbie Boles, Ayla Burns, Nuthawin Charoensri, Oren Cohen, Susan Countryman, Mary Ann Cristobal, Bobbi Croy, Suzanne Dale, Hrushikesh Deshmukh, Amanda Douglas, Vincent Drouillon, Marcia Eisenberg, Howard Engler, Rama Ghatti, Prashant Gupta, Susan Hicks, Jake Humphrey, Lax Iyer, Lisa Pfefferle, Manoj Jain, Matthew Robinson, Mohan Kolli, Brian Krueger, Tim Kuphal, Stanley Letovsky, Michael Levandoski, Craig Lukasik, Jonathan Meltzer, Brian Norvell, Mindy Nye, Scott Parker, Christos Petropoulos, John Pruitt, Steven Ragan, Scott Ryan, Mike Sapeta, Jana Schroth, Suresh Babu Selvaraju, Goran Stevovic, Amanda Suchanek, Andrea Throop, Lyndon Tilson, Thomas Urban, Joe Voshell, Kimberly Wagner, Jonathan Williams, Mary Williamson, Qian Zeng, Tricia Zwiefelhofer, Clinton R. Paden, Duncan MacCannell |
| EPI_ISL_3325833 | TGen North | TGen North | Jolene Bowers, Heather Centner, Brett Van Tassel, Chris French, Hayley Yaglom, Darrin Lemmer, Dave Engelthaler |
| EPI_ISL_3325834, EPI_ISL_3325836 | Laboratory Corporation of America | Centers for Disease Control and Prevention Division of Viral Diseases, Pathogen Discovery | Dakota Howard, Dhwani Batra, Peter W. Cook, Kara Moser, Adrian Paskey, Jason Caravas, Benjamin Rambo-Martin, Shatavia Morrison, Christopher Gulvick, Scott Sammons, Yvette Unoarumhi, Darlene Wagner, Matthew Schmerer, Minoo Agarwal, Eyad Almasri, Debbie Boles, Ayla Burns, Nuthawin Charoensri, Oren Cohen, Susan Countryman, Mary Ann Cristobal, Bobbi Croy, Suzanne Dale, Hrushikesh Deshmukh, Amanda Douglas, Vincent Drouillon, Marcia Eisenberg, Howard Engler, Rama Ghatti, Prashant Gupta, Susan Hicks, Jake Humphrey, Lax Iyer, Lisa Pfefferle, Manoj Jain, Matthew Robinson, Mohan Kolli, Brian Krueger, Tim Kuphal, Stanley Letovsky, Michael Levandoski, Craig Lukasik, Jonathan Meltzer, Brian Norvell, Mindy Nye, Scott Parker, Christos Petropoulos, John Pruitt, Steven Ragan, Scott Ryan, Mike Sapeta, Jana Schroth, Suresh Babu Selvaraju, Goran Stevovic, Amanda Suchanek, Andrea Throop, Lyndon Tilson, Thomas Urban, Joe Voshell, Kimberly Wagner, Jonathan Williams, Mary Williamson, Qian Zeng, Tricia Zwiefelhofer, Clinton R. Paden, Duncan MacCannell |
| EPI_ISL_3325837 | TGen North | TGen North | Jolene Bowers, Heather Centner, Brett Van Tassel, Chris French, Hayley Yaglom, Darrin Lemmer, Dave Engelthaler |
| EPI_ISL_3325838 | Laboratory Corporation of America | Centers for Disease Control and Prevention Division of Viral Diseases, Pathogen Discovery | Dakota Howard, Dhwani Batra, Peter W. Cook, Kara Moser, Adrian Paskey, Jason Caravas, Benjamin Rambo-Martin, Shatavia Morrison, Christopher Gulvick, Scott Sammons, Yvette Unoarumhi, Darlene Wagner, Matthew Schmerer, Minoo Agarwal, Eyad Almasri, Debbie Boles, Ayla Burns, Nuthawin Charoensri, Oren Cohen, Susan Countryman, Mary Ann Cristobal, Bobbi Croy, Suzanne Dale, Hrushikesh Deshmukh, Amanda Douglas, Vincent Drouillon, Marcia Eisenberg, Howard Engler, Rama Ghatti, Prashant Gupta, Susan Hicks, Jake Humphrey, Lax Iyer, Lisa Pfefferle, Manoj Jain, Matthew Robinson, Mohan Kolli, Brian Krueger, Tim Kuphal, Stanley Letovsky, Michael Levandoski, Craig Lukasik, Jonathan Meltzer, Brian Norvell, Mindy Nye, Scott Parker, Christos Petropoulos, John Pruitt, Steven Ragan, Scott Ryan, Mike Sapeta, Jana Schroth, Suresh Babu Selvaraju, Goran Stevovic, Amanda Suchanek, Andrea Throop, Lyndon Tilson, Thomas Urban, Joe Voshell, Kimberly Wagner, Jonathan Williams, Mary Williamson, Qian Zeng, Tricia Zwiefelhofer, Clinton R. Paden, Duncan MacCannell |

| | | | |
|---|---|---|---|
| EPI_ISL_3325839 | TGen North | TGen North | Jolene Bowers, Heather Centner, Brett Van Tassel, Chris French, Hayley Yaglom, Darrin Lemmer, Dave Engelthaler |
| EPI_ISL_3325840, EPI_ISL_3325841, EPI_ISL_3325843 | Laboratory Corporation of America | Centers for Disease Control and Prevention Division of Viral Diseases, Pathogen Discovery | Dakota Howard, Dhwani Batra, Peter W. Cook, Kara Moser, Adrian Paskey, Jason Caravas, Benjamin Rambo-Martin, Shatavia Morrison, Christopher Gulvick, Scott Sammons, Yvette Unoarumhi, Darlene Wagner, Matthew Schmerer, Minoo Agarwal, Eyad Almasri, Debbie Boles, Ayla Burns, Nuthawin Charoensri, Oren Cohen, Susan Countryman, Mary Ann Cristobal, Bobbi Croy, Suzanne Dale, Hrushikesh Deshmukh, Amanda Douglas, Vincent Drouillon, Marcia Eisenberg, Howard Engler, Rama Ghatti, Prashant Gupta, Susan Hicks, Jake Humphrey, Lax Iyer, Lisa Pfefferle, Manoj Jain, Matthew Robinson, Mohan Kolli, Brian Krueger, Tim Kuphal, Stanley Letovsky, Michael Levandoski, Craig Lukasik, Jonathan Meltzer, Brian Norvell, Mindy Nye, Scott Parker, Christos Petropoulos, John Pruitt, Steven Ragan, Scott Ryan, Mike Sapeta, Jana Schroth, Suresh Babu Selvaraju, Goran Stevovic, Amanda Suchanek, Andrea Throop, Lyndon Tilson, Thomas Urban, Joe Voshell, Kimberly Wagner, Jonathan Williams, Mary Williamson, Qian Zeng, Tricia Zwiefelhofer, Clinton R. Paden, Duncan MacCannell |
| EPI_ISL_3325844 | TGen North | TGen North | Jolene Bowers, Heather Centner, Brett Van Tassel, Chris French, Hayley Yaglom, Darrin Lemmer, Dave Engelthaler |
| EPI_ISL_3325845 | Laboratory Corporation of America | Centers for Disease Control and Prevention Division of Viral Diseases, Pathogen Discovery | Dakota Howard, Dhwani Batra, Peter W. Cook, Kara Moser, Adrian Paskey, Jason Caravas, Benjamin Rambo-Martin, Shatavia Morrison, Christopher Gulvick, Scott Sammons, Yvette Unoarumhi, Darlene Wagner, Matthew Schmerer, Minoo Agarwal, Eyad Almasri, Debbie Boles, Ayla Burns, Nuthawin Charoensri, Oren Cohen, Susan Countryman, Mary Ann Cristobal, Bobbi Croy, Suzanne Dale, Hrushikesh Deshmukh, Amanda Douglas, Vincent Drouillon, Marcia Eisenberg, Howard Engler, Rama Ghatti, Prashant Gupta, Susan Hicks, Jake Humphrey, Lax Iyer, Lisa Pfefferle, Manoj Jain, Matthew Robinson, Mohan Kolli, Brian Krueger, Tim Kuphal, Stanley Letovsky, Michael Levandoski, Craig Lukasik, Jonathan Meltzer, Brian Norvell, Mindy Nye, Scott Parker, Christos Petropoulos, John Pruitt, Steven Ragan, Scott Ryan, Mike Sapeta, Jana Schroth, Suresh Babu Selvaraju, Goran Stevovic, Amanda Suchanek, Andrea Throop, Lyndon Tilson, Thomas Urban, Joe Voshell, Kimberly Wagner, Jonathan Williams, Mary Williamson, Qian Zeng, Tricia Zwiefelhofer, Clinton R. Paden, Duncan MacCannell |
| EPI_ISL_3325846 | TGen North | TGen North | Jolene Bowers, Heather Centner, Brett Van Tassel, Chris French, Hayley Yaglom, Darrin Lemmer, Dave Engelthaler |
| EPI_ISL_3325848 | Laboratory Corporation of America | Centers for Disease Control and Prevention Division of Viral Diseases, Pathogen Discovery | Dakota Howard, Dhwani Batra, Peter W. Cook, Kara Moser, Adrian Paskey, Jason Caravas, Benjamin Rambo-Martin, Shatavia Morrison, Christopher Gulvick, Scott Sammons, Yvette Unoarumhi, Darlene Wagner, Matthew Schmerer, Minoo Agarwal, Eyad Almasri, Debbie Boles, Ayla Burns, Nuthawin Charoensri, Oren Cohen, Susan Countryman, Mary Ann Cristobal, Bobbi Croy, Suzanne Dale, Hrushikesh Deshmukh, Amanda Douglas, Vincent Drouillon, Marcia Eisenberg, Howard Engler, Rama Ghatti, Prashant Gupta, Susan Hicks, Jake Humphrey, Lax Iyer, Lisa Pfefferle, Manoj Jain, Matthew Robinson, Mohan Kolli, Brian Krueger, Tim Kuphal, Stanley Letovsky, Michael Levandoski, Craig Lukasik, Jonathan Meltzer, Brian Norvell, Mindy Nye, Scott Parker, Christos Petropoulos, John Pruitt, Steven Ragan, Scott Ryan, Mike Sapeta, Jana Schroth, Suresh Babu Selvaraju, Goran Stevovic, Amanda Suchanek, Andrea Throop, Lyndon Tilson, Thomas Urban, Joe Voshell, Kimberly Wagner, Jonathan Williams, Mary Williamson, Qian Zeng, Tricia Zwiefelhofer, Clinton R. Paden, Duncan MacCannell |
| EPI_ISL_3325850 | TGen North | TGen North | Jolene Bowers, Heather Centner, Brett Van Tassel, Chris French, Hayley Yaglom, Darrin Lemmer, Dave Engelthaler |
| EPI_ISL_3325851, EPI_ISL_3325852 | Laboratory Corporation of America | Centers for Disease Control and Prevention Division of Viral Diseases, Pathogen Discovery | Dakota Howard, Dhwani Batra, Peter W. Cook, Kara Moser, Adrian Paskey, Jason Caravas, Benjamin Rambo-Martin, Shatavia Morrison, Christopher Gulvick, Scott Sammons, Yvette Unoarumhi, Darlene Wagner, Matthew Schmerer, Minoo Agarwal, Eyad Almasri, Debbie Boles, Ayla Burns, Nuthawin Charoensri, Oren Cohen, Susan Countryman, Mary Ann Cristobal, Bobbi Croy, Suzanne Dale, Hrushikesh Deshmukh, Amanda Douglas, Vincent Drouillon, Marcia Eisenberg, Howard Engler, Rama Ghatti, Prashant Gupta, Susan Hicks, Jake Humphrey, Lax Iyer, Lisa Pfefferle, Manoj Jain, Matthew Robinson, Mohan Kolli, Brian Krueger, Tim Kuphal, Stanley Letovsky, Michael Levandoski, Craig Lukasik, Jonathan Meltzer, Brian Norvell, Mindy Nye, Scott Parker, Christos Petropoulos, John Pruitt, Steven Ragan, Scott Ryan, Mike Sapeta, Jana Schroth, Suresh Babu Selvaraju, Goran Stevovic, Amanda Suchanek, Andrea Throop, Lyndon Tilson, Thomas Urban, Joe Voshell, Kimberly Wagner, Jonathan Williams, Mary Williamson, Qian Zeng, Tricia Zwiefelhofer, Clinton R. Paden, Duncan MacCannell |
| EPI_ISL_3325853, EPI_ISL_3325855 | TGen North | TGen North | Jolene Bowers, Heather Centner, Brett Van Tassel, Chris French, Hayley Yaglom, Darrin Lemmer, Dave Engelthaler |
| EPI_ISL_3325856, EPI_ISL_3325858 | Laboratory Corporation of America | Centers for Disease Control and Prevention Division of Viral Diseases, Pathogen Discovery | Dakota Howard, Dhwani Batra, Peter W. Cook, Kara Moser, Adrian Paskey, Jason Caravas, Benjamin Rambo-Martin, Shatavia Morrison, Christopher Gulvick, Scott Sammons, Yvette Unoarumhi, Darlene Wagner, Matthew Schmerer, Minoo Agarwal, Eyad Almasri, Debbie Boles, Ayla Burns, Nuthawin Charoensri, Oren Cohen, Susan Countryman, Mary Ann Cristobal, Bobbi Croy, Suzanne Dale, Hrushikesh Deshmukh, Amanda Douglas, Vincent Drouillon, Marcia Eisenberg, Howard Engler, Rama Ghatti, Prashant Gupta, Susan Hicks, Jake Humphrey, Lax Iyer, Lisa Pfefferle, Manoj Jain, Matthew Robinson, Mohan Kolli, Brian Krueger, Tim Kuphal, Stanley Letovsky, Michael Levandoski, Craig Lukasik, Jonathan Meltzer, Brian Norvell, Mindy Nye, Scott Parker, Christos Petropoulos, John Pruitt, Steven Ragan, Scott Ryan, Mike Sapeta, Jana Schroth, Suresh Babu Selvaraju, Goran Stevovic, Amanda Suchanek, Andrea Throop, Lyndon Tilson, Thomas Urban, Joe Voshell, Kimberly Wagner, Jonathan Williams, Mary Williamson, Qian Zeng, Tricia Zwiefelhofer, Clinton R. Paden, Duncan MacCannell |
| EPI_ISL_3325859 | TGen North | TGen North | Jolene Bowers, Heather Centner, Brett Van Tassel, Chris French, Hayley Yaglom, Darrin Lemmer, Dave Engelthaler |
| EPI_ISL_3325860, EPI_ISL_3325861 | Laboratory Corporation of America | Centers for Disease Control and Prevention Division of Viral Diseases, Pathogen Discovery | Dakota Howard, Dhwani Batra, Peter W. Cook, Kara Moser, Adrian Paskey, Jason Caravas, Benjamin Rambo-Martin, Shatavia Morrison, Christopher Gulvick, Scott Sammons, Yvette Unoarumhi, Darlene Wagner, Matthew Schmerer, Minoo Agarwal, Eyad Almasri, Debbie Boles, Ayla Burns, Nuthawin Charoensri, Oren Cohen, Susan Countryman, Mary Ann Cristobal, Bobbi Croy, Suzanne Dale, Hrushikesh Deshmukh, Amanda Douglas, Vincent Drouillon, Marcia Eisenberg, Howard Engler, Rama Ghatti, Prashant Gupta, Susan Hicks, Jake Humphrey, Lax Iyer, Lisa Pfefferle, Manoj Jain, Matthew Robinson, Mohan Kolli, Brian Krueger, Tim Kuphal, Stanley Letovsky, Michael Levandoski, Craig Lukasik, Jonathan Meltzer, Brian Norvell, Mindy Nye, Scott Parker, Christos Petropoulos, John Pruitt, Steven Ragan, Scott Ryan, Mike Sapeta, Jana Schroth, Suresh Babu Selvaraju, Goran Stevovic, Amanda Suchanek, Andrea Throop, Lyndon Tilson, Thomas Urban, Joe Voshell, Kimberly Wagner, Jonathan Williams, Mary Williamson, Qian Zeng, Tricia Zwiefelhofer, Clinton R. Paden, Duncan MacCannell |
| EPI_ISL_3325862, EPI_ISL_3325863, EPI_ISL_3325865 | TGen North | TGen North | Jolene Bowers, Heather Centner, Brett Van Tassel, Chris French, Hayley Yaglom, Darrin Lemmer, Dave Engelthaler |
| EPI_ISL_3325866 | Laboratory Corporation of America | Centers for Disease Control and Prevention Division of Viral Diseases, Pathogen Discovery | Dakota Howard, Dhwani Batra, Peter W. Cook, Kara Moser, Adrian Paskey, Jason Caravas, Benjamin Rambo-Martin, Shatavia Morrison, Christopher Gulvick, Scott Sammons, Yvette Unoarumhi, Darlene Wagner, Matthew Schmerer, Minoo Agarwal, Eyad Almasri, Debbie Boles, Ayla Burns, Nuthawin Charoensri, Oren Cohen, Susan Countryman, Mary Ann Cristobal, Bobbi Croy, Suzanne Dale, Hrushikesh Deshmukh, Amanda Douglas, Vincent Drouillon, Marcia Eisenberg, Howard Engler, Rama Ghatti, Prashant Gupta, Susan Hicks, Jake Humphrey, Lax Iyer, Lisa Pfefferle, Manoj Jain, Matthew Robinson, Mohan Kolli, Brian Krueger, Tim Kuphal, Stanley Letovsky, Michael Levandoski, Craig Lukasik, Jonathan Meltzer, Brian Norvell, Mindy Nye, Scott Parker, Christos Petropoulos, John Pruitt, Steven Ragan, Scott Ryan, Mike Sapeta, Jana Schroth, Suresh Babu Selvaraju, Goran Stevovic, Amanda Suchanek, Andrea Throop, Lyndon Tilson, Thomas Urban, Joe Voshell, Kimberly Wagner, Jonathan Williams, Mary Williamson, Qian Zeng, Tricia Zwiefelhofer, Clinton R. Paden, Duncan MacCannell |
| EPI_ISL_3325867 | TGen North | TGen North | Jolene Bowers, Heather Centner, Brett Van Tassel, Chris French, Hayley Yaglom, Darrin Lemmer, Dave Engelthaler |
| EPI_ISL_3325868 | Laboratory Corporation of America | Centers for Disease Control and Prevention Division of Viral Diseases, Pathogen Discovery | Dakota Howard, Dhwani Batra, Peter W. Cook, Kara Moser, Adrian Paskey, Jason Caravas, Benjamin Rambo-Martin, Shatavia Morrison, Christopher Gulvick, Scott Sammons, Yvette Unoarumhi, Darlene Wagner, Matthew Schmerer, Minoo Agarwal, Eyad Almasri, Debbie Boles, Ayla Burns, Nuthawin Charoensri, Oren Cohen, Susan Countryman, Mary Ann Cristobal, Bobbi Croy, Suzanne Dale, Hrushikesh Deshmukh, Amanda Douglas, Vincent Drouillon, Marcia Eisenberg, Howard Engler, Rama Ghatti, Prashant Gupta, Susan Hicks, Jake Humphrey, Lax Iyer, Lisa Pfefferle, Manoj Jain, Matthew Robinson, Mohan Kolli, Brian Krueger, Tim Kuphal, Stanley Letovsky, Michael Levandoski, Craig Lukasik, Jonathan Meltzer, Brian Norvell, Mindy Nye, Scott Parker, Christos Petropoulos, John Pruitt, Steven Ragan, Scott Ryan, Mike Sapeta, Jana Schroth, Suresh Babu Selvaraju, Goran Stevovic, Amanda Suchanek, Andrea Throop, Lyndon Tilson, Thomas Urban, Joe Voshell, Kimberly Wagner, Jonathan Williams, Mary Williamson, Qian Zeng, Tricia Zwiefelhofer, Clinton R. Paden, Duncan MacCannell |
| EPI_ISL_3325869 | TGen North | TGen North | Jolene Bowers, Heather Centner, Brett Van Tassel, Chris French, Hayley Yaglom, Darrin Lemmer, Dave Engelthaler |
| EPI_ISL_3325870, EPI_ISL_3325873 | Laboratory Corporation of America | Centers for Disease Control and Prevention Division of Viral Diseases, Pathogen Discovery | Dakota Howard, Dhwani Batra, Peter W. Cook, Kara Moser, Adrian Paskey, Jason Caravas, Benjamin Rambo-Martin, Shatavia Morrison, Christopher Gulvick, Scott Sammons, Yvette Unoarumhi, Darlene Wagner, Matthew Schmerer, Minoo Agarwal, Eyad Almasri, Debbie Boles, Ayla Burns, Nuthawin Charoensri, Oren Cohen, Susan Countryman, Mary Ann Cristobal, Bobbi Croy, Suzanne Dale, Hrushikesh Deshmukh, Amanda Douglas, Vincent Drouillon, Marcia Eisenberg, Howard Engler, Rama Ghatti, Prashant Gupta, Susan Hicks, Jake Humphrey, Lax Iyer, Lisa Pfefferle, Manoj Jain, Matthew Robinson, Mohan Kolli, Brian Krueger, Tim Kuphal, Stanley Letovsky, Michael Levandoski, Craig Lukasik, Jonathan Meltzer, Brian Norvell, Mindy Nye, Scott Parker, Christos Petropoulos, John Pruitt, Steven Ragan, Scott Ryan, Mike Sapeta, Jana Schroth, Suresh Babu Selvaraju, Goran Stevovic, Amanda Suchanek, Andrea Throop, Lyndon Tilson, Thomas Urban, Joe Voshell, Kimberly Wagner, Jonathan Williams, Mary Williamson, Qian Zeng, Tricia Zwiefelhofer, Clinton R. Paden, Duncan MacCannell |
| EPI_ISL_3325874 | TGen North | TGen North | Jolene Bowers, Heather Centner, Brett Van Tassel, Chris French, Hayley Yaglom, Darrin Lemmer, Dave Engelthaler |
| EPI_ISL_3325875, EPI_ISL_3325877, EPI_ISL_3325879, EPI_ISL_3325880 | Laboratory Corporation of America | Centers for Disease Control and Prevention Division of Viral Diseases, Pathogen Discovery | Dakota Howard, Dhwani Batra, Peter W. Cook, Kara Moser, Adrian Paskey, Jason Caravas, Benjamin Rambo-Martin, Shatavia Morrison, Christopher Gulvick, Scott Sammons, Yvette Unoarumhi, Darlene Wagner, Matthew Schmerer, Minoo Agarwal, Eyad Almasri, Debbie Boles, Ayla Burns, Nuthawin Charoensri, Oren Cohen, Susan Countryman, Mary Ann Cristobal, Bobbi Croy, Suzanne Dale, Hrushikesh Deshmukh, Amanda Douglas, Vincent Drouillon, Marcia Eisenberg, Howard Engler, Rama Ghatti, Prashant Gupta, Susan Hicks, Jake Humphrey, Lax Iyer, Lisa Pfefferle, Manoj Jain, Matthew Robinson, Mohan Kolli, Brian Krueger, Tim Kuphal, Stanley Letovsky, Michael Levandoski, Craig Lukasik, Jonathan Meltzer, Brian Norvell, Mindy Nye, Scott Parker, Christos Petropoulos, John Pruitt, Steven Ragan, Scott Ryan, Mike Sapeta, Jana Schroth, Suresh Babu Selvaraju, Goran Stevovic, Amanda Suchanek, Andrea Throop, Lyndon Tilson, Thomas Urban, Joe Voshell, Kimberly Wagner, Jonathan Williams, Mary Williamson, Qian Zeng, Tricia Zwiefelhofer, Clinton R. Paden, Duncan MacCannell |
| EPI_ISL_3325881 | TGen North | TGen North | Jolene Bowers, Heather Centner, Brett Van Tassel, Chris French, Hayley Yaglom, Darrin Lemmer, Dave Engelthaler |
| EPI_ISL_3325882 | Laboratory Corporation of America | Centers for Disease Control and Prevention Division of Viral Diseases, Pathogen Discovery | Dakota Howard, Dhwani Batra, Peter W. Cook, Kara Moser, Adrian Paskey, Jason Caravas, Benjamin Rambo-Martin, Shatavia Morrison, Christopher Gulvick, Scott Sammons, Yvette Unoarumhi, Darlene Wagner, Matthew Schmerer, Minoo Agarwal, Eyad Almasri, Debbie Boles, Ayla Burns, Nuthawin Charoensri, Oren Cohen, Susan Countryman, Mary Ann Cristobal, Bobbi Croy, Suzanne Dale, Hrushikesh Deshmukh, Amanda Douglas, Vincent Drouillon, Marcia Eisenberg, Howard Engler, Rama Ghatti, Prashant Gupta, Susan Hicks, Jake Humphrey, Lax Iyer, Lisa Pfefferle, Manoj Jain, Matthew Robinson, Mohan Kolli, Brian Krueger, Tim Kuphal, Stanley Letovsky, Michael Levandoski, Craig Lukasik, Jonathan Meltzer, Brian Norvell, Mindy Nye, Scott Parker, Christos Petropoulos, John Pruitt, Steven Ragan, Scott Ryan, Mike Sapeta, Jana Schroth, Suresh Babu Selvaraju, Goran Stevovic, Amanda Suchanek, Andrea Throop, Lyndon Tilson, Thomas Urban, Joe Voshell, Kimberly Wagner, Jonathan Williams, Mary Williamson, Qian Zeng, Tricia Zwiefelhofer, Clinton R. Paden, Duncan MacCannell |
| EPI_ISL_3325883 | TGen North | TGen North | Jolene Bowers, Heather Centner, Brett Van Tassel, Chris French, Hayley Yaglom, Darrin Lemmer, Dave Engelthaler |
| EPI_ISL_3325884 | Laboratory Corporation of America | Centers for Disease Control and Prevention Division of Viral Diseases, Pathogen Discovery | Dakota Howard, Dhwani Batra, Peter W. Cook, Kara Moser, Adrian Paskey, Jason Caravas, Benjamin Rambo-Martin, Shatavia Morrison, Christopher Gulvick, Scott Sammons, Yvette Unoarumhi, Darlene Wagner, Matthew Schmerer, Minoo Agarwal, Eyad Almasri, Debbie Boles, Ayla Burns, Nuthawin Charoensri, Oren Cohen, Susan Countryman, Mary Ann Cristobal, Bobbi Croy, Suzanne Dale, Hrushikesh Deshmukh, Amanda Douglas, Vincent Drouillon, Marcia Eisenberg, Howard Engler, Rama Ghatti, Prashant Gupta, Susan Hicks, Jake Humphrey, Lax Iyer, Lisa Pfefferle, Manoj Jain, Matthew Robinson, Mohan Kolli, Brian Krueger, Tim Kuphal, Stanley Letovsky, Michael Levandoski, Craig Lukasik, Jonathan Meltzer, Brian Norvell, Mindy Nye, Scott Parker, Christos Petropoulos, John Pruitt, Steven Ragan, Scott Ryan, Mike Sapeta, Jana Schroth, Suresh Babu Selvaraju, Goran Stevovic, Amanda Suchanek, Andrea Throop, Lyndon Tilson, Thomas Urban, Joe Voshell, Kimberly Wagner, Jonathan Williams, Mary Williamson, Qian Zeng, Tricia Zwiefelhofer, Clinton R. Paden, Duncan MacCannell |
| EPI_ISL_3325885 | TGen North | TGen North | Jolene Bowers, Heather Centner, Brett Van Tassel, Chris French, Hayley Yaglom, Darrin Lemmer, Dave Engelthaler |
| EPI_ISL_3325886, EPI_ISL_3325888 | Laboratory Corporation of America | Centers for Disease Control and Prevention Division of Viral Diseases, Pathogen Discovery | Dakota Howard, Dhwani Batra, Peter W. Cook, Kara Moser, Adrian Paskey, Jason Caravas, Benjamin Rambo-Martin, Shatavia Morrison, Christopher Gulvick, Scott Sammons, Yvette Unoarumhi, Darlene Wagner, Matthew Schmerer, Minoo Agarwal, Eyad Almasri, Debbie Boles, Ayla Burns, Nuthawin Charoensri, Oren Cohen, Susan Countryman, Mary Ann Cristobal, Bobbi Croy, Suzanne Dale, Hrushikesh Deshmukh, Amanda Douglas, Vincent Drouillon, Marcia Eisenberg, Howard Engler, Rama Ghatti, Prashant Gupta, Susan Hicks, Jake Humphrey, Lax Iyer, Lisa Pfefferle, Manoj Jain, Matthew Robinson, Mohan Kolli, Brian Krueger, Tim Kuphal, Stanley Letovsky, Michael Levandoski, Craig Lukasik, Jonathan Meltzer, Brian Norvell, Mindy Nye, Scott Parker, Christos Petropoulos, John Pruitt, Steven Ragan, Scott Ryan, Mike Sapeta, Jana Schroth, Suresh Babu Selvaraju, Goran Stevovic, Amanda Suchanek, Andrea Throop, Lyndon Tilson, Thomas Urban, Joe Voshell, Kimberly Wagner, Jonathan Williams, Mary Williamson, Qian Zeng, Tricia Zwiefelhofer, Clinton R. Paden, Duncan MacCannell |
| EPI_ISL_3325889 | TGen North | TGen North | Jolene Bowers, Heather Centner, Brett Van Tassel, Chris French, Hayley Yaglom, Darrin Lemmer, Dave Engelthaler |
| EPI_ISL_3325890, EPI_ISL_3325891, EPI_ISL_3325893 | Laboratory Corporation of America | Centers for Disease Control and Prevention Division of Viral Diseases, Pathogen Discovery | Dakota Howard, Dhwani Batra, Peter W. Cook, Kara Moser, Adrian Paskey, Jason Caravas, Benjamin Rambo-Martin, Shatavia Morrison, Christopher Gulvick, Scott Sammons, Yvette Unoarumhi, Darlene Wagner, Matthew Schmerer, Minoo Agarwal, Eyad Almasri, Debbie Boles, Ayla Burns, Nuthawin Charoensri, Oren Cohen, Susan Countryman, Mary Ann Cristobal, Bobbi Croy, Suzanne Dale, Hrushikesh Deshmukh, Amanda Douglas, Vincent Drouillon, Marcia Eisenberg, Howard Engler, Rama Ghatti, Prashant Gupta, Susan Hicks, Jake Humphrey, Lax Iyer, Lisa Pfefferle, Manoj Jain, Matthew Robinson, Mohan Kolli, Brian Krueger, Tim Kuphal, Stanley Letovsky, Michael Levandoski, Craig Lukasik, Jonathan Meltzer, Brian Norvell, Mindy Nye, Scott Parker, Christos Petropoulos, John Pruitt, Steven Ragan, Scott Ryan, Mike Sapeta, Jana Schroth, Suresh Babu Selvaraju, Goran Stevovic, Amanda Suchanek, Andrea Throop, Lyndon Tilson, Thomas Urban, Joe Voshell, Kimberly Wagner, Jonathan Williams, Mary Williamson, Qian Zeng, Tricia Zwiefelhofer, Clinton R. Paden, Duncan MacCannell |
| EPI_ISL_3325894 | TGen North | TGen North | Jolene Bowers, Heather Centner, Brett Van Tassel, Chris French, Hayley Yaglom, Darrin Lemmer, Dave Engelthaler |
| EPI_ISL_3325899, EPI_ISL_3325900 | Laboratory Corporation of America | Centers for Disease Control and Prevention Division of Viral Diseases, Pathogen Discovery | Dakota Howard, Dhwani Batra, Peter W. Cook, Kara Moser, Adrian Paskey, Jason Caravas, Benjamin Rambo-Martin, Shatavia Morrison, Christopher Gulvick, Scott Sammons, Yvette Unoarumhi, Darlene Wagner, Matthew Schmerer, Minoo Agarwal, Eyad Almasri, Debbie Boles, Ayla Burns, Nuthawin Charoensri, Oren Cohen, Susan Countryman, Mary Ann Cristobal, Bobbi Croy, Suzanne Dale, Hrushikesh Deshmukh, Amanda Douglas, Vincent Drouillon, Marcia Eisenberg, Howard Engler, Rama Ghatti, Prashant Gupta, Susan Hicks, Jake Humphrey, Lax Iyer, Lisa Pfefferle, Manoj Jain, Matthew Robinson, Mohan Kolli, Brian Krueger, Tim Kuphal, Stanley Letovsky, Michael Levandoski, Craig Lukasik, Jonathan Meltzer, Brian Norvell, Mindy Nye, Scott Parker, Christos Petropoulos, John Pruitt, Steven Ragan, Scott Ryan, Mike Sapeta, Jana Schroth, Suresh Babu Selvaraju, Goran Stevovic, Amanda Suchanek, Andrea Throop, Lyndon Tilson, Thomas Urban, Joe Voshell, Kimberly Wagner, Jonathan Williams, Mary Williamson, Qian Zeng, Tricia Zwiefelhofer, Clinton R. Paden, Duncan MacCannell |
| EPI_ISL_3325901 | TGen North | TGen North | Jolene Bowers, Heather Centner, Brett Van Tassel, Chris French, Hayley Yaglom, Darrin Lemmer, Dave Engelthaler |
| EPI_ISL_3325902, EPI_ISL_3325904 | Laboratory Corporation of America | Centers for Disease Control and Prevention Division of Viral Diseases, Pathogen Discovery | Dakota Howard, Dhwani Batra, Peter W. Cook, Kara Moser, Adrian Paskey, Jason Caravas, Benjamin Rambo-Martin, Shatavia Morrison, Christopher Gulvick, Scott Sammons, Yvette Unoarumhi, Darlene Wagner, Matthew Schmerer, Minoo Agarwal, Eyad Almasri, Debbie Boles, Ayla Burns, Nuthawin Charoensri, Oren Cohen, Susan Countryman, Mary Ann Cristobal, Bobbi Croy, Suzanne Dale, Hrushikesh Deshmukh, Amanda Douglas, Vincent Drouillon, Marcia Eisenberg, Howard |

| | | | |
|---|---|---|---|
| | | | Engler, Rama Ghatti, Prashant Gupta, Susan Hicks, Jake Humphrey, Lax Iyer, Lisa Pfefferle, Manoj Jain, Matthew Robinson, Mohan Kolli, Brian Krueger, Tim Kuphal, Stanley Letovsky, Michael Levandoski, Craig Lukasik, Jonathan Meltzer, Brian Norvell, Mindy Nye, Scott Parker, Christos Petropoulos, John Pruitt, Steven Ragan, Scott Ryan, Mike Sapeta, Jana Schroth, Suresh Babu Selvaraju, Goran Stevovic, Amanda Suchanek, Andrea Throop, Lyndon Tilson, Thomas Urban, Joe Voshell, Kimberly Wagner, Jonathan Williams, Mary Williamson, Qian Zeng, Tricia Zwiefelhofer, Clinton R. Paden, Duncan MacCannell |
| EPI_ISL_3325905 | TGen North | TGen North | Jolene Bowers, Heather Centner, Brett Van Tassel, Chris French, Hayley Yaglom, Darrin Lemmer, Dave Engelthaler |
| EPI_ISL_3325906 | Laboratory Corporation of America | Centers for Disease Control and Prevention Division of Viral Diseases, Pathogen Discovery | Dakota Howard, Dhwani Batra, Peter W. Cook, Kara Moser, Adrian Paskey, Jason Caravas, Benjamin Rambo-Martin, Shatavia Morrison, Christopher Gulvick, Scott Sammons, Yvette Unoarumhi, Darlene Wagner, Matthew Schmerer, Minoo Agarwal, Eyad Almasri, Debbie Boles, Ayla Burns, Nuthawin Charoensri, Oren Cohen, Susan Countryman, Mary Ann Cristobal, Bobbi Croy, Suzanne Dale, Hrushikesh Deshmukh, Amanda Douglas, Vincent Drouillon, Marcia Eisenberg, Howard Engler, Rama Ghatti, Prashant Gupta, Susan Hicks, Jake Humphrey, Lax Iyer, Lisa Pfefferle, Manoj Jain, Matthew Robinson, Mohan Kolli, Brian Krueger, Tim Kuphal, Stanley Letovsky, Michael Levandoski, Craig Lukasik, Jonathan Meltzer, Brian Norvell, Mindy Nye, Scott Parker, Christos Petropoulos, John Pruitt, Steven Ragan, Scott Ryan, Mike Sapeta, Jana Schroth, Suresh Babu Selvaraju, Goran Stevovic, Amanda Suchanek, Andrea Throop, Lyndon Tilson, Thomas Urban, Joe Voshell, Kimberly Wagner, Jonathan Williams, Mary Williamson, Qian Zeng, Tricia Zwiefelhofer, Clinton R. Paden, Duncan MacCannell |
| EPI_ISL_3325907 | TGen North | TGen North | Jolene Bowers, Heather Centner, Brett Van Tassel, Chris French, Hayley Yaglom, Darrin Lemmer, Dave Engelthaler |
| EPI_ISL_3325908 | Laboratory Corporation of America | Centers for Disease Control and Prevention Division of Viral Diseases, Pathogen Discovery | Dakota Howard, Dhwani Batra, Peter W. Cook, Kara Moser, Adrian Paskey, Jason Caravas, Benjamin Rambo-Martin, Shatavia Morrison, Christopher Gulvick, Scott Sammons, Yvette Unoarumhi, Darlene Wagner, Matthew Schmerer, Minoo Agarwal, Eyad Almasri, Debbie Boles, Ayla Burns, Nuthawin Charoensri, Oren Cohen, Susan Countryman, Mary Ann Cristobal, Bobbi Croy, Suzanne Dale, Hrushikesh Deshmukh, Amanda Douglas, Vincent Drouillon, Marcia Eisenberg, Howard Engler, Rama Ghatti, Prashant Gupta, Susan Hicks, Jake Humphrey, Lax Iyer, Lisa Pfefferle, Manoj Jain, Matthew Robinson, Mohan Kolli, Brian Krueger, Tim Kuphal, Stanley Letovsky, Michael Levandoski, Craig Lukasik, Jonathan Meltzer, Brian Norvell, Mindy Nye, Scott Parker, Christos Petropoulos, John Pruitt, Steven Ragan, Scott Ryan, Mike Sapeta, Jana Schroth, Suresh Babu Selvaraju, Goran Stevovic, Amanda Suchanek, Andrea Throop, Lyndon Tilson, Thomas Urban, Joe Voshell, Kimberly Wagner, Jonathan Williams, Mary Williamson, Qian Zeng, Tricia Zwiefelhofer, Clinton R. Paden, Duncan MacCannell |
| EPI_ISL_3325909 | TGen North | TGen North | Jolene Bowers, Heather Centner, Brett Van Tassel, Chris French, Hayley Yaglom, Darrin Lemmer, Dave Engelthaler |
| EPI_ISL_3325910, EPI_ISL_3325911 | Laboratory Corporation of America | Centers for Disease Control and Prevention Division of Viral Diseases, Pathogen Discovery | Dakota Howard, Dhwani Batra, Peter W. Cook, Kara Moser, Adrian Paskey, Jason Caravas, Benjamin Rambo-Martin, Shatavia Morrison, Christopher Gulvick, Scott Sammons, Yvette Unoarumhi, Darlene Wagner, Matthew Schmerer, Minoo Agarwal, Eyad Almasri, Debbie Boles, Ayla Burns, Nuthawin Charoensri, Oren Cohen, Susan Countryman, Mary Ann Cristobal, Bobbi Croy, Suzanne Dale, Hrushikesh Deshmukh, Amanda Douglas, Vincent Drouillon, Marcia Eisenberg, Howard Engler, Rama Ghatti, Prashant Gupta, Susan Hicks, Jake Humphrey, Lax Iyer, Lisa Pfefferle, Manoj Jain, Matthew Robinson, Mohan Kolli, Brian Krueger, Tim Kuphal, Stanley Letovsky, Michael Levandoski, Craig Lukasik, Jonathan Meltzer, Brian Norvell, Mindy Nye, Scott Parker, Christos Petropoulos, John Pruitt, Steven Ragan, Scott Ryan, Mike Sapeta, Jana Schroth, Suresh Babu Selvaraju, Goran Stevovic, Amanda Suchanek, Andrea Throop, Lyndon Tilson, Thomas Urban, Joe Voshell, Kimberly Wagner, Jonathan Williams, Mary Williamson, Qian Zeng, Tricia Zwiefelhofer, Clinton R. Paden, Duncan MacCannell |
| EPI_ISL_3325912 | TGen North | TGen North | Jolene Bowers, Heather Centner, Brett Van Tassel, Chris French, Hayley Yaglom, Darrin Lemmer, Dave Engelthaler |
| EPI_ISL_3325913 | Laboratory Corporation of America | Centers for Disease Control and Prevention Division of Viral Diseases, Pathogen Discovery | Dakota Howard, Dhwani Batra, Peter W. Cook, Kara Moser, Adrian Paskey, Jason Caravas, Benjamin Rambo-Martin, Shatavia Morrison, Christopher Gulvick, Scott Sammons, Yvette Unoarumhi, Darlene Wagner, Matthew Schmerer, Minoo Agarwal, Eyad Almasri, Debbie Boles, Ayla Burns, Nuthawin Charoensri, Oren Cohen, Susan Countryman, Mary Ann Cristobal, Bobbi Croy, Suzanne Dale, Hrushikesh Deshmukh, Amanda Douglas, Vincent Drouillon, Marcia Eisenberg, Howard Engler, Rama Ghatti, Prashant Gupta, Susan Hicks, Jake Humphrey, Lax Iyer, Lisa Pfefferle, Manoj Jain, Matthew Robinson, Mohan Kolli, Brian Krueger, Tim Kuphal, Stanley Letovsky, Michael Levandoski, Craig Lukasik, Jonathan Meltzer, Brian Norvell, Mindy Nye, Scott Parker, Christos Petropoulos, John Pruitt, Steven Ragan, Scott Ryan, Mike Sapeta, Jana Schroth, Suresh Babu Selvaraju, Goran Stevovic, Amanda Suchanek, Andrea Throop, Lyndon Tilson, Thomas Urban, Joe Voshell, Kimberly Wagner, Jonathan Williams, Mary Williamson, Qian Zeng, Tricia Zwiefelhofer, Clinton R. Paden, Duncan MacCannell |
| EPI_ISL_3325914 | TGen North | TGen North | Jolene Bowers, Heather Centner, Brett Van Tassel, Chris French, Hayley Yaglom, Darrin Lemmer, Dave Engelthaler |
| EPI_ISL_3325915 | Laboratory Corporation of America | Centers for Disease Control and Prevention Division of Viral Diseases, Pathogen Discovery | Dakota Howard, Dhwani Batra, Peter W. Cook, Kara Moser, Adrian Paskey, Jason Caravas, Benjamin Rambo-Martin, Shatavia Morrison, Christopher Gulvick, Scott Sammons, Yvette Unoarumhi, Darlene Wagner, Matthew Schmerer, Minoo Agarwal, Eyad Almasri, Debbie Boles, Ayla Burns, Nuthawin Charoensri, Oren Cohen, Susan Countryman, Mary Ann Cristobal, Bobbi Croy, Suzanne Dale, Hrushikesh Deshmukh, Amanda Douglas, Vincent Drouillon, Marcia Eisenberg, Howard Engler, Rama Ghatti, Prashant Gupta, Susan Hicks, Jake Humphrey, Lax Iyer, Lisa Pfefferle, Manoj Jain, Matthew Robinson, Mohan Kolli, Brian Krueger, Tim Kuphal, Stanley Letovsky, Michael Levandoski, Craig Lukasik, Jonathan Meltzer, Brian Norvell, Mindy Nye, Scott Parker, Christos Petropoulos, John Pruitt, Steven Ragan, Scott Ryan, Mike Sapeta, Jana Schroth, Suresh Babu Selvaraju, Goran Stevovic, Amanda Suchanek, Andrea Throop, Lyndon Tilson, Thomas Urban, Joe Voshell, Kimberly Wagner, Jonathan Williams, Mary Williamson, Qian Zeng, Tricia Zwiefelhofer, Clinton R. Paden, Duncan MacCannell |
| EPI_ISL_3325916 | TGen North | TGen North | Jolene Bowers, Heather Centner, Brett Van Tassel, Chris French, Hayley Yaglom, Darrin Lemmer, Dave Engelthaler |
| EPI_ISL_3325917 | Laboratory Corporation of America | Centers for Disease Control and Prevention Division of Viral Diseases, Pathogen Discovery | Dakota Howard, Dhwani Batra, Peter W. Cook, Kara Moser, Adrian Paskey, Jason Caravas, Benjamin Rambo-Martin, Shatavia Morrison, Christopher Gulvick, Scott Sammons, Yvette Unoarumhi, Darlene Wagner, Matthew Schmerer, Minoo Agarwal, Eyad Almasri, Debbie Boles, Ayla Burns, Nuthawin Charoensri, Oren Cohen, Susan Countryman, Mary Ann Cristobal, Bobbi Croy, Suzanne Dale, Hrushikesh Deshmukh, Amanda Douglas, Vincent Drouillon, Marcia Eisenberg, Howard Engler, Rama Ghatti, Prashant Gupta, Susan Hicks, Jake Humphrey, Lax Iyer, Lisa Pfefferle, Manoj Jain, Matthew Robinson, Mohan Kolli, Brian Krueger, Tim Kuphal, Stanley Letovsky, Michael Levandoski, Craig Lukasik, Jonathan Meltzer, Brian Norvell, Mindy Nye, Scott Parker, Christos Petropoulos, John Pruitt, Steven Ragan, Scott Ryan, Mike Sapeta, Jana Schroth, Suresh Babu Selvaraju, Goran Stevovic, Amanda Suchanek, Andrea Throop, Lyndon Tilson, Thomas Urban, Joe Voshell, Kimberly Wagner, Jonathan Williams, Mary Williamson, Qian Zeng, Tricia Zwiefelhofer, Clinton R. Paden, Duncan MacCannell |
| EPI_ISL_3325918 | TGen North | TGen North | Jolene Bowers, Heather Centner, Brett Van Tassel, Chris French, Hayley Yaglom, Darrin Lemmer, Dave Engelthaler |
| EPI_ISL_3325919 | Laboratory Corporation of America | Centers for Disease Control and Prevention Division of Viral Diseases, Pathogen Discovery | Dakota Howard, Dhwani Batra, Peter W. Cook, Kara Moser, Adrian Paskey, Jason Caravas, Benjamin Rambo-Martin, Shatavia Morrison, Christopher Gulvick, Scott Sammons, Yvette Unoarumhi, Darlene Wagner, Matthew Schmerer, Minoo Agarwal, Eyad Almasri, Debbie Boles, Ayla Burns, Nuthawin Charoensri, Oren Cohen, Susan Countryman, Mary Ann Cristobal, Bobbi Croy, Suzanne Dale, Hrushikesh Deshmukh, Amanda Douglas, Vincent Drouillon, Marcia Eisenberg, Howard Engler, Rama Ghatti, Prashant Gupta, Susan Hicks, Jake Humphrey, Lax Iyer, Lisa Pfefferle, Manoj Jain, Matthew Robinson, Mohan Kolli, Brian Krueger, Tim Kuphal, Stanley Letovsky, Michael Levandoski, Craig Lukasik, Jonathan Meltzer, Brian Norvell, Mindy Nye, Scott Parker, Christos Petropoulos, John Pruitt, Steven Ragan, Scott Ryan, Mike Sapeta, Jana Schroth, Suresh Babu Selvaraju, Goran Stevovic, Amanda Suchanek, Andrea Throop, Lyndon Tilson, Thomas Urban, Joe Voshell, Kimberly Wagner, Jonathan Williams, Mary Williamson, Qian Zeng, Tricia Zwiefelhofer, Clinton R. Paden, Duncan MacCannell |
| EPI_ISL_3325920 | TGen North | TGen North | Jolene Bowers, Heather Centner, Brett Van Tassel, Chris French, Hayley Yaglom, Darrin Lemmer, Dave Engelthaler |
| EPI_ISL_3325921, EPI_ISL_3325923 | Laboratory Corporation of America | Centers for Disease Control and Prevention Division of Viral Diseases, Pathogen Discovery | Dakota Howard, Dhwani Batra, Peter W. Cook, Kara Moser, Adrian Paskey, Jason Caravas, Benjamin Rambo-Martin, Shatavia Morrison, Christopher Gulvick, Scott Sammons, Yvette Unoarumhi, Darlene Wagner, Matthew Schmerer, Minoo Agarwal, Eyad Almasri, Debbie Boles, Ayla Burns, Nuthawin Charoensri, Oren Cohen, Susan Countryman, Mary Ann Cristobal, Bobbi Croy, Suzanne Dale, Hrushikesh Deshmukh, Amanda Douglas, Vincent Drouillon, Marcia Eisenberg, Howard Engler, Rama Ghatti, Prashant Gupta, Susan Hicks, Jake Humphrey, Lax Iyer, Lisa Pfefferle, Manoj Jain, Matthew Robinson, Mohan Kolli, Brian Krueger, Tim Kuphal, Stanley Letovsky, Michael Levandoski, Craig Lukasik, Jonathan Meltzer, Brian Norvell, Mindy Nye, Scott Parker, Christos Petropoulos, John Pruitt, Steven Ragan, Scott Ryan, Mike Sapeta, Jana Schroth, Suresh Babu Selvaraju, Goran Stevovic, Amanda Suchanek, Andrea Throop, Lyndon Tilson, Thomas Urban, Joe Voshell, Kimberly Wagner, Jonathan Williams, Mary Williamson, Qian Zeng, Tricia Zwiefelhofer, Clinton R. Paden, Duncan MacCannell |
| EPI_ISL_3325924 | TGen North | TGen North | Jolene Bowers, Heather Centner, Brett Van Tassel, Chris French, Hayley Yaglom, Darrin Lemmer, Dave Engelthaler |
| EPI_ISL_3325925 | Laboratory Corporation of America | Centers for Disease Control and Prevention Division of Viral Diseases, Pathogen Discovery | Dakota Howard, Dhwani Batra, Peter W. Cook, Kara Moser, Adrian Paskey, Jason Caravas, Benjamin Rambo-Martin, Shatavia Morrison, Christopher Gulvick, Scott Sammons, Yvette Unoarumhi, Darlene Wagner, Matthew Schmerer, Minoo Agarwal, Eyad Almasri, Debbie Boles, Ayla Burns, Nuthawin Charoensri, Oren Cohen, Susan Countryman, Mary Ann Cristobal, Bobbi Croy, Suzanne Dale, Hrushikesh Deshmukh, Amanda Douglas, Vincent Drouillon, Marcia Eisenberg, Howard Engler, Rama Ghatti, Prashant Gupta, Susan Hicks, Jake Humphrey, Lax Iyer, Lisa Pfefferle, Manoj Jain, Matthew Robinson, Mohan Kolli, Brian Krueger, Tim Kuphal, Stanley Letovsky, Michael Levandoski, Craig Lukasik, Jonathan Meltzer, Brian Norvell, Mindy Nye, Scott Parker, Christos Petropoulos, John Pruitt, Steven Ragan, Scott Ryan, Mike Sapeta, Jana Schroth, Suresh Babu Selvaraju, Goran Stevovic, Amanda Suchanek, Andrea Throop, Lyndon Tilson, Thomas Urban, Joe Voshell, Kimberly Wagner, Jonathan Williams, Mary Williamson, Qian Zeng, Tricia Zwiefelhofer, Clinton R. Paden, Duncan MacCannell |
| EPI_ISL_3325926 | TGen North | TGen North | Jolene Bowers, Heather Centner, Brett Van Tassel, Chris French, Hayley Yaglom, Darrin Lemmer, Dave Engelthaler |
| EPI_ISL_3325927 | Laboratory Corporation of America | Centers for Disease Control and Prevention Division of Viral Diseases, Pathogen Discovery | Dakota Howard, Dhwani Batra, Peter W. Cook, Kara Moser, Adrian Paskey, Jason Caravas, Benjamin Rambo-Martin, Shatavia Morrison, Christopher Gulvick, Scott Sammons, Yvette Unoarumhi, Darlene Wagner, Matthew Schmerer, Minoo Agarwal, Eyad Almasri, Debbie Boles, Ayla Burns, Nuthawin Charoensri, Oren Cohen, Susan Countryman, Mary Ann Cristobal, Bobbi Croy, Suzanne Dale, Hrushikesh Deshmukh, Amanda Douglas, Vincent Drouillon, Marcia Eisenberg, Howard Engler, Rama Ghatti, Prashant Gupta, Susan Hicks, Jake Humphrey, Lax Iyer, Lisa Pfefferle, Manoj Jain, Matthew Robinson, Mohan Kolli, Brian Krueger, Tim Kuphal, Stanley Letovsky, Michael Levandoski, Craig Lukasik, Jonathan Meltzer, Brian Norvell, Mindy Nye, Scott Parker, Christos Petropoulos, John Pruitt, Steven Ragan, Scott Ryan, Mike Sapeta, Jana Schroth, Suresh Babu Selvaraju, Goran Stevovic, Amanda Suchanek, Andrea Throop, Lyndon Tilson, Thomas Urban, Joe Voshell, Kimberly Wagner, Jonathan Williams, Mary Williamson, Qian Zeng, Tricia Zwiefelhofer, Clinton R. Paden, Duncan MacCannell |
| EPI_ISL_3325928 | TGen North | TGen North | Jolene Bowers, Heather Centner, Brett Van Tassel, Chris French, Hayley Yaglom, Darrin Lemmer, Dave Engelthaler |
| EPI_ISL_3325929 | Laboratory Corporation of America | Centers for Disease Control and Prevention Division of Viral Diseases, Pathogen Discovery | Dakota Howard, Dhwani Batra, Peter W. Cook, Kara Moser, Adrian Paskey, Jason Caravas, Benjamin Rambo-Martin, Shatavia Morrison, Christopher Gulvick, Scott Sammons, Yvette Unoarumhi, Darlene Wagner, Matthew Schmerer, Minoo Agarwal, Eyad Almasri, Debbie Boles, Ayla Burns, Nuthawin Charoensri, Oren Cohen, Susan Countryman, Mary Ann Cristobal, Bobbi Croy, Suzanne Dale, Hrushikesh Deshmukh, Amanda Douglas, Vincent Drouillon, Marcia Eisenberg, Howard Engler, Rama Ghatti, Prashant Gupta, Susan Hicks, Jake Humphrey, Lax Iyer, Lisa Pfefferle, Manoj Jain, Matthew Robinson, Mohan Kolli, Brian Krueger, Tim Kuphal, Stanley Letovsky, Michael Levandoski, Craig Lukasik, Jonathan Meltzer, Brian Norvell, Mindy Nye, Scott Parker, Christos Petropoulos, John Pruitt, Steven Ragan, Scott Ryan, Mike Sapeta, Jana Schroth, Suresh Babu Selvaraju, Goran Stevovic, Amanda Suchanek, Andrea Throop, Lyndon Tilson, Thomas Urban, Joe Voshell, Kimberly Wagner, Jonathan Williams, Mary Williamson, Qian Zeng, Tricia Zwiefelhofer, Clinton R. Paden, Duncan MacCannell |
| EPI_ISL_3325930 | TGen North | TGen North | Jolene Bowers, Heather Centner, Brett Van Tassel, Chris French, Hayley Yaglom, Darrin Lemmer, Dave Engelthaler |
| EPI_ISL_3325931 | Laboratory Corporation of America | Centers for Disease Control and Prevention Division of Viral Diseases, Pathogen Discovery | Dakota Howard, Dhwani Batra, Peter W. Cook, Kara Moser, Adrian Paskey, Jason Caravas, Benjamin Rambo-Martin, Shatavia Morrison, Christopher Gulvick, Scott Sammons, Yvette Unoarumhi, Darlene Wagner, Matthew Schmerer, Minoo Agarwal, Eyad Almasri, Debbie Boles, Ayla Burns, Nuthawin Charoensri, Oren Cohen, Susan Countryman, Mary Ann Cristobal, Bobbi Croy, Suzanne Dale, Hrushikesh Deshmukh, Amanda Douglas, Vincent Drouillon, Marcia Eisenberg, Howard Engler, Rama Ghatti, Prashant Gupta, Susan Hicks, Jake Humphrey, Lax Iyer, Lisa Pfefferle, Manoj Jain, Matthew Robinson, Mohan Kolli, Brian Krueger, Tim Kuphal, Stanley Letovsky, Michael Levandoski, Craig Lukasik, Jonathan Meltzer, Brian Norvell, Mindy Nye, Scott Parker, Christos Petropoulos, John Pruitt, Steven Ragan, Scott Ryan, Mike Sapeta, Jana Schroth, Suresh Babu Selvaraju, Goran Stevovic, Amanda Suchanek, Andrea Throop, Lyndon Tilson, Thomas Urban, Joe Voshell, Kimberly Wagner, Jonathan Williams, Mary Williamson, Qian Zeng, Tricia Zwiefelhofer, Clinton R. Paden, Duncan MacCannell |
| EPI_ISL_3325932 | TGen North | TGen North | Jolene Bowers, Heather Centner, Brett Van Tassel, Chris French, Hayley Yaglom, Darrin Lemmer, Dave Engelthaler |
| EPI_ISL_3325933 | Laboratory Corporation of America | Centers for Disease Control and Prevention Division of Viral Diseases, Pathogen Discovery | Dakota Howard, Dhwani Batra, Peter W. Cook, Kara Moser, Adrian Paskey, Jason Caravas, Benjamin Rambo-Martin, Shatavia Morrison, Christopher Gulvick, Scott Sammons, Yvette Unoarumhi, Darlene Wagner, Matthew Schmerer, Minoo Agarwal, Eyad Almasri, Debbie Boles, Ayla Burns, Nuthawin Charoensri, Oren Cohen, Susan Countryman, Mary Ann Cristobal, Bobbi Croy, Suzanne Dale, Hrushikesh Deshmukh, Amanda Douglas, Vincent Drouillon, Marcia Eisenberg, Howard Engler, Rama Ghatti, Prashant Gupta, Susan Hicks, Jake Humphrey, Lax Iyer, Lisa Pfefferle, Manoj Jain, Matthew Robinson, Mohan Kolli, Brian Krueger, Tim Kuphal, Stanley Letovsky, Michael Levandoski, Craig Lukasik, Jonathan Meltzer, Brian Norvell, Mindy Nye, Scott Parker, Christos Petropoulos, John Pruitt, Steven Ragan, Scott Ryan, Mike Sapeta, Jana Schroth, Suresh Babu Selvaraju, Goran Stevovic, Amanda Suchanek, Andrea Throop, Lyndon Tilson, Thomas Urban, Joe Voshell, Kimberly Wagner, Jonathan Williams, Mary Williamson, Qian Zeng, Tricia Zwiefelhofer, Clinton R. Paden, Duncan MacCannell |
| EPI_ISL_3325934 | TGen North | TGen North | Jolene Bowers, Heather Centner, Brett Van Tassel, Chris French, Hayley Yaglom, Darrin Lemmer, Dave Engelthaler |
| EPI_ISL_3325935 | Laboratory Corporation of America | Centers for Disease Control and Prevention Division of Viral Diseases, Pathogen Discovery | Dakota Howard, Dhwani Batra, Peter W. Cook, Kara Moser, Adrian Paskey, Jason Caravas, Benjamin Rambo-Martin, Shatavia Morrison, Christopher Gulvick, Scott Sammons, Yvette Unoarumhi, Darlene Wagner, Matthew Schmerer, Minoo Agarwal, Eyad Almasri, Debbie Boles, Ayla Burns, Nuthawin Charoensri, Oren Cohen, Susan Countryman, Mary Ann Cristobal, Bobbi Croy, Suzanne Dale, Hrushikesh Deshmukh, Amanda Douglas, Vincent Drouillon, Marcia Eisenberg, Howard Engler, Rama Ghatti, Prashant Gupta, Susan Hicks, Jake Humphrey, Lax Iyer, Lisa Pfefferle, Manoj Jain, Matthew Robinson, Mohan Kolli, Brian Krueger, Tim Kuphal, Stanley Letovsky, Michael Levandoski, Craig Lukasik, Jonathan Meltzer, Brian Norvell, Mindy Nye, Scott Parker, Christos Petropoulos, John Pruitt, Steven Ragan, Scott Ryan, Mike Sapeta, Jana Schroth, Suresh Babu Selvaraju, Goran Stevovic, Amanda Suchanek, Andrea Throop, Lyndon Tilson, Thomas Urban, Joe Voshell, Kimberly Wagner, Jonathan Williams, Mary Williamson, Qian Zeng, Tricia Zwiefelhofer, Clinton R. Paden, Duncan MacCannell |
| EPI_ISL_3325937 | TGen North | TGen North | Jolene Bowers, Heather Centner, Brett Van Tassel, Chris French, Hayley Yaglom, Darrin Lemmer, Dave Engelthaler |
| EPI_ISL_3325938 | Laboratory Corporation of America | Centers for Disease Control and Prevention Division of Viral Diseases, Pathogen Discovery | Dakota Howard, Dhwani Batra, Peter W. Cook, Kara Moser, Adrian Paskey, Jason Caravas, Benjamin Rambo-Martin, Shatavia Morrison, Christopher Gulvick, Scott Sammons, Yvette Unoarumhi, Darlene Wagner, Matthew Schmerer, Minoo Agarwal, Eyad Almasri, Debbie Boles, Ayla Burns, Nuthawin Charoensri, Oren Cohen, Susan Countryman, Mary Ann Cristobal, Bobbi Croy, Suzanne Dale, Hrushikesh Deshmukh, Amanda Douglas, Vincent Drouillon, Marcia Eisenberg, Howard Engler, Rama Ghatti, Prashant Gupta, Susan Hicks, Jake Humphrey, Lax Iyer, Lisa Pfefferle, Manoj Jain, Matthew Robinson, Mohan Kolli, Brian Krueger, Tim Kuphal, Stanley Letovsky, Michael Levandoski, Craig Lukasik, Jonathan Meltzer, Brian Norvell, Mindy Nye, Scott Parker, Christos Petropoulos, John Pruitt, Steven Ragan, Scott Ryan, Mike Sapeta, Jana Schroth, Suresh Babu Selvaraju, Goran Stevovic, Amanda Suchanek, Andrea Throop, Lyndon Tilson, Thomas Urban, Joe Voshell, Kimberly Wagner, Jonathan Williams, Mary Williamson, Qian Zeng, Tricia Zwiefelhofer, Clinton R. Paden, Duncan MacCannell |
| EPI_ISL_3325939 | TGen North | TGen North | Jolene Bowers, Heather Centner, Brett Van Tassel, Chris French, Hayley Yaglom, Darrin Lemmer, Dave Engelthaler |
| EPI_ISL_3325940, | Laboratory Corporation of America | Centers for Disease Control and Prevention Division of Viral | Dakota Howard, Dhwani Batra, Peter W. Cook, Kara Moser, Adrian Paskey, Jason Caravas, Benjamin Rambo-Martin, Shatavia Morrison, Christopher Gulvick, Scott Sammons, Yvette Unoarumhi, Darlene Wagner, Matthew Schmerer, Minoo |

| | | | |
|---|---|---|---|
| EPI_ISL_3325942 | | Diseases, Pathogen Discovery | Agarwal, Eyad Almasri, Debbie Boles, Ayla Burns, Nuthawin Charoensri, Oren Cohen, Susan Countryman, Mary Ann Cristobal, Bobbi Croy, Suzanne Dale, Hrushikesh Deshmukh, Amanda Douglas, Vincent Drouillon, Marcia Eisenberg, Howard Engler, Rama Ghatti, Prashant Gupta, Susan Hicks, Jake Humphrey, Lax Iyer, Lisa Pfefferle, Manoj Jain, Matthew Robinson, Mohan Kolli, Brian Krueger, Tim Kuphal, Stanley Letovsky, Michael Levandoski, Craig Lukasik, Jonathan Meltzer, Brian Norvell, Mindy Nye, Scott Parker, Christos Petropoulos, John Pruitt, Steven Ragan, Scott Ryan, Mike Sapeta, Jana Schroth, Suresh Babu Selvaraju, Goran Stevovic, Amanda Suchanek, Andrea Throop, Lyndon Tilson, Thomas Urban, Joe Voshell, Kimberly Wagner, Jonathan Williams, Mary Williamson, Qian Zeng, Tricia Zwiefelhofer, Clinton R. Paden, Duncan MacCannell |
| EPI_ISL_3325943 | TGen North | TGen North | Jolene Bowers, Heather Centner, Brett Van Tassel, Chris French, Hayley Yaglom, Darrin Lemmer, Dave Engelthaler |
| EPI_ISL_3325944, EPI_ISL_3325945 | Laboratory Corporation of America | Centers for Disease Control and Prevention Division of Viral Diseases, Pathogen Discovery | Dakota Howard, Dhwani Batra, Peter W. Cook, Kara Moser, Adrian Paskey, Jason Caravas, Benjamin Rambo-Martin, Shatavia Morrison, Christopher Gulvick, Scott Sammons, Yvette Unoarumhi, Darlene Wagner, Matthew Schmerer, Minoo Agarwal, Eyad Almasri, Debbie Boles, Ayla Burns, Nuthawin Charoensri, Oren Cohen, Susan Countryman, Mary Ann Cristobal, Bobbi Croy, Suzanne Dale, Hrushikesh Deshmukh, Amanda Douglas, Vincent Drouillon, Marcia Eisenberg, Howard Engler, Rama Ghatti, Prashant Gupta, Susan Hicks, Jake Humphrey, Lax Iyer, Lisa Pfefferle, Manoj Jain, Matthew Robinson, Mohan Kolli, Brian Krueger, Tim Kuphal, Stanley Letovsky, Michael Levandoski, Craig Lukasik, Jonathan Meltzer, Brian Norvell, Mindy Nye, Scott Parker, Christos Petropoulos, John Pruitt, Steven Ragan, Scott Ryan, Mike Sapeta, Jana Schroth, Suresh Babu Selvaraju, Goran Stevovic, Amanda Suchanek, Andrea Throop, Lyndon Tilson, Thomas Urban, Joe Voshell, Kimberly Wagner, Jonathan Williams, Mary Williamson, Qian Zeng, Tricia Zwiefelhofer, Clinton R. Paden, Duncan MacCannell |
| EPI_ISL_3325946 | TGen North | TGen North | Jolene Bowers, Heather Centner, Brett Van Tassel, Chris French, Hayley Yaglom, Darrin Lemmer, Dave Engelthaler |
| EPI_ISL_3325947 | Laboratory Corporation of America | Centers for Disease Control and Prevention Division of Viral Diseases, Pathogen Discovery | Dakota Howard, Dhwani Batra, Peter W. Cook, Kara Moser, Adrian Paskey, Jason Caravas, Benjamin Rambo-Martin, Shatavia Morrison, Christopher Gulvick, Scott Sammons, Yvette Unoarumhi, Darlene Wagner, Matthew Schmerer, Minoo Agarwal, Eyad Almasri, Debbie Boles, Ayla Burns, Nuthawin Charoensri, Oren Cohen, Susan Countryman, Mary Ann Cristobal, Bobbi Croy, Suzanne Dale, Hrushikesh Deshmukh, Amanda Douglas, Vincent Drouillon, Marcia Eisenberg, Howard Engler, Rama Ghatti, Prashant Gupta, Susan Hicks, Jake Humphrey, Lax Iyer, Lisa Pfefferle, Manoj Jain, Matthew Robinson, Mohan Kolli, Brian Krueger, Tim Kuphal, Stanley Letovsky, Michael Levandoski, Craig Lukasik, Jonathan Meltzer, Brian Norvell, Mindy Nye, Scott Parker, Christos Petropoulos, John Pruitt, Steven Ragan, Scott Ryan, Mike Sapeta, Jana Schroth, Suresh Babu Selvaraju, Goran Stevovic, Amanda Suchanek, Andrea Throop, Lyndon Tilson, Thomas Urban, Joe Voshell, Kimberly Wagner, Jonathan Williams, Mary Williamson, Qian Zeng, Tricia Zwiefelhofer, Clinton R. Paden, Duncan MacCannell |
| EPI_ISL_3325948 | TGen North | TGen North | Jolene Bowers, Heather Centner, Brett Van Tassel, Chris French, Hayley Yaglom, Darrin Lemmer, Dave Engelthaler |
| EPI_ISL_3325949, EPI_ISL_3325951 | Laboratory Corporation of America | Centers for Disease Control and Prevention Division of Viral Diseases, Pathogen Discovery | Dakota Howard, Dhwani Batra, Peter W. Cook, Kara Moser, Adrian Paskey, Jason Caravas, Benjamin Rambo-Martin, Shatavia Morrison, Christopher Gulvick, Scott Sammons, Yvette Unoarumhi, Darlene Wagner, Matthew Schmerer, Minoo Agarwal, Eyad Almasri, Debbie Boles, Ayla Burns, Nuthawin Charoensri, Oren Cohen, Susan Countryman, Mary Ann Cristobal, Bobbi Croy, Suzanne Dale, Hrushikesh Deshmukh, Amanda Douglas, Vincent Drouillon, Marcia Eisenberg, Howard Engler, Rama Ghatti, Prashant Gupta, Susan Hicks, Jake Humphrey, Lax Iyer, Lisa Pfefferle, Manoj Jain, Matthew Robinson, Mohan Kolli, Brian Krueger, Tim Kuphal, Stanley Letovsky, Michael Levandoski, Craig Lukasik, Jonathan Meltzer, Brian Norvell, Mindy Nye, Scott Parker, Christos Petropoulos, John Pruitt, Steven Ragan, Scott Ryan, Mike Sapeta, Jana Schroth, Suresh Babu Selvaraju, Goran Stevovic, Amanda Suchanek, Andrea Throop, Lyndon Tilson, Thomas Urban, Joe Voshell, Kimberly Wagner, Jonathan Williams, Mary Williamson, Qian Zeng, Tricia Zwiefelhofer, Clinton R. Paden, Duncan MacCannell |
| EPI_ISL_3325952 | TGen North | TGen North | Jolene Bowers, Heather Centner, Brett Van Tassel, Chris French, Hayley Yaglom, Darrin Lemmer, Dave Engelthaler |
| EPI_ISL_3325953, EPI_ISL_3325954, EPI_ISL_3325956, EPI_ISL_3325958 | Laboratory Corporation of America | Centers for Disease Control and Prevention Division of Viral Diseases, Pathogen Discovery | Dakota Howard, Dhwani Batra, Peter W. Cook, Kara Moser, Adrian Paskey, Jason Caravas, Benjamin Rambo-Martin, Shatavia Morrison, Christopher Gulvick, Scott Sammons, Yvette Unoarumhi, Darlene Wagner, Matthew Schmerer, Minoo Agarwal, Eyad Almasri, Debbie Boles, Ayla Burns, Nuthawin Charoensri, Oren Cohen, Susan Countryman, Mary Ann Cristobal, Bobbi Croy, Suzanne Dale, Hrushikesh Deshmukh, Amanda Douglas, Vincent Drouillon, Marcia Eisenberg, Howard Engler, Rama Ghatti, Prashant Gupta, Susan Hicks, Jake Humphrey, Lax Iyer, Lisa Pfefferle, Manoj Jain, Matthew Robinson, Mohan Kolli, Brian Krueger, Tim Kuphal, Stanley Letovsky, Michael Levandoski, Craig Lukasik, Jonathan Meltzer, Brian Norvell, Mindy Nye, Scott Parker, Christos Petropoulos, John Pruitt, Steven Ragan, Scott Ryan, Mike Sapeta, Jana Schroth, Suresh Babu Selvaraju, Goran Stevovic, Amanda Suchanek, Andrea Throop, Lyndon Tilson, Thomas Urban, Joe Voshell, Kimberly Wagner, Jonathan Williams, Mary Williamson, Qian Zeng, Tricia Zwiefelhofer, Clinton R. Paden, Duncan MacCannell |
| EPI_ISL_3325959 | TGen North | TGen North | Jolene Bowers, Heather Centner, Brett Van Tassel, Chris French, Hayley Yaglom, Darrin Lemmer, Dave Engelthaler |
| EPI_ISL_3325960, EPI_ISL_3325962, EPI_ISL_3325964, EPI_ISL_3325965 | Laboratory Corporation of America | Centers for Disease Control and Prevention Division of Viral Diseases, Pathogen Discovery | Dakota Howard, Dhwani Batra, Peter W. Cook, Kara Moser, Adrian Paskey, Jason Caravas, Benjamin Rambo-Martin, Shatavia Morrison, Christopher Gulvick, Scott Sammons, Yvette Unoarumhi, Darlene Wagner, Matthew Schmerer, Minoo Agarwal, Eyad Almasri, Debbie Boles, Ayla Burns, Nuthawin Charoensri, Oren Cohen, Susan Countryman, Mary Ann Cristobal, Bobbi Croy, Suzanne Dale, Hrushikesh Deshmukh, Amanda Douglas, Vincent Drouillon, Marcia Eisenberg, Howard Engler, Rama Ghatti, Prashant Gupta, Susan Hicks, Jake Humphrey, Lax Iyer, Lisa Pfefferle, Manoj Jain, Matthew Robinson, Mohan Kolli, Brian Krueger, Tim Kuphal, Stanley Letovsky, Michael Levandoski, Craig Lukasik, Jonathan Meltzer, Brian Norvell, Mindy Nye, Scott Parker, Christos Petropoulos, John Pruitt, Steven Ragan, Scott Ryan, Mike Sapeta, Jana Schroth, Suresh Babu Selvaraju, Goran Stevovic, Amanda Suchanek, Andrea Throop, Lyndon Tilson, Thomas Urban, Joe Voshell, Kimberly Wagner, Jonathan Williams, Mary Williamson, Qian Zeng, Tricia Zwiefelhofer, Clinton R. Paden, Duncan MacCannell |
| EPI_ISL_3325966 | TGen North | TGen North | Jolene Bowers, Heather Centner, Brett Van Tassel, Chris French, Hayley Yaglom, Darrin Lemmer, Dave Engelthaler |
| EPI_ISL_3325967, EPI_ISL_3325969 | Laboratory Corporation of America | Centers for Disease Control and Prevention Division of Viral Diseases, Pathogen Discovery | Dakota Howard, Dhwani Batra, Peter W. Cook, Kara Moser, Adrian Paskey, Jason Caravas, Benjamin Rambo-Martin, Shatavia Morrison, Christopher Gulvick, Scott Sammons, Yvette Unoarumhi, Darlene Wagner, Matthew Schmerer, Minoo Agarwal, Eyad Almasri, Debbie Boles, Ayla Burns, Nuthawin Charoensri, Oren Cohen, Susan Countryman, Mary Ann Cristobal, Bobbi Croy, Suzanne Dale, Hrushikesh Deshmukh, Amanda Douglas, Vincent Drouillon, Marcia Eisenberg, Howard Engler, Rama Ghatti, Prashant Gupta, Susan Hicks, Jake Humphrey, Lax Iyer, Lisa Pfefferle, Manoj Jain, Matthew Robinson, Mohan Kolli, Brian Krueger, Tim Kuphal, Stanley Letovsky, Michael Levandoski, Craig Lukasik, Jonathan Meltzer, Brian Norvell, Mindy Nye, Scott Parker, Christos Petropoulos, John Pruitt, Steven Ragan, Scott Ryan, Mike Sapeta, Jana Schroth, Suresh Babu Selvaraju, Goran Stevovic, Amanda Suchanek, Andrea Throop, Lyndon Tilson, Thomas Urban, Joe Voshell, Kimberly Wagner, Jonathan Williams, Mary Williamson, Qian Zeng, Tricia Zwiefelhofer, Clinton R. Paden, Duncan MacCannell |
| EPI_ISL_3325970 | TGen North | TGen North | Jolene Bowers, Heather Centner, Brett Van Tassel, Chris French, Hayley Yaglom, Darrin Lemmer, Dave Engelthaler |
| EPI_ISL_3325971 | Laboratory Corporation of America | Centers for Disease Control and Prevention Division of Viral Diseases, Pathogen Discovery | Dakota Howard, Dhwani Batra, Peter W. Cook, Kara Moser, Adrian Paskey, Jason Caravas, Benjamin Rambo-Martin, Shatavia Morrison, Christopher Gulvick, Scott Sammons, Yvette Unoarumhi, Darlene Wagner, Matthew Schmerer, Minoo Agarwal, Eyad Almasri, Debbie Boles, Ayla Burns, Nuthawin Charoensri, Oren Cohen, Susan Countryman, Mary Ann Cristobal, Bobbi Croy, Suzanne Dale, Hrushikesh Deshmukh, Amanda Douglas, Vincent Drouillon, Marcia Eisenberg, Howard Engler, Rama Ghatti, Prashant Gupta, Susan Hicks, Jake Humphrey, Lax Iyer, Lisa Pfefferle, Manoj Jain, Matthew Robinson, Mohan Kolli, Brian Krueger, Tim Kuphal, Stanley Letovsky, Michael Levandoski, Craig Lukasik, Jonathan Meltzer, Brian Norvell, Mindy Nye, Scott Parker, Christos Petropoulos, John Pruitt, Steven Ragan, Scott Ryan, Mike Sapeta, Jana Schroth, Suresh Babu Selvaraju, Goran Stevovic, Amanda Suchanek, Andrea Throop, Lyndon Tilson, Thomas Urban, Joe Voshell, Kimberly Wagner, Jonathan Williams, Mary Williamson, Qian Zeng, Tricia Zwiefelhofer, Clinton R. Paden, Duncan MacCannell |
| EPI_ISL_3325972 | TGen North | TGen North | Jolene Bowers, Heather Centner, Brett Van Tassel, Chris French, Hayley Yaglom, Darrin Lemmer, Dave Engelthaler |
| EPI_ISL_3325973 | Laboratory Corporation of America | Centers for Disease Control and Prevention Division of Viral Diseases, Pathogen Discovery | Dakota Howard, Dhwani Batra, Peter W. Cook, Kara Moser, Adrian Paskey, Jason Caravas, Benjamin Rambo-Martin, Shatavia Morrison, Christopher Gulvick, Scott Sammons, Yvette Unoarumhi, Darlene Wagner, Matthew Schmerer, Minoo Agarwal, Eyad Almasri, Debbie Boles, Ayla Burns, Nuthawin Charoensri, Oren Cohen, Susan Countryman, Mary Ann Cristobal, Bobbi Croy, Suzanne Dale, Hrushikesh Deshmukh, Amanda Douglas, Vincent Drouillon, Marcia Eisenberg, Howard Engler, Rama Ghatti, Prashant Gupta, Susan Hicks, Jake Humphrey, Lax Iyer, Lisa Pfefferle, Manoj Jain, Matthew Robinson, Mohan Kolli, Brian Krueger, Tim Kuphal, Stanley Letovsky, Michael Levandoski, Craig Lukasik, Jonathan Meltzer, Brian Norvell, Mindy Nye, Scott Parker, Christos Petropoulos, John Pruitt, Steven Ragan, Scott Ryan, Mike Sapeta, Jana Schroth, Suresh Babu Selvaraju, Goran Stevovic, Amanda Suchanek, Andrea Throop, Lyndon Tilson, Thomas Urban, Joe Voshell, Kimberly Wagner, Jonathan Williams, Mary Williamson, Qian Zeng, Tricia Zwiefelhofer, Clinton R. Paden, Duncan MacCannell |
| EPI_ISL_3325974 | TGen North | TGen North | Jolene Bowers, Heather Centner, Brett Van Tassel, Chris French, Hayley Yaglom, Darrin Lemmer, Dave Engelthaler |
| EPI_ISL_3325977, EPI_ISL_3325978 | Laboratory Corporation of America | Centers for Disease Control and Prevention Division of Viral Diseases, Pathogen Discovery | Dakota Howard, Dhwani Batra, Peter W. Cook, Kara Moser, Adrian Paskey, Jason Caravas, Benjamin Rambo-Martin, Shatavia Morrison, Christopher Gulvick, Scott Sammons, Yvette Unoarumhi, Darlene Wagner, Matthew Schmerer, Minoo Agarwal, Eyad Almasri, Debbie Boles, Ayla Burns, Nuthawin Charoensri, Oren Cohen, Susan Countryman, Mary Ann Cristobal, Bobbi Croy, Suzanne Dale, Hrushikesh Deshmukh, Amanda Douglas, Vincent Drouillon, Marcia Eisenberg, Howard Engler, Rama Ghatti, Prashant Gupta, Susan Hicks, Jake Humphrey, Lax Iyer, Lisa Pfefferle, Manoj Jain, Matthew Robinson, Mohan Kolli, Brian Krueger, Tim Kuphal, Stanley Letovsky, Michael Levandoski, Craig Lukasik, Jonathan Meltzer, Brian Norvell, Mindy Nye, Scott Parker, Christos Petropoulos, John Pruitt, Steven Ragan, Scott Ryan, Mike Sapeta, Jana Schroth, Suresh Babu Selvaraju, Goran Stevovic, Amanda Suchanek, Andrea Throop, Lyndon Tilson, Thomas Urban, Joe Voshell, Kimberly Wagner, Jonathan Williams, Mary Williamson, Qian Zeng, Tricia Zwiefelhofer, Clinton R. Paden, Duncan MacCannell |
| EPI_ISL_3325979 | TGen North | TGen North | Jolene Bowers, Heather Centner, Brett Van Tassel, Chris French, Hayley Yaglom, Darrin Lemmer, Dave Engelthaler |
| EPI_ISL_3325980 | Laboratory Corporation of America | Centers for Disease Control and Prevention Division of Viral Diseases, Pathogen Discovery | Dakota Howard, Dhwani Batra, Peter W. Cook, Kara Moser, Adrian Paskey, Jason Caravas, Benjamin Rambo-Martin, Shatavia Morrison, Christopher Gulvick, Scott Sammons, Yvette Unoarumhi, Darlene Wagner, Matthew Schmerer, Minoo Agarwal, Eyad Almasri, Debbie Boles, Ayla Burns, Nuthawin Charoensri, Oren Cohen, Susan Countryman, Mary Ann Cristobal, Bobbi Croy, Suzanne Dale, Hrushikesh Deshmukh, Amanda Douglas, Vincent Drouillon, Marcia Eisenberg, Howard Engler, Rama Ghatti, Prashant Gupta, Susan Hicks, Jake Humphrey, Lax Iyer, Lisa Pfefferle, Manoj Jain, Matthew Robinson, Mohan Kolli, Brian Krueger, Tim Kuphal, Stanley Letovsky, Michael Levandoski, Craig Lukasik, Jonathan Meltzer, Brian Norvell, Mindy Nye, Scott Parker, Christos Petropoulos, John Pruitt, Steven Ragan, Scott Ryan, Mike Sapeta, Jana Schroth, Suresh Babu Selvaraju, Goran Stevovic, Amanda Suchanek, Andrea Throop, Lyndon Tilson, Thomas Urban, Joe Voshell, Kimberly Wagner, Jonathan Williams, Mary Williamson, Qian Zeng, Tricia Zwiefelhofer, Clinton R. Paden, Duncan MacCannell |
| EPI_ISL_3325981 | TGen North | TGen North | Jolene Bowers, Heather Centner, Brett Van Tassel, Chris French, Hayley Yaglom, Darrin Lemmer, Dave Engelthaler |
| EPI_ISL_3325982 | Laboratory Corporation of America | Centers for Disease Control and Prevention Division of Viral Diseases, Pathogen Discovery | Dakota Howard, Dhwani Batra, Peter W. Cook, Kara Moser, Adrian Paskey, Jason Caravas, Benjamin Rambo-Martin, Shatavia Morrison, Christopher Gulvick, Scott Sammons, Yvette Unoarumhi, Darlene Wagner, Matthew Schmerer, Minoo Agarwal, Eyad Almasri, Debbie Boles, Ayla Burns, Nuthawin Charoensri, Oren Cohen, Susan Countryman, Mary Ann Cristobal, Bobbi Croy, Suzanne Dale, Hrushikesh Deshmukh, Amanda Douglas, Vincent Drouillon, Marcia Eisenberg, Howard Engler, Rama Ghatti, Prashant Gupta, Susan Hicks, Jake Humphrey, Lax Iyer, Lisa Pfefferle, Manoj Jain, Matthew Robinson, Mohan Kolli, Brian Krueger, Tim Kuphal, Stanley Letovsky, Michael Levandoski, Craig Lukasik, Jonathan Meltzer, Brian Norvell, Mindy Nye, Scott Parker, Christos Petropoulos, John Pruitt, Steven Ragan, Scott Ryan, Mike Sapeta, Jana Schroth, Suresh Babu Selvaraju, Goran Stevovic, Amanda Suchanek, Andrea Throop, Lyndon Tilson, Thomas Urban, Joe Voshell, Kimberly Wagner, Jonathan Williams, Mary Williamson, Qian Zeng, Tricia Zwiefelhofer, Clinton R. Paden, Duncan MacCannell |
| EPI_ISL_3325983 | TGen North | TGen North | Jolene Bowers, Heather Centner, Brett Van Tassel, Chris French, Hayley Yaglom, Darrin Lemmer, Dave Engelthaler |
| EPI_ISL_3325984, EPI_ISL_3325985, EPI_ISL_3325987 | Laboratory Corporation of America | Centers for Disease Control and Prevention Division of Viral Diseases, Pathogen Discovery | Dakota Howard, Dhwani Batra, Peter W. Cook, Kara Moser, Adrian Paskey, Jason Caravas, Benjamin Rambo-Martin, Shatavia Morrison, Christopher Gulvick, Scott Sammons, Yvette Unoarumhi, Darlene Wagner, Matthew Schmerer, Minoo Agarwal, Eyad Almasri, Debbie Boles, Ayla Burns, Nuthawin Charoensri, Oren Cohen, Susan Countryman, Mary Ann Cristobal, Bobbi Croy, Suzanne Dale, Hrushikesh Deshmukh, Amanda Douglas, Vincent Drouillon, Marcia Eisenberg, Howard Engler, Rama Ghatti, Prashant Gupta, Susan Hicks, Jake Humphrey, Lax Iyer, Lisa Pfefferle, Manoj Jain, Matthew Robinson, Mohan Kolli, Brian Krueger, Tim Kuphal, Stanley Letovsky, Michael Levandoski, Craig Lukasik, Jonathan Meltzer, Brian Norvell, Mindy Nye, Scott Parker, Christos Petropoulos, John Pruitt, Steven Ragan, Scott Ryan, Mike Sapeta, Jana Schroth, Suresh Babu Selvaraju, Goran Stevovic, Amanda Suchanek, Andrea Throop, Lyndon Tilson, Thomas Urban, Joe Voshell, Kimberly Wagner, Jonathan Williams, Mary Williamson, Qian Zeng, Tricia Zwiefelhofer, Clinton R. Paden, Duncan MacCannell |
| EPI_ISL_3325988 | TGen North | TGen North | Jolene Bowers, Heather Centner, Brett Van Tassel, Chris French, Hayley Yaglom, Darrin Lemmer, Dave Engelthaler |
| EPI_ISL_3325989 | Laboratory Corporation of America | Centers for Disease Control and Prevention Division of Viral Diseases, Pathogen Discovery | Dakota Howard, Dhwani Batra, Peter W. Cook, Kara Moser, Adrian Paskey, Jason Caravas, Benjamin Rambo-Martin, Shatavia Morrison, Christopher Gulvick, Scott Sammons, Yvette Unoarumhi, Darlene Wagner, Matthew Schmerer, Minoo Agarwal, Eyad Almasri, Debbie Boles, Ayla Burns, Nuthawin Charoensri, Oren Cohen, Susan Countryman, Mary Ann Cristobal, Bobbi Croy, Suzanne Dale, Hrushikesh Deshmukh, Amanda Douglas, Vincent Drouillon, Marcia Eisenberg, Howard Engler, Rama Ghatti, Prashant Gupta, Susan Hicks, Jake Humphrey, Lax Iyer, Lisa Pfefferle, Manoj Jain, Matthew Robinson, Mohan Kolli, Brian Krueger, Tim Kuphal, Stanley Letovsky, Michael Levandoski, Craig Lukasik, Jonathan Meltzer, Brian Norvell, Mindy Nye, Scott Parker, Christos Petropoulos, John Pruitt, Steven Ragan, Scott Ryan, Mike Sapeta, Jana Schroth, Suresh Babu Selvaraju, Goran Stevovic, Amanda Suchanek, Andrea Throop, Lyndon Tilson, Thomas Urban, Joe Voshell, Kimberly Wagner, Jonathan Williams, Mary Williamson, Qian Zeng, Tricia Zwiefelhofer, Clinton R. Paden, Duncan MacCannell |
| EPI_ISL_3325990 | TGen North | TGen North | Jolene Bowers, Heather Centner, Brett Van Tassel, Chris French, Hayley Yaglom, Darrin Lemmer, Dave Engelthaler |
| EPI_ISL_3325991, EPI_ISL_3325993, EPI_ISL_3325995, EPI_ISL_3325996 | Laboratory Corporation of America | Centers for Disease Control and Prevention Division of Viral Diseases, Pathogen Discovery | Dakota Howard, Dhwani Batra, Peter W. Cook, Kara Moser, Adrian Paskey, Jason Caravas, Benjamin Rambo-Martin, Shatavia Morrison, Christopher Gulvick, Scott Sammons, Yvette Unoarumhi, Darlene Wagner, Matthew Schmerer, Minoo Agarwal, Eyad Almasri, Debbie Boles, Ayla Burns, Nuthawin Charoensri, Oren Cohen, Susan Countryman, Mary Ann Cristobal, Bobbi Croy, Suzanne Dale, Hrushikesh Deshmukh, Amanda Douglas, Vincent Drouillon, Marcia Eisenberg, Howard Engler, Rama Ghatti, Prashant Gupta, Susan Hicks, Jake Humphrey, Lax Iyer, Lisa Pfefferle, Manoj Jain, Matthew Robinson, Mohan Kolli, Brian Krueger, Tim Kuphal, Stanley Letovsky, Michael Levandoski, Craig Lukasik, Jonathan Meltzer, Brian Norvell, Mindy Nye, Scott Parker, Christos Petropoulos, John Pruitt, Steven Ragan, Scott Ryan, Mike Sapeta, Jana Schroth, Suresh Babu Selvaraju, Goran Stevovic, Amanda Suchanek, Andrea Throop, Lyndon Tilson, Thomas Urban, Joe Voshell, Kimberly Wagner, Jonathan Williams, Mary Williamson, Qian Zeng, Tricia Zwiefelhofer, Clinton R. Paden, Duncan MacCannell |
| EPI_ISL_3325997 | TGen North | TGen North | Jolene Bowers, Heather Centner, Brett Van Tassel, Chris French, Hayley Yaglom, Darrin Lemmer, Dave Engelthaler |
| EPI_ISL_3325998, EPI_ISL_3326000 | Laboratory Corporation of America | Centers for Disease Control and Prevention Division of Viral Diseases, Pathogen Discovery | Dakota Howard, Dhwani Batra, Peter W. Cook, Kara Moser, Adrian Paskey, Jason Caravas, Benjamin Rambo-Martin, Shatavia Morrison, Christopher Gulvick, Scott Sammons, Yvette Unoarumhi, Darlene Wagner, Matthew Schmerer, Minoo Agarwal, Eyad Almasri, Debbie Boles, Ayla Burns, Nuthawin Charoensri, Oren Cohen, Susan Countryman, Mary Ann Cristobal, Bobbi Croy, Suzanne Dale, Hrushikesh Deshmukh, Amanda Douglas, Vincent Drouillon, Marcia Eisenberg, Howard Engler, Rama Ghatti, Prashant Gupta, Susan Hicks, Jake Humphrey, Lax Iyer, Lisa Pfefferle, Manoj Jain, Matthew Robinson, Mohan Kolli, Brian Krueger, Tim Kuphal, Stanley Letovsky, Michael Levandoski, Craig Lukasik, Jonathan Meltzer, Brian Norvell, Mindy Nye, Scott Parker, Christos Petropoulos, John Pruitt, Steven Ragan, Scott Ryan, Mike Sapeta, Jana Schroth, Suresh Babu Selvaraju, Goran Stevovic, Amanda Suchanek, Andrea Throop, Lyndon Tilson, Thomas Urban, Joe Voshell, Kimberly Wagner, Jonathan Williams, Mary Williamson, Qian Zeng, Tricia Zwiefelhofer, Clinton R. Paden, Duncan MacCannell |
| EPI_ISL_3326001 | TGen North | TGen North | Jolene Bowers, Heather Centner, Brett Van Tassel, Chris French, Hayley Yaglom, Darrin Lemmer, Dave Engelthaler |
| EPI_ISL_3326002 | Laboratory Corporation of America | Centers for Disease Control and Prevention Division of Viral | Dakota Howard, Dhwani Batra, Peter W. Cook, Kara Moser, Adrian Paskey, Jason Caravas, Benjamin Rambo-Martin, Shatavia Morrison, Christopher Gulvick, Scott Sammons, Yvette Unoarumhi, Darlene Wagner, Matthew Schmerer, Minoo |

| | | | |
|---|---|---|---|
| | | Diseases, Pathogen Discovery | Agarwal, Eyad Almasri, Debbie Boles, Ayla Burns, Nuthawin Charoensri, Oren Cohen, Susan Countryman, Mary Ann Cristobal, Bobbi Croy, Suzanne Dale, Hrushikesh Deshmukh, Amanda Douglas, Vincent Drouillon, Marcia Eisenberg, Howard Engler, Rama Ghatti, Prashant Gupta, Susan Hicks, Jake Humphrey, Lax Iyer, Lisa Pfefferle, Manoj Jain, Matthew Robinson, Mohan Kolli, Brian Krueger, Tim Kuphal, Stanley Letovsky, Michael Levandoski, Craig Lukasik, Jonathan Meltzer, Brian Norvell, Mindy Nye, Scott Parker, Christos Petropoulos, John Pruitt, Steven Ragan, Scott Ryan, Mike Sapeta, Jana Schroth, Suresh Babu Selvaraju, Goran Stevovic, Amanda Suchanek, Andrea Throop, Lyndon Tilson, Thomas Urban, Joe Voshell, Kimberly Wagner, Jonathan Williams, Mary Williamson, Qian Zeng, Tricia Zwiefelhofer, Clinton R. Paden, Duncan MacCannell |
| EPI_ISL_3326003 | TGen North | TGen North | Jolene Bowers, Heather Centner, Brett Van Tassel, Chris French, Hayley Yaglom, Darrin Lemmer, Dave Engelthaler |
| EPI_ISL_3326004 | Laboratory Corporation of America | Centers for Disease Control and Prevention Division of Viral Diseases, Pathogen Discovery | Dakota Howard, Dhwani Batra, Peter W. Cook, Kara Moser, Adrian Paskey, Jason Caravas, Benjamin Rambo-Martin, Shatavia Morrison, Christopher Gulvick, Scott Sammons, Yvette Unoarumhi, Darlene Wagner, Matthew Schmerer, Minoo Agarwal, Eyad Almasri, Debbie Boles, Ayla Burns, Nuthawin Charoensri, Oren Cohen, Susan Countryman, Mary Ann Cristobal, Bobbi Croy, Suzanne Dale, Hrushikesh Deshmukh, Amanda Douglas, Vincent Drouillon, Marcia Eisenberg, Howard Engler, Rama Ghatti, Prashant Gupta, Susan Hicks, Jake Humphrey, Lax Iyer, Lisa Pfefferle, Manoj Jain, Matthew Robinson, Mohan Kolli, Brian Krueger, Tim Kuphal, Stanley Letovsky, Michael Levandoski, Craig Lukasik, Jonathan Meltzer, Brian Norvell, Mindy Nye, Scott Parker, Christos Petropoulos, John Pruitt, Steven Ragan, Scott Ryan, Mike Sapeta, Jana Schroth, Suresh Babu Selvaraju, Goran Stevovic, Amanda Suchanek, Andrea Throop, Lyndon Tilson, Thomas Urban, Joe Voshell, Kimberly Wagner, Jonathan Williams, Mary Williamson, Qian Zeng, Tricia Zwiefelhofer, Clinton R. Paden, Duncan MacCannell |
| EPI_ISL_3326005 | TGen North | TGen North | Jolene Bowers, Heather Centner, Brett Van Tassel, Chris French, Hayley Yaglom, Darrin Lemmer, Dave Engelthaler |
| EPI_ISL_3326006, EPI_ISL_3326007 | Laboratory Corporation of America | Centers for Disease Control and Prevention Division of Viral Diseases, Pathogen Discovery | Dakota Howard, Dhwani Batra, Peter W. Cook, Kara Moser, Adrian Paskey, Jason Caravas, Benjamin Rambo-Martin, Shatavia Morrison, Christopher Gulvick, Scott Sammons, Yvette Unoarumhi, Darlene Wagner, Matthew Schmerer, Minoo Agarwal, Eyad Almasri, Debbie Boles, Ayla Burns, Nuthawin Charoensri, Oren Cohen, Susan Countryman, Mary Ann Cristobal, Bobbi Croy, Suzanne Dale, Hrushikesh Deshmukh, Amanda Douglas, Vincent Drouillon, Marcia Eisenberg, Howard Engler, Rama Ghatti, Prashant Gupta, Susan Hicks, Jake Humphrey, Lax Iyer, Lisa Pfefferle, Manoj Jain, Matthew Robinson, Mohan Kolli, Brian Krueger, Tim Kuphal, Stanley Letovsky, Michael Levandoski, Craig Lukasik, Jonathan Meltzer, Brian Norvell, Mindy Nye, Scott Parker, Christos Petropoulos, John Pruitt, Steven Ragan, Scott Ryan, Mike Sapeta, Jana Schroth, Suresh Babu Selvaraju, Goran Stevovic, Amanda Suchanek, Andrea Throop, Lyndon Tilson, Thomas Urban, Joe Voshell, Kimberly Wagner, Jonathan Williams, Mary Williamson, Qian Zeng, Tricia Zwiefelhofer, Clinton R. Paden, Duncan MacCannell |
| EPI_ISL_3326008 | TGen North | TGen North | Jolene Bowers, Heather Centner, Brett Van Tassel, Chris French, Hayley Yaglom, Darrin Lemmer, Dave Engelthaler |
| EPI_ISL_3326009 | Laboratory Corporation of America | Centers for Disease Control and Prevention Division of Viral Diseases, Pathogen Discovery | Dakota Howard, Dhwani Batra, Peter W. Cook, Kara Moser, Adrian Paskey, Jason Caravas, Benjamin Rambo-Martin, Shatavia Morrison, Christopher Gulvick, Scott Sammons, Yvette Unoarumhi, Darlene Wagner, Matthew Schmerer, Minoo Agarwal, Eyad Almasri, Debbie Boles, Ayla Burns, Nuthawin Charoensri, Oren Cohen, Susan Countryman, Mary Ann Cristobal, Bobbi Croy, Suzanne Dale, Hrushikesh Deshmukh, Amanda Douglas, Vincent Drouillon, Marcia Eisenberg, Howard Engler, Rama Ghatti, Prashant Gupta, Susan Hicks, Jake Humphrey, Lax Iyer, Lisa Pfefferle, Manoj Jain, Matthew Robinson, Mohan Kolli, Brian Krueger, Tim Kuphal, Stanley Letovsky, Michael Levandoski, Craig Lukasik, Jonathan Meltzer, Brian Norvell, Mindy Nye, Scott Parker, Christos Petropoulos, John Pruitt, Steven Ragan, Scott Ryan, Mike Sapeta, Jana Schroth, Suresh Babu Selvaraju, Goran Stevovic, Amanda Suchanek, Andrea Throop, Lyndon Tilson, Thomas Urban, Joe Voshell, Kimberly Wagner, Jonathan Williams, Mary Williamson, Qian Zeng, Tricia Zwiefelhofer, Clinton R. Paden, Duncan MacCannell |
| EPI_ISL_3326010 | TGen North | TGen North | Jolene Bowers, Heather Centner, Brett Van Tassel, Chris French, Hayley Yaglom, Darrin Lemmer, Dave Engelthaler |
| EPI_ISL_3326011 | Laboratory Corporation of America | Centers for Disease Control and Prevention Division of Viral Diseases, Pathogen Discovery | Dakota Howard, Dhwani Batra, Peter W. Cook, Kara Moser, Adrian Paskey, Jason Caravas, Benjamin Rambo-Martin, Shatavia Morrison, Christopher Gulvick, Scott Sammons, Yvette Unoarumhi, Darlene Wagner, Matthew Schmerer, Minoo Agarwal, Eyad Almasri, Debbie Boles, Ayla Burns, Nuthawin Charoensri, Oren Cohen, Susan Countryman, Mary Ann Cristobal, Bobbi Croy, Suzanne Dale, Hrushikesh Deshmukh, Amanda Douglas, Vincent Drouillon, Marcia Eisenberg, Howard Engler, Rama Ghatti, Prashant Gupta, Susan Hicks, Jake Humphrey, Lax Iyer, Lisa Pfefferle, Manoj Jain, Matthew Robinson, Mohan Kolli, Brian Krueger, Tim Kuphal, Stanley Letovsky, Michael Levandoski, Craig Lukasik, Jonathan Meltzer, Brian Norvell, Mindy Nye, Scott Parker, Christos Petropoulos, John Pruitt, Steven Ragan, Scott Ryan, Mike Sapeta, Jana Schroth, Suresh Babu Selvaraju, Goran Stevovic, Amanda Suchanek, Andrea Throop, Lyndon Tilson, Thomas Urban, Joe Voshell, Kimberly Wagner, Jonathan Williams, Mary Williamson, Qian Zeng, Tricia Zwiefelhofer, Clinton R. Paden, Duncan MacCannell |
| EPI_ISL_3326012 | TGen North | TGen North | Jolene Bowers, Heather Centner, Brett Van Tassel, Chris French, Hayley Yaglom, Darrin Lemmer, Dave Engelthaler |
| EPI_ISL_3326013, EPI_ISL_3326015, EPI_ISL_3326017, EPI_ISL_3326018 | Laboratory Corporation of America | Centers for Disease Control and Prevention Division of Viral Diseases, Pathogen Discovery | Dakota Howard, Dhwani Batra, Peter W. Cook, Kara Moser, Adrian Paskey, Jason Caravas, Benjamin Rambo-Martin, Shatavia Morrison, Christopher Gulvick, Scott Sammons, Yvette Unoarumhi, Darlene Wagner, Matthew Schmerer, Minoo Agarwal, Eyad Almasri, Debbie Boles, Ayla Burns, Nuthawin Charoensri, Oren Cohen, Susan Countryman, Mary Ann Cristobal, Bobbi Croy, Suzanne Dale, Hrushikesh Deshmukh, Amanda Douglas, Vincent Drouillon, Marcia Eisenberg, Howard Engler, Rama Ghatti, Prashant Gupta, Susan Hicks, Jake Humphrey, Lax Iyer, Lisa Pfefferle, Manoj Jain, Matthew Robinson, Mohan Kolli, Brian Krueger, Tim Kuphal, Stanley Letovsky, Michael Levandoski, Craig Lukasik, Jonathan Meltzer, Brian Norvell, Mindy Nye, Scott Parker, Christos Petropoulos, John Pruitt, Steven Ragan, Scott Ryan, Mike Sapeta, Jana Schroth, Suresh Babu Selvaraju, Goran Stevovic, Amanda Suchanek, Andrea Throop, Lyndon Tilson, Thomas Urban, Joe Voshell, Kimberly Wagner, Jonathan Williams, Mary Williamson, Qian Zeng, Tricia Zwiefelhofer, Clinton R. Paden, Duncan MacCannell |
| EPI_ISL_3326019 | TGen North | TGen North | Jolene Bowers, Heather Centner, Brett Van Tassel, Chris French, Hayley Yaglom, Darrin Lemmer, Dave Engelthaler |
| EPI_ISL_3326020 | Laboratory Corporation of America | Centers for Disease Control and Prevention Division of Viral Diseases, Pathogen Discovery | Dakota Howard, Dhwani Batra, Peter W. Cook, Kara Moser, Adrian Paskey, Jason Caravas, Benjamin Rambo-Martin, Shatavia Morrison, Christopher Gulvick, Scott Sammons, Yvette Unoarumhi, Darlene Wagner, Matthew Schmerer, Minoo Agarwal, Eyad Almasri, Debbie Boles, Ayla Burns, Nuthawin Charoensri, Oren Cohen, Susan Countryman, Mary Ann Cristobal, Bobbi Croy, Suzanne Dale, Hrushikesh Deshmukh, Amanda Douglas, Vincent Drouillon, Marcia Eisenberg, Howard Engler, Rama Ghatti, Prashant Gupta, Susan Hicks, Jake Humphrey, Lax Iyer, Lisa Pfefferle, Manoj Jain, Matthew Robinson, Mohan Kolli, Brian Krueger, Tim Kuphal, Stanley Letovsky, Michael Levandoski, Craig Lukasik, Jonathan Meltzer, Brian Norvell, Mindy Nye, Scott Parker, Christos Petropoulos, John Pruitt, Steven Ragan, Scott Ryan, Mike Sapeta, Jana Schroth, Suresh Babu Selvaraju, Goran Stevovic, Amanda Suchanek, Andrea Throop, Lyndon Tilson, Thomas Urban, Joe Voshell, Kimberly Wagner, Jonathan Williams, Mary Williamson, Qian Zeng, Tricia Zwiefelhofer, Clinton R. Paden, Duncan MacCannell |
| EPI_ISL_3326021 | TGen North | TGen North | Jolene Bowers, Heather Centner, Brett Van Tassel, Chris French, Hayley Yaglom, Darrin Lemmer, Dave Engelthaler |
| EPI_ISL_3326022 | Laboratory Corporation of America | Centers for Disease Control and Prevention Division of Viral Diseases, Pathogen Discovery | Dakota Howard, Dhwani Batra, Peter W. Cook, Kara Moser, Adrian Paskey, Jason Caravas, Benjamin Rambo-Martin, Shatavia Morrison, Christopher Gulvick, Scott Sammons, Yvette Unoarumhi, Darlene Wagner, Matthew Schmerer, Minoo Agarwal, Eyad Almasri, Debbie Boles, Ayla Burns, Nuthawin Charoensri, Oren Cohen, Susan Countryman, Mary Ann Cristobal, Bobbi Croy, Suzanne Dale, Hrushikesh Deshmukh, Amanda Douglas, Vincent Drouillon, Marcia Eisenberg, Howard Engler, Rama Ghatti, Prashant Gupta, Susan Hicks, Jake Humphrey, Lax Iyer, Lisa Pfefferle, Manoj Jain, Matthew Robinson, Mohan Kolli, Brian Krueger, Tim Kuphal, Stanley Letovsky, Michael Levandoski, Craig Lukasik, Jonathan Meltzer, Brian Norvell, Mindy Nye, Scott Parker, Christos Petropoulos, John Pruitt, Steven Ragan, Scott Ryan, Mike Sapeta, Jana Schroth, Suresh Babu Selvaraju, Goran Stevovic, Amanda Suchanek, Andrea Throop, Lyndon Tilson, Thomas Urban, Joe Voshell, Kimberly Wagner, Jonathan Williams, Mary Williamson, Qian Zeng, Tricia Zwiefelhofer, Clinton R. Paden, Duncan MacCannell |
| EPI_ISL_3326023 | TGen North | TGen North | Jolene Bowers, Heather Centner, Brett Van Tassel, Chris French, Hayley Yaglom, Darrin Lemmer, Dave Engelthaler |
| EPI_ISL_3326024, EPI_ISL_3326026, EPI_ISL_3326028, EPI_ISL_3326029 | Laboratory Corporation of America | Centers for Disease Control and Prevention Division of Viral Diseases, Pathogen Discovery | Dakota Howard, Dhwani Batra, Peter W. Cook, Kara Moser, Adrian Paskey, Jason Caravas, Benjamin Rambo-Martin, Shatavia Morrison, Christopher Gulvick, Scott Sammons, Yvette Unoarumhi, Darlene Wagner, Matthew Schmerer, Minoo Agarwal, Eyad Almasri, Debbie Boles, Ayla Burns, Nuthawin Charoensri, Oren Cohen, Susan Countryman, Mary Ann Cristobal, Bobbi Croy, Suzanne Dale, Hrushikesh Deshmukh, Amanda Douglas, Vincent Drouillon, Marcia Eisenberg, Howard Engler, Rama Ghatti, Prashant Gupta, Susan Hicks, Jake Humphrey, Lax Iyer, Lisa Pfefferle, Manoj Jain, Matthew Robinson, Mohan Kolli, Brian Krueger, Tim Kuphal, Stanley Letovsky, Michael Levandoski, Craig Lukasik, Jonathan Meltzer, Brian Norvell, Mindy Nye, Scott Parker, Christos Petropoulos, John Pruitt, Steven Ragan, Scott Ryan, Mike Sapeta, Jana Schroth, Suresh Babu Selvaraju, Goran Stevovic, Amanda Suchanek, Andrea Throop, Lyndon Tilson, Thomas Urban, Joe Voshell, Kimberly Wagner, Jonathan Williams, Mary Williamson, Qian Zeng, Tricia Zwiefelhofer, Clinton R. Paden, Duncan MacCannell |
| EPI_ISL_3326030, EPI_ISL_3326031 | TGen North | TGen North | Jolene Bowers, Heather Centner, Brett Van Tassel, Chris French, Hayley Yaglom, Darrin Lemmer, Dave Engelthaler |
| EPI_ISL_3326032, EPI_ISL_3326033 | Laboratory Corporation of America | Centers for Disease Control and Prevention Division of Viral Diseases, Pathogen Discovery | Dakota Howard, Dhwani Batra, Peter W. Cook, Kara Moser, Adrian Paskey, Jason Caravas, Benjamin Rambo-Martin, Shatavia Morrison, Christopher Gulvick, Scott Sammons, Yvette Unoarumhi, Darlene Wagner, Matthew Schmerer, Minoo Agarwal, Eyad Almasri, Debbie Boles, Ayla Burns, Nuthawin Charoensri, Oren Cohen, Susan Countryman, Mary Ann Cristobal, Bobbi Croy, Suzanne Dale, Hrushikesh Deshmukh, Amanda Douglas, Vincent Drouillon, Marcia Eisenberg, Howard Engler, Rama Ghatti, Prashant Gupta, Susan Hicks, Jake Humphrey, Lax Iyer, Lisa Pfefferle, Manoj Jain, Matthew Robinson, Mohan Kolli, Brian Krueger, Tim Kuphal, Stanley Letovsky, Michael Levandoski, Craig Lukasik, Jonathan Meltzer, Brian Norvell, Mindy Nye, Scott Parker, Christos Petropoulos, John Pruitt, Steven Ragan, Scott Ryan, Mike Sapeta, Jana Schroth, Suresh Babu Selvaraju, Goran Stevovic, Amanda Suchanek, Andrea Throop, Lyndon Tilson, Thomas Urban, Joe Voshell, Kimberly Wagner, Jonathan Williams, Mary Williamson, Qian Zeng, Tricia Zwiefelhofer, Clinton R. Paden, Duncan MacCannell |
| EPI_ISL_3326034 | TGen North | TGen North | Jolene Bowers, Heather Centner, Brett Van Tassel, Chris French, Hayley Yaglom, Darrin Lemmer, Dave Engelthaler |
| EPI_ISL_3326035 | Laboratory Corporation of America | Centers for Disease Control and Prevention Division of Viral Diseases, Pathogen Discovery | Dakota Howard, Dhwani Batra, Peter W. Cook, Kara Moser, Adrian Paskey, Jason Caravas, Benjamin Rambo-Martin, Shatavia Morrison, Christopher Gulvick, Scott Sammons, Yvette Unoarumhi, Darlene Wagner, Matthew Schmerer, Minoo Agarwal, Eyad Almasri, Debbie Boles, Ayla Burns, Nuthawin Charoensri, Oren Cohen, Susan Countryman, Mary Ann Cristobal, Bobbi Croy, Suzanne Dale, Hrushikesh Deshmukh, Amanda Douglas, Vincent Drouillon, Marcia Eisenberg, Howard Engler, Rama Ghatti, Prashant Gupta, Susan Hicks, Jake Humphrey, Lax Iyer, Lisa Pfefferle, Manoj Jain, Matthew Robinson, Mohan Kolli, Brian Krueger, Tim Kuphal, Stanley Letovsky, Michael Levandoski, Craig Lukasik, Jonathan Meltzer, Brian Norvell, Mindy Nye, Scott Parker, Christos Petropoulos, John Pruitt, Steven Ragan, Scott Ryan, Mike Sapeta, Jana Schroth, Suresh Babu Selvaraju, Goran Stevovic, Amanda Suchanek, Andrea Throop, Lyndon Tilson, Thomas Urban, Joe Voshell, Kimberly Wagner, Jonathan Williams, Mary Williamson, Qian Zeng, Tricia Zwiefelhofer, Clinton R. Paden, Duncan MacCannell |
| EPI_ISL_3326036 | TGen North | TGen North | Jolene Bowers, Heather Centner, Brett Van Tassel, Chris French, Hayley Yaglom, Darrin Lemmer, Dave Engelthaler |
| EPI_ISL_3326037 | Laboratory Corporation of America | Centers for Disease Control and Prevention Division of Viral Diseases, Pathogen Discovery | Dakota Howard, Dhwani Batra, Peter W. Cook, Kara Moser, Adrian Paskey, Jason Caravas, Benjamin Rambo-Martin, Shatavia Morrison, Christopher Gulvick, Scott Sammons, Yvette Unoarumhi, Darlene Wagner, Matthew Schmerer, Minoo Agarwal, Eyad Almasri, Debbie Boles, Ayla Burns, Nuthawin Charoensri, Oren Cohen, Susan Countryman, Mary Ann Cristobal, Bobbi Croy, Suzanne Dale, Hrushikesh Deshmukh, Amanda Douglas, Vincent Drouillon, Marcia Eisenberg, Howard Engler, Rama Ghatti, Prashant Gupta, Susan Hicks, Jake Humphrey, Lax Iyer, Lisa Pfefferle, Manoj Jain, Matthew Robinson, Mohan Kolli, Brian Krueger, Tim Kuphal, Stanley Letovsky, Michael Levandoski, Craig Lukasik, Jonathan Meltzer, Brian Norvell, Mindy Nye, Scott Parker, Christos Petropoulos, John Pruitt, Steven Ragan, Scott Ryan, Mike Sapeta, Jana Schroth, Suresh Babu Selvaraju, Goran Stevovic, Amanda Suchanek, Andrea Throop, Lyndon Tilson, Thomas Urban, Joe Voshell, Kimberly Wagner, Jonathan Williams, Mary Williamson, Qian Zeng, Tricia Zwiefelhofer, Clinton R. Paden, Duncan MacCannell |
| EPI_ISL_3326038 | TGen North | TGen North | Jolene Bowers, Heather Centner, Brett Van Tassel, Chris French, Hayley Yaglom, Darrin Lemmer, Dave Engelthaler |
| EPI_ISL_3326039, EPI_ISL_3326040 | Laboratory Corporation of America | Centers for Disease Control and Prevention Division of Viral Diseases, Pathogen Discovery | Dakota Howard, Dhwani Batra, Peter W. Cook, Kara Moser, Adrian Paskey, Jason Caravas, Benjamin Rambo-Martin, Shatavia Morrison, Christopher Gulvick, Scott Sammons, Yvette Unoarumhi, Darlene Wagner, Matthew Schmerer, Minoo Agarwal, Eyad Almasri, Debbie Boles, Ayla Burns, Nuthawin Charoensri, Oren Cohen, Susan Countryman, Mary Ann Cristobal, Bobbi Croy, Suzanne Dale, Hrushikesh Deshmukh, Amanda Douglas, Vincent Drouillon, Marcia Eisenberg, Howard Engler, Rama Ghatti, Prashant Gupta, Susan Hicks, Jake Humphrey, Lax Iyer, Lisa Pfefferle, Manoj Jain, Matthew Robinson, Mohan Kolli, Brian Krueger, Tim Kuphal, Stanley Letovsky, Michael Levandoski, Craig Lukasik, Jonathan Meltzer, Brian Norvell, Mindy Nye, Scott Parker, Christos Petropoulos, John Pruitt, Steven Ragan, Scott Ryan, Mike Sapeta, Jana Schroth, Suresh Babu Selvaraju, Goran Stevovic, Amanda Suchanek, Andrea Throop, Lyndon Tilson, Thomas Urban, Joe Voshell, Kimberly Wagner, Jonathan Williams, Mary Williamson, Qian Zeng, Tricia Zwiefelhofer, Clinton R. Paden, Duncan MacCannell |
| EPI_ISL_3326041, EPI_ISL_3326042, EPI_ISL_3326043, EPI_ISL_3326044 | TGen North | TGen North | Jolene Bowers, Heather Centner, Brett Van Tassel, Chris French, Hayley Yaglom, Darrin Lemmer, Dave Engelthaler |
| EPI_ISL_3326045, EPI_ISL_3326047, EPI_ISL_3326048, EPI_ISL_3326052, EPI_ISL_3326054 | Laboratory Corporation of America | Centers for Disease Control and Prevention Division of Viral Diseases, Pathogen Discovery | Dakota Howard, Dhwani Batra, Peter W. Cook, Kara Moser, Adrian Paskey, Jason Caravas, Benjamin Rambo-Martin, Shatavia Morrison, Christopher Gulvick, Scott Sammons, Yvette Unoarumhi, Darlene Wagner, Matthew Schmerer, Minoo Agarwal, Eyad Almasri, Debbie Boles, Ayla Burns, Nuthawin Charoensri, Oren Cohen, Susan Countryman, Mary Ann Cristobal, Bobbi Croy, Suzanne Dale, Hrushikesh Deshmukh, Amanda Douglas, Vincent Drouillon, Marcia Eisenberg, Howard Engler, Rama Ghatti, Prashant Gupta, Susan Hicks, Jake Humphrey, Lax Iyer, Lisa Pfefferle, Manoj Jain, Matthew Robinson, Mohan Kolli, Brian Krueger, Tim Kuphal, Stanley Letovsky, Michael Levandoski, Craig Lukasik, Jonathan Meltzer, Brian Norvell, Mindy Nye, Scott Parker, Christos Petropoulos, John Pruitt, Steven Ragan, Scott Ryan, Mike Sapeta, Jana Schroth, Suresh Babu Selvaraju, Goran Stevovic, Amanda Suchanek, Andrea Throop, Lyndon Tilson, Thomas Urban, Joe Voshell, Kimberly Wagner, Jonathan Williams, Mary Williamson, Qian Zeng, Tricia Zwiefelhofer, Clinton R. Paden, Duncan MacCannell |
| EPI_ISL_3326055 | TGen North | TGen North | Jolene Bowers, Heather Centner, Brett Van Tassel, Chris French, Hayley Yaglom, Darrin Lemmer, Dave Engelthaler |
| EPI_ISL_3326056, EPI_ISL_3326059, EPI_ISL_3326061 | Laboratory Corporation of America | Centers for Disease Control and Prevention Division of Viral Diseases, Pathogen Discovery | Dakota Howard, Dhwani Batra, Peter W. Cook, Kara Moser, Adrian Paskey, Jason Caravas, Benjamin Rambo-Martin, Shatavia Morrison, Christopher Gulvick, Scott Sammons, Yvette Unoarumhi, Darlene Wagner, Matthew Schmerer, Minoo Agarwal, Eyad Almasri, Debbie Boles, Ayla Burns, Nuthawin Charoensri, Oren Cohen, Susan Countryman, Mary Ann Cristobal, Bobbi Croy, Suzanne Dale, Hrushikesh Deshmukh, Amanda Douglas, Vincent Drouillon, Marcia Eisenberg, Howard Engler, Rama Ghatti, Prashant Gupta, Susan Hicks, Jake Humphrey, Lax Iyer, Lisa Pfefferle, Manoj Jain, Matthew Robinson, Mohan Kolli, Brian Krueger, Tim Kuphal, Stanley Letovsky, Michael Levandoski, Craig Lukasik, Jonathan Meltzer, Brian Norvell, Mindy Nye, Scott Parker, Christos Petropoulos, John Pruitt, Steven Ragan, Scott Ryan, Mike Sapeta, Jana Schroth, Suresh Babu Selvaraju, Goran Stevovic, Amanda Suchanek, Andrea Throop, Lyndon Tilson, Thomas Urban, Joe Voshell, Kimberly Wagner, Jonathan Williams, Mary Williamson, Qian Zeng, Tricia Zwiefelhofer, Clinton R. Paden, Duncan MacCannell |
| EPI_ISL_3326062 | TGen North | TGen North | Jolene Bowers, Heather Centner, Brett Van Tassel, Chris French, Hayley Yaglom, Darrin Lemmer, Dave Engelthaler |
| EPI_ISL_3326063 | Laboratory Corporation of America | Centers for Disease Control and Prevention Division of Viral Diseases, Pathogen Discovery | Dakota Howard, Dhwani Batra, Peter W. Cook, Kara Moser, Adrian Paskey, Jason Caravas, Benjamin Rambo-Martin, Shatavia Morrison, Christopher Gulvick, Scott Sammons, Yvette Unoarumhi, Darlene Wagner, Matthew Schmerer, Minoo Agarwal, Eyad Almasri, Debbie Boles, Ayla Burns, Nuthawin Charoensri, Oren Cohen, Susan Countryman, Mary Ann Cristobal, Bobbi Croy, Suzanne Dale, Hrushikesh Deshmukh, Amanda Douglas, Vincent Drouillon, Marcia Eisenberg, Howard Engler, Rama Ghatti, Prashant Gupta, Susan Hicks, Jake Humphrey, Lax Iyer, Lisa Pfefferle, Manoj Jain, Matthew Robinson, Mohan Kolli, Brian Krueger, Tim Kuphal, Stanley Letovsky, Michael Levandoski, Craig Lukasik, Jonathan Meltzer, Brian |

| | | | |
|---|---|---|---|
| | | | Norvell, Mindy Nye, Scott Parker, Christos Petropoulos, John Pruitt, Steven Ragan, Scott Ryan, Mike Sapeta, Jana Schroth, Suresh Babu Selvaraju, Goran Stevovic, Amanda Suchanek, Andrea Throop, Lyndon Tilson, Thomas Urban, Joe Voshell, Kimberly Wagner, Jonathan Williams, Mary Williamson, Qian Zeng, Tricia Zwiefelhofer, Clinton R. Paden, Duncan MacCannell |
| EPI_ISL_3326064 | TGen North | TGen North | Jolene Bowers, Heather Centner, Brett Van Tassel, Chris French, Hayley Yaglom, Darrin Lemmer, Dave Engelthaler |
| EPI_ISL_3326065 | Laboratory Corporation of America | Centers for Disease Control and Prevention Division of Viral Diseases, Pathogen Discovery | Dakota Howard, Dhwani Batra, Peter W. Cook, Kara Moser, Adrian Paskey, Jason Caravas, Benjamin Rambo-Martin, Shatavia Morrison, Christopher Gulvick, Scott Sammons, Yvette Unoarumhi, Darlene Wagner, Matthew Schmerer, Minoo Agarwal, Eyad Almasri, Debbie Boles, Ayla Burns, Nuthawin Charoensri, Oren Cohen, Susan Countryman, Mary Ann Cristobal, Bobbi Croy, Suzanne Dale, Hrushikesh Deshmukh, Amanda Douglas, Vincent Drouillon, Marcia Eisenberg, Howard Engler, Rama Ghatti, Prashant Gupta, Susan Hicks, Jake Humphrey, Lax Iyer, Lisa Pfefferle, Manoj Jain, Matthew Robinson, Mohan Kolli, Brian Krueger, Tim Kuphal, Stanley Letovsky, Michael Levandoski, Craig Lukasik, Jonathan Meltzer, Brian Norvell, Mindy Nye, Scott Parker, Christos Petropoulos, John Pruitt, Steven Ragan, Scott Ryan, Mike Sapeta, Jana Schroth, Suresh Babu Selvaraju, Goran Stevovic, Amanda Suchanek, Andrea Throop, Lyndon Tilson, Thomas Urban, Joe Voshell, Kimberly Wagner, Jonathan Williams, Mary Williamson, Qian Zeng, Tricia Zwiefelhofer, Clinton R. Paden, Duncan MacCannell |
| EPI_ISL_3326066 | TGen North | TGen North | Jolene Bowers, Heather Centner, Brett Van Tassel, Chris French, Hayley Yaglom, Darrin Lemmer, Dave Engelthaler |
| EPI_ISL_3326067, EPI_ISL_3326068 | Laboratory Corporation of America | Centers for Disease Control and Prevention Division of Viral Diseases, Pathogen Discovery | Dakota Howard, Dhwani Batra, Peter W. Cook, Kara Moser, Adrian Paskey, Jason Caravas, Benjamin Rambo-Martin, Shatavia Morrison, Christopher Gulvick, Scott Sammons, Yvette Unoarumhi, Darlene Wagner, Matthew Schmerer, Minoo Agarwal, Eyad Almasri, Debbie Boles, Ayla Burns, Nuthawin Charoensri, Oren Cohen, Susan Countryman, Mary Ann Cristobal, Bobbi Croy, Suzanne Dale, Hrushikesh Deshmukh, Amanda Douglas, Vincent Drouillon, Marcia Eisenberg, Howard Engler, Rama Ghatti, Prashant Gupta, Susan Hicks, Jake Humphrey, Lax Iyer, Lisa Pfefferle, Manoj Jain, Matthew Robinson, Mohan Kolli, Brian Krueger, Tim Kuphal, Stanley Letovsky, Michael Levandoski, Craig Lukasik, Jonathan Meltzer, Brian Norvell, Mindy Nye, Scott Parker, Christos Petropoulos, John Pruitt, Steven Ragan, Scott Ryan, Mike Sapeta, Jana Schroth, Suresh Babu Selvaraju, Goran Stevovic, Amanda Suchanek, Andrea Throop, Lyndon Tilson, Thomas Urban, Joe Voshell, Kimberly Wagner, Jonathan Williams, Mary Williamson, Qian Zeng, Tricia Zwiefelhofer, Clinton R. Paden, Duncan MacCannell |
| EPI_ISL_3326069 | TGen North | TGen North | Jolene Bowers, Heather Centner, Brett Van Tassel, Chris French, Hayley Yaglom, Darrin Lemmer, Dave Engelthaler |
| EPI_ISL_3326070 | Laboratory Corporation of America | Centers for Disease Control and Prevention Division of Viral Diseases, Pathogen Discovery | Dakota Howard, Dhwani Batra, Peter W. Cook, Kara Moser, Adrian Paskey, Jason Caravas, Benjamin Rambo-Martin, Shatavia Morrison, Christopher Gulvick, Scott Sammons, Yvette Unoarumhi, Darlene Wagner, Matthew Schmerer, Minoo Agarwal, Eyad Almasri, Debbie Boles, Ayla Burns, Nuthawin Charoensri, Oren Cohen, Susan Countryman, Mary Ann Cristobal, Bobbi Croy, Suzanne Dale, Hrushikesh Deshmukh, Amanda Douglas, Vincent Drouillon, Marcia Eisenberg, Howard Engler, Rama Ghatti, Prashant Gupta, Susan Hicks, Jake Humphrey, Lax Iyer, Lisa Pfefferle, Manoj Jain, Matthew Robinson, Mohan Kolli, Brian Krueger, Tim Kuphal, Stanley Letovsky, Michael Levandoski, Craig Lukasik, Jonathan Meltzer, Brian Norvell, Mindy Nye, Scott Parker, Christos Petropoulos, John Pruitt, Steven Ragan, Scott Ryan, Mike Sapeta, Jana Schroth, Suresh Babu Selvaraju, Goran Stevovic, Amanda Suchanek, Andrea Throop, Lyndon Tilson, Thomas Urban, Joe Voshell, Kimberly Wagner, Jonathan Williams, Mary Williamson, Qian Zeng, Tricia Zwiefelhofer, Clinton R. Paden, Duncan MacCannell |
| EPI_ISL_3326071 | TGen North | TGen North | Jolene Bowers, Heather Centner, Brett Van Tassel, Chris French, Hayley Yaglom, Darrin Lemmer, Dave Engelthaler |
| EPI_ISL_3326072 | Laboratory Corporation of America | Centers for Disease Control and Prevention Division of Viral Diseases, Pathogen Discovery | Dakota Howard, Dhwani Batra, Peter W. Cook, Kara Moser, Adrian Paskey, Jason Caravas, Benjamin Rambo-Martin, Shatavia Morrison, Christopher Gulvick, Scott Sammons, Yvette Unoarumhi, Darlene Wagner, Matthew Schmerer, Minoo Agarwal, Eyad Almasri, Debbie Boles, Ayla Burns, Nuthawin Charoensri, Oren Cohen, Susan Countryman, Mary Ann Cristobal, Bobbi Croy, Suzanne Dale, Hrushikesh Deshmukh, Amanda Douglas, Vincent Drouillon, Marcia Eisenberg, Howard Engler, Rama Ghatti, Prashant Gupta, Susan Hicks, Jake Humphrey, Lax Iyer, Lisa Pfefferle, Manoj Jain, Matthew Robinson, Mohan Kolli, Brian Krueger, Tim Kuphal, Stanley Letovsky, Michael Levandoski, Craig Lukasik, Jonathan Meltzer, Brian Norvell, Mindy Nye, Scott Parker, Christos Petropoulos, John Pruitt, Steven Ragan, Scott Ryan, Mike Sapeta, Jana Schroth, Suresh Babu Selvaraju, Goran Stevovic, Amanda Suchanek, Andrea Throop, Lyndon Tilson, Thomas Urban, Joe Voshell, Kimberly Wagner, Jonathan Williams, Mary Williamson, Qian Zeng, Tricia Zwiefelhofer, Clinton R. Paden, Duncan MacCannell |
| EPI_ISL_3326073 | TGen North | TGen North | Jolene Bowers, Heather Centner, Brett Van Tassel, Chris French, Hayley Yaglom, Darrin Lemmer, Dave Engelthaler |
| EPI_ISL_3326074, EPI_ISL_3326076, EPI_ISL_3326077 | Laboratory Corporation of America | Centers for Disease Control and Prevention Division of Viral Diseases, Pathogen Discovery | Dakota Howard, Dhwani Batra, Peter W. Cook, Kara Moser, Adrian Paskey, Jason Caravas, Benjamin Rambo-Martin, Shatavia Morrison, Christopher Gulvick, Scott Sammons, Yvette Unoarumhi, Darlene Wagner, Matthew Schmerer, Minoo Agarwal, Eyad Almasri, Debbie Boles, Ayla Burns, Nuthawin Charoensri, Oren Cohen, Susan Countryman, Mary Ann Cristobal, Bobbi Croy, Suzanne Dale, Hrushikesh Deshmukh, Amanda Douglas, Vincent Drouillon, Marcia Eisenberg, Howard Engler, Rama Ghatti, Prashant Gupta, Susan Hicks, Jake Humphrey, Lax Iyer, Lisa Pfefferle, Manoj Jain, Matthew Robinson, Mohan Kolli, Brian Krueger, Tim Kuphal, Stanley Letovsky, Michael Levandoski, Craig Lukasik, Jonathan Meltzer, Brian Norvell, Mindy Nye, Scott Parker, Christos Petropoulos, John Pruitt, Steven Ragan, Scott Ryan, Mike Sapeta, Jana Schroth, Suresh Babu Selvaraju, Goran Stevovic, Amanda Suchanek, Andrea Throop, Lyndon Tilson, Thomas Urban, Joe Voshell, Kimberly Wagner, Jonathan Williams, Mary Williamson, Qian Zeng, Tricia Zwiefelhofer, Clinton R. Paden, Duncan MacCannell |
| EPI_ISL_3326078 | TGen North | TGen North | Jolene Bowers, Heather Centner, Brett Van Tassel, Chris French, Hayley Yaglom, Darrin Lemmer, Dave Engelthaler |
| EPI_ISL_3326079 | Laboratory Corporation of America | Centers for Disease Control and Prevention Division of Viral Diseases, Pathogen Discovery | Dakota Howard, Dhwani Batra, Peter W. Cook, Kara Moser, Adrian Paskey, Jason Caravas, Benjamin Rambo-Martin, Shatavia Morrison, Christopher Gulvick, Scott Sammons, Yvette Unoarumhi, Darlene Wagner, Matthew Schmerer, Minoo Agarwal, Eyad Almasri, Debbie Boles, Ayla Burns, Nuthawin Charoensri, Oren Cohen, Susan Countryman, Mary Ann Cristobal, Bobbi Croy, Suzanne Dale, Hrushikesh Deshmukh, Amanda Douglas, Vincent Drouillon, Marcia Eisenberg, Howard Engler, Rama Ghatti, Prashant Gupta, Susan Hicks, Jake Humphrey, Lax Iyer, Lisa Pfefferle, Manoj Jain, Matthew Robinson, Mohan Kolli, Brian Krueger, Tim Kuphal, Stanley Letovsky, Michael Levandoski, Craig Lukasik, Jonathan Meltzer, Brian Norvell, Mindy Nye, Scott Parker, Christos Petropoulos, John Pruitt, Steven Ragan, Scott Ryan, Mike Sapeta, Jana Schroth, Suresh Babu Selvaraju, Goran Stevovic, Amanda Suchanek, Andrea Throop, Lyndon Tilson, Thomas Urban, Joe Voshell, Kimberly Wagner, Jonathan Williams, Mary Williamson, Qian Zeng, Tricia Zwiefelhofer, Clinton R. Paden, Duncan MacCannell |
| EPI_ISL_3326080 | TGen North | TGen North | Jolene Bowers, Heather Centner, Brett Van Tassel, Chris French, Hayley Yaglom, Darrin Lemmer, Dave Engelthaler |
| EPI_ISL_3326081, EPI_ISL_3326082 | Laboratory Corporation of America | Centers for Disease Control and Prevention Division of Viral Diseases, Pathogen Discovery | Dakota Howard, Dhwani Batra, Peter W. Cook, Kara Moser, Adrian Paskey, Jason Caravas, Benjamin Rambo-Martin, Shatavia Morrison, Christopher Gulvick, Scott Sammons, Yvette Unoarumhi, Darlene Wagner, Matthew Schmerer, Minoo Agarwal, Eyad Almasri, Debbie Boles, Ayla Burns, Nuthawin Charoensri, Oren Cohen, Susan Countryman, Mary Ann Cristobal, Bobbi Croy, Suzanne Dale, Hrushikesh Deshmukh, Amanda Douglas, Vincent Drouillon, Marcia Eisenberg, Howard Engler, Rama Ghatti, Prashant Gupta, Susan Hicks, Jake Humphrey, Lax Iyer, Lisa Pfefferle, Manoj Jain, Matthew Robinson, Mohan Kolli, Brian Krueger, Tim Kuphal, Stanley Letovsky, Michael Levandoski, Craig Lukasik, Jonathan Meltzer, Brian Norvell, Mindy Nye, Scott Parker, Christos Petropoulos, John Pruitt, Steven Ragan, Scott Ryan, Mike Sapeta, Jana Schroth, Suresh Babu Selvaraju, Goran Stevovic, Amanda Suchanek, Andrea Throop, Lyndon Tilson, Thomas Urban, Joe Voshell, Kimberly Wagner, Jonathan Williams, Mary Williamson, Qian Zeng, Tricia Zwiefelhofer, Clinton R. Paden, Duncan MacCannell |
| EPI_ISL_3326083 | TGen North | TGen North | Jolene Bowers, Heather Centner, Brett Van Tassel, Chris French, Hayley Yaglom, Darrin Lemmer, Dave Engelthaler |
| EPI_ISL_3326084 | Laboratory Corporation of America | Centers for Disease Control and Prevention Division of Viral Diseases, Pathogen Discovery | Dakota Howard, Dhwani Batra, Peter W. Cook, Kara Moser, Adrian Paskey, Jason Caravas, Benjamin Rambo-Martin, Shatavia Morrison, Christopher Gulvick, Scott Sammons, Yvette Unoarumhi, Darlene Wagner, Matthew Schmerer, Minoo Agarwal, Eyad Almasri, Debbie Boles, Ayla Burns, Nuthawin Charoensri, Oren Cohen, Susan Countryman, Mary Ann Cristobal, Bobbi Croy, Suzanne Dale, Hrushikesh Deshmukh, Amanda Douglas, Vincent Drouillon, Marcia Eisenberg, Howard Engler, Rama Ghatti, Prashant Gupta, Susan Hicks, Jake Humphrey, Lax Iyer, Lisa Pfefferle, Manoj Jain, Matthew Robinson, Mohan Kolli, Brian Krueger, Tim Kuphal, Stanley Letovsky, Michael Levandoski, Craig Lukasik, Jonathan Meltzer, Brian Norvell, Mindy Nye, Scott Parker, Christos Petropoulos, John Pruitt, Steven Ragan, Scott Ryan, Mike Sapeta, Jana Schroth, Suresh Babu Selvaraju, Goran Stevovic, Amanda Suchanek, Andrea Throop, Lyndon Tilson, Thomas Urban, Joe Voshell, Kimberly Wagner, Jonathan Williams, Mary Williamson, Qian Zeng, Tricia Zwiefelhofer, Clinton R. Paden, Duncan MacCannell |
| EPI_ISL_3326085 | TGen North | TGen North | Jolene Bowers, Heather Centner, Brett Van Tassel, Chris French, Hayley Yaglom, Darrin Lemmer, Dave Engelthaler |
| EPI_ISL_3326086 | Laboratory Corporation of America | Centers for Disease Control and Prevention Division of Viral Diseases, Pathogen Discovery | Dakota Howard, Dhwani Batra, Peter W. Cook, Kara Moser, Adrian Paskey, Jason Caravas, Benjamin Rambo-Martin, Shatavia Morrison, Christopher Gulvick, Scott Sammons, Yvette Unoarumhi, Darlene Wagner, Matthew Schmerer, Minoo Agarwal, Eyad Almasri, Debbie Boles, Ayla Burns, Nuthawin Charoensri, Oren Cohen, Susan Countryman, Mary Ann Cristobal, Bobbi Croy, Suzanne Dale, Hrushikesh Deshmukh, Amanda Douglas, Vincent Drouillon, Marcia Eisenberg, Howard Engler, Rama Ghatti, Prashant Gupta, Susan Hicks, Jake Humphrey, Lax Iyer, Lisa Pfefferle, Manoj Jain, Matthew Robinson, Mohan Kolli, Brian Krueger, Tim Kuphal, Stanley Letovsky, Michael Levandoski, Craig Lukasik, Jonathan Meltzer, Brian Norvell, Mindy Nye, Scott Parker, Christos Petropoulos, John Pruitt, Steven Ragan, Scott Ryan, Mike Sapeta, Jana Schroth, Suresh Babu Selvaraju, Goran Stevovic, Amanda Suchanek, Andrea Throop, Lyndon Tilson, Thomas Urban, Joe Voshell, Kimberly Wagner, Jonathan Williams, Mary Williamson, Qian Zeng, Tricia Zwiefelhofer, Clinton R. Paden, Duncan MacCannell |
| EPI_ISL_3326087 | TGen North | TGen North | Jolene Bowers, Heather Centner, Brett Van Tassel, Chris French, Hayley Yaglom, Darrin Lemmer, Dave Engelthaler |
| EPI_ISL_3326088 | Laboratory Corporation of America | Centers for Disease Control and Prevention Division of Viral Diseases, Pathogen Discovery | Dakota Howard, Dhwani Batra, Peter W. Cook, Kara Moser, Adrian Paskey, Jason Caravas, Benjamin Rambo-Martin, Shatavia Morrison, Christopher Gulvick, Scott Sammons, Yvette Unoarumhi, Darlene Wagner, Matthew Schmerer, Minoo Agarwal, Eyad Almasri, Debbie Boles, Ayla Burns, Nuthawin Charoensri, Oren Cohen, Susan Countryman, Mary Ann Cristobal, Bobbi Croy, Suzanne Dale, Hrushikesh Deshmukh, Amanda Douglas, Vincent Drouillon, Marcia Eisenberg, Howard Engler, Rama Ghatti, Prashant Gupta, Susan Hicks, Jake Humphrey, Lax Iyer, Lisa Pfefferle, Manoj Jain, Matthew Robinson, Mohan Kolli, Brian Krueger, Tim Kuphal, Stanley Letovsky, Michael Levandoski, Craig Lukasik, Jonathan Meltzer, Brian Norvell, Mindy Nye, Scott Parker, Christos Petropoulos, John Pruitt, Steven Ragan, Scott Ryan, Mike Sapeta, Jana Schroth, Suresh Babu Selvaraju, Goran Stevovic, Amanda Suchanek, Andrea Throop, Lyndon Tilson, Thomas Urban, Joe Voshell, Kimberly Wagner, Jonathan Williams, Mary Williamson, Qian Zeng, Tricia Zwiefelhofer, Clinton R. Paden, Duncan MacCannell |
| EPI_ISL_3326089 | TGen North | TGen North | Jolene Bowers, Heather Centner, Brett Van Tassel, Chris French, Hayley Yaglom, Darrin Lemmer, Dave Engelthaler |
| EPI_ISL_3326091 | Laboratory Corporation of America | Centers for Disease Control and Prevention Division of Viral Diseases, Pathogen Discovery | Dakota Howard, Dhwani Batra, Peter W. Cook, Kara Moser, Adrian Paskey, Jason Caravas, Benjamin Rambo-Martin, Shatavia Morrison, Christopher Gulvick, Scott Sammons, Yvette Unoarumhi, Darlene Wagner, Matthew Schmerer, Minoo Agarwal, Eyad Almasri, Debbie Boles, Ayla Burns, Nuthawin Charoensri, Oren Cohen, Susan Countryman, Mary Ann Cristobal, Bobbi Croy, Suzanne Dale, Hrushikesh Deshmukh, Amanda Douglas, Vincent Drouillon, Marcia Eisenberg, Howard Engler, Rama Ghatti, Prashant Gupta, Susan Hicks, Jake Humphrey, Lax Iyer, Lisa Pfefferle, Manoj Jain, Matthew Robinson, Mohan Kolli, Brian Krueger, Tim Kuphal, Stanley Letovsky, Michael Levandoski, Craig Lukasik, Jonathan Meltzer, Brian Norvell, Mindy Nye, Scott Parker, Christos Petropoulos, John Pruitt, Steven Ragan, Scott Ryan, Mike Sapeta, Jana Schroth, Suresh Babu Selvaraju, Goran Stevovic, Amanda Suchanek, Andrea Throop, Lyndon Tilson, Thomas Urban, Joe Voshell, Kimberly Wagner, Jonathan Williams, Mary Williamson, Qian Zeng, Tricia Zwiefelhofer, Clinton R. Paden, Duncan MacCannell |
| EPI_ISL_3326092 | TGen North | TGen North | Jolene Bowers, Heather Centner, Brett Van Tassel, Chris French, Hayley Yaglom, Darrin Lemmer, Dave Engelthaler |
| EPI_ISL_3326093, EPI_ISL_3326095 | Laboratory Corporation of America | Centers for Disease Control and Prevention Division of Viral Diseases, Pathogen Discovery | Dakota Howard, Dhwani Batra, Peter W. Cook, Kara Moser, Adrian Paskey, Jason Caravas, Benjamin Rambo-Martin, Shatavia Morrison, Christopher Gulvick, Scott Sammons, Yvette Unoarumhi, Darlene Wagner, Matthew Schmerer, Minoo Agarwal, Eyad Almasri, Debbie Boles, Ayla Burns, Nuthawin Charoensri, Oren Cohen, Susan Countryman, Mary Ann Cristobal, Bobbi Croy, Suzanne Dale, Hrushikesh Deshmukh, Amanda Douglas, Vincent Drouillon, Marcia Eisenberg, Howard Engler, Rama Ghatti, Prashant Gupta, Susan Hicks, Jake Humphrey, Lax Iyer, Lisa Pfefferle, Manoj Jain, Matthew Robinson, Mohan Kolli, Brian Krueger, Tim Kuphal, Stanley Letovsky, Michael Levandoski, Craig Lukasik, Jonathan Meltzer, Brian Norvell, Mindy Nye, Scott Parker, Christos Petropoulos, John Pruitt, Steven Ragan, Scott Ryan, Mike Sapeta, Jana Schroth, Suresh Babu Selvaraju, Goran Stevovic, Amanda Suchanek, Andrea Throop, Lyndon Tilson, Thomas Urban, Joe Voshell, Kimberly Wagner, Jonathan Williams, Mary Williamson, Qian Zeng, Tricia Zwiefelhofer, Clinton R. Paden, Duncan MacCannell |
| EPI_ISL_3326096 | TGen North | TGen North | Jolene Bowers, Heather Centner, Brett Van Tassel, Chris French, Hayley Yaglom, Darrin Lemmer, Dave Engelthaler |
| EPI_ISL_3326097 | Laboratory Corporation of America | Centers for Disease Control and Prevention Division of Viral Diseases, Pathogen Discovery | Dakota Howard, Dhwani Batra, Peter W. Cook, Kara Moser, Adrian Paskey, Jason Caravas, Benjamin Rambo-Martin, Shatavia Morrison, Christopher Gulvick, Scott Sammons, Yvette Unoarumhi, Darlene Wagner, Matthew Schmerer, Minoo Agarwal, Eyad Almasri, Debbie Boles, Ayla Burns, Nuthawin Charoensri, Oren Cohen, Susan Countryman, Mary Ann Cristobal, Bobbi Croy, Suzanne Dale, Hrushikesh Deshmukh, Amanda Douglas, Vincent Drouillon, Marcia Eisenberg, Howard Engler, Rama Ghatti, Prashant Gupta, Susan Hicks, Jake Humphrey, Lax Iyer, Lisa Pfefferle, Manoj Jain, Matthew Robinson, Mohan Kolli, Brian Krueger, Tim Kuphal, Stanley Letovsky, Michael Levandoski, Craig Lukasik, Jonathan Meltzer, Brian Norvell, Mindy Nye, Scott Parker, Christos Petropoulos, John Pruitt, Steven Ragan, Scott Ryan, Mike Sapeta, Jana Schroth, Suresh Babu Selvaraju, Goran Stevovic, Amanda Suchanek, Andrea Throop, Lyndon Tilson, Thomas Urban, Joe Voshell, Kimberly Wagner, Jonathan Williams, Mary Williamson, Qian Zeng, Tricia Zwiefelhofer, Clinton R. Paden, Duncan MacCannell |
| EPI_ISL_3326098 | TGen North | TGen North | Jolene Bowers, Heather Centner, Brett Van Tassel, Chris French, Hayley Yaglom, Darrin Lemmer, Dave Engelthaler |
| EPI_ISL_3326099 | Laboratory Corporation of America | Centers for Disease Control and Prevention Division of Viral Diseases, Pathogen Discovery | Dakota Howard, Dhwani Batra, Peter W. Cook, Kara Moser, Adrian Paskey, Jason Caravas, Benjamin Rambo-Martin, Shatavia Morrison, Christopher Gulvick, Scott Sammons, Yvette Unoarumhi, Darlene Wagner, Matthew Schmerer, Minoo Agarwal, Eyad Almasri, Debbie Boles, Ayla Burns, Nuthawin Charoensri, Oren Cohen, Susan Countryman, Mary Ann Cristobal, Bobbi Croy, Suzanne Dale, Hrushikesh Deshmukh, Amanda Douglas, Vincent Drouillon, Marcia Eisenberg, Howard Engler, Rama Ghatti, Prashant Gupta, Susan Hicks, Jake Humphrey, Lax Iyer, Lisa Pfefferle, Manoj Jain, Matthew Robinson, Mohan Kolli, Brian Krueger, Tim Kuphal, Stanley Letovsky, Michael Levandoski, Craig Lukasik, Jonathan Meltzer, Brian Norvell, Mindy Nye, Scott Parker, Christos Petropoulos, John Pruitt, Steven Ragan, Scott Ryan, Mike Sapeta, Jana Schroth, Suresh Babu Selvaraju, Goran Stevovic, Amanda Suchanek, Andrea Throop, Lyndon Tilson, Thomas Urban, Joe Voshell, Kimberly Wagner, Jonathan Williams, Mary Williamson, Qian Zeng, Tricia Zwiefelhofer, Clinton R. Paden, Duncan MacCannell |
| EPI_ISL_3326100 | TGen North | TGen North | Jolene Bowers, Heather Centner, Brett Van Tassel, Chris French, Hayley Yaglom, Darrin Lemmer, Dave Engelthaler |
| EPI_ISL_3326101 | Laboratory Corporation of America | Centers for Disease Control and Prevention Division of Viral Diseases, Pathogen Discovery | Dakota Howard, Dhwani Batra, Peter W. Cook, Kara Moser, Adrian Paskey, Jason Caravas, Benjamin Rambo-Martin, Shatavia Morrison, Christopher Gulvick, Scott Sammons, Yvette Unoarumhi, Darlene Wagner, Matthew Schmerer, Minoo Agarwal, Eyad Almasri, Debbie Boles, Ayla Burns, Nuthawin Charoensri, Oren Cohen, Susan Countryman, Mary Ann Cristobal, Bobbi Croy, Suzanne Dale, Hrushikesh Deshmukh, Amanda Douglas, Vincent Drouillon, Marcia Eisenberg, Howard Engler, Rama Ghatti, Prashant Gupta, Susan Hicks, Jake Humphrey, Lax Iyer, Lisa Pfefferle, Manoj Jain, Matthew Robinson, Mohan Kolli, Brian Krueger, Tim Kuphal, Stanley Letovsky, Michael Levandoski, Craig Lukasik, Jonathan Meltzer, Brian Norvell, Mindy Nye, Scott Parker, Christos Petropoulos, John Pruitt, Steven Ragan, Scott Ryan, Mike Sapeta, Jana Schroth, Suresh Babu Selvaraju, Goran Stevovic, Amanda Suchanek, Andrea Throop, Lyndon Tilson, Thomas Urban, Joe Voshell, Kimberly Wagner, Jonathan Williams, Mary Williamson, Qian Zeng, Tricia Zwiefelhofer, Clinton R. Paden, Duncan MacCannell |
| EPI_ISL_3326102 | TGen North | TGen North | Jolene Bowers, Heather Centner, Brett Van Tassel, Chris French, Hayley Yaglom, Darrin Lemmer, Dave Engelthaler |
| EPI_ISL_3326103 | Laboratory Corporation of America | Centers for Disease Control and Prevention Division of Viral Diseases, Pathogen Discovery | Dakota Howard, Dhwani Batra, Peter W. Cook, Kara Moser, Adrian Paskey, Jason Caravas, Benjamin Rambo-Martin, Shatavia Morrison, Christopher Gulvick, Scott Sammons, Yvette Unoarumhi, Darlene Wagner, Matthew Schmerer, Minoo Agarwal, Eyad Almasri, Debbie Boles, Ayla Burns, Nuthawin Charoensri, Oren Cohen, Susan Countryman, Mary Ann Cristobal, Bobbi Croy, Suzanne Dale, Hrushikesh Deshmukh, Amanda Douglas, Vincent Drouillon, Marcia Eisenberg, Howard |

| | | | |
|---|---|---|---|
| | | | Engler, Rama Ghatti, Prashant Gupta, Susan Hicks, Jake Humphrey, Lax Iyer, Lisa Pfefferle, Manoj Jain, Matthew Robinson, Mohan Kolli, Brian Krueger, Tim Kuphal, Stanley Letovsky, Michael Levandoski, Craig Lukasik, Jonathan Meltzer, Brian Norvell, Mindy Nye, Scott Parker, Christos Petropoulos, John Pruitt, Steven Ragan, Scott Ryan, Mike Sapeta, Jana Schroth, Suresh Babu Selvaraju, Goran Stevovic, Amanda Suchanek, Andrea Throop, Lyndon Tilson, Thomas Urban, Joe Voshell, Kimberly Wagner, Jonathan Williams, Mary Williamson, Qian Zeng, Tricia Zwiefelhofer, Clinton R. Paden, Duncan MacCannell |
| EPI_ISL_3326104 | TGen North | TGen North | Jolene Bowers, Heather Centner, Brett Van Tassel, Chris French, Hayley Yaglom, Darrin Lemmer, Dave Engelthaler |
| EPI_ISL_3326105, EPI_ISL_3326106, EPI_ISL_3326108 | Laboratory Corporation of America | Centers for Disease Control and Prevention Division of Viral Diseases, Pathogen Discovery | Dakota Howard, Dhwani Batra, Peter W. Cook, Kara Moser, Adrian Paskey, Jason Caravas, Benjamin Rambo-Martin, Shatavia Morrison, Christopher Gulvick, Scott Sammons, Yvette Unoarumhi, Darlene Wagner, Matthew Schmerer, Minoo Agarwal, Eyad Almasri, Debbie Boles, Ayla Burns, Nuthawin Charoensri, Oren Cohen, Susan Countryman, Mary Ann Cristobal, Bobbi Croy, Suzanne Dale, Hrushikesh Deshmukh, Amanda Douglas, Vincent Drouillon, Marcia Eisenberg, Howard Engler, Rama Ghatti, Prashant Gupta, Susan Hicks, Jake Humphrey, Lax Iyer, Lisa Pfefferle, Manoj Jain, Matthew Robinson, Mohan Kolli, Brian Krueger, Tim Kuphal, Stanley Letovsky, Michael Levandoski, Craig Lukasik, Jonathan Meltzer, Brian Norvell, Mindy Nye, Scott Parker, Christos Petropoulos, John Pruitt, Steven Ragan, Scott Ryan, Mike Sapeta, Jana Schroth, Suresh Babu Selvaraju, Goran Stevovic, Amanda Suchanek, Andrea Throop, Lyndon Tilson, Thomas Urban, Joe Voshell, Kimberly Wagner, Jonathan Williams, Mary Williamson, Qian Zeng, Tricia Zwiefelhofer, Clinton R. Paden, Duncan MacCannell |
| EPI_ISL_3326109 | TGen North | TGen North | Jolene Bowers, Heather Centner, Brett Van Tassel, Chris French, Hayley Yaglom, Darrin Lemmer, Dave Engelthaler |
| EPI_ISL_3326110 | Hospital Universitari Vall d'Hebron - Vall d'Hebron Institut de Recerca | Hospital Universitari Vall d'Hebron - Vall d'Hebron Institut de Recerca | Cristina Andrés, Maria Piñana, Alejandra González-Sánchez, Damir Garcia-Cehic, Ariadna Rando, Juliana Esperalba, Maria Gema Codina, Carla Castillo, Karen García, Rodrigo Vásquez, Maria Carmen Martin, Tomàs Pumarola, Josep Quer, Andrés Antón |
| EPI_ISL_3326111 | Laboratory Corporation of America | Centers for Disease Control and Prevention Division of Viral Diseases, Pathogen Discovery | Dakota Howard, Dhwani Batra, Peter W. Cook, Kara Moser, Adrian Paskey, Jason Caravas, Benjamin Rambo-Martin, Shatavia Morrison, Christopher Gulvick, Scott Sammons, Yvette Unoarumhi, Darlene Wagner, Matthew Schmerer, Minoo Agarwal, Eyad Almasri, Debbie Boles, Ayla Burns, Nuthawin Charoensri, Oren Cohen, Susan Countryman, Mary Ann Cristobal, Bobbi Croy, Suzanne Dale, Hrushikesh Deshmukh, Amanda Douglas, Vincent Drouillon, Marcia Eisenberg, Howard Engler, Rama Ghatti, Prashant Gupta, Susan Hicks, Jake Humphrey, Lax Iyer, Lisa Pfefferle, Manoj Jain, Matthew Robinson, Mohan Kolli, Brian Krueger, Tim Kuphal, Stanley Letovsky, Michael Levandoski, Craig Lukasik, Jonathan Meltzer, Brian Norvell, Mindy Nye, Scott Parker, Christos Petropoulos, John Pruitt, Steven Ragan, Scott Ryan, Mike Sapeta, Jana Schroth, Suresh Babu Selvaraju, Goran Stevovic, Amanda Suchanek, Andrea Throop, Lyndon Tilson, Thomas Urban, Joe Voshell, Kimberly Wagner, Jonathan Williams, Mary Williamson, Qian Zeng, Tricia Zwiefelhofer, Clinton R. Paden, Duncan MacCannell |
| EPI_ISL_3326112 | TGen North | TGen North | Jolene Bowers, Heather Centner, Brett Van Tassel, Chris French, Hayley Yaglom, Darrin Lemmer, Dave Engelthaler |
| EPI_ISL_3326113, EPI_ISL_3326114 | Laboratory Corporation of America | Centers for Disease Control and Prevention Division of Viral Diseases, Pathogen Discovery | Dakota Howard, Dhwani Batra, Peter W. Cook, Kara Moser, Adrian Paskey, Jason Caravas, Benjamin Rambo-Martin, Shatavia Morrison, Christopher Gulvick, Scott Sammons, Yvette Unoarumhi, Darlene Wagner, Matthew Schmerer, Minoo Agarwal, Eyad Almasri, Debbie Boles, Ayla Burns, Nuthawin Charoensri, Oren Cohen, Susan Countryman, Mary Ann Cristobal, Bobbi Croy, Suzanne Dale, Hrushikesh Deshmukh, Amanda Douglas, Vincent Drouillon, Marcia Eisenberg, Howard Engler, Rama Ghatti, Prashant Gupta, Susan Hicks, Jake Humphrey, Lax Iyer, Lisa Pfefferle, Manoj Jain, Matthew Robinson, Mohan Kolli, Brian Krueger, Tim Kuphal, Stanley Letovsky, Michael Levandoski, Craig Lukasik, Jonathan Meltzer, Brian Norvell, Mindy Nye, Scott Parker, Christos Petropoulos, John Pruitt, Steven Ragan, Scott Ryan, Mike Sapeta, Jana Schroth, Suresh Babu Selvaraju, Goran Stevovic, Amanda Suchanek, Andrea Throop, Lyndon Tilson, Thomas Urban, Joe Voshell, Kimberly Wagner, Jonathan Williams, Mary Williamson, Qian Zeng, Tricia Zwiefelhofer, Clinton R. Paden, Duncan MacCannell |
| EPI_ISL_3326115 | TGen North | TGen North | Jolene Bowers, Heather Centner, Brett Van Tassel, Chris French, Hayley Yaglom, Darrin Lemmer, Dave Engelthaler |
| EPI_ISL_3326116 | Laboratory Corporation of America | Centers for Disease Control and Prevention Division of Viral Diseases, Pathogen Discovery | Dakota Howard, Dhwani Batra, Peter W. Cook, Kara Moser, Adrian Paskey, Jason Caravas, Benjamin Rambo-Martin, Shatavia Morrison, Christopher Gulvick, Scott Sammons, Yvette Unoarumhi, Darlene Wagner, Matthew Schmerer, Minoo Agarwal, Eyad Almasri, Debbie Boles, Ayla Burns, Nuthawin Charoensri, Oren Cohen, Susan Countryman, Mary Ann Cristobal, Bobbi Croy, Suzanne Dale, Hrushikesh Deshmukh, Amanda Douglas, Vincent Drouillon, Marcia Eisenberg, Howard Engler, Rama Ghatti, Prashant Gupta, Susan Hicks, Jake Humphrey, Lax Iyer, Lisa Pfefferle, Manoj Jain, Matthew Robinson, Mohan Kolli, Brian Krueger, Tim Kuphal, Stanley Letovsky, Michael Levandoski, Craig Lukasik, Jonathan Meltzer, Brian Norvell, Mindy Nye, Scott Parker, Christos Petropoulos, John Pruitt, Steven Ragan, Scott Ryan, Mike Sapeta, Jana Schroth, Suresh Babu Selvaraju, Goran Stevovic, Amanda Suchanek, Andrea Throop, Lyndon Tilson, Thomas Urban, Joe Voshell, Kimberly Wagner, Jonathan Williams, Mary Williamson, Qian Zeng, Tricia Zwiefelhofer, Clinton R. Paden, Duncan MacCannell |
| EPI_ISL_3326117 | TGen North | TGen North | Jolene Bowers, Heather Centner, Brett Van Tassel, Chris French, Hayley Yaglom, Darrin Lemmer, Dave Engelthaler |
| EPI_ISL_3326118 | Laboratory Corporation of America | Centers for Disease Control and Prevention Division of Viral Diseases, Pathogen Discovery | Dakota Howard, Dhwani Batra, Peter W. Cook, Kara Moser, Adrian Paskey, Jason Caravas, Benjamin Rambo-Martin, Shatavia Morrison, Christopher Gulvick, Scott Sammons, Yvette Unoarumhi, Darlene Wagner, Matthew Schmerer, Minoo Agarwal, Eyad Almasri, Debbie Boles, Ayla Burns, Nuthawin Charoensri, Oren Cohen, Susan Countryman, Mary Ann Cristobal, Bobbi Croy, Suzanne Dale, Hrushikesh Deshmukh, Amanda Douglas, Vincent Drouillon, Marcia Eisenberg, Howard Engler, Rama Ghatti, Prashant Gupta, Susan Hicks, Jake Humphrey, Lax Iyer, Lisa Pfefferle, Manoj Jain, Matthew Robinson, Mohan Kolli, Brian Krueger, Tim Kuphal, Stanley Letovsky, Michael Levandoski, Craig Lukasik, Jonathan Meltzer, Brian Norvell, Mindy Nye, Scott Parker, Christos Petropoulos, John Pruitt, Steven Ragan, Scott Ryan, Mike Sapeta, Jana Schroth, Suresh Babu Selvaraju, Goran Stevovic, Amanda Suchanek, Andrea Throop, Lyndon Tilson, Thomas Urban, Joe Voshell, Kimberly Wagner, Jonathan Williams, Mary Williamson, Qian Zeng, Tricia Zwiefelhofer, Clinton R. Paden, Duncan MacCannell |
| EPI_ISL_3326119 | Hospital Universitari Vall d'Hebron - Vall d'Hebron Institut de Recerca | Hospital Universitari Vall d'Hebron - Vall d'Hebron Institut de Recerca | Cristina Andrés, Maria Piñana, Alejandra González-Sánchez, Damir Garcia-Cehic, Ariadna Rando, Juliana Esperalba, Maria Gema Codina, Carla Castillo, Karen García, Rodrigo Vásquez, Maria Carmen Martin, Tomàs Pumarola, Josep Quer, Andrés Antón |
| EPI_ISL_3326120 | TGen North | TGen North | Jolene Bowers, Heather Centner, Brett Van Tassel, Chris French, Hayley Yaglom, Darrin Lemmer, Dave Engelthaler |
| EPI_ISL_3326121 | Laboratory Corporation of America | Centers for Disease Control and Prevention Division of Viral Diseases, Pathogen Discovery | Dakota Howard, Dhwani Batra, Peter W. Cook, Kara Moser, Adrian Paskey, Jason Caravas, Benjamin Rambo-Martin, Shatavia Morrison, Christopher Gulvick, Scott Sammons, Yvette Unoarumhi, Darlene Wagner, Matthew Schmerer, Minoo Agarwal, Eyad Almasri, Debbie Boles, Ayla Burns, Nuthawin Charoensri, Oren Cohen, Susan Countryman, Mary Ann Cristobal, Bobbi Croy, Suzanne Dale, Hrushikesh Deshmukh, Amanda Douglas, Vincent Drouillon, Marcia Eisenberg, Howard Engler, Rama Ghatti, Prashant Gupta, Susan Hicks, Jake Humphrey, Lax Iyer, Lisa Pfefferle, Manoj Jain, Matthew Robinson, Mohan Kolli, Brian Krueger, Tim Kuphal, Stanley Letovsky, Michael Levandoski, Craig Lukasik, Jonathan Meltzer, Brian Norvell, Mindy Nye, Scott Parker, Christos Petropoulos, John Pruitt, Steven Ragan, Scott Ryan, Mike Sapeta, Jana Schroth, Suresh Babu Selvaraju, Goran Stevovic, Amanda Suchanek, Andrea Throop, Lyndon Tilson, Thomas Urban, Joe Voshell, Kimberly Wagner, Jonathan Williams, Mary Williamson, Qian Zeng, Tricia Zwiefelhofer, Clinton R. Paden, Duncan MacCannell |
| EPI_ISL_3326122, EPI_ISL_3326123 | Hospital Universitari Vall d'Hebron - Vall d'Hebron Institut de Recerca | Hospital Universitari Vall d'Hebron - Vall d'Hebron Institut de Recerca | Cristina Andrés, Maria Piñana, Alejandra González-Sánchez, Damir Garcia-Cehic, Ariadna Rando, Juliana Esperalba, Maria Gema Codina, Carla Castillo, Karen García, Rodrigo Vásquez, Maria Carmen Martin, Tomàs Pumarola, Josep Quer, Andrés Antón |
| EPI_ISL_3326125 | Laboratory Corporation of America | Centers for Disease Control and Prevention Division of Viral Diseases, Pathogen Discovery | Dakota Howard, Dhwani Batra, Peter W. Cook, Kara Moser, Adrian Paskey, Jason Caravas, Benjamin Rambo-Martin, Shatavia Morrison, Christopher Gulvick, Scott Sammons, Yvette Unoarumhi, Darlene Wagner, Matthew Schmerer, Minoo Agarwal, Eyad Almasri, Debbie Boles, Ayla Burns, Nuthawin Charoensri, Oren Cohen, Susan Countryman, Mary Ann Cristobal, Bobbi Croy, Suzanne Dale, Hrushikesh Deshmukh, Amanda Douglas, Vincent Drouillon, Marcia Eisenberg, Howard Engler, Rama Ghatti, Prashant Gupta, Susan Hicks, Jake Humphrey, Lax Iyer, Lisa Pfefferle, Manoj Jain, Matthew Robinson, Mohan Kolli, Brian Krueger, Tim Kuphal, Stanley Letovsky, Michael Levandoski, Craig Lukasik, Jonathan Meltzer, Brian Norvell, Mindy Nye, Scott Parker, Christos Petropoulos, John Pruitt, Steven Ragan, Scott Ryan, Mike Sapeta, Jana Schroth, Suresh Babu Selvaraju, Goran Stevovic, Amanda Suchanek, Andrea Throop, Lyndon Tilson, Thomas Urban, Joe Voshell, Kimberly Wagner, Jonathan Williams, Mary Williamson, Qian Zeng, Tricia Zwiefelhofer, Clinton R. Paden, Duncan MacCannell |
| EPI_ISL_3326126, EPI_ISL_3326127 | Hospital Universitari Vall d'Hebron - Vall d'Hebron Institut de Recerca | Hospital Universitari Vall d'Hebron - Vall d'Hebron Institut de Recerca | Cristina Andrés, Maria Piñana, Alejandra González-Sánchez, Damir Garcia-Cehic, Ariadna Rando, Juliana Esperalba, Maria Gema Codina, Carla Castillo, Karen García, Rodrigo Vásquez, Maria Carmen Martin, Tomàs Pumarola, Josep Quer, Andrés Antón |
| EPI_ISL_3326128 | Laboratory Corporation of America | Centers for Disease Control and Prevention Division of Viral Diseases, Pathogen Discovery | Dakota Howard, Dhwani Batra, Peter W. Cook, Kara Moser, Adrian Paskey, Jason Caravas, Benjamin Rambo-Martin, Shatavia Morrison, Christopher Gulvick, Scott Sammons, Yvette Unoarumhi, Darlene Wagner, Matthew Schmerer, Minoo Agarwal, Eyad Almasri, Debbie Boles, Ayla Burns, Nuthawin Charoensri, Oren Cohen, Susan Countryman, Mary Ann Cristobal, Bobbi Croy, Suzanne Dale, Hrushikesh Deshmukh, Amanda Douglas, Vincent Drouillon, Marcia Eisenberg, Howard Engler, Rama Ghatti, Prashant Gupta, Susan Hicks, Jake Humphrey, Lax Iyer, Lisa Pfefferle, Manoj Jain, Matthew Robinson, Mohan Kolli, Brian Krueger, Tim Kuphal, Stanley Letovsky, Michael Levandoski, Craig Lukasik, Jonathan Meltzer, Brian Norvell, Mindy Nye, Scott Parker, Christos Petropoulos, John Pruitt, Steven Ragan, Scott Ryan, Mike Sapeta, Jana Schroth, Suresh Babu Selvaraju, Goran Stevovic, Amanda Suchanek, Andrea Throop, Lyndon Tilson, Thomas Urban, Joe Voshell, Kimberly Wagner, Jonathan Williams, Mary Williamson, Qian Zeng, Tricia Zwiefelhofer, Clinton R. Paden, Duncan MacCannell |
| EPI_ISL_3326129 | TGen North | TGen North | Jolene Bowers, Heather Centner, Brett Van Tassel, Chris French, Hayley Yaglom, Darrin Lemmer, Dave Engelthaler |
| EPI_ISL_3326130 | Hospital Universitari Vall d'Hebron - Vall d'Hebron Institut de Recerca | Hospital Universitari Vall d'Hebron - Vall d'Hebron Institut de Recerca | Cristina Andrés, Maria Piñana, Alejandra González-Sánchez, Damir Garcia-Cehic, Ariadna Rando, Juliana Esperalba, Maria Gema Codina, Carla Castillo, Karen García, Rodrigo Vásquez, Maria Carmen Martin, Tomàs Pumarola, Josep Quer, Andrés Antón |
| EPI_ISL_3326131 | Laboratory Corporation of America | Centers for Disease Control and Prevention Division of Viral Diseases, Pathogen Discovery | Dakota Howard, Dhwani Batra, Peter W. Cook, Kara Moser, Adrian Paskey, Jason Caravas, Benjamin Rambo-Martin, Shatavia Morrison, Christopher Gulvick, Scott Sammons, Yvette Unoarumhi, Darlene Wagner, Matthew Schmerer, Minoo Agarwal, Eyad Almasri, Debbie Boles, Ayla Burns, Nuthawin Charoensri, Oren Cohen, Susan Countryman, Mary Ann Cristobal, Bobbi Croy, Suzanne Dale, Hrushikesh Deshmukh, Amanda Douglas, Vincent Drouillon, Marcia Eisenberg, Howard Engler, Rama Ghatti, Prashant Gupta, Susan Hicks, Jake Humphrey, Lax Iyer, Lisa Pfefferle, Manoj Jain, Matthew Robinson, Mohan Kolli, Brian Krueger, Tim Kuphal, Stanley Letovsky, Michael Levandoski, Craig Lukasik, Jonathan Meltzer, Brian Norvell, Mindy Nye, Scott Parker, Christos Petropoulos, John Pruitt, Steven Ragan, Scott Ryan, Mike Sapeta, Jana Schroth, Suresh Babu Selvaraju, Goran Stevovic, Amanda Suchanek, Andrea Throop, Lyndon Tilson, Thomas Urban, Joe Voshell, Kimberly Wagner, Jonathan Williams, Mary Williamson, Qian Zeng, Tricia Zwiefelhofer, Clinton R. Paden, Duncan MacCannell |
| EPI_ISL_3326132 | TGen North | TGen North | Jolene Bowers, Heather Centner, Brett Van Tassel, Chris French, Hayley Yaglom, Darrin Lemmer, Dave Engelthaler |
| EPI_ISL_3326133, EPI_ISL_3326134 | Hospital Universitari Vall d'Hebron - Vall d'Hebron Institut de Recerca | Hospital Universitari Vall d'Hebron - Vall d'Hebron Institut de Recerca | Cristina Andrés, Maria Piñana, Alejandra González-Sánchez, Damir Garcia-Cehic, Ariadna Rando, Juliana Esperalba, Maria Gema Codina, Carla Castillo, Karen García, Rodrigo Vásquez, Maria Carmen Martin, Tomàs Pumarola, Josep Quer, Andrés Antón |
| EPI_ISL_3326135 | Laboratory Corporation of America | Centers for Disease Control and Prevention Division of Viral Diseases, Pathogen Discovery | Dakota Howard, Dhwani Batra, Peter W. Cook, Kara Moser, Adrian Paskey, Jason Caravas, Benjamin Rambo-Martin, Shatavia Morrison, Christopher Gulvick, Scott Sammons, Yvette Unoarumhi, Darlene Wagner, Matthew Schmerer, Minoo Agarwal, Eyad Almasri, Debbie Boles, Ayla Burns, Nuthawin Charoensri, Oren Cohen, Susan Countryman, Mary Ann Cristobal, Bobbi Croy, Suzanne Dale, Hrushikesh Deshmukh, Amanda Douglas, Vincent Drouillon, Marcia Eisenberg, Howard Engler, Rama Ghatti, Prashant Gupta, Susan Hicks, Jake Humphrey, Lax Iyer, Lisa Pfefferle, Manoj Jain, Matthew Robinson, Mohan Kolli, Brian Krueger, Tim Kuphal, Stanley Letovsky, Michael Levandoski, Craig Lukasik, Jonathan Meltzer, Brian Norvell, Mindy Nye, Scott Parker, Christos Petropoulos, John Pruitt, Steven Ragan, Scott Ryan, Mike Sapeta, Jana Schroth, Suresh Babu Selvaraju, Goran Stevovic, Amanda Suchanek, Andrea Throop, Lyndon Tilson, Thomas Urban, Joe Voshell, Kimberly Wagner, Jonathan Williams, Mary Williamson, Qian Zeng, Tricia Zwiefelhofer, Clinton R. Paden, Duncan MacCannell |
| EPI_ISL_3326137 | Hospital Universitari Vall d'Hebron - Vall d'Hebron Institut de Recerca | Hospital Universitari Vall d'Hebron - Vall d'Hebron Institut de Recerca | Cristina Andrés, Maria Piñana, Alejandra González-Sánchez, Damir Garcia-Cehic, Ariadna Rando, Juliana Esperalba, Maria Gema Codina, Carla Castillo, Karen García, Rodrigo Vásquez, Maria Carmen Martin, Tomàs Pumarola, Josep Quer, Andrés Antón |
| EPI_ISL_3326138 | Laboratory Corporation of America | Centers for Disease Control and Prevention Division of Viral Diseases, Pathogen Discovery | Dakota Howard, Dhwani Batra, Peter W. Cook, Kara Moser, Adrian Paskey, Jason Caravas, Benjamin Rambo-Martin, Shatavia Morrison, Christopher Gulvick, Scott Sammons, Yvette Unoarumhi, Darlene Wagner, Matthew Schmerer, Minoo Agarwal, Eyad Almasri, Debbie Boles, Ayla Burns, Nuthawin Charoensri, Oren Cohen, Susan Countryman, Mary Ann Cristobal, Bobbi Croy, Suzanne Dale, Hrushikesh Deshmukh, Amanda Douglas, Vincent Drouillon, Marcia Eisenberg, Howard Engler, Rama Ghatti, Prashant Gupta, Susan Hicks, Jake Humphrey, Lax Iyer, Lisa Pfefferle, Manoj Jain, Matthew Robinson, Mohan Kolli, Brian Krueger, Tim Kuphal, Stanley Letovsky, Michael Levandoski, Craig Lukasik, Jonathan Meltzer, Brian Norvell, Mindy Nye, Scott Parker, Christos Petropoulos, John Pruitt, Steven Ragan, Scott Ryan, Mike Sapeta, Jana Schroth, Suresh Babu Selvaraju, Goran Stevovic, Amanda Suchanek, Andrea Throop, Lyndon Tilson, Thomas Urban, Joe Voshell, Kimberly Wagner, Jonathan Williams, Mary Williamson, Qian Zeng, Tricia Zwiefelhofer, Clinton R. Paden, Duncan MacCannell |
| EPI_ISL_3326139, EPI_ISL_3326140 | Hospital Universitari Vall d'Hebron - Vall d'Hebron Institut de Recerca | Hospital Universitari Vall d'Hebron - Vall d'Hebron Institut de Recerca | Cristina Andrés, Maria Piñana, Alejandra González-Sánchez, Damir Garcia-Cehic, Ariadna Rando, Juliana Esperalba, Maria Gema Codina, Carla Castillo, Karen García, Rodrigo Vásquez, Maria Carmen Martin, Tomàs Pumarola, Josep Quer, Andrés Antón |
| EPI_ISL_3326141 | TGen North | TGen North | Jolene Bowers, Heather Centner, Brett Van Tassel, Chris French, Hayley Yaglom, Darrin Lemmer, Dave Engelthaler |
| EPI_ISL_3326142 | Laboratory Corporation of America | Centers for Disease Control and Prevention Division of Viral Diseases, Pathogen Discovery | Dakota Howard, Dhwani Batra, Peter W. Cook, Kara Moser, Adrian Paskey, Jason Caravas, Benjamin Rambo-Martin, Shatavia Morrison, Christopher Gulvick, Scott Sammons, Yvette Unoarumhi, Darlene Wagner, Matthew Schmerer, Minoo Agarwal, Eyad Almasri, Debbie Boles, Ayla Burns, Nuthawin Charoensri, Oren Cohen, Susan Countryman, Mary Ann Cristobal, Bobbi Croy, Suzanne Dale, Hrushikesh Deshmukh, Amanda Douglas, Vincent Drouillon, Marcia Eisenberg, Howard Engler, Rama Ghatti, Prashant Gupta, Susan Hicks, Jake Humphrey, Lax Iyer, Lisa Pfefferle, Manoj Jain, Matthew Robinson, Mohan Kolli, Brian Krueger, Tim Kuphal, Stanley Letovsky, Michael Levandoski, Craig Lukasik, Jonathan Meltzer, Brian Norvell, Mindy Nye, Scott Parker, Christos Petropoulos, John Pruitt, Steven Ragan, Scott Ryan, Mike Sapeta, Jana Schroth, Suresh Babu Selvaraju, Goran Stevovic, Amanda Suchanek, Andrea Throop, Lyndon Tilson, Thomas Urban, Joe Voshell, Kimberly Wagner, Jonathan Williams, Mary Williamson, Qian Zeng, Tricia Zwiefelhofer, Clinton R. Paden, Duncan MacCannell |
| EPI_ISL_3326143, EPI_ISL_3326144 | Hospital Universitari Vall d'Hebron - Vall d'Hebron Institut de Recerca | Hospital Universitari Vall d'Hebron - Vall d'Hebron Institut de Recerca | Cristina Andrés, Maria Piñana, Alejandra González-Sánchez, Damir Garcia-Cehic, Ariadna Rando, Juliana Esperalba, Maria Gema Codina, Carla Castillo, Karen García, Rodrigo Vásquez, Maria Carmen Martin, Tomàs Pumarola, Josep Quer, Andrés Antón |
| EPI_ISL_3326145 | TGen North | TGen North | Jolene Bowers, Heather Centner, Brett Van Tassel, Chris French, Hayley Yaglom, Darrin Lemmer, Dave Engelthaler |
| EPI_ISL_3326146 | Laboratory Corporation of America | Centers for Disease Control and Prevention Division of Viral Diseases, Pathogen Discovery | Dakota Howard, Dhwani Batra, Peter W. Cook, Kara Moser, Adrian Paskey, Jason Caravas, Benjamin Rambo-Martin, Shatavia Morrison, Christopher Gulvick, Scott Sammons, Yvette Unoarumhi, Darlene Wagner, Matthew Schmerer, Minoo Agarwal, Eyad Almasri, Debbie Boles, Ayla Burns, Nuthawin Charoensri, Oren Cohen, Susan Countryman, Mary Ann Cristobal, Bobbi Croy, Suzanne Dale, Hrushikesh Deshmukh, Amanda Douglas, Vincent Drouillon, Marcia Eisenberg, Howard Engler, Rama Ghatti, Prashant Gupta, Susan Hicks, Jake Humphrey, Lax Iyer, Lisa Pfefferle, Manoj Jain, Matthew Robinson, Mohan Kolli, Brian Krueger, Tim Kuphal, Stanley Letovsky, Michael Levandoski, Craig Lukasik, Jonathan Meltzer, Brian Norvell, Mindy Nye, Scott Parker, Christos Petropoulos, John Pruitt, Steven Ragan, Scott Ryan, Mike Sapeta, Jana Schroth, Suresh Babu Selvaraju, Goran Stevovic, Amanda Suchanek, Andrea Throop, Lyndon Tilson, Thomas Urban, Joe Voshell, Kimberly Wagner, Jonathan Williams, Mary Williamson, Qian Zeng, Tricia Zwiefelhofer, Clinton R. Paden, Duncan MacCannell |

| | | | |
|---|---|---|---|
| EPI_ISL_3326147 | Hospital Universitari Vall d'Hebron - Vall d'Hebron Institut de Recerca | Hospital Universitari Vall d'Hebron - Vall d'Hebron Institut de Recerca | Cristina Andrés, Maria Piñana, Alejandra González-Sánchez, Damir Garcia-Cehic, Ariadna Rando, Juliana Esperalba, Maria Gema Codina, Carla Castillo, Karen García, Rodrigo Vásquez, Maria Carmen Martin, Tomàs Pumarola, Josep Quer, Andrés Antón |
| EPI_ISL_3326148 | Laboratory Corporation of America | Centers for Disease Control and Prevention Division of Viral Diseases, Pathogen Discovery | Dakota Howard, Dhwani Batra, Peter W. Cook, Kara Moser, Adrian Paskey, Jason Caravas, Benjamin Rambo-Martin, Shatavia Morrison, Christopher Gulvick, Scott Sammons, Yvette Unoarumhi, Darlene Wagner, Matthew Schmerer, Minoo Agarwal, Eyad Almasri, Debbie Boles, Ayla Burns, Nuthawin Charoensri, Oren Cohen, Susan Countryman, Mary Ann Cristobal, Bobbi Croy, Suzanne Dale, Hrushikesh Deshmukh, Amanda Douglas, Vincent Drouillon, Marcia Eisenberg, Howard Engler, Rama Ghatti, Prashant Gupta, Susan Hicks, Jake Humphrey, Lax Iyer, Lisa Pfefferle, Manoj Jain, Matthew Robinson, Mohan Kolli, Brian Krueger, Tim Kuphal, Stanley Letovsky, Michael Levandoski, Craig Lukasik, Jonathan Meltzer, Brian Norvell, Mindy Nye, Scott Parker, Christos Petropoulos, John Pruitt, Steven Ragan, Scott Ryan, Mike Sapeta, Jana Schroth, Suresh Babu Selvaraju, Goran Stevovic, Amanda Suchanek, Andrew Throop, Lyndon Tilson, Thomas Urban, Joe Voshell, Kimberly Wagner, Jonathan Williams, Mary Williamson, Qian Zeng, Tricia Zwiefelhofer, Clinton R. Paden, Duncan MacCannell |
| EPI_ISL_3326149, EPI_ISL_3326151 | Hospital Universitari Vall d'Hebron - Vall d'Hebron Institut de Recerca | Hospital Universitari Vall d'Hebron - Vall d'Hebron Institut de Recerca | Cristina Andrés, Maria Piñana, Alejandra González-Sánchez, Damir Garcia-Cehic, Ariadna Rando, Juliana Esperalba, Maria Gema Codina, Carla Castillo, Karen García, Rodrigo Vásquez, Maria Carmen Martin, Tomàs Pumarola, Josep Quer, Andrés Antón |
| EPI_ISL_3326152 | Laboratory Corporation of America | Centers for Disease Control and Prevention Division of Viral Diseases, Pathogen Discovery | Dakota Howard, Dhwani Batra, Peter W. Cook, Kara Moser, Adrian Paskey, Jason Caravas, Benjamin Rambo-Martin, Shatavia Morrison, Christopher Gulvick, Scott Sammons, Yvette Unoarumhi, Darlene Wagner, Matthew Schmerer, Minoo Agarwal, Eyad Almasri, Debbie Boles, Ayla Burns, Nuthawin Charoensri, Oren Cohen, Susan Countryman, Mary Ann Cristobal, Bobbi Croy, Suzanne Dale, Hrushikesh Deshmukh, Amanda Douglas, Vincent Drouillon, Marcia Eisenberg, Howard Engler, Rama Ghatti, Prashant Gupta, Susan Hicks, Jake Humphrey, Lax Iyer, Lisa Pfefferle, Manoj Jain, Matthew Robinson, Mohan Kolli, Brian Krueger, Tim Kuphal, Stanley Letovsky, Michael Levandoski, Craig Lukasik, Jonathan Meltzer, Brian Norvell, Mindy Nye, Scott Parker, Christos Petropoulos, John Pruitt, Steven Ragan, Scott Ryan, Mike Sapeta, Jana Schroth, Suresh Babu Selvaraju, Goran Stevovic, Amanda Suchanek, Andrew Throop, Lyndon Tilson, Thomas Urban, Joe Voshell, Kimberly Wagner, Jonathan Williams, Mary Williamson, Qian Zeng, Tricia Zwiefelhofer, Clinton R. Paden, Duncan MacCannell |
| EPI_ISL_3326153 | TGen North | TGen North | Jolene Bowers, Heather Centner, Brett Van Tassel, Chris French, Hayley Yaglom, Darrin Lemmer, Dave Engelthaler |
| EPI_ISL_3326154, EPI_ISL_3326155 | Hospital Universitari Vall d'Hebron - Vall d'Hebron Institut de Recerca | Hospital Universitari Vall d'Hebron - Vall d'Hebron Institut de Recerca | Cristina Andrés, Maria Piñana, Alejandra González-Sánchez, Damir Garcia-Cehic, Ariadna Rando, Juliana Esperalba, Maria Gema Codina, Carla Castillo, Karen García, Rodrigo Vásquez, Maria Carmen Martin, Tomàs Pumarola, Josep Quer, Andrés Antón |
| EPI_ISL_3326156 | Laboratory Corporation of America | Centers for Disease Control and Prevention Division of Viral Diseases, Pathogen Discovery | Dakota Howard, Dhwani Batra, Peter W. Cook, Kara Moser, Adrian Paskey, Jason Caravas, Benjamin Rambo-Martin, Shatavia Morrison, Christopher Gulvick, Scott Sammons, Yvette Unoarumhi, Darlene Wagner, Matthew Schmerer, Minoo Agarwal, Eyad Almasri, Debbie Boles, Ayla Burns, Nuthawin Charoensri, Oren Cohen, Susan Countryman, Mary Ann Cristobal, Bobbi Croy, Suzanne Dale, Hrushikesh Deshmukh, Amanda Douglas, Vincent Drouillon, Marcia Eisenberg, Howard Engler, Rama Ghatti, Prashant Gupta, Susan Hicks, Jake Humphrey, Lax Iyer, Lisa Pfefferle, Manoj Jain, Matthew Robinson, Mohan Kolli, Brian Krueger, Tim Kuphal, Stanley Letovsky, Michael Levandoski, Craig Lukasik, Jonathan Meltzer, Brian Norvell, Mindy Nye, Scott Parker, Christos Petropoulos, John Pruitt, Steven Ragan, Scott Ryan, Mike Sapeta, Jana Schroth, Suresh Babu Selvaraju, Goran Stevovic, Amanda Suchanek, Andrew Throop, Lyndon Tilson, Thomas Urban, Joe Voshell, Kimberly Wagner, Jonathan Williams, Mary Williamson, Qian Zeng, Tricia Zwiefelhofer, Clinton R. Paden, Duncan MacCannell |
| EPI_ISL_3326158 | Hospital Universitari Vall d'Hebron - Vall d'Hebron Institut de Recerca | Hospital Universitari Vall d'Hebron - Vall d'Hebron Institut de Recerca | Cristina Andrés, Maria Piñana, Alejandra González-Sánchez, Damir Garcia-Cehic, Ariadna Rando, Juliana Esperalba, Maria Gema Codina, Carla Castillo, Karen García, Rodrigo Vásquez, Maria Carmen Martin, Tomàs Pumarola, Josep Quer, Andrés Antón |
| EPI_ISL_3326159 | Laboratory Corporation of America | Centers for Disease Control and Prevention Division of Viral Diseases, Pathogen Discovery | Dakota Howard, Dhwani Batra, Peter W. Cook, Kara Moser, Adrian Paskey, Jason Caravas, Benjamin Rambo-Martin, Shatavia Morrison, Christopher Gulvick, Scott Sammons, Yvette Unoarumhi, Darlene Wagner, Matthew Schmerer, Minoo Agarwal, Eyad Almasri, Debbie Boles, Ayla Burns, Nuthawin Charoensri, Oren Cohen, Susan Countryman, Mary Ann Cristobal, Bobbi Croy, Suzanne Dale, Hrushikesh Deshmukh, Amanda Douglas, Vincent Drouillon, Marcia Eisenberg, Howard Engler, Rama Ghatti, Prashant Gupta, Susan Hicks, Jake Humphrey, Lax Iyer, Lisa Pfefferle, Manoj Jain, Matthew Robinson, Mohan Kolli, Brian Krueger, Tim Kuphal, Stanley Letovsky, Michael Levandoski, Craig Lukasik, Jonathan Meltzer, Brian Norvell, Mindy Nye, Scott Parker, Christos Petropoulos, John Pruitt, Steven Ragan, Scott Ryan, Mike Sapeta, Jana Schroth, Suresh Babu Selvaraju, Goran Stevovic, Amanda Suchanek, Andrew Throop, Lyndon Tilson, Thomas Urban, Joe Voshell, Kimberly Wagner, Jonathan Williams, Mary Williamson, Qian Zeng, Tricia Zwiefelhofer, Clinton R. Paden, Duncan MacCannell |
| EPI_ISL_3326160 | Hospital Universitari Vall d'Hebron - Vall d'Hebron Institut de Recerca | Hospital Universitari Vall d'Hebron - Vall d'Hebron Institut de Recerca | Cristina Andrés, Maria Piñana, Alejandra González-Sánchez, Damir Garcia-Cehic, Ariadna Rando, Juliana Esperalba, Maria Gema Codina, Carla Castillo, Karen García, Rodrigo Vásquez, Maria Carmen Martin, Tomàs Pumarola, Josep Quer, Andrés Antón |
| EPI_ISL_3326161 | TGen North | TGen North | Jolene Bowers, Heather Centner, Brett Van Tassel, Chris French, Hayley Yaglom, Darrin Lemmer, Dave Engelthaler |
| EPI_ISL_3326162 | Hospital Universitari Vall d'Hebron - Vall d'Hebron Institut de Recerca | Hospital Universitari Vall d'Hebron - Vall d'Hebron Institut de Recerca | Cristina Andrés, Maria Piñana, Alejandra González-Sánchez, Damir Garcia-Cehic, Ariadna Rando, Juliana Esperalba, Maria Gema Codina, Carla Castillo, Karen García, Rodrigo Vásquez, Maria Carmen Martin, Tomàs Pumarola, Josep Quer, Andrés Antón |
| EPI_ISL_3326163 | Laboratory Corporation of America | Centers for Disease Control and Prevention Division of Viral Diseases, Pathogen Discovery | Dakota Howard, Dhwani Batra, Peter W. Cook, Kara Moser, Adrian Paskey, Jason Caravas, Benjamin Rambo-Martin, Shatavia Morrison, Christopher Gulvick, Scott Sammons, Yvette Unoarumhi, Darlene Wagner, Matthew Schmerer, Minoo Agarwal, Eyad Almasri, Debbie Boles, Ayla Burns, Nuthawin Charoensri, Oren Cohen, Susan Countryman, Mary Ann Cristobal, Bobbi Croy, Suzanne Dale, Hrushikesh Deshmukh, Amanda Douglas, Vincent Drouillon, Marcia Eisenberg, Howard Engler, Rama Ghatti, Prashant Gupta, Susan Hicks, Jake Humphrey, Lax Iyer, Lisa Pfefferle, Manoj Jain, Matthew Robinson, Mohan Kolli, Brian Krueger, Tim Kuphal, Stanley Letovsky, Michael Levandoski, Craig Lukasik, Jonathan Meltzer, Brian Norvell, Mindy Nye, Scott Parker, Christos Petropoulos, John Pruitt, Steven Ragan, Scott Ryan, Mike Sapeta, Jana Schroth, Suresh Babu Selvaraju, Goran Stevovic, Amanda Suchanek, Andrew Throop, Lyndon Tilson, Thomas Urban, Joe Voshell, Kimberly Wagner, Jonathan Williams, Mary Williamson, Qian Zeng, Tricia Zwiefelhofer, Clinton R. Paden, Duncan MacCannell |
| EPI_ISL_3326164 | Hospital Universitari Vall d'Hebron - Vall d'Hebron Institut de Recerca | Hospital Universitari Vall d'Hebron - Vall d'Hebron Institut de Recerca | Cristina Andrés, Maria Piñana, Alejandra González-Sánchez, Damir Garcia-Cehic, Ariadna Rando, Juliana Esperalba, Maria Gema Codina, Carla Castillo, Karen García, Rodrigo Vásquez, Maria Carmen Martin, Tomàs Pumarola, Josep Quer, Andrés Antón |
| EPI_ISL_3326166 | Laboratory Corporation of America | Centers for Disease Control and Prevention Division of Viral Diseases, Pathogen Discovery | Dakota Howard, Dhwani Batra, Peter W. Cook, Kara Moser, Adrian Paskey, Jason Caravas, Benjamin Rambo-Martin, Shatavia Morrison, Christopher Gulvick, Scott Sammons, Yvette Unoarumhi, Darlene Wagner, Matthew Schmerer, Minoo Agarwal, Eyad Almasri, Debbie Boles, Ayla Burns, Nuthawin Charoensri, Oren Cohen, Susan Countryman, Mary Ann Cristobal, Bobbi Croy, Suzanne Dale, Hrushikesh Deshmukh, Amanda Douglas, Vincent Drouillon, Marcia Eisenberg, Howard Engler, Rama Ghatti, Prashant Gupta, Susan Hicks, Jake Humphrey, Lax Iyer, Lisa Pfefferle, Manoj Jain, Matthew Robinson, Mohan Kolli, Brian Krueger, Tim Kuphal, Stanley Letovsky, Michael Levandoski, Craig Lukasik, Jonathan Meltzer, Brian Norvell, Mindy Nye, Scott Parker, Christos Petropoulos, John Pruitt, Steven Ragan, Scott Ryan, Mike Sapeta, Jana Schroth, Suresh Babu Selvaraju, Goran Stevovic, Amanda Suchanek, Andrew Throop, Lyndon Tilson, Thomas Urban, Joe Voshell, Kimberly Wagner, Jonathan Williams, Mary Williamson, Qian Zeng, Tricia Zwiefelhofer, Clinton R. Paden, Duncan MacCannell |
| EPI_ISL_3326167, EPI_ISL_3326168 | Hospital Universitari Vall d'Hebron - Vall d'Hebron Institut de Recerca | Hospital Universitari Vall d'Hebron - Vall d'Hebron Institut de Recerca | Cristina Andrés, Maria Piñana, Alejandra González-Sánchez, Damir Garcia-Cehic, Ariadna Rando, Juliana Esperalba, Maria Gema Codina, Carla Castillo, Karen García, Rodrigo Vásquez, Maria Carmen Martin, Tomàs Pumarola, Josep Quer, Andrés Antón |
| EPI_ISL_3326169 | Laboratory Corporation of America | Centers for Disease Control and Prevention Division of Viral Diseases, Pathogen Discovery | Dakota Howard, Dhwani Batra, Peter W. Cook, Kara Moser, Adrian Paskey, Jason Caravas, Benjamin Rambo-Martin, Shatavia Morrison, Christopher Gulvick, Scott Sammons, Yvette Unoarumhi, Darlene Wagner, Matthew Schmerer, Minoo Agarwal, Eyad Almasri, Debbie Boles, Ayla Burns, Nuthawin Charoensri, Oren Cohen, Susan Countryman, Mary Ann Cristobal, Bobbi Croy, Suzanne Dale, Hrushikesh Deshmukh, Amanda Douglas, Vincent Drouillon, Marcia Eisenberg, Howard Engler, Rama Ghatti, Prashant Gupta, Susan Hicks, Jake Humphrey, Lax Iyer, Lisa Pfefferle, Manoj Jain, Matthew Robinson, Mohan Kolli, Brian Krueger, Tim Kuphal, Stanley Letovsky, Michael Levandoski, Craig Lukasik, Jonathan Meltzer, Brian Norvell, Mindy Nye, Scott Parker, Christos Petropoulos, John Pruitt, Steven Ragan, Scott Ryan, Mike Sapeta, Jana Schroth, Suresh Babu Selvaraju, Goran Stevovic, Amanda Suchanek, Andrew Throop, Lyndon Tilson, Thomas Urban, Joe Voshell, Kimberly Wagner, Jonathan Williams, Mary Williamson, Qian Zeng, Tricia Zwiefelhofer, Clinton R. Paden, Duncan MacCannell |
| EPI_ISL_3326171, EPI_ISL_3326172 | Hospital Universitari Vall d'Hebron - Vall d'Hebron Institut de Recerca | Hospital Universitari Vall d'Hebron - Vall d'Hebron Institut de Recerca | Cristina Andrés, Maria Piñana, Alejandra González-Sánchez, Damir Garcia-Cehic, Ariadna Rando, Juliana Esperalba, Maria Gema Codina, Carla Castillo, Karen García, Rodrigo Vásquez, Maria Carmen Martin, Tomàs Pumarola, Josep Quer, Andrés Antón |
| EPI_ISL_3326173 | Laboratory Corporation of America | Centers for Disease Control and Prevention Division of Viral Diseases, Pathogen Discovery | Dakota Howard, Dhwani Batra, Peter W. Cook, Kara Moser, Adrian Paskey, Jason Caravas, Benjamin Rambo-Martin, Shatavia Morrison, Christopher Gulvick, Scott Sammons, Yvette Unoarumhi, Darlene Wagner, Matthew Schmerer, Minoo Agarwal, Eyad Almasri, Debbie Boles, Ayla Burns, Nuthawin Charoensri, Oren Cohen, Susan Countryman, Mary Ann Cristobal, Bobbi Croy, Suzanne Dale, Hrushikesh Deshmukh, Amanda Douglas, Vincent Drouillon, Marcia Eisenberg, Howard Engler, Rama Ghatti, Prashant Gupta, Susan Hicks, Jake Humphrey, Lax Iyer, Lisa Pfefferle, Manoj Jain, Matthew Robinson, Mohan Kolli, Brian Krueger, Tim Kuphal, Stanley Letovsky, Michael Levandoski, Craig Lukasik, Jonathan Meltzer, Brian Norvell, Mindy Nye, Scott Parker, Christos Petropoulos, John Pruitt, Steven Ragan, Scott Ryan, Mike Sapeta, Jana Schroth, Suresh Babu Selvaraju, Goran Stevovic, Amanda Suchanek, Andrew Throop, Lyndon Tilson, Thomas Urban, Joe Voshell, Kimberly Wagner, Jonathan Williams, Mary Williamson, Qian Zeng, Tricia Zwiefelhofer, Clinton R. Paden, Duncan MacCannell |
| EPI_ISL_3326174 | TGen North | TGen North | Jolene Bowers, Heather Centner, Brett Van Tassel, Chris French, Hayley Yaglom, Darrin Lemmer, Dave Engelthaler |
| EPI_ISL_3326175 | Hospital Universitari Vall d'Hebron - Vall d'Hebron Institut de Recerca | Hospital Universitari Vall d'Hebron - Vall d'Hebron Institut de Recerca | Cristina Andrés, Maria Piñana, Alejandra González-Sánchez, Damir Garcia-Cehic, Ariadna Rando, Juliana Esperalba, Maria Gema Codina, Carla Castillo, Karen García, Rodrigo Vásquez, Maria Carmen Martin, Tomàs Pumarola, Josep Quer, Andrés Antón |
| EPI_ISL_3326176 | Laboratory Corporation of America | Centers for Disease Control and Prevention Division of Viral Diseases, Pathogen Discovery | Dakota Howard, Dhwani Batra, Peter W. Cook, Kara Moser, Adrian Paskey, Jason Caravas, Benjamin Rambo-Martin, Shatavia Morrison, Christopher Gulvick, Scott Sammons, Yvette Unoarumhi, Darlene Wagner, Matthew Schmerer, Minoo Agarwal, Eyad Almasri, Debbie Boles, Ayla Burns, Nuthawin Charoensri, Oren Cohen, Susan Countryman, Mary Ann Cristobal, Bobbi Croy, Suzanne Dale, Hrushikesh Deshmukh, Amanda Douglas, Vincent Drouillon, Marcia Eisenberg, Howard Engler, Rama Ghatti, Prashant Gupta, Susan Hicks, Jake Humphrey, Lax Iyer, Lisa Pfefferle, Manoj Jain, Matthew Robinson, Mohan Kolli, Brian Krueger, Tim Kuphal, Stanley Letovsky, Michael Levandoski, Craig Lukasik, Jonathan Meltzer, Brian Norvell, Mindy Nye, Scott Parker, Christos Petropoulos, John Pruitt, Steven Ragan, Scott Ryan, Mike Sapeta, Jana Schroth, Suresh Babu Selvaraju, Goran Stevovic, Amanda Suchanek, Andrew Throop, Lyndon Tilson, Thomas Urban, Joe Voshell, Kimberly Wagner, Jonathan Williams, Mary Williamson, Qian Zeng, Tricia Zwiefelhofer, Clinton R. Paden, Duncan MacCannell |
| EPI_ISL_3326177, EPI_ISL_3326178 | Hospital Universitari Vall d'Hebron - Vall d'Hebron Institut de Recerca | Hospital Universitari Vall d'Hebron - Vall d'Hebron Institut de Recerca | Cristina Andrés, Maria Piñana, Alejandra González-Sánchez, Damir Garcia-Cehic, Ariadna Rando, Juliana Esperalba, Maria Gema Codina, Carla Castillo, Karen García, Rodrigo Vásquez, Maria Carmen Martin, Tomàs Pumarola, Josep Quer, Andrés Antón |
| EPI_ISL_3326180 | Laboratory Corporation of America | Centers for Disease Control and Prevention Division of Viral Diseases, Pathogen Discovery | Dakota Howard, Dhwani Batra, Peter W. Cook, Kara Moser, Adrian Paskey, Jason Caravas, Benjamin Rambo-Martin, Shatavia Morrison, Christopher Gulvick, Scott Sammons, Yvette Unoarumhi, Darlene Wagner, Matthew Schmerer, Minoo Agarwal, Eyad Almasri, Debbie Boles, Ayla Burns, Nuthawin Charoensri, Oren Cohen, Susan Countryman, Mary Ann Cristobal, Bobbi Croy, Suzanne Dale, Hrushikesh Deshmukh, Amanda Douglas, Vincent Drouillon, Marcia Eisenberg, Howard Engler, Rama Ghatti, Prashant Gupta, Susan Hicks, Jake Humphrey, Lax Iyer, Lisa Pfefferle, Manoj Jain, Matthew Robinson, Mohan Kolli, Brian Krueger, Tim Kuphal, Stanley Letovsky, Michael Levandoski, Craig Lukasik, Jonathan Meltzer, Brian Norvell, Mindy Nye, Scott Parker, Christos Petropoulos, John Pruitt, Steven Ragan, Scott Ryan, Mike Sapeta, Jana Schroth, Suresh Babu Selvaraju, Goran Stevovic, Amanda Suchanek, Andrew Throop, Lyndon Tilson, Thomas Urban, Joe Voshell, Kimberly Wagner, Jonathan Williams, Mary Williamson, Qian Zeng, Tricia Zwiefelhofer, Clinton R. Paden, Duncan MacCannell |
| EPI_ISL_3326181, EPI_ISL_3326182 | Hospital Universitari Vall d'Hebron - Vall d'Hebron Institut de Recerca | Hospital Universitari Vall d'Hebron - Vall d'Hebron Institut de Recerca | Cristina Andrés, Maria Piñana, Alejandra González-Sánchez, Damir Garcia-Cehic, Ariadna Rando, Juliana Esperalba, Maria Gema Codina, Carla Castillo, Karen García, Rodrigo Vásquez, Maria Carmen Martin, Tomàs Pumarola, Josep Quer, Andrés Antón |
| EPI_ISL_3326184 | Laboratory Corporation of America | Centers for Disease Control and Prevention Division of Viral Diseases, Pathogen Discovery | Dakota Howard, Dhwani Batra, Peter W. Cook, Kara Moser, Adrian Paskey, Jason Caravas, Benjamin Rambo-Martin, Shatavia Morrison, Christopher Gulvick, Scott Sammons, Yvette Unoarumhi, Darlene Wagner, Matthew Schmerer, Minoo Agarwal, Eyad Almasri, Debbie Boles, Ayla Burns, Nuthawin Charoensri, Oren Cohen, Susan Countryman, Mary Ann Cristobal, Bobbi Croy, Suzanne Dale, Hrushikesh Deshmukh, Amanda Douglas, Vincent Drouillon, Marcia Eisenberg, Howard Engler, Rama Ghatti, Prashant Gupta, Susan Hicks, Jake Humphrey, Lax Iyer, Lisa Pfefferle, Manoj Jain, Matthew Robinson, Mohan Kolli, Brian Krueger, Tim Kuphal, Stanley Letovsky, Michael Levandoski, Craig Lukasik, Jonathan Meltzer, Brian Norvell, Mindy Nye, Scott Parker, Christos Petropoulos, John Pruitt, Steven Ragan, Scott Ryan, Mike Sapeta, Jana Schroth, Suresh Babu Selvaraju, Goran Stevovic, Amanda Suchanek, Andrew Throop, Lyndon Tilson, Thomas Urban, Joe Voshell, Kimberly Wagner, Jonathan Williams, Mary Williamson, Qian Zeng, Tricia Zwiefelhofer, Clinton R. Paden, Duncan MacCannell |
| EPI_ISL_3326185, EPI_ISL_3326186 | Hospital Universitari Vall d'Hebron - Vall d'Hebron Institut de Recerca | Hospital Universitari Vall d'Hebron - Vall d'Hebron Institut de Recerca | Cristina Andrés, Maria Piñana, Alejandra González-Sánchez, Damir Garcia-Cehic, Ariadna Rando, Juliana Esperalba, Maria Gema Codina, Carla Castillo, Karen García, Rodrigo Vásquez, Maria Carmen Martin, Tomàs Pumarola, Josep Quer, Andrés Antón |
| EPI_ISL_3326187 | Laboratory Corporation of America | Centers for Disease Control and Prevention Division of Viral Diseases, Pathogen Discovery | Dakota Howard, Dhwani Batra, Peter W. Cook, Kara Moser, Adrian Paskey, Jason Caravas, Benjamin Rambo-Martin, Shatavia Morrison, Christopher Gulvick, Scott Sammons, Yvette Unoarumhi, Darlene Wagner, Matthew Schmerer, Minoo Agarwal, Eyad Almasri, Debbie Boles, Ayla Burns, Nuthawin Charoensri, Oren Cohen, Susan Countryman, Mary Ann Cristobal, Bobbi Croy, Suzanne Dale, Hrushikesh Deshmukh, Amanda Douglas, Vincent Drouillon, Marcia Eisenberg, Howard Engler, Rama Ghatti, Prashant Gupta, Susan Hicks, Jake Humphrey, Lax Iyer, Lisa Pfefferle, Manoj Jain, Matthew Robinson, Mohan Kolli, Brian Krueger, Tim Kuphal, Stanley Letovsky, Michael Levandoski, Craig Lukasik, Jonathan Meltzer, Brian Norvell, Mindy Nye, Scott Parker, Christos Petropoulos, John Pruitt, Steven Ragan, Scott Ryan, Mike Sapeta, Jana Schroth, Suresh Babu Selvaraju, Goran Stevovic, Amanda Suchanek, Andrew Throop, Lyndon Tilson, Thomas Urban, Joe Voshell, Kimberly Wagner, Jonathan Williams, Mary Williamson, Qian Zeng, Tricia Zwiefelhofer, Clinton R. Paden, Duncan MacCannell |
| EPI_ISL_3326189 | Hospital Universitari Vall d'Hebron - Vall d'Hebron Institut de Recerca | Hospital Universitari Vall d'Hebron - Vall d'Hebron Institut de Recerca | Cristina Andrés, Maria Piñana, Alejandra González-Sánchez, Damir Garcia-Cehic, Ariadna Rando, Juliana Esperalba, Maria Gema Codina, Carla Castillo, Karen García, Rodrigo Vásquez, Maria Carmen Martin, Tomàs Pumarola, Josep Quer, Andrés Antón |
| EPI_ISL_3326190 | Laboratory Corporation of America | Centers for Disease Control and Prevention Division of Viral Diseases, Pathogen Discovery | Dakota Howard, Dhwani Batra, Peter W. Cook, Kara Moser, Adrian Paskey, Jason Caravas, Benjamin Rambo-Martin, Shatavia Morrison, Christopher Gulvick, Scott Sammons, Yvette Unoarumhi, Darlene Wagner, Matthew Schmerer, Minoo Agarwal, Eyad Almasri, Debbie Boles, Ayla Burns, Nuthawin Charoensri, Oren Cohen, Susan Countryman, Mary Ann Cristobal, Bobbi Croy, Suzanne Dale, Hrushikesh Deshmukh, Amanda Douglas, Vincent Drouillon, Marcia Eisenberg, Howard Engler, Rama Ghatti, Prashant Gupta, Susan Hicks, Jake Humphrey, Lax Iyer, Lisa Pfefferle, Manoj Jain, Matthew Robinson, Mohan Kolli, Brian Krueger, Tim Kuphal, Stanley Letovsky, Michael Levandoski, Craig Lukasik, Jonathan Meltzer, Brian Norvell, Mindy Nye, Scott Parker, Christos Petropoulos, John Pruitt, Steven Ragan, Scott Ryan, Mike Sapeta, Jana Schroth, Suresh Babu Selvaraju, Goran Stevovic, Amanda Suchanek, Andrew Throop, Lyndon Tilson, Thomas Urban, Joe Voshell, Kimberly Wagner, Jonathan Williams, Mary Williamson, Qian Zeng, Tricia Zwiefelhofer, Clinton R. Paden, Duncan MacCannell |
| EPI_ISL_3326191 | Hospital Universitari Vall d'Hebron - Vall d'Hebron Institut de | Hospital Universitari Vall d'Hebron - Vall d'Hebron Institut de | Cristina Andrés, Maria Piñana, Alejandra González-Sánchez, Damir Garcia-Cehic, Ariadna Rando, Juliana Esperalba, Maria Gema Codina, Carla Castillo, Karen García, Rodrigo Vásquez, Maria Carmen Martin, Tomàs Pumarola, Josep Quer, Andrés |

| | | | |
|---|---|---|---|
| | Recerca | Recerca | Antón |
| EPI_ISL_3326192 | TGen North | TGen North | Jolene Bowers, Heather Centner, Brett Van Tassel, Chris French, Hayley Yaglom, Darrin Lemmer, Dave Engelthaler |
| EPI_ISL_3326193 | Hospital Universitari Vall d'Hebron - Vall d'Hebron Institut de Recerca | Hospital Universitari Vall d'Hebron - Vall d'Hebron Institut de Recerca | Cristina Andrés, Maria Piñana, Alejandra González-Sánchez, Damir Garcia-Cehic, Ariadna Rando, Juliana Esperalba, Maria Gema Codina, Carla Castillo, Karen García, Rodrigo Vásquez, Maria Carmen Martin, Tomàs Pumarola, Josep Quer, Andrés Antón |
| EPI_ISL_3326194 | Laboratory Corporation of America | Centers for Disease Control and Prevention Division of Viral Diseases, Pathogen Discovery | Dakota Howard, Dhwani Batra, Peter W. Cook, Kara Moser, Adrian Paskey, Jason Caravas, Benjamin Rambo-Martin, Shatavia Morrison, Christopher Gulvick, Scott Sammons, Yvette Unoarumhi, Darlene Wagner, Matthew Schmerer, Minoo Agarwal, Eyad Almasri, Debbie Boles, Ayla Burns, Nuthawin Charoensri, Oren Cohen, Susan Countryman, Mary Ann Cristobal, Bobbi Croy, Suzanne Dale, Hrushikesh Deshmukh, Amanda Douglas, Vincent Drouillon, Marcia Eisenberg, Howard Engler, Rama Ghatti, Prashant Gupta, Susan Hicks, Jake Humphrey, Lax Iyer, Lisa Pfefferle, Manoj Jain, Matthew Robinson, Mohan Kolli, Brian Krueger, Tim Kuphal, Stanley Letovsky, Michael Levandoski, Craig Lukasik, Jonathan Meltzer, Brian Norvell, Mindy Nye, Scott Parker, Christos Petropoulos, John Pruitt, Steven Ragan, Scott Ryan, Mike Sapeta, Jana Schroth, Suresh Babu Selvaraju, Goran Stevovic, Amanda Suchanek, Andrea Throop, Lyndon Tilson, Thomas Urban, Joe Voshell, Kimberly Wagner, Jonathan Williams, Mary Williamson, Qian Zeng, Tricia Zwiefelhofer, Clinton R. Paden, Duncan MacCannell |
| EPI_ISL_3326195 | Hospital Universitari Vall d'Hebron - Vall d'Hebron Institut de Recerca | Hospital Universitari Vall d'Hebron - Vall d'Hebron Institut de Recerca | Cristina Andrés, Maria Piñana, Alejandra González-Sánchez, Damir Garcia-Cehic, Ariadna Rando, Juliana Esperalba, Maria Gema Codina, Carla Castillo, Karen García, Rodrigo Vásquez, Maria Carmen Martin, Tomàs Pumarola, Josep Quer, Andrés Antón |
| EPI_ISL_3326196 | TGen North | TGen North | Jolene Bowers, Heather Centner, Brett Van Tassel, Chris French, Hayley Yaglom, Darrin Lemmer, Dave Engelthaler |
| EPI_ISL_3326197 | Hospital Universitari Vall d'Hebron - Vall d'Hebron Institut de Recerca | Hospital Universitari Vall d'Hebron - Vall d'Hebron Institut de Recerca | Cristina Andrés, Maria Piñana, Alejandra González-Sánchez, Damir Garcia-Cehic, Ariadna Rando, Juliana Esperalba, Maria Gema Codina, Carla Castillo, Karen García, Rodrigo Vásquez, Maria Carmen Martin, Tomàs Pumarola, Josep Quer, Andrés Antón |
| EPI_ISL_3326198 | Laboratory Corporation of America | Centers for Disease Control and Prevention Division of Viral Diseases, Pathogen Discovery | Dakota Howard, Dhwani Batra, Peter W. Cook, Kara Moser, Adrian Paskey, Jason Caravas, Benjamin Rambo-Martin, Shatavia Morrison, Christopher Gulvick, Scott Sammons, Yvette Unoarumhi, Darlene Wagner, Matthew Schmerer, Minoo Agarwal, Eyad Almasri, Debbie Boles, Ayla Burns, Nuthawin Charoensri, Oren Cohen, Susan Countryman, Mary Ann Cristobal, Bobbi Croy, Suzanne Dale, Hrushikesh Deshmukh, Amanda Douglas, Vincent Drouillon, Marcia Eisenberg, Howard Engler, Rama Ghatti, Prashant Gupta, Susan Hicks, Jake Humphrey, Lax Iyer, Lisa Pfefferle, Manoj Jain, Matthew Robinson, Mohan Kolli, Brian Krueger, Tim Kuphal, Stanley Letovsky, Michael Levandoski, Craig Lukasik, Jonathan Meltzer, Brian Norvell, Mindy Nye, Scott Parker, Christos Petropoulos, John Pruitt, Steven Ragan, Scott Ryan, Mike Sapeta, Jana Schroth, Suresh Babu Selvaraju, Goran Stevovic, Amanda Suchanek, Andrea Throop, Lyndon Tilson, Thomas Urban, Joe Voshell, Kimberly Wagner, Jonathan Williams, Mary Williamson, Qian Zeng, Tricia Zwiefelhofer, Clinton R. Paden, Duncan MacCannell |
| EPI_ISL_3326199 | TGen North | TGen North | Jolene Bowers, Heather Centner, Brett Van Tassel, Chris French, Hayley Yaglom, Darrin Lemmer, Dave Engelthaler |
| EPI_ISL_3326200 | Hospital Universitari Vall d'Hebron - Vall d'Hebron Institut de Recerca | Hospital Universitari Vall d'Hebron - Vall d'Hebron Institut de Recerca | Cristina Andrés, Maria Piñana, Alejandra González-Sánchez, Damir Garcia-Cehic, Ariadna Rando, Juliana Esperalba, Maria Gema Codina, Carla Castillo, Karen García, Rodrigo Vásquez, Maria Carmen Martin, Tomàs Pumarola, Josep Quer, Andrés Antón |
| EPI_ISL_3326201 | Laboratory Corporation of America | Centers for Disease Control and Prevention Division of Viral Diseases, Pathogen Discovery | Dakota Howard, Dhwani Batra, Peter W. Cook, Kara Moser, Adrian Paskey, Jason Caravas, Benjamin Rambo-Martin, Shatavia Morrison, Christopher Gulvick, Scott Sammons, Yvette Unoarumhi, Darlene Wagner, Matthew Schmerer, Minoo Agarwal, Eyad Almasri, Debbie Boles, Ayla Burns, Nuthawin Charoensri, Oren Cohen, Susan Countryman, Mary Ann Cristobal, Bobbi Croy, Suzanne Dale, Hrushikesh Deshmukh, Amanda Douglas, Vincent Drouillon, Marcia Eisenberg, Howard Engler, Rama Ghatti, Prashant Gupta, Susan Hicks, Jake Humphrey, Lax Iyer, Lisa Pfefferle, Manoj Jain, Matthew Robinson, Mohan Kolli, Brian Krueger, Tim Kuphal, Stanley Letovsky, Michael Levandoski, Craig Lukasik, Jonathan Meltzer, Brian Norvell, Mindy Nye, Scott Parker, Christos Petropoulos, John Pruitt, Steven Ragan, Scott Ryan, Mike Sapeta, Jana Schroth, Suresh Babu Selvaraju, Goran Stevovic, Amanda Suchanek, Andrea Throop, Lyndon Tilson, Thomas Urban, Joe Voshell, Kimberly Wagner, Jonathan Williams, Mary Williamson, Qian Zeng, Tricia Zwiefelhofer, Clinton R. Paden, Duncan MacCannell |
| EPI_ISL_3326202 | TGen North | TGen North | Jolene Bowers, Heather Centner, Brett Van Tassel, Chris French, Hayley Yaglom, Darrin Lemmer, Dave Engelthaler |
| EPI_ISL_3326203, EPI_ISL_3326204 | Hospital Universitari Vall d'Hebron - Vall d'Hebron Institut de Recerca | Hospital Universitari Vall d'Hebron - Vall d'Hebron Institut de Recerca | Cristina Andrés, Maria Piñana, Alejandra González-Sánchez, Damir Garcia-Cehic, Ariadna Rando, Juliana Esperalba, Maria Gema Codina, Carla Castillo, Karen García, Rodrigo Vásquez, Maria Carmen Martin, Tomàs Pumarola, Josep Quer, Andrés Antón |
| EPI_ISL_3326205 | Laboratory Corporation of America | Centers for Disease Control and Prevention Division of Viral Diseases, Pathogen Discovery | Dakota Howard, Dhwani Batra, Peter W. Cook, Kara Moser, Adrian Paskey, Jason Caravas, Benjamin Rambo-Martin, Shatavia Morrison, Christopher Gulvick, Scott Sammons, Yvette Unoarumhi, Darlene Wagner, Matthew Schmerer, Minoo Agarwal, Eyad Almasri, Debbie Boles, Ayla Burns, Nuthawin Charoensri, Oren Cohen, Susan Countryman, Mary Ann Cristobal, Bobbi Croy, Suzanne Dale, Hrushikesh Deshmukh, Amanda Douglas, Vincent Drouillon, Marcia Eisenberg, Howard Engler, Rama Ghatti, Prashant Gupta, Susan Hicks, Jake Humphrey, Lax Iyer, Lisa Pfefferle, Manoj Jain, Matthew Robinson, Mohan Kolli, Brian Krueger, Tim Kuphal, Stanley Letovsky, Michael Levandoski, Craig Lukasik, Jonathan Meltzer, Brian Norvell, Mindy Nye, Scott Parker, Christos Petropoulos, John Pruitt, Steven Ragan, Scott Ryan, Mike Sapeta, Jana Schroth, Suresh Babu Selvaraju, Goran Stevovic, Amanda Suchanek, Andrea Throop, Lyndon Tilson, Thomas Urban, Joe Voshell, Kimberly Wagner, Jonathan Williams, Mary Williamson, Qian Zeng, Tricia Zwiefelhofer, Clinton R. Paden, Duncan MacCannell |
| EPI_ISL_3326206 | TGen North | TGen North | Jolene Bowers, Heather Centner, Brett Van Tassel, Chris French, Hayley Yaglom, Darrin Lemmer, Dave Engelthaler |
| EPI_ISL_3326207, EPI_ISL_3326208 | Hospital Universitari Vall d'Hebron - Vall d'Hebron Institut de Recerca | Hospital Universitari Vall d'Hebron - Vall d'Hebron Institut de Recerca | Cristina Andrés, Maria Piñana, Alejandra González-Sánchez, Damir Garcia-Cehic, Ariadna Rando, Juliana Esperalba, Maria Gema Codina, Carla Castillo, Karen García, Rodrigo Vásquez, Maria Carmen Martin, Tomàs Pumarola, Josep Quer, Andrés Antón |
| EPI_ISL_3326209 | Laboratory Corporation of America | Centers for Disease Control and Prevention Division of Viral Diseases, Pathogen Discovery | Dakota Howard, Dhwani Batra, Peter W. Cook, Kara Moser, Adrian Paskey, Jason Caravas, Benjamin Rambo-Martin, Shatavia Morrison, Christopher Gulvick, Scott Sammons, Yvette Unoarumhi, Darlene Wagner, Matthew Schmerer, Minoo Agarwal, Eyad Almasri, Debbie Boles, Ayla Burns, Nuthawin Charoensri, Oren Cohen, Susan Countryman, Mary Ann Cristobal, Bobbi Croy, Suzanne Dale, Hrushikesh Deshmukh, Amanda Douglas, Vincent Drouillon, Marcia Eisenberg, Howard Engler, Rama Ghatti, Prashant Gupta, Susan Hicks, Jake Humphrey, Lax Iyer, Lisa Pfefferle, Manoj Jain, Matthew Robinson, Mohan Kolli, Brian Krueger, Tim Kuphal, Stanley Letovsky, Michael Levandoski, Craig Lukasik, Jonathan Meltzer, Brian Norvell, Mindy Nye, Scott Parker, Christos Petropoulos, John Pruitt, Steven Ragan, Scott Ryan, Mike Sapeta, Jana Schroth, Suresh Babu Selvaraju, Goran Stevovic, Amanda Suchanek, Andrea Throop, Lyndon Tilson, Thomas Urban, Joe Voshell, Kimberly Wagner, Jonathan Williams, Mary Williamson, Qian Zeng, Tricia Zwiefelhofer, Clinton R. Paden, Duncan MacCannell |
| EPI_ISL_3326210 | TGen North | TGen North | Jolene Bowers, Heather Centner, Brett Van Tassel, Chris French, Hayley Yaglom, Darrin Lemmer, Dave Engelthaler |
| EPI_ISL_3326211 | Hospital Universitari Vall d'Hebron - Vall d'Hebron Institut de Recerca | Hospital Universitari Vall d'Hebron - Vall d'Hebron Institut de Recerca | Cristina Andrés, Maria Piñana, Alejandra González-Sánchez, Damir Garcia-Cehic, Ariadna Rando, Juliana Esperalba, Maria Gema Codina, Carla Castillo, Karen García, Rodrigo Vásquez, Maria Carmen Martin, Tomàs Pumarola, Josep Quer, Andrés Antón |
| EPI_ISL_3326212 | Laboratory Corporation of America | Centers for Disease Control and Prevention Division of Viral Diseases, Pathogen Discovery | Dakota Howard, Dhwani Batra, Peter W. Cook, Kara Moser, Adrian Paskey, Jason Caravas, Benjamin Rambo-Martin, Shatavia Morrison, Christopher Gulvick, Scott Sammons, Yvette Unoarumhi, Darlene Wagner, Matthew Schmerer, Minoo Agarwal, Eyad Almasri, Debbie Boles, Ayla Burns, Nuthawin Charoensri, Oren Cohen, Susan Countryman, Mary Ann Cristobal, Bobbi Croy, Suzanne Dale, Hrushikesh Deshmukh, Amanda Douglas, Vincent Drouillon, Marcia Eisenberg, Howard Engler, Rama Ghatti, Prashant Gupta, Susan Hicks, Jake Humphrey, Lax Iyer, Lisa Pfefferle, Manoj Jain, Matthew Robinson, Mohan Kolli, Brian Krueger, Tim Kuphal, Stanley Letovsky, Michael Levandoski, Craig Lukasik, Jonathan Meltzer, Brian Norvell, Mindy Nye, Scott Parker, Christos Petropoulos, John Pruitt, Steven Ragan, Scott Ryan, Mike Sapeta, Jana Schroth, Suresh Babu Selvaraju, Goran Stevovic, Amanda Suchanek, Andrea Throop, Lyndon Tilson, Thomas Urban, Joe Voshell, Kimberly Wagner, Jonathan Williams, Mary Williamson, Qian Zeng, Tricia Zwiefelhofer, Clinton R. Paden, Duncan MacCannell |
| EPI_ISL_3326213 | Hospital Universitari Vall d'Hebron - Vall d'Hebron Institut de Recerca | Hospital Universitari Vall d'Hebron - Vall d'Hebron Institut de Recerca | Cristina Andrés, Maria Piñana, Alejandra González-Sánchez, Damir Garcia-Cehic, Ariadna Rando, Juliana Esperalba, Maria Gema Codina, Carla Castillo, Karen García, Rodrigo Vásquez, Maria Carmen Martin, Tomàs Pumarola, Josep Quer, Andrés Antón |
| EPI_ISL_3326214 | TGen North | TGen North | Jolene Bowers, Heather Centner, Brett Van Tassel, Chris French, Hayley Yaglom, Darrin Lemmer, Dave Engelthaler |
| EPI_ISL_3326215 | Hospital Universitari Vall d'Hebron - Vall d'Hebron Institut de Recerca | Hospital Universitari Vall d'Hebron - Vall d'Hebron Institut de Recerca | Cristina Andrés, Maria Piñana, Alejandra González-Sánchez, Damir Garcia-Cehic, Ariadna Rando, Juliana Esperalba, Maria Gema Codina, Carla Castillo, Karen García, Rodrigo Vásquez, Maria Carmen Martin, Tomàs Pumarola, Josep Quer, Andrés Antón |
| EPI_ISL_3326216 | Laboratory Corporation of America | Centers for Disease Control and Prevention Division of Viral Diseases, Pathogen Discovery | Dakota Howard, Dhwani Batra, Peter W. Cook, Kara Moser, Adrian Paskey, Jason Caravas, Benjamin Rambo-Martin, Shatavia Morrison, Christopher Gulvick, Scott Sammons, Yvette Unoarumhi, Darlene Wagner, Matthew Schmerer, Minoo Agarwal, Eyad Almasri, Debbie Boles, Ayla Burns, Nuthawin Charoensri, Oren Cohen, Susan Countryman, Mary Ann Cristobal, Bobbi Croy, Suzanne Dale, Hrushikesh Deshmukh, Amanda Douglas, Vincent Drouillon, Marcia Eisenberg, Howard Engler, Rama Ghatti, Prashant Gupta, Susan Hicks, Jake Humphrey, Lax Iyer, Lisa Pfefferle, Manoj Jain, Matthew Robinson, Mohan Kolli, Brian Krueger, Tim Kuphal, Stanley Letovsky, Michael Levandoski, Craig Lukasik, Jonathan Meltzer, Brian Norvell, Mindy Nye, Scott Parker, Christos Petropoulos, John Pruitt, Steven Ragan, Scott Ryan, Mike Sapeta, Jana Schroth, Suresh Babu Selvaraju, Goran Stevovic, Amanda Suchanek, Andrea Throop, Lyndon Tilson, Thomas Urban, Joe Voshell, Kimberly Wagner, Jonathan Williams, Mary Williamson, Qian Zeng, Tricia Zwiefelhofer, Clinton R. Paden, Duncan MacCannell |
| EPI_ISL_3326217 | Hospital Universitari Vall d'Hebron - Vall d'Hebron Institut de Recerca | Hospital Universitari Vall d'Hebron - Vall d'Hebron Institut de Recerca | Cristina Andrés, Maria Piñana, Alejandra González-Sánchez, Damir Garcia-Cehic, Ariadna Rando, Juliana Esperalba, Maria Gema Codina, Carla Castillo, Karen García, Rodrigo Vásquez, Maria Carmen Martin, Tomàs Pumarola, Josep Quer, Andrés Antón |
| EPI_ISL_3326218 | TGen North | TGen North | Jolene Bowers, Heather Centner, Brett Van Tassel, Chris French, Hayley Yaglom, Darrin Lemmer, Dave Engelthaler |
| EPI_ISL_3326219, EPI_ISL_3326221, EPI_ISL_3326222 | Hospital Universitari Vall d'Hebron - Vall d'Hebron Institut de Recerca | Hospital Universitari Vall d'Hebron - Vall d'Hebron Institut de Recerca | Cristina Andrés, Maria Piñana, Alejandra González-Sánchez, Damir Garcia-Cehic, Ariadna Rando, Juliana Esperalba, Maria Gema Codina, Carla Castillo, Karen García, Rodrigo Vásquez, Maria Carmen Martin, Tomàs Pumarola, Josep Quer, Andrés Antón |
| EPI_ISL_3326223 | Laboratory Corporation of America | Centers for Disease Control and Prevention Division of Viral Diseases, Pathogen Discovery | Dakota Howard, Dhwani Batra, Peter W. Cook, Kara Moser, Adrian Paskey, Jason Caravas, Benjamin Rambo-Martin, Shatavia Morrison, Christopher Gulvick, Scott Sammons, Yvette Unoarumhi, Darlene Wagner, Matthew Schmerer, Minoo Agarwal, Eyad Almasri, Debbie Boles, Ayla Burns, Nuthawin Charoensri, Oren Cohen, Susan Countryman, Mary Ann Cristobal, Bobbi Croy, Suzanne Dale, Hrushikesh Deshmukh, Amanda Douglas, Vincent Drouillon, Marcia Eisenberg, Howard Engler, Rama Ghatti, Prashant Gupta, Susan Hicks, Jake Humphrey, Lax Iyer, Lisa Pfefferle, Manoj Jain, Matthew Robinson, Mohan Kolli, Brian Krueger, Tim Kuphal, Stanley Letovsky, Michael Levandoski, Craig Lukasik, Jonathan Meltzer, Brian Norvell, Mindy Nye, Scott Parker, Christos Petropoulos, John Pruitt, Steven Ragan, Scott Ryan, Mike Sapeta, Jana Schroth, Suresh Babu Selvaraju, Goran Stevovic, Amanda Suchanek, Andrea Throop, Lyndon Tilson, Thomas Urban, Joe Voshell, Kimberly Wagner, Jonathan Williams, Mary Williamson, Qian Zeng, Tricia Zwiefelhofer, Clinton R. Paden, Duncan MacCannell |
| EPI_ISL_3326224 | TGen North | TGen North | Jolene Bowers, Heather Centner, Brett Van Tassel, Chris French, Hayley Yaglom, Darrin Lemmer, Dave Engelthaler |
| EPI_ISL_3326225 | Hospital Universitari Vall d'Hebron - Vall d'Hebron Institut de Recerca | Hospital Universitari Vall d'Hebron - Vall d'Hebron Institut de Recerca | Cristina Andrés, Maria Piñana, Alejandra González-Sánchez, Damir Garcia-Cehic, Ariadna Rando, Juliana Esperalba, Maria Gema Codina, Carla Castillo, Karen García, Rodrigo Vásquez, Maria Carmen Martin, Tomàs Pumarola, Josep Quer, Andrés Antón |
| EPI_ISL_3326226 | Laboratory Corporation of America | Centers for Disease Control and Prevention Division of Viral Diseases, Pathogen Discovery | Dakota Howard, Dhwani Batra, Peter W. Cook, Kara Moser, Adrian Paskey, Jason Caravas, Benjamin Rambo-Martin, Shatavia Morrison, Christopher Gulvick, Scott Sammons, Yvette Unoarumhi, Darlene Wagner, Matthew Schmerer, Minoo Agarwal, Eyad Almasri, Debbie Boles, Ayla Burns, Nuthawin Charoensri, Oren Cohen, Susan Countryman, Mary Ann Cristobal, Bobbi Croy, Suzanne Dale, Hrushikesh Deshmukh, Amanda Douglas, Vincent Drouillon, Marcia Eisenberg, Howard Engler, Rama Ghatti, Prashant Gupta, Susan Hicks, Jake Humphrey, Lax Iyer, Lisa Pfefferle, Manoj Jain, Matthew Robinson, Mohan Kolli, Brian Krueger, Tim Kuphal, Stanley Letovsky, Michael Levandoski, Craig Lukasik, Jonathan Meltzer, Brian Norvell, Mindy Nye, Scott Parker, Christos Petropoulos, John Pruitt, Steven Ragan, Scott Ryan, Mike Sapeta, Jana Schroth, Suresh Babu Selvaraju, Goran Stevovic, Amanda Suchanek, Andrea Throop, Lyndon Tilson, Thomas Urban, Joe Voshell, Kimberly Wagner, Jonathan Williams, Mary Williamson, Qian Zeng, Tricia Zwiefelhofer, Clinton R. Paden, Duncan MacCannell |
| EPI_ISL_3326227, EPI_ISL_3326229 | Hospital Universitari Vall d'Hebron - Vall d'Hebron Institut de Recerca | Hospital Universitari Vall d'Hebron - Vall d'Hebron Institut de Recerca | Cristina Andrés, Maria Piñana, Alejandra González-Sánchez, Damir Garcia-Cehic, Ariadna Rando, Juliana Esperalba, Maria Gema Codina, Carla Castillo, Karen García, Rodrigo Vásquez, Maria Carmen Martin, Tomàs Pumarola, Josep Quer, Andrés Antón |
| EPI_ISL_3326230 | Laboratory Corporation of America | Centers for Disease Control and Prevention Division of Viral Diseases, Pathogen Discovery | Dakota Howard, Dhwani Batra, Peter W. Cook, Kara Moser, Adrian Paskey, Jason Caravas, Benjamin Rambo-Martin, Shatavia Morrison, Christopher Gulvick, Scott Sammons, Yvette Unoarumhi, Darlene Wagner, Matthew Schmerer, Minoo Agarwal, Eyad Almasri, Debbie Boles, Ayla Burns, Nuthawin Charoensri, Oren Cohen, Susan Countryman, Mary Ann Cristobal, Bobbi Croy, Suzanne Dale, Hrushikesh Deshmukh, Amanda Douglas, Vincent Drouillon, Marcia Eisenberg, Howard Engler, Rama Ghatti, Prashant Gupta, Susan Hicks, Jake Humphrey, Lax Iyer, Lisa Pfefferle, Manoj Jain, Matthew Robinson, Mohan Kolli, Brian Krueger, Tim Kuphal, Stanley Letovsky, Michael Levandoski, Craig Lukasik, Jonathan Meltzer, Brian Norvell, Mindy Nye, Scott Parker, Christos Petropoulos, John Pruitt, Steven Ragan, Scott Ryan, Mike Sapeta, Jana Schroth, Suresh Babu Selvaraju, Goran Stevovic, Amanda Suchanek, Andrea Throop, Lyndon Tilson, Thomas Urban, Joe Voshell, Kimberly Wagner, Jonathan Williams, Mary Williamson, Qian Zeng, Tricia Zwiefelhofer, Clinton R. Paden, Duncan MacCannell |
| EPI_ISL_3326231 | Hospital Universitari Vall d'Hebron - Vall d'Hebron Institut de Recerca | Hospital Universitari Vall d'Hebron - Vall d'Hebron Institut de Recerca | Cristina Andrés, Maria Piñana, Alejandra González-Sánchez, Damir Garcia-Cehic, Ariadna Rando, Juliana Esperalba, Maria Gema Codina, Carla Castillo, Karen García, Rodrigo Vásquez, Maria Carmen Martin, Tomàs Pumarola, Josep Quer, Andrés Antón |
| EPI_ISL_3326232 | TGen North | TGen North | Jolene Bowers, Heather Centner, Brett Van Tassel, Chris French, Hayley Yaglom, Darrin Lemmer, Dave Engelthaler |
| EPI_ISL_3326233 | Laboratory Corporation of America | Centers for Disease Control and Prevention Division of Viral Diseases, Pathogen Discovery | Dakota Howard, Dhwani Batra, Peter W. Cook, Kara Moser, Adrian Paskey, Jason Caravas, Benjamin Rambo-Martin, Shatavia Morrison, Christopher Gulvick, Scott Sammons, Yvette Unoarumhi, Darlene Wagner, Matthew Schmerer, Minoo Agarwal, Eyad Almasri, Debbie Boles, Ayla Burns, Nuthawin Charoensri, Oren Cohen, Susan Countryman, Mary Ann Cristobal, Bobbi Croy, Suzanne Dale, Hrushikesh Deshmukh, Amanda Douglas, Vincent Drouillon, Marcia Eisenberg, Howard Engler, Rama Ghatti, Prashant Gupta, Susan Hicks, Jake Humphrey, Lax Iyer, Lisa Pfefferle, Manoj Jain, Matthew Robinson, Mohan Kolli, Brian Krueger, Tim Kuphal, Stanley Letovsky, Michael Levandoski, Craig Lukasik, Jonathan Meltzer, Brian Norvell, Mindy Nye, Scott Parker, Christos Petropoulos, John Pruitt, Steven Ragan, Scott Ryan, Mike Sapeta, Jana Schroth, Suresh Babu Selvaraju, Goran Stevovic, Amanda Suchanek, Andrea Throop, Lyndon Tilson, Thomas Urban, Joe Voshell, |

| | | | |
|---|---|---|---|
| | | | Kimberly Wagner, Jonathan Williams, Mary Williamson, Qian Zeng, Tricia Zwiefelhofer, Clinton R. Paden, Duncan MacCannell |
| EPI_ISL_3326234, EPI_ISL_3326235 | Hospital Universitari Vall d'Hebron - Vall d'Hebron Institut de Recerca | Hospital Universitari Vall d'Hebron - Vall d'Hebron Institut de Recerca | Cristina Andrés, Maria Piñana, Alejandra González-Sánchez, Damir Garcia-Cehic, Ariadna Rando, Juliana Esperalba, Maria Gema Codina, Carla Castillo, Karen García, Rodrigo Vásquez, Maria Carmen Martin, Tomàs Pumarola, Josep Quer, Andrés Antón |
| EPI_ISL_3326236 | TGen North | TGen North | Jolene Bowers, Heather Centner, Brett Van Tassel, Chris French, Hayley Yaglom, Darrin Lemmer, Dave Engelthaler |
| EPI_ISL_3326237 | Laboratory Corporation of America | Centers for Disease Control and Prevention Division of Viral Diseases, Pathogen Discovery | Dakota Howard, Dhwani Batra, Peter W. Cook, Kara Moser, Adrian Paskey, Jason Caravas, Benjamin Rambo-Martin, Shatavia Morrison, Christopher Gulvick, Scott Sammons, Yvette Unoarumhi, Darlene Wagner, Matthew Schmerer, Minoo Agarwal, Eyad Almasri, Debbie Boles, Ayla Burns, Nuthawin Charoensri, Oren Cohen, Susan Countryman, Mary Ann Cristobal, Bobbi Croy, Suzanne Dale, Hrushikesh Deshmukh, Amanda Douglas, Vincent Drouillon, Marcia Eisenberg, Howard Engler, Rama Ghatti, Prashant Gupta, Susan Hicks, Jake Humphrey, Lax Iyer, Lisa Pfefferle, Manoj Jain, Matthew Robinson, Mohan Kolli, Brian Krueger, Tim Kuphal, Stanley Letovsky, Michael Levandoski, Craig Lukasik, Jonathan Meltzer, Brian Norvell, Mindy Nye, Scott Parker, Christos Petropoulos, John Pruitt, Steven Ragan, Scott Ryan, Mike Sapeta, Jana Schroth, Suresh Babu Selvaraju, Goran Stevovic, Amanda Suchanek, Andrea Throop, Lyndon Tilson, Thomas Urban, Joe Voshell, Kimberly Wagner, Jonathan Williams, Mary Williamson, Qian Zeng, Tricia Zwiefelhofer, Clinton R. Paden, Duncan MacCannell |
| EPI_ISL_3326238, EPI_ISL_3326240 | Hospital Universitari Vall d'Hebron - Vall d'Hebron Institut de Recerca | Hospital Universitari Vall d'Hebron - Vall d'Hebron Institut de Recerca | Cristina Andrés, Maria Piñana, Alejandra González-Sánchez, Damir Garcia-Cehic, Ariadna Rando, Juliana Esperalba, Maria Gema Codina, Carla Castillo, Karen García, Rodrigo Vásquez, Maria Carmen Martin, Tomàs Pumarola, Josep Quer, Andrés Antón |
| EPI_ISL_3326241 | TGen North | TGen North | Jolene Bowers, Heather Centner, Brett Van Tassel, Chris French, Hayley Yaglom, Darrin Lemmer, Dave Engelthaler |
| EPI_ISL_3326242 | Hospital Universitari Vall d'Hebron - Vall d'Hebron Institut de Recerca | Hospital Universitari Vall d'Hebron - Vall d'Hebron Institut de Recerca | Cristina Andrés, Maria Piñana, Alejandra González-Sánchez, Damir Garcia-Cehic, Ariadna Rando, Juliana Esperalba, Maria Gema Codina, Carla Castillo, Karen García, Rodrigo Vásquez, Maria Carmen Martin, Tomàs Pumarola, Josep Quer, Andrés Antón |
| EPI_ISL_3326243 | Laboratory Corporation of America | Centers for Disease Control and Prevention Division of Viral Diseases, Pathogen Discovery | Dakota Howard, Dhwani Batra, Peter W. Cook, Kara Moser, Adrian Paskey, Jason Caravas, Benjamin Rambo-Martin, Shatavia Morrison, Christopher Gulvick, Scott Sammons, Yvette Unoarumhi, Darlene Wagner, Matthew Schmerer, Minoo Agarwal, Eyad Almasri, Debbie Boles, Ayla Burns, Nuthawin Charoensri, Oren Cohen, Susan Countryman, Mary Ann Cristobal, Bobbi Croy, Suzanne Dale, Hrushikesh Deshmukh, Amanda Douglas, Vincent Drouillon, Marcia Eisenberg, Howard Engler, Rama Ghatti, Prashant Gupta, Susan Hicks, Jake Humphrey, Lax Iyer, Lisa Pfefferle, Manoj Jain, Matthew Robinson, Mohan Kolli, Brian Krueger, Tim Kuphal, Stanley Letovsky, Michael Levandoski, Craig Lukasik, Jonathan Meltzer, Brian Norvell, Mindy Nye, Scott Parker, Christos Petropoulos, John Pruitt, Steven Ragan, Scott Ryan, Mike Sapeta, Jana Schroth, Suresh Babu Selvaraju, Goran Stevovic, Amanda Suchanek, Andrea Throop, Lyndon Tilson, Thomas Urban, Joe Voshell, Kimberly Wagner, Jonathan Williams, Mary Williamson, Qian Zeng, Tricia Zwiefelhofer, Clinton R. Paden, Duncan MacCannell |
| EPI_ISL_3326244 | Hospital Universitari Vall d'Hebron - Vall d'Hebron Institut de Recerca | Hospital Universitari Vall d'Hebron - Vall d'Hebron Institut de Recerca | Cristina Andrés, Maria Piñana, Alejandra González-Sánchez, Damir Garcia-Cehic, Ariadna Rando, Juliana Esperalba, Maria Gema Codina, Carla Castillo, Karen García, Rodrigo Vásquez, Maria Carmen Martin, Tomàs Pumarola, Josep Quer, Andrés Antón |
| EPI_ISL_3326245 | TGen North | TGen North | Jolene Bowers, Heather Centner, Brett Van Tassel, Chris French, Hayley Yaglom, Darrin Lemmer, Dave Engelthaler |
| EPI_ISL_3326246 | Hospital Universitari Vall d'Hebron - Vall d'Hebron Institut de Recerca | Hospital Universitari Vall d'Hebron - Vall d'Hebron Institut de Recerca | Cristina Andrés, Maria Piñana, Alejandra González-Sánchez, Damir Garcia-Cehic, Ariadna Rando, Juliana Esperalba, Maria Gema Codina, Carla Castillo, Karen García, Rodrigo Vásquez, Maria Carmen Martin, Tomàs Pumarola, Josep Quer, Andrés Antón |
| EPI_ISL_3326247 | Laboratory Corporation of America | Centers for Disease Control and Prevention Division of Viral Diseases, Pathogen Discovery | Dakota Howard, Dhwani Batra, Peter W. Cook, Kara Moser, Adrian Paskey, Jason Caravas, Benjamin Rambo-Martin, Shatavia Morrison, Christopher Gulvick, Scott Sammons, Yvette Unoarumhi, Darlene Wagner, Matthew Schmerer, Minoo Agarwal, Eyad Almasri, Debbie Boles, Ayla Burns, Nuthawin Charoensri, Oren Cohen, Susan Countryman, Mary Ann Cristobal, Bobbi Croy, Suzanne Dale, Hrushikesh Deshmukh, Amanda Douglas, Vincent Drouillon, Marcia Eisenberg, Howard Engler, Rama Ghatti, Prashant Gupta, Susan Hicks, Jake Humphrey, Lax Iyer, Lisa Pfefferle, Manoj Jain, Matthew Robinson, Mohan Kolli, Brian Krueger, Tim Kuphal, Stanley Letovsky, Michael Levandoski, Craig Lukasik, Jonathan Meltzer, Brian Norvell, Mindy Nye, Scott Parker, Christos Petropoulos, John Pruitt, Steven Ragan, Scott Ryan, Mike Sapeta, Jana Schroth, Suresh Babu Selvaraju, Goran Stevovic, Amanda Suchanek, Andrea Throop, Lyndon Tilson, Thomas Urban, Joe Voshell, Kimberly Wagner, Jonathan Williams, Mary Williamson, Qian Zeng, Tricia Zwiefelhofer, Clinton R. Paden, Duncan MacCannell |
| EPI_ISL_3326249 | Hospital Universitari Vall d'Hebron - Vall d'Hebron Institut de Recerca | Hospital Universitari Vall d'Hebron - Vall d'Hebron Institut de Recerca | Cristina Andrés, Maria Piñana, Alejandra González-Sánchez, Damir Garcia-Cehic, Ariadna Rando, Juliana Esperalba, Maria Gema Codina, Carla Castillo, Karen García, Rodrigo Vásquez, Maria Carmen Martin, Tomàs Pumarola, Josep Quer, Andrés Antón |
| EPI_ISL_3326250 | Laboratory Corporation of America | Centers for Disease Control and Prevention Division of Viral Diseases, Pathogen Discovery | Dakota Howard, Dhwani Batra, Peter W. Cook, Kara Moser, Adrian Paskey, Jason Caravas, Benjamin Rambo-Martin, Shatavia Morrison, Christopher Gulvick, Scott Sammons, Yvette Unoarumhi, Darlene Wagner, Matthew Schmerer, Minoo Agarwal, Eyad Almasri, Debbie Boles, Ayla Burns, Nuthawin Charoensri, Oren Cohen, Susan Countryman, Mary Ann Cristobal, Bobbi Croy, Suzanne Dale, Hrushikesh Deshmukh, Amanda Douglas, Vincent Drouillon, Marcia Eisenberg, Howard Engler, Rama Ghatti, Prashant Gupta, Susan Hicks, Jake Humphrey, Lax Iyer, Lisa Pfefferle, Manoj Jain, Matthew Robinson, Mohan Kolli, Brian Krueger, Tim Kuphal, Stanley Letovsky, Michael Levandoski, Craig Lukasik, Jonathan Meltzer, Brian Norvell, Mindy Nye, Scott Parker, Christos Petropoulos, John Pruitt, Steven Ragan, Scott Ryan, Mike Sapeta, Jana Schroth, Suresh Babu Selvaraju, Goran Stevovic, Amanda Suchanek, Andrea Throop, Lyndon Tilson, Thomas Urban, Joe Voshell, Kimberly Wagner, Jonathan Williams, Mary Williamson, Qian Zeng, Tricia Zwiefelhofer, Clinton R. Paden, Duncan MacCannell |
| EPI_ISL_3326251 | Hospital Universitari Vall d'Hebron - Vall d'Hebron Institut de Recerca | Hospital Universitari Vall d'Hebron - Vall d'Hebron Institut de Recerca | Cristina Andrés, Maria Piñana, Alejandra González-Sánchez, Damir Garcia-Cehic, Ariadna Rando, Juliana Esperalba, Maria Gema Codina, Carla Castillo, Karen García, Rodrigo Vásquez, Maria Carmen Martin, Tomàs Pumarola, Josep Quer, Andrés Antón |
| EPI_ISL_3326252 | TGen North | TGen North | Jolene Bowers, Heather Centner, Brett Van Tassel, Chris French, Hayley Yaglom, Darrin Lemmer, Dave Engelthaler |
| EPI_ISL_3326253, EPI_ISL_3326255 | Hospital Universitari Vall d'Hebron - Vall d'Hebron Institut de Recerca | Hospital Universitari Vall d'Hebron - Vall d'Hebron Institut de Recerca | Cristina Andrés, Maria Piñana, Alejandra González-Sánchez, Damir Garcia-Cehic, Ariadna Rando, Juliana Esperalba, Maria Gema Codina, Carla Castillo, Karen García, Rodrigo Vásquez, Maria Carmen Martin, Tomàs Pumarola, Josep Quer, Andrés Antón |
| EPI_ISL_3326256 | TGen North | TGen North | Jolene Bowers, Heather Centner, Brett Van Tassel, Chris French, Hayley Yaglom, Darrin Lemmer, Dave Engelthaler |
| EPI_ISL_3326257 | Hospital Universitari Vall d'Hebron - Vall d'Hebron Institut de Recerca | Hospital Universitari Vall d'Hebron - Vall d'Hebron Institut de Recerca | Cristina Andrés, Maria Piñana, Alejandra González-Sánchez, Damir Garcia-Cehic, Ariadna Rando, Juliana Esperalba, Maria Gema Codina, Carla Castillo, Karen García, Rodrigo Vásquez, Maria Carmen Martin, Tomàs Pumarola, Josep Quer, Andrés Antón |
| EPI_ISL_3326258 | Laboratory Corporation of America | Centers for Disease Control and Prevention Division of Viral Diseases, Pathogen Discovery | Dakota Howard, Dhwani Batra, Peter W. Cook, Kara Moser, Adrian Paskey, Jason Caravas, Benjamin Rambo-Martin, Shatavia Morrison, Christopher Gulvick, Scott Sammons, Yvette Unoarumhi, Darlene Wagner, Matthew Schmerer, Minoo Agarwal, Eyad Almasri, Debbie Boles, Ayla Burns, Nuthawin Charoensri, Oren Cohen, Susan Countryman, Mary Ann Cristobal, Bobbi Croy, Suzanne Dale, Hrushikesh Deshmukh, Amanda Douglas, Vincent Drouillon, Marcia Eisenberg, Howard Engler, Rama Ghatti, Prashant Gupta, Susan Hicks, Jake Humphrey, Lax Iyer, Lisa Pfefferle, Manoj Jain, Matthew Robinson, Mohan Kolli, Brian Krueger, Tim Kuphal, Stanley Letovsky, Michael Levandoski, Craig Lukasik, Jonathan Meltzer, Brian Norvell, Mindy Nye, Scott Parker, Christos Petropoulos, John Pruitt, Steven Ragan, Scott Ryan, Mike Sapeta, Jana Schroth, Suresh Babu Selvaraju, Goran Stevovic, Amanda Suchanek, Andrea Throop, Lyndon Tilson, Thomas Urban, Joe Voshell, Kimberly Wagner, Jonathan Williams, Mary Williamson, Qian Zeng, Tricia Zwiefelhofer, Clinton R. Paden, Duncan MacCannell |
| EPI_ISL_3326259 | Hospital Universitari Vall d'Hebron - Vall d'Hebron Institut de Recerca | Hospital Universitari Vall d'Hebron - Vall d'Hebron Institut de Recerca | Cristina Andrés, Maria Piñana, Alejandra González-Sánchez, Damir Garcia-Cehic, Ariadna Rando, Juliana Esperalba, Maria Gema Codina, Carla Castillo, Karen García, Rodrigo Vásquez, Maria Carmen Martin, Tomàs Pumarola, Josep Quer, Andrés Antón |
| EPI_ISL_3326260 | TGen North | TGen North | Jolene Bowers, Heather Centner, Brett Van Tassel, Chris French, Hayley Yaglom, Darrin Lemmer, Dave Engelthaler |
| EPI_ISL_3326261 | Hospital Universitari Vall d'Hebron - Vall d'Hebron Institut de Recerca | Hospital Universitari Vall d'Hebron - Vall d'Hebron Institut de Recerca | Cristina Andrés, Maria Piñana, Alejandra González-Sánchez, Damir Garcia-Cehic, Ariadna Rando, Juliana Esperalba, Maria Gema Codina, Carla Castillo, Karen García, Rodrigo Vásquez, Maria Carmen Martin, Tomàs Pumarola, Josep Quer, Andrés Antón |
| EPI_ISL_3326262 | Laboratory Corporation of America | Centers for Disease Control and Prevention Division of Viral Diseases, Pathogen Discovery | Dakota Howard, Dhwani Batra, Peter W. Cook, Kara Moser, Adrian Paskey, Jason Caravas, Benjamin Rambo-Martin, Shatavia Morrison, Christopher Gulvick, Scott Sammons, Yvette Unoarumhi, Darlene Wagner, Matthew Schmerer, Minoo Agarwal, Eyad Almasri, Debbie Boles, Ayla Burns, Nuthawin Charoensri, Oren Cohen, Susan Countryman, Mary Ann Cristobal, Bobbi Croy, Suzanne Dale, Hrushikesh Deshmukh, Amanda Douglas, Vincent Drouillon, Marcia Eisenberg, Howard Engler, Rama Ghatti, Prashant Gupta, Susan Hicks, Jake Humphrey, Lax Iyer, Lisa Pfefferle, Manoj Jain, Matthew Robinson, Mohan Kolli, Brian Krueger, Tim Kuphal, Stanley Letovsky, Michael Levandoski, Craig Lukasik, Jonathan Meltzer, Brian Norvell, Mindy Nye, Scott Parker, Christos Petropoulos, John Pruitt, Steven Ragan, Scott Ryan, Mike Sapeta, Jana Schroth, Suresh Babu Selvaraju, Goran Stevovic, Amanda Suchanek, Andrea Throop, Lyndon Tilson, Thomas Urban, Joe Voshell, Kimberly Wagner, Jonathan Williams, Mary Williamson, Qian Zeng, Tricia Zwiefelhofer, Clinton R. Paden, Duncan MacCannell |
| EPI_ISL_3326264 | TGen North | TGen North | Jolene Bowers, Heather Centner, Brett Van Tassel, Chris French, Hayley Yaglom, Darrin Lemmer, Dave Engelthaler |
| EPI_ISL_3326265 | Laboratory Corporation of America | Centers for Disease Control and Prevention Division of Viral Diseases, Pathogen Discovery | Dakota Howard, Dhwani Batra, Peter W. Cook, Kara Moser, Adrian Paskey, Jason Caravas, Benjamin Rambo-Martin, Shatavia Morrison, Christopher Gulvick, Scott Sammons, Yvette Unoarumhi, Darlene Wagner, Matthew Schmerer, Minoo Agarwal, Eyad Almasri, Debbie Boles, Ayla Burns, Nuthawin Charoensri, Oren Cohen, Susan Countryman, Mary Ann Cristobal, Bobbi Croy, Suzanne Dale, Hrushikesh Deshmukh, Amanda Douglas, Vincent Drouillon, Marcia Eisenberg, Howard Engler, Rama Ghatti, Prashant Gupta, Susan Hicks, Jake Humphrey, Lax Iyer, Lisa Pfefferle, Manoj Jain, Matthew Robinson, Mohan Kolli, Brian Krueger, Tim Kuphal, Stanley Letovsky, Michael Levandoski, Craig Lukasik, Jonathan Meltzer, Brian Norvell, Mindy Nye, Scott Parker, Christos Petropoulos, John Pruitt, Steven Ragan, Scott Ryan, Mike Sapeta, Jana Schroth, Suresh Babu Selvaraju, Goran Stevovic, Amanda Suchanek, Andrea Throop, Lyndon Tilson, Thomas Urban, Joe Voshell, Kimberly Wagner, Jonathan Williams, Mary Williamson, Qian Zeng, Tricia Zwiefelhofer, Clinton R. Paden, Duncan MacCannell |
| EPI_ISL_3326266, EPI_ISL_3326267 | Hospital Universitari Vall d'Hebron - Vall d'Hebron Institut de Recerca | Hospital Universitari Vall d'Hebron - Vall d'Hebron Institut de Recerca | Cristina Andrés, Maria Piñana, Alejandra González-Sánchez, Damir Garcia-Cehic, Ariadna Rando, Juliana Esperalba, Maria Gema Codina, Carla Castillo, Karen García, Rodrigo Vásquez, Maria Carmen Martin, Tomàs Pumarola, Josep Quer, Andrés Antón |
| EPI_ISL_3326268 | Laboratory Corporation of America | Centers for Disease Control and Prevention Division of Viral Diseases, Pathogen Discovery | Dakota Howard, Dhwani Batra, Peter W. Cook, Kara Moser, Adrian Paskey, Jason Caravas, Benjamin Rambo-Martin, Shatavia Morrison, Christopher Gulvick, Scott Sammons, Yvette Unoarumhi, Darlene Wagner, Matthew Schmerer, Minoo Agarwal, Eyad Almasri, Debbie Boles, Ayla Burns, Nuthawin Charoensri, Oren Cohen, Susan Countryman, Mary Ann Cristobal, Bobbi Croy, Suzanne Dale, Hrushikesh Deshmukh, Amanda Douglas, Vincent Drouillon, Marcia Eisenberg, Howard Engler, Rama Ghatti, Prashant Gupta, Susan Hicks, Jake Humphrey, Lax Iyer, Lisa Pfefferle, Manoj Jain, Matthew Robinson, Mohan Kolli, Brian Krueger, Tim Kuphal, Stanley Letovsky, Michael Levandoski, Craig Lukasik, Jonathan Meltzer, Brian Norvell, Mindy Nye, Scott Parker, Christos Petropoulos, John Pruitt, Steven Ragan, Scott Ryan, Mike Sapeta, Jana Schroth, Suresh Babu Selvaraju, Goran Stevovic, Amanda Suchanek, Andrea Throop, Lyndon Tilson, Thomas Urban, Joe Voshell, Kimberly Wagner, Jonathan Williams, Mary Williamson, Qian Zeng, Tricia Zwiefelhofer, Clinton R. Paden, Duncan MacCannell |
| EPI_ISL_3326269 | TGen North | TGen North | Jolene Bowers, Heather Centner, Brett Van Tassel, Chris French, Hayley Yaglom, Darrin Lemmer, Dave Engelthaler |
| EPI_ISL_3326270, EPI_ISL_3326271 | Hospital Universitari Vall d'Hebron - Vall d'Hebron Institut de Recerca | Hospital Universitari Vall d'Hebron - Vall d'Hebron Institut de Recerca | Cristina Andrés, Maria Piñana, Alejandra González-Sánchez, Damir Garcia-Cehic, Ariadna Rando, Juliana Esperalba, Maria Gema Codina, Carla Castillo, Karen García, Rodrigo Vásquez, Maria Carmen Martin, Tomàs Pumarola, Josep Quer, Andrés Antón |
| EPI_ISL_3326272 | Laboratory Corporation of America | Centers for Disease Control and Prevention Division of Viral Diseases, Pathogen Discovery | Dakota Howard, Dhwani Batra, Peter W. Cook, Kara Moser, Adrian Paskey, Jason Caravas, Benjamin Rambo-Martin, Shatavia Morrison, Christopher Gulvick, Scott Sammons, Yvette Unoarumhi, Darlene Wagner, Matthew Schmerer, Minoo Agarwal, Eyad Almasri, Debbie Boles, Ayla Burns, Nuthawin Charoensri, Oren Cohen, Susan Countryman, Mary Ann Cristobal, Bobbi Croy, Suzanne Dale, Hrushikesh Deshmukh, Amanda Douglas, Vincent Drouillon, Marcia Eisenberg, Howard Engler, Rama Ghatti, Prashant Gupta, Susan Hicks, Jake Humphrey, Lax Iyer, Lisa Pfefferle, Manoj Jain, Matthew Robinson, Mohan Kolli, Brian Krueger, Tim Kuphal, Stanley Letovsky, Michael Levandoski, Craig Lukasik, Jonathan Meltzer, Brian Norvell, Mindy Nye, Scott Parker, Christos Petropoulos, John Pruitt, Steven Ragan, Scott Ryan, Mike Sapeta, Jana Schroth, Suresh Babu Selvaraju, Goran Stevovic, Amanda Suchanek, Andrea Throop, Lyndon Tilson, Thomas Urban, Joe Voshell, Kimberly Wagner, Jonathan Williams, Mary Williamson, Qian Zeng, Tricia Zwiefelhofer, Clinton R. Paden, Duncan MacCannell |
| EPI_ISL_3326273 | TGen North | TGen North | Jolene Bowers, Heather Centner, Brett Van Tassel, Chris French, Hayley Yaglom, Darrin Lemmer, Dave Engelthaler |
| EPI_ISL_3326274 | Hospital Universitari Vall d'Hebron - Vall d'Hebron Institut de Recerca | Hospital Universitari Vall d'Hebron - Vall d'Hebron Institut de Recerca | Cristina Andrés, Maria Piñana, Alejandra González-Sánchez, Damir Garcia-Cehic, Ariadna Rando, Juliana Esperalba, Maria Gema Codina, Carla Castillo, Karen García, Rodrigo Vásquez, Maria Carmen Martin, Tomàs Pumarola, Josep Quer, Andrés Antón |
| EPI_ISL_3326275 | Laboratory Corporation of America | Centers for Disease Control and Prevention Division of Viral Diseases, Pathogen Discovery | Dakota Howard, Dhwani Batra, Peter W. Cook, Kara Moser, Adrian Paskey, Jason Caravas, Benjamin Rambo-Martin, Shatavia Morrison, Christopher Gulvick, Scott Sammons, Yvette Unoarumhi, Darlene Wagner, Matthew Schmerer, Minoo Agarwal, Eyad Almasri, Debbie Boles, Ayla Burns, Nuthawin Charoensri, Oren Cohen, Susan Countryman, Mary Ann Cristobal, Bobbi Croy, Suzanne Dale, Hrushikesh Deshmukh, Amanda Douglas, Vincent Drouillon, Marcia Eisenberg, Howard Engler, Rama Ghatti, Prashant Gupta, Susan Hicks, Jake Humphrey, Lax Iyer, Lisa Pfefferle, Manoj Jain, Matthew Robinson, Mohan Kolli, Brian Krueger, Tim Kuphal, Stanley Letovsky, Michael Levandoski, Craig Lukasik, Jonathan Meltzer, Brian Norvell, Mindy Nye, Scott Parker, Christos Petropoulos, John Pruitt, Steven Ragan, Scott Ryan, Mike Sapeta, Jana Schroth, Suresh Babu Selvaraju, Goran Stevovic, Amanda Suchanek, Andrea Throop, Lyndon Tilson, Thomas Urban, Joe Voshell, Kimberly Wagner, Jonathan Williams, Mary Williamson, Qian Zeng, Tricia Zwiefelhofer, Clinton R. Paden, Duncan MacCannell |
| EPI_ISL_3326276 | Hospital Universitari Vall d'Hebron - Vall d'Hebron Institut de Recerca | Hospital Universitari Vall d'Hebron - Vall d'Hebron Institut de Recerca | Cristina Andrés, Maria Piñana, Alejandra González-Sánchez, Damir Garcia-Cehic, Ariadna Rando, Juliana Esperalba, Maria Gema Codina, Carla Castillo, Karen García, Rodrigo Vásquez, Maria Carmen Martin, Tomàs Pumarola, Josep Quer, Andrés Antón |
| EPI_ISL_3326277 | TGen North | TGen North | Jolene Bowers, Heather Centner, Brett Van Tassel, Chris French, Hayley Yaglom, Darrin Lemmer, Dave Engelthaler |
| EPI_ISL_3326278 | Hospital Universitari Vall d'Hebron - Vall d'Hebron Institut de Recerca | Hospital Universitari Vall d'Hebron - Vall d'Hebron Institut de Recerca | Cristina Andrés, Maria Piñana, Alejandra González-Sánchez, Damir Garcia-Cehic, Ariadna Rando, Juliana Esperalba, Maria Gema Codina, Carla Castillo, Karen García, Rodrigo Vásquez, Maria Carmen Martin, Tomàs Pumarola, Josep Quer, Andrés Antón |

| | | | |
|---|---|---|---|
| EPI_ISL_3326279 | Laboratory Corporation of America | Centers for Disease Control and Prevention Division of Viral Diseases, Pathogen Discovery | Dakota Howard, Dhwani Batra, Peter W. Cook, Kara Moser, Adrian Paskey, Jason Caravas, Benjamin Rambo-Martin, Shatavia Morrison, Christopher Gulvick, Scott Sammons, Yvette Unoarumhi, Darlene Wagner, Matthew Schmerer, Minoo Agarwal, Eyad Almasri, Debbie Boles, Ayla Burns, Nuthawin Charoensri, Oren Cohen, Susan Countryman, Mary Ann Cristobal, Bobbi Croy, Suzanne Dale, Hrushikesh Deshmukh, Amanda Douglas, Vincent Drouillon, Marcia Eisenberg, Howard Engler, Rama Ghatti, Prashant Gupta, Susan Hicks, Jake Humphrey, Lax Iyer, Lisa Pfefferle, Manoj Jain, Matthew Robinson, Mohan Kolli, Brian Krueger, Tim Kuphal, Stanley Letovsky, Michael Levandoski, Craig Lukasik, Jonathan Meltzer, Brian Norvell, Mindy Nye, Scott Parker, Christos Petropoulos, John Pruitt, Steven Ragan, Scott Ryan, Mike Sapeta, Jana Schroth, Suresh Babu Selvaraju, Goran Stevovic, Amanda Suchanek, Andrea Throop, Lyndon Tilson, Thomas Urban, Joe Voshell, Kimberly Wagner, Jonathan Williams, Mary Williamson, Qian Zeng, Tricia Zwiefelhofer, Clinton R. Paden, Duncan MacCannell |
| EPI_ISL_3326280 | Hospital Universitari Vall d'Hebron - Vall d'Hebron Institut de Recerca | Hospital Universitari Vall d'Hebron - Vall d'Hebron Institut de Recerca | Cristina Andrés, Maria Piñana, Alejandra González-Sánchez, Damir Garcia-Cehic, Ariadna Rando, Juliana Esperalba, Maria Gema Codina, Carla Castillo, Karen García, Rodrigo Vásquez, Maria Carmen Martin, Tomàs Pumarola, Josep Quer, Andrés Antón |
| EPI_ISL_3326281 | TGen North | TGen North | Jolene Bowers, Heather Centner, Brett Van Tassel, Chris French, Hayley Yaglom, Darrin Lemmer, Dave Engelthaler |
| EPI_ISL_3326283, EPI_ISL_3326284 | Hospital Universitari Vall d'Hebron - Vall d'Hebron Institut de Recerca | Hospital Universitari Vall d'Hebron - Vall d'Hebron Institut de Recerca | Cristina Andrés, Maria Piñana, Alejandra González-Sánchez, Damir Garcia-Cehic, Ariadna Rando, Juliana Esperalba, Maria Gema Codina, Carla Castillo, Karen García, Rodrigo Vásquez, Maria Carmen Martin, Tomàs Pumarola, Josep Quer, Andrés Antón |
| EPI_ISL_3326285 | Laboratory Corporation of America | Centers for Disease Control and Prevention Division of Viral Diseases, Pathogen Discovery | Dakota Howard, Dhwani Batra, Peter W. Cook, Kara Moser, Adrian Paskey, Jason Caravas, Benjamin Rambo-Martin, Shatavia Morrison, Christopher Gulvick, Scott Sammons, Yvette Unoarumhi, Darlene Wagner, Matthew Schmerer, Minoo Agarwal, Eyad Almasri, Debbie Boles, Ayla Burns, Nuthawin Charoensri, Oren Cohen, Susan Countryman, Mary Ann Cristobal, Bobbi Croy, Suzanne Dale, Hrushikesh Deshmukh, Amanda Douglas, Vincent Drouillon, Marcia Eisenberg, Howard Engler, Rama Ghatti, Prashant Gupta, Susan Hicks, Jake Humphrey, Lax Iyer, Lisa Pfefferle, Manoj Jain, Matthew Robinson, Mohan Kolli, Brian Krueger, Tim Kuphal, Stanley Letovsky, Michael Levandoski, Craig Lukasik, Jonathan Meltzer, Brian Norvell, Mindy Nye, Scott Parker, Christos Petropoulos, John Pruitt, Steven Ragan, Scott Ryan, Mike Sapeta, Jana Schroth, Suresh Babu Selvaraju, Goran Stevovic, Amanda Suchanek, Andrea Throop, Lyndon Tilson, Thomas Urban, Joe Voshell, Kimberly Wagner, Jonathan Williams, Mary Williamson, Qian Zeng, Tricia Zwiefelhofer, Clinton R. Paden, Duncan MacCannell |
| EPI_ISL_3326286 | TGen North | TGen North | Jolene Bowers, Heather Centner, Brett Van Tassel, Chris French, Hayley Yaglom, Darrin Lemmer, Dave Engelthaler |
| EPI_ISL_3326287, EPI_ISL_3326288 | Hospital Universitari Vall d'Hebron - Vall d'Hebron Institut de Recerca | Hospital Universitari Vall d'Hebron - Vall d'Hebron Institut de Recerca | Cristina Andrés, Maria Piñana, Alejandra González-Sánchez, Damir Garcia-Cehic, Ariadna Rando, Juliana Esperalba, Maria Gema Codina, Carla Castillo, Karen García, Rodrigo Vásquez, Maria Carmen Martin, Tomàs Pumarola, Josep Quer, Andrés Antón |
| EPI_ISL_3326289 | Laboratory Corporation of America | Centers for Disease Control and Prevention Division of Viral Diseases, Pathogen Discovery | Dakota Howard, Dhwani Batra, Peter W. Cook, Kara Moser, Adrian Paskey, Jason Caravas, Benjamin Rambo-Martin, Shatavia Morrison, Christopher Gulvick, Scott Sammons, Yvette Unoarumhi, Darlene Wagner, Matthew Schmerer, Minoo Agarwal, Eyad Almasri, Debbie Boles, Ayla Burns, Nuthawin Charoensri, Oren Cohen, Susan Countryman, Mary Ann Cristobal, Bobbi Croy, Suzanne Dale, Hrushikesh Deshmukh, Amanda Douglas, Vincent Drouillon, Marcia Eisenberg, Howard Engler, Rama Ghatti, Prashant Gupta, Susan Hicks, Jake Humphrey, Lax Iyer, Lisa Pfefferle, Manoj Jain, Matthew Robinson, Mohan Kolli, Brian Krueger, Tim Kuphal, Stanley Letovsky, Michael Levandoski, Craig Lukasik, Jonathan Meltzer, Brian Norvell, Mindy Nye, Scott Parker, Christos Petropoulos, John Pruitt, Steven Ragan, Scott Ryan, Mike Sapeta, Jana Schroth, Suresh Babu Selvaraju, Goran Stevovic, Amanda Suchanek, Andrea Throop, Lyndon Tilson, Thomas Urban, Joe Voshell, Kimberly Wagner, Jonathan Williams, Mary Williamson, Qian Zeng, Tricia Zwiefelhofer, Clinton R. Paden, Duncan MacCannell |
| EPI_ISL_3326290 | TGen North | TGen North | Jolene Bowers, Heather Centner, Brett Van Tassel, Chris French, Hayley Yaglom, Darrin Lemmer, Dave Engelthaler |
| EPI_ISL_3326291 | Hospital Universitari Vall d'Hebron - Vall d'Hebron Institut de Recerca | Hospital Universitari Vall d'Hebron - Vall d'Hebron Institut de Recerca | Cristina Andrés, Maria Piñana, Alejandra González-Sánchez, Damir Garcia-Cehic, Ariadna Rando, Juliana Esperalba, Maria Gema Codina, Carla Castillo, Karen García, Rodrigo Vásquez, Maria Carmen Martin, Tomàs Pumarola, Josep Quer, Andrés Antón |
| EPI_ISL_3326292 | Laboratory Corporation of America | Centers for Disease Control and Prevention Division of Viral Diseases, Pathogen Discovery | Dakota Howard, Dhwani Batra, Peter W. Cook, Kara Moser, Adrian Paskey, Jason Caravas, Benjamin Rambo-Martin, Shatavia Morrison, Christopher Gulvick, Scott Sammons, Yvette Unoarumhi, Darlene Wagner, Matthew Schmerer, Minoo Agarwal, Eyad Almasri, Debbie Boles, Ayla Burns, Nuthawin Charoensri, Oren Cohen, Susan Countryman, Mary Ann Cristobal, Bobbi Croy, Suzanne Dale, Hrushikesh Deshmukh, Amanda Douglas, Vincent Drouillon, Marcia Eisenberg, Howard Engler, Rama Ghatti, Prashant Gupta, Susan Hicks, Jake Humphrey, Lax Iyer, Lisa Pfefferle, Manoj Jain, Matthew Robinson, Mohan Kolli, Brian Krueger, Tim Kuphal, Stanley Letovsky, Michael Levandoski, Craig Lukasik, Jonathan Meltzer, Brian Norvell, Mindy Nye, Scott Parker, Christos Petropoulos, John Pruitt, Steven Ragan, Scott Ryan, Mike Sapeta, Jana Schroth, Suresh Babu Selvaraju, Goran Stevovic, Amanda Suchanek, Andrea Throop, Lyndon Tilson, Thomas Urban, Joe Voshell, Kimberly Wagner, Jonathan Williams, Mary Williamson, Qian Zeng, Tricia Zwiefelhofer, Clinton R. Paden, Duncan MacCannell |
| EPI_ISL_3326293, EPI_ISL_3326295 | Hospital Universitari Vall d'Hebron - Vall d'Hebron Institut de Recerca | Hospital Universitari Vall d'Hebron - Vall d'Hebron Institut de Recerca | Cristina Andrés, Maria Piñana, Alejandra González-Sánchez, Damir Garcia-Cehic, Ariadna Rando, Juliana Esperalba, Maria Gema Codina, Carla Castillo, Karen García, Rodrigo Vásquez, Maria Carmen Martin, Tomàs Pumarola, Josep Quer, Andrés Antón |
| EPI_ISL_3326296 | Laboratory Corporation of America | Centers for Disease Control and Prevention Division of Viral Diseases, Pathogen Discovery | Dakota Howard, Dhwani Batra, Peter W. Cook, Kara Moser, Adrian Paskey, Jason Caravas, Benjamin Rambo-Martin, Shatavia Morrison, Christopher Gulvick, Scott Sammons, Yvette Unoarumhi, Darlene Wagner, Matthew Schmerer, Minoo Agarwal, Eyad Almasri, Debbie Boles, Ayla Burns, Nuthawin Charoensri, Oren Cohen, Susan Countryman, Mary Ann Cristobal, Bobbi Croy, Suzanne Dale, Hrushikesh Deshmukh, Amanda Douglas, Vincent Drouillon, Marcia Eisenberg, Howard Engler, Rama Ghatti, Prashant Gupta, Susan Hicks, Jake Humphrey, Lax Iyer, Lisa Pfefferle, Manoj Jain, Matthew Robinson, Mohan Kolli, Brian Krueger, Tim Kuphal, Stanley Letovsky, Michael Levandoski, Craig Lukasik, Jonathan Meltzer, Brian Norvell, Mindy Nye, Scott Parker, Christos Petropoulos, John Pruitt, Steven Ragan, Scott Ryan, Mike Sapeta, Jana Schroth, Suresh Babu Selvaraju, Goran Stevovic, Amanda Suchanek, Andrea Throop, Lyndon Tilson, Thomas Urban, Joe Voshell, Kimberly Wagner, Jonathan Williams, Mary Williamson, Qian Zeng, Tricia Zwiefelhofer, Clinton R. Paden, Duncan MacCannell |
| EPI_ISL_3326297, EPI_ISL_3326299 | Hospital Universitari Vall d'Hebron - Vall d'Hebron Institut de Recerca | Hospital Universitari Vall d'Hebron - Vall d'Hebron Institut de Recerca | Cristina Andrés, Maria Piñana, Alejandra González-Sánchez, Damir Garcia-Cehic, Ariadna Rando, Juliana Esperalba, Maria Gema Codina, Carla Castillo, Karen García, Rodrigo Vásquez, Maria Carmen Martin, Tomàs Pumarola, Josep Quer, Andrés Antón |
| EPI_ISL_3326300, EPI_ISL_3326302, EPI_ISL_3326303, EPI_ISL_3326305 | Laboratory Corporation of America | Centers for Disease Control and Prevention Division of Viral Diseases, Pathogen Discovery | Dakota Howard, Dhwani Batra, Peter W. Cook, Kara Moser, Adrian Paskey, Jason Caravas, Benjamin Rambo-Martin, Shatavia Morrison, Christopher Gulvick, Scott Sammons, Yvette Unoarumhi, Darlene Wagner, Matthew Schmerer, Minoo Agarwal, Eyad Almasri, Debbie Boles, Ayla Burns, Nuthawin Charoensri, Oren Cohen, Susan Countryman, Mary Ann Cristobal, Bobbi Croy, Suzanne Dale, Hrushikesh Deshmukh, Amanda Douglas, Vincent Drouillon, Marcia Eisenberg, Howard Engler, Rama Ghatti, Prashant Gupta, Susan Hicks, Jake Humphrey, Lax Iyer, Lisa Pfefferle, Manoj Jain, Matthew Robinson, Mohan Kolli, Brian Krueger, Tim Kuphal, Stanley Letovsky, Michael Levandoski, Craig Lukasik, Jonathan Meltzer, Brian Norvell, Mindy Nye, Scott Parker, Christos Petropoulos, John Pruitt, Steven Ragan, Scott Ryan, Mike Sapeta, Jana Schroth, Suresh Babu Selvaraju, Goran Stevovic, Amanda Suchanek, Andrea Throop, Lyndon Tilson, Thomas Urban, Joe Voshell, Kimberly Wagner, Jonathan Williams, Mary Williamson, Qian Zeng, Tricia Zwiefelhofer, Clinton R. Paden, Duncan MacCannell |
| EPI_ISL_3326306 | TGen North | TGen North | Jolene Bowers, Heather Centner, Brett Van Tassel, Chris French, Hayley Yaglom, Darrin Lemmer, Dave Engelthaler |
| EPI_ISL_3326307, EPI_ISL_3326309 | Laboratory Corporation of America | Centers for Disease Control and Prevention Division of Viral Diseases, Pathogen Discovery | Dakota Howard, Dhwani Batra, Peter W. Cook, Kara Moser, Adrian Paskey, Jason Caravas, Benjamin Rambo-Martin, Shatavia Morrison, Christopher Gulvick, Scott Sammons, Yvette Unoarumhi, Darlene Wagner, Matthew Schmerer, Minoo Agarwal, Eyad Almasri, Debbie Boles, Ayla Burns, Nuthawin Charoensri, Oren Cohen, Susan Countryman, Mary Ann Cristobal, Bobbi Croy, Suzanne Dale, Hrushikesh Deshmukh, Amanda Douglas, Vincent Drouillon, Marcia Eisenberg, Howard Engler, Rama Ghatti, Prashant Gupta, Susan Hicks, Jake Humphrey, Lax Iyer, Lisa Pfefferle, Manoj Jain, Matthew Robinson, Mohan Kolli, Brian Krueger, Tim Kuphal, Stanley Letovsky, Michael Levandoski, Craig Lukasik, Jonathan Meltzer, Brian Norvell, Mindy Nye, Scott Parker, Christos Petropoulos, John Pruitt, Steven Ragan, Scott Ryan, Mike Sapeta, Jana Schroth, Suresh Babu Selvaraju, Goran Stevovic, Amanda Suchanek, Andrea Throop, Lyndon Tilson, Thomas Urban, Joe Voshell, Kimberly Wagner, Jonathan Williams, Mary Williamson, Qian Zeng, Tricia Zwiefelhofer, Clinton R. Paden, Duncan MacCannell |
| EPI_ISL_3326310 | TGen North | TGen North | Jolene Bowers, Heather Centner, Brett Van Tassel, Chris French, Hayley Yaglom, Darrin Lemmer, Dave Engelthaler |
| EPI_ISL_3326311, EPI_ISL_3326312, EPI_ISL_3326314 | Laboratory Corporation of America | Centers for Disease Control and Prevention Division of Viral Diseases, Pathogen Discovery | Dakota Howard, Dhwani Batra, Peter W. Cook, Kara Moser, Adrian Paskey, Jason Caravas, Benjamin Rambo-Martin, Shatavia Morrison, Christopher Gulvick, Scott Sammons, Yvette Unoarumhi, Darlene Wagner, Matthew Schmerer, Minoo Agarwal, Eyad Almasri, Debbie Boles, Ayla Burns, Nuthawin Charoensri, Oren Cohen, Susan Countryman, Mary Ann Cristobal, Bobbi Croy, Suzanne Dale, Hrushikesh Deshmukh, Amanda Douglas, Vincent Drouillon, Marcia Eisenberg, Howard Engler, Rama Ghatti, Prashant Gupta, Susan Hicks, Jake Humphrey, Lax Iyer, Lisa Pfefferle, Manoj Jain, Matthew Robinson, Mohan Kolli, Brian Krueger, Tim Kuphal, Stanley Letovsky, Michael Levandoski, Craig Lukasik, Jonathan Meltzer, Brian Norvell, Mindy Nye, Scott Parker, Christos Petropoulos, John Pruitt, Steven Ragan, Scott Ryan, Mike Sapeta, Jana Schroth, Suresh Babu Selvaraju, Goran Stevovic, Amanda Suchanek, Andrea Throop, Lyndon Tilson, Thomas Urban, Joe Voshell, Kimberly Wagner, Jonathan Williams, Mary Williamson, Qian Zeng, Tricia Zwiefelhofer, Clinton R. Paden, Duncan MacCannell |
| EPI_ISL_3326315 | TGen North | TGen North | Jolene Bowers, Heather Centner, Brett Van Tassel, Chris French, Hayley Yaglom, Darrin Lemmer, Dave Engelthaler |
| EPI_ISL_3326316 | Laboratory Corporation of America | Centers for Disease Control and Prevention Division of Viral Diseases, Pathogen Discovery | Dakota Howard, Dhwani Batra, Peter W. Cook, Kara Moser, Adrian Paskey, Jason Caravas, Benjamin Rambo-Martin, Shatavia Morrison, Christopher Gulvick, Scott Sammons, Yvette Unoarumhi, Darlene Wagner, Matthew Schmerer, Minoo Agarwal, Eyad Almasri, Debbie Boles, Ayla Burns, Nuthawin Charoensri, Oren Cohen, Susan Countryman, Mary Ann Cristobal, Bobbi Croy, Suzanne Dale, Hrushikesh Deshmukh, Amanda Douglas, Vincent Drouillon, Marcia Eisenberg, Howard Engler, Rama Ghatti, Prashant Gupta, Susan Hicks, Jake Humphrey, Lax Iyer, Lisa Pfefferle, Manoj Jain, Matthew Robinson, Mohan Kolli, Brian Krueger, Tim Kuphal, Stanley Letovsky, Michael Levandoski, Craig Lukasik, Jonathan Meltzer, Brian Norvell, Mindy Nye, Scott Parker, Christos Petropoulos, John Pruitt, Steven Ragan, Scott Ryan, Mike Sapeta, Jana Schroth, Suresh Babu Selvaraju, Goran Stevovic, Amanda Suchanek, Andrea Throop, Lyndon Tilson, Thomas Urban, Joe Voshell, Kimberly Wagner, Jonathan Williams, Mary Williamson, Qian Zeng, Tricia Zwiefelhofer, Clinton R. Paden, Duncan MacCannell |
| EPI_ISL_3326317 | TGen North | TGen North | Jolene Bowers, Heather Centner, Brett Van Tassel, Chris French, Hayley Yaglom, Darrin Lemmer, Dave Engelthaler |
| EPI_ISL_3326318 | Laboratory Corporation of America | Centers for Disease Control and Prevention Division of Viral Diseases, Pathogen Discovery | Dakota Howard, Dhwani Batra, Peter W. Cook, Kara Moser, Adrian Paskey, Jason Caravas, Benjamin Rambo-Martin, Shatavia Morrison, Christopher Gulvick, Scott Sammons, Yvette Unoarumhi, Darlene Wagner, Matthew Schmerer, Minoo Agarwal, Eyad Almasri, Debbie Boles, Ayla Burns, Nuthawin Charoensri, Oren Cohen, Susan Countryman, Mary Ann Cristobal, Bobbi Croy, Suzanne Dale, Hrushikesh Deshmukh, Amanda Douglas, Vincent Drouillon, Marcia Eisenberg, Howard Engler, Rama Ghatti, Prashant Gupta, Susan Hicks, Jake Humphrey, Lax Iyer, Lisa Pfefferle, Manoj Jain, Matthew Robinson, Mohan Kolli, Brian Krueger, Tim Kuphal, Stanley Letovsky, Michael Levandoski, Craig Lukasik, Jonathan Meltzer, Brian Norvell, Mindy Nye, Scott Parker, Christos Petropoulos, John Pruitt, Steven Ragan, Scott Ryan, Mike Sapeta, Jana Schroth, Suresh Babu Selvaraju, Goran Stevovic, Amanda Suchanek, Andrea Throop, Lyndon Tilson, Thomas Urban, Joe Voshell, Kimberly Wagner, Jonathan Williams, Mary Williamson, Qian Zeng, Tricia Zwiefelhofer, Clinton R. Paden, Duncan MacCannell |
| EPI_ISL_3326319 | TGen North | TGen North | Jolene Bowers, Heather Centner, Brett Van Tassel, Chris French, Hayley Yaglom, Darrin Lemmer, Dave Engelthaler |
| EPI_ISL_3326320 | Laboratory Corporation of America | Centers for Disease Control and Prevention Division of Viral Diseases, Pathogen Discovery | Dakota Howard, Dhwani Batra, Peter W. Cook, Kara Moser, Adrian Paskey, Jason Caravas, Benjamin Rambo-Martin, Shatavia Morrison, Christopher Gulvick, Scott Sammons, Yvette Unoarumhi, Darlene Wagner, Matthew Schmerer, Minoo Agarwal, Eyad Almasri, Debbie Boles, Ayla Burns, Nuthawin Charoensri, Oren Cohen, Susan Countryman, Mary Ann Cristobal, Bobbi Croy, Suzanne Dale, Hrushikesh Deshmukh, Amanda Douglas, Vincent Drouillon, Marcia Eisenberg, Howard Engler, Rama Ghatti, Prashant Gupta, Susan Hicks, Jake Humphrey, Lax Iyer, Lisa Pfefferle, Manoj Jain, Matthew Robinson, Mohan Kolli, Brian Krueger, Tim Kuphal, Stanley Letovsky, Michael Levandoski, Craig Lukasik, Jonathan Meltzer, Brian Norvell, Mindy Nye, Scott Parker, Christos Petropoulos, John Pruitt, Steven Ragan, Scott Ryan, Mike Sapeta, Jana Schroth, Suresh Babu Selvaraju, Goran Stevovic, Amanda Suchanek, Andrea Throop, Lyndon Tilson, Thomas Urban, Joe Voshell, Kimberly Wagner, Jonathan Williams, Mary Williamson, Qian Zeng, Tricia Zwiefelhofer, Clinton R. Paden, Duncan MacCannell |
| EPI_ISL_3326321 | TGen North | TGen North | Jolene Bowers, Heather Centner, Brett Van Tassel, Chris French, Hayley Yaglom, Darrin Lemmer, Dave Engelthaler |
| EPI_ISL_3326322 | Laboratory Corporation of America | Centers for Disease Control and Prevention Division of Viral Diseases, Pathogen Discovery | Dakota Howard, Dhwani Batra, Peter W. Cook, Kara Moser, Adrian Paskey, Jason Caravas, Benjamin Rambo-Martin, Shatavia Morrison, Christopher Gulvick, Scott Sammons, Yvette Unoarumhi, Darlene Wagner, Matthew Schmerer, Minoo Agarwal, Eyad Almasri, Debbie Boles, Ayla Burns, Nuthawin Charoensri, Oren Cohen, Susan Countryman, Mary Ann Cristobal, Bobbi Croy, Suzanne Dale, Hrushikesh Deshmukh, Amanda Douglas, Vincent Drouillon, Marcia Eisenberg, Howard Engler, Rama Ghatti, Prashant Gupta, Susan Hicks, Jake Humphrey, Lax Iyer, Lisa Pfefferle, Manoj Jain, Matthew Robinson, Mohan Kolli, Brian Krueger, Tim Kuphal, Stanley Letovsky, Michael Levandoski, Craig Lukasik, Jonathan Meltzer, Brian Norvell, Mindy Nye, Scott Parker, Christos Petropoulos, John Pruitt, Steven Ragan, Scott Ryan, Mike Sapeta, Jana Schroth, Suresh Babu Selvaraju, Goran Stevovic, Amanda Suchanek, Andrea Throop, Lyndon Tilson, Thomas Urban, Joe Voshell, Kimberly Wagner, Jonathan Williams, Mary Williamson, Qian Zeng, Tricia Zwiefelhofer, Clinton R. Paden, Duncan MacCannell |
| EPI_ISL_3326323 | TGen North | TGen North | Jolene Bowers, Heather Centner, Brett Van Tassel, Chris French, Hayley Yaglom, Darrin Lemmer, Dave Engelthaler |
| EPI_ISL_3326324 | Laboratory Corporation of America | Centers for Disease Control and Prevention Division of Viral Diseases, Pathogen Discovery | Dakota Howard, Dhwani Batra, Peter W. Cook, Kara Moser, Adrian Paskey, Jason Caravas, Benjamin Rambo-Martin, Shatavia Morrison, Christopher Gulvick, Scott Sammons, Yvette Unoarumhi, Darlene Wagner, Matthew Schmerer, Minoo Agarwal, Eyad Almasri, Debbie Boles, Ayla Burns, Nuthawin Charoensri, Oren Cohen, Susan Countryman, Mary Ann Cristobal, Bobbi Croy, Suzanne Dale, Hrushikesh Deshmukh, Amanda Douglas, Vincent Drouillon, Marcia Eisenberg, Howard Engler, Rama Ghatti, Prashant Gupta, Susan Hicks, Jake Humphrey, Lax Iyer, Lisa Pfefferle, Manoj Jain, Matthew Robinson, Mohan Kolli, Brian Krueger, Tim Kuphal, Stanley Letovsky, Michael Levandoski, Craig Lukasik, Jonathan Meltzer, Brian Norvell, Mindy Nye, Scott Parker, Christos Petropoulos, John Pruitt, Steven Ragan, Scott Ryan, Mike Sapeta, Jana Schroth, Suresh Babu Selvaraju, Goran Stevovic, Amanda Suchanek, Andrea Throop, Lyndon Tilson, Thomas Urban, Joe Voshell, Kimberly Wagner, Jonathan Williams, Mary Williamson, Qian Zeng, Tricia Zwiefelhofer, Clinton R. Paden, Duncan MacCannell |
| EPI_ISL_3326325 | TGen North | TGen North | Jolene Bowers, Heather Centner, Brett Van Tassel, Chris French, Hayley Yaglom, Darrin Lemmer, Dave Engelthaler |
| EPI_ISL_3326326 | Laboratory Corporation of America | Centers for Disease Control and Prevention Division of Viral Diseases, Pathogen Discovery | Dakota Howard, Dhwani Batra, Peter W. Cook, Kara Moser, Adrian Paskey, Jason Caravas, Benjamin Rambo-Martin, Shatavia Morrison, Christopher Gulvick, Scott Sammons, Yvette Unoarumhi, Darlene Wagner, Matthew Schmerer, Minoo Agarwal, Eyad Almasri, Debbie Boles, Ayla Burns, Nuthawin Charoensri, Oren Cohen, Susan Countryman, Mary Ann Cristobal, Bobbi Croy, Suzanne Dale, Hrushikesh Deshmukh, Amanda Douglas, Vincent Drouillon, Marcia Eisenberg, Howard Engler, Rama Ghatti, Prashant Gupta, Susan Hicks, Jake Humphrey, Lax Iyer, Lisa Pfefferle, Manoj Jain, Matthew Robinson, Mohan Kolli, Brian Krueger, Tim Kuphal, Stanley Letovsky, Michael Levandoski, Craig Lukasik, Jonathan Meltzer, Brian Norvell, Mindy Nye, Scott Parker, Christos Petropoulos, John Pruitt, Steven Ragan, Scott Ryan, Mike Sapeta, Jana Schroth, Suresh Babu Selvaraju, Goran Stevovic, Amanda Suchanek, Andrea Throop, Lyndon Tilson, Thomas Urban, Joe Voshell, Kimberly Wagner, Jonathan Williams, Mary Williamson, Qian Zeng, Tricia Zwiefelhofer, Clinton R. Paden, Duncan MacCannell |
| EPI_ISL_3326327, EPI_ISL_3326329 | TGen North | TGen North | Jolene Bowers, Heather Centner, Brett Van Tassel, Chris French, Hayley Yaglom, Darrin Lemmer, Dave Engelthaler |
| EPI_ISL_3326330 | Laboratory Corporation of America | Centers for Disease Control and Prevention Division of Viral | Dakota Howard, Dhwani Batra, Peter W. Cook, Kara Moser, Adrian Paskey, Jason Caravas, Benjamin Rambo-Martin, Shatavia Morrison, Christopher Gulvick, Scott Sammons, Yvette Unoarumhi, Darlene Wagner, Matthew Schmerer, Minoo |

| | | | |
|---|---|---|---|
| | | Diseases, Pathogen Discovery | Agarwal, Eyad Almasri, Debbie Boles, Ayla Burns, Nuthawin Charoensri, Oren Cohen, Susan Countryman, Mary Ann Cristobal, Bobbi Croy, Suzanne Dale, Hrushikesh Deshmukh, Amanda Douglas, Vincent Drouillon, Marcia Eisenberg, Howard Engler, Rama Ghatti, Prashant Gupta, Susan Hicks, Jake Humphrey, Lax Iyer, Lisa Pfefferle, Manoj Jain, Matthew Robinson, Mohan Kolli, Brian Krueger, Tim Kuphal, Stanley Letovsky, Michael Levandoski, Craig Lukasik, Jonathan Meltzer, Brian Norvell, Mindy Nye, Scott Parker, Christos Petropoulos, John Pruitt, Steven Ragan, Scott Ryan, Mike Sapeta, Jana Schroth, Suresh Babu Selvaraju, Goran Stevovic, Amanda Suchanek, Andrea Throop, Lyndon Tilson, Thomas Urban, Joe Voshell, Kimberly Wagner, Jonathan Williams, Mary Williamson, Qian Zeng, Tricia Zwiefelhofer, Clinton R. Paden, Duncan MacCannell |
| EPI_ISL_3326331 | TGen North | TGen North | Jolene Bowers, Heather Centner, Brett Van Tassel, Chris French, Hayley Yaglom, Darrin Lemmer, Dave Engelthaler |
| EPI_ISL_3326332, EPI_ISL_3326333, EPI_ISL_3326337 | Laboratory Corporation of America | Centers for Disease Control and Prevention Division of Viral Diseases, Pathogen Discovery | Dakota Howard, Dhwani Batra, Peter W. Cook, Kara Moser, Adrian Paskey, Jason Caravas, Benjamin Rambo-Martin, Shatavia Morrison, Christopher Gulvick, Scott Sammons, Yvette Unoarumhi, Darlene Wagner, Matthew Schmerer, Minoo Agarwal, Eyad Almasri, Debbie Boles, Ayla Burns, Nuthawin Charoensri, Oren Cohen, Susan Countryman, Mary Ann Cristobal, Bobbi Croy, Suzanne Dale, Hrushikesh Deshmukh, Amanda Douglas, Vincent Drouillon, Marcia Eisenberg, Howard Engler, Rama Ghatti, Prashant Gupta, Susan Hicks, Jake Humphrey, Lax Iyer, Lisa Pfefferle, Manoj Jain, Matthew Robinson, Mohan Kolli, Brian Krueger, Tim Kuphal, Stanley Letovsky, Michael Levandoski, Craig Lukasik, Jonathan Meltzer, Brian Norvell, Mindy Nye, Scott Parker, Christos Petropoulos, John Pruitt, Steven Ragan, Scott Ryan, Mike Sapeta, Jana Schroth, Suresh Babu Selvaraju, Goran Stevovic, Amanda Suchanek, Andrea Throop, Lyndon Tilson, Thomas Urban, Joe Voshell, Kimberly Wagner, Jonathan Williams, Mary Williamson, Qian Zeng, Tricia Zwiefelhofer, Clinton R. Paden, Duncan MacCannell |
| EPI_ISL_3326338 | TGen North | TGen North | Jolene Bowers, Heather Centner, Brett Van Tassel, Chris French, Hayley Yaglom, Darrin Lemmer, Dave Engelthaler |
| EPI_ISL_3326339 | Laboratory Corporation of America | Centers for Disease Control and Prevention Division of Viral Diseases, Pathogen Discovery | Dakota Howard, Dhwani Batra, Peter W. Cook, Kara Moser, Adrian Paskey, Jason Caravas, Benjamin Rambo-Martin, Shatavia Morrison, Christopher Gulvick, Scott Sammons, Yvette Unoarumhi, Darlene Wagner, Matthew Schmerer, Minoo Agarwal, Eyad Almasri, Debbie Boles, Ayla Burns, Nuthawin Charoensri, Oren Cohen, Susan Countryman, Mary Ann Cristobal, Bobbi Croy, Suzanne Dale, Hrushikesh Deshmukh, Amanda Douglas, Vincent Drouillon, Marcia Eisenberg, Howard Engler, Rama Ghatti, Prashant Gupta, Susan Hicks, Jake Humphrey, Lax Iyer, Lisa Pfefferle, Manoj Jain, Matthew Robinson, Mohan Kolli, Brian Krueger, Tim Kuphal, Stanley Letovsky, Michael Levandoski, Craig Lukasik, Jonathan Meltzer, Brian Norvell, Mindy Nye, Scott Parker, Christos Petropoulos, John Pruitt, Steven Ragan, Scott Ryan, Mike Sapeta, Jana Schroth, Suresh Babu Selvaraju, Goran Stevovic, Amanda Suchanek, Andrea Throop, Lyndon Tilson, Thomas Urban, Joe Voshell, Kimberly Wagner, Jonathan Williams, Mary Williamson, Qian Zeng, Tricia Zwiefelhofer, Clinton R. Paden, Duncan MacCannell |
| EPI_ISL_3326340 | TGen North | TGen North | Jolene Bowers, Heather Centner, Brett Van Tassel, Chris French, Hayley Yaglom, Darrin Lemmer, Dave Engelthaler |
| EPI_ISL_3326341, EPI_ISL_3326345 | Laboratory Corporation of America | Centers for Disease Control and Prevention Division of Viral Diseases, Pathogen Discovery | Dakota Howard, Dhwani Batra, Peter W. Cook, Kara Moser, Adrian Paskey, Jason Caravas, Benjamin Rambo-Martin, Shatavia Morrison, Christopher Gulvick, Scott Sammons, Yvette Unoarumhi, Darlene Wagner, Matthew Schmerer, Minoo Agarwal, Eyad Almasri, Debbie Boles, Ayla Burns, Nuthawin Charoensri, Oren Cohen, Susan Countryman, Mary Ann Cristobal, Bobbi Croy, Suzanne Dale, Hrushikesh Deshmukh, Amanda Douglas, Vincent Drouillon, Marcia Eisenberg, Howard Engler, Rama Ghatti, Prashant Gupta, Susan Hicks, Jake Humphrey, Lax Iyer, Lisa Pfefferle, Manoj Jain, Matthew Robinson, Mohan Kolli, Brian Krueger, Tim Kuphal, Stanley Letovsky, Michael Levandoski, Craig Lukasik, Jonathan Meltzer, Brian Norvell, Mindy Nye, Scott Parker, Christos Petropoulos, John Pruitt, Steven Ragan, Scott Ryan, Mike Sapeta, Jana Schroth, Suresh Babu Selvaraju, Goran Stevovic, Amanda Suchanek, Andrea Throop, Lyndon Tilson, Thomas Urban, Joe Voshell, Kimberly Wagner, Jonathan Williams, Mary Williamson, Qian Zeng, Tricia Zwiefelhofer, Clinton R. Paden, Duncan MacCannell |
| EPI_ISL_3326346 | TGen North | TGen North | Jolene Bowers, Heather Centner, Brett Van Tassel, Chris French, Hayley Yaglom, Darrin Lemmer, Dave Engelthaler |
| EPI_ISL_3326347, EPI_ISL_3326349, EPI_ISL_3326350, EPI_ISL_3326352 | Laboratory Corporation of America | Centers for Disease Control and Prevention Division of Viral Diseases, Pathogen Discovery | Dakota Howard, Dhwani Batra, Peter W. Cook, Kara Moser, Adrian Paskey, Jason Caravas, Benjamin Rambo-Martin, Shatavia Morrison, Christopher Gulvick, Scott Sammons, Yvette Unoarumhi, Darlene Wagner, Matthew Schmerer, Minoo Agarwal, Eyad Almasri, Debbie Boles, Ayla Burns, Nuthawin Charoensri, Oren Cohen, Susan Countryman, Mary Ann Cristobal, Bobbi Croy, Suzanne Dale, Hrushikesh Deshmukh, Amanda Douglas, Vincent Drouillon, Marcia Eisenberg, Howard Engler, Rama Ghatti, Prashant Gupta, Susan Hicks, Jake Humphrey, Lax Iyer, Lisa Pfefferle, Manoj Jain, Matthew Robinson, Mohan Kolli, Brian Krueger, Tim Kuphal, Stanley Letovsky, Michael Levandoski, Craig Lukasik, Jonathan Meltzer, Brian Norvell, Mindy Nye, Scott Parker, Christos Petropoulos, John Pruitt, Steven Ragan, Scott Ryan, Mike Sapeta, Jana Schroth, Suresh Babu Selvaraju, Goran Stevovic, Amanda Suchanek, Andrea Throop, Lyndon Tilson, Thomas Urban, Joe Voshell, Kimberly Wagner, Jonathan Williams, Mary Williamson, Qian Zeng, Tricia Zwiefelhofer, Clinton R. Paden, Duncan MacCannell |
| EPI_ISL_3326353, EPI_ISL_3326355 | TGen North | TGen North | Jolene Bowers, Heather Centner, Brett Van Tassel, Chris French, Hayley Yaglom, Darrin Lemmer, Dave Engelthaler |
| EPI_ISL_3326356 | Laboratory Corporation of America | Centers for Disease Control and Prevention Division of Viral Diseases, Pathogen Discovery | Dakota Howard, Dhwani Batra, Peter W. Cook, Kara Moser, Adrian Paskey, Jason Caravas, Benjamin Rambo-Martin, Shatavia Morrison, Christopher Gulvick, Scott Sammons, Yvette Unoarumhi, Darlene Wagner, Matthew Schmerer, Minoo Agarwal, Eyad Almasri, Debbie Boles, Ayla Burns, Nuthawin Charoensri, Oren Cohen, Susan Countryman, Mary Ann Cristobal, Bobbi Croy, Suzanne Dale, Hrushikesh Deshmukh, Amanda Douglas, Vincent Drouillon, Marcia Eisenberg, Howard Engler, Rama Ghatti, Prashant Gupta, Susan Hicks, Jake Humphrey, Lax Iyer, Lisa Pfefferle, Manoj Jain, Matthew Robinson, Mohan Kolli, Brian Krueger, Tim Kuphal, Stanley Letovsky, Michael Levandoski, Craig Lukasik, Jonathan Meltzer, Brian Norvell, Mindy Nye, Scott Parker, Christos Petropoulos, John Pruitt, Steven Ragan, Scott Ryan, Mike Sapeta, Jana Schroth, Suresh Babu Selvaraju, Goran Stevovic, Amanda Suchanek, Andrea Throop, Lyndon Tilson, Thomas Urban, Joe Voshell, Kimberly Wagner, Jonathan Williams, Mary Williamson, Qian Zeng, Tricia Zwiefelhofer, Clinton R. Paden, Duncan MacCannell |
| EPI_ISL_3326357 | TGen North | TGen North | Jolene Bowers, Heather Centner, Brett Van Tassel, Chris French, Hayley Yaglom, Darrin Lemmer, Dave Engelthaler |
| EPI_ISL_3326358, EPI_ISL_3326359 | Laboratory Corporation of America | Centers for Disease Control and Prevention Division of Viral Diseases, Pathogen Discovery | Dakota Howard, Dhwani Batra, Peter W. Cook, Kara Moser, Adrian Paskey, Jason Caravas, Benjamin Rambo-Martin, Shatavia Morrison, Christopher Gulvick, Scott Sammons, Yvette Unoarumhi, Darlene Wagner, Matthew Schmerer, Minoo Agarwal, Eyad Almasri, Debbie Boles, Ayla Burns, Nuthawin Charoensri, Oren Cohen, Susan Countryman, Mary Ann Cristobal, Bobbi Croy, Suzanne Dale, Hrushikesh Deshmukh, Amanda Douglas, Vincent Drouillon, Marcia Eisenberg, Howard Engler, Rama Ghatti, Prashant Gupta, Susan Hicks, Jake Humphrey, Lax Iyer, Lisa Pfefferle, Manoj Jain, Matthew Robinson, Mohan Kolli, Brian Krueger, Tim Kuphal, Stanley Letovsky, Michael Levandoski, Craig Lukasik, Jonathan Meltzer, Brian Norvell, Mindy Nye, Scott Parker, Christos Petropoulos, John Pruitt, Steven Ragan, Scott Ryan, Mike Sapeta, Jana Schroth, Suresh Babu Selvaraju, Goran Stevovic, Amanda Suchanek, Andrea Throop, Lyndon Tilson, Thomas Urban, Joe Voshell, Kimberly Wagner, Jonathan Williams, Mary Williamson, Qian Zeng, Tricia Zwiefelhofer, Clinton R. Paden, Duncan MacCannell |
| EPI_ISL_3326360 | TGen North | TGen North | Jolene Bowers, Heather Centner, Brett Van Tassel, Chris French, Hayley Yaglom, Darrin Lemmer, Dave Engelthaler |
| EPI_ISL_3326361, EPI_ISL_3326363 | Laboratory Corporation of America | Centers for Disease Control and Prevention Division of Viral Diseases, Pathogen Discovery | Dakota Howard, Dhwani Batra, Peter W. Cook, Kara Moser, Adrian Paskey, Jason Caravas, Benjamin Rambo-Martin, Shatavia Morrison, Christopher Gulvick, Scott Sammons, Yvette Unoarumhi, Darlene Wagner, Matthew Schmerer, Minoo Agarwal, Eyad Almasri, Debbie Boles, Ayla Burns, Nuthawin Charoensri, Oren Cohen, Susan Countryman, Mary Ann Cristobal, Bobbi Croy, Suzanne Dale, Hrushikesh Deshmukh, Amanda Douglas, Vincent Drouillon, Marcia Eisenberg, Howard Engler, Rama Ghatti, Prashant Gupta, Susan Hicks, Jake Humphrey, Lax Iyer, Lisa Pfefferle, Manoj Jain, Matthew Robinson, Mohan Kolli, Brian Krueger, Tim Kuphal, Stanley Letovsky, Michael Levandoski, Craig Lukasik, Jonathan Meltzer, Brian Norvell, Mindy Nye, Scott Parker, Christos Petropoulos, John Pruitt, Steven Ragan, Scott Ryan, Mike Sapeta, Jana Schroth, Suresh Babu Selvaraju, Goran Stevovic, Amanda Suchanek, Andrea Throop, Lyndon Tilson, Thomas Urban, Joe Voshell, Kimberly Wagner, Jonathan Williams, Mary Williamson, Qian Zeng, Tricia Zwiefelhofer, Clinton R. Paden, Duncan MacCannell |
| EPI_ISL_3326364 | TGen North | TGen North | Jolene Bowers, Heather Centner, Brett Van Tassel, Chris French, Hayley Yaglom, Darrin Lemmer, Dave Engelthaler |
| EPI_ISL_3326365, EPI_ISL_3326366 | Laboratory Corporation of America | Centers for Disease Control and Prevention Division of Viral Diseases, Pathogen Discovery | Dakota Howard, Dhwani Batra, Peter W. Cook, Kara Moser, Adrian Paskey, Jason Caravas, Benjamin Rambo-Martin, Shatavia Morrison, Christopher Gulvick, Scott Sammons, Yvette Unoarumhi, Darlene Wagner, Matthew Schmerer, Minoo Agarwal, Eyad Almasri, Debbie Boles, Ayla Burns, Nuthawin Charoensri, Oren Cohen, Susan Countryman, Mary Ann Cristobal, Bobbi Croy, Suzanne Dale, Hrushikesh Deshmukh, Amanda Douglas, Vincent Drouillon, Marcia Eisenberg, Howard Engler, Rama Ghatti, Prashant Gupta, Susan Hicks, Jake Humphrey, Lax Iyer, Lisa Pfefferle, Manoj Jain, Matthew Robinson, Mohan Kolli, Brian Krueger, Tim Kuphal, Stanley Letovsky, Michael Levandoski, Craig Lukasik, Jonathan Meltzer, Brian Norvell, Mindy Nye, Scott Parker, Christos Petropoulos, John Pruitt, Steven Ragan, Scott Ryan, Mike Sapeta, Jana Schroth, Suresh Babu Selvaraju, Goran Stevovic, Amanda Suchanek, Andrea Throop, Lyndon Tilson, Thomas Urban, Joe Voshell, Kimberly Wagner, Jonathan Williams, Mary Williamson, Qian Zeng, Tricia Zwiefelhofer, Clinton R. Paden, Duncan MacCannell |
| EPI_ISL_3326367 | TGen North | TGen North | Jolene Bowers, Heather Centner, Brett Van Tassel, Chris French, Hayley Yaglom, Darrin Lemmer, Dave Engelthaler |
| EPI_ISL_3326368 | Laboratory Corporation of America | Centers for Disease Control and Prevention Division of Viral Diseases, Pathogen Discovery | Dakota Howard, Dhwani Batra, Peter W. Cook, Kara Moser, Adrian Paskey, Jason Caravas, Benjamin Rambo-Martin, Shatavia Morrison, Christopher Gulvick, Scott Sammons, Yvette Unoarumhi, Darlene Wagner, Matthew Schmerer, Minoo Agarwal, Eyad Almasri, Debbie Boles, Ayla Burns, Nuthawin Charoensri, Oren Cohen, Susan Countryman, Mary Ann Cristobal, Bobbi Croy, Suzanne Dale, Hrushikesh Deshmukh, Amanda Douglas, Vincent Drouillon, Marcia Eisenberg, Howard Engler, Rama Ghatti, Prashant Gupta, Susan Hicks, Jake Humphrey, Lax Iyer, Lisa Pfefferle, Manoj Jain, Matthew Robinson, Mohan Kolli, Brian Krueger, Tim Kuphal, Stanley Letovsky, Michael Levandoski, Craig Lukasik, Jonathan Meltzer, Brian Norvell, Mindy Nye, Scott Parker, Christos Petropoulos, John Pruitt, Steven Ragan, Scott Ryan, Mike Sapeta, Jana Schroth, Suresh Babu Selvaraju, Goran Stevovic, Amanda Suchanek, Andrea Throop, Lyndon Tilson, Thomas Urban, Joe Voshell, Kimberly Wagner, Jonathan Williams, Mary Williamson, Qian Zeng, Tricia Zwiefelhofer, Clinton R. Paden, Duncan MacCannell |
| EPI_ISL_3326369 | TGen North | TGen North | Jolene Bowers, Heather Centner, Brett Van Tassel, Chris French, Hayley Yaglom, Darrin Lemmer, Dave Engelthaler |
| EPI_ISL_3326370, EPI_ISL_3326372, EPI_ISL_3326374 | Laboratory Corporation of America | Centers for Disease Control and Prevention Division of Viral Diseases, Pathogen Discovery | Dakota Howard, Dhwani Batra, Peter W. Cook, Kara Moser, Adrian Paskey, Jason Caravas, Benjamin Rambo-Martin, Shatavia Morrison, Christopher Gulvick, Scott Sammons, Yvette Unoarumhi, Darlene Wagner, Matthew Schmerer, Minoo Agarwal, Eyad Almasri, Debbie Boles, Ayla Burns, Nuthawin Charoensri, Oren Cohen, Susan Countryman, Mary Ann Cristobal, Bobbi Croy, Suzanne Dale, Hrushikesh Deshmukh, Amanda Douglas, Vincent Drouillon, Marcia Eisenberg, Howard Engler, Rama Ghatti, Prashant Gupta, Susan Hicks, Jake Humphrey, Lax Iyer, Lisa Pfefferle, Manoj Jain, Matthew Robinson, Mohan Kolli, Brian Krueger, Tim Kuphal, Stanley Letovsky, Michael Levandoski, Craig Lukasik, Jonathan Meltzer, Brian Norvell, Mindy Nye, Scott Parker, Christos Petropoulos, John Pruitt, Steven Ragan, Scott Ryan, Mike Sapeta, Jana Schroth, Suresh Babu Selvaraju, Goran Stevovic, Amanda Suchanek, Andrea Throop, Lyndon Tilson, Thomas Urban, Joe Voshell, Kimberly Wagner, Jonathan Williams, Mary Williamson, Qian Zeng, Tricia Zwiefelhofer, Clinton R. Paden, Duncan MacCannell |
| EPI_ISL_3326377 | TGen North | TGen North | Jolene Bowers, Heather Centner, Brett Van Tassel, Chris French, Hayley Yaglom, Darrin Lemmer, Dave Engelthaler |
| EPI_ISL_3326378 | Laboratory Corporation of America | Centers for Disease Control and Prevention Division of Viral Diseases, Pathogen Discovery | Dakota Howard, Dhwani Batra, Peter W. Cook, Kara Moser, Adrian Paskey, Jason Caravas, Benjamin Rambo-Martin, Shatavia Morrison, Christopher Gulvick, Scott Sammons, Yvette Unoarumhi, Darlene Wagner, Matthew Schmerer, Minoo Agarwal, Eyad Almasri, Debbie Boles, Ayla Burns, Nuthawin Charoensri, Oren Cohen, Susan Countryman, Mary Ann Cristobal, Bobbi Croy, Suzanne Dale, Hrushikesh Deshmukh, Amanda Douglas, Vincent Drouillon, Marcia Eisenberg, Howard Engler, Rama Ghatti, Prashant Gupta, Susan Hicks, Jake Humphrey, Lax Iyer, Lisa Pfefferle, Manoj Jain, Matthew Robinson, Mohan Kolli, Brian Krueger, Tim Kuphal, Stanley Letovsky, Michael Levandoski, Craig Lukasik, Jonathan Meltzer, Brian Norvell, Mindy Nye, Scott Parker, Christos Petropoulos, John Pruitt, Steven Ragan, Scott Ryan, Mike Sapeta, Jana Schroth, Suresh Babu Selvaraju, Goran Stevovic, Amanda Suchanek, Andrea Throop, Lyndon Tilson, Thomas Urban, Joe Voshell, Kimberly Wagner, Jonathan Williams, Mary Williamson, Qian Zeng, Tricia Zwiefelhofer, Clinton R. Paden, Duncan MacCannell |
| EPI_ISL_3326379 | TGen North | TGen North | Jolene Bowers, Heather Centner, Brett Van Tassel, Chris French, Hayley Yaglom, Darrin Lemmer, Dave Engelthaler |
| EPI_ISL_3326380, EPI_ISL_3326381 | Laboratory Corporation of America | Centers for Disease Control and Prevention Division of Viral Diseases, Pathogen Discovery | Dakota Howard, Dhwani Batra, Peter W. Cook, Kara Moser, Adrian Paskey, Jason Caravas, Benjamin Rambo-Martin, Shatavia Morrison, Christopher Gulvick, Scott Sammons, Yvette Unoarumhi, Darlene Wagner, Matthew Schmerer, Minoo Agarwal, Eyad Almasri, Debbie Boles, Ayla Burns, Nuthawin Charoensri, Oren Cohen, Susan Countryman, Mary Ann Cristobal, Bobbi Croy, Suzanne Dale, Hrushikesh Deshmukh, Amanda Douglas, Vincent Drouillon, Marcia Eisenberg, Howard Engler, Rama Ghatti, Prashant Gupta, Susan Hicks, Jake Humphrey, Lax Iyer, Lisa Pfefferle, Manoj Jain, Matthew Robinson, Mohan Kolli, Brian Krueger, Tim Kuphal, Stanley Letovsky, Michael Levandoski, Craig Lukasik, Jonathan Meltzer, Brian Norvell, Mindy Nye, Scott Parker, Christos Petropoulos, John Pruitt, Steven Ragan, Scott Ryan, Mike Sapeta, Jana Schroth, Suresh Babu Selvaraju, Goran Stevovic, Amanda Suchanek, Andrea Throop, Lyndon Tilson, Thomas Urban, Joe Voshell, Kimberly Wagner, Jonathan Williams, Mary Williamson, Qian Zeng, Tricia Zwiefelhofer, Clinton R. Paden, Duncan MacCannell |
| EPI_ISL_3326382 | TGen North | TGen North | Jolene Bowers, Heather Centner, Brett Van Tassel, Chris French, Hayley Yaglom, Darrin Lemmer, Dave Engelthaler |
| EPI_ISL_3326383, EPI_ISL_3326385 | Laboratory Corporation of America | Centers for Disease Control and Prevention Division of Viral Diseases, Pathogen Discovery | Dakota Howard, Dhwani Batra, Peter W. Cook, Kara Moser, Adrian Paskey, Jason Caravas, Benjamin Rambo-Martin, Shatavia Morrison, Christopher Gulvick, Scott Sammons, Yvette Unoarumhi, Darlene Wagner, Matthew Schmerer, Minoo Agarwal, Eyad Almasri, Debbie Boles, Ayla Burns, Nuthawin Charoensri, Oren Cohen, Susan Countryman, Mary Ann Cristobal, Bobbi Croy, Suzanne Dale, Hrushikesh Deshmukh, Amanda Douglas, Vincent Drouillon, Marcia Eisenberg, Howard Engler, Rama Ghatti, Prashant Gupta, Susan Hicks, Jake Humphrey, Lax Iyer, Lisa Pfefferle, Manoj Jain, Matthew Robinson, Mohan Kolli, Brian Krueger, Tim Kuphal, Stanley Letovsky, Michael Levandoski, Craig Lukasik, Jonathan Meltzer, Brian Norvell, Mindy Nye, Scott Parker, Christos Petropoulos, John Pruitt, Steven Ragan, Scott Ryan, Mike Sapeta, Jana Schroth, Suresh Babu Selvaraju, Goran Stevovic, Amanda Suchanek, Andrea Throop, Lyndon Tilson, Thomas Urban, Joe Voshell, Kimberly Wagner, Jonathan Williams, Mary Williamson, Qian Zeng, Tricia Zwiefelhofer, Clinton R. Paden, Duncan MacCannell |
| EPI_ISL_3326386 | TGen North | TGen North | Jolene Bowers, Heather Centner, Brett Van Tassel, Chris French, Hayley Yaglom, Darrin Lemmer, Dave Engelthaler |
| EPI_ISL_3326387 | Laboratory Corporation of America | Centers for Disease Control and Prevention Division of Viral Diseases, Pathogen Discovery | Dakota Howard, Dhwani Batra, Peter W. Cook, Kara Moser, Adrian Paskey, Jason Caravas, Benjamin Rambo-Martin, Shatavia Morrison, Christopher Gulvick, Scott Sammons, Yvette Unoarumhi, Darlene Wagner, Matthew Schmerer, Minoo Agarwal, Eyad Almasri, Debbie Boles, Ayla Burns, Nuthawin Charoensri, Oren Cohen, Susan Countryman, Mary Ann Cristobal, Bobbi Croy, Suzanne Dale, Hrushikesh Deshmukh, Amanda Douglas, Vincent Drouillon, Marcia Eisenberg, Howard Engler, Rama Ghatti, Prashant Gupta, Susan Hicks, Jake Humphrey, Lax Iyer, Lisa Pfefferle, Manoj Jain, Matthew Robinson, Mohan Kolli, Brian Krueger, Tim Kuphal, Stanley Letovsky, Michael Levandoski, Craig Lukasik, Jonathan Meltzer, Brian Norvell, Mindy Nye, Scott Parker, Christos Petropoulos, John Pruitt, Steven Ragan, Scott Ryan, Mike Sapeta, Jana Schroth, Suresh Babu Selvaraju, Goran Stevovic, Amanda Suchanek, Andrea Throop, Lyndon Tilson, Thomas Urban, Joe Voshell, Kimberly Wagner, Jonathan Williams, Mary Williamson, Qian Zeng, Tricia Zwiefelhofer, Clinton R. Paden, Duncan MacCannell |
| EPI_ISL_3326388 | TGen North | TGen North | Jolene Bowers, Heather Centner, Brett Van Tassel, Chris French, Hayley Yaglom, Darrin Lemmer, Dave Engelthaler |
| EPI_ISL_3326391, EPI_ISL_3326392, EPI_ISL_3326394, EPI_ISL_3326398, EPI_ISL_3326400, EPI_ISL_3326401 | Laboratory Corporation of America | Centers for Disease Control and Prevention Division of Viral Diseases, Pathogen Discovery | Dakota Howard, Dhwani Batra, Peter W. Cook, Kara Moser, Adrian Paskey, Jason Caravas, Benjamin Rambo-Martin, Shatavia Morrison, Christopher Gulvick, Scott Sammons, Yvette Unoarumhi, Darlene Wagner, Matthew Schmerer, Minoo Agarwal, Eyad Almasri, Debbie Boles, Ayla Burns, Nuthawin Charoensri, Oren Cohen, Susan Countryman, Mary Ann Cristobal, Bobbi Croy, Suzanne Dale, Hrushikesh Deshmukh, Amanda Douglas, Vincent Drouillon, Marcia Eisenberg, Howard Engler, Rama Ghatti, Prashant Gupta, Susan Hicks, Jake Humphrey, Lax Iyer, Lisa Pfefferle, Manoj Jain, Matthew Robinson, Mohan Kolli, Brian Krueger, Tim Kuphal, Stanley Letovsky, Michael Levandoski, Craig Lukasik, Jonathan Meltzer, Brian Norvell, Mindy Nye, Scott Parker, Christos Petropoulos, John Pruitt, Steven Ragan, Scott Ryan, Mike Sapeta, Jana Schroth, Suresh Babu Selvaraju, Goran Stevovic, Amanda Suchanek, Andrea Throop, Lyndon Tilson, Thomas Urban, Joe Voshell, Kimberly Wagner, Jonathan Williams, Mary Williamson, Qian Zeng, Tricia Zwiefelhofer, Clinton R. Paden, Duncan MacCannell |
| EPI_ISL_3326402 | TGen North | TGen North | Jolene Bowers, Heather Centner, Brett Van Tassel, Chris French, Hayley Yaglom, Darrin Lemmer, Dave Engelthaler |

| | | | |
|---|---|---|---|
| EPI_ISL_3326403 | Laboratory Corporation of America | Centers for Disease Control and Prevention Division of Viral Diseases, Pathogen Discovery | Dakota Howard, Dhwani Batra, Peter W. Cook, Kara Moser, Adrian Paskey, Jason Caravas, Benjamin Rambo-Martin, Shatavia Morrison, Christopher Gulvick, Scott Sammons, Yvette Unoarumhi, Darlene Wagner, Matthew Schmerer, Minoo Agarwal, Eyad Almasri, Debbie Boles, Ayla Burns, Nuthawin Charoensri, Oren Cohen, Susan Countryman, Mary Ann Cristobal, Bobbi Croy, Suzanne Dale, Hrushikesh Deshmukh, Amanda Douglas, Vincent Drouillon, Marcia Eisenberg, Howard Engler, Rama Ghatti, Prashant Gupta, Susan Hicks, Jake Humphrey, Lax Iyer, Lisa Pfefferle, Manoj Jain, Matthew Robinson, Mohan Kolli, Brian Krueger, Tim Kuphal, Stanley Letovsky, Michael Levandoski, Craig Lukasik, Jonathan Meltzer, Brian Norvell, Mindy Nye, Scott Parker, Christos Petropoulos, John Pruitt, Steven Ragan, Scott Ryan, Mike Sapeta, Jana Schroth, Suresh Babu Selvaraju, Goran Stevovic, Amanda Suchanek, Andrea Throop, Lyndon Tilson, Thomas Urban, Joe Voshell, Kimberly Wagner, Jonathan Williams, Mary Williamson, Qian Zeng, Tricia Zwiefelhofer, Clinton R. Paden, Duncan MacCannell |
| EPI_ISL_3326404 | TGen North | TGen North | Jolene Bowers, Heather Centner, Brett Van Tassel, Chris French, Hayley Yaglom, Darrin Lemmer, Dave Engelthaler |
| EPI_ISL_3326405 | Laboratory Corporation of America | Centers for Disease Control and Prevention Division of Viral Diseases, Pathogen Discovery | Dakota Howard, Dhwani Batra, Peter W. Cook, Kara Moser, Adrian Paskey, Jason Caravas, Benjamin Rambo-Martin, Shatavia Morrison, Christopher Gulvick, Scott Sammons, Yvette Unoarumhi, Darlene Wagner, Matthew Schmerer, Minoo Agarwal, Eyad Almasri, Debbie Boles, Ayla Burns, Nuthawin Charoensri, Oren Cohen, Susan Countryman, Mary Ann Cristobal, Bobbi Croy, Suzanne Dale, Hrushikesh Deshmukh, Amanda Douglas, Vincent Drouillon, Marcia Eisenberg, Howard Engler, Rama Ghatti, Prashant Gupta, Susan Hicks, Jake Humphrey, Lax Iyer, Lisa Pfefferle, Manoj Jain, Matthew Robinson, Mohan Kolli, Brian Krueger, Tim Kuphal, Stanley Letovsky, Michael Levandoski, Craig Lukasik, Jonathan Meltzer, Brian Norvell, Mindy Nye, Scott Parker, Christos Petropoulos, John Pruitt, Steven Ragan, Scott Ryan, Mike Sapeta, Jana Schroth, Suresh Babu Selvaraju, Goran Stevovic, Amanda Suchanek, Andrea Throop, Lyndon Tilson, Thomas Urban, Joe Voshell, Kimberly Wagner, Jonathan Williams, Mary Williamson, Qian Zeng, Tricia Zwiefelhofer, Clinton R. Paden, Duncan MacCannell |
| EPI_ISL_3326406 | TGen North | TGen North | Jolene Bowers, Heather Centner, Brett Van Tassel, Chris French, Hayley Yaglom, Darrin Lemmer, Dave Engelthaler |
| EPI_ISL_3326407 | Laboratory Corporation of America | Centers for Disease Control and Prevention Division of Viral Diseases, Pathogen Discovery | Dakota Howard, Dhwani Batra, Peter W. Cook, Kara Moser, Adrian Paskey, Jason Caravas, Benjamin Rambo-Martin, Shatavia Morrison, Christopher Gulvick, Scott Sammons, Yvette Unoarumhi, Darlene Wagner, Matthew Schmerer, Minoo Agarwal, Eyad Almasri, Debbie Boles, Ayla Burns, Nuthawin Charoensri, Oren Cohen, Susan Countryman, Mary Ann Cristobal, Bobbi Croy, Suzanne Dale, Hrushikesh Deshmukh, Amanda Douglas, Vincent Drouillon, Marcia Eisenberg, Howard Engler, Rama Ghatti, Prashant Gupta, Susan Hicks, Jake Humphrey, Lax Iyer, Lisa Pfefferle, Manoj Jain, Matthew Robinson, Mohan Kolli, Brian Krueger, Tim Kuphal, Stanley Letovsky, Michael Levandoski, Craig Lukasik, Jonathan Meltzer, Brian Norvell, Mindy Nye, Scott Parker, Christos Petropoulos, John Pruitt, Steven Ragan, Scott Ryan, Mike Sapeta, Jana Schroth, Suresh Babu Selvaraju, Goran Stevovic, Amanda Suchanek, Andrea Throop, Lyndon Tilson, Thomas Urban, Joe Voshell, Kimberly Wagner, Jonathan Williams, Mary Williamson, Qian Zeng, Tricia Zwiefelhofer, Clinton R. Paden, Duncan MacCannell |
| EPI_ISL_3326408 | TGen North | TGen North | Jolene Bowers, Heather Centner, Brett Van Tassel, Chris French, Hayley Yaglom, Darrin Lemmer, Dave Engelthaler |
| EPI_ISL_3326409 | Laboratory Corporation of America | Centers for Disease Control and Prevention Division of Viral Diseases, Pathogen Discovery | Dakota Howard, Dhwani Batra, Peter W. Cook, Kara Moser, Adrian Paskey, Jason Caravas, Benjamin Rambo-Martin, Shatavia Morrison, Christopher Gulvick, Scott Sammons, Yvette Unoarumhi, Darlene Wagner, Matthew Schmerer, Minoo Agarwal, Eyad Almasri, Debbie Boles, Ayla Burns, Nuthawin Charoensri, Oren Cohen, Susan Countryman, Mary Ann Cristobal, Bobbi Croy, Suzanne Dale, Hrushikesh Deshmukh, Amanda Douglas, Vincent Drouillon, Marcia Eisenberg, Howard Engler, Rama Ghatti, Prashant Gupta, Susan Hicks, Jake Humphrey, Lax Iyer, Lisa Pfefferle, Manoj Jain, Matthew Robinson, Mohan Kolli, Brian Krueger, Tim Kuphal, Stanley Letovsky, Michael Levandoski, Craig Lukasik, Jonathan Meltzer, Brian Norvell, Mindy Nye, Scott Parker, Christos Petropoulos, John Pruitt, Steven Ragan, Scott Ryan, Mike Sapeta, Jana Schroth, Suresh Babu Selvaraju, Goran Stevovic, Amanda Suchanek, Andrea Throop, Lyndon Tilson, Thomas Urban, Joe Voshell, Kimberly Wagner, Jonathan Williams, Mary Williamson, Qian Zeng, Tricia Zwiefelhofer, Clinton R. Paden, Duncan MacCannell |
| EPI_ISL_3326410 | TGen North | TGen North | Jolene Bowers, Heather Centner, Brett Van Tassel, Chris French, Hayley Yaglom, Darrin Lemmer, Dave Engelthaler |
| EPI_ISL_3326411, EPI_ISL_3326412 | Laboratory Corporation of America | Centers for Disease Control and Prevention Division of Viral Diseases, Pathogen Discovery | Dakota Howard, Dhwani Batra, Peter W. Cook, Kara Moser, Adrian Paskey, Jason Caravas, Benjamin Rambo-Martin, Shatavia Morrison, Christopher Gulvick, Scott Sammons, Yvette Unoarumhi, Darlene Wagner, Matthew Schmerer, Minoo Agarwal, Eyad Almasri, Debbie Boles, Ayla Burns, Nuthawin Charoensri, Oren Cohen, Susan Countryman, Mary Ann Cristobal, Bobbi Croy, Suzanne Dale, Hrushikesh Deshmukh, Amanda Douglas, Vincent Drouillon, Marcia Eisenberg, Howard Engler, Rama Ghatti, Prashant Gupta, Susan Hicks, Jake Humphrey, Lax Iyer, Lisa Pfefferle, Manoj Jain, Matthew Robinson, Mohan Kolli, Brian Krueger, Tim Kuphal, Stanley Letovsky, Michael Levandoski, Craig Lukasik, Jonathan Meltzer, Brian Norvell, Mindy Nye, Scott Parker, Christos Petropoulos, John Pruitt, Steven Ragan, Scott Ryan, Mike Sapeta, Jana Schroth, Suresh Babu Selvaraju, Goran Stevovic, Amanda Suchanek, Andrea Throop, Lyndon Tilson, Thomas Urban, Joe Voshell, Kimberly Wagner, Jonathan Williams, Mary Williamson, Qian Zeng, Tricia Zwiefelhofer, Clinton R. Paden, Duncan MacCannell |
| EPI_ISL_3326413 | TGen North | TGen North | Jolene Bowers, Heather Centner, Brett Van Tassel, Chris French, Hayley Yaglom, Darrin Lemmer, Dave Engelthaler |
| EPI_ISL_3326414 | Laboratory Corporation of America | Centers for Disease Control and Prevention Division of Viral Diseases, Pathogen Discovery | Dakota Howard, Dhwani Batra, Peter W. Cook, Kara Moser, Adrian Paskey, Jason Caravas, Benjamin Rambo-Martin, Shatavia Morrison, Christopher Gulvick, Scott Sammons, Yvette Unoarumhi, Darlene Wagner, Matthew Schmerer, Minoo Agarwal, Eyad Almasri, Debbie Boles, Ayla Burns, Nuthawin Charoensri, Oren Cohen, Susan Countryman, Mary Ann Cristobal, Bobbi Croy, Suzanne Dale, Hrushikesh Deshmukh, Amanda Douglas, Vincent Drouillon, Marcia Eisenberg, Howard Engler, Rama Ghatti, Prashant Gupta, Susan Hicks, Jake Humphrey, Lax Iyer, Lisa Pfefferle, Manoj Jain, Matthew Robinson, Mohan Kolli, Brian Krueger, Tim Kuphal, Stanley Letovsky, Michael Levandoski, Craig Lukasik, Jonathan Meltzer, Brian Norvell, Mindy Nye, Scott Parker, Christos Petropoulos, John Pruitt, Steven Ragan, Scott Ryan, Mike Sapeta, Jana Schroth, Suresh Babu Selvaraju, Goran Stevovic, Amanda Suchanek, Andrea Throop, Lyndon Tilson, Thomas Urban, Joe Voshell, Kimberly Wagner, Jonathan Williams, Mary Williamson, Qian Zeng, Tricia Zwiefelhofer, Clinton R. Paden, Duncan MacCannell |
| EPI_ISL_3326415 | TGen North | TGen North | Jolene Bowers, Heather Centner, Brett Van Tassel, Chris French, Hayley Yaglom, Darrin Lemmer, Dave Engelthaler |
| EPI_ISL_3326416, EPI_ISL_3326418 | Laboratory Corporation of America | Centers for Disease Control and Prevention Division of Viral Diseases, Pathogen Discovery | Dakota Howard, Dhwani Batra, Peter W. Cook, Kara Moser, Adrian Paskey, Jason Caravas, Benjamin Rambo-Martin, Shatavia Morrison, Christopher Gulvick, Scott Sammons, Yvette Unoarumhi, Darlene Wagner, Matthew Schmerer, Minoo Agarwal, Eyad Almasri, Debbie Boles, Ayla Burns, Nuthawin Charoensri, Oren Cohen, Susan Countryman, Mary Ann Cristobal, Bobbi Croy, Suzanne Dale, Hrushikesh Deshmukh, Amanda Douglas, Vincent Drouillon, Marcia Eisenberg, Howard Engler, Rama Ghatti, Prashant Gupta, Susan Hicks, Jake Humphrey, Lax Iyer, Lisa Pfefferle, Manoj Jain, Matthew Robinson, Mohan Kolli, Brian Krueger, Tim Kuphal, Stanley Letovsky, Michael Levandoski, Craig Lukasik, Jonathan Meltzer, Brian Norvell, Mindy Nye, Scott Parker, Christos Petropoulos, John Pruitt, Steven Ragan, Scott Ryan, Mike Sapeta, Jana Schroth, Suresh Babu Selvaraju, Goran Stevovic, Amanda Suchanek, Andrea Throop, Lyndon Tilson, Thomas Urban, Joe Voshell, Kimberly Wagner, Jonathan Williams, Mary Williamson, Qian Zeng, Tricia Zwiefelhofer, Clinton R. Paden, Duncan MacCannell |
| EPI_ISL_3326420 | TGen North | TGen North | Jolene Bowers, Heather Centner, Brett Van Tassel, Chris French, Hayley Yaglom, Darrin Lemmer, Dave Engelthaler |
| EPI_ISL_3326421 | Laboratory Corporation of America | Centers for Disease Control and Prevention Division of Viral Diseases, Pathogen Discovery | Dakota Howard, Dhwani Batra, Peter W. Cook, Kara Moser, Adrian Paskey, Jason Caravas, Benjamin Rambo-Martin, Shatavia Morrison, Christopher Gulvick, Scott Sammons, Yvette Unoarumhi, Darlene Wagner, Matthew Schmerer, Minoo Agarwal, Eyad Almasri, Debbie Boles, Ayla Burns, Nuthawin Charoensri, Oren Cohen, Susan Countryman, Mary Ann Cristobal, Bobbi Croy, Suzanne Dale, Hrushikesh Deshmukh, Amanda Douglas, Vincent Drouillon, Marcia Eisenberg, Howard Engler, Rama Ghatti, Prashant Gupta, Susan Hicks, Jake Humphrey, Lax Iyer, Lisa Pfefferle, Manoj Jain, Matthew Robinson, Mohan Kolli, Brian Krueger, Tim Kuphal, Stanley Letovsky, Michael Levandoski, Craig Lukasik, Jonathan Meltzer, Brian Norvell, Mindy Nye, Scott Parker, Christos Petropoulos, John Pruitt, Steven Ragan, Scott Ryan, Mike Sapeta, Jana Schroth, Suresh Babu Selvaraju, Goran Stevovic, Amanda Suchanek, Andrea Throop, Lyndon Tilson, Thomas Urban, Joe Voshell, Kimberly Wagner, Jonathan Williams, Mary Williamson, Qian Zeng, Tricia Zwiefelhofer, Clinton R. Paden, Duncan MacCannell |
| EPI_ISL_3326422 | TGen North | TGen North | Jolene Bowers, Heather Centner, Brett Van Tassel, Chris French, Hayley Yaglom, Darrin Lemmer, Dave Engelthaler |
| EPI_ISL_3326423 | Laboratory Corporation of America | Centers for Disease Control and Prevention Division of Viral Diseases, Pathogen Discovery | Dakota Howard, Dhwani Batra, Peter W. Cook, Kara Moser, Adrian Paskey, Jason Caravas, Benjamin Rambo-Martin, Shatavia Morrison, Christopher Gulvick, Scott Sammons, Yvette Unoarumhi, Darlene Wagner, Matthew Schmerer, Minoo Agarwal, Eyad Almasri, Debbie Boles, Ayla Burns, Nuthawin Charoensri, Oren Cohen, Susan Countryman, Mary Ann Cristobal, Bobbi Croy, Suzanne Dale, Hrushikesh Deshmukh, Amanda Douglas, Vincent Drouillon, Marcia Eisenberg, Howard Engler, Rama Ghatti, Prashant Gupta, Susan Hicks, Jake Humphrey, Lax Iyer, Lisa Pfefferle, Manoj Jain, Matthew Robinson, Mohan Kolli, Brian Krueger, Tim Kuphal, Stanley Letovsky, Michael Levandoski, Craig Lukasik, Jonathan Meltzer, Brian Norvell, Mindy Nye, Scott Parker, Christos Petropoulos, John Pruitt, Steven Ragan, Scott Ryan, Mike Sapeta, Jana Schroth, Suresh Babu Selvaraju, Goran Stevovic, Amanda Suchanek, Andrea Throop, Lyndon Tilson, Thomas Urban, Joe Voshell, Kimberly Wagner, Jonathan Williams, Mary Williamson, Qian Zeng, Tricia Zwiefelhofer, Clinton R. Paden, Duncan MacCannell |
| EPI_ISL_3326424 | TGen North | TGen North | Jolene Bowers, Heather Centner, Brett Van Tassel, Chris French, Hayley Yaglom, Darrin Lemmer, Dave Engelthaler |
| EPI_ISL_3326425, EPI_ISL_3326427 | Laboratory Corporation of America | Centers for Disease Control and Prevention Division of Viral Diseases, Pathogen Discovery | Dakota Howard, Dhwani Batra, Peter W. Cook, Kara Moser, Adrian Paskey, Jason Caravas, Benjamin Rambo-Martin, Shatavia Morrison, Christopher Gulvick, Scott Sammons, Yvette Unoarumhi, Darlene Wagner, Matthew Schmerer, Minoo Agarwal, Eyad Almasri, Debbie Boles, Ayla Burns, Nuthawin Charoensri, Oren Cohen, Susan Countryman, Mary Ann Cristobal, Bobbi Croy, Suzanne Dale, Hrushikesh Deshmukh, Amanda Douglas, Vincent Drouillon, Marcia Eisenberg, Howard Engler, Rama Ghatti, Prashant Gupta, Susan Hicks, Jake Humphrey, Lax Iyer, Lisa Pfefferle, Manoj Jain, Matthew Robinson, Mohan Kolli, Brian Krueger, Tim Kuphal, Stanley Letovsky, Michael Levandoski, Craig Lukasik, Jonathan Meltzer, Brian Norvell, Mindy Nye, Scott Parker, Christos Petropoulos, John Pruitt, Steven Ragan, Scott Ryan, Mike Sapeta, Jana Schroth, Suresh Babu Selvaraju, Goran Stevovic, Amanda Suchanek, Andrea Throop, Lyndon Tilson, Thomas Urban, Joe Voshell, Kimberly Wagner, Jonathan Williams, Mary Williamson, Qian Zeng, Tricia Zwiefelhofer, Clinton R. Paden, Duncan MacCannell |
| EPI_ISL_3326428 | TGen North | TGen North | Jolene Bowers, Heather Centner, Brett Van Tassel, Chris French, Hayley Yaglom, Darrin Lemmer, Dave Engelthaler |
| EPI_ISL_3326429 | Laboratory Corporation of America | Centers for Disease Control and Prevention Division of Viral Diseases, Pathogen Discovery | Dakota Howard, Dhwani Batra, Peter W. Cook, Kara Moser, Adrian Paskey, Jason Caravas, Benjamin Rambo-Martin, Shatavia Morrison, Christopher Gulvick, Scott Sammons, Yvette Unoarumhi, Darlene Wagner, Matthew Schmerer, Minoo Agarwal, Eyad Almasri, Debbie Boles, Ayla Burns, Nuthawin Charoensri, Oren Cohen, Susan Countryman, Mary Ann Cristobal, Bobbi Croy, Suzanne Dale, Hrushikesh Deshmukh, Amanda Douglas, Vincent Drouillon, Marcia Eisenberg, Howard Engler, Rama Ghatti, Prashant Gupta, Susan Hicks, Jake Humphrey, Lax Iyer, Lisa Pfefferle, Manoj Jain, Matthew Robinson, Mohan Kolli, Brian Krueger, Tim Kuphal, Stanley Letovsky, Michael Levandoski, Craig Lukasik, Jonathan Meltzer, Brian Norvell, Mindy Nye, Scott Parker, Christos Petropoulos, John Pruitt, Steven Ragan, Scott Ryan, Mike Sapeta, Jana Schroth, Suresh Babu Selvaraju, Goran Stevovic, Amanda Suchanek, Andrea Throop, Lyndon Tilson, Thomas Urban, Joe Voshell, Kimberly Wagner, Jonathan Williams, Mary Williamson, Qian Zeng, Tricia Zwiefelhofer, Clinton R. Paden, Duncan MacCannell |
| EPI_ISL_3326430 | TGen North | TGen North | Jolene Bowers, Heather Centner, Brett Van Tassel, Chris French, Hayley Yaglom, Darrin Lemmer, Dave Engelthaler |
| EPI_ISL_3326431 | Laboratory Corporation of America | Centers for Disease Control and Prevention Division of Viral Diseases, Pathogen Discovery | Dakota Howard, Dhwani Batra, Peter W. Cook, Kara Moser, Adrian Paskey, Jason Caravas, Benjamin Rambo-Martin, Shatavia Morrison, Christopher Gulvick, Scott Sammons, Yvette Unoarumhi, Darlene Wagner, Matthew Schmerer, Minoo Agarwal, Eyad Almasri, Debbie Boles, Ayla Burns, Nuthawin Charoensri, Oren Cohen, Susan Countryman, Mary Ann Cristobal, Bobbi Croy, Suzanne Dale, Hrushikesh Deshmukh, Amanda Douglas, Vincent Drouillon, Marcia Eisenberg, Howard Engler, Rama Ghatti, Prashant Gupta, Susan Hicks, Jake Humphrey, Lax Iyer, Lisa Pfefferle, Manoj Jain, Matthew Robinson, Mohan Kolli, Brian Krueger, Tim Kuphal, Stanley Letovsky, Michael Levandoski, Craig Lukasik, Jonathan Meltzer, Brian Norvell, Mindy Nye, Scott Parker, Christos Petropoulos, John Pruitt, Steven Ragan, Scott Ryan, Mike Sapeta, Jana Schroth, Suresh Babu Selvaraju, Goran Stevovic, Amanda Suchanek, Andrea Throop, Lyndon Tilson, Thomas Urban, Joe Voshell, Kimberly Wagner, Jonathan Williams, Mary Williamson, Qian Zeng, Tricia Zwiefelhofer, Clinton R. Paden, Duncan MacCannell |
| EPI_ISL_3326432 | TGen North | TGen North | Jolene Bowers, Heather Centner, Brett Van Tassel, Chris French, Hayley Yaglom, Darrin Lemmer, Dave Engelthaler |
| EPI_ISL_3326433, EPI_ISL_3326434 | Laboratory Corporation of America | Centers for Disease Control and Prevention Division of Viral Diseases, Pathogen Discovery | Dakota Howard, Dhwani Batra, Peter W. Cook, Kara Moser, Adrian Paskey, Jason Caravas, Benjamin Rambo-Martin, Shatavia Morrison, Christopher Gulvick, Scott Sammons, Yvette Unoarumhi, Darlene Wagner, Matthew Schmerer, Minoo Agarwal, Eyad Almasri, Debbie Boles, Ayla Burns, Nuthawin Charoensri, Oren Cohen, Susan Countryman, Mary Ann Cristobal, Bobbi Croy, Suzanne Dale, Hrushikesh Deshmukh, Amanda Douglas, Vincent Drouillon, Marcia Eisenberg, Howard Engler, Rama Ghatti, Prashant Gupta, Susan Hicks, Jake Humphrey, Lax Iyer, Lisa Pfefferle, Manoj Jain, Matthew Robinson, Mohan Kolli, Brian Krueger, Tim Kuphal, Stanley Letovsky, Michael Levandoski, Craig Lukasik, Jonathan Meltzer, Brian Norvell, Mindy Nye, Scott Parker, Christos Petropoulos, John Pruitt, Steven Ragan, Scott Ryan, Mike Sapeta, Jana Schroth, Suresh Babu Selvaraju, Goran Stevovic, Amanda Suchanek, Andrea Throop, Lyndon Tilson, Thomas Urban, Joe Voshell, Kimberly Wagner, Jonathan Williams, Mary Williamson, Qian Zeng, Tricia Zwiefelhofer, Clinton R. Paden, Duncan MacCannell |
| EPI_ISL_3326435 | TGen North | TGen North | Jolene Bowers, Heather Centner, Brett Van Tassel, Chris French, Hayley Yaglom, Darrin Lemmer, Dave Engelthaler |
| EPI_ISL_3326436, EPI_ISL_3326438 | Laboratory Corporation of America | Centers for Disease Control and Prevention Division of Viral Diseases, Pathogen Discovery | Dakota Howard, Dhwani Batra, Peter W. Cook, Kara Moser, Adrian Paskey, Jason Caravas, Benjamin Rambo-Martin, Shatavia Morrison, Christopher Gulvick, Scott Sammons, Yvette Unoarumhi, Darlene Wagner, Matthew Schmerer, Minoo Agarwal, Eyad Almasri, Debbie Boles, Ayla Burns, Nuthawin Charoensri, Oren Cohen, Susan Countryman, Mary Ann Cristobal, Bobbi Croy, Suzanne Dale, Hrushikesh Deshmukh, Amanda Douglas, Vincent Drouillon, Marcia Eisenberg, Howard Engler, Rama Ghatti, Prashant Gupta, Susan Hicks, Jake Humphrey, Lax Iyer, Lisa Pfefferle, Manoj Jain, Matthew Robinson, Mohan Kolli, Brian Krueger, Tim Kuphal, Stanley Letovsky, Michael Levandoski, Craig Lukasik, Jonathan Meltzer, Brian Norvell, Mindy Nye, Scott Parker, Christos Petropoulos, John Pruitt, Steven Ragan, Scott Ryan, Mike Sapeta, Jana Schroth, Suresh Babu Selvaraju, Goran Stevovic, Amanda Suchanek, Andrea Throop, Lyndon Tilson, Thomas Urban, Joe Voshell, Kimberly Wagner, Jonathan Williams, Mary Williamson, Qian Zeng, Tricia Zwiefelhofer, Clinton R. Paden, Duncan MacCannell |
| EPI_ISL_3326439 | TGen North | TGen North | Jolene Bowers, Heather Centner, Brett Van Tassel, Chris French, Hayley Yaglom, Darrin Lemmer, Dave Engelthaler |
| EPI_ISL_3326440 | Laboratory Corporation of America | Centers for Disease Control and Prevention Division of Viral Diseases, Pathogen Discovery | Dakota Howard, Dhwani Batra, Peter W. Cook, Kara Moser, Adrian Paskey, Jason Caravas, Benjamin Rambo-Martin, Shatavia Morrison, Christopher Gulvick, Scott Sammons, Yvette Unoarumhi, Darlene Wagner, Matthew Schmerer, Minoo Agarwal, Eyad Almasri, Debbie Boles, Ayla Burns, Nuthawin Charoensri, Oren Cohen, Susan Countryman, Mary Ann Cristobal, Bobbi Croy, Suzanne Dale, Hrushikesh Deshmukh, Amanda Douglas, Vincent Drouillon, Marcia Eisenberg, Howard Engler, Rama Ghatti, Prashant Gupta, Susan Hicks, Jake Humphrey, Lax Iyer, Lisa Pfefferle, Manoj Jain, Matthew Robinson, Mohan Kolli, Brian Krueger, Tim Kuphal, Stanley Letovsky, Michael Levandoski, Craig Lukasik, Jonathan Meltzer, Brian Norvell, Mindy Nye, Scott Parker, Christos Petropoulos, John Pruitt, Steven Ragan, Scott Ryan, Mike Sapeta, Jana Schroth, Suresh Babu Selvaraju, Goran Stevovic, Amanda Suchanek, Andrea Throop, Lyndon Tilson, Thomas Urban, Joe Voshell, Kimberly Wagner, Jonathan Williams, Mary Williamson, Qian Zeng, Tricia Zwiefelhofer, Clinton R. Paden, Duncan MacCannell |
| EPI_ISL_3326441 | TGen North | TGen North | Jolene Bowers, Heather Centner, Brett Van Tassel, Chris French, Hayley Yaglom, Darrin Lemmer, Dave Engelthaler |
| EPI_ISL_3326442, EPI_ISL_3326444, EPI_ISL_3326445 | Laboratory Corporation of America | Centers for Disease Control and Prevention Division of Viral Diseases, Pathogen Discovery | Dakota Howard, Dhwani Batra, Peter W. Cook, Kara Moser, Adrian Paskey, Jason Caravas, Benjamin Rambo-Martin, Shatavia Morrison, Christopher Gulvick, Scott Sammons, Yvette Unoarumhi, Darlene Wagner, Matthew Schmerer, Minoo Agarwal, Eyad Almasri, Debbie Boles, Ayla Burns, Nuthawin Charoensri, Oren Cohen, Susan Countryman, Mary Ann Cristobal, Bobbi Croy, Suzanne Dale, Hrushikesh Deshmukh, Amanda Douglas, Vincent Drouillon, Marcia Eisenberg, Howard Engler, Rama Ghatti, Prashant Gupta, Susan Hicks, Jake Humphrey, Lax Iyer, Lisa Pfefferle, Manoj Jain, Matthew Robinson, Mohan Kolli, Brian Krueger, Tim Kuphal, Stanley Letovsky, Michael Levandoski, Craig Lukasik, Jonathan Meltzer, Brian Norvell, Mindy Nye, Scott Parker, Christos Petropoulos, John Pruitt, Steven Ragan, Scott Ryan, Mike Sapeta, Jana Schroth, Suresh Babu Selvaraju, Goran Stevovic, Amanda Suchanek, Andrea Throop, Lyndon Tilson, Thomas Urban, Joe Voshell, Kimberly Wagner, Jonathan Williams, Mary Williamson, Qian Zeng, Tricia Zwiefelhofer, Clinton R. Paden, Duncan MacCannell |

| | | | |
|---|---|---|---|
| EPI_ISL_3326446 | TGen North | TGen North | Jolene Bowers, Heather Centner, Brett Van Tassel, Chris French, Hayley Yaglom, Darrin Lemmer, Dave Engelthaler |
| EPI_ISL_3326447, EPI_ISL_3326449 | Laboratory Corporation of America | Centers for Disease Control and Prevention Division of Viral Diseases, Pathogen Discovery | Dakota Howard, Dhwani Batra, Peter W. Cook, Kara Moser, Adrian Paskey, Jason Caravas, Benjamin Rambo-Martin, Shatavia Morrison, Christopher Gulvick, Scott Sammons, Yvette Unoarumhi, Darlene Wagner, Matthew Schmerer, Minoo Agarwal, Eyad Almasri, Debbie Boles, Ayla Burns, Nuthawin Charoensri, Oren Cohen, Susan Countryman, Mary Ann Cristobal, Bobbi Croy, Suzanne Dale, Hrushikesh Deshmukh, Amanda Douglas, Vincent Drouillon, Marcia Eisenberg, Howard Engler, Rama Ghatti, Prashant Gupta, Susan Hicks, Jake Humphrey, Lax Iyer, Lisa Pfefferle, Manoj Jain, Matthew Robinson, Mohan Kolli, Brian Krueger, Tim Kuphal, Stanley Letovsky, Michael Levandoski, Craig Lukasik, Jonathan Meltzer, Brian Norvell, Mindy Nye, Scott Parker, Christos Petropoulos, John Pruitt, Steven Ragan, Scott Ryan, Mike Sapeta, Jana Schroth, Suresh Babu Selvaraju, Goran Stevovic, Amanda Suchanek, Andrea Throop, Lyndon Tilson, Thomas Urban, Joe Voshell, Kimberly Wagner, Jonathan Williams, Mary Williamson, Qian Zeng, Tricia Zwiefelhofer, Clinton R. Paden, Duncan MacCannell |
| EPI_ISL_3326450 | TGen North | TGen North | Jolene Bowers, Heather Centner, Brett Van Tassel, Chris French, Hayley Yaglom, Darrin Lemmer, Dave Engelthaler |
| EPI_ISL_3326451 | Laboratory Corporation of America | Centers for Disease Control and Prevention Division of Viral Diseases, Pathogen Discovery | Dakota Howard, Dhwani Batra, Peter W. Cook, Kara Moser, Adrian Paskey, Jason Caravas, Benjamin Rambo-Martin, Shatavia Morrison, Christopher Gulvick, Scott Sammons, Yvette Unoarumhi, Darlene Wagner, Matthew Schmerer, Minoo Agarwal, Eyad Almasri, Debbie Boles, Ayla Burns, Nuthawin Charoensri, Oren Cohen, Susan Countryman, Mary Ann Cristobal, Bobbi Croy, Suzanne Dale, Hrushikesh Deshmukh, Amanda Douglas, Vincent Drouillon, Marcia Eisenberg, Howard Engler, Rama Ghatti, Prashant Gupta, Susan Hicks, Jake Humphrey, Lax Iyer, Lisa Pfefferle, Manoj Jain, Matthew Robinson, Mohan Kolli, Brian Krueger, Tim Kuphal, Stanley Letovsky, Michael Levandoski, Craig Lukasik, Jonathan Meltzer, Brian Norvell, Mindy Nye, Scott Parker, Christos Petropoulos, John Pruitt, Steven Ragan, Scott Ryan, Mike Sapeta, Jana Schroth, Suresh Babu Selvaraju, Goran Stevovic, Amanda Suchanek, Andrea Throop, Lyndon Tilson, Thomas Urban, Joe Voshell, Kimberly Wagner, Jonathan Williams, Mary Williamson, Qian Zeng, Tricia Zwiefelhofer, Clinton R. Paden, Duncan MacCannell |
| EPI_ISL_3326452 | TGen North | TGen North | Jolene Bowers, Heather Centner, Brett Van Tassel, Chris French, Hayley Yaglom, Darrin Lemmer, Dave Engelthaler |
| EPI_ISL_3326453 | Laboratory Corporation of America | Centers for Disease Control and Prevention Division of Viral Diseases, Pathogen Discovery | Dakota Howard, Dhwani Batra, Peter W. Cook, Kara Moser, Adrian Paskey, Jason Caravas, Benjamin Rambo-Martin, Shatavia Morrison, Christopher Gulvick, Scott Sammons, Yvette Unoarumhi, Darlene Wagner, Matthew Schmerer, Minoo Agarwal, Eyad Almasri, Debbie Boles, Ayla Burns, Nuthawin Charoensri, Oren Cohen, Susan Countryman, Mary Ann Cristobal, Bobbi Croy, Suzanne Dale, Hrushikesh Deshmukh, Amanda Douglas, Vincent Drouillon, Marcia Eisenberg, Howard Engler, Rama Ghatti, Prashant Gupta, Susan Hicks, Jake Humphrey, Lax Iyer, Lisa Pfefferle, Manoj Jain, Matthew Robinson, Mohan Kolli, Brian Krueger, Tim Kuphal, Stanley Letovsky, Michael Levandoski, Craig Lukasik, Jonathan Meltzer, Brian Norvell, Mindy Nye, Scott Parker, Christos Petropoulos, John Pruitt, Steven Ragan, Scott Ryan, Mike Sapeta, Jana Schroth, Suresh Babu Selvaraju, Goran Stevovic, Amanda Suchanek, Andrea Throop, Lyndon Tilson, Thomas Urban, Joe Voshell, Kimberly Wagner, Jonathan Williams, Mary Williamson, Qian Zeng, Tricia Zwiefelhofer, Clinton R. Paden, Duncan MacCannell |
| EPI_ISL_3326454 | TGen North | TGen North | Jolene Bowers, Heather Centner, Brett Van Tassel, Chris French, Hayley Yaglom, Darrin Lemmer, Dave Engelthaler |
| EPI_ISL_3326455 | Laboratory Corporation of America | Centers for Disease Control and Prevention Division of Viral Diseases, Pathogen Discovery | Dakota Howard, Dhwani Batra, Peter W. Cook, Kara Moser, Adrian Paskey, Jason Caravas, Benjamin Rambo-Martin, Shatavia Morrison, Christopher Gulvick, Scott Sammons, Yvette Unoarumhi, Darlene Wagner, Matthew Schmerer, Minoo Agarwal, Eyad Almasri, Debbie Boles, Ayla Burns, Nuthawin Charoensri, Oren Cohen, Susan Countryman, Mary Ann Cristobal, Bobbi Croy, Suzanne Dale, Hrushikesh Deshmukh, Amanda Douglas, Vincent Drouillon, Marcia Eisenberg, Howard Engler, Rama Ghatti, Prashant Gupta, Susan Hicks, Jake Humphrey, Lax Iyer, Lisa Pfefferle, Manoj Jain, Matthew Robinson, Mohan Kolli, Brian Krueger, Tim Kuphal, Stanley Letovsky, Michael Levandoski, Craig Lukasik, Jonathan Meltzer, Brian Norvell, Mindy Nye, Scott Parker, Christos Petropoulos, John Pruitt, Steven Ragan, Scott Ryan, Mike Sapeta, Jana Schroth, Suresh Babu Selvaraju, Goran Stevovic, Amanda Suchanek, Andrea Throop, Lyndon Tilson, Thomas Urban, Joe Voshell, Kimberly Wagner, Jonathan Williams, Mary Williamson, Qian Zeng, Tricia Zwiefelhofer, Clinton R. Paden, Duncan MacCannell |
| EPI_ISL_3326456 | TGen North | TGen North | Jolene Bowers, Heather Centner, Brett Van Tassel, Chris French, Hayley Yaglom, Darrin Lemmer, Dave Engelthaler |
| EPI_ISL_3326457 | Laboratory Corporation of America | Centers for Disease Control and Prevention Division of Viral Diseases, Pathogen Discovery | Dakota Howard, Dhwani Batra, Peter W. Cook, Kara Moser, Adrian Paskey, Jason Caravas, Benjamin Rambo-Martin, Shatavia Morrison, Christopher Gulvick, Scott Sammons, Yvette Unoarumhi, Darlene Wagner, Matthew Schmerer, Minoo Agarwal, Eyad Almasri, Debbie Boles, Ayla Burns, Nuthawin Charoensri, Oren Cohen, Susan Countryman, Mary Ann Cristobal, Bobbi Croy, Suzanne Dale, Hrushikesh Deshmukh, Amanda Douglas, Vincent Drouillon, Marcia Eisenberg, Howard Engler, Rama Ghatti, Prashant Gupta, Susan Hicks, Jake Humphrey, Lax Iyer, Lisa Pfefferle, Manoj Jain, Matthew Robinson, Mohan Kolli, Brian Krueger, Tim Kuphal, Stanley Letovsky, Michael Levandoski, Craig Lukasik, Jonathan Meltzer, Brian Norvell, Mindy Nye, Scott Parker, Christos Petropoulos, John Pruitt, Steven Ragan, Scott Ryan, Mike Sapeta, Jana Schroth, Suresh Babu Selvaraju, Goran Stevovic, Amanda Suchanek, Andrea Throop, Lyndon Tilson, Thomas Urban, Joe Voshell, Kimberly Wagner, Jonathan Williams, Mary Williamson, Qian Zeng, Tricia Zwiefelhofer, Clinton R. Paden, Duncan MacCannell |
| EPI_ISL_3326458 | TGen North | TGen North | Jolene Bowers, Heather Centner, Brett Van Tassel, Chris French, Hayley Yaglom, Darrin Lemmer, Dave Engelthaler |
| EPI_ISL_3326459 | Laboratory Corporation of America | Centers for Disease Control and Prevention Division of Viral Diseases, Pathogen Discovery | Dakota Howard, Dhwani Batra, Peter W. Cook, Kara Moser, Adrian Paskey, Jason Caravas, Benjamin Rambo-Martin, Shatavia Morrison, Christopher Gulvick, Scott Sammons, Yvette Unoarumhi, Darlene Wagner, Matthew Schmerer, Minoo Agarwal, Eyad Almasri, Debbie Boles, Ayla Burns, Nuthawin Charoensri, Oren Cohen, Susan Countryman, Mary Ann Cristobal, Bobbi Croy, Suzanne Dale, Hrushikesh Deshmukh, Amanda Douglas, Vincent Drouillon, Marcia Eisenberg, Howard Engler, Rama Ghatti, Prashant Gupta, Susan Hicks, Jake Humphrey, Lax Iyer, Lisa Pfefferle, Manoj Jain, Matthew Robinson, Mohan Kolli, Brian Krueger, Tim Kuphal, Stanley Letovsky, Michael Levandoski, Craig Lukasik, Jonathan Meltzer, Brian Norvell, Mindy Nye, Scott Parker, Christos Petropoulos, John Pruitt, Steven Ragan, Scott Ryan, Mike Sapeta, Jana Schroth, Suresh Babu Selvaraju, Goran Stevovic, Amanda Suchanek, Andrea Throop, Lyndon Tilson, Thomas Urban, Joe Voshell, Kimberly Wagner, Jonathan Williams, Mary Williamson, Qian Zeng, Tricia Zwiefelhofer, Clinton R. Paden, Duncan MacCannell |
| EPI_ISL_3326460 | TGen North | TGen North | Jolene Bowers, Heather Centner, Brett Van Tassel, Chris French, Hayley Yaglom, Darrin Lemmer, Dave Engelthaler |
| EPI_ISL_3326461, EPI_ISL_3326463, EPI_ISL_3326464 | Laboratory Corporation of America | Centers for Disease Control and Prevention Division of Viral Diseases, Pathogen Discovery | Dakota Howard, Dhwani Batra, Peter W. Cook, Kara Moser, Adrian Paskey, Jason Caravas, Benjamin Rambo-Martin, Shatavia Morrison, Christopher Gulvick, Scott Sammons, Yvette Unoarumhi, Darlene Wagner, Matthew Schmerer, Minoo Agarwal, Eyad Almasri, Debbie Boles, Ayla Burns, Nuthawin Charoensri, Oren Cohen, Susan Countryman, Mary Ann Cristobal, Bobbi Croy, Suzanne Dale, Hrushikesh Deshmukh, Amanda Douglas, Vincent Drouillon, Marcia Eisenberg, Howard Engler, Rama Ghatti, Prashant Gupta, Susan Hicks, Jake Humphrey, Lax Iyer, Lisa Pfefferle, Manoj Jain, Matthew Robinson, Mohan Kolli, Brian Krueger, Tim Kuphal, Stanley Letovsky, Michael Levandoski, Craig Lukasik, Jonathan Meltzer, Brian Norvell, Mindy Nye, Scott Parker, Christos Petropoulos, John Pruitt, Steven Ragan, Scott Ryan, Mike Sapeta, Jana Schroth, Suresh Babu Selvaraju, Goran Stevovic, Amanda Suchanek, Andrea Throop, Lyndon Tilson, Thomas Urban, Joe Voshell, Kimberly Wagner, Jonathan Williams, Mary Williamson, Qian Zeng, Tricia Zwiefelhofer, Clinton R. Paden, Duncan MacCannell |
| EPI_ISL_3326465 | TGen North | TGen North | Jolene Bowers, Heather Centner, Brett Van Tassel, Chris French, Hayley Yaglom, Darrin Lemmer, Dave Engelthaler |
| EPI_ISL_3326466 | Laboratory Corporation of America | Centers for Disease Control and Prevention Division of Viral Diseases, Pathogen Discovery | Dakota Howard, Dhwani Batra, Peter W. Cook, Kara Moser, Adrian Paskey, Jason Caravas, Benjamin Rambo-Martin, Shatavia Morrison, Christopher Gulvick, Scott Sammons, Yvette Unoarumhi, Darlene Wagner, Matthew Schmerer, Minoo Agarwal, Eyad Almasri, Debbie Boles, Ayla Burns, Nuthawin Charoensri, Oren Cohen, Susan Countryman, Mary Ann Cristobal, Bobbi Croy, Suzanne Dale, Hrushikesh Deshmukh, Amanda Douglas, Vincent Drouillon, Marcia Eisenberg, Howard Engler, Rama Ghatti, Prashant Gupta, Susan Hicks, Jake Humphrey, Lax Iyer, Lisa Pfefferle, Manoj Jain, Matthew Robinson, Mohan Kolli, Brian Krueger, Tim Kuphal, Stanley Letovsky, Michael Levandoski, Craig Lukasik, Jonathan Meltzer, Brian Norvell, Mindy Nye, Scott Parker, Christos Petropoulos, John Pruitt, Steven Ragan, Scott Ryan, Mike Sapeta, Jana Schroth, Suresh Babu Selvaraju, Goran Stevovic, Amanda Suchanek, Andrea Throop, Lyndon Tilson, Thomas Urban, Joe Voshell, Kimberly Wagner, Jonathan Williams, Mary Williamson, Qian Zeng, Tricia Zwiefelhofer, Clinton R. Paden, Duncan MacCannell |
| EPI_ISL_3326467 | TGen North | TGen North | Jolene Bowers, Heather Centner, Brett Van Tassel, Chris French, Hayley Yaglom, Darrin Lemmer, Dave Engelthaler |
| EPI_ISL_3326468, EPI_ISL_3326469, EPI_ISL_3326471 | Laboratory Corporation of America | Centers for Disease Control and Prevention Division of Viral Diseases, Pathogen Discovery | Dakota Howard, Dhwani Batra, Peter W. Cook, Kara Moser, Adrian Paskey, Jason Caravas, Benjamin Rambo-Martin, Shatavia Morrison, Christopher Gulvick, Scott Sammons, Yvette Unoarumhi, Darlene Wagner, Matthew Schmerer, Minoo Agarwal, Eyad Almasri, Debbie Boles, Ayla Burns, Nuthawin Charoensri, Oren Cohen, Susan Countryman, Mary Ann Cristobal, Bobbi Croy, Suzanne Dale, Hrushikesh Deshmukh, Amanda Douglas, Vincent Drouillon, Marcia Eisenberg, Howard Engler, Rama Ghatti, Prashant Gupta, Susan Hicks, Jake Humphrey, Lax Iyer, Lisa Pfefferle, Manoj Jain, Matthew Robinson, Mohan Kolli, Brian Krueger, Tim Kuphal, Stanley Letovsky, Michael Levandoski, Craig Lukasik, Jonathan Meltzer, Brian Norvell, Mindy Nye, Scott Parker, Christos Petropoulos, John Pruitt, Steven Ragan, Scott Ryan, Mike Sapeta, Jana Schroth, Suresh Babu Selvaraju, Goran Stevovic, Amanda Suchanek, Andrea Throop, Lyndon Tilson, Thomas Urban, Joe Voshell, Kimberly Wagner, Jonathan Williams, Mary Williamson, Qian Zeng, Tricia Zwiefelhofer, Clinton R. Paden, Duncan MacCannell |
| EPI_ISL_3326472 | TGen North | TGen North | Jolene Bowers, Heather Centner, Brett Van Tassel, Chris French, Hayley Yaglom, Darrin Lemmer, Dave Engelthaler |
| EPI_ISL_3326473 | Laboratory Corporation of America | Centers for Disease Control and Prevention Division of Viral Diseases, Pathogen Discovery | Dakota Howard, Dhwani Batra, Peter W. Cook, Kara Moser, Adrian Paskey, Jason Caravas, Benjamin Rambo-Martin, Shatavia Morrison, Christopher Gulvick, Scott Sammons, Yvette Unoarumhi, Darlene Wagner, Matthew Schmerer, Minoo Agarwal, Eyad Almasri, Debbie Boles, Ayla Burns, Nuthawin Charoensri, Oren Cohen, Susan Countryman, Mary Ann Cristobal, Bobbi Croy, Suzanne Dale, Hrushikesh Deshmukh, Amanda Douglas, Vincent Drouillon, Marcia Eisenberg, Howard Engler, Rama Ghatti, Prashant Gupta, Susan Hicks, Jake Humphrey, Lax Iyer, Lisa Pfefferle, Manoj Jain, Matthew Robinson, Mohan Kolli, Brian Krueger, Tim Kuphal, Stanley Letovsky, Michael Levandoski, Craig Lukasik, Jonathan Meltzer, Brian Norvell, Mindy Nye, Scott Parker, Christos Petropoulos, John Pruitt, Steven Ragan, Scott Ryan, Mike Sapeta, Jana Schroth, Suresh Babu Selvaraju, Goran Stevovic, Amanda Suchanek, Andrea Throop, Lyndon Tilson, Thomas Urban, Joe Voshell, Kimberly Wagner, Jonathan Williams, Mary Williamson, Qian Zeng, Tricia Zwiefelhofer, Clinton R. Paden, Duncan MacCannell |
| EPI_ISL_3326474 | TGen North | TGen North | Jolene Bowers, Heather Centner, Brett Van Tassel, Chris French, Hayley Yaglom, Darrin Lemmer, Dave Engelthaler |
| EPI_ISL_3326475, EPI_ISL_3326477 | Laboratory Corporation of America | Centers for Disease Control and Prevention Division of Viral Diseases, Pathogen Discovery | Dakota Howard, Dhwani Batra, Peter W. Cook, Kara Moser, Adrian Paskey, Jason Caravas, Benjamin Rambo-Martin, Shatavia Morrison, Christopher Gulvick, Scott Sammons, Yvette Unoarumhi, Darlene Wagner, Matthew Schmerer, Minoo Agarwal, Eyad Almasri, Debbie Boles, Ayla Burns, Nuthawin Charoensri, Oren Cohen, Susan Countryman, Mary Ann Cristobal, Bobbi Croy, Suzanne Dale, Hrushikesh Deshmukh, Amanda Douglas, Vincent Drouillon, Marcia Eisenberg, Howard Engler, Rama Ghatti, Prashant Gupta, Susan Hicks, Jake Humphrey, Lax Iyer, Lisa Pfefferle, Manoj Jain, Matthew Robinson, Mohan Kolli, Brian Krueger, Tim Kuphal, Stanley Letovsky, Michael Levandoski, Craig Lukasik, Jonathan Meltzer, Brian Norvell, Mindy Nye, Scott Parker, Christos Petropoulos, John Pruitt, Steven Ragan, Scott Ryan, Mike Sapeta, Jana Schroth, Suresh Babu Selvaraju, Goran Stevovic, Amanda Suchanek, Andrea Throop, Lyndon Tilson, Thomas Urban, Joe Voshell, Kimberly Wagner, Jonathan Williams, Mary Williamson, Qian Zeng, Tricia Zwiefelhofer, Clinton R. Paden, Duncan MacCannell |
| EPI_ISL_3326478 | TGen North | TGen North | Jolene Bowers, Heather Centner, Brett Van Tassel, Chris French, Hayley Yaglom, Darrin Lemmer, Dave Engelthaler |
| EPI_ISL_3326479 | Laboratory Corporation of America | Centers for Disease Control and Prevention Division of Viral Diseases, Pathogen Discovery | Dakota Howard, Dhwani Batra, Peter W. Cook, Kara Moser, Adrian Paskey, Jason Caravas, Benjamin Rambo-Martin, Shatavia Morrison, Christopher Gulvick, Scott Sammons, Yvette Unoarumhi, Darlene Wagner, Matthew Schmerer, Minoo Agarwal, Eyad Almasri, Debbie Boles, Ayla Burns, Nuthawin Charoensri, Oren Cohen, Susan Countryman, Mary Ann Cristobal, Bobbi Croy, Suzanne Dale, Hrushikesh Deshmukh, Amanda Douglas, Vincent Drouillon, Marcia Eisenberg, Howard Engler, Rama Ghatti, Prashant Gupta, Susan Hicks, Jake Humphrey, Lax Iyer, Lisa Pfefferle, Manoj Jain, Matthew Robinson, Mohan Kolli, Brian Krueger, Tim Kuphal, Stanley Letovsky, Michael Levandoski, Craig Lukasik, Jonathan Meltzer, Brian Norvell, Mindy Nye, Scott Parker, Christos Petropoulos, John Pruitt, Steven Ragan, Scott Ryan, Mike Sapeta, Jana Schroth, Suresh Babu Selvaraju, Goran Stevovic, Amanda Suchanek, Andrea Throop, Lyndon Tilson, Thomas Urban, Joe Voshell, Kimberly Wagner, Jonathan Williams, Mary Williamson, Qian Zeng, Tricia Zwiefelhofer, Clinton R. Paden, Duncan MacCannell |
| EPI_ISL_3326480 | TGen North | TGen North | Jolene Bowers, Heather Centner, Brett Van Tassel, Chris French, Hayley Yaglom, Darrin Lemmer, Dave Engelthaler |
| EPI_ISL_3326481 | Laboratory Corporation of America | Centers for Disease Control and Prevention Division of Viral Diseases, Pathogen Discovery | Dakota Howard, Dhwani Batra, Peter W. Cook, Kara Moser, Adrian Paskey, Jason Caravas, Benjamin Rambo-Martin, Shatavia Morrison, Christopher Gulvick, Scott Sammons, Yvette Unoarumhi, Darlene Wagner, Matthew Schmerer, Minoo Agarwal, Eyad Almasri, Debbie Boles, Ayla Burns, Nuthawin Charoensri, Oren Cohen, Susan Countryman, Mary Ann Cristobal, Bobbi Croy, Suzanne Dale, Hrushikesh Deshmukh, Amanda Douglas, Vincent Drouillon, Marcia Eisenberg, Howard Engler, Rama Ghatti, Prashant Gupta, Susan Hicks, Jake Humphrey, Lax Iyer, Lisa Pfefferle, Manoj Jain, Matthew Robinson, Mohan Kolli, Brian Krueger, Tim Kuphal, Stanley Letovsky, Michael Levandoski, Craig Lukasik, Jonathan Meltzer, Brian Norvell, Mindy Nye, Scott Parker, Christos Petropoulos, John Pruitt, Steven Ragan, Scott Ryan, Mike Sapeta, Jana Schroth, Suresh Babu Selvaraju, Goran Stevovic, Amanda Suchanek, Andrea Throop, Lyndon Tilson, Thomas Urban, Joe Voshell, Kimberly Wagner, Jonathan Williams, Mary Williamson, Qian Zeng, Tricia Zwiefelhofer, Clinton R. Paden, Duncan MacCannell |
| EPI_ISL_3326482 | TGen North | TGen North | Jolene Bowers, Heather Centner, Brett Van Tassel, Chris French, Hayley Yaglom, Darrin Lemmer, Dave Engelthaler |
| EPI_ISL_3326483 | Laboratory Corporation of America | Centers for Disease Control and Prevention Division of Viral Diseases, Pathogen Discovery | Dakota Howard, Dhwani Batra, Peter W. Cook, Kara Moser, Adrian Paskey, Jason Caravas, Benjamin Rambo-Martin, Shatavia Morrison, Christopher Gulvick, Scott Sammons, Yvette Unoarumhi, Darlene Wagner, Matthew Schmerer, Minoo Agarwal, Eyad Almasri, Debbie Boles, Ayla Burns, Nuthawin Charoensri, Oren Cohen, Susan Countryman, Mary Ann Cristobal, Bobbi Croy, Suzanne Dale, Hrushikesh Deshmukh, Amanda Douglas, Vincent Drouillon, Marcia Eisenberg, Howard Engler, Rama Ghatti, Prashant Gupta, Susan Hicks, Jake Humphrey, Lax Iyer, Lisa Pfefferle, Manoj Jain, Matthew Robinson, Mohan Kolli, Brian Krueger, Tim Kuphal, Stanley Letovsky, Michael Levandoski, Craig Lukasik, Jonathan Meltzer, Brian Norvell, Mindy Nye, Scott Parker, Christos Petropoulos, John Pruitt, Steven Ragan, Scott Ryan, Mike Sapeta, Jana Schroth, Suresh Babu Selvaraju, Goran Stevovic, Amanda Suchanek, Andrea Throop, Lyndon Tilson, Thomas Urban, Joe Voshell, Kimberly Wagner, Jonathan Williams, Mary Williamson, Qian Zeng, Tricia Zwiefelhofer, Clinton R. Paden, Duncan MacCannell |
| EPI_ISL_3326486 | TGen North | TGen North | Jolene Bowers, Heather Centner, Brett Van Tassel, Chris French, Hayley Yaglom, Darrin Lemmer, Dave Engelthaler |
| EPI_ISL_3326487, EPI_ISL_3326488 | Laboratory Corporation of America | Centers for Disease Control and Prevention Division of Viral Diseases, Pathogen Discovery | Dakota Howard, Dhwani Batra, Peter W. Cook, Kara Moser, Adrian Paskey, Jason Caravas, Benjamin Rambo-Martin, Shatavia Morrison, Christopher Gulvick, Scott Sammons, Yvette Unoarumhi, Darlene Wagner, Matthew Schmerer, Minoo Agarwal, Eyad Almasri, Debbie Boles, Ayla Burns, Nuthawin Charoensri, Oren Cohen, Susan Countryman, Mary Ann Cristobal, Bobbi Croy, Suzanne Dale, Hrushikesh Deshmukh, Amanda Douglas, Vincent Drouillon, Marcia Eisenberg, Howard Engler, Rama Ghatti, Prashant Gupta, Susan Hicks, Jake Humphrey, Lax Iyer, Lisa Pfefferle, Manoj Jain, Matthew Robinson, Mohan Kolli, Brian Krueger, Tim Kuphal, Stanley Letovsky, Michael Levandoski, Craig Lukasik, Jonathan Meltzer, Brian Norvell, Mindy Nye, Scott Parker, Christos Petropoulos, John Pruitt, Steven Ragan, Scott Ryan, Mike Sapeta, Jana Schroth, Suresh Babu Selvaraju, Goran Stevovic, Amanda Suchanek, Andrea Throop, Lyndon Tilson, Thomas Urban, Joe Voshell, Kimberly Wagner, Jonathan Williams, Mary Williamson, Qian Zeng, Tricia Zwiefelhofer, Clinton R. Paden, Duncan MacCannell |
| EPI_ISL_3326489 | TGen North | TGen North | Jolene Bowers, Heather Centner, Brett Van Tassel, Chris French, Hayley Yaglom, Darrin Lemmer, Dave Engelthaler |
| EPI_ISL_3326490 | Laboratory Corporation of America | Centers for Disease Control and Prevention Division of Viral Diseases, Pathogen Discovery | Dakota Howard, Dhwani Batra, Peter W. Cook, Kara Moser, Adrian Paskey, Jason Caravas, Benjamin Rambo-Martin, Shatavia Morrison, Christopher Gulvick, Scott Sammons, Yvette Unoarumhi, Darlene Wagner, Matthew Schmerer, Minoo Agarwal, Eyad Almasri, Debbie Boles, Ayla Burns, Nuthawin Charoensri, Oren Cohen, Susan Countryman, Mary Ann Cristobal, Bobbi Croy, Suzanne Dale, Hrushikesh Deshmukh, Amanda Douglas, Vincent Drouillon, Marcia Eisenberg, Howard Engler, Rama Ghatti, Prashant Gupta, Susan Hicks, Jake Humphrey, Lax Iyer, Lisa Pfefferle, Manoj Jain, Matthew Robinson, Mohan Kolli, Brian Krueger, Tim Kuphal, Stanley Letovsky, Michael Levandoski, Craig Lukasik, Jonathan Meltzer, Brian Norvell, Mindy Nye, Scott Parker, Christos Petropoulos, John Pruitt, Steven Ragan, Scott Ryan, Mike Sapeta, Jana Schroth, Suresh Babu Selvaraju, Goran Stevovic, Amanda Suchanek, Andrea Throop, Lyndon Tilson, Thomas Urban, Joe Voshell, Kimberly Wagner, Jonathan Williams, Mary Williamson, Qian Zeng, Tricia Zwiefelhofer, Clinton R. Paden, Duncan MacCannell |

| | | | |
|---|---|---|---|
| EPI_ISL_3326491 | TGen North | TGen North | Jolene Bowers, Heather Centner, Brett Van Tassel, Chris French, Hayley Yaglom, Darrin Lemmer, Dave Engelthaler |
| EPI_ISL_3326492 | Laboratory Corporation of America | Centers for Disease Control and Prevention Division of Viral Diseases, Pathogen Discovery | Dakota Howard, Dhwani Batra, Peter W. Cook, Kara Moser, Adrian Paskey, Jason Caravas, Benjamin Rambo-Martin, Shatavia Morrison, Christopher Gulvick, Scott Sammons, Yvette Unoarumhi, Darlene Wagner, Matthew Schmerer, Minoo Agarwal, Eyad Almasri, Debbie Boles, Ayla Burns, Nuthawin Charoensri, Oren Cohen, Susan Countryman, Mary Ann Cristobal, Bobbi Croy, Suzanne Dale, Hrushikesh Deshmukh, Amanda Douglas, Vincent Drouillon, Marcia Eisenberg, Howard Engler, Rama Ghatti, Prashant Gupta, Susan Hicks, Jake Humphrey, Lax Iyer, Lisa Pfefferle, Manoj Jain, Matthew Robinson, Mohan Kolli, Brian Krueger, Tim Kuphal, Stanley Letovsky, Michael Levandoski, Craig Lukasik, Jonathan Meltzer, Brian Norvell, Mindy Nye, Scott Parker, Christos Petropoulos, John Pruitt, Steven Ragan, Scott Ryan, Mike Sapeta, Jana Schroth, Suresh Babu Selvaraju, Goran Stevovic, Amanda Suchanek, Andrea Throop, Lyndon Tilson, Thomas Urban, Joe Voshell, Kimberly Wagner, Jonathan Williams, Mary Williamson, Qian Zeng, Tricia Zwiefelhofer, Clinton R. Paden, Duncan MacCannell |
| EPI_ISL_3326493 | TGen North | TGen North | Jolene Bowers, Heather Centner, Brett Van Tassel, Chris French, Hayley Yaglom, Darrin Lemmer, Dave Engelthaler |
| EPI_ISL_3326494 | Laboratory Corporation of America | Centers for Disease Control and Prevention Division of Viral Diseases, Pathogen Discovery | Dakota Howard, Dhwani Batra, Peter W. Cook, Kara Moser, Adrian Paskey, Jason Caravas, Benjamin Rambo-Martin, Shatavia Morrison, Christopher Gulvick, Scott Sammons, Yvette Unoarumhi, Darlene Wagner, Matthew Schmerer, Minoo Agarwal, Eyad Almasri, Debbie Boles, Ayla Burns, Nuthawin Charoensri, Oren Cohen, Susan Countryman, Mary Ann Cristobal, Bobbi Croy, Suzanne Dale, Hrushikesh Deshmukh, Amanda Douglas, Vincent Drouillon, Marcia Eisenberg, Howard Engler, Rama Ghatti, Prashant Gupta, Susan Hicks, Jake Humphrey, Lax Iyer, Lisa Pfefferle, Manoj Jain, Matthew Robinson, Mohan Kolli, Brian Krueger, Tim Kuphal, Stanley Letovsky, Michael Levandoski, Craig Lukasik, Jonathan Meltzer, Brian Norvell, Mindy Nye, Scott Parker, Christos Petropoulos, John Pruitt, Steven Ragan, Scott Ryan, Mike Sapeta, Jana Schroth, Suresh Babu Selvaraju, Goran Stevovic, Amanda Suchanek, Andrea Throop, Lyndon Tilson, Thomas Urban, Joe Voshell, Kimberly Wagner, Jonathan Williams, Mary Williamson, Qian Zeng, Tricia Zwiefelhofer, Clinton R. Paden, Duncan MacCannell |
| EPI_ISL_3326495 | TGen North | TGen North | Jolene Bowers, Heather Centner, Brett Van Tassel, Chris French, Hayley Yaglom, Darrin Lemmer, Dave Engelthaler |
| EPI_ISL_3326496, EPI_ISL_3326498 | Laboratory Corporation of America | Centers for Disease Control and Prevention Division of Viral Diseases, Pathogen Discovery | Dakota Howard, Dhwani Batra, Peter W. Cook, Kara Moser, Adrian Paskey, Jason Caravas, Benjamin Rambo-Martin, Shatavia Morrison, Christopher Gulvick, Scott Sammons, Yvette Unoarumhi, Darlene Wagner, Matthew Schmerer, Minoo Agarwal, Eyad Almasri, Debbie Boles, Ayla Burns, Nuthawin Charoensri, Oren Cohen, Susan Countryman, Mary Ann Cristobal, Bobbi Croy, Suzanne Dale, Hrushikesh Deshmukh, Amanda Douglas, Vincent Drouillon, Marcia Eisenberg, Howard Engler, Rama Ghatti, Prashant Gupta, Susan Hicks, Jake Humphrey, Lax Iyer, Lisa Pfefferle, Manoj Jain, Matthew Robinson, Mohan Kolli, Brian Krueger, Tim Kuphal, Stanley Letovsky, Michael Levandoski, Craig Lukasik, Jonathan Meltzer, Brian Norvell, Mindy Nye, Scott Parker, Christos Petropoulos, John Pruitt, Steven Ragan, Scott Ryan, Mike Sapeta, Jana Schroth, Suresh Babu Selvaraju, Goran Stevovic, Amanda Suchanek, Andrea Throop, Lyndon Tilson, Thomas Urban, Joe Voshell, Kimberly Wagner, Jonathan Williams, Mary Williamson, Qian Zeng, Tricia Zwiefelhofer, Clinton R. Paden, Duncan MacCannell |
| EPI_ISL_3326499 | TGen North | TGen North | Jolene Bowers, Heather Centner, Brett Van Tassel, Chris French, Hayley Yaglom, Darrin Lemmer, Dave Engelthaler |
| EPI_ISL_3326500 | Laboratory Corporation of America | Centers for Disease Control and Prevention Division of Viral Diseases, Pathogen Discovery | Dakota Howard, Dhwani Batra, Peter W. Cook, Kara Moser, Adrian Paskey, Jason Caravas, Benjamin Rambo-Martin, Shatavia Morrison, Christopher Gulvick, Scott Sammons, Yvette Unoarumhi, Darlene Wagner, Matthew Schmerer, Minoo Agarwal, Eyad Almasri, Debbie Boles, Ayla Burns, Nuthawin Charoensri, Oren Cohen, Susan Countryman, Mary Ann Cristobal, Bobbi Croy, Suzanne Dale, Hrushikesh Deshmukh, Amanda Douglas, Vincent Drouillon, Marcia Eisenberg, Howard Engler, Rama Ghatti, Prashant Gupta, Susan Hicks, Jake Humphrey, Lax Iyer, Lisa Pfefferle, Manoj Jain, Matthew Robinson, Mohan Kolli, Brian Krueger, Tim Kuphal, Stanley Letovsky, Michael Levandoski, Craig Lukasik, Jonathan Meltzer, Brian Norvell, Mindy Nye, Scott Parker, Christos Petropoulos, John Pruitt, Steven Ragan, Scott Ryan, Mike Sapeta, Jana Schroth, Suresh Babu Selvaraju, Goran Stevovic, Amanda Suchanek, Andrea Throop, Lyndon Tilson, Thomas Urban, Joe Voshell, Kimberly Wagner, Jonathan Williams, Mary Williamson, Qian Zeng, Tricia Zwiefelhofer, Clinton R. Paden, Duncan MacCannell |
| EPI_ISL_3326502, EPI_ISL_3326504, EPI_ISL_3326508 | TGen North | TGen North | Jolene Bowers, Heather Centner, Brett Van Tassel, Chris French, Hayley Yaglom, Darrin Lemmer, Dave Engelthaler |
| EPI_ISL_3326509, EPI_ISL_3326510 | Laboratory Corporation of America | Centers for Disease Control and Prevention Division of Viral Diseases, Pathogen Discovery | Dakota Howard, Dhwani Batra, Peter W. Cook, Kara Moser, Adrian Paskey, Jason Caravas, Benjamin Rambo-Martin, Shatavia Morrison, Christopher Gulvick, Scott Sammons, Yvette Unoarumhi, Darlene Wagner, Matthew Schmerer, Minoo Agarwal, Eyad Almasri, Debbie Boles, Ayla Burns, Nuthawin Charoensri, Oren Cohen, Susan Countryman, Mary Ann Cristobal, Bobbi Croy, Suzanne Dale, Hrushikesh Deshmukh, Amanda Douglas, Vincent Drouillon, Marcia Eisenberg, Howard Engler, Rama Ghatti, Prashant Gupta, Susan Hicks, Jake Humphrey, Lax Iyer, Lisa Pfefferle, Manoj Jain, Matthew Robinson, Mohan Kolli, Brian Krueger, Tim Kuphal, Stanley Letovsky, Michael Levandoski, Craig Lukasik, Jonathan Meltzer, Brian Norvell, Mindy Nye, Scott Parker, Christos Petropoulos, John Pruitt, Steven Ragan, Scott Ryan, Mike Sapeta, Jana Schroth, Suresh Babu Selvaraju, Goran Stevovic, Amanda Suchanek, Andrea Throop, Lyndon Tilson, Thomas Urban, Joe Voshell, Kimberly Wagner, Jonathan Williams, Mary Williamson, Qian Zeng, Tricia Zwiefelhofer, Clinton R. Paden, Duncan MacCannell |
| EPI_ISL_3326511 | TGen North | TGen North | Jolene Bowers, Heather Centner, Brett Van Tassel, Chris French, Hayley Yaglom, Darrin Lemmer, Dave Engelthaler |
| EPI_ISL_3326512 | Laboratory Corporation of America | Centers for Disease Control and Prevention Division of Viral Diseases, Pathogen Discovery | Dakota Howard, Dhwani Batra, Peter W. Cook, Kara Moser, Adrian Paskey, Jason Caravas, Benjamin Rambo-Martin, Shatavia Morrison, Christopher Gulvick, Scott Sammons, Yvette Unoarumhi, Darlene Wagner, Matthew Schmerer, Minoo Agarwal, Eyad Almasri, Debbie Boles, Ayla Burns, Nuthawin Charoensri, Oren Cohen, Susan Countryman, Mary Ann Cristobal, Bobbi Croy, Suzanne Dale, Hrushikesh Deshmukh, Amanda Douglas, Vincent Drouillon, Marcia Eisenberg, Howard Engler, Rama Ghatti, Prashant Gupta, Susan Hicks, Jake Humphrey, Lax Iyer, Lisa Pfefferle, Manoj Jain, Matthew Robinson, Mohan Kolli, Brian Krueger, Tim Kuphal, Stanley Letovsky, Michael Levandoski, Craig Lukasik, Jonathan Meltzer, Brian Norvell, Mindy Nye, Scott Parker, Christos Petropoulos, John Pruitt, Steven Ragan, Scott Ryan, Mike Sapeta, Jana Schroth, Suresh Babu Selvaraju, Goran Stevovic, Amanda Suchanek, Andrea Throop, Lyndon Tilson, Thomas Urban, Joe Voshell, Kimberly Wagner, Jonathan Williams, Mary Williamson, Qian Zeng, Tricia Zwiefelhofer, Clinton R. Paden, Duncan MacCannell |
| EPI_ISL_3326513 | TGen North | TGen North | Jolene Bowers, Heather Centner, Brett Van Tassel, Chris French, Hayley Yaglom, Darrin Lemmer, Dave Engelthaler |
| EPI_ISL_3326514 | Laboratory Corporation of America | Centers for Disease Control and Prevention Division of Viral Diseases, Pathogen Discovery | Dakota Howard, Dhwani Batra, Peter W. Cook, Kara Moser, Adrian Paskey, Jason Caravas, Benjamin Rambo-Martin, Shatavia Morrison, Christopher Gulvick, Scott Sammons, Yvette Unoarumhi, Darlene Wagner, Matthew Schmerer, Minoo Agarwal, Eyad Almasri, Debbie Boles, Ayla Burns, Nuthawin Charoensri, Oren Cohen, Susan Countryman, Mary Ann Cristobal, Bobbi Croy, Suzanne Dale, Hrushikesh Deshmukh, Amanda Douglas, Vincent Drouillon, Marcia Eisenberg, Howard Engler, Rama Ghatti, Prashant Gupta, Susan Hicks, Jake Humphrey, Lax Iyer, Lisa Pfefferle, Manoj Jain, Matthew Robinson, Mohan Kolli, Brian Krueger, Tim Kuphal, Stanley Letovsky, Michael Levandoski, Craig Lukasik, Jonathan Meltzer, Brian Norvell, Mindy Nye, Scott Parker, Christos Petropoulos, John Pruitt, Steven Ragan, Scott Ryan, Mike Sapeta, Jana Schroth, Suresh Babu Selvaraju, Goran Stevovic, Amanda Suchanek, Andrea Throop, Lyndon Tilson, Thomas Urban, Joe Voshell, Kimberly Wagner, Jonathan Williams, Mary Williamson, Qian Zeng, Tricia Zwiefelhofer, Clinton R. Paden, Duncan MacCannell |
| EPI_ISL_3326515 | TGen North | TGen North | Jolene Bowers, Heather Centner, Brett Van Tassel, Chris French, Hayley Yaglom, Darrin Lemmer, Dave Engelthaler |
| EPI_ISL_3326516 | Laboratory Corporation of America | Centers for Disease Control and Prevention Division of Viral Diseases, Pathogen Discovery | Dakota Howard, Dhwani Batra, Peter W. Cook, Kara Moser, Adrian Paskey, Jason Caravas, Benjamin Rambo-Martin, Shatavia Morrison, Christopher Gulvick, Scott Sammons, Yvette Unoarumhi, Darlene Wagner, Matthew Schmerer, Minoo Agarwal, Eyad Almasri, Debbie Boles, Ayla Burns, Nuthawin Charoensri, Oren Cohen, Susan Countryman, Mary Ann Cristobal, Bobbi Croy, Suzanne Dale, Hrushikesh Deshmukh, Amanda Douglas, Vincent Drouillon, Marcia Eisenberg, Howard Engler, Rama Ghatti, Prashant Gupta, Susan Hicks, Jake Humphrey, Lax Iyer, Lisa Pfefferle, Manoj Jain, Matthew Robinson, Mohan Kolli, Brian Krueger, Tim Kuphal, Stanley Letovsky, Michael Levandoski, Craig Lukasik, Jonathan Meltzer, Brian Norvell, Mindy Nye, Scott Parker, Christos Petropoulos, John Pruitt, Steven Ragan, Scott Ryan, Mike Sapeta, Jana Schroth, Suresh Babu Selvaraju, Goran Stevovic, Amanda Suchanek, Andrea Throop, Lyndon Tilson, Thomas Urban, Joe Voshell, Kimberly Wagner, Jonathan Williams, Mary Williamson, Qian Zeng, Tricia Zwiefelhofer, Clinton R. Paden, Duncan MacCannell |
| EPI_ISL_3326517 | TGen North | TGen North | Jolene Bowers, Heather Centner, Brett Van Tassel, Chris French, Hayley Yaglom, Darrin Lemmer, Dave Engelthaler |
| EPI_ISL_3326518 | Laboratory Corporation of America | Centers for Disease Control and Prevention Division of Viral Diseases, Pathogen Discovery | Dakota Howard, Dhwani Batra, Peter W. Cook, Kara Moser, Adrian Paskey, Jason Caravas, Benjamin Rambo-Martin, Shatavia Morrison, Christopher Gulvick, Scott Sammons, Yvette Unoarumhi, Darlene Wagner, Matthew Schmerer, Minoo Agarwal, Eyad Almasri, Debbie Boles, Ayla Burns, Nuthawin Charoensri, Oren Cohen, Susan Countryman, Mary Ann Cristobal, Bobbi Croy, Suzanne Dale, Hrushikesh Deshmukh, Amanda Douglas, Vincent Drouillon, Marcia Eisenberg, Howard Engler, Rama Ghatti, Prashant Gupta, Susan Hicks, Jake Humphrey, Lax Iyer, Lisa Pfefferle, Manoj Jain, Matthew Robinson, Mohan Kolli, Brian Krueger, Tim Kuphal, Stanley Letovsky, Michael Levandoski, Craig Lukasik, Jonathan Meltzer, Brian Norvell, Mindy Nye, Scott Parker, Christos Petropoulos, John Pruitt, Steven Ragan, Scott Ryan, Mike Sapeta, Jana Schroth, Suresh Babu Selvaraju, Goran Stevovic, Amanda Suchanek, Andrea Throop, Lyndon Tilson, Thomas Urban, Joe Voshell, Kimberly Wagner, Jonathan Williams, Mary Williamson, Qian Zeng, Tricia Zwiefelhofer, Clinton R. Paden, Duncan MacCannell |
| EPI_ISL_3326519 | TGen North | TGen North | Jolene Bowers, Heather Centner, Brett Van Tassel, Chris French, Hayley Yaglom, Darrin Lemmer, Dave Engelthaler |
| EPI_ISL_3326520 | Laboratory Corporation of America | Centers for Disease Control and Prevention Division of Viral Diseases, Pathogen Discovery | Dakota Howard, Dhwani Batra, Peter W. Cook, Kara Moser, Adrian Paskey, Jason Caravas, Benjamin Rambo-Martin, Shatavia Morrison, Christopher Gulvick, Scott Sammons, Yvette Unoarumhi, Darlene Wagner, Matthew Schmerer, Minoo Agarwal, Eyad Almasri, Debbie Boles, Ayla Burns, Nuthawin Charoensri, Oren Cohen, Susan Countryman, Mary Ann Cristobal, Bobbi Croy, Suzanne Dale, Hrushikesh Deshmukh, Amanda Douglas, Vincent Drouillon, Marcia Eisenberg, Howard Engler, Rama Ghatti, Prashant Gupta, Susan Hicks, Jake Humphrey, Lax Iyer, Lisa Pfefferle, Manoj Jain, Matthew Robinson, Mohan Kolli, Brian Krueger, Tim Kuphal, Stanley Letovsky, Michael Levandoski, Craig Lukasik, Jonathan Meltzer, Brian Norvell, Mindy Nye, Scott Parker, Christos Petropoulos, John Pruitt, Steven Ragan, Scott Ryan, Mike Sapeta, Jana Schroth, Suresh Babu Selvaraju, Goran Stevovic, Amanda Suchanek, Andrea Throop, Lyndon Tilson, Thomas Urban, Joe Voshell, Kimberly Wagner, Jonathan Williams, Mary Williamson, Qian Zeng, Tricia Zwiefelhofer, Clinton R. Paden, Duncan MacCannell |
| EPI_ISL_3326521 | TGen North | TGen North | Jolene Bowers, Heather Centner, Brett Van Tassel, Chris French, Hayley Yaglom, Darrin Lemmer, Dave Engelthaler |
| EPI_ISL_3326522, EPI_ISL_3326523, EPI_ISL_3326525 | Laboratory Corporation of America | Centers for Disease Control and Prevention Division of Viral Diseases, Pathogen Discovery | Dakota Howard, Dhwani Batra, Peter W. Cook, Kara Moser, Adrian Paskey, Jason Caravas, Benjamin Rambo-Martin, Shatavia Morrison, Christopher Gulvick, Scott Sammons, Yvette Unoarumhi, Darlene Wagner, Matthew Schmerer, Minoo Agarwal, Eyad Almasri, Debbie Boles, Ayla Burns, Nuthawin Charoensri, Oren Cohen, Susan Countryman, Mary Ann Cristobal, Bobbi Croy, Suzanne Dale, Hrushikesh Deshmukh, Amanda Douglas, Vincent Drouillon, Marcia Eisenberg, Howard Engler, Rama Ghatti, Prashant Gupta, Susan Hicks, Jake Humphrey, Lax Iyer, Lisa Pfefferle, Manoj Jain, Matthew Robinson, Mohan Kolli, Brian Krueger, Tim Kuphal, Stanley Letovsky, Michael Levandoski, Craig Lukasik, Jonathan Meltzer, Brian Norvell, Mindy Nye, Scott Parker, Christos Petropoulos, John Pruitt, Steven Ragan, Scott Ryan, Mike Sapeta, Jana Schroth, Suresh Babu Selvaraju, Goran Stevovic, Amanda Suchanek, Andrea Throop, Lyndon Tilson, Thomas Urban, Joe Voshell, Kimberly Wagner, Jonathan Williams, Mary Williamson, Qian Zeng, Tricia Zwiefelhofer, Clinton R. Paden, Duncan MacCannell |
| EPI_ISL_3326526 | TGen North | TGen North | Jolene Bowers, Heather Centner, Brett Van Tassel, Chris French, Hayley Yaglom, Darrin Lemmer, Dave Engelthaler |
| EPI_ISL_3326527 | Laboratory Corporation of America | Centers for Disease Control and Prevention Division of Viral Diseases, Pathogen Discovery | Dakota Howard, Dhwani Batra, Peter W. Cook, Kara Moser, Adrian Paskey, Jason Caravas, Benjamin Rambo-Martin, Shatavia Morrison, Christopher Gulvick, Scott Sammons, Yvette Unoarumhi, Darlene Wagner, Matthew Schmerer, Minoo Agarwal, Eyad Almasri, Debbie Boles, Ayla Burns, Nuthawin Charoensri, Oren Cohen, Susan Countryman, Mary Ann Cristobal, Bobbi Croy, Suzanne Dale, Hrushikesh Deshmukh, Amanda Douglas, Vincent Drouillon, Marcia Eisenberg, Howard Engler, Rama Ghatti, Prashant Gupta, Susan Hicks, Jake Humphrey, Lax Iyer, Lisa Pfefferle, Manoj Jain, Matthew Robinson, Mohan Kolli, Brian Krueger, Tim Kuphal, Stanley Letovsky, Michael Levandoski, Craig Lukasik, Jonathan Meltzer, Brian Norvell, Mindy Nye, Scott Parker, Christos Petropoulos, John Pruitt, Steven Ragan, Scott Ryan, Mike Sapeta, Jana Schroth, Suresh Babu Selvaraju, Goran Stevovic, Amanda Suchanek, Andrea Throop, Lyndon Tilson, Thomas Urban, Joe Voshell, Kimberly Wagner, Jonathan Williams, Mary Williamson, Qian Zeng, Tricia Zwiefelhofer, Clinton R. Paden, Duncan MacCannell |
| EPI_ISL_3326528, EPI_ISL_3326530, EPI_ISL_3326532, EPI_ISL_3326536 | TGen North | TGen North | Jolene Bowers, Heather Centner, Brett Van Tassel, Chris French, Hayley Yaglom, Darrin Lemmer, Dave Engelthaler |
| EPI_ISL_3326537 | Laboratory Corporation of America | Centers for Disease Control and Prevention Division of Viral Diseases, Pathogen Discovery | Dakota Howard, Dhwani Batra, Peter W. Cook, Kara Moser, Adrian Paskey, Jason Caravas, Benjamin Rambo-Martin, Shatavia Morrison, Christopher Gulvick, Scott Sammons, Yvette Unoarumhi, Darlene Wagner, Matthew Schmerer, Minoo Agarwal, Eyad Almasri, Debbie Boles, Ayla Burns, Nuthawin Charoensri, Oren Cohen, Susan Countryman, Mary Ann Cristobal, Bobbi Croy, Suzanne Dale, Hrushikesh Deshmukh, Amanda Douglas, Vincent Drouillon, Marcia Eisenberg, Howard Engler, Rama Ghatti, Prashant Gupta, Susan Hicks, Jake Humphrey, Lax Iyer, Lisa Pfefferle, Manoj Jain, Matthew Robinson, Mohan Kolli, Brian Krueger, Tim Kuphal, Stanley Letovsky, Michael Levandoski, Craig Lukasik, Jonathan Meltzer, Brian Norvell, Mindy Nye, Scott Parker, Christos Petropoulos, John Pruitt, Steven Ragan, Scott Ryan, Mike Sapeta, Jana Schroth, Suresh Babu Selvaraju, Goran Stevovic, Amanda Suchanek, Andrea Throop, Lyndon Tilson, Thomas Urban, Joe Voshell, Kimberly Wagner, Jonathan Williams, Mary Williamson, Qian Zeng, Tricia Zwiefelhofer, Clinton R. Paden, Duncan MacCannell |
| EPI_ISL_3326538 | TGen North | TGen North | Jolene Bowers, Heather Centner, Brett Van Tassel, Chris French, Hayley Yaglom, Darrin Lemmer, Dave Engelthaler |
| EPI_ISL_3326540, EPI_ISL_3326542, EPI_ISL_3326544 | Laboratory Corporation of America | Centers for Disease Control and Prevention Division of Viral Diseases, Pathogen Discovery | Dakota Howard, Dhwani Batra, Peter W. Cook, Kara Moser, Adrian Paskey, Jason Caravas, Benjamin Rambo-Martin, Shatavia Morrison, Christopher Gulvick, Scott Sammons, Yvette Unoarumhi, Darlene Wagner, Matthew Schmerer, Minoo Agarwal, Eyad Almasri, Debbie Boles, Ayla Burns, Nuthawin Charoensri, Oren Cohen, Susan Countryman, Mary Ann Cristobal, Bobbi Croy, Suzanne Dale, Hrushikesh Deshmukh, Amanda Douglas, Vincent Drouillon, Marcia Eisenberg, Howard Engler, Rama Ghatti, Prashant Gupta, Susan Hicks, Jake Humphrey, Lax Iyer, Lisa Pfefferle, Manoj Jain, Matthew Robinson, Mohan Kolli, Brian Krueger, Tim Kuphal, Stanley Letovsky, Michael Levandoski, Craig Lukasik, Jonathan Meltzer, Brian Norvell, Mindy Nye, Scott Parker, Christos Petropoulos, John Pruitt, Steven Ragan, Scott Ryan, Mike Sapeta, Jana Schroth, Suresh Babu Selvaraju, Goran Stevovic, Amanda Suchanek, Andrea Throop, Lyndon Tilson, Thomas Urban, Joe Voshell, Kimberly Wagner, Jonathan Williams, Mary Williamson, Qian Zeng, Tricia Zwiefelhofer, Clinton R. Paden, Duncan MacCannell |
| EPI_ISL_3326545, EPI_ISL_3326547 | TGen North | TGen North | Jolene Bowers, Heather Centner, Brett Van Tassel, Chris French, Hayley Yaglom, Darrin Lemmer, Dave Engelthaler |
| EPI_ISL_3326548, EPI_ISL_3326549 | Laboratory Corporation of America | Centers for Disease Control and Prevention Division of Viral Diseases, Pathogen Discovery | Dakota Howard, Dhwani Batra, Peter W. Cook, Kara Moser, Adrian Paskey, Jason Caravas, Benjamin Rambo-Martin, Shatavia Morrison, Christopher Gulvick, Scott Sammons, Yvette Unoarumhi, Darlene Wagner, Matthew Schmerer, Minoo Agarwal, Eyad Almasri, Debbie Boles, Ayla Burns, Nuthawin Charoensri, Oren Cohen, Susan Countryman, Mary Ann Cristobal, Bobbi Croy, Suzanne Dale, Hrushikesh Deshmukh, Amanda Douglas, Vincent Drouillon, Marcia Eisenberg, Howard Engler, Rama Ghatti, Prashant Gupta, Susan Hicks, Jake Humphrey, Lax Iyer, Lisa Pfefferle, Manoj Jain, Matthew Robinson, Mohan Kolli, Brian Krueger, Tim Kuphal, Stanley Letovsky, Michael Levandoski, Craig Lukasik, Jonathan Meltzer, Brian Norvell, Mindy Nye, Scott Parker, Christos Petropoulos, John Pruitt, Steven Ragan, Scott Ryan, Mike Sapeta, Jana Schroth, Suresh Babu Selvaraju, Goran Stevovic, Amanda Suchanek, Andrea Throop, Lyndon Tilson, Thomas Urban, Joe Voshell, Kimberly Wagner, Jonathan Williams, Mary Williamson, Qian Zeng, Tricia Zwiefelhofer, Clinton R. Paden, Duncan MacCannell |
| EPI_ISL_3326550, | TGen North | TGen North | Jolene Bowers, Heather Centner, Brett Van Tassel, Chris French, Hayley Yaglom, Darrin Lemmer, Dave Engelthaler |

| | | | |
|---|---|---|---|
| EPI_ISL_3326552 | | | |
| EPI_ISL_3326553, EPI_ISL_3326555 | Laboratory Corporation of America | Centers for Disease Control and Prevention Division of Viral Diseases, Pathogen Discovery | Dakota Howard, Dhwani Batra, Peter W. Cook, Kara Moser, Adrian Paskey, Jason Caravas, Benjamin Rambo-Martin, Shatavia Morrison, Christopher Gulvick, Scott Sammons, Yvette Unoarumhi, Darlene Wagner, Matthew Schmerer, Minoo Agarwal, Eyad Almasri, Debbie Boles, Ayla Burns, Nuthawin Charoensri, Oren Cohen, Susan Countryman, Mary Ann Cristobal, Bobbi Croy, Suzanne Dale, Hrushikesh Deshmukh, Amanda Douglas, Vincent Drouillon, Marcia Eisenberg, Howard Engler, Rama Ghatti, Prashant Gupta, Susan Hicks, Jake Humphrey, Lax Iyer, Lisa Pfefferle, Manoj Jain, Matthew Robinson, Mohan Kolli, Brian Krueger, Tim Kuphal, Stanley Letovsky, Michael Levandoski, Craig Lukasik, Jonathan Meltzer, Brian Norvell, Mindy Nye, Scott Parker, Christos Petropoulos, John Pruitt, Steven Ragan, Scott Ryan, Mike Sapeta, Jana Schroth, Suresh Babu Selvaraju, Goran Stevovic, Amanda Suchanek, Andrea Throop, Lyndon Tilson, Thomas Urban, Joe Voshell, Kimberly Wagner, Jonathan Williams, Mary Williamson, Qian Zeng, Tricia Zwiefelhofer, Clinton R. Paden, Duncan MacCannell |
| EPI_ISL_3326556 | TGen North | TGen North | Jolene Bowers, Heather Centner, Brett Van Tassel, Chris French, Hayley Yaglom, Darrin Lemmer, Dave Engelthaler |
| EPI_ISL_3326559 | Laboratory Corporation of America | Centers for Disease Control and Prevention Division of Viral Diseases, Pathogen Discovery | Dakota Howard, Dhwani Batra, Peter W. Cook, Kara Moser, Adrian Paskey, Jason Caravas, Benjamin Rambo-Martin, Shatavia Morrison, Christopher Gulvick, Scott Sammons, Yvette Unoarumhi, Darlene Wagner, Matthew Schmerer, Minoo Agarwal, Eyad Almasri, Debbie Boles, Ayla Burns, Nuthawin Charoensri, Oren Cohen, Susan Countryman, Mary Ann Cristobal, Bobbi Croy, Suzanne Dale, Hrushikesh Deshmukh, Amanda Douglas, Vincent Drouillon, Marcia Eisenberg, Howard Engler, Rama Ghatti, Prashant Gupta, Susan Hicks, Jake Humphrey, Lax Iyer, Lisa Pfefferle, Manoj Jain, Matthew Robinson, Mohan Kolli, Brian Krueger, Tim Kuphal, Stanley Letovsky, Michael Levandoski, Craig Lukasik, Jonathan Meltzer, Brian Norvell, Mindy Nye, Scott Parker, Christos Petropoulos, John Pruitt, Steven Ragan, Scott Ryan, Mike Sapeta, Jana Schroth, Suresh Babu Selvaraju, Goran Stevovic, Amanda Suchanek, Andrea Throop, Lyndon Tilson, Thomas Urban, Joe Voshell, Kimberly Wagner, Jonathan Williams, Mary Williamson, Qian Zeng, Tricia Zwiefelhofer, Clinton R. Paden, Duncan MacCannell |
| EPI_ISL_3326563 | TGen North | TGen North | Jolene Bowers, Heather Centner, Brett Van Tassel, Chris French, Hayley Yaglom, Darrin Lemmer, Dave Engelthaler |
| EPI_ISL_3326564 | Laboratory Corporation of America | Centers for Disease Control and Prevention Division of Viral Diseases, Pathogen Discovery | Dakota Howard, Dhwani Batra, Peter W. Cook, Kara Moser, Adrian Paskey, Jason Caravas, Benjamin Rambo-Martin, Shatavia Morrison, Christopher Gulvick, Scott Sammons, Yvette Unoarumhi, Darlene Wagner, Matthew Schmerer, Minoo Agarwal, Eyad Almasri, Debbie Boles, Ayla Burns, Nuthawin Charoensri, Oren Cohen, Susan Countryman, Mary Ann Cristobal, Bobbi Croy, Suzanne Dale, Hrushikesh Deshmukh, Amanda Douglas, Vincent Drouillon, Marcia Eisenberg, Howard Engler, Rama Ghatti, Prashant Gupta, Susan Hicks, Jake Humphrey, Lax Iyer, Lisa Pfefferle, Manoj Jain, Matthew Robinson, Mohan Kolli, Brian Krueger, Tim Kuphal, Stanley Letovsky, Michael Levandoski, Craig Lukasik, Jonathan Meltzer, Brian Norvell, Mindy Nye, Scott Parker, Christos Petropoulos, John Pruitt, Steven Ragan, Scott Ryan, Mike Sapeta, Jana Schroth, Suresh Babu Selvaraju, Goran Stevovic, Amanda Suchanek, Andrea Throop, Lyndon Tilson, Thomas Urban, Joe Voshell, Kimberly Wagner, Jonathan Williams, Mary Williamson, Qian Zeng, Tricia Zwiefelhofer, Clinton R. Paden, Duncan MacCannell |
| EPI_ISL_3326565 | TGen North | TGen North | Jolene Bowers, Heather Centner, Brett Van Tassel, Chris French, Hayley Yaglom, Darrin Lemmer, Dave Engelthaler |
| EPI_ISL_3326566 | Laboratory Corporation of America | Centers for Disease Control and Prevention Division of Viral Diseases, Pathogen Discovery | Dakota Howard, Dhwani Batra, Peter W. Cook, Kara Moser, Adrian Paskey, Jason Caravas, Benjamin Rambo-Martin, Shatavia Morrison, Christopher Gulvick, Scott Sammons, Yvette Unoarumhi, Darlene Wagner, Matthew Schmerer, Minoo Agarwal, Eyad Almasri, Debbie Boles, Ayla Burns, Nuthawin Charoensri, Oren Cohen, Susan Countryman, Mary Ann Cristobal, Bobbi Croy, Suzanne Dale, Hrushikesh Deshmukh, Amanda Douglas, Vincent Drouillon, Marcia Eisenberg, Howard Engler, Rama Ghatti, Prashant Gupta, Susan Hicks, Jake Humphrey, Lax Iyer, Lisa Pfefferle, Manoj Jain, Matthew Robinson, Mohan Kolli, Brian Krueger, Tim Kuphal, Stanley Letovsky, Michael Levandoski, Craig Lukasik, Jonathan Meltzer, Brian Norvell, Mindy Nye, Scott Parker, Christos Petropoulos, John Pruitt, Steven Ragan, Scott Ryan, Mike Sapeta, Jana Schroth, Suresh Babu Selvaraju, Goran Stevovic, Amanda Suchanek, Andrea Throop, Lyndon Tilson, Thomas Urban, Joe Voshell, Kimberly Wagner, Jonathan Williams, Mary Williamson, Qian Zeng, Tricia Zwiefelhofer, Clinton R. Paden, Duncan MacCannell |
| EPI_ISL_3326567 | TGen North | TGen North | Jolene Bowers, Heather Centner, Brett Van Tassel, Chris French, Hayley Yaglom, Darrin Lemmer, Dave Engelthaler |
| EPI_ISL_3326568 | Laboratory Corporation of America | Centers for Disease Control and Prevention Division of Viral Diseases, Pathogen Discovery | Dakota Howard, Dhwani Batra, Peter W. Cook, Kara Moser, Adrian Paskey, Jason Caravas, Benjamin Rambo-Martin, Shatavia Morrison, Christopher Gulvick, Scott Sammons, Yvette Unoarumhi, Darlene Wagner, Matthew Schmerer, Minoo Agarwal, Eyad Almasri, Debbie Boles, Ayla Burns, Nuthawin Charoensri, Oren Cohen, Susan Countryman, Mary Ann Cristobal, Bobbi Croy, Suzanne Dale, Hrushikesh Deshmukh, Amanda Douglas, Vincent Drouillon, Marcia Eisenberg, Howard Engler, Rama Ghatti, Prashant Gupta, Susan Hicks, Jake Humphrey, Lax Iyer, Lisa Pfefferle, Manoj Jain, Matthew Robinson, Mohan Kolli, Brian Krueger, Tim Kuphal, Stanley Letovsky, Michael Levandoski, Craig Lukasik, Jonathan Meltzer, Brian Norvell, Mindy Nye, Scott Parker, Christos Petropoulos, John Pruitt, Steven Ragan, Scott Ryan, Mike Sapeta, Jana Schroth, Suresh Babu Selvaraju, Goran Stevovic, Amanda Suchanek, Andrea Throop, Lyndon Tilson, Thomas Urban, Joe Voshell, Kimberly Wagner, Jonathan Williams, Mary Williamson, Qian Zeng, Tricia Zwiefelhofer, Clinton R. Paden, Duncan MacCannell |
| EPI_ISL_3326571 | TGen North | TGen North | Jolene Bowers, Heather Centner, Brett Van Tassel, Chris French, Hayley Yaglom, Darrin Lemmer, Dave Engelthaler |
| EPI_ISL_3326573 | Laboratory Corporation of America | Centers for Disease Control and Prevention Division of Viral Diseases, Pathogen Discovery | Dakota Howard, Dhwani Batra, Peter W. Cook, Kara Moser, Adrian Paskey, Jason Caravas, Benjamin Rambo-Martin, Shatavia Morrison, Christopher Gulvick, Scott Sammons, Yvette Unoarumhi, Darlene Wagner, Matthew Schmerer, Minoo Agarwal, Eyad Almasri, Debbie Boles, Ayla Burns, Nuthawin Charoensri, Oren Cohen, Susan Countryman, Mary Ann Cristobal, Bobbi Croy, Suzanne Dale, Hrushikesh Deshmukh, Amanda Douglas, Vincent Drouillon, Marcia Eisenberg, Howard Engler, Rama Ghatti, Prashant Gupta, Susan Hicks, Jake Humphrey, Lax Iyer, Lisa Pfefferle, Manoj Jain, Matthew Robinson, Mohan Kolli, Brian Krueger, Tim Kuphal, Stanley Letovsky, Michael Levandoski, Craig Lukasik, Jonathan Meltzer, Brian Norvell, Mindy Nye, Scott Parker, Christos Petropoulos, John Pruitt, Steven Ragan, Scott Ryan, Mike Sapeta, Jana Schroth, Suresh Babu Selvaraju, Goran Stevovic, Amanda Suchanek, Andrea Throop, Lyndon Tilson, Thomas Urban, Joe Voshell, Kimberly Wagner, Jonathan Williams, Mary Williamson, Qian Zeng, Tricia Zwiefelhofer, Clinton R. Paden, Duncan MacCannell |
| EPI_ISL_3326574 | TGen North | TGen North | Jolene Bowers, Heather Centner, Brett Van Tassel, Chris French, Hayley Yaglom, Darrin Lemmer, Dave Engelthaler |
| EPI_ISL_3326575 | Laboratory Corporation of America | Centers for Disease Control and Prevention Division of Viral Diseases, Pathogen Discovery | Dakota Howard, Dhwani Batra, Peter W. Cook, Kara Moser, Adrian Paskey, Jason Caravas, Benjamin Rambo-Martin, Shatavia Morrison, Christopher Gulvick, Scott Sammons, Yvette Unoarumhi, Darlene Wagner, Matthew Schmerer, Minoo Agarwal, Eyad Almasri, Debbie Boles, Ayla Burns, Nuthawin Charoensri, Oren Cohen, Susan Countryman, Mary Ann Cristobal, Bobbi Croy, Suzanne Dale, Hrushikesh Deshmukh, Amanda Douglas, Vincent Drouillon, Marcia Eisenberg, Howard Engler, Rama Ghatti, Prashant Gupta, Susan Hicks, Jake Humphrey, Lax Iyer, Lisa Pfefferle, Manoj Jain, Matthew Robinson, Mohan Kolli, Brian Krueger, Tim Kuphal, Stanley Letovsky, Michael Levandoski, Craig Lukasik, Jonathan Meltzer, Brian Norvell, Mindy Nye, Scott Parker, Christos Petropoulos, John Pruitt, Steven Ragan, Scott Ryan, Mike Sapeta, Jana Schroth, Suresh Babu Selvaraju, Goran Stevovic, Amanda Suchanek, Andrea Throop, Lyndon Tilson, Thomas Urban, Joe Voshell, Kimberly Wagner, Jonathan Williams, Mary Williamson, Qian Zeng, Tricia Zwiefelhofer, Clinton R. Paden, Duncan MacCannell |
| EPI_ISL_3326576 | TGen North | TGen North | Jolene Bowers, Heather Centner, Brett Van Tassel, Chris French, Hayley Yaglom, Darrin Lemmer, Dave Engelthaler |
| EPI_ISL_3326577 | Laboratory Corporation of America | Centers for Disease Control and Prevention Division of Viral Diseases, Pathogen Discovery | Dakota Howard, Dhwani Batra, Peter W. Cook, Kara Moser, Adrian Paskey, Jason Caravas, Benjamin Rambo-Martin, Shatavia Morrison, Christopher Gulvick, Scott Sammons, Yvette Unoarumhi, Darlene Wagner, Matthew Schmerer, Minoo Agarwal, Eyad Almasri, Debbie Boles, Ayla Burns, Nuthawin Charoensri, Oren Cohen, Susan Countryman, Mary Ann Cristobal, Bobbi Croy, Suzanne Dale, Hrushikesh Deshmukh, Amanda Douglas, Vincent Drouillon, Marcia Eisenberg, Howard Engler, Rama Ghatti, Prashant Gupta, Susan Hicks, Jake Humphrey, Lax Iyer, Lisa Pfefferle, Manoj Jain, Matthew Robinson, Mohan Kolli, Brian Krueger, Tim Kuphal, Stanley Letovsky, Michael Levandoski, Craig Lukasik, Jonathan Meltzer, Brian Norvell, Mindy Nye, Scott Parker, Christos Petropoulos, John Pruitt, Steven Ragan, Scott Ryan, Mike Sapeta, Jana Schroth, Suresh Babu Selvaraju, Goran Stevovic, Amanda Suchanek, Andrea Throop, Lyndon Tilson, Thomas Urban, Joe Voshell, Kimberly Wagner, Jonathan Williams, Mary Williamson, Qian Zeng, Tricia Zwiefelhofer, Clinton R. Paden, Duncan MacCannell |
| EPI_ISL_3326578 | TGen North | TGen North | Jolene Bowers, Heather Centner, Brett Van Tassel, Chris French, Hayley Yaglom, Darrin Lemmer, Dave Engelthaler |
| EPI_ISL_3326579 | Laboratory Corporation of America | Centers for Disease Control and Prevention Division of Viral Diseases, Pathogen Discovery | Dakota Howard, Dhwani Batra, Peter W. Cook, Kara Moser, Adrian Paskey, Jason Caravas, Benjamin Rambo-Martin, Shatavia Morrison, Christopher Gulvick, Scott Sammons, Yvette Unoarumhi, Darlene Wagner, Matthew Schmerer, Minoo Agarwal, Eyad Almasri, Debbie Boles, Ayla Burns, Nuthawin Charoensri, Oren Cohen, Susan Countryman, Mary Ann Cristobal, Bobbi Croy, Suzanne Dale, Hrushikesh Deshmukh, Amanda Douglas, Vincent Drouillon, Marcia Eisenberg, Howard Engler, Rama Ghatti, Prashant Gupta, Susan Hicks, Jake Humphrey, Lax Iyer, Lisa Pfefferle, Manoj Jain, Matthew Robinson, Mohan Kolli, Brian Krueger, Tim Kuphal, Stanley Letovsky, Michael Levandoski, Craig Lukasik, Jonathan Meltzer, Brian Norvell, Mindy Nye, Scott Parker, Christos Petropoulos, John Pruitt, Steven Ragan, Scott Ryan, Mike Sapeta, Jana Schroth, Suresh Babu Selvaraju, Goran Stevovic, Amanda Suchanek, Andrea Throop, Lyndon Tilson, Thomas Urban, Joe Voshell, Kimberly Wagner, Jonathan Williams, Mary Williamson, Qian Zeng, Tricia Zwiefelhofer, Clinton R. Paden, Duncan MacCannell |
| EPI_ISL_3326580 | TGen North | TGen North | Jolene Bowers, Heather Centner, Brett Van Tassel, Chris French, Hayley Yaglom, Darrin Lemmer, Dave Engelthaler |
| EPI_ISL_3326581, EPI_ISL_3326582 | Laboratory Corporation of America | Centers for Disease Control and Prevention Division of Viral Diseases, Pathogen Discovery | Dakota Howard, Dhwani Batra, Peter W. Cook, Kara Moser, Adrian Paskey, Jason Caravas, Benjamin Rambo-Martin, Shatavia Morrison, Christopher Gulvick, Scott Sammons, Yvette Unoarumhi, Darlene Wagner, Matthew Schmerer, Minoo Agarwal, Eyad Almasri, Debbie Boles, Ayla Burns, Nuthawin Charoensri, Oren Cohen, Susan Countryman, Mary Ann Cristobal, Bobbi Croy, Suzanne Dale, Hrushikesh Deshmukh, Amanda Douglas, Vincent Drouillon, Marcia Eisenberg, Howard Engler, Rama Ghatti, Prashant Gupta, Susan Hicks, Jake Humphrey, Lax Iyer, Lisa Pfefferle, Manoj Jain, Matthew Robinson, Mohan Kolli, Brian Krueger, Tim Kuphal, Stanley Letovsky, Michael Levandoski, Craig Lukasik, Jonathan Meltzer, Brian Norvell, Mindy Nye, Scott Parker, Christos Petropoulos, John Pruitt, Steven Ragan, Scott Ryan, Mike Sapeta, Jana Schroth, Suresh Babu Selvaraju, Goran Stevovic, Amanda Suchanek, Andrea Throop, Lyndon Tilson, Thomas Urban, Joe Voshell, Kimberly Wagner, Jonathan Williams, Mary Williamson, Qian Zeng, Tricia Zwiefelhofer, Clinton R. Paden, Duncan MacCannell |
| EPI_ISL_3326583 | TGen North | TGen North | Jolene Bowers, Heather Centner, Brett Van Tassel, Chris French, Hayley Yaglom, Darrin Lemmer, Dave Engelthaler |
| EPI_ISL_3326584 | Laboratory Corporation of America | Centers for Disease Control and Prevention Division of Viral Diseases, Pathogen Discovery | Dakota Howard, Dhwani Batra, Peter W. Cook, Kara Moser, Adrian Paskey, Jason Caravas, Benjamin Rambo-Martin, Shatavia Morrison, Christopher Gulvick, Scott Sammons, Yvette Unoarumhi, Darlene Wagner, Matthew Schmerer, Minoo Agarwal, Eyad Almasri, Debbie Boles, Ayla Burns, Nuthawin Charoensri, Oren Cohen, Susan Countryman, Mary Ann Cristobal, Bobbi Croy, Suzanne Dale, Hrushikesh Deshmukh, Amanda Douglas, Vincent Drouillon, Marcia Eisenberg, Howard Engler, Rama Ghatti, Prashant Gupta, Susan Hicks, Jake Humphrey, Lax Iyer, Lisa Pfefferle, Manoj Jain, Matthew Robinson, Mohan Kolli, Brian Krueger, Tim Kuphal, Stanley Letovsky, Michael Levandoski, Craig Lukasik, Jonathan Meltzer, Brian Norvell, Mindy Nye, Scott Parker, Christos Petropoulos, John Pruitt, Steven Ragan, Scott Ryan, Mike Sapeta, Jana Schroth, Suresh Babu Selvaraju, Goran Stevovic, Amanda Suchanek, Andrea Throop, Lyndon Tilson, Thomas Urban, Joe Voshell, Kimberly Wagner, Jonathan Williams, Mary Williamson, Qian Zeng, Tricia Zwiefelhofer, Clinton R. Paden, Duncan MacCannell |
| EPI_ISL_3326585 | TGen North | TGen North | Jolene Bowers, Heather Centner, Brett Van Tassel, Chris French, Hayley Yaglom, Darrin Lemmer, Dave Engelthaler |
| EPI_ISL_3326586 | Laboratory Corporation of America | Centers for Disease Control and Prevention Division of Viral Diseases, Pathogen Discovery | Dakota Howard, Dhwani Batra, Peter W. Cook, Kara Moser, Adrian Paskey, Jason Caravas, Benjamin Rambo-Martin, Shatavia Morrison, Christopher Gulvick, Scott Sammons, Yvette Unoarumhi, Darlene Wagner, Matthew Schmerer, Minoo Agarwal, Eyad Almasri, Debbie Boles, Ayla Burns, Nuthawin Charoensri, Oren Cohen, Susan Countryman, Mary Ann Cristobal, Bobbi Croy, Suzanne Dale, Hrushikesh Deshmukh, Amanda Douglas, Vincent Drouillon, Marcia Eisenberg, Howard Engler, Rama Ghatti, Prashant Gupta, Susan Hicks, Jake Humphrey, Lax Iyer, Lisa Pfefferle, Manoj Jain, Matthew Robinson, Mohan Kolli, Brian Krueger, Tim Kuphal, Stanley Letovsky, Michael Levandoski, Craig Lukasik, Jonathan Meltzer, Brian Norvell, Mindy Nye, Scott Parker, Christos Petropoulos, John Pruitt, Steven Ragan, Scott Ryan, Mike Sapeta, Jana Schroth, Suresh Babu Selvaraju, Goran Stevovic, Amanda Suchanek, Andrea Throop, Lyndon Tilson, Thomas Urban, Joe Voshell, Kimberly Wagner, Jonathan Williams, Mary Williamson, Qian Zeng, Tricia Zwiefelhofer, Clinton R. Paden, Duncan MacCannell |
| EPI_ISL_3326587 | TGen North | TGen North | Jolene Bowers, Heather Centner, Brett Van Tassel, Chris French, Hayley Yaglom, Darrin Lemmer, Dave Engelthaler |
| EPI_ISL_3326588, EPI_ISL_3326589, EPI_ISL_3326591, EPI_ISL_3326593, EPI_ISL_3326595, EPI_ISL_3326596 | Laboratory Corporation of America | Centers for Disease Control and Prevention Division of Viral Diseases, Pathogen Discovery | Dakota Howard, Dhwani Batra, Peter W. Cook, Kara Moser, Adrian Paskey, Jason Caravas, Benjamin Rambo-Martin, Shatavia Morrison, Christopher Gulvick, Scott Sammons, Yvette Unoarumhi, Darlene Wagner, Matthew Schmerer, Minoo Agarwal, Eyad Almasri, Debbie Boles, Ayla Burns, Nuthawin Charoensri, Oren Cohen, Susan Countryman, Mary Ann Cristobal, Bobbi Croy, Suzanne Dale, Hrushikesh Deshmukh, Amanda Douglas, Vincent Drouillon, Marcia Eisenberg, Howard Engler, Rama Ghatti, Prashant Gupta, Susan Hicks, Jake Humphrey, Lax Iyer, Lisa Pfefferle, Manoj Jain, Matthew Robinson, Mohan Kolli, Brian Krueger, Tim Kuphal, Stanley Letovsky, Michael Levandoski, Craig Lukasik, Jonathan Meltzer, Brian Norvell, Mindy Nye, Scott Parker, Christos Petropoulos, John Pruitt, Steven Ragan, Scott Ryan, Mike Sapeta, Jana Schroth, Suresh Babu Selvaraju, Goran Stevovic, Amanda Suchanek, Andrea Throop, Lyndon Tilson, Thomas Urban, Joe Voshell, Kimberly Wagner, Jonathan Williams, Mary Williamson, Qian Zeng, Tricia Zwiefelhofer, Clinton R. Paden, Duncan MacCannell |
| EPI_ISL_3326597 | TGen North | TGen North | Jolene Bowers, Heather Centner, Brett Van Tassel, Chris French, Hayley Yaglom, Darrin Lemmer, Dave Engelthaler |
| EPI_ISL_3326598 | Laboratory Corporation of America | Centers for Disease Control and Prevention Division of Viral Diseases, Pathogen Discovery | Dakota Howard, Dhwani Batra, Peter W. Cook, Kara Moser, Adrian Paskey, Jason Caravas, Benjamin Rambo-Martin, Shatavia Morrison, Christopher Gulvick, Scott Sammons, Yvette Unoarumhi, Darlene Wagner, Matthew Schmerer, Minoo Agarwal, Eyad Almasri, Debbie Boles, Ayla Burns, Nuthawin Charoensri, Oren Cohen, Susan Countryman, Mary Ann Cristobal, Bobbi Croy, Suzanne Dale, Hrushikesh Deshmukh, Amanda Douglas, Vincent Drouillon, Marcia Eisenberg, Howard Engler, Rama Ghatti, Prashant Gupta, Susan Hicks, Jake Humphrey, Lax Iyer, Lisa Pfefferle, Manoj Jain, Matthew Robinson, Mohan Kolli, Brian Krueger, Tim Kuphal, Stanley Letovsky, Michael Levandoski, Craig Lukasik, Jonathan Meltzer, Brian Norvell, Mindy Nye, Scott Parker, Christos Petropoulos, John Pruitt, Steven Ragan, Scott Ryan, Mike Sapeta, Jana Schroth, Suresh Babu Selvaraju, Goran Stevovic, Amanda Suchanek, Andrea Throop, Lyndon Tilson, Thomas Urban, Joe Voshell, Kimberly Wagner, Jonathan Williams, Mary Williamson, Qian Zeng, Tricia Zwiefelhofer, Clinton R. Paden, Duncan MacCannell |
| EPI_ISL_3326599 | TGen North | TGen North | Jolene Bowers, Heather Centner, Brett Van Tassel, Chris French, Hayley Yaglom, Darrin Lemmer, Dave Engelthaler |
| EPI_ISL_3326600 | Laboratory Corporation of America | Centers for Disease Control and Prevention Division of Viral Diseases, Pathogen Discovery | Dakota Howard, Dhwani Batra, Peter W. Cook, Kara Moser, Adrian Paskey, Jason Caravas, Benjamin Rambo-Martin, Shatavia Morrison, Christopher Gulvick, Scott Sammons, Yvette Unoarumhi, Darlene Wagner, Matthew Schmerer, Minoo Agarwal, Eyad Almasri, Debbie Boles, Ayla Burns, Nuthawin Charoensri, Oren Cohen, Susan Countryman, Mary Ann Cristobal, Bobbi Croy, Suzanne Dale, Hrushikesh Deshmukh, Amanda Douglas, Vincent Drouillon, Marcia Eisenberg, Howard Engler, Rama Ghatti, Prashant Gupta, Susan Hicks, Jake Humphrey, Lax Iyer, Lisa Pfefferle, Manoj Jain, Matthew Robinson, Mohan Kolli, Brian Krueger, Tim Kuphal, Stanley Letovsky, Michael Levandoski, Craig Lukasik, Jonathan Meltzer, Brian Norvell, Mindy Nye, Scott Parker, Christos Petropoulos, John Pruitt, Steven Ragan, Scott Ryan, Mike Sapeta, Jana Schroth, Suresh Babu Selvaraju, Goran Stevovic, Amanda Suchanek, Andrea Throop, Lyndon Tilson, Thomas Urban, Joe Voshell, Kimberly Wagner, Jonathan Williams, Mary Williamson, Qian Zeng, Tricia Zwiefelhofer, Clinton R. Paden, Duncan MacCannell |
| EPI_ISL_3326601 | TGen North | TGen North | Jolene Bowers, Heather Centner, Brett Van Tassel, Chris French, Hayley Yaglom, Darrin Lemmer, Dave Engelthaler |
| EPI_ISL_3326602 | Laboratory Corporation of America | Centers for Disease Control and Prevention Division of Viral Diseases, Pathogen Discovery | Dakota Howard, Dhwani Batra, Peter W. Cook, Kara Moser, Adrian Paskey, Jason Caravas, Benjamin Rambo-Martin, Shatavia Morrison, Christopher Gulvick, Scott Sammons, Yvette Unoarumhi, Darlene Wagner, Matthew Schmerer, Minoo Agarwal, Eyad Almasri, Debbie Boles, Ayla Burns, Nuthawin Charoensri, Oren Cohen, Susan Countryman, Mary Ann Cristobal, Bobbi Croy, Suzanne Dale, Hrushikesh Deshmukh, Amanda Douglas, Vincent Drouillon, Marcia Eisenberg, Howard Engler, Rama Ghatti, Prashant Gupta, Susan Hicks, Jake Humphrey, Lax Iyer, Lisa Pfefferle, Manoj Jain, Matthew Robinson, Mohan Kolli, Brian Krueger, Tim Kuphal, Stanley Letovsky, Michael Levandoski, Craig Lukasik, Jonathan Meltzer, Brian Norvell, Mindy Nye, Scott Parker, Christos Petropoulos, John Pruitt, Steven Ragan, Scott Ryan, Mike Sapeta, Jana Schroth, Suresh Babu Selvaraju, Goran Stevovic, Amanda Suchanek, Andrea Throop, Lyndon Tilson, Thomas Urban, Joe Voshell, |

| | | | |
|---|---|---|---|
| | | | Kimberly Wagner, Jonathan Williams, Mary Williamson, Qian Zeng, Tricia Zwiefelhofer, Clinton R. Paden, Duncan MacCannell |
| EPI_ISL_3326603 | TGen North | TGen North | Jolene Bowers, Heather Centner, Brett Van Tassel, Chris French, Hayley Yaglom, Darrin Lemmer, Dave Engelthaler |
| EPI_ISL_3326604, EPI_ISL_3326606, EPI_ISL_3326607 | Laboratory Corporation of America | Centers for Disease Control and Prevention Division of Viral Diseases, Pathogen Discovery | Dakota Howard, Dhwani Batra, Peter W. Cook, Kara Moser, Adrian Paskey, Jason Caravas, Benjamin Rambo-Martin, Shatavia Morrison, Christopher Gulvick, Scott Sammons, Yvette Unoarumhi, Darlene Wagner, Matthew Schmerer, Minoo Agarwal, Eyad Almasri, Debbie Boles, Ayla Burns, Nuthawin Charoensri, Oren Cohen, Susan Countryman, Mary Ann Cristobal, Bobbi Croy, Suzanne Dale, Hrushikesh Deshmukh, Amanda Douglas, Vincent Drouillon, Marcia Eisenberg, Howard Engler, Rama Ghatti, Prashant Gupta, Susan Hicks, Jake Humphrey, Lax Iyer, Lisa Pfefferle, Manoj Jain, Matthew Robinson, Mohan Kolli, Brian Krueger, Tim Kuphal, Stanley Letovsky, Michael Levandoski, Craig Lukasik, Jonathan Meltzer, Brian Norvell, Mindy Nye, Scott Parker, Christos Petropoulos, John Pruitt, Steven Ragan, Scott Ryan, Mike Sapeta, Jana Schroth, Suresh Babu Selvaraju, Goran Stevovic, Amanda Suchanek, Andrea Throop, Lyndon Tilson, Thomas Urban, Joe Voshell, Kimberly Wagner, Jonathan Williams, Mary Williamson, Qian Zeng, Tricia Zwiefelhofer, Clinton R. Paden, Duncan MacCannell |
| EPI_ISL_3326608 | TGen North | TGen North | Jolene Bowers, Heather Centner, Brett Van Tassel, Chris French, Hayley Yaglom, Darrin Lemmer, Dave Engelthaler |
| EPI_ISL_3326609, EPI_ISL_3326611 | Laboratory Corporation of America | Centers for Disease Control and Prevention Division of Viral Diseases, Pathogen Discovery | Dakota Howard, Dhwani Batra, Peter W. Cook, Kara Moser, Adrian Paskey, Jason Caravas, Benjamin Rambo-Martin, Shatavia Morrison, Christopher Gulvick, Scott Sammons, Yvette Unoarumhi, Darlene Wagner, Matthew Schmerer, Minoo Agarwal, Eyad Almasri, Debbie Boles, Ayla Burns, Nuthawin Charoensri, Oren Cohen, Susan Countryman, Mary Ann Cristobal, Bobbi Croy, Suzanne Dale, Hrushikesh Deshmukh, Amanda Douglas, Vincent Drouillon, Marcia Eisenberg, Howard Engler, Rama Ghatti, Prashant Gupta, Susan Hicks, Jake Humphrey, Lax Iyer, Lisa Pfefferle, Manoj Jain, Matthew Robinson, Mohan Kolli, Brian Krueger, Tim Kuphal, Stanley Letovsky, Michael Levandoski, Craig Lukasik, Jonathan Meltzer, Brian Norvell, Mindy Nye, Scott Parker, Christos Petropoulos, John Pruitt, Steven Ragan, Scott Ryan, Mike Sapeta, Jana Schroth, Suresh Babu Selvaraju, Goran Stevovic, Amanda Suchanek, Andrea Throop, Lyndon Tilson, Thomas Urban, Joe Voshell, Kimberly Wagner, Jonathan Williams, Mary Williamson, Qian Zeng, Tricia Zwiefelhofer, Clinton R. Paden, Duncan MacCannell |
| EPI_ISL_3326612 | TGen North | TGen North | Jolene Bowers, Heather Centner, Brett Van Tassel, Chris French, Hayley Yaglom, Darrin Lemmer, Dave Engelthaler |
| EPI_ISL_3326614 | Laboratory Corporation of America | Centers for Disease Control and Prevention Division of Viral Diseases, Pathogen Discovery | Dakota Howard, Dhwani Batra, Peter W. Cook, Kara Moser, Adrian Paskey, Jason Caravas, Benjamin Rambo-Martin, Shatavia Morrison, Christopher Gulvick, Scott Sammons, Yvette Unoarumhi, Darlene Wagner, Matthew Schmerer, Minoo Agarwal, Eyad Almasri, Debbie Boles, Ayla Burns, Nuthawin Charoensri, Oren Cohen, Susan Countryman, Mary Ann Cristobal, Bobbi Croy, Suzanne Dale, Hrushikesh Deshmukh, Amanda Douglas, Vincent Drouillon, Marcia Eisenberg, Howard Engler, Rama Ghatti, Prashant Gupta, Susan Hicks, Jake Humphrey, Lax Iyer, Lisa Pfefferle, Manoj Jain, Matthew Robinson, Mohan Kolli, Brian Krueger, Tim Kuphal, Stanley Letovsky, Michael Levandoski, Craig Lukasik, Jonathan Meltzer, Brian Norvell, Mindy Nye, Scott Parker, Christos Petropoulos, John Pruitt, Steven Ragan, Scott Ryan, Mike Sapeta, Jana Schroth, Suresh Babu Selvaraju, Goran Stevovic, Amanda Suchanek, Andrea Throop, Lyndon Tilson, Thomas Urban, Joe Voshell, Kimberly Wagner, Jonathan Williams, Mary Williamson, Qian Zeng, Tricia Zwiefelhofer, Clinton R. Paden, Duncan MacCannell |
| EPI_ISL_3326615 | TGen North | TGen North | Jolene Bowers, Heather Centner, Brett Van Tassel, Chris French, Hayley Yaglom, Darrin Lemmer, Dave Engelthaler |
| EPI_ISL_3326616 | Laboratory Corporation of America | Centers for Disease Control and Prevention Division of Viral Diseases, Pathogen Discovery | Dakota Howard, Dhwani Batra, Peter W. Cook, Kara Moser, Adrian Paskey, Jason Caravas, Benjamin Rambo-Martin, Shatavia Morrison, Christopher Gulvick, Scott Sammons, Yvette Unoarumhi, Darlene Wagner, Matthew Schmerer, Minoo Agarwal, Eyad Almasri, Debbie Boles, Ayla Burns, Nuthawin Charoensri, Oren Cohen, Susan Countryman, Mary Ann Cristobal, Bobbi Croy, Suzanne Dale, Hrushikesh Deshmukh, Amanda Douglas, Vincent Drouillon, Marcia Eisenberg, Howard Engler, Rama Ghatti, Prashant Gupta, Susan Hicks, Jake Humphrey, Lax Iyer, Lisa Pfefferle, Manoj Jain, Matthew Robinson, Mohan Kolli, Brian Krueger, Tim Kuphal, Stanley Letovsky, Michael Levandoski, Craig Lukasik, Jonathan Meltzer, Brian Norvell, Mindy Nye, Scott Parker, Christos Petropoulos, John Pruitt, Steven Ragan, Scott Ryan, Mike Sapeta, Jana Schroth, Suresh Babu Selvaraju, Goran Stevovic, Amanda Suchanek, Andrea Throop, Lyndon Tilson, Thomas Urban, Joe Voshell, Kimberly Wagner, Jonathan Williams, Mary Williamson, Qian Zeng, Tricia Zwiefelhofer, Clinton R. Paden, Duncan MacCannell |
| EPI_ISL_3326617 | TGen North | TGen North | Jolene Bowers, Heather Centner, Brett Van Tassel, Chris French, Hayley Yaglom, Darrin Lemmer, Dave Engelthaler |
| EPI_ISL_3326618 | Laboratory Corporation of America | Centers for Disease Control and Prevention Division of Viral Diseases, Pathogen Discovery | Dakota Howard, Dhwani Batra, Peter W. Cook, Kara Moser, Adrian Paskey, Jason Caravas, Benjamin Rambo-Martin, Shatavia Morrison, Christopher Gulvick, Scott Sammons, Yvette Unoarumhi, Darlene Wagner, Matthew Schmerer, Minoo Agarwal, Eyad Almasri, Debbie Boles, Ayla Burns, Nuthawin Charoensri, Oren Cohen, Susan Countryman, Mary Ann Cristobal, Bobbi Croy, Suzanne Dale, Hrushikesh Deshmukh, Amanda Douglas, Vincent Drouillon, Marcia Eisenberg, Howard Engler, Rama Ghatti, Prashant Gupta, Susan Hicks, Jake Humphrey, Lax Iyer, Lisa Pfefferle, Manoj Jain, Matthew Robinson, Mohan Kolli, Brian Krueger, Tim Kuphal, Stanley Letovsky, Michael Levandoski, Craig Lukasik, Jonathan Meltzer, Brian Norvell, Mindy Nye, Scott Parker, Christos Petropoulos, John Pruitt, Steven Ragan, Scott Ryan, Mike Sapeta, Jana Schroth, Suresh Babu Selvaraju, Goran Stevovic, Amanda Suchanek, Andrea Throop, Lyndon Tilson, Thomas Urban, Joe Voshell, Kimberly Wagner, Jonathan Williams, Mary Williamson, Qian Zeng, Tricia Zwiefelhofer, Clinton R. Paden, Duncan MacCannell |
| EPI_ISL_3326619 | TGen North | TGen North | Jolene Bowers, Heather Centner, Brett Van Tassel, Chris French, Hayley Yaglom, Darrin Lemmer, Dave Engelthaler |
| EPI_ISL_3326620 | Laboratory Corporation of America | Centers for Disease Control and Prevention Division of Viral Diseases, Pathogen Discovery | Dakota Howard, Dhwani Batra, Peter W. Cook, Kara Moser, Adrian Paskey, Jason Caravas, Benjamin Rambo-Martin, Shatavia Morrison, Christopher Gulvick, Scott Sammons, Yvette Unoarumhi, Darlene Wagner, Matthew Schmerer, Minoo Agarwal, Eyad Almasri, Debbie Boles, Ayla Burns, Nuthawin Charoensri, Oren Cohen, Susan Countryman, Mary Ann Cristobal, Bobbi Croy, Suzanne Dale, Hrushikesh Deshmukh, Amanda Douglas, Vincent Drouillon, Marcia Eisenberg, Howard Engler, Rama Ghatti, Prashant Gupta, Susan Hicks, Jake Humphrey, Lax Iyer, Lisa Pfefferle, Manoj Jain, Matthew Robinson, Mohan Kolli, Brian Krueger, Tim Kuphal, Stanley Letovsky, Michael Levandoski, Craig Lukasik, Jonathan Meltzer, Brian Norvell, Mindy Nye, Scott Parker, Christos Petropoulos, John Pruitt, Steven Ragan, Scott Ryan, Mike Sapeta, Jana Schroth, Suresh Babu Selvaraju, Goran Stevovic, Amanda Suchanek, Andrea Throop, Lyndon Tilson, Thomas Urban, Joe Voshell, Kimberly Wagner, Jonathan Williams, Mary Williamson, Qian Zeng, Tricia Zwiefelhofer, Clinton R. Paden, Duncan MacCannell |
| EPI_ISL_3326621 | TGen North | TGen North | Jolene Bowers, Heather Centner, Brett Van Tassel, Chris French, Hayley Yaglom, Darrin Lemmer, Dave Engelthaler |
| EPI_ISL_3326622 | Laboratory Corporation of America | Centers for Disease Control and Prevention Division of Viral Diseases, Pathogen Discovery | Dakota Howard, Dhwani Batra, Peter W. Cook, Kara Moser, Adrian Paskey, Jason Caravas, Benjamin Rambo-Martin, Shatavia Morrison, Christopher Gulvick, Scott Sammons, Yvette Unoarumhi, Darlene Wagner, Matthew Schmerer, Minoo Agarwal, Eyad Almasri, Debbie Boles, Ayla Burns, Nuthawin Charoensri, Oren Cohen, Susan Countryman, Mary Ann Cristobal, Bobbi Croy, Suzanne Dale, Hrushikesh Deshmukh, Amanda Douglas, Vincent Drouillon, Marcia Eisenberg, Howard Engler, Rama Ghatti, Prashant Gupta, Susan Hicks, Jake Humphrey, Lax Iyer, Lisa Pfefferle, Manoj Jain, Matthew Robinson, Mohan Kolli, Brian Krueger, Tim Kuphal, Stanley Letovsky, Michael Levandoski, Craig Lukasik, Jonathan Meltzer, Brian Norvell, Mindy Nye, Scott Parker, Christos Petropoulos, John Pruitt, Steven Ragan, Scott Ryan, Mike Sapeta, Jana Schroth, Suresh Babu Selvaraju, Goran Stevovic, Amanda Suchanek, Andrea Throop, Lyndon Tilson, Thomas Urban, Joe Voshell, Kimberly Wagner, Jonathan Williams, Mary Williamson, Qian Zeng, Tricia Zwiefelhofer, Clinton R. Paden, Duncan MacCannell |
| EPI_ISL_3326623 | TGen North | TGen North | Jolene Bowers, Heather Centner, Brett Van Tassel, Chris French, Hayley Yaglom, Darrin Lemmer, Dave Engelthaler |
| EPI_ISL_3326624, EPI_ISL_3326625, EPI_ISL_3326627 | Laboratory Corporation of America | Centers for Disease Control and Prevention Division of Viral Diseases, Pathogen Discovery | Dakota Howard, Dhwani Batra, Peter W. Cook, Kara Moser, Adrian Paskey, Jason Caravas, Benjamin Rambo-Martin, Shatavia Morrison, Christopher Gulvick, Scott Sammons, Yvette Unoarumhi, Darlene Wagner, Matthew Schmerer, Minoo Agarwal, Eyad Almasri, Debbie Boles, Ayla Burns, Nuthawin Charoensri, Oren Cohen, Susan Countryman, Mary Ann Cristobal, Bobbi Croy, Suzanne Dale, Hrushikesh Deshmukh, Amanda Douglas, Vincent Drouillon, Marcia Eisenberg, Howard Engler, Rama Ghatti, Prashant Gupta, Susan Hicks, Jake Humphrey, Lax Iyer, Lisa Pfefferle, Manoj Jain, Matthew Robinson, Mohan Kolli, Brian Krueger, Tim Kuphal, Stanley Letovsky, Michael Levandoski, Craig Lukasik, Jonathan Meltzer, Brian Norvell, Mindy Nye, Scott Parker, Christos Petropoulos, John Pruitt, Steven Ragan, Scott Ryan, Mike Sapeta, Jana Schroth, Suresh Babu Selvaraju, Goran Stevovic, Amanda Suchanek, Andrea Throop, Lyndon Tilson, Thomas Urban, Joe Voshell, Kimberly Wagner, Jonathan Williams, Mary Williamson, Qian Zeng, Tricia Zwiefelhofer, Clinton R. Paden, Duncan MacCannell |
| EPI_ISL_3326628 | TGen North | TGen North | Jolene Bowers, Heather Centner, Brett Van Tassel, Chris French, Hayley Yaglom, Darrin Lemmer, Dave Engelthaler |
| EPI_ISL_3326629, EPI_ISL_3326630 | Laboratory Corporation of America | Centers for Disease Control and Prevention Division of Viral Diseases, Pathogen Discovery | Dakota Howard, Dhwani Batra, Peter W. Cook, Kara Moser, Adrian Paskey, Jason Caravas, Benjamin Rambo-Martin, Shatavia Morrison, Christopher Gulvick, Scott Sammons, Yvette Unoarumhi, Darlene Wagner, Matthew Schmerer, Minoo Agarwal, Eyad Almasri, Debbie Boles, Ayla Burns, Nuthawin Charoensri, Oren Cohen, Susan Countryman, Mary Ann Cristobal, Bobbi Croy, Suzanne Dale, Hrushikesh Deshmukh, Amanda Douglas, Vincent Drouillon, Marcia Eisenberg, Howard Engler, Rama Ghatti, Prashant Gupta, Susan Hicks, Jake Humphrey, Lax Iyer, Lisa Pfefferle, Manoj Jain, Matthew Robinson, Mohan Kolli, Brian Krueger, Tim Kuphal, Stanley Letovsky, Michael Levandoski, Craig Lukasik, Jonathan Meltzer, Brian Norvell, Mindy Nye, Scott Parker, Christos Petropoulos, John Pruitt, Steven Ragan, Scott Ryan, Mike Sapeta, Jana Schroth, Suresh Babu Selvaraju, Goran Stevovic, Amanda Suchanek, Andrea Throop, Lyndon Tilson, Thomas Urban, Joe Voshell, Kimberly Wagner, Jonathan Williams, Mary Williamson, Qian Zeng, Tricia Zwiefelhofer, Clinton R. Paden, Duncan MacCannell |
| EPI_ISL_3326631 | TGen North | TGen North | Jolene Bowers, Heather Centner, Brett Van Tassel, Chris French, Hayley Yaglom, Darrin Lemmer, Dave Engelthaler |
| EPI_ISL_3326632 | Laboratory Corporation of America | Centers for Disease Control and Prevention Division of Viral Diseases, Pathogen Discovery | Dakota Howard, Dhwani Batra, Peter W. Cook, Kara Moser, Adrian Paskey, Jason Caravas, Benjamin Rambo-Martin, Shatavia Morrison, Christopher Gulvick, Scott Sammons, Yvette Unoarumhi, Darlene Wagner, Matthew Schmerer, Minoo Agarwal, Eyad Almasri, Debbie Boles, Ayla Burns, Nuthawin Charoensri, Oren Cohen, Susan Countryman, Mary Ann Cristobal, Bobbi Croy, Suzanne Dale, Hrushikesh Deshmukh, Amanda Douglas, Vincent Drouillon, Marcia Eisenberg, Howard Engler, Rama Ghatti, Prashant Gupta, Susan Hicks, Jake Humphrey, Lax Iyer, Lisa Pfefferle, Manoj Jain, Matthew Robinson, Mohan Kolli, Brian Krueger, Tim Kuphal, Stanley Letovsky, Michael Levandoski, Craig Lukasik, Jonathan Meltzer, Brian Norvell, Mindy Nye, Scott Parker, Christos Petropoulos, John Pruitt, Steven Ragan, Scott Ryan, Mike Sapeta, Jana Schroth, Suresh Babu Selvaraju, Goran Stevovic, Amanda Suchanek, Andrea Throop, Lyndon Tilson, Thomas Urban, Joe Voshell, Kimberly Wagner, Jonathan Williams, Mary Williamson, Qian Zeng, Tricia Zwiefelhofer, Clinton R. Paden, Duncan MacCannell |
| EPI_ISL_3326633 | TGen North | TGen North | Jolene Bowers, Heather Centner, Brett Van Tassel, Chris French, Hayley Yaglom, Darrin Lemmer, Dave Engelthaler |
| EPI_ISL_3326634 | Laboratory Corporation of America | Centers for Disease Control and Prevention Division of Viral Diseases, Pathogen Discovery | Dakota Howard, Dhwani Batra, Peter W. Cook, Kara Moser, Adrian Paskey, Jason Caravas, Benjamin Rambo-Martin, Shatavia Morrison, Christopher Gulvick, Scott Sammons, Yvette Unoarumhi, Darlene Wagner, Matthew Schmerer, Minoo Agarwal, Eyad Almasri, Debbie Boles, Ayla Burns, Nuthawin Charoensri, Oren Cohen, Susan Countryman, Mary Ann Cristobal, Bobbi Croy, Suzanne Dale, Hrushikesh Deshmukh, Amanda Douglas, Vincent Drouillon, Marcia Eisenberg, Howard Engler, Rama Ghatti, Prashant Gupta, Susan Hicks, Jake Humphrey, Lax Iyer, Lisa Pfefferle, Manoj Jain, Matthew Robinson, Mohan Kolli, Brian Krueger, Tim Kuphal, Stanley Letovsky, Michael Levandoski, Craig Lukasik, Jonathan Meltzer, Brian Norvell, Mindy Nye, Scott Parker, Christos Petropoulos, John Pruitt, Steven Ragan, Scott Ryan, Mike Sapeta, Jana Schroth, Suresh Babu Selvaraju, Goran Stevovic, Amanda Suchanek, Andrea Throop, Lyndon Tilson, Thomas Urban, Joe Voshell, Kimberly Wagner, Jonathan Williams, Mary Williamson, Qian Zeng, Tricia Zwiefelhofer, Clinton R. Paden, Duncan MacCannell |
| EPI_ISL_3326635 | TGen North | TGen North | Jolene Bowers, Heather Centner, Brett Van Tassel, Chris French, Hayley Yaglom, Darrin Lemmer, Dave Engelthaler |
| EPI_ISL_3326636 | Laboratory Corporation of America | Centers for Disease Control and Prevention Division of Viral Diseases, Pathogen Discovery | Dakota Howard, Dhwani Batra, Peter W. Cook, Kara Moser, Adrian Paskey, Jason Caravas, Benjamin Rambo-Martin, Shatavia Morrison, Christopher Gulvick, Scott Sammons, Yvette Unoarumhi, Darlene Wagner, Matthew Schmerer, Minoo Agarwal, Eyad Almasri, Debbie Boles, Ayla Burns, Nuthawin Charoensri, Oren Cohen, Susan Countryman, Mary Ann Cristobal, Bobbi Croy, Suzanne Dale, Hrushikesh Deshmukh, Amanda Douglas, Vincent Drouillon, Marcia Eisenberg, Howard Engler, Rama Ghatti, Prashant Gupta, Susan Hicks, Jake Humphrey, Lax Iyer, Lisa Pfefferle, Manoj Jain, Matthew Robinson, Mohan Kolli, Brian Krueger, Tim Kuphal, Stanley Letovsky, Michael Levandoski, Craig Lukasik, Jonathan Meltzer, Brian Norvell, Mindy Nye, Scott Parker, Christos Petropoulos, John Pruitt, Steven Ragan, Scott Ryan, Mike Sapeta, Jana Schroth, Suresh Babu Selvaraju, Goran Stevovic, Amanda Suchanek, Andrea Throop, Lyndon Tilson, Thomas Urban, Joe Voshell, Kimberly Wagner, Jonathan Williams, Mary Williamson, Qian Zeng, Tricia Zwiefelhofer, Clinton R. Paden, Duncan MacCannell |
| EPI_ISL_3326637 | TGen North | TGen North | Jolene Bowers, Heather Centner, Brett Van Tassel, Chris French, Hayley Yaglom, Darrin Lemmer, Dave Engelthaler |
| EPI_ISL_3326638 | Laboratory Corporation of America | Centers for Disease Control and Prevention Division of Viral Diseases, Pathogen Discovery | Dakota Howard, Dhwani Batra, Peter W. Cook, Kara Moser, Adrian Paskey, Jason Caravas, Benjamin Rambo-Martin, Shatavia Morrison, Christopher Gulvick, Scott Sammons, Yvette Unoarumhi, Darlene Wagner, Matthew Schmerer, Minoo Agarwal, Eyad Almasri, Debbie Boles, Ayla Burns, Nuthawin Charoensri, Oren Cohen, Susan Countryman, Mary Ann Cristobal, Bobbi Croy, Suzanne Dale, Hrushikesh Deshmukh, Amanda Douglas, Vincent Drouillon, Marcia Eisenberg, Howard Engler, Rama Ghatti, Prashant Gupta, Susan Hicks, Jake Humphrey, Lax Iyer, Lisa Pfefferle, Manoj Jain, Matthew Robinson, Mohan Kolli, Brian Krueger, Tim Kuphal, Stanley Letovsky, Michael Levandoski, Craig Lukasik, Jonathan Meltzer, Brian Norvell, Mindy Nye, Scott Parker, Christos Petropoulos, John Pruitt, Steven Ragan, Scott Ryan, Mike Sapeta, Jana Schroth, Suresh Babu Selvaraju, Goran Stevovic, Amanda Suchanek, Andrea Throop, Lyndon Tilson, Thomas Urban, Joe Voshell, Kimberly Wagner, Jonathan Williams, Mary Williamson, Qian Zeng, Tricia Zwiefelhofer, Clinton R. Paden, Duncan MacCannell |
| EPI_ISL_3326639 | TGen North | TGen North | Jolene Bowers, Heather Centner, Brett Van Tassel, Chris French, Hayley Yaglom, Darrin Lemmer, Dave Engelthaler |
| EPI_ISL_3326640 | Laboratory Corporation of America | Centers for Disease Control and Prevention Division of Viral Diseases, Pathogen Discovery | Dakota Howard, Dhwani Batra, Peter W. Cook, Kara Moser, Adrian Paskey, Jason Caravas, Benjamin Rambo-Martin, Shatavia Morrison, Christopher Gulvick, Scott Sammons, Yvette Unoarumhi, Darlene Wagner, Matthew Schmerer, Minoo Agarwal, Eyad Almasri, Debbie Boles, Ayla Burns, Nuthawin Charoensri, Oren Cohen, Susan Countryman, Mary Ann Cristobal, Bobbi Croy, Suzanne Dale, Hrushikesh Deshmukh, Amanda Douglas, Vincent Drouillon, Marcia Eisenberg, Howard Engler, Rama Ghatti, Prashant Gupta, Susan Hicks, Jake Humphrey, Lax Iyer, Lisa Pfefferle, Manoj Jain, Matthew Robinson, Mohan Kolli, Brian Krueger, Tim Kuphal, Stanley Letovsky, Michael Levandoski, Craig Lukasik, Jonathan Meltzer, Brian Norvell, Mindy Nye, Scott Parker, Christos Petropoulos, John Pruitt, Steven Ragan, Scott Ryan, Mike Sapeta, Jana Schroth, Suresh Babu Selvaraju, Goran Stevovic, Amanda Suchanek, Andrea Throop, Lyndon Tilson, Thomas Urban, Joe Voshell, Kimberly Wagner, Jonathan Williams, Mary Williamson, Qian Zeng, Tricia Zwiefelhofer, Clinton R. Paden, Duncan MacCannell |
| EPI_ISL_3326641 | TGen North | TGen North | Jolene Bowers, Heather Centner, Brett Van Tassel, Chris French, Hayley Yaglom, Darrin Lemmer, Dave Engelthaler |
| EPI_ISL_3326642, EPI_ISL_3326643 | Laboratory Corporation of America | Centers for Disease Control and Prevention Division of Viral Diseases, Pathogen Discovery | Dakota Howard, Dhwani Batra, Peter W. Cook, Kara Moser, Adrian Paskey, Jason Caravas, Benjamin Rambo-Martin, Shatavia Morrison, Christopher Gulvick, Scott Sammons, Yvette Unoarumhi, Darlene Wagner, Matthew Schmerer, Minoo Agarwal, Eyad Almasri, Debbie Boles, Ayla Burns, Nuthawin Charoensri, Oren Cohen, Susan Countryman, Mary Ann Cristobal, Bobbi Croy, Suzanne Dale, Hrushikesh Deshmukh, Amanda Douglas, Vincent Drouillon, Marcia Eisenberg, Howard Engler, Rama Ghatti, Prashant Gupta, Susan Hicks, Jake Humphrey, Lax Iyer, Lisa Pfefferle, Manoj Jain, Matthew Robinson, Mohan Kolli, Brian Krueger, Tim Kuphal, Stanley Letovsky, Michael Levandoski, Craig Lukasik, Jonathan Meltzer, Brian Norvell, Mindy Nye, Scott Parker, Christos Petropoulos, John Pruitt, Steven Ragan, Scott Ryan, Mike Sapeta, Jana Schroth, Suresh Babu Selvaraju, Goran Stevovic, Amanda Suchanek, Andrea Throop, Lyndon Tilson, Thomas Urban, Joe Voshell, Kimberly Wagner, Jonathan Williams, Mary Williamson, Qian Zeng, Tricia Zwiefelhofer, Clinton R. Paden, Duncan MacCannell |
| EPI_ISL_3326644 | TGen North | TGen North | Jolene Bowers, Heather Centner, Brett Van Tassel, Chris French, Hayley Yaglom, Darrin Lemmer, Dave Engelthaler |
| EPI_ISL_3326645, EPI_ISL_3326647 | Laboratory Corporation of America | Centers for Disease Control and Prevention Division of Viral Diseases, Pathogen Discovery | Dakota Howard, Dhwani Batra, Peter W. Cook, Kara Moser, Adrian Paskey, Jason Caravas, Benjamin Rambo-Martin, Shatavia Morrison, Christopher Gulvick, Scott Sammons, Yvette Unoarumhi, Darlene Wagner, Matthew Schmerer, Minoo Agarwal, Eyad Almasri, Debbie Boles, Ayla Burns, Nuthawin Charoensri, Oren Cohen, Susan Countryman, Mary Ann Cristobal, Bobbi Croy, Suzanne Dale, Hrushikesh Deshmukh, Amanda Douglas, Vincent Drouillon, Marcia Eisenberg, Howard Engler, Rama Ghatti, Prashant Gupta, Susan Hicks, Jake Humphrey, Lax Iyer, Lisa Pfefferle, Manoj Jain, Matthew Robinson, Mohan Kolli, Brian Krueger, Tim Kuphal, Stanley Letovsky, Michael Levandoski, Craig Lukasik, Jonathan Meltzer, Brian Norvell, Mindy Nye, Scott Parker, Christos Petropoulos, John Pruitt, Steven Ragan, Scott Ryan, Mike Sapeta, Jana Schroth, Suresh Babu Selvaraju, Goran Stevovic, Amanda Suchanek, Andrea Throop, Lyndon Tilson, Thomas Urban, Joe Voshell, |

| | | | |
|---|---|---|---|
| | | | Kimberly Wagner, Jonathan Williams, Mary Williamson, Qian Zeng, Tricia Zwiefelhofer, Clinton R. Paden, Duncan MacCannell |
| EPI_ISL_3326648 | TGen North | TGen North | Jolene Bowers, Heather Centner, Brett Van Tassel, Chris French, Hayley Yaglom, Darrin Lemmer, Dave Engelthaler |
| EPI_ISL_3326651 | Laboratory Corporation of America | Centers for Disease Control and Prevention Division of Viral Diseases, Pathogen Discovery | Dakota Howard, Dhwani Batra, Peter W. Cook, Kara Moser, Adrian Paskey, Jason Caravas, Benjamin Rambo-Martin, Shatavia Morrison, Christopher Gulvick, Scott Sammons, Yvette Unoarumhi, Darlene Wagner, Matthew Schmerer, Minoo Agarwal, Eyad Almasri, Debbie Boles, Ayla Burns, Nuthawin Charoensri, Oren Cohen, Susan Countryman, Mary Ann Cristobal, Bobbi Croy, Suzanne Dale, Hrushikesh Deshmukh, Amanda Douglas, Vincent Drouillon, Marcia Eisenberg, Howard Engler, Rama Ghatti, Prashant Gupta, Susan Hicks, Jake Humphrey, Lax Iyer, Lisa Pfefferle, Manoj Jain, Matthew Robinson, Mohan Kolli, Brian Krueger, Tim Kuphal, Stanley Letovsky, Michael Levandoski, Craig Lukasik, Jonathan Meltzer, Brian Norvell, Mindy Nye, Scott Parker, Christos Petropoulos, John Pruitt, Steven Ragan, Scott Ryan, Mike Sapeta, Jana Schroth, Suresh Babu Selvaraju, Goran Stevovic, Amanda Suchanek, Andrea Throop, Lyndon Tilson, Thomas Urban, Joe Voshell, Kimberly Wagner, Jonathan Williams, Mary Williamson, Qian Zeng, Tricia Zwiefelhofer, Clinton R. Paden, Duncan MacCannell |
| EPI_ISL_3326652 | TGen North | TGen North | Jolene Bowers, Heather Centner, Brett Van Tassel, Chris French, Hayley Yaglom, Darrin Lemmer, Dave Engelthaler |
| EPI_ISL_3326653 | Laboratory Corporation of America | Centers for Disease Control and Prevention Division of Viral Diseases, Pathogen Discovery | Dakota Howard, Dhwani Batra, Peter W. Cook, Kara Moser, Adrian Paskey, Jason Caravas, Benjamin Rambo-Martin, Shatavia Morrison, Christopher Gulvick, Scott Sammons, Yvette Unoarumhi, Darlene Wagner, Matthew Schmerer, Minoo Agarwal, Eyad Almasri, Debbie Boles, Ayla Burns, Nuthawin Charoensri, Oren Cohen, Susan Countryman, Mary Ann Cristobal, Bobbi Croy, Suzanne Dale, Hrushikesh Deshmukh, Amanda Douglas, Vincent Drouillon, Marcia Eisenberg, Howard Engler, Rama Ghatti, Prashant Gupta, Susan Hicks, Jake Humphrey, Lax Iyer, Lisa Pfefferle, Manoj Jain, Matthew Robinson, Mohan Kolli, Brian Krueger, Tim Kuphal, Stanley Letovsky, Michael Levandoski, Craig Lukasik, Jonathan Meltzer, Brian Norvell, Mindy Nye, Scott Parker, Christos Petropoulos, John Pruitt, Steven Ragan, Scott Ryan, Mike Sapeta, Jana Schroth, Suresh Babu Selvaraju, Goran Stevovic, Amanda Suchanek, Andrea Throop, Lyndon Tilson, Thomas Urban, Joe Voshell, Kimberly Wagner, Jonathan Williams, Mary Williamson, Qian Zeng, Tricia Zwiefelhofer, Clinton R. Paden, Duncan MacCannell |
| EPI_ISL_3326654, EPI_ISL_3326656 | TGen North | TGen North | Jolene Bowers, Heather Centner, Brett Van Tassel, Chris French, Hayley Yaglom, Darrin Lemmer, Dave Engelthaler |
| EPI_ISL_3326657, EPI_ISL_3326660 | Laboratory Corporation of America | Centers for Disease Control and Prevention Division of Viral Diseases, Pathogen Discovery | Dakota Howard, Dhwani Batra, Peter W. Cook, Kara Moser, Adrian Paskey, Jason Caravas, Benjamin Rambo-Martin, Shatavia Morrison, Christopher Gulvick, Scott Sammons, Yvette Unoarumhi, Darlene Wagner, Matthew Schmerer, Minoo Agarwal, Eyad Almasri, Debbie Boles, Ayla Burns, Nuthawin Charoensri, Oren Cohen, Susan Countryman, Mary Ann Cristobal, Bobbi Croy, Suzanne Dale, Hrushikesh Deshmukh, Amanda Douglas, Vincent Drouillon, Marcia Eisenberg, Howard Engler, Rama Ghatti, Prashant Gupta, Susan Hicks, Jake Humphrey, Lax Iyer, Lisa Pfefferle, Manoj Jain, Matthew Robinson, Mohan Kolli, Brian Krueger, Tim Kuphal, Stanley Letovsky, Michael Levandoski, Craig Lukasik, Jonathan Meltzer, Brian Norvell, Mindy Nye, Scott Parker, Christos Petropoulos, John Pruitt, Steven Ragan, Scott Ryan, Mike Sapeta, Jana Schroth, Suresh Babu Selvaraju, Goran Stevovic, Amanda Suchanek, Andrea Throop, Lyndon Tilson, Thomas Urban, Joe Voshell, Kimberly Wagner, Jonathan Williams, Mary Williamson, Qian Zeng, Tricia Zwiefelhofer, Clinton R. Paden, Duncan MacCannell |
| EPI_ISL_3326661 | TGen North | TGen North | Jolene Bowers, Heather Centner, Brett Van Tassel, Chris French, Hayley Yaglom, Darrin Lemmer, Dave Engelthaler |
| EPI_ISL_3326666, EPI_ISL_3326668 | Laboratory Corporation of America | Centers for Disease Control and Prevention Division of Viral Diseases, Pathogen Discovery | Dakota Howard, Dhwani Batra, Peter W. Cook, Kara Moser, Adrian Paskey, Jason Caravas, Benjamin Rambo-Martin, Shatavia Morrison, Christopher Gulvick, Scott Sammons, Yvette Unoarumhi, Darlene Wagner, Matthew Schmerer, Minoo Agarwal, Eyad Almasri, Debbie Boles, Ayla Burns, Nuthawin Charoensri, Oren Cohen, Susan Countryman, Mary Ann Cristobal, Bobbi Croy, Suzanne Dale, Hrushikesh Deshmukh, Amanda Douglas, Vincent Drouillon, Marcia Eisenberg, Howard Engler, Rama Ghatti, Prashant Gupta, Susan Hicks, Jake Humphrey, Lax Iyer, Lisa Pfefferle, Manoj Jain, Matthew Robinson, Mohan Kolli, Brian Krueger, Tim Kuphal, Stanley Letovsky, Michael Levandoski, Craig Lukasik, Jonathan Meltzer, Brian Norvell, Mindy Nye, Scott Parker, Christos Petropoulos, John Pruitt, Steven Ragan, Scott Ryan, Mike Sapeta, Jana Schroth, Suresh Babu Selvaraju, Goran Stevovic, Amanda Suchanek, Andrea Throop, Lyndon Tilson, Thomas Urban, Joe Voshell, Kimberly Wagner, Jonathan Williams, Mary Williamson, Qian Zeng, Tricia Zwiefelhofer, Clinton R. Paden, Duncan MacCannell |
| EPI_ISL_3326669 | TGen North | TGen North | Jolene Bowers, Heather Centner, Brett Van Tassel, Chris French, Hayley Yaglom, Darrin Lemmer, Dave Engelthaler |
| EPI_ISL_3326670, EPI_ISL_3326673 | Laboratory Corporation of America | Centers for Disease Control and Prevention Division of Viral Diseases, Pathogen Discovery | Dakota Howard, Dhwani Batra, Peter W. Cook, Kara Moser, Adrian Paskey, Jason Caravas, Benjamin Rambo-Martin, Shatavia Morrison, Christopher Gulvick, Scott Sammons, Yvette Unoarumhi, Darlene Wagner, Matthew Schmerer, Minoo Agarwal, Eyad Almasri, Debbie Boles, Ayla Burns, Nuthawin Charoensri, Oren Cohen, Susan Countryman, Mary Ann Cristobal, Bobbi Croy, Suzanne Dale, Hrushikesh Deshmukh, Amanda Douglas, Vincent Drouillon, Marcia Eisenberg, Howard Engler, Rama Ghatti, Prashant Gupta, Susan Hicks, Jake Humphrey, Lax Iyer, Lisa Pfefferle, Manoj Jain, Matthew Robinson, Mohan Kolli, Brian Krueger, Tim Kuphal, Stanley Letovsky, Michael Levandoski, Craig Lukasik, Jonathan Meltzer, Brian Norvell, Mindy Nye, Scott Parker, Christos Petropoulos, John Pruitt, Steven Ragan, Scott Ryan, Mike Sapeta, Jana Schroth, Suresh Babu Selvaraju, Goran Stevovic, Amanda Suchanek, Andrea Throop, Lyndon Tilson, Thomas Urban, Joe Voshell, Kimberly Wagner, Jonathan Williams, Mary Williamson, Qian Zeng, Tricia Zwiefelhofer, Clinton R. Paden, Duncan MacCannell |
| EPI_ISL_3326674 | TGen North | TGen North | Jolene Bowers, Heather Centner, Brett Van Tassel, Chris French, Hayley Yaglom, Darrin Lemmer, Dave Engelthaler |
| EPI_ISL_3326675 | Laboratory Corporation of America | Centers for Disease Control and Prevention Division of Viral Diseases, Pathogen Discovery | Dakota Howard, Dhwani Batra, Peter W. Cook, Kara Moser, Adrian Paskey, Jason Caravas, Benjamin Rambo-Martin, Shatavia Morrison, Christopher Gulvick, Scott Sammons, Yvette Unoarumhi, Darlene Wagner, Matthew Schmerer, Minoo Agarwal, Eyad Almasri, Debbie Boles, Ayla Burns, Nuthawin Charoensri, Oren Cohen, Susan Countryman, Mary Ann Cristobal, Bobbi Croy, Suzanne Dale, Hrushikesh Deshmukh, Amanda Douglas, Vincent Drouillon, Marcia Eisenberg, Howard Engler, Rama Ghatti, Prashant Gupta, Susan Hicks, Jake Humphrey, Lax Iyer, Lisa Pfefferle, Manoj Jain, Matthew Robinson, Mohan Kolli, Brian Krueger, Tim Kuphal, Stanley Letovsky, Michael Levandoski, Craig Lukasik, Jonathan Meltzer, Brian Norvell, Mindy Nye, Scott Parker, Christos Petropoulos, John Pruitt, Steven Ragan, Scott Ryan, Mike Sapeta, Jana Schroth, Suresh Babu Selvaraju, Goran Stevovic, Amanda Suchanek, Andrea Throop, Lyndon Tilson, Thomas Urban, Joe Voshell, Kimberly Wagner, Jonathan Williams, Mary Williamson, Qian Zeng, Tricia Zwiefelhofer, Clinton R. Paden, Duncan MacCannell |
| EPI_ISL_3326676 | TGen North | TGen North | Jolene Bowers, Heather Centner, Brett Van Tassel, Chris French, Hayley Yaglom, Darrin Lemmer, Dave Engelthaler |
| EPI_ISL_3326677 | Laboratory Corporation of America | Centers for Disease Control and Prevention Division of Viral Diseases, Pathogen Discovery | Dakota Howard, Dhwani Batra, Peter W. Cook, Kara Moser, Adrian Paskey, Jason Caravas, Benjamin Rambo-Martin, Shatavia Morrison, Christopher Gulvick, Scott Sammons, Yvette Unoarumhi, Darlene Wagner, Matthew Schmerer, Minoo Agarwal, Eyad Almasri, Debbie Boles, Ayla Burns, Nuthawin Charoensri, Oren Cohen, Susan Countryman, Mary Ann Cristobal, Bobbi Croy, Suzanne Dale, Hrushikesh Deshmukh, Amanda Douglas, Vincent Drouillon, Marcia Eisenberg, Howard Engler, Rama Ghatti, Prashant Gupta, Susan Hicks, Jake Humphrey, Lax Iyer, Lisa Pfefferle, Manoj Jain, Matthew Robinson, Mohan Kolli, Brian Krueger, Tim Kuphal, Stanley Letovsky, Michael Levandoski, Craig Lukasik, Jonathan Meltzer, Brian Norvell, Mindy Nye, Scott Parker, Christos Petropoulos, John Pruitt, Steven Ragan, Scott Ryan, Mike Sapeta, Jana Schroth, Suresh Babu Selvaraju, Goran Stevovic, Amanda Suchanek, Andrea Throop, Lyndon Tilson, Thomas Urban, Joe Voshell, Kimberly Wagner, Jonathan Williams, Mary Williamson, Qian Zeng, Tricia Zwiefelhofer, Clinton R. Paden, Duncan MacCannell |
| EPI_ISL_3326678 | TGen North | TGen North | Jolene Bowers, Heather Centner, Brett Van Tassel, Chris French, Hayley Yaglom, Darrin Lemmer, Dave Engelthaler |
| EPI_ISL_3326679 | Laboratory Corporation of America | Centers for Disease Control and Prevention Division of Viral Diseases, Pathogen Discovery | Dakota Howard, Dhwani Batra, Peter W. Cook, Kara Moser, Adrian Paskey, Jason Caravas, Benjamin Rambo-Martin, Shatavia Morrison, Christopher Gulvick, Scott Sammons, Yvette Unoarumhi, Darlene Wagner, Matthew Schmerer, Minoo Agarwal, Eyad Almasri, Debbie Boles, Ayla Burns, Nuthawin Charoensri, Oren Cohen, Susan Countryman, Mary Ann Cristobal, Bobbi Croy, Suzanne Dale, Hrushikesh Deshmukh, Amanda Douglas, Vincent Drouillon, Marcia Eisenberg, Howard Engler, Rama Ghatti, Prashant Gupta, Susan Hicks, Jake Humphrey, Lax Iyer, Lisa Pfefferle, Manoj Jain, Matthew Robinson, Mohan Kolli, Brian Krueger, Tim Kuphal, Stanley Letovsky, Michael Levandoski, Craig Lukasik, Jonathan Meltzer, Brian Norvell, Mindy Nye, Scott Parker, Christos Petropoulos, John Pruitt, Steven Ragan, Scott Ryan, Mike Sapeta, Jana Schroth, Suresh Babu Selvaraju, Goran Stevovic, Amanda Suchanek, Andrea Throop, Lyndon Tilson, Thomas Urban, Joe Voshell, Kimberly Wagner, Jonathan Williams, Mary Williamson, Qian Zeng, Tricia Zwiefelhofer, Clinton R. Paden, Duncan MacCannell |
| EPI_ISL_3326680 | TGen North | TGen North | Jolene Bowers, Heather Centner, Brett Van Tassel, Chris French, Hayley Yaglom, Darrin Lemmer, Dave Engelthaler |
| EPI_ISL_3326681 | Laboratory Corporation of America | Centers for Disease Control and Prevention Division of Viral Diseases, Pathogen Discovery | Dakota Howard, Dhwani Batra, Peter W. Cook, Kara Moser, Adrian Paskey, Jason Caravas, Benjamin Rambo-Martin, Shatavia Morrison, Christopher Gulvick, Scott Sammons, Yvette Unoarumhi, Darlene Wagner, Matthew Schmerer, Minoo Agarwal, Eyad Almasri, Debbie Boles, Ayla Burns, Nuthawin Charoensri, Oren Cohen, Susan Countryman, Mary Ann Cristobal, Bobbi Croy, Suzanne Dale, Hrushikesh Deshmukh, Amanda Douglas, Vincent Drouillon, Marcia Eisenberg, Howard Engler, Rama Ghatti, Prashant Gupta, Susan Hicks, Jake Humphrey, Lax Iyer, Lisa Pfefferle, Manoj Jain, Matthew Robinson, Mohan Kolli, Brian Krueger, Tim Kuphal, Stanley Letovsky, Michael Levandoski, Craig Lukasik, Jonathan Meltzer, Brian Norvell, Mindy Nye, Scott Parker, Christos Petropoulos, John Pruitt, Steven Ragan, Scott Ryan, Mike Sapeta, Jana Schroth, Suresh Babu Selvaraju, Goran Stevovic, Amanda Suchanek, Andrea Throop, Lyndon Tilson, Thomas Urban, Joe Voshell, Kimberly Wagner, Jonathan Williams, Mary Williamson, Qian Zeng, Tricia Zwiefelhofer, Clinton R. Paden, Duncan MacCannell |
| EPI_ISL_3326682 | TGen North | TGen North | Jolene Bowers, Heather Centner, Brett Van Tassel, Chris French, Hayley Yaglom, Darrin Lemmer, Dave Engelthaler |
| EPI_ISL_3326683, EPI_ISL_3326684 | Laboratory Corporation of America | Centers for Disease Control and Prevention Division of Viral Diseases, Pathogen Discovery | Dakota Howard, Dhwani Batra, Peter W. Cook, Kara Moser, Adrian Paskey, Jason Caravas, Benjamin Rambo-Martin, Shatavia Morrison, Christopher Gulvick, Scott Sammons, Yvette Unoarumhi, Darlene Wagner, Matthew Schmerer, Minoo Agarwal, Eyad Almasri, Debbie Boles, Ayla Burns, Nuthawin Charoensri, Oren Cohen, Susan Countryman, Mary Ann Cristobal, Bobbi Croy, Suzanne Dale, Hrushikesh Deshmukh, Amanda Douglas, Vincent Drouillon, Marcia Eisenberg, Howard Engler, Rama Ghatti, Prashant Gupta, Susan Hicks, Jake Humphrey, Lax Iyer, Lisa Pfefferle, Manoj Jain, Matthew Robinson, Mohan Kolli, Brian Krueger, Tim Kuphal, Stanley Letovsky, Michael Levandoski, Craig Lukasik, Jonathan Meltzer, Brian Norvell, Mindy Nye, Scott Parker, Christos Petropoulos, John Pruitt, Steven Ragan, Scott Ryan, Mike Sapeta, Jana Schroth, Suresh Babu Selvaraju, Goran Stevovic, Amanda Suchanek, Andrea Throop, Lyndon Tilson, Thomas Urban, Joe Voshell, Kimberly Wagner, Jonathan Williams, Mary Williamson, Qian Zeng, Tricia Zwiefelhofer, Clinton R. Paden, Duncan MacCannell |
| EPI_ISL_3326685 | TGen North | TGen North | Jolene Bowers, Heather Centner, Brett Van Tassel, Chris French, Hayley Yaglom, Darrin Lemmer, Dave Engelthaler |
| EPI_ISL_3326686 | Laboratory Corporation of America | Centers for Disease Control and Prevention Division of Viral Diseases, Pathogen Discovery | Dakota Howard, Dhwani Batra, Peter W. Cook, Kara Moser, Adrian Paskey, Jason Caravas, Benjamin Rambo-Martin, Shatavia Morrison, Christopher Gulvick, Scott Sammons, Yvette Unoarumhi, Darlene Wagner, Matthew Schmerer, Minoo Agarwal, Eyad Almasri, Debbie Boles, Ayla Burns, Nuthawin Charoensri, Oren Cohen, Susan Countryman, Mary Ann Cristobal, Bobbi Croy, Suzanne Dale, Hrushikesh Deshmukh, Amanda Douglas, Vincent Drouillon, Marcia Eisenberg, Howard Engler, Rama Ghatti, Prashant Gupta, Susan Hicks, Jake Humphrey, Lax Iyer, Lisa Pfefferle, Manoj Jain, Matthew Robinson, Mohan Kolli, Brian Krueger, Tim Kuphal, Stanley Letovsky, Michael Levandoski, Craig Lukasik, Jonathan Meltzer, Brian Norvell, Mindy Nye, Scott Parker, Christos Petropoulos, John Pruitt, Steven Ragan, Scott Ryan, Mike Sapeta, Jana Schroth, Suresh Babu Selvaraju, Goran Stevovic, Amanda Suchanek, Andrea Throop, Lyndon Tilson, Thomas Urban, Joe Voshell, Kimberly Wagner, Jonathan Williams, Mary Williamson, Qian Zeng, Tricia Zwiefelhofer, Clinton R. Paden, Duncan MacCannell |
| EPI_ISL_3326687 | TGen North | TGen North | Jolene Bowers, Heather Centner, Brett Van Tassel, Chris French, Hayley Yaglom, Darrin Lemmer, Dave Engelthaler |
| EPI_ISL_3326688, EPI_ISL_3326690 | Laboratory Corporation of America | Centers for Disease Control and Prevention Division of Viral Diseases, Pathogen Discovery | Dakota Howard, Dhwani Batra, Peter W. Cook, Kara Moser, Adrian Paskey, Jason Caravas, Benjamin Rambo-Martin, Shatavia Morrison, Christopher Gulvick, Scott Sammons, Yvette Unoarumhi, Darlene Wagner, Matthew Schmerer, Minoo Agarwal, Eyad Almasri, Debbie Boles, Ayla Burns, Nuthawin Charoensri, Oren Cohen, Susan Countryman, Mary Ann Cristobal, Bobbi Croy, Suzanne Dale, Hrushikesh Deshmukh, Amanda Douglas, Vincent Drouillon, Marcia Eisenberg, Howard Engler, Rama Ghatti, Prashant Gupta, Susan Hicks, Jake Humphrey, Lax Iyer, Lisa Pfefferle, Manoj Jain, Matthew Robinson, Mohan Kolli, Brian Krueger, Tim Kuphal, Stanley Letovsky, Michael Levandoski, Craig Lukasik, Jonathan Meltzer, Brian Norvell, Mindy Nye, Scott Parker, Christos Petropoulos, John Pruitt, Steven Ragan, Scott Ryan, Mike Sapeta, Jana Schroth, Suresh Babu Selvaraju, Goran Stevovic, Amanda Suchanek, Andrea Throop, Lyndon Tilson, Thomas Urban, Joe Voshell, Kimberly Wagner, Jonathan Williams, Mary Williamson, Qian Zeng, Tricia Zwiefelhofer, Clinton R. Paden, Duncan MacCannell |
| EPI_ISL_3326691 | TGen North | TGen North | Jolene Bowers, Heather Centner, Brett Van Tassel, Chris French, Hayley Yaglom, Darrin Lemmer, Dave Engelthaler |
| EPI_ISL_3326692, EPI_ISL_3326694, EPI_ISL_3326695 | Laboratory Corporation of America | Centers for Disease Control and Prevention Division of Viral Diseases, Pathogen Discovery | Dakota Howard, Dhwani Batra, Peter W. Cook, Kara Moser, Adrian Paskey, Jason Caravas, Benjamin Rambo-Martin, Shatavia Morrison, Christopher Gulvick, Scott Sammons, Yvette Unoarumhi, Darlene Wagner, Matthew Schmerer, Minoo Agarwal, Eyad Almasri, Debbie Boles, Ayla Burns, Nuthawin Charoensri, Oren Cohen, Susan Countryman, Mary Ann Cristobal, Bobbi Croy, Suzanne Dale, Hrushikesh Deshmukh, Amanda Douglas, Vincent Drouillon, Marcia Eisenberg, Howard Engler, Rama Ghatti, Prashant Gupta, Susan Hicks, Jake Humphrey, Lax Iyer, Lisa Pfefferle, Manoj Jain, Matthew Robinson, Mohan Kolli, Brian Krueger, Tim Kuphal, Stanley Letovsky, Michael Levandoski, Craig Lukasik, Jonathan Meltzer, Brian Norvell, Mindy Nye, Scott Parker, Christos Petropoulos, John Pruitt, Steven Ragan, Scott Ryan, Mike Sapeta, Jana Schroth, Suresh Babu Selvaraju, Goran Stevovic, Amanda Suchanek, Andrea Throop, Lyndon Tilson, Thomas Urban, Joe Voshell, Kimberly Wagner, Jonathan Williams, Mary Williamson, Qian Zeng, Tricia Zwiefelhofer, Clinton R. Paden, Duncan MacCannell |
| EPI_ISL_3326696, EPI_ISL_3326698 | TGen North | TGen North | Jolene Bowers, Heather Centner, Brett Van Tassel, Chris French, Hayley Yaglom, Darrin Lemmer, Dave Engelthaler |
| EPI_ISL_3326699, EPI_ISL_3326701 | Laboratory Corporation of America | Centers for Disease Control and Prevention Division of Viral Diseases, Pathogen Discovery | Dakota Howard, Dhwani Batra, Peter W. Cook, Kara Moser, Adrian Paskey, Jason Caravas, Benjamin Rambo-Martin, Shatavia Morrison, Christopher Gulvick, Scott Sammons, Yvette Unoarumhi, Darlene Wagner, Matthew Schmerer, Minoo Agarwal, Eyad Almasri, Debbie Boles, Ayla Burns, Nuthawin Charoensri, Oren Cohen, Susan Countryman, Mary Ann Cristobal, Bobbi Croy, Suzanne Dale, Hrushikesh Deshmukh, Amanda Douglas, Vincent Drouillon, Marcia Eisenberg, Howard Engler, Rama Ghatti, Prashant Gupta, Susan Hicks, Jake Humphrey, Lax Iyer, Lisa Pfefferle, Manoj Jain, Matthew Robinson, Mohan Kolli, Brian Krueger, Tim Kuphal, Stanley Letovsky, Michael Levandoski, Craig Lukasik, Jonathan Meltzer, Brian Norvell, Mindy Nye, Scott Parker, Christos Petropoulos, John Pruitt, Steven Ragan, Scott Ryan, Mike Sapeta, Jana Schroth, Suresh Babu Selvaraju, Goran Stevovic, Amanda Suchanek, Andrea Throop, Lyndon Tilson, Thomas Urban, Joe Voshell, Kimberly Wagner, Jonathan Williams, Mary Williamson, Qian Zeng, Tricia Zwiefelhofer, Clinton R. Paden, Duncan MacCannell |
| EPI_ISL_3326702 | TGen North | TGen North | Jolene Bowers, Heather Centner, Brett Van Tassel, Chris French, Hayley Yaglom, Darrin Lemmer, Dave Engelthaler |
| EPI_ISL_3326703, EPI_ISL_3326705, EPI_ISL_3326707, EPI_ISL_3326708 | Laboratory Corporation of America | Centers for Disease Control and Prevention Division of Viral Diseases, Pathogen Discovery | Dakota Howard, Dhwani Batra, Peter W. Cook, Kara Moser, Adrian Paskey, Jason Caravas, Benjamin Rambo-Martin, Shatavia Morrison, Christopher Gulvick, Scott Sammons, Yvette Unoarumhi, Darlene Wagner, Matthew Schmerer, Minoo Agarwal, Eyad Almasri, Debbie Boles, Ayla Burns, Nuthawin Charoensri, Oren Cohen, Susan Countryman, Mary Ann Cristobal, Bobbi Croy, Suzanne Dale, Hrushikesh Deshmukh, Amanda Douglas, Vincent Drouillon, Marcia Eisenberg, Howard Engler, Rama Ghatti, Prashant Gupta, Susan Hicks, Jake Humphrey, Lax Iyer, Lisa Pfefferle, Manoj Jain, Matthew Robinson, Mohan Kolli, Brian Krueger, Tim Kuphal, Stanley Letovsky, Michael Levandoski, Craig Lukasik, Jonathan Meltzer, Brian Norvell, Mindy Nye, Scott Parker, Christos Petropoulos, John Pruitt, Steven Ragan, Scott Ryan, Mike Sapeta, Jana Schroth, Suresh Babu Selvaraju, Goran Stevovic, Amanda Suchanek, Andrea Throop, Lyndon Tilson, Thomas Urban, Joe Voshell, Kimberly Wagner, Jonathan Williams, Mary Williamson, Qian Zeng, Tricia Zwiefelhofer, Clinton R. Paden, Duncan MacCannell |
| EPI_ISL_3326709 | TGen North | TGen North | Jolene Bowers, Heather Centner, Brett Van Tassel, Chris French, Hayley Yaglom, Darrin Lemmer, Dave Engelthaler |
| EPI_ISL_3326710, EPI_ISL_3326712 | Laboratory Corporation of America | Centers for Disease Control and Prevention Division of Viral Diseases, Pathogen Discovery | Dakota Howard, Dhwani Batra, Peter W. Cook, Kara Moser, Adrian Paskey, Jason Caravas, Benjamin Rambo-Martin, Shatavia Morrison, Christopher Gulvick, Scott Sammons, Yvette Unoarumhi, Darlene Wagner, Matthew Schmerer, Minoo Agarwal, Eyad Almasri, Debbie Boles, Ayla Burns, Nuthawin Charoensri, Oren Cohen, Susan Countryman, Mary Ann Cristobal, Bobbi Croy, Suzanne Dale, Hrushikesh Deshmukh, Amanda Douglas, Vincent Drouillon, Marcia Eisenberg, Howard |

| | | | |
|---|---|---|---|
| | | | Engler, Rama Ghatti, Prashant Gupta, Susan Hicks, Jake Humphrey, Lax Iyer, Lisa Pfefferle, Manoj Jain, Matthew Robinson, Mohan Kolli, Brian Krueger, Tim Kuphal, Stanley Letovsky, Michael Levandoski, Craig Lukasik, Jonathan Meltzer, Brian Norvell, Mindy Nye, Scott Parker, Christos Petropoulos, John Pruitt, Steven Ragan, Scott Ryan, Mike Sapeta, Jana Schroth, Suresh Babu Selvaraju, Goran Stevovic, Amanda Suchanek, Andrea Throop, Lyndon Tilson, Thomas Urban, Joe Voshell, Kimberly Wagner, Jonathan Williams, Mary Williamson, Qian Zeng, Tricia Zwiefelhofer, Clinton R. Paden, Duncan MacCannell |
| EPI_ISL_3326714 | TGen North | TGen North | Jolene Bowers, Heather Centner, Brett Van Tassel, Chris French, Hayley Yaglom, Darrin Lemmer, Dave Engelthaler |
| EPI_ISL_3326715 | Laboratory Corporation of America | Centers for Disease Control and Prevention Division of Viral Diseases, Pathogen Discovery | Dakota Howard, Dhwani Batra, Peter W. Cook, Kara Moser, Adrian Paskey, Jason Caravas, Benjamin Rambo-Martin, Shatavia Morrison, Christopher Gulvick, Scott Sammons, Yvette Unoarumhi, Darlene Wagner, Matthew Schmerer, Minoo Agarwal, Eyad Almasri, Debbie Boles, Ayla Burns, Nuthawin Charoensri, Oren Cohen, Susan Countryman, Mary Ann Cristobal, Bobbi Croy, Suzanne Dale, Hrushikesh Deshmukh, Amanda Douglas, Vincent Drouillon, Marcia Eisenberg, Howard Engler, Rama Ghatti, Prashant Gupta, Susan Hicks, Jake Humphrey, Lax Iyer, Lisa Pfefferle, Manoj Jain, Matthew Robinson, Mohan Kolli, Brian Krueger, Tim Kuphal, Stanley Letovsky, Michael Levandoski, Craig Lukasik, Jonathan Meltzer, Brian Norvell, Mindy Nye, Scott Parker, Christos Petropoulos, John Pruitt, Steven Ragan, Scott Ryan, Mike Sapeta, Jana Schroth, Suresh Babu Selvaraju, Goran Stevovic, Amanda Suchanek, Andrea Throop, Lyndon Tilson, Thomas Urban, Joe Voshell, Kimberly Wagner, Jonathan Williams, Mary Williamson, Qian Zeng, Tricia Zwiefelhofer, Clinton R. Paden, Duncan MacCannell |
| EPI_ISL_3326716 | TGen North | TGen North | Jolene Bowers, Heather Centner, Brett Van Tassel, Chris French, Hayley Yaglom, Darrin Lemmer, Dave Engelthaler |
| EPI_ISL_3326717 | Laboratory Corporation of America | Centers for Disease Control and Prevention Division of Viral Diseases, Pathogen Discovery | Dakota Howard, Dhwani Batra, Peter W. Cook, Kara Moser, Adrian Paskey, Jason Caravas, Benjamin Rambo-Martin, Shatavia Morrison, Christopher Gulvick, Scott Sammons, Yvette Unoarumhi, Darlene Wagner, Matthew Schmerer, Minoo Agarwal, Eyad Almasri, Debbie Boles, Ayla Burns, Nuthawin Charoensri, Oren Cohen, Susan Countryman, Mary Ann Cristobal, Bobbi Croy, Suzanne Dale, Hrushikesh Deshmukh, Amanda Douglas, Vincent Drouillon, Marcia Eisenberg, Howard Engler, Rama Ghatti, Prashant Gupta, Susan Hicks, Jake Humphrey, Lax Iyer, Lisa Pfefferle, Manoj Jain, Matthew Robinson, Mohan Kolli, Brian Krueger, Tim Kuphal, Stanley Letovsky, Michael Levandoski, Craig Lukasik, Jonathan Meltzer, Brian Norvell, Mindy Nye, Scott Parker, Christos Petropoulos, John Pruitt, Steven Ragan, Scott Ryan, Mike Sapeta, Jana Schroth, Suresh Babu Selvaraju, Goran Stevovic, Amanda Suchanek, Andrea Throop, Lyndon Tilson, Thomas Urban, Joe Voshell, Kimberly Wagner, Jonathan Williams, Mary Williamson, Qian Zeng, Tricia Zwiefelhofer, Clinton R. Paden, Duncan MacCannell |
| EPI_ISL_3326718 | TGen North | TGen North | Jolene Bowers, Heather Centner, Brett Van Tassel, Chris French, Hayley Yaglom, Darrin Lemmer, Dave Engelthaler |
| EPI_ISL_3326719 | Laboratory Corporation of America | Centers for Disease Control and Prevention Division of Viral Diseases, Pathogen Discovery | Dakota Howard, Dhwani Batra, Peter W. Cook, Kara Moser, Adrian Paskey, Jason Caravas, Benjamin Rambo-Martin, Shatavia Morrison, Christopher Gulvick, Scott Sammons, Yvette Unoarumhi, Darlene Wagner, Matthew Schmerer, Minoo Agarwal, Eyad Almasri, Debbie Boles, Ayla Burns, Nuthawin Charoensri, Oren Cohen, Susan Countryman, Mary Ann Cristobal, Bobbi Croy, Suzanne Dale, Hrushikesh Deshmukh, Amanda Douglas, Vincent Drouillon, Marcia Eisenberg, Howard Engler, Rama Ghatti, Prashant Gupta, Susan Hicks, Jake Humphrey, Lax Iyer, Lisa Pfefferle, Manoj Jain, Matthew Robinson, Mohan Kolli, Brian Krueger, Tim Kuphal, Stanley Letovsky, Michael Levandoski, Craig Lukasik, Jonathan Meltzer, Brian Norvell, Mindy Nye, Scott Parker, Christos Petropoulos, John Pruitt, Steven Ragan, Scott Ryan, Mike Sapeta, Jana Schroth, Suresh Babu Selvaraju, Goran Stevovic, Amanda Suchanek, Andrea Throop, Lyndon Tilson, Thomas Urban, Joe Voshell, Kimberly Wagner, Jonathan Williams, Mary Williamson, Qian Zeng, Tricia Zwiefelhofer, Clinton R. Paden, Duncan MacCannell |
| EPI_ISL_3326720 | TGen North | TGen North | Jolene Bowers, Heather Centner, Brett Van Tassel, Chris French, Hayley Yaglom, Darrin Lemmer, Dave Engelthaler |
| EPI_ISL_3326721 | Laboratory Corporation of America | Centers for Disease Control and Prevention Division of Viral Diseases, Pathogen Discovery | Dakota Howard, Dhwani Batra, Peter W. Cook, Kara Moser, Adrian Paskey, Jason Caravas, Benjamin Rambo-Martin, Shatavia Morrison, Christopher Gulvick, Scott Sammons, Yvette Unoarumhi, Darlene Wagner, Matthew Schmerer, Minoo Agarwal, Eyad Almasri, Debbie Boles, Ayla Burns, Nuthawin Charoensri, Oren Cohen, Susan Countryman, Mary Ann Cristobal, Bobbi Croy, Suzanne Dale, Hrushikesh Deshmukh, Amanda Douglas, Vincent Drouillon, Marcia Eisenberg, Howard Engler, Rama Ghatti, Prashant Gupta, Susan Hicks, Jake Humphrey, Lax Iyer, Lisa Pfefferle, Manoj Jain, Matthew Robinson, Mohan Kolli, Brian Krueger, Tim Kuphal, Stanley Letovsky, Michael Levandoski, Craig Lukasik, Jonathan Meltzer, Brian Norvell, Mindy Nye, Scott Parker, Christos Petropoulos, John Pruitt, Steven Ragan, Scott Ryan, Mike Sapeta, Jana Schroth, Suresh Babu Selvaraju, Goran Stevovic, Amanda Suchanek, Andrea Throop, Lyndon Tilson, Thomas Urban, Joe Voshell, Kimberly Wagner, Jonathan Williams, Mary Williamson, Qian Zeng, Tricia Zwiefelhofer, Clinton R. Paden, Duncan MacCannell |
| EPI_ISL_3326722 | TGen North | TGen North | Jolene Bowers, Heather Centner, Brett Van Tassel, Chris French, Hayley Yaglom, Darrin Lemmer, Dave Engelthaler |
| EPI_ISL_3326723 | Laboratory Corporation of America | Centers for Disease Control and Prevention Division of Viral Diseases, Pathogen Discovery | Dakota Howard, Dhwani Batra, Peter W. Cook, Kara Moser, Adrian Paskey, Jason Caravas, Benjamin Rambo-Martin, Shatavia Morrison, Christopher Gulvick, Scott Sammons, Yvette Unoarumhi, Darlene Wagner, Matthew Schmerer, Minoo Agarwal, Eyad Almasri, Debbie Boles, Ayla Burns, Nuthawin Charoensri, Oren Cohen, Susan Countryman, Mary Ann Cristobal, Bobbi Croy, Suzanne Dale, Hrushikesh Deshmukh, Amanda Douglas, Vincent Drouillon, Marcia Eisenberg, Howard Engler, Rama Ghatti, Prashant Gupta, Susan Hicks, Jake Humphrey, Lax Iyer, Lisa Pfefferle, Manoj Jain, Matthew Robinson, Mohan Kolli, Brian Krueger, Tim Kuphal, Stanley Letovsky, Michael Levandoski, Craig Lukasik, Jonathan Meltzer, Brian Norvell, Mindy Nye, Scott Parker, Christos Petropoulos, John Pruitt, Steven Ragan, Scott Ryan, Mike Sapeta, Jana Schroth, Suresh Babu Selvaraju, Goran Stevovic, Amanda Suchanek, Andrea Throop, Lyndon Tilson, Thomas Urban, Joe Voshell, Kimberly Wagner, Jonathan Williams, Mary Williamson, Qian Zeng, Tricia Zwiefelhofer, Clinton R. Paden, Duncan MacCannell |
| EPI_ISL_3326724 | TGen North | TGen North | Jolene Bowers, Heather Centner, Brett Van Tassel, Chris French, Hayley Yaglom, Darrin Lemmer, Dave Engelthaler |
| EPI_ISL_3326725 | Laboratory Corporation of America | Centers for Disease Control and Prevention Division of Viral Diseases, Pathogen Discovery | Dakota Howard, Dhwani Batra, Peter W. Cook, Kara Moser, Adrian Paskey, Jason Caravas, Benjamin Rambo-Martin, Shatavia Morrison, Christopher Gulvick, Scott Sammons, Yvette Unoarumhi, Darlene Wagner, Matthew Schmerer, Minoo Agarwal, Eyad Almasri, Debbie Boles, Ayla Burns, Nuthawin Charoensri, Oren Cohen, Susan Countryman, Mary Ann Cristobal, Bobbi Croy, Suzanne Dale, Hrushikesh Deshmukh, Amanda Douglas, Vincent Drouillon, Marcia Eisenberg, Howard Engler, Rama Ghatti, Prashant Gupta, Susan Hicks, Jake Humphrey, Lax Iyer, Lisa Pfefferle, Manoj Jain, Matthew Robinson, Mohan Kolli, Brian Krueger, Tim Kuphal, Stanley Letovsky, Michael Levandoski, Craig Lukasik, Jonathan Meltzer, Brian Norvell, Mindy Nye, Scott Parker, Christos Petropoulos, John Pruitt, Steven Ragan, Scott Ryan, Mike Sapeta, Jana Schroth, Suresh Babu Selvaraju, Goran Stevovic, Amanda Suchanek, Andrea Throop, Lyndon Tilson, Thomas Urban, Joe Voshell, Kimberly Wagner, Jonathan Williams, Mary Williamson, Qian Zeng, Tricia Zwiefelhofer, Clinton R. Paden, Duncan MacCannell |
| EPI_ISL_3326726 | TGen North | TGen North | Jolene Bowers, Heather Centner, Brett Van Tassel, Chris French, Hayley Yaglom, Darrin Lemmer, Dave Engelthaler |
| EPI_ISL_3326727, EPI_ISL_3326728 | Laboratory Corporation of America | Centers for Disease Control and Prevention Division of Viral Diseases, Pathogen Discovery | Dakota Howard, Dhwani Batra, Peter W. Cook, Kara Moser, Adrian Paskey, Jason Caravas, Benjamin Rambo-Martin, Shatavia Morrison, Christopher Gulvick, Scott Sammons, Yvette Unoarumhi, Darlene Wagner, Matthew Schmerer, Minoo Agarwal, Eyad Almasri, Debbie Boles, Ayla Burns, Nuthawin Charoensri, Oren Cohen, Susan Countryman, Mary Ann Cristobal, Bobbi Croy, Suzanne Dale, Hrushikesh Deshmukh, Amanda Douglas, Vincent Drouillon, Marcia Eisenberg, Howard Engler, Rama Ghatti, Prashant Gupta, Susan Hicks, Jake Humphrey, Lax Iyer, Lisa Pfefferle, Manoj Jain, Matthew Robinson, Mohan Kolli, Brian Krueger, Tim Kuphal, Stanley Letovsky, Michael Levandoski, Craig Lukasik, Jonathan Meltzer, Brian Norvell, Mindy Nye, Scott Parker, Christos Petropoulos, John Pruitt, Steven Ragan, Scott Ryan, Mike Sapeta, Jana Schroth, Suresh Babu Selvaraju, Goran Stevovic, Amanda Suchanek, Andrea Throop, Lyndon Tilson, Thomas Urban, Joe Voshell, Kimberly Wagner, Jonathan Williams, Mary Williamson, Qian Zeng, Tricia Zwiefelhofer, Clinton R. Paden, Duncan MacCannell |
| EPI_ISL_3326729 | TGen North | TGen North | Jolene Bowers, Heather Centner, Brett Van Tassel, Chris French, Hayley Yaglom, Darrin Lemmer, Dave Engelthaler |
| EPI_ISL_3326730 | Laboratory Corporation of America | Centers for Disease Control and Prevention Division of Viral Diseases, Pathogen Discovery | Dakota Howard, Dhwani Batra, Peter W. Cook, Kara Moser, Adrian Paskey, Jason Caravas, Benjamin Rambo-Martin, Shatavia Morrison, Christopher Gulvick, Scott Sammons, Yvette Unoarumhi, Darlene Wagner, Matthew Schmerer, Minoo Agarwal, Eyad Almasri, Debbie Boles, Ayla Burns, Nuthawin Charoensri, Oren Cohen, Susan Countryman, Mary Ann Cristobal, Bobbi Croy, Suzanne Dale, Hrushikesh Deshmukh, Amanda Douglas, Vincent Drouillon, Marcia Eisenberg, Howard Engler, Rama Ghatti, Prashant Gupta, Susan Hicks, Jake Humphrey, Lax Iyer, Lisa Pfefferle, Manoj Jain, Matthew Robinson, Mohan Kolli, Brian Krueger, Tim Kuphal, Stanley Letovsky, Michael Levandoski, Craig Lukasik, Jonathan Meltzer, Brian Norvell, Mindy Nye, Scott Parker, Christos Petropoulos, John Pruitt, Steven Ragan, Scott Ryan, Mike Sapeta, Jana Schroth, Suresh Babu Selvaraju, Goran Stevovic, Amanda Suchanek, Andrea Throop, Lyndon Tilson, Thomas Urban, Joe Voshell, Kimberly Wagner, Jonathan Williams, Mary Williamson, Qian Zeng, Tricia Zwiefelhofer, Clinton R. Paden, Duncan MacCannell |
| EPI_ISL_3326731 | TGen North | TGen North | Jolene Bowers, Heather Centner, Brett Van Tassel, Chris French, Hayley Yaglom, Darrin Lemmer, Dave Engelthaler |
| EPI_ISL_3326732, EPI_ISL_3326734 | Laboratory Corporation of America | Centers for Disease Control and Prevention Division of Viral Diseases, Pathogen Discovery | Dakota Howard, Dhwani Batra, Peter W. Cook, Kara Moser, Adrian Paskey, Jason Caravas, Benjamin Rambo-Martin, Shatavia Morrison, Christopher Gulvick, Scott Sammons, Yvette Unoarumhi, Darlene Wagner, Matthew Schmerer, Minoo Agarwal, Eyad Almasri, Debbie Boles, Ayla Burns, Nuthawin Charoensri, Oren Cohen, Susan Countryman, Mary Ann Cristobal, Bobbi Croy, Suzanne Dale, Hrushikesh Deshmukh, Amanda Douglas, Vincent Drouillon, Marcia Eisenberg, Howard Engler, Rama Ghatti, Prashant Gupta, Susan Hicks, Jake Humphrey, Lax Iyer, Lisa Pfefferle, Manoj Jain, Matthew Robinson, Mohan Kolli, Brian Krueger, Tim Kuphal, Stanley Letovsky, Michael Levandoski, Craig Lukasik, Jonathan Meltzer, Brian Norvell, Mindy Nye, Scott Parker, Christos Petropoulos, John Pruitt, Steven Ragan, Scott Ryan, Mike Sapeta, Jana Schroth, Suresh Babu Selvaraju, Goran Stevovic, Amanda Suchanek, Andrea Throop, Lyndon Tilson, Thomas Urban, Joe Voshell, Kimberly Wagner, Jonathan Williams, Mary Williamson, Qian Zeng, Tricia Zwiefelhofer, Clinton R. Paden, Duncan MacCannell |
| EPI_ISL_3326735 | TGen North | TGen North | Jolene Bowers, Heather Centner, Brett Van Tassel, Chris French, Hayley Yaglom, Darrin Lemmer, Dave Engelthaler |
| EPI_ISL_3326738, EPI_ISL_3326739 | Laboratory Corporation of America | Centers for Disease Control and Prevention Division of Viral Diseases, Pathogen Discovery | Dakota Howard, Dhwani Batra, Peter W. Cook, Kara Moser, Adrian Paskey, Jason Caravas, Benjamin Rambo-Martin, Shatavia Morrison, Christopher Gulvick, Scott Sammons, Yvette Unoarumhi, Darlene Wagner, Matthew Schmerer, Minoo Agarwal, Eyad Almasri, Debbie Boles, Ayla Burns, Nuthawin Charoensri, Oren Cohen, Susan Countryman, Mary Ann Cristobal, Bobbi Croy, Suzanne Dale, Hrushikesh Deshmukh, Amanda Douglas, Vincent Drouillon, Marcia Eisenberg, Howard Engler, Rama Ghatti, Prashant Gupta, Susan Hicks, Jake Humphrey, Lax Iyer, Lisa Pfefferle, Manoj Jain, Matthew Robinson, Mohan Kolli, Brian Krueger, Tim Kuphal, Stanley Letovsky, Michael Levandoski, Craig Lukasik, Jonathan Meltzer, Brian Norvell, Mindy Nye, Scott Parker, Christos Petropoulos, John Pruitt, Steven Ragan, Scott Ryan, Mike Sapeta, Jana Schroth, Suresh Babu Selvaraju, Goran Stevovic, Amanda Suchanek, Andrea Throop, Lyndon Tilson, Thomas Urban, Joe Voshell, Kimberly Wagner, Jonathan Williams, Mary Williamson, Qian Zeng, Tricia Zwiefelhofer, Clinton R. Paden, Duncan MacCannell |
| EPI_ISL_3326740 | TGen North | TGen North | Jolene Bowers, Heather Centner, Brett Van Tassel, Chris French, Hayley Yaglom, Darrin Lemmer, Dave Engelthaler |
| EPI_ISL_3326741, EPI_ISL_3326743 | Laboratory Corporation of America | Centers for Disease Control and Prevention Division of Viral Diseases, Pathogen Discovery | Dakota Howard, Dhwani Batra, Peter W. Cook, Kara Moser, Adrian Paskey, Jason Caravas, Benjamin Rambo-Martin, Shatavia Morrison, Christopher Gulvick, Scott Sammons, Yvette Unoarumhi, Darlene Wagner, Matthew Schmerer, Minoo Agarwal, Eyad Almasri, Debbie Boles, Ayla Burns, Nuthawin Charoensri, Oren Cohen, Susan Countryman, Mary Ann Cristobal, Bobbi Croy, Suzanne Dale, Hrushikesh Deshmukh, Amanda Douglas, Vincent Drouillon, Marcia Eisenberg, Howard Engler, Rama Ghatti, Prashant Gupta, Susan Hicks, Jake Humphrey, Lax Iyer, Lisa Pfefferle, Manoj Jain, Matthew Robinson, Mohan Kolli, Brian Krueger, Tim Kuphal, Stanley Letovsky, Michael Levandoski, Craig Lukasik, Jonathan Meltzer, Brian Norvell, Mindy Nye, Scott Parker, Christos Petropoulos, John Pruitt, Steven Ragan, Scott Ryan, Mike Sapeta, Jana Schroth, Suresh Babu Selvaraju, Goran Stevovic, Amanda Suchanek, Andrea Throop, Lyndon Tilson, Thomas Urban, Joe Voshell, Kimberly Wagner, Jonathan Williams, Mary Williamson, Qian Zeng, Tricia Zwiefelhofer, Clinton R. Paden, Duncan MacCannell |
| EPI_ISL_3326744 | TGen North | TGen North | Jolene Bowers, Heather Centner, Brett Van Tassel, Chris French, Hayley Yaglom, Darrin Lemmer, Dave Engelthaler |
| EPI_ISL_3326745 | Laboratory Corporation of America | Centers for Disease Control and Prevention Division of Viral Diseases, Pathogen Discovery | Dakota Howard, Dhwani Batra, Peter W. Cook, Kara Moser, Adrian Paskey, Jason Caravas, Benjamin Rambo-Martin, Shatavia Morrison, Christopher Gulvick, Scott Sammons, Yvette Unoarumhi, Darlene Wagner, Matthew Schmerer, Minoo Agarwal, Eyad Almasri, Debbie Boles, Ayla Burns, Nuthawin Charoensri, Oren Cohen, Susan Countryman, Mary Ann Cristobal, Bobbi Croy, Suzanne Dale, Hrushikesh Deshmukh, Amanda Douglas, Vincent Drouillon, Marcia Eisenberg, Howard Engler, Rama Ghatti, Prashant Gupta, Susan Hicks, Jake Humphrey, Lax Iyer, Lisa Pfefferle, Manoj Jain, Matthew Robinson, Mohan Kolli, Brian Krueger, Tim Kuphal, Stanley Letovsky, Michael Levandoski, Craig Lukasik, Jonathan Meltzer, Brian Norvell, Mindy Nye, Scott Parker, Christos Petropoulos, John Pruitt, Steven Ragan, Scott Ryan, Mike Sapeta, Jana Schroth, Suresh Babu Selvaraju, Goran Stevovic, Amanda Suchanek, Andrea Throop, Lyndon Tilson, Thomas Urban, Joe Voshell, Kimberly Wagner, Jonathan Williams, Mary Williamson, Qian Zeng, Tricia Zwiefelhofer, Clinton R. Paden, Duncan MacCannell |
| EPI_ISL_3326746 | TGen North | TGen North | Jolene Bowers, Heather Centner, Brett Van Tassel, Chris French, Hayley Yaglom, Darrin Lemmer, Dave Engelthaler |
| EPI_ISL_3326747, EPI_ISL_3326749, EPI_ISL_3326750 | Laboratory Corporation of America | Centers for Disease Control and Prevention Division of Viral Diseases, Pathogen Discovery | Dakota Howard, Dhwani Batra, Peter W. Cook, Kara Moser, Adrian Paskey, Jason Caravas, Benjamin Rambo-Martin, Shatavia Morrison, Christopher Gulvick, Scott Sammons, Yvette Unoarumhi, Darlene Wagner, Matthew Schmerer, Minoo Agarwal, Eyad Almasri, Debbie Boles, Ayla Burns, Nuthawin Charoensri, Oren Cohen, Susan Countryman, Mary Ann Cristobal, Bobbi Croy, Suzanne Dale, Hrushikesh Deshmukh, Amanda Douglas, Vincent Drouillon, Marcia Eisenberg, Howard Engler, Rama Ghatti, Prashant Gupta, Susan Hicks, Jake Humphrey, Lax Iyer, Lisa Pfefferle, Manoj Jain, Matthew Robinson, Mohan Kolli, Brian Krueger, Tim Kuphal, Stanley Letovsky, Michael Levandoski, Craig Lukasik, Jonathan Meltzer, Brian Norvell, Mindy Nye, Scott Parker, Christos Petropoulos, John Pruitt, Steven Ragan, Scott Ryan, Mike Sapeta, Jana Schroth, Suresh Babu Selvaraju, Goran Stevovic, Amanda Suchanek, Andrea Throop, Lyndon Tilson, Thomas Urban, Joe Voshell, Kimberly Wagner, Jonathan Williams, Mary Williamson, Qian Zeng, Tricia Zwiefelhofer, Clinton R. Paden, Duncan MacCannell |
| EPI_ISL_3326751 | TGen North | TGen North | Jolene Bowers, Heather Centner, Brett Van Tassel, Chris French, Hayley Yaglom, Darrin Lemmer, Dave Engelthaler |
| EPI_ISL_3326752 | Laboratory Corporation of America | Centers for Disease Control and Prevention Division of Viral Diseases, Pathogen Discovery | Dakota Howard, Dhwani Batra, Peter W. Cook, Kara Moser, Adrian Paskey, Jason Caravas, Benjamin Rambo-Martin, Shatavia Morrison, Christopher Gulvick, Scott Sammons, Yvette Unoarumhi, Darlene Wagner, Matthew Schmerer, Minoo Agarwal, Eyad Almasri, Debbie Boles, Ayla Burns, Nuthawin Charoensri, Oren Cohen, Susan Countryman, Mary Ann Cristobal, Bobbi Croy, Suzanne Dale, Hrushikesh Deshmukh, Amanda Douglas, Vincent Drouillon, Marcia Eisenberg, Howard Engler, Rama Ghatti, Prashant Gupta, Susan Hicks, Jake Humphrey, Lax Iyer, Lisa Pfefferle, Manoj Jain, Matthew Robinson, Mohan Kolli, Brian Krueger, Tim Kuphal, Stanley Letovsky, Michael Levandoski, Craig Lukasik, Jonathan Meltzer, Brian Norvell, Mindy Nye, Scott Parker, Christos Petropoulos, John Pruitt, Steven Ragan, Scott Ryan, Mike Sapeta, Jana Schroth, Suresh Babu Selvaraju, Goran Stevovic, Amanda Suchanek, Andrea Throop, Lyndon Tilson, Thomas Urban, Joe Voshell, Kimberly Wagner, Jonathan Williams, Mary Williamson, Qian Zeng, Tricia Zwiefelhofer, Clinton R. Paden, Duncan MacCannell |
| EPI_ISL_3326753 | TGen North | TGen North | Jolene Bowers, Heather Centner, Brett Van Tassel, Chris French, Hayley Yaglom, Darrin Lemmer, Dave Engelthaler |
| EPI_ISL_3326754, EPI_ISL_3326756 | Laboratory Corporation of America | Centers for Disease Control and Prevention Division of Viral Diseases, Pathogen Discovery | Dakota Howard, Dhwani Batra, Peter W. Cook, Kara Moser, Adrian Paskey, Jason Caravas, Benjamin Rambo-Martin, Shatavia Morrison, Christopher Gulvick, Scott Sammons, Yvette Unoarumhi, Darlene Wagner, Matthew Schmerer, Minoo Agarwal, Eyad Almasri, Debbie Boles, Ayla Burns, Nuthawin Charoensri, Oren Cohen, Susan Countryman, Mary Ann Cristobal, Bobbi Croy, Suzanne Dale, Hrushikesh Deshmukh, Amanda Douglas, Vincent Drouillon, Marcia Eisenberg, Howard Engler, Rama Ghatti, Prashant Gupta, Susan Hicks, Jake Humphrey, Lax Iyer, Lisa Pfefferle, Manoj Jain, Matthew Robinson, Mohan Kolli, Brian Krueger, Tim Kuphal, Stanley Letovsky, Michael Levandoski, Craig Lukasik, Jonathan Meltzer, Brian Norvell, Mindy Nye, Scott Parker, Christos Petropoulos, John Pruitt, Steven Ragan, Scott Ryan, Mike Sapeta, Jana Schroth, Suresh Babu Selvaraju, Goran Stevovic, Amanda Suchanek, Andrea Throop, Lyndon Tilson, Thomas Urban, Joe Voshell, Kimberly Wagner, Jonathan Williams, Mary Williamson, Qian Zeng, Tricia Zwiefelhofer, Clinton R. Paden, Duncan MacCannell |
| EPI_ISL_3326757 | TGen North | TGen North | Jolene Bowers, Heather Centner, Brett Van Tassel, Chris French, Hayley Yaglom, Darrin Lemmer, Dave Engelthaler |
| EPI_ISL_3326758, EPI_ISL_3326760 | Laboratory Corporation of America | Centers for Disease Control and Prevention Division of Viral Diseases, Pathogen Discovery | Dakota Howard, Dhwani Batra, Peter W. Cook, Kara Moser, Adrian Paskey, Jason Caravas, Benjamin Rambo-Martin, Shatavia Morrison, Christopher Gulvick, Scott Sammons, Yvette Unoarumhi, Darlene Wagner, Matthew Schmerer, Minoo Agarwal, Eyad Almasri, Debbie Boles, Ayla Burns, Nuthawin Charoensri, Oren Cohen, Susan Countryman, Mary Ann Cristobal, Bobbi Croy, Suzanne Dale, Hrushikesh Deshmukh, Amanda Douglas, Vincent Drouillon, Marcia Eisenberg, Howard |

| | | | |
|---|---|---|---|
| | | | Engler, Rama Ghatti, Prashant Gupta, Susan Hicks, Jake Humphrey, Lax Iyer, Lisa Pfefferle, Manoj Jain, Matthew Robinson, Mohan Kolli, Brian Krueger, Tim Kuphal, Stanley Letovsky, Michael Levandoski, Craig Lukasik, Jonathan Meltzer, Brian Norvell, Mindy Nye, Scott Parker, Christos Petropoulos, John Pruitt, Steven Ragan, Scott Ryan, Mike Sapeta, Jana Schroth, Suresh Babu Selvaraju, Goran Stevovic, Amanda Suchanek, Andrea Throop, Lyndon Tilson, Thomas Urban, Joe Voshell, Kimberly Wagner, Jonathan Williams, Mary Williamson, Qian Zeng, Tricia Zwiefelhofer, Clinton R. Paden, Duncan MacCannell |
| EPI_ISL_3326761, EPI_ISL_3326763 | TGen North | TGen North | Jolene Bowers, Heather Centner, Brett Van Tassel, Chris French, Hayley Yaglom, Darrin Lemmer, Dave Engelthaler |
| EPI_ISL_3326764, EPI_ISL_3326765, EPI_ISL_3326767 | Laboratory Corporation of America | Centers for Disease Control and Prevention Division of Viral Diseases, Pathogen Discovery | Dakota Howard, Dhwani Batra, Peter W. Cook, Kara Moser, Adrian Paskey, Jason Caravas, Benjamin Rambo-Martin, Shatavia Morrison, Christopher Gulvick, Scott Sammons, Yvette Unoarumhi, Darlene Wagner, Matthew Schmerer, Minoo Agarwal, Eyad Almasri, Debbie Boles, Ayla Burns, Nuthawin Charoensri, Oren Cohen, Susan Countryman, Mary Ann Cristobal, Bobbi Croy, Suzanne Dale, Hrushikesh Deshmukh, Amanda Douglas, Vincent Drouillon, Marcia Eisenberg, Howard Engler, Rama Ghatti, Prashant Gupta, Susan Hicks, Jake Humphrey, Lax Iyer, Lisa Pfefferle, Manoj Jain, Matthew Robinson, Mohan Kolli, Brian Krueger, Tim Kuphal, Stanley Letovsky, Michael Levandoski, Craig Lukasik, Jonathan Meltzer, Brian Norvell, Mindy Nye, Scott Parker, Christos Petropoulos, John Pruitt, Steven Ragan, Scott Ryan, Mike Sapeta, Jana Schroth, Suresh Babu Selvaraju, Goran Stevovic, Amanda Suchanek, Andrea Throop, Lyndon Tilson, Thomas Urban, Joe Voshell, Kimberly Wagner, Jonathan Williams, Mary Williamson, Qian Zeng, Tricia Zwiefelhofer, Clinton R. Paden, Duncan MacCannell |
| EPI_ISL_3326768 | TGen North | TGen North | Jolene Bowers, Heather Centner, Brett Van Tassel, Chris French, Hayley Yaglom, Darrin Lemmer, Dave Engelthaler |
| EPI_ISL_3326769 | Laboratory Corporation of America | Centers for Disease Control and Prevention Division of Viral Diseases, Pathogen Discovery | Dakota Howard, Dhwani Batra, Peter W. Cook, Kara Moser, Adrian Paskey, Jason Caravas, Benjamin Rambo-Martin, Shatavia Morrison, Christopher Gulvick, Scott Sammons, Yvette Unoarumhi, Darlene Wagner, Matthew Schmerer, Minoo Agarwal, Eyad Almasri, Debbie Boles, Ayla Burns, Nuthawin Charoensri, Oren Cohen, Susan Countryman, Mary Ann Cristobal, Bobbi Croy, Suzanne Dale, Hrushikesh Deshmukh, Amanda Douglas, Vincent Drouillon, Marcia Eisenberg, Howard Engler, Rama Ghatti, Prashant Gupta, Susan Hicks, Jake Humphrey, Lax Iyer, Lisa Pfefferle, Manoj Jain, Matthew Robinson, Mohan Kolli, Brian Krueger, Tim Kuphal, Stanley Letovsky, Michael Levandoski, Craig Lukasik, Jonathan Meltzer, Brian Norvell, Mindy Nye, Scott Parker, Christos Petropoulos, John Pruitt, Steven Ragan, Scott Ryan, Mike Sapeta, Jana Schroth, Suresh Babu Selvaraju, Goran Stevovic, Amanda Suchanek, Andrea Throop, Lyndon Tilson, Thomas Urban, Joe Voshell, Kimberly Wagner, Jonathan Williams, Mary Williamson, Qian Zeng, Tricia Zwiefelhofer, Clinton R. Paden, Duncan MacCannell |
| EPI_ISL_3326770 | TGen North | TGen North | Jolene Bowers, Heather Centner, Brett Van Tassel, Chris French, Hayley Yaglom, Darrin Lemmer, Dave Engelthaler |
| EPI_ISL_3326771, EPI_ISL_3326773, EPI_ISL_3326775, EPI_ISL_3326776 | Laboratory Corporation of America | Centers for Disease Control and Prevention Division of Viral Diseases, Pathogen Discovery | Dakota Howard, Dhwani Batra, Peter W. Cook, Kara Moser, Adrian Paskey, Jason Caravas, Benjamin Rambo-Martin, Shatavia Morrison, Christopher Gulvick, Scott Sammons, Yvette Unoarumhi, Darlene Wagner, Matthew Schmerer, Minoo Agarwal, Eyad Almasri, Debbie Boles, Ayla Burns, Nuthawin Charoensri, Oren Cohen, Susan Countryman, Mary Ann Cristobal, Bobbi Croy, Suzanne Dale, Hrushikesh Deshmukh, Amanda Douglas, Vincent Drouillon, Marcia Eisenberg, Howard Engler, Rama Ghatti, Prashant Gupta, Susan Hicks, Jake Humphrey, Lax Iyer, Lisa Pfefferle, Manoj Jain, Matthew Robinson, Mohan Kolli, Brian Krueger, Tim Kuphal, Stanley Letovsky, Michael Levandoski, Craig Lukasik, Jonathan Meltzer, Brian Norvell, Mindy Nye, Scott Parker, Christos Petropoulos, John Pruitt, Steven Ragan, Scott Ryan, Mike Sapeta, Jana Schroth, Suresh Babu Selvaraju, Goran Stevovic, Amanda Suchanek, Andrea Throop, Lyndon Tilson, Thomas Urban, Joe Voshell, Kimberly Wagner, Jonathan Williams, Mary Williamson, Qian Zeng, Tricia Zwiefelhofer, Clinton R. Paden, Duncan MacCannell |
| EPI_ISL_3326777 | TGen North | TGen North | Jolene Bowers, Heather Centner, Brett Van Tassel, Chris French, Hayley Yaglom, Darrin Lemmer, Dave Engelthaler |
| EPI_ISL_3326778 | Laboratory Corporation of America | Centers for Disease Control and Prevention Division of Viral Diseases, Pathogen Discovery | Dakota Howard, Dhwani Batra, Peter W. Cook, Kara Moser, Adrian Paskey, Jason Caravas, Benjamin Rambo-Martin, Shatavia Morrison, Christopher Gulvick, Scott Sammons, Yvette Unoarumhi, Darlene Wagner, Matthew Schmerer, Minoo Agarwal, Eyad Almasri, Debbie Boles, Ayla Burns, Nuthawin Charoensri, Oren Cohen, Susan Countryman, Mary Ann Cristobal, Bobbi Croy, Suzanne Dale, Hrushikesh Deshmukh, Amanda Douglas, Vincent Drouillon, Marcia Eisenberg, Howard Engler, Rama Ghatti, Prashant Gupta, Susan Hicks, Jake Humphrey, Lax Iyer, Lisa Pfefferle, Manoj Jain, Matthew Robinson, Mohan Kolli, Brian Krueger, Tim Kuphal, Stanley Letovsky, Michael Levandoski, Craig Lukasik, Jonathan Meltzer, Brian Norvell, Mindy Nye, Scott Parker, Christos Petropoulos, John Pruitt, Steven Ragan, Scott Ryan, Mike Sapeta, Jana Schroth, Suresh Babu Selvaraju, Goran Stevovic, Amanda Suchanek, Andrea Throop, Lyndon Tilson, Thomas Urban, Joe Voshell, Kimberly Wagner, Jonathan Williams, Mary Williamson, Qian Zeng, Tricia Zwiefelhofer, Clinton R. Paden, Duncan MacCannell |
| EPI_ISL_3326779 | TGen North | TGen North | Jolene Bowers, Heather Centner, Brett Van Tassel, Chris French, Hayley Yaglom, Darrin Lemmer, Dave Engelthaler |
| EPI_ISL_3326780 | Laboratory Corporation of America | Centers for Disease Control and Prevention Division of Viral Diseases, Pathogen Discovery | Dakota Howard, Dhwani Batra, Peter W. Cook, Kara Moser, Adrian Paskey, Jason Caravas, Benjamin Rambo-Martin, Shatavia Morrison, Christopher Gulvick, Scott Sammons, Yvette Unoarumhi, Darlene Wagner, Matthew Schmerer, Minoo Agarwal, Eyad Almasri, Debbie Boles, Ayla Burns, Nuthawin Charoensri, Oren Cohen, Susan Countryman, Mary Ann Cristobal, Bobbi Croy, Suzanne Dale, Hrushikesh Deshmukh, Amanda Douglas, Vincent Drouillon, Marcia Eisenberg, Howard Engler, Rama Ghatti, Prashant Gupta, Susan Hicks, Jake Humphrey, Lax Iyer, Lisa Pfefferle, Manoj Jain, Matthew Robinson, Mohan Kolli, Brian Krueger, Tim Kuphal, Stanley Letovsky, Michael Levandoski, Craig Lukasik, Jonathan Meltzer, Brian Norvell, Mindy Nye, Scott Parker, Christos Petropoulos, John Pruitt, Steven Ragan, Scott Ryan, Mike Sapeta, Jana Schroth, Suresh Babu Selvaraju, Goran Stevovic, Amanda Suchanek, Andrea Throop, Lyndon Tilson, Thomas Urban, Joe Voshell, Kimberly Wagner, Jonathan Williams, Mary Williamson, Qian Zeng, Tricia Zwiefelhofer, Clinton R. Paden, Duncan MacCannell |
| EPI_ISL_3326781 | TGen North | TGen North | Jolene Bowers, Heather Centner, Brett Van Tassel, Chris French, Hayley Yaglom, Darrin Lemmer, Dave Engelthaler |
| EPI_ISL_3326782 | Laboratory Corporation of America | Centers for Disease Control and Prevention Division of Viral Diseases, Pathogen Discovery | Dakota Howard, Dhwani Batra, Peter W. Cook, Kara Moser, Adrian Paskey, Jason Caravas, Benjamin Rambo-Martin, Shatavia Morrison, Christopher Gulvick, Scott Sammons, Yvette Unoarumhi, Darlene Wagner, Matthew Schmerer, Minoo Agarwal, Eyad Almasri, Debbie Boles, Ayla Burns, Nuthawin Charoensri, Oren Cohen, Susan Countryman, Mary Ann Cristobal, Bobbi Croy, Suzanne Dale, Hrushikesh Deshmukh, Amanda Douglas, Vincent Drouillon, Marcia Eisenberg, Howard Engler, Rama Ghatti, Prashant Gupta, Susan Hicks, Jake Humphrey, Lax Iyer, Lisa Pfefferle, Manoj Jain, Matthew Robinson, Mohan Kolli, Brian Krueger, Tim Kuphal, Stanley Letovsky, Michael Levandoski, Craig Lukasik, Jonathan Meltzer, Brian Norvell, Mindy Nye, Scott Parker, Christos Petropoulos, John Pruitt, Steven Ragan, Scott Ryan, Mike Sapeta, Jana Schroth, Suresh Babu Selvaraju, Goran Stevovic, Amanda Suchanek, Andrea Throop, Lyndon Tilson, Thomas Urban, Joe Voshell, Kimberly Wagner, Jonathan Williams, Mary Williamson, Qian Zeng, Tricia Zwiefelhofer, Clinton R. Paden, Duncan MacCannell |
| EPI_ISL_3326783 | TGen North | TGen North | Jolene Bowers, Heather Centner, Brett Van Tassel, Chris French, Hayley Yaglom, Darrin Lemmer, Dave Engelthaler |
| EPI_ISL_3326784 | Laboratory Corporation of America | Centers for Disease Control and Prevention Division of Viral Diseases, Pathogen Discovery | Dakota Howard, Dhwani Batra, Peter W. Cook, Kara Moser, Adrian Paskey, Jason Caravas, Benjamin Rambo-Martin, Shatavia Morrison, Christopher Gulvick, Scott Sammons, Yvette Unoarumhi, Darlene Wagner, Matthew Schmerer, Minoo Agarwal, Eyad Almasri, Debbie Boles, Ayla Burns, Nuthawin Charoensri, Oren Cohen, Susan Countryman, Mary Ann Cristobal, Bobbi Croy, Suzanne Dale, Hrushikesh Deshmukh, Amanda Douglas, Vincent Drouillon, Marcia Eisenberg, Howard Engler, Rama Ghatti, Prashant Gupta, Susan Hicks, Jake Humphrey, Lax Iyer, Lisa Pfefferle, Manoj Jain, Matthew Robinson, Mohan Kolli, Brian Krueger, Tim Kuphal, Stanley Letovsky, Michael Levandoski, Craig Lukasik, Jonathan Meltzer, Brian Norvell, Mindy Nye, Scott Parker, Christos Petropoulos, John Pruitt, Steven Ragan, Scott Ryan, Mike Sapeta, Jana Schroth, Suresh Babu Selvaraju, Goran Stevovic, Amanda Suchanek, Andrea Throop, Lyndon Tilson, Thomas Urban, Joe Voshell, Kimberly Wagner, Jonathan Williams, Mary Williamson, Qian Zeng, Tricia Zwiefelhofer, Clinton R. Paden, Duncan MacCannell |
| EPI_ISL_3326785 | TGen North | TGen North | Jolene Bowers, Heather Centner, Brett Van Tassel, Chris French, Hayley Yaglom, Darrin Lemmer, Dave Engelthaler |
| EPI_ISL_3326786, EPI_ISL_3326788 | Laboratory Corporation of America | Centers for Disease Control and Prevention Division of Viral Diseases, Pathogen Discovery | Dakota Howard, Dhwani Batra, Peter W. Cook, Kara Moser, Adrian Paskey, Jason Caravas, Benjamin Rambo-Martin, Shatavia Morrison, Christopher Gulvick, Scott Sammons, Yvette Unoarumhi, Darlene Wagner, Matthew Schmerer, Minoo Agarwal, Eyad Almasri, Debbie Boles, Ayla Burns, Nuthawin Charoensri, Oren Cohen, Susan Countryman, Mary Ann Cristobal, Bobbi Croy, Suzanne Dale, Hrushikesh Deshmukh, Amanda Douglas, Vincent Drouillon, Marcia Eisenberg, Howard Engler, Rama Ghatti, Prashant Gupta, Susan Hicks, Jake Humphrey, Lax Iyer, Lisa Pfefferle, Manoj Jain, Matthew Robinson, Mohan Kolli, Brian Krueger, Tim Kuphal, Stanley Letovsky, Michael Levandoski, Craig Lukasik, Jonathan Meltzer, Brian Norvell, Mindy Nye, Scott Parker, Christos Petropoulos, John Pruitt, Steven Ragan, Scott Ryan, Mike Sapeta, Jana Schroth, Suresh Babu Selvaraju, Goran Stevovic, Amanda Suchanek, Andrea Throop, Lyndon Tilson, Thomas Urban, Joe Voshell, Kimberly Wagner, Jonathan Williams, Mary Williamson, Qian Zeng, Tricia Zwiefelhofer, Clinton R. Paden, Duncan MacCannell |
| EPI_ISL_3326789 | TGen North | TGen North | Jolene Bowers, Heather Centner, Brett Van Tassel, Chris French, Hayley Yaglom, Darrin Lemmer, Dave Engelthaler |
| EPI_ISL_3326790, EPI_ISL_3326791 | Laboratory Corporation of America | Centers for Disease Control and Prevention Division of Viral Diseases, Pathogen Discovery | Dakota Howard, Dhwani Batra, Peter W. Cook, Kara Moser, Adrian Paskey, Jason Caravas, Benjamin Rambo-Martin, Shatavia Morrison, Christopher Gulvick, Scott Sammons, Yvette Unoarumhi, Darlene Wagner, Matthew Schmerer, Minoo Agarwal, Eyad Almasri, Debbie Boles, Ayla Burns, Nuthawin Charoensri, Oren Cohen, Susan Countryman, Mary Ann Cristobal, Bobbi Croy, Suzanne Dale, Hrushikesh Deshmukh, Amanda Douglas, Vincent Drouillon, Marcia Eisenberg, Howard Engler, Rama Ghatti, Prashant Gupta, Susan Hicks, Jake Humphrey, Lax Iyer, Lisa Pfefferle, Manoj Jain, Matthew Robinson, Mohan Kolli, Brian Krueger, Tim Kuphal, Stanley Letovsky, Michael Levandoski, Craig Lukasik, Jonathan Meltzer, Brian Norvell, Mindy Nye, Scott Parker, Christos Petropoulos, John Pruitt, Steven Ragan, Scott Ryan, Mike Sapeta, Jana Schroth, Suresh Babu Selvaraju, Goran Stevovic, Amanda Suchanek, Andrea Throop, Lyndon Tilson, Thomas Urban, Joe Voshell, Kimberly Wagner, Jonathan Williams, Mary Williamson, Qian Zeng, Tricia Zwiefelhofer, Clinton R. Paden, Duncan MacCannell |
| EPI_ISL_3326792 | TGen North | TGen North | Jolene Bowers, Heather Centner, Brett Van Tassel, Chris French, Hayley Yaglom, Darrin Lemmer, Dave Engelthaler |
| EPI_ISL_3326793, EPI_ISL_3326795 | Laboratory Corporation of America | Centers for Disease Control and Prevention Division of Viral Diseases, Pathogen Discovery | Dakota Howard, Dhwani Batra, Peter W. Cook, Kara Moser, Adrian Paskey, Jason Caravas, Benjamin Rambo-Martin, Shatavia Morrison, Christopher Gulvick, Scott Sammons, Yvette Unoarumhi, Darlene Wagner, Matthew Schmerer, Minoo Agarwal, Eyad Almasri, Debbie Boles, Ayla Burns, Nuthawin Charoensri, Oren Cohen, Susan Countryman, Mary Ann Cristobal, Bobbi Croy, Suzanne Dale, Hrushikesh Deshmukh, Amanda Douglas, Vincent Drouillon, Marcia Eisenberg, Howard Engler, Rama Ghatti, Prashant Gupta, Susan Hicks, Jake Humphrey, Lax Iyer, Lisa Pfefferle, Manoj Jain, Matthew Robinson, Mohan Kolli, Brian Krueger, Tim Kuphal, Stanley Letovsky, Michael Levandoski, Craig Lukasik, Jonathan Meltzer, Brian Norvell, Mindy Nye, Scott Parker, Christos Petropoulos, John Pruitt, Steven Ragan, Scott Ryan, Mike Sapeta, Jana Schroth, Suresh Babu Selvaraju, Goran Stevovic, Amanda Suchanek, Andrea Throop, Lyndon Tilson, Thomas Urban, Joe Voshell, Kimberly Wagner, Jonathan Williams, Mary Williamson, Qian Zeng, Tricia Zwiefelhofer, Clinton R. Paden, Duncan MacCannell |
| EPI_ISL_3326796 | TGen North | TGen North | Jolene Bowers, Heather Centner, Brett Van Tassel, Chris French, Hayley Yaglom, Darrin Lemmer, Dave Engelthaler |
| EPI_ISL_3326797, EPI_ISL_3326799 | Laboratory Corporation of America | Centers for Disease Control and Prevention Division of Viral Diseases, Pathogen Discovery | Dakota Howard, Dhwani Batra, Peter W. Cook, Kara Moser, Adrian Paskey, Jason Caravas, Benjamin Rambo-Martin, Shatavia Morrison, Christopher Gulvick, Scott Sammons, Yvette Unoarumhi, Darlene Wagner, Matthew Schmerer, Minoo Agarwal, Eyad Almasri, Debbie Boles, Ayla Burns, Nuthawin Charoensri, Oren Cohen, Susan Countryman, Mary Ann Cristobal, Bobbi Croy, Suzanne Dale, Hrushikesh Deshmukh, Amanda Douglas, Vincent Drouillon, Marcia Eisenberg, Howard Engler, Rama Ghatti, Prashant Gupta, Susan Hicks, Jake Humphrey, Lax Iyer, Lisa Pfefferle, Manoj Jain, Matthew Robinson, Mohan Kolli, Brian Krueger, Tim Kuphal, Stanley Letovsky, Michael Levandoski, Craig Lukasik, Jonathan Meltzer, Brian Norvell, Mindy Nye, Scott Parker, Christos Petropoulos, John Pruitt, Steven Ragan, Scott Ryan, Mike Sapeta, Jana Schroth, Suresh Babu Selvaraju, Goran Stevovic, Amanda Suchanek, Andrea Throop, Lyndon Tilson, Thomas Urban, Joe Voshell, Kimberly Wagner, Jonathan Williams, Mary Williamson, Qian Zeng, Tricia Zwiefelhofer, Clinton R. Paden, Duncan MacCannell |
| EPI_ISL_3326800 | TGen North | TGen North | Jolene Bowers, Heather Centner, Brett Van Tassel, Chris French, Hayley Yaglom, Darrin Lemmer, Dave Engelthaler |
| EPI_ISL_3326801 | Laboratory Corporation of America | Centers for Disease Control and Prevention Division of Viral Diseases, Pathogen Discovery | Dakota Howard, Dhwani Batra, Peter W. Cook, Kara Moser, Adrian Paskey, Jason Caravas, Benjamin Rambo-Martin, Shatavia Morrison, Christopher Gulvick, Scott Sammons, Yvette Unoarumhi, Darlene Wagner, Matthew Schmerer, Minoo Agarwal, Eyad Almasri, Debbie Boles, Ayla Burns, Nuthawin Charoensri, Oren Cohen, Susan Countryman, Mary Ann Cristobal, Bobbi Croy, Suzanne Dale, Hrushikesh Deshmukh, Amanda Douglas, Vincent Drouillon, Marcia Eisenberg, Howard Engler, Rama Ghatti, Prashant Gupta, Susan Hicks, Jake Humphrey, Lax Iyer, Lisa Pfefferle, Manoj Jain, Matthew Robinson, Mohan Kolli, Brian Krueger, Tim Kuphal, Stanley Letovsky, Michael Levandoski, Craig Lukasik, Jonathan Meltzer, Brian Norvell, Mindy Nye, Scott Parker, Christos Petropoulos, John Pruitt, Steven Ragan, Scott Ryan, Mike Sapeta, Jana Schroth, Suresh Babu Selvaraju, Goran Stevovic, Amanda Suchanek, Andrea Throop, Lyndon Tilson, Thomas Urban, Joe Voshell, Kimberly Wagner, Jonathan Williams, Mary Williamson, Qian Zeng, Tricia Zwiefelhofer, Clinton R. Paden, Duncan MacCannell |
| EPI_ISL_3326802 | TGen North | TGen North | Jolene Bowers, Heather Centner, Brett Van Tassel, Chris French, Hayley Yaglom, Darrin Lemmer, Dave Engelthaler |
| EPI_ISL_3326803 | Laboratory Corporation of America | Centers for Disease Control and Prevention Division of Viral Diseases, Pathogen Discovery | Dakota Howard, Dhwani Batra, Peter W. Cook, Kara Moser, Adrian Paskey, Jason Caravas, Benjamin Rambo-Martin, Shatavia Morrison, Christopher Gulvick, Scott Sammons, Yvette Unoarumhi, Darlene Wagner, Matthew Schmerer, Minoo Agarwal, Eyad Almasri, Debbie Boles, Ayla Burns, Nuthawin Charoensri, Oren Cohen, Susan Countryman, Mary Ann Cristobal, Bobbi Croy, Suzanne Dale, Hrushikesh Deshmukh, Amanda Douglas, Vincent Drouillon, Marcia Eisenberg, Howard Engler, Rama Ghatti, Prashant Gupta, Susan Hicks, Jake Humphrey, Lax Iyer, Lisa Pfefferle, Manoj Jain, Matthew Robinson, Mohan Kolli, Brian Krueger, Tim Kuphal, Stanley Letovsky, Michael Levandoski, Craig Lukasik, Jonathan Meltzer, Brian Norvell, Mindy Nye, Scott Parker, Christos Petropoulos, John Pruitt, Steven Ragan, Scott Ryan, Mike Sapeta, Jana Schroth, Suresh Babu Selvaraju, Goran Stevovic, Amanda Suchanek, Andrea Throop, Lyndon Tilson, Thomas Urban, Joe Voshell, Kimberly Wagner, Jonathan Williams, Mary Williamson, Qian Zeng, Tricia Zwiefelhofer, Clinton R. Paden, Duncan MacCannell |
| EPI_ISL_3326804 | TGen North | TGen North | Jolene Bowers, Heather Centner, Brett Van Tassel, Chris French, Hayley Yaglom, Darrin Lemmer, Dave Engelthaler |
| EPI_ISL_3326805, EPI_ISL_3326806 | Laboratory Corporation of America | Centers for Disease Control and Prevention Division of Viral Diseases, Pathogen Discovery | Dakota Howard, Dhwani Batra, Peter W. Cook, Kara Moser, Adrian Paskey, Jason Caravas, Benjamin Rambo-Martin, Shatavia Morrison, Christopher Gulvick, Scott Sammons, Yvette Unoarumhi, Darlene Wagner, Matthew Schmerer, Minoo Agarwal, Eyad Almasri, Debbie Boles, Ayla Burns, Nuthawin Charoensri, Oren Cohen, Susan Countryman, Mary Ann Cristobal, Bobbi Croy, Suzanne Dale, Hrushikesh Deshmukh, Amanda Douglas, Vincent Drouillon, Marcia Eisenberg, Howard Engler, Rama Ghatti, Prashant Gupta, Susan Hicks, Jake Humphrey, Lax Iyer, Lisa Pfefferle, Manoj Jain, Matthew Robinson, Mohan Kolli, Brian Krueger, Tim Kuphal, Stanley Letovsky, Michael Levandoski, Craig Lukasik, Jonathan Meltzer, Brian Norvell, Mindy Nye, Scott Parker, Christos Petropoulos, John Pruitt, Steven Ragan, Scott Ryan, Mike Sapeta, Jana Schroth, Suresh Babu Selvaraju, Goran Stevovic, Amanda Suchanek, Andrea Throop, Lyndon Tilson, Thomas Urban, Joe Voshell, Kimberly Wagner, Jonathan Williams, Mary Williamson, Qian Zeng, Tricia Zwiefelhofer, Clinton R. Paden, Duncan MacCannell |
| EPI_ISL_3326807 | TGen North | TGen North | Jolene Bowers, Heather Centner, Brett Van Tassel, Chris French, Hayley Yaglom, Darrin Lemmer, Dave Engelthaler |
| EPI_ISL_3326808 | Laboratory Corporation of America | Centers for Disease Control and Prevention Division of Viral Diseases, Pathogen Discovery | Dakota Howard, Dhwani Batra, Peter W. Cook, Kara Moser, Adrian Paskey, Jason Caravas, Benjamin Rambo-Martin, Shatavia Morrison, Christopher Gulvick, Scott Sammons, Yvette Unoarumhi, Darlene Wagner, Matthew Schmerer, Minoo Agarwal, Eyad Almasri, Debbie Boles, Ayla Burns, Nuthawin Charoensri, Oren Cohen, Susan Countryman, Mary Ann Cristobal, Bobbi Croy, Suzanne Dale, Hrushikesh Deshmukh, Amanda Douglas, Vincent Drouillon, Marcia Eisenberg, Howard Engler, Rama Ghatti, Prashant Gupta, Susan Hicks, Jake Humphrey, Lax Iyer, Lisa Pfefferle, Manoj Jain, Matthew Robinson, Mohan Kolli, Brian Krueger, Tim Kuphal, Stanley Letovsky, Michael Levandoski, Craig Lukasik, Jonathan Meltzer, Brian Norvell, Mindy Nye, Scott Parker, Christos Petropoulos, John Pruitt, Steven Ragan, Scott Ryan, Mike Sapeta, Jana Schroth, Suresh Babu Selvaraju, Goran Stevovic, Amanda Suchanek, Andrea Throop, Lyndon Tilson, Thomas Urban, Joe Voshell, Kimberly Wagner, Jonathan Williams, Mary Williamson, Qian Zeng, Tricia Zwiefelhofer, Clinton R. Paden, Duncan MacCannell |
| EPI_ISL_3326809 | TGen North | TGen North | Jolene Bowers, Heather Centner, Brett Van Tassel, Chris French, Hayley Yaglom, Darrin Lemmer, Dave Engelthaler |
| EPI_ISL_3326810 | Laboratory Corporation of America | Centers for Disease Control and Prevention Division of Viral | Dakota Howard, Dhwani Batra, Peter W. Cook, Kara Moser, Adrian Paskey, Jason Caravas, Benjamin Rambo-Martin, Shatavia Morrison, Christopher Gulvick, Scott Sammons, Yvette Unoarumhi, Darlene Wagner, Matthew Schmerer, Minoo |

| | | | |
|---|---|---|---|
| | | Diseases, Pathogen Discovery | Agarwal, Eyad Almasri, Debbie Boles, Ayla Burns, Nuthawin Charoensri, Oren Cohen, Susan Countryman, Mary Ann Cristobal, Bobbi Croy, Suzanne Dale, Hrushikesh Deshmukh, Amanda Douglas, Vincent Drouillon, Marcia Eisenberg, Howard Engler, Rama Ghatti, Prashant Gupta, Susan Hicks, Jake Humphrey, Lax Iyer, Lisa Pfefferle, Manoj Jain, Matthew Robinson, Mohan Kolli, Brian Krueger, Tim Kuphal, Stanley Letovsky, Michael Levandoski, Craig Lukasik, Jonathan Meltzer, Brian Norvell, Mindy Nye, Scott Parker, Christos Petropoulos, John Pruitt, Steven Ragan, Scott Ryan, Mike Sapeta, Jana Schroth, Suresh Babu Selvaraju, Goran Stevovic, Amanda Suchanek, Andrea Throop, Lyndon Tilson, Thomas Urban, Joe Voshell, Kimberly Wagner, Jonathan Williams, Mary Williamson, Qian Zeng, Tricia Zwiefelhofer, Clinton R. Paden, Duncan MacCannell |
| EPI_ISL_3326811 | TGen North | TGen North | Jolene Bowers, Heather Centner, Brett Van Tassel, Chris French, Hayley Yaglom, Darrin Lemmer, Dave Engelthaler |
| EPI_ISL_3326812, EPI_ISL_3326814, EPI_ISL_3326815 | Laboratory Corporation of America | Centers for Disease Control and Prevention Division of Viral Diseases, Pathogen Discovery | Dakota Howard, Dhwani Batra, Peter W. Cook, Kara Moser, Adrian Paskey, Jason Caravas, Benjamin Rambo-Martin, Shatavia Morrison, Christopher Gulvick, Scott Sammons, Yvette Unoarumhi, Darlene Wagner, Matthew Schmerer, Minoo Agarwal, Eyad Almasri, Debbie Boles, Ayla Burns, Nuthawin Charoensri, Oren Cohen, Susan Countryman, Mary Ann Cristobal, Bobbi Croy, Suzanne Dale, Hrushikesh Deshmukh, Amanda Douglas, Vincent Drouillon, Marcia Eisenberg, Howard Engler, Rama Ghatti, Prashant Gupta, Susan Hicks, Jake Humphrey, Lax Iyer, Lisa Pfefferle, Manoj Jain, Matthew Robinson, Mohan Kolli, Brian Krueger, Tim Kuphal, Stanley Letovsky, Michael Levandoski, Craig Lukasik, Jonathan Meltzer, Brian Norvell, Mindy Nye, Scott Parker, Christos Petropoulos, John Pruitt, Steven Ragan, Scott Ryan, Mike Sapeta, Jana Schroth, Suresh Babu Selvaraju, Goran Stevovic, Amanda Suchanek, Andrea Throop, Lyndon Tilson, Thomas Urban, Joe Voshell, Kimberly Wagner, Jonathan Williams, Mary Williamson, Qian Zeng, Tricia Zwiefelhofer, Clinton R. Paden, Duncan MacCannell |
| EPI_ISL_3326816 | TGen North | TGen North | Jolene Bowers, Heather Centner, Brett Van Tassel, Chris French, Hayley Yaglom, Darrin Lemmer, Dave Engelthaler |
| EPI_ISL_3326817 | Laboratory Corporation of America | Centers for Disease Control and Prevention Division of Viral Diseases, Pathogen Discovery | Dakota Howard, Dhwani Batra, Peter W. Cook, Kara Moser, Adrian Paskey, Jason Caravas, Benjamin Rambo-Martin, Shatavia Morrison, Christopher Gulvick, Scott Sammons, Yvette Unoarumhi, Darlene Wagner, Matthew Schmerer, Minoo Agarwal, Eyad Almasri, Debbie Boles, Ayla Burns, Nuthawin Charoensri, Oren Cohen, Susan Countryman, Mary Ann Cristobal, Bobbi Croy, Suzanne Dale, Hrushikesh Deshmukh, Amanda Douglas, Vincent Drouillon, Marcia Eisenberg, Howard Engler, Rama Ghatti, Prashant Gupta, Susan Hicks, Jake Humphrey, Lax Iyer, Lisa Pfefferle, Manoj Jain, Matthew Robinson, Mohan Kolli, Brian Krueger, Tim Kuphal, Stanley Letovsky, Michael Levandoski, Craig Lukasik, Jonathan Meltzer, Brian Norvell, Mindy Nye, Scott Parker, Christos Petropoulos, John Pruitt, Steven Ragan, Scott Ryan, Mike Sapeta, Jana Schroth, Suresh Babu Selvaraju, Goran Stevovic, Amanda Suchanek, Andrea Throop, Lyndon Tilson, Thomas Urban, Joe Voshell, Kimberly Wagner, Jonathan Williams, Mary Williamson, Qian Zeng, Tricia Zwiefelhofer, Clinton R. Paden, Duncan MacCannell |
| EPI_ISL_3326818 | TGen North | TGen North | Jolene Bowers, Heather Centner, Brett Van Tassel, Chris French, Hayley Yaglom, Darrin Lemmer, Dave Engelthaler |
| EPI_ISL_3326819 | Laboratory Corporation of America | Centers for Disease Control and Prevention Division of Viral Diseases, Pathogen Discovery | Dakota Howard, Dhwani Batra, Peter W. Cook, Kara Moser, Adrian Paskey, Jason Caravas, Benjamin Rambo-Martin, Shatavia Morrison, Christopher Gulvick, Scott Sammons, Yvette Unoarumhi, Darlene Wagner, Matthew Schmerer, Minoo Agarwal, Eyad Almasri, Debbie Boles, Ayla Burns, Nuthawin Charoensri, Oren Cohen, Susan Countryman, Mary Ann Cristobal, Bobbi Croy, Suzanne Dale, Hrushikesh Deshmukh, Amanda Douglas, Vincent Drouillon, Marcia Eisenberg, Howard Engler, Rama Ghatti, Prashant Gupta, Susan Hicks, Jake Humphrey, Lax Iyer, Lisa Pfefferle, Manoj Jain, Matthew Robinson, Mohan Kolli, Brian Krueger, Tim Kuphal, Stanley Letovsky, Michael Levandoski, Craig Lukasik, Jonathan Meltzer, Brian Norvell, Mindy Nye, Scott Parker, Christos Petropoulos, John Pruitt, Steven Ragan, Scott Ryan, Mike Sapeta, Jana Schroth, Suresh Babu Selvaraju, Goran Stevovic, Amanda Suchanek, Andrea Throop, Lyndon Tilson, Thomas Urban, Joe Voshell, Kimberly Wagner, Jonathan Williams, Mary Williamson, Qian Zeng, Tricia Zwiefelhofer, Clinton R. Paden, Duncan MacCannell |
| EPI_ISL_3326820 | TGen North | TGen North | Jolene Bowers, Heather Centner, Brett Van Tassel, Chris French, Hayley Yaglom, Darrin Lemmer, Dave Engelthaler |
| EPI_ISL_3326821 | Laboratory Corporation of America | Centers for Disease Control and Prevention Division of Viral Diseases, Pathogen Discovery | Dakota Howard, Dhwani Batra, Peter W. Cook, Kara Moser, Adrian Paskey, Jason Caravas, Benjamin Rambo-Martin, Shatavia Morrison, Christopher Gulvick, Scott Sammons, Yvette Unoarumhi, Darlene Wagner, Matthew Schmerer, Minoo Agarwal, Eyad Almasri, Debbie Boles, Ayla Burns, Nuthawin Charoensri, Oren Cohen, Susan Countryman, Mary Ann Cristobal, Bobbi Croy, Suzanne Dale, Hrushikesh Deshmukh, Amanda Douglas, Vincent Drouillon, Marcia Eisenberg, Howard Engler, Rama Ghatti, Prashant Gupta, Susan Hicks, Jake Humphrey, Lax Iyer, Lisa Pfefferle, Manoj Jain, Matthew Robinson, Mohan Kolli, Brian Krueger, Tim Kuphal, Stanley Letovsky, Michael Levandoski, Craig Lukasik, Jonathan Meltzer, Brian Norvell, Mindy Nye, Scott Parker, Christos Petropoulos, John Pruitt, Steven Ragan, Scott Ryan, Mike Sapeta, Jana Schroth, Suresh Babu Selvaraju, Goran Stevovic, Amanda Suchanek, Andrea Throop, Lyndon Tilson, Thomas Urban, Joe Voshell, Kimberly Wagner, Jonathan Williams, Mary Williamson, Qian Zeng, Tricia Zwiefelhofer, Clinton R. Paden, Duncan MacCannell |
| EPI_ISL_3326822 | TGen North | TGen North | Jolene Bowers, Heather Centner, Brett Van Tassel, Chris French, Hayley Yaglom, Darrin Lemmer, Dave Engelthaler |
| EPI_ISL_3326823 | Laboratory Corporation of America | Centers for Disease Control and Prevention Division of Viral Diseases, Pathogen Discovery | Dakota Howard, Dhwani Batra, Peter W. Cook, Kara Moser, Adrian Paskey, Jason Caravas, Benjamin Rambo-Martin, Shatavia Morrison, Christopher Gulvick, Scott Sammons, Yvette Unoarumhi, Darlene Wagner, Matthew Schmerer, Minoo Agarwal, Eyad Almasri, Debbie Boles, Ayla Burns, Nuthawin Charoensri, Oren Cohen, Susan Countryman, Mary Ann Cristobal, Bobbi Croy, Suzanne Dale, Hrushikesh Deshmukh, Amanda Douglas, Vincent Drouillon, Marcia Eisenberg, Howard Engler, Rama Ghatti, Prashant Gupta, Susan Hicks, Jake Humphrey, Lax Iyer, Lisa Pfefferle, Manoj Jain, Matthew Robinson, Mohan Kolli, Brian Krueger, Tim Kuphal, Stanley Letovsky, Michael Levandoski, Craig Lukasik, Jonathan Meltzer, Brian Norvell, Mindy Nye, Scott Parker, Christos Petropoulos, John Pruitt, Steven Ragan, Scott Ryan, Mike Sapeta, Jana Schroth, Suresh Babu Selvaraju, Goran Stevovic, Amanda Suchanek, Andrea Throop, Lyndon Tilson, Thomas Urban, Joe Voshell, Kimberly Wagner, Jonathan Williams, Mary Williamson, Qian Zeng, Tricia Zwiefelhofer, Clinton R. Paden, Duncan MacCannell |
| EPI_ISL_3326824 | TGen North | TGen North | Jolene Bowers, Heather Centner, Brett Van Tassel, Chris French, Hayley Yaglom, Darrin Lemmer, Dave Engelthaler |
| EPI_ISL_3326825 | Laboratory Corporation of America | Centers for Disease Control and Prevention Division of Viral Diseases, Pathogen Discovery | Dakota Howard, Dhwani Batra, Peter W. Cook, Kara Moser, Adrian Paskey, Jason Caravas, Benjamin Rambo-Martin, Shatavia Morrison, Christopher Gulvick, Scott Sammons, Yvette Unoarumhi, Darlene Wagner, Matthew Schmerer, Minoo Agarwal, Eyad Almasri, Debbie Boles, Ayla Burns, Nuthawin Charoensri, Oren Cohen, Susan Countryman, Mary Ann Cristobal, Bobbi Croy, Suzanne Dale, Hrushikesh Deshmukh, Amanda Douglas, Vincent Drouillon, Marcia Eisenberg, Howard Engler, Rama Ghatti, Prashant Gupta, Susan Hicks, Jake Humphrey, Lax Iyer, Lisa Pfefferle, Manoj Jain, Matthew Robinson, Mohan Kolli, Brian Krueger, Tim Kuphal, Stanley Letovsky, Michael Levandoski, Craig Lukasik, Jonathan Meltzer, Brian Norvell, Mindy Nye, Scott Parker, Christos Petropoulos, John Pruitt, Steven Ragan, Scott Ryan, Mike Sapeta, Jana Schroth, Suresh Babu Selvaraju, Goran Stevovic, Amanda Suchanek, Andrea Throop, Lyndon Tilson, Thomas Urban, Joe Voshell, Kimberly Wagner, Jonathan Williams, Mary Williamson, Qian Zeng, Tricia Zwiefelhofer, Clinton R. Paden, Duncan MacCannell |
| EPI_ISL_3326826 | TGen North | TGen North | Jolene Bowers, Heather Centner, Brett Van Tassel, Chris French, Hayley Yaglom, Darrin Lemmer, Dave Engelthaler |
| EPI_ISL_3326827 | Laboratory Corporation of America | Centers for Disease Control and Prevention Division of Viral Diseases, Pathogen Discovery | Dakota Howard, Dhwani Batra, Peter W. Cook, Kara Moser, Adrian Paskey, Jason Caravas, Benjamin Rambo-Martin, Shatavia Morrison, Christopher Gulvick, Scott Sammons, Yvette Unoarumhi, Darlene Wagner, Matthew Schmerer, Minoo Agarwal, Eyad Almasri, Debbie Boles, Ayla Burns, Nuthawin Charoensri, Oren Cohen, Susan Countryman, Mary Ann Cristobal, Bobbi Croy, Suzanne Dale, Hrushikesh Deshmukh, Amanda Douglas, Vincent Drouillon, Marcia Eisenberg, Howard Engler, Rama Ghatti, Prashant Gupta, Susan Hicks, Jake Humphrey, Lax Iyer, Lisa Pfefferle, Manoj Jain, Matthew Robinson, Mohan Kolli, Brian Krueger, Tim Kuphal, Stanley Letovsky, Michael Levandoski, Craig Lukasik, Jonathan Meltzer, Brian Norvell, Mindy Nye, Scott Parker, Christos Petropoulos, John Pruitt, Steven Ragan, Scott Ryan, Mike Sapeta, Jana Schroth, Suresh Babu Selvaraju, Goran Stevovic, Amanda Suchanek, Andrea Throop, Lyndon Tilson, Thomas Urban, Joe Voshell, Kimberly Wagner, Jonathan Williams, Mary Williamson, Qian Zeng, Tricia Zwiefelhofer, Clinton R. Paden, Duncan MacCannell |
| EPI_ISL_3326828 | TGen North | TGen North | Jolene Bowers, Heather Centner, Brett Van Tassel, Chris French, Hayley Yaglom, Darrin Lemmer, Dave Engelthaler |
| EPI_ISL_3326829, EPI_ISL_3326831, EPI_ISL_3326832, EPI_ISL_3326834 | Laboratory Corporation of America | Centers for Disease Control and Prevention Division of Viral Diseases, Pathogen Discovery | Dakota Howard, Dhwani Batra, Peter W. Cook, Kara Moser, Adrian Paskey, Jason Caravas, Benjamin Rambo-Martin, Shatavia Morrison, Christopher Gulvick, Scott Sammons, Yvette Unoarumhi, Darlene Wagner, Matthew Schmerer, Minoo Agarwal, Eyad Almasri, Debbie Boles, Ayla Burns, Nuthawin Charoensri, Oren Cohen, Susan Countryman, Mary Ann Cristobal, Bobbi Croy, Suzanne Dale, Hrushikesh Deshmukh, Amanda Douglas, Vincent Drouillon, Marcia Eisenberg, Howard Engler, Rama Ghatti, Prashant Gupta, Susan Hicks, Jake Humphrey, Lax Iyer, Lisa Pfefferle, Manoj Jain, Matthew Robinson, Mohan Kolli, Brian Krueger, Tim Kuphal, Stanley Letovsky, Michael Levandoski, Craig Lukasik, Jonathan Meltzer, Brian Norvell, Mindy Nye, Scott Parker, Christos Petropoulos, John Pruitt, Steven Ragan, Scott Ryan, Mike Sapeta, Jana Schroth, Suresh Babu Selvaraju, Goran Stevovic, Amanda Suchanek, Andrea Throop, Lyndon Tilson, Thomas Urban, Joe Voshell, Kimberly Wagner, Jonathan Williams, Mary Williamson, Qian Zeng, Tricia Zwiefelhofer, Clinton R. Paden, Duncan MacCannell |
| EPI_ISL_3326835, EPI_ISL_3326836 | TGen North | TGen North | Jolene Bowers, Heather Centner, Brett Van Tassel, Chris French, Hayley Yaglom, Darrin Lemmer, Dave Engelthaler |
| EPI_ISL_3326837, EPI_ISL_3326838 | Laboratory Corporation of America | Centers for Disease Control and Prevention Division of Viral Diseases, Pathogen Discovery | Dakota Howard, Dhwani Batra, Peter W. Cook, Kara Moser, Adrian Paskey, Jason Caravas, Benjamin Rambo-Martin, Shatavia Morrison, Christopher Gulvick, Scott Sammons, Yvette Unoarumhi, Darlene Wagner, Matthew Schmerer, Minoo Agarwal, Eyad Almasri, Debbie Boles, Ayla Burns, Nuthawin Charoensri, Oren Cohen, Susan Countryman, Mary Ann Cristobal, Bobbi Croy, Suzanne Dale, Hrushikesh Deshmukh, Amanda Douglas, Vincent Drouillon, Marcia Eisenberg, Howard Engler, Rama Ghatti, Prashant Gupta, Susan Hicks, Jake Humphrey, Lax Iyer, Lisa Pfefferle, Manoj Jain, Matthew Robinson, Mohan Kolli, Brian Krueger, Tim Kuphal, Stanley Letovsky, Michael Levandoski, Craig Lukasik, Jonathan Meltzer, Brian Norvell, Mindy Nye, Scott Parker, Christos Petropoulos, John Pruitt, Steven Ragan, Scott Ryan, Mike Sapeta, Jana Schroth, Suresh Babu Selvaraju, Goran Stevovic, Amanda Suchanek, Andrea Throop, Lyndon Tilson, Thomas Urban, Joe Voshell, Kimberly Wagner, Jonathan Williams, Mary Williamson, Qian Zeng, Tricia Zwiefelhofer, Clinton R. Paden, Duncan MacCannell |
| EPI_ISL_3326839 | TGen North | TGen North | Jolene Bowers, Heather Centner, Brett Van Tassel, Chris French, Hayley Yaglom, Darrin Lemmer, Dave Engelthaler |
| EPI_ISL_3326840 | Laboratory Corporation of America | Centers for Disease Control and Prevention Division of Viral Diseases, Pathogen Discovery | Dakota Howard, Dhwani Batra, Peter W. Cook, Kara Moser, Adrian Paskey, Jason Caravas, Benjamin Rambo-Martin, Shatavia Morrison, Christopher Gulvick, Scott Sammons, Yvette Unoarumhi, Darlene Wagner, Matthew Schmerer, Minoo Agarwal, Eyad Almasri, Debbie Boles, Ayla Burns, Nuthawin Charoensri, Oren Cohen, Susan Countryman, Mary Ann Cristobal, Bobbi Croy, Suzanne Dale, Hrushikesh Deshmukh, Amanda Douglas, Vincent Drouillon, Marcia Eisenberg, Howard Engler, Rama Ghatti, Prashant Gupta, Susan Hicks, Jake Humphrey, Lax Iyer, Lisa Pfefferle, Manoj Jain, Matthew Robinson, Mohan Kolli, Brian Krueger, Tim Kuphal, Stanley Letovsky, Michael Levandoski, Craig Lukasik, Jonathan Meltzer, Brian Norvell, Mindy Nye, Scott Parker, Christos Petropoulos, John Pruitt, Steven Ragan, Scott Ryan, Mike Sapeta, Jana Schroth, Suresh Babu Selvaraju, Goran Stevovic, Amanda Suchanek, Andrea Throop, Lyndon Tilson, Thomas Urban, Joe Voshell, Kimberly Wagner, Jonathan Williams, Mary Williamson, Qian Zeng, Tricia Zwiefelhofer, Clinton R. Paden, Duncan MacCannell |
| EPI_ISL_3326841 | TGen North | TGen North | Jolene Bowers, Heather Centner, Brett Van Tassel, Chris French, Hayley Yaglom, Darrin Lemmer, Dave Engelthaler |
| EPI_ISL_3326842 | Laboratory Corporation of America | Centers for Disease Control and Prevention Division of Viral Diseases, Pathogen Discovery | Dakota Howard, Dhwani Batra, Peter W. Cook, Kara Moser, Adrian Paskey, Jason Caravas, Benjamin Rambo-Martin, Shatavia Morrison, Christopher Gulvick, Scott Sammons, Yvette Unoarumhi, Darlene Wagner, Matthew Schmerer, Minoo Agarwal, Eyad Almasri, Debbie Boles, Ayla Burns, Nuthawin Charoensri, Oren Cohen, Susan Countryman, Mary Ann Cristobal, Bobbi Croy, Suzanne Dale, Hrushikesh Deshmukh, Amanda Douglas, Vincent Drouillon, Marcia Eisenberg, Howard Engler, Rama Ghatti, Prashant Gupta, Susan Hicks, Jake Humphrey, Lax Iyer, Lisa Pfefferle, Manoj Jain, Matthew Robinson, Mohan Kolli, Brian Krueger, Tim Kuphal, Stanley Letovsky, Michael Levandoski, Craig Lukasik, Jonathan Meltzer, Brian Norvell, Mindy Nye, Scott Parker, Christos Petropoulos, John Pruitt, Steven Ragan, Scott Ryan, Mike Sapeta, Jana Schroth, Suresh Babu Selvaraju, Goran Stevovic, Amanda Suchanek, Andrea Throop, Lyndon Tilson, Thomas Urban, Joe Voshell, Kimberly Wagner, Jonathan Williams, Mary Williamson, Qian Zeng, Tricia Zwiefelhofer, Clinton R. Paden, Duncan MacCannell |
| EPI_ISL_3326843 | TGen North | TGen North | Jolene Bowers, Heather Centner, Brett Van Tassel, Chris French, Hayley Yaglom, Darrin Lemmer, Dave Engelthaler |
| EPI_ISL_3326844, EPI_ISL_3326845 | Laboratory Corporation of America | Centers for Disease Control and Prevention Division of Viral Diseases, Pathogen Discovery | Dakota Howard, Dhwani Batra, Peter W. Cook, Kara Moser, Adrian Paskey, Jason Caravas, Benjamin Rambo-Martin, Shatavia Morrison, Christopher Gulvick, Scott Sammons, Yvette Unoarumhi, Darlene Wagner, Matthew Schmerer, Minoo Agarwal, Eyad Almasri, Debbie Boles, Ayla Burns, Nuthawin Charoensri, Oren Cohen, Susan Countryman, Mary Ann Cristobal, Bobbi Croy, Suzanne Dale, Hrushikesh Deshmukh, Amanda Douglas, Vincent Drouillon, Marcia Eisenberg, Howard Engler, Rama Ghatti, Prashant Gupta, Susan Hicks, Jake Humphrey, Lax Iyer, Lisa Pfefferle, Manoj Jain, Matthew Robinson, Mohan Kolli, Brian Krueger, Tim Kuphal, Stanley Letovsky, Michael Levandoski, Craig Lukasik, Jonathan Meltzer, Brian Norvell, Mindy Nye, Scott Parker, Christos Petropoulos, John Pruitt, Steven Ragan, Scott Ryan, Mike Sapeta, Jana Schroth, Suresh Babu Selvaraju, Goran Stevovic, Amanda Suchanek, Andrea Throop, Lyndon Tilson, Thomas Urban, Joe Voshell, Kimberly Wagner, Jonathan Williams, Mary Williamson, Qian Zeng, Tricia Zwiefelhofer, Clinton R. Paden, Duncan MacCannell |
| EPI_ISL_3326846 | TGen North | TGen North | Jolene Bowers, Heather Centner, Brett Van Tassel, Chris French, Hayley Yaglom, Darrin Lemmer, Dave Engelthaler |
| EPI_ISL_3326847, EPI_ISL_3326849 | Laboratory Corporation of America | Centers for Disease Control and Prevention Division of Viral Diseases, Pathogen Discovery | Dakota Howard, Dhwani Batra, Peter W. Cook, Kara Moser, Adrian Paskey, Jason Caravas, Benjamin Rambo-Martin, Shatavia Morrison, Christopher Gulvick, Scott Sammons, Yvette Unoarumhi, Darlene Wagner, Matthew Schmerer, Minoo Agarwal, Eyad Almasri, Debbie Boles, Ayla Burns, Nuthawin Charoensri, Oren Cohen, Susan Countryman, Mary Ann Cristobal, Bobbi Croy, Suzanne Dale, Hrushikesh Deshmukh, Amanda Douglas, Vincent Drouillon, Marcia Eisenberg, Howard Engler, Rama Ghatti, Prashant Gupta, Susan Hicks, Jake Humphrey, Lax Iyer, Lisa Pfefferle, Manoj Jain, Matthew Robinson, Mohan Kolli, Brian Krueger, Tim Kuphal, Stanley Letovsky, Michael Levandoski, Craig Lukasik, Jonathan Meltzer, Brian Norvell, Mindy Nye, Scott Parker, Christos Petropoulos, John Pruitt, Steven Ragan, Scott Ryan, Mike Sapeta, Jana Schroth, Suresh Babu Selvaraju, Goran Stevovic, Amanda Suchanek, Andrea Throop, Lyndon Tilson, Thomas Urban, Joe Voshell, Kimberly Wagner, Jonathan Williams, Mary Williamson, Qian Zeng, Tricia Zwiefelhofer, Clinton R. Paden, Duncan MacCannell |
| EPI_ISL_3326850 | TGen North | TGen North | Jolene Bowers, Heather Centner, Brett Van Tassel, Chris French, Hayley Yaglom, Darrin Lemmer, Dave Engelthaler |
| EPI_ISL_3326851, EPI_ISL_3326853, EPI_ISL_3326854 | Laboratory Corporation of America | Centers for Disease Control and Prevention Division of Viral Diseases, Pathogen Discovery | Dakota Howard, Dhwani Batra, Peter W. Cook, Kara Moser, Adrian Paskey, Jason Caravas, Benjamin Rambo-Martin, Shatavia Morrison, Christopher Gulvick, Scott Sammons, Yvette Unoarumhi, Darlene Wagner, Matthew Schmerer, Minoo Agarwal, Eyad Almasri, Debbie Boles, Ayla Burns, Nuthawin Charoensri, Oren Cohen, Susan Countryman, Mary Ann Cristobal, Bobbi Croy, Suzanne Dale, Hrushikesh Deshmukh, Amanda Douglas, Vincent Drouillon, Marcia Eisenberg, Howard Engler, Rama Ghatti, Prashant Gupta, Susan Hicks, Jake Humphrey, Lax Iyer, Lisa Pfefferle, Manoj Jain, Matthew Robinson, Mohan Kolli, Brian Krueger, Tim Kuphal, Stanley Letovsky, Michael Levandoski, Craig Lukasik, Jonathan Meltzer, Brian Norvell, Mindy Nye, Scott Parker, Christos Petropoulos, John Pruitt, Steven Ragan, Scott Ryan, Mike Sapeta, Jana Schroth, Suresh Babu Selvaraju, Goran Stevovic, Amanda Suchanek, Andrea Throop, Lyndon Tilson, Thomas Urban, Joe Voshell, Kimberly Wagner, Jonathan Williams, Mary Williamson, Qian Zeng, Tricia Zwiefelhofer, Clinton R. Paden, Duncan MacCannell |
| EPI_ISL_3326855 | TGen North | TGen North | Jolene Bowers, Heather Centner, Brett Van Tassel, Chris French, Hayley Yaglom, Darrin Lemmer, Dave Engelthaler |
| EPI_ISL_3326856, EPI_ISL_3326857, EPI_ISL_3326858, EPI_ISL_3326859, EPI_ISL_3326861, EPI_ISL_3326862, EPI_ISL_3326863, EPI_ISL_3326864, EPI_ISL_3326865, EPI_ISL_3326866, EPI_ISL_3326867, EPI_ISL_3326869, EPI_ISL_3326871, EPI_ISL_3326872, EPI_ISL_3326873, EPI_ISL_3326874, EPI_ISL_3326875, EPI_ISL_3326876, EPI_ISL_3326877, EPI_ISL_3326878, EPI_ISL_3326879, EPI_ISL_3326880, EPI_ISL_3326881, EPI_ISL_3326882, EPI_ISL_3326883, EPI_ISL_3326884, EPI_ISL_3326885, EPI_ISL_3326886, EPI_ISL_3326887, EPI_ISL_3326888, EPI_ISL_3326889, EPI_ISL_3326890, EPI_ISL_3326891, EPI_ISL_3326892, EPI_ISL_3326894, EPI_ISL_3326895, EPI_ISL_3326896, EPI_ISL_3326897, EPI_ISL_3326900, EPI_ISL_3326901, EPI_ISL_3326902, EPI_ISL_3326903, EPI_ISL_3326904, EPI_ISL_3326905, EPI_ISL_3326906, EPI_ISL_3326908, EPI_ISL_3326909, EPI_ISL_3326910, EPI_ISL_3326912, EPI_ISL_3326913, EPI_ISL_3326914, EPI_ISL_3326915, EPI_ISL_3326916, EPI_ISL_3326917, EPI_ISL_3326918, EPI_ISL_3326920, EPI_ISL_3326921, EPI_ISL_3326922, EPI_ISL_3326923, EPI_ISL_3326924, EPI_ISL_3326925, EPI_ISL_3326926, EPI_ISL_3326927, EPI_ISL_3326928, EPI_ISL_3326929, EPI_ISL_3326930, EPI_ISL_3326931, EPI_ISL_3326932, EPI_ISL_3326933, EPI_ISL_3326934, EPI_ISL_3326935, EPI_ISL_3326936, EPI_ISL_3326937, EPI_ISL_3326938, EPI_ISL_3326939, EPI_ISL_3326940, EPI_ISL_3326941, EPI_ISL_3326943, EPI_ISL_3326944, EPI_ISL_3326946, EPI_ISL_3326949, EPI_ISL_3326950, EPI_ISL_3326951, EPI_ISL_3326952, EPI_ISL_3326953, EPI_ISL_3326954, EPI_ISL_3326955, EPI_ISL_3326956, EPI_ISL_3326957, EPI_ISL_3326959, EPI_ISL_3326960, EPI_ISL_3326961, EPI_ISL_3326962, EPI_ISL_3326963, EPI_ISL_3326964, EPI_ISL_3326965, EPI_ISL_3326966, EPI_ISL_3326967, EPI_ISL_3326968, EPI_ISL_3326969, EPI_ISL_3326970, EPI_ISL_3326971, EPI_ISL_3326972, EPI_ISL_3326973, EPI_ISL_3326974, EPI_ISL_3326975, EPI_ISL_3326976, EPI_ISL_3326977, EPI_ISL_3326978, EPI_ISL_3326979, EPI_ISL_3326980, EPI_ISL_3326981, EPI_ISL_3326982, EPI_ISL_3326983, EPI_ISL_3326984, EPI_ISL_3326985, EPI_ISL_3326986, EPI_ISL_3326987, EPI_ISL_3326988, EPI_ISL_3326989, EPI_ISL_3326990, EPI_ISL_3326991, EPI_ISL_3326992, EPI_ISL_3326993, EPI_ISL_3326994, EPI_ISL_3326995, EPI_ISL_3326996, EPI_ISL_3326997, EPI_ISL_3326998, EPI_ISL_3326999, EPI_ISL_3327000, EPI_ISL_3327001, EPI_ISL_3327002, EPI_ISL_3327003, EPI_ISL_3327004, EPI_ISL_3327005, EPI_ISL_3327006, EPI_ISL_3327008, EPI_ISL_3327009, EPI_ISL_3327010, EPI_ISL_3327011, EPI_ISL_3327012, EPI_ISL_3327013, EPI_ISL_3327014, EPI_ISL_3327015, EPI_ISL_3327016, EPI_ISL_3327017, EPI_ISL_3327018, EPI_ISL_3327019, EPI_ISL_3327020, EPI_ISL_3327021, EPI_ISL_3327022, EPI_ISL_3327023, EPI_ISL_3327024, | | | |

| | | | |
|---|---|---|---|
| EPI_ISL_3327025, EPI_ISL_3327026, EPI_ISL_3327027, EPI_ISL_3327028, EPI_ISL_3327029, EPI_ISL_3327030, EPI_ISL_3327031, EPI_ISL_3327032, EPI_ISL_3327033, EPI_ISL_3327034, EPI_ISL_3327035, EPI_ISL_3327036, EPI_ISL_3327037, EPI_ISL_3327038, EPI_ISL_3327039, EPI_ISL_3327040, EPI_ISL_3327041, EPI_ISL_3327042, EPI_ISL_3327043, EPI_ISL_3327044, EPI_ISL_3327045, EPI_ISL_3327046, EPI_ISL_3327048, EPI_ISL_3327049, EPI_ISL_3327050, EPI_ISL_3327051, EPI_ISL_3327052, EPI_ISL_3327053, EPI_ISL_3327054, EPI_ISL_3327055, EPI_ISL_3327056, EPI_ISL_3327057, EPI_ISL_3327058, EPI_ISL_3327059, EPI_ISL_3327060, EPI_ISL_3327061, EPI_ISL_3327062, EPI_ISL_3327063, EPI_ISL_3327064, EPI_ISL_3327065, EPI_ISL_3327066, EPI_ISL_3327067, EPI_ISL_3327068, EPI_ISL_3327069, EPI_ISL_3327070, EPI_ISL_3327071, EPI_ISL_3327072, EPI_ISL_3327074, EPI_ISL_3327076, EPI_ISL_3327077, EPI_ISL_3327078, EPI_ISL_3327080, EPI_ISL_3327082, EPI_ISL_3327083, EPI_ISL_3327084, EPI_ISL_3327085, EPI_ISL_3327086, EPI_ISL_3327087, EPI_ISL_3327088, EPI_ISL_3327089, EPI_ISL_3327090, EPI_ISL_3327092, EPI_ISL_3327093, EPI_ISL_3327094, EPI_ISL_3327095, EPI_ISL_3327096, EPI_ISL_3327097, EPI_ISL_3327098, EPI_ISL_3327100, EPI_ISL_3327101, EPI_ISL_3327102, EPI_ISL_3327103, EPI_ISL_3327104, EPI_ISL_3327105, EPI_ISL_3327107, EPI_ISL_3327108, EPI_ISL_3327109, EPI_ISL_3327110, EPI_ISL_3327111, EPI_ISL_3327112, EPI_ISL_3327113, EPI_ISL_3327115, EPI_ISL_3327116, EPI_ISL_3327117, EPI_ISL_3327118, EPI_ISL_3327119, EPI_ISL_3327120, EPI_ISL_3327121, EPI_ISL_3327122, EPI_ISL_3327123, EPI_ISL_3327124, EPI_ISL_3327125, EPI_ISL_3327126, EPI_ISL_3327127, EPI_ISL_3327128, EPI_ISL_3327129, EPI_ISL_3327130, EPI_ISL_3327132, EPI_ISL_3327133, EPI_ISL_3327134, EPI_ISL_3327135, EPI_ISL_3327136, EPI_ISL_3327137, EPI_ISL_3327139, EPI_ISL_3327140, EPI_ISL_3327141, EPI_ISL_3327142, EPI_ISL_3327143, EPI_ISL_3327144, EPI_ISL_3327145, EPI_ISL_3327146, EPI_ISL_3327147, EPI_ISL_3327148, EPI_ISL_3327149, EPI_ISL_3327150, EPI_ISL_3327151, EPI_ISL_3327152, EPI_ISL_3327153, EPI_ISL_3327154, EPI_ISL_3327155, EPI_ISL_3327156, EPI_ISL_3327157, EPI_ISL_3327158, EPI_ISL_3327159, EPI_ISL_3327160, EPI_ISL_3327161, EPI_ISL_3327162, EPI_ISL_3327163, EPI_ISL_3327164, EPI_ISL_3327165, EPI_ISL_3327166, EPI_ISL_3327167, EPI_ISL_3327168, EPI_ISL_3327171, EPI_ISL_3327172, EPI_ISL_3327173, EPI_ISL_3327174, EPI_ISL_3327175, EPI_ISL_3327176, EPI_ISL_3327178, EPI_ISL_3327179, EPI_ISL_3327180, EPI_ISL_3327181, EPI_ISL_3327182, EPI_ISL_3327184, EPI_ISL_3327185, EPI_ISL_3327187, EPI_ISL_3327188, EPI_ISL_3327189, EPI_ISL_3327190, EPI_ISL_3327191, EPI_ISL_3327192, EPI_ISL_3327193, EPI_ISL_3327194, EPI_ISL_3327195, EPI_ISL_3327196, EPI_ISL_3327197, EPI_ISL_3327198, EPI_ISL_3327199, EPI_ISL_3327200, EPI_ISL_3327201, EPI_ISL_3327202, EPI_ISL_3327203, EPI_ISL_3327204, EPI_ISL_3327205, EPI_ISL_3327206, EPI_ISL_3327208, EPI_ISL_3327209, EPI_ISL_3327210, EPI_ISL_3327211, EPI_ISL_3327212, EPI_ISL_3327213, EPI_ISL_3327214, EPI_ISL_3327215, EPI_ISL_3327217, EPI_ISL_3327218, EPI_ISL_3327219, EPI_ISL_3327220, EPI_ISL_3327221, EPI_ISL_3327222, EPI_ISL_3327223, EPI_ISL_3327224, EPI_ISL_3327225, EPI_ISL_3327226, EPI_ISL_3327227, EPI_ISL_3327228, EPI_ISL_3327229, EPI_ISL_3327230, EPI_ISL_3327231, EPI_ISL_3327233, EPI_ISL_3327234, EPI_ISL_3327235, EPI_ISL_3327236, EPI_ISL_3327237, EPI_ISL_3327238, EPI_ISL_3327239, EPI_ISL_3327240, EPI_ISL_3327241, EPI_ISL_3327242, EPI_ISL_3327243, EPI_ISL_3327244, EPI_ISL_3327245, EPI_ISL_3327246, EPI_ISL_3327247, EPI_ISL_3327248, EPI_ISL_3327249, EPI_ISL_3327250, EPI_ISL_3327251, EPI_ISL_3327252, EPI_ISL_3327253, EPI_ISL_3327254, EPI_ISL_3327255, EPI_ISL_3327256, EPI_ISL_3327257, EPI_ISL_3327258, EPI_ISL_3327259, EPI_ISL_3327260, EPI_ISL_3327261, EPI_ISL_3327262, EPI_ISL_3327263, EPI_ISL_3327264, EPI_ISL_3327265, EPI_ISL_3327266, EPI_ISL_3327267, EPI_ISL_3327269, EPI_ISL_3327271, EPI_ISL_3327272, EPI_ISL_3327273, EPI_ISL_3327274, EPI_ISL_3327276, EPI_ISL_3327277, EPI_ISL_3327278, EPI_ISL_3327279, EPI_ISL_3327280, EPI_ISL_3327281, EPI_ISL_3327282, EPI_ISL_3327284, EPI_ISL_3327285, EPI_ISL_3327289, EPI_ISL_3327290, EPI_ISL_3327291, EPI_ISL_3327293, EPI_ISL_3327294, EPI_ISL_3327295, EPI_ISL_3327296, EPI_ISL_3327297, EPI_ISL_3327298, EPI_ISL_3327299, EPI_ISL_3327300, EPI_ISL_3327301, EPI_ISL_3327302, EPI_ISL_3327303, EPI_ISL_3327304, EPI_ISL_3327305, EPI_ISL_3327306, EPI_ISL_3327307, EPI_ISL_3327308, EPI_ISL_3327309, EPI_ISL_3327310, EPI_ISL_3327311, EPI_ISL_3327312, EPI_ISL_3327313, EPI_ISL_3327314, EPI_ISL_3327315, EPI_ISL_3327317, EPI_ISL_3327318, EPI_ISL_3327319, EPI_ISL_3327320, EPI_ISL_3327321, EPI_ISL_3327322, EPI_ISL_3327323, EPI_ISL_3327324, EPI_ISL_3327327, EPI_ISL_3327328, EPI_ISL_3327329, EPI_ISL_3327330, EPI_ISL_3327331, EPI_ISL_3327332, EPI_ISL_3327333, EPI_ISL_3327334, EPI_ISL_3327335, EPI_ISL_3327336, EPI_ISL_3327337, EPI_ISL_3327338, EPI_ISL_3327339, EPI_ISL_3327340, EPI_ISL_3327341, EPI_ISL_3327342, EPI_ISL_3327343, EPI_ISL_3327344, EPI_ISL_3327345, EPI_ISL_3327346, EPI_ISL_3327347, EPI_ISL_3327348, EPI_ISL_3327349, EPI_ISL_3327350, EPI_ISL_3327351, EPI_ISL_3327352, EPI_ISL_3327353, EPI_ISL_3327354, EPI_ISL_3327355, EPI_ISL_3327356, EPI_ISL_3327357, EPI_ISL_3327358, EPI_ISL_3327359, EPI_ISL_3327360, EPI_ISL_3327361, EPI_ISL_3327362, EPI_ISL_3327363, EPI_ISL_3327364, EPI_ISL_3327365, EPI_ISL_3327367, EPI_ISL_3327368, EPI_ISL_3327369, EPI_ISL_3327370, EPI_ISL_3327372, EPI_ISL_3327373, EPI_ISL_3327374, EPI_ISL_3327375, EPI_ISL_3327377, EPI_ISL_3327378, EPI_ISL_3327379, EPI_ISL_3327380, EPI_ISL_3327381, EPI_ISL_3327382, EPI_ISL_3327383, EPI_ISL_3327384, EPI_ISL_3327386, EPI_ISL_3327387, EPI_ISL_3327388, EPI_ISL_3327389, EPI_ISL_3327390, EPI_ISL_3327391, EPI_ISL_3327392, EPI_ISL_3327393, EPI_ISL_3327394, EPI_ISL_3327395, EPI_ISL_3327396, EPI_ISL_3327397, EPI_ISL_3327398, EPI_ISL_3327399, EPI_ISL_3327400, EPI_ISL_3327402, EPI_ISL_3327403, EPI_ISL_3327404, EPI_ISL_3327405, EPI_ISL_3327406, EPI_ISL_3327407, EPI_ISL_3327408, EPI_ISL_3327409, EPI_ISL_3327410, EPI_ISL_3327411, EPI_ISL_3327412, EPI_ISL_3327413, EPI_ISL_3327414, EPI_ISL_3327415, EPI_ISL_3327416, EPI_ISL_3327417, EPI_ISL_3327418, EPI_ISL_3327419, EPI_ISL_3327420, EPI_ISL_3327421, EPI_ISL_3327422, EPI_ISL_3327423, EPI_ISL_3327424, EPI_ISL_3327425, EPI_ISL_3327426, EPI_ISL_3327428, EPI_ISL_3327429, EPI_ISL_3327430, EPI_ISL_3327431, EPI_ISL_3327432, EPI_ISL_3327433, EPI_ISL_3327434, EPI_ISL_3327435, EPI_ISL_3327436, EPI_ISL_3327437, EPI_ISL_3327438, EPI_ISL_3327439, EPI_ISL_3327440, EPI_ISL_3327441, EPI_ISL_3327442, EPI_ISL_3327443, EPI_ISL_3327444, EPI_ISL_3327445, EPI_ISL_3327446, EPI_ISL_3327447, EPI_ISL_3327448, EPI_ISL_3327449, EPI_ISL_3327451, EPI_ISL_3327452, EPI_ISL_3327453, EPI_ISL_3327454, EPI_ISL_3327455, EPI_ISL_3327456, EPI_ISL_3327457, EPI_ISL_3327458, EPI_ISL_3327459, EPI_ISL_3327460, EPI_ISL_3327461, EPI_ISL_3327462, EPI_ISL_3327463, EPI_ISL_3327464, EPI_ISL_3327465, EPI_ISL_3327466, EPI_ISL_3327467, EPI_ISL_3327468, EPI_ISL_3327469, EPI_ISL_3327470, EPI_ISL_3327471, EPI_ISL_3327472, EPI_ISL_3327473, EPI_ISL_3327474, EPI_ISL_3327475, EPI_ISL_3327476, EPI_ISL_3327477, EPI_ISL_3327478, EPI_ISL_3327479, EPI_ISL_3327480, EPI_ISL_3327482, EPI_ISL_3327483, EPI_ISL_3327484, EPI_ISL_3327485, EPI_ISL_3327486, EPI_ISL_3327487, EPI_ISL_3327488, EPI_ISL_3327489, EPI_ISL_3327490, EPI_ISL_3327491, EPI_ISL_3327492, EPI_ISL_3327493, EPI_ISL_3327494, EPI_ISL_3327495, EPI_ISL_3327498, EPI_ISL_3327500, EPI_ISL_3327501, EPI_ISL_3327502, EPI_ISL_3327503, EPI_ISL_3327504, EPI_ISL_3327505, EPI_ISL_3327506, EPI_ISL_3327507, EPI_ISL_3327508, EPI_ISL_3327509, EPI_ISL_3327510, EPI_ISL_3327512, EPI_ISL_3327513, EPI_ISL_3327514, EPI_ISL_3327516, EPI_ISL_3327517, EPI_ISL_3327518, EPI_ISL_3327519, EPI_ISL_3327520, EPI_ISL_3327521, EPI_ISL_3327522, EPI_ISL_3327523, EPI_ISL_3327524, EPI_ISL_3327525, EPI_ISL_3327526, EPI_ISL_3327527, EPI_ISL_3327528, EPI_ISL_3327529, EPI_ISL_3327530, EPI_ISL_3327531, EPI_ISL_3327532, EPI_ISL_3327533, EPI_ISL_3327534, EPI_ISL_3327536, EPI_ISL_3327537, EPI_ISL_3327538, EPI_ISL_3327539, EPI_ISL_3327540, EPI_ISL_3327541, EPI_ISL_3327542, EPI_ISL_3327543, EPI_ISL_3327544, EPI_ISL_3327545, EPI_ISL_3327546, EPI_ISL_3327547, EPI_ISL_3327548, EPI_ISL_3327549, EPI_ISL_3327550, EPI_ISL_3327551, EPI_ISL_3327552, EPI_ISL_3327553, EPI_ISL_3327554, EPI_ISL_3327556, EPI_ISL_3327557, EPI_ISL_3327558, EPI_ISL_3327559, EPI_ISL_3327560, EPI_ISL_3327561, EPI_ISL_3327562, EPI_ISL_3327563, EPI_ISL_3327564, EPI_ISL_3327565, EPI_ISL_3327566, EPI_ISL_3327567, EPI_ISL_3327568, EPI_ISL_3327569, EPI_ISL_3327570, EPI_ISL_3327571, EPI_ISL_3327572, EPI_ISL_3327573, EPI_ISL_3327574, EPI_ISL_3327575, EPI_ISL_3327576, EPI_ISL_3327577, EPI_ISL_3327578, EPI_ISL_3327579, EPI_ISL_3327580, EPI_ISL_3327581, EPI_ISL_3327582, EPI_ISL_3327583, EPI_ISL_3327584, EPI_ISL_3327585, EPI_ISL_3327586, EPI_ISL_3327587, EPI_ISL_3327589, EPI_ISL_3327590, EPI_ISL_3327591, EPI_ISL_3327592, EPI_ISL_3327593, EPI_ISL_3327594, EPI_ISL_3327595, EPI_ISL_3327596, EPI_ISL_3327597, EPI_ISL_3327598, EPI_ISL_3327599, EPI_ISL_3327600, EPI_ISL_3327601, EPI_ISL_3327602, EPI_ISL_3327603, EPI_ISL_3327604, EPI_ISL_3327605, EPI_ISL_3327606, EPI_ISL_3327607, EPI_ISL_3327608, EPI_ISL_3327609, EPI_ISL_3327610, EPI_ISL_3327611, EPI_ISL_3327612, EPI_ISL_3327613, EPI_ISL_3327614, EPI_ISL_3327615, EPI_ISL_3327616, EPI_ISL_3327617, EPI_ISL_3327618, EPI_ISL_3327619, EPI_ISL_3327620, EPI_ISL_3327621, EPI_ISL_3327622, EPI_ISL_3327623, EPI_ISL_3327624, EPI_ISL_3327625, EPI_ISL_3327626, EPI_ISL_3327627, EPI_ISL_3327628, EPI_ISL_3327629, EPI_ISL_3327630, EPI_ISL_3327631, EPI_ISL_3327632, EPI_ISL_3327633, EPI_ISL_3327634, EPI_ISL_3327635, EPI_ISL_3327636, EPI_ISL_3327637, EPI_ISL_3327638, EPI_ISL_3327639, EPI_ISL_3327640, EPI_ISL_3327641, EPI_ISL_3327642, EPI_ISL_3327644, EPI_ISL_3327646, EPI_ISL_3327647, EPI_ISL_3327648, EPI_ISL_3327649, EPI_ISL_3327650, EPI_ISL_3327652, EPI_ISL_3327653, EPI_ISL_3327654, EPI_ISL_3327655, EPI_ISL_3327656, EPI_ISL_3327657, EPI_ISL_3327658, EPI_ISL_3327659, EPI_ISL_3327660, EPI_ISL_3327661, EPI_ISL_3327664, EPI_ISL_3327665, EPI_ISL_3327666, EPI_ISL_3327667, EPI_ISL_3327668, EPI_ISL_3327669, EPI_ISL_3327670, EPI_ISL_3327671, EPI_ISL_3327672, EPI_ISL_3327673, EPI_ISL_3327674, EPI_ISL_3327675, EPI_ISL_3327676, EPI_ISL_3327677, EPI_ISL_3327678, EPI_ISL_3327679, EPI_ISL_3327680, EPI_ISL_3327681, EPI_ISL_3327682, EPI_ISL_3327683, EPI_ISL_3327684, EPI_ISL_3327685, EPI_ISL_3327686, EPI_ISL_3327687, EPI_ISL_3327688, EPI_ISL_3327689, EPI_ISL_3327690, EPI_ISL_3327691, EPI_ISL_3327693, EPI_ISL_3327694, EPI_ISL_3327695, EPI_ISL_3327696, EPI_ISL_3327697, EPI_ISL_3327698, EPI_ISL_3327699, EPI_ISL_3327700, EPI_ISL_3327701, EPI_ISL_3327703, EPI_ISL_3327704, EPI_ISL_3327705, EPI_ISL_3327706, EPI_ISL_3327707, EPI_ISL_3327708, EPI_ISL_3327709, EPI_ISL_3327710, EPI_ISL_3327711, EPI_ISL_3327712, EPI_ISL_3327713, EPI_ISL_3327714, EPI_ISL_3327715, EPI_ISL_3327716, EPI_ISL_3327717, EPI_ISL_3327718, EPI_ISL_3327719, EPI_ISL_3327720, EPI_ISL_3327721, EPI_ISL_3327722, EPI_ISL_3327723, EPI_ISL_3327724, EPI_ISL_3327725, EPI_ISL_3327726, EPI_ISL_3327727, EPI_ISL_3327728, EPI_ISL_3327729, EPI_ISL_3327730, EPI_ISL_3327731, EPI_ISL_3327732, EPI_ISL_3327733, EPI_ISL_3327734, EPI_ISL_3327735, EPI_ISL_3327736, EPI_ISL_3327737, EPI_ISL_3327738, EPI_ISL_3327739, EPI_ISL_3327740, EPI_ISL_3327742, EPI_ISL_3327743, EPI_ISL_3327744, EPI_ISL_3327745, EPI_ISL_3327747, EPI_ISL_3327748, EPI_ISL_3327749, EPI_ISL_3327750, EPI_ISL_3327751, EPI_ISL_3327752, EPI_ISL_3327753, EPI_ISL_3327755, EPI_ISL_3327756, EPI_ISL_3327757, EPI_ISL_3327758, EPI_ISL_3327759, EPI_ISL_3327760, EPI_ISL_3327761, EPI_ISL_3327762, EPI_ISL_3327764, EPI_ISL_3327765, EPI_ISL_3327766, EPI_ISL_3327767, EPI_ISL_3327768, EPI_ISL_3327769, EPI_ISL_3327770, EPI_ISL_3327771, EPI_ISL_3327772, EPI_ISL_3327773, EPI_ISL_3327774, EPI_ISL_3327775, EPI_ISL_3327776, EPI_ISL_3327777, EPI_ISL_3327778, EPI_ISL_3327779, EPI_ISL_3327780, EPI_ISL_3327781, EPI_ISL_3327783, EPI_ISL_3327784, EPI_ISL_3327786, EPI_ISL_3327787, EPI_ISL_3327788, EPI_ISL_3327789, EPI_ISL_3327790, EPI_ISL_3327791, EPI_ISL_3327793, EPI_ISL_3327794, EPI_ISL_3327795, EPI_ISL_3327796, EPI_ISL_3327798, EPI_ISL_3327799, EPI_ISL_3327800, EPI_ISL_3327801, EPI_ISL_3327802, EPI_ISL_3327803, EPI_ISL_3327804, EPI_ISL_3327805, EPI_ISL_3327806, EPI_ISL_3327807, EPI_ISL_3327809, EPI_ISL_3327810, EPI_ISL_3327813, EPI_ISL_3327814, EPI_ISL_3327815, EPI_ISL_3327816, EPI_ISL_3327817, EPI_ISL_3327819, EPI_ISL_3327820, EPI_ISL_3327822, EPI_ISL_3327823, EPI_ISL_3327824, EPI_ISL_3327825, EPI_ISL_3327826, EPI_ISL_3327827, EPI_ISL_3327828, EPI_ISL_3327829, EPI_ISL_3327830, EPI_ISL_3327831, EPI_ISL_3327832, EPI_ISL_3327833, EPI_ISL_3327834, EPI_ISL_3327835, EPI_ISL_3327836, EPI_ISL_3327837, EPI_ISL_3327838, EPI_ISL_3327839, EPI_ISL_3327840, EPI_ISL_3327841, EPI_ISL_3327842, EPI_ISL_3327843, EPI_ISL_3327844, EPI_ISL_3327845, EPI_ISL_3327846, EPI_ISL_3327847, EPI_ISL_3327848, EPI_ISL_3327849, EPI_ISL_3327850, EPI_ISL_3327851, EPI_ISL_3327852, EPI_ISL_3327853, EPI_ISL_3327854, EPI_ISL_3327855, EPI_ISL_3327856, EPI_ISL_3327857, EPI_ISL_3327858, EPI_ISL_3327859, EPI_ISL_3327860, EPI_ISL_3327861, EPI_ISL_3327862, EPI_ISL_3327863, EPI_ISL_3327864, EPI_ISL_3327865, EPI_ISL_3327866, EPI_ISL_3327867, EPI_ISL_3327868, EPI_ISL_3327869, EPI_ISL_3327870, EPI_ISL_3327871, EPI_ISL_3327872, EPI_ISL_3327873, EPI_ISL_3327874, EPI_ISL_3327875 | | | |
| see above | Laboratory Corporation of America | Centers for Disease Control and Prevention Division of Viral Diseases, Pathogen Discovery | Dakota Howard, Dhwani Batra, Peter W. Cook, Kara Moser, Adrian Paskey, Jason Caravas, Benjamin Rambo-Martin, Shatavia Morrison, Christopher Gulvick, Scott Sammons, Yvette Unoarumhi, Darlene Wagner, Matthew Schmerer, Minoo Agarwal, Eyad Almasri, Debbie Boles, Ayla Burns, Nuthawin Charoensri, Oren Cohen, Susan Countryman, Mary Ann Cristobal, Bobbi Croy, Suzanne Dale, Hrushikesh Deshmukh, Amanda Douglas, Vincent Drouillon, Marcia Eisenberg, Howard Engler, Rama Ghatti, Prashant Gupta, Susan Hicks, Jake Humphrey, Lax Iyer, Lisa Pfefferle, Manoj Jain, Matthew Robinson, Mohan Kolli, Brian Krueger, Tim Kuphal, Stanley Letovsky, Michael Levandoski, Craig Lukasik, Jonathan Meltzer, Brian Norvell, Mindy Nye, Scott Parker, Christos Petropoulos, John Pruitt, Steven Ragan, Scott Ryan, Mike Sapeta, Jana Schroth, Suresh Babu Selvaraju, Goran Stevovic, Amanda Suchanek, Andrea Throop, Lyndon Tilson, Thomas Urban, Joe Voshell, Kimberly Wagner, Jonathan Williams, Mary Williamson, Qian Zeng, Tricia Zwiefelhofer, Clinton R. Paden, Duncan MacCannell |
| EPI_ISL_3327876 | NJDOH, Public Health and Environmental Laboratories | NJ_PHEL | Lindsey Bodnar, Shiv K. Verma, Ryan Pachucki, Chelsea San Filippo, Jacquelyn Deverell, Dana Woell, Allison Roder, Byeong Jeong |
| EPI_ISL_3327877 | Laboratory Corporation of America | Centers for Disease Control and Prevention Division of Viral Diseases, Pathogen Discovery | Dakota Howard, Dhwani Batra, Peter W. Cook, Kara Moser, Adrian Paskey, Jason Caravas, Benjamin Rambo-Martin, Shatavia Morrison, Christopher Gulvick, Scott Sammons, Yvette Unoarumhi, Darlene Wagner, Matthew Schmerer, Minoo Agarwal, Eyad Almasri, Debbie Boles, Ayla Burns, Nuthawin Charoensri, Oren Cohen, Susan Countryman, Mary Ann Cristobal, Bobbi Croy, Suzanne Dale, Hrushikesh Deshmukh, Amanda Douglas, Vincent Drouillon, Marcia Eisenberg, Howard Engler, Rama Ghatti, Prashant Gupta, Susan Hicks, Jake Humphrey, Lax Iyer, Lisa Pfefferle, Manoj Jain, Matthew Robinson, Mohan Kolli, Brian Krueger, Tim Kuphal, Stanley Letovsky, Michael Levandoski, Craig Lukasik, Jonathan Meltzer, Brian Norvell, Mindy Nye, Scott Parker, Christos Petropoulos, John Pruitt, Steven Ragan, Scott Ryan, Mike Sapeta, Jana Schroth, Suresh Babu Selvaraju, Goran Stevovic, Amanda Suchanek, Andrea Throop, Lyndon Tilson, Thomas Urban, Joe Voshell, Kimberly Wagner, Jonathan Williams, Mary Williamson, Qian Zeng, Tricia Zwiefelhofer, Clinton R. Paden, Duncan MacCannell |
| EPI_ISL_3327878, EPI_ISL_3327879 | NJDOH, Public Health and Environmental Laboratories | NJ_PHEL | Lindsey Bodnar, Shiv K. Verma, Ryan Pachucki, Chelsea San Filippo, Jacquelyn Deverell, Dana Woell, Allison Roder, Byeong Jeong |
| EPI_ISL_3327880 | Laboratory Corporation of America | Centers for Disease Control and Prevention Division of Viral Diseases, Pathogen Discovery | Dakota Howard, Dhwani Batra, Peter W. Cook, Kara Moser, Adrian Paskey, Jason Caravas, Benjamin Rambo-Martin, Shatavia Morrison, Christopher Gulvick, Scott Sammons, Yvette Unoarumhi, Darlene Wagner, Matthew Schmerer, Minoo Agarwal, Eyad Almasri, Debbie Boles, Ayla Burns, Nuthawin Charoensri, Oren Cohen, Susan Countryman, Mary Ann Cristobal, Bobbi Croy, Suzanne Dale, Hrushikesh Deshmukh, Amanda Douglas, Vincent Drouillon, Marcia Eisenberg, Howard Engler, Rama Ghatti, Prashant Gupta, Susan Hicks, Jake Humphrey, Lax Iyer, Lisa Pfefferle, Manoj Jain, Matthew Robinson, Mohan Kolli, Brian Krueger, Tim Kuphal, Stanley Letovsky, Michael Levandoski, Craig Lukasik, Jonathan Meltzer, Brian Norvell, Mindy Nye, Scott Parker, Christos Petropoulos, John Pruitt, Steven Ragan, Scott Ryan, Mike Sapeta, Jana Schroth, Suresh Babu Selvaraju, Goran Stevovic, Amanda Suchanek, Andrea Throop, Lyndon Tilson, Thomas Urban, Joe Voshell, Kimberly Wagner, Jonathan Williams, Mary Williamson, Qian Zeng, Tricia Zwiefelhofer, Clinton R. Paden, Duncan MacCannell |
| EPI_ISL_3327881 | NJDOH, Public Health and Environmental Laboratories | NJ_PHEL | Lindsey Bodnar, Shiv K. Verma, Ryan Pachucki, Chelsea San Filippo, Jacquelyn Deverell, Dana Woell, Allison Roder, Byeong Jeong |
| EPI_ISL_3327882 | Laboratory Corporation of America | Centers for Disease Control and Prevention Division of Viral Diseases, Pathogen Discovery | Dakota Howard, Dhwani Batra, Peter W. Cook, Kara Moser, Adrian Paskey, Jason Caravas, Benjamin Rambo-Martin, Shatavia Morrison, Christopher Gulvick, Scott Sammons, Yvette Unoarumhi, Darlene Wagner, Matthew Schmerer, Minoo Agarwal, Eyad Almasri, Debbie Boles, Ayla Burns, Nuthawin Charoensri, Oren Cohen, Susan Countryman, Mary Ann Cristobal, Bobbi Croy, Suzanne Dale, Hrushikesh Deshmukh, Amanda Douglas, Vincent Drouillon, Marcia Eisenberg, Howard Engler, Rama Ghatti, Prashant Gupta, Susan Hicks, Jake Humphrey, Lax Iyer, Lisa Pfefferle, Manoj Jain, Matthew Robinson, Mohan Kolli, Brian Krueger, Tim Kuphal, Stanley Letovsky, Michael Levandoski, Craig Lukasik, Jonathan Meltzer, Brian Norvell, Mindy Nye, Scott Parker, Christos Petropoulos, John Pruitt, Steven Ragan, Scott Ryan, Mike Sapeta, Jana Schroth, Suresh Babu Selvaraju, Goran Stevovic, Amanda Suchanek, Andrea Throop, Lyndon Tilson, Thomas Urban, Joe Voshell, Kimberly Wagner, Jonathan Williams, Mary Williamson, Qian Zeng, Tricia Zwiefelhofer, Clinton R. Paden, Duncan MacCannell |
| EPI_ISL_3327883 | NJDOH, Public Health and Environmental Laboratories | NJ_PHEL | Lindsey Bodnar, Shiv K. Verma, Ryan Pachucki, Chelsea San Filippo, Jacquelyn Deverell, Dana Woell, Allison Roder, Byeong Jeong |
| EPI_ISL_3327884, EPI_ISL_3327885 | Laboratory Corporation of America | Centers for Disease Control and Prevention Division of Viral Diseases, Pathogen Discovery | Dakota Howard, Dhwani Batra, Peter W. Cook, Kara Moser, Adrian Paskey, Jason Caravas, Benjamin Rambo-Martin, Shatavia Morrison, Christopher Gulvick, Scott Sammons, Yvette Unoarumhi, Darlene Wagner, Matthew Schmerer, Minoo Agarwal, Eyad Almasri, Debbie Boles, Ayla Burns, Nuthawin Charoensri, Oren Cohen, Susan Countryman, Mary Ann Cristobal, Bobbi Croy, Suzanne Dale, Hrushikesh Deshmukh, Amanda Douglas, Vincent Drouillon, Marcia Eisenberg, Howard Engler, Rama Ghatti, Prashant Gupta, Susan Hicks, Jake Humphrey, Lax Iyer, Lisa Pfefferle, Manoj Jain, Matthew Robinson, Mohan Kolli, Brian Krueger, Tim Kuphal, Stanley Letovsky, Michael Levandoski, Craig Lukasik, Jonathan Meltzer, Brian Norvell, Mindy Nye, Scott Parker, Christos Petropoulos, John Pruitt, Steven Ragan, Scott Ryan, Mike Sapeta, Jana Schroth, Suresh Babu Selvaraju, Goran Stevovic, Amanda Suchanek, Andrea Throop, Lyndon Tilson, Thomas Urban, Joe Voshell, Kimberly Wagner, Jonathan Williams, Mary Williamson, Qian Zeng, Tricia Zwiefelhofer, Clinton R. Paden, Duncan MacCannell |
| EPI_ISL_3327886, EPI_ISL_3327888, EPI_ISL_3327889 | NJDOH, Public Health and Environmental Laboratories | NJ_PHEL | Lindsey Bodnar, Shiv K. Verma, Ryan Pachucki, Chelsea San Filippo, Jacquelyn Deverell, Dana Woell, Allison Roder, Byeong Jeong |
| EPI_ISL_3327890 | Laboratory Corporation of America | Centers for Disease Control and Prevention Division of Viral Diseases, Pathogen Discovery | Dakota Howard, Dhwani Batra, Peter W. Cook, Kara Moser, Adrian Paskey, Jason Caravas, Benjamin Rambo-Martin, Shatavia Morrison, Christopher Gulvick, Scott Sammons, Yvette Unoarumhi, Darlene Wagner, Matthew Schmerer, Minoo Agarwal, Eyad Almasri, Debbie Boles, Ayla Burns, Nuthawin Charoensri, Oren Cohen, Susan Countryman, Mary Ann Cristobal, Bobbi Croy, Suzanne Dale, Hrushikesh Deshmukh, Amanda Douglas, Vincent Drouillon, Marcia Eisenberg, Howard Engler, Rama Ghatti, Prashant Gupta, Susan Hicks, Jake Humphrey, Lax Iyer, Lisa Pfefferle, Manoj Jain, Matthew Robinson, Mohan Kolli, Brian Krueger, Tim Kuphal, Stanley Letovsky, Michael Levandoski, Craig Lukasik, Jonathan Meltzer, Brian Norvell, Mindy Nye, Scott Parker, Christos Petropoulos, John Pruitt, Steven Ragan, Scott Ryan, Mike Sapeta, Jana Schroth, Suresh Babu Selvaraju, Goran Stevovic, Amanda Suchanek, Andrea Throop, Lyndon Tilson, Thomas Urban, Joe Voshell, Kimberly Wagner, Jonathan Williams, Mary Williamson, Qian Zeng, Tricia Zwiefelhofer, Clinton R. Paden, Duncan MacCannell |
| EPI_ISL_3327891 | NJDOH, Public Health and Environmental Laboratories | NJ_PHEL | Lindsey Bodnar, Shiv K. Verma, Ryan Pachucki, Chelsea San Filippo, Jacquelyn Deverell, Dana Woell, Allison Roder, Byeong Jeong |
| EPI_ISL_3327892, EPI_ISL_3327893, EPI_ISL_3327894 | Laboratory Corporation of America | Centers for Disease Control and Prevention Division of Viral Diseases, Pathogen Discovery | Dakota Howard, Dhwani Batra, Peter W. Cook, Kara Moser, Adrian Paskey, Jason Caravas, Benjamin Rambo-Martin, Shatavia Morrison, Christopher Gulvick, Scott Sammons, Yvette Unoarumhi, Darlene Wagner, Matthew Schmerer, Minoo Agarwal, Eyad Almasri, Debbie Boles, Ayla Burns, Nuthawin Charoensri, Oren Cohen, Susan Countryman, Mary Ann Cristobal, Bobbi Croy, Suzanne Dale, Hrushikesh Deshmukh, Amanda Douglas, Vincent Drouillon, Marcia Eisenberg, Howard Engler, Rama Ghatti, Prashant Gupta, Susan Hicks, Jake Humphrey, Lax Iyer, Lisa Pfefferle, Manoj Jain, Matthew Robinson, Mohan Kolli, Brian Krueger, Tim Kuphal, Stanley Letovsky, Michael Levandoski, Craig Lukasik, Jonathan Meltzer, Brian Norvell, Mindy Nye, Scott Parker, Christos Petropoulos, John Pruitt, Steven Ragan, Scott Ryan, Mike Sapeta, Jana Schroth, Suresh Babu Selvaraju, Goran Stevovic, Amanda Suchanek, Andrea Throop, Lyndon Tilson, Thomas Urban, Joe Voshell, Kimberly Wagner, Jonathan Williams, Mary Williamson, Qian Zeng, Tricia Zwiefelhofer, Clinton R. Paden, Duncan MacCannell |
| EPI_ISL_3327895 | NJDOH, Public Health and Environmental Laboratories | NJ_PHEL | Lindsey Bodnar, Shiv K. Verma, Ryan Pachucki, Chelsea San Filippo, Jacquelyn Deverell, Dana Woell, Allison Roder, Byeong Jeong |
| EPI_ISL_3327896, EPI_ISL_3327897 | Laboratory Corporation of America | Centers for Disease Control and Prevention Division of Viral Diseases, Pathogen Discovery | Dakota Howard, Dhwani Batra, Peter W. Cook, Kara Moser, Adrian Paskey, Jason Caravas, Benjamin Rambo-Martin, Shatavia Morrison, Christopher Gulvick, Scott Sammons, Yvette Unoarumhi, Darlene Wagner, Matthew Schmerer, Minoo Agarwal, Eyad Almasri, Debbie Boles, Ayla Burns, Nuthawin Charoensri, Oren Cohen, Susan Countryman, Mary Ann Cristobal, Bobbi Croy, Suzanne Dale, Hrushikesh Deshmukh, Amanda Douglas, Vincent Drouillon, Marcia Eisenberg, Howard Engler, Rama Ghatti, Prashant Gupta, Susan Hicks, Jake Humphrey, Lax Iyer, Lisa Pfefferle, Manoj Jain, Matthew Robinson, Mohan Kolli, Brian Krueger, Tim Kuphal, Stanley Letovsky, Michael Levandoski, Craig Lukasik, Jonathan Meltzer, Brian Norvell, Mindy Nye, Scott Parker, Christos Petropoulos, John Pruitt, Steven Ragan, Scott Ryan, Mike Sapeta, Jana Schroth, Suresh Babu Selvaraju, Goran Stevovic, Amanda Suchanek, Andrea Throop, Lyndon Tilson, Thomas Urban, Joe Voshell, Kimberly Wagner, Jonathan Williams, Mary Williamson, Qian Zeng, Tricia Zwiefelhofer, Clinton R. Paden, Duncan MacCannell |
| EPI_ISL_3327898 | NJDOH, Public Health and Environmental Laboratories | NJ_PHEL | Lindsey Bodnar, Shiv K. Verma, Ryan Pachucki, Chelsea San Filippo, Jacquelyn Deverell, Dana Woell, Allison Roder, Byeong Jeong |
| EPI_ISL_3327899, EPI_ISL_3327900 | Laboratory Corporation of America | Centers for Disease Control and Prevention Division of Viral Diseases, Pathogen Discovery | Dakota Howard, Dhwani Batra, Peter W. Cook, Kara Moser, Adrian Paskey, Jason Caravas, Benjamin Rambo-Martin, Shatavia Morrison, Christopher Gulvick, Scott Sammons, Yvette Unoarumhi, Darlene Wagner, Matthew Schmerer, Minoo Agarwal, Eyad Almasri, Debbie Boles, Ayla Burns, Nuthawin Charoensri, Oren Cohen, Susan Countryman, Mary Ann Cristobal, Bobbi Croy, Suzanne Dale, Hrushikesh Deshmukh, Amanda Douglas, Vincent Drouillon, Marcia Eisenberg, Howard Engler, Rama Ghatti, Prashant Gupta, Susan Hicks, Jake Humphrey, Lax Iyer, Lisa Pfefferle, Manoj Jain, Matthew Robinson, Mohan Kolli, Brian Krueger, Tim Kuphal, Stanley Letovsky, Michael Levandoski, Craig Lukasik, Jonathan Meltzer, Brian Norvell, Mindy Nye, Scott Parker, Christos Petropoulos, John Pruitt, Steven Ragan, Scott Ryan, Mike Sapeta, Jana Schroth, Suresh Babu Selvaraju, Goran Stevovic, Amanda Suchanek, Andrea Throop, Lyndon Tilson, Thomas Urban, Joe Voshell, Kimberly Wagner, Jonathan Williams, Mary Williamson, Qian Zeng, Tricia Zwiefelhofer, Clinton R. Paden, Duncan MacCannell |
| EPI_ISL_3327901 | NJDOH, Public Health and Environmental Laboratories | NJ_PHEL | Lindsey Bodnar, Shiv K. Verma, Ryan Pachucki, Chelsea San Filippo, Jacquelyn Deverell, Dana Woell, Allison Roder, Byeong Jeong |
| EPI_ISL_3327902, EPI_ISL_3327903 | Laboratory Corporation of America | Centers for Disease Control and Prevention Division of Viral Diseases, Pathogen Discovery | Dakota Howard, Dhwani Batra, Peter W. Cook, Kara Moser, Adrian Paskey, Jason Caravas, Benjamin Rambo-Martin, Shatavia Morrison, Christopher Gulvick, Scott Sammons, Yvette Unoarumhi, Darlene Wagner, Matthew Schmerer, Minoo Agarwal, Eyad Almasri, Debbie Boles, Ayla Burns, Nuthawin Charoensri, Oren Cohen, Susan Countryman, Mary Ann Cristobal, Bobbi Croy, Suzanne Dale, Hrushikesh Deshmukh, Amanda Douglas, Vincent Drouillon, Marcia Eisenberg, Howard Engler, Rama Ghatti, Prashant Gupta, Susan Hicks, Jake Humphrey, Lax Iyer, Lisa Pfefferle, Manoj Jain, Matthew Robinson, Mohan Kolli, Brian Krueger, Tim Kuphal, Stanley Letovsky, Michael Levandoski, Craig Lukasik, Jonathan Meltzer, Brian Norvell, Mindy Nye, Scott Parker, Christos Petropoulos, John Pruitt, Steven Ragan, Scott Ryan, Mike Sapeta, Jana Schroth, Suresh Babu Selvaraju, Goran Stevovic, Amanda Suchanek, Andrea Throop, Lyndon Tilson, Thomas Urban, Joe Voshell, Kimberly Wagner, Jonathan Williams, Mary Williamson, Qian Zeng, Tricia Zwiefelhofer, Clinton R. Paden, Duncan MacCannell |
| EPI_ISL_3327904 | NJDOH, Public Health and Environmental Laboratories | NJ_PHEL | Lindsey Bodnar, Shiv K. Verma, Ryan Pachucki, Chelsea San Filippo, Jacquelyn Deverell, Dana Woell, Allison Roder, Byeong Jeong |
| EPI_ISL_3327905, EPI_ISL_3327906 | Laboratory Corporation of America | Centers for Disease Control and Prevention Division of Viral Diseases, Pathogen Discovery | Dakota Howard, Dhwani Batra, Peter W. Cook, Kara Moser, Adrian Paskey, Jason Caravas, Benjamin Rambo-Martin, Shatavia Morrison, Christopher Gulvick, Scott Sammons, Yvette Unoarumhi, Darlene Wagner, Matthew Schmerer, Minoo Agarwal, Eyad Almasri, Debbie Boles, Ayla Burns, Nuthawin Charoensri, Oren Cohen, Susan Countryman, Mary Ann Cristobal, Bobbi Croy, Suzanne Dale, Hrushikesh Deshmukh, Amanda Douglas, Vincent Drouillon, Marcia Eisenberg, Howard |

| | | | |
|---|---|---|---|
| | | | Engler, Rama Ghatti, Prashant Gupta, Susan Hicks, Jake Humphrey, Lax Iyer, Lisa Pfefferle, Manoj Jain, Matthew Robinson, Mohan Kolli, Brian Krueger, Tim Kuphal, Stanley Letovsky, Michael Levandoski, Craig Lukasik, Jonathan Meltzer, Brian Norvell, Mindy Nye, Scott Parker, Christos Petropoulos, John Pruitt, Steven Ragan, Scott Ryan, Mike Sapeta, Jana Schroth, Suresh Babu Selvaraju, Goran Stevovic, Amanda Suchanek, Andrea Throop, Lyndon Tilson, Thomas Urban, Joe Voshell, Kimberly Wagner, Jonathan Williams, Mary Williamson, Qian Zeng, Tricia Zwiefelhofer, Clinton R. Paden, Duncan MacCannell |
| EPI_ISL_3327907 | NJDOH, Public Health and Environmental Laboratories | NJ_PHEL | Lindsey Bodnar, Shiv K. Verma, Ryan Pachucki, Chelsea San Filippo, Jacquelyn Deverell, Dana Woell, Allison Roder, Byeong Jeong |
| EPI_ISL_3327908 | Laboratory Corporation of America | Centers for Disease Control and Prevention Division of Viral Diseases, Pathogen Discovery | Dakota Howard, Dhwani Batra, Peter W. Cook, Kara Moser, Adrian Paskey, Jason Caravas, Benjamin Rambo-Martin, Shatavia Morrison, Christopher Gulvick, Scott Sammons, Yvette Unoarumhi, Darlene Wagner, Matthew Schmerer, Minoo Agarwal, Eyad Almasri, Debbie Boles, Ayla Burns, Nuthawin Charoensri, Oren Cohen, Susan Countryman, Mary Ann Cristobal, Bobbi Croy, Suzanne Dale, Hrushikesh Deshmukh, Amanda Douglas, Vincent Drouillon, Marcia Eisenberg, Howard Engler, Rama Ghatti, Prashant Gupta, Susan Hicks, Jake Humphrey, Lax Iyer, Lisa Pfefferle, Manoj Jain, Matthew Robinson, Mohan Kolli, Brian Krueger, Tim Kuphal, Stanley Letovsky, Michael Levandoski, Craig Lukasik, Jonathan Meltzer, Brian Norvell, Mindy Nye, Scott Parker, Christos Petropoulos, John Pruitt, Steven Ragan, Scott Ryan, Mike Sapeta, Jana Schroth, Suresh Babu Selvaraju, Goran Stevovic, Amanda Suchanek, Andrea Throop, Lyndon Tilson, Thomas Urban, Joe Voshell, Kimberly Wagner, Jonathan Williams, Mary Williamson, Qian Zeng, Tricia Zwiefelhofer, Clinton R. Paden, Duncan MacCannell |
| EPI_ISL_3327909 | NJDOH, Public Health and Environmental Laboratories | NJ_PHEL | Lindsey Bodnar, Shiv K. Verma, Ryan Pachucki, Chelsea San Filippo, Jacquelyn Deverell, Dana Woell, Allison Roder, Byeong Jeong |
| EPI_ISL_3327910 | Laboratory Corporation of America | Centers for Disease Control and Prevention Division of Viral Diseases, Pathogen Discovery | Dakota Howard, Dhwani Batra, Peter W. Cook, Kara Moser, Adrian Paskey, Jason Caravas, Benjamin Rambo-Martin, Shatavia Morrison, Christopher Gulvick, Scott Sammons, Yvette Unoarumhi, Darlene Wagner, Matthew Schmerer, Minoo Agarwal, Eyad Almasri, Debbie Boles, Ayla Burns, Nuthawin Charoensri, Oren Cohen, Susan Countryman, Mary Ann Cristobal, Bobbi Croy, Suzanne Dale, Hrushikesh Deshmukh, Amanda Douglas, Vincent Drouillon, Marcia Eisenberg, Howard Engler, Rama Ghatti, Prashant Gupta, Susan Hicks, Jake Humphrey, Lax Iyer, Lisa Pfefferle, Manoj Jain, Matthew Robinson, Mohan Kolli, Brian Krueger, Tim Kuphal, Stanley Letovsky, Michael Levandoski, Craig Lukasik, Jonathan Meltzer, Brian Norvell, Mindy Nye, Scott Parker, Christos Petropoulos, John Pruitt, Steven Ragan, Scott Ryan, Mike Sapeta, Jana Schroth, Suresh Babu Selvaraju, Goran Stevovic, Amanda Suchanek, Andrea Throop, Lyndon Tilson, Thomas Urban, Joe Voshell, Kimberly Wagner, Jonathan Williams, Mary Williamson, Qian Zeng, Tricia Zwiefelhofer, Clinton R. Paden, Duncan MacCannell |
| EPI_ISL_3327911 | NJDOH, Public Health and Environmental Laboratories | NJ_PHEL | Lindsey Bodnar, Shiv K. Verma, Ryan Pachucki, Chelsea San Filippo, Jacquelyn Deverell, Dana Woell, Allison Roder, Byeong Jeong |
| EPI_ISL_3327912, EPI_ISL_3327914 | Laboratory Corporation of America | Centers for Disease Control and Prevention Division of Viral Diseases, Pathogen Discovery | Dakota Howard, Dhwani Batra, Peter W. Cook, Kara Moser, Adrian Paskey, Jason Caravas, Benjamin Rambo-Martin, Shatavia Morrison, Christopher Gulvick, Scott Sammons, Yvette Unoarumhi, Darlene Wagner, Matthew Schmerer, Minoo Agarwal, Eyad Almasri, Debbie Boles, Ayla Burns, Nuthawin Charoensri, Oren Cohen, Susan Countryman, Mary Ann Cristobal, Bobbi Croy, Suzanne Dale, Hrushikesh Deshmukh, Amanda Douglas, Vincent Drouillon, Marcia Eisenberg, Howard Engler, Rama Ghatti, Prashant Gupta, Susan Hicks, Jake Humphrey, Lax Iyer, Lisa Pfefferle, Manoj Jain, Matthew Robinson, Mohan Kolli, Brian Krueger, Tim Kuphal, Stanley Letovsky, Michael Levandoski, Craig Lukasik, Jonathan Meltzer, Brian Norvell, Mindy Nye, Scott Parker, Christos Petropoulos, John Pruitt, Steven Ragan, Scott Ryan, Mike Sapeta, Jana Schroth, Suresh Babu Selvaraju, Goran Stevovic, Amanda Suchanek, Andrea Throop, Lyndon Tilson, Thomas Urban, Joe Voshell, Kimberly Wagner, Jonathan Williams, Mary Williamson, Qian Zeng, Tricia Zwiefelhofer, Clinton R. Paden, Duncan MacCannell |
| EPI_ISL_3327916 | NJDOH, Public Health and Environmental Laboratories | NJ_PHEL | Lindsey Bodnar, Shiv K. Verma, Ryan Pachucki, Chelsea San Filippo, Jacquelyn Deverell, Dana Woell, Allison Roder, Byeong Jeong |
| EPI_ISL_3327917, EPI_ISL_3327918 | Laboratory Corporation of America | Centers for Disease Control and Prevention Division of Viral Diseases, Pathogen Discovery | Dakota Howard, Dhwani Batra, Peter W. Cook, Kara Moser, Adrian Paskey, Jason Caravas, Benjamin Rambo-Martin, Shatavia Morrison, Christopher Gulvick, Scott Sammons, Yvette Unoarumhi, Darlene Wagner, Matthew Schmerer, Minoo Agarwal, Eyad Almasri, Debbie Boles, Ayla Burns, Nuthawin Charoensri, Oren Cohen, Susan Countryman, Mary Ann Cristobal, Bobbi Croy, Suzanne Dale, Hrushikesh Deshmukh, Amanda Douglas, Vincent Drouillon, Marcia Eisenberg, Howard Engler, Rama Ghatti, Prashant Gupta, Susan Hicks, Jake Humphrey, Lax Iyer, Lisa Pfefferle, Manoj Jain, Matthew Robinson, Mohan Kolli, Brian Krueger, Tim Kuphal, Stanley Letovsky, Michael Levandoski, Craig Lukasik, Jonathan Meltzer, Brian Norvell, Mindy Nye, Scott Parker, Christos Petropoulos, John Pruitt, Steven Ragan, Scott Ryan, Mike Sapeta, Jana Schroth, Suresh Babu Selvaraju, Goran Stevovic, Amanda Suchanek, Andrea Throop, Lyndon Tilson, Thomas Urban, Joe Voshell, Kimberly Wagner, Jonathan Williams, Mary Williamson, Qian Zeng, Tricia Zwiefelhofer, Clinton R. Paden, Duncan MacCannell |
| EPI_ISL_3327919 | NJDOH, Public Health and Environmental Laboratories | NJ_PHEL | Lindsey Bodnar, Shiv K. Verma, Ryan Pachucki, Chelsea San Filippo, Jacquelyn Deverell, Dana Woell, Allison Roder, Byeong Jeong |
| EPI_ISL_3327920, EPI_ISL_3327921 | Laboratory Corporation of America | Centers for Disease Control and Prevention Division of Viral Diseases, Pathogen Discovery | Dakota Howard, Dhwani Batra, Peter W. Cook, Kara Moser, Adrian Paskey, Jason Caravas, Benjamin Rambo-Martin, Shatavia Morrison, Christopher Gulvick, Scott Sammons, Yvette Unoarumhi, Darlene Wagner, Matthew Schmerer, Minoo Agarwal, Eyad Almasri, Debbie Boles, Ayla Burns, Nuthawin Charoensri, Oren Cohen, Susan Countryman, Mary Ann Cristobal, Bobbi Croy, Suzanne Dale, Hrushikesh Deshmukh, Amanda Douglas, Vincent Drouillon, Marcia Eisenberg, Howard Engler, Rama Ghatti, Prashant Gupta, Susan Hicks, Jake Humphrey, Lax Iyer, Lisa Pfefferle, Manoj Jain, Matthew Robinson, Mohan Kolli, Brian Krueger, Tim Kuphal, Stanley Letovsky, Michael Levandoski, Craig Lukasik, Jonathan Meltzer, Brian Norvell, Mindy Nye, Scott Parker, Christos Petropoulos, John Pruitt, Steven Ragan, Scott Ryan, Mike Sapeta, Jana Schroth, Suresh Babu Selvaraju, Goran Stevovic, Amanda Suchanek, Andrea Throop, Lyndon Tilson, Thomas Urban, Joe Voshell, Kimberly Wagner, Jonathan Williams, Mary Williamson, Qian Zeng, Tricia Zwiefelhofer, Clinton R. Paden, Duncan MacCannell |
| EPI_ISL_3327922, EPI_ISL_3327925 | NJDOH, Public Health and Environmental Laboratories | NJ_PHEL | Lindsey Bodnar, Shiv K. Verma, Ryan Pachucki, Chelsea San Filippo, Jacquelyn Deverell, Dana Woell, Allison Roder, Byeong Jeong |
| EPI_ISL_3327926 | Laboratory Corporation of America | Centers for Disease Control and Prevention Division of Viral Diseases, Pathogen Discovery | Dakota Howard, Dhwani Batra, Peter W. Cook, Kara Moser, Adrian Paskey, Jason Caravas, Benjamin Rambo-Martin, Shatavia Morrison, Christopher Gulvick, Scott Sammons, Yvette Unoarumhi, Darlene Wagner, Matthew Schmerer, Minoo Agarwal, Eyad Almasri, Debbie Boles, Ayla Burns, Nuthawin Charoensri, Oren Cohen, Susan Countryman, Mary Ann Cristobal, Bobbi Croy, Suzanne Dale, Hrushikesh Deshmukh, Amanda Douglas, Vincent Drouillon, Marcia Eisenberg, Howard Engler, Rama Ghatti, Prashant Gupta, Susan Hicks, Jake Humphrey, Lax Iyer, Lisa Pfefferle, Manoj Jain, Matthew Robinson, Mohan Kolli, Brian Krueger, Tim Kuphal, Stanley Letovsky, Michael Levandoski, Craig Lukasik, Jonathan Meltzer, Brian Norvell, Mindy Nye, Scott Parker, Christos Petropoulos, John Pruitt, Steven Ragan, Scott Ryan, Mike Sapeta, Jana Schroth, Suresh Babu Selvaraju, Goran Stevovic, Amanda Suchanek, Andrea Throop, Lyndon Tilson, Thomas Urban, Joe Voshell, Kimberly Wagner, Jonathan Williams, Mary Williamson, Qian Zeng, Tricia Zwiefelhofer, Clinton R. Paden, Duncan MacCannell |
| EPI_ISL_3327927, EPI_ISL_3327928 | NJDOH, Public Health and Environmental Laboratories | NJ_PHEL | Lindsey Bodnar, Shiv K. Verma, Ryan Pachucki, Chelsea San Filippo, Jacquelyn Deverell, Dana Woell, Allison Roder, Byeong Jeong |
| EPI_ISL_3327929 | Laboratory Corporation of America | Centers for Disease Control and Prevention Division of Viral Diseases, Pathogen Discovery | Dakota Howard, Dhwani Batra, Peter W. Cook, Kara Moser, Adrian Paskey, Jason Caravas, Benjamin Rambo-Martin, Shatavia Morrison, Christopher Gulvick, Scott Sammons, Yvette Unoarumhi, Darlene Wagner, Matthew Schmerer, Minoo Agarwal, Eyad Almasri, Debbie Boles, Ayla Burns, Nuthawin Charoensri, Oren Cohen, Susan Countryman, Mary Ann Cristobal, Bobbi Croy, Suzanne Dale, Hrushikesh Deshmukh, Amanda Douglas, Vincent Drouillon, Marcia Eisenberg, Howard Engler, Rama Ghatti, Prashant Gupta, Susan Hicks, Jake Humphrey, Lax Iyer, Lisa Pfefferle, Manoj Jain, Matthew Robinson, Mohan Kolli, Brian Krueger, Tim Kuphal, Stanley Letovsky, Michael Levandoski, Craig Lukasik, Jonathan Meltzer, Brian Norvell, Mindy Nye, Scott Parker, Christos Petropoulos, John Pruitt, Steven Ragan, Scott Ryan, Mike Sapeta, Jana Schroth, Suresh Babu Selvaraju, Goran Stevovic, Amanda Suchanek, Andrea Throop, Lyndon Tilson, Thomas Urban, Joe Voshell, Kimberly Wagner, Jonathan Williams, Mary Williamson, Qian Zeng, Tricia Zwiefelhofer, Clinton R. Paden, Duncan MacCannell |
| EPI_ISL_3327930 | NJDOH, Public Health and Environmental Laboratories | NJ_PHEL | Lindsey Bodnar, Shiv K. Verma, Ryan Pachucki, Chelsea San Filippo, Jacquelyn Deverell, Dana Woell, Allison Roder, Byeong Jeong |
| EPI_ISL_3327931 | Laboratory Corporation of America | Centers for Disease Control and Prevention Division of Viral Diseases, Pathogen Discovery | Dakota Howard, Dhwani Batra, Peter W. Cook, Kara Moser, Adrian Paskey, Jason Caravas, Benjamin Rambo-Martin, Shatavia Morrison, Christopher Gulvick, Scott Sammons, Yvette Unoarumhi, Darlene Wagner, Matthew Schmerer, Minoo Agarwal, Eyad Almasri, Debbie Boles, Ayla Burns, Nuthawin Charoensri, Oren Cohen, Susan Countryman, Mary Ann Cristobal, Bobbi Croy, Suzanne Dale, Hrushikesh Deshmukh, Amanda Douglas, Vincent Drouillon, Marcia Eisenberg, Howard Engler, Rama Ghatti, Prashant Gupta, Susan Hicks, Jake Humphrey, Lax Iyer, Lisa Pfefferle, Manoj Jain, Matthew Robinson, Mohan Kolli, Brian Krueger, Tim Kuphal, Stanley Letovsky, Michael Levandoski, Craig Lukasik, Jonathan Meltzer, Brian Norvell, Mindy Nye, Scott Parker, Christos Petropoulos, John Pruitt, Steven Ragan, Scott Ryan, Mike Sapeta, Jana Schroth, Suresh Babu Selvaraju, Goran Stevovic, Amanda Suchanek, Andrea Throop, Lyndon Tilson, Thomas Urban, Joe Voshell, Kimberly Wagner, Jonathan Williams, Mary Williamson, Qian Zeng, Tricia Zwiefelhofer, Clinton R. Paden, Duncan MacCannell |
| EPI_ISL_3327932, EPI_ISL_3327933 | NJDOH, Public Health and Environmental Laboratories | NJ_PHEL | Lindsey Bodnar, Shiv K. Verma, Ryan Pachucki, Chelsea San Filippo, Jacquelyn Deverell, Dana Woell, Allison Roder, Byeong Jeong |
| EPI_ISL_3327934, EPI_ISL_3327935 | Laboratory Corporation of America | Centers for Disease Control and Prevention Division of Viral Diseases, Pathogen Discovery | Dakota Howard, Dhwani Batra, Peter W. Cook, Kara Moser, Adrian Paskey, Jason Caravas, Benjamin Rambo-Martin, Shatavia Morrison, Christopher Gulvick, Scott Sammons, Yvette Unoarumhi, Darlene Wagner, Matthew Schmerer, Minoo Agarwal, Eyad Almasri, Debbie Boles, Ayla Burns, Nuthawin Charoensri, Oren Cohen, Susan Countryman, Mary Ann Cristobal, Bobbi Croy, Suzanne Dale, Hrushikesh Deshmukh, Amanda Douglas, Vincent Drouillon, Marcia Eisenberg, Howard Engler, Rama Ghatti, Prashant Gupta, Susan Hicks, Jake Humphrey, Lax Iyer, Lisa Pfefferle, Manoj Jain, Matthew Robinson, Mohan Kolli, Brian Krueger, Tim Kuphal, Stanley Letovsky, Michael Levandoski, Craig Lukasik, Jonathan Meltzer, Brian Norvell, Mindy Nye, Scott Parker, Christos Petropoulos, John Pruitt, Steven Ragan, Scott Ryan, Mike Sapeta, Jana Schroth, Suresh Babu Selvaraju, Goran Stevovic, Amanda Suchanek, Andrea Throop, Lyndon Tilson, Thomas Urban, Joe Voshell, Kimberly Wagner, Jonathan Williams, Mary Williamson, Qian Zeng, Tricia Zwiefelhofer, Clinton R. Paden, Duncan MacCannell |
| EPI_ISL_3327936 | NJDOH, Public Health and Environmental Laboratories | NJ_PHEL | Lindsey Bodnar, Shiv K. Verma, Ryan Pachucki, Chelsea San Filippo, Jacquelyn Deverell, Dana Woell, Allison Roder, Byeong Jeong |
| EPI_ISL_3327937, EPI_ISL_3327938 | Laboratory Corporation of America | Centers for Disease Control and Prevention Division of Viral Diseases, Pathogen Discovery | Dakota Howard, Dhwani Batra, Peter W. Cook, Kara Moser, Adrian Paskey, Jason Caravas, Benjamin Rambo-Martin, Shatavia Morrison, Christopher Gulvick, Scott Sammons, Yvette Unoarumhi, Darlene Wagner, Matthew Schmerer, Minoo Agarwal, Eyad Almasri, Debbie Boles, Ayla Burns, Nuthawin Charoensri, Oren Cohen, Susan Countryman, Mary Ann Cristobal, Bobbi Croy, Suzanne Dale, Hrushikesh Deshmukh, Amanda Douglas, Vincent Drouillon, Marcia Eisenberg, Howard Engler, Rama Ghatti, Prashant Gupta, Susan Hicks, Jake Humphrey, Lax Iyer, Lisa Pfefferle, Manoj Jain, Matthew Robinson, Mohan Kolli, Brian Krueger, Tim Kuphal, Stanley Letovsky, Michael Levandoski, Craig Lukasik, Jonathan Meltzer, Brian Norvell, Mindy Nye, Scott Parker, Christos Petropoulos, John Pruitt, Steven Ragan, Scott Ryan, Mike Sapeta, Jana Schroth, Suresh Babu Selvaraju, Goran Stevovic, Amanda Suchanek, Andrea Throop, Lyndon Tilson, Thomas Urban, Joe Voshell, Kimberly Wagner, Jonathan Williams, Mary Williamson, Qian Zeng, Tricia Zwiefelhofer, Clinton R. Paden, Duncan MacCannell |
| EPI_ISL_3327939 | NJDOH, Public Health and Environmental Laboratories | NJ_PHEL | Lindsey Bodnar, Shiv K. Verma, Ryan Pachucki, Chelsea San Filippo, Jacquelyn Deverell, Dana Woell, Allison Roder, Byeong Jeong |
| EPI_ISL_3327940, EPI_ISL_3327941, EPI_ISL_3327942 | Laboratory Corporation of America | Centers for Disease Control and Prevention Division of Viral Diseases, Pathogen Discovery | Dakota Howard, Dhwani Batra, Peter W. Cook, Kara Moser, Adrian Paskey, Jason Caravas, Benjamin Rambo-Martin, Shatavia Morrison, Christopher Gulvick, Scott Sammons, Yvette Unoarumhi, Darlene Wagner, Matthew Schmerer, Minoo Agarwal, Eyad Almasri, Debbie Boles, Ayla Burns, Nuthawin Charoensri, Oren Cohen, Susan Countryman, Mary Ann Cristobal, Bobbi Croy, Suzanne Dale, Hrushikesh Deshmukh, Amanda Douglas, Vincent Drouillon, Marcia Eisenberg, Howard Engler, Rama Ghatti, Prashant Gupta, Susan Hicks, Jake Humphrey, Lax Iyer, Lisa Pfefferle, Manoj Jain, Matthew Robinson, Mohan Kolli, Brian Krueger, Tim Kuphal, Stanley Letovsky, Michael Levandoski, Craig Lukasik, Jonathan Meltzer, Brian Norvell, Mindy Nye, Scott Parker, Christos Petropoulos, John Pruitt, Steven Ragan, Scott Ryan, Mike Sapeta, Jana Schroth, Suresh Babu Selvaraju, Goran Stevovic, Amanda Suchanek, Andrea Throop, Lyndon Tilson, Thomas Urban, Joe Voshell, Kimberly Wagner, Jonathan Williams, Mary Williamson, Qian Zeng, Tricia Zwiefelhofer, Clinton R. Paden, Duncan MacCannell |
| EPI_ISL_3327943 | NJDOH, Public Health and Environmental Laboratories | NJ_PHEL | Lindsey Bodnar, Shiv K. Verma, Ryan Pachucki, Chelsea San Filippo, Jacquelyn Deverell, Dana Woell, Allison Roder, Byeong Jeong |
| EPI_ISL_3327944 | Laboratory Corporation of America | Centers for Disease Control and Prevention Division of Viral Diseases, Pathogen Discovery | Dakota Howard, Dhwani Batra, Peter W. Cook, Kara Moser, Adrian Paskey, Jason Caravas, Benjamin Rambo-Martin, Shatavia Morrison, Christopher Gulvick, Scott Sammons, Yvette Unoarumhi, Darlene Wagner, Matthew Schmerer, Minoo Agarwal, Eyad Almasri, Debbie Boles, Ayla Burns, Nuthawin Charoensri, Oren Cohen, Susan Countryman, Mary Ann Cristobal, Bobbi Croy, Suzanne Dale, Hrushikesh Deshmukh, Amanda Douglas, Vincent Drouillon, Marcia Eisenberg, Howard Engler, Rama Ghatti, Prashant Gupta, Susan Hicks, Jake Humphrey, Lax Iyer, Lisa Pfefferle, Manoj Jain, Matthew Robinson, Mohan Kolli, Brian Krueger, Tim Kuphal, Stanley Letovsky, Michael Levandoski, Craig Lukasik, Jonathan Meltzer, Brian Norvell, Mindy Nye, Scott Parker, Christos Petropoulos, John Pruitt, Steven Ragan, Scott Ryan, Mike Sapeta, Jana Schroth, Suresh Babu Selvaraju, Goran Stevovic, Amanda Suchanek, Andrea Throop, Lyndon Tilson, Thomas Urban, Joe Voshell, Kimberly Wagner, Jonathan Williams, Mary Williamson, Qian Zeng, Tricia Zwiefelhofer, Clinton R. Paden, Duncan MacCannell |
| EPI_ISL_3327945, EPI_ISL_3327946 | NJDOH, Public Health and Environmental Laboratories | NJ_PHEL | Lindsey Bodnar, Shiv K. Verma, Ryan Pachucki, Chelsea San Filippo, Jacquelyn Deverell, Dana Woell, Allison Roder, Byeong Jeong |
| EPI_ISL_3327947 | Laboratory Corporation of America | Centers for Disease Control and Prevention Division of Viral Diseases, Pathogen Discovery | Dakota Howard, Dhwani Batra, Peter W. Cook, Kara Moser, Adrian Paskey, Jason Caravas, Benjamin Rambo-Martin, Shatavia Morrison, Christopher Gulvick, Scott Sammons, Yvette Unoarumhi, Darlene Wagner, Matthew Schmerer, Minoo Agarwal, Eyad Almasri, Debbie Boles, Ayla Burns, Nuthawin Charoensri, Oren Cohen, Susan Countryman, Mary Ann Cristobal, Bobbi Croy, Suzanne Dale, Hrushikesh Deshmukh, Amanda Douglas, Vincent Drouillon, Marcia Eisenberg, Howard Engler, Rama Ghatti, Prashant Gupta, Susan Hicks, Jake Humphrey, Lax Iyer, Lisa Pfefferle, Manoj Jain, Matthew Robinson, Mohan Kolli, Brian Krueger, Tim Kuphal, Stanley Letovsky, Michael Levandoski, Craig Lukasik, Jonathan Meltzer, Brian Norvell, Mindy Nye, Scott Parker, Christos Petropoulos, John Pruitt, Steven Ragan, Scott Ryan, Mike Sapeta, Jana Schroth, Suresh Babu Selvaraju, Goran Stevovic, Amanda Suchanek, Andrea Throop, Lyndon Tilson, Thomas Urban, Joe Voshell, Kimberly Wagner, Jonathan Williams, Mary Williamson, Qian Zeng, Tricia Zwiefelhofer, Clinton R. Paden, Duncan MacCannell |
| EPI_ISL_3327948 | NJDOH, Public Health and Environmental Laboratories | NJ_PHEL | Lindsey Bodnar, Shiv K. Verma, Ryan Pachucki, Chelsea San Filippo, Jacquelyn Deverell, Dana Woell, Allison Roder, Byeong Jeong |
| EPI_ISL_3327949, EPI_ISL_3327950, EPI_ISL_3327951 | Laboratory Corporation of America | Centers for Disease Control and Prevention Division of Viral Diseases, Pathogen Discovery | Dakota Howard, Dhwani Batra, Peter W. Cook, Kara Moser, Adrian Paskey, Jason Caravas, Benjamin Rambo-Martin, Shatavia Morrison, Christopher Gulvick, Scott Sammons, Yvette Unoarumhi, Darlene Wagner, Matthew Schmerer, Minoo Agarwal, Eyad Almasri, Debbie Boles, Ayla Burns, Nuthawin Charoensri, Oren Cohen, Susan Countryman, Mary Ann Cristobal, Bobbi Croy, Suzanne Dale, Hrushikesh Deshmukh, Amanda Douglas, Vincent Drouillon, Marcia Eisenberg, Howard Engler, Rama Ghatti, Prashant Gupta, Susan Hicks, Jake Humphrey, Lax Iyer, Lisa Pfefferle, Manoj Jain, Matthew Robinson, Mohan Kolli, Brian Krueger, Tim Kuphal, Stanley Letovsky, Michael Levandoski, Craig Lukasik, Jonathan Meltzer, Brian Norvell, Mindy Nye, Scott Parker, Christos Petropoulos, John Pruitt, Steven Ragan, Scott Ryan, Mike Sapeta, Jana Schroth, Suresh Babu Selvaraju, Goran Stevovic, Amanda Suchanek, Andrea Throop, Lyndon Tilson, Thomas Urban, Joe Voshell, Kimberly Wagner, Jonathan Williams, Mary Williamson, Qian Zeng, Tricia Zwiefelhofer, Clinton R. Paden, Duncan MacCannell |
| EPI_ISL_3327952 | NJDOH, Public Health and Environmental Laboratories | NJ_PHEL | Lindsey Bodnar, Shiv K. Verma, Ryan Pachucki, Chelsea San Filippo, Jacquelyn Deverell, Dana Woell, Allison Roder, Byeong Jeong |
| EPI_ISL_3327953, EPI_ISL_3327954 | Laboratory Corporation of America | Centers for Disease Control and Prevention Division of Viral Diseases, Pathogen Discovery | Dakota Howard, Dhwani Batra, Peter W. Cook, Kara Moser, Adrian Paskey, Jason Caravas, Benjamin Rambo-Martin, Shatavia Morrison, Christopher Gulvick, Scott Sammons, Yvette Unoarumhi, Darlene Wagner, Matthew Schmerer, Minoo Agarwal, Eyad Almasri, Debbie Boles, Ayla Burns, Nuthawin Charoensri, Oren Cohen, Susan Countryman, Mary Ann Cristobal, Bobbi Croy, Suzanne Dale, Hrushikesh Deshmukh, Amanda Douglas, Vincent Drouillon, Marcia Eisenberg, Howard Engler, Rama Ghatti, Prashant Gupta, Susan Hicks, Jake Humphrey, Lax Iyer, Lisa Pfefferle, Manoj Jain, Matthew Robinson, Mohan Kolli, Brian Krueger, Tim Kuphal, Stanley Letovsky, Michael Levandoski, Craig Lukasik, Jonathan Meltzer, Brian Norvell, Mindy Nye, Scott Parker, Christos Petropoulos, John Pruitt, Steven Ragan, Scott Ryan, Mike Sapeta, Jana Schroth, Suresh Babu Selvaraju, Goran Stevovic, Amanda Suchanek, Andrea Throop, Lyndon Tilson, Thomas Urban, Joe Voshell, |

| | | | |
|---|---|---|---|
| | | | Kimberly Wagner, Jonathan Williams, Mary Williamson, Qian Zeng, Tricia Zwiefelhofer, Clinton R. Paden, Duncan MacCannell |
| EPI_ISL_3327955 | NJDOH, Public Health and Environmental Laboratories | NJ_PHEL | Lindsey Bodnar, Shiv K. Verma, Ryan Pachucki, Chelsea San Filippo, Jacquelyn Deverell, Dana Woell, Allison Roder, Byeong Jeong |
| EPI_ISL_3327956, EPI_ISL_3327957 | Laboratory Corporation of America | Centers for Disease Control and Prevention Division of Viral Diseases, Pathogen Discovery | Dakota Howard, Dhwani Batra, Peter W. Cook, Kara Moser, Adrian Paskey, Jason Caravas, Benjamin Rambo-Martin, Shatavia Morrison, Christopher Gulvick, Scott Sammons, Yvette Unoarumhi, Darlene Wagner, Matthew Schmerer, Minoo Agarwal, Eyad Almasri, Debbie Boles, Ayla Burns, Nuthawin Charoensri, Oren Cohen, Susan Countryman, Mary Ann Cristobal, Bobbi Croy, Suzanne Dale, Hrushikesh Deshmukh, Amanda Douglas, Vincent Drouillon, Marcia Eisenberg, Howard Engler, Rama Ghatti, Prashant Gupta, Susan Hicks, Jake Humphrey, Lax Iyer, Lisa Pfefferle, Manoj Jain, Matthew Robinson, Mohan Kolli, Brian Krueger, Tim Kuphal, Stanley Letovsky, Michael Levandoski, Craig Lukasik, Jonathan Meltzer, Brian Norvell, Mindy Nye, Scott Parker, Christos Petropoulos, John Pruitt, Steven Ragan, Scott Ryan, Mike Sapeta, Jana Schroth, Suresh Babu Selvaraju, Goran Stevovic, Amanda Suchanek, Andrea Throop, Lyndon Tilson, Thomas Urban, Joe Voshell, Kimberly Wagner, Jonathan Williams, Mary Williamson, Qian Zeng, Tricia Zwiefelhofer, Clinton R. Paden, Duncan MacCannell |
| EPI_ISL_3327958 | NJDOH, Public Health and Environmental Laboratories | NJ_PHEL | Lindsey Bodnar, Shiv K. Verma, Ryan Pachucki, Chelsea San Filippo, Jacquelyn Deverell, Dana Woell, Allison Roder, Byeong Jeong |
| EPI_ISL_3327959, EPI_ISL_3327960, EPI_ISL_3327961 | Laboratory Corporation of America | Centers for Disease Control and Prevention Division of Viral Diseases, Pathogen Discovery | Dakota Howard, Dhwani Batra, Peter W. Cook, Kara Moser, Adrian Paskey, Jason Caravas, Benjamin Rambo-Martin, Shatavia Morrison, Christopher Gulvick, Scott Sammons, Yvette Unoarumhi, Darlene Wagner, Matthew Schmerer, Minoo Agarwal, Eyad Almasri, Debbie Boles, Ayla Burns, Nuthawin Charoensri, Oren Cohen, Susan Countryman, Mary Ann Cristobal, Bobbi Croy, Suzanne Dale, Hrushikesh Deshmukh, Amanda Douglas, Vincent Drouillon, Marcia Eisenberg, Howard Engler, Rama Ghatti, Prashant Gupta, Susan Hicks, Jake Humphrey, Lax Iyer, Lisa Pfefferle, Manoj Jain, Matthew Robinson, Mohan Kolli, Brian Krueger, Tim Kuphal, Stanley Letovsky, Michael Levandoski, Craig Lukasik, Jonathan Meltzer, Brian Norvell, Mindy Nye, Scott Parker, Christos Petropoulos, John Pruitt, Steven Ragan, Scott Ryan, Mike Sapeta, Jana Schroth, Suresh Babu Selvaraju, Goran Stevovic, Amanda Suchanek, Andrea Throop, Lyndon Tilson, Thomas Urban, Joe Voshell, Kimberly Wagner, Jonathan Williams, Mary Williamson, Qian Zeng, Tricia Zwiefelhofer, Clinton R. Paden, Duncan MacCannell |
| EPI_ISL_3327962 | NJDOH, Public Health and Environmental Laboratories | NJ_PHEL | Lindsey Bodnar, Shiv K. Verma, Ryan Pachucki, Chelsea San Filippo, Jacquelyn Deverell, Dana Woell, Allison Roder, Byeong Jeong |
| EPI_ISL_3327963, EPI_ISL_3327964 | Laboratory Corporation of America | Centers for Disease Control and Prevention Division of Viral Diseases, Pathogen Discovery | Dakota Howard, Dhwani Batra, Peter W. Cook, Kara Moser, Adrian Paskey, Jason Caravas, Benjamin Rambo-Martin, Shatavia Morrison, Christopher Gulvick, Scott Sammons, Yvette Unoarumhi, Darlene Wagner, Matthew Schmerer, Minoo Agarwal, Eyad Almasri, Debbie Boles, Ayla Burns, Nuthawin Charoensri, Oren Cohen, Susan Countryman, Mary Ann Cristobal, Bobbi Croy, Suzanne Dale, Hrushikesh Deshmukh, Amanda Douglas, Vincent Drouillon, Marcia Eisenberg, Howard Engler, Rama Ghatti, Prashant Gupta, Susan Hicks, Jake Humphrey, Lax Iyer, Lisa Pfefferle, Manoj Jain, Matthew Robinson, Mohan Kolli, Brian Krueger, Tim Kuphal, Stanley Letovsky, Michael Levandoski, Craig Lukasik, Jonathan Meltzer, Brian Norvell, Mindy Nye, Scott Parker, Christos Petropoulos, John Pruitt, Steven Ragan, Scott Ryan, Mike Sapeta, Jana Schroth, Suresh Babu Selvaraju, Goran Stevovic, Amanda Suchanek, Andrea Throop, Lyndon Tilson, Thomas Urban, Joe Voshell, Kimberly Wagner, Jonathan Williams, Mary Williamson, Qian Zeng, Tricia Zwiefelhofer, Clinton R. Paden, Duncan MacCannell |
| EPI_ISL_3327965 | NJDOH, Public Health and Environmental Laboratories | NJ_PHEL | Lindsey Bodnar, Shiv K. Verma, Ryan Pachucki, Chelsea San Filippo, Jacquelyn Deverell, Dana Woell, Allison Roder, Byeong Jeong |
| EPI_ISL_3327966, EPI_ISL_3327967 | Laboratory Corporation of America | Centers for Disease Control and Prevention Division of Viral Diseases, Pathogen Discovery | Dakota Howard, Dhwani Batra, Peter W. Cook, Kara Moser, Adrian Paskey, Jason Caravas, Benjamin Rambo-Martin, Shatavia Morrison, Christopher Gulvick, Scott Sammons, Yvette Unoarumhi, Darlene Wagner, Matthew Schmerer, Minoo Agarwal, Eyad Almasri, Debbie Boles, Ayla Burns, Nuthawin Charoensri, Oren Cohen, Susan Countryman, Mary Ann Cristobal, Bobbi Croy, Suzanne Dale, Hrushikesh Deshmukh, Amanda Douglas, Vincent Drouillon, Marcia Eisenberg, Howard Engler, Rama Ghatti, Prashant Gupta, Susan Hicks, Jake Humphrey, Lax Iyer, Lisa Pfefferle, Manoj Jain, Matthew Robinson, Mohan Kolli, Brian Krueger, Tim Kuphal, Stanley Letovsky, Michael Levandoski, Craig Lukasik, Jonathan Meltzer, Brian Norvell, Mindy Nye, Scott Parker, Christos Petropoulos, John Pruitt, Steven Ragan, Scott Ryan, Mike Sapeta, Jana Schroth, Suresh Babu Selvaraju, Goran Stevovic, Amanda Suchanek, Andrea Throop, Lyndon Tilson, Thomas Urban, Joe Voshell, Kimberly Wagner, Jonathan Williams, Mary Williamson, Qian Zeng, Tricia Zwiefelhofer, Clinton R. Paden, Duncan MacCannell |
| EPI_ISL_3327968 | NJDOH, Public Health and Environmental Laboratories | NJ_PHEL | Lindsey Bodnar, Shiv K. Verma, Ryan Pachucki, Chelsea San Filippo, Jacquelyn Deverell, Dana Woell, Allison Roder, Byeong Jeong |
| EPI_ISL_3327969, EPI_ISL_3327970 | Laboratory Corporation of America | Centers for Disease Control and Prevention Division of Viral Diseases, Pathogen Discovery | Dakota Howard, Dhwani Batra, Peter W. Cook, Kara Moser, Adrian Paskey, Jason Caravas, Benjamin Rambo-Martin, Shatavia Morrison, Christopher Gulvick, Scott Sammons, Yvette Unoarumhi, Darlene Wagner, Matthew Schmerer, Minoo Agarwal, Eyad Almasri, Debbie Boles, Ayla Burns, Nuthawin Charoensri, Oren Cohen, Susan Countryman, Mary Ann Cristobal, Bobbi Croy, Suzanne Dale, Hrushikesh Deshmukh, Amanda Douglas, Vincent Drouillon, Marcia Eisenberg, Howard Engler, Rama Ghatti, Prashant Gupta, Susan Hicks, Jake Humphrey, Lax Iyer, Lisa Pfefferle, Manoj Jain, Matthew Robinson, Mohan Kolli, Brian Krueger, Tim Kuphal, Stanley Letovsky, Michael Levandoski, Craig Lukasik, Jonathan Meltzer, Brian Norvell, Mindy Nye, Scott Parker, Christos Petropoulos, John Pruitt, Steven Ragan, Scott Ryan, Mike Sapeta, Jana Schroth, Suresh Babu Selvaraju, Goran Stevovic, Amanda Suchanek, Andrea Throop, Lyndon Tilson, Thomas Urban, Joe Voshell, Kimberly Wagner, Jonathan Williams, Mary Williamson, Qian Zeng, Tricia Zwiefelhofer, Clinton R. Paden, Duncan MacCannell |
| EPI_ISL_3327971 | NJDOH, Public Health and Environmental Laboratories | NJ_PHEL | Lindsey Bodnar, Shiv K. Verma, Ryan Pachucki, Chelsea San Filippo, Jacquelyn Deverell, Dana Woell, Allison Roder, Byeong Jeong |
| EPI_ISL_3327972, EPI_ISL_3327973, EPI_ISL_3327974 | Laboratory Corporation of America | Centers for Disease Control and Prevention Division of Viral Diseases, Pathogen Discovery | Dakota Howard, Dhwani Batra, Peter W. Cook, Kara Moser, Adrian Paskey, Jason Caravas, Benjamin Rambo-Martin, Shatavia Morrison, Christopher Gulvick, Scott Sammons, Yvette Unoarumhi, Darlene Wagner, Matthew Schmerer, Minoo Agarwal, Eyad Almasri, Debbie Boles, Ayla Burns, Nuthawin Charoensri, Oren Cohen, Susan Countryman, Mary Ann Cristobal, Bobbi Croy, Suzanne Dale, Hrushikesh Deshmukh, Amanda Douglas, Vincent Drouillon, Marcia Eisenberg, Howard Engler, Rama Ghatti, Prashant Gupta, Susan Hicks, Jake Humphrey, Lax Iyer, Lisa Pfefferle, Manoj Jain, Matthew Robinson, Mohan Kolli, Brian Krueger, Tim Kuphal, Stanley Letovsky, Michael Levandoski, Craig Lukasik, Jonathan Meltzer, Brian Norvell, Mindy Nye, Scott Parker, Christos Petropoulos, John Pruitt, Steven Ragan, Scott Ryan, Mike Sapeta, Jana Schroth, Suresh Babu Selvaraju, Goran Stevovic, Amanda Suchanek, Andrea Throop, Lyndon Tilson, Thomas Urban, Joe Voshell, Kimberly Wagner, Jonathan Williams, Mary Williamson, Qian Zeng, Tricia Zwiefelhofer, Clinton R. Paden, Duncan MacCannell |
| EPI_ISL_3327975 | NJDOH, Public Health and Environmental Laboratories | NJ_PHEL | Lindsey Bodnar, Shiv K. Verma, Ryan Pachucki, Chelsea San Filippo, Jacquelyn Deverell, Dana Woell, Allison Roder, Byeong Jeong |
| EPI_ISL_3327976, EPI_ISL_3327977 | Laboratory Corporation of America | Centers for Disease Control and Prevention Division of Viral Diseases, Pathogen Discovery | Dakota Howard, Dhwani Batra, Peter W. Cook, Kara Moser, Adrian Paskey, Jason Caravas, Benjamin Rambo-Martin, Shatavia Morrison, Christopher Gulvick, Scott Sammons, Yvette Unoarumhi, Darlene Wagner, Matthew Schmerer, Minoo Agarwal, Eyad Almasri, Debbie Boles, Ayla Burns, Nuthawin Charoensri, Oren Cohen, Susan Countryman, Mary Ann Cristobal, Bobbi Croy, Suzanne Dale, Hrushikesh Deshmukh, Amanda Douglas, Vincent Drouillon, Marcia Eisenberg, Howard Engler, Rama Ghatti, Prashant Gupta, Susan Hicks, Jake Humphrey, Lax Iyer, Lisa Pfefferle, Manoj Jain, Matthew Robinson, Mohan Kolli, Brian Krueger, Tim Kuphal, Stanley Letovsky, Michael Levandoski, Craig Lukasik, Jonathan Meltzer, Brian Norvell, Mindy Nye, Scott Parker, Christos Petropoulos, John Pruitt, Steven Ragan, Scott Ryan, Mike Sapeta, Jana Schroth, Suresh Babu Selvaraju, Goran Stevovic, Amanda Suchanek, Andrea Throop, Lyndon Tilson, Thomas Urban, Joe Voshell, Kimberly Wagner, Jonathan Williams, Mary Williamson, Qian Zeng, Tricia Zwiefelhofer, Clinton R. Paden, Duncan MacCannell |
| EPI_ISL_3327978 | NJDOH, Public Health and Environmental Laboratories | NJ_PHEL | Lindsey Bodnar, Shiv K. Verma, Ryan Pachucki, Chelsea San Filippo, Jacquelyn Deverell, Dana Woell, Allison Roder, Byeong Jeong |
| EPI_ISL_3327979, EPI_ISL_3327980 | Laboratory Corporation of America | Centers for Disease Control and Prevention Division of Viral Diseases, Pathogen Discovery | Dakota Howard, Dhwani Batra, Peter W. Cook, Kara Moser, Adrian Paskey, Jason Caravas, Benjamin Rambo-Martin, Shatavia Morrison, Christopher Gulvick, Scott Sammons, Yvette Unoarumhi, Darlene Wagner, Matthew Schmerer, Minoo Agarwal, Eyad Almasri, Debbie Boles, Ayla Burns, Nuthawin Charoensri, Oren Cohen, Susan Countryman, Mary Ann Cristobal, Bobbi Croy, Suzanne Dale, Hrushikesh Deshmukh, Amanda Douglas, Vincent Drouillon, Marcia Eisenberg, Howard Engler, Rama Ghatti, Prashant Gupta, Susan Hicks, Jake Humphrey, Lax Iyer, Lisa Pfefferle, Manoj Jain, Matthew Robinson, Mohan Kolli, Brian Krueger, Tim Kuphal, Stanley Letovsky, Michael Levandoski, Craig Lukasik, Jonathan Meltzer, Brian Norvell, Mindy Nye, Scott Parker, Christos Petropoulos, John Pruitt, Steven Ragan, Scott Ryan, Mike Sapeta, Jana Schroth, Suresh Babu Selvaraju, Goran Stevovic, Amanda Suchanek, Andrea Throop, Lyndon Tilson, Thomas Urban, Joe Voshell, Kimberly Wagner, Jonathan Williams, Mary Williamson, Qian Zeng, Tricia Zwiefelhofer, Clinton R. Paden, Duncan MacCannell |
| EPI_ISL_3327981 | NJDOH, Public Health and Environmental Laboratories | NJ_PHEL | Lindsey Bodnar, Shiv K. Verma, Ryan Pachucki, Chelsea San Filippo, Jacquelyn Deverell, Dana Woell, Allison Roder, Byeong Jeong |
| EPI_ISL_3327982, EPI_ISL_3327983, EPI_ISL_3327984, EPI_ISL_3327986, EPI_ISL_3327987 | Laboratory Corporation of America | Centers for Disease Control and Prevention Division of Viral Diseases, Pathogen Discovery | Dakota Howard, Dhwani Batra, Peter W. Cook, Kara Moser, Adrian Paskey, Jason Caravas, Benjamin Rambo-Martin, Shatavia Morrison, Christopher Gulvick, Scott Sammons, Yvette Unoarumhi, Darlene Wagner, Matthew Schmerer, Minoo Agarwal, Eyad Almasri, Debbie Boles, Ayla Burns, Nuthawin Charoensri, Oren Cohen, Susan Countryman, Mary Ann Cristobal, Bobbi Croy, Suzanne Dale, Hrushikesh Deshmukh, Amanda Douglas, Vincent Drouillon, Marcia Eisenberg, Howard Engler, Rama Ghatti, Prashant Gupta, Susan Hicks, Jake Humphrey, Lax Iyer, Lisa Pfefferle, Manoj Jain, Matthew Robinson, Mohan Kolli, Brian Krueger, Tim Kuphal, Stanley Letovsky, Michael Levandoski, Craig Lukasik, Jonathan Meltzer, Brian Norvell, Mindy Nye, Scott Parker, Christos Petropoulos, John Pruitt, Steven Ragan, Scott Ryan, Mike Sapeta, Jana Schroth, Suresh Babu Selvaraju, Goran Stevovic, Amanda Suchanek, Andrea Throop, Lyndon Tilson, Thomas Urban, Joe Voshell, Kimberly Wagner, Jonathan Williams, Mary Williamson, Qian Zeng, Tricia Zwiefelhofer, Clinton R. Paden, Duncan MacCannell |
| EPI_ISL_3327988 | NJDOH, Public Health and Environmental Laboratories | NJ_PHEL | Lindsey Bodnar, Shiv K. Verma, Ryan Pachucki, Chelsea San Filippo, Jacquelyn Deverell, Dana Woell, Allison Roder, Byeong Jeong |
| EPI_ISL_3327989, EPI_ISL_3327990 | Laboratory Corporation of America | Centers for Disease Control and Prevention Division of Viral Diseases, Pathogen Discovery | Dakota Howard, Dhwani Batra, Peter W. Cook, Kara Moser, Adrian Paskey, Jason Caravas, Benjamin Rambo-Martin, Shatavia Morrison, Christopher Gulvick, Scott Sammons, Yvette Unoarumhi, Darlene Wagner, Matthew Schmerer, Minoo Agarwal, Eyad Almasri, Debbie Boles, Ayla Burns, Nuthawin Charoensri, Oren Cohen, Susan Countryman, Mary Ann Cristobal, Bobbi Croy, Suzanne Dale, Hrushikesh Deshmukh, Amanda Douglas, Vincent Drouillon, Marcia Eisenberg, Howard Engler, Rama Ghatti, Prashant Gupta, Susan Hicks, Jake Humphrey, Lax Iyer, Lisa Pfefferle, Manoj Jain, Matthew Robinson, Mohan Kolli, Brian Krueger, Tim Kuphal, Stanley Letovsky, Michael Levandoski, Craig Lukasik, Jonathan Meltzer, Brian Norvell, Mindy Nye, Scott Parker, Christos Petropoulos, John Pruitt, Steven Ragan, Scott Ryan, Mike Sapeta, Jana Schroth, Suresh Babu Selvaraju, Goran Stevovic, Amanda Suchanek, Andrea Throop, Lyndon Tilson, Thomas Urban, Joe Voshell, Kimberly Wagner, Jonathan Williams, Mary Williamson, Qian Zeng, Tricia Zwiefelhofer, Clinton R. Paden, Duncan MacCannell |
| EPI_ISL_3327991 | NJDOH, Public Health and Environmental Laboratories | NJ_PHEL | Lindsey Bodnar, Shiv K. Verma, Ryan Pachucki, Chelsea San Filippo, Jacquelyn Deverell, Dana Woell, Allison Roder, Byeong Jeong |
| EPI_ISL_3327992, EPI_ISL_3327993, EPI_ISL_3327994 | Laboratory Corporation of America | Centers for Disease Control and Prevention Division of Viral Diseases, Pathogen Discovery | Dakota Howard, Dhwani Batra, Peter W. Cook, Kara Moser, Adrian Paskey, Jason Caravas, Benjamin Rambo-Martin, Shatavia Morrison, Christopher Gulvick, Scott Sammons, Yvette Unoarumhi, Darlene Wagner, Matthew Schmerer, Minoo Agarwal, Eyad Almasri, Debbie Boles, Ayla Burns, Nuthawin Charoensri, Oren Cohen, Susan Countryman, Mary Ann Cristobal, Bobbi Croy, Suzanne Dale, Hrushikesh Deshmukh, Amanda Douglas, Vincent Drouillon, Marcia Eisenberg, Howard Engler, Rama Ghatti, Prashant Gupta, Susan Hicks, Jake Humphrey, Lax Iyer, Lisa Pfefferle, Manoj Jain, Matthew Robinson, Mohan Kolli, Brian Krueger, Tim Kuphal, Stanley Letovsky, Michael Levandoski, Craig Lukasik, Jonathan Meltzer, Brian Norvell, Mindy Nye, Scott Parker, Christos Petropoulos, John Pruitt, Steven Ragan, Scott Ryan, Mike Sapeta, Jana Schroth, Suresh Babu Selvaraju, Goran Stevovic, Amanda Suchanek, Andrea Throop, Lyndon Tilson, Thomas Urban, Joe Voshell, Kimberly Wagner, Jonathan Williams, Mary Williamson, Qian Zeng, Tricia Zwiefelhofer, Clinton R. Paden, Duncan MacCannell |
| EPI_ISL_3327995 | NJDOH, Public Health and Environmental Laboratories | NJ_PHEL | Lindsey Bodnar, Shiv K. Verma, Ryan Pachucki, Chelsea San Filippo, Jacquelyn Deverell, Dana Woell, Allison Roder, Byeong Jeong |
| EPI_ISL_3327996, EPI_ISL_3327997 | Laboratory Corporation of America | Centers for Disease Control and Prevention Division of Viral Diseases, Pathogen Discovery | Dakota Howard, Dhwani Batra, Peter W. Cook, Kara Moser, Adrian Paskey, Jason Caravas, Benjamin Rambo-Martin, Shatavia Morrison, Christopher Gulvick, Scott Sammons, Yvette Unoarumhi, Darlene Wagner, Matthew Schmerer, Minoo Agarwal, Eyad Almasri, Debbie Boles, Ayla Burns, Nuthawin Charoensri, Oren Cohen, Susan Countryman, Mary Ann Cristobal, Bobbi Croy, Suzanne Dale, Hrushikesh Deshmukh, Amanda Douglas, Vincent Drouillon, Marcia Eisenberg, Howard Engler, Rama Ghatti, Prashant Gupta, Susan Hicks, Jake Humphrey, Lax Iyer, Lisa Pfefferle, Manoj Jain, Matthew Robinson, Mohan Kolli, Brian Krueger, Tim Kuphal, Stanley Letovsky, Michael Levandoski, Craig Lukasik, Jonathan Meltzer, Brian Norvell, Mindy Nye, Scott Parker, Christos Petropoulos, John Pruitt, Steven Ragan, Scott Ryan, Mike Sapeta, Jana Schroth, Suresh Babu Selvaraju, Goran Stevovic, Amanda Suchanek, Andrea Throop, Lyndon Tilson, Thomas Urban, Joe Voshell, Kimberly Wagner, Jonathan Williams, Mary Williamson, Qian Zeng, Tricia Zwiefelhofer, Clinton R. Paden, Duncan MacCannell |
| EPI_ISL_3327998 | NJDOH, Public Health and Environmental Laboratories | NJ_PHEL | Lindsey Bodnar, Shiv K. Verma, Ryan Pachucki, Chelsea San Filippo, Jacquelyn Deverell, Dana Woell, Allison Roder, Byeong Jeong |
| EPI_ISL_3327999, EPI_ISL_3328000 | Laboratory Corporation of America | Centers for Disease Control and Prevention Division of Viral Diseases, Pathogen Discovery | Dakota Howard, Dhwani Batra, Peter W. Cook, Kara Moser, Adrian Paskey, Jason Caravas, Benjamin Rambo-Martin, Shatavia Morrison, Christopher Gulvick, Scott Sammons, Yvette Unoarumhi, Darlene Wagner, Matthew Schmerer, Minoo Agarwal, Eyad Almasri, Debbie Boles, Ayla Burns, Nuthawin Charoensri, Oren Cohen, Susan Countryman, Mary Ann Cristobal, Bobbi Croy, Suzanne Dale, Hrushikesh Deshmukh, Amanda Douglas, Vincent Drouillon, Marcia Eisenberg, Howard Engler, Rama Ghatti, Prashant Gupta, Susan Hicks, Jake Humphrey, Lax Iyer, Lisa Pfefferle, Manoj Jain, Matthew Robinson, Mohan Kolli, Brian Krueger, Tim Kuphal, Stanley Letovsky, Michael Levandoski, Craig Lukasik, Jonathan Meltzer, Brian Norvell, Mindy Nye, Scott Parker, Christos Petropoulos, John Pruitt, Steven Ragan, Scott Ryan, Mike Sapeta, Jana Schroth, Suresh Babu Selvaraju, Goran Stevovic, Amanda Suchanek, Andrea Throop, Lyndon Tilson, Thomas Urban, Joe Voshell, Kimberly Wagner, Jonathan Williams, Mary Williamson, Qian Zeng, Tricia Zwiefelhofer, Clinton R. Paden, Duncan MacCannell |
| EPI_ISL_3328001 | NJDOH, Public Health and Environmental Laboratories | NJ_PHEL | Lindsey Bodnar, Shiv K. Verma, Ryan Pachucki, Chelsea San Filippo, Jacquelyn Deverell, Dana Woell, Allison Roder, Byeong Jeong |
| EPI_ISL_3328002, EPI_ISL_3328003 | Laboratory Corporation of America | Centers for Disease Control and Prevention Division of Viral Diseases, Pathogen Discovery | Dakota Howard, Dhwani Batra, Peter W. Cook, Kara Moser, Adrian Paskey, Jason Caravas, Benjamin Rambo-Martin, Shatavia Morrison, Christopher Gulvick, Scott Sammons, Yvette Unoarumhi, Darlene Wagner, Matthew Schmerer, Minoo Agarwal, Eyad Almasri, Debbie Boles, Ayla Burns, Nuthawin Charoensri, Oren Cohen, Susan Countryman, Mary Ann Cristobal, Bobbi Croy, Suzanne Dale, Hrushikesh Deshmukh, Amanda Douglas, Vincent Drouillon, Marcia Eisenberg, Howard Engler, Rama Ghatti, Prashant Gupta, Susan Hicks, Jake Humphrey, Lax Iyer, Lisa Pfefferle, Manoj Jain, Matthew Robinson, Mohan Kolli, Brian Krueger, Tim Kuphal, Stanley Letovsky, Michael Levandoski, Craig Lukasik, Jonathan Meltzer, Brian Norvell, Mindy Nye, Scott Parker, Christos Petropoulos, John Pruitt, Steven Ragan, Scott Ryan, Mike Sapeta, Jana Schroth, Suresh Babu Selvaraju, Goran Stevovic, Amanda Suchanek, Andrea Throop, Lyndon Tilson, Thomas Urban, Joe Voshell, Kimberly Wagner, Jonathan Williams, Mary Williamson, Qian Zeng, Tricia Zwiefelhofer, Clinton R. Paden, Duncan MacCannell |
| EPI_ISL_3328004 | NJDOH, Public Health and Environmental Laboratories | NJ_PHEL | Lindsey Bodnar, Shiv K. Verma, Ryan Pachucki, Chelsea San Filippo, Jacquelyn Deverell, Dana Woell, Allison Roder, Byeong Jeong |

EPI_ISL_3328005, EPI_ISL_3328006, EPI_ISL_3328007, EPI_ISL_3328009, EPI_ISL_3328010, EPI_ISL_3328011, EPI_ISL_3328012, EPI_ISL_3328014, EPI_ISL_3328015, EPI_ISL_3328016, EPI_ISL_3328017, EPI_ISL_3328018, EPI_ISL_3328019, EPI_ISL_3328020, EPI_ISL_3328021, EPI_ISL_3328022, EPI_ISL_3328023, EPI_ISL_3328024, EPI_ISL_3328027, EPI_ISL_3328029, EPI_ISL_3328030, EPI_ISL_3328031, EPI_ISL_3328032, EPI_ISL_3328033, EPI_ISL_3328034, EPI_ISL_3328035, EPI_ISL_3328037, EPI_ISL_3328038, EPI_ISL_3328039, EPI_ISL_3328040, EPI_ISL_3328041, EPI_ISL_3328042, EPI_ISL_3328043, EPI_ISL_3328045, EPI_ISL_3328046, EPI_ISL_3328048, EPI_ISL_3328049, EPI_ISL_3328050, EPI_ISL_3328051, EPI_ISL_3328052, EPI_ISL_3328053, EPI_ISL_3328054, EPI_ISL_3328055, EPI_ISL_3328056, EPI_ISL_3328057, EPI_ISL_3328058, EPI_ISL_3328059, EPI_ISL_3328060, EPI_ISL_3328061, EPI_ISL_3328062, EPI_ISL_3328063, EPI_ISL_3328064, EPI_ISL_3328065, EPI_ISL_3328066, EPI_ISL_3328068, EPI_ISL_3328069, EPI_ISL_3328070, EPI_ISL_3328071, EPI_ISL_3328072, EPI_ISL_3328073, EPI_ISL_3328074, EPI_ISL_3328075, EPI_ISL_3328076, EPI_ISL_3328077, EPI_ISL_3328078, EPI_ISL_3328079, EPI_ISL_3328080, EPI_ISL_3328081, EPI_ISL_3328082, EPI_ISL_3328083, EPI_ISL_3328084, EPI_ISL_3328085, EPI_ISL_3328086, EPI_ISL_3328087, EPI_ISL_3328089, EPI_ISL_3328091, EPI_ISL_3328092, EPI_ISL_3328093, EPI_ISL_3328094, EPI_ISL_3328095, EPI_ISL_3328096, EPI_ISL_3328099, EPI_ISL_3328100, EPI_ISL_3328102, EPI_ISL_3328103, EPI_ISL_3328104, EPI_ISL_3328105, EPI_ISL_3328106, EPI_ISL_3328107, EPI_ISL_3328109, EPI_ISL_3328110, EPI_ISL_3328111, EPI_ISL_3328112, EPI_ISL_3328113, EPI_ISL_3328114, EPI_ISL_3328115, EPI_ISL_3328116, EPI_ISL_3328117, EPI_ISL_3328118, EPI_ISL_3328119, EPI_ISL_3328120, EPI_ISL_3328121, EPI_ISL_3328122, EPI_ISL_3328123, EPI_ISL_3328124, EPI_ISL_3328127, EPI_ISL_3328128, EPI_ISL_3328129, EPI_ISL_3328131, EPI_ISL_3328132, EPI_ISL_3328133, EPI_ISL_3328134, EPI_ISL_3328135, EPI_ISL_3328137, EPI_ISL_3328139, EPI_ISL_3328141, EPI_ISL_3328142, EPI_ISL_3328144, EPI_ISL_3328145, EPI_ISL_3328146, EPI_ISL_3328147, EPI_ISL_3328148, EPI_ISL_3328149, EPI_ISL_3328150, EPI_ISL_3328151, EPI_ISL_3328152, EPI_ISL_3328153, EPI_ISL_3328154, EPI_ISL_3328155, EPI_ISL_3328156, EPI_ISL_3328157, EPI_ISL_3328158, EPI_ISL_3328159, EPI_ISL_3328161, EPI_ISL_3328162, EPI_ISL_3328163, EPI_ISL_3328164, EPI_ISL_3328165, EPI_ISL_3328168, EPI_ISL_3328169, EPI_ISL_3328171, EPI_ISL_3328172, EPI_ISL_3328173, EPI_ISL_3328174, EPI_ISL_3328175, EPI_ISL_3328176, EPI_ISL_3328177, EPI_ISL_3328178, EPI_ISL_3328179, EPI_ISL_3328180, EPI_ISL_3328181, EPI_ISL_3328182, EPI_ISL_3328185, EPI_ISL_3328186, EPI_ISL_3328187, EPI_ISL_3328188, EPI_ISL_3328189, EPI_ISL_3328190, EPI_ISL_3328191, EPI_ISL_3328192, EPI_ISL_3328193, EPI_ISL_3328194, EPI_ISL_3328195, EPI_ISL_3328196, EPI_ISL_3328197, EPI_ISL_3328198, EPI_ISL_3328199, EPI_ISL_3328200, EPI_ISL_3328203, EPI_ISL_3328204, EPI_ISL_3328205, EPI_ISL_3328206, EPI_ISL_3328207, EPI_ISL_3328208, EPI_ISL_3328209, EPI_ISL_3328210, EPI_ISL_3328211, EPI_ISL_3328212, EPI_ISL_3328213, EPI_ISL_3328214, EPI_ISL_3328215, EPI_ISL_3328216, EPI_ISL_3328218, EPI_ISL_3328219, EPI_ISL_3328220, EPI_ISL_3328221, EPI_ISL_3328222, EPI_ISL_3328223, EPI_ISL_3328224, EPI_ISL_3328225, EPI_ISL_3328226, EPI_ISL_3328227, EPI_ISL_3328228, EPI_ISL_3328229, EPI_ISL_3328230, EPI_ISL_3328231, EPI_ISL_3328232, EPI_ISL_3328233, EPI_ISL_3328234, EPI_ISL_3328235, EPI_ISL_3328236, EPI_ISL_3328237, EPI_ISL_3328238, EPI_ISL_3328239, EPI_ISL_3328240, EPI_ISL_3328241, EPI_ISL_3328242, EPI_ISL_3328243, EPI_ISL_3328244, EPI_ISL_3328245, EPI_ISL_3328246, EPI_ISL_3328248, EPI_ISL_3328249, EPI_ISL_3328250, EPI_ISL_3328251, EPI_ISL_3328253, EPI_ISL_3328254, EPI_ISL_3328255, EPI_ISL_3328256, EPI_ISL_3328257,

EPI_ISL_3328258, EPI_ISL_3328259, EPI_ISL_3328260, EPI_ISL_3328261, EPI_ISL_3328262, EPI_ISL_3328263, EPI_ISL_3328264, EPI_ISL_3328267, EPI_ISL_3328268, EPI_ISL_3328269, EPI_ISL_3328270, EPI_ISL_3328271, EPI_ISL_3328272, EPI_ISL_3328273, EPI_ISL_3328274, EPI_ISL_3328276, EPI_ISL_3328277, EPI_ISL_3328278, EPI_ISL_3328279, EPI_ISL_3328280, EPI_ISL_3328282, EPI_ISL_3328283, EPI_ISL_3328284, EPI_ISL_3328285, EPI_ISL_3328286, EPI_ISL_3328287, EPI_ISL_3328288, EPI_ISL_3328289, EPI_ISL_3328291, EPI_ISL_3328292, EPI_ISL_3328293, EPI_ISL_3328294, EPI_ISL_3328296, EPI_ISL_3328297, EPI_ISL_3328298, EPI_ISL_3328299, EPI_ISL_3328300, EPI_ISL_3328301, EPI_ISL_3328302, EPI_ISL_3328303, EPI_ISL_3328304, EPI_ISL_3328305, EPI_ISL_3328306, EPI_ISL_3328309, EPI_ISL_3328310, EPI_ISL_3328311, EPI_ISL_3328313, EPI_ISL_3328314, EPI_ISL_3328315, EPI_ISL_3328316, EPI_ISL_3328317, EPI_ISL_3328318, EPI_ISL_3328319, EPI_ISL_3328320, EPI_ISL_3328321, EPI_ISL_3328323, EPI_ISL_3328324, EPI_ISL_3328325, EPI_ISL_3328327, EPI_ISL_3328328, EPI_ISL_3328329, EPI_ISL_3328330, EPI_ISL_3328331, EPI_ISL_3328332, EPI_ISL_3328333, EPI_ISL_3328334, EPI_ISL_3328335, EPI_ISL_3328336, EPI_ISL_3328337, EPI_ISL_3328338, EPI_ISL_3328340, EPI_ISL_3328341, EPI_ISL_3328342, EPI_ISL_3328343, EPI_ISL_3328344, EPI_ISL_3328345, EPI_ISL_3328346, EPI_ISL_3328347, EPI_ISL_3328348, EPI_ISL_3328349, EPI_ISL_3328350, EPI_ISL_3328351, EPI_ISL_3328352, EPI_ISL_3328353, EPI_ISL_3328354, EPI_ISL_3328355, EPI_ISL_3328356, EPI_ISL_3328357, EPI_ISL_3328358, EPI_ISL_3328359, EPI_ISL_3328360, EPI_ISL_3328361, EPI_ISL_3328362, EPI_ISL_3328363, EPI_ISL_3328364, EPI_ISL_3328365, EPI_ISL_3328366, EPI_ISL_3328367, EPI_ISL_3328368, EPI_ISL_3328369, EPI_ISL_3328370, EPI_ISL_3328371, EPI_ISL_3328372, EPI_ISL_3328373, EPI_ISL_3328374, EPI_ISL_3328375, EPI_ISL_3328376, EPI_ISL_3328377, EPI_ISL_3328378, EPI_ISL_3328379, EPI_ISL_3328380, EPI_ISL_3328381, EPI_ISL_3328382, EPI_ISL_3328383, EPI_ISL_3328384, EPI_ISL_3328385, EPI_ISL_3328386, EPI_ISL_3328387, EPI_ISL_3328388, EPI_ISL_3328389, EPI_ISL_3328390, EPI_ISL_3328391, EPI_ISL_3328392, EPI_ISL_3328393, EPI_ISL_3328394, EPI_ISL_3328395, EPI_ISL_3328396, EPI_ISL_3328397, EPI_ISL_3328398, EPI_ISL_3328399, EPI_ISL_3328400, EPI_ISL_3328401, EPI_ISL_3328402, EPI_ISL_3328403, EPI_ISL_3328404, EPI_ISL_3328405, EPI_ISL_3328406, EPI_ISL_3328407, EPI_ISL_3328408, EPI_ISL_3328409, EPI_ISL_3328410, EPI_ISL_3328411, EPI_ISL_3328412, EPI_ISL_3328413, EPI_ISL_3328414, EPI_ISL_3328415, EPI_ISL_3328416, EPI_ISL_3328417, EPI_ISL_3328418, EPI_ISL_3328419, EPI_ISL_3328420, EPI_ISL_3328421, EPI_ISL_3328422, EPI_ISL_3328424, EPI_ISL_3328425, EPI_ISL_3328426, EPI_ISL_3328427, EPI_ISL_3328428, EPI_ISL_3328429, EPI_ISL_3328430, EPI_ISL_3328431, EPI_ISL_3328432, EPI_ISL_3328433, EPI_ISL_3328434, EPI_ISL_3328435, EPI_ISL_3328436, EPI_ISL_3328437, EPI_ISL_3328438, EPI_ISL_3328439, EPI_ISL_3328440, EPI_ISL_3328441, EPI_ISL_3328445, EPI_ISL_3328446, EPI_ISL_3328447, EPI_ISL_3328448, EPI_ISL_3328449, EPI_ISL_3328450, EPI_ISL_3328454, EPI_ISL_3328456, EPI_ISL_3328457, EPI_ISL_3328461, EPI_ISL_3328462, EPI_ISL_3328463, EPI_ISL_3328465, EPI_ISL_3328466, EPI_ISL_3328469, EPI_ISL_3328470, EPI_ISL_3328471, EPI_ISL_3328472, EPI_ISL_3328473, EPI_ISL_3328474, EPI_ISL_3328475, EPI_ISL_3328477, EPI_ISL_3328478, EPI_ISL_3328479, EPI_ISL_3328482, EPI_ISL_3328484, EPI_ISL_3328485, EPI_ISL_3328486, EPI_ISL_3328487, EPI_ISL_3328488, EPI_ISL_3328489, EPI_ISL_3328490, EPI_ISL_3328491, EPI_ISL_3328492, EPI_ISL_3328493, EPI_ISL_3328494, EPI_ISL_3328495, EPI_ISL_3328497, EPI_ISL_3328498, EPI_ISL_3328500, EPI_ISL_3328501, EPI_ISL_3328502, EPI_ISL_3328503, EPI_ISL_3328504, EPI_ISL_3328505, EPI_ISL_3328506, EPI_ISL_3328508, EPI_ISL_3328509, EPI_ISL_3328510, EPI_ISL_3328511, EPI_ISL_3328512, EPI_ISL_3328513, EPI_ISL_3328515, EPI_ISL_3328516, EPI_ISL_3328517, EPI_ISL_3328518, EPI_ISL_3328520, EPI_ISL_3328521, EPI_ISL_3328522, EPI_ISL_3328523, EPI_ISL_3328524, EPI_ISL_3328525, EPI_ISL_3328526, EPI_ISL_3328528, EPI_ISL_3328529, EPI_ISL_3328530, EPI_ISL_3328531, EPI_ISL_3328533, EPI_ISL_3328534, EPI_ISL_3328535, EPI_ISL_3328536, EPI_ISL_3328537, EPI_ISL_3328538, EPI_ISL_3328539, EPI_ISL_3328540, EPI_ISL_3328541, EPI_ISL_3328542, EPI_ISL_3328543, EPI_ISL_3328544, EPI_ISL_3328545, EPI_ISL_3328546, EPI_ISL_3328547, EPI_ISL_3328548, EPI_ISL_3328549, EPI_ISL_3328550, EPI_ISL_3328551, EPI_ISL_3328553, EPI_ISL_3328554, EPI_ISL_3328556, EPI_ISL_3328557, EPI_ISL_3328558, EPI_ISL_3328559, EPI_ISL_3328560, EPI_ISL_3328561, EPI_ISL_3328562, EPI_ISL_3328563, EPI_ISL_3328564, EPI_ISL_3328565, EPI_ISL_3328566, EPI_ISL_3328567, EPI_ISL_3328568, EPI_ISL_3328569, EPI_ISL_3328570, EPI_ISL_3328571, EPI_ISL_3328572, EPI_ISL_3328573, EPI_ISL_3328574, EPI_ISL_3328575, EPI_ISL_3328576, EPI_ISL_3328577, EPI_ISL_3328579, EPI_ISL_3328580, EPI_ISL_3328581, EPI_ISL_3328582, EPI_ISL_3328583, EPI_ISL_3328584, EPI_ISL_3328585, EPI_ISL_3328586, EPI_ISL_3328587, EPI_ISL_3328588, EPI_ISL_3328589, EPI_ISL_3328590, EPI_ISL_3328591, EPI_ISL_3328593, EPI_ISL_3328594, EPI_ISL_3328595, EPI_ISL_3328596, EPI_ISL_3328597, EPI_ISL_3328598, EPI_ISL_3328601, EPI_ISL_3328603, EPI_ISL_3328604, EPI_ISL_3328605, EPI_ISL_3328606, EPI_ISL_3328607, EPI_ISL_3328608, EPI_ISL_3328609, EPI_ISL_3328611, EPI_ISL_3328612, EPI_ISL_3328613, EPI_ISL_3328614, EPI_ISL_3328615, EPI_ISL_3328616, EPI_ISL_3328617, EPI_ISL_3328618, EPI_ISL_3328619, EPI_ISL_3328620, EPI_ISL_3328621, EPI_ISL_3328623, EPI_ISL_3328624, EPI_ISL_3328625, EPI_ISL_3328626, EPI_ISL_3328627, EPI_ISL_3328628, EPI_ISL_3328629, EPI_ISL_3328630, EPI_ISL_3328631, EPI_ISL_3328632, EPI_ISL_3328633, EPI_ISL_3328634, EPI_ISL_3328635, EPI_ISL_3328636, EPI_ISL_3328637, EPI_ISL_3328638, EPI_ISL_3328639, EPI_ISL_3328641, EPI_ISL_3328642, EPI_ISL_3328643, EPI_ISL_3328644, EPI_ISL_3328645, EPI_ISL_3328646, EPI_ISL_3328647, EPI_ISL_3328648, EPI_ISL_3328649, EPI_ISL_3328650, EPI_ISL_3328651, EPI_ISL_3328652, EPI_ISL_3328653, EPI_ISL_3328654, EPI_ISL_3328655, EPI_ISL_3328656, EPI_ISL_3328658, EPI_ISL_3328659, EPI_ISL_3328660, EPI_ISL_3328661, EPI_ISL_3328662, EPI_ISL_3328663, EPI_ISL_3328664, EPI_ISL_3328665, EPI_ISL_3328666, EPI_ISL_3328667, EPI_ISL_3328669, EPI_ISL_3328670, EPI_ISL_3328671, EPI_ISL_3328672, EPI_ISL_3328673, EPI_ISL_3328674, EPI_ISL_3328675, EPI_ISL_3328676, EPI_ISL_3328677, EPI_ISL_3328678, EPI_ISL_3328679, EPI_ISL_3328680, EPI_ISL_3328681, EPI_ISL_3328683, EPI_ISL_3328684, EPI_ISL_3328685, EPI_ISL_3328686, EPI_ISL_3328687, EPI_ISL_3328688, EPI_ISL_3328689, EPI_ISL_3328690, EPI_ISL_3328691, EPI_ISL_3328692, EPI_ISL_3328693, EPI_ISL_3328694, EPI_ISL_3328695, EPI_ISL_3328696, EPI_ISL_3328697, EPI_ISL_3328698, EPI_ISL_3328699, EPI_ISL_3328700, EPI_ISL_3328701, EPI_ISL_3328702, EPI_ISL_3328704, EPI_ISL_3328705, EPI_ISL_3328707, EPI_ISL_3328708, EPI_ISL_3328709, EPI_ISL_3328710, EPI_ISL_3328711, EPI_ISL_3328712, EPI_ISL_3328713, EPI_ISL_3328714, EPI_ISL_3328715, EPI_ISL_3328716, EPI_ISL_3328717, EPI_ISL_3328718, EPI_ISL_3328719, EPI_ISL_3328720, EPI_ISL_3328721, EPI_ISL_3328722, EPI_ISL_3328723, EPI_ISL_3328724, EPI_ISL_3328725, EPI_ISL_3328726, EPI_ISL_3328727, EPI_ISL_3328728, EPI_ISL_3328729, EPI_ISL_3328730, EPI_ISL_3328731, EPI_ISL_3328732, EPI_ISL_3328733, EPI_ISL_3328734, EPI_ISL_3328735, EPI_ISL_3328736, EPI_ISL_3328737, EPI_ISL_3328738, EPI_ISL_3328739, EPI_ISL_3328740, EPI_ISL_3328741, EPI_ISL_3328742, EPI_ISL_3328743, EPI_ISL_3328744, EPI_ISL_3328745, EPI_ISL_3328746, EPI_ISL_3328747, EPI_ISL_3328748, EPI_ISL_3328749, EPI_ISL_3328750, EPI_ISL_3328751, EPI_ISL_3328752, EPI_ISL_3328753, EPI_ISL_3328754, EPI_ISL_3328755, EPI_ISL_3328757, EPI_ISL_3328758, EPI_ISL_3328759, EPI_ISL_3328760, EPI_ISL_3328761, EPI_ISL_3328762, EPI_ISL_3328763, EPI_ISL_3328764, EPI_ISL_3328765, EPI_ISL_3328766, EPI_ISL_3328767, EPI_ISL_3328768, EPI_ISL_3328769, EPI_ISL_3328770, EPI_ISL_3328771, EPI_ISL_3328772, EPI_ISL_3328773, EPI_ISL_3328774, EPI_ISL_3328775, EPI_ISL_3328776, EPI_ISL_3328777, EPI_ISL_3328778, EPI_ISL_3328779, EPI_ISL_3328780, EPI_ISL_3328781, EPI_ISL_3328782, EPI_ISL_3328783, EPI_ISL_3328784, EPI_ISL_3328785, EPI_ISL_3328786, EPI_ISL_3328787, EPI_ISL_3328788, EPI_ISL_3328789, EPI_ISL_3328790, EPI_ISL_3328791, EPI_ISL_3328793, EPI_ISL_3328794, EPI_ISL_3328795, EPI_ISL_3328796, EPI_ISL_3328797, EPI_ISL_3328798, EPI_ISL_3328799, EPI_ISL_3328801, EPI_ISL_3328802, EPI_ISL_3328803, EPI_ISL_3328804, EPI_ISL_3328805, EPI_ISL_3328807, EPI_ISL_3328808, EPI_ISL_3328809, EPI_ISL_3328810, EPI_ISL_3328811, EPI_ISL_3328812, EPI_ISL_3328813, EPI_ISL_3328814, EPI_ISL_3328815, EPI_ISL_3328816, EPI_ISL_3328817, EPI_ISL_3328818, EPI_ISL_3328819, EPI_ISL_3328820, EPI_ISL_3328821, EPI_ISL_3328822, EPI_ISL_3328823, EPI_ISL_3328824, EPI_ISL_3328825, EPI_ISL_3328826, EPI_ISL_3328827, EPI_ISL_3328829, EPI_ISL_3328831, EPI_ISL_3328832, EPI_ISL_3328833, EPI_ISL_3328834, EPI_ISL_3328835, EPI_ISL_3328836, EPI_ISL_3328837, EPI_ISL_3328838, EPI_ISL_3328839, EPI_ISL_3328840, EPI_ISL_3328841, EPI_ISL_3328842, EPI_ISL_3328843, EPI_ISL_3328844, EPI_ISL_3328845, EPI_ISL_3328846, EPI_ISL_3328847, EPI_ISL_3328848, EPI_ISL_3328849, EPI_ISL_3328850, EPI_ISL_3328853, EPI_ISL_3328854, EPI_ISL_3328856, EPI_ISL_3328857, EPI_ISL_3328858, EPI_ISL_3328859, EPI_ISL_3328860, EPI_ISL_3328861, EPI_ISL_3328862, EPI_ISL_3328863, EPI_ISL_3328864, EPI_ISL_3328866, EPI_ISL_3328867, EPI_ISL_3328868, EPI_ISL_3328869, EPI_ISL_3328870, EPI_ISL_3328871, EPI_ISL_3328872, EPI_ISL_3328873, EPI_ISL_3328874, EPI_ISL_3328875, EPI_ISL_3328876, EPI_ISL_3328877, EPI_ISL_3328878, EPI_ISL_3328879, EPI_ISL_3328880, EPI_ISL_3328881, EPI_ISL_3328882, EPI_ISL_3328883, EPI_ISL_3328884, EPI_ISL_3328885, EPI_ISL_3328886, EPI_ISL_3328887, EPI_ISL_3328888, EPI_ISL_3328889, EPI_ISL_3328890, EPI_ISL_3328891, EPI_ISL_3328892, EPI_ISL_3328893, EPI_ISL_3328894, EPI_ISL_3328895, EPI_ISL_3328896, EPI_ISL_3328897, EPI_ISL_3328898, EPI_ISL_3328900, EPI_ISL_3328901, EPI_ISL_3328902, EPI_ISL_3328903, EPI_ISL_3328904, EPI_ISL_3328905, EPI_ISL_3328906, EPI_ISL_3328907, EPI_ISL_3328908, EPI_ISL_3328909, EPI_ISL_3328911, EPI_ISL_3328912, EPI_ISL_3328913, EPI_ISL_3328914, EPI_ISL_3328915, EPI_ISL_3328916, EPI_ISL_3328917, EPI_ISL_3328918, EPI_ISL_3328919, EPI_ISL_3328920, EPI_ISL_3328921, EPI_ISL_3328922, EPI_ISL_3328923, EPI_ISL_3328924, EPI_ISL_3328925, EPI_ISL_3328926, EPI_ISL_3328927, EPI_ISL_3328928, EPI_ISL_3328929, EPI_ISL_3328930, EPI_ISL_3328931, EPI_ISL_3328932, EPI_ISL_3328934, EPI_ISL_3328935, EPI_ISL_3328936, EPI_ISL_3328938, EPI_ISL_3328939, EPI_ISL_3328940, EPI_ISL_3328941, EPI_ISL_3328942, EPI_ISL_3328943, EPI_ISL_3328944, EPI_ISL_3328945, EPI_ISL_3328946, EPI_ISL_3328947, EPI_ISL_3328948, EPI_ISL_3328949, EPI_ISL_3328950, EPI_ISL_3328951, EPI_ISL_3328952, EPI_ISL_3328953, EPI_ISL_3328954, EPI_ISL_3328955, EPI_ISL_3328956, EPI_ISL_3328957, EPI_ISL_3328958, EPI_ISL_3328959, EPI_ISL_3328960, EPI_ISL_3328961, EPI_ISL_3328962, EPI_ISL_3328963, EPI_ISL_3328965, EPI_ISL_3328966, EPI_ISL_3328967, EPI_ISL_3328968, EPI_ISL_3328969, EPI_ISL_3328970, EPI_ISL_3328971, EPI_ISL_3328972, EPI_ISL_3328973, EPI_ISL_3328974, EPI_ISL_3328975, EPI_ISL_3328976, EPI_ISL_3328977, EPI_ISL_3328978, EPI_ISL_3328979, EPI_ISL_3328980, EPI_ISL_3328981, EPI_ISL_3328982, EPI_ISL_3328983, EPI_ISL_3328984, EPI_ISL_3328985, EPI_ISL_3328986, EPI_ISL_3328987, EPI_ISL_3328988, EPI_ISL_3328989, EPI_ISL_3328990, EPI_ISL_3328991, EPI_ISL_3328992, EPI_ISL_3328993, EPI_ISL_3328994, EPI_ISL_3328995, EPI_ISL_3328996, EPI_ISL_3328997, EPI_ISL_3328998, EPI_ISL_3328999, EPI_ISL_3329000, EPI_ISL_3329001, EPI_ISL_3329002, EPI_ISL_3329003, EPI_ISL_3329004, EPI_ISL_3329005, EPI_ISL_3329006, EPI_ISL_3329007, EPI_ISL_3329008, EPI_ISL_3329009, EPI_ISL_3329010, EPI_ISL_3329011, EPI_ISL_3329012, EPI_ISL_3329013, EPI_ISL_3329014, EPI_ISL_3329015, EPI_ISL_3329016, EPI_ISL_3329017, EPI_ISL_3329018, EPI_ISL_3329019, EPI_ISL_3329020, EPI_ISL_3329021, EPI_ISL_3329022, EPI_ISL_3329023, EPI_ISL_3329024, EPI_ISL_3329025, EPI_ISL_3329026, EPI_ISL_3329027, EPI_ISL_3329028, EPI_ISL_3329029, EPI_ISL_3329031, EPI_ISL_3329032, EPI_ISL_3329033, EPI_ISL_3329034, EPI_ISL_3329035, EPI_ISL_3329036, EPI_ISL_3329037, EPI_ISL_3329038, EPI_ISL_3329039, EPI_ISL_3329040, EPI_ISL_3329041, EPI_ISL_3329042, EPI_ISL_3329043, EPI_ISL_3329044, EPI_ISL_3329045, EPI_ISL_3329046, EPI_ISL_3329047, EPI_ISL_3329049, EPI_ISL_3329050, EPI_ISL_3329051, EPI_ISL_3329052, EPI_ISL_3329053, EPI_ISL_3329054, EPI_ISL_3329055, EPI_ISL_3329056, EPI_ISL_3329057, EPI_ISL_3329058, EPI_ISL_3329060, EPI_ISL_3329061, EPI_ISL_3329062, EPI_ISL_3329063, EPI_ISL_3329064, EPI_ISL_3329065, EPI_ISL_3329066, EPI_ISL_3329067, EPI_ISL_3329068, EPI_ISL_3329069, EPI_ISL_3329070, EPI_ISL_3329071, EPI_ISL_3329072, EPI_ISL_3329073, EPI_ISL_3329074, EPI_ISL_3329075, EPI_ISL_3329076, EPI_ISL_3329077, EPI_ISL_3329078, EPI_ISL_3329079, EPI_ISL_3329080, EPI_ISL_3329081, EPI_ISL_3329082, EPI_ISL_3329083, EPI_ISL_3329084, EPI_ISL_3329087, EPI_ISL_3329088, EPI_ISL_3329089, EPI_ISL_3329090, EPI_ISL_3329091, EPI_ISL_3329092, EPI_ISL_3329093, EPI_ISL_3329094, EPI_ISL_3329095, EPI_ISL_3329096, EPI_ISL_3329097, EPI_ISL_3329098, EPI_ISL_3329099, EPI_ISL_3329100, EPI_ISL_3329101, EPI_ISL_3329102, EPI_ISL_3329103, EPI_ISL_3329104, EPI_ISL_3329105, EPI_ISL_3329106, EPI_ISL_3329107, EPI_ISL_3329108, EPI_ISL_3329109, EPI_ISL_3329110, EPI_ISL_3329111, EPI_ISL_3329112, EPI_ISL_3329113, EPI_ISL_3329114, EPI_ISL_3329116, EPI_ISL_3329118, EPI_ISL_3329119, EPI_ISL_3329120, EPI_ISL_3329121, EPI_ISL_3329122, EPI_ISL_3329123, EPI_ISL_3329124, EPI_ISL_3329125, EPI_ISL_3329126, EPI_ISL_3329127, EPI_ISL_3329128, EPI_ISL_3329129, EPI_ISL_3329130, EPI_ISL_3329131, EPI_ISL_3329132, EPI_ISL_3329133, EPI_ISL_3329134, EPI_ISL_3329135, EPI_ISL_3329136, EPI_ISL_3329138, EPI_ISL_3329139, EPI_ISL_3329140, EPI_ISL_3329141, EPI_ISL_3329142, EPI_ISL_3329143, EPI_ISL_3329145, EPI_ISL_3329146, EPI_ISL_3329148, EPI_ISL_3329149, EPI_ISL_3329150

| | | | |
|---|---|---|---|
| see above | Laboratory Corporation of America | Centers for Disease Control and Prevention Division of Viral Diseases, Pathogen Discovery | Dakota Howard, Dhwani Batra, Peter W. Cook, Kara Moser, Adrian Paskey, Jason Caravas, Benjamin Rambo-Martin, Shatavia Morrison, Christopher Gulvick, Scott Sammons, Yvette Unoarumhi, Darlene Wagner, Matthew Schmerer, Minoo Agarwal, Eyad Almasri, Debbie Boles, Ayla Burns, Nuthawin Charoensri, Oren Cohen, Susan Countryman, Mary Ann Cristobal, Bobbi Croy, Suzanne Dale, Hrushikesh Deshmukh, Amanda Douglas, Vincent Drouillon, Marcia Eisenberg, Howard Engler, Rama Ghatti, Prashant Gupta, Susan Hicks, Jake Humphrey, Lax Iyer, Lisa Pfefferle, Manoj Jain, Matthew Robinson, Mohan Kolli, Brian Krueger, Tim Kuphal, Stanley Letovsky, Michael Levandoski, Craig Lukasik, Jonathan Meltzer, Brian Norvell, Mindy Nye, Scott Parker, Christos Petropoulos, John Pruitt, Steven Ragan, Scott Ryan, Mike Sapeta, Jana Schroth, Suresh Babu Selvaraju, Goran Stevovic, Amanda Suchanek, Andrea Throop, Lyndon Tilson, Thomas Urban, Joe Voshell, Kimberly Wagner, Jonathan Williams, Mary Williamson, Qian Zeng, Tricia Zwiefelhofer, Clinton R. Paden, Duncan MacCannell |
| EPI_ISL_3329151 | New Mexico Department of Health Scientific Laboratory | New Mexico Department of Health Scientific Laboratory | Ellie Johnson, D'eldra Malone, Jennifer Benoit, Ratheesh Rajan, Linda Salazar, Anastacia Griego-Fisher |
| EPI_ISL_3329152 | Laboratory Corporation of America | Centers for Disease Control and Prevention Division of Viral Diseases, Pathogen Discovery | Dakota Howard, Dhwani Batra, Peter W. Cook, Kara Moser, Adrian Paskey, Jason Caravas, Benjamin Rambo-Martin, Shatavia Morrison, Christopher Gulvick, Scott Sammons, Yvette Unoarumhi, Darlene Wagner, Matthew Schmerer, Minoo Agarwal, Eyad Almasri, Debbie Boles, Ayla Burns, Nuthawin Charoensri, Oren Cohen, Susan Countryman, Mary Ann Cristobal, Bobbi Croy, Suzanne Dale, Hrushikesh Deshmukh, Amanda Douglas, Vincent Drouillon, Marcia Eisenberg, Howard Engler, Rama Ghatti, Prashant Gupta, Susan Hicks, Jake Humphrey, Lax Iyer, Lisa Pfefferle, Manoj Jain, Matthew Robinson, Mohan Kolli, Brian Krueger, Tim Kuphal, Stanley Letovsky, Michael Levandoski, Craig Lukasik, Jonathan Meltzer, Brian Norvell, Mindy Nye, Scott Parker, Christos Petropoulos, John Pruitt, Steven Ragan, Scott Ryan, Mike Sapeta, Jana Schroth, Suresh Babu Selvaraju, Goran Stevovic, Amanda Suchanek, Andrea Throop, Lyndon Tilson, Thomas Urban, Joe Voshell, Kimberly Wagner, Jonathan Williams, Mary Williamson, Qian Zeng, Tricia Zwiefelhofer, Clinton R. Paden, Duncan MacCannell |
| EPI_ISL_3329153 | New Mexico Department of Health Scientific Laboratory | New Mexico Department of Health Scientific Laboratory | Ellie Johnson, D'eldra Malone, Jennifer Benoit, Ratheesh Rajan, Linda Salazar, Anastacia Griego-Fisher |
| EPI_ISL_3329155, EPI_ISL_3329157 | Laboratory Corporation of America | Centers for Disease Control and Prevention Division of Viral Diseases, Pathogen Discovery | Dakota Howard, Dhwani Batra, Peter W. Cook, Kara Moser, Adrian Paskey, Jason Caravas, Benjamin Rambo-Martin, Shatavia Morrison, Christopher Gulvick, Scott Sammons, Yvette Unoarumhi, Darlene Wagner, Matthew Schmerer, Minoo Agarwal, Eyad Almasri, Debbie Boles, Ayla Burns, Nuthawin Charoensri, Oren Cohen, Susan Countryman, Mary Ann Cristobal, Bobbi Croy, Suzanne Dale, Hrushikesh Deshmukh, Amanda Douglas, Vincent Drouillon, Marcia Eisenberg, Howard Engler, Rama Ghatti, Prashant Gupta, Susan Hicks, Jake Humphrey, Lax Iyer, Lisa Pfefferle, Manoj Jain, Matthew Robinson, Mohan Kolli, Brian Krueger, Tim Kuphal, Stanley Letovsky, Michael Levandoski, Craig Lukasik, Jonathan Meltzer, Brian Norvell, Mindy Nye, Scott Parker, Christos Petropoulos, John Pruitt, Steven Ragan, Scott Ryan, Mike Sapeta, Jana Schroth, Suresh Babu Selvaraju, Goran Stevovic, Amanda Suchanek, Andrea Throop, Lyndon Tilson, Thomas Urban, Joe Voshell, Kimberly Wagner, Jonathan Williams, Mary Williamson, Qian Zeng, Tricia Zwiefelhofer, Clinton R. Paden, Duncan MacCannell |
| EPI_ISL_3329158 | New Mexico Department of Health Scientific Laboratory | New Mexico Department of Health Scientific Laboratory | Ellie Johnson, D'eldra Malone, Jennifer Benoit, Ratheesh Rajan, Linda Salazar, Anastacia Griego-Fisher |
| EPI_ISL_3329159 | Laboratory Corporation of America | Centers for Disease Control and Prevention Division of Viral Diseases, Pathogen Discovery | Dakota Howard, Dhwani Batra, Peter W. Cook, Kara Moser, Adrian Paskey, Jason Caravas, Benjamin Rambo-Martin, Shatavia Morrison, Christopher Gulvick, Scott Sammons, Yvette Unoarumhi, Darlene Wagner, Matthew Schmerer, Minoo Agarwal, Eyad Almasri, Debbie Boles, Ayla Burns, Nuthawin Charoensri, Oren Cohen, Susan Countryman, Mary Ann Cristobal, Bobbi Croy, Suzanne Dale, Hrushikesh Deshmukh, Amanda Douglas, Vincent Drouillon, Marcia Eisenberg, Howard Engler, Rama Ghatti, Prashant Gupta, Susan Hicks, Jake Humphrey, Lax Iyer, Lisa Pfefferle, Manoj Jain, Matthew Robinson, Mohan Kolli, Brian Krueger, Tim Kuphal, Stanley Letovsky, Michael Levandoski, Craig Lukasik, Jonathan Meltzer, Brian Norvell, Mindy Nye, Scott Parker, Christos Petropoulos, John Pruitt, Steven Ragan, Scott Ryan, Mike Sapeta, Jana Schroth, Suresh Babu Selvaraju, Goran Stevovic, Amanda Suchanek, Andrea Throop, Lyndon Tilson, Thomas Urban, Joe Voshell, Kimberly Wagner, Jonathan Williams, Mary Williamson, Qian Zeng, Tricia Zwiefelhofer, Clinton R. Paden, Duncan MacCannell |
| EPI_ISL_3329160 | New Mexico Department of Health Scientific Laboratory | New Mexico Department of Health Scientific Laboratory | Ellie Johnson, D'eldra Malone, Jennifer Benoit, Ratheesh Rajan, Linda Salazar, Anastacia Griego-Fisher |
| EPI_ISL_3329161, EPI_ISL_3329162 | Laboratory Corporation of America | Centers for Disease Control and Prevention Division of Viral Diseases, Pathogen Discovery | Dakota Howard, Dhwani Batra, Peter W. Cook, Kara Moser, Adrian Paskey, Jason Caravas, Benjamin Rambo-Martin, Shatavia Morrison, Christopher Gulvick, Scott Sammons, Yvette Unoarumhi, Darlene Wagner, Matthew Schmerer, Minoo Agarwal, Eyad Almasri, Debbie Boles, Ayla Burns, Nuthawin Charoensri, Oren Cohen, Susan Countryman, Mary Ann Cristobal, Bobbi Croy, Suzanne Dale, Hrushikesh Deshmukh, Amanda Douglas, Vincent Drouillon, Marcia Eisenberg, Howard Engler, Rama Ghatti, Prashant Gupta, Susan Hicks, Jake Humphrey, Lax Iyer, Lisa Pfefferle, Manoj Jain, Matthew Robinson, Mohan Kolli, Brian Krueger, Tim Kuphal, Stanley Letovsky, Michael Levandoski, Craig Lukasik, Jonathan Meltzer, Brian Norvell, Mindy Nye, Scott Parker, Christos Petropoulos, John Pruitt, Steven Ragan, Scott Ryan, Mike Sapeta, Jana Schroth, Suresh Babu Selvaraju, Goran Stevovic, Amanda Suchanek, Andrea Throop, Lyndon Tilson, Thomas Urban, Joe Voshell, Kimberly Wagner, Jonathan Williams, Mary Williamson, Qian Zeng, Tricia Zwiefelhofer, Clinton R. Paden, Duncan MacCannell |
| EPI_ISL_3329163 | New Mexico Department of Health Scientific Laboratory | New Mexico Department of Health Scientific Laboratory | Ellie Johnson, D'eldra Malone, Jennifer Benoit, Ratheesh Rajan, Linda Salazar, Anastacia Griego-Fisher |
| EPI_ISL_3329164 | Laboratory Corporation of America | Centers for Disease Control and Prevention Division of Viral Diseases, Pathogen Discovery | Dakota Howard, Dhwani Batra, Peter W. Cook, Kara Moser, Adrian Paskey, Jason Caravas, Benjamin Rambo-Martin, Shatavia Morrison, Christopher Gulvick, Scott Sammons, Yvette Unoarumhi, Darlene Wagner, Matthew Schmerer, Minoo Agarwal, Eyad Almasri, Debbie Boles, Ayla Burns, Nuthawin Charoensri, Oren Cohen, Susan Countryman, Mary Ann Cristobal, Bobbi Croy, Suzanne Dale, Hrushikesh Deshmukh, Amanda Douglas, Vincent Drouillon, Marcia Eisenberg, Howard Engler, Rama Ghatti, Prashant Gupta, Susan Hicks, Jake Humphrey, Lax Iyer, Lisa Pfefferle, Manoj Jain, Matthew Robinson, Mohan Kolli, Brian Krueger, Tim Kuphal, Stanley Letovsky, Michael Levandoski, Craig Lukasik, Jonathan Meltzer, Brian Norvell, Mindy Nye, Scott Parker, Christos Petropoulos, John Pruitt, Steven Ragan, Scott Ryan, Mike Sapeta, Jana Schroth, Suresh Babu Selvaraju, Goran Stevovic, Amanda Suchanek, Andrea Throop, Lyndon Tilson, Thomas Urban, Joe Voshell, Kimberly Wagner, Jonathan Williams, Mary Williamson, Qian Zeng, Tricia Zwiefelhofer, Clinton R. Paden, Duncan MacCannell |
| EPI_ISL_3329165 | New Mexico Department of Health Scientific Laboratory | New Mexico Department of Health Scientific Laboratory | Ellie Johnson, D'eldra Malone, Jennifer Benoit, Ratheesh Rajan, Linda Salazar, Anastacia Griego-Fisher |
| EPI_ISL_3329166 | Laboratory Corporation of America | Centers for Disease Control and Prevention Division of Viral Diseases, Pathogen Discovery | Dakota Howard, Dhwani Batra, Peter W. Cook, Kara Moser, Adrian Paskey, Jason Caravas, Benjamin Rambo-Martin, Shatavia Morrison, Christopher Gulvick, Scott Sammons, Yvette Unoarumhi, Darlene Wagner, Matthew Schmerer, Minoo Agarwal, Eyad Almasri, Debbie Boles, Ayla Burns, Nuthawin Charoensri, Oren Cohen, Susan Countryman, Mary Ann Cristobal, Bobbi Croy, Suzanne Dale, Hrushikesh Deshmukh, Amanda Douglas, Vincent Drouillon, Marcia Eisenberg, Howard Engler, Rama Ghatti, Prashant Gupta, Susan Hicks, Jake Humphrey, Lax Iyer, Lisa Pfefferle, Manoj Jain, Matthew Robinson, Mohan Kolli, Brian Krueger, Tim Kuphal, Stanley Letovsky, Michael Levandoski, Craig Lukasik, Jonathan Meltzer, Brian Norvell, Mindy Nye, Scott Parker, Christos Petropoulos, John Pruitt, Steven Ragan, Scott Ryan, Mike Sapeta, Jana Schroth, Suresh Babu Selvaraju, Goran Stevovic, Amanda Suchanek, Andrea Throop, Lyndon Tilson, Thomas Urban, Joe Voshell, Kimberly Wagner, Jonathan Williams, Mary Williamson, Qian Zeng, Tricia Zwiefelhofer, Clinton R. Paden, Duncan MacCannell |
| EPI_ISL_3329167 | New Mexico Department of Health Scientific Laboratory | New Mexico Department of Health Scientific Laboratory | Ellie Johnson, D'eldra Malone, Jennifer Benoit, Ratheesh Rajan, Linda Salazar, Anastacia Griego-Fisher |
| EPI_ISL_3329168 | Laboratory Corporation of America | Centers for Disease Control and Prevention Division of Viral Diseases, Pathogen Discovery | Dakota Howard, Dhwani Batra, Peter W. Cook, Kara Moser, Adrian Paskey, Jason Caravas, Benjamin Rambo-Martin, Shatavia Morrison, Christopher Gulvick, Scott Sammons, Yvette Unoarumhi, Darlene Wagner, Matthew Schmerer, Minoo Agarwal, Eyad Almasri, Debbie Boles, Ayla Burns, Nuthawin Charoensri, Oren Cohen, Susan Countryman, Mary Ann Cristobal, Bobbi Croy, Suzanne Dale, Hrushikesh Deshmukh, Amanda Douglas, Vincent Drouillon, Marcia Eisenberg, Howard Engler, Rama Ghatti, Prashant Gupta, Susan Hicks, Jake Humphrey, Lax Iyer, Lisa Pfefferle, Manoj Jain, Matthew Robinson, Mohan Kolli, Brian Krueger, Tim Kuphal, Stanley Letovsky, Michael Levandoski, Craig Lukasik, Jonathan Meltzer, Brian Norvell, Mindy Nye, Scott Parker, Christos Petropoulos, John Pruitt, Steven Ragan, Scott Ryan, Mike Sapeta, Jana Schroth, Suresh Babu Selvaraju, Goran Stevovic, Amanda Suchanek, Andrea Throop, Lyndon Tilson, Thomas Urban, Joe Voshell, Kimberly Wagner, Jonathan Williams, Mary Williamson, Qian Zeng, Tricia Zwiefelhofer, Clinton R. Paden, Duncan MacCannell |
| EPI_ISL_3329169 | New Mexico Department of Health Scientific Laboratory | New Mexico Department of Health Scientific Laboratory | Ellie Johnson, D'eldra Malone, Jennifer Benoit, Ratheesh Rajan, Linda Salazar, Anastacia Griego-Fisher |
| EPI_ISL_3329170 | Laboratory Corporation of America | Centers for Disease Control and Prevention Division of Viral Diseases, Pathogen Discovery | Dakota Howard, Dhwani Batra, Peter W. Cook, Kara Moser, Adrian Paskey, Jason Caravas, Benjamin Rambo-Martin, Shatavia Morrison, Christopher Gulvick, Scott Sammons, Yvette Unoarumhi, Darlene Wagner, Matthew Schmerer, Minoo Agarwal, Eyad Almasri, Debbie Boles, Ayla Burns, Nuthawin Charoensri, Oren Cohen, Susan Countryman, Mary Ann Cristobal, Bobbi Croy, Suzanne Dale, Hrushikesh Deshmukh, Amanda Douglas, Vincent Drouillon, Marcia Eisenberg, Howard Engler, Rama Ghatti, Prashant Gupta, Susan Hicks, Jake Humphrey, Lax Iyer, Lisa Pfefferle, Manoj Jain, Matthew Robinson, Mohan Kolli, Brian Krueger, Tim Kuphal, Stanley Letovsky, Michael Levandoski, Craig Lukasik, Jonathan Meltzer, Brian Norvell, Mindy Nye, Scott Parker, Christos Petropoulos, John Pruitt, Steven Ragan, Scott Ryan, Mike Sapeta, Jana Schroth, Suresh Babu Selvaraju, Goran Stevovic, Amanda Suchanek, Andrea Throop, Lyndon Tilson, Thomas Urban, Joe Voshell, Kimberly Wagner, Jonathan Williams, Mary Williamson, Qian Zeng, Tricia Zwiefelhofer, Clinton R. Paden, Duncan MacCannell |
| EPI_ISL_3329172 | New Mexico Department of Health Scientific Laboratory | New Mexico Department of Health Scientific Laboratory | Ellie Johnson, D'eldra Malone, Jennifer Benoit, Ratheesh Rajan, Linda Salazar, Anastacia Griego-Fisher |
| EPI_ISL_3329173 | Laboratory Corporation of America | Centers for Disease Control and Prevention Division of Viral Diseases, Pathogen Discovery | Dakota Howard, Dhwani Batra, Peter W. Cook, Kara Moser, Adrian Paskey, Jason Caravas, Benjamin Rambo-Martin, Shatavia Morrison, Christopher Gulvick, Scott Sammons, Yvette Unoarumhi, Darlene Wagner, Matthew Schmerer, Minoo Agarwal, Eyad Almasri, Debbie Boles, Ayla Burns, Nuthawin Charoensri, Oren Cohen, Susan Countryman, Mary Ann Cristobal, Bobbi Croy, Suzanne Dale, Hrushikesh Deshmukh, Amanda Douglas, Vincent Drouillon, Marcia Eisenberg, Howard Engler, Rama Ghatti, Prashant Gupta, Susan Hicks, Jake Humphrey, Lax Iyer, Lisa Pfefferle, Manoj Jain, Matthew Robinson, Mohan Kolli, Brian Krueger, Tim Kuphal, Stanley Letovsky, Michael Levandoski, Craig Lukasik, Jonathan Meltzer, Brian Norvell, Mindy Nye, Scott Parker, Christos Petropoulos, John Pruitt, Steven Ragan, Scott Ryan, Mike Sapeta, Jana Schroth, Suresh Babu Selvaraju, Goran Stevovic, Amanda Suchanek, Andrea Throop, Lyndon Tilson, Thomas Urban, Joe Voshell, Kimberly Wagner, Jonathan Williams, Mary Williamson, Qian Zeng, Tricia Zwiefelhofer, Clinton R. Paden, Duncan MacCannell |
| EPI_ISL_3329174 | New Mexico Department of Health Scientific Laboratory | New Mexico Department of Health Scientific Laboratory | Ellie Johnson, D'eldra Malone, Jennifer Benoit, Ratheesh Rajan, Linda Salazar, Anastacia Griego-Fisher |
| EPI_ISL_3329175, EPI_ISL_3329176 | Laboratory Corporation of America | Centers for Disease Control and Prevention Division of Viral Diseases, Pathogen Discovery | Dakota Howard, Dhwani Batra, Peter W. Cook, Kara Moser, Adrian Paskey, Jason Caravas, Benjamin Rambo-Martin, Shatavia Morrison, Christopher Gulvick, Scott Sammons, Yvette Unoarumhi, Darlene Wagner, Matthew Schmerer, Minoo Agarwal, Eyad Almasri, Debbie Boles, Ayla Burns, Nuthawin Charoensri, Oren Cohen, Susan Countryman, Mary Ann Cristobal, Bobbi Croy, Suzanne Dale, Hrushikesh Deshmukh, Amanda Douglas, Vincent Drouillon, Marcia Eisenberg, Howard Engler, Rama Ghatti, Prashant Gupta, Susan Hicks, Jake Humphrey, Lax Iyer, Lisa Pfefferle, Manoj Jain, Matthew Robinson, Mohan Kolli, Brian Krueger, Tim Kuphal, Stanley Letovsky, Michael Levandoski, Craig Lukasik, Jonathan Meltzer, Brian |

| | | | |
|---|---|---|---|
| | | | Norvell, Mindy Nye, Scott Parker, Christos Petropoulos, John Pruitt, Steven Ragan, Scott Ryan, Mike Sapeta, Jana Schroth, Suresh Babu Selvaraju, Goran Stevovic, Amanda Suchanek, Andrea Throop, Lyndon Tilson, Thomas Urban, Joe Voshell, Kimberly Wagner, Jonathan Williams, Mary Williamson, Qian Zeng, Tricia Zwiefelhofer, Clinton R. Paden, Duncan MacCannell |
| EPI_ISL_3329177 | New Mexico Department of Health Scientific Laboratory | New Mexico Department of Health Scientific Laboratory | Ellie Johnson, D'eldra Malone, Jennifer Benoit, Ratheesh Rajan, Linda Salazar, Anastacia Griego-Fisher |
| EPI_ISL_3329178 | Laboratory Corporation of America | Centers for Disease Control and Prevention Division of Viral Diseases, Pathogen Discovery | Dakota Howard, Dhwani Batra, Peter W. Cook, Kara Moser, Adrian Paskey, Jason Caravas, Benjamin Rambo-Martin, Shatavia Morrison, Christopher Gulvick, Scott Sammons, Yvette Unoarumhi, Darlene Wagner, Matthew Schmerer, Minoo Agarwal, Eyad Almasri, Debbie Boles, Ayla Burns, Nuthawin Charoensri, Oren Cohen, Susan Countryman, Mary Ann Cristobal, Bobbi Croy, Suzanne Dale, Hrushikesh Deshmukh, Amanda Douglas, Vincent Drouillon, Marcia Eisenberg, Howard Engler, Rama Ghatti, Prashant Gupta, Susan Hicks, Jake Humphrey, Lax Iyer, Lisa Pfefferle, Manoj Jain, Matthew Robinson, Mohan Kolli, Brian Krueger, Tim Kuphal, Stanley Letovsky, Michael Levandoski, Craig Lukasik, Jonathan Meltzer, Brian Norvell, Mindy Nye, Scott Parker, Christos Petropoulos, John Pruitt, Steven Ragan, Scott Ryan, Mike Sapeta, Jana Schroth, Suresh Babu Selvaraju, Goran Stevovic, Amanda Suchanek, Andrea Throop, Lyndon Tilson, Thomas Urban, Joe Voshell, Kimberly Wagner, Jonathan Williams, Mary Williamson, Qian Zeng, Tricia Zwiefelhofer, Clinton R. Paden, Duncan MacCannell |
| EPI_ISL_3329179, EPI_ISL_3329181 | New Mexico Department of Health Scientific Laboratory | New Mexico Department of Health Scientific Laboratory | Ellie Johnson, D'eldra Malone, Jennifer Benoit, Ratheesh Rajan, Linda Salazar, Anastacia Griego-Fisher |
| EPI_ISL_3329182, EPI_ISL_3329183 | Laboratory Corporation of America | Centers for Disease Control and Prevention Division of Viral Diseases, Pathogen Discovery | Dakota Howard, Dhwani Batra, Peter W. Cook, Kara Moser, Adrian Paskey, Jason Caravas, Benjamin Rambo-Martin, Shatavia Morrison, Christopher Gulvick, Scott Sammons, Yvette Unoarumhi, Darlene Wagner, Matthew Schmerer, Minoo Agarwal, Eyad Almasri, Debbie Boles, Ayla Burns, Nuthawin Charoensri, Oren Cohen, Susan Countryman, Mary Ann Cristobal, Bobbi Croy, Suzanne Dale, Hrushikesh Deshmukh, Amanda Douglas, Vincent Drouillon, Marcia Eisenberg, Howard Engler, Rama Ghatti, Prashant Gupta, Susan Hicks, Jake Humphrey, Lax Iyer, Lisa Pfefferle, Manoj Jain, Matthew Robinson, Mohan Kolli, Brian Krueger, Tim Kuphal, Stanley Letovsky, Michael Levandoski, Craig Lukasik, Jonathan Meltzer, Brian Norvell, Mindy Nye, Scott Parker, Christos Petropoulos, John Pruitt, Steven Ragan, Scott Ryan, Mike Sapeta, Jana Schroth, Suresh Babu Selvaraju, Goran Stevovic, Amanda Suchanek, Andrea Throop, Lyndon Tilson, Thomas Urban, Joe Voshell, Kimberly Wagner, Jonathan Williams, Mary Williamson, Qian Zeng, Tricia Zwiefelhofer, Clinton R. Paden, Duncan MacCannell |
| EPI_ISL_3329184 | New Mexico Department of Health Scientific Laboratory | New Mexico Department of Health Scientific Laboratory | Ellie Johnson, D'eldra Malone, Jennifer Benoit, Ratheesh Rajan, Linda Salazar, Anastacia Griego-Fisher |
| EPI_ISL_3329185 | Laboratory Corporation of America | Centers for Disease Control and Prevention Division of Viral Diseases, Pathogen Discovery | Dakota Howard, Dhwani Batra, Peter W. Cook, Kara Moser, Adrian Paskey, Jason Caravas, Benjamin Rambo-Martin, Shatavia Morrison, Christopher Gulvick, Scott Sammons, Yvette Unoarumhi, Darlene Wagner, Matthew Schmerer, Minoo Agarwal, Eyad Almasri, Debbie Boles, Ayla Burns, Nuthawin Charoensri, Oren Cohen, Susan Countryman, Mary Ann Cristobal, Bobbi Croy, Suzanne Dale, Hrushikesh Deshmukh, Amanda Douglas, Vincent Drouillon, Marcia Eisenberg, Howard Engler, Rama Ghatti, Prashant Gupta, Susan Hicks, Jake Humphrey, Lax Iyer, Lisa Pfefferle, Manoj Jain, Matthew Robinson, Mohan Kolli, Brian Krueger, Tim Kuphal, Stanley Letovsky, Michael Levandoski, Craig Lukasik, Jonathan Meltzer, Brian Norvell, Mindy Nye, Scott Parker, Christos Petropoulos, John Pruitt, Steven Ragan, Scott Ryan, Mike Sapeta, Jana Schroth, Suresh Babu Selvaraju, Goran Stevovic, Amanda Suchanek, Andrea Throop, Lyndon Tilson, Thomas Urban, Joe Voshell, Kimberly Wagner, Jonathan Williams, Mary Williamson, Qian Zeng, Tricia Zwiefelhofer, Clinton R. Paden, Duncan MacCannell |
| EPI_ISL_3329186 | New Mexico Department of Health Scientific Laboratory | New Mexico Department of Health Scientific Laboratory | Ellie Johnson, D'eldra Malone, Jennifer Benoit, Ratheesh Rajan, Linda Salazar, Anastacia Griego-Fisher |
| EPI_ISL_3329187 | Laboratory Corporation of America | Centers for Disease Control and Prevention Division of Viral Diseases, Pathogen Discovery | Dakota Howard, Dhwani Batra, Peter W. Cook, Kara Moser, Adrian Paskey, Jason Caravas, Benjamin Rambo-Martin, Shatavia Morrison, Christopher Gulvick, Scott Sammons, Yvette Unoarumhi, Darlene Wagner, Matthew Schmerer, Minoo Agarwal, Eyad Almasri, Debbie Boles, Ayla Burns, Nuthawin Charoensri, Oren Cohen, Susan Countryman, Mary Ann Cristobal, Bobbi Croy, Suzanne Dale, Hrushikesh Deshmukh, Amanda Douglas, Vincent Drouillon, Marcia Eisenberg, Howard Engler, Rama Ghatti, Prashant Gupta, Susan Hicks, Jake Humphrey, Lax Iyer, Lisa Pfefferle, Manoj Jain, Matthew Robinson, Mohan Kolli, Brian Krueger, Tim Kuphal, Stanley Letovsky, Michael Levandoski, Craig Lukasik, Jonathan Meltzer, Brian Norvell, Mindy Nye, Scott Parker, Christos Petropoulos, John Pruitt, Steven Ragan, Scott Ryan, Mike Sapeta, Jana Schroth, Suresh Babu Selvaraju, Goran Stevovic, Amanda Suchanek, Andrea Throop, Lyndon Tilson, Thomas Urban, Joe Voshell, Kimberly Wagner, Jonathan Williams, Mary Williamson, Qian Zeng, Tricia Zwiefelhofer, Clinton R. Paden, Duncan MacCannell |
| EPI_ISL_3329188 | New Mexico Department of Health Scientific Laboratory | New Mexico Department of Health Scientific Laboratory | Ellie Johnson, D'eldra Malone, Jennifer Benoit, Ratheesh Rajan, Linda Salazar, Anastacia Griego-Fisher |
| EPI_ISL_3329189 | Laboratory Corporation of America | Centers for Disease Control and Prevention Division of Viral Diseases, Pathogen Discovery | Dakota Howard, Dhwani Batra, Peter W. Cook, Kara Moser, Adrian Paskey, Jason Caravas, Benjamin Rambo-Martin, Shatavia Morrison, Christopher Gulvick, Scott Sammons, Yvette Unoarumhi, Darlene Wagner, Matthew Schmerer, Minoo Agarwal, Eyad Almasri, Debbie Boles, Ayla Burns, Nuthawin Charoensri, Oren Cohen, Susan Countryman, Mary Ann Cristobal, Bobbi Croy, Suzanne Dale, Hrushikesh Deshmukh, Amanda Douglas, Vincent Drouillon, Marcia Eisenberg, Howard Engler, Rama Ghatti, Prashant Gupta, Susan Hicks, Jake Humphrey, Lax Iyer, Lisa Pfefferle, Manoj Jain, Matthew Robinson, Mohan Kolli, Brian Krueger, Tim Kuphal, Stanley Letovsky, Michael Levandoski, Craig Lukasik, Jonathan Meltzer, Brian Norvell, Mindy Nye, Scott Parker, Christos Petropoulos, John Pruitt, Steven Ragan, Scott Ryan, Mike Sapeta, Jana Schroth, Suresh Babu Selvaraju, Goran Stevovic, Amanda Suchanek, Andrea Throop, Lyndon Tilson, Thomas Urban, Joe Voshell, Kimberly Wagner, Jonathan Williams, Mary Williamson, Qian Zeng, Tricia Zwiefelhofer, Clinton R. Paden, Duncan MacCannell |
| EPI_ISL_3329190 | New Mexico Department of Health Scientific Laboratory | New Mexico Department of Health Scientific Laboratory | Ellie Johnson, D'eldra Malone, Jennifer Benoit, Ratheesh Rajan, Linda Salazar, Anastacia Griego-Fisher |
| EPI_ISL_3329191 | Laboratory Corporation of America | Centers for Disease Control and Prevention Division of Viral Diseases, Pathogen Discovery | Dakota Howard, Dhwani Batra, Peter W. Cook, Kara Moser, Adrian Paskey, Jason Caravas, Benjamin Rambo-Martin, Shatavia Morrison, Christopher Gulvick, Scott Sammons, Yvette Unoarumhi, Darlene Wagner, Matthew Schmerer, Minoo Agarwal, Eyad Almasri, Debbie Boles, Ayla Burns, Nuthawin Charoensri, Oren Cohen, Susan Countryman, Mary Ann Cristobal, Bobbi Croy, Suzanne Dale, Hrushikesh Deshmukh, Amanda Douglas, Vincent Drouillon, Marcia Eisenberg, Howard Engler, Rama Ghatti, Prashant Gupta, Susan Hicks, Jake Humphrey, Lax Iyer, Lisa Pfefferle, Manoj Jain, Matthew Robinson, Mohan Kolli, Brian Krueger, Tim Kuphal, Stanley Letovsky, Michael Levandoski, Craig Lukasik, Jonathan Meltzer, Brian Norvell, Mindy Nye, Scott Parker, Christos Petropoulos, John Pruitt, Steven Ragan, Scott Ryan, Mike Sapeta, Jana Schroth, Suresh Babu Selvaraju, Goran Stevovic, Amanda Suchanek, Andrea Throop, Lyndon Tilson, Thomas Urban, Joe Voshell, Kimberly Wagner, Jonathan Williams, Mary Williamson, Qian Zeng, Tricia Zwiefelhofer, Clinton R. Paden, Duncan MacCannell |
| EPI_ISL_3329192 | New Mexico Department of Health Scientific Laboratory | New Mexico Department of Health Scientific Laboratory | Ellie Johnson, D'eldra Malone, Jennifer Benoit, Ratheesh Rajan, Linda Salazar, Anastacia Griego-Fisher |
| EPI_ISL_3329193 | Laboratory Corporation of America | Centers for Disease Control and Prevention Division of Viral Diseases, Pathogen Discovery | Dakota Howard, Dhwani Batra, Peter W. Cook, Kara Moser, Adrian Paskey, Jason Caravas, Benjamin Rambo-Martin, Shatavia Morrison, Christopher Gulvick, Scott Sammons, Yvette Unoarumhi, Darlene Wagner, Matthew Schmerer, Minoo Agarwal, Eyad Almasri, Debbie Boles, Ayla Burns, Nuthawin Charoensri, Oren Cohen, Susan Countryman, Mary Ann Cristobal, Bobbi Croy, Suzanne Dale, Hrushikesh Deshmukh, Amanda Douglas, Vincent Drouillon, Marcia Eisenberg, Howard Engler, Rama Ghatti, Prashant Gupta, Susan Hicks, Jake Humphrey, Lax Iyer, Lisa Pfefferle, Manoj Jain, Matthew Robinson, Mohan Kolli, Brian Krueger, Tim Kuphal, Stanley Letovsky, Michael Levandoski, Craig Lukasik, Jonathan Meltzer, Brian Norvell, Mindy Nye, Scott Parker, Christos Petropoulos, John Pruitt, Steven Ragan, Scott Ryan, Mike Sapeta, Jana Schroth, Suresh Babu Selvaraju, Goran Stevovic, Amanda Suchanek, Andrea Throop, Lyndon Tilson, Thomas Urban, Joe Voshell, Kimberly Wagner, Jonathan Williams, Mary Williamson, Qian Zeng, Tricia Zwiefelhofer, Clinton R. Paden, Duncan MacCannell |
| EPI_ISL_3329194, EPI_ISL_3329196 | New Mexico Department of Health Scientific Laboratory | New Mexico Department of Health Scientific Laboratory | Ellie Johnson, D'eldra Malone, Jennifer Benoit, Ratheesh Rajan, Linda Salazar, Anastacia Griego-Fisher |
| EPI_ISL_3329197, EPI_ISL_3329198 | Laboratory Corporation of America | Centers for Disease Control and Prevention Division of Viral Diseases, Pathogen Discovery | Dakota Howard, Dhwani Batra, Peter W. Cook, Kara Moser, Adrian Paskey, Jason Caravas, Benjamin Rambo-Martin, Shatavia Morrison, Christopher Gulvick, Scott Sammons, Yvette Unoarumhi, Darlene Wagner, Matthew Schmerer, Minoo Agarwal, Eyad Almasri, Debbie Boles, Ayla Burns, Nuthawin Charoensri, Oren Cohen, Susan Countryman, Mary Ann Cristobal, Bobbi Croy, Suzanne Dale, Hrushikesh Deshmukh, Amanda Douglas, Vincent Drouillon, Marcia Eisenberg, Howard Engler, Rama Ghatti, Prashant Gupta, Susan Hicks, Jake Humphrey, Lax Iyer, Lisa Pfefferle, Manoj Jain, Matthew Robinson, Mohan Kolli, Brian Krueger, Tim Kuphal, Stanley Letovsky, Michael Levandoski, Craig Lukasik, Jonathan Meltzer, Brian Norvell, Mindy Nye, Scott Parker, Christos Petropoulos, John Pruitt, Steven Ragan, Scott Ryan, Mike Sapeta, Jana Schroth, Suresh Babu Selvaraju, Goran Stevovic, Amanda Suchanek, Andrea Throop, Lyndon Tilson, Thomas Urban, Joe Voshell, Kimberly Wagner, Jonathan Williams, Mary Williamson, Qian Zeng, Tricia Zwiefelhofer, Clinton R. Paden, Duncan MacCannell |
| EPI_ISL_3329199 | New Mexico Department of Health Scientific Laboratory | New Mexico Department of Health Scientific Laboratory | Ellie Johnson, D'eldra Malone, Jennifer Benoit, Ratheesh Rajan, Linda Salazar, Anastacia Griego-Fisher |
| EPI_ISL_3329200 | Laboratory Corporation of America | Centers for Disease Control and Prevention Division of Viral Diseases, Pathogen Discovery | Dakota Howard, Dhwani Batra, Peter W. Cook, Kara Moser, Adrian Paskey, Jason Caravas, Benjamin Rambo-Martin, Shatavia Morrison, Christopher Gulvick, Scott Sammons, Yvette Unoarumhi, Darlene Wagner, Matthew Schmerer, Minoo Agarwal, Eyad Almasri, Debbie Boles, Ayla Burns, Nuthawin Charoensri, Oren Cohen, Susan Countryman, Mary Ann Cristobal, Bobbi Croy, Suzanne Dale, Hrushikesh Deshmukh, Amanda Douglas, Vincent Drouillon, Marcia Eisenberg, Howard Engler, Rama Ghatti, Prashant Gupta, Susan Hicks, Jake Humphrey, Lax Iyer, Lisa Pfefferle, Manoj Jain, Matthew Robinson, Mohan Kolli, Brian Krueger, Tim Kuphal, Stanley Letovsky, Michael Levandoski, Craig Lukasik, Jonathan Meltzer, Brian Norvell, Mindy Nye, Scott Parker, Christos Petropoulos, John Pruitt, Steven Ragan, Scott Ryan, Mike Sapeta, Jana Schroth, Suresh Babu Selvaraju, Goran Stevovic, Amanda Suchanek, Andrea Throop, Lyndon Tilson, Thomas Urban, Joe Voshell, Kimberly Wagner, Jonathan Williams, Mary Williamson, Qian Zeng, Tricia Zwiefelhofer, Clinton R. Paden, Duncan MacCannell |
| EPI_ISL_3329201 | New Mexico Department of Health Scientific Laboratory | New Mexico Department of Health Scientific Laboratory | Ellie Johnson, D'eldra Malone, Jennifer Benoit, Ratheesh Rajan, Linda Salazar, Anastacia Griego-Fisher |
| EPI_ISL_3329202 | Laboratory Corporation of America | Centers for Disease Control and Prevention Division of Viral Diseases, Pathogen Discovery | Dakota Howard, Dhwani Batra, Peter W. Cook, Kara Moser, Adrian Paskey, Jason Caravas, Benjamin Rambo-Martin, Shatavia Morrison, Christopher Gulvick, Scott Sammons, Yvette Unoarumhi, Darlene Wagner, Matthew Schmerer, Minoo Agarwal, Eyad Almasri, Debbie Boles, Ayla Burns, Nuthawin Charoensri, Oren Cohen, Susan Countryman, Mary Ann Cristobal, Bobbi Croy, Suzanne Dale, Hrushikesh Deshmukh, Amanda Douglas, Vincent Drouillon, Marcia Eisenberg, Howard Engler, Rama Ghatti, Prashant Gupta, Susan Hicks, Jake Humphrey, Lax Iyer, Lisa Pfefferle, Manoj Jain, Matthew Robinson, Mohan Kolli, Brian Krueger, Tim Kuphal, Stanley Letovsky, Michael Levandoski, Craig Lukasik, Jonathan Meltzer, Brian Norvell, Mindy Nye, Scott Parker, Christos Petropoulos, John Pruitt, Steven Ragan, Scott Ryan, Mike Sapeta, Jana Schroth, Suresh Babu Selvaraju, Goran Stevovic, Amanda Suchanek, Andrea Throop, Lyndon Tilson, Thomas Urban, Joe Voshell, Kimberly Wagner, Jonathan Williams, Mary Williamson, Qian Zeng, Tricia Zwiefelhofer, Clinton R. Paden, Duncan MacCannell |
| EPI_ISL_3329203 | New Mexico Department of Health Scientific Laboratory | New Mexico Department of Health Scientific Laboratory | Ellie Johnson, D'eldra Malone, Jennifer Benoit, Ratheesh Rajan, Linda Salazar, Anastacia Griego-Fisher |
| EPI_ISL_3329204 | Laboratory Corporation of America | Centers for Disease Control and Prevention Division of Viral Diseases, Pathogen Discovery | Dakota Howard, Dhwani Batra, Peter W. Cook, Kara Moser, Adrian Paskey, Jason Caravas, Benjamin Rambo-Martin, Shatavia Morrison, Christopher Gulvick, Scott Sammons, Yvette Unoarumhi, Darlene Wagner, Matthew Schmerer, Minoo Agarwal, Eyad Almasri, Debbie Boles, Ayla Burns, Nuthawin Charoensri, Oren Cohen, Susan Countryman, Mary Ann Cristobal, Bobbi Croy, Suzanne Dale, Hrushikesh Deshmukh, Amanda Douglas, Vincent Drouillon, Marcia Eisenberg, Howard Engler, Rama Ghatti, Prashant Gupta, Susan Hicks, Jake Humphrey, Lax Iyer, Lisa Pfefferle, Manoj Jain, Matthew Robinson, Mohan Kolli, Brian Krueger, Tim Kuphal, Stanley Letovsky, Michael Levandoski, Craig Lukasik, Jonathan Meltzer, Brian Norvell, Mindy Nye, Scott Parker, Christos Petropoulos, John Pruitt, Steven Ragan, Scott Ryan, Mike Sapeta, Jana Schroth, Suresh Babu Selvaraju, Goran Stevovic, Amanda Suchanek, Andrea Throop, Lyndon Tilson, Thomas Urban, Joe Voshell, Kimberly Wagner, Jonathan Williams, Mary Williamson, Qian Zeng, Tricia Zwiefelhofer, Clinton R. Paden, Duncan MacCannell |
| EPI_ISL_3329205 | New Mexico Department of Health Scientific Laboratory | New Mexico Department of Health Scientific Laboratory | Ellie Johnson, D'eldra Malone, Jennifer Benoit, Ratheesh Rajan, Linda Salazar, Anastacia Griego-Fisher |
| EPI_ISL_3329206, EPI_ISL_3329207 | Laboratory Corporation of America | Centers for Disease Control and Prevention Division of Viral Diseases, Pathogen Discovery | Dakota Howard, Dhwani Batra, Peter W. Cook, Kara Moser, Adrian Paskey, Jason Caravas, Benjamin Rambo-Martin, Shatavia Morrison, Christopher Gulvick, Scott Sammons, Yvette Unoarumhi, Darlene Wagner, Matthew Schmerer, Minoo Agarwal, Eyad Almasri, Debbie Boles, Ayla Burns, Nuthawin Charoensri, Oren Cohen, Susan Countryman, Mary Ann Cristobal, Bobbi Croy, Suzanne Dale, Hrushikesh Deshmukh, Amanda Douglas, Vincent Drouillon, Marcia Eisenberg, Howard Engler, Rama Ghatti, Prashant Gupta, Susan Hicks, Jake Humphrey, Lax Iyer, Lisa Pfefferle, Manoj Jain, Matthew Robinson, Mohan Kolli, Brian Krueger, Tim Kuphal, Stanley Letovsky, Michael Levandoski, Craig Lukasik, Jonathan Meltzer, Brian Norvell, Mindy Nye, Scott Parker, Christos Petropoulos, John Pruitt, Steven Ragan, Scott Ryan, Mike Sapeta, Jana Schroth, Suresh Babu Selvaraju, Goran Stevovic, Amanda Suchanek, Andrea Throop, Lyndon Tilson, Thomas Urban, Joe Voshell, Kimberly Wagner, Jonathan Williams, Mary Williamson, Qian Zeng, Tricia Zwiefelhofer, Clinton R. Paden, Duncan MacCannell |
| EPI_ISL_3329208 | New Mexico Department of Health Scientific Laboratory | New Mexico Department of Health Scientific Laboratory | Ellie Johnson, D'eldra Malone, Jennifer Benoit, Ratheesh Rajan, Linda Salazar, Anastacia Griego-Fisher |
| EPI_ISL_3329209 | Laboratory Corporation of America | Centers for Disease Control and Prevention Division of Viral Diseases, Pathogen Discovery | Dakota Howard, Dhwani Batra, Peter W. Cook, Kara Moser, Adrian Paskey, Jason Caravas, Benjamin Rambo-Martin, Shatavia Morrison, Christopher Gulvick, Scott Sammons, Yvette Unoarumhi, Darlene Wagner, Matthew Schmerer, Minoo Agarwal, Eyad Almasri, Debbie Boles, Ayla Burns, Nuthawin Charoensri, Oren Cohen, Susan Countryman, Mary Ann Cristobal, Bobbi Croy, Suzanne Dale, Hrushikesh Deshmukh, Amanda Douglas, Vincent Drouillon, Marcia Eisenberg, Howard Engler, Rama Ghatti, Prashant Gupta, Susan Hicks, Jake Humphrey, Lax Iyer, Lisa Pfefferle, Manoj Jain, Matthew Robinson, Mohan Kolli, Brian Krueger, Tim Kuphal, Stanley Letovsky, Michael Levandoski, Craig Lukasik, Jonathan Meltzer, Brian Norvell, Mindy Nye, Scott Parker, Christos Petropoulos, John Pruitt, Steven Ragan, Scott Ryan, Mike Sapeta, Jana Schroth, Suresh Babu Selvaraju, Goran Stevovic, Amanda Suchanek, Andrea Throop, Lyndon Tilson, Thomas Urban, Joe Voshell, Kimberly Wagner, Jonathan Williams, Mary Williamson, Qian Zeng, Tricia Zwiefelhofer, Clinton R. Paden, Duncan MacCannell |
| EPI_ISL_3329210, EPI_ISL_3329212 | New Mexico Department of Health Scientific Laboratory | New Mexico Department of Health Scientific Laboratory | Ellie Johnson, D'eldra Malone, Jennifer Benoit, Ratheesh Rajan, Linda Salazar, Anastacia Griego-Fisher |
| EPI_ISL_3329213, EPI_ISL_3329214 | Laboratory Corporation of America | Centers for Disease Control and Prevention Division of Viral Diseases, Pathogen Discovery | Dakota Howard, Dhwani Batra, Peter W. Cook, Kara Moser, Adrian Paskey, Jason Caravas, Benjamin Rambo-Martin, Shatavia Morrison, Christopher Gulvick, Scott Sammons, Yvette Unoarumhi, Darlene Wagner, Matthew Schmerer, Minoo Agarwal, Eyad Almasri, Debbie Boles, Ayla Burns, Nuthawin Charoensri, Oren Cohen, Susan Countryman, Mary Ann Cristobal, Bobbi Croy, Suzanne Dale, Hrushikesh Deshmukh, Amanda Douglas, Vincent Drouillon, Marcia Eisenberg, Howard Engler, Rama Ghatti, Prashant Gupta, Susan Hicks, Jake Humphrey, Lax Iyer, Lisa Pfefferle, Manoj Jain, Matthew Robinson, Mohan Kolli, Brian Krueger, Tim Kuphal, Stanley Letovsky, Michael Levandoski, Craig Lukasik, Jonathan Meltzer, Brian Norvell, Mindy Nye, Scott Parker, Christos Petropoulos, John Pruitt, Steven Ragan, Scott Ryan, Mike Sapeta, Jana Schroth, Suresh Babu Selvaraju, Goran Stevovic, Amanda Suchanek, Andrea Throop, Lyndon Tilson, Thomas Urban, Joe Voshell, Kimberly Wagner, Jonathan Williams, Mary Williamson, Qian Zeng, Tricia Zwiefelhofer, Clinton R. Paden, Duncan MacCannell |
| EPI_ISL_3329215, EPI_ISL_3329216 | New Mexico Department of Health Scientific Laboratory | New Mexico Department of Health Scientific Laboratory | Ellie Johnson, D'eldra Malone, Jennifer Benoit, Ratheesh Rajan, Linda Salazar, Anastacia Griego-Fisher |
| EPI_ISL_3329217, EPI_ISL_3329218 | Laboratory Corporation of America | Centers for Disease Control and Prevention Division of Viral Diseases, Pathogen Discovery | Dakota Howard, Dhwani Batra, Peter W. Cook, Kara Moser, Adrian Paskey, Jason Caravas, Benjamin Rambo-Martin, Shatavia Morrison, Christopher Gulvick, Scott Sammons, Yvette Unoarumhi, Darlene Wagner, Matthew Schmerer, Minoo Agarwal, Eyad Almasri, Debbie Boles, Ayla Burns, Nuthawin Charoensri, Oren Cohen, Susan Countryman, Mary Ann Cristobal, Bobbi Croy, Suzanne Dale, Hrushikesh Deshmukh, Amanda Douglas, Vincent Drouillon, Marcia Eisenberg, Howard Engler, Rama Ghatti, Prashant Gupta, Susan Hicks, Jake Humphrey, Lax Iyer, Lisa Pfefferle, Manoj Jain, Matthew Robinson, Mohan Kolli, Brian Krueger, Tim Kuphal, Stanley Letovsky, Michael Levandoski, Craig Lukasik, Jonathan Meltzer, Brian Norvell, Mindy Nye, Scott Parker, Christos Petropoulos, John Pruitt, Steven Ragan, Scott Ryan, Mike Sapeta, Jana Schroth, Suresh Babu Selvaraju, Goran Stevovic, Amanda Suchanek, Andrea Throop, Lyndon Tilson, Thomas Urban, Joe Voshell, Kimberly Wagner, Jonathan Williams, Mary Williamson, Qian Zeng, Tricia Zwiefelhofer, Clinton R. Paden, Duncan MacCannell |

| | | | |
|---|---|---|---|
| EPI_ISL_3329219 | New Mexico Department of Health Scientific Laboratory | New Mexico Department of Health Scientific Laboratory | Ellie Johnson, D'eldra Malone, Jennifer Benoit, Ratheesh Rajan, Linda Salazar, Anastacia Griego-Fisher |
| EPI_ISL_3329220, EPI_ISL_3329222 | Laboratory Corporation of America | Centers for Disease Control and Prevention Division of Viral Diseases, Pathogen Discovery | Dakota Howard, Dhwani Batra, Peter W. Cook, Kara Moser, Adrian Paskey, Jason Caravas, Benjamin Rambo-Martin, Shatavia Morrison, Christopher Gulvick, Scott Sammons, Yvette Unoarumhi, Darlene Wagner, Matthew Schmerer, Minoo Agarwal, Eyad Almasri, Debbie Boles, Ayla Burns, Nuthawin Charoensri, Oren Cohen, Susan Countryman, Mary Ann Cristobal, Bobbi Croy, Suzanne Dale, Hrushikesh Deshmukh, Amanda Douglas, Vincent Drouillon, Marcia Eisenberg, Howard Engler, Rama Ghatti, Prashant Gupta, Susan Hicks, Jake Humphrey, Lax Iyer, Lisa Pfefferle, Manoj Jain, Matthew Robinson, Mohan Kolli, Brian Krueger, Tim Kuphal, Stanley Letovsky, Michael Levandoski, Craig Lukasik, Jonathan Meltzer, Brian Norvell, Mindy Nye, Scott Parker, Christos Petropoulos, John Pruitt, Steven Ragan, Scott Ryan, Mike Sapeta, Jana Schroth, Suresh Babu Selvaraju, Goran Stevovic, Amanda Suchanek, Andrea Throop, Lyndon Tilson, Thomas Urban, Joe Voshell, Kimberly Wagner, Jonathan Williams, Mary Williamson, Qian Zeng, Tricia Zwiefelhofer, Clinton R. Paden, Duncan MacCannell |
| EPI_ISL_3329223 | New Mexico Department of Health Scientific Laboratory | New Mexico Department of Health Scientific Laboratory | Ellie Johnson, D'eldra Malone, Jennifer Benoit, Ratheesh Rajan, Linda Salazar, Anastacia Griego-Fisher |
| EPI_ISL_3329225 | Laboratory Corporation of America | Centers for Disease Control and Prevention Division of Viral Diseases, Pathogen Discovery | Dakota Howard, Dhwani Batra, Peter W. Cook, Kara Moser, Adrian Paskey, Jason Caravas, Benjamin Rambo-Martin, Shatavia Morrison, Christopher Gulvick, Scott Sammons, Yvette Unoarumhi, Darlene Wagner, Matthew Schmerer, Minoo Agarwal, Eyad Almasri, Debbie Boles, Ayla Burns, Nuthawin Charoensri, Oren Cohen, Susan Countryman, Mary Ann Cristobal, Bobbi Croy, Suzanne Dale, Hrushikesh Deshmukh, Amanda Douglas, Vincent Drouillon, Marcia Eisenberg, Howard Engler, Rama Ghatti, Prashant Gupta, Susan Hicks, Jake Humphrey, Lax Iyer, Lisa Pfefferle, Manoj Jain, Matthew Robinson, Mohan Kolli, Brian Krueger, Tim Kuphal, Stanley Letovsky, Michael Levandoski, Craig Lukasik, Jonathan Meltzer, Brian Norvell, Mindy Nye, Scott Parker, Christos Petropoulos, John Pruitt, Steven Ragan, Scott Ryan, Mike Sapeta, Jana Schroth, Suresh Babu Selvaraju, Goran Stevovic, Amanda Suchanek, Andrea Throop, Lyndon Tilson, Thomas Urban, Joe Voshell, Kimberly Wagner, Jonathan Williams, Mary Williamson, Qian Zeng, Tricia Zwiefelhofer, Clinton R. Paden, Duncan MacCannell |
| EPI_ISL_3329226 | New Mexico Department of Health Scientific Laboratory | New Mexico Department of Health Scientific Laboratory | Ellie Johnson, D'eldra Malone, Jennifer Benoit, Ratheesh Rajan, Linda Salazar, Anastacia Griego-Fisher |
| EPI_ISL_3329227, EPI_ISL_3329229, EPI_ISL_3329231, EPI_ISL_3329232 | Laboratory Corporation of America | Centers for Disease Control and Prevention Division of Viral Diseases, Pathogen Discovery | Dakota Howard, Dhwani Batra, Peter W. Cook, Kara Moser, Adrian Paskey, Jason Caravas, Benjamin Rambo-Martin, Shatavia Morrison, Christopher Gulvick, Scott Sammons, Yvette Unoarumhi, Darlene Wagner, Matthew Schmerer, Minoo Agarwal, Eyad Almasri, Debbie Boles, Ayla Burns, Nuthawin Charoensri, Oren Cohen, Susan Countryman, Mary Ann Cristobal, Bobbi Croy, Suzanne Dale, Hrushikesh Deshmukh, Amanda Douglas, Vincent Drouillon, Marcia Eisenberg, Howard Engler, Rama Ghatti, Prashant Gupta, Susan Hicks, Jake Humphrey, Lax Iyer, Lisa Pfefferle, Manoj Jain, Matthew Robinson, Mohan Kolli, Brian Krueger, Tim Kuphal, Stanley Letovsky, Michael Levandoski, Craig Lukasik, Jonathan Meltzer, Brian Norvell, Mindy Nye, Scott Parker, Christos Petropoulos, John Pruitt, Steven Ragan, Scott Ryan, Mike Sapeta, Jana Schroth, Suresh Babu Selvaraju, Goran Stevovic, Amanda Suchanek, Andrea Throop, Lyndon Tilson, Thomas Urban, Joe Voshell, Kimberly Wagner, Jonathan Williams, Mary Williamson, Qian Zeng, Tricia Zwiefelhofer, Clinton R. Paden, Duncan MacCannell |
| EPI_ISL_3329233 | New Mexico Department of Health Scientific Laboratory | New Mexico Department of Health Scientific Laboratory | Ellie Johnson, D'eldra Malone, Jennifer Benoit, Ratheesh Rajan, Linda Salazar, Anastacia Griego-Fisher |
| EPI_ISL_3329234 | Laboratory Corporation of America | Centers for Disease Control and Prevention Division of Viral Diseases, Pathogen Discovery | Dakota Howard, Dhwani Batra, Peter W. Cook, Kara Moser, Adrian Paskey, Jason Caravas, Benjamin Rambo-Martin, Shatavia Morrison, Christopher Gulvick, Scott Sammons, Yvette Unoarumhi, Darlene Wagner, Matthew Schmerer, Minoo Agarwal, Eyad Almasri, Debbie Boles, Ayla Burns, Nuthawin Charoensri, Oren Cohen, Susan Countryman, Mary Ann Cristobal, Bobbi Croy, Suzanne Dale, Hrushikesh Deshmukh, Amanda Douglas, Vincent Drouillon, Marcia Eisenberg, Howard Engler, Rama Ghatti, Prashant Gupta, Susan Hicks, Jake Humphrey, Lax Iyer, Lisa Pfefferle, Manoj Jain, Matthew Robinson, Mohan Kolli, Brian Krueger, Tim Kuphal, Stanley Letovsky, Michael Levandoski, Craig Lukasik, Jonathan Meltzer, Brian Norvell, Mindy Nye, Scott Parker, Christos Petropoulos, John Pruitt, Steven Ragan, Scott Ryan, Mike Sapeta, Jana Schroth, Suresh Babu Selvaraju, Goran Stevovic, Amanda Suchanek, Andrea Throop, Lyndon Tilson, Thomas Urban, Joe Voshell, Kimberly Wagner, Jonathan Williams, Mary Williamson, Qian Zeng, Tricia Zwiefelhofer, Clinton R. Paden, Duncan MacCannell |
| EPI_ISL_3329235 | New Mexico Department of Health Scientific Laboratory | New Mexico Department of Health Scientific Laboratory | Ellie Johnson, D'eldra Malone, Jennifer Benoit, Ratheesh Rajan, Linda Salazar, Anastacia Griego-Fisher |
| EPI_ISL_3329236 | Laboratory Corporation of America | Centers for Disease Control and Prevention Division of Viral Diseases, Pathogen Discovery | Dakota Howard, Dhwani Batra, Peter W. Cook, Kara Moser, Adrian Paskey, Jason Caravas, Benjamin Rambo-Martin, Shatavia Morrison, Christopher Gulvick, Scott Sammons, Yvette Unoarumhi, Darlene Wagner, Matthew Schmerer, Minoo Agarwal, Eyad Almasri, Debbie Boles, Ayla Burns, Nuthawin Charoensri, Oren Cohen, Susan Countryman, Mary Ann Cristobal, Bobbi Croy, Suzanne Dale, Hrushikesh Deshmukh, Amanda Douglas, Vincent Drouillon, Marcia Eisenberg, Howard Engler, Rama Ghatti, Prashant Gupta, Susan Hicks, Jake Humphrey, Lax Iyer, Lisa Pfefferle, Manoj Jain, Matthew Robinson, Mohan Kolli, Brian Krueger, Tim Kuphal, Stanley Letovsky, Michael Levandoski, Craig Lukasik, Jonathan Meltzer, Brian Norvell, Mindy Nye, Scott Parker, Christos Petropoulos, John Pruitt, Steven Ragan, Scott Ryan, Mike Sapeta, Jana Schroth, Suresh Babu Selvaraju, Goran Stevovic, Amanda Suchanek, Andrea Throop, Lyndon Tilson, Thomas Urban, Joe Voshell, Kimberly Wagner, Jonathan Williams, Mary Williamson, Qian Zeng, Tricia Zwiefelhofer, Clinton R. Paden, Duncan MacCannell |
| EPI_ISL_3329237 | New Mexico Department of Health Scientific Laboratory | New Mexico Department of Health Scientific Laboratory | Ellie Johnson, D'eldra Malone, Jennifer Benoit, Ratheesh Rajan, Linda Salazar, Anastacia Griego-Fisher |
| EPI_ISL_3329238, EPI_ISL_3329239 | Laboratory Corporation of America | Centers for Disease Control and Prevention Division of Viral Diseases, Pathogen Discovery | Dakota Howard, Dhwani Batra, Peter W. Cook, Kara Moser, Adrian Paskey, Jason Caravas, Benjamin Rambo-Martin, Shatavia Morrison, Christopher Gulvick, Scott Sammons, Yvette Unoarumhi, Darlene Wagner, Matthew Schmerer, Minoo Agarwal, Eyad Almasri, Debbie Boles, Ayla Burns, Nuthawin Charoensri, Oren Cohen, Susan Countryman, Mary Ann Cristobal, Bobbi Croy, Suzanne Dale, Hrushikesh Deshmukh, Amanda Douglas, Vincent Drouillon, Marcia Eisenberg, Howard Engler, Rama Ghatti, Prashant Gupta, Susan Hicks, Jake Humphrey, Lax Iyer, Lisa Pfefferle, Manoj Jain, Matthew Robinson, Mohan Kolli, Brian Krueger, Tim Kuphal, Stanley Letovsky, Michael Levandoski, Craig Lukasik, Jonathan Meltzer, Brian Norvell, Mindy Nye, Scott Parker, Christos Petropoulos, John Pruitt, Steven Ragan, Scott Ryan, Mike Sapeta, Jana Schroth, Suresh Babu Selvaraju, Goran Stevovic, Amanda Suchanek, Andrea Throop, Lyndon Tilson, Thomas Urban, Joe Voshell, Kimberly Wagner, Jonathan Williams, Mary Williamson, Qian Zeng, Tricia Zwiefelhofer, Clinton R. Paden, Duncan MacCannell |
| EPI_ISL_3329240 | New Mexico Department of Health Scientific Laboratory | New Mexico Department of Health Scientific Laboratory | Ellie Johnson, D'eldra Malone, Jennifer Benoit, Ratheesh Rajan, Linda Salazar, Anastacia Griego-Fisher |
| EPI_ISL_3329241 | Laboratory Corporation of America | Centers for Disease Control and Prevention Division of Viral Diseases, Pathogen Discovery | Dakota Howard, Dhwani Batra, Peter W. Cook, Kara Moser, Adrian Paskey, Jason Caravas, Benjamin Rambo-Martin, Shatavia Morrison, Christopher Gulvick, Scott Sammons, Yvette Unoarumhi, Darlene Wagner, Matthew Schmerer, Minoo Agarwal, Eyad Almasri, Debbie Boles, Ayla Burns, Nuthawin Charoensri, Oren Cohen, Susan Countryman, Mary Ann Cristobal, Bobbi Croy, Suzanne Dale, Hrushikesh Deshmukh, Amanda Douglas, Vincent Drouillon, Marcia Eisenberg, Howard Engler, Rama Ghatti, Prashant Gupta, Susan Hicks, Jake Humphrey, Lax Iyer, Lisa Pfefferle, Manoj Jain, Matthew Robinson, Mohan Kolli, Brian Krueger, Tim Kuphal, Stanley Letovsky, Michael Levandoski, Craig Lukasik, Jonathan Meltzer, Brian Norvell, Mindy Nye, Scott Parker, Christos Petropoulos, John Pruitt, Steven Ragan, Scott Ryan, Mike Sapeta, Jana Schroth, Suresh Babu Selvaraju, Goran Stevovic, Amanda Suchanek, Andrea Throop, Lyndon Tilson, Thomas Urban, Joe Voshell, Kimberly Wagner, Jonathan Williams, Mary Williamson, Qian Zeng, Tricia Zwiefelhofer, Clinton R. Paden, Duncan MacCannell |
| EPI_ISL_3329242 | New Mexico Department of Health Scientific Laboratory | New Mexico Department of Health Scientific Laboratory | Ellie Johnson, D'eldra Malone, Jennifer Benoit, Ratheesh Rajan, Linda Salazar, Anastacia Griego-Fisher |
| EPI_ISL_3329243 | Laboratory Corporation of America | Centers for Disease Control and Prevention Division of Viral Diseases, Pathogen Discovery | Dakota Howard, Dhwani Batra, Peter W. Cook, Kara Moser, Adrian Paskey, Jason Caravas, Benjamin Rambo-Martin, Shatavia Morrison, Christopher Gulvick, Scott Sammons, Yvette Unoarumhi, Darlene Wagner, Matthew Schmerer, Minoo Agarwal, Eyad Almasri, Debbie Boles, Ayla Burns, Nuthawin Charoensri, Oren Cohen, Susan Countryman, Mary Ann Cristobal, Bobbi Croy, Suzanne Dale, Hrushikesh Deshmukh, Amanda Douglas, Vincent Drouillon, Marcia Eisenberg, Howard Engler, Rama Ghatti, Prashant Gupta, Susan Hicks, Jake Humphrey, Lax Iyer, Lisa Pfefferle, Manoj Jain, Matthew Robinson, Mohan Kolli, Brian Krueger, Tim Kuphal, Stanley Letovsky, Michael Levandoski, Craig Lukasik, Jonathan Meltzer, Brian Norvell, Mindy Nye, Scott Parker, Christos Petropoulos, John Pruitt, Steven Ragan, Scott Ryan, Mike Sapeta, Jana Schroth, Suresh Babu Selvaraju, Goran Stevovic, Amanda Suchanek, Andrea Throop, Lyndon Tilson, Thomas Urban, Joe Voshell, Kimberly Wagner, Jonathan Williams, Mary Williamson, Qian Zeng, Tricia Zwiefelhofer, Clinton R. Paden, Duncan MacCannell |
| EPI_ISL_3329244 | New Mexico Department of Health Scientific Laboratory | New Mexico Department of Health Scientific Laboratory | Ellie Johnson, D'eldra Malone, Jennifer Benoit, Ratheesh Rajan, Linda Salazar, Anastacia Griego-Fisher |
| EPI_ISL_3329245, EPI_ISL_3329247, EPI_ISL_3329248 | Laboratory Corporation of America | Centers for Disease Control and Prevention Division of Viral Diseases, Pathogen Discovery | Dakota Howard, Dhwani Batra, Peter W. Cook, Kara Moser, Adrian Paskey, Jason Caravas, Benjamin Rambo-Martin, Shatavia Morrison, Christopher Gulvick, Scott Sammons, Yvette Unoarumhi, Darlene Wagner, Matthew Schmerer, Minoo Agarwal, Eyad Almasri, Debbie Boles, Ayla Burns, Nuthawin Charoensri, Oren Cohen, Susan Countryman, Mary Ann Cristobal, Bobbi Croy, Suzanne Dale, Hrushikesh Deshmukh, Amanda Douglas, Vincent Drouillon, Marcia Eisenberg, Howard Engler, Rama Ghatti, Prashant Gupta, Susan Hicks, Jake Humphrey, Lax Iyer, Lisa Pfefferle, Manoj Jain, Matthew Robinson, Mohan Kolli, Brian Krueger, Tim Kuphal, Stanley Letovsky, Michael Levandoski, Craig Lukasik, Jonathan Meltzer, Brian Norvell, Mindy Nye, Scott Parker, Christos Petropoulos, John Pruitt, Steven Ragan, Scott Ryan, Mike Sapeta, Jana Schroth, Suresh Babu Selvaraju, Goran Stevovic, Amanda Suchanek, Andrea Throop, Lyndon Tilson, Thomas Urban, Joe Voshell, Kimberly Wagner, Jonathan Williams, Mary Williamson, Qian Zeng, Tricia Zwiefelhofer, Clinton R. Paden, Duncan MacCannell |
| EPI_ISL_3329249 | New Mexico Department of Health Scientific Laboratory | New Mexico Department of Health Scientific Laboratory | Ellie Johnson, D'eldra Malone, Jennifer Benoit, Ratheesh Rajan, Linda Salazar, Anastacia Griego-Fisher |
| EPI_ISL_3329250 | Laboratory Corporation of America | Centers for Disease Control and Prevention Division of Viral Diseases, Pathogen Discovery | Dakota Howard, Dhwani Batra, Peter W. Cook, Kara Moser, Adrian Paskey, Jason Caravas, Benjamin Rambo-Martin, Shatavia Morrison, Christopher Gulvick, Scott Sammons, Yvette Unoarumhi, Darlene Wagner, Matthew Schmerer, Minoo Agarwal, Eyad Almasri, Debbie Boles, Ayla Burns, Nuthawin Charoensri, Oren Cohen, Susan Countryman, Mary Ann Cristobal, Bobbi Croy, Suzanne Dale, Hrushikesh Deshmukh, Amanda Douglas, Vincent Drouillon, Marcia Eisenberg, Howard Engler, Rama Ghatti, Prashant Gupta, Susan Hicks, Jake Humphrey, Lax Iyer, Lisa Pfefferle, Manoj Jain, Matthew Robinson, Mohan Kolli, Brian Krueger, Tim Kuphal, Stanley Letovsky, Michael Levandoski, Craig Lukasik, Jonathan Meltzer, Brian Norvell, Mindy Nye, Scott Parker, Christos Petropoulos, John Pruitt, Steven Ragan, Scott Ryan, Mike Sapeta, Jana Schroth, Suresh Babu Selvaraju, Goran Stevovic, Amanda Suchanek, Andrea Throop, Lyndon Tilson, Thomas Urban, Joe Voshell, Kimberly Wagner, Jonathan Williams, Mary Williamson, Qian Zeng, Tricia Zwiefelhofer, Clinton R. Paden, Duncan MacCannell |
| EPI_ISL_3329251 | New Mexico Department of Health Scientific Laboratory | New Mexico Department of Health Scientific Laboratory | Ellie Johnson, D'eldra Malone, Jennifer Benoit, Ratheesh Rajan, Linda Salazar, Anastacia Griego-Fisher |
| EPI_ISL_3329252 | Laboratory Corporation of America | Centers for Disease Control and Prevention Division of Viral Diseases, Pathogen Discovery | Dakota Howard, Dhwani Batra, Peter W. Cook, Kara Moser, Adrian Paskey, Jason Caravas, Benjamin Rambo-Martin, Shatavia Morrison, Christopher Gulvick, Scott Sammons, Yvette Unoarumhi, Darlene Wagner, Matthew Schmerer, Minoo Agarwal, Eyad Almasri, Debbie Boles, Ayla Burns, Nuthawin Charoensri, Oren Cohen, Susan Countryman, Mary Ann Cristobal, Bobbi Croy, Suzanne Dale, Hrushikesh Deshmukh, Amanda Douglas, Vincent Drouillon, Marcia Eisenberg, Howard Engler, Rama Ghatti, Prashant Gupta, Susan Hicks, Jake Humphrey, Lax Iyer, Lisa Pfefferle, Manoj Jain, Matthew Robinson, Mohan Kolli, Brian Krueger, Tim Kuphal, Stanley Letovsky, Michael Levandoski, Craig Lukasik, Jonathan Meltzer, Brian Norvell, Mindy Nye, Scott Parker, Christos Petropoulos, John Pruitt, Steven Ragan, Scott Ryan, Mike Sapeta, Jana Schroth, Suresh Babu Selvaraju, Goran Stevovic, Amanda Suchanek, Andrea Throop, Lyndon Tilson, Thomas Urban, Joe Voshell, Kimberly Wagner, Jonathan Williams, Mary Williamson, Qian Zeng, Tricia Zwiefelhofer, Clinton R. Paden, Duncan MacCannell |
| EPI_ISL_3329253, EPI_ISL_3329255 | New Mexico Department of Health Scientific Laboratory | New Mexico Department of Health Scientific Laboratory | Ellie Johnson, D'eldra Malone, Jennifer Benoit, Ratheesh Rajan, Linda Salazar, Anastacia Griego-Fisher |
| EPI_ISL_3329256 | Laboratory Corporation of America | Centers for Disease Control and Prevention Division of Viral Diseases, Pathogen Discovery | Dakota Howard, Dhwani Batra, Peter W. Cook, Kara Moser, Adrian Paskey, Jason Caravas, Benjamin Rambo-Martin, Shatavia Morrison, Christopher Gulvick, Scott Sammons, Yvette Unoarumhi, Darlene Wagner, Matthew Schmerer, Minoo Agarwal, Eyad Almasri, Debbie Boles, Ayla Burns, Nuthawin Charoensri, Oren Cohen, Susan Countryman, Mary Ann Cristobal, Bobbi Croy, Suzanne Dale, Hrushikesh Deshmukh, Amanda Douglas, Vincent Drouillon, Marcia Eisenberg, Howard Engler, Rama Ghatti, Prashant Gupta, Susan Hicks, Jake Humphrey, Lax Iyer, Lisa Pfefferle, Manoj Jain, Matthew Robinson, Mohan Kolli, Brian Krueger, Tim Kuphal, Stanley Letovsky, Michael Levandoski, Craig Lukasik, Jonathan Meltzer, Brian Norvell, Mindy Nye, Scott Parker, Christos Petropoulos, John Pruitt, Steven Ragan, Scott Ryan, Mike Sapeta, Jana Schroth, Suresh Babu Selvaraju, Goran Stevovic, Amanda Suchanek, Andrea Throop, Lyndon Tilson, Thomas Urban, Joe Voshell, Kimberly Wagner, Jonathan Williams, Mary Williamson, Qian Zeng, Tricia Zwiefelhofer, Clinton R. Paden, Duncan MacCannell |
| EPI_ISL_3329257 | New Mexico Department of Health Scientific Laboratory | New Mexico Department of Health Scientific Laboratory | Ellie Johnson, D'eldra Malone, Jennifer Benoit, Ratheesh Rajan, Linda Salazar, Anastacia Griego-Fisher |
| EPI_ISL_3329259 | Laboratory Corporation of America | Centers for Disease Control and Prevention Division of Viral Diseases, Pathogen Discovery | Dakota Howard, Dhwani Batra, Peter W. Cook, Kara Moser, Adrian Paskey, Jason Caravas, Benjamin Rambo-Martin, Shatavia Morrison, Christopher Gulvick, Scott Sammons, Yvette Unoarumhi, Darlene Wagner, Matthew Schmerer, Minoo Agarwal, Eyad Almasri, Debbie Boles, Ayla Burns, Nuthawin Charoensri, Oren Cohen, Susan Countryman, Mary Ann Cristobal, Bobbi Croy, Suzanne Dale, Hrushikesh Deshmukh, Amanda Douglas, Vincent Drouillon, Marcia Eisenberg, Howard Engler, Rama Ghatti, Prashant Gupta, Susan Hicks, Jake Humphrey, Lax Iyer, Lisa Pfefferle, Manoj Jain, Matthew Robinson, Mohan Kolli, Brian Krueger, Tim Kuphal, Stanley Letovsky, Michael Levandoski, Craig Lukasik, Jonathan Meltzer, Brian Norvell, Mindy Nye, Scott Parker, Christos Petropoulos, John Pruitt, Steven Ragan, Scott Ryan, Mike Sapeta, Jana Schroth, Suresh Babu Selvaraju, Goran Stevovic, Amanda Suchanek, Andrea Throop, Lyndon Tilson, Thomas Urban, Joe Voshell, Kimberly Wagner, Jonathan Williams, Mary Williamson, Qian Zeng, Tricia Zwiefelhofer, Clinton R. Paden, Duncan MacCannell |
| EPI_ISL_3329260 | New Mexico Department of Health Scientific Laboratory | New Mexico Department of Health Scientific Laboratory | Ellie Johnson, D'eldra Malone, Jennifer Benoit, Ratheesh Rajan, Linda Salazar, Anastacia Griego-Fisher |
| EPI_ISL_3329261 | Laboratory Corporation of America | Centers for Disease Control and Prevention Division of Viral Diseases, Pathogen Discovery | Dakota Howard, Dhwani Batra, Peter W. Cook, Kara Moser, Adrian Paskey, Jason Caravas, Benjamin Rambo-Martin, Shatavia Morrison, Christopher Gulvick, Scott Sammons, Yvette Unoarumhi, Darlene Wagner, Matthew Schmerer, Minoo Agarwal, Eyad Almasri, Debbie Boles, Ayla Burns, Nuthawin Charoensri, Oren Cohen, Susan Countryman, Mary Ann Cristobal, Bobbi Croy, Suzanne Dale, Hrushikesh Deshmukh, Amanda Douglas, Vincent Drouillon, Marcia Eisenberg, Howard Engler, Rama Ghatti, Prashant Gupta, Susan Hicks, Jake Humphrey, Lax Iyer, Lisa Pfefferle, Manoj Jain, Matthew Robinson, Mohan Kolli, Brian Krueger, Tim Kuphal, Stanley Letovsky, Michael Levandoski, Craig Lukasik, Jonathan Meltzer, Brian Norvell, Mindy Nye, Scott Parker, Christos Petropoulos, John Pruitt, Steven Ragan, Scott Ryan, Mike Sapeta, Jana Schroth, Suresh Babu Selvaraju, Goran Stevovic, Amanda Suchanek, Andrea Throop, Lyndon Tilson, Thomas Urban, Joe Voshell, Kimberly Wagner, Jonathan Williams, Mary Williamson, Qian Zeng, Tricia Zwiefelhofer, Clinton R. Paden, Duncan MacCannell |
| EPI_ISL_3329262 | New Mexico Department of Health Scientific Laboratory | New Mexico Department of Health Scientific Laboratory | Ellie Johnson, D'eldra Malone, Jennifer Benoit, Ratheesh Rajan, Linda Salazar, Anastacia Griego-Fisher |
| EPI_ISL_3329263 | Laboratory Corporation of America | Centers for Disease Control and Prevention Division of Viral Diseases, Pathogen Discovery | Dakota Howard, Dhwani Batra, Peter W. Cook, Kara Moser, Adrian Paskey, Jason Caravas, Benjamin Rambo-Martin, Shatavia Morrison, Christopher Gulvick, Scott Sammons, Yvette Unoarumhi, Darlene Wagner, Matthew Schmerer, Minoo Agarwal, Eyad Almasri, Debbie Boles, Ayla Burns, Nuthawin Charoensri, Oren Cohen, Susan Countryman, Mary Ann Cristobal, Bobbi Croy, Suzanne Dale, Hrushikesh Deshmukh, Amanda Douglas, Vincent Drouillon, Marcia Eisenberg, Howard Engler, Rama Ghatti, Prashant Gupta, Susan Hicks, Jake Humphrey, Lax Iyer, Lisa Pfefferle, Manoj Jain, Matthew Robinson, Mohan Kolli, Brian Krueger, Tim Kuphal, Stanley Letovsky, Michael Levandoski, Craig Lukasik, Jonathan Meltzer, Brian Norvell, Mindy Nye, Scott Parker, Christos Petropoulos, John Pruitt, Steven Ragan, Scott Ryan, Mike Sapeta, Jana Schroth, Suresh Babu Selvaraju, Goran Stevovic, Amanda Suchanek, Andrea Throop, Lyndon Tilson, Thomas Urban, Joe Voshell, Kimberly Wagner, Jonathan Williams, Mary Williamson, Qian Zeng, Tricia Zwiefelhofer, Clinton R. Paden, Duncan MacCannell |
| EPI_ISL_3329264 | New Mexico Department of Health Scientific Laboratory | New Mexico Department of Health Scientific Laboratory | Ellie Johnson, D'eldra Malone, Jennifer Benoit, Ratheesh Rajan, Linda Salazar, Anastacia Griego-Fisher |
| EPI_ISL_3329265 | Laboratory Corporation of America | Centers for Disease Control and Prevention Division of Viral Diseases, Pathogen Discovery | Dakota Howard, Dhwani Batra, Peter W. Cook, Kara Moser, Adrian Paskey, Jason Caravas, Benjamin Rambo-Martin, Shatavia Morrison, Christopher Gulvick, Scott Sammons, Yvette Unoarumhi, Darlene Wagner, Matthew Schmerer, Minoo Agarwal, Eyad Almasri, Debbie Boles, Ayla Burns, Nuthawin Charoensri, Oren Cohen, Susan Countryman, Mary Ann Cristobal, Bobbi Croy, Suzanne Dale, Hrushikesh Deshmukh, Amanda Douglas, Vincent Drouillon, Marcia Eisenberg, Howard Engler, Rama Ghatti, Prashant Gupta, Susan Hicks, Jake Humphrey, Lax Iyer, Lisa Pfefferle, Manoj Jain, Matthew Robinson, Mohan Kolli, Brian Krueger, Tim Kuphal, Stanley Letovsky, Michael Levandoski, Craig Lukasik, Jonathan Meltzer, Brian Norvell, Mindy Nye, Scott Parker, Christos Petropoulos, John Pruitt, Steven Ragan, Scott Ryan, Mike Sapeta, Jana Schroth, Suresh Babu Selvaraju, Goran Stevovic, Amanda Suchanek, Andrea Throop, Lyndon Tilson, Thomas Urban, Joe Voshell, |

| | | | |
|---|---|---|---|
| | | | Kimberly Wagner, Jonathan Williams, Mary Williamson, Qian Zeng, Tricia Zwiefelhofer, Clinton R. Paden, Duncan MacCannell |
| EPI_ISL_3329266 | New Mexico Department of Health Scientific Laboratory | New Mexico Department of Health Scientific Laboratory | Ellie Johnson, D'eldra Malone, Jennifer Benoit, Ratheesh Rajan, Linda Salazar, Anastacia Griego-Fisher |
| EPI_ISL_3329267, EPI_ISL_3329268 | Laboratory Corporation of America | Centers for Disease Control and Prevention Division of Viral Diseases, Pathogen Discovery | Dakota Howard, Dhwani Batra, Peter W. Cook, Kara Moser, Adrian Paskey, Jason Caravas, Benjamin Rambo-Martin, Shatavia Morrison, Christopher Gulvick, Scott Sammons, Yvette Unoarumhi, Darlene Wagner, Matthew Schmerer, Minoo Agarwal, Eyad Almasri, Debbie Boles, Ayla Burns, Nuthawin Charoensri, Oren Cohen, Susan Countryman, Mary Ann Cristobal, Bobbi Croy, Suzanne Dale, Hrushikesh Deshmukh, Amanda Douglas, Vincent Drouillon, Marcia Eisenberg, Howard Engler, Rama Ghatti, Prashant Gupta, Susan Hicks, Jake Humphrey, Lax Iyer, Lisa Pfefferle, Manoj Jain, Matthew Robinson, Mohan Kolli, Brian Krueger, Tim Kuphal, Stanley Letovsky, Michael Levandoski, Craig Lukasik, Jonathan Meltzer, Brian Norvell, Mindy Nye, Scott Parker, Christos Petropoulos, John Pruitt, Steven Ragan, Scott Ryan, Mike Sapeta, Jana Schroth, Suresh Babu Selvaraju, Goran Stevovic, Amanda Suchanek, Andrea Throop, Lyndon Tilson, Thomas Urban, Joe Voshell, Kimberly Wagner, Jonathan Williams, Mary Williamson, Qian Zeng, Tricia Zwiefelhofer, Clinton R. Paden, Duncan MacCannell |
| EPI_ISL_3329269 | New Mexico Department of Health Scientific Laboratory | New Mexico Department of Health Scientific Laboratory | Ellie Johnson, D'eldra Malone, Jennifer Benoit, Ratheesh Rajan, Linda Salazar, Anastacia Griego-Fisher |
| EPI_ISL_3329270 | Laboratory Corporation of America | Centers for Disease Control and Prevention Division of Viral Diseases, Pathogen Discovery | Dakota Howard, Dhwani Batra, Peter W. Cook, Kara Moser, Adrian Paskey, Jason Caravas, Benjamin Rambo-Martin, Shatavia Morrison, Christopher Gulvick, Scott Sammons, Yvette Unoarumhi, Darlene Wagner, Matthew Schmerer, Minoo Agarwal, Eyad Almasri, Debbie Boles, Ayla Burns, Nuthawin Charoensri, Oren Cohen, Susan Countryman, Mary Ann Cristobal, Bobbi Croy, Suzanne Dale, Hrushikesh Deshmukh, Amanda Douglas, Vincent Drouillon, Marcia Eisenberg, Howard Engler, Rama Ghatti, Prashant Gupta, Susan Hicks, Jake Humphrey, Lax Iyer, Lisa Pfefferle, Manoj Jain, Matthew Robinson, Mohan Kolli, Brian Krueger, Tim Kuphal, Stanley Letovsky, Michael Levandoski, Craig Lukasik, Jonathan Meltzer, Brian Norvell, Mindy Nye, Scott Parker, Christos Petropoulos, John Pruitt, Steven Ragan, Scott Ryan, Mike Sapeta, Jana Schroth, Suresh Babu Selvaraju, Goran Stevovic, Amanda Suchanek, Andrea Throop, Lyndon Tilson, Thomas Urban, Joe Voshell, Kimberly Wagner, Jonathan Williams, Mary Williamson, Qian Zeng, Tricia Zwiefelhofer, Clinton R. Paden, Duncan MacCannell |
| EPI_ISL_3329271 | New Mexico Department of Health Scientific Laboratory | New Mexico Department of Health Scientific Laboratory | Ellie Johnson, D'eldra Malone, Jennifer Benoit, Ratheesh Rajan, Linda Salazar, Anastacia Griego-Fisher |
| EPI_ISL_3329272 | Laboratory Corporation of America | Centers for Disease Control and Prevention Division of Viral Diseases, Pathogen Discovery | Dakota Howard, Dhwani Batra, Peter W. Cook, Kara Moser, Adrian Paskey, Jason Caravas, Benjamin Rambo-Martin, Shatavia Morrison, Christopher Gulvick, Scott Sammons, Yvette Unoarumhi, Darlene Wagner, Matthew Schmerer, Minoo Agarwal, Eyad Almasri, Debbie Boles, Ayla Burns, Nuthawin Charoensri, Oren Cohen, Susan Countryman, Mary Ann Cristobal, Bobbi Croy, Suzanne Dale, Hrushikesh Deshmukh, Amanda Douglas, Vincent Drouillon, Marcia Eisenberg, Howard Engler, Rama Ghatti, Prashant Gupta, Susan Hicks, Jake Humphrey, Lax Iyer, Lisa Pfefferle, Manoj Jain, Matthew Robinson, Mohan Kolli, Brian Krueger, Tim Kuphal, Stanley Letovsky, Michael Levandoski, Craig Lukasik, Jonathan Meltzer, Brian Norvell, Mindy Nye, Scott Parker, Christos Petropoulos, John Pruitt, Steven Ragan, Scott Ryan, Mike Sapeta, Jana Schroth, Suresh Babu Selvaraju, Goran Stevovic, Amanda Suchanek, Andrea Throop, Lyndon Tilson, Thomas Urban, Joe Voshell, Kimberly Wagner, Jonathan Williams, Mary Williamson, Qian Zeng, Tricia Zwiefelhofer, Clinton R. Paden, Duncan MacCannell |
| EPI_ISL_3329273 | New Mexico Department of Health Scientific Laboratory | New Mexico Department of Health Scientific Laboratory | Ellie Johnson, D'eldra Malone, Jennifer Benoit, Ratheesh Rajan, Linda Salazar, Anastacia Griego-Fisher |
| EPI_ISL_3329274, EPI_ISL_3329275 | Laboratory Corporation of America | Centers for Disease Control and Prevention Division of Viral Diseases, Pathogen Discovery | Dakota Howard, Dhwani Batra, Peter W. Cook, Kara Moser, Adrian Paskey, Jason Caravas, Benjamin Rambo-Martin, Shatavia Morrison, Christopher Gulvick, Scott Sammons, Yvette Unoarumhi, Darlene Wagner, Matthew Schmerer, Minoo Agarwal, Eyad Almasri, Debbie Boles, Ayla Burns, Nuthawin Charoensri, Oren Cohen, Susan Countryman, Mary Ann Cristobal, Bobbi Croy, Suzanne Dale, Hrushikesh Deshmukh, Amanda Douglas, Vincent Drouillon, Marcia Eisenberg, Howard Engler, Rama Ghatti, Prashant Gupta, Susan Hicks, Jake Humphrey, Lax Iyer, Lisa Pfefferle, Manoj Jain, Matthew Robinson, Mohan Kolli, Brian Krueger, Tim Kuphal, Stanley Letovsky, Michael Levandoski, Craig Lukasik, Jonathan Meltzer, Brian Norvell, Mindy Nye, Scott Parker, Christos Petropoulos, John Pruitt, Steven Ragan, Scott Ryan, Mike Sapeta, Jana Schroth, Suresh Babu Selvaraju, Goran Stevovic, Amanda Suchanek, Andrea Throop, Lyndon Tilson, Thomas Urban, Joe Voshell, Kimberly Wagner, Jonathan Williams, Mary Williamson, Qian Zeng, Tricia Zwiefelhofer, Clinton R. Paden, Duncan MacCannell |
| EPI_ISL_3329276 | New Mexico Department of Health Scientific Laboratory | New Mexico Department of Health Scientific Laboratory | Ellie Johnson, D'eldra Malone, Jennifer Benoit, Ratheesh Rajan, Linda Salazar, Anastacia Griego-Fisher |
| EPI_ISL_3329277 | Laboratory Corporation of America | Centers for Disease Control and Prevention Division of Viral Diseases, Pathogen Discovery | Dakota Howard, Dhwani Batra, Peter W. Cook, Kara Moser, Adrian Paskey, Jason Caravas, Benjamin Rambo-Martin, Shatavia Morrison, Christopher Gulvick, Scott Sammons, Yvette Unoarumhi, Darlene Wagner, Matthew Schmerer, Minoo Agarwal, Eyad Almasri, Debbie Boles, Ayla Burns, Nuthawin Charoensri, Oren Cohen, Susan Countryman, Mary Ann Cristobal, Bobbi Croy, Suzanne Dale, Hrushikesh Deshmukh, Amanda Douglas, Vincent Drouillon, Marcia Eisenberg, Howard Engler, Rama Ghatti, Prashant Gupta, Susan Hicks, Jake Humphrey, Lax Iyer, Lisa Pfefferle, Manoj Jain, Matthew Robinson, Mohan Kolli, Brian Krueger, Tim Kuphal, Stanley Letovsky, Michael Levandoski, Craig Lukasik, Jonathan Meltzer, Brian Norvell, Mindy Nye, Scott Parker, Christos Petropoulos, John Pruitt, Steven Ragan, Scott Ryan, Mike Sapeta, Jana Schroth, Suresh Babu Selvaraju, Goran Stevovic, Amanda Suchanek, Andrea Throop, Lyndon Tilson, Thomas Urban, Joe Voshell, Kimberly Wagner, Jonathan Williams, Mary Williamson, Qian Zeng, Tricia Zwiefelhofer, Clinton R. Paden, Duncan MacCannell |
| EPI_ISL_3329278 | New Mexico Department of Health Scientific Laboratory | New Mexico Department of Health Scientific Laboratory | Ellie Johnson, D'eldra Malone, Jennifer Benoit, Ratheesh Rajan, Linda Salazar, Anastacia Griego-Fisher |
| EPI_ISL_3329279 | Laboratory Corporation of America | Centers for Disease Control and Prevention Division of Viral Diseases, Pathogen Discovery | Dakota Howard, Dhwani Batra, Peter W. Cook, Kara Moser, Adrian Paskey, Jason Caravas, Benjamin Rambo-Martin, Shatavia Morrison, Christopher Gulvick, Scott Sammons, Yvette Unoarumhi, Darlene Wagner, Matthew Schmerer, Minoo Agarwal, Eyad Almasri, Debbie Boles, Ayla Burns, Nuthawin Charoensri, Oren Cohen, Susan Countryman, Mary Ann Cristobal, Bobbi Croy, Suzanne Dale, Hrushikesh Deshmukh, Amanda Douglas, Vincent Drouillon, Marcia Eisenberg, Howard Engler, Rama Ghatti, Prashant Gupta, Susan Hicks, Jake Humphrey, Lax Iyer, Lisa Pfefferle, Manoj Jain, Matthew Robinson, Mohan Kolli, Brian Krueger, Tim Kuphal, Stanley Letovsky, Michael Levandoski, Craig Lukasik, Jonathan Meltzer, Brian Norvell, Mindy Nye, Scott Parker, Christos Petropoulos, John Pruitt, Steven Ragan, Scott Ryan, Mike Sapeta, Jana Schroth, Suresh Babu Selvaraju, Goran Stevovic, Amanda Suchanek, Andrea Throop, Lyndon Tilson, Thomas Urban, Joe Voshell, Kimberly Wagner, Jonathan Williams, Mary Williamson, Qian Zeng, Tricia Zwiefelhofer, Clinton R. Paden, Duncan MacCannell |
| EPI_ISL_3329280 | New Mexico Department of Health Scientific Laboratory | New Mexico Department of Health Scientific Laboratory | Ellie Johnson, D'eldra Malone, Jennifer Benoit, Ratheesh Rajan, Linda Salazar, Anastacia Griego-Fisher |
| EPI_ISL_3329282 | Laboratory Corporation of America | Centers for Disease Control and Prevention Division of Viral Diseases, Pathogen Discovery | Dakota Howard, Dhwani Batra, Peter W. Cook, Kara Moser, Adrian Paskey, Jason Caravas, Benjamin Rambo-Martin, Shatavia Morrison, Christopher Gulvick, Scott Sammons, Yvette Unoarumhi, Darlene Wagner, Matthew Schmerer, Minoo Agarwal, Eyad Almasri, Debbie Boles, Ayla Burns, Nuthawin Charoensri, Oren Cohen, Susan Countryman, Mary Ann Cristobal, Bobbi Croy, Suzanne Dale, Hrushikesh Deshmukh, Amanda Douglas, Vincent Drouillon, Marcia Eisenberg, Howard Engler, Rama Ghatti, Prashant Gupta, Susan Hicks, Jake Humphrey, Lax Iyer, Lisa Pfefferle, Manoj Jain, Matthew Robinson, Mohan Kolli, Brian Krueger, Tim Kuphal, Stanley Letovsky, Michael Levandoski, Craig Lukasik, Jonathan Meltzer, Brian Norvell, Mindy Nye, Scott Parker, Christos Petropoulos, John Pruitt, Steven Ragan, Scott Ryan, Mike Sapeta, Jana Schroth, Suresh Babu Selvaraju, Goran Stevovic, Amanda Suchanek, Andrea Throop, Lyndon Tilson, Thomas Urban, Joe Voshell, Kimberly Wagner, Jonathan Williams, Mary Williamson, Qian Zeng, Tricia Zwiefelhofer, Clinton R. Paden, Duncan MacCannell |
| EPI_ISL_3329283 | New Mexico Department of Health Scientific Laboratory | New Mexico Department of Health Scientific Laboratory | Ellie Johnson, D'eldra Malone, Jennifer Benoit, Ratheesh Rajan, Linda Salazar, Anastacia Griego-Fisher |
| EPI_ISL_3329284 | Laboratory Corporation of America | Centers for Disease Control and Prevention Division of Viral Diseases, Pathogen Discovery | Dakota Howard, Dhwani Batra, Peter W. Cook, Kara Moser, Adrian Paskey, Jason Caravas, Benjamin Rambo-Martin, Shatavia Morrison, Christopher Gulvick, Scott Sammons, Yvette Unoarumhi, Darlene Wagner, Matthew Schmerer, Minoo Agarwal, Eyad Almasri, Debbie Boles, Ayla Burns, Nuthawin Charoensri, Oren Cohen, Susan Countryman, Mary Ann Cristobal, Bobbi Croy, Suzanne Dale, Hrushikesh Deshmukh, Amanda Douglas, Vincent Drouillon, Marcia Eisenberg, Howard Engler, Rama Ghatti, Prashant Gupta, Susan Hicks, Jake Humphrey, Lax Iyer, Lisa Pfefferle, Manoj Jain, Matthew Robinson, Mohan Kolli, Brian Krueger, Tim Kuphal, Stanley Letovsky, Michael Levandoski, Craig Lukasik, Jonathan Meltzer, Brian Norvell, Mindy Nye, Scott Parker, Christos Petropoulos, John Pruitt, Steven Ragan, Scott Ryan, Mike Sapeta, Jana Schroth, Suresh Babu Selvaraju, Goran Stevovic, Amanda Suchanek, Andrea Throop, Lyndon Tilson, Thomas Urban, Joe Voshell, Kimberly Wagner, Jonathan Williams, Mary Williamson, Qian Zeng, Tricia Zwiefelhofer, Clinton R. Paden, Duncan MacCannell |
| EPI_ISL_3329285 | New Mexico Department of Health Scientific Laboratory | New Mexico Department of Health Scientific Laboratory | Ellie Johnson, D'eldra Malone, Jennifer Benoit, Ratheesh Rajan, Linda Salazar, Anastacia Griego-Fisher |
| EPI_ISL_3329286, EPI_ISL_3329287 | Laboratory Corporation of America | Centers for Disease Control and Prevention Division of Viral Diseases, Pathogen Discovery | Dakota Howard, Dhwani Batra, Peter W. Cook, Kara Moser, Adrian Paskey, Jason Caravas, Benjamin Rambo-Martin, Shatavia Morrison, Christopher Gulvick, Scott Sammons, Yvette Unoarumhi, Darlene Wagner, Matthew Schmerer, Minoo Agarwal, Eyad Almasri, Debbie Boles, Ayla Burns, Nuthawin Charoensri, Oren Cohen, Susan Countryman, Mary Ann Cristobal, Bobbi Croy, Suzanne Dale, Hrushikesh Deshmukh, Amanda Douglas, Vincent Drouillon, Marcia Eisenberg, Howard Engler, Rama Ghatti, Prashant Gupta, Susan Hicks, Jake Humphrey, Lax Iyer, Lisa Pfefferle, Manoj Jain, Matthew Robinson, Mohan Kolli, Brian Krueger, Tim Kuphal, Stanley Letovsky, Michael Levandoski, Craig Lukasik, Jonathan Meltzer, Brian Norvell, Mindy Nye, Scott Parker, Christos Petropoulos, John Pruitt, Steven Ragan, Scott Ryan, Mike Sapeta, Jana Schroth, Suresh Babu Selvaraju, Goran Stevovic, Amanda Suchanek, Andrea Throop, Lyndon Tilson, Thomas Urban, Joe Voshell, Kimberly Wagner, Jonathan Williams, Mary Williamson, Qian Zeng, Tricia Zwiefelhofer, Clinton R. Paden, Duncan MacCannell |
| EPI_ISL_3329288 | New Mexico Department of Health Scientific Laboratory | New Mexico Department of Health Scientific Laboratory | Ellie Johnson, D'eldra Malone, Jennifer Benoit, Ratheesh Rajan, Linda Salazar, Anastacia Griego-Fisher |
| EPI_ISL_3329289 | Laboratory Corporation of America | Centers for Disease Control and Prevention Division of Viral Diseases, Pathogen Discovery | Dakota Howard, Dhwani Batra, Peter W. Cook, Kara Moser, Adrian Paskey, Jason Caravas, Benjamin Rambo-Martin, Shatavia Morrison, Christopher Gulvick, Scott Sammons, Yvette Unoarumhi, Darlene Wagner, Matthew Schmerer, Minoo Agarwal, Eyad Almasri, Debbie Boles, Ayla Burns, Nuthawin Charoensri, Oren Cohen, Susan Countryman, Mary Ann Cristobal, Bobbi Croy, Suzanne Dale, Hrushikesh Deshmukh, Amanda Douglas, Vincent Drouillon, Marcia Eisenberg, Howard Engler, Rama Ghatti, Prashant Gupta, Susan Hicks, Jake Humphrey, Lax Iyer, Lisa Pfefferle, Manoj Jain, Matthew Robinson, Mohan Kolli, Brian Krueger, Tim Kuphal, Stanley Letovsky, Michael Levandoski, Craig Lukasik, Jonathan Meltzer, Brian Norvell, Mindy Nye, Scott Parker, Christos Petropoulos, John Pruitt, Steven Ragan, Scott Ryan, Mike Sapeta, Jana Schroth, Suresh Babu Selvaraju, Goran Stevovic, Amanda Suchanek, Andrea Throop, Lyndon Tilson, Thomas Urban, Joe Voshell, Kimberly Wagner, Jonathan Williams, Mary Williamson, Qian Zeng, Tricia Zwiefelhofer, Clinton R. Paden, Duncan MacCannell |
| EPI_ISL_3329290, EPI_ISL_3329291, EPI_ISL_3329292 | New Mexico Department of Health Scientific Laboratory | New Mexico Department of Health Scientific Laboratory | Ellie Johnson, D'eldra Malone, Jennifer Benoit, Ratheesh Rajan, Linda Salazar, Anastacia Griego-Fisher |
| EPI_ISL_3329293 | Laboratory Corporation of America | Centers for Disease Control and Prevention Division of Viral Diseases, Pathogen Discovery | Dakota Howard, Dhwani Batra, Peter W. Cook, Kara Moser, Adrian Paskey, Jason Caravas, Benjamin Rambo-Martin, Shatavia Morrison, Christopher Gulvick, Scott Sammons, Yvette Unoarumhi, Darlene Wagner, Matthew Schmerer, Minoo Agarwal, Eyad Almasri, Debbie Boles, Ayla Burns, Nuthawin Charoensri, Oren Cohen, Susan Countryman, Mary Ann Cristobal, Bobbi Croy, Suzanne Dale, Hrushikesh Deshmukh, Amanda Douglas, Vincent Drouillon, Marcia Eisenberg, Howard Engler, Rama Ghatti, Prashant Gupta, Susan Hicks, Jake Humphrey, Lax Iyer, Lisa Pfefferle, Manoj Jain, Matthew Robinson, Mohan Kolli, Brian Krueger, Tim Kuphal, Stanley Letovsky, Michael Levandoski, Craig Lukasik, Jonathan Meltzer, Brian Norvell, Mindy Nye, Scott Parker, Christos Petropoulos, John Pruitt, Steven Ragan, Scott Ryan, Mike Sapeta, Jana Schroth, Suresh Babu Selvaraju, Goran Stevovic, Amanda Suchanek, Andrea Throop, Lyndon Tilson, Thomas Urban, Joe Voshell, Kimberly Wagner, Jonathan Williams, Mary Williamson, Qian Zeng, Tricia Zwiefelhofer, Clinton R. Paden, Duncan MacCannell |
| EPI_ISL_3329294 | New Mexico Department of Health Scientific Laboratory | New Mexico Department of Health Scientific Laboratory | Ellie Johnson, D'eldra Malone, Jennifer Benoit, Ratheesh Rajan, Linda Salazar, Anastacia Griego-Fisher |
| EPI_ISL_3329295, EPI_ISL_3329296 | Laboratory Corporation of America | Centers for Disease Control and Prevention Division of Viral Diseases, Pathogen Discovery | Dakota Howard, Dhwani Batra, Peter W. Cook, Kara Moser, Adrian Paskey, Jason Caravas, Benjamin Rambo-Martin, Shatavia Morrison, Christopher Gulvick, Scott Sammons, Yvette Unoarumhi, Darlene Wagner, Matthew Schmerer, Minoo Agarwal, Eyad Almasri, Debbie Boles, Ayla Burns, Nuthawin Charoensri, Oren Cohen, Susan Countryman, Mary Ann Cristobal, Bobbi Croy, Suzanne Dale, Hrushikesh Deshmukh, Amanda Douglas, Vincent Drouillon, Marcia Eisenberg, Howard Engler, Rama Ghatti, Prashant Gupta, Susan Hicks, Jake Humphrey, Lax Iyer, Lisa Pfefferle, Manoj Jain, Matthew Robinson, Mohan Kolli, Brian Krueger, Tim Kuphal, Stanley Letovsky, Michael Levandoski, Craig Lukasik, Jonathan Meltzer, Brian Norvell, Mindy Nye, Scott Parker, Christos Petropoulos, John Pruitt, Steven Ragan, Scott Ryan, Mike Sapeta, Jana Schroth, Suresh Babu Selvaraju, Goran Stevovic, Amanda Suchanek, Andrea Throop, Lyndon Tilson, Thomas Urban, Joe Voshell, Kimberly Wagner, Jonathan Williams, Mary Williamson, Qian Zeng, Tricia Zwiefelhofer, Clinton R. Paden, Duncan MacCannell |
| EPI_ISL_3329297 | New Mexico Department of Health Scientific Laboratory | New Mexico Department of Health Scientific Laboratory | Ellie Johnson, D'eldra Malone, Jennifer Benoit, Ratheesh Rajan, Linda Salazar, Anastacia Griego-Fisher |
| EPI_ISL_3329298 | Laboratory Corporation of America | Centers for Disease Control and Prevention Division of Viral Diseases, Pathogen Discovery | Dakota Howard, Dhwani Batra, Peter W. Cook, Kara Moser, Adrian Paskey, Jason Caravas, Benjamin Rambo-Martin, Shatavia Morrison, Christopher Gulvick, Scott Sammons, Yvette Unoarumhi, Darlene Wagner, Matthew Schmerer, Minoo Agarwal, Eyad Almasri, Debbie Boles, Ayla Burns, Nuthawin Charoensri, Oren Cohen, Susan Countryman, Mary Ann Cristobal, Bobbi Croy, Suzanne Dale, Hrushikesh Deshmukh, Amanda Douglas, Vincent Drouillon, Marcia Eisenberg, Howard Engler, Rama Ghatti, Prashant Gupta, Susan Hicks, Jake Humphrey, Lax Iyer, Lisa Pfefferle, Manoj Jain, Matthew Robinson, Mohan Kolli, Brian Krueger, Tim Kuphal, Stanley Letovsky, Michael Levandoski, Craig Lukasik, Jonathan Meltzer, Brian Norvell, Mindy Nye, Scott Parker, Christos Petropoulos, John Pruitt, Steven Ragan, Scott Ryan, Mike Sapeta, Jana Schroth, Suresh Babu Selvaraju, Goran Stevovic, Amanda Suchanek, Andrea Throop, Lyndon Tilson, Thomas Urban, Joe Voshell, Kimberly Wagner, Jonathan Williams, Mary Williamson, Qian Zeng, Tricia Zwiefelhofer, Clinton R. Paden, Duncan MacCannell |
| EPI_ISL_3329299 | New Mexico Department of Health Scientific Laboratory | New Mexico Department of Health Scientific Laboratory | Ellie Johnson, D'eldra Malone, Jennifer Benoit, Ratheesh Rajan, Linda Salazar, Anastacia Griego-Fisher |
| EPI_ISL_3329300, EPI_ISL_3329301 | Laboratory Corporation of America | Centers for Disease Control and Prevention Division of Viral Diseases, Pathogen Discovery | Dakota Howard, Dhwani Batra, Peter W. Cook, Kara Moser, Adrian Paskey, Jason Caravas, Benjamin Rambo-Martin, Shatavia Morrison, Christopher Gulvick, Scott Sammons, Yvette Unoarumhi, Darlene Wagner, Matthew Schmerer, Minoo Agarwal, Eyad Almasri, Debbie Boles, Ayla Burns, Nuthawin Charoensri, Oren Cohen, Susan Countryman, Mary Ann Cristobal, Bobbi Croy, Suzanne Dale, Hrushikesh Deshmukh, Amanda Douglas, Vincent Drouillon, Marcia Eisenberg, Howard Engler, Rama Ghatti, Prashant Gupta, Susan Hicks, Jake Humphrey, Lax Iyer, Lisa Pfefferle, Manoj Jain, Matthew Robinson, Mohan Kolli, Brian Krueger, Tim Kuphal, Stanley Letovsky, Michael Levandoski, Craig Lukasik, Jonathan Meltzer, Brian Norvell, Mindy Nye, Scott Parker, Christos Petropoulos, John Pruitt, Steven Ragan, Scott Ryan, Mike Sapeta, Jana Schroth, Suresh Babu Selvaraju, Goran Stevovic, Amanda Suchanek, Andrea Throop, Lyndon Tilson, Thomas Urban, Joe Voshell, Kimberly Wagner, Jonathan Williams, Mary Williamson, Qian Zeng, Tricia Zwiefelhofer, Clinton R. Paden, Duncan MacCannell |
| EPI_ISL_3329302 | New Mexico Department of Health Scientific Laboratory | New Mexico Department of Health Scientific Laboratory | Ellie Johnson, D'eldra Malone, Jennifer Benoit, Ratheesh Rajan, Linda Salazar, Anastacia Griego-Fisher |
| EPI_ISL_3329303 | Laboratory Corporation of America | Centers for Disease Control and Prevention Division of Viral Diseases, Pathogen Discovery | Dakota Howard, Dhwani Batra, Peter W. Cook, Kara Moser, Adrian Paskey, Jason Caravas, Benjamin Rambo-Martin, Shatavia Morrison, Christopher Gulvick, Scott Sammons, Yvette Unoarumhi, Darlene Wagner, Matthew Schmerer, Minoo Agarwal, Eyad Almasri, Debbie Boles, Ayla Burns, Nuthawin Charoensri, Oren Cohen, Susan Countryman, Mary Ann Cristobal, Bobbi Croy, Suzanne Dale, Hrushikesh Deshmukh, Amanda Douglas, Vincent Drouillon, Marcia Eisenberg, Howard Engler, Rama Ghatti, Prashant Gupta, Susan Hicks, Jake Humphrey, Lax Iyer, Lisa Pfefferle, Manoj Jain, Matthew Robinson, Mohan Kolli, Brian Krueger, Tim Kuphal, Stanley Letovsky, Michael Levandoski, Craig Lukasik, Jonathan Meltzer, Brian Norvell, Mindy Nye, Scott Parker, Christos Petropoulos, John Pruitt, Steven Ragan, Scott Ryan, Mike Sapeta, Jana Schroth, Suresh Babu Selvaraju, Goran Stevovic, Amanda Suchanek, Andrea Throop, Lyndon Tilson, Thomas Urban, Joe Voshell, Kimberly Wagner, Jonathan Williams, Mary Williamson, Qian Zeng, Tricia Zwiefelhofer, Clinton R. Paden, Duncan MacCannell |
| EPI_ISL_3329304 | New Mexico Department of Health Scientific Laboratory | New Mexico Department of Health Scientific Laboratory | Ellie Johnson, D'eldra Malone, Jennifer Benoit, Ratheesh Rajan, Linda Salazar, Anastacia Griego-Fisher |
| EPI_ISL_3329305, EPI_ISL_3329306 | Laboratory Corporation of America | Centers for Disease Control and Prevention Division of Viral Diseases, Pathogen Discovery | Dakota Howard, Dhwani Batra, Peter W. Cook, Kara Moser, Adrian Paskey, Jason Caravas, Benjamin Rambo-Martin, Shatavia Morrison, Christopher Gulvick, Scott Sammons, Yvette Unoarumhi, Darlene Wagner, Matthew Schmerer, Minoo Agarwal, Eyad Almasri, Debbie Boles, Ayla Burns, Nuthawin Charoensri, Oren Cohen, Susan Countryman, Mary Ann Cristobal, Bobbi Croy, Suzanne Dale, Hrushikesh Deshmukh, Amanda Douglas, Vincent Drouillon, Marcia Eisenberg, Howard |

| | | | |
|---|---|---|---|
| | | | Engler, Rama Ghatti, Prashant Gupta, Susan Hicks, Jake Humphrey, Lax Iyer, Lisa Pfefferle, Manoj Jain, Matthew Robinson, Mohan Kolli, Brian Krueger, Tim Kuphal, Stanley Letovsky, Michael Levandoski, Craig Lukasik, Jonathan Meltzer, Brian Norvell, Mindy Nye, Scott Parker, Christos Petropoulos, John Pruitt, Steven Ragan, Scott Ryan, Mike Sapeta, Jana Schroth, Suresh Babu Selvaraju, Goran Stevovic, Amanda Suchanek, Andrea Throop, Lyndon Tilson, Thomas Urban, Joe Voshell, Kimberly Wagner, Jonathan Williams, Mary Williamson, Qian Zeng, Tricia Zwiefelhofer, Clinton R. Paden, Duncan MacCannell |
| EPI_ISL_3329307 | New Mexico Department of Health Scientific Laboratory | New Mexico Department of Health Scientific Laboratory | Ellie Johnson, D'eldra Malone, Jennifer Benoit, Ratheesh Rajan, Linda Salazar, Anastacia Griego-Fisher |
| EPI_ISL_3329308, EPI_ISL_3329310 | Laboratory Corporation of America | Centers for Disease Control and Prevention Division of Viral Diseases, Pathogen Discovery | Dakota Howard, Dhwani Batra, Peter W. Cook, Kara Moser, Adrian Paskey, Jason Caravas, Benjamin Rambo-Martin, Shatavia Morrison, Christopher Gulvick, Scott Sammons, Yvette Unoarumhi, Darlene Wagner, Matthew Schmerer, Minoo Agarwal, Eyad Almasri, Debbie Boles, Ayla Burns, Nuthawin Charoensri, Oren Cohen, Susan Countryman, Mary Ann Cristobal, Bobbi Croy, Suzanne Dale, Hrushikesh Deshmukh, Amanda Douglas, Vincent Drouillon, Marcia Eisenberg, Howard Engler, Rama Ghatti, Prashant Gupta, Susan Hicks, Jake Humphrey, Lax Iyer, Lisa Pfefferle, Manoj Jain, Matthew Robinson, Mohan Kolli, Brian Krueger, Tim Kuphal, Stanley Letovsky, Michael Levandoski, Craig Lukasik, Jonathan Meltzer, Brian Norvell, Mindy Nye, Scott Parker, Christos Petropoulos, John Pruitt, Steven Ragan, Scott Ryan, Mike Sapeta, Jana Schroth, Suresh Babu Selvaraju, Goran Stevovic, Amanda Suchanek, Andrea Throop, Lyndon Tilson, Thomas Urban, Joe Voshell, Kimberly Wagner, Jonathan Williams, Mary Williamson, Qian Zeng, Tricia Zwiefelhofer, Clinton R. Paden, Duncan MacCannell |
| EPI_ISL_3329311 | New Mexico Department of Health Scientific Laboratory | New Mexico Department of Health Scientific Laboratory | Ellie Johnson, D'eldra Malone, Jennifer Benoit, Ratheesh Rajan, Linda Salazar, Anastacia Griego-Fisher |
| EPI_ISL_3329312, EPI_ISL_3329313 | Laboratory Corporation of America | Centers for Disease Control and Prevention Division of Viral Diseases, Pathogen Discovery | Dakota Howard, Dhwani Batra, Peter W. Cook, Kara Moser, Adrian Paskey, Jason Caravas, Benjamin Rambo-Martin, Shatavia Morrison, Christopher Gulvick, Scott Sammons, Yvette Unoarumhi, Darlene Wagner, Matthew Schmerer, Minoo Agarwal, Eyad Almasri, Debbie Boles, Ayla Burns, Nuthawin Charoensri, Oren Cohen, Susan Countryman, Mary Ann Cristobal, Bobbi Croy, Suzanne Dale, Hrushikesh Deshmukh, Amanda Douglas, Vincent Drouillon, Marcia Eisenberg, Howard Engler, Rama Ghatti, Prashant Gupta, Susan Hicks, Jake Humphrey, Lax Iyer, Lisa Pfefferle, Manoj Jain, Matthew Robinson, Mohan Kolli, Brian Krueger, Tim Kuphal, Stanley Letovsky, Michael Levandoski, Craig Lukasik, Jonathan Meltzer, Brian Norvell, Mindy Nye, Scott Parker, Christos Petropoulos, John Pruitt, Steven Ragan, Scott Ryan, Mike Sapeta, Jana Schroth, Suresh Babu Selvaraju, Goran Stevovic, Amanda Suchanek, Andrea Throop, Lyndon Tilson, Thomas Urban, Joe Voshell, Kimberly Wagner, Jonathan Williams, Mary Williamson, Qian Zeng, Tricia Zwiefelhofer, Clinton R. Paden, Duncan MacCannell |
| EPI_ISL_3329314 | New Mexico Department of Health Scientific Laboratory | New Mexico Department of Health Scientific Laboratory | Ellie Johnson, D'eldra Malone, Jennifer Benoit, Ratheesh Rajan, Linda Salazar, Anastacia Griego-Fisher |
| EPI_ISL_3329315 | Laboratory Corporation of America | Centers for Disease Control and Prevention Division of Viral Diseases, Pathogen Discovery | Dakota Howard, Dhwani Batra, Peter W. Cook, Kara Moser, Adrian Paskey, Jason Caravas, Benjamin Rambo-Martin, Shatavia Morrison, Christopher Gulvick, Scott Sammons, Yvette Unoarumhi, Darlene Wagner, Matthew Schmerer, Minoo Agarwal, Eyad Almasri, Debbie Boles, Ayla Burns, Nuthawin Charoensri, Oren Cohen, Susan Countryman, Mary Ann Cristobal, Bobbi Croy, Suzanne Dale, Hrushikesh Deshmukh, Amanda Douglas, Vincent Drouillon, Marcia Eisenberg, Howard Engler, Rama Ghatti, Prashant Gupta, Susan Hicks, Jake Humphrey, Lax Iyer, Lisa Pfefferle, Manoj Jain, Matthew Robinson, Mohan Kolli, Brian Krueger, Tim Kuphal, Stanley Letovsky, Michael Levandoski, Craig Lukasik, Jonathan Meltzer, Brian Norvell, Mindy Nye, Scott Parker, Christos Petropoulos, John Pruitt, Steven Ragan, Scott Ryan, Mike Sapeta, Jana Schroth, Suresh Babu Selvaraju, Goran Stevovic, Amanda Suchanek, Andrea Throop, Lyndon Tilson, Thomas Urban, Joe Voshell, Kimberly Wagner, Jonathan Williams, Mary Williamson, Qian Zeng, Tricia Zwiefelhofer, Clinton R. Paden, Duncan MacCannell |
| EPI_ISL_3329316 | New Mexico Department of Health Scientific Laboratory | New Mexico Department of Health Scientific Laboratory | Ellie Johnson, D'eldra Malone, Jennifer Benoit, Ratheesh Rajan, Linda Salazar, Anastacia Griego-Fisher |
| EPI_ISL_3329317 | Laboratory Corporation of America | Centers for Disease Control and Prevention Division of Viral Diseases, Pathogen Discovery | Dakota Howard, Dhwani Batra, Peter W. Cook, Kara Moser, Adrian Paskey, Jason Caravas, Benjamin Rambo-Martin, Shatavia Morrison, Christopher Gulvick, Scott Sammons, Yvette Unoarumhi, Darlene Wagner, Matthew Schmerer, Minoo Agarwal, Eyad Almasri, Debbie Boles, Ayla Burns, Nuthawin Charoensri, Oren Cohen, Susan Countryman, Mary Ann Cristobal, Bobbi Croy, Suzanne Dale, Hrushikesh Deshmukh, Amanda Douglas, Vincent Drouillon, Marcia Eisenberg, Howard Engler, Rama Ghatti, Prashant Gupta, Susan Hicks, Jake Humphrey, Lax Iyer, Lisa Pfefferle, Manoj Jain, Matthew Robinson, Mohan Kolli, Brian Krueger, Tim Kuphal, Stanley Letovsky, Michael Levandoski, Craig Lukasik, Jonathan Meltzer, Brian Norvell, Mindy Nye, Scott Parker, Christos Petropoulos, John Pruitt, Steven Ragan, Scott Ryan, Mike Sapeta, Jana Schroth, Suresh Babu Selvaraju, Goran Stevovic, Amanda Suchanek, Andrea Throop, Lyndon Tilson, Thomas Urban, Joe Voshell, Kimberly Wagner, Jonathan Williams, Mary Williamson, Qian Zeng, Tricia Zwiefelhofer, Clinton R. Paden, Duncan MacCannell |
| EPI_ISL_3329318 | New Mexico Department of Health Scientific Laboratory | New Mexico Department of Health Scientific Laboratory | Ellie Johnson, D'eldra Malone, Jennifer Benoit, Ratheesh Rajan, Linda Salazar, Anastacia Griego-Fisher |
| EPI_ISL_3329319 | Laboratory Corporation of America | Centers for Disease Control and Prevention Division of Viral Diseases, Pathogen Discovery | Dakota Howard, Dhwani Batra, Peter W. Cook, Kara Moser, Adrian Paskey, Jason Caravas, Benjamin Rambo-Martin, Shatavia Morrison, Christopher Gulvick, Scott Sammons, Yvette Unoarumhi, Darlene Wagner, Matthew Schmerer, Minoo Agarwal, Eyad Almasri, Debbie Boles, Ayla Burns, Nuthawin Charoensri, Oren Cohen, Susan Countryman, Mary Ann Cristobal, Bobbi Croy, Suzanne Dale, Hrushikesh Deshmukh, Amanda Douglas, Vincent Drouillon, Marcia Eisenberg, Howard Engler, Rama Ghatti, Prashant Gupta, Susan Hicks, Jake Humphrey, Lax Iyer, Lisa Pfefferle, Manoj Jain, Matthew Robinson, Mohan Kolli, Brian Krueger, Tim Kuphal, Stanley Letovsky, Michael Levandoski, Craig Lukasik, Jonathan Meltzer, Brian Norvell, Mindy Nye, Scott Parker, Christos Petropoulos, John Pruitt, Steven Ragan, Scott Ryan, Mike Sapeta, Jana Schroth, Suresh Babu Selvaraju, Goran Stevovic, Amanda Suchanek, Andrea Throop, Lyndon Tilson, Thomas Urban, Joe Voshell, Kimberly Wagner, Jonathan Williams, Mary Williamson, Qian Zeng, Tricia Zwiefelhofer, Clinton R. Paden, Duncan MacCannell |
| EPI_ISL_3329320 | New Mexico Department of Health Scientific Laboratory | New Mexico Department of Health Scientific Laboratory | Ellie Johnson, D'eldra Malone, Jennifer Benoit, Ratheesh Rajan, Linda Salazar, Anastacia Griego-Fisher |
| EPI_ISL_3329321, EPI_ISL_3329322 | Laboratory Corporation of America | Centers for Disease Control and Prevention Division of Viral Diseases, Pathogen Discovery | Dakota Howard, Dhwani Batra, Peter W. Cook, Kara Moser, Adrian Paskey, Jason Caravas, Benjamin Rambo-Martin, Shatavia Morrison, Christopher Gulvick, Scott Sammons, Yvette Unoarumhi, Darlene Wagner, Matthew Schmerer, Minoo Agarwal, Eyad Almasri, Debbie Boles, Ayla Burns, Nuthawin Charoensri, Oren Cohen, Susan Countryman, Mary Ann Cristobal, Bobbi Croy, Suzanne Dale, Hrushikesh Deshmukh, Amanda Douglas, Vincent Drouillon, Marcia Eisenberg, Howard Engler, Rama Ghatti, Prashant Gupta, Susan Hicks, Jake Humphrey, Lax Iyer, Lisa Pfefferle, Manoj Jain, Matthew Robinson, Mohan Kolli, Brian Krueger, Tim Kuphal, Stanley Letovsky, Michael Levandoski, Craig Lukasik, Jonathan Meltzer, Brian Norvell, Mindy Nye, Scott Parker, Christos Petropoulos, John Pruitt, Steven Ragan, Scott Ryan, Mike Sapeta, Jana Schroth, Suresh Babu Selvaraju, Goran Stevovic, Amanda Suchanek, Andrea Throop, Lyndon Tilson, Thomas Urban, Joe Voshell, Kimberly Wagner, Jonathan Williams, Mary Williamson, Qian Zeng, Tricia Zwiefelhofer, Clinton R. Paden, Duncan MacCannell |
| EPI_ISL_3329323 | New Mexico Department of Health Scientific Laboratory | New Mexico Department of Health Scientific Laboratory | Ellie Johnson, D'eldra Malone, Jennifer Benoit, Ratheesh Rajan, Linda Salazar, Anastacia Griego-Fisher |
| EPI_ISL_3329324 | Laboratory Corporation of America | Centers for Disease Control and Prevention Division of Viral Diseases, Pathogen Discovery | Dakota Howard, Dhwani Batra, Peter W. Cook, Kara Moser, Adrian Paskey, Jason Caravas, Benjamin Rambo-Martin, Shatavia Morrison, Christopher Gulvick, Scott Sammons, Yvette Unoarumhi, Darlene Wagner, Matthew Schmerer, Minoo Agarwal, Eyad Almasri, Debbie Boles, Ayla Burns, Nuthawin Charoensri, Oren Cohen, Susan Countryman, Mary Ann Cristobal, Bobbi Croy, Suzanne Dale, Hrushikesh Deshmukh, Amanda Douglas, Vincent Drouillon, Marcia Eisenberg, Howard Engler, Rama Ghatti, Prashant Gupta, Susan Hicks, Jake Humphrey, Lax Iyer, Lisa Pfefferle, Manoj Jain, Matthew Robinson, Mohan Kolli, Brian Krueger, Tim Kuphal, Stanley Letovsky, Michael Levandoski, Craig Lukasik, Jonathan Meltzer, Brian Norvell, Mindy Nye, Scott Parker, Christos Petropoulos, John Pruitt, Steven Ragan, Scott Ryan, Mike Sapeta, Jana Schroth, Suresh Babu Selvaraju, Goran Stevovic, Amanda Suchanek, Andrea Throop, Lyndon Tilson, Thomas Urban, Joe Voshell, Kimberly Wagner, Jonathan Williams, Mary Williamson, Qian Zeng, Tricia Zwiefelhofer, Clinton R. Paden, Duncan MacCannell |
| EPI_ISL_3329325 | New Mexico Department of Health Scientific Laboratory | New Mexico Department of Health Scientific Laboratory | Ellie Johnson, D'eldra Malone, Jennifer Benoit, Ratheesh Rajan, Linda Salazar, Anastacia Griego-Fisher |
| EPI_ISL_3329326, EPI_ISL_3329328, EPI_ISL_3329329 | Laboratory Corporation of America | Centers for Disease Control and Prevention Division of Viral Diseases, Pathogen Discovery | Dakota Howard, Dhwani Batra, Peter W. Cook, Kara Moser, Adrian Paskey, Jason Caravas, Benjamin Rambo-Martin, Shatavia Morrison, Christopher Gulvick, Scott Sammons, Yvette Unoarumhi, Darlene Wagner, Matthew Schmerer, Minoo Agarwal, Eyad Almasri, Debbie Boles, Ayla Burns, Nuthawin Charoensri, Oren Cohen, Susan Countryman, Mary Ann Cristobal, Bobbi Croy, Suzanne Dale, Hrushikesh Deshmukh, Amanda Douglas, Vincent Drouillon, Marcia Eisenberg, Howard Engler, Rama Ghatti, Prashant Gupta, Susan Hicks, Jake Humphrey, Lax Iyer, Lisa Pfefferle, Manoj Jain, Matthew Robinson, Mohan Kolli, Brian Krueger, Tim Kuphal, Stanley Letovsky, Michael Levandoski, Craig Lukasik, Jonathan Meltzer, Brian Norvell, Mindy Nye, Scott Parker, Christos Petropoulos, John Pruitt, Steven Ragan, Scott Ryan, Mike Sapeta, Jana Schroth, Suresh Babu Selvaraju, Goran Stevovic, Amanda Suchanek, Andrea Throop, Lyndon Tilson, Thomas Urban, Joe Voshell, Kimberly Wagner, Jonathan Williams, Mary Williamson, Qian Zeng, Tricia Zwiefelhofer, Clinton R. Paden, Duncan MacCannell |
| EPI_ISL_3329330 | New Mexico Department of Health Scientific Laboratory | New Mexico Department of Health Scientific Laboratory | Ellie Johnson, D'eldra Malone, Jennifer Benoit, Ratheesh Rajan, Linda Salazar, Anastacia Griego-Fisher |
| EPI_ISL_3329331 | Laboratory Corporation of America | Centers for Disease Control and Prevention Division of Viral Diseases, Pathogen Discovery | Dakota Howard, Dhwani Batra, Peter W. Cook, Kara Moser, Adrian Paskey, Jason Caravas, Benjamin Rambo-Martin, Shatavia Morrison, Christopher Gulvick, Scott Sammons, Yvette Unoarumhi, Darlene Wagner, Matthew Schmerer, Minoo Agarwal, Eyad Almasri, Debbie Boles, Ayla Burns, Nuthawin Charoensri, Oren Cohen, Susan Countryman, Mary Ann Cristobal, Bobbi Croy, Suzanne Dale, Hrushikesh Deshmukh, Amanda Douglas, Vincent Drouillon, Marcia Eisenberg, Howard Engler, Rama Ghatti, Prashant Gupta, Susan Hicks, Jake Humphrey, Lax Iyer, Lisa Pfefferle, Manoj Jain, Matthew Robinson, Mohan Kolli, Brian Krueger, Tim Kuphal, Stanley Letovsky, Michael Levandoski, Craig Lukasik, Jonathan Meltzer, Brian Norvell, Mindy Nye, Scott Parker, Christos Petropoulos, John Pruitt, Steven Ragan, Scott Ryan, Mike Sapeta, Jana Schroth, Suresh Babu Selvaraju, Goran Stevovic, Amanda Suchanek, Andrea Throop, Lyndon Tilson, Thomas Urban, Joe Voshell, Kimberly Wagner, Jonathan Williams, Mary Williamson, Qian Zeng, Tricia Zwiefelhofer, Clinton R. Paden, Duncan MacCannell |
| EPI_ISL_3329332 | New Mexico Department of Health Scientific Laboratory | New Mexico Department of Health Scientific Laboratory | Ellie Johnson, D'eldra Malone, Jennifer Benoit, Ratheesh Rajan, Linda Salazar, Anastacia Griego-Fisher |
| EPI_ISL_3329333 | Laboratory Corporation of America | Centers for Disease Control and Prevention Division of Viral Diseases, Pathogen Discovery | Dakota Howard, Dhwani Batra, Peter W. Cook, Kara Moser, Adrian Paskey, Jason Caravas, Benjamin Rambo-Martin, Shatavia Morrison, Christopher Gulvick, Scott Sammons, Yvette Unoarumhi, Darlene Wagner, Matthew Schmerer, Minoo Agarwal, Eyad Almasri, Debbie Boles, Ayla Burns, Nuthawin Charoensri, Oren Cohen, Susan Countryman, Mary Ann Cristobal, Bobbi Croy, Suzanne Dale, Hrushikesh Deshmukh, Amanda Douglas, Vincent Drouillon, Marcia Eisenberg, Howard Engler, Rama Ghatti, Prashant Gupta, Susan Hicks, Jake Humphrey, Lax Iyer, Lisa Pfefferle, Manoj Jain, Matthew Robinson, Mohan Kolli, Brian Krueger, Tim Kuphal, Stanley Letovsky, Michael Levandoski, Craig Lukasik, Jonathan Meltzer, Brian Norvell, Mindy Nye, Scott Parker, Christos Petropoulos, John Pruitt, Steven Ragan, Scott Ryan, Mike Sapeta, Jana Schroth, Suresh Babu Selvaraju, Goran Stevovic, Amanda Suchanek, Andrea Throop, Lyndon Tilson, Thomas Urban, Joe Voshell, Kimberly Wagner, Jonathan Williams, Mary Williamson, Qian Zeng, Tricia Zwiefelhofer, Clinton R. Paden, Duncan MacCannell |
| EPI_ISL_3329334 | New Mexico Department of Health Scientific Laboratory | New Mexico Department of Health Scientific Laboratory | Ellie Johnson, D'eldra Malone, Jennifer Benoit, Ratheesh Rajan, Linda Salazar, Anastacia Griego-Fisher |
| EPI_ISL_3329335, EPI_ISL_3329336 | Laboratory Corporation of America | Centers for Disease Control and Prevention Division of Viral Diseases, Pathogen Discovery | Dakota Howard, Dhwani Batra, Peter W. Cook, Kara Moser, Adrian Paskey, Jason Caravas, Benjamin Rambo-Martin, Shatavia Morrison, Christopher Gulvick, Scott Sammons, Yvette Unoarumhi, Darlene Wagner, Matthew Schmerer, Minoo Agarwal, Eyad Almasri, Debbie Boles, Ayla Burns, Nuthawin Charoensri, Oren Cohen, Susan Countryman, Mary Ann Cristobal, Bobbi Croy, Suzanne Dale, Hrushikesh Deshmukh, Amanda Douglas, Vincent Drouillon, Marcia Eisenberg, Howard Engler, Rama Ghatti, Prashant Gupta, Susan Hicks, Jake Humphrey, Lax Iyer, Lisa Pfefferle, Manoj Jain, Matthew Robinson, Mohan Kolli, Brian Krueger, Tim Kuphal, Stanley Letovsky, Michael Levandoski, Craig Lukasik, Jonathan Meltzer, Brian Norvell, Mindy Nye, Scott Parker, Christos Petropoulos, John Pruitt, Steven Ragan, Scott Ryan, Mike Sapeta, Jana Schroth, Suresh Babu Selvaraju, Goran Stevovic, Amanda Suchanek, Andrea Throop, Lyndon Tilson, Thomas Urban, Joe Voshell, Kimberly Wagner, Jonathan Williams, Mary Williamson, Qian Zeng, Tricia Zwiefelhofer, Clinton R. Paden, Duncan MacCannell |
| EPI_ISL_3329337, EPI_ISL_3329338 | New Mexico Department of Health Scientific Laboratory | New Mexico Department of Health Scientific Laboratory | Ellie Johnson, D'eldra Malone, Jennifer Benoit, Ratheesh Rajan, Linda Salazar, Anastacia Griego-Fisher |
| EPI_ISL_3329339, EPI_ISL_3329340 | Laboratory Corporation of America | Centers for Disease Control and Prevention Division of Viral Diseases, Pathogen Discovery | Dakota Howard, Dhwani Batra, Peter W. Cook, Kara Moser, Adrian Paskey, Jason Caravas, Benjamin Rambo-Martin, Shatavia Morrison, Christopher Gulvick, Scott Sammons, Yvette Unoarumhi, Darlene Wagner, Matthew Schmerer, Minoo Agarwal, Eyad Almasri, Debbie Boles, Ayla Burns, Nuthawin Charoensri, Oren Cohen, Susan Countryman, Mary Ann Cristobal, Bobbi Croy, Suzanne Dale, Hrushikesh Deshmukh, Amanda Douglas, Vincent Drouillon, Marcia Eisenberg, Howard Engler, Rama Ghatti, Prashant Gupta, Susan Hicks, Jake Humphrey, Lax Iyer, Lisa Pfefferle, Manoj Jain, Matthew Robinson, Mohan Kolli, Brian Krueger, Tim Kuphal, Stanley Letovsky, Michael Levandoski, Craig Lukasik, Jonathan Meltzer, Brian Norvell, Mindy Nye, Scott Parker, Christos Petropoulos, John Pruitt, Steven Ragan, Scott Ryan, Mike Sapeta, Jana Schroth, Suresh Babu Selvaraju, Goran Stevovic, Amanda Suchanek, Andrea Throop, Lyndon Tilson, Thomas Urban, Joe Voshell, Kimberly Wagner, Jonathan Williams, Mary Williamson, Qian Zeng, Tricia Zwiefelhofer, Clinton R. Paden, Duncan MacCannell |
| EPI_ISL_3329341 | New Mexico Department of Health Scientific Laboratory | New Mexico Department of Health Scientific Laboratory | Ellie Johnson, D'eldra Malone, Jennifer Benoit, Ratheesh Rajan, Linda Salazar, Anastacia Griego-Fisher |
| EPI_ISL_3329342 | Laboratory Corporation of America | Centers for Disease Control and Prevention Division of Viral Diseases, Pathogen Discovery | Dakota Howard, Dhwani Batra, Peter W. Cook, Kara Moser, Adrian Paskey, Jason Caravas, Benjamin Rambo-Martin, Shatavia Morrison, Christopher Gulvick, Scott Sammons, Yvette Unoarumhi, Darlene Wagner, Matthew Schmerer, Minoo Agarwal, Eyad Almasri, Debbie Boles, Ayla Burns, Nuthawin Charoensri, Oren Cohen, Susan Countryman, Mary Ann Cristobal, Bobbi Croy, Suzanne Dale, Hrushikesh Deshmukh, Amanda Douglas, Vincent Drouillon, Marcia Eisenberg, Howard Engler, Rama Ghatti, Prashant Gupta, Susan Hicks, Jake Humphrey, Lax Iyer, Lisa Pfefferle, Manoj Jain, Matthew Robinson, Mohan Kolli, Brian Krueger, Tim Kuphal, Stanley Letovsky, Michael Levandoski, Craig Lukasik, Jonathan Meltzer, Brian Norvell, Mindy Nye, Scott Parker, Christos Petropoulos, John Pruitt, Steven Ragan, Scott Ryan, Mike Sapeta, Jana Schroth, Suresh Babu Selvaraju, Goran Stevovic, Amanda Suchanek, Andrea Throop, Lyndon Tilson, Thomas Urban, Joe Voshell, Kimberly Wagner, Jonathan Williams, Mary Williamson, Qian Zeng, Tricia Zwiefelhofer, Clinton R. Paden, Duncan MacCannell |
| EPI_ISL_3329343, EPI_ISL_3329345 | New Mexico Department of Health Scientific Laboratory | New Mexico Department of Health Scientific Laboratory | Ellie Johnson, D'eldra Malone, Jennifer Benoit, Ratheesh Rajan, Linda Salazar, Anastacia Griego-Fisher |
| EPI_ISL_3329346 | Laboratory Corporation of America | Centers for Disease Control and Prevention Division of Viral Diseases, Pathogen Discovery | Dakota Howard, Dhwani Batra, Peter W. Cook, Kara Moser, Adrian Paskey, Jason Caravas, Benjamin Rambo-Martin, Shatavia Morrison, Christopher Gulvick, Scott Sammons, Yvette Unoarumhi, Darlene Wagner, Matthew Schmerer, Minoo Agarwal, Eyad Almasri, Debbie Boles, Ayla Burns, Nuthawin Charoensri, Oren Cohen, Susan Countryman, Mary Ann Cristobal, Bobbi Croy, Suzanne Dale, Hrushikesh Deshmukh, Amanda Douglas, Vincent Drouillon, Marcia Eisenberg, Howard Engler, Rama Ghatti, Prashant Gupta, Susan Hicks, Jake Humphrey, Lax Iyer, Lisa Pfefferle, Manoj Jain, Matthew Robinson, Mohan Kolli, Brian Krueger, Tim Kuphal, Stanley Letovsky, Michael Levandoski, Craig Lukasik, Jonathan Meltzer, Brian Norvell, Mindy Nye, Scott Parker, Christos Petropoulos, John Pruitt, Steven Ragan, Scott Ryan, Mike Sapeta, Jana Schroth, Suresh Babu Selvaraju, Goran Stevovic, Amanda Suchanek, Andrea Throop, Lyndon Tilson, Thomas Urban, Joe Voshell, Kimberly Wagner, Jonathan Williams, Mary Williamson, Qian Zeng, Tricia Zwiefelhofer, Clinton R. Paden, Duncan MacCannell |
| EPI_ISL_3329347, EPI_ISL_3329349 | New Mexico Department of Health Scientific Laboratory | New Mexico Department of Health Scientific Laboratory | Ellie Johnson, D'eldra Malone, Jennifer Benoit, Ratheesh Rajan, Linda Salazar, Anastacia Griego-Fisher |
| EPI_ISL_3329350 | Laboratory Corporation of America | Centers for Disease Control and Prevention Division of Viral Diseases, Pathogen Discovery | Dakota Howard, Dhwani Batra, Peter W. Cook, Kara Moser, Adrian Paskey, Jason Caravas, Benjamin Rambo-Martin, Shatavia Morrison, Christopher Gulvick, Scott Sammons, Yvette Unoarumhi, Darlene Wagner, Matthew Schmerer, Minoo Agarwal, Eyad Almasri, Debbie Boles, Ayla Burns, Nuthawin Charoensri, Oren Cohen, Susan Countryman, Mary Ann Cristobal, Bobbi Croy, Suzanne Dale, Hrushikesh Deshmukh, Amanda Douglas, Vincent Drouillon, Marcia Eisenberg, Howard Engler, Rama Ghatti, Prashant Gupta, Susan Hicks, Jake Humphrey, Lax Iyer, Lisa Pfefferle, Manoj Jain, Matthew Robinson, Mohan Kolli, Brian Krueger, Tim Kuphal, Stanley Letovsky, Michael Levandoski, Craig Lukasik, Jonathan Meltzer, Brian Norvell, Mindy Nye, Scott Parker, Christos Petropoulos, John Pruitt, Steven Ragan, Scott Ryan, Mike Sapeta, Jana Schroth, Suresh Babu Selvaraju, Goran Stevovic, Amanda Suchanek, Andrea Throop, Lyndon Tilson, Thomas Urban, Joe Voshell, Kimberly Wagner, Jonathan Williams, Mary Williamson, Qian Zeng, Tricia Zwiefelhofer, Clinton R. Paden, Duncan MacCannell |
| EPI_ISL_3329351 | New Mexico Department of Health Scientific Laboratory | New Mexico Department of Health Scientific Laboratory | Ellie Johnson, D'eldra Malone, Jennifer Benoit, Ratheesh Rajan, Linda Salazar, Anastacia Griego-Fisher |

| | | | |
|---|---|---|---|
| EPI_ISL_3329352 | Laboratory Corporation of America | Centers for Disease Control and Prevention Division of Viral Diseases, Pathogen Discovery | Dakota Howard, Dhwani Batra, Peter W. Cook, Kara Moser, Adrian Paskey, Jason Caravas, Benjamin Rambo-Martin, Shatavia Morrison, Christopher Gulvick, Scott Sammons, Yvette Unoarumhi, Darlene Wagner, Matthew Schmerer, Minoo Agarwal, Eyad Almasri, Debbie Boles, Ayla Burns, Nuthawin Charoensri, Oren Cohen, Susan Countryman, Mary Ann Cristobal, Bobbi Croy, Suzanne Dale, Hrushikesh Deshmukh, Amanda Douglas, Vincent Drouillon, Marcia Eisenberg, Howard Engler, Rama Ghatti, Prashant Gupta, Susan Hicks, Jake Humphrey, Lax Iyer, Lisa Pfefferle, Manoj Jain, Matthew Robinson, Mohan Kolli, Brian Krueger, Tim Kuphal, Stanley Letovsky, Michael Levandoski, Craig Lukasik, Jonathan Meltzer, Brian Norvell, Mindy Nye, Scott Parker, Christos Petropoulos, John Pruitt, Steven Ragan, Scott Ryan, Mike Sapeta, Jana Schroth, Suresh Babu Selvaraju, Goran Stevovic, Amanda Suchanek, Andrea Throop, Lyndon Tilson, Thomas Urban, Joe Voshell, Kimberly Wagner, Jonathan Williams, Mary Williamson, Qian Zeng, Tricia Zwiefelhofer, Clinton R. Paden, Duncan MacCannell |
| EPI_ISL_3329353 | New Mexico Department of Health Scientific Laboratory | New Mexico Department of Health Scientific Laboratory | Ellie Johnson, D'eldra Malone, Jennifer Benoit, Ratheesh Rajan, Linda Salazar, Anastacia Griego-Fisher |
| EPI_ISL_3329354, EPI_ISL_3329355 | Laboratory Corporation of America | Centers for Disease Control and Prevention Division of Viral Diseases, Pathogen Discovery | Dakota Howard, Dhwani Batra, Peter W. Cook, Kara Moser, Adrian Paskey, Jason Caravas, Benjamin Rambo-Martin, Shatavia Morrison, Christopher Gulvick, Scott Sammons, Yvette Unoarumhi, Darlene Wagner, Matthew Schmerer, Minoo Agarwal, Eyad Almasri, Debbie Boles, Ayla Burns, Nuthawin Charoensri, Oren Cohen, Susan Countryman, Mary Ann Cristobal, Bobbi Croy, Suzanne Dale, Hrushikesh Deshmukh, Amanda Douglas, Vincent Drouillon, Marcia Eisenberg, Howard Engler, Rama Ghatti, Prashant Gupta, Susan Hicks, Jake Humphrey, Lax Iyer, Lisa Pfefferle, Manoj Jain, Matthew Robinson, Mohan Kolli, Brian Krueger, Tim Kuphal, Stanley Letovsky, Michael Levandoski, Craig Lukasik, Jonathan Meltzer, Brian Norvell, Mindy Nye, Scott Parker, Christos Petropoulos, John Pruitt, Steven Ragan, Scott Ryan, Mike Sapeta, Jana Schroth, Suresh Babu Selvaraju, Goran Stevovic, Amanda Suchanek, Andrea Throop, Lyndon Tilson, Thomas Urban, Joe Voshell, Kimberly Wagner, Jonathan Williams, Mary Williamson, Qian Zeng, Tricia Zwiefelhofer, Clinton R. Paden, Duncan MacCannell |
| EPI_ISL_3329356 | New Mexico Department of Health Scientific Laboratory | New Mexico Department of Health Scientific Laboratory | Ellie Johnson, D'eldra Malone, Jennifer Benoit, Ratheesh Rajan, Linda Salazar, Anastacia Griego-Fisher |
| EPI_ISL_3329357 | Laboratory Corporation of America | Centers for Disease Control and Prevention Division of Viral Diseases, Pathogen Discovery | Dakota Howard, Dhwani Batra, Peter W. Cook, Kara Moser, Adrian Paskey, Jason Caravas, Benjamin Rambo-Martin, Shatavia Morrison, Christopher Gulvick, Scott Sammons, Yvette Unoarumhi, Darlene Wagner, Matthew Schmerer, Minoo Agarwal, Eyad Almasri, Debbie Boles, Ayla Burns, Nuthawin Charoensri, Oren Cohen, Susan Countryman, Mary Ann Cristobal, Bobbi Croy, Suzanne Dale, Hrushikesh Deshmukh, Amanda Douglas, Vincent Drouillon, Marcia Eisenberg, Howard Engler, Rama Ghatti, Prashant Gupta, Susan Hicks, Jake Humphrey, Lax Iyer, Lisa Pfefferle, Manoj Jain, Matthew Robinson, Mohan Kolli, Brian Krueger, Tim Kuphal, Stanley Letovsky, Michael Levandoski, Craig Lukasik, Jonathan Meltzer, Brian Norvell, Mindy Nye, Scott Parker, Christos Petropoulos, John Pruitt, Steven Ragan, Scott Ryan, Mike Sapeta, Jana Schroth, Suresh Babu Selvaraju, Goran Stevovic, Amanda Suchanek, Andrea Throop, Lyndon Tilson, Thomas Urban, Joe Voshell, Kimberly Wagner, Jonathan Williams, Mary Williamson, Qian Zeng, Tricia Zwiefelhofer, Clinton R. Paden, Duncan MacCannell |
| EPI_ISL_3329358 | New Mexico Department of Health Scientific Laboratory | New Mexico Department of Health Scientific Laboratory | Ellie Johnson, D'eldra Malone, Jennifer Benoit, Ratheesh Rajan, Linda Salazar, Anastacia Griego-Fisher |
| EPI_ISL_3329359, EPI_ISL_3329360, EPI_ISL_3329361, EPI_ISL_3329362, EPI_ISL_3329363, EPI_ISL_3329364, EPI_ISL_3329365, EPI_ISL_3329366, EPI_ISL_3329367, EPI_ISL_3329368, EPI_ISL_3329369, EPI_ISL_3329370, EPI_ISL_3329372, EPI_ISL_3329373, EPI_ISL_3329374, EPI_ISL_3329375, EPI_ISL_3329376, EPI_ISL_3329377, EPI_ISL_3329378, EPI_ISL_3329379, EPI_ISL_3329381, EPI_ISL_3329382, EPI_ISL_3329383, EPI_ISL_3329384, EPI_ISL_3329385, EPI_ISL_3329387, EPI_ISL_3329388, EPI_ISL_3329389, EPI_ISL_3329390, EPI_ISL_3329391, EPI_ISL_3329392, EPI_ISL_3329393, EPI_ISL_3329394, EPI_ISL_3329395, EPI_ISL_3329396, EPI_ISL_3329397, EPI_ISL_3329398, EPI_ISL_3329399, EPI_ISL_3329400, EPI_ISL_3329401, EPI_ISL_3329402, EPI_ISL_3329403, EPI_ISL_3329404, EPI_ISL_3329405, EPI_ISL_3329406, EPI_ISL_3329407, EPI_ISL_3329408, EPI_ISL_3329409, EPI_ISL_3329410, EPI_ISL_3329411, EPI_ISL_3329412, EPI_ISL_3329413, EPI_ISL_3329414, EPI_ISL_3329415, EPI_ISL_3329417, EPI_ISL_3329420, EPI_ISL_3329421, EPI_ISL_3329422, EPI_ISL_3329423, EPI_ISL_3329424, EPI_ISL_3329426, EPI_ISL_3329427, EPI_ISL_3329428, EPI_ISL_3329429, EPI_ISL_3329430, EPI_ISL_3329431, EPI_ISL_3329432, EPI_ISL_3329433, EPI_ISL_3329434, EPI_ISL_3329435, EPI_ISL_3329436, EPI_ISL_3329437, EPI_ISL_3329438, EPI_ISL_3329439, EPI_ISL_3329440, EPI_ISL_3329441, EPI_ISL_3329442, EPI_ISL_3329443, EPI_ISL_3329444, EPI_ISL_3329445, EPI_ISL_3329446, EPI_ISL_3329447, EPI_ISL_3329448, EPI_ISL_3329449, EPI_ISL_3329450, EPI_ISL_3329451, EPI_ISL_3329452, EPI_ISL_3329454, EPI_ISL_3329455, EPI_ISL_3329456, EPI_ISL_3329458, EPI_ISL_3329459, EPI_ISL_3329460, EPI_ISL_3329461, EPI_ISL_3329463, EPI_ISL_3329464, EPI_ISL_3329465, EPI_ISL_3329466, EPI_ISL_3329467, EPI_ISL_3329468, EPI_ISL_3329469, EPI_ISL_3329470, EPI_ISL_3329471, EPI_ISL_3329472, EPI_ISL_3329473, EPI_ISL_3329475, EPI_ISL_3329476, EPI_ISL_3329477, EPI_ISL_3329478, EPI_ISL_3329479, EPI_ISL_3329480, EPI_ISL_3329481, EPI_ISL_3329482, EPI_ISL_3329483, EPI_ISL_3329484, EPI_ISL_3329485, EPI_ISL_3329486, EPI_ISL_3329488, EPI_ISL_3329489, EPI_ISL_3329490, EPI_ISL_3329492, EPI_ISL_3329493, EPI_ISL_3329494, EPI_ISL_3329495, EPI_ISL_3329496, EPI_ISL_3329497, EPI_ISL_3329498, EPI_ISL_3329499, EPI_ISL_3329500, EPI_ISL_3329501, EPI_ISL_3329502, EPI_ISL_3329503, EPI_ISL_3329504, EPI_ISL_3329505, EPI_ISL_3329506, EPI_ISL_3329507, EPI_ISL_3329508, EPI_ISL_3329509, EPI_ISL_3329510, EPI_ISL_3329511, EPI_ISL_3329512, EPI_ISL_3329513, EPI_ISL_3329514, EPI_ISL_3329515, EPI_ISL_3329516, EPI_ISL_3329518, EPI_ISL_3329519, EPI_ISL_3329520, EPI_ISL_3329521, EPI_ISL_3329522, EPI_ISL_3329523, EPI_ISL_3329524, EPI_ISL_3329525, EPI_ISL_3329526, EPI_ISL_3329527, EPI_ISL_3329528, EPI_ISL_3329529, EPI_ISL_3329530, EPI_ISL_3329531, EPI_ISL_3329533, EPI_ISL_3329534, EPI_ISL_3329535, EPI_ISL_3329536, EPI_ISL_3329537, EPI_ISL_3329538, EPI_ISL_3329539, EPI_ISL_3329540, EPI_ISL_3329541, EPI_ISL_3329543, EPI_ISL_3329544, EPI_ISL_3329545, EPI_ISL_3329546, EPI_ISL_3329547, EPI_ISL_3329548, EPI_ISL_3329549, EPI_ISL_3329550, EPI_ISL_3329551, EPI_ISL_3329552, EPI_ISL_3329553, EPI_ISL_3329554, EPI_ISL_3329555, EPI_ISL_3329556, EPI_ISL_3329557, EPI_ISL_3329558, EPI_ISL_3329559, EPI_ISL_3329560, EPI_ISL_3329561, EPI_ISL_3329562, EPI_ISL_3329563, EPI_ISL_3329564, EPI_ISL_3329565, EPI_ISL_3329566, EPI_ISL_3329567, EPI_ISL_3329568, EPI_ISL_3329569, EPI_ISL_3329570, EPI_ISL_3329571, EPI_ISL_3329573, EPI_ISL_3329574, EPI_ISL_3329575, EPI_ISL_3329576, EPI_ISL_3329577, EPI_ISL_3329578, EPI_ISL_3329579, EPI_ISL_3329580, EPI_ISL_3329581, EPI_ISL_3329582, EPI_ISL_3329583, EPI_ISL_3329584, EPI_ISL_3329585, EPI_ISL_3329586, EPI_ISL_3329587, EPI_ISL_3329588, EPI_ISL_3329589, EPI_ISL_3329590, EPI_ISL_3329591, EPI_ISL_3329592, EPI_ISL_3329593, EPI_ISL_3329594, EPI_ISL_3329595, EPI_ISL_3329596, EPI_ISL_3329597, EPI_ISL_3329598, EPI_ISL_3329599, EPI_ISL_3329600, EPI_ISL_3329601, EPI_ISL_3329602, EPI_ISL_3329603, EPI_ISL_3329604, EPI_ISL_3329605, EPI_ISL_3329607, EPI_ISL_3329608, EPI_ISL_3329609, EPI_ISL_3329610, EPI_ISL_3329611, EPI_ISL_3329612, EPI_ISL_3329613, EPI_ISL_3329614, EPI_ISL_3329616, EPI_ISL_3329617, EPI_ISL_3329618, EPI_ISL_3329619, EPI_ISL_3329620, EPI_ISL_3329621, EPI_ISL_3329622, EPI_ISL_3329623, EPI_ISL_3329624, EPI_ISL_3329625, EPI_ISL_3329626, EPI_ISL_3329627, EPI_ISL_3329628, EPI_ISL_3329629, EPI_ISL_3329630, EPI_ISL_3329631, EPI_ISL_3329632, EPI_ISL_3329633, EPI_ISL_3329634, EPI_ISL_3329635, EPI_ISL_3329636, EPI_ISL_3329637, EPI_ISL_3329638, EPI_ISL_3329639, EPI_ISL_3329640, EPI_ISL_3329641, EPI_ISL_3329642, EPI_ISL_3329643, EPI_ISL_3329644, EPI_ISL_3329645, EPI_ISL_3329646, EPI_ISL_3329647, EPI_ISL_3329648, EPI_ISL_3329649, EPI_ISL_3329650, EPI_ISL_3329651, EPI_ISL_3329652, EPI_ISL_3329653, EPI_ISL_3329654, EPI_ISL_3329655, EPI_ISL_3329656, EPI_ISL_3329657, EPI_ISL_3329658, EPI_ISL_3329659, EPI_ISL_3329660, EPI_ISL_3329661, EPI_ISL_3329662, EPI_ISL_3329663, EPI_ISL_3329664, EPI_ISL_3329665, EPI_ISL_3329666, EPI_ISL_3329667, EPI_ISL_3329668, EPI_ISL_3329669, EPI_ISL_3329671, EPI_ISL_3329672, EPI_ISL_3329674, EPI_ISL_3329675, EPI_ISL_3329676, EPI_ISL_3329677, EPI_ISL_3329678, EPI_ISL_3329679, EPI_ISL_3329680, EPI_ISL_3329681, EPI_ISL_3329682, EPI_ISL_3329683, EPI_ISL_3329684, EPI_ISL_3329685, EPI_ISL_3329686, EPI_ISL_3329687, EPI_ISL_3329688, EPI_ISL_3329689, EPI_ISL_3329690, EPI_ISL_3329691, EPI_ISL_3329692, EPI_ISL_3329693, EPI_ISL_3329694, EPI_ISL_3329695, EPI_ISL_3329696, EPI_ISL_3329697, EPI_ISL_3329698, EPI_ISL_3329699, EPI_ISL_3329700, EPI_ISL_3329701, EPI_ISL_3329702, EPI_ISL_3329703, EPI_ISL_3329704, EPI_ISL_3329705, EPI_ISL_3329706, EPI_ISL_3329707, EPI_ISL_3329708, EPI_ISL_3329709, EPI_ISL_3329710, EPI_ISL_3329711, EPI_ISL_3329712, EPI_ISL_3329714, EPI_ISL_3329715, EPI_ISL_3329716, EPI_ISL_3329717, EPI_ISL_3329718, EPI_ISL_3329719, EPI_ISL_3329721, EPI_ISL_3329722, EPI_ISL_3329723, EPI_ISL_3329724, EPI_ISL_3329725, EPI_ISL_3329727, EPI_ISL_3329728, EPI_ISL_3329730, EPI_ISL_3329731, EPI_ISL_3329733, EPI_ISL_3329734, EPI_ISL_3329735, EPI_ISL_3329736, EPI_ISL_3329737, EPI_ISL_3329738, EPI_ISL_3329739, EPI_ISL_3329740, EPI_ISL_3329741, EPI_ISL_3329742, EPI_ISL_3329743, EPI_ISL_3329744, EPI_ISL_3329745, EPI_ISL_3329746, EPI_ISL_3329747, EPI_ISL_3329748, EPI_ISL_3329749, EPI_ISL_3329750, EPI_ISL_3329753, EPI_ISL_3329754, EPI_ISL_3329755, EPI_ISL_3329756, EPI_ISL_3329757, EPI_ISL_3329758, EPI_ISL_3329759, EPI_ISL_3329760, EPI_ISL_3329761, EPI_ISL_3329762, EPI_ISL_3329763, EPI_ISL_3329764, EPI_ISL_3329765, EPI_ISL_3329766, EPI_ISL_3329767, EPI_ISL_3329768, EPI_ISL_3329769, EPI_ISL_3329770, EPI_ISL_3329771, EPI_ISL_3329772, EPI_ISL_3329773, EPI_ISL_3329774, EPI_ISL_3329775, EPI_ISL_3329776, EPI_ISL_3329777, EPI_ISL_3329778, EPI_ISL_3329779, EPI_ISL_3329780, EPI_ISL_3329781, EPI_ISL_3329782, EPI_ISL_3329783, EPI_ISL_3329784, EPI_ISL_3329785, EPI_ISL_3329786, EPI_ISL_3329787, EPI_ISL_3329788, EPI_ISL_3329789, EPI_ISL_3329790, EPI_ISL_3329791, EPI_ISL_3329792, EPI_ISL_3329793, EPI_ISL_3329794, EPI_ISL_3329795, EPI_ISL_3329796, EPI_ISL_3329797, EPI_ISL_3329798, EPI_ISL_3329799, EPI_ISL_3329800, EPI_ISL_3329801, EPI_ISL_3329802, EPI_ISL_3329803, EPI_ISL_3329805, EPI_ISL_3329806, EPI_ISL_3329807, EPI_ISL_3329808, EPI_ISL_3329809, EPI_ISL_3329811, EPI_ISL_3329812, EPI_ISL_3329813, EPI_ISL_3329814, EPI_ISL_3329815, EPI_ISL_3329816, EPI_ISL_3329817, EPI_ISL_3329818, EPI_ISL_3329819, EPI_ISL_3329820, EPI_ISL_3329821, EPI_ISL_3329822, EPI_ISL_3329823, EPI_ISL_3329824, EPI_ISL_3329825, EPI_ISL_3329826, EPI_ISL_3329827, EPI_ISL_3329828, EPI_ISL_3329829, EPI_ISL_3329830, EPI_ISL_3329831, EPI_ISL_3329832, EPI_ISL_3329833, EPI_ISL_3329834, EPI_ISL_3329835, EPI_ISL_3329836, EPI_ISL_3329837, EPI_ISL_3329839, EPI_ISL_3329840, EPI_ISL_3329841, EPI_ISL_3329842, EPI_ISL_3329843, EPI_ISL_3329844, EPI_ISL_3329845, EPI_ISL_3329846, EPI_ISL_3329847, EPI_ISL_3329848, EPI_ISL_3329849, EPI_ISL_3329850, EPI_ISL_3329851, EPI_ISL_3329852, EPI_ISL_3329853, EPI_ISL_3329854, EPI_ISL_3329855, EPI_ISL_3329856, EPI_ISL_3329857, EPI_ISL_3329858, EPI_ISL_3329859, EPI_ISL_3329860, EPI_ISL_3329861, EPI_ISL_3329862, EPI_ISL_3329863, EPI_ISL_3329864, EPI_ISL_3329865, EPI_ISL_3329866, EPI_ISL_3329867, EPI_ISL_3329868, EPI_ISL_3329869, EPI_ISL_3329870, EPI_ISL_3329871, EPI_ISL_3329872, EPI_ISL_3329873, EPI_ISL_3329875, EPI_ISL_3329876, EPI_ISL_3329877, EPI_ISL_3329878, EPI_ISL_3329879, EPI_ISL_3329880, EPI_ISL_3329881, EPI_ISL_3329882, EPI_ISL_3329883, EPI_ISL_3329884, EPI_ISL_3329885, EPI_ISL_3329886, EPI_ISL_3329887, EPI_ISL_3329888, EPI_ISL_3329889, EPI_ISL_3329890, EPI_ISL_3329891, EPI_ISL_3329892, EPI_ISL_3329893, EPI_ISL_3329894, EPI_ISL_3329895, EPI_ISL_3329896, EPI_ISL_3329897, EPI_ISL_3329898, EPI_ISL_3329899, EPI_ISL_3329900, EPI_ISL_3329901, EPI_ISL_3329902, EPI_ISL_3329903, EPI_ISL_3329904, EPI_ISL_3329905, EPI_ISL_3329906, EPI_ISL_3329907, EPI_ISL_3329908, EPI_ISL_3329909, EPI_ISL_3329910, EPI_ISL_3329911, EPI_ISL_3329912, EPI_ISL_3329913, EPI_ISL_3329914, EPI_ISL_3329915, EPI_ISL_3329916, EPI_ISL_3329917, EPI_ISL_3329918, EPI_ISL_3329919, EPI_ISL_3329920, EPI_ISL_3329922, EPI_ISL_3329923, EPI_ISL_3329925, EPI_ISL_3329926, EPI_ISL_3329927, EPI_ISL_3329928, EPI_ISL_3329929, EPI_ISL_3329930, EPI_ISL_3329932, EPI_ISL_3329933, EPI_ISL_3329934, EPI_ISL_3329935, EPI_ISL_3329936, EPI_ISL_3329937, EPI_ISL_3329938 | | | |
| see above | Laboratory Corporation of America | Centers for Disease Control and Prevention Division of Viral Diseases, Pathogen Discovery | Dakota Howard, Dhwani Batra, Peter W. Cook, Kara Moser, Adrian Paskey, Jason Caravas, Benjamin Rambo-Martin, Shatavia Morrison, Christopher Gulvick, Scott Sammons, Yvette Unoarumhi, Darlene Wagner, Matthew Schmerer, Minoo Agarwal, Eyad Almasri, Debbie Boles, Ayla Burns, Nuthawin Charoensri, Oren Cohen, Susan Countryman, Mary Ann Cristobal, Bobbi Croy, Suzanne Dale, Hrushikesh Deshmukh, Amanda Douglas, Vincent Drouillon, Marcia Eisenberg, Howard Engler, Rama Ghatti, Prashant Gupta, Susan Hicks, Jake Humphrey, Lax Iyer, Lisa Pfefferle, Manoj Jain, Matthew Robinson, Mohan Kolli, Brian Krueger, Tim Kuphal, Stanley Letovsky, Michael Levandoski, Craig Lukasik, Jonathan Meltzer, Brian Norvell, Mindy Nye, Scott Parker, Christos Petropoulos, John Pruitt, Steven Ragan, Scott Ryan, Mike Sapeta, Jana Schroth, Suresh Babu Selvaraju, Goran Stevovic, Amanda Suchanek, Andrea Throop, Lyndon Tilson, Thomas Urban, Joe Voshell, Kimberly Wagner, Jonathan Williams, Mary Williamson, Qian Zeng, Tricia Zwiefelhofer, Clinton R. Paden, Duncan MacCannell |
| EPI_ISL_3329939 | Illinois Department of Public Health - Springfield Lab | Illinois Department of Public Health - Springfield Lab | Bryan Sim, Gordon McCall, Marshall Smock |
| EPI_ISL_3329940 | Laboratory Corporation of America | Centers for Disease Control and Prevention Division of Viral Diseases, Pathogen Discovery | Dakota Howard, Dhwani Batra, Peter W. Cook, Kara Moser, Adrian Paskey, Jason Caravas, Benjamin Rambo-Martin, Shatavia Morrison, Christopher Gulvick, Scott Sammons, Yvette Unoarumhi, Darlene Wagner, Matthew Schmerer, Minoo Agarwal, Eyad Almasri, Debbie Boles, Ayla Burns, Nuthawin Charoensri, Oren Cohen, Susan Countryman, Mary Ann Cristobal, Bobbi Croy, Suzanne Dale, Hrushikesh Deshmukh, Amanda Douglas, Vincent Drouillon, Marcia Eisenberg, Howard Engler, Rama Ghatti, Prashant Gupta, Susan Hicks, Jake Humphrey, Lax Iyer, Lisa Pfefferle, Manoj Jain, Matthew Robinson, Mohan Kolli, Brian Krueger, Tim Kuphal, Stanley Letovsky, Michael Levandoski, Craig Lukasik, Jonathan Meltzer, Brian Norvell, Mindy Nye, Scott Parker, Christos Petropoulos, John Pruitt, Steven Ragan, Scott Ryan, Mike Sapeta, Jana Schroth, Suresh Babu Selvaraju, Goran Stevovic, Amanda Suchanek, Andrea Throop, Lyndon Tilson, Thomas Urban, Joe Voshell, Kimberly Wagner, Jonathan Williams, Mary Williamson, Qian Zeng, Tricia Zwiefelhofer, Clinton R. Paden, Duncan MacCannell |
| EPI_ISL_3329941 | Illinois Department of Public Health - Springfield Lab | Illinois Department of Public Health - Springfield Lab | Bryan Sim, Gordon McCall, Marshall Smock |
| EPI_ISL_3329942 | Laboratory Corporation of America | Centers for Disease Control and Prevention Division of Viral Diseases, Pathogen Discovery | Dakota Howard, Dhwani Batra, Peter W. Cook, Kara Moser, Adrian Paskey, Jason Caravas, Benjamin Rambo-Martin, Shatavia Morrison, Christopher Gulvick, Scott Sammons, Yvette Unoarumhi, Darlene Wagner, Matthew Schmerer, Minoo Agarwal, Eyad Almasri, Debbie Boles, Ayla Burns, Nuthawin Charoensri, Oren Cohen, Susan Countryman, Mary Ann Cristobal, Bobbi Croy, Suzanne Dale, Hrushikesh Deshmukh, Amanda Douglas, Vincent Drouillon, Marcia Eisenberg, Howard Engler, Rama Ghatti, Prashant Gupta, Susan Hicks, Jake Humphrey, Lax Iyer, Lisa Pfefferle, Manoj Jain, Matthew Robinson, Mohan Kolli, Brian Krueger, Tim Kuphal, Stanley Letovsky, Michael Levandoski, Craig Lukasik, Jonathan Meltzer, Brian Norvell, Mindy Nye, Scott Parker, Christos Petropoulos, John Pruitt, Steven Ragan, Scott Ryan, Mike Sapeta, Jana Schroth, Suresh Babu Selvaraju, Goran Stevovic, Amanda Suchanek, Andrea Throop, Lyndon Tilson, Thomas Urban, Joe Voshell, Kimberly Wagner, Jonathan Williams, Mary Williamson, Qian Zeng, Tricia Zwiefelhofer, Clinton R. Paden, Duncan MacCannell |
| EPI_ISL_3329943 | Illinois Department of Public Health - Springfield Lab | Illinois Department of Public Health - Springfield Lab | Bryan Sim, Gordon McCall, Marshall Smock |
| EPI_ISL_3329944 | Laboratory Corporation of America | Centers for Disease Control and Prevention Division of Viral Diseases, Pathogen Discovery | Dakota Howard, Dhwani Batra, Peter W. Cook, Kara Moser, Adrian Paskey, Jason Caravas, Benjamin Rambo-Martin, Shatavia Morrison, Christopher Gulvick, Scott Sammons, Yvette Unoarumhi, Darlene Wagner, Matthew Schmerer, Minoo Agarwal, Eyad Almasri, Debbie Boles, Ayla Burns, Nuthawin Charoensri, Oren Cohen, Susan Countryman, Mary Ann Cristobal, Bobbi Croy, Suzanne Dale, Hrushikesh Deshmukh, Amanda Douglas, Vincent Drouillon, Marcia Eisenberg, Howard Engler, Rama Ghatti, Prashant Gupta, Susan Hicks, Jake Humphrey, Lax Iyer, Lisa Pfefferle, Manoj Jain, Matthew Robinson, Mohan Kolli, Brian Krueger, Tim Kuphal, Stanley Letovsky, Michael Levandoski, Craig Lukasik, Jonathan Meltzer, Brian Norvell, Mindy Nye, Scott Parker, Christos Petropoulos, John Pruitt, Steven Ragan, Scott Ryan, Mike Sapeta, Jana Schroth, Suresh Babu Selvaraju, Goran Stevovic, Amanda Suchanek, Andrea Throop, Lyndon Tilson, Thomas Urban, Joe Voshell, Kimberly Wagner, Jonathan Williams, Mary Williamson, Qian Zeng, Tricia Zwiefelhofer, Clinton R. Paden, Duncan MacCannell |
| EPI_ISL_3329945, EPI_ISL_3329946 | Illinois Department of Public Health - Springfield Lab | Illinois Department of Public Health - Springfield Lab | Bryan Sim, Gordon McCall, Marshall Smock |
| EPI_ISL_3329947 | Laboratory Corporation of America | Centers for Disease Control and Prevention Division of Viral Diseases, Pathogen Discovery | Dakota Howard, Dhwani Batra, Peter W. Cook, Kara Moser, Adrian Paskey, Jason Caravas, Benjamin Rambo-Martin, Shatavia Morrison, Christopher Gulvick, Scott Sammons, Yvette Unoarumhi, Darlene Wagner, Matthew Schmerer, Minoo Agarwal, Eyad Almasri, Debbie Boles, Ayla Burns, Nuthawin Charoensri, Oren Cohen, Susan Countryman, Mary Ann Cristobal, Bobbi Croy, Suzanne Dale, Hrushikesh Deshmukh, Amanda Douglas, Vincent Drouillon, Marcia Eisenberg, Howard Engler, Rama Ghatti, Prashant Gupta, Susan Hicks, Jake Humphrey, Lax Iyer, Lisa Pfefferle, Manoj Jain, Matthew Robinson, Mohan Kolli, Brian Krueger, Tim Kuphal, Stanley Letovsky, Michael Levandoski, Craig Lukasik, Jonathan Meltzer, Brian Norvell, Mindy Nye, Scott Parker, Christos Petropoulos, John Pruitt, Steven Ragan, Scott Ryan, Mike Sapeta, Jana Schroth, Suresh Babu Selvaraju, Goran Stevovic, Amanda Suchanek, Andrea Throop, Lyndon Tilson, Thomas Urban, Joe Voshell, Kimberly Wagner, Jonathan Williams, Mary Williamson, Qian Zeng, Tricia Zwiefelhofer, Clinton R. Paden, Duncan MacCannell |
| EPI_ISL_3329948 | Illinois Department of Public Health - Springfield Lab | Illinois Department of Public Health - Springfield Lab | Bryan Sim, Gordon McCall, Marshall Smock |
| EPI_ISL_3329949 | Laboratory Corporation of America | Centers for Disease Control and Prevention Division of Viral Diseases, Pathogen Discovery | Dakota Howard, Dhwani Batra, Peter W. Cook, Kara Moser, Adrian Paskey, Jason Caravas, Benjamin Rambo-Martin, Shatavia Morrison, Christopher Gulvick, Scott Sammons, Yvette Unoarumhi, Darlene Wagner, Matthew Schmerer, Minoo Agarwal, Eyad Almasri, Debbie Boles, Ayla Burns, Nuthawin Charoensri, Oren Cohen, Susan Countryman, Mary Ann Cristobal, Bobbi Croy, Suzanne Dale, Hrushikesh Deshmukh, Amanda Douglas, Vincent Drouillon, Marcia Eisenberg, Howard Engler, Rama Ghatti, Prashant Gupta, Susan Hicks, Jake Humphrey, Lax Iyer, Lisa Pfefferle, Manoj Jain, Matthew Robinson, Mohan Kolli, Brian Krueger, Tim Kuphal, Stanley Letovsky, Michael Levandoski, Craig Lukasik, Jonathan Meltzer, Brian Norvell, Mindy Nye, Scott Parker, Christos Petropoulos, John Pruitt, Steven Ragan, Scott Ryan, Mike Sapeta, Jana Schroth, Suresh Babu Selvaraju, Goran Stevovic, Amanda Suchanek, Andrea Throop, Lyndon Tilson, Thomas Urban, Joe Voshell, Kimberly Wagner, Jonathan Williams, Mary Williamson, Qian Zeng, Tricia Zwiefelhofer, Clinton R. Paden, Duncan MacCannell |
| EPI_ISL_3329950 | Illinois Department of Public Health - Springfield Lab | Illinois Department of Public Health - Springfield Lab | Bryan Sim, Gordon McCall, Marshall Smock |
| EPI_ISL_3329951 | Laboratory Corporation of America | Centers for Disease Control and Prevention Division of Viral Diseases, Pathogen Discovery | Dakota Howard, Dhwani Batra, Peter W. Cook, Kara Moser, Adrian Paskey, Jason Caravas, Benjamin Rambo-Martin, Shatavia Morrison, Christopher Gulvick, Scott Sammons, Yvette Unoarumhi, Darlene Wagner, Matthew Schmerer, Minoo Agarwal, Eyad Almasri, Debbie Boles, Ayla Burns, Nuthawin Charoensri, Oren Cohen, Susan Countryman, Mary Ann Cristobal, Bobbi Croy, Suzanne Dale, Hrushikesh Deshmukh, Amanda Douglas, Vincent Drouillon, Marcia Eisenberg, Howard Engler, Rama Ghatti, Prashant Gupta, Susan Hicks, Jake Humphrey, Lax Iyer, Lisa Pfefferle, Manoj Jain, Matthew Robinson, Mohan Kolli, Brian Krueger, Tim Kuphal, Stanley Letovsky, Michael Levandoski, Craig Lukasik, Jonathan Meltzer, Brian Norvell, Mindy Nye, Scott Parker, Christos Petropoulos, John Pruitt, Steven Ragan, Scott Ryan, Mike Sapeta, Jana Schroth, Suresh Babu Selvaraju, Goran Stevovic, Amanda Suchanek, Andrea Throop, Lyndon Tilson, Thomas Urban, Joe Voshell, Kimberly Wagner, Jonathan Williams, Mary Williamson, Qian Zeng, Tricia Zwiefelhofer, Clinton R. Paden, Duncan MacCannell |
| EPI_ISL_3329952, EPI_ISL_3329953, EPI_ISL_3329955 | Illinois Department of Public Health - Springfield Lab | Illinois Department of Public Health - Springfield Lab | Bryan Sim, Gordon McCall, Marshall Smock |
| EPI_ISL_3329956 | Laboratory Corporation of America | Centers for Disease Control and Prevention Division of Viral Diseases, Pathogen Discovery | Dakota Howard, Dhwani Batra, Peter W. Cook, Kara Moser, Adrian Paskey, Jason Caravas, Benjamin Rambo-Martin, Shatavia Morrison, Christopher Gulvick, Scott Sammons, Yvette Unoarumhi, Darlene Wagner, Matthew Schmerer, Minoo Agarwal, Eyad Almasri, Debbie Boles, Ayla Burns, Nuthawin Charoensri, Oren Cohen, Susan Countryman, Mary Ann Cristobal, Bobbi Croy, Suzanne Dale, Hrushikesh Deshmukh, Amanda Douglas, Vincent Drouillon, Marcia Eisenberg, Howard Engler, Rama Ghatti, Prashant Gupta, Susan Hicks, Jake Humphrey, Lax Iyer, Lisa Pfefferle, Manoj Jain, Matthew Robinson, Mohan Kolli, Brian Krueger, Tim Kuphal, Stanley Letovsky, Michael Levandoski, Craig Lukasik, Jonathan Meltzer, Brian Norvell, Mindy Nye, Scott Parker, Christos Petropoulos, John Pruitt, Steven Ragan, Scott Ryan, Mike Sapeta, Jana Schroth, Suresh Babu Selvaraju, Goran Stevovic, Amanda Suchanek, Andrea Throop, Lyndon Tilson, Thomas Urban, Joe Voshell, Kimberly Wagner, Jonathan Williams, Mary Williamson, Qian Zeng, Tricia Zwiefelhofer, Clinton R. Paden, Duncan MacCannell |
| EPI_ISL_3329957, EPI_ISL_3329958 | Illinois Department of Public Health - Springfield Lab | Illinois Department of Public Health - Springfield Lab | Bryan Sim, Gordon McCall, Marshall Smock |
| EPI_ISL_3329959 | Laboratory Corporation of America | Centers for Disease Control and Prevention Division of Viral Diseases, Pathogen Discovery | Dakota Howard, Dhwani Batra, Peter W. Cook, Kara Moser, Adrian Paskey, Jason Caravas, Benjamin Rambo-Martin, Shatavia Morrison, Christopher Gulvick, Scott Sammons, Yvette Unoarumhi, Darlene Wagner, Matthew Schmerer, Minoo Agarwal, Eyad Almasri, Debbie Boles, Ayla Burns, Nuthawin Charoensri, Oren Cohen, Susan Countryman, Mary Ann Cristobal, Bobbi Croy, Suzanne Dale, Hrushikesh Deshmukh, Amanda Douglas, Vincent Drouillon, Marcia Eisenberg, Howard Engler, Rama Ghatti, Prashant Gupta, Susan Hicks, Jake Humphrey, Lax Iyer, Lisa Pfefferle, Manoj Jain, Matthew Robinson, Mohan Kolli, Brian Krueger, Tim Kuphal, Stanley Letovsky, Michael Levandoski, Craig Lukasik, Jonathan Meltzer, Brian Norvell, Mindy Nye, Scott Parker, Christos Petropoulos, John Pruitt, Steven Ragan, Scott Ryan, Mike Sapeta, Jana Schroth, Suresh Babu Selvaraju, Goran Stevovic, Amanda Suchanek, Andrea Throop, Lyndon Tilson, Thomas Urban, Joe Voshell, |

| | | | |
|---|---|---|---|
| | | | Kimberly Wagner, Jonathan Williams, Mary Williamson, Qian Zeng, Tricia Zwiefelhofer, Clinton R. Paden, Duncan MacCannell |
| EPI_ISL_3329960 | Illinois Department of Public Health - Springfield Lab | Illinois Department of Public Health - Springfield Lab | Bryan Sim, Gordon McCall, Marshall Smock |
| EPI_ISL_3329961 | Laboratory Corporation of America | Centers for Disease Control and Prevention Division of Viral Diseases, Pathogen Discovery | Dakota Howard, Dhwani Batra, Peter W. Cook, Kara Moser, Adrian Paskey, Jason Caravas, Benjamin Rambo-Martin, Shatavia Morrison, Christopher Gulvick, Scott Sammons, Yvette Unoarumhi, Darlene Wagner, Matthew Schmerer, Minoo Agarwal, Eyad Almasri, Debbie Boles, Ayla Burns, Nuthawin Charoensri, Oren Cohen, Susan Countryman, Mary Ann Cristobal, Bobbi Croy, Suzanne Dale, Hrushikesh Deshmukh, Amanda Douglas, Vincent Drouillon, Marcia Eisenberg, Howard Engler, Rama Ghatti, Prashant Gupta, Susan Hicks, Jake Humphrey, Lax Iyer, Lisa Pfefferle, Manoj Jain, Matthew Robinson, Mohan Kolli, Brian Krueger, Tim Kuphal, Stanley Letovsky, Michael Levandoski, Craig Lukasik, Jonathan Meltzer, Brian Norvell, Mindy Nye, Scott Parker, Christos Petropoulos, John Pruitt, Steven Ragan, Scott Ryan, Mike Sapeta, Jana Schroth, Suresh Babu Selvaraju, Goran Stevovic, Amanda Suchanek, Andrea Throop, Lyndon Tilson, Thomas Urban, Joe Voshell, Kimberly Wagner, Jonathan Williams, Mary Williamson, Qian Zeng, Tricia Zwiefelhofer, Clinton R. Paden, Duncan MacCannell |
| EPI_ISL_3329962, EPI_ISL_3329963 | Illinois Department of Public Health - Springfield Lab | Illinois Department of Public Health - Springfield Lab | Bryan Sim, Gordon McCall, Marshall Smock |
| EPI_ISL_3329964 | Laboratory Corporation of America | Centers for Disease Control and Prevention Division of Viral Diseases, Pathogen Discovery | Dakota Howard, Dhwani Batra, Peter W. Cook, Kara Moser, Adrian Paskey, Jason Caravas, Benjamin Rambo-Martin, Shatavia Morrison, Christopher Gulvick, Scott Sammons, Yvette Unoarumhi, Darlene Wagner, Matthew Schmerer, Minoo Agarwal, Eyad Almasri, Debbie Boles, Ayla Burns, Nuthawin Charoensri, Oren Cohen, Susan Countryman, Mary Ann Cristobal, Bobbi Croy, Suzanne Dale, Hrushikesh Deshmukh, Amanda Douglas, Vincent Drouillon, Marcia Eisenberg, Howard Engler, Rama Ghatti, Prashant Gupta, Susan Hicks, Jake Humphrey, Lax Iyer, Lisa Pfefferle, Manoj Jain, Matthew Robinson, Mohan Kolli, Brian Krueger, Tim Kuphal, Stanley Letovsky, Michael Levandoski, Craig Lukasik, Jonathan Meltzer, Brian Norvell, Mindy Nye, Scott Parker, Christos Petropoulos, John Pruitt, Steven Ragan, Scott Ryan, Mike Sapeta, Jana Schroth, Suresh Babu Selvaraju, Goran Stevovic, Amanda Suchanek, Andrea Throop, Lyndon Tilson, Thomas Urban, Joe Voshell, Kimberly Wagner, Jonathan Williams, Mary Williamson, Qian Zeng, Tricia Zwiefelhofer, Clinton R. Paden, Duncan MacCannell |
| EPI_ISL_3329965, EPI_ISL_3329966 | Illinois Department of Public Health - Springfield Lab | Illinois Department of Public Health - Springfield Lab | Bryan Sim, Gordon McCall, Marshall Smock |
| EPI_ISL_3329967 | Laboratory Corporation of America | Centers for Disease Control and Prevention Division of Viral Diseases, Pathogen Discovery | Dakota Howard, Dhwani Batra, Peter W. Cook, Kara Moser, Adrian Paskey, Jason Caravas, Benjamin Rambo-Martin, Shatavia Morrison, Christopher Gulvick, Scott Sammons, Yvette Unoarumhi, Darlene Wagner, Matthew Schmerer, Minoo Agarwal, Eyad Almasri, Debbie Boles, Ayla Burns, Nuthawin Charoensri, Oren Cohen, Susan Countryman, Mary Ann Cristobal, Bobbi Croy, Suzanne Dale, Hrushikesh Deshmukh, Amanda Douglas, Vincent Drouillon, Marcia Eisenberg, Howard Engler, Rama Ghatti, Prashant Gupta, Susan Hicks, Jake Humphrey, Lax Iyer, Lisa Pfefferle, Manoj Jain, Matthew Robinson, Mohan Kolli, Brian Krueger, Tim Kuphal, Stanley Letovsky, Michael Levandoski, Craig Lukasik, Jonathan Meltzer, Brian Norvell, Mindy Nye, Scott Parker, Christos Petropoulos, John Pruitt, Steven Ragan, Scott Ryan, Mike Sapeta, Jana Schroth, Suresh Babu Selvaraju, Goran Stevovic, Amanda Suchanek, Andrea Throop, Lyndon Tilson, Thomas Urban, Joe Voshell, Kimberly Wagner, Jonathan Williams, Mary Williamson, Qian Zeng, Tricia Zwiefelhofer, Clinton R. Paden, Duncan MacCannell |
| EPI_ISL_3329968 | Illinois Department of Public Health - Springfield Lab | Illinois Department of Public Health - Springfield Lab | Bryan Sim, Gordon McCall, Marshall Smock |
| EPI_ISL_3329969 | Laboratory Corporation of America | Centers for Disease Control and Prevention Division of Viral Diseases, Pathogen Discovery | Dakota Howard, Dhwani Batra, Peter W. Cook, Kara Moser, Adrian Paskey, Jason Caravas, Benjamin Rambo-Martin, Shatavia Morrison, Christopher Gulvick, Scott Sammons, Yvette Unoarumhi, Darlene Wagner, Matthew Schmerer, Minoo Agarwal, Eyad Almasri, Debbie Boles, Ayla Burns, Nuthawin Charoensri, Oren Cohen, Susan Countryman, Mary Ann Cristobal, Bobbi Croy, Suzanne Dale, Hrushikesh Deshmukh, Amanda Douglas, Vincent Drouillon, Marcia Eisenberg, Howard Engler, Rama Ghatti, Prashant Gupta, Susan Hicks, Jake Humphrey, Lax Iyer, Lisa Pfefferle, Manoj Jain, Matthew Robinson, Mohan Kolli, Brian Krueger, Tim Kuphal, Stanley Letovsky, Michael Levandoski, Craig Lukasik, Jonathan Meltzer, Brian Norvell, Mindy Nye, Scott Parker, Christos Petropoulos, John Pruitt, Steven Ragan, Scott Ryan, Mike Sapeta, Jana Schroth, Suresh Babu Selvaraju, Goran Stevovic, Amanda Suchanek, Andrea Throop, Lyndon Tilson, Thomas Urban, Joe Voshell, Kimberly Wagner, Jonathan Williams, Mary Williamson, Qian Zeng, Tricia Zwiefelhofer, Clinton R. Paden, Duncan MacCannell |
| EPI_ISL_3329970, EPI_ISL_3329971 | Illinois Department of Public Health - Springfield Lab | Illinois Department of Public Health - Springfield Lab | Bryan Sim, Gordon McCall, Marshall Smock |
| EPI_ISL_3329972 | Laboratory Corporation of America | Centers for Disease Control and Prevention Division of Viral Diseases, Pathogen Discovery | Dakota Howard, Dhwani Batra, Peter W. Cook, Kara Moser, Adrian Paskey, Jason Caravas, Benjamin Rambo-Martin, Shatavia Morrison, Christopher Gulvick, Scott Sammons, Yvette Unoarumhi, Darlene Wagner, Matthew Schmerer, Minoo Agarwal, Eyad Almasri, Debbie Boles, Ayla Burns, Nuthawin Charoensri, Oren Cohen, Susan Countryman, Mary Ann Cristobal, Bobbi Croy, Suzanne Dale, Hrushikesh Deshmukh, Amanda Douglas, Vincent Drouillon, Marcia Eisenberg, Howard Engler, Rama Ghatti, Prashant Gupta, Susan Hicks, Jake Humphrey, Lax Iyer, Lisa Pfefferle, Manoj Jain, Matthew Robinson, Mohan Kolli, Brian Krueger, Tim Kuphal, Stanley Letovsky, Michael Levandoski, Craig Lukasik, Jonathan Meltzer, Brian Norvell, Mindy Nye, Scott Parker, Christos Petropoulos, John Pruitt, Steven Ragan, Scott Ryan, Mike Sapeta, Jana Schroth, Suresh Babu Selvaraju, Goran Stevovic, Amanda Suchanek, Andrea Throop, Lyndon Tilson, Thomas Urban, Joe Voshell, Kimberly Wagner, Jonathan Williams, Mary Williamson, Qian Zeng, Tricia Zwiefelhofer, Clinton R. Paden, Duncan MacCannell |
| EPI_ISL_3329973 | Illinois Department of Public Health - Springfield Lab | Illinois Department of Public Health - Springfield Lab | Bryan Sim, Gordon McCall, Marshall Smock |
| EPI_ISL_3329974 | Laboratory Corporation of America | Centers for Disease Control and Prevention Division of Viral Diseases, Pathogen Discovery | Dakota Howard, Dhwani Batra, Peter W. Cook, Kara Moser, Adrian Paskey, Jason Caravas, Benjamin Rambo-Martin, Shatavia Morrison, Christopher Gulvick, Scott Sammons, Yvette Unoarumhi, Darlene Wagner, Matthew Schmerer, Minoo Agarwal, Eyad Almasri, Debbie Boles, Ayla Burns, Nuthawin Charoensri, Oren Cohen, Susan Countryman, Mary Ann Cristobal, Bobbi Croy, Suzanne Dale, Hrushikesh Deshmukh, Amanda Douglas, Vincent Drouillon, Marcia Eisenberg, Howard Engler, Rama Ghatti, Prashant Gupta, Susan Hicks, Jake Humphrey, Lax Iyer, Lisa Pfefferle, Manoj Jain, Matthew Robinson, Mohan Kolli, Brian Krueger, Tim Kuphal, Stanley Letovsky, Michael Levandoski, Craig Lukasik, Jonathan Meltzer, Brian Norvell, Mindy Nye, Scott Parker, Christos Petropoulos, John Pruitt, Steven Ragan, Scott Ryan, Mike Sapeta, Jana Schroth, Suresh Babu Selvaraju, Goran Stevovic, Amanda Suchanek, Andrea Throop, Lyndon Tilson, Thomas Urban, Joe Voshell, Kimberly Wagner, Jonathan Williams, Mary Williamson, Qian Zeng, Tricia Zwiefelhofer, Clinton R. Paden, Duncan MacCannell |
| EPI_ISL_3329975, EPI_ISL_3329976 | Illinois Department of Public Health - Springfield Lab | Illinois Department of Public Health - Springfield Lab | Bryan Sim, Gordon McCall, Marshall Smock |
| EPI_ISL_3329977 | Laboratory Corporation of America | Centers for Disease Control and Prevention Division of Viral Diseases, Pathogen Discovery | Dakota Howard, Dhwani Batra, Peter W. Cook, Kara Moser, Adrian Paskey, Jason Caravas, Benjamin Rambo-Martin, Shatavia Morrison, Christopher Gulvick, Scott Sammons, Yvette Unoarumhi, Darlene Wagner, Matthew Schmerer, Minoo Agarwal, Eyad Almasri, Debbie Boles, Ayla Burns, Nuthawin Charoensri, Oren Cohen, Susan Countryman, Mary Ann Cristobal, Bobbi Croy, Suzanne Dale, Hrushikesh Deshmukh, Amanda Douglas, Vincent Drouillon, Marcia Eisenberg, Howard Engler, Rama Ghatti, Prashant Gupta, Susan Hicks, Jake Humphrey, Lax Iyer, Lisa Pfefferle, Manoj Jain, Matthew Robinson, Mohan Kolli, Brian Krueger, Tim Kuphal, Stanley Letovsky, Michael Levandoski, Craig Lukasik, Jonathan Meltzer, Brian Norvell, Mindy Nye, Scott Parker, Christos Petropoulos, John Pruitt, Steven Ragan, Scott Ryan, Mike Sapeta, Jana Schroth, Suresh Babu Selvaraju, Goran Stevovic, Amanda Suchanek, Andrea Throop, Lyndon Tilson, Thomas Urban, Joe Voshell, Kimberly Wagner, Jonathan Williams, Mary Williamson, Qian Zeng, Tricia Zwiefelhofer, Clinton R. Paden, Duncan MacCannell |
| EPI_ISL_3329978, EPI_ISL_3329979 | Illinois Department of Public Health - Springfield Lab | Illinois Department of Public Health - Springfield Lab | Bryan Sim, Gordon McCall, Marshall Smock |
| EPI_ISL_3329980 | Laboratory Corporation of America | Centers for Disease Control and Prevention Division of Viral Diseases, Pathogen Discovery | Dakota Howard, Dhwani Batra, Peter W. Cook, Kara Moser, Adrian Paskey, Jason Caravas, Benjamin Rambo-Martin, Shatavia Morrison, Christopher Gulvick, Scott Sammons, Yvette Unoarumhi, Darlene Wagner, Matthew Schmerer, Minoo Agarwal, Eyad Almasri, Debbie Boles, Ayla Burns, Nuthawin Charoensri, Oren Cohen, Susan Countryman, Mary Ann Cristobal, Bobbi Croy, Suzanne Dale, Hrushikesh Deshmukh, Amanda Douglas, Vincent Drouillon, Marcia Eisenberg, Howard Engler, Rama Ghatti, Prashant Gupta, Susan Hicks, Jake Humphrey, Lax Iyer, Lisa Pfefferle, Manoj Jain, Matthew Robinson, Mohan Kolli, Brian Krueger, Tim Kuphal, Stanley Letovsky, Michael Levandoski, Craig Lukasik, Jonathan Meltzer, Brian Norvell, Mindy Nye, Scott Parker, Christos Petropoulos, John Pruitt, Steven Ragan, Scott Ryan, Mike Sapeta, Jana Schroth, Suresh Babu Selvaraju, Goran Stevovic, Amanda Suchanek, Andrea Throop, Lyndon Tilson, Thomas Urban, Joe Voshell, Kimberly Wagner, Jonathan Williams, Mary Williamson, Qian Zeng, Tricia Zwiefelhofer, Clinton R. Paden, Duncan MacCannell |
| EPI_ISL_3329981 | Illinois Department of Public Health - Springfield Lab | Illinois Department of Public Health - Springfield Lab | Bryan Sim, Gordon McCall, Marshall Smock |
| EPI_ISL_3329982 | Laboratory Corporation of America | Centers for Disease Control and Prevention Division of Viral Diseases, Pathogen Discovery | Dakota Howard, Dhwani Batra, Peter W. Cook, Kara Moser, Adrian Paskey, Jason Caravas, Benjamin Rambo-Martin, Shatavia Morrison, Christopher Gulvick, Scott Sammons, Yvette Unoarumhi, Darlene Wagner, Matthew Schmerer, Minoo Agarwal, Eyad Almasri, Debbie Boles, Ayla Burns, Nuthawin Charoensri, Oren Cohen, Susan Countryman, Mary Ann Cristobal, Bobbi Croy, Suzanne Dale, Hrushikesh Deshmukh, Amanda Douglas, Vincent Drouillon, Marcia Eisenberg, Howard Engler, Rama Ghatti, Prashant Gupta, Susan Hicks, Jake Humphrey, Lax Iyer, Lisa Pfefferle, Manoj Jain, Matthew Robinson, Mohan Kolli, Brian Krueger, Tim Kuphal, Stanley Letovsky, Michael Levandoski, Craig Lukasik, Jonathan Meltzer, Brian Norvell, Mindy Nye, Scott Parker, Christos Petropoulos, John Pruitt, Steven Ragan, Scott Ryan, Mike Sapeta, Jana Schroth, Suresh Babu Selvaraju, Goran Stevovic, Amanda Suchanek, Andrea Throop, Lyndon Tilson, Thomas Urban, Joe Voshell, Kimberly Wagner, Jonathan Williams, Mary Williamson, Qian Zeng, Tricia Zwiefelhofer, Clinton R. Paden, Duncan MacCannell |
| EPI_ISL_3329983, EPI_ISL_3329984 | Illinois Department of Public Health - Springfield Lab | Illinois Department of Public Health - Springfield Lab | Bryan Sim, Gordon McCall, Marshall Smock |
| EPI_ISL_3329985 | Laboratory Corporation of America | Centers for Disease Control and Prevention Division of Viral Diseases, Pathogen Discovery | Dakota Howard, Dhwani Batra, Peter W. Cook, Kara Moser, Adrian Paskey, Jason Caravas, Benjamin Rambo-Martin, Shatavia Morrison, Christopher Gulvick, Scott Sammons, Yvette Unoarumhi, Darlene Wagner, Matthew Schmerer, Minoo Agarwal, Eyad Almasri, Debbie Boles, Ayla Burns, Nuthawin Charoensri, Oren Cohen, Susan Countryman, Mary Ann Cristobal, Bobbi Croy, Suzanne Dale, Hrushikesh Deshmukh, Amanda Douglas, Vincent Drouillon, Marcia Eisenberg, Howard Engler, Rama Ghatti, Prashant Gupta, Susan Hicks, Jake Humphrey, Lax Iyer, Lisa Pfefferle, Manoj Jain, Matthew Robinson, Mohan Kolli, Brian Krueger, Tim Kuphal, Stanley Letovsky, Michael Levandoski, Craig Lukasik, Jonathan Meltzer, Brian Norvell, Mindy Nye, Scott Parker, Christos Petropoulos, John Pruitt, Steven Ragan, Scott Ryan, Mike Sapeta, Jana Schroth, Suresh Babu Selvaraju, Goran Stevovic, Amanda Suchanek, Andrea Throop, Lyndon Tilson, Thomas Urban, Joe Voshell, Kimberly Wagner, Jonathan Williams, Mary Williamson, Qian Zeng, Tricia Zwiefelhofer, Clinton R. Paden, Duncan MacCannell |
| EPI_ISL_3329986 | Illinois Department of Public Health - Springfield Lab | Illinois Department of Public Health - Springfield Lab | Bryan Sim, Gordon McCall, Marshall Smock |
| EPI_ISL_3329987, EPI_ISL_3329988 | Laboratory Corporation of America | Centers for Disease Control and Prevention Division of Viral Diseases, Pathogen Discovery | Dakota Howard, Dhwani Batra, Peter W. Cook, Kara Moser, Adrian Paskey, Jason Caravas, Benjamin Rambo-Martin, Shatavia Morrison, Christopher Gulvick, Scott Sammons, Yvette Unoarumhi, Darlene Wagner, Matthew Schmerer, Minoo Agarwal, Eyad Almasri, Debbie Boles, Ayla Burns, Nuthawin Charoensri, Oren Cohen, Susan Countryman, Mary Ann Cristobal, Bobbi Croy, Suzanne Dale, Hrushikesh Deshmukh, Amanda Douglas, Vincent Drouillon, Marcia Eisenberg, Howard Engler, Rama Ghatti, Prashant Gupta, Susan Hicks, Jake Humphrey, Lax Iyer, Lisa Pfefferle, Manoj Jain, Matthew Robinson, Mohan Kolli, Brian Krueger, Tim Kuphal, Stanley Letovsky, Michael Levandoski, Craig Lukasik, Jonathan Meltzer, Brian Norvell, Mindy Nye, Scott Parker, Christos Petropoulos, John Pruitt, Steven Ragan, Scott Ryan, Mike Sapeta, Jana Schroth, Suresh Babu Selvaraju, Goran Stevovic, Amanda Suchanek, Andrea Throop, Lyndon Tilson, Thomas Urban, Joe Voshell, Kimberly Wagner, Jonathan Williams, Mary Williamson, Qian Zeng, Tricia Zwiefelhofer, Clinton R. Paden, Duncan MacCannell |
| EPI_ISL_3329989 | Illinois Department of Public Health - Springfield Lab | Illinois Department of Public Health - Springfield Lab | Bryan Sim, Gordon McCall, Marshall Smock |
| EPI_ISL_3329990, EPI_ISL_3329991 | Laboratory Corporation of America | Centers for Disease Control and Prevention Division of Viral Diseases, Pathogen Discovery | Dakota Howard, Dhwani Batra, Peter W. Cook, Kara Moser, Adrian Paskey, Jason Caravas, Benjamin Rambo-Martin, Shatavia Morrison, Christopher Gulvick, Scott Sammons, Yvette Unoarumhi, Darlene Wagner, Matthew Schmerer, Minoo Agarwal, Eyad Almasri, Debbie Boles, Ayla Burns, Nuthawin Charoensri, Oren Cohen, Susan Countryman, Mary Ann Cristobal, Bobbi Croy, Suzanne Dale, Hrushikesh Deshmukh, Amanda Douglas, Vincent Drouillon, Marcia Eisenberg, Howard Engler, Rama Ghatti, Prashant Gupta, Susan Hicks, Jake Humphrey, Lax Iyer, Lisa Pfefferle, Manoj Jain, Matthew Robinson, Mohan Kolli, Brian Krueger, Tim Kuphal, Stanley Letovsky, Michael Levandoski, Craig Lukasik, Jonathan Meltzer, Brian Norvell, Mindy Nye, Scott Parker, Christos Petropoulos, John Pruitt, Steven Ragan, Scott Ryan, Mike Sapeta, Jana Schroth, Suresh Babu Selvaraju, Goran Stevovic, Amanda Suchanek, Andrea Throop, Lyndon Tilson, Thomas Urban, Joe Voshell, Kimberly Wagner, Jonathan Williams, Mary Williamson, Qian Zeng, Tricia Zwiefelhofer, Clinton R. Paden, Duncan MacCannell |
| EPI_ISL_3329992 | Illinois Department of Public Health - Springfield Lab | Illinois Department of Public Health - Springfield Lab | Bryan Sim, Gordon McCall, Marshall Smock |
| EPI_ISL_3329993, EPI_ISL_3329994, EPI_ISL_3329995 | Laboratory Corporation of America | Centers for Disease Control and Prevention Division of Viral Diseases, Pathogen Discovery | Dakota Howard, Dhwani Batra, Peter W. Cook, Kara Moser, Adrian Paskey, Jason Caravas, Benjamin Rambo-Martin, Shatavia Morrison, Christopher Gulvick, Scott Sammons, Yvette Unoarumhi, Darlene Wagner, Matthew Schmerer, Minoo Agarwal, Eyad Almasri, Debbie Boles, Ayla Burns, Nuthawin Charoensri, Oren Cohen, Susan Countryman, Mary Ann Cristobal, Bobbi Croy, Suzanne Dale, Hrushikesh Deshmukh, Amanda Douglas, Vincent Drouillon, Marcia Eisenberg, Howard Engler, Rama Ghatti, Prashant Gupta, Susan Hicks, Jake Humphrey, Lax Iyer, Lisa Pfefferle, Manoj Jain, Matthew Robinson, Mohan Kolli, Brian Krueger, Tim Kuphal, Stanley Letovsky, Michael Levandoski, Craig Lukasik, Jonathan Meltzer, Brian Norvell, Mindy Nye, Scott Parker, Christos Petropoulos, John Pruitt, Steven Ragan, Scott Ryan, Mike Sapeta, Jana Schroth, Suresh Babu Selvaraju, Goran Stevovic, Amanda Suchanek, Andrea Throop, Lyndon Tilson, Thomas Urban, Joe Voshell, Kimberly Wagner, Jonathan Williams, Mary Williamson, Qian Zeng, Tricia Zwiefelhofer, Clinton R. Paden, Duncan MacCannell |
| EPI_ISL_3329996, EPI_ISL_3329997 | Illinois Department of Public Health - Springfield Lab | Illinois Department of Public Health - Springfield Lab | Bryan Sim, Gordon McCall, Marshall Smock |
| EPI_ISL_3329998 | Laboratory Corporation of America | Centers for Disease Control and Prevention Division of Viral Diseases, Pathogen Discovery | Dakota Howard, Dhwani Batra, Peter W. Cook, Kara Moser, Adrian Paskey, Jason Caravas, Benjamin Rambo-Martin, Shatavia Morrison, Christopher Gulvick, Scott Sammons, Yvette Unoarumhi, Darlene Wagner, Matthew Schmerer, Minoo Agarwal, Eyad Almasri, Debbie Boles, Ayla Burns, Nuthawin Charoensri, Oren Cohen, Susan Countryman, Mary Ann Cristobal, Bobbi Croy, Suzanne Dale, Hrushikesh Deshmukh, Amanda Douglas, Vincent Drouillon, Marcia Eisenberg, Howard Engler, Rama Ghatti, Prashant Gupta, Susan Hicks, Jake Humphrey, Lax Iyer, Lisa Pfefferle, Manoj Jain, Matthew Robinson, Mohan Kolli, Brian Krueger, Tim Kuphal, Stanley Letovsky, Michael Levandoski, Craig Lukasik, Jonathan Meltzer, Brian Norvell, Mindy Nye, Scott Parker, Christos Petropoulos, John Pruitt, Steven Ragan, Scott Ryan, Mike Sapeta, Jana Schroth, Suresh Babu Selvaraju, Goran Stevovic, Amanda Suchanek, Andrea Throop, Lyndon Tilson, Thomas Urban, Joe Voshell, Kimberly Wagner, Jonathan Williams, Mary Williamson, Qian Zeng, Tricia Zwiefelhofer, Clinton R. Paden, Duncan MacCannell |
| EPI_ISL_3329999 | Illinois Department of Public Health - Springfield Lab | Illinois Department of Public Health - Springfield Lab | Bryan Sim, Gordon McCall, Marshall Smock |
| EPI_ISL_3330000, EPI_ISL_3330003 | Laboratory Corporation of America | Centers for Disease Control and Prevention Division of Viral Diseases, Pathogen Discovery | Dakota Howard, Dhwani Batra, Peter W. Cook, Kara Moser, Adrian Paskey, Jason Caravas, Benjamin Rambo-Martin, Shatavia Morrison, Christopher Gulvick, Scott Sammons, Yvette Unoarumhi, Darlene Wagner, Matthew Schmerer, Minoo Agarwal, Eyad Almasri, Debbie Boles, Ayla Burns, Nuthawin Charoensri, Oren Cohen, Susan Countryman, Mary Ann Cristobal, Bobbi Croy, Suzanne Dale, Hrushikesh Deshmukh, Amanda Douglas, Vincent Drouillon, Marcia Eisenberg, Howard Engler, Rama Ghatti, Prashant Gupta, Susan Hicks, Jake Humphrey, Lax Iyer, Lisa Pfefferle, Manoj Jain, Matthew Robinson, Mohan Kolli, Brian Krueger, Tim Kuphal, Stanley Letovsky, Michael Levandoski, Craig Lukasik, Jonathan Meltzer, Brian Norvell, Mindy Nye, Scott Parker, Christos Petropoulos, John Pruitt, Steven Ragan, Scott Ryan, Mike Sapeta, Jana Schroth, Suresh Babu Selvaraju, Goran Stevovic, Amanda Suchanek, Andrea Throop, Lyndon Tilson, Thomas Urban, Joe Voshell, |

| | | | |
|---|---|---|---|
| | | | Kimberly Wagner, Jonathan Williams, Mary Williamson, Qian Zeng, Tricia Zwiefelhofer, Clinton R. Paden, Duncan MacCannell |
| EPI_ISL_3330004 | Illinois Department of Public Health - Springfield Lab | Illinois Department of Public Health - Springfield Lab | Bryan Sim, Gordon McCall, Marshall Smock |
| EPI_ISL_3330005 | Laboratory Corporation of America | Centers for Disease Control and Prevention Division of Viral Diseases, Pathogen Discovery | Dakota Howard, Dhwani Batra, Peter W. Cook, Kara Moser, Adrian Paskey, Jason Caravas, Benjamin Rambo-Martin, Shatavia Morrison, Christopher Gulvick, Scott Sammons, Yvette Unoarumhi, Darlene Wagner, Matthew Schmerer, Minoo Agarwal, Eyad Almasri, Debbie Boles, Ayla Burns, Nuthawin Charoensri, Oren Cohen, Susan Countryman, Mary Ann Cristobal, Bobbi Croy, Suzanne Dale, Hrushikesh Deshmukh, Amanda Douglas, Vincent Drouillon, Marcia Eisenberg, Howard Engler, Rama Ghatti, Prashant Gupta, Susan Hicks, Jake Humphrey, Lax Iyer, Lisa Pfefferle, Manoj Jain, Matthew Robinson, Mohan Kolli, Brian Krueger, Tim Kuphal, Stanley Letovsky, Michael Levandoski, Craig Lukasik, Jonathan Meltzer, Brian Norvell, Mindy Nye, Scott Parker, Christos Petropoulos, John Pruitt, Steven Ragan, Scott Ryan, Mike Sapeta, Jana Schroth, Suresh Babu Selvaraju, Goran Stevovic, Amanda Suchanek, Andrea Throop, Lyndon Tilson, Thomas Urban, Joe Voshell, Kimberly Wagner, Jonathan Williams, Mary Williamson, Qian Zeng, Tricia Zwiefelhofer, Clinton R. Paden, Duncan MacCannell |
| EPI_ISL_3330006, EPI_ISL_3330007 | Illinois Department of Public Health - Springfield Lab | Illinois Department of Public Health - Springfield Lab | Bryan Sim, Gordon McCall, Marshall Smock |
| EPI_ISL_3330008, EPI_ISL_3330009, EPI_ISL_3330010 | Laboratory Corporation of America | Centers for Disease Control and Prevention Division of Viral Diseases, Pathogen Discovery | Dakota Howard, Dhwani Batra, Peter W. Cook, Kara Moser, Adrian Paskey, Jason Caravas, Benjamin Rambo-Martin, Shatavia Morrison, Christopher Gulvick, Scott Sammons, Yvette Unoarumhi, Darlene Wagner, Matthew Schmerer, Minoo Agarwal, Eyad Almasri, Debbie Boles, Ayla Burns, Nuthawin Charoensri, Oren Cohen, Susan Countryman, Mary Ann Cristobal, Bobbi Croy, Suzanne Dale, Hrushikesh Deshmukh, Amanda Douglas, Vincent Drouillon, Marcia Eisenberg, Howard Engler, Rama Ghatti, Prashant Gupta, Susan Hicks, Jake Humphrey, Lax Iyer, Lisa Pfefferle, Manoj Jain, Matthew Robinson, Mohan Kolli, Brian Krueger, Tim Kuphal, Stanley Letovsky, Michael Levandoski, Craig Lukasik, Jonathan Meltzer, Brian Norvell, Mindy Nye, Scott Parker, Christos Petropoulos, John Pruitt, Steven Ragan, Scott Ryan, Mike Sapeta, Jana Schroth, Suresh Babu Selvaraju, Goran Stevovic, Amanda Suchanek, Andrea Throop, Lyndon Tilson, Thomas Urban, Joe Voshell, Kimberly Wagner, Jonathan Williams, Mary Williamson, Qian Zeng, Tricia Zwiefelhofer, Clinton R. Paden, Duncan MacCannell |
| EPI_ISL_3330011 | Illinois Department of Public Health - Springfield Lab | Illinois Department of Public Health - Springfield Lab | Bryan Sim, Gordon McCall, Marshall Smock |
| EPI_ISL_3330012 | Laboratory Corporation of America | Centers for Disease Control and Prevention Division of Viral Diseases, Pathogen Discovery | Dakota Howard, Dhwani Batra, Peter W. Cook, Kara Moser, Adrian Paskey, Jason Caravas, Benjamin Rambo-Martin, Shatavia Morrison, Christopher Gulvick, Scott Sammons, Yvette Unoarumhi, Darlene Wagner, Matthew Schmerer, Minoo Agarwal, Eyad Almasri, Debbie Boles, Ayla Burns, Nuthawin Charoensri, Oren Cohen, Susan Countryman, Mary Ann Cristobal, Bobbi Croy, Suzanne Dale, Hrushikesh Deshmukh, Amanda Douglas, Vincent Drouillon, Marcia Eisenberg, Howard Engler, Rama Ghatti, Prashant Gupta, Susan Hicks, Jake Humphrey, Lax Iyer, Lisa Pfefferle, Manoj Jain, Matthew Robinson, Mohan Kolli, Brian Krueger, Tim Kuphal, Stanley Letovsky, Michael Levandoski, Craig Lukasik, Jonathan Meltzer, Brian Norvell, Mindy Nye, Scott Parker, Christos Petropoulos, John Pruitt, Steven Ragan, Scott Ryan, Mike Sapeta, Jana Schroth, Suresh Babu Selvaraju, Goran Stevovic, Amanda Suchanek, Andrea Throop, Lyndon Tilson, Thomas Urban, Joe Voshell, Kimberly Wagner, Jonathan Williams, Mary Williamson, Qian Zeng, Tricia Zwiefelhofer, Clinton R. Paden, Duncan MacCannell |
| EPI_ISL_3330013, EPI_ISL_3330014, EPI_ISL_3330016, EPI_ISL_3330017 | Illinois Department of Public Health - Springfield Lab | Illinois Department of Public Health - Springfield Lab | Bryan Sim, Gordon McCall, Marshall Smock |
| EPI_ISL_3330018 | Laboratory Corporation of America | Centers for Disease Control and Prevention Division of Viral Diseases, Pathogen Discovery | Dakota Howard, Dhwani Batra, Peter W. Cook, Kara Moser, Adrian Paskey, Jason Caravas, Benjamin Rambo-Martin, Shatavia Morrison, Christopher Gulvick, Scott Sammons, Yvette Unoarumhi, Darlene Wagner, Matthew Schmerer, Minoo Agarwal, Eyad Almasri, Debbie Boles, Ayla Burns, Nuthawin Charoensri, Oren Cohen, Susan Countryman, Mary Ann Cristobal, Bobbi Croy, Suzanne Dale, Hrushikesh Deshmukh, Amanda Douglas, Vincent Drouillon, Marcia Eisenberg, Howard Engler, Rama Ghatti, Prashant Gupta, Susan Hicks, Jake Humphrey, Lax Iyer, Lisa Pfefferle, Manoj Jain, Matthew Robinson, Mohan Kolli, Brian Krueger, Tim Kuphal, Stanley Letovsky, Michael Levandoski, Craig Lukasik, Jonathan Meltzer, Brian Norvell, Mindy Nye, Scott Parker, Christos Petropoulos, John Pruitt, Steven Ragan, Scott Ryan, Mike Sapeta, Jana Schroth, Suresh Babu Selvaraju, Goran Stevovic, Amanda Suchanek, Andrea Throop, Lyndon Tilson, Thomas Urban, Joe Voshell, Kimberly Wagner, Jonathan Williams, Mary Williamson, Qian Zeng, Tricia Zwiefelhofer, Clinton R. Paden, Duncan MacCannell |
| EPI_ISL_3330019 | Illinois Department of Public Health - Springfield Lab | Illinois Department of Public Health - Springfield Lab | Bryan Sim, Gordon McCall, Marshall Smock |
| EPI_ISL_3330020 | Laboratory Corporation of America | Centers for Disease Control and Prevention Division of Viral Diseases, Pathogen Discovery | Dakota Howard, Dhwani Batra, Peter W. Cook, Kara Moser, Adrian Paskey, Jason Caravas, Benjamin Rambo-Martin, Shatavia Morrison, Christopher Gulvick, Scott Sammons, Yvette Unoarumhi, Darlene Wagner, Matthew Schmerer, Minoo Agarwal, Eyad Almasri, Debbie Boles, Ayla Burns, Nuthawin Charoensri, Oren Cohen, Susan Countryman, Mary Ann Cristobal, Bobbi Croy, Suzanne Dale, Hrushikesh Deshmukh, Amanda Douglas, Vincent Drouillon, Marcia Eisenberg, Howard Engler, Rama Ghatti, Prashant Gupta, Susan Hicks, Jake Humphrey, Lax Iyer, Lisa Pfefferle, Manoj Jain, Matthew Robinson, Mohan Kolli, Brian Krueger, Tim Kuphal, Stanley Letovsky, Michael Levandoski, Craig Lukasik, Jonathan Meltzer, Brian Norvell, Mindy Nye, Scott Parker, Christos Petropoulos, John Pruitt, Steven Ragan, Scott Ryan, Mike Sapeta, Jana Schroth, Suresh Babu Selvaraju, Goran Stevovic, Amanda Suchanek, Andrea Throop, Lyndon Tilson, Thomas Urban, Joe Voshell, Kimberly Wagner, Jonathan Williams, Mary Williamson, Qian Zeng, Tricia Zwiefelhofer, Clinton R. Paden, Duncan MacCannell |
| EPI_ISL_3330021, EPI_ISL_3330022 | Illinois Department of Public Health - Springfield Lab | Illinois Department of Public Health - Springfield Lab | Bryan Sim, Gordon McCall, Marshall Smock |
| EPI_ISL_3330023 | Laboratory Corporation of America | Centers for Disease Control and Prevention Division of Viral Diseases, Pathogen Discovery | Dakota Howard, Dhwani Batra, Peter W. Cook, Kara Moser, Adrian Paskey, Jason Caravas, Benjamin Rambo-Martin, Shatavia Morrison, Christopher Gulvick, Scott Sammons, Yvette Unoarumhi, Darlene Wagner, Matthew Schmerer, Minoo Agarwal, Eyad Almasri, Debbie Boles, Ayla Burns, Nuthawin Charoensri, Oren Cohen, Susan Countryman, Mary Ann Cristobal, Bobbi Croy, Suzanne Dale, Hrushikesh Deshmukh, Amanda Douglas, Vincent Drouillon, Marcia Eisenberg, Howard Engler, Rama Ghatti, Prashant Gupta, Susan Hicks, Jake Humphrey, Lax Iyer, Lisa Pfefferle, Manoj Jain, Matthew Robinson, Mohan Kolli, Brian Krueger, Tim Kuphal, Stanley Letovsky, Michael Levandoski, Craig Lukasik, Jonathan Meltzer, Brian Norvell, Mindy Nye, Scott Parker, Christos Petropoulos, John Pruitt, Steven Ragan, Scott Ryan, Mike Sapeta, Jana Schroth, Suresh Babu Selvaraju, Goran Stevovic, Amanda Suchanek, Andrea Throop, Lyndon Tilson, Thomas Urban, Joe Voshell, Kimberly Wagner, Jonathan Williams, Mary Williamson, Qian Zeng, Tricia Zwiefelhofer, Clinton R. Paden, Duncan MacCannell |
| EPI_ISL_3330024 | Illinois Department of Public Health - Springfield Lab | Illinois Department of Public Health - Springfield Lab | Bryan Sim, Gordon McCall, Marshall Smock |
| EPI_ISL_3330025 | Laboratory Corporation of America | Centers for Disease Control and Prevention Division of Viral Diseases, Pathogen Discovery | Dakota Howard, Dhwani Batra, Peter W. Cook, Kara Moser, Adrian Paskey, Jason Caravas, Benjamin Rambo-Martin, Shatavia Morrison, Christopher Gulvick, Scott Sammons, Yvette Unoarumhi, Darlene Wagner, Matthew Schmerer, Minoo Agarwal, Eyad Almasri, Debbie Boles, Ayla Burns, Nuthawin Charoensri, Oren Cohen, Susan Countryman, Mary Ann Cristobal, Bobbi Croy, Suzanne Dale, Hrushikesh Deshmukh, Amanda Douglas, Vincent Drouillon, Marcia Eisenberg, Howard Engler, Rama Ghatti, Prashant Gupta, Susan Hicks, Jake Humphrey, Lax Iyer, Lisa Pfefferle, Manoj Jain, Matthew Robinson, Mohan Kolli, Brian Krueger, Tim Kuphal, Stanley Letovsky, Michael Levandoski, Craig Lukasik, Jonathan Meltzer, Brian Norvell, Mindy Nye, Scott Parker, Christos Petropoulos, John Pruitt, Steven Ragan, Scott Ryan, Mike Sapeta, Jana Schroth, Suresh Babu Selvaraju, Goran Stevovic, Amanda Suchanek, Andrea Throop, Lyndon Tilson, Thomas Urban, Joe Voshell, Kimberly Wagner, Jonathan Williams, Mary Williamson, Qian Zeng, Tricia Zwiefelhofer, Clinton R. Paden, Duncan MacCannell |
| EPI_ISL_3330026, EPI_ISL_3330027 | Illinois Department of Public Health - Springfield Lab | Illinois Department of Public Health - Springfield Lab | Bryan Sim, Gordon McCall, Marshall Smock |
| EPI_ISL_3330028 | Laboratory Corporation of America | Centers for Disease Control and Prevention Division of Viral Diseases, Pathogen Discovery | Dakota Howard, Dhwani Batra, Peter W. Cook, Kara Moser, Adrian Paskey, Jason Caravas, Benjamin Rambo-Martin, Shatavia Morrison, Christopher Gulvick, Scott Sammons, Yvette Unoarumhi, Darlene Wagner, Matthew Schmerer, Minoo Agarwal, Eyad Almasri, Debbie Boles, Ayla Burns, Nuthawin Charoensri, Oren Cohen, Susan Countryman, Mary Ann Cristobal, Bobbi Croy, Suzanne Dale, Hrushikesh Deshmukh, Amanda Douglas, Vincent Drouillon, Marcia Eisenberg, Howard Engler, Rama Ghatti, Prashant Gupta, Susan Hicks, Jake Humphrey, Lax Iyer, Lisa Pfefferle, Manoj Jain, Matthew Robinson, Mohan Kolli, Brian Krueger, Tim Kuphal, Stanley Letovsky, Michael Levandoski, Craig Lukasik, Jonathan Meltzer, Brian Norvell, Mindy Nye, Scott Parker, Christos Petropoulos, John Pruitt, Steven Ragan, Scott Ryan, Mike Sapeta, Jana Schroth, Suresh Babu Selvaraju, Goran Stevovic, Amanda Suchanek, Andrea Throop, Lyndon Tilson, Thomas Urban, Joe Voshell, Kimberly Wagner, Jonathan Williams, Mary Williamson, Qian Zeng, Tricia Zwiefelhofer, Clinton R. Paden, Duncan MacCannell |
| EPI_ISL_3330029, EPI_ISL_3330030 | Illinois Department of Public Health - Springfield Lab | Illinois Department of Public Health - Springfield Lab | Bryan Sim, Gordon McCall, Marshall Smock |
| EPI_ISL_3330031 | Laboratory Corporation of America | Centers for Disease Control and Prevention Division of Viral Diseases, Pathogen Discovery | Dakota Howard, Dhwani Batra, Peter W. Cook, Kara Moser, Adrian Paskey, Jason Caravas, Benjamin Rambo-Martin, Shatavia Morrison, Christopher Gulvick, Scott Sammons, Yvette Unoarumhi, Darlene Wagner, Matthew Schmerer, Minoo Agarwal, Eyad Almasri, Debbie Boles, Ayla Burns, Nuthawin Charoensri, Oren Cohen, Susan Countryman, Mary Ann Cristobal, Bobbi Croy, Suzanne Dale, Hrushikesh Deshmukh, Amanda Douglas, Vincent Drouillon, Marcia Eisenberg, Howard Engler, Rama Ghatti, Prashant Gupta, Susan Hicks, Jake Humphrey, Lax Iyer, Lisa Pfefferle, Manoj Jain, Matthew Robinson, Mohan Kolli, Brian Krueger, Tim Kuphal, Stanley Letovsky, Michael Levandoski, Craig Lukasik, Jonathan Meltzer, Brian Norvell, Mindy Nye, Scott Parker, Christos Petropoulos, John Pruitt, Steven Ragan, Scott Ryan, Mike Sapeta, Jana Schroth, Suresh Babu Selvaraju, Goran Stevovic, Amanda Suchanek, Andrea Throop, Lyndon Tilson, Thomas Urban, Joe Voshell, Kimberly Wagner, Jonathan Williams, Mary Williamson, Qian Zeng, Tricia Zwiefelhofer, Clinton R. Paden, Duncan MacCannell |
| EPI_ISL_3330032, EPI_ISL_3330034, EPI_ISL_3330035, EPI_ISL_3330037 | Illinois Department of Public Health - Springfield Lab | Illinois Department of Public Health - Springfield Lab | Bryan Sim, Gordon McCall, Marshall Smock |
| EPI_ISL_3330038 | Laboratory Corporation of America | Centers for Disease Control and Prevention Division of Viral Diseases, Pathogen Discovery | Dakota Howard, Dhwani Batra, Peter W. Cook, Kara Moser, Adrian Paskey, Jason Caravas, Benjamin Rambo-Martin, Shatavia Morrison, Christopher Gulvick, Scott Sammons, Yvette Unoarumhi, Darlene Wagner, Matthew Schmerer, Minoo Agarwal, Eyad Almasri, Debbie Boles, Ayla Burns, Nuthawin Charoensri, Oren Cohen, Susan Countryman, Mary Ann Cristobal, Bobbi Croy, Suzanne Dale, Hrushikesh Deshmukh, Amanda Douglas, Vincent Drouillon, Marcia Eisenberg, Howard Engler, Rama Ghatti, Prashant Gupta, Susan Hicks, Jake Humphrey, Lax Iyer, Lisa Pfefferle, Manoj Jain, Matthew Robinson, Mohan Kolli, Brian Krueger, Tim Kuphal, Stanley Letovsky, Michael Levandoski, Craig Lukasik, Jonathan Meltzer, Brian Norvell, Mindy Nye, Scott Parker, Christos Petropoulos, John Pruitt, Steven Ragan, Scott Ryan, Mike Sapeta, Jana Schroth, Suresh Babu Selvaraju, Goran Stevovic, Amanda Suchanek, Andrea Throop, Lyndon Tilson, Thomas Urban, Joe Voshell, Kimberly Wagner, Jonathan Williams, Mary Williamson, Qian Zeng, Tricia Zwiefelhofer, Clinton R. Paden, Duncan MacCannell |
| EPI_ISL_3330039, EPI_ISL_3330040 | Illinois Department of Public Health - Springfield Lab | Illinois Department of Public Health - Springfield Lab | Bryan Sim, Gordon McCall, Marshall Smock |
| EPI_ISL_3330041, EPI_ISL_3330042, EPI_ISL_3330043 | Laboratory Corporation of America | Centers for Disease Control and Prevention Division of Viral Diseases, Pathogen Discovery | Dakota Howard, Dhwani Batra, Peter W. Cook, Kara Moser, Adrian Paskey, Jason Caravas, Benjamin Rambo-Martin, Shatavia Morrison, Christopher Gulvick, Scott Sammons, Yvette Unoarumhi, Darlene Wagner, Matthew Schmerer, Minoo Agarwal, Eyad Almasri, Debbie Boles, Ayla Burns, Nuthawin Charoensri, Oren Cohen, Susan Countryman, Mary Ann Cristobal, Bobbi Croy, Suzanne Dale, Hrushikesh Deshmukh, Amanda Douglas, Vincent Drouillon, Marcia Eisenberg, Howard Engler, Rama Ghatti, Prashant Gupta, Susan Hicks, Jake Humphrey, Lax Iyer, Lisa Pfefferle, Manoj Jain, Matthew Robinson, Mohan Kolli, Brian Krueger, Tim Kuphal, Stanley Letovsky, Michael Levandoski, Craig Lukasik, Jonathan Meltzer, Brian Norvell, Mindy Nye, Scott Parker, Christos Petropoulos, John Pruitt, Steven Ragan, Scott Ryan, Mike Sapeta, Jana Schroth, Suresh Babu Selvaraju, Goran Stevovic, Amanda Suchanek, Andrea Throop, Lyndon Tilson, Thomas Urban, Joe Voshell, Kimberly Wagner, Jonathan Williams, Mary Williamson, Qian Zeng, Tricia Zwiefelhofer, Clinton R. Paden, Duncan MacCannell |
| EPI_ISL_3330044 | Illinois Department of Public Health - Springfield Lab | Illinois Department of Public Health - Springfield Lab | Bryan Sim, Gordon McCall, Marshall Smock |
| EPI_ISL_3330045, EPI_ISL_3330046 | Laboratory Corporation of America | Centers for Disease Control and Prevention Division of Viral Diseases, Pathogen Discovery | Dakota Howard, Dhwani Batra, Peter W. Cook, Kara Moser, Adrian Paskey, Jason Caravas, Benjamin Rambo-Martin, Shatavia Morrison, Christopher Gulvick, Scott Sammons, Yvette Unoarumhi, Darlene Wagner, Matthew Schmerer, Minoo Agarwal, Eyad Almasri, Debbie Boles, Ayla Burns, Nuthawin Charoensri, Oren Cohen, Susan Countryman, Mary Ann Cristobal, Bobbi Croy, Suzanne Dale, Hrushikesh Deshmukh, Amanda Douglas, Vincent Drouillon, Marcia Eisenberg, Howard Engler, Rama Ghatti, Prashant Gupta, Susan Hicks, Jake Humphrey, Lax Iyer, Lisa Pfefferle, Manoj Jain, Matthew Robinson, Mohan Kolli, Brian Krueger, Tim Kuphal, Stanley Letovsky, Michael Levandoski, Craig Lukasik, Jonathan Meltzer, Brian Norvell, Mindy Nye, Scott Parker, Christos Petropoulos, John Pruitt, Steven Ragan, Scott Ryan, Mike Sapeta, Jana Schroth, Suresh Babu Selvaraju, Goran Stevovic, Amanda Suchanek, Andrea Throop, Lyndon Tilson, Thomas Urban, Joe Voshell, Kimberly Wagner, Jonathan Williams, Mary Williamson, Qian Zeng, Tricia Zwiefelhofer, Clinton R. Paden, Duncan MacCannell |
| EPI_ISL_3330047 | Illinois Department of Public Health - Springfield Lab | Illinois Department of Public Health - Springfield Lab | Bryan Sim, Gordon McCall, Marshall Smock |
| EPI_ISL_3330049 | Laboratory Corporation of America | Centers for Disease Control and Prevention Division of Viral Diseases, Pathogen Discovery | Dakota Howard, Dhwani Batra, Peter W. Cook, Kara Moser, Adrian Paskey, Jason Caravas, Benjamin Rambo-Martin, Shatavia Morrison, Christopher Gulvick, Scott Sammons, Yvette Unoarumhi, Darlene Wagner, Matthew Schmerer, Minoo Agarwal, Eyad Almasri, Debbie Boles, Ayla Burns, Nuthawin Charoensri, Oren Cohen, Susan Countryman, Mary Ann Cristobal, Bobbi Croy, Suzanne Dale, Hrushikesh Deshmukh, Amanda Douglas, Vincent Drouillon, Marcia Eisenberg, Howard Engler, Rama Ghatti, Prashant Gupta, Susan Hicks, Jake Humphrey, Lax Iyer, Lisa Pfefferle, Manoj Jain, Matthew Robinson, Mohan Kolli, Brian Krueger, Tim Kuphal, Stanley Letovsky, Michael Levandoski, Craig Lukasik, Jonathan Meltzer, Brian Norvell, Mindy Nye, Scott Parker, Christos Petropoulos, John Pruitt, Steven Ragan, Scott Ryan, Mike Sapeta, Jana Schroth, Suresh Babu Selvaraju, Goran Stevovic, Amanda Suchanek, Andrea Throop, Lyndon Tilson, Thomas Urban, Joe Voshell, Kimberly Wagner, Jonathan Williams, Mary Williamson, Qian Zeng, Tricia Zwiefelhofer, Clinton R. Paden, Duncan MacCannell |
| EPI_ISL_3330050 | Illinois Department of Public Health - Springfield Lab | Illinois Department of Public Health - Springfield Lab | Bryan Sim, Gordon McCall, Marshall Smock |
| EPI_ISL_3330051, EPI_ISL_3330052, EPI_ISL_3330053 | Laboratory Corporation of America | Centers for Disease Control and Prevention Division of Viral Diseases, Pathogen Discovery | Dakota Howard, Dhwani Batra, Peter W. Cook, Kara Moser, Adrian Paskey, Jason Caravas, Benjamin Rambo-Martin, Shatavia Morrison, Christopher Gulvick, Scott Sammons, Yvette Unoarumhi, Darlene Wagner, Matthew Schmerer, Minoo Agarwal, Eyad Almasri, Debbie Boles, Ayla Burns, Nuthawin Charoensri, Oren Cohen, Susan Countryman, Mary Ann Cristobal, Bobbi Croy, Suzanne Dale, Hrushikesh Deshmukh, Amanda Douglas, Vincent Drouillon, Marcia Eisenberg, Howard Engler, Rama Ghatti, Prashant Gupta, Susan Hicks, Jake Humphrey, Lax Iyer, Lisa Pfefferle, Manoj Jain, Matthew Robinson, Mohan Kolli, Brian Krueger, Tim Kuphal, Stanley Letovsky, Michael Levandoski, Craig Lukasik, Jonathan Meltzer, Brian Norvell, Mindy Nye, Scott Parker, Christos Petropoulos, John Pruitt, Steven Ragan, Scott Ryan, Mike Sapeta, Jana Schroth, Suresh Babu Selvaraju, Goran Stevovic, Amanda Suchanek, Andrea Throop, Lyndon Tilson, Thomas Urban, Joe Voshell, Kimberly Wagner, Jonathan Williams, Mary Williamson, Qian Zeng, Tricia Zwiefelhofer, Clinton R. Paden, Duncan MacCannell |
| EPI_ISL_3330054 | Illinois Department of Public Health - Springfield Lab | Illinois Department of Public Health - Springfield Lab | Bryan Sim, Gordon McCall, Marshall Smock |

| | | | |
|---|---|---|---|
| EPI_ISL_3330055, EPI_ISL_3330056 | Laboratory Corporation of America | Centers for Disease Control and Prevention Division of Viral Diseases, Pathogen Discovery | Dakota Howard, Dhwani Batra, Peter W. Cook, Kara Moser, Adrian Paskey, Jason Caravas, Benjamin Rambo-Martin, Shatavia Morrison, Christopher Gulvick, Scott Sammons, Yvette Unoarumhi, Darlene Wagner, Matthew Schmerer, Minoo Agarwal, Eyad Almasri, Debbie Boles, Ayla Burns, Nuthawin Charoensri, Oren Cohen, Susan Countryman, Mary Ann Cristobal, Bobbi Croy, Suzanne Dale, Hrushikesh Deshmukh, Amanda Douglas, Vincent Drouillon, Marcia Eisenberg, Howard Engler, Rama Ghatti, Prashant Gupta, Susan Hicks, Jake Humphrey, Lax Iyer, Lisa Pfefferle, Manoj Jain, Matthew Robinson, Mohan Kolli, Brian Krueger, Tim Kuphal, Stanley Letovsky, Michael Levandoski, Craig Lukasik, Jonathan Meltzer, Brian Norvell, Mindy Nye, Scott Parker, Christos Petropoulos, John Pruitt, Steven Ragan, Scott Ryan, Mike Sapeta, Jana Schroth, Suresh Babu Selvaraju, Goran Stevovic, Amanda Suchanek, Andrea Throop, Lyndon Tilson, Thomas Urban, Joe Voshell, Kimberly Wagner, Jonathan Williams, Qian Zeng, Tricia Zwiefelhofer, Clinton R. Paden, Duncan MacCannell |
| EPI_ISL_3330057 | Illinois Department of Public Health - Springfield Lab | Illinois Department of Public Health - Springfield Lab | Bryan Sim, Gordon McCall, Marshall Smock |
| EPI_ISL_3330059 | Laboratory Corporation of America | Centers for Disease Control and Prevention Division of Viral Diseases, Pathogen Discovery | Dakota Howard, Dhwani Batra, Peter W. Cook, Kara Moser, Adrian Paskey, Jason Caravas, Benjamin Rambo-Martin, Shatavia Morrison, Christopher Gulvick, Scott Sammons, Yvette Unoarumhi, Darlene Wagner, Matthew Schmerer, Minoo Agarwal, Eyad Almasri, Debbie Boles, Ayla Burns, Nuthawin Charoensri, Oren Cohen, Susan Countryman, Mary Ann Cristobal, Bobbi Croy, Suzanne Dale, Hrushikesh Deshmukh, Amanda Douglas, Vincent Drouillon, Marcia Eisenberg, Howard Engler, Rama Ghatti, Prashant Gupta, Susan Hicks, Jake Humphrey, Lax Iyer, Lisa Pfefferle, Manoj Jain, Matthew Robinson, Mohan Kolli, Brian Krueger, Tim Kuphal, Stanley Letovsky, Michael Levandoski, Craig Lukasik, Jonathan Meltzer, Brian Norvell, Mindy Nye, Scott Parker, Christos Petropoulos, John Pruitt, Steven Ragan, Scott Ryan, Mike Sapeta, Jana Schroth, Suresh Babu Selvaraju, Goran Stevovic, Amanda Suchanek, Andrea Throop, Lyndon Tilson, Thomas Urban, Joe Voshell, Kimberly Wagner, Jonathan Williams, Mary Williamson, Qian Zeng, Tricia Zwiefelhofer, Clinton R. Paden, Duncan MacCannell |
| EPI_ISL_3330060 | Illinois Department of Public Health - Springfield Lab | Illinois Department of Public Health - Springfield Lab | Bryan Sim, Gordon McCall, Marshall Smock |
| EPI_ISL_3330062 | Laboratory Corporation of America | Centers for Disease Control and Prevention Division of Viral Diseases, Pathogen Discovery | Dakota Howard, Dhwani Batra, Peter W. Cook, Kara Moser, Adrian Paskey, Jason Caravas, Benjamin Rambo-Martin, Shatavia Morrison, Christopher Gulvick, Scott Sammons, Yvette Unoarumhi, Darlene Wagner, Matthew Schmerer, Minoo Agarwal, Eyad Almasri, Debbie Boles, Ayla Burns, Nuthawin Charoensri, Oren Cohen, Susan Countryman, Mary Ann Cristobal, Bobbi Croy, Suzanne Dale, Hrushikesh Deshmukh, Amanda Douglas, Vincent Drouillon, Marcia Eisenberg, Howard Engler, Rama Ghatti, Prashant Gupta, Susan Hicks, Jake Humphrey, Lax Iyer, Lisa Pfefferle, Manoj Jain, Matthew Robinson, Mohan Kolli, Brian Krueger, Tim Kuphal, Stanley Letovsky, Michael Levandoski, Craig Lukasik, Jonathan Meltzer, Brian Norvell, Mindy Nye, Scott Parker, Christos Petropoulos, John Pruitt, Steven Ragan, Scott Ryan, Mike Sapeta, Jana Schroth, Suresh Babu Selvaraju, Goran Stevovic, Amanda Suchanek, Andrea Throop, Lyndon Tilson, Thomas Urban, Joe Voshell, Kimberly Wagner, Jonathan Williams, Mary Williamson, Qian Zeng, Tricia Zwiefelhofer, Clinton R. Paden, Duncan MacCannell |
| EPI_ISL_3330063 | Illinois Department of Public Health - Springfield Lab | Illinois Department of Public Health - Springfield Lab | Bryan Sim, Gordon McCall, Marshall Smock |
| EPI_ISL_3330064 | Laboratory Corporation of America | Centers for Disease Control and Prevention Division of Viral Diseases, Pathogen Discovery | Dakota Howard, Dhwani Batra, Peter W. Cook, Kara Moser, Adrian Paskey, Jason Caravas, Benjamin Rambo-Martin, Shatavia Morrison, Christopher Gulvick, Scott Sammons, Yvette Unoarumhi, Darlene Wagner, Matthew Schmerer, Minoo Agarwal, Eyad Almasri, Debbie Boles, Ayla Burns, Nuthawin Charoensri, Oren Cohen, Susan Countryman, Mary Ann Cristobal, Bobbi Croy, Suzanne Dale, Hrushikesh Deshmukh, Amanda Douglas, Vincent Drouillon, Marcia Eisenberg, Howard Engler, Rama Ghatti, Prashant Gupta, Susan Hicks, Jake Humphrey, Lax Iyer, Lisa Pfefferle, Manoj Jain, Matthew Robinson, Mohan Kolli, Brian Krueger, Tim Kuphal, Stanley Letovsky, Michael Levandoski, Craig Lukasik, Jonathan Meltzer, Brian Norvell, Mindy Nye, Scott Parker, Christos Petropoulos, John Pruitt, Steven Ragan, Scott Ryan, Mike Sapeta, Jana Schroth, Suresh Babu Selvaraju, Goran Stevovic, Amanda Suchanek, Andrea Throop, Lyndon Tilson, Thomas Urban, Joe Voshell, Kimberly Wagner, Jonathan Williams, Mary Williamson, Qian Zeng, Tricia Zwiefelhofer, Clinton R. Paden, Duncan MacCannell |
| EPI_ISL_3330065 | Illinois Department of Public Health - Springfield Lab | Illinois Department of Public Health - Springfield Lab | Bryan Sim, Gordon McCall, Marshall Smock |
| EPI_ISL_3330066 | Laboratory Corporation of America | Centers for Disease Control and Prevention Division of Viral Diseases, Pathogen Discovery | Dakota Howard, Dhwani Batra, Peter W. Cook, Kara Moser, Adrian Paskey, Jason Caravas, Benjamin Rambo-Martin, Shatavia Morrison, Christopher Gulvick, Scott Sammons, Yvette Unoarumhi, Darlene Wagner, Matthew Schmerer, Minoo Agarwal, Eyad Almasri, Debbie Boles, Ayla Burns, Nuthawin Charoensri, Oren Cohen, Susan Countryman, Mary Ann Cristobal, Bobbi Croy, Suzanne Dale, Hrushikesh Deshmukh, Amanda Douglas, Vincent Drouillon, Marcia Eisenberg, Howard Engler, Rama Ghatti, Prashant Gupta, Susan Hicks, Jake Humphrey, Lax Iyer, Lisa Pfefferle, Manoj Jain, Matthew Robinson, Mohan Kolli, Brian Krueger, Tim Kuphal, Stanley Letovsky, Michael Levandoski, Craig Lukasik, Jonathan Meltzer, Brian Norvell, Mindy Nye, Scott Parker, Christos Petropoulos, John Pruitt, Steven Ragan, Scott Ryan, Mike Sapeta, Jana Schroth, Suresh Babu Selvaraju, Goran Stevovic, Amanda Suchanek, Andrea Throop, Lyndon Tilson, Thomas Urban, Joe Voshell, Kimberly Wagner, Jonathan Williams, Mary Williamson, Qian Zeng, Tricia Zwiefelhofer, Clinton R. Paden, Duncan MacCannell |
| EPI_ISL_3330067, EPI_ISL_3330068 | Illinois Department of Public Health - Springfield Lab | Illinois Department of Public Health - Springfield Lab | Bryan Sim, Gordon McCall, Marshall Smock |
| EPI_ISL_3330069, EPI_ISL_3330070, EPI_ISL_3330071 | Laboratory Corporation of America | Centers for Disease Control and Prevention Division of Viral Diseases, Pathogen Discovery | Dakota Howard, Dhwani Batra, Peter W. Cook, Kara Moser, Adrian Paskey, Jason Caravas, Benjamin Rambo-Martin, Shatavia Morrison, Christopher Gulvick, Scott Sammons, Yvette Unoarumhi, Darlene Wagner, Matthew Schmerer, Minoo Agarwal, Eyad Almasri, Debbie Boles, Ayla Burns, Nuthawin Charoensri, Oren Cohen, Susan Countryman, Mary Ann Cristobal, Bobbi Croy, Suzanne Dale, Hrushikesh Deshmukh, Amanda Douglas, Vincent Drouillon, Marcia Eisenberg, Howard Engler, Rama Ghatti, Prashant Gupta, Susan Hicks, Jake Humphrey, Lax Iyer, Lisa Pfefferle, Manoj Jain, Matthew Robinson, Mohan Kolli, Brian Krueger, Tim Kuphal, Stanley Letovsky, Michael Levandoski, Craig Lukasik, Jonathan Meltzer, Brian Norvell, Mindy Nye, Scott Parker, Christos Petropoulos, John Pruitt, Steven Ragan, Scott Ryan, Mike Sapeta, Jana Schroth, Suresh Babu Selvaraju, Goran Stevovic, Amanda Suchanek, Andrea Throop, Lyndon Tilson, Thomas Urban, Joe Voshell, Kimberly Wagner, Jonathan Williams, Mary Williamson, Qian Zeng, Tricia Zwiefelhofer, Clinton R. Paden, Duncan MacCannell |
| EPI_ISL_3330072 | Illinois Department of Public Health - Springfield Lab | Illinois Department of Public Health - Springfield Lab | Bryan Sim, Gordon McCall, Marshall Smock |
| EPI_ISL_3330073, EPI_ISL_3330074 | Laboratory Corporation of America | Centers for Disease Control and Prevention Division of Viral Diseases, Pathogen Discovery | Dakota Howard, Dhwani Batra, Peter W. Cook, Kara Moser, Adrian Paskey, Jason Caravas, Benjamin Rambo-Martin, Shatavia Morrison, Christopher Gulvick, Scott Sammons, Yvette Unoarumhi, Darlene Wagner, Matthew Schmerer, Minoo Agarwal, Eyad Almasri, Debbie Boles, Ayla Burns, Nuthawin Charoensri, Oren Cohen, Susan Countryman, Mary Ann Cristobal, Bobbi Croy, Suzanne Dale, Hrushikesh Deshmukh, Amanda Douglas, Vincent Drouillon, Marcia Eisenberg, Howard Engler, Rama Ghatti, Prashant Gupta, Susan Hicks, Jake Humphrey, Lax Iyer, Lisa Pfefferle, Manoj Jain, Matthew Robinson, Mohan Kolli, Brian Krueger, Tim Kuphal, Stanley Letovsky, Michael Levandoski, Craig Lukasik, Jonathan Meltzer, Brian Norvell, Mindy Nye, Scott Parker, Christos Petropoulos, John Pruitt, Steven Ragan, Scott Ryan, Mike Sapeta, Jana Schroth, Suresh Babu Selvaraju, Goran Stevovic, Amanda Suchanek, Andrea Throop, Lyndon Tilson, Thomas Urban, Joe Voshell, Kimberly Wagner, Jonathan Williams, Mary Williamson, Qian Zeng, Tricia Zwiefelhofer, Clinton R. Paden, Duncan MacCannell |
| EPI_ISL_3330075 | Illinois Department of Public Health - Springfield Lab | Illinois Department of Public Health - Springfield Lab | Bryan Sim, Gordon McCall, Marshall Smock |
| EPI_ISL_3330076, EPI_ISL_3330077, EPI_ISL_3330078 | Laboratory Corporation of America | Centers for Disease Control and Prevention Division of Viral Diseases, Pathogen Discovery | Dakota Howard, Dhwani Batra, Peter W. Cook, Kara Moser, Adrian Paskey, Jason Caravas, Benjamin Rambo-Martin, Shatavia Morrison, Christopher Gulvick, Scott Sammons, Yvette Unoarumhi, Darlene Wagner, Matthew Schmerer, Minoo Agarwal, Eyad Almasri, Debbie Boles, Ayla Burns, Nuthawin Charoensri, Oren Cohen, Susan Countryman, Mary Ann Cristobal, Bobbi Croy, Suzanne Dale, Hrushikesh Deshmukh, Amanda Douglas, Vincent Drouillon, Marcia Eisenberg, Howard Engler, Rama Ghatti, Prashant Gupta, Susan Hicks, Jake Humphrey, Lax Iyer, Lisa Pfefferle, Manoj Jain, Matthew Robinson, Mohan Kolli, Brian Krueger, Tim Kuphal, Stanley Letovsky, Michael Levandoski, Craig Lukasik, Jonathan Meltzer, Brian Norvell, Mindy Nye, Scott Parker, Christos Petropoulos, John Pruitt, Steven Ragan, Scott Ryan, Mike Sapeta, Jana Schroth, Suresh Babu Selvaraju, Goran Stevovic, Amanda Suchanek, Andrea Throop, Lyndon Tilson, Thomas Urban, Joe Voshell, Kimberly Wagner, Jonathan Williams, Mary Williamson, Qian Zeng, Tricia Zwiefelhofer, Clinton R. Paden, Duncan MacCannell |
| EPI_ISL_3330079 | Illinois Department of Public Health - Springfield Lab | Illinois Department of Public Health - Springfield Lab | Bryan Sim, Gordon McCall, Marshall Smock |
| EPI_ISL_3330080, EPI_ISL_3330081 | Laboratory Corporation of America | Centers for Disease Control and Prevention Division of Viral Diseases, Pathogen Discovery | Dakota Howard, Dhwani Batra, Peter W. Cook, Kara Moser, Adrian Paskey, Jason Caravas, Benjamin Rambo-Martin, Shatavia Morrison, Christopher Gulvick, Scott Sammons, Yvette Unoarumhi, Darlene Wagner, Matthew Schmerer, Minoo Agarwal, Eyad Almasri, Debbie Boles, Ayla Burns, Nuthawin Charoensri, Oren Cohen, Susan Countryman, Mary Ann Cristobal, Bobbi Croy, Suzanne Dale, Hrushikesh Deshmukh, Amanda Douglas, Vincent Drouillon, Marcia Eisenberg, Howard Engler, Rama Ghatti, Prashant Gupta, Susan Hicks, Jake Humphrey, Lax Iyer, Lisa Pfefferle, Manoj Jain, Matthew Robinson, Mohan Kolli, Brian Krueger, Tim Kuphal, Stanley Letovsky, Michael Levandoski, Craig Lukasik, Jonathan Meltzer, Brian Norvell, Mindy Nye, Scott Parker, Christos Petropoulos, John Pruitt, Steven Ragan, Scott Ryan, Mike Sapeta, Jana Schroth, Suresh Babu Selvaraju, Goran Stevovic, Amanda Suchanek, Andrea Throop, Lyndon Tilson, Thomas Urban, Joe Voshell, Kimberly Wagner, Jonathan Williams, Mary Williamson, Qian Zeng, Tricia Zwiefelhofer, Clinton R. Paden, Duncan MacCannell |
| EPI_ISL_3330082 | Illinois Department of Public Health - Springfield Lab | Illinois Department of Public Health - Springfield Lab | Bryan Sim, Gordon McCall, Marshall Smock |
| EPI_ISL_3330083, EPI_ISL_3330084, EPI_ISL_3330085 | Laboratory Corporation of America | Centers for Disease Control and Prevention Division of Viral Diseases, Pathogen Discovery | Dakota Howard, Dhwani Batra, Peter W. Cook, Kara Moser, Adrian Paskey, Jason Caravas, Benjamin Rambo-Martin, Shatavia Morrison, Christopher Gulvick, Scott Sammons, Yvette Unoarumhi, Darlene Wagner, Matthew Schmerer, Minoo Agarwal, Eyad Almasri, Debbie Boles, Ayla Burns, Nuthawin Charoensri, Oren Cohen, Susan Countryman, Mary Ann Cristobal, Bobbi Croy, Suzanne Dale, Hrushikesh Deshmukh, Amanda Douglas, Vincent Drouillon, Marcia Eisenberg, Howard Engler, Rama Ghatti, Prashant Gupta, Susan Hicks, Jake Humphrey, Lax Iyer, Lisa Pfefferle, Manoj Jain, Matthew Robinson, Mohan Kolli, Brian Krueger, Tim Kuphal, Stanley Letovsky, Michael Levandoski, Craig Lukasik, Jonathan Meltzer, Brian Norvell, Mindy Nye, Scott Parker, Christos Petropoulos, John Pruitt, Steven Ragan, Scott Ryan, Mike Sapeta, Jana Schroth, Suresh Babu Selvaraju, Goran Stevovic, Amanda Suchanek, Andrea Throop, Lyndon Tilson, Thomas Urban, Joe Voshell, Kimberly Wagner, Jonathan Williams, Mary Williamson, Qian Zeng, Tricia Zwiefelhofer, Clinton R. Paden, Duncan MacCannell |
| EPI_ISL_3330086 | Illinois Department of Public Health - Springfield Lab | Illinois Department of Public Health - Springfield Lab | Bryan Sim, Gordon McCall, Marshall Smock |
| EPI_ISL_3330087, EPI_ISL_3330088 | Laboratory Corporation of America | Centers for Disease Control and Prevention Division of Viral Diseases, Pathogen Discovery | Dakota Howard, Dhwani Batra, Peter W. Cook, Kara Moser, Adrian Paskey, Jason Caravas, Benjamin Rambo-Martin, Shatavia Morrison, Christopher Gulvick, Scott Sammons, Yvette Unoarumhi, Darlene Wagner, Matthew Schmerer, Minoo Agarwal, Eyad Almasri, Debbie Boles, Ayla Burns, Nuthawin Charoensri, Oren Cohen, Susan Countryman, Mary Ann Cristobal, Bobbi Croy, Suzanne Dale, Hrushikesh Deshmukh, Amanda Douglas, Vincent Drouillon, Marcia Eisenberg, Howard Engler, Rama Ghatti, Prashant Gupta, Susan Hicks, Jake Humphrey, Lax Iyer, Lisa Pfefferle, Manoj Jain, Matthew Robinson, Mohan Kolli, Brian Krueger, Tim Kuphal, Stanley Letovsky, Michael Levandoski, Craig Lukasik, Jonathan Meltzer, Brian Norvell, Mindy Nye, Scott Parker, Christos Petropoulos, John Pruitt, Steven Ragan, Scott Ryan, Mike Sapeta, Jana Schroth, Suresh Babu Selvaraju, Goran Stevovic, Amanda Suchanek, Andrea Throop, Lyndon Tilson, Thomas Urban, Joe Voshell, Kimberly Wagner, Jonathan Williams, Mary Williamson, Qian Zeng, Tricia Zwiefelhofer, Clinton R. Paden, Duncan MacCannell |
| EPI_ISL_3330089 | Illinois Department of Public Health - Springfield Lab | Illinois Department of Public Health - Springfield Lab | Bryan Sim, Gordon McCall, Marshall Smock |
| EPI_ISL_3330090, EPI_ISL_3330091, EPI_ISL_3330092 | Laboratory Corporation of America | Centers for Disease Control and Prevention Division of Viral Diseases, Pathogen Discovery | Dakota Howard, Dhwani Batra, Peter W. Cook, Kara Moser, Adrian Paskey, Jason Caravas, Benjamin Rambo-Martin, Shatavia Morrison, Christopher Gulvick, Scott Sammons, Yvette Unoarumhi, Darlene Wagner, Matthew Schmerer, Minoo Agarwal, Eyad Almasri, Debbie Boles, Ayla Burns, Nuthawin Charoensri, Oren Cohen, Susan Countryman, Mary Ann Cristobal, Bobbi Croy, Suzanne Dale, Hrushikesh Deshmukh, Amanda Douglas, Vincent Drouillon, Marcia Eisenberg, Howard Engler, Rama Ghatti, Prashant Gupta, Susan Hicks, Jake Humphrey, Lax Iyer, Lisa Pfefferle, Manoj Jain, Matthew Robinson, Mohan Kolli, Brian Krueger, Tim Kuphal, Stanley Letovsky, Michael Levandoski, Craig Lukasik, Jonathan Meltzer, Brian Norvell, Mindy Nye, Scott Parker, Christos Petropoulos, John Pruitt, Steven Ragan, Scott Ryan, Mike Sapeta, Jana Schroth, Suresh Babu Selvaraju, Goran Stevovic, Amanda Suchanek, Andrea Throop, Lyndon Tilson, Thomas Urban, Joe Voshell, Kimberly Wagner, Jonathan Williams, Mary Williamson, Qian Zeng, Tricia Zwiefelhofer, Clinton R. Paden, Duncan MacCannell |
| EPI_ISL_3330093 | Illinois Department of Public Health - Springfield Lab | Illinois Department of Public Health - Springfield Lab | Bryan Sim, Gordon McCall, Marshall Smock |
| EPI_ISL_3330094, EPI_ISL_3330095 | Laboratory Corporation of America | Centers for Disease Control and Prevention Division of Viral Diseases, Pathogen Discovery | Dakota Howard, Dhwani Batra, Peter W. Cook, Kara Moser, Adrian Paskey, Jason Caravas, Benjamin Rambo-Martin, Shatavia Morrison, Christopher Gulvick, Scott Sammons, Yvette Unoarumhi, Darlene Wagner, Matthew Schmerer, Minoo Agarwal, Eyad Almasri, Debbie Boles, Ayla Burns, Nuthawin Charoensri, Oren Cohen, Susan Countryman, Mary Ann Cristobal, Bobbi Croy, Suzanne Dale, Hrushikesh Deshmukh, Amanda Douglas, Vincent Drouillon, Marcia Eisenberg, Howard Engler, Rama Ghatti, Prashant Gupta, Susan Hicks, Jake Humphrey, Lax Iyer, Lisa Pfefferle, Manoj Jain, Matthew Robinson, Mohan Kolli, Brian Krueger, Tim Kuphal, Stanley Letovsky, Michael Levandoski, Craig Lukasik, Jonathan Meltzer, Brian Norvell, Mindy Nye, Scott Parker, Christos Petropoulos, John Pruitt, Steven Ragan, Scott Ryan, Mike Sapeta, Jana Schroth, Suresh Babu Selvaraju, Goran Stevovic, Amanda Suchanek, Andrea Throop, Lyndon Tilson, Thomas Urban, Joe Voshell, Kimberly Wagner, Jonathan Williams, Mary Williamson, Qian Zeng, Tricia Zwiefelhofer, Clinton R. Paden, Duncan MacCannell |
| EPI_ISL_3330096 | Illinois Department of Public Health - Springfield Lab | Illinois Department of Public Health - Springfield Lab | Bryan Sim, Gordon McCall, Marshall Smock |
| EPI_ISL_3330097, EPI_ISL_3330098, EPI_ISL_3330099 | Laboratory Corporation of America | Centers for Disease Control and Prevention Division of Viral Diseases, Pathogen Discovery | Dakota Howard, Dhwani Batra, Peter W. Cook, Kara Moser, Adrian Paskey, Jason Caravas, Benjamin Rambo-Martin, Shatavia Morrison, Christopher Gulvick, Scott Sammons, Yvette Unoarumhi, Darlene Wagner, Matthew Schmerer, Minoo Agarwal, Eyad Almasri, Debbie Boles, Ayla Burns, Nuthawin Charoensri, Oren Cohen, Susan Countryman, Mary Ann Cristobal, Bobbi Croy, Suzanne Dale, Hrushikesh Deshmukh, Amanda Douglas, Vincent Drouillon, Marcia Eisenberg, Howard Engler, Rama Ghatti, Prashant Gupta, Susan Hicks, Jake Humphrey, Lax Iyer, Lisa Pfefferle, Manoj Jain, Matthew Robinson, Mohan Kolli, Brian Krueger, Tim Kuphal, Stanley Letovsky, Michael Levandoski, Craig Lukasik, Jonathan Meltzer, Brian Norvell, Mindy Nye, Scott Parker, Christos Petropoulos, John Pruitt, Steven Ragan, Scott Ryan, Mike Sapeta, Jana Schroth, Suresh Babu Selvaraju, Goran Stevovic, Amanda Suchanek, Andrea Throop, Lyndon Tilson, Thomas Urban, Joe Voshell, Kimberly Wagner, Jonathan Williams, Mary Williamson, Qian Zeng, Tricia Zwiefelhofer, Clinton R. Paden, Duncan MacCannell |
| EPI_ISL_3330100 | Illinois Department of Public Health - Springfield Lab | Illinois Department of Public Health - Springfield Lab | Bryan Sim, Gordon McCall, Marshall Smock |
| EPI_ISL_3330101, EPI_ISL_3330102 | Laboratory Corporation of America | Centers for Disease Control and Prevention Division of Viral Diseases, Pathogen Discovery | Dakota Howard, Dhwani Batra, Peter W. Cook, Kara Moser, Adrian Paskey, Jason Caravas, Benjamin Rambo-Martin, Shatavia Morrison, Christopher Gulvick, Scott Sammons, Yvette Unoarumhi, Darlene Wagner, Matthew Schmerer, Minoo Agarwal, Eyad Almasri, Debbie Boles, Ayla Burns, Nuthawin Charoensri, Oren Cohen, Susan Countryman, Mary Ann Cristobal, Bobbi Croy, Suzanne Dale, Hrushikesh Deshmukh, Amanda Douglas, Vincent Drouillon, Marcia Eisenberg, Howard Engler, Rama Ghatti, Prashant Gupta, Susan Hicks, Jake Humphrey, Lax Iyer, Lisa Pfefferle, Manoj Jain, Matthew Robinson, Mohan Kolli, Brian Krueger, Tim Kuphal, Stanley Letovsky, Michael Levandoski, Craig Lukasik, Jonathan Meltzer, Brian Norvell, Mindy Nye, Scott Parker, Christos Petropoulos, John Pruitt, Steven Ragan, Scott Ryan, Mike Sapeta, Jana Schroth, Suresh Babu Selvaraju, Goran Stevovic, Amanda Suchanek, Andrea Throop, Lyndon Tilson, Thomas Urban, Joe Voshell, Kimberly Wagner, Jonathan Williams, Mary Williamson, Qian Zeng, Tricia Zwiefelhofer, Clinton R. Paden, Duncan MacCannell |
| EPI_ISL_3330104 | Illinois Department of Public Health - Springfield Lab | Illinois Department of Public Health - Springfield Lab | Bryan Sim, Gordon McCall, Marshall Smock |
| EPI_ISL_3330106 | Laboratory Corporation of America | Centers for Disease Control and Prevention Division of Viral Diseases, Pathogen Discovery | Dakota Howard, Dhwani Batra, Peter W. Cook, Kara Moser, Adrian Paskey, Jason Caravas, Benjamin Rambo-Martin, Shatavia Morrison, Christopher Gulvick, Scott Sammons, Yvette Unoarumhi, Darlene Wagner, Matthew Schmerer, Minoo Agarwal, Eyad Almasri, Debbie Boles, Ayla Burns, Nuthawin Charoensri, Oren Cohen, Susan Countryman, Mary Ann Cristobal, Bobbi Croy, Suzanne Dale, Hrushikesh Deshmukh, Amanda Douglas, Vincent Drouillon, Marcia Eisenberg, Howard Engler, Rama Ghatti, Prashant Gupta, Susan Hicks, Jake Humphrey, Lax Iyer, Lisa Pfefferle, Manoj Jain, Matthew Robinson, Mohan Kolli, Brian Krueger, Tim Kuphal, Stanley Letovsky, Michael Levandoski, Craig Lukasik, Jonathan Meltzer, Brian Norvell, Mindy Nye, Scott Parker, Christos Petropoulos, John Pruitt, Steven Ragan, Scott Ryan, Mike Sapeta, Jana Schroth, Suresh Babu Selvaraju, Goran Stevovic, Amanda Suchanek, Andrea Throop, Lyndon Tilson, Thomas Urban, Joe Voshell, Kimberly Wagner, Jonathan Williams, Mary Williamson, Qian Zeng, Tricia Zwiefelhofer, Clinton R. Paden, Duncan MacCannell |
| EPI_ISL_3330107 | Illinois Department of Public Health - Springfield Lab | Illinois Department of Public Health - Springfield Lab | Bryan Sim, Gordon McCall, Marshall Smock |

| | | | |
|---|---|---|---|
| EPI_ISL_3330108 | Laboratory Corporation of America | Centers for Disease Control and Prevention Division of Viral Diseases, Pathogen Discovery | Dakota Howard, Dhwani Batra, Peter W. Cook, Kara Moser, Adrian Paskey, Jason Caravas, Benjamin Rambo-Martin, Shatavia Morrison, Christopher Gulvick, Scott Sammons, Yvette Unoarumhi, Darlene Wagner, Matthew Schmerer, Minoo Agarwal, Eyad Almasri, Debbie Boles, Ayla Burns, Nuthawin Charoensri, Oren Cohen, Susan Countryman, Mary Ann Cristobal, Bobbi Croy, Suzanne Dale, Hrushikesh Deshmukh, Amanda Douglas, Vincent Drouillon, Marcia Eisenberg, Howard Engler, Rama Ghatti, Prashant Gupta, Susan Hicks, Jake Humphrey, Lax Iyer, Lisa Pfefferle, Manoj Jain, Matthew Robinson, Mohan Kolli, Brian Krueger, Tim Kuphal, Stanley Letovsky, Michael Levandoski, Craig Lukasik, Jonathan Meltzer, Brian Norvell, Mindy Nye, Scott Parker, Christos Petropoulos, John Pruitt, Steven Ragan, Scott Ryan, Mike Sapeta, Jana Schroth, Suresh Babu Selvaraju, Goran Stevovic, Amanda Suchanek, Andrea Throop, Lyndon Tilson, Thomas Urban, Joe Voshell, Kimberly Wagner, Jonathan Williams, Mary Williamson, Qian Zeng, Tricia Zwiefelhofer, Clinton R. Paden, Duncan MacCannell |
| EPI_ISL_3330110 | Illinois Department of Public Health - Springfield Lab | Illinois Department of Public Health - Springfield Lab | Bryan Sim, Gordon McCall, Marshall Smock |
| EPI_ISL_3330111, EPI_ISL_3330112 | Laboratory Corporation of America | Centers for Disease Control and Prevention Division of Viral Diseases, Pathogen Discovery | Dakota Howard, Dhwani Batra, Peter W. Cook, Kara Moser, Adrian Paskey, Jason Caravas, Benjamin Rambo-Martin, Shatavia Morrison, Christopher Gulvick, Scott Sammons, Yvette Unoarumhi, Darlene Wagner, Matthew Schmerer, Minoo Agarwal, Eyad Almasri, Debbie Boles, Ayla Burns, Nuthawin Charoensri, Oren Cohen, Susan Countryman, Mary Ann Cristobal, Bobbi Croy, Suzanne Dale, Hrushikesh Deshmukh, Amanda Douglas, Vincent Drouillon, Marcia Eisenberg, Howard Engler, Rama Ghatti, Prashant Gupta, Susan Hicks, Jake Humphrey, Lax Iyer, Lisa Pfefferle, Manoj Jain, Matthew Robinson, Mohan Kolli, Brian Krueger, Tim Kuphal, Stanley Letovsky, Michael Levandoski, Craig Lukasik, Jonathan Meltzer, Brian Norvell, Mindy Nye, Scott Parker, Christos Petropoulos, John Pruitt, Steven Ragan, Scott Ryan, Mike Sapeta, Jana Schroth, Suresh Babu Selvaraju, Goran Stevovic, Amanda Suchanek, Andrea Throop, Lyndon Tilson, Thomas Urban, Joe Voshell, Kimberly Wagner, Jonathan Williams, Mary Williamson, Qian Zeng, Tricia Zwiefelhofer, Clinton R. Paden, Duncan MacCannell |
| EPI_ISL_3330113 | Illinois Department of Public Health - Springfield Lab | Illinois Department of Public Health - Springfield Lab | Bryan Sim, Gordon McCall, Marshall Smock |
| EPI_ISL_3330114, EPI_ISL_3330115, EPI_ISL_3330116 | Laboratory Corporation of America | Centers for Disease Control and Prevention Division of Viral Diseases, Pathogen Discovery | Dakota Howard, Dhwani Batra, Peter W. Cook, Kara Moser, Adrian Paskey, Jason Caravas, Benjamin Rambo-Martin, Shatavia Morrison, Christopher Gulvick, Scott Sammons, Yvette Unoarumhi, Darlene Wagner, Matthew Schmerer, Minoo Agarwal, Eyad Almasri, Debbie Boles, Ayla Burns, Nuthawin Charoensri, Oren Cohen, Susan Countryman, Mary Ann Cristobal, Bobbi Croy, Suzanne Dale, Hrushikesh Deshmukh, Amanda Douglas, Vincent Drouillon, Marcia Eisenberg, Howard Engler, Rama Ghatti, Prashant Gupta, Susan Hicks, Jake Humphrey, Lax Iyer, Lisa Pfefferle, Manoj Jain, Matthew Robinson, Mohan Kolli, Brian Krueger, Tim Kuphal, Stanley Letovsky, Michael Levandoski, Craig Lukasik, Jonathan Meltzer, Brian Norvell, Mindy Nye, Scott Parker, Christos Petropoulos, John Pruitt, Steven Ragan, Scott Ryan, Mike Sapeta, Jana Schroth, Suresh Babu Selvaraju, Goran Stevovic, Amanda Suchanek, Andrea Throop, Lyndon Tilson, Thomas Urban, Joe Voshell, Kimberly Wagner, Jonathan Williams, Mary Williamson, Qian Zeng, Tricia Zwiefelhofer, Clinton R. Paden, Duncan MacCannell |
| EPI_ISL_3330117, EPI_ISL_3330118 | Illinois Department of Public Health - Springfield Lab | Illinois Department of Public Health - Springfield Lab | Bryan Sim, Gordon McCall, Marshall Smock |
| EPI_ISL_3330119 | Laboratory Corporation of America | Centers for Disease Control and Prevention Division of Viral Diseases, Pathogen Discovery | Dakota Howard, Dhwani Batra, Peter W. Cook, Kara Moser, Adrian Paskey, Jason Caravas, Benjamin Rambo-Martin, Shatavia Morrison, Christopher Gulvick, Scott Sammons, Yvette Unoarumhi, Darlene Wagner, Matthew Schmerer, Minoo Agarwal, Eyad Almasri, Debbie Boles, Ayla Burns, Nuthawin Charoensri, Oren Cohen, Susan Countryman, Mary Ann Cristobal, Bobbi Croy, Suzanne Dale, Hrushikesh Deshmukh, Amanda Douglas, Vincent Drouillon, Marcia Eisenberg, Howard Engler, Rama Ghatti, Prashant Gupta, Susan Hicks, Jake Humphrey, Lax Iyer, Lisa Pfefferle, Manoj Jain, Matthew Robinson, Mohan Kolli, Brian Krueger, Tim Kuphal, Stanley Letovsky, Michael Levandoski, Craig Lukasik, Jonathan Meltzer, Brian Norvell, Mindy Nye, Scott Parker, Christos Petropoulos, John Pruitt, Steven Ragan, Scott Ryan, Mike Sapeta, Jana Schroth, Suresh Babu Selvaraju, Goran Stevovic, Amanda Suchanek, Andrea Throop, Lyndon Tilson, Thomas Urban, Joe Voshell, Kimberly Wagner, Jonathan Williams, Mary Williamson, Qian Zeng, Tricia Zwiefelhofer, Clinton R. Paden, Duncan MacCannell |
| EPI_ISL_3330120, EPI_ISL_3330121 | Illinois Department of Public Health - Springfield Lab | Illinois Department of Public Health - Springfield Lab | Bryan Sim, Gordon McCall, Marshall Smock |
| EPI_ISL_3330122 | Laboratory Corporation of America | Centers for Disease Control and Prevention Division of Viral Diseases, Pathogen Discovery | Dakota Howard, Dhwani Batra, Peter W. Cook, Kara Moser, Adrian Paskey, Jason Caravas, Benjamin Rambo-Martin, Shatavia Morrison, Christopher Gulvick, Scott Sammons, Yvette Unoarumhi, Darlene Wagner, Matthew Schmerer, Minoo Agarwal, Eyad Almasri, Debbie Boles, Ayla Burns, Nuthawin Charoensri, Oren Cohen, Susan Countryman, Mary Ann Cristobal, Bobbi Croy, Suzanne Dale, Hrushikesh Deshmukh, Amanda Douglas, Vincent Drouillon, Marcia Eisenberg, Howard Engler, Rama Ghatti, Prashant Gupta, Susan Hicks, Jake Humphrey, Lax Iyer, Lisa Pfefferle, Manoj Jain, Matthew Robinson, Mohan Kolli, Brian Krueger, Tim Kuphal, Stanley Letovsky, Michael Levandoski, Craig Lukasik, Jonathan Meltzer, Brian Norvell, Mindy Nye, Scott Parker, Christos Petropoulos, John Pruitt, Steven Ragan, Scott Ryan, Mike Sapeta, Jana Schroth, Suresh Babu Selvaraju, Goran Stevovic, Amanda Suchanek, Andrea Throop, Lyndon Tilson, Thomas Urban, Joe Voshell, Kimberly Wagner, Jonathan Williams, Mary Williamson, Qian Zeng, Tricia Zwiefelhofer, Clinton R. Paden, Duncan MacCannell |
| EPI_ISL_3330123 | Illinois Department of Public Health - Springfield Lab | Illinois Department of Public Health - Springfield Lab | Bryan Sim, Gordon McCall, Marshall Smock |
| EPI_ISL_3330124, EPI_ISL_3330125, EPI_ISL_3330126 | Laboratory Corporation of America | Centers for Disease Control and Prevention Division of Viral Diseases, Pathogen Discovery | Dakota Howard, Dhwani Batra, Peter W. Cook, Kara Moser, Adrian Paskey, Jason Caravas, Benjamin Rambo-Martin, Shatavia Morrison, Christopher Gulvick, Scott Sammons, Yvette Unoarumhi, Darlene Wagner, Matthew Schmerer, Minoo Agarwal, Eyad Almasri, Debbie Boles, Ayla Burns, Nuthawin Charoensri, Oren Cohen, Susan Countryman, Mary Ann Cristobal, Bobbi Croy, Suzanne Dale, Hrushikesh Deshmukh, Amanda Douglas, Vincent Drouillon, Marcia Eisenberg, Howard Engler, Rama Ghatti, Prashant Gupta, Susan Hicks, Jake Humphrey, Lax Iyer, Lisa Pfefferle, Manoj Jain, Matthew Robinson, Mohan Kolli, Brian Krueger, Tim Kuphal, Stanley Letovsky, Michael Levandoski, Craig Lukasik, Jonathan Meltzer, Brian Norvell, Mindy Nye, Scott Parker, Christos Petropoulos, John Pruitt, Steven Ragan, Scott Ryan, Mike Sapeta, Jana Schroth, Suresh Babu Selvaraju, Goran Stevovic, Amanda Suchanek, Andrea Throop, Lyndon Tilson, Thomas Urban, Joe Voshell, Kimberly Wagner, Jonathan Williams, Mary Williamson, Qian Zeng, Tricia Zwiefelhofer, Clinton R. Paden, Duncan MacCannell |
| EPI_ISL_3330127 | Illinois Department of Public Health - Springfield Lab | Illinois Department of Public Health - Springfield Lab | Bryan Sim, Gordon McCall, Marshall Smock |
| EPI_ISL_3330128, EPI_ISL_3330129 | Laboratory Corporation of America | Centers for Disease Control and Prevention Division of Viral Diseases, Pathogen Discovery | Dakota Howard, Dhwani Batra, Peter W. Cook, Kara Moser, Adrian Paskey, Jason Caravas, Benjamin Rambo-Martin, Shatavia Morrison, Christopher Gulvick, Scott Sammons, Yvette Unoarumhi, Darlene Wagner, Matthew Schmerer, Minoo Agarwal, Eyad Almasri, Debbie Boles, Ayla Burns, Nuthawin Charoensri, Oren Cohen, Susan Countryman, Mary Ann Cristobal, Bobbi Croy, Suzanne Dale, Hrushikesh Deshmukh, Amanda Douglas, Vincent Drouillon, Marcia Eisenberg, Howard Engler, Rama Ghatti, Prashant Gupta, Susan Hicks, Jake Humphrey, Lax Iyer, Lisa Pfefferle, Manoj Jain, Matthew Robinson, Mohan Kolli, Brian Krueger, Tim Kuphal, Stanley Letovsky, Michael Levandoski, Craig Lukasik, Jonathan Meltzer, Brian Norvell, Mindy Nye, Scott Parker, Christos Petropoulos, John Pruitt, Steven Ragan, Scott Ryan, Mike Sapeta, Jana Schroth, Suresh Babu Selvaraju, Goran Stevovic, Amanda Suchanek, Andrea Throop, Lyndon Tilson, Thomas Urban, Joe Voshell, Kimberly Wagner, Jonathan Williams, Mary Williamson, Qian Zeng, Tricia Zwiefelhofer, Clinton R. Paden, Duncan MacCannell |
| EPI_ISL_3330130 | Illinois Department of Public Health - Springfield Lab | Illinois Department of Public Health - Springfield Lab | Bryan Sim, Gordon McCall, Marshall Smock |
| EPI_ISL_3330131, EPI_ISL_3330132, EPI_ISL_3330133 | Laboratory Corporation of America | Centers for Disease Control and Prevention Division of Viral Diseases, Pathogen Discovery | Dakota Howard, Dhwani Batra, Peter W. Cook, Kara Moser, Adrian Paskey, Jason Caravas, Benjamin Rambo-Martin, Shatavia Morrison, Christopher Gulvick, Scott Sammons, Yvette Unoarumhi, Darlene Wagner, Matthew Schmerer, Minoo Agarwal, Eyad Almasri, Debbie Boles, Ayla Burns, Nuthawin Charoensri, Oren Cohen, Susan Countryman, Mary Ann Cristobal, Bobbi Croy, Suzanne Dale, Hrushikesh Deshmukh, Amanda Douglas, Vincent Drouillon, Marcia Eisenberg, Howard Engler, Rama Ghatti, Prashant Gupta, Susan Hicks, Jake Humphrey, Lax Iyer, Lisa Pfefferle, Manoj Jain, Matthew Robinson, Mohan Kolli, Brian Krueger, Tim Kuphal, Stanley Letovsky, Michael Levandoski, Craig Lukasik, Jonathan Meltzer, Brian Norvell, Mindy Nye, Scott Parker, Christos Petropoulos, John Pruitt, Steven Ragan, Scott Ryan, Mike Sapeta, Jana Schroth, Suresh Babu Selvaraju, Goran Stevovic, Amanda Suchanek, Andrea Throop, Lyndon Tilson, Thomas Urban, Joe Voshell, Kimberly Wagner, Jonathan Williams, Mary Williamson, Qian Zeng, Tricia Zwiefelhofer, Clinton R. Paden, Duncan MacCannell |
| EPI_ISL_3330134 | Illinois Department of Public Health - Springfield Lab | Illinois Department of Public Health - Springfield Lab | Bryan Sim, Gordon McCall, Marshall Smock |
| EPI_ISL_3330135, EPI_ISL_3330136 | Laboratory Corporation of America | Centers for Disease Control and Prevention Division of Viral Diseases, Pathogen Discovery | Dakota Howard, Dhwani Batra, Peter W. Cook, Kara Moser, Adrian Paskey, Jason Caravas, Benjamin Rambo-Martin, Shatavia Morrison, Christopher Gulvick, Scott Sammons, Yvette Unoarumhi, Darlene Wagner, Matthew Schmerer, Minoo Agarwal, Eyad Almasri, Debbie Boles, Ayla Burns, Nuthawin Charoensri, Oren Cohen, Susan Countryman, Mary Ann Cristobal, Bobbi Croy, Suzanne Dale, Hrushikesh Deshmukh, Amanda Douglas, Vincent Drouillon, Marcia Eisenberg, Howard Engler, Rama Ghatti, Prashant Gupta, Susan Hicks, Jake Humphrey, Lax Iyer, Lisa Pfefferle, Manoj Jain, Matthew Robinson, Mohan Kolli, Brian Krueger, Tim Kuphal, Stanley Letovsky, Michael Levandoski, Craig Lukasik, Jonathan Meltzer, Brian Norvell, Mindy Nye, Scott Parker, Christos Petropoulos, John Pruitt, Steven Ragan, Scott Ryan, Mike Sapeta, Jana Schroth, Suresh Babu Selvaraju, Goran Stevovic, Amanda Suchanek, Andrea Throop, Lyndon Tilson, Thomas Urban, Joe Voshell, Kimberly Wagner, Jonathan Williams, Mary Williamson, Qian Zeng, Tricia Zwiefelhofer, Clinton R. Paden, Duncan MacCannell |
| EPI_ISL_3330137 | Illinois Department of Public Health - Springfield Lab | Illinois Department of Public Health - Springfield Lab | Bryan Sim, Gordon McCall, Marshall Smock |
| EPI_ISL_3330138, EPI_ISL_3330139, EPI_ISL_3330140 | Laboratory Corporation of America | Centers for Disease Control and Prevention Division of Viral Diseases, Pathogen Discovery | Dakota Howard, Dhwani Batra, Peter W. Cook, Kara Moser, Adrian Paskey, Jason Caravas, Benjamin Rambo-Martin, Shatavia Morrison, Christopher Gulvick, Scott Sammons, Yvette Unoarumhi, Darlene Wagner, Matthew Schmerer, Minoo Agarwal, Eyad Almasri, Debbie Boles, Ayla Burns, Nuthawin Charoensri, Oren Cohen, Susan Countryman, Mary Ann Cristobal, Bobbi Croy, Suzanne Dale, Hrushikesh Deshmukh, Amanda Douglas, Vincent Drouillon, Marcia Eisenberg, Howard Engler, Rama Ghatti, Prashant Gupta, Susan Hicks, Jake Humphrey, Lax Iyer, Lisa Pfefferle, Manoj Jain, Matthew Robinson, Mohan Kolli, Brian Krueger, Tim Kuphal, Stanley Letovsky, Michael Levandoski, Craig Lukasik, Jonathan Meltzer, Brian Norvell, Mindy Nye, Scott Parker, Christos Petropoulos, John Pruitt, Steven Ragan, Scott Ryan, Mike Sapeta, Jana Schroth, Suresh Babu Selvaraju, Goran Stevovic, Amanda Suchanek, Andrea Throop, Lyndon Tilson, Thomas Urban, Joe Voshell, Kimberly Wagner, Jonathan Williams, Mary Williamson, Qian Zeng, Tricia Zwiefelhofer, Clinton R. Paden, Duncan MacCannell |
| EPI_ISL_3330141 | Illinois Department of Public Health - Springfield Lab | Illinois Department of Public Health - Springfield Lab | Bryan Sim, Gordon McCall, Marshall Smock |
| EPI_ISL_3330142, EPI_ISL_3330143 | Laboratory Corporation of America | Centers for Disease Control and Prevention Division of Viral Diseases, Pathogen Discovery | Dakota Howard, Dhwani Batra, Peter W. Cook, Kara Moser, Adrian Paskey, Jason Caravas, Benjamin Rambo-Martin, Shatavia Morrison, Christopher Gulvick, Scott Sammons, Yvette Unoarumhi, Darlene Wagner, Matthew Schmerer, Minoo Agarwal, Eyad Almasri, Debbie Boles, Ayla Burns, Nuthawin Charoensri, Oren Cohen, Susan Countryman, Mary Ann Cristobal, Bobbi Croy, Suzanne Dale, Hrushikesh Deshmukh, Amanda Douglas, Vincent Drouillon, Marcia Eisenberg, Howard Engler, Rama Ghatti, Prashant Gupta, Susan Hicks, Jake Humphrey, Lax Iyer, Lisa Pfefferle, Manoj Jain, Matthew Robinson, Mohan Kolli, Brian Krueger, Tim Kuphal, Stanley Letovsky, Michael Levandoski, Craig Lukasik, Jonathan Meltzer, Brian Norvell, Mindy Nye, Scott Parker, Christos Petropoulos, John Pruitt, Steven Ragan, Scott Ryan, Mike Sapeta, Jana Schroth, Suresh Babu Selvaraju, Goran Stevovic, Amanda Suchanek, Andrea Throop, Lyndon Tilson, Thomas Urban, Joe Voshell, Kimberly Wagner, Jonathan Williams, Mary Williamson, Qian Zeng, Tricia Zwiefelhofer, Clinton R. Paden, Duncan MacCannell |
| EPI_ISL_3330144 | Illinois Department of Public Health - Springfield Lab | Illinois Department of Public Health - Springfield Lab | Bryan Sim, Gordon McCall, Marshall Smock |
| EPI_ISL_3330145, EPI_ISL_3330147 | Laboratory Corporation of America | Centers for Disease Control and Prevention Division of Viral Diseases, Pathogen Discovery | Dakota Howard, Dhwani Batra, Peter W. Cook, Kara Moser, Adrian Paskey, Jason Caravas, Benjamin Rambo-Martin, Shatavia Morrison, Christopher Gulvick, Scott Sammons, Yvette Unoarumhi, Darlene Wagner, Matthew Schmerer, Minoo Agarwal, Eyad Almasri, Debbie Boles, Ayla Burns, Nuthawin Charoensri, Oren Cohen, Susan Countryman, Mary Ann Cristobal, Bobbi Croy, Suzanne Dale, Hrushikesh Deshmukh, Amanda Douglas, Vincent Drouillon, Marcia Eisenberg, Howard Engler, Rama Ghatti, Prashant Gupta, Susan Hicks, Jake Humphrey, Lax Iyer, Lisa Pfefferle, Manoj Jain, Matthew Robinson, Mohan Kolli, Brian Krueger, Tim Kuphal, Stanley Letovsky, Michael Levandoski, Craig Lukasik, Jonathan Meltzer, Brian Norvell, Mindy Nye, Scott Parker, Christos Petropoulos, John Pruitt, Steven Ragan, Scott Ryan, Mike Sapeta, Jana Schroth, Suresh Babu Selvaraju, Goran Stevovic, Amanda Suchanek, Andrea Throop, Lyndon Tilson, Thomas Urban, Joe Voshell, Kimberly Wagner, Jonathan Williams, Mary Williamson, Qian Zeng, Tricia Zwiefelhofer, Clinton R. Paden, Duncan MacCannell |
| EPI_ISL_3330148 | Illinois Department of Public Health - Springfield Lab | Illinois Department of Public Health - Springfield Lab | Bryan Sim, Gordon McCall, Marshall Smock |
| EPI_ISL_3330149, EPI_ISL_3330150 | Laboratory Corporation of America | Centers for Disease Control and Prevention Division of Viral Diseases, Pathogen Discovery | Dakota Howard, Dhwani Batra, Peter W. Cook, Kara Moser, Adrian Paskey, Jason Caravas, Benjamin Rambo-Martin, Shatavia Morrison, Christopher Gulvick, Scott Sammons, Yvette Unoarumhi, Darlene Wagner, Matthew Schmerer, Minoo Agarwal, Eyad Almasri, Debbie Boles, Ayla Burns, Nuthawin Charoensri, Oren Cohen, Susan Countryman, Mary Ann Cristobal, Bobbi Croy, Suzanne Dale, Hrushikesh Deshmukh, Amanda Douglas, Vincent Drouillon, Marcia Eisenberg, Howard Engler, Rama Ghatti, Prashant Gupta, Susan Hicks, Jake Humphrey, Lax Iyer, Lisa Pfefferle, Manoj Jain, Matthew Robinson, Mohan Kolli, Brian Krueger, Tim Kuphal, Stanley Letovsky, Michael Levandoski, Craig Lukasik, Jonathan Meltzer, Brian Norvell, Mindy Nye, Scott Parker, Christos Petropoulos, John Pruitt, Steven Ragan, Scott Ryan, Mike Sapeta, Jana Schroth, Suresh Babu Selvaraju, Goran Stevovic, Amanda Suchanek, Andrea Throop, Lyndon Tilson, Thomas Urban, Joe Voshell, Kimberly Wagner, Jonathan Williams, Mary Williamson, Qian Zeng, Tricia Zwiefelhofer, Clinton R. Paden, Duncan MacCannell |
| EPI_ISL_3330151 | Illinois Department of Public Health - Springfield Lab | Illinois Department of Public Health - Springfield Lab | Bryan Sim, Gordon McCall, Marshall Smock |
| EPI_ISL_3330152, EPI_ISL_3330153, EPI_ISL_3330154 | Laboratory Corporation of America | Centers for Disease Control and Prevention Division of Viral Diseases, Pathogen Discovery | Dakota Howard, Dhwani Batra, Peter W. Cook, Kara Moser, Adrian Paskey, Jason Caravas, Benjamin Rambo-Martin, Shatavia Morrison, Christopher Gulvick, Scott Sammons, Yvette Unoarumhi, Darlene Wagner, Matthew Schmerer, Minoo Agarwal, Eyad Almasri, Debbie Boles, Ayla Burns, Nuthawin Charoensri, Oren Cohen, Susan Countryman, Mary Ann Cristobal, Bobbi Croy, Suzanne Dale, Hrushikesh Deshmukh, Amanda Douglas, Vincent Drouillon, Marcia Eisenberg, Howard Engler, Rama Ghatti, Prashant Gupta, Susan Hicks, Jake Humphrey, Lax Iyer, Lisa Pfefferle, Manoj Jain, Matthew Robinson, Mohan Kolli, Brian Krueger, Tim Kuphal, Stanley Letovsky, Michael Levandoski, Craig Lukasik, Jonathan Meltzer, Brian Norvell, Mindy Nye, Scott Parker, Christos Petropoulos, John Pruitt, Steven Ragan, Scott Ryan, Mike Sapeta, Jana Schroth, Suresh Babu Selvaraju, Goran Stevovic, Amanda Suchanek, Andrea Throop, Lyndon Tilson, Thomas Urban, Joe Voshell, Kimberly Wagner, Jonathan Williams, Mary Williamson, Qian Zeng, Tricia Zwiefelhofer, Clinton R. Paden, Duncan MacCannell |
| EPI_ISL_3330155 | Illinois Department of Public Health - Springfield Lab | Illinois Department of Public Health - Springfield Lab | Bryan Sim, Gordon McCall, Marshall Smock |
| EPI_ISL_3330156, EPI_ISL_3330157 | Laboratory Corporation of America | Centers for Disease Control and Prevention Division of Viral Diseases, Pathogen Discovery | Dakota Howard, Dhwani Batra, Peter W. Cook, Kara Moser, Adrian Paskey, Jason Caravas, Benjamin Rambo-Martin, Shatavia Morrison, Christopher Gulvick, Scott Sammons, Yvette Unoarumhi, Darlene Wagner, Matthew Schmerer, Minoo Agarwal, Eyad Almasri, Debbie Boles, Ayla Burns, Nuthawin Charoensri, Oren Cohen, Susan Countryman, Mary Ann Cristobal, Bobbi Croy, Suzanne Dale, Hrushikesh Deshmukh, Amanda Douglas, Vincent Drouillon, Marcia Eisenberg, Howard Engler, Rama Ghatti, Prashant Gupta, Susan Hicks, Jake Humphrey, Lax Iyer, Lisa Pfefferle, Manoj Jain, Matthew Robinson, Mohan Kolli, Brian Krueger, Tim Kuphal, Stanley Letovsky, Michael Levandoski, Craig Lukasik, Jonathan Meltzer, Brian Norvell, Mindy Nye, Scott Parker, Christos Petropoulos, John Pruitt, Steven Ragan, Scott Ryan, Mike Sapeta, Jana Schroth, Suresh Babu Selvaraju, Goran Stevovic, Amanda Suchanek, Andrea Throop, Lyndon Tilson, Thomas Urban, Joe Voshell, Kimberly Wagner, Jonathan Williams, Mary Williamson, Qian Zeng, Tricia Zwiefelhofer, Clinton R. Paden, Duncan MacCannell |
| EPI_ISL_3330158 | Illinois Department of Public Health - Springfield Lab | Illinois Department of Public Health - Springfield Lab | Bryan Sim, Gordon McCall, Marshall Smock |
| EPI_ISL_3330160 | Laboratory Corporation of America | Centers for Disease Control and Prevention Division of Viral Diseases, Pathogen Discovery | Dakota Howard, Dhwani Batra, Peter W. Cook, Kara Moser, Adrian Paskey, Jason Caravas, Benjamin Rambo-Martin, Shatavia Morrison, Christopher Gulvick, Scott Sammons, Yvette Unoarumhi, Darlene Wagner, Matthew Schmerer, Minoo Agarwal, Eyad Almasri, Debbie Boles, Ayla Burns, Nuthawin Charoensri, Oren Cohen, Susan Countryman, Mary Ann Cristobal, Bobbi Croy, Suzanne Dale, Hrushikesh Deshmukh, Amanda Douglas, Vincent Drouillon, Marcia Eisenberg, Howard Engler, Rama Ghatti, Prashant Gupta, Susan Hicks, Jake Humphrey, Lax Iyer, Lisa Pfefferle, Manoj Jain, Matthew Robinson, Mohan Kolli, Brian Krueger, Tim Kuphal, Stanley Letovsky, Michael Levandoski, Craig Lukasik, Jonathan Meltzer, Brian Norvell, Mindy Nye, Scott Parker, Christos Petropoulos, John Pruitt, Steven Ragan, Scott Ryan, Mike Sapeta, Jana Schroth, Suresh Babu Selvaraju, Goran Stevovic, Amanda Suchanek, Andrea Throop, Lyndon Tilson, Thomas Urban, Joe Voshell, Kimberly Wagner, Jonathan Williams, Mary Williamson, Qian Zeng, Tricia Zwiefelhofer, Clinton R. Paden, Duncan MacCannell |

| | | | |
|---|---|---|---|
| EPI_ISL_3330161 | Illinois Department of Public Health - Springfield Lab | Illinois Department of Public Health - Springfield Lab | Bryan Sim, Gordon McCall, Marshall Smock |
| EPI_ISL_3330162, EPI_ISL_3330163 | Laboratory Corporation of America | Centers for Disease Control and Prevention Division of Viral Diseases, Pathogen Discovery | Dakota Howard, Dhwani Batra, Peter W. Cook, Kara Moser, Adrian Paskey, Jason Caravas, Benjamin Rambo-Martin, Shatavia Morrison, Christopher Gulvick, Scott Sammons, Yvette Unoarumhi, Darlene Wagner, Matthew Schmerer, Minoo Agarwal, Eyad Almasri, Debbie Boles, Ayla Burns, Nuthawin Charoensri, Oren Cohen, Susan Countryman, Mary Ann Cristobal, Bobbi Croy, Suzanne Dale, Hrushikesh Deshmukh, Amanda Douglas, Vincent Drouillon, Marcia Eisenberg, Howard Engler, Rama Ghatti, Prashant Gupta, Susan Hicks, Jake Humphrey, Lax Iyer, Lisa Pfefferle, Manoj Jain, Matthew Robinson, Mohan Kolli, Brian Krueger, Tim Kuphal, Stanley Letovsky, Michael Levandoski, Craig Lukasik, Jonathan Meltzer, Brian Norvell, Mindy Nye, Scott Parker, Christos Petropoulos, John Pruitt, Steven Ragan, Scott Ryan, Mike Sapeta, Jana Schroth, Suresh Babu Selvaraju, Goran Stevovic, Amanda Suchanek, Andrea Throop, Lyndon Tilson, Thomas Urban, Joe Voshell, Kimberly Wagner, Jonathan Williams, Mary Williamson, Qian Zeng, Tricia Zwiefelhofer, Clinton R. Paden, Duncan MacCannell |
| EPI_ISL_3330164 | Illinois Department of Public Health - Springfield Lab | Illinois Department of Public Health - Springfield Lab | Bryan Sim, Gordon McCall, Marshall Smock |
| EPI_ISL_3330165, EPI_ISL_3330166, EPI_ISL_3330167 | Laboratory Corporation of America | Centers for Disease Control and Prevention Division of Viral Diseases, Pathogen Discovery | Dakota Howard, Dhwani Batra, Peter W. Cook, Kara Moser, Adrian Paskey, Jason Caravas, Benjamin Rambo-Martin, Shatavia Morrison, Christopher Gulvick, Scott Sammons, Yvette Unoarumhi, Darlene Wagner, Matthew Schmerer, Minoo Agarwal, Eyad Almasri, Debbie Boles, Ayla Burns, Nuthawin Charoensri, Oren Cohen, Susan Countryman, Mary Ann Cristobal, Bobbi Croy, Suzanne Dale, Hrushikesh Deshmukh, Amanda Douglas, Vincent Drouillon, Marcia Eisenberg, Howard Engler, Rama Ghatti, Prashant Gupta, Susan Hicks, Jake Humphrey, Lax Iyer, Lisa Pfefferle, Manoj Jain, Matthew Robinson, Mohan Kolli, Brian Krueger, Tim Kuphal, Stanley Letovsky, Michael Levandoski, Craig Lukasik, Jonathan Meltzer, Brian Norvell, Mindy Nye, Scott Parker, Christos Petropoulos, John Pruitt, Steven Ragan, Scott Ryan, Mike Sapeta, Jana Schroth, Suresh Babu Selvaraju, Goran Stevovic, Amanda Suchanek, Andrea Throop, Lyndon Tilson, Thomas Urban, Joe Voshell, Kimberly Wagner, Jonathan Williams, Mary Williamson, Qian Zeng, Tricia Zwiefelhofer, Clinton R. Paden, Duncan MacCannell |
| EPI_ISL_3330168 | Illinois Department of Public Health - Springfield Lab | Illinois Department of Public Health - Springfield Lab | Bryan Sim, Gordon McCall, Marshall Smock |
| EPI_ISL_3330169, EPI_ISL_3330170 | Laboratory Corporation of America | Centers for Disease Control and Prevention Division of Viral Diseases, Pathogen Discovery | Dakota Howard, Dhwani Batra, Peter W. Cook, Kara Moser, Adrian Paskey, Jason Caravas, Benjamin Rambo-Martin, Shatavia Morrison, Christopher Gulvick, Scott Sammons, Yvette Unoarumhi, Darlene Wagner, Matthew Schmerer, Minoo Agarwal, Eyad Almasri, Debbie Boles, Ayla Burns, Nuthawin Charoensri, Oren Cohen, Susan Countryman, Mary Ann Cristobal, Bobbi Croy, Suzanne Dale, Hrushikesh Deshmukh, Amanda Douglas, Vincent Drouillon, Marcia Eisenberg, Howard Engler, Rama Ghatti, Prashant Gupta, Susan Hicks, Jake Humphrey, Lax Iyer, Lisa Pfefferle, Manoj Jain, Matthew Robinson, Mohan Kolli, Brian Krueger, Tim Kuphal, Stanley Letovsky, Michael Levandoski, Craig Lukasik, Jonathan Meltzer, Brian Norvell, Mindy Nye, Scott Parker, Christos Petropoulos, John Pruitt, Steven Ragan, Scott Ryan, Mike Sapeta, Jana Schroth, Suresh Babu Selvaraju, Goran Stevovic, Amanda Suchanek, Andrea Throop, Lyndon Tilson, Thomas Urban, Joe Voshell, Kimberly Wagner, Jonathan Williams, Mary Williamson, Qian Zeng, Tricia Zwiefelhofer, Clinton R. Paden, Duncan MacCannell |
| EPI_ISL_3330171 | Illinois Department of Public Health - Springfield Lab | Illinois Department of Public Health - Springfield Lab | Bryan Sim, Gordon McCall, Marshall Smock |
| EPI_ISL_3330172, EPI_ISL_3330173, EPI_ISL_3330174 | Laboratory Corporation of America | Centers for Disease Control and Prevention Division of Viral Diseases, Pathogen Discovery | Dakota Howard, Dhwani Batra, Peter W. Cook, Kara Moser, Adrian Paskey, Jason Caravas, Benjamin Rambo-Martin, Shatavia Morrison, Christopher Gulvick, Scott Sammons, Yvette Unoarumhi, Darlene Wagner, Matthew Schmerer, Minoo Agarwal, Eyad Almasri, Debbie Boles, Ayla Burns, Nuthawin Charoensri, Oren Cohen, Susan Countryman, Mary Ann Cristobal, Bobbi Croy, Suzanne Dale, Hrushikesh Deshmukh, Amanda Douglas, Vincent Drouillon, Marcia Eisenberg, Howard Engler, Rama Ghatti, Prashant Gupta, Susan Hicks, Jake Humphrey, Lax Iyer, Lisa Pfefferle, Manoj Jain, Matthew Robinson, Mohan Kolli, Brian Krueger, Tim Kuphal, Stanley Letovsky, Michael Levandoski, Craig Lukasik, Jonathan Meltzer, Brian Norvell, Mindy Nye, Scott Parker, Christos Petropoulos, John Pruitt, Steven Ragan, Scott Ryan, Mike Sapeta, Jana Schroth, Suresh Babu Selvaraju, Goran Stevovic, Amanda Suchanek, Andrea Throop, Lyndon Tilson, Thomas Urban, Joe Voshell, Kimberly Wagner, Jonathan Williams, Mary Williamson, Qian Zeng, Tricia Zwiefelhofer, Clinton R. Paden, Duncan MacCannell |
| EPI_ISL_3330175 | Illinois Department of Public Health - Springfield Lab | Illinois Department of Public Health - Springfield Lab | Bryan Sim, Gordon McCall, Marshall Smock |
| EPI_ISL_3330176, EPI_ISL_3330177 | Laboratory Corporation of America | Centers for Disease Control and Prevention Division of Viral Diseases, Pathogen Discovery | Dakota Howard, Dhwani Batra, Peter W. Cook, Kara Moser, Adrian Paskey, Jason Caravas, Benjamin Rambo-Martin, Shatavia Morrison, Christopher Gulvick, Scott Sammons, Yvette Unoarumhi, Darlene Wagner, Matthew Schmerer, Minoo Agarwal, Eyad Almasri, Debbie Boles, Ayla Burns, Nuthawin Charoensri, Oren Cohen, Susan Countryman, Mary Ann Cristobal, Bobbi Croy, Suzanne Dale, Hrushikesh Deshmukh, Amanda Douglas, Vincent Drouillon, Marcia Eisenberg, Howard Engler, Rama Ghatti, Prashant Gupta, Susan Hicks, Jake Humphrey, Lax Iyer, Lisa Pfefferle, Manoj Jain, Matthew Robinson, Mohan Kolli, Brian Krueger, Tim Kuphal, Stanley Letovsky, Michael Levandoski, Craig Lukasik, Jonathan Meltzer, Brian Norvell, Mindy Nye, Scott Parker, Christos Petropoulos, John Pruitt, Steven Ragan, Scott Ryan, Mike Sapeta, Jana Schroth, Suresh Babu Selvaraju, Goran Stevovic, Amanda Suchanek, Andrea Throop, Lyndon Tilson, Thomas Urban, Joe Voshell, Kimberly Wagner, Jonathan Williams, Mary Williamson, Qian Zeng, Tricia Zwiefelhofer, Clinton R. Paden, Duncan MacCannell |
| EPI_ISL_3330178 | Illinois Department of Public Health - Springfield Lab | Illinois Department of Public Health - Springfield Lab | Bryan Sim, Gordon McCall, Marshall Smock |
| EPI_ISL_3330179, EPI_ISL_3330180 | Laboratory Corporation of America | Centers for Disease Control and Prevention Division of Viral Diseases, Pathogen Discovery | Dakota Howard, Dhwani Batra, Peter W. Cook, Kara Moser, Adrian Paskey, Jason Caravas, Benjamin Rambo-Martin, Shatavia Morrison, Christopher Gulvick, Scott Sammons, Yvette Unoarumhi, Darlene Wagner, Matthew Schmerer, Minoo Agarwal, Eyad Almasri, Debbie Boles, Ayla Burns, Nuthawin Charoensri, Oren Cohen, Susan Countryman, Mary Ann Cristobal, Bobbi Croy, Suzanne Dale, Hrushikesh Deshmukh, Amanda Douglas, Vincent Drouillon, Marcia Eisenberg, Howard Engler, Rama Ghatti, Prashant Gupta, Susan Hicks, Jake Humphrey, Lax Iyer, Lisa Pfefferle, Manoj Jain, Matthew Robinson, Mohan Kolli, Brian Krueger, Tim Kuphal, Stanley Letovsky, Michael Levandoski, Craig Lukasik, Jonathan Meltzer, Brian Norvell, Mindy Nye, Scott Parker, Christos Petropoulos, John Pruitt, Steven Ragan, Scott Ryan, Mike Sapeta, Jana Schroth, Suresh Babu Selvaraju, Goran Stevovic, Amanda Suchanek, Andrea Throop, Lyndon Tilson, Thomas Urban, Joe Voshell, Kimberly Wagner, Jonathan Williams, Mary Williamson, Qian Zeng, Tricia Zwiefelhofer, Clinton R. Paden, Duncan MacCannell |
| EPI_ISL_3330181 | Illinois Department of Public Health - Springfield Lab | Illinois Department of Public Health - Springfield Lab | Bryan Sim, Gordon McCall, Marshall Smock |
| EPI_ISL_3330182, EPI_ISL_3330183 | Laboratory Corporation of America | Centers for Disease Control and Prevention Division of Viral Diseases, Pathogen Discovery | Dakota Howard, Dhwani Batra, Peter W. Cook, Kara Moser, Adrian Paskey, Jason Caravas, Benjamin Rambo-Martin, Shatavia Morrison, Christopher Gulvick, Scott Sammons, Yvette Unoarumhi, Darlene Wagner, Matthew Schmerer, Minoo Agarwal, Eyad Almasri, Debbie Boles, Ayla Burns, Nuthawin Charoensri, Oren Cohen, Susan Countryman, Mary Ann Cristobal, Bobbi Croy, Suzanne Dale, Hrushikesh Deshmukh, Amanda Douglas, Vincent Drouillon, Marcia Eisenberg, Howard Engler, Rama Ghatti, Prashant Gupta, Susan Hicks, Jake Humphrey, Lax Iyer, Lisa Pfefferle, Manoj Jain, Matthew Robinson, Mohan Kolli, Brian Krueger, Tim Kuphal, Stanley Letovsky, Michael Levandoski, Craig Lukasik, Jonathan Meltzer, Brian Norvell, Mindy Nye, Scott Parker, Christos Petropoulos, John Pruitt, Steven Ragan, Scott Ryan, Mike Sapeta, Jana Schroth, Suresh Babu Selvaraju, Goran Stevovic, Amanda Suchanek, Andrea Throop, Lyndon Tilson, Thomas Urban, Joe Voshell, Kimberly Wagner, Jonathan Williams, Mary Williamson, Qian Zeng, Tricia Zwiefelhofer, Clinton R. Paden, Duncan MacCannell |
| EPI_ISL_3330184 | Illinois Department of Public Health - Springfield Lab | Illinois Department of Public Health - Springfield Lab | Bryan Sim, Gordon McCall, Marshall Smock |
| EPI_ISL_3330185, EPI_ISL_3330186 | Laboratory Corporation of America | Centers for Disease Control and Prevention Division of Viral Diseases, Pathogen Discovery | Dakota Howard, Dhwani Batra, Peter W. Cook, Kara Moser, Adrian Paskey, Jason Caravas, Benjamin Rambo-Martin, Shatavia Morrison, Christopher Gulvick, Scott Sammons, Yvette Unoarumhi, Darlene Wagner, Matthew Schmerer, Minoo Agarwal, Eyad Almasri, Debbie Boles, Ayla Burns, Nuthawin Charoensri, Oren Cohen, Susan Countryman, Mary Ann Cristobal, Bobbi Croy, Suzanne Dale, Hrushikesh Deshmukh, Amanda Douglas, Vincent Drouillon, Marcia Eisenberg, Howard Engler, Rama Ghatti, Prashant Gupta, Susan Hicks, Jake Humphrey, Lax Iyer, Lisa Pfefferle, Manoj Jain, Matthew Robinson, Mohan Kolli, Brian Krueger, Tim Kuphal, Stanley Letovsky, Michael Levandoski, Craig Lukasik, Jonathan Meltzer, Brian Norvell, Mindy Nye, Scott Parker, Christos Petropoulos, John Pruitt, Steven Ragan, Scott Ryan, Mike Sapeta, Jana Schroth, Suresh Babu Selvaraju, Goran Stevovic, Amanda Suchanek, Andrea Throop, Lyndon Tilson, Thomas Urban, Joe Voshell, Kimberly Wagner, Jonathan Williams, Mary Williamson, Qian Zeng, Tricia Zwiefelhofer, Clinton R. Paden, Duncan MacCannell |
| EPI_ISL_3330187 | Illinois Department of Public Health - Springfield Lab | Illinois Department of Public Health - Springfield Lab | Bryan Sim, Gordon McCall, Marshall Smock |
| EPI_ISL_3330188, EPI_ISL_3330189, EPI_ISL_3330190 | Laboratory Corporation of America | Centers for Disease Control and Prevention Division of Viral Diseases, Pathogen Discovery | Dakota Howard, Dhwani Batra, Peter W. Cook, Kara Moser, Adrian Paskey, Jason Caravas, Benjamin Rambo-Martin, Shatavia Morrison, Christopher Gulvick, Scott Sammons, Yvette Unoarumhi, Darlene Wagner, Matthew Schmerer, Minoo Agarwal, Eyad Almasri, Debbie Boles, Ayla Burns, Nuthawin Charoensri, Oren Cohen, Susan Countryman, Mary Ann Cristobal, Bobbi Croy, Suzanne Dale, Hrushikesh Deshmukh, Amanda Douglas, Vincent Drouillon, Marcia Eisenberg, Howard Engler, Rama Ghatti, Prashant Gupta, Susan Hicks, Jake Humphrey, Lax Iyer, Lisa Pfefferle, Manoj Jain, Matthew Robinson, Mohan Kolli, Brian Krueger, Tim Kuphal, Stanley Letovsky, Michael Levandoski, Craig Lukasik, Jonathan Meltzer, Brian Norvell, Mindy Nye, Scott Parker, Christos Petropoulos, John Pruitt, Steven Ragan, Scott Ryan, Mike Sapeta, Jana Schroth, Suresh Babu Selvaraju, Goran Stevovic, Amanda Suchanek, Andrea Throop, Lyndon Tilson, Thomas Urban, Joe Voshell, Kimberly Wagner, Jonathan Williams, Mary Williamson, Qian Zeng, Tricia Zwiefelhofer, Clinton R. Paden, Duncan MacCannell |
| EPI_ISL_3330191 | Illinois Department of Public Health - Springfield Lab | Illinois Department of Public Health - Springfield Lab | Bryan Sim, Gordon McCall, Marshall Smock |
| EPI_ISL_3330192, EPI_ISL_3330193 | Laboratory Corporation of America | Centers for Disease Control and Prevention Division of Viral Diseases, Pathogen Discovery | Dakota Howard, Dhwani Batra, Peter W. Cook, Kara Moser, Adrian Paskey, Jason Caravas, Benjamin Rambo-Martin, Shatavia Morrison, Christopher Gulvick, Scott Sammons, Yvette Unoarumhi, Darlene Wagner, Matthew Schmerer, Minoo Agarwal, Eyad Almasri, Debbie Boles, Ayla Burns, Nuthawin Charoensri, Oren Cohen, Susan Countryman, Mary Ann Cristobal, Bobbi Croy, Suzanne Dale, Hrushikesh Deshmukh, Amanda Douglas, Vincent Drouillon, Marcia Eisenberg, Howard Engler, Rama Ghatti, Prashant Gupta, Susan Hicks, Jake Humphrey, Lax Iyer, Lisa Pfefferle, Manoj Jain, Matthew Robinson, Mohan Kolli, Brian Krueger, Tim Kuphal, Stanley Letovsky, Michael Levandoski, Craig Lukasik, Jonathan Meltzer, Brian Norvell, Mindy Nye, Scott Parker, Christos Petropoulos, John Pruitt, Steven Ragan, Scott Ryan, Mike Sapeta, Jana Schroth, Suresh Babu Selvaraju, Goran Stevovic, Amanda Suchanek, Andrea Throop, Lyndon Tilson, Thomas Urban, Joe Voshell, Kimberly Wagner, Jonathan Williams, Mary Williamson, Qian Zeng, Tricia Zwiefelhofer, Clinton R. Paden, Duncan MacCannell |
| EPI_ISL_3330194 | Illinois Department of Public Health - Springfield Lab | Illinois Department of Public Health - Springfield Lab | Bryan Sim, Gordon McCall, Marshall Smock |
| EPI_ISL_3330195, EPI_ISL_3330196, EPI_ISL_3330197 | Laboratory Corporation of America | Centers for Disease Control and Prevention Division of Viral Diseases, Pathogen Discovery | Dakota Howard, Dhwani Batra, Peter W. Cook, Kara Moser, Adrian Paskey, Jason Caravas, Benjamin Rambo-Martin, Shatavia Morrison, Christopher Gulvick, Scott Sammons, Yvette Unoarumhi, Darlene Wagner, Matthew Schmerer, Minoo Agarwal, Eyad Almasri, Debbie Boles, Ayla Burns, Nuthawin Charoensri, Oren Cohen, Susan Countryman, Mary Ann Cristobal, Bobbi Croy, Suzanne Dale, Hrushikesh Deshmukh, Amanda Douglas, Vincent Drouillon, Marcia Eisenberg, Howard Engler, Rama Ghatti, Prashant Gupta, Susan Hicks, Jake Humphrey, Lax Iyer, Lisa Pfefferle, Manoj Jain, Matthew Robinson, Mohan Kolli, Brian Krueger, Tim Kuphal, Stanley Letovsky, Michael Levandoski, Craig Lukasik, Jonathan Meltzer, Brian Norvell, Mindy Nye, Scott Parker, Christos Petropoulos, John Pruitt, Steven Ragan, Scott Ryan, Mike Sapeta, Jana Schroth, Suresh Babu Selvaraju, Goran Stevovic, Amanda Suchanek, Andrea Throop, Lyndon Tilson, Thomas Urban, Joe Voshell, Kimberly Wagner, Jonathan Williams, Mary Williamson, Qian Zeng, Tricia Zwiefelhofer, Clinton R. Paden, Duncan MacCannell |
| EPI_ISL_3330198 | Illinois Department of Public Health - Springfield Lab | Illinois Department of Public Health - Springfield Lab | Bryan Sim, Gordon McCall, Marshall Smock |
| EPI_ISL_3330199, EPI_ISL_3330200, EPI_ISL_3330201 | Laboratory Corporation of America | Centers for Disease Control and Prevention Division of Viral Diseases, Pathogen Discovery | Dakota Howard, Dhwani Batra, Peter W. Cook, Kara Moser, Adrian Paskey, Jason Caravas, Benjamin Rambo-Martin, Shatavia Morrison, Christopher Gulvick, Scott Sammons, Yvette Unoarumhi, Darlene Wagner, Matthew Schmerer, Minoo Agarwal, Eyad Almasri, Debbie Boles, Ayla Burns, Nuthawin Charoensri, Oren Cohen, Susan Countryman, Mary Ann Cristobal, Bobbi Croy, Suzanne Dale, Hrushikesh Deshmukh, Amanda Douglas, Vincent Drouillon, Marcia Eisenberg, Howard Engler, Rama Ghatti, Prashant Gupta, Susan Hicks, Jake Humphrey, Lax Iyer, Lisa Pfefferle, Manoj Jain, Matthew Robinson, Mohan Kolli, Brian Krueger, Tim Kuphal, Stanley Letovsky, Michael Levandoski, Craig Lukasik, Jonathan Meltzer, Brian Norvell, Mindy Nye, Scott Parker, Christos Petropoulos, John Pruitt, Steven Ragan, Scott Ryan, Mike Sapeta, Jana Schroth, Suresh Babu Selvaraju, Goran Stevovic, Amanda Suchanek, Andrea Throop, Lyndon Tilson, Thomas Urban, Joe Voshell, Kimberly Wagner, Jonathan Williams, Mary Williamson, Qian Zeng, Tricia Zwiefelhofer, Clinton R. Paden, Duncan MacCannell |
| EPI_ISL_3330202 | Illinois Department of Public Health - Springfield Lab | Illinois Department of Public Health - Springfield Lab | Bryan Sim, Gordon McCall, Marshall Smock |
| EPI_ISL_3330203, EPI_ISL_3330204 | Laboratory Corporation of America | Centers for Disease Control and Prevention Division of Viral Diseases, Pathogen Discovery | Dakota Howard, Dhwani Batra, Peter W. Cook, Kara Moser, Adrian Paskey, Jason Caravas, Benjamin Rambo-Martin, Shatavia Morrison, Christopher Gulvick, Scott Sammons, Yvette Unoarumhi, Darlene Wagner, Matthew Schmerer, Minoo Agarwal, Eyad Almasri, Debbie Boles, Ayla Burns, Nuthawin Charoensri, Oren Cohen, Susan Countryman, Mary Ann Cristobal, Bobbi Croy, Suzanne Dale, Hrushikesh Deshmukh, Amanda Douglas, Vincent Drouillon, Marcia Eisenberg, Howard Engler, Rama Ghatti, Prashant Gupta, Susan Hicks, Jake Humphrey, Lax Iyer, Lisa Pfefferle, Manoj Jain, Matthew Robinson, Mohan Kolli, Brian Krueger, Tim Kuphal, Stanley Letovsky, Michael Levandoski, Craig Lukasik, Jonathan Meltzer, Brian Norvell, Mindy Nye, Scott Parker, Christos Petropoulos, John Pruitt, Steven Ragan, Scott Ryan, Mike Sapeta, Jana Schroth, Suresh Babu Selvaraju, Goran Stevovic, Amanda Suchanek, Andrea Throop, Lyndon Tilson, Thomas Urban, Joe Voshell, Kimberly Wagner, Jonathan Williams, Mary Williamson, Qian Zeng, Tricia Zwiefelhofer, Clinton R. Paden, Duncan MacCannell |
| EPI_ISL_3330205 | Illinois Department of Public Health - Springfield Lab | Illinois Department of Public Health - Springfield Lab | Bryan Sim, Gordon McCall, Marshall Smock |
| EPI_ISL_3330206, EPI_ISL_3330207 | Laboratory Corporation of America | Centers for Disease Control and Prevention Division of Viral Diseases, Pathogen Discovery | Dakota Howard, Dhwani Batra, Peter W. Cook, Kara Moser, Adrian Paskey, Jason Caravas, Benjamin Rambo-Martin, Shatavia Morrison, Christopher Gulvick, Scott Sammons, Yvette Unoarumhi, Darlene Wagner, Matthew Schmerer, Minoo Agarwal, Eyad Almasri, Debbie Boles, Ayla Burns, Nuthawin Charoensri, Oren Cohen, Susan Countryman, Mary Ann Cristobal, Bobbi Croy, Suzanne Dale, Hrushikesh Deshmukh, Amanda Douglas, Vincent Drouillon, Marcia Eisenberg, Howard Engler, Rama Ghatti, Prashant Gupta, Susan Hicks, Jake Humphrey, Lax Iyer, Lisa Pfefferle, Manoj Jain, Matthew Robinson, Mohan Kolli, Brian Krueger, Tim Kuphal, Stanley Letovsky, Michael Levandoski, Craig Lukasik, Jonathan Meltzer, Brian Norvell, Mindy Nye, Scott Parker, Christos Petropoulos, John Pruitt, Steven Ragan, Scott Ryan, Mike Sapeta, Jana Schroth, Suresh Babu Selvaraju, Goran Stevovic, Amanda Suchanek, Andrea Throop, Lyndon Tilson, Thomas Urban, Joe Voshell, Kimberly Wagner, Jonathan Williams, Mary Williamson, Qian Zeng, Tricia Zwiefelhofer, Clinton R. Paden, Duncan MacCannell |
| EPI_ISL_3330208 | Illinois Department of Public Health - Springfield Lab | Illinois Department of Public Health - Springfield Lab | Bryan Sim, Gordon McCall, Marshall Smock |
| EPI_ISL_3330209, EPI_ISL_3330211, EPI_ISL_3330213 | Laboratory Corporation of America | Centers for Disease Control and Prevention Division of Viral Diseases, Pathogen Discovery | Dakota Howard, Dhwani Batra, Peter W. Cook, Kara Moser, Adrian Paskey, Jason Caravas, Benjamin Rambo-Martin, Shatavia Morrison, Christopher Gulvick, Scott Sammons, Yvette Unoarumhi, Darlene Wagner, Matthew Schmerer, Minoo Agarwal, Eyad Almasri, Debbie Boles, Ayla Burns, Nuthawin Charoensri, Oren Cohen, Susan Countryman, Mary Ann Cristobal, Bobbi Croy, Suzanne Dale, Hrushikesh Deshmukh, Amanda Douglas, Vincent Drouillon, Marcia Eisenberg, Howard Engler, Rama Ghatti, Prashant Gupta, Susan Hicks, Jake Humphrey, Lax Iyer, Lisa Pfefferle, Manoj Jain, Matthew Robinson, Mohan Kolli, Brian Krueger, Tim Kuphal, Stanley Letovsky, Michael Levandoski, Craig Lukasik, Jonathan Meltzer, Brian Norvell, Mindy Nye, Scott Parker, Christos Petropoulos, John Pruitt, Steven Ragan, Scott Ryan, Mike Sapeta, Jana Schroth, Suresh Babu Selvaraju, Goran Stevovic, Amanda Suchanek, Andrea Throop, Lyndon Tilson, Thomas Urban, Joe Voshell, Kimberly Wagner, Jonathan Williams, Mary Williamson, Qian Zeng, Tricia Zwiefelhofer, Clinton R. Paden, Duncan MacCannell |
| EPI_ISL_3330214 | Illinois Department of Public Health - Springfield Lab | Illinois Department of Public Health - Springfield Lab | Bryan Sim, Gordon McCall, Marshall Smock |
| EPI_ISL_3330215, EPI_ISL_3330217 | Laboratory Corporation of America | Centers for Disease Control and Prevention Division of Viral Diseases, Pathogen Discovery | Dakota Howard, Dhwani Batra, Peter W. Cook, Kara Moser, Adrian Paskey, Jason Caravas, Benjamin Rambo-Martin, Shatavia Morrison, Christopher Gulvick, Scott Sammons, Yvette Unoarumhi, Darlene Wagner, Matthew Schmerer, Minoo Agarwal, Eyad Almasri, Debbie Boles, Ayla Burns, Nuthawin Charoensri, Oren Cohen, Susan Countryman, Mary Ann Cristobal, Bobbi Croy, Suzanne Dale, Hrushikesh Deshmukh, Amanda Douglas, Vincent Drouillon, Marcia Eisenberg, Howard Engler, Rama Ghatti, Prashant Gupta, Susan Hicks, Jake Humphrey, Lax Iyer, Lisa Pfefferle, Manoj Jain, Matthew Robinson, Mohan Kolli, Brian Krueger, Tim Kuphal, Stanley Letovsky, Michael Levandoski, Craig Lukasik, Jonathan Meltzer, Brian Norvell, Mindy Nye, Scott Parker, Christos Petropoulos, John Pruitt, Steven Ragan, Scott Ryan, Mike Sapeta, Jana Schroth, Suresh Babu Selvaraju, Goran Stevovic, Amanda Suchanek, Andrea Throop, Lyndon Tilson, Thomas Urban, Joe Voshell, Kimberly Wagner, Jonathan Williams, Mary Williamson, Qian Zeng, Tricia Zwiefelhofer, Clinton R. Paden, Duncan MacCannell |

| | | | |
|---|---|---|---|
| EPI_ISL_3330218 | Illinois Department of Public Health - Springfield Lab | Illinois Department of Public Health - Springfield Lab | Bryan Sim, Gordon McCall, Marshall Smock |
| EPI_ISL_3330220, EPI_ISL_3330222 | Laboratory Corporation of America | Centers for Disease Control and Prevention Division of Viral Diseases, Pathogen Discovery | Dakota Howard, Dhwani Batra, Peter W. Cook, Kara Moser, Adrian Paskey, Jason Caravas, Benjamin Rambo-Martin, Shatavia Morrison, Christopher Gulvick, Scott Sammons, Yvette Unoarumhi, Darlene Wagner, Matthew Schmerer, Minoo Agarwal, Eyad Almasri, Debbie Boles, Ayla Burns, Nuthawin Charoensri, Oren Cohen, Susan Countryman, Mary Ann Cristobal, Bobbi Croy, Suzanne Dale, Hrushikesh Deshmukh, Amanda Douglas, Vincent Drouillon, Marcia Eisenberg, Howard Engler, Rama Ghatti, Prashant Gupta, Susan Hicks, Jake Humphrey, Lax Iyer, Lisa Pfefferle, Manoj Jain, Matthew Robinson, Mohan Kolli, Brian Krueger, Tim Kuphal, Stanley Letovsky, Michael Levandoski, Craig Lukasik, Jonathan Meltzer, Brian Norvell, Mindy Nye, Scott Parker, Christos Petropoulos, John Pruitt, Steven Ragan, Scott Ryan, Mike Sapeta, Jana Schroth, Suresh Babu Selvaraju, Goran Stevovic, Amanda Suchanek, Andrea Throop, Lyndon Tilson, Thomas Urban, Joe Voshell, Kimberly Wagner, Jonathan Williams, Mary Williamson, Qian Zeng, Tricia Zwiefelhofer, Clinton R. Paden, Duncan MacCannell |
| EPI_ISL_3330224 | Illinois Department of Public Health - Springfield Lab | Illinois Department of Public Health - Springfield Lab | Bryan Sim, Gordon McCall, Marshall Smock |
| EPI_ISL_3330225 | Laboratory Corporation of America | Centers for Disease Control and Prevention Division of Viral Diseases, Pathogen Discovery | Dakota Howard, Dhwani Batra, Peter W. Cook, Kara Moser, Adrian Paskey, Jason Caravas, Benjamin Rambo-Martin, Shatavia Morrison, Christopher Gulvick, Scott Sammons, Yvette Unoarumhi, Darlene Wagner, Matthew Schmerer, Minoo Agarwal, Eyad Almasri, Debbie Boles, Ayla Burns, Nuthawin Charoensri, Oren Cohen, Susan Countryman, Mary Ann Cristobal, Bobbi Croy, Suzanne Dale, Hrushikesh Deshmukh, Amanda Douglas, Vincent Drouillon, Marcia Eisenberg, Howard Engler, Rama Ghatti, Prashant Gupta, Susan Hicks, Jake Humphrey, Lax Iyer, Lisa Pfefferle, Manoj Jain, Matthew Robinson, Mohan Kolli, Brian Krueger, Tim Kuphal, Stanley Letovsky, Michael Levandoski, Craig Lukasik, Jonathan Meltzer, Brian Norvell, Mindy Nye, Scott Parker, Christos Petropoulos, John Pruitt, Steven Ragan, Scott Ryan, Mike Sapeta, Jana Schroth, Suresh Babu Selvaraju, Goran Stevovic, Amanda Suchanek, Andrea Throop, Lyndon Tilson, Thomas Urban, Joe Voshell, Kimberly Wagner, Jonathan Williams, Mary Williamson, Qian Zeng, Tricia Zwiefelhofer, Clinton R. Paden, Duncan MacCannell |
| EPI_ISL_3330226, EPI_ISL_3330228, EPI_ISL_3330229, EPI_ISL_3330231 | Illinois Department of Public Health - Springfield Lab | Illinois Department of Public Health - Springfield Lab | Bryan Sim, Gordon McCall, Marshall Smock |
| EPI_ISL_3330233 | Laboratory Corporation of America | Centers for Disease Control and Prevention Division of Viral Diseases, Pathogen Discovery | Dakota Howard, Dhwani Batra, Peter W. Cook, Kara Moser, Adrian Paskey, Jason Caravas, Benjamin Rambo-Martin, Shatavia Morrison, Christopher Gulvick, Scott Sammons, Yvette Unoarumhi, Darlene Wagner, Matthew Schmerer, Minoo Agarwal, Eyad Almasri, Debbie Boles, Ayla Burns, Nuthawin Charoensri, Oren Cohen, Susan Countryman, Mary Ann Cristobal, Bobbi Croy, Suzanne Dale, Hrushikesh Deshmukh, Amanda Douglas, Vincent Drouillon, Marcia Eisenberg, Howard Engler, Rama Ghatti, Prashant Gupta, Susan Hicks, Jake Humphrey, Lax Iyer, Lisa Pfefferle, Manoj Jain, Matthew Robinson, Mohan Kolli, Brian Krueger, Tim Kuphal, Stanley Letovsky, Michael Levandoski, Craig Lukasik, Jonathan Meltzer, Brian Norvell, Mindy Nye, Scott Parker, Christos Petropoulos, John Pruitt, Steven Ragan, Scott Ryan, Mike Sapeta, Jana Schroth, Suresh Babu Selvaraju, Goran Stevovic, Amanda Suchanek, Andrea Throop, Lyndon Tilson, Thomas Urban, Joe Voshell, Kimberly Wagner, Jonathan Williams, Mary Williamson, Qian Zeng, Tricia Zwiefelhofer, Clinton R. Paden, Duncan MacCannell |
| EPI_ISL_3330234 | Illinois Department of Public Health - Springfield Lab | Illinois Department of Public Health - Springfield Lab | Bryan Sim, Gordon McCall, Marshall Smock |
| EPI_ISL_3330236, EPI_ISL_3330237, EPI_ISL_3330238 | Laboratory Corporation of America | Centers for Disease Control and Prevention Division of Viral Diseases, Pathogen Discovery | Dakota Howard, Dhwani Batra, Peter W. Cook, Kara Moser, Adrian Paskey, Jason Caravas, Benjamin Rambo-Martin, Shatavia Morrison, Christopher Gulvick, Scott Sammons, Yvette Unoarumhi, Darlene Wagner, Matthew Schmerer, Minoo Agarwal, Eyad Almasri, Debbie Boles, Ayla Burns, Nuthawin Charoensri, Oren Cohen, Susan Countryman, Mary Ann Cristobal, Bobbi Croy, Suzanne Dale, Hrushikesh Deshmukh, Amanda Douglas, Vincent Drouillon, Marcia Eisenberg, Howard Engler, Rama Ghatti, Prashant Gupta, Susan Hicks, Jake Humphrey, Lax Iyer, Lisa Pfefferle, Manoj Jain, Matthew Robinson, Mohan Kolli, Brian Krueger, Tim Kuphal, Stanley Letovsky, Michael Levandoski, Craig Lukasik, Jonathan Meltzer, Brian Norvell, Mindy Nye, Scott Parker, Christos Petropoulos, John Pruitt, Steven Ragan, Scott Ryan, Mike Sapeta, Jana Schroth, Suresh Babu Selvaraju, Goran Stevovic, Amanda Suchanek, Andrea Throop, Lyndon Tilson, Thomas Urban, Joe Voshell, Kimberly Wagner, Jonathan Williams, Mary Williamson, Qian Zeng, Tricia Zwiefelhofer, Clinton R. Paden, Duncan MacCannell |
| EPI_ISL_3330239 | Illinois Department of Public Health - Springfield Lab | Illinois Department of Public Health - Springfield Lab | Bryan Sim, Gordon McCall, Marshall Smock |
| EPI_ISL_3330240, EPI_ISL_3330241, EPI_ISL_3330242 | Laboratory Corporation of America | Centers for Disease Control and Prevention Division of Viral Diseases, Pathogen Discovery | Dakota Howard, Dhwani Batra, Peter W. Cook, Kara Moser, Adrian Paskey, Jason Caravas, Benjamin Rambo-Martin, Shatavia Morrison, Christopher Gulvick, Scott Sammons, Yvette Unoarumhi, Darlene Wagner, Matthew Schmerer, Minoo Agarwal, Eyad Almasri, Debbie Boles, Ayla Burns, Nuthawin Charoensri, Oren Cohen, Susan Countryman, Mary Ann Cristobal, Bobbi Croy, Suzanne Dale, Hrushikesh Deshmukh, Amanda Douglas, Vincent Drouillon, Marcia Eisenberg, Howard Engler, Rama Ghatti, Prashant Gupta, Susan Hicks, Jake Humphrey, Lax Iyer, Lisa Pfefferle, Manoj Jain, Matthew Robinson, Mohan Kolli, Brian Krueger, Tim Kuphal, Stanley Letovsky, Michael Levandoski, Craig Lukasik, Jonathan Meltzer, Brian Norvell, Mindy Nye, Scott Parker, Christos Petropoulos, John Pruitt, Steven Ragan, Scott Ryan, Mike Sapeta, Jana Schroth, Suresh Babu Selvaraju, Goran Stevovic, Amanda Suchanek, Andrea Throop, Lyndon Tilson, Thomas Urban, Joe Voshell, Kimberly Wagner, Jonathan Williams, Mary Williamson, Qian Zeng, Tricia Zwiefelhofer, Clinton R. Paden, Duncan MacCannell |
| EPI_ISL_3330243 | Illinois Department of Public Health - Springfield Lab | Illinois Department of Public Health - Springfield Lab | Bryan Sim, Gordon McCall, Marshall Smock |
| EPI_ISL_3330244, EPI_ISL_3330245 | Laboratory Corporation of America | Centers for Disease Control and Prevention Division of Viral Diseases, Pathogen Discovery | Dakota Howard, Dhwani Batra, Peter W. Cook, Kara Moser, Adrian Paskey, Jason Caravas, Benjamin Rambo-Martin, Shatavia Morrison, Christopher Gulvick, Scott Sammons, Yvette Unoarumhi, Darlene Wagner, Matthew Schmerer, Minoo Agarwal, Eyad Almasri, Debbie Boles, Ayla Burns, Nuthawin Charoensri, Oren Cohen, Susan Countryman, Mary Ann Cristobal, Bobbi Croy, Suzanne Dale, Hrushikesh Deshmukh, Amanda Douglas, Vincent Drouillon, Marcia Eisenberg, Howard Engler, Rama Ghatti, Prashant Gupta, Susan Hicks, Jake Humphrey, Lax Iyer, Lisa Pfefferle, Manoj Jain, Matthew Robinson, Mohan Kolli, Brian Krueger, Tim Kuphal, Stanley Letovsky, Michael Levandoski, Craig Lukasik, Jonathan Meltzer, Brian Norvell, Mindy Nye, Scott Parker, Christos Petropoulos, John Pruitt, Steven Ragan, Scott Ryan, Mike Sapeta, Jana Schroth, Suresh Babu Selvaraju, Goran Stevovic, Amanda Suchanek, Andrea Throop, Lyndon Tilson, Thomas Urban, Joe Voshell, Kimberly Wagner, Jonathan Williams, Mary Williamson, Qian Zeng, Tricia Zwiefelhofer, Clinton R. Paden, Duncan MacCannell |
| EPI_ISL_3330246 | Illinois Department of Public Health - Springfield Lab | Illinois Department of Public Health - Springfield Lab | Bryan Sim, Gordon McCall, Marshall Smock |
| EPI_ISL_3330247, EPI_ISL_3330248 | Laboratory Corporation of America | Centers for Disease Control and Prevention Division of Viral Diseases, Pathogen Discovery | Dakota Howard, Dhwani Batra, Peter W. Cook, Kara Moser, Adrian Paskey, Jason Caravas, Benjamin Rambo-Martin, Shatavia Morrison, Christopher Gulvick, Scott Sammons, Yvette Unoarumhi, Darlene Wagner, Matthew Schmerer, Minoo Agarwal, Eyad Almasri, Debbie Boles, Ayla Burns, Nuthawin Charoensri, Oren Cohen, Susan Countryman, Mary Ann Cristobal, Bobbi Croy, Suzanne Dale, Hrushikesh Deshmukh, Amanda Douglas, Vincent Drouillon, Marcia Eisenberg, Howard Engler, Rama Ghatti, Prashant Gupta, Susan Hicks, Jake Humphrey, Lax Iyer, Lisa Pfefferle, Manoj Jain, Matthew Robinson, Mohan Kolli, Brian Krueger, Tim Kuphal, Stanley Letovsky, Michael Levandoski, Craig Lukasik, Jonathan Meltzer, Brian Norvell, Mindy Nye, Scott Parker, Christos Petropoulos, John Pruitt, Steven Ragan, Scott Ryan, Mike Sapeta, Jana Schroth, Suresh Babu Selvaraju, Goran Stevovic, Amanda Suchanek, Andrea Throop, Lyndon Tilson, Thomas Urban, Joe Voshell, Kimberly Wagner, Jonathan Williams, Mary Williamson, Qian Zeng, Tricia Zwiefelhofer, Clinton R. Paden, Duncan MacCannell |
| EPI_ISL_3330249 | Illinois Department of Public Health - Springfield Lab | Illinois Department of Public Health - Springfield Lab | Bryan Sim, Gordon McCall, Marshall Smock |
| EPI_ISL_3330250, EPI_ISL_3330251 | Laboratory Corporation of America | Centers for Disease Control and Prevention Division of Viral Diseases, Pathogen Discovery | Dakota Howard, Dhwani Batra, Peter W. Cook, Kara Moser, Adrian Paskey, Jason Caravas, Benjamin Rambo-Martin, Shatavia Morrison, Christopher Gulvick, Scott Sammons, Yvette Unoarumhi, Darlene Wagner, Matthew Schmerer, Minoo Agarwal, Eyad Almasri, Debbie Boles, Ayla Burns, Nuthawin Charoensri, Oren Cohen, Susan Countryman, Mary Ann Cristobal, Bobbi Croy, Suzanne Dale, Hrushikesh Deshmukh, Amanda Douglas, Vincent Drouillon, Marcia Eisenberg, Howard Engler, Rama Ghatti, Prashant Gupta, Susan Hicks, Jake Humphrey, Lax Iyer, Lisa Pfefferle, Manoj Jain, Matthew Robinson, Mohan Kolli, Brian Krueger, Tim Kuphal, Stanley Letovsky, Michael Levandoski, Craig Lukasik, Jonathan Meltzer, Brian Norvell, Mindy Nye, Scott Parker, Christos Petropoulos, John Pruitt, Steven Ragan, Scott Ryan, Mike Sapeta, Jana Schroth, Suresh Babu Selvaraju, Goran Stevovic, Amanda Suchanek, Andrea Throop, Lyndon Tilson, Thomas Urban, Joe Voshell, Kimberly Wagner, Jonathan Williams, Mary Williamson, Qian Zeng, Tricia Zwiefelhofer, Clinton R. Paden, Duncan MacCannell |
| EPI_ISL_3330252 | Illinois Department of Public Health - Springfield Lab | Illinois Department of Public Health - Springfield Lab | Bryan Sim, Gordon McCall, Marshall Smock |
| EPI_ISL_3330253, EPI_ISL_3330254, EPI_ISL_3330255 | Laboratory Corporation of America | Centers for Disease Control and Prevention Division of Viral Diseases, Pathogen Discovery | Dakota Howard, Dhwani Batra, Peter W. Cook, Kara Moser, Adrian Paskey, Jason Caravas, Benjamin Rambo-Martin, Shatavia Morrison, Christopher Gulvick, Scott Sammons, Yvette Unoarumhi, Darlene Wagner, Matthew Schmerer, Minoo Agarwal, Eyad Almasri, Debbie Boles, Ayla Burns, Nuthawin Charoensri, Oren Cohen, Susan Countryman, Mary Ann Cristobal, Bobbi Croy, Suzanne Dale, Hrushikesh Deshmukh, Amanda Douglas, Vincent Drouillon, Marcia Eisenberg, Howard Engler, Rama Ghatti, Prashant Gupta, Susan Hicks, Jake Humphrey, Lax Iyer, Lisa Pfefferle, Manoj Jain, Matthew Robinson, Mohan Kolli, Brian Krueger, Tim Kuphal, Stanley Letovsky, Michael Levandoski, Craig Lukasik, Jonathan Meltzer, Brian Norvell, Mindy Nye, Scott Parker, Christos Petropoulos, John Pruitt, Steven Ragan, Scott Ryan, Mike Sapeta, Jana Schroth, Suresh Babu Selvaraju, Goran Stevovic, Amanda Suchanek, Andrea Throop, Lyndon Tilson, Thomas Urban, Joe Voshell, Kimberly Wagner, Jonathan Williams, Mary Williamson, Qian Zeng, Tricia Zwiefelhofer, Clinton R. Paden, Duncan MacCannell |
| EPI_ISL_3330256 | Illinois Department of Public Health - Springfield Lab | Illinois Department of Public Health - Springfield Lab | Bryan Sim, Gordon McCall, Marshall Smock |
| EPI_ISL_3330257, EPI_ISL_3330258, EPI_ISL_3330259 | Laboratory Corporation of America | Centers for Disease Control and Prevention Division of Viral Diseases, Pathogen Discovery | Dakota Howard, Dhwani Batra, Peter W. Cook, Kara Moser, Adrian Paskey, Jason Caravas, Benjamin Rambo-Martin, Shatavia Morrison, Christopher Gulvick, Scott Sammons, Yvette Unoarumhi, Darlene Wagner, Matthew Schmerer, Minoo Agarwal, Eyad Almasri, Debbie Boles, Ayla Burns, Nuthawin Charoensri, Oren Cohen, Susan Countryman, Mary Ann Cristobal, Bobbi Croy, Suzanne Dale, Hrushikesh Deshmukh, Amanda Douglas, Vincent Drouillon, Marcia Eisenberg, Howard Engler, Rama Ghatti, Prashant Gupta, Susan Hicks, Jake Humphrey, Lax Iyer, Lisa Pfefferle, Manoj Jain, Matthew Robinson, Mohan Kolli, Brian Krueger, Tim Kuphal, Stanley Letovsky, Michael Levandoski, Craig Lukasik, Jonathan Meltzer, Brian Norvell, Mindy Nye, Scott Parker, Christos Petropoulos, John Pruitt, Steven Ragan, Scott Ryan, Mike Sapeta, Jana Schroth, Suresh Babu Selvaraju, Goran Stevovic, Amanda Suchanek, Andrea Throop, Lyndon Tilson, Thomas Urban, Joe Voshell, Kimberly Wagner, Jonathan Williams, Mary Williamson, Qian Zeng, Tricia Zwiefelhofer, Clinton R. Paden, Duncan MacCannell |
| EPI_ISL_3330260 | Illinois Department of Public Health - Springfield Lab | Illinois Department of Public Health - Springfield Lab | Bryan Sim, Gordon McCall, Marshall Smock |
| EPI_ISL_3330261, EPI_ISL_3330262, EPI_ISL_3330263 | Laboratory Corporation of America | Centers for Disease Control and Prevention Division of Viral Diseases, Pathogen Discovery | Dakota Howard, Dhwani Batra, Peter W. Cook, Kara Moser, Adrian Paskey, Jason Caravas, Benjamin Rambo-Martin, Shatavia Morrison, Christopher Gulvick, Scott Sammons, Yvette Unoarumhi, Darlene Wagner, Matthew Schmerer, Minoo Agarwal, Eyad Almasri, Debbie Boles, Ayla Burns, Nuthawin Charoensri, Oren Cohen, Susan Countryman, Mary Ann Cristobal, Bobbi Croy, Suzanne Dale, Hrushikesh Deshmukh, Amanda Douglas, Vincent Drouillon, Marcia Eisenberg, Howard Engler, Rama Ghatti, Prashant Gupta, Susan Hicks, Jake Humphrey, Lax Iyer, Lisa Pfefferle, Manoj Jain, Matthew Robinson, Mohan Kolli, Brian Krueger, Tim Kuphal, Stanley Letovsky, Michael Levandoski, Craig Lukasik, Jonathan Meltzer, Brian Norvell, Mindy Nye, Scott Parker, Christos Petropoulos, John Pruitt, Steven Ragan, Scott Ryan, Mike Sapeta, Jana Schroth, Suresh Babu Selvaraju, Goran Stevovic, Amanda Suchanek, Andrea Throop, Lyndon Tilson, Thomas Urban, Joe Voshell, Kimberly Wagner, Jonathan Williams, Mary Williamson, Qian Zeng, Tricia Zwiefelhofer, Clinton R. Paden, Duncan MacCannell |
| EPI_ISL_3330264 | Illinois Department of Public Health - Springfield Lab | Illinois Department of Public Health - Springfield Lab | Bryan Sim, Gordon McCall, Marshall Smock |
| EPI_ISL_3330265, EPI_ISL_3330266, EPI_ISL_3330267 | Laboratory Corporation of America | Centers for Disease Control and Prevention Division of Viral Diseases, Pathogen Discovery | Dakota Howard, Dhwani Batra, Peter W. Cook, Kara Moser, Adrian Paskey, Jason Caravas, Benjamin Rambo-Martin, Shatavia Morrison, Christopher Gulvick, Scott Sammons, Yvette Unoarumhi, Darlene Wagner, Matthew Schmerer, Minoo Agarwal, Eyad Almasri, Debbie Boles, Ayla Burns, Nuthawin Charoensri, Oren Cohen, Susan Countryman, Mary Ann Cristobal, Bobbi Croy, Suzanne Dale, Hrushikesh Deshmukh, Amanda Douglas, Vincent Drouillon, Marcia Eisenberg, Howard Engler, Rama Ghatti, Prashant Gupta, Susan Hicks, Jake Humphrey, Lax Iyer, Lisa Pfefferle, Manoj Jain, Matthew Robinson, Mohan Kolli, Brian Krueger, Tim Kuphal, Stanley Letovsky, Michael Levandoski, Craig Lukasik, Jonathan Meltzer, Brian Norvell, Mindy Nye, Scott Parker, Christos Petropoulos, John Pruitt, Steven Ragan, Scott Ryan, Mike Sapeta, Jana Schroth, Suresh Babu Selvaraju, Goran Stevovic, Amanda Suchanek, Andrea Throop, Lyndon Tilson, Thomas Urban, Joe Voshell, Kimberly Wagner, Jonathan Williams, Mary Williamson, Qian Zeng, Tricia Zwiefelhofer, Clinton R. Paden, Duncan MacCannell |
| EPI_ISL_3330268, EPI_ISL_3330269 | Illinois Department of Public Health - Springfield Lab | Illinois Department of Public Health - Springfield Lab | Bryan Sim, Gordon McCall, Marshall Smock |
| EPI_ISL_3330270 | Laboratory Corporation of America | Centers for Disease Control and Prevention Division of Viral Diseases, Pathogen Discovery | Dakota Howard, Dhwani Batra, Peter W. Cook, Kara Moser, Adrian Paskey, Jason Caravas, Benjamin Rambo-Martin, Shatavia Morrison, Christopher Gulvick, Scott Sammons, Yvette Unoarumhi, Darlene Wagner, Matthew Schmerer, Minoo Agarwal, Eyad Almasri, Debbie Boles, Ayla Burns, Nuthawin Charoensri, Oren Cohen, Susan Countryman, Mary Ann Cristobal, Bobbi Croy, Suzanne Dale, Hrushikesh Deshmukh, Amanda Douglas, Vincent Drouillon, Marcia Eisenberg, Howard Engler, Rama Ghatti, Prashant Gupta, Susan Hicks, Jake Humphrey, Lax Iyer, Lisa Pfefferle, Manoj Jain, Matthew Robinson, Mohan Kolli, Brian Krueger, Tim Kuphal, Stanley Letovsky, Michael Levandoski, Craig Lukasik, Jonathan Meltzer, Brian Norvell, Mindy Nye, Scott Parker, Christos Petropoulos, John Pruitt, Steven Ragan, Scott Ryan, Mike Sapeta, Jana Schroth, Suresh Babu Selvaraju, Goran Stevovic, Amanda Suchanek, Andrea Throop, Lyndon Tilson, Thomas Urban, Joe Voshell, Kimberly Wagner, Jonathan Williams, Mary Williamson, Qian Zeng, Tricia Zwiefelhofer, Clinton R. Paden, Duncan MacCannell |
| EPI_ISL_3330271 | Illinois Department of Public Health - Springfield Lab | Illinois Department of Public Health - Springfield Lab | Bryan Sim, Gordon McCall, Marshall Smock |
| EPI_ISL_3330272 | Laboratory Corporation of America | Centers for Disease Control and Prevention Division of Viral Diseases, Pathogen Discovery | Dakota Howard, Dhwani Batra, Peter W. Cook, Kara Moser, Adrian Paskey, Jason Caravas, Benjamin Rambo-Martin, Shatavia Morrison, Christopher Gulvick, Scott Sammons, Yvette Unoarumhi, Darlene Wagner, Matthew Schmerer, Minoo Agarwal, Eyad Almasri, Debbie Boles, Ayla Burns, Nuthawin Charoensri, Oren Cohen, Susan Countryman, Mary Ann Cristobal, Bobbi Croy, Suzanne Dale, Hrushikesh Deshmukh, Amanda Douglas, Vincent Drouillon, Marcia Eisenberg, Howard Engler, Rama Ghatti, Prashant Gupta, Susan Hicks, Jake Humphrey, Lax Iyer, Lisa Pfefferle, Manoj Jain, Matthew Robinson, Mohan Kolli, Brian Krueger, Tim Kuphal, Stanley Letovsky, Michael Levandoski, Craig Lukasik, Jonathan Meltzer, Brian Norvell, Mindy Nye, Scott Parker, Christos Petropoulos, John Pruitt, Steven Ragan, Scott Ryan, Mike Sapeta, Jana Schroth, Suresh Babu Selvaraju, Goran Stevovic, Amanda Suchanek, Andrea Throop, Lyndon Tilson, Thomas Urban, Joe Voshell, Kimberly Wagner, Jonathan Williams, Mary Williamson, Qian Zeng, Tricia Zwiefelhofer, Clinton R. Paden, Duncan MacCannell |
| EPI_ISL_3330273 | Illinois Department of Public Health - Springfield Lab | Illinois Department of Public Health - Springfield Lab | Bryan Sim, Gordon McCall, Marshall Smock |
| EPI_ISL_3330274 | Laboratory Corporation of America | Centers for Disease Control and Prevention Division of Viral Diseases, Pathogen Discovery | Dakota Howard, Dhwani Batra, Peter W. Cook, Kara Moser, Adrian Paskey, Jason Caravas, Benjamin Rambo-Martin, Shatavia Morrison, Christopher Gulvick, Scott Sammons, Yvette Unoarumhi, Darlene Wagner, Matthew Schmerer, Minoo Agarwal, Eyad Almasri, Debbie Boles, Ayla Burns, Nuthawin Charoensri, Oren Cohen, Susan Countryman, Mary Ann Cristobal, Bobbi Croy, Suzanne Dale, Hrushikesh Deshmukh, Amanda Douglas, Vincent Drouillon, Marcia Eisenberg, Howard Engler, Rama Ghatti, Prashant Gupta, Susan Hicks, Jake Humphrey, Lax Iyer, Lisa Pfefferle, Manoj Jain, Matthew Robinson, Mohan Kolli, Brian Krueger, Tim Kuphal, Stanley Letovsky, Michael Levandoski, Craig Lukasik, Jonathan Meltzer, Brian Norvell, Mindy Nye, Scott Parker, Christos Petropoulos, John Pruitt, Steven Ragan, Scott Ryan, Mike Sapeta, Jana Schroth, Suresh Babu Selvaraju, Goran Stevovic, Amanda Suchanek, Andrea Throop, Lyndon Tilson, Thomas Urban, Joe Voshell, Kimberly Wagner, Jonathan Williams, Mary Williamson, Qian Zeng, Tricia Zwiefelhofer, Clinton R. Paden, Duncan MacCannell |
| EPI_ISL_3330275 | Illinois Department of Public Health - Springfield Lab | Illinois Department of Public Health - Springfield Lab | Bryan Sim, Gordon McCall, Marshall Smock |
| EPI_ISL_3330276, | Laboratory Corporation of America | Centers for Disease Control and Prevention Division of Viral | Dakota Howard, Dhwani Batra, Peter W. Cook, Kara Moser, Adrian Paskey, Jason Caravas, Benjamin Rambo-Martin, Shatavia Morrison, Christopher Gulvick, Scott Sammons, Yvette Unoarumhi, Darlene Wagner, Matthew Schmerer, Minoo |

| | | | |
|---|---|---|---|
| EPI_ISL_3330277 | | Diseases, Pathogen Discovery | Agarwal, Eyad Almasri, Debbie Boles, Ayla Burns, Nuthawin Charoensri, Oren Cohen, Susan Countryman, Mary Ann Cristobal, Bobbi Croy, Suzanne Dale, Hrushikesh Deshmukh, Amanda Douglas, Vincent Drouillon, Marcia Eisenberg, Howard Engler, Rama Ghatti, Prashant Gupta, Susan Hicks, Jake Humphrey, Lax Iyer, Lisa Pfefferle, Manoj Jain, Matthew Robinson, Mohan Kolli, Brian Krueger, Tim Kuphal, Stanley Letovsky, Michael Levandoski, Craig Lukasik, Jonathan Meltzer, Brian Norvell, Mindy Nye, Scott Parker, Christos Petropoulos, John Pruitt, Steven Ragan, Scott Ryan, Mike Sapeta, Jana Schroth, Suresh Babu Selvaraju, Goran Stevovic, Amanda Suchanek, Andrea Throop, Lyndon Tilson, Thomas Urban, Joe Voshell, Kimberly Wagner, Jonathan Williams, Mary Williamson, Qian Zeng, Tricia Zwiefelhofer, Clinton R. Paden, Duncan MacCannell |
| EPI_ISL_3330278 | Illinois Department of Public Health - Springfield Lab | Illinois Department of Public Health - Springfield Lab | Bryan Sim, Gordon McCall, Marshall Smock |
| EPI_ISL_3330279, EPI_ISL_3330280 | Laboratory Corporation of America | Centers for Disease Control and Prevention Division of Viral Diseases, Pathogen Discovery | Dakota Howard, Dhwani Batra, Peter W. Cook, Kara Moser, Adrian Paskey, Jason Caravas, Benjamin Rambo-Martin, Shatavia Morrison, Christopher Gulvick, Scott Sammons, Yvette Unoarumhi, Darlene Wagner, Matthew Schmerer, Minoo Agarwal, Eyad Almasri, Debbie Boles, Ayla Burns, Nuthawin Charoensri, Oren Cohen, Susan Countryman, Mary Ann Cristobal, Bobbi Croy, Suzanne Dale, Hrushikesh Deshmukh, Amanda Douglas, Vincent Drouillon, Marcia Eisenberg, Howard Engler, Rama Ghatti, Prashant Gupta, Susan Hicks, Jake Humphrey, Lax Iyer, Lisa Pfefferle, Manoj Jain, Matthew Robinson, Mohan Kolli, Brian Krueger, Tim Kuphal, Stanley Letovsky, Michael Levandoski, Craig Lukasik, Jonathan Meltzer, Brian Norvell, Mindy Nye, Scott Parker, Christos Petropoulos, John Pruitt, Steven Ragan, Scott Ryan, Mike Sapeta, Jana Schroth, Suresh Babu Selvaraju, Goran Stevovic, Amanda Suchanek, Andrea Throop, Lyndon Tilson, Thomas Urban, Joe Voshell, Kimberly Wagner, Jonathan Williams, Mary Williamson, Qian Zeng, Tricia Zwiefelhofer, Clinton R. Paden, Duncan MacCannell |
| EPI_ISL_3330281 | Illinois Department of Public Health - Springfield Lab | Illinois Department of Public Health - Springfield Lab | Bryan Sim, Gordon McCall, Marshall Smock |
| EPI_ISL_3330282, EPI_ISL_3330283, EPI_ISL_3330284 | Laboratory Corporation of America | Centers for Disease Control and Prevention Division of Viral Diseases, Pathogen Discovery | Dakota Howard, Dhwani Batra, Peter W. Cook, Kara Moser, Adrian Paskey, Jason Caravas, Benjamin Rambo-Martin, Shatavia Morrison, Christopher Gulvick, Scott Sammons, Yvette Unoarumhi, Darlene Wagner, Matthew Schmerer, Minoo Agarwal, Eyad Almasri, Debbie Boles, Ayla Burns, Nuthawin Charoensri, Oren Cohen, Susan Countryman, Mary Ann Cristobal, Bobbi Croy, Suzanne Dale, Hrushikesh Deshmukh, Amanda Douglas, Vincent Drouillon, Marcia Eisenberg, Howard Engler, Rama Ghatti, Prashant Gupta, Susan Hicks, Jake Humphrey, Lax Iyer, Lisa Pfefferle, Manoj Jain, Matthew Robinson, Mohan Kolli, Brian Krueger, Tim Kuphal, Stanley Letovsky, Michael Levandoski, Craig Lukasik, Jonathan Meltzer, Brian Norvell, Mindy Nye, Scott Parker, Christos Petropoulos, John Pruitt, Steven Ragan, Scott Ryan, Mike Sapeta, Jana Schroth, Suresh Babu Selvaraju, Goran Stevovic, Amanda Suchanek, Andrea Throop, Lyndon Tilson, Thomas Urban, Joe Voshell, Kimberly Wagner, Jonathan Williams, Mary Williamson, Qian Zeng, Tricia Zwiefelhofer, Clinton R. Paden, Duncan MacCannell |
| EPI_ISL_3330285 | Illinois Department of Public Health - Springfield Lab | Illinois Department of Public Health - Springfield Lab | Bryan Sim, Gordon McCall, Marshall Smock |
| EPI_ISL_3330286 | Laboratory Corporation of America | Centers for Disease Control and Prevention Division of Viral Diseases, Pathogen Discovery | Dakota Howard, Dhwani Batra, Peter W. Cook, Kara Moser, Adrian Paskey, Jason Caravas, Benjamin Rambo-Martin, Shatavia Morrison, Christopher Gulvick, Scott Sammons, Yvette Unoarumhi, Darlene Wagner, Matthew Schmerer, Minoo Agarwal, Eyad Almasri, Debbie Boles, Ayla Burns, Nuthawin Charoensri, Oren Cohen, Susan Countryman, Mary Ann Cristobal, Bobbi Croy, Suzanne Dale, Hrushikesh Deshmukh, Amanda Douglas, Vincent Drouillon, Marcia Eisenberg, Howard Engler, Rama Ghatti, Prashant Gupta, Susan Hicks, Jake Humphrey, Lax Iyer, Lisa Pfefferle, Manoj Jain, Matthew Robinson, Mohan Kolli, Brian Krueger, Tim Kuphal, Stanley Letovsky, Michael Levandoski, Craig Lukasik, Jonathan Meltzer, Brian Norvell, Mindy Nye, Scott Parker, Christos Petropoulos, John Pruitt, Steven Ragan, Scott Ryan, Mike Sapeta, Jana Schroth, Suresh Babu Selvaraju, Goran Stevovic, Amanda Suchanek, Andrea Throop, Lyndon Tilson, Thomas Urban, Joe Voshell, Kimberly Wagner, Jonathan Williams, Mary Williamson, Qian Zeng, Tricia Zwiefelhofer, Clinton R. Paden, Duncan MacCannell |
| EPI_ISL_3330287 | Illinois Department of Public Health - Springfield Lab | Illinois Department of Public Health - Springfield Lab | Bryan Sim, Gordon McCall, Marshall Smock |
| EPI_ISL_3330288 | Laboratory Corporation of America | Centers for Disease Control and Prevention Division of Viral Diseases, Pathogen Discovery | Dakota Howard, Dhwani Batra, Peter W. Cook, Kara Moser, Adrian Paskey, Jason Caravas, Benjamin Rambo-Martin, Shatavia Morrison, Christopher Gulvick, Scott Sammons, Yvette Unoarumhi, Darlene Wagner, Matthew Schmerer, Minoo Agarwal, Eyad Almasri, Debbie Boles, Ayla Burns, Nuthawin Charoensri, Oren Cohen, Susan Countryman, Mary Ann Cristobal, Bobbi Croy, Suzanne Dale, Hrushikesh Deshmukh, Amanda Douglas, Vincent Drouillon, Marcia Eisenberg, Howard Engler, Rama Ghatti, Prashant Gupta, Susan Hicks, Jake Humphrey, Lax Iyer, Lisa Pfefferle, Manoj Jain, Matthew Robinson, Mohan Kolli, Brian Krueger, Tim Kuphal, Stanley Letovsky, Michael Levandoski, Craig Lukasik, Jonathan Meltzer, Brian Norvell, Mindy Nye, Scott Parker, Christos Petropoulos, John Pruitt, Steven Ragan, Scott Ryan, Mike Sapeta, Jana Schroth, Suresh Babu Selvaraju, Goran Stevovic, Amanda Suchanek, Andrea Throop, Lyndon Tilson, Thomas Urban, Joe Voshell, Kimberly Wagner, Jonathan Williams, Mary Williamson, Qian Zeng, Tricia Zwiefelhofer, Clinton R. Paden, Duncan MacCannell |
| EPI_ISL_3330289, EPI_ISL_3330290 | Illinois Department of Public Health - Springfield Lab | Illinois Department of Public Health - Springfield Lab | Bryan Sim, Gordon McCall, Marshall Smock |
| EPI_ISL_3330291, EPI_ISL_3330292, EPI_ISL_3330293, EPI_ISL_3330294 | Laboratory Corporation of America | Centers for Disease Control and Prevention Division of Viral Diseases, Pathogen Discovery | Dakota Howard, Dhwani Batra, Peter W. Cook, Kara Moser, Adrian Paskey, Jason Caravas, Benjamin Rambo-Martin, Shatavia Morrison, Christopher Gulvick, Scott Sammons, Yvette Unoarumhi, Darlene Wagner, Matthew Schmerer, Minoo Agarwal, Eyad Almasri, Debbie Boles, Ayla Burns, Nuthawin Charoensri, Oren Cohen, Susan Countryman, Mary Ann Cristobal, Bobbi Croy, Suzanne Dale, Hrushikesh Deshmukh, Amanda Douglas, Vincent Drouillon, Marcia Eisenberg, Howard Engler, Rama Ghatti, Prashant Gupta, Susan Hicks, Jake Humphrey, Lax Iyer, Lisa Pfefferle, Manoj Jain, Matthew Robinson, Mohan Kolli, Brian Krueger, Tim Kuphal, Stanley Letovsky, Michael Levandoski, Craig Lukasik, Jonathan Meltzer, Brian Norvell, Mindy Nye, Scott Parker, Christos Petropoulos, John Pruitt, Steven Ragan, Scott Ryan, Mike Sapeta, Jana Schroth, Suresh Babu Selvaraju, Goran Stevovic, Amanda Suchanek, Andrea Throop, Lyndon Tilson, Thomas Urban, Joe Voshell, Kimberly Wagner, Jonathan Williams, Mary Williamson, Qian Zeng, Tricia Zwiefelhofer, Clinton R. Paden, Duncan MacCannell |
| EPI_ISL_3330295 | Illinois Department of Public Health - Springfield Lab | Illinois Department of Public Health - Springfield Lab | Bryan Sim, Gordon McCall, Marshall Smock |
| EPI_ISL_3330296, EPI_ISL_3330297, EPI_ISL_3330298 | Laboratory Corporation of America | Centers for Disease Control and Prevention Division of Viral Diseases, Pathogen Discovery | Dakota Howard, Dhwani Batra, Peter W. Cook, Kara Moser, Adrian Paskey, Jason Caravas, Benjamin Rambo-Martin, Shatavia Morrison, Christopher Gulvick, Scott Sammons, Yvette Unoarumhi, Darlene Wagner, Matthew Schmerer, Minoo Agarwal, Eyad Almasri, Debbie Boles, Ayla Burns, Nuthawin Charoensri, Oren Cohen, Susan Countryman, Mary Ann Cristobal, Bobbi Croy, Suzanne Dale, Hrushikesh Deshmukh, Amanda Douglas, Vincent Drouillon, Marcia Eisenberg, Howard Engler, Rama Ghatti, Prashant Gupta, Susan Hicks, Jake Humphrey, Lax Iyer, Lisa Pfefferle, Manoj Jain, Matthew Robinson, Mohan Kolli, Brian Krueger, Tim Kuphal, Stanley Letovsky, Michael Levandoski, Craig Lukasik, Jonathan Meltzer, Brian Norvell, Mindy Nye, Scott Parker, Christos Petropoulos, John Pruitt, Steven Ragan, Scott Ryan, Mike Sapeta, Jana Schroth, Suresh Babu Selvaraju, Goran Stevovic, Amanda Suchanek, Andrea Throop, Lyndon Tilson, Thomas Urban, Joe Voshell, Kimberly Wagner, Jonathan Williams, Mary Williamson, Qian Zeng, Tricia Zwiefelhofer, Clinton R. Paden, Duncan MacCannell |
| EPI_ISL_3330299 | Illinois Department of Public Health - Springfield Lab | Illinois Department of Public Health - Springfield Lab | Bryan Sim, Gordon McCall, Marshall Smock |
| EPI_ISL_3330300, EPI_ISL_3330301 | Laboratory Corporation of America | Centers for Disease Control and Prevention Division of Viral Diseases, Pathogen Discovery | Dakota Howard, Dhwani Batra, Peter W. Cook, Kara Moser, Adrian Paskey, Jason Caravas, Benjamin Rambo-Martin, Shatavia Morrison, Christopher Gulvick, Scott Sammons, Yvette Unoarumhi, Darlene Wagner, Matthew Schmerer, Minoo Agarwal, Eyad Almasri, Debbie Boles, Ayla Burns, Nuthawin Charoensri, Oren Cohen, Susan Countryman, Mary Ann Cristobal, Bobbi Croy, Suzanne Dale, Hrushikesh Deshmukh, Amanda Douglas, Vincent Drouillon, Marcia Eisenberg, Howard Engler, Rama Ghatti, Prashant Gupta, Susan Hicks, Jake Humphrey, Lax Iyer, Lisa Pfefferle, Manoj Jain, Matthew Robinson, Mohan Kolli, Brian Krueger, Tim Kuphal, Stanley Letovsky, Michael Levandoski, Craig Lukasik, Jonathan Meltzer, Brian Norvell, Mindy Nye, Scott Parker, Christos Petropoulos, John Pruitt, Steven Ragan, Scott Ryan, Mike Sapeta, Jana Schroth, Suresh Babu Selvaraju, Goran Stevovic, Amanda Suchanek, Andrea Throop, Lyndon Tilson, Thomas Urban, Joe Voshell, Kimberly Wagner, Jonathan Williams, Mary Williamson, Qian Zeng, Tricia Zwiefelhofer, Clinton R. Paden, Duncan MacCannell |
| EPI_ISL_3330302 | Illinois Department of Public Health - Springfield Lab | Illinois Department of Public Health - Springfield Lab | Bryan Sim, Gordon McCall, Marshall Smock |
| EPI_ISL_3330303, EPI_ISL_3330304, EPI_ISL_3330305 | Laboratory Corporation of America | Centers for Disease Control and Prevention Division of Viral Diseases, Pathogen Discovery | Dakota Howard, Dhwani Batra, Peter W. Cook, Kara Moser, Adrian Paskey, Jason Caravas, Benjamin Rambo-Martin, Shatavia Morrison, Christopher Gulvick, Scott Sammons, Yvette Unoarumhi, Darlene Wagner, Matthew Schmerer, Minoo Agarwal, Eyad Almasri, Debbie Boles, Ayla Burns, Nuthawin Charoensri, Oren Cohen, Susan Countryman, Mary Ann Cristobal, Bobbi Croy, Suzanne Dale, Hrushikesh Deshmukh, Amanda Douglas, Vincent Drouillon, Marcia Eisenberg, Howard Engler, Rama Ghatti, Prashant Gupta, Susan Hicks, Jake Humphrey, Lax Iyer, Lisa Pfefferle, Manoj Jain, Matthew Robinson, Mohan Kolli, Brian Krueger, Tim Kuphal, Stanley Letovsky, Michael Levandoski, Craig Lukasik, Jonathan Meltzer, Brian Norvell, Mindy Nye, Scott Parker, Christos Petropoulos, John Pruitt, Steven Ragan, Scott Ryan, Mike Sapeta, Jana Schroth, Suresh Babu Selvaraju, Goran Stevovic, Amanda Suchanek, Andrea Throop, Lyndon Tilson, Thomas Urban, Joe Voshell, Kimberly Wagner, Jonathan Williams, Mary Williamson, Qian Zeng, Tricia Zwiefelhofer, Clinton R. Paden, Duncan MacCannell |
| EPI_ISL_3330306, EPI_ISL_3330307 | Illinois Department of Public Health - Springfield Lab | Illinois Department of Public Health - Springfield Lab | Bryan Sim, Gordon McCall, Marshall Smock |
| EPI_ISL_3330308 | Laboratory Corporation of America | Centers for Disease Control and Prevention Division of Viral Diseases, Pathogen Discovery | Dakota Howard, Dhwani Batra, Peter W. Cook, Kara Moser, Adrian Paskey, Jason Caravas, Benjamin Rambo-Martin, Shatavia Morrison, Christopher Gulvick, Scott Sammons, Yvette Unoarumhi, Darlene Wagner, Matthew Schmerer, Minoo Agarwal, Eyad Almasri, Debbie Boles, Ayla Burns, Nuthawin Charoensri, Oren Cohen, Susan Countryman, Mary Ann Cristobal, Bobbi Croy, Suzanne Dale, Hrushikesh Deshmukh, Amanda Douglas, Vincent Drouillon, Marcia Eisenberg, Howard Engler, Rama Ghatti, Prashant Gupta, Susan Hicks, Jake Humphrey, Lax Iyer, Lisa Pfefferle, Manoj Jain, Matthew Robinson, Mohan Kolli, Brian Krueger, Tim Kuphal, Stanley Letovsky, Michael Levandoski, Craig Lukasik, Jonathan Meltzer, Brian Norvell, Mindy Nye, Scott Parker, Christos Petropoulos, John Pruitt, Steven Ragan, Scott Ryan, Mike Sapeta, Jana Schroth, Suresh Babu Selvaraju, Goran Stevovic, Amanda Suchanek, Andrea Throop, Lyndon Tilson, Thomas Urban, Joe Voshell, Kimberly Wagner, Jonathan Williams, Mary Williamson, Qian Zeng, Tricia Zwiefelhofer, Clinton R. Paden, Duncan MacCannell |
| EPI_ISL_3330309 | Illinois Department of Public Health - Springfield Lab | Illinois Department of Public Health - Springfield Lab | Bryan Sim, Gordon McCall, Marshall Smock |
| EPI_ISL_3330310, EPI_ISL_3330312, EPI_ISL_3330313, EPI_ISL_3330314 | Laboratory Corporation of America | Centers for Disease Control and Prevention Division of Viral Diseases, Pathogen Discovery | Dakota Howard, Dhwani Batra, Peter W. Cook, Kara Moser, Adrian Paskey, Jason Caravas, Benjamin Rambo-Martin, Shatavia Morrison, Christopher Gulvick, Scott Sammons, Yvette Unoarumhi, Darlene Wagner, Matthew Schmerer, Minoo Agarwal, Eyad Almasri, Debbie Boles, Ayla Burns, Nuthawin Charoensri, Oren Cohen, Susan Countryman, Mary Ann Cristobal, Bobbi Croy, Suzanne Dale, Hrushikesh Deshmukh, Amanda Douglas, Vincent Drouillon, Marcia Eisenberg, Howard Engler, Rama Ghatti, Prashant Gupta, Susan Hicks, Jake Humphrey, Lax Iyer, Lisa Pfefferle, Manoj Jain, Matthew Robinson, Mohan Kolli, Brian Krueger, Tim Kuphal, Stanley Letovsky, Michael Levandoski, Craig Lukasik, Jonathan Meltzer, Brian Norvell, Mindy Nye, Scott Parker, Christos Petropoulos, John Pruitt, Steven Ragan, Scott Ryan, Mike Sapeta, Jana Schroth, Suresh Babu Selvaraju, Goran Stevovic, Amanda Suchanek, Andrea Throop, Lyndon Tilson, Thomas Urban, Joe Voshell, Kimberly Wagner, Jonathan Williams, Mary Williamson, Qian Zeng, Tricia Zwiefelhofer, Clinton R. Paden, Duncan MacCannell |
| EPI_ISL_3330315 | Illinois Department of Public Health - Springfield Lab | Illinois Department of Public Health - Springfield Lab | Bryan Sim, Gordon McCall, Marshall Smock |
| EPI_ISL_3330316 | Laboratory Corporation of America | Centers for Disease Control and Prevention Division of Viral Diseases, Pathogen Discovery | Dakota Howard, Dhwani Batra, Peter W. Cook, Kara Moser, Adrian Paskey, Jason Caravas, Benjamin Rambo-Martin, Shatavia Morrison, Christopher Gulvick, Scott Sammons, Yvette Unoarumhi, Darlene Wagner, Matthew Schmerer, Minoo Agarwal, Eyad Almasri, Debbie Boles, Ayla Burns, Nuthawin Charoensri, Oren Cohen, Susan Countryman, Mary Ann Cristobal, Bobbi Croy, Suzanne Dale, Hrushikesh Deshmukh, Amanda Douglas, Vincent Drouillon, Marcia Eisenberg, Howard Engler, Rama Ghatti, Prashant Gupta, Susan Hicks, Jake Humphrey, Lax Iyer, Lisa Pfefferle, Manoj Jain, Matthew Robinson, Mohan Kolli, Brian Krueger, Tim Kuphal, Stanley Letovsky, Michael Levandoski, Craig Lukasik, Jonathan Meltzer, Brian Norvell, Mindy Nye, Scott Parker, Christos Petropoulos, John Pruitt, Steven Ragan, Scott Ryan, Mike Sapeta, Jana Schroth, Suresh Babu Selvaraju, Goran Stevovic, Amanda Suchanek, Andrea Throop, Lyndon Tilson, Thomas Urban, Joe Voshell, Kimberly Wagner, Jonathan Williams, Mary Williamson, Qian Zeng, Tricia Zwiefelhofer, Clinton R. Paden, Duncan MacCannell |
| EPI_ISL_3330317, EPI_ISL_3330318 | Illinois Department of Public Health - Springfield Lab | Illinois Department of Public Health - Springfield Lab | Bryan Sim, Gordon McCall, Marshall Smock |
| EPI_ISL_3330319 | Laboratory Corporation of America | Centers for Disease Control and Prevention Division of Viral Diseases, Pathogen Discovery | Dakota Howard, Dhwani Batra, Peter W. Cook, Kara Moser, Adrian Paskey, Jason Caravas, Benjamin Rambo-Martin, Shatavia Morrison, Christopher Gulvick, Scott Sammons, Yvette Unoarumhi, Darlene Wagner, Matthew Schmerer, Minoo Agarwal, Eyad Almasri, Debbie Boles, Ayla Burns, Nuthawin Charoensri, Oren Cohen, Susan Countryman, Mary Ann Cristobal, Bobbi Croy, Suzanne Dale, Hrushikesh Deshmukh, Amanda Douglas, Vincent Drouillon, Marcia Eisenberg, Howard Engler, Rama Ghatti, Prashant Gupta, Susan Hicks, Jake Humphrey, Lax Iyer, Lisa Pfefferle, Manoj Jain, Matthew Robinson, Mohan Kolli, Brian Krueger, Tim Kuphal, Stanley Letovsky, Michael Levandoski, Craig Lukasik, Jonathan Meltzer, Brian Norvell, Mindy Nye, Scott Parker, Christos Petropoulos, John Pruitt, Steven Ragan, Scott Ryan, Mike Sapeta, Jana Schroth, Suresh Babu Selvaraju, Goran Stevovic, Amanda Suchanek, Andrea Throop, Lyndon Tilson, Thomas Urban, Joe Voshell, Kimberly Wagner, Jonathan Williams, Mary Williamson, Qian Zeng, Tricia Zwiefelhofer, Clinton R. Paden, Duncan MacCannell |
| EPI_ISL_3330320 | Illinois Department of Public Health - Springfield Lab | Illinois Department of Public Health - Springfield Lab | Bryan Sim, Gordon McCall, Marshall Smock |
| EPI_ISL_3330321 | Laboratory Corporation of America | Centers for Disease Control and Prevention Division of Viral Diseases, Pathogen Discovery | Dakota Howard, Dhwani Batra, Peter W. Cook, Kara Moser, Adrian Paskey, Jason Caravas, Benjamin Rambo-Martin, Shatavia Morrison, Christopher Gulvick, Scott Sammons, Yvette Unoarumhi, Darlene Wagner, Matthew Schmerer, Minoo Agarwal, Eyad Almasri, Debbie Boles, Ayla Burns, Nuthawin Charoensri, Oren Cohen, Susan Countryman, Mary Ann Cristobal, Bobbi Croy, Suzanne Dale, Hrushikesh Deshmukh, Amanda Douglas, Vincent Drouillon, Marcia Eisenberg, Howard Engler, Rama Ghatti, Prashant Gupta, Susan Hicks, Jake Humphrey, Lax Iyer, Lisa Pfefferle, Manoj Jain, Matthew Robinson, Mohan Kolli, Brian Krueger, Tim Kuphal, Stanley Letovsky, Michael Levandoski, Craig Lukasik, Jonathan Meltzer, Brian Norvell, Mindy Nye, Scott Parker, Christos Petropoulos, John Pruitt, Steven Ragan, Scott Ryan, Mike Sapeta, Jana Schroth, Suresh Babu Selvaraju, Goran Stevovic, Amanda Suchanek, Andrea Throop, Lyndon Tilson, Thomas Urban, Joe Voshell, Kimberly Wagner, Jonathan Williams, Mary Williamson, Qian Zeng, Tricia Zwiefelhofer, Clinton R. Paden, Duncan MacCannell |
| EPI_ISL_3330322, EPI_ISL_3330323 | Illinois Department of Public Health - Springfield Lab | Illinois Department of Public Health - Springfield Lab | Bryan Sim, Gordon McCall, Marshall Smock |
| EPI_ISL_3330324 | Laboratory Corporation of America | Centers for Disease Control and Prevention Division of Viral Diseases, Pathogen Discovery | Dakota Howard, Dhwani Batra, Peter W. Cook, Kara Moser, Adrian Paskey, Jason Caravas, Benjamin Rambo-Martin, Shatavia Morrison, Christopher Gulvick, Scott Sammons, Yvette Unoarumhi, Darlene Wagner, Matthew Schmerer, Minoo Agarwal, Eyad Almasri, Debbie Boles, Ayla Burns, Nuthawin Charoensri, Oren Cohen, Susan Countryman, Mary Ann Cristobal, Bobbi Croy, Suzanne Dale, Hrushikesh Deshmukh, Amanda Douglas, Vincent Drouillon, Marcia Eisenberg, Howard Engler, Rama Ghatti, Prashant Gupta, Susan Hicks, Jake Humphrey, Lax Iyer, Lisa Pfefferle, Manoj Jain, Matthew Robinson, Mohan Kolli, Brian Krueger, Tim Kuphal, Stanley Letovsky, Michael Levandoski, Craig Lukasik, Jonathan Meltzer, Brian Norvell, Mindy Nye, Scott Parker, Christos Petropoulos, John Pruitt, Steven Ragan, Scott Ryan, Mike Sapeta, Jana Schroth, Suresh Babu Selvaraju, Goran Stevovic, Amanda Suchanek, Andrea Throop, Lyndon Tilson, Thomas Urban, Joe Voshell, Kimberly Wagner, Jonathan Williams, Mary Williamson, Qian Zeng, Tricia Zwiefelhofer, Clinton R. Paden, Duncan MacCannell |
| EPI_ISL_3330325 | Illinois Department of Public Health - Springfield Lab | Illinois Department of Public Health - Springfield Lab | Bryan Sim, Gordon McCall, Marshall Smock |
| EPI_ISL_3330327 | Laboratory Corporation of America | Centers for Disease Control and Prevention Division of Viral Diseases, Pathogen Discovery | Dakota Howard, Dhwani Batra, Peter W. Cook, Kara Moser, Adrian Paskey, Jason Caravas, Benjamin Rambo-Martin, Shatavia Morrison, Christopher Gulvick, Scott Sammons, Yvette Unoarumhi, Darlene Wagner, Matthew Schmerer, Minoo Agarwal, Eyad Almasri, Debbie Boles, Ayla Burns, Nuthawin Charoensri, Oren Cohen, Susan Countryman, Mary Ann Cristobal, Bobbi Croy, Suzanne Dale, Hrushikesh Deshmukh, Amanda Douglas, Vincent Drouillon, Marcia Eisenberg, Howard Engler, Rama Ghatti, Prashant Gupta, Susan Hicks, Jake Humphrey, Lax Iyer, Lisa Pfefferle, Manoj Jain, Matthew Robinson, Mohan Kolli, Brian Krueger, Tim Kuphal, Stanley Letovsky, Michael Levandoski, Craig Lukasik, Jonathan Meltzer, Brian Norvell, Mindy Nye, Scott Parker, Christos Petropoulos, John Pruitt, Steven Ragan, Scott Ryan, Mike Sapeta, Jana Schroth, Suresh Babu Selvaraju, Goran Stevovic, Amanda Suchanek, Andrea Throop, Lyndon Tilson, Thomas Urban, Joe Voshell, |

| | | | |
|---|---|---|---|
| | | | Kimberly Wagner, Jonathan Williams, Mary Williamson, Qian Zeng, Tricia Zwiefelhofer, Clinton R. Paden, Duncan MacCannell |
| EPI_ISL_3330328, EPI_ISL_3330330 | Illinois Department of Public Health - Springfield Lab | Illinois Department of Public Health - Springfield Lab | Bryan Sim, Gordon McCall, Marshall Smock |
| EPI_ISL_3330331 | Laboratory Corporation of America | Centers for Disease Control and Prevention Division of Viral Diseases, Pathogen Discovery | Dakota Howard, Dhwani Batra, Peter W. Cook, Kara Moser, Adrian Paskey, Jason Caravas, Benjamin Rambo-Martin, Shatavia Morrison, Christopher Gulvick, Scott Sammons, Yvette Unoarumhi, Darlene Wagner, Matthew Schmerer, Minoo Agarwal, Eyad Almasri, Debbie Boles, Ayla Burns, Nuthawin Charoensri, Oren Cohen, Susan Countryman, Mary Ann Cristobal, Bobbi Croy, Suzanne Dale, Hrushikesh Deshmukh, Amanda Douglas, Vincent Drouillon, Marcia Eisenberg, Howard Engler, Rama Ghatti, Prashant Gupta, Susan Hicks, Jake Humphrey, Lax Iyer, Lisa Pfefferle, Manoj Jain, Matthew Robinson, Mohan Kolli, Brian Krueger, Tim Kuphal, Stanley Letovsky, Michael Levandoski, Craig Lukasik, Jonathan Meltzer, Brian Norvell, Mindy Nye, Scott Parker, Christos Petropoulos, John Pruitt, Steven Ragan, Scott Ryan, Mike Sapeta, Jana Schroth, Suresh Babu Selvaraju, Goran Stevovic, Amanda Suchanek, Andrea Throop, Lyndon Tilson, Thomas Urban, Joe Voshell, Kimberly Wagner, Jonathan Williams, Mary Williamson, Qian Zeng, Tricia Zwiefelhofer, Clinton R. Paden, Duncan MacCannell |
| EPI_ISL_3330332 | Illinois Department of Public Health - Springfield Lab | Illinois Department of Public Health - Springfield Lab | Bryan Sim, Gordon McCall, Marshall Smock |
| EPI_ISL_3330334 | Laboratory Corporation of America | Centers for Disease Control and Prevention Division of Viral Diseases, Pathogen Discovery | Dakota Howard, Dhwani Batra, Peter W. Cook, Kara Moser, Adrian Paskey, Jason Caravas, Benjamin Rambo-Martin, Shatavia Morrison, Christopher Gulvick, Scott Sammons, Yvette Unoarumhi, Darlene Wagner, Matthew Schmerer, Minoo Agarwal, Eyad Almasri, Debbie Boles, Ayla Burns, Nuthawin Charoensri, Oren Cohen, Susan Countryman, Mary Ann Cristobal, Bobbi Croy, Suzanne Dale, Hrushikesh Deshmukh, Amanda Douglas, Vincent Drouillon, Marcia Eisenberg, Howard Engler, Rama Ghatti, Prashant Gupta, Susan Hicks, Jake Humphrey, Lax Iyer, Lisa Pfefferle, Manoj Jain, Matthew Robinson, Mohan Kolli, Brian Krueger, Tim Kuphal, Stanley Letovsky, Michael Levandoski, Craig Lukasik, Jonathan Meltzer, Brian Norvell, Mindy Nye, Scott Parker, Christos Petropoulos, John Pruitt, Steven Ragan, Scott Ryan, Mike Sapeta, Jana Schroth, Suresh Babu Selvaraju, Goran Stevovic, Amanda Suchanek, Andrea Throop, Lyndon Tilson, Thomas Urban, Joe Voshell, Kimberly Wagner, Jonathan Williams, Mary Williamson, Qian Zeng, Tricia Zwiefelhofer, Clinton R. Paden, Duncan MacCannell |
| EPI_ISL_3330335 | Illinois Department of Public Health - Springfield Lab | Illinois Department of Public Health - Springfield Lab | Bryan Sim, Gordon McCall, Marshall Smock |
| EPI_ISL_3330336, EPI_ISL_3330338, EPI_ISL_3330340 | Laboratory Corporation of America | Centers for Disease Control and Prevention Division of Viral Diseases, Pathogen Discovery | Dakota Howard, Dhwani Batra, Peter W. Cook, Kara Moser, Adrian Paskey, Jason Caravas, Benjamin Rambo-Martin, Shatavia Morrison, Christopher Gulvick, Scott Sammons, Yvette Unoarumhi, Darlene Wagner, Matthew Schmerer, Minoo Agarwal, Eyad Almasri, Debbie Boles, Ayla Burns, Nuthawin Charoensri, Oren Cohen, Susan Countryman, Mary Ann Cristobal, Bobbi Croy, Suzanne Dale, Hrushikesh Deshmukh, Amanda Douglas, Vincent Drouillon, Marcia Eisenberg, Howard Engler, Rama Ghatti, Prashant Gupta, Susan Hicks, Jake Humphrey, Lax Iyer, Lisa Pfefferle, Manoj Jain, Matthew Robinson, Mohan Kolli, Brian Krueger, Tim Kuphal, Stanley Letovsky, Michael Levandoski, Craig Lukasik, Jonathan Meltzer, Brian Norvell, Mindy Nye, Scott Parker, Christos Petropoulos, John Pruitt, Steven Ragan, Scott Ryan, Mike Sapeta, Jana Schroth, Suresh Babu Selvaraju, Goran Stevovic, Amanda Suchanek, Andrea Throop, Lyndon Tilson, Thomas Urban, Joe Voshell, Kimberly Wagner, Jonathan Williams, Mary Williamson, Qian Zeng, Tricia Zwiefelhofer, Clinton R. Paden, Duncan MacCannell |
| EPI_ISL_3330342 | Illinois Department of Public Health - Springfield Lab | Illinois Department of Public Health - Springfield Lab | Bryan Sim, Gordon McCall, Marshall Smock |
| EPI_ISL_3330343, EPI_ISL_3330345, EPI_ISL_3330348, EPI_ISL_3330349, EPI_ISL_3330351 | Laboratory Corporation of America | Centers for Disease Control and Prevention Division of Viral Diseases, Pathogen Discovery | Dakota Howard, Dhwani Batra, Peter W. Cook, Kara Moser, Adrian Paskey, Jason Caravas, Benjamin Rambo-Martin, Shatavia Morrison, Christopher Gulvick, Scott Sammons, Yvette Unoarumhi, Darlene Wagner, Matthew Schmerer, Minoo Agarwal, Eyad Almasri, Debbie Boles, Ayla Burns, Nuthawin Charoensri, Oren Cohen, Susan Countryman, Mary Ann Cristobal, Bobbi Croy, Suzanne Dale, Hrushikesh Deshmukh, Amanda Douglas, Vincent Drouillon, Marcia Eisenberg, Howard Engler, Rama Ghatti, Prashant Gupta, Susan Hicks, Jake Humphrey, Lax Iyer, Lisa Pfefferle, Manoj Jain, Matthew Robinson, Mohan Kolli, Brian Krueger, Tim Kuphal, Stanley Letovsky, Michael Levandoski, Craig Lukasik, Jonathan Meltzer, Brian Norvell, Mindy Nye, Scott Parker, Christos Petropoulos, John Pruitt, Steven Ragan, Scott Ryan, Mike Sapeta, Jana Schroth, Suresh Babu Selvaraju, Goran Stevovic, Amanda Suchanek, Andrea Throop, Lyndon Tilson, Thomas Urban, Joe Voshell, Kimberly Wagner, Jonathan Williams, Mary Williamson, Qian Zeng, Tricia Zwiefelhofer, Clinton R. Paden, Duncan MacCannell |
| EPI_ISL_3330352 | Illinois Department of Public Health - Springfield Lab | Illinois Department of Public Health - Springfield Lab | Bryan Sim, Gordon McCall, Marshall Smock |
| EPI_ISL_3330354, EPI_ISL_3330355, EPI_ISL_3330356 | Laboratory Corporation of America | Centers for Disease Control and Prevention Division of Viral Diseases, Pathogen Discovery | Dakota Howard, Dhwani Batra, Peter W. Cook, Kara Moser, Adrian Paskey, Jason Caravas, Benjamin Rambo-Martin, Shatavia Morrison, Christopher Gulvick, Scott Sammons, Yvette Unoarumhi, Darlene Wagner, Matthew Schmerer, Minoo Agarwal, Eyad Almasri, Debbie Boles, Ayla Burns, Nuthawin Charoensri, Oren Cohen, Susan Countryman, Mary Ann Cristobal, Bobbi Croy, Suzanne Dale, Hrushikesh Deshmukh, Amanda Douglas, Vincent Drouillon, Marcia Eisenberg, Howard Engler, Rama Ghatti, Prashant Gupta, Susan Hicks, Jake Humphrey, Lax Iyer, Lisa Pfefferle, Manoj Jain, Matthew Robinson, Mohan Kolli, Brian Krueger, Tim Kuphal, Stanley Letovsky, Michael Levandoski, Craig Lukasik, Jonathan Meltzer, Brian Norvell, Mindy Nye, Scott Parker, Christos Petropoulos, John Pruitt, Steven Ragan, Scott Ryan, Mike Sapeta, Jana Schroth, Suresh Babu Selvaraju, Goran Stevovic, Amanda Suchanek, Andrea Throop, Lyndon Tilson, Thomas Urban, Joe Voshell, Kimberly Wagner, Jonathan Williams, Mary Williamson, Qian Zeng, Tricia Zwiefelhofer, Clinton R. Paden, Duncan MacCannell |
| EPI_ISL_3330357 | Illinois Department of Public Health - Springfield Lab | Illinois Department of Public Health - Springfield Lab | Bryan Sim, Gordon McCall, Marshall Smock |
| EPI_ISL_3330358, EPI_ISL_3330359 | Laboratory Corporation of America | Centers for Disease Control and Prevention Division of Viral Diseases, Pathogen Discovery | Dakota Howard, Dhwani Batra, Peter W. Cook, Kara Moser, Adrian Paskey, Jason Caravas, Benjamin Rambo-Martin, Shatavia Morrison, Christopher Gulvick, Scott Sammons, Yvette Unoarumhi, Darlene Wagner, Matthew Schmerer, Minoo Agarwal, Eyad Almasri, Debbie Boles, Ayla Burns, Nuthawin Charoensri, Oren Cohen, Susan Countryman, Mary Ann Cristobal, Bobbi Croy, Suzanne Dale, Hrushikesh Deshmukh, Amanda Douglas, Vincent Drouillon, Marcia Eisenberg, Howard Engler, Rama Ghatti, Prashant Gupta, Susan Hicks, Jake Humphrey, Lax Iyer, Lisa Pfefferle, Manoj Jain, Matthew Robinson, Mohan Kolli, Brian Krueger, Tim Kuphal, Stanley Letovsky, Michael Levandoski, Craig Lukasik, Jonathan Meltzer, Brian Norvell, Mindy Nye, Scott Parker, Christos Petropoulos, John Pruitt, Steven Ragan, Scott Ryan, Mike Sapeta, Jana Schroth, Suresh Babu Selvaraju, Goran Stevovic, Amanda Suchanek, Andrea Throop, Lyndon Tilson, Thomas Urban, Joe Voshell, Kimberly Wagner, Jonathan Williams, Mary Williamson, Qian Zeng, Tricia Zwiefelhofer, Clinton R. Paden, Duncan MacCannell |
| EPI_ISL_3330360 | Illinois Department of Public Health - Springfield Lab | Illinois Department of Public Health - Springfield Lab | Bryan Sim, Gordon McCall, Marshall Smock |
| EPI_ISL_3330361, EPI_ISL_3330363, EPI_ISL_3330365, EPI_ISL_3330366, EPI_ISL_3330367 | Laboratory Corporation of America | Centers for Disease Control and Prevention Division of Viral Diseases, Pathogen Discovery | Dakota Howard, Dhwani Batra, Peter W. Cook, Kara Moser, Adrian Paskey, Jason Caravas, Benjamin Rambo-Martin, Shatavia Morrison, Christopher Gulvick, Scott Sammons, Yvette Unoarumhi, Darlene Wagner, Matthew Schmerer, Minoo Agarwal, Eyad Almasri, Debbie Boles, Ayla Burns, Nuthawin Charoensri, Oren Cohen, Susan Countryman, Mary Ann Cristobal, Bobbi Croy, Suzanne Dale, Hrushikesh Deshmukh, Amanda Douglas, Vincent Drouillon, Marcia Eisenberg, Howard Engler, Rama Ghatti, Prashant Gupta, Susan Hicks, Jake Humphrey, Lax Iyer, Lisa Pfefferle, Manoj Jain, Matthew Robinson, Mohan Kolli, Brian Krueger, Tim Kuphal, Stanley Letovsky, Michael Levandoski, Craig Lukasik, Jonathan Meltzer, Brian Norvell, Mindy Nye, Scott Parker, Christos Petropoulos, John Pruitt, Steven Ragan, Scott Ryan, Mike Sapeta, Jana Schroth, Suresh Babu Selvaraju, Goran Stevovic, Amanda Suchanek, Andrea Throop, Lyndon Tilson, Thomas Urban, Joe Voshell, Kimberly Wagner, Jonathan Williams, Mary Williamson, Qian Zeng, Tricia Zwiefelhofer, Clinton R. Paden, Duncan MacCannell |
| EPI_ISL_3330368 | Illinois Department of Public Health - Springfield Lab | Illinois Department of Public Health - Springfield Lab | Bryan Sim, Gordon McCall, Marshall Smock |
| EPI_ISL_3330369 | Laboratory Corporation of America | Centers for Disease Control and Prevention Division of Viral Diseases, Pathogen Discovery | Dakota Howard, Dhwani Batra, Peter W. Cook, Kara Moser, Adrian Paskey, Jason Caravas, Benjamin Rambo-Martin, Shatavia Morrison, Christopher Gulvick, Scott Sammons, Yvette Unoarumhi, Darlene Wagner, Matthew Schmerer, Minoo Agarwal, Eyad Almasri, Debbie Boles, Ayla Burns, Nuthawin Charoensri, Oren Cohen, Susan Countryman, Mary Ann Cristobal, Bobbi Croy, Suzanne Dale, Hrushikesh Deshmukh, Amanda Douglas, Vincent Drouillon, Marcia Eisenberg, Howard Engler, Rama Ghatti, Prashant Gupta, Susan Hicks, Jake Humphrey, Lax Iyer, Lisa Pfefferle, Manoj Jain, Matthew Robinson, Mohan Kolli, Brian Krueger, Tim Kuphal, Stanley Letovsky, Michael Levandoski, Craig Lukasik, Jonathan Meltzer, Brian Norvell, Mindy Nye, Scott Parker, Christos Petropoulos, John Pruitt, Steven Ragan, Scott Ryan, Mike Sapeta, Jana Schroth, Suresh Babu Selvaraju, Goran Stevovic, Amanda Suchanek, Andrea Throop, Lyndon Tilson, Thomas Urban, Joe Voshell, Kimberly Wagner, Jonathan Williams, Mary Williamson, Qian Zeng, Tricia Zwiefelhofer, Clinton R. Paden, Duncan MacCannell |
| EPI_ISL_3330370, EPI_ISL_3330371, EPI_ISL_3330373 | Illinois Department of Public Health - Springfield Lab | Illinois Department of Public Health - Springfield Lab | Bryan Sim, Gordon McCall, Marshall Smock |
| EPI_ISL_3330374 | Laboratory Corporation of America | Centers for Disease Control and Prevention Division of Viral Diseases, Pathogen Discovery | Dakota Howard, Dhwani Batra, Peter W. Cook, Kara Moser, Adrian Paskey, Jason Caravas, Benjamin Rambo-Martin, Shatavia Morrison, Christopher Gulvick, Scott Sammons, Yvette Unoarumhi, Darlene Wagner, Matthew Schmerer, Minoo Agarwal, Eyad Almasri, Debbie Boles, Ayla Burns, Nuthawin Charoensri, Oren Cohen, Susan Countryman, Mary Ann Cristobal, Bobbi Croy, Suzanne Dale, Hrushikesh Deshmukh, Amanda Douglas, Vincent Drouillon, Marcia Eisenberg, Howard Engler, Rama Ghatti, Prashant Gupta, Susan Hicks, Jake Humphrey, Lax Iyer, Lisa Pfefferle, Manoj Jain, Matthew Robinson, Mohan Kolli, Brian Krueger, Tim Kuphal, Stanley Letovsky, Michael Levandoski, Craig Lukasik, Jonathan Meltzer, Brian Norvell, Mindy Nye, Scott Parker, Christos Petropoulos, John Pruitt, Steven Ragan, Scott Ryan, Mike Sapeta, Jana Schroth, Suresh Babu Selvaraju, Goran Stevovic, Amanda Suchanek, Andrea Throop, Lyndon Tilson, Thomas Urban, Joe Voshell, Kimberly Wagner, Jonathan Williams, Mary Williamson, Qian Zeng, Tricia Zwiefelhofer, Clinton R. Paden, Duncan MacCannell |
| EPI_ISL_3330375 | Illinois Department of Public Health - Springfield Lab | Illinois Department of Public Health - Springfield Lab | Bryan Sim, Gordon McCall, Marshall Smock |
| EPI_ISL_3330376 | Laboratory Corporation of America | Centers for Disease Control and Prevention Division of Viral Diseases, Pathogen Discovery | Dakota Howard, Dhwani Batra, Peter W. Cook, Kara Moser, Adrian Paskey, Jason Caravas, Benjamin Rambo-Martin, Shatavia Morrison, Christopher Gulvick, Scott Sammons, Yvette Unoarumhi, Darlene Wagner, Matthew Schmerer, Minoo Agarwal, Eyad Almasri, Debbie Boles, Ayla Burns, Nuthawin Charoensri, Oren Cohen, Susan Countryman, Mary Ann Cristobal, Bobbi Croy, Suzanne Dale, Hrushikesh Deshmukh, Amanda Douglas, Vincent Drouillon, Marcia Eisenberg, Howard Engler, Rama Ghatti, Prashant Gupta, Susan Hicks, Jake Humphrey, Lax Iyer, Lisa Pfefferle, Manoj Jain, Matthew Robinson, Mohan Kolli, Brian Krueger, Tim Kuphal, Stanley Letovsky, Michael Levandoski, Craig Lukasik, Jonathan Meltzer, Brian Norvell, Mindy Nye, Scott Parker, Christos Petropoulos, John Pruitt, Steven Ragan, Scott Ryan, Mike Sapeta, Jana Schroth, Suresh Babu Selvaraju, Goran Stevovic, Amanda Suchanek, Andrea Throop, Lyndon Tilson, Thomas Urban, Joe Voshell, Kimberly Wagner, Jonathan Williams, Mary Williamson, Qian Zeng, Tricia Zwiefelhofer, Clinton R. Paden, Duncan MacCannell |
| EPI_ISL_3330377, EPI_ISL_3330378 | Illinois Department of Public Health - Springfield Lab | Illinois Department of Public Health - Springfield Lab | Bryan Sim, Gordon McCall, Marshall Smock |
| EPI_ISL_3330379 | Laboratory Corporation of America | Centers for Disease Control and Prevention Division of Viral Diseases, Pathogen Discovery | Dakota Howard, Dhwani Batra, Peter W. Cook, Kara Moser, Adrian Paskey, Jason Caravas, Benjamin Rambo-Martin, Shatavia Morrison, Christopher Gulvick, Scott Sammons, Yvette Unoarumhi, Darlene Wagner, Matthew Schmerer, Minoo Agarwal, Eyad Almasri, Debbie Boles, Ayla Burns, Nuthawin Charoensri, Oren Cohen, Susan Countryman, Mary Ann Cristobal, Bobbi Croy, Suzanne Dale, Hrushikesh Deshmukh, Amanda Douglas, Vincent Drouillon, Marcia Eisenberg, Howard Engler, Rama Ghatti, Prashant Gupta, Susan Hicks, Jake Humphrey, Lax Iyer, Lisa Pfefferle, Manoj Jain, Matthew Robinson, Mohan Kolli, Brian Krueger, Tim Kuphal, Stanley Letovsky, Michael Levandoski, Craig Lukasik, Jonathan Meltzer, Brian Norvell, Mindy Nye, Scott Parker, Christos Petropoulos, John Pruitt, Steven Ragan, Scott Ryan, Mike Sapeta, Jana Schroth, Suresh Babu Selvaraju, Goran Stevovic, Amanda Suchanek, Andrea Throop, Lyndon Tilson, Thomas Urban, Joe Voshell, Kimberly Wagner, Jonathan Williams, Mary Williamson, Qian Zeng, Tricia Zwiefelhofer, Clinton R. Paden, Duncan MacCannell |
| EPI_ISL_3330380 | Illinois Department of Public Health - Springfield Lab | Illinois Department of Public Health - Springfield Lab | Bryan Sim, Gordon McCall, Marshall Smock |
| EPI_ISL_3330381 | Laboratory Corporation of America | Centers for Disease Control and Prevention Division of Viral Diseases, Pathogen Discovery | Dakota Howard, Dhwani Batra, Peter W. Cook, Kara Moser, Adrian Paskey, Jason Caravas, Benjamin Rambo-Martin, Shatavia Morrison, Christopher Gulvick, Scott Sammons, Yvette Unoarumhi, Darlene Wagner, Matthew Schmerer, Minoo Agarwal, Eyad Almasri, Debbie Boles, Ayla Burns, Nuthawin Charoensri, Oren Cohen, Susan Countryman, Mary Ann Cristobal, Bobbi Croy, Suzanne Dale, Hrushikesh Deshmukh, Amanda Douglas, Vincent Drouillon, Marcia Eisenberg, Howard Engler, Rama Ghatti, Prashant Gupta, Susan Hicks, Jake Humphrey, Lax Iyer, Lisa Pfefferle, Manoj Jain, Matthew Robinson, Mohan Kolli, Brian Krueger, Tim Kuphal, Stanley Letovsky, Michael Levandoski, Craig Lukasik, Jonathan Meltzer, Brian Norvell, Mindy Nye, Scott Parker, Christos Petropoulos, John Pruitt, Steven Ragan, Scott Ryan, Mike Sapeta, Jana Schroth, Suresh Babu Selvaraju, Goran Stevovic, Amanda Suchanek, Andrea Throop, Lyndon Tilson, Thomas Urban, Joe Voshell, Kimberly Wagner, Jonathan Williams, Mary Williamson, Qian Zeng, Tricia Zwiefelhofer, Clinton R. Paden, Duncan MacCannell |
| EPI_ISL_3330382 | Illinois Department of Public Health - Springfield Lab | Illinois Department of Public Health - Springfield Lab | Bryan Sim, Gordon McCall, Marshall Smock |
| EPI_ISL_3330383 | Laboratory Corporation of America | Centers for Disease Control and Prevention Division of Viral Diseases, Pathogen Discovery | Dakota Howard, Dhwani Batra, Peter W. Cook, Kara Moser, Adrian Paskey, Jason Caravas, Benjamin Rambo-Martin, Shatavia Morrison, Christopher Gulvick, Scott Sammons, Yvette Unoarumhi, Darlene Wagner, Matthew Schmerer, Minoo Agarwal, Eyad Almasri, Debbie Boles, Ayla Burns, Nuthawin Charoensri, Oren Cohen, Susan Countryman, Mary Ann Cristobal, Bobbi Croy, Suzanne Dale, Hrushikesh Deshmukh, Amanda Douglas, Vincent Drouillon, Marcia Eisenberg, Howard Engler, Rama Ghatti, Prashant Gupta, Susan Hicks, Jake Humphrey, Lax Iyer, Lisa Pfefferle, Manoj Jain, Matthew Robinson, Mohan Kolli, Brian Krueger, Tim Kuphal, Stanley Letovsky, Michael Levandoski, Craig Lukasik, Jonathan Meltzer, Brian Norvell, Mindy Nye, Scott Parker, Christos Petropoulos, John Pruitt, Steven Ragan, Scott Ryan, Mike Sapeta, Jana Schroth, Suresh Babu Selvaraju, Goran Stevovic, Amanda Suchanek, Andrea Throop, Lyndon Tilson, Thomas Urban, Joe Voshell, Kimberly Wagner, Jonathan Williams, Mary Williamson, Qian Zeng, Tricia Zwiefelhofer, Clinton R. Paden, Duncan MacCannell |
| EPI_ISL_3330384, EPI_ISL_3330385 | Illinois Department of Public Health - Springfield Lab | Illinois Department of Public Health - Springfield Lab | Bryan Sim, Gordon McCall, Marshall Smock |
| EPI_ISL_3330386 | Laboratory Corporation of America | Centers for Disease Control and Prevention Division of Viral Diseases, Pathogen Discovery | Dakota Howard, Dhwani Batra, Peter W. Cook, Kara Moser, Adrian Paskey, Jason Caravas, Benjamin Rambo-Martin, Shatavia Morrison, Christopher Gulvick, Scott Sammons, Yvette Unoarumhi, Darlene Wagner, Matthew Schmerer, Minoo Agarwal, Eyad Almasri, Debbie Boles, Ayla Burns, Nuthawin Charoensri, Oren Cohen, Susan Countryman, Mary Ann Cristobal, Bobbi Croy, Suzanne Dale, Hrushikesh Deshmukh, Amanda Douglas, Vincent Drouillon, Marcia Eisenberg, Howard Engler, Rama Ghatti, Prashant Gupta, Susan Hicks, Jake Humphrey, Lax Iyer, Lisa Pfefferle, Manoj Jain, Matthew Robinson, Mohan Kolli, Brian Krueger, Tim Kuphal, Stanley Letovsky, Michael Levandoski, Craig Lukasik, Jonathan Meltzer, Brian Norvell, Mindy Nye, Scott Parker, Christos Petropoulos, John Pruitt, Steven Ragan, Scott Ryan, Mike Sapeta, Jana Schroth, Suresh Babu Selvaraju, Goran Stevovic, Amanda Suchanek, Andrea Throop, Lyndon Tilson, Thomas Urban, Joe Voshell, Kimberly Wagner, Jonathan Williams, Mary Williamson, Qian Zeng, Tricia Zwiefelhofer, Clinton R. Paden, Duncan MacCannell |
| EPI_ISL_3330387 | Illinois Department of Public Health - Springfield Lab | Illinois Department of Public Health - Springfield Lab | Bryan Sim, Gordon McCall, Marshall Smock |
| EPI_ISL_3330388 | Laboratory Corporation of America | Centers for Disease Control and Prevention Division of Viral Diseases, Pathogen Discovery | Dakota Howard, Dhwani Batra, Peter W. Cook, Kara Moser, Adrian Paskey, Jason Caravas, Benjamin Rambo-Martin, Shatavia Morrison, Christopher Gulvick, Scott Sammons, Yvette Unoarumhi, Darlene Wagner, Matthew Schmerer, Minoo Agarwal, Eyad Almasri, Debbie Boles, Ayla Burns, Nuthawin Charoensri, Oren Cohen, Susan Countryman, Mary Ann Cristobal, Bobbi Croy, Suzanne Dale, Hrushikesh Deshmukh, Amanda Douglas, Vincent Drouillon, Marcia Eisenberg, Howard Engler, Rama Ghatti, Prashant Gupta, Susan Hicks, Jake Humphrey, Lax Iyer, Lisa Pfefferle, Manoj Jain, Matthew Robinson, Mohan Kolli, Brian Krueger, Tim Kuphal, Stanley Letovsky, Michael Levandoski, Craig Lukasik, Jonathan Meltzer, Brian Norvell, Mindy Nye, Scott Parker, Christos Petropoulos, John Pruitt, Steven Ragan, Scott Ryan, Mike Sapeta, Jana Schroth, Suresh Babu Selvaraju, Goran Stevovic, Amanda Suchanek, Andrea Throop, Lyndon Tilson, Thomas Urban, Joe Voshell, Kimberly Wagner, Jonathan Williams, Mary Williamson, Qian Zeng, Tricia Zwiefelhofer, Clinton R. Paden, Duncan MacCannell |

| | | | |
|---|---|---|---|
| EPI_ISL_3330389, EPI_ISL_3330390 | Illinois Department of Public Health - Springfield Lab | Illinois Department of Public Health - Springfield Lab | Bryan Sim, Gordon McCall, Marshall Smock |
| EPI_ISL_3330391 | Laboratory Corporation of America | Centers for Disease Control and Prevention Division of Viral Diseases, Pathogen Discovery | Dakota Howard, Dhwani Batra, Peter W. Cook, Kara Moser, Adrian Paskey, Jason Caravas, Benjamin Rambo-Martin, Shatavia Morrison, Christopher Gulvick, Scott Sammons, Yvette Unoarumhi, Darlene Wagner, Matthew Schmerer, Minoo Agarwal, Eyad Almasri, Debbie Boles, Ayla Burns, Nuthawin Charoensri, Oren Cohen, Susan Countryman, Mary Ann Cristobal, Bobbi Croy, Suzanne Dale, Hrushikesh Deshmukh, Amanda Douglas, Vincent Drouillon, Marcia Eisenberg, Howard Engler, Rama Ghatti, Prashant Gupta, Susan Hicks, Jake Humphrey, Lax Iyer, Lisa Pfefferle, Manoj Jain, Matthew Robinson, Mohan Kolli, Brian Krueger, Tim Kuphal, Stanley Letovsky, Michael Levandoski, Craig Lukasik, Jonathan Meltzer, Brian Norvell, Mindy Nye, Scott Parker, Christos Petropoulos, John Pruitt, Steven Ragan, Scott Ryan, Mike Sapeta, Jana Schroth, Suresh Babu Selvaraju, Goran Stevovic, Amanda Suchanek, Andrea Throop, Lyndon Tilson, Thomas Urban, Joe Voshell, Kimberly Wagner, Jonathan Williams, Mary Williamson, Qian Zeng, Tricia Zwiefelhofer, Clinton R. Paden, Duncan MacCannell |
| EPI_ISL_3330392 | Illinois Department of Public Health - Springfield Lab | Illinois Department of Public Health - Springfield Lab | Bryan Sim, Gordon McCall, Marshall Smock |
| EPI_ISL_3330393 | Laboratory Corporation of America | Centers for Disease Control and Prevention Division of Viral Diseases, Pathogen Discovery | Dakota Howard, Dhwani Batra, Peter W. Cook, Kara Moser, Adrian Paskey, Jason Caravas, Benjamin Rambo-Martin, Shatavia Morrison, Christopher Gulvick, Scott Sammons, Yvette Unoarumhi, Darlene Wagner, Matthew Schmerer, Minoo Agarwal, Eyad Almasri, Debbie Boles, Ayla Burns, Nuthawin Charoensri, Oren Cohen, Susan Countryman, Mary Ann Cristobal, Bobbi Croy, Suzanne Dale, Hrushikesh Deshmukh, Amanda Douglas, Vincent Drouillon, Marcia Eisenberg, Howard Engler, Rama Ghatti, Prashant Gupta, Susan Hicks, Jake Humphrey, Lax Iyer, Lisa Pfefferle, Manoj Jain, Matthew Robinson, Mohan Kolli, Brian Krueger, Tim Kuphal, Stanley Letovsky, Michael Levandoski, Craig Lukasik, Jonathan Meltzer, Brian Norvell, Mindy Nye, Scott Parker, Christos Petropoulos, John Pruitt, Steven Ragan, Scott Ryan, Mike Sapeta, Jana Schroth, Suresh Babu Selvaraju, Goran Stevovic, Amanda Suchanek, Andrea Throop, Lyndon Tilson, Thomas Urban, Joe Voshell, Kimberly Wagner, Jonathan Williams, Mary Williamson, Qian Zeng, Tricia Zwiefelhofer, Clinton R. Paden, Duncan MacCannell |
| EPI_ISL_3330394, EPI_ISL_3330395 | Illinois Department of Public Health - Springfield Lab | Illinois Department of Public Health - Springfield Lab | Bryan Sim, Gordon McCall, Marshall Smock |
| EPI_ISL_3330396, EPI_ISL_3330397, EPI_ISL_3330398, EPI_ISL_3330399, EPI_ISL_3330400, EPI_ISL_3330401, EPI_ISL_3330402, EPI_ISL_3330403, EPI_ISL_3330404, EPI_ISL_3330406, EPI_ISL_3330407, EPI_ISL_3330409, EPI_ISL_3330411, EPI_ISL_3330413, EPI_ISL_3330415, EPI_ISL_3330417, EPI_ISL_3330418, EPI_ISL_3330420, EPI_ISL_3330422, EPI_ISL_3330424, EPI_ISL_3330426, EPI_ISL_3330427, EPI_ISL_3330428, EPI_ISL_3330429, EPI_ISL_3330430, EPI_ISL_3330431, EPI_ISL_3330432, EPI_ISL_3330433, EPI_ISL_3330434, EPI_ISL_3330435, EPI_ISL_3330436, EPI_ISL_3330437, EPI_ISL_3330438, EPI_ISL_3330439, EPI_ISL_3330440, EPI_ISL_3330441, EPI_ISL_3330442, EPI_ISL_3330443, EPI_ISL_3330444, EPI_ISL_3330445, EPI_ISL_3330446, EPI_ISL_3330447, EPI_ISL_3330448, EPI_ISL_3330449, EPI_ISL_3330450, EPI_ISL_3330451, EPI_ISL_3330453, EPI_ISL_3330454, EPI_ISL_3330455, EPI_ISL_3330456, EPI_ISL_3330457, EPI_ISL_3330458, EPI_ISL_3330459, EPI_ISL_3330460, EPI_ISL_3330461, EPI_ISL_3330462, EPI_ISL_3330463, EPI_ISL_3330465, EPI_ISL_3330467, EPI_ISL_3330468, EPI_ISL_3330470, EPI_ISL_3330471, EPI_ISL_3330473, EPI_ISL_3330475, EPI_ISL_3330477, EPI_ISL_3330478, EPI_ISL_3330480, EPI_ISL_3330482, EPI_ISL_3330484, EPI_ISL_3330485, EPI_ISL_3330486, EPI_ISL_3330487, EPI_ISL_3330488, EPI_ISL_3330489, EPI_ISL_3330490, EPI_ISL_3330492, EPI_ISL_3330493, EPI_ISL_3330494, EPI_ISL_3330495, EPI_ISL_3330496, EPI_ISL_3330497, EPI_ISL_3330498, EPI_ISL_3330499, EPI_ISL_3330500, EPI_ISL_3330501, EPI_ISL_3330502, EPI_ISL_3330503, EPI_ISL_3330504, EPI_ISL_3330505, EPI_ISL_3330506, EPI_ISL_3330507, EPI_ISL_3330508, EPI_ISL_3330509, EPI_ISL_3330510, EPI_ISL_3330511, EPI_ISL_3330513, EPI_ISL_3330514, EPI_ISL_3330515, EPI_ISL_3330516, EPI_ISL_3330517, EPI_ISL_3330518, EPI_ISL_3330519, EPI_ISL_3330520, EPI_ISL_3330521, EPI_ISL_3330522, EPI_ISL_3330523, EPI_ISL_3330524, EPI_ISL_3330525, EPI_ISL_3330526, EPI_ISL_3330527, EPI_ISL_3330528, EPI_ISL_3330529, EPI_ISL_3330530, EPI_ISL_3330531, EPI_ISL_3330532, EPI_ISL_3330533, EPI_ISL_3330535, EPI_ISL_3330537, EPI_ISL_3330539, EPI_ISL_3330541, EPI_ISL_3330542, EPI_ISL_3330544, EPI_ISL_3330546, EPI_ISL_3330548, EPI_ISL_3330550, EPI_ISL_3330551, EPI_ISL_3330553, EPI_ISL_3330554, EPI_ISL_3330555, EPI_ISL_3330556, EPI_ISL_3330557, EPI_ISL_3330558, EPI_ISL_3330559, EPI_ISL_3330560, EPI_ISL_3330561, EPI_ISL_3330562, EPI_ISL_3330565, EPI_ISL_3330566, EPI_ISL_3330567, EPI_ISL_3330568, EPI_ISL_3330569, EPI_ISL_3330571, EPI_ISL_3330572, EPI_ISL_3330573, EPI_ISL_3330574, EPI_ISL_3330576, EPI_ISL_3330577, EPI_ISL_3330578, EPI_ISL_3330579, EPI_ISL_3330580, EPI_ISL_3330581, EPI_ISL_3330582, EPI_ISL_3330583, EPI_ISL_3330584, EPI_ISL_3330585, EPI_ISL_3330586, EPI_ISL_3330587, EPI_ISL_3330588, EPI_ISL_3330589, EPI_ISL_3330590, EPI_ISL_3330591, EPI_ISL_3330592, EPI_ISL_3330593, EPI_ISL_3330594, EPI_ISL_3330595, EPI_ISL_3330596, EPI_ISL_3330597, EPI_ISL_3330598, EPI_ISL_3330600, EPI_ISL_3330601, EPI_ISL_3330602, EPI_ISL_3330603, EPI_ISL_3330604, EPI_ISL_3330605, EPI_ISL_3330606, EPI_ISL_3330607, EPI_ISL_3330608, EPI_ISL_3330609, EPI_ISL_3330611, EPI_ISL_3330612, EPI_ISL_3330614, EPI_ISL_3330615, EPI_ISL_3330616, EPI_ISL_3330617, EPI_ISL_3330618, EPI_ISL_3330619, EPI_ISL_3330620, EPI_ISL_3330621, EPI_ISL_3330622, EPI_ISL_3330623, EPI_ISL_3330624, EPI_ISL_3330625, EPI_ISL_3330626, EPI_ISL_3330627, EPI_ISL_3330628, EPI_ISL_3330629, EPI_ISL_3330631, EPI_ISL_3330632, EPI_ISL_3330635, EPI_ISL_3330639, EPI_ISL_3330643, EPI_ISL_3330644, EPI_ISL_3330645, EPI_ISL_3330646, EPI_ISL_3330647, EPI_ISL_3330648, EPI_ISL_3330649, EPI_ISL_3330650, EPI_ISL_3330651, EPI_ISL_3330652, EPI_ISL_3330653, EPI_ISL_3330654, EPI_ISL_3330655, EPI_ISL_3330656, EPI_ISL_3330658, EPI_ISL_3330659, EPI_ISL_3330660, EPI_ISL_3330661, EPI_ISL_3330662, EPI_ISL_3330663, EPI_ISL_3330664, EPI_ISL_3330665, EPI_ISL_3330666, EPI_ISL_3330667, EPI_ISL_3330668, EPI_ISL_3330669, EPI_ISL_3330670, EPI_ISL_3330671, EPI_ISL_3330672, EPI_ISL_3330675, EPI_ISL_3330676, EPI_ISL_3330677, EPI_ISL_3330678, EPI_ISL_3330679, EPI_ISL_3330680, EPI_ISL_3330681, EPI_ISL_3330682, EPI_ISL_3330683, EPI_ISL_3330684, EPI_ISL_3330685, EPI_ISL_3330686, EPI_ISL_3330687, EPI_ISL_3330688, EPI_ISL_3330689, EPI_ISL_3330690, EPI_ISL_3330691, EPI_ISL_3330694, EPI_ISL_3330695, EPI_ISL_3330696, EPI_ISL_3330699, EPI_ISL_3330700, EPI_ISL_3330701, EPI_ISL_3330702, EPI_ISL_3330703, EPI_ISL_3330707, EPI_ISL_3330708, EPI_ISL_3330709, EPI_ISL_3330710, EPI_ISL_3330711, EPI_ISL_3330712, EPI_ISL_3330713, EPI_ISL_3330714, EPI_ISL_3330715, EPI_ISL_3330716, EPI_ISL_3330717, EPI_ISL_3330718, EPI_ISL_3330719, EPI_ISL_3330720, EPI_ISL_3330721, EPI_ISL_3330722, EPI_ISL_3330723, EPI_ISL_3330725, EPI_ISL_3330726, EPI_ISL_3330727, EPI_ISL_3330728, EPI_ISL_3330729, EPI_ISL_3330730, EPI_ISL_3330731 | | | |
| see above | Laboratory Corporation of America | Centers for Disease Control and Prevention Division of Viral Diseases, Pathogen Discovery | Dakota Howard, Dhwani Batra, Peter W. Cook, Kara Moser, Adrian Paskey, Jason Caravas, Benjamin Rambo-Martin, Shatavia Morrison, Christopher Gulvick, Scott Sammons, Yvette Unoarumhi, Darlene Wagner, Matthew Schmerer, Minoo Agarwal, Eyad Almasri, Debbie Boles, Ayla Burns, Nuthawin Charoensri, Oren Cohen, Susan Countryman, Mary Ann Cristobal, Bobbi Croy, Suzanne Dale, Hrushikesh Deshmukh, Amanda Douglas, Vincent Drouillon, Marcia Eisenberg, Howard Engler, Rama Ghatti, Prashant Gupta, Susan Hicks, Jake Humphrey, Lax Iyer, Lisa Pfefferle, Manoj Jain, Matthew Robinson, Mohan Kolli, Brian Krueger, Tim Kuphal, Stanley Letovsky, Michael Levandoski, Craig Lukasik, Jonathan Meltzer, Brian Norvell, Mindy Nye, Scott Parker, Christos Petropoulos, John Pruitt, Steven Ragan, Scott Ryan, Mike Sapeta, Jana Schroth, Suresh Babu Selvaraju, Goran Stevovic, Amanda Suchanek, Andrea Throop, Lyndon Tilson, Thomas Urban, Joe Voshell, Kimberly Wagner, Jonathan Williams, Mary Williamson, Qian Zeng, Tricia Zwiefelhofer, Clinton R. Paden, Duncan MacCannell |
| EPI_ISL_3330732, EPI_ISL_3330733, EPI_ISL_3330734, EPI_ISL_3330735, EPI_ISL_3330736, EPI_ISL_3330737, EPI_ISL_3330738, EPI_ISL_3330739, EPI_ISL_3330740, EPI_ISL_3330741, EPI_ISL_3330742, EPI_ISL_3330743, EPI_ISL_3330744, EPI_ISL_3330745, EPI_ISL_3330746, EPI_ISL_3330747, EPI_ISL_3330748, EPI_ISL_3330749, EPI_ISL_3330750, EPI_ISL_3330751, EPI_ISL_3330752, EPI_ISL_3330753, EPI_ISL_3330754, EPI_ISL_3330755, EPI_ISL_3330756, EPI_ISL_3330757, EPI_ISL_3330758, EPI_ISL_3330759, EPI_ISL_3330760, EPI_ISL_3330761, EPI_ISL_3330762, EPI_ISL_3330763, EPI_ISL_3330764, EPI_ISL_3330765, EPI_ISL_3330766, EPI_ISL_3330767, EPI_ISL_3330768, EPI_ISL_3330769, EPI_ISL_3330770, EPI_ISL_3330771, EPI_ISL_3330772, EPI_ISL_3330773, EPI_ISL_3330774, EPI_ISL_3330775, EPI_ISL_3330776, EPI_ISL_3330777, EPI_ISL_3330778, EPI_ISL_3330779, EPI_ISL_3330780, EPI_ISL_3330781, EPI_ISL_3330782 | | | |
| see above | Quest Diagnostics Incorporated | Centers for Disease Control and Prevention Division of Viral Diseases, Pathogen Discovery | Dakota Howard, Dhwani Batra, Peter W. Cook, Kara Moser, Adrian Paskey, Jason Caravas, Benjamin Rambo-Martin, Shatavia Morrison, Christopher Gulvick, Scott Sammons, Yvette Unoarumhi, Darlene Wagner, Matthew Schmerer, S. H. Rosenthal, A. Gerasimova, R. M. Kagan, B. Anderson, M. Hua, Y. Liu, L.E. Bernstein, K.E. Livingston, A. Perez, I. A. Shlyakhter, R. V. Rolando, R. Owen, P. Tanpaiboon, F. Lacbawan, Clinton R. Paden, Duncan MacCannell |
| EPI_ISL_3330783 | TXDSHS | TXDSHS | Rashmi Tuladhar, Bonnie Oh, Jenny Zhang, Maliha Rahman, Mayela Pedrueza, Anita Pokharel, Karen Bobier, Lorraine Rodriguez, Myong Koag, Chun Wang, Rachel Lee, Grace Kubin |
| EPI_ISL_3330784, EPI_ISL_3330785, EPI_ISL_3330786, EPI_ISL_3330787 | Quest Diagnostics Incorporated | Centers for Disease Control and Prevention Division of Viral Diseases, Pathogen Discovery | Dakota Howard, Dhwani Batra, Peter W. Cook, Kara Moser, Adrian Paskey, Jason Caravas, Benjamin Rambo-Martin, Shatavia Morrison, Christopher Gulvick, Scott Sammons, Yvette Unoarumhi, Darlene Wagner, Matthew Schmerer, S. H. Rosenthal, A. Gerasimova, R. M. Kagan, B. Anderson, M. Hua, Y. Liu, L.E. Bernstein, K.E. Livingston, A. Perez, I. A. Shlyakhter, R. V. Rolando, R. Owen, P. Tanpaiboon, F. Lacbawan, Clinton R. Paden, Duncan MacCannell |
| EPI_ISL_3330788 | TXDSHS | TXDSHS | Rashmi Tuladhar, Bonnie Oh, Jenny Zhang, Maliha Rahman, Mayela Pedrueza, Anita Pokharel, Karen Bobier, Lorraine Rodriguez, Myong Koag, Chun Wang, Rachel Lee, Grace Kubin |
| EPI_ISL_3330789, EPI_ISL_3330790 | Quest Diagnostics Incorporated | Centers for Disease Control and Prevention Division of Viral Diseases, Pathogen Discovery | Dakota Howard, Dhwani Batra, Peter W. Cook, Kara Moser, Adrian Paskey, Jason Caravas, Benjamin Rambo-Martin, Shatavia Morrison, Christopher Gulvick, Scott Sammons, Yvette Unoarumhi, Darlene Wagner, Matthew Schmerer, S. H. Rosenthal, A. Gerasimova, R. M. Kagan, B. Anderson, M. Hua, Y. Liu, L.E. Bernstein, K.E. Livingston, A. Perez, I. A. Shlyakhter, R. V. Rolando, R. Owen, P. Tanpaiboon, F. Lacbawan, Clinton R. Paden, Duncan MacCannell |
| EPI_ISL_3330791 | TXDSHS | TXDSHS | Rashmi Tuladhar, Bonnie Oh, Jenny Zhang, Maliha Rahman, Mayela Pedrueza, Anita Pokharel, Karen Bobier, Lorraine Rodriguez, Myong Koag, Chun Wang, Rachel Lee, Grace Kubin |
| EPI_ISL_3330792, EPI_ISL_3330793, EPI_ISL_3330794 | Quest Diagnostics Incorporated | Centers for Disease Control and Prevention Division of Viral Diseases, Pathogen Discovery | Dakota Howard, Dhwani Batra, Peter W. Cook, Kara Moser, Adrian Paskey, Jason Caravas, Benjamin Rambo-Martin, Shatavia Morrison, Christopher Gulvick, Scott Sammons, Yvette Unoarumhi, Darlene Wagner, Matthew Schmerer, S. H. Rosenthal, A. Gerasimova, R. M. Kagan, B. Anderson, M. Hua, Y. Liu, L.E. Bernstein, K.E. Livingston, A. Perez, I. A. Shlyakhter, R. V. Rolando, R. Owen, P. Tanpaiboon, F. Lacbawan, Clinton R. Paden, Duncan MacCannell |
| EPI_ISL_3330795 | TXDSHS | TXDSHS | Rashmi Tuladhar, Bonnie Oh, Jenny Zhang, Maliha Rahman, Mayela Pedrueza, Anita Pokharel, Karen Bobier, Lorraine Rodriguez, Myong Koag, Chun Wang, Rachel Lee, Grace Kubin |
| EPI_ISL_3330796, EPI_ISL_3330797, EPI_ISL_3330798 | Quest Diagnostics Incorporated | Centers for Disease Control and Prevention Division of Viral Diseases, Pathogen Discovery | Dakota Howard, Dhwani Batra, Peter W. Cook, Kara Moser, Adrian Paskey, Jason Caravas, Benjamin Rambo-Martin, Shatavia Morrison, Christopher Gulvick, Scott Sammons, Yvette Unoarumhi, Darlene Wagner, Matthew Schmerer, S. H. Rosenthal, A. Gerasimova, R. M. Kagan, B. Anderson, M. Hua, Y. Liu, L.E. Bernstein, K.E. Livingston, A. Perez, I. A. Shlyakhter, R. V. Rolando, R. Owen, P. Tanpaiboon, F. Lacbawan, Clinton R. Paden, Duncan MacCannell |
| EPI_ISL_3330799, EPI_ISL_3330800 | TXDSHS | TXDSHS | Rashmi Tuladhar, Bonnie Oh, Jenny Zhang, Maliha Rahman, Mayela Pedrueza, Anita Pokharel, Karen Bobier, Lorraine Rodriguez, Myong Koag, Chun Wang, Rachel Lee, Grace Kubin |
| EPI_ISL_3330801 | Quest Diagnostics Incorporated | Centers for Disease Control and Prevention Division of Viral Diseases, Pathogen Discovery | Dakota Howard, Dhwani Batra, Peter W. Cook, Kara Moser, Adrian Paskey, Jason Caravas, Benjamin Rambo-Martin, Shatavia Morrison, Christopher Gulvick, Scott Sammons, Yvette Unoarumhi, Darlene Wagner, Matthew Schmerer, S. H. Rosenthal, A. Gerasimova, R. M. Kagan, B. Anderson, M. Hua, Y. Liu, L.E. Bernstein, K.E. Livingston, A. Perez, I. A. Shlyakhter, R. V. Rolando, R. Owen, P. Tanpaiboon, F. Lacbawan, Clinton R. Paden, Duncan MacCannell |
| EPI_ISL_3330802 | TXDSHS | TXDSHS | Rashmi Tuladhar, Bonnie Oh, Jenny Zhang, Maliha Rahman, Mayela Pedrueza, Anita Pokharel, Karen Bobier, Lorraine Rodriguez, Myong Koag, Chun Wang, Rachel Lee, Grace Kubin |
| EPI_ISL_3330803 | Quest Diagnostics Incorporated | Centers for Disease Control and Prevention Division of Viral Diseases, Pathogen Discovery | Dakota Howard, Dhwani Batra, Peter W. Cook, Kara Moser, Adrian Paskey, Jason Caravas, Benjamin Rambo-Martin, Shatavia Morrison, Christopher Gulvick, Scott Sammons, Yvette Unoarumhi, Darlene Wagner, Matthew Schmerer, S. H. Rosenthal, A. Gerasimova, R. M. Kagan, B. Anderson, M. Hua, Y. Liu, L.E. Bernstein, K.E. Livingston, A. Perez, I. A. Shlyakhter, R. V. Rolando, R. Owen, P. Tanpaiboon, F. Lacbawan, Clinton R. Paden, Duncan MacCannell |
| EPI_ISL_3330804, EPI_ISL_3330805 | TXDSHS | TXDSHS | Rashmi Tuladhar, Bonnie Oh, Jenny Zhang, Maliha Rahman, Mayela Pedrueza, Anita Pokharel, Karen Bobier, Lorraine Rodriguez, Myong Koag, Chun Wang, Rachel Lee, Grace Kubin |
| EPI_ISL_3330806, EPI_ISL_3330807, EPI_ISL_3330808, EPI_ISL_3330809 | Quest Diagnostics Incorporated | Centers for Disease Control and Prevention Division of Viral Diseases, Pathogen Discovery | Dakota Howard, Dhwani Batra, Peter W. Cook, Kara Moser, Adrian Paskey, Jason Caravas, Benjamin Rambo-Martin, Shatavia Morrison, Christopher Gulvick, Scott Sammons, Yvette Unoarumhi, Darlene Wagner, Matthew Schmerer, S. H. Rosenthal, A. Gerasimova, R. M. Kagan, B. Anderson, M. Hua, Y. Liu, L.E. Bernstein, K.E. Livingston, A. Perez, I. A. Shlyakhter, R. V. Rolando, R. Owen, P. Tanpaiboon, F. Lacbawan, Clinton R. Paden, Duncan MacCannell |
| EPI_ISL_3330810 | TXDSHS | TXDSHS | Rashmi Tuladhar, Bonnie Oh, Jenny Zhang, Maliha Rahman, Mayela Pedrueza, Anita Pokharel, Karen Bobier, Lorraine Rodriguez, Myong Koag, Chun Wang, Rachel Lee, Grace Kubin |
| EPI_ISL_3330811, EPI_ISL_3330812, EPI_ISL_3330813 | Quest Diagnostics Incorporated | Centers for Disease Control and Prevention Division of Viral Diseases, Pathogen Discovery | Dakota Howard, Dhwani Batra, Peter W. Cook, Kara Moser, Adrian Paskey, Jason Caravas, Benjamin Rambo-Martin, Shatavia Morrison, Christopher Gulvick, Scott Sammons, Yvette Unoarumhi, Darlene Wagner, Matthew Schmerer, S. H. Rosenthal, A. Gerasimova, R. M. Kagan, B. Anderson, M. Hua, Y. Liu, L.E. Bernstein, K.E. Livingston, A. Perez, I. A. Shlyakhter, R. V. Rolando, R. Owen, P. Tanpaiboon, F. Lacbawan, Clinton R. Paden, Duncan MacCannell |
| EPI_ISL_3330814 | TXDSHS | TXDSHS | Rashmi Tuladhar, Bonnie Oh, Jenny Zhang, Maliha Rahman, Mayela Pedrueza, Anita Pokharel, Karen Bobier, Lorraine Rodriguez, Myong Koag, Chun Wang, Rachel Lee, Grace Kubin |
| EPI_ISL_3330815, EPI_ISL_3330816 | Quest Diagnostics Incorporated | Centers for Disease Control and Prevention Division of Viral Diseases, Pathogen Discovery | Dakota Howard, Dhwani Batra, Peter W. Cook, Kara Moser, Adrian Paskey, Jason Caravas, Benjamin Rambo-Martin, Shatavia Morrison, Christopher Gulvick, Scott Sammons, Yvette Unoarumhi, Darlene Wagner, Matthew Schmerer, S. H. Rosenthal, A. Gerasimova, R. M. Kagan, B. Anderson, M. Hua, Y. Liu, L.E. Bernstein, K.E. Livingston, A. Perez, I. A. Shlyakhter, R. V. Rolando, R. Owen, P. Tanpaiboon, F. Lacbawan, Clinton R. Paden, Duncan MacCannell |
| EPI_ISL_3330817 | TXDSHS | TXDSHS | Rashmi Tuladhar, Bonnie Oh, Jenny Zhang, Maliha Rahman, Mayela Pedrueza, Anita Pokharel, Karen Bobier, Lorraine Rodriguez, Myong Koag, Chun Wang, Rachel Lee, Grace Kubin |
| EPI_ISL_3330818 | Quest Diagnostics Incorporated | Centers for Disease Control and Prevention Division of Viral Diseases, Pathogen Discovery | Dakota Howard, Dhwani Batra, Peter W. Cook, Kara Moser, Adrian Paskey, Jason Caravas, Benjamin Rambo-Martin, Shatavia Morrison, Christopher Gulvick, Scott Sammons, Yvette Unoarumhi, Darlene Wagner, Matthew Schmerer, S. H. Rosenthal, A. Gerasimova, R. M. Kagan, B. Anderson, M. Hua, Y. Liu, L.E. Bernstein, K.E. Livingston, A. Perez, I. A. Shlyakhter, R. V. Rolando, R. Owen, P. Tanpaiboon, F. Lacbawan, Clinton R. Paden, Duncan MacCannell |
| EPI_ISL_3330819 | TXDSHS | TXDSHS | Rashmi Tuladhar, Bonnie Oh, Jenny Zhang, Maliha Rahman, Mayela Pedrueza, Anita Pokharel, Karen Bobier, Lorraine Rodriguez, Myong Koag, Chun Wang, Rachel Lee, Grace Kubin |
| EPI_ISL_3330820 | Quest Diagnostics Incorporated | Centers for Disease Control and Prevention Division of Viral Diseases, Pathogen Discovery | Dakota Howard, Dhwani Batra, Peter W. Cook, Kara Moser, Adrian Paskey, Jason Caravas, Benjamin Rambo-Martin, Shatavia Morrison, Christopher Gulvick, Scott Sammons, Yvette Unoarumhi, Darlene Wagner, Matthew Schmerer, S. H. Rosenthal, A. Gerasimova, R. M. Kagan, B. Anderson, M. Hua, Y. Liu, L.E. Bernstein, K.E. Livingston, A. Perez, I. A. Shlyakhter, R. V. Rolando, R. Owen, P. Tanpaiboon, F. Lacbawan, Clinton R. Paden, Duncan MacCannell |
| EPI_ISL_3330821 | TXDSHS | TXDSHS | Rashmi Tuladhar, Bonnie Oh, Jenny Zhang, Maliha Rahman, Mayela Pedrueza, Anita Pokharel, Karen Bobier, Lorraine Rodriguez, Myong Koag, Chun Wang, Rachel Lee, Grace Kubin |
| EPI_ISL_3330822 | Quest Diagnostics Incorporated | Centers for Disease Control and Prevention Division of Viral Diseases, Pathogen Discovery | Dakota Howard, Dhwani Batra, Peter W. Cook, Kara Moser, Adrian Paskey, Jason Caravas, Benjamin Rambo-Martin, Shatavia Morrison, Christopher Gulvick, Scott Sammons, Yvette Unoarumhi, Darlene Wagner, Matthew Schmerer, S. H. Rosenthal, A. Gerasimova, R. M. Kagan, B. Anderson, M. Hua, Y. Liu, L.E. Bernstein, K.E. Livingston, A. Perez, I. A. Shlyakhter, R. V. Rolando, R. Owen, P. Tanpaiboon, F. Lacbawan, Clinton R. Paden, Duncan MacCannell |
| EPI_ISL_3330823 | TXDSHS | TXDSHS | Rashmi Tuladhar, Bonnie Oh, Jenny Zhang, Maliha Rahman, Mayela Pedrueza, Anita Pokharel, Karen Bobier, Lorraine Rodriguez, Myong Koag, Chun Wang, Rachel Lee, Grace Kubin |
| EPI_ISL_3330824 | Quest Diagnostics Incorporated | Centers for Disease Control and Prevention Division of Viral Diseases, Pathogen Discovery | Dakota Howard, Dhwani Batra, Peter W. Cook, Kara Moser, Adrian Paskey, Jason Caravas, Benjamin Rambo-Martin, Shatavia Morrison, Christopher Gulvick, Scott Sammons, Yvette Unoarumhi, Darlene Wagner, Matthew Schmerer, S. H. Rosenthal, A. Gerasimova, R. M. Kagan, B. Anderson, M. Hua, Y. Liu, L.E. Bernstein, K.E. Livingston, A. Perez, I. A. Shlyakhter, R. V. Rolando, R. Owen, P. Tanpaiboon, F. Lacbawan, Clinton R. Paden, Duncan MacCannell |
| EPI_ISL_3330825, EPI_ISL_3330826 | TXDSHS | TXDSHS | Rashmi Tuladhar, Bonnie Oh, Jenny Zhang, Maliha Rahman, Mayela Pedrueza, Anita Pokharel, Karen Bobier, Lorraine Rodriguez, Myong Koag, Chun Wang, Rachel Lee, Grace Kubin |
| EPI_ISL_3330827 | Quest Diagnostics Incorporated | Centers for Disease Control and Prevention Division of Viral Diseases, Pathogen Discovery | Dakota Howard, Dhwani Batra, Peter W. Cook, Kara Moser, Adrian Paskey, Jason Caravas, Benjamin Rambo-Martin, Shatavia Morrison, Christopher Gulvick, Scott Sammons, Yvette Unoarumhi, Darlene Wagner, Matthew Schmerer, S. H. Rosenthal, A. Gerasimova, R. M. Kagan, B. Anderson, M. Hua, Y. Liu, L.E. Bernstein, K.E. Livingston, A. Perez, I. A. Shlyakhter, R. V. Rolando, R. Owen, P. Tanpaiboon, F. Lacbawan, Clinton R. Paden, Duncan MacCannell |
| EPI_ISL_3330828 | TXDSHS | TXDSHS | Rashmi Tuladhar, Bonnie Oh, Jenny Zhang, Maliha Rahman, Mayela Pedrueza, Anita Pokharel, Karen Bobier, Lorraine Rodriguez, Myong Koag, Chun Wang, Rachel Lee, Grace Kubin |
| EPI_ISL_3330829 | Quest Diagnostics Incorporated | Centers for Disease Control and Prevention Division of Viral Diseases, Pathogen Discovery | Dakota Howard, Dhwani Batra, Peter W. Cook, Kara Moser, Adrian Paskey, Jason Caravas, Benjamin Rambo-Martin, Shatavia Morrison, Christopher Gulvick, Scott Sammons, Yvette Unoarumhi, Darlene Wagner, Matthew Schmerer, S. H. Rosenthal, A. Gerasimova, R. M. Kagan, B. Anderson, M. Hua, Y. Liu, L.E. Bernstein, K.E. Livingston, A. Perez, I. A. Shlyakhter, R. V. Rolando, R. Owen, P. Tanpaiboon, F. Lacbawan, Clinton R. Paden, Duncan MacCannell |

| | | | |
|---|---|---|---|
| EPI_ISL_3330830 | TXDSHS | TXDSHS | Rashmi Tuladhar, Bonnie Oh, Jenny Zhang, Maliha Rahman, Mayela Pedrueza, Anita Pokharel, Karen Bobier, Lorraine Rodriguez, Myong Koag, Chun Wang, Rachel Lee, Grace Kubin |
| EPI_ISL_3330832, EPI_ISL_3330833 | Quest Diagnostics Incorporated | Centers for Disease Control and Prevention Division of Viral Diseases, Pathogen Discovery | Dakota Howard, Dhwani Batra, Peter W. Cook, Kara Moser, Adrian Paskey, Jason Caravas, Benjamin Rambo-Martin, Shatavia Morrison, Christopher Gulvick, Scott Sammons, Yvette Unoarumhi, Darlene Wagner, Matthew Schmerer, S. H. Rosenthal, A. Gerasimova, R. M. Kagan, B. Anderson, M. Hua, Y. Liu, L.E. Bernstein, K.E. Livingston, A. Perez, I. A. Shlyakhter, R. V. Rolando, R. Owen, P. Tanpaiboon, F. Lacbawan, Clinton R. Paden, Duncan MacCannell |
| EPI_ISL_3330834 | TXDSHS | TXDSHS | Rashmi Tuladhar, Bonnie Oh, Jenny Zhang, Maliha Rahman, Mayela Pedrueza, Anita Pokharel, Karen Bobier, Lorraine Rodriguez, Myong Koag, Chun Wang, Rachel Lee, Grace Kubin |
| EPI_ISL_3330835, EPI_ISL_3330836 | Quest Diagnostics Incorporated | Centers for Disease Control and Prevention Division of Viral Diseases, Pathogen Discovery | Dakota Howard, Dhwani Batra, Peter W. Cook, Kara Moser, Adrian Paskey, Jason Caravas, Benjamin Rambo-Martin, Shatavia Morrison, Christopher Gulvick, Scott Sammons, Yvette Unoarumhi, Darlene Wagner, Matthew Schmerer, S. H. Rosenthal, A. Gerasimova, R. M. Kagan, B. Anderson, M. Hua, Y. Liu, L.E. Bernstein, K.E. Livingston, A. Perez, I. A. Shlyakhter, R. V. Rolando, R. Owen, P. Tanpaiboon, F. Lacbawan, Clinton R. Paden, Duncan MacCannell |
| EPI_ISL_3330837 | TXDSHS | TXDSHS | Rashmi Tuladhar, Bonnie Oh, Jenny Zhang, Maliha Rahman, Mayela Pedrueza, Anita Pokharel, Karen Bobier, Lorraine Rodriguez, Myong Koag, Chun Wang, Rachel Lee, Grace Kubin |
| EPI_ISL_3330838 | Quest Diagnostics Incorporated | Centers for Disease Control and Prevention Division of Viral Diseases, Pathogen Discovery | Dakota Howard, Dhwani Batra, Peter W. Cook, Kara Moser, Adrian Paskey, Jason Caravas, Benjamin Rambo-Martin, Shatavia Morrison, Christopher Gulvick, Scott Sammons, Yvette Unoarumhi, Darlene Wagner, Matthew Schmerer, S. H. Rosenthal, A. Gerasimova, R. M. Kagan, B. Anderson, M. Hua, Y. Liu, L.E. Bernstein, K.E. Livingston, A. Perez, I. A. Shlyakhter, R. V. Rolando, R. Owen, P. Tanpaiboon, F. Lacbawan, Clinton R. Paden, Duncan MacCannell |
| EPI_ISL_3330839 | TXDSHS | TXDSHS | Rashmi Tuladhar, Bonnie Oh, Jenny Zhang, Maliha Rahman, Mayela Pedrueza, Anita Pokharel, Karen Bobier, Lorraine Rodriguez, Myong Koag, Chun Wang, Rachel Lee, Grace Kubin |
| EPI_ISL_3330840 | Quest Diagnostics Incorporated | Centers for Disease Control and Prevention Division of Viral Diseases, Pathogen Discovery | Dakota Howard, Dhwani Batra, Peter W. Cook, Kara Moser, Adrian Paskey, Jason Caravas, Benjamin Rambo-Martin, Shatavia Morrison, Christopher Gulvick, Scott Sammons, Yvette Unoarumhi, Darlene Wagner, Matthew Schmerer, S. H. Rosenthal, A. Gerasimova, R. M. Kagan, B. Anderson, M. Hua, Y. Liu, L.E. Bernstein, K.E. Livingston, A. Perez, I. A. Shlyakhter, R. V. Rolando, R. Owen, P. Tanpaiboon, F. Lacbawan, Clinton R. Paden, Duncan MacCannell |
| EPI_ISL_3330841 | TXDSHS | TXDSHS | Rashmi Tuladhar, Bonnie Oh, Jenny Zhang, Maliha Rahman, Mayela Pedrueza, Anita Pokharel, Karen Bobier, Lorraine Rodriguez, Myong Koag, Chun Wang, Rachel Lee, Grace Kubin |
| EPI_ISL_3330842 | Quest Diagnostics Incorporated | Centers for Disease Control and Prevention Division of Viral Diseases, Pathogen Discovery | Dakota Howard, Dhwani Batra, Peter W. Cook, Kara Moser, Adrian Paskey, Jason Caravas, Benjamin Rambo-Martin, Shatavia Morrison, Christopher Gulvick, Scott Sammons, Yvette Unoarumhi, Darlene Wagner, Matthew Schmerer, S. H. Rosenthal, A. Gerasimova, R. M. Kagan, B. Anderson, M. Hua, Y. Liu, L.E. Bernstein, K.E. Livingston, A. Perez, I. A. Shlyakhter, R. V. Rolando, R. Owen, P. Tanpaiboon, F. Lacbawan, Clinton R. Paden, Duncan MacCannell |
| EPI_ISL_3330843, EPI_ISL_3330844 | TXDSHS | TXDSHS | Rashmi Tuladhar, Bonnie Oh, Jenny Zhang, Maliha Rahman, Mayela Pedrueza, Anita Pokharel, Karen Bobier, Lorraine Rodriguez, Myong Koag, Chun Wang, Rachel Lee, Grace Kubin |
| EPI_ISL_3330845 | Quest Diagnostics Incorporated | Centers for Disease Control and Prevention Division of Viral Diseases, Pathogen Discovery | Dakota Howard, Dhwani Batra, Peter W. Cook, Kara Moser, Adrian Paskey, Jason Caravas, Benjamin Rambo-Martin, Shatavia Morrison, Christopher Gulvick, Scott Sammons, Yvette Unoarumhi, Darlene Wagner, Matthew Schmerer, S. H. Rosenthal, A. Gerasimova, R. M. Kagan, B. Anderson, M. Hua, Y. Liu, L.E. Bernstein, K.E. Livingston, A. Perez, I. A. Shlyakhter, R. V. Rolando, R. Owen, P. Tanpaiboon, F. Lacbawan, Clinton R. Paden, Duncan MacCannell |
| EPI_ISL_3330846, EPI_ISL_3330847 | TXDSHS | TXDSHS | Rashmi Tuladhar, Bonnie Oh, Jenny Zhang, Maliha Rahman, Mayela Pedrueza, Anita Pokharel, Karen Bobier, Lorraine Rodriguez, Myong Koag, Chun Wang, Rachel Lee, Grace Kubin |
| EPI_ISL_3330848 | Quest Diagnostics Incorporated | Centers for Disease Control and Prevention Division of Viral Diseases, Pathogen Discovery | Dakota Howard, Dhwani Batra, Peter W. Cook, Kara Moser, Adrian Paskey, Jason Caravas, Benjamin Rambo-Martin, Shatavia Morrison, Christopher Gulvick, Scott Sammons, Yvette Unoarumhi, Darlene Wagner, Matthew Schmerer, S. H. Rosenthal, A. Gerasimova, R. M. Kagan, B. Anderson, M. Hua, Y. Liu, L.E. Bernstein, K.E. Livingston, A. Perez, I. A. Shlyakhter, R. V. Rolando, R. Owen, P. Tanpaiboon, F. Lacbawan, Clinton R. Paden, Duncan MacCannell |
| EPI_ISL_3330849 | TXDSHS | TXDSHS | Rashmi Tuladhar, Bonnie Oh, Jenny Zhang, Maliha Rahman, Mayela Pedrueza, Anita Pokharel, Karen Bobier, Lorraine Rodriguez, Myong Koag, Chun Wang, Rachel Lee, Grace Kubin |
| EPI_ISL_3330850 | Quest Diagnostics Incorporated | Centers for Disease Control and Prevention Division of Viral Diseases, Pathogen Discovery | Dakota Howard, Dhwani Batra, Peter W. Cook, Kara Moser, Adrian Paskey, Jason Caravas, Benjamin Rambo-Martin, Shatavia Morrison, Christopher Gulvick, Scott Sammons, Yvette Unoarumhi, Darlene Wagner, Matthew Schmerer, S. H. Rosenthal, A. Gerasimova, R. M. Kagan, B. Anderson, M. Hua, Y. Liu, L.E. Bernstein, K.E. Livingston, A. Perez, I. A. Shlyakhter, R. V. Rolando, R. Owen, P. Tanpaiboon, F. Lacbawan, Clinton R. Paden, Duncan MacCannell |
| EPI_ISL_3330851, EPI_ISL_3330852 | TXDSHS | TXDSHS | Rashmi Tuladhar, Bonnie Oh, Jenny Zhang, Maliha Rahman, Mayela Pedrueza, Anita Pokharel, Karen Bobier, Lorraine Rodriguez, Myong Koag, Chun Wang, Rachel Lee, Grace Kubin |
| EPI_ISL_3330853, EPI_ISL_3330854, EPI_ISL_3330855, EPI_ISL_3330856, EPI_ISL_3330857, EPI_ISL_3330858, EPI_ISL_3330859, EPI_ISL_3330860, EPI_ISL_3330861, EPI_ISL_3330862, EPI_ISL_3330864, EPI_ISL_3330865, EPI_ISL_3330866, EPI_ISL_3330867, EPI_ISL_3330868, EPI_ISL_3330869, EPI_ISL_3330870, EPI_ISL_3330871, EPI_ISL_3330872, EPI_ISL_3330873, EPI_ISL_3330874, EPI_ISL_3330875, EPI_ISL_3330876, EPI_ISL_3330877, EPI_ISL_3330878, EPI_ISL_3330879, EPI_ISL_3330880, EPI_ISL_3330881, EPI_ISL_3330883, EPI_ISL_3330884, EPI_ISL_3330885, EPI_ISL_3330886, EPI_ISL_3330887, EPI_ISL_3330888, EPI_ISL_3330889, EPI_ISL_3330890, EPI_ISL_3330891, EPI_ISL_3330892, EPI_ISL_3330893, EPI_ISL_3330894, EPI_ISL_3330895, EPI_ISL_3330896, EPI_ISL_3330897, EPI_ISL_3330898, EPI_ISL_3330899, EPI_ISL_3330900, EPI_ISL_3330901, EPI_ISL_3330902, EPI_ISL_3330903, EPI_ISL_3330904, EPI_ISL_3330905, EPI_ISL_3330906, EPI_ISL_3330907, EPI_ISL_3330908, EPI_ISL_3330909, EPI_ISL_3330910, EPI_ISL_3330911, EPI_ISL_3330912, EPI_ISL_3330913, EPI_ISL_3330914, EPI_ISL_3330915, EPI_ISL_3330916, EPI_ISL_3330918, EPI_ISL_3330919, EPI_ISL_3330920, EPI_ISL_3330921, EPI_ISL_3330922, EPI_ISL_3330923, EPI_ISL_3330924, EPI_ISL_3330925, EPI_ISL_3330926, EPI_ISL_3330927, EPI_ISL_3330928, EPI_ISL_3330929, EPI_ISL_3330930, EPI_ISL_3330931, EPI_ISL_3330932, EPI_ISL_3330933, EPI_ISL_3330934, EPI_ISL_3330935, EPI_ISL_3330936, EPI_ISL_3330937, EPI_ISL_3330938, EPI_ISL_3330939, EPI_ISL_3330940, EPI_ISL_3330941, EPI_ISL_3330942, EPI_ISL_3330943, EPI_ISL_3330944, EPI_ISL_3330946, EPI_ISL_3330947, EPI_ISL_3330948, EPI_ISL_3330949, EPI_ISL_3330950, EPI_ISL_3330951, EPI_ISL_3330952, EPI_ISL_3330953, EPI_ISL_3330954, EPI_ISL_3330955, EPI_ISL_3330956, EPI_ISL_3330957, EPI_ISL_3330958, EPI_ISL_3330959, EPI_ISL_3330960, EPI_ISL_3330961, EPI_ISL_3330962, EPI_ISL_3330963, EPI_ISL_3330964, EPI_ISL_3330965, EPI_ISL_3330966, EPI_ISL_3330967, EPI_ISL_3330968, EPI_ISL_3330969, EPI_ISL_3330970, EPI_ISL_3330971, EPI_ISL_3330972, EPI_ISL_3330973, EPI_ISL_3330974, EPI_ISL_3330975, EPI_ISL_3330976, EPI_ISL_3330977, EPI_ISL_3330978, EPI_ISL_3330979, EPI_ISL_3330980, EPI_ISL_3330981, EPI_ISL_3330982, EPI_ISL_3330983, EPI_ISL_3330984, EPI_ISL_3330985, EPI_ISL_3330986, EPI_ISL_3330987, EPI_ISL_3330988, EPI_ISL_3330989, EPI_ISL_3330990, EPI_ISL_3330991, EPI_ISL_3330992, EPI_ISL_3330993, EPI_ISL_3330994, EPI_ISL_3330995, EPI_ISL_3330996, EPI_ISL_3330997, EPI_ISL_3330998, EPI_ISL_3330999, EPI_ISL_3331000, EPI_ISL_3331001, EPI_ISL_3331002, EPI_ISL_3331003, EPI_ISL_3331004, EPI_ISL_3331005, EPI_ISL_3331006, EPI_ISL_3331007, EPI_ISL_3331008, EPI_ISL_3331009, EPI_ISL_3331010, EPI_ISL_3331011, EPI_ISL_3331012, EPI_ISL_3331013, EPI_ISL_3331014, EPI_ISL_3331016, EPI_ISL_3331017, EPI_ISL_3331018, EPI_ISL_3331019, EPI_ISL_3331020, EPI_ISL_3331021, EPI_ISL_3331022, EPI_ISL_3331023, EPI_ISL_3331024, EPI_ISL_3331025, EPI_ISL_3331026, EPI_ISL_3331027, EPI_ISL_3331029, EPI_ISL_3331030, EPI_ISL_3331031, EPI_ISL_3331032, EPI_ISL_3331033, EPI_ISL_3331034, EPI_ISL_3331035, EPI_ISL_3331036, EPI_ISL_3331037, EPI_ISL_3331038, EPI_ISL_3331039, EPI_ISL_3331040, EPI_ISL_3331041, EPI_ISL_3331042, EPI_ISL_3331043, EPI_ISL_3331044, EPI_ISL_3331045, EPI_ISL_3331046, EPI_ISL_3331047, EPI_ISL_3331048, EPI_ISL_3331049, EPI_ISL_3331050, EPI_ISL_3331051, EPI_ISL_3331052, EPI_ISL_3331053, EPI_ISL_3331054, EPI_ISL_3331055, EPI_ISL_3331056, EPI_ISL_3331057, EPI_ISL_3331058, EPI_ISL_3331059, EPI_ISL_3331060, EPI_ISL_3331061, EPI_ISL_3331062, EPI_ISL_3331063, EPI_ISL_3331064, EPI_ISL_3331065, EPI_ISL_3331066, EPI_ISL_3331067, EPI_ISL_3331068, EPI_ISL_3331069, EPI_ISL_3331070, EPI_ISL_3331071, EPI_ISL_3331072, EPI_ISL_3331073, EPI_ISL_3331074, EPI_ISL_3331075, EPI_ISL_3331076, EPI_ISL_3331077, EPI_ISL_3331078, EPI_ISL_3331079, EPI_ISL_3331080, EPI_ISL_3331081, EPI_ISL_3331083, EPI_ISL_3331084, EPI_ISL_3331085, EPI_ISL_3331086, EPI_ISL_3331087, EPI_ISL_3331088, EPI_ISL_3331089, EPI_ISL_3331090, EPI_ISL_3331091, EPI_ISL_3331092, EPI_ISL_3331093, EPI_ISL_3331094, EPI_ISL_3331095, EPI_ISL_3331096, EPI_ISL_3331097, EPI_ISL_3331098, EPI_ISL_3331099, EPI_ISL_3331100, EPI_ISL_3331101, EPI_ISL_3331102, EPI_ISL_3331103, EPI_ISL_3331104, EPI_ISL_3331105, EPI_ISL_3331106, EPI_ISL_3331107, EPI_ISL_3331108, EPI_ISL_3331109, EPI_ISL_3331110, EPI_ISL_3331111, EPI_ISL_3331112, EPI_ISL_3331113, EPI_ISL_3331114, EPI_ISL_3331115, EPI_ISL_3331116, EPI_ISL_3331117, EPI_ISL_3331118, EPI_ISL_3331119, EPI_ISL_3331120, EPI_ISL_3331121, EPI_ISL_3331122, EPI_ISL_3331123, EPI_ISL_3331124, EPI_ISL_3331125, EPI_ISL_3331126, EPI_ISL_3331127, EPI_ISL_3331128, EPI_ISL_3331129, EPI_ISL_3331130, EPI_ISL_3331131, EPI_ISL_3331132, EPI_ISL_3331133, EPI_ISL_3331134, EPI_ISL_3331135, EPI_ISL_3331136, EPI_ISL_3331138, EPI_ISL_3331139, EPI_ISL_3331140, EPI_ISL_3331141, EPI_ISL_3331142, EPI_ISL_3331143, EPI_ISL_3331144, EPI_ISL_3331145, EPI_ISL_3331146, EPI_ISL_3331147, EPI_ISL_3331148, EPI_ISL_3331149, EPI_ISL_3331150, EPI_ISL_3331151, EPI_ISL_3331152, EPI_ISL_3331153, EPI_ISL_3331154, EPI_ISL_3331155, EPI_ISL_3331156, EPI_ISL_3331157, EPI_ISL_3331158, EPI_ISL_3331159, EPI_ISL_3331160, EPI_ISL_3331161, EPI_ISL_3331162, EPI_ISL_3331163, EPI_ISL_3331164, EPI_ISL_3331165, EPI_ISL_3331166, EPI_ISL_3331167, EPI_ISL_3331168, EPI_ISL_3331169, EPI_ISL_3331170, EPI_ISL_3331171, EPI_ISL_3331172, EPI_ISL_3331173, EPI_ISL_3331174, EPI_ISL_3331176, EPI_ISL_3331177, EPI_ISL_3331178, EPI_ISL_3331179, EPI_ISL_3331181, EPI_ISL_3331182, EPI_ISL_3331183, EPI_ISL_3331184, EPI_ISL_3331185, EPI_ISL_3331186, EPI_ISL_3331187, EPI_ISL_3331188, EPI_ISL_3331189, EPI_ISL_3331190, EPI_ISL_3331191, EPI_ISL_3331192, EPI_ISL_3331193, EPI_ISL_3331194, EPI_ISL_3331195, EPI_ISL_3331196, EPI_ISL_3331197, EPI_ISL_3331198, EPI_ISL_3331199, EPI_ISL_3331200, EPI_ISL_3331201, EPI_ISL_3331202, EPI_ISL_3331203, EPI_ISL_3331204, EPI_ISL_3331205, EPI_ISL_3331206, EPI_ISL_3331207, EPI_ISL_3331208, EPI_ISL_3331209, EPI_ISL_3331210, EPI_ISL_3331211, EPI_ISL_3331212, EPI_ISL_3331213, EPI_ISL_3331214, EPI_ISL_3331215, EPI_ISL_3331216, EPI_ISL_3331217, EPI_ISL_3331218, EPI_ISL_3331219, EPI_ISL_3331220, EPI_ISL_3331221, EPI_ISL_3331222, EPI_ISL_3331223, EPI_ISL_3331224, EPI_ISL_3331226, EPI_ISL_3331228, EPI_ISL_3331229, EPI_ISL_3331230, EPI_ISL_3331231, EPI_ISL_3331232, EPI_ISL_3331233, EPI_ISL_3331234, EPI_ISL_3331235, EPI_ISL_3331236, EPI_ISL_3331237, EPI_ISL_3331238, EPI_ISL_3331239, EPI_ISL_3331240, EPI_ISL_3331241, EPI_ISL_3331242, EPI_ISL_3331243, EPI_ISL_3331244, EPI_ISL_3331245, EPI_ISL_3331246, EPI_ISL_3331248, EPI_ISL_3331249, EPI_ISL_3331250, EPI_ISL_3331251 | | | |
| see above | Quest Diagnostics Incorporated | Centers for Disease Control and Prevention Division of Viral Diseases, Pathogen Discovery | Dakota Howard, Dhwani Batra, Peter W. Cook, Kara Moser, Adrian Paskey, Jason Caravas, Benjamin Rambo-Martin, Shatavia Morrison, Christopher Gulvick, Scott Sammons, Yvette Unoarumhi, Darlene Wagner, Matthew Schmerer, S. H. Rosenthal, A. Gerasimova, R. M. Kagan, B. Anderson, M. Hua, Y. Liu, L.E. Bernstein, K.E. Livingston, A. Perez, I. A. Shlyakhter, R. V. Rolando, R. Owen, P. Tanpaiboon, F. Lacbawan, Clinton R. Paden, Duncan MacCannell |
| EPI_ISL_3331252, EPI_ISL_3331253, EPI_ISL_3331254, EPI_ISL_3331255, EPI_ISL_3331256, EPI_ISL_3331257, EPI_ISL_3331258, EPI_ISL_3331259, EPI_ISL_3331260, EPI_ISL_3331261, EPI_ISL_3331262, EPI_ISL_3331263, EPI_ISL_3331264, EPI_ISL_3331265, EPI_ISL_3331266, EPI_ISL_3331267, EPI_ISL_3331268, EPI_ISL_3331269, EPI_ISL_3331270, EPI_ISL_3331272, EPI_ISL_3331273, EPI_ISL_3331274, EPI_ISL_3331275, EPI_ISL_3331276, EPI_ISL_3331277, EPI_ISL_3331278, EPI_ISL_3331279, EPI_ISL_3331280, EPI_ISL_3331281, EPI_ISL_3331282, EPI_ISL_3331283, EPI_ISL_3331284, EPI_ISL_3331285, EPI_ISL_3331286, EPI_ISL_3331287, EPI_ISL_3331288, EPI_ISL_3331289, EPI_ISL_3331290, EPI_ISL_3331291, EPI_ISL_3331292, EPI_ISL_3331293, EPI_ISL_3331294, EPI_ISL_3331295, EPI_ISL_3331296, EPI_ISL_3331297, EPI_ISL_3331298, EPI_ISL_3331299, EPI_ISL_3331300, EPI_ISL_3331301, EPI_ISL_3331302, EPI_ISL_3331303, EPI_ISL_3331304, EPI_ISL_3331305, EPI_ISL_3331306, EPI_ISL_3331307, EPI_ISL_3331308, EPI_ISL_3331309, EPI_ISL_3331310, EPI_ISL_3331311, EPI_ISL_3331312, EPI_ISL_3331313, EPI_ISL_3331315, EPI_ISL_3331316, EPI_ISL_3331317, EPI_ISL_3331318, EPI_ISL_3331319, EPI_ISL_3331320, EPI_ISL_3331321, EPI_ISL_3331322, EPI_ISL_3331323, EPI_ISL_3331324, EPI_ISL_3331325, EPI_ISL_3331326, EPI_ISL_3331327, EPI_ISL_3331328, EPI_ISL_3331329, EPI_ISL_3331330, EPI_ISL_3331331, EPI_ISL_3331332, EPI_ISL_3331333, EPI_ISL_3331334, EPI_ISL_3331335, EPI_ISL_3331336, EPI_ISL_3331337, EPI_ISL_3331338, EPI_ISL_3331340, EPI_ISL_3331341, EPI_ISL_3331342, EPI_ISL_3331343, EPI_ISL_3331344, EPI_ISL_3331345, EPI_ISL_3331346, EPI_ISL_3331347, EPI_ISL_3331348, EPI_ISL_3331349, EPI_ISL_3331350, EPI_ISL_3331351, EPI_ISL_3331352, EPI_ISL_3331353, EPI_ISL_3331354, EPI_ISL_3331355, EPI_ISL_3331356, EPI_ISL_3331357, EPI_ISL_3331358, EPI_ISL_3331359, EPI_ISL_3331360, EPI_ISL_3331361, EPI_ISL_3331362, EPI_ISL_3331363, EPI_ISL_3331364, EPI_ISL_3331365, EPI_ISL_3331366, EPI_ISL_3331367, EPI_ISL_3331368, EPI_ISL_3331369, EPI_ISL_3331370, EPI_ISL_3331371, EPI_ISL_3331372, EPI_ISL_3331373, EPI_ISL_3331374, EPI_ISL_3331375, EPI_ISL_3331376, EPI_ISL_3331377, EPI_ISL_3331378, EPI_ISL_3331379, EPI_ISL_3331380, EPI_ISL_3331381, EPI_ISL_3331382, EPI_ISL_3331383, EPI_ISL_3331384, EPI_ISL_3331385, EPI_ISL_3331386, EPI_ISL_3331387, EPI_ISL_3331388, EPI_ISL_3331390, EPI_ISL_3331391, EPI_ISL_3331392, EPI_ISL_3331393, EPI_ISL_3331394, EPI_ISL_3331395, EPI_ISL_3331396, EPI_ISL_3331397, EPI_ISL_3331398, EPI_ISL_3331399, EPI_ISL_3331400, EPI_ISL_3331401, EPI_ISL_3331402, EPI_ISL_3331403, EPI_ISL_3331404, EPI_ISL_3331405, EPI_ISL_3331406, EPI_ISL_3331407, EPI_ISL_3331408, EPI_ISL_3331409, EPI_ISL_3331410, EPI_ISL_3331412, EPI_ISL_3331413, EPI_ISL_3331414, EPI_ISL_3331415, EPI_ISL_3331416, EPI_ISL_3331417, EPI_ISL_3331418, EPI_ISL_3331419, EPI_ISL_3331420, EPI_ISL_3331421, EPI_ISL_3331422, EPI_ISL_3331423, EPI_ISL_3331424, EPI_ISL_3331425, EPI_ISL_3331426, EPI_ISL_3331427, EPI_ISL_3331428, EPI_ISL_3331429, EPI_ISL_3331430, EPI_ISL_3331431, EPI_ISL_3331432, EPI_ISL_3331433, EPI_ISL_3331434, EPI_ISL_3331435, EPI_ISL_3331436, EPI_ISL_3331437, EPI_ISL_3331438, EPI_ISL_3331439, EPI_ISL_3331440, EPI_ISL_3331441, EPI_ISL_3331442, EPI_ISL_3331443, EPI_ISL_3331444, EPI_ISL_3331445, EPI_ISL_3331446, EPI_ISL_3331447, EPI_ISL_3331448, EPI_ISL_3331449, EPI_ISL_3331450, EPI_ISL_3331451, EPI_ISL_3331452, EPI_ISL_3331453, EPI_ISL_3331454, EPI_ISL_3331455, EPI_ISL_3331456, EPI_ISL_3331457, EPI_ISL_3331458, EPI_ISL_3331459, EPI_ISL_3331460, EPI_ISL_3331461, EPI_ISL_3331462, EPI_ISL_3331463, EPI_ISL_3331464, EPI_ISL_3331465 | | | |
| see above | Fulgent Genetics | Centers for Disease Control and Prevention Division of Viral Diseases, Pathogen Discovery | Dakota Howard, Dhwani Batra, Peter W. Cook, Kara Moser, Adrian Paskey, Jason Caravas, Benjamin Rambo-Martin, Shatavia Morrison, Christopher Gulvick, Scott Sammons, Yvette Unoarumhi, Darlene Wagner, Matthew Schmerer, Harry Gao, Mickey Li, John Gao, Joseph Fierro, Benafsh Sapra, Becky Tsai, Yan Meng, Doreen Ng, James Xie, Clinton R. Paden, Duncan MacCannell |
| EPI_ISL_3331466 | Pandemic Response Lab - NYC | Pandemic Response Lab, R&D | Henry Lee, Michael Hammerling, Melissa Hopkins, Cybill del Castillo, Shinyoung Clair Kang, William Ward, Pradeep Bugga, Sol Rey, Dylan Law, Alex Carpio, Isabel Fernandez Escapa, Haiping Hao, Jon Laurent |
| EPI_ISL_3331467, EPI_ISL_3331468 | Fulgent Genetics | Centers for Disease Control and Prevention Division of Viral Diseases, Pathogen Discovery | Dakota Howard, Dhwani Batra, Peter W. Cook, Kara Moser, Adrian Paskey, Jason Caravas, Benjamin Rambo-Martin, Shatavia Morrison, Christopher Gulvick, Scott Sammons, Yvette Unoarumhi, Darlene Wagner, Matthew Schmerer, Harry Gao, Mickey Li, John Gao, Joseph Fierro, Benafsh Sapra, Becky Tsai, Yan Meng, Doreen Ng, James Xie, Clinton R. Paden, Duncan MacCannell |
| EPI_ISL_3331469, EPI_ISL_3331470 | Pandemic Response Lab - NYC | Pandemic Response Lab, R&D | Henry Lee, Michael Hammerling, Melissa Hopkins, Cybill del Castillo, Shinyoung Clair Kang, William Ward, Pradeep Bugga, Sol Rey, Dylan Law, Alex Carpio, Isabel Fernandez Escapa, Haiping Hao, Jon Laurent |
| EPI_ISL_3331471 | Fulgent Genetics | Centers for Disease Control and Prevention Division of Viral Diseases, Pathogen Discovery | Dakota Howard, Dhwani Batra, Peter W. Cook, Kara Moser, Adrian Paskey, Jason Caravas, Benjamin Rambo-Martin, Shatavia Morrison, Christopher Gulvick, Scott Sammons, Yvette Unoarumhi, Darlene Wagner, Matthew Schmerer, Harry Gao, Mickey Li, John Gao, Joseph Fierro, Benafsh Sapra, Becky Tsai, Yan Meng, Doreen Ng, James Xie, Clinton R. Paden, Duncan MacCannell |
| EPI_ISL_3331472 | Pandemic Response Lab - NYC | Pandemic Response Lab, R&D | Henry Lee, Michael Hammerling, Melissa Hopkins, Cybill del Castillo, Shinyoung Clair Kang, William Ward, Pradeep Bugga, Sol Rey, Dylan Law, Alex Carpio, Isabel Fernandez Escapa, Haiping Hao, Jon Laurent |
| EPI_ISL_3331473 | Fulgent Genetics | Centers for Disease Control and Prevention Division of Viral Diseases, Pathogen Discovery | Dakota Howard, Dhwani Batra, Peter W. Cook, Kara Moser, Adrian Paskey, Jason Caravas, Benjamin Rambo-Martin, Shatavia Morrison, Christopher Gulvick, Scott Sammons, Yvette Unoarumhi, Darlene Wagner, Matthew Schmerer, Harry Gao, Mickey Li, John Gao, Joseph Fierro, Benafsh Sapra, Becky Tsai, Yan Meng, Doreen Ng, James Xie, Clinton R. Paden, Duncan MacCannell |
| EPI_ISL_3331474 | Pandemic Response Lab - NYC | Pandemic Response Lab, R&D | Henry Lee, Michael Hammerling, Melissa Hopkins, Cybill del Castillo, Shinyoung Clair Kang, William Ward, Pradeep Bugga, Sol Rey, Dylan Law, Alex Carpio, Isabel Fernandez Escapa, Haiping Hao, Jon Laurent |
| EPI_ISL_3331475 | Fulgent Genetics | Centers for Disease Control and Prevention Division of Viral Diseases, Pathogen Discovery | Dakota Howard, Dhwani Batra, Peter W. Cook, Kara Moser, Adrian Paskey, Jason Caravas, Benjamin Rambo-Martin, Shatavia Morrison, Christopher Gulvick, Scott Sammons, Yvette Unoarumhi, Darlene Wagner, Matthew Schmerer, Harry Gao, Mickey Li, John Gao, Joseph Fierro, Benafsh Sapra, Becky Tsai, Yan Meng, Doreen Ng, James Xie, Clinton R. Paden, Duncan MacCannell |
| EPI_ISL_3331476, EPI_ISL_3331477 | Pandemic Response Lab - NYC | Pandemic Response Lab, R&D | Henry Lee, Michael Hammerling, Melissa Hopkins, Cybill del Castillo, Shinyoung Clair Kang, William Ward, Pradeep Bugga, Sol Rey, Dylan Law, Alex Carpio, Isabel Fernandez Escapa, Haiping Hao, Jon Laurent |
| EPI_ISL_3331478 | Fulgent Genetics | Centers for Disease Control and Prevention Division of Viral Diseases, Pathogen Discovery | Dakota Howard, Dhwani Batra, Peter W. Cook, Kara Moser, Adrian Paskey, Jason Caravas, Benjamin Rambo-Martin, Shatavia Morrison, Christopher Gulvick, Scott Sammons, Yvette Unoarumhi, Darlene Wagner, Matthew Schmerer, Harry Gao, Mickey Li, John Gao, Joseph Fierro, Benafsh Sapra, Becky Tsai, Yan Meng, Doreen Ng, James Xie, Clinton R. Paden, Duncan MacCannell |
| EPI_ISL_3331479 | Pandemic Response Lab - NYC | Pandemic Response Lab, R&D | Henry Lee, Michael Hammerling, Melissa Hopkins, Cybill del Castillo, Shinyoung Clair Kang, William Ward, Pradeep Bugga, Sol Rey, Dylan Law, Alex Carpio, Isabel Fernandez Escapa, Haiping Hao, Jon Laurent |
| EPI_ISL_3331480 | Fulgent Genetics | Centers for Disease Control and Prevention Division of Viral Diseases, Pathogen Discovery | Dakota Howard, Dhwani Batra, Peter W. Cook, Kara Moser, Adrian Paskey, Jason Caravas, Benjamin Rambo-Martin, Shatavia Morrison, Christopher Gulvick, Scott Sammons, Yvette Unoarumhi, Darlene Wagner, Matthew Schmerer, Harry Gao, Mickey Li, John Gao, Joseph Fierro, Benafsh Sapra, Becky Tsai, Yan Meng, Doreen Ng, James Xie, Clinton R. Paden, Duncan MacCannell |
| EPI_ISL_3331481, EPI_ISL_3331482 | Pandemic Response Lab - NYC | Pandemic Response Lab, R&D | Henry Lee, Michael Hammerling, Melissa Hopkins, Cybill del Castillo, Shinyoung Clair Kang, William Ward, Pradeep Bugga, Sol Rey, Dylan Law, Alex Carpio, Isabel Fernandez Escapa, Haiping Hao, Jon Laurent |
| EPI_ISL_3331483 | Fulgent Genetics | Centers for Disease Control and Prevention Division of Viral Diseases, Pathogen Discovery | Dakota Howard, Dhwani Batra, Peter W. Cook, Kara Moser, Adrian Paskey, Jason Caravas, Benjamin Rambo-Martin, Shatavia Morrison, Christopher Gulvick, Scott Sammons, Yvette Unoarumhi, Darlene Wagner, Matthew Schmerer, Harry Gao, Mickey Li, John Gao, Joseph Fierro, Benafsh Sapra, Becky Tsai, Yan Meng, Doreen Ng, James Xie, Clinton R. Paden, Duncan MacCannell |
| EPI_ISL_3331484 | Pandemic Response Lab - NYC | Pandemic Response Lab, R&D | Henry Lee, Michael Hammerling, Melissa Hopkins, Cybill del Castillo, Shinyoung Clair Kang, William Ward, Pradeep Bugga, Sol Rey, Dylan Law, Alex Carpio, Isabel Fernandez Escapa, Haiping Hao, Jon Laurent |
| EPI_ISL_3331485 | Fulgent Genetics | Centers for Disease Control and Prevention Division of Viral Diseases, Pathogen Discovery | Dakota Howard, Dhwani Batra, Peter W. Cook, Kara Moser, Adrian Paskey, Jason Caravas, Benjamin Rambo-Martin, Shatavia Morrison, Christopher Gulvick, Scott Sammons, Yvette Unoarumhi, Darlene Wagner, Matthew Schmerer, Harry Gao, Mickey Li, John Gao, Joseph Fierro, Benafsh Sapra, Becky Tsai, Yan Meng, Doreen Ng, James Xie, Clinton R. Paden, Duncan MacCannell |
| EPI_ISL_3331486, EPI_ISL_3331487 | Pandemic Response Lab - NYC | Pandemic Response Lab, R&D | Henry Lee, Michael Hammerling, Melissa Hopkins, Cybill del Castillo, Shinyoung Clair Kang, William Ward, Pradeep Bugga, Sol Rey, Dylan Law, Alex Carpio, Isabel Fernandez Escapa, Haiping Hao, Jon Laurent |
| EPI_ISL_3331488 | Fulgent Genetics | Centers for Disease Control and Prevention Division of Viral Diseases, Pathogen Discovery | Dakota Howard, Dhwani Batra, Peter W. Cook, Kara Moser, Adrian Paskey, Jason Caravas, Benjamin Rambo-Martin, Shatavia Morrison, Christopher Gulvick, Scott Sammons, Yvette Unoarumhi, Darlene Wagner, Matthew Schmerer, Harry Gao, Mickey Li, John Gao, Joseph Fierro, Benafsh Sapra, Becky Tsai, Yan Meng, Doreen Ng, James Xie, Clinton R. Paden, Duncan MacCannell |
| EPI_ISL_3331489 | Pandemic Response Lab - NYC | Pandemic Response Lab, R&D | Henry Lee, Michael Hammerling, Melissa Hopkins, Cybill del Castillo, Shinyoung Clair Kang, William Ward, Pradeep Bugga, Sol Rey, Dylan Law, Alex Carpio, Isabel Fernandez Escapa, Haiping Hao, Jon Laurent |
| EPI_ISL_3331490 | Fulgent Genetics | Centers for Disease Control and Prevention Division of Viral | Dakota Howard, Dhwani Batra, Peter W. Cook, Kara Moser, Adrian Paskey, Jason Caravas, Benjamin Rambo-Martin, Shatavia Morrison, Christopher Gulvick, Scott Sammons, Yvette Unoarumhi, Darlene Wagner, Matthew Schmerer, Harry Gao, |

| | | | |
|---|---|---|---|
| | | Diseases, Pathogen Discovery | Mickey Li, John Gao, Joseph Fierro, Benafsh Sapra, Becky Tsai, Yan Meng, Doreen Ng, James Xie, Clinton R. Paden, Duncan MacCannell |
| EPI_ISL_3331491, EPI_ISL_3331492 | Pandemic Response Lab - NYC | Pandemic Response Lab, R&D | Henry Lee, Michael Hammerling, Melissa Hopkins, Cybill del Castillo, Shinyoung Clair Kang, William Ward, Pradeep Bugga, Sol Rey, Dylan Law, Alex Carpio, Isabel Fernandez Escapa, Haiping Hao, Jon Laurent |
| EPI_ISL_3331493 | Fulgent Genetics | Centers for Disease Control and Prevention Division of Viral Diseases, Pathogen Discovery | Dakota Howard, Dhwani Batra, Peter W. Cook, Kara Moser, Adrian Paskey, Jason Caravas, Benjamin Rambo-Martin, Shatavia Morrison, Christopher Gulvick, Scott Sammons, Yvette Unoarumhi, Darlene Wagner, Matthew Schmerer, Harry Gao, Mickey Li, John Gao, Joseph Fierro, Benafsh Sapra, Becky Tsai, Yan Meng, Doreen Ng, James Xie, Clinton R. Paden, Duncan MacCannell |
| EPI_ISL_3331494, EPI_ISL_3331495 | Pandemic Response Lab - NYC | Pandemic Response Lab, R&D | Henry Lee, Michael Hammerling, Melissa Hopkins, Cybill del Castillo, Shinyoung Clair Kang, William Ward, Pradeep Bugga, Sol Rey, Dylan Law, Alex Carpio, Isabel Fernandez Escapa, Haiping Hao, Jon Laurent |
| EPI_ISL_3331496 | Fulgent Genetics | Centers for Disease Control and Prevention Division of Viral Diseases, Pathogen Discovery | Dakota Howard, Dhwani Batra, Peter W. Cook, Kara Moser, Adrian Paskey, Jason Caravas, Benjamin Rambo-Martin, Shatavia Morrison, Christopher Gulvick, Scott Sammons, Yvette Unoarumhi, Darlene Wagner, Matthew Schmerer, Harry Gao, Mickey Li, John Gao, Joseph Fierro, Benafsh Sapra, Becky Tsai, Yan Meng, Doreen Ng, James Xie, Clinton R. Paden, Duncan MacCannell |
| EPI_ISL_3331497 | Pandemic Response Lab - NYC | Pandemic Response Lab, R&D | Henry Lee, Michael Hammerling, Melissa Hopkins, Cybill del Castillo, Shinyoung Clair Kang, William Ward, Pradeep Bugga, Sol Rey, Dylan Law, Alex Carpio, Isabel Fernandez Escapa, Haiping Hao, Jon Laurent |
| EPI_ISL_3331498 | Fulgent Genetics | Centers for Disease Control and Prevention Division of Viral Diseases, Pathogen Discovery | Dakota Howard, Dhwani Batra, Peter W. Cook, Kara Moser, Adrian Paskey, Jason Caravas, Benjamin Rambo-Martin, Shatavia Morrison, Christopher Gulvick, Scott Sammons, Yvette Unoarumhi, Darlene Wagner, Matthew Schmerer, Harry Gao, Mickey Li, John Gao, Joseph Fierro, Benafsh Sapra, Becky Tsai, Yan Meng, Doreen Ng, James Xie, Clinton R. Paden, Duncan MacCannell |
| EPI_ISL_3331499 | Pandemic Response Lab - NYC | Pandemic Response Lab, R&D | Henry Lee, Michael Hammerling, Melissa Hopkins, Cybill del Castillo, Shinyoung Clair Kang, William Ward, Pradeep Bugga, Sol Rey, Dylan Law, Alex Carpio, Isabel Fernandez Escapa, Haiping Hao, Jon Laurent |
| EPI_ISL_3331500 | Fulgent Genetics | Centers for Disease Control and Prevention Division of Viral Diseases, Pathogen Discovery | Dakota Howard, Dhwani Batra, Peter W. Cook, Kara Moser, Adrian Paskey, Jason Caravas, Benjamin Rambo-Martin, Shatavia Morrison, Christopher Gulvick, Scott Sammons, Yvette Unoarumhi, Darlene Wagner, Matthew Schmerer, Harry Gao, Mickey Li, John Gao, Joseph Fierro, Benafsh Sapra, Becky Tsai, Yan Meng, Doreen Ng, James Xie, Clinton R. Paden, Duncan MacCannell |
| EPI_ISL_3331501, EPI_ISL_3331502 | Pandemic Response Lab - NYC | Pandemic Response Lab, R&D | Henry Lee, Michael Hammerling, Melissa Hopkins, Cybill del Castillo, Shinyoung Clair Kang, William Ward, Pradeep Bugga, Sol Rey, Dylan Law, Alex Carpio, Isabel Fernandez Escapa, Haiping Hao, Jon Laurent |
| EPI_ISL_3331503 | Fulgent Genetics | Centers for Disease Control and Prevention Division of Viral Diseases, Pathogen Discovery | Dakota Howard, Dhwani Batra, Peter W. Cook, Kara Moser, Adrian Paskey, Jason Caravas, Benjamin Rambo-Martin, Shatavia Morrison, Christopher Gulvick, Scott Sammons, Yvette Unoarumhi, Darlene Wagner, Matthew Schmerer, Harry Gao, Mickey Li, John Gao, Joseph Fierro, Benafsh Sapra, Becky Tsai, Yan Meng, Doreen Ng, James Xie, Clinton R. Paden, Duncan MacCannell |
| EPI_ISL_3331504, EPI_ISL_3331505 | Pandemic Response Lab - NYC | Pandemic Response Lab, R&D | Henry Lee, Michael Hammerling, Melissa Hopkins, Cybill del Castillo, Shinyoung Clair Kang, William Ward, Pradeep Bugga, Sol Rey, Dylan Law, Alex Carpio, Isabel Fernandez Escapa, Haiping Hao, Jon Laurent |
| EPI_ISL_3331506 | Fulgent Genetics | Centers for Disease Control and Prevention Division of Viral Diseases, Pathogen Discovery | Dakota Howard, Dhwani Batra, Peter W. Cook, Kara Moser, Adrian Paskey, Jason Caravas, Benjamin Rambo-Martin, Shatavia Morrison, Christopher Gulvick, Scott Sammons, Yvette Unoarumhi, Darlene Wagner, Matthew Schmerer, Harry Gao, Mickey Li, John Gao, Joseph Fierro, Benafsh Sapra, Becky Tsai, Yan Meng, Doreen Ng, James Xie, Clinton R. Paden, Duncan MacCannell |
| EPI_ISL_3331507 | Pandemic Response Lab - NYC | Pandemic Response Lab, R&D | Henry Lee, Michael Hammerling, Melissa Hopkins, Cybill del Castillo, Shinyoung Clair Kang, William Ward, Pradeep Bugga, Sol Rey, Dylan Law, Alex Carpio, Isabel Fernandez Escapa, Haiping Hao, Jon Laurent |
| EPI_ISL_3331508 | Fulgent Genetics | Centers for Disease Control and Prevention Division of Viral Diseases, Pathogen Discovery | Dakota Howard, Dhwani Batra, Peter W. Cook, Kara Moser, Adrian Paskey, Jason Caravas, Benjamin Rambo-Martin, Shatavia Morrison, Christopher Gulvick, Scott Sammons, Yvette Unoarumhi, Darlene Wagner, Matthew Schmerer, Harry Gao, Mickey Li, John Gao, Joseph Fierro, Benafsh Sapra, Becky Tsai, Yan Meng, Doreen Ng, James Xie, Clinton R. Paden, Duncan MacCannell |
| EPI_ISL_3331509 | Pandemic Response Lab - NYC | Pandemic Response Lab, R&D | Henry Lee, Michael Hammerling, Melissa Hopkins, Cybill del Castillo, Shinyoung Clair Kang, William Ward, Pradeep Bugga, Sol Rey, Dylan Law, Alex Carpio, Isabel Fernandez Escapa, Haiping Hao, Jon Laurent |
| EPI_ISL_3331510 | Fulgent Genetics | Centers for Disease Control and Prevention Division of Viral Diseases, Pathogen Discovery | Dakota Howard, Dhwani Batra, Peter W. Cook, Kara Moser, Adrian Paskey, Jason Caravas, Benjamin Rambo-Martin, Shatavia Morrison, Christopher Gulvick, Scott Sammons, Yvette Unoarumhi, Darlene Wagner, Matthew Schmerer, Harry Gao, Mickey Li, John Gao, Joseph Fierro, Benafsh Sapra, Becky Tsai, Yan Meng, Doreen Ng, James Xie, Clinton R. Paden, Duncan MacCannell |
| EPI_ISL_3331511, EPI_ISL_3331512 | Pandemic Response Lab - NYC | Pandemic Response Lab, R&D | Henry Lee, Michael Hammerling, Melissa Hopkins, Cybill del Castillo, Shinyoung Clair Kang, William Ward, Pradeep Bugga, Sol Rey, Dylan Law, Alex Carpio, Isabel Fernandez Escapa, Haiping Hao, Jon Laurent |
| EPI_ISL_3331513 | Fulgent Genetics | Centers for Disease Control and Prevention Division of Viral Diseases, Pathogen Discovery | Dakota Howard, Dhwani Batra, Peter W. Cook, Kara Moser, Adrian Paskey, Jason Caravas, Benjamin Rambo-Martin, Shatavia Morrison, Christopher Gulvick, Scott Sammons, Yvette Unoarumhi, Darlene Wagner, Matthew Schmerer, Harry Gao, Mickey Li, John Gao, Joseph Fierro, Benafsh Sapra, Becky Tsai, Yan Meng, Doreen Ng, James Xie, Clinton R. Paden, Duncan MacCannell |
| EPI_ISL_3331514 | Pandemic Response Lab - NYC | Pandemic Response Lab, R&D | Henry Lee, Michael Hammerling, Melissa Hopkins, Cybill del Castillo, Shinyoung Clair Kang, William Ward, Pradeep Bugga, Sol Rey, Dylan Law, Alex Carpio, Isabel Fernandez Escapa, Haiping Hao, Jon Laurent |
| EPI_ISL_3331515 | Fulgent Genetics | Centers for Disease Control and Prevention Division of Viral Diseases, Pathogen Discovery | Dakota Howard, Dhwani Batra, Peter W. Cook, Kara Moser, Adrian Paskey, Jason Caravas, Benjamin Rambo-Martin, Shatavia Morrison, Christopher Gulvick, Scott Sammons, Yvette Unoarumhi, Darlene Wagner, Matthew Schmerer, Harry Gao, Mickey Li, John Gao, Joseph Fierro, Benafsh Sapra, Becky Tsai, Yan Meng, Doreen Ng, James Xie, Clinton R. Paden, Duncan MacCannell |
| EPI_ISL_3331516, EPI_ISL_3331517 | Pandemic Response Lab - NYC | Pandemic Response Lab, R&D | Henry Lee, Michael Hammerling, Melissa Hopkins, Cybill del Castillo, Shinyoung Clair Kang, William Ward, Pradeep Bugga, Sol Rey, Dylan Law, Alex Carpio, Isabel Fernandez Escapa, Haiping Hao, Jon Laurent |
| EPI_ISL_3331518 | Fulgent Genetics | Centers for Disease Control and Prevention Division of Viral Diseases, Pathogen Discovery | Dakota Howard, Dhwani Batra, Peter W. Cook, Kara Moser, Adrian Paskey, Jason Caravas, Benjamin Rambo-Martin, Shatavia Morrison, Christopher Gulvick, Scott Sammons, Yvette Unoarumhi, Darlene Wagner, Matthew Schmerer, Harry Gao, Mickey Li, John Gao, Joseph Fierro, Benafsh Sapra, Becky Tsai, Yan Meng, Doreen Ng, James Xie, Clinton R. Paden, Duncan MacCannell |
| EPI_ISL_3331519 | Pandemic Response Lab - NYC | Pandemic Response Lab, R&D | Henry Lee, Michael Hammerling, Melissa Hopkins, Cybill del Castillo, Shinyoung Clair Kang, William Ward, Pradeep Bugga, Sol Rey, Dylan Law, Alex Carpio, Isabel Fernandez Escapa, Haiping Hao, Jon Laurent |
| EPI_ISL_3331520 | Fulgent Genetics | Centers for Disease Control and Prevention Division of Viral Diseases, Pathogen Discovery | Dakota Howard, Dhwani Batra, Peter W. Cook, Kara Moser, Adrian Paskey, Jason Caravas, Benjamin Rambo-Martin, Shatavia Morrison, Christopher Gulvick, Scott Sammons, Yvette Unoarumhi, Darlene Wagner, Matthew Schmerer, Harry Gao, Mickey Li, John Gao, Joseph Fierro, Benafsh Sapra, Becky Tsai, Yan Meng, Doreen Ng, James Xie, Clinton R. Paden, Duncan MacCannell |
| EPI_ISL_3331521 | Pandemic Response Lab - NYC | Pandemic Response Lab, R&D | Henry Lee, Michael Hammerling, Melissa Hopkins, Cybill del Castillo, Shinyoung Clair Kang, William Ward, Pradeep Bugga, Sol Rey, Dylan Law, Alex Carpio, Isabel Fernandez Escapa, Haiping Hao, Jon Laurent |
| EPI_ISL_3331522, EPI_ISL_3331523, EPI_ISL_3331524 | Fulgent Genetics | Centers for Disease Control and Prevention Division of Viral Diseases, Pathogen Discovery | Dakota Howard, Dhwani Batra, Peter W. Cook, Kara Moser, Adrian Paskey, Jason Caravas, Benjamin Rambo-Martin, Shatavia Morrison, Christopher Gulvick, Scott Sammons, Yvette Unoarumhi, Darlene Wagner, Matthew Schmerer, Harry Gao, Mickey Li, John Gao, Joseph Fierro, Benafsh Sapra, Becky Tsai, Yan Meng, Doreen Ng, James Xie, Clinton R. Paden, Duncan MacCannell |
| EPI_ISL_3331525 | Pandemic Response Lab - NYC | Pandemic Response Lab, R&D | Henry Lee, Michael Hammerling, Melissa Hopkins, Cybill del Castillo, Shinyoung Clair Kang, William Ward, Pradeep Bugga, Sol Rey, Dylan Law, Alex Carpio, Isabel Fernandez Escapa, Haiping Hao, Jon Laurent |
| EPI_ISL_3331526, EPI_ISL_3331527 | Fulgent Genetics | Centers for Disease Control and Prevention Division of Viral Diseases, Pathogen Discovery | Dakota Howard, Dhwani Batra, Peter W. Cook, Kara Moser, Adrian Paskey, Jason Caravas, Benjamin Rambo-Martin, Shatavia Morrison, Christopher Gulvick, Scott Sammons, Yvette Unoarumhi, Darlene Wagner, Matthew Schmerer, Harry Gao, Mickey Li, John Gao, Joseph Fierro, Benafsh Sapra, Becky Tsai, Yan Meng, Doreen Ng, James Xie, Clinton R. Paden, Duncan MacCannell |
| EPI_ISL_3331528 | Pandemic Response Lab - NYC | Pandemic Response Lab, R&D | Henry Lee, Michael Hammerling, Melissa Hopkins, Cybill del Castillo, Shinyoung Clair Kang, William Ward, Pradeep Bugga, Sol Rey, Dylan Law, Alex Carpio, Isabel Fernandez Escapa, Haiping Hao, Jon Laurent |
| EPI_ISL_3331529 | Fulgent Genetics | Centers for Disease Control and Prevention Division of Viral Diseases, Pathogen Discovery | Dakota Howard, Dhwani Batra, Peter W. Cook, Kara Moser, Adrian Paskey, Jason Caravas, Benjamin Rambo-Martin, Shatavia Morrison, Christopher Gulvick, Scott Sammons, Yvette Unoarumhi, Darlene Wagner, Matthew Schmerer, Harry Gao, Mickey Li, John Gao, Joseph Fierro, Benafsh Sapra, Becky Tsai, Yan Meng, Doreen Ng, James Xie, Clinton R. Paden, Duncan MacCannell |
| EPI_ISL_3331530, EPI_ISL_3331531 | Pandemic Response Lab - NYC | Pandemic Response Lab, R&D | Henry Lee, Michael Hammerling, Melissa Hopkins, Cybill del Castillo, Shinyoung Clair Kang, William Ward, Pradeep Bugga, Sol Rey, Dylan Law, Alex Carpio, Isabel Fernandez Escapa, Haiping Hao, Jon Laurent |
| EPI_ISL_3331532 | Fulgent Genetics | Centers for Disease Control and Prevention Division of Viral Diseases, Pathogen Discovery | Dakota Howard, Dhwani Batra, Peter W. Cook, Kara Moser, Adrian Paskey, Jason Caravas, Benjamin Rambo-Martin, Shatavia Morrison, Christopher Gulvick, Scott Sammons, Yvette Unoarumhi, Darlene Wagner, Matthew Schmerer, Harry Gao, Mickey Li, John Gao, Joseph Fierro, Benafsh Sapra, Becky Tsai, Yan Meng, Doreen Ng, James Xie, Clinton R. Paden, Duncan MacCannell |
| EPI_ISL_3331533 | Pandemic Response Lab - NYC | Pandemic Response Lab, R&D | Henry Lee, Michael Hammerling, Melissa Hopkins, Cybill del Castillo, Shinyoung Clair Kang, William Ward, Pradeep Bugga, Sol Rey, Dylan Law, Alex Carpio, Isabel Fernandez Escapa, Haiping Hao, Jon Laurent |
| EPI_ISL_3331534 | Fulgent Genetics | Centers for Disease Control and Prevention Division of Viral Diseases, Pathogen Discovery | Dakota Howard, Dhwani Batra, Peter W. Cook, Kara Moser, Adrian Paskey, Jason Caravas, Benjamin Rambo-Martin, Shatavia Morrison, Christopher Gulvick, Scott Sammons, Yvette Unoarumhi, Darlene Wagner, Matthew Schmerer, Harry Gao, Mickey Li, John Gao, Joseph Fierro, Benafsh Sapra, Becky Tsai, Yan Meng, Doreen Ng, James Xie, Clinton R. Paden, Duncan MacCannell |
| EPI_ISL_3331535 | Pandemic Response Lab - NYC | Pandemic Response Lab, R&D | Henry Lee, Michael Hammerling, Melissa Hopkins, Cybill del Castillo, Shinyoung Clair Kang, William Ward, Pradeep Bugga, Sol Rey, Dylan Law, Alex Carpio, Isabel Fernandez Escapa, Haiping Hao, Jon Laurent |
| EPI_ISL_3331536, EPI_ISL_3331537, EPI_ISL_3331538 | Fulgent Genetics | Centers for Disease Control and Prevention Division of Viral Diseases, Pathogen Discovery | Dakota Howard, Dhwani Batra, Peter W. Cook, Kara Moser, Adrian Paskey, Jason Caravas, Benjamin Rambo-Martin, Shatavia Morrison, Christopher Gulvick, Scott Sammons, Yvette Unoarumhi, Darlene Wagner, Matthew Schmerer, Harry Gao, Mickey Li, John Gao, Joseph Fierro, Benafsh Sapra, Becky Tsai, Yan Meng, Doreen Ng, James Xie, Clinton R. Paden, Duncan MacCannell |
| EPI_ISL_3331539 | Pandemic Response Lab - NYC | Pandemic Response Lab, R&D | Henry Lee, Michael Hammerling, Melissa Hopkins, Cybill del Castillo, Shinyoung Clair Kang, William Ward, Pradeep Bugga, Sol Rey, Dylan Law, Alex Carpio, Isabel Fernandez Escapa, Haiping Hao, Jon Laurent |
| EPI_ISL_3331540 | Fulgent Genetics | Centers for Disease Control and Prevention Division of Viral Diseases, Pathogen Discovery | Dakota Howard, Dhwani Batra, Peter W. Cook, Kara Moser, Adrian Paskey, Jason Caravas, Benjamin Rambo-Martin, Shatavia Morrison, Christopher Gulvick, Scott Sammons, Yvette Unoarumhi, Darlene Wagner, Matthew Schmerer, Harry Gao, Mickey Li, John Gao, Joseph Fierro, Benafsh Sapra, Becky Tsai, Yan Meng, Doreen Ng, James Xie, Clinton R. Paden, Duncan MacCannell |
| EPI_ISL_3331541 | Pandemic Response Lab - NYC | Pandemic Response Lab, R&D | Henry Lee, Michael Hammerling, Melissa Hopkins, Cybill del Castillo, Shinyoung Clair Kang, William Ward, Pradeep Bugga, Sol Rey, Dylan Law, Alex Carpio, Isabel Fernandez Escapa, Haiping Hao, Jon Laurent |
| EPI_ISL_3331542 | Fulgent Genetics | Centers for Disease Control and Prevention Division of Viral Diseases, Pathogen Discovery | Dakota Howard, Dhwani Batra, Peter W. Cook, Kara Moser, Adrian Paskey, Jason Caravas, Benjamin Rambo-Martin, Shatavia Morrison, Christopher Gulvick, Scott Sammons, Yvette Unoarumhi, Darlene Wagner, Matthew Schmerer, Harry Gao, Mickey Li, John Gao, Joseph Fierro, Benafsh Sapra, Becky Tsai, Yan Meng, Doreen Ng, James Xie, Clinton R. Paden, Duncan MacCannell |
| EPI_ISL_3331543, EPI_ISL_3331544 | Pandemic Response Lab - NYC | Pandemic Response Lab, R&D | Henry Lee, Michael Hammerling, Melissa Hopkins, Cybill del Castillo, Shinyoung Clair Kang, William Ward, Pradeep Bugga, Sol Rey, Dylan Law, Alex Carpio, Isabel Fernandez Escapa, Haiping Hao, Jon Laurent |
| EPI_ISL_3331545 | Fulgent Genetics | Centers for Disease Control and Prevention Division of Viral Diseases, Pathogen Discovery | Dakota Howard, Dhwani Batra, Peter W. Cook, Kara Moser, Adrian Paskey, Jason Caravas, Benjamin Rambo-Martin, Shatavia Morrison, Christopher Gulvick, Scott Sammons, Yvette Unoarumhi, Darlene Wagner, Matthew Schmerer, Harry Gao, Mickey Li, John Gao, Joseph Fierro, Benafsh Sapra, Becky Tsai, Yan Meng, Doreen Ng, James Xie, Clinton R. Paden, Duncan MacCannell |
| EPI_ISL_3331546 | Pandemic Response Lab - NYC | Pandemic Response Lab, R&D | Henry Lee, Michael Hammerling, Melissa Hopkins, Cybill del Castillo, Shinyoung Clair Kang, William Ward, Pradeep Bugga, Sol Rey, Dylan Law, Alex Carpio, Isabel Fernandez Escapa, Haiping Hao, Jon Laurent |
| EPI_ISL_3331547 | Fulgent Genetics | Centers for Disease Control and Prevention Division of Viral Diseases, Pathogen Discovery | Dakota Howard, Dhwani Batra, Peter W. Cook, Kara Moser, Adrian Paskey, Jason Caravas, Benjamin Rambo-Martin, Shatavia Morrison, Christopher Gulvick, Scott Sammons, Yvette Unoarumhi, Darlene Wagner, Matthew Schmerer, Harry Gao, Mickey Li, John Gao, Joseph Fierro, Benafsh Sapra, Becky Tsai, Yan Meng, Doreen Ng, James Xie, Clinton R. Paden, Duncan MacCannell |
| EPI_ISL_3331548, EPI_ISL_3331549 | Pandemic Response Lab - NYC | Pandemic Response Lab, R&D | Henry Lee, Michael Hammerling, Melissa Hopkins, Cybill del Castillo, Shinyoung Clair Kang, William Ward, Pradeep Bugga, Sol Rey, Dylan Law, Alex Carpio, Isabel Fernandez Escapa, Haiping Hao, Jon Laurent |
| EPI_ISL_3331550 | Fulgent Genetics | Centers for Disease Control and Prevention Division of Viral Diseases, Pathogen Discovery | Dakota Howard, Dhwani Batra, Peter W. Cook, Kara Moser, Adrian Paskey, Jason Caravas, Benjamin Rambo-Martin, Shatavia Morrison, Christopher Gulvick, Scott Sammons, Yvette Unoarumhi, Darlene Wagner, Matthew Schmerer, Harry Gao, Mickey Li, John Gao, Joseph Fierro, Benafsh Sapra, Becky Tsai, Yan Meng, Doreen Ng, James Xie, Clinton R. Paden, Duncan MacCannell |
| EPI_ISL_3331551 | Pandemic Response Lab - NYC | Pandemic Response Lab, R&D | Henry Lee, Michael Hammerling, Melissa Hopkins, Cybill del Castillo, Shinyoung Clair Kang, William Ward, Pradeep Bugga, Sol Rey, Dylan Law, Alex Carpio, Isabel Fernandez Escapa, Haiping Hao, Jon Laurent |
| EPI_ISL_3331552 | Fulgent Genetics | Centers for Disease Control and Prevention Division of Viral Diseases, Pathogen Discovery | Dakota Howard, Dhwani Batra, Peter W. Cook, Kara Moser, Adrian Paskey, Jason Caravas, Benjamin Rambo-Martin, Shatavia Morrison, Christopher Gulvick, Scott Sammons, Yvette Unoarumhi, Darlene Wagner, Matthew Schmerer, Harry Gao, Mickey Li, John Gao, Joseph Fierro, Benafsh Sapra, Becky Tsai, Yan Meng, Doreen Ng, James Xie, Clinton R. Paden, Duncan MacCannell |
| EPI_ISL_3331553 | Pandemic Response Lab - NYC | Pandemic Response Lab, R&D | Henry Lee, Michael Hammerling, Melissa Hopkins, Cybill del Castillo, Shinyoung Clair Kang, William Ward, Pradeep Bugga, Sol Rey, Dylan Law, Alex Carpio, Isabel Fernandez Escapa, Haiping Hao, Jon Laurent |
| EPI_ISL_3331554 | Fulgent Genetics | Centers for Disease Control and Prevention Division of Viral Diseases, Pathogen Discovery | Dakota Howard, Dhwani Batra, Peter W. Cook, Kara Moser, Adrian Paskey, Jason Caravas, Benjamin Rambo-Martin, Shatavia Morrison, Christopher Gulvick, Scott Sammons, Yvette Unoarumhi, Darlene Wagner, Matthew Schmerer, Harry Gao, Mickey Li, John Gao, Joseph Fierro, Benafsh Sapra, Becky Tsai, Yan Meng, Doreen Ng, James Xie, Clinton R. Paden, Duncan MacCannell |
| EPI_ISL_3331555 | Pandemic Response Lab - NYC | Pandemic Response Lab, R&D | Henry Lee, Michael Hammerling, Melissa Hopkins, Cybill del Castillo, Shinyoung Clair Kang, William Ward, Pradeep Bugga, Sol Rey, Dylan Law, Alex Carpio, Isabel Fernandez Escapa, Haiping Hao, Jon Laurent |

| | | | |
|---|---|---|---|
| EPI_ISL_3331556, EPI_ISL_3331557, EPI_ISL_3331558, EPI_ISL_3331559 | Fulgent Genetics | Centers for Disease Control and Prevention Division of Viral Diseases, Pathogen Discovery | Dakota Howard, Dhwani Batra, Peter W. Cook, Kara Moser, Adrian Paskey, Jason Caravas, Benjamin Rambo-Martin, Shatavia Morrison, Christopher Gulvick, Scott Sammons, Yvette Unoarumhi, Darlene Wagner, Matthew Schmerer, Harry Gao, Mickey Li, John Gao, Joseph Fierro, Benafsh Sapra, Becky Tsai, Yan Meng, Doreen Ng, James Xie, Clinton R. Paden, Duncan MacCannell |
| EPI_ISL_3331560, EPI_ISL_3331561 | Pandemic Response Lab - NYC | Pandemic Response Lab, R&D | Henry Lee, Michael Hammerling, Melissa Hopkins, Cybill del Castillo, Shinyoung Clair Kang, William Ward, Pradeep Bugga, Sol Rey, Dylan Law, Alex Carpio, Isabel Fernandez Escapa, Haiping Hao, Jon Laurent |
| EPI_ISL_3331562 | Fulgent Genetics | Centers for Disease Control and Prevention Division of Viral Diseases, Pathogen Discovery | Dakota Howard, Dhwani Batra, Peter W. Cook, Kara Moser, Adrian Paskey, Jason Caravas, Benjamin Rambo-Martin, Shatavia Morrison, Christopher Gulvick, Scott Sammons, Yvette Unoarumhi, Darlene Wagner, Matthew Schmerer, Harry Gao, Mickey Li, John Gao, Joseph Fierro, Benafsh Sapra, Becky Tsai, Yan Meng, Doreen Ng, James Xie, Clinton R. Paden, Duncan MacCannell |
| EPI_ISL_3331563 | Pandemic Response Lab - NYC | Pandemic Response Lab, R&D | Henry Lee, Michael Hammerling, Melissa Hopkins, Cybill del Castillo, Shinyoung Clair Kang, William Ward, Pradeep Bugga, Sol Rey, Dylan Law, Alex Carpio, Isabel Fernandez Escapa, Haiping Hao, Jon Laurent |
| EPI_ISL_3331564, EPI_ISL_3331565 | Fulgent Genetics | Centers for Disease Control and Prevention Division of Viral Diseases, Pathogen Discovery | Dakota Howard, Dhwani Batra, Peter W. Cook, Kara Moser, Adrian Paskey, Jason Caravas, Benjamin Rambo-Martin, Shatavia Morrison, Christopher Gulvick, Scott Sammons, Yvette Unoarumhi, Darlene Wagner, Matthew Schmerer, Harry Gao, Mickey Li, John Gao, Joseph Fierro, Benafsh Sapra, Becky Tsai, Yan Meng, Doreen Ng, James Xie, Clinton R. Paden, Duncan MacCannell |
| EPI_ISL_3331566, EPI_ISL_3331567 | Pandemic Response Lab - NYC | Pandemic Response Lab, R&D | Henry Lee, Michael Hammerling, Melissa Hopkins, Cybill del Castillo, Shinyoung Clair Kang, William Ward, Pradeep Bugga, Sol Rey, Dylan Law, Alex Carpio, Isabel Fernandez Escapa, Haiping Hao, Jon Laurent |
| EPI_ISL_3331568 | Fulgent Genetics | Centers for Disease Control and Prevention Division of Viral Diseases, Pathogen Discovery | Dakota Howard, Dhwani Batra, Peter W. Cook, Kara Moser, Adrian Paskey, Jason Caravas, Benjamin Rambo-Martin, Shatavia Morrison, Christopher Gulvick, Scott Sammons, Yvette Unoarumhi, Darlene Wagner, Matthew Schmerer, Harry Gao, Mickey Li, John Gao, Joseph Fierro, Benafsh Sapra, Becky Tsai, Yan Meng, Doreen Ng, James Xie, Clinton R. Paden, Duncan MacCannell |
| EPI_ISL_3331569 | Pandemic Response Lab - NYC | Pandemic Response Lab, R&D | Henry Lee, Michael Hammerling, Melissa Hopkins, Cybill del Castillo, Shinyoung Clair Kang, William Ward, Pradeep Bugga, Sol Rey, Dylan Law, Alex Carpio, Isabel Fernandez Escapa, Haiping Hao, Jon Laurent |
| EPI_ISL_3331570 | Fulgent Genetics | Centers for Disease Control and Prevention Division of Viral Diseases, Pathogen Discovery | Dakota Howard, Dhwani Batra, Peter W. Cook, Kara Moser, Adrian Paskey, Jason Caravas, Benjamin Rambo-Martin, Shatavia Morrison, Christopher Gulvick, Scott Sammons, Yvette Unoarumhi, Darlene Wagner, Matthew Schmerer, Harry Gao, Mickey Li, John Gao, Joseph Fierro, Benafsh Sapra, Becky Tsai, Yan Meng, Doreen Ng, James Xie, Clinton R. Paden, Duncan MacCannell |
| EPI_ISL_3331571 | Pandemic Response Lab - NYC | Pandemic Response Lab, R&D | Henry Lee, Michael Hammerling, Melissa Hopkins, Cybill del Castillo, Shinyoung Clair Kang, William Ward, Pradeep Bugga, Sol Rey, Dylan Law, Alex Carpio, Isabel Fernandez Escapa, Haiping Hao, Jon Laurent |
| EPI_ISL_3331572 | Fulgent Genetics | Centers for Disease Control and Prevention Division of Viral Diseases, Pathogen Discovery | Dakota Howard, Dhwani Batra, Peter W. Cook, Kara Moser, Adrian Paskey, Jason Caravas, Benjamin Rambo-Martin, Shatavia Morrison, Christopher Gulvick, Scott Sammons, Yvette Unoarumhi, Darlene Wagner, Matthew Schmerer, Harry Gao, Mickey Li, John Gao, Joseph Fierro, Benafsh Sapra, Becky Tsai, Yan Meng, Doreen Ng, James Xie, Clinton R. Paden, Duncan MacCannell |
| EPI_ISL_3331573, EPI_ISL_3331574 | Pandemic Response Lab - NYC | Pandemic Response Lab, R&D | Henry Lee, Michael Hammerling, Melissa Hopkins, Cybill del Castillo, Shinyoung Clair Kang, William Ward, Pradeep Bugga, Sol Rey, Dylan Law, Alex Carpio, Isabel Fernandez Escapa, Haiping Hao, Jon Laurent |
| EPI_ISL_3331575 | Fulgent Genetics | Centers for Disease Control and Prevention Division of Viral Diseases, Pathogen Discovery | Dakota Howard, Dhwani Batra, Peter W. Cook, Kara Moser, Adrian Paskey, Jason Caravas, Benjamin Rambo-Martin, Shatavia Morrison, Christopher Gulvick, Scott Sammons, Yvette Unoarumhi, Darlene Wagner, Matthew Schmerer, Harry Gao, Mickey Li, John Gao, Joseph Fierro, Benafsh Sapra, Becky Tsai, Yan Meng, Doreen Ng, James Xie, Clinton R. Paden, Duncan MacCannell |
| EPI_ISL_3331576 | Pandemic Response Lab - NYC | Pandemic Response Lab, R&D | Henry Lee, Michael Hammerling, Melissa Hopkins, Cybill del Castillo, Shinyoung Clair Kang, William Ward, Pradeep Bugga, Sol Rey, Dylan Law, Alex Carpio, Isabel Fernandez Escapa, Haiping Hao, Jon Laurent |
| EPI_ISL_3331577 | Fulgent Genetics | Centers for Disease Control and Prevention Division of Viral Diseases, Pathogen Discovery | Dakota Howard, Dhwani Batra, Peter W. Cook, Kara Moser, Adrian Paskey, Jason Caravas, Benjamin Rambo-Martin, Shatavia Morrison, Christopher Gulvick, Scott Sammons, Yvette Unoarumhi, Darlene Wagner, Matthew Schmerer, Harry Gao, Mickey Li, John Gao, Joseph Fierro, Benafsh Sapra, Becky Tsai, Yan Meng, Doreen Ng, James Xie, Clinton R. Paden, Duncan MacCannell |
| EPI_ISL_3331578, EPI_ISL_3331579 | Pandemic Response Lab - NYC | Pandemic Response Lab, R&D | Henry Lee, Michael Hammerling, Melissa Hopkins, Cybill del Castillo, Shinyoung Clair Kang, William Ward, Pradeep Bugga, Sol Rey, Dylan Law, Alex Carpio, Isabel Fernandez Escapa, Haiping Hao, Jon Laurent |
| EPI_ISL_3331580 | Fulgent Genetics | Centers for Disease Control and Prevention Division of Viral Diseases, Pathogen Discovery | Dakota Howard, Dhwani Batra, Peter W. Cook, Kara Moser, Adrian Paskey, Jason Caravas, Benjamin Rambo-Martin, Shatavia Morrison, Christopher Gulvick, Scott Sammons, Yvette Unoarumhi, Darlene Wagner, Matthew Schmerer, Harry Gao, Mickey Li, John Gao, Joseph Fierro, Benafsh Sapra, Becky Tsai, Yan Meng, Doreen Ng, James Xie, Clinton R. Paden, Duncan MacCannell |
| EPI_ISL_3331581 | Pandemic Response Lab - NYC | Pandemic Response Lab, R&D | Henry Lee, Michael Hammerling, Melissa Hopkins, Cybill del Castillo, Shinyoung Clair Kang, William Ward, Pradeep Bugga, Sol Rey, Dylan Law, Alex Carpio, Isabel Fernandez Escapa, Haiping Hao, Jon Laurent |
| EPI_ISL_3331582 | Fulgent Genetics | Centers for Disease Control and Prevention Division of Viral Diseases, Pathogen Discovery | Dakota Howard, Dhwani Batra, Peter W. Cook, Kara Moser, Adrian Paskey, Jason Caravas, Benjamin Rambo-Martin, Shatavia Morrison, Christopher Gulvick, Scott Sammons, Yvette Unoarumhi, Darlene Wagner, Matthew Schmerer, Harry Gao, Mickey Li, John Gao, Joseph Fierro, Benafsh Sapra, Becky Tsai, Yan Meng, Doreen Ng, James Xie, Clinton R. Paden, Duncan MacCannell |
| EPI_ISL_3331583 | Pandemic Response Lab - NYC | Pandemic Response Lab, R&D | Henry Lee, Michael Hammerling, Melissa Hopkins, Cybill del Castillo, Shinyoung Clair Kang, William Ward, Pradeep Bugga, Sol Rey, Dylan Law, Alex Carpio, Isabel Fernandez Escapa, Haiping Hao, Jon Laurent |
| EPI_ISL_3331584 | Fulgent Genetics | Centers for Disease Control and Prevention Division of Viral Diseases, Pathogen Discovery | Dakota Howard, Dhwani Batra, Peter W. Cook, Kara Moser, Adrian Paskey, Jason Caravas, Benjamin Rambo-Martin, Shatavia Morrison, Christopher Gulvick, Scott Sammons, Yvette Unoarumhi, Darlene Wagner, Matthew Schmerer, Harry Gao, Mickey Li, John Gao, Joseph Fierro, Benafsh Sapra, Becky Tsai, Yan Meng, Doreen Ng, James Xie, Clinton R. Paden, Duncan MacCannell |
| EPI_ISL_3331585 | Pandemic Response Lab - NYC | Pandemic Response Lab, R&D | Henry Lee, Michael Hammerling, Melissa Hopkins, Cybill del Castillo, Shinyoung Clair Kang, William Ward, Pradeep Bugga, Sol Rey, Dylan Law, Alex Carpio, Isabel Fernandez Escapa, Haiping Hao, Jon Laurent |
| EPI_ISL_3331586 | Fulgent Genetics | Centers for Disease Control and Prevention Division of Viral Diseases, Pathogen Discovery | Dakota Howard, Dhwani Batra, Peter W. Cook, Kara Moser, Adrian Paskey, Jason Caravas, Benjamin Rambo-Martin, Shatavia Morrison, Christopher Gulvick, Scott Sammons, Yvette Unoarumhi, Darlene Wagner, Matthew Schmerer, Harry Gao, Mickey Li, John Gao, Joseph Fierro, Benafsh Sapra, Becky Tsai, Yan Meng, Doreen Ng, James Xie, Clinton R. Paden, Duncan MacCannell |
| EPI_ISL_3331587, EPI_ISL_3331588 | Pandemic Response Lab - NYC | Pandemic Response Lab, R&D | Henry Lee, Michael Hammerling, Melissa Hopkins, Cybill del Castillo, Shinyoung Clair Kang, William Ward, Pradeep Bugga, Sol Rey, Dylan Law, Alex Carpio, Isabel Fernandez Escapa, Haiping Hao, Jon Laurent |
| EPI_ISL_3331589 | Fulgent Genetics | Centers for Disease Control and Prevention Division of Viral Diseases, Pathogen Discovery | Dakota Howard, Dhwani Batra, Peter W. Cook, Kara Moser, Adrian Paskey, Jason Caravas, Benjamin Rambo-Martin, Shatavia Morrison, Christopher Gulvick, Scott Sammons, Yvette Unoarumhi, Darlene Wagner, Matthew Schmerer, Harry Gao, Mickey Li, John Gao, Joseph Fierro, Benafsh Sapra, Becky Tsai, Yan Meng, Doreen Ng, James Xie, Clinton R. Paden, Duncan MacCannell |
| EPI_ISL_3331590 | Pandemic Response Lab - NYC | Pandemic Response Lab, R&D | Henry Lee, Michael Hammerling, Melissa Hopkins, Cybill del Castillo, Shinyoung Clair Kang, William Ward, Pradeep Bugga, Sol Rey, Dylan Law, Alex Carpio, Isabel Fernandez Escapa, Haiping Hao, Jon Laurent |
| EPI_ISL_3331591 | Fulgent Genetics | Centers for Disease Control and Prevention Division of Viral Diseases, Pathogen Discovery | Dakota Howard, Dhwani Batra, Peter W. Cook, Kara Moser, Adrian Paskey, Jason Caravas, Benjamin Rambo-Martin, Shatavia Morrison, Christopher Gulvick, Scott Sammons, Yvette Unoarumhi, Darlene Wagner, Matthew Schmerer, Harry Gao, Mickey Li, John Gao, Joseph Fierro, Benafsh Sapra, Becky Tsai, Yan Meng, Doreen Ng, James Xie, Clinton R. Paden, Duncan MacCannell |
| EPI_ISL_3331592 | Pandemic Response Lab - NYC | Pandemic Response Lab, R&D | Henry Lee, Michael Hammerling, Melissa Hopkins, Cybill del Castillo, Shinyoung Clair Kang, William Ward, Pradeep Bugga, Sol Rey, Dylan Law, Alex Carpio, Isabel Fernandez Escapa, Haiping Hao, Jon Laurent |
| EPI_ISL_3331593 | Fulgent Genetics | Centers for Disease Control and Prevention Division of Viral Diseases, Pathogen Discovery | Dakota Howard, Dhwani Batra, Peter W. Cook, Kara Moser, Adrian Paskey, Jason Caravas, Benjamin Rambo-Martin, Shatavia Morrison, Christopher Gulvick, Scott Sammons, Yvette Unoarumhi, Darlene Wagner, Matthew Schmerer, Harry Gao, Mickey Li, John Gao, Joseph Fierro, Benafsh Sapra, Becky Tsai, Yan Meng, Doreen Ng, James Xie, Clinton R. Paden, Duncan MacCannell |
| EPI_ISL_3331594, EPI_ISL_3331595 | Pandemic Response Lab - NYC | Pandemic Response Lab, R&D | Henry Lee, Michael Hammerling, Melissa Hopkins, Cybill del Castillo, Shinyoung Clair Kang, William Ward, Pradeep Bugga, Sol Rey, Dylan Law, Alex Carpio, Isabel Fernandez Escapa, Haiping Hao, Jon Laurent |
| EPI_ISL_3331596 | Fulgent Genetics | Centers for Disease Control and Prevention Division of Viral Diseases, Pathogen Discovery | Dakota Howard, Dhwani Batra, Peter W. Cook, Kara Moser, Adrian Paskey, Jason Caravas, Benjamin Rambo-Martin, Shatavia Morrison, Christopher Gulvick, Scott Sammons, Yvette Unoarumhi, Darlene Wagner, Matthew Schmerer, Harry Gao, Mickey Li, John Gao, Joseph Fierro, Benafsh Sapra, Becky Tsai, Yan Meng, Doreen Ng, James Xie, Clinton R. Paden, Duncan MacCannell |
| EPI_ISL_3331597 | Pandemic Response Lab - NYC | Pandemic Response Lab, R&D | Henry Lee, Michael Hammerling, Melissa Hopkins, Cybill del Castillo, Shinyoung Clair Kang, William Ward, Pradeep Bugga, Sol Rey, Dylan Law, Alex Carpio, Isabel Fernandez Escapa, Haiping Hao, Jon Laurent |
| EPI_ISL_3331598 | Fulgent Genetics | Centers for Disease Control and Prevention Division of Viral Diseases, Pathogen Discovery | Dakota Howard, Dhwani Batra, Peter W. Cook, Kara Moser, Adrian Paskey, Jason Caravas, Benjamin Rambo-Martin, Shatavia Morrison, Christopher Gulvick, Scott Sammons, Yvette Unoarumhi, Darlene Wagner, Matthew Schmerer, Harry Gao, Mickey Li, John Gao, Joseph Fierro, Benafsh Sapra, Becky Tsai, Yan Meng, Doreen Ng, James Xie, Clinton R. Paden, Duncan MacCannell |
| EPI_ISL_3331599 | Pandemic Response Lab - NYC | Pandemic Response Lab, R&D | Henry Lee, Michael Hammerling, Melissa Hopkins, Cybill del Castillo, Shinyoung Clair Kang, William Ward, Pradeep Bugga, Sol Rey, Dylan Law, Alex Carpio, Isabel Fernandez Escapa, Haiping Hao, Jon Laurent |
| EPI_ISL_3331600, EPI_ISL_3331601, EPI_ISL_3331602, EPI_ISL_3331603 | Fulgent Genetics | Centers for Disease Control and Prevention Division of Viral Diseases, Pathogen Discovery | Dakota Howard, Dhwani Batra, Peter W. Cook, Kara Moser, Adrian Paskey, Jason Caravas, Benjamin Rambo-Martin, Shatavia Morrison, Christopher Gulvick, Scott Sammons, Yvette Unoarumhi, Darlene Wagner, Matthew Schmerer, Harry Gao, Mickey Li, John Gao, Joseph Fierro, Benafsh Sapra, Becky Tsai, Yan Meng, Doreen Ng, James Xie, Clinton R. Paden, Duncan MacCannell |
| EPI_ISL_3331604 | Pandemic Response Lab - NYC | Pandemic Response Lab, R&D | Henry Lee, Michael Hammerling, Melissa Hopkins, Cybill del Castillo, Shinyoung Clair Kang, William Ward, Pradeep Bugga, Sol Rey, Dylan Law, Alex Carpio, Isabel Fernandez Escapa, Haiping Hao, Jon Laurent |
| EPI_ISL_3331605, EPI_ISL_3331606, EPI_ISL_3331607, EPI_ISL_3331608 | Fulgent Genetics | Centers for Disease Control and Prevention Division of Viral Diseases, Pathogen Discovery | Dakota Howard, Dhwani Batra, Peter W. Cook, Kara Moser, Adrian Paskey, Jason Caravas, Benjamin Rambo-Martin, Shatavia Morrison, Christopher Gulvick, Scott Sammons, Yvette Unoarumhi, Darlene Wagner, Matthew Schmerer, Harry Gao, Mickey Li, John Gao, Joseph Fierro, Benafsh Sapra, Becky Tsai, Yan Meng, Doreen Ng, James Xie, Clinton R. Paden, Duncan MacCannell |
| EPI_ISL_3331610 | Pandemic Response Lab - NYC | Pandemic Response Lab, R&D | Henry Lee, Michael Hammerling, Melissa Hopkins, Cybill del Castillo, Shinyoung Clair Kang, William Ward, Pradeep Bugga, Sol Rey, Dylan Law, Alex Carpio, Isabel Fernandez Escapa, Haiping Hao, Jon Laurent |
| EPI_ISL_3331611 | Fulgent Genetics | Centers for Disease Control and Prevention Division of Viral Diseases, Pathogen Discovery | Dakota Howard, Dhwani Batra, Peter W. Cook, Kara Moser, Adrian Paskey, Jason Caravas, Benjamin Rambo-Martin, Shatavia Morrison, Christopher Gulvick, Scott Sammons, Yvette Unoarumhi, Darlene Wagner, Matthew Schmerer, Harry Gao, Mickey Li, John Gao, Joseph Fierro, Benafsh Sapra, Becky Tsai, Yan Meng, Doreen Ng, James Xie, Clinton R. Paden, Duncan MacCannell |
| EPI_ISL_3331612 | Pandemic Response Lab - NYC | Pandemic Response Lab, R&D | Henry Lee, Michael Hammerling, Melissa Hopkins, Cybill del Castillo, Shinyoung Clair Kang, William Ward, Pradeep Bugga, Sol Rey, Dylan Law, Alex Carpio, Isabel Fernandez Escapa, Haiping Hao, Jon Laurent |
| EPI_ISL_3331613 | Fulgent Genetics | Centers for Disease Control and Prevention Division of Viral Diseases, Pathogen Discovery | Dakota Howard, Dhwani Batra, Peter W. Cook, Kara Moser, Adrian Paskey, Jason Caravas, Benjamin Rambo-Martin, Shatavia Morrison, Christopher Gulvick, Scott Sammons, Yvette Unoarumhi, Darlene Wagner, Matthew Schmerer, Harry Gao, Mickey Li, John Gao, Joseph Fierro, Benafsh Sapra, Becky Tsai, Yan Meng, Doreen Ng, James Xie, Clinton R. Paden, Duncan MacCannell |
| EPI_ISL_3331614 | Pandemic Response Lab - NYC | Pandemic Response Lab, R&D | Henry Lee, Michael Hammerling, Melissa Hopkins, Cybill del Castillo, Shinyoung Clair Kang, William Ward, Pradeep Bugga, Sol Rey, Dylan Law, Alex Carpio, Isabel Fernandez Escapa, Haiping Hao, Jon Laurent |
| EPI_ISL_3331615 | Fulgent Genetics | Centers for Disease Control and Prevention Division of Viral Diseases, Pathogen Discovery | Dakota Howard, Dhwani Batra, Peter W. Cook, Kara Moser, Adrian Paskey, Jason Caravas, Benjamin Rambo-Martin, Shatavia Morrison, Christopher Gulvick, Scott Sammons, Yvette Unoarumhi, Darlene Wagner, Matthew Schmerer, Harry Gao, Mickey Li, John Gao, Joseph Fierro, Benafsh Sapra, Becky Tsai, Yan Meng, Doreen Ng, James Xie, Clinton R. Paden, Duncan MacCannell |
| EPI_ISL_3331616, EPI_ISL_3331617 | Pandemic Response Lab - NYC | Pandemic Response Lab, R&D | Henry Lee, Michael Hammerling, Melissa Hopkins, Cybill del Castillo, Shinyoung Clair Kang, William Ward, Pradeep Bugga, Sol Rey, Dylan Law, Alex Carpio, Isabel Fernandez Escapa, Haiping Hao, Jon Laurent |
| EPI_ISL_3331618 | Fulgent Genetics | Centers for Disease Control and Prevention Division of Viral Diseases, Pathogen Discovery | Dakota Howard, Dhwani Batra, Peter W. Cook, Kara Moser, Adrian Paskey, Jason Caravas, Benjamin Rambo-Martin, Shatavia Morrison, Christopher Gulvick, Scott Sammons, Yvette Unoarumhi, Darlene Wagner, Matthew Schmerer, Harry Gao, Mickey Li, John Gao, Joseph Fierro, Benafsh Sapra, Becky Tsai, Yan Meng, Doreen Ng, James Xie, Clinton R. Paden, Duncan MacCannell |
| EPI_ISL_3331619 | Pandemic Response Lab - NYC | Pandemic Response Lab, R&D | Henry Lee, Michael Hammerling, Melissa Hopkins, Cybill del Castillo, Shinyoung Clair Kang, William Ward, Pradeep Bugga, Sol Rey, Dylan Law, Alex Carpio, Isabel Fernandez Escapa, Haiping Hao, Jon Laurent |
| EPI_ISL_3331620 | Valais Hospital, Central Institute | Valais Hospital, Central Institute | Alexis Dumoulin, Lorenzo Cerutti, Cedric Howald, Henri Pegeot, Melyssa Elies, Deborah Penet, Keith Harshman, Ioannis Xenarios |
| EPI_ISL_3331621 | Fulgent Genetics | Centers for Disease Control and Prevention Division of Viral Diseases, Pathogen Discovery | Dakota Howard, Dhwani Batra, Peter W. Cook, Kara Moser, Adrian Paskey, Jason Caravas, Benjamin Rambo-Martin, Shatavia Morrison, Christopher Gulvick, Scott Sammons, Yvette Unoarumhi, Darlene Wagner, Matthew Schmerer, Harry Gao, Mickey Li, John Gao, Joseph Fierro, Benafsh Sapra, Becky Tsai, Yan Meng, Doreen Ng, James Xie, Clinton R. Paden, Duncan MacCannell |
| EPI_ISL_3331622 | Valais Hospital, Central Institute | Valais Hospital, Central Institute | Alexis Dumoulin, Lorenzo Cerutti, Cedric Howald, Henri Pegeot, Melyssa Elies, Deborah Penet, Keith Harshman, Ioannis Xenarios |
| EPI_ISL_3331623 | Pandemic Response Lab - NYC | Pandemic Response Lab, R&D | Henry Lee, Michael Hammerling, Melissa Hopkins, Cybill del Castillo, Shinyoung Clair Kang, William Ward, Pradeep Bugga, Sol Rey, Dylan Law, Alex Carpio, Isabel Fernandez Escapa, Haiping Hao, Jon Laurent |

| | | | |
|---|---|---|---|
| EPI_ISL_3331624 | Valais Hospital, Central Institute | Valais Hospital, Central Institute | Alexis Dumoulin, Lorenzo Cerutti, Cedric Howald, Henri Pegeot, Melyssa Elies, Deborah Penet, Keith Harshman, Ioannis Xenarios |
| EPI_ISL_3331625 | Pandemic Response Lab - NYC | Pandemic Response Lab, R&D | Henry Lee, Michael Hammerling, Melissa Hopkins, Cybill del Castillo, Shinyoung Clair Kang, William Ward, Pradeep Bugga, Sol Rey, Dylan Law, Alex Carpio, Isabel Fernandez Escapa, Haiping Hao, Jon Laurent |
| EPI_ISL_3331626 | Fulgent Genetics | Centers for Disease Control and Prevention Division of Viral Diseases, Pathogen Discovery | Dakota Howard, Dhwani Batra, Peter W. Cook, Kara Moser, Adrian Paskey, Jason Caravas, Benjamin Rambo-Martin, Shatavia Morrison, Christopher Gulvick, Scott Sammons, Yvette Unoarumhi, Darlene Wagner, Matthew Schmerer, Harry Gao, Mickey Li, John Gao, Joseph Fierro, Benafsh Sapra, Becky Tsai, Yan Meng, Doreen Ng, James Xie, Clinton R. Paden, Duncan MacCannell |
| EPI_ISL_3331628 | Pandemic Response Lab - NYC | Pandemic Response Lab, R&D | Henry Lee, Michael Hammerling, Melissa Hopkins, Cybill del Castillo, Shinyoung Clair Kang, William Ward, Pradeep Bugga, Sol Rey, Dylan Law, Alex Carpio, Isabel Fernandez Escapa, Haiping Hao, Jon Laurent |
| EPI_ISL_3331629 | Fulgent Genetics | Centers for Disease Control and Prevention Division of Viral Diseases, Pathogen Discovery | Dakota Howard, Dhwani Batra, Peter W. Cook, Kara Moser, Adrian Paskey, Jason Caravas, Benjamin Rambo-Martin, Shatavia Morrison, Christopher Gulvick, Scott Sammons, Yvette Unoarumhi, Darlene Wagner, Matthew Schmerer, Harry Gao, Mickey Li, John Gao, Joseph Fierro, Benafsh Sapra, Becky Tsai, Yan Meng, Doreen Ng, James Xie, Clinton R. Paden, Duncan MacCannell |
| EPI_ISL_3331630, EPI_ISL_3331631 | Valais Hospital, Central Institute | Valais Hospital, Central Institute | Alexis Dumoulin, Lorenzo Cerutti, Cedric Howald, Henri Pegeot, Melyssa Elies, Deborah Penet, Keith Harshman, Ioannis Xenarios |
| EPI_ISL_3331632 | Fulgent Genetics | Centers for Disease Control and Prevention Division of Viral Diseases, Pathogen Discovery | Dakota Howard, Dhwani Batra, Peter W. Cook, Kara Moser, Adrian Paskey, Jason Caravas, Benjamin Rambo-Martin, Shatavia Morrison, Christopher Gulvick, Scott Sammons, Yvette Unoarumhi, Darlene Wagner, Matthew Schmerer, Harry Gao, Mickey Li, John Gao, Joseph Fierro, Benafsh Sapra, Becky Tsai, Yan Meng, Doreen Ng, James Xie, Clinton R. Paden, Duncan MacCannell |
| EPI_ISL_3331633 | Laboratoire d'analyses médicales Dr. Etienne Kessler | Valais Hospital, Central Institute | Alexis Dumoulin, Lorenzo Cerutti, Cedric Howald, Henri Pegeot, Melyssa Elies, Deborah Penet, Keith Harshman, Ioannis Xenarios |
| EPI_ISL_3331634 | Fulgent Genetics | Centers for Disease Control and Prevention Division of Viral Diseases, Pathogen Discovery | Dakota Howard, Dhwani Batra, Peter W. Cook, Kara Moser, Adrian Paskey, Jason Caravas, Benjamin Rambo-Martin, Shatavia Morrison, Christopher Gulvick, Scott Sammons, Yvette Unoarumhi, Darlene Wagner, Matthew Schmerer, Harry Gao, Mickey Li, John Gao, Joseph Fierro, Benafsh Sapra, Becky Tsai, Yan Meng, Doreen Ng, James Xie, Clinton R. Paden, Duncan MacCannell |
| EPI_ISL_3331635, EPI_ISL_3331636 | Valais Hospital, Central Institute | Valais Hospital, Central Institute | Alexis Dumoulin, Lorenzo Cerutti, Cedric Howald, Henri Pegeot, Melyssa Elies, Deborah Penet, Keith Harshman, Ioannis Xenarios |
| EPI_ISL_3331637 | Fulgent Genetics | Centers for Disease Control and Prevention Division of Viral Diseases, Pathogen Discovery | Dakota Howard, Dhwani Batra, Peter W. Cook, Kara Moser, Adrian Paskey, Jason Caravas, Benjamin Rambo-Martin, Shatavia Morrison, Christopher Gulvick, Scott Sammons, Yvette Unoarumhi, Darlene Wagner, Matthew Schmerer, Harry Gao, Mickey Li, John Gao, Joseph Fierro, Benafsh Sapra, Becky Tsai, Yan Meng, Doreen Ng, James Xie, Clinton R. Paden, Duncan MacCannell |
| EPI_ISL_3331638 | Valais Hospital, Central Institute | Valais Hospital, Central Institute | Alexis Dumoulin, Lorenzo Cerutti, Cedric Howald, Henri Pegeot, Melyssa Elies, Deborah Penet, Keith Harshman, Ioannis Xenarios |
| EPI_ISL_3331639 | Fulgent Genetics | Centers for Disease Control and Prevention Division of Viral Diseases, Pathogen Discovery | Dakota Howard, Dhwani Batra, Peter W. Cook, Kara Moser, Adrian Paskey, Jason Caravas, Benjamin Rambo-Martin, Shatavia Morrison, Christopher Gulvick, Scott Sammons, Yvette Unoarumhi, Darlene Wagner, Matthew Schmerer, Harry Gao, Mickey Li, John Gao, Joseph Fierro, Benafsh Sapra, Becky Tsai, Yan Meng, Doreen Ng, James Xie, Clinton R. Paden, Duncan MacCannell |
| EPI_ISL_3331640 | Pandemic Response Lab - NYC | Pandemic Response Lab, R&D | Henry Lee, Michael Hammerling, Melissa Hopkins, Cybill del Castillo, Shinyoung Clair Kang, William Ward, Pradeep Bugga, Sol Rey, Dylan Law, Alex Carpio, Isabel Fernandez Escapa, Haiping Hao, Jon Laurent |
| EPI_ISL_3331641, EPI_ISL_3331642 | Valais Hospital, Central Institute | Valais Hospital, Central Institute | Alexis Dumoulin, Lorenzo Cerutti, Cedric Howald, Henri Pegeot, Melyssa Elies, Deborah Penet, Keith Harshman, Ioannis Xenarios |
| EPI_ISL_3331643 | Pandemic Response Lab - NYC | Pandemic Response Lab, R&D | Henry Lee, Michael Hammerling, Melissa Hopkins, Cybill del Castillo, Shinyoung Clair Kang, William Ward, Pradeep Bugga, Sol Rey, Dylan Law, Alex Carpio, Isabel Fernandez Escapa, Haiping Hao, Jon Laurent |
| EPI_ISL_3331644 | Fulgent Genetics | Centers for Disease Control and Prevention Division of Viral Diseases, Pathogen Discovery | Dakota Howard, Dhwani Batra, Peter W. Cook, Kara Moser, Adrian Paskey, Jason Caravas, Benjamin Rambo-Martin, Shatavia Morrison, Christopher Gulvick, Scott Sammons, Yvette Unoarumhi, Darlene Wagner, Matthew Schmerer, Harry Gao, Mickey Li, John Gao, Joseph Fierro, Benafsh Sapra, Becky Tsai, Yan Meng, Doreen Ng, James Xie, Clinton R. Paden, Duncan MacCannell |
| EPI_ISL_3331645 | Valais Hospital, Central Institute | Valais Hospital, Central Institute | Alexis Dumoulin, Lorenzo Cerutti, Cedric Howald, Henri Pegeot, Melyssa Elies, Deborah Penet, Keith Harshman, Ioannis Xenarios |
| EPI_ISL_3331646 | Pandemic Response Lab - NYC | Pandemic Response Lab, R&D | Henry Lee, Michael Hammerling, Melissa Hopkins, Cybill del Castillo, Shinyoung Clair Kang, William Ward, Pradeep Bugga, Sol Rey, Dylan Law, Alex Carpio, Isabel Fernandez Escapa, Haiping Hao, Jon Laurent |
| EPI_ISL_3331647 | Fulgent Genetics | Centers for Disease Control and Prevention Division of Viral Diseases, Pathogen Discovery | Dakota Howard, Dhwani Batra, Peter W. Cook, Kara Moser, Adrian Paskey, Jason Caravas, Benjamin Rambo-Martin, Shatavia Morrison, Christopher Gulvick, Scott Sammons, Yvette Unoarumhi, Darlene Wagner, Matthew Schmerer, Harry Gao, Mickey Li, John Gao, Joseph Fierro, Benafsh Sapra, Becky Tsai, Yan Meng, Doreen Ng, James Xie, Clinton R. Paden, Duncan MacCannell |
| EPI_ISL_3331648 | Valais Hospital, Central Institute | Valais Hospital, Central Institute | Alexis Dumoulin, Lorenzo Cerutti, Cedric Howald, Henri Pegeot, Melyssa Elies, Deborah Penet, Keith Harshman, Ioannis Xenarios |
| EPI_ISL_3331649 | Pandemic Response Lab - NYC | Pandemic Response Lab, R&D | Henry Lee, Michael Hammerling, Melissa Hopkins, Cybill del Castillo, Shinyoung Clair Kang, William Ward, Pradeep Bugga, Sol Rey, Dylan Law, Alex Carpio, Isabel Fernandez Escapa, Haiping Hao, Jon Laurent |
| EPI_ISL_3331651 | Fulgent Genetics | Centers for Disease Control and Prevention Division of Viral Diseases, Pathogen Discovery | Dakota Howard, Dhwani Batra, Peter W. Cook, Kara Moser, Adrian Paskey, Jason Caravas, Benjamin Rambo-Martin, Shatavia Morrison, Christopher Gulvick, Scott Sammons, Yvette Unoarumhi, Darlene Wagner, Matthew Schmerer, Harry Gao, Mickey Li, John Gao, Joseph Fierro, Benafsh Sapra, Becky Tsai, Yan Meng, Doreen Ng, James Xie, Clinton R. Paden, Duncan MacCannell |
| EPI_ISL_3331652 | Valais Hospital, Central Institute | Valais Hospital, Central Institute | Alexis Dumoulin, Lorenzo Cerutti, Cedric Howald, Henri Pegeot, Melyssa Elies, Deborah Penet, Keith Harshman, Ioannis Xenarios |
| EPI_ISL_3331653 | Pandemic Response Lab - NYC | Pandemic Response Lab, R&D | Henry Lee, Michael Hammerling, Melissa Hopkins, Cybill del Castillo, Shinyoung Clair Kang, William Ward, Pradeep Bugga, Sol Rey, Dylan Law, Alex Carpio, Isabel Fernandez Escapa, Haiping Hao, Jon Laurent |
| EPI_ISL_3331655 | Fulgent Genetics | Centers for Disease Control and Prevention Division of Viral Diseases, Pathogen Discovery | Dakota Howard, Dhwani Batra, Peter W. Cook, Kara Moser, Adrian Paskey, Jason Caravas, Benjamin Rambo-Martin, Shatavia Morrison, Christopher Gulvick, Scott Sammons, Yvette Unoarumhi, Darlene Wagner, Matthew Schmerer, Harry Gao, Mickey Li, John Gao, Joseph Fierro, Benafsh Sapra, Becky Tsai, Yan Meng, Doreen Ng, James Xie, Clinton R. Paden, Duncan MacCannell |
| EPI_ISL_3331657 | Valais Hospital, Central Institute | Valais Hospital, Central Institute | Alexis Dumoulin, Lorenzo Cerutti, Cedric Howald, Henri Pegeot, Melyssa Elies, Deborah Penet, Keith Harshman, Ioannis Xenarios |
| EPI_ISL_3331658 | Fulgent Genetics | Centers for Disease Control and Prevention Division of Viral Diseases, Pathogen Discovery | Dakota Howard, Dhwani Batra, Peter W. Cook, Kara Moser, Adrian Paskey, Jason Caravas, Benjamin Rambo-Martin, Shatavia Morrison, Christopher Gulvick, Scott Sammons, Yvette Unoarumhi, Darlene Wagner, Matthew Schmerer, Harry Gao, Mickey Li, John Gao, Joseph Fierro, Benafsh Sapra, Becky Tsai, Yan Meng, Doreen Ng, James Xie, Clinton R. Paden, Duncan MacCannell |
| EPI_ISL_3331659, EPI_ISL_3331661 | Pandemic Response Lab - NYC | Pandemic Response Lab, R&D | Henry Lee, Michael Hammerling, Melissa Hopkins, Cybill del Castillo, Shinyoung Clair Kang, William Ward, Pradeep Bugga, Sol Rey, Dylan Law, Alex Carpio, Isabel Fernandez Escapa, Haiping Hao, Jon Laurent |
| EPI_ISL_3331662 | Valais Hospital, Central Institute | Valais Hospital, Central Institute | Alexis Dumoulin, Lorenzo Cerutti, Cedric Howald, Henri Pegeot, Melyssa Elies, Deborah Penet, Keith Harshman, Ioannis Xenarios |
| EPI_ISL_3331663 | Fulgent Genetics | Centers for Disease Control and Prevention Division of Viral Diseases, Pathogen Discovery | Dakota Howard, Dhwani Batra, Peter W. Cook, Kara Moser, Adrian Paskey, Jason Caravas, Benjamin Rambo-Martin, Shatavia Morrison, Christopher Gulvick, Scott Sammons, Yvette Unoarumhi, Darlene Wagner, Matthew Schmerer, Harry Gao, Mickey Li, John Gao, Joseph Fierro, Benafsh Sapra, Becky Tsai, Yan Meng, Doreen Ng, James Xie, Clinton R. Paden, Duncan MacCannell |
| EPI_ISL_3331664 | Valais Hospital, Central Institute | Valais Hospital, Central Institute | Alexis Dumoulin, Lorenzo Cerutti, Cedric Howald, Henri Pegeot, Melyssa Elies, Deborah Penet, Keith Harshman, Ioannis Xenarios |
| EPI_ISL_3331665 | Pandemic Response Lab - NYC | Pandemic Response Lab, R&D | Henry Lee, Michael Hammerling, Melissa Hopkins, Cybill del Castillo, Shinyoung Clair Kang, William Ward, Pradeep Bugga, Sol Rey, Dylan Law, Alex Carpio, Isabel Fernandez Escapa, Haiping Hao, Jon Laurent |
| EPI_ISL_3331666 | Fulgent Genetics | Centers for Disease Control and Prevention Division of Viral Diseases, Pathogen Discovery | Dakota Howard, Dhwani Batra, Peter W. Cook, Kara Moser, Adrian Paskey, Jason Caravas, Benjamin Rambo-Martin, Shatavia Morrison, Christopher Gulvick, Scott Sammons, Yvette Unoarumhi, Darlene Wagner, Matthew Schmerer, Harry Gao, Mickey Li, John Gao, Joseph Fierro, Benafsh Sapra, Becky Tsai, Yan Meng, Doreen Ng, James Xie, Clinton R. Paden, Duncan MacCannell |
| EPI_ISL_3331668 | Valais Hospital, Central Institute | Valais Hospital, Central Institute | Alexis Dumoulin, Lorenzo Cerutti, Cedric Howald, Henri Pegeot, Melyssa Elies, Deborah Penet, Keith Harshman, Ioannis Xenarios |
| EPI_ISL_3331669 | Pandemic Response Lab - NYC | Pandemic Response Lab, R&D | Henry Lee, Michael Hammerling, Melissa Hopkins, Cybill del Castillo, Shinyoung Clair Kang, William Ward, Pradeep Bugga, Sol Rey, Dylan Law, Alex Carpio, Isabel Fernandez Escapa, Haiping Hao, Jon Laurent |
| EPI_ISL_3331670 | Fulgent Genetics | Centers for Disease Control and Prevention Division of Viral Diseases, Pathogen Discovery | Dakota Howard, Dhwani Batra, Peter W. Cook, Kara Moser, Adrian Paskey, Jason Caravas, Benjamin Rambo-Martin, Shatavia Morrison, Christopher Gulvick, Scott Sammons, Yvette Unoarumhi, Darlene Wagner, Matthew Schmerer, Harry Gao, Mickey Li, John Gao, Joseph Fierro, Benafsh Sapra, Becky Tsai, Yan Meng, Doreen Ng, James Xie, Clinton R. Paden, Duncan MacCannell |
| EPI_ISL_3331671 | Valais Hospital, Central Institute | Valais Hospital, Central Institute | Alexis Dumoulin, Lorenzo Cerutti, Cedric Howald, Henri Pegeot, Melyssa Elies, Deborah Penet, Keith Harshman, Ioannis Xenarios |
| EPI_ISL_3331672 | Pandemic Response Lab - NYC | Pandemic Response Lab, R&D | Henry Lee, Michael Hammerling, Melissa Hopkins, Cybill del Castillo, Shinyoung Clair Kang, William Ward, Pradeep Bugga, Sol Rey, Dylan Law, Alex Carpio, Isabel Fernandez Escapa, Haiping Hao, Jon Laurent |
| EPI_ISL_3331673 | Valais Hospital, Central Institute | Valais Hospital, Central Institute | Alexis Dumoulin, Lorenzo Cerutti, Cedric Howald, Henri Pegeot, Melyssa Elies, Deborah Penet, Keith Harshman, Ioannis Xenarios |
| EPI_ISL_3331674 | Fulgent Genetics | Centers for Disease Control and Prevention Division of Viral Diseases, Pathogen Discovery | Dakota Howard, Dhwani Batra, Peter W. Cook, Kara Moser, Adrian Paskey, Jason Caravas, Benjamin Rambo-Martin, Shatavia Morrison, Christopher Gulvick, Scott Sammons, Yvette Unoarumhi, Darlene Wagner, Matthew Schmerer, Harry Gao, Mickey Li, John Gao, Joseph Fierro, Benafsh Sapra, Becky Tsai, Yan Meng, Doreen Ng, James Xie, Clinton R. Paden, Duncan MacCannell |
| EPI_ISL_3331675 | Pandemic Response Lab - NYC | Pandemic Response Lab, R&D | Henry Lee, Michael Hammerling, Melissa Hopkins, Cybill del Castillo, Shinyoung Clair Kang, William Ward, Pradeep Bugga, Sol Rey, Dylan Law, Alex Carpio, Isabel Fernandez Escapa, Haiping Hao, Jon Laurent |
| EPI_ISL_3331676 | Valais Hospital, Central Institute | Valais Hospital, Central Institute | Alexis Dumoulin, Lorenzo Cerutti, Cedric Howald, Henri Pegeot, Melyssa Elies, Deborah Penet, Keith Harshman, Ioannis Xenarios |
| EPI_ISL_3331677 | Pandemic Response Lab - NYC | Pandemic Response Lab, R&D | Henry Lee, Michael Hammerling, Melissa Hopkins, Cybill del Castillo, Shinyoung Clair Kang, William Ward, Pradeep Bugga, Sol Rey, Dylan Law, Alex Carpio, Isabel Fernandez Escapa, Haiping Hao, Jon Laurent |
| EPI_ISL_3331678 | Valais Hospital, Central Institute | Valais Hospital, Central Institute | Alexis Dumoulin, Lorenzo Cerutti, Cedric Howald, Henri Pegeot, Melyssa Elies, Deborah Penet, Keith Harshman, Ioannis Xenarios |
| EPI_ISL_3331679 | Fulgent Genetics | Centers for Disease Control and Prevention Division of Viral Diseases, Pathogen Discovery | Dakota Howard, Dhwani Batra, Peter W. Cook, Kara Moser, Adrian Paskey, Jason Caravas, Benjamin Rambo-Martin, Shatavia Morrison, Christopher Gulvick, Scott Sammons, Yvette Unoarumhi, Darlene Wagner, Matthew Schmerer, Harry Gao, Mickey Li, John Gao, Joseph Fierro, Benafsh Sapra, Becky Tsai, Yan Meng, Doreen Ng, James Xie, Clinton R. Paden, Duncan MacCannell |
| EPI_ISL_3331680 | Valais Hospital, Central Institute | Valais Hospital, Central Institute | Alexis Dumoulin, Lorenzo Cerutti, Cedric Howald, Henri Pegeot, Melyssa Elies, Deborah Penet, Keith Harshman, Ioannis Xenarios |
| EPI_ISL_3331681 | Pandemic Response Lab - NYC | Pandemic Response Lab, R&D | Henry Lee, Michael Hammerling, Melissa Hopkins, Cybill del Castillo, Shinyoung Clair Kang, William Ward, Pradeep Bugga, Sol Rey, Dylan Law, Alex Carpio, Isabel Fernandez Escapa, Haiping Hao, Jon Laurent |
| EPI_ISL_3331682 | Fulgent Genetics | Centers for Disease Control and Prevention Division of Viral Diseases, Pathogen Discovery | Dakota Howard, Dhwani Batra, Peter W. Cook, Kara Moser, Adrian Paskey, Jason Caravas, Benjamin Rambo-Martin, Shatavia Morrison, Christopher Gulvick, Scott Sammons, Yvette Unoarumhi, Darlene Wagner, Matthew Schmerer, Harry Gao, Mickey Li, John Gao, Joseph Fierro, Benafsh Sapra, Becky Tsai, Yan Meng, Doreen Ng, James Xie, Clinton R. Paden, Duncan MacCannell |
| EPI_ISL_3331683, EPI_ISL_3331684 | Valais Hospital, Central Institute | Valais Hospital, Central Institute | Alexis Dumoulin, Lorenzo Cerutti, Cedric Howald, Henri Pegeot, Melyssa Elies, Deborah Penet, Keith Harshman, Ioannis Xenarios |
| EPI_ISL_3331685 | Pandemic Response Lab - NYC | Pandemic Response Lab, R&D | Henry Lee, Michael Hammerling, Melissa Hopkins, Cybill del Castillo, Shinyoung Clair Kang, William Ward, Pradeep Bugga, Sol Rey, Dylan Law, Alex Carpio, Isabel Fernandez Escapa, Haiping Hao, Jon Laurent |
| EPI_ISL_3331686 | Fulgent Genetics | Centers for Disease Control and Prevention Division of Viral Diseases, Pathogen Discovery | Dakota Howard, Dhwani Batra, Peter W. Cook, Kara Moser, Adrian Paskey, Jason Caravas, Benjamin Rambo-Martin, Shatavia Morrison, Christopher Gulvick, Scott Sammons, Yvette Unoarumhi, Darlene Wagner, Matthew Schmerer, Harry Gao, Mickey Li, John Gao, Joseph Fierro, Benafsh Sapra, Becky Tsai, Yan Meng, Doreen Ng, James Xie, Clinton R. Paden, Duncan MacCannell |
| EPI_ISL_3331687 | Valais Hospital, Central Institute | Valais Hospital, Central Institute | Alexis Dumoulin, Lorenzo Cerutti, Cedric Howald, Henri Pegeot, Melyssa Elies, Deborah Penet, Keith Harshman, Ioannis Xenarios |
| EPI_ISL_3331688 | Pandemic Response Lab - NYC | Pandemic Response Lab, R&D | Henry Lee, Michael Hammerling, Melissa Hopkins, Cybill del Castillo, Shinyoung Clair Kang, William Ward, Pradeep Bugga, Sol Rey, Dylan Law, Alex Carpio, Isabel Fernandez Escapa, Haiping Hao, Jon Laurent |
| EPI_ISL_3331689 | Valais Hospital, Central Institute | Valais Hospital, Central Institute | Alexis Dumoulin, Lorenzo Cerutti, Cedric Howald, Henri Pegeot, Melyssa Elies, Deborah Penet, Keith Harshman, Ioannis Xenarios |
| EPI_ISL_3331690 | Fulgent Genetics | Centers for Disease Control and Prevention Division of Viral Diseases, Pathogen Discovery | Dakota Howard, Dhwani Batra, Peter W. Cook, Kara Moser, Adrian Paskey, Jason Caravas, Benjamin Rambo-Martin, Shatavia Morrison, Christopher Gulvick, Scott Sammons, Yvette Unoarumhi, Darlene Wagner, Matthew Schmerer, Harry Gao, Mickey Li, John Gao, Joseph Fierro, Benafsh Sapra, Becky Tsai, Yan Meng, Doreen Ng, James Xie, Clinton R. Paden, Duncan MacCannell |
| EPI_ISL_3331691 | Pandemic Response Lab - NYC | Pandemic Response Lab, R&D | Henry Lee, Michael Hammerling, Melissa Hopkins, Cybill del Castillo, Shinyoung Clair Kang, William Ward, Pradeep Bugga, Sol Rey, Dylan Law, Alex Carpio, Isabel Fernandez Escapa, Haiping Hao, Jon Laurent |
| EPI_ISL_3331692 | Valais Hospital, Central Institute | Valais Hospital, Central Institute | Alexis Dumoulin, Lorenzo Cerutti, Cedric Howald, Henri Pegeot, Melyssa Elies, Deborah Penet, Keith Harshman, Ioannis Xenarios |
| EPI_ISL_3331693 | Fulgent Genetics | Centers for Disease Control and Prevention Division of Viral Diseases, Pathogen Discovery | Dakota Howard, Dhwani Batra, Peter W. Cook, Kara Moser, Adrian Paskey, Jason Caravas, Benjamin Rambo-Martin, Shatavia Morrison, Christopher Gulvick, Scott Sammons, Yvette Unoarumhi, Darlene Wagner, Matthew Schmerer, Harry Gao, Mickey Li, John Gao, Joseph Fierro, Benafsh Sapra, Becky Tsai, Yan Meng, Doreen Ng, James Xie, Clinton R. Paden, Duncan MacCannell |

| | | | |
|---|---|---|---|
| EPI_ISL_3331694 | Valais Hospital, Central Institute | Valais Hospital, Central Institute | Alexis Dumoulin, Lorenzo Cerutti, Cedric Howald, Henri Pegeot, Melyssa Elies, Deborah Penet, Keith Harshman, Ioannis Xenarios |
| EPI_ISL_3331696 | Fulgent Genetics | Centers for Disease Control and Prevention Division of Viral Diseases, Pathogen Discovery | Dakota Howard, Dhwani Batra, Peter W. Cook, Kara Moser, Adrian Paskey, Jason Caravas, Benjamin Rambo-Martin, Shatavia Morrison, Christopher Gulvick, Scott Sammons, Yvette Unoarumhi, Darlene Wagner, Matthew Schmerer, Harry Gao, Mickey Li, John Gao, Joseph Fierro, Benafsh Sapra, Becky Tsai, Yan Meng, Doreen Ng, James Xie, Clinton R. Paden, Duncan MacCannell |
| EPI_ISL_3331697, EPI_ISL_3331698 | Valais Hospital, Central Institute | Valais Hospital, Central Institute | Alexis Dumoulin, Lorenzo Cerutti, Cedric Howald, Henri Pegeot, Melyssa Elies, Deborah Penet, Keith Harshman, Ioannis Xenarios |
| EPI_ISL_3331699 | Fulgent Genetics | Centers for Disease Control and Prevention Division of Viral Diseases, Pathogen Discovery | Dakota Howard, Dhwani Batra, Peter W. Cook, Kara Moser, Adrian Paskey, Jason Caravas, Benjamin Rambo-Martin, Shatavia Morrison, Christopher Gulvick, Scott Sammons, Yvette Unoarumhi, Darlene Wagner, Matthew Schmerer, Harry Gao, Mickey Li, John Gao, Joseph Fierro, Benafsh Sapra, Becky Tsai, Yan Meng, Doreen Ng, James Xie, Clinton R. Paden, Duncan MacCannell |
| EPI_ISL_3331700 | Valais Hospital, Central Institute | Valais Hospital, Central Institute | Alexis Dumoulin, Lorenzo Cerutti, Cedric Howald, Henri Pegeot, Melyssa Elies, Deborah Penet, Keith Harshman, Ioannis Xenarios |
| EPI_ISL_3331701 | Pandemic Response Lab - NYC | Pandemic Response Lab, R&D | Henry Lee, Michael Hammerling, Melissa Hopkins, Cybill del Castillo, Shinyoung Clair Kang, William Ward, Pradeep Bugga, Sol Rey, Dylan Law, Alex Carpio, Isabel Fernandez Escapa, Haiping Hao, Jon Laurent |
| EPI_ISL_3331702 | Valais Hospital, Central Institute | Valais Hospital, Central Institute | Alexis Dumoulin, Lorenzo Cerutti, Cedric Howald, Henri Pegeot, Melyssa Elies, Deborah Penet, Keith Harshman, Ioannis Xenarios |
| EPI_ISL_3331703 | Fulgent Genetics | Centers for Disease Control and Prevention Division of Viral Diseases, Pathogen Discovery | Dakota Howard, Dhwani Batra, Peter W. Cook, Kara Moser, Adrian Paskey, Jason Caravas, Benjamin Rambo-Martin, Shatavia Morrison, Christopher Gulvick, Scott Sammons, Yvette Unoarumhi, Darlene Wagner, Matthew Schmerer, Harry Gao, Mickey Li, John Gao, Joseph Fierro, Benafsh Sapra, Becky Tsai, Yan Meng, Doreen Ng, James Xie, Clinton R. Paden, Duncan MacCannell |
| EPI_ISL_3331704 | Valais Hospital, Central Institute | Valais Hospital, Central Institute | Alexis Dumoulin, Lorenzo Cerutti, Cedric Howald, Henri Pegeot, Melyssa Elies, Deborah Penet, Keith Harshman, Ioannis Xenarios |
| EPI_ISL_3331705 | Pandemic Response Lab - NYC | Pandemic Response Lab, R&D | Henry Lee, Michael Hammerling, Melissa Hopkins, Cybill del Castillo, Shinyoung Clair Kang, William Ward, Pradeep Bugga, Sol Rey, Dylan Law, Alex Carpio, Isabel Fernandez Escapa, Haiping Hao, Jon Laurent |
| EPI_ISL_3331706 | Fulgent Genetics | Centers for Disease Control and Prevention Division of Viral Diseases, Pathogen Discovery | Dakota Howard, Dhwani Batra, Peter W. Cook, Kara Moser, Adrian Paskey, Jason Caravas, Benjamin Rambo-Martin, Shatavia Morrison, Christopher Gulvick, Scott Sammons, Yvette Unoarumhi, Darlene Wagner, Matthew Schmerer, Harry Gao, Mickey Li, John Gao, Joseph Fierro, Benafsh Sapra, Becky Tsai, Yan Meng, Doreen Ng, James Xie, Clinton R. Paden, Duncan MacCannell |
| EPI_ISL_3331707 | Valais Hospital, Central Institute | Valais Hospital, Central Institute | Alexis Dumoulin, Lorenzo Cerutti, Cedric Howald, Henri Pegeot, Melyssa Elies, Deborah Penet, Keith Harshman, Ioannis Xenarios |
| EPI_ISL_3331708 | Pandemic Response Lab - NYC | Pandemic Response Lab, R&D | Henry Lee, Michael Hammerling, Melissa Hopkins, Cybill del Castillo, Shinyoung Clair Kang, William Ward, Pradeep Bugga, Sol Rey, Dylan Law, Alex Carpio, Isabel Fernandez Escapa, Haiping Hao, Jon Laurent |
| EPI_ISL_3331709 | Valais Hospital, Central Institute | Valais Hospital, Central Institute | Alexis Dumoulin, Lorenzo Cerutti, Cedric Howald, Henri Pegeot, Melyssa Elies, Deborah Penet, Keith Harshman, Ioannis Xenarios |
| EPI_ISL_3331710 | Fulgent Genetics | Centers for Disease Control and Prevention Division of Viral Diseases, Pathogen Discovery | Dakota Howard, Dhwani Batra, Peter W. Cook, Kara Moser, Adrian Paskey, Jason Caravas, Benjamin Rambo-Martin, Shatavia Morrison, Christopher Gulvick, Scott Sammons, Yvette Unoarumhi, Darlene Wagner, Matthew Schmerer, Harry Gao, Mickey Li, John Gao, Joseph Fierro, Benafsh Sapra, Becky Tsai, Yan Meng, Doreen Ng, James Xie, Clinton R. Paden, Duncan MacCannell |
| EPI_ISL_3331711 | Valais Hospital, Central Institute | Valais Hospital, Central Institute | Alexis Dumoulin, Lorenzo Cerutti, Cedric Howald, Henri Pegeot, Melyssa Elies, Deborah Penet, Keith Harshman, Ioannis Xenarios |
| EPI_ISL_3331712 | Pandemic Response Lab - NYC | Pandemic Response Lab, R&D | Henry Lee, Michael Hammerling, Melissa Hopkins, Cybill del Castillo, Shinyoung Clair Kang, William Ward, Pradeep Bugga, Sol Rey, Dylan Law, Alex Carpio, Isabel Fernandez Escapa, Haiping Hao, Jon Laurent |
| EPI_ISL_3331713 | Fulgent Genetics | Centers for Disease Control and Prevention Division of Viral Diseases, Pathogen Discovery | Dakota Howard, Dhwani Batra, Peter W. Cook, Kara Moser, Adrian Paskey, Jason Caravas, Benjamin Rambo-Martin, Shatavia Morrison, Christopher Gulvick, Scott Sammons, Yvette Unoarumhi, Darlene Wagner, Matthew Schmerer, Harry Gao, Mickey Li, John Gao, Joseph Fierro, Benafsh Sapra, Becky Tsai, Yan Meng, Doreen Ng, James Xie, Clinton R. Paden, Duncan MacCannell |
| EPI_ISL_3331714 | Valais Hospital, Central Institute | Valais Hospital, Central Institute | Alexis Dumoulin, Lorenzo Cerutti, Cedric Howald, Henri Pegeot, Melyssa Elies, Deborah Penet, Keith Harshman, Ioannis Xenarios |
| EPI_ISL_3331715 | Pandemic Response Lab - NYC | Pandemic Response Lab, R&D | Henry Lee, Michael Hammerling, Melissa Hopkins, Cybill del Castillo, Shinyoung Clair Kang, William Ward, Pradeep Bugga, Sol Rey, Dylan Law, Alex Carpio, Isabel Fernandez Escapa, Haiping Hao, Jon Laurent |
| EPI_ISL_3331716 | Valais Hospital, Central Institute | Valais Hospital, Central Institute | Alexis Dumoulin, Lorenzo Cerutti, Cedric Howald, Henri Pegeot, Melyssa Elies, Deborah Penet, Keith Harshman, Ioannis Xenarios |
| EPI_ISL_3331717 | Fulgent Genetics | Centers for Disease Control and Prevention Division of Viral Diseases, Pathogen Discovery | Dakota Howard, Dhwani Batra, Peter W. Cook, Kara Moser, Adrian Paskey, Jason Caravas, Benjamin Rambo-Martin, Shatavia Morrison, Christopher Gulvick, Scott Sammons, Yvette Unoarumhi, Darlene Wagner, Matthew Schmerer, Harry Gao, Mickey Li, John Gao, Joseph Fierro, Benafsh Sapra, Becky Tsai, Yan Meng, Doreen Ng, James Xie, Clinton R. Paden, Duncan MacCannell |
| EPI_ISL_3331718, EPI_ISL_3331720 | Pandemic Response Lab - NYC | Pandemic Response Lab, R&D | Henry Lee, Michael Hammerling, Melissa Hopkins, Cybill del Castillo, Shinyoung Clair Kang, William Ward, Pradeep Bugga, Sol Rey, Dylan Law, Alex Carpio, Isabel Fernandez Escapa, Haiping Hao, Jon Laurent |
| EPI_ISL_3331721, EPI_ISL_3331722 | Valais Hospital, Central Institute | Valais Hospital, Central Institute | Alexis Dumoulin, Lorenzo Cerutti, Cedric Howald, Henri Pegeot, Melyssa Elies, Deborah Penet, Keith Harshman, Ioannis Xenarios |
| EPI_ISL_3331723 | Pandemic Response Lab - NYC | Pandemic Response Lab, R&D | Henry Lee, Michael Hammerling, Melissa Hopkins, Cybill del Castillo, Shinyoung Clair Kang, William Ward, Pradeep Bugga, Sol Rey, Dylan Law, Alex Carpio, Isabel Fernandez Escapa, Haiping Hao, Jon Laurent |
| EPI_ISL_3331724 | Valais Hospital, Central Institute | Valais Hospital, Central Institute | Alexis Dumoulin, Lorenzo Cerutti, Cedric Howald, Henri Pegeot, Melyssa Elies, Deborah Penet, Keith Harshman, Ioannis Xenarios |
| EPI_ISL_3331725 | Pandemic Response Lab - NYC | Pandemic Response Lab, R&D | Henry Lee, Michael Hammerling, Melissa Hopkins, Cybill del Castillo, Shinyoung Clair Kang, William Ward, Pradeep Bugga, Sol Rey, Dylan Law, Alex Carpio, Isabel Fernandez Escapa, Haiping Hao, Jon Laurent |
| EPI_ISL_3331726 | Valais Hospital, Central Institute | Valais Hospital, Central Institute | Alexis Dumoulin, Lorenzo Cerutti, Cedric Howald, Henri Pegeot, Melyssa Elies, Deborah Penet, Keith Harshman, Ioannis Xenarios |
| EPI_ISL_3331727 | Pandemic Response Lab - NYC | Pandemic Response Lab, R&D | Henry Lee, Michael Hammerling, Melissa Hopkins, Cybill del Castillo, Shinyoung Clair Kang, William Ward, Pradeep Bugga, Sol Rey, Dylan Law, Alex Carpio, Isabel Fernandez Escapa, Haiping Hao, Jon Laurent |
| EPI_ISL_3331728 | Valais Hospital, Central Institute | Valais Hospital, Central Institute | Alexis Dumoulin, Lorenzo Cerutti, Cedric Howald, Henri Pegeot, Melyssa Elies, Deborah Penet, Keith Harshman, Ioannis Xenarios |
| EPI_ISL_3331729 | Pandemic Response Lab - NYC | Pandemic Response Lab, R&D | Henry Lee, Michael Hammerling, Melissa Hopkins, Cybill del Castillo, Shinyoung Clair Kang, William Ward, Pradeep Bugga, Sol Rey, Dylan Law, Alex Carpio, Isabel Fernandez Escapa, Haiping Hao, Jon Laurent |
| EPI_ISL_3331730 | Valais Hospital, Central Institute | Valais Hospital, Central Institute | Alexis Dumoulin, Lorenzo Cerutti, Cedric Howald, Henri Pegeot, Melyssa Elies, Deborah Penet, Keith Harshman, Ioannis Xenarios |
| EPI_ISL_3331731 | Pandemic Response Lab - NYC | Pandemic Response Lab, R&D | Henry Lee, Michael Hammerling, Melissa Hopkins, Cybill del Castillo, Shinyoung Clair Kang, William Ward, Pradeep Bugga, Sol Rey, Dylan Law, Alex Carpio, Isabel Fernandez Escapa, Haiping Hao, Jon Laurent |
| EPI_ISL_3331732, EPI_ISL_3331733 | Valais Hospital, Central Institute | Valais Hospital, Central Institute | Alexis Dumoulin, Lorenzo Cerutti, Cedric Howald, Henri Pegeot, Melyssa Elies, Deborah Penet, Keith Harshman, Ioannis Xenarios |
| EPI_ISL_3331734, EPI_ISL_3331736, EPI_ISL_3331738 | Pandemic Response Lab - NYC | Pandemic Response Lab, R&D | Henry Lee, Michael Hammerling, Melissa Hopkins, Cybill del Castillo, Shinyoung Clair Kang, William Ward, Pradeep Bugga, Sol Rey, Dylan Law, Alex Carpio, Isabel Fernandez Escapa, Haiping Hao, Jon Laurent |
| EPI_ISL_3331739, EPI_ISL_3331740 | Valais Hospital, Central Institute | Valais Hospital, Central Institute | Alexis Dumoulin, Lorenzo Cerutti, Cedric Howald, Henri Pegeot, Melyssa Elies, Deborah Penet, Keith Harshman, Ioannis Xenarios |
| EPI_ISL_3331741 | Pandemic Response Lab - NYC | Pandemic Response Lab, R&D | Henry Lee, Michael Hammerling, Melissa Hopkins, Cybill del Castillo, Shinyoung Clair Kang, William Ward, Pradeep Bugga, Sol Rey, Dylan Law, Alex Carpio, Isabel Fernandez Escapa, Haiping Hao, Jon Laurent |
| EPI_ISL_3331742 | Valais Hospital, Central Institute | Valais Hospital, Central Institute | Alexis Dumoulin, Lorenzo Cerutti, Cedric Howald, Henri Pegeot, Melyssa Elies, Deborah Penet, Keith Harshman, Ioannis Xenarios |
| EPI_ISL_3331743 | Pandemic Response Lab - NYC | Pandemic Response Lab, R&D | Henry Lee, Michael Hammerling, Melissa Hopkins, Cybill del Castillo, Shinyoung Clair Kang, William Ward, Pradeep Bugga, Sol Rey, Dylan Law, Alex Carpio, Isabel Fernandez Escapa, Haiping Hao, Jon Laurent |
| EPI_ISL_3331744 | Valais Hospital, Central Institute | Valais Hospital, Central Institute | Alexis Dumoulin, Lorenzo Cerutti, Cedric Howald, Henri Pegeot, Melyssa Elies, Deborah Penet, Keith Harshman, Ioannis Xenarios |
| EPI_ISL_3331745, EPI_ISL_3331746, EPI_ISL_3331747, EPI_ISL_3331748, EPI_ISL_3331749, EPI_ISL_3331750, EPI_ISL_3331751, EPI_ISL_3331752, EPI_ISL_3331753, EPI_ISL_3331754, EPI_ISL_3331755, EPI_ISL_3331756, EPI_ISL_3331757, EPI_ISL_3331758, EPI_ISL_3331759, EPI_ISL_3331760, EPI_ISL_3331761, EPI_ISL_3331762, EPI_ISL_3331763, EPI_ISL_3331764, EPI_ISL_3331765, EPI_ISL_3331766, EPI_ISL_3331767, EPI_ISL_3331768, EPI_ISL_3331769, EPI_ISL_3331770, EPI_ISL_3331771, EPI_ISL_3331772, EPI_ISL_3331773, EPI_ISL_3331774, EPI_ISL_3331775, EPI_ISL_3331776, EPI_ISL_3331777, EPI_ISL_3331778, EPI_ISL_3331779, EPI_ISL_3331780, EPI_ISL_3331781, EPI_ISL_3331782, EPI_ISL_3331783, EPI_ISL_3331784, EPI_ISL_3331785, EPI_ISL_3331786, EPI_ISL_3331787, EPI_ISL_3331788, EPI_ISL_3331789, EPI_ISL_3331790, EPI_ISL_3331791, EPI_ISL_3331792, EPI_ISL_3331793, EPI_ISL_3331794, EPI_ISL_3331795, EPI_ISL_3331796, EPI_ISL_3331797, EPI_ISL_3331798, EPI_ISL_3331799, EPI_ISL_3331800, EPI_ISL_3331801, EPI_ISL_3331802, EPI_ISL_3331803, EPI_ISL_3331804, EPI_ISL_3331805, EPI_ISL_3331806, EPI_ISL_3331807, EPI_ISL_3331808, EPI_ISL_3331809, EPI_ISL_3331810, EPI_ISL_3331811, EPI_ISL_3331812, EPI_ISL_3331813, EPI_ISL_3331814, EPI_ISL_3331815, EPI_ISL_3331816, EPI_ISL_3331817, EPI_ISL_3331818, EPI_ISL_3331819, EPI_ISL_3331820, EPI_ISL_3331821, EPI_ISL_3331822, EPI_ISL_3331823, EPI_ISL_3331824, EPI_ISL_3331825, EPI_ISL_3331826, EPI_ISL_3331827, EPI_ISL_3331828, EPI_ISL_3331829, EPI_ISL_3331830, EPI_ISL_3331831, EPI_ISL_3331832, EPI_ISL_3331833, EPI_ISL_3331834, EPI_ISL_3331835, EPI_ISL_3331836, EPI_ISL_3331837, EPI_ISL_3331838, EPI_ISL_3331839, EPI_ISL_3331840, EPI_ISL_3331841, EPI_ISL_3331842, EPI_ISL_3331843, EPI_ISL_3331844, EPI_ISL_3331845, EPI_ISL_3331846, EPI_ISL_3331847, EPI_ISL_3331848, EPI_ISL_3331849, EPI_ISL_3331850, EPI_ISL_3331851, EPI_ISL_3331852, EPI_ISL_3331853, EPI_ISL_3331854, EPI_ISL_3331855, EPI_ISL_3331856, EPI_ISL_3331857, EPI_ISL_3331858, EPI_ISL_3331859, EPI_ISL_3331860, EPI_ISL_3331861, EPI_ISL_3331862, EPI_ISL_3331863, EPI_ISL_3331864, EPI_ISL_3331865, EPI_ISL_3331866, EPI_ISL_3331867, EPI_ISL_3331868, EPI_ISL_3331869, EPI_ISL_3331870, EPI_ISL_3331871, EPI_ISL_3331872, EPI_ISL_3331873, EPI_ISL_3331874, EPI_ISL_3331875, EPI_ISL_3331876, EPI_ISL_3331877, EPI_ISL_3331878, EPI_ISL_3331880, EPI_ISL_3331881, EPI_ISL_3331882, EPI_ISL_3331883, EPI_ISL_3331884, EPI_ISL_3331885, EPI_ISL_3331886, EPI_ISL_3331887, EPI_ISL_3331888, EPI_ISL_3331889, EPI_ISL_3331890, EPI_ISL_3331891, EPI_ISL_3331892, EPI_ISL_3331893, EPI_ISL_3331894, EPI_ISL_3331895, EPI_ISL_3331896, EPI_ISL_3331897, EPI_ISL_3331898, EPI_ISL_3331899, EPI_ISL_3331900, EPI_ISL_3331901, EPI_ISL_3331902, EPI_ISL_3331903, EPI_ISL_3331904, EPI_ISL_3331905, EPI_ISL_3331906, EPI_ISL_3331907, EPI_ISL_3331908, EPI_ISL_3331909, EPI_ISL_3331910, EPI_ISL_3331911, EPI_ISL_3331912, EPI_ISL_3331913, EPI_ISL_3331914, EPI_ISL_3331915, EPI_ISL_3331916, EPI_ISL_3331917, EPI_ISL_3331918, EPI_ISL_3331919, EPI_ISL_3331920, EPI_ISL_3331921, EPI_ISL_3331922, EPI_ISL_3331923, EPI_ISL_3331924, EPI_ISL_3331925, EPI_ISL_3331926, EPI_ISL_3331927, EPI_ISL_3331928, EPI_ISL_3331929, EPI_ISL_3331930, EPI_ISL_3331931, EPI_ISL_3331932, EPI_ISL_3331933, EPI_ISL_3331934, EPI_ISL_3331935, EPI_ISL_3331936, EPI_ISL_3331937, EPI_ISL_3331938, EPI_ISL_3331939, EPI_ISL_3331940, EPI_ISL_3331941, EPI_ISL_3331942, EPI_ISL_3331943, EPI_ISL_3331944, EPI_ISL_3331945, EPI_ISL_3331946, EPI_ISL_3331947, EPI_ISL_3331948, EPI_ISL_3331949, EPI_ISL_3331950, EPI_ISL_3331951, EPI_ISL_3331952, EPI_ISL_3331953, EPI_ISL_3331954, EPI_ISL_3331955, EPI_ISL_3331956, EPI_ISL_3331957, EPI_ISL_3331958, EPI_ISL_3331959, EPI_ISL_3331960, EPI_ISL_3331961, EPI_ISL_3331962, EPI_ISL_3331963, EPI_ISL_3331965, EPI_ISL_3331966, EPI_ISL_3331967, EPI_ISL_3331968, EPI_ISL_3331969, EPI_ISL_3331970, EPI_ISL_3331971, EPI_ISL_3331972, EPI_ISL_3331973, EPI_ISL_3331974, EPI_ISL_3331975, EPI_ISL_3331976, EPI_ISL_3331977, EPI_ISL_3331978, EPI_ISL_3331979, EPI_ISL_3331980, EPI_ISL_3331981, EPI_ISL_3331982, EPI_ISL_3331983, EPI_ISL_3331984, EPI_ISL_3331985, EPI_ISL_3331986, EPI_ISL_3331987, EPI_ISL_3331988, EPI_ISL_3331989, EPI_ISL_3331990, EPI_ISL_3331991, EPI_ISL_3331992, EPI_ISL_3331993, EPI_ISL_3331994, EPI_ISL_3331995, EPI_ISL_3331996, EPI_ISL_3331997, EPI_ISL_3331998, EPI_ISL_3331999, EPI_ISL_3332000, EPI_ISL_3332001, EPI_ISL_3332002, EPI_ISL_3332003, EPI_ISL_3332004, EPI_ISL_3332005, EPI_ISL_3332006, EPI_ISL_3332007, EPI_ISL_3332008, EPI_ISL_3332009, EPI_ISL_3332010, EPI_ISL_3332011, EPI_ISL_3332012, EPI_ISL_3332013, EPI_ISL_3332014, EPI_ISL_3332015, EPI_ISL_3332016, EPI_ISL_3332017, EPI_ISL_3332018, EPI_ISL_3332019, EPI_ISL_3332020, EPI_ISL_3332021, EPI_ISL_3332022, EPI_ISL_3332023, EPI_ISL_3332024, EPI_ISL_3332025, EPI_ISL_3332026, EPI_ISL_3332027, EPI_ISL_3332028, EPI_ISL_3332029, EPI_ISL_3332030, EPI_ISL_3332031, EPI_ISL_3332032, EPI_ISL_3332033, EPI_ISL_3332034, EPI_ISL_3332035, EPI_ISL_3332037, EPI_ISL_3332038, EPI_ISL_3332039, EPI_ISL_3332040, EPI_ISL_3332041, EPI_ISL_3332042, EPI_ISL_3332043, EPI_ISL_3332044, EPI_ISL_3332045, EPI_ISL_3332046, EPI_ISL_3332047, EPI_ISL_3332048, EPI_ISL_3332049, EPI_ISL_3332050, EPI_ISL_3332051, EPI_ISL_3332052, EPI_ISL_3332053, EPI_ISL_3332054, EPI_ISL_3332055, EPI_ISL_3332056, EPI_ISL_3332057, EPI_ISL_3332058, EPI_ISL_3332059, EPI_ISL_3332060, EPI_ISL_3332061, EPI_ISL_3332062, EPI_ISL_3332063, EPI_ISL_3332064, EPI_ISL_3332065, EPI_ISL_3332066, EPI_ISL_3332067, EPI_ISL_3332068, EPI_ISL_3332069, EPI_ISL_3332070, EPI_ISL_3332071, EPI_ISL_3332072, EPI_ISL_3332073, EPI_ISL_3332074, EPI_ISL_3332077, EPI_ISL_3332078, EPI_ISL_3332079, EPI_ISL_3332080, EPI_ISL_3332081, EPI_ISL_3332082, EPI_ISL_3332083, EPI_ISL_3332084, EPI_ISL_3332085, EPI_ISL_3332086, EPI_ISL_3332087, EPI_ISL_3332088, EPI_ISL_3332089, EPI_ISL_3332090, EPI_ISL_3332091, EPI_ISL_3332092, EPI_ISL_3332093, EPI_ISL_3332095, EPI_ISL_3332096, EPI_ISL_3332097, EPI_ISL_3332098, EPI_ISL_3332099, EPI_ISL_3332100, EPI_ISL_3332101, EPI_ISL_3332102, EPI_ISL_3332103, EPI_ISL_3332104, EPI_ISL_3332105, EPI_ISL_3332106, EPI_ISL_3332107, EPI_ISL_3332109, EPI_ISL_3332110, EPI_ISL_3332111, EPI_ISL_3332112, EPI_ISL_3332113, EPI_ISL_3332114, EPI_ISL_3332115, EPI_ISL_3332116, EPI_ISL_3332117, EPI_ISL_3332118, EPI_ISL_3332119, EPI_ISL_3332120, EPI_ISL_3332122, EPI_ISL_3332123, EPI_ISL_3332124, EPI_ISL_3332125, EPI_ISL_3332126, EPI_ISL_3332127, EPI_ISL_3332128, EPI_ISL_3332129, EPI_ISL_3332130, EPI_ISL_3332131, EPI_ISL_3332132, EPI_ISL_3332133, EPI_ISL_3332134, EPI_ISL_3332135, EPI_ISL_3332136, EPI_ISL_3332137, EPI_ISL_3332140, EPI_ISL_3332141, EPI_ISL_3332142, EPI_ISL_3332143, EPI_ISL_3332144, EPI_ISL_3332145, EPI_ISL_3332146, EPI_ISL_3332148, EPI_ISL_3332149, EPI_ISL_3332150, EPI_ISL_3332151, EPI_ISL_3332152, EPI_ISL_3332153, EPI_ISL_3332154, EPI_ISL_3332155, EPI_ISL_3332157, EPI_ISL_3332158, EPI_ISL_3332160, EPI_ISL_3332161, EPI_ISL_3332162, EPI_ISL_3332163, EPI_ISL_3332164, EPI_ISL_3332165, EPI_ISL_3332166, EPI_ISL_3332167, EPI_ISL_3332168, EPI_ISL_3332169, EPI_ISL_3332170, EPI_ISL_3332171, EPI_ISL_3332172, EPI_ISL_3332173, EPI_ISL_3332174, EPI_ISL_3332175, EPI_ISL_3332176, EPI_ISL_3332177, EPI_ISL_3332178, EPI_ISL_3332179, EPI_ISL_3332180, EPI_ISL_3332181, EPI_ISL_3332182, EPI_ISL_3332183, EPI_ISL_3332184, EPI_ISL_3332185, EPI_ISL_3332186, EPI_ISL_3332187, EPI_ISL_3332188, EPI_ISL_3332189, EPI_ISL_3332190, EPI_ISL_3332191, EPI_ISL_3332192, EPI_ISL_3332193, EPI_ISL_3332194 | | | |
| see above | Pandemic Response Lab - NYC | Pandemic Response Lab, R&D | Henry Lee, Michael Hammerling, Melissa Hopkins, Cybill del Castillo, Shinyoung Clair Kang, William Ward, Pradeep Bugga, Sol Rey, Dylan Law, Alex Carpio, Isabel Fernandez Escapa, Haiping Hao, Jon Laurent |
| EPI_ISL_3332196, EPI_ISL_3332197, EPI_ISL_3332198, EPI_ISL_3332199, EPI_ISL_3332200, EPI_ISL_3332201, EPI_ISL_3332202, EPI_ISL_3332204, EPI_ISL_3332205, EPI_ISL_3332206, EPI_ISL_3332207, EPI_ISL_3332208, EPI_ISL_3332209, EPI_ISL_3332210, EPI_ISL_3332211, EPI_ISL_3332212, EPI_ISL_3332213, EPI_ISL_3332214, EPI_ISL_3332215, EPI_ISL_3332217, EPI_ISL_3332219, EPI_ISL_3332222 | | | |
| see above | Sonic Reference Laboratory | Sonic Reference Laboratory | Robert Moller, Bijaya Dhakal, Krishna Maddula, John Woody, Cora Lahey, Ellen Pizarek |
| EPI_ISL_3332223, EPI_ISL_3332225, EPI_ISL_3332226, EPI_ISL_3332227, EPI_ISL_3332228, EPI_ISL_3332229, EPI_ISL_3332230, EPI_ISL_3332231, EPI_ISL_3332232, EPI_ISL_3332233, EPI_ISL_3332234, EPI_ISL_3332235, EPI_ISL_3332236, EPI_ISL_3332237, EPI_ISL_3332238, EPI_ISL_3332239, EPI_ISL_3332240, EPI_ISL_3332241, EPI_ISL_3332242, EPI_ISL_3332243, EPI_ISL_3332244, EPI_ISL_3332245, EPI_ISL_3332246, EPI_ISL_3332247, EPI_ISL_3332248, EPI_ISL_3332249, EPI_ISL_3332250, EPI_ISL_3332251, EPI_ISL_3332252, EPI_ISL_3332253 | | | |
| see above | IN State Department of Health Laboratory Services | IN State Department of Health Laboratory Services | Cassandra Campion, Jamie Yeadon, Brian Pope, Lixia Liu, Kyle Brownlee, Melissa Hindenlang, Mark Glazier |
| EPI_ISL_3332254 | University Hospital Brno, OKMI | University Hospital Brno, CMBG | Matej Bezdicek, Jan Svaton, Martina Lengerova, Kristyna Dufkova, Pavlina Volfova |
| EPI_ISL_3332255, EPI_ISL_3332256, EPI_ISL_3332257 | SYNLAB | University Hospital Brno, CMBG | Matej Bezdicek, Jan Svaton, Martina Lengerova, Kristyna Dufkova, Pavlina Volfova |
| EPI_ISL_3332258, EPI_ISL_3332259, EPI_ISL_3332260, EPI_ISL_3332261, EPI_ISL_3332262 | St. Anne´s University Hospital Brno | University Hospital Brno, CMBG | Matej Bezdicek, Jan Svaton, Martina Lengerova, Kristyna Dufkova, Pavlina Volfova |

| | | | |
|---|---|---|---|
| EPI_ISL_3332263, EPI_ISL_3332264, EPI_ISL_3332265, EPI_ISL_3332266 | Cytogeneticka laborator Brno | University Hospital Brno, CMBG | Matej Bezdicek, Jan Svaton, Martina Lengerova, Kristyna Dufkova, Pavlina Volfova |
| EPI_ISL_3332267, EPI_ISL_3332268, EPI_ISL_3332269, EPI_ISL_3332270, EPI_ISL_3332271, EPI_ISL_3332272, EPI_ISL_3332273, EPI_ISL_3332274 | University Hospital Brno, OKMI | University Hospital Brno, CMBG | Matej Bezdicek, Jan Svaton, Martina Lengerova, Kristyna Dufkova, Pavlina Volfova |
| EPI_ISL_3332275 | Cytogeneticka laborator Brno | University Hospital Brno, CMBG | Matej Bezdicek, Jan Svaton, Martina Lengerova, Kristyna Dufkova, Pavlina Volfova |
| EPI_ISL_3332276, EPI_ISL_3332277, EPI_ISL_3332278 | SYNLAB | University Hospital Brno, CMBG | Matej Bezdicek, Jan Svaton, Martina Lengerova, Kristyna Dufkova, Pavlina Volfova |
| EPI_ISL_3332279, EPI_ISL_3332280 | Nemocnice Vyskov | University Hospital Brno, CMBG | Matej Bezdicek, Jan Svaton, Martina Lengerova, Kristyna Dufkova, Pavlina Volfova |
| EPI_ISL_3332281, EPI_ISL_3332283, EPI_ISL_3332284, EPI_ISL_3332285, EPI_ISL_3332286 | St. Anne´s University Hospital Brno | University Hospital Brno, CMBG | Matej Bezdicek, Jan Svaton, Martina Lengerova, Kristyna Dufkova, Pavlina Volfova |
| EPI_ISL_3332287, EPI_ISL_3332288, EPI_ISL_3332289, EPI_ISL_3332290, EPI_ISL_3332291 | Nemocnice Havlickuv Brod | University Hospital Brno, CMBG | Matej Bezdicek, Jan Svaton, Martina Lengerova, Kristyna Dufkova, Pavlina Volfova |
| EPI_ISL_3332292, EPI_ISL_3332293, EPI_ISL_3332294, EPI_ISL_3332295, EPI_ISL_3332296, EPI_ISL_3332297, EPI_ISL_3332298, EPI_ISL_3332299, EPI_ISL_3332300, EPI_ISL_3332301, EPI_ISL_3332302 | | | |
| see above | Nemocnice Jihlava | University Hospital Brno, CMBG | Matej Bezdicek, Jan Svaton, Martina Lengerova, Kristyna Dufkova, Pavlina Volfova |
| EPI_ISL_3332303, EPI_ISL_3332304, EPI_ISL_3332305, EPI_ISL_3332306, EPI_ISL_3332307, EPI_ISL_3332309, EPI_ISL_3332310, EPI_ISL_3332311, EPI_ISL_3332312, EPI_ISL_3332313, EPI_ISL_3332314, EPI_ISL_3332315, EPI_ISL_3332316, EPI_ISL_3332317, EPI_ISL_3332318, EPI_ISL_3332320, EPI_ISL_3332321, EPI_ISL_3332322, EPI_ISL_3332323, EPI_ISL_3332324, EPI_ISL_3332325, EPI_ISL_3332326, EPI_ISL_3332327, EPI_ISL_3332328 | | | |
| see above | Sonic Reference Laboratory | Sonic Reference Laboratory | Robert Moller, Bijaya Dhakal, Krishna Maddula, John Woody, Cora Lahey, Ellen Pizarek |
| EPI_ISL_3332330, EPI_ISL_3332331, EPI_ISL_3332333, EPI_ISL_3332334, EPI_ISL_3332335, EPI_ISL_3332336, EPI_ISL_3332337, EPI_ISL_3332338, EPI_ISL_3332339, EPI_ISL_3332340, EPI_ISL_3332341, EPI_ISL_3332342, EPI_ISL_3332343, EPI_ISL_3332344, EPI_ISL_3332345, EPI_ISL_3332346, EPI_ISL_3332347, EPI_ISL_3332350, EPI_ISL_3332351, EPI_ISL_3332352, EPI_ISL_3332353, EPI_ISL_3332354, EPI_ISL_3332355, EPI_ISL_3332356, EPI_ISL_3332357, EPI_ISL_3332358, EPI_ISL_3332359, EPI_ISL_3332360, EPI_ISL_3332361, EPI_ISL_3332362, EPI_ISL_3332363, EPI_ISL_3332364, EPI_ISL_3332365, EPI_ISL_3332366, EPI_ISL_3332375, EPI_ISL_3332376 | | | |
| see above | MD PHL | Maryland Department of Health Laboratories Administration | Eric N. Keller, Jillian Loomis, Kwang Low, Terence L. Moore, Ami Patel, and Robert Myers |
| EPI_ISL_3332719, EPI_ISL_3332729, EPI_ISL_3332730 | Tokyo Medical and Dental University | Tokyo Medical and Dental University | Kousuke Tanimoto, Yukie Tanaka, Yoko Nukui, Akinori Kimura, Hiroaki Takeuchi |
| EPI_ISL_3332743, EPI_ISL_3332744, EPI_ISL_3332746, EPI_ISL_3332747, EPI_ISL_3332748, EPI_ISL_3332749, EPI_ISL_3332750, EPI_ISL_3332751, EPI_ISL_3332752, EPI_ISL_3332753, EPI_ISL_3332754, EPI_ISL_3332755, EPI_ISL_3332756, EPI_ISL_3332758, EPI_ISL_3332760, EPI_ISL_3332761, EPI_ISL_3332762, EPI_ISL_3332763, EPI_ISL_3332764, EPI_ISL_3332765, EPI_ISL_3332766, EPI_ISL_3332767, EPI_ISL_3332768, EPI_ISL_3332769, EPI_ISL_3332770, EPI_ISL_3332771, EPI_ISL_3332772, EPI_ISL_3332773, EPI_ISL_3332774, EPI_ISL_3332775, EPI_ISL_3332776, EPI_ISL_3332777, EPI_ISL_3332778, EPI_ISL_3332779, EPI_ISL_3332780, EPI_ISL_3332781, EPI_ISL_3332782, EPI_ISL_3332783, EPI_ISL_3332784, EPI_ISL_3332785, EPI_ISL_3332786, EPI_ISL_3332787, EPI_ISL_3332788, EPI_ISL_3332789, EPI_ISL_3332790, EPI_ISL_3332791, EPI_ISL_3332792, EPI_ISL_3332794, EPI_ISL_3332795, EPI_ISL_3332796, EPI_ISL_3332797, EPI_ISL_3332798, EPI_ISL_3332799, EPI_ISL_3332800, EPI_ISL_3332801, EPI_ISL_3332802, EPI_ISL_3332803, EPI_ISL_3332804 | | | |
| see above | Aegis Sciences Corporation | Centers for Disease Control and Prevention Division of Viral Diseases, Pathogen Discovery | Dakota Howard, Dhwani Batra, Peter W. Cook, Kara Moser, Adrian Paskey, Jason Caravas, Benjamin Rambo-Martin, Shatavia Morrison, Christopher Gulvick, Scott Sammons, Yvette Unoarumhi, Darlene Wagner, Matthew Schmerer, Cyndi Clark, Patrick Campbell, Rob Case, Vikramsinha Ghorpade, Holly Houdeshell, Ola Kvalvaag, Dillon Nall, Ethan Sanders, Alec Vest, Shaun Westlund, Matthew Hardison, Clinton R. Paden, Duncan MacCannell |
| EPI_ISL_3332806 | Vitalabo lab.medyczne Sp. Z o.o. | Wojewodzka Stacja Sanitarno-Epidemiologiczna w Lodzi, Oddzial Laboratoryjny Mikrobiologii i Parazytologii | Elian Terpo, Szymon Konkolewski, Marta Gawronska, Martyna Matera, Agnieszka Karnowska, Danuta Bartczak, Katarzyna Kosowska, Lucyna Kochanowska, Aleksandra Krawczyk |
| EPI_ISL_3332807, EPI_ISL_3332808 | Zaklad Diagnostyki Mikrobiologicznej | Wojewodzka Stacja Sanitarno-Epidemiologiczna w Lodzi, Oddzial Laboratoryjny Mikrobiologii i Parazytologii | Elian Terpo, Szymon Konkolewski, Marta Gawronska, Martyna Matera, Agnieszka Karnowska, Danuta Bartczak, Katarzyna Kosowska, Lucyna Kochanowska, Aleksandra Krawczyk |
| EPI_ISL_3332809 | Laboratorium Mikrobiologiczne RSS w Grudziadzu | Wojewodzka Stacja Sanitarno-Epidemiologiczna w Lodzi, Oddzial Laboratoryjny Mikrobiologii i Parazytologii | Elian Terpo, Szymon Konkolewski, Marta Gawronska, Martyna Matera, Agnieszka Karnowska, Danuta Bartczak, Katarzyna Kosowska, Lucyna Kochanowska, Aleksandra Krawczyk |
| EPI_ISL_3332810 | Laboratorium Badan Epidemiologiczno-Klinicznych WSSE w Olsztynie | Wojewodzka Stacja Sanitarno-Epidemiologiczna w Lodzi, Oddzial Laboratoryjny Mikrobiologii i Parazytologii | Elian Terpo, Szymon Konkolewski, Marta Gawronska, Martyna Matera, Agnieszka Karnowska, Danuta Bartczak, Katarzyna Kosowska, Lucyna Kochanowska, Aleksandra Krawczyk |
| EPI_ISL_3332811, EPI_ISL_3332813, EPI_ISL_3332814 | Laboratoria Medyczne Optimed | Wojewodzka Stacja Sanitarno-Epidemiologiczna w Lodzi, Oddzial Laboratoryjny Mikrobiologii i Parazytologii | Elian Terpo, Szymon Konkolewski, Marta Gawronska, Martyna Matera, Agnieszka Karnowska, Danuta Bartczak, Katarzyna Kosowska, Lucyna Kochanowska, Aleksandra Krawczyk |
| EPI_ISL_3332815, EPI_ISL_3332816, EPI_ISL_3332817 | Medyczne Laboratorium Mikrobiologiczne Pro-Medica w Elku Sp. z o.o. | Wojewodzka Stacja Sanitarno-Epidemiologiczna w Lodzi, Oddzial Laboratoryjny Mikrobiologii i Parazytologii | Elian Terpo, Szymon Konkolewski, Marta Gawronska, Martyna Matera, Agnieszka Karnowska, Danuta Bartczak, Katarzyna Kosowska, Lucyna Kochanowska, Aleksandra Krawczyk |
| EPI_ISL_3332818, EPI_ISL_3332819, EPI_ISL_3332820 | Pracownia Genetyki i Biologii Molekularnej WSSD w Olsztynie | Wojewodzka Stacja Sanitarno-Epidemiologiczna w Lodzi, Oddzial Laboratoryjny Mikrobiologii i Parazytologii | Elian Terpo, Szymon Konkolewski, Marta Gawronska, Martyna Matera, Agnieszka Karnowska, Danuta Bartczak, Katarzyna Kosowska, Lucyna Kochanowska, Aleksandra Krawczyk |
| EPI_ISL_3332821, EPI_ISL_3332822 | GEN LAB Laboratorium Medyczne | Wojewodzka Stacja Sanitarno-Epidemiologiczna w Lodzi, Oddzial Laboratoryjny Mikrobiologii i Parazytologii | Elian Terpo, Szymon Konkolewski, Marta Gawronska, Martyna Matera, Agnieszka Karnowska, Danuta Bartczak, Katarzyna Kosowska, Lucyna Kochanowska, Aleksandra Krawczyk |
| EPI_ISL_3332823 | Laboratorium Badan Klinicznych WSSE w Opolu | Wojewodzka Stacja Sanitarno-Epidemiologiczna w Lodzi, Oddzial Laboratoryjny Mikrobiologii i Parazytologii | Elian Terpo, Szymon Konkolewski, Marta Gawronska, Martyna Matera, Agnieszka Karnowska, Danuta Bartczak, Katarzyna Kosowska, Lucyna Kochanowska, Aleksandra Krawczyk |
| EPI_ISL_3332824, EPI_ISL_3332825, EPI_ISL_3332826, EPI_ISL_3332827 | Cellis Laboratorium | Wojewodzka Stacja Sanitarno-Epidemiologiczna w Lodzi, Oddzial Laboratoryjny Mikrobiologii i Parazytologii | Elian Terpo, Szymon Konkolewski, Marta Gawronska, Martyna Matera, Agnieszka Karnowska, Danuta Bartczak, Katarzyna Kosowska, Lucyna Kochanowska, Aleksandra Krawczyk |
| EPI_ISL_3332828 | ZDL CSK MSWiA | Wojewodzka Stacja Sanitarno-Epidemiologiczna w Lodzi, Oddzial Laboratoryjny Mikrobiologii i Parazytologii | Elian Terpo, Szymon Konkolewski, Marta Gawronska, Martyna Matera, Agnieszka Karnowska, Danuta Bartczak, Katarzyna Kosowska, Lucyna Kochanowska, Aleksandra Krawczyk |
| EPI_ISL_3332829, EPI_ISL_3332830, EPI_ISL_3332831 | NZOZ Medigen Diagnostyka Molekularna | Wojewodzka Stacja Sanitarno-Epidemiologiczna w Lodzi, Oddzial Laboratoryjny Mikrobiologii i Parazytologii | Elian Terpo, Szymon Konkolewski, Marta Gawronska, Martyna Matera, Agnieszka Karnowska, Danuta Bartczak, Katarzyna Kosowska, Lucyna Kochanowska, Aleksandra Krawczyk |
| EPI_ISL_3332832, EPI_ISL_3332833 | Laboratorium Zakladu Wirusologii NIZP-PZH-PIB | Wojewodzka Stacja Sanitarno-Epidemiologiczna w Lodzi, Oddzial Laboratoryjny Mikrobiologii i Parazytologii | Elian Terpo, Szymon Konkolewski, Marta Gawronska, Martyna Matera, Agnieszka Karnowska, Danuta Bartczak, Katarzyna Kosowska, Lucyna Kochanowska, Aleksandra Krawczyk |
| EPI_ISL_3332834 | Laboratorium Diagnostyczne Centrum Medycznego Luxmed | Wojewodzka Stacja Sanitarno-Epidemiologiczna w Lodzi, Oddzial Laboratoryjny Mikrobiologii i Parazytologii | Elian Terpo, Szymon Konkolewski, Marta Gawronska, Martyna Matera, Agnieszka Karnowska, Danuta Bartczak, Katarzyna Kosowska, Lucyna Kochanowska, Aleksandra Krawczyk |
| EPI_ISL_3332835 | Laboratorium Diagnostyczne Maklab | Wojewodzka Stacja Sanitarno-Epidemiologiczna w Lodzi, Oddzial Laboratoryjny Mikrobiologii i Parazytologii | Elian Terpo, Szymon Konkolewski, Marta Gawronska, Martyna Matera, Agnieszka Karnowska, Danuta Bartczak, Katarzyna Kosowska, Lucyna Kochanowska, Aleksandra Krawczyk |
| EPI_ISL_3332836 | Laboratorium Arion Szpitale Sp. z o.o. | Wojewodzka Stacja Sanitarno-Epidemiologiczna w Lodzi, Oddzial Laboratoryjny Mikrobiologii i Parazytologii | Elian Terpo, Szymon Konkolewski, Marta Gawronska, Martyna Matera, Agnieszka Karnowska, Danuta Bartczak, Katarzyna Kosowska, Lucyna Kochanowska, Aleksandra Krawczyk |
| EPI_ISL_3332837, EPI_ISL_3332838 | Laboratorium Biol. Molek. USK | Wojewodzka Stacja Sanitarno-Epidemiologiczna w Lodzi, Oddzial Laboratoryjny Mikrobiologii i Parazytologii | Elian Terpo, Szymon Konkolewski, Marta Gawronska, Martyna Matera, Agnieszka Karnowska, Danuta Bartczak, Katarzyna Kosowska, Lucyna Kochanowska, Aleksandra Krawczyk |
| EPI_ISL_3332839 | Laboratorium WSSE w Lublinie | Wojewodzka Stacja Sanitarno-Epidemiologiczna w Lodzi, Oddzial Laboratoryjny Mikrobiologii i Parazytologii | Elian Terpo, Szymon Konkolewski, Marta Gawronska, Martyna Matera, Agnieszka Karnowska, Danuta Bartczak, Katarzyna Kosowska, Lucyna Kochanowska, Aleksandra Krawczyk |
| EPI_ISL_3332840 | Zaklad Diagnostyki Laboratoryjnej | Wojewodzka Stacja Sanitarno-Epidemiologiczna w Lodzi, Oddzial Laboratoryjny Mikrobiologii i Parazytologii | Elian Terpo, Szymon Konkolewski, Marta Gawronska, Martyna Matera, Agnieszka Karnowska, Danuta Bartczak, Katarzyna Kosowska, Lucyna Kochanowska, Aleksandra Krawczyk |
| EPI_ISL_3332841 | Laboratorium Diagnostyczne Maklab | Wojewodzka Stacja Sanitarno-Epidemiologiczna w Lodzi, Oddzial Laboratoryjny Mikrobiologii i Parazytologii | Elian Terpo, Szymon Konkolewski, Marta Gawronska, Martyna Matera, Agnieszka Karnowska, Danuta Bartczak, Katarzyna Kosowska, Lucyna Kochanowska, Aleksandra Krawczyk |
| EPI_ISL_3332842, EPI_ISL_3332843 | Laboratorium Diagnostyczne Centrum Medycznego Luxmed | Wojewodzka Stacja Sanitarno-Epidemiologiczna w Lodzi, Oddzial Laboratoryjny Mikrobiologii i Parazytologii | Elian Terpo, Szymon Konkolewski, Marta Gawronska, Martyna Matera, Agnieszka Karnowska, Danuta Bartczak, Katarzyna Kosowska, Lucyna Kochanowska, Aleksandra Krawczyk |
| EPI_ISL_3332844 | Laboratorium Biol. Molek. USK | Wojewodzka Stacja Sanitarno-Epidemiologiczna w Lodzi, Oddzial | Elian Terpo, Szymon Konkolewski, Marta Gawronska, Martyna Matera, Agnieszka Karnowska, Danuta Bartczak, Katarzyna Kosowska, Lucyna Kochanowska, Aleksandra Krawczyk |

| | | | |
|---|---|---|---|
| | | Laboratoryjny Mikrobiologii i Parazytologii | |
| EPI_ISL_3332845 | Laboratorium WSSE w Lublinie | Wojewodzka Stacja Sanitarno-Epidemiologiczna w Lodzi, Oddzial Laboratoryjny Mikrobiologii i Parazytologii | Elian Terpo, Szymon Konkolewski, Marta Gawronska, Martyna Matera, Agnieszka Karnowska, Danuta Bartczak, Katarzyna Kosowska, Lucyna Kochanowska, Aleksandra Krawczyk |
| EPI_ISL_3332846 | Laboratorium ALAB Lublin | Wojewodzka Stacja Sanitarno-Epidemiologiczna w Lodzi, Oddzial Laboratoryjny Mikrobiologii i Parazytologii | Elian Terpo, Szymon Konkolewski, Marta Gawronska, Martyna Matera, Agnieszka Karnowska, Danuta Bartczak, Katarzyna Kosowska, Lucyna Kochanowska, Aleksandra Krawczyk |
| EPI_ISL_3332847 | Laboratorium Diagnostyki Mikrobiologicznej z Pracownia Pratka Gruzlicy SP SzW im. Jana Bozego w Lublinie | Wojewodzka Stacja Sanitarno-Epidemiologiczna w Lodzi, Oddzial Laboratoryjny Mikrobiologii i Parazytologii | Elian Terpo, Szymon Konkolewski, Marta Gawronska, Martyna Matera, Agnieszka Karnowska, Danuta Bartczak, Katarzyna Kosowska, Lucyna Kochanowska, Aleksandra Krawczyk |
| EPI_ISL_3332848 | Laboratorium Diagnostyczne Centrum Medycznego Luxmed | Wojewodzka Stacja Sanitarno-Epidemiologiczna w Lodzi, Oddzial Laboratoryjny Mikrobiologii i Parazytologii | Elian Terpo, Szymon Konkolewski, Marta Gawronska, Martyna Matera, Agnieszka Karnowska, Danuta Bartczak, Katarzyna Kosowska, Lucyna Kochanowska, Aleksandra Krawczyk |
| EPI_ISL_3332849 | MLDiM Szpital Wojewodzki w Suwalkach | Wojewodzka Stacja Sanitarno-Epidemiologiczna w Lodzi, Oddzial Laboratoryjny Mikrobiologii i Parazytologii | Elian Terpo, Szymon Konkolewski, Marta Gawronska, Martyna Matera, Agnieszka Karnowska, Danuta Bartczak, Katarzyna Kosowska, Lucyna Kochanowska, Aleksandra Krawczyk |
| EPI_ISL_3332850 | Laboratorium Swietokrzyskie | Wojewodzka Stacja Sanitarno-Epidemiologiczna w Lodzi, Oddzial Laboratoryjny Mikrobiologii i Parazytologii | Elian Terpo, Szymon Konkolewski, Marta Gawronska, Martyna Matera, Agnieszka Karnowska, Danuta Bartczak, Katarzyna Kosowska, Lucyna Kochanowska, Aleksandra Krawczyk |
| EPI_ISL_3332851 | Zaklad Mikrobiologii Klinicznej WSZ im. dr R. Ostrzyckiego w Koninie | Wojewodzka Stacja Sanitarno-Epidemiologiczna w Lodzi, Oddzial Laboratoryjny Mikrobiologii i Parazytologii | Elian Terpo, Szymon Konkolewski, Marta Gawronska, Martyna Matera, Agnieszka Karnowska, Danuta Bartczak, Katarzyna Kosowska, Lucyna Kochanowska, Aleksandra Krawczyk |
| EPI_ISL_3332852 | Centrum Badan DNA | Wojewodzka Stacja Sanitarno-Epidemiologiczna w Lodzi, Oddzial Laboratoryjny Mikrobiologii i Parazytologii | Elian Terpo, Szymon Konkolewski, Marta Gawronska, Martyna Matera, Agnieszka Karnowska, Danuta Bartczak, Katarzyna Kosowska, Lucyna Kochanowska, Aleksandra Krawczyk |
| EPI_ISL_3332853 | Instytut Mikroekologii Sp. z o. o. & Co. Sp.k. | Wojewodzka Stacja Sanitarno-Epidemiologiczna w Lodzi, Oddzial Laboratoryjny Mikrobiologii i Parazytologii | Elian Terpo, Szymon Konkolewski, Marta Gawronska, Martyna Matera, Agnieszka Karnowska, Danuta Bartczak, Katarzyna Kosowska, Lucyna Kochanowska, Aleksandra Krawczyk |
| EPI_ISL_3332854 | GenXOne SA | Wojewodzka Stacja Sanitarno-Epidemiologiczna w Lodzi, Oddzial Laboratoryjny Mikrobiologii i Parazytologii | Elian Terpo, Szymon Konkolewski, Marta Gawronska, Martyna Matera, Agnieszka Karnowska, Danuta Bartczak, Katarzyna Kosowska, Lucyna Kochanowska, Aleksandra Krawczyk |
| EPI_ISL_3332855 | Wojewodzki Specjalistyczny Szpital im. Dr. WL. Bieganskiego | Wojewodzka Stacja Sanitarno-Epidemiologiczna w Lodzi, Oddzial Laboratoryjny Mikrobiologii i Parazytologii | Elian Terpo, Szymon Konkolewski, Marta Gawronska, Martyna Matera, Agnieszka Karnowska, Danuta Bartczak, Katarzyna Kosowska, Lucyna Kochanowska, Aleksandra Krawczyk |
| EPI_ISL_3332856 | Centrum Medyczne MEDYK Sp. z o.o. Sp. k. Zaklad Diagnostyki Medycznej | Wojewodzka Stacja Sanitarno-Epidemiologiczna w Lodzi, Oddzial Laboratoryjny Mikrobiologii i Parazytologii | Elian Terpo, Szymon Konkolewski, Marta Gawronska, Martyna Matera, Agnieszka Karnowska, Danuta Bartczak, Katarzyna Kosowska, Lucyna Kochanowska, Aleksandra Krawczyk |
| EPI_ISL_3332857 | Laboratorium Mikrobiologiczne Synevo Lodz | Wojewodzka Stacja Sanitarno-Epidemiologiczna w Lodzi, Oddzial Laboratoryjny Mikrobiologii i Parazytologii | Elian Terpo, Szymon Konkolewski, Marta Gawronska, Martyna Matera, Agnieszka Karnowska, Danuta Bartczak, Katarzyna Kosowska, Lucyna Kochanowska, Aleksandra Krawczyk |
| EPI_ISL_3332858, EPI_ISL_3332859 | Centrum Medyczne MEDYK Sp. z o.o. Sp. k. Zaklad Diagnostyki Medycznej | Wojewodzka Stacja Sanitarno-Epidemiologiczna w Lodzi, Oddzial Laboratoryjny Mikrobiologii i Parazytologii | Elian Terpo, Szymon Konkolewski, Marta Gawronska, Martyna Matera, Agnieszka Karnowska, Danuta Bartczak, Katarzyna Kosowska, Lucyna Kochanowska, Aleksandra Krawczyk |
| EPI_ISL_3332860 | Laboratoria Medyczne Bruss grupa Alab sp. z o.o. Medyczne Laboratorium Diagnostyczne Pracownia Genetyki | Wojewodzka Stacja Sanitarno-Epidemiologiczna w Lodzi, Oddzial Laboratoryjny Mikrobiologii i Parazytologii | Elian Terpo, Szymon Konkolewski, Marta Gawronska, Martyna Matera, Agnieszka Karnowska, Danuta Bartczak, Katarzyna Kosowska, Lucyna Kochanowska, Aleksandra Krawczyk |
| EPI_ISL_3332861, EPI_ISL_3332863, EPI_ISL_3332865, EPI_ISL_3332866, EPI_ISL_3332867 | Medyczne Laboratoria Diagnostyczne INVICTA | Wojewodzka Stacja Sanitarno-Epidemiologiczna w Lodzi, Oddzial Laboratoryjny Mikrobiologii i Parazytologii | Elian Terpo, Szymon Konkolewski, Marta Gawronska, Martyna Matera, Agnieszka Karnowska, Danuta Bartczak, Katarzyna Kosowska, Lucyna Kochanowska, Aleksandra Krawczyk |
| EPI_ISL_3332868, EPI_ISL_3332869, EPI_ISL_3332870, EPI_ISL_3332871, EPI_ISL_3332872, EPI_ISL_3332873, EPI_ISL_3332874, EPI_ISL_3332875, EPI_ISL_3332876 | Uniwersyteckie Centrum Kliniczne | Wojewodzka Stacja Sanitarno-Epidemiologiczna w Lodzi, Oddzial Laboratoryjny Mikrobiologii i Parazytologii | Elian Terpo, Szymon Konkolewski, Marta Gawronska, Martyna Matera, Agnieszka Karnowska, Danuta Bartczak, Katarzyna Kosowska, Lucyna Kochanowska, Aleksandra Krawczyk |
| EPI_ISL_3332877 | WSSE w Gdansku - Laboratorium | Wojewodzka Stacja Sanitarno-Epidemiologiczna w Lodzi, Oddzial Laboratoryjny Mikrobiologii i Parazytologii | Elian Terpo, Szymon Konkolewski, Marta Gawronska, Martyna Matera, Agnieszka Karnowska, Danuta Bartczak, Katarzyna Kosowska, Lucyna Kochanowska, Aleksandra Krawczyk |
| EPI_ISL_3332878 | Uniwersyteckie Centrum Kliniczne | Wojewodzka Stacja Sanitarno-Epidemiologiczna w Lodzi, Oddzial Laboratoryjny Mikrobiologii i Parazytologii | Elian Terpo, Szymon Konkolewski, Marta Gawronska, Martyna Matera, Agnieszka Karnowska, Danuta Bartczak, Katarzyna Kosowska, Lucyna Kochanowska, Aleksandra Krawczyk |
| EPI_ISL_3332880 | WSSE w Gdansku - Laboratorium | Wojewodzka Stacja Sanitarno-Epidemiologiczna w Lodzi, Oddzial Laboratoryjny Mikrobiologii i Parazytologii | Elian Terpo, Szymon Konkolewski, Marta Gawronska, Martyna Matera, Agnieszka Karnowska, Danuta Bartczak, Katarzyna Kosowska, Lucyna Kochanowska, Aleksandra Krawczyk |
| EPI_ISL_3332881 | Diagnostyka Sp. z o.o. | Wojewodzka Stacja Sanitarno-Epidemiologiczna w Lodzi, Oddzial Laboratoryjny Mikrobiologii i Parazytologii | Elian Terpo, Szymon Konkolewski, Marta Gawronska, Martyna Matera, Agnieszka Karnowska, Danuta Bartczak, Katarzyna Kosowska, Lucyna Kochanowska, Aleksandra Krawczyk |
| EPI_ISL_3332882, EPI_ISL_3332883 | Dzial Diagnostyki Laboratoryjnej Wojewodzkie Centrum Szpitalne Kotliny Jeleniogorskiej | Wojewodzka Stacja Sanitarno-Epidemiologiczna w Lodzi, Oddzial Laboratoryjny Mikrobiologii i Parazytologii | Elian Terpo, Szymon Konkolewski, Marta Gawronska, Martyna Matera, Agnieszka Karnowska, Danuta Bartczak, Katarzyna Kosowska, Lucyna Kochanowska, Aleksandra Krawczyk |
| EPI_ISL_3332884, EPI_ISL_3332885 | Laboratorium Biol. Molek. USK | Wojewodzka Stacja Sanitarno-Epidemiologiczna w Lodzi, Oddzial Laboratoryjny Mikrobiologii i Parazytologii | Elian Terpo, Szymon Konkolewski, Marta Gawronska, Martyna Matera, Agnieszka Karnowska, Danuta Bartczak, Katarzyna Kosowska, Lucyna Kochanowska, Aleksandra Krawczyk |
| EPI_ISL_3332886 | Area of Virology, Serology and Virology Division (SAViD), New South Wales Health Pathology Randwick | Virology Research Laboratory; Area of Virology, Serology and Virology Division (SAViD), New South Wales Health Pathology Randwick | Foster, C.; Au, J.; Ruiz Silva, M.; Deveson, I.; Bull, R.; Van Hal, S.; Rawlinson, W. |
| EPI_ISL_3332898, EPI_ISL_3332899, EPI_ISL_3332900 | Lafene Health Center | Kansas State Veterinary Diagnostic Laboratory | Tyler Doerksen, Andreea Lu, Lance Noll, Elizabeth Porter, Emily Cox, Jianfa Bai, Jamie Henningson, Rachel Palinski |
| EPI_ISL_3332901, EPI_ISL_3332902, EPI_ISL_3332903, EPI_ISL_3332904, EPI_ISL_3332905, EPI_ISL_3332906, EPI_ISL_3332907, EPI_ISL_3332909, EPI_ISL_3332910, EPI_ISL_3332911, EPI_ISL_3332912, EPI_ISL_3332931, EPI_ISL_3332957, EPI_ISL_3332991, EPI_ISL_3332992, EPI_ISL_3332993, EPI_ISL_3332994, EPI_ISL_3332995, EPI_ISL_3332996, EPI_ISL_3332997, EPI_ISL_3332998, EPI_ISL_3332999, EPI_ISL_3333000, EPI_ISL_3333001, EPI_ISL_3333006, EPI_ISL_3333010, EPI_ISL_3333016, EPI_ISL_3333017, EPI_ISL_3333018, EPI_ISL_3333019, EPI_ISL_3333020, EPI_ISL_3333021, EPI_ISL_3333022, EPI_ISL_3333023, EPI_ISL_3333024, EPI_ISL_3333025, EPI_ISL_3333026, EPI_ISL_3333029, EPI_ISL_3333032, EPI_ISL_3333033, EPI_ISL_3333034, EPI_ISL_3333036, EPI_ISL_3333037, EPI_ISL_3333040, EPI_ISL_3333043, EPI_ISL_3333045 | | | |
| see above | Wyoming Public Health Laboratory | Wyoming Public Health Laboratory | Jim Mildenberger, Taylor Fearing, Channing Weber, Ashley Norberg, Chayse Rowley, Marley Goetz, Brian Dominguez, Elliot Thomasson, Sam Britz, Cari Sloma, Robert Petit, and Rob Christensen |
| EPI_ISL_3333062 | University of Iowa Hospitals and Clinics | State Hygienic Laboratory at the University of Iowa | Valerie Reeb, Erik Twait, Anna Yakos, Mariah Knutson, Kris Eveland, Kristen Zanon, Jeff, Benfer, Davis Rieckenberg, Wes Hottel, Alankar Kampoowale |
| EPI_ISL_3333063, EPI_ISL_3333064, EPI_ISL_3333065, EPI_ISL_3333066, EPI_ISL_3333067, EPI_ISL_3333068, EPI_ISL_3333070, EPI_ISL_3333071, EPI_ISL_3333072, EPI_ISL_3333073, EPI_ISL_3333074, EPI_ISL_3333075, EPI_ISL_3333076, EPI_ISL_3333077, EPI_ISL_3333078, EPI_ISL_3333080, EPI_ISL_3333081, EPI_ISL_3333082, EPI_ISL_3333083, EPI_ISL_3333085, EPI_ISL_3333086, EPI_ISL_3333087, EPI_ISL_3333088, EPI_ISL_3333090, EPI_ISL_3333092, EPI_ISL_3333093, EPI_ISL_3333094, EPI_ISL_3333095, EPI_ISL_3333096, EPI_ISL_3333097 | | | |
| see above | State Hygienic Laboratory at the University of Iowa | State Hygienic Laboratory at the University of Iowa | Valerie Reeb, Erik Twait, Anna Yakos, Mariah Knutson, Kris Eveland, Kristen Zanon, Jeff, Benfer, Davis Rieckenberg, Wes Hottel, Alankar Kampoowale |
| EPI_ISL_3333099 | University of Iowa Hospitals and Clinics | State Hygienic Laboratory at the University of Iowa | Valerie Reeb, Erik Twait, Anna Yakos, Mariah Knutson, Kris Eveland, Kristen Zanon, Jeff, Benfer, Davis Rieckenberg, Wes Hottel, Alankar Kampoowale |
| EPI_ISL_3333100, EPI_ISL_3333101, EPI_ISL_3333102, EPI_ISL_3333103 | State Hygienic Laboratory at the University of Iowa | State Hygienic Laboratory at the University of Iowa | Valerie Reeb, Erik Twait, Anna Yakos, Mariah Knutson, Kris Eveland, Kristen Zanon, Jeff, Benfer, Davis Rieckenberg, Wes Hottel, Alankar Kampoowale |
| EPI_ISL_3333104 | University of Iowa Hospitals and Clinics | State Hygienic Laboratory at the University of Iowa | Valerie Reeb, Erik Twait, Anna Yakos, Mariah Knutson, Kris Eveland, Kristen Zanon, Jeff, Benfer, Davis Rieckenberg, Wes Hottel, Alankar Kampoowale |
| EPI_ISL_3333105, EPI_ISL_3333106, EPI_ISL_3333107, EPI_ISL_3333108 | State Hygienic Laboratory at the University of Iowa | State Hygienic Laboratory at the University of Iowa | Valerie Reeb, Erik Twait, Anna Yakos, Mariah Knutson, Kris Eveland, Kristen Zanon, Jeff, Benfer, Davis Rieckenberg, Wes Hottel, Alankar Kampoowale |
| EPI_ISL_3333109 | University of Iowa Hospitals and Clinics | State Hygienic Laboratory at the University of Iowa | Valerie Reeb, Erik Twait, Anna Yakos, Mariah Knutson, Kris Eveland, Kristen Zanon, Jeff, Benfer, Davis Rieckenberg, Wes Hottel, Alankar Kampoowale |
| EPI_ISL_3333111, EPI_ISL_3333112, EPI_ISL_3333113, EPI_ISL_3333114, EPI_ISL_3333115, EPI_ISL_3333116, EPI_ISL_3333117, EPI_ISL_3333118, EPI_ISL_3333119, EPI_ISL_3333120, EPI_ISL_3333122, EPI_ISL_3333124, EPI_ISL_3333125, EPI_ISL_3333126, EPI_ISL_3333127, EPI_ISL_3333128, EPI_ISL_3333129, EPI_ISL_3333130, EPI_ISL_3333131, EPI_ISL_3333132, EPI_ISL_3333133, EPI_ISL_3333134, EPI_ISL_3333135, EPI_ISL_3333136, EPI_ISL_3333137, EPI_ISL_3333138, EPI_ISL_3333139, EPI_ISL_3333140, EPI_ISL_3333141, EPI_ISL_3333142, EPI_ISL_3333143, EPI_ISL_3333144, EPI_ISL_3333145, EPI_ISL_3333146, EPI_ISL_3333147, EPI_ISL_3333148, EPI_ISL_3333149, EPI_ISL_3333150, EPI_ISL_3333151, EPI_ISL_3333152, EPI_ISL_3333153 | | | |
| see above | State Hygienic Laboratory at the University of Iowa | State Hygienic Laboratory at the University of Iowa | Valerie Reeb, Erik Twait, Anna Yakos, Mariah Knutson, Kris Eveland, Kristen Zanon, Jeff, Benfer, Davis Rieckenberg, Wes Hottel, Alankar Kampoowale |
| EPI_ISL_3333154, EPI_ISL_3333155, EPI_ISL_3333156, EPI_ISL_3333157 | University of Iowa Hospitals and Clinics | State Hygienic Laboratory at the University of Iowa | Valerie Reeb, Erik Twait, Anna Yakos, Mariah Knutson, Kris Eveland, Kristen Zanon, Jeff, Benfer, Davis Rieckenberg, Wes Hottel, Alankar Kampoowale |
| EPI_ISL_3333158 | State Hygienic Laboratory at the University of Iowa | State Hygienic Laboratory at the University of Iowa | Valerie Reeb, Erik Twait, Anna Yakos, Mariah Knutson, Kris Eveland, Kristen Zanon, Jeff, Benfer, Davis Rieckenberg, Wes Hottel, Alankar Kampoowale |
| EPI_ISL_3333159, EPI_ISL_3333160 | University of Iowa Hospitals and Clinics | State Hygienic Laboratory at the University of Iowa | Valerie Reeb, Erik Twait, Anna Yakos, Mariah Knutson, Kris Eveland, Kristen Zanon, Jeff, Benfer, Davis Rieckenberg, Wes Hottel, Alankar Kampoowale |

| | | | |
|---|---|---|---|
| EPI_ISL_3333161 | State Hygienic Laboratory at the University of Iowa | State Hygienic Laboratory at the University of Iowa | Valerie Reeb, Erik Twait, Anna Yakos, Mariah Knutson, Kris Eveland, Kristen Zanon, Jeff, Benfer, Davis Rieckenberg, Wes Hottel, Alankar Kampoowale |
| EPI_ISL_3333162, EPI_ISL_3333164 | University of Iowa Hospitals and Clinics | State Hygienic Laboratory at the University of Iowa | Valerie Reeb, Erik Twait, Anna Yakos, Mariah Knutson, Kris Eveland, Kristen Zanon, Jeff, Benfer, Davis Rieckenberg, Wes Hottel, Alankar Kampoowale |
| EPI_ISL_3333165, EPI_ISL_3333166 | State Hygienic Laboratory at the University of Iowa | State Hygienic Laboratory at the University of Iowa | Valerie Reeb, Erik Twait, Anna Yakos, Mariah Knutson, Kris Eveland, Kristen Zanon, Jeff, Benfer, Davis Rieckenberg, Wes Hottel, Alankar Kampoowale |
| EPI_ISL_3333167, EPI_ISL_3333168 | University of Iowa Hospitals and Clinics | State Hygienic Laboratory at the University of Iowa | Valerie Reeb, Erik Twait, Anna Yakos, Mariah Knutson, Kris Eveland, Kristen Zanon, Jeff, Benfer, Davis Rieckenberg, Wes Hottel, Alankar Kampoowale |
| EPI_ISL_3333169, EPI_ISL_3333170, EPI_ISL_3333171, EPI_ISL_3333172 | State Hygienic Laboratory at the University of Iowa | State Hygienic Laboratory at the University of Iowa | Valerie Reeb, Erik Twait, Anna Yakos, Mariah Knutson, Kris Eveland, Kristen Zanon, Jeff, Benfer, Davis Rieckenberg, Wes Hottel, Alankar Kampoowale |
| EPI_ISL_3333173, EPI_ISL_3333174 | University of Iowa Hospitals and Clinics | State Hygienic Laboratory at the University of Iowa | Valerie Reeb, Erik Twait, Anna Yakos, Mariah Knutson, Kris Eveland, Kristen Zanon, Jeff, Benfer, Davis Rieckenberg, Wes Hottel, Alankar Kampoowale |
| EPI_ISL_3333177 | Cellis Laboratorium | Wojewodzka Stacja Sanitarno-Epidemiologiczna w Rzeszowie, Laboratorium Diagnostyki Medycznej | Marzena Baranowska, Katarzyna Wilk, Karolina Ostrowska, Anna Nowakowska |
| EPI_ISL_3333178 | Centrum Medyczne MEDYK Zaklad Diagnostyki Medycznej | Wojewodzka Stacja Sanitarno-Epidemiologiczna w Rzeszowie, Laboratorium Diagnostyki Medycznej | Marzena Baranowska, Katarzyna Wilk, Karolina Ostrowska, Anna Nowakowska |
| EPI_ISL_3333179, EPI_ISL_3333180, EPI_ISL_3333181, EPI_ISL_3333182, EPI_ISL_3333183, EPI_ISL_3333184, EPI_ISL_3333185, EPI_ISL_3333186, EPI_ISL_3333187, EPI_ISL_3333188 | New South Wales Health Pathology Royal Prince Alfred Hospital | Microbiology RPAH | Foster, C.; Au, J.; Ruiz Silva, M.; Deveson, I.; Bull, R.; Van Hal, S.; Rawlinson, W. |
| EPI_ISL_3333189, EPI_ISL_3333190, EPI_ISL_3333191, EPI_ISL_3333192, EPI_ISL_3333193, EPI_ISL_3333194, EPI_ISL_3333195, EPI_ISL_3333196, EPI_ISL_3333197, EPI_ISL_3333198, EPI_ISL_3333199, EPI_ISL_3333200, EPI_ISL_3333201, EPI_ISL_3333202, EPI_ISL_3333203, EPI_ISL_3333204 | | | |
| see above | Queensland Health Forensic and Scientific Services | Queensland Health Forensic and Scientific Services | Chenwei Wang on behalf of Q-PHIRE Genomics |
| EPI_ISL_3333205 | PHV-FSS | PHV-FSS | Chenwei Wang on behalf of Q-PHIRE Genomics |
| EPI_ISL_3333206, EPI_ISL_3333207, EPI_ISL_3333208, EPI_ISL_3333209, EPI_ISL_3333210, EPI_ISL_3333211, EPI_ISL_3333212, EPI_ISL_3333213, EPI_ISL_3333214, EPI_ISL_3333215, EPI_ISL_3333216, EPI_ISL_3333217, EPI_ISL_3333218, EPI_ISL_3333219, EPI_ISL_3333220, EPI_ISL_3333221, EPI_ISL_3333222, EPI_ISL_3333223, EPI_ISL_3333224, EPI_ISL_3333225, EPI_ISL_3333226, EPI_ISL_3333227 | | | |
| see above | Queensland Health Forensic and Scientific Services | Queensland Health Forensic and Scientific Services | Chenwei Wang on behalf of Q-PHIRE Genomics |
| EPI_ISL_3333228 | PHV-FSS | PHV-FSS | Chenwei Wang on behalf of Q-PHIRE Genomics |
| EPI_ISL_3333229, EPI_ISL_3333230, EPI_ISL_3333231 | Queensland Health Forensic and Scientific Services | Queensland Health Forensic and Scientific Services | Chenwei Wang on behalf of Q-PHIRE Genomics |
| EPI_ISL_3333232 | Gribbles Pathology | Institute for Medical Research, Infectious Disease Research Centre, National Institutes of Health, Ministry of Health Malaysia | Suppiah J, Kamel K, Mohd Zawawi Z, Azizan MA, Ramly N, Robert F, Anasir MI, Thayan R |
| EPI_ISL_3333233 | Gribbles Pathology (M) Sdn Bhd | Institute for Medical Research, Infectious Disease Research Centre, National Institutes of Health, Ministry of Health Malaysia | Suppiah J, Kamel K, Mohd Zawawi Z, Azizan MA, Ramly N, Robert F, Anasir MI, Thayan R |
| EPI_ISL_3333234 | Kajang Hospital | Institute for Medical Research, Infectious Disease Research Centre, National Institutes of Health, Ministry of Health Malaysia | Suppiah J, Kamel K, Mohd Zawawi Z, Azizan MA, Ramly N, Robert F, Anasir MI, Thayan R |
| EPI_ISL_3333235 | Serdang Hospital | Institute for Medical Research, Infectious Disease Research Centre, National Institutes of Health, Ministry of Health Malaysia | Suppiah J, Kamel K, Mohd Zawawi Z, Azizan MA, Ramly N, Robert F, Anasir MI, Thayan R |
| EPI_ISL_3333236 | Kota Bharu Public Health Laboratory | Institute for Medical Research, Infectious Disease Research Centre, National Institutes of Health, Ministry of Health Malaysia | Suppiah J, Kamel K, Mohd Zawawi Z, Azizan MA, Ramly N, Robert F, Anasir MI, Thayan R |
| EPI_ISL_3333237 | Sultanah Bahiyah Hospital, Alor Setar | Institute for Medical Research, Infectious Disease Research Centre, National Institutes of Health, Ministry of Health Malaysia | Suppiah J, Kamel K, Mohd Zawawi Z, Azizan MA, Ramly N, Robert F, Anasir MI, Thayan R |
| EPI_ISL_3333238 | RSPI SULIANTI SAROSO | Genomik Solidaritas Indonesia Laboratorium | Reinhart Gabriel, Carissa Sintca Wijaya, Vania Gavrila Wikasa, Dhahlia Agustina Cahyono, Gracia Felias Enos Korompis, Annisa Muthiah Sukirman, Anuraj Shankar, Louisa Markus, Meutia Ayuputeri Kumaheri, Ariel Pradipta |
| EPI_ISL_3333239 | Genomik Solidaritas Indonesia Laboratorium | Genomik Solidaritas Indonesia Laboratorium | Reinhart Gabriel, Carissa Sintca Wijaya, Vania Gavrila Wikasa, Dhahlia Agustina Cahyono, Gracia Felias Enos Korompis, Annisa Muthiah Sukirman, Anuraj Shankar, Louisa Markus, Meutia Ayuputeri Kumaheri, Ariel Pradipta |
| EPI_ISL_3333240, EPI_ISL_3333241, EPI_ISL_3333242, EPI_ISL_3333243 | RSPI SULIANTI SAROSO | Genomik Solidaritas Indonesia Laboratorium | Carissa Sintca Wijaya, Vania Gavrila Wikasa, Dhahlia Agustina Cahyono, Reinhart Gabriel, Gracia Felias Enos Korompis, Annisa Muthiah Sukirman, Anuraj Shankar, Louisa Markus, Meutia Ayuputeri Kumaheri, Ariel Pradipta |
| EPI_ISL_3333244, EPI_ISL_3333245, EPI_ISL_3333246 | RSPI SULIANTI SAROSO | Genomik Solidaritas Indonesia Laboratorium | Vania Gavrila Wikasa, Dhahlia Agustina Cahyono, Reinhart Gabriel, Carissa Sintca Wijaya, Annisa Muthiah Sukirman, Gracia Felias Enos Korompis, Anuraj Shankar, Louisa Markus, Meutia Ayuputeri Kumaheri, Ariel Pradipta |
| EPI_ISL_3333247, EPI_ISL_3333248 | RSPI SULIANTI SAROSO | Genomik Solidaritas Indonesia Laboratorium | Dhahlia Agustina Cahyono, Reinhart Gabriel, Carissa Sintca Wijaya, Vania Gavrila Wikasa, Annisa Muthiah Sukirman, Gracia Felias Enos Korompis, Anuraj Shankar, Louisa Markus, Meutia Ayuputeri Kumaheri, Ariel Pradipta |
| EPI_ISL_3333249 | Genomik Solidaritas Indonesia Laboratorium | Genomik Solidaritas Indonesia Laboratorium | Dhahlia Agustina Cahyono, Reinhart Gabriel, Carissa Sintca Wijaya, Vania Gavrila Wikasa, Annisa Muthiah Sukirman, Gracia Felias Enos Korompis, Anuraj Shankar, Louisa Markus, Meutia Ayuputeri Kumaheri, Ariel Pradipta |
| EPI_ISL_3333250 | Genomik Solidaritas Indonesia Laboratorium | Genomik Solidaritas Indonesia Laboratorium | Reinhart Gabriel, Carissa Sintca Wijaya, Vania Gavrila Wikasa, Dhahlia Agustina Cahyono, Gracia Felias Enos Korompis, Annisa Muthiah Sukirman, Anuraj Shankar, Louisa Markus, Meutia Ayuputeri Kumaheri, Ariel Pradipta |
| EPI_ISL_3333251 | Genomik Solidaritas Indonesia Laboratorium | Genomik Solidaritas Indonesia Laboratorium | Vania Gavrila Wikasa, Dhahlia Agustina Cahyono, Reinhart Gabriel, Carissa Sintca Wijaya, Annisa Muthiah Sukirman, Gracia Felias Enos Korompis, Anuraj Shankar, Louisa Markus, Meutia Ayuputeri Kumaheri, Ariel Pradipta |
| EPI_ISL_3333562 | Tuanku Fauziah Hospital | Institute for Medical Research, Infectious Disease Research Centre, National Institutes of Health, Ministry of Health Malaysia | Suppiah J, Kamel K, Mohd Zawawi Z, Azizan MA, Ramly N, Robert F, Anasir MI, Thayan R |
| EPI_ISL_3333563 | Slim River Hospital | Institute for Medical Research, Infectious Disease Research Centre, National Institutes of Health, Ministry of Health Malaysia | Suppiah J, Kamel K, Mohd Zawawi Z, Azizan MA, Ramly N, Robert F, Anasir MI, Thayan R |
| EPI_ISL_3333581 | Kota Bharu Public Health Laboratory | Institute for Medical Research, Infectious Disease Research Centre, National Institutes of Health, Ministry of Health Malaysia | Suppiah J, Kamel K, Mohd Zawawi Z, Azizan MA, Ramly N, Robert F, Anasir MI, Thayan R |
| EPI_ISL_3333594, EPI_ISL_3333595, EPI_ISL_3333596, EPI_ISL_3333597 | SC (UCO) Igiene e Sanità Pubblica, ASUGI, Trieste | SC (UCO) Igiene e Sanità Pubblica, ASUGI, Trieste | Licastro D, Dal Monego S, Degasperi M, Marcello A, Segat L, Piscianz E, Busetti M, Fontana F, Breda C, Barbone F, D'Agaro P |
| EPI_ISL_3333598, EPI_ISL_3333599, EPI_ISL_3333600, EPI_ISL_3333601, EPI_ISL_3333602, EPI_ISL_3333603, EPI_ISL_3333604, EPI_ISL_3333605, EPI_ISL_3333606, EPI_ISL_3333607 | SC (UCO) Igiene e Sanità Pubblica, ASUGI, Trieste | SC (UCO) Igiene e Sanità Pubblica, ASUGI, Trieste | Licastro D, Dal Monego S, Degasperi M, Marcello A, Segat L, De Rosa R, Breda C, Barbone F, D'Agaro P |
| EPI_ISL_3333608, EPI_ISL_3333609, EPI_ISL_3333610, EPI_ISL_3333611, EPI_ISL_3333612, EPI_ISL_3333613, EPI_ISL_3333614, EPI_ISL_3333615, EPI_ISL_3333616, EPI_ISL_3333617 | SC (UCO) Igiene e Sanità Pubblica, ASUGI, Trieste | SC (UCO) Igiene e Sanità Pubblica, ASUGI, Trieste | Licastro D, Dal Monego S, Degasperi M, Marcello A, Segat L, Piscianz E, Busetti M, Fontana F, Breda C, Barbone F, D'Agaro P |
| EPI_ISL_3333618, | SC (UCO) Igiene e Sanità Pubblica, ASUGI, Trieste | SC (UCO) Igiene e Sanità Pubblica, ASUGI, Trieste | Licastro D, Dal Monego S, Degasperi M, Marcello A, Segat L, Pipan C, Breda C, Barbone F, D'Agaro P |

| | | | |
|---|---|---|---|
| EPI_ISL_3333619, EPI_ISL_3333620, EPI_ISL_3333621, EPI_ISL_3333622, EPI_ISL_3333623 | | | |
| EPI_ISL_3333626, EPI_ISL_3333627, EPI_ISL_3333628 | AULSS 3 Venezia | UOSD Genetica e Citogenetica - Azienda ULSS 3 Serenissima; Istituto Zooprofilattico Sperimentale delle Venezie | Mosé Favarato, Elisa Squarcina, Laura Squarzon, Claudia Perini, Laura Bevilacqua, Adelaide Milani, Alessia Schivo, Annalisa Salviato, Elisa Palumbo, Erika Giorgia Quaranta, Luca Tassoni, Ambra Pastori, Edoardo Giussani, Alice Fusaro, Isabella Monne, Calogero Terregino, Antonia Ricci |
| EPI_ISL_3333629, EPI_ISL_3333630 | AULSS 2 Marca Trevigiana | UOSD Genetica e Citogenetica - Azienda ULSS 3 Serenissima; Istituto Zooprofilattico Sperimentale delle Venezie | Mosé Favarato, Elisa Squarcina, Laura Squarzon, Claudia Perini, Laura Bevilacqua, Adelaide Milani, Alessia Schivo, Annalisa Salviato, Elisa Palumbo, Erika Giorgia Quaranta, Luca Tassoni, Ambra Pastori, Edoardo Giussani, Alice Fusaro, Isabella Monne, Calogero Terregino, Antonia Ricci |
| EPI_ISL_3333631 | AULSS 9 Scaligera - Villafranca | UOSD Genetica e Citogenetica - Azienda ULSS 3 Serenissima; Istituto Zooprofilattico Sperimentale delle Venezie | Mosé Favarato, Elisa Squarcina, Laura Squarzon, Claudia Perini, Laura Bevilacqua, Adelaide Milani, Alessia Schivo, Annalisa Salviato, Elisa Palumbo, Erika Giorgia Quaranta, Luca Tassoni, Ambra Pastori, Edoardo Giussani, Alice Fusaro, Isabella Monne, Calogero Terregino, Antonia Ricci |
| EPI_ISL_3333632, EPI_ISL_3333633, EPI_ISL_3333634, EPI_ISL_3333635, EPI_ISL_3333636, EPI_ISL_3333637 | AULSS 2 Marca Trevigiana | UOSD Genetica e Citogenetica - Azienda ULSS 3 Serenissima; Istituto Zooprofilattico Sperimentale delle Venezie | Mosé Favarato, Elisa Squarcina, Laura Squarzon, Claudia Perini, Laura Bevilacqua, Adelaide Milani, Alessia Schivo, Annalisa Salviato, Elisa Palumbo, Erika Giorgia Quaranta, Luca Tassoni, Ambra Pastori, Edoardo Giussani, Alice Fusaro, Isabella Monne, Calogero Terregino, Antonia Ricci |
| EPI_ISL_3333638, EPI_ISL_3333639, EPI_ISL_3333640, EPI_ISL_3333642, EPI_ISL_3333643, EPI_ISL_3333644, EPI_ISL_3333645, EPI_ISL_3333646, EPI_ISL_3333647, EPI_ISL_3333648 | AULSS 9 Scaligera - Villafranca | UOSD Genetica e Citogenetica - Azienda ULSS 3 Serenissima; Istituto Zooprofilattico Sperimentale delle Venezie | Mosé Favarato, Elisa Squarcina, Laura Squarzon, Claudia Perini, Laura Bevilacqua, Adelaide Milani, Alessia Schivo, Annalisa Salviato, Elisa Palumbo, Erika Giorgia Quaranta, Luca Tassoni, Ambra Pastori, Edoardo Giussani, Alice Fusaro, Isabella Monne, Calogero Terregino, Antonia Ricci |
| EPI_ISL_3333649, EPI_ISL_3333650, EPI_ISL_3333651, EPI_ISL_3333652 | AULSS 4 Veneto Orientale | UOSD Genetica e Citogenetica - Azienda ULSS 3 Serenissima; Istituto Zooprofilattico Sperimentale delle Venezie | Mosé Favarato, Elisa Squarcina, Laura Squarzon, Claudia Perini, Laura Bevilacqua, Adelaide Milani, Alessia Schivo, Annalisa Salviato, Elisa Palumbo, Erika Giorgia Quaranta, Luca Tassoni, Ambra Pastori, Edoardo Giussani, Alice Fusaro, Isabella Monne, Calogero Terregino, Antonia Ricci |
| EPI_ISL_3333653, EPI_ISL_3333654, EPI_ISL_3333655 | AULSS 3 Venezia | UOSD Genetica e Citogenetica - Azienda ULSS 3 Serenissima; Istituto Zooprofilattico Sperimentale delle Venezie | Mosé Favarato, Elisa Squarcina, Laura Squarzon, Claudia Perini, Laura Bevilacqua, Adelaide Milani, Alessia Schivo, Annalisa Salviato, Elisa Palumbo, Erika Giorgia Quaranta, Luca Tassoni, Ambra Pastori, Edoardo Giussani, Alice Fusaro, Isabella Monne, Calogero Terregino, Antonia Ricci |
| EPI_ISL_3333656, EPI_ISL_3333657, EPI_ISL_3333929, EPI_ISL_3333930, EPI_ISL_3333931 | Sultanah Bahiyah Hospital, Alor Setar | Institute for Medical Research, Infectious Disease Research Centre, National Institutes of Health, Ministry of Health Malaysia | Suppiah J, Kamel K, Mohd Zawawi Z, Azizan MA, Ramly N, Robert F, Anasir MI, Thayan R |
| EPI_ISL_3333932, EPI_ISL_3333933, EPI_ISL_3333934, EPI_ISL_3333935, EPI_ISL_3333936, EPI_ISL_3333937, EPI_ISL_3333938, EPI_ISL_3333939, EPI_ISL_3333940, EPI_ISL_3333941, EPI_ISL_3333942, EPI_ISL_3333943, EPI_ISL_3333944, EPI_ISL_3333945, EPI_ISL_3333946, EPI_ISL_3333947, EPI_ISL_3333948, EPI_ISL_3333949, EPI_ISL_3333950, EPI_ISL_3333951, EPI_ISL_3333952, EPI_ISL_3333953, EPI_ISL_3333954, EPI_ISL_3333955, EPI_ISL_3333956, EPI_ISL_3333957, EPI_ISL_3333958, EPI_ISL_3333959, EPI_ISL_3333960, EPI_ISL_3333961, EPI_ISL_3333962, EPI_ISL_3333963, EPI_ISL_3333964, EPI_ISL_3333966, EPI_ISL_3333967, EPI_ISL_3333968, EPI_ISL_3333969, EPI_ISL_3333970, EPI_ISL_3333971, EPI_ISL_3333972, EPI_ISL_3333973, EPI_ISL_3333974, EPI_ISL_3333975, EPI_ISL_3333976, EPI_ISL_3333977, EPI_ISL_3333978, EPI_ISL_3333979, EPI_ISL_3333980, EPI_ISL_3333981, EPI_ISL_3333982, EPI_ISL_3333983, EPI_ISL_3333984, EPI_ISL_3333985, EPI_ISL_3333986, EPI_ISL_3333987, EPI_ISL_3333988, EPI_ISL_3333991, EPI_ISL_3333992, EPI_ISL_3333993, EPI_ISL_3333994, EPI_ISL_3333995, EPI_ISL_3333996, EPI_ISL_3333997, EPI_ISL_3333998, EPI_ISL_3333999, EPI_ISL_3334000, EPI_ISL_3334001, EPI_ISL_3334002, EPI_ISL_3334003, EPI_ISL_3334004, EPI_ISL_3334005, EPI_ISL_3334006, EPI_ISL_3334007, EPI_ISL_3334008, EPI_ISL_3334009, EPI_ISL_3334010, EPI_ISL_3334011, EPI_ISL_3334012, EPI_ISL_3334013, EPI_ISL_3334014, EPI_ISL_3334015, EPI_ISL_3334016, EPI_ISL_3334017, EPI_ISL_3334018, EPI_ISL_3334019, EPI_ISL_3334020 | | | |
| see above | Santa Clara Valley Medical Center | Santa Clara County Public Health Laboratory | Santa Clara County Public Health Department |
| EPI_ISL_3334021 | Ipoh Public Health Laboratory (MKAI), Ministry of Health Malaysia | Institute for Medical Research, Infectious Disease Research Centre, National Institutes of Health, Ministry of Health Malaysia | Suppiah J, Kamel K, Mohd Zawawi Z, Azizan MA, Ramly N, Robert F, Anasir MI, Thayan R |
| EPI_ISL_3334022 | Santa Clara County Public Health Laboratory | Santa Clara County Public Health Laboratory | Santa Clara County Public Health Department |
| EPI_ISL_3334023, EPI_ISL_3334024, EPI_ISL_3334025, EPI_ISL_3334026, EPI_ISL_3334027, EPI_ISL_3334028, EPI_ISL_3334029, EPI_ISL_3334030, EPI_ISL_3334031, EPI_ISL_3334032, EPI_ISL_3334034, EPI_ISL_3334035, EPI_ISL_3334036, EPI_ISL_3334037, EPI_ISL_3334038, EPI_ISL_3334039, EPI_ISL_3334040, EPI_ISL_3334042, EPI_ISL_3334043, EPI_ISL_3334044, EPI_ISL_3334046, EPI_ISL_3334047, EPI_ISL_3334048, EPI_ISL_3334049, EPI_ISL_3334050, EPI_ISL_3334052, EPI_ISL_3334053, EPI_ISL_3334054, EPI_ISL_3334055, EPI_ISL_3334056, EPI_ISL_3334057, EPI_ISL_3334058, EPI_ISL_3334059, EPI_ISL_3334060, EPI_ISL_3334061, EPI_ISL_3334062, EPI_ISL_3334063, EPI_ISL_3334064, EPI_ISL_3334065, EPI_ISL_3334066, EPI_ISL_3334067, EPI_ISL_3334068, EPI_ISL_3334069, EPI_ISL_3334070, EPI_ISL_3334071, EPI_ISL_3334072, EPI_ISL_3334073, EPI_ISL_3334074, EPI_ISL_3334075, EPI_ISL_3334076, EPI_ISL_3334077, EPI_ISL_3334078, EPI_ISL_3334079, EPI_ISL_3334080, EPI_ISL_3334082, EPI_ISL_3334083, EPI_ISL_3334085, EPI_ISL_3334086, EPI_ISL_3334087, EPI_ISL_3334088, EPI_ISL_3334089, EPI_ISL_3334090, EPI_ISL_3334091, EPI_ISL_3334092, EPI_ISL_3334093, EPI_ISL_3334094, EPI_ISL_3334095, EPI_ISL_3334096, EPI_ISL_3334097, EPI_ISL_3334098, EPI_ISL_3334099, EPI_ISL_3334100, EPI_ISL_3334101, EPI_ISL_3334102, EPI_ISL_3334103, EPI_ISL_3334104, EPI_ISL_3334105, EPI_ISL_3334106, EPI_ISL_3334107, EPI_ISL_3334108, EPI_ISL_3334109, EPI_ISL_3334110, EPI_ISL_3334113, EPI_ISL_3334114, EPI_ISL_3334115, EPI_ISL_3334116, EPI_ISL_3334117, EPI_ISL_3334118, EPI_ISL_3334119 | | | |
| see above | Santa Clara Valley Medical Center | Santa Clara County Public Health Laboratory | Santa Clara County Public Health Department |
| EPI_ISL_3334120 | Santa Clara County Public Health Laboratory | Santa Clara County Public Health Laboratory | Santa Clara County Public Health Department |
| EPI_ISL_3334121, EPI_ISL_3334122, EPI_ISL_3334123, EPI_ISL_3334124, EPI_ISL_3334125, EPI_ISL_3334126, EPI_ISL_3334127, EPI_ISL_3334128, EPI_ISL_3334129, EPI_ISL_3334130, EPI_ISL_3334131, EPI_ISL_3334132, EPI_ISL_3334133, EPI_ISL_3334134, EPI_ISL_3334135, EPI_ISL_3334136, EPI_ISL_3334137, EPI_ISL_3334138, EPI_ISL_3334139, EPI_ISL_3334140, EPI_ISL_3334141, EPI_ISL_3334142, EPI_ISL_3334143, EPI_ISL_3334144, EPI_ISL_3334145, EPI_ISL_3334146, EPI_ISL_3334148, EPI_ISL_3334149, EPI_ISL_3334150, EPI_ISL_3334151, EPI_ISL_3334152, EPI_ISL_3334153, EPI_ISL_3334154, EPI_ISL_3334155, EPI_ISL_3334156, EPI_ISL_3334157, EPI_ISL_3334158, EPI_ISL_3334159, EPI_ISL_3334160, EPI_ISL_3334161, EPI_ISL_3334162, EPI_ISL_3334163, EPI_ISL_3334164, EPI_ISL_3334165, EPI_ISL_3334166, EPI_ISL_3334167, EPI_ISL_3334168, EPI_ISL_3334169, EPI_ISL_3334170, EPI_ISL_3334171, EPI_ISL_3334172, EPI_ISL_3334173, EPI_ISL_3334175, EPI_ISL_3334176, EPI_ISL_3334177, EPI_ISL_3334178, EPI_ISL_3334179, EPI_ISL_3334180, EPI_ISL_3334181 | | | |
| see above | Santa Clara Valley Medical Center | Santa Clara County Public Health Laboratory | Santa Clara County Public Health Department |
| EPI_ISL_3334182 | Ipoh Public Health Laboratory (MKAI), Ministry of Health Malaysia | Institute for Medical Research, Infectious Disease Research Centre, National Institutes of Health, Ministry of Health Malaysia | Suppiah J, Kamel K, Mohd Zawawi Z, Azizan MA, Ramly N, Robert F, Anasir MI, Thayan R |
| EPI_ISL_3334184, EPI_ISL_3334185, EPI_ISL_3334186 | UCSC Genomics Institute | UCSC Genomics Institute | Mark Akeson, Ikenna Anigbogu, Eric Beraut, Isabel Bjork, Molly Cassatt-Johnstone, Terren Chang, Russell Corbett-Detig, Namrita Dhillon, Maximilian Haeussler, David Haussler, Angie Hinrichs, Miten Jain, Joshua Kapp, A. Marm Kilpatrick, Jakob McBroome, Hugh Olsen, Jeremy Sanford, Beth Shapiro, Michael Stone, Bryan Thornlow, Yatish Turakhia, Ciara Wanket |
| EPI_ISL_3334189 | Valais Hospital, Central Institute | Valais Hospital, Central Institute | Alexis Dumoulin, Lorenzo Cerutti, Cedric Howald, Henri Pegeot, Melyssa Elies, Deborah Penet, Keith Harshman, Ioannis Xenarios |
| EPI_ISL_3334190 | National Public Health Laboratory, Sungai Buloh | Institute for Medical Research, Infectious Disease Research Centre, National Institutes of Health, Ministry of Health Malaysia | Suppiah J, Kamel K, Mohd Zawawi Z, Azizan MA, Ramly N, Robert F, Anasir MI, Thayan R |
| EPI_ISL_3334210, EPI_ISL_3334211, EPI_ISL_3334212, EPI_ISL_3334213, EPI_ISL_3334214 | NB-Hôpital Georges L. Dumont | New Brunswick – Vitalité Health Network | Chacko S., Shaw W., Lacroix J., Allain E., Lyons P., Garceau R., Desnoyers G., Crapoulet N. |
| EPI_ISL_3334217 | National Public Health Laboratory, Sungai Buloh | Institute for Medical Research, Infectious Disease Research Centre, National Institutes of Health, Ministry of Health Malaysia | Suppiah J, Kamel K, Mohd Zawawi Z, Azizan MA, Ramly N, Robert F, Anasir MI, Thayan R |
| EPI_ISL_3334218 | Gleneagles Hospital Medini Johor | Institute for Medical Research, Infectious Disease Research Centre, National Institutes of Health, Ministry of Health Malaysia | Suppiah J, Kamel K, Mohd Zawawi Z, Azizan MA, Ramly N, Robert F, Anasir MI, Thayan R |
| EPI_ISL_3334236, EPI_ISL_3334237, EPI_ISL_3334238, EPI_ISL_3334240, EPI_ISL_3334241, EPI_ISL_3334242, EPI_ISL_3334243, EPI_ISL_3334244, EPI_ISL_3334246, EPI_ISL_3334247, EPI_ISL_3334249, EPI_ISL_3334250, EPI_ISL_3334256, EPI_ISL_3334257, EPI_ISL_3334266, EPI_ISL_3334267, EPI_ISL_3334268, EPI_ISL_3334270, EPI_ISL_3334271, EPI_ISL_3334272, EPI_ISL_3334273, EPI_ISL_3334277, EPI_ISL_3334279, EPI_ISL_3334280, EPI_ISL_3334285, EPI_ISL_3334286, EPI_ISL_3334289, EPI_ISL_3334290, EPI_ISL_3334294, EPI_ISL_3334295, EPI_ISL_3334296, EPI_ISL_3334297, EPI_ISL_3334299, EPI_ISL_3334301, EPI_ISL_3334302, EPI_ISL_3334303, EPI_ISL_3334305, EPI_ISL_3334306, EPI_ISL_3334307, EPI_ISL_3334309, EPI_ISL_3334311, EPI_ISL_3334312, EPI_ISL_3334314, EPI_ISL_3334315, EPI_ISL_3334316, EPI_ISL_3334317, EPI_ISL_3334318, EPI_ISL_3334319, EPI_ISL_3334320, EPI_ISL_3334322, EPI_ISL_3334325, EPI_ISL_3334327, EPI_ISL_3334328, EPI_ISL_3334330, EPI_ISL_3334331, EPI_ISL_3334332, EPI_ISL_3334333, EPI_ISL_3334334, EPI_ISL_3334335, EPI_ISL_3334337, EPI_ISL_3334338, EPI_ISL_3334339, EPI_ISL_3334340, EPI_ISL_3334341, EPI_ISL_3334342, EPI_ISL_3334343, EPI_ISL_3334344, EPI_ISL_3334345, EPI_ISL_3334346, EPI_ISL_3334347, EPI_ISL_3334348, EPI_ISL_3334349, EPI_ISL_3334350, EPI_ISL_3334352, EPI_ISL_3334353, EPI_ISL_3334354, EPI_ISL_3334355, EPI_ISL_3334356, EPI_ISL_3334358, EPI_ISL_3334359, EPI_ISL_3334360, EPI_ISL_3334361, EPI_ISL_3334362, EPI_ISL_3334363, EPI_ISL_3334364, EPI_ISL_3334365, EPI_ISL_3334366, EPI_ISL_3334367, EPI_ISL_3334368, EPI_ISL_3334369, EPI_ISL_3334370, EPI_ISL_3334371, EPI_ISL_3334372, EPI_ISL_3334373, EPI_ISL_3334374, EPI_ISL_3334375, EPI_ISL_3334376, EPI_ISL_3334377, EPI_ISL_3334378, EPI_ISL_3334379, EPI_ISL_3334380, EPI_ISL_3334381, EPI_ISL_3334382, EPI_ISL_3334387, EPI_ISL_3334389, EPI_ISL_3334390, EPI_ISL_3334391, EPI_ISL_3334392, EPI_ISL_3334394, EPI_ISL_3334395, EPI_ISL_3334396, EPI_ISL_3334397, EPI_ISL_3334398, EPI_ISL_3334399, EPI_ISL_3334400, EPI_ISL_3334401, EPI_ISL_3334402, EPI_ISL_3334404, EPI_ISL_3334405, EPI_ISL_3334406, EPI_ISL_3334407, EPI_ISL_3334408, EPI_ISL_3334409, EPI_ISL_3334410, EPI_ISL_3334411, EPI_ISL_3334412, EPI_ISL_3334413, EPI_ISL_3334414, EPI_ISL_3334415, EPI_ISL_3334416, EPI_ISL_3334419, EPI_ISL_3334420, EPI_ISL_3334421, EPI_ISL_3334422, EPI_ISL_3334423, EPI_ISL_3334424, EPI_ISL_3334425, EPI_ISL_3334426, EPI_ISL_3334427, EPI_ISL_3334428, EPI_ISL_3334430, EPI_ISL_3334431, EPI_ISL_3334433, EPI_ISL_3334434, EPI_ISL_3334435, EPI_ISL_3334436, EPI_ISL_3334437, EPI_ISL_3334438, EPI_ISL_3334439, EPI_ISL_3334441, EPI_ISL_3334442, EPI_ISL_3334443, EPI_ISL_3334444, EPI_ISL_3334445, EPI_ISL_3334446, EPI_ISL_3334447, EPI_ISL_3334448, EPI_ISL_3334451, EPI_ISL_3334452, EPI_ISL_3334453, EPI_ISL_3334454, EPI_ISL_3334455, EPI_ISL_3334456, EPI_ISL_3334458, EPI_ISL_3334459, EPI_ISL_3334460, EPI_ISL_3334461, EPI_ISL_3334463, EPI_ISL_3334464, EPI_ISL_3334465, EPI_ISL_3334467, EPI_ISL_3334469, EPI_ISL_3334470, EPI_ISL_3334471, EPI_ISL_3334472, EPI_ISL_3334473, EPI_ISL_3334474, EPI_ISL_3334475, EPI_ISL_3334477, EPI_ISL_3334478, EPI_ISL_3334479, EPI_ISL_3334480, EPI_ISL_3334481, EPI_ISL_3334483, EPI_ISL_3334484, EPI_ISL_3334485, EPI_ISL_3334486, EPI_ISL_3334487, EPI_ISL_3334488, EPI_ISL_3334489, EPI_ISL_3334490, EPI_ISL_3334492, EPI_ISL_3334493, EPI_ISL_3334494, EPI_ISL_3334495, EPI_ISL_3334496, EPI_ISL_3334497, EPI_ISL_3334499, EPI_ISL_3334500, EPI_ISL_3334501, EPI_ISL_3334502, EPI_ISL_3334504, EPI_ISL_3334505, EPI_ISL_3334506, EPI_ISL_3334507, EPI_ISL_3334508, EPI_ISL_3334510, EPI_ISL_3334511, EPI_ISL_3334512, EPI_ISL_3334513, EPI_ISL_3334514, EPI_ISL_3334515, EPI_ISL_3334517, EPI_ISL_3334518, EPI_ISL_3334519, EPI_ISL_3334520, EPI_ISL_3334521, EPI_ISL_3334523, EPI_ISL_3334524, EPI_ISL_3334525, EPI_ISL_3334526, EPI_ISL_3334528, EPI_ISL_3334529, EPI_ISL_3334530, EPI_ISL_3334531, EPI_ISL_3334532, EPI_ISL_3334533, EPI_ISL_3334534, EPI_ISL_3334535, EPI_ISL_3334536, EPI_ISL_3334537, EPI_ISL_3334538, EPI_ISL_3334539, EPI_ISL_3334540, EPI_ISL_3334541, EPI_ISL_3334542, EPI_ISL_3334543, EPI_ISL_3334544, EPI_ISL_3334545, EPI_ISL_3334546, EPI_ISL_3334548, EPI_ISL_3334550, EPI_ISL_3334551, EPI_ISL_3334552, EPI_ISL_3334553, EPI_ISL_3334554, EPI_ISL_3334555 | | | |
| see above | BioneXt Lab | Laboratoire national de sante, Microbiology, Microbial Genomics Platform | Anke Wienecke-Baldacchino, Catherine Ragimbeau,Jessica Tapp, Fatu Djabi, Lise Pignon, Raoul Salmon,Virginie Jover, Elodie Solarino, Thibault Ferrandon, Tamir Abdelrahman |
| EPI_ISL_3334556, EPI_ISL_3334557, EPI_ISL_3334558, EPI_ISL_3334559, EPI_ISL_3334561, EPI_ISL_3334562, EPI_ISL_3334563 | Centre Hospitalier du Nord | Laboratoire national de sante, Microbiology, Microbial Genomics Platform | Anke Wienecke-Baldacchino, Catherine Ragimbeau,Jessica Tapp, Fatu Djabi, Lise Pignon, Raoul Salmon,Virginie Jover, Elodie Solarino, Fatiha Boulmerka, Jacqueline Parmentier, Tamir Abdelrahman |
| EPI_ISL_3334564, | Hopitaux Robert Schuman | Laboratoire national de sante, Microbiology, Microbial Genomics | Anke Wienecke-Baldacchino, Catherine Ragimbeau,Jessica Tapp, Fatu Djabi, Lise Pignon, Raoul Salmon,Virginie Jover, Elodie Solarino, Alain Hakim, Nawfal Faik, Tamir Abdelrahman |

| | | | |
|---|---|---|---|
| EPI_ISL_3334565, EPI_ISL_3334566, EPI_ISL_3334567, EPI_ISL_3334569, EPI_ISL_3334570, EPI_ISL_3334571 | | Platform | |
| EPI_ISL_3334572, EPI_ISL_3334574, EPI_ISL_3334575, EPI_ISL_3334577, EPI_ISL_3334578, EPI_ISL_3334580, EPI_ISL_3334582, EPI_ISL_3334584, EPI_ISL_3334585, EPI_ISL_3334586, EPI_ISL_3334587, EPI_ISL_3334588, EPI_ISL_3334589, EPI_ISL_3334591, EPI_ISL_3334592, EPI_ISL_3334593 | | | |
| see above | Hospital Center Emile Mayrisch | Laboratoire national de sante, Microbiology, Microbial Genomics Platform | Anke Wienecke-Baldacchino, Catherine Ragimbeau,Jessica Tapp, Fatu Djabi, Lise Pignon, Raoul Salmon,Virginie Jover, Elodie Solarino, Cynthia Oxacelay, Tamir Abdelrahman |
| EPI_ISL_3334594, EPI_ISL_3334598, EPI_ISL_3334601, EPI_ISL_3334602, EPI_ISL_3334603, EPI_ISL_3334604, EPI_ISL_3334605, EPI_ISL_3334607, EPI_ISL_3334608, EPI_ISL_3334609, EPI_ISL_3334610, EPI_ISL_3334611, EPI_ISL_3334613, EPI_ISL_3334614, EPI_ISL_3334616, EPI_ISL_3334618, EPI_ISL_3334619, EPI_ISL_3334620, EPI_ISL_3334621, EPI_ISL_3334622, EPI_ISL_3334623, EPI_ISL_3334627, EPI_ISL_3334628, EPI_ISL_3334629, EPI_ISL_3334630, EPI_ISL_3334631, EPI_ISL_3334632, EPI_ISL_3334633 | | | |
| see above | Hospital Center Luxembourg | Laboratoire national de sante, Microbiology, Microbial Genomics Platform | Anke Wienecke-Baldacchino, Catherine Ragimbeau,Jessica Tapp, Fatu Djabi, Lise Pignon, Raoul Salmon,Virginie Jover, Elodie Solarino, Michel Kohnen, Jean-Hugues Francois, Tamir Abdelrahman |
| EPI_ISL_3334635, EPI_ISL_3334636, EPI_ISL_3334637, EPI_ISL_3334638, EPI_ISL_3334639, EPI_ISL_3334640, EPI_ISL_3334642, EPI_ISL_3334643, EPI_ISL_3334644, EPI_ISL_3334645, EPI_ISL_3334648, EPI_ISL_3334649, EPI_ISL_3334650, EPI_ISL_3334651, EPI_ISL_3334652, EPI_ISL_3334654, EPI_ISL_3334655, EPI_ISL_3334656, EPI_ISL_3334657 | | | |
| see above | Laboratoire national de sante, 1 rue Louis Rech, L-3555 Dudelange, Luxembourg | Laboratoire national de sante, Microbiology, Microbial Genomics Platform | Anke Wienecke-Baldacchino, Catherine Ragimbeau, Jessica Tapp, Fatu Djabi, Lise Pignon, Raoul Salmon,Virginie Jover, Elodie Solarino, Trung Nguyen Nguyen, Tamir Abdelrahman |
| EPI_ISL_3334666, EPI_ISL_3334667, EPI_ISL_3334668, EPI_ISL_3334669, EPI_ISL_3334670, EPI_ISL_3334672, EPI_ISL_3334673, EPI_ISL_3334675, EPI_ISL_3334676, EPI_ISL_3334677, EPI_ISL_3334678, EPI_ISL_3334681, EPI_ISL_3334682, EPI_ISL_3334683, EPI_ISL_3334684, EPI_ISL_3334686, EPI_ISL_3334687, EPI_ISL_3334688, EPI_ISL_3334689, EPI_ISL_3334691, EPI_ISL_3334692, EPI_ISL_3334695, EPI_ISL_3334696, EPI_ISL_3334697, EPI_ISL_3334698, EPI_ISL_3334699, EPI_ISL_3334700, EPI_ISL_3334701, EPI_ISL_3334702, EPI_ISL_3334703, EPI_ISL_3334704, EPI_ISL_3334705, EPI_ISL_3334706, EPI_ISL_3334707, EPI_ISL_3334708, EPI_ISL_3334709, EPI_ISL_3334710, EPI_ISL_3334711, EPI_ISL_3334712, EPI_ISL_3334713, EPI_ISL_3334714, EPI_ISL_3334715, EPI_ISL_3334716, EPI_ISL_3334717, EPI_ISL_3334721, EPI_ISL_3334722, EPI_ISL_3334723, EPI_ISL_3334724, EPI_ISL_3334725, EPI_ISL_3334726, EPI_ISL_3334727, EPI_ISL_3334729, EPI_ISL_3334730, EPI_ISL_3334731, EPI_ISL_3334732, EPI_ISL_3334734, EPI_ISL_3334735, EPI_ISL_3334736, EPI_ISL_3334737, EPI_ISL_3334738, EPI_ISL_3334741, EPI_ISL_3334742, EPI_ISL_3334744, EPI_ISL_3334745, EPI_ISL_3334746, EPI_ISL_3334747, EPI_ISL_3334748, EPI_ISL_3334749, EPI_ISL_3334751, EPI_ISL_3334753, EPI_ISL_3334754, EPI_ISL_3334755, EPI_ISL_3334757, EPI_ISL_3334758, EPI_ISL_3334759, EPI_ISL_3334760, EPI_ISL_3334761, EPI_ISL_3334763, EPI_ISL_3334764, EPI_ISL_3334765, EPI_ISL_3334767, EPI_ISL_3334768, EPI_ISL_3334769, EPI_ISL_3334770, EPI_ISL_3334771 | | | |
| see above | Laboratoires d'analyses medicales - Ketterthill | Laboratoire national de sante, Microbiology, Microbial Genomics Platform | Anke Wienecke-Baldacchino, Catherine Ragimbeau,Jessica Tapp, Fatu Djabi, Lise Pignon, Raoul Salmon,Virginie Jover, Elodie Solarino, Serge Vedy, Caroline Scheiber, Tamir Abdelrahman |
| EPI_ISL_3334793, EPI_ISL_3334794, EPI_ISL_3334795, EPI_ISL_3334797, EPI_ISL_3334799, EPI_ISL_3334800, EPI_ISL_3334801, EPI_ISL_3334802, EPI_ISL_3334803, EPI_ISL_3334804, EPI_ISL_3334805, EPI_ISL_3334807, EPI_ISL_3334808, EPI_ISL_3334809, EPI_ISL_3334810, EPI_ISL_3334813, EPI_ISL_3334818, EPI_ISL_3334820, EPI_ISL_3334821, EPI_ISL_3334822, EPI_ISL_3334823, EPI_ISL_3334824, EPI_ISL_3334825, EPI_ISL_3334827, EPI_ISL_3334828, EPI_ISL_3334829, EPI_ISL_3334830, EPI_ISL_3334831, EPI_ISL_3334832, EPI_ISL_3334833, EPI_ISL_3334834, EPI_ISL_3334835, EPI_ISL_3334837, EPI_ISL_3334838, EPI_ISL_3334839, EPI_ISL_3334840, EPI_ISL_3334841, EPI_ISL_3334842 | | | |
| see above | Laboratoires Reunis, 38 Rue Hiel, 6131 Junglinster, Luxembourg | Laboratoire national de sante, Microbiology, Microbial Genomics Platform | Anke Wienecke-Baldacchino, Catherine Ragimbeau,Jessica Tapp, Fatu Djabi, Lise Pignon, Raoul Salmon,Virginie Jover, Elodie Solarino, Bernard Weber, Tamir Abdelrahman |
| EPI_ISL_3334843 | Lablink (M) Sdn. Bhd | Institute for Medical Research, Infectious Disease Research Centre, National Institutes of Health, Ministry of Health Malaysia | Suppiah J, Kamel K, Mohd Zawawi Z, Azizan MA, Ramly N, Robert F, Anasir MI, Thayan R |
| EPI_ISL_3334857 | Lighthouse Lab in Milton Keynes | Wellcome Sanger Institute for the COVID-19 Genomics UK (COG-UK) Consortium | The Lighthouse Lab in Milton Keynes and Alex Alderton, Roberto Amato, Jeffrey Barrett, Sonia Goncalves, Ewan Harrison, David K. Jackson, Ian Johnston, Dominic Kwiatkowski, Cordelia Langford, John Sillitoe on behalf of the Wellcome Sanger Institute COVID-19 Surveillance Team |
| EPI_ISL_3334858, EPI_ISL_3334859 | Lighthouse Laboratory Plymouth | Wellcome Sanger Institute for the COVID-19 Genomics UK (COG-UK) Consortium | Lighthouse Laboratory Plymouth and Alex Alderton, Roberto Amato, Jeffrey Barrett, Sonia Goncalves, Ewan Harrison, David K. Jackson, Ian Johnston, Dominic Kwiatkowski, Cordelia Langford, John Sillitoe on behalf of the Wellcome Sanger Institute COVID-19 Surveillance Team |
| EPI_ISL_3334860, EPI_ISL_3334861 | Lighthouse Lab in Milton Keynes | Wellcome Sanger Institute for the COVID-19 Genomics UK (COG-UK) Consortium | The Lighthouse Lab in Milton Keynes and Alex Alderton, Roberto Amato, Jeffrey Barrett, Sonia Goncalves, Ewan Harrison, David K. Jackson, Ian Johnston, Dominic Kwiatkowski, Cordelia Langford, John Sillitoe on behalf of the Wellcome Sanger Institute COVID-19 Surveillance Team |
| EPI_ISL_3334862, EPI_ISL_3334863 | Lighthouse Laboratory Plymouth | Wellcome Sanger Institute for the COVID-19 Genomics UK (COG-UK) Consortium | Lighthouse Laboratory Plymouth and Alex Alderton, Roberto Amato, Jeffrey Barrett, Sonia Goncalves, Ewan Harrison, David K. Jackson, Ian Johnston, Dominic Kwiatkowski, Cordelia Langford, John Sillitoe on behalf of the Wellcome Sanger Institute COVID-19 Surveillance Team |
| EPI_ISL_3334864, EPI_ISL_3334865 | Lighthouse Lab in Milton Keynes | Wellcome Sanger Institute for the COVID-19 Genomics UK (COG-UK) Consortium | The Lighthouse Lab in Milton Keynes and Alex Alderton, Roberto Amato, Jeffrey Barrett, Sonia Goncalves, Ewan Harrison, David K. Jackson, Ian Johnston, Dominic Kwiatkowski, Cordelia Langford, John Sillitoe on behalf of the Wellcome Sanger Institute COVID-19 Surveillance Team |
| EPI_ISL_3334866, EPI_ISL_3334867 | Lighthouse Laboratory Plymouth | Wellcome Sanger Institute for the COVID-19 Genomics UK (COG-UK) Consortium | Lighthouse Laboratory Plymouth and Alex Alderton, Roberto Amato, Jeffrey Barrett, Sonia Goncalves, Ewan Harrison, David K. Jackson, Ian Johnston, Dominic Kwiatkowski, Cordelia Langford, John Sillitoe on behalf of the Wellcome Sanger Institute COVID-19 Surveillance Team |
| EPI_ISL_3334868 | Berkshire and Surrey Pathology Services Lighthouse Laboratory | Wellcome Sanger Institute for the COVID-19 Genomics UK (COG-UK) Consortium | Berkshire and Surrey Pathology Services Lighthouse Laboratory and Alex Alderton, Roberto Amato, Jeffrey Barrett, Sonia Goncalves, Ewan Harrison, David K. Jackson, Ian Johnston, Dominic Kwiatkowski, Cordelia Langford, John Sillitoe on behalf of the Wellcome Sanger Institute COVID-19 Surveillance Team |
| EPI_ISL_3334869, EPI_ISL_3334870 | Lighthouse Lab in Milton Keynes | Wellcome Sanger Institute for the COVID-19 Genomics UK (COG-UK) Consortium | The Lighthouse Lab in Milton Keynes and Alex Alderton, Roberto Amato, Jeffrey Barrett, Sonia Goncalves, Ewan Harrison, David K. Jackson, Ian Johnston, Dominic Kwiatkowski, Cordelia Langford, John Sillitoe on behalf of the Wellcome Sanger Institute COVID-19 Surveillance Team |
| EPI_ISL_3334871 | Lighthouse Laboratory Plymouth | Wellcome Sanger Institute for the COVID-19 Genomics UK (COG-UK) Consortium | Lighthouse Laboratory Plymouth and Alex Alderton, Roberto Amato, Jeffrey Barrett, Sonia Goncalves, Ewan Harrison, David K. Jackson, Ian Johnston, Dominic Kwiatkowski, Cordelia Langford, John Sillitoe on behalf of the Wellcome Sanger Institute COVID-19 Surveillance Team |
| EPI_ISL_3334872, EPI_ISL_3334873 | Lighthouse Lab in Milton Keynes | Wellcome Sanger Institute for the COVID-19 Genomics UK (COG-UK) Consortium | The Lighthouse Lab in Milton Keynes and Alex Alderton, Roberto Amato, Jeffrey Barrett, Sonia Goncalves, Ewan Harrison, David K. Jackson, Ian Johnston, Dominic Kwiatkowski, Cordelia Langford, John Sillitoe on behalf of the Wellcome Sanger Institute COVID-19 Surveillance Team |
| EPI_ISL_3334874 | Lighthouse Laboratory Plymouth | Wellcome Sanger Institute for the COVID-19 Genomics UK (COG-UK) Consortium | Lighthouse Laboratory Plymouth and Alex Alderton, Roberto Amato, Jeffrey Barrett, Sonia Goncalves, Ewan Harrison, David K. Jackson, Ian Johnston, Dominic Kwiatkowski, Cordelia Langford, John Sillitoe on behalf of the Wellcome Sanger Institute COVID-19 Surveillance Team |
| EPI_ISL_3334875 | Berkshire and Surrey Pathology Services Lighthouse Laboratory | Wellcome Sanger Institute for the COVID-19 Genomics UK (COG-UK) Consortium | Berkshire and Surrey Pathology Services Lighthouse Laboratory and Alex Alderton, Roberto Amato, Jeffrey Barrett, Sonia Goncalves, Ewan Harrison, David K. Jackson, Ian Johnston, Dominic Kwiatkowski, Cordelia Langford, John Sillitoe on behalf of the Wellcome Sanger Institute COVID-19 Surveillance Team |
| EPI_ISL_3334876 | Lighthouse Lab in Milton Keynes | Wellcome Sanger Institute for the COVID-19 Genomics UK (COG-UK) Consortium | The Lighthouse Lab in Milton Keynes and Alex Alderton, Roberto Amato, Jeffrey Barrett, Sonia Goncalves, Ewan Harrison, David K. Jackson, Ian Johnston, Dominic Kwiatkowski, Cordelia Langford, John Sillitoe on behalf of the Wellcome Sanger Institute COVID-19 Surveillance Team |
| EPI_ISL_3334877, EPI_ISL_3334878 | Berkshire and Surrey Pathology Services Lighthouse Laboratory | Wellcome Sanger Institute for the COVID-19 Genomics UK (COG-UK) Consortium | Berkshire and Surrey Pathology Services Lighthouse Laboratory and Alex Alderton, Roberto Amato, Jeffrey Barrett, Sonia Goncalves, Ewan Harrison, David K. Jackson, Ian Johnston, Dominic Kwiatkowski, Cordelia Langford, John Sillitoe on behalf of the Wellcome Sanger Institute COVID-19 Surveillance Team |
| EPI_ISL_3334879, EPI_ISL_3334880, EPI_ISL_3334881, EPI_ISL_3334882 | Lighthouse Lab in Milton Keynes | Wellcome Sanger Institute for the COVID-19 Genomics UK (COG-UK) Consortium | The Lighthouse Lab in Milton Keynes and Alex Alderton, Roberto Amato, Jeffrey Barrett, Sonia Goncalves, Ewan Harrison, David K. Jackson, Ian Johnston, Dominic Kwiatkowski, Cordelia Langford, John Sillitoe on behalf of the Wellcome Sanger Institute COVID-19 Surveillance Team |
| EPI_ISL_3334883 | Lighthouse Laboratory Plymouth | Wellcome Sanger Institute for the COVID-19 Genomics UK (COG-UK) Consortium | Lighthouse Laboratory Plymouth and Alex Alderton, Roberto Amato, Jeffrey Barrett, Sonia Goncalves, Ewan Harrison, David K. Jackson, Ian Johnston, Dominic Kwiatkowski, Cordelia Langford, John Sillitoe on behalf of the Wellcome Sanger Institute COVID-19 Surveillance Team |
| EPI_ISL_3334884 | Berkshire and Surrey Pathology Services Lighthouse Laboratory | Wellcome Sanger Institute for the COVID-19 Genomics UK (COG-UK) Consortium | Berkshire and Surrey Pathology Services Lighthouse Laboratory and Alex Alderton, Roberto Amato, Jeffrey Barrett, Sonia Goncalves, Ewan Harrison, David K. Jackson, Ian Johnston, Dominic Kwiatkowski, Cordelia Langford, John Sillitoe on behalf of the Wellcome Sanger Institute COVID-19 Surveillance Team |
| EPI_ISL_3334885, EPI_ISL_3334886, EPI_ISL_3334887, EPI_ISL_3334888 | Lighthouse Lab in Milton Keynes | Wellcome Sanger Institute for the COVID-19 Genomics UK (COG-UK) Consortium | The Lighthouse Lab in Milton Keynes and Alex Alderton, Roberto Amato, Jeffrey Barrett, Sonia Goncalves, Ewan Harrison, David K. Jackson, Ian Johnston, Dominic Kwiatkowski, Cordelia Langford, John Sillitoe on behalf of the Wellcome Sanger Institute COVID-19 Surveillance Team |
| EPI_ISL_3334889 | Lighthouse Laboratory Plymouth | Wellcome Sanger Institute for the COVID-19 Genomics UK (COG-UK) Consortium | Lighthouse Laboratory Plymouth and Alex Alderton, Roberto Amato, Jeffrey Barrett, Sonia Goncalves, Ewan Harrison, David K. Jackson, Ian Johnston, Dominic Kwiatkowski, Cordelia Langford, John Sillitoe on behalf of the Wellcome Sanger Institute COVID-19 Surveillance Team |
| EPI_ISL_3334890 | Lighthouse Lab in Milton Keynes | Wellcome Sanger Institute for the COVID-19 Genomics UK (COG-UK) Consortium | The Lighthouse Lab in Milton Keynes and Alex Alderton, Roberto Amato, Jeffrey Barrett, Sonia Goncalves, Ewan Harrison, David K. Jackson, Ian Johnston, Dominic Kwiatkowski, Cordelia Langford, John Sillitoe on behalf of the Wellcome Sanger Institute COVID-19 Surveillance Team |
| EPI_ISL_3334891 | Lighthouse Laboratory Plymouth | Wellcome Sanger Institute for the COVID-19 Genomics UK (COG-UK) Consortium | Lighthouse Laboratory Plymouth and Alex Alderton, Roberto Amato, Jeffrey Barrett, Sonia Goncalves, Ewan Harrison, David K. Jackson, Ian Johnston, Dominic Kwiatkowski, Cordelia Langford, John Sillitoe on behalf of the Wellcome Sanger Institute COVID-19 Surveillance Team |
| EPI_ISL_3334892, EPI_ISL_3334893, EPI_ISL_3334894, EPI_ISL_3334895, EPI_ISL_3334896 | Lighthouse Lab in Milton Keynes | Wellcome Sanger Institute for the COVID-19 Genomics UK (COG-UK) Consortium | The Lighthouse Lab in Milton Keynes and Alex Alderton, Roberto Amato, Jeffrey Barrett, Sonia Goncalves, Ewan Harrison, David K. Jackson, Ian Johnston, Dominic Kwiatkowski, Cordelia Langford, John Sillitoe on behalf of the Wellcome Sanger Institute COVID-19 Surveillance Team |
| EPI_ISL_3334897 | Berkshire and Surrey Pathology Services Lighthouse Laboratory | Wellcome Sanger Institute for the COVID-19 Genomics UK (COG-UK) Consortium | Berkshire and Surrey Pathology Services Lighthouse Laboratory and Alex Alderton, Roberto Amato, Jeffrey Barrett, Sonia Goncalves, Ewan Harrison, David K. Jackson, Ian Johnston, Dominic Kwiatkowski, Cordelia Langford, John Sillitoe on behalf of the Wellcome Sanger Institute COVID-19 Surveillance Team |
| EPI_ISL_3334898 | Lighthouse Lab in Milton Keynes | Wellcome Sanger Institute for the COVID-19 Genomics UK (COG-UK) Consortium | The Lighthouse Lab in Milton Keynes and Alex Alderton, Roberto Amato, Jeffrey Barrett, Sonia Goncalves, Ewan Harrison, David K. Jackson, Ian Johnston, Dominic Kwiatkowski, Cordelia Langford, John Sillitoe on behalf of the Wellcome Sanger Institute COVID-19 Surveillance Team |
| EPI_ISL_3334899 | Berkshire and Surrey Pathology Services Lighthouse Laboratory | Wellcome Sanger Institute for the COVID-19 Genomics UK (COG-UK) Consortium | Berkshire and Surrey Pathology Services Lighthouse Laboratory and Alex Alderton, Roberto Amato, Jeffrey Barrett, Sonia Goncalves, Ewan Harrison, David K. Jackson, Ian Johnston, Dominic Kwiatkowski, Cordelia Langford, John Sillitoe on behalf of the Wellcome Sanger Institute COVID-19 Surveillance Team |
| EPI_ISL_3334900, EPI_ISL_3334901, EPI_ISL_3334902, EPI_ISL_3334903 | Lighthouse Lab in Milton Keynes | Wellcome Sanger Institute for the COVID-19 Genomics UK (COG-UK) Consortium | The Lighthouse Lab in Milton Keynes and Alex Alderton, Roberto Amato, Jeffrey Barrett, Sonia Goncalves, Ewan Harrison, David K. Jackson, Ian Johnston, Dominic Kwiatkowski, Cordelia Langford, John Sillitoe on behalf of the Wellcome Sanger Institute COVID-19 Surveillance Team |
| EPI_ISL_3334904, EPI_ISL_3334905 | Berkshire and Surrey Pathology Services Lighthouse Laboratory | Wellcome Sanger Institute for the COVID-19 Genomics UK (COG-UK) Consortium | Berkshire and Surrey Pathology Services Lighthouse Laboratory and Alex Alderton, Roberto Amato, Jeffrey Barrett, Sonia Goncalves, Ewan Harrison, David K. Jackson, Ian Johnston, Dominic Kwiatkowski, Cordelia Langford, John Sillitoe on behalf of the Wellcome Sanger Institute COVID-19 Surveillance Team |
| EPI_ISL_3334906 | Lighthouse Laboratory Plymouth | Wellcome Sanger Institute for the COVID-19 Genomics UK (COG-UK) Consortium | Lighthouse Laboratory Plymouth and Alex Alderton, Roberto Amato, Jeffrey Barrett, Sonia Goncalves, Ewan Harrison, David K. Jackson, Ian Johnston, Dominic Kwiatkowski, Cordelia Langford, John Sillitoe on behalf of the Wellcome Sanger Institute COVID-19 Surveillance Team |

| | | | |
|---|---|---|---|
| EPI_ISL_3334907, EPI_ISL_3334908 | Lighthouse Lab in Milton Keynes | Wellcome Sanger Institute for the COVID-19 Genomics UK (COG-UK) Consortium | The Lighthouse Lab in Milton Keynes and Alex Alderton, Roberto Amato, Jeffrey Barrett, Sonia Goncalves, Ewan Harrison, David K. Jackson, Ian Johnston, Dominic Kwiatkowski, Cordelia Langford, John Sillitoe on behalf of the Wellcome Sanger Institute COVID-19 Surveillance Team |
| EPI_ISL_3334909, EPI_ISL_3334910 | Lighthouse Laboratory Plymouth | Wellcome Sanger Institute for the COVID-19 Genomics UK (COG-UK) Consortium | Lighthouse Laboratory Plymouth and Alex Alderton, Roberto Amato, Jeffrey Barrett, Sonia Goncalves, Ewan Harrison, David K. Jackson, Ian Johnston, Dominic Kwiatkowski, Cordelia Langford, John Sillitoe on behalf of the Wellcome Sanger Institute COVID-19 Surveillance Team |
| EPI_ISL_3334911 | Lighthouse Lab in Milton Keynes | Wellcome Sanger Institute for the COVID-19 Genomics UK (COG-UK) Consortium | The Lighthouse Lab in Milton Keynes and Alex Alderton, Roberto Amato, Jeffrey Barrett, Sonia Goncalves, Ewan Harrison, David K. Jackson, Ian Johnston, Dominic Kwiatkowski, Cordelia Langford, John Sillitoe on behalf of the Wellcome Sanger Institute COVID-19 Surveillance Team |
| EPI_ISL_3334912 | Berkshire and Surrey Pathology Services Lighthouse Laboratory | Wellcome Sanger Institute for the COVID-19 Genomics UK (COG-UK) Consortium | Berkshire and Surrey Pathology Services Lighthouse Laboratory and Alex Alderton, Roberto Amato, Jeffrey Barrett, Sonia Goncalves, Ewan Harrison, David K. Jackson, Ian Johnston, Dominic Kwiatkowski, Cordelia Langford, John Sillitoe on behalf of the Wellcome Sanger Institute COVID-19 Surveillance Team |
| EPI_ISL_3334913 | Lighthouse Laboratory Plymouth | Wellcome Sanger Institute for the COVID-19 Genomics UK (COG-UK) Consortium | Lighthouse Laboratory Plymouth and Alex Alderton, Roberto Amato, Jeffrey Barrett, Sonia Goncalves, Ewan Harrison, David K. Jackson, Ian Johnston, Dominic Kwiatkowski, Cordelia Langford, John Sillitoe on behalf of the Wellcome Sanger Institute COVID-19 Surveillance Team |
| EPI_ISL_3334914 | Berkshire and Surrey Pathology Services Lighthouse Laboratory | Wellcome Sanger Institute for the COVID-19 Genomics UK (COG-UK) Consortium | Berkshire and Surrey Pathology Services Lighthouse Laboratory and Alex Alderton, Roberto Amato, Jeffrey Barrett, Sonia Goncalves, Ewan Harrison, David K. Jackson, Ian Johnston, Dominic Kwiatkowski, Cordelia Langford, John Sillitoe on behalf of the Wellcome Sanger Institute COVID-19 Surveillance Team |
| EPI_ISL_3334915, EPI_ISL_3334916, EPI_ISL_3334917, EPI_ISL_3334918, EPI_ISL_3334919 | Lighthouse Lab in Milton Keynes | Wellcome Sanger Institute for the COVID-19 Genomics UK (COG-UK) Consortium | The Lighthouse Lab in Milton Keynes and Alex Alderton, Roberto Amato, Jeffrey Barrett, Sonia Goncalves, Ewan Harrison, David K. Jackson, Ian Johnston, Dominic Kwiatkowski, Cordelia Langford, John Sillitoe on behalf of the Wellcome Sanger Institute COVID-19 Surveillance Team |
| EPI_ISL_3334920 | Berkshire and Surrey Pathology Services Lighthouse Laboratory | Wellcome Sanger Institute for the COVID-19 Genomics UK (COG-UK) Consortium | Berkshire and Surrey Pathology Services Lighthouse Laboratory and Alex Alderton, Roberto Amato, Jeffrey Barrett, Sonia Goncalves, Ewan Harrison, David K. Jackson, Ian Johnston, Dominic Kwiatkowski, Cordelia Langford, John Sillitoe on behalf of the Wellcome Sanger Institute COVID-19 Surveillance Team |
| EPI_ISL_3334921 | Lighthouse Lab in Milton Keynes | Wellcome Sanger Institute for the COVID-19 Genomics UK (COG-UK) Consortium | The Lighthouse Lab in Milton Keynes and Alex Alderton, Roberto Amato, Jeffrey Barrett, Sonia Goncalves, Ewan Harrison, David K. Jackson, Ian Johnston, Dominic Kwiatkowski, Cordelia Langford, John Sillitoe on behalf of the Wellcome Sanger Institute COVID-19 Surveillance Team |
| EPI_ISL_3334922 | Lighthouse Laboratory Plymouth | Wellcome Sanger Institute for the COVID-19 Genomics UK (COG-UK) Consortium | Lighthouse Laboratory Plymouth and Alex Alderton, Roberto Amato, Jeffrey Barrett, Sonia Goncalves, Ewan Harrison, David K. Jackson, Ian Johnston, Dominic Kwiatkowski, Cordelia Langford, John Sillitoe on behalf of the Wellcome Sanger Institute COVID-19 Surveillance Team |
| EPI_ISL_3334923 | Lighthouse Lab in Milton Keynes | Wellcome Sanger Institute for the COVID-19 Genomics UK (COG-UK) Consortium | The Lighthouse Lab in Milton Keynes and Alex Alderton, Roberto Amato, Jeffrey Barrett, Sonia Goncalves, Ewan Harrison, David K. Jackson, Ian Johnston, Dominic Kwiatkowski, Cordelia Langford, John Sillitoe on behalf of the Wellcome Sanger Institute COVID-19 Surveillance Team |
| EPI_ISL_3334924 | Lighthouse Laboratory Plymouth | Wellcome Sanger Institute for the COVID-19 Genomics UK (COG-UK) Consortium | Lighthouse Laboratory Plymouth and Alex Alderton, Roberto Amato, Jeffrey Barrett, Sonia Goncalves, Ewan Harrison, David K. Jackson, Ian Johnston, Dominic Kwiatkowski, Cordelia Langford, John Sillitoe on behalf of the Wellcome Sanger Institute COVID-19 Surveillance Team |
| EPI_ISL_3334925 | Berkshire and Surrey Pathology Services Lighthouse Laboratory | Wellcome Sanger Institute for the COVID-19 Genomics UK (COG-UK) Consortium | Berkshire and Surrey Pathology Services Lighthouse Laboratory and Alex Alderton, Roberto Amato, Jeffrey Barrett, Sonia Goncalves, Ewan Harrison, David K. Jackson, Ian Johnston, Dominic Kwiatkowski, Cordelia Langford, John Sillitoe on behalf of the Wellcome Sanger Institute COVID-19 Surveillance Team |
| EPI_ISL_3334926 | Lighthouse Laboratory Plymouth | Wellcome Sanger Institute for the COVID-19 Genomics UK (COG-UK) Consortium | Lighthouse Laboratory Plymouth and Alex Alderton, Roberto Amato, Jeffrey Barrett, Sonia Goncalves, Ewan Harrison, David K. Jackson, Ian Johnston, Dominic Kwiatkowski, Cordelia Langford, John Sillitoe on behalf of the Wellcome Sanger Institute COVID-19 Surveillance Team |
| EPI_ISL_3334927 | Berkshire and Surrey Pathology Services Lighthouse Laboratory | Wellcome Sanger Institute for the COVID-19 Genomics UK (COG-UK) Consortium | Berkshire and Surrey Pathology Services Lighthouse Laboratory and Alex Alderton, Roberto Amato, Jeffrey Barrett, Sonia Goncalves, Ewan Harrison, David K. Jackson, Ian Johnston, Dominic Kwiatkowski, Cordelia Langford, John Sillitoe on behalf of the Wellcome Sanger Institute COVID-19 Surveillance Team |
| EPI_ISL_3334928, EPI_ISL_3334929, EPI_ISL_3334930, EPI_ISL_3334931 | Lighthouse Lab in Milton Keynes | Wellcome Sanger Institute for the COVID-19 Genomics UK (COG-UK) Consortium | The Lighthouse Lab in Milton Keynes and Alex Alderton, Roberto Amato, Jeffrey Barrett, Sonia Goncalves, Ewan Harrison, David K. Jackson, Ian Johnston, Dominic Kwiatkowski, Cordelia Langford, John Sillitoe on behalf of the Wellcome Sanger Institute COVID-19 Surveillance Team |
| EPI_ISL_3334932, EPI_ISL_3334933 | Berkshire and Surrey Pathology Services Lighthouse Laboratory | Wellcome Sanger Institute for the COVID-19 Genomics UK (COG-UK) Consortium | Berkshire and Surrey Pathology Services Lighthouse Laboratory and Alex Alderton, Roberto Amato, Jeffrey Barrett, Sonia Goncalves, Ewan Harrison, David K. Jackson, Ian Johnston, Dominic Kwiatkowski, Cordelia Langford, John Sillitoe on behalf of the Wellcome Sanger Institute COVID-19 Surveillance Team |
| EPI_ISL_3334934 | Lighthouse Lab in Milton Keynes | Wellcome Sanger Institute for the COVID-19 Genomics UK (COG-UK) Consortium | The Lighthouse Lab in Milton Keynes and Alex Alderton, Roberto Amato, Jeffrey Barrett, Sonia Goncalves, Ewan Harrison, David K. Jackson, Ian Johnston, Dominic Kwiatkowski, Cordelia Langford, John Sillitoe on behalf of the Wellcome Sanger Institute COVID-19 Surveillance Team |
| EPI_ISL_3334935 | Lighthouse Laboratory Plymouth | Wellcome Sanger Institute for the COVID-19 Genomics UK (COG-UK) Consortium | Lighthouse Laboratory Plymouth and Alex Alderton, Roberto Amato, Jeffrey Barrett, Sonia Goncalves, Ewan Harrison, David K. Jackson, Ian Johnston, Dominic Kwiatkowski, Cordelia Langford, John Sillitoe on behalf of the Wellcome Sanger Institute COVID-19 Surveillance Team |
| EPI_ISL_3334936 | Lighthouse Lab in Milton Keynes | Wellcome Sanger Institute for the COVID-19 Genomics UK (COG-UK) Consortium | The Lighthouse Lab in Milton Keynes and Alex Alderton, Roberto Amato, Jeffrey Barrett, Sonia Goncalves, Ewan Harrison, David K. Jackson, Ian Johnston, Dominic Kwiatkowski, Cordelia Langford, John Sillitoe on behalf of the Wellcome Sanger Institute COVID-19 Surveillance Team |
| EPI_ISL_3334937, EPI_ISL_3334938, EPI_ISL_3334939 | Lighthouse Laboratory Plymouth | Wellcome Sanger Institute for the COVID-19 Genomics UK (COG-UK) Consortium | Lighthouse Laboratory Plymouth and Alex Alderton, Roberto Amato, Jeffrey Barrett, Sonia Goncalves, Ewan Harrison, David K. Jackson, Ian Johnston, Dominic Kwiatkowski, Cordelia Langford, John Sillitoe on behalf of the Wellcome Sanger Institute COVID-19 Surveillance Team |
| EPI_ISL_3334940 | Lighthouse Lab in Milton Keynes | Wellcome Sanger Institute for the COVID-19 Genomics UK (COG-UK) Consortium | The Lighthouse Lab in Milton Keynes and Alex Alderton, Roberto Amato, Jeffrey Barrett, Sonia Goncalves, Ewan Harrison, David K. Jackson, Ian Johnston, Dominic Kwiatkowski, Cordelia Langford, John Sillitoe on behalf of the Wellcome Sanger Institute COVID-19 Surveillance Team |
| EPI_ISL_3334941 | Lighthouse Laboratory Plymouth | Wellcome Sanger Institute for the COVID-19 Genomics UK (COG-UK) Consortium | Lighthouse Laboratory Plymouth and Alex Alderton, Roberto Amato, Jeffrey Barrett, Sonia Goncalves, Ewan Harrison, David K. Jackson, Ian Johnston, Dominic Kwiatkowski, Cordelia Langford, John Sillitoe on behalf of the Wellcome Sanger Institute COVID-19 Surveillance Team |
| EPI_ISL_3334942 | Lighthouse Lab in Milton Keynes | Wellcome Sanger Institute for the COVID-19 Genomics UK (COG-UK) Consortium | The Lighthouse Lab in Milton Keynes and Alex Alderton, Roberto Amato, Jeffrey Barrett, Sonia Goncalves, Ewan Harrison, David K. Jackson, Ian Johnston, Dominic Kwiatkowski, Cordelia Langford, John Sillitoe on behalf of the Wellcome Sanger Institute COVID-19 Surveillance Team |
| EPI_ISL_3334943 | Berkshire and Surrey Pathology Services Lighthouse Laboratory | Wellcome Sanger Institute for the COVID-19 Genomics UK (COG-UK) Consortium | Berkshire and Surrey Pathology Services Lighthouse Laboratory and Alex Alderton, Roberto Amato, Jeffrey Barrett, Sonia Goncalves, Ewan Harrison, David K. Jackson, Ian Johnston, Dominic Kwiatkowski, Cordelia Langford, John Sillitoe on behalf of the Wellcome Sanger Institute COVID-19 Surveillance Team |
| EPI_ISL_3334944 | Lighthouse Lab in Milton Keynes | Wellcome Sanger Institute for the COVID-19 Genomics UK (COG-UK) Consortium | The Lighthouse Lab in Milton Keynes and Alex Alderton, Roberto Amato, Jeffrey Barrett, Sonia Goncalves, Ewan Harrison, David K. Jackson, Ian Johnston, Dominic Kwiatkowski, Cordelia Langford, John Sillitoe on behalf of the Wellcome Sanger Institute COVID-19 Surveillance Team |
| EPI_ISL_3334945 | Lighthouse Laboratory Plymouth | Wellcome Sanger Institute for the COVID-19 Genomics UK (COG-UK) Consortium | Lighthouse Laboratory Plymouth and Alex Alderton, Roberto Amato, Jeffrey Barrett, Sonia Goncalves, Ewan Harrison, David K. Jackson, Ian Johnston, Dominic Kwiatkowski, Cordelia Langford, John Sillitoe on behalf of the Wellcome Sanger Institute COVID-19 Surveillance Team |
| EPI_ISL_3334946, EPI_ISL_3334947, EPI_ISL_3334948 | Lighthouse Lab in Milton Keynes | Wellcome Sanger Institute for the COVID-19 Genomics UK (COG-UK) Consortium | The Lighthouse Lab in Milton Keynes and Alex Alderton, Roberto Amato, Jeffrey Barrett, Sonia Goncalves, Ewan Harrison, David K. Jackson, Ian Johnston, Dominic Kwiatkowski, Cordelia Langford, John Sillitoe on behalf of the Wellcome Sanger Institute COVID-19 Surveillance Team |
| EPI_ISL_3334949, EPI_ISL_3334950 | Lighthouse Laboratory Plymouth | Wellcome Sanger Institute for the COVID-19 Genomics UK (COG-UK) Consortium | Lighthouse Laboratory Plymouth and Alex Alderton, Roberto Amato, Jeffrey Barrett, Sonia Goncalves, Ewan Harrison, David K. Jackson, Ian Johnston, Dominic Kwiatkowski, Cordelia Langford, John Sillitoe on behalf of the Wellcome Sanger Institute COVID-19 Surveillance Team |
| EPI_ISL_3334951, EPI_ISL_3334952 | Lighthouse Lab in Milton Keynes | Wellcome Sanger Institute for the COVID-19 Genomics UK (COG-UK) Consortium | The Lighthouse Lab in Milton Keynes and Alex Alderton, Roberto Amato, Jeffrey Barrett, Sonia Goncalves, Ewan Harrison, David K. Jackson, Ian Johnston, Dominic Kwiatkowski, Cordelia Langford, John Sillitoe on behalf of the Wellcome Sanger Institute COVID-19 Surveillance Team |
| EPI_ISL_3334953 | Lighthouse Laboratory Plymouth | Wellcome Sanger Institute for the COVID-19 Genomics UK (COG-UK) Consortium | Lighthouse Laboratory Plymouth and Alex Alderton, Roberto Amato, Jeffrey Barrett, Sonia Goncalves, Ewan Harrison, David K. Jackson, Ian Johnston, Dominic Kwiatkowski, Cordelia Langford, John Sillitoe on behalf of the Wellcome Sanger Institute COVID-19 Surveillance Team |
| EPI_ISL_3334954, EPI_ISL_3334955, EPI_ISL_3334956, EPI_ISL_3334957 | Lighthouse Lab in Milton Keynes | Wellcome Sanger Institute for the COVID-19 Genomics UK (COG-UK) Consortium | The Lighthouse Lab in Milton Keynes and Alex Alderton, Roberto Amato, Jeffrey Barrett, Sonia Goncalves, Ewan Harrison, David K. Jackson, Ian Johnston, Dominic Kwiatkowski, Cordelia Langford, John Sillitoe on behalf of the Wellcome Sanger Institute COVID-19 Surveillance Team |
| EPI_ISL_3334958 | Lighthouse Laboratory Plymouth | Wellcome Sanger Institute for the COVID-19 Genomics UK (COG-UK) Consortium | Lighthouse Laboratory Plymouth and Alex Alderton, Roberto Amato, Jeffrey Barrett, Sonia Goncalves, Ewan Harrison, David K. Jackson, Ian Johnston, Dominic Kwiatkowski, Cordelia Langford, John Sillitoe on behalf of the Wellcome Sanger Institute COVID-19 Surveillance Team |
| EPI_ISL_3334959, EPI_ISL_3334960 | Lighthouse Lab in Milton Keynes | Wellcome Sanger Institute for the COVID-19 Genomics UK (COG-UK) Consortium | The Lighthouse Lab in Milton Keynes and Alex Alderton, Roberto Amato, Jeffrey Barrett, Sonia Goncalves, Ewan Harrison, David K. Jackson, Ian Johnston, Dominic Kwiatkowski, Cordelia Langford, John Sillitoe on behalf of the Wellcome Sanger Institute COVID-19 Surveillance Team |
| EPI_ISL_3334961, EPI_ISL_3334962 | Lighthouse Laboratory Plymouth | Wellcome Sanger Institute for the COVID-19 Genomics UK (COG-UK) Consortium | Lighthouse Laboratory Plymouth and Alex Alderton, Roberto Amato, Jeffrey Barrett, Sonia Goncalves, Ewan Harrison, David K. Jackson, Ian Johnston, Dominic Kwiatkowski, Cordelia Langford, John Sillitoe on behalf of the Wellcome Sanger Institute COVID-19 Surveillance Team |
| EPI_ISL_3334963 | Lighthouse Lab in Milton Keynes | Wellcome Sanger Institute for the COVID-19 Genomics UK (COG-UK) Consortium | The Lighthouse Lab in Milton Keynes and Alex Alderton, Roberto Amato, Jeffrey Barrett, Sonia Goncalves, Ewan Harrison, David K. Jackson, Ian Johnston, Dominic Kwiatkowski, Cordelia Langford, John Sillitoe on behalf of the Wellcome Sanger Institute COVID-19 Surveillance Team |
| EPI_ISL_3334964 | Lighthouse Laboratory Plymouth | Wellcome Sanger Institute for the COVID-19 Genomics UK (COG-UK) Consortium | Lighthouse Laboratory Plymouth and Alex Alderton, Roberto Amato, Jeffrey Barrett, Sonia Goncalves, Ewan Harrison, David K. Jackson, Ian Johnston, Dominic Kwiatkowski, Cordelia Langford, John Sillitoe on behalf of the Wellcome Sanger Institute COVID-19 Surveillance Team |
| EPI_ISL_3334965 | Lighthouse Lab in Milton Keynes | Wellcome Sanger Institute for the COVID-19 Genomics UK (COG-UK) Consortium | The Lighthouse Lab in Milton Keynes and Alex Alderton, Roberto Amato, Jeffrey Barrett, Sonia Goncalves, Ewan Harrison, David K. Jackson, Ian Johnston, Dominic Kwiatkowski, Cordelia Langford, John Sillitoe on behalf of the Wellcome Sanger Institute COVID-19 Surveillance Team |
| EPI_ISL_3334966 | Lighthouse Laboratory Plymouth | Wellcome Sanger Institute for the COVID-19 Genomics UK (COG-UK) Consortium | Lighthouse Laboratory Plymouth and Alex Alderton, Roberto Amato, Jeffrey Barrett, Sonia Goncalves, Ewan Harrison, David K. Jackson, Ian Johnston, Dominic Kwiatkowski, Cordelia Langford, John Sillitoe on behalf of the Wellcome Sanger Institute COVID-19 Surveillance Team |
| EPI_ISL_3334967, EPI_ISL_3334968, EPI_ISL_3334969, EPI_ISL_3334970, EPI_ISL_3334971 | Lighthouse Lab in Milton Keynes | Wellcome Sanger Institute for the COVID-19 Genomics UK (COG-UK) Consortium | The Lighthouse Lab in Milton Keynes and Alex Alderton, Roberto Amato, Jeffrey Barrett, Sonia Goncalves, Ewan Harrison, David K. Jackson, Ian Johnston, Dominic Kwiatkowski, Cordelia Langford, John Sillitoe on behalf of the Wellcome Sanger Institute COVID-19 Surveillance Team |

| | | | |
|---|---|---|---|
| EPI_ISL_3334972 | Lighthouse Laboratory Plymouth | Wellcome Sanger Institute for the COVID-19 Genomics UK (COG-UK) Consortium | Lighthouse Laboratory Plymouth and Alex Alderton, Roberto Amato, Jeffrey Barrett, Sonia Goncalves, Ewan Harrison, David K. Jackson, Ian Johnston, Dominic Kwiatkowski, Cordelia Langford, John Sillitoe on behalf of the Wellcome Sanger Institute COVID-19 Surveillance Team |
| EPI_ISL_3334973 | Lighthouse Lab in Milton Keynes | Wellcome Sanger Institute for the COVID-19 Genomics UK (COG-UK) Consortium | The Lighthouse Lab in Milton Keynes and Alex Alderton, Roberto Amato, Jeffrey Barrett, Sonia Goncalves, Ewan Harrison, David K. Jackson, Ian Johnston, Dominic Kwiatkowski, Cordelia Langford, John Sillitoe on behalf of the Wellcome Sanger Institute COVID-19 Surveillance Team |
| EPI_ISL_3334974 | Berkshire and Surrey Pathology Services Lighthouse Laboratory | Wellcome Sanger Institute for the COVID-19 Genomics UK (COG-UK) Consortium | Berkshire and Surrey Pathology Services Lighthouse Laboratory and Alex Alderton, Roberto Amato, Jeffrey Barrett, Sonia Goncalves, Ewan Harrison, David K. Jackson, Ian Johnston, Dominic Kwiatkowski, Cordelia Langford, John Sillitoe on behalf of the Wellcome Sanger Institute COVID-19 Surveillance Team |
| EPI_ISL_3334975 | Lighthouse Laboratory Plymouth | Wellcome Sanger Institute for the COVID-19 Genomics UK (COG-UK) Consortium | Lighthouse Laboratory Plymouth and Alex Alderton, Roberto Amato, Jeffrey Barrett, Sonia Goncalves, Ewan Harrison, David K. Jackson, Ian Johnston, Dominic Kwiatkowski, Cordelia Langford, John Sillitoe on behalf of the Wellcome Sanger Institute COVID-19 Surveillance Team |
| EPI_ISL_3334976 | Berkshire and Surrey Pathology Services Lighthouse Laboratory | Wellcome Sanger Institute for the COVID-19 Genomics UK (COG-UK) Consortium | Berkshire and Surrey Pathology Services Lighthouse Laboratory and Alex Alderton, Roberto Amato, Jeffrey Barrett, Sonia Goncalves, Ewan Harrison, David K. Jackson, Ian Johnston, Dominic Kwiatkowski, Cordelia Langford, John Sillitoe on behalf of the Wellcome Sanger Institute COVID-19 Surveillance Team |
| EPI_ISL_3334977 | Lighthouse Laboratory Plymouth | Wellcome Sanger Institute for the COVID-19 Genomics UK (COG-UK) Consortium | Lighthouse Laboratory Plymouth and Alex Alderton, Roberto Amato, Jeffrey Barrett, Sonia Goncalves, Ewan Harrison, David K. Jackson, Ian Johnston, Dominic Kwiatkowski, Cordelia Langford, John Sillitoe on behalf of the Wellcome Sanger Institute COVID-19 Surveillance Team |
| EPI_ISL_3334978 | Lighthouse Lab in Milton Keynes | Wellcome Sanger Institute for the COVID-19 Genomics UK (COG-UK) Consortium | The Lighthouse Lab in Milton Keynes and Alex Alderton, Roberto Amato, Jeffrey Barrett, Sonia Goncalves, Ewan Harrison, David K. Jackson, Ian Johnston, Dominic Kwiatkowski, Cordelia Langford, John Sillitoe on behalf of the Wellcome Sanger Institute COVID-19 Surveillance Team |
| EPI_ISL_3334979, EPI_ISL_3334980 | Lighthouse Laboratory Plymouth | Wellcome Sanger Institute for the COVID-19 Genomics UK (COG-UK) Consortium | Lighthouse Laboratory Plymouth and Alex Alderton, Roberto Amato, Jeffrey Barrett, Sonia Goncalves, Ewan Harrison, David K. Jackson, Ian Johnston, Dominic Kwiatkowski, Cordelia Langford, John Sillitoe on behalf of the Wellcome Sanger Institute COVID-19 Surveillance Team |
| EPI_ISL_3334981, EPI_ISL_3334982, EPI_ISL_3334983, EPI_ISL_3334984 | Lighthouse Lab in Milton Keynes | Wellcome Sanger Institute for the COVID-19 Genomics UK (COG-UK) Consortium | The Lighthouse Lab in Milton Keynes and Alex Alderton, Roberto Amato, Jeffrey Barrett, Sonia Goncalves, Ewan Harrison, David K. Jackson, Ian Johnston, Dominic Kwiatkowski, Cordelia Langford, John Sillitoe on behalf of the Wellcome Sanger Institute COVID-19 Surveillance Team |
| EPI_ISL_3334985 | Berkshire and Surrey Pathology Services Lighthouse Laboratory | Wellcome Sanger Institute for the COVID-19 Genomics UK (COG-UK) Consortium | Berkshire and Surrey Pathology Services Lighthouse Laboratory and Alex Alderton, Roberto Amato, Jeffrey Barrett, Sonia Goncalves, Ewan Harrison, David K. Jackson, Ian Johnston, Dominic Kwiatkowski, Cordelia Langford, John Sillitoe on behalf of the Wellcome Sanger Institute COVID-19 Surveillance Team |
| EPI_ISL_3334986, EPI_ISL_3334987, EPI_ISL_3334988 | Lighthouse Lab in Milton Keynes | Wellcome Sanger Institute for the COVID-19 Genomics UK (COG-UK) Consortium | The Lighthouse Lab in Milton Keynes and Alex Alderton, Roberto Amato, Jeffrey Barrett, Sonia Goncalves, Ewan Harrison, David K. Jackson, Ian Johnston, Dominic Kwiatkowski, Cordelia Langford, John Sillitoe on behalf of the Wellcome Sanger Institute COVID-19 Surveillance Team |
| EPI_ISL_3334989, EPI_ISL_3334990 | Lighthouse Laboratory Plymouth | Wellcome Sanger Institute for the COVID-19 Genomics UK (COG-UK) Consortium | Lighthouse Laboratory Plymouth and Alex Alderton, Roberto Amato, Jeffrey Barrett, Sonia Goncalves, Ewan Harrison, David K. Jackson, Ian Johnston, Dominic Kwiatkowski, Cordelia Langford, John Sillitoe on behalf of the Wellcome Sanger Institute COVID-19 Surveillance Team |
| EPI_ISL_3334991 | Lighthouse Lab in Milton Keynes | Wellcome Sanger Institute for the COVID-19 Genomics UK (COG-UK) Consortium | The Lighthouse Lab in Milton Keynes and Alex Alderton, Roberto Amato, Jeffrey Barrett, Sonia Goncalves, Ewan Harrison, David K. Jackson, Ian Johnston, Dominic Kwiatkowski, Cordelia Langford, John Sillitoe on behalf of the Wellcome Sanger Institute COVID-19 Surveillance Team |
| EPI_ISL_3334992, EPI_ISL_3334993 | Lighthouse Laboratory Plymouth | Wellcome Sanger Institute for the COVID-19 Genomics UK (COG-UK) Consortium | Lighthouse Laboratory Plymouth and Alex Alderton, Roberto Amato, Jeffrey Barrett, Sonia Goncalves, Ewan Harrison, David K. Jackson, Ian Johnston, Dominic Kwiatkowski, Cordelia Langford, John Sillitoe on behalf of the Wellcome Sanger Institute COVID-19 Surveillance Team |
| EPI_ISL_3334994, EPI_ISL_3334995 | Berkshire and Surrey Pathology Services Lighthouse Laboratory | Wellcome Sanger Institute for the COVID-19 Genomics UK (COG-UK) Consortium | Berkshire and Surrey Pathology Services Lighthouse Laboratory and Alex Alderton, Roberto Amato, Jeffrey Barrett, Sonia Goncalves, Ewan Harrison, David K. Jackson, Ian Johnston, Dominic Kwiatkowski, Cordelia Langford, John Sillitoe on behalf of the Wellcome Sanger Institute COVID-19 Surveillance Team |
| EPI_ISL_3334996 | Lighthouse Laboratory Plymouth | Wellcome Sanger Institute for the COVID-19 Genomics UK (COG-UK) Consortium | Lighthouse Laboratory Plymouth and Alex Alderton, Roberto Amato, Jeffrey Barrett, Sonia Goncalves, Ewan Harrison, David K. Jackson, Ian Johnston, Dominic Kwiatkowski, Cordelia Langford, John Sillitoe on behalf of the Wellcome Sanger Institute COVID-19 Surveillance Team |
| EPI_ISL_3334997 | Lighthouse Lab in Milton Keynes | Wellcome Sanger Institute for the COVID-19 Genomics UK (COG-UK) Consortium | The Lighthouse Lab in Milton Keynes and Alex Alderton, Roberto Amato, Jeffrey Barrett, Sonia Goncalves, Ewan Harrison, David K. Jackson, Ian Johnston, Dominic Kwiatkowski, Cordelia Langford, John Sillitoe on behalf of the Wellcome Sanger Institute COVID-19 Surveillance Team |
| EPI_ISL_3334998 | Lighthouse Laboratory Plymouth | Wellcome Sanger Institute for the COVID-19 Genomics UK (COG-UK) Consortium | Lighthouse Laboratory Plymouth and Alex Alderton, Roberto Amato, Jeffrey Barrett, Sonia Goncalves, Ewan Harrison, David K. Jackson, Ian Johnston, Dominic Kwiatkowski, Cordelia Langford, John Sillitoe on behalf of the Wellcome Sanger Institute COVID-19 Surveillance Team |
| EPI_ISL_3334999, EPI_ISL_3335000 | Lighthouse Lab in Milton Keynes | Wellcome Sanger Institute for the COVID-19 Genomics UK (COG-UK) Consortium | The Lighthouse Lab in Milton Keynes and Alex Alderton, Roberto Amato, Jeffrey Barrett, Sonia Goncalves, Ewan Harrison, David K. Jackson, Ian Johnston, Dominic Kwiatkowski, Cordelia Langford, John Sillitoe on behalf of the Wellcome Sanger Institute COVID-19 Surveillance Team |
| EPI_ISL_3335001, EPI_ISL_3335002, EPI_ISL_3335003, EPI_ISL_3335004 | Berkshire and Surrey Pathology Services Lighthouse Laboratory | Wellcome Sanger Institute for the COVID-19 Genomics UK (COG-UK) Consortium | Berkshire and Surrey Pathology Services Lighthouse Laboratory and Alex Alderton, Roberto Amato, Jeffrey Barrett, Sonia Goncalves, Ewan Harrison, David K. Jackson, Ian Johnston, Dominic Kwiatkowski, Cordelia Langford, John Sillitoe on behalf of the Wellcome Sanger Institute COVID-19 Surveillance Team |
| EPI_ISL_3335005, EPI_ISL_3335006 | Lighthouse Lab in Milton Keynes | Wellcome Sanger Institute for the COVID-19 Genomics UK (COG-UK) Consortium | The Lighthouse Lab in Milton Keynes and Alex Alderton, Roberto Amato, Jeffrey Barrett, Sonia Goncalves, Ewan Harrison, David K. Jackson, Ian Johnston, Dominic Kwiatkowski, Cordelia Langford, John Sillitoe on behalf of the Wellcome Sanger Institute COVID-19 Surveillance Team |
| EPI_ISL_3335007 | Berkshire and Surrey Pathology Services Lighthouse Laboratory | Wellcome Sanger Institute for the COVID-19 Genomics UK (COG-UK) Consortium | Berkshire and Surrey Pathology Services Lighthouse Laboratory and Alex Alderton, Roberto Amato, Jeffrey Barrett, Sonia Goncalves, Ewan Harrison, David K. Jackson, Ian Johnston, Dominic Kwiatkowski, Cordelia Langford, John Sillitoe on behalf of the Wellcome Sanger Institute COVID-19 Surveillance Team |
| EPI_ISL_3335008, EPI_ISL_3335009 | Lighthouse Lab in Milton Keynes | Wellcome Sanger Institute for the COVID-19 Genomics UK (COG-UK) Consortium | The Lighthouse Lab in Milton Keynes and Alex Alderton, Roberto Amato, Jeffrey Barrett, Sonia Goncalves, Ewan Harrison, David K. Jackson, Ian Johnston, Dominic Kwiatkowski, Cordelia Langford, John Sillitoe on behalf of the Wellcome Sanger Institute COVID-19 Surveillance Team |
| EPI_ISL_3335010 | Lighthouse Laboratory Plymouth | Wellcome Sanger Institute for the COVID-19 Genomics UK (COG-UK) Consortium | Lighthouse Laboratory Plymouth and Alex Alderton, Roberto Amato, Jeffrey Barrett, Sonia Goncalves, Ewan Harrison, David K. Jackson, Ian Johnston, Dominic Kwiatkowski, Cordelia Langford, John Sillitoe on behalf of the Wellcome Sanger Institute COVID-19 Surveillance Team |
| EPI_ISL_3335011, EPI_ISL_3335012 | Lighthouse Lab in Milton Keynes | Wellcome Sanger Institute for the COVID-19 Genomics UK (COG-UK) Consortium | The Lighthouse Lab in Milton Keynes and Alex Alderton, Roberto Amato, Jeffrey Barrett, Sonia Goncalves, Ewan Harrison, David K. Jackson, Ian Johnston, Dominic Kwiatkowski, Cordelia Langford, John Sillitoe on behalf of the Wellcome Sanger Institute COVID-19 Surveillance Team |
| EPI_ISL_3335013, EPI_ISL_3335014 | Berkshire and Surrey Pathology Services Lighthouse Laboratory | Wellcome Sanger Institute for the COVID-19 Genomics UK (COG-UK) Consortium | Berkshire and Surrey Pathology Services Lighthouse Laboratory and Alex Alderton, Roberto Amato, Jeffrey Barrett, Sonia Goncalves, Ewan Harrison, David K. Jackson, Ian Johnston, Dominic Kwiatkowski, Cordelia Langford, John Sillitoe on behalf of the Wellcome Sanger Institute COVID-19 Surveillance Team |
| EPI_ISL_3335015 | Lighthouse Laboratory Plymouth | Wellcome Sanger Institute for the COVID-19 Genomics UK (COG-UK) Consortium | Lighthouse Laboratory Plymouth and Alex Alderton, Roberto Amato, Jeffrey Barrett, Sonia Goncalves, Ewan Harrison, David K. Jackson, Ian Johnston, Dominic Kwiatkowski, Cordelia Langford, John Sillitoe on behalf of the Wellcome Sanger Institute COVID-19 Surveillance Team |
| EPI_ISL_3335016, EPI_ISL_3335017, EPI_ISL_3335018 | Lighthouse Lab in Milton Keynes | Wellcome Sanger Institute for the COVID-19 Genomics UK (COG-UK) Consortium | The Lighthouse Lab in Milton Keynes and Alex Alderton, Roberto Amato, Jeffrey Barrett, Sonia Goncalves, Ewan Harrison, David K. Jackson, Ian Johnston, Dominic Kwiatkowski, Cordelia Langford, John Sillitoe on behalf of the Wellcome Sanger Institute COVID-19 Surveillance Team |
| EPI_ISL_3335019 | Lighthouse Laboratory Plymouth | Wellcome Sanger Institute for the COVID-19 Genomics UK (COG-UK) Consortium | Lighthouse Laboratory Plymouth and Alex Alderton, Roberto Amato, Jeffrey Barrett, Sonia Goncalves, Ewan Harrison, David K. Jackson, Ian Johnston, Dominic Kwiatkowski, Cordelia Langford, John Sillitoe on behalf of the Wellcome Sanger Institute COVID-19 Surveillance Team |
| EPI_ISL_3335020, EPI_ISL_3335021 | Lighthouse Lab in Milton Keynes | Wellcome Sanger Institute for the COVID-19 Genomics UK (COG-UK) Consortium | The Lighthouse Lab in Milton Keynes and Alex Alderton, Roberto Amato, Jeffrey Barrett, Sonia Goncalves, Ewan Harrison, David K. Jackson, Ian Johnston, Dominic Kwiatkowski, Cordelia Langford, John Sillitoe on behalf of the Wellcome Sanger Institute COVID-19 Surveillance Team |
| EPI_ISL_3335022, EPI_ISL_3335023, EPI_ISL_3335024 | Lighthouse Laboratory Plymouth | Wellcome Sanger Institute for the COVID-19 Genomics UK (COG-UK) Consortium | Lighthouse Laboratory Plymouth and Alex Alderton, Roberto Amato, Jeffrey Barrett, Sonia Goncalves, Ewan Harrison, David K. Jackson, Ian Johnston, Dominic Kwiatkowski, Cordelia Langford, John Sillitoe on behalf of the Wellcome Sanger Institute COVID-19 Surveillance Team |
| EPI_ISL_3335025 | Berkshire and Surrey Pathology Services Lighthouse Laboratory | Wellcome Sanger Institute for the COVID-19 Genomics UK (COG-UK) Consortium | Berkshire and Surrey Pathology Services Lighthouse Laboratory and Alex Alderton, Roberto Amato, Jeffrey Barrett, Sonia Goncalves, Ewan Harrison, David K. Jackson, Ian Johnston, Dominic Kwiatkowski, Cordelia Langford, John Sillitoe on behalf of the Wellcome Sanger Institute COVID-19 Surveillance Team |
| EPI_ISL_3335026, EPI_ISL_3335027 | Lighthouse Lab in Milton Keynes | Wellcome Sanger Institute for the COVID-19 Genomics UK (COG-UK) Consortium | The Lighthouse Lab in Milton Keynes and Alex Alderton, Roberto Amato, Jeffrey Barrett, Sonia Goncalves, Ewan Harrison, David K. Jackson, Ian Johnston, Dominic Kwiatkowski, Cordelia Langford, John Sillitoe on behalf of the Wellcome Sanger Institute COVID-19 Surveillance Team |
| EPI_ISL_3335028 | Lighthouse Laboratory Plymouth | Wellcome Sanger Institute for the COVID-19 Genomics UK (COG-UK) Consortium | Lighthouse Laboratory Plymouth and Alex Alderton, Roberto Amato, Jeffrey Barrett, Sonia Goncalves, Ewan Harrison, David K. Jackson, Ian Johnston, Dominic Kwiatkowski, Cordelia Langford, John Sillitoe on behalf of the Wellcome Sanger Institute COVID-19 Surveillance Team |
| EPI_ISL_3335029 | Berkshire and Surrey Pathology Services Lighthouse Laboratory | Wellcome Sanger Institute for the COVID-19 Genomics UK (COG-UK) Consortium | Berkshire and Surrey Pathology Services Lighthouse Laboratory and Alex Alderton, Roberto Amato, Jeffrey Barrett, Sonia Goncalves, Ewan Harrison, David K. Jackson, Ian Johnston, Dominic Kwiatkowski, Cordelia Langford, John Sillitoe on behalf of the Wellcome Sanger Institute COVID-19 Surveillance Team |
| EPI_ISL_3335030, EPI_ISL_3335031 | Lighthouse Laboratory Plymouth | Wellcome Sanger Institute for the COVID-19 Genomics UK (COG-UK) Consortium | Lighthouse Laboratory Plymouth and Alex Alderton, Roberto Amato, Jeffrey Barrett, Sonia Goncalves, Ewan Harrison, David K. Jackson, Ian Johnston, Dominic Kwiatkowski, Cordelia Langford, John Sillitoe on behalf of the Wellcome Sanger Institute COVID-19 Surveillance Team |
| EPI_ISL_3335032 | Lighthouse Lab in Milton Keynes | Wellcome Sanger Institute for the COVID-19 Genomics UK (COG-UK) Consortium | The Lighthouse Lab in Milton Keynes and Alex Alderton, Roberto Amato, Jeffrey Barrett, Sonia Goncalves, Ewan Harrison, David K. Jackson, Ian Johnston, Dominic Kwiatkowski, Cordelia Langford, John Sillitoe on behalf of the Wellcome Sanger Institute COVID-19 Surveillance Team |
| EPI_ISL_3335033, EPI_ISL_3335034 | Lighthouse Laboratory Plymouth | Wellcome Sanger Institute for the COVID-19 Genomics UK (COG-UK) Consortium | Lighthouse Laboratory Plymouth and Alex Alderton, Roberto Amato, Jeffrey Barrett, Sonia Goncalves, Ewan Harrison, David K. Jackson, Ian Johnston, Dominic Kwiatkowski, Cordelia Langford, John Sillitoe on behalf of the Wellcome Sanger Institute COVID-19 Surveillance Team |
| EPI_ISL_3335035 | Lighthouse Lab in Milton Keynes | Wellcome Sanger Institute for the COVID-19 Genomics UK (COG-UK) Consortium | The Lighthouse Lab in Milton Keynes and Alex Alderton, Roberto Amato, Jeffrey Barrett, Sonia Goncalves, Ewan Harrison, David K. Jackson, Ian Johnston, Dominic Kwiatkowski, Cordelia Langford, John Sillitoe on behalf of the Wellcome Sanger Institute COVID-19 Surveillance Team |
| EPI_ISL_3335036 | Lighthouse Laboratory Plymouth | Wellcome Sanger Institute for the COVID-19 Genomics UK (COG-UK) Consortium | Lighthouse Laboratory Plymouth and Alex Alderton, Roberto Amato, Jeffrey Barrett, Sonia Goncalves, Ewan Harrison, David K. Jackson, Ian Johnston, Dominic Kwiatkowski, Cordelia Langford, John Sillitoe on behalf of the Wellcome Sanger Institute COVID-19 Surveillance Team |
| EPI_ISL_3335037, EPI_ISL_3335038 | Lighthouse Lab in Milton Keynes | Wellcome Sanger Institute for the COVID-19 Genomics UK (COG-UK) Consortium | The Lighthouse Lab in Milton Keynes and Alex Alderton, Roberto Amato, Jeffrey Barrett, Sonia Goncalves, Ewan Harrison, David K. Jackson, Ian Johnston, Dominic Kwiatkowski, Cordelia Langford, John Sillitoe on behalf of the Wellcome Sanger Institute COVID-19 Surveillance Team |
| EPI_ISL_3335039 | Lighthouse Laboratory Plymouth | Wellcome Sanger Institute for the COVID-19 Genomics UK (COG-UK) Consortium | Lighthouse Laboratory Plymouth and Alex Alderton, Roberto Amato, Jeffrey Barrett, Sonia Goncalves, Ewan Harrison, David K. Jackson, Ian Johnston, Dominic Kwiatkowski, Cordelia Langford, John Sillitoe on behalf of the Wellcome Sanger Institute COVID-19 Surveillance Team |
| EPI_ISL_3335040, | Lighthouse Lab in Milton Keynes | Wellcome Sanger Institute for the COVID-19 Genomics UK (COG-UK) | The Lighthouse Lab in Milton Keynes and Alex Alderton, Roberto Amato, Jeffrey Barrett, Sonia Goncalves, Ewan Harrison, David K. Jackson, Ian Johnston, Dominic Kwiatkowski, Cordelia Langford, John Sillitoe on behalf of the Wellcome Sanger |

| | | | |
|---|---|---|---|
| EPI_ISL_3335041, EPI_ISL_3335042, EPI_ISL_3335043 | | Consortium | Institute COVID-19 Surveillance Team |
| EPI_ISL_3335044 | Berkshire and Surrey Pathology Services Lighthouse Laboratory | Wellcome Sanger Institute for the COVID-19 Genomics UK (COG-UK) Consortium | Berkshire and Surrey Pathology Services Lighthouse Laboratory and Alex Alderton, Roberto Amato, Jeffrey Barrett, Sonia Goncalves, Ewan Harrison, David K. Jackson, Ian Johnston, Dominic Kwiatkowski, Cordelia Langford, John Sillitoe on behalf of the Wellcome Sanger Institute COVID-19 Surveillance Team |
| EPI_ISL_3335045 | Lighthouse Lab in Milton Keynes | Wellcome Sanger Institute for the COVID-19 Genomics UK (COG-UK) Consortium | The Lighthouse Lab in Milton Keynes and Alex Alderton, Roberto Amato, Jeffrey Barrett, Sonia Goncalves, Ewan Harrison, David K. Jackson, Ian Johnston, Dominic Kwiatkowski, Cordelia Langford, John Sillitoe on behalf of the Wellcome Sanger Institute COVID-19 Surveillance Team |
| EPI_ISL_3335046, EPI_ISL_3335047 | Lighthouse Laboratory Plymouth | Wellcome Sanger Institute for the COVID-19 Genomics UK (COG-UK) Consortium | Lighthouse Laboratory Plymouth and Alex Alderton, Roberto Amato, Jeffrey Barrett, Sonia Goncalves, Ewan Harrison, David K. Jackson, Ian Johnston, Dominic Kwiatkowski, Cordelia Langford, John Sillitoe on behalf of the Wellcome Sanger Institute COVID-19 Surveillance Team |
| EPI_ISL_3335048, EPI_ISL_3335049 | Lighthouse Lab in Milton Keynes | Wellcome Sanger Institute for the COVID-19 Genomics UK (COG-UK) Consortium | The Lighthouse Lab in Milton Keynes and Alex Alderton, Roberto Amato, Jeffrey Barrett, Sonia Goncalves, Ewan Harrison, David K. Jackson, Ian Johnston, Dominic Kwiatkowski, Cordelia Langford, John Sillitoe on behalf of the Wellcome Sanger Institute COVID-19 Surveillance Team |
| EPI_ISL_3335050 | Lighthouse Laboratory Plymouth | Wellcome Sanger Institute for the COVID-19 Genomics UK (COG-UK) Consortium | Lighthouse Laboratory Plymouth and Alex Alderton, Roberto Amato, Jeffrey Barrett, Sonia Goncalves, Ewan Harrison, David K. Jackson, Ian Johnston, Dominic Kwiatkowski, Cordelia Langford, John Sillitoe on behalf of the Wellcome Sanger Institute COVID-19 Surveillance Team |
| EPI_ISL_3335051 | Lighthouse Lab in Milton Keynes | Wellcome Sanger Institute for the COVID-19 Genomics UK (COG-UK) Consortium | The Lighthouse Lab in Milton Keynes and Alex Alderton, Roberto Amato, Jeffrey Barrett, Sonia Goncalves, Ewan Harrison, David K. Jackson, Ian Johnston, Dominic Kwiatkowski, Cordelia Langford, John Sillitoe on behalf of the Wellcome Sanger Institute COVID-19 Surveillance Team |
| EPI_ISL_3335052 | Lighthouse Laboratory Plymouth | Wellcome Sanger Institute for the COVID-19 Genomics UK (COG-UK) Consortium | Lighthouse Laboratory Plymouth and Alex Alderton, Roberto Amato, Jeffrey Barrett, Sonia Goncalves, Ewan Harrison, David K. Jackson, Ian Johnston, Dominic Kwiatkowski, Cordelia Langford, John Sillitoe on behalf of the Wellcome Sanger Institute COVID-19 Surveillance Team |
| EPI_ISL_3335053, EPI_ISL_3335054 | Lighthouse Lab in Milton Keynes | Wellcome Sanger Institute for the COVID-19 Genomics UK (COG-UK) Consortium | The Lighthouse Lab in Milton Keynes and Alex Alderton, Roberto Amato, Jeffrey Barrett, Sonia Goncalves, Ewan Harrison, David K. Jackson, Ian Johnston, Dominic Kwiatkowski, Cordelia Langford, John Sillitoe on behalf of the Wellcome Sanger Institute COVID-19 Surveillance Team |
| EPI_ISL_3335055 | Berkshire and Surrey Pathology Services Lighthouse Laboratory | Wellcome Sanger Institute for the COVID-19 Genomics UK (COG-UK) Consortium | Berkshire and Surrey Pathology Services Lighthouse Laboratory and Alex Alderton, Roberto Amato, Jeffrey Barrett, Sonia Goncalves, Ewan Harrison, David K. Jackson, Ian Johnston, Dominic Kwiatkowski, Cordelia Langford, John Sillitoe on behalf of the Wellcome Sanger Institute COVID-19 Surveillance Team |
| EPI_ISL_3335056, EPI_ISL_3335057, EPI_ISL_3335058, EPI_ISL_3335059 | Lighthouse Lab in Milton Keynes | Wellcome Sanger Institute for the COVID-19 Genomics UK (COG-UK) Consortium | The Lighthouse Lab in Milton Keynes and Alex Alderton, Roberto Amato, Jeffrey Barrett, Sonia Goncalves, Ewan Harrison, David K. Jackson, Ian Johnston, Dominic Kwiatkowski, Cordelia Langford, John Sillitoe on behalf of the Wellcome Sanger Institute COVID-19 Surveillance Team |
| EPI_ISL_3335060, EPI_ISL_3335061 | Berkshire and Surrey Pathology Services Lighthouse Laboratory | Wellcome Sanger Institute for the COVID-19 Genomics UK (COG-UK) Consortium | Berkshire and Surrey Pathology Services Lighthouse Laboratory and Alex Alderton, Roberto Amato, Jeffrey Barrett, Sonia Goncalves, Ewan Harrison, David K. Jackson, Ian Johnston, Dominic Kwiatkowski, Cordelia Langford, John Sillitoe on behalf of the Wellcome Sanger Institute COVID-19 Surveillance Team |
| EPI_ISL_3335062, EPI_ISL_3335063 | Lighthouse Lab in Milton Keynes | Wellcome Sanger Institute for the COVID-19 Genomics UK (COG-UK) Consortium | The Lighthouse Lab in Milton Keynes and Alex Alderton, Roberto Amato, Jeffrey Barrett, Sonia Goncalves, Ewan Harrison, David K. Jackson, Ian Johnston, Dominic Kwiatkowski, Cordelia Langford, John Sillitoe on behalf of the Wellcome Sanger Institute COVID-19 Surveillance Team |
| EPI_ISL_3335064 | Lighthouse Laboratory Plymouth | Wellcome Sanger Institute for the COVID-19 Genomics UK (COG-UK) Consortium | Lighthouse Laboratory Plymouth and Alex Alderton, Roberto Amato, Jeffrey Barrett, Sonia Goncalves, Ewan Harrison, David K. Jackson, Ian Johnston, Dominic Kwiatkowski, Cordelia Langford, John Sillitoe on behalf of the Wellcome Sanger Institute COVID-19 Surveillance Team |
| EPI_ISL_3335065, EPI_ISL_3335066, EPI_ISL_3335067, EPI_ISL_3335068, EPI_ISL_3335069, EPI_ISL_3335070, EPI_ISL_3335071 | Lighthouse Lab in Milton Keynes | Wellcome Sanger Institute for the COVID-19 Genomics UK (COG-UK) Consortium | The Lighthouse Lab in Milton Keynes and Alex Alderton, Roberto Amato, Jeffrey Barrett, Sonia Goncalves, Ewan Harrison, David K. Jackson, Ian Johnston, Dominic Kwiatkowski, Cordelia Langford, John Sillitoe on behalf of the Wellcome Sanger Institute COVID-19 Surveillance Team |
| EPI_ISL_3335072 | Lighthouse Laboratory Plymouth | Wellcome Sanger Institute for the COVID-19 Genomics UK (COG-UK) Consortium | Lighthouse Laboratory Plymouth and Alex Alderton, Roberto Amato, Jeffrey Barrett, Sonia Goncalves, Ewan Harrison, David K. Jackson, Ian Johnston, Dominic Kwiatkowski, Cordelia Langford, John Sillitoe on behalf of the Wellcome Sanger Institute COVID-19 Surveillance Team |
| EPI_ISL_3335073, EPI_ISL_3335074, EPI_ISL_3335075, EPI_ISL_3335076 | Lighthouse Lab in Milton Keynes | Wellcome Sanger Institute for the COVID-19 Genomics UK (COG-UK) Consortium | The Lighthouse Lab in Milton Keynes and Alex Alderton, Roberto Amato, Jeffrey Barrett, Sonia Goncalves, Ewan Harrison, David K. Jackson, Ian Johnston, Dominic Kwiatkowski, Cordelia Langford, John Sillitoe on behalf of the Wellcome Sanger Institute COVID-19 Surveillance Team |
| EPI_ISL_3335077 | Lighthouse Laboratory Plymouth | Wellcome Sanger Institute for the COVID-19 Genomics UK (COG-UK) Consortium | Lighthouse Laboratory Plymouth and Alex Alderton, Roberto Amato, Jeffrey Barrett, Sonia Goncalves, Ewan Harrison, David K. Jackson, Ian Johnston, Dominic Kwiatkowski, Cordelia Langford, John Sillitoe on behalf of the Wellcome Sanger Institute COVID-19 Surveillance Team |
| EPI_ISL_3335078, EPI_ISL_3335079, EPI_ISL_3335080 | Lighthouse Lab in Milton Keynes | Wellcome Sanger Institute for the COVID-19 Genomics UK (COG-UK) Consortium | The Lighthouse Lab in Milton Keynes and Alex Alderton, Roberto Amato, Jeffrey Barrett, Sonia Goncalves, Ewan Harrison, David K. Jackson, Ian Johnston, Dominic Kwiatkowski, Cordelia Langford, John Sillitoe on behalf of the Wellcome Sanger Institute COVID-19 Surveillance Team |
| EPI_ISL_3335081, EPI_ISL_3335082 | Lighthouse Laboratory Plymouth | Wellcome Sanger Institute for the COVID-19 Genomics UK (COG-UK) Consortium | Lighthouse Laboratory Plymouth and Alex Alderton, Roberto Amato, Jeffrey Barrett, Sonia Goncalves, Ewan Harrison, David K. Jackson, Ian Johnston, Dominic Kwiatkowski, Cordelia Langford, John Sillitoe on behalf of the Wellcome Sanger Institute COVID-19 Surveillance Team |
| EPI_ISL_3335083 | Lighthouse Lab in Milton Keynes | Wellcome Sanger Institute for the COVID-19 Genomics UK (COG-UK) Consortium | The Lighthouse Lab in Milton Keynes and Alex Alderton, Roberto Amato, Jeffrey Barrett, Sonia Goncalves, Ewan Harrison, David K. Jackson, Ian Johnston, Dominic Kwiatkowski, Cordelia Langford, John Sillitoe on behalf of the Wellcome Sanger Institute COVID-19 Surveillance Team |
| EPI_ISL_3335084 | Lighthouse Laboratory Plymouth | Wellcome Sanger Institute for the COVID-19 Genomics UK (COG-UK) Consortium | Lighthouse Laboratory Plymouth and Alex Alderton, Roberto Amato, Jeffrey Barrett, Sonia Goncalves, Ewan Harrison, David K. Jackson, Ian Johnston, Dominic Kwiatkowski, Cordelia Langford, John Sillitoe on behalf of the Wellcome Sanger Institute COVID-19 Surveillance Team |
| EPI_ISL_3335085, EPI_ISL_3335086, EPI_ISL_3335087, EPI_ISL_3335088, EPI_ISL_3335089 | Lighthouse Lab in Milton Keynes | Wellcome Sanger Institute for the COVID-19 Genomics UK (COG-UK) Consortium | The Lighthouse Lab in Milton Keynes and Alex Alderton, Roberto Amato, Jeffrey Barrett, Sonia Goncalves, Ewan Harrison, David K. Jackson, Ian Johnston, Dominic Kwiatkowski, Cordelia Langford, John Sillitoe on behalf of the Wellcome Sanger Institute COVID-19 Surveillance Team |
| EPI_ISL_3335090 | Lighthouse Laboratory Plymouth | Wellcome Sanger Institute for the COVID-19 Genomics UK (COG-UK) Consortium | Lighthouse Laboratory Plymouth and Alex Alderton, Roberto Amato, Jeffrey Barrett, Sonia Goncalves, Ewan Harrison, David K. Jackson, Ian Johnston, Dominic Kwiatkowski, Cordelia Langford, John Sillitoe on behalf of the Wellcome Sanger Institute COVID-19 Surveillance Team |
| EPI_ISL_3335091 | Lighthouse Lab in Milton Keynes | Wellcome Sanger Institute for the COVID-19 Genomics UK (COG-UK) Consortium | The Lighthouse Lab in Milton Keynes and Alex Alderton, Roberto Amato, Jeffrey Barrett, Sonia Goncalves, Ewan Harrison, David K. Jackson, Ian Johnston, Dominic Kwiatkowski, Cordelia Langford, John Sillitoe on behalf of the Wellcome Sanger Institute COVID-19 Surveillance Team |
| EPI_ISL_3335092 | Lighthouse Laboratory Plymouth | Wellcome Sanger Institute for the COVID-19 Genomics UK (COG-UK) Consortium | Lighthouse Laboratory Plymouth and Alex Alderton, Roberto Amato, Jeffrey Barrett, Sonia Goncalves, Ewan Harrison, David K. Jackson, Ian Johnston, Dominic Kwiatkowski, Cordelia Langford, John Sillitoe on behalf of the Wellcome Sanger Institute COVID-19 Surveillance Team |
| EPI_ISL_3335093 | Lighthouse Lab in Milton Keynes | Wellcome Sanger Institute for the COVID-19 Genomics UK (COG-UK) Consortium | The Lighthouse Lab in Milton Keynes and Alex Alderton, Roberto Amato, Jeffrey Barrett, Sonia Goncalves, Ewan Harrison, David K. Jackson, Ian Johnston, Dominic Kwiatkowski, Cordelia Langford, John Sillitoe on behalf of the Wellcome Sanger Institute COVID-19 Surveillance Team |
| EPI_ISL_3335094 | Lighthouse Laboratory Plymouth | Wellcome Sanger Institute for the COVID-19 Genomics UK (COG-UK) Consortium | Lighthouse Laboratory Plymouth and Alex Alderton, Roberto Amato, Jeffrey Barrett, Sonia Goncalves, Ewan Harrison, David K. Jackson, Ian Johnston, Dominic Kwiatkowski, Cordelia Langford, John Sillitoe on behalf of the Wellcome Sanger Institute COVID-19 Surveillance Team |
| EPI_ISL_3335095, EPI_ISL_3335096 | Lighthouse Lab in Milton Keynes | Wellcome Sanger Institute for the COVID-19 Genomics UK (COG-UK) Consortium | The Lighthouse Lab in Milton Keynes and Alex Alderton, Roberto Amato, Jeffrey Barrett, Sonia Goncalves, Ewan Harrison, David K. Jackson, Ian Johnston, Dominic Kwiatkowski, Cordelia Langford, John Sillitoe on behalf of the Wellcome Sanger Institute COVID-19 Surveillance Team |
| EPI_ISL_3335097 | Berkshire and Surrey Pathology Services Lighthouse Laboratory | Wellcome Sanger Institute for the COVID-19 Genomics UK (COG-UK) Consortium | Berkshire and Surrey Pathology Services Lighthouse Laboratory and Alex Alderton, Roberto Amato, Jeffrey Barrett, Sonia Goncalves, Ewan Harrison, David K. Jackson, Ian Johnston, Dominic Kwiatkowski, Cordelia Langford, John Sillitoe on behalf of the Wellcome Sanger Institute COVID-19 Surveillance Team |
| EPI_ISL_3335098 | Lighthouse Laboratory Plymouth | Wellcome Sanger Institute for the COVID-19 Genomics UK (COG-UK) Consortium | Lighthouse Laboratory Plymouth and Alex Alderton, Roberto Amato, Jeffrey Barrett, Sonia Goncalves, Ewan Harrison, David K. Jackson, Ian Johnston, Dominic Kwiatkowski, Cordelia Langford, John Sillitoe on behalf of the Wellcome Sanger Institute COVID-19 Surveillance Team |
| EPI_ISL_3335099 | Lighthouse Lab in Milton Keynes | Wellcome Sanger Institute for the COVID-19 Genomics UK (COG-UK) Consortium | The Lighthouse Lab in Milton Keynes and Alex Alderton, Roberto Amato, Jeffrey Barrett, Sonia Goncalves, Ewan Harrison, David K. Jackson, Ian Johnston, Dominic Kwiatkowski, Cordelia Langford, John Sillitoe on behalf of the Wellcome Sanger Institute COVID-19 Surveillance Team |
| EPI_ISL_3335100 | Lighthouse Laboratory Plymouth | Wellcome Sanger Institute for the COVID-19 Genomics UK (COG-UK) Consortium | Lighthouse Laboratory Plymouth and Alex Alderton, Roberto Amato, Jeffrey Barrett, Sonia Goncalves, Ewan Harrison, David K. Jackson, Ian Johnston, Dominic Kwiatkowski, Cordelia Langford, John Sillitoe on behalf of the Wellcome Sanger Institute COVID-19 Surveillance Team |
| EPI_ISL_3335101 | Lighthouse Lab in Milton Keynes | Wellcome Sanger Institute for the COVID-19 Genomics UK (COG-UK) Consortium | The Lighthouse Lab in Milton Keynes and Alex Alderton, Roberto Amato, Jeffrey Barrett, Sonia Goncalves, Ewan Harrison, David K. Jackson, Ian Johnston, Dominic Kwiatkowski, Cordelia Langford, John Sillitoe on behalf of the Wellcome Sanger Institute COVID-19 Surveillance Team |
| EPI_ISL_3335102 | Lighthouse Laboratory Plymouth | Wellcome Sanger Institute for the COVID-19 Genomics UK (COG-UK) Consortium | Lighthouse Laboratory Plymouth and Alex Alderton, Roberto Amato, Jeffrey Barrett, Sonia Goncalves, Ewan Harrison, David K. Jackson, Ian Johnston, Dominic Kwiatkowski, Cordelia Langford, John Sillitoe on behalf of the Wellcome Sanger Institute COVID-19 Surveillance Team |
| EPI_ISL_3335103 | Berkshire and Surrey Pathology Services Lighthouse Laboratory | Wellcome Sanger Institute for the COVID-19 Genomics UK (COG-UK) Consortium | Berkshire and Surrey Pathology Services Lighthouse Laboratory and Alex Alderton, Roberto Amato, Jeffrey Barrett, Sonia Goncalves, Ewan Harrison, David K. Jackson, Ian Johnston, Dominic Kwiatkowski, Cordelia Langford, John Sillitoe on behalf of the Wellcome Sanger Institute COVID-19 Surveillance Team |
| EPI_ISL_3335104 | Lighthouse Lab in Milton Keynes | Wellcome Sanger Institute for the COVID-19 Genomics UK (COG-UK) Consortium | The Lighthouse Lab in Milton Keynes and Alex Alderton, Roberto Amato, Jeffrey Barrett, Sonia Goncalves, Ewan Harrison, David K. Jackson, Ian Johnston, Dominic Kwiatkowski, Cordelia Langford, John Sillitoe on behalf of the Wellcome Sanger Institute COVID-19 Surveillance Team |
| EPI_ISL_3335105 | Lighthouse Laboratory Plymouth | Wellcome Sanger Institute for the COVID-19 Genomics UK (COG-UK) Consortium | Lighthouse Laboratory Plymouth and Alex Alderton, Roberto Amato, Jeffrey Barrett, Sonia Goncalves, Ewan Harrison, David K. Jackson, Ian Johnston, Dominic Kwiatkowski, Cordelia Langford, John Sillitoe on behalf of the Wellcome Sanger Institute COVID-19 Surveillance Team |
| EPI_ISL_3335106 | Berkshire and Surrey Pathology Services Lighthouse Laboratory | Wellcome Sanger Institute for the COVID-19 Genomics UK (COG-UK) Consortium | Berkshire and Surrey Pathology Services Lighthouse Laboratory and Alex Alderton, Roberto Amato, Jeffrey Barrett, Sonia Goncalves, Ewan Harrison, David K. Jackson, Ian Johnston, Dominic Kwiatkowski, Cordelia Langford, John Sillitoe on behalf of the Wellcome Sanger Institute COVID-19 Surveillance Team |

| | | | |
|---|---|---|---|
| EPI_ISL_3335107 | Lighthouse Lab in Milton Keynes | Wellcome Sanger Institute for the COVID-19 Genomics UK (COG-UK) Consortium | The Lighthouse Lab in Milton Keynes and Alex Alderton, Roberto Amato, Jeffrey Barrett, Sonia Goncalves, Ewan Harrison, David K. Jackson, Ian Johnston, Dominic Kwiatkowski, Cordelia Langford, John Sillitoe on behalf of the Wellcome Sanger Institute COVID-19 Surveillance Team |
| EPI_ISL_3335108 | Berkshire and Surrey Pathology Services Lighthouse Laboratory | Wellcome Sanger Institute for the COVID-19 Genomics UK (COG-UK) Consortium | Berkshire and Surrey Pathology Services Lighthouse Laboratory and Alex Alderton, Roberto Amato, Jeffrey Barrett, Sonia Goncalves, Ewan Harrison, David K. Jackson, Ian Johnston, Dominic Kwiatkowski, Cordelia Langford, John Sillitoe on behalf of the Wellcome Sanger Institute COVID-19 Surveillance Team |
| EPI_ISL_3335109 | Lighthouse Lab in Milton Keynes | Wellcome Sanger Institute for the COVID-19 Genomics UK (COG-UK) Consortium | The Lighthouse Lab in Milton Keynes and Alex Alderton, Roberto Amato, Jeffrey Barrett, Sonia Goncalves, Ewan Harrison, David K. Jackson, Ian Johnston, Dominic Kwiatkowski, Cordelia Langford, John Sillitoe on behalf of the Wellcome Sanger Institute COVID-19 Surveillance Team |
| EPI_ISL_3335110 | Berkshire and Surrey Pathology Services Lighthouse Laboratory | Wellcome Sanger Institute for the COVID-19 Genomics UK (COG-UK) Consortium | Berkshire and Surrey Pathology Services Lighthouse Laboratory and Alex Alderton, Roberto Amato, Jeffrey Barrett, Sonia Goncalves, Ewan Harrison, David K. Jackson, Ian Johnston, Dominic Kwiatkowski, Cordelia Langford, John Sillitoe on behalf of the Wellcome Sanger Institute COVID-19 Surveillance Team |
| EPI_ISL_3335111 | Lighthouse Lab in Milton Keynes | Wellcome Sanger Institute for the COVID-19 Genomics UK (COG-UK) Consortium | The Lighthouse Lab in Milton Keynes and Alex Alderton, Roberto Amato, Jeffrey Barrett, Sonia Goncalves, Ewan Harrison, David K. Jackson, Ian Johnston, Dominic Kwiatkowski, Cordelia Langford, John Sillitoe on behalf of the Wellcome Sanger Institute COVID-19 Surveillance Team |
| EPI_ISL_3335112 | Berkshire and Surrey Pathology Services Lighthouse Laboratory | Wellcome Sanger Institute for the COVID-19 Genomics UK (COG-UK) Consortium | Berkshire and Surrey Pathology Services Lighthouse Laboratory and Alex Alderton, Roberto Amato, Jeffrey Barrett, Sonia Goncalves, Ewan Harrison, David K. Jackson, Ian Johnston, Dominic Kwiatkowski, Cordelia Langford, John Sillitoe on behalf of the Wellcome Sanger Institute COVID-19 Surveillance Team |
| EPI_ISL_3335113, EPI_ISL_3335114 | Lighthouse Laboratory Plymouth | Wellcome Sanger Institute for the COVID-19 Genomics UK (COG-UK) Consortium | Lighthouse Laboratory Plymouth and Alex Alderton, Roberto Amato, Jeffrey Barrett, Sonia Goncalves, Ewan Harrison, David K. Jackson, Ian Johnston, Dominic Kwiatkowski, Cordelia Langford, John Sillitoe on behalf of the Wellcome Sanger Institute COVID-19 Surveillance Team |
| EPI_ISL_3335115, EPI_ISL_3335116 | Lighthouse Lab in Milton Keynes | Wellcome Sanger Institute for the COVID-19 Genomics UK (COG-UK) Consortium | The Lighthouse Lab in Milton Keynes and Alex Alderton, Roberto Amato, Jeffrey Barrett, Sonia Goncalves, Ewan Harrison, David K. Jackson, Ian Johnston, Dominic Kwiatkowski, Cordelia Langford, John Sillitoe on behalf of the Wellcome Sanger Institute COVID-19 Surveillance Team |
| EPI_ISL_3335117, EPI_ISL_3335118, EPI_ISL_3335119 | Lighthouse Laboratory Plymouth | Wellcome Sanger Institute for the COVID-19 Genomics UK (COG-UK) Consortium | Lighthouse Laboratory Plymouth and Alex Alderton, Roberto Amato, Jeffrey Barrett, Sonia Goncalves, Ewan Harrison, David K. Jackson, Ian Johnston, Dominic Kwiatkowski, Cordelia Langford, John Sillitoe on behalf of the Wellcome Sanger Institute COVID-19 Surveillance Team |
| EPI_ISL_3335120, EPI_ISL_3335121 | Lighthouse Lab in Milton Keynes | Wellcome Sanger Institute for the COVID-19 Genomics UK (COG-UK) Consortium | The Lighthouse Lab in Milton Keynes and Alex Alderton, Roberto Amato, Jeffrey Barrett, Sonia Goncalves, Ewan Harrison, David K. Jackson, Ian Johnston, Dominic Kwiatkowski, Cordelia Langford, John Sillitoe on behalf of the Wellcome Sanger Institute COVID-19 Surveillance Team |
| EPI_ISL_3335122, EPI_ISL_3335123, EPI_ISL_3335124 | Berkshire and Surrey Pathology Services Lighthouse Laboratory | Wellcome Sanger Institute for the COVID-19 Genomics UK (COG-UK) Consortium | Berkshire and Surrey Pathology Services Lighthouse Laboratory and Alex Alderton, Roberto Amato, Jeffrey Barrett, Sonia Goncalves, Ewan Harrison, David K. Jackson, Ian Johnston, Dominic Kwiatkowski, Cordelia Langford, John Sillitoe on behalf of the Wellcome Sanger Institute COVID-19 Surveillance Team |
| EPI_ISL_3335125 | Lighthouse Lab in Milton Keynes | Wellcome Sanger Institute for the COVID-19 Genomics UK (COG-UK) Consortium | The Lighthouse Lab in Milton Keynes and Alex Alderton, Roberto Amato, Jeffrey Barrett, Sonia Goncalves, Ewan Harrison, David K. Jackson, Ian Johnston, Dominic Kwiatkowski, Cordelia Langford, John Sillitoe on behalf of the Wellcome Sanger Institute COVID-19 Surveillance Team |
| EPI_ISL_3335126 | Lighthouse Laboratory Plymouth | Wellcome Sanger Institute for the COVID-19 Genomics UK (COG-UK) Consortium | Lighthouse Laboratory Plymouth and Alex Alderton, Roberto Amato, Jeffrey Barrett, Sonia Goncalves, Ewan Harrison, David K. Jackson, Ian Johnston, Dominic Kwiatkowski, Cordelia Langford, John Sillitoe on behalf of the Wellcome Sanger Institute COVID-19 Surveillance Team |
| EPI_ISL_3335127, EPI_ISL_3335128, EPI_ISL_3335129, EPI_ISL_3335130 | Lighthouse Lab in Milton Keynes | Wellcome Sanger Institute for the COVID-19 Genomics UK (COG-UK) Consortium | The Lighthouse Lab in Milton Keynes and Alex Alderton, Roberto Amato, Jeffrey Barrett, Sonia Goncalves, Ewan Harrison, David K. Jackson, Ian Johnston, Dominic Kwiatkowski, Cordelia Langford, John Sillitoe on behalf of the Wellcome Sanger Institute COVID-19 Surveillance Team |
| EPI_ISL_3335131 | Lighthouse Laboratory Plymouth | Wellcome Sanger Institute for the COVID-19 Genomics UK (COG-UK) Consortium | Lighthouse Laboratory Plymouth and Alex Alderton, Roberto Amato, Jeffrey Barrett, Sonia Goncalves, Ewan Harrison, David K. Jackson, Ian Johnston, Dominic Kwiatkowski, Cordelia Langford, John Sillitoe on behalf of the Wellcome Sanger Institute COVID-19 Surveillance Team |
| EPI_ISL_3335132 | Lighthouse Lab in Alderley Park | Wellcome Sanger Institute for the COVID-19 Genomics UK (COG-UK) Consortium | Jacquelyn Wynn, Mairead Hyland, The Lighthouse Lab in Alderley Park and Alex Alderton, Roberto Amato, Jeffrey Barrett, Sonia Goncalves, Ewan Harrison, David K. Jackson, Ian Johnston, Dominic Kwiatkowski, Cordelia Langford, John Sillitoe on behalf of the Wellcome Sanger Institute COVID-19 Surveillance Team |
| EPI_ISL_3335133, EPI_ISL_3335134 | Lighthouse Lab in Milton Keynes | Wellcome Sanger Institute for the COVID-19 Genomics UK (COG-UK) Consortium | The Lighthouse Lab in Milton Keynes and Alex Alderton, Roberto Amato, Jeffrey Barrett, Sonia Goncalves, Ewan Harrison, David K. Jackson, Ian Johnston, Dominic Kwiatkowski, Cordelia Langford, John Sillitoe on behalf of the Wellcome Sanger Institute COVID-19 Surveillance Team |
| EPI_ISL_3335135 | Health Services Laboratories | Wellcome Sanger Institute for the COVID-19 Genomics UK (COG-UK) Consortium | Health Services Laboratories and Alex Alderton, Roberto Amato, Jeffrey Barrett, Sonia Goncalves, Ewan Harrison, David K. Jackson, Ian Johnston, Dominic Kwiatkowski, Cordelia Langford, John Sillitoe on behalf of the Wellcome Sanger Institute COVID-19 Surveillance Team |
| EPI_ISL_3335136 | Lighthouse Lab in Milton Keynes | Wellcome Sanger Institute for the COVID-19 Genomics UK (COG-UK) Consortium | The Lighthouse Lab in Milton Keynes and Alex Alderton, Roberto Amato, Jeffrey Barrett, Sonia Goncalves, Ewan Harrison, David K. Jackson, Ian Johnston, Dominic Kwiatkowski, Cordelia Langford, John Sillitoe on behalf of the Wellcome Sanger Institute COVID-19 Surveillance Team |
| EPI_ISL_3335137 | Lighthouse Lab in Alderley Park | Wellcome Sanger Institute for the COVID-19 Genomics UK (COG-UK) Consortium | Jacquelyn Wynn, Mairead Hyland, The Lighthouse Lab in Alderley Park and Alex Alderton, Roberto Amato, Jeffrey Barrett, Sonia Goncalves, Ewan Harrison, David K. Jackson, Ian Johnston, Dominic Kwiatkowski, Cordelia Langford, John Sillitoe on behalf of the Wellcome Sanger Institute COVID-19 Surveillance Team |
| EPI_ISL_3335138 | Lighthouse Lab in Milton Keynes | Wellcome Sanger Institute for the COVID-19 Genomics UK (COG-UK) Consortium | The Lighthouse Lab in Milton Keynes and Alex Alderton, Roberto Amato, Jeffrey Barrett, Sonia Goncalves, Ewan Harrison, David K. Jackson, Ian Johnston, Dominic Kwiatkowski, Cordelia Langford, John Sillitoe on behalf of the Wellcome Sanger Institute COVID-19 Surveillance Team |
| EPI_ISL_3335139 | Lighthouse Lab in Alderley Park | Wellcome Sanger Institute for the COVID-19 Genomics UK (COG-UK) Consortium | Jacquelyn Wynn, Mairead Hyland, The Lighthouse Lab in Alderley Park and Alex Alderton, Roberto Amato, Jeffrey Barrett, Sonia Goncalves, Ewan Harrison, David K. Jackson, Ian Johnston, Dominic Kwiatkowski, Cordelia Langford, John Sillitoe on behalf of the Wellcome Sanger Institute COVID-19 Surveillance Team |
| EPI_ISL_3335140 | Lighthouse Lab in Milton Keynes | Wellcome Sanger Institute for the COVID-19 Genomics UK (COG-UK) Consortium | The Lighthouse Lab in Milton Keynes and Alex Alderton, Roberto Amato, Jeffrey Barrett, Sonia Goncalves, Ewan Harrison, David K. Jackson, Ian Johnston, Dominic Kwiatkowski, Cordelia Langford, John Sillitoe on behalf of the Wellcome Sanger Institute COVID-19 Surveillance Team |
| EPI_ISL_3335141 | Lighthouse Laboratory Plymouth | Wellcome Sanger Institute for the COVID-19 Genomics UK (COG-UK) Consortium | Lighthouse Laboratory Plymouth and Alex Alderton, Roberto Amato, Jeffrey Barrett, Sonia Goncalves, Ewan Harrison, David K. Jackson, Ian Johnston, Dominic Kwiatkowski, Cordelia Langford, John Sillitoe on behalf of the Wellcome Sanger Institute COVID-19 Surveillance Team |
| EPI_ISL_3335142 | Lighthouse Lab in Alderley Park | Wellcome Sanger Institute for the COVID-19 Genomics UK (COG-UK) Consortium | Jacquelyn Wynn, Mairead Hyland, The Lighthouse Lab in Alderley Park and Alex Alderton, Roberto Amato, Jeffrey Barrett, Sonia Goncalves, Ewan Harrison, David K. Jackson, Ian Johnston, Dominic Kwiatkowski, Cordelia Langford, John Sillitoe on behalf of the Wellcome Sanger Institute COVID-19 Surveillance Team |
| EPI_ISL_3335143, EPI_ISL_3335144 | Lighthouse Laboratory Plymouth | Wellcome Sanger Institute for the COVID-19 Genomics UK (COG-UK) Consortium | Lighthouse Laboratory Plymouth and Alex Alderton, Roberto Amato, Jeffrey Barrett, Sonia Goncalves, Ewan Harrison, David K. Jackson, Ian Johnston, Dominic Kwiatkowski, Cordelia Langford, John Sillitoe on behalf of the Wellcome Sanger Institute COVID-19 Surveillance Team |
| EPI_ISL_3335145 | Lighthouse Lab in Alderley Park | Wellcome Sanger Institute for the COVID-19 Genomics UK (COG-UK) Consortium | Jacquelyn Wynn, Mairead Hyland, The Lighthouse Lab in Alderley Park and Alex Alderton, Roberto Amato, Jeffrey Barrett, Sonia Goncalves, Ewan Harrison, David K. Jackson, Ian Johnston, Dominic Kwiatkowski, Cordelia Langford, John Sillitoe on behalf of the Wellcome Sanger Institute COVID-19 Surveillance Team |
| EPI_ISL_3335146 | Lighthouse Laboratory Plymouth | Wellcome Sanger Institute for the COVID-19 Genomics UK (COG-UK) Consortium | Lighthouse Laboratory Plymouth and Alex Alderton, Roberto Amato, Jeffrey Barrett, Sonia Goncalves, Ewan Harrison, David K. Jackson, Ian Johnston, Dominic Kwiatkowski, Cordelia Langford, John Sillitoe on behalf of the Wellcome Sanger Institute COVID-19 Surveillance Team |
| EPI_ISL_3335147, EPI_ISL_3335148 | Health Services Laboratories | Wellcome Sanger Institute for the COVID-19 Genomics UK (COG-UK) Consortium | Health Services Laboratories and Alex Alderton, Roberto Amato, Jeffrey Barrett, Sonia Goncalves, Ewan Harrison, David K. Jackson, Ian Johnston, Dominic Kwiatkowski, Cordelia Langford, John Sillitoe on behalf of the Wellcome Sanger Institute COVID-19 Surveillance Team |
| EPI_ISL_3335149 | Lighthouse Lab in Milton Keynes | Wellcome Sanger Institute for the COVID-19 Genomics UK (COG-UK) Consortium | The Lighthouse Lab in Milton Keynes and Alex Alderton, Roberto Amato, Jeffrey Barrett, Sonia Goncalves, Ewan Harrison, David K. Jackson, Ian Johnston, Dominic Kwiatkowski, Cordelia Langford, John Sillitoe on behalf of the Wellcome Sanger Institute COVID-19 Surveillance Team |
| EPI_ISL_3335150 | Lighthouse Lab in Alderley Park | Wellcome Sanger Institute for the COVID-19 Genomics UK (COG-UK) Consortium | Jacquelyn Wynn, Mairead Hyland, The Lighthouse Lab in Alderley Park and Alex Alderton, Roberto Amato, Jeffrey Barrett, Sonia Goncalves, Ewan Harrison, David K. Jackson, Ian Johnston, Dominic Kwiatkowski, Cordelia Langford, John Sillitoe on behalf of the Wellcome Sanger Institute COVID-19 Surveillance Team |
| EPI_ISL_3335151, EPI_ISL_3335152 | Lighthouse Laboratory Plymouth | Wellcome Sanger Institute for the COVID-19 Genomics UK (COG-UK) Consortium | Lighthouse Laboratory Plymouth and Alex Alderton, Roberto Amato, Jeffrey Barrett, Sonia Goncalves, Ewan Harrison, David K. Jackson, Ian Johnston, Dominic Kwiatkowski, Cordelia Langford, John Sillitoe on behalf of the Wellcome Sanger Institute COVID-19 Surveillance Team |
| EPI_ISL_3335153 | Health Services Laboratories | Wellcome Sanger Institute for the COVID-19 Genomics UK (COG-UK) Consortium | Health Services Laboratories and Alex Alderton, Roberto Amato, Jeffrey Barrett, Sonia Goncalves, Ewan Harrison, David K. Jackson, Ian Johnston, Dominic Kwiatkowski, Cordelia Langford, John Sillitoe on behalf of the Wellcome Sanger Institute COVID-19 Surveillance Team |
| EPI_ISL_3335154, EPI_ISL_3335155 | Lighthouse Lab in Milton Keynes | Wellcome Sanger Institute for the COVID-19 Genomics UK (COG-UK) Consortium | The Lighthouse Lab in Milton Keynes and Alex Alderton, Roberto Amato, Jeffrey Barrett, Sonia Goncalves, Ewan Harrison, David K. Jackson, Ian Johnston, Dominic Kwiatkowski, Cordelia Langford, John Sillitoe on behalf of the Wellcome Sanger Institute COVID-19 Surveillance Team |
| EPI_ISL_3335156, EPI_ISL_3335157 | Health Services Laboratories | Wellcome Sanger Institute for the COVID-19 Genomics UK (COG-UK) Consortium | Health Services Laboratories and Alex Alderton, Roberto Amato, Jeffrey Barrett, Sonia Goncalves, Ewan Harrison, David K. Jackson, Ian Johnston, Dominic Kwiatkowski, Cordelia Langford, John Sillitoe on behalf of the Wellcome Sanger Institute COVID-19 Surveillance Team |
| EPI_ISL_3335158, EPI_ISL_3335159 | Lighthouse Lab in Milton Keynes | Wellcome Sanger Institute for the COVID-19 Genomics UK (COG-UK) Consortium | The Lighthouse Lab in Milton Keynes and Alex Alderton, Roberto Amato, Jeffrey Barrett, Sonia Goncalves, Ewan Harrison, David K. Jackson, Ian Johnston, Dominic Kwiatkowski, Cordelia Langford, John Sillitoe on behalf of the Wellcome Sanger Institute COVID-19 Surveillance Team |
| EPI_ISL_3335160 | Health Services Laboratories | Wellcome Sanger Institute for the COVID-19 Genomics UK (COG-UK) Consortium | Health Services Laboratories and Alex Alderton, Roberto Amato, Jeffrey Barrett, Sonia Goncalves, Ewan Harrison, David K. Jackson, Ian Johnston, Dominic Kwiatkowski, Cordelia Langford, John Sillitoe on behalf of the Wellcome Sanger Institute COVID-19 Surveillance Team |
| EPI_ISL_3335161 | Lighthouse Laboratory Plymouth | Wellcome Sanger Institute for the COVID-19 Genomics UK (COG-UK) Consortium | Lighthouse Laboratory Plymouth and Alex Alderton, Roberto Amato, Jeffrey Barrett, Sonia Goncalves, Ewan Harrison, David K. Jackson, Ian Johnston, Dominic Kwiatkowski, Cordelia Langford, John Sillitoe on behalf of the Wellcome Sanger Institute COVID-19 Surveillance Team |
| EPI_ISL_3335162 | Lighthouse Lab in Alderley Park | Wellcome Sanger Institute for the COVID-19 Genomics UK (COG-UK) Consortium | Jacquelyn Wynn, Mairead Hyland, The Lighthouse Lab in Alderley Park and Alex Alderton, Roberto Amato, Jeffrey Barrett, Sonia Goncalves, Ewan Harrison, David K. Jackson, Ian Johnston, Dominic Kwiatkowski, Cordelia Langford, John Sillitoe on behalf of the Wellcome Sanger Institute COVID-19 Surveillance Team |
| EPI_ISL_3335163, EPI_ISL_3335164, EPI_ISL_3335165, EPI_ISL_3335166 | Lighthouse Laboratory Plymouth | Wellcome Sanger Institute for the COVID-19 Genomics UK (COG-UK) Consortium | Lighthouse Laboratory Plymouth and Alex Alderton, Roberto Amato, Jeffrey Barrett, Sonia Goncalves, Ewan Harrison, David K. Jackson, Ian Johnston, Dominic Kwiatkowski, Cordelia Langford, John Sillitoe on behalf of the Wellcome Sanger Institute COVID-19 Surveillance Team |
| EPI_ISL_3335167, EPI_ISL_3335168 | Lighthouse Lab in Milton Keynes | Wellcome Sanger Institute for the COVID-19 Genomics UK (COG-UK) Consortium | The Lighthouse Lab in Milton Keynes and Alex Alderton, Roberto Amato, Jeffrey Barrett, Sonia Goncalves, Ewan Harrison, David K. Jackson, Ian Johnston, Dominic Kwiatkowski, Cordelia Langford, John Sillitoe on behalf of the Wellcome Sanger Institute COVID-19 Surveillance Team |
| EPI_ISL_3335169 | Lighthouse Lab in Alderley Park | Wellcome Sanger Institute for the COVID-19 Genomics UK (COG-UK) Consortium | Jacquelyn Wynn, Mairead Hyland, The Lighthouse Lab in Alderley Park and Alex Alderton, Roberto Amato, Jeffrey Barrett, Sonia Goncalves, Ewan Harrison, David K. Jackson, Ian Johnston, Dominic Kwiatkowski, Cordelia Langford, John Sillitoe on behalf of the Wellcome Sanger Institute COVID-19 Surveillance Team |
| EPI_ISL_3335170, | Lighthouse Lab in Milton Keynes | Wellcome Sanger Institute for the COVID-19 Genomics UK (COG-UK) | The Lighthouse Lab in Milton Keynes and Alex Alderton, Roberto Amato, Jeffrey Barrett, Sonia Goncalves, Ewan Harrison, David K. Jackson, Ian Johnston, Dominic Kwiatkowski, Cordelia Langford, John Sillitoe on behalf of the Wellcome Sanger |

| | | | |
|---|---|---|---|
| EPI_ISL_3335171 | | Consortium | Institute COVID-19 Surveillance Team |
| EPI_ISL_3335172, EPI_ISL_3335173 | Health Services Laboratories | Wellcome Sanger Institute for the COVID-19 Genomics UK (COG-UK) Consortium | Health Services Laboratories and Alex Alderton, Roberto Amato, Jeffrey Barrett, Sonia Goncalves, Ewan Harrison, David K. Jackson, Ian Johnston, Dominic Kwiatkowski, Cordelia Langford, John Sillitoe on behalf of the Wellcome Sanger Institute COVID-19 Surveillance Team |
| EPI_ISL_3335174 | Lighthouse Lab in Alderley Park | Wellcome Sanger Institute for the COVID-19 Genomics UK (COG-UK) Consortium | Jacquelyn Wynn, Mairead Hyland, The Lighthouse Lab in Alderley Park and Alex Alderton, Roberto Amato, Jeffrey Barrett, Sonia Goncalves, Ewan Harrison, David K. Jackson, Ian Johnston, Dominic Kwiatkowski, Cordelia Langford, John Sillitoe on behalf of the Wellcome Sanger Institute COVID-19 Surveillance Team |
| EPI_ISL_3335175 | Lighthouse Laboratory Plymouth | Wellcome Sanger Institute for the COVID-19 Genomics UK (COG-UK) Consortium | Lighthouse Laboratory Plymouth and Alex Alderton, Roberto Amato, Jeffrey Barrett, Sonia Goncalves, Ewan Harrison, David K. Jackson, Ian Johnston, Dominic Kwiatkowski, Cordelia Langford, John Sillitoe on behalf of the Wellcome Sanger Institute COVID-19 Surveillance Team |
| EPI_ISL_3335176 | Health Services Laboratories | Wellcome Sanger Institute for the COVID-19 Genomics UK (COG-UK) Consortium | Health Services Laboratories and Alex Alderton, Roberto Amato, Jeffrey Barrett, Sonia Goncalves, Ewan Harrison, David K. Jackson, Ian Johnston, Dominic Kwiatkowski, Cordelia Langford, John Sillitoe on behalf of the Wellcome Sanger Institute COVID-19 Surveillance Team |
| EPI_ISL_3335177 | Lighthouse Lab in Milton Keynes | Wellcome Sanger Institute for the COVID-19 Genomics UK (COG-UK) Consortium | The Lighthouse Lab in Milton Keynes and Alex Alderton, Roberto Amato, Jeffrey Barrett, Sonia Goncalves, Ewan Harrison, David K. Jackson, Ian Johnston, Dominic Kwiatkowski, Cordelia Langford, John Sillitoe on behalf of the Wellcome Sanger Institute COVID-19 Surveillance Team |
| EPI_ISL_3335178 | Lighthouse Laboratory Plymouth | Wellcome Sanger Institute for the COVID-19 Genomics UK (COG-UK) Consortium | Lighthouse Laboratory Plymouth and Alex Alderton, Roberto Amato, Jeffrey Barrett, Sonia Goncalves, Ewan Harrison, David K. Jackson, Ian Johnston, Dominic Kwiatkowski, Cordelia Langford, John Sillitoe on behalf of the Wellcome Sanger Institute COVID-19 Surveillance Team |
| EPI_ISL_3335179, EPI_ISL_3335180 | Lighthouse Lab in Milton Keynes | Wellcome Sanger Institute for the COVID-19 Genomics UK (COG-UK) Consortium | The Lighthouse Lab in Milton Keynes and Alex Alderton, Roberto Amato, Jeffrey Barrett, Sonia Goncalves, Ewan Harrison, David K. Jackson, Ian Johnston, Dominic Kwiatkowski, Cordelia Langford, John Sillitoe on behalf of the Wellcome Sanger Institute COVID-19 Surveillance Team |
| EPI_ISL_3335181 | Health Services Laboratories | Wellcome Sanger Institute for the COVID-19 Genomics UK (COG-UK) Consortium | Health Services Laboratories and Alex Alderton, Roberto Amato, Jeffrey Barrett, Sonia Goncalves, Ewan Harrison, David K. Jackson, Ian Johnston, Dominic Kwiatkowski, Cordelia Langford, John Sillitoe on behalf of the Wellcome Sanger Institute COVID-19 Surveillance Team |
| EPI_ISL_3335182 | Lighthouse Lab in Milton Keynes | Wellcome Sanger Institute for the COVID-19 Genomics UK (COG-UK) Consortium | The Lighthouse Lab in Milton Keynes and Alex Alderton, Roberto Amato, Jeffrey Barrett, Sonia Goncalves, Ewan Harrison, David K. Jackson, Ian Johnston, Dominic Kwiatkowski, Cordelia Langford, John Sillitoe on behalf of the Wellcome Sanger Institute COVID-19 Surveillance Team |
| EPI_ISL_3335183 | Lighthouse Lab in Alderley Park | Wellcome Sanger Institute for the COVID-19 Genomics UK (COG-UK) Consortium | Jacquelyn Wynn, Mairead Hyland, The Lighthouse Lab in Alderley Park and Alex Alderton, Roberto Amato, Jeffrey Barrett, Sonia Goncalves, Ewan Harrison, David K. Jackson, Ian Johnston, Dominic Kwiatkowski, Cordelia Langford, John Sillitoe on behalf of the Wellcome Sanger Institute COVID-19 Surveillance Team |
| EPI_ISL_3335184, EPI_ISL_3335185 | Lighthouse Laboratory Plymouth | Wellcome Sanger Institute for the COVID-19 Genomics UK (COG-UK) Consortium | Lighthouse Laboratory Plymouth and Alex Alderton, Roberto Amato, Jeffrey Barrett, Sonia Goncalves, Ewan Harrison, David K. Jackson, Ian Johnston, Dominic Kwiatkowski, Cordelia Langford, John Sillitoe on behalf of the Wellcome Sanger Institute COVID-19 Surveillance Team |
| EPI_ISL_3335186, EPI_ISL_3335187, EPI_ISL_3335188 | Lighthouse Lab in Milton Keynes | Wellcome Sanger Institute for the COVID-19 Genomics UK (COG-UK) Consortium | The Lighthouse Lab in Milton Keynes and Alex Alderton, Roberto Amato, Jeffrey Barrett, Sonia Goncalves, Ewan Harrison, David K. Jackson, Ian Johnston, Dominic Kwiatkowski, Cordelia Langford, John Sillitoe on behalf of the Wellcome Sanger Institute COVID-19 Surveillance Team |
| EPI_ISL_3335189 | Lighthouse Lab in Alderley Park | Wellcome Sanger Institute for the COVID-19 Genomics UK (COG-UK) Consortium | Jacquelyn Wynn, Mairead Hyland, The Lighthouse Lab in Alderley Park and Alex Alderton, Roberto Amato, Jeffrey Barrett, Sonia Goncalves, Ewan Harrison, David K. Jackson, Ian Johnston, Dominic Kwiatkowski, Cordelia Langford, John Sillitoe on behalf of the Wellcome Sanger Institute COVID-19 Surveillance Team |
| EPI_ISL_3335190 | Lighthouse Lab in Milton Keynes | Wellcome Sanger Institute for the COVID-19 Genomics UK (COG-UK) Consortium | The Lighthouse Lab in Milton Keynes and Alex Alderton, Roberto Amato, Jeffrey Barrett, Sonia Goncalves, Ewan Harrison, David K. Jackson, Ian Johnston, Dominic Kwiatkowski, Cordelia Langford, John Sillitoe on behalf of the Wellcome Sanger Institute COVID-19 Surveillance Team |
| EPI_ISL_3335191, EPI_ISL_3335192 | Lighthouse Laboratory Plymouth | Wellcome Sanger Institute for the COVID-19 Genomics UK (COG-UK) Consortium | Lighthouse Laboratory Plymouth and Alex Alderton, Roberto Amato, Jeffrey Barrett, Sonia Goncalves, Ewan Harrison, David K. Jackson, Ian Johnston, Dominic Kwiatkowski, Cordelia Langford, John Sillitoe on behalf of the Wellcome Sanger Institute COVID-19 Surveillance Team |
| EPI_ISL_3335193, EPI_ISL_3335194 | Lighthouse Lab in Milton Keynes | Wellcome Sanger Institute for the COVID-19 Genomics UK (COG-UK) Consortium | The Lighthouse Lab in Milton Keynes and Alex Alderton, Roberto Amato, Jeffrey Barrett, Sonia Goncalves, Ewan Harrison, David K. Jackson, Ian Johnston, Dominic Kwiatkowski, Cordelia Langford, John Sillitoe on behalf of the Wellcome Sanger Institute COVID-19 Surveillance Team |
| EPI_ISL_3335195, EPI_ISL_3335196 | Lighthouse Laboratory Plymouth | Wellcome Sanger Institute for the COVID-19 Genomics UK (COG-UK) Consortium | Lighthouse Laboratory Plymouth and Alex Alderton, Roberto Amato, Jeffrey Barrett, Sonia Goncalves, Ewan Harrison, David K. Jackson, Ian Johnston, Dominic Kwiatkowski, Cordelia Langford, John Sillitoe on behalf of the Wellcome Sanger Institute COVID-19 Surveillance Team |
| EPI_ISL_3335197 | Lighthouse Lab in Alderley Park | Wellcome Sanger Institute for the COVID-19 Genomics UK (COG-UK) Consortium | Jacquelyn Wynn, Mairead Hyland, The Lighthouse Lab in Alderley Park and Alex Alderton, Roberto Amato, Jeffrey Barrett, Sonia Goncalves, Ewan Harrison, David K. Jackson, Ian Johnston, Dominic Kwiatkowski, Cordelia Langford, John Sillitoe on behalf of the Wellcome Sanger Institute COVID-19 Surveillance Team |
| EPI_ISL_3335198, EPI_ISL_3335199, EPI_ISL_3335200 | Lighthouse Lab in Milton Keynes | Wellcome Sanger Institute for the COVID-19 Genomics UK (COG-UK) Consortium | The Lighthouse Lab in Milton Keynes and Alex Alderton, Roberto Amato, Jeffrey Barrett, Sonia Goncalves, Ewan Harrison, David K. Jackson, Ian Johnston, Dominic Kwiatkowski, Cordelia Langford, John Sillitoe on behalf of the Wellcome Sanger Institute COVID-19 Surveillance Team |
| EPI_ISL_3335201, EPI_ISL_3335202, EPI_ISL_3335203 | Lighthouse Laboratory Plymouth | Wellcome Sanger Institute for the COVID-19 Genomics UK (COG-UK) Consortium | Lighthouse Laboratory Plymouth and Alex Alderton, Roberto Amato, Jeffrey Barrett, Sonia Goncalves, Ewan Harrison, David K. Jackson, Ian Johnston, Dominic Kwiatkowski, Cordelia Langford, John Sillitoe on behalf of the Wellcome Sanger Institute COVID-19 Surveillance Team |
| EPI_ISL_3335204, EPI_ISL_3335205 | Lighthouse Lab in Milton Keynes | Wellcome Sanger Institute for the COVID-19 Genomics UK (COG-UK) Consortium | The Lighthouse Lab in Milton Keynes and Alex Alderton, Roberto Amato, Jeffrey Barrett, Sonia Goncalves, Ewan Harrison, David K. Jackson, Ian Johnston, Dominic Kwiatkowski, Cordelia Langford, John Sillitoe on behalf of the Wellcome Sanger Institute COVID-19 Surveillance Team |
| EPI_ISL_3335206 | Health Services Laboratories | Wellcome Sanger Institute for the COVID-19 Genomics UK (COG-UK) Consortium | Health Services Laboratories and Alex Alderton, Roberto Amato, Jeffrey Barrett, Sonia Goncalves, Ewan Harrison, David K. Jackson, Ian Johnston, Dominic Kwiatkowski, Cordelia Langford, John Sillitoe on behalf of the Wellcome Sanger Institute COVID-19 Surveillance Team |
| EPI_ISL_3335207, EPI_ISL_3335208 | Lighthouse Lab in Alderley Park | Wellcome Sanger Institute for the COVID-19 Genomics UK (COG-UK) Consortium | Jacquelyn Wynn, Mairead Hyland, The Lighthouse Lab in Alderley Park and Alex Alderton, Roberto Amato, Jeffrey Barrett, Sonia Goncalves, Ewan Harrison, David K. Jackson, Ian Johnston, Dominic Kwiatkowski, Cordelia Langford, John Sillitoe on behalf of the Wellcome Sanger Institute COVID-19 Surveillance Team |
| EPI_ISL_3335209, EPI_ISL_3335210 | Lighthouse Laboratory Plymouth | Wellcome Sanger Institute for the COVID-19 Genomics UK (COG-UK) Consortium | Lighthouse Laboratory Plymouth and Alex Alderton, Roberto Amato, Jeffrey Barrett, Sonia Goncalves, Ewan Harrison, David K. Jackson, Ian Johnston, Dominic Kwiatkowski, Cordelia Langford, John Sillitoe on behalf of the Wellcome Sanger Institute COVID-19 Surveillance Team |
| EPI_ISL_3335211 | Lighthouse Lab in Milton Keynes | Wellcome Sanger Institute for the COVID-19 Genomics UK (COG-UK) Consortium | The Lighthouse Lab in Milton Keynes and Alex Alderton, Roberto Amato, Jeffrey Barrett, Sonia Goncalves, Ewan Harrison, David K. Jackson, Ian Johnston, Dominic Kwiatkowski, Cordelia Langford, John Sillitoe on behalf of the Wellcome Sanger Institute COVID-19 Surveillance Team |
| EPI_ISL_3335212 | Lighthouse Lab in Alderley Park | Wellcome Sanger Institute for the COVID-19 Genomics UK (COG-UK) Consortium | Jacquelyn Wynn, Mairead Hyland, The Lighthouse Lab in Alderley Park and Alex Alderton, Roberto Amato, Jeffrey Barrett, Sonia Goncalves, Ewan Harrison, David K. Jackson, Ian Johnston, Dominic Kwiatkowski, Cordelia Langford, John Sillitoe on behalf of the Wellcome Sanger Institute COVID-19 Surveillance Team |
| EPI_ISL_3335213 | Health Services Laboratories | Wellcome Sanger Institute for the COVID-19 Genomics UK (COG-UK) Consortium | Health Services Laboratories and Alex Alderton, Roberto Amato, Jeffrey Barrett, Sonia Goncalves, Ewan Harrison, David K. Jackson, Ian Johnston, Dominic Kwiatkowski, Cordelia Langford, John Sillitoe on behalf of the Wellcome Sanger Institute COVID-19 Surveillance Team |
| EPI_ISL_3335214 | Lighthouse Lab in Alderley Park | Wellcome Sanger Institute for the COVID-19 Genomics UK (COG-UK) Consortium | Jacquelyn Wynn, Mairead Hyland, The Lighthouse Lab in Alderley Park and Alex Alderton, Roberto Amato, Jeffrey Barrett, Sonia Goncalves, Ewan Harrison, David K. Jackson, Ian Johnston, Dominic Kwiatkowski, Cordelia Langford, John Sillitoe on behalf of the Wellcome Sanger Institute COVID-19 Surveillance Team |
| EPI_ISL_3335215 | Lighthouse Lab in Milton Keynes | Wellcome Sanger Institute for the COVID-19 Genomics UK (COG-UK) Consortium | The Lighthouse Lab in Milton Keynes and Alex Alderton, Roberto Amato, Jeffrey Barrett, Sonia Goncalves, Ewan Harrison, David K. Jackson, Ian Johnston, Dominic Kwiatkowski, Cordelia Langford, John Sillitoe on behalf of the Wellcome Sanger Institute COVID-19 Surveillance Team |
| EPI_ISL_3335216 | Lighthouse Lab in Alderley Park | Wellcome Sanger Institute for the COVID-19 Genomics UK (COG-UK) Consortium | Jacquelyn Wynn, Mairead Hyland, The Lighthouse Lab in Alderley Park and Alex Alderton, Roberto Amato, Jeffrey Barrett, Sonia Goncalves, Ewan Harrison, David K. Jackson, Ian Johnston, Dominic Kwiatkowski, Cordelia Langford, John Sillitoe on behalf of the Wellcome Sanger Institute COVID-19 Surveillance Team |
| EPI_ISL_3335217 | Lighthouse Laboratory Plymouth | Wellcome Sanger Institute for the COVID-19 Genomics UK (COG-UK) Consortium | Lighthouse Laboratory Plymouth and Alex Alderton, Roberto Amato, Jeffrey Barrett, Sonia Goncalves, Ewan Harrison, David K. Jackson, Ian Johnston, Dominic Kwiatkowski, Cordelia Langford, John Sillitoe on behalf of the Wellcome Sanger Institute COVID-19 Surveillance Team |
| EPI_ISL_3335218, EPI_ISL_3335219 | Lighthouse Lab in Alderley Park | Wellcome Sanger Institute for the COVID-19 Genomics UK (COG-UK) Consortium | Jacquelyn Wynn, Mairead Hyland, The Lighthouse Lab in Alderley Park and Alex Alderton, Roberto Amato, Jeffrey Barrett, Sonia Goncalves, Ewan Harrison, David K. Jackson, Ian Johnston, Dominic Kwiatkowski, Cordelia Langford, John Sillitoe on behalf of the Wellcome Sanger Institute COVID-19 Surveillance Team |
| EPI_ISL_3335220 | Health Services Laboratories | Wellcome Sanger Institute for the COVID-19 Genomics UK (COG-UK) Consortium | Health Services Laboratories and Alex Alderton, Roberto Amato, Jeffrey Barrett, Sonia Goncalves, Ewan Harrison, David K. Jackson, Ian Johnston, Dominic Kwiatkowski, Cordelia Langford, John Sillitoe on behalf of the Wellcome Sanger Institute COVID-19 Surveillance Team |
| EPI_ISL_3335221, EPI_ISL_3335222 | Lighthouse Lab in Alderley Park | Wellcome Sanger Institute for the COVID-19 Genomics UK (COG-UK) Consortium | Jacquelyn Wynn, Mairead Hyland, The Lighthouse Lab in Alderley Park and Alex Alderton, Roberto Amato, Jeffrey Barrett, Sonia Goncalves, Ewan Harrison, David K. Jackson, Ian Johnston, Dominic Kwiatkowski, Cordelia Langford, John Sillitoe on behalf of the Wellcome Sanger Institute COVID-19 Surveillance Team |
| EPI_ISL_3335223 | Lighthouse Lab in Milton Keynes | Wellcome Sanger Institute for the COVID-19 Genomics UK (COG-UK) Consortium | The Lighthouse Lab in Milton Keynes and Alex Alderton, Roberto Amato, Jeffrey Barrett, Sonia Goncalves, Ewan Harrison, David K. Jackson, Ian Johnston, Dominic Kwiatkowski, Cordelia Langford, John Sillitoe on behalf of the Wellcome Sanger Institute COVID-19 Surveillance Team |
| EPI_ISL_3335224, EPI_ISL_3335225 | Health Services Laboratories | Wellcome Sanger Institute for the COVID-19 Genomics UK (COG-UK) Consortium | Health Services Laboratories and Alex Alderton, Roberto Amato, Jeffrey Barrett, Sonia Goncalves, Ewan Harrison, David K. Jackson, Ian Johnston, Dominic Kwiatkowski, Cordelia Langford, John Sillitoe on behalf of the Wellcome Sanger Institute COVID-19 Surveillance Team |
| EPI_ISL_3335226 | Lighthouse Lab in Alderley Park | Wellcome Sanger Institute for the COVID-19 Genomics UK (COG-UK) Consortium | Jacquelyn Wynn, Mairead Hyland, The Lighthouse Lab in Alderley Park and Alex Alderton, Roberto Amato, Jeffrey Barrett, Sonia Goncalves, Ewan Harrison, David K. Jackson, Ian Johnston, Dominic Kwiatkowski, Cordelia Langford, John Sillitoe on behalf of the Wellcome Sanger Institute COVID-19 Surveillance Team |
| EPI_ISL_3335227 | Lighthouse Lab in Milton Keynes | Wellcome Sanger Institute for the COVID-19 Genomics UK (COG-UK) Consortium | The Lighthouse Lab in Milton Keynes and Alex Alderton, Roberto Amato, Jeffrey Barrett, Sonia Goncalves, Ewan Harrison, David K. Jackson, Ian Johnston, Dominic Kwiatkowski, Cordelia Langford, John Sillitoe on behalf of the Wellcome Sanger Institute COVID-19 Surveillance Team |
| EPI_ISL_3335228, EPI_ISL_3335229 | Health Services Laboratories | Wellcome Sanger Institute for the COVID-19 Genomics UK (COG-UK) Consortium | Health Services Laboratories and Alex Alderton, Roberto Amato, Jeffrey Barrett, Sonia Goncalves, Ewan Harrison, David K. Jackson, Ian Johnston, Dominic Kwiatkowski, Cordelia Langford, John Sillitoe on behalf of the Wellcome Sanger Institute COVID-19 Surveillance Team |
| EPI_ISL_3335230, EPI_ISL_3335231 | Lighthouse Laboratory Plymouth | Wellcome Sanger Institute for the COVID-19 Genomics UK (COG-UK) Consortium | Lighthouse Laboratory Plymouth and Alex Alderton, Roberto Amato, Jeffrey Barrett, Sonia Goncalves, Ewan Harrison, David K. Jackson, Ian Johnston, Dominic Kwiatkowski, Cordelia Langford, John Sillitoe on behalf of the Wellcome Sanger Institute COVID-19 Surveillance Team |
| EPI_ISL_3335232 | Lighthouse Lab in Alderley Park | Wellcome Sanger Institute for the COVID-19 Genomics UK (COG-UK) Consortium | Jacquelyn Wynn, Mairead Hyland, The Lighthouse Lab in Alderley Park and Alex Alderton, Roberto Amato, Jeffrey Barrett, Sonia Goncalves, Ewan Harrison, David K. Jackson, Ian Johnston, Dominic Kwiatkowski, Cordelia Langford, John Sillitoe on behalf of the Wellcome Sanger Institute COVID-19 Surveillance Team |
| EPI_ISL_3335233 | Lighthouse Laboratory Plymouth | Wellcome Sanger Institute for the COVID-19 Genomics UK (COG-UK) Consortium | Lighthouse Laboratory Plymouth and Alex Alderton, Roberto Amato, Jeffrey Barrett, Sonia Goncalves, Ewan Harrison, David K. Jackson, Ian Johnston, Dominic Kwiatkowski, Cordelia Langford, John Sillitoe on behalf of the Wellcome Sanger Institute COVID-19 Surveillance Team |
| EPI_ISL_3335234 | Lighthouse Lab in Alderley Park | Wellcome Sanger Institute for the COVID-19 Genomics UK (COG-UK) Consortium | Jacquelyn Wynn, Mairead Hyland, The Lighthouse Lab in Alderley Park and Alex Alderton, Roberto Amato, Jeffrey Barrett, Sonia Goncalves, Ewan Harrison, David K. Jackson, Ian Johnston, Dominic Kwiatkowski, Cordelia Langford, John Sillitoe on behalf of the Wellcome Sanger Institute COVID-19 Surveillance Team |
| EPI_ISL_3335235 | Lighthouse Laboratory Plymouth | Wellcome Sanger Institute for the COVID-19 Genomics UK (COG-UK) | Lighthouse Laboratory Plymouth and Alex Alderton, Roberto Amato, Jeffrey Barrett, Sonia Goncalves, Ewan Harrison, David K. Jackson, Ian Johnston, Dominic Kwiatkowski, Cordelia Langford, John Sillitoe on behalf of the Wellcome Sanger |

| | | | |
|---|---|---|---|
| | | Consortium | Institute COVID-19 Surveillance Team |
| EPI_ISL_3335236 | Health Services Laboratories | Wellcome Sanger Institute for the COVID-19 Genomics UK (COG-UK) Consortium | Health Services Laboratories and Alex Alderton, Roberto Amato, Jeffrey Barrett, Sonia Goncalves, Ewan Harrison, David K. Jackson, Ian Johnston, Dominic Kwiatkowski, Cordelia Langford, John Sillitoe on behalf of the Wellcome Sanger Institute COVID-19 Surveillance Team |
| EPI_ISL_3335237 | Lighthouse Laboratory Plymouth | Wellcome Sanger Institute for the COVID-19 Genomics UK (COG-UK) Consortium | Lighthouse Laboratory Plymouth and Alex Alderton, Roberto Amato, Jeffrey Barrett, Sonia Goncalves, Ewan Harrison, David K. Jackson, Ian Johnston, Dominic Kwiatkowski, Cordelia Langford, John Sillitoe on behalf of the Wellcome Sanger Institute COVID-19 Surveillance Team |
| EPI_ISL_3335238 | Lighthouse Lab in Milton Keynes | Wellcome Sanger Institute for the COVID-19 Genomics UK (COG-UK) Consortium | The Lighthouse Lab in Milton Keynes and Alex Alderton, Roberto Amato, Jeffrey Barrett, Sonia Goncalves, Ewan Harrison, David K. Jackson, Ian Johnston, Dominic Kwiatkowski, Cordelia Langford, John Sillitoe on behalf of the Wellcome Sanger Institute COVID-19 Surveillance Team |
| EPI_ISL_3335239, EPI_ISL_3335240 | Lighthouse Lab in Alderley Park | Wellcome Sanger Institute for the COVID-19 Genomics UK (COG-UK) Consortium | Jacquelyn Wynn, Mairead Hyland, The Lighthouse Lab in Alderley Park and Alex Alderton, Roberto Amato, Jeffrey Barrett, Sonia Goncalves, Ewan Harrison, David K. Jackson, Ian Johnston, Dominic Kwiatkowski, Cordelia Langford, John Sillitoe on behalf of the Wellcome Sanger Institute COVID-19 Surveillance Team |
| EPI_ISL_3335241, EPI_ISL_3335242 | Lighthouse Laboratory Plymouth | Wellcome Sanger Institute for the COVID-19 Genomics UK (COG-UK) Consortium | Lighthouse Laboratory Plymouth and Alex Alderton, Roberto Amato, Jeffrey Barrett, Sonia Goncalves, Ewan Harrison, David K. Jackson, Ian Johnston, Dominic Kwiatkowski, Cordelia Langford, John Sillitoe on behalf of the Wellcome Sanger Institute COVID-19 Surveillance Team |
| EPI_ISL_3335243 | Lighthouse Lab in Milton Keynes | Wellcome Sanger Institute for the COVID-19 Genomics UK (COG-UK) Consortium | The Lighthouse Lab in Milton Keynes and Alex Alderton, Roberto Amato, Jeffrey Barrett, Sonia Goncalves, Ewan Harrison, David K. Jackson, Ian Johnston, Dominic Kwiatkowski, Cordelia Langford, John Sillitoe on behalf of the Wellcome Sanger Institute COVID-19 Surveillance Team |
| EPI_ISL_3335244 | Lighthouse Lab in Alderley Park | Wellcome Sanger Institute for the COVID-19 Genomics UK (COG-UK) Consortium | Jacquelyn Wynn, Mairead Hyland, The Lighthouse Lab in Alderley Park and Alex Alderton, Roberto Amato, Jeffrey Barrett, Sonia Goncalves, Ewan Harrison, David K. Jackson, Ian Johnston, Dominic Kwiatkowski, Cordelia Langford, John Sillitoe on behalf of the Wellcome Sanger Institute COVID-19 Surveillance Team |
| EPI_ISL_3335245 | Lighthouse Lab in Milton Keynes | Wellcome Sanger Institute for the COVID-19 Genomics UK (COG-UK) Consortium | The Lighthouse Lab in Milton Keynes and Alex Alderton, Roberto Amato, Jeffrey Barrett, Sonia Goncalves, Ewan Harrison, David K. Jackson, Ian Johnston, Dominic Kwiatkowski, Cordelia Langford, John Sillitoe on behalf of the Wellcome Sanger Institute COVID-19 Surveillance Team |
| EPI_ISL_3335246 | Lighthouse Lab in Alderley Park | Wellcome Sanger Institute for the COVID-19 Genomics UK (COG-UK) Consortium | Jacquelyn Wynn, Mairead Hyland, The Lighthouse Lab in Alderley Park and Alex Alderton, Roberto Amato, Jeffrey Barrett, Sonia Goncalves, Ewan Harrison, David K. Jackson, Ian Johnston, Dominic Kwiatkowski, Cordelia Langford, John Sillitoe on behalf of the Wellcome Sanger Institute COVID-19 Surveillance Team |
| EPI_ISL_3335247 | Health Services Laboratories | Wellcome Sanger Institute for the COVID-19 Genomics UK (COG-UK) Consortium | Health Services Laboratories and Alex Alderton, Roberto Amato, Jeffrey Barrett, Sonia Goncalves, Ewan Harrison, David K. Jackson, Ian Johnston, Dominic Kwiatkowski, Cordelia Langford, John Sillitoe on behalf of the Wellcome Sanger Institute COVID-19 Surveillance Team |
| EPI_ISL_3335248, EPI_ISL_3335249 | Lighthouse Lab in Alderley Park | Wellcome Sanger Institute for the COVID-19 Genomics UK (COG-UK) Consortium | Jacquelyn Wynn, Mairead Hyland, The Lighthouse Lab in Alderley Park and Alex Alderton, Roberto Amato, Jeffrey Barrett, Sonia Goncalves, Ewan Harrison, David K. Jackson, Ian Johnston, Dominic Kwiatkowski, Cordelia Langford, John Sillitoe on behalf of the Wellcome Sanger Institute COVID-19 Surveillance Team |
| EPI_ISL_3335250 | Lighthouse Lab in Milton Keynes | Wellcome Sanger Institute for the COVID-19 Genomics UK (COG-UK) Consortium | The Lighthouse Lab in Milton Keynes and Alex Alderton, Roberto Amato, Jeffrey Barrett, Sonia Goncalves, Ewan Harrison, David K. Jackson, Ian Johnston, Dominic Kwiatkowski, Cordelia Langford, John Sillitoe on behalf of the Wellcome Sanger Institute COVID-19 Surveillance Team |
| EPI_ISL_3335251 | Lighthouse Laboratory Plymouth | Wellcome Sanger Institute for the COVID-19 Genomics UK (COG-UK) Consortium | Lighthouse Laboratory Plymouth and Alex Alderton, Roberto Amato, Jeffrey Barrett, Sonia Goncalves, Ewan Harrison, David K. Jackson, Ian Johnston, Dominic Kwiatkowski, Cordelia Langford, John Sillitoe on behalf of the Wellcome Sanger Institute COVID-19 Surveillance Team |
| EPI_ISL_3335252 | Lighthouse Lab in Alderley Park | Wellcome Sanger Institute for the COVID-19 Genomics UK (COG-UK) Consortium | Jacquelyn Wynn, Mairead Hyland, The Lighthouse Lab in Alderley Park and Alex Alderton, Roberto Amato, Jeffrey Barrett, Sonia Goncalves, Ewan Harrison, David K. Jackson, Ian Johnston, Dominic Kwiatkowski, Cordelia Langford, John Sillitoe on behalf of the Wellcome Sanger Institute COVID-19 Surveillance Team |
| EPI_ISL_3335253, EPI_ISL_3335254 | Lighthouse Laboratory Plymouth | Wellcome Sanger Institute for the COVID-19 Genomics UK (COG-UK) Consortium | Lighthouse Laboratory Plymouth and Alex Alderton, Roberto Amato, Jeffrey Barrett, Sonia Goncalves, Ewan Harrison, David K. Jackson, Ian Johnston, Dominic Kwiatkowski, Cordelia Langford, John Sillitoe on behalf of the Wellcome Sanger Institute COVID-19 Surveillance Team |
| EPI_ISL_3335255 | Lighthouse Lab in Milton Keynes | Wellcome Sanger Institute for the COVID-19 Genomics UK (COG-UK) Consortium | The Lighthouse Lab in Milton Keynes and Alex Alderton, Roberto Amato, Jeffrey Barrett, Sonia Goncalves, Ewan Harrison, David K. Jackson, Ian Johnston, Dominic Kwiatkowski, Cordelia Langford, John Sillitoe on behalf of the Wellcome Sanger Institute COVID-19 Surveillance Team |
| EPI_ISL_3335256 | Health Services Laboratories | Wellcome Sanger Institute for the COVID-19 Genomics UK (COG-UK) Consortium | Health Services Laboratories and Alex Alderton, Roberto Amato, Jeffrey Barrett, Sonia Goncalves, Ewan Harrison, David K. Jackson, Ian Johnston, Dominic Kwiatkowski, Cordelia Langford, John Sillitoe on behalf of the Wellcome Sanger Institute COVID-19 Surveillance Team |
| EPI_ISL_3335257, EPI_ISL_3335258 | Lighthouse Lab in Milton Keynes | Wellcome Sanger Institute for the COVID-19 Genomics UK (COG-UK) Consortium | The Lighthouse Lab in Milton Keynes and Alex Alderton, Roberto Amato, Jeffrey Barrett, Sonia Goncalves, Ewan Harrison, David K. Jackson, Ian Johnston, Dominic Kwiatkowski, Cordelia Langford, John Sillitoe on behalf of the Wellcome Sanger Institute COVID-19 Surveillance Team |
| EPI_ISL_3335259 | Lighthouse Lab in Alderley Park | Wellcome Sanger Institute for the COVID-19 Genomics UK (COG-UK) Consortium | Jacquelyn Wynn, Mairead Hyland, The Lighthouse Lab in Alderley Park and Alex Alderton, Roberto Amato, Jeffrey Barrett, Sonia Goncalves, Ewan Harrison, David K. Jackson, Ian Johnston, Dominic Kwiatkowski, Cordelia Langford, John Sillitoe on behalf of the Wellcome Sanger Institute COVID-19 Surveillance Team |
| EPI_ISL_3335260, EPI_ISL_3335261, EPI_ISL_3335262 | Lighthouse Laboratory Plymouth | Wellcome Sanger Institute for the COVID-19 Genomics UK (COG-UK) Consortium | Lighthouse Laboratory Plymouth and Alex Alderton, Roberto Amato, Jeffrey Barrett, Sonia Goncalves, Ewan Harrison, David K. Jackson, Ian Johnston, Dominic Kwiatkowski, Cordelia Langford, John Sillitoe on behalf of the Wellcome Sanger Institute COVID-19 Surveillance Team |
| EPI_ISL_3335263 | Health Services Laboratories | Wellcome Sanger Institute for the COVID-19 Genomics UK (COG-UK) Consortium | Health Services Laboratories and Alex Alderton, Roberto Amato, Jeffrey Barrett, Sonia Goncalves, Ewan Harrison, David K. Jackson, Ian Johnston, Dominic Kwiatkowski, Cordelia Langford, John Sillitoe on behalf of the Wellcome Sanger Institute COVID-19 Surveillance Team |
| EPI_ISL_3335264 | Lighthouse Laboratory Plymouth | Wellcome Sanger Institute for the COVID-19 Genomics UK (COG-UK) Consortium | Lighthouse Laboratory Plymouth and Alex Alderton, Roberto Amato, Jeffrey Barrett, Sonia Goncalves, Ewan Harrison, David K. Jackson, Ian Johnston, Dominic Kwiatkowski, Cordelia Langford, John Sillitoe on behalf of the Wellcome Sanger Institute COVID-19 Surveillance Team |
| EPI_ISL_3335265 | Lighthouse Lab in Alderley Park | Wellcome Sanger Institute for the COVID-19 Genomics UK (COG-UK) Consortium | Jacquelyn Wynn, Mairead Hyland, The Lighthouse Lab in Alderley Park and Alex Alderton, Roberto Amato, Jeffrey Barrett, Sonia Goncalves, Ewan Harrison, David K. Jackson, Ian Johnston, Dominic Kwiatkowski, Cordelia Langford, John Sillitoe on behalf of the Wellcome Sanger Institute COVID-19 Surveillance Team |
| EPI_ISL_3335266, EPI_ISL_3335267 | Lighthouse Laboratory Plymouth | Wellcome Sanger Institute for the COVID-19 Genomics UK (COG-UK) Consortium | Lighthouse Laboratory Plymouth and Alex Alderton, Roberto Amato, Jeffrey Barrett, Sonia Goncalves, Ewan Harrison, David K. Jackson, Ian Johnston, Dominic Kwiatkowski, Cordelia Langford, John Sillitoe on behalf of the Wellcome Sanger Institute COVID-19 Surveillance Team |
| EPI_ISL_3335268 | Lighthouse Lab in Alderley Park | Wellcome Sanger Institute for the COVID-19 Genomics UK (COG-UK) Consortium | Jacquelyn Wynn, Mairead Hyland, The Lighthouse Lab in Alderley Park and Alex Alderton, Roberto Amato, Jeffrey Barrett, Sonia Goncalves, Ewan Harrison, David K. Jackson, Ian Johnston, Dominic Kwiatkowski, Cordelia Langford, John Sillitoe on behalf of the Wellcome Sanger Institute COVID-19 Surveillance Team |
| EPI_ISL_3335269 | Health Services Laboratories | Wellcome Sanger Institute for the COVID-19 Genomics UK (COG-UK) Consortium | Health Services Laboratories and Alex Alderton, Roberto Amato, Jeffrey Barrett, Sonia Goncalves, Ewan Harrison, David K. Jackson, Ian Johnston, Dominic Kwiatkowski, Cordelia Langford, John Sillitoe on behalf of the Wellcome Sanger Institute COVID-19 Surveillance Team |
| EPI_ISL_3335270 | Lighthouse Lab in Alderley Park | Wellcome Sanger Institute for the COVID-19 Genomics UK (COG-UK) Consortium | Jacquelyn Wynn, Mairead Hyland, The Lighthouse Lab in Alderley Park and Alex Alderton, Roberto Amato, Jeffrey Barrett, Sonia Goncalves, Ewan Harrison, David K. Jackson, Ian Johnston, Dominic Kwiatkowski, Cordelia Langford, John Sillitoe on behalf of the Wellcome Sanger Institute COVID-19 Surveillance Team |
| EPI_ISL_3335271 | Health Services Laboratories | Wellcome Sanger Institute for the COVID-19 Genomics UK (COG-UK) Consortium | Health Services Laboratories and Alex Alderton, Roberto Amato, Jeffrey Barrett, Sonia Goncalves, Ewan Harrison, David K. Jackson, Ian Johnston, Dominic Kwiatkowski, Cordelia Langford, John Sillitoe on behalf of the Wellcome Sanger Institute COVID-19 Surveillance Team |
| EPI_ISL_3335272 | Lighthouse Lab in Milton Keynes | Wellcome Sanger Institute for the COVID-19 Genomics UK (COG-UK) Consortium | The Lighthouse Lab in Milton Keynes and Alex Alderton, Roberto Amato, Jeffrey Barrett, Sonia Goncalves, Ewan Harrison, David K. Jackson, Ian Johnston, Dominic Kwiatkowski, Cordelia Langford, John Sillitoe on behalf of the Wellcome Sanger Institute COVID-19 Surveillance Team |
| EPI_ISL_3335273 | Lighthouse Lab in Alderley Park | Wellcome Sanger Institute for the COVID-19 Genomics UK (COG-UK) Consortium | Jacquelyn Wynn, Mairead Hyland, The Lighthouse Lab in Alderley Park and Alex Alderton, Roberto Amato, Jeffrey Barrett, Sonia Goncalves, Ewan Harrison, David K. Jackson, Ian Johnston, Dominic Kwiatkowski, Cordelia Langford, John Sillitoe on behalf of the Wellcome Sanger Institute COVID-19 Surveillance Team |
| EPI_ISL_3335274, EPI_ISL_3335275 | Lighthouse Lab in Milton Keynes | Wellcome Sanger Institute for the COVID-19 Genomics UK (COG-UK) Consortium | The Lighthouse Lab in Milton Keynes and Alex Alderton, Roberto Amato, Jeffrey Barrett, Sonia Goncalves, Ewan Harrison, David K. Jackson, Ian Johnston, Dominic Kwiatkowski, Cordelia Langford, John Sillitoe on behalf of the Wellcome Sanger Institute COVID-19 Surveillance Team |
| EPI_ISL_3335276 | Health Services Laboratories | Wellcome Sanger Institute for the COVID-19 Genomics UK (COG-UK) Consortium | Health Services Laboratories and Alex Alderton, Roberto Amato, Jeffrey Barrett, Sonia Goncalves, Ewan Harrison, David K. Jackson, Ian Johnston, Dominic Kwiatkowski, Cordelia Langford, John Sillitoe on behalf of the Wellcome Sanger Institute COVID-19 Surveillance Team |
| EPI_ISL_3335277 | Lighthouse Laboratory Plymouth | Wellcome Sanger Institute for the COVID-19 Genomics UK (COG-UK) Consortium | Lighthouse Laboratory Plymouth and Alex Alderton, Roberto Amato, Jeffrey Barrett, Sonia Goncalves, Ewan Harrison, David K. Jackson, Ian Johnston, Dominic Kwiatkowski, Cordelia Langford, John Sillitoe on behalf of the Wellcome Sanger Institute COVID-19 Surveillance Team |
| EPI_ISL_3335278, EPI_ISL_3335279, EPI_ISL_3335280 | Lighthouse Lab in Milton Keynes | Wellcome Sanger Institute for the COVID-19 Genomics UK (COG-UK) Consortium | The Lighthouse Lab in Milton Keynes and Alex Alderton, Roberto Amato, Jeffrey Barrett, Sonia Goncalves, Ewan Harrison, David K. Jackson, Ian Johnston, Dominic Kwiatkowski, Cordelia Langford, John Sillitoe on behalf of the Wellcome Sanger Institute COVID-19 Surveillance Team |
| EPI_ISL_3335281 | Lighthouse Laboratory Plymouth | Wellcome Sanger Institute for the COVID-19 Genomics UK (COG-UK) Consortium | Lighthouse Laboratory Plymouth and Alex Alderton, Roberto Amato, Jeffrey Barrett, Sonia Goncalves, Ewan Harrison, David K. Jackson, Ian Johnston, Dominic Kwiatkowski, Cordelia Langford, John Sillitoe on behalf of the Wellcome Sanger Institute COVID-19 Surveillance Team |
| EPI_ISL_3335282, EPI_ISL_3335283 | Lighthouse Lab in Alderley Park | Wellcome Sanger Institute for the COVID-19 Genomics UK (COG-UK) Consortium | Jacquelyn Wynn, Mairead Hyland, The Lighthouse Lab in Alderley Park and Alex Alderton, Roberto Amato, Jeffrey Barrett, Sonia Goncalves, Ewan Harrison, David K. Jackson, Ian Johnston, Dominic Kwiatkowski, Cordelia Langford, John Sillitoe on behalf of the Wellcome Sanger Institute COVID-19 Surveillance Team |
| EPI_ISL_3335284 | Lighthouse Laboratory Plymouth | Wellcome Sanger Institute for the COVID-19 Genomics UK (COG-UK) Consortium | Lighthouse Laboratory Plymouth and Alex Alderton, Roberto Amato, Jeffrey Barrett, Sonia Goncalves, Ewan Harrison, David K. Jackson, Ian Johnston, Dominic Kwiatkowski, Cordelia Langford, John Sillitoe on behalf of the Wellcome Sanger Institute COVID-19 Surveillance Team |
| EPI_ISL_3335285, EPI_ISL_3335286, EPI_ISL_3335287 | Lighthouse Lab in Alderley Park | Wellcome Sanger Institute for the COVID-19 Genomics UK (COG-UK) Consortium | Jacquelyn Wynn, Mairead Hyland, The Lighthouse Lab in Alderley Park and Alex Alderton, Roberto Amato, Jeffrey Barrett, Sonia Goncalves, Ewan Harrison, David K. Jackson, Ian Johnston, Dominic Kwiatkowski, Cordelia Langford, John Sillitoe on behalf of the Wellcome Sanger Institute COVID-19 Surveillance Team |
| EPI_ISL_3335288 | Health Services Laboratories | Wellcome Sanger Institute for the COVID-19 Genomics UK (COG-UK) Consortium | Health Services Laboratories and Alex Alderton, Roberto Amato, Jeffrey Barrett, Sonia Goncalves, Ewan Harrison, David K. Jackson, Ian Johnston, Dominic Kwiatkowski, Cordelia Langford, John Sillitoe on behalf of the Wellcome Sanger Institute COVID-19 Surveillance Team |
| EPI_ISL_3335289, EPI_ISL_3335290 | Lighthouse Lab in Alderley Park | Wellcome Sanger Institute for the COVID-19 Genomics UK (COG-UK) Consortium | Jacquelyn Wynn, Mairead Hyland, The Lighthouse Lab in Alderley Park and Alex Alderton, Roberto Amato, Jeffrey Barrett, Sonia Goncalves, Ewan Harrison, David K. Jackson, Ian Johnston, Dominic Kwiatkowski, Cordelia Langford, John Sillitoe on behalf of the Wellcome Sanger Institute COVID-19 Surveillance Team |
| EPI_ISL_3335291, EPI_ISL_3335292 | Lighthouse Lab in Milton Keynes | Wellcome Sanger Institute for the COVID-19 Genomics UK (COG-UK) Consortium | The Lighthouse Lab in Milton Keynes and Alex Alderton, Roberto Amato, Jeffrey Barrett, Sonia Goncalves, Ewan Harrison, David K. Jackson, Ian Johnston, Dominic Kwiatkowski, Cordelia Langford, John Sillitoe on behalf of the Wellcome Sanger Institute COVID-19 Surveillance Team |
| EPI_ISL_3335293 | Health Services Laboratories | Wellcome Sanger Institute for the COVID-19 Genomics UK (COG-UK) Consortium | Health Services Laboratories and Alex Alderton, Roberto Amato, Jeffrey Barrett, Sonia Goncalves, Ewan Harrison, David K. Jackson, Ian Johnston, Dominic Kwiatkowski, Cordelia Langford, John Sillitoe on behalf of the Wellcome Sanger Institute COVID-19 Surveillance Team |
| EPI_ISL_3335294 | Lighthouse Lab in Milton Keynes | Wellcome Sanger Institute for the COVID-19 Genomics UK (COG-UK) Consortium | The Lighthouse Lab in Milton Keynes and Alex Alderton, Roberto Amato, Jeffrey Barrett, Sonia Goncalves, Ewan Harrison, David K. Jackson, Ian Johnston, Dominic Kwiatkowski, Cordelia Langford, John Sillitoe on behalf of the Wellcome Sanger Institute COVID-19 Surveillance Team |
| EPI_ISL_3335295 | Lighthouse Lab in Alderley Park | Wellcome Sanger Institute for the COVID-19 Genomics UK (COG-UK) | Jacquelyn Wynn, Mairead Hyland, The Lighthouse Lab in Alderley Park and Alex Alderton, Roberto Amato, Jeffrey Barrett, Sonia Goncalves, Ewan Harrison, David K. Jackson, Ian Johnston, Dominic Kwiatkowski, Cordelia Langford, John Sillitoe on |

| | | | |
|---|---|---|---|
| | | Consortium | behalf of the Wellcome Sanger Institute COVID-19 Surveillance Team |
| EPI_ISL_3335296 | Lighthouse Laboratory Plymouth | Wellcome Sanger Institute for the COVID-19 Genomics UK (COG-UK) Consortium | Lighthouse Laboratory Plymouth and Alex Alderton, Roberto Amato, Jeffrey Barrett, Sonia Goncalves, Ewan Harrison, David K. Jackson, Ian Johnston, Dominic Kwiatkowski, Cordelia Langford, John Sillitoe on behalf of the Wellcome Sanger Institute COVID-19 Surveillance Team |
| EPI_ISL_3335297 | Lighthouse Lab in Milton Keynes | Wellcome Sanger Institute for the COVID-19 Genomics UK (COG-UK) Consortium | The Lighthouse Lab in Milton Keynes and Alex Alderton, Roberto Amato, Jeffrey Barrett, Sonia Goncalves, Ewan Harrison, David K. Jackson, Ian Johnston, Dominic Kwiatkowski, Cordelia Langford, John Sillitoe on behalf of the Wellcome Sanger Institute COVID-19 Surveillance Team |
| EPI_ISL_3335298, EPI_ISL_3335299 | Lighthouse Lab in Alderley Park | Wellcome Sanger Institute for the COVID-19 Genomics UK (COG-UK) Consortium | Jacquelyn Wynn, Mairead Hyland, The Lighthouse Lab in Alderley Park and Alex Alderton, Roberto Amato, Jeffrey Barrett, Sonia Goncalves, Ewan Harrison, David K. Jackson, Ian Johnston, Dominic Kwiatkowski, Cordelia Langford, John Sillitoe on behalf of the Wellcome Sanger Institute COVID-19 Surveillance Team |
| EPI_ISL_3335300, EPI_ISL_3335301 | Health Services Laboratories | Wellcome Sanger Institute for the COVID-19 Genomics UK (COG-UK) Consortium | Health Services Laboratories and Alex Alderton, Roberto Amato, Jeffrey Barrett, Sonia Goncalves, Ewan Harrison, David K. Jackson, Ian Johnston, Dominic Kwiatkowski, Cordelia Langford, John Sillitoe on behalf of the Wellcome Sanger Institute COVID-19 Surveillance Team |
| EPI_ISL_3335302, EPI_ISL_3335303 | Lighthouse Lab in Alderley Park | Wellcome Sanger Institute for the COVID-19 Genomics UK (COG-UK) Consortium | Jacquelyn Wynn, Mairead Hyland, The Lighthouse Lab in Alderley Park and Alex Alderton, Roberto Amato, Jeffrey Barrett, Sonia Goncalves, Ewan Harrison, David K. Jackson, Ian Johnston, Dominic Kwiatkowski, Cordelia Langford, John Sillitoe on behalf of the Wellcome Sanger Institute COVID-19 Surveillance Team |
| EPI_ISL_3335304 | Health Services Laboratories | Wellcome Sanger Institute for the COVID-19 Genomics UK (COG-UK) Consortium | Health Services Laboratories and Alex Alderton, Roberto Amato, Jeffrey Barrett, Sonia Goncalves, Ewan Harrison, David K. Jackson, Ian Johnston, Dominic Kwiatkowski, Cordelia Langford, John Sillitoe on behalf of the Wellcome Sanger Institute COVID-19 Surveillance Team |
| EPI_ISL_3335305 | Lighthouse Lab in Alderley Park | Wellcome Sanger Institute for the COVID-19 Genomics UK (COG-UK) Consortium | Jacquelyn Wynn, Mairead Hyland, The Lighthouse Lab in Alderley Park and Alex Alderton, Roberto Amato, Jeffrey Barrett, Sonia Goncalves, Ewan Harrison, David K. Jackson, Ian Johnston, Dominic Kwiatkowski, Cordelia Langford, John Sillitoe on behalf of the Wellcome Sanger Institute COVID-19 Surveillance Team |
| EPI_ISL_3335306 | Health Services Laboratories | Wellcome Sanger Institute for the COVID-19 Genomics UK (COG-UK) Consortium | Health Services Laboratories and Alex Alderton, Roberto Amato, Jeffrey Barrett, Sonia Goncalves, Ewan Harrison, David K. Jackson, Ian Johnston, Dominic Kwiatkowski, Cordelia Langford, John Sillitoe on behalf of the Wellcome Sanger Institute COVID-19 Surveillance Team |
| EPI_ISL_3335307 | Lighthouse Lab in Milton Keynes | Wellcome Sanger Institute for the COVID-19 Genomics UK (COG-UK) Consortium | The Lighthouse Lab in Milton Keynes and Alex Alderton, Roberto Amato, Jeffrey Barrett, Sonia Goncalves, Ewan Harrison, David K. Jackson, Ian Johnston, Dominic Kwiatkowski, Cordelia Langford, John Sillitoe on behalf of the Wellcome Sanger Institute COVID-19 Surveillance Team |
| EPI_ISL_3335308, EPI_ISL_3335309 | Lighthouse Laboratory Plymouth | Wellcome Sanger Institute for the COVID-19 Genomics UK (COG-UK) Consortium | Lighthouse Laboratory Plymouth and Alex Alderton, Roberto Amato, Jeffrey Barrett, Sonia Goncalves, Ewan Harrison, David K. Jackson, Ian Johnston, Dominic Kwiatkowski, Cordelia Langford, John Sillitoe on behalf of the Wellcome Sanger Institute COVID-19 Surveillance Team |
| EPI_ISL_3335310 | Lighthouse Lab in Milton Keynes | Wellcome Sanger Institute for the COVID-19 Genomics UK (COG-UK) Consortium | The Lighthouse Lab in Milton Keynes and Alex Alderton, Roberto Amato, Jeffrey Barrett, Sonia Goncalves, Ewan Harrison, David K. Jackson, Ian Johnston, Dominic Kwiatkowski, Cordelia Langford, John Sillitoe on behalf of the Wellcome Sanger Institute COVID-19 Surveillance Team |
| EPI_ISL_3335311 | Health Services Laboratories | Wellcome Sanger Institute for the COVID-19 Genomics UK (COG-UK) Consortium | Health Services Laboratories and Alex Alderton, Roberto Amato, Jeffrey Barrett, Sonia Goncalves, Ewan Harrison, David K. Jackson, Ian Johnston, Dominic Kwiatkowski, Cordelia Langford, John Sillitoe on behalf of the Wellcome Sanger Institute COVID-19 Surveillance Team |
| EPI_ISL_3335312 | Lighthouse Lab in Milton Keynes | Wellcome Sanger Institute for the COVID-19 Genomics UK (COG-UK) Consortium | The Lighthouse Lab in Milton Keynes and Alex Alderton, Roberto Amato, Jeffrey Barrett, Sonia Goncalves, Ewan Harrison, David K. Jackson, Ian Johnston, Dominic Kwiatkowski, Cordelia Langford, John Sillitoe on behalf of the Wellcome Sanger Institute COVID-19 Surveillance Team |
| EPI_ISL_3335313 | Health Services Laboratories | Wellcome Sanger Institute for the COVID-19 Genomics UK (COG-UK) Consortium | Health Services Laboratories and Alex Alderton, Roberto Amato, Jeffrey Barrett, Sonia Goncalves, Ewan Harrison, David K. Jackson, Ian Johnston, Dominic Kwiatkowski, Cordelia Langford, John Sillitoe on behalf of the Wellcome Sanger Institute COVID-19 Surveillance Team |
| EPI_ISL_3335314, EPI_ISL_3335315 | Lighthouse Lab in Alderley Park | Wellcome Sanger Institute for the COVID-19 Genomics UK (COG-UK) Consortium | Jacquelyn Wynn, Mairead Hyland, The Lighthouse Lab in Alderley Park and Alex Alderton, Roberto Amato, Jeffrey Barrett, Sonia Goncalves, Ewan Harrison, David K. Jackson, Ian Johnston, Dominic Kwiatkowski, Cordelia Langford, John Sillitoe on behalf of the Wellcome Sanger Institute COVID-19 Surveillance Team |
| EPI_ISL_3335316 | Lighthouse Laboratory Plymouth | Wellcome Sanger Institute for the COVID-19 Genomics UK (COG-UK) Consortium | Lighthouse Laboratory Plymouth and Alex Alderton, Roberto Amato, Jeffrey Barrett, Sonia Goncalves, Ewan Harrison, David K. Jackson, Ian Johnston, Dominic Kwiatkowski, Cordelia Langford, John Sillitoe on behalf of the Wellcome Sanger Institute COVID-19 Surveillance Team |
| EPI_ISL_3335317 | Health Services Laboratories | Wellcome Sanger Institute for the COVID-19 Genomics UK (COG-UK) Consortium | Health Services Laboratories and Alex Alderton, Roberto Amato, Jeffrey Barrett, Sonia Goncalves, Ewan Harrison, David K. Jackson, Ian Johnston, Dominic Kwiatkowski, Cordelia Langford, John Sillitoe on behalf of the Wellcome Sanger Institute COVID-19 Surveillance Team |
| EPI_ISL_3335318, EPI_ISL_3335319, EPI_ISL_3335320 | Lighthouse Lab in Alderley Park | Wellcome Sanger Institute for the COVID-19 Genomics UK (COG-UK) Consortium | Jacquelyn Wynn, Mairead Hyland, The Lighthouse Lab in Alderley Park and Alex Alderton, Roberto Amato, Jeffrey Barrett, Sonia Goncalves, Ewan Harrison, David K. Jackson, Ian Johnston, Dominic Kwiatkowski, Cordelia Langford, John Sillitoe on behalf of the Wellcome Sanger Institute COVID-19 Surveillance Team |
| EPI_ISL_3335321, EPI_ISL_3335322, EPI_ISL_3335323 | Health Services Laboratories | Wellcome Sanger Institute for the COVID-19 Genomics UK (COG-UK) Consortium | Health Services Laboratories and Alex Alderton, Roberto Amato, Jeffrey Barrett, Sonia Goncalves, Ewan Harrison, David K. Jackson, Ian Johnston, Dominic Kwiatkowski, Cordelia Langford, John Sillitoe on behalf of the Wellcome Sanger Institute COVID-19 Surveillance Team |
| EPI_ISL_3335324, EPI_ISL_3335325 | Lighthouse Laboratory Plymouth | Wellcome Sanger Institute for the COVID-19 Genomics UK (COG-UK) Consortium | Lighthouse Laboratory Plymouth and Alex Alderton, Roberto Amato, Jeffrey Barrett, Sonia Goncalves, Ewan Harrison, David K. Jackson, Ian Johnston, Dominic Kwiatkowski, Cordelia Langford, John Sillitoe on behalf of the Wellcome Sanger Institute COVID-19 Surveillance Team |
| EPI_ISL_3335326 | Lighthouse Lab in Alderley Park | Wellcome Sanger Institute for the COVID-19 Genomics UK (COG-UK) Consortium | Jacquelyn Wynn, Mairead Hyland, The Lighthouse Lab in Alderley Park and Alex Alderton, Roberto Amato, Jeffrey Barrett, Sonia Goncalves, Ewan Harrison, David K. Jackson, Ian Johnston, Dominic Kwiatkowski, Cordelia Langford, John Sillitoe on behalf of the Wellcome Sanger Institute COVID-19 Surveillance Team |
| EPI_ISL_3335327 | Health Services Laboratories | Wellcome Sanger Institute for the COVID-19 Genomics UK (COG-UK) Consortium | Health Services Laboratories and Alex Alderton, Roberto Amato, Jeffrey Barrett, Sonia Goncalves, Ewan Harrison, David K. Jackson, Ian Johnston, Dominic Kwiatkowski, Cordelia Langford, John Sillitoe on behalf of the Wellcome Sanger Institute COVID-19 Surveillance Team |
| EPI_ISL_3335328 | Lighthouse Lab in Milton Keynes | Wellcome Sanger Institute for the COVID-19 Genomics UK (COG-UK) Consortium | The Lighthouse Lab in Milton Keynes and Alex Alderton, Roberto Amato, Jeffrey Barrett, Sonia Goncalves, Ewan Harrison, David K. Jackson, Ian Johnston, Dominic Kwiatkowski, Cordelia Langford, John Sillitoe on behalf of the Wellcome Sanger Institute COVID-19 Surveillance Team |
| EPI_ISL_3335329 | Health Services Laboratories | Wellcome Sanger Institute for the COVID-19 Genomics UK (COG-UK) Consortium | Health Services Laboratories and Alex Alderton, Roberto Amato, Jeffrey Barrett, Sonia Goncalves, Ewan Harrison, David K. Jackson, Ian Johnston, Dominic Kwiatkowski, Cordelia Langford, John Sillitoe on behalf of the Wellcome Sanger Institute COVID-19 Surveillance Team |
| EPI_ISL_3335330 | Lighthouse Lab in Alderley Park | Wellcome Sanger Institute for the COVID-19 Genomics UK (COG-UK) Consortium | Jacquelyn Wynn, Mairead Hyland, The Lighthouse Lab in Alderley Park and Alex Alderton, Roberto Amato, Jeffrey Barrett, Sonia Goncalves, Ewan Harrison, David K. Jackson, Ian Johnston, Dominic Kwiatkowski, Cordelia Langford, John Sillitoe on behalf of the Wellcome Sanger Institute COVID-19 Surveillance Team |
| EPI_ISL_3335331 | Lighthouse Lab in Milton Keynes | Wellcome Sanger Institute for the COVID-19 Genomics UK (COG-UK) Consortium | The Lighthouse Lab in Milton Keynes and Alex Alderton, Roberto Amato, Jeffrey Barrett, Sonia Goncalves, Ewan Harrison, David K. Jackson, Ian Johnston, Dominic Kwiatkowski, Cordelia Langford, John Sillitoe on behalf of the Wellcome Sanger Institute COVID-19 Surveillance Team |
| EPI_ISL_3335332, EPI_ISL_3335333 | Lighthouse Lab in Alderley Park | Wellcome Sanger Institute for the COVID-19 Genomics UK (COG-UK) Consortium | Jacquelyn Wynn, Mairead Hyland, The Lighthouse Lab in Alderley Park and Alex Alderton, Roberto Amato, Jeffrey Barrett, Sonia Goncalves, Ewan Harrison, David K. Jackson, Ian Johnston, Dominic Kwiatkowski, Cordelia Langford, John Sillitoe on behalf of the Wellcome Sanger Institute COVID-19 Surveillance Team |
| EPI_ISL_3335334 | Lighthouse Lab in Milton Keynes | Wellcome Sanger Institute for the COVID-19 Genomics UK (COG-UK) Consortium | The Lighthouse Lab in Milton Keynes and Alex Alderton, Roberto Amato, Jeffrey Barrett, Sonia Goncalves, Ewan Harrison, David K. Jackson, Ian Johnston, Dominic Kwiatkowski, Cordelia Langford, John Sillitoe on behalf of the Wellcome Sanger Institute COVID-19 Surveillance Team |
| EPI_ISL_3335335 | Lighthouse Lab in Alderley Park | Wellcome Sanger Institute for the COVID-19 Genomics UK (COG-UK) Consortium | Jacquelyn Wynn, Mairead Hyland, The Lighthouse Lab in Alderley Park and Alex Alderton, Roberto Amato, Jeffrey Barrett, Sonia Goncalves, Ewan Harrison, David K. Jackson, Ian Johnston, Dominic Kwiatkowski, Cordelia Langford, John Sillitoe on behalf of the Wellcome Sanger Institute COVID-19 Surveillance Team |
| EPI_ISL_3335336 | Lighthouse Lab in Milton Keynes | Wellcome Sanger Institute for the COVID-19 Genomics UK (COG-UK) Consortium | The Lighthouse Lab in Milton Keynes and Alex Alderton, Roberto Amato, Jeffrey Barrett, Sonia Goncalves, Ewan Harrison, David K. Jackson, Ian Johnston, Dominic Kwiatkowski, Cordelia Langford, John Sillitoe on behalf of the Wellcome Sanger Institute COVID-19 Surveillance Team |
| EPI_ISL_3335337 | Lighthouse Lab in Alderley Park | Wellcome Sanger Institute for the COVID-19 Genomics UK (COG-UK) Consortium | Jacquelyn Wynn, Mairead Hyland, The Lighthouse Lab in Alderley Park and Alex Alderton, Roberto Amato, Jeffrey Barrett, Sonia Goncalves, Ewan Harrison, David K. Jackson, Ian Johnston, Dominic Kwiatkowski, Cordelia Langford, John Sillitoe on behalf of the Wellcome Sanger Institute COVID-19 Surveillance Team |
| EPI_ISL_3335338, EPI_ISL_3335339 | Lighthouse Laboratory Plymouth | Wellcome Sanger Institute for the COVID-19 Genomics UK (COG-UK) Consortium | Lighthouse Laboratory Plymouth and Alex Alderton, Roberto Amato, Jeffrey Barrett, Sonia Goncalves, Ewan Harrison, David K. Jackson, Ian Johnston, Dominic Kwiatkowski, Cordelia Langford, John Sillitoe on behalf of the Wellcome Sanger Institute COVID-19 Surveillance Team |
| EPI_ISL_3335340 | Lighthouse Lab in Milton Keynes | Wellcome Sanger Institute for the COVID-19 Genomics UK (COG-UK) Consortium | The Lighthouse Lab in Milton Keynes and Alex Alderton, Roberto Amato, Jeffrey Barrett, Sonia Goncalves, Ewan Harrison, David K. Jackson, Ian Johnston, Dominic Kwiatkowski, Cordelia Langford, John Sillitoe on behalf of the Wellcome Sanger Institute COVID-19 Surveillance Team |
| EPI_ISL_3335341 | Health Services Laboratories | Wellcome Sanger Institute for the COVID-19 Genomics UK (COG-UK) Consortium | Health Services Laboratories and Alex Alderton, Roberto Amato, Jeffrey Barrett, Sonia Goncalves, Ewan Harrison, David K. Jackson, Ian Johnston, Dominic Kwiatkowski, Cordelia Langford, John Sillitoe on behalf of the Wellcome Sanger Institute COVID-19 Surveillance Team |
| EPI_ISL_3335342 | Lighthouse Lab in Milton Keynes | Wellcome Sanger Institute for the COVID-19 Genomics UK (COG-UK) Consortium | The Lighthouse Lab in Milton Keynes and Alex Alderton, Roberto Amato, Jeffrey Barrett, Sonia Goncalves, Ewan Harrison, David K. Jackson, Ian Johnston, Dominic Kwiatkowski, Cordelia Langford, John Sillitoe on behalf of the Wellcome Sanger Institute COVID-19 Surveillance Team |
| EPI_ISL_3335343 | Lighthouse Lab in Alderley Park | Wellcome Sanger Institute for the COVID-19 Genomics UK (COG-UK) Consortium | Jacquelyn Wynn, Mairead Hyland, The Lighthouse Lab in Alderley Park and Alex Alderton, Roberto Amato, Jeffrey Barrett, Sonia Goncalves, Ewan Harrison, David K. Jackson, Ian Johnston, Dominic Kwiatkowski, Cordelia Langford, John Sillitoe on behalf of the Wellcome Sanger Institute COVID-19 Surveillance Team |
| EPI_ISL_3335344, EPI_ISL_3335345 | Lighthouse Laboratory Plymouth | Wellcome Sanger Institute for the COVID-19 Genomics UK (COG-UK) Consortium | Lighthouse Laboratory Plymouth and Alex Alderton, Roberto Amato, Jeffrey Barrett, Sonia Goncalves, Ewan Harrison, David K. Jackson, Ian Johnston, Dominic Kwiatkowski, Cordelia Langford, John Sillitoe on behalf of the Wellcome Sanger Institute COVID-19 Surveillance Team |
| EPI_ISL_3335346 | Lighthouse Lab in Alderley Park | Wellcome Sanger Institute for the COVID-19 Genomics UK (COG-UK) Consortium | Jacquelyn Wynn, Mairead Hyland, The Lighthouse Lab in Alderley Park and Alex Alderton, Roberto Amato, Jeffrey Barrett, Sonia Goncalves, Ewan Harrison, David K. Jackson, Ian Johnston, Dominic Kwiatkowski, Cordelia Langford, John Sillitoe on behalf of the Wellcome Sanger Institute COVID-19 Surveillance Team |
| EPI_ISL_3335347 | Lighthouse Laboratory Plymouth | Wellcome Sanger Institute for the COVID-19 Genomics UK (COG-UK) Consortium | Lighthouse Laboratory Plymouth and Alex Alderton, Roberto Amato, Jeffrey Barrett, Sonia Goncalves, Ewan Harrison, David K. Jackson, Ian Johnston, Dominic Kwiatkowski, Cordelia Langford, John Sillitoe on behalf of the Wellcome Sanger Institute COVID-19 Surveillance Team |
| EPI_ISL_3335348, EPI_ISL_3335349 | Lighthouse Lab in Milton Keynes | Wellcome Sanger Institute for the COVID-19 Genomics UK (COG-UK) Consortium | The Lighthouse Lab in Milton Keynes and Alex Alderton, Roberto Amato, Jeffrey Barrett, Sonia Goncalves, Ewan Harrison, David K. Jackson, Ian Johnston, Dominic Kwiatkowski, Cordelia Langford, John Sillitoe on behalf of the Wellcome Sanger Institute COVID-19 Surveillance Team |
| EPI_ISL_3335350 | Lighthouse Laboratory Plymouth | Wellcome Sanger Institute for the COVID-19 Genomics UK (COG-UK) Consortium | Lighthouse Laboratory Plymouth and Alex Alderton, Roberto Amato, Jeffrey Barrett, Sonia Goncalves, Ewan Harrison, David K. Jackson, Ian Johnston, Dominic Kwiatkowski, Cordelia Langford, John Sillitoe on behalf of the Wellcome Sanger Institute COVID-19 Surveillance Team |
| EPI_ISL_3335351 | Lighthouse Lab in Milton Keynes | Wellcome Sanger Institute for the COVID-19 Genomics UK (COG-UK) Consortium | The Lighthouse Lab in Milton Keynes and Alex Alderton, Roberto Amato, Jeffrey Barrett, Sonia Goncalves, Ewan Harrison, David K. Jackson, Ian Johnston, Dominic Kwiatkowski, Cordelia Langford, John Sillitoe on behalf of the Wellcome Sanger Institute COVID-19 Surveillance Team |
| EPI_ISL_3335352 | Lighthouse Laboratory Plymouth | Wellcome Sanger Institute for the COVID-19 Genomics UK (COG-UK) Consortium | Lighthouse Laboratory Plymouth and Alex Alderton, Roberto Amato, Jeffrey Barrett, Sonia Goncalves, Ewan Harrison, David K. Jackson, Ian Johnston, Dominic Kwiatkowski, Cordelia Langford, John Sillitoe on behalf of the Wellcome Sanger Institute COVID-19 Surveillance Team |
| EPI_ISL_3335353 | Lighthouse Lab in Milton Keynes | Wellcome Sanger Institute for the COVID-19 Genomics UK (COG-UK) | The Lighthouse Lab in Milton Keynes and Alex Alderton, Roberto Amato, Jeffrey Barrett, Sonia Goncalves, Ewan Harrison, David K. Jackson, Ian Johnston, Dominic Kwiatkowski, Cordelia Langford, John Sillitoe on behalf of the Wellcome Sanger |

| | | | |
|---|---|---|---|
| | | Consortium | Institute COVID-19 Surveillance Team |
| EPI_ISL_3335354 | Lighthouse Laboratory Plymouth | Wellcome Sanger Institute for the COVID-19 Genomics UK (COG-UK) Consortium | Lighthouse Laboratory Plymouth and Alex Alderton, Roberto Amato, Jeffrey Barrett, Sonia Goncalves, Ewan Harrison, David K. Jackson, Ian Johnston, Dominic Kwiatkowski, Cordelia Langford, John Sillitoe on behalf of the Wellcome Sanger Institute COVID-19 Surveillance Team |
| EPI_ISL_3335355, EPI_ISL_3335356 | Lighthouse Lab in Alderley Park | Wellcome Sanger Institute for the COVID-19 Genomics UK (COG-UK) Consortium | Jacquelyn Wynn, Mairead Hyland, The Lighthouse Lab in Alderley Park and Alex Alderton, Roberto Amato, Jeffrey Barrett, Sonia Goncalves, Ewan Harrison, David K. Jackson, Ian Johnston, Dominic Kwiatkowski, Cordelia Langford, John Sillitoe on behalf of the Wellcome Sanger Institute COVID-19 Surveillance Team |
| EPI_ISL_3335357 | Lighthouse Laboratory Plymouth | Wellcome Sanger Institute for the COVID-19 Genomics UK (COG-UK) Consortium | Lighthouse Laboratory Plymouth and Alex Alderton, Roberto Amato, Jeffrey Barrett, Sonia Goncalves, Ewan Harrison, David K. Jackson, Ian Johnston, Dominic Kwiatkowski, Cordelia Langford, John Sillitoe on behalf of the Wellcome Sanger Institute COVID-19 Surveillance Team |
| EPI_ISL_3335358, EPI_ISL_3335359 | Lighthouse Lab in Milton Keynes | Wellcome Sanger Institute for the COVID-19 Genomics UK (COG-UK) Consortium | The Lighthouse Lab in Milton Keynes and Alex Alderton, Roberto Amato, Jeffrey Barrett, Sonia Goncalves, Ewan Harrison, David K. Jackson, Ian Johnston, Dominic Kwiatkowski, Cordelia Langford, John Sillitoe on behalf of the Wellcome Sanger Institute COVID-19 Surveillance Team |
| EPI_ISL_3335360 | Health Services Laboratories | Wellcome Sanger Institute for the COVID-19 Genomics UK (COG-UK) Consortium | Health Services Laboratories and Alex Alderton, Roberto Amato, Jeffrey Barrett, Sonia Goncalves, Ewan Harrison, David K. Jackson, Ian Johnston, Dominic Kwiatkowski, Cordelia Langford, John Sillitoe on behalf of the Wellcome Sanger Institute COVID-19 Surveillance Team |
| EPI_ISL_3335361 | Lighthouse Laboratory Plymouth | Wellcome Sanger Institute for the COVID-19 Genomics UK (COG-UK) Consortium | Lighthouse Laboratory Plymouth and Alex Alderton, Roberto Amato, Jeffrey Barrett, Sonia Goncalves, Ewan Harrison, David K. Jackson, Ian Johnston, Dominic Kwiatkowski, Cordelia Langford, John Sillitoe on behalf of the Wellcome Sanger Institute COVID-19 Surveillance Team |
| EPI_ISL_3335362 | Health Services Laboratories | Wellcome Sanger Institute for the COVID-19 Genomics UK (COG-UK) Consortium | Health Services Laboratories and Alex Alderton, Roberto Amato, Jeffrey Barrett, Sonia Goncalves, Ewan Harrison, David K. Jackson, Ian Johnston, Dominic Kwiatkowski, Cordelia Langford, John Sillitoe on behalf of the Wellcome Sanger Institute COVID-19 Surveillance Team |
| EPI_ISL_3335363, EPI_ISL_3335364 | Lighthouse Lab in Alderley Park | Wellcome Sanger Institute for the COVID-19 Genomics UK (COG-UK) Consortium | Jacquelyn Wynn, Mairead Hyland, The Lighthouse Lab in Alderley Park and Alex Alderton, Roberto Amato, Jeffrey Barrett, Sonia Goncalves, Ewan Harrison, David K. Jackson, Ian Johnston, Dominic Kwiatkowski, Cordelia Langford, John Sillitoe on behalf of the Wellcome Sanger Institute COVID-19 Surveillance Team |
| EPI_ISL_3335365, EPI_ISL_3335366 | Lighthouse Laboratory Plymouth | Wellcome Sanger Institute for the COVID-19 Genomics UK (COG-UK) Consortium | Lighthouse Laboratory Plymouth and Alex Alderton, Roberto Amato, Jeffrey Barrett, Sonia Goncalves, Ewan Harrison, David K. Jackson, Ian Johnston, Dominic Kwiatkowski, Cordelia Langford, John Sillitoe on behalf of the Wellcome Sanger Institute COVID-19 Surveillance Team |
| EPI_ISL_3335367 | Health Services Laboratories | Wellcome Sanger Institute for the COVID-19 Genomics UK (COG-UK) Consortium | Health Services Laboratories and Alex Alderton, Roberto Amato, Jeffrey Barrett, Sonia Goncalves, Ewan Harrison, David K. Jackson, Ian Johnston, Dominic Kwiatkowski, Cordelia Langford, John Sillitoe on behalf of the Wellcome Sanger Institute COVID-19 Surveillance Team |
| EPI_ISL_3335368, EPI_ISL_3335369 | Lighthouse Laboratory Plymouth | Wellcome Sanger Institute for the COVID-19 Genomics UK (COG-UK) Consortium | Lighthouse Laboratory Plymouth and Alex Alderton, Roberto Amato, Jeffrey Barrett, Sonia Goncalves, Ewan Harrison, David K. Jackson, Ian Johnston, Dominic Kwiatkowski, Cordelia Langford, John Sillitoe on behalf of the Wellcome Sanger Institute COVID-19 Surveillance Team |
| EPI_ISL_3335370, EPI_ISL_3335371, EPI_ISL_3335372 | Lighthouse Lab in Alderley Park | Wellcome Sanger Institute for the COVID-19 Genomics UK (COG-UK) Consortium | Jacquelyn Wynn, Mairead Hyland, The Lighthouse Lab in Alderley Park and Alex Alderton, Roberto Amato, Jeffrey Barrett, Sonia Goncalves, Ewan Harrison, David K. Jackson, Ian Johnston, Dominic Kwiatkowski, Cordelia Langford, John Sillitoe on behalf of the Wellcome Sanger Institute COVID-19 Surveillance Team |
| EPI_ISL_3335373 | Health Services Laboratories | Wellcome Sanger Institute for the COVID-19 Genomics UK (COG-UK) Consortium | Health Services Laboratories and Alex Alderton, Roberto Amato, Jeffrey Barrett, Sonia Goncalves, Ewan Harrison, David K. Jackson, Ian Johnston, Dominic Kwiatkowski, Cordelia Langford, John Sillitoe on behalf of the Wellcome Sanger Institute COVID-19 Surveillance Team |
| EPI_ISL_3335374 | Lighthouse Lab in Alderley Park | Wellcome Sanger Institute for the COVID-19 Genomics UK (COG-UK) Consortium | Jacquelyn Wynn, Mairead Hyland, The Lighthouse Lab in Alderley Park and Alex Alderton, Roberto Amato, Jeffrey Barrett, Sonia Goncalves, Ewan Harrison, David K. Jackson, Ian Johnston, Dominic Kwiatkowski, Cordelia Langford, John Sillitoe on behalf of the Wellcome Sanger Institute COVID-19 Surveillance Team |
| EPI_ISL_3335375 | Health Services Laboratories | Wellcome Sanger Institute for the COVID-19 Genomics UK (COG-UK) Consortium | Health Services Laboratories and Alex Alderton, Roberto Amato, Jeffrey Barrett, Sonia Goncalves, Ewan Harrison, David K. Jackson, Ian Johnston, Dominic Kwiatkowski, Cordelia Langford, John Sillitoe on behalf of the Wellcome Sanger Institute COVID-19 Surveillance Team |
| EPI_ISL_3335376, EPI_ISL_3335377, EPI_ISL_3335378, EPI_ISL_3335379 | Lighthouse Laboratory Plymouth | Wellcome Sanger Institute for the COVID-19 Genomics UK (COG-UK) Consortium | Lighthouse Laboratory Plymouth and Alex Alderton, Roberto Amato, Jeffrey Barrett, Sonia Goncalves, Ewan Harrison, David K. Jackson, Ian Johnston, Dominic Kwiatkowski, Cordelia Langford, John Sillitoe on behalf of the Wellcome Sanger Institute COVID-19 Surveillance Team |
| EPI_ISL_3335380, EPI_ISL_3335381 | Lighthouse Lab in Alderley Park | Wellcome Sanger Institute for the COVID-19 Genomics UK (COG-UK) Consortium | Jacquelyn Wynn, Mairead Hyland, The Lighthouse Lab in Alderley Park and Alex Alderton, Roberto Amato, Jeffrey Barrett, Sonia Goncalves, Ewan Harrison, David K. Jackson, Ian Johnston, Dominic Kwiatkowski, Cordelia Langford, John Sillitoe on behalf of the Wellcome Sanger Institute COVID-19 Surveillance Team |
| EPI_ISL_3335382, EPI_ISL_3335383 | Lighthouse Laboratory Plymouth | Wellcome Sanger Institute for the COVID-19 Genomics UK (COG-UK) Consortium | Lighthouse Laboratory Plymouth and Alex Alderton, Roberto Amato, Jeffrey Barrett, Sonia Goncalves, Ewan Harrison, David K. Jackson, Ian Johnston, Dominic Kwiatkowski, Cordelia Langford, John Sillitoe on behalf of the Wellcome Sanger Institute COVID-19 Surveillance Team |
| EPI_ISL_3335384 | Health Services Laboratories | Wellcome Sanger Institute for the COVID-19 Genomics UK (COG-UK) Consortium | Health Services Laboratories and Alex Alderton, Roberto Amato, Jeffrey Barrett, Sonia Goncalves, Ewan Harrison, David K. Jackson, Ian Johnston, Dominic Kwiatkowski, Cordelia Langford, John Sillitoe on behalf of the Wellcome Sanger Institute COVID-19 Surveillance Team |
| EPI_ISL_3335385 | Lighthouse Laboratory Plymouth | Wellcome Sanger Institute for the COVID-19 Genomics UK (COG-UK) Consortium | Lighthouse Laboratory Plymouth and Alex Alderton, Roberto Amato, Jeffrey Barrett, Sonia Goncalves, Ewan Harrison, David K. Jackson, Ian Johnston, Dominic Kwiatkowski, Cordelia Langford, John Sillitoe on behalf of the Wellcome Sanger Institute COVID-19 Surveillance Team |
| EPI_ISL_3335386 | Health Services Laboratories | Wellcome Sanger Institute for the COVID-19 Genomics UK (COG-UK) Consortium | Health Services Laboratories and Alex Alderton, Roberto Amato, Jeffrey Barrett, Sonia Goncalves, Ewan Harrison, David K. Jackson, Ian Johnston, Dominic Kwiatkowski, Cordelia Langford, John Sillitoe on behalf of the Wellcome Sanger Institute COVID-19 Surveillance Team |
| EPI_ISL_3335387 | Lighthouse Lab in Milton Keynes | Wellcome Sanger Institute for the COVID-19 Genomics UK (COG-UK) Consortium | The Lighthouse Lab in Milton Keynes and Alex Alderton, Roberto Amato, Jeffrey Barrett, Sonia Goncalves, Ewan Harrison, David K. Jackson, Ian Johnston, Dominic Kwiatkowski, Cordelia Langford, John Sillitoe on behalf of the Wellcome Sanger Institute COVID-19 Surveillance Team |
| EPI_ISL_3335388, EPI_ISL_3335389, EPI_ISL_3335390 | Health Services Laboratories | Wellcome Sanger Institute for the COVID-19 Genomics UK (COG-UK) Consortium | Health Services Laboratories and Alex Alderton, Roberto Amato, Jeffrey Barrett, Sonia Goncalves, Ewan Harrison, David K. Jackson, Ian Johnston, Dominic Kwiatkowski, Cordelia Langford, John Sillitoe on behalf of the Wellcome Sanger Institute COVID-19 Surveillance Team |
| EPI_ISL_3335391 | Lighthouse Lab in Milton Keynes | Wellcome Sanger Institute for the COVID-19 Genomics UK (COG-UK) Consortium | The Lighthouse Lab in Milton Keynes and Alex Alderton, Roberto Amato, Jeffrey Barrett, Sonia Goncalves, Ewan Harrison, David K. Jackson, Ian Johnston, Dominic Kwiatkowski, Cordelia Langford, John Sillitoe on behalf of the Wellcome Sanger Institute COVID-19 Surveillance Team |
| EPI_ISL_3335392, EPI_ISL_3335393 | Health Services Laboratories | Wellcome Sanger Institute for the COVID-19 Genomics UK (COG-UK) Consortium | Health Services Laboratories and Alex Alderton, Roberto Amato, Jeffrey Barrett, Sonia Goncalves, Ewan Harrison, David K. Jackson, Ian Johnston, Dominic Kwiatkowski, Cordelia Langford, John Sillitoe on behalf of the Wellcome Sanger Institute COVID-19 Surveillance Team |
| EPI_ISL_3335394 | Lighthouse Laboratory Plymouth | Wellcome Sanger Institute for the COVID-19 Genomics UK (COG-UK) Consortium | Lighthouse Laboratory Plymouth and Alex Alderton, Roberto Amato, Jeffrey Barrett, Sonia Goncalves, Ewan Harrison, David K. Jackson, Ian Johnston, Dominic Kwiatkowski, Cordelia Langford, John Sillitoe on behalf of the Wellcome Sanger Institute COVID-19 Surveillance Team |
| EPI_ISL_3335395, EPI_ISL_3335396 | Lighthouse Lab in Alderley Park | Wellcome Sanger Institute for the COVID-19 Genomics UK (COG-UK) Consortium | Jacquelyn Wynn, Mairead Hyland, The Lighthouse Lab in Alderley Park and Alex Alderton, Roberto Amato, Jeffrey Barrett, Sonia Goncalves, Ewan Harrison, David K. Jackson, Ian Johnston, Dominic Kwiatkowski, Cordelia Langford, John Sillitoe on behalf of the Wellcome Sanger Institute COVID-19 Surveillance Team |
| EPI_ISL_3335397 | Lighthouse Lab in Milton Keynes | Wellcome Sanger Institute for the COVID-19 Genomics UK (COG-UK) Consortium | The Lighthouse Lab in Milton Keynes and Alex Alderton, Roberto Amato, Jeffrey Barrett, Sonia Goncalves, Ewan Harrison, David K. Jackson, Ian Johnston, Dominic Kwiatkowski, Cordelia Langford, John Sillitoe on behalf of the Wellcome Sanger Institute COVID-19 Surveillance Team |
| EPI_ISL_3335398 | Lighthouse Laboratory Plymouth | Wellcome Sanger Institute for the COVID-19 Genomics UK (COG-UK) Consortium | Lighthouse Laboratory Plymouth and Alex Alderton, Roberto Amato, Jeffrey Barrett, Sonia Goncalves, Ewan Harrison, David K. Jackson, Ian Johnston, Dominic Kwiatkowski, Cordelia Langford, John Sillitoe on behalf of the Wellcome Sanger Institute COVID-19 Surveillance Team |
| EPI_ISL_3335399 | Lighthouse Lab in Alderley Park | Wellcome Sanger Institute for the COVID-19 Genomics UK (COG-UK) Consortium | Jacquelyn Wynn, Mairead Hyland, The Lighthouse Lab in Alderley Park and Alex Alderton, Roberto Amato, Jeffrey Barrett, Sonia Goncalves, Ewan Harrison, David K. Jackson, Ian Johnston, Dominic Kwiatkowski, Cordelia Langford, John Sillitoe on behalf of the Wellcome Sanger Institute COVID-19 Surveillance Team |
| EPI_ISL_3335400 | Lighthouse Lab in Milton Keynes | Wellcome Sanger Institute for the COVID-19 Genomics UK (COG-UK) Consortium | The Lighthouse Lab in Milton Keynes and Alex Alderton, Roberto Amato, Jeffrey Barrett, Sonia Goncalves, Ewan Harrison, David K. Jackson, Ian Johnston, Dominic Kwiatkowski, Cordelia Langford, John Sillitoe on behalf of the Wellcome Sanger Institute COVID-19 Surveillance Team |
| EPI_ISL_3335401, EPI_ISL_3335402 | Lighthouse Lab in Alderley Park | Wellcome Sanger Institute for the COVID-19 Genomics UK (COG-UK) Consortium | Jacquelyn Wynn, Mairead Hyland, The Lighthouse Lab in Alderley Park and Alex Alderton, Roberto Amato, Jeffrey Barrett, Sonia Goncalves, Ewan Harrison, David K. Jackson, Ian Johnston, Dominic Kwiatkowski, Cordelia Langford, John Sillitoe on behalf of the Wellcome Sanger Institute COVID-19 Surveillance Team |
| EPI_ISL_3335403 | Lighthouse Laboratory Plymouth | Wellcome Sanger Institute for the COVID-19 Genomics UK (COG-UK) Consortium | Lighthouse Laboratory Plymouth and Alex Alderton, Roberto Amato, Jeffrey Barrett, Sonia Goncalves, Ewan Harrison, David K. Jackson, Ian Johnston, Dominic Kwiatkowski, Cordelia Langford, John Sillitoe on behalf of the Wellcome Sanger Institute COVID-19 Surveillance Team |
| EPI_ISL_3335404 | Lighthouse Lab in Alderley Park | Wellcome Sanger Institute for the COVID-19 Genomics UK (COG-UK) Consortium | Jacquelyn Wynn, Mairead Hyland, The Lighthouse Lab in Alderley Park and Alex Alderton, Roberto Amato, Jeffrey Barrett, Sonia Goncalves, Ewan Harrison, David K. Jackson, Ian Johnston, Dominic Kwiatkowski, Cordelia Langford, John Sillitoe on behalf of the Wellcome Sanger Institute COVID-19 Surveillance Team |
| EPI_ISL_3335405 | Health Services Laboratories | Wellcome Sanger Institute for the COVID-19 Genomics UK (COG-UK) Consortium | Health Services Laboratories and Alex Alderton, Roberto Amato, Jeffrey Barrett, Sonia Goncalves, Ewan Harrison, David K. Jackson, Ian Johnston, Dominic Kwiatkowski, Cordelia Langford, John Sillitoe on behalf of the Wellcome Sanger Institute COVID-19 Surveillance Team |
| EPI_ISL_3335406 | Lighthouse Lab in Milton Keynes | Wellcome Sanger Institute for the COVID-19 Genomics UK (COG-UK) Consortium | The Lighthouse Lab in Milton Keynes and Alex Alderton, Roberto Amato, Jeffrey Barrett, Sonia Goncalves, Ewan Harrison, David K. Jackson, Ian Johnston, Dominic Kwiatkowski, Cordelia Langford, John Sillitoe on behalf of the Wellcome Sanger Institute COVID-19 Surveillance Team |
| EPI_ISL_3335407 | Health Services Laboratories | Wellcome Sanger Institute for the COVID-19 Genomics UK (COG-UK) Consortium | Health Services Laboratories and Alex Alderton, Roberto Amato, Jeffrey Barrett, Sonia Goncalves, Ewan Harrison, David K. Jackson, Ian Johnston, Dominic Kwiatkowski, Cordelia Langford, John Sillitoe on behalf of the Wellcome Sanger Institute COVID-19 Surveillance Team |
| EPI_ISL_3335408 | Lighthouse Lab in Milton Keynes | Wellcome Sanger Institute for the COVID-19 Genomics UK (COG-UK) Consortium | The Lighthouse Lab in Milton Keynes and Alex Alderton, Roberto Amato, Jeffrey Barrett, Sonia Goncalves, Ewan Harrison, David K. Jackson, Ian Johnston, Dominic Kwiatkowski, Cordelia Langford, John Sillitoe on behalf of the Wellcome Sanger Institute COVID-19 Surveillance Team |
| EPI_ISL_3335409 | Lighthouse Laboratory Plymouth | Wellcome Sanger Institute for the COVID-19 Genomics UK (COG-UK) Consortium | Lighthouse Laboratory Plymouth and Alex Alderton, Roberto Amato, Jeffrey Barrett, Sonia Goncalves, Ewan Harrison, David K. Jackson, Ian Johnston, Dominic Kwiatkowski, Cordelia Langford, John Sillitoe on behalf of the Wellcome Sanger Institute COVID-19 Surveillance Team |
| EPI_ISL_3335410 | Health Services Laboratories | Wellcome Sanger Institute for the COVID-19 Genomics UK (COG-UK) Consortium | Health Services Laboratories and Alex Alderton, Roberto Amato, Jeffrey Barrett, Sonia Goncalves, Ewan Harrison, David K. Jackson, Ian Johnston, Dominic Kwiatkowski, Cordelia Langford, John Sillitoe on behalf of the Wellcome Sanger Institute COVID-19 Surveillance Team |
| EPI_ISL_3335411, EPI_ISL_3335412 | Lighthouse Lab in Milton Keynes | Wellcome Sanger Institute for the COVID-19 Genomics UK (COG-UK) Consortium | The Lighthouse Lab in Milton Keynes and Alex Alderton, Roberto Amato, Jeffrey Barrett, Sonia Goncalves, Ewan Harrison, David K. Jackson, Ian Johnston, Dominic Kwiatkowski, Cordelia Langford, John Sillitoe on behalf of the Wellcome Sanger Institute COVID-19 Surveillance Team |
| EPI_ISL_3335413, EPI_ISL_3335414, EPI_ISL_3335415 | Lighthouse Laboratory Plymouth | Wellcome Sanger Institute for the COVID-19 Genomics UK (COG-UK) Consortium | Lighthouse Laboratory Plymouth and Alex Alderton, Roberto Amato, Jeffrey Barrett, Sonia Goncalves, Ewan Harrison, David K. Jackson, Ian Johnston, Dominic Kwiatkowski, Cordelia Langford, John Sillitoe on behalf of the Wellcome Sanger Institute COVID-19 Surveillance Team |
| EPI_ISL_3335416 | Lighthouse Lab in Milton Keynes | Wellcome Sanger Institute for the COVID-19 Genomics UK (COG-UK) | The Lighthouse Lab in Milton Keynes and Alex Alderton, Roberto Amato, Jeffrey Barrett, Sonia Goncalves, Ewan Harrison, David K. Jackson, Ian Johnston, Dominic Kwiatkowski, Cordelia Langford, John Sillitoe on behalf of the Wellcome Sanger |

| | | | |
|---|---|---|---|
| | | Consortium | Institute COVID-19 Surveillance Team |
| EPI_ISL_3335417, EPI_ISL_3335418 | Lighthouse Lab in Alderley Park | Wellcome Sanger Institute for the COVID-19 Genomics UK (COG-UK) Consortium | Jacquelyn Wynn, Mairead Hyland, The Lighthouse Lab in Alderley Park and Alex Alderton, Roberto Amato, Jeffrey Barrett, Sonia Goncalves, Ewan Harrison, David K. Jackson, Ian Johnston, Dominic Kwiatkowski, Cordelia Langford, John Sillitoe on behalf of the Wellcome Sanger Institute COVID-19 Surveillance Team |
| EPI_ISL_3335419, EPI_ISL_3335420 | Lighthouse Laboratory Plymouth | Wellcome Sanger Institute for the COVID-19 Genomics UK (COG-UK) Consortium | Lighthouse Laboratory Plymouth and Alex Alderton, Roberto Amato, Jeffrey Barrett, Sonia Goncalves, Ewan Harrison, David K. Jackson, Ian Johnston, Dominic Kwiatkowski, Cordelia Langford, John Sillitoe on behalf of the Wellcome Sanger Institute COVID-19 Surveillance Team |
| EPI_ISL_3335421 | Lighthouse Lab in Alderley Park | Wellcome Sanger Institute for the COVID-19 Genomics UK (COG-UK) Consortium | Jacquelyn Wynn, Mairead Hyland, The Lighthouse Lab in Alderley Park and Alex Alderton, Roberto Amato, Jeffrey Barrett, Sonia Goncalves, Ewan Harrison, David K. Jackson, Ian Johnston, Dominic Kwiatkowski, Cordelia Langford, John Sillitoe on behalf of the Wellcome Sanger Institute COVID-19 Surveillance Team |
| EPI_ISL_3335422 | Lighthouse Lab in Milton Keynes | Wellcome Sanger Institute for the COVID-19 Genomics UK (COG-UK) Consortium | The Lighthouse Lab in Milton Keynes and Alex Alderton, Roberto Amato, Jeffrey Barrett, Sonia Goncalves, Ewan Harrison, David K. Jackson, Ian Johnston, Dominic Kwiatkowski, Cordelia Langford, John Sillitoe on behalf of the Wellcome Sanger Institute COVID-19 Surveillance Team |
| EPI_ISL_3335423 | Lighthouse Laboratory Plymouth | Wellcome Sanger Institute for the COVID-19 Genomics UK (COG-UK) Consortium | Lighthouse Laboratory Plymouth and Alex Alderton, Roberto Amato, Jeffrey Barrett, Sonia Goncalves, Ewan Harrison, David K. Jackson, Ian Johnston, Dominic Kwiatkowski, Cordelia Langford, John Sillitoe on behalf of the Wellcome Sanger Institute COVID-19 Surveillance Team |
| EPI_ISL_3335424 | Lighthouse Lab in Milton Keynes | Wellcome Sanger Institute for the COVID-19 Genomics UK (COG-UK) Consortium | The Lighthouse Lab in Milton Keynes and Alex Alderton, Roberto Amato, Jeffrey Barrett, Sonia Goncalves, Ewan Harrison, David K. Jackson, Ian Johnston, Dominic Kwiatkowski, Cordelia Langford, John Sillitoe on behalf of the Wellcome Sanger Institute COVID-19 Surveillance Team |
| EPI_ISL_3335425 | Health Services Laboratories | Wellcome Sanger Institute for the COVID-19 Genomics UK (COG-UK) Consortium | Health Services Laboratories and Alex Alderton, Roberto Amato, Jeffrey Barrett, Sonia Goncalves, Ewan Harrison, David K. Jackson, Ian Johnston, Dominic Kwiatkowski, Cordelia Langford, John Sillitoe on behalf of the Wellcome Sanger Institute COVID-19 Surveillance Team |
| EPI_ISL_3335426 | Lighthouse Lab in Alderley Park | Wellcome Sanger Institute for the COVID-19 Genomics UK (COG-UK) Consortium | Jacquelyn Wynn, Mairead Hyland, The Lighthouse Lab in Alderley Park and Alex Alderton, Roberto Amato, Jeffrey Barrett, Sonia Goncalves, Ewan Harrison, David K. Jackson, Ian Johnston, Dominic Kwiatkowski, Cordelia Langford, John Sillitoe on behalf of the Wellcome Sanger Institute COVID-19 Surveillance Team |
| EPI_ISL_3335427 | Health Services Laboratories | Wellcome Sanger Institute for the COVID-19 Genomics UK (COG-UK) Consortium | Health Services Laboratories and Alex Alderton, Roberto Amato, Jeffrey Barrett, Sonia Goncalves, Ewan Harrison, David K. Jackson, Ian Johnston, Dominic Kwiatkowski, Cordelia Langford, John Sillitoe on behalf of the Wellcome Sanger Institute COVID-19 Surveillance Team |
| EPI_ISL_3335429 | Lighthouse Laboratory Plymouth | Wellcome Sanger Institute for the COVID-19 Genomics UK (COG-UK) Consortium | Lighthouse Laboratory Plymouth and Alex Alderton, Roberto Amato, Jeffrey Barrett, Sonia Goncalves, Ewan Harrison, David K. Jackson, Ian Johnston, Dominic Kwiatkowski, Cordelia Langford, John Sillitoe on behalf of the Wellcome Sanger Institute COVID-19 Surveillance Team |
| EPI_ISL_3335430 | Health Services Laboratories | Wellcome Sanger Institute for the COVID-19 Genomics UK (COG-UK) Consortium | Health Services Laboratories and Alex Alderton, Roberto Amato, Jeffrey Barrett, Sonia Goncalves, Ewan Harrison, David K. Jackson, Ian Johnston, Dominic Kwiatkowski, Cordelia Langford, John Sillitoe on behalf of the Wellcome Sanger Institute COVID-19 Surveillance Team |
| EPI_ISL_3335431 | Lighthouse Laboratory Plymouth | Wellcome Sanger Institute for the COVID-19 Genomics UK (COG-UK) Consortium | Lighthouse Laboratory Plymouth and Alex Alderton, Roberto Amato, Jeffrey Barrett, Sonia Goncalves, Ewan Harrison, David K. Jackson, Ian Johnston, Dominic Kwiatkowski, Cordelia Langford, John Sillitoe on behalf of the Wellcome Sanger Institute COVID-19 Surveillance Team |
| EPI_ISL_3335432, EPI_ISL_3335433, EPI_ISL_3335434, EPI_ISL_3335435, EPI_ISL_3335436, EPI_ISL_3335437 | Lighthouse Lab in Alderley Park | Wellcome Sanger Institute for the COVID-19 Genomics UK (COG-UK) Consortium | Jacquelyn Wynn, Mairead Hyland, The Lighthouse Lab in Alderley Park and Alex Alderton, Roberto Amato, Jeffrey Barrett, Sonia Goncalves, Ewan Harrison, David K. Jackson, Ian Johnston, Dominic Kwiatkowski, Cordelia Langford, John Sillitoe on behalf of the Wellcome Sanger Institute COVID-19 Surveillance Team |
| EPI_ISL_3335438, EPI_ISL_3335439 | Lighthouse Laboratory Plymouth | Wellcome Sanger Institute for the COVID-19 Genomics UK (COG-UK) Consortium | Lighthouse Laboratory Plymouth and Alex Alderton, Roberto Amato, Jeffrey Barrett, Sonia Goncalves, Ewan Harrison, David K. Jackson, Ian Johnston, Dominic Kwiatkowski, Cordelia Langford, John Sillitoe on behalf of the Wellcome Sanger Institute COVID-19 Surveillance Team |
| EPI_ISL_3335440 | Lighthouse Lab in Milton Keynes | Wellcome Sanger Institute for the COVID-19 Genomics UK (COG-UK) Consortium | The Lighthouse Lab in Milton Keynes and Alex Alderton, Roberto Amato, Jeffrey Barrett, Sonia Goncalves, Ewan Harrison, David K. Jackson, Ian Johnston, Dominic Kwiatkowski, Cordelia Langford, John Sillitoe on behalf of the Wellcome Sanger Institute COVID-19 Surveillance Team |
| EPI_ISL_3335441 | Lighthouse Lab in Alderley Park | Wellcome Sanger Institute for the COVID-19 Genomics UK (COG-UK) Consortium | Jacquelyn Wynn, Mairead Hyland, The Lighthouse Lab in Alderley Park and Alex Alderton, Roberto Amato, Jeffrey Barrett, Sonia Goncalves, Ewan Harrison, David K. Jackson, Ian Johnston, Dominic Kwiatkowski, Cordelia Langford, John Sillitoe on behalf of the Wellcome Sanger Institute COVID-19 Surveillance Team |
| EPI_ISL_3335442 | Lighthouse Lab in Milton Keynes | Wellcome Sanger Institute for the COVID-19 Genomics UK (COG-UK) Consortium | The Lighthouse Lab in Milton Keynes and Alex Alderton, Roberto Amato, Jeffrey Barrett, Sonia Goncalves, Ewan Harrison, David K. Jackson, Ian Johnston, Dominic Kwiatkowski, Cordelia Langford, John Sillitoe on behalf of the Wellcome Sanger Institute COVID-19 Surveillance Team |
| EPI_ISL_3335443 | Lighthouse Lab in Alderley Park | Wellcome Sanger Institute for the COVID-19 Genomics UK (COG-UK) Consortium | Jacquelyn Wynn, Mairead Hyland, The Lighthouse Lab in Alderley Park and Alex Alderton, Roberto Amato, Jeffrey Barrett, Sonia Goncalves, Ewan Harrison, David K. Jackson, Ian Johnston, Dominic Kwiatkowski, Cordelia Langford, John Sillitoe on behalf of the Wellcome Sanger Institute COVID-19 Surveillance Team |
| EPI_ISL_3335444 | Health Services Laboratories | Wellcome Sanger Institute for the COVID-19 Genomics UK (COG-UK) Consortium | Health Services Laboratories and Alex Alderton, Roberto Amato, Jeffrey Barrett, Sonia Goncalves, Ewan Harrison, David K. Jackson, Ian Johnston, Dominic Kwiatkowski, Cordelia Langford, John Sillitoe on behalf of the Wellcome Sanger Institute COVID-19 Surveillance Team |
| EPI_ISL_3335445, EPI_ISL_3335446 | Lighthouse Laboratory Plymouth | Wellcome Sanger Institute for the COVID-19 Genomics UK (COG-UK) Consortium | Lighthouse Laboratory Plymouth and Alex Alderton, Roberto Amato, Jeffrey Barrett, Sonia Goncalves, Ewan Harrison, David K. Jackson, Ian Johnston, Dominic Kwiatkowski, Cordelia Langford, John Sillitoe on behalf of the Wellcome Sanger Institute COVID-19 Surveillance Team |
| EPI_ISL_3335447 | Lighthouse Lab in Milton Keynes | Wellcome Sanger Institute for the COVID-19 Genomics UK (COG-UK) Consortium | The Lighthouse Lab in Milton Keynes and Alex Alderton, Roberto Amato, Jeffrey Barrett, Sonia Goncalves, Ewan Harrison, David K. Jackson, Ian Johnston, Dominic Kwiatkowski, Cordelia Langford, John Sillitoe on behalf of the Wellcome Sanger Institute COVID-19 Surveillance Team |
| EPI_ISL_3335448, EPI_ISL_3335449 | Health Services Laboratories | Wellcome Sanger Institute for the COVID-19 Genomics UK (COG-UK) Consortium | Health Services Laboratories and Alex Alderton, Roberto Amato, Jeffrey Barrett, Sonia Goncalves, Ewan Harrison, David K. Jackson, Ian Johnston, Dominic Kwiatkowski, Cordelia Langford, John Sillitoe on behalf of the Wellcome Sanger Institute COVID-19 Surveillance Team |
| EPI_ISL_3335450 | Lighthouse Lab in Milton Keynes | Wellcome Sanger Institute for the COVID-19 Genomics UK (COG-UK) Consortium | The Lighthouse Lab in Milton Keynes and Alex Alderton, Roberto Amato, Jeffrey Barrett, Sonia Goncalves, Ewan Harrison, David K. Jackson, Ian Johnston, Dominic Kwiatkowski, Cordelia Langford, John Sillitoe on behalf of the Wellcome Sanger Institute COVID-19 Surveillance Team |
| EPI_ISL_3335451 | Lighthouse Laboratory Plymouth | Wellcome Sanger Institute for the COVID-19 Genomics UK (COG-UK) Consortium | Lighthouse Laboratory Plymouth and Alex Alderton, Roberto Amato, Jeffrey Barrett, Sonia Goncalves, Ewan Harrison, David K. Jackson, Ian Johnston, Dominic Kwiatkowski, Cordelia Langford, John Sillitoe on behalf of the Wellcome Sanger Institute COVID-19 Surveillance Team |
| EPI_ISL_3335452, EPI_ISL_3335453 | Lighthouse Lab in Alderley Park | Wellcome Sanger Institute for the COVID-19 Genomics UK (COG-UK) Consortium | Jacquelyn Wynn, Mairead Hyland, The Lighthouse Lab in Alderley Park and Alex Alderton, Roberto Amato, Jeffrey Barrett, Sonia Goncalves, Ewan Harrison, David K. Jackson, Ian Johnston, Dominic Kwiatkowski, Cordelia Langford, John Sillitoe on behalf of the Wellcome Sanger Institute COVID-19 Surveillance Team |
| EPI_ISL_3335454 | Lighthouse Laboratory Plymouth | Wellcome Sanger Institute for the COVID-19 Genomics UK (COG-UK) Consortium | Lighthouse Laboratory Plymouth and Alex Alderton, Roberto Amato, Jeffrey Barrett, Sonia Goncalves, Ewan Harrison, David K. Jackson, Ian Johnston, Dominic Kwiatkowski, Cordelia Langford, John Sillitoe on behalf of the Wellcome Sanger Institute COVID-19 Surveillance Team |
| EPI_ISL_3335455, EPI_ISL_3335456, EPI_ISL_3335457 | Lighthouse Lab in Alderley Park | Wellcome Sanger Institute for the COVID-19 Genomics UK (COG-UK) Consortium | Jacquelyn Wynn, Mairead Hyland, The Lighthouse Lab in Alderley Park and Alex Alderton, Roberto Amato, Jeffrey Barrett, Sonia Goncalves, Ewan Harrison, David K. Jackson, Ian Johnston, Dominic Kwiatkowski, Cordelia Langford, John Sillitoe on behalf of the Wellcome Sanger Institute COVID-19 Surveillance Team |
| EPI_ISL_3335458 | Lighthouse Laboratory Plymouth | Wellcome Sanger Institute for the COVID-19 Genomics UK (COG-UK) Consortium | Lighthouse Laboratory Plymouth and Alex Alderton, Roberto Amato, Jeffrey Barrett, Sonia Goncalves, Ewan Harrison, David K. Jackson, Ian Johnston, Dominic Kwiatkowski, Cordelia Langford, John Sillitoe on behalf of the Wellcome Sanger Institute COVID-19 Surveillance Team |
| EPI_ISL_3335459 | Lighthouse Lab in Alderley Park | Wellcome Sanger Institute for the COVID-19 Genomics UK (COG-UK) Consortium | Jacquelyn Wynn, Mairead Hyland, The Lighthouse Lab in Alderley Park and Alex Alderton, Roberto Amato, Jeffrey Barrett, Sonia Goncalves, Ewan Harrison, David K. Jackson, Ian Johnston, Dominic Kwiatkowski, Cordelia Langford, John Sillitoe on behalf of the Wellcome Sanger Institute COVID-19 Surveillance Team |
| EPI_ISL_3335460 | Lighthouse Laboratory Plymouth | Wellcome Sanger Institute for the COVID-19 Genomics UK (COG-UK) Consortium | Lighthouse Laboratory Plymouth and Alex Alderton, Roberto Amato, Jeffrey Barrett, Sonia Goncalves, Ewan Harrison, David K. Jackson, Ian Johnston, Dominic Kwiatkowski, Cordelia Langford, John Sillitoe on behalf of the Wellcome Sanger Institute COVID-19 Surveillance Team |
| EPI_ISL_3335461, EPI_ISL_3335462, EPI_ISL_3335463, EPI_ISL_3335464 | Lighthouse Lab in Alderley Park | Wellcome Sanger Institute for the COVID-19 Genomics UK (COG-UK) Consortium | Jacquelyn Wynn, Mairead Hyland, The Lighthouse Lab in Alderley Park and Alex Alderton, Roberto Amato, Jeffrey Barrett, Sonia Goncalves, Ewan Harrison, David K. Jackson, Ian Johnston, Dominic Kwiatkowski, Cordelia Langford, John Sillitoe on behalf of the Wellcome Sanger Institute COVID-19 Surveillance Team |
| EPI_ISL_3335465 | Lighthouse Laboratory Plymouth | Wellcome Sanger Institute for the COVID-19 Genomics UK (COG-UK) Consortium | Lighthouse Laboratory Plymouth and Alex Alderton, Roberto Amato, Jeffrey Barrett, Sonia Goncalves, Ewan Harrison, David K. Jackson, Ian Johnston, Dominic Kwiatkowski, Cordelia Langford, John Sillitoe on behalf of the Wellcome Sanger Institute COVID-19 Surveillance Team |
| EPI_ISL_3335466, EPI_ISL_3335467, EPI_ISL_3335468, EPI_ISL_3335469 | Lighthouse Lab in Alderley Park | Wellcome Sanger Institute for the COVID-19 Genomics UK (COG-UK) Consortium | Jacquelyn Wynn, Mairead Hyland, The Lighthouse Lab in Alderley Park and Alex Alderton, Roberto Amato, Jeffrey Barrett, Sonia Goncalves, Ewan Harrison, David K. Jackson, Ian Johnston, Dominic Kwiatkowski, Cordelia Langford, John Sillitoe on behalf of the Wellcome Sanger Institute COVID-19 Surveillance Team |
| EPI_ISL_3335470 | Lighthouse Laboratory Plymouth | Wellcome Sanger Institute for the COVID-19 Genomics UK (COG-UK) Consortium | Lighthouse Laboratory Plymouth and Alex Alderton, Roberto Amato, Jeffrey Barrett, Sonia Goncalves, Ewan Harrison, David K. Jackson, Ian Johnston, Dominic Kwiatkowski, Cordelia Langford, John Sillitoe on behalf of the Wellcome Sanger Institute COVID-19 Surveillance Team |
| EPI_ISL_3335471 | Lighthouse Lab in Alderley Park | Wellcome Sanger Institute for the COVID-19 Genomics UK (COG-UK) Consortium | Jacquelyn Wynn, Mairead Hyland, The Lighthouse Lab in Alderley Park and Alex Alderton, Roberto Amato, Jeffrey Barrett, Sonia Goncalves, Ewan Harrison, David K. Jackson, Ian Johnston, Dominic Kwiatkowski, Cordelia Langford, John Sillitoe on behalf of the Wellcome Sanger Institute COVID-19 Surveillance Team |
| EPI_ISL_3335472 | Lighthouse Laboratory Plymouth | Wellcome Sanger Institute for the COVID-19 Genomics UK (COG-UK) Consortium | Lighthouse Laboratory Plymouth and Alex Alderton, Roberto Amato, Jeffrey Barrett, Sonia Goncalves, Ewan Harrison, David K. Jackson, Ian Johnston, Dominic Kwiatkowski, Cordelia Langford, John Sillitoe on behalf of the Wellcome Sanger Institute COVID-19 Surveillance Team |
| EPI_ISL_3335473, EPI_ISL_3335474, EPI_ISL_3335475, EPI_ISL_3335476, EPI_ISL_3335477, EPI_ISL_3335478 | Lighthouse Lab in Alderley Park | Wellcome Sanger Institute for the COVID-19 Genomics UK (COG-UK) Consortium | Jacquelyn Wynn, Mairead Hyland, The Lighthouse Lab in Alderley Park and Alex Alderton, Roberto Amato, Jeffrey Barrett, Sonia Goncalves, Ewan Harrison, David K. Jackson, Ian Johnston, Dominic Kwiatkowski, Cordelia Langford, John Sillitoe on behalf of the Wellcome Sanger Institute COVID-19 Surveillance Team |
| EPI_ISL_3335479 | Lighthouse Laboratory Plymouth | Wellcome Sanger Institute for the COVID-19 Genomics UK (COG-UK) Consortium | Lighthouse Laboratory Plymouth and Alex Alderton, Roberto Amato, Jeffrey Barrett, Sonia Goncalves, Ewan Harrison, David K. Jackson, Ian Johnston, Dominic Kwiatkowski, Cordelia Langford, John Sillitoe on behalf of the Wellcome Sanger Institute COVID-19 Surveillance Team |
| EPI_ISL_3335480 | Lighthouse Lab in Alderley Park | Wellcome Sanger Institute for the COVID-19 Genomics UK (COG-UK) Consortium | Jacquelyn Wynn, Mairead Hyland, The Lighthouse Lab in Alderley Park and Alex Alderton, Roberto Amato, Jeffrey Barrett, Sonia Goncalves, Ewan Harrison, David K. Jackson, Ian Johnston, Dominic Kwiatkowski, Cordelia Langford, John Sillitoe on behalf of the Wellcome Sanger Institute COVID-19 Surveillance Team |
| EPI_ISL_3335481 | Lighthouse Laboratory Plymouth | Wellcome Sanger Institute for the COVID-19 Genomics UK (COG-UK) | Lighthouse Laboratory Plymouth and Alex Alderton, Roberto Amato, Jeffrey Barrett, Sonia Goncalves, Ewan Harrison, David K. Jackson, Ian Johnston, Dominic Kwiatkowski, Cordelia Langford, John Sillitoe on behalf of the Wellcome Sanger |

| | | | |
|---|---|---|---|
| | | Consortium | Institute COVID-19 Surveillance Team |
| EPI_ISL_3335482, EPI_ISL_3335483, EPI_ISL_3335484, EPI_ISL_3335485, EPI_ISL_3335486, EPI_ISL_3335487 | Lighthouse Lab in Alderley Park | Wellcome Sanger Institute for the COVID-19 Genomics UK (COG-UK) Consortium | Jacquelyn Wynn, Mairead Hyland, The Lighthouse Lab in Alderley Park and Alex Alderton, Roberto Amato, Jeffrey Barrett, Sonia Goncalves, Ewan Harrison, David K. Jackson, Ian Johnston, Dominic Kwiatkowski, Cordelia Langford, John Sillitoe on behalf of the Wellcome Sanger Institute COVID-19 Surveillance Team |
| EPI_ISL_3335488 | Lighthouse Laboratory Plymouth | Wellcome Sanger Institute for the COVID-19 Genomics UK (COG-UK) Consortium | Lighthouse Laboratory Plymouth and Alex Alderton, Roberto Amato, Jeffrey Barrett, Sonia Goncalves, Ewan Harrison, David K. Jackson, Ian Johnston, Dominic Kwiatkowski, Cordelia Langford, John Sillitoe on behalf of the Wellcome Sanger Institute COVID-19 Surveillance Team |
| EPI_ISL_3335489, EPI_ISL_3335490, EPI_ISL_3335491 | Lighthouse Lab in Alderley Park | Wellcome Sanger Institute for the COVID-19 Genomics UK (COG-UK) Consortium | Jacquelyn Wynn, Mairead Hyland, The Lighthouse Lab in Alderley Park and Alex Alderton, Roberto Amato, Jeffrey Barrett, Sonia Goncalves, Ewan Harrison, David K. Jackson, Ian Johnston, Dominic Kwiatkowski, Cordelia Langford, John Sillitoe on behalf of the Wellcome Sanger Institute COVID-19 Surveillance Team |
| EPI_ISL_3335492 | Lighthouse Laboratory Plymouth | Wellcome Sanger Institute for the COVID-19 Genomics UK (COG-UK) Consortium | Lighthouse Laboratory Plymouth and Alex Alderton, Roberto Amato, Jeffrey Barrett, Sonia Goncalves, Ewan Harrison, David K. Jackson, Ian Johnston, Dominic Kwiatkowski, Cordelia Langford, John Sillitoe on behalf of the Wellcome Sanger Institute COVID-19 Surveillance Team |
| EPI_ISL_3335493 | Lighthouse Lab in Alderley Park | Wellcome Sanger Institute for the COVID-19 Genomics UK (COG-UK) Consortium | Jacquelyn Wynn, Mairead Hyland, The Lighthouse Lab in Alderley Park and Alex Alderton, Roberto Amato, Jeffrey Barrett, Sonia Goncalves, Ewan Harrison, David K. Jackson, Ian Johnston, Dominic Kwiatkowski, Cordelia Langford, John Sillitoe on behalf of the Wellcome Sanger Institute COVID-19 Surveillance Team |
| EPI_ISL_3335494 | Lighthouse Laboratory Plymouth | Wellcome Sanger Institute for the COVID-19 Genomics UK (COG-UK) Consortium | Lighthouse Laboratory Plymouth and Alex Alderton, Roberto Amato, Jeffrey Barrett, Sonia Goncalves, Ewan Harrison, David K. Jackson, Ian Johnston, Dominic Kwiatkowski, Cordelia Langford, John Sillitoe on behalf of the Wellcome Sanger Institute COVID-19 Surveillance Team |
| EPI_ISL_3335495, EPI_ISL_3335496 | Lighthouse Lab in Alderley Park | Wellcome Sanger Institute for the COVID-19 Genomics UK (COG-UK) Consortium | Jacquelyn Wynn, Mairead Hyland, The Lighthouse Lab in Alderley Park and Alex Alderton, Roberto Amato, Jeffrey Barrett, Sonia Goncalves, Ewan Harrison, David K. Jackson, Ian Johnston, Dominic Kwiatkowski, Cordelia Langford, John Sillitoe on behalf of the Wellcome Sanger Institute COVID-19 Surveillance Team |
| EPI_ISL_3335497 | Lighthouse Lab in Milton Keynes | Wellcome Sanger Institute for the COVID-19 Genomics UK (COG-UK) Consortium | The Lighthouse Lab in Milton Keynes and Alex Alderton, Roberto Amato, Jeffrey Barrett, Sonia Goncalves, Ewan Harrison, David K. Jackson, Ian Johnston, Dominic Kwiatkowski, Cordelia Langford, John Sillitoe on behalf of the Wellcome Sanger Institute COVID-19 Surveillance Team |
| EPI_ISL_3335498 | Lighthouse Lab in Alderley Park | Wellcome Sanger Institute for the COVID-19 Genomics UK (COG-UK) Consortium | Jacquelyn Wynn, Mairead Hyland, The Lighthouse Lab in Alderley Park and Alex Alderton, Roberto Amato, Jeffrey Barrett, Sonia Goncalves, Ewan Harrison, David K. Jackson, Ian Johnston, Dominic Kwiatkowski, Cordelia Langford, John Sillitoe on behalf of the Wellcome Sanger Institute COVID-19 Surveillance Team |
| EPI_ISL_3335499 | Lighthouse Laboratory Plymouth | Wellcome Sanger Institute for the COVID-19 Genomics UK (COG-UK) Consortium | Lighthouse Laboratory Plymouth and Alex Alderton, Roberto Amato, Jeffrey Barrett, Sonia Goncalves, Ewan Harrison, David K. Jackson, Ian Johnston, Dominic Kwiatkowski, Cordelia Langford, John Sillitoe on behalf of the Wellcome Sanger Institute COVID-19 Surveillance Team |
| EPI_ISL_3335500, EPI_ISL_3335501, EPI_ISL_3335502, EPI_ISL_3335503, EPI_ISL_3335504, EPI_ISL_3335505, EPI_ISL_3335506, EPI_ISL_3335507, EPI_ISL_3335508, EPI_ISL_3335509, EPI_ISL_3335510, EPI_ISL_3335511, EPI_ISL_3335512, EPI_ISL_3335513 | | | |
| see above | Lighthouse Lab in Alderley Park | Wellcome Sanger Institute for the COVID-19 Genomics UK (COG-UK) Consortium | Jacquelyn Wynn, Mairead Hyland, The Lighthouse Lab in Alderley Park and Alex Alderton, Roberto Amato, Jeffrey Barrett, Sonia Goncalves, Ewan Harrison, David K. Jackson, Ian Johnston, Dominic Kwiatkowski, Cordelia Langford, John Sillitoe on behalf of the Wellcome Sanger Institute COVID-19 Surveillance Team |
| EPI_ISL_3335514 | Lighthouse Laboratory Plymouth | Wellcome Sanger Institute for the COVID-19 Genomics UK (COG-UK) Consortium | Lighthouse Laboratory Plymouth and Alex Alderton, Roberto Amato, Jeffrey Barrett, Sonia Goncalves, Ewan Harrison, David K. Jackson, Ian Johnston, Dominic Kwiatkowski, Cordelia Langford, John Sillitoe on behalf of the Wellcome Sanger Institute COVID-19 Surveillance Team |
| EPI_ISL_3335515, EPI_ISL_3335516, EPI_ISL_3335517, EPI_ISL_3335518, EPI_ISL_3335519 | Lighthouse Lab in Alderley Park | Wellcome Sanger Institute for the COVID-19 Genomics UK (COG-UK) Consortium | Jacquelyn Wynn, Mairead Hyland, The Lighthouse Lab in Alderley Park and Alex Alderton, Roberto Amato, Jeffrey Barrett, Sonia Goncalves, Ewan Harrison, David K. Jackson, Ian Johnston, Dominic Kwiatkowski, Cordelia Langford, John Sillitoe on behalf of the Wellcome Sanger Institute COVID-19 Surveillance Team |
| EPI_ISL_3335520, EPI_ISL_3335521 | Lighthouse Laboratory Plymouth | Wellcome Sanger Institute for the COVID-19 Genomics UK (COG-UK) Consortium | Lighthouse Laboratory Plymouth and Alex Alderton, Roberto Amato, Jeffrey Barrett, Sonia Goncalves, Ewan Harrison, David K. Jackson, Ian Johnston, Dominic Kwiatkowski, Cordelia Langford, John Sillitoe on behalf of the Wellcome Sanger Institute COVID-19 Surveillance Team |
| EPI_ISL_3335522 | Lighthouse Lab in Alderley Park | Wellcome Sanger Institute for the COVID-19 Genomics UK (COG-UK) Consortium | Jacquelyn Wynn, Mairead Hyland, The Lighthouse Lab in Alderley Park and Alex Alderton, Roberto Amato, Jeffrey Barrett, Sonia Goncalves, Ewan Harrison, David K. Jackson, Ian Johnston, Dominic Kwiatkowski, Cordelia Langford, John Sillitoe on behalf of the Wellcome Sanger Institute COVID-19 Surveillance Team |
| EPI_ISL_3335523 | Lighthouse Laboratory Plymouth | Wellcome Sanger Institute for the COVID-19 Genomics UK (COG-UK) Consortium | Lighthouse Laboratory Plymouth and Alex Alderton, Roberto Amato, Jeffrey Barrett, Sonia Goncalves, Ewan Harrison, David K. Jackson, Ian Johnston, Dominic Kwiatkowski, Cordelia Langford, John Sillitoe on behalf of the Wellcome Sanger Institute COVID-19 Surveillance Team |
| EPI_ISL_3335524 | Lighthouse Lab in Alderley Park | Wellcome Sanger Institute for the COVID-19 Genomics UK (COG-UK) Consortium | Jacquelyn Wynn, Mairead Hyland, The Lighthouse Lab in Alderley Park and Alex Alderton, Roberto Amato, Jeffrey Barrett, Sonia Goncalves, Ewan Harrison, David K. Jackson, Ian Johnston, Dominic Kwiatkowski, Cordelia Langford, John Sillitoe on behalf of the Wellcome Sanger Institute COVID-19 Surveillance Team |
| EPI_ISL_3335525 | Lighthouse Laboratory Plymouth | Wellcome Sanger Institute for the COVID-19 Genomics UK (COG-UK) Consortium | Lighthouse Laboratory Plymouth and Alex Alderton, Roberto Amato, Jeffrey Barrett, Sonia Goncalves, Ewan Harrison, David K. Jackson, Ian Johnston, Dominic Kwiatkowski, Cordelia Langford, John Sillitoe on behalf of the Wellcome Sanger Institute COVID-19 Surveillance Team |
| EPI_ISL_3335526, EPI_ISL_3335527 | Lighthouse Lab in Alderley Park | Wellcome Sanger Institute for the COVID-19 Genomics UK (COG-UK) Consortium | Jacquelyn Wynn, Mairead Hyland, The Lighthouse Lab in Alderley Park and Alex Alderton, Roberto Amato, Jeffrey Barrett, Sonia Goncalves, Ewan Harrison, David K. Jackson, Ian Johnston, Dominic Kwiatkowski, Cordelia Langford, John Sillitoe on behalf of the Wellcome Sanger Institute COVID-19 Surveillance Team |
| EPI_ISL_3335528 | Lighthouse Laboratory Plymouth | Wellcome Sanger Institute for the COVID-19 Genomics UK (COG-UK) Consortium | Lighthouse Laboratory Plymouth and Alex Alderton, Roberto Amato, Jeffrey Barrett, Sonia Goncalves, Ewan Harrison, David K. Jackson, Ian Johnston, Dominic Kwiatkowski, Cordelia Langford, John Sillitoe on behalf of the Wellcome Sanger Institute COVID-19 Surveillance Team |
| EPI_ISL_3335529, EPI_ISL_3335530, EPI_ISL_3335531, EPI_ISL_3335532 | Lighthouse Lab in Alderley Park | Wellcome Sanger Institute for the COVID-19 Genomics UK (COG-UK) Consortium | Jacquelyn Wynn, Mairead Hyland, The Lighthouse Lab in Alderley Park and Alex Alderton, Roberto Amato, Jeffrey Barrett, Sonia Goncalves, Ewan Harrison, David K. Jackson, Ian Johnston, Dominic Kwiatkowski, Cordelia Langford, John Sillitoe on behalf of the Wellcome Sanger Institute COVID-19 Surveillance Team |
| EPI_ISL_3335533, EPI_ISL_3335534 | Lighthouse Laboratory Plymouth | Wellcome Sanger Institute for the COVID-19 Genomics UK (COG-UK) Consortium | Lighthouse Laboratory Plymouth and Alex Alderton, Roberto Amato, Jeffrey Barrett, Sonia Goncalves, Ewan Harrison, David K. Jackson, Ian Johnston, Dominic Kwiatkowski, Cordelia Langford, John Sillitoe on behalf of the Wellcome Sanger Institute COVID-19 Surveillance Team |
| EPI_ISL_3335535 | Lighthouse Lab in Alderley Park | Wellcome Sanger Institute for the COVID-19 Genomics UK (COG-UK) Consortium | Jacquelyn Wynn, Mairead Hyland, The Lighthouse Lab in Alderley Park and Alex Alderton, Roberto Amato, Jeffrey Barrett, Sonia Goncalves, Ewan Harrison, David K. Jackson, Ian Johnston, Dominic Kwiatkowski, Cordelia Langford, John Sillitoe on behalf of the Wellcome Sanger Institute COVID-19 Surveillance Team |
| EPI_ISL_3335536, EPI_ISL_3335537, EPI_ISL_3335538 | Lighthouse Laboratory Plymouth | Wellcome Sanger Institute for the COVID-19 Genomics UK (COG-UK) Consortium | Lighthouse Laboratory Plymouth and Alex Alderton, Roberto Amato, Jeffrey Barrett, Sonia Goncalves, Ewan Harrison, David K. Jackson, Ian Johnston, Dominic Kwiatkowski, Cordelia Langford, John Sillitoe on behalf of the Wellcome Sanger Institute COVID-19 Surveillance Team |
| EPI_ISL_3335539 | Lighthouse Lab in Alderley Park | Wellcome Sanger Institute for the COVID-19 Genomics UK (COG-UK) Consortium | Jacquelyn Wynn, Mairead Hyland, The Lighthouse Lab in Alderley Park and Alex Alderton, Roberto Amato, Jeffrey Barrett, Sonia Goncalves, Ewan Harrison, David K. Jackson, Ian Johnston, Dominic Kwiatkowski, Cordelia Langford, John Sillitoe on behalf of the Wellcome Sanger Institute COVID-19 Surveillance Team |
| EPI_ISL_3335540 | Lighthouse Laboratory Plymouth | Wellcome Sanger Institute for the COVID-19 Genomics UK (COG-UK) Consortium | Lighthouse Laboratory Plymouth and Alex Alderton, Roberto Amato, Jeffrey Barrett, Sonia Goncalves, Ewan Harrison, David K. Jackson, Ian Johnston, Dominic Kwiatkowski, Cordelia Langford, John Sillitoe on behalf of the Wellcome Sanger Institute COVID-19 Surveillance Team |
| EPI_ISL_3335541, EPI_ISL_3335542, EPI_ISL_3335543 | Lighthouse Lab in Alderley Park | Wellcome Sanger Institute for the COVID-19 Genomics UK (COG-UK) Consortium | Jacquelyn Wynn, Mairead Hyland, The Lighthouse Lab in Alderley Park and Alex Alderton, Roberto Amato, Jeffrey Barrett, Sonia Goncalves, Ewan Harrison, David K. Jackson, Ian Johnston, Dominic Kwiatkowski, Cordelia Langford, John Sillitoe on behalf of the Wellcome Sanger Institute COVID-19 Surveillance Team |
| EPI_ISL_3335544 | Lighthouse Laboratory Plymouth | Wellcome Sanger Institute for the COVID-19 Genomics UK (COG-UK) Consortium | Lighthouse Laboratory Plymouth and Alex Alderton, Roberto Amato, Jeffrey Barrett, Sonia Goncalves, Ewan Harrison, David K. Jackson, Ian Johnston, Dominic Kwiatkowski, Cordelia Langford, John Sillitoe on behalf of the Wellcome Sanger Institute COVID-19 Surveillance Team |
| EPI_ISL_3335545 | Lighthouse Lab in Alderley Park | Wellcome Sanger Institute for the COVID-19 Genomics UK (COG-UK) Consortium | Jacquelyn Wynn, Mairead Hyland, The Lighthouse Lab in Alderley Park and Alex Alderton, Roberto Amato, Jeffrey Barrett, Sonia Goncalves, Ewan Harrison, David K. Jackson, Ian Johnston, Dominic Kwiatkowski, Cordelia Langford, John Sillitoe on behalf of the Wellcome Sanger Institute COVID-19 Surveillance Team |
| EPI_ISL_3335546, EPI_ISL_3335547 | Lighthouse Laboratory Plymouth | Wellcome Sanger Institute for the COVID-19 Genomics UK (COG-UK) Consortium | Lighthouse Laboratory Plymouth and Alex Alderton, Roberto Amato, Jeffrey Barrett, Sonia Goncalves, Ewan Harrison, David K. Jackson, Ian Johnston, Dominic Kwiatkowski, Cordelia Langford, John Sillitoe on behalf of the Wellcome Sanger Institute COVID-19 Surveillance Team |
| EPI_ISL_3335548, EPI_ISL_3335549, EPI_ISL_3335550, EPI_ISL_3335551 | Lighthouse Lab in Alderley Park | Wellcome Sanger Institute for the COVID-19 Genomics UK (COG-UK) Consortium | Jacquelyn Wynn, Mairead Hyland, The Lighthouse Lab in Alderley Park and Alex Alderton, Roberto Amato, Jeffrey Barrett, Sonia Goncalves, Ewan Harrison, David K. Jackson, Ian Johnston, Dominic Kwiatkowski, Cordelia Langford, John Sillitoe on behalf of the Wellcome Sanger Institute COVID-19 Surveillance Team |
| EPI_ISL_3335552, EPI_ISL_3335553 | Lighthouse Lab in Milton Keynes | Wellcome Sanger Institute for the COVID-19 Genomics UK (COG-UK) Consortium | The Lighthouse Lab in Milton Keynes and Alex Alderton, Roberto Amato, Jeffrey Barrett, Sonia Goncalves, Ewan Harrison, David K. Jackson, Ian Johnston, Dominic Kwiatkowski, Cordelia Langford, John Sillitoe on behalf of the Wellcome Sanger Institute COVID-19 Surveillance Team |
| EPI_ISL_3335554 | Lighthouse Lab in Alderley Park | Wellcome Sanger Institute for the COVID-19 Genomics UK (COG-UK) Consortium | Jacquelyn Wynn, Mairead Hyland, The Lighthouse Lab in Alderley Park and Alex Alderton, Roberto Amato, Jeffrey Barrett, Sonia Goncalves, Ewan Harrison, David K. Jackson, Ian Johnston, Dominic Kwiatkowski, Cordelia Langford, John Sillitoe on behalf of the Wellcome Sanger Institute COVID-19 Surveillance Team |
| EPI_ISL_3335555 | Lighthouse Laboratory Plymouth | Wellcome Sanger Institute for the COVID-19 Genomics UK (COG-UK) Consortium | Lighthouse Laboratory Plymouth and Alex Alderton, Roberto Amato, Jeffrey Barrett, Sonia Goncalves, Ewan Harrison, David K. Jackson, Ian Johnston, Dominic Kwiatkowski, Cordelia Langford, John Sillitoe on behalf of the Wellcome Sanger Institute COVID-19 Surveillance Team |
| EPI_ISL_3335556, EPI_ISL_3335557 | Lighthouse Lab in Alderley Park | Wellcome Sanger Institute for the COVID-19 Genomics UK (COG-UK) Consortium | Jacquelyn Wynn, Mairead Hyland, The Lighthouse Lab in Alderley Park and Alex Alderton, Roberto Amato, Jeffrey Barrett, Sonia Goncalves, Ewan Harrison, David K. Jackson, Ian Johnston, Dominic Kwiatkowski, Cordelia Langford, John Sillitoe on behalf of the Wellcome Sanger Institute COVID-19 Surveillance Team |
| EPI_ISL_3335558 | Lighthouse Laboratory Plymouth | Wellcome Sanger Institute for the COVID-19 Genomics UK (COG-UK) Consortium | Lighthouse Laboratory Plymouth and Alex Alderton, Roberto Amato, Jeffrey Barrett, Sonia Goncalves, Ewan Harrison, David K. Jackson, Ian Johnston, Dominic Kwiatkowski, Cordelia Langford, John Sillitoe on behalf of the Wellcome Sanger Institute COVID-19 Surveillance Team |
| EPI_ISL_3335559, EPI_ISL_3335560 | Lighthouse Lab in Alderley Park | Wellcome Sanger Institute for the COVID-19 Genomics UK (COG-UK) Consortium | Jacquelyn Wynn, Mairead Hyland, The Lighthouse Lab in Alderley Park and Alex Alderton, Roberto Amato, Jeffrey Barrett, Sonia Goncalves, Ewan Harrison, David K. Jackson, Ian Johnston, Dominic Kwiatkowski, Cordelia Langford, John Sillitoe on behalf of the Wellcome Sanger Institute COVID-19 Surveillance Team |
| EPI_ISL_3335561 | Lighthouse Laboratory Plymouth | Wellcome Sanger Institute for the COVID-19 Genomics UK (COG-UK) Consortium | Lighthouse Laboratory Plymouth and Alex Alderton, Roberto Amato, Jeffrey Barrett, Sonia Goncalves, Ewan Harrison, David K. Jackson, Ian Johnston, Dominic Kwiatkowski, Cordelia Langford, John Sillitoe on behalf of the Wellcome Sanger Institute COVID-19 Surveillance Team |
| EPI_ISL_3335562, | Lighthouse Lab in Alderley Park | Wellcome Sanger Institute for the COVID-19 Genomics UK (COG-UK) | Jacquelyn Wynn, Mairead Hyland, The Lighthouse Lab in Alderley Park and Alex Alderton, Roberto Amato, Jeffrey Barrett, Sonia Goncalves, Ewan Harrison, David K. Jackson, Ian Johnston, Dominic Kwiatkowski, Cordelia Langford, John Sillitoe on |

| | | | |
|---|---|---|---|
| EPI_ISL_3335563, EPI_ISL_3335564, EPI_ISL_3335565, EPI_ISL_3335566 | | Consortium | behalf of the Wellcome Sanger Institute COVID-19 Surveillance Team |
| EPI_ISL_3335567, EPI_ISL_3335568 | Lighthouse Laboratory Plymouth | Wellcome Sanger Institute for the COVID-19 Genomics UK (COG-UK) Consortium | Lighthouse Laboratory Plymouth and Alex Alderton, Roberto Amato, Jeffrey Barrett, Sonia Goncalves, Ewan Harrison, David K. Jackson, Ian Johnston, Dominic Kwiatkowski, Cordelia Langford, John Sillitoe on behalf of the Wellcome Sanger Institute COVID-19 Surveillance Team |
| EPI_ISL_3335569 | Lighthouse Lab in Alderley Park | Wellcome Sanger Institute for the COVID-19 Genomics UK (COG-UK) Consortium | Jacquelyn Wynn, Mairead Hyland, The Lighthouse Lab in Alderley Park and Alex Alderton, Roberto Amato, Jeffrey Barrett, Sonia Goncalves, Ewan Harrison, David K. Jackson, Ian Johnston, Dominic Kwiatkowski, Cordelia Langford, John Sillitoe on behalf of the Wellcome Sanger Institute COVID-19 Surveillance Team |
| EPI_ISL_3335570, EPI_ISL_3335571, EPI_ISL_3335572 | Lighthouse Laboratory Plymouth | Wellcome Sanger Institute for the COVID-19 Genomics UK (COG-UK) Consortium | Lighthouse Laboratory Plymouth and Alex Alderton, Roberto Amato, Jeffrey Barrett, Sonia Goncalves, Ewan Harrison, David K. Jackson, Ian Johnston, Dominic Kwiatkowski, Cordelia Langford, John Sillitoe on behalf of the Wellcome Sanger Institute COVID-19 Surveillance Team |
| EPI_ISL_3335573, EPI_ISL_3335574, EPI_ISL_3335575, EPI_ISL_3335576, EPI_ISL_3335577, EPI_ISL_3335578, EPI_ISL_3335579 | Lighthouse Lab in Alderley Park | Wellcome Sanger Institute for the COVID-19 Genomics UK (COG-UK) Consortium | Jacquelyn Wynn, Mairead Hyland, The Lighthouse Lab in Alderley Park and Alex Alderton, Roberto Amato, Jeffrey Barrett, Sonia Goncalves, Ewan Harrison, David K. Jackson, Ian Johnston, Dominic Kwiatkowski, Cordelia Langford, John Sillitoe on behalf of the Wellcome Sanger Institute COVID-19 Surveillance Team |
| EPI_ISL_3335580 | Lighthouse Laboratory Plymouth | Wellcome Sanger Institute for the COVID-19 Genomics UK (COG-UK) Consortium | Lighthouse Laboratory Plymouth and Alex Alderton, Roberto Amato, Jeffrey Barrett, Sonia Goncalves, Ewan Harrison, David K. Jackson, Ian Johnston, Dominic Kwiatkowski, Cordelia Langford, John Sillitoe on behalf of the Wellcome Sanger Institute COVID-19 Surveillance Team |
| EPI_ISL_3335581, EPI_ISL_3335582 | Lighthouse Lab in Alderley Park | Wellcome Sanger Institute for the COVID-19 Genomics UK (COG-UK) Consortium | Jacquelyn Wynn, Mairead Hyland, The Lighthouse Lab in Alderley Park and Alex Alderton, Roberto Amato, Jeffrey Barrett, Sonia Goncalves, Ewan Harrison, David K. Jackson, Ian Johnston, Dominic Kwiatkowski, Cordelia Langford, John Sillitoe on behalf of the Wellcome Sanger Institute COVID-19 Surveillance Team |
| EPI_ISL_3335583 | Lighthouse Laboratory Plymouth | Wellcome Sanger Institute for the COVID-19 Genomics UK (COG-UK) Consortium | Lighthouse Laboratory Plymouth and Alex Alderton, Roberto Amato, Jeffrey Barrett, Sonia Goncalves, Ewan Harrison, David K. Jackson, Ian Johnston, Dominic Kwiatkowski, Cordelia Langford, John Sillitoe on behalf of the Wellcome Sanger Institute COVID-19 Surveillance Team |
| EPI_ISL_3335584, EPI_ISL_3335585 | Lighthouse Lab in Alderley Park | Wellcome Sanger Institute for the COVID-19 Genomics UK (COG-UK) Consortium | Jacquelyn Wynn, Mairead Hyland, The Lighthouse Lab in Alderley Park and Alex Alderton, Roberto Amato, Jeffrey Barrett, Sonia Goncalves, Ewan Harrison, David K. Jackson, Ian Johnston, Dominic Kwiatkowski, Cordelia Langford, John Sillitoe on behalf of the Wellcome Sanger Institute COVID-19 Surveillance Team |
| EPI_ISL_3335586 | Lighthouse Laboratory Plymouth | Wellcome Sanger Institute for the COVID-19 Genomics UK (COG-UK) Consortium | Lighthouse Laboratory Plymouth and Alex Alderton, Roberto Amato, Jeffrey Barrett, Sonia Goncalves, Ewan Harrison, David K. Jackson, Ian Johnston, Dominic Kwiatkowski, Cordelia Langford, John Sillitoe on behalf of the Wellcome Sanger Institute COVID-19 Surveillance Team |
| EPI_ISL_3335587, EPI_ISL_3335588, EPI_ISL_3335589, EPI_ISL_3335590, EPI_ISL_3335591 | Lighthouse Lab in Alderley Park | Wellcome Sanger Institute for the COVID-19 Genomics UK (COG-UK) Consortium | Jacquelyn Wynn, Mairead Hyland, The Lighthouse Lab in Alderley Park and Alex Alderton, Roberto Amato, Jeffrey Barrett, Sonia Goncalves, Ewan Harrison, David K. Jackson, Ian Johnston, Dominic Kwiatkowski, Cordelia Langford, John Sillitoe on behalf of the Wellcome Sanger Institute COVID-19 Surveillance Team |
| EPI_ISL_3335592, EPI_ISL_3335593, EPI_ISL_3335594 | Lighthouse Laboratory Plymouth | Wellcome Sanger Institute for the COVID-19 Genomics UK (COG-UK) Consortium | Lighthouse Laboratory Plymouth and Alex Alderton, Roberto Amato, Jeffrey Barrett, Sonia Goncalves, Ewan Harrison, David K. Jackson, Ian Johnston, Dominic Kwiatkowski, Cordelia Langford, John Sillitoe on behalf of the Wellcome Sanger Institute COVID-19 Surveillance Team |
| EPI_ISL_3335595, EPI_ISL_3335596, EPI_ISL_3335597 | Lighthouse Lab in Alderley Park | Wellcome Sanger Institute for the COVID-19 Genomics UK (COG-UK) Consortium | Jacquelyn Wynn, Mairead Hyland, The Lighthouse Lab in Alderley Park and Alex Alderton, Roberto Amato, Jeffrey Barrett, Sonia Goncalves, Ewan Harrison, David K. Jackson, Ian Johnston, Dominic Kwiatkowski, Cordelia Langford, John Sillitoe on behalf of the Wellcome Sanger Institute COVID-19 Surveillance Team |
| EPI_ISL_3335598 | Lighthouse Laboratory Plymouth | Wellcome Sanger Institute for the COVID-19 Genomics UK (COG-UK) Consortium | Lighthouse Laboratory Plymouth and Alex Alderton, Roberto Amato, Jeffrey Barrett, Sonia Goncalves, Ewan Harrison, David K. Jackson, Ian Johnston, Dominic Kwiatkowski, Cordelia Langford, John Sillitoe on behalf of the Wellcome Sanger Institute COVID-19 Surveillance Team |
| EPI_ISL_3335599, EPI_ISL_3335600, EPI_ISL_3335601 | Lighthouse Lab in Alderley Park | Wellcome Sanger Institute for the COVID-19 Genomics UK (COG-UK) Consortium | Jacquelyn Wynn, Mairead Hyland, The Lighthouse Lab in Alderley Park and Alex Alderton, Roberto Amato, Jeffrey Barrett, Sonia Goncalves, Ewan Harrison, David K. Jackson, Ian Johnston, Dominic Kwiatkowski, Cordelia Langford, John Sillitoe on behalf of the Wellcome Sanger Institute COVID-19 Surveillance Team |
| EPI_ISL_3335602 | Lighthouse Laboratory Plymouth | Wellcome Sanger Institute for the COVID-19 Genomics UK (COG-UK) Consortium | Lighthouse Laboratory Plymouth and Alex Alderton, Roberto Amato, Jeffrey Barrett, Sonia Goncalves, Ewan Harrison, David K. Jackson, Ian Johnston, Dominic Kwiatkowski, Cordelia Langford, John Sillitoe on behalf of the Wellcome Sanger Institute COVID-19 Surveillance Team |
| EPI_ISL_3335603, EPI_ISL_3335604 | Lighthouse Lab in Alderley Park | Wellcome Sanger Institute for the COVID-19 Genomics UK (COG-UK) Consortium | Jacquelyn Wynn, Mairead Hyland, The Lighthouse Lab in Alderley Park and Alex Alderton, Roberto Amato, Jeffrey Barrett, Sonia Goncalves, Ewan Harrison, David K. Jackson, Ian Johnston, Dominic Kwiatkowski, Cordelia Langford, John Sillitoe on behalf of the Wellcome Sanger Institute COVID-19 Surveillance Team |
| EPI_ISL_3335605, EPI_ISL_3335606, EPI_ISL_3335607, EPI_ISL_3335608, EPI_ISL_3335609, EPI_ISL_3335610, EPI_ISL_3335611, EPI_ISL_3335612 | Lighthouse Laboratory Plymouth | Wellcome Sanger Institute for the COVID-19 Genomics UK (COG-UK) Consortium | Lighthouse Laboratory Plymouth and Alex Alderton, Roberto Amato, Jeffrey Barrett, Sonia Goncalves, Ewan Harrison, David K. Jackson, Ian Johnston, Dominic Kwiatkowski, Cordelia Langford, John Sillitoe on behalf of the Wellcome Sanger Institute COVID-19 Surveillance Team |
| EPI_ISL_3335613, EPI_ISL_3335614 | Lighthouse Lab in Alderley Park | Wellcome Sanger Institute for the COVID-19 Genomics UK (COG-UK) Consortium | Jacquelyn Wynn, Mairead Hyland, The Lighthouse Lab in Alderley Park and Alex Alderton, Roberto Amato, Jeffrey Barrett, Sonia Goncalves, Ewan Harrison, David K. Jackson, Ian Johnston, Dominic Kwiatkowski, Cordelia Langford, John Sillitoe on behalf of the Wellcome Sanger Institute COVID-19 Surveillance Team |
| EPI_ISL_3335615 | Lighthouse Laboratory Plymouth | Wellcome Sanger Institute for the COVID-19 Genomics UK (COG-UK) Consortium | Lighthouse Laboratory Plymouth and Alex Alderton, Roberto Amato, Jeffrey Barrett, Sonia Goncalves, Ewan Harrison, David K. Jackson, Ian Johnston, Dominic Kwiatkowski, Cordelia Langford, John Sillitoe on behalf of the Wellcome Sanger Institute COVID-19 Surveillance Team |
| EPI_ISL_3335616, EPI_ISL_3335617, EPI_ISL_3335618, EPI_ISL_3335619, EPI_ISL_3335620 | Lighthouse Lab in Alderley Park | Wellcome Sanger Institute for the COVID-19 Genomics UK (COG-UK) Consortium | Jacquelyn Wynn, Mairead Hyland, The Lighthouse Lab in Alderley Park and Alex Alderton, Roberto Amato, Jeffrey Barrett, Sonia Goncalves, Ewan Harrison, David K. Jackson, Ian Johnston, Dominic Kwiatkowski, Cordelia Langford, John Sillitoe on behalf of the Wellcome Sanger Institute COVID-19 Surveillance Team |
| EPI_ISL_3335621 | Lighthouse Laboratory Plymouth | Wellcome Sanger Institute for the COVID-19 Genomics UK (COG-UK) Consortium | Lighthouse Laboratory Plymouth and Alex Alderton, Roberto Amato, Jeffrey Barrett, Sonia Goncalves, Ewan Harrison, David K. Jackson, Ian Johnston, Dominic Kwiatkowski, Cordelia Langford, John Sillitoe on behalf of the Wellcome Sanger Institute COVID-19 Surveillance Team |
| EPI_ISL_3335622 | Lighthouse Lab in Alderley Park | Wellcome Sanger Institute for the COVID-19 Genomics UK (COG-UK) Consortium | Jacquelyn Wynn, Mairead Hyland, The Lighthouse Lab in Alderley Park and Alex Alderton, Roberto Amato, Jeffrey Barrett, Sonia Goncalves, Ewan Harrison, David K. Jackson, Ian Johnston, Dominic Kwiatkowski, Cordelia Langford, John Sillitoe on behalf of the Wellcome Sanger Institute COVID-19 Surveillance Team |
| EPI_ISL_3335623 | Lighthouse Laboratory Plymouth | Wellcome Sanger Institute for the COVID-19 Genomics UK (COG-UK) Consortium | Lighthouse Laboratory Plymouth and Alex Alderton, Roberto Amato, Jeffrey Barrett, Sonia Goncalves, Ewan Harrison, David K. Jackson, Ian Johnston, Dominic Kwiatkowski, Cordelia Langford, John Sillitoe on behalf of the Wellcome Sanger Institute COVID-19 Surveillance Team |
| EPI_ISL_3335624, EPI_ISL_3335625, EPI_ISL_3335626 | Lighthouse Lab in Alderley Park | Wellcome Sanger Institute for the COVID-19 Genomics UK (COG-UK) Consortium | Jacquelyn Wynn, Mairead Hyland, The Lighthouse Lab in Alderley Park and Alex Alderton, Roberto Amato, Jeffrey Barrett, Sonia Goncalves, Ewan Harrison, David K. Jackson, Ian Johnston, Dominic Kwiatkowski, Cordelia Langford, John Sillitoe on behalf of the Wellcome Sanger Institute COVID-19 Surveillance Team |
| EPI_ISL_3335627, EPI_ISL_3335628, EPI_ISL_3335629 | Lighthouse Laboratory Plymouth | Wellcome Sanger Institute for the COVID-19 Genomics UK (COG-UK) Consortium | Lighthouse Laboratory Plymouth and Alex Alderton, Roberto Amato, Jeffrey Barrett, Sonia Goncalves, Ewan Harrison, David K. Jackson, Ian Johnston, Dominic Kwiatkowski, Cordelia Langford, John Sillitoe on behalf of the Wellcome Sanger Institute COVID-19 Surveillance Team |
| EPI_ISL_3335630 | Lighthouse Lab in Alderley Park | Wellcome Sanger Institute for the COVID-19 Genomics UK (COG-UK) Consortium | Jacquelyn Wynn, Mairead Hyland, The Lighthouse Lab in Alderley Park and Alex Alderton, Roberto Amato, Jeffrey Barrett, Sonia Goncalves, Ewan Harrison, David K. Jackson, Ian Johnston, Dominic Kwiatkowski, Cordelia Langford, John Sillitoe on behalf of the Wellcome Sanger Institute COVID-19 Surveillance Team |
| EPI_ISL_3335631 | Lighthouse Laboratory Plymouth | Wellcome Sanger Institute for the COVID-19 Genomics UK (COG-UK) Consortium | Lighthouse Laboratory Plymouth and Alex Alderton, Roberto Amato, Jeffrey Barrett, Sonia Goncalves, Ewan Harrison, David K. Jackson, Ian Johnston, Dominic Kwiatkowski, Cordelia Langford, John Sillitoe on behalf of the Wellcome Sanger Institute COVID-19 Surveillance Team |
| EPI_ISL_3335632 | Lighthouse Lab in Alderley Park | Wellcome Sanger Institute for the COVID-19 Genomics UK (COG-UK) Consortium | Jacquelyn Wynn, Mairead Hyland, The Lighthouse Lab in Alderley Park and Alex Alderton, Roberto Amato, Jeffrey Barrett, Sonia Goncalves, Ewan Harrison, David K. Jackson, Ian Johnston, Dominic Kwiatkowski, Cordelia Langford, John Sillitoe on behalf of the Wellcome Sanger Institute COVID-19 Surveillance Team |
| EPI_ISL_3335633 | Lighthouse Laboratory Plymouth | Wellcome Sanger Institute for the COVID-19 Genomics UK (COG-UK) Consortium | Lighthouse Laboratory Plymouth and Alex Alderton, Roberto Amato, Jeffrey Barrett, Sonia Goncalves, Ewan Harrison, David K. Jackson, Ian Johnston, Dominic Kwiatkowski, Cordelia Langford, John Sillitoe on behalf of the Wellcome Sanger Institute COVID-19 Surveillance Team |
| EPI_ISL_3335634, EPI_ISL_3335635 | Lighthouse Lab in Alderley Park | Wellcome Sanger Institute for the COVID-19 Genomics UK (COG-UK) Consortium | Jacquelyn Wynn, Mairead Hyland, The Lighthouse Lab in Alderley Park and Alex Alderton, Roberto Amato, Jeffrey Barrett, Sonia Goncalves, Ewan Harrison, David K. Jackson, Ian Johnston, Dominic Kwiatkowski, Cordelia Langford, John Sillitoe on behalf of the Wellcome Sanger Institute COVID-19 Surveillance Team |
| EPI_ISL_3335636, EPI_ISL_3335637 | Lighthouse Laboratory Plymouth | Wellcome Sanger Institute for the COVID-19 Genomics UK (COG-UK) Consortium | Lighthouse Laboratory Plymouth and Alex Alderton, Roberto Amato, Jeffrey Barrett, Sonia Goncalves, Ewan Harrison, David K. Jackson, Ian Johnston, Dominic Kwiatkowski, Cordelia Langford, John Sillitoe on behalf of the Wellcome Sanger Institute COVID-19 Surveillance Team |
| EPI_ISL_3335638 | Lighthouse Lab in Alderley Park | Wellcome Sanger Institute for the COVID-19 Genomics UK (COG-UK) Consortium | Jacquelyn Wynn, Mairead Hyland, The Lighthouse Lab in Alderley Park and Alex Alderton, Roberto Amato, Jeffrey Barrett, Sonia Goncalves, Ewan Harrison, David K. Jackson, Ian Johnston, Dominic Kwiatkowski, Cordelia Langford, John Sillitoe on behalf of the Wellcome Sanger Institute COVID-19 Surveillance Team |
| EPI_ISL_3335639 | Lighthouse Laboratory Plymouth | Wellcome Sanger Institute for the COVID-19 Genomics UK (COG-UK) Consortium | Lighthouse Laboratory Plymouth and Alex Alderton, Roberto Amato, Jeffrey Barrett, Sonia Goncalves, Ewan Harrison, David K. Jackson, Ian Johnston, Dominic Kwiatkowski, Cordelia Langford, John Sillitoe on behalf of the Wellcome Sanger Institute COVID-19 Surveillance Team |
| EPI_ISL_3335640, | Lighthouse Lab in Alderley Park | Wellcome Sanger Institute for the COVID-19 Genomics UK (COG-UK) | Jacquelyn Wynn, Mairead Hyland, The Lighthouse Lab in Alderley Park and Alex Alderton, Roberto Amato, Jeffrey Barrett, Sonia Goncalves, Ewan Harrison, David K. Jackson, Ian Johnston, Dominic Kwiatkowski, Cordelia Langford, John Sillitoe on |

| | | | |
|---|---|---|---|
| EPI_ISL_3335641, EPI_ISL_3335642, EPI_ISL_3335643, EPI_ISL_3335644, EPI_ISL_3335645 | | Consortium | behalf of the Wellcome Sanger Institute COVID-19 Surveillance Team |
| EPI_ISL_3335646 | Lighthouse Laboratory Plymouth | Wellcome Sanger Institute for the COVID-19 Genomics UK (COG-UK) Consortium | Lighthouse Laboratory Plymouth and Alex Alderton, Roberto Amato, Jeffrey Barrett, Sonia Goncalves, Ewan Harrison, David K. Jackson, Ian Johnston, Dominic Kwiatkowski, Cordelia Langford, John Sillitoe on behalf of the Wellcome Sanger Institute COVID-19 Surveillance Team |
| EPI_ISL_3335647 | Lighthouse Lab in Alderley Park | Wellcome Sanger Institute for the COVID-19 Genomics UK (COG-UK) Consortium | Jacquelyn Wynn, Mairead Hyland, The Lighthouse Lab in Alderley Park and Alex Alderton, Roberto Amato, Jeffrey Barrett, Sonia Goncalves, Ewan Harrison, David K. Jackson, Ian Johnston, Dominic Kwiatkowski, Cordelia Langford, John Sillitoe on behalf of the Wellcome Sanger Institute COVID-19 Surveillance Team |
| EPI_ISL_3335648 | Lighthouse Laboratory Plymouth | Wellcome Sanger Institute for the COVID-19 Genomics UK (COG-UK) Consortium | Lighthouse Laboratory Plymouth and Alex Alderton, Roberto Amato, Jeffrey Barrett, Sonia Goncalves, Ewan Harrison, David K. Jackson, Ian Johnston, Dominic Kwiatkowski, Cordelia Langford, John Sillitoe on behalf of the Wellcome Sanger Institute COVID-19 Surveillance Team |
| EPI_ISL_3335649, EPI_ISL_3335650 | Lighthouse Lab in Alderley Park | Wellcome Sanger Institute for the COVID-19 Genomics UK (COG-UK) Consortium | Jacquelyn Wynn, Mairead Hyland, The Lighthouse Lab in Alderley Park and Alex Alderton, Roberto Amato, Jeffrey Barrett, Sonia Goncalves, Ewan Harrison, David K. Jackson, Ian Johnston, Dominic Kwiatkowski, Cordelia Langford, John Sillitoe on behalf of the Wellcome Sanger Institute COVID-19 Surveillance Team |
| EPI_ISL_3335651 | Lighthouse Laboratory Plymouth | Wellcome Sanger Institute for the COVID-19 Genomics UK (COG-UK) Consortium | Lighthouse Laboratory Plymouth and Alex Alderton, Roberto Amato, Jeffrey Barrett, Sonia Goncalves, Ewan Harrison, David K. Jackson, Ian Johnston, Dominic Kwiatkowski, Cordelia Langford, John Sillitoe on behalf of the Wellcome Sanger Institute COVID-19 Surveillance Team |
| EPI_ISL_3335652, EPI_ISL_3335653, EPI_ISL_3335654, EPI_ISL_3335655, EPI_ISL_3335656, EPI_ISL_3335657 | Lighthouse Lab in Alderley Park | Wellcome Sanger Institute for the COVID-19 Genomics UK (COG-UK) Consortium | Jacquelyn Wynn, Mairead Hyland, The Lighthouse Lab in Alderley Park and Alex Alderton, Roberto Amato, Jeffrey Barrett, Sonia Goncalves, Ewan Harrison, David K. Jackson, Ian Johnston, Dominic Kwiatkowski, Cordelia Langford, John Sillitoe on behalf of the Wellcome Sanger Institute COVID-19 Surveillance Team |
| EPI_ISL_3335658 | Lighthouse Laboratory Plymouth | Wellcome Sanger Institute for the COVID-19 Genomics UK (COG-UK) Consortium | Lighthouse Laboratory Plymouth and Alex Alderton, Roberto Amato, Jeffrey Barrett, Sonia Goncalves, Ewan Harrison, David K. Jackson, Ian Johnston, Dominic Kwiatkowski, Cordelia Langford, John Sillitoe on behalf of the Wellcome Sanger Institute COVID-19 Surveillance Team |
| EPI_ISL_3335659 | Lighthouse Lab in Alderley Park | Wellcome Sanger Institute for the COVID-19 Genomics UK (COG-UK) Consortium | Jacquelyn Wynn, Mairead Hyland, The Lighthouse Lab in Alderley Park and Alex Alderton, Roberto Amato, Jeffrey Barrett, Sonia Goncalves, Ewan Harrison, David K. Jackson, Ian Johnston, Dominic Kwiatkowski, Cordelia Langford, John Sillitoe on behalf of the Wellcome Sanger Institute COVID-19 Surveillance Team |
| EPI_ISL_3335660, EPI_ISL_3335661 | Lighthouse Laboratory Plymouth | Wellcome Sanger Institute for the COVID-19 Genomics UK (COG-UK) Consortium | Lighthouse Laboratory Plymouth and Alex Alderton, Roberto Amato, Jeffrey Barrett, Sonia Goncalves, Ewan Harrison, David K. Jackson, Ian Johnston, Dominic Kwiatkowski, Cordelia Langford, John Sillitoe on behalf of the Wellcome Sanger Institute COVID-19 Surveillance Team |
| EPI_ISL_3335662, EPI_ISL_3335663 | Lighthouse Lab in Alderley Park | Wellcome Sanger Institute for the COVID-19 Genomics UK (COG-UK) Consortium | Jacquelyn Wynn, Mairead Hyland, The Lighthouse Lab in Alderley Park and Alex Alderton, Roberto Amato, Jeffrey Barrett, Sonia Goncalves, Ewan Harrison, David K. Jackson, Ian Johnston, Dominic Kwiatkowski, Cordelia Langford, John Sillitoe on behalf of the Wellcome Sanger Institute COVID-19 Surveillance Team |
| EPI_ISL_3335664 | Lighthouse Lab in Milton Keynes | Wellcome Sanger Institute for the COVID-19 Genomics UK (COG-UK) Consortium | The Lighthouse Lab in Milton Keynes and Alex Alderton, Roberto Amato, Jeffrey Barrett, Sonia Goncalves, Ewan Harrison, David K. Jackson, Ian Johnston, Dominic Kwiatkowski, Cordelia Langford, John Sillitoe on behalf of the Wellcome Sanger Institute COVID-19 Surveillance Team |
| EPI_ISL_3335665 | Lighthouse Laboratory Plymouth | Wellcome Sanger Institute for the COVID-19 Genomics UK (COG-UK) Consortium | Lighthouse Laboratory Plymouth and Alex Alderton, Roberto Amato, Jeffrey Barrett, Sonia Goncalves, Ewan Harrison, David K. Jackson, Ian Johnston, Dominic Kwiatkowski, Cordelia Langford, John Sillitoe on behalf of the Wellcome Sanger Institute COVID-19 Surveillance Team |
| EPI_ISL_3335666, EPI_ISL_3335667, EPI_ISL_3335668, EPI_ISL_3335669 | Lighthouse Lab in Alderley Park | Wellcome Sanger Institute for the COVID-19 Genomics UK (COG-UK) Consortium | Jacquelyn Wynn, Mairead Hyland, The Lighthouse Lab in Alderley Park and Alex Alderton, Roberto Amato, Jeffrey Barrett, Sonia Goncalves, Ewan Harrison, David K. Jackson, Ian Johnston, Dominic Kwiatkowski, Cordelia Langford, John Sillitoe on behalf of the Wellcome Sanger Institute COVID-19 Surveillance Team |
| EPI_ISL_3335670 | Lighthouse Laboratory Plymouth | Wellcome Sanger Institute for the COVID-19 Genomics UK (COG-UK) Consortium | Lighthouse Laboratory Plymouth and Alex Alderton, Roberto Amato, Jeffrey Barrett, Sonia Goncalves, Ewan Harrison, David K. Jackson, Ian Johnston, Dominic Kwiatkowski, Cordelia Langford, John Sillitoe on behalf of the Wellcome Sanger Institute COVID-19 Surveillance Team |
| EPI_ISL_3335671, EPI_ISL_3335672 | Lighthouse Lab in Alderley Park | Wellcome Sanger Institute for the COVID-19 Genomics UK (COG-UK) Consortium | Jacquelyn Wynn, Mairead Hyland, The Lighthouse Lab in Alderley Park and Alex Alderton, Roberto Amato, Jeffrey Barrett, Sonia Goncalves, Ewan Harrison, David K. Jackson, Ian Johnston, Dominic Kwiatkowski, Cordelia Langford, John Sillitoe on behalf of the Wellcome Sanger Institute COVID-19 Surveillance Team |
| EPI_ISL_3335673, EPI_ISL_3335674 | Lighthouse Laboratory Plymouth | Wellcome Sanger Institute for the COVID-19 Genomics UK (COG-UK) Consortium | Lighthouse Laboratory Plymouth and Alex Alderton, Roberto Amato, Jeffrey Barrett, Sonia Goncalves, Ewan Harrison, David K. Jackson, Ian Johnston, Dominic Kwiatkowski, Cordelia Langford, John Sillitoe on behalf of the Wellcome Sanger Institute COVID-19 Surveillance Team |
| EPI_ISL_3335675, EPI_ISL_3335676, EPI_ISL_3335677, EPI_ISL_3335678, EPI_ISL_3335679 | Lighthouse Lab in Alderley Park | Wellcome Sanger Institute for the COVID-19 Genomics UK (COG-UK) Consortium | Jacquelyn Wynn, Mairead Hyland, The Lighthouse Lab in Alderley Park and Alex Alderton, Roberto Amato, Jeffrey Barrett, Sonia Goncalves, Ewan Harrison, David K. Jackson, Ian Johnston, Dominic Kwiatkowski, Cordelia Langford, John Sillitoe on behalf of the Wellcome Sanger Institute COVID-19 Surveillance Team |
| EPI_ISL_3335680, EPI_ISL_3335681, EPI_ISL_3335682 | Lighthouse Laboratory Plymouth | Wellcome Sanger Institute for the COVID-19 Genomics UK (COG-UK) Consortium | Lighthouse Laboratory Plymouth and Alex Alderton, Roberto Amato, Jeffrey Barrett, Sonia Goncalves, Ewan Harrison, David K. Jackson, Ian Johnston, Dominic Kwiatkowski, Cordelia Langford, John Sillitoe on behalf of the Wellcome Sanger Institute COVID-19 Surveillance Team |
| EPI_ISL_3335683, EPI_ISL_3335684 | Lighthouse Lab in Alderley Park | Wellcome Sanger Institute for the COVID-19 Genomics UK (COG-UK) Consortium | Jacquelyn Wynn, Mairead Hyland, The Lighthouse Lab in Alderley Park and Alex Alderton, Roberto Amato, Jeffrey Barrett, Sonia Goncalves, Ewan Harrison, David K. Jackson, Ian Johnston, Dominic Kwiatkowski, Cordelia Langford, John Sillitoe on behalf of the Wellcome Sanger Institute COVID-19 Surveillance Team |
| EPI_ISL_3335685 | Lighthouse Laboratory Plymouth | Wellcome Sanger Institute for the COVID-19 Genomics UK (COG-UK) Consortium | Lighthouse Laboratory Plymouth and Alex Alderton, Roberto Amato, Jeffrey Barrett, Sonia Goncalves, Ewan Harrison, David K. Jackson, Ian Johnston, Dominic Kwiatkowski, Cordelia Langford, John Sillitoe on behalf of the Wellcome Sanger Institute COVID-19 Surveillance Team |
| EPI_ISL_3335686 | Lighthouse Lab in Alderley Park | Wellcome Sanger Institute for the COVID-19 Genomics UK (COG-UK) Consortium | Jacquelyn Wynn, Mairead Hyland, The Lighthouse Lab in Alderley Park and Alex Alderton, Roberto Amato, Jeffrey Barrett, Sonia Goncalves, Ewan Harrison, David K. Jackson, Ian Johnston, Dominic Kwiatkowski, Cordelia Langford, John Sillitoe on behalf of the Wellcome Sanger Institute COVID-19 Surveillance Team |
| EPI_ISL_3335687 | Lighthouse Laboratory Plymouth | Wellcome Sanger Institute for the COVID-19 Genomics UK (COG-UK) Consortium | Lighthouse Laboratory Plymouth and Alex Alderton, Roberto Amato, Jeffrey Barrett, Sonia Goncalves, Ewan Harrison, David K. Jackson, Ian Johnston, Dominic Kwiatkowski, Cordelia Langford, John Sillitoe on behalf of the Wellcome Sanger Institute COVID-19 Surveillance Team |
| EPI_ISL_3335688, EPI_ISL_3335689 | Lighthouse Lab in Alderley Park | Wellcome Sanger Institute for the COVID-19 Genomics UK (COG-UK) Consortium | Jacquelyn Wynn, Mairead Hyland, The Lighthouse Lab in Alderley Park and Alex Alderton, Roberto Amato, Jeffrey Barrett, Sonia Goncalves, Ewan Harrison, David K. Jackson, Ian Johnston, Dominic Kwiatkowski, Cordelia Langford, John Sillitoe on behalf of the Wellcome Sanger Institute COVID-19 Surveillance Team |
| EPI_ISL_3335690, EPI_ISL_3335691, EPI_ISL_3335692 | Lighthouse Laboratory Plymouth | Wellcome Sanger Institute for the COVID-19 Genomics UK (COG-UK) Consortium | Lighthouse Laboratory Plymouth and Alex Alderton, Roberto Amato, Jeffrey Barrett, Sonia Goncalves, Ewan Harrison, David K. Jackson, Ian Johnston, Dominic Kwiatkowski, Cordelia Langford, John Sillitoe on behalf of the Wellcome Sanger Institute COVID-19 Surveillance Team |
| EPI_ISL_3335693, EPI_ISL_3335694, EPI_ISL_3335695, EPI_ISL_3335696 | Lighthouse Lab in Alderley Park | Wellcome Sanger Institute for the COVID-19 Genomics UK (COG-UK) Consortium | Jacquelyn Wynn, Mairead Hyland, The Lighthouse Lab in Alderley Park and Alex Alderton, Roberto Amato, Jeffrey Barrett, Sonia Goncalves, Ewan Harrison, David K. Jackson, Ian Johnston, Dominic Kwiatkowski, Cordelia Langford, John Sillitoe on behalf of the Wellcome Sanger Institute COVID-19 Surveillance Team |
| EPI_ISL_3335697 | Lighthouse Laboratory Plymouth | Wellcome Sanger Institute for the COVID-19 Genomics UK (COG-UK) Consortium | Lighthouse Laboratory Plymouth and Alex Alderton, Roberto Amato, Jeffrey Barrett, Sonia Goncalves, Ewan Harrison, David K. Jackson, Ian Johnston, Dominic Kwiatkowski, Cordelia Langford, John Sillitoe on behalf of the Wellcome Sanger Institute COVID-19 Surveillance Team |
| EPI_ISL_3335698, EPI_ISL_3335699, EPI_ISL_3335700, EPI_ISL_3335701, EPI_ISL_3335702, EPI_ISL_3335703, EPI_ISL_3335704 | Lighthouse Lab in Alderley Park | Wellcome Sanger Institute for the COVID-19 Genomics UK (COG-UK) Consortium | Jacquelyn Wynn, Mairead Hyland, The Lighthouse Lab in Alderley Park and Alex Alderton, Roberto Amato, Jeffrey Barrett, Sonia Goncalves, Ewan Harrison, David K. Jackson, Ian Johnston, Dominic Kwiatkowski, Cordelia Langford, John Sillitoe on behalf of the Wellcome Sanger Institute COVID-19 Surveillance Team |
| EPI_ISL_3335705, EPI_ISL_3335706 | Lighthouse Laboratory Plymouth | Wellcome Sanger Institute for the COVID-19 Genomics UK (COG-UK) Consortium | Lighthouse Laboratory Plymouth and Alex Alderton, Roberto Amato, Jeffrey Barrett, Sonia Goncalves, Ewan Harrison, David K. Jackson, Ian Johnston, Dominic Kwiatkowski, Cordelia Langford, John Sillitoe on behalf of the Wellcome Sanger Institute COVID-19 Surveillance Team |
| EPI_ISL_3335707 | Lighthouse Lab in Milton Keynes | Wellcome Sanger Institute for the COVID-19 Genomics UK (COG-UK) Consortium | The Lighthouse Lab in Milton Keynes and Alex Alderton, Roberto Amato, Jeffrey Barrett, Sonia Goncalves, Ewan Harrison, David K. Jackson, Ian Johnston, Dominic Kwiatkowski, Cordelia Langford, John Sillitoe on behalf of the Wellcome Sanger Institute COVID-19 Surveillance Team |
| EPI_ISL_3335708 | Lighthouse Laboratory Plymouth | Wellcome Sanger Institute for the COVID-19 Genomics UK (COG-UK) Consortium | Lighthouse Laboratory Plymouth and Alex Alderton, Roberto Amato, Jeffrey Barrett, Sonia Goncalves, Ewan Harrison, David K. Jackson, Ian Johnston, Dominic Kwiatkowski, Cordelia Langford, John Sillitoe on behalf of the Wellcome Sanger Institute COVID-19 Surveillance Team |
| EPI_ISL_3335709, EPI_ISL_3335710 | Lighthouse Lab in Milton Keynes | Wellcome Sanger Institute for the COVID-19 Genomics UK (COG-UK) Consortium | The Lighthouse Lab in Milton Keynes and Alex Alderton, Roberto Amato, Jeffrey Barrett, Sonia Goncalves, Ewan Harrison, David K. Jackson, Ian Johnston, Dominic Kwiatkowski, Cordelia Langford, John Sillitoe on behalf of the Wellcome Sanger Institute COVID-19 Surveillance Team |
| EPI_ISL_3335711 | Health Services Laboratories | Wellcome Sanger Institute for the COVID-19 Genomics UK (COG-UK) Consortium | Health Services Laboratories and Alex Alderton, Roberto Amato, Jeffrey Barrett, Sonia Goncalves, Ewan Harrison, David K. Jackson, Ian Johnston, Dominic Kwiatkowski, Cordelia Langford, John Sillitoe on behalf of the Wellcome Sanger Institute COVID-19 Surveillance Team |
| EPI_ISL_3335712, EPI_ISL_3335713, EPI_ISL_3335714 | Lighthouse Laboratory Plymouth | Wellcome Sanger Institute for the COVID-19 Genomics UK (COG-UK) Consortium | Lighthouse Laboratory Plymouth and Alex Alderton, Roberto Amato, Jeffrey Barrett, Sonia Goncalves, Ewan Harrison, David K. Jackson, Ian Johnston, Dominic Kwiatkowski, Cordelia Langford, John Sillitoe on behalf of the Wellcome Sanger Institute COVID-19 Surveillance Team |
| EPI_ISL_3335715, EPI_ISL_3335716, EPI_ISL_3335717, EPI_ISL_3335718 | Lighthouse Lab in Milton Keynes | Wellcome Sanger Institute for the COVID-19 Genomics UK (COG-UK) Consortium | The Lighthouse Lab in Milton Keynes and Alex Alderton, Roberto Amato, Jeffrey Barrett, Sonia Goncalves, Ewan Harrison, David K. Jackson, Ian Johnston, Dominic Kwiatkowski, Cordelia Langford, John Sillitoe on behalf of the Wellcome Sanger Institute COVID-19 Surveillance Team |

| | | | |
|---|---|---|---|
| EPI_ISL_3335719 | Health Services Laboratories | Wellcome Sanger Institute for the COVID-19 Genomics UK (COG-UK) Consortium | Health Services Laboratories and Alex Alderton, Roberto Amato, Jeffrey Barrett, Sonia Goncalves, Ewan Harrison, David K. Jackson, Ian Johnston, Dominic Kwiatkowski, Cordelia Langford, John Sillitoe on behalf of the Wellcome Sanger Institute COVID-19 Surveillance Team |
| EPI_ISL_3335720 | Lighthouse Lab in Milton Keynes | Wellcome Sanger Institute for the COVID-19 Genomics UK (COG-UK) Consortium | The Lighthouse Lab in Milton Keynes and Alex Alderton, Roberto Amato, Jeffrey Barrett, Sonia Goncalves, Ewan Harrison, David K. Jackson, Ian Johnston, Dominic Kwiatkowski, Cordelia Langford, John Sillitoe on behalf of the Wellcome Sanger Institute COVID-19 Surveillance Team |
| EPI_ISL_3335721 | Lighthouse Laboratory Plymouth | Wellcome Sanger Institute for the COVID-19 Genomics UK (COG-UK) Consortium | Lighthouse Laboratory Plymouth and Alex Alderton, Roberto Amato, Jeffrey Barrett, Sonia Goncalves, Ewan Harrison, David K. Jackson, Ian Johnston, Dominic Kwiatkowski, Cordelia Langford, John Sillitoe on behalf of the Wellcome Sanger Institute COVID-19 Surveillance Team |
| EPI_ISL_3335722 | Health Services Laboratories | Wellcome Sanger Institute for the COVID-19 Genomics UK (COG-UK) Consortium | Health Services Laboratories and Alex Alderton, Roberto Amato, Jeffrey Barrett, Sonia Goncalves, Ewan Harrison, David K. Jackson, Ian Johnston, Dominic Kwiatkowski, Cordelia Langford, John Sillitoe on behalf of the Wellcome Sanger Institute COVID-19 Surveillance Team |
| EPI_ISL_3335723, EPI_ISL_3335724, EPI_ISL_3335725 | Lighthouse Lab in Milton Keynes | Wellcome Sanger Institute for the COVID-19 Genomics UK (COG-UK) Consortium | The Lighthouse Lab in Milton Keynes and Alex Alderton, Roberto Amato, Jeffrey Barrett, Sonia Goncalves, Ewan Harrison, David K. Jackson, Ian Johnston, Dominic Kwiatkowski, Cordelia Langford, John Sillitoe on behalf of the Wellcome Sanger Institute COVID-19 Surveillance Team |
| EPI_ISL_3335726 | Lighthouse Laboratory Plymouth | Wellcome Sanger Institute for the COVID-19 Genomics UK (COG-UK) Consortium | Lighthouse Laboratory Plymouth and Alex Alderton, Roberto Amato, Jeffrey Barrett, Sonia Goncalves, Ewan Harrison, David K. Jackson, Ian Johnston, Dominic Kwiatkowski, Cordelia Langford, John Sillitoe on behalf of the Wellcome Sanger Institute COVID-19 Surveillance Team |
| EPI_ISL_3335727, EPI_ISL_3335728, EPI_ISL_3335729, EPI_ISL_3335730 | Lighthouse Lab in Milton Keynes | Wellcome Sanger Institute for the COVID-19 Genomics UK (COG-UK) Consortium | The Lighthouse Lab in Milton Keynes and Alex Alderton, Roberto Amato, Jeffrey Barrett, Sonia Goncalves, Ewan Harrison, David K. Jackson, Ian Johnston, Dominic Kwiatkowski, Cordelia Langford, John Sillitoe on behalf of the Wellcome Sanger Institute COVID-19 Surveillance Team |
| EPI_ISL_3335731 | Lighthouse Laboratory Plymouth | Wellcome Sanger Institute for the COVID-19 Genomics UK (COG-UK) Consortium | Lighthouse Laboratory Plymouth and Alex Alderton, Roberto Amato, Jeffrey Barrett, Sonia Goncalves, Ewan Harrison, David K. Jackson, Ian Johnston, Dominic Kwiatkowski, Cordelia Langford, John Sillitoe on behalf of the Wellcome Sanger Institute COVID-19 Surveillance Team |
| EPI_ISL_3335732, EPI_ISL_3335733 | Lighthouse Lab in Milton Keynes | Wellcome Sanger Institute for the COVID-19 Genomics UK (COG-UK) Consortium | The Lighthouse Lab in Milton Keynes and Alex Alderton, Roberto Amato, Jeffrey Barrett, Sonia Goncalves, Ewan Harrison, David K. Jackson, Ian Johnston, Dominic Kwiatkowski, Cordelia Langford, John Sillitoe on behalf of the Wellcome Sanger Institute COVID-19 Surveillance Team |
| EPI_ISL_3335734, EPI_ISL_3335735 | Health Services Laboratories | Wellcome Sanger Institute for the COVID-19 Genomics UK (COG-UK) Consortium | Health Services Laboratories and Alex Alderton, Roberto Amato, Jeffrey Barrett, Sonia Goncalves, Ewan Harrison, David K. Jackson, Ian Johnston, Dominic Kwiatkowski, Cordelia Langford, John Sillitoe on behalf of the Wellcome Sanger Institute COVID-19 Surveillance Team |
| EPI_ISL_3335736 | Lighthouse Lab in Milton Keynes | Wellcome Sanger Institute for the COVID-19 Genomics UK (COG-UK) Consortium | The Lighthouse Lab in Milton Keynes and Alex Alderton, Roberto Amato, Jeffrey Barrett, Sonia Goncalves, Ewan Harrison, David K. Jackson, Ian Johnston, Dominic Kwiatkowski, Cordelia Langford, John Sillitoe on behalf of the Wellcome Sanger Institute COVID-19 Surveillance Team |
| EPI_ISL_3335737 | Health Services Laboratories | Wellcome Sanger Institute for the COVID-19 Genomics UK (COG-UK) Consortium | Health Services Laboratories and Alex Alderton, Roberto Amato, Jeffrey Barrett, Sonia Goncalves, Ewan Harrison, David K. Jackson, Ian Johnston, Dominic Kwiatkowski, Cordelia Langford, John Sillitoe on behalf of the Wellcome Sanger Institute COVID-19 Surveillance Team |
| EPI_ISL_3335738 | Lighthouse Lab in Milton Keynes | Wellcome Sanger Institute for the COVID-19 Genomics UK (COG-UK) Consortium | The Lighthouse Lab in Milton Keynes and Alex Alderton, Roberto Amato, Jeffrey Barrett, Sonia Goncalves, Ewan Harrison, David K. Jackson, Ian Johnston, Dominic Kwiatkowski, Cordelia Langford, John Sillitoe on behalf of the Wellcome Sanger Institute COVID-19 Surveillance Team |
| EPI_ISL_3335739 | Lighthouse Laboratory Plymouth | Wellcome Sanger Institute for the COVID-19 Genomics UK (COG-UK) Consortium | Lighthouse Laboratory Plymouth and Alex Alderton, Roberto Amato, Jeffrey Barrett, Sonia Goncalves, Ewan Harrison, David K. Jackson, Ian Johnston, Dominic Kwiatkowski, Cordelia Langford, John Sillitoe on behalf of the Wellcome Sanger Institute COVID-19 Surveillance Team |
| EPI_ISL_3335740 | Lighthouse Lab in Milton Keynes | Wellcome Sanger Institute for the COVID-19 Genomics UK (COG-UK) Consortium | The Lighthouse Lab in Milton Keynes and Alex Alderton, Roberto Amato, Jeffrey Barrett, Sonia Goncalves, Ewan Harrison, David K. Jackson, Ian Johnston, Dominic Kwiatkowski, Cordelia Langford, John Sillitoe on behalf of the Wellcome Sanger Institute COVID-19 Surveillance Team |
| EPI_ISL_3335741, EPI_ISL_3335742 | Health Services Laboratories | Wellcome Sanger Institute for the COVID-19 Genomics UK (COG-UK) Consortium | Health Services Laboratories and Alex Alderton, Roberto Amato, Jeffrey Barrett, Sonia Goncalves, Ewan Harrison, David K. Jackson, Ian Johnston, Dominic Kwiatkowski, Cordelia Langford, John Sillitoe on behalf of the Wellcome Sanger Institute COVID-19 Surveillance Team |
| EPI_ISL_3335743 | Lighthouse Laboratory Plymouth | Wellcome Sanger Institute for the COVID-19 Genomics UK (COG-UK) Consortium | Lighthouse Laboratory Plymouth and Alex Alderton, Roberto Amato, Jeffrey Barrett, Sonia Goncalves, Ewan Harrison, David K. Jackson, Ian Johnston, Dominic Kwiatkowski, Cordelia Langford, John Sillitoe on behalf of the Wellcome Sanger Institute COVID-19 Surveillance Team |
| EPI_ISL_3335744 | Health Services Laboratories | Wellcome Sanger Institute for the COVID-19 Genomics UK (COG-UK) Consortium | Health Services Laboratories and Alex Alderton, Roberto Amato, Jeffrey Barrett, Sonia Goncalves, Ewan Harrison, David K. Jackson, Ian Johnston, Dominic Kwiatkowski, Cordelia Langford, John Sillitoe on behalf of the Wellcome Sanger Institute COVID-19 Surveillance Team |
| EPI_ISL_3335745 | Lighthouse Laboratory Plymouth | Wellcome Sanger Institute for the COVID-19 Genomics UK (COG-UK) Consortium | Lighthouse Laboratory Plymouth and Alex Alderton, Roberto Amato, Jeffrey Barrett, Sonia Goncalves, Ewan Harrison, David K. Jackson, Ian Johnston, Dominic Kwiatkowski, Cordelia Langford, John Sillitoe on behalf of the Wellcome Sanger Institute COVID-19 Surveillance Team |
| EPI_ISL_3335746 | Lighthouse Lab in Milton Keynes | Wellcome Sanger Institute for the COVID-19 Genomics UK (COG-UK) Consortium | The Lighthouse Lab in Milton Keynes and Alex Alderton, Roberto Amato, Jeffrey Barrett, Sonia Goncalves, Ewan Harrison, David K. Jackson, Ian Johnston, Dominic Kwiatkowski, Cordelia Langford, John Sillitoe on behalf of the Wellcome Sanger Institute COVID-19 Surveillance Team |
| EPI_ISL_3335747, EPI_ISL_3335748 | Lighthouse Laboratory Plymouth | Wellcome Sanger Institute for the COVID-19 Genomics UK (COG-UK) Consortium | Lighthouse Laboratory Plymouth and Alex Alderton, Roberto Amato, Jeffrey Barrett, Sonia Goncalves, Ewan Harrison, David K. Jackson, Ian Johnston, Dominic Kwiatkowski, Cordelia Langford, John Sillitoe on behalf of the Wellcome Sanger Institute COVID-19 Surveillance Team |
| EPI_ISL_3335749 | Health Services Laboratories | Wellcome Sanger Institute for the COVID-19 Genomics UK (COG-UK) Consortium | Health Services Laboratories and Alex Alderton, Roberto Amato, Jeffrey Barrett, Sonia Goncalves, Ewan Harrison, David K. Jackson, Ian Johnston, Dominic Kwiatkowski, Cordelia Langford, John Sillitoe on behalf of the Wellcome Sanger Institute COVID-19 Surveillance Team |
| EPI_ISL_3335750 | Lighthouse Lab in Milton Keynes | Wellcome Sanger Institute for the COVID-19 Genomics UK (COG-UK) Consortium | The Lighthouse Lab in Milton Keynes and Alex Alderton, Roberto Amato, Jeffrey Barrett, Sonia Goncalves, Ewan Harrison, David K. Jackson, Ian Johnston, Dominic Kwiatkowski, Cordelia Langford, John Sillitoe on behalf of the Wellcome Sanger Institute COVID-19 Surveillance Team |
| EPI_ISL_3335751 | Lighthouse Laboratory Plymouth | Wellcome Sanger Institute for the COVID-19 Genomics UK (COG-UK) Consortium | Lighthouse Laboratory Plymouth and Alex Alderton, Roberto Amato, Jeffrey Barrett, Sonia Goncalves, Ewan Harrison, David K. Jackson, Ian Johnston, Dominic Kwiatkowski, Cordelia Langford, John Sillitoe on behalf of the Wellcome Sanger Institute COVID-19 Surveillance Team |
| EPI_ISL_3335752, EPI_ISL_3335753 | Lighthouse Lab in Milton Keynes | Wellcome Sanger Institute for the COVID-19 Genomics UK (COG-UK) Consortium | The Lighthouse Lab in Milton Keynes and Alex Alderton, Roberto Amato, Jeffrey Barrett, Sonia Goncalves, Ewan Harrison, David K. Jackson, Ian Johnston, Dominic Kwiatkowski, Cordelia Langford, John Sillitoe on behalf of the Wellcome Sanger Institute COVID-19 Surveillance Team |
| EPI_ISL_3335754 | Health Services Laboratories | Wellcome Sanger Institute for the COVID-19 Genomics UK (COG-UK) Consortium | Health Services Laboratories and Alex Alderton, Roberto Amato, Jeffrey Barrett, Sonia Goncalves, Ewan Harrison, David K. Jackson, Ian Johnston, Dominic Kwiatkowski, Cordelia Langford, John Sillitoe on behalf of the Wellcome Sanger Institute COVID-19 Surveillance Team |
| EPI_ISL_3335755, EPI_ISL_3335756 | Lighthouse Lab in Milton Keynes | Wellcome Sanger Institute for the COVID-19 Genomics UK (COG-UK) Consortium | The Lighthouse Lab in Milton Keynes and Alex Alderton, Roberto Amato, Jeffrey Barrett, Sonia Goncalves, Ewan Harrison, David K. Jackson, Ian Johnston, Dominic Kwiatkowski, Cordelia Langford, John Sillitoe on behalf of the Wellcome Sanger Institute COVID-19 Surveillance Team |
| EPI_ISL_3335757 | Health Services Laboratories | Wellcome Sanger Institute for the COVID-19 Genomics UK (COG-UK) Consortium | Health Services Laboratories and Alex Alderton, Roberto Amato, Jeffrey Barrett, Sonia Goncalves, Ewan Harrison, David K. Jackson, Ian Johnston, Dominic Kwiatkowski, Cordelia Langford, John Sillitoe on behalf of the Wellcome Sanger Institute COVID-19 Surveillance Team |
| EPI_ISL_3335758 | Lighthouse Lab in Milton Keynes | Wellcome Sanger Institute for the COVID-19 Genomics UK (COG-UK) Consortium | The Lighthouse Lab in Milton Keynes and Alex Alderton, Roberto Amato, Jeffrey Barrett, Sonia Goncalves, Ewan Harrison, David K. Jackson, Ian Johnston, Dominic Kwiatkowski, Cordelia Langford, John Sillitoe on behalf of the Wellcome Sanger Institute COVID-19 Surveillance Team |
| EPI_ISL_3335759, EPI_ISL_3335760 | Health Services Laboratories | Wellcome Sanger Institute for the COVID-19 Genomics UK (COG-UK) Consortium | Health Services Laboratories and Alex Alderton, Roberto Amato, Jeffrey Barrett, Sonia Goncalves, Ewan Harrison, David K. Jackson, Ian Johnston, Dominic Kwiatkowski, Cordelia Langford, John Sillitoe on behalf of the Wellcome Sanger Institute COVID-19 Surveillance Team |
| EPI_ISL_3335761 | Lighthouse Laboratory Plymouth | Wellcome Sanger Institute for the COVID-19 Genomics UK (COG-UK) Consortium | Lighthouse Laboratory Plymouth and Alex Alderton, Roberto Amato, Jeffrey Barrett, Sonia Goncalves, Ewan Harrison, David K. Jackson, Ian Johnston, Dominic Kwiatkowski, Cordelia Langford, John Sillitoe on behalf of the Wellcome Sanger Institute COVID-19 Surveillance Team |
| EPI_ISL_3335762, EPI_ISL_3335763, EPI_ISL_3335764, EPI_ISL_3335765, EPI_ISL_3335766, EPI_ISL_3335767, EPI_ISL_3335768 | Lighthouse Lab in Milton Keynes | Wellcome Sanger Institute for the COVID-19 Genomics UK (COG-UK) Consortium | The Lighthouse Lab in Milton Keynes and Alex Alderton, Roberto Amato, Jeffrey Barrett, Sonia Goncalves, Ewan Harrison, David K. Jackson, Ian Johnston, Dominic Kwiatkowski, Cordelia Langford, John Sillitoe on behalf of the Wellcome Sanger Institute COVID-19 Surveillance Team |
| EPI_ISL_3335769 | Lighthouse Laboratory Plymouth | Wellcome Sanger Institute for the COVID-19 Genomics UK (COG-UK) Consortium | Lighthouse Laboratory Plymouth and Alex Alderton, Roberto Amato, Jeffrey Barrett, Sonia Goncalves, Ewan Harrison, David K. Jackson, Ian Johnston, Dominic Kwiatkowski, Cordelia Langford, John Sillitoe on behalf of the Wellcome Sanger Institute COVID-19 Surveillance Team |
| EPI_ISL_3335770, EPI_ISL_3335771 | Health Services Laboratories | Wellcome Sanger Institute for the COVID-19 Genomics UK (COG-UK) Consortium | Health Services Laboratories and Alex Alderton, Roberto Amato, Jeffrey Barrett, Sonia Goncalves, Ewan Harrison, David K. Jackson, Ian Johnston, Dominic Kwiatkowski, Cordelia Langford, John Sillitoe on behalf of the Wellcome Sanger Institute COVID-19 Surveillance Team |
| EPI_ISL_3335772 | Lighthouse Lab in Milton Keynes | Wellcome Sanger Institute for the COVID-19 Genomics UK (COG-UK) Consortium | The Lighthouse Lab in Milton Keynes and Alex Alderton, Roberto Amato, Jeffrey Barrett, Sonia Goncalves, Ewan Harrison, David K. Jackson, Ian Johnston, Dominic Kwiatkowski, Cordelia Langford, John Sillitoe on behalf of the Wellcome Sanger Institute COVID-19 Surveillance Team |
| EPI_ISL_3335773 | Health Services Laboratories | Wellcome Sanger Institute for the COVID-19 Genomics UK (COG-UK) Consortium | Health Services Laboratories and Alex Alderton, Roberto Amato, Jeffrey Barrett, Sonia Goncalves, Ewan Harrison, David K. Jackson, Ian Johnston, Dominic Kwiatkowski, Cordelia Langford, John Sillitoe on behalf of the Wellcome Sanger Institute COVID-19 Surveillance Team |
| EPI_ISL_3335774, EPI_ISL_3335775, EPI_ISL_3335776 | Lighthouse Lab in Milton Keynes | Wellcome Sanger Institute for the COVID-19 Genomics UK (COG-UK) Consortium | The Lighthouse Lab in Milton Keynes and Alex Alderton, Roberto Amato, Jeffrey Barrett, Sonia Goncalves, Ewan Harrison, David K. Jackson, Ian Johnston, Dominic Kwiatkowski, Cordelia Langford, John Sillitoe on behalf of the Wellcome Sanger Institute COVID-19 Surveillance Team |
| EPI_ISL_3335777, EPI_ISL_3335778 | Lighthouse Laboratory Plymouth | Wellcome Sanger Institute for the COVID-19 Genomics UK (COG-UK) Consortium | Lighthouse Laboratory Plymouth and Alex Alderton, Roberto Amato, Jeffrey Barrett, Sonia Goncalves, Ewan Harrison, David K. Jackson, Ian Johnston, Dominic Kwiatkowski, Cordelia Langford, John Sillitoe on behalf of the Wellcome Sanger Institute COVID-19 Surveillance Team |
| EPI_ISL_3335779, EPI_ISL_3335780 | Lighthouse Lab in Milton Keynes | Wellcome Sanger Institute for the COVID-19 Genomics UK (COG-UK) Consortium | The Lighthouse Lab in Milton Keynes and Alex Alderton, Roberto Amato, Jeffrey Barrett, Sonia Goncalves, Ewan Harrison, David K. Jackson, Ian Johnston, Dominic Kwiatkowski, Cordelia Langford, John Sillitoe on behalf of the Wellcome Sanger Institute COVID-19 Surveillance Team |
| EPI_ISL_3335781 | Lighthouse Laboratory Plymouth | Wellcome Sanger Institute for the COVID-19 Genomics UK (COG-UK) Consortium | Lighthouse Laboratory Plymouth and Alex Alderton, Roberto Amato, Jeffrey Barrett, Sonia Goncalves, Ewan Harrison, David K. Jackson, Ian Johnston, Dominic Kwiatkowski, Cordelia Langford, John Sillitoe on behalf of the Wellcome Sanger Institute COVID-19 Surveillance Team |
| EPI_ISL_3335782, EPI_ISL_3335783 | Lighthouse Lab in Milton Keynes | Wellcome Sanger Institute for the COVID-19 Genomics UK (COG-UK) Consortium | The Lighthouse Lab in Milton Keynes and Alex Alderton, Roberto Amato, Jeffrey Barrett, Sonia Goncalves, Ewan Harrison, David K. Jackson, Ian Johnston, Dominic Kwiatkowski, Cordelia Langford, John Sillitoe on behalf of the Wellcome Sanger Institute COVID-19 Surveillance Team |
| EPI_ISL_3335784, | Lighthouse Laboratory Plymouth | Wellcome Sanger Institute for the COVID-19 Genomics UK (COG-UK) | Lighthouse Laboratory Plymouth and Alex Alderton, Roberto Amato, Jeffrey Barrett, Sonia Goncalves, Ewan Harrison, David K. Jackson, Ian Johnston, Dominic Kwiatkowski, Cordelia Langford, John Sillitoe on behalf of the Wellcome Sanger |

| | | | |
|---|---|---|---|
| EPI_ISL_3335785 | | Consortium | Institute COVID-19 Surveillance Team |
| EPI_ISL_3335786, EPI_ISL_3335787, EPI_ISL_3335788, EPI_ISL_3335789 | Lighthouse Lab in Milton Keynes | Wellcome Sanger Institute for the COVID-19 Genomics UK (COG-UK) Consortium | The Lighthouse Lab in Milton Keynes and Alex Alderton, Roberto Amato, Jeffrey Barrett, Sonia Goncalves, Ewan Harrison, David K. Jackson, Ian Johnston, Dominic Kwiatkowski, Cordelia Langford, John Sillitoe on behalf of the Wellcome Sanger Institute COVID-19 Surveillance Team |
| EPI_ISL_3335790 | Lighthouse Laboratory Plymouth | Wellcome Sanger Institute for the COVID-19 Genomics UK (COG-UK) Consortium | Lighthouse Laboratory Plymouth and Alex Alderton, Roberto Amato, Jeffrey Barrett, Sonia Goncalves, Ewan Harrison, David K. Jackson, Ian Johnston, Dominic Kwiatkowski, Cordelia Langford, John Sillitoe on behalf of the Wellcome Sanger Institute COVID-19 Surveillance Team |
| EPI_ISL_3335791, EPI_ISL_3335792, EPI_ISL_3335793 | Lighthouse Lab in Milton Keynes | Wellcome Sanger Institute for the COVID-19 Genomics UK (COG-UK) Consortium | The Lighthouse Lab in Milton Keynes and Alex Alderton, Roberto Amato, Jeffrey Barrett, Sonia Goncalves, Ewan Harrison, David K. Jackson, Ian Johnston, Dominic Kwiatkowski, Cordelia Langford, John Sillitoe on behalf of the Wellcome Sanger Institute COVID-19 Surveillance Team |
| EPI_ISL_3335794 | Lighthouse Laboratory Plymouth | Wellcome Sanger Institute for the COVID-19 Genomics UK (COG-UK) Consortium | Lighthouse Laboratory Plymouth and Alex Alderton, Roberto Amato, Jeffrey Barrett, Sonia Goncalves, Ewan Harrison, David K. Jackson, Ian Johnston, Dominic Kwiatkowski, Cordelia Langford, John Sillitoe on behalf of the Wellcome Sanger Institute COVID-19 Surveillance Team |
| EPI_ISL_3335795, EPI_ISL_3335796 | Health Services Laboratories | Wellcome Sanger Institute for the COVID-19 Genomics UK (COG-UK) Consortium | Health Services Laboratories and Alex Alderton, Roberto Amato, Jeffrey Barrett, Sonia Goncalves, Ewan Harrison, David K. Jackson, Ian Johnston, Dominic Kwiatkowski, Cordelia Langford, John Sillitoe on behalf of the Wellcome Sanger Institute COVID-19 Surveillance Team |
| EPI_ISL_3335797 | Lighthouse Laboratory Plymouth | Wellcome Sanger Institute for the COVID-19 Genomics UK (COG-UK) Consortium | Lighthouse Laboratory Plymouth and Alex Alderton, Roberto Amato, Jeffrey Barrett, Sonia Goncalves, Ewan Harrison, David K. Jackson, Ian Johnston, Dominic Kwiatkowski, Cordelia Langford, John Sillitoe on behalf of the Wellcome Sanger Institute COVID-19 Surveillance Team |
| EPI_ISL_3335798 | Lighthouse Lab in Milton Keynes | Wellcome Sanger Institute for the COVID-19 Genomics UK (COG-UK) Consortium | The Lighthouse Lab in Milton Keynes and Alex Alderton, Roberto Amato, Jeffrey Barrett, Sonia Goncalves, Ewan Harrison, David K. Jackson, Ian Johnston, Dominic Kwiatkowski, Cordelia Langford, John Sillitoe on behalf of the Wellcome Sanger Institute COVID-19 Surveillance Team |
| EPI_ISL_3335799 | Lighthouse Laboratory Plymouth | Wellcome Sanger Institute for the COVID-19 Genomics UK (COG-UK) Consortium | Lighthouse Laboratory Plymouth and Alex Alderton, Roberto Amato, Jeffrey Barrett, Sonia Goncalves, Ewan Harrison, David K. Jackson, Ian Johnston, Dominic Kwiatkowski, Cordelia Langford, John Sillitoe on behalf of the Wellcome Sanger Institute COVID-19 Surveillance Team |
| EPI_ISL_3335800 | Lighthouse Lab in Milton Keynes | Wellcome Sanger Institute for the COVID-19 Genomics UK (COG-UK) Consortium | The Lighthouse Lab in Milton Keynes and Alex Alderton, Roberto Amato, Jeffrey Barrett, Sonia Goncalves, Ewan Harrison, David K. Jackson, Ian Johnston, Dominic Kwiatkowski, Cordelia Langford, John Sillitoe on behalf of the Wellcome Sanger Institute COVID-19 Surveillance Team |
| EPI_ISL_3335801, EPI_ISL_3335802 | Lighthouse Laboratory Plymouth | Wellcome Sanger Institute for the COVID-19 Genomics UK (COG-UK) Consortium | Lighthouse Laboratory Plymouth and Alex Alderton, Roberto Amato, Jeffrey Barrett, Sonia Goncalves, Ewan Harrison, David K. Jackson, Ian Johnston, Dominic Kwiatkowski, Cordelia Langford, John Sillitoe on behalf of the Wellcome Sanger Institute COVID-19 Surveillance Team |
| EPI_ISL_3335803, EPI_ISL_3335804, EPI_ISL_3335805, EPI_ISL_3335806 | Lighthouse Lab in Milton Keynes | Wellcome Sanger Institute for the COVID-19 Genomics UK (COG-UK) Consortium | The Lighthouse Lab in Milton Keynes and Alex Alderton, Roberto Amato, Jeffrey Barrett, Sonia Goncalves, Ewan Harrison, David K. Jackson, Ian Johnston, Dominic Kwiatkowski, Cordelia Langford, John Sillitoe on behalf of the Wellcome Sanger Institute COVID-19 Surveillance Team |
| EPI_ISL_3335807, EPI_ISL_3335808 | Lighthouse Laboratory Plymouth | Wellcome Sanger Institute for the COVID-19 Genomics UK (COG-UK) Consortium | Lighthouse Laboratory Plymouth and Alex Alderton, Roberto Amato, Jeffrey Barrett, Sonia Goncalves, Ewan Harrison, David K. Jackson, Ian Johnston, Dominic Kwiatkowski, Cordelia Langford, John Sillitoe on behalf of the Wellcome Sanger Institute COVID-19 Surveillance Team |
| EPI_ISL_3335809 | Lighthouse Lab in Milton Keynes | Wellcome Sanger Institute for the COVID-19 Genomics UK (COG-UK) Consortium | The Lighthouse Lab in Milton Keynes and Alex Alderton, Roberto Amato, Jeffrey Barrett, Sonia Goncalves, Ewan Harrison, David K. Jackson, Ian Johnston, Dominic Kwiatkowski, Cordelia Langford, John Sillitoe on behalf of the Wellcome Sanger Institute COVID-19 Surveillance Team |
| EPI_ISL_3335810, EPI_ISL_3335811, EPI_ISL_3335812 | Lighthouse Laboratory Plymouth | Wellcome Sanger Institute for the COVID-19 Genomics UK (COG-UK) Consortium | Lighthouse Laboratory Plymouth and Alex Alderton, Roberto Amato, Jeffrey Barrett, Sonia Goncalves, Ewan Harrison, David K. Jackson, Ian Johnston, Dominic Kwiatkowski, Cordelia Langford, John Sillitoe on behalf of the Wellcome Sanger Institute COVID-19 Surveillance Team |
| EPI_ISL_3335813 | Lighthouse Lab in Milton Keynes | Wellcome Sanger Institute for the COVID-19 Genomics UK (COG-UK) Consortium | The Lighthouse Lab in Milton Keynes and Alex Alderton, Roberto Amato, Jeffrey Barrett, Sonia Goncalves, Ewan Harrison, David K. Jackson, Ian Johnston, Dominic Kwiatkowski, Cordelia Langford, John Sillitoe on behalf of the Wellcome Sanger Institute COVID-19 Surveillance Team |
| EPI_ISL_3335814 | Lighthouse Laboratory Plymouth | Wellcome Sanger Institute for the COVID-19 Genomics UK (COG-UK) Consortium | Lighthouse Laboratory Plymouth and Alex Alderton, Roberto Amato, Jeffrey Barrett, Sonia Goncalves, Ewan Harrison, David K. Jackson, Ian Johnston, Dominic Kwiatkowski, Cordelia Langford, John Sillitoe on behalf of the Wellcome Sanger Institute COVID-19 Surveillance Team |
| EPI_ISL_3335815, EPI_ISL_3335816, EPI_ISL_3335817, EPI_ISL_3335818, EPI_ISL_3335819 | Lighthouse Lab in Milton Keynes | Wellcome Sanger Institute for the COVID-19 Genomics UK (COG-UK) Consortium | The Lighthouse Lab in Milton Keynes and Alex Alderton, Roberto Amato, Jeffrey Barrett, Sonia Goncalves, Ewan Harrison, David K. Jackson, Ian Johnston, Dominic Kwiatkowski, Cordelia Langford, John Sillitoe on behalf of the Wellcome Sanger Institute COVID-19 Surveillance Team |
| EPI_ISL_3335820 | Lighthouse Laboratory Plymouth | Wellcome Sanger Institute for the COVID-19 Genomics UK (COG-UK) Consortium | Lighthouse Laboratory Plymouth and Alex Alderton, Roberto Amato, Jeffrey Barrett, Sonia Goncalves, Ewan Harrison, David K. Jackson, Ian Johnston, Dominic Kwiatkowski, Cordelia Langford, John Sillitoe on behalf of the Wellcome Sanger Institute COVID-19 Surveillance Team |
| EPI_ISL_3335821 | Health Services Laboratories | Wellcome Sanger Institute for the COVID-19 Genomics UK (COG-UK) Consortium | Health Services Laboratories and Alex Alderton, Roberto Amato, Jeffrey Barrett, Sonia Goncalves, Ewan Harrison, David K. Jackson, Ian Johnston, Dominic Kwiatkowski, Cordelia Langford, John Sillitoe on behalf of the Wellcome Sanger Institute COVID-19 Surveillance Team |
| EPI_ISL_3335822 | Lighthouse Lab in Milton Keynes | Wellcome Sanger Institute for the COVID-19 Genomics UK (COG-UK) Consortium | The Lighthouse Lab in Milton Keynes and Alex Alderton, Roberto Amato, Jeffrey Barrett, Sonia Goncalves, Ewan Harrison, David K. Jackson, Ian Johnston, Dominic Kwiatkowski, Cordelia Langford, John Sillitoe on behalf of the Wellcome Sanger Institute COVID-19 Surveillance Team |
| EPI_ISL_3335823 | Lighthouse Laboratory Plymouth | Wellcome Sanger Institute for the COVID-19 Genomics UK (COG-UK) Consortium | Lighthouse Laboratory Plymouth and Alex Alderton, Roberto Amato, Jeffrey Barrett, Sonia Goncalves, Ewan Harrison, David K. Jackson, Ian Johnston, Dominic Kwiatkowski, Cordelia Langford, John Sillitoe on behalf of the Wellcome Sanger Institute COVID-19 Surveillance Team |
| EPI_ISL_3335824 | Lighthouse Lab in Milton Keynes | Wellcome Sanger Institute for the COVID-19 Genomics UK (COG-UK) Consortium | The Lighthouse Lab in Milton Keynes and Alex Alderton, Roberto Amato, Jeffrey Barrett, Sonia Goncalves, Ewan Harrison, David K. Jackson, Ian Johnston, Dominic Kwiatkowski, Cordelia Langford, John Sillitoe on behalf of the Wellcome Sanger Institute COVID-19 Surveillance Team |
| EPI_ISL_3335825, EPI_ISL_3335826 | Lighthouse Laboratory Plymouth | Wellcome Sanger Institute for the COVID-19 Genomics UK (COG-UK) Consortium | Lighthouse Laboratory Plymouth and Alex Alderton, Roberto Amato, Jeffrey Barrett, Sonia Goncalves, Ewan Harrison, David K. Jackson, Ian Johnston, Dominic Kwiatkowski, Cordelia Langford, John Sillitoe on behalf of the Wellcome Sanger Institute COVID-19 Surveillance Team |
| EPI_ISL_3335827, EPI_ISL_3335828 | Lighthouse Lab in Milton Keynes | Wellcome Sanger Institute for the COVID-19 Genomics UK (COG-UK) Consortium | The Lighthouse Lab in Milton Keynes and Alex Alderton, Roberto Amato, Jeffrey Barrett, Sonia Goncalves, Ewan Harrison, David K. Jackson, Ian Johnston, Dominic Kwiatkowski, Cordelia Langford, John Sillitoe on behalf of the Wellcome Sanger Institute COVID-19 Surveillance Team |
| EPI_ISL_3335829 | Lighthouse Laboratory Plymouth | Wellcome Sanger Institute for the COVID-19 Genomics UK (COG-UK) Consortium | Lighthouse Laboratory Plymouth and Alex Alderton, Roberto Amato, Jeffrey Barrett, Sonia Goncalves, Ewan Harrison, David K. Jackson, Ian Johnston, Dominic Kwiatkowski, Cordelia Langford, John Sillitoe on behalf of the Wellcome Sanger Institute COVID-19 Surveillance Team |
| EPI_ISL_3335830 | Lighthouse Lab in Milton Keynes | Wellcome Sanger Institute for the COVID-19 Genomics UK (COG-UK) Consortium | The Lighthouse Lab in Milton Keynes and Alex Alderton, Roberto Amato, Jeffrey Barrett, Sonia Goncalves, Ewan Harrison, David K. Jackson, Ian Johnston, Dominic Kwiatkowski, Cordelia Langford, John Sillitoe on behalf of the Wellcome Sanger Institute COVID-19 Surveillance Team |
| EPI_ISL_3335831 | Lighthouse Laboratory Plymouth | Wellcome Sanger Institute for the COVID-19 Genomics UK (COG-UK) Consortium | Lighthouse Laboratory Plymouth and Alex Alderton, Roberto Amato, Jeffrey Barrett, Sonia Goncalves, Ewan Harrison, David K. Jackson, Ian Johnston, Dominic Kwiatkowski, Cordelia Langford, John Sillitoe on behalf of the Wellcome Sanger Institute COVID-19 Surveillance Team |
| EPI_ISL_3335832 | Lighthouse Lab in Milton Keynes | Wellcome Sanger Institute for the COVID-19 Genomics UK (COG-UK) Consortium | The Lighthouse Lab in Milton Keynes and Alex Alderton, Roberto Amato, Jeffrey Barrett, Sonia Goncalves, Ewan Harrison, David K. Jackson, Ian Johnston, Dominic Kwiatkowski, Cordelia Langford, John Sillitoe on behalf of the Wellcome Sanger Institute COVID-19 Surveillance Team |
| EPI_ISL_3335833, EPI_ISL_3335834, EPI_ISL_3335835 | Health Services Laboratories | Wellcome Sanger Institute for the COVID-19 Genomics UK (COG-UK) Consortium | Health Services Laboratories and Alex Alderton, Roberto Amato, Jeffrey Barrett, Sonia Goncalves, Ewan Harrison, David K. Jackson, Ian Johnston, Dominic Kwiatkowski, Cordelia Langford, John Sillitoe on behalf of the Wellcome Sanger Institute COVID-19 Surveillance Team |
| EPI_ISL_3335836, EPI_ISL_3335837 | Lighthouse Lab in Milton Keynes | Wellcome Sanger Institute for the COVID-19 Genomics UK (COG-UK) Consortium | The Lighthouse Lab in Milton Keynes and Alex Alderton, Roberto Amato, Jeffrey Barrett, Sonia Goncalves, Ewan Harrison, David K. Jackson, Ian Johnston, Dominic Kwiatkowski, Cordelia Langford, John Sillitoe on behalf of the Wellcome Sanger Institute COVID-19 Surveillance Team |
| EPI_ISL_3335838, EPI_ISL_3335839 | Lighthouse Laboratory Plymouth | Wellcome Sanger Institute for the COVID-19 Genomics UK (COG-UK) Consortium | Lighthouse Laboratory Plymouth and Alex Alderton, Roberto Amato, Jeffrey Barrett, Sonia Goncalves, Ewan Harrison, David K. Jackson, Ian Johnston, Dominic Kwiatkowski, Cordelia Langford, John Sillitoe on behalf of the Wellcome Sanger Institute COVID-19 Surveillance Team |
| EPI_ISL_3335840 | Lighthouse Lab in Milton Keynes | Wellcome Sanger Institute for the COVID-19 Genomics UK (COG-UK) Consortium | The Lighthouse Lab in Milton Keynes and Alex Alderton, Roberto Amato, Jeffrey Barrett, Sonia Goncalves, Ewan Harrison, David K. Jackson, Ian Johnston, Dominic Kwiatkowski, Cordelia Langford, John Sillitoe on behalf of the Wellcome Sanger Institute COVID-19 Surveillance Team |
| EPI_ISL_3335841, EPI_ISL_3335842 | Health Services Laboratories | Wellcome Sanger Institute for the COVID-19 Genomics UK (COG-UK) Consortium | Health Services Laboratories and Alex Alderton, Roberto Amato, Jeffrey Barrett, Sonia Goncalves, Ewan Harrison, David K. Jackson, Ian Johnston, Dominic Kwiatkowski, Cordelia Langford, John Sillitoe on behalf of the Wellcome Sanger Institute COVID-19 Surveillance Team |
| EPI_ISL_3335843 | Lighthouse Lab in Milton Keynes | Wellcome Sanger Institute for the COVID-19 Genomics UK (COG-UK) Consortium | The Lighthouse Lab in Milton Keynes and Alex Alderton, Roberto Amato, Jeffrey Barrett, Sonia Goncalves, Ewan Harrison, David K. Jackson, Ian Johnston, Dominic Kwiatkowski, Cordelia Langford, John Sillitoe on behalf of the Wellcome Sanger Institute COVID-19 Surveillance Team |
| EPI_ISL_3335844 | Lighthouse Laboratory Plymouth | Wellcome Sanger Institute for the COVID-19 Genomics UK (COG-UK) Consortium | Lighthouse Laboratory Plymouth and Alex Alderton, Roberto Amato, Jeffrey Barrett, Sonia Goncalves, Ewan Harrison, David K. Jackson, Ian Johnston, Dominic Kwiatkowski, Cordelia Langford, John Sillitoe on behalf of the Wellcome Sanger Institute COVID-19 Surveillance Team |
| EPI_ISL_3335845 | Lighthouse Lab in Milton Keynes | Wellcome Sanger Institute for the COVID-19 Genomics UK (COG-UK) Consortium | The Lighthouse Lab in Milton Keynes and Alex Alderton, Roberto Amato, Jeffrey Barrett, Sonia Goncalves, Ewan Harrison, David K. Jackson, Ian Johnston, Dominic Kwiatkowski, Cordelia Langford, John Sillitoe on behalf of the Wellcome Sanger Institute COVID-19 Surveillance Team |
| EPI_ISL_3335846, EPI_ISL_3335847 | Health Services Laboratories | Wellcome Sanger Institute for the COVID-19 Genomics UK (COG-UK) Consortium | Health Services Laboratories and Alex Alderton, Roberto Amato, Jeffrey Barrett, Sonia Goncalves, Ewan Harrison, David K. Jackson, Ian Johnston, Dominic Kwiatkowski, Cordelia Langford, John Sillitoe on behalf of the Wellcome Sanger Institute COVID-19 Surveillance Team |
| EPI_ISL_3335848 | Lighthouse Laboratory Plymouth | Wellcome Sanger Institute for the COVID-19 Genomics UK (COG-UK) Consortium | Lighthouse Laboratory Plymouth and Alex Alderton, Roberto Amato, Jeffrey Barrett, Sonia Goncalves, Ewan Harrison, David K. Jackson, Ian Johnston, Dominic Kwiatkowski, Cordelia Langford, John Sillitoe on behalf of the Wellcome Sanger Institute COVID-19 Surveillance Team |
| EPI_ISL_3335849, EPI_ISL_3335850, EPI_ISL_3335851 | Lighthouse Lab in Milton Keynes | Wellcome Sanger Institute for the COVID-19 Genomics UK (COG-UK) Consortium | The Lighthouse Lab in Milton Keynes and Alex Alderton, Roberto Amato, Jeffrey Barrett, Sonia Goncalves, Ewan Harrison, David K. Jackson, Ian Johnston, Dominic Kwiatkowski, Cordelia Langford, John Sillitoe on behalf of the Wellcome Sanger Institute COVID-19 Surveillance Team |
| EPI_ISL_3335852 | Lighthouse Laboratory Plymouth | Wellcome Sanger Institute for the COVID-19 Genomics UK (COG-UK) Consortium | Lighthouse Laboratory Plymouth and Alex Alderton, Roberto Amato, Jeffrey Barrett, Sonia Goncalves, Ewan Harrison, David K. Jackson, Ian Johnston, Dominic Kwiatkowski, Cordelia Langford, John Sillitoe on behalf of the Wellcome Sanger Institute COVID-19 Surveillance Team |

| | | | |
|---|---|---|---|
| EPI_ISL_3335853, EPI_ISL_3335854 | Lighthouse Lab in Milton Keynes | Wellcome Sanger Institute for the COVID-19 Genomics UK (COG-UK) Consortium | The Lighthouse Lab in Milton Keynes and Alex Alderton, Roberto Amato, Jeffrey Barrett, Sonia Goncalves, Ewan Harrison, David K. Jackson, Ian Johnston, Dominic Kwiatkowski, Cordelia Langford, John Sillitoe on behalf of the Wellcome Sanger Institute COVID-19 Surveillance Team |
| EPI_ISL_3335855 | Lighthouse Laboratory Plymouth | Wellcome Sanger Institute for the COVID-19 Genomics UK (COG-UK) Consortium | Lighthouse Laboratory Plymouth and Alex Alderton, Roberto Amato, Jeffrey Barrett, Sonia Goncalves, Ewan Harrison, David K. Jackson, Ian Johnston, Dominic Kwiatkowski, Cordelia Langford, John Sillitoe on behalf of the Wellcome Sanger Institute COVID-19 Surveillance Team |
| EPI_ISL_3335856 | Lighthouse Lab in Milton Keynes | Wellcome Sanger Institute for the COVID-19 Genomics UK (COG-UK) Consortium | The Lighthouse Lab in Milton Keynes and Alex Alderton, Roberto Amato, Jeffrey Barrett, Sonia Goncalves, Ewan Harrison, David K. Jackson, Ian Johnston, Dominic Kwiatkowski, Cordelia Langford, John Sillitoe on behalf of the Wellcome Sanger Institute COVID-19 Surveillance Team |
| EPI_ISL_3335857 | Health Services Laboratories | Wellcome Sanger Institute for the COVID-19 Genomics UK (COG-UK) Consortium | Health Services Laboratories and Alex Alderton, Roberto Amato, Jeffrey Barrett, Sonia Goncalves, Ewan Harrison, David K. Jackson, Ian Johnston, Dominic Kwiatkowski, Cordelia Langford, John Sillitoe on behalf of the Wellcome Sanger Institute COVID-19 Surveillance Team |
| EPI_ISL_3335858 | Lighthouse Laboratory Plymouth | Wellcome Sanger Institute for the COVID-19 Genomics UK (COG-UK) Consortium | Lighthouse Laboratory Plymouth and Alex Alderton, Roberto Amato, Jeffrey Barrett, Sonia Goncalves, Ewan Harrison, David K. Jackson, Ian Johnston, Dominic Kwiatkowski, Cordelia Langford, John Sillitoe on behalf of the Wellcome Sanger Institute COVID-19 Surveillance Team |
| EPI_ISL_3335859, EPI_ISL_3335860, EPI_ISL_3335861 | Lighthouse Lab in Milton Keynes | Wellcome Sanger Institute for the COVID-19 Genomics UK (COG-UK) Consortium | The Lighthouse Lab in Milton Keynes and Alex Alderton, Roberto Amato, Jeffrey Barrett, Sonia Goncalves, Ewan Harrison, David K. Jackson, Ian Johnston, Dominic Kwiatkowski, Cordelia Langford, John Sillitoe on behalf of the Wellcome Sanger Institute COVID-19 Surveillance Team |
| EPI_ISL_3335862 | Health Services Laboratories | Wellcome Sanger Institute for the COVID-19 Genomics UK (COG-UK) Consortium | Health Services Laboratories and Alex Alderton, Roberto Amato, Jeffrey Barrett, Sonia Goncalves, Ewan Harrison, David K. Jackson, Ian Johnston, Dominic Kwiatkowski, Cordelia Langford, John Sillitoe on behalf of the Wellcome Sanger Institute COVID-19 Surveillance Team |
| EPI_ISL_3335863, EPI_ISL_3335864 | Lighthouse Lab in Milton Keynes | Wellcome Sanger Institute for the COVID-19 Genomics UK (COG-UK) Consortium | The Lighthouse Lab in Milton Keynes and Alex Alderton, Roberto Amato, Jeffrey Barrett, Sonia Goncalves, Ewan Harrison, David K. Jackson, Ian Johnston, Dominic Kwiatkowski, Cordelia Langford, John Sillitoe on behalf of the Wellcome Sanger Institute COVID-19 Surveillance Team |
| EPI_ISL_3335865 | Lighthouse Laboratory Plymouth | Wellcome Sanger Institute for the COVID-19 Genomics UK (COG-UK) Consortium | Lighthouse Laboratory Plymouth and Alex Alderton, Roberto Amato, Jeffrey Barrett, Sonia Goncalves, Ewan Harrison, David K. Jackson, Ian Johnston, Dominic Kwiatkowski, Cordelia Langford, John Sillitoe on behalf of the Wellcome Sanger Institute COVID-19 Surveillance Team |
| EPI_ISL_3335866, EPI_ISL_3335867, EPI_ISL_3335868 | Lighthouse Lab in Milton Keynes | Wellcome Sanger Institute for the COVID-19 Genomics UK (COG-UK) Consortium | The Lighthouse Lab in Milton Keynes and Alex Alderton, Roberto Amato, Jeffrey Barrett, Sonia Goncalves, Ewan Harrison, David K. Jackson, Ian Johnston, Dominic Kwiatkowski, Cordelia Langford, John Sillitoe on behalf of the Wellcome Sanger Institute COVID-19 Surveillance Team |
| EPI_ISL_3335869 | Health Services Laboratories | Wellcome Sanger Institute for the COVID-19 Genomics UK (COG-UK) Consortium | Health Services Laboratories and Alex Alderton, Roberto Amato, Jeffrey Barrett, Sonia Goncalves, Ewan Harrison, David K. Jackson, Ian Johnston, Dominic Kwiatkowski, Cordelia Langford, John Sillitoe on behalf of the Wellcome Sanger Institute COVID-19 Surveillance Team |
| EPI_ISL_3335870, EPI_ISL_3335871, EPI_ISL_3335872, EPI_ISL_3335873 | Lighthouse Lab in Milton Keynes | Wellcome Sanger Institute for the COVID-19 Genomics UK (COG-UK) Consortium | The Lighthouse Lab in Milton Keynes and Alex Alderton, Roberto Amato, Jeffrey Barrett, Sonia Goncalves, Ewan Harrison, David K. Jackson, Ian Johnston, Dominic Kwiatkowski, Cordelia Langford, John Sillitoe on behalf of the Wellcome Sanger Institute COVID-19 Surveillance Team |
| EPI_ISL_3335874, EPI_ISL_3335875 | Lighthouse Laboratory Plymouth | Wellcome Sanger Institute for the COVID-19 Genomics UK (COG-UK) Consortium | Lighthouse Laboratory Plymouth and Alex Alderton, Roberto Amato, Jeffrey Barrett, Sonia Goncalves, Ewan Harrison, David K. Jackson, Ian Johnston, Dominic Kwiatkowski, Cordelia Langford, John Sillitoe on behalf of the Wellcome Sanger Institute COVID-19 Surveillance Team |
| EPI_ISL_3335876 | Lighthouse Lab in Milton Keynes | Wellcome Sanger Institute for the COVID-19 Genomics UK (COG-UK) Consortium | The Lighthouse Lab in Milton Keynes and Alex Alderton, Roberto Amato, Jeffrey Barrett, Sonia Goncalves, Ewan Harrison, David K. Jackson, Ian Johnston, Dominic Kwiatkowski, Cordelia Langford, John Sillitoe on behalf of the Wellcome Sanger Institute COVID-19 Surveillance Team |
| EPI_ISL_3335877 | Health Services Laboratories | Wellcome Sanger Institute for the COVID-19 Genomics UK (COG-UK) Consortium | Health Services Laboratories and Alex Alderton, Roberto Amato, Jeffrey Barrett, Sonia Goncalves, Ewan Harrison, David K. Jackson, Ian Johnston, Dominic Kwiatkowski, Cordelia Langford, John Sillitoe on behalf of the Wellcome Sanger Institute COVID-19 Surveillance Team |
| EPI_ISL_3335878 | Lighthouse Lab in Milton Keynes | Wellcome Sanger Institute for the COVID-19 Genomics UK (COG-UK) Consortium | The Lighthouse Lab in Milton Keynes and Alex Alderton, Roberto Amato, Jeffrey Barrett, Sonia Goncalves, Ewan Harrison, David K. Jackson, Ian Johnston, Dominic Kwiatkowski, Cordelia Langford, John Sillitoe on behalf of the Wellcome Sanger Institute COVID-19 Surveillance Team |
| EPI_ISL_3335879 | Health Services Laboratories | Wellcome Sanger Institute for the COVID-19 Genomics UK (COG-UK) Consortium | Health Services Laboratories and Alex Alderton, Roberto Amato, Jeffrey Barrett, Sonia Goncalves, Ewan Harrison, David K. Jackson, Ian Johnston, Dominic Kwiatkowski, Cordelia Langford, John Sillitoe on behalf of the Wellcome Sanger Institute COVID-19 Surveillance Team |
| EPI_ISL_3335880 | Lighthouse Laboratory Plymouth | Wellcome Sanger Institute for the COVID-19 Genomics UK (COG-UK) Consortium | Lighthouse Laboratory Plymouth and Alex Alderton, Roberto Amato, Jeffrey Barrett, Sonia Goncalves, Ewan Harrison, David K. Jackson, Ian Johnston, Dominic Kwiatkowski, Cordelia Langford, John Sillitoe on behalf of the Wellcome Sanger Institute COVID-19 Surveillance Team |
| EPI_ISL_3335881 | Lighthouse Lab in Milton Keynes | Wellcome Sanger Institute for the COVID-19 Genomics UK (COG-UK) Consortium | The Lighthouse Lab in Milton Keynes and Alex Alderton, Roberto Amato, Jeffrey Barrett, Sonia Goncalves, Ewan Harrison, David K. Jackson, Ian Johnston, Dominic Kwiatkowski, Cordelia Langford, John Sillitoe on behalf of the Wellcome Sanger Institute COVID-19 Surveillance Team |
| EPI_ISL_3335882, EPI_ISL_3335883 | Lighthouse Laboratory Plymouth | Wellcome Sanger Institute for the COVID-19 Genomics UK (COG-UK) Consortium | Lighthouse Laboratory Plymouth and Alex Alderton, Roberto Amato, Jeffrey Barrett, Sonia Goncalves, Ewan Harrison, David K. Jackson, Ian Johnston, Dominic Kwiatkowski, Cordelia Langford, John Sillitoe on behalf of the Wellcome Sanger Institute COVID-19 Surveillance Team |
| EPI_ISL_3335884, EPI_ISL_3335885, EPI_ISL_3335886 | Lighthouse Lab in Milton Keynes | Wellcome Sanger Institute for the COVID-19 Genomics UK (COG-UK) Consortium | The Lighthouse Lab in Milton Keynes and Alex Alderton, Roberto Amato, Jeffrey Barrett, Sonia Goncalves, Ewan Harrison, David K. Jackson, Ian Johnston, Dominic Kwiatkowski, Cordelia Langford, John Sillitoe on behalf of the Wellcome Sanger Institute COVID-19 Surveillance Team |
| EPI_ISL_3335887, EPI_ISL_3335888, EPI_ISL_3335889 | Health Services Laboratories | Wellcome Sanger Institute for the COVID-19 Genomics UK (COG-UK) Consortium | Health Services Laboratories and Alex Alderton, Roberto Amato, Jeffrey Barrett, Sonia Goncalves, Ewan Harrison, David K. Jackson, Ian Johnston, Dominic Kwiatkowski, Cordelia Langford, John Sillitoe on behalf of the Wellcome Sanger Institute COVID-19 Surveillance Team |
| EPI_ISL_3335890, EPI_ISL_3335891 | Lighthouse Lab in Milton Keynes | Wellcome Sanger Institute for the COVID-19 Genomics UK (COG-UK) Consortium | The Lighthouse Lab in Milton Keynes and Alex Alderton, Roberto Amato, Jeffrey Barrett, Sonia Goncalves, Ewan Harrison, David K. Jackson, Ian Johnston, Dominic Kwiatkowski, Cordelia Langford, John Sillitoe on behalf of the Wellcome Sanger Institute COVID-19 Surveillance Team |
| EPI_ISL_3335892 | Health Services Laboratories | Wellcome Sanger Institute for the COVID-19 Genomics UK (COG-UK) Consortium | Health Services Laboratories and Alex Alderton, Roberto Amato, Jeffrey Barrett, Sonia Goncalves, Ewan Harrison, David K. Jackson, Ian Johnston, Dominic Kwiatkowski, Cordelia Langford, John Sillitoe on behalf of the Wellcome Sanger Institute COVID-19 Surveillance Team |
| EPI_ISL_3335893 | Lighthouse Laboratory Plymouth | Wellcome Sanger Institute for the COVID-19 Genomics UK (COG-UK) Consortium | Lighthouse Laboratory Plymouth and Alex Alderton, Roberto Amato, Jeffrey Barrett, Sonia Goncalves, Ewan Harrison, David K. Jackson, Ian Johnston, Dominic Kwiatkowski, Cordelia Langford, John Sillitoe on behalf of the Wellcome Sanger Institute COVID-19 Surveillance Team |
| EPI_ISL_3335894, EPI_ISL_3335895 | Lighthouse Lab in Milton Keynes | Wellcome Sanger Institute for the COVID-19 Genomics UK (COG-UK) Consortium | The Lighthouse Lab in Milton Keynes and Alex Alderton, Roberto Amato, Jeffrey Barrett, Sonia Goncalves, Ewan Harrison, David K. Jackson, Ian Johnston, Dominic Kwiatkowski, Cordelia Langford, John Sillitoe on behalf of the Wellcome Sanger Institute COVID-19 Surveillance Team |
| EPI_ISL_3335896 | Lighthouse Laboratory Plymouth | Wellcome Sanger Institute for the COVID-19 Genomics UK (COG-UK) Consortium | Lighthouse Laboratory Plymouth and Alex Alderton, Roberto Amato, Jeffrey Barrett, Sonia Goncalves, Ewan Harrison, David K. Jackson, Ian Johnston, Dominic Kwiatkowski, Cordelia Langford, John Sillitoe on behalf of the Wellcome Sanger Institute COVID-19 Surveillance Team |
| EPI_ISL_3335897, EPI_ISL_3335898, EPI_ISL_3335899, EPI_ISL_3335900, EPI_ISL_3335901 | Lighthouse Lab in Milton Keynes | Wellcome Sanger Institute for the COVID-19 Genomics UK (COG-UK) Consortium | The Lighthouse Lab in Milton Keynes and Alex Alderton, Roberto Amato, Jeffrey Barrett, Sonia Goncalves, Ewan Harrison, David K. Jackson, Ian Johnston, Dominic Kwiatkowski, Cordelia Langford, John Sillitoe on behalf of the Wellcome Sanger Institute COVID-19 Surveillance Team |
| EPI_ISL_3335902, EPI_ISL_3335903 | Health Services Laboratories | Wellcome Sanger Institute for the COVID-19 Genomics UK (COG-UK) Consortium | Health Services Laboratories and Alex Alderton, Roberto Amato, Jeffrey Barrett, Sonia Goncalves, Ewan Harrison, David K. Jackson, Ian Johnston, Dominic Kwiatkowski, Cordelia Langford, John Sillitoe on behalf of the Wellcome Sanger Institute COVID-19 Surveillance Team |
| EPI_ISL_3335904 | Lighthouse Laboratory Plymouth | Wellcome Sanger Institute for the COVID-19 Genomics UK (COG-UK) Consortium | Lighthouse Laboratory Plymouth and Alex Alderton, Roberto Amato, Jeffrey Barrett, Sonia Goncalves, Ewan Harrison, David K. Jackson, Ian Johnston, Dominic Kwiatkowski, Cordelia Langford, John Sillitoe on behalf of the Wellcome Sanger Institute COVID-19 Surveillance Team |
| EPI_ISL_3335905 | Lighthouse Lab in Milton Keynes | Wellcome Sanger Institute for the COVID-19 Genomics UK (COG-UK) Consortium | The Lighthouse Lab in Milton Keynes and Alex Alderton, Roberto Amato, Jeffrey Barrett, Sonia Goncalves, Ewan Harrison, David K. Jackson, Ian Johnston, Dominic Kwiatkowski, Cordelia Langford, John Sillitoe on behalf of the Wellcome Sanger Institute COVID-19 Surveillance Team |
| EPI_ISL_3335906 | Lighthouse Laboratory Plymouth | Wellcome Sanger Institute for the COVID-19 Genomics UK (COG-UK) Consortium | Lighthouse Laboratory Plymouth and Alex Alderton, Roberto Amato, Jeffrey Barrett, Sonia Goncalves, Ewan Harrison, David K. Jackson, Ian Johnston, Dominic Kwiatkowski, Cordelia Langford, John Sillitoe on behalf of the Wellcome Sanger Institute COVID-19 Surveillance Team |
| EPI_ISL_3335907 | Health Services Laboratories | Wellcome Sanger Institute for the COVID-19 Genomics UK (COG-UK) Consortium | Health Services Laboratories and Alex Alderton, Roberto Amato, Jeffrey Barrett, Sonia Goncalves, Ewan Harrison, David K. Jackson, Ian Johnston, Dominic Kwiatkowski, Cordelia Langford, John Sillitoe on behalf of the Wellcome Sanger Institute COVID-19 Surveillance Team |
| EPI_ISL_3335908 | Lighthouse Lab in Milton Keynes | Wellcome Sanger Institute for the COVID-19 Genomics UK (COG-UK) Consortium | The Lighthouse Lab in Milton Keynes and Alex Alderton, Roberto Amato, Jeffrey Barrett, Sonia Goncalves, Ewan Harrison, David K. Jackson, Ian Johnston, Dominic Kwiatkowski, Cordelia Langford, John Sillitoe on behalf of the Wellcome Sanger Institute COVID-19 Surveillance Team |
| EPI_ISL_3335909, EPI_ISL_3335910 | Lighthouse Laboratory Plymouth | Wellcome Sanger Institute for the COVID-19 Genomics UK (COG-UK) Consortium | Lighthouse Laboratory Plymouth and Alex Alderton, Roberto Amato, Jeffrey Barrett, Sonia Goncalves, Ewan Harrison, David K. Jackson, Ian Johnston, Dominic Kwiatkowski, Cordelia Langford, John Sillitoe on behalf of the Wellcome Sanger Institute COVID-19 Surveillance Team |
| EPI_ISL_3335911 | Health Services Laboratories | Wellcome Sanger Institute for the COVID-19 Genomics UK (COG-UK) Consortium | Health Services Laboratories and Alex Alderton, Roberto Amato, Jeffrey Barrett, Sonia Goncalves, Ewan Harrison, David K. Jackson, Ian Johnston, Dominic Kwiatkowski, Cordelia Langford, John Sillitoe on behalf of the Wellcome Sanger Institute COVID-19 Surveillance Team |
| EPI_ISL_3335912 | Lighthouse Laboratory Plymouth | Wellcome Sanger Institute for the COVID-19 Genomics UK (COG-UK) Consortium | Lighthouse Laboratory Plymouth and Alex Alderton, Roberto Amato, Jeffrey Barrett, Sonia Goncalves, Ewan Harrison, David K. Jackson, Ian Johnston, Dominic Kwiatkowski, Cordelia Langford, John Sillitoe on behalf of the Wellcome Sanger Institute COVID-19 Surveillance Team |
| EPI_ISL_3335913 | Health Services Laboratories | Wellcome Sanger Institute for the COVID-19 Genomics UK (COG-UK) Consortium | Health Services Laboratories and Alex Alderton, Roberto Amato, Jeffrey Barrett, Sonia Goncalves, Ewan Harrison, David K. Jackson, Ian Johnston, Dominic Kwiatkowski, Cordelia Langford, John Sillitoe on behalf of the Wellcome Sanger Institute COVID-19 Surveillance Team |
| EPI_ISL_3335914 | Lighthouse Laboratory Plymouth | Wellcome Sanger Institute for the COVID-19 Genomics UK (COG-UK) Consortium | Lighthouse Laboratory Plymouth and Alex Alderton, Roberto Amato, Jeffrey Barrett, Sonia Goncalves, Ewan Harrison, David K. Jackson, Ian Johnston, Dominic Kwiatkowski, Cordelia Langford, John Sillitoe on behalf of the Wellcome Sanger Institute COVID-19 Surveillance Team |
| EPI_ISL_3335915 | Lighthouse Lab in Milton Keynes | Wellcome Sanger Institute for the COVID-19 Genomics UK (COG-UK) Consortium | The Lighthouse Lab in Milton Keynes and Alex Alderton, Roberto Amato, Jeffrey Barrett, Sonia Goncalves, Ewan Harrison, David K. Jackson, Ian Johnston, Dominic Kwiatkowski, Cordelia Langford, John Sillitoe on behalf of the Wellcome Sanger Institute COVID-19 Surveillance Team |
| EPI_ISL_3335916, EPI_ISL_3335917, EPI_ISL_3335918 | Lighthouse Laboratory Plymouth | Wellcome Sanger Institute for the COVID-19 Genomics UK (COG-UK) Consortium | Lighthouse Laboratory Plymouth and Alex Alderton, Roberto Amato, Jeffrey Barrett, Sonia Goncalves, Ewan Harrison, David K. Jackson, Ian Johnston, Dominic Kwiatkowski, Cordelia Langford, John Sillitoe on behalf of the Wellcome Sanger Institute COVID-19 Surveillance Team |

| | | | |
|---|---|---|---|
| EPI_ISL_3335919, EPI_ISL_3335920, EPI_ISL_3335921, EPI_ISL_3335922, EPI_ISL_3335923, EPI_ISL_3335924, EPI_ISL_3335925, EPI_ISL_3335926, EPI_ISL_3335927 | Lighthouse Lab in Milton Keynes | Wellcome Sanger Institute for the COVID-19 Genomics UK (COG-UK) Consortium | The Lighthouse Lab in Milton Keynes and Alex Alderton, Roberto Amato, Jeffrey Barrett, Sonia Goncalves, Ewan Harrison, David K. Jackson, Ian Johnston, Dominic Kwiatkowski, Cordelia Langford, John Sillitoe on behalf of the Wellcome Sanger Institute COVID-19 Surveillance Team |
| EPI_ISL_3335928, EPI_ISL_3335929 | Health Services Laboratories | Wellcome Sanger Institute for the COVID-19 Genomics UK (COG-UK) Consortium | Health Services Laboratories and Alex Alderton, Roberto Amato, Jeffrey Barrett, Sonia Goncalves, Ewan Harrison, David K. Jackson, Ian Johnston, Dominic Kwiatkowski, Cordelia Langford, John Sillitoe on behalf of the Wellcome Sanger Institute COVID-19 Surveillance Team |
| EPI_ISL_3335930, EPI_ISL_3335931 | Lighthouse Lab in Milton Keynes | Wellcome Sanger Institute for the COVID-19 Genomics UK (COG-UK) Consortium | The Lighthouse Lab in Milton Keynes and Alex Alderton, Roberto Amato, Jeffrey Barrett, Sonia Goncalves, Ewan Harrison, David K. Jackson, Ian Johnston, Dominic Kwiatkowski, Cordelia Langford, John Sillitoe on behalf of the Wellcome Sanger Institute COVID-19 Surveillance Team |
| EPI_ISL_3335932 | Health Services Laboratories | Wellcome Sanger Institute for the COVID-19 Genomics UK (COG-UK) Consortium | Health Services Laboratories and Alex Alderton, Roberto Amato, Jeffrey Barrett, Sonia Goncalves, Ewan Harrison, David K. Jackson, Ian Johnston, Dominic Kwiatkowski, Cordelia Langford, John Sillitoe on behalf of the Wellcome Sanger Institute COVID-19 Surveillance Team |
| EPI_ISL_3335933, EPI_ISL_3335934 | Lighthouse Lab in Milton Keynes | Wellcome Sanger Institute for the COVID-19 Genomics UK (COG-UK) Consortium | The Lighthouse Lab in Milton Keynes and Alex Alderton, Roberto Amato, Jeffrey Barrett, Sonia Goncalves, Ewan Harrison, David K. Jackson, Ian Johnston, Dominic Kwiatkowski, Cordelia Langford, John Sillitoe on behalf of the Wellcome Sanger Institute COVID-19 Surveillance Team |
| EPI_ISL_3335935 | Lighthouse Laboratory Plymouth | Wellcome Sanger Institute for the COVID-19 Genomics UK (COG-UK) Consortium | Lighthouse Laboratory Plymouth and Alex Alderton, Roberto Amato, Jeffrey Barrett, Sonia Goncalves, Ewan Harrison, David K. Jackson, Ian Johnston, Dominic Kwiatkowski, Cordelia Langford, John Sillitoe on behalf of the Wellcome Sanger Institute COVID-19 Surveillance Team |
| EPI_ISL_3335936, EPI_ISL_3335937, EPI_ISL_3335938, EPI_ISL_3335939 | Lighthouse Lab in Milton Keynes | Wellcome Sanger Institute for the COVID-19 Genomics UK (COG-UK) Consortium | The Lighthouse Lab in Milton Keynes and Alex Alderton, Roberto Amato, Jeffrey Barrett, Sonia Goncalves, Ewan Harrison, David K. Jackson, Ian Johnston, Dominic Kwiatkowski, Cordelia Langford, John Sillitoe on behalf of the Wellcome Sanger Institute COVID-19 Surveillance Team |
| EPI_ISL_3335940, EPI_ISL_3335941 | Health Services Laboratories | Wellcome Sanger Institute for the COVID-19 Genomics UK (COG-UK) Consortium | Health Services Laboratories and Alex Alderton, Roberto Amato, Jeffrey Barrett, Sonia Goncalves, Ewan Harrison, David K. Jackson, Ian Johnston, Dominic Kwiatkowski, Cordelia Langford, John Sillitoe on behalf of the Wellcome Sanger Institute COVID-19 Surveillance Team |
| EPI_ISL_3335942, EPI_ISL_3335943 | Lighthouse Lab in Milton Keynes | Wellcome Sanger Institute for the COVID-19 Genomics UK (COG-UK) Consortium | The Lighthouse Lab in Milton Keynes and Alex Alderton, Roberto Amato, Jeffrey Barrett, Sonia Goncalves, Ewan Harrison, David K. Jackson, Ian Johnston, Dominic Kwiatkowski, Cordelia Langford, John Sillitoe on behalf of the Wellcome Sanger Institute COVID-19 Surveillance Team |
| EPI_ISL_3335944 | Lighthouse Laboratory Plymouth | Wellcome Sanger Institute for the COVID-19 Genomics UK (COG-UK) Consortium | Lighthouse Laboratory Plymouth and Alex Alderton, Roberto Amato, Jeffrey Barrett, Sonia Goncalves, Ewan Harrison, David K. Jackson, Ian Johnston, Dominic Kwiatkowski, Cordelia Langford, John Sillitoe on behalf of the Wellcome Sanger Institute COVID-19 Surveillance Team |
| EPI_ISL_3335945, EPI_ISL_3335946, EPI_ISL_3335947 | Lighthouse Lab in Milton Keynes | Wellcome Sanger Institute for the COVID-19 Genomics UK (COG-UK) Consortium | The Lighthouse Lab in Milton Keynes and Alex Alderton, Roberto Amato, Jeffrey Barrett, Sonia Goncalves, Ewan Harrison, David K. Jackson, Ian Johnston, Dominic Kwiatkowski, Cordelia Langford, John Sillitoe on behalf of the Wellcome Sanger Institute COVID-19 Surveillance Team |
| EPI_ISL_3335948, EPI_ISL_3335949 | Health Services Laboratories | Wellcome Sanger Institute for the COVID-19 Genomics UK (COG-UK) Consortium | Health Services Laboratories and Alex Alderton, Roberto Amato, Jeffrey Barrett, Sonia Goncalves, Ewan Harrison, David K. Jackson, Ian Johnston, Dominic Kwiatkowski, Cordelia Langford, John Sillitoe on behalf of the Wellcome Sanger Institute COVID-19 Surveillance Team |
| EPI_ISL_3335950 | Lighthouse Lab in Milton Keynes | Wellcome Sanger Institute for the COVID-19 Genomics UK (COG-UK) Consortium | The Lighthouse Lab in Milton Keynes and Alex Alderton, Roberto Amato, Jeffrey Barrett, Sonia Goncalves, Ewan Harrison, David K. Jackson, Ian Johnston, Dominic Kwiatkowski, Cordelia Langford, John Sillitoe on behalf of the Wellcome Sanger Institute COVID-19 Surveillance Team |
| EPI_ISL_3335951 | Lighthouse Laboratory Plymouth | Wellcome Sanger Institute for the COVID-19 Genomics UK (COG-UK) Consortium | Lighthouse Laboratory Plymouth and Alex Alderton, Roberto Amato, Jeffrey Barrett, Sonia Goncalves, Ewan Harrison, David K. Jackson, Ian Johnston, Dominic Kwiatkowski, Cordelia Langford, John Sillitoe on behalf of the Wellcome Sanger Institute COVID-19 Surveillance Team |
| EPI_ISL_3335952 | Health Services Laboratories | Wellcome Sanger Institute for the COVID-19 Genomics UK (COG-UK) Consortium | Health Services Laboratories and Alex Alderton, Roberto Amato, Jeffrey Barrett, Sonia Goncalves, Ewan Harrison, David K. Jackson, Ian Johnston, Dominic Kwiatkowski, Cordelia Langford, John Sillitoe on behalf of the Wellcome Sanger Institute COVID-19 Surveillance Team |
| EPI_ISL_3335953, EPI_ISL_3335954 | Lighthouse Lab in Milton Keynes | Wellcome Sanger Institute for the COVID-19 Genomics UK (COG-UK) Consortium | The Lighthouse Lab in Milton Keynes and Alex Alderton, Roberto Amato, Jeffrey Barrett, Sonia Goncalves, Ewan Harrison, David K. Jackson, Ian Johnston, Dominic Kwiatkowski, Cordelia Langford, John Sillitoe on behalf of the Wellcome Sanger Institute COVID-19 Surveillance Team |
| EPI_ISL_3335955 | Health Services Laboratories | Wellcome Sanger Institute for the COVID-19 Genomics UK (COG-UK) Consortium | Health Services Laboratories and Alex Alderton, Roberto Amato, Jeffrey Barrett, Sonia Goncalves, Ewan Harrison, David K. Jackson, Ian Johnston, Dominic Kwiatkowski, Cordelia Langford, John Sillitoe on behalf of the Wellcome Sanger Institute COVID-19 Surveillance Team |
| EPI_ISL_3335956 | Lighthouse Laboratory Plymouth | Wellcome Sanger Institute for the COVID-19 Genomics UK (COG-UK) Consortium | Lighthouse Laboratory Plymouth and Alex Alderton, Roberto Amato, Jeffrey Barrett, Sonia Goncalves, Ewan Harrison, David K. Jackson, Ian Johnston, Dominic Kwiatkowski, Cordelia Langford, John Sillitoe on behalf of the Wellcome Sanger Institute COVID-19 Surveillance Team |
| EPI_ISL_3335957, EPI_ISL_3335958 | Lighthouse Lab in Milton Keynes | Wellcome Sanger Institute for the COVID-19 Genomics UK (COG-UK) Consortium | The Lighthouse Lab in Milton Keynes and Alex Alderton, Roberto Amato, Jeffrey Barrett, Sonia Goncalves, Ewan Harrison, David K. Jackson, Ian Johnston, Dominic Kwiatkowski, Cordelia Langford, John Sillitoe on behalf of the Wellcome Sanger Institute COVID-19 Surveillance Team |
| EPI_ISL_3335959, EPI_ISL_3335960 | Lighthouse Laboratory Plymouth | Wellcome Sanger Institute for the COVID-19 Genomics UK (COG-UK) Consortium | Lighthouse Laboratory Plymouth and Alex Alderton, Roberto Amato, Jeffrey Barrett, Sonia Goncalves, Ewan Harrison, David K. Jackson, Ian Johnston, Dominic Kwiatkowski, Cordelia Langford, John Sillitoe on behalf of the Wellcome Sanger Institute COVID-19 Surveillance Team |
| EPI_ISL_3335961, EPI_ISL_3335962, EPI_ISL_3335963 | Lighthouse Lab in Milton Keynes | Wellcome Sanger Institute for the COVID-19 Genomics UK (COG-UK) Consortium | The Lighthouse Lab in Milton Keynes and Alex Alderton, Roberto Amato, Jeffrey Barrett, Sonia Goncalves, Ewan Harrison, David K. Jackson, Ian Johnston, Dominic Kwiatkowski, Cordelia Langford, John Sillitoe on behalf of the Wellcome Sanger Institute COVID-19 Surveillance Team |
| EPI_ISL_3335964 | Health Services Laboratories | Wellcome Sanger Institute for the COVID-19 Genomics UK (COG-UK) Consortium | Health Services Laboratories and Alex Alderton, Roberto Amato, Jeffrey Barrett, Sonia Goncalves, Ewan Harrison, David K. Jackson, Ian Johnston, Dominic Kwiatkowski, Cordelia Langford, John Sillitoe on behalf of the Wellcome Sanger Institute COVID-19 Surveillance Team |
| EPI_ISL_3335965 | Lighthouse Laboratory Plymouth | Wellcome Sanger Institute for the COVID-19 Genomics UK (COG-UK) Consortium | Lighthouse Laboratory Plymouth and Alex Alderton, Roberto Amato, Jeffrey Barrett, Sonia Goncalves, Ewan Harrison, David K. Jackson, Ian Johnston, Dominic Kwiatkowski, Cordelia Langford, John Sillitoe on behalf of the Wellcome Sanger Institute COVID-19 Surveillance Team |
| EPI_ISL_3335966 | Health Services Laboratories | Wellcome Sanger Institute for the COVID-19 Genomics UK (COG-UK) Consortium | Health Services Laboratories and Alex Alderton, Roberto Amato, Jeffrey Barrett, Sonia Goncalves, Ewan Harrison, David K. Jackson, Ian Johnston, Dominic Kwiatkowski, Cordelia Langford, John Sillitoe on behalf of the Wellcome Sanger Institute COVID-19 Surveillance Team |
| EPI_ISL_3335967 | Lighthouse Lab in Milton Keynes | Wellcome Sanger Institute for the COVID-19 Genomics UK (COG-UK) Consortium | The Lighthouse Lab in Milton Keynes and Alex Alderton, Roberto Amato, Jeffrey Barrett, Sonia Goncalves, Ewan Harrison, David K. Jackson, Ian Johnston, Dominic Kwiatkowski, Cordelia Langford, John Sillitoe on behalf of the Wellcome Sanger Institute COVID-19 Surveillance Team |
| EPI_ISL_3335968 | Lighthouse Laboratory Plymouth | Wellcome Sanger Institute for the COVID-19 Genomics UK (COG-UK) Consortium | Lighthouse Laboratory Plymouth and Alex Alderton, Roberto Amato, Jeffrey Barrett, Sonia Goncalves, Ewan Harrison, David K. Jackson, Ian Johnston, Dominic Kwiatkowski, Cordelia Langford, John Sillitoe on behalf of the Wellcome Sanger Institute COVID-19 Surveillance Team |
| EPI_ISL_3335969 | Lighthouse Lab in Milton Keynes | Wellcome Sanger Institute for the COVID-19 Genomics UK (COG-UK) Consortium | The Lighthouse Lab in Milton Keynes and Alex Alderton, Roberto Amato, Jeffrey Barrett, Sonia Goncalves, Ewan Harrison, David K. Jackson, Ian Johnston, Dominic Kwiatkowski, Cordelia Langford, John Sillitoe on behalf of the Wellcome Sanger Institute COVID-19 Surveillance Team |
| EPI_ISL_3335970 | Lighthouse Laboratory Plymouth | Wellcome Sanger Institute for the COVID-19 Genomics UK (COG-UK) Consortium | Lighthouse Laboratory Plymouth and Alex Alderton, Roberto Amato, Jeffrey Barrett, Sonia Goncalves, Ewan Harrison, David K. Jackson, Ian Johnston, Dominic Kwiatkowski, Cordelia Langford, John Sillitoe on behalf of the Wellcome Sanger Institute COVID-19 Surveillance Team |
| EPI_ISL_3335971 | Lighthouse Lab in Milton Keynes | Wellcome Sanger Institute for the COVID-19 Genomics UK (COG-UK) Consortium | The Lighthouse Lab in Milton Keynes and Alex Alderton, Roberto Amato, Jeffrey Barrett, Sonia Goncalves, Ewan Harrison, David K. Jackson, Ian Johnston, Dominic Kwiatkowski, Cordelia Langford, John Sillitoe on behalf of the Wellcome Sanger Institute COVID-19 Surveillance Team |
| EPI_ISL_3335972 | Health Services Laboratories | Wellcome Sanger Institute for the COVID-19 Genomics UK (COG-UK) Consortium | Health Services Laboratories and Alex Alderton, Roberto Amato, Jeffrey Barrett, Sonia Goncalves, Ewan Harrison, David K. Jackson, Ian Johnston, Dominic Kwiatkowski, Cordelia Langford, John Sillitoe on behalf of the Wellcome Sanger Institute COVID-19 Surveillance Team |
| EPI_ISL_3335973, EPI_ISL_3335974, EPI_ISL_3335975 | Lighthouse Lab in Milton Keynes | Wellcome Sanger Institute for the COVID-19 Genomics UK (COG-UK) Consortium | The Lighthouse Lab in Milton Keynes and Alex Alderton, Roberto Amato, Jeffrey Barrett, Sonia Goncalves, Ewan Harrison, David K. Jackson, Ian Johnston, Dominic Kwiatkowski, Cordelia Langford, John Sillitoe on behalf of the Wellcome Sanger Institute COVID-19 Surveillance Team |
| EPI_ISL_3335976, EPI_ISL_3335977 | Lighthouse Laboratory Plymouth | Wellcome Sanger Institute for the COVID-19 Genomics UK (COG-UK) Consortium | Lighthouse Laboratory Plymouth and Alex Alderton, Roberto Amato, Jeffrey Barrett, Sonia Goncalves, Ewan Harrison, David K. Jackson, Ian Johnston, Dominic Kwiatkowski, Cordelia Langford, John Sillitoe on behalf of the Wellcome Sanger Institute COVID-19 Surveillance Team |
| EPI_ISL_3335978 | Lighthouse Lab in Milton Keynes | Wellcome Sanger Institute for the COVID-19 Genomics UK (COG-UK) Consortium | The Lighthouse Lab in Milton Keynes and Alex Alderton, Roberto Amato, Jeffrey Barrett, Sonia Goncalves, Ewan Harrison, David K. Jackson, Ian Johnston, Dominic Kwiatkowski, Cordelia Langford, John Sillitoe on behalf of the Wellcome Sanger Institute COVID-19 Surveillance Team |
| EPI_ISL_3335979 | Health Services Laboratories | Wellcome Sanger Institute for the COVID-19 Genomics UK (COG-UK) Consortium | Health Services Laboratories and Alex Alderton, Roberto Amato, Jeffrey Barrett, Sonia Goncalves, Ewan Harrison, David K. Jackson, Ian Johnston, Dominic Kwiatkowski, Cordelia Langford, John Sillitoe on behalf of the Wellcome Sanger Institute COVID-19 Surveillance Team |
| EPI_ISL_3335980, EPI_ISL_3335981 | Lighthouse Lab in Milton Keynes | Wellcome Sanger Institute for the COVID-19 Genomics UK (COG-UK) Consortium | The Lighthouse Lab in Milton Keynes and Alex Alderton, Roberto Amato, Jeffrey Barrett, Sonia Goncalves, Ewan Harrison, David K. Jackson, Ian Johnston, Dominic Kwiatkowski, Cordelia Langford, John Sillitoe on behalf of the Wellcome Sanger Institute COVID-19 Surveillance Team |
| EPI_ISL_3335982, EPI_ISL_3335983 | Health Services Laboratories | Wellcome Sanger Institute for the COVID-19 Genomics UK (COG-UK) Consortium | Health Services Laboratories and Alex Alderton, Roberto Amato, Jeffrey Barrett, Sonia Goncalves, Ewan Harrison, David K. Jackson, Ian Johnston, Dominic Kwiatkowski, Cordelia Langford, John Sillitoe on behalf of the Wellcome Sanger Institute COVID-19 Surveillance Team |
| EPI_ISL_3335984 | Lighthouse Laboratory Plymouth | Wellcome Sanger Institute for the COVID-19 Genomics UK (COG-UK) Consortium | Lighthouse Laboratory Plymouth and Alex Alderton, Roberto Amato, Jeffrey Barrett, Sonia Goncalves, Ewan Harrison, David K. Jackson, Ian Johnston, Dominic Kwiatkowski, Cordelia Langford, John Sillitoe on behalf of the Wellcome Sanger Institute COVID-19 Surveillance Team |
| EPI_ISL_3335985 | Lighthouse Lab in Milton Keynes | Wellcome Sanger Institute for the COVID-19 Genomics UK (COG-UK) Consortium | The Lighthouse Lab in Milton Keynes and Alex Alderton, Roberto Amato, Jeffrey Barrett, Sonia Goncalves, Ewan Harrison, David K. Jackson, Ian Johnston, Dominic Kwiatkowski, Cordelia Langford, John Sillitoe on behalf of the Wellcome Sanger Institute COVID-19 Surveillance Team |
| EPI_ISL_3335986 | Lighthouse Laboratory Plymouth | Wellcome Sanger Institute for the COVID-19 Genomics UK (COG-UK) Consortium | Lighthouse Laboratory Plymouth and Alex Alderton, Roberto Amato, Jeffrey Barrett, Sonia Goncalves, Ewan Harrison, David K. Jackson, Ian Johnston, Dominic Kwiatkowski, Cordelia Langford, John Sillitoe on behalf of the Wellcome Sanger Institute COVID-19 Surveillance Team |
| EPI_ISL_3335987, EPI_ISL_3335988 | Health Services Laboratories | Wellcome Sanger Institute for the COVID-19 Genomics UK (COG-UK) Consortium | Health Services Laboratories and Alex Alderton, Roberto Amato, Jeffrey Barrett, Sonia Goncalves, Ewan Harrison, David K. Jackson, Ian Johnston, Dominic Kwiatkowski, Cordelia Langford, John Sillitoe on behalf of the Wellcome Sanger Institute COVID-19 Surveillance Team |

| | | | |
|---|---|---|---|
| EPI_ISL_3335989, EPI_ISL_3335990, EPI_ISL_3335991 | Lighthouse Lab in Milton Keynes | Wellcome Sanger Institute for the COVID-19 Genomics UK (COG-UK) Consortium | The Lighthouse Lab in Milton Keynes and Alex Alderton, Roberto Amato, Jeffrey Barrett, Sonia Goncalves, Ewan Harrison, David K. Jackson, Ian Johnston, Dominic Kwiatkowski, Cordelia Langford, John Sillitoe on behalf of the Wellcome Sanger Institute COVID-19 Surveillance Team |
| EPI_ISL_3335992 | Lighthouse Laboratory Plymouth | Wellcome Sanger Institute for the COVID-19 Genomics UK (COG-UK) Consortium | Lighthouse Laboratory Plymouth and Alex Alderton, Roberto Amato, Jeffrey Barrett, Sonia Goncalves, Ewan Harrison, David K. Jackson, Ian Johnston, Dominic Kwiatkowski, Cordelia Langford, John Sillitoe on behalf of the Wellcome Sanger Institute COVID-19 Surveillance Team |
| EPI_ISL_3335993 | Health Services Laboratories | Wellcome Sanger Institute for the COVID-19 Genomics UK (COG-UK) Consortium | Health Services Laboratories and Alex Alderton, Roberto Amato, Jeffrey Barrett, Sonia Goncalves, Ewan Harrison, David K. Jackson, Ian Johnston, Dominic Kwiatkowski, Cordelia Langford, John Sillitoe on behalf of the Wellcome Sanger Institute COVID-19 Surveillance Team |
| EPI_ISL_3335994, EPI_ISL_3335995 | Lighthouse Lab in Milton Keynes | Wellcome Sanger Institute for the COVID-19 Genomics UK (COG-UK) Consortium | The Lighthouse Lab in Milton Keynes and Alex Alderton, Roberto Amato, Jeffrey Barrett, Sonia Goncalves, Ewan Harrison, David K. Jackson, Ian Johnston, Dominic Kwiatkowski, Cordelia Langford, John Sillitoe on behalf of the Wellcome Sanger Institute COVID-19 Surveillance Team |
| EPI_ISL_3335996 | Lighthouse Laboratory Plymouth | Wellcome Sanger Institute for the COVID-19 Genomics UK (COG-UK) Consortium | Lighthouse Laboratory Plymouth and Alex Alderton, Roberto Amato, Jeffrey Barrett, Sonia Goncalves, Ewan Harrison, David K. Jackson, Ian Johnston, Dominic Kwiatkowski, Cordelia Langford, John Sillitoe on behalf of the Wellcome Sanger Institute COVID-19 Surveillance Team |
| EPI_ISL_3335997 | Health Services Laboratories | Wellcome Sanger Institute for the COVID-19 Genomics UK (COG-UK) Consortium | Health Services Laboratories and Alex Alderton, Roberto Amato, Jeffrey Barrett, Sonia Goncalves, Ewan Harrison, David K. Jackson, Ian Johnston, Dominic Kwiatkowski, Cordelia Langford, John Sillitoe on behalf of the Wellcome Sanger Institute COVID-19 Surveillance Team |
| EPI_ISL_3335998, EPI_ISL_3335999 | Lighthouse Laboratory Plymouth | Wellcome Sanger Institute for the COVID-19 Genomics UK (COG-UK) Consortium | Lighthouse Laboratory Plymouth and Alex Alderton, Roberto Amato, Jeffrey Barrett, Sonia Goncalves, Ewan Harrison, David K. Jackson, Ian Johnston, Dominic Kwiatkowski, Cordelia Langford, John Sillitoe on behalf of the Wellcome Sanger Institute COVID-19 Surveillance Team |
| EPI_ISL_3336000 | Health Services Laboratories | Wellcome Sanger Institute for the COVID-19 Genomics UK (COG-UK) Consortium | Health Services Laboratories and Alex Alderton, Roberto Amato, Jeffrey Barrett, Sonia Goncalves, Ewan Harrison, David K. Jackson, Ian Johnston, Dominic Kwiatkowski, Cordelia Langford, John Sillitoe on behalf of the Wellcome Sanger Institute COVID-19 Surveillance Team |
| EPI_ISL_3336001, EPI_ISL_3336002 | Lighthouse Laboratory Plymouth | Wellcome Sanger Institute for the COVID-19 Genomics UK (COG-UK) Consortium | Lighthouse Laboratory Plymouth and Alex Alderton, Roberto Amato, Jeffrey Barrett, Sonia Goncalves, Ewan Harrison, David K. Jackson, Ian Johnston, Dominic Kwiatkowski, Cordelia Langford, John Sillitoe on behalf of the Wellcome Sanger Institute COVID-19 Surveillance Team |
| EPI_ISL_3336003, EPI_ISL_3336004, EPI_ISL_3336005, EPI_ISL_3336006, EPI_ISL_3336007, EPI_ISL_3336008, EPI_ISL_3336009, EPI_ISL_3336010, EPI_ISL_3336011, EPI_ISL_3336012, EPI_ISL_3336013, EPI_ISL_3336014, EPI_ISL_3336015, EPI_ISL_3336016, EPI_ISL_3336017, EPI_ISL_3336018, EPI_ISL_3336019, EPI_ISL_3336020, EPI_ISL_3336021, EPI_ISL_3336022, EPI_ISL_3336023, EPI_ISL_3336024, EPI_ISL_3336025, EPI_ISL_3336026, EPI_ISL_3336027, EPI_ISL_3336028, EPI_ISL_3336029, EPI_ISL_3336030, EPI_ISL_3336031, EPI_ISL_3336032, EPI_ISL_3336033, EPI_ISL_3336034, EPI_ISL_3336035, EPI_ISL_3336036, EPI_ISL_3336037, EPI_ISL_3336038, EPI_ISL_3336039, EPI_ISL_3336040, EPI_ISL_3336041, EPI_ISL_3336042, EPI_ISL_3336043, EPI_ISL_3336044, EPI_ISL_3336045, EPI_ISL_3336046, EPI_ISL_3336047, EPI_ISL_3336048, EPI_ISL_3336049, EPI_ISL_3336050, EPI_ISL_3336051, EPI_ISL_3336052, EPI_ISL_3336053, EPI_ISL_3336054, EPI_ISL_3336055, EPI_ISL_3336056, EPI_ISL_3336057, EPI_ISL_3336058, EPI_ISL_3336059, EPI_ISL_3336060, EPI_ISL_3336061, EPI_ISL_3336062, EPI_ISL_3336063, EPI_ISL_3336064, EPI_ISL_3336065, EPI_ISL_3336066, EPI_ISL_3336067, EPI_ISL_3336068, EPI_ISL_3336069, EPI_ISL_3336070, EPI_ISL_3336071, EPI_ISL_3336072, EPI_ISL_3336073, EPI_ISL_3336074, EPI_ISL_3336075, EPI_ISL_3336076, EPI_ISL_3336077, EPI_ISL_3336078, EPI_ISL_3336079, EPI_ISL_3336080, EPI_ISL_3336081, EPI_ISL_3336082, EPI_ISL_3336083, EPI_ISL_3336084, EPI_ISL_3336085, EPI_ISL_3336086, EPI_ISL_3336087, EPI_ISL_3336088, EPI_ISL_3336089, EPI_ISL_3336090, EPI_ISL_3336091, EPI_ISL_3336092, EPI_ISL_3336093, EPI_ISL_3336094, EPI_ISL_3336095, EPI_ISL_3336096, EPI_ISL_3336097, EPI_ISL_3336098, EPI_ISL_3336099, EPI_ISL_3336100, EPI_ISL_3336101, EPI_ISL_3336102, EPI_ISL_3336103, EPI_ISL_3336104, EPI_ISL_3336105, EPI_ISL_3336106, EPI_ISL_3336107, EPI_ISL_3336108, EPI_ISL_3336109, EPI_ISL_3336110, EPI_ISL_3336111, EPI_ISL_3336112, EPI_ISL_3336113, EPI_ISL_3336114, EPI_ISL_3336115, EPI_ISL_3336116, EPI_ISL_3336117, EPI_ISL_3336118, EPI_ISL_3336119, EPI_ISL_3336120, EPI_ISL_3336121, EPI_ISL_3336122, EPI_ISL_3336123, EPI_ISL_3336124, EPI_ISL_3336125, EPI_ISL_3336126, EPI_ISL_3336127, EPI_ISL_3336128, EPI_ISL_3336129, EPI_ISL_3336130, EPI_ISL_3336131, EPI_ISL_3336132, EPI_ISL_3336133, EPI_ISL_3336134, EPI_ISL_3336135, EPI_ISL_3336136, EPI_ISL_3336137, EPI_ISL_3336138, EPI_ISL_3336139, EPI_ISL_3336140, EPI_ISL_3336141, EPI_ISL_3336142 | | | |
| see above | Lighthouse Lab in Milton Keynes | Wellcome Sanger Institute for the COVID-19 Genomics UK (COG-UK) Consortium | The Lighthouse Lab in Milton Keynes and Alex Alderton, Roberto Amato, Jeffrey Barrett, Sonia Goncalves, Ewan Harrison, David K. Jackson, Ian Johnston, Dominic Kwiatkowski, Cordelia Langford, John Sillitoe on behalf of the Wellcome Sanger Institute COVID-19 Surveillance Team |
| EPI_ISL_3336143 | Berkshire and Surrey Pathology Services Lighthouse Laboratory | Wellcome Sanger Institute for the COVID-19 Genomics UK (COG-UK) Consortium | Berkshire and Surrey Pathology Services Lighthouse Laboratory and Alex Alderton, Roberto Amato, Jeffrey Barrett, Sonia Goncalves, Ewan Harrison, David K. Jackson, Ian Johnston, Dominic Kwiatkowski, Cordelia Langford, John Sillitoe on behalf of the Wellcome Sanger Institute COVID-19 Surveillance Team |
| EPI_ISL_3336144, EPI_ISL_3336145, EPI_ISL_3336146 | Lighthouse Laboratory Plymouth | Wellcome Sanger Institute for the COVID-19 Genomics UK (COG-UK) Consortium | Lighthouse Laboratory Plymouth and Alex Alderton, Roberto Amato, Jeffrey Barrett, Sonia Goncalves, Ewan Harrison, David K. Jackson, Ian Johnston, Dominic Kwiatkowski, Cordelia Langford, John Sillitoe on behalf of the Wellcome Sanger Institute COVID-19 Surveillance Team |
| EPI_ISL_3336147, EPI_ISL_3336148 | Lighthouse Lab in Milton Keynes | Wellcome Sanger Institute for the COVID-19 Genomics UK (COG-UK) Consortium | The Lighthouse Lab in Milton Keynes and Alex Alderton, Roberto Amato, Jeffrey Barrett, Sonia Goncalves, Ewan Harrison, David K. Jackson, Ian Johnston, Dominic Kwiatkowski, Cordelia Langford, John Sillitoe on behalf of the Wellcome Sanger Institute COVID-19 Surveillance Team |
| EPI_ISL_3336149 | Lighthouse Laboratory Plymouth | Wellcome Sanger Institute for the COVID-19 Genomics UK (COG-UK) Consortium | Lighthouse Laboratory Plymouth and Alex Alderton, Roberto Amato, Jeffrey Barrett, Sonia Goncalves, Ewan Harrison, David K. Jackson, Ian Johnston, Dominic Kwiatkowski, Cordelia Langford, John Sillitoe on behalf of the Wellcome Sanger Institute COVID-19 Surveillance Team |
| EPI_ISL_3336150, EPI_ISL_3336151 | Lighthouse Lab in Glasgow | Wellcome Sanger Institute for the COVID-19 Genomics UK (COG-UK) Consortium | Harper VanSteenhouse, Yumi Kasai, David Gray, Carol Clugston, Anna Dominiczak and Alex Alderton, Roberto Amato, Jeffrey Barrett, Sonia Goncalves, Ewan Harrison, David K. Jackson, Ian Johnston, Dominic Kwiatkowski, Cordelia Langford, John Sillitoe on behalf of the Wellcome Sanger Institute COVID-19 Surveillance Team |
| EPI_ISL_3336152, EPI_ISL_3336153, EPI_ISL_3336154 | Lighthouse Laboratory Plymouth | Wellcome Sanger Institute for the COVID-19 Genomics UK (COG-UK) Consortium | Lighthouse Laboratory Plymouth and Alex Alderton, Roberto Amato, Jeffrey Barrett, Sonia Goncalves, Ewan Harrison, David K. Jackson, Ian Johnston, Dominic Kwiatkowski, Cordelia Langford, John Sillitoe on behalf of the Wellcome Sanger Institute COVID-19 Surveillance Team |
| EPI_ISL_3336155 | Lighthouse Lab in Milton Keynes | Wellcome Sanger Institute for the COVID-19 Genomics UK (COG-UK) Consortium | The Lighthouse Lab in Milton Keynes and Alex Alderton, Roberto Amato, Jeffrey Barrett, Sonia Goncalves, Ewan Harrison, David K. Jackson, Ian Johnston, Dominic Kwiatkowski, Cordelia Langford, John Sillitoe on behalf of the Wellcome Sanger Institute COVID-19 Surveillance Team |
| EPI_ISL_3336156 | Berkshire and Surrey Pathology Services Lighthouse Laboratory | Wellcome Sanger Institute for the COVID-19 Genomics UK (COG-UK) Consortium | Berkshire and Surrey Pathology Services Lighthouse Laboratory and Alex Alderton, Roberto Amato, Jeffrey Barrett, Sonia Goncalves, Ewan Harrison, David K. Jackson, Ian Johnston, Dominic Kwiatkowski, Cordelia Langford, John Sillitoe on behalf of the Wellcome Sanger Institute COVID-19 Surveillance Team |
| EPI_ISL_3336157 | Lighthouse Lab in Glasgow | Wellcome Sanger Institute for the COVID-19 Genomics UK (COG-UK) Consortium | Harper VanSteenhouse, Yumi Kasai, David Gray, Carol Clugston, Anna Dominiczak and Alex Alderton, Roberto Amato, Jeffrey Barrett, Sonia Goncalves, Ewan Harrison, David K. Jackson, Ian Johnston, Dominic Kwiatkowski, Cordelia Langford, John Sillitoe on behalf of the Wellcome Sanger Institute COVID-19 Surveillance Team |
| EPI_ISL_3336158 | Lighthouse Lab in Milton Keynes | Wellcome Sanger Institute for the COVID-19 Genomics UK (COG-UK) Consortium | The Lighthouse Lab in Milton Keynes and Alex Alderton, Roberto Amato, Jeffrey Barrett, Sonia Goncalves, Ewan Harrison, David K. Jackson, Ian Johnston, Dominic Kwiatkowski, Cordelia Langford, John Sillitoe on behalf of the Wellcome Sanger Institute COVID-19 Surveillance Team |
| EPI_ISL_3336159 | Lighthouse Laboratory Plymouth | Wellcome Sanger Institute for the COVID-19 Genomics UK (COG-UK) Consortium | Lighthouse Laboratory Plymouth and Alex Alderton, Roberto Amato, Jeffrey Barrett, Sonia Goncalves, Ewan Harrison, David K. Jackson, Ian Johnston, Dominic Kwiatkowski, Cordelia Langford, John Sillitoe on behalf of the Wellcome Sanger Institute COVID-19 Surveillance Team |
| EPI_ISL_3336160 | Lighthouse Lab in Glasgow | Wellcome Sanger Institute for the COVID-19 Genomics UK (COG-UK) Consortium | Harper VanSteenhouse, Yumi Kasai, David Gray, Carol Clugston, Anna Dominiczak and Alex Alderton, Roberto Amato, Jeffrey Barrett, Sonia Goncalves, Ewan Harrison, David K. Jackson, Ian Johnston, Dominic Kwiatkowski, Cordelia Langford, John Sillitoe on behalf of the Wellcome Sanger Institute COVID-19 Surveillance Team |
| EPI_ISL_3336161, EPI_ISL_3336162 | Lighthouse Laboratory Plymouth | Wellcome Sanger Institute for the COVID-19 Genomics UK (COG-UK) Consortium | Lighthouse Laboratory Plymouth and Alex Alderton, Roberto Amato, Jeffrey Barrett, Sonia Goncalves, Ewan Harrison, David K. Jackson, Ian Johnston, Dominic Kwiatkowski, Cordelia Langford, John Sillitoe on behalf of the Wellcome Sanger Institute COVID-19 Surveillance Team |
| EPI_ISL_3336163, EPI_ISL_3336164 | Lighthouse Lab in Milton Keynes | Wellcome Sanger Institute for the COVID-19 Genomics UK (COG-UK) Consortium | The Lighthouse Lab in Milton Keynes and Alex Alderton, Roberto Amato, Jeffrey Barrett, Sonia Goncalves, Ewan Harrison, David K. Jackson, Ian Johnston, Dominic Kwiatkowski, Cordelia Langford, John Sillitoe on behalf of the Wellcome Sanger Institute COVID-19 Surveillance Team |
| EPI_ISL_3336165 | Berkshire and Surrey Pathology Services Lighthouse Laboratory | Wellcome Sanger Institute for the COVID-19 Genomics UK (COG-UK) Consortium | Berkshire and Surrey Pathology Services Lighthouse Laboratory and Alex Alderton, Roberto Amato, Jeffrey Barrett, Sonia Goncalves, Ewan Harrison, David K. Jackson, Ian Johnston, Dominic Kwiatkowski, Cordelia Langford, John Sillitoe on behalf of the Wellcome Sanger Institute COVID-19 Surveillance Team |
| EPI_ISL_3336166 | Lighthouse Lab in Glasgow | Wellcome Sanger Institute for the COVID-19 Genomics UK (COG-UK) Consortium | Harper VanSteenhouse, Yumi Kasai, David Gray, Carol Clugston, Anna Dominiczak and Alex Alderton, Roberto Amato, Jeffrey Barrett, Sonia Goncalves, Ewan Harrison, David K. Jackson, Ian Johnston, Dominic Kwiatkowski, Cordelia Langford, John Sillitoe on behalf of the Wellcome Sanger Institute COVID-19 Surveillance Team |
| EPI_ISL_3336167, EPI_ISL_3336168 | Lighthouse Lab in Milton Keynes | Wellcome Sanger Institute for the COVID-19 Genomics UK (COG-UK) Consortium | The Lighthouse Lab in Milton Keynes and Alex Alderton, Roberto Amato, Jeffrey Barrett, Sonia Goncalves, Ewan Harrison, David K. Jackson, Ian Johnston, Dominic Kwiatkowski, Cordelia Langford, John Sillitoe on behalf of the Wellcome Sanger Institute COVID-19 Surveillance Team |
| EPI_ISL_3336169, EPI_ISL_3336170 | Lighthouse Laboratory Plymouth | Wellcome Sanger Institute for the COVID-19 Genomics UK (COG-UK) Consortium | Lighthouse Laboratory Plymouth and Alex Alderton, Roberto Amato, Jeffrey Barrett, Sonia Goncalves, Ewan Harrison, David K. Jackson, Ian Johnston, Dominic Kwiatkowski, Cordelia Langford, John Sillitoe on behalf of the Wellcome Sanger Institute COVID-19 Surveillance Team |
| EPI_ISL_3336171 | Lighthouse Lab in Glasgow | Wellcome Sanger Institute for the COVID-19 Genomics UK (COG-UK) Consortium | Harper VanSteenhouse, Yumi Kasai, David Gray, Carol Clugston, Anna Dominiczak and Alex Alderton, Roberto Amato, Jeffrey Barrett, Sonia Goncalves, Ewan Harrison, David K. Jackson, Ian Johnston, Dominic Kwiatkowski, Cordelia Langford, John Sillitoe on behalf of the Wellcome Sanger Institute COVID-19 Surveillance Team |
| EPI_ISL_3336172, EPI_ISL_3336173 | Lighthouse Laboratory Plymouth | Wellcome Sanger Institute for the COVID-19 Genomics UK (COG-UK) Consortium | Lighthouse Laboratory Plymouth and Alex Alderton, Roberto Amato, Jeffrey Barrett, Sonia Goncalves, Ewan Harrison, David K. Jackson, Ian Johnston, Dominic Kwiatkowski, Cordelia Langford, John Sillitoe on behalf of the Wellcome Sanger Institute COVID-19 Surveillance Team |
| EPI_ISL_3336174 | Lighthouse Lab in Milton Keynes | Wellcome Sanger Institute for the COVID-19 Genomics UK (COG-UK) Consortium | The Lighthouse Lab in Milton Keynes and Alex Alderton, Roberto Amato, Jeffrey Barrett, Sonia Goncalves, Ewan Harrison, David K. Jackson, Ian Johnston, Dominic Kwiatkowski, Cordelia Langford, John Sillitoe on behalf of the Wellcome Sanger Institute COVID-19 Surveillance Team |
| EPI_ISL_3336175 | Lighthouse Lab in Glasgow | Wellcome Sanger Institute for the COVID-19 Genomics UK (COG-UK) Consortium | Harper VanSteenhouse, Yumi Kasai, David Gray, Carol Clugston, Anna Dominiczak and Alex Alderton, Roberto Amato, Jeffrey Barrett, Sonia Goncalves, Ewan Harrison, David K. Jackson, Ian Johnston, Dominic Kwiatkowski, Cordelia Langford, John Sillitoe on behalf of the Wellcome Sanger Institute COVID-19 Surveillance Team |
| EPI_ISL_3336176 | Berkshire and Surrey Pathology Services Lighthouse Laboratory | Wellcome Sanger Institute for the COVID-19 Genomics UK (COG-UK) Consortium | Berkshire and Surrey Pathology Services Lighthouse Laboratory and Alex Alderton, Roberto Amato, Jeffrey Barrett, Sonia Goncalves, Ewan Harrison, David K. Jackson, Ian Johnston, Dominic Kwiatkowski, Cordelia Langford, John Sillitoe on behalf of the Wellcome Sanger Institute COVID-19 Surveillance Team |
| EPI_ISL_3336177 | Lighthouse Lab in Milton Keynes | Wellcome Sanger Institute for the COVID-19 Genomics UK (COG-UK) Consortium | The Lighthouse Lab in Milton Keynes and Alex Alderton, Roberto Amato, Jeffrey Barrett, Sonia Goncalves, Ewan Harrison, David K. Jackson, Ian Johnston, Dominic Kwiatkowski, Cordelia Langford, John Sillitoe on behalf of the Wellcome Sanger Institute COVID-19 Surveillance Team |
| EPI_ISL_3336178 | Berkshire and Surrey Pathology Services Lighthouse Laboratory | Wellcome Sanger Institute for the COVID-19 Genomics UK (COG-UK) Consortium | Berkshire and Surrey Pathology Services Lighthouse Laboratory and Alex Alderton, Roberto Amato, Jeffrey Barrett, Sonia Goncalves, Ewan Harrison, David K. Jackson, Ian Johnston, Dominic Kwiatkowski, Cordelia Langford, John Sillitoe on behalf of the Wellcome Sanger Institute COVID-19 Surveillance Team |
| EPI_ISL_3336179 | Lighthouse Lab in Milton Keynes | Wellcome Sanger Institute for the COVID-19 Genomics UK (COG-UK) Consortium | The Lighthouse Lab in Milton Keynes and Alex Alderton, Roberto Amato, Jeffrey Barrett, Sonia Goncalves, Ewan Harrison, David K. Jackson, Ian Johnston, Dominic Kwiatkowski, Cordelia Langford, John Sillitoe on behalf of the Wellcome Sanger Institute COVID-19 Surveillance Team |
| EPI_ISL_3336180 | Lighthouse Lab in Glasgow | Wellcome Sanger Institute for the COVID-19 Genomics UK (COG-UK) Consortium | Harper VanSteenhouse, Yumi Kasai, David Gray, Carol Clugston, Anna Dominiczak and Alex Alderton, Roberto Amato, Jeffrey Barrett, Sonia Goncalves, Ewan Harrison, David K. Jackson, Ian Johnston, Dominic Kwiatkowski, Cordelia Langford, John Sillitoe on behalf of the Wellcome Sanger Institute COVID-19 Surveillance Team |
| EPI_ISL_3336181 | Berkshire and Surrey Pathology Services Lighthouse Laboratory | Wellcome Sanger Institute for the COVID-19 Genomics UK (COG-UK) Consortium | Berkshire and Surrey Pathology Services Lighthouse Laboratory and Alex Alderton, Roberto Amato, Jeffrey Barrett, Sonia Goncalves, Ewan Harrison, David K. Jackson, Ian Johnston, Dominic Kwiatkowski, Cordelia Langford, John Sillitoe on behalf of the Wellcome Sanger Institute COVID-19 Surveillance Team |
| EPI_ISL_3336182 | Lighthouse Laboratory Plymouth | Wellcome Sanger Institute for the COVID-19 Genomics UK (COG-UK) Consortium | Lighthouse Laboratory Plymouth and Alex Alderton, Roberto Amato, Jeffrey Barrett, Sonia Goncalves, Ewan Harrison, David K. Jackson, Ian Johnston, Dominic Kwiatkowski, Cordelia Langford, John Sillitoe on behalf of the Wellcome Sanger Institute COVID-19 Surveillance Team |
| EPI_ISL_3336183 | Berkshire and Surrey Pathology Services Lighthouse Laboratory | Wellcome Sanger Institute for the COVID-19 Genomics UK (COG-UK) Consortium | Berkshire and Surrey Pathology Services Lighthouse Laboratory and Alex Alderton, Roberto Amato, Jeffrey Barrett, Sonia Goncalves, Ewan Harrison, David K. Jackson, Ian Johnston, Dominic Kwiatkowski, Cordelia Langford, John Sillitoe on behalf of the Wellcome Sanger Institute COVID-19 Surveillance Team |
| EPI_ISL_3336184 | Lighthouse Laboratory Plymouth | Wellcome Sanger Institute for the COVID-19 Genomics UK (COG-UK) Consortium | Lighthouse Laboratory Plymouth and Alex Alderton, Roberto Amato, Jeffrey Barrett, Sonia Goncalves, Ewan Harrison, David K. Jackson, Ian Johnston, Dominic Kwiatkowski, Cordelia Langford, John Sillitoe on behalf of the Wellcome Sanger Institute COVID-19 Surveillance Team |

| | | | |
|---|---|---|---|
| EPI_ISL_3336185 | Lighthouse Lab in Glasgow | Wellcome Sanger Institute for the COVID-19 Genomics UK (COG-UK) Consortium | Harper VanSteenhouse, Yumi Kasai, David Gray, Carol Clugston, Anna Dominiczak and Alex Alderton, Roberto Amato, Jeffrey Barrett, Sonia Goncalves, Ewan Harrison, David K. Jackson, Ian Johnston, Dominic Kwiatkowski, Cordelia Langford, John Sillitoe on behalf of the Wellcome Sanger Institute COVID-19 Surveillance Team |
| EPI_ISL_3336186 | Lighthouse Laboratory Plymouth | Wellcome Sanger Institute for the COVID-19 Genomics UK (COG-UK) Consortium | Lighthouse Laboratory Plymouth and Alex Alderton, Roberto Amato, Jeffrey Barrett, Sonia Goncalves, Ewan Harrison, David K. Jackson, Ian Johnston, Dominic Kwiatkowski, Cordelia Langford, John Sillitoe on behalf of the Wellcome Sanger Institute COVID-19 Surveillance Team |
| EPI_ISL_3336187, EPI_ISL_3336188, EPI_ISL_3336189, EPI_ISL_3336190 | Lighthouse Lab in Milton Keynes | Wellcome Sanger Institute for the COVID-19 Genomics UK (COG-UK) Consortium | The Lighthouse Lab in Milton Keynes and Alex Alderton, Roberto Amato, Jeffrey Barrett, Sonia Goncalves, Ewan Harrison, David K. Jackson, Ian Johnston, Dominic Kwiatkowski, Cordelia Langford, John Sillitoe on behalf of the Wellcome Sanger Institute COVID-19 Surveillance Team |
| EPI_ISL_3336191, EPI_ISL_3336192 | Lighthouse Laboratory Plymouth | Wellcome Sanger Institute for the COVID-19 Genomics UK (COG-UK) Consortium | Lighthouse Laboratory Plymouth and Alex Alderton, Roberto Amato, Jeffrey Barrett, Sonia Goncalves, Ewan Harrison, David K. Jackson, Ian Johnston, Dominic Kwiatkowski, Cordelia Langford, John Sillitoe on behalf of the Wellcome Sanger Institute COVID-19 Surveillance Team |
| EPI_ISL_3336193 | Lighthouse Lab in Milton Keynes | Wellcome Sanger Institute for the COVID-19 Genomics UK (COG-UK) Consortium | The Lighthouse Lab in Milton Keynes and Alex Alderton, Roberto Amato, Jeffrey Barrett, Sonia Goncalves, Ewan Harrison, David K. Jackson, Ian Johnston, Dominic Kwiatkowski, Cordelia Langford, John Sillitoe on behalf of the Wellcome Sanger Institute COVID-19 Surveillance Team |
| EPI_ISL_3336194 | Lighthouse Lab in Glasgow | Wellcome Sanger Institute for the COVID-19 Genomics UK (COG-UK) Consortium | Harper VanSteenhouse, Yumi Kasai, David Gray, Carol Clugston, Anna Dominiczak and Alex Alderton, Roberto Amato, Jeffrey Barrett, Sonia Goncalves, Ewan Harrison, David K. Jackson, Ian Johnston, Dominic Kwiatkowski, Cordelia Langford, John Sillitoe on behalf of the Wellcome Sanger Institute COVID-19 Surveillance Team |
| EPI_ISL_3336195, EPI_ISL_3336196 | Lighthouse Laboratory Plymouth | Wellcome Sanger Institute for the COVID-19 Genomics UK (COG-UK) Consortium | Lighthouse Laboratory Plymouth and Alex Alderton, Roberto Amato, Jeffrey Barrett, Sonia Goncalves, Ewan Harrison, David K. Jackson, Ian Johnston, Dominic Kwiatkowski, Cordelia Langford, John Sillitoe on behalf of the Wellcome Sanger Institute COVID-19 Surveillance Team |
| EPI_ISL_3336197, EPI_ISL_3336198 | Lighthouse Lab in Glasgow | Wellcome Sanger Institute for the COVID-19 Genomics UK (COG-UK) Consortium | Harper VanSteenhouse, Yumi Kasai, David Gray, Carol Clugston, Anna Dominiczak and Alex Alderton, Roberto Amato, Jeffrey Barrett, Sonia Goncalves, Ewan Harrison, David K. Jackson, Ian Johnston, Dominic Kwiatkowski, Cordelia Langford, John Sillitoe on behalf of the Wellcome Sanger Institute COVID-19 Surveillance Team |
| EPI_ISL_3336199 | Lighthouse Laboratory Plymouth | Wellcome Sanger Institute for the COVID-19 Genomics UK (COG-UK) Consortium | Lighthouse Laboratory Plymouth and Alex Alderton, Roberto Amato, Jeffrey Barrett, Sonia Goncalves, Ewan Harrison, David K. Jackson, Ian Johnston, Dominic Kwiatkowski, Cordelia Langford, John Sillitoe on behalf of the Wellcome Sanger Institute COVID-19 Surveillance Team |
| EPI_ISL_3336200, EPI_ISL_3336201, EPI_ISL_3336202 | Berkshire and Surrey Pathology Services Lighthouse Laboratory | Wellcome Sanger Institute for the COVID-19 Genomics UK (COG-UK) Consortium | Berkshire and Surrey Pathology Services Lighthouse Laboratory and Alex Alderton, Roberto Amato, Jeffrey Barrett, Sonia Goncalves, Ewan Harrison, David K. Jackson, Ian Johnston, Dominic Kwiatkowski, Cordelia Langford, John Sillitoe on behalf of the Wellcome Sanger Institute COVID-19 Surveillance Team |
| EPI_ISL_3336203 | Lighthouse Lab in Glasgow | Wellcome Sanger Institute for the COVID-19 Genomics UK (COG-UK) Consortium | Harper VanSteenhouse, Yumi Kasai, David Gray, Carol Clugston, Anna Dominiczak and Alex Alderton, Roberto Amato, Jeffrey Barrett, Sonia Goncalves, Ewan Harrison, David K. Jackson, Ian Johnston, Dominic Kwiatkowski, Cordelia Langford, John Sillitoe on behalf of the Wellcome Sanger Institute COVID-19 Surveillance Team |
| EPI_ISL_3336204, EPI_ISL_3336205 | Lighthouse Lab in Milton Keynes | Wellcome Sanger Institute for the COVID-19 Genomics UK (COG-UK) Consortium | The Lighthouse Lab in Milton Keynes and Alex Alderton, Roberto Amato, Jeffrey Barrett, Sonia Goncalves, Ewan Harrison, David K. Jackson, Ian Johnston, Dominic Kwiatkowski, Cordelia Langford, John Sillitoe on behalf of the Wellcome Sanger Institute COVID-19 Surveillance Team |
| EPI_ISL_3336206, EPI_ISL_3336207 | Lighthouse Lab in Glasgow | Wellcome Sanger Institute for the COVID-19 Genomics UK (COG-UK) Consortium | Harper VanSteenhouse, Yumi Kasai, David Gray, Carol Clugston, Anna Dominiczak and Alex Alderton, Roberto Amato, Jeffrey Barrett, Sonia Goncalves, Ewan Harrison, David K. Jackson, Ian Johnston, Dominic Kwiatkowski, Cordelia Langford, John Sillitoe on behalf of the Wellcome Sanger Institute COVID-19 Surveillance Team |
| EPI_ISL_3336208 | Lighthouse Laboratory Plymouth | Wellcome Sanger Institute for the COVID-19 Genomics UK (COG-UK) Consortium | Lighthouse Laboratory Plymouth and Alex Alderton, Roberto Amato, Jeffrey Barrett, Sonia Goncalves, Ewan Harrison, David K. Jackson, Ian Johnston, Dominic Kwiatkowski, Cordelia Langford, John Sillitoe on behalf of the Wellcome Sanger Institute COVID-19 Surveillance Team |
| EPI_ISL_3336209, EPI_ISL_3336210 | Berkshire and Surrey Pathology Services Lighthouse Laboratory | Wellcome Sanger Institute for the COVID-19 Genomics UK (COG-UK) Consortium | Berkshire and Surrey Pathology Services Lighthouse Laboratory and Alex Alderton, Roberto Amato, Jeffrey Barrett, Sonia Goncalves, Ewan Harrison, David K. Jackson, Ian Johnston, Dominic Kwiatkowski, Cordelia Langford, John Sillitoe on behalf of the Wellcome Sanger Institute COVID-19 Surveillance Team |
| EPI_ISL_3336211 | Lighthouse Lab in Milton Keynes | Wellcome Sanger Institute for the COVID-19 Genomics UK (COG-UK) Consortium | The Lighthouse Lab in Milton Keynes and Alex Alderton, Roberto Amato, Jeffrey Barrett, Sonia Goncalves, Ewan Harrison, David K. Jackson, Ian Johnston, Dominic Kwiatkowski, Cordelia Langford, John Sillitoe on behalf of the Wellcome Sanger Institute COVID-19 Surveillance Team |
| EPI_ISL_3336212 | Berkshire and Surrey Pathology Services Lighthouse Laboratory | Wellcome Sanger Institute for the COVID-19 Genomics UK (COG-UK) Consortium | Berkshire and Surrey Pathology Services Lighthouse Laboratory and Alex Alderton, Roberto Amato, Jeffrey Barrett, Sonia Goncalves, Ewan Harrison, David K. Jackson, Ian Johnston, Dominic Kwiatkowski, Cordelia Langford, John Sillitoe on behalf of the Wellcome Sanger Institute COVID-19 Surveillance Team |
| EPI_ISL_3336213 | Lighthouse Lab in Glasgow | Wellcome Sanger Institute for the COVID-19 Genomics UK (COG-UK) Consortium | Harper VanSteenhouse, Yumi Kasai, David Gray, Carol Clugston, Anna Dominiczak and Alex Alderton, Roberto Amato, Jeffrey Barrett, Sonia Goncalves, Ewan Harrison, David K. Jackson, Ian Johnston, Dominic Kwiatkowski, Cordelia Langford, John Sillitoe on behalf of the Wellcome Sanger Institute COVID-19 Surveillance Team |
| EPI_ISL_3336214, EPI_ISL_3336215 | Lighthouse Lab in Milton Keynes | Wellcome Sanger Institute for the COVID-19 Genomics UK (COG-UK) Consortium | The Lighthouse Lab in Milton Keynes and Alex Alderton, Roberto Amato, Jeffrey Barrett, Sonia Goncalves, Ewan Harrison, David K. Jackson, Ian Johnston, Dominic Kwiatkowski, Cordelia Langford, John Sillitoe on behalf of the Wellcome Sanger Institute COVID-19 Surveillance Team |
| EPI_ISL_3336216 | Berkshire and Surrey Pathology Services Lighthouse Laboratory | Wellcome Sanger Institute for the COVID-19 Genomics UK (COG-UK) Consortium | Berkshire and Surrey Pathology Services Lighthouse Laboratory and Alex Alderton, Roberto Amato, Jeffrey Barrett, Sonia Goncalves, Ewan Harrison, David K. Jackson, Ian Johnston, Dominic Kwiatkowski, Cordelia Langford, John Sillitoe on behalf of the Wellcome Sanger Institute COVID-19 Surveillance Team |
| EPI_ISL_3336217 | Lighthouse Lab in Milton Keynes | Wellcome Sanger Institute for the COVID-19 Genomics UK (COG-UK) Consortium | The Lighthouse Lab in Milton Keynes and Alex Alderton, Roberto Amato, Jeffrey Barrett, Sonia Goncalves, Ewan Harrison, David K. Jackson, Ian Johnston, Dominic Kwiatkowski, Cordelia Langford, John Sillitoe on behalf of the Wellcome Sanger Institute COVID-19 Surveillance Team |
| EPI_ISL_3336218 | Berkshire and Surrey Pathology Services Lighthouse Laboratory | Wellcome Sanger Institute for the COVID-19 Genomics UK (COG-UK) Consortium | Berkshire and Surrey Pathology Services Lighthouse Laboratory and Alex Alderton, Roberto Amato, Jeffrey Barrett, Sonia Goncalves, Ewan Harrison, David K. Jackson, Ian Johnston, Dominic Kwiatkowski, Cordelia Langford, John Sillitoe on behalf of the Wellcome Sanger Institute COVID-19 Surveillance Team |
| EPI_ISL_3336219 | Lighthouse Lab in Milton Keynes | Wellcome Sanger Institute for the COVID-19 Genomics UK (COG-UK) Consortium | The Lighthouse Lab in Milton Keynes and Alex Alderton, Roberto Amato, Jeffrey Barrett, Sonia Goncalves, Ewan Harrison, David K. Jackson, Ian Johnston, Dominic Kwiatkowski, Cordelia Langford, John Sillitoe on behalf of the Wellcome Sanger Institute COVID-19 Surveillance Team |
| EPI_ISL_3336220 | Lighthouse Lab in Glasgow | Wellcome Sanger Institute for the COVID-19 Genomics UK (COG-UK) Consortium | Harper VanSteenhouse, Yumi Kasai, David Gray, Carol Clugston, Anna Dominiczak and Alex Alderton, Roberto Amato, Jeffrey Barrett, Sonia Goncalves, Ewan Harrison, David K. Jackson, Ian Johnston, Dominic Kwiatkowski, Cordelia Langford, John Sillitoe on behalf of the Wellcome Sanger Institute COVID-19 Surveillance Team |
| EPI_ISL_3336221 | Lighthouse Laboratory Plymouth | Wellcome Sanger Institute for the COVID-19 Genomics UK (COG-UK) Consortium | Lighthouse Laboratory Plymouth and Alex Alderton, Roberto Amato, Jeffrey Barrett, Sonia Goncalves, Ewan Harrison, David K. Jackson, Ian Johnston, Dominic Kwiatkowski, Cordelia Langford, John Sillitoe on behalf of the Wellcome Sanger Institute COVID-19 Surveillance Team |
| EPI_ISL_3336222 | Lighthouse Lab in Milton Keynes | Wellcome Sanger Institute for the COVID-19 Genomics UK (COG-UK) Consortium | The Lighthouse Lab in Milton Keynes and Alex Alderton, Roberto Amato, Jeffrey Barrett, Sonia Goncalves, Ewan Harrison, David K. Jackson, Ian Johnston, Dominic Kwiatkowski, Cordelia Langford, John Sillitoe on behalf of the Wellcome Sanger Institute COVID-19 Surveillance Team |
| EPI_ISL_3336223 | Lighthouse Laboratory Plymouth | Wellcome Sanger Institute for the COVID-19 Genomics UK (COG-UK) Consortium | Lighthouse Laboratory Plymouth and Alex Alderton, Roberto Amato, Jeffrey Barrett, Sonia Goncalves, Ewan Harrison, David K. Jackson, Ian Johnston, Dominic Kwiatkowski, Cordelia Langford, John Sillitoe on behalf of the Wellcome Sanger Institute COVID-19 Surveillance Team |
| EPI_ISL_3336224, EPI_ISL_3336225 | Lighthouse Lab in Milton Keynes | Wellcome Sanger Institute for the COVID-19 Genomics UK (COG-UK) Consortium | The Lighthouse Lab in Milton Keynes and Alex Alderton, Roberto Amato, Jeffrey Barrett, Sonia Goncalves, Ewan Harrison, David K. Jackson, Ian Johnston, Dominic Kwiatkowski, Cordelia Langford, John Sillitoe on behalf of the Wellcome Sanger Institute COVID-19 Surveillance Team |
| EPI_ISL_3336226, EPI_ISL_3336227 | Berkshire and Surrey Pathology Services Lighthouse Laboratory | Wellcome Sanger Institute for the COVID-19 Genomics UK (COG-UK) Consortium | Berkshire and Surrey Pathology Services Lighthouse Laboratory and Alex Alderton, Roberto Amato, Jeffrey Barrett, Sonia Goncalves, Ewan Harrison, David K. Jackson, Ian Johnston, Dominic Kwiatkowski, Cordelia Langford, John Sillitoe on behalf of the Wellcome Sanger Institute COVID-19 Surveillance Team |
| EPI_ISL_3336228 | Lighthouse Lab in Glasgow | Wellcome Sanger Institute for the COVID-19 Genomics UK (COG-UK) Consortium | Harper VanSteenhouse, Yumi Kasai, David Gray, Carol Clugston, Anna Dominiczak and Alex Alderton, Roberto Amato, Jeffrey Barrett, Sonia Goncalves, Ewan Harrison, David K. Jackson, Ian Johnston, Dominic Kwiatkowski, Cordelia Langford, John Sillitoe on behalf of the Wellcome Sanger Institute COVID-19 Surveillance Team |
| EPI_ISL_3336229 | Lighthouse Lab in Milton Keynes | Wellcome Sanger Institute for the COVID-19 Genomics UK (COG-UK) Consortium | The Lighthouse Lab in Milton Keynes and Alex Alderton, Roberto Amato, Jeffrey Barrett, Sonia Goncalves, Ewan Harrison, David K. Jackson, Ian Johnston, Dominic Kwiatkowski, Cordelia Langford, John Sillitoe on behalf of the Wellcome Sanger Institute COVID-19 Surveillance Team |
| EPI_ISL_3336230 | Lighthouse Lab in Glasgow | Wellcome Sanger Institute for the COVID-19 Genomics UK (COG-UK) Consortium | Harper VanSteenhouse, Yumi Kasai, David Gray, Carol Clugston, Anna Dominiczak and Alex Alderton, Roberto Amato, Jeffrey Barrett, Sonia Goncalves, Ewan Harrison, David K. Jackson, Ian Johnston, Dominic Kwiatkowski, Cordelia Langford, John Sillitoe on behalf of the Wellcome Sanger Institute COVID-19 Surveillance Team |
| EPI_ISL_3336231 | Lighthouse Lab in Milton Keynes | Wellcome Sanger Institute for the COVID-19 Genomics UK (COG-UK) Consortium | The Lighthouse Lab in Milton Keynes and Alex Alderton, Roberto Amato, Jeffrey Barrett, Sonia Goncalves, Ewan Harrison, David K. Jackson, Ian Johnston, Dominic Kwiatkowski, Cordelia Langford, John Sillitoe on behalf of the Wellcome Sanger Institute COVID-19 Surveillance Team |
| EPI_ISL_3336232, EPI_ISL_3336233 | Lighthouse Laboratory Plymouth | Wellcome Sanger Institute for the COVID-19 Genomics UK (COG-UK) Consortium | Lighthouse Laboratory Plymouth and Alex Alderton, Roberto Amato, Jeffrey Barrett, Sonia Goncalves, Ewan Harrison, David K. Jackson, Ian Johnston, Dominic Kwiatkowski, Cordelia Langford, John Sillitoe on behalf of the Wellcome Sanger Institute COVID-19 Surveillance Team |
| EPI_ISL_3336234 | Lighthouse Lab in Milton Keynes | Wellcome Sanger Institute for the COVID-19 Genomics UK (COG-UK) Consortium | The Lighthouse Lab in Milton Keynes and Alex Alderton, Roberto Amato, Jeffrey Barrett, Sonia Goncalves, Ewan Harrison, David K. Jackson, Ian Johnston, Dominic Kwiatkowski, Cordelia Langford, John Sillitoe on behalf of the Wellcome Sanger Institute COVID-19 Surveillance Team |
| EPI_ISL_3336235 | Berkshire and Surrey Pathology Services Lighthouse Laboratory | Wellcome Sanger Institute for the COVID-19 Genomics UK (COG-UK) Consortium | Berkshire and Surrey Pathology Services Lighthouse Laboratory and Alex Alderton, Roberto Amato, Jeffrey Barrett, Sonia Goncalves, Ewan Harrison, David K. Jackson, Ian Johnston, Dominic Kwiatkowski, Cordelia Langford, John Sillitoe on behalf of the Wellcome Sanger Institute COVID-19 Surveillance Team |
| EPI_ISL_3336236 | Lighthouse Laboratory Plymouth | Wellcome Sanger Institute for the COVID-19 Genomics UK (COG-UK) Consortium | Lighthouse Laboratory Plymouth and Alex Alderton, Roberto Amato, Jeffrey Barrett, Sonia Goncalves, Ewan Harrison, David K. Jackson, Ian Johnston, Dominic Kwiatkowski, Cordelia Langford, John Sillitoe on behalf of the Wellcome Sanger Institute COVID-19 Surveillance Team |
| EPI_ISL_3336237 | Lighthouse Lab in Milton Keynes | Wellcome Sanger Institute for the COVID-19 Genomics UK (COG-UK) Consortium | The Lighthouse Lab in Milton Keynes and Alex Alderton, Roberto Amato, Jeffrey Barrett, Sonia Goncalves, Ewan Harrison, David K. Jackson, Ian Johnston, Dominic Kwiatkowski, Cordelia Langford, John Sillitoe on behalf of the Wellcome Sanger Institute COVID-19 Surveillance Team |
| EPI_ISL_3336238 | Berkshire and Surrey Pathology Services Lighthouse Laboratory | Wellcome Sanger Institute for the COVID-19 Genomics UK (COG-UK) Consortium | Berkshire and Surrey Pathology Services Lighthouse Laboratory and Alex Alderton, Roberto Amato, Jeffrey Barrett, Sonia Goncalves, Ewan Harrison, David K. Jackson, Ian Johnston, Dominic Kwiatkowski, Cordelia Langford, John Sillitoe on behalf of the Wellcome Sanger Institute COVID-19 Surveillance Team |
| EPI_ISL_3336239 | Lighthouse Laboratory Plymouth | Wellcome Sanger Institute for the COVID-19 Genomics UK (COG-UK) Consortium | Lighthouse Laboratory Plymouth and Alex Alderton, Roberto Amato, Jeffrey Barrett, Sonia Goncalves, Ewan Harrison, David K. Jackson, Ian Johnston, Dominic Kwiatkowski, Cordelia Langford, John Sillitoe on behalf of the Wellcome Sanger Institute COVID-19 Surveillance Team |
| EPI_ISL_3336240 | Lighthouse Lab in Glasgow | Wellcome Sanger Institute for the COVID-19 Genomics UK (COG-UK) Consortium | Harper VanSteenhouse, Yumi Kasai, David Gray, Carol Clugston, Anna Dominiczak and Alex Alderton, Roberto Amato, Jeffrey Barrett, Sonia Goncalves, Ewan Harrison, David K. Jackson, Ian Johnston, Dominic Kwiatkowski, Cordelia Langford, John Sillitoe on behalf of the Wellcome Sanger Institute COVID-19 Surveillance Team |
| EPI_ISL_3336241 | Lighthouse Laboratory Plymouth | Wellcome Sanger Institute for the COVID-19 Genomics UK (COG-UK) Consortium | Lighthouse Laboratory Plymouth and Alex Alderton, Roberto Amato, Jeffrey Barrett, Sonia Goncalves, Ewan Harrison, David K. Jackson, Ian Johnston, Dominic Kwiatkowski, Cordelia Langford, John Sillitoe on behalf of the Wellcome Sanger Institute COVID-19 Surveillance Team |
| EPI_ISL_3336242 | Lighthouse Lab in Milton Keynes | Wellcome Sanger Institute for the COVID-19 Genomics UK (COG-UK) Consortium | The Lighthouse Lab in Milton Keynes and Alex Alderton, Roberto Amato, Jeffrey Barrett, Sonia Goncalves, Ewan Harrison, David K. Jackson, Ian Johnston, Dominic Kwiatkowski, Cordelia Langford, John Sillitoe on behalf of the Wellcome Sanger Institute COVID-19 Surveillance Team |
| EPI_ISL_3336243 | Lighthouse Laboratory Plymouth | Wellcome Sanger Institute for the COVID-19 Genomics UK (COG-UK) Consortium | Lighthouse Laboratory Plymouth and Alex Alderton, Roberto Amato, Jeffrey Barrett, Sonia Goncalves, Ewan Harrison, David K. Jackson, Ian Johnston, Dominic Kwiatkowski, Cordelia Langford, John Sillitoe on behalf of the Wellcome Sanger Institute COVID-19 Surveillance Team |

| | | | |
|---|---|---|---|
| EPI_ISL_3336244 | Berkshire and Surrey Pathology Services Lighthouse Laboratory | Wellcome Sanger Institute for the COVID-19 Genomics UK (COG-UK) Consortium | Berkshire and Surrey Pathology Services Lighthouse Laboratory and Alex Alderton, Roberto Amato, Jeffrey Barrett, Sonia Goncalves, Ewan Harrison, David K. Jackson, Ian Johnston, Dominic Kwiatkowski, Cordelia Langford, John Sillitoe on behalf of the Wellcome Sanger Institute COVID-19 Surveillance Team |
| EPI_ISL_3336245, EPI_ISL_3336246 | Lighthouse Lab in Glasgow | Wellcome Sanger Institute for the COVID-19 Genomics UK (COG-UK) Consortium | Harper VanSteenhouse, Yumi Kasai, David Gray, Carol Clugston, Anna Dominiczak and Alex Alderton, Roberto Amato, Jeffrey Barrett, Sonia Goncalves, Ewan Harrison, David K. Jackson, Ian Johnston, Dominic Kwiatkowski, Cordelia Langford, John Sillitoe on behalf of the Wellcome Sanger Institute COVID-19 Surveillance Team |
| EPI_ISL_3336247 | Lighthouse Laboratory Plymouth | Wellcome Sanger Institute for the COVID-19 Genomics UK (COG-UK) Consortium | Lighthouse Laboratory Plymouth and Alex Alderton, Roberto Amato, Jeffrey Barrett, Sonia Goncalves, Ewan Harrison, David K. Jackson, Ian Johnston, Dominic Kwiatkowski, Cordelia Langford, John Sillitoe on behalf of the Wellcome Sanger Institute COVID-19 Surveillance Team |
| EPI_ISL_3336248, EPI_ISL_3336249 | Lighthouse Lab in Glasgow | Wellcome Sanger Institute for the COVID-19 Genomics UK (COG-UK) Consortium | Harper VanSteenhouse, Yumi Kasai, David Gray, Carol Clugston, Anna Dominiczak and Alex Alderton, Roberto Amato, Jeffrey Barrett, Sonia Goncalves, Ewan Harrison, David K. Jackson, Ian Johnston, Dominic Kwiatkowski, Cordelia Langford, John Sillitoe on behalf of the Wellcome Sanger Institute COVID-19 Surveillance Team |
| EPI_ISL_3336250 | Lighthouse Laboratory Plymouth | Wellcome Sanger Institute for the COVID-19 Genomics UK (COG-UK) Consortium | Lighthouse Laboratory Plymouth and Alex Alderton, Roberto Amato, Jeffrey Barrett, Sonia Goncalves, Ewan Harrison, David K. Jackson, Ian Johnston, Dominic Kwiatkowski, Cordelia Langford, John Sillitoe on behalf of the Wellcome Sanger Institute COVID-19 Surveillance Team |
| EPI_ISL_3336251 | Berkshire and Surrey Pathology Services Lighthouse Laboratory | Wellcome Sanger Institute for the COVID-19 Genomics UK (COG-UK) Consortium | Berkshire and Surrey Pathology Services Lighthouse Laboratory and Alex Alderton, Roberto Amato, Jeffrey Barrett, Sonia Goncalves, Ewan Harrison, David K. Jackson, Ian Johnston, Dominic Kwiatkowski, Cordelia Langford, John Sillitoe on behalf of the Wellcome Sanger Institute COVID-19 Surveillance Team |
| EPI_ISL_3336252 | Lighthouse Laboratory Plymouth | Wellcome Sanger Institute for the COVID-19 Genomics UK (COG-UK) Consortium | Lighthouse Laboratory Plymouth and Alex Alderton, Roberto Amato, Jeffrey Barrett, Sonia Goncalves, Ewan Harrison, David K. Jackson, Ian Johnston, Dominic Kwiatkowski, Cordelia Langford, John Sillitoe on behalf of the Wellcome Sanger Institute COVID-19 Surveillance Team |
| EPI_ISL_3336253 | Berkshire and Surrey Pathology Services Lighthouse Laboratory | Wellcome Sanger Institute for the COVID-19 Genomics UK (COG-UK) Consortium | Berkshire and Surrey Pathology Services Lighthouse Laboratory and Alex Alderton, Roberto Amato, Jeffrey Barrett, Sonia Goncalves, Ewan Harrison, David K. Jackson, Ian Johnston, Dominic Kwiatkowski, Cordelia Langford, John Sillitoe on behalf of the Wellcome Sanger Institute COVID-19 Surveillance Team |
| EPI_ISL_3336254 | Lighthouse Lab in Milton Keynes | Wellcome Sanger Institute for the COVID-19 Genomics UK (COG-UK) Consortium | The Lighthouse Lab in Milton Keynes and Alex Alderton, Roberto Amato, Jeffrey Barrett, Sonia Goncalves, Ewan Harrison, David K. Jackson, Ian Johnston, Dominic Kwiatkowski, Cordelia Langford, John Sillitoe on behalf of the Wellcome Sanger Institute COVID-19 Surveillance Team |
| EPI_ISL_3336255 | Lighthouse Laboratory Plymouth | Wellcome Sanger Institute for the COVID-19 Genomics UK (COG-UK) Consortium | Lighthouse Laboratory Plymouth and Alex Alderton, Roberto Amato, Jeffrey Barrett, Sonia Goncalves, Ewan Harrison, David K. Jackson, Ian Johnston, Dominic Kwiatkowski, Cordelia Langford, John Sillitoe on behalf of the Wellcome Sanger Institute COVID-19 Surveillance Team |
| EPI_ISL_3336256, EPI_ISL_3336257 | Lighthouse Lab in Milton Keynes | Wellcome Sanger Institute for the COVID-19 Genomics UK (COG-UK) Consortium | The Lighthouse Lab in Milton Keynes and Alex Alderton, Roberto Amato, Jeffrey Barrett, Sonia Goncalves, Ewan Harrison, David K. Jackson, Ian Johnston, Dominic Kwiatkowski, Cordelia Langford, John Sillitoe on behalf of the Wellcome Sanger Institute COVID-19 Surveillance Team |
| EPI_ISL_3336258 | Berkshire and Surrey Pathology Services Lighthouse Laboratory | Wellcome Sanger Institute for the COVID-19 Genomics UK (COG-UK) Consortium | Berkshire and Surrey Pathology Services Lighthouse Laboratory and Alex Alderton, Roberto Amato, Jeffrey Barrett, Sonia Goncalves, Ewan Harrison, David K. Jackson, Ian Johnston, Dominic Kwiatkowski, Cordelia Langford, John Sillitoe on behalf of the Wellcome Sanger Institute COVID-19 Surveillance Team |
| EPI_ISL_3336259, EPI_ISL_3336260, EPI_ISL_3336261 | Lighthouse Lab in Glasgow | Wellcome Sanger Institute for the COVID-19 Genomics UK (COG-UK) Consortium | Harper VanSteenhouse, Yumi Kasai, David Gray, Carol Clugston, Anna Dominiczak and Alex Alderton, Roberto Amato, Jeffrey Barrett, Sonia Goncalves, Ewan Harrison, David K. Jackson, Ian Johnston, Dominic Kwiatkowski, Cordelia Langford, John Sillitoe on behalf of the Wellcome Sanger Institute COVID-19 Surveillance Team |
| EPI_ISL_3336262 | Lighthouse Laboratory Plymouth | Wellcome Sanger Institute for the COVID-19 Genomics UK (COG-UK) Consortium | Lighthouse Laboratory Plymouth and Alex Alderton, Roberto Amato, Jeffrey Barrett, Sonia Goncalves, Ewan Harrison, David K. Jackson, Ian Johnston, Dominic Kwiatkowski, Cordelia Langford, John Sillitoe on behalf of the Wellcome Sanger Institute COVID-19 Surveillance Team |
| EPI_ISL_3336263 | Berkshire and Surrey Pathology Services Lighthouse Laboratory | Wellcome Sanger Institute for the COVID-19 Genomics UK (COG-UK) Consortium | Berkshire and Surrey Pathology Services Lighthouse Laboratory and Alex Alderton, Roberto Amato, Jeffrey Barrett, Sonia Goncalves, Ewan Harrison, David K. Jackson, Ian Johnston, Dominic Kwiatkowski, Cordelia Langford, John Sillitoe on behalf of the Wellcome Sanger Institute COVID-19 Surveillance Team |
| EPI_ISL_3336264 | Lighthouse Lab in Glasgow | Wellcome Sanger Institute for the COVID-19 Genomics UK (COG-UK) Consortium | Harper VanSteenhouse, Yumi Kasai, David Gray, Carol Clugston, Anna Dominiczak and Alex Alderton, Roberto Amato, Jeffrey Barrett, Sonia Goncalves, Ewan Harrison, David K. Jackson, Ian Johnston, Dominic Kwiatkowski, Cordelia Langford, John Sillitoe on behalf of the Wellcome Sanger Institute COVID-19 Surveillance Team |
| EPI_ISL_3336265, EPI_ISL_3336266 | Lighthouse Laboratory Plymouth | Wellcome Sanger Institute for the COVID-19 Genomics UK (COG-UK) Consortium | Lighthouse Laboratory Plymouth and Alex Alderton, Roberto Amato, Jeffrey Barrett, Sonia Goncalves, Ewan Harrison, David K. Jackson, Ian Johnston, Dominic Kwiatkowski, Cordelia Langford, John Sillitoe on behalf of the Wellcome Sanger Institute COVID-19 Surveillance Team |
| EPI_ISL_3336267, EPI_ISL_3336268 | Berkshire and Surrey Pathology Services Lighthouse Laboratory | Wellcome Sanger Institute for the COVID-19 Genomics UK (COG-UK) Consortium | Berkshire and Surrey Pathology Services Lighthouse Laboratory and Alex Alderton, Roberto Amato, Jeffrey Barrett, Sonia Goncalves, Ewan Harrison, David K. Jackson, Ian Johnston, Dominic Kwiatkowski, Cordelia Langford, John Sillitoe on behalf of the Wellcome Sanger Institute COVID-19 Surveillance Team |
| EPI_ISL_3336269 | Lighthouse Lab in Glasgow | Wellcome Sanger Institute for the COVID-19 Genomics UK (COG-UK) Consortium | Harper VanSteenhouse, Yumi Kasai, David Gray, Carol Clugston, Anna Dominiczak and Alex Alderton, Roberto Amato, Jeffrey Barrett, Sonia Goncalves, Ewan Harrison, David K. Jackson, Ian Johnston, Dominic Kwiatkowski, Cordelia Langford, John Sillitoe on behalf of the Wellcome Sanger Institute COVID-19 Surveillance Team |
| EPI_ISL_3336270 | Berkshire and Surrey Pathology Services Lighthouse Laboratory | Wellcome Sanger Institute for the COVID-19 Genomics UK (COG-UK) Consortium | Berkshire and Surrey Pathology Services Lighthouse Laboratory and Alex Alderton, Roberto Amato, Jeffrey Barrett, Sonia Goncalves, Ewan Harrison, David K. Jackson, Ian Johnston, Dominic Kwiatkowski, Cordelia Langford, John Sillitoe on behalf of the Wellcome Sanger Institute COVID-19 Surveillance Team |
| EPI_ISL_3336271, EPI_ISL_3336272 | Lighthouse Laboratory Plymouth | Wellcome Sanger Institute for the COVID-19 Genomics UK (COG-UK) Consortium | Lighthouse Laboratory Plymouth and Alex Alderton, Roberto Amato, Jeffrey Barrett, Sonia Goncalves, Ewan Harrison, David K. Jackson, Ian Johnston, Dominic Kwiatkowski, Cordelia Langford, John Sillitoe on behalf of the Wellcome Sanger Institute COVID-19 Surveillance Team |
| EPI_ISL_3336273, EPI_ISL_3336274 | Lighthouse Lab in Milton Keynes | Wellcome Sanger Institute for the COVID-19 Genomics UK (COG-UK) Consortium | The Lighthouse Lab in Milton Keynes and Alex Alderton, Roberto Amato, Jeffrey Barrett, Sonia Goncalves, Ewan Harrison, David K. Jackson, Ian Johnston, Dominic Kwiatkowski, Cordelia Langford, John Sillitoe on behalf of the Wellcome Sanger Institute COVID-19 Surveillance Team |
| EPI_ISL_3336275 | Berkshire and Surrey Pathology Services Lighthouse Laboratory | Wellcome Sanger Institute for the COVID-19 Genomics UK (COG-UK) Consortium | Berkshire and Surrey Pathology Services Lighthouse Laboratory and Alex Alderton, Roberto Amato, Jeffrey Barrett, Sonia Goncalves, Ewan Harrison, David K. Jackson, Ian Johnston, Dominic Kwiatkowski, Cordelia Langford, John Sillitoe on behalf of the Wellcome Sanger Institute COVID-19 Surveillance Team |
| EPI_ISL_3336276 | Lighthouse Lab in Milton Keynes | Wellcome Sanger Institute for the COVID-19 Genomics UK (COG-UK) Consortium | The Lighthouse Lab in Milton Keynes and Alex Alderton, Roberto Amato, Jeffrey Barrett, Sonia Goncalves, Ewan Harrison, David K. Jackson, Ian Johnston, Dominic Kwiatkowski, Cordelia Langford, John Sillitoe on behalf of the Wellcome Sanger Institute COVID-19 Surveillance Team |
| EPI_ISL_3336277 | Lighthouse Lab in Glasgow | Wellcome Sanger Institute for the COVID-19 Genomics UK (COG-UK) Consortium | Harper VanSteenhouse, Yumi Kasai, David Gray, Carol Clugston, Anna Dominiczak and Alex Alderton, Roberto Amato, Jeffrey Barrett, Sonia Goncalves, Ewan Harrison, David K. Jackson, Ian Johnston, Dominic Kwiatkowski, Cordelia Langford, John Sillitoe on behalf of the Wellcome Sanger Institute COVID-19 Surveillance Team |
| EPI_ISL_3336278, EPI_ISL_3336279 | Lighthouse Lab in Milton Keynes | Wellcome Sanger Institute for the COVID-19 Genomics UK (COG-UK) Consortium | The Lighthouse Lab in Milton Keynes and Alex Alderton, Roberto Amato, Jeffrey Barrett, Sonia Goncalves, Ewan Harrison, David K. Jackson, Ian Johnston, Dominic Kwiatkowski, Cordelia Langford, John Sillitoe on behalf of the Wellcome Sanger Institute COVID-19 Surveillance Team |
| EPI_ISL_3336280, EPI_ISL_3336281 | Lighthouse Laboratory Plymouth | Wellcome Sanger Institute for the COVID-19 Genomics UK (COG-UK) Consortium | Lighthouse Laboratory Plymouth and Alex Alderton, Roberto Amato, Jeffrey Barrett, Sonia Goncalves, Ewan Harrison, David K. Jackson, Ian Johnston, Dominic Kwiatkowski, Cordelia Langford, John Sillitoe on behalf of the Wellcome Sanger Institute COVID-19 Surveillance Team |
| EPI_ISL_3336282, EPI_ISL_3336283 | Berkshire and Surrey Pathology Services Lighthouse Laboratory | Wellcome Sanger Institute for the COVID-19 Genomics UK (COG-UK) Consortium | Berkshire and Surrey Pathology Services Lighthouse Laboratory and Alex Alderton, Roberto Amato, Jeffrey Barrett, Sonia Goncalves, Ewan Harrison, David K. Jackson, Ian Johnston, Dominic Kwiatkowski, Cordelia Langford, John Sillitoe on behalf of the Wellcome Sanger Institute COVID-19 Surveillance Team |
| EPI_ISL_3336284, EPI_ISL_3336285 | Lighthouse Lab in Milton Keynes | Wellcome Sanger Institute for the COVID-19 Genomics UK (COG-UK) Consortium | The Lighthouse Lab in Milton Keynes and Alex Alderton, Roberto Amato, Jeffrey Barrett, Sonia Goncalves, Ewan Harrison, David K. Jackson, Ian Johnston, Dominic Kwiatkowski, Cordelia Langford, John Sillitoe on behalf of the Wellcome Sanger Institute COVID-19 Surveillance Team |
| EPI_ISL_3336286 | Lighthouse Laboratory Plymouth | Wellcome Sanger Institute for the COVID-19 Genomics UK (COG-UK) Consortium | Lighthouse Laboratory Plymouth and Alex Alderton, Roberto Amato, Jeffrey Barrett, Sonia Goncalves, Ewan Harrison, David K. Jackson, Ian Johnston, Dominic Kwiatkowski, Cordelia Langford, John Sillitoe on behalf of the Wellcome Sanger Institute COVID-19 Surveillance Team |
| EPI_ISL_3336287, EPI_ISL_3336288 | Lighthouse Lab in Milton Keynes | Wellcome Sanger Institute for the COVID-19 Genomics UK (COG-UK) Consortium | The Lighthouse Lab in Milton Keynes and Alex Alderton, Roberto Amato, Jeffrey Barrett, Sonia Goncalves, Ewan Harrison, David K. Jackson, Ian Johnston, Dominic Kwiatkowski, Cordelia Langford, John Sillitoe on behalf of the Wellcome Sanger Institute COVID-19 Surveillance Team |
| EPI_ISL_3336289, EPI_ISL_3336290 | Lighthouse Laboratory Plymouth | Wellcome Sanger Institute for the COVID-19 Genomics UK (COG-UK) Consortium | Lighthouse Laboratory Plymouth and Alex Alderton, Roberto Amato, Jeffrey Barrett, Sonia Goncalves, Ewan Harrison, David K. Jackson, Ian Johnston, Dominic Kwiatkowski, Cordelia Langford, John Sillitoe on behalf of the Wellcome Sanger Institute COVID-19 Surveillance Team |
| EPI_ISL_3336291, EPI_ISL_3336292, EPI_ISL_3336293, EPI_ISL_3336294 | Lighthouse Lab in Milton Keynes | Wellcome Sanger Institute for the COVID-19 Genomics UK (COG-UK) Consortium | The Lighthouse Lab in Milton Keynes and Alex Alderton, Roberto Amato, Jeffrey Barrett, Sonia Goncalves, Ewan Harrison, David K. Jackson, Ian Johnston, Dominic Kwiatkowski, Cordelia Langford, John Sillitoe on behalf of the Wellcome Sanger Institute COVID-19 Surveillance Team |
| EPI_ISL_3336295 | Lighthouse Laboratory Plymouth | Wellcome Sanger Institute for the COVID-19 Genomics UK (COG-UK) Consortium | Lighthouse Laboratory Plymouth and Alex Alderton, Roberto Amato, Jeffrey Barrett, Sonia Goncalves, Ewan Harrison, David K. Jackson, Ian Johnston, Dominic Kwiatkowski, Cordelia Langford, John Sillitoe on behalf of the Wellcome Sanger Institute COVID-19 Surveillance Team |
| EPI_ISL_3336296 | Lighthouse Lab in Milton Keynes | Wellcome Sanger Institute for the COVID-19 Genomics UK (COG-UK) Consortium | The Lighthouse Lab in Milton Keynes and Alex Alderton, Roberto Amato, Jeffrey Barrett, Sonia Goncalves, Ewan Harrison, David K. Jackson, Ian Johnston, Dominic Kwiatkowski, Cordelia Langford, John Sillitoe on behalf of the Wellcome Sanger Institute COVID-19 Surveillance Team |
| EPI_ISL_3336297, EPI_ISL_3336298 | Lighthouse Laboratory Plymouth | Wellcome Sanger Institute for the COVID-19 Genomics UK (COG-UK) Consortium | Lighthouse Laboratory Plymouth and Alex Alderton, Roberto Amato, Jeffrey Barrett, Sonia Goncalves, Ewan Harrison, David K. Jackson, Ian Johnston, Dominic Kwiatkowski, Cordelia Langford, John Sillitoe on behalf of the Wellcome Sanger Institute COVID-19 Surveillance Team |
| EPI_ISL_3336299 | Lighthouse Lab in Glasgow | Wellcome Sanger Institute for the COVID-19 Genomics UK (COG-UK) Consortium | Harper VanSteenhouse, Yumi Kasai, David Gray, Carol Clugston, Anna Dominiczak and Alex Alderton, Roberto Amato, Jeffrey Barrett, Sonia Goncalves, Ewan Harrison, David K. Jackson, Ian Johnston, Dominic Kwiatkowski, Cordelia Langford, John Sillitoe on behalf of the Wellcome Sanger Institute COVID-19 Surveillance Team |
| EPI_ISL_3336300, EPI_ISL_3336301 | Lighthouse Laboratory Plymouth | Wellcome Sanger Institute for the COVID-19 Genomics UK (COG-UK) Consortium | Lighthouse Laboratory Plymouth and Alex Alderton, Roberto Amato, Jeffrey Barrett, Sonia Goncalves, Ewan Harrison, David K. Jackson, Ian Johnston, Dominic Kwiatkowski, Cordelia Langford, John Sillitoe on behalf of the Wellcome Sanger Institute COVID-19 Surveillance Team |
| EPI_ISL_3336302, EPI_ISL_3336303 | Lighthouse Lab in Milton Keynes | Wellcome Sanger Institute for the COVID-19 Genomics UK (COG-UK) Consortium | The Lighthouse Lab in Milton Keynes and Alex Alderton, Roberto Amato, Jeffrey Barrett, Sonia Goncalves, Ewan Harrison, David K. Jackson, Ian Johnston, Dominic Kwiatkowski, Cordelia Langford, John Sillitoe on behalf of the Wellcome Sanger Institute COVID-19 Surveillance Team |
| EPI_ISL_3336304, EPI_ISL_3336305 | Berkshire and Surrey Pathology Services Lighthouse Laboratory | Wellcome Sanger Institute for the COVID-19 Genomics UK (COG-UK) Consortium | Berkshire and Surrey Pathology Services Lighthouse Laboratory and Alex Alderton, Roberto Amato, Jeffrey Barrett, Sonia Goncalves, Ewan Harrison, David K. Jackson, Ian Johnston, Dominic Kwiatkowski, Cordelia Langford, John Sillitoe on behalf of the Wellcome Sanger Institute COVID-19 Surveillance Team |
| EPI_ISL_3336306 | Lighthouse Lab in Glasgow | Wellcome Sanger Institute for the COVID-19 Genomics UK (COG-UK) Consortium | Harper VanSteenhouse, Yumi Kasai, David Gray, Carol Clugston, Anna Dominiczak and Alex Alderton, Roberto Amato, Jeffrey Barrett, Sonia Goncalves, Ewan Harrison, David K. Jackson, Ian Johnston, Dominic Kwiatkowski, Cordelia Langford, John Sillitoe on behalf of the Wellcome Sanger Institute COVID-19 Surveillance Team |
| EPI_ISL_3336307 | Berkshire and Surrey Pathology Services Lighthouse Laboratory | Wellcome Sanger Institute for the COVID-19 Genomics UK (COG-UK) Consortium | Berkshire and Surrey Pathology Services Lighthouse Laboratory and Alex Alderton, Roberto Amato, Jeffrey Barrett, Sonia Goncalves, Ewan Harrison, David K. Jackson, Ian Johnston, Dominic Kwiatkowski, Cordelia Langford, John Sillitoe on behalf of the Wellcome Sanger Institute COVID-19 Surveillance Team |
| EPI_ISL_3336308, EPI_ISL_3336309 | Lighthouse Laboratory Plymouth | Wellcome Sanger Institute for the COVID-19 Genomics UK (COG-UK) Consortium | Lighthouse Laboratory Plymouth and Alex Alderton, Roberto Amato, Jeffrey Barrett, Sonia Goncalves, Ewan Harrison, David K. Jackson, Ian Johnston, Dominic Kwiatkowski, Cordelia Langford, John Sillitoe on behalf of the Wellcome Sanger Institute COVID-19 Surveillance Team |
| EPI_ISL_3336310 | Berkshire and Surrey Pathology Services Lighthouse Laboratory | Wellcome Sanger Institute for the COVID-19 Genomics UK (COG-UK) Consortium | Berkshire and Surrey Pathology Services Lighthouse Laboratory and Alex Alderton, Roberto Amato, Jeffrey Barrett, Sonia Goncalves, Ewan Harrison, David K. Jackson, Ian Johnston, Dominic Kwiatkowski, Cordelia Langford, John Sillitoe on behalf of the Wellcome Sanger Institute COVID-19 Surveillance Team |

| | | | |
|---|---|---|---|
| EPI_ISL_3336311 | Lighthouse Lab in Milton Keynes | Wellcome Sanger Institute for the COVID-19 Genomics UK (COG-UK) Consortium | The Lighthouse Lab in Milton Keynes and Alex Alderton, Roberto Amato, Jeffrey Barrett, Sonia Goncalves, Ewan Harrison, David K. Jackson, Ian Johnston, Dominic Kwiatkowski, Cordelia Langford, John Sillitoe on behalf of the Wellcome Sanger Institute COVID-19 Surveillance Team |
| EPI_ISL_3336312, EPI_ISL_3336313 | Berkshire and Surrey Pathology Services Lighthouse Laboratory | Wellcome Sanger Institute for the COVID-19 Genomics UK (COG-UK) Consortium | Berkshire and Surrey Pathology Services Lighthouse Laboratory and Alex Alderton, Roberto Amato, Jeffrey Barrett, Sonia Goncalves, Ewan Harrison, David K. Jackson, Ian Johnston, Dominic Kwiatkowski, Cordelia Langford, John Sillitoe on behalf of the Wellcome Sanger Institute COVID-19 Surveillance Team |
| EPI_ISL_3336314 | Lighthouse Lab in Glasgow | Wellcome Sanger Institute for the COVID-19 Genomics UK (COG-UK) Consortium | Harper VanSteenhouse, Yumi Kasai, David Gray, Carol Clugston, Anna Dominiczak and Alex Alderton, Roberto Amato, Jeffrey Barrett, Sonia Goncalves, Ewan Harrison, David K. Jackson, Ian Johnston, Dominic Kwiatkowski, Cordelia Langford, John Sillitoe on behalf of the Wellcome Sanger Institute COVID-19 Surveillance Team |
| EPI_ISL_3336315, EPI_ISL_3336316, EPI_ISL_3336317 | Lighthouse Lab in Milton Keynes | Wellcome Sanger Institute for the COVID-19 Genomics UK (COG-UK) Consortium | The Lighthouse Lab in Milton Keynes and Alex Alderton, Roberto Amato, Jeffrey Barrett, Sonia Goncalves, Ewan Harrison, David K. Jackson, Ian Johnston, Dominic Kwiatkowski, Cordelia Langford, John Sillitoe on behalf of the Wellcome Sanger Institute COVID-19 Surveillance Team |
| EPI_ISL_3336318 | Berkshire and Surrey Pathology Services Lighthouse Laboratory | Wellcome Sanger Institute for the COVID-19 Genomics UK (COG-UK) Consortium | Berkshire and Surrey Pathology Services Lighthouse Laboratory and Alex Alderton, Roberto Amato, Jeffrey Barrett, Sonia Goncalves, Ewan Harrison, David K. Jackson, Ian Johnston, Dominic Kwiatkowski, Cordelia Langford, John Sillitoe on behalf of the Wellcome Sanger Institute COVID-19 Surveillance Team |
| EPI_ISL_3336319 | Lighthouse Laboratory Plymouth | Wellcome Sanger Institute for the COVID-19 Genomics UK (COG-UK) Consortium | Lighthouse Laboratory Plymouth and Alex Alderton, Roberto Amato, Jeffrey Barrett, Sonia Goncalves, Ewan Harrison, David K. Jackson, Ian Johnston, Dominic Kwiatkowski, Cordelia Langford, John Sillitoe on behalf of the Wellcome Sanger Institute COVID-19 Surveillance Team |
| EPI_ISL_3336320 | Lighthouse Lab in Milton Keynes | Wellcome Sanger Institute for the COVID-19 Genomics UK (COG-UK) Consortium | The Lighthouse Lab in Milton Keynes and Alex Alderton, Roberto Amato, Jeffrey Barrett, Sonia Goncalves, Ewan Harrison, David K. Jackson, Ian Johnston, Dominic Kwiatkowski, Cordelia Langford, John Sillitoe on behalf of the Wellcome Sanger Institute COVID-19 Surveillance Team |
| EPI_ISL_3336321 | Lighthouse Laboratory Plymouth | Wellcome Sanger Institute for the COVID-19 Genomics UK (COG-UK) Consortium | Lighthouse Laboratory Plymouth and Alex Alderton, Roberto Amato, Jeffrey Barrett, Sonia Goncalves, Ewan Harrison, David K. Jackson, Ian Johnston, Dominic Kwiatkowski, Cordelia Langford, John Sillitoe on behalf of the Wellcome Sanger Institute COVID-19 Surveillance Team |
| EPI_ISL_3336322 | Berkshire and Surrey Pathology Services Lighthouse Laboratory | Wellcome Sanger Institute for the COVID-19 Genomics UK (COG-UK) Consortium | Berkshire and Surrey Pathology Services Lighthouse Laboratory and Alex Alderton, Roberto Amato, Jeffrey Barrett, Sonia Goncalves, Ewan Harrison, David K. Jackson, Ian Johnston, Dominic Kwiatkowski, Cordelia Langford, John Sillitoe on behalf of the Wellcome Sanger Institute COVID-19 Surveillance Team |
| EPI_ISL_3336323 | Lighthouse Laboratory Plymouth | Wellcome Sanger Institute for the COVID-19 Genomics UK (COG-UK) Consortium | Lighthouse Laboratory Plymouth and Alex Alderton, Roberto Amato, Jeffrey Barrett, Sonia Goncalves, Ewan Harrison, David K. Jackson, Ian Johnston, Dominic Kwiatkowski, Cordelia Langford, John Sillitoe on behalf of the Wellcome Sanger Institute COVID-19 Surveillance Team |
| EPI_ISL_3336324, EPI_ISL_3336325 | Lighthouse Lab in Milton Keynes | Wellcome Sanger Institute for the COVID-19 Genomics UK (COG-UK) Consortium | The Lighthouse Lab in Milton Keynes and Alex Alderton, Roberto Amato, Jeffrey Barrett, Sonia Goncalves, Ewan Harrison, David K. Jackson, Ian Johnston, Dominic Kwiatkowski, Cordelia Langford, John Sillitoe on behalf of the Wellcome Sanger Institute COVID-19 Surveillance Team |
| EPI_ISL_3336326 | Berkshire and Surrey Pathology Services Lighthouse Laboratory | Wellcome Sanger Institute for the COVID-19 Genomics UK (COG-UK) Consortium | Berkshire and Surrey Pathology Services Lighthouse Laboratory and Alex Alderton, Roberto Amato, Jeffrey Barrett, Sonia Goncalves, Ewan Harrison, David K. Jackson, Ian Johnston, Dominic Kwiatkowski, Cordelia Langford, John Sillitoe on behalf of the Wellcome Sanger Institute COVID-19 Surveillance Team |
| EPI_ISL_3336327, EPI_ISL_3336328 | Lighthouse Laboratory Plymouth | Wellcome Sanger Institute for the COVID-19 Genomics UK (COG-UK) Consortium | Lighthouse Laboratory Plymouth and Alex Alderton, Roberto Amato, Jeffrey Barrett, Sonia Goncalves, Ewan Harrison, David K. Jackson, Ian Johnston, Dominic Kwiatkowski, Cordelia Langford, John Sillitoe on behalf of the Wellcome Sanger Institute COVID-19 Surveillance Team |
| EPI_ISL_3336329 | Lighthouse Lab in Glasgow | Wellcome Sanger Institute for the COVID-19 Genomics UK (COG-UK) Consortium | Harper VanSteenhouse, Yumi Kasai, David Gray, Carol Clugston, Anna Dominiczak and Alex Alderton, Roberto Amato, Jeffrey Barrett, Sonia Goncalves, Ewan Harrison, David K. Jackson, Ian Johnston, Dominic Kwiatkowski, Cordelia Langford, John Sillitoe on behalf of the Wellcome Sanger Institute COVID-19 Surveillance Team |
| EPI_ISL_3336330 | Lighthouse Laboratory Plymouth | Wellcome Sanger Institute for the COVID-19 Genomics UK (COG-UK) Consortium | Lighthouse Laboratory Plymouth and Alex Alderton, Roberto Amato, Jeffrey Barrett, Sonia Goncalves, Ewan Harrison, David K. Jackson, Ian Johnston, Dominic Kwiatkowski, Cordelia Langford, John Sillitoe on behalf of the Wellcome Sanger Institute COVID-19 Surveillance Team |
| EPI_ISL_3336331 | Berkshire and Surrey Pathology Services Lighthouse Laboratory | Wellcome Sanger Institute for the COVID-19 Genomics UK (COG-UK) Consortium | Berkshire and Surrey Pathology Services Lighthouse Laboratory and Alex Alderton, Roberto Amato, Jeffrey Barrett, Sonia Goncalves, Ewan Harrison, David K. Jackson, Ian Johnston, Dominic Kwiatkowski, Cordelia Langford, John Sillitoe on behalf of the Wellcome Sanger Institute COVID-19 Surveillance Team |
| EPI_ISL_3336332 | Lighthouse Lab in Glasgow | Wellcome Sanger Institute for the COVID-19 Genomics UK (COG-UK) Consortium | Harper VanSteenhouse, Yumi Kasai, David Gray, Carol Clugston, Anna Dominiczak and Alex Alderton, Roberto Amato, Jeffrey Barrett, Sonia Goncalves, Ewan Harrison, David K. Jackson, Ian Johnston, Dominic Kwiatkowski, Cordelia Langford, John Sillitoe on behalf of the Wellcome Sanger Institute COVID-19 Surveillance Team |
| EPI_ISL_3336333, EPI_ISL_3336334 | Lighthouse Laboratory Plymouth | Wellcome Sanger Institute for the COVID-19 Genomics UK (COG-UK) Consortium | Lighthouse Laboratory Plymouth and Alex Alderton, Roberto Amato, Jeffrey Barrett, Sonia Goncalves, Ewan Harrison, David K. Jackson, Ian Johnston, Dominic Kwiatkowski, Cordelia Langford, John Sillitoe on behalf of the Wellcome Sanger Institute COVID-19 Surveillance Team |
| EPI_ISL_3336335 | Lighthouse Lab in Glasgow | Wellcome Sanger Institute for the COVID-19 Genomics UK (COG-UK) Consortium | Harper VanSteenhouse, Yumi Kasai, David Gray, Carol Clugston, Anna Dominiczak and Alex Alderton, Roberto Amato, Jeffrey Barrett, Sonia Goncalves, Ewan Harrison, David K. Jackson, Ian Johnston, Dominic Kwiatkowski, Cordelia Langford, John Sillitoe on behalf of the Wellcome Sanger Institute COVID-19 Surveillance Team |
| EPI_ISL_3336336 | Berkshire and Surrey Pathology Services Lighthouse Laboratory | Wellcome Sanger Institute for the COVID-19 Genomics UK (COG-UK) Consortium | Berkshire and Surrey Pathology Services Lighthouse Laboratory and Alex Alderton, Roberto Amato, Jeffrey Barrett, Sonia Goncalves, Ewan Harrison, David K. Jackson, Ian Johnston, Dominic Kwiatkowski, Cordelia Langford, John Sillitoe on behalf of the Wellcome Sanger Institute COVID-19 Surveillance Team |
| EPI_ISL_3336337 | Lighthouse Lab in Glasgow | Wellcome Sanger Institute for the COVID-19 Genomics UK (COG-UK) Consortium | Harper VanSteenhouse, Yumi Kasai, David Gray, Carol Clugston, Anna Dominiczak and Alex Alderton, Roberto Amato, Jeffrey Barrett, Sonia Goncalves, Ewan Harrison, David K. Jackson, Ian Johnston, Dominic Kwiatkowski, Cordelia Langford, John Sillitoe on behalf of the Wellcome Sanger Institute COVID-19 Surveillance Team |
| EPI_ISL_3336338 | Lighthouse Laboratory Plymouth | Wellcome Sanger Institute for the COVID-19 Genomics UK (COG-UK) Consortium | Lighthouse Laboratory Plymouth and Alex Alderton, Roberto Amato, Jeffrey Barrett, Sonia Goncalves, Ewan Harrison, David K. Jackson, Ian Johnston, Dominic Kwiatkowski, Cordelia Langford, John Sillitoe on behalf of the Wellcome Sanger Institute COVID-19 Surveillance Team |
| EPI_ISL_3336339 | Berkshire and Surrey Pathology Services Lighthouse Laboratory | Wellcome Sanger Institute for the COVID-19 Genomics UK (COG-UK) Consortium | Berkshire and Surrey Pathology Services Lighthouse Laboratory and Alex Alderton, Roberto Amato, Jeffrey Barrett, Sonia Goncalves, Ewan Harrison, David K. Jackson, Ian Johnston, Dominic Kwiatkowski, Cordelia Langford, John Sillitoe on behalf of the Wellcome Sanger Institute COVID-19 Surveillance Team |
| EPI_ISL_3336340 | Lighthouse Lab in Glasgow | Wellcome Sanger Institute for the COVID-19 Genomics UK (COG-UK) Consortium | Harper VanSteenhouse, Yumi Kasai, David Gray, Carol Clugston, Anna Dominiczak and Alex Alderton, Roberto Amato, Jeffrey Barrett, Sonia Goncalves, Ewan Harrison, David K. Jackson, Ian Johnston, Dominic Kwiatkowski, Cordelia Langford, John Sillitoe on behalf of the Wellcome Sanger Institute COVID-19 Surveillance Team |
| EPI_ISL_3336341 | Lighthouse Lab in Milton Keynes | Wellcome Sanger Institute for the COVID-19 Genomics UK (COG-UK) Consortium | The Lighthouse Lab in Milton Keynes and Alex Alderton, Roberto Amato, Jeffrey Barrett, Sonia Goncalves, Ewan Harrison, David K. Jackson, Ian Johnston, Dominic Kwiatkowski, Cordelia Langford, John Sillitoe on behalf of the Wellcome Sanger Institute COVID-19 Surveillance Team |
| EPI_ISL_3336342 | Lighthouse Laboratory Plymouth | Wellcome Sanger Institute for the COVID-19 Genomics UK (COG-UK) Consortium | Lighthouse Laboratory Plymouth and Alex Alderton, Roberto Amato, Jeffrey Barrett, Sonia Goncalves, Ewan Harrison, David K. Jackson, Ian Johnston, Dominic Kwiatkowski, Cordelia Langford, John Sillitoe on behalf of the Wellcome Sanger Institute COVID-19 Surveillance Team |
| EPI_ISL_3336343 | Lighthouse Lab in Glasgow | Wellcome Sanger Institute for the COVID-19 Genomics UK (COG-UK) Consortium | Harper VanSteenhouse, Yumi Kasai, David Gray, Carol Clugston, Anna Dominiczak and Alex Alderton, Roberto Amato, Jeffrey Barrett, Sonia Goncalves, Ewan Harrison, David K. Jackson, Ian Johnston, Dominic Kwiatkowski, Cordelia Langford, John Sillitoe on behalf of the Wellcome Sanger Institute COVID-19 Surveillance Team |
| EPI_ISL_3336344 | Berkshire and Surrey Pathology Services Lighthouse Laboratory | Wellcome Sanger Institute for the COVID-19 Genomics UK (COG-UK) Consortium | Berkshire and Surrey Pathology Services Lighthouse Laboratory and Alex Alderton, Roberto Amato, Jeffrey Barrett, Sonia Goncalves, Ewan Harrison, David K. Jackson, Ian Johnston, Dominic Kwiatkowski, Cordelia Langford, John Sillitoe on behalf of the Wellcome Sanger Institute COVID-19 Surveillance Team |
| EPI_ISL_3336345 | Lighthouse Lab in Glasgow | Wellcome Sanger Institute for the COVID-19 Genomics UK (COG-UK) Consortium | Harper VanSteenhouse, Yumi Kasai, David Gray, Carol Clugston, Anna Dominiczak and Alex Alderton, Roberto Amato, Jeffrey Barrett, Sonia Goncalves, Ewan Harrison, David K. Jackson, Ian Johnston, Dominic Kwiatkowski, Cordelia Langford, John Sillitoe on behalf of the Wellcome Sanger Institute COVID-19 Surveillance Team |
| EPI_ISL_3336346 | Berkshire and Surrey Pathology Services Lighthouse Laboratory | Wellcome Sanger Institute for the COVID-19 Genomics UK (COG-UK) Consortium | Berkshire and Surrey Pathology Services Lighthouse Laboratory and Alex Alderton, Roberto Amato, Jeffrey Barrett, Sonia Goncalves, Ewan Harrison, David K. Jackson, Ian Johnston, Dominic Kwiatkowski, Cordelia Langford, John Sillitoe on behalf of the Wellcome Sanger Institute COVID-19 Surveillance Team |
| EPI_ISL_3336347, EPI_ISL_3336348, EPI_ISL_3336349 | Lighthouse Laboratory Plymouth | Wellcome Sanger Institute for the COVID-19 Genomics UK (COG-UK) Consortium | Lighthouse Laboratory Plymouth and Alex Alderton, Roberto Amato, Jeffrey Barrett, Sonia Goncalves, Ewan Harrison, David K. Jackson, Ian Johnston, Dominic Kwiatkowski, Cordelia Langford, John Sillitoe on behalf of the Wellcome Sanger Institute COVID-19 Surveillance Team |
| EPI_ISL_3336350 | Berkshire and Surrey Pathology Services Lighthouse Laboratory | Wellcome Sanger Institute for the COVID-19 Genomics UK (COG-UK) Consortium | Berkshire and Surrey Pathology Services Lighthouse Laboratory and Alex Alderton, Roberto Amato, Jeffrey Barrett, Sonia Goncalves, Ewan Harrison, David K. Jackson, Ian Johnston, Dominic Kwiatkowski, Cordelia Langford, John Sillitoe on behalf of the Wellcome Sanger Institute COVID-19 Surveillance Team |
| EPI_ISL_3336351 | Lighthouse Lab in Glasgow | Wellcome Sanger Institute for the COVID-19 Genomics UK (COG-UK) Consortium | Harper VanSteenhouse, Yumi Kasai, David Gray, Carol Clugston, Anna Dominiczak and Alex Alderton, Roberto Amato, Jeffrey Barrett, Sonia Goncalves, Ewan Harrison, David K. Jackson, Ian Johnston, Dominic Kwiatkowski, Cordelia Langford, John Sillitoe on behalf of the Wellcome Sanger Institute COVID-19 Surveillance Team |
| EPI_ISL_3336352, EPI_ISL_3336353, EPI_ISL_3336354 | Lighthouse Lab in Milton Keynes | Wellcome Sanger Institute for the COVID-19 Genomics UK (COG-UK) Consortium | The Lighthouse Lab in Milton Keynes and Alex Alderton, Roberto Amato, Jeffrey Barrett, Sonia Goncalves, Ewan Harrison, David K. Jackson, Ian Johnston, Dominic Kwiatkowski, Cordelia Langford, John Sillitoe on behalf of the Wellcome Sanger Institute COVID-19 Surveillance Team |
| EPI_ISL_3336355, EPI_ISL_3336356 | Lighthouse Lab in Glasgow | Wellcome Sanger Institute for the COVID-19 Genomics UK (COG-UK) Consortium | Harper VanSteenhouse, Yumi Kasai, David Gray, Carol Clugston, Anna Dominiczak and Alex Alderton, Roberto Amato, Jeffrey Barrett, Sonia Goncalves, Ewan Harrison, David K. Jackson, Ian Johnston, Dominic Kwiatkowski, Cordelia Langford, John Sillitoe on behalf of the Wellcome Sanger Institute COVID-19 Surveillance Team |
| EPI_ISL_3336357 | Dipartimento di Medicina di Laboratorio, Azienda sanitaria universitaria Friuli Centrale (ASU FC) | Dipartimento di Medicina di Laboratorio, Azienda sanitaria universitaria Friuli Centrale (ASU FC) | Corrado Pipan, Catia Mio, Chiara Dal Secco, Stefania Marzinotto, Francesco Curcio |
| EPI_ISL_3336358 | Berkshire and Surrey Pathology Services Lighthouse Laboratory | Wellcome Sanger Institute for the COVID-19 Genomics UK (COG-UK) Consortium | Berkshire and Surrey Pathology Services Lighthouse Laboratory and Alex Alderton, Roberto Amato, Jeffrey Barrett, Sonia Goncalves, Ewan Harrison, David K. Jackson, Ian Johnston, Dominic Kwiatkowski, Cordelia Langford, John Sillitoe on behalf of the Wellcome Sanger Institute COVID-19 Surveillance Team |
| EPI_ISL_3336359 | Lighthouse Lab in Milton Keynes | Wellcome Sanger Institute for the COVID-19 Genomics UK (COG-UK) Consortium | The Lighthouse Lab in Milton Keynes and Alex Alderton, Roberto Amato, Jeffrey Barrett, Sonia Goncalves, Ewan Harrison, David K. Jackson, Ian Johnston, Dominic Kwiatkowski, Cordelia Langford, John Sillitoe on behalf of the Wellcome Sanger Institute COVID-19 Surveillance Team |
| EPI_ISL_3336360 | Dipartimento di Medicina di Laboratorio, Azienda sanitaria universitaria Friuli Centrale (ASU FC) | Dipartimento di Medicina di Laboratorio, Azienda sanitaria universitaria Friuli Centrale (ASU FC) | Corrado Pipan, Catia Mio, Chiara Dal Secco, Stefania Marzinotto, Francesco Curcio |
| EPI_ISL_3336361 | Lighthouse Lab in Glasgow | Wellcome Sanger Institute for the COVID-19 Genomics UK (COG-UK) Consortium | Harper VanSteenhouse, Yumi Kasai, David Gray, Carol Clugston, Anna Dominiczak and Alex Alderton, Roberto Amato, Jeffrey Barrett, Sonia Goncalves, Ewan Harrison, David K. Jackson, Ian Johnston, Dominic Kwiatkowski, Cordelia Langford, John Sillitoe on behalf of the Wellcome Sanger Institute COVID-19 Surveillance Team |
| EPI_ISL_3336362 | Lighthouse Laboratory Plymouth | Wellcome Sanger Institute for the COVID-19 Genomics UK (COG-UK) Consortium | Lighthouse Laboratory Plymouth and Alex Alderton, Roberto Amato, Jeffrey Barrett, Sonia Goncalves, Ewan Harrison, David K. Jackson, Ian Johnston, Dominic Kwiatkowski, Cordelia Langford, John Sillitoe on behalf of the Wellcome Sanger Institute COVID-19 Surveillance Team |
| EPI_ISL_3336363 | Dipartimento di Medicina di Laboratorio, Azienda sanitaria universitaria Friuli Centrale (ASU FC) | Dipartimento di Medicina di Laboratorio, Azienda sanitaria universitaria Friuli Centrale (ASU FC) | Corrado Pipan, Catia Mio, Chiara Dal Secco, Stefania Marzinotto, Francesco Curcio |
| EPI_ISL_3336364 | Lighthouse Lab in Glasgow | Wellcome Sanger Institute for the COVID-19 Genomics UK (COG-UK) Consortium | Harper VanSteenhouse, Yumi Kasai, David Gray, Carol Clugston, Anna Dominiczak and Alex Alderton, Roberto Amato, Jeffrey Barrett, Sonia Goncalves, Ewan Harrison, David K. Jackson, Ian Johnston, Dominic Kwiatkowski, Cordelia Langford, John Sillitoe on behalf of the Wellcome Sanger Institute COVID-19 Surveillance Team |
| EPI_ISL_3336365 | Berkshire and Surrey Pathology Services Lighthouse Laboratory | Wellcome Sanger Institute for the COVID-19 Genomics UK (COG-UK) Consortium | Berkshire and Surrey Pathology Services Lighthouse Laboratory and Alex Alderton, Roberto Amato, Jeffrey Barrett, Sonia Goncalves, Ewan Harrison, David K. Jackson, Ian Johnston, Dominic Kwiatkowski, Cordelia Langford, John Sillitoe on behalf of the Wellcome Sanger Institute COVID-19 Surveillance Team |

| | | | |
|---|---|---|---|
| EPI_ISL_3336366 | Dipartimento di Medicina di Laboratorio, Azienda sanitaria universitaria Friuli Centrale (ASU FC) | Dipartimento di Medicina di Laboratorio, Azienda sanitaria universitaria Friuli Centrale (ASU FC) | Corrado Pipan, Catia Mio, Chiara Dal Secco, Stefania Marzinotto, Francesco Curcio |
| EPI_ISL_3336367, EPI_ISL_3336368 | Berkshire and Surrey Pathology Services Lighthouse Laboratory | Wellcome Sanger Institute for the COVID-19 Genomics UK (COG-UK) Consortium | Berkshire and Surrey Pathology Services Lighthouse Laboratory and Alex Alderton, Roberto Amato, Jeffrey Barrett, Sonia Goncalves, Ewan Harrison, David K. Jackson, Ian Johnston, Dominic Kwiatkowski, Cordelia Langford, John Sillitoe on behalf of the Wellcome Sanger Institute COVID-19 Surveillance Team |
| EPI_ISL_3336369 | Lighthouse Lab in Milton Keynes | Wellcome Sanger Institute for the COVID-19 Genomics UK (COG-UK) Consortium | The Lighthouse Lab in Milton Keynes and Alex Alderton, Roberto Amato, Jeffrey Barrett, Sonia Goncalves, Ewan Harrison, David K. Jackson, Ian Johnston, Dominic Kwiatkowski, Cordelia Langford, John Sillitoe on behalf of the Wellcome Sanger Institute COVID-19 Surveillance Team |
| EPI_ISL_3336370 | Berkshire and Surrey Pathology Services Lighthouse Laboratory | Wellcome Sanger Institute for the COVID-19 Genomics UK (COG-UK) Consortium | Berkshire and Surrey Pathology Services Lighthouse Laboratory and Alex Alderton, Roberto Amato, Jeffrey Barrett, Sonia Goncalves, Ewan Harrison, David K. Jackson, Ian Johnston, Dominic Kwiatkowski, Cordelia Langford, John Sillitoe on behalf of the Wellcome Sanger Institute COVID-19 Surveillance Team |
| EPI_ISL_3336371 | Lighthouse Lab in Milton Keynes | Wellcome Sanger Institute for the COVID-19 Genomics UK (COG-UK) Consortium | The Lighthouse Lab in Milton Keynes and Alex Alderton, Roberto Amato, Jeffrey Barrett, Sonia Goncalves, Ewan Harrison, David K. Jackson, Ian Johnston, Dominic Kwiatkowski, Cordelia Langford, John Sillitoe on behalf of the Wellcome Sanger Institute COVID-19 Surveillance Team |
| EPI_ISL_3336372 | Lighthouse Laboratory Plymouth | Wellcome Sanger Institute for the COVID-19 Genomics UK (COG-UK) Consortium | Lighthouse Laboratory Plymouth and Alex Alderton, Roberto Amato, Jeffrey Barrett, Sonia Goncalves, Ewan Harrison, David K. Jackson, Ian Johnston, Dominic Kwiatkowski, Cordelia Langford, John Sillitoe on behalf of the Wellcome Sanger Institute COVID-19 Surveillance Team |
| EPI_ISL_3336373 | Lighthouse Lab in Glasgow | Wellcome Sanger Institute for the COVID-19 Genomics UK (COG-UK) Consortium | Harper VanSteenhouse, Yumi Kasai, David Gray, Carol Clugston, Anna Dominiczak and Alex Alderton, Roberto Amato, Jeffrey Barrett, Sonia Goncalves, Ewan Harrison, David K. Jackson, Ian Johnston, Dominic Kwiatkowski, Cordelia Langford, John Sillitoe on behalf of the Wellcome Sanger Institute COVID-19 Surveillance Team |
| EPI_ISL_3336374, EPI_ISL_3336375, EPI_ISL_3336376 | Lighthouse Laboratory Plymouth | Wellcome Sanger Institute for the COVID-19 Genomics UK (COG-UK) Consortium | Lighthouse Laboratory Plymouth and Alex Alderton, Roberto Amato, Jeffrey Barrett, Sonia Goncalves, Ewan Harrison, David K. Jackson, Ian Johnston, Dominic Kwiatkowski, Cordelia Langford, John Sillitoe on behalf of the Wellcome Sanger Institute COVID-19 Surveillance Team |
| EPI_ISL_3336377 | Lighthouse Lab in Glasgow | Wellcome Sanger Institute for the COVID-19 Genomics UK (COG-UK) Consortium | Harper VanSteenhouse, Yumi Kasai, David Gray, Carol Clugston, Anna Dominiczak and Alex Alderton, Roberto Amato, Jeffrey Barrett, Sonia Goncalves, Ewan Harrison, David K. Jackson, Ian Johnston, Dominic Kwiatkowski, Cordelia Langford, John Sillitoe on behalf of the Wellcome Sanger Institute COVID-19 Surveillance Team |
| EPI_ISL_3336378 | Berkshire and Surrey Pathology Services Lighthouse Laboratory | Wellcome Sanger Institute for the COVID-19 Genomics UK (COG-UK) Consortium | Berkshire and Surrey Pathology Services Lighthouse Laboratory and Alex Alderton, Roberto Amato, Jeffrey Barrett, Sonia Goncalves, Ewan Harrison, David K. Jackson, Ian Johnston, Dominic Kwiatkowski, Cordelia Langford, John Sillitoe on behalf of the Wellcome Sanger Institute COVID-19 Surveillance Team |
| EPI_ISL_3336379 | Lighthouse Lab in Glasgow | Wellcome Sanger Institute for the COVID-19 Genomics UK (COG-UK) Consortium | Harper VanSteenhouse, Yumi Kasai, David Gray, Carol Clugston, Anna Dominiczak and Alex Alderton, Roberto Amato, Jeffrey Barrett, Sonia Goncalves, Ewan Harrison, David K. Jackson, Ian Johnston, Dominic Kwiatkowski, Cordelia Langford, John Sillitoe on behalf of the Wellcome Sanger Institute COVID-19 Surveillance Team |
| EPI_ISL_3336380 | Berkshire and Surrey Pathology Services Lighthouse Laboratory | Wellcome Sanger Institute for the COVID-19 Genomics UK (COG-UK) Consortium | Berkshire and Surrey Pathology Services Lighthouse Laboratory and Alex Alderton, Roberto Amato, Jeffrey Barrett, Sonia Goncalves, Ewan Harrison, David K. Jackson, Ian Johnston, Dominic Kwiatkowski, Cordelia Langford, John Sillitoe on behalf of the Wellcome Sanger Institute COVID-19 Surveillance Team |
| EPI_ISL_3336381 | Lighthouse Laboratory Plymouth | Wellcome Sanger Institute for the COVID-19 Genomics UK (COG-UK) Consortium | Lighthouse Laboratory Plymouth and Alex Alderton, Roberto Amato, Jeffrey Barrett, Sonia Goncalves, Ewan Harrison, David K. Jackson, Ian Johnston, Dominic Kwiatkowski, Cordelia Langford, John Sillitoe on behalf of the Wellcome Sanger Institute COVID-19 Surveillance Team |
| EPI_ISL_3336382, EPI_ISL_3336383 | Berkshire and Surrey Pathology Services Lighthouse Laboratory | Wellcome Sanger Institute for the COVID-19 Genomics UK (COG-UK) Consortium | Berkshire and Surrey Pathology Services Lighthouse Laboratory and Alex Alderton, Roberto Amato, Jeffrey Barrett, Sonia Goncalves, Ewan Harrison, David K. Jackson, Ian Johnston, Dominic Kwiatkowski, Cordelia Langford, John Sillitoe on behalf of the Wellcome Sanger Institute COVID-19 Surveillance Team |
| EPI_ISL_3336384 | Lighthouse Lab in Glasgow | Wellcome Sanger Institute for the COVID-19 Genomics UK (COG-UK) Consortium | Harper VanSteenhouse, Yumi Kasai, David Gray, Carol Clugston, Anna Dominiczak and Alex Alderton, Roberto Amato, Jeffrey Barrett, Sonia Goncalves, Ewan Harrison, David K. Jackson, Ian Johnston, Dominic Kwiatkowski, Cordelia Langford, John Sillitoe on behalf of the Wellcome Sanger Institute COVID-19 Surveillance Team |
| EPI_ISL_3336385 | Lighthouse Lab in Milton Keynes | Wellcome Sanger Institute for the COVID-19 Genomics UK (COG-UK) Consortium | The Lighthouse Lab in Milton Keynes and Alex Alderton, Roberto Amato, Jeffrey Barrett, Sonia Goncalves, Ewan Harrison, David K. Jackson, Ian Johnston, Dominic Kwiatkowski, Cordelia Langford, John Sillitoe on behalf of the Wellcome Sanger Institute COVID-19 Surveillance Team |
| EPI_ISL_3336386 | Berkshire and Surrey Pathology Services Lighthouse Laboratory | Wellcome Sanger Institute for the COVID-19 Genomics UK (COG-UK) Consortium | Berkshire and Surrey Pathology Services Lighthouse Laboratory and Alex Alderton, Roberto Amato, Jeffrey Barrett, Sonia Goncalves, Ewan Harrison, David K. Jackson, Ian Johnston, Dominic Kwiatkowski, Cordelia Langford, John Sillitoe on behalf of the Wellcome Sanger Institute COVID-19 Surveillance Team |
| EPI_ISL_3336387 | Lighthouse Lab in Glasgow | Wellcome Sanger Institute for the COVID-19 Genomics UK (COG-UK) Consortium | Harper VanSteenhouse, Yumi Kasai, David Gray, Carol Clugston, Anna Dominiczak and Alex Alderton, Roberto Amato, Jeffrey Barrett, Sonia Goncalves, Ewan Harrison, David K. Jackson, Ian Johnston, Dominic Kwiatkowski, Cordelia Langford, John Sillitoe on behalf of the Wellcome Sanger Institute COVID-19 Surveillance Team |
| EPI_ISL_3336388, EPI_ISL_3336389 | Lighthouse Laboratory Plymouth | Wellcome Sanger Institute for the COVID-19 Genomics UK (COG-UK) Consortium | Lighthouse Laboratory Plymouth and Alex Alderton, Roberto Amato, Jeffrey Barrett, Sonia Goncalves, Ewan Harrison, David K. Jackson, Ian Johnston, Dominic Kwiatkowski, Cordelia Langford, John Sillitoe on behalf of the Wellcome Sanger Institute COVID-19 Surveillance Team |
| EPI_ISL_3336390, EPI_ISL_3336391 | Lighthouse Lab in Milton Keynes | Wellcome Sanger Institute for the COVID-19 Genomics UK (COG-UK) Consortium | The Lighthouse Lab in Milton Keynes and Alex Alderton, Roberto Amato, Jeffrey Barrett, Sonia Goncalves, Ewan Harrison, David K. Jackson, Ian Johnston, Dominic Kwiatkowski, Cordelia Langford, John Sillitoe on behalf of the Wellcome Sanger Institute COVID-19 Surveillance Team |
| EPI_ISL_3336392, EPI_ISL_3336393 | Lighthouse Laboratory Plymouth | Wellcome Sanger Institute for the COVID-19 Genomics UK (COG-UK) Consortium | Lighthouse Laboratory Plymouth and Alex Alderton, Roberto Amato, Jeffrey Barrett, Sonia Goncalves, Ewan Harrison, David K. Jackson, Ian Johnston, Dominic Kwiatkowski, Cordelia Langford, John Sillitoe on behalf of the Wellcome Sanger Institute COVID-19 Surveillance Team |
| EPI_ISL_3336394 | Berkshire and Surrey Pathology Services Lighthouse Laboratory | Wellcome Sanger Institute for the COVID-19 Genomics UK (COG-UK) Consortium | Berkshire and Surrey Pathology Services Lighthouse Laboratory and Alex Alderton, Roberto Amato, Jeffrey Barrett, Sonia Goncalves, Ewan Harrison, David K. Jackson, Ian Johnston, Dominic Kwiatkowski, Cordelia Langford, John Sillitoe on behalf of the Wellcome Sanger Institute COVID-19 Surveillance Team |
| EPI_ISL_3336395 | Lighthouse Lab in Glasgow | Wellcome Sanger Institute for the COVID-19 Genomics UK (COG-UK) Consortium | Harper VanSteenhouse, Yumi Kasai, David Gray, Carol Clugston, Anna Dominiczak and Alex Alderton, Roberto Amato, Jeffrey Barrett, Sonia Goncalves, Ewan Harrison, David K. Jackson, Ian Johnston, Dominic Kwiatkowski, Cordelia Langford, John Sillitoe on behalf of the Wellcome Sanger Institute COVID-19 Surveillance Team |
| EPI_ISL_3336396 | Lighthouse Lab in Milton Keynes | Wellcome Sanger Institute for the COVID-19 Genomics UK (COG-UK) Consortium | The Lighthouse Lab in Milton Keynes and Alex Alderton, Roberto Amato, Jeffrey Barrett, Sonia Goncalves, Ewan Harrison, David K. Jackson, Ian Johnston, Dominic Kwiatkowski, Cordelia Langford, John Sillitoe on behalf of the Wellcome Sanger Institute COVID-19 Surveillance Team |
| EPI_ISL_3336397 | Berkshire and Surrey Pathology Services Lighthouse Laboratory | Wellcome Sanger Institute for the COVID-19 Genomics UK (COG-UK) Consortium | Berkshire and Surrey Pathology Services Lighthouse Laboratory and Alex Alderton, Roberto Amato, Jeffrey Barrett, Sonia Goncalves, Ewan Harrison, David K. Jackson, Ian Johnston, Dominic Kwiatkowski, Cordelia Langford, John Sillitoe on behalf of the Wellcome Sanger Institute COVID-19 Surveillance Team |
| EPI_ISL_3336398 | Lighthouse Laboratory Plymouth | Wellcome Sanger Institute for the COVID-19 Genomics UK (COG-UK) Consortium | Lighthouse Laboratory Plymouth and Alex Alderton, Roberto Amato, Jeffrey Barrett, Sonia Goncalves, Ewan Harrison, David K. Jackson, Ian Johnston, Dominic Kwiatkowski, Cordelia Langford, John Sillitoe on behalf of the Wellcome Sanger Institute COVID-19 Surveillance Team |
| EPI_ISL_3336399, EPI_ISL_3336400 | Lighthouse Lab in Milton Keynes | Wellcome Sanger Institute for the COVID-19 Genomics UK (COG-UK) Consortium | The Lighthouse Lab in Milton Keynes and Alex Alderton, Roberto Amato, Jeffrey Barrett, Sonia Goncalves, Ewan Harrison, David K. Jackson, Ian Johnston, Dominic Kwiatkowski, Cordelia Langford, John Sillitoe on behalf of the Wellcome Sanger Institute COVID-19 Surveillance Team |
| EPI_ISL_3336401 | Berkshire and Surrey Pathology Services Lighthouse Laboratory | Wellcome Sanger Institute for the COVID-19 Genomics UK (COG-UK) Consortium | Berkshire and Surrey Pathology Services Lighthouse Laboratory and Alex Alderton, Roberto Amato, Jeffrey Barrett, Sonia Goncalves, Ewan Harrison, David K. Jackson, Ian Johnston, Dominic Kwiatkowski, Cordelia Langford, John Sillitoe on behalf of the Wellcome Sanger Institute COVID-19 Surveillance Team |
| EPI_ISL_3336402 | Lighthouse Lab in Milton Keynes | Wellcome Sanger Institute for the COVID-19 Genomics UK (COG-UK) Consortium | The Lighthouse Lab in Milton Keynes and Alex Alderton, Roberto Amato, Jeffrey Barrett, Sonia Goncalves, Ewan Harrison, David K. Jackson, Ian Johnston, Dominic Kwiatkowski, Cordelia Langford, John Sillitoe on behalf of the Wellcome Sanger Institute COVID-19 Surveillance Team |
| EPI_ISL_3336403 | Lighthouse Lab in Alderley Park | Wellcome Sanger Institute for the COVID-19 Genomics UK (COG-UK) Consortium | Jacquelyn Wynn, Mairead Hyland, The Lighthouse Lab in Alderley Park and Alex Alderton, Roberto Amato, Jeffrey Barrett, Sonia Goncalves, Ewan Harrison, David K. Jackson, Ian Johnston, Dominic Kwiatkowski, Cordelia Langford, John Sillitoe on behalf of the Wellcome Sanger Institute COVID-19 Surveillance Team |
| EPI_ISL_3336404, EPI_ISL_3336405 | Lighthouse Lab in Milton Keynes | Wellcome Sanger Institute for the COVID-19 Genomics UK (COG-UK) Consortium | The Lighthouse Lab in Milton Keynes and Alex Alderton, Roberto Amato, Jeffrey Barrett, Sonia Goncalves, Ewan Harrison, David K. Jackson, Ian Johnston, Dominic Kwiatkowski, Cordelia Langford, John Sillitoe on behalf of the Wellcome Sanger Institute COVID-19 Surveillance Team |
| EPI_ISL_3336406, EPI_ISL_3336407 | Lighthouse Laboratory Plymouth | Wellcome Sanger Institute for the COVID-19 Genomics UK (COG-UK) Consortium | Lighthouse Laboratory Plymouth and Alex Alderton, Roberto Amato, Jeffrey Barrett, Sonia Goncalves, Ewan Harrison, David K. Jackson, Ian Johnston, Dominic Kwiatkowski, Cordelia Langford, John Sillitoe on behalf of the Wellcome Sanger Institute COVID-19 Surveillance Team |
| EPI_ISL_3336408 | Berkshire and Surrey Pathology Services Lighthouse Laboratory | Wellcome Sanger Institute for the COVID-19 Genomics UK (COG-UK) Consortium | Berkshire and Surrey Pathology Services Lighthouse Laboratory and Alex Alderton, Roberto Amato, Jeffrey Barrett, Sonia Goncalves, Ewan Harrison, David K. Jackson, Ian Johnston, Dominic Kwiatkowski, Cordelia Langford, John Sillitoe on behalf of the Wellcome Sanger Institute COVID-19 Surveillance Team |
| EPI_ISL_3336409 | Lighthouse Lab in Alderley Park | Wellcome Sanger Institute for the COVID-19 Genomics UK (COG-UK) Consortium | Jacquelyn Wynn, Mairead Hyland, The Lighthouse Lab in Alderley Park and Alex Alderton, Roberto Amato, Jeffrey Barrett, Sonia Goncalves, Ewan Harrison, David K. Jackson, Ian Johnston, Dominic Kwiatkowski, Cordelia Langford, John Sillitoe on behalf of the Wellcome Sanger Institute COVID-19 Surveillance Team |
| EPI_ISL_3336410 | Berkshire and Surrey Pathology Services Lighthouse Laboratory | Wellcome Sanger Institute for the COVID-19 Genomics UK (COG-UK) Consortium | Berkshire and Surrey Pathology Services Lighthouse Laboratory and Alex Alderton, Roberto Amato, Jeffrey Barrett, Sonia Goncalves, Ewan Harrison, David K. Jackson, Ian Johnston, Dominic Kwiatkowski, Cordelia Langford, John Sillitoe on behalf of the Wellcome Sanger Institute COVID-19 Surveillance Team |
| EPI_ISL_3336411 | Lighthouse Laboratory Plymouth | Wellcome Sanger Institute for the COVID-19 Genomics UK (COG-UK) Consortium | Lighthouse Laboratory Plymouth and Alex Alderton, Roberto Amato, Jeffrey Barrett, Sonia Goncalves, Ewan Harrison, David K. Jackson, Ian Johnston, Dominic Kwiatkowski, Cordelia Langford, John Sillitoe on behalf of the Wellcome Sanger Institute COVID-19 Surveillance Team |
| EPI_ISL_3336412 | Berkshire and Surrey Pathology Services Lighthouse Laboratory | Wellcome Sanger Institute for the COVID-19 Genomics UK (COG-UK) Consortium | Berkshire and Surrey Pathology Services Lighthouse Laboratory and Alex Alderton, Roberto Amato, Jeffrey Barrett, Sonia Goncalves, Ewan Harrison, David K. Jackson, Ian Johnston, Dominic Kwiatkowski, Cordelia Langford, John Sillitoe on behalf of the Wellcome Sanger Institute COVID-19 Surveillance Team |
| EPI_ISL_3336413 | Lighthouse Laboratory Plymouth | Wellcome Sanger Institute for the COVID-19 Genomics UK (COG-UK) Consortium | Lighthouse Laboratory Plymouth and Alex Alderton, Roberto Amato, Jeffrey Barrett, Sonia Goncalves, Ewan Harrison, David K. Jackson, Ian Johnston, Dominic Kwiatkowski, Cordelia Langford, John Sillitoe on behalf of the Wellcome Sanger Institute COVID-19 Surveillance Team |
| EPI_ISL_3336414 | Lighthouse Lab in Alderley Park | Wellcome Sanger Institute for the COVID-19 Genomics UK (COG-UK) Consortium | Jacquelyn Wynn, Mairead Hyland, The Lighthouse Lab in Alderley Park and Alex Alderton, Roberto Amato, Jeffrey Barrett, Sonia Goncalves, Ewan Harrison, David K. Jackson, Ian Johnston, Dominic Kwiatkowski, Cordelia Langford, John Sillitoe on behalf of the Wellcome Sanger Institute COVID-19 Surveillance Team |
| EPI_ISL_3336415, EPI_ISL_3336416, EPI_ISL_3336417 | Lighthouse Laboratory Plymouth | Wellcome Sanger Institute for the COVID-19 Genomics UK (COG-UK) Consortium | Lighthouse Laboratory Plymouth and Alex Alderton, Roberto Amato, Jeffrey Barrett, Sonia Goncalves, Ewan Harrison, David K. Jackson, Ian Johnston, Dominic Kwiatkowski, Cordelia Langford, John Sillitoe on behalf of the Wellcome Sanger Institute COVID-19 Surveillance Team |
| EPI_ISL_3336418 | Lighthouse Lab in Alderley Park | Wellcome Sanger Institute for the COVID-19 Genomics UK (COG-UK) Consortium | Jacquelyn Wynn, Mairead Hyland, The Lighthouse Lab in Alderley Park and Alex Alderton, Roberto Amato, Jeffrey Barrett, Sonia Goncalves, Ewan Harrison, David K. Jackson, Ian Johnston, Dominic Kwiatkowski, Cordelia Langford, John Sillitoe on behalf of the Wellcome Sanger Institute COVID-19 Surveillance Team |
| EPI_ISL_3336419 | Berkshire and Surrey Pathology Services Lighthouse Laboratory | Wellcome Sanger Institute for the COVID-19 Genomics UK (COG-UK) Consortium | Berkshire and Surrey Pathology Services Lighthouse Laboratory and Alex Alderton, Roberto Amato, Jeffrey Barrett, Sonia Goncalves, Ewan Harrison, David K. Jackson, Ian Johnston, Dominic Kwiatkowski, Cordelia Langford, John Sillitoe on behalf of the Wellcome Sanger Institute COVID-19 Surveillance Team |
| EPI_ISL_3336420 | Lighthouse Lab in Alderley Park | Wellcome Sanger Institute for the COVID-19 Genomics UK (COG-UK) Consortium | Jacquelyn Wynn, Mairead Hyland, The Lighthouse Lab in Alderley Park and Alex Alderton, Roberto Amato, Jeffrey Barrett, Sonia Goncalves, Ewan Harrison, David K. Jackson, Ian Johnston, Dominic Kwiatkowski, Cordelia Langford, John Sillitoe on behalf of the Wellcome Sanger Institute COVID-19 Surveillance Team |
| EPI_ISL_3336421, EPI_ISL_3336422, EPI_ISL_3336423, | Berkshire and Surrey Pathology Services Lighthouse Laboratory | Wellcome Sanger Institute for the COVID-19 Genomics UK (COG-UK) Consortium | Berkshire and Surrey Pathology Services Lighthouse Laboratory and Alex Alderton, Roberto Amato, Jeffrey Barrett, Sonia Goncalves, Ewan Harrison, David K. Jackson, Ian Johnston, Dominic Kwiatkowski, Cordelia Langford, John Sillitoe on behalf of the Wellcome Sanger Institute COVID-19 Surveillance Team |

| | | | |
|---|---|---|---|
| EPI_ISL_3336424 | | | |
| EPI_ISL_3336425, EPI_ISL_3336426, EPI_ISL_3336427 | Lighthouse Laboratory Plymouth | Wellcome Sanger Institute for the COVID-19 Genomics UK (COG-UK) Consortium | Lighthouse Laboratory Plymouth and Alex Alderton, Roberto Amato, Jeffrey Barrett, Sonia Goncalves, Ewan Harrison, David K. Jackson, Ian Johnston, Dominic Kwiatkowski, Cordelia Langford, John Sillitoe on behalf of the Wellcome Sanger Institute COVID-19 Surveillance Team |
| EPI_ISL_3336428 | Lighthouse Lab in Milton Keynes | Wellcome Sanger Institute for the COVID-19 Genomics UK (COG-UK) Consortium | The Lighthouse Lab in Milton Keynes and Alex Alderton, Roberto Amato, Jeffrey Barrett, Sonia Goncalves, Ewan Harrison, David K. Jackson, Ian Johnston, Dominic Kwiatkowski, Cordelia Langford, John Sillitoe on behalf of the Wellcome Sanger Institute COVID-19 Surveillance Team |
| EPI_ISL_3336429 | Lighthouse Laboratory Plymouth | Wellcome Sanger Institute for the COVID-19 Genomics UK (COG-UK) Consortium | Lighthouse Laboratory Plymouth and Alex Alderton, Roberto Amato, Jeffrey Barrett, Sonia Goncalves, Ewan Harrison, David K. Jackson, Ian Johnston, Dominic Kwiatkowski, Cordelia Langford, John Sillitoe on behalf of the Wellcome Sanger Institute COVID-19 Surveillance Team |
| EPI_ISL_3336430 | Berkshire and Surrey Pathology Services Lighthouse Laboratory | Wellcome Sanger Institute for the COVID-19 Genomics UK (COG-UK) Consortium | Berkshire and Surrey Pathology Services Lighthouse Laboratory and Alex Alderton, Roberto Amato, Jeffrey Barrett, Sonia Goncalves, Ewan Harrison, David K. Jackson, Ian Johnston, Dominic Kwiatkowski, Cordelia Langford, John Sillitoe on behalf of the Wellcome Sanger Institute COVID-19 Surveillance Team |
| EPI_ISL_3336431 | Lighthouse Laboratory Plymouth | Wellcome Sanger Institute for the COVID-19 Genomics UK (COG-UK) Consortium | Lighthouse Laboratory Plymouth and Alex Alderton, Roberto Amato, Jeffrey Barrett, Sonia Goncalves, Ewan Harrison, David K. Jackson, Ian Johnston, Dominic Kwiatkowski, Cordelia Langford, John Sillitoe on behalf of the Wellcome Sanger Institute COVID-19 Surveillance Team |
| EPI_ISL_3336432 | Lighthouse Lab in Milton Keynes | Wellcome Sanger Institute for the COVID-19 Genomics UK (COG-UK) Consortium | The Lighthouse Lab in Milton Keynes and Alex Alderton, Roberto Amato, Jeffrey Barrett, Sonia Goncalves, Ewan Harrison, David K. Jackson, Ian Johnston, Dominic Kwiatkowski, Cordelia Langford, John Sillitoe on behalf of the Wellcome Sanger Institute COVID-19 Surveillance Team |
| EPI_ISL_3336433 | Berkshire and Surrey Pathology Services Lighthouse Laboratory | Wellcome Sanger Institute for the COVID-19 Genomics UK (COG-UK) Consortium | Berkshire and Surrey Pathology Services Lighthouse Laboratory and Alex Alderton, Roberto Amato, Jeffrey Barrett, Sonia Goncalves, Ewan Harrison, David K. Jackson, Ian Johnston, Dominic Kwiatkowski, Cordelia Langford, John Sillitoe on behalf of the Wellcome Sanger Institute COVID-19 Surveillance Team |
| EPI_ISL_3336434 | Lighthouse Laboratory Plymouth | Wellcome Sanger Institute for the COVID-19 Genomics UK (COG-UK) Consortium | Lighthouse Laboratory Plymouth and Alex Alderton, Roberto Amato, Jeffrey Barrett, Sonia Goncalves, Ewan Harrison, David K. Jackson, Ian Johnston, Dominic Kwiatkowski, Cordelia Langford, John Sillitoe on behalf of the Wellcome Sanger Institute COVID-19 Surveillance Team |
| EPI_ISL_3336435 | Berkshire and Surrey Pathology Services Lighthouse Laboratory | Wellcome Sanger Institute for the COVID-19 Genomics UK (COG-UK) Consortium | Berkshire and Surrey Pathology Services Lighthouse Laboratory and Alex Alderton, Roberto Amato, Jeffrey Barrett, Sonia Goncalves, Ewan Harrison, David K. Jackson, Ian Johnston, Dominic Kwiatkowski, Cordelia Langford, John Sillitoe on behalf of the Wellcome Sanger Institute COVID-19 Surveillance Team |
| EPI_ISL_3336436 | Lighthouse Lab in Alderley Park | Wellcome Sanger Institute for the COVID-19 Genomics UK (COG-UK) Consortium | Jacquelyn Wynn, Mairead Hyland, The Lighthouse Lab in Alderley Park and Alex Alderton, Roberto Amato, Jeffrey Barrett, Sonia Goncalves, Ewan Harrison, David K. Jackson, Ian Johnston, Dominic Kwiatkowski, Cordelia Langford, John Sillitoe on behalf of the Wellcome Sanger Institute COVID-19 Surveillance Team |
| EPI_ISL_3336437 | Lighthouse Laboratory Plymouth | Wellcome Sanger Institute for the COVID-19 Genomics UK (COG-UK) Consortium | Lighthouse Laboratory Plymouth and Alex Alderton, Roberto Amato, Jeffrey Barrett, Sonia Goncalves, Ewan Harrison, David K. Jackson, Ian Johnston, Dominic Kwiatkowski, Cordelia Langford, John Sillitoe on behalf of the Wellcome Sanger Institute COVID-19 Surveillance Team |
| EPI_ISL_3336438 | Lighthouse Lab in Milton Keynes | Wellcome Sanger Institute for the COVID-19 Genomics UK (COG-UK) Consortium | The Lighthouse Lab in Milton Keynes and Alex Alderton, Roberto Amato, Jeffrey Barrett, Sonia Goncalves, Ewan Harrison, David K. Jackson, Ian Johnston, Dominic Kwiatkowski, Cordelia Langford, John Sillitoe on behalf of the Wellcome Sanger Institute COVID-19 Surveillance Team |
| EPI_ISL_3336439, EPI_ISL_3336440 | Lighthouse Laboratory Plymouth | Wellcome Sanger Institute for the COVID-19 Genomics UK (COG-UK) Consortium | Lighthouse Laboratory Plymouth and Alex Alderton, Roberto Amato, Jeffrey Barrett, Sonia Goncalves, Ewan Harrison, David K. Jackson, Ian Johnston, Dominic Kwiatkowski, Cordelia Langford, John Sillitoe on behalf of the Wellcome Sanger Institute COVID-19 Surveillance Team |
| EPI_ISL_3336441 | Lighthouse Lab in Milton Keynes | Wellcome Sanger Institute for the COVID-19 Genomics UK (COG-UK) Consortium | The Lighthouse Lab in Milton Keynes and Alex Alderton, Roberto Amato, Jeffrey Barrett, Sonia Goncalves, Ewan Harrison, David K. Jackson, Ian Johnston, Dominic Kwiatkowski, Cordelia Langford, John Sillitoe on behalf of the Wellcome Sanger Institute COVID-19 Surveillance Team |
| EPI_ISL_3336443 | Lighthouse Laboratory Plymouth | Wellcome Sanger Institute for the COVID-19 Genomics UK (COG-UK) Consortium | Lighthouse Laboratory Plymouth and Alex Alderton, Roberto Amato, Jeffrey Barrett, Sonia Goncalves, Ewan Harrison, David K. Jackson, Ian Johnston, Dominic Kwiatkowski, Cordelia Langford, John Sillitoe on behalf of the Wellcome Sanger Institute COVID-19 Surveillance Team |
| EPI_ISL_3336444, EPI_ISL_3336445 | Lighthouse Lab in Alderley Park | Wellcome Sanger Institute for the COVID-19 Genomics UK (COG-UK) Consortium | Jacquelyn Wynn, Mairead Hyland, The Lighthouse Lab in Alderley Park and Alex Alderton, Roberto Amato, Jeffrey Barrett, Sonia Goncalves, Ewan Harrison, David K. Jackson, Ian Johnston, Dominic Kwiatkowski, Cordelia Langford, John Sillitoe on behalf of the Wellcome Sanger Institute COVID-19 Surveillance Team |
| EPI_ISL_3336446, EPI_ISL_3336447 | Lighthouse Laboratory Plymouth | Wellcome Sanger Institute for the COVID-19 Genomics UK (COG-UK) Consortium | Lighthouse Laboratory Plymouth and Alex Alderton, Roberto Amato, Jeffrey Barrett, Sonia Goncalves, Ewan Harrison, David K. Jackson, Ian Johnston, Dominic Kwiatkowski, Cordelia Langford, John Sillitoe on behalf of the Wellcome Sanger Institute COVID-19 Surveillance Team |
| EPI_ISL_3336448, EPI_ISL_3336449 | Lighthouse Lab in Milton Keynes | Wellcome Sanger Institute for the COVID-19 Genomics UK (COG-UK) Consortium | The Lighthouse Lab in Milton Keynes and Alex Alderton, Roberto Amato, Jeffrey Barrett, Sonia Goncalves, Ewan Harrison, David K. Jackson, Ian Johnston, Dominic Kwiatkowski, Cordelia Langford, John Sillitoe on behalf of the Wellcome Sanger Institute COVID-19 Surveillance Team |
| EPI_ISL_3336450 | Lighthouse Lab in Alderley Park | Wellcome Sanger Institute for the COVID-19 Genomics UK (COG-UK) Consortium | Jacquelyn Wynn, Mairead Hyland, The Lighthouse Lab in Alderley Park and Alex Alderton, Roberto Amato, Jeffrey Barrett, Sonia Goncalves, Ewan Harrison, David K. Jackson, Ian Johnston, Dominic Kwiatkowski, Cordelia Langford, John Sillitoe on behalf of the Wellcome Sanger Institute COVID-19 Surveillance Team |
| EPI_ISL_3336451 | Lighthouse Lab in Milton Keynes | Wellcome Sanger Institute for the COVID-19 Genomics UK (COG-UK) Consortium | The Lighthouse Lab in Milton Keynes and Alex Alderton, Roberto Amato, Jeffrey Barrett, Sonia Goncalves, Ewan Harrison, David K. Jackson, Ian Johnston, Dominic Kwiatkowski, Cordelia Langford, John Sillitoe on behalf of the Wellcome Sanger Institute COVID-19 Surveillance Team |
| EPI_ISL_3336452 | Lighthouse Laboratory Plymouth | Wellcome Sanger Institute for the COVID-19 Genomics UK (COG-UK) Consortium | Lighthouse Laboratory Plymouth and Alex Alderton, Roberto Amato, Jeffrey Barrett, Sonia Goncalves, Ewan Harrison, David K. Jackson, Ian Johnston, Dominic Kwiatkowski, Cordelia Langford, John Sillitoe on behalf of the Wellcome Sanger Institute COVID-19 Surveillance Team |
| EPI_ISL_3336453 | Berkshire and Surrey Pathology Services Lighthouse Laboratory | Wellcome Sanger Institute for the COVID-19 Genomics UK (COG-UK) Consortium | Berkshire and Surrey Pathology Services Lighthouse Laboratory and Alex Alderton, Roberto Amato, Jeffrey Barrett, Sonia Goncalves, Ewan Harrison, David K. Jackson, Ian Johnston, Dominic Kwiatkowski, Cordelia Langford, John Sillitoe on behalf of the Wellcome Sanger Institute COVID-19 Surveillance Team |
| EPI_ISL_3336454, EPI_ISL_3336455 | Lighthouse Laboratory Plymouth | Wellcome Sanger Institute for the COVID-19 Genomics UK (COG-UK) Consortium | Lighthouse Laboratory Plymouth and Alex Alderton, Roberto Amato, Jeffrey Barrett, Sonia Goncalves, Ewan Harrison, David K. Jackson, Ian Johnston, Dominic Kwiatkowski, Cordelia Langford, John Sillitoe on behalf of the Wellcome Sanger Institute COVID-19 Surveillance Team |
| EPI_ISL_3336456 | Lighthouse Lab in Milton Keynes | Wellcome Sanger Institute for the COVID-19 Genomics UK (COG-UK) Consortium | The Lighthouse Lab in Milton Keynes and Alex Alderton, Roberto Amato, Jeffrey Barrett, Sonia Goncalves, Ewan Harrison, David K. Jackson, Ian Johnston, Dominic Kwiatkowski, Cordelia Langford, John Sillitoe on behalf of the Wellcome Sanger Institute COVID-19 Surveillance Team |
| EPI_ISL_3336457, EPI_ISL_3336458 | Lighthouse Lab in Alderley Park | Wellcome Sanger Institute for the COVID-19 Genomics UK (COG-UK) Consortium | Jacquelyn Wynn, Mairead Hyland, The Lighthouse Lab in Alderley Park and Alex Alderton, Roberto Amato, Jeffrey Barrett, Sonia Goncalves, Ewan Harrison, David K. Jackson, Ian Johnston, Dominic Kwiatkowski, Cordelia Langford, John Sillitoe on behalf of the Wellcome Sanger Institute COVID-19 Surveillance Team |
| EPI_ISL_3336459 | Berkshire and Surrey Pathology Services Lighthouse Laboratory | Wellcome Sanger Institute for the COVID-19 Genomics UK (COG-UK) Consortium | Berkshire and Surrey Pathology Services Lighthouse Laboratory and Alex Alderton, Roberto Amato, Jeffrey Barrett, Sonia Goncalves, Ewan Harrison, David K. Jackson, Ian Johnston, Dominic Kwiatkowski, Cordelia Langford, John Sillitoe on behalf of the Wellcome Sanger Institute COVID-19 Surveillance Team |
| EPI_ISL_3336460, EPI_ISL_3336461 | Lighthouse Lab in Alderley Park | Wellcome Sanger Institute for the COVID-19 Genomics UK (COG-UK) Consortium | Jacquelyn Wynn, Mairead Hyland, The Lighthouse Lab in Alderley Park and Alex Alderton, Roberto Amato, Jeffrey Barrett, Sonia Goncalves, Ewan Harrison, David K. Jackson, Ian Johnston, Dominic Kwiatkowski, Cordelia Langford, John Sillitoe on behalf of the Wellcome Sanger Institute COVID-19 Surveillance Team |
| EPI_ISL_3336462 | Lighthouse Laboratory Plymouth | Wellcome Sanger Institute for the COVID-19 Genomics UK (COG-UK) Consortium | Lighthouse Laboratory Plymouth and Alex Alderton, Roberto Amato, Jeffrey Barrett, Sonia Goncalves, Ewan Harrison, David K. Jackson, Ian Johnston, Dominic Kwiatkowski, Cordelia Langford, John Sillitoe on behalf of the Wellcome Sanger Institute COVID-19 Surveillance Team |
| EPI_ISL_3336463 | Berkshire and Surrey Pathology Services Lighthouse Laboratory | Wellcome Sanger Institute for the COVID-19 Genomics UK (COG-UK) Consortium | Berkshire and Surrey Pathology Services Lighthouse Laboratory and Alex Alderton, Roberto Amato, Jeffrey Barrett, Sonia Goncalves, Ewan Harrison, David K. Jackson, Ian Johnston, Dominic Kwiatkowski, Cordelia Langford, John Sillitoe on behalf of the Wellcome Sanger Institute COVID-19 Surveillance Team |
| EPI_ISL_3336464 | Lighthouse Lab in Alderley Park | Wellcome Sanger Institute for the COVID-19 Genomics UK (COG-UK) Consortium | Jacquelyn Wynn, Mairead Hyland, The Lighthouse Lab in Alderley Park and Alex Alderton, Roberto Amato, Jeffrey Barrett, Sonia Goncalves, Ewan Harrison, David K. Jackson, Ian Johnston, Dominic Kwiatkowski, Cordelia Langford, John Sillitoe on behalf of the Wellcome Sanger Institute COVID-19 Surveillance Team |
| EPI_ISL_3336465, EPI_ISL_3336466 | Lighthouse Laboratory Plymouth | Wellcome Sanger Institute for the COVID-19 Genomics UK (COG-UK) Consortium | Lighthouse Laboratory Plymouth and Alex Alderton, Roberto Amato, Jeffrey Barrett, Sonia Goncalves, Ewan Harrison, David K. Jackson, Ian Johnston, Dominic Kwiatkowski, Cordelia Langford, John Sillitoe on behalf of the Wellcome Sanger Institute COVID-19 Surveillance Team |
| EPI_ISL_3336467 | Lighthouse Lab in Milton Keynes | Wellcome Sanger Institute for the COVID-19 Genomics UK (COG-UK) Consortium | The Lighthouse Lab in Milton Keynes and Alex Alderton, Roberto Amato, Jeffrey Barrett, Sonia Goncalves, Ewan Harrison, David K. Jackson, Ian Johnston, Dominic Kwiatkowski, Cordelia Langford, John Sillitoe on behalf of the Wellcome Sanger Institute COVID-19 Surveillance Team |
| EPI_ISL_3336468 | Berkshire and Surrey Pathology Services Lighthouse Laboratory | Wellcome Sanger Institute for the COVID-19 Genomics UK (COG-UK) Consortium | Berkshire and Surrey Pathology Services Lighthouse Laboratory and Alex Alderton, Roberto Amato, Jeffrey Barrett, Sonia Goncalves, Ewan Harrison, David K. Jackson, Ian Johnston, Dominic Kwiatkowski, Cordelia Langford, John Sillitoe on behalf of the Wellcome Sanger Institute COVID-19 Surveillance Team |
| EPI_ISL_3336469 | Lighthouse Laboratory Plymouth | Wellcome Sanger Institute for the COVID-19 Genomics UK (COG-UK) Consortium | Lighthouse Laboratory Plymouth and Alex Alderton, Roberto Amato, Jeffrey Barrett, Sonia Goncalves, Ewan Harrison, David K. Jackson, Ian Johnston, Dominic Kwiatkowski, Cordelia Langford, John Sillitoe on behalf of the Wellcome Sanger Institute COVID-19 Surveillance Team |
| EPI_ISL_3336470 | Lighthouse Lab in Alderley Park | Wellcome Sanger Institute for the COVID-19 Genomics UK (COG-UK) Consortium | Jacquelyn Wynn, Mairead Hyland, The Lighthouse Lab in Alderley Park and Alex Alderton, Roberto Amato, Jeffrey Barrett, Sonia Goncalves, Ewan Harrison, David K. Jackson, Ian Johnston, Dominic Kwiatkowski, Cordelia Langford, John Sillitoe on behalf of the Wellcome Sanger Institute COVID-19 Surveillance Team |
| EPI_ISL_3336471 | Lighthouse Lab in Milton Keynes | Wellcome Sanger Institute for the COVID-19 Genomics UK (COG-UK) Consortium | The Lighthouse Lab in Milton Keynes and Alex Alderton, Roberto Amato, Jeffrey Barrett, Sonia Goncalves, Ewan Harrison, David K. Jackson, Ian Johnston, Dominic Kwiatkowski, Cordelia Langford, John Sillitoe on behalf of the Wellcome Sanger Institute COVID-19 Surveillance Team |
| EPI_ISL_3336472 | Lighthouse Laboratory Plymouth | Wellcome Sanger Institute for the COVID-19 Genomics UK (COG-UK) Consortium | Lighthouse Laboratory Plymouth and Alex Alderton, Roberto Amato, Jeffrey Barrett, Sonia Goncalves, Ewan Harrison, David K. Jackson, Ian Johnston, Dominic Kwiatkowski, Cordelia Langford, John Sillitoe on behalf of the Wellcome Sanger Institute COVID-19 Surveillance Team |
| EPI_ISL_3336473 | Lighthouse Lab in Milton Keynes | Wellcome Sanger Institute for the COVID-19 Genomics UK (COG-UK) Consortium | The Lighthouse Lab in Milton Keynes and Alex Alderton, Roberto Amato, Jeffrey Barrett, Sonia Goncalves, Ewan Harrison, David K. Jackson, Ian Johnston, Dominic Kwiatkowski, Cordelia Langford, John Sillitoe on behalf of the Wellcome Sanger Institute COVID-19 Surveillance Team |
| EPI_ISL_3336474 | Lighthouse Lab in Alderley Park | Wellcome Sanger Institute for the COVID-19 Genomics UK (COG-UK) Consortium | Jacquelyn Wynn, Mairead Hyland, The Lighthouse Lab in Alderley Park and Alex Alderton, Roberto Amato, Jeffrey Barrett, Sonia Goncalves, Ewan Harrison, David K. Jackson, Ian Johnston, Dominic Kwiatkowski, Cordelia Langford, John Sillitoe on behalf of the Wellcome Sanger Institute COVID-19 Surveillance Team |
| EPI_ISL_3336475 | Berkshire and Surrey Pathology Services Lighthouse Laboratory | Wellcome Sanger Institute for the COVID-19 Genomics UK (COG-UK) Consortium | Berkshire and Surrey Pathology Services Lighthouse Laboratory and Alex Alderton, Roberto Amato, Jeffrey Barrett, Sonia Goncalves, Ewan Harrison, David K. Jackson, Ian Johnston, Dominic Kwiatkowski, Cordelia Langford, John Sillitoe on behalf of the Wellcome Sanger Institute COVID-19 Surveillance Team |
| EPI_ISL_3336476 | Lighthouse Lab in Alderley Park | Wellcome Sanger Institute for the COVID-19 Genomics UK (COG-UK) Consortium | Jacquelyn Wynn, Mairead Hyland, The Lighthouse Lab in Alderley Park and Alex Alderton, Roberto Amato, Jeffrey Barrett, Sonia Goncalves, Ewan Harrison, David K. Jackson, Ian Johnston, Dominic Kwiatkowski, Cordelia Langford, John Sillitoe on behalf of the Wellcome Sanger Institute COVID-19 Surveillance Team |
| EPI_ISL_3336477 | Lighthouse Laboratory Plymouth | Wellcome Sanger Institute for the COVID-19 Genomics UK (COG-UK) Consortium | Lighthouse Laboratory Plymouth and Alex Alderton, Roberto Amato, Jeffrey Barrett, Sonia Goncalves, Ewan Harrison, David K. Jackson, Ian Johnston, Dominic Kwiatkowski, Cordelia Langford, John Sillitoe on behalf of the Wellcome Sanger Institute COVID-19 Surveillance Team |
| EPI_ISL_3336478 | Berkshire and Surrey Pathology Services Lighthouse Laboratory | Wellcome Sanger Institute for the COVID-19 Genomics UK (COG-UK) Consortium | Berkshire and Surrey Pathology Services Lighthouse Laboratory and Alex Alderton, Roberto Amato, Jeffrey Barrett, Sonia Goncalves, Ewan Harrison, David K. Jackson, Ian Johnston, Dominic Kwiatkowski, Cordelia Langford, John Sillitoe on behalf of the Wellcome Sanger Institute COVID-19 Surveillance Team |
| EPI_ISL_3336479, EPI_ISL_3336480 | Lighthouse Lab in Milton Keynes | Wellcome Sanger Institute for the COVID-19 Genomics UK (COG-UK) Consortium | The Lighthouse Lab in Milton Keynes and Alex Alderton, Roberto Amato, Jeffrey Barrett, Sonia Goncalves, Ewan Harrison, David K. Jackson, Ian Johnston, Dominic Kwiatkowski, Cordelia Langford, John Sillitoe on behalf of the Wellcome Sanger Institute COVID-19 Surveillance Team |

| | | | |
|---|---|---|---|
| EPI_ISL_3336481 | Lighthouse Lab in Alderley Park | Wellcome Sanger Institute for the COVID-19 Genomics UK (COG-UK) Consortium | Jacquelyn Wynn, Mairead Hyland, The Lighthouse Lab in Alderley Park and Alex Alderton, Roberto Amato, Jeffrey Barrett, Sonia Goncalves, Ewan Harrison, David K. Jackson, Ian Johnston, Dominic Kwiatkowski, Cordelia Langford, John Sillitoe on behalf of the Wellcome Sanger Institute COVID-19 Surveillance Team |
| EPI_ISL_3336482, EPI_ISL_3336483, EPI_ISL_3336484 | Lighthouse Laboratory Plymouth | Wellcome Sanger Institute for the COVID-19 Genomics UK (COG-UK) Consortium | Lighthouse Laboratory Plymouth and Alex Alderton, Roberto Amato, Jeffrey Barrett, Sonia Goncalves, Ewan Harrison, David K. Jackson, Ian Johnston, Dominic Kwiatkowski, Cordelia Langford, John Sillitoe on behalf of the Wellcome Sanger Institute COVID-19 Surveillance Team |
| EPI_ISL_3336485 | Lighthouse Lab in Milton Keynes | Wellcome Sanger Institute for the COVID-19 Genomics UK (COG-UK) Consortium | The Lighthouse Lab in Milton Keynes and Alex Alderton, Roberto Amato, Jeffrey Barrett, Sonia Goncalves, Ewan Harrison, David K. Jackson, Ian Johnston, Dominic Kwiatkowski, Cordelia Langford, John Sillitoe on behalf of the Wellcome Sanger Institute COVID-19 Surveillance Team |
| EPI_ISL_3336486, EPI_ISL_3336487 | Berkshire and Surrey Pathology Services Lighthouse Laboratory | Wellcome Sanger Institute for the COVID-19 Genomics UK (COG-UK) Consortium | Berkshire and Surrey Pathology Services Lighthouse Laboratory and Alex Alderton, Roberto Amato, Jeffrey Barrett, Sonia Goncalves, Ewan Harrison, David K. Jackson, Ian Johnston, Dominic Kwiatkowski, Cordelia Langford, John Sillitoe on behalf of the Wellcome Sanger Institute COVID-19 Surveillance Team |
| EPI_ISL_3336488 | Lighthouse Lab in Alderley Park | Wellcome Sanger Institute for the COVID-19 Genomics UK (COG-UK) Consortium | Jacquelyn Wynn, Mairead Hyland, The Lighthouse Lab in Alderley Park and Alex Alderton, Roberto Amato, Jeffrey Barrett, Sonia Goncalves, Ewan Harrison, David K. Jackson, Ian Johnston, Dominic Kwiatkowski, Cordelia Langford, John Sillitoe on behalf of the Wellcome Sanger Institute COVID-19 Surveillance Team |
| EPI_ISL_3336489 | Lighthouse Lab in Milton Keynes | Wellcome Sanger Institute for the COVID-19 Genomics UK (COG-UK) Consortium | The Lighthouse Lab in Milton Keynes and Alex Alderton, Roberto Amato, Jeffrey Barrett, Sonia Goncalves, Ewan Harrison, David K. Jackson, Ian Johnston, Dominic Kwiatkowski, Cordelia Langford, John Sillitoe on behalf of the Wellcome Sanger Institute COVID-19 Surveillance Team |
| EPI_ISL_3336490 | Berkshire and Surrey Pathology Services Lighthouse Laboratory | Wellcome Sanger Institute for the COVID-19 Genomics UK (COG-UK) Consortium | Berkshire and Surrey Pathology Services Lighthouse Laboratory and Alex Alderton, Roberto Amato, Jeffrey Barrett, Sonia Goncalves, Ewan Harrison, David K. Jackson, Ian Johnston, Dominic Kwiatkowski, Cordelia Langford, John Sillitoe on behalf of the Wellcome Sanger Institute COVID-19 Surveillance Team |
| EPI_ISL_3336491 | Lighthouse Lab in Alderley Park | Wellcome Sanger Institute for the COVID-19 Genomics UK (COG-UK) Consortium | Jacquelyn Wynn, Mairead Hyland, The Lighthouse Lab in Alderley Park and Alex Alderton, Roberto Amato, Jeffrey Barrett, Sonia Goncalves, Ewan Harrison, David K. Jackson, Ian Johnston, Dominic Kwiatkowski, Cordelia Langford, John Sillitoe on behalf of the Wellcome Sanger Institute COVID-19 Surveillance Team |
| EPI_ISL_3336492 | Lighthouse Lab in Milton Keynes | Wellcome Sanger Institute for the COVID-19 Genomics UK (COG-UK) Consortium | The Lighthouse Lab in Milton Keynes and Alex Alderton, Roberto Amato, Jeffrey Barrett, Sonia Goncalves, Ewan Harrison, David K. Jackson, Ian Johnston, Dominic Kwiatkowski, Cordelia Langford, John Sillitoe on behalf of the Wellcome Sanger Institute COVID-19 Surveillance Team |
| EPI_ISL_3336493 | Lighthouse Lab in Alderley Park | Wellcome Sanger Institute for the COVID-19 Genomics UK (COG-UK) Consortium | Jacquelyn Wynn, Mairead Hyland, The Lighthouse Lab in Alderley Park and Alex Alderton, Roberto Amato, Jeffrey Barrett, Sonia Goncalves, Ewan Harrison, David K. Jackson, Ian Johnston, Dominic Kwiatkowski, Cordelia Langford, John Sillitoe on behalf of the Wellcome Sanger Institute COVID-19 Surveillance Team |
| EPI_ISL_3336494 | Lighthouse Lab in Milton Keynes | Wellcome Sanger Institute for the COVID-19 Genomics UK (COG-UK) Consortium | The Lighthouse Lab in Milton Keynes and Alex Alderton, Roberto Amato, Jeffrey Barrett, Sonia Goncalves, Ewan Harrison, David K. Jackson, Ian Johnston, Dominic Kwiatkowski, Cordelia Langford, John Sillitoe on behalf of the Wellcome Sanger Institute COVID-19 Surveillance Team |
| EPI_ISL_3336495 | Lighthouse Laboratory Plymouth | Wellcome Sanger Institute for the COVID-19 Genomics UK (COG-UK) Consortium | Lighthouse Laboratory Plymouth and Alex Alderton, Roberto Amato, Jeffrey Barrett, Sonia Goncalves, Ewan Harrison, David K. Jackson, Ian Johnston, Dominic Kwiatkowski, Cordelia Langford, John Sillitoe on behalf of the Wellcome Sanger Institute COVID-19 Surveillance Team |
| EPI_ISL_3336496 | Lighthouse Lab in Alderley Park | Wellcome Sanger Institute for the COVID-19 Genomics UK (COG-UK) Consortium | Jacquelyn Wynn, Mairead Hyland, The Lighthouse Lab in Alderley Park and Alex Alderton, Roberto Amato, Jeffrey Barrett, Sonia Goncalves, Ewan Harrison, David K. Jackson, Ian Johnston, Dominic Kwiatkowski, Cordelia Langford, John Sillitoe on behalf of the Wellcome Sanger Institute COVID-19 Surveillance Team |
| EPI_ISL_3336497 | Berkshire and Surrey Pathology Services Lighthouse Laboratory | Wellcome Sanger Institute for the COVID-19 Genomics UK (COG-UK) Consortium | Berkshire and Surrey Pathology Services Lighthouse Laboratory and Alex Alderton, Roberto Amato, Jeffrey Barrett, Sonia Goncalves, Ewan Harrison, David K. Jackson, Ian Johnston, Dominic Kwiatkowski, Cordelia Langford, John Sillitoe on behalf of the Wellcome Sanger Institute COVID-19 Surveillance Team |
| EPI_ISL_3336498, EPI_ISL_3336499 | Lighthouse Lab in Milton Keynes | Wellcome Sanger Institute for the COVID-19 Genomics UK (COG-UK) Consortium | The Lighthouse Lab in Milton Keynes and Alex Alderton, Roberto Amato, Jeffrey Barrett, Sonia Goncalves, Ewan Harrison, David K. Jackson, Ian Johnston, Dominic Kwiatkowski, Cordelia Langford, John Sillitoe on behalf of the Wellcome Sanger Institute COVID-19 Surveillance Team |
| EPI_ISL_3336500, EPI_ISL_3336501 | Lighthouse Laboratory Plymouth | Wellcome Sanger Institute for the COVID-19 Genomics UK (COG-UK) Consortium | Lighthouse Laboratory Plymouth and Alex Alderton, Roberto Amato, Jeffrey Barrett, Sonia Goncalves, Ewan Harrison, David K. Jackson, Ian Johnston, Dominic Kwiatkowski, Cordelia Langford, John Sillitoe on behalf of the Wellcome Sanger Institute COVID-19 Surveillance Team |
| EPI_ISL_3336502 | Lighthouse Lab in Alderley Park | Wellcome Sanger Institute for the COVID-19 Genomics UK (COG-UK) Consortium | Jacquelyn Wynn, Mairead Hyland, The Lighthouse Lab in Alderley Park and Alex Alderton, Roberto Amato, Jeffrey Barrett, Sonia Goncalves, Ewan Harrison, David K. Jackson, Ian Johnston, Dominic Kwiatkowski, Cordelia Langford, John Sillitoe on behalf of the Wellcome Sanger Institute COVID-19 Surveillance Team |
| EPI_ISL_3336503 | Lighthouse Laboratory Plymouth | Wellcome Sanger Institute for the COVID-19 Genomics UK (COG-UK) Consortium | Lighthouse Laboratory Plymouth and Alex Alderton, Roberto Amato, Jeffrey Barrett, Sonia Goncalves, Ewan Harrison, David K. Jackson, Ian Johnston, Dominic Kwiatkowski, Cordelia Langford, John Sillitoe on behalf of the Wellcome Sanger Institute COVID-19 Surveillance Team |
| EPI_ISL_3336504, EPI_ISL_3336505 | Lighthouse Lab in Milton Keynes | Wellcome Sanger Institute for the COVID-19 Genomics UK (COG-UK) Consortium | The Lighthouse Lab in Milton Keynes and Alex Alderton, Roberto Amato, Jeffrey Barrett, Sonia Goncalves, Ewan Harrison, David K. Jackson, Ian Johnston, Dominic Kwiatkowski, Cordelia Langford, John Sillitoe on behalf of the Wellcome Sanger Institute COVID-19 Surveillance Team |
| EPI_ISL_3336506, EPI_ISL_3336507 | Lighthouse Lab in Alderley Park | Wellcome Sanger Institute for the COVID-19 Genomics UK (COG-UK) Consortium | Jacquelyn Wynn, Mairead Hyland, The Lighthouse Lab in Alderley Park and Alex Alderton, Roberto Amato, Jeffrey Barrett, Sonia Goncalves, Ewan Harrison, David K. Jackson, Ian Johnston, Dominic Kwiatkowski, Cordelia Langford, John Sillitoe on behalf of the Wellcome Sanger Institute COVID-19 Surveillance Team |
| EPI_ISL_3336508 | Lighthouse Laboratory Plymouth | Wellcome Sanger Institute for the COVID-19 Genomics UK (COG-UK) Consortium | Lighthouse Laboratory Plymouth and Alex Alderton, Roberto Amato, Jeffrey Barrett, Sonia Goncalves, Ewan Harrison, David K. Jackson, Ian Johnston, Dominic Kwiatkowski, Cordelia Langford, John Sillitoe on behalf of the Wellcome Sanger Institute COVID-19 Surveillance Team |
| EPI_ISL_3336509 | Lighthouse Lab in Alderley Park | Wellcome Sanger Institute for the COVID-19 Genomics UK (COG-UK) Consortium | Jacquelyn Wynn, Mairead Hyland, The Lighthouse Lab in Alderley Park and Alex Alderton, Roberto Amato, Jeffrey Barrett, Sonia Goncalves, Ewan Harrison, David K. Jackson, Ian Johnston, Dominic Kwiatkowski, Cordelia Langford, John Sillitoe on behalf of the Wellcome Sanger Institute COVID-19 Surveillance Team |
| EPI_ISL_3336510, EPI_ISL_3336511 | Lighthouse Laboratory Plymouth | Wellcome Sanger Institute for the COVID-19 Genomics UK (COG-UK) Consortium | Lighthouse Laboratory Plymouth and Alex Alderton, Roberto Amato, Jeffrey Barrett, Sonia Goncalves, Ewan Harrison, David K. Jackson, Ian Johnston, Dominic Kwiatkowski, Cordelia Langford, John Sillitoe on behalf of the Wellcome Sanger Institute COVID-19 Surveillance Team |
| EPI_ISL_3336512 | Berkshire and Surrey Pathology Services Lighthouse Laboratory | Wellcome Sanger Institute for the COVID-19 Genomics UK (COG-UK) Consortium | Berkshire and Surrey Pathology Services Lighthouse Laboratory and Alex Alderton, Roberto Amato, Jeffrey Barrett, Sonia Goncalves, Ewan Harrison, David K. Jackson, Ian Johnston, Dominic Kwiatkowski, Cordelia Langford, John Sillitoe on behalf of the Wellcome Sanger Institute COVID-19 Surveillance Team |
| EPI_ISL_3336513, EPI_ISL_3336514 | Lighthouse Laboratory Plymouth | Wellcome Sanger Institute for the COVID-19 Genomics UK (COG-UK) Consortium | Lighthouse Laboratory Plymouth and Alex Alderton, Roberto Amato, Jeffrey Barrett, Sonia Goncalves, Ewan Harrison, David K. Jackson, Ian Johnston, Dominic Kwiatkowski, Cordelia Langford, John Sillitoe on behalf of the Wellcome Sanger Institute COVID-19 Surveillance Team |
| EPI_ISL_3336515 | Lighthouse Lab in Milton Keynes | Wellcome Sanger Institute for the COVID-19 Genomics UK (COG-UK) Consortium | The Lighthouse Lab in Milton Keynes and Alex Alderton, Roberto Amato, Jeffrey Barrett, Sonia Goncalves, Ewan Harrison, David K. Jackson, Ian Johnston, Dominic Kwiatkowski, Cordelia Langford, John Sillitoe on behalf of the Wellcome Sanger Institute COVID-19 Surveillance Team |
| EPI_ISL_3336516, EPI_ISL_3336517 | Lighthouse Lab in Alderley Park | Wellcome Sanger Institute for the COVID-19 Genomics UK (COG-UK) Consortium | Jacquelyn Wynn, Mairead Hyland, The Lighthouse Lab in Alderley Park and Alex Alderton, Roberto Amato, Jeffrey Barrett, Sonia Goncalves, Ewan Harrison, David K. Jackson, Ian Johnston, Dominic Kwiatkowski, Cordelia Langford, John Sillitoe on behalf of the Wellcome Sanger Institute COVID-19 Surveillance Team |
| EPI_ISL_3336518 | Lighthouse Laboratory Plymouth | Wellcome Sanger Institute for the COVID-19 Genomics UK (COG-UK) Consortium | Lighthouse Laboratory Plymouth and Alex Alderton, Roberto Amato, Jeffrey Barrett, Sonia Goncalves, Ewan Harrison, David K. Jackson, Ian Johnston, Dominic Kwiatkowski, Cordelia Langford, John Sillitoe on behalf of the Wellcome Sanger Institute COVID-19 Surveillance Team |
| EPI_ISL_3336519 | Berkshire and Surrey Pathology Services Lighthouse Laboratory | Wellcome Sanger Institute for the COVID-19 Genomics UK (COG-UK) Consortium | Berkshire and Surrey Pathology Services Lighthouse Laboratory and Alex Alderton, Roberto Amato, Jeffrey Barrett, Sonia Goncalves, Ewan Harrison, David K. Jackson, Ian Johnston, Dominic Kwiatkowski, Cordelia Langford, John Sillitoe on behalf of the Wellcome Sanger Institute COVID-19 Surveillance Team |
| EPI_ISL_3336520 | Lighthouse Lab in Milton Keynes | Wellcome Sanger Institute for the COVID-19 Genomics UK (COG-UK) Consortium | The Lighthouse Lab in Milton Keynes and Alex Alderton, Roberto Amato, Jeffrey Barrett, Sonia Goncalves, Ewan Harrison, David K. Jackson, Ian Johnston, Dominic Kwiatkowski, Cordelia Langford, John Sillitoe on behalf of the Wellcome Sanger Institute COVID-19 Surveillance Team |
| EPI_ISL_3336521, EPI_ISL_3336522, EPI_ISL_3336524 | Lighthouse Lab in Alderley Park | Wellcome Sanger Institute for the COVID-19 Genomics UK (COG-UK) Consortium | Jacquelyn Wynn, Mairead Hyland, The Lighthouse Lab in Alderley Park and Alex Alderton, Roberto Amato, Jeffrey Barrett, Sonia Goncalves, Ewan Harrison, David K. Jackson, Ian Johnston, Dominic Kwiatkowski, Cordelia Langford, John Sillitoe on behalf of the Wellcome Sanger Institute COVID-19 Surveillance Team |
| EPI_ISL_3336525 | Berkshire and Surrey Pathology Services Lighthouse Laboratory | Wellcome Sanger Institute for the COVID-19 Genomics UK (COG-UK) Consortium | Berkshire and Surrey Pathology Services Lighthouse Laboratory and Alex Alderton, Roberto Amato, Jeffrey Barrett, Sonia Goncalves, Ewan Harrison, David K. Jackson, Ian Johnston, Dominic Kwiatkowski, Cordelia Langford, John Sillitoe on behalf of the Wellcome Sanger Institute COVID-19 Surveillance Team |
| EPI_ISL_3336526, EPI_ISL_3336527, EPI_ISL_3336528, EPI_ISL_3336529, EPI_ISL_3336530, EPI_ISL_3336531 | Lighthouse Lab in Alderley Park | Wellcome Sanger Institute for the COVID-19 Genomics UK (COG-UK) Consortium | Jacquelyn Wynn, Mairead Hyland, The Lighthouse Lab in Alderley Park and Alex Alderton, Roberto Amato, Jeffrey Barrett, Sonia Goncalves, Ewan Harrison, David K. Jackson, Ian Johnston, Dominic Kwiatkowski, Cordelia Langford, John Sillitoe on behalf of the Wellcome Sanger Institute COVID-19 Surveillance Team |
| EPI_ISL_3336532 | Berkshire and Surrey Pathology Services Lighthouse Laboratory | Wellcome Sanger Institute for the COVID-19 Genomics UK (COG-UK) Consortium | Berkshire and Surrey Pathology Services Lighthouse Laboratory and Alex Alderton, Roberto Amato, Jeffrey Barrett, Sonia Goncalves, Ewan Harrison, David K. Jackson, Ian Johnston, Dominic Kwiatkowski, Cordelia Langford, John Sillitoe on behalf of the Wellcome Sanger Institute COVID-19 Surveillance Team |
| EPI_ISL_3336533 | Lighthouse Lab in Alderley Park | Wellcome Sanger Institute for the COVID-19 Genomics UK (COG-UK) Consortium | Jacquelyn Wynn, Mairead Hyland, The Lighthouse Lab in Alderley Park and Alex Alderton, Roberto Amato, Jeffrey Barrett, Sonia Goncalves, Ewan Harrison, David K. Jackson, Ian Johnston, Dominic Kwiatkowski, Cordelia Langford, John Sillitoe on behalf of the Wellcome Sanger Institute COVID-19 Surveillance Team |
| EPI_ISL_3336534 | Lighthouse Lab in Milton Keynes | Wellcome Sanger Institute for the COVID-19 Genomics UK (COG-UK) Consortium | The Lighthouse Lab in Milton Keynes and Alex Alderton, Roberto Amato, Jeffrey Barrett, Sonia Goncalves, Ewan Harrison, David K. Jackson, Ian Johnston, Dominic Kwiatkowski, Cordelia Langford, John Sillitoe on behalf of the Wellcome Sanger Institute COVID-19 Surveillance Team |
| EPI_ISL_3336535 | Berkshire and Surrey Pathology Services Lighthouse Laboratory | Wellcome Sanger Institute for the COVID-19 Genomics UK (COG-UK) Consortium | Berkshire and Surrey Pathology Services Lighthouse Laboratory and Alex Alderton, Roberto Amato, Jeffrey Barrett, Sonia Goncalves, Ewan Harrison, David K. Jackson, Ian Johnston, Dominic Kwiatkowski, Cordelia Langford, John Sillitoe on behalf of the Wellcome Sanger Institute COVID-19 Surveillance Team |
| EPI_ISL_3336536 | Lighthouse Lab in Milton Keynes | Wellcome Sanger Institute for the COVID-19 Genomics UK (COG-UK) Consortium | The Lighthouse Lab in Milton Keynes and Alex Alderton, Roberto Amato, Jeffrey Barrett, Sonia Goncalves, Ewan Harrison, David K. Jackson, Ian Johnston, Dominic Kwiatkowski, Cordelia Langford, John Sillitoe on behalf of the Wellcome Sanger Institute COVID-19 Surveillance Team |
| EPI_ISL_3336537 | Berkshire and Surrey Pathology Services Lighthouse Laboratory | Wellcome Sanger Institute for the COVID-19 Genomics UK (COG-UK) Consortium | Berkshire and Surrey Pathology Services Lighthouse Laboratory and Alex Alderton, Roberto Amato, Jeffrey Barrett, Sonia Goncalves, Ewan Harrison, David K. Jackson, Ian Johnston, Dominic Kwiatkowski, Cordelia Langford, John Sillitoe on behalf of the Wellcome Sanger Institute COVID-19 Surveillance Team |
| EPI_ISL_3336538 | Lighthouse Laboratory Plymouth | Wellcome Sanger Institute for the COVID-19 Genomics UK (COG-UK) Consortium | Lighthouse Laboratory Plymouth and Alex Alderton, Roberto Amato, Jeffrey Barrett, Sonia Goncalves, Ewan Harrison, David K. Jackson, Ian Johnston, Dominic Kwiatkowski, Cordelia Langford, John Sillitoe on behalf of the Wellcome Sanger Institute COVID-19 Surveillance Team |
| EPI_ISL_3336539 | Lighthouse Lab in Alderley Park | Wellcome Sanger Institute for the COVID-19 Genomics UK (COG-UK) Consortium | Jacquelyn Wynn, Mairead Hyland, The Lighthouse Lab in Alderley Park and Alex Alderton, Roberto Amato, Jeffrey Barrett, Sonia Goncalves, Ewan Harrison, David K. Jackson, Ian Johnston, Dominic Kwiatkowski, Cordelia Langford, John Sillitoe on behalf of the Wellcome Sanger Institute COVID-19 Surveillance Team |
| EPI_ISL_3336540, EPI_ISL_3336541, EPI_ISL_3336542 | Lighthouse Lab in Milton Keynes | Wellcome Sanger Institute for the COVID-19 Genomics UK (COG-UK) Consortium | The Lighthouse Lab in Milton Keynes and Alex Alderton, Roberto Amato, Jeffrey Barrett, Sonia Goncalves, Ewan Harrison, David K. Jackson, Ian Johnston, Dominic Kwiatkowski, Cordelia Langford, John Sillitoe on behalf of the Wellcome Sanger Institute COVID-19 Surveillance Team |
| EPI_ISL_3336543 | Lighthouse Laboratory Plymouth | Wellcome Sanger Institute for the COVID-19 Genomics UK (COG-UK) | Lighthouse Laboratory Plymouth and Alex Alderton, Roberto Amato, Jeffrey Barrett, Sonia Goncalves, Ewan Harrison, David K. Jackson, Ian Johnston, Dominic Kwiatkowski, Cordelia Langford, John Sillitoe on behalf of the Wellcome Sanger |

| | | | |
|---|---|---|---|
| | | Consortium | Institute COVID-19 Surveillance Team |
| EPI_ISL_3336544 | Berkshire and Surrey Pathology Services Lighthouse Laboratory | Wellcome Sanger Institute for the COVID-19 Genomics UK (COG-UK) Consortium | Berkshire and Surrey Pathology Services Lighthouse Laboratory and Alex Alderton, Roberto Amato, Jeffrey Barrett, Sonia Goncalves, Ewan Harrison, David K. Jackson, Ian Johnston, Dominic Kwiatkowski, Cordelia Langford, John Sillitoe on behalf of the Wellcome Sanger Institute COVID-19 Surveillance Team |
| EPI_ISL_3336545 | Lighthouse Lab in Alderley Park | Wellcome Sanger Institute for the COVID-19 Genomics UK (COG-UK) Consortium | Jacquelyn Wynn, Mairead Hyland, The Lighthouse Lab in Alderley Park and Alex Alderton, Roberto Amato, Jeffrey Barrett, Sonia Goncalves, Ewan Harrison, David K. Jackson, Ian Johnston, Dominic Kwiatkowski, Cordelia Langford, John Sillitoe on behalf of the Wellcome Sanger Institute COVID-19 Surveillance Team |
| EPI_ISL_3336546 | Lighthouse Lab in Milton Keynes | Wellcome Sanger Institute for the COVID-19 Genomics UK (COG-UK) Consortium | The Lighthouse Lab in Milton Keynes and Alex Alderton, Roberto Amato, Jeffrey Barrett, Sonia Goncalves, Ewan Harrison, David K. Jackson, Ian Johnston, Dominic Kwiatkowski, Cordelia Langford, John Sillitoe on behalf of the Wellcome Sanger Institute COVID-19 Surveillance Team |
| EPI_ISL_3336547 | Berkshire and Surrey Pathology Services Lighthouse Laboratory | Wellcome Sanger Institute for the COVID-19 Genomics UK (COG-UK) Consortium | Berkshire and Surrey Pathology Services Lighthouse Laboratory and Alex Alderton, Roberto Amato, Jeffrey Barrett, Sonia Goncalves, Ewan Harrison, David K. Jackson, Ian Johnston, Dominic Kwiatkowski, Cordelia Langford, John Sillitoe on behalf of the Wellcome Sanger Institute COVID-19 Surveillance Team |
| EPI_ISL_3336548, EPI_ISL_3336549, EPI_ISL_3336550 | Lighthouse Lab in Milton Keynes | Wellcome Sanger Institute for the COVID-19 Genomics UK (COG-UK) Consortium | The Lighthouse Lab in Milton Keynes and Alex Alderton, Roberto Amato, Jeffrey Barrett, Sonia Goncalves, Ewan Harrison, David K. Jackson, Ian Johnston, Dominic Kwiatkowski, Cordelia Langford, John Sillitoe on behalf of the Wellcome Sanger Institute COVID-19 Surveillance Team |
| EPI_ISL_3336551 | Berkshire and Surrey Pathology Services Lighthouse Laboratory | Wellcome Sanger Institute for the COVID-19 Genomics UK (COG-UK) Consortium | Berkshire and Surrey Pathology Services Lighthouse Laboratory and Alex Alderton, Roberto Amato, Jeffrey Barrett, Sonia Goncalves, Ewan Harrison, David K. Jackson, Ian Johnston, Dominic Kwiatkowski, Cordelia Langford, John Sillitoe on behalf of the Wellcome Sanger Institute COVID-19 Surveillance Team |
| EPI_ISL_3336552 | Lighthouse Lab in Alderley Park | Wellcome Sanger Institute for the COVID-19 Genomics UK (COG-UK) Consortium | Jacquelyn Wynn, Mairead Hyland, The Lighthouse Lab in Alderley Park and Alex Alderton, Roberto Amato, Jeffrey Barrett, Sonia Goncalves, Ewan Harrison, David K. Jackson, Ian Johnston, Dominic Kwiatkowski, Cordelia Langford, John Sillitoe on behalf of the Wellcome Sanger Institute COVID-19 Surveillance Team |
| EPI_ISL_3336553 | Lighthouse Lab in Milton Keynes | Wellcome Sanger Institute for the COVID-19 Genomics UK (COG-UK) Consortium | The Lighthouse Lab in Milton Keynes and Alex Alderton, Roberto Amato, Jeffrey Barrett, Sonia Goncalves, Ewan Harrison, David K. Jackson, Ian Johnston, Dominic Kwiatkowski, Cordelia Langford, John Sillitoe on behalf of the Wellcome Sanger Institute COVID-19 Surveillance Team |
| EPI_ISL_3336554, EPI_ISL_3336555 | Lighthouse Laboratory Plymouth | Wellcome Sanger Institute for the COVID-19 Genomics UK (COG-UK) Consortium | Lighthouse Laboratory Plymouth and Alex Alderton, Roberto Amato, Jeffrey Barrett, Sonia Goncalves, Ewan Harrison, David K. Jackson, Ian Johnston, Dominic Kwiatkowski, Cordelia Langford, John Sillitoe on behalf of the Wellcome Sanger Institute COVID-19 Surveillance Team |
| EPI_ISL_3336556 | Lighthouse Lab in Alderley Park | Wellcome Sanger Institute for the COVID-19 Genomics UK (COG-UK) Consortium | Jacquelyn Wynn, Mairead Hyland, The Lighthouse Lab in Alderley Park and Alex Alderton, Roberto Amato, Jeffrey Barrett, Sonia Goncalves, Ewan Harrison, David K. Jackson, Ian Johnston, Dominic Kwiatkowski, Cordelia Langford, John Sillitoe on behalf of the Wellcome Sanger Institute COVID-19 Surveillance Team |
| EPI_ISL_3336557 | Berkshire and Surrey Pathology Services Lighthouse Laboratory | Wellcome Sanger Institute for the COVID-19 Genomics UK (COG-UK) Consortium | Berkshire and Surrey Pathology Services Lighthouse Laboratory and Alex Alderton, Roberto Amato, Jeffrey Barrett, Sonia Goncalves, Ewan Harrison, David K. Jackson, Ian Johnston, Dominic Kwiatkowski, Cordelia Langford, John Sillitoe on behalf of the Wellcome Sanger Institute COVID-19 Surveillance Team |
| EPI_ISL_3336558 | Lighthouse Lab in Alderley Park | Wellcome Sanger Institute for the COVID-19 Genomics UK (COG-UK) Consortium | Jacquelyn Wynn, Mairead Hyland, The Lighthouse Lab in Alderley Park and Alex Alderton, Roberto Amato, Jeffrey Barrett, Sonia Goncalves, Ewan Harrison, David K. Jackson, Ian Johnston, Dominic Kwiatkowski, Cordelia Langford, John Sillitoe on behalf of the Wellcome Sanger Institute COVID-19 Surveillance Team |
| EPI_ISL_3336559 | Lighthouse Lab in Milton Keynes | Wellcome Sanger Institute for the COVID-19 Genomics UK (COG-UK) Consortium | The Lighthouse Lab in Milton Keynes and Alex Alderton, Roberto Amato, Jeffrey Barrett, Sonia Goncalves, Ewan Harrison, David K. Jackson, Ian Johnston, Dominic Kwiatkowski, Cordelia Langford, John Sillitoe on behalf of the Wellcome Sanger Institute COVID-19 Surveillance Team |
| EPI_ISL_3336560 | Lighthouse Laboratory Plymouth | Wellcome Sanger Institute for the COVID-19 Genomics UK (COG-UK) Consortium | Lighthouse Laboratory Plymouth and Alex Alderton, Roberto Amato, Jeffrey Barrett, Sonia Goncalves, Ewan Harrison, David K. Jackson, Ian Johnston, Dominic Kwiatkowski, Cordelia Langford, John Sillitoe on behalf of the Wellcome Sanger Institute COVID-19 Surveillance Team |
| EPI_ISL_3336561, EPI_ISL_3336562 | Berkshire and Surrey Pathology Services Lighthouse Laboratory | Wellcome Sanger Institute for the COVID-19 Genomics UK (COG-UK) Consortium | Berkshire and Surrey Pathology Services Lighthouse Laboratory and Alex Alderton, Roberto Amato, Jeffrey Barrett, Sonia Goncalves, Ewan Harrison, David K. Jackson, Ian Johnston, Dominic Kwiatkowski, Cordelia Langford, John Sillitoe on behalf of the Wellcome Sanger Institute COVID-19 Surveillance Team |
| EPI_ISL_3336563, EPI_ISL_3336564 | Lighthouse Laboratory Plymouth | Wellcome Sanger Institute for the COVID-19 Genomics UK (COG-UK) Consortium | Lighthouse Laboratory Plymouth and Alex Alderton, Roberto Amato, Jeffrey Barrett, Sonia Goncalves, Ewan Harrison, David K. Jackson, Ian Johnston, Dominic Kwiatkowski, Cordelia Langford, John Sillitoe on behalf of the Wellcome Sanger Institute COVID-19 Surveillance Team |
| EPI_ISL_3336565 | Berkshire and Surrey Pathology Services Lighthouse Laboratory | Wellcome Sanger Institute for the COVID-19 Genomics UK (COG-UK) Consortium | Berkshire and Surrey Pathology Services Lighthouse Laboratory and Alex Alderton, Roberto Amato, Jeffrey Barrett, Sonia Goncalves, Ewan Harrison, David K. Jackson, Ian Johnston, Dominic Kwiatkowski, Cordelia Langford, John Sillitoe on behalf of the Wellcome Sanger Institute COVID-19 Surveillance Team |
| EPI_ISL_3336566 | Lighthouse Lab in Milton Keynes | Wellcome Sanger Institute for the COVID-19 Genomics UK (COG-UK) Consortium | The Lighthouse Lab in Milton Keynes and Alex Alderton, Roberto Amato, Jeffrey Barrett, Sonia Goncalves, Ewan Harrison, David K. Jackson, Ian Johnston, Dominic Kwiatkowski, Cordelia Langford, John Sillitoe on behalf of the Wellcome Sanger Institute COVID-19 Surveillance Team |
| EPI_ISL_3336567 | Lighthouse Lab in Alderley Park | Wellcome Sanger Institute for the COVID-19 Genomics UK (COG-UK) Consortium | Jacquelyn Wynn, Mairead Hyland, The Lighthouse Lab in Alderley Park and Alex Alderton, Roberto Amato, Jeffrey Barrett, Sonia Goncalves, Ewan Harrison, David K. Jackson, Ian Johnston, Dominic Kwiatkowski, Cordelia Langford, John Sillitoe on behalf of the Wellcome Sanger Institute COVID-19 Surveillance Team |
| EPI_ISL_3336568 | Lighthouse Lab in Milton Keynes | Wellcome Sanger Institute for the COVID-19 Genomics UK (COG-UK) Consortium | The Lighthouse Lab in Milton Keynes and Alex Alderton, Roberto Amato, Jeffrey Barrett, Sonia Goncalves, Ewan Harrison, David K. Jackson, Ian Johnston, Dominic Kwiatkowski, Cordelia Langford, John Sillitoe on behalf of the Wellcome Sanger Institute COVID-19 Surveillance Team |
| EPI_ISL_3336569 | Lighthouse Laboratory Plymouth | Wellcome Sanger Institute for the COVID-19 Genomics UK (COG-UK) Consortium | Lighthouse Laboratory Plymouth and Alex Alderton, Roberto Amato, Jeffrey Barrett, Sonia Goncalves, Ewan Harrison, David K. Jackson, Ian Johnston, Dominic Kwiatkowski, Cordelia Langford, John Sillitoe on behalf of the Wellcome Sanger Institute COVID-19 Surveillance Team |
| EPI_ISL_3336570 | Lighthouse Lab in Milton Keynes | Wellcome Sanger Institute for the COVID-19 Genomics UK (COG-UK) Consortium | The Lighthouse Lab in Milton Keynes and Alex Alderton, Roberto Amato, Jeffrey Barrett, Sonia Goncalves, Ewan Harrison, David K. Jackson, Ian Johnston, Dominic Kwiatkowski, Cordelia Langford, John Sillitoe on behalf of the Wellcome Sanger Institute COVID-19 Surveillance Team |
| EPI_ISL_3336571 | Berkshire and Surrey Pathology Services Lighthouse Laboratory | Wellcome Sanger Institute for the COVID-19 Genomics UK (COG-UK) Consortium | Berkshire and Surrey Pathology Services Lighthouse Laboratory and Alex Alderton, Roberto Amato, Jeffrey Barrett, Sonia Goncalves, Ewan Harrison, David K. Jackson, Ian Johnston, Dominic Kwiatkowski, Cordelia Langford, John Sillitoe on behalf of the Wellcome Sanger Institute COVID-19 Surveillance Team |
| EPI_ISL_3336572 | Lighthouse Lab in Alderley Park | Wellcome Sanger Institute for the COVID-19 Genomics UK (COG-UK) Consortium | Jacquelyn Wynn, Mairead Hyland, The Lighthouse Lab in Alderley Park and Alex Alderton, Roberto Amato, Jeffrey Barrett, Sonia Goncalves, Ewan Harrison, David K. Jackson, Ian Johnston, Dominic Kwiatkowski, Cordelia Langford, John Sillitoe on behalf of the Wellcome Sanger Institute COVID-19 Surveillance Team |
| EPI_ISL_3336573 | Berkshire and Surrey Pathology Services Lighthouse Laboratory | Wellcome Sanger Institute for the COVID-19 Genomics UK (COG-UK) Consortium | Berkshire and Surrey Pathology Services Lighthouse Laboratory and Alex Alderton, Roberto Amato, Jeffrey Barrett, Sonia Goncalves, Ewan Harrison, David K. Jackson, Ian Johnston, Dominic Kwiatkowski, Cordelia Langford, John Sillitoe on behalf of the Wellcome Sanger Institute COVID-19 Surveillance Team |
| EPI_ISL_3336574 | Lighthouse Laboratory Plymouth | Wellcome Sanger Institute for the COVID-19 Genomics UK (COG-UK) Consortium | Lighthouse Laboratory Plymouth and Alex Alderton, Roberto Amato, Jeffrey Barrett, Sonia Goncalves, Ewan Harrison, David K. Jackson, Ian Johnston, Dominic Kwiatkowski, Cordelia Langford, John Sillitoe on behalf of the Wellcome Sanger Institute COVID-19 Surveillance Team |
| EPI_ISL_3336575 | Lighthouse Lab in Milton Keynes | Wellcome Sanger Institute for the COVID-19 Genomics UK (COG-UK) Consortium | The Lighthouse Lab in Milton Keynes and Alex Alderton, Roberto Amato, Jeffrey Barrett, Sonia Goncalves, Ewan Harrison, David K. Jackson, Ian Johnston, Dominic Kwiatkowski, Cordelia Langford, John Sillitoe on behalf of the Wellcome Sanger Institute COVID-19 Surveillance Team |
| EPI_ISL_3336576, EPI_ISL_3336577 | Lighthouse Laboratory Plymouth | Wellcome Sanger Institute for the COVID-19 Genomics UK (COG-UK) Consortium | Lighthouse Laboratory Plymouth and Alex Alderton, Roberto Amato, Jeffrey Barrett, Sonia Goncalves, Ewan Harrison, David K. Jackson, Ian Johnston, Dominic Kwiatkowski, Cordelia Langford, John Sillitoe on behalf of the Wellcome Sanger Institute COVID-19 Surveillance Team |
| EPI_ISL_3336578, EPI_ISL_3336579 | Berkshire and Surrey Pathology Services Lighthouse Laboratory | Wellcome Sanger Institute for the COVID-19 Genomics UK (COG-UK) Consortium | Berkshire and Surrey Pathology Services Lighthouse Laboratory and Alex Alderton, Roberto Amato, Jeffrey Barrett, Sonia Goncalves, Ewan Harrison, David K. Jackson, Ian Johnston, Dominic Kwiatkowski, Cordelia Langford, John Sillitoe on behalf of the Wellcome Sanger Institute COVID-19 Surveillance Team |
| EPI_ISL_3336580, EPI_ISL_3336581, EPI_ISL_3336582 | Lighthouse Lab in Alderley Park | Wellcome Sanger Institute for the COVID-19 Genomics UK (COG-UK) Consortium | Jacquelyn Wynn, Mairead Hyland, The Lighthouse Lab in Alderley Park and Alex Alderton, Roberto Amato, Jeffrey Barrett, Sonia Goncalves, Ewan Harrison, David K. Jackson, Ian Johnston, Dominic Kwiatkowski, Cordelia Langford, John Sillitoe on behalf of the Wellcome Sanger Institute COVID-19 Surveillance Team |
| EPI_ISL_3336583, EPI_ISL_3336584 | Lighthouse Laboratory Plymouth | Wellcome Sanger Institute for the COVID-19 Genomics UK (COG-UK) Consortium | Lighthouse Laboratory Plymouth and Alex Alderton, Roberto Amato, Jeffrey Barrett, Sonia Goncalves, Ewan Harrison, David K. Jackson, Ian Johnston, Dominic Kwiatkowski, Cordelia Langford, John Sillitoe on behalf of the Wellcome Sanger Institute COVID-19 Surveillance Team |
| EPI_ISL_3336585 | Berkshire and Surrey Pathology Services Lighthouse Laboratory | Wellcome Sanger Institute for the COVID-19 Genomics UK (COG-UK) Consortium | Berkshire and Surrey Pathology Services Lighthouse Laboratory and Alex Alderton, Roberto Amato, Jeffrey Barrett, Sonia Goncalves, Ewan Harrison, David K. Jackson, Ian Johnston, Dominic Kwiatkowski, Cordelia Langford, John Sillitoe on behalf of the Wellcome Sanger Institute COVID-19 Surveillance Team |
| EPI_ISL_3336586 | Lighthouse Laboratory Plymouth | Wellcome Sanger Institute for the COVID-19 Genomics UK (COG-UK) Consortium | Lighthouse Laboratory Plymouth and Alex Alderton, Roberto Amato, Jeffrey Barrett, Sonia Goncalves, Ewan Harrison, David K. Jackson, Ian Johnston, Dominic Kwiatkowski, Cordelia Langford, John Sillitoe on behalf of the Wellcome Sanger Institute COVID-19 Surveillance Team |
| EPI_ISL_3336587, EPI_ISL_3336588 | Berkshire and Surrey Pathology Services Lighthouse Laboratory | Wellcome Sanger Institute for the COVID-19 Genomics UK (COG-UK) Consortium | Berkshire and Surrey Pathology Services Lighthouse Laboratory and Alex Alderton, Roberto Amato, Jeffrey Barrett, Sonia Goncalves, Ewan Harrison, David K. Jackson, Ian Johnston, Dominic Kwiatkowski, Cordelia Langford, John Sillitoe on behalf of the Wellcome Sanger Institute COVID-19 Surveillance Team |
| EPI_ISL_3336589, EPI_ISL_3336590 | Lighthouse Laboratory Plymouth | Wellcome Sanger Institute for the COVID-19 Genomics UK (COG-UK) Consortium | Lighthouse Laboratory Plymouth and Alex Alderton, Roberto Amato, Jeffrey Barrett, Sonia Goncalves, Ewan Harrison, David K. Jackson, Ian Johnston, Dominic Kwiatkowski, Cordelia Langford, John Sillitoe on behalf of the Wellcome Sanger Institute COVID-19 Surveillance Team |
| EPI_ISL_3336591, EPI_ISL_3336592, EPI_ISL_3336593 | Lighthouse Lab in Alderley Park | Wellcome Sanger Institute for the COVID-19 Genomics UK (COG-UK) Consortium | Jacquelyn Wynn, Mairead Hyland, The Lighthouse Lab in Alderley Park and Alex Alderton, Roberto Amato, Jeffrey Barrett, Sonia Goncalves, Ewan Harrison, David K. Jackson, Ian Johnston, Dominic Kwiatkowski, Cordelia Langford, John Sillitoe on behalf of the Wellcome Sanger Institute COVID-19 Surveillance Team |
| EPI_ISL_3336594, EPI_ISL_3336595 | Lighthouse Lab in Milton Keynes | Wellcome Sanger Institute for the COVID-19 Genomics UK (COG-UK) Consortium | The Lighthouse Lab in Milton Keynes and Alex Alderton, Roberto Amato, Jeffrey Barrett, Sonia Goncalves, Ewan Harrison, David K. Jackson, Ian Johnston, Dominic Kwiatkowski, Cordelia Langford, John Sillitoe on behalf of the Wellcome Sanger Institute COVID-19 Surveillance Team |
| EPI_ISL_3336596, EPI_ISL_3336597, EPI_ISL_3336598 | Lighthouse Lab in Alderley Park | Wellcome Sanger Institute for the COVID-19 Genomics UK (COG-UK) Consortium | Jacquelyn Wynn, Mairead Hyland, The Lighthouse Lab in Alderley Park and Alex Alderton, Roberto Amato, Jeffrey Barrett, Sonia Goncalves, Ewan Harrison, David K. Jackson, Ian Johnston, Dominic Kwiatkowski, Cordelia Langford, John Sillitoe on behalf of the Wellcome Sanger Institute COVID-19 Surveillance Team |
| EPI_ISL_3336599 | Lighthouse Lab in Milton Keynes | Wellcome Sanger Institute for the COVID-19 Genomics UK (COG-UK) Consortium | The Lighthouse Lab in Milton Keynes and Alex Alderton, Roberto Amato, Jeffrey Barrett, Sonia Goncalves, Ewan Harrison, David K. Jackson, Ian Johnston, Dominic Kwiatkowski, Cordelia Langford, John Sillitoe on behalf of the Wellcome Sanger Institute COVID-19 Surveillance Team |
| EPI_ISL_3336600 | Berkshire and Surrey Pathology Services Lighthouse Laboratory | Wellcome Sanger Institute for the COVID-19 Genomics UK (COG-UK) Consortium | Berkshire and Surrey Pathology Services Lighthouse Laboratory and Alex Alderton, Roberto Amato, Jeffrey Barrett, Sonia Goncalves, Ewan Harrison, David K. Jackson, Ian Johnston, Dominic Kwiatkowski, Cordelia Langford, John Sillitoe on behalf of the Wellcome Sanger Institute COVID-19 Surveillance Team |
| EPI_ISL_3336601 | Lighthouse Lab in Milton Keynes | Wellcome Sanger Institute for the COVID-19 Genomics UK (COG-UK) Consortium | The Lighthouse Lab in Milton Keynes and Alex Alderton, Roberto Amato, Jeffrey Barrett, Sonia Goncalves, Ewan Harrison, David K. Jackson, Ian Johnston, Dominic Kwiatkowski, Cordelia Langford, John Sillitoe on behalf of the Wellcome Sanger Institute COVID-19 Surveillance Team |
| EPI_ISL_3336602 | Lighthouse Lab in Alderley Park | Wellcome Sanger Institute for the COVID-19 Genomics UK (COG-UK) Consortium | Jacquelyn Wynn, Mairead Hyland, The Lighthouse Lab in Alderley Park and Alex Alderton, Roberto Amato, Jeffrey Barrett, Sonia Goncalves, Ewan Harrison, David K. Jackson, Ian Johnston, Dominic Kwiatkowski, Cordelia Langford, John Sillitoe on behalf of the Wellcome Sanger Institute COVID-19 Surveillance Team |
| EPI_ISL_3336603, EPI_ISL_3336604 | Lighthouse Lab in Milton Keynes | Wellcome Sanger Institute for the COVID-19 Genomics UK (COG-UK) Consortium | The Lighthouse Lab in Milton Keynes and Alex Alderton, Roberto Amato, Jeffrey Barrett, Sonia Goncalves, Ewan Harrison, David K. Jackson, Ian Johnston, Dominic Kwiatkowski, Cordelia Langford, John Sillitoe on behalf of the Wellcome Sanger Institute COVID-19 Surveillance Team |

| | | | |
|---|---|---|---|
| EPI_ISL_3336605, EPI_ISL_3336606 | Berkshire and Surrey Pathology Services Lighthouse Laboratory | Wellcome Sanger Institute for the COVID-19 Genomics UK (COG-UK) Consortium | Berkshire and Surrey Pathology Services Lighthouse Laboratory and Alex Alderton, Roberto Amato, Jeffrey Barrett, Sonia Goncalves, Ewan Harrison, David K. Jackson, Ian Johnston, Dominic Kwiatkowski, Cordelia Langford, John Sillitoe on behalf of the Wellcome Sanger Institute COVID-19 Surveillance Team |
| EPI_ISL_3336607 | Lighthouse Lab in Milton Keynes | Wellcome Sanger Institute for the COVID-19 Genomics UK (COG-UK) Consortium | The Lighthouse Lab in Milton Keynes and Alex Alderton, Roberto Amato, Jeffrey Barrett, Sonia Goncalves, Ewan Harrison, David K. Jackson, Ian Johnston, Dominic Kwiatkowski, Cordelia Langford, John Sillitoe on behalf of the Wellcome Sanger Institute COVID-19 Surveillance Team |
| EPI_ISL_3336608 | Berkshire and Surrey Pathology Services Lighthouse Laboratory | Wellcome Sanger Institute for the COVID-19 Genomics UK (COG-UK) Consortium | Berkshire and Surrey Pathology Services Lighthouse Laboratory and Alex Alderton, Roberto Amato, Jeffrey Barrett, Sonia Goncalves, Ewan Harrison, David K. Jackson, Ian Johnston, Dominic Kwiatkowski, Cordelia Langford, John Sillitoe on behalf of the Wellcome Sanger Institute COVID-19 Surveillance Team |
| EPI_ISL_3336609, EPI_ISL_3336610 | Lighthouse Lab in Milton Keynes | Wellcome Sanger Institute for the COVID-19 Genomics UK (COG-UK) Consortium | The Lighthouse Lab in Milton Keynes and Alex Alderton, Roberto Amato, Jeffrey Barrett, Sonia Goncalves, Ewan Harrison, David K. Jackson, Ian Johnston, Dominic Kwiatkowski, Cordelia Langford, John Sillitoe on behalf of the Wellcome Sanger Institute COVID-19 Surveillance Team |
| EPI_ISL_3336611 | Berkshire and Surrey Pathology Services Lighthouse Laboratory | Wellcome Sanger Institute for the COVID-19 Genomics UK (COG-UK) Consortium | Berkshire and Surrey Pathology Services Lighthouse Laboratory and Alex Alderton, Roberto Amato, Jeffrey Barrett, Sonia Goncalves, Ewan Harrison, David K. Jackson, Ian Johnston, Dominic Kwiatkowski, Cordelia Langford, John Sillitoe on behalf of the Wellcome Sanger Institute COVID-19 Surveillance Team |
| EPI_ISL_3336612 | Lighthouse Lab in Alderley Park | Wellcome Sanger Institute for the COVID-19 Genomics UK (COG-UK) Consortium | Jacquelyn Wynn, Mairead Hyland, The Lighthouse Lab in Alderley Park and Alex Alderton, Roberto Amato, Jeffrey Barrett, Sonia Goncalves, Ewan Harrison, David K. Jackson, Ian Johnston, Dominic Kwiatkowski, Cordelia Langford, John Sillitoe on behalf of the Wellcome Sanger Institute COVID-19 Surveillance Team |
| EPI_ISL_3336613 | Lighthouse Laboratory Plymouth | Wellcome Sanger Institute for the COVID-19 Genomics UK (COG-UK) Consortium | Lighthouse Laboratory Plymouth and Alex Alderton, Roberto Amato, Jeffrey Barrett, Sonia Goncalves, Ewan Harrison, David K. Jackson, Ian Johnston, Dominic Kwiatkowski, Cordelia Langford, John Sillitoe on behalf of the Wellcome Sanger Institute COVID-19 Surveillance Team |
| EPI_ISL_3336614 | Berkshire and Surrey Pathology Services Lighthouse Laboratory | Wellcome Sanger Institute for the COVID-19 Genomics UK (COG-UK) Consortium | Berkshire and Surrey Pathology Services Lighthouse Laboratory and Alex Alderton, Roberto Amato, Jeffrey Barrett, Sonia Goncalves, Ewan Harrison, David K. Jackson, Ian Johnston, Dominic Kwiatkowski, Cordelia Langford, John Sillitoe on behalf of the Wellcome Sanger Institute COVID-19 Surveillance Team |
| EPI_ISL_3336615 | Lighthouse Lab in Milton Keynes | Wellcome Sanger Institute for the COVID-19 Genomics UK (COG-UK) Consortium | The Lighthouse Lab in Milton Keynes and Alex Alderton, Roberto Amato, Jeffrey Barrett, Sonia Goncalves, Ewan Harrison, David K. Jackson, Ian Johnston, Dominic Kwiatkowski, Cordelia Langford, John Sillitoe on behalf of the Wellcome Sanger Institute COVID-19 Surveillance Team |
| EPI_ISL_3336616 | Berkshire and Surrey Pathology Services Lighthouse Laboratory | Wellcome Sanger Institute for the COVID-19 Genomics UK (COG-UK) Consortium | Berkshire and Surrey Pathology Services Lighthouse Laboratory and Alex Alderton, Roberto Amato, Jeffrey Barrett, Sonia Goncalves, Ewan Harrison, David K. Jackson, Ian Johnston, Dominic Kwiatkowski, Cordelia Langford, John Sillitoe on behalf of the Wellcome Sanger Institute COVID-19 Surveillance Team |
| EPI_ISL_3336617, EPI_ISL_3336618 | Lighthouse Lab in Alderley Park | Wellcome Sanger Institute for the COVID-19 Genomics UK (COG-UK) Consortium | Jacquelyn Wynn, Mairead Hyland, The Lighthouse Lab in Alderley Park and Alex Alderton, Roberto Amato, Jeffrey Barrett, Sonia Goncalves, Ewan Harrison, David K. Jackson, Ian Johnston, Dominic Kwiatkowski, Cordelia Langford, John Sillitoe on behalf of the Wellcome Sanger Institute COVID-19 Surveillance Team |
| EPI_ISL_3336619 | Lighthouse Laboratory Plymouth | Wellcome Sanger Institute for the COVID-19 Genomics UK (COG-UK) Consortium | Lighthouse Laboratory Plymouth and Alex Alderton, Roberto Amato, Jeffrey Barrett, Sonia Goncalves, Ewan Harrison, David K. Jackson, Ian Johnston, Dominic Kwiatkowski, Cordelia Langford, John Sillitoe on behalf of the Wellcome Sanger Institute COVID-19 Surveillance Team |
| EPI_ISL_3336620, EPI_ISL_3336621, EPI_ISL_3336622, EPI_ISL_3336623 | Lighthouse Lab in Milton Keynes | Wellcome Sanger Institute for the COVID-19 Genomics UK (COG-UK) Consortium | The Lighthouse Lab in Milton Keynes and Alex Alderton, Roberto Amato, Jeffrey Barrett, Sonia Goncalves, Ewan Harrison, David K. Jackson, Ian Johnston, Dominic Kwiatkowski, Cordelia Langford, John Sillitoe on behalf of the Wellcome Sanger Institute COVID-19 Surveillance Team |
| EPI_ISL_3336624, EPI_ISL_3336625 | Lighthouse Lab in Alderley Park | Wellcome Sanger Institute for the COVID-19 Genomics UK (COG-UK) Consortium | Jacquelyn Wynn, Mairead Hyland, The Lighthouse Lab in Alderley Park and Alex Alderton, Roberto Amato, Jeffrey Barrett, Sonia Goncalves, Ewan Harrison, David K. Jackson, Ian Johnston, Dominic Kwiatkowski, Cordelia Langford, John Sillitoe on behalf of the Wellcome Sanger Institute COVID-19 Surveillance Team |
| EPI_ISL_3336626 | Lighthouse Lab in Milton Keynes | Wellcome Sanger Institute for the COVID-19 Genomics UK (COG-UK) Consortium | The Lighthouse Lab in Milton Keynes and Alex Alderton, Roberto Amato, Jeffrey Barrett, Sonia Goncalves, Ewan Harrison, David K. Jackson, Ian Johnston, Dominic Kwiatkowski, Cordelia Langford, John Sillitoe on behalf of the Wellcome Sanger Institute COVID-19 Surveillance Team |
| EPI_ISL_3336627 | Berkshire and Surrey Pathology Services Lighthouse Laboratory | Wellcome Sanger Institute for the COVID-19 Genomics UK (COG-UK) Consortium | Berkshire and Surrey Pathology Services Lighthouse Laboratory and Alex Alderton, Roberto Amato, Jeffrey Barrett, Sonia Goncalves, Ewan Harrison, David K. Jackson, Ian Johnston, Dominic Kwiatkowski, Cordelia Langford, John Sillitoe on behalf of the Wellcome Sanger Institute COVID-19 Surveillance Team |
| EPI_ISL_3336628, EPI_ISL_3336629 | Lighthouse Lab in Milton Keynes | Wellcome Sanger Institute for the COVID-19 Genomics UK (COG-UK) Consortium | The Lighthouse Lab in Milton Keynes and Alex Alderton, Roberto Amato, Jeffrey Barrett, Sonia Goncalves, Ewan Harrison, David K. Jackson, Ian Johnston, Dominic Kwiatkowski, Cordelia Langford, John Sillitoe on behalf of the Wellcome Sanger Institute COVID-19 Surveillance Team |
| EPI_ISL_3336630 | Lighthouse Lab in Alderley Park | Wellcome Sanger Institute for the COVID-19 Genomics UK (COG-UK) Consortium | Jacquelyn Wynn, Mairead Hyland, The Lighthouse Lab in Alderley Park and Alex Alderton, Roberto Amato, Jeffrey Barrett, Sonia Goncalves, Ewan Harrison, David K. Jackson, Ian Johnston, Dominic Kwiatkowski, Cordelia Langford, John Sillitoe on behalf of the Wellcome Sanger Institute COVID-19 Surveillance Team |
| EPI_ISL_3336631 | Berkshire and Surrey Pathology Services Lighthouse Laboratory | Wellcome Sanger Institute for the COVID-19 Genomics UK (COG-UK) Consortium | Berkshire and Surrey Pathology Services Lighthouse Laboratory and Alex Alderton, Roberto Amato, Jeffrey Barrett, Sonia Goncalves, Ewan Harrison, David K. Jackson, Ian Johnston, Dominic Kwiatkowski, Cordelia Langford, John Sillitoe on behalf of the Wellcome Sanger Institute COVID-19 Surveillance Team |
| EPI_ISL_3336632 | Lighthouse Lab in Alderley Park | Wellcome Sanger Institute for the COVID-19 Genomics UK (COG-UK) Consortium | Jacquelyn Wynn, Mairead Hyland, The Lighthouse Lab in Alderley Park and Alex Alderton, Roberto Amato, Jeffrey Barrett, Sonia Goncalves, Ewan Harrison, David K. Jackson, Ian Johnston, Dominic Kwiatkowski, Cordelia Langford, John Sillitoe on behalf of the Wellcome Sanger Institute COVID-19 Surveillance Team |
| EPI_ISL_3336633 | Berkshire and Surrey Pathology Services Lighthouse Laboratory | Wellcome Sanger Institute for the COVID-19 Genomics UK (COG-UK) Consortium | Berkshire and Surrey Pathology Services Lighthouse Laboratory and Alex Alderton, Roberto Amato, Jeffrey Barrett, Sonia Goncalves, Ewan Harrison, David K. Jackson, Ian Johnston, Dominic Kwiatkowski, Cordelia Langford, John Sillitoe on behalf of the Wellcome Sanger Institute COVID-19 Surveillance Team |
| EPI_ISL_3336634 | Lighthouse Lab in Alderley Park | Wellcome Sanger Institute for the COVID-19 Genomics UK (COG-UK) Consortium | Jacquelyn Wynn, Mairead Hyland, The Lighthouse Lab in Alderley Park and Alex Alderton, Roberto Amato, Jeffrey Barrett, Sonia Goncalves, Ewan Harrison, David K. Jackson, Ian Johnston, Dominic Kwiatkowski, Cordelia Langford, John Sillitoe on behalf of the Wellcome Sanger Institute COVID-19 Surveillance Team |
| EPI_ISL_3336635, EPI_ISL_3336636 | Lighthouse Lab in Milton Keynes | Wellcome Sanger Institute for the COVID-19 Genomics UK (COG-UK) Consortium | The Lighthouse Lab in Milton Keynes and Alex Alderton, Roberto Amato, Jeffrey Barrett, Sonia Goncalves, Ewan Harrison, David K. Jackson, Ian Johnston, Dominic Kwiatkowski, Cordelia Langford, John Sillitoe on behalf of the Wellcome Sanger Institute COVID-19 Surveillance Team |
| EPI_ISL_3336637 | Lighthouse Laboratory Plymouth | Wellcome Sanger Institute for the COVID-19 Genomics UK (COG-UK) Consortium | Lighthouse Laboratory Plymouth and Alex Alderton, Roberto Amato, Jeffrey Barrett, Sonia Goncalves, Ewan Harrison, David K. Jackson, Ian Johnston, Dominic Kwiatkowski, Cordelia Langford, John Sillitoe on behalf of the Wellcome Sanger Institute COVID-19 Surveillance Team |
| EPI_ISL_3336638 | Berkshire and Surrey Pathology Services Lighthouse Laboratory | Wellcome Sanger Institute for the COVID-19 Genomics UK (COG-UK) Consortium | Berkshire and Surrey Pathology Services Lighthouse Laboratory and Alex Alderton, Roberto Amato, Jeffrey Barrett, Sonia Goncalves, Ewan Harrison, David K. Jackson, Ian Johnston, Dominic Kwiatkowski, Cordelia Langford, John Sillitoe on behalf of the Wellcome Sanger Institute COVID-19 Surveillance Team |
| EPI_ISL_3336639 | Lighthouse Lab in Alderley Park | Wellcome Sanger Institute for the COVID-19 Genomics UK (COG-UK) Consortium | Jacquelyn Wynn, Mairead Hyland, The Lighthouse Lab in Alderley Park and Alex Alderton, Roberto Amato, Jeffrey Barrett, Sonia Goncalves, Ewan Harrison, David K. Jackson, Ian Johnston, Dominic Kwiatkowski, Cordelia Langford, John Sillitoe on behalf of the Wellcome Sanger Institute COVID-19 Surveillance Team |
| EPI_ISL_3336640, EPI_ISL_3336641 | Lighthouse Laboratory Plymouth | Wellcome Sanger Institute for the COVID-19 Genomics UK (COG-UK) Consortium | Lighthouse Laboratory Plymouth and Alex Alderton, Roberto Amato, Jeffrey Barrett, Sonia Goncalves, Ewan Harrison, David K. Jackson, Ian Johnston, Dominic Kwiatkowski, Cordelia Langford, John Sillitoe on behalf of the Wellcome Sanger Institute COVID-19 Surveillance Team |
| EPI_ISL_3336642 | Berkshire and Surrey Pathology Services Lighthouse Laboratory | Wellcome Sanger Institute for the COVID-19 Genomics UK (COG-UK) Consortium | Berkshire and Surrey Pathology Services Lighthouse Laboratory and Alex Alderton, Roberto Amato, Jeffrey Barrett, Sonia Goncalves, Ewan Harrison, David K. Jackson, Ian Johnston, Dominic Kwiatkowski, Cordelia Langford, John Sillitoe on behalf of the Wellcome Sanger Institute COVID-19 Surveillance Team |
| EPI_ISL_3336643 | Lighthouse Lab in Milton Keynes | Wellcome Sanger Institute for the COVID-19 Genomics UK (COG-UK) Consortium | The Lighthouse Lab in Milton Keynes and Alex Alderton, Roberto Amato, Jeffrey Barrett, Sonia Goncalves, Ewan Harrison, David K. Jackson, Ian Johnston, Dominic Kwiatkowski, Cordelia Langford, John Sillitoe on behalf of the Wellcome Sanger Institute COVID-19 Surveillance Team |
| EPI_ISL_3336644 | Lighthouse Lab in Alderley Park | Wellcome Sanger Institute for the COVID-19 Genomics UK (COG-UK) Consortium | Jacquelyn Wynn, Mairead Hyland, The Lighthouse Lab in Alderley Park and Alex Alderton, Roberto Amato, Jeffrey Barrett, Sonia Goncalves, Ewan Harrison, David K. Jackson, Ian Johnston, Dominic Kwiatkowski, Cordelia Langford, John Sillitoe on behalf of the Wellcome Sanger Institute COVID-19 Surveillance Team |
| EPI_ISL_3336645 | Lighthouse Lab in Milton Keynes | Wellcome Sanger Institute for the COVID-19 Genomics UK (COG-UK) Consortium | The Lighthouse Lab in Milton Keynes and Alex Alderton, Roberto Amato, Jeffrey Barrett, Sonia Goncalves, Ewan Harrison, David K. Jackson, Ian Johnston, Dominic Kwiatkowski, Cordelia Langford, John Sillitoe on behalf of the Wellcome Sanger Institute COVID-19 Surveillance Team |
| EPI_ISL_3336646 | Lighthouse Laboratory Plymouth | Wellcome Sanger Institute for the COVID-19 Genomics UK (COG-UK) Consortium | Lighthouse Laboratory Plymouth and Alex Alderton, Roberto Amato, Jeffrey Barrett, Sonia Goncalves, Ewan Harrison, David K. Jackson, Ian Johnston, Dominic Kwiatkowski, Cordelia Langford, John Sillitoe on behalf of the Wellcome Sanger Institute COVID-19 Surveillance Team |
| EPI_ISL_3336647 | Berkshire and Surrey Pathology Services Lighthouse Laboratory | Wellcome Sanger Institute for the COVID-19 Genomics UK (COG-UK) Consortium | Berkshire and Surrey Pathology Services Lighthouse Laboratory and Alex Alderton, Roberto Amato, Jeffrey Barrett, Sonia Goncalves, Ewan Harrison, David K. Jackson, Ian Johnston, Dominic Kwiatkowski, Cordelia Langford, John Sillitoe on behalf of the Wellcome Sanger Institute COVID-19 Surveillance Team |
| EPI_ISL_3336648 | Lighthouse Laboratory Plymouth | Wellcome Sanger Institute for the COVID-19 Genomics UK (COG-UK) Consortium | Lighthouse Laboratory Plymouth and Alex Alderton, Roberto Amato, Jeffrey Barrett, Sonia Goncalves, Ewan Harrison, David K. Jackson, Ian Johnston, Dominic Kwiatkowski, Cordelia Langford, John Sillitoe on behalf of the Wellcome Sanger Institute COVID-19 Surveillance Team |
| EPI_ISL_3336649 | Berkshire and Surrey Pathology Services Lighthouse Laboratory | Wellcome Sanger Institute for the COVID-19 Genomics UK (COG-UK) Consortium | Berkshire and Surrey Pathology Services Lighthouse Laboratory and Alex Alderton, Roberto Amato, Jeffrey Barrett, Sonia Goncalves, Ewan Harrison, David K. Jackson, Ian Johnston, Dominic Kwiatkowski, Cordelia Langford, John Sillitoe on behalf of the Wellcome Sanger Institute COVID-19 Surveillance Team |
| EPI_ISL_3336650 | Lighthouse Lab in Milton Keynes | Wellcome Sanger Institute for the COVID-19 Genomics UK (COG-UK) Consortium | The Lighthouse Lab in Milton Keynes and Alex Alderton, Roberto Amato, Jeffrey Barrett, Sonia Goncalves, Ewan Harrison, David K. Jackson, Ian Johnston, Dominic Kwiatkowski, Cordelia Langford, John Sillitoe on behalf of the Wellcome Sanger Institute COVID-19 Surveillance Team |
| EPI_ISL_3336651 | Berkshire and Surrey Pathology Services Lighthouse Laboratory | Wellcome Sanger Institute for the COVID-19 Genomics UK (COG-UK) Consortium | Berkshire and Surrey Pathology Services Lighthouse Laboratory and Alex Alderton, Roberto Amato, Jeffrey Barrett, Sonia Goncalves, Ewan Harrison, David K. Jackson, Ian Johnston, Dominic Kwiatkowski, Cordelia Langford, John Sillitoe on behalf of the Wellcome Sanger Institute COVID-19 Surveillance Team |
| EPI_ISL_3336652 | Lighthouse Lab in Alderley Park | Wellcome Sanger Institute for the COVID-19 Genomics UK (COG-UK) Consortium | Jacquelyn Wynn, Mairead Hyland, The Lighthouse Lab in Alderley Park and Alex Alderton, Roberto Amato, Jeffrey Barrett, Sonia Goncalves, Ewan Harrison, David K. Jackson, Ian Johnston, Dominic Kwiatkowski, Cordelia Langford, John Sillitoe on behalf of the Wellcome Sanger Institute COVID-19 Surveillance Team |
| EPI_ISL_3336653 | Lighthouse Lab in Milton Keynes | Wellcome Sanger Institute for the COVID-19 Genomics UK (COG-UK) Consortium | The Lighthouse Lab in Milton Keynes and Alex Alderton, Roberto Amato, Jeffrey Barrett, Sonia Goncalves, Ewan Harrison, David K. Jackson, Ian Johnston, Dominic Kwiatkowski, Cordelia Langford, John Sillitoe on behalf of the Wellcome Sanger Institute COVID-19 Surveillance Team |
| EPI_ISL_3336654 | Berkshire and Surrey Pathology Services Lighthouse Laboratory | Wellcome Sanger Institute for the COVID-19 Genomics UK (COG-UK) Consortium | Berkshire and Surrey Pathology Services Lighthouse Laboratory and Alex Alderton, Roberto Amato, Jeffrey Barrett, Sonia Goncalves, Ewan Harrison, David K. Jackson, Ian Johnston, Dominic Kwiatkowski, Cordelia Langford, John Sillitoe on behalf of the Wellcome Sanger Institute COVID-19 Surveillance Team |
| EPI_ISL_3336655 | Lighthouse Laboratory Plymouth | Wellcome Sanger Institute for the COVID-19 Genomics UK (COG-UK) Consortium | Lighthouse Laboratory Plymouth and Alex Alderton, Roberto Amato, Jeffrey Barrett, Sonia Goncalves, Ewan Harrison, David K. Jackson, Ian Johnston, Dominic Kwiatkowski, Cordelia Langford, John Sillitoe on behalf of the Wellcome Sanger Institute COVID-19 Surveillance Team |
| EPI_ISL_3336656 | Lighthouse Lab in Milton Keynes | Wellcome Sanger Institute for the COVID-19 Genomics UK (COG-UK) Consortium | The Lighthouse Lab in Milton Keynes and Alex Alderton, Roberto Amato, Jeffrey Barrett, Sonia Goncalves, Ewan Harrison, David K. Jackson, Ian Johnston, Dominic Kwiatkowski, Cordelia Langford, John Sillitoe on behalf of the Wellcome Sanger Institute COVID-19 Surveillance Team |
| EPI_ISL_3336657 | Berkshire and Surrey Pathology Services Lighthouse Laboratory | Wellcome Sanger Institute for the COVID-19 Genomics UK (COG-UK) Consortium | Berkshire and Surrey Pathology Services Lighthouse Laboratory and Alex Alderton, Roberto Amato, Jeffrey Barrett, Sonia Goncalves, Ewan Harrison, David K. Jackson, Ian Johnston, Dominic Kwiatkowski, Cordelia Langford, John Sillitoe on behalf of the Wellcome Sanger Institute COVID-19 Surveillance Team |
| EPI_ISL_3336658 | Lighthouse Laboratory Plymouth | Wellcome Sanger Institute for the COVID-19 Genomics UK (COG-UK) Consortium | Lighthouse Laboratory Plymouth and Alex Alderton, Roberto Amato, Jeffrey Barrett, Sonia Goncalves, Ewan Harrison, David K. Jackson, Ian Johnston, Dominic Kwiatkowski, Cordelia Langford, John Sillitoe on behalf of the Wellcome Sanger Institute COVID-19 Surveillance Team |
| EPI_ISL_3336659, | Lighthouse Lab in Alderley Park | Wellcome Sanger Institute for the COVID-19 Genomics UK (COG-UK) | Jacquelyn Wynn, Mairead Hyland, The Lighthouse Lab in Alderley Park and Alex Alderton, Roberto Amato, Jeffrey Barrett, Sonia Goncalves, Ewan Harrison, David K. Jackson, Ian Johnston, Dominic Kwiatkowski, Cordelia Langford, John Sillitoe on |

| | | | |
|---|---|---|---|
| EPI_ISL_3336660 | | Consortium | behalf of the Wellcome Sanger Institute COVID-19 Surveillance Team |
| EPI_ISL_3336661, EPI_ISL_3336662 | Lighthouse Lab in Milton Keynes | Wellcome Sanger Institute for the COVID-19 Genomics UK (COG-UK) Consortium | The Lighthouse Lab in Milton Keynes and Alex Alderton, Roberto Amato, Jeffrey Barrett, Sonia Goncalves, Ewan Harrison, David K. Jackson, Ian Johnston, Dominic Kwiatkowski, Cordelia Langford, John Sillitoe on behalf of the Wellcome Sanger Institute COVID-19 Surveillance Team |
| EPI_ISL_3336663 | Berkshire and Surrey Pathology Services Lighthouse Laboratory | Wellcome Sanger Institute for the COVID-19 Genomics UK (COG-UK) Consortium | Berkshire and Surrey Pathology Services Lighthouse Laboratory and Alex Alderton, Roberto Amato, Jeffrey Barrett, Sonia Goncalves, Ewan Harrison, David K. Jackson, Ian Johnston, Dominic Kwiatkowski, Cordelia Langford, John Sillitoe on behalf of the Wellcome Sanger Institute COVID-19 Surveillance Team |
| EPI_ISL_3336664 | Lighthouse Lab in Alderley Park | Wellcome Sanger Institute for the COVID-19 Genomics UK (COG-UK) Consortium | Jacquelyn Wynn, Mairead Hyland, The Lighthouse Lab in Alderley Park and Alex Alderton, Roberto Amato, Jeffrey Barrett, Sonia Goncalves, Ewan Harrison, David K. Jackson, Ian Johnston, Dominic Kwiatkowski, Cordelia Langford, John Sillitoe on behalf of the Wellcome Sanger Institute COVID-19 Surveillance Team |
| EPI_ISL_3336665 | Lighthouse Lab in Milton Keynes | Wellcome Sanger Institute for the COVID-19 Genomics UK (COG-UK) Consortium | The Lighthouse Lab in Milton Keynes and Alex Alderton, Roberto Amato, Jeffrey Barrett, Sonia Goncalves, Ewan Harrison, David K. Jackson, Ian Johnston, Dominic Kwiatkowski, Cordelia Langford, John Sillitoe on behalf of the Wellcome Sanger Institute COVID-19 Surveillance Team |
| EPI_ISL_3336666 | Lighthouse Laboratory Plymouth | Wellcome Sanger Institute for the COVID-19 Genomics UK (COG-UK) Consortium | Lighthouse Laboratory Plymouth and Alex Alderton, Roberto Amato, Jeffrey Barrett, Sonia Goncalves, Ewan Harrison, David K. Jackson, Ian Johnston, Dominic Kwiatkowski, Cordelia Langford, John Sillitoe on behalf of the Wellcome Sanger Institute COVID-19 Surveillance Team |
| EPI_ISL_3336667 | Berkshire and Surrey Pathology Services Lighthouse Laboratory | Wellcome Sanger Institute for the COVID-19 Genomics UK (COG-UK) Consortium | Berkshire and Surrey Pathology Services Lighthouse Laboratory and Alex Alderton, Roberto Amato, Jeffrey Barrett, Sonia Goncalves, Ewan Harrison, David K. Jackson, Ian Johnston, Dominic Kwiatkowski, Cordelia Langford, John Sillitoe on behalf of the Wellcome Sanger Institute COVID-19 Surveillance Team |
| EPI_ISL_3336668 | Lighthouse Laboratory Plymouth | Wellcome Sanger Institute for the COVID-19 Genomics UK (COG-UK) Consortium | Lighthouse Laboratory Plymouth and Alex Alderton, Roberto Amato, Jeffrey Barrett, Sonia Goncalves, Ewan Harrison, David K. Jackson, Ian Johnston, Dominic Kwiatkowski, Cordelia Langford, John Sillitoe on behalf of the Wellcome Sanger Institute COVID-19 Surveillance Team |
| EPI_ISL_3336669, EPI_ISL_3336670 | Lighthouse Lab in Milton Keynes | Wellcome Sanger Institute for the COVID-19 Genomics UK (COG-UK) Consortium | The Lighthouse Lab in Milton Keynes and Alex Alderton, Roberto Amato, Jeffrey Barrett, Sonia Goncalves, Ewan Harrison, David K. Jackson, Ian Johnston, Dominic Kwiatkowski, Cordelia Langford, John Sillitoe on behalf of the Wellcome Sanger Institute COVID-19 Surveillance Team |
| EPI_ISL_3336671, EPI_ISL_3336672 | Lighthouse Lab in Alderley Park | Wellcome Sanger Institute for the COVID-19 Genomics UK (COG-UK) Consortium | Jacquelyn Wynn, Mairead Hyland, The Lighthouse Lab in Alderley Park and Alex Alderton, Roberto Amato, Jeffrey Barrett, Sonia Goncalves, Ewan Harrison, David K. Jackson, Ian Johnston, Dominic Kwiatkowski, Cordelia Langford, John Sillitoe on behalf of the Wellcome Sanger Institute COVID-19 Surveillance Team |
| EPI_ISL_3336673 | Lighthouse Laboratory Plymouth | Wellcome Sanger Institute for the COVID-19 Genomics UK (COG-UK) Consortium | Lighthouse Laboratory Plymouth and Alex Alderton, Roberto Amato, Jeffrey Barrett, Sonia Goncalves, Ewan Harrison, David K. Jackson, Ian Johnston, Dominic Kwiatkowski, Cordelia Langford, John Sillitoe on behalf of the Wellcome Sanger Institute COVID-19 Surveillance Team |
| EPI_ISL_3336674, EPI_ISL_3336675 | Lighthouse Lab in Milton Keynes | Wellcome Sanger Institute for the COVID-19 Genomics UK (COG-UK) Consortium | The Lighthouse Lab in Milton Keynes and Alex Alderton, Roberto Amato, Jeffrey Barrett, Sonia Goncalves, Ewan Harrison, David K. Jackson, Ian Johnston, Dominic Kwiatkowski, Cordelia Langford, John Sillitoe on behalf of the Wellcome Sanger Institute COVID-19 Surveillance Team |
| EPI_ISL_3336676, EPI_ISL_3336677 | Lighthouse Laboratory Plymouth | Wellcome Sanger Institute for the COVID-19 Genomics UK (COG-UK) Consortium | Lighthouse Laboratory Plymouth and Alex Alderton, Roberto Amato, Jeffrey Barrett, Sonia Goncalves, Ewan Harrison, David K. Jackson, Ian Johnston, Dominic Kwiatkowski, Cordelia Langford, John Sillitoe on behalf of the Wellcome Sanger Institute COVID-19 Surveillance Team |
| EPI_ISL_3336678 | Berkshire and Surrey Pathology Services Lighthouse Laboratory | Wellcome Sanger Institute for the COVID-19 Genomics UK (COG-UK) Consortium | Berkshire and Surrey Pathology Services Lighthouse Laboratory and Alex Alderton, Roberto Amato, Jeffrey Barrett, Sonia Goncalves, Ewan Harrison, David K. Jackson, Ian Johnston, Dominic Kwiatkowski, Cordelia Langford, John Sillitoe on behalf of the Wellcome Sanger Institute COVID-19 Surveillance Team |
| EPI_ISL_3336679 | Lighthouse Lab in Alderley Park | Wellcome Sanger Institute for the COVID-19 Genomics UK (COG-UK) Consortium | Jacquelyn Wynn, Mairead Hyland, The Lighthouse Lab in Alderley Park and Alex Alderton, Roberto Amato, Jeffrey Barrett, Sonia Goncalves, Ewan Harrison, David K. Jackson, Ian Johnston, Dominic Kwiatkowski, Cordelia Langford, John Sillitoe on behalf of the Wellcome Sanger Institute COVID-19 Surveillance Team |
| EPI_ISL_3336680 | Lighthouse Laboratory Plymouth | Wellcome Sanger Institute for the COVID-19 Genomics UK (COG-UK) Consortium | Lighthouse Laboratory Plymouth and Alex Alderton, Roberto Amato, Jeffrey Barrett, Sonia Goncalves, Ewan Harrison, David K. Jackson, Ian Johnston, Dominic Kwiatkowski, Cordelia Langford, John Sillitoe on behalf of the Wellcome Sanger Institute COVID-19 Surveillance Team |
| EPI_ISL_3336681 | Lighthouse Lab in Alderley Park | Wellcome Sanger Institute for the COVID-19 Genomics UK (COG-UK) Consortium | Jacquelyn Wynn, Mairead Hyland, The Lighthouse Lab in Alderley Park and Alex Alderton, Roberto Amato, Jeffrey Barrett, Sonia Goncalves, Ewan Harrison, David K. Jackson, Ian Johnston, Dominic Kwiatkowski, Cordelia Langford, John Sillitoe on behalf of the Wellcome Sanger Institute COVID-19 Surveillance Team |
| EPI_ISL_3336682 | Lighthouse Laboratory Plymouth | Wellcome Sanger Institute for the COVID-19 Genomics UK (COG-UK) Consortium | Lighthouse Laboratory Plymouth and Alex Alderton, Roberto Amato, Jeffrey Barrett, Sonia Goncalves, Ewan Harrison, David K. Jackson, Ian Johnston, Dominic Kwiatkowski, Cordelia Langford, John Sillitoe on behalf of the Wellcome Sanger Institute COVID-19 Surveillance Team |
| EPI_ISL_3336683 | Berkshire and Surrey Pathology Services Lighthouse Laboratory | Wellcome Sanger Institute for the COVID-19 Genomics UK (COG-UK) Consortium | Berkshire and Surrey Pathology Services Lighthouse Laboratory and Alex Alderton, Roberto Amato, Jeffrey Barrett, Sonia Goncalves, Ewan Harrison, David K. Jackson, Ian Johnston, Dominic Kwiatkowski, Cordelia Langford, John Sillitoe on behalf of the Wellcome Sanger Institute COVID-19 Surveillance Team |
| EPI_ISL_3336684 | Lighthouse Lab in Alderley Park | Wellcome Sanger Institute for the COVID-19 Genomics UK (COG-UK) Consortium | Jacquelyn Wynn, Mairead Hyland, The Lighthouse Lab in Alderley Park and Alex Alderton, Roberto Amato, Jeffrey Barrett, Sonia Goncalves, Ewan Harrison, David K. Jackson, Ian Johnston, Dominic Kwiatkowski, Cordelia Langford, John Sillitoe on behalf of the Wellcome Sanger Institute COVID-19 Surveillance Team |
| EPI_ISL_3336685 | Lighthouse Laboratory Plymouth | Wellcome Sanger Institute for the COVID-19 Genomics UK (COG-UK) Consortium | Lighthouse Laboratory Plymouth and Alex Alderton, Roberto Amato, Jeffrey Barrett, Sonia Goncalves, Ewan Harrison, David K. Jackson, Ian Johnston, Dominic Kwiatkowski, Cordelia Langford, John Sillitoe on behalf of the Wellcome Sanger Institute COVID-19 Surveillance Team |
| EPI_ISL_3336686, EPI_ISL_3336687 | Lighthouse Lab in Milton Keynes | Wellcome Sanger Institute for the COVID-19 Genomics UK (COG-UK) Consortium | The Lighthouse Lab in Milton Keynes and Alex Alderton, Roberto Amato, Jeffrey Barrett, Sonia Goncalves, Ewan Harrison, David K. Jackson, Ian Johnston, Dominic Kwiatkowski, Cordelia Langford, John Sillitoe on behalf of the Wellcome Sanger Institute COVID-19 Surveillance Team |
| EPI_ISL_3336688 | Lighthouse Lab in Alderley Park | Wellcome Sanger Institute for the COVID-19 Genomics UK (COG-UK) Consortium | Jacquelyn Wynn, Mairead Hyland, The Lighthouse Lab in Alderley Park and Alex Alderton, Roberto Amato, Jeffrey Barrett, Sonia Goncalves, Ewan Harrison, David K. Jackson, Ian Johnston, Dominic Kwiatkowski, Cordelia Langford, John Sillitoe on behalf of the Wellcome Sanger Institute COVID-19 Surveillance Team |
| EPI_ISL_3336689 | Lighthouse Lab in Milton Keynes | Wellcome Sanger Institute for the COVID-19 Genomics UK (COG-UK) Consortium | The Lighthouse Lab in Milton Keynes and Alex Alderton, Roberto Amato, Jeffrey Barrett, Sonia Goncalves, Ewan Harrison, David K. Jackson, Ian Johnston, Dominic Kwiatkowski, Cordelia Langford, John Sillitoe on behalf of the Wellcome Sanger Institute COVID-19 Surveillance Team |
| EPI_ISL_3336690 | Lighthouse Lab in Alderley Park | Wellcome Sanger Institute for the COVID-19 Genomics UK (COG-UK) Consortium | Jacquelyn Wynn, Mairead Hyland, The Lighthouse Lab in Alderley Park and Alex Alderton, Roberto Amato, Jeffrey Barrett, Sonia Goncalves, Ewan Harrison, David K. Jackson, Ian Johnston, Dominic Kwiatkowski, Cordelia Langford, John Sillitoe on behalf of the Wellcome Sanger Institute COVID-19 Surveillance Team |
| EPI_ISL_3336691 | Lighthouse Laboratory Plymouth | Wellcome Sanger Institute for the COVID-19 Genomics UK (COG-UK) Consortium | Lighthouse Laboratory Plymouth and Alex Alderton, Roberto Amato, Jeffrey Barrett, Sonia Goncalves, Ewan Harrison, David K. Jackson, Ian Johnston, Dominic Kwiatkowski, Cordelia Langford, John Sillitoe on behalf of the Wellcome Sanger Institute COVID-19 Surveillance Team |
| EPI_ISL_3336692 | Lighthouse Lab in Milton Keynes | Wellcome Sanger Institute for the COVID-19 Genomics UK (COG-UK) Consortium | The Lighthouse Lab in Milton Keynes and Alex Alderton, Roberto Amato, Jeffrey Barrett, Sonia Goncalves, Ewan Harrison, David K. Jackson, Ian Johnston, Dominic Kwiatkowski, Cordelia Langford, John Sillitoe on behalf of the Wellcome Sanger Institute COVID-19 Surveillance Team |
| EPI_ISL_3336693 | Berkshire and Surrey Pathology Services Lighthouse Laboratory | Wellcome Sanger Institute for the COVID-19 Genomics UK (COG-UK) Consortium | Berkshire and Surrey Pathology Services Lighthouse Laboratory and Alex Alderton, Roberto Amato, Jeffrey Barrett, Sonia Goncalves, Ewan Harrison, David K. Jackson, Ian Johnston, Dominic Kwiatkowski, Cordelia Langford, John Sillitoe on behalf of the Wellcome Sanger Institute COVID-19 Surveillance Team |
| EPI_ISL_3336694 | Lighthouse Lab in Milton Keynes | Wellcome Sanger Institute for the COVID-19 Genomics UK (COG-UK) Consortium | The Lighthouse Lab in Milton Keynes and Alex Alderton, Roberto Amato, Jeffrey Barrett, Sonia Goncalves, Ewan Harrison, David K. Jackson, Ian Johnston, Dominic Kwiatkowski, Cordelia Langford, John Sillitoe on behalf of the Wellcome Sanger Institute COVID-19 Surveillance Team |
| EPI_ISL_3336695 | Lighthouse Laboratory Plymouth | Wellcome Sanger Institute for the COVID-19 Genomics UK (COG-UK) Consortium | Lighthouse Laboratory Plymouth and Alex Alderton, Roberto Amato, Jeffrey Barrett, Sonia Goncalves, Ewan Harrison, David K. Jackson, Ian Johnston, Dominic Kwiatkowski, Cordelia Langford, John Sillitoe on behalf of the Wellcome Sanger Institute COVID-19 Surveillance Team |
| EPI_ISL_3336696 | Berkshire and Surrey Pathology Services Lighthouse Laboratory | Wellcome Sanger Institute for the COVID-19 Genomics UK (COG-UK) Consortium | Berkshire and Surrey Pathology Services Lighthouse Laboratory and Alex Alderton, Roberto Amato, Jeffrey Barrett, Sonia Goncalves, Ewan Harrison, David K. Jackson, Ian Johnston, Dominic Kwiatkowski, Cordelia Langford, John Sillitoe on behalf of the Wellcome Sanger Institute COVID-19 Surveillance Team |
| EPI_ISL_3336697, EPI_ISL_3336698, EPI_ISL_3336699 | Lighthouse Lab in Milton Keynes | Wellcome Sanger Institute for the COVID-19 Genomics UK (COG-UK) Consortium | The Lighthouse Lab in Milton Keynes and Alex Alderton, Roberto Amato, Jeffrey Barrett, Sonia Goncalves, Ewan Harrison, David K. Jackson, Ian Johnston, Dominic Kwiatkowski, Cordelia Langford, John Sillitoe on behalf of the Wellcome Sanger Institute COVID-19 Surveillance Team |
| EPI_ISL_3336700 | Berkshire and Surrey Pathology Services Lighthouse Laboratory | Wellcome Sanger Institute for the COVID-19 Genomics UK (COG-UK) Consortium | Berkshire and Surrey Pathology Services Lighthouse Laboratory and Alex Alderton, Roberto Amato, Jeffrey Barrett, Sonia Goncalves, Ewan Harrison, David K. Jackson, Ian Johnston, Dominic Kwiatkowski, Cordelia Langford, John Sillitoe on behalf of the Wellcome Sanger Institute COVID-19 Surveillance Team |
| EPI_ISL_3336701 | Lighthouse Lab in Alderley Park | Wellcome Sanger Institute for the COVID-19 Genomics UK (COG-UK) Consortium | Jacquelyn Wynn, Mairead Hyland, The Lighthouse Lab in Alderley Park and Alex Alderton, Roberto Amato, Jeffrey Barrett, Sonia Goncalves, Ewan Harrison, David K. Jackson, Ian Johnston, Dominic Kwiatkowski, Cordelia Langford, John Sillitoe on behalf of the Wellcome Sanger Institute COVID-19 Surveillance Team |
| EPI_ISL_3336702, EPI_ISL_3336703 | Lighthouse Lab in Milton Keynes | Wellcome Sanger Institute for the COVID-19 Genomics UK (COG-UK) Consortium | The Lighthouse Lab in Milton Keynes and Alex Alderton, Roberto Amato, Jeffrey Barrett, Sonia Goncalves, Ewan Harrison, David K. Jackson, Ian Johnston, Dominic Kwiatkowski, Cordelia Langford, John Sillitoe on behalf of the Wellcome Sanger Institute COVID-19 Surveillance Team |
| EPI_ISL_3336704 | Lighthouse Laboratory Plymouth | Wellcome Sanger Institute for the COVID-19 Genomics UK (COG-UK) Consortium | Lighthouse Laboratory Plymouth and Alex Alderton, Roberto Amato, Jeffrey Barrett, Sonia Goncalves, Ewan Harrison, David K. Jackson, Ian Johnston, Dominic Kwiatkowski, Cordelia Langford, John Sillitoe on behalf of the Wellcome Sanger Institute COVID-19 Surveillance Team |
| EPI_ISL_3336705 | Lighthouse Lab in Milton Keynes | Wellcome Sanger Institute for the COVID-19 Genomics UK (COG-UK) Consortium | The Lighthouse Lab in Milton Keynes and Alex Alderton, Roberto Amato, Jeffrey Barrett, Sonia Goncalves, Ewan Harrison, David K. Jackson, Ian Johnston, Dominic Kwiatkowski, Cordelia Langford, John Sillitoe on behalf of the Wellcome Sanger Institute COVID-19 Surveillance Team |
| EPI_ISL_3336706, EPI_ISL_3336707 | Lighthouse Lab in Alderley Park | Wellcome Sanger Institute for the COVID-19 Genomics UK (COG-UK) Consortium | Jacquelyn Wynn, Mairead Hyland, The Lighthouse Lab in Alderley Park and Alex Alderton, Roberto Amato, Jeffrey Barrett, Sonia Goncalves, Ewan Harrison, David K. Jackson, Ian Johnston, Dominic Kwiatkowski, Cordelia Langford, John Sillitoe on behalf of the Wellcome Sanger Institute COVID-19 Surveillance Team |
| EPI_ISL_3336708 | Berkshire and Surrey Pathology Services Lighthouse Laboratory | Wellcome Sanger Institute for the COVID-19 Genomics UK (COG-UK) Consortium | Berkshire and Surrey Pathology Services Lighthouse Laboratory and Alex Alderton, Roberto Amato, Jeffrey Barrett, Sonia Goncalves, Ewan Harrison, David K. Jackson, Ian Johnston, Dominic Kwiatkowski, Cordelia Langford, John Sillitoe on behalf of the Wellcome Sanger Institute COVID-19 Surveillance Team |
| EPI_ISL_3336709 | Lighthouse Lab in Milton Keynes | Wellcome Sanger Institute for the COVID-19 Genomics UK (COG-UK) Consortium | The Lighthouse Lab in Milton Keynes and Alex Alderton, Roberto Amato, Jeffrey Barrett, Sonia Goncalves, Ewan Harrison, David K. Jackson, Ian Johnston, Dominic Kwiatkowski, Cordelia Langford, John Sillitoe on behalf of the Wellcome Sanger Institute COVID-19 Surveillance Team |
| EPI_ISL_3336710 | Berkshire and Surrey Pathology Services Lighthouse Laboratory | Wellcome Sanger Institute for the COVID-19 Genomics UK (COG-UK) Consortium | Berkshire and Surrey Pathology Services Lighthouse Laboratory and Alex Alderton, Roberto Amato, Jeffrey Barrett, Sonia Goncalves, Ewan Harrison, David K. Jackson, Ian Johnston, Dominic Kwiatkowski, Cordelia Langford, John Sillitoe on behalf of the Wellcome Sanger Institute COVID-19 Surveillance Team |
| EPI_ISL_3336711 | Lighthouse Lab in Milton Keynes | Wellcome Sanger Institute for the COVID-19 Genomics UK (COG-UK) Consortium | The Lighthouse Lab in Milton Keynes and Alex Alderton, Roberto Amato, Jeffrey Barrett, Sonia Goncalves, Ewan Harrison, David K. Jackson, Ian Johnston, Dominic Kwiatkowski, Cordelia Langford, John Sillitoe on behalf of the Wellcome Sanger Institute COVID-19 Surveillance Team |
| EPI_ISL_3336712 | Berkshire and Surrey Pathology Services Lighthouse Laboratory | Wellcome Sanger Institute for the COVID-19 Genomics UK (COG-UK) Consortium | Berkshire and Surrey Pathology Services Lighthouse Laboratory and Alex Alderton, Roberto Amato, Jeffrey Barrett, Sonia Goncalves, Ewan Harrison, David K. Jackson, Ian Johnston, Dominic Kwiatkowski, Cordelia Langford, John Sillitoe on behalf of the Wellcome Sanger Institute COVID-19 Surveillance Team |
| EPI_ISL_3336713 | Lighthouse Lab in Alderley Park | Wellcome Sanger Institute for the COVID-19 Genomics UK (COG-UK) Consortium | Jacquelyn Wynn, Mairead Hyland, The Lighthouse Lab in Alderley Park and Alex Alderton, Roberto Amato, Jeffrey Barrett, Sonia Goncalves, Ewan Harrison, David K. Jackson, Ian Johnston, Dominic Kwiatkowski, Cordelia Langford, John Sillitoe on behalf of the Wellcome Sanger Institute COVID-19 Surveillance Team |
| EPI_ISL_3336714 | Lighthouse Lab in Milton Keynes | Wellcome Sanger Institute for the COVID-19 Genomics UK (COG-UK) Consortium | The Lighthouse Lab in Milton Keynes and Alex Alderton, Roberto Amato, Jeffrey Barrett, Sonia Goncalves, Ewan Harrison, David K. Jackson, Ian Johnston, Dominic Kwiatkowski, Cordelia Langford, John Sillitoe on behalf of the Wellcome Sanger Institute COVID-19 Surveillance Team |

| | | | |
|---|---|---|---|
| EPI_ISL_3336715 | Lighthouse Laboratory Plymouth | Wellcome Sanger Institute for the COVID-19 Genomics UK (COG-UK) Consortium | Lighthouse Laboratory Plymouth and Alex Alderton, Roberto Amato, Jeffrey Barrett, Sonia Goncalves, Ewan Harrison, David K. Jackson, Ian Johnston, Dominic Kwiatkowski, Cordelia Langford, John Sillitoe on behalf of the Wellcome Sanger Institute COVID-19 Surveillance Team |
| EPI_ISL_3336716 | Lighthouse Lab in Milton Keynes | Wellcome Sanger Institute for the COVID-19 Genomics UK (COG-UK) Consortium | The Lighthouse Lab in Milton Keynes and Alex Alderton, Roberto Amato, Jeffrey Barrett, Sonia Goncalves, Ewan Harrison, David K. Jackson, Ian Johnston, Dominic Kwiatkowski, Cordelia Langford, John Sillitoe on behalf of the Wellcome Sanger Institute COVID-19 Surveillance Team |
| EPI_ISL_3336717 | Berkshire and Surrey Pathology Services Lighthouse Laboratory | Wellcome Sanger Institute for the COVID-19 Genomics UK (COG-UK) Consortium | Berkshire and Surrey Pathology Services Lighthouse Laboratory and Alex Alderton, Roberto Amato, Jeffrey Barrett, Sonia Goncalves, Ewan Harrison, David K. Jackson, Ian Johnston, Dominic Kwiatkowski, Cordelia Langford, John Sillitoe on behalf of the Wellcome Sanger Institute COVID-19 Surveillance Team |
| EPI_ISL_3336718, EPI_ISL_3336719 | Lighthouse Lab in Milton Keynes | Wellcome Sanger Institute for the COVID-19 Genomics UK (COG-UK) Consortium | The Lighthouse Lab in Milton Keynes and Alex Alderton, Roberto Amato, Jeffrey Barrett, Sonia Goncalves, Ewan Harrison, David K. Jackson, Ian Johnston, Dominic Kwiatkowski, Cordelia Langford, John Sillitoe on behalf of the Wellcome Sanger Institute COVID-19 Surveillance Team |
| EPI_ISL_3336720, EPI_ISL_3336721 | Lighthouse Laboratory Plymouth | Wellcome Sanger Institute for the COVID-19 Genomics UK (COG-UK) Consortium | Lighthouse Laboratory Plymouth and Alex Alderton, Roberto Amato, Jeffrey Barrett, Sonia Goncalves, Ewan Harrison, David K. Jackson, Ian Johnston, Dominic Kwiatkowski, Cordelia Langford, John Sillitoe on behalf of the Wellcome Sanger Institute COVID-19 Surveillance Team |
| EPI_ISL_3336722 | Lighthouse Lab in Milton Keynes | Wellcome Sanger Institute for the COVID-19 Genomics UK (COG-UK) Consortium | The Lighthouse Lab in Milton Keynes and Alex Alderton, Roberto Amato, Jeffrey Barrett, Sonia Goncalves, Ewan Harrison, David K. Jackson, Ian Johnston, Dominic Kwiatkowski, Cordelia Langford, John Sillitoe on behalf of the Wellcome Sanger Institute COVID-19 Surveillance Team |
| EPI_ISL_3336723 | Berkshire and Surrey Pathology Services Lighthouse Laboratory | Wellcome Sanger Institute for the COVID-19 Genomics UK (COG-UK) Consortium | Berkshire and Surrey Pathology Services Lighthouse Laboratory and Alex Alderton, Roberto Amato, Jeffrey Barrett, Sonia Goncalves, Ewan Harrison, David K. Jackson, Ian Johnston, Dominic Kwiatkowski, Cordelia Langford, John Sillitoe on behalf of the Wellcome Sanger Institute COVID-19 Surveillance Team |
| EPI_ISL_3336724 | Lighthouse Lab in Alderley Park | Wellcome Sanger Institute for the COVID-19 Genomics UK (COG-UK) Consortium | Jacquelyn Wynn, Mairead Hyland, The Lighthouse Lab in Alderley Park and Alex Alderton, Roberto Amato, Jeffrey Barrett, Sonia Goncalves, Ewan Harrison, David K. Jackson, Ian Johnston, Dominic Kwiatkowski, Cordelia Langford, John Sillitoe on behalf of the Wellcome Sanger Institute COVID-19 Surveillance Team |
| EPI_ISL_3336725 | Berkshire and Surrey Pathology Services Lighthouse Laboratory | Wellcome Sanger Institute for the COVID-19 Genomics UK (COG-UK) Consortium | Berkshire and Surrey Pathology Services Lighthouse Laboratory and Alex Alderton, Roberto Amato, Jeffrey Barrett, Sonia Goncalves, Ewan Harrison, David K. Jackson, Ian Johnston, Dominic Kwiatkowski, Cordelia Langford, John Sillitoe on behalf of the Wellcome Sanger Institute COVID-19 Surveillance Team |
| EPI_ISL_3336726 | Lighthouse Laboratory Plymouth | Wellcome Sanger Institute for the COVID-19 Genomics UK (COG-UK) Consortium | Lighthouse Laboratory Plymouth and Alex Alderton, Roberto Amato, Jeffrey Barrett, Sonia Goncalves, Ewan Harrison, David K. Jackson, Ian Johnston, Dominic Kwiatkowski, Cordelia Langford, John Sillitoe on behalf of the Wellcome Sanger Institute COVID-19 Surveillance Team |
| EPI_ISL_3336727 | Lighthouse Lab in Milton Keynes | Wellcome Sanger Institute for the COVID-19 Genomics UK (COG-UK) Consortium | The Lighthouse Lab in Milton Keynes and Alex Alderton, Roberto Amato, Jeffrey Barrett, Sonia Goncalves, Ewan Harrison, David K. Jackson, Ian Johnston, Dominic Kwiatkowski, Cordelia Langford, John Sillitoe on behalf of the Wellcome Sanger Institute COVID-19 Surveillance Team |
| EPI_ISL_3336728 | Lighthouse Laboratory Plymouth | Wellcome Sanger Institute for the COVID-19 Genomics UK (COG-UK) Consortium | Lighthouse Laboratory Plymouth and Alex Alderton, Roberto Amato, Jeffrey Barrett, Sonia Goncalves, Ewan Harrison, David K. Jackson, Ian Johnston, Dominic Kwiatkowski, Cordelia Langford, John Sillitoe on behalf of the Wellcome Sanger Institute COVID-19 Surveillance Team |
| EPI_ISL_3336729 | Lighthouse Lab in Alderley Park | Wellcome Sanger Institute for the COVID-19 Genomics UK (COG-UK) Consortium | Jacquelyn Wynn, Mairead Hyland, The Lighthouse Lab in Alderley Park and Alex Alderton, Roberto Amato, Jeffrey Barrett, Sonia Goncalves, Ewan Harrison, David K. Jackson, Ian Johnston, Dominic Kwiatkowski, Cordelia Langford, John Sillitoe on behalf of the Wellcome Sanger Institute COVID-19 Surveillance Team |
| EPI_ISL_3336730 | Lighthouse Laboratory Plymouth | Wellcome Sanger Institute for the COVID-19 Genomics UK (COG-UK) Consortium | Lighthouse Laboratory Plymouth and Alex Alderton, Roberto Amato, Jeffrey Barrett, Sonia Goncalves, Ewan Harrison, David K. Jackson, Ian Johnston, Dominic Kwiatkowski, Cordelia Langford, John Sillitoe on behalf of the Wellcome Sanger Institute COVID-19 Surveillance Team |
| EPI_ISL_3336731 | Lighthouse Lab in Alderley Park | Wellcome Sanger Institute for the COVID-19 Genomics UK (COG-UK) Consortium | Jacquelyn Wynn, Mairead Hyland, The Lighthouse Lab in Alderley Park and Alex Alderton, Roberto Amato, Jeffrey Barrett, Sonia Goncalves, Ewan Harrison, David K. Jackson, Ian Johnston, Dominic Kwiatkowski, Cordelia Langford, John Sillitoe on behalf of the Wellcome Sanger Institute COVID-19 Surveillance Team |
| EPI_ISL_3336732 | Lighthouse Laboratory Plymouth | Wellcome Sanger Institute for the COVID-19 Genomics UK (COG-UK) Consortium | Lighthouse Laboratory Plymouth and Alex Alderton, Roberto Amato, Jeffrey Barrett, Sonia Goncalves, Ewan Harrison, David K. Jackson, Ian Johnston, Dominic Kwiatkowski, Cordelia Langford, John Sillitoe on behalf of the Wellcome Sanger Institute COVID-19 Surveillance Team |
| EPI_ISL_3336733 | Lighthouse Lab in Alderley Park | Wellcome Sanger Institute for the COVID-19 Genomics UK (COG-UK) Consortium | Jacquelyn Wynn, Mairead Hyland, The Lighthouse Lab in Alderley Park and Alex Alderton, Roberto Amato, Jeffrey Barrett, Sonia Goncalves, Ewan Harrison, David K. Jackson, Ian Johnston, Dominic Kwiatkowski, Cordelia Langford, John Sillitoe on behalf of the Wellcome Sanger Institute COVID-19 Surveillance Team |
| EPI_ISL_3336734 | Lighthouse Lab in Milton Keynes | Wellcome Sanger Institute for the COVID-19 Genomics UK (COG-UK) Consortium | The Lighthouse Lab in Milton Keynes and Alex Alderton, Roberto Amato, Jeffrey Barrett, Sonia Goncalves, Ewan Harrison, David K. Jackson, Ian Johnston, Dominic Kwiatkowski, Cordelia Langford, John Sillitoe on behalf of the Wellcome Sanger Institute COVID-19 Surveillance Team |
| EPI_ISL_3336735 | Lighthouse Lab in Alderley Park | Wellcome Sanger Institute for the COVID-19 Genomics UK (COG-UK) Consortium | Jacquelyn Wynn, Mairead Hyland, The Lighthouse Lab in Alderley Park and Alex Alderton, Roberto Amato, Jeffrey Barrett, Sonia Goncalves, Ewan Harrison, David K. Jackson, Ian Johnston, Dominic Kwiatkowski, Cordelia Langford, John Sillitoe on behalf of the Wellcome Sanger Institute COVID-19 Surveillance Team |
| EPI_ISL_3336736 | Lighthouse Lab in Milton Keynes | Wellcome Sanger Institute for the COVID-19 Genomics UK (COG-UK) Consortium | The Lighthouse Lab in Milton Keynes and Alex Alderton, Roberto Amato, Jeffrey Barrett, Sonia Goncalves, Ewan Harrison, David K. Jackson, Ian Johnston, Dominic Kwiatkowski, Cordelia Langford, John Sillitoe on behalf of the Wellcome Sanger Institute COVID-19 Surveillance Team |
| EPI_ISL_3336737 | Lighthouse Lab in Alderley Park | Wellcome Sanger Institute for the COVID-19 Genomics UK (COG-UK) Consortium | Jacquelyn Wynn, Mairead Hyland, The Lighthouse Lab in Alderley Park and Alex Alderton, Roberto Amato, Jeffrey Barrett, Sonia Goncalves, Ewan Harrison, David K. Jackson, Ian Johnston, Dominic Kwiatkowski, Cordelia Langford, John Sillitoe on behalf of the Wellcome Sanger Institute COVID-19 Surveillance Team |
| EPI_ISL_3336738 | Lighthouse Lab in Milton Keynes | Wellcome Sanger Institute for the COVID-19 Genomics UK (COG-UK) Consortium | The Lighthouse Lab in Milton Keynes and Alex Alderton, Roberto Amato, Jeffrey Barrett, Sonia Goncalves, Ewan Harrison, David K. Jackson, Ian Johnston, Dominic Kwiatkowski, Cordelia Langford, John Sillitoe on behalf of the Wellcome Sanger Institute COVID-19 Surveillance Team |
| EPI_ISL_3336739 | Lighthouse Laboratory Plymouth | Wellcome Sanger Institute for the COVID-19 Genomics UK (COG-UK) Consortium | Lighthouse Laboratory Plymouth and Alex Alderton, Roberto Amato, Jeffrey Barrett, Sonia Goncalves, Ewan Harrison, David K. Jackson, Ian Johnston, Dominic Kwiatkowski, Cordelia Langford, John Sillitoe on behalf of the Wellcome Sanger Institute COVID-19 Surveillance Team |
| EPI_ISL_3336740 | Lighthouse Lab in Alderley Park | Wellcome Sanger Institute for the COVID-19 Genomics UK (COG-UK) Consortium | Jacquelyn Wynn, Mairead Hyland, The Lighthouse Lab in Alderley Park and Alex Alderton, Roberto Amato, Jeffrey Barrett, Sonia Goncalves, Ewan Harrison, David K. Jackson, Ian Johnston, Dominic Kwiatkowski, Cordelia Langford, John Sillitoe on behalf of the Wellcome Sanger Institute COVID-19 Surveillance Team |
| EPI_ISL_3336741 | Lighthouse Lab in Milton Keynes | Wellcome Sanger Institute for the COVID-19 Genomics UK (COG-UK) Consortium | The Lighthouse Lab in Milton Keynes and Alex Alderton, Roberto Amato, Jeffrey Barrett, Sonia Goncalves, Ewan Harrison, David K. Jackson, Ian Johnston, Dominic Kwiatkowski, Cordelia Langford, John Sillitoe on behalf of the Wellcome Sanger Institute COVID-19 Surveillance Team |
| EPI_ISL_3336742 | Lighthouse Laboratory Plymouth | Wellcome Sanger Institute for the COVID-19 Genomics UK (COG-UK) Consortium | Lighthouse Laboratory Plymouth and Alex Alderton, Roberto Amato, Jeffrey Barrett, Sonia Goncalves, Ewan Harrison, David K. Jackson, Ian Johnston, Dominic Kwiatkowski, Cordelia Langford, John Sillitoe on behalf of the Wellcome Sanger Institute COVID-19 Surveillance Team |
| EPI_ISL_3336743 | Berkshire and Surrey Pathology Services Lighthouse Laboratory | Wellcome Sanger Institute for the COVID-19 Genomics UK (COG-UK) Consortium | Berkshire and Surrey Pathology Services Lighthouse Laboratory and Alex Alderton, Roberto Amato, Jeffrey Barrett, Sonia Goncalves, Ewan Harrison, David K. Jackson, Ian Johnston, Dominic Kwiatkowski, Cordelia Langford, John Sillitoe on behalf of the Wellcome Sanger Institute COVID-19 Surveillance Team |
| EPI_ISL_3336744, EPI_ISL_3336745 | Lighthouse Laboratory Plymouth | Wellcome Sanger Institute for the COVID-19 Genomics UK (COG-UK) Consortium | Lighthouse Laboratory Plymouth and Alex Alderton, Roberto Amato, Jeffrey Barrett, Sonia Goncalves, Ewan Harrison, David K. Jackson, Ian Johnston, Dominic Kwiatkowski, Cordelia Langford, John Sillitoe on behalf of the Wellcome Sanger Institute COVID-19 Surveillance Team |
| EPI_ISL_3336746 | Berkshire and Surrey Pathology Services Lighthouse Laboratory | Wellcome Sanger Institute for the COVID-19 Genomics UK (COG-UK) Consortium | Berkshire and Surrey Pathology Services Lighthouse Laboratory and Alex Alderton, Roberto Amato, Jeffrey Barrett, Sonia Goncalves, Ewan Harrison, David K. Jackson, Ian Johnston, Dominic Kwiatkowski, Cordelia Langford, John Sillitoe on behalf of the Wellcome Sanger Institute COVID-19 Surveillance Team |
| EPI_ISL_3336747, EPI_ISL_3336748, EPI_ISL_3336749 | Lighthouse Laboratory Plymouth | Wellcome Sanger Institute for the COVID-19 Genomics UK (COG-UK) Consortium | Lighthouse Laboratory Plymouth and Alex Alderton, Roberto Amato, Jeffrey Barrett, Sonia Goncalves, Ewan Harrison, David K. Jackson, Ian Johnston, Dominic Kwiatkowski, Cordelia Langford, John Sillitoe on behalf of the Wellcome Sanger Institute COVID-19 Surveillance Team |
| EPI_ISL_3336750 | Lighthouse Lab in Alderley Park | Wellcome Sanger Institute for the COVID-19 Genomics UK (COG-UK) Consortium | Jacquelyn Wynn, Mairead Hyland, The Lighthouse Lab in Alderley Park and Alex Alderton, Roberto Amato, Jeffrey Barrett, Sonia Goncalves, Ewan Harrison, David K. Jackson, Ian Johnston, Dominic Kwiatkowski, Cordelia Langford, John Sillitoe on behalf of the Wellcome Sanger Institute COVID-19 Surveillance Team |
| EPI_ISL_3336751 | Lighthouse Lab in Milton Keynes | Wellcome Sanger Institute for the COVID-19 Genomics UK (COG-UK) Consortium | The Lighthouse Lab in Milton Keynes and Alex Alderton, Roberto Amato, Jeffrey Barrett, Sonia Goncalves, Ewan Harrison, David K. Jackson, Ian Johnston, Dominic Kwiatkowski, Cordelia Langford, John Sillitoe on behalf of the Wellcome Sanger Institute COVID-19 Surveillance Team |
| EPI_ISL_3336752 | Lighthouse Lab in Alderley Park | Wellcome Sanger Institute for the COVID-19 Genomics UK (COG-UK) Consortium | Jacquelyn Wynn, Mairead Hyland, The Lighthouse Lab in Alderley Park and Alex Alderton, Roberto Amato, Jeffrey Barrett, Sonia Goncalves, Ewan Harrison, David K. Jackson, Ian Johnston, Dominic Kwiatkowski, Cordelia Langford, John Sillitoe on behalf of the Wellcome Sanger Institute COVID-19 Surveillance Team |
| EPI_ISL_3336753, EPI_ISL_3336754 | Lighthouse Lab in Milton Keynes | Wellcome Sanger Institute for the COVID-19 Genomics UK (COG-UK) Consortium | The Lighthouse Lab in Milton Keynes and Alex Alderton, Roberto Amato, Jeffrey Barrett, Sonia Goncalves, Ewan Harrison, David K. Jackson, Ian Johnston, Dominic Kwiatkowski, Cordelia Langford, John Sillitoe on behalf of the Wellcome Sanger Institute COVID-19 Surveillance Team |
| EPI_ISL_3336755, EPI_ISL_3336756 | Lighthouse Laboratory Plymouth | Wellcome Sanger Institute for the COVID-19 Genomics UK (COG-UK) Consortium | Lighthouse Laboratory Plymouth and Alex Alderton, Roberto Amato, Jeffrey Barrett, Sonia Goncalves, Ewan Harrison, David K. Jackson, Ian Johnston, Dominic Kwiatkowski, Cordelia Langford, John Sillitoe on behalf of the Wellcome Sanger Institute COVID-19 Surveillance Team |
| EPI_ISL_3336757 | Berkshire and Surrey Pathology Services Lighthouse Laboratory | Wellcome Sanger Institute for the COVID-19 Genomics UK (COG-UK) Consortium | Berkshire and Surrey Pathology Services Lighthouse Laboratory and Alex Alderton, Roberto Amato, Jeffrey Barrett, Sonia Goncalves, Ewan Harrison, David K. Jackson, Ian Johnston, Dominic Kwiatkowski, Cordelia Langford, John Sillitoe on behalf of the Wellcome Sanger Institute COVID-19 Surveillance Team |
| EPI_ISL_3336758, EPI_ISL_3336759 | Lighthouse Lab in Milton Keynes | Wellcome Sanger Institute for the COVID-19 Genomics UK (COG-UK) Consortium | The Lighthouse Lab in Milton Keynes and Alex Alderton, Roberto Amato, Jeffrey Barrett, Sonia Goncalves, Ewan Harrison, David K. Jackson, Ian Johnston, Dominic Kwiatkowski, Cordelia Langford, John Sillitoe on behalf of the Wellcome Sanger Institute COVID-19 Surveillance Team |
| EPI_ISL_3336760, EPI_ISL_3336761 | Berkshire and Surrey Pathology Services Lighthouse Laboratory | Wellcome Sanger Institute for the COVID-19 Genomics UK (COG-UK) Consortium | Berkshire and Surrey Pathology Services Lighthouse Laboratory and Alex Alderton, Roberto Amato, Jeffrey Barrett, Sonia Goncalves, Ewan Harrison, David K. Jackson, Ian Johnston, Dominic Kwiatkowski, Cordelia Langford, John Sillitoe on behalf of the Wellcome Sanger Institute COVID-19 Surveillance Team |
| EPI_ISL_3336762, EPI_ISL_3336763, EPI_ISL_3336764 | Lighthouse Laboratory Plymouth | Wellcome Sanger Institute for the COVID-19 Genomics UK (COG-UK) Consortium | Lighthouse Laboratory Plymouth and Alex Alderton, Roberto Amato, Jeffrey Barrett, Sonia Goncalves, Ewan Harrison, David K. Jackson, Ian Johnston, Dominic Kwiatkowski, Cordelia Langford, John Sillitoe on behalf of the Wellcome Sanger Institute COVID-19 Surveillance Team |
| EPI_ISL_3336765 | Berkshire and Surrey Pathology Services Lighthouse Laboratory | Wellcome Sanger Institute for the COVID-19 Genomics UK (COG-UK) Consortium | Berkshire and Surrey Pathology Services Lighthouse Laboratory and Alex Alderton, Roberto Amato, Jeffrey Barrett, Sonia Goncalves, Ewan Harrison, David K. Jackson, Ian Johnston, Dominic Kwiatkowski, Cordelia Langford, John Sillitoe on behalf of the Wellcome Sanger Institute COVID-19 Surveillance Team |
| EPI_ISL_3336766 | Lighthouse Laboratory Plymouth | Wellcome Sanger Institute for the COVID-19 Genomics UK (COG-UK) Consortium | Lighthouse Laboratory Plymouth and Alex Alderton, Roberto Amato, Jeffrey Barrett, Sonia Goncalves, Ewan Harrison, David K. Jackson, Ian Johnston, Dominic Kwiatkowski, Cordelia Langford, John Sillitoe on behalf of the Wellcome Sanger Institute COVID-19 Surveillance Team |
| EPI_ISL_3336767, EPI_ISL_3336768 | Lighthouse Lab in Alderley Park | Wellcome Sanger Institute for the COVID-19 Genomics UK (COG-UK) Consortium | Jacquelyn Wynn, Mairead Hyland, The Lighthouse Lab in Alderley Park and Alex Alderton, Roberto Amato, Jeffrey Barrett, Sonia Goncalves, Ewan Harrison, David K. Jackson, Ian Johnston, Dominic Kwiatkowski, Cordelia Langford, John Sillitoe on behalf of the Wellcome Sanger Institute COVID-19 Surveillance Team |
| EPI_ISL_3336769 | Lighthouse Lab in Milton Keynes | Wellcome Sanger Institute for the COVID-19 Genomics UK (COG-UK) Consortium | The Lighthouse Lab in Milton Keynes and Alex Alderton, Roberto Amato, Jeffrey Barrett, Sonia Goncalves, Ewan Harrison, David K. Jackson, Ian Johnston, Dominic Kwiatkowski, Cordelia Langford, John Sillitoe on behalf of the Wellcome Sanger Institute COVID-19 Surveillance Team |
| EPI_ISL_3336770 | Lighthouse Laboratory Plymouth | Wellcome Sanger Institute for the COVID-19 Genomics UK (COG-UK) Consortium | Lighthouse Laboratory Plymouth and Alex Alderton, Roberto Amato, Jeffrey Barrett, Sonia Goncalves, Ewan Harrison, David K. Jackson, Ian Johnston, Dominic Kwiatkowski, Cordelia Langford, John Sillitoe on behalf of the Wellcome Sanger Institute COVID-19 Surveillance Team |

| | | | |
|---|---|---|---|
| EPI_ISL_3336771 | Lighthouse Lab in Alderley Park | Wellcome Sanger Institute for the COVID-19 Genomics UK (COG-UK) Consortium | Jacquelyn Wynn, Mairead Hyland, The Lighthouse Lab in Alderley Park and Alex Alderton, Roberto Amato, Jeffrey Barrett, Sonia Goncalves, Ewan Harrison, David K. Jackson, Ian Johnston, Dominic Kwiatkowski, Cordelia Langford, John Sillitoe on behalf of the Wellcome Sanger Institute COVID-19 Surveillance Team |
| EPI_ISL_3336772, EPI_ISL_3336773 | Lighthouse Lab in Milton Keynes | Wellcome Sanger Institute for the COVID-19 Genomics UK (COG-UK) Consortium | The Lighthouse Lab in Milton Keynes and Alex Alderton, Roberto Amato, Jeffrey Barrett, Sonia Goncalves, Ewan Harrison, David K. Jackson, Ian Johnston, Dominic Kwiatkowski, Cordelia Langford, John Sillitoe on behalf of the Wellcome Sanger Institute COVID-19 Surveillance Team |
| EPI_ISL_3336774 | Berkshire and Surrey Pathology Services Lighthouse Laboratory | Wellcome Sanger Institute for the COVID-19 Genomics UK (COG-UK) Consortium | Berkshire and Surrey Pathology Services Lighthouse Laboratory and Alex Alderton, Roberto Amato, Jeffrey Barrett, Sonia Goncalves, Ewan Harrison, David K. Jackson, Ian Johnston, Dominic Kwiatkowski, Cordelia Langford, John Sillitoe on behalf of the Wellcome Sanger Institute COVID-19 Surveillance Team |
| EPI_ISL_3336775 | Lighthouse Lab in Alderley Park | Wellcome Sanger Institute for the COVID-19 Genomics UK (COG-UK) Consortium | Jacquelyn Wynn, Mairead Hyland, The Lighthouse Lab in Alderley Park and Alex Alderton, Roberto Amato, Jeffrey Barrett, Sonia Goncalves, Ewan Harrison, David K. Jackson, Ian Johnston, Dominic Kwiatkowski, Cordelia Langford, John Sillitoe on behalf of the Wellcome Sanger Institute COVID-19 Surveillance Team |
| EPI_ISL_3336776 | Lighthouse Lab in Milton Keynes | Wellcome Sanger Institute for the COVID-19 Genomics UK (COG-UK) Consortium | The Lighthouse Lab in Milton Keynes and Alex Alderton, Roberto Amato, Jeffrey Barrett, Sonia Goncalves, Ewan Harrison, David K. Jackson, Ian Johnston, Dominic Kwiatkowski, Cordelia Langford, John Sillitoe on behalf of the Wellcome Sanger Institute COVID-19 Surveillance Team |
| EPI_ISL_3336777 | Lighthouse Lab in Alderley Park | Wellcome Sanger Institute for the COVID-19 Genomics UK (COG-UK) Consortium | Jacquelyn Wynn, Mairead Hyland, The Lighthouse Lab in Alderley Park and Alex Alderton, Roberto Amato, Jeffrey Barrett, Sonia Goncalves, Ewan Harrison, David K. Jackson, Ian Johnston, Dominic Kwiatkowski, Cordelia Langford, John Sillitoe on behalf of the Wellcome Sanger Institute COVID-19 Surveillance Team |
| EPI_ISL_3336778, EPI_ISL_3336779 | Berkshire and Surrey Pathology Services Lighthouse Laboratory | Wellcome Sanger Institute for the COVID-19 Genomics UK (COG-UK) Consortium | Berkshire and Surrey Pathology Services Lighthouse Laboratory and Alex Alderton, Roberto Amato, Jeffrey Barrett, Sonia Goncalves, Ewan Harrison, David K. Jackson, Ian Johnston, Dominic Kwiatkowski, Cordelia Langford, John Sillitoe on behalf of the Wellcome Sanger Institute COVID-19 Surveillance Team |
| EPI_ISL_3336780 | Lighthouse Laboratory Plymouth | Wellcome Sanger Institute for the COVID-19 Genomics UK (COG-UK) Consortium | Lighthouse Laboratory Plymouth and Alex Alderton, Roberto Amato, Jeffrey Barrett, Sonia Goncalves, Ewan Harrison, David K. Jackson, Ian Johnston, Dominic Kwiatkowski, Cordelia Langford, John Sillitoe on behalf of the Wellcome Sanger Institute COVID-19 Surveillance Team |
| EPI_ISL_3336781 | Lighthouse Lab in Alderley Park | Wellcome Sanger Institute for the COVID-19 Genomics UK (COG-UK) Consortium | Jacquelyn Wynn, Mairead Hyland, The Lighthouse Lab in Alderley Park and Alex Alderton, Roberto Amato, Jeffrey Barrett, Sonia Goncalves, Ewan Harrison, David K. Jackson, Ian Johnston, Dominic Kwiatkowski, Cordelia Langford, John Sillitoe on behalf of the Wellcome Sanger Institute COVID-19 Surveillance Team |
| EPI_ISL_3336782 | Berkshire and Surrey Pathology Services Lighthouse Laboratory | Wellcome Sanger Institute for the COVID-19 Genomics UK (COG-UK) Consortium | Berkshire and Surrey Pathology Services Lighthouse Laboratory and Alex Alderton, Roberto Amato, Jeffrey Barrett, Sonia Goncalves, Ewan Harrison, David K. Jackson, Ian Johnston, Dominic Kwiatkowski, Cordelia Langford, John Sillitoe on behalf of the Wellcome Sanger Institute COVID-19 Surveillance Team |
| EPI_ISL_3336783 | Lighthouse Lab in Alderley Park | Wellcome Sanger Institute for the COVID-19 Genomics UK (COG-UK) Consortium | Jacquelyn Wynn, Mairead Hyland, The Lighthouse Lab in Alderley Park and Alex Alderton, Roberto Amato, Jeffrey Barrett, Sonia Goncalves, Ewan Harrison, David K. Jackson, Ian Johnston, Dominic Kwiatkowski, Cordelia Langford, John Sillitoe on behalf of the Wellcome Sanger Institute COVID-19 Surveillance Team |
| EPI_ISL_3336784, EPI_ISL_3336785 | Lighthouse Laboratory Plymouth | Wellcome Sanger Institute for the COVID-19 Genomics UK (COG-UK) Consortium | Lighthouse Laboratory Plymouth and Alex Alderton, Roberto Amato, Jeffrey Barrett, Sonia Goncalves, Ewan Harrison, David K. Jackson, Ian Johnston, Dominic Kwiatkowski, Cordelia Langford, John Sillitoe on behalf of the Wellcome Sanger Institute COVID-19 Surveillance Team |
| EPI_ISL_3336786 | Lighthouse Lab in Milton Keynes | Wellcome Sanger Institute for the COVID-19 Genomics UK (COG-UK) Consortium | The Lighthouse Lab in Milton Keynes and Alex Alderton, Roberto Amato, Jeffrey Barrett, Sonia Goncalves, Ewan Harrison, David K. Jackson, Ian Johnston, Dominic Kwiatkowski, Cordelia Langford, John Sillitoe on behalf of the Wellcome Sanger Institute COVID-19 Surveillance Team |
| EPI_ISL_3336787, EPI_ISL_3336788 | Lighthouse Laboratory Plymouth | Wellcome Sanger Institute for the COVID-19 Genomics UK (COG-UK) Consortium | Lighthouse Laboratory Plymouth and Alex Alderton, Roberto Amato, Jeffrey Barrett, Sonia Goncalves, Ewan Harrison, David K. Jackson, Ian Johnston, Dominic Kwiatkowski, Cordelia Langford, John Sillitoe on behalf of the Wellcome Sanger Institute COVID-19 Surveillance Team |
| EPI_ISL_3336789 | Lighthouse Lab in Milton Keynes | Wellcome Sanger Institute for the COVID-19 Genomics UK (COG-UK) Consortium | The Lighthouse Lab in Milton Keynes and Alex Alderton, Roberto Amato, Jeffrey Barrett, Sonia Goncalves, Ewan Harrison, David K. Jackson, Ian Johnston, Dominic Kwiatkowski, Cordelia Langford, John Sillitoe on behalf of the Wellcome Sanger Institute COVID-19 Surveillance Team |
| EPI_ISL_3336790 | Lighthouse Lab in Alderley Park | Wellcome Sanger Institute for the COVID-19 Genomics UK (COG-UK) Consortium | Jacquelyn Wynn, Mairead Hyland, The Lighthouse Lab in Alderley Park and Alex Alderton, Roberto Amato, Jeffrey Barrett, Sonia Goncalves, Ewan Harrison, David K. Jackson, Ian Johnston, Dominic Kwiatkowski, Cordelia Langford, John Sillitoe on behalf of the Wellcome Sanger Institute COVID-19 Surveillance Team |
| EPI_ISL_3336791 | Lighthouse Lab in Milton Keynes | Wellcome Sanger Institute for the COVID-19 Genomics UK (COG-UK) Consortium | The Lighthouse Lab in Milton Keynes and Alex Alderton, Roberto Amato, Jeffrey Barrett, Sonia Goncalves, Ewan Harrison, David K. Jackson, Ian Johnston, Dominic Kwiatkowski, Cordelia Langford, John Sillitoe on behalf of the Wellcome Sanger Institute COVID-19 Surveillance Team |
| EPI_ISL_3336792, EPI_ISL_3336793 | Lighthouse Lab in Alderley Park | Wellcome Sanger Institute for the COVID-19 Genomics UK (COG-UK) Consortium | Jacquelyn Wynn, Mairead Hyland, The Lighthouse Lab in Alderley Park and Alex Alderton, Roberto Amato, Jeffrey Barrett, Sonia Goncalves, Ewan Harrison, David K. Jackson, Ian Johnston, Dominic Kwiatkowski, Cordelia Langford, John Sillitoe on behalf of the Wellcome Sanger Institute COVID-19 Surveillance Team |
| EPI_ISL_3336794 | Lighthouse Lab in Milton Keynes | Wellcome Sanger Institute for the COVID-19 Genomics UK (COG-UK) Consortium | The Lighthouse Lab in Milton Keynes and Alex Alderton, Roberto Amato, Jeffrey Barrett, Sonia Goncalves, Ewan Harrison, David K. Jackson, Ian Johnston, Dominic Kwiatkowski, Cordelia Langford, John Sillitoe on behalf of the Wellcome Sanger Institute COVID-19 Surveillance Team |
| EPI_ISL_3336795 | Lighthouse Lab in Alderley Park | Wellcome Sanger Institute for the COVID-19 Genomics UK (COG-UK) Consortium | Jacquelyn Wynn, Mairead Hyland, The Lighthouse Lab in Alderley Park and Alex Alderton, Roberto Amato, Jeffrey Barrett, Sonia Goncalves, Ewan Harrison, David K. Jackson, Ian Johnston, Dominic Kwiatkowski, Cordelia Langford, John Sillitoe on behalf of the Wellcome Sanger Institute COVID-19 Surveillance Team |
| EPI_ISL_3336796 | Berkshire and Surrey Pathology Services Lighthouse Laboratory | Wellcome Sanger Institute for the COVID-19 Genomics UK (COG-UK) Consortium | Berkshire and Surrey Pathology Services Lighthouse Laboratory and Alex Alderton, Roberto Amato, Jeffrey Barrett, Sonia Goncalves, Ewan Harrison, David K. Jackson, Ian Johnston, Dominic Kwiatkowski, Cordelia Langford, John Sillitoe on behalf of the Wellcome Sanger Institute COVID-19 Surveillance Team |
| EPI_ISL_3336797 | Lighthouse Lab in Alderley Park | Wellcome Sanger Institute for the COVID-19 Genomics UK (COG-UK) Consortium | Jacquelyn Wynn, Mairead Hyland, The Lighthouse Lab in Alderley Park and Alex Alderton, Roberto Amato, Jeffrey Barrett, Sonia Goncalves, Ewan Harrison, David K. Jackson, Ian Johnston, Dominic Kwiatkowski, Cordelia Langford, John Sillitoe on behalf of the Wellcome Sanger Institute COVID-19 Surveillance Team |
| EPI_ISL_3336798 | Berkshire and Surrey Pathology Services Lighthouse Laboratory | Wellcome Sanger Institute for the COVID-19 Genomics UK (COG-UK) Consortium | Berkshire and Surrey Pathology Services Lighthouse Laboratory and Alex Alderton, Roberto Amato, Jeffrey Barrett, Sonia Goncalves, Ewan Harrison, David K. Jackson, Ian Johnston, Dominic Kwiatkowski, Cordelia Langford, John Sillitoe on behalf of the Wellcome Sanger Institute COVID-19 Surveillance Team |
| EPI_ISL_3336799 | Lighthouse Laboratory Plymouth | Wellcome Sanger Institute for the COVID-19 Genomics UK (COG-UK) Consortium | Lighthouse Laboratory Plymouth and Alex Alderton, Roberto Amato, Jeffrey Barrett, Sonia Goncalves, Ewan Harrison, David K. Jackson, Ian Johnston, Dominic Kwiatkowski, Cordelia Langford, John Sillitoe on behalf of the Wellcome Sanger Institute COVID-19 Surveillance Team |
| EPI_ISL_3336800 | Lighthouse Lab in Milton Keynes | Wellcome Sanger Institute for the COVID-19 Genomics UK (COG-UK) Consortium | The Lighthouse Lab in Milton Keynes and Alex Alderton, Roberto Amato, Jeffrey Barrett, Sonia Goncalves, Ewan Harrison, David K. Jackson, Ian Johnston, Dominic Kwiatkowski, Cordelia Langford, John Sillitoe on behalf of the Wellcome Sanger Institute COVID-19 Surveillance Team |
| EPI_ISL_3336801 | Berkshire and Surrey Pathology Services Lighthouse Laboratory | Wellcome Sanger Institute for the COVID-19 Genomics UK (COG-UK) Consortium | Berkshire and Surrey Pathology Services Lighthouse Laboratory and Alex Alderton, Roberto Amato, Jeffrey Barrett, Sonia Goncalves, Ewan Harrison, David K. Jackson, Ian Johnston, Dominic Kwiatkowski, Cordelia Langford, John Sillitoe on behalf of the Wellcome Sanger Institute COVID-19 Surveillance Team |
| EPI_ISL_3336802, EPI_ISL_3336803 | Lighthouse Lab in Milton Keynes | Wellcome Sanger Institute for the COVID-19 Genomics UK (COG-UK) Consortium | The Lighthouse Lab in Milton Keynes and Alex Alderton, Roberto Amato, Jeffrey Barrett, Sonia Goncalves, Ewan Harrison, David K. Jackson, Ian Johnston, Dominic Kwiatkowski, Cordelia Langford, John Sillitoe on behalf of the Wellcome Sanger Institute COVID-19 Surveillance Team |
| EPI_ISL_3336804 | Lighthouse Laboratory Plymouth | Wellcome Sanger Institute for the COVID-19 Genomics UK (COG-UK) Consortium | Lighthouse Laboratory Plymouth and Alex Alderton, Roberto Amato, Jeffrey Barrett, Sonia Goncalves, Ewan Harrison, David K. Jackson, Ian Johnston, Dominic Kwiatkowski, Cordelia Langford, John Sillitoe on behalf of the Wellcome Sanger Institute COVID-19 Surveillance Team |
| EPI_ISL_3336805 | Lighthouse Lab in Alderley Park | Wellcome Sanger Institute for the COVID-19 Genomics UK (COG-UK) Consortium | Jacquelyn Wynn, Mairead Hyland, The Lighthouse Lab in Alderley Park and Alex Alderton, Roberto Amato, Jeffrey Barrett, Sonia Goncalves, Ewan Harrison, David K. Jackson, Ian Johnston, Dominic Kwiatkowski, Cordelia Langford, John Sillitoe on behalf of the Wellcome Sanger Institute COVID-19 Surveillance Team |
| EPI_ISL_3336806 | Lighthouse Lab in Milton Keynes | Wellcome Sanger Institute for the COVID-19 Genomics UK (COG-UK) Consortium | The Lighthouse Lab in Milton Keynes and Alex Alderton, Roberto Amato, Jeffrey Barrett, Sonia Goncalves, Ewan Harrison, David K. Jackson, Ian Johnston, Dominic Kwiatkowski, Cordelia Langford, John Sillitoe on behalf of the Wellcome Sanger Institute COVID-19 Surveillance Team |
| EPI_ISL_3336807 | Berkshire and Surrey Pathology Services Lighthouse Laboratory | Wellcome Sanger Institute for the COVID-19 Genomics UK (COG-UK) Consortium | Berkshire and Surrey Pathology Services Lighthouse Laboratory and Alex Alderton, Roberto Amato, Jeffrey Barrett, Sonia Goncalves, Ewan Harrison, David K. Jackson, Ian Johnston, Dominic Kwiatkowski, Cordelia Langford, John Sillitoe on behalf of the Wellcome Sanger Institute COVID-19 Surveillance Team |
| EPI_ISL_3336808, EPI_ISL_3336809 | Lighthouse Laboratory Plymouth | Wellcome Sanger Institute for the COVID-19 Genomics UK (COG-UK) Consortium | Lighthouse Laboratory Plymouth and Alex Alderton, Roberto Amato, Jeffrey Barrett, Sonia Goncalves, Ewan Harrison, David K. Jackson, Ian Johnston, Dominic Kwiatkowski, Cordelia Langford, John Sillitoe on behalf of the Wellcome Sanger Institute COVID-19 Surveillance Team |
| EPI_ISL_3336810 | Lighthouse Lab in Milton Keynes | Wellcome Sanger Institute for the COVID-19 Genomics UK (COG-UK) Consortium | The Lighthouse Lab in Milton Keynes and Alex Alderton, Roberto Amato, Jeffrey Barrett, Sonia Goncalves, Ewan Harrison, David K. Jackson, Ian Johnston, Dominic Kwiatkowski, Cordelia Langford, John Sillitoe on behalf of the Wellcome Sanger Institute COVID-19 Surveillance Team |
| EPI_ISL_3336811 | Lighthouse Lab in Alderley Park | Wellcome Sanger Institute for the COVID-19 Genomics UK (COG-UK) Consortium | Jacquelyn Wynn, Mairead Hyland, The Lighthouse Lab in Alderley Park and Alex Alderton, Roberto Amato, Jeffrey Barrett, Sonia Goncalves, Ewan Harrison, David K. Jackson, Ian Johnston, Dominic Kwiatkowski, Cordelia Langford, John Sillitoe on behalf of the Wellcome Sanger Institute COVID-19 Surveillance Team |
| EPI_ISL_3336812, EPI_ISL_3336813 | Berkshire and Surrey Pathology Services Lighthouse Laboratory | Wellcome Sanger Institute for the COVID-19 Genomics UK (COG-UK) Consortium | Berkshire and Surrey Pathology Services Lighthouse Laboratory and Alex Alderton, Roberto Amato, Jeffrey Barrett, Sonia Goncalves, Ewan Harrison, David K. Jackson, Ian Johnston, Dominic Kwiatkowski, Cordelia Langford, John Sillitoe on behalf of the Wellcome Sanger Institute COVID-19 Surveillance Team |
| EPI_ISL_3336814 | Lighthouse Lab in Milton Keynes | Wellcome Sanger Institute for the COVID-19 Genomics UK (COG-UK) Consortium | The Lighthouse Lab in Milton Keynes and Alex Alderton, Roberto Amato, Jeffrey Barrett, Sonia Goncalves, Ewan Harrison, David K. Jackson, Ian Johnston, Dominic Kwiatkowski, Cordelia Langford, John Sillitoe on behalf of the Wellcome Sanger Institute COVID-19 Surveillance Team |
| EPI_ISL_3336815 | Berkshire and Surrey Pathology Services Lighthouse Laboratory | Wellcome Sanger Institute for the COVID-19 Genomics UK (COG-UK) Consortium | Berkshire and Surrey Pathology Services Lighthouse Laboratory and Alex Alderton, Roberto Amato, Jeffrey Barrett, Sonia Goncalves, Ewan Harrison, David K. Jackson, Ian Johnston, Dominic Kwiatkowski, Cordelia Langford, John Sillitoe on behalf of the Wellcome Sanger Institute COVID-19 Surveillance Team |
| EPI_ISL_3336816 | Lighthouse Lab in Alderley Park | Wellcome Sanger Institute for the COVID-19 Genomics UK (COG-UK) Consortium | Jacquelyn Wynn, Mairead Hyland, The Lighthouse Lab in Alderley Park and Alex Alderton, Roberto Amato, Jeffrey Barrett, Sonia Goncalves, Ewan Harrison, David K. Jackson, Ian Johnston, Dominic Kwiatkowski, Cordelia Langford, John Sillitoe on behalf of the Wellcome Sanger Institute COVID-19 Surveillance Team |
| EPI_ISL_3336817 | Lighthouse Laboratory Plymouth | Wellcome Sanger Institute for the COVID-19 Genomics UK (COG-UK) Consortium | Lighthouse Laboratory Plymouth and Alex Alderton, Roberto Amato, Jeffrey Barrett, Sonia Goncalves, Ewan Harrison, David K. Jackson, Ian Johnston, Dominic Kwiatkowski, Cordelia Langford, John Sillitoe on behalf of the Wellcome Sanger Institute COVID-19 Surveillance Team |
| EPI_ISL_3336818, EPI_ISL_3336819 | Lighthouse Lab in Alderley Park | Wellcome Sanger Institute for the COVID-19 Genomics UK (COG-UK) Consortium | Jacquelyn Wynn, Mairead Hyland, The Lighthouse Lab in Alderley Park and Alex Alderton, Roberto Amato, Jeffrey Barrett, Sonia Goncalves, Ewan Harrison, David K. Jackson, Ian Johnston, Dominic Kwiatkowski, Cordelia Langford, John Sillitoe on behalf of the Wellcome Sanger Institute COVID-19 Surveillance Team |
| EPI_ISL_3336820, EPI_ISL_3336821 | Lighthouse Lab in Milton Keynes | Wellcome Sanger Institute for the COVID-19 Genomics UK (COG-UK) Consortium | The Lighthouse Lab in Milton Keynes and Alex Alderton, Roberto Amato, Jeffrey Barrett, Sonia Goncalves, Ewan Harrison, David K. Jackson, Ian Johnston, Dominic Kwiatkowski, Cordelia Langford, John Sillitoe on behalf of the Wellcome Sanger Institute COVID-19 Surveillance Team |
| EPI_ISL_3336822 | Lighthouse Laboratory Plymouth | Wellcome Sanger Institute for the COVID-19 Genomics UK (COG-UK) Consortium | Lighthouse Laboratory Plymouth and Alex Alderton, Roberto Amato, Jeffrey Barrett, Sonia Goncalves, Ewan Harrison, David K. Jackson, Ian Johnston, Dominic Kwiatkowski, Cordelia Langford, John Sillitoe on behalf of the Wellcome Sanger Institute COVID-19 Surveillance Team |
| EPI_ISL_3336823 | Berkshire and Surrey Pathology Services Lighthouse Laboratory | Wellcome Sanger Institute for the COVID-19 Genomics UK (COG-UK) Consortium | Berkshire and Surrey Pathology Services Lighthouse Laboratory and Alex Alderton, Roberto Amato, Jeffrey Barrett, Sonia Goncalves, Ewan Harrison, David K. Jackson, Ian Johnston, Dominic Kwiatkowski, Cordelia Langford, John Sillitoe on behalf of the Wellcome Sanger Institute COVID-19 Surveillance Team |
| EPI_ISL_3336824, EPI_ISL_3336825 | Lighthouse Laboratory Plymouth | Wellcome Sanger Institute for the COVID-19 Genomics UK (COG-UK) Consortium | Lighthouse Laboratory Plymouth and Alex Alderton, Roberto Amato, Jeffrey Barrett, Sonia Goncalves, Ewan Harrison, David K. Jackson, Ian Johnston, Dominic Kwiatkowski, Cordelia Langford, John Sillitoe on behalf of the Wellcome Sanger Institute COVID-19 Surveillance Team |
| EPI_ISL_3336826 | Berkshire and Surrey Pathology Services Lighthouse Laboratory | Wellcome Sanger Institute for the COVID-19 Genomics UK (COG-UK) Consortium | Berkshire and Surrey Pathology Services Lighthouse Laboratory and Alex Alderton, Roberto Amato, Jeffrey Barrett, Sonia Goncalves, Ewan Harrison, David K. Jackson, Ian Johnston, Dominic Kwiatkowski, Cordelia Langford, John Sillitoe on behalf of the Wellcome Sanger Institute COVID-19 Surveillance Team |

| | | | |
|---|---|---|---|
| EPI_ISL_3336827, EPI_ISL_3336828 | Lighthouse Lab in Alderley Park | Wellcome Sanger Institute for the COVID-19 Genomics UK (COG-UK) Consortium | Jacquelyn Wynn, Mairead Hyland, The Lighthouse Lab in Alderley Park and Alex Alderton, Roberto Amato, Jeffrey Barrett, Sonia Goncalves, Ewan Harrison, David K. Jackson, Ian Johnston, Dominic Kwiatkowski, Cordelia Langford, John Sillitoe on behalf of the Wellcome Sanger Institute COVID-19 Surveillance Team |
| EPI_ISL_3336829, EPI_ISL_3336830 | Lighthouse Lab in Milton Keynes | Wellcome Sanger Institute for the COVID-19 Genomics UK (COG-UK) Consortium | The Lighthouse Lab in Milton Keynes and Alex Alderton, Roberto Amato, Jeffrey Barrett, Sonia Goncalves, Ewan Harrison, David K. Jackson, Ian Johnston, Dominic Kwiatkowski, Cordelia Langford, John Sillitoe on behalf of the Wellcome Sanger Institute COVID-19 Surveillance Team |
| EPI_ISL_3336831 | Berkshire and Surrey Pathology Services Lighthouse Laboratory | Wellcome Sanger Institute for the COVID-19 Genomics UK (COG-UK) Consortium | Berkshire and Surrey Pathology Services Lighthouse Laboratory and Alex Alderton, Roberto Amato, Jeffrey Barrett, Sonia Goncalves, Ewan Harrison, David K. Jackson, Ian Johnston, Dominic Kwiatkowski, Cordelia Langford, John Sillitoe on behalf of the Wellcome Sanger Institute COVID-19 Surveillance Team |
| EPI_ISL_3336832 | Lighthouse Lab in Milton Keynes | Wellcome Sanger Institute for the COVID-19 Genomics UK (COG-UK) Consortium | The Lighthouse Lab in Milton Keynes and Alex Alderton, Roberto Amato, Jeffrey Barrett, Sonia Goncalves, Ewan Harrison, David K. Jackson, Ian Johnston, Dominic Kwiatkowski, Cordelia Langford, John Sillitoe on behalf of the Wellcome Sanger Institute COVID-19 Surveillance Team |
| EPI_ISL_3336833 | Lighthouse Lab in Alderley Park | Wellcome Sanger Institute for the COVID-19 Genomics UK (COG-UK) Consortium | Jacquelyn Wynn, Mairead Hyland, The Lighthouse Lab in Alderley Park and Alex Alderton, Roberto Amato, Jeffrey Barrett, Sonia Goncalves, Ewan Harrison, David K. Jackson, Ian Johnston, Dominic Kwiatkowski, Cordelia Langford, John Sillitoe on behalf of the Wellcome Sanger Institute COVID-19 Surveillance Team |
| EPI_ISL_3336834 | Lighthouse Laboratory Plymouth | Wellcome Sanger Institute for the COVID-19 Genomics UK (COG-UK) Consortium | Lighthouse Laboratory Plymouth and Alex Alderton, Roberto Amato, Jeffrey Barrett, Sonia Goncalves, Ewan Harrison, David K. Jackson, Ian Johnston, Dominic Kwiatkowski, Cordelia Langford, John Sillitoe on behalf of the Wellcome Sanger Institute COVID-19 Surveillance Team |
| EPI_ISL_3336835 | Lighthouse Lab in Alderley Park | Wellcome Sanger Institute for the COVID-19 Genomics UK (COG-UK) Consortium | Jacquelyn Wynn, Mairead Hyland, The Lighthouse Lab in Alderley Park and Alex Alderton, Roberto Amato, Jeffrey Barrett, Sonia Goncalves, Ewan Harrison, David K. Jackson, Ian Johnston, Dominic Kwiatkowski, Cordelia Langford, John Sillitoe on behalf of the Wellcome Sanger Institute COVID-19 Surveillance Team |
| EPI_ISL_3336836 | Berkshire and Surrey Pathology Services Lighthouse Laboratory | Wellcome Sanger Institute for the COVID-19 Genomics UK (COG-UK) Consortium | Berkshire and Surrey Pathology Services Lighthouse Laboratory and Alex Alderton, Roberto Amato, Jeffrey Barrett, Sonia Goncalves, Ewan Harrison, David K. Jackson, Ian Johnston, Dominic Kwiatkowski, Cordelia Langford, John Sillitoe on behalf of the Wellcome Sanger Institute COVID-19 Surveillance Team |
| EPI_ISL_3336837 | Lighthouse Lab in Milton Keynes | Wellcome Sanger Institute for the COVID-19 Genomics UK (COG-UK) Consortium | The Lighthouse Lab in Milton Keynes and Alex Alderton, Roberto Amato, Jeffrey Barrett, Sonia Goncalves, Ewan Harrison, David K. Jackson, Ian Johnston, Dominic Kwiatkowski, Cordelia Langford, John Sillitoe on behalf of the Wellcome Sanger Institute COVID-19 Surveillance Team |
| EPI_ISL_3336838, EPI_ISL_3336839, EPI_ISL_3336840 | Berkshire and Surrey Pathology Services Lighthouse Laboratory | Wellcome Sanger Institute for the COVID-19 Genomics UK (COG-UK) Consortium | Berkshire and Surrey Pathology Services Lighthouse Laboratory and Alex Alderton, Roberto Amato, Jeffrey Barrett, Sonia Goncalves, Ewan Harrison, David K. Jackson, Ian Johnston, Dominic Kwiatkowski, Cordelia Langford, John Sillitoe on behalf of the Wellcome Sanger Institute COVID-19 Surveillance Team |
| EPI_ISL_3336841 | Lighthouse Laboratory Plymouth | Wellcome Sanger Institute for the COVID-19 Genomics UK (COG-UK) Consortium | Lighthouse Laboratory Plymouth and Alex Alderton, Roberto Amato, Jeffrey Barrett, Sonia Goncalves, Ewan Harrison, David K. Jackson, Ian Johnston, Dominic Kwiatkowski, Cordelia Langford, John Sillitoe on behalf of the Wellcome Sanger Institute COVID-19 Surveillance Team |
| EPI_ISL_3336842 | Lighthouse Lab in Alderley Park | Wellcome Sanger Institute for the COVID-19 Genomics UK (COG-UK) Consortium | Jacquelyn Wynn, Mairead Hyland, The Lighthouse Lab in Alderley Park and Alex Alderton, Roberto Amato, Jeffrey Barrett, Sonia Goncalves, Ewan Harrison, David K. Jackson, Ian Johnston, Dominic Kwiatkowski, Cordelia Langford, John Sillitoe on behalf of the Wellcome Sanger Institute COVID-19 Surveillance Team |
| EPI_ISL_3336843 | Berkshire and Surrey Pathology Services Lighthouse Laboratory | Wellcome Sanger Institute for the COVID-19 Genomics UK (COG-UK) Consortium | Berkshire and Surrey Pathology Services Lighthouse Laboratory and Alex Alderton, Roberto Amato, Jeffrey Barrett, Sonia Goncalves, Ewan Harrison, David K. Jackson, Ian Johnston, Dominic Kwiatkowski, Cordelia Langford, John Sillitoe on behalf of the Wellcome Sanger Institute COVID-19 Surveillance Team |
| EPI_ISL_3336844 | Lighthouse Lab in Alderley Park | Wellcome Sanger Institute for the COVID-19 Genomics UK (COG-UK) Consortium | Jacquelyn Wynn, Mairead Hyland, The Lighthouse Lab in Alderley Park and Alex Alderton, Roberto Amato, Jeffrey Barrett, Sonia Goncalves, Ewan Harrison, David K. Jackson, Ian Johnston, Dominic Kwiatkowski, Cordelia Langford, John Sillitoe on behalf of the Wellcome Sanger Institute COVID-19 Surveillance Team |
| EPI_ISL_3336845, EPI_ISL_3336846 | Lighthouse Laboratory Plymouth | Wellcome Sanger Institute for the COVID-19 Genomics UK (COG-UK) Consortium | Lighthouse Laboratory Plymouth and Alex Alderton, Roberto Amato, Jeffrey Barrett, Sonia Goncalves, Ewan Harrison, David K. Jackson, Ian Johnston, Dominic Kwiatkowski, Cordelia Langford, John Sillitoe on behalf of the Wellcome Sanger Institute COVID-19 Surveillance Team |
| EPI_ISL_3336847 | Lighthouse Lab in Milton Keynes | Wellcome Sanger Institute for the COVID-19 Genomics UK (COG-UK) Consortium | The Lighthouse Lab in Milton Keynes and Alex Alderton, Roberto Amato, Jeffrey Barrett, Sonia Goncalves, Ewan Harrison, David K. Jackson, Ian Johnston, Dominic Kwiatkowski, Cordelia Langford, John Sillitoe on behalf of the Wellcome Sanger Institute COVID-19 Surveillance Team |
| EPI_ISL_3336848 | Lighthouse Lab in Alderley Park | Wellcome Sanger Institute for the COVID-19 Genomics UK (COG-UK) Consortium | Jacquelyn Wynn, Mairead Hyland, The Lighthouse Lab in Alderley Park and Alex Alderton, Roberto Amato, Jeffrey Barrett, Sonia Goncalves, Ewan Harrison, David K. Jackson, Ian Johnston, Dominic Kwiatkowski, Cordelia Langford, John Sillitoe on behalf of the Wellcome Sanger Institute COVID-19 Surveillance Team |
| EPI_ISL_3336849 | Berkshire and Surrey Pathology Services Lighthouse Laboratory | Wellcome Sanger Institute for the COVID-19 Genomics UK (COG-UK) Consortium | Berkshire and Surrey Pathology Services Lighthouse Laboratory and Alex Alderton, Roberto Amato, Jeffrey Barrett, Sonia Goncalves, Ewan Harrison, David K. Jackson, Ian Johnston, Dominic Kwiatkowski, Cordelia Langford, John Sillitoe on behalf of the Wellcome Sanger Institute COVID-19 Surveillance Team |
| EPI_ISL_3336850 | Lighthouse Laboratory Plymouth | Wellcome Sanger Institute for the COVID-19 Genomics UK (COG-UK) Consortium | Lighthouse Laboratory Plymouth and Alex Alderton, Roberto Amato, Jeffrey Barrett, Sonia Goncalves, Ewan Harrison, David K. Jackson, Ian Johnston, Dominic Kwiatkowski, Cordelia Langford, John Sillitoe on behalf of the Wellcome Sanger Institute COVID-19 Surveillance Team |
| EPI_ISL_3336851 | Lighthouse Lab in Alderley Park | Wellcome Sanger Institute for the COVID-19 Genomics UK (COG-UK) Consortium | Jacquelyn Wynn, Mairead Hyland, The Lighthouse Lab in Alderley Park and Alex Alderton, Roberto Amato, Jeffrey Barrett, Sonia Goncalves, Ewan Harrison, David K. Jackson, Ian Johnston, Dominic Kwiatkowski, Cordelia Langford, John Sillitoe on behalf of the Wellcome Sanger Institute COVID-19 Surveillance Team |
| EPI_ISL_3336852 | Lighthouse Lab in Milton Keynes | Wellcome Sanger Institute for the COVID-19 Genomics UK (COG-UK) Consortium | The Lighthouse Lab in Milton Keynes and Alex Alderton, Roberto Amato, Jeffrey Barrett, Sonia Goncalves, Ewan Harrison, David K. Jackson, Ian Johnston, Dominic Kwiatkowski, Cordelia Langford, John Sillitoe on behalf of the Wellcome Sanger Institute COVID-19 Surveillance Team |
| EPI_ISL_3336853 | Berkshire and Surrey Pathology Services Lighthouse Laboratory | Wellcome Sanger Institute for the COVID-19 Genomics UK (COG-UK) Consortium | Berkshire and Surrey Pathology Services Lighthouse Laboratory and Alex Alderton, Roberto Amato, Jeffrey Barrett, Sonia Goncalves, Ewan Harrison, David K. Jackson, Ian Johnston, Dominic Kwiatkowski, Cordelia Langford, John Sillitoe on behalf of the Wellcome Sanger Institute COVID-19 Surveillance Team |
| EPI_ISL_3336854 | Lighthouse Lab in Milton Keynes | Wellcome Sanger Institute for the COVID-19 Genomics UK (COG-UK) Consortium | The Lighthouse Lab in Milton Keynes and Alex Alderton, Roberto Amato, Jeffrey Barrett, Sonia Goncalves, Ewan Harrison, David K. Jackson, Ian Johnston, Dominic Kwiatkowski, Cordelia Langford, John Sillitoe on behalf of the Wellcome Sanger Institute COVID-19 Surveillance Team |
| EPI_ISL_3336855 | Lighthouse Lab in Alderley Park | Wellcome Sanger Institute for the COVID-19 Genomics UK (COG-UK) Consortium | Jacquelyn Wynn, Mairead Hyland, The Lighthouse Lab in Alderley Park and Alex Alderton, Roberto Amato, Jeffrey Barrett, Sonia Goncalves, Ewan Harrison, David K. Jackson, Ian Johnston, Dominic Kwiatkowski, Cordelia Langford, John Sillitoe on behalf of the Wellcome Sanger Institute COVID-19 Surveillance Team |
| EPI_ISL_3336856, EPI_ISL_3336857 | Berkshire and Surrey Pathology Services Lighthouse Laboratory | Wellcome Sanger Institute for the COVID-19 Genomics UK (COG-UK) Consortium | Berkshire and Surrey Pathology Services Lighthouse Laboratory and Alex Alderton, Roberto Amato, Jeffrey Barrett, Sonia Goncalves, Ewan Harrison, David K. Jackson, Ian Johnston, Dominic Kwiatkowski, Cordelia Langford, John Sillitoe on behalf of the Wellcome Sanger Institute COVID-19 Surveillance Team |
| EPI_ISL_3336858 | Lighthouse Laboratory Plymouth | Wellcome Sanger Institute for the COVID-19 Genomics UK (COG-UK) Consortium | Lighthouse Laboratory Plymouth and Alex Alderton, Roberto Amato, Jeffrey Barrett, Sonia Goncalves, Ewan Harrison, David K. Jackson, Ian Johnston, Dominic Kwiatkowski, Cordelia Langford, John Sillitoe on behalf of the Wellcome Sanger Institute COVID-19 Surveillance Team |
| EPI_ISL_3336859, EPI_ISL_3336860 | Lighthouse Lab in Alderley Park | Wellcome Sanger Institute for the COVID-19 Genomics UK (COG-UK) Consortium | Jacquelyn Wynn, Mairead Hyland, The Lighthouse Lab in Alderley Park and Alex Alderton, Roberto Amato, Jeffrey Barrett, Sonia Goncalves, Ewan Harrison, David K. Jackson, Ian Johnston, Dominic Kwiatkowski, Cordelia Langford, John Sillitoe on behalf of the Wellcome Sanger Institute COVID-19 Surveillance Team |
| EPI_ISL_3336861 | Berkshire and Surrey Pathology Services Lighthouse Laboratory | Wellcome Sanger Institute for the COVID-19 Genomics UK (COG-UK) Consortium | Berkshire and Surrey Pathology Services Lighthouse Laboratory and Alex Alderton, Roberto Amato, Jeffrey Barrett, Sonia Goncalves, Ewan Harrison, David K. Jackson, Ian Johnston, Dominic Kwiatkowski, Cordelia Langford, John Sillitoe on behalf of the Wellcome Sanger Institute COVID-19 Surveillance Team |
| EPI_ISL_3336862 | Lighthouse Lab in Milton Keynes | Wellcome Sanger Institute for the COVID-19 Genomics UK (COG-UK) Consortium | The Lighthouse Lab in Milton Keynes and Alex Alderton, Roberto Amato, Jeffrey Barrett, Sonia Goncalves, Ewan Harrison, David K. Jackson, Ian Johnston, Dominic Kwiatkowski, Cordelia Langford, John Sillitoe on behalf of the Wellcome Sanger Institute COVID-19 Surveillance Team |
| EPI_ISL_3336863 | Berkshire and Surrey Pathology Services Lighthouse Laboratory | Wellcome Sanger Institute for the COVID-19 Genomics UK (COG-UK) Consortium | Berkshire and Surrey Pathology Services Lighthouse Laboratory and Alex Alderton, Roberto Amato, Jeffrey Barrett, Sonia Goncalves, Ewan Harrison, David K. Jackson, Ian Johnston, Dominic Kwiatkowski, Cordelia Langford, John Sillitoe on behalf of the Wellcome Sanger Institute COVID-19 Surveillance Team |
| EPI_ISL_3336864 | Lighthouse Lab in Alderley Park | Wellcome Sanger Institute for the COVID-19 Genomics UK (COG-UK) Consortium | Jacquelyn Wynn, Mairead Hyland, The Lighthouse Lab in Alderley Park and Alex Alderton, Roberto Amato, Jeffrey Barrett, Sonia Goncalves, Ewan Harrison, David K. Jackson, Ian Johnston, Dominic Kwiatkowski, Cordelia Langford, John Sillitoe on behalf of the Wellcome Sanger Institute COVID-19 Surveillance Team |
| EPI_ISL_3336865 | Lighthouse Lab in Milton Keynes | Wellcome Sanger Institute for the COVID-19 Genomics UK (COG-UK) Consortium | The Lighthouse Lab in Milton Keynes and Alex Alderton, Roberto Amato, Jeffrey Barrett, Sonia Goncalves, Ewan Harrison, David K. Jackson, Ian Johnston, Dominic Kwiatkowski, Cordelia Langford, John Sillitoe on behalf of the Wellcome Sanger Institute COVID-19 Surveillance Team |
| EPI_ISL_3336866, EPI_ISL_3336867 | Berkshire and Surrey Pathology Services Lighthouse Laboratory | Wellcome Sanger Institute for the COVID-19 Genomics UK (COG-UK) Consortium | Berkshire and Surrey Pathology Services Lighthouse Laboratory and Alex Alderton, Roberto Amato, Jeffrey Barrett, Sonia Goncalves, Ewan Harrison, David K. Jackson, Ian Johnston, Dominic Kwiatkowski, Cordelia Langford, John Sillitoe on behalf of the Wellcome Sanger Institute COVID-19 Surveillance Team |
| EPI_ISL_3336868 | Lighthouse Lab in Milton Keynes | Wellcome Sanger Institute for the COVID-19 Genomics UK (COG-UK) Consortium | The Lighthouse Lab in Milton Keynes and Alex Alderton, Roberto Amato, Jeffrey Barrett, Sonia Goncalves, Ewan Harrison, David K. Jackson, Ian Johnston, Dominic Kwiatkowski, Cordelia Langford, John Sillitoe on behalf of the Wellcome Sanger Institute COVID-19 Surveillance Team |
| EPI_ISL_3336869 | Lighthouse Lab in Alderley Park | Wellcome Sanger Institute for the COVID-19 Genomics UK (COG-UK) Consortium | Jacquelyn Wynn, Mairead Hyland, The Lighthouse Lab in Alderley Park and Alex Alderton, Roberto Amato, Jeffrey Barrett, Sonia Goncalves, Ewan Harrison, David K. Jackson, Ian Johnston, Dominic Kwiatkowski, Cordelia Langford, John Sillitoe on behalf of the Wellcome Sanger Institute COVID-19 Surveillance Team |
| EPI_ISL_3336870 | Lighthouse Lab in Milton Keynes | Wellcome Sanger Institute for the COVID-19 Genomics UK (COG-UK) Consortium | The Lighthouse Lab in Milton Keynes and Alex Alderton, Roberto Amato, Jeffrey Barrett, Sonia Goncalves, Ewan Harrison, David K. Jackson, Ian Johnston, Dominic Kwiatkowski, Cordelia Langford, John Sillitoe on behalf of the Wellcome Sanger Institute COVID-19 Surveillance Team |
| EPI_ISL_3336871, EPI_ISL_3336872, EPI_ISL_3336873, EPI_ISL_3336874 | Lighthouse Laboratory Plymouth | Wellcome Sanger Institute for the COVID-19 Genomics UK (COG-UK) Consortium | Lighthouse Laboratory Plymouth and Alex Alderton, Roberto Amato, Jeffrey Barrett, Sonia Goncalves, Ewan Harrison, David K. Jackson, Ian Johnston, Dominic Kwiatkowski, Cordelia Langford, John Sillitoe on behalf of the Wellcome Sanger Institute COVID-19 Surveillance Team |
| EPI_ISL_3336875 | Lighthouse Lab in Milton Keynes | Wellcome Sanger Institute for the COVID-19 Genomics UK (COG-UK) Consortium | The Lighthouse Lab in Milton Keynes and Alex Alderton, Roberto Amato, Jeffrey Barrett, Sonia Goncalves, Ewan Harrison, David K. Jackson, Ian Johnston, Dominic Kwiatkowski, Cordelia Langford, John Sillitoe on behalf of the Wellcome Sanger Institute COVID-19 Surveillance Team |
| EPI_ISL_3336876 | Lighthouse Lab in Alderley Park | Wellcome Sanger Institute for the COVID-19 Genomics UK (COG-UK) Consortium | Jacquelyn Wynn, Mairead Hyland, The Lighthouse Lab in Alderley Park and Alex Alderton, Roberto Amato, Jeffrey Barrett, Sonia Goncalves, Ewan Harrison, David K. Jackson, Ian Johnston, Dominic Kwiatkowski, Cordelia Langford, John Sillitoe on behalf of the Wellcome Sanger Institute COVID-19 Surveillance Team |
| EPI_ISL_3336877 | Lighthouse Lab in Milton Keynes | Wellcome Sanger Institute for the COVID-19 Genomics UK (COG-UK) Consortium | The Lighthouse Lab in Milton Keynes and Alex Alderton, Roberto Amato, Jeffrey Barrett, Sonia Goncalves, Ewan Harrison, David K. Jackson, Ian Johnston, Dominic Kwiatkowski, Cordelia Langford, John Sillitoe on behalf of the Wellcome Sanger Institute COVID-19 Surveillance Team |
| EPI_ISL_3336878 | Lighthouse Laboratory Plymouth | Wellcome Sanger Institute for the COVID-19 Genomics UK (COG-UK) Consortium | Lighthouse Laboratory Plymouth and Alex Alderton, Roberto Amato, Jeffrey Barrett, Sonia Goncalves, Ewan Harrison, David K. Jackson, Ian Johnston, Dominic Kwiatkowski, Cordelia Langford, John Sillitoe on behalf of the Wellcome Sanger Institute COVID-19 Surveillance Team |
| EPI_ISL_3336879 | Department of Virology, Istituto Zooprofilattico Sperimentale del Lazio e della Toscana (IZSLT) | Department of General Diagnostics; Department of Virology; Istituto Zooprofilattico Sperimentale del Lazio e della Toscana (IZSLT) | Patricia Alba, Giuseppe Manna, Elena L. Diaconu, Fabiola Feltrin, Raffaela Conti, Teresa Scicluna, Virginia Carfora, Antonella Cersini, Alessia Franco, Antonio Battisti. |
| EPI_ISL_3336880, EPI_ISL_3336881 | Berkshire and Surrey Pathology Services Lighthouse Laboratory | Wellcome Sanger Institute for the COVID-19 Genomics UK (COG-UK) Consortium | Berkshire and Surrey Pathology Services Lighthouse Laboratory and Alex Alderton, Roberto Amato, Jeffrey Barrett, Sonia Goncalves, Ewan Harrison, David K. Jackson, Ian Johnston, Dominic Kwiatkowski, Cordelia Langford, John Sillitoe on behalf of the Wellcome Sanger Institute COVID-19 Surveillance Team |
| EPI_ISL_3336882 | Department of Virology, Istituto Zooprofilattico Sperimentale del Lazio e della Toscana (IZSLT) | Department of General Diagnostics; Department of Virology; Istituto Zooprofilattico Sperimentale del Lazio e della Toscana (IZSLT) | Patricia Alba, Giuseppe Manna, Elena L. Diaconu, Fabiola Feltrin, Raffaela Conti, Teresa Scicluna, Virginia Carfora, Antonella Cersini, Alessia Franco, Antonio Battisti. |

| | | | |
|---|---|---|---|
| EPI_ISL_3336883, EPI_ISL_3336884 | Lighthouse Lab in Milton Keynes | Wellcome Sanger Institute for the COVID-19 Genomics UK (COG-UK) Consortium | The Lighthouse Lab in Milton Keynes and Alex Alderton, Roberto Amato, Jeffrey Barrett, Sonia Goncalves, Ewan Harrison, David K. Jackson, Ian Johnston, Dominic Kwiatkowski, Cordelia Langford, John Sillitoe on behalf of the Wellcome Sanger Institute COVID-19 Surveillance Team |
| EPI_ISL_3336885 | Lighthouse Laboratory Plymouth | Wellcome Sanger Institute for the COVID-19 Genomics UK (COG-UK) Consortium | Lighthouse Laboratory Plymouth and Alex Alderton, Roberto Amato, Jeffrey Barrett, Sonia Goncalves, Ewan Harrison, David K. Jackson, Ian Johnston, Dominic Kwiatkowski, Cordelia Langford, John Sillitoe on behalf of the Wellcome Sanger Institute COVID-19 Surveillance Team |
| EPI_ISL_3336886 | University of Rome Tor Vergata: Departm Experim Medicine Chair of Virology | Department of General Diagnostics; Department of Virology; Istituto Zooprofilattico Sperimentale del Lazio e della Toscana (IZSLT) | Patricia Alba, Giuseppe Manna, Elena L. Diaconu, Fabiola Feltrin, Raffaela Conti, Teresa Scicluna, Virginia Carfora, Antonella Cersini, Alessia Franco, Antonio Battisti. |
| EPI_ISL_3336887, EPI_ISL_3336888 | Lighthouse Lab in Milton Keynes | Wellcome Sanger Institute for the COVID-19 Genomics UK (COG-UK) Consortium | The Lighthouse Lab in Milton Keynes and Alex Alderton, Roberto Amato, Jeffrey Barrett, Sonia Goncalves, Ewan Harrison, David K. Jackson, Ian Johnston, Dominic Kwiatkowski, Cordelia Langford, John Sillitoe on behalf of the Wellcome Sanger Institute COVID-19 Surveillance Team |
| EPI_ISL_3336889 | Department of Virology, Istituto Zooprofilattico Sperimentale del Lazio e della Toscana (IZSLT) | Department of General Diagnostics; Department of Virology; Istituto Zooprofilattico Sperimentale del Lazio e della Toscana (IZSLT) | Patricia Alba, Giuseppe Manna, Elena L. Diaconu, Fabiola Feltrin, Raffaela Conti, Teresa Scicluna, Virginia Carfora, Antonella Cersini, Alessia Franco, Antonio Battisti. |
| EPI_ISL_3336890 | Lighthouse Lab in Milton Keynes | Wellcome Sanger Institute for the COVID-19 Genomics UK (COG-UK) Consortium | The Lighthouse Lab in Milton Keynes and Alex Alderton, Roberto Amato, Jeffrey Barrett, Sonia Goncalves, Ewan Harrison, David K. Jackson, Ian Johnston, Dominic Kwiatkowski, Cordelia Langford, John Sillitoe on behalf of the Wellcome Sanger Institute COVID-19 Surveillance Team |
| EPI_ISL_3336891, EPI_ISL_3336892 | Lighthouse Lab in Alderley Park | Wellcome Sanger Institute for the COVID-19 Genomics UK (COG-UK) Consortium | Jacquelyn Wynn, Mairead Hyland, The Lighthouse Lab in Alderley Park and Alex Alderton, Roberto Amato, Jeffrey Barrett, Sonia Goncalves, Ewan Harrison, David K. Jackson, Ian Johnston, Dominic Kwiatkowski, Cordelia Langford, John Sillitoe on behalf of the Wellcome Sanger Institute COVID-19 Surveillance Team |
| EPI_ISL_3336893 | Department of Virology, Istituto Zooprofilattico Sperimentale del Lazio e della Toscana (IZSLT) | Department of General Diagnostics; Department of Virology; Istituto Zooprofilattico Sperimentale del Lazio e della Toscana (IZSLT) | Patricia Alba, Giuseppe Manna, Elena L. Diaconu, Fabiola Feltrin, Raffaela Conti, Teresa Scicluna, Virginia Carfora, Antonella Cersini, Alessia Franco, Antonio Battisti. |
| EPI_ISL_3336894 | Lighthouse Lab in Alderley Park | Wellcome Sanger Institute for the COVID-19 Genomics UK (COG-UK) Consortium | Jacquelyn Wynn, Mairead Hyland, The Lighthouse Lab in Alderley Park and Alex Alderton, Roberto Amato, Jeffrey Barrett, Sonia Goncalves, Ewan Harrison, David K. Jackson, Ian Johnston, Dominic Kwiatkowski, Cordelia Langford, John Sillitoe on behalf of the Wellcome Sanger Institute COVID-19 Surveillance Team |
| EPI_ISL_3336895 | Lighthouse Laboratory Plymouth | Wellcome Sanger Institute for the COVID-19 Genomics UK (COG-UK) Consortium | Lighthouse Laboratory Plymouth and Alex Alderton, Roberto Amato, Jeffrey Barrett, Sonia Goncalves, Ewan Harrison, David K. Jackson, Ian Johnston, Dominic Kwiatkowski, Cordelia Langford, John Sillitoe on behalf of the Wellcome Sanger Institute COVID-19 Surveillance Team |
| EPI_ISL_3336896 | Lighthouse Lab in Alderley Park | Wellcome Sanger Institute for the COVID-19 Genomics UK (COG-UK) Consortium | Jacquelyn Wynn, Mairead Hyland, The Lighthouse Lab in Alderley Park and Alex Alderton, Roberto Amato, Jeffrey Barrett, Sonia Goncalves, Ewan Harrison, David K. Jackson, Ian Johnston, Dominic Kwiatkowski, Cordelia Langford, John Sillitoe on behalf of the Wellcome Sanger Institute COVID-19 Surveillance Team |
| EPI_ISL_3336897 | Department of Virology, Istituto Zooprofilattico Sperimentale del Lazio e della Toscana (IZSLT) | Department of General Diagnostics; Department of Virology; Istituto Zooprofilattico Sperimentale del Lazio e della Toscana (IZSLT) | Patricia Alba, Giuseppe Manna, Elena L. Diaconu, Fabiola Feltrin, Raffaela Conti, Teresa Scicluna, Virginia Carfora, Antonella Cersini, Alessia Franco, Antonio Battisti. |
| EPI_ISL_3336898 | Lighthouse Lab in Milton Keynes | Wellcome Sanger Institute for the COVID-19 Genomics UK (COG-UK) Consortium | The Lighthouse Lab in Milton Keynes and Alex Alderton, Roberto Amato, Jeffrey Barrett, Sonia Goncalves, Ewan Harrison, David K. Jackson, Ian Johnston, Dominic Kwiatkowski, Cordelia Langford, John Sillitoe on behalf of the Wellcome Sanger Institute COVID-19 Surveillance Team |
| EPI_ISL_3336899 | Lighthouse Lab in Alderley Park | Wellcome Sanger Institute for the COVID-19 Genomics UK (COG-UK) Consortium | Jacquelyn Wynn, Mairead Hyland, The Lighthouse Lab in Alderley Park and Alex Alderton, Roberto Amato, Jeffrey Barrett, Sonia Goncalves, Ewan Harrison, David K. Jackson, Ian Johnston, Dominic Kwiatkowski, Cordelia Langford, John Sillitoe on behalf of the Wellcome Sanger Institute COVID-19 Surveillance Team |
| EPI_ISL_3336900 | Department of Virology, Istituto Zooprofilattico Sperimentale del Lazio e della Toscana (IZSLT) | Department of General Diagnostics; Department of Virology; Istituto Zooprofilattico Sperimentale del Lazio e della Toscana (IZSLT) | Patricia Alba, Giuseppe Manna, Elena L. Diaconu, Fabiola Feltrin, Raffaela Conti, Teresa Scicluna, Virginia Carfora, Antonella Cersini, Alessia Franco, Antonio Battisti. |
| EPI_ISL_3336901 | Lighthouse Lab in Milton Keynes | Wellcome Sanger Institute for the COVID-19 Genomics UK (COG-UK) Consortium | The Lighthouse Lab in Milton Keynes and Alex Alderton, Roberto Amato, Jeffrey Barrett, Sonia Goncalves, Ewan Harrison, David K. Jackson, Ian Johnston, Dominic Kwiatkowski, Cordelia Langford, John Sillitoe on behalf of the Wellcome Sanger Institute COVID-19 Surveillance Team |
| EPI_ISL_3336902, EPI_ISL_3336903 | Berkshire and Surrey Pathology Services Lighthouse Laboratory | Wellcome Sanger Institute for the COVID-19 Genomics UK (COG-UK) Consortium | Berkshire and Surrey Pathology Services Lighthouse Laboratory and Alex Alderton, Roberto Amato, Jeffrey Barrett, Sonia Goncalves, Ewan Harrison, David K. Jackson, Ian Johnston, Dominic Kwiatkowski, Cordelia Langford, John Sillitoe on behalf of the Wellcome Sanger Institute COVID-19 Surveillance Team |
| EPI_ISL_3336904 | Department of Virology, Istituto Zooprofilattico Sperimentale del Lazio e della Toscana (IZSLT) | Department of General Diagnostics; Department of Virology; Istituto Zooprofilattico Sperimentale del Lazio e della Toscana (IZSLT) | Patricia Alba, Giuseppe Manna, Elena L. Diaconu, Fabiola Feltrin, Raffaela Conti, Teresa Scicluna, Virginia Carfora, Antonella Cersini, Alessia Franco, Antonio Battisti. |
| EPI_ISL_3336905 | Lighthouse Lab in Alderley Park | Wellcome Sanger Institute for the COVID-19 Genomics UK (COG-UK) Consortium | Jacquelyn Wynn, Mairead Hyland, The Lighthouse Lab in Alderley Park and Alex Alderton, Roberto Amato, Jeffrey Barrett, Sonia Goncalves, Ewan Harrison, David K. Jackson, Ian Johnston, Dominic Kwiatkowski, Cordelia Langford, John Sillitoe on behalf of the Wellcome Sanger Institute COVID-19 Surveillance Team |
| EPI_ISL_3336906 | Lighthouse Laboratory Plymouth | Wellcome Sanger Institute for the COVID-19 Genomics UK (COG-UK) Consortium | Lighthouse Laboratory Plymouth and Alex Alderton, Roberto Amato, Jeffrey Barrett, Sonia Goncalves, Ewan Harrison, David K. Jackson, Ian Johnston, Dominic Kwiatkowski, Cordelia Langford, John Sillitoe on behalf of the Wellcome Sanger Institute COVID-19 Surveillance Team |
| EPI_ISL_3336907 | University of Rome Tor Vergata: Departm Experim Medicine Chair of Virology | Department of General Diagnostics; Department of Virology; Istituto Zooprofilattico Sperimentale del Lazio e della Toscana (IZSLT) | Patricia Alba, Giuseppe Manna, Elena L. Diaconu, Fabiola Feltrin, Raffaela Conti, Teresa Scicluna, Virginia Carfora, Antonella Cersini, Alessia Franco, Antonio Battisti. |
| EPI_ISL_3336908 | Lighthouse Lab in Alderley Park | Wellcome Sanger Institute for the COVID-19 Genomics UK (COG-UK) Consortium | Jacquelyn Wynn, Mairead Hyland, The Lighthouse Lab in Alderley Park and Alex Alderton, Roberto Amato, Jeffrey Barrett, Sonia Goncalves, Ewan Harrison, David K. Jackson, Ian Johnston, Dominic Kwiatkowski, Cordelia Langford, John Sillitoe on behalf of the Wellcome Sanger Institute COVID-19 Surveillance Team |
| EPI_ISL_3336909, EPI_ISL_3336910 | Berkshire and Surrey Pathology Services Lighthouse Laboratory | Wellcome Sanger Institute for the COVID-19 Genomics UK (COG-UK) Consortium | Berkshire and Surrey Pathology Services Lighthouse Laboratory and Alex Alderton, Roberto Amato, Jeffrey Barrett, Sonia Goncalves, Ewan Harrison, David K. Jackson, Ian Johnston, Dominic Kwiatkowski, Cordelia Langford, John Sillitoe on behalf of the Wellcome Sanger Institute COVID-19 Surveillance Team |
| EPI_ISL_3336911 | Department of Virology, Istituto Zooprofilattico Sperimentale del Lazio e della Toscana (IZSLT) | Department of General Diagnostics; Department of Virology; Istituto Zooprofilattico Sperimentale del Lazio e della Toscana (IZSLT) | Patricia Alba, Giuseppe Manna, Elena L. Diaconu, Fabiola Feltrin, Raffaela Conti, Teresa Scicluna, Virginia Carfora, Antonella Cersini, Alessia Franco, Antonio Battisti. |
| EPI_ISL_3336912, EPI_ISL_3336913 | Lighthouse Lab in Milton Keynes | Wellcome Sanger Institute for the COVID-19 Genomics UK (COG-UK) Consortium | The Lighthouse Lab in Milton Keynes and Alex Alderton, Roberto Amato, Jeffrey Barrett, Sonia Goncalves, Ewan Harrison, David K. Jackson, Ian Johnston, Dominic Kwiatkowski, Cordelia Langford, John Sillitoe on behalf of the Wellcome Sanger Institute COVID-19 Surveillance Team |
| EPI_ISL_3336914 | University of Rome Tor Vergata: Departm Experim Medicine Chair of Virology | Department of General Diagnostics; Department of Virology; Istituto Zooprofilattico Sperimentale del Lazio e della Toscana (IZSLT) | Patricia Alba, Giuseppe Manna, Elena L. Diaconu, Fabiola Feltrin, Raffaela Conti, Teresa Scicluna, Virginia Carfora, Antonella Cersini, Alessia Franco, Antonio Battisti. |
| EPI_ISL_3336915, EPI_ISL_3336916 | Lighthouse Lab in Milton Keynes | Wellcome Sanger Institute for the COVID-19 Genomics UK (COG-UK) Consortium | The Lighthouse Lab in Milton Keynes and Alex Alderton, Roberto Amato, Jeffrey Barrett, Sonia Goncalves, Ewan Harrison, David K. Jackson, Ian Johnston, Dominic Kwiatkowski, Cordelia Langford, John Sillitoe on behalf of the Wellcome Sanger Institute COVID-19 Surveillance Team |
| EPI_ISL_3336917 | Berkshire and Surrey Pathology Services Lighthouse Laboratory | Wellcome Sanger Institute for the COVID-19 Genomics UK (COG-UK) Consortium | Berkshire and Surrey Pathology Services Lighthouse Laboratory and Alex Alderton, Roberto Amato, Jeffrey Barrett, Sonia Goncalves, Ewan Harrison, David K. Jackson, Ian Johnston, Dominic Kwiatkowski, Cordelia Langford, John Sillitoe on behalf of the Wellcome Sanger Institute COVID-19 Surveillance Team |
| EPI_ISL_3336918 | University of Rome Tor Vergata: Departm Experim Medicine Chair of Virology | Department of General Diagnostics; Department of Virology; Istituto Zooprofilattico Sperimentale del Lazio e della Toscana (IZSLT) | Patricia Alba, Giuseppe Manna, Elena L. Diaconu, Fabiola Feltrin, Raffaela Conti, Teresa Scicluna, Virginia Carfora, Antonella Cersini, Alessia Franco, Antonio Battisti. |
| EPI_ISL_3336919 | Lighthouse Lab in Alderley Park | Wellcome Sanger Institute for the COVID-19 Genomics UK (COG-UK) Consortium | Jacquelyn Wynn, Mairead Hyland, The Lighthouse Lab in Alderley Park and Alex Alderton, Roberto Amato, Jeffrey Barrett, Sonia Goncalves, Ewan Harrison, David K. Jackson, Ian Johnston, Dominic Kwiatkowski, Cordelia Langford, John Sillitoe on behalf of the Wellcome Sanger Institute COVID-19 Surveillance Team |
| EPI_ISL_3336920, EPI_ISL_3336921 | Lighthouse Lab in Milton Keynes | Wellcome Sanger Institute for the COVID-19 Genomics UK (COG-UK) Consortium | The Lighthouse Lab in Milton Keynes and Alex Alderton, Roberto Amato, Jeffrey Barrett, Sonia Goncalves, Ewan Harrison, David K. Jackson, Ian Johnston, Dominic Kwiatkowski, Cordelia Langford, John Sillitoe on behalf of the Wellcome Sanger Institute COVID-19 Surveillance Team |
| EPI_ISL_3336922 | Department of Virology, Istituto Zooprofilattico Sperimentale del Lazio e della Toscana (IZSLT) | Department of General Diagnostics; Department of Virology; Istituto Zooprofilattico Sperimentale del Lazio e della Toscana (IZSLT) | Patricia Alba, Giuseppe Manna, Elena L. Diaconu, Fabiola Feltrin, Raffaela Conti, Teresa Scicluna, Virginia Carfora, Antonella Cersini, Alessia Franco, Antonio Battisti. |
| EPI_ISL_3336923, EPI_ISL_3336924 | Lighthouse Lab in Milton Keynes | Wellcome Sanger Institute for the COVID-19 Genomics UK (COG-UK) Consortium | The Lighthouse Lab in Milton Keynes and Alex Alderton, Roberto Amato, Jeffrey Barrett, Sonia Goncalves, Ewan Harrison, David K. Jackson, Ian Johnston, Dominic Kwiatkowski, Cordelia Langford, John Sillitoe on behalf of the Wellcome Sanger Institute COVID-19 Surveillance Team |
| EPI_ISL_3336925 | Lighthouse Lab in Alderley Park | Wellcome Sanger Institute for the COVID-19 Genomics UK (COG-UK) Consortium | Jacquelyn Wynn, Mairead Hyland, The Lighthouse Lab in Alderley Park and Alex Alderton, Roberto Amato, Jeffrey Barrett, Sonia Goncalves, Ewan Harrison, David K. Jackson, Ian Johnston, Dominic Kwiatkowski, Cordelia Langford, John Sillitoe on behalf of the Wellcome Sanger Institute COVID-19 Surveillance Team |
| EPI_ISL_3336926, EPI_ISL_3336927 | Berkshire and Surrey Pathology Services Lighthouse Laboratory | Wellcome Sanger Institute for the COVID-19 Genomics UK (COG-UK) Consortium | Berkshire and Surrey Pathology Services Lighthouse Laboratory and Alex Alderton, Roberto Amato, Jeffrey Barrett, Sonia Goncalves, Ewan Harrison, David K. Jackson, Ian Johnston, Dominic Kwiatkowski, Cordelia Langford, John Sillitoe on behalf of the Wellcome Sanger Institute COVID-19 Surveillance Team |
| EPI_ISL_3336928 | Lighthouse Lab in Milton Keynes | Wellcome Sanger Institute for the COVID-19 Genomics UK (COG-UK) Consortium | The Lighthouse Lab in Milton Keynes and Alex Alderton, Roberto Amato, Jeffrey Barrett, Sonia Goncalves, Ewan Harrison, David K. Jackson, Ian Johnston, Dominic Kwiatkowski, Cordelia Langford, John Sillitoe on behalf of the Wellcome Sanger Institute COVID-19 Surveillance Team |
| EPI_ISL_3336929, EPI_ISL_3336930 | Berkshire and Surrey Pathology Services Lighthouse Laboratory | Wellcome Sanger Institute for the COVID-19 Genomics UK (COG-UK) Consortium | Berkshire and Surrey Pathology Services Lighthouse Laboratory and Alex Alderton, Roberto Amato, Jeffrey Barrett, Sonia Goncalves, Ewan Harrison, David K. Jackson, Ian Johnston, Dominic Kwiatkowski, Cordelia Langford, John Sillitoe on behalf of the Wellcome Sanger Institute COVID-19 Surveillance Team |
| EPI_ISL_3336931 | Lighthouse Laboratory Plymouth | Wellcome Sanger Institute for the COVID-19 Genomics UK (COG-UK) Consortium | Lighthouse Laboratory Plymouth and Alex Alderton, Roberto Amato, Jeffrey Barrett, Sonia Goncalves, Ewan Harrison, David K. Jackson, Ian Johnston, Dominic Kwiatkowski, Cordelia Langford, John Sillitoe on behalf of the Wellcome Sanger Institute COVID-19 Surveillance Team |
| EPI_ISL_3336932 | Berkshire and Surrey Pathology Services Lighthouse Laboratory | Wellcome Sanger Institute for the COVID-19 Genomics UK (COG-UK) Consortium | Berkshire and Surrey Pathology Services Lighthouse Laboratory and Alex Alderton, Roberto Amato, Jeffrey Barrett, Sonia Goncalves, Ewan Harrison, David K. Jackson, Ian Johnston, Dominic Kwiatkowski, Cordelia Langford, John Sillitoe on behalf of the Wellcome Sanger Institute COVID-19 Surveillance Team |
| EPI_ISL_3336933 | Lighthouse Laboratory Plymouth | Wellcome Sanger Institute for the COVID-19 Genomics UK (COG-UK) Consortium | Lighthouse Laboratory Plymouth and Alex Alderton, Roberto Amato, Jeffrey Barrett, Sonia Goncalves, Ewan Harrison, David K. Jackson, Ian Johnston, Dominic Kwiatkowski, Cordelia Langford, John Sillitoe on behalf of the Wellcome Sanger Institute COVID-19 Surveillance Team |
| EPI_ISL_3336934 | Lighthouse Lab in Alderley Park | Wellcome Sanger Institute for the COVID-19 Genomics UK (COG-UK) Consortium | Jacquelyn Wynn, Mairead Hyland, The Lighthouse Lab in Alderley Park and Alex Alderton, Roberto Amato, Jeffrey Barrett, Sonia Goncalves, Ewan Harrison, David K. Jackson, Ian Johnston, Dominic Kwiatkowski, Cordelia Langford, John Sillitoe on behalf of the Wellcome Sanger Institute COVID-19 Surveillance Team |
| EPI_ISL_3336935 | Lighthouse Lab in Milton Keynes | Wellcome Sanger Institute for the COVID-19 Genomics UK (COG-UK) Consortium | The Lighthouse Lab in Milton Keynes and Alex Alderton, Roberto Amato, Jeffrey Barrett, Sonia Goncalves, Ewan Harrison, David K. Jackson, Ian Johnston, Dominic Kwiatkowski, Cordelia Langford, John Sillitoe on behalf of the Wellcome Sanger Institute COVID-19 Surveillance Team |
| EPI_ISL_3336936, EPI_ISL_3336937, EPI_ISL_3336938, EPI_ISL_3336939 | Lighthouse Lab in Alderley Park | Wellcome Sanger Institute for the COVID-19 Genomics UK (COG-UK) Consortium | Jacquelyn Wynn, Mairead Hyland, The Lighthouse Lab in Alderley Park and Alex Alderton, Roberto Amato, Jeffrey Barrett, Sonia Goncalves, Ewan Harrison, David K. Jackson, Ian Johnston, Dominic Kwiatkowski, Cordelia Langford, John Sillitoe on behalf of the Wellcome Sanger Institute COVID-19 Surveillance Team |
| EPI_ISL_3336940 | Lighthouse Laboratory Plymouth | Wellcome Sanger Institute for the COVID-19 Genomics UK (COG-UK) Consortium | Lighthouse Laboratory Plymouth and Alex Alderton, Roberto Amato, Jeffrey Barrett, Sonia Goncalves, Ewan Harrison, David K. Jackson, Ian Johnston, Dominic Kwiatkowski, Cordelia Langford, John Sillitoe on behalf of the Wellcome Sanger Institute COVID-19 Surveillance Team |

| | | | |
|---|---|---|---|
| EPI_ISL_3336941, EPI_ISL_3336942 | Lighthouse Lab in Milton Keynes | Wellcome Sanger Institute for the COVID-19 Genomics UK (COG-UK) Consortium | The Lighthouse Lab in Milton Keynes and Alex Alderton, Roberto Amato, Jeffrey Barrett, Sonia Goncalves, Ewan Harrison, David K. Jackson, Ian Johnston, Dominic Kwiatkowski, Cordelia Langford, John Sillitoe on behalf of the Wellcome Sanger Institute COVID-19 Surveillance Team |
| EPI_ISL_3336943 | Lighthouse Laboratory Plymouth | Wellcome Sanger Institute for the COVID-19 Genomics UK (COG-UK) Consortium | Lighthouse Laboratory Plymouth and Alex Alderton, Roberto Amato, Jeffrey Barrett, Sonia Goncalves, Ewan Harrison, David K. Jackson, Ian Johnston, Dominic Kwiatkowski, Cordelia Langford, John Sillitoe on behalf of the Wellcome Sanger Institute COVID-19 Surveillance Team |
| EPI_ISL_3336944 | Lighthouse Lab in Milton Keynes | Wellcome Sanger Institute for the COVID-19 Genomics UK (COG-UK) Consortium | The Lighthouse Lab in Milton Keynes and Alex Alderton, Roberto Amato, Jeffrey Barrett, Sonia Goncalves, Ewan Harrison, David K. Jackson, Ian Johnston, Dominic Kwiatkowski, Cordelia Langford, John Sillitoe on behalf of the Wellcome Sanger Institute COVID-19 Surveillance Team |
| EPI_ISL_3336945 | Lighthouse Lab in Alderley Park | Wellcome Sanger Institute for the COVID-19 Genomics UK (COG-UK) Consortium | Jacquelyn Wynn, Mairead Hyland, The Lighthouse Lab in Alderley Park and Alex Alderton, Roberto Amato, Jeffrey Barrett, Sonia Goncalves, Ewan Harrison, David K. Jackson, Ian Johnston, Dominic Kwiatkowski, Cordelia Langford, John Sillitoe on behalf of the Wellcome Sanger Institute COVID-19 Surveillance Team |
| EPI_ISL_3336946 | Lighthouse Laboratory Plymouth | Wellcome Sanger Institute for the COVID-19 Genomics UK (COG-UK) Consortium | Lighthouse Laboratory Plymouth and Alex Alderton, Roberto Amato, Jeffrey Barrett, Sonia Goncalves, Ewan Harrison, David K. Jackson, Ian Johnston, Dominic Kwiatkowski, Cordelia Langford, John Sillitoe on behalf of the Wellcome Sanger Institute COVID-19 Surveillance Team |
| EPI_ISL_3336947 | Lighthouse Lab in Milton Keynes | Wellcome Sanger Institute for the COVID-19 Genomics UK (COG-UK) Consortium | The Lighthouse Lab in Milton Keynes and Alex Alderton, Roberto Amato, Jeffrey Barrett, Sonia Goncalves, Ewan Harrison, David K. Jackson, Ian Johnston, Dominic Kwiatkowski, Cordelia Langford, John Sillitoe on behalf of the Wellcome Sanger Institute COVID-19 Surveillance Team |
| EPI_ISL_3336948 | Lighthouse Lab in Alderley Park | Wellcome Sanger Institute for the COVID-19 Genomics UK (COG-UK) Consortium | Jacquelyn Wynn, Mairead Hyland, The Lighthouse Lab in Alderley Park and Alex Alderton, Roberto Amato, Jeffrey Barrett, Sonia Goncalves, Ewan Harrison, David K. Jackson, Ian Johnston, Dominic Kwiatkowski, Cordelia Langford, John Sillitoe on behalf of the Wellcome Sanger Institute COVID-19 Surveillance Team |
| EPI_ISL_3336949 | Lighthouse Lab in Milton Keynes | Wellcome Sanger Institute for the COVID-19 Genomics UK (COG-UK) Consortium | The Lighthouse Lab in Milton Keynes and Alex Alderton, Roberto Amato, Jeffrey Barrett, Sonia Goncalves, Ewan Harrison, David K. Jackson, Ian Johnston, Dominic Kwiatkowski, Cordelia Langford, John Sillitoe on behalf of the Wellcome Sanger Institute COVID-19 Surveillance Team |
| EPI_ISL_3336950, EPI_ISL_3336951 | Lighthouse Lab in Alderley Park | Wellcome Sanger Institute for the COVID-19 Genomics UK (COG-UK) Consortium | Jacquelyn Wynn, Mairead Hyland, The Lighthouse Lab in Alderley Park and Alex Alderton, Roberto Amato, Jeffrey Barrett, Sonia Goncalves, Ewan Harrison, David K. Jackson, Ian Johnston, Dominic Kwiatkowski, Cordelia Langford, John Sillitoe on behalf of the Wellcome Sanger Institute COVID-19 Surveillance Team |
| EPI_ISL_3336952, EPI_ISL_3336953 | Lighthouse Lab in Milton Keynes | Wellcome Sanger Institute for the COVID-19 Genomics UK (COG-UK) Consortium | The Lighthouse Lab in Milton Keynes and Alex Alderton, Roberto Amato, Jeffrey Barrett, Sonia Goncalves, Ewan Harrison, David K. Jackson, Ian Johnston, Dominic Kwiatkowski, Cordelia Langford, John Sillitoe on behalf of the Wellcome Sanger Institute COVID-19 Surveillance Team |
| EPI_ISL_3336954 | Lighthouse Lab in Alderley Park | Wellcome Sanger Institute for the COVID-19 Genomics UK (COG-UK) Consortium | Jacquelyn Wynn, Mairead Hyland, The Lighthouse Lab in Alderley Park and Alex Alderton, Roberto Amato, Jeffrey Barrett, Sonia Goncalves, Ewan Harrison, David K. Jackson, Ian Johnston, Dominic Kwiatkowski, Cordelia Langford, John Sillitoe on behalf of the Wellcome Sanger Institute COVID-19 Surveillance Team |
| EPI_ISL_3336955 | Lighthouse Lab in Milton Keynes | Wellcome Sanger Institute for the COVID-19 Genomics UK (COG-UK) Consortium | The Lighthouse Lab in Milton Keynes and Alex Alderton, Roberto Amato, Jeffrey Barrett, Sonia Goncalves, Ewan Harrison, David K. Jackson, Ian Johnston, Dominic Kwiatkowski, Cordelia Langford, John Sillitoe on behalf of the Wellcome Sanger Institute COVID-19 Surveillance Team |
| EPI_ISL_3336956, EPI_ISL_3336957 | Berkshire and Surrey Pathology Services Lighthouse Laboratory | Wellcome Sanger Institute for the COVID-19 Genomics UK (COG-UK) Consortium | Berkshire and Surrey Pathology Services Lighthouse Laboratory and Alex Alderton, Roberto Amato, Jeffrey Barrett, Sonia Goncalves, Ewan Harrison, David K. Jackson, Ian Johnston, Dominic Kwiatkowski, Cordelia Langford, John Sillitoe on behalf of the Wellcome Sanger Institute COVID-19 Surveillance Team |
| EPI_ISL_3336958 | Lighthouse Lab in Alderley Park | Wellcome Sanger Institute for the COVID-19 Genomics UK (COG-UK) Consortium | Jacquelyn Wynn, Mairead Hyland, The Lighthouse Lab in Alderley Park and Alex Alderton, Roberto Amato, Jeffrey Barrett, Sonia Goncalves, Ewan Harrison, David K. Jackson, Ian Johnston, Dominic Kwiatkowski, Cordelia Langford, John Sillitoe on behalf of the Wellcome Sanger Institute COVID-19 Surveillance Team |
| EPI_ISL_3336959 | Berkshire and Surrey Pathology Services Lighthouse Laboratory | Wellcome Sanger Institute for the COVID-19 Genomics UK (COG-UK) Consortium | Berkshire and Surrey Pathology Services Lighthouse Laboratory and Alex Alderton, Roberto Amato, Jeffrey Barrett, Sonia Goncalves, Ewan Harrison, David K. Jackson, Ian Johnston, Dominic Kwiatkowski, Cordelia Langford, John Sillitoe on behalf of the Wellcome Sanger Institute COVID-19 Surveillance Team |
| EPI_ISL_3336960 | Lighthouse Lab in Alderley Park | Wellcome Sanger Institute for the COVID-19 Genomics UK (COG-UK) Consortium | Jacquelyn Wynn, Mairead Hyland, The Lighthouse Lab in Alderley Park and Alex Alderton, Roberto Amato, Jeffrey Barrett, Sonia Goncalves, Ewan Harrison, David K. Jackson, Ian Johnston, Dominic Kwiatkowski, Cordelia Langford, John Sillitoe on behalf of the Wellcome Sanger Institute COVID-19 Surveillance Team |
| EPI_ISL_3336961, EPI_ISL_3336962 | Berkshire and Surrey Pathology Services Lighthouse Laboratory | Wellcome Sanger Institute for the COVID-19 Genomics UK (COG-UK) Consortium | Berkshire and Surrey Pathology Services Lighthouse Laboratory and Alex Alderton, Roberto Amato, Jeffrey Barrett, Sonia Goncalves, Ewan Harrison, David K. Jackson, Ian Johnston, Dominic Kwiatkowski, Cordelia Langford, John Sillitoe on behalf of the Wellcome Sanger Institute COVID-19 Surveillance Team |
| EPI_ISL_3336963 | Lighthouse Lab in Milton Keynes | Wellcome Sanger Institute for the COVID-19 Genomics UK (COG-UK) Consortium | The Lighthouse Lab in Milton Keynes and Alex Alderton, Roberto Amato, Jeffrey Barrett, Sonia Goncalves, Ewan Harrison, David K. Jackson, Ian Johnston, Dominic Kwiatkowski, Cordelia Langford, John Sillitoe on behalf of the Wellcome Sanger Institute COVID-19 Surveillance Team |
| EPI_ISL_3336964, EPI_ISL_3336965 | Berkshire and Surrey Pathology Services Lighthouse Laboratory | Wellcome Sanger Institute for the COVID-19 Genomics UK (COG-UK) Consortium | Berkshire and Surrey Pathology Services Lighthouse Laboratory and Alex Alderton, Roberto Amato, Jeffrey Barrett, Sonia Goncalves, Ewan Harrison, David K. Jackson, Ian Johnston, Dominic Kwiatkowski, Cordelia Langford, John Sillitoe on behalf of the Wellcome Sanger Institute COVID-19 Surveillance Team |
| EPI_ISL_3336966, EPI_ISL_3336967, EPI_ISL_3336968 | Lighthouse Lab in Milton Keynes | Wellcome Sanger Institute for the COVID-19 Genomics UK (COG-UK) Consortium | The Lighthouse Lab in Milton Keynes and Alex Alderton, Roberto Amato, Jeffrey Barrett, Sonia Goncalves, Ewan Harrison, David K. Jackson, Ian Johnston, Dominic Kwiatkowski, Cordelia Langford, John Sillitoe on behalf of the Wellcome Sanger Institute COVID-19 Surveillance Team |
| EPI_ISL_3336969, EPI_ISL_3336970 | Lighthouse Lab in Alderley Park | Wellcome Sanger Institute for the COVID-19 Genomics UK (COG-UK) Consortium | Jacquelyn Wynn, Mairead Hyland, The Lighthouse Lab in Alderley Park and Alex Alderton, Roberto Amato, Jeffrey Barrett, Sonia Goncalves, Ewan Harrison, David K. Jackson, Ian Johnston, Dominic Kwiatkowski, Cordelia Langford, John Sillitoe on behalf of the Wellcome Sanger Institute COVID-19 Surveillance Team |
| EPI_ISL_3336971, EPI_ISL_3336972 | Lighthouse Lab in Milton Keynes | Wellcome Sanger Institute for the COVID-19 Genomics UK (COG-UK) Consortium | The Lighthouse Lab in Milton Keynes and Alex Alderton, Roberto Amato, Jeffrey Barrett, Sonia Goncalves, Ewan Harrison, David K. Jackson, Ian Johnston, Dominic Kwiatkowski, Cordelia Langford, John Sillitoe on behalf of the Wellcome Sanger Institute COVID-19 Surveillance Team |
| EPI_ISL_3336973 | Berkshire and Surrey Pathology Services Lighthouse Laboratory | Wellcome Sanger Institute for the COVID-19 Genomics UK (COG-UK) Consortium | Berkshire and Surrey Pathology Services Lighthouse Laboratory and Alex Alderton, Roberto Amato, Jeffrey Barrett, Sonia Goncalves, Ewan Harrison, David K. Jackson, Ian Johnston, Dominic Kwiatkowski, Cordelia Langford, John Sillitoe on behalf of the Wellcome Sanger Institute COVID-19 Surveillance Team |
| EPI_ISL_3336974 | Lighthouse Lab in Alderley Park | Wellcome Sanger Institute for the COVID-19 Genomics UK (COG-UK) Consortium | Jacquelyn Wynn, Mairead Hyland, The Lighthouse Lab in Alderley Park and Alex Alderton, Roberto Amato, Jeffrey Barrett, Sonia Goncalves, Ewan Harrison, David K. Jackson, Ian Johnston, Dominic Kwiatkowski, Cordelia Langford, John Sillitoe on behalf of the Wellcome Sanger Institute COVID-19 Surveillance Team |
| EPI_ISL_3336975 | Berkshire and Surrey Pathology Services Lighthouse Laboratory | Wellcome Sanger Institute for the COVID-19 Genomics UK (COG-UK) Consortium | Berkshire and Surrey Pathology Services Lighthouse Laboratory and Alex Alderton, Roberto Amato, Jeffrey Barrett, Sonia Goncalves, Ewan Harrison, David K. Jackson, Ian Johnston, Dominic Kwiatkowski, Cordelia Langford, John Sillitoe on behalf of the Wellcome Sanger Institute COVID-19 Surveillance Team |
| EPI_ISL_3336976 | Lighthouse Laboratory Plymouth | Wellcome Sanger Institute for the COVID-19 Genomics UK (COG-UK) Consortium | Lighthouse Laboratory Plymouth and Alex Alderton, Roberto Amato, Jeffrey Barrett, Sonia Goncalves, Ewan Harrison, David K. Jackson, Ian Johnston, Dominic Kwiatkowski, Cordelia Langford, John Sillitoe on behalf of the Wellcome Sanger Institute COVID-19 Surveillance Team |
| EPI_ISL_3336977, EPI_ISL_3336978 | Lighthouse Lab in Alderley Park | Wellcome Sanger Institute for the COVID-19 Genomics UK (COG-UK) Consortium | Jacquelyn Wynn, Mairead Hyland, The Lighthouse Lab in Alderley Park and Alex Alderton, Roberto Amato, Jeffrey Barrett, Sonia Goncalves, Ewan Harrison, David K. Jackson, Ian Johnston, Dominic Kwiatkowski, Cordelia Langford, John Sillitoe on behalf of the Wellcome Sanger Institute COVID-19 Surveillance Team |
| EPI_ISL_3336979 | Berkshire and Surrey Pathology Services Lighthouse Laboratory | Wellcome Sanger Institute for the COVID-19 Genomics UK (COG-UK) Consortium | Berkshire and Surrey Pathology Services Lighthouse Laboratory and Alex Alderton, Roberto Amato, Jeffrey Barrett, Sonia Goncalves, Ewan Harrison, David K. Jackson, Ian Johnston, Dominic Kwiatkowski, Cordelia Langford, John Sillitoe on behalf of the Wellcome Sanger Institute COVID-19 Surveillance Team |
| EPI_ISL_3336980 | Lighthouse Lab in Alderley Park | Wellcome Sanger Institute for the COVID-19 Genomics UK (COG-UK) Consortium | Jacquelyn Wynn, Mairead Hyland, The Lighthouse Lab in Alderley Park and Alex Alderton, Roberto Amato, Jeffrey Barrett, Sonia Goncalves, Ewan Harrison, David K. Jackson, Ian Johnston, Dominic Kwiatkowski, Cordelia Langford, John Sillitoe on behalf of the Wellcome Sanger Institute COVID-19 Surveillance Team |
| EPI_ISL_3336981, EPI_ISL_3336982 | Lighthouse Lab in Milton Keynes | Wellcome Sanger Institute for the COVID-19 Genomics UK (COG-UK) Consortium | The Lighthouse Lab in Milton Keynes and Alex Alderton, Roberto Amato, Jeffrey Barrett, Sonia Goncalves, Ewan Harrison, David K. Jackson, Ian Johnston, Dominic Kwiatkowski, Cordelia Langford, John Sillitoe on behalf of the Wellcome Sanger Institute COVID-19 Surveillance Team |
| EPI_ISL_3336983 | Lighthouse Lab in Alderley Park | Wellcome Sanger Institute for the COVID-19 Genomics UK (COG-UK) Consortium | Jacquelyn Wynn, Mairead Hyland, The Lighthouse Lab in Alderley Park and Alex Alderton, Roberto Amato, Jeffrey Barrett, Sonia Goncalves, Ewan Harrison, David K. Jackson, Ian Johnston, Dominic Kwiatkowski, Cordelia Langford, John Sillitoe on behalf of the Wellcome Sanger Institute COVID-19 Surveillance Team |
| EPI_ISL_3336984 | Lighthouse Laboratory Plymouth | Wellcome Sanger Institute for the COVID-19 Genomics UK (COG-UK) Consortium | Lighthouse Laboratory Plymouth and Alex Alderton, Roberto Amato, Jeffrey Barrett, Sonia Goncalves, Ewan Harrison, David K. Jackson, Ian Johnston, Dominic Kwiatkowski, Cordelia Langford, John Sillitoe on behalf of the Wellcome Sanger Institute COVID-19 Surveillance Team |
| EPI_ISL_3336985 | Lighthouse Lab in Milton Keynes | Wellcome Sanger Institute for the COVID-19 Genomics UK (COG-UK) Consortium | The Lighthouse Lab in Milton Keynes and Alex Alderton, Roberto Amato, Jeffrey Barrett, Sonia Goncalves, Ewan Harrison, David K. Jackson, Ian Johnston, Dominic Kwiatkowski, Cordelia Langford, John Sillitoe on behalf of the Wellcome Sanger Institute COVID-19 Surveillance Team |
| EPI_ISL_3336986 | Berkshire and Surrey Pathology Services Lighthouse Laboratory | Wellcome Sanger Institute for the COVID-19 Genomics UK (COG-UK) Consortium | Berkshire and Surrey Pathology Services Lighthouse Laboratory and Alex Alderton, Roberto Amato, Jeffrey Barrett, Sonia Goncalves, Ewan Harrison, David K. Jackson, Ian Johnston, Dominic Kwiatkowski, Cordelia Langford, John Sillitoe on behalf of the Wellcome Sanger Institute COVID-19 Surveillance Team |
| EPI_ISL_3336987 | Lighthouse Lab in Alderley Park | Wellcome Sanger Institute for the COVID-19 Genomics UK (COG-UK) Consortium | Jacquelyn Wynn, Mairead Hyland, The Lighthouse Lab in Alderley Park and Alex Alderton, Roberto Amato, Jeffrey Barrett, Sonia Goncalves, Ewan Harrison, David K. Jackson, Ian Johnston, Dominic Kwiatkowski, Cordelia Langford, John Sillitoe on behalf of the Wellcome Sanger Institute COVID-19 Surveillance Team |
| EPI_ISL_3336988 | Berkshire and Surrey Pathology Services Lighthouse Laboratory | Wellcome Sanger Institute for the COVID-19 Genomics UK (COG-UK) Consortium | Berkshire and Surrey Pathology Services Lighthouse Laboratory and Alex Alderton, Roberto Amato, Jeffrey Barrett, Sonia Goncalves, Ewan Harrison, David K. Jackson, Ian Johnston, Dominic Kwiatkowski, Cordelia Langford, John Sillitoe on behalf of the Wellcome Sanger Institute COVID-19 Surveillance Team |
| EPI_ISL_3336989 | Lighthouse Lab in Milton Keynes | Wellcome Sanger Institute for the COVID-19 Genomics UK (COG-UK) Consortium | The Lighthouse Lab in Milton Keynes and Alex Alderton, Roberto Amato, Jeffrey Barrett, Sonia Goncalves, Ewan Harrison, David K. Jackson, Ian Johnston, Dominic Kwiatkowski, Cordelia Langford, John Sillitoe on behalf of the Wellcome Sanger Institute COVID-19 Surveillance Team |
| EPI_ISL_3336990 | Lighthouse Lab in Alderley Park | Wellcome Sanger Institute for the COVID-19 Genomics UK (COG-UK) Consortium | Jacquelyn Wynn, Mairead Hyland, The Lighthouse Lab in Alderley Park and Alex Alderton, Roberto Amato, Jeffrey Barrett, Sonia Goncalves, Ewan Harrison, David K. Jackson, Ian Johnston, Dominic Kwiatkowski, Cordelia Langford, John Sillitoe on behalf of the Wellcome Sanger Institute COVID-19 Surveillance Team |
| EPI_ISL_3336991 | Berkshire and Surrey Pathology Services Lighthouse Laboratory | Wellcome Sanger Institute for the COVID-19 Genomics UK (COG-UK) Consortium | Berkshire and Surrey Pathology Services Lighthouse Laboratory and Alex Alderton, Roberto Amato, Jeffrey Barrett, Sonia Goncalves, Ewan Harrison, David K. Jackson, Ian Johnston, Dominic Kwiatkowski, Cordelia Langford, John Sillitoe on behalf of the Wellcome Sanger Institute COVID-19 Surveillance Team |
| EPI_ISL_3336992 | Lighthouse Lab in Milton Keynes | Wellcome Sanger Institute for the COVID-19 Genomics UK (COG-UK) Consortium | The Lighthouse Lab in Milton Keynes and Alex Alderton, Roberto Amato, Jeffrey Barrett, Sonia Goncalves, Ewan Harrison, David K. Jackson, Ian Johnston, Dominic Kwiatkowski, Cordelia Langford, John Sillitoe on behalf of the Wellcome Sanger Institute COVID-19 Surveillance Team |
| EPI_ISL_3336993 | Lighthouse Lab in Alderley Park | Wellcome Sanger Institute for the COVID-19 Genomics UK (COG-UK) Consortium | Jacquelyn Wynn, Mairead Hyland, The Lighthouse Lab in Alderley Park and Alex Alderton, Roberto Amato, Jeffrey Barrett, Sonia Goncalves, Ewan Harrison, David K. Jackson, Ian Johnston, Dominic Kwiatkowski, Cordelia Langford, John Sillitoe on behalf of the Wellcome Sanger Institute COVID-19 Surveillance Team |
| EPI_ISL_3336994 | Lighthouse Lab in Milton Keynes | Wellcome Sanger Institute for the COVID-19 Genomics UK (COG-UK) Consortium | The Lighthouse Lab in Milton Keynes and Alex Alderton, Roberto Amato, Jeffrey Barrett, Sonia Goncalves, Ewan Harrison, David K. Jackson, Ian Johnston, Dominic Kwiatkowski, Cordelia Langford, John Sillitoe on behalf of the Wellcome Sanger Institute COVID-19 Surveillance Team |
| EPI_ISL_3336995 | Lighthouse Laboratory Plymouth | Wellcome Sanger Institute for the COVID-19 Genomics UK (COG-UK) Consortium | Lighthouse Laboratory Plymouth and Alex Alderton, Roberto Amato, Jeffrey Barrett, Sonia Goncalves, Ewan Harrison, David K. Jackson, Ian Johnston, Dominic Kwiatkowski, Cordelia Langford, John Sillitoe on behalf of the Wellcome Sanger Institute COVID-19 Surveillance Team |
| EPI_ISL_3336996 | Lighthouse Lab in Alderley Park | Wellcome Sanger Institute for the COVID-19 Genomics UK (COG-UK) Consortium | Jacquelyn Wynn, Mairead Hyland, The Lighthouse Lab in Alderley Park and Alex Alderton, Roberto Amato, Jeffrey Barrett, Sonia Goncalves, Ewan Harrison, David K. Jackson, Ian Johnston, Dominic Kwiatkowski, Cordelia Langford, John Sillitoe on behalf of the Wellcome Sanger Institute COVID-19 Surveillance Team |

| | | | |
|---|---|---|---|
| EPI_ISL_3336997 | Berkshire and Surrey Pathology Services Lighthouse Laboratory | Wellcome Sanger Institute for the COVID-19 Genomics UK (COG-UK) Consortium | Berkshire and Surrey Pathology Services Lighthouse Laboratory and Alex Alderton, Roberto Amato, Jeffrey Barrett, Sonia Goncalves, Ewan Harrison, David K. Jackson, Ian Johnston, Dominic Kwiatkowski, Cordelia Langford, John Sillitoe on behalf of the Wellcome Sanger Institute COVID-19 Surveillance Team |
| EPI_ISL_3336998, EPI_ISL_3336999, EPI_ISL_3337000 | Lighthouse Laboratory Plymouth | Wellcome Sanger Institute for the COVID-19 Genomics UK (COG-UK) Consortium | Lighthouse Laboratory Plymouth and Alex Alderton, Roberto Amato, Jeffrey Barrett, Sonia Goncalves, Ewan Harrison, David K. Jackson, Ian Johnston, Dominic Kwiatkowski, Cordelia Langford, John Sillitoe on behalf of the Wellcome Sanger Institute COVID-19 Surveillance Team |
| EPI_ISL_3337001, EPI_ISL_3337002 | Berkshire and Surrey Pathology Services Lighthouse Laboratory | Wellcome Sanger Institute for the COVID-19 Genomics UK (COG-UK) Consortium | Berkshire and Surrey Pathology Services Lighthouse Laboratory and Alex Alderton, Roberto Amato, Jeffrey Barrett, Sonia Goncalves, Ewan Harrison, David K. Jackson, Ian Johnston, Dominic Kwiatkowski, Cordelia Langford, John Sillitoe on behalf of the Wellcome Sanger Institute COVID-19 Surveillance Team |
| EPI_ISL_3337003 | Lighthouse Lab in Alderley Park | Wellcome Sanger Institute for the COVID-19 Genomics UK (COG-UK) Consortium | Jacquelyn Wynn, Mairead Hyland, The Lighthouse Lab in Alderley Park and Alex Alderton, Roberto Amato, Jeffrey Barrett, Sonia Goncalves, Ewan Harrison, David K. Jackson, Ian Johnston, Dominic Kwiatkowski, Cordelia Langford, John Sillitoe on behalf of the Wellcome Sanger Institute COVID-19 Surveillance Team |
| EPI_ISL_3337004 | Lighthouse Lab in Milton Keynes | Wellcome Sanger Institute for the COVID-19 Genomics UK (COG-UK) Consortium | The Lighthouse Lab in Milton Keynes and Alex Alderton, Roberto Amato, Jeffrey Barrett, Sonia Goncalves, Ewan Harrison, David K. Jackson, Ian Johnston, Dominic Kwiatkowski, Cordelia Langford, John Sillitoe on behalf of the Wellcome Sanger Institute COVID-19 Surveillance Team |
| EPI_ISL_3337005 | Lighthouse Lab in Alderley Park | Wellcome Sanger Institute for the COVID-19 Genomics UK (COG-UK) Consortium | Jacquelyn Wynn, Mairead Hyland, The Lighthouse Lab in Alderley Park and Alex Alderton, Roberto Amato, Jeffrey Barrett, Sonia Goncalves, Ewan Harrison, David K. Jackson, Ian Johnston, Dominic Kwiatkowski, Cordelia Langford, John Sillitoe on behalf of the Wellcome Sanger Institute COVID-19 Surveillance Team |
| EPI_ISL_3337006 | Berkshire and Surrey Pathology Services Lighthouse Laboratory | Wellcome Sanger Institute for the COVID-19 Genomics UK (COG-UK) Consortium | Berkshire and Surrey Pathology Services Lighthouse Laboratory and Alex Alderton, Roberto Amato, Jeffrey Barrett, Sonia Goncalves, Ewan Harrison, David K. Jackson, Ian Johnston, Dominic Kwiatkowski, Cordelia Langford, John Sillitoe on behalf of the Wellcome Sanger Institute COVID-19 Surveillance Team |
| EPI_ISL_3337007 | Lighthouse Laboratory Plymouth | Wellcome Sanger Institute for the COVID-19 Genomics UK (COG-UK) Consortium | Lighthouse Laboratory Plymouth and Alex Alderton, Roberto Amato, Jeffrey Barrett, Sonia Goncalves, Ewan Harrison, David K. Jackson, Ian Johnston, Dominic Kwiatkowski, Cordelia Langford, John Sillitoe on behalf of the Wellcome Sanger Institute COVID-19 Surveillance Team |
| EPI_ISL_3337008, EPI_ISL_3337009, EPI_ISL_3337010 | Lighthouse Lab in Milton Keynes | Wellcome Sanger Institute for the COVID-19 Genomics UK (COG-UK) Consortium | The Lighthouse Lab in Milton Keynes and Alex Alderton, Roberto Amato, Jeffrey Barrett, Sonia Goncalves, Ewan Harrison, David K. Jackson, Ian Johnston, Dominic Kwiatkowski, Cordelia Langford, John Sillitoe on behalf of the Wellcome Sanger Institute COVID-19 Surveillance Team |
| EPI_ISL_3337011 | Lighthouse Laboratory Plymouth | Wellcome Sanger Institute for the COVID-19 Genomics UK (COG-UK) Consortium | Lighthouse Laboratory Plymouth and Alex Alderton, Roberto Amato, Jeffrey Barrett, Sonia Goncalves, Ewan Harrison, David K. Jackson, Ian Johnston, Dominic Kwiatkowski, Cordelia Langford, John Sillitoe on behalf of the Wellcome Sanger Institute COVID-19 Surveillance Team |
| EPI_ISL_3337012 | Berkshire and Surrey Pathology Services Lighthouse Laboratory | Wellcome Sanger Institute for the COVID-19 Genomics UK (COG-UK) Consortium | Berkshire and Surrey Pathology Services Lighthouse Laboratory and Alex Alderton, Roberto Amato, Jeffrey Barrett, Sonia Goncalves, Ewan Harrison, David K. Jackson, Ian Johnston, Dominic Kwiatkowski, Cordelia Langford, John Sillitoe on behalf of the Wellcome Sanger Institute COVID-19 Surveillance Team |
| EPI_ISL_3337013 | Lighthouse Laboratory Plymouth | Wellcome Sanger Institute for the COVID-19 Genomics UK (COG-UK) Consortium | Lighthouse Laboratory Plymouth and Alex Alderton, Roberto Amato, Jeffrey Barrett, Sonia Goncalves, Ewan Harrison, David K. Jackson, Ian Johnston, Dominic Kwiatkowski, Cordelia Langford, John Sillitoe on behalf of the Wellcome Sanger Institute COVID-19 Surveillance Team |
| EPI_ISL_3337014 | Lighthouse Lab in Alderley Park | Wellcome Sanger Institute for the COVID-19 Genomics UK (COG-UK) Consortium | Jacquelyn Wynn, Mairead Hyland, The Lighthouse Lab in Alderley Park and Alex Alderton, Roberto Amato, Jeffrey Barrett, Sonia Goncalves, Ewan Harrison, David K. Jackson, Ian Johnston, Dominic Kwiatkowski, Cordelia Langford, John Sillitoe on behalf of the Wellcome Sanger Institute COVID-19 Surveillance Team |
| EPI_ISL_3337015, EPI_ISL_3337016 | Berkshire and Surrey Pathology Services Lighthouse Laboratory | Wellcome Sanger Institute for the COVID-19 Genomics UK (COG-UK) Consortium | Berkshire and Surrey Pathology Services Lighthouse Laboratory and Alex Alderton, Roberto Amato, Jeffrey Barrett, Sonia Goncalves, Ewan Harrison, David K. Jackson, Ian Johnston, Dominic Kwiatkowski, Cordelia Langford, John Sillitoe on behalf of the Wellcome Sanger Institute COVID-19 Surveillance Team |
| EPI_ISL_3337017 | Lighthouse Lab in Alderley Park | Wellcome Sanger Institute for the COVID-19 Genomics UK (COG-UK) Consortium | Jacquelyn Wynn, Mairead Hyland, The Lighthouse Lab in Alderley Park and Alex Alderton, Roberto Amato, Jeffrey Barrett, Sonia Goncalves, Ewan Harrison, David K. Jackson, Ian Johnston, Dominic Kwiatkowski, Cordelia Langford, John Sillitoe on behalf of the Wellcome Sanger Institute COVID-19 Surveillance Team |
| EPI_ISL_3337018, EPI_ISL_3337019, EPI_ISL_3337020 | Lighthouse Laboratory Plymouth | Wellcome Sanger Institute for the COVID-19 Genomics UK (COG-UK) Consortium | Lighthouse Laboratory Plymouth and Alex Alderton, Roberto Amato, Jeffrey Barrett, Sonia Goncalves, Ewan Harrison, David K. Jackson, Ian Johnston, Dominic Kwiatkowski, Cordelia Langford, John Sillitoe on behalf of the Wellcome Sanger Institute COVID-19 Surveillance Team |
| EPI_ISL_3337021 | Lighthouse Lab in Alderley Park | Wellcome Sanger Institute for the COVID-19 Genomics UK (COG-UK) Consortium | Jacquelyn Wynn, Mairead Hyland, The Lighthouse Lab in Alderley Park and Alex Alderton, Roberto Amato, Jeffrey Barrett, Sonia Goncalves, Ewan Harrison, David K. Jackson, Ian Johnston, Dominic Kwiatkowski, Cordelia Langford, John Sillitoe on behalf of the Wellcome Sanger Institute COVID-19 Surveillance Team |
| EPI_ISL_3337022, EPI_ISL_3337023, EPI_ISL_3337024 | Berkshire and Surrey Pathology Services Lighthouse Laboratory | Wellcome Sanger Institute for the COVID-19 Genomics UK (COG-UK) Consortium | Berkshire and Surrey Pathology Services Lighthouse Laboratory and Alex Alderton, Roberto Amato, Jeffrey Barrett, Sonia Goncalves, Ewan Harrison, David K. Jackson, Ian Johnston, Dominic Kwiatkowski, Cordelia Langford, John Sillitoe on behalf of the Wellcome Sanger Institute COVID-19 Surveillance Team |
| EPI_ISL_3337025, EPI_ISL_3337026, EPI_ISL_3337027 | Lighthouse Laboratory Plymouth | Wellcome Sanger Institute for the COVID-19 Genomics UK (COG-UK) Consortium | Lighthouse Laboratory Plymouth and Alex Alderton, Roberto Amato, Jeffrey Barrett, Sonia Goncalves, Ewan Harrison, David K. Jackson, Ian Johnston, Dominic Kwiatkowski, Cordelia Langford, John Sillitoe on behalf of the Wellcome Sanger Institute COVID-19 Surveillance Team |
| EPI_ISL_3337028 | Lighthouse Lab in Milton Keynes | Wellcome Sanger Institute for the COVID-19 Genomics UK (COG-UK) Consortium | The Lighthouse Lab in Milton Keynes and Alex Alderton, Roberto Amato, Jeffrey Barrett, Sonia Goncalves, Ewan Harrison, David K. Jackson, Ian Johnston, Dominic Kwiatkowski, Cordelia Langford, John Sillitoe on behalf of the Wellcome Sanger Institute COVID-19 Surveillance Team |
| EPI_ISL_3337029 | Lighthouse Lab in Alderley Park | Wellcome Sanger Institute for the COVID-19 Genomics UK (COG-UK) Consortium | Jacquelyn Wynn, Mairead Hyland, The Lighthouse Lab in Alderley Park and Alex Alderton, Roberto Amato, Jeffrey Barrett, Sonia Goncalves, Ewan Harrison, David K. Jackson, Ian Johnston, Dominic Kwiatkowski, Cordelia Langford, John Sillitoe on behalf of the Wellcome Sanger Institute COVID-19 Surveillance Team |
| EPI_ISL_3337030, EPI_ISL_3337031, EPI_ISL_3337032 | Berkshire and Surrey Pathology Services Lighthouse Laboratory | Wellcome Sanger Institute for the COVID-19 Genomics UK (COG-UK) Consortium | Berkshire and Surrey Pathology Services Lighthouse Laboratory and Alex Alderton, Roberto Amato, Jeffrey Barrett, Sonia Goncalves, Ewan Harrison, David K. Jackson, Ian Johnston, Dominic Kwiatkowski, Cordelia Langford, John Sillitoe on behalf of the Wellcome Sanger Institute COVID-19 Surveillance Team |
| EPI_ISL_3337033 | Lighthouse Lab in Alderley Park | Wellcome Sanger Institute for the COVID-19 Genomics UK (COG-UK) Consortium | Jacquelyn Wynn, Mairead Hyland, The Lighthouse Lab in Alderley Park and Alex Alderton, Roberto Amato, Jeffrey Barrett, Sonia Goncalves, Ewan Harrison, David K. Jackson, Ian Johnston, Dominic Kwiatkowski, Cordelia Langford, John Sillitoe on behalf of the Wellcome Sanger Institute COVID-19 Surveillance Team |
| EPI_ISL_3337034 | Lighthouse Laboratory Plymouth | Wellcome Sanger Institute for the COVID-19 Genomics UK (COG-UK) Consortium | Lighthouse Laboratory Plymouth and Alex Alderton, Roberto Amato, Jeffrey Barrett, Sonia Goncalves, Ewan Harrison, David K. Jackson, Ian Johnston, Dominic Kwiatkowski, Cordelia Langford, John Sillitoe on behalf of the Wellcome Sanger Institute COVID-19 Surveillance Team |
| EPI_ISL_3337035, EPI_ISL_3337036 | Lighthouse Lab in Milton Keynes | Wellcome Sanger Institute for the COVID-19 Genomics UK (COG-UK) Consortium | The Lighthouse Lab in Milton Keynes and Alex Alderton, Roberto Amato, Jeffrey Barrett, Sonia Goncalves, Ewan Harrison, David K. Jackson, Ian Johnston, Dominic Kwiatkowski, Cordelia Langford, John Sillitoe on behalf of the Wellcome Sanger Institute COVID-19 Surveillance Team |
| EPI_ISL_3337037 | Berkshire and Surrey Pathology Services Lighthouse Laboratory | Wellcome Sanger Institute for the COVID-19 Genomics UK (COG-UK) Consortium | Berkshire and Surrey Pathology Services Lighthouse Laboratory and Alex Alderton, Roberto Amato, Jeffrey Barrett, Sonia Goncalves, Ewan Harrison, David K. Jackson, Ian Johnston, Dominic Kwiatkowski, Cordelia Langford, John Sillitoe on behalf of the Wellcome Sanger Institute COVID-19 Surveillance Team |
| EPI_ISL_3337038, EPI_ISL_3337039, EPI_ISL_3337040, EPI_ISL_3337041 | Lighthouse Lab in Milton Keynes | Wellcome Sanger Institute for the COVID-19 Genomics UK (COG-UK) Consortium | The Lighthouse Lab in Milton Keynes and Alex Alderton, Roberto Amato, Jeffrey Barrett, Sonia Goncalves, Ewan Harrison, David K. Jackson, Ian Johnston, Dominic Kwiatkowski, Cordelia Langford, John Sillitoe on behalf of the Wellcome Sanger Institute COVID-19 Surveillance Team |
| EPI_ISL_3337042 | Lighthouse Lab in Alderley Park | Wellcome Sanger Institute for the COVID-19 Genomics UK (COG-UK) Consortium | Jacquelyn Wynn, Mairead Hyland, The Lighthouse Lab in Alderley Park and Alex Alderton, Roberto Amato, Jeffrey Barrett, Sonia Goncalves, Ewan Harrison, David K. Jackson, Ian Johnston, Dominic Kwiatkowski, Cordelia Langford, John Sillitoe on behalf of the Wellcome Sanger Institute COVID-19 Surveillance Team |
| EPI_ISL_3337043 | Lighthouse Lab in Milton Keynes | Wellcome Sanger Institute for the COVID-19 Genomics UK (COG-UK) Consortium | The Lighthouse Lab in Milton Keynes and Alex Alderton, Roberto Amato, Jeffrey Barrett, Sonia Goncalves, Ewan Harrison, David K. Jackson, Ian Johnston, Dominic Kwiatkowski, Cordelia Langford, John Sillitoe on behalf of the Wellcome Sanger Institute COVID-19 Surveillance Team |
| EPI_ISL_3337044 | Lighthouse Lab in Alderley Park | Wellcome Sanger Institute for the COVID-19 Genomics UK (COG-UK) Consortium | Jacquelyn Wynn, Mairead Hyland, The Lighthouse Lab in Alderley Park and Alex Alderton, Roberto Amato, Jeffrey Barrett, Sonia Goncalves, Ewan Harrison, David K. Jackson, Ian Johnston, Dominic Kwiatkowski, Cordelia Langford, John Sillitoe on behalf of the Wellcome Sanger Institute COVID-19 Surveillance Team |
| EPI_ISL_3337045 | Lighthouse Laboratory Plymouth | Wellcome Sanger Institute for the COVID-19 Genomics UK (COG-UK) Consortium | Lighthouse Laboratory Plymouth and Alex Alderton, Roberto Amato, Jeffrey Barrett, Sonia Goncalves, Ewan Harrison, David K. Jackson, Ian Johnston, Dominic Kwiatkowski, Cordelia Langford, John Sillitoe on behalf of the Wellcome Sanger Institute COVID-19 Surveillance Team |
| EPI_ISL_3337046 | Berkshire and Surrey Pathology Services Lighthouse Laboratory | Wellcome Sanger Institute for the COVID-19 Genomics UK (COG-UK) Consortium | Berkshire and Surrey Pathology Services Lighthouse Laboratory and Alex Alderton, Roberto Amato, Jeffrey Barrett, Sonia Goncalves, Ewan Harrison, David K. Jackson, Ian Johnston, Dominic Kwiatkowski, Cordelia Langford, John Sillitoe on behalf of the Wellcome Sanger Institute COVID-19 Surveillance Team |
| EPI_ISL_3337047 | Lighthouse Lab in Alderley Park | Wellcome Sanger Institute for the COVID-19 Genomics UK (COG-UK) Consortium | Jacquelyn Wynn, Mairead Hyland, The Lighthouse Lab in Alderley Park and Alex Alderton, Roberto Amato, Jeffrey Barrett, Sonia Goncalves, Ewan Harrison, David K. Jackson, Ian Johnston, Dominic Kwiatkowski, Cordelia Langford, John Sillitoe on behalf of the Wellcome Sanger Institute COVID-19 Surveillance Team |
| EPI_ISL_3337048 | Berkshire and Surrey Pathology Services Lighthouse Laboratory | Wellcome Sanger Institute for the COVID-19 Genomics UK (COG-UK) Consortium | Berkshire and Surrey Pathology Services Lighthouse Laboratory and Alex Alderton, Roberto Amato, Jeffrey Barrett, Sonia Goncalves, Ewan Harrison, David K. Jackson, Ian Johnston, Dominic Kwiatkowski, Cordelia Langford, John Sillitoe on behalf of the Wellcome Sanger Institute COVID-19 Surveillance Team |
| EPI_ISL_3337049 | Lighthouse Laboratory Plymouth | Wellcome Sanger Institute for the COVID-19 Genomics UK (COG-UK) Consortium | Lighthouse Laboratory Plymouth and Alex Alderton, Roberto Amato, Jeffrey Barrett, Sonia Goncalves, Ewan Harrison, David K. Jackson, Ian Johnston, Dominic Kwiatkowski, Cordelia Langford, John Sillitoe on behalf of the Wellcome Sanger Institute COVID-19 Surveillance Team |
| EPI_ISL_3337050 | Lighthouse Lab in Milton Keynes | Wellcome Sanger Institute for the COVID-19 Genomics UK (COG-UK) Consortium | The Lighthouse Lab in Milton Keynes and Alex Alderton, Roberto Amato, Jeffrey Barrett, Sonia Goncalves, Ewan Harrison, David K. Jackson, Ian Johnston, Dominic Kwiatkowski, Cordelia Langford, John Sillitoe on behalf of the Wellcome Sanger Institute COVID-19 Surveillance Team |
| EPI_ISL_3337051 | Lighthouse Laboratory Plymouth | Wellcome Sanger Institute for the COVID-19 Genomics UK (COG-UK) Consortium | Lighthouse Laboratory Plymouth and Alex Alderton, Roberto Amato, Jeffrey Barrett, Sonia Goncalves, Ewan Harrison, David K. Jackson, Ian Johnston, Dominic Kwiatkowski, Cordelia Langford, John Sillitoe on behalf of the Wellcome Sanger Institute COVID-19 Surveillance Team |
| EPI_ISL_3337052 | Lighthouse Lab in Alderley Park | Wellcome Sanger Institute for the COVID-19 Genomics UK (COG-UK) Consortium | Jacquelyn Wynn, Mairead Hyland, The Lighthouse Lab in Alderley Park and Alex Alderton, Roberto Amato, Jeffrey Barrett, Sonia Goncalves, Ewan Harrison, David K. Jackson, Ian Johnston, Dominic Kwiatkowski, Cordelia Langford, John Sillitoe on behalf of the Wellcome Sanger Institute COVID-19 Surveillance Team |
| EPI_ISL_3337053 | Lighthouse Lab in Milton Keynes | Wellcome Sanger Institute for the COVID-19 Genomics UK (COG-UK) Consortium | The Lighthouse Lab in Milton Keynes and Alex Alderton, Roberto Amato, Jeffrey Barrett, Sonia Goncalves, Ewan Harrison, David K. Jackson, Ian Johnston, Dominic Kwiatkowski, Cordelia Langford, John Sillitoe on behalf of the Wellcome Sanger Institute COVID-19 Surveillance Team |
| EPI_ISL_3337054 | Lighthouse Laboratory Plymouth | Wellcome Sanger Institute for the COVID-19 Genomics UK (COG-UK) Consortium | Lighthouse Laboratory Plymouth and Alex Alderton, Roberto Amato, Jeffrey Barrett, Sonia Goncalves, Ewan Harrison, David K. Jackson, Ian Johnston, Dominic Kwiatkowski, Cordelia Langford, John Sillitoe on behalf of the Wellcome Sanger Institute COVID-19 Surveillance Team |
| EPI_ISL_3337055 | Berkshire and Surrey Pathology Services Lighthouse Laboratory | Wellcome Sanger Institute for the COVID-19 Genomics UK (COG-UK) Consortium | Berkshire and Surrey Pathology Services Lighthouse Laboratory and Alex Alderton, Roberto Amato, Jeffrey Barrett, Sonia Goncalves, Ewan Harrison, David K. Jackson, Ian Johnston, Dominic Kwiatkowski, Cordelia Langford, John Sillitoe on behalf of the Wellcome Sanger Institute COVID-19 Surveillance Team |
| EPI_ISL_3337056 | Lighthouse Lab in Alderley Park | Wellcome Sanger Institute for the COVID-19 Genomics UK (COG-UK) Consortium | Jacquelyn Wynn, Mairead Hyland, The Lighthouse Lab in Alderley Park and Alex Alderton, Roberto Amato, Jeffrey Barrett, Sonia Goncalves, Ewan Harrison, David K. Jackson, Ian Johnston, Dominic Kwiatkowski, Cordelia Langford, John Sillitoe on behalf of the Wellcome Sanger Institute COVID-19 Surveillance Team |

| | | | |
|---|---|---|---|
| EPI_ISL_3337057 | Berkshire and Surrey Pathology Services Lighthouse Laboratory | Wellcome Sanger Institute for the COVID-19 Genomics UK (COG-UK) Consortium | Berkshire and Surrey Pathology Services Lighthouse Laboratory and Alex Alderton, Roberto Amato, Jeffrey Barrett, Sonia Goncalves, Ewan Harrison, David K. Jackson, Ian Johnston, Dominic Kwiatkowski, Cordelia Langford, John Sillitoe on behalf of the Wellcome Sanger Institute COVID-19 Surveillance Team |
| EPI_ISL_3337058 | Lighthouse Lab in Alderley Park | Wellcome Sanger Institute for the COVID-19 Genomics UK (COG-UK) Consortium | Jacquelyn Wynn, Mairead Hyland, The Lighthouse Lab in Alderley Park and Alex Alderton, Roberto Amato, Jeffrey Barrett, Sonia Goncalves, Ewan Harrison, David K. Jackson, Ian Johnston, Dominic Kwiatkowski, Cordelia Langford, John Sillitoe on behalf of the Wellcome Sanger Institute COVID-19 Surveillance Team |
| EPI_ISL_3337059 | Lighthouse Laboratory Plymouth | Wellcome Sanger Institute for the COVID-19 Genomics UK (COG-UK) Consortium | Lighthouse Laboratory Plymouth and Alex Alderton, Roberto Amato, Jeffrey Barrett, Sonia Goncalves, Ewan Harrison, David K. Jackson, Ian Johnston, Dominic Kwiatkowski, Cordelia Langford, John Sillitoe on behalf of the Wellcome Sanger Institute COVID-19 Surveillance Team |
| EPI_ISL_3337060 | Berkshire and Surrey Pathology Services Lighthouse Laboratory | Wellcome Sanger Institute for the COVID-19 Genomics UK (COG-UK) Consortium | Berkshire and Surrey Pathology Services Lighthouse Laboratory and Alex Alderton, Roberto Amato, Jeffrey Barrett, Sonia Goncalves, Ewan Harrison, David K. Jackson, Ian Johnston, Dominic Kwiatkowski, Cordelia Langford, John Sillitoe on behalf of the Wellcome Sanger Institute COVID-19 Surveillance Team |
| EPI_ISL_3337061 | Lighthouse Lab in Alderley Park | Wellcome Sanger Institute for the COVID-19 Genomics UK (COG-UK) Consortium | Jacquelyn Wynn, Mairead Hyland, The Lighthouse Lab in Alderley Park and Alex Alderton, Roberto Amato, Jeffrey Barrett, Sonia Goncalves, Ewan Harrison, David K. Jackson, Ian Johnston, Dominic Kwiatkowski, Cordelia Langford, John Sillitoe on behalf of the Wellcome Sanger Institute COVID-19 Surveillance Team |
| EPI_ISL_3337062 | Lighthouse Laboratory Plymouth | Wellcome Sanger Institute for the COVID-19 Genomics UK (COG-UK) Consortium | Lighthouse Laboratory Plymouth and Alex Alderton, Roberto Amato, Jeffrey Barrett, Sonia Goncalves, Ewan Harrison, David K. Jackson, Ian Johnston, Dominic Kwiatkowski, Cordelia Langford, John Sillitoe on behalf of the Wellcome Sanger Institute COVID-19 Surveillance Team |
| EPI_ISL_3337063 | Berkshire and Surrey Pathology Services Lighthouse Laboratory | Wellcome Sanger Institute for the COVID-19 Genomics UK (COG-UK) Consortium | Berkshire and Surrey Pathology Services Lighthouse Laboratory and Alex Alderton, Roberto Amato, Jeffrey Barrett, Sonia Goncalves, Ewan Harrison, David K. Jackson, Ian Johnston, Dominic Kwiatkowski, Cordelia Langford, John Sillitoe on behalf of the Wellcome Sanger Institute COVID-19 Surveillance Team |
| EPI_ISL_3337064, EPI_ISL_3337065, EPI_ISL_3337066, EPI_ISL_3337067, EPI_ISL_3337068, EPI_ISL_3337069, EPI_ISL_3337070, EPI_ISL_3337071, EPI_ISL_3337072, EPI_ISL_3337073 | Lighthouse Lab in Milton Keynes | Wellcome Sanger Institute for the COVID-19 Genomics UK (COG-UK) Consortium | The Lighthouse Lab in Milton Keynes and Alex Alderton, Roberto Amato, Jeffrey Barrett, Sonia Goncalves, Ewan Harrison, David K. Jackson, Ian Johnston, Dominic Kwiatkowski, Cordelia Langford, John Sillitoe on behalf of the Wellcome Sanger Institute COVID-19 Surveillance Team |
| EPI_ISL_3337074 | Berkshire and Surrey Pathology Services Lighthouse Laboratory | Wellcome Sanger Institute for the COVID-19 Genomics UK (COG-UK) Consortium | Berkshire and Surrey Pathology Services Lighthouse Laboratory and Alex Alderton, Roberto Amato, Jeffrey Barrett, Sonia Goncalves, Ewan Harrison, David K. Jackson, Ian Johnston, Dominic Kwiatkowski, Cordelia Langford, John Sillitoe on behalf of the Wellcome Sanger Institute COVID-19 Surveillance Team |
| EPI_ISL_3337075, EPI_ISL_3337076, EPI_ISL_3337077, EPI_ISL_3337078, EPI_ISL_3337079 | Lighthouse Lab in Milton Keynes | Wellcome Sanger Institute for the COVID-19 Genomics UK (COG-UK) Consortium | The Lighthouse Lab in Milton Keynes and Alex Alderton, Roberto Amato, Jeffrey Barrett, Sonia Goncalves, Ewan Harrison, David K. Jackson, Ian Johnston, Dominic Kwiatkowski, Cordelia Langford, John Sillitoe on behalf of the Wellcome Sanger Institute COVID-19 Surveillance Team |
| EPI_ISL_3337080 | Berkshire and Surrey Pathology Services Lighthouse Laboratory | Wellcome Sanger Institute for the COVID-19 Genomics UK (COG-UK) Consortium | Berkshire and Surrey Pathology Services Lighthouse Laboratory and Alex Alderton, Roberto Amato, Jeffrey Barrett, Sonia Goncalves, Ewan Harrison, David K. Jackson, Ian Johnston, Dominic Kwiatkowski, Cordelia Langford, John Sillitoe on behalf of the Wellcome Sanger Institute COVID-19 Surveillance Team |
| EPI_ISL_3337081, EPI_ISL_3337082, EPI_ISL_3337083, EPI_ISL_3337084, EPI_ISL_3337085 | Lighthouse Lab in Milton Keynes | Wellcome Sanger Institute for the COVID-19 Genomics UK (COG-UK) Consortium | The Lighthouse Lab in Milton Keynes and Alex Alderton, Roberto Amato, Jeffrey Barrett, Sonia Goncalves, Ewan Harrison, David K. Jackson, Ian Johnston, Dominic Kwiatkowski, Cordelia Langford, John Sillitoe on behalf of the Wellcome Sanger Institute COVID-19 Surveillance Team |
| EPI_ISL_3337086 | Berkshire and Surrey Pathology Services Lighthouse Laboratory | Wellcome Sanger Institute for the COVID-19 Genomics UK (COG-UK) Consortium | Berkshire and Surrey Pathology Services Lighthouse Laboratory and Alex Alderton, Roberto Amato, Jeffrey Barrett, Sonia Goncalves, Ewan Harrison, David K. Jackson, Ian Johnston, Dominic Kwiatkowski, Cordelia Langford, John Sillitoe on behalf of the Wellcome Sanger Institute COVID-19 Surveillance Team |
| EPI_ISL_3337087, EPI_ISL_3337088, EPI_ISL_3337089, EPI_ISL_3337090, EPI_ISL_3337091, EPI_ISL_3337092, EPI_ISL_3337093, EPI_ISL_3337094 | Lighthouse Lab in Milton Keynes | Wellcome Sanger Institute for the COVID-19 Genomics UK (COG-UK) Consortium | The Lighthouse Lab in Milton Keynes and Alex Alderton, Roberto Amato, Jeffrey Barrett, Sonia Goncalves, Ewan Harrison, David K. Jackson, Ian Johnston, Dominic Kwiatkowski, Cordelia Langford, John Sillitoe on behalf of the Wellcome Sanger Institute COVID-19 Surveillance Team |
| EPI_ISL_3337095, EPI_ISL_3337096 | Berkshire and Surrey Pathology Services Lighthouse Laboratory | Wellcome Sanger Institute for the COVID-19 Genomics UK (COG-UK) Consortium | Berkshire and Surrey Pathology Services Lighthouse Laboratory and Alex Alderton, Roberto Amato, Jeffrey Barrett, Sonia Goncalves, Ewan Harrison, David K. Jackson, Ian Johnston, Dominic Kwiatkowski, Cordelia Langford, John Sillitoe on behalf of the Wellcome Sanger Institute COVID-19 Surveillance Team |
| EPI_ISL_3337097, EPI_ISL_3337098, EPI_ISL_3337099, EPI_ISL_3337100, EPI_ISL_3337101, EPI_ISL_3337102, EPI_ISL_3337103, EPI_ISL_3337104 | Lighthouse Lab in Milton Keynes | Wellcome Sanger Institute for the COVID-19 Genomics UK (COG-UK) Consortium | The Lighthouse Lab in Milton Keynes and Alex Alderton, Roberto Amato, Jeffrey Barrett, Sonia Goncalves, Ewan Harrison, David K. Jackson, Ian Johnston, Dominic Kwiatkowski, Cordelia Langford, John Sillitoe on behalf of the Wellcome Sanger Institute COVID-19 Surveillance Team |
| EPI_ISL_3337105 | Berkshire and Surrey Pathology Services Lighthouse Laboratory | Wellcome Sanger Institute for the COVID-19 Genomics UK (COG-UK) Consortium | Berkshire and Surrey Pathology Services Lighthouse Laboratory and Alex Alderton, Roberto Amato, Jeffrey Barrett, Sonia Goncalves, Ewan Harrison, David K. Jackson, Ian Johnston, Dominic Kwiatkowski, Cordelia Langford, John Sillitoe on behalf of the Wellcome Sanger Institute COVID-19 Surveillance Team |
| EPI_ISL_3337106 | Lighthouse Lab in Milton Keynes | Wellcome Sanger Institute for the COVID-19 Genomics UK (COG-UK) Consortium | The Lighthouse Lab in Milton Keynes and Alex Alderton, Roberto Amato, Jeffrey Barrett, Sonia Goncalves, Ewan Harrison, David K. Jackson, Ian Johnston, Dominic Kwiatkowski, Cordelia Langford, John Sillitoe on behalf of the Wellcome Sanger Institute COVID-19 Surveillance Team |
| EPI_ISL_3337107 | Berkshire and Surrey Pathology Services Lighthouse Laboratory | Wellcome Sanger Institute for the COVID-19 Genomics UK (COG-UK) Consortium | Berkshire and Surrey Pathology Services Lighthouse Laboratory and Alex Alderton, Roberto Amato, Jeffrey Barrett, Sonia Goncalves, Ewan Harrison, David K. Jackson, Ian Johnston, Dominic Kwiatkowski, Cordelia Langford, John Sillitoe on behalf of the Wellcome Sanger Institute COVID-19 Surveillance Team |
| EPI_ISL_3337108, EPI_ISL_3337109, EPI_ISL_3337110 | Lighthouse Lab in Milton Keynes | Wellcome Sanger Institute for the COVID-19 Genomics UK (COG-UK) Consortium | The Lighthouse Lab in Milton Keynes and Alex Alderton, Roberto Amato, Jeffrey Barrett, Sonia Goncalves, Ewan Harrison, David K. Jackson, Ian Johnston, Dominic Kwiatkowski, Cordelia Langford, John Sillitoe on behalf of the Wellcome Sanger Institute COVID-19 Surveillance Team |
| EPI_ISL_3337111 | Berkshire and Surrey Pathology Services Lighthouse Laboratory | Wellcome Sanger Institute for the COVID-19 Genomics UK (COG-UK) Consortium | Berkshire and Surrey Pathology Services Lighthouse Laboratory and Alex Alderton, Roberto Amato, Jeffrey Barrett, Sonia Goncalves, Ewan Harrison, David K. Jackson, Ian Johnston, Dominic Kwiatkowski, Cordelia Langford, John Sillitoe on behalf of the Wellcome Sanger Institute COVID-19 Surveillance Team |
| EPI_ISL_3337112, EPI_ISL_3337113, EPI_ISL_3337114, EPI_ISL_3337115, EPI_ISL_3337116, EPI_ISL_3337117, EPI_ISL_3337118 | Lighthouse Lab in Milton Keynes | Wellcome Sanger Institute for the COVID-19 Genomics UK (COG-UK) Consortium | The Lighthouse Lab in Milton Keynes and Alex Alderton, Roberto Amato, Jeffrey Barrett, Sonia Goncalves, Ewan Harrison, David K. Jackson, Ian Johnston, Dominic Kwiatkowski, Cordelia Langford, John Sillitoe on behalf of the Wellcome Sanger Institute COVID-19 Surveillance Team |
| EPI_ISL_3337119 | Berkshire and Surrey Pathology Services Lighthouse Laboratory | Wellcome Sanger Institute for the COVID-19 Genomics UK (COG-UK) Consortium | Berkshire and Surrey Pathology Services Lighthouse Laboratory and Alex Alderton, Roberto Amato, Jeffrey Barrett, Sonia Goncalves, Ewan Harrison, David K. Jackson, Ian Johnston, Dominic Kwiatkowski, Cordelia Langford, John Sillitoe on behalf of the Wellcome Sanger Institute COVID-19 Surveillance Team |
| EPI_ISL_3337120, EPI_ISL_3337121, EPI_ISL_3337122, EPI_ISL_3337123, EPI_ISL_3337124, EPI_ISL_3337125, EPI_ISL_3337126, EPI_ISL_3337127, EPI_ISL_3337128, EPI_ISL_3337129, EPI_ISL_3337130, EPI_ISL_3337131, EPI_ISL_3337132, EPI_ISL_3337133 | | | |
| see above | Lighthouse Lab in Milton Keynes | Wellcome Sanger Institute for the COVID-19 Genomics UK (COG-UK) Consortium | The Lighthouse Lab in Milton Keynes and Alex Alderton, Roberto Amato, Jeffrey Barrett, Sonia Goncalves, Ewan Harrison, David K. Jackson, Ian Johnston, Dominic Kwiatkowski, Cordelia Langford, John Sillitoe on behalf of the Wellcome Sanger Institute COVID-19 Surveillance Team |
| EPI_ISL_3337134 | Berkshire and Surrey Pathology Services Lighthouse Laboratory | Wellcome Sanger Institute for the COVID-19 Genomics UK (COG-UK) Consortium | Berkshire and Surrey Pathology Services Lighthouse Laboratory and Alex Alderton, Roberto Amato, Jeffrey Barrett, Sonia Goncalves, Ewan Harrison, David K. Jackson, Ian Johnston, Dominic Kwiatkowski, Cordelia Langford, John Sillitoe on behalf of the Wellcome Sanger Institute COVID-19 Surveillance Team |
| EPI_ISL_3337135, EPI_ISL_3337136, EPI_ISL_3337137, EPI_ISL_3337138, EPI_ISL_3337139 | Lighthouse Lab in Milton Keynes | Wellcome Sanger Institute for the COVID-19 Genomics UK (COG-UK) Consortium | The Lighthouse Lab in Milton Keynes and Alex Alderton, Roberto Amato, Jeffrey Barrett, Sonia Goncalves, Ewan Harrison, David K. Jackson, Ian Johnston, Dominic Kwiatkowski, Cordelia Langford, John Sillitoe on behalf of the Wellcome Sanger Institute COVID-19 Surveillance Team |
| EPI_ISL_3337140 | Berkshire and Surrey Pathology Services Lighthouse Laboratory | Wellcome Sanger Institute for the COVID-19 Genomics UK (COG-UK) Consortium | Berkshire and Surrey Pathology Services Lighthouse Laboratory and Alex Alderton, Roberto Amato, Jeffrey Barrett, Sonia Goncalves, Ewan Harrison, David K. Jackson, Ian Johnston, Dominic Kwiatkowski, Cordelia Langford, John Sillitoe on behalf of the Wellcome Sanger Institute COVID-19 Surveillance Team |
| EPI_ISL_3337141, EPI_ISL_3337142, EPI_ISL_3337143, EPI_ISL_3337144, EPI_ISL_3337145, EPI_ISL_3337146, EPI_ISL_3337147, EPI_ISL_3337148, EPI_ISL_3337149, EPI_ISL_3337150, EPI_ISL_3337151, EPI_ISL_3337152 | | | |
| see above | Lighthouse Lab in Milton Keynes | Wellcome Sanger Institute for the COVID-19 Genomics UK (COG-UK) Consortium | The Lighthouse Lab in Milton Keynes and Alex Alderton, Roberto Amato, Jeffrey Barrett, Sonia Goncalves, Ewan Harrison, David K. Jackson, Ian Johnston, Dominic Kwiatkowski, Cordelia Langford, John Sillitoe on behalf of the Wellcome Sanger Institute COVID-19 Surveillance Team |
| EPI_ISL_3337153 | Berkshire and Surrey Pathology Services Lighthouse Laboratory | Wellcome Sanger Institute for the COVID-19 Genomics UK (COG-UK) Consortium | Berkshire and Surrey Pathology Services Lighthouse Laboratory and Alex Alderton, Roberto Amato, Jeffrey Barrett, Sonia Goncalves, Ewan Harrison, David K. Jackson, Ian Johnston, Dominic Kwiatkowski, Cordelia Langford, John Sillitoe on behalf of the Wellcome Sanger Institute COVID-19 Surveillance Team |
| EPI_ISL_3337154, | Lighthouse Lab in Milton Keynes | Wellcome Sanger Institute for the COVID-19 Genomics UK (COG-UK) | The Lighthouse Lab in Milton Keynes and Alex Alderton, Roberto Amato, Jeffrey Barrett, Sonia Goncalves, Ewan Harrison, David K. Jackson, Ian Johnston, Dominic Kwiatkowski, Cordelia Langford, John Sillitoe on behalf of the Wellcome Sanger |

| | | | |
|---|---|---|---|
| EPI_ISL_3337155 | | Consortium | Institute COVID-19 Surveillance Team |
| EPI_ISL_3337156 | Berkshire and Surrey Pathology Services Lighthouse Laboratory | Wellcome Sanger Institute for the COVID-19 Genomics UK (COG-UK) Consortium | Berkshire and Surrey Pathology Services Lighthouse Laboratory and Alex Alderton, Roberto Amato, Jeffrey Barrett, Sonia Goncalves, Ewan Harrison, David K. Jackson, Ian Johnston, Dominic Kwiatkowski, Cordelia Langford, John Sillitoe on behalf of the Wellcome Sanger Institute COVID-19 Surveillance Team |
| EPI_ISL_3337157 | Lighthouse Lab in Milton Keynes | Wellcome Sanger Institute for the COVID-19 Genomics UK (COG-UK) Consortium | The Lighthouse Lab in Milton Keynes and Alex Alderton, Roberto Amato, Jeffrey Barrett, Sonia Goncalves, Ewan Harrison, David K. Jackson, Ian Johnston, Dominic Kwiatkowski, Cordelia Langford, John Sillitoe on behalf of the Wellcome Sanger Institute COVID-19 Surveillance Team |
| EPI_ISL_3337158 | Berkshire and Surrey Pathology Services Lighthouse Laboratory | Wellcome Sanger Institute for the COVID-19 Genomics UK (COG-UK) Consortium | Berkshire and Surrey Pathology Services Lighthouse Laboratory and Alex Alderton, Roberto Amato, Jeffrey Barrett, Sonia Goncalves, Ewan Harrison, David K. Jackson, Ian Johnston, Dominic Kwiatkowski, Cordelia Langford, John Sillitoe on behalf of the Wellcome Sanger Institute COVID-19 Surveillance Team |
| EPI_ISL_3337159, EPI_ISL_3337160, EPI_ISL_3337161, EPI_ISL_3337162, EPI_ISL_3337163, EPI_ISL_3337164, EPI_ISL_3337165, EPI_ISL_3337166, EPI_ISL_3337167, EPI_ISL_3337168, EPI_ISL_3337169, EPI_ISL_3337170 | | | |
| see above | Lighthouse Lab in Milton Keynes | Wellcome Sanger Institute for the COVID-19 Genomics UK (COG-UK) Consortium | The Lighthouse Lab in Milton Keynes and Alex Alderton, Roberto Amato, Jeffrey Barrett, Sonia Goncalves, Ewan Harrison, David K. Jackson, Ian Johnston, Dominic Kwiatkowski, Cordelia Langford, John Sillitoe on behalf of the Wellcome Sanger Institute COVID-19 Surveillance Team |
| EPI_ISL_3337171 | Berkshire and Surrey Pathology Services Lighthouse Laboratory | Wellcome Sanger Institute for the COVID-19 Genomics UK (COG-UK) Consortium | Berkshire and Surrey Pathology Services Lighthouse Laboratory and Alex Alderton, Roberto Amato, Jeffrey Barrett, Sonia Goncalves, Ewan Harrison, David K. Jackson, Ian Johnston, Dominic Kwiatkowski, Cordelia Langford, John Sillitoe on behalf of the Wellcome Sanger Institute COVID-19 Surveillance Team |
| EPI_ISL_3337172, EPI_ISL_3337173, EPI_ISL_3337174, EPI_ISL_3337175, EPI_ISL_3337176, EPI_ISL_3337177, EPI_ISL_3337178, EPI_ISL_3337179, EPI_ISL_3337180, EPI_ISL_3337181, EPI_ISL_3337182, EPI_ISL_3337183, EPI_ISL_3337184, EPI_ISL_3337185, EPI_ISL_3337186, EPI_ISL_3337187 | | | |
| see above | Lighthouse Lab in Milton Keynes | Wellcome Sanger Institute for the COVID-19 Genomics UK (COG-UK) Consortium | The Lighthouse Lab in Milton Keynes and Alex Alderton, Roberto Amato, Jeffrey Barrett, Sonia Goncalves, Ewan Harrison, David K. Jackson, Ian Johnston, Dominic Kwiatkowski, Cordelia Langford, John Sillitoe on behalf of the Wellcome Sanger Institute COVID-19 Surveillance Team |
| EPI_ISL_3337188, EPI_ISL_3337189 | Berkshire and Surrey Pathology Services Lighthouse Laboratory | Wellcome Sanger Institute for the COVID-19 Genomics UK (COG-UK) Consortium | Berkshire and Surrey Pathology Services Lighthouse Laboratory and Alex Alderton, Roberto Amato, Jeffrey Barrett, Sonia Goncalves, Ewan Harrison, David K. Jackson, Ian Johnston, Dominic Kwiatkowski, Cordelia Langford, John Sillitoe on behalf of the Wellcome Sanger Institute COVID-19 Surveillance Team |
| EPI_ISL_3337190, EPI_ISL_3337191 | Lighthouse Lab in Milton Keynes | Wellcome Sanger Institute for the COVID-19 Genomics UK (COG-UK) Consortium | The Lighthouse Lab in Milton Keynes and Alex Alderton, Roberto Amato, Jeffrey Barrett, Sonia Goncalves, Ewan Harrison, David K. Jackson, Ian Johnston, Dominic Kwiatkowski, Cordelia Langford, John Sillitoe on behalf of the Wellcome Sanger Institute COVID-19 Surveillance Team |
| EPI_ISL_3337192 | Berkshire and Surrey Pathology Services Lighthouse Laboratory | Wellcome Sanger Institute for the COVID-19 Genomics UK (COG-UK) Consortium | Berkshire and Surrey Pathology Services Lighthouse Laboratory and Alex Alderton, Roberto Amato, Jeffrey Barrett, Sonia Goncalves, Ewan Harrison, David K. Jackson, Ian Johnston, Dominic Kwiatkowski, Cordelia Langford, John Sillitoe on behalf of the Wellcome Sanger Institute COVID-19 Surveillance Team |
| EPI_ISL_3337193, EPI_ISL_3337194, EPI_ISL_3337195, EPI_ISL_3337196 | Lighthouse Lab in Milton Keynes | Wellcome Sanger Institute for the COVID-19 Genomics UK (COG-UK) Consortium | The Lighthouse Lab in Milton Keynes and Alex Alderton, Roberto Amato, Jeffrey Barrett, Sonia Goncalves, Ewan Harrison, David K. Jackson, Ian Johnston, Dominic Kwiatkowski, Cordelia Langford, John Sillitoe on behalf of the Wellcome Sanger Institute COVID-19 Surveillance Team |
| EPI_ISL_3337197 | Berkshire and Surrey Pathology Services Lighthouse Laboratory | Wellcome Sanger Institute for the COVID-19 Genomics UK (COG-UK) Consortium | Berkshire and Surrey Pathology Services Lighthouse Laboratory and Alex Alderton, Roberto Amato, Jeffrey Barrett, Sonia Goncalves, Ewan Harrison, David K. Jackson, Ian Johnston, Dominic Kwiatkowski, Cordelia Langford, John Sillitoe on behalf of the Wellcome Sanger Institute COVID-19 Surveillance Team |
| EPI_ISL_3337198 | Lighthouse Lab in Milton Keynes | Wellcome Sanger Institute for the COVID-19 Genomics UK (COG-UK) Consortium | The Lighthouse Lab in Milton Keynes and Alex Alderton, Roberto Amato, Jeffrey Barrett, Sonia Goncalves, Ewan Harrison, David K. Jackson, Ian Johnston, Dominic Kwiatkowski, Cordelia Langford, John Sillitoe on behalf of the Wellcome Sanger Institute COVID-19 Surveillance Team |
| EPI_ISL_3337199 | Berkshire and Surrey Pathology Services Lighthouse Laboratory | Wellcome Sanger Institute for the COVID-19 Genomics UK (COG-UK) Consortium | Berkshire and Surrey Pathology Services Lighthouse Laboratory and Alex Alderton, Roberto Amato, Jeffrey Barrett, Sonia Goncalves, Ewan Harrison, David K. Jackson, Ian Johnston, Dominic Kwiatkowski, Cordelia Langford, John Sillitoe on behalf of the Wellcome Sanger Institute COVID-19 Surveillance Team |
| EPI_ISL_3337200, EPI_ISL_3337201 | Lighthouse Lab in Milton Keynes | Wellcome Sanger Institute for the COVID-19 Genomics UK (COG-UK) Consortium | The Lighthouse Lab in Milton Keynes and Alex Alderton, Roberto Amato, Jeffrey Barrett, Sonia Goncalves, Ewan Harrison, David K. Jackson, Ian Johnston, Dominic Kwiatkowski, Cordelia Langford, John Sillitoe on behalf of the Wellcome Sanger Institute COVID-19 Surveillance Team |
| EPI_ISL_3337202 | Berkshire and Surrey Pathology Services Lighthouse Laboratory | Wellcome Sanger Institute for the COVID-19 Genomics UK (COG-UK) Consortium | Berkshire and Surrey Pathology Services Lighthouse Laboratory and Alex Alderton, Roberto Amato, Jeffrey Barrett, Sonia Goncalves, Ewan Harrison, David K. Jackson, Ian Johnston, Dominic Kwiatkowski, Cordelia Langford, John Sillitoe on behalf of the Wellcome Sanger Institute COVID-19 Surveillance Team |
| EPI_ISL_3337203 | Lighthouse Lab in Milton Keynes | Wellcome Sanger Institute for the COVID-19 Genomics UK (COG-UK) Consortium | The Lighthouse Lab in Milton Keynes and Alex Alderton, Roberto Amato, Jeffrey Barrett, Sonia Goncalves, Ewan Harrison, David K. Jackson, Ian Johnston, Dominic Kwiatkowski, Cordelia Langford, John Sillitoe on behalf of the Wellcome Sanger Institute COVID-19 Surveillance Team |
| EPI_ISL_3337204 | Berkshire and Surrey Pathology Services Lighthouse Laboratory | Wellcome Sanger Institute for the COVID-19 Genomics UK (COG-UK) Consortium | Berkshire and Surrey Pathology Services Lighthouse Laboratory and Alex Alderton, Roberto Amato, Jeffrey Barrett, Sonia Goncalves, Ewan Harrison, David K. Jackson, Ian Johnston, Dominic Kwiatkowski, Cordelia Langford, John Sillitoe on behalf of the Wellcome Sanger Institute COVID-19 Surveillance Team |
| EPI_ISL_3337205, EPI_ISL_3337206, EPI_ISL_3337207, EPI_ISL_3337208, EPI_ISL_3337209 | Lighthouse Lab in Milton Keynes | Wellcome Sanger Institute for the COVID-19 Genomics UK (COG-UK) Consortium | The Lighthouse Lab in Milton Keynes and Alex Alderton, Roberto Amato, Jeffrey Barrett, Sonia Goncalves, Ewan Harrison, David K. Jackson, Ian Johnston, Dominic Kwiatkowski, Cordelia Langford, John Sillitoe on behalf of the Wellcome Sanger Institute COVID-19 Surveillance Team |
| EPI_ISL_3337210 | Berkshire and Surrey Pathology Services Lighthouse Laboratory | Wellcome Sanger Institute for the COVID-19 Genomics UK (COG-UK) Consortium | Berkshire and Surrey Pathology Services Lighthouse Laboratory and Alex Alderton, Roberto Amato, Jeffrey Barrett, Sonia Goncalves, Ewan Harrison, David K. Jackson, Ian Johnston, Dominic Kwiatkowski, Cordelia Langford, John Sillitoe on behalf of the Wellcome Sanger Institute COVID-19 Surveillance Team |
| EPI_ISL_3337211, EPI_ISL_3337212, EPI_ISL_3337213 | Lighthouse Lab in Milton Keynes | Wellcome Sanger Institute for the COVID-19 Genomics UK (COG-UK) Consortium | The Lighthouse Lab in Milton Keynes and Alex Alderton, Roberto Amato, Jeffrey Barrett, Sonia Goncalves, Ewan Harrison, David K. Jackson, Ian Johnston, Dominic Kwiatkowski, Cordelia Langford, John Sillitoe on behalf of the Wellcome Sanger Institute COVID-19 Surveillance Team |
| EPI_ISL_3337214 | Berkshire and Surrey Pathology Services Lighthouse Laboratory | Wellcome Sanger Institute for the COVID-19 Genomics UK (COG-UK) Consortium | Berkshire and Surrey Pathology Services Lighthouse Laboratory and Alex Alderton, Roberto Amato, Jeffrey Barrett, Sonia Goncalves, Ewan Harrison, David K. Jackson, Ian Johnston, Dominic Kwiatkowski, Cordelia Langford, John Sillitoe on behalf of the Wellcome Sanger Institute COVID-19 Surveillance Team |
| EPI_ISL_3337215, EPI_ISL_3337216, EPI_ISL_3337217, EPI_ISL_3337218, EPI_ISL_3337219, EPI_ISL_3337220, EPI_ISL_3337221, EPI_ISL_3337222 | Lighthouse Lab in Milton Keynes | Wellcome Sanger Institute for the COVID-19 Genomics UK (COG-UK) Consortium | The Lighthouse Lab in Milton Keynes and Alex Alderton, Roberto Amato, Jeffrey Barrett, Sonia Goncalves, Ewan Harrison, David K. Jackson, Ian Johnston, Dominic Kwiatkowski, Cordelia Langford, John Sillitoe on behalf of the Wellcome Sanger Institute COVID-19 Surveillance Team |
| EPI_ISL_3337223, EPI_ISL_3337224, EPI_ISL_3337225 | Berkshire and Surrey Pathology Services Lighthouse Laboratory | Wellcome Sanger Institute for the COVID-19 Genomics UK (COG-UK) Consortium | Berkshire and Surrey Pathology Services Lighthouse Laboratory and Alex Alderton, Roberto Amato, Jeffrey Barrett, Sonia Goncalves, Ewan Harrison, David K. Jackson, Ian Johnston, Dominic Kwiatkowski, Cordelia Langford, John Sillitoe on behalf of the Wellcome Sanger Institute COVID-19 Surveillance Team |
| EPI_ISL_3337226, EPI_ISL_3337227, EPI_ISL_3337228, EPI_ISL_3337229, EPI_ISL_3337230 | Lighthouse Lab in Milton Keynes | Wellcome Sanger Institute for the COVID-19 Genomics UK (COG-UK) Consortium | The Lighthouse Lab in Milton Keynes and Alex Alderton, Roberto Amato, Jeffrey Barrett, Sonia Goncalves, Ewan Harrison, David K. Jackson, Ian Johnston, Dominic Kwiatkowski, Cordelia Langford, John Sillitoe on behalf of the Wellcome Sanger Institute COVID-19 Surveillance Team |
| EPI_ISL_3337231, EPI_ISL_3337232 | Berkshire and Surrey Pathology Services Lighthouse Laboratory | Wellcome Sanger Institute for the COVID-19 Genomics UK (COG-UK) Consortium | Berkshire and Surrey Pathology Services Lighthouse Laboratory and Alex Alderton, Roberto Amato, Jeffrey Barrett, Sonia Goncalves, Ewan Harrison, David K. Jackson, Ian Johnston, Dominic Kwiatkowski, Cordelia Langford, John Sillitoe on behalf of the Wellcome Sanger Institute COVID-19 Surveillance Team |
| EPI_ISL_3337233, EPI_ISL_3337234 | Lighthouse Lab in Milton Keynes | Wellcome Sanger Institute for the COVID-19 Genomics UK (COG-UK) Consortium | The Lighthouse Lab in Milton Keynes and Alex Alderton, Roberto Amato, Jeffrey Barrett, Sonia Goncalves, Ewan Harrison, David K. Jackson, Ian Johnston, Dominic Kwiatkowski, Cordelia Langford, John Sillitoe on behalf of the Wellcome Sanger Institute COVID-19 Surveillance Team |
| EPI_ISL_3337235, EPI_ISL_3337236 | Berkshire and Surrey Pathology Services Lighthouse Laboratory | Wellcome Sanger Institute for the COVID-19 Genomics UK (COG-UK) Consortium | Berkshire and Surrey Pathology Services Lighthouse Laboratory and Alex Alderton, Roberto Amato, Jeffrey Barrett, Sonia Goncalves, Ewan Harrison, David K. Jackson, Ian Johnston, Dominic Kwiatkowski, Cordelia Langford, John Sillitoe on behalf of the Wellcome Sanger Institute COVID-19 Surveillance Team |
| EPI_ISL_3337237, EPI_ISL_3337238, EPI_ISL_3337239 | Lighthouse Lab in Milton Keynes | Wellcome Sanger Institute for the COVID-19 Genomics UK (COG-UK) Consortium | The Lighthouse Lab in Milton Keynes and Alex Alderton, Roberto Amato, Jeffrey Barrett, Sonia Goncalves, Ewan Harrison, David K. Jackson, Ian Johnston, Dominic Kwiatkowski, Cordelia Langford, John Sillitoe on behalf of the Wellcome Sanger Institute COVID-19 Surveillance Team |
| EPI_ISL_3337240 | Berkshire and Surrey Pathology Services Lighthouse Laboratory | Wellcome Sanger Institute for the COVID-19 Genomics UK (COG-UK) Consortium | Berkshire and Surrey Pathology Services Lighthouse Laboratory and Alex Alderton, Roberto Amato, Jeffrey Barrett, Sonia Goncalves, Ewan Harrison, David K. Jackson, Ian Johnston, Dominic Kwiatkowski, Cordelia Langford, John Sillitoe on behalf of the Wellcome Sanger Institute COVID-19 Surveillance Team |
| EPI_ISL_3337241, EPI_ISL_3337242, EPI_ISL_3337243, EPI_ISL_3337244, EPI_ISL_3337245, EPI_ISL_3337246, EPI_ISL_3337247, EPI_ISL_3337248, EPI_ISL_3337249 | Lighthouse Lab in Milton Keynes | Wellcome Sanger Institute for the COVID-19 Genomics UK (COG-UK) Consortium | The Lighthouse Lab in Milton Keynes and Alex Alderton, Roberto Amato, Jeffrey Barrett, Sonia Goncalves, Ewan Harrison, David K. Jackson, Ian Johnston, Dominic Kwiatkowski, Cordelia Langford, John Sillitoe on behalf of the Wellcome Sanger Institute COVID-19 Surveillance Team |
| EPI_ISL_3337250 | Berkshire and Surrey Pathology Services Lighthouse Laboratory | Wellcome Sanger Institute for the COVID-19 Genomics UK (COG-UK) Consortium | Berkshire and Surrey Pathology Services Lighthouse Laboratory and Alex Alderton, Roberto Amato, Jeffrey Barrett, Sonia Goncalves, Ewan Harrison, David K. Jackson, Ian Johnston, Dominic Kwiatkowski, Cordelia Langford, John Sillitoe on behalf of the Wellcome Sanger Institute COVID-19 Surveillance Team |
| EPI_ISL_3337251 | Lighthouse Lab in Milton Keynes | Wellcome Sanger Institute for the COVID-19 Genomics UK (COG-UK) Consortium | The Lighthouse Lab in Milton Keynes and Alex Alderton, Roberto Amato, Jeffrey Barrett, Sonia Goncalves, Ewan Harrison, David K. Jackson, Ian Johnston, Dominic Kwiatkowski, Cordelia Langford, John Sillitoe on behalf of the Wellcome Sanger Institute COVID-19 Surveillance Team |
| EPI_ISL_3337252 | Berkshire and Surrey Pathology Services Lighthouse Laboratory | Wellcome Sanger Institute for the COVID-19 Genomics UK (COG-UK) Consortium | Berkshire and Surrey Pathology Services Lighthouse Laboratory and Alex Alderton, Roberto Amato, Jeffrey Barrett, Sonia Goncalves, Ewan Harrison, David K. Jackson, Ian Johnston, Dominic Kwiatkowski, Cordelia Langford, John Sillitoe on behalf of the Wellcome Sanger Institute COVID-19 Surveillance Team |
| EPI_ISL_3337253, | Lighthouse Lab in Milton Keynes | Wellcome Sanger Institute for the COVID-19 Genomics UK (COG-UK) | The Lighthouse Lab in Milton Keynes and Alex Alderton, Roberto Amato, Jeffrey Barrett, Sonia Goncalves, Ewan Harrison, David K. Jackson, Ian Johnston, Dominic Kwiatkowski, Cordelia Langford, John Sillitoe on behalf of the Wellcome Sanger |

| | | | |
|---|---|---|---|
| EPI_ISL_3337254, EPI_ISL_3337255 | | Consortium | Institute COVID-19 Surveillance Team |
| EPI_ISL_3337256 | Berkshire and Surrey Pathology Services Lighthouse Laboratory | Wellcome Sanger Institute for the COVID-19 Genomics UK (COG-UK) Consortium | Berkshire and Surrey Pathology Services Lighthouse Laboratory and Alex Alderton, Roberto Amato, Jeffrey Barrett, Sonia Goncalves, Ewan Harrison, David K. Jackson, Ian Johnston, Dominic Kwiatkowski, Cordelia Langford, John Sillitoe on behalf of the Wellcome Sanger Institute COVID-19 Surveillance Team |
| EPI_ISL_3337257, EPI_ISL_3337258 | Lighthouse Lab in Milton Keynes | Wellcome Sanger Institute for the COVID-19 Genomics UK (COG-UK) Consortium | The Lighthouse Lab in Milton Keynes and Alex Alderton, Roberto Amato, Jeffrey Barrett, Sonia Goncalves, Ewan Harrison, David K. Jackson, Ian Johnston, Dominic Kwiatkowski, Cordelia Langford, John Sillitoe on behalf of the Wellcome Sanger Institute COVID-19 Surveillance Team |
| EPI_ISL_3337259, EPI_ISL_3337260 | Berkshire and Surrey Pathology Services Lighthouse Laboratory | Wellcome Sanger Institute for the COVID-19 Genomics UK (COG-UK) Consortium | Berkshire and Surrey Pathology Services Lighthouse Laboratory and Alex Alderton, Roberto Amato, Jeffrey Barrett, Sonia Goncalves, Ewan Harrison, David K. Jackson, Ian Johnston, Dominic Kwiatkowski, Cordelia Langford, John Sillitoe on behalf of the Wellcome Sanger Institute COVID-19 Surveillance Team |
| EPI_ISL_3337261, EPI_ISL_3337262 | Lighthouse Lab in Milton Keynes | Wellcome Sanger Institute for the COVID-19 Genomics UK (COG-UK) Consortium | The Lighthouse Lab in Milton Keynes and Alex Alderton, Roberto Amato, Jeffrey Barrett, Sonia Goncalves, Ewan Harrison, David K. Jackson, Ian Johnston, Dominic Kwiatkowski, Cordelia Langford, John Sillitoe on behalf of the Wellcome Sanger Institute COVID-19 Surveillance Team |
| EPI_ISL_3337263 | Berkshire and Surrey Pathology Services Lighthouse Laboratory | Wellcome Sanger Institute for the COVID-19 Genomics UK (COG-UK) Consortium | Berkshire and Surrey Pathology Services Lighthouse Laboratory and Alex Alderton, Roberto Amato, Jeffrey Barrett, Sonia Goncalves, Ewan Harrison, David K. Jackson, Ian Johnston, Dominic Kwiatkowski, Cordelia Langford, John Sillitoe on behalf of the Wellcome Sanger Institute COVID-19 Surveillance Team |
| EPI_ISL_3337264 | Lighthouse Lab in Milton Keynes | Wellcome Sanger Institute for the COVID-19 Genomics UK (COG-UK) Consortium | The Lighthouse Lab in Milton Keynes and Alex Alderton, Roberto Amato, Jeffrey Barrett, Sonia Goncalves, Ewan Harrison, David K. Jackson, Ian Johnston, Dominic Kwiatkowski, Cordelia Langford, John Sillitoe on behalf of the Wellcome Sanger Institute COVID-19 Surveillance Team |
| EPI_ISL_3337265 | Berkshire and Surrey Pathology Services Lighthouse Laboratory | Wellcome Sanger Institute for the COVID-19 Genomics UK (COG-UK) Consortium | Berkshire and Surrey Pathology Services Lighthouse Laboratory and Alex Alderton, Roberto Amato, Jeffrey Barrett, Sonia Goncalves, Ewan Harrison, David K. Jackson, Ian Johnston, Dominic Kwiatkowski, Cordelia Langford, John Sillitoe on behalf of the Wellcome Sanger Institute COVID-19 Surveillance Team |
| EPI_ISL_3337266 | Lighthouse Lab in Milton Keynes | Wellcome Sanger Institute for the COVID-19 Genomics UK (COG-UK) Consortium | The Lighthouse Lab in Milton Keynes and Alex Alderton, Roberto Amato, Jeffrey Barrett, Sonia Goncalves, Ewan Harrison, David K. Jackson, Ian Johnston, Dominic Kwiatkowski, Cordelia Langford, John Sillitoe on behalf of the Wellcome Sanger Institute COVID-19 Surveillance Team |
| EPI_ISL_3337267 | Berkshire and Surrey Pathology Services Lighthouse Laboratory | Wellcome Sanger Institute for the COVID-19 Genomics UK (COG-UK) Consortium | Berkshire and Surrey Pathology Services Lighthouse Laboratory and Alex Alderton, Roberto Amato, Jeffrey Barrett, Sonia Goncalves, Ewan Harrison, David K. Jackson, Ian Johnston, Dominic Kwiatkowski, Cordelia Langford, John Sillitoe on behalf of the Wellcome Sanger Institute COVID-19 Surveillance Team |
| EPI_ISL_3337268, EPI_ISL_3337269, EPI_ISL_3337270 | Lighthouse Lab in Milton Keynes | Wellcome Sanger Institute for the COVID-19 Genomics UK (COG-UK) Consortium | The Lighthouse Lab in Milton Keynes and Alex Alderton, Roberto Amato, Jeffrey Barrett, Sonia Goncalves, Ewan Harrison, David K. Jackson, Ian Johnston, Dominic Kwiatkowski, Cordelia Langford, John Sillitoe on behalf of the Wellcome Sanger Institute COVID-19 Surveillance Team |
| EPI_ISL_3337271 | Berkshire and Surrey Pathology Services Lighthouse Laboratory | Wellcome Sanger Institute for the COVID-19 Genomics UK (COG-UK) Consortium | Berkshire and Surrey Pathology Services Lighthouse Laboratory and Alex Alderton, Roberto Amato, Jeffrey Barrett, Sonia Goncalves, Ewan Harrison, David K. Jackson, Ian Johnston, Dominic Kwiatkowski, Cordelia Langford, John Sillitoe on behalf of the Wellcome Sanger Institute COVID-19 Surveillance Team |
| EPI_ISL_3337272, EPI_ISL_3337273, EPI_ISL_3337274 | Lighthouse Lab in Milton Keynes | Wellcome Sanger Institute for the COVID-19 Genomics UK (COG-UK) Consortium | The Lighthouse Lab in Milton Keynes and Alex Alderton, Roberto Amato, Jeffrey Barrett, Sonia Goncalves, Ewan Harrison, David K. Jackson, Ian Johnston, Dominic Kwiatkowski, Cordelia Langford, John Sillitoe on behalf of the Wellcome Sanger Institute COVID-19 Surveillance Team |
| EPI_ISL_3337275 | Berkshire and Surrey Pathology Services Lighthouse Laboratory | Wellcome Sanger Institute for the COVID-19 Genomics UK (COG-UK) Consortium | Berkshire and Surrey Pathology Services Lighthouse Laboratory and Alex Alderton, Roberto Amato, Jeffrey Barrett, Sonia Goncalves, Ewan Harrison, David K. Jackson, Ian Johnston, Dominic Kwiatkowski, Cordelia Langford, John Sillitoe on behalf of the Wellcome Sanger Institute COVID-19 Surveillance Team |
| EPI_ISL_3337276 | Lighthouse Lab in Milton Keynes | Wellcome Sanger Institute for the COVID-19 Genomics UK (COG-UK) Consortium | The Lighthouse Lab in Milton Keynes and Alex Alderton, Roberto Amato, Jeffrey Barrett, Sonia Goncalves, Ewan Harrison, David K. Jackson, Ian Johnston, Dominic Kwiatkowski, Cordelia Langford, John Sillitoe on behalf of the Wellcome Sanger Institute COVID-19 Surveillance Team |
| EPI_ISL_3337277, EPI_ISL_3337278 | Berkshire and Surrey Pathology Services Lighthouse Laboratory | Wellcome Sanger Institute for the COVID-19 Genomics UK (COG-UK) Consortium | Berkshire and Surrey Pathology Services Lighthouse Laboratory and Alex Alderton, Roberto Amato, Jeffrey Barrett, Sonia Goncalves, Ewan Harrison, David K. Jackson, Ian Johnston, Dominic Kwiatkowski, Cordelia Langford, John Sillitoe on behalf of the Wellcome Sanger Institute COVID-19 Surveillance Team |
| EPI_ISL_3337279, EPI_ISL_3337280 | Lighthouse Lab in Milton Keynes | Wellcome Sanger Institute for the COVID-19 Genomics UK (COG-UK) Consortium | The Lighthouse Lab in Milton Keynes and Alex Alderton, Roberto Amato, Jeffrey Barrett, Sonia Goncalves, Ewan Harrison, David K. Jackson, Ian Johnston, Dominic Kwiatkowski, Cordelia Langford, John Sillitoe on behalf of the Wellcome Sanger Institute COVID-19 Surveillance Team |
| EPI_ISL_3337281 | Berkshire and Surrey Pathology Services Lighthouse Laboratory | Wellcome Sanger Institute for the COVID-19 Genomics UK (COG-UK) Consortium | Berkshire and Surrey Pathology Services Lighthouse Laboratory and Alex Alderton, Roberto Amato, Jeffrey Barrett, Sonia Goncalves, Ewan Harrison, David K. Jackson, Ian Johnston, Dominic Kwiatkowski, Cordelia Langford, John Sillitoe on behalf of the Wellcome Sanger Institute COVID-19 Surveillance Team |
| EPI_ISL_3337282, EPI_ISL_3337283 | Lighthouse Lab in Milton Keynes | Wellcome Sanger Institute for the COVID-19 Genomics UK (COG-UK) Consortium | The Lighthouse Lab in Milton Keynes and Alex Alderton, Roberto Amato, Jeffrey Barrett, Sonia Goncalves, Ewan Harrison, David K. Jackson, Ian Johnston, Dominic Kwiatkowski, Cordelia Langford, John Sillitoe on behalf of the Wellcome Sanger Institute COVID-19 Surveillance Team |
| EPI_ISL_3337284 | Berkshire and Surrey Pathology Services Lighthouse Laboratory | Wellcome Sanger Institute for the COVID-19 Genomics UK (COG-UK) Consortium | Berkshire and Surrey Pathology Services Lighthouse Laboratory and Alex Alderton, Roberto Amato, Jeffrey Barrett, Sonia Goncalves, Ewan Harrison, David K. Jackson, Ian Johnston, Dominic Kwiatkowski, Cordelia Langford, John Sillitoe on behalf of the Wellcome Sanger Institute COVID-19 Surveillance Team |
| EPI_ISL_3337285, EPI_ISL_3337286, EPI_ISL_3337287 | Lighthouse Lab in Milton Keynes | Wellcome Sanger Institute for the COVID-19 Genomics UK (COG-UK) Consortium | The Lighthouse Lab in Milton Keynes and Alex Alderton, Roberto Amato, Jeffrey Barrett, Sonia Goncalves, Ewan Harrison, David K. Jackson, Ian Johnston, Dominic Kwiatkowski, Cordelia Langford, John Sillitoe on behalf of the Wellcome Sanger Institute COVID-19 Surveillance Team |
| EPI_ISL_3337288 | Berkshire and Surrey Pathology Services Lighthouse Laboratory | Wellcome Sanger Institute for the COVID-19 Genomics UK (COG-UK) Consortium | Berkshire and Surrey Pathology Services Lighthouse Laboratory and Alex Alderton, Roberto Amato, Jeffrey Barrett, Sonia Goncalves, Ewan Harrison, David K. Jackson, Ian Johnston, Dominic Kwiatkowski, Cordelia Langford, John Sillitoe on behalf of the Wellcome Sanger Institute COVID-19 Surveillance Team |
| EPI_ISL_3337289 | Lighthouse Lab in Milton Keynes | Wellcome Sanger Institute for the COVID-19 Genomics UK (COG-UK) Consortium | The Lighthouse Lab in Milton Keynes and Alex Alderton, Roberto Amato, Jeffrey Barrett, Sonia Goncalves, Ewan Harrison, David K. Jackson, Ian Johnston, Dominic Kwiatkowski, Cordelia Langford, John Sillitoe on behalf of the Wellcome Sanger Institute COVID-19 Surveillance Team |
| EPI_ISL_3337290 | Berkshire and Surrey Pathology Services Lighthouse Laboratory | Wellcome Sanger Institute for the COVID-19 Genomics UK (COG-UK) Consortium | Berkshire and Surrey Pathology Services Lighthouse Laboratory and Alex Alderton, Roberto Amato, Jeffrey Barrett, Sonia Goncalves, Ewan Harrison, David K. Jackson, Ian Johnston, Dominic Kwiatkowski, Cordelia Langford, John Sillitoe on behalf of the Wellcome Sanger Institute COVID-19 Surveillance Team |
| EPI_ISL_3337291, EPI_ISL_3337292, EPI_ISL_3337293, EPI_ISL_3337294, EPI_ISL_3337295, EPI_ISL_3337296, EPI_ISL_3337297, EPI_ISL_3337298, EPI_ISL_3337299, EPI_ISL_3337300, EPI_ISL_3337301, EPI_ISL_3337302 | | | |
| see above | Lighthouse Lab in Milton Keynes | Wellcome Sanger Institute for the COVID-19 Genomics UK (COG-UK) Consortium | The Lighthouse Lab in Milton Keynes and Alex Alderton, Roberto Amato, Jeffrey Barrett, Sonia Goncalves, Ewan Harrison, David K. Jackson, Ian Johnston, Dominic Kwiatkowski, Cordelia Langford, John Sillitoe on behalf of the Wellcome Sanger Institute COVID-19 Surveillance Team |
| EPI_ISL_3337303 | Berkshire and Surrey Pathology Services Lighthouse Laboratory | Wellcome Sanger Institute for the COVID-19 Genomics UK (COG-UK) Consortium | Berkshire and Surrey Pathology Services Lighthouse Laboratory and Alex Alderton, Roberto Amato, Jeffrey Barrett, Sonia Goncalves, Ewan Harrison, David K. Jackson, Ian Johnston, Dominic Kwiatkowski, Cordelia Langford, John Sillitoe on behalf of the Wellcome Sanger Institute COVID-19 Surveillance Team |
| EPI_ISL_3337304 | Lighthouse Lab in Milton Keynes | Wellcome Sanger Institute for the COVID-19 Genomics UK (COG-UK) Consortium | The Lighthouse Lab in Milton Keynes and Alex Alderton, Roberto Amato, Jeffrey Barrett, Sonia Goncalves, Ewan Harrison, David K. Jackson, Ian Johnston, Dominic Kwiatkowski, Cordelia Langford, John Sillitoe on behalf of the Wellcome Sanger Institute COVID-19 Surveillance Team |
| EPI_ISL_3337305 | Berkshire and Surrey Pathology Services Lighthouse Laboratory | Wellcome Sanger Institute for the COVID-19 Genomics UK (COG-UK) Consortium | Berkshire and Surrey Pathology Services Lighthouse Laboratory and Alex Alderton, Roberto Amato, Jeffrey Barrett, Sonia Goncalves, Ewan Harrison, David K. Jackson, Ian Johnston, Dominic Kwiatkowski, Cordelia Langford, John Sillitoe on behalf of the Wellcome Sanger Institute COVID-19 Surveillance Team |
| EPI_ISL_3337306 | Lighthouse Lab in Milton Keynes | Wellcome Sanger Institute for the COVID-19 Genomics UK (COG-UK) Consortium | The Lighthouse Lab in Milton Keynes and Alex Alderton, Roberto Amato, Jeffrey Barrett, Sonia Goncalves, Ewan Harrison, David K. Jackson, Ian Johnston, Dominic Kwiatkowski, Cordelia Langford, John Sillitoe on behalf of the Wellcome Sanger Institute COVID-19 Surveillance Team |
| EPI_ISL_3337307 | Berkshire and Surrey Pathology Services Lighthouse Laboratory | Wellcome Sanger Institute for the COVID-19 Genomics UK (COG-UK) Consortium | Berkshire and Surrey Pathology Services Lighthouse Laboratory and Alex Alderton, Roberto Amato, Jeffrey Barrett, Sonia Goncalves, Ewan Harrison, David K. Jackson, Ian Johnston, Dominic Kwiatkowski, Cordelia Langford, John Sillitoe on behalf of the Wellcome Sanger Institute COVID-19 Surveillance Team |
| EPI_ISL_3337308, EPI_ISL_3337309, EPI_ISL_3337310, EPI_ISL_3337311, EPI_ISL_3337312 | Lighthouse Lab in Milton Keynes | Wellcome Sanger Institute for the COVID-19 Genomics UK (COG-UK) Consortium | The Lighthouse Lab in Milton Keynes and Alex Alderton, Roberto Amato, Jeffrey Barrett, Sonia Goncalves, Ewan Harrison, David K. Jackson, Ian Johnston, Dominic Kwiatkowski, Cordelia Langford, John Sillitoe on behalf of the Wellcome Sanger Institute COVID-19 Surveillance Team |
| EPI_ISL_3337313 | Berkshire and Surrey Pathology Services Lighthouse Laboratory | Wellcome Sanger Institute for the COVID-19 Genomics UK (COG-UK) Consortium | Berkshire and Surrey Pathology Services Lighthouse Laboratory and Alex Alderton, Roberto Amato, Jeffrey Barrett, Sonia Goncalves, Ewan Harrison, David K. Jackson, Ian Johnston, Dominic Kwiatkowski, Cordelia Langford, John Sillitoe on behalf of the Wellcome Sanger Institute COVID-19 Surveillance Team |
| EPI_ISL_3337314, EPI_ISL_3337315, EPI_ISL_3337316, EPI_ISL_3337317, EPI_ISL_3337318 | Lighthouse Lab in Milton Keynes | Wellcome Sanger Institute for the COVID-19 Genomics UK (COG-UK) Consortium | The Lighthouse Lab in Milton Keynes and Alex Alderton, Roberto Amato, Jeffrey Barrett, Sonia Goncalves, Ewan Harrison, David K. Jackson, Ian Johnston, Dominic Kwiatkowski, Cordelia Langford, John Sillitoe on behalf of the Wellcome Sanger Institute COVID-19 Surveillance Team |
| EPI_ISL_3337319 | Berkshire and Surrey Pathology Services Lighthouse Laboratory | Wellcome Sanger Institute for the COVID-19 Genomics UK (COG-UK) Consortium | Berkshire and Surrey Pathology Services Lighthouse Laboratory and Alex Alderton, Roberto Amato, Jeffrey Barrett, Sonia Goncalves, Ewan Harrison, David K. Jackson, Ian Johnston, Dominic Kwiatkowski, Cordelia Langford, John Sillitoe on behalf of the Wellcome Sanger Institute COVID-19 Surveillance Team |
| EPI_ISL_3337320, EPI_ISL_3337321, EPI_ISL_3337322, EPI_ISL_3337323, EPI_ISL_3337324, EPI_ISL_3337325, EPI_ISL_3337326, EPI_ISL_3337327, EPI_ISL_3337328, EPI_ISL_3337329, EPI_ISL_3337330, EPI_ISL_3337331, EPI_ISL_3337332, EPI_ISL_3337333, EPI_ISL_3337334, EPI_ISL_3337335, EPI_ISL_3337336, EPI_ISL_3337337 | | | |
| see above | Lighthouse Lab in Milton Keynes | Wellcome Sanger Institute for the COVID-19 Genomics UK (COG-UK) Consortium | The Lighthouse Lab in Milton Keynes and Alex Alderton, Roberto Amato, Jeffrey Barrett, Sonia Goncalves, Ewan Harrison, David K. Jackson, Ian Johnston, Dominic Kwiatkowski, Cordelia Langford, John Sillitoe on behalf of the Wellcome Sanger Institute COVID-19 Surveillance Team |
| EPI_ISL_3337338 | Berkshire and Surrey Pathology Services Lighthouse Laboratory | Wellcome Sanger Institute for the COVID-19 Genomics UK (COG-UK) Consortium | Berkshire and Surrey Pathology Services Lighthouse Laboratory and Alex Alderton, Roberto Amato, Jeffrey Barrett, Sonia Goncalves, Ewan Harrison, David K. Jackson, Ian Johnston, Dominic Kwiatkowski, Cordelia Langford, John Sillitoe on behalf of the Wellcome Sanger Institute COVID-19 Surveillance Team |
| EPI_ISL_3337339, EPI_ISL_3337340, EPI_ISL_3337341, EPI_ISL_3337342, EPI_ISL_3337343, EPI_ISL_3337344, EPI_ISL_3337345 | Lighthouse Lab in Milton Keynes | Wellcome Sanger Institute for the COVID-19 Genomics UK (COG-UK) Consortium | The Lighthouse Lab in Milton Keynes and Alex Alderton, Roberto Amato, Jeffrey Barrett, Sonia Goncalves, Ewan Harrison, David K. Jackson, Ian Johnston, Dominic Kwiatkowski, Cordelia Langford, John Sillitoe on behalf of the Wellcome Sanger Institute COVID-19 Surveillance Team |
| EPI_ISL_3337346 | Berkshire and Surrey Pathology Services Lighthouse Laboratory | Wellcome Sanger Institute for the COVID-19 Genomics UK (COG-UK) Consortium | Berkshire and Surrey Pathology Services Lighthouse Laboratory and Alex Alderton, Roberto Amato, Jeffrey Barrett, Sonia Goncalves, Ewan Harrison, David K. Jackson, Ian Johnston, Dominic Kwiatkowski, Cordelia Langford, John Sillitoe on behalf of the Wellcome Sanger Institute COVID-19 Surveillance Team |
| EPI_ISL_3337347, | Lighthouse Lab in Milton Keynes | Wellcome Sanger Institute for the COVID-19 Genomics UK (COG-UK) | The Lighthouse Lab in Milton Keynes and Alex Alderton, Roberto Amato, Jeffrey Barrett, Sonia Goncalves, Ewan Harrison, David K. Jackson, Ian Johnston, Dominic Kwiatkowski, Cordelia Langford, John Sillitoe on behalf of the Wellcome Sanger |

| | | | |
|---|---|---|---|
| EPI_ISL_3337348, EPI_ISL_3337349, EPI_ISL_3337350, EPI_ISL_3337351, EPI_ISL_3337352, EPI_ISL_3337353 | | Consortium | Institute COVID-19 Surveillance Team |
| EPI_ISL_3337354 | Berkshire and Surrey Pathology Services Lighthouse Laboratory | Wellcome Sanger Institute for the COVID-19 Genomics UK (COG-UK) Consortium | Berkshire and Surrey Pathology Services Lighthouse Laboratory and Alex Alderton, Roberto Amato, Jeffrey Barrett, Sonia Goncalves, Ewan Harrison, David K. Jackson, Ian Johnston, Dominic Kwiatkowski, Cordelia Langford, John Sillitoe on behalf of the Wellcome Sanger Institute COVID-19 Surveillance Team |
| EPI_ISL_3337355, EPI_ISL_3337356, EPI_ISL_3337357, EPI_ISL_3337358, EPI_ISL_3337359 | Lighthouse Lab in Milton Keynes | Wellcome Sanger Institute for the COVID-19 Genomics UK (COG-UK) Consortium | The Lighthouse Lab in Milton Keynes and Alex Alderton, Roberto Amato, Jeffrey Barrett, Sonia Goncalves, Ewan Harrison, David K. Jackson, Ian Johnston, Dominic Kwiatkowski, Cordelia Langford, John Sillitoe on behalf of the Wellcome Sanger Institute COVID-19 Surveillance Team |
| EPI_ISL_3337360 | Berkshire and Surrey Pathology Services Lighthouse Laboratory | Wellcome Sanger Institute for the COVID-19 Genomics UK (COG-UK) Consortium | Berkshire and Surrey Pathology Services Lighthouse Laboratory and Alex Alderton, Roberto Amato, Jeffrey Barrett, Sonia Goncalves, Ewan Harrison, David K. Jackson, Ian Johnston, Dominic Kwiatkowski, Cordelia Langford, John Sillitoe on behalf of the Wellcome Sanger Institute COVID-19 Surveillance Team |
| EPI_ISL_3337361, EPI_ISL_3337362, EPI_ISL_3337363, EPI_ISL_3337364, EPI_ISL_3337365 | Lighthouse Lab in Milton Keynes | Wellcome Sanger Institute for the COVID-19 Genomics UK (COG-UK) Consortium | The Lighthouse Lab in Milton Keynes and Alex Alderton, Roberto Amato, Jeffrey Barrett, Sonia Goncalves, Ewan Harrison, David K. Jackson, Ian Johnston, Dominic Kwiatkowski, Cordelia Langford, John Sillitoe on behalf of the Wellcome Sanger Institute COVID-19 Surveillance Team |
| EPI_ISL_3337366, EPI_ISL_3337367 | Berkshire and Surrey Pathology Services Lighthouse Laboratory | Wellcome Sanger Institute for the COVID-19 Genomics UK (COG-UK) Consortium | Berkshire and Surrey Pathology Services Lighthouse Laboratory and Alex Alderton, Roberto Amato, Jeffrey Barrett, Sonia Goncalves, Ewan Harrison, David K. Jackson, Ian Johnston, Dominic Kwiatkowski, Cordelia Langford, John Sillitoe on behalf of the Wellcome Sanger Institute COVID-19 Surveillance Team |
| EPI_ISL_3337368, EPI_ISL_3337369, EPI_ISL_3337370, EPI_ISL_3337371, EPI_ISL_3337372, EPI_ISL_3337373, EPI_ISL_3337374, EPI_ISL_3337375, EPI_ISL_3337376 | Lighthouse Lab in Milton Keynes | Wellcome Sanger Institute for the COVID-19 Genomics UK (COG-UK) Consortium | The Lighthouse Lab in Milton Keynes and Alex Alderton, Roberto Amato, Jeffrey Barrett, Sonia Goncalves, Ewan Harrison, David K. Jackson, Ian Johnston, Dominic Kwiatkowski, Cordelia Langford, John Sillitoe on behalf of the Wellcome Sanger Institute COVID-19 Surveillance Team |
| EPI_ISL_3337377 | Berkshire and Surrey Pathology Services Lighthouse Laboratory | Wellcome Sanger Institute for the COVID-19 Genomics UK (COG-UK) Consortium | Berkshire and Surrey Pathology Services Lighthouse Laboratory and Alex Alderton, Roberto Amato, Jeffrey Barrett, Sonia Goncalves, Ewan Harrison, David K. Jackson, Ian Johnston, Dominic Kwiatkowski, Cordelia Langford, John Sillitoe on behalf of the Wellcome Sanger Institute COVID-19 Surveillance Team |
| EPI_ISL_3337378 | Lighthouse Lab in Milton Keynes | Wellcome Sanger Institute for the COVID-19 Genomics UK (COG-UK) Consortium | The Lighthouse Lab in Milton Keynes and Alex Alderton, Roberto Amato, Jeffrey Barrett, Sonia Goncalves, Ewan Harrison, David K. Jackson, Ian Johnston, Dominic Kwiatkowski, Cordelia Langford, John Sillitoe on behalf of the Wellcome Sanger Institute COVID-19 Surveillance Team |
| EPI_ISL_3337379 | Berkshire and Surrey Pathology Services Lighthouse Laboratory | Wellcome Sanger Institute for the COVID-19 Genomics UK (COG-UK) Consortium | Berkshire and Surrey Pathology Services Lighthouse Laboratory and Alex Alderton, Roberto Amato, Jeffrey Barrett, Sonia Goncalves, Ewan Harrison, David K. Jackson, Ian Johnston, Dominic Kwiatkowski, Cordelia Langford, John Sillitoe on behalf of the Wellcome Sanger Institute COVID-19 Surveillance Team |
| EPI_ISL_3337380 | Health Services Laboratories | Wellcome Sanger Institute for the COVID-19 Genomics UK (COG-UK) Consortium | Health Services Laboratories and Alex Alderton, Roberto Amato, Jeffrey Barrett, Sonia Goncalves, Ewan Harrison, David K. Jackson, Ian Johnston, Dominic Kwiatkowski, Cordelia Langford, John Sillitoe on behalf of the Wellcome Sanger Institute COVID-19 Surveillance Team |
| EPI_ISL_3337381 | Berkshire and Surrey Pathology Services Lighthouse Laboratory | Wellcome Sanger Institute for the COVID-19 Genomics UK (COG-UK) Consortium | Berkshire and Surrey Pathology Services Lighthouse Laboratory and Alex Alderton, Roberto Amato, Jeffrey Barrett, Sonia Goncalves, Ewan Harrison, David K. Jackson, Ian Johnston, Dominic Kwiatkowski, Cordelia Langford, John Sillitoe on behalf of the Wellcome Sanger Institute COVID-19 Surveillance Team |
| EPI_ISL_3337382 | Health Services Laboratories | Wellcome Sanger Institute for the COVID-19 Genomics UK (COG-UK) Consortium | Health Services Laboratories and Alex Alderton, Roberto Amato, Jeffrey Barrett, Sonia Goncalves, Ewan Harrison, David K. Jackson, Ian Johnston, Dominic Kwiatkowski, Cordelia Langford, John Sillitoe on behalf of the Wellcome Sanger Institute COVID-19 Surveillance Team |
| EPI_ISL_3337383, EPI_ISL_3337384 | Lighthouse Lab in Milton Keynes | Wellcome Sanger Institute for the COVID-19 Genomics UK (COG-UK) Consortium | The Lighthouse Lab in Milton Keynes and Alex Alderton, Roberto Amato, Jeffrey Barrett, Sonia Goncalves, Ewan Harrison, David K. Jackson, Ian Johnston, Dominic Kwiatkowski, Cordelia Langford, John Sillitoe on behalf of the Wellcome Sanger Institute COVID-19 Surveillance Team |
| EPI_ISL_3337385 | Berkshire and Surrey Pathology Services Lighthouse Laboratory | Wellcome Sanger Institute for the COVID-19 Genomics UK (COG-UK) Consortium | Berkshire and Surrey Pathology Services Lighthouse Laboratory and Alex Alderton, Roberto Amato, Jeffrey Barrett, Sonia Goncalves, Ewan Harrison, David K. Jackson, Ian Johnston, Dominic Kwiatkowski, Cordelia Langford, John Sillitoe on behalf of the Wellcome Sanger Institute COVID-19 Surveillance Team |
| EPI_ISL_3337386, EPI_ISL_3337387, EPI_ISL_3337388, EPI_ISL_3337389, EPI_ISL_3337390, EPI_ISL_3337391 | Lighthouse Lab in Milton Keynes | Wellcome Sanger Institute for the COVID-19 Genomics UK (COG-UK) Consortium | The Lighthouse Lab in Milton Keynes and Alex Alderton, Roberto Amato, Jeffrey Barrett, Sonia Goncalves, Ewan Harrison, David K. Jackson, Ian Johnston, Dominic Kwiatkowski, Cordelia Langford, John Sillitoe on behalf of the Wellcome Sanger Institute COVID-19 Surveillance Team |
| EPI_ISL_3337392 | Berkshire and Surrey Pathology Services Lighthouse Laboratory | Wellcome Sanger Institute for the COVID-19 Genomics UK (COG-UK) Consortium | Berkshire and Surrey Pathology Services Lighthouse Laboratory and Alex Alderton, Roberto Amato, Jeffrey Barrett, Sonia Goncalves, Ewan Harrison, David K. Jackson, Ian Johnston, Dominic Kwiatkowski, Cordelia Langford, John Sillitoe on behalf of the Wellcome Sanger Institute COVID-19 Surveillance Team |
| EPI_ISL_3337393, EPI_ISL_3337394, EPI_ISL_3337395, EPI_ISL_3337396, EPI_ISL_3337397, EPI_ISL_3337398, EPI_ISL_3337399, EPI_ISL_3337400, EPI_ISL_3337401, EPI_ISL_3337402, EPI_ISL_3337403 | | | |
| see above | Lighthouse Lab in Milton Keynes | Wellcome Sanger Institute for the COVID-19 Genomics UK (COG-UK) Consortium | The Lighthouse Lab in Milton Keynes and Alex Alderton, Roberto Amato, Jeffrey Barrett, Sonia Goncalves, Ewan Harrison, David K. Jackson, Ian Johnston, Dominic Kwiatkowski, Cordelia Langford, John Sillitoe on behalf of the Wellcome Sanger Institute COVID-19 Surveillance Team |
| EPI_ISL_3337404 | Berkshire and Surrey Pathology Services Lighthouse Laboratory | Wellcome Sanger Institute for the COVID-19 Genomics UK (COG-UK) Consortium | Berkshire and Surrey Pathology Services Lighthouse Laboratory and Alex Alderton, Roberto Amato, Jeffrey Barrett, Sonia Goncalves, Ewan Harrison, David K. Jackson, Ian Johnston, Dominic Kwiatkowski, Cordelia Langford, John Sillitoe on behalf of the Wellcome Sanger Institute COVID-19 Surveillance Team |
| EPI_ISL_3337405, EPI_ISL_3337406 | Lighthouse Lab in Milton Keynes | Wellcome Sanger Institute for the COVID-19 Genomics UK (COG-UK) Consortium | The Lighthouse Lab in Milton Keynes and Alex Alderton, Roberto Amato, Jeffrey Barrett, Sonia Goncalves, Ewan Harrison, David K. Jackson, Ian Johnston, Dominic Kwiatkowski, Cordelia Langford, John Sillitoe on behalf of the Wellcome Sanger Institute COVID-19 Surveillance Team |
| EPI_ISL_3337407, EPI_ISL_3337408 | Berkshire and Surrey Pathology Services Lighthouse Laboratory | Wellcome Sanger Institute for the COVID-19 Genomics UK (COG-UK) Consortium | Berkshire and Surrey Pathology Services Lighthouse Laboratory and Alex Alderton, Roberto Amato, Jeffrey Barrett, Sonia Goncalves, Ewan Harrison, David K. Jackson, Ian Johnston, Dominic Kwiatkowski, Cordelia Langford, John Sillitoe on behalf of the Wellcome Sanger Institute COVID-19 Surveillance Team |
| EPI_ISL_3337409, EPI_ISL_3337410, EPI_ISL_3337411 | Lighthouse Lab in Milton Keynes | Wellcome Sanger Institute for the COVID-19 Genomics UK (COG-UK) Consortium | The Lighthouse Lab in Milton Keynes and Alex Alderton, Roberto Amato, Jeffrey Barrett, Sonia Goncalves, Ewan Harrison, David K. Jackson, Ian Johnston, Dominic Kwiatkowski, Cordelia Langford, John Sillitoe on behalf of the Wellcome Sanger Institute COVID-19 Surveillance Team |
| EPI_ISL_3337412 | Berkshire and Surrey Pathology Services Lighthouse Laboratory | Wellcome Sanger Institute for the COVID-19 Genomics UK (COG-UK) Consortium | Berkshire and Surrey Pathology Services Lighthouse Laboratory and Alex Alderton, Roberto Amato, Jeffrey Barrett, Sonia Goncalves, Ewan Harrison, David K. Jackson, Ian Johnston, Dominic Kwiatkowski, Cordelia Langford, John Sillitoe on behalf of the Wellcome Sanger Institute COVID-19 Surveillance Team |
| EPI_ISL_3337413 | Lighthouse Lab in Milton Keynes | Wellcome Sanger Institute for the COVID-19 Genomics UK (COG-UK) Consortium | The Lighthouse Lab in Milton Keynes and Alex Alderton, Roberto Amato, Jeffrey Barrett, Sonia Goncalves, Ewan Harrison, David K. Jackson, Ian Johnston, Dominic Kwiatkowski, Cordelia Langford, John Sillitoe on behalf of the Wellcome Sanger Institute COVID-19 Surveillance Team |
| EPI_ISL_3337414 | Health Services Laboratories | Wellcome Sanger Institute for the COVID-19 Genomics UK (COG-UK) Consortium | Health Services Laboratories and Alex Alderton, Roberto Amato, Jeffrey Barrett, Sonia Goncalves, Ewan Harrison, David K. Jackson, Ian Johnston, Dominic Kwiatkowski, Cordelia Langford, John Sillitoe on behalf of the Wellcome Sanger Institute COVID-19 Surveillance Team |
| EPI_ISL_3337415 | Berkshire and Surrey Pathology Services Lighthouse Laboratory | Wellcome Sanger Institute for the COVID-19 Genomics UK (COG-UK) Consortium | Berkshire and Surrey Pathology Services Lighthouse Laboratory and Alex Alderton, Roberto Amato, Jeffrey Barrett, Sonia Goncalves, Ewan Harrison, David K. Jackson, Ian Johnston, Dominic Kwiatkowski, Cordelia Langford, John Sillitoe on behalf of the Wellcome Sanger Institute COVID-19 Surveillance Team |
| EPI_ISL_3337416 | Lighthouse Lab in Milton Keynes | Wellcome Sanger Institute for the COVID-19 Genomics UK (COG-UK) Consortium | The Lighthouse Lab in Milton Keynes and Alex Alderton, Roberto Amato, Jeffrey Barrett, Sonia Goncalves, Ewan Harrison, David K. Jackson, Ian Johnston, Dominic Kwiatkowski, Cordelia Langford, John Sillitoe on behalf of the Wellcome Sanger Institute COVID-19 Surveillance Team |
| EPI_ISL_3337417 | Berkshire and Surrey Pathology Services Lighthouse Laboratory | Wellcome Sanger Institute for the COVID-19 Genomics UK (COG-UK) Consortium | Berkshire and Surrey Pathology Services Lighthouse Laboratory and Alex Alderton, Roberto Amato, Jeffrey Barrett, Sonia Goncalves, Ewan Harrison, David K. Jackson, Ian Johnston, Dominic Kwiatkowski, Cordelia Langford, John Sillitoe on behalf of the Wellcome Sanger Institute COVID-19 Surveillance Team |
| EPI_ISL_3337418, EPI_ISL_3337419 | Lighthouse Lab in Milton Keynes | Wellcome Sanger Institute for the COVID-19 Genomics UK (COG-UK) Consortium | The Lighthouse Lab in Milton Keynes and Alex Alderton, Roberto Amato, Jeffrey Barrett, Sonia Goncalves, Ewan Harrison, David K. Jackson, Ian Johnston, Dominic Kwiatkowski, Cordelia Langford, John Sillitoe on behalf of the Wellcome Sanger Institute COVID-19 Surveillance Team |
| EPI_ISL_3337420 | Berkshire and Surrey Pathology Services Lighthouse Laboratory | Wellcome Sanger Institute for the COVID-19 Genomics UK (COG-UK) Consortium | Berkshire and Surrey Pathology Services Lighthouse Laboratory and Alex Alderton, Roberto Amato, Jeffrey Barrett, Sonia Goncalves, Ewan Harrison, David K. Jackson, Ian Johnston, Dominic Kwiatkowski, Cordelia Langford, John Sillitoe on behalf of the Wellcome Sanger Institute COVID-19 Surveillance Team |
| EPI_ISL_3337421, EPI_ISL_3337422, EPI_ISL_3337423 | Lighthouse Lab in Milton Keynes | Wellcome Sanger Institute for the COVID-19 Genomics UK (COG-UK) Consortium | The Lighthouse Lab in Milton Keynes and Alex Alderton, Roberto Amato, Jeffrey Barrett, Sonia Goncalves, Ewan Harrison, David K. Jackson, Ian Johnston, Dominic Kwiatkowski, Cordelia Langford, John Sillitoe on behalf of the Wellcome Sanger Institute COVID-19 Surveillance Team |
| EPI_ISL_3337424 | Health Services Laboratories | Wellcome Sanger Institute for the COVID-19 Genomics UK (COG-UK) Consortium | Health Services Laboratories and Alex Alderton, Roberto Amato, Jeffrey Barrett, Sonia Goncalves, Ewan Harrison, David K. Jackson, Ian Johnston, Dominic Kwiatkowski, Cordelia Langford, John Sillitoe on behalf of the Wellcome Sanger Institute COVID-19 Surveillance Team |
| EPI_ISL_3337425 | Lighthouse Lab in Milton Keynes | Wellcome Sanger Institute for the COVID-19 Genomics UK (COG-UK) Consortium | The Lighthouse Lab in Milton Keynes and Alex Alderton, Roberto Amato, Jeffrey Barrett, Sonia Goncalves, Ewan Harrison, David K. Jackson, Ian Johnston, Dominic Kwiatkowski, Cordelia Langford, John Sillitoe on behalf of the Wellcome Sanger Institute COVID-19 Surveillance Team |
| EPI_ISL_3337426, EPI_ISL_3337427 | Health Services Laboratories | Wellcome Sanger Institute for the COVID-19 Genomics UK (COG-UK) Consortium | Health Services Laboratories and Alex Alderton, Roberto Amato, Jeffrey Barrett, Sonia Goncalves, Ewan Harrison, David K. Jackson, Ian Johnston, Dominic Kwiatkowski, Cordelia Langford, John Sillitoe on behalf of the Wellcome Sanger Institute COVID-19 Surveillance Team |
| EPI_ISL_3337428 | Berkshire and Surrey Pathology Services Lighthouse Laboratory | Wellcome Sanger Institute for the COVID-19 Genomics UK (COG-UK) Consortium | Berkshire and Surrey Pathology Services Lighthouse Laboratory and Alex Alderton, Roberto Amato, Jeffrey Barrett, Sonia Goncalves, Ewan Harrison, David K. Jackson, Ian Johnston, Dominic Kwiatkowski, Cordelia Langford, John Sillitoe on behalf of the Wellcome Sanger Institute COVID-19 Surveillance Team |

| | | | |
|---|---|---|---|
| EPI_ISL_3337429, EPI_ISL_3337430 | Health Services Laboratories | Wellcome Sanger Institute for the COVID-19 Genomics UK (COG-UK) Consortium | Health Services Laboratories and Alex Alderton, Roberto Amato, Jeffrey Barrett, Sonia Goncalves, Ewan Harrison, David K. Jackson, Ian Johnston, Dominic Kwiatkowski, Cordelia Langford, John Sillitoe on behalf of the Wellcome Sanger Institute COVID-19 Surveillance Team |
| EPI_ISL_3337431, EPI_ISL_3337432 | Lighthouse Lab in Milton Keynes | Wellcome Sanger Institute for the COVID-19 Genomics UK (COG-UK) Consortium | The Lighthouse Lab in Milton Keynes and Alex Alderton, Roberto Amato, Jeffrey Barrett, Sonia Goncalves, Ewan Harrison, David K. Jackson, Ian Johnston, Dominic Kwiatkowski, Cordelia Langford, John Sillitoe on behalf of the Wellcome Sanger Institute COVID-19 Surveillance Team |
| EPI_ISL_3337433, EPI_ISL_3337434 | Health Services Laboratories | Wellcome Sanger Institute for the COVID-19 Genomics UK (COG-UK) Consortium | Health Services Laboratories and Alex Alderton, Roberto Amato, Jeffrey Barrett, Sonia Goncalves, Ewan Harrison, David K. Jackson, Ian Johnston, Dominic Kwiatkowski, Cordelia Langford, John Sillitoe on behalf of the Wellcome Sanger Institute COVID-19 Surveillance Team |
| EPI_ISL_3337435 | Berkshire and Surrey Pathology Services Lighthouse Laboratory | Wellcome Sanger Institute for the COVID-19 Genomics UK (COG-UK) Consortium | Berkshire and Surrey Pathology Services Lighthouse Laboratory and Alex Alderton, Roberto Amato, Jeffrey Barrett, Sonia Goncalves, Ewan Harrison, David K. Jackson, Ian Johnston, Dominic Kwiatkowski, Cordelia Langford, John Sillitoe on behalf of the Wellcome Sanger Institute COVID-19 Surveillance Team |
| EPI_ISL_3337436 | Lighthouse Lab in Milton Keynes | Wellcome Sanger Institute for the COVID-19 Genomics UK (COG-UK) Consortium | The Lighthouse Lab in Milton Keynes and Alex Alderton, Roberto Amato, Jeffrey Barrett, Sonia Goncalves, Ewan Harrison, David K. Jackson, Ian Johnston, Dominic Kwiatkowski, Cordelia Langford, John Sillitoe on behalf of the Wellcome Sanger Institute COVID-19 Surveillance Team |
| EPI_ISL_3337437 | Berkshire and Surrey Pathology Services Lighthouse Laboratory | Wellcome Sanger Institute for the COVID-19 Genomics UK (COG-UK) Consortium | Berkshire and Surrey Pathology Services Lighthouse Laboratory and Alex Alderton, Roberto Amato, Jeffrey Barrett, Sonia Goncalves, Ewan Harrison, David K. Jackson, Ian Johnston, Dominic Kwiatkowski, Cordelia Langford, John Sillitoe on behalf of the Wellcome Sanger Institute COVID-19 Surveillance Team |
| EPI_ISL_3337438 | Lighthouse Lab in Milton Keynes | Wellcome Sanger Institute for the COVID-19 Genomics UK (COG-UK) Consortium | The Lighthouse Lab in Milton Keynes and Alex Alderton, Roberto Amato, Jeffrey Barrett, Sonia Goncalves, Ewan Harrison, David K. Jackson, Ian Johnston, Dominic Kwiatkowski, Cordelia Langford, John Sillitoe on behalf of the Wellcome Sanger Institute COVID-19 Surveillance Team |
| EPI_ISL_3337439 | Berkshire and Surrey Pathology Services Lighthouse Laboratory | Wellcome Sanger Institute for the COVID-19 Genomics UK (COG-UK) Consortium | Berkshire and Surrey Pathology Services Lighthouse Laboratory and Alex Alderton, Roberto Amato, Jeffrey Barrett, Sonia Goncalves, Ewan Harrison, David K. Jackson, Ian Johnston, Dominic Kwiatkowski, Cordelia Langford, John Sillitoe on behalf of the Wellcome Sanger Institute COVID-19 Surveillance Team |
| EPI_ISL_3337440 | Lighthouse Lab in Milton Keynes | Wellcome Sanger Institute for the COVID-19 Genomics UK (COG-UK) Consortium | The Lighthouse Lab in Milton Keynes and Alex Alderton, Roberto Amato, Jeffrey Barrett, Sonia Goncalves, Ewan Harrison, David K. Jackson, Ian Johnston, Dominic Kwiatkowski, Cordelia Langford, John Sillitoe on behalf of the Wellcome Sanger Institute COVID-19 Surveillance Team |
| EPI_ISL_3337441 | Berkshire and Surrey Pathology Services Lighthouse Laboratory | Wellcome Sanger Institute for the COVID-19 Genomics UK (COG-UK) Consortium | Berkshire and Surrey Pathology Services Lighthouse Laboratory and Alex Alderton, Roberto Amato, Jeffrey Barrett, Sonia Goncalves, Ewan Harrison, David K. Jackson, Ian Johnston, Dominic Kwiatkowski, Cordelia Langford, John Sillitoe on behalf of the Wellcome Sanger Institute COVID-19 Surveillance Team |
| EPI_ISL_3337442, EPI_ISL_3337443, EPI_ISL_3337444, EPI_ISL_3337445, EPI_ISL_3337446, EPI_ISL_3337447, EPI_ISL_3337448 | Lighthouse Lab in Milton Keynes | Wellcome Sanger Institute for the COVID-19 Genomics UK (COG-UK) Consortium | The Lighthouse Lab in Milton Keynes and Alex Alderton, Roberto Amato, Jeffrey Barrett, Sonia Goncalves, Ewan Harrison, David K. Jackson, Ian Johnston, Dominic Kwiatkowski, Cordelia Langford, John Sillitoe on behalf of the Wellcome Sanger Institute COVID-19 Surveillance Team |
| EPI_ISL_3337449 | Berkshire and Surrey Pathology Services Lighthouse Laboratory | Wellcome Sanger Institute for the COVID-19 Genomics UK (COG-UK) Consortium | Berkshire and Surrey Pathology Services Lighthouse Laboratory and Alex Alderton, Roberto Amato, Jeffrey Barrett, Sonia Goncalves, Ewan Harrison, David K. Jackson, Ian Johnston, Dominic Kwiatkowski, Cordelia Langford, John Sillitoe on behalf of the Wellcome Sanger Institute COVID-19 Surveillance Team |
| EPI_ISL_3337450, EPI_ISL_3337451, EPI_ISL_3337452, EPI_ISL_3337453 | Lighthouse Lab in Milton Keynes | Wellcome Sanger Institute for the COVID-19 Genomics UK (COG-UK) Consortium | The Lighthouse Lab in Milton Keynes and Alex Alderton, Roberto Amato, Jeffrey Barrett, Sonia Goncalves, Ewan Harrison, David K. Jackson, Ian Johnston, Dominic Kwiatkowski, Cordelia Langford, John Sillitoe on behalf of the Wellcome Sanger Institute COVID-19 Surveillance Team |
| EPI_ISL_3337454 | Berkshire and Surrey Pathology Services Lighthouse Laboratory | Wellcome Sanger Institute for the COVID-19 Genomics UK (COG-UK) Consortium | Berkshire and Surrey Pathology Services Lighthouse Laboratory and Alex Alderton, Roberto Amato, Jeffrey Barrett, Sonia Goncalves, Ewan Harrison, David K. Jackson, Ian Johnston, Dominic Kwiatkowski, Cordelia Langford, John Sillitoe on behalf of the Wellcome Sanger Institute COVID-19 Surveillance Team |
| EPI_ISL_3337455 | Lighthouse Lab in Milton Keynes | Wellcome Sanger Institute for the COVID-19 Genomics UK (COG-UK) Consortium | The Lighthouse Lab in Milton Keynes and Alex Alderton, Roberto Amato, Jeffrey Barrett, Sonia Goncalves, Ewan Harrison, David K. Jackson, Ian Johnston, Dominic Kwiatkowski, Cordelia Langford, John Sillitoe on behalf of the Wellcome Sanger Institute COVID-19 Surveillance Team |
| EPI_ISL_3337456 | Health Services Laboratories | Wellcome Sanger Institute for the COVID-19 Genomics UK (COG-UK) Consortium | Health Services Laboratories and Alex Alderton, Roberto Amato, Jeffrey Barrett, Sonia Goncalves, Ewan Harrison, David K. Jackson, Ian Johnston, Dominic Kwiatkowski, Cordelia Langford, John Sillitoe on behalf of the Wellcome Sanger Institute COVID-19 Surveillance Team |
| EPI_ISL_3337457 | Berkshire and Surrey Pathology Services Lighthouse Laboratory | Wellcome Sanger Institute for the COVID-19 Genomics UK (COG-UK) Consortium | Berkshire and Surrey Pathology Services Lighthouse Laboratory and Alex Alderton, Roberto Amato, Jeffrey Barrett, Sonia Goncalves, Ewan Harrison, David K. Jackson, Ian Johnston, Dominic Kwiatkowski, Cordelia Langford, John Sillitoe on behalf of the Wellcome Sanger Institute COVID-19 Surveillance Team |
| EPI_ISL_3337458, EPI_ISL_3337459, EPI_ISL_3337460, EPI_ISL_3337461 | Lighthouse Lab in Milton Keynes | Wellcome Sanger Institute for the COVID-19 Genomics UK (COG-UK) Consortium | The Lighthouse Lab in Milton Keynes and Alex Alderton, Roberto Amato, Jeffrey Barrett, Sonia Goncalves, Ewan Harrison, David K. Jackson, Ian Johnston, Dominic Kwiatkowski, Cordelia Langford, John Sillitoe on behalf of the Wellcome Sanger Institute COVID-19 Surveillance Team |
| EPI_ISL_3337462, EPI_ISL_3337463 | Health Services Laboratories | Wellcome Sanger Institute for the COVID-19 Genomics UK (COG-UK) Consortium | Health Services Laboratories and Alex Alderton, Roberto Amato, Jeffrey Barrett, Sonia Goncalves, Ewan Harrison, David K. Jackson, Ian Johnston, Dominic Kwiatkowski, Cordelia Langford, John Sillitoe on behalf of the Wellcome Sanger Institute COVID-19 Surveillance Team |
| EPI_ISL_3337464 | Berkshire and Surrey Pathology Services Lighthouse Laboratory | Wellcome Sanger Institute for the COVID-19 Genomics UK (COG-UK) Consortium | Berkshire and Surrey Pathology Services Lighthouse Laboratory and Alex Alderton, Roberto Amato, Jeffrey Barrett, Sonia Goncalves, Ewan Harrison, David K. Jackson, Ian Johnston, Dominic Kwiatkowski, Cordelia Langford, John Sillitoe on behalf of the Wellcome Sanger Institute COVID-19 Surveillance Team |
| EPI_ISL_3337465, EPI_ISL_3337466, EPI_ISL_3337467, EPI_ISL_3337468, EPI_ISL_3337469, EPI_ISL_3337470, EPI_ISL_3337471, EPI_ISL_3337472, EPI_ISL_3337473 | Lighthouse Lab in Milton Keynes | Wellcome Sanger Institute for the COVID-19 Genomics UK (COG-UK) Consortium | The Lighthouse Lab in Milton Keynes and Alex Alderton, Roberto Amato, Jeffrey Barrett, Sonia Goncalves, Ewan Harrison, David K. Jackson, Ian Johnston, Dominic Kwiatkowski, Cordelia Langford, John Sillitoe on behalf of the Wellcome Sanger Institute COVID-19 Surveillance Team |
| EPI_ISL_3337474 | Berkshire and Surrey Pathology Services Lighthouse Laboratory | Wellcome Sanger Institute for the COVID-19 Genomics UK (COG-UK) Consortium | Berkshire and Surrey Pathology Services Lighthouse Laboratory and Alex Alderton, Roberto Amato, Jeffrey Barrett, Sonia Goncalves, Ewan Harrison, David K. Jackson, Ian Johnston, Dominic Kwiatkowski, Cordelia Langford, John Sillitoe on behalf of the Wellcome Sanger Institute COVID-19 Surveillance Team |
| EPI_ISL_3337475, EPI_ISL_3337476 | Lighthouse Lab in Milton Keynes | Wellcome Sanger Institute for the COVID-19 Genomics UK (COG-UK) Consortium | The Lighthouse Lab in Milton Keynes and Alex Alderton, Roberto Amato, Jeffrey Barrett, Sonia Goncalves, Ewan Harrison, David K. Jackson, Ian Johnston, Dominic Kwiatkowski, Cordelia Langford, John Sillitoe on behalf of the Wellcome Sanger Institute COVID-19 Surveillance Team |
| EPI_ISL_3337477, EPI_ISL_3337478 | Health Services Laboratories | Wellcome Sanger Institute for the COVID-19 Genomics UK (COG-UK) Consortium | Health Services Laboratories and Alex Alderton, Roberto Amato, Jeffrey Barrett, Sonia Goncalves, Ewan Harrison, David K. Jackson, Ian Johnston, Dominic Kwiatkowski, Cordelia Langford, John Sillitoe on behalf of the Wellcome Sanger Institute COVID-19 Surveillance Team |
| EPI_ISL_3337479 | Berkshire and Surrey Pathology Services Lighthouse Laboratory | Wellcome Sanger Institute for the COVID-19 Genomics UK (COG-UK) Consortium | Berkshire and Surrey Pathology Services Lighthouse Laboratory and Alex Alderton, Roberto Amato, Jeffrey Barrett, Sonia Goncalves, Ewan Harrison, David K. Jackson, Ian Johnston, Dominic Kwiatkowski, Cordelia Langford, John Sillitoe on behalf of the Wellcome Sanger Institute COVID-19 Surveillance Team |
| EPI_ISL_3337480, EPI_ISL_3337481 | Health Services Laboratories | Wellcome Sanger Institute for the COVID-19 Genomics UK (COG-UK) Consortium | Health Services Laboratories and Alex Alderton, Roberto Amato, Jeffrey Barrett, Sonia Goncalves, Ewan Harrison, David K. Jackson, Ian Johnston, Dominic Kwiatkowski, Cordelia Langford, John Sillitoe on behalf of the Wellcome Sanger Institute COVID-19 Surveillance Team |
| EPI_ISL_3337482, EPI_ISL_3337483, EPI_ISL_3337484, EPI_ISL_3337485 | Lighthouse Lab in Milton Keynes | Wellcome Sanger Institute for the COVID-19 Genomics UK (COG-UK) Consortium | The Lighthouse Lab in Milton Keynes and Alex Alderton, Roberto Amato, Jeffrey Barrett, Sonia Goncalves, Ewan Harrison, David K. Jackson, Ian Johnston, Dominic Kwiatkowski, Cordelia Langford, John Sillitoe on behalf of the Wellcome Sanger Institute COVID-19 Surveillance Team |
| EPI_ISL_3337486, EPI_ISL_3337487 | Health Services Laboratories | Wellcome Sanger Institute for the COVID-19 Genomics UK (COG-UK) Consortium | Health Services Laboratories and Alex Alderton, Roberto Amato, Jeffrey Barrett, Sonia Goncalves, Ewan Harrison, David K. Jackson, Ian Johnston, Dominic Kwiatkowski, Cordelia Langford, John Sillitoe on behalf of the Wellcome Sanger Institute COVID-19 Surveillance Team |
| EPI_ISL_3337488 | Lighthouse Lab in Milton Keynes | Wellcome Sanger Institute for the COVID-19 Genomics UK (COG-UK) Consortium | The Lighthouse Lab in Milton Keynes and Alex Alderton, Roberto Amato, Jeffrey Barrett, Sonia Goncalves, Ewan Harrison, David K. Jackson, Ian Johnston, Dominic Kwiatkowski, Cordelia Langford, John Sillitoe on behalf of the Wellcome Sanger Institute COVID-19 Surveillance Team |
| EPI_ISL_3337489 | Health Services Laboratories | Wellcome Sanger Institute for the COVID-19 Genomics UK (COG-UK) Consortium | Health Services Laboratories and Alex Alderton, Roberto Amato, Jeffrey Barrett, Sonia Goncalves, Ewan Harrison, David K. Jackson, Ian Johnston, Dominic Kwiatkowski, Cordelia Langford, John Sillitoe on behalf of the Wellcome Sanger Institute COVID-19 Surveillance Team |
| EPI_ISL_3337490 | Lighthouse Lab in Milton Keynes | Wellcome Sanger Institute for the COVID-19 Genomics UK (COG-UK) Consortium | The Lighthouse Lab in Milton Keynes and Alex Alderton, Roberto Amato, Jeffrey Barrett, Sonia Goncalves, Ewan Harrison, David K. Jackson, Ian Johnston, Dominic Kwiatkowski, Cordelia Langford, John Sillitoe on behalf of the Wellcome Sanger Institute COVID-19 Surveillance Team |
| EPI_ISL_3337491 | Health Services Laboratories | Wellcome Sanger Institute for the COVID-19 Genomics UK (COG-UK) Consortium | Health Services Laboratories and Alex Alderton, Roberto Amato, Jeffrey Barrett, Sonia Goncalves, Ewan Harrison, David K. Jackson, Ian Johnston, Dominic Kwiatkowski, Cordelia Langford, John Sillitoe on behalf of the Wellcome Sanger Institute COVID-19 Surveillance Team |
| EPI_ISL_3337492 | Lighthouse Lab in Milton Keynes | Wellcome Sanger Institute for the COVID-19 Genomics UK (COG-UK) Consortium | The Lighthouse Lab in Milton Keynes and Alex Alderton, Roberto Amato, Jeffrey Barrett, Sonia Goncalves, Ewan Harrison, David K. Jackson, Ian Johnston, Dominic Kwiatkowski, Cordelia Langford, John Sillitoe on behalf of the Wellcome Sanger Institute COVID-19 Surveillance Team |
| EPI_ISL_3337493 | Berkshire and Surrey Pathology Services Lighthouse Laboratory | Wellcome Sanger Institute for the COVID-19 Genomics UK (COG-UK) Consortium | Berkshire and Surrey Pathology Services Lighthouse Laboratory and Alex Alderton, Roberto Amato, Jeffrey Barrett, Sonia Goncalves, Ewan Harrison, David K. Jackson, Ian Johnston, Dominic Kwiatkowski, Cordelia Langford, John Sillitoe on behalf of the Wellcome Sanger Institute COVID-19 Surveillance Team |
| EPI_ISL_3337494 | Lighthouse Lab in Milton Keynes | Wellcome Sanger Institute for the COVID-19 Genomics UK (COG-UK) Consortium | The Lighthouse Lab in Milton Keynes and Alex Alderton, Roberto Amato, Jeffrey Barrett, Sonia Goncalves, Ewan Harrison, David K. Jackson, Ian Johnston, Dominic Kwiatkowski, Cordelia Langford, John Sillitoe on behalf of the Wellcome Sanger Institute COVID-19 Surveillance Team |
| EPI_ISL_3337495 | Health Services Laboratories | Wellcome Sanger Institute for the COVID-19 Genomics UK (COG-UK) Consortium | Health Services Laboratories and Alex Alderton, Roberto Amato, Jeffrey Barrett, Sonia Goncalves, Ewan Harrison, David K. Jackson, Ian Johnston, Dominic Kwiatkowski, Cordelia Langford, John Sillitoe on behalf of the Wellcome Sanger Institute COVID-19 Surveillance Team |
| EPI_ISL_3337496 | Lighthouse Lab in Milton Keynes | Wellcome Sanger Institute for the COVID-19 Genomics UK (COG-UK) Consortium | The Lighthouse Lab in Milton Keynes and Alex Alderton, Roberto Amato, Jeffrey Barrett, Sonia Goncalves, Ewan Harrison, David K. Jackson, Ian Johnston, Dominic Kwiatkowski, Cordelia Langford, John Sillitoe on behalf of the Wellcome Sanger Institute COVID-19 Surveillance Team |
| EPI_ISL_3337497 | Berkshire and Surrey Pathology Services Lighthouse Laboratory | Wellcome Sanger Institute for the COVID-19 Genomics UK (COG-UK) | Berkshire and Surrey Pathology Services Lighthouse Laboratory and Alex Alderton, Roberto Amato, Jeffrey Barrett, Sonia Goncalves, Ewan Harrison, David K. Jackson, Ian Johnston, Dominic Kwiatkowski, Cordelia Langford, John Sillitoe on behalf |

| | | | |
|---|---|---|---|
| | | Consortium | of the Wellcome Sanger Institute COVID-19 Surveillance Team |
| EPI_ISL_3337498 | Lighthouse Lab in Milton Keynes | Wellcome Sanger Institute for the COVID-19 Genomics UK (COG-UK) Consortium | The Lighthouse Lab in Milton Keynes and Alex Alderton, Roberto Amato, Jeffrey Barrett, Sonia Goncalves, Ewan Harrison, David K. Jackson, Ian Johnston, Dominic Kwiatkowski, Cordelia Langford, John Sillitoe on behalf of the Wellcome Sanger Institute COVID-19 Surveillance Team |
| EPI_ISL_3337499 | Health Services Laboratories | Wellcome Sanger Institute for the COVID-19 Genomics UK (COG-UK) Consortium | Health Services Laboratories and Alex Alderton, Roberto Amato, Jeffrey Barrett, Sonia Goncalves, Ewan Harrison, David K. Jackson, Ian Johnston, Dominic Kwiatkowski, Cordelia Langford, John Sillitoe on behalf of the Wellcome Sanger Institute COVID-19 Surveillance Team |
| EPI_ISL_3337500 | Berkshire and Surrey Pathology Services Lighthouse Laboratory | Wellcome Sanger Institute for the COVID-19 Genomics UK (COG-UK) Consortium | Berkshire and Surrey Pathology Services Lighthouse Laboratory and Alex Alderton, Roberto Amato, Jeffrey Barrett, Sonia Goncalves, Ewan Harrison, David K. Jackson, Ian Johnston, Dominic Kwiatkowski, Cordelia Langford, John Sillitoe on behalf of the Wellcome Sanger Institute COVID-19 Surveillance Team |
| EPI_ISL_3337501, EPI_ISL_3337502 | Lighthouse Lab in Milton Keynes | Wellcome Sanger Institute for the COVID-19 Genomics UK (COG-UK) Consortium | The Lighthouse Lab in Milton Keynes and Alex Alderton, Roberto Amato, Jeffrey Barrett, Sonia Goncalves, Ewan Harrison, David K. Jackson, Ian Johnston, Dominic Kwiatkowski, Cordelia Langford, John Sillitoe on behalf of the Wellcome Sanger Institute COVID-19 Surveillance Team |
| EPI_ISL_3337503 | Berkshire and Surrey Pathology Services Lighthouse Laboratory | Wellcome Sanger Institute for the COVID-19 Genomics UK (COG-UK) Consortium | Berkshire and Surrey Pathology Services Lighthouse Laboratory and Alex Alderton, Roberto Amato, Jeffrey Barrett, Sonia Goncalves, Ewan Harrison, David K. Jackson, Ian Johnston, Dominic Kwiatkowski, Cordelia Langford, John Sillitoe on behalf of the Wellcome Sanger Institute COVID-19 Surveillance Team |
| EPI_ISL_3337504 | Health Services Laboratories | Wellcome Sanger Institute for the COVID-19 Genomics UK (COG-UK) Consortium | Health Services Laboratories and Alex Alderton, Roberto Amato, Jeffrey Barrett, Sonia Goncalves, Ewan Harrison, David K. Jackson, Ian Johnston, Dominic Kwiatkowski, Cordelia Langford, John Sillitoe on behalf of the Wellcome Sanger Institute COVID-19 Surveillance Team |
| EPI_ISL_3337505, EPI_ISL_3337506, EPI_ISL_3337507 | Lighthouse Lab in Milton Keynes | Wellcome Sanger Institute for the COVID-19 Genomics UK (COG-UK) Consortium | The Lighthouse Lab in Milton Keynes and Alex Alderton, Roberto Amato, Jeffrey Barrett, Sonia Goncalves, Ewan Harrison, David K. Jackson, Ian Johnston, Dominic Kwiatkowski, Cordelia Langford, John Sillitoe on behalf of the Wellcome Sanger Institute COVID-19 Surveillance Team |
| EPI_ISL_3337508 | Health Services Laboratories | Wellcome Sanger Institute for the COVID-19 Genomics UK (COG-UK) Consortium | Health Services Laboratories and Alex Alderton, Roberto Amato, Jeffrey Barrett, Sonia Goncalves, Ewan Harrison, David K. Jackson, Ian Johnston, Dominic Kwiatkowski, Cordelia Langford, John Sillitoe on behalf of the Wellcome Sanger Institute COVID-19 Surveillance Team |
| EPI_ISL_3337509 | Berkshire and Surrey Pathology Services Lighthouse Laboratory | Wellcome Sanger Institute for the COVID-19 Genomics UK (COG-UK) Consortium | Berkshire and Surrey Pathology Services Lighthouse Laboratory and Alex Alderton, Roberto Amato, Jeffrey Barrett, Sonia Goncalves, Ewan Harrison, David K. Jackson, Ian Johnston, Dominic Kwiatkowski, Cordelia Langford, John Sillitoe on behalf of the Wellcome Sanger Institute COVID-19 Surveillance Team |
| EPI_ISL_3337510 | Lighthouse Lab in Milton Keynes | Wellcome Sanger Institute for the COVID-19 Genomics UK (COG-UK) Consortium | The Lighthouse Lab in Milton Keynes and Alex Alderton, Roberto Amato, Jeffrey Barrett, Sonia Goncalves, Ewan Harrison, David K. Jackson, Ian Johnston, Dominic Kwiatkowski, Cordelia Langford, John Sillitoe on behalf of the Wellcome Sanger Institute COVID-19 Surveillance Team |
| EPI_ISL_3337511 | Berkshire and Surrey Pathology Services Lighthouse Laboratory | Wellcome Sanger Institute for the COVID-19 Genomics UK (COG-UK) Consortium | Berkshire and Surrey Pathology Services Lighthouse Laboratory and Alex Alderton, Roberto Amato, Jeffrey Barrett, Sonia Goncalves, Ewan Harrison, David K. Jackson, Ian Johnston, Dominic Kwiatkowski, Cordelia Langford, John Sillitoe on behalf of the Wellcome Sanger Institute COVID-19 Surveillance Team |
| EPI_ISL_3337512 | Lighthouse Lab in Milton Keynes | Wellcome Sanger Institute for the COVID-19 Genomics UK (COG-UK) Consortium | The Lighthouse Lab in Milton Keynes and Alex Alderton, Roberto Amato, Jeffrey Barrett, Sonia Goncalves, Ewan Harrison, David K. Jackson, Ian Johnston, Dominic Kwiatkowski, Cordelia Langford, John Sillitoe on behalf of the Wellcome Sanger Institute COVID-19 Surveillance Team |
| EPI_ISL_3337513 | Health Services Laboratories | Wellcome Sanger Institute for the COVID-19 Genomics UK (COG-UK) Consortium | Health Services Laboratories and Alex Alderton, Roberto Amato, Jeffrey Barrett, Sonia Goncalves, Ewan Harrison, David K. Jackson, Ian Johnston, Dominic Kwiatkowski, Cordelia Langford, John Sillitoe on behalf of the Wellcome Sanger Institute COVID-19 Surveillance Team |
| EPI_ISL_3337514, EPI_ISL_3337515 | Lighthouse Lab in Milton Keynes | Wellcome Sanger Institute for the COVID-19 Genomics UK (COG-UK) Consortium | The Lighthouse Lab in Milton Keynes and Alex Alderton, Roberto Amato, Jeffrey Barrett, Sonia Goncalves, Ewan Harrison, David K. Jackson, Ian Johnston, Dominic Kwiatkowski, Cordelia Langford, John Sillitoe on behalf of the Wellcome Sanger Institute COVID-19 Surveillance Team |
| EPI_ISL_3337516 | Berkshire and Surrey Pathology Services Lighthouse Laboratory | Wellcome Sanger Institute for the COVID-19 Genomics UK (COG-UK) Consortium | Berkshire and Surrey Pathology Services Lighthouse Laboratory and Alex Alderton, Roberto Amato, Jeffrey Barrett, Sonia Goncalves, Ewan Harrison, David K. Jackson, Ian Johnston, Dominic Kwiatkowski, Cordelia Langford, John Sillitoe on behalf of the Wellcome Sanger Institute COVID-19 Surveillance Team |
| EPI_ISL_3337517 | Lighthouse Lab in Milton Keynes | Wellcome Sanger Institute for the COVID-19 Genomics UK (COG-UK) Consortium | The Lighthouse Lab in Milton Keynes and Alex Alderton, Roberto Amato, Jeffrey Barrett, Sonia Goncalves, Ewan Harrison, David K. Jackson, Ian Johnston, Dominic Kwiatkowski, Cordelia Langford, John Sillitoe on behalf of the Wellcome Sanger Institute COVID-19 Surveillance Team |
| EPI_ISL_3337518, EPI_ISL_3337519, EPI_ISL_3337520, EPI_ISL_3337521, EPI_ISL_3337522, EPI_ISL_3337523 | Berkshire and Surrey Pathology Services Lighthouse Laboratory | Wellcome Sanger Institute for the COVID-19 Genomics UK (COG-UK) Consortium | Berkshire and Surrey Pathology Services Lighthouse Laboratory and Alex Alderton, Roberto Amato, Jeffrey Barrett, Sonia Goncalves, Ewan Harrison, David K. Jackson, Ian Johnston, Dominic Kwiatkowski, Cordelia Langford, John Sillitoe on behalf of the Wellcome Sanger Institute COVID-19 Surveillance Team |
| EPI_ISL_3337524, EPI_ISL_3337525, EPI_ISL_3337526, EPI_ISL_3337527, EPI_ISL_3337528, EPI_ISL_3337529, EPI_ISL_3337530, EPI_ISL_3337531, EPI_ISL_3337532 | Lighthouse Lab in Milton Keynes | Wellcome Sanger Institute for the COVID-19 Genomics UK (COG-UK) Consortium | The Lighthouse Lab in Milton Keynes and Alex Alderton, Roberto Amato, Jeffrey Barrett, Sonia Goncalves, Ewan Harrison, David K. Jackson, Ian Johnston, Dominic Kwiatkowski, Cordelia Langford, John Sillitoe on behalf of the Wellcome Sanger Institute COVID-19 Surveillance Team |
| EPI_ISL_3337533 | Health Services Laboratories | Wellcome Sanger Institute for the COVID-19 Genomics UK (COG-UK) Consortium | Health Services Laboratories and Alex Alderton, Roberto Amato, Jeffrey Barrett, Sonia Goncalves, Ewan Harrison, David K. Jackson, Ian Johnston, Dominic Kwiatkowski, Cordelia Langford, John Sillitoe on behalf of the Wellcome Sanger Institute COVID-19 Surveillance Team |
| EPI_ISL_3337534, EPI_ISL_3337535 | Lighthouse Lab in Milton Keynes | Wellcome Sanger Institute for the COVID-19 Genomics UK (COG-UK) Consortium | The Lighthouse Lab in Milton Keynes and Alex Alderton, Roberto Amato, Jeffrey Barrett, Sonia Goncalves, Ewan Harrison, David K. Jackson, Ian Johnston, Dominic Kwiatkowski, Cordelia Langford, John Sillitoe on behalf of the Wellcome Sanger Institute COVID-19 Surveillance Team |
| EPI_ISL_3337536 | Berkshire and Surrey Pathology Services Lighthouse Laboratory | Wellcome Sanger Institute for the COVID-19 Genomics UK (COG-UK) Consortium | Berkshire and Surrey Pathology Services Lighthouse Laboratory and Alex Alderton, Roberto Amato, Jeffrey Barrett, Sonia Goncalves, Ewan Harrison, David K. Jackson, Ian Johnston, Dominic Kwiatkowski, Cordelia Langford, John Sillitoe on behalf of the Wellcome Sanger Institute COVID-19 Surveillance Team |
| EPI_ISL_3337537, EPI_ISL_3337538, EPI_ISL_3337539 | Lighthouse Lab in Milton Keynes | Wellcome Sanger Institute for the COVID-19 Genomics UK (COG-UK) Consortium | The Lighthouse Lab in Milton Keynes and Alex Alderton, Roberto Amato, Jeffrey Barrett, Sonia Goncalves, Ewan Harrison, David K. Jackson, Ian Johnston, Dominic Kwiatkowski, Cordelia Langford, John Sillitoe on behalf of the Wellcome Sanger Institute COVID-19 Surveillance Team |
| EPI_ISL_3337540 | Berkshire and Surrey Pathology Services Lighthouse Laboratory | Wellcome Sanger Institute for the COVID-19 Genomics UK (COG-UK) Consortium | Berkshire and Surrey Pathology Services Lighthouse Laboratory and Alex Alderton, Roberto Amato, Jeffrey Barrett, Sonia Goncalves, Ewan Harrison, David K. Jackson, Ian Johnston, Dominic Kwiatkowski, Cordelia Langford, John Sillitoe on behalf of the Wellcome Sanger Institute COVID-19 Surveillance Team |
| EPI_ISL_3337541 | Health Services Laboratories | Wellcome Sanger Institute for the COVID-19 Genomics UK (COG-UK) Consortium | Health Services Laboratories and Alex Alderton, Roberto Amato, Jeffrey Barrett, Sonia Goncalves, Ewan Harrison, David K. Jackson, Ian Johnston, Dominic Kwiatkowski, Cordelia Langford, John Sillitoe on behalf of the Wellcome Sanger Institute COVID-19 Surveillance Team |
| EPI_ISL_3337542 | Berkshire and Surrey Pathology Services Lighthouse Laboratory | Wellcome Sanger Institute for the COVID-19 Genomics UK (COG-UK) Consortium | Berkshire and Surrey Pathology Services Lighthouse Laboratory and Alex Alderton, Roberto Amato, Jeffrey Barrett, Sonia Goncalves, Ewan Harrison, David K. Jackson, Ian Johnston, Dominic Kwiatkowski, Cordelia Langford, John Sillitoe on behalf of the Wellcome Sanger Institute COVID-19 Surveillance Team |
| EPI_ISL_3337543, EPI_ISL_3337544, EPI_ISL_3337545 | Lighthouse Lab in Milton Keynes | Wellcome Sanger Institute for the COVID-19 Genomics UK (COG-UK) Consortium | The Lighthouse Lab in Milton Keynes and Alex Alderton, Roberto Amato, Jeffrey Barrett, Sonia Goncalves, Ewan Harrison, David K. Jackson, Ian Johnston, Dominic Kwiatkowski, Cordelia Langford, John Sillitoe on behalf of the Wellcome Sanger Institute COVID-19 Surveillance Team |
| EPI_ISL_3337546 | Berkshire and Surrey Pathology Services Lighthouse Laboratory | Wellcome Sanger Institute for the COVID-19 Genomics UK (COG-UK) Consortium | Berkshire and Surrey Pathology Services Lighthouse Laboratory and Alex Alderton, Roberto Amato, Jeffrey Barrett, Sonia Goncalves, Ewan Harrison, David K. Jackson, Ian Johnston, Dominic Kwiatkowski, Cordelia Langford, John Sillitoe on behalf of the Wellcome Sanger Institute COVID-19 Surveillance Team |
| EPI_ISL_3337547 | Lighthouse Lab in Milton Keynes | Wellcome Sanger Institute for the COVID-19 Genomics UK (COG-UK) Consortium | The Lighthouse Lab in Milton Keynes and Alex Alderton, Roberto Amato, Jeffrey Barrett, Sonia Goncalves, Ewan Harrison, David K. Jackson, Ian Johnston, Dominic Kwiatkowski, Cordelia Langford, John Sillitoe on behalf of the Wellcome Sanger Institute COVID-19 Surveillance Team |
| EPI_ISL_3337548, EPI_ISL_3337549 | Berkshire and Surrey Pathology Services Lighthouse Laboratory | Wellcome Sanger Institute for the COVID-19 Genomics UK (COG-UK) Consortium | Berkshire and Surrey Pathology Services Lighthouse Laboratory and Alex Alderton, Roberto Amato, Jeffrey Barrett, Sonia Goncalves, Ewan Harrison, David K. Jackson, Ian Johnston, Dominic Kwiatkowski, Cordelia Langford, John Sillitoe on behalf of the Wellcome Sanger Institute COVID-19 Surveillance Team |
| EPI_ISL_3337550, EPI_ISL_3337551 | Lighthouse Lab in Milton Keynes | Wellcome Sanger Institute for the COVID-19 Genomics UK (COG-UK) Consortium | The Lighthouse Lab in Milton Keynes and Alex Alderton, Roberto Amato, Jeffrey Barrett, Sonia Goncalves, Ewan Harrison, David K. Jackson, Ian Johnston, Dominic Kwiatkowski, Cordelia Langford, John Sillitoe on behalf of the Wellcome Sanger Institute COVID-19 Surveillance Team |
| EPI_ISL_3337552 | Berkshire and Surrey Pathology Services Lighthouse Laboratory | Wellcome Sanger Institute for the COVID-19 Genomics UK (COG-UK) Consortium | Berkshire and Surrey Pathology Services Lighthouse Laboratory and Alex Alderton, Roberto Amato, Jeffrey Barrett, Sonia Goncalves, Ewan Harrison, David K. Jackson, Ian Johnston, Dominic Kwiatkowski, Cordelia Langford, John Sillitoe on behalf of the Wellcome Sanger Institute COVID-19 Surveillance Team |
| EPI_ISL_3337553 | Lighthouse Lab in Milton Keynes | Wellcome Sanger Institute for the COVID-19 Genomics UK (COG-UK) Consortium | The Lighthouse Lab in Milton Keynes and Alex Alderton, Roberto Amato, Jeffrey Barrett, Sonia Goncalves, Ewan Harrison, David K. Jackson, Ian Johnston, Dominic Kwiatkowski, Cordelia Langford, John Sillitoe on behalf of the Wellcome Sanger Institute COVID-19 Surveillance Team |
| EPI_ISL_3337554 | Health Services Laboratories | Wellcome Sanger Institute for the COVID-19 Genomics UK (COG-UK) Consortium | Health Services Laboratories and Alex Alderton, Roberto Amato, Jeffrey Barrett, Sonia Goncalves, Ewan Harrison, David K. Jackson, Ian Johnston, Dominic Kwiatkowski, Cordelia Langford, John Sillitoe on behalf of the Wellcome Sanger Institute COVID-19 Surveillance Team |
| EPI_ISL_3337555 | Berkshire and Surrey Pathology Services Lighthouse Laboratory | Wellcome Sanger Institute for the COVID-19 Genomics UK (COG-UK) Consortium | Berkshire and Surrey Pathology Services Lighthouse Laboratory and Alex Alderton, Roberto Amato, Jeffrey Barrett, Sonia Goncalves, Ewan Harrison, David K. Jackson, Ian Johnston, Dominic Kwiatkowski, Cordelia Langford, John Sillitoe on behalf of the Wellcome Sanger Institute COVID-19 Surveillance Team |
| EPI_ISL_3337556 | Lighthouse Lab in Milton Keynes | Wellcome Sanger Institute for the COVID-19 Genomics UK (COG-UK) Consortium | The Lighthouse Lab in Milton Keynes and Alex Alderton, Roberto Amato, Jeffrey Barrett, Sonia Goncalves, Ewan Harrison, David K. Jackson, Ian Johnston, Dominic Kwiatkowski, Cordelia Langford, John Sillitoe on behalf of the Wellcome Sanger Institute COVID-19 Surveillance Team |
| EPI_ISL_3337557, EPI_ISL_3337558, EPI_ISL_3337559 | Berkshire and Surrey Pathology Services Lighthouse Laboratory | Wellcome Sanger Institute for the COVID-19 Genomics UK (COG-UK) Consortium | Berkshire and Surrey Pathology Services Lighthouse Laboratory and Alex Alderton, Roberto Amato, Jeffrey Barrett, Sonia Goncalves, Ewan Harrison, David K. Jackson, Ian Johnston, Dominic Kwiatkowski, Cordelia Langford, John Sillitoe on behalf of the Wellcome Sanger Institute COVID-19 Surveillance Team |
| EPI_ISL_3337560 | Lighthouse Lab in Milton Keynes | Wellcome Sanger Institute for the COVID-19 Genomics UK (COG-UK) Consortium | The Lighthouse Lab in Milton Keynes and Alex Alderton, Roberto Amato, Jeffrey Barrett, Sonia Goncalves, Ewan Harrison, David K. Jackson, Ian Johnston, Dominic Kwiatkowski, Cordelia Langford, John Sillitoe on behalf of the Wellcome Sanger Institute COVID-19 Surveillance Team |
| EPI_ISL_3337561 | Health Services Laboratories | Wellcome Sanger Institute for the COVID-19 Genomics UK (COG-UK) Consortium | Health Services Laboratories and Alex Alderton, Roberto Amato, Jeffrey Barrett, Sonia Goncalves, Ewan Harrison, David K. Jackson, Ian Johnston, Dominic Kwiatkowski, Cordelia Langford, John Sillitoe on behalf of the Wellcome Sanger Institute COVID-19 Surveillance Team |
| EPI_ISL_3337562, | Lighthouse Lab in Milton Keynes | Wellcome Sanger Institute for the COVID-19 Genomics UK (COG-UK) | The Lighthouse Lab in Milton Keynes and Alex Alderton, Roberto Amato, Jeffrey Barrett, Sonia Goncalves, Ewan Harrison, David K. Jackson, Ian Johnston, Dominic Kwiatkowski, Cordelia Langford, John Sillitoe on behalf of the Wellcome Sanger |

| | | | |
|---|---|---|---|
| EPI_ISL_3337563 | | Consortium | Institute COVID-19 Surveillance Team |
| EPI_ISL_3337564 | Health Services Laboratories | Wellcome Sanger Institute for the COVID-19 Genomics UK (COG-UK) Consortium | Health Services Laboratories and Alex Alderton, Roberto Amato, Jeffrey Barrett, Sonia Goncalves, Ewan Harrison, David K. Jackson, Ian Johnston, Dominic Kwiatkowski, Cordelia Langford, John Sillitoe on behalf of the Wellcome Sanger Institute COVID-19 Surveillance Team |
| EPI_ISL_3337565 | Berkshire and Surrey Pathology Services Lighthouse Laboratory | Wellcome Sanger Institute for the COVID-19 Genomics UK (COG-UK) Consortium | Berkshire and Surrey Pathology Services Lighthouse Laboratory and Alex Alderton, Roberto Amato, Jeffrey Barrett, Sonia Goncalves, Ewan Harrison, David K. Jackson, Ian Johnston, Dominic Kwiatkowski, Cordelia Langford, John Sillitoe on behalf of the Wellcome Sanger Institute COVID-19 Surveillance Team |
| EPI_ISL_3337566, EPI_ISL_3337567 | Health Services Laboratories | Wellcome Sanger Institute for the COVID-19 Genomics UK (COG-UK) Consortium | Health Services Laboratories and Alex Alderton, Roberto Amato, Jeffrey Barrett, Sonia Goncalves, Ewan Harrison, David K. Jackson, Ian Johnston, Dominic Kwiatkowski, Cordelia Langford, John Sillitoe on behalf of the Wellcome Sanger Institute COVID-19 Surveillance Team |
| EPI_ISL_3337568 | Berkshire and Surrey Pathology Services Lighthouse Laboratory | Wellcome Sanger Institute for the COVID-19 Genomics UK (COG-UK) Consortium | Berkshire and Surrey Pathology Services Lighthouse Laboratory and Alex Alderton, Roberto Amato, Jeffrey Barrett, Sonia Goncalves, Ewan Harrison, David K. Jackson, Ian Johnston, Dominic Kwiatkowski, Cordelia Langford, John Sillitoe on behalf of the Wellcome Sanger Institute COVID-19 Surveillance Team |
| EPI_ISL_3337569, EPI_ISL_3337570, EPI_ISL_3337571, EPI_ISL_3337572, EPI_ISL_3337573 | Lighthouse Lab in Milton Keynes | Wellcome Sanger Institute for the COVID-19 Genomics UK (COG-UK) Consortium | The Lighthouse Lab in Milton Keynes and Alex Alderton, Roberto Amato, Jeffrey Barrett, Sonia Goncalves, Ewan Harrison, David K. Jackson, Ian Johnston, Dominic Kwiatkowski, Cordelia Langford, John Sillitoe on behalf of the Wellcome Sanger Institute COVID-19 Surveillance Team |
| EPI_ISL_3337574 | Berkshire and Surrey Pathology Services Lighthouse Laboratory | Wellcome Sanger Institute for the COVID-19 Genomics UK (COG-UK) Consortium | Berkshire and Surrey Pathology Services Lighthouse Laboratory and Alex Alderton, Roberto Amato, Jeffrey Barrett, Sonia Goncalves, Ewan Harrison, David K. Jackson, Ian Johnston, Dominic Kwiatkowski, Cordelia Langford, John Sillitoe on behalf of the Wellcome Sanger Institute COVID-19 Surveillance Team |
| EPI_ISL_3337575 | Health Services Laboratories | Wellcome Sanger Institute for the COVID-19 Genomics UK (COG-UK) Consortium | Health Services Laboratories and Alex Alderton, Roberto Amato, Jeffrey Barrett, Sonia Goncalves, Ewan Harrison, David K. Jackson, Ian Johnston, Dominic Kwiatkowski, Cordelia Langford, John Sillitoe on behalf of the Wellcome Sanger Institute COVID-19 Surveillance Team |
| EPI_ISL_3337576 | Lighthouse Lab in Milton Keynes | Wellcome Sanger Institute for the COVID-19 Genomics UK (COG-UK) Consortium | The Lighthouse Lab in Milton Keynes and Alex Alderton, Roberto Amato, Jeffrey Barrett, Sonia Goncalves, Ewan Harrison, David K. Jackson, Ian Johnston, Dominic Kwiatkowski, Cordelia Langford, John Sillitoe on behalf of the Wellcome Sanger Institute COVID-19 Surveillance Team |
| EPI_ISL_3337577, EPI_ISL_3337578 | Berkshire and Surrey Pathology Services Lighthouse Laboratory | Wellcome Sanger Institute for the COVID-19 Genomics UK (COG-UK) Consortium | Berkshire and Surrey Pathology Services Lighthouse Laboratory and Alex Alderton, Roberto Amato, Jeffrey Barrett, Sonia Goncalves, Ewan Harrison, David K. Jackson, Ian Johnston, Dominic Kwiatkowski, Cordelia Langford, John Sillitoe on behalf of the Wellcome Sanger Institute COVID-19 Surveillance Team |
| EPI_ISL_3337579 | Health Services Laboratories | Wellcome Sanger Institute for the COVID-19 Genomics UK (COG-UK) Consortium | Health Services Laboratories and Alex Alderton, Roberto Amato, Jeffrey Barrett, Sonia Goncalves, Ewan Harrison, David K. Jackson, Ian Johnston, Dominic Kwiatkowski, Cordelia Langford, John Sillitoe on behalf of the Wellcome Sanger Institute COVID-19 Surveillance Team |
| EPI_ISL_3337580 | Lighthouse Lab in Milton Keynes | Wellcome Sanger Institute for the COVID-19 Genomics UK (COG-UK) Consortium | The Lighthouse Lab in Milton Keynes and Alex Alderton, Roberto Amato, Jeffrey Barrett, Sonia Goncalves, Ewan Harrison, David K. Jackson, Ian Johnston, Dominic Kwiatkowski, Cordelia Langford, John Sillitoe on behalf of the Wellcome Sanger Institute COVID-19 Surveillance Team |
| EPI_ISL_3337581 | Health Services Laboratories | Wellcome Sanger Institute for the COVID-19 Genomics UK (COG-UK) Consortium | Health Services Laboratories and Alex Alderton, Roberto Amato, Jeffrey Barrett, Sonia Goncalves, Ewan Harrison, David K. Jackson, Ian Johnston, Dominic Kwiatkowski, Cordelia Langford, John Sillitoe on behalf of the Wellcome Sanger Institute COVID-19 Surveillance Team |
| EPI_ISL_3337582 | Berkshire and Surrey Pathology Services Lighthouse Laboratory | Wellcome Sanger Institute for the COVID-19 Genomics UK (COG-UK) Consortium | Berkshire and Surrey Pathology Services Lighthouse Laboratory and Alex Alderton, Roberto Amato, Jeffrey Barrett, Sonia Goncalves, Ewan Harrison, David K. Jackson, Ian Johnston, Dominic Kwiatkowski, Cordelia Langford, John Sillitoe on behalf of the Wellcome Sanger Institute COVID-19 Surveillance Team |
| EPI_ISL_3337583 | Lighthouse Lab in Milton Keynes | Wellcome Sanger Institute for the COVID-19 Genomics UK (COG-UK) Consortium | The Lighthouse Lab in Milton Keynes and Alex Alderton, Roberto Amato, Jeffrey Barrett, Sonia Goncalves, Ewan Harrison, David K. Jackson, Ian Johnston, Dominic Kwiatkowski, Cordelia Langford, John Sillitoe on behalf of the Wellcome Sanger Institute COVID-19 Surveillance Team |
| EPI_ISL_3337584 | Health Services Laboratories | Wellcome Sanger Institute for the COVID-19 Genomics UK (COG-UK) Consortium | Health Services Laboratories and Alex Alderton, Roberto Amato, Jeffrey Barrett, Sonia Goncalves, Ewan Harrison, David K. Jackson, Ian Johnston, Dominic Kwiatkowski, Cordelia Langford, John Sillitoe on behalf of the Wellcome Sanger Institute COVID-19 Surveillance Team |
| EPI_ISL_3337585, EPI_ISL_3337586 | Berkshire and Surrey Pathology Services Lighthouse Laboratory | Wellcome Sanger Institute for the COVID-19 Genomics UK (COG-UK) Consortium | Berkshire and Surrey Pathology Services Lighthouse Laboratory and Alex Alderton, Roberto Amato, Jeffrey Barrett, Sonia Goncalves, Ewan Harrison, David K. Jackson, Ian Johnston, Dominic Kwiatkowski, Cordelia Langford, John Sillitoe on behalf of the Wellcome Sanger Institute COVID-19 Surveillance Team |
| EPI_ISL_3337587, EPI_ISL_3337588 | Lighthouse Lab in Milton Keynes | Wellcome Sanger Institute for the COVID-19 Genomics UK (COG-UK) Consortium | The Lighthouse Lab in Milton Keynes and Alex Alderton, Roberto Amato, Jeffrey Barrett, Sonia Goncalves, Ewan Harrison, David K. Jackson, Ian Johnston, Dominic Kwiatkowski, Cordelia Langford, John Sillitoe on behalf of the Wellcome Sanger Institute COVID-19 Surveillance Team |
| EPI_ISL_3337589 | Health Services Laboratories | Wellcome Sanger Institute for the COVID-19 Genomics UK (COG-UK) Consortium | Health Services Laboratories and Alex Alderton, Roberto Amato, Jeffrey Barrett, Sonia Goncalves, Ewan Harrison, David K. Jackson, Ian Johnston, Dominic Kwiatkowski, Cordelia Langford, John Sillitoe on behalf of the Wellcome Sanger Institute COVID-19 Surveillance Team |
| EPI_ISL_3337590 | Berkshire and Surrey Pathology Services Lighthouse Laboratory | Wellcome Sanger Institute for the COVID-19 Genomics UK (COG-UK) Consortium | Berkshire and Surrey Pathology Services Lighthouse Laboratory and Alex Alderton, Roberto Amato, Jeffrey Barrett, Sonia Goncalves, Ewan Harrison, David K. Jackson, Ian Johnston, Dominic Kwiatkowski, Cordelia Langford, John Sillitoe on behalf of the Wellcome Sanger Institute COVID-19 Surveillance Team |
| EPI_ISL_3337591, EPI_ISL_3337592, EPI_ISL_3337593 | Lighthouse Lab in Milton Keynes | Wellcome Sanger Institute for the COVID-19 Genomics UK (COG-UK) Consortium | The Lighthouse Lab in Milton Keynes and Alex Alderton, Roberto Amato, Jeffrey Barrett, Sonia Goncalves, Ewan Harrison, David K. Jackson, Ian Johnston, Dominic Kwiatkowski, Cordelia Langford, John Sillitoe on behalf of the Wellcome Sanger Institute COVID-19 Surveillance Team |
| EPI_ISL_3337594 | Health Services Laboratories | Wellcome Sanger Institute for the COVID-19 Genomics UK (COG-UK) Consortium | Health Services Laboratories and Alex Alderton, Roberto Amato, Jeffrey Barrett, Sonia Goncalves, Ewan Harrison, David K. Jackson, Ian Johnston, Dominic Kwiatkowski, Cordelia Langford, John Sillitoe on behalf of the Wellcome Sanger Institute COVID-19 Surveillance Team |
| EPI_ISL_3337595, EPI_ISL_3337596 | Lighthouse Lab in Milton Keynes | Wellcome Sanger Institute for the COVID-19 Genomics UK (COG-UK) Consortium | The Lighthouse Lab in Milton Keynes and Alex Alderton, Roberto Amato, Jeffrey Barrett, Sonia Goncalves, Ewan Harrison, David K. Jackson, Ian Johnston, Dominic Kwiatkowski, Cordelia Langford, John Sillitoe on behalf of the Wellcome Sanger Institute COVID-19 Surveillance Team |
| EPI_ISL_3337597, EPI_ISL_3337598 | Berkshire and Surrey Pathology Services Lighthouse Laboratory | Wellcome Sanger Institute for the COVID-19 Genomics UK (COG-UK) Consortium | Berkshire and Surrey Pathology Services Lighthouse Laboratory and Alex Alderton, Roberto Amato, Jeffrey Barrett, Sonia Goncalves, Ewan Harrison, David K. Jackson, Ian Johnston, Dominic Kwiatkowski, Cordelia Langford, John Sillitoe on behalf of the Wellcome Sanger Institute COVID-19 Surveillance Team |
| EPI_ISL_3337599 | Lighthouse Lab in Milton Keynes | Wellcome Sanger Institute for the COVID-19 Genomics UK (COG-UK) Consortium | The Lighthouse Lab in Milton Keynes and Alex Alderton, Roberto Amato, Jeffrey Barrett, Sonia Goncalves, Ewan Harrison, David K. Jackson, Ian Johnston, Dominic Kwiatkowski, Cordelia Langford, John Sillitoe on behalf of the Wellcome Sanger Institute COVID-19 Surveillance Team |
| EPI_ISL_3337600 | Health Services Laboratories | Wellcome Sanger Institute for the COVID-19 Genomics UK (COG-UK) Consortium | Health Services Laboratories and Alex Alderton, Roberto Amato, Jeffrey Barrett, Sonia Goncalves, Ewan Harrison, David K. Jackson, Ian Johnston, Dominic Kwiatkowski, Cordelia Langford, John Sillitoe on behalf of the Wellcome Sanger Institute COVID-19 Surveillance Team |
| EPI_ISL_3337601, EPI_ISL_3337602 | Lighthouse Lab in Milton Keynes | Wellcome Sanger Institute for the COVID-19 Genomics UK (COG-UK) Consortium | The Lighthouse Lab in Milton Keynes and Alex Alderton, Roberto Amato, Jeffrey Barrett, Sonia Goncalves, Ewan Harrison, David K. Jackson, Ian Johnston, Dominic Kwiatkowski, Cordelia Langford, John Sillitoe on behalf of the Wellcome Sanger Institute COVID-19 Surveillance Team |
| EPI_ISL_3337603 | Health Services Laboratories | Wellcome Sanger Institute for the COVID-19 Genomics UK (COG-UK) Consortium | Health Services Laboratories and Alex Alderton, Roberto Amato, Jeffrey Barrett, Sonia Goncalves, Ewan Harrison, David K. Jackson, Ian Johnston, Dominic Kwiatkowski, Cordelia Langford, John Sillitoe on behalf of the Wellcome Sanger Institute COVID-19 Surveillance Team |
| EPI_ISL_3337604, EPI_ISL_3337605, EPI_ISL_3337606 | Lighthouse Lab in Milton Keynes | Wellcome Sanger Institute for the COVID-19 Genomics UK (COG-UK) Consortium | The Lighthouse Lab in Milton Keynes and Alex Alderton, Roberto Amato, Jeffrey Barrett, Sonia Goncalves, Ewan Harrison, David K. Jackson, Ian Johnston, Dominic Kwiatkowski, Cordelia Langford, John Sillitoe on behalf of the Wellcome Sanger Institute COVID-19 Surveillance Team |
| EPI_ISL_3337607, EPI_ISL_3337608 | Berkshire and Surrey Pathology Services Lighthouse Laboratory | Wellcome Sanger Institute for the COVID-19 Genomics UK (COG-UK) Consortium | Berkshire and Surrey Pathology Services Lighthouse Laboratory and Alex Alderton, Roberto Amato, Jeffrey Barrett, Sonia Goncalves, Ewan Harrison, David K. Jackson, Ian Johnston, Dominic Kwiatkowski, Cordelia Langford, John Sillitoe on behalf of the Wellcome Sanger Institute COVID-19 Surveillance Team |
| EPI_ISL_3337609 | Health Services Laboratories | Wellcome Sanger Institute for the COVID-19 Genomics UK (COG-UK) Consortium | Health Services Laboratories and Alex Alderton, Roberto Amato, Jeffrey Barrett, Sonia Goncalves, Ewan Harrison, David K. Jackson, Ian Johnston, Dominic Kwiatkowski, Cordelia Langford, John Sillitoe on behalf of the Wellcome Sanger Institute COVID-19 Surveillance Team |
| EPI_ISL_3337610 | Lighthouse Lab in Milton Keynes | Wellcome Sanger Institute for the COVID-19 Genomics UK (COG-UK) Consortium | The Lighthouse Lab in Milton Keynes and Alex Alderton, Roberto Amato, Jeffrey Barrett, Sonia Goncalves, Ewan Harrison, David K. Jackson, Ian Johnston, Dominic Kwiatkowski, Cordelia Langford, John Sillitoe on behalf of the Wellcome Sanger Institute COVID-19 Surveillance Team |
| EPI_ISL_3337611 | Health Services Laboratories | Wellcome Sanger Institute for the COVID-19 Genomics UK (COG-UK) Consortium | Health Services Laboratories and Alex Alderton, Roberto Amato, Jeffrey Barrett, Sonia Goncalves, Ewan Harrison, David K. Jackson, Ian Johnston, Dominic Kwiatkowski, Cordelia Langford, John Sillitoe on behalf of the Wellcome Sanger Institute COVID-19 Surveillance Team |
| EPI_ISL_3337612, EPI_ISL_3337613 | Berkshire and Surrey Pathology Services Lighthouse Laboratory | Wellcome Sanger Institute for the COVID-19 Genomics UK (COG-UK) Consortium | Berkshire and Surrey Pathology Services Lighthouse Laboratory and Alex Alderton, Roberto Amato, Jeffrey Barrett, Sonia Goncalves, Ewan Harrison, David K. Jackson, Ian Johnston, Dominic Kwiatkowski, Cordelia Langford, John Sillitoe on behalf of the Wellcome Sanger Institute COVID-19 Surveillance Team |
| EPI_ISL_3337614, EPI_ISL_3337615 | Lighthouse Lab in Milton Keynes | Wellcome Sanger Institute for the COVID-19 Genomics UK (COG-UK) Consortium | The Lighthouse Lab in Milton Keynes and Alex Alderton, Roberto Amato, Jeffrey Barrett, Sonia Goncalves, Ewan Harrison, David K. Jackson, Ian Johnston, Dominic Kwiatkowski, Cordelia Langford, John Sillitoe on behalf of the Wellcome Sanger Institute COVID-19 Surveillance Team |
| EPI_ISL_3337616, EPI_ISL_3337617 | Berkshire and Surrey Pathology Services Lighthouse Laboratory | Wellcome Sanger Institute for the COVID-19 Genomics UK (COG-UK) Consortium | Berkshire and Surrey Pathology Services Lighthouse Laboratory and Alex Alderton, Roberto Amato, Jeffrey Barrett, Sonia Goncalves, Ewan Harrison, David K. Jackson, Ian Johnston, Dominic Kwiatkowski, Cordelia Langford, John Sillitoe on behalf of the Wellcome Sanger Institute COVID-19 Surveillance Team |
| EPI_ISL_3337618, EPI_ISL_3337619, EPI_ISL_3337620 | Lighthouse Lab in Milton Keynes | Wellcome Sanger Institute for the COVID-19 Genomics UK (COG-UK) Consortium | The Lighthouse Lab in Milton Keynes and Alex Alderton, Roberto Amato, Jeffrey Barrett, Sonia Goncalves, Ewan Harrison, David K. Jackson, Ian Johnston, Dominic Kwiatkowski, Cordelia Langford, John Sillitoe on behalf of the Wellcome Sanger Institute COVID-19 Surveillance Team |
| EPI_ISL_3337621, EPI_ISL_3337622 | Health Services Laboratories | Wellcome Sanger Institute for the COVID-19 Genomics UK (COG-UK) Consortium | Health Services Laboratories and Alex Alderton, Roberto Amato, Jeffrey Barrett, Sonia Goncalves, Ewan Harrison, David K. Jackson, Ian Johnston, Dominic Kwiatkowski, Cordelia Langford, John Sillitoe on behalf of the Wellcome Sanger Institute COVID-19 Surveillance Team |
| EPI_ISL_3337623, EPI_ISL_3337624, EPI_ISL_3337625, EPI_ISL_3337626 | Lighthouse Lab in Milton Keynes | Wellcome Sanger Institute for the COVID-19 Genomics UK (COG-UK) Consortium | The Lighthouse Lab in Milton Keynes and Alex Alderton, Roberto Amato, Jeffrey Barrett, Sonia Goncalves, Ewan Harrison, David K. Jackson, Ian Johnston, Dominic Kwiatkowski, Cordelia Langford, John Sillitoe on behalf of the Wellcome Sanger Institute COVID-19 Surveillance Team |
| EPI_ISL_3337627, EPI_ISL_3337628 | Health Services Laboratories | Wellcome Sanger Institute for the COVID-19 Genomics UK (COG-UK) Consortium | Health Services Laboratories and Alex Alderton, Roberto Amato, Jeffrey Barrett, Sonia Goncalves, Ewan Harrison, David K. Jackson, Ian Johnston, Dominic Kwiatkowski, Cordelia Langford, John Sillitoe on behalf of the Wellcome Sanger Institute COVID-19 Surveillance Team |
| EPI_ISL_3337629 | Lighthouse Lab in Milton Keynes | Wellcome Sanger Institute for the COVID-19 Genomics UK (COG-UK) Consortium | The Lighthouse Lab in Milton Keynes and Alex Alderton, Roberto Amato, Jeffrey Barrett, Sonia Goncalves, Ewan Harrison, David K. Jackson, Ian Johnston, Dominic Kwiatkowski, Cordelia Langford, John Sillitoe on behalf of the Wellcome Sanger Institute COVID-19 Surveillance Team |
| EPI_ISL_3337630 | Berkshire and Surrey Pathology Services Lighthouse Laboratory | Wellcome Sanger Institute for the COVID-19 Genomics UK (COG-UK) Consortium | Berkshire and Surrey Pathology Services Lighthouse Laboratory and Alex Alderton, Roberto Amato, Jeffrey Barrett, Sonia Goncalves, Ewan Harrison, David K. Jackson, Ian Johnston, Dominic Kwiatkowski, Cordelia Langford, John Sillitoe on behalf of the Wellcome Sanger Institute COVID-19 Surveillance Team |
| EPI_ISL_3337631, | Lighthouse Lab in Milton Keynes | Wellcome Sanger Institute for the COVID-19 Genomics UK (COG-UK) | The Lighthouse Lab in Milton Keynes and Alex Alderton, Roberto Amato, Jeffrey Barrett, Sonia Goncalves, Ewan Harrison, David K. Jackson, Ian Johnston, Dominic Kwiatkowski, Cordelia Langford, John Sillitoe on behalf of the Wellcome Sanger |

| | | | |
|---|---|---|---|
| EPI_ISL_3337632, EPI_ISL_3337633, EPI_ISL_3337634, EPI_ISL_3337635, EPI_ISL_3337636, EPI_ISL_3337637 | | Consortium | Institute COVID-19 Surveillance Team |
| EPI_ISL_3337638 | Berkshire and Surrey Pathology Services Lighthouse Laboratory | Wellcome Sanger Institute for the COVID-19 Genomics UK (COG-UK) Consortium | Berkshire and Surrey Pathology Services Lighthouse Laboratory and Alex Alderton, Roberto Amato, Jeffrey Barrett, Sonia Goncalves, Ewan Harrison, David K. Jackson, Ian Johnston, Dominic Kwiatkowski, Cordelia Langford, John Sillitoe on behalf of the Wellcome Sanger Institute COVID-19 Surveillance Team |
| EPI_ISL_3337639 | Health Services Laboratories | Wellcome Sanger Institute for the COVID-19 Genomics UK (COG-UK) Consortium | Health Services Laboratories and Alex Alderton, Roberto Amato, Jeffrey Barrett, Sonia Goncalves, Ewan Harrison, David K. Jackson, Ian Johnston, Dominic Kwiatkowski, Cordelia Langford, John Sillitoe on behalf of the Wellcome Sanger Institute COVID-19 Surveillance Team |
| EPI_ISL_3337640 | Berkshire and Surrey Pathology Services Lighthouse Laboratory | Wellcome Sanger Institute for the COVID-19 Genomics UK (COG-UK) Consortium | Berkshire and Surrey Pathology Services Lighthouse Laboratory and Alex Alderton, Roberto Amato, Jeffrey Barrett, Sonia Goncalves, Ewan Harrison, David K. Jackson, Ian Johnston, Dominic Kwiatkowski, Cordelia Langford, John Sillitoe on behalf of the Wellcome Sanger Institute COVID-19 Surveillance Team |
| EPI_ISL_3337641 | Lighthouse Lab in Milton Keynes | Wellcome Sanger Institute for the COVID-19 Genomics UK (COG-UK) Consortium | The Lighthouse Lab in Milton Keynes and Alex Alderton, Roberto Amato, Jeffrey Barrett, Sonia Goncalves, Ewan Harrison, David K. Jackson, Ian Johnston, Dominic Kwiatkowski, Cordelia Langford, John Sillitoe on behalf of the Wellcome Sanger Institute COVID-19 Surveillance Team |
| EPI_ISL_3337642 | Health Services Laboratories | Wellcome Sanger Institute for the COVID-19 Genomics UK (COG-UK) Consortium | Health Services Laboratories and Alex Alderton, Roberto Amato, Jeffrey Barrett, Sonia Goncalves, Ewan Harrison, David K. Jackson, Ian Johnston, Dominic Kwiatkowski, Cordelia Langford, John Sillitoe on behalf of the Wellcome Sanger Institute COVID-19 Surveillance Team |
| EPI_ISL_3337643, EPI_ISL_3337644, EPI_ISL_3337645 | Lighthouse Lab in Milton Keynes | Wellcome Sanger Institute for the COVID-19 Genomics UK (COG-UK) Consortium | The Lighthouse Lab in Milton Keynes and Alex Alderton, Roberto Amato, Jeffrey Barrett, Sonia Goncalves, Ewan Harrison, David K. Jackson, Ian Johnston, Dominic Kwiatkowski, Cordelia Langford, John Sillitoe on behalf of the Wellcome Sanger Institute COVID-19 Surveillance Team |
| EPI_ISL_3337646 | Health Services Laboratories | Wellcome Sanger Institute for the COVID-19 Genomics UK (COG-UK) Consortium | Health Services Laboratories and Alex Alderton, Roberto Amato, Jeffrey Barrett, Sonia Goncalves, Ewan Harrison, David K. Jackson, Ian Johnston, Dominic Kwiatkowski, Cordelia Langford, John Sillitoe on behalf of the Wellcome Sanger Institute COVID-19 Surveillance Team |
| EPI_ISL_3337647, EPI_ISL_3337648 | Lighthouse Lab in Milton Keynes | Wellcome Sanger Institute for the COVID-19 Genomics UK (COG-UK) Consortium | The Lighthouse Lab in Milton Keynes and Alex Alderton, Roberto Amato, Jeffrey Barrett, Sonia Goncalves, Ewan Harrison, David K. Jackson, Ian Johnston, Dominic Kwiatkowski, Cordelia Langford, John Sillitoe on behalf of the Wellcome Sanger Institute COVID-19 Surveillance Team |
| EPI_ISL_3337649 | Health Services Laboratories | Wellcome Sanger Institute for the COVID-19 Genomics UK (COG-UK) Consortium | Health Services Laboratories and Alex Alderton, Roberto Amato, Jeffrey Barrett, Sonia Goncalves, Ewan Harrison, David K. Jackson, Ian Johnston, Dominic Kwiatkowski, Cordelia Langford, John Sillitoe on behalf of the Wellcome Sanger Institute COVID-19 Surveillance Team |
| EPI_ISL_3337650 | Berkshire and Surrey Pathology Services Lighthouse Laboratory | Wellcome Sanger Institute for the COVID-19 Genomics UK (COG-UK) Consortium | Berkshire and Surrey Pathology Services Lighthouse Laboratory and Alex Alderton, Roberto Amato, Jeffrey Barrett, Sonia Goncalves, Ewan Harrison, David K. Jackson, Ian Johnston, Dominic Kwiatkowski, Cordelia Langford, John Sillitoe on behalf of the Wellcome Sanger Institute COVID-19 Surveillance Team |
| EPI_ISL_3337651, EPI_ISL_3337652, EPI_ISL_3337653, EPI_ISL_3337654 | Lighthouse Lab in Milton Keynes | Wellcome Sanger Institute for the COVID-19 Genomics UK (COG-UK) Consortium | The Lighthouse Lab in Milton Keynes and Alex Alderton, Roberto Amato, Jeffrey Barrett, Sonia Goncalves, Ewan Harrison, David K. Jackson, Ian Johnston, Dominic Kwiatkowski, Cordelia Langford, John Sillitoe on behalf of the Wellcome Sanger Institute COVID-19 Surveillance Team |
| EPI_ISL_3337655 | Berkshire and Surrey Pathology Services Lighthouse Laboratory | Wellcome Sanger Institute for the COVID-19 Genomics UK (COG-UK) Consortium | Berkshire and Surrey Pathology Services Lighthouse Laboratory and Alex Alderton, Roberto Amato, Jeffrey Barrett, Sonia Goncalves, Ewan Harrison, David K. Jackson, Ian Johnston, Dominic Kwiatkowski, Cordelia Langford, John Sillitoe on behalf of the Wellcome Sanger Institute COVID-19 Surveillance Team |
| EPI_ISL_3337656, EPI_ISL_3337657 | Lighthouse Lab in Alderley Park | Wellcome Sanger Institute for the COVID-19 Genomics UK (COG-UK) Consortium | Jacquelyn Wynn, Mairead Hyland, The Lighthouse Lab in Alderley Park and Alex Alderton, Roberto Amato, Jeffrey Barrett, Sonia Goncalves, Ewan Harrison, David K. Jackson, Ian Johnston, Dominic Kwiatkowski, Cordelia Langford, John Sillitoe on behalf of the Wellcome Sanger Institute COVID-19 Surveillance Team |
| EPI_ISL_3337658 | Lighthouse Lab in Milton Keynes | Wellcome Sanger Institute for the COVID-19 Genomics UK (COG-UK) Consortium | The Lighthouse Lab in Milton Keynes and Alex Alderton, Roberto Amato, Jeffrey Barrett, Sonia Goncalves, Ewan Harrison, David K. Jackson, Ian Johnston, Dominic Kwiatkowski, Cordelia Langford, John Sillitoe on behalf of the Wellcome Sanger Institute COVID-19 Surveillance Team |
| EPI_ISL_3337659 | Lighthouse Lab in Alderley Park | Wellcome Sanger Institute for the COVID-19 Genomics UK (COG-UK) Consortium | Jacquelyn Wynn, Mairead Hyland, The Lighthouse Lab in Alderley Park and Alex Alderton, Roberto Amato, Jeffrey Barrett, Sonia Goncalves, Ewan Harrison, David K. Jackson, Ian Johnston, Dominic Kwiatkowski, Cordelia Langford, John Sillitoe on behalf of the Wellcome Sanger Institute COVID-19 Surveillance Team |
| EPI_ISL_3337660 | Lighthouse Laboratory Plymouth | Wellcome Sanger Institute for the COVID-19 Genomics UK (COG-UK) Consortium | Lighthouse Laboratory Plymouth and Alex Alderton, Roberto Amato, Jeffrey Barrett, Sonia Goncalves, Ewan Harrison, David K. Jackson, Ian Johnston, Dominic Kwiatkowski, Cordelia Langford, John Sillitoe on behalf of the Wellcome Sanger Institute COVID-19 Surveillance Team |
| EPI_ISL_3337661 | Lighthouse Lab in Alderley Park | Wellcome Sanger Institute for the COVID-19 Genomics UK (COG-UK) Consortium | Jacquelyn Wynn, Mairead Hyland, The Lighthouse Lab in Alderley Park and Alex Alderton, Roberto Amato, Jeffrey Barrett, Sonia Goncalves, Ewan Harrison, David K. Jackson, Ian Johnston, Dominic Kwiatkowski, Cordelia Langford, John Sillitoe on behalf of the Wellcome Sanger Institute COVID-19 Surveillance Team |
| EPI_ISL_3337662 | Lighthouse Lab in Milton Keynes | Wellcome Sanger Institute for the COVID-19 Genomics UK (COG-UK) Consortium | The Lighthouse Lab in Milton Keynes and Alex Alderton, Roberto Amato, Jeffrey Barrett, Sonia Goncalves, Ewan Harrison, David K. Jackson, Ian Johnston, Dominic Kwiatkowski, Cordelia Langford, John Sillitoe on behalf of the Wellcome Sanger Institute COVID-19 Surveillance Team |
| EPI_ISL_3337663 | Lighthouse Laboratory Plymouth | Wellcome Sanger Institute for the COVID-19 Genomics UK (COG-UK) Consortium | Lighthouse Laboratory Plymouth and Alex Alderton, Roberto Amato, Jeffrey Barrett, Sonia Goncalves, Ewan Harrison, David K. Jackson, Ian Johnston, Dominic Kwiatkowski, Cordelia Langford, John Sillitoe on behalf of the Wellcome Sanger Institute COVID-19 Surveillance Team |
| EPI_ISL_3337664, EPI_ISL_3337665 | Lighthouse Lab in Alderley Park | Wellcome Sanger Institute for the COVID-19 Genomics UK (COG-UK) Consortium | Jacquelyn Wynn, Mairead Hyland, The Lighthouse Lab in Alderley Park and Alex Alderton, Roberto Amato, Jeffrey Barrett, Sonia Goncalves, Ewan Harrison, David K. Jackson, Ian Johnston, Dominic Kwiatkowski, Cordelia Langford, John Sillitoe on behalf of the Wellcome Sanger Institute COVID-19 Surveillance Team |
| EPI_ISL_3337666 | Lighthouse Lab in Milton Keynes | Wellcome Sanger Institute for the COVID-19 Genomics UK (COG-UK) Consortium | The Lighthouse Lab in Milton Keynes and Alex Alderton, Roberto Amato, Jeffrey Barrett, Sonia Goncalves, Ewan Harrison, David K. Jackson, Ian Johnston, Dominic Kwiatkowski, Cordelia Langford, John Sillitoe on behalf of the Wellcome Sanger Institute COVID-19 Surveillance Team |
| EPI_ISL_3337667 | Lighthouse Lab in Alderley Park | Wellcome Sanger Institute for the COVID-19 Genomics UK (COG-UK) Consortium | Jacquelyn Wynn, Mairead Hyland, The Lighthouse Lab in Alderley Park and Alex Alderton, Roberto Amato, Jeffrey Barrett, Sonia Goncalves, Ewan Harrison, David K. Jackson, Ian Johnston, Dominic Kwiatkowski, Cordelia Langford, John Sillitoe on behalf of the Wellcome Sanger Institute COVID-19 Surveillance Team |
| EPI_ISL_3337668 | Lighthouse Laboratory Plymouth | Wellcome Sanger Institute for the COVID-19 Genomics UK (COG-UK) Consortium | Lighthouse Laboratory Plymouth and Alex Alderton, Roberto Amato, Jeffrey Barrett, Sonia Goncalves, Ewan Harrison, David K. Jackson, Ian Johnston, Dominic Kwiatkowski, Cordelia Langford, John Sillitoe on behalf of the Wellcome Sanger Institute COVID-19 Surveillance Team |
| EPI_ISL_3337669 | Lighthouse Lab in Milton Keynes | Wellcome Sanger Institute for the COVID-19 Genomics UK (COG-UK) Consortium | The Lighthouse Lab in Milton Keynes and Alex Alderton, Roberto Amato, Jeffrey Barrett, Sonia Goncalves, Ewan Harrison, David K. Jackson, Ian Johnston, Dominic Kwiatkowski, Cordelia Langford, John Sillitoe on behalf of the Wellcome Sanger Institute COVID-19 Surveillance Team |
| EPI_ISL_3337670, EPI_ISL_3337671, EPI_ISL_3337672 | Lighthouse Lab in Alderley Park | Wellcome Sanger Institute for the COVID-19 Genomics UK (COG-UK) Consortium | Jacquelyn Wynn, Mairead Hyland, The Lighthouse Lab in Alderley Park and Alex Alderton, Roberto Amato, Jeffrey Barrett, Sonia Goncalves, Ewan Harrison, David K. Jackson, Ian Johnston, Dominic Kwiatkowski, Cordelia Langford, John Sillitoe on behalf of the Wellcome Sanger Institute COVID-19 Surveillance Team |
| EPI_ISL_3337673 | Lighthouse Lab in Milton Keynes | Wellcome Sanger Institute for the COVID-19 Genomics UK (COG-UK) Consortium | The Lighthouse Lab in Milton Keynes and Alex Alderton, Roberto Amato, Jeffrey Barrett, Sonia Goncalves, Ewan Harrison, David K. Jackson, Ian Johnston, Dominic Kwiatkowski, Cordelia Langford, John Sillitoe on behalf of the Wellcome Sanger Institute COVID-19 Surveillance Team |
| EPI_ISL_3337674 | Lighthouse Lab in Alderley Park | Wellcome Sanger Institute for the COVID-19 Genomics UK (COG-UK) Consortium | Jacquelyn Wynn, Mairead Hyland, The Lighthouse Lab in Alderley Park and Alex Alderton, Roberto Amato, Jeffrey Barrett, Sonia Goncalves, Ewan Harrison, David K. Jackson, Ian Johnston, Dominic Kwiatkowski, Cordelia Langford, John Sillitoe on behalf of the Wellcome Sanger Institute COVID-19 Surveillance Team |
| EPI_ISL_3337675 | Lighthouse Laboratory Plymouth | Wellcome Sanger Institute for the COVID-19 Genomics UK (COG-UK) Consortium | Lighthouse Laboratory Plymouth and Alex Alderton, Roberto Amato, Jeffrey Barrett, Sonia Goncalves, Ewan Harrison, David K. Jackson, Ian Johnston, Dominic Kwiatkowski, Cordelia Langford, John Sillitoe on behalf of the Wellcome Sanger Institute COVID-19 Surveillance Team |
| EPI_ISL_3337676, EPI_ISL_3337677, EPI_ISL_3337678, EPI_ISL_3337679 | Lighthouse Lab in Alderley Park | Wellcome Sanger Institute for the COVID-19 Genomics UK (COG-UK) Consortium | Jacquelyn Wynn, Mairead Hyland, The Lighthouse Lab in Alderley Park and Alex Alderton, Roberto Amato, Jeffrey Barrett, Sonia Goncalves, Ewan Harrison, David K. Jackson, Ian Johnston, Dominic Kwiatkowski, Cordelia Langford, John Sillitoe on behalf of the Wellcome Sanger Institute COVID-19 Surveillance Team |
| EPI_ISL_3337680 | Lighthouse Lab in Milton Keynes | Wellcome Sanger Institute for the COVID-19 Genomics UK (COG-UK) Consortium | The Lighthouse Lab in Milton Keynes and Alex Alderton, Roberto Amato, Jeffrey Barrett, Sonia Goncalves, Ewan Harrison, David K. Jackson, Ian Johnston, Dominic Kwiatkowski, Cordelia Langford, John Sillitoe on behalf of the Wellcome Sanger Institute COVID-19 Surveillance Team |
| EPI_ISL_3337681 | Lighthouse Lab in Alderley Park | Wellcome Sanger Institute for the COVID-19 Genomics UK (COG-UK) Consortium | Jacquelyn Wynn, Mairead Hyland, The Lighthouse Lab in Alderley Park and Alex Alderton, Roberto Amato, Jeffrey Barrett, Sonia Goncalves, Ewan Harrison, David K. Jackson, Ian Johnston, Dominic Kwiatkowski, Cordelia Langford, John Sillitoe on behalf of the Wellcome Sanger Institute COVID-19 Surveillance Team |
| EPI_ISL_3337682, EPI_ISL_3337683 | Lighthouse Laboratory Plymouth | Wellcome Sanger Institute for the COVID-19 Genomics UK (COG-UK) Consortium | Lighthouse Laboratory Plymouth and Alex Alderton, Roberto Amato, Jeffrey Barrett, Sonia Goncalves, Ewan Harrison, David K. Jackson, Ian Johnston, Dominic Kwiatkowski, Cordelia Langford, John Sillitoe on behalf of the Wellcome Sanger Institute COVID-19 Surveillance Team |
| EPI_ISL_3337684 | Lighthouse Lab in Alderley Park | Wellcome Sanger Institute for the COVID-19 Genomics UK (COG-UK) Consortium | Jacquelyn Wynn, Mairead Hyland, The Lighthouse Lab in Alderley Park and Alex Alderton, Roberto Amato, Jeffrey Barrett, Sonia Goncalves, Ewan Harrison, David K. Jackson, Ian Johnston, Dominic Kwiatkowski, Cordelia Langford, John Sillitoe on behalf of the Wellcome Sanger Institute COVID-19 Surveillance Team |
| EPI_ISL_3337685, EPI_ISL_3337686 | Lighthouse Laboratory Plymouth | Wellcome Sanger Institute for the COVID-19 Genomics UK (COG-UK) Consortium | Lighthouse Laboratory Plymouth and Alex Alderton, Roberto Amato, Jeffrey Barrett, Sonia Goncalves, Ewan Harrison, David K. Jackson, Ian Johnston, Dominic Kwiatkowski, Cordelia Langford, John Sillitoe on behalf of the Wellcome Sanger Institute COVID-19 Surveillance Team |
| EPI_ISL_3337687 | Lighthouse Lab in Alderley Park | Wellcome Sanger Institute for the COVID-19 Genomics UK (COG-UK) Consortium | Jacquelyn Wynn, Mairead Hyland, The Lighthouse Lab in Alderley Park and Alex Alderton, Roberto Amato, Jeffrey Barrett, Sonia Goncalves, Ewan Harrison, David K. Jackson, Ian Johnston, Dominic Kwiatkowski, Cordelia Langford, John Sillitoe on behalf of the Wellcome Sanger Institute COVID-19 Surveillance Team |
| EPI_ISL_3337688 | Lighthouse Lab in Milton Keynes | Wellcome Sanger Institute for the COVID-19 Genomics UK (COG-UK) Consortium | The Lighthouse Lab in Milton Keynes and Alex Alderton, Roberto Amato, Jeffrey Barrett, Sonia Goncalves, Ewan Harrison, David K. Jackson, Ian Johnston, Dominic Kwiatkowski, Cordelia Langford, John Sillitoe on behalf of the Wellcome Sanger Institute COVID-19 Surveillance Team |
| EPI_ISL_3337689 | Lighthouse Lab in Alderley Park | Wellcome Sanger Institute for the COVID-19 Genomics UK (COG-UK) Consortium | Jacquelyn Wynn, Mairead Hyland, The Lighthouse Lab in Alderley Park and Alex Alderton, Roberto Amato, Jeffrey Barrett, Sonia Goncalves, Ewan Harrison, David K. Jackson, Ian Johnston, Dominic Kwiatkowski, Cordelia Langford, John Sillitoe on behalf of the Wellcome Sanger Institute COVID-19 Surveillance Team |
| EPI_ISL_3337690, EPI_ISL_3337691 | Lighthouse Lab in Milton Keynes | Wellcome Sanger Institute for the COVID-19 Genomics UK (COG-UK) Consortium | The Lighthouse Lab in Milton Keynes and Alex Alderton, Roberto Amato, Jeffrey Barrett, Sonia Goncalves, Ewan Harrison, David K. Jackson, Ian Johnston, Dominic Kwiatkowski, Cordelia Langford, John Sillitoe on behalf of the Wellcome Sanger Institute COVID-19 Surveillance Team |
| EPI_ISL_3337692 | Lighthouse Lab in Alderley Park | Wellcome Sanger Institute for the COVID-19 Genomics UK (COG-UK) Consortium | Jacquelyn Wynn, Mairead Hyland, The Lighthouse Lab in Alderley Park and Alex Alderton, Roberto Amato, Jeffrey Barrett, Sonia Goncalves, Ewan Harrison, David K. Jackson, Ian Johnston, Dominic Kwiatkowski, Cordelia Langford, John Sillitoe on behalf of the Wellcome Sanger Institute COVID-19 Surveillance Team |
| EPI_ISL_3337693 | Lighthouse Laboratory Plymouth | Wellcome Sanger Institute for the COVID-19 Genomics UK (COG-UK) Consortium | Lighthouse Laboratory Plymouth and Alex Alderton, Roberto Amato, Jeffrey Barrett, Sonia Goncalves, Ewan Harrison, David K. Jackson, Ian Johnston, Dominic Kwiatkowski, Cordelia Langford, John Sillitoe on behalf of the Wellcome Sanger Institute COVID-19 Surveillance Team |

| | | | |
|---|---|---|---|
| EPI_ISL_3337694, EPI_ISL_3337695 | Lighthouse Lab in Milton Keynes | Wellcome Sanger Institute for the COVID-19 Genomics UK (COG-UK) Consortium | The Lighthouse Lab in Milton Keynes and Alex Alderton, Roberto Amato, Jeffrey Barrett, Sonia Goncalves, Ewan Harrison, David K. Jackson, Ian Johnston, Dominic Kwiatkowski, Cordelia Langford, John Sillitoe on behalf of the Wellcome Sanger Institute COVID-19 Surveillance Team |
| EPI_ISL_3337696, EPI_ISL_3337697, EPI_ISL_3337698 | Lighthouse Lab in Alderley Park | Wellcome Sanger Institute for the COVID-19 Genomics UK (COG-UK) Consortium | Jacquelyn Wynn, Mairead Hyland, The Lighthouse Lab in Alderley Park and Alex Alderton, Roberto Amato, Jeffrey Barrett, Sonia Goncalves, Ewan Harrison, David K. Jackson, Ian Johnston, Dominic Kwiatkowski, Cordelia Langford, John Sillitoe on behalf of the Wellcome Sanger Institute COVID-19 Surveillance Team |
| EPI_ISL_3337699 | Lighthouse Laboratory Plymouth | Wellcome Sanger Institute for the COVID-19 Genomics UK (COG-UK) Consortium | Lighthouse Laboratory Plymouth and Alex Alderton, Roberto Amato, Jeffrey Barrett, Sonia Goncalves, Ewan Harrison, David K. Jackson, Ian Johnston, Dominic Kwiatkowski, Cordelia Langford, John Sillitoe on behalf of the Wellcome Sanger Institute COVID-19 Surveillance Team |
| EPI_ISL_3337700 | Lighthouse Lab in Milton Keynes | Wellcome Sanger Institute for the COVID-19 Genomics UK (COG-UK) Consortium | The Lighthouse Lab in Milton Keynes and Alex Alderton, Roberto Amato, Jeffrey Barrett, Sonia Goncalves, Ewan Harrison, David K. Jackson, Ian Johnston, Dominic Kwiatkowski, Cordelia Langford, John Sillitoe on behalf of the Wellcome Sanger Institute COVID-19 Surveillance Team |
| EPI_ISL_3337701, EPI_ISL_3337702 | Lighthouse Lab in Alderley Park | Wellcome Sanger Institute for the COVID-19 Genomics UK (COG-UK) Consortium | Jacquelyn Wynn, Mairead Hyland, The Lighthouse Lab in Alderley Park and Alex Alderton, Roberto Amato, Jeffrey Barrett, Sonia Goncalves, Ewan Harrison, David K. Jackson, Ian Johnston, Dominic Kwiatkowski, Cordelia Langford, John Sillitoe on behalf of the Wellcome Sanger Institute COVID-19 Surveillance Team |
| EPI_ISL_3337703, EPI_ISL_3337704 | Lighthouse Lab in Milton Keynes | Wellcome Sanger Institute for the COVID-19 Genomics UK (COG-UK) Consortium | The Lighthouse Lab in Milton Keynes and Alex Alderton, Roberto Amato, Jeffrey Barrett, Sonia Goncalves, Ewan Harrison, David K. Jackson, Ian Johnston, Dominic Kwiatkowski, Cordelia Langford, John Sillitoe on behalf of the Wellcome Sanger Institute COVID-19 Surveillance Team |
| EPI_ISL_3337705, EPI_ISL_3337706, EPI_ISL_3337707 | Lighthouse Laboratory Plymouth | Wellcome Sanger Institute for the COVID-19 Genomics UK (COG-UK) Consortium | Lighthouse Laboratory Plymouth and Alex Alderton, Roberto Amato, Jeffrey Barrett, Sonia Goncalves, Ewan Harrison, David K. Jackson, Ian Johnston, Dominic Kwiatkowski, Cordelia Langford, John Sillitoe on behalf of the Wellcome Sanger Institute COVID-19 Surveillance Team |
| EPI_ISL_3337708, EPI_ISL_3337709 | Lighthouse Lab in Alderley Park | Wellcome Sanger Institute for the COVID-19 Genomics UK (COG-UK) Consortium | Jacquelyn Wynn, Mairead Hyland, The Lighthouse Lab in Alderley Park and Alex Alderton, Roberto Amato, Jeffrey Barrett, Sonia Goncalves, Ewan Harrison, David K. Jackson, Ian Johnston, Dominic Kwiatkowski, Cordelia Langford, John Sillitoe on behalf of the Wellcome Sanger Institute COVID-19 Surveillance Team |
| EPI_ISL_3337710 | Lighthouse Laboratory Plymouth | Wellcome Sanger Institute for the COVID-19 Genomics UK (COG-UK) Consortium | Lighthouse Laboratory Plymouth and Alex Alderton, Roberto Amato, Jeffrey Barrett, Sonia Goncalves, Ewan Harrison, David K. Jackson, Ian Johnston, Dominic Kwiatkowski, Cordelia Langford, John Sillitoe on behalf of the Wellcome Sanger Institute COVID-19 Surveillance Team |
| EPI_ISL_3337711 | Lighthouse Lab in Alderley Park | Wellcome Sanger Institute for the COVID-19 Genomics UK (COG-UK) Consortium | Jacquelyn Wynn, Mairead Hyland, The Lighthouse Lab in Alderley Park and Alex Alderton, Roberto Amato, Jeffrey Barrett, Sonia Goncalves, Ewan Harrison, David K. Jackson, Ian Johnston, Dominic Kwiatkowski, Cordelia Langford, John Sillitoe on behalf of the Wellcome Sanger Institute COVID-19 Surveillance Team |
| EPI_ISL_3337712 | Lighthouse Laboratory Plymouth | Wellcome Sanger Institute for the COVID-19 Genomics UK (COG-UK) Consortium | Lighthouse Laboratory Plymouth and Alex Alderton, Roberto Amato, Jeffrey Barrett, Sonia Goncalves, Ewan Harrison, David K. Jackson, Ian Johnston, Dominic Kwiatkowski, Cordelia Langford, John Sillitoe on behalf of the Wellcome Sanger Institute COVID-19 Surveillance Team |
| EPI_ISL_3337713 | Lighthouse Lab in Alderley Park | Wellcome Sanger Institute for the COVID-19 Genomics UK (COG-UK) Consortium | Jacquelyn Wynn, Mairead Hyland, The Lighthouse Lab in Alderley Park and Alex Alderton, Roberto Amato, Jeffrey Barrett, Sonia Goncalves, Ewan Harrison, David K. Jackson, Ian Johnston, Dominic Kwiatkowski, Cordelia Langford, John Sillitoe on behalf of the Wellcome Sanger Institute COVID-19 Surveillance Team |
| EPI_ISL_3337714 | Lighthouse Laboratory Plymouth | Wellcome Sanger Institute for the COVID-19 Genomics UK (COG-UK) Consortium | Lighthouse Laboratory Plymouth and Alex Alderton, Roberto Amato, Jeffrey Barrett, Sonia Goncalves, Ewan Harrison, David K. Jackson, Ian Johnston, Dominic Kwiatkowski, Cordelia Langford, John Sillitoe on behalf of the Wellcome Sanger Institute COVID-19 Surveillance Team |
| EPI_ISL_3337715 | Lighthouse Lab in Alderley Park | Wellcome Sanger Institute for the COVID-19 Genomics UK (COG-UK) Consortium | Jacquelyn Wynn, Mairead Hyland, The Lighthouse Lab in Alderley Park and Alex Alderton, Roberto Amato, Jeffrey Barrett, Sonia Goncalves, Ewan Harrison, David K. Jackson, Ian Johnston, Dominic Kwiatkowski, Cordelia Langford, John Sillitoe on behalf of the Wellcome Sanger Institute COVID-19 Surveillance Team |
| EPI_ISL_3337716 | Lighthouse Laboratory Plymouth | Wellcome Sanger Institute for the COVID-19 Genomics UK (COG-UK) Consortium | Lighthouse Laboratory Plymouth and Alex Alderton, Roberto Amato, Jeffrey Barrett, Sonia Goncalves, Ewan Harrison, David K. Jackson, Ian Johnston, Dominic Kwiatkowski, Cordelia Langford, John Sillitoe on behalf of the Wellcome Sanger Institute COVID-19 Surveillance Team |
| EPI_ISL_3337717, EPI_ISL_3337718, EPI_ISL_3337719, EPI_ISL_3337720, EPI_ISL_3337721, EPI_ISL_3337722 | Lighthouse Lab in Alderley Park | Wellcome Sanger Institute for the COVID-19 Genomics UK (COG-UK) Consortium | Jacquelyn Wynn, Mairead Hyland, The Lighthouse Lab in Alderley Park and Alex Alderton, Roberto Amato, Jeffrey Barrett, Sonia Goncalves, Ewan Harrison, David K. Jackson, Ian Johnston, Dominic Kwiatkowski, Cordelia Langford, John Sillitoe on behalf of the Wellcome Sanger Institute COVID-19 Surveillance Team |
| EPI_ISL_3337723 | Lighthouse Laboratory Plymouth | Wellcome Sanger Institute for the COVID-19 Genomics UK (COG-UK) Consortium | Lighthouse Laboratory Plymouth and Alex Alderton, Roberto Amato, Jeffrey Barrett, Sonia Goncalves, Ewan Harrison, David K. Jackson, Ian Johnston, Dominic Kwiatkowski, Cordelia Langford, John Sillitoe on behalf of the Wellcome Sanger Institute COVID-19 Surveillance Team |
| EPI_ISL_3337724 | Lighthouse Lab in Alderley Park | Wellcome Sanger Institute for the COVID-19 Genomics UK (COG-UK) Consortium | Jacquelyn Wynn, Mairead Hyland, The Lighthouse Lab in Alderley Park and Alex Alderton, Roberto Amato, Jeffrey Barrett, Sonia Goncalves, Ewan Harrison, David K. Jackson, Ian Johnston, Dominic Kwiatkowski, Cordelia Langford, John Sillitoe on behalf of the Wellcome Sanger Institute COVID-19 Surveillance Team |
| EPI_ISL_3337725 | Lighthouse Laboratory Plymouth | Wellcome Sanger Institute for the COVID-19 Genomics UK (COG-UK) Consortium | Lighthouse Laboratory Plymouth and Alex Alderton, Roberto Amato, Jeffrey Barrett, Sonia Goncalves, Ewan Harrison, David K. Jackson, Ian Johnston, Dominic Kwiatkowski, Cordelia Langford, John Sillitoe on behalf of the Wellcome Sanger Institute COVID-19 Surveillance Team |
| EPI_ISL_3337726 | Lighthouse Lab in Milton Keynes | Wellcome Sanger Institute for the COVID-19 Genomics UK (COG-UK) Consortium | The Lighthouse Lab in Milton Keynes and Alex Alderton, Roberto Amato, Jeffrey Barrett, Sonia Goncalves, Ewan Harrison, David K. Jackson, Ian Johnston, Dominic Kwiatkowski, Cordelia Langford, John Sillitoe on behalf of the Wellcome Sanger Institute COVID-19 Surveillance Team |
| EPI_ISL_3337727 | Lighthouse Lab in Alderley Park | Wellcome Sanger Institute for the COVID-19 Genomics UK (COG-UK) Consortium | Jacquelyn Wynn, Mairead Hyland, The Lighthouse Lab in Alderley Park and Alex Alderton, Roberto Amato, Jeffrey Barrett, Sonia Goncalves, Ewan Harrison, David K. Jackson, Ian Johnston, Dominic Kwiatkowski, Cordelia Langford, John Sillitoe on behalf of the Wellcome Sanger Institute COVID-19 Surveillance Team |
| EPI_ISL_3337728 | Lighthouse Laboratory Plymouth | Wellcome Sanger Institute for the COVID-19 Genomics UK (COG-UK) Consortium | Lighthouse Laboratory Plymouth and Alex Alderton, Roberto Amato, Jeffrey Barrett, Sonia Goncalves, Ewan Harrison, David K. Jackson, Ian Johnston, Dominic Kwiatkowski, Cordelia Langford, John Sillitoe on behalf of the Wellcome Sanger Institute COVID-19 Surveillance Team |
| EPI_ISL_3337729, EPI_ISL_3337730, EPI_ISL_3337731 | Lighthouse Lab in Alderley Park | Wellcome Sanger Institute for the COVID-19 Genomics UK (COG-UK) Consortium | Jacquelyn Wynn, Mairead Hyland, The Lighthouse Lab in Alderley Park and Alex Alderton, Roberto Amato, Jeffrey Barrett, Sonia Goncalves, Ewan Harrison, David K. Jackson, Ian Johnston, Dominic Kwiatkowski, Cordelia Langford, John Sillitoe on behalf of the Wellcome Sanger Institute COVID-19 Surveillance Team |
| EPI_ISL_3337732 | Lighthouse Lab in Milton Keynes | Wellcome Sanger Institute for the COVID-19 Genomics UK (COG-UK) Consortium | The Lighthouse Lab in Milton Keynes and Alex Alderton, Roberto Amato, Jeffrey Barrett, Sonia Goncalves, Ewan Harrison, David K. Jackson, Ian Johnston, Dominic Kwiatkowski, Cordelia Langford, John Sillitoe on behalf of the Wellcome Sanger Institute COVID-19 Surveillance Team |
| EPI_ISL_3337733 | Lighthouse Lab in Alderley Park | Wellcome Sanger Institute for the COVID-19 Genomics UK (COG-UK) Consortium | Jacquelyn Wynn, Mairead Hyland, The Lighthouse Lab in Alderley Park and Alex Alderton, Roberto Amato, Jeffrey Barrett, Sonia Goncalves, Ewan Harrison, David K. Jackson, Ian Johnston, Dominic Kwiatkowski, Cordelia Langford, John Sillitoe on behalf of the Wellcome Sanger Institute COVID-19 Surveillance Team |
| EPI_ISL_3337734 | Lighthouse Lab in Milton Keynes | Wellcome Sanger Institute for the COVID-19 Genomics UK (COG-UK) Consortium | The Lighthouse Lab in Milton Keynes and Alex Alderton, Roberto Amato, Jeffrey Barrett, Sonia Goncalves, Ewan Harrison, David K. Jackson, Ian Johnston, Dominic Kwiatkowski, Cordelia Langford, John Sillitoe on behalf of the Wellcome Sanger Institute COVID-19 Surveillance Team |
| EPI_ISL_3337735 | Lighthouse Laboratory Plymouth | Wellcome Sanger Institute for the COVID-19 Genomics UK (COG-UK) Consortium | Lighthouse Laboratory Plymouth and Alex Alderton, Roberto Amato, Jeffrey Barrett, Sonia Goncalves, Ewan Harrison, David K. Jackson, Ian Johnston, Dominic Kwiatkowski, Cordelia Langford, John Sillitoe on behalf of the Wellcome Sanger Institute COVID-19 Surveillance Team |
| EPI_ISL_3337736 | Lighthouse Lab in Milton Keynes | Wellcome Sanger Institute for the COVID-19 Genomics UK (COG-UK) Consortium | The Lighthouse Lab in Milton Keynes and Alex Alderton, Roberto Amato, Jeffrey Barrett, Sonia Goncalves, Ewan Harrison, David K. Jackson, Ian Johnston, Dominic Kwiatkowski, Cordelia Langford, John Sillitoe on behalf of the Wellcome Sanger Institute COVID-19 Surveillance Team |
| EPI_ISL_3337737, EPI_ISL_3337738 | Lighthouse Lab in Alderley Park | Wellcome Sanger Institute for the COVID-19 Genomics UK (COG-UK) Consortium | Jacquelyn Wynn, Mairead Hyland, The Lighthouse Lab in Alderley Park and Alex Alderton, Roberto Amato, Jeffrey Barrett, Sonia Goncalves, Ewan Harrison, David K. Jackson, Ian Johnston, Dominic Kwiatkowski, Cordelia Langford, John Sillitoe on behalf of the Wellcome Sanger Institute COVID-19 Surveillance Team |
| EPI_ISL_3337739 | Lighthouse Laboratory Plymouth | Wellcome Sanger Institute for the COVID-19 Genomics UK (COG-UK) Consortium | Lighthouse Laboratory Plymouth and Alex Alderton, Roberto Amato, Jeffrey Barrett, Sonia Goncalves, Ewan Harrison, David K. Jackson, Ian Johnston, Dominic Kwiatkowski, Cordelia Langford, John Sillitoe on behalf of the Wellcome Sanger Institute COVID-19 Surveillance Team |
| EPI_ISL_3337740 | Lighthouse Lab in Alderley Park | Wellcome Sanger Institute for the COVID-19 Genomics UK (COG-UK) Consortium | Jacquelyn Wynn, Mairead Hyland, The Lighthouse Lab in Alderley Park and Alex Alderton, Roberto Amato, Jeffrey Barrett, Sonia Goncalves, Ewan Harrison, David K. Jackson, Ian Johnston, Dominic Kwiatkowski, Cordelia Langford, John Sillitoe on behalf of the Wellcome Sanger Institute COVID-19 Surveillance Team |
| EPI_ISL_3337741, EPI_ISL_3337742 | Lighthouse Lab in Milton Keynes | Wellcome Sanger Institute for the COVID-19 Genomics UK (COG-UK) Consortium | The Lighthouse Lab in Milton Keynes and Alex Alderton, Roberto Amato, Jeffrey Barrett, Sonia Goncalves, Ewan Harrison, David K. Jackson, Ian Johnston, Dominic Kwiatkowski, Cordelia Langford, John Sillitoe on behalf of the Wellcome Sanger Institute COVID-19 Surveillance Team |
| EPI_ISL_3337743, EPI_ISL_3337744, EPI_ISL_3337745 | Lighthouse Laboratory Plymouth | Wellcome Sanger Institute for the COVID-19 Genomics UK (COG-UK) Consortium | Lighthouse Laboratory Plymouth and Alex Alderton, Roberto Amato, Jeffrey Barrett, Sonia Goncalves, Ewan Harrison, David K. Jackson, Ian Johnston, Dominic Kwiatkowski, Cordelia Langford, John Sillitoe on behalf of the Wellcome Sanger Institute COVID-19 Surveillance Team |
| EPI_ISL_3337746 | Lighthouse Lab in Alderley Park | Wellcome Sanger Institute for the COVID-19 Genomics UK (COG-UK) Consortium | Jacquelyn Wynn, Mairead Hyland, The Lighthouse Lab in Alderley Park and Alex Alderton, Roberto Amato, Jeffrey Barrett, Sonia Goncalves, Ewan Harrison, David K. Jackson, Ian Johnston, Dominic Kwiatkowski, Cordelia Langford, John Sillitoe on behalf of the Wellcome Sanger Institute COVID-19 Surveillance Team |
| EPI_ISL_3337747 | Lighthouse Lab in Milton Keynes | Wellcome Sanger Institute for the COVID-19 Genomics UK (COG-UK) Consortium | The Lighthouse Lab in Milton Keynes and Alex Alderton, Roberto Amato, Jeffrey Barrett, Sonia Goncalves, Ewan Harrison, David K. Jackson, Ian Johnston, Dominic Kwiatkowski, Cordelia Langford, John Sillitoe on behalf of the Wellcome Sanger Institute COVID-19 Surveillance Team |
| EPI_ISL_3337748 | Lighthouse Lab in Alderley Park | Wellcome Sanger Institute for the COVID-19 Genomics UK (COG-UK) Consortium | Jacquelyn Wynn, Mairead Hyland, The Lighthouse Lab in Alderley Park and Alex Alderton, Roberto Amato, Jeffrey Barrett, Sonia Goncalves, Ewan Harrison, David K. Jackson, Ian Johnston, Dominic Kwiatkowski, Cordelia Langford, John Sillitoe on behalf of the Wellcome Sanger Institute COVID-19 Surveillance Team |
| EPI_ISL_3337749 | Lighthouse Lab in Milton Keynes | Wellcome Sanger Institute for the COVID-19 Genomics UK (COG-UK) Consortium | The Lighthouse Lab in Milton Keynes and Alex Alderton, Roberto Amato, Jeffrey Barrett, Sonia Goncalves, Ewan Harrison, David K. Jackson, Ian Johnston, Dominic Kwiatkowski, Cordelia Langford, John Sillitoe on behalf of the Wellcome Sanger Institute COVID-19 Surveillance Team |
| EPI_ISL_3337750 | Lighthouse Laboratory Plymouth | Wellcome Sanger Institute for the COVID-19 Genomics UK (COG-UK) Consortium | Lighthouse Laboratory Plymouth and Alex Alderton, Roberto Amato, Jeffrey Barrett, Sonia Goncalves, Ewan Harrison, David K. Jackson, Ian Johnston, Dominic Kwiatkowski, Cordelia Langford, John Sillitoe on behalf of the Wellcome Sanger Institute COVID-19 Surveillance Team |
| EPI_ISL_3337751 | Lighthouse Lab in Alderley Park | Wellcome Sanger Institute for the COVID-19 Genomics UK (COG-UK) Consortium | Jacquelyn Wynn, Mairead Hyland, The Lighthouse Lab in Alderley Park and Alex Alderton, Roberto Amato, Jeffrey Barrett, Sonia Goncalves, Ewan Harrison, David K. Jackson, Ian Johnston, Dominic Kwiatkowski, Cordelia Langford, John Sillitoe on behalf of the Wellcome Sanger Institute COVID-19 Surveillance Team |
| EPI_ISL_3337752, EPI_ISL_3337753 | Lighthouse Lab in Milton Keynes | Wellcome Sanger Institute for the COVID-19 Genomics UK (COG-UK) Consortium | The Lighthouse Lab in Milton Keynes and Alex Alderton, Roberto Amato, Jeffrey Barrett, Sonia Goncalves, Ewan Harrison, David K. Jackson, Ian Johnston, Dominic Kwiatkowski, Cordelia Langford, John Sillitoe on behalf of the Wellcome Sanger Institute COVID-19 Surveillance Team |
| EPI_ISL_3337754, EPI_ISL_3337755, EPI_ISL_3337756 | Lighthouse Lab in Alderley Park | Wellcome Sanger Institute for the COVID-19 Genomics UK (COG-UK) Consortium | Jacquelyn Wynn, Mairead Hyland, The Lighthouse Lab in Alderley Park and Alex Alderton, Roberto Amato, Jeffrey Barrett, Sonia Goncalves, Ewan Harrison, David K. Jackson, Ian Johnston, Dominic Kwiatkowski, Cordelia Langford, John Sillitoe on behalf of the Wellcome Sanger Institute COVID-19 Surveillance Team |
| EPI_ISL_3337757, | Lighthouse Lab in Milton Keynes | Wellcome Sanger Institute for the COVID-19 Genomics UK (COG-UK) | The Lighthouse Lab in Milton Keynes and Alex Alderton, Roberto Amato, Jeffrey Barrett, Sonia Goncalves, Ewan Harrison, David K. Jackson, Ian Johnston, Dominic Kwiatkowski, Cordelia Langford, John Sillitoe on behalf of the Wellcome Sanger |

| | | | |
|---|---|---|---|
| EPI_ISL_3337758 | | Consortium | Institute COVID-19 Surveillance Team |
| EPI_ISL_3337759, EPI_ISL_3337760 | Lighthouse Lab in Alderley Park | Wellcome Sanger Institute for the COVID-19 Genomics UK (COG-UK) Consortium | Jacquelyn Wynn, Mairead Hyland, The Lighthouse Lab in Alderley Park and Alex Alderton, Roberto Amato, Jeffrey Barrett, Sonia Goncalves, Ewan Harrison, David K. Jackson, Ian Johnston, Dominic Kwiatkowski, Cordelia Langford, John Sillitoe on behalf of the Wellcome Sanger Institute COVID-19 Surveillance Team |
| EPI_ISL_3337761 | Lighthouse Lab in Milton Keynes | Wellcome Sanger Institute for the COVID-19 Genomics UK (COG-UK) Consortium | The Lighthouse Lab in Milton Keynes and Alex Alderton, Roberto Amato, Jeffrey Barrett, Sonia Goncalves, Ewan Harrison, David K. Jackson, Ian Johnston, Dominic Kwiatkowski, Cordelia Langford, John Sillitoe on behalf of the Wellcome Sanger Institute COVID-19 Surveillance Team |
| EPI_ISL_3337762 | Lighthouse Lab in Alderley Park | Wellcome Sanger Institute for the COVID-19 Genomics UK (COG-UK) Consortium | Jacquelyn Wynn, Mairead Hyland, The Lighthouse Lab in Alderley Park and Alex Alderton, Roberto Amato, Jeffrey Barrett, Sonia Goncalves, Ewan Harrison, David K. Jackson, Ian Johnston, Dominic Kwiatkowski, Cordelia Langford, John Sillitoe on behalf of the Wellcome Sanger Institute COVID-19 Surveillance Team |
| EPI_ISL_3337763 | Lighthouse Lab in Milton Keynes | Wellcome Sanger Institute for the COVID-19 Genomics UK (COG-UK) Consortium | The Lighthouse Lab in Milton Keynes and Alex Alderton, Roberto Amato, Jeffrey Barrett, Sonia Goncalves, Ewan Harrison, David K. Jackson, Ian Johnston, Dominic Kwiatkowski, Cordelia Langford, John Sillitoe on behalf of the Wellcome Sanger Institute COVID-19 Surveillance Team |
| EPI_ISL_3337764 | Lighthouse Lab in Alderley Park | Wellcome Sanger Institute for the COVID-19 Genomics UK (COG-UK) Consortium | Jacquelyn Wynn, Mairead Hyland, The Lighthouse Lab in Alderley Park and Alex Alderton, Roberto Amato, Jeffrey Barrett, Sonia Goncalves, Ewan Harrison, David K. Jackson, Ian Johnston, Dominic Kwiatkowski, Cordelia Langford, John Sillitoe on behalf of the Wellcome Sanger Institute COVID-19 Surveillance Team |
| EPI_ISL_3337765 | Lighthouse Laboratory Plymouth | Wellcome Sanger Institute for the COVID-19 Genomics UK (COG-UK) Consortium | Lighthouse Laboratory Plymouth and Alex Alderton, Roberto Amato, Jeffrey Barrett, Sonia Goncalves, Ewan Harrison, David K. Jackson, Ian Johnston, Dominic Kwiatkowski, Cordelia Langford, John Sillitoe on behalf of the Wellcome Sanger Institute COVID-19 Surveillance Team |
| EPI_ISL_3337766, EPI_ISL_3337767, EPI_ISL_3337768, EPI_ISL_3337769 | Lighthouse Lab in Alderley Park | Wellcome Sanger Institute for the COVID-19 Genomics UK (COG-UK) Consortium | Jacquelyn Wynn, Mairead Hyland, The Lighthouse Lab in Alderley Park and Alex Alderton, Roberto Amato, Jeffrey Barrett, Sonia Goncalves, Ewan Harrison, David K. Jackson, Ian Johnston, Dominic Kwiatkowski, Cordelia Langford, John Sillitoe on behalf of the Wellcome Sanger Institute COVID-19 Surveillance Team |
| EPI_ISL_3337770 | Lighthouse Laboratory Plymouth | Wellcome Sanger Institute for the COVID-19 Genomics UK (COG-UK) Consortium | Lighthouse Laboratory Plymouth and Alex Alderton, Roberto Amato, Jeffrey Barrett, Sonia Goncalves, Ewan Harrison, David K. Jackson, Ian Johnston, Dominic Kwiatkowski, Cordelia Langford, John Sillitoe on behalf of the Wellcome Sanger Institute COVID-19 Surveillance Team |
| EPI_ISL_3337771, EPI_ISL_3337772 | Lighthouse Lab in Milton Keynes | Wellcome Sanger Institute for the COVID-19 Genomics UK (COG-UK) Consortium | The Lighthouse Lab in Milton Keynes and Alex Alderton, Roberto Amato, Jeffrey Barrett, Sonia Goncalves, Ewan Harrison, David K. Jackson, Ian Johnston, Dominic Kwiatkowski, Cordelia Langford, John Sillitoe on behalf of the Wellcome Sanger Institute COVID-19 Surveillance Team |
| EPI_ISL_3337773 | Lighthouse Laboratory Plymouth | Wellcome Sanger Institute for the COVID-19 Genomics UK (COG-UK) Consortium | Lighthouse Laboratory Plymouth and Alex Alderton, Roberto Amato, Jeffrey Barrett, Sonia Goncalves, Ewan Harrison, David K. Jackson, Ian Johnston, Dominic Kwiatkowski, Cordelia Langford, John Sillitoe on behalf of the Wellcome Sanger Institute COVID-19 Surveillance Team |
| EPI_ISL_3337774 | Lighthouse Lab in Alderley Park | Wellcome Sanger Institute for the COVID-19 Genomics UK (COG-UK) Consortium | Jacquelyn Wynn, Mairead Hyland, The Lighthouse Lab in Alderley Park and Alex Alderton, Roberto Amato, Jeffrey Barrett, Sonia Goncalves, Ewan Harrison, David K. Jackson, Ian Johnston, Dominic Kwiatkowski, Cordelia Langford, John Sillitoe on behalf of the Wellcome Sanger Institute COVID-19 Surveillance Team |
| EPI_ISL_3337775 | Lighthouse Lab in Milton Keynes | Wellcome Sanger Institute for the COVID-19 Genomics UK (COG-UK) Consortium | The Lighthouse Lab in Milton Keynes and Alex Alderton, Roberto Amato, Jeffrey Barrett, Sonia Goncalves, Ewan Harrison, David K. Jackson, Ian Johnston, Dominic Kwiatkowski, Cordelia Langford, John Sillitoe on behalf of the Wellcome Sanger Institute COVID-19 Surveillance Team |
| EPI_ISL_3337776 | Lighthouse Lab in Alderley Park | Wellcome Sanger Institute for the COVID-19 Genomics UK (COG-UK) Consortium | Jacquelyn Wynn, Mairead Hyland, The Lighthouse Lab in Alderley Park and Alex Alderton, Roberto Amato, Jeffrey Barrett, Sonia Goncalves, Ewan Harrison, David K. Jackson, Ian Johnston, Dominic Kwiatkowski, Cordelia Langford, John Sillitoe on behalf of the Wellcome Sanger Institute COVID-19 Surveillance Team |
| EPI_ISL_3337777, EPI_ISL_3337778 | Lighthouse Lab in Milton Keynes | Wellcome Sanger Institute for the COVID-19 Genomics UK (COG-UK) Consortium | The Lighthouse Lab in Milton Keynes and Alex Alderton, Roberto Amato, Jeffrey Barrett, Sonia Goncalves, Ewan Harrison, David K. Jackson, Ian Johnston, Dominic Kwiatkowski, Cordelia Langford, John Sillitoe on behalf of the Wellcome Sanger Institute COVID-19 Surveillance Team |
| EPI_ISL_3337779 | Lighthouse Laboratory Plymouth | Wellcome Sanger Institute for the COVID-19 Genomics UK (COG-UK) Consortium | Lighthouse Laboratory Plymouth and Alex Alderton, Roberto Amato, Jeffrey Barrett, Sonia Goncalves, Ewan Harrison, David K. Jackson, Ian Johnston, Dominic Kwiatkowski, Cordelia Langford, John Sillitoe on behalf of the Wellcome Sanger Institute COVID-19 Surveillance Team |
| EPI_ISL_3337780, EPI_ISL_3337781, EPI_ISL_3337782 | Lighthouse Lab in Alderley Park | Wellcome Sanger Institute for the COVID-19 Genomics UK (COG-UK) Consortium | Jacquelyn Wynn, Mairead Hyland, The Lighthouse Lab in Alderley Park and Alex Alderton, Roberto Amato, Jeffrey Barrett, Sonia Goncalves, Ewan Harrison, David K. Jackson, Ian Johnston, Dominic Kwiatkowski, Cordelia Langford, John Sillitoe on behalf of the Wellcome Sanger Institute COVID-19 Surveillance Team |
| EPI_ISL_3337783 | Lighthouse Laboratory Plymouth | Wellcome Sanger Institute for the COVID-19 Genomics UK (COG-UK) Consortium | Lighthouse Laboratory Plymouth and Alex Alderton, Roberto Amato, Jeffrey Barrett, Sonia Goncalves, Ewan Harrison, David K. Jackson, Ian Johnston, Dominic Kwiatkowski, Cordelia Langford, John Sillitoe on behalf of the Wellcome Sanger Institute COVID-19 Surveillance Team |
| EPI_ISL_3337784 | Lighthouse Lab in Milton Keynes | Wellcome Sanger Institute for the COVID-19 Genomics UK (COG-UK) Consortium | The Lighthouse Lab in Milton Keynes and Alex Alderton, Roberto Amato, Jeffrey Barrett, Sonia Goncalves, Ewan Harrison, David K. Jackson, Ian Johnston, Dominic Kwiatkowski, Cordelia Langford, John Sillitoe on behalf of the Wellcome Sanger Institute COVID-19 Surveillance Team |
| EPI_ISL_3337785, EPI_ISL_3337786, EPI_ISL_3337787 | Lighthouse Lab in Alderley Park | Wellcome Sanger Institute for the COVID-19 Genomics UK (COG-UK) Consortium | Jacquelyn Wynn, Mairead Hyland, The Lighthouse Lab in Alderley Park and Alex Alderton, Roberto Amato, Jeffrey Barrett, Sonia Goncalves, Ewan Harrison, David K. Jackson, Ian Johnston, Dominic Kwiatkowski, Cordelia Langford, John Sillitoe on behalf of the Wellcome Sanger Institute COVID-19 Surveillance Team |
| EPI_ISL_3337788 | Lighthouse Laboratory Plymouth | Wellcome Sanger Institute for the COVID-19 Genomics UK (COG-UK) Consortium | Lighthouse Laboratory Plymouth and Alex Alderton, Roberto Amato, Jeffrey Barrett, Sonia Goncalves, Ewan Harrison, David K. Jackson, Ian Johnston, Dominic Kwiatkowski, Cordelia Langford, John Sillitoe on behalf of the Wellcome Sanger Institute COVID-19 Surveillance Team |
| EPI_ISL_3337789 | Lighthouse Lab in Milton Keynes | Wellcome Sanger Institute for the COVID-19 Genomics UK (COG-UK) Consortium | The Lighthouse Lab in Milton Keynes and Alex Alderton, Roberto Amato, Jeffrey Barrett, Sonia Goncalves, Ewan Harrison, David K. Jackson, Ian Johnston, Dominic Kwiatkowski, Cordelia Langford, John Sillitoe on behalf of the Wellcome Sanger Institute COVID-19 Surveillance Team |
| EPI_ISL_3337790 | Lighthouse Lab in Alderley Park | Wellcome Sanger Institute for the COVID-19 Genomics UK (COG-UK) Consortium | Jacquelyn Wynn, Mairead Hyland, The Lighthouse Lab in Alderley Park and Alex Alderton, Roberto Amato, Jeffrey Barrett, Sonia Goncalves, Ewan Harrison, David K. Jackson, Ian Johnston, Dominic Kwiatkowski, Cordelia Langford, John Sillitoe on behalf of the Wellcome Sanger Institute COVID-19 Surveillance Team |
| EPI_ISL_3337791 | Lighthouse Laboratory Plymouth | Wellcome Sanger Institute for the COVID-19 Genomics UK (COG-UK) Consortium | Lighthouse Laboratory Plymouth and Alex Alderton, Roberto Amato, Jeffrey Barrett, Sonia Goncalves, Ewan Harrison, David K. Jackson, Ian Johnston, Dominic Kwiatkowski, Cordelia Langford, John Sillitoe on behalf of the Wellcome Sanger Institute COVID-19 Surveillance Team |
| EPI_ISL_3337792, EPI_ISL_3337793 | Lighthouse Lab in Alderley Park | Wellcome Sanger Institute for the COVID-19 Genomics UK (COG-UK) Consortium | Jacquelyn Wynn, Mairead Hyland, The Lighthouse Lab in Alderley Park and Alex Alderton, Roberto Amato, Jeffrey Barrett, Sonia Goncalves, Ewan Harrison, David K. Jackson, Ian Johnston, Dominic Kwiatkowski, Cordelia Langford, John Sillitoe on behalf of the Wellcome Sanger Institute COVID-19 Surveillance Team |
| EPI_ISL_3337794 | Lighthouse Laboratory Plymouth | Wellcome Sanger Institute for the COVID-19 Genomics UK (COG-UK) Consortium | Lighthouse Laboratory Plymouth and Alex Alderton, Roberto Amato, Jeffrey Barrett, Sonia Goncalves, Ewan Harrison, David K. Jackson, Ian Johnston, Dominic Kwiatkowski, Cordelia Langford, John Sillitoe on behalf of the Wellcome Sanger Institute COVID-19 Surveillance Team |
| EPI_ISL_3337795 | Lighthouse Lab in Alderley Park | Wellcome Sanger Institute for the COVID-19 Genomics UK (COG-UK) Consortium | Jacquelyn Wynn, Mairead Hyland, The Lighthouse Lab in Alderley Park and Alex Alderton, Roberto Amato, Jeffrey Barrett, Sonia Goncalves, Ewan Harrison, David K. Jackson, Ian Johnston, Dominic Kwiatkowski, Cordelia Langford, John Sillitoe on behalf of the Wellcome Sanger Institute COVID-19 Surveillance Team |
| EPI_ISL_3337796 | Lighthouse Laboratory Plymouth | Wellcome Sanger Institute for the COVID-19 Genomics UK (COG-UK) Consortium | Lighthouse Laboratory Plymouth and Alex Alderton, Roberto Amato, Jeffrey Barrett, Sonia Goncalves, Ewan Harrison, David K. Jackson, Ian Johnston, Dominic Kwiatkowski, Cordelia Langford, John Sillitoe on behalf of the Wellcome Sanger Institute COVID-19 Surveillance Team |
| EPI_ISL_3337797 | Lighthouse Lab in Milton Keynes | Wellcome Sanger Institute for the COVID-19 Genomics UK (COG-UK) Consortium | The Lighthouse Lab in Milton Keynes and Alex Alderton, Roberto Amato, Jeffrey Barrett, Sonia Goncalves, Ewan Harrison, David K. Jackson, Ian Johnston, Dominic Kwiatkowski, Cordelia Langford, John Sillitoe on behalf of the Wellcome Sanger Institute COVID-19 Surveillance Team |
| EPI_ISL_3337798 | Lighthouse Laboratory Plymouth | Wellcome Sanger Institute for the COVID-19 Genomics UK (COG-UK) Consortium | Lighthouse Laboratory Plymouth and Alex Alderton, Roberto Amato, Jeffrey Barrett, Sonia Goncalves, Ewan Harrison, David K. Jackson, Ian Johnston, Dominic Kwiatkowski, Cordelia Langford, John Sillitoe on behalf of the Wellcome Sanger Institute COVID-19 Surveillance Team |
| EPI_ISL_3337799, EPI_ISL_3337800 | Lighthouse Lab in Alderley Park | Wellcome Sanger Institute for the COVID-19 Genomics UK (COG-UK) Consortium | Jacquelyn Wynn, Mairead Hyland, The Lighthouse Lab in Alderley Park and Alex Alderton, Roberto Amato, Jeffrey Barrett, Sonia Goncalves, Ewan Harrison, David K. Jackson, Ian Johnston, Dominic Kwiatkowski, Cordelia Langford, John Sillitoe on behalf of the Wellcome Sanger Institute COVID-19 Surveillance Team |
| EPI_ISL_3337801, EPI_ISL_3337802 | Lighthouse Laboratory Plymouth | Wellcome Sanger Institute for the COVID-19 Genomics UK (COG-UK) Consortium | Lighthouse Laboratory Plymouth and Alex Alderton, Roberto Amato, Jeffrey Barrett, Sonia Goncalves, Ewan Harrison, David K. Jackson, Ian Johnston, Dominic Kwiatkowski, Cordelia Langford, John Sillitoe on behalf of the Wellcome Sanger Institute COVID-19 Surveillance Team |
| EPI_ISL_3337803, EPI_ISL_3337804, EPI_ISL_3337805, EPI_ISL_3337806 | Lighthouse Lab in Alderley Park | Wellcome Sanger Institute for the COVID-19 Genomics UK (COG-UK) Consortium | Jacquelyn Wynn, Mairead Hyland, The Lighthouse Lab in Alderley Park and Alex Alderton, Roberto Amato, Jeffrey Barrett, Sonia Goncalves, Ewan Harrison, David K. Jackson, Ian Johnston, Dominic Kwiatkowski, Cordelia Langford, John Sillitoe on behalf of the Wellcome Sanger Institute COVID-19 Surveillance Team |
| EPI_ISL_3337807, EPI_ISL_3337808 | Lighthouse Lab in Milton Keynes | Wellcome Sanger Institute for the COVID-19 Genomics UK (COG-UK) Consortium | The Lighthouse Lab in Milton Keynes and Alex Alderton, Roberto Amato, Jeffrey Barrett, Sonia Goncalves, Ewan Harrison, David K. Jackson, Ian Johnston, Dominic Kwiatkowski, Cordelia Langford, John Sillitoe on behalf of the Wellcome Sanger Institute COVID-19 Surveillance Team |
| EPI_ISL_3337809 | Lighthouse Laboratory Plymouth | Wellcome Sanger Institute for the COVID-19 Genomics UK (COG-UK) Consortium | Lighthouse Laboratory Plymouth and Alex Alderton, Roberto Amato, Jeffrey Barrett, Sonia Goncalves, Ewan Harrison, David K. Jackson, Ian Johnston, Dominic Kwiatkowski, Cordelia Langford, John Sillitoe on behalf of the Wellcome Sanger Institute COVID-19 Surveillance Team |
| EPI_ISL_3337810 | Lighthouse Lab in Alderley Park | Wellcome Sanger Institute for the COVID-19 Genomics UK (COG-UK) Consortium | Jacquelyn Wynn, Mairead Hyland, The Lighthouse Lab in Alderley Park and Alex Alderton, Roberto Amato, Jeffrey Barrett, Sonia Goncalves, Ewan Harrison, David K. Jackson, Ian Johnston, Dominic Kwiatkowski, Cordelia Langford, John Sillitoe on behalf of the Wellcome Sanger Institute COVID-19 Surveillance Team |
| EPI_ISL_3337811 | Lighthouse Laboratory Plymouth | Wellcome Sanger Institute for the COVID-19 Genomics UK (COG-UK) Consortium | Lighthouse Laboratory Plymouth and Alex Alderton, Roberto Amato, Jeffrey Barrett, Sonia Goncalves, Ewan Harrison, David K. Jackson, Ian Johnston, Dominic Kwiatkowski, Cordelia Langford, John Sillitoe on behalf of the Wellcome Sanger Institute COVID-19 Surveillance Team |
| EPI_ISL_3337812 | Lighthouse Lab in Alderley Park | Wellcome Sanger Institute for the COVID-19 Genomics UK (COG-UK) Consortium | Jacquelyn Wynn, Mairead Hyland, The Lighthouse Lab in Alderley Park and Alex Alderton, Roberto Amato, Jeffrey Barrett, Sonia Goncalves, Ewan Harrison, David K. Jackson, Ian Johnston, Dominic Kwiatkowski, Cordelia Langford, John Sillitoe on behalf of the Wellcome Sanger Institute COVID-19 Surveillance Team |
| EPI_ISL_3337813 | Lighthouse Laboratory Plymouth | Wellcome Sanger Institute for the COVID-19 Genomics UK (COG-UK) Consortium | Lighthouse Laboratory Plymouth and Alex Alderton, Roberto Amato, Jeffrey Barrett, Sonia Goncalves, Ewan Harrison, David K. Jackson, Ian Johnston, Dominic Kwiatkowski, Cordelia Langford, John Sillitoe on behalf of the Wellcome Sanger Institute COVID-19 Surveillance Team |
| EPI_ISL_3337814, EPI_ISL_3337815 | Lighthouse Lab in Alderley Park | Wellcome Sanger Institute for the COVID-19 Genomics UK (COG-UK) Consortium | Jacquelyn Wynn, Mairead Hyland, The Lighthouse Lab in Alderley Park and Alex Alderton, Roberto Amato, Jeffrey Barrett, Sonia Goncalves, Ewan Harrison, David K. Jackson, Ian Johnston, Dominic Kwiatkowski, Cordelia Langford, John Sillitoe on behalf of the Wellcome Sanger Institute COVID-19 Surveillance Team |
| EPI_ISL_3337816 | Lighthouse Lab in Milton Keynes | Wellcome Sanger Institute for the COVID-19 Genomics UK (COG-UK) Consortium | The Lighthouse Lab in Milton Keynes and Alex Alderton, Roberto Amato, Jeffrey Barrett, Sonia Goncalves, Ewan Harrison, David K. Jackson, Ian Johnston, Dominic Kwiatkowski, Cordelia Langford, John Sillitoe on behalf of the Wellcome Sanger Institute COVID-19 Surveillance Team |
| EPI_ISL_3337817 | Lighthouse Lab in Alderley Park | Wellcome Sanger Institute for the COVID-19 Genomics UK (COG-UK) Consortium | Jacquelyn Wynn, Mairead Hyland, The Lighthouse Lab in Alderley Park and Alex Alderton, Roberto Amato, Jeffrey Barrett, Sonia Goncalves, Ewan Harrison, David K. Jackson, Ian Johnston, Dominic Kwiatkowski, Cordelia Langford, John Sillitoe on behalf of the Wellcome Sanger Institute COVID-19 Surveillance Team |
| EPI_ISL_3337818 | Lighthouse Laboratory Plymouth | Wellcome Sanger Institute for the COVID-19 Genomics UK (COG-UK) Consortium | Lighthouse Laboratory Plymouth and Alex Alderton, Roberto Amato, Jeffrey Barrett, Sonia Goncalves, Ewan Harrison, David K. Jackson, Ian Johnston, Dominic Kwiatkowski, Cordelia Langford, John Sillitoe on behalf of the Wellcome Sanger Institute COVID-19 Surveillance Team |

| | | | |
|---|---|---|---|
| EPI_ISL_3337819, EPI_ISL_3337820 | Lighthouse Lab in Alderley Park | Wellcome Sanger Institute for the COVID-19 Genomics UK (COG-UK) Consortium | Jacquelyn Wynn, Mairead Hyland, The Lighthouse Lab in Alderley Park and Alex Alderton, Roberto Amato, Jeffrey Barrett, Sonia Goncalves, Ewan Harrison, David K. Jackson, Ian Johnston, Dominic Kwiatkowski, Cordelia Langford, John Sillitoe on behalf of the Wellcome Sanger Institute COVID-19 Surveillance Team |
| EPI_ISL_3337821 | Lighthouse Laboratory Plymouth | Wellcome Sanger Institute for the COVID-19 Genomics UK (COG-UK) Consortium | Lighthouse Laboratory Plymouth and Alex Alderton, Roberto Amato, Jeffrey Barrett, Sonia Goncalves, Ewan Harrison, David K. Jackson, Ian Johnston, Dominic Kwiatkowski, Cordelia Langford, John Sillitoe on behalf of the Wellcome Sanger Institute COVID-19 Surveillance Team |
| EPI_ISL_3337822, EPI_ISL_3337823 | Lighthouse Lab in Milton Keynes | Wellcome Sanger Institute for the COVID-19 Genomics UK (COG-UK) Consortium | The Lighthouse Lab in Milton Keynes and Alex Alderton, Roberto Amato, Jeffrey Barrett, Sonia Goncalves, Ewan Harrison, David K. Jackson, Ian Johnston, Dominic Kwiatkowski, Cordelia Langford, John Sillitoe on behalf of the Wellcome Sanger Institute COVID-19 Surveillance Team |
| EPI_ISL_3337824 | Lighthouse Lab in Alderley Park | Wellcome Sanger Institute for the COVID-19 Genomics UK (COG-UK) Consortium | Jacquelyn Wynn, Mairead Hyland, The Lighthouse Lab in Alderley Park and Alex Alderton, Roberto Amato, Jeffrey Barrett, Sonia Goncalves, Ewan Harrison, David K. Jackson, Ian Johnston, Dominic Kwiatkowski, Cordelia Langford, John Sillitoe on behalf of the Wellcome Sanger Institute COVID-19 Surveillance Team |
| EPI_ISL_3337825 | Lighthouse Laboratory Plymouth | Wellcome Sanger Institute for the COVID-19 Genomics UK (COG-UK) Consortium | Lighthouse Laboratory Plymouth and Alex Alderton, Roberto Amato, Jeffrey Barrett, Sonia Goncalves, Ewan Harrison, David K. Jackson, Ian Johnston, Dominic Kwiatkowski, Cordelia Langford, John Sillitoe on behalf of the Wellcome Sanger Institute COVID-19 Surveillance Team |
| EPI_ISL_3337826, EPI_ISL_3337827, EPI_ISL_3337828 | Lighthouse Lab in Alderley Park | Wellcome Sanger Institute for the COVID-19 Genomics UK (COG-UK) Consortium | Jacquelyn Wynn, Mairead Hyland, The Lighthouse Lab in Alderley Park and Alex Alderton, Roberto Amato, Jeffrey Barrett, Sonia Goncalves, Ewan Harrison, David K. Jackson, Ian Johnston, Dominic Kwiatkowski, Cordelia Langford, John Sillitoe on behalf of the Wellcome Sanger Institute COVID-19 Surveillance Team |
| EPI_ISL_3337829, EPI_ISL_3337830, EPI_ISL_3337831 | Lighthouse Lab in Milton Keynes | Wellcome Sanger Institute for the COVID-19 Genomics UK (COG-UK) Consortium | The Lighthouse Lab in Milton Keynes and Alex Alderton, Roberto Amato, Jeffrey Barrett, Sonia Goncalves, Ewan Harrison, David K. Jackson, Ian Johnston, Dominic Kwiatkowski, Cordelia Langford, John Sillitoe on behalf of the Wellcome Sanger Institute COVID-19 Surveillance Team |
| EPI_ISL_3337832, EPI_ISL_3337833 | Lighthouse Lab in Alderley Park | Wellcome Sanger Institute for the COVID-19 Genomics UK (COG-UK) Consortium | Jacquelyn Wynn, Mairead Hyland, The Lighthouse Lab in Alderley Park and Alex Alderton, Roberto Amato, Jeffrey Barrett, Sonia Goncalves, Ewan Harrison, David K. Jackson, Ian Johnston, Dominic Kwiatkowski, Cordelia Langford, John Sillitoe on behalf of the Wellcome Sanger Institute COVID-19 Surveillance Team |
| EPI_ISL_3337834 | Lighthouse Laboratory Plymouth | Wellcome Sanger Institute for the COVID-19 Genomics UK (COG-UK) Consortium | Lighthouse Laboratory Plymouth and Alex Alderton, Roberto Amato, Jeffrey Barrett, Sonia Goncalves, Ewan Harrison, David K. Jackson, Ian Johnston, Dominic Kwiatkowski, Cordelia Langford, John Sillitoe on behalf of the Wellcome Sanger Institute COVID-19 Surveillance Team |
| EPI_ISL_3337835 | Lighthouse Lab in Alderley Park | Wellcome Sanger Institute for the COVID-19 Genomics UK (COG-UK) Consortium | Jacquelyn Wynn, Mairead Hyland, The Lighthouse Lab in Alderley Park and Alex Alderton, Roberto Amato, Jeffrey Barrett, Sonia Goncalves, Ewan Harrison, David K. Jackson, Ian Johnston, Dominic Kwiatkowski, Cordelia Langford, John Sillitoe on behalf of the Wellcome Sanger Institute COVID-19 Surveillance Team |
| EPI_ISL_3337836 | Lighthouse Lab in Milton Keynes | Wellcome Sanger Institute for the COVID-19 Genomics UK (COG-UK) Consortium | The Lighthouse Lab in Milton Keynes and Alex Alderton, Roberto Amato, Jeffrey Barrett, Sonia Goncalves, Ewan Harrison, David K. Jackson, Ian Johnston, Dominic Kwiatkowski, Cordelia Langford, John Sillitoe on behalf of the Wellcome Sanger Institute COVID-19 Surveillance Team |
| EPI_ISL_3337837 | Lighthouse Laboratory Plymouth | Wellcome Sanger Institute for the COVID-19 Genomics UK (COG-UK) Consortium | Lighthouse Laboratory Plymouth and Alex Alderton, Roberto Amato, Jeffrey Barrett, Sonia Goncalves, Ewan Harrison, David K. Jackson, Ian Johnston, Dominic Kwiatkowski, Cordelia Langford, John Sillitoe on behalf of the Wellcome Sanger Institute COVID-19 Surveillance Team |
| EPI_ISL_3337838 | Lighthouse Lab in Alderley Park | Wellcome Sanger Institute for the COVID-19 Genomics UK (COG-UK) Consortium | Jacquelyn Wynn, Mairead Hyland, The Lighthouse Lab in Alderley Park and Alex Alderton, Roberto Amato, Jeffrey Barrett, Sonia Goncalves, Ewan Harrison, David K. Jackson, Ian Johnston, Dominic Kwiatkowski, Cordelia Langford, John Sillitoe on behalf of the Wellcome Sanger Institute COVID-19 Surveillance Team |
| EPI_ISL_3337839 | Lighthouse Laboratory Plymouth | Wellcome Sanger Institute for the COVID-19 Genomics UK (COG-UK) Consortium | Lighthouse Laboratory Plymouth and Alex Alderton, Roberto Amato, Jeffrey Barrett, Sonia Goncalves, Ewan Harrison, David K. Jackson, Ian Johnston, Dominic Kwiatkowski, Cordelia Langford, John Sillitoe on behalf of the Wellcome Sanger Institute COVID-19 Surveillance Team |
| EPI_ISL_3337840 | Lighthouse Lab in Milton Keynes | Wellcome Sanger Institute for the COVID-19 Genomics UK (COG-UK) Consortium | The Lighthouse Lab in Milton Keynes and Alex Alderton, Roberto Amato, Jeffrey Barrett, Sonia Goncalves, Ewan Harrison, David K. Jackson, Ian Johnston, Dominic Kwiatkowski, Cordelia Langford, John Sillitoe on behalf of the Wellcome Sanger Institute COVID-19 Surveillance Team |
| EPI_ISL_3337841 | Lighthouse Lab in Alderley Park | Wellcome Sanger Institute for the COVID-19 Genomics UK (COG-UK) Consortium | Jacquelyn Wynn, Mairead Hyland, The Lighthouse Lab in Alderley Park and Alex Alderton, Roberto Amato, Jeffrey Barrett, Sonia Goncalves, Ewan Harrison, David K. Jackson, Ian Johnston, Dominic Kwiatkowski, Cordelia Langford, John Sillitoe on behalf of the Wellcome Sanger Institute COVID-19 Surveillance Team |
| EPI_ISL_3337842 | Lighthouse Lab in Milton Keynes | Wellcome Sanger Institute for the COVID-19 Genomics UK (COG-UK) Consortium | The Lighthouse Lab in Milton Keynes and Alex Alderton, Roberto Amato, Jeffrey Barrett, Sonia Goncalves, Ewan Harrison, David K. Jackson, Ian Johnston, Dominic Kwiatkowski, Cordelia Langford, John Sillitoe on behalf of the Wellcome Sanger Institute COVID-19 Surveillance Team |
| EPI_ISL_3337843, EPI_ISL_3337844 | Lighthouse Laboratory Plymouth | Wellcome Sanger Institute for the COVID-19 Genomics UK (COG-UK) Consortium | Lighthouse Laboratory Plymouth and Alex Alderton, Roberto Amato, Jeffrey Barrett, Sonia Goncalves, Ewan Harrison, David K. Jackson, Ian Johnston, Dominic Kwiatkowski, Cordelia Langford, John Sillitoe on behalf of the Wellcome Sanger Institute COVID-19 Surveillance Team |
| EPI_ISL_3337845 | Lighthouse Lab in Milton Keynes | Wellcome Sanger Institute for the COVID-19 Genomics UK (COG-UK) Consortium | The Lighthouse Lab in Milton Keynes and Alex Alderton, Roberto Amato, Jeffrey Barrett, Sonia Goncalves, Ewan Harrison, David K. Jackson, Ian Johnston, Dominic Kwiatkowski, Cordelia Langford, John Sillitoe on behalf of the Wellcome Sanger Institute COVID-19 Surveillance Team |
| EPI_ISL_3337846 | Lighthouse Laboratory Plymouth | Wellcome Sanger Institute for the COVID-19 Genomics UK (COG-UK) Consortium | Lighthouse Laboratory Plymouth and Alex Alderton, Roberto Amato, Jeffrey Barrett, Sonia Goncalves, Ewan Harrison, David K. Jackson, Ian Johnston, Dominic Kwiatkowski, Cordelia Langford, John Sillitoe on behalf of the Wellcome Sanger Institute COVID-19 Surveillance Team |
| EPI_ISL_3337847 | Lighthouse Lab in Alderley Park | Wellcome Sanger Institute for the COVID-19 Genomics UK (COG-UK) Consortium | Jacquelyn Wynn, Mairead Hyland, The Lighthouse Lab in Alderley Park and Alex Alderton, Roberto Amato, Jeffrey Barrett, Sonia Goncalves, Ewan Harrison, David K. Jackson, Ian Johnston, Dominic Kwiatkowski, Cordelia Langford, John Sillitoe on behalf of the Wellcome Sanger Institute COVID-19 Surveillance Team |
| EPI_ISL_3337848 | Lighthouse Lab in Milton Keynes | Wellcome Sanger Institute for the COVID-19 Genomics UK (COG-UK) Consortium | The Lighthouse Lab in Milton Keynes and Alex Alderton, Roberto Amato, Jeffrey Barrett, Sonia Goncalves, Ewan Harrison, David K. Jackson, Ian Johnston, Dominic Kwiatkowski, Cordelia Langford, John Sillitoe on behalf of the Wellcome Sanger Institute COVID-19 Surveillance Team |
| EPI_ISL_3337849 | Lighthouse Lab in Alderley Park | Wellcome Sanger Institute for the COVID-19 Genomics UK (COG-UK) Consortium | Jacquelyn Wynn, Mairead Hyland, The Lighthouse Lab in Alderley Park and Alex Alderton, Roberto Amato, Jeffrey Barrett, Sonia Goncalves, Ewan Harrison, David K. Jackson, Ian Johnston, Dominic Kwiatkowski, Cordelia Langford, John Sillitoe on behalf of the Wellcome Sanger Institute COVID-19 Surveillance Team |
| EPI_ISL_3337850 | Lighthouse Laboratory Plymouth | Wellcome Sanger Institute for the COVID-19 Genomics UK (COG-UK) Consortium | Lighthouse Laboratory Plymouth and Alex Alderton, Roberto Amato, Jeffrey Barrett, Sonia Goncalves, Ewan Harrison, David K. Jackson, Ian Johnston, Dominic Kwiatkowski, Cordelia Langford, John Sillitoe on behalf of the Wellcome Sanger Institute COVID-19 Surveillance Team |
| EPI_ISL_3337851 | Lighthouse Lab in Alderley Park | Wellcome Sanger Institute for the COVID-19 Genomics UK (COG-UK) Consortium | Jacquelyn Wynn, Mairead Hyland, The Lighthouse Lab in Alderley Park and Alex Alderton, Roberto Amato, Jeffrey Barrett, Sonia Goncalves, Ewan Harrison, David K. Jackson, Ian Johnston, Dominic Kwiatkowski, Cordelia Langford, John Sillitoe on behalf of the Wellcome Sanger Institute COVID-19 Surveillance Team |
| EPI_ISL_3337852 | Lighthouse Laboratory Plymouth | Wellcome Sanger Institute for the COVID-19 Genomics UK (COG-UK) Consortium | Lighthouse Laboratory Plymouth and Alex Alderton, Roberto Amato, Jeffrey Barrett, Sonia Goncalves, Ewan Harrison, David K. Jackson, Ian Johnston, Dominic Kwiatkowski, Cordelia Langford, John Sillitoe on behalf of the Wellcome Sanger Institute COVID-19 Surveillance Team |
| EPI_ISL_3337853 | Lighthouse Lab in Milton Keynes | Wellcome Sanger Institute for the COVID-19 Genomics UK (COG-UK) Consortium | The Lighthouse Lab in Milton Keynes and Alex Alderton, Roberto Amato, Jeffrey Barrett, Sonia Goncalves, Ewan Harrison, David K. Jackson, Ian Johnston, Dominic Kwiatkowski, Cordelia Langford, John Sillitoe on behalf of the Wellcome Sanger Institute COVID-19 Surveillance Team |
| EPI_ISL_3337854, EPI_ISL_3337855 | Lighthouse Lab in Alderley Park | Wellcome Sanger Institute for the COVID-19 Genomics UK (COG-UK) Consortium | Jacquelyn Wynn, Mairead Hyland, The Lighthouse Lab in Alderley Park and Alex Alderton, Roberto Amato, Jeffrey Barrett, Sonia Goncalves, Ewan Harrison, David K. Jackson, Ian Johnston, Dominic Kwiatkowski, Cordelia Langford, John Sillitoe on behalf of the Wellcome Sanger Institute COVID-19 Surveillance Team |
| EPI_ISL_3337856 | Lighthouse Laboratory Plymouth | Wellcome Sanger Institute for the COVID-19 Genomics UK (COG-UK) Consortium | Lighthouse Laboratory Plymouth and Alex Alderton, Roberto Amato, Jeffrey Barrett, Sonia Goncalves, Ewan Harrison, David K. Jackson, Ian Johnston, Dominic Kwiatkowski, Cordelia Langford, John Sillitoe on behalf of the Wellcome Sanger Institute COVID-19 Surveillance Team |
| EPI_ISL_3337857, EPI_ISL_3337858, EPI_ISL_3337859 | Lighthouse Lab in Alderley Park | Wellcome Sanger Institute for the COVID-19 Genomics UK (COG-UK) Consortium | Jacquelyn Wynn, Mairead Hyland, The Lighthouse Lab in Alderley Park and Alex Alderton, Roberto Amato, Jeffrey Barrett, Sonia Goncalves, Ewan Harrison, David K. Jackson, Ian Johnston, Dominic Kwiatkowski, Cordelia Langford, John Sillitoe on behalf of the Wellcome Sanger Institute COVID-19 Surveillance Team |
| EPI_ISL_3337860, EPI_ISL_3337861 | Lighthouse Lab in Milton Keynes | Wellcome Sanger Institute for the COVID-19 Genomics UK (COG-UK) Consortium | The Lighthouse Lab in Milton Keynes and Alex Alderton, Roberto Amato, Jeffrey Barrett, Sonia Goncalves, Ewan Harrison, David K. Jackson, Ian Johnston, Dominic Kwiatkowski, Cordelia Langford, John Sillitoe on behalf of the Wellcome Sanger Institute COVID-19 Surveillance Team |
| EPI_ISL_3337862, EPI_ISL_3337863 | Lighthouse Lab in Alderley Park | Wellcome Sanger Institute for the COVID-19 Genomics UK (COG-UK) Consortium | Jacquelyn Wynn, Mairead Hyland, The Lighthouse Lab in Alderley Park and Alex Alderton, Roberto Amato, Jeffrey Barrett, Sonia Goncalves, Ewan Harrison, David K. Jackson, Ian Johnston, Dominic Kwiatkowski, Cordelia Langford, John Sillitoe on behalf of the Wellcome Sanger Institute COVID-19 Surveillance Team |
| EPI_ISL_3337864 | Lighthouse Lab in Milton Keynes | Wellcome Sanger Institute for the COVID-19 Genomics UK (COG-UK) Consortium | The Lighthouse Lab in Milton Keynes and Alex Alderton, Roberto Amato, Jeffrey Barrett, Sonia Goncalves, Ewan Harrison, David K. Jackson, Ian Johnston, Dominic Kwiatkowski, Cordelia Langford, John Sillitoe on behalf of the Wellcome Sanger Institute COVID-19 Surveillance Team |
| EPI_ISL_3337865, EPI_ISL_3337866 | Lighthouse Lab in Alderley Park | Wellcome Sanger Institute for the COVID-19 Genomics UK (COG-UK) Consortium | Jacquelyn Wynn, Mairead Hyland, The Lighthouse Lab in Alderley Park and Alex Alderton, Roberto Amato, Jeffrey Barrett, Sonia Goncalves, Ewan Harrison, David K. Jackson, Ian Johnston, Dominic Kwiatkowski, Cordelia Langford, John Sillitoe on behalf of the Wellcome Sanger Institute COVID-19 Surveillance Team |
| EPI_ISL_3337867, EPI_ISL_3337868 | Lighthouse Lab in Milton Keynes | Wellcome Sanger Institute for the COVID-19 Genomics UK (COG-UK) Consortium | The Lighthouse Lab in Milton Keynes and Alex Alderton, Roberto Amato, Jeffrey Barrett, Sonia Goncalves, Ewan Harrison, David K. Jackson, Ian Johnston, Dominic Kwiatkowski, Cordelia Langford, John Sillitoe on behalf of the Wellcome Sanger Institute COVID-19 Surveillance Team |
| EPI_ISL_3337869, EPI_ISL_3337870, EPI_ISL_3337871 | Lighthouse Lab in Alderley Park | Wellcome Sanger Institute for the COVID-19 Genomics UK (COG-UK) Consortium | Jacquelyn Wynn, Mairead Hyland, The Lighthouse Lab in Alderley Park and Alex Alderton, Roberto Amato, Jeffrey Barrett, Sonia Goncalves, Ewan Harrison, David K. Jackson, Ian Johnston, Dominic Kwiatkowski, Cordelia Langford, John Sillitoe on behalf of the Wellcome Sanger Institute COVID-19 Surveillance Team |
| EPI_ISL_3337872, EPI_ISL_3337873 | Lighthouse Laboratory Plymouth | Wellcome Sanger Institute for the COVID-19 Genomics UK (COG-UK) Consortium | Lighthouse Laboratory Plymouth and Alex Alderton, Roberto Amato, Jeffrey Barrett, Sonia Goncalves, Ewan Harrison, David K. Jackson, Ian Johnston, Dominic Kwiatkowski, Cordelia Langford, John Sillitoe on behalf of the Wellcome Sanger Institute COVID-19 Surveillance Team |
| EPI_ISL_3337874 | Lighthouse Lab in Milton Keynes | Wellcome Sanger Institute for the COVID-19 Genomics UK (COG-UK) Consortium | The Lighthouse Lab in Milton Keynes and Alex Alderton, Roberto Amato, Jeffrey Barrett, Sonia Goncalves, Ewan Harrison, David K. Jackson, Ian Johnston, Dominic Kwiatkowski, Cordelia Langford, John Sillitoe on behalf of the Wellcome Sanger Institute COVID-19 Surveillance Team |
| EPI_ISL_3337875 | Lighthouse Lab in Alderley Park | Wellcome Sanger Institute for the COVID-19 Genomics UK (COG-UK) Consortium | Jacquelyn Wynn, Mairead Hyland, The Lighthouse Lab in Alderley Park and Alex Alderton, Roberto Amato, Jeffrey Barrett, Sonia Goncalves, Ewan Harrison, David K. Jackson, Ian Johnston, Dominic Kwiatkowski, Cordelia Langford, John Sillitoe on behalf of the Wellcome Sanger Institute COVID-19 Surveillance Team |
| EPI_ISL_3337876 | Lighthouse Laboratory Plymouth | Wellcome Sanger Institute for the COVID-19 Genomics UK (COG-UK) Consortium | Lighthouse Laboratory Plymouth and Alex Alderton, Roberto Amato, Jeffrey Barrett, Sonia Goncalves, Ewan Harrison, David K. Jackson, Ian Johnston, Dominic Kwiatkowski, Cordelia Langford, John Sillitoe on behalf of the Wellcome Sanger Institute COVID-19 Surveillance Team |
| EPI_ISL_3337877, EPI_ISL_3337878, EPI_ISL_3337879, EPI_ISL_3337880 | Lighthouse Lab in Alderley Park | Wellcome Sanger Institute for the COVID-19 Genomics UK (COG-UK) Consortium | Jacquelyn Wynn, Mairead Hyland, The Lighthouse Lab in Alderley Park and Alex Alderton, Roberto Amato, Jeffrey Barrett, Sonia Goncalves, Ewan Harrison, David K. Jackson, Ian Johnston, Dominic Kwiatkowski, Cordelia Langford, John Sillitoe on behalf of the Wellcome Sanger Institute COVID-19 Surveillance Team |
| EPI_ISL_3337881 | Lighthouse Laboratory Plymouth | Wellcome Sanger Institute for the COVID-19 Genomics UK (COG-UK) Consortium | Lighthouse Laboratory Plymouth and Alex Alderton, Roberto Amato, Jeffrey Barrett, Sonia Goncalves, Ewan Harrison, David K. Jackson, Ian Johnston, Dominic Kwiatkowski, Cordelia Langford, John Sillitoe on behalf of the Wellcome Sanger Institute COVID-19 Surveillance Team |
| EPI_ISL_3337882, | Lighthouse Lab in Milton Keynes | Wellcome Sanger Institute for the COVID-19 Genomics UK (COG-UK) | The Lighthouse Lab in Milton Keynes and Alex Alderton, Roberto Amato, Jeffrey Barrett, Sonia Goncalves, Ewan Harrison, David K. Jackson, Ian Johnston, Dominic Kwiatkowski, Cordelia Langford, John Sillitoe on behalf of the Wellcome Sanger |

| | | | |
|---|---|---|---|
| EPI_ISL_3337883 | | Consortium | Institute COVID-19 Surveillance Team |
| EPI_ISL_3337884 | Lighthouse Lab in Alderley Park | Wellcome Sanger Institute for the COVID-19 Genomics UK (COG-UK) Consortium | Jacquelyn Wynn, Mairead Hyland, The Lighthouse Lab in Alderley Park and Alex Alderton, Roberto Amato, Jeffrey Barrett, Sonia Goncalves, Ewan Harrison, David K. Jackson, Ian Johnston, Dominic Kwiatkowski, Cordelia Langford, John Sillitoe on behalf of the Wellcome Sanger Institute COVID-19 Surveillance Team |
| EPI_ISL_3337885 | Lighthouse Lab in Milton Keynes | Wellcome Sanger Institute for the COVID-19 Genomics UK (COG-UK) Consortium | The Lighthouse Lab in Milton Keynes and Alex Alderton, Roberto Amato, Jeffrey Barrett, Sonia Goncalves, Ewan Harrison, David K. Jackson, Ian Johnston, Dominic Kwiatkowski, Cordelia Langford, John Sillitoe on behalf of the Wellcome Sanger Institute COVID-19 Surveillance Team |
| EPI_ISL_3337886 | Lighthouse Laboratory Plymouth | Wellcome Sanger Institute for the COVID-19 Genomics UK (COG-UK) Consortium | Lighthouse Laboratory Plymouth and Alex Alderton, Roberto Amato, Jeffrey Barrett, Sonia Goncalves, Ewan Harrison, David K. Jackson, Ian Johnston, Dominic Kwiatkowski, Cordelia Langford, John Sillitoe on behalf of the Wellcome Sanger Institute COVID-19 Surveillance Team |
| EPI_ISL_3337887, EPI_ISL_3337888 | Lighthouse Lab in Alderley Park | Wellcome Sanger Institute for the COVID-19 Genomics UK (COG-UK) Consortium | Jacquelyn Wynn, Mairead Hyland, The Lighthouse Lab in Alderley Park and Alex Alderton, Roberto Amato, Jeffrey Barrett, Sonia Goncalves, Ewan Harrison, David K. Jackson, Ian Johnston, Dominic Kwiatkowski, Cordelia Langford, John Sillitoe on behalf of the Wellcome Sanger Institute COVID-19 Surveillance Team |
| EPI_ISL_3337889 | Lighthouse Lab in Milton Keynes | Wellcome Sanger Institute for the COVID-19 Genomics UK (COG-UK) Consortium | The Lighthouse Lab in Milton Keynes and Alex Alderton, Roberto Amato, Jeffrey Barrett, Sonia Goncalves, Ewan Harrison, David K. Jackson, Ian Johnston, Dominic Kwiatkowski, Cordelia Langford, John Sillitoe on behalf of the Wellcome Sanger Institute COVID-19 Surveillance Team |
| EPI_ISL_3337890, EPI_ISL_3337891, EPI_ISL_3337892, EPI_ISL_3337893 | Lighthouse Laboratory Plymouth | Wellcome Sanger Institute for the COVID-19 Genomics UK (COG-UK) Consortium | Lighthouse Laboratory Plymouth and Alex Alderton, Roberto Amato, Jeffrey Barrett, Sonia Goncalves, Ewan Harrison, David K. Jackson, Ian Johnston, Dominic Kwiatkowski, Cordelia Langford, John Sillitoe on behalf of the Wellcome Sanger Institute COVID-19 Surveillance Team |
| EPI_ISL_3337894 | Lighthouse Lab in Alderley Park | Wellcome Sanger Institute for the COVID-19 Genomics UK (COG-UK) Consortium | Jacquelyn Wynn, Mairead Hyland, The Lighthouse Lab in Alderley Park and Alex Alderton, Roberto Amato, Jeffrey Barrett, Sonia Goncalves, Ewan Harrison, David K. Jackson, Ian Johnston, Dominic Kwiatkowski, Cordelia Langford, John Sillitoe on behalf of the Wellcome Sanger Institute COVID-19 Surveillance Team |
| EPI_ISL_3337895 | Lighthouse Lab in Milton Keynes | Wellcome Sanger Institute for the COVID-19 Genomics UK (COG-UK) Consortium | The Lighthouse Lab in Milton Keynes and Alex Alderton, Roberto Amato, Jeffrey Barrett, Sonia Goncalves, Ewan Harrison, David K. Jackson, Ian Johnston, Dominic Kwiatkowski, Cordelia Langford, John Sillitoe on behalf of the Wellcome Sanger Institute COVID-19 Surveillance Team |
| EPI_ISL_3337896 | Lighthouse Laboratory Plymouth | Wellcome Sanger Institute for the COVID-19 Genomics UK (COG-UK) Consortium | Lighthouse Laboratory Plymouth and Alex Alderton, Roberto Amato, Jeffrey Barrett, Sonia Goncalves, Ewan Harrison, David K. Jackson, Ian Johnston, Dominic Kwiatkowski, Cordelia Langford, John Sillitoe on behalf of the Wellcome Sanger Institute COVID-19 Surveillance Team |
| EPI_ISL_3337897, EPI_ISL_3337898 | Lighthouse Lab in Alderley Park | Wellcome Sanger Institute for the COVID-19 Genomics UK (COG-UK) Consortium | Jacquelyn Wynn, Mairead Hyland, The Lighthouse Lab in Alderley Park and Alex Alderton, Roberto Amato, Jeffrey Barrett, Sonia Goncalves, Ewan Harrison, David K. Jackson, Ian Johnston, Dominic Kwiatkowski, Cordelia Langford, John Sillitoe on behalf of the Wellcome Sanger Institute COVID-19 Surveillance Team |
| EPI_ISL_3337899 | Lighthouse Laboratory Plymouth | Wellcome Sanger Institute for the COVID-19 Genomics UK (COG-UK) Consortium | Lighthouse Laboratory Plymouth and Alex Alderton, Roberto Amato, Jeffrey Barrett, Sonia Goncalves, Ewan Harrison, David K. Jackson, Ian Johnston, Dominic Kwiatkowski, Cordelia Langford, John Sillitoe on behalf of the Wellcome Sanger Institute COVID-19 Surveillance Team |
| EPI_ISL_3337900, EPI_ISL_3337901, EPI_ISL_3337902 | Lighthouse Lab in Alderley Park | Wellcome Sanger Institute for the COVID-19 Genomics UK (COG-UK) Consortium | Jacquelyn Wynn, Mairead Hyland, The Lighthouse Lab in Alderley Park and Alex Alderton, Roberto Amato, Jeffrey Barrett, Sonia Goncalves, Ewan Harrison, David K. Jackson, Ian Johnston, Dominic Kwiatkowski, Cordelia Langford, John Sillitoe on behalf of the Wellcome Sanger Institute COVID-19 Surveillance Team |
| EPI_ISL_3337903 | Lighthouse Lab in Milton Keynes | Wellcome Sanger Institute for the COVID-19 Genomics UK (COG-UK) Consortium | The Lighthouse Lab in Milton Keynes and Alex Alderton, Roberto Amato, Jeffrey Barrett, Sonia Goncalves, Ewan Harrison, David K. Jackson, Ian Johnston, Dominic Kwiatkowski, Cordelia Langford, John Sillitoe on behalf of the Wellcome Sanger Institute COVID-19 Surveillance Team |
| EPI_ISL_3337904, EPI_ISL_3337905, EPI_ISL_3337906, EPI_ISL_3337907 | Lighthouse Lab in Alderley Park | Wellcome Sanger Institute for the COVID-19 Genomics UK (COG-UK) Consortium | Jacquelyn Wynn, Mairead Hyland, The Lighthouse Lab in Alderley Park and Alex Alderton, Roberto Amato, Jeffrey Barrett, Sonia Goncalves, Ewan Harrison, David K. Jackson, Ian Johnston, Dominic Kwiatkowski, Cordelia Langford, John Sillitoe on behalf of the Wellcome Sanger Institute COVID-19 Surveillance Team |
| EPI_ISL_3337908 | Lighthouse Lab in Milton Keynes | Wellcome Sanger Institute for the COVID-19 Genomics UK (COG-UK) Consortium | The Lighthouse Lab in Milton Keynes and Alex Alderton, Roberto Amato, Jeffrey Barrett, Sonia Goncalves, Ewan Harrison, David K. Jackson, Ian Johnston, Dominic Kwiatkowski, Cordelia Langford, John Sillitoe on behalf of the Wellcome Sanger Institute COVID-19 Surveillance Team |
| EPI_ISL_3337909 | Lighthouse Laboratory Plymouth | Wellcome Sanger Institute for the COVID-19 Genomics UK (COG-UK) Consortium | Lighthouse Laboratory Plymouth and Alex Alderton, Roberto Amato, Jeffrey Barrett, Sonia Goncalves, Ewan Harrison, David K. Jackson, Ian Johnston, Dominic Kwiatkowski, Cordelia Langford, John Sillitoe on behalf of the Wellcome Sanger Institute COVID-19 Surveillance Team |
| EPI_ISL_3337910 | Lighthouse Lab in Alderley Park | Wellcome Sanger Institute for the COVID-19 Genomics UK (COG-UK) Consortium | Jacquelyn Wynn, Mairead Hyland, The Lighthouse Lab in Alderley Park and Alex Alderton, Roberto Amato, Jeffrey Barrett, Sonia Goncalves, Ewan Harrison, David K. Jackson, Ian Johnston, Dominic Kwiatkowski, Cordelia Langford, John Sillitoe on behalf of the Wellcome Sanger Institute COVID-19 Surveillance Team |
| EPI_ISL_3337911, EPI_ISL_3337912 | Lighthouse Laboratory Plymouth | Wellcome Sanger Institute for the COVID-19 Genomics UK (COG-UK) Consortium | Lighthouse Laboratory Plymouth and Alex Alderton, Roberto Amato, Jeffrey Barrett, Sonia Goncalves, Ewan Harrison, David K. Jackson, Ian Johnston, Dominic Kwiatkowski, Cordelia Langford, John Sillitoe on behalf of the Wellcome Sanger Institute COVID-19 Surveillance Team |
| EPI_ISL_3337913, EPI_ISL_3337914 | Lighthouse Lab in Alderley Park | Wellcome Sanger Institute for the COVID-19 Genomics UK (COG-UK) Consortium | Jacquelyn Wynn, Mairead Hyland, The Lighthouse Lab in Alderley Park and Alex Alderton, Roberto Amato, Jeffrey Barrett, Sonia Goncalves, Ewan Harrison, David K. Jackson, Ian Johnston, Dominic Kwiatkowski, Cordelia Langford, John Sillitoe on behalf of the Wellcome Sanger Institute COVID-19 Surveillance Team |
| EPI_ISL_3337915 | Lighthouse Lab in Milton Keynes | Wellcome Sanger Institute for the COVID-19 Genomics UK (COG-UK) Consortium | The Lighthouse Lab in Milton Keynes and Alex Alderton, Roberto Amato, Jeffrey Barrett, Sonia Goncalves, Ewan Harrison, David K. Jackson, Ian Johnston, Dominic Kwiatkowski, Cordelia Langford, John Sillitoe on behalf of the Wellcome Sanger Institute COVID-19 Surveillance Team |
| EPI_ISL_3337916 | Lighthouse Lab in Alderley Park | Wellcome Sanger Institute for the COVID-19 Genomics UK (COG-UK) Consortium | Jacquelyn Wynn, Mairead Hyland, The Lighthouse Lab in Alderley Park and Alex Alderton, Roberto Amato, Jeffrey Barrett, Sonia Goncalves, Ewan Harrison, David K. Jackson, Ian Johnston, Dominic Kwiatkowski, Cordelia Langford, John Sillitoe on behalf of the Wellcome Sanger Institute COVID-19 Surveillance Team |
| EPI_ISL_3337917 | Lighthouse Laboratory Plymouth | Wellcome Sanger Institute for the COVID-19 Genomics UK (COG-UK) Consortium | Lighthouse Laboratory Plymouth and Alex Alderton, Roberto Amato, Jeffrey Barrett, Sonia Goncalves, Ewan Harrison, David K. Jackson, Ian Johnston, Dominic Kwiatkowski, Cordelia Langford, John Sillitoe on behalf of the Wellcome Sanger Institute COVID-19 Surveillance Team |
| EPI_ISL_3337918, EPI_ISL_3337919, EPI_ISL_3337920, EPI_ISL_3337921, EPI_ISL_3337922, EPI_ISL_3337923 | Lighthouse Lab in Alderley Park | Wellcome Sanger Institute for the COVID-19 Genomics UK (COG-UK) Consortium | Jacquelyn Wynn, Mairead Hyland, The Lighthouse Lab in Alderley Park and Alex Alderton, Roberto Amato, Jeffrey Barrett, Sonia Goncalves, Ewan Harrison, David K. Jackson, Ian Johnston, Dominic Kwiatkowski, Cordelia Langford, John Sillitoe on behalf of the Wellcome Sanger Institute COVID-19 Surveillance Team |
| EPI_ISL_3337924 | Lighthouse Lab in Milton Keynes | Wellcome Sanger Institute for the COVID-19 Genomics UK (COG-UK) Consortium | The Lighthouse Lab in Milton Keynes and Alex Alderton, Roberto Amato, Jeffrey Barrett, Sonia Goncalves, Ewan Harrison, David K. Jackson, Ian Johnston, Dominic Kwiatkowski, Cordelia Langford, John Sillitoe on behalf of the Wellcome Sanger Institute COVID-19 Surveillance Team |
| EPI_ISL_3337925, EPI_ISL_3337926, EPI_ISL_3337927 | Lighthouse Laboratory Plymouth | Wellcome Sanger Institute for the COVID-19 Genomics UK (COG-UK) Consortium | Lighthouse Laboratory Plymouth and Alex Alderton, Roberto Amato, Jeffrey Barrett, Sonia Goncalves, Ewan Harrison, David K. Jackson, Ian Johnston, Dominic Kwiatkowski, Cordelia Langford, John Sillitoe on behalf of the Wellcome Sanger Institute COVID-19 Surveillance Team |
| EPI_ISL_3337928, EPI_ISL_3337929 | Lighthouse Lab in Milton Keynes | Wellcome Sanger Institute for the COVID-19 Genomics UK (COG-UK) Consortium | The Lighthouse Lab in Milton Keynes and Alex Alderton, Roberto Amato, Jeffrey Barrett, Sonia Goncalves, Ewan Harrison, David K. Jackson, Ian Johnston, Dominic Kwiatkowski, Cordelia Langford, John Sillitoe on behalf of the Wellcome Sanger Institute COVID-19 Surveillance Team |
| EPI_ISL_3337930 | Lighthouse Laboratory Plymouth | Wellcome Sanger Institute for the COVID-19 Genomics UK (COG-UK) Consortium | Lighthouse Laboratory Plymouth and Alex Alderton, Roberto Amato, Jeffrey Barrett, Sonia Goncalves, Ewan Harrison, David K. Jackson, Ian Johnston, Dominic Kwiatkowski, Cordelia Langford, John Sillitoe on behalf of the Wellcome Sanger Institute COVID-19 Surveillance Team |
| EPI_ISL_3337931 | Lighthouse Lab in Milton Keynes | Wellcome Sanger Institute for the COVID-19 Genomics UK (COG-UK) Consortium | The Lighthouse Lab in Milton Keynes and Alex Alderton, Roberto Amato, Jeffrey Barrett, Sonia Goncalves, Ewan Harrison, David K. Jackson, Ian Johnston, Dominic Kwiatkowski, Cordelia Langford, John Sillitoe on behalf of the Wellcome Sanger Institute COVID-19 Surveillance Team |
| EPI_ISL_3337932, EPI_ISL_3337933 | Lighthouse Lab in Alderley Park | Wellcome Sanger Institute for the COVID-19 Genomics UK (COG-UK) Consortium | Jacquelyn Wynn, Mairead Hyland, The Lighthouse Lab in Alderley Park and Alex Alderton, Roberto Amato, Jeffrey Barrett, Sonia Goncalves, Ewan Harrison, David K. Jackson, Ian Johnston, Dominic Kwiatkowski, Cordelia Langford, John Sillitoe on behalf of the Wellcome Sanger Institute COVID-19 Surveillance Team |
| EPI_ISL_3337934 | Lighthouse Lab in Milton Keynes | Wellcome Sanger Institute for the COVID-19 Genomics UK (COG-UK) Consortium | The Lighthouse Lab in Milton Keynes and Alex Alderton, Roberto Amato, Jeffrey Barrett, Sonia Goncalves, Ewan Harrison, David K. Jackson, Ian Johnston, Dominic Kwiatkowski, Cordelia Langford, John Sillitoe on behalf of the Wellcome Sanger Institute COVID-19 Surveillance Team |
| EPI_ISL_3337935, EPI_ISL_3337936 | Lighthouse Lab in Alderley Park | Wellcome Sanger Institute for the COVID-19 Genomics UK (COG-UK) Consortium | Jacquelyn Wynn, Mairead Hyland, The Lighthouse Lab in Alderley Park and Alex Alderton, Roberto Amato, Jeffrey Barrett, Sonia Goncalves, Ewan Harrison, David K. Jackson, Ian Johnston, Dominic Kwiatkowski, Cordelia Langford, John Sillitoe on behalf of the Wellcome Sanger Institute COVID-19 Surveillance Team |
| EPI_ISL_3337937 | Lighthouse Laboratory Plymouth | Wellcome Sanger Institute for the COVID-19 Genomics UK (COG-UK) Consortium | Lighthouse Laboratory Plymouth and Alex Alderton, Roberto Amato, Jeffrey Barrett, Sonia Goncalves, Ewan Harrison, David K. Jackson, Ian Johnston, Dominic Kwiatkowski, Cordelia Langford, John Sillitoe on behalf of the Wellcome Sanger Institute COVID-19 Surveillance Team |
| EPI_ISL_3337938 | Lighthouse Lab in Milton Keynes | Wellcome Sanger Institute for the COVID-19 Genomics UK (COG-UK) Consortium | The Lighthouse Lab in Milton Keynes and Alex Alderton, Roberto Amato, Jeffrey Barrett, Sonia Goncalves, Ewan Harrison, David K. Jackson, Ian Johnston, Dominic Kwiatkowski, Cordelia Langford, John Sillitoe on behalf of the Wellcome Sanger Institute COVID-19 Surveillance Team |
| EPI_ISL_3337939 | Lighthouse Laboratory Plymouth | Wellcome Sanger Institute for the COVID-19 Genomics UK (COG-UK) Consortium | Lighthouse Laboratory Plymouth and Alex Alderton, Roberto Amato, Jeffrey Barrett, Sonia Goncalves, Ewan Harrison, David K. Jackson, Ian Johnston, Dominic Kwiatkowski, Cordelia Langford, John Sillitoe on behalf of the Wellcome Sanger Institute COVID-19 Surveillance Team |
| EPI_ISL_3337940, EPI_ISL_3337941, EPI_ISL_3337942 | Lighthouse Lab in Alderley Park | Wellcome Sanger Institute for the COVID-19 Genomics UK (COG-UK) Consortium | Jacquelyn Wynn, Mairead Hyland, The Lighthouse Lab in Alderley Park and Alex Alderton, Roberto Amato, Jeffrey Barrett, Sonia Goncalves, Ewan Harrison, David K. Jackson, Ian Johnston, Dominic Kwiatkowski, Cordelia Langford, John Sillitoe on behalf of the Wellcome Sanger Institute COVID-19 Surveillance Team |
| EPI_ISL_3337943 | Lighthouse Laboratory Plymouth | Wellcome Sanger Institute for the COVID-19 Genomics UK (COG-UK) Consortium | Lighthouse Laboratory Plymouth and Alex Alderton, Roberto Amato, Jeffrey Barrett, Sonia Goncalves, Ewan Harrison, David K. Jackson, Ian Johnston, Dominic Kwiatkowski, Cordelia Langford, John Sillitoe on behalf of the Wellcome Sanger Institute COVID-19 Surveillance Team |
| EPI_ISL_3337944, EPI_ISL_3337945 | Lighthouse Lab in Alderley Park | Wellcome Sanger Institute for the COVID-19 Genomics UK (COG-UK) Consortium | Jacquelyn Wynn, Mairead Hyland, The Lighthouse Lab in Alderley Park and Alex Alderton, Roberto Amato, Jeffrey Barrett, Sonia Goncalves, Ewan Harrison, David K. Jackson, Ian Johnston, Dominic Kwiatkowski, Cordelia Langford, John Sillitoe on behalf of the Wellcome Sanger Institute COVID-19 Surveillance Team |
| EPI_ISL_3337946 | Lighthouse Laboratory Plymouth | Wellcome Sanger Institute for the COVID-19 Genomics UK (COG-UK) Consortium | Lighthouse Laboratory Plymouth and Alex Alderton, Roberto Amato, Jeffrey Barrett, Sonia Goncalves, Ewan Harrison, David K. Jackson, Ian Johnston, Dominic Kwiatkowski, Cordelia Langford, John Sillitoe on behalf of the Wellcome Sanger Institute COVID-19 Surveillance Team |
| EPI_ISL_3337947, EPI_ISL_3337948, EPI_ISL_3337949 | Lighthouse Lab in Alderley Park | Wellcome Sanger Institute for the COVID-19 Genomics UK (COG-UK) Consortium | Jacquelyn Wynn, Mairead Hyland, The Lighthouse Lab in Alderley Park and Alex Alderton, Roberto Amato, Jeffrey Barrett, Sonia Goncalves, Ewan Harrison, David K. Jackson, Ian Johnston, Dominic Kwiatkowski, Cordelia Langford, John Sillitoe on behalf of the Wellcome Sanger Institute COVID-19 Surveillance Team |
| EPI_ISL_3337950 | Lighthouse Lab in Milton Keynes | Wellcome Sanger Institute for the COVID-19 Genomics UK (COG-UK) Consortium | The Lighthouse Lab in Milton Keynes and Alex Alderton, Roberto Amato, Jeffrey Barrett, Sonia Goncalves, Ewan Harrison, David K. Jackson, Ian Johnston, Dominic Kwiatkowski, Cordelia Langford, John Sillitoe on behalf of the Wellcome Sanger Institute COVID-19 Surveillance Team |

| | | | |
|---|---|---|---|
| EPI_ISL_3337951 | Lighthouse Lab in Alderley Park | Wellcome Sanger Institute for the COVID-19 Genomics UK (COG-UK) Consortium | Jacquelyn Wynn, Mairead Hyland, The Lighthouse Lab in Alderley Park and Alex Alderton, Roberto Amato, Jeffrey Barrett, Sonia Goncalves, Ewan Harrison, David K. Jackson, Ian Johnston, Dominic Kwiatkowski, Cordelia Langford, John Sillitoe on behalf of the Wellcome Sanger Institute COVID-19 Surveillance Team |
| EPI_ISL_3337952 | Lighthouse Lab in Milton Keynes | Wellcome Sanger Institute for the COVID-19 Genomics UK (COG-UK) Consortium | The Lighthouse Lab in Milton Keynes and Alex Alderton, Roberto Amato, Jeffrey Barrett, Sonia Goncalves, Ewan Harrison, David K. Jackson, Ian Johnston, Dominic Kwiatkowski, Cordelia Langford, John Sillitoe on behalf of the Wellcome Sanger Institute COVID-19 Surveillance Team |
| EPI_ISL_3337953 | Lighthouse Laboratory Plymouth | Wellcome Sanger Institute for the COVID-19 Genomics UK (COG-UK) Consortium | Lighthouse Laboratory Plymouth and Alex Alderton, Roberto Amato, Jeffrey Barrett, Sonia Goncalves, Ewan Harrison, David K. Jackson, Ian Johnston, Dominic Kwiatkowski, Cordelia Langford, John Sillitoe on behalf of the Wellcome Sanger Institute COVID-19 Surveillance Team |
| EPI_ISL_3337954, EPI_ISL_3337955, EPI_ISL_3337956, EPI_ISL_3337957 | Lighthouse Lab in Alderley Park | Wellcome Sanger Institute for the COVID-19 Genomics UK (COG-UK) Consortium | Jacquelyn Wynn, Mairead Hyland, The Lighthouse Lab in Alderley Park and Alex Alderton, Roberto Amato, Jeffrey Barrett, Sonia Goncalves, Ewan Harrison, David K. Jackson, Ian Johnston, Dominic Kwiatkowski, Cordelia Langford, John Sillitoe on behalf of the Wellcome Sanger Institute COVID-19 Surveillance Team |
| EPI_ISL_3337958 | Lighthouse Lab in Milton Keynes | Wellcome Sanger Institute for the COVID-19 Genomics UK (COG-UK) Consortium | The Lighthouse Lab in Milton Keynes and Alex Alderton, Roberto Amato, Jeffrey Barrett, Sonia Goncalves, Ewan Harrison, David K. Jackson, Ian Johnston, Dominic Kwiatkowski, Cordelia Langford, John Sillitoe on behalf of the Wellcome Sanger Institute COVID-19 Surveillance Team |
| EPI_ISL_3337959, EPI_ISL_3337960 | Lighthouse Lab in Alderley Park | Wellcome Sanger Institute for the COVID-19 Genomics UK (COG-UK) Consortium | Jacquelyn Wynn, Mairead Hyland, The Lighthouse Lab in Alderley Park and Alex Alderton, Roberto Amato, Jeffrey Barrett, Sonia Goncalves, Ewan Harrison, David K. Jackson, Ian Johnston, Dominic Kwiatkowski, Cordelia Langford, John Sillitoe on behalf of the Wellcome Sanger Institute COVID-19 Surveillance Team |
| EPI_ISL_3337961 | Lighthouse Lab in Milton Keynes | Wellcome Sanger Institute for the COVID-19 Genomics UK (COG-UK) Consortium | The Lighthouse Lab in Milton Keynes and Alex Alderton, Roberto Amato, Jeffrey Barrett, Sonia Goncalves, Ewan Harrison, David K. Jackson, Ian Johnston, Dominic Kwiatkowski, Cordelia Langford, John Sillitoe on behalf of the Wellcome Sanger Institute COVID-19 Surveillance Team |
| EPI_ISL_3337962 | Lighthouse Laboratory Plymouth | Wellcome Sanger Institute for the COVID-19 Genomics UK (COG-UK) Consortium | Lighthouse Laboratory Plymouth and Alex Alderton, Roberto Amato, Jeffrey Barrett, Sonia Goncalves, Ewan Harrison, David K. Jackson, Ian Johnston, Dominic Kwiatkowski, Cordelia Langford, John Sillitoe on behalf of the Wellcome Sanger Institute COVID-19 Surveillance Team |
| EPI_ISL_3337963, EPI_ISL_3337964, EPI_ISL_3337965, EPI_ISL_3337966 | Lighthouse Lab in Alderley Park | Wellcome Sanger Institute for the COVID-19 Genomics UK (COG-UK) Consortium | Jacquelyn Wynn, Mairead Hyland, The Lighthouse Lab in Alderley Park and Alex Alderton, Roberto Amato, Jeffrey Barrett, Sonia Goncalves, Ewan Harrison, David K. Jackson, Ian Johnston, Dominic Kwiatkowski, Cordelia Langford, John Sillitoe on behalf of the Wellcome Sanger Institute COVID-19 Surveillance Team |
| EPI_ISL_3337967 | Lighthouse Lab in Milton Keynes | Wellcome Sanger Institute for the COVID-19 Genomics UK (COG-UK) Consortium | The Lighthouse Lab in Milton Keynes and Alex Alderton, Roberto Amato, Jeffrey Barrett, Sonia Goncalves, Ewan Harrison, David K. Jackson, Ian Johnston, Dominic Kwiatkowski, Cordelia Langford, John Sillitoe on behalf of the Wellcome Sanger Institute COVID-19 Surveillance Team |
| EPI_ISL_3337968 | Lighthouse Lab in Alderley Park | Wellcome Sanger Institute for the COVID-19 Genomics UK (COG-UK) Consortium | Jacquelyn Wynn, Mairead Hyland, The Lighthouse Lab in Alderley Park and Alex Alderton, Roberto Amato, Jeffrey Barrett, Sonia Goncalves, Ewan Harrison, David K. Jackson, Ian Johnston, Dominic Kwiatkowski, Cordelia Langford, John Sillitoe on behalf of the Wellcome Sanger Institute COVID-19 Surveillance Team |
| EPI_ISL_3337969 | Lighthouse Lab in Milton Keynes | Wellcome Sanger Institute for the COVID-19 Genomics UK (COG-UK) Consortium | The Lighthouse Lab in Milton Keynes and Alex Alderton, Roberto Amato, Jeffrey Barrett, Sonia Goncalves, Ewan Harrison, David K. Jackson, Ian Johnston, Dominic Kwiatkowski, Cordelia Langford, John Sillitoe on behalf of the Wellcome Sanger Institute COVID-19 Surveillance Team |
| EPI_ISL_3337970 | Lighthouse Lab in Alderley Park | Wellcome Sanger Institute for the COVID-19 Genomics UK (COG-UK) Consortium | Jacquelyn Wynn, Mairead Hyland, The Lighthouse Lab in Alderley Park and Alex Alderton, Roberto Amato, Jeffrey Barrett, Sonia Goncalves, Ewan Harrison, David K. Jackson, Ian Johnston, Dominic Kwiatkowski, Cordelia Langford, John Sillitoe on behalf of the Wellcome Sanger Institute COVID-19 Surveillance Team |
| EPI_ISL_3337971 | Lighthouse Lab in Milton Keynes | Wellcome Sanger Institute for the COVID-19 Genomics UK (COG-UK) Consortium | The Lighthouse Lab in Milton Keynes and Alex Alderton, Roberto Amato, Jeffrey Barrett, Sonia Goncalves, Ewan Harrison, David K. Jackson, Ian Johnston, Dominic Kwiatkowski, Cordelia Langford, John Sillitoe on behalf of the Wellcome Sanger Institute COVID-19 Surveillance Team |
| EPI_ISL_3337972, EPI_ISL_3337973, EPI_ISL_3337974, EPI_ISL_3337975 | Lighthouse Lab in Alderley Park | Wellcome Sanger Institute for the COVID-19 Genomics UK (COG-UK) Consortium | Jacquelyn Wynn, Mairead Hyland, The Lighthouse Lab in Alderley Park and Alex Alderton, Roberto Amato, Jeffrey Barrett, Sonia Goncalves, Ewan Harrison, David K. Jackson, Ian Johnston, Dominic Kwiatkowski, Cordelia Langford, John Sillitoe on behalf of the Wellcome Sanger Institute COVID-19 Surveillance Team |
| EPI_ISL_3337976 | Lighthouse Laboratory Plymouth | Wellcome Sanger Institute for the COVID-19 Genomics UK (COG-UK) Consortium | Lighthouse Laboratory Plymouth and Alex Alderton, Roberto Amato, Jeffrey Barrett, Sonia Goncalves, Ewan Harrison, David K. Jackson, Ian Johnston, Dominic Kwiatkowski, Cordelia Langford, John Sillitoe on behalf of the Wellcome Sanger Institute COVID-19 Surveillance Team |
| EPI_ISL_3337977, EPI_ISL_3337978 | Lighthouse Lab in Alderley Park | Wellcome Sanger Institute for the COVID-19 Genomics UK (COG-UK) Consortium | Jacquelyn Wynn, Mairead Hyland, The Lighthouse Lab in Alderley Park and Alex Alderton, Roberto Amato, Jeffrey Barrett, Sonia Goncalves, Ewan Harrison, David K. Jackson, Ian Johnston, Dominic Kwiatkowski, Cordelia Langford, John Sillitoe on behalf of the Wellcome Sanger Institute COVID-19 Surveillance Team |
| EPI_ISL_3337979 | Lighthouse Lab in Milton Keynes | Wellcome Sanger Institute for the COVID-19 Genomics UK (COG-UK) Consortium | The Lighthouse Lab in Milton Keynes and Alex Alderton, Roberto Amato, Jeffrey Barrett, Sonia Goncalves, Ewan Harrison, David K. Jackson, Ian Johnston, Dominic Kwiatkowski, Cordelia Langford, John Sillitoe on behalf of the Wellcome Sanger Institute COVID-19 Surveillance Team |
| EPI_ISL_3337980 | Lighthouse Laboratory Plymouth | Wellcome Sanger Institute for the COVID-19 Genomics UK (COG-UK) Consortium | Lighthouse Laboratory Plymouth and Alex Alderton, Roberto Amato, Jeffrey Barrett, Sonia Goncalves, Ewan Harrison, David K. Jackson, Ian Johnston, Dominic Kwiatkowski, Cordelia Langford, John Sillitoe on behalf of the Wellcome Sanger Institute COVID-19 Surveillance Team |
| EPI_ISL_3337981 | Lighthouse Lab in Milton Keynes | Wellcome Sanger Institute for the COVID-19 Genomics UK (COG-UK) Consortium | The Lighthouse Lab in Milton Keynes and Alex Alderton, Roberto Amato, Jeffrey Barrett, Sonia Goncalves, Ewan Harrison, David K. Jackson, Ian Johnston, Dominic Kwiatkowski, Cordelia Langford, John Sillitoe on behalf of the Wellcome Sanger Institute COVID-19 Surveillance Team |
| EPI_ISL_3337982, EPI_ISL_3337983 | Health Services Laboratories | Wellcome Sanger Institute for the COVID-19 Genomics UK (COG-UK) Consortium | Health Services Laboratories and Alex Alderton, Roberto Amato, Jeffrey Barrett, Sonia Goncalves, Ewan Harrison, David K. Jackson, Ian Johnston, Dominic Kwiatkowski, Cordelia Langford, John Sillitoe on behalf of the Wellcome Sanger Institute COVID-19 Surveillance Team |
| EPI_ISL_3337984 | Lighthouse Lab in Milton Keynes | Wellcome Sanger Institute for the COVID-19 Genomics UK (COG-UK) Consortium | The Lighthouse Lab in Milton Keynes and Alex Alderton, Roberto Amato, Jeffrey Barrett, Sonia Goncalves, Ewan Harrison, David K. Jackson, Ian Johnston, Dominic Kwiatkowski, Cordelia Langford, John Sillitoe on behalf of the Wellcome Sanger Institute COVID-19 Surveillance Team |
| EPI_ISL_3337985 | Lighthouse Laboratory Plymouth | Wellcome Sanger Institute for the COVID-19 Genomics UK (COG-UK) Consortium | Lighthouse Laboratory Plymouth and Alex Alderton, Roberto Amato, Jeffrey Barrett, Sonia Goncalves, Ewan Harrison, David K. Jackson, Ian Johnston, Dominic Kwiatkowski, Cordelia Langford, John Sillitoe on behalf of the Wellcome Sanger Institute COVID-19 Surveillance Team |
| EPI_ISL_3337986 | Lighthouse Lab in Milton Keynes | Wellcome Sanger Institute for the COVID-19 Genomics UK (COG-UK) Consortium | The Lighthouse Lab in Milton Keynes and Alex Alderton, Roberto Amato, Jeffrey Barrett, Sonia Goncalves, Ewan Harrison, David K. Jackson, Ian Johnston, Dominic Kwiatkowski, Cordelia Langford, John Sillitoe on behalf of the Wellcome Sanger Institute COVID-19 Surveillance Team |
| EPI_ISL_3337987, EPI_ISL_3337988 | Health Services Laboratories | Wellcome Sanger Institute for the COVID-19 Genomics UK (COG-UK) Consortium | Health Services Laboratories and Alex Alderton, Roberto Amato, Jeffrey Barrett, Sonia Goncalves, Ewan Harrison, David K. Jackson, Ian Johnston, Dominic Kwiatkowski, Cordelia Langford, John Sillitoe on behalf of the Wellcome Sanger Institute COVID-19 Surveillance Team |
| EPI_ISL_3337989, EPI_ISL_3337990 | Berkshire and Surrey Pathology Services Lighthouse Laboratory | Wellcome Sanger Institute for the COVID-19 Genomics UK (COG-UK) Consortium | Berkshire and Surrey Pathology Services Lighthouse Laboratory and Alex Alderton, Roberto Amato, Jeffrey Barrett, Sonia Goncalves, Ewan Harrison, David K. Jackson, Ian Johnston, Dominic Kwiatkowski, Cordelia Langford, John Sillitoe on behalf of the Wellcome Sanger Institute COVID-19 Surveillance Team |
| EPI_ISL_3337991 | Lighthouse Laboratory Plymouth | Wellcome Sanger Institute for the COVID-19 Genomics UK (COG-UK) Consortium | Lighthouse Laboratory Plymouth and Alex Alderton, Roberto Amato, Jeffrey Barrett, Sonia Goncalves, Ewan Harrison, David K. Jackson, Ian Johnston, Dominic Kwiatkowski, Cordelia Langford, John Sillitoe on behalf of the Wellcome Sanger Institute COVID-19 Surveillance Team |
| EPI_ISL_3337992 | Berkshire and Surrey Pathology Services Lighthouse Laboratory | Wellcome Sanger Institute for the COVID-19 Genomics UK (COG-UK) Consortium | Berkshire and Surrey Pathology Services Lighthouse Laboratory and Alex Alderton, Roberto Amato, Jeffrey Barrett, Sonia Goncalves, Ewan Harrison, David K. Jackson, Ian Johnston, Dominic Kwiatkowski, Cordelia Langford, John Sillitoe on behalf of the Wellcome Sanger Institute COVID-19 Surveillance Team |
| EPI_ISL_3337993 | Health Services Laboratories | Wellcome Sanger Institute for the COVID-19 Genomics UK (COG-UK) Consortium | Health Services Laboratories and Alex Alderton, Roberto Amato, Jeffrey Barrett, Sonia Goncalves, Ewan Harrison, David K. Jackson, Ian Johnston, Dominic Kwiatkowski, Cordelia Langford, John Sillitoe on behalf of the Wellcome Sanger Institute COVID-19 Surveillance Team |
| EPI_ISL_3337994 | Berkshire and Surrey Pathology Services Lighthouse Laboratory | Wellcome Sanger Institute for the COVID-19 Genomics UK (COG-UK) Consortium | Berkshire and Surrey Pathology Services Lighthouse Laboratory and Alex Alderton, Roberto Amato, Jeffrey Barrett, Sonia Goncalves, Ewan Harrison, David K. Jackson, Ian Johnston, Dominic Kwiatkowski, Cordelia Langford, John Sillitoe on behalf of the Wellcome Sanger Institute COVID-19 Surveillance Team |
| EPI_ISL_3337995, EPI_ISL_3337996 | Health Services Laboratories | Wellcome Sanger Institute for the COVID-19 Genomics UK (COG-UK) Consortium | Health Services Laboratories and Alex Alderton, Roberto Amato, Jeffrey Barrett, Sonia Goncalves, Ewan Harrison, David K. Jackson, Ian Johnston, Dominic Kwiatkowski, Cordelia Langford, John Sillitoe on behalf of the Wellcome Sanger Institute COVID-19 Surveillance Team |
| EPI_ISL_3337997, EPI_ISL_3337998 | Lighthouse Lab in Milton Keynes | Wellcome Sanger Institute for the COVID-19 Genomics UK (COG-UK) Consortium | The Lighthouse Lab in Milton Keynes and Alex Alderton, Roberto Amato, Jeffrey Barrett, Sonia Goncalves, Ewan Harrison, David K. Jackson, Ian Johnston, Dominic Kwiatkowski, Cordelia Langford, John Sillitoe on behalf of the Wellcome Sanger Institute COVID-19 Surveillance Team |
| EPI_ISL_3337999 | Lighthouse Laboratory Plymouth | Wellcome Sanger Institute for the COVID-19 Genomics UK (COG-UK) Consortium | Lighthouse Laboratory Plymouth and Alex Alderton, Roberto Amato, Jeffrey Barrett, Sonia Goncalves, Ewan Harrison, David K. Jackson, Ian Johnston, Dominic Kwiatkowski, Cordelia Langford, John Sillitoe on behalf of the Wellcome Sanger Institute COVID-19 Surveillance Team |
| EPI_ISL_3338000 | Health Services Laboratories | Wellcome Sanger Institute for the COVID-19 Genomics UK (COG-UK) Consortium | Health Services Laboratories and Alex Alderton, Roberto Amato, Jeffrey Barrett, Sonia Goncalves, Ewan Harrison, David K. Jackson, Ian Johnston, Dominic Kwiatkowski, Cordelia Langford, John Sillitoe on behalf of the Wellcome Sanger Institute COVID-19 Surveillance Team |
| EPI_ISL_3338001 | Berkshire and Surrey Pathology Services Lighthouse Laboratory | Wellcome Sanger Institute for the COVID-19 Genomics UK (COG-UK) Consortium | Berkshire and Surrey Pathology Services Lighthouse Laboratory and Alex Alderton, Roberto Amato, Jeffrey Barrett, Sonia Goncalves, Ewan Harrison, David K. Jackson, Ian Johnston, Dominic Kwiatkowski, Cordelia Langford, John Sillitoe on behalf of the Wellcome Sanger Institute COVID-19 Surveillance Team |
| EPI_ISL_3338002 | Lighthouse Laboratory Plymouth | Wellcome Sanger Institute for the COVID-19 Genomics UK (COG-UK) Consortium | Lighthouse Laboratory Plymouth and Alex Alderton, Roberto Amato, Jeffrey Barrett, Sonia Goncalves, Ewan Harrison, David K. Jackson, Ian Johnston, Dominic Kwiatkowski, Cordelia Langford, John Sillitoe on behalf of the Wellcome Sanger Institute COVID-19 Surveillance Team |
| EPI_ISL_3338003, EPI_ISL_3338004, EPI_ISL_3338005 | Berkshire and Surrey Pathology Services Lighthouse Laboratory | Wellcome Sanger Institute for the COVID-19 Genomics UK (COG-UK) Consortium | Berkshire and Surrey Pathology Services Lighthouse Laboratory and Alex Alderton, Roberto Amato, Jeffrey Barrett, Sonia Goncalves, Ewan Harrison, David K. Jackson, Ian Johnston, Dominic Kwiatkowski, Cordelia Langford, John Sillitoe on behalf of the Wellcome Sanger Institute COVID-19 Surveillance Team |
| EPI_ISL_3338006, EPI_ISL_3338007 | Health Services Laboratories | Wellcome Sanger Institute for the COVID-19 Genomics UK (COG-UK) Consortium | Health Services Laboratories and Alex Alderton, Roberto Amato, Jeffrey Barrett, Sonia Goncalves, Ewan Harrison, David K. Jackson, Ian Johnston, Dominic Kwiatkowski, Cordelia Langford, John Sillitoe on behalf of the Wellcome Sanger Institute COVID-19 Surveillance Team |
| EPI_ISL_3338008, EPI_ISL_3338009 | Lighthouse Laboratory Plymouth | Wellcome Sanger Institute for the COVID-19 Genomics UK (COG-UK) Consortium | Lighthouse Laboratory Plymouth and Alex Alderton, Roberto Amato, Jeffrey Barrett, Sonia Goncalves, Ewan Harrison, David K. Jackson, Ian Johnston, Dominic Kwiatkowski, Cordelia Langford, John Sillitoe on behalf of the Wellcome Sanger Institute COVID-19 Surveillance Team |
| EPI_ISL_3338010 | Berkshire and Surrey Pathology Services Lighthouse Laboratory | Wellcome Sanger Institute for the COVID-19 Genomics UK (COG-UK) Consortium | Berkshire and Surrey Pathology Services Lighthouse Laboratory and Alex Alderton, Roberto Amato, Jeffrey Barrett, Sonia Goncalves, Ewan Harrison, David K. Jackson, Ian Johnston, Dominic Kwiatkowski, Cordelia Langford, John Sillitoe on behalf of the Wellcome Sanger Institute COVID-19 Surveillance Team |
| EPI_ISL_3338011 | Health Services Laboratories | Wellcome Sanger Institute for the COVID-19 Genomics UK (COG-UK) Consortium | Health Services Laboratories and Alex Alderton, Roberto Amato, Jeffrey Barrett, Sonia Goncalves, Ewan Harrison, David K. Jackson, Ian Johnston, Dominic Kwiatkowski, Cordelia Langford, John Sillitoe on behalf of the Wellcome Sanger Institute COVID-19 Surveillance Team |
| EPI_ISL_3338012, EPI_ISL_3338013 | Lighthouse Laboratory Plymouth | Wellcome Sanger Institute for the COVID-19 Genomics UK (COG-UK) Consortium | Lighthouse Laboratory Plymouth and Alex Alderton, Roberto Amato, Jeffrey Barrett, Sonia Goncalves, Ewan Harrison, David K. Jackson, Ian Johnston, Dominic Kwiatkowski, Cordelia Langford, John Sillitoe on behalf of the Wellcome Sanger Institute COVID-19 Surveillance Team |

| | | | |
|---|---|---|---|
| EPI_ISL_3338014 | Lighthouse Lab in Milton Keynes | Wellcome Sanger Institute for the COVID-19 Genomics UK (COG-UK) Consortium | The Lighthouse Lab in Milton Keynes and Alex Alderton, Roberto Amato, Jeffrey Barrett, Sonia Goncalves, Ewan Harrison, David K. Jackson, Ian Johnston, Dominic Kwiatkowski, Cordelia Langford, John Sillitoe on behalf of the Wellcome Sanger Institute COVID-19 Surveillance Team |
| EPI_ISL_3338015 | Health Services Laboratories | Wellcome Sanger Institute for the COVID-19 Genomics UK (COG-UK) Consortium | Health Services Laboratories and Alex Alderton, Roberto Amato, Jeffrey Barrett, Sonia Goncalves, Ewan Harrison, David K. Jackson, Ian Johnston, Dominic Kwiatkowski, Cordelia Langford, John Sillitoe on behalf of the Wellcome Sanger Institute COVID-19 Surveillance Team |
| EPI_ISL_3338016 | Lighthouse Laboratory Plymouth | Wellcome Sanger Institute for the COVID-19 Genomics UK (COG-UK) Consortium | Lighthouse Laboratory Plymouth and Alex Alderton, Roberto Amato, Jeffrey Barrett, Sonia Goncalves, Ewan Harrison, David K. Jackson, Ian Johnston, Dominic Kwiatkowski, Cordelia Langford, John Sillitoe on behalf of the Wellcome Sanger Institute COVID-19 Surveillance Team |
| EPI_ISL_3338017, EPI_ISL_3338018, EPI_ISL_3338019 | Health Services Laboratories | Wellcome Sanger Institute for the COVID-19 Genomics UK (COG-UK) Consortium | Health Services Laboratories and Alex Alderton, Roberto Amato, Jeffrey Barrett, Sonia Goncalves, Ewan Harrison, David K. Jackson, Ian Johnston, Dominic Kwiatkowski, Cordelia Langford, John Sillitoe on behalf of the Wellcome Sanger Institute COVID-19 Surveillance Team |
| EPI_ISL_3338020, EPI_ISL_3338021 | Lighthouse Laboratory Plymouth | Wellcome Sanger Institute for the COVID-19 Genomics UK (COG-UK) Consortium | Lighthouse Laboratory Plymouth and Alex Alderton, Roberto Amato, Jeffrey Barrett, Sonia Goncalves, Ewan Harrison, David K. Jackson, Ian Johnston, Dominic Kwiatkowski, Cordelia Langford, John Sillitoe on behalf of the Wellcome Sanger Institute COVID-19 Surveillance Team |
| EPI_ISL_3338022 | Health Services Laboratories | Wellcome Sanger Institute for the COVID-19 Genomics UK (COG-UK) Consortium | Health Services Laboratories and Alex Alderton, Roberto Amato, Jeffrey Barrett, Sonia Goncalves, Ewan Harrison, David K. Jackson, Ian Johnston, Dominic Kwiatkowski, Cordelia Langford, John Sillitoe on behalf of the Wellcome Sanger Institute COVID-19 Surveillance Team |
| EPI_ISL_3338023, EPI_ISL_3338024 | Lighthouse Lab in Milton Keynes | Wellcome Sanger Institute for the COVID-19 Genomics UK (COG-UK) Consortium | The Lighthouse Lab in Milton Keynes and Alex Alderton, Roberto Amato, Jeffrey Barrett, Sonia Goncalves, Ewan Harrison, David K. Jackson, Ian Johnston, Dominic Kwiatkowski, Cordelia Langford, John Sillitoe on behalf of the Wellcome Sanger Institute COVID-19 Surveillance Team |
| EPI_ISL_3338025 | Lighthouse Laboratory Plymouth | Wellcome Sanger Institute for the COVID-19 Genomics UK (COG-UK) Consortium | Lighthouse Laboratory Plymouth and Alex Alderton, Roberto Amato, Jeffrey Barrett, Sonia Goncalves, Ewan Harrison, David K. Jackson, Ian Johnston, Dominic Kwiatkowski, Cordelia Langford, John Sillitoe on behalf of the Wellcome Sanger Institute COVID-19 Surveillance Team |
| EPI_ISL_3338026, EPI_ISL_3338027, EPI_ISL_3338028 | Health Services Laboratories | Wellcome Sanger Institute for the COVID-19 Genomics UK (COG-UK) Consortium | Health Services Laboratories and Alex Alderton, Roberto Amato, Jeffrey Barrett, Sonia Goncalves, Ewan Harrison, David K. Jackson, Ian Johnston, Dominic Kwiatkowski, Cordelia Langford, John Sillitoe on behalf of the Wellcome Sanger Institute COVID-19 Surveillance Team |
| EPI_ISL_3338029 | Lighthouse Laboratory Plymouth | Wellcome Sanger Institute for the COVID-19 Genomics UK (COG-UK) Consortium | Lighthouse Laboratory Plymouth and Alex Alderton, Roberto Amato, Jeffrey Barrett, Sonia Goncalves, Ewan Harrison, David K. Jackson, Ian Johnston, Dominic Kwiatkowski, Cordelia Langford, John Sillitoe on behalf of the Wellcome Sanger Institute COVID-19 Surveillance Team |
| EPI_ISL_3338030 | Health Services Laboratories | Wellcome Sanger Institute for the COVID-19 Genomics UK (COG-UK) Consortium | Health Services Laboratories and Alex Alderton, Roberto Amato, Jeffrey Barrett, Sonia Goncalves, Ewan Harrison, David K. Jackson, Ian Johnston, Dominic Kwiatkowski, Cordelia Langford, John Sillitoe on behalf of the Wellcome Sanger Institute COVID-19 Surveillance Team |
| EPI_ISL_3338031 | Lighthouse Laboratory Plymouth | Wellcome Sanger Institute for the COVID-19 Genomics UK (COG-UK) Consortium | Lighthouse Laboratory Plymouth and Alex Alderton, Roberto Amato, Jeffrey Barrett, Sonia Goncalves, Ewan Harrison, David K. Jackson, Ian Johnston, Dominic Kwiatkowski, Cordelia Langford, John Sillitoe on behalf of the Wellcome Sanger Institute COVID-19 Surveillance Team |
| EPI_ISL_3338032 | Health Services Laboratories | Wellcome Sanger Institute for the COVID-19 Genomics UK (COG-UK) Consortium | Health Services Laboratories and Alex Alderton, Roberto Amato, Jeffrey Barrett, Sonia Goncalves, Ewan Harrison, David K. Jackson, Ian Johnston, Dominic Kwiatkowski, Cordelia Langford, John Sillitoe on behalf of the Wellcome Sanger Institute COVID-19 Surveillance Team |
| EPI_ISL_3338033 | Lighthouse Lab in Milton Keynes | Wellcome Sanger Institute for the COVID-19 Genomics UK (COG-UK) Consortium | The Lighthouse Lab in Milton Keynes and Alex Alderton, Roberto Amato, Jeffrey Barrett, Sonia Goncalves, Ewan Harrison, David K. Jackson, Ian Johnston, Dominic Kwiatkowski, Cordelia Langford, John Sillitoe on behalf of the Wellcome Sanger Institute COVID-19 Surveillance Team |
| EPI_ISL_3338034 | Lighthouse Laboratory Plymouth | Wellcome Sanger Institute for the COVID-19 Genomics UK (COG-UK) Consortium | Lighthouse Laboratory Plymouth and Alex Alderton, Roberto Amato, Jeffrey Barrett, Sonia Goncalves, Ewan Harrison, David K. Jackson, Ian Johnston, Dominic Kwiatkowski, Cordelia Langford, John Sillitoe on behalf of the Wellcome Sanger Institute COVID-19 Surveillance Team |
| EPI_ISL_3338035 | Health Services Laboratories | Wellcome Sanger Institute for the COVID-19 Genomics UK (COG-UK) Consortium | Health Services Laboratories and Alex Alderton, Roberto Amato, Jeffrey Barrett, Sonia Goncalves, Ewan Harrison, David K. Jackson, Ian Johnston, Dominic Kwiatkowski, Cordelia Langford, John Sillitoe on behalf of the Wellcome Sanger Institute COVID-19 Surveillance Team |
| EPI_ISL_3338036, EPI_ISL_3338037 | Lighthouse Lab in Milton Keynes | Wellcome Sanger Institute for the COVID-19 Genomics UK (COG-UK) Consortium | The Lighthouse Lab in Milton Keynes and Alex Alderton, Roberto Amato, Jeffrey Barrett, Sonia Goncalves, Ewan Harrison, David K. Jackson, Ian Johnston, Dominic Kwiatkowski, Cordelia Langford, John Sillitoe on behalf of the Wellcome Sanger Institute COVID-19 Surveillance Team |
| EPI_ISL_3338038 | Lighthouse Laboratory Plymouth | Wellcome Sanger Institute for the COVID-19 Genomics UK (COG-UK) Consortium | Lighthouse Laboratory Plymouth and Alex Alderton, Roberto Amato, Jeffrey Barrett, Sonia Goncalves, Ewan Harrison, David K. Jackson, Ian Johnston, Dominic Kwiatkowski, Cordelia Langford, John Sillitoe on behalf of the Wellcome Sanger Institute COVID-19 Surveillance Team |
| EPI_ISL_3338039, EPI_ISL_3338040 | Lighthouse Lab in Milton Keynes | Wellcome Sanger Institute for the COVID-19 Genomics UK (COG-UK) Consortium | The Lighthouse Lab in Milton Keynes and Alex Alderton, Roberto Amato, Jeffrey Barrett, Sonia Goncalves, Ewan Harrison, David K. Jackson, Ian Johnston, Dominic Kwiatkowski, Cordelia Langford, John Sillitoe on behalf of the Wellcome Sanger Institute COVID-19 Surveillance Team |
| EPI_ISL_3338041, EPI_ISL_3338042 | Health Services Laboratories | Wellcome Sanger Institute for the COVID-19 Genomics UK (COG-UK) Consortium | Health Services Laboratories and Alex Alderton, Roberto Amato, Jeffrey Barrett, Sonia Goncalves, Ewan Harrison, David K. Jackson, Ian Johnston, Dominic Kwiatkowski, Cordelia Langford, John Sillitoe on behalf of the Wellcome Sanger Institute COVID-19 Surveillance Team |
| EPI_ISL_3338043 | Lighthouse Laboratory Plymouth | Wellcome Sanger Institute for the COVID-19 Genomics UK (COG-UK) Consortium | Lighthouse Laboratory Plymouth and Alex Alderton, Roberto Amato, Jeffrey Barrett, Sonia Goncalves, Ewan Harrison, David K. Jackson, Ian Johnston, Dominic Kwiatkowski, Cordelia Langford, John Sillitoe on behalf of the Wellcome Sanger Institute COVID-19 Surveillance Team |
| EPI_ISL_3338044, EPI_ISL_3338045, EPI_ISL_3338046 | Lighthouse Lab in Milton Keynes | Wellcome Sanger Institute for the COVID-19 Genomics UK (COG-UK) Consortium | The Lighthouse Lab in Milton Keynes and Alex Alderton, Roberto Amato, Jeffrey Barrett, Sonia Goncalves, Ewan Harrison, David K. Jackson, Ian Johnston, Dominic Kwiatkowski, Cordelia Langford, John Sillitoe on behalf of the Wellcome Sanger Institute COVID-19 Surveillance Team |
| EPI_ISL_3338047, EPI_ISL_3338048, EPI_ISL_3338049, EPI_ISL_3338050, EPI_ISL_3338051 | Lighthouse Laboratory Plymouth | Wellcome Sanger Institute for the COVID-19 Genomics UK (COG-UK) Consortium | Lighthouse Laboratory Plymouth and Alex Alderton, Roberto Amato, Jeffrey Barrett, Sonia Goncalves, Ewan Harrison, David K. Jackson, Ian Johnston, Dominic Kwiatkowski, Cordelia Langford, John Sillitoe on behalf of the Wellcome Sanger Institute COVID-19 Surveillance Team |
| EPI_ISL_3338052 | Berkshire and Surrey Pathology Services Lighthouse Laboratory | Wellcome Sanger Institute for the COVID-19 Genomics UK (COG-UK) Consortium | Berkshire and Surrey Pathology Services Lighthouse Laboratory and Alex Alderton, Roberto Amato, Jeffrey Barrett, Sonia Goncalves, Ewan Harrison, David K. Jackson, Ian Johnston, Dominic Kwiatkowski, Cordelia Langford, John Sillitoe on behalf of the Wellcome Sanger Institute COVID-19 Surveillance Team |
| EPI_ISL_3338053, EPI_ISL_3338054 | Lighthouse Lab in Milton Keynes | Wellcome Sanger Institute for the COVID-19 Genomics UK (COG-UK) Consortium | The Lighthouse Lab in Milton Keynes and Alex Alderton, Roberto Amato, Jeffrey Barrett, Sonia Goncalves, Ewan Harrison, David K. Jackson, Ian Johnston, Dominic Kwiatkowski, Cordelia Langford, John Sillitoe on behalf of the Wellcome Sanger Institute COVID-19 Surveillance Team |
| EPI_ISL_3338055 | Health Services Laboratories | Wellcome Sanger Institute for the COVID-19 Genomics UK (COG-UK) Consortium | Health Services Laboratories and Alex Alderton, Roberto Amato, Jeffrey Barrett, Sonia Goncalves, Ewan Harrison, David K. Jackson, Ian Johnston, Dominic Kwiatkowski, Cordelia Langford, John Sillitoe on behalf of the Wellcome Sanger Institute COVID-19 Surveillance Team |
| EPI_ISL_3338056 | Lighthouse Laboratory Plymouth | Wellcome Sanger Institute for the COVID-19 Genomics UK (COG-UK) Consortium | Lighthouse Laboratory Plymouth and Alex Alderton, Roberto Amato, Jeffrey Barrett, Sonia Goncalves, Ewan Harrison, David K. Jackson, Ian Johnston, Dominic Kwiatkowski, Cordelia Langford, John Sillitoe on behalf of the Wellcome Sanger Institute COVID-19 Surveillance Team |
| EPI_ISL_3338057 | Lighthouse Lab in Milton Keynes | Wellcome Sanger Institute for the COVID-19 Genomics UK (COG-UK) Consortium | The Lighthouse Lab in Milton Keynes and Alex Alderton, Roberto Amato, Jeffrey Barrett, Sonia Goncalves, Ewan Harrison, David K. Jackson, Ian Johnston, Dominic Kwiatkowski, Cordelia Langford, John Sillitoe on behalf of the Wellcome Sanger Institute COVID-19 Surveillance Team |
| EPI_ISL_3338058 | Berkshire and Surrey Pathology Services Lighthouse Laboratory | Wellcome Sanger Institute for the COVID-19 Genomics UK (COG-UK) Consortium | Berkshire and Surrey Pathology Services Lighthouse Laboratory and Alex Alderton, Roberto Amato, Jeffrey Barrett, Sonia Goncalves, Ewan Harrison, David K. Jackson, Ian Johnston, Dominic Kwiatkowski, Cordelia Langford, John Sillitoe on behalf of the Wellcome Sanger Institute COVID-19 Surveillance Team |
| EPI_ISL_3338059, EPI_ISL_3338060, EPI_ISL_3338061, EPI_ISL_3338062 | Health Services Laboratories | Wellcome Sanger Institute for the COVID-19 Genomics UK (COG-UK) Consortium | Health Services Laboratories and Alex Alderton, Roberto Amato, Jeffrey Barrett, Sonia Goncalves, Ewan Harrison, David K. Jackson, Ian Johnston, Dominic Kwiatkowski, Cordelia Langford, John Sillitoe on behalf of the Wellcome Sanger Institute COVID-19 Surveillance Team |
| EPI_ISL_3338063 | Lighthouse Lab in Milton Keynes | Wellcome Sanger Institute for the COVID-19 Genomics UK (COG-UK) Consortium | The Lighthouse Lab in Milton Keynes and Alex Alderton, Roberto Amato, Jeffrey Barrett, Sonia Goncalves, Ewan Harrison, David K. Jackson, Ian Johnston, Dominic Kwiatkowski, Cordelia Langford, John Sillitoe on behalf of the Wellcome Sanger Institute COVID-19 Surveillance Team |
| EPI_ISL_3338064 | Health Services Laboratories | Wellcome Sanger Institute for the COVID-19 Genomics UK (COG-UK) Consortium | Health Services Laboratories and Alex Alderton, Roberto Amato, Jeffrey Barrett, Sonia Goncalves, Ewan Harrison, David K. Jackson, Ian Johnston, Dominic Kwiatkowski, Cordelia Langford, John Sillitoe on behalf of the Wellcome Sanger Institute COVID-19 Surveillance Team |
| EPI_ISL_3338065, EPI_ISL_3338066, EPI_ISL_3338067 | Lighthouse Laboratory Plymouth | Wellcome Sanger Institute for the COVID-19 Genomics UK (COG-UK) Consortium | Lighthouse Laboratory Plymouth and Alex Alderton, Roberto Amato, Jeffrey Barrett, Sonia Goncalves, Ewan Harrison, David K. Jackson, Ian Johnston, Dominic Kwiatkowski, Cordelia Langford, John Sillitoe on behalf of the Wellcome Sanger Institute COVID-19 Surveillance Team |
| EPI_ISL_3338068 | Berkshire and Surrey Pathology Services Lighthouse Laboratory | Wellcome Sanger Institute for the COVID-19 Genomics UK (COG-UK) Consortium | Berkshire and Surrey Pathology Services Lighthouse Laboratory and Alex Alderton, Roberto Amato, Jeffrey Barrett, Sonia Goncalves, Ewan Harrison, David K. Jackson, Ian Johnston, Dominic Kwiatkowski, Cordelia Langford, John Sillitoe on behalf of the Wellcome Sanger Institute COVID-19 Surveillance Team |
| EPI_ISL_3338069 | Lighthouse Laboratory Plymouth | Wellcome Sanger Institute for the COVID-19 Genomics UK (COG-UK) Consortium | Lighthouse Laboratory Plymouth and Alex Alderton, Roberto Amato, Jeffrey Barrett, Sonia Goncalves, Ewan Harrison, David K. Jackson, Ian Johnston, Dominic Kwiatkowski, Cordelia Langford, John Sillitoe on behalf of the Wellcome Sanger Institute COVID-19 Surveillance Team |
| EPI_ISL_3338070 | Berkshire and Surrey Pathology Services Lighthouse Laboratory | Wellcome Sanger Institute for the COVID-19 Genomics UK (COG-UK) Consortium | Berkshire and Surrey Pathology Services Lighthouse Laboratory and Alex Alderton, Roberto Amato, Jeffrey Barrett, Sonia Goncalves, Ewan Harrison, David K. Jackson, Ian Johnston, Dominic Kwiatkowski, Cordelia Langford, John Sillitoe on behalf of the Wellcome Sanger Institute COVID-19 Surveillance Team |
| EPI_ISL_3338071 | Health Services Laboratories | Wellcome Sanger Institute for the COVID-19 Genomics UK (COG-UK) Consortium | Health Services Laboratories and Alex Alderton, Roberto Amato, Jeffrey Barrett, Sonia Goncalves, Ewan Harrison, David K. Jackson, Ian Johnston, Dominic Kwiatkowski, Cordelia Langford, John Sillitoe on behalf of the Wellcome Sanger Institute COVID-19 Surveillance Team |
| EPI_ISL_3338072 | Berkshire and Surrey Pathology Services Lighthouse Laboratory | Wellcome Sanger Institute for the COVID-19 Genomics UK (COG-UK) Consortium | Berkshire and Surrey Pathology Services Lighthouse Laboratory and Alex Alderton, Roberto Amato, Jeffrey Barrett, Sonia Goncalves, Ewan Harrison, David K. Jackson, Ian Johnston, Dominic Kwiatkowski, Cordelia Langford, John Sillitoe on behalf of the Wellcome Sanger Institute COVID-19 Surveillance Team |
| EPI_ISL_3338073 | Health Services Laboratories | Wellcome Sanger Institute for the COVID-19 Genomics UK (COG-UK) Consortium | Health Services Laboratories and Alex Alderton, Roberto Amato, Jeffrey Barrett, Sonia Goncalves, Ewan Harrison, David K. Jackson, Ian Johnston, Dominic Kwiatkowski, Cordelia Langford, John Sillitoe on behalf of the Wellcome Sanger Institute COVID-19 Surveillance Team |
| EPI_ISL_3338074 | Lighthouse Laboratory Plymouth | Wellcome Sanger Institute for the COVID-19 Genomics UK (COG-UK) Consortium | Lighthouse Laboratory Plymouth and Alex Alderton, Roberto Amato, Jeffrey Barrett, Sonia Goncalves, Ewan Harrison, David K. Jackson, Ian Johnston, Dominic Kwiatkowski, Cordelia Langford, John Sillitoe on behalf of the Wellcome Sanger Institute COVID-19 Surveillance Team |
| EPI_ISL_3338075 | Berkshire and Surrey Pathology Services Lighthouse Laboratory | Wellcome Sanger Institute for the COVID-19 Genomics UK (COG-UK) Consortium | Berkshire and Surrey Pathology Services Lighthouse Laboratory and Alex Alderton, Roberto Amato, Jeffrey Barrett, Sonia Goncalves, Ewan Harrison, David K. Jackson, Ian Johnston, Dominic Kwiatkowski, Cordelia Langford, John Sillitoe on behalf of the Wellcome Sanger Institute COVID-19 Surveillance Team |
| EPI_ISL_3338076, | Lighthouse Laboratory Plymouth | Wellcome Sanger Institute for the COVID-19 Genomics UK (COG-UK) | Lighthouse Laboratory Plymouth and Alex Alderton, Roberto Amato, Jeffrey Barrett, Sonia Goncalves, Ewan Harrison, David K. Jackson, Ian Johnston, Dominic Kwiatkowski, Cordelia Langford, John Sillitoe on behalf of the Wellcome Sanger |

| | | | |
|---|---|---|---|
| EPI_ISL_3338077 | | Consortium | Institute COVID-19 Surveillance Team |
| EPI_ISL_3338078, EPI_ISL_3338079 | Lighthouse Lab in Milton Keynes | Wellcome Sanger Institute for the COVID-19 Genomics UK (COG-UK) Consortium | The Lighthouse Lab in Milton Keynes and Alex Alderton, Roberto Amato, Jeffrey Barrett, Sonia Goncalves, Ewan Harrison, David K. Jackson, Ian Johnston, Dominic Kwiatkowski, Cordelia Langford, John Sillitoe on behalf of the Wellcome Sanger Institute COVID-19 Surveillance Team |
| EPI_ISL_3338080 | Health Services Laboratories | Wellcome Sanger Institute for the COVID-19 Genomics UK (COG-UK) Consortium | Health Services Laboratories and Alex Alderton, Roberto Amato, Jeffrey Barrett, Sonia Goncalves, Ewan Harrison, David K. Jackson, Ian Johnston, Dominic Kwiatkowski, Cordelia Langford, John Sillitoe on behalf of the Wellcome Sanger Institute COVID-19 Surveillance Team |
| EPI_ISL_3338081 | Lighthouse Lab in Milton Keynes | Wellcome Sanger Institute for the COVID-19 Genomics UK (COG-UK) Consortium | The Lighthouse Lab in Milton Keynes and Alex Alderton, Roberto Amato, Jeffrey Barrett, Sonia Goncalves, Ewan Harrison, David K. Jackson, Ian Johnston, Dominic Kwiatkowski, Cordelia Langford, John Sillitoe on behalf of the Wellcome Sanger Institute COVID-19 Surveillance Team |
| EPI_ISL_3338082, EPI_ISL_3338083 | Health Services Laboratories | Wellcome Sanger Institute for the COVID-19 Genomics UK (COG-UK) Consortium | Health Services Laboratories and Alex Alderton, Roberto Amato, Jeffrey Barrett, Sonia Goncalves, Ewan Harrison, David K. Jackson, Ian Johnston, Dominic Kwiatkowski, Cordelia Langford, John Sillitoe on behalf of the Wellcome Sanger Institute COVID-19 Surveillance Team |
| EPI_ISL_3338084, EPI_ISL_3338085, EPI_ISL_3338086 | Lighthouse Lab in Milton Keynes | Wellcome Sanger Institute for the COVID-19 Genomics UK (COG-UK) Consortium | The Lighthouse Lab in Milton Keynes and Alex Alderton, Roberto Amato, Jeffrey Barrett, Sonia Goncalves, Ewan Harrison, David K. Jackson, Ian Johnston, Dominic Kwiatkowski, Cordelia Langford, John Sillitoe on behalf of the Wellcome Sanger Institute COVID-19 Surveillance Team |
| EPI_ISL_3338087 | Health Services Laboratories | Wellcome Sanger Institute for the COVID-19 Genomics UK (COG-UK) Consortium | Health Services Laboratories and Alex Alderton, Roberto Amato, Jeffrey Barrett, Sonia Goncalves, Ewan Harrison, David K. Jackson, Ian Johnston, Dominic Kwiatkowski, Cordelia Langford, John Sillitoe on behalf of the Wellcome Sanger Institute COVID-19 Surveillance Team |
| EPI_ISL_3338088, EPI_ISL_3338089, EPI_ISL_3338090 | Lighthouse Lab in Milton Keynes | Wellcome Sanger Institute for the COVID-19 Genomics UK (COG-UK) Consortium | The Lighthouse Lab in Milton Keynes and Alex Alderton, Roberto Amato, Jeffrey Barrett, Sonia Goncalves, Ewan Harrison, David K. Jackson, Ian Johnston, Dominic Kwiatkowski, Cordelia Langford, John Sillitoe on behalf of the Wellcome Sanger Institute COVID-19 Surveillance Team |
| EPI_ISL_3338091 | Lighthouse Laboratory Plymouth | Wellcome Sanger Institute for the COVID-19 Genomics UK (COG-UK) Consortium | Lighthouse Laboratory Plymouth and Alex Alderton, Roberto Amato, Jeffrey Barrett, Sonia Goncalves, Ewan Harrison, David K. Jackson, Ian Johnston, Dominic Kwiatkowski, Cordelia Langford, John Sillitoe on behalf of the Wellcome Sanger Institute COVID-19 Surveillance Team |
| EPI_ISL_3338092 | Lighthouse Lab in Milton Keynes | Wellcome Sanger Institute for the COVID-19 Genomics UK (COG-UK) Consortium | The Lighthouse Lab in Milton Keynes and Alex Alderton, Roberto Amato, Jeffrey Barrett, Sonia Goncalves, Ewan Harrison, David K. Jackson, Ian Johnston, Dominic Kwiatkowski, Cordelia Langford, John Sillitoe on behalf of the Wellcome Sanger Institute COVID-19 Surveillance Team |
| EPI_ISL_3338093 | Health Services Laboratories | Wellcome Sanger Institute for the COVID-19 Genomics UK (COG-UK) Consortium | Health Services Laboratories and Alex Alderton, Roberto Amato, Jeffrey Barrett, Sonia Goncalves, Ewan Harrison, David K. Jackson, Ian Johnston, Dominic Kwiatkowski, Cordelia Langford, John Sillitoe on behalf of the Wellcome Sanger Institute COVID-19 Surveillance Team |
| EPI_ISL_3338094 | Berkshire and Surrey Pathology Services Lighthouse Laboratory | Wellcome Sanger Institute for the COVID-19 Genomics UK (COG-UK) Consortium | Berkshire and Surrey Pathology Services Lighthouse Laboratory and Alex Alderton, Roberto Amato, Jeffrey Barrett, Sonia Goncalves, Ewan Harrison, David K. Jackson, Ian Johnston, Dominic Kwiatkowski, Cordelia Langford, John Sillitoe on behalf of the Wellcome Sanger Institute COVID-19 Surveillance Team |
| EPI_ISL_3338095, EPI_ISL_3338096 | Lighthouse Laboratory Plymouth | Wellcome Sanger Institute for the COVID-19 Genomics UK (COG-UK) Consortium | Lighthouse Laboratory Plymouth and Alex Alderton, Roberto Amato, Jeffrey Barrett, Sonia Goncalves, Ewan Harrison, David K. Jackson, Ian Johnston, Dominic Kwiatkowski, Cordelia Langford, John Sillitoe on behalf of the Wellcome Sanger Institute COVID-19 Surveillance Team |
| EPI_ISL_3338097 | Lighthouse Lab in Milton Keynes | Wellcome Sanger Institute for the COVID-19 Genomics UK (COG-UK) Consortium | The Lighthouse Lab in Milton Keynes and Alex Alderton, Roberto Amato, Jeffrey Barrett, Sonia Goncalves, Ewan Harrison, David K. Jackson, Ian Johnston, Dominic Kwiatkowski, Cordelia Langford, John Sillitoe on behalf of the Wellcome Sanger Institute COVID-19 Surveillance Team |
| EPI_ISL_3338098, EPI_ISL_3338099 | Lighthouse Laboratory Plymouth | Wellcome Sanger Institute for the COVID-19 Genomics UK (COG-UK) Consortium | Lighthouse Laboratory Plymouth and Alex Alderton, Roberto Amato, Jeffrey Barrett, Sonia Goncalves, Ewan Harrison, David K. Jackson, Ian Johnston, Dominic Kwiatkowski, Cordelia Langford, John Sillitoe on behalf of the Wellcome Sanger Institute COVID-19 Surveillance Team |
| EPI_ISL_3338100 | Lighthouse Lab in Milton Keynes | Wellcome Sanger Institute for the COVID-19 Genomics UK (COG-UK) Consortium | The Lighthouse Lab in Milton Keynes and Alex Alderton, Roberto Amato, Jeffrey Barrett, Sonia Goncalves, Ewan Harrison, David K. Jackson, Ian Johnston, Dominic Kwiatkowski, Cordelia Langford, John Sillitoe on behalf of the Wellcome Sanger Institute COVID-19 Surveillance Team |
| EPI_ISL_3338101 | Lighthouse Laboratory Plymouth | Wellcome Sanger Institute for the COVID-19 Genomics UK (COG-UK) Consortium | Lighthouse Laboratory Plymouth and Alex Alderton, Roberto Amato, Jeffrey Barrett, Sonia Goncalves, Ewan Harrison, David K. Jackson, Ian Johnston, Dominic Kwiatkowski, Cordelia Langford, John Sillitoe on behalf of the Wellcome Sanger Institute COVID-19 Surveillance Team |
| EPI_ISL_3338102 | Lighthouse Lab in Milton Keynes | Wellcome Sanger Institute for the COVID-19 Genomics UK (COG-UK) Consortium | The Lighthouse Lab in Milton Keynes and Alex Alderton, Roberto Amato, Jeffrey Barrett, Sonia Goncalves, Ewan Harrison, David K. Jackson, Ian Johnston, Dominic Kwiatkowski, Cordelia Langford, John Sillitoe on behalf of the Wellcome Sanger Institute COVID-19 Surveillance Team |
| EPI_ISL_3338103, EPI_ISL_3338104 | Berkshire and Surrey Pathology Services Lighthouse Laboratory | Wellcome Sanger Institute for the COVID-19 Genomics UK (COG-UK) Consortium | Berkshire and Surrey Pathology Services Lighthouse Laboratory and Alex Alderton, Roberto Amato, Jeffrey Barrett, Sonia Goncalves, Ewan Harrison, David K. Jackson, Ian Johnston, Dominic Kwiatkowski, Cordelia Langford, John Sillitoe on behalf of the Wellcome Sanger Institute COVID-19 Surveillance Team |
| EPI_ISL_3338105 | Lighthouse Laboratory Plymouth | Wellcome Sanger Institute for the COVID-19 Genomics UK (COG-UK) Consortium | Lighthouse Laboratory Plymouth and Alex Alderton, Roberto Amato, Jeffrey Barrett, Sonia Goncalves, Ewan Harrison, David K. Jackson, Ian Johnston, Dominic Kwiatkowski, Cordelia Langford, John Sillitoe on behalf of the Wellcome Sanger Institute COVID-19 Surveillance Team |
| EPI_ISL_3338106, EPI_ISL_3338107 | Health Services Laboratories | Wellcome Sanger Institute for the COVID-19 Genomics UK (COG-UK) Consortium | Health Services Laboratories and Alex Alderton, Roberto Amato, Jeffrey Barrett, Sonia Goncalves, Ewan Harrison, David K. Jackson, Ian Johnston, Dominic Kwiatkowski, Cordelia Langford, John Sillitoe on behalf of the Wellcome Sanger Institute COVID-19 Surveillance Team |
| EPI_ISL_3338108, EPI_ISL_3338109, EPI_ISL_3338110 | Lighthouse Laboratory Plymouth | Wellcome Sanger Institute for the COVID-19 Genomics UK (COG-UK) Consortium | Lighthouse Laboratory Plymouth and Alex Alderton, Roberto Amato, Jeffrey Barrett, Sonia Goncalves, Ewan Harrison, David K. Jackson, Ian Johnston, Dominic Kwiatkowski, Cordelia Langford, John Sillitoe on behalf of the Wellcome Sanger Institute COVID-19 Surveillance Team |
| EPI_ISL_3338111 | Health Services Laboratories | Wellcome Sanger Institute for the COVID-19 Genomics UK (COG-UK) Consortium | Health Services Laboratories and Alex Alderton, Roberto Amato, Jeffrey Barrett, Sonia Goncalves, Ewan Harrison, David K. Jackson, Ian Johnston, Dominic Kwiatkowski, Cordelia Langford, John Sillitoe on behalf of the Wellcome Sanger Institute COVID-19 Surveillance Team |
| EPI_ISL_3338112 | Lighthouse Lab in Milton Keynes | Wellcome Sanger Institute for the COVID-19 Genomics UK (COG-UK) Consortium | The Lighthouse Lab in Milton Keynes and Alex Alderton, Roberto Amato, Jeffrey Barrett, Sonia Goncalves, Ewan Harrison, David K. Jackson, Ian Johnston, Dominic Kwiatkowski, Cordelia Langford, John Sillitoe on behalf of the Wellcome Sanger Institute COVID-19 Surveillance Team |
| EPI_ISL_3338113 | Health Services Laboratories | Wellcome Sanger Institute for the COVID-19 Genomics UK (COG-UK) Consortium | Health Services Laboratories and Alex Alderton, Roberto Amato, Jeffrey Barrett, Sonia Goncalves, Ewan Harrison, David K. Jackson, Ian Johnston, Dominic Kwiatkowski, Cordelia Langford, John Sillitoe on behalf of the Wellcome Sanger Institute COVID-19 Surveillance Team |
| EPI_ISL_3338114, EPI_ISL_3338115, EPI_ISL_3338116 | Lighthouse Laboratory Plymouth | Wellcome Sanger Institute for the COVID-19 Genomics UK (COG-UK) Consortium | Lighthouse Laboratory Plymouth and Alex Alderton, Roberto Amato, Jeffrey Barrett, Sonia Goncalves, Ewan Harrison, David K. Jackson, Ian Johnston, Dominic Kwiatkowski, Cordelia Langford, John Sillitoe on behalf of the Wellcome Sanger Institute COVID-19 Surveillance Team |
| EPI_ISL_3338117 | Health Services Laboratories | Wellcome Sanger Institute for the COVID-19 Genomics UK (COG-UK) Consortium | Health Services Laboratories and Alex Alderton, Roberto Amato, Jeffrey Barrett, Sonia Goncalves, Ewan Harrison, David K. Jackson, Ian Johnston, Dominic Kwiatkowski, Cordelia Langford, John Sillitoe on behalf of the Wellcome Sanger Institute COVID-19 Surveillance Team |
| EPI_ISL_3338118 | Lighthouse Lab in Milton Keynes | Wellcome Sanger Institute for the COVID-19 Genomics UK (COG-UK) Consortium | The Lighthouse Lab in Milton Keynes and Alex Alderton, Roberto Amato, Jeffrey Barrett, Sonia Goncalves, Ewan Harrison, David K. Jackson, Ian Johnston, Dominic Kwiatkowski, Cordelia Langford, John Sillitoe on behalf of the Wellcome Sanger Institute COVID-19 Surveillance Team |
| EPI_ISL_3338119, EPI_ISL_3338120, EPI_ISL_3338121 | Lighthouse Laboratory Plymouth | Wellcome Sanger Institute for the COVID-19 Genomics UK (COG-UK) Consortium | Lighthouse Laboratory Plymouth and Alex Alderton, Roberto Amato, Jeffrey Barrett, Sonia Goncalves, Ewan Harrison, David K. Jackson, Ian Johnston, Dominic Kwiatkowski, Cordelia Langford, John Sillitoe on behalf of the Wellcome Sanger Institute COVID-19 Surveillance Team |
| EPI_ISL_3338122 | Lighthouse Lab in Milton Keynes | Wellcome Sanger Institute for the COVID-19 Genomics UK (COG-UK) Consortium | The Lighthouse Lab in Milton Keynes and Alex Alderton, Roberto Amato, Jeffrey Barrett, Sonia Goncalves, Ewan Harrison, David K. Jackson, Ian Johnston, Dominic Kwiatkowski, Cordelia Langford, John Sillitoe on behalf of the Wellcome Sanger Institute COVID-19 Surveillance Team |
| EPI_ISL_3338123 | Berkshire and Surrey Pathology Services Lighthouse Laboratory | Wellcome Sanger Institute for the COVID-19 Genomics UK (COG-UK) Consortium | Berkshire and Surrey Pathology Services Lighthouse Laboratory and Alex Alderton, Roberto Amato, Jeffrey Barrett, Sonia Goncalves, Ewan Harrison, David K. Jackson, Ian Johnston, Dominic Kwiatkowski, Cordelia Langford, John Sillitoe on behalf of the Wellcome Sanger Institute COVID-19 Surveillance Team |
| EPI_ISL_3338124, EPI_ISL_3338125 | Lighthouse Laboratory Plymouth | Wellcome Sanger Institute for the COVID-19 Genomics UK (COG-UK) Consortium | Lighthouse Laboratory Plymouth and Alex Alderton, Roberto Amato, Jeffrey Barrett, Sonia Goncalves, Ewan Harrison, David K. Jackson, Ian Johnston, Dominic Kwiatkowski, Cordelia Langford, John Sillitoe on behalf of the Wellcome Sanger Institute COVID-19 Surveillance Team |
| EPI_ISL_3338126 | Lighthouse Lab in Milton Keynes | Wellcome Sanger Institute for the COVID-19 Genomics UK (COG-UK) Consortium | The Lighthouse Lab in Milton Keynes and Alex Alderton, Roberto Amato, Jeffrey Barrett, Sonia Goncalves, Ewan Harrison, David K. Jackson, Ian Johnston, Dominic Kwiatkowski, Cordelia Langford, John Sillitoe on behalf of the Wellcome Sanger Institute COVID-19 Surveillance Team |
| EPI_ISL_3338127 | Health Services Laboratories | Wellcome Sanger Institute for the COVID-19 Genomics UK (COG-UK) Consortium | Health Services Laboratories and Alex Alderton, Roberto Amato, Jeffrey Barrett, Sonia Goncalves, Ewan Harrison, David K. Jackson, Ian Johnston, Dominic Kwiatkowski, Cordelia Langford, John Sillitoe on behalf of the Wellcome Sanger Institute COVID-19 Surveillance Team |
| EPI_ISL_3338128 | Lighthouse Lab in Milton Keynes | Wellcome Sanger Institute for the COVID-19 Genomics UK (COG-UK) Consortium | The Lighthouse Lab in Milton Keynes and Alex Alderton, Roberto Amato, Jeffrey Barrett, Sonia Goncalves, Ewan Harrison, David K. Jackson, Ian Johnston, Dominic Kwiatkowski, Cordelia Langford, John Sillitoe on behalf of the Wellcome Sanger Institute COVID-19 Surveillance Team |
| EPI_ISL_3338129 | Health Services Laboratories | Wellcome Sanger Institute for the COVID-19 Genomics UK (COG-UK) Consortium | Health Services Laboratories and Alex Alderton, Roberto Amato, Jeffrey Barrett, Sonia Goncalves, Ewan Harrison, David K. Jackson, Ian Johnston, Dominic Kwiatkowski, Cordelia Langford, John Sillitoe on behalf of the Wellcome Sanger Institute COVID-19 Surveillance Team |
| EPI_ISL_3338130 | Lighthouse Laboratory Plymouth | Wellcome Sanger Institute for the COVID-19 Genomics UK (COG-UK) Consortium | Lighthouse Laboratory Plymouth and Alex Alderton, Roberto Amato, Jeffrey Barrett, Sonia Goncalves, Ewan Harrison, David K. Jackson, Ian Johnston, Dominic Kwiatkowski, Cordelia Langford, John Sillitoe on behalf of the Wellcome Sanger Institute COVID-19 Surveillance Team |
| EPI_ISL_3338131 | Health Services Laboratories | Wellcome Sanger Institute for the COVID-19 Genomics UK (COG-UK) Consortium | Health Services Laboratories and Alex Alderton, Roberto Amato, Jeffrey Barrett, Sonia Goncalves, Ewan Harrison, David K. Jackson, Ian Johnston, Dominic Kwiatkowski, Cordelia Langford, John Sillitoe on behalf of the Wellcome Sanger Institute COVID-19 Surveillance Team |
| EPI_ISL_3338132 | Berkshire and Surrey Pathology Services Lighthouse Laboratory | Wellcome Sanger Institute for the COVID-19 Genomics UK (COG-UK) Consortium | Berkshire and Surrey Pathology Services Lighthouse Laboratory and Alex Alderton, Roberto Amato, Jeffrey Barrett, Sonia Goncalves, Ewan Harrison, David K. Jackson, Ian Johnston, Dominic Kwiatkowski, Cordelia Langford, John Sillitoe on behalf of the Wellcome Sanger Institute COVID-19 Surveillance Team |
| EPI_ISL_3338133, EPI_ISL_3338134 | Lighthouse Laboratory Plymouth | Wellcome Sanger Institute for the COVID-19 Genomics UK (COG-UK) Consortium | Lighthouse Laboratory Plymouth and Alex Alderton, Roberto Amato, Jeffrey Barrett, Sonia Goncalves, Ewan Harrison, David K. Jackson, Ian Johnston, Dominic Kwiatkowski, Cordelia Langford, John Sillitoe on behalf of the Wellcome Sanger Institute COVID-19 Surveillance Team |
| EPI_ISL_3338135 | Health Services Laboratories | Wellcome Sanger Institute for the COVID-19 Genomics UK (COG-UK) Consortium | Health Services Laboratories and Alex Alderton, Roberto Amato, Jeffrey Barrett, Sonia Goncalves, Ewan Harrison, David K. Jackson, Ian Johnston, Dominic Kwiatkowski, Cordelia Langford, John Sillitoe on behalf of the Wellcome Sanger Institute COVID-19 Surveillance Team |
| EPI_ISL_3338136 | Lighthouse Lab in Milton Keynes | Wellcome Sanger Institute for the COVID-19 Genomics UK (COG-UK) Consortium | The Lighthouse Lab in Milton Keynes and Alex Alderton, Roberto Amato, Jeffrey Barrett, Sonia Goncalves, Ewan Harrison, David K. Jackson, Ian Johnston, Dominic Kwiatkowski, Cordelia Langford, John Sillitoe on behalf of the Wellcome Sanger Institute COVID-19 Surveillance Team |
| EPI_ISL_3338137 | Berkshire and Surrey Pathology Services Lighthouse Laboratory | Wellcome Sanger Institute for the COVID-19 Genomics UK (COG-UK) Consortium | Berkshire and Surrey Pathology Services Lighthouse Laboratory and Alex Alderton, Roberto Amato, Jeffrey Barrett, Sonia Goncalves, Ewan Harrison, David K. Jackson, Ian Johnston, Dominic Kwiatkowski, Cordelia Langford, John Sillitoe on behalf of the Wellcome Sanger Institute COVID-19 Surveillance Team |
| EPI_ISL_3338138, | Lighthouse Laboratory Plymouth | Wellcome Sanger Institute for the COVID-19 Genomics UK (COG-UK) | Lighthouse Laboratory Plymouth and Alex Alderton, Roberto Amato, Jeffrey Barrett, Sonia Goncalves, Ewan Harrison, David K. Jackson, Ian Johnston, Dominic Kwiatkowski, Cordelia Langford, John Sillitoe on behalf of the Wellcome Sanger |

| | | | |
|---|---|---|---|
| EPI_ISL_3338139 | | Consortium | Institute COVID-19 Surveillance Team |
| EPI_ISL_3338140 | Lighthouse Lab in Milton Keynes | Wellcome Sanger Institute for the COVID-19 Genomics UK (COG-UK) Consortium | The Lighthouse Lab in Milton Keynes and Alex Alderton, Roberto Amato, Jeffrey Barrett, Sonia Goncalves, Ewan Harrison, David K. Jackson, Ian Johnston, Dominic Kwiatkowski, Cordelia Langford, John Sillitoe on behalf of the Wellcome Sanger Institute COVID-19 Surveillance Team |
| EPI_ISL_3338141 | Health Services Laboratories | Wellcome Sanger Institute for the COVID-19 Genomics UK (COG-UK) Consortium | Health Services Laboratories and Alex Alderton, Roberto Amato, Jeffrey Barrett, Sonia Goncalves, Ewan Harrison, David K. Jackson, Ian Johnston, Dominic Kwiatkowski, Cordelia Langford, John Sillitoe on behalf of the Wellcome Sanger Institute COVID-19 Surveillance Team |
| EPI_ISL_3338142 | Lighthouse Lab in Milton Keynes | Wellcome Sanger Institute for the COVID-19 Genomics UK (COG-UK) Consortium | The Lighthouse Lab in Milton Keynes and Alex Alderton, Roberto Amato, Jeffrey Barrett, Sonia Goncalves, Ewan Harrison, David K. Jackson, Ian Johnston, Dominic Kwiatkowski, Cordelia Langford, John Sillitoe on behalf of the Wellcome Sanger Institute COVID-19 Surveillance Team |
| EPI_ISL_3338143 | Lighthouse Laboratory Plymouth | Wellcome Sanger Institute for the COVID-19 Genomics UK (COG-UK) Consortium | Lighthouse Laboratory Plymouth and Alex Alderton, Roberto Amato, Jeffrey Barrett, Sonia Goncalves, Ewan Harrison, David K. Jackson, Ian Johnston, Dominic Kwiatkowski, Cordelia Langford, John Sillitoe on behalf of the Wellcome Sanger Institute COVID-19 Surveillance Team |
| EPI_ISL_3338144 | Lighthouse Lab in Milton Keynes | Wellcome Sanger Institute for the COVID-19 Genomics UK (COG-UK) Consortium | The Lighthouse Lab in Milton Keynes and Alex Alderton, Roberto Amato, Jeffrey Barrett, Sonia Goncalves, Ewan Harrison, David K. Jackson, Ian Johnston, Dominic Kwiatkowski, Cordelia Langford, John Sillitoe on behalf of the Wellcome Sanger Institute COVID-19 Surveillance Team |
| EPI_ISL_3338145 | Lighthouse Laboratory Plymouth | Wellcome Sanger Institute for the COVID-19 Genomics UK (COG-UK) Consortium | Lighthouse Laboratory Plymouth and Alex Alderton, Roberto Amato, Jeffrey Barrett, Sonia Goncalves, Ewan Harrison, David K. Jackson, Ian Johnston, Dominic Kwiatkowski, Cordelia Langford, John Sillitoe on behalf of the Wellcome Sanger Institute COVID-19 Surveillance Team |
| EPI_ISL_3338146, EPI_ISL_3338147, EPI_ISL_3338148, EPI_ISL_3338149, EPI_ISL_3338150, EPI_ISL_3338151 | Lighthouse Lab in Milton Keynes | Wellcome Sanger Institute for the COVID-19 Genomics UK (COG-UK) Consortium | The Lighthouse Lab in Milton Keynes and Alex Alderton, Roberto Amato, Jeffrey Barrett, Sonia Goncalves, Ewan Harrison, David K. Jackson, Ian Johnston, Dominic Kwiatkowski, Cordelia Langford, John Sillitoe on behalf of the Wellcome Sanger Institute COVID-19 Surveillance Team |
| EPI_ISL_3338152 | Health Services Laboratories | Wellcome Sanger Institute for the COVID-19 Genomics UK (COG-UK) Consortium | Health Services Laboratories and Alex Alderton, Roberto Amato, Jeffrey Barrett, Sonia Goncalves, Ewan Harrison, David K. Jackson, Ian Johnston, Dominic Kwiatkowski, Cordelia Langford, John Sillitoe on behalf of the Wellcome Sanger Institute COVID-19 Surveillance Team |
| EPI_ISL_3338153 | Lighthouse Lab in Milton Keynes | Wellcome Sanger Institute for the COVID-19 Genomics UK (COG-UK) Consortium | The Lighthouse Lab in Milton Keynes and Alex Alderton, Roberto Amato, Jeffrey Barrett, Sonia Goncalves, Ewan Harrison, David K. Jackson, Ian Johnston, Dominic Kwiatkowski, Cordelia Langford, John Sillitoe on behalf of the Wellcome Sanger Institute COVID-19 Surveillance Team |
| EPI_ISL_3338154 | Berkshire and Surrey Pathology Services Lighthouse Laboratory | Wellcome Sanger Institute for the COVID-19 Genomics UK (COG-UK) Consortium | Berkshire and Surrey Pathology Services Lighthouse Laboratory and Alex Alderton, Roberto Amato, Jeffrey Barrett, Sonia Goncalves, Ewan Harrison, David K. Jackson, Ian Johnston, Dominic Kwiatkowski, Cordelia Langford, John Sillitoe on behalf of the Wellcome Sanger Institute COVID-19 Surveillance Team |
| EPI_ISL_3338155 | Lighthouse Laboratory Plymouth | Wellcome Sanger Institute for the COVID-19 Genomics UK (COG-UK) Consortium | Lighthouse Laboratory Plymouth and Alex Alderton, Roberto Amato, Jeffrey Barrett, Sonia Goncalves, Ewan Harrison, David K. Jackson, Ian Johnston, Dominic Kwiatkowski, Cordelia Langford, John Sillitoe on behalf of the Wellcome Sanger Institute COVID-19 Surveillance Team |
| EPI_ISL_3338156 | Health Services Laboratories | Wellcome Sanger Institute for the COVID-19 Genomics UK (COG-UK) Consortium | Health Services Laboratories and Alex Alderton, Roberto Amato, Jeffrey Barrett, Sonia Goncalves, Ewan Harrison, David K. Jackson, Ian Johnston, Dominic Kwiatkowski, Cordelia Langford, John Sillitoe on behalf of the Wellcome Sanger Institute COVID-19 Surveillance Team |
| EPI_ISL_3338157, EPI_ISL_3338158 | Lighthouse Lab in Milton Keynes | Wellcome Sanger Institute for the COVID-19 Genomics UK (COG-UK) Consortium | The Lighthouse Lab in Milton Keynes and Alex Alderton, Roberto Amato, Jeffrey Barrett, Sonia Goncalves, Ewan Harrison, David K. Jackson, Ian Johnston, Dominic Kwiatkowski, Cordelia Langford, John Sillitoe on behalf of the Wellcome Sanger Institute COVID-19 Surveillance Team |
| EPI_ISL_3338159 | Lighthouse Laboratory Plymouth | Wellcome Sanger Institute for the COVID-19 Genomics UK (COG-UK) Consortium | Lighthouse Laboratory Plymouth and Alex Alderton, Roberto Amato, Jeffrey Barrett, Sonia Goncalves, Ewan Harrison, David K. Jackson, Ian Johnston, Dominic Kwiatkowski, Cordelia Langford, John Sillitoe on behalf of the Wellcome Sanger Institute COVID-19 Surveillance Team |
| EPI_ISL_3338160 | Lighthouse Lab in Milton Keynes | Wellcome Sanger Institute for the COVID-19 Genomics UK (COG-UK) Consortium | The Lighthouse Lab in Milton Keynes and Alex Alderton, Roberto Amato, Jeffrey Barrett, Sonia Goncalves, Ewan Harrison, David K. Jackson, Ian Johnston, Dominic Kwiatkowski, Cordelia Langford, John Sillitoe on behalf of the Wellcome Sanger Institute COVID-19 Surveillance Team |
| EPI_ISL_3338161, EPI_ISL_3338162 | Health Services Laboratories | Wellcome Sanger Institute for the COVID-19 Genomics UK (COG-UK) Consortium | Health Services Laboratories and Alex Alderton, Roberto Amato, Jeffrey Barrett, Sonia Goncalves, Ewan Harrison, David K. Jackson, Ian Johnston, Dominic Kwiatkowski, Cordelia Langford, John Sillitoe on behalf of the Wellcome Sanger Institute COVID-19 Surveillance Team |
| EPI_ISL_3338163 | Berkshire and Surrey Pathology Services Lighthouse Laboratory | Wellcome Sanger Institute for the COVID-19 Genomics UK (COG-UK) Consortium | Berkshire and Surrey Pathology Services Lighthouse Laboratory and Alex Alderton, Roberto Amato, Jeffrey Barrett, Sonia Goncalves, Ewan Harrison, David K. Jackson, Ian Johnston, Dominic Kwiatkowski, Cordelia Langford, John Sillitoe on behalf of the Wellcome Sanger Institute COVID-19 Surveillance Team |
| EPI_ISL_3338164 | Health Services Laboratories | Wellcome Sanger Institute for the COVID-19 Genomics UK (COG-UK) Consortium | Health Services Laboratories and Alex Alderton, Roberto Amato, Jeffrey Barrett, Sonia Goncalves, Ewan Harrison, David K. Jackson, Ian Johnston, Dominic Kwiatkowski, Cordelia Langford, John Sillitoe on behalf of the Wellcome Sanger Institute COVID-19 Surveillance Team |
| EPI_ISL_3338165 | Lighthouse Lab in Milton Keynes | Wellcome Sanger Institute for the COVID-19 Genomics UK (COG-UK) Consortium | The Lighthouse Lab in Milton Keynes and Alex Alderton, Roberto Amato, Jeffrey Barrett, Sonia Goncalves, Ewan Harrison, David K. Jackson, Ian Johnston, Dominic Kwiatkowski, Cordelia Langford, John Sillitoe on behalf of the Wellcome Sanger Institute COVID-19 Surveillance Team |
| EPI_ISL_3338166 | Lighthouse Laboratory Plymouth | Wellcome Sanger Institute for the COVID-19 Genomics UK (COG-UK) Consortium | Lighthouse Laboratory Plymouth and Alex Alderton, Roberto Amato, Jeffrey Barrett, Sonia Goncalves, Ewan Harrison, David K. Jackson, Ian Johnston, Dominic Kwiatkowski, Cordelia Langford, John Sillitoe on behalf of the Wellcome Sanger Institute COVID-19 Surveillance Team |
| EPI_ISL_3338167 | Health Services Laboratories | Wellcome Sanger Institute for the COVID-19 Genomics UK (COG-UK) Consortium | Health Services Laboratories and Alex Alderton, Roberto Amato, Jeffrey Barrett, Sonia Goncalves, Ewan Harrison, David K. Jackson, Ian Johnston, Dominic Kwiatkowski, Cordelia Langford, John Sillitoe on behalf of the Wellcome Sanger Institute COVID-19 Surveillance Team |
| EPI_ISL_3338168 | Lighthouse Laboratory Plymouth | Wellcome Sanger Institute for the COVID-19 Genomics UK (COG-UK) Consortium | Lighthouse Laboratory Plymouth and Alex Alderton, Roberto Amato, Jeffrey Barrett, Sonia Goncalves, Ewan Harrison, David K. Jackson, Ian Johnston, Dominic Kwiatkowski, Cordelia Langford, John Sillitoe on behalf of the Wellcome Sanger Institute COVID-19 Surveillance Team |
| EPI_ISL_3338169, EPI_ISL_3338170 | Health Services Laboratories | Wellcome Sanger Institute for the COVID-19 Genomics UK (COG-UK) Consortium | Health Services Laboratories and Alex Alderton, Roberto Amato, Jeffrey Barrett, Sonia Goncalves, Ewan Harrison, David K. Jackson, Ian Johnston, Dominic Kwiatkowski, Cordelia Langford, John Sillitoe on behalf of the Wellcome Sanger Institute COVID-19 Surveillance Team |
| EPI_ISL_3338171 | Lighthouse Laboratory Plymouth | Wellcome Sanger Institute for the COVID-19 Genomics UK (COG-UK) Consortium | Lighthouse Laboratory Plymouth and Alex Alderton, Roberto Amato, Jeffrey Barrett, Sonia Goncalves, Ewan Harrison, David K. Jackson, Ian Johnston, Dominic Kwiatkowski, Cordelia Langford, John Sillitoe on behalf of the Wellcome Sanger Institute COVID-19 Surveillance Team |
| EPI_ISL_3338172 | Health Services Laboratories | Wellcome Sanger Institute for the COVID-19 Genomics UK (COG-UK) Consortium | Health Services Laboratories and Alex Alderton, Roberto Amato, Jeffrey Barrett, Sonia Goncalves, Ewan Harrison, David K. Jackson, Ian Johnston, Dominic Kwiatkowski, Cordelia Langford, John Sillitoe on behalf of the Wellcome Sanger Institute COVID-19 Surveillance Team |
| EPI_ISL_3338173, EPI_ISL_3338174 | Lighthouse Lab in Milton Keynes | Wellcome Sanger Institute for the COVID-19 Genomics UK (COG-UK) Consortium | The Lighthouse Lab in Milton Keynes and Alex Alderton, Roberto Amato, Jeffrey Barrett, Sonia Goncalves, Ewan Harrison, David K. Jackson, Ian Johnston, Dominic Kwiatkowski, Cordelia Langford, John Sillitoe on behalf of the Wellcome Sanger Institute COVID-19 Surveillance Team |
| EPI_ISL_3338175 | Lighthouse Laboratory Plymouth | Wellcome Sanger Institute for the COVID-19 Genomics UK (COG-UK) Consortium | Lighthouse Laboratory Plymouth and Alex Alderton, Roberto Amato, Jeffrey Barrett, Sonia Goncalves, Ewan Harrison, David K. Jackson, Ian Johnston, Dominic Kwiatkowski, Cordelia Langford, John Sillitoe on behalf of the Wellcome Sanger Institute COVID-19 Surveillance Team |
| EPI_ISL_3338176 | Lighthouse Lab in Milton Keynes | Wellcome Sanger Institute for the COVID-19 Genomics UK (COG-UK) Consortium | The Lighthouse Lab in Milton Keynes and Alex Alderton, Roberto Amato, Jeffrey Barrett, Sonia Goncalves, Ewan Harrison, David K. Jackson, Ian Johnston, Dominic Kwiatkowski, Cordelia Langford, John Sillitoe on behalf of the Wellcome Sanger Institute COVID-19 Surveillance Team |
| EPI_ISL_3338177 | Lighthouse Laboratory Plymouth | Wellcome Sanger Institute for the COVID-19 Genomics UK (COG-UK) Consortium | Lighthouse Laboratory Plymouth and Alex Alderton, Roberto Amato, Jeffrey Barrett, Sonia Goncalves, Ewan Harrison, David K. Jackson, Ian Johnston, Dominic Kwiatkowski, Cordelia Langford, John Sillitoe on behalf of the Wellcome Sanger Institute COVID-19 Surveillance Team |
| EPI_ISL_3338178 | Health Services Laboratories | Wellcome Sanger Institute for the COVID-19 Genomics UK (COG-UK) Consortium | Health Services Laboratories and Alex Alderton, Roberto Amato, Jeffrey Barrett, Sonia Goncalves, Ewan Harrison, David K. Jackson, Ian Johnston, Dominic Kwiatkowski, Cordelia Langford, John Sillitoe on behalf of the Wellcome Sanger Institute COVID-19 Surveillance Team |
| EPI_ISL_3338179 | Lighthouse Laboratory Plymouth | Wellcome Sanger Institute for the COVID-19 Genomics UK (COG-UK) Consortium | Lighthouse Laboratory Plymouth and Alex Alderton, Roberto Amato, Jeffrey Barrett, Sonia Goncalves, Ewan Harrison, David K. Jackson, Ian Johnston, Dominic Kwiatkowski, Cordelia Langford, John Sillitoe on behalf of the Wellcome Sanger Institute COVID-19 Surveillance Team |
| EPI_ISL_3338180 | Health Services Laboratories | Wellcome Sanger Institute for the COVID-19 Genomics UK (COG-UK) Consortium | Health Services Laboratories and Alex Alderton, Roberto Amato, Jeffrey Barrett, Sonia Goncalves, Ewan Harrison, David K. Jackson, Ian Johnston, Dominic Kwiatkowski, Cordelia Langford, John Sillitoe on behalf of the Wellcome Sanger Institute COVID-19 Surveillance Team |
| EPI_ISL_3338181 | Lighthouse Lab in Milton Keynes | Wellcome Sanger Institute for the COVID-19 Genomics UK (COG-UK) Consortium | The Lighthouse Lab in Milton Keynes and Alex Alderton, Roberto Amato, Jeffrey Barrett, Sonia Goncalves, Ewan Harrison, David K. Jackson, Ian Johnston, Dominic Kwiatkowski, Cordelia Langford, John Sillitoe on behalf of the Wellcome Sanger Institute COVID-19 Surveillance Team |
| EPI_ISL_3338182 | Berkshire and Surrey Pathology Services Lighthouse Laboratory | Wellcome Sanger Institute for the COVID-19 Genomics UK (COG-UK) Consortium | Berkshire and Surrey Pathology Services Lighthouse Laboratory and Alex Alderton, Roberto Amato, Jeffrey Barrett, Sonia Goncalves, Ewan Harrison, David K. Jackson, Ian Johnston, Dominic Kwiatkowski, Cordelia Langford, John Sillitoe on behalf of the Wellcome Sanger Institute COVID-19 Surveillance Team |
| EPI_ISL_3338183 | Health Services Laboratories | Wellcome Sanger Institute for the COVID-19 Genomics UK (COG-UK) Consortium | Health Services Laboratories and Alex Alderton, Roberto Amato, Jeffrey Barrett, Sonia Goncalves, Ewan Harrison, David K. Jackson, Ian Johnston, Dominic Kwiatkowski, Cordelia Langford, John Sillitoe on behalf of the Wellcome Sanger Institute COVID-19 Surveillance Team |
| EPI_ISL_3338184, EPI_ISL_3338185 | Lighthouse Lab in Milton Keynes | Wellcome Sanger Institute for the COVID-19 Genomics UK (COG-UK) Consortium | The Lighthouse Lab in Milton Keynes and Alex Alderton, Roberto Amato, Jeffrey Barrett, Sonia Goncalves, Ewan Harrison, David K. Jackson, Ian Johnston, Dominic Kwiatkowski, Cordelia Langford, John Sillitoe on behalf of the Wellcome Sanger Institute COVID-19 Surveillance Team |
| EPI_ISL_3338186 | Lighthouse Laboratory Plymouth | Wellcome Sanger Institute for the COVID-19 Genomics UK (COG-UK) Consortium | Lighthouse Laboratory Plymouth and Alex Alderton, Roberto Amato, Jeffrey Barrett, Sonia Goncalves, Ewan Harrison, David K. Jackson, Ian Johnston, Dominic Kwiatkowski, Cordelia Langford, John Sillitoe on behalf of the Wellcome Sanger Institute COVID-19 Surveillance Team |
| EPI_ISL_3338187 | Lighthouse Lab in Milton Keynes | Wellcome Sanger Institute for the COVID-19 Genomics UK (COG-UK) Consortium | The Lighthouse Lab in Milton Keynes and Alex Alderton, Roberto Amato, Jeffrey Barrett, Sonia Goncalves, Ewan Harrison, David K. Jackson, Ian Johnston, Dominic Kwiatkowski, Cordelia Langford, John Sillitoe on behalf of the Wellcome Sanger Institute COVID-19 Surveillance Team |
| EPI_ISL_3338188, EPI_ISL_3338189 | Lighthouse Laboratory Plymouth | Wellcome Sanger Institute for the COVID-19 Genomics UK (COG-UK) Consortium | Lighthouse Laboratory Plymouth and Alex Alderton, Roberto Amato, Jeffrey Barrett, Sonia Goncalves, Ewan Harrison, David K. Jackson, Ian Johnston, Dominic Kwiatkowski, Cordelia Langford, John Sillitoe on behalf of the Wellcome Sanger Institute COVID-19 Surveillance Team |
| EPI_ISL_3338190 | Lighthouse Lab in Milton Keynes | Wellcome Sanger Institute for the COVID-19 Genomics UK (COG-UK) Consortium | The Lighthouse Lab in Milton Keynes and Alex Alderton, Roberto Amato, Jeffrey Barrett, Sonia Goncalves, Ewan Harrison, David K. Jackson, Ian Johnston, Dominic Kwiatkowski, Cordelia Langford, John Sillitoe on behalf of the Wellcome Sanger Institute COVID-19 Surveillance Team |
| EPI_ISL_3338191 | Lighthouse Laboratory Plymouth | Wellcome Sanger Institute for the COVID-19 Genomics UK (COG-UK) Consortium | Lighthouse Laboratory Plymouth and Alex Alderton, Roberto Amato, Jeffrey Barrett, Sonia Goncalves, Ewan Harrison, David K. Jackson, Ian Johnston, Dominic Kwiatkowski, Cordelia Langford, John Sillitoe on behalf of the Wellcome Sanger Institute COVID-19 Surveillance Team |
| EPI_ISL_3338192 | Health Services Laboratories | Wellcome Sanger Institute for the COVID-19 Genomics UK (COG-UK) Consortium | Health Services Laboratories and Alex Alderton, Roberto Amato, Jeffrey Barrett, Sonia Goncalves, Ewan Harrison, David K. Jackson, Ian Johnston, Dominic Kwiatkowski, Cordelia Langford, John Sillitoe on behalf of the Wellcome Sanger Institute COVID-19 Surveillance Team |
| EPI_ISL_3338193 | Berkshire and Surrey Pathology Services Lighthouse Laboratory | Wellcome Sanger Institute for the COVID-19 Genomics UK (COG-UK) Consortium | Berkshire and Surrey Pathology Services Lighthouse Laboratory and Alex Alderton, Roberto Amato, Jeffrey Barrett, Sonia Goncalves, Ewan Harrison, David K. Jackson, Ian Johnston, Dominic Kwiatkowski, Cordelia Langford, John Sillitoe on behalf of the Wellcome Sanger Institute COVID-19 Surveillance Team |

| | | | |
|---|---|---|---|
| EPI_ISL_3338194, EPI_ISL_3338195 | Lighthouse Laboratory Plymouth | Wellcome Sanger Institute for the COVID-19 Genomics UK (COG-UK) Consortium | Lighthouse Laboratory Plymouth and Alex Alderton, Roberto Amato, Jeffrey Barrett, Sonia Goncalves, Ewan Harrison, David K. Jackson, Ian Johnston, Dominic Kwiatkowski, Cordelia Langford, John Sillitoe on behalf of the Wellcome Sanger Institute COVID-19 Surveillance Team |
| EPI_ISL_3338196 | Lighthouse Lab in Milton Keynes | Wellcome Sanger Institute for the COVID-19 Genomics UK (COG-UK) Consortium | The Lighthouse Lab in Milton Keynes and Alex Alderton, Roberto Amato, Jeffrey Barrett, Sonia Goncalves, Ewan Harrison, David K. Jackson, Ian Johnston, Dominic Kwiatkowski, Cordelia Langford, John Sillitoe on behalf of the Wellcome Sanger Institute COVID-19 Surveillance Team |
| EPI_ISL_3338197 | Lighthouse Laboratory Plymouth | Wellcome Sanger Institute for the COVID-19 Genomics UK (COG-UK) Consortium | Lighthouse Laboratory Plymouth and Alex Alderton, Roberto Amato, Jeffrey Barrett, Sonia Goncalves, Ewan Harrison, David K. Jackson, Ian Johnston, Dominic Kwiatkowski, Cordelia Langford, John Sillitoe on behalf of the Wellcome Sanger Institute COVID-19 Surveillance Team |
| EPI_ISL_3338198 | Berkshire and Surrey Pathology Services Lighthouse Laboratory | Wellcome Sanger Institute for the COVID-19 Genomics UK (COG-UK) Consortium | Berkshire and Surrey Pathology Services Lighthouse Laboratory and Alex Alderton, Roberto Amato, Jeffrey Barrett, Sonia Goncalves, Ewan Harrison, David K. Jackson, Ian Johnston, Dominic Kwiatkowski, Cordelia Langford, John Sillitoe on behalf of the Wellcome Sanger Institute COVID-19 Surveillance Team |
| EPI_ISL_3338199 | Health Services Laboratories | Wellcome Sanger Institute for the COVID-19 Genomics UK (COG-UK) Consortium | Health Services Laboratories and Alex Alderton, Roberto Amato, Jeffrey Barrett, Sonia Goncalves, Ewan Harrison, David K. Jackson, Ian Johnston, Dominic Kwiatkowski, Cordelia Langford, John Sillitoe on behalf of the Wellcome Sanger Institute COVID-19 Surveillance Team |
| EPI_ISL_3338200 | Lighthouse Laboratory Plymouth | Wellcome Sanger Institute for the COVID-19 Genomics UK (COG-UK) Consortium | Lighthouse Laboratory Plymouth and Alex Alderton, Roberto Amato, Jeffrey Barrett, Sonia Goncalves, Ewan Harrison, David K. Jackson, Ian Johnston, Dominic Kwiatkowski, Cordelia Langford, John Sillitoe on behalf of the Wellcome Sanger Institute COVID-19 Surveillance Team |
| EPI_ISL_3338201, EPI_ISL_3338202 | Health Services Laboratories | Wellcome Sanger Institute for the COVID-19 Genomics UK (COG-UK) Consortium | Health Services Laboratories and Alex Alderton, Roberto Amato, Jeffrey Barrett, Sonia Goncalves, Ewan Harrison, David K. Jackson, Ian Johnston, Dominic Kwiatkowski, Cordelia Langford, John Sillitoe on behalf of the Wellcome Sanger Institute COVID-19 Surveillance Team |
| EPI_ISL_3338203 | Lighthouse Laboratory Plymouth | Wellcome Sanger Institute for the COVID-19 Genomics UK (COG-UK) Consortium | Lighthouse Laboratory Plymouth and Alex Alderton, Roberto Amato, Jeffrey Barrett, Sonia Goncalves, Ewan Harrison, David K. Jackson, Ian Johnston, Dominic Kwiatkowski, Cordelia Langford, John Sillitoe on behalf of the Wellcome Sanger Institute COVID-19 Surveillance Team |
| EPI_ISL_3338204 | Lighthouse Lab in Milton Keynes | Wellcome Sanger Institute for the COVID-19 Genomics UK (COG-UK) Consortium | The Lighthouse Lab in Milton Keynes and Alex Alderton, Roberto Amato, Jeffrey Barrett, Sonia Goncalves, Ewan Harrison, David K. Jackson, Ian Johnston, Dominic Kwiatkowski, Cordelia Langford, John Sillitoe on behalf of the Wellcome Sanger Institute COVID-19 Surveillance Team |
| EPI_ISL_3338205 | Health Services Laboratories | Wellcome Sanger Institute for the COVID-19 Genomics UK (COG-UK) Consortium | Health Services Laboratories and Alex Alderton, Roberto Amato, Jeffrey Barrett, Sonia Goncalves, Ewan Harrison, David K. Jackson, Ian Johnston, Dominic Kwiatkowski, Cordelia Langford, John Sillitoe on behalf of the Wellcome Sanger Institute COVID-19 Surveillance Team |
| EPI_ISL_3338206 | Lighthouse Lab in Milton Keynes | Wellcome Sanger Institute for the COVID-19 Genomics UK (COG-UK) Consortium | The Lighthouse Lab in Milton Keynes and Alex Alderton, Roberto Amato, Jeffrey Barrett, Sonia Goncalves, Ewan Harrison, David K. Jackson, Ian Johnston, Dominic Kwiatkowski, Cordelia Langford, John Sillitoe on behalf of the Wellcome Sanger Institute COVID-19 Surveillance Team |
| EPI_ISL_3338207 | Lighthouse Laboratory Plymouth | Wellcome Sanger Institute for the COVID-19 Genomics UK (COG-UK) Consortium | Lighthouse Laboratory Plymouth and Alex Alderton, Roberto Amato, Jeffrey Barrett, Sonia Goncalves, Ewan Harrison, David K. Jackson, Ian Johnston, Dominic Kwiatkowski, Cordelia Langford, John Sillitoe on behalf of the Wellcome Sanger Institute COVID-19 Surveillance Team |
| EPI_ISL_3338208, EPI_ISL_3338209 | Lighthouse Lab in Milton Keynes | Wellcome Sanger Institute for the COVID-19 Genomics UK (COG-UK) Consortium | The Lighthouse Lab in Milton Keynes and Alex Alderton, Roberto Amato, Jeffrey Barrett, Sonia Goncalves, Ewan Harrison, David K. Jackson, Ian Johnston, Dominic Kwiatkowski, Cordelia Langford, John Sillitoe on behalf of the Wellcome Sanger Institute COVID-19 Surveillance Team |
| EPI_ISL_3338210 | Health Services Laboratories | Wellcome Sanger Institute for the COVID-19 Genomics UK (COG-UK) Consortium | Health Services Laboratories and Alex Alderton, Roberto Amato, Jeffrey Barrett, Sonia Goncalves, Ewan Harrison, David K. Jackson, Ian Johnston, Dominic Kwiatkowski, Cordelia Langford, John Sillitoe on behalf of the Wellcome Sanger Institute COVID-19 Surveillance Team |
| EPI_ISL_3338211 | Lighthouse Laboratory Plymouth | Wellcome Sanger Institute for the COVID-19 Genomics UK (COG-UK) Consortium | Lighthouse Laboratory Plymouth and Alex Alderton, Roberto Amato, Jeffrey Barrett, Sonia Goncalves, Ewan Harrison, David K. Jackson, Ian Johnston, Dominic Kwiatkowski, Cordelia Langford, John Sillitoe on behalf of the Wellcome Sanger Institute COVID-19 Surveillance Team |
| EPI_ISL_3338212 | Lighthouse Lab in Milton Keynes | Wellcome Sanger Institute for the COVID-19 Genomics UK (COG-UK) Consortium | The Lighthouse Lab in Milton Keynes and Alex Alderton, Roberto Amato, Jeffrey Barrett, Sonia Goncalves, Ewan Harrison, David K. Jackson, Ian Johnston, Dominic Kwiatkowski, Cordelia Langford, John Sillitoe on behalf of the Wellcome Sanger Institute COVID-19 Surveillance Team |
| EPI_ISL_3338213, EPI_ISL_3338214, EPI_ISL_3338215 | Health Services Laboratories | Wellcome Sanger Institute for the COVID-19 Genomics UK (COG-UK) Consortium | Health Services Laboratories and Alex Alderton, Roberto Amato, Jeffrey Barrett, Sonia Goncalves, Ewan Harrison, David K. Jackson, Ian Johnston, Dominic Kwiatkowski, Cordelia Langford, John Sillitoe on behalf of the Wellcome Sanger Institute COVID-19 Surveillance Team |
| EPI_ISL_3338216 | Lighthouse Lab in Milton Keynes | Wellcome Sanger Institute for the COVID-19 Genomics UK (COG-UK) Consortium | The Lighthouse Lab in Milton Keynes and Alex Alderton, Roberto Amato, Jeffrey Barrett, Sonia Goncalves, Ewan Harrison, David K. Jackson, Ian Johnston, Dominic Kwiatkowski, Cordelia Langford, John Sillitoe on behalf of the Wellcome Sanger Institute COVID-19 Surveillance Team |
| EPI_ISL_3338217, EPI_ISL_3338218 | Lighthouse Laboratory Plymouth | Wellcome Sanger Institute for the COVID-19 Genomics UK (COG-UK) Consortium | Lighthouse Laboratory Plymouth and Alex Alderton, Roberto Amato, Jeffrey Barrett, Sonia Goncalves, Ewan Harrison, David K. Jackson, Ian Johnston, Dominic Kwiatkowski, Cordelia Langford, John Sillitoe on behalf of the Wellcome Sanger Institute COVID-19 Surveillance Team |
| EPI_ISL_3338219 | Lighthouse Lab in Milton Keynes | Wellcome Sanger Institute for the COVID-19 Genomics UK (COG-UK) Consortium | The Lighthouse Lab in Milton Keynes and Alex Alderton, Roberto Amato, Jeffrey Barrett, Sonia Goncalves, Ewan Harrison, David K. Jackson, Ian Johnston, Dominic Kwiatkowski, Cordelia Langford, John Sillitoe on behalf of the Wellcome Sanger Institute COVID-19 Surveillance Team |
| EPI_ISL_3338220 | Lighthouse Laboratory Plymouth | Wellcome Sanger Institute for the COVID-19 Genomics UK (COG-UK) Consortium | Lighthouse Laboratory Plymouth and Alex Alderton, Roberto Amato, Jeffrey Barrett, Sonia Goncalves, Ewan Harrison, David K. Jackson, Ian Johnston, Dominic Kwiatkowski, Cordelia Langford, John Sillitoe on behalf of the Wellcome Sanger Institute COVID-19 Surveillance Team |
| EPI_ISL_3338221, EPI_ISL_3338222, EPI_ISL_3338223 | Health Services Laboratories | Wellcome Sanger Institute for the COVID-19 Genomics UK (COG-UK) Consortium | Health Services Laboratories and Alex Alderton, Roberto Amato, Jeffrey Barrett, Sonia Goncalves, Ewan Harrison, David K. Jackson, Ian Johnston, Dominic Kwiatkowski, Cordelia Langford, John Sillitoe on behalf of the Wellcome Sanger Institute COVID-19 Surveillance Team |
| EPI_ISL_3338224, EPI_ISL_3338225 | Lighthouse Lab in Milton Keynes | Wellcome Sanger Institute for the COVID-19 Genomics UK (COG-UK) Consortium | The Lighthouse Lab in Milton Keynes and Alex Alderton, Roberto Amato, Jeffrey Barrett, Sonia Goncalves, Ewan Harrison, David K. Jackson, Ian Johnston, Dominic Kwiatkowski, Cordelia Langford, John Sillitoe on behalf of the Wellcome Sanger Institute COVID-19 Surveillance Team |
| EPI_ISL_3338226 | Health Services Laboratories | Wellcome Sanger Institute for the COVID-19 Genomics UK (COG-UK) Consortium | Health Services Laboratories and Alex Alderton, Roberto Amato, Jeffrey Barrett, Sonia Goncalves, Ewan Harrison, David K. Jackson, Ian Johnston, Dominic Kwiatkowski, Cordelia Langford, John Sillitoe on behalf of the Wellcome Sanger Institute COVID-19 Surveillance Team |
| EPI_ISL_3338227, EPI_ISL_3338228, EPI_ISL_3338229 | Lighthouse Lab in Milton Keynes | Wellcome Sanger Institute for the COVID-19 Genomics UK (COG-UK) Consortium | The Lighthouse Lab in Milton Keynes and Alex Alderton, Roberto Amato, Jeffrey Barrett, Sonia Goncalves, Ewan Harrison, David K. Jackson, Ian Johnston, Dominic Kwiatkowski, Cordelia Langford, John Sillitoe on behalf of the Wellcome Sanger Institute COVID-19 Surveillance Team |
| EPI_ISL_3338230 | Lighthouse Laboratory Plymouth | Wellcome Sanger Institute for the COVID-19 Genomics UK (COG-UK) Consortium | Lighthouse Laboratory Plymouth and Alex Alderton, Roberto Amato, Jeffrey Barrett, Sonia Goncalves, Ewan Harrison, David K. Jackson, Ian Johnston, Dominic Kwiatkowski, Cordelia Langford, John Sillitoe on behalf of the Wellcome Sanger Institute COVID-19 Surveillance Team |
| EPI_ISL_3338231, EPI_ISL_3338232 | Lighthouse Lab in Milton Keynes | Wellcome Sanger Institute for the COVID-19 Genomics UK (COG-UK) Consortium | The Lighthouse Lab in Milton Keynes and Alex Alderton, Roberto Amato, Jeffrey Barrett, Sonia Goncalves, Ewan Harrison, David K. Jackson, Ian Johnston, Dominic Kwiatkowski, Cordelia Langford, John Sillitoe on behalf of the Wellcome Sanger Institute COVID-19 Surveillance Team |
| EPI_ISL_3338233 | Lighthouse Laboratory Plymouth | Wellcome Sanger Institute for the COVID-19 Genomics UK (COG-UK) Consortium | Lighthouse Laboratory Plymouth and Alex Alderton, Roberto Amato, Jeffrey Barrett, Sonia Goncalves, Ewan Harrison, David K. Jackson, Ian Johnston, Dominic Kwiatkowski, Cordelia Langford, John Sillitoe on behalf of the Wellcome Sanger Institute COVID-19 Surveillance Team |
| EPI_ISL_3338234 | Lighthouse Lab in Milton Keynes | Wellcome Sanger Institute for the COVID-19 Genomics UK (COG-UK) Consortium | The Lighthouse Lab in Milton Keynes and Alex Alderton, Roberto Amato, Jeffrey Barrett, Sonia Goncalves, Ewan Harrison, David K. Jackson, Ian Johnston, Dominic Kwiatkowski, Cordelia Langford, John Sillitoe on behalf of the Wellcome Sanger Institute COVID-19 Surveillance Team |
| EPI_ISL_3338235 | Health Services Laboratories | Wellcome Sanger Institute for the COVID-19 Genomics UK (COG-UK) Consortium | Health Services Laboratories and Alex Alderton, Roberto Amato, Jeffrey Barrett, Sonia Goncalves, Ewan Harrison, David K. Jackson, Ian Johnston, Dominic Kwiatkowski, Cordelia Langford, John Sillitoe on behalf of the Wellcome Sanger Institute COVID-19 Surveillance Team |
| EPI_ISL_3338236, EPI_ISL_3338237 | Lighthouse Laboratory Plymouth | Wellcome Sanger Institute for the COVID-19 Genomics UK (COG-UK) Consortium | Lighthouse Laboratory Plymouth and Alex Alderton, Roberto Amato, Jeffrey Barrett, Sonia Goncalves, Ewan Harrison, David K. Jackson, Ian Johnston, Dominic Kwiatkowski, Cordelia Langford, John Sillitoe on behalf of the Wellcome Sanger Institute COVID-19 Surveillance Team |
| EPI_ISL_3338238 | Health Services Laboratories | Wellcome Sanger Institute for the COVID-19 Genomics UK (COG-UK) Consortium | Health Services Laboratories and Alex Alderton, Roberto Amato, Jeffrey Barrett, Sonia Goncalves, Ewan Harrison, David K. Jackson, Ian Johnston, Dominic Kwiatkowski, Cordelia Langford, John Sillitoe on behalf of the Wellcome Sanger Institute COVID-19 Surveillance Team |
| EPI_ISL_3338239 | Lighthouse Laboratory Plymouth | Wellcome Sanger Institute for the COVID-19 Genomics UK (COG-UK) Consortium | Lighthouse Laboratory Plymouth and Alex Alderton, Roberto Amato, Jeffrey Barrett, Sonia Goncalves, Ewan Harrison, David K. Jackson, Ian Johnston, Dominic Kwiatkowski, Cordelia Langford, John Sillitoe on behalf of the Wellcome Sanger Institute COVID-19 Surveillance Team |
| EPI_ISL_3338240 | Health Services Laboratories | Wellcome Sanger Institute for the COVID-19 Genomics UK (COG-UK) Consortium | Health Services Laboratories and Alex Alderton, Roberto Amato, Jeffrey Barrett, Sonia Goncalves, Ewan Harrison, David K. Jackson, Ian Johnston, Dominic Kwiatkowski, Cordelia Langford, John Sillitoe on behalf of the Wellcome Sanger Institute COVID-19 Surveillance Team |
| EPI_ISL_3338241, EPI_ISL_3338242, EPI_ISL_3338243, EPI_ISL_3338244 | Lighthouse Lab in Milton Keynes | Wellcome Sanger Institute for the COVID-19 Genomics UK (COG-UK) Consortium | The Lighthouse Lab in Milton Keynes and Alex Alderton, Roberto Amato, Jeffrey Barrett, Sonia Goncalves, Ewan Harrison, David K. Jackson, Ian Johnston, Dominic Kwiatkowski, Cordelia Langford, John Sillitoe on behalf of the Wellcome Sanger Institute COVID-19 Surveillance Team |
| EPI_ISL_3338245, EPI_ISL_3338246, EPI_ISL_3338247 | Lighthouse Laboratory Plymouth | Wellcome Sanger Institute for the COVID-19 Genomics UK (COG-UK) Consortium | Lighthouse Laboratory Plymouth and Alex Alderton, Roberto Amato, Jeffrey Barrett, Sonia Goncalves, Ewan Harrison, David K. Jackson, Ian Johnston, Dominic Kwiatkowski, Cordelia Langford, John Sillitoe on behalf of the Wellcome Sanger Institute COVID-19 Surveillance Team |
| EPI_ISL_3338248 | Health Services Laboratories | Wellcome Sanger Institute for the COVID-19 Genomics UK (COG-UK) Consortium | Health Services Laboratories and Alex Alderton, Roberto Amato, Jeffrey Barrett, Sonia Goncalves, Ewan Harrison, David K. Jackson, Ian Johnston, Dominic Kwiatkowski, Cordelia Langford, John Sillitoe on behalf of the Wellcome Sanger Institute COVID-19 Surveillance Team |
| EPI_ISL_3338249 | Lighthouse Lab in Alderley Park | Wellcome Sanger Institute for the COVID-19 Genomics UK (COG-UK) Consortium | Jacquelyn Wynn, Mairead Hyland, The Lighthouse Lab in Alderley Park and Alex Alderton, Roberto Amato, Jeffrey Barrett, Sonia Goncalves, Ewan Harrison, David K. Jackson, Ian Johnston, Dominic Kwiatkowski, Cordelia Langford, John Sillitoe on behalf of the Wellcome Sanger Institute COVID-19 Surveillance Team |
| EPI_ISL_3338250, EPI_ISL_3338251, EPI_ISL_3338252 | Lighthouse Lab in Glasgow | Wellcome Sanger Institute for the COVID-19 Genomics UK (COG-UK) Consortium | Harper VanSteenhouse, Yumi Kasai, David Gray, Carol Clugston, Anna Dominiczak and Alex Alderton, Roberto Amato, Jeffrey Barrett, Sonia Goncalves, Ewan Harrison, David K. Jackson, Ian Johnston, Dominic Kwiatkowski, Cordelia Langford, John Sillitoe on behalf of the Wellcome Sanger Institute COVID-19 Surveillance Team |
| EPI_ISL_3338253, EPI_ISL_3338254, EPI_ISL_3338255 | Lighthouse Lab in Milton Keynes | Wellcome Sanger Institute for the COVID-19 Genomics UK (COG-UK) Consortium | The Lighthouse Lab in Milton Keynes and Alex Alderton, Roberto Amato, Jeffrey Barrett, Sonia Goncalves, Ewan Harrison, David K. Jackson, Ian Johnston, Dominic Kwiatkowski, Cordelia Langford, John Sillitoe on behalf of the Wellcome Sanger Institute COVID-19 Surveillance Team |
| EPI_ISL_3338256, EPI_ISL_3338257 | Lighthouse Laboratory Plymouth | Wellcome Sanger Institute for the COVID-19 Genomics UK (COG-UK) Consortium | Lighthouse Laboratory Plymouth and Alex Alderton, Roberto Amato, Jeffrey Barrett, Sonia Goncalves, Ewan Harrison, David K. Jackson, Ian Johnston, Dominic Kwiatkowski, Cordelia Langford, John Sillitoe on behalf of the Wellcome Sanger Institute COVID-19 Surveillance Team |
| EPI_ISL_3338258 | Lighthouse Lab in Glasgow | Wellcome Sanger Institute for the COVID-19 Genomics UK (COG-UK) | Harper VanSteenhouse, Yumi Kasai, David Gray, Carol Clugston, Anna Dominiczak and Alex Alderton, Roberto Amato, Jeffrey Barrett, Sonia Goncalves, Ewan Harrison, David K. Jackson, Ian Johnston, Dominic Kwiatkowski, Cordelia Langford, |

| | | | |
|---|---|---|---|
| | | Consortium | John Sillitoe on behalf of the Wellcome Sanger Institute COVID-19 Surveillance Team |
| EPI_ISL_3338259 | Lighthouse Lab in Milton Keynes | Wellcome Sanger Institute for the COVID-19 Genomics UK (COG-UK) Consortium | The Lighthouse Lab in Milton Keynes and Alex Alderton, Roberto Amato, Jeffrey Barrett, Sonia Goncalves, Ewan Harrison, David K. Jackson, Ian Johnston, Dominic Kwiatkowski, Cordelia Langford, John Sillitoe on behalf of the Wellcome Sanger Institute COVID-19 Surveillance Team |
| EPI_ISL_3338260 | Lighthouse Lab in Glasgow | Wellcome Sanger Institute for the COVID-19 Genomics UK (COG-UK) Consortium | Harper VanSteenhouse, Yumi Kasai, David Gray, Carol Clugston, Anna Dominiczak and Alex Alderton, Roberto Amato, Jeffrey Barrett, Sonia Goncalves, Ewan Harrison, David K. Jackson, Ian Johnston, Dominic Kwiatkowski, Cordelia Langford, John Sillitoe on behalf of the Wellcome Sanger Institute COVID-19 Surveillance Team |
| EPI_ISL_3338261, EPI_ISL_3338262 | Lighthouse Lab in Milton Keynes | Wellcome Sanger Institute for the COVID-19 Genomics UK (COG-UK) Consortium | The Lighthouse Lab in Milton Keynes and Alex Alderton, Roberto Amato, Jeffrey Barrett, Sonia Goncalves, Ewan Harrison, David K. Jackson, Ian Johnston, Dominic Kwiatkowski, Cordelia Langford, John Sillitoe on behalf of the Wellcome Sanger Institute COVID-19 Surveillance Team |
| EPI_ISL_3338263 | Lighthouse Lab in Glasgow | Wellcome Sanger Institute for the COVID-19 Genomics UK (COG-UK) Consortium | Harper VanSteenhouse, Yumi Kasai, David Gray, Carol Clugston, Anna Dominiczak and Alex Alderton, Roberto Amato, Jeffrey Barrett, Sonia Goncalves, Ewan Harrison, David K. Jackson, Ian Johnston, Dominic Kwiatkowski, Cordelia Langford, John Sillitoe on behalf of the Wellcome Sanger Institute COVID-19 Surveillance Team |
| EPI_ISL_3338264, EPI_ISL_3338265 | Lighthouse Lab in Alderley Park | Wellcome Sanger Institute for the COVID-19 Genomics UK (COG-UK) Consortium | Jacquelyn Wynn, Mairead Hyland, The Lighthouse Lab in Alderley Park and Alex Alderton, Roberto Amato, Jeffrey Barrett, Sonia Goncalves, Ewan Harrison, David K. Jackson, Ian Johnston, Dominic Kwiatkowski, Cordelia Langford, John Sillitoe on behalf of the Wellcome Sanger Institute COVID-19 Surveillance Team |
| EPI_ISL_3338266, EPI_ISL_3338267 | Lighthouse Lab in Glasgow | Wellcome Sanger Institute for the COVID-19 Genomics UK (COG-UK) Consortium | Harper VanSteenhouse, Yumi Kasai, David Gray, Carol Clugston, Anna Dominiczak and Alex Alderton, Roberto Amato, Jeffrey Barrett, Sonia Goncalves, Ewan Harrison, David K. Jackson, Ian Johnston, Dominic Kwiatkowski, Cordelia Langford, John Sillitoe on behalf of the Wellcome Sanger Institute COVID-19 Surveillance Team |
| EPI_ISL_3338268, EPI_ISL_3338269 | Lighthouse Lab in Milton Keynes | Wellcome Sanger Institute for the COVID-19 Genomics UK (COG-UK) Consortium | The Lighthouse Lab in Milton Keynes and Alex Alderton, Roberto Amato, Jeffrey Barrett, Sonia Goncalves, Ewan Harrison, David K. Jackson, Ian Johnston, Dominic Kwiatkowski, Cordelia Langford, John Sillitoe on behalf of the Wellcome Sanger Institute COVID-19 Surveillance Team |
| EPI_ISL_3338270, EPI_ISL_3338271, EPI_ISL_3338272 | Lighthouse Lab in Alderley Park | Wellcome Sanger Institute for the COVID-19 Genomics UK (COG-UK) Consortium | Jacquelyn Wynn, Mairead Hyland, The Lighthouse Lab in Alderley Park and Alex Alderton, Roberto Amato, Jeffrey Barrett, Sonia Goncalves, Ewan Harrison, David K. Jackson, Ian Johnston, Dominic Kwiatkowski, Cordelia Langford, John Sillitoe on behalf of the Wellcome Sanger Institute COVID-19 Surveillance Team |
| EPI_ISL_3338273 | Berkshire and Surrey Pathology Services Lighthouse Laboratory | Wellcome Sanger Institute for the COVID-19 Genomics UK (COG-UK) Consortium | Berkshire and Surrey Pathology Services Lighthouse Laboratory and Alex Alderton, Roberto Amato, Jeffrey Barrett, Sonia Goncalves, Ewan Harrison, David K. Jackson, Ian Johnston, Dominic Kwiatkowski, Cordelia Langford, John Sillitoe on behalf of the Wellcome Sanger Institute COVID-19 Surveillance Team |
| EPI_ISL_3338274 | Lighthouse Lab in Alderley Park | Wellcome Sanger Institute for the COVID-19 Genomics UK (COG-UK) Consortium | Jacquelyn Wynn, Mairead Hyland, The Lighthouse Lab in Alderley Park and Alex Alderton, Roberto Amato, Jeffrey Barrett, Sonia Goncalves, Ewan Harrison, David K. Jackson, Ian Johnston, Dominic Kwiatkowski, Cordelia Langford, John Sillitoe on behalf of the Wellcome Sanger Institute COVID-19 Surveillance Team |
| EPI_ISL_3338275 | Berkshire and Surrey Pathology Services Lighthouse Laboratory | Wellcome Sanger Institute for the COVID-19 Genomics UK (COG-UK) Consortium | Berkshire and Surrey Pathology Services Lighthouse Laboratory and Alex Alderton, Roberto Amato, Jeffrey Barrett, Sonia Goncalves, Ewan Harrison, David K. Jackson, Ian Johnston, Dominic Kwiatkowski, Cordelia Langford, John Sillitoe on behalf of the Wellcome Sanger Institute COVID-19 Surveillance Team |
| EPI_ISL_3338276, EPI_ISL_3338277, EPI_ISL_3338278, EPI_ISL_3338279, EPI_ISL_3338280 | Lighthouse Lab in Alderley Park | Wellcome Sanger Institute for the COVID-19 Genomics UK (COG-UK) Consortium | Jacquelyn Wynn, Mairead Hyland, The Lighthouse Lab in Alderley Park and Alex Alderton, Roberto Amato, Jeffrey Barrett, Sonia Goncalves, Ewan Harrison, David K. Jackson, Ian Johnston, Dominic Kwiatkowski, Cordelia Langford, John Sillitoe on behalf of the Wellcome Sanger Institute COVID-19 Surveillance Team |
| EPI_ISL_3338281 | Lighthouse Laboratory Plymouth | Wellcome Sanger Institute for the COVID-19 Genomics UK (COG-UK) Consortium | Lighthouse Laboratory Plymouth and Alex Alderton, Roberto Amato, Jeffrey Barrett, Sonia Goncalves, Ewan Harrison, David K. Jackson, Ian Johnston, Dominic Kwiatkowski, Cordelia Langford, John Sillitoe on behalf of the Wellcome Sanger Institute COVID-19 Surveillance Team |
| EPI_ISL_3338282, EPI_ISL_3338283 | Lighthouse Lab in Alderley Park | Wellcome Sanger Institute for the COVID-19 Genomics UK (COG-UK) Consortium | Jacquelyn Wynn, Mairead Hyland, The Lighthouse Lab in Alderley Park and Alex Alderton, Roberto Amato, Jeffrey Barrett, Sonia Goncalves, Ewan Harrison, David K. Jackson, Ian Johnston, Dominic Kwiatkowski, Cordelia Langford, John Sillitoe on behalf of the Wellcome Sanger Institute COVID-19 Surveillance Team |
| EPI_ISL_3338284 | Lighthouse Laboratory Plymouth | Wellcome Sanger Institute for the COVID-19 Genomics UK (COG-UK) Consortium | Lighthouse Laboratory Plymouth and Alex Alderton, Roberto Amato, Jeffrey Barrett, Sonia Goncalves, Ewan Harrison, David K. Jackson, Ian Johnston, Dominic Kwiatkowski, Cordelia Langford, John Sillitoe on behalf of the Wellcome Sanger Institute COVID-19 Surveillance Team |
| EPI_ISL_3338285, EPI_ISL_3338286, EPI_ISL_3338287 | Lighthouse Lab in Alderley Park | Wellcome Sanger Institute for the COVID-19 Genomics UK (COG-UK) Consortium | Jacquelyn Wynn, Mairead Hyland, The Lighthouse Lab in Alderley Park and Alex Alderton, Roberto Amato, Jeffrey Barrett, Sonia Goncalves, Ewan Harrison, David K. Jackson, Ian Johnston, Dominic Kwiatkowski, Cordelia Langford, John Sillitoe on behalf of the Wellcome Sanger Institute COVID-19 Surveillance Team |
| EPI_ISL_3338289 | Lighthouse Lab in Glasgow | Wellcome Sanger Institute for the COVID-19 Genomics UK (COG-UK) Consortium | Harper VanSteenhouse, Yumi Kasai, David Gray, Carol Clugston, Anna Dominiczak and Alex Alderton, Roberto Amato, Jeffrey Barrett, Sonia Goncalves, Ewan Harrison, David K. Jackson, Ian Johnston, Dominic Kwiatkowski, Cordelia Langford, John Sillitoe on behalf of the Wellcome Sanger Institute COVID-19 Surveillance Team |
| EPI_ISL_3338290 | Lighthouse Lab in Alderley Park | Wellcome Sanger Institute for the COVID-19 Genomics UK (COG-UK) Consortium | Jacquelyn Wynn, Mairead Hyland, The Lighthouse Lab in Alderley Park and Alex Alderton, Roberto Amato, Jeffrey Barrett, Sonia Goncalves, Ewan Harrison, David K. Jackson, Ian Johnston, Dominic Kwiatkowski, Cordelia Langford, John Sillitoe on behalf of the Wellcome Sanger Institute COVID-19 Surveillance Team |
| EPI_ISL_3338291 | Lighthouse Laboratory Plymouth | Wellcome Sanger Institute for the COVID-19 Genomics UK (COG-UK) Consortium | Lighthouse Laboratory Plymouth and Alex Alderton, Roberto Amato, Jeffrey Barrett, Sonia Goncalves, Ewan Harrison, David K. Jackson, Ian Johnston, Dominic Kwiatkowski, Cordelia Langford, John Sillitoe on behalf of the Wellcome Sanger Institute COVID-19 Surveillance Team |
| EPI_ISL_3338292 | Lighthouse Lab in Glasgow | Wellcome Sanger Institute for the COVID-19 Genomics UK (COG-UK) Consortium | Harper VanSteenhouse, Yumi Kasai, David Gray, Carol Clugston, Anna Dominiczak and Alex Alderton, Roberto Amato, Jeffrey Barrett, Sonia Goncalves, Ewan Harrison, David K. Jackson, Ian Johnston, Dominic Kwiatkowski, Cordelia Langford, John Sillitoe on behalf of the Wellcome Sanger Institute COVID-19 Surveillance Team |
| EPI_ISL_3338293, EPI_ISL_3338294, EPI_ISL_3338295, EPI_ISL_3338296, EPI_ISL_3338297 | Lighthouse Lab in Alderley Park | Wellcome Sanger Institute for the COVID-19 Genomics UK (COG-UK) Consortium | Jacquelyn Wynn, Mairead Hyland, The Lighthouse Lab in Alderley Park and Alex Alderton, Roberto Amato, Jeffrey Barrett, Sonia Goncalves, Ewan Harrison, David K. Jackson, Ian Johnston, Dominic Kwiatkowski, Cordelia Langford, John Sillitoe on behalf of the Wellcome Sanger Institute COVID-19 Surveillance Team |
| EPI_ISL_3338298, EPI_ISL_3338299 | Lighthouse Laboratory Plymouth | Wellcome Sanger Institute for the COVID-19 Genomics UK (COG-UK) Consortium | Lighthouse Laboratory Plymouth and Alex Alderton, Roberto Amato, Jeffrey Barrett, Sonia Goncalves, Ewan Harrison, David K. Jackson, Ian Johnston, Dominic Kwiatkowski, Cordelia Langford, John Sillitoe on behalf of the Wellcome Sanger Institute COVID-19 Surveillance Team |
| EPI_ISL_3338300, EPI_ISL_3338301, EPI_ISL_3338302, EPI_ISL_3338303, EPI_ISL_3338304, EPI_ISL_3338305 | Lighthouse Lab in Alderley Park | Wellcome Sanger Institute for the COVID-19 Genomics UK (COG-UK) Consortium | Jacquelyn Wynn, Mairead Hyland, The Lighthouse Lab in Alderley Park and Alex Alderton, Roberto Amato, Jeffrey Barrett, Sonia Goncalves, Ewan Harrison, David K. Jackson, Ian Johnston, Dominic Kwiatkowski, Cordelia Langford, John Sillitoe on behalf of the Wellcome Sanger Institute COVID-19 Surveillance Team |
| EPI_ISL_3338306 | Lighthouse Lab in Glasgow | Wellcome Sanger Institute for the COVID-19 Genomics UK (COG-UK) Consortium | Harper VanSteenhouse, Yumi Kasai, David Gray, Carol Clugston, Anna Dominiczak and Alex Alderton, Roberto Amato, Jeffrey Barrett, Sonia Goncalves, Ewan Harrison, David K. Jackson, Ian Johnston, Dominic Kwiatkowski, Cordelia Langford, John Sillitoe on behalf of the Wellcome Sanger Institute COVID-19 Surveillance Team |
| EPI_ISL_3338307, EPI_ISL_3338308 | Lighthouse Laboratory Plymouth | Wellcome Sanger Institute for the COVID-19 Genomics UK (COG-UK) Consortium | Lighthouse Laboratory Plymouth and Alex Alderton, Roberto Amato, Jeffrey Barrett, Sonia Goncalves, Ewan Harrison, David K. Jackson, Ian Johnston, Dominic Kwiatkowski, Cordelia Langford, John Sillitoe on behalf of the Wellcome Sanger Institute COVID-19 Surveillance Team |
| EPI_ISL_3338309, EPI_ISL_3338310 | Lighthouse Lab in Alderley Park | Wellcome Sanger Institute for the COVID-19 Genomics UK (COG-UK) Consortium | Jacquelyn Wynn, Mairead Hyland, The Lighthouse Lab in Alderley Park and Alex Alderton, Roberto Amato, Jeffrey Barrett, Sonia Goncalves, Ewan Harrison, David K. Jackson, Ian Johnston, Dominic Kwiatkowski, Cordelia Langford, John Sillitoe on behalf of the Wellcome Sanger Institute COVID-19 Surveillance Team |
| EPI_ISL_3338311 | Lighthouse Lab in Glasgow | Wellcome Sanger Institute for the COVID-19 Genomics UK (COG-UK) Consortium | Harper VanSteenhouse, Yumi Kasai, David Gray, Carol Clugston, Anna Dominiczak and Alex Alderton, Roberto Amato, Jeffrey Barrett, Sonia Goncalves, Ewan Harrison, David K. Jackson, Ian Johnston, Dominic Kwiatkowski, Cordelia Langford, John Sillitoe on behalf of the Wellcome Sanger Institute COVID-19 Surveillance Team |
| EPI_ISL_3338312, EPI_ISL_3338313 | Lighthouse Lab in Alderley Park | Wellcome Sanger Institute for the COVID-19 Genomics UK (COG-UK) Consortium | Jacquelyn Wynn, Mairead Hyland, The Lighthouse Lab in Alderley Park and Alex Alderton, Roberto Amato, Jeffrey Barrett, Sonia Goncalves, Ewan Harrison, David K. Jackson, Ian Johnston, Dominic Kwiatkowski, Cordelia Langford, John Sillitoe on behalf of the Wellcome Sanger Institute COVID-19 Surveillance Team |
| EPI_ISL_3338314 | Lighthouse Lab in Glasgow | Wellcome Sanger Institute for the COVID-19 Genomics UK (COG-UK) Consortium | Harper VanSteenhouse, Yumi Kasai, David Gray, Carol Clugston, Anna Dominiczak and Alex Alderton, Roberto Amato, Jeffrey Barrett, Sonia Goncalves, Ewan Harrison, David K. Jackson, Ian Johnston, Dominic Kwiatkowski, Cordelia Langford, John Sillitoe on behalf of the Wellcome Sanger Institute COVID-19 Surveillance Team |
| EPI_ISL_3338315 | Lighthouse Lab in Alderley Park | Wellcome Sanger Institute for the COVID-19 Genomics UK (COG-UK) Consortium | Jacquelyn Wynn, Mairead Hyland, The Lighthouse Lab in Alderley Park and Alex Alderton, Roberto Amato, Jeffrey Barrett, Sonia Goncalves, Ewan Harrison, David K. Jackson, Ian Johnston, Dominic Kwiatkowski, Cordelia Langford, John Sillitoe on behalf of the Wellcome Sanger Institute COVID-19 Surveillance Team |
| EPI_ISL_3338316, EPI_ISL_3338317, EPI_ISL_3338318 | Lighthouse Laboratory Plymouth | Wellcome Sanger Institute for the COVID-19 Genomics UK (COG-UK) Consortium | Lighthouse Laboratory Plymouth and Alex Alderton, Roberto Amato, Jeffrey Barrett, Sonia Goncalves, Ewan Harrison, David K. Jackson, Ian Johnston, Dominic Kwiatkowski, Cordelia Langford, John Sillitoe on behalf of the Wellcome Sanger Institute COVID-19 Surveillance Team |
| EPI_ISL_3338319 | Lighthouse Lab in Alderley Park | Wellcome Sanger Institute for the COVID-19 Genomics UK (COG-UK) Consortium | Jacquelyn Wynn, Mairead Hyland, The Lighthouse Lab in Alderley Park and Alex Alderton, Roberto Amato, Jeffrey Barrett, Sonia Goncalves, Ewan Harrison, David K. Jackson, Ian Johnston, Dominic Kwiatkowski, Cordelia Langford, John Sillitoe on behalf of the Wellcome Sanger Institute COVID-19 Surveillance Team |
| EPI_ISL_3338320 | Lighthouse Lab in Glasgow | Wellcome Sanger Institute for the COVID-19 Genomics UK (COG-UK) Consortium | Harper VanSteenhouse, Yumi Kasai, David Gray, Carol Clugston, Anna Dominiczak and Alex Alderton, Roberto Amato, Jeffrey Barrett, Sonia Goncalves, Ewan Harrison, David K. Jackson, Ian Johnston, Dominic Kwiatkowski, Cordelia Langford, John Sillitoe on behalf of the Wellcome Sanger Institute COVID-19 Surveillance Team |
| EPI_ISL_3338321 | Lighthouse Lab in Alderley Park | Wellcome Sanger Institute for the COVID-19 Genomics UK (COG-UK) Consortium | Jacquelyn Wynn, Mairead Hyland, The Lighthouse Lab in Alderley Park and Alex Alderton, Roberto Amato, Jeffrey Barrett, Sonia Goncalves, Ewan Harrison, David K. Jackson, Ian Johnston, Dominic Kwiatkowski, Cordelia Langford, John Sillitoe on behalf of the Wellcome Sanger Institute COVID-19 Surveillance Team |
| EPI_ISL_3338322 | Lighthouse Lab in Glasgow | Wellcome Sanger Institute for the COVID-19 Genomics UK (COG-UK) Consortium | Harper VanSteenhouse, Yumi Kasai, David Gray, Carol Clugston, Anna Dominiczak and Alex Alderton, Roberto Amato, Jeffrey Barrett, Sonia Goncalves, Ewan Harrison, David K. Jackson, Ian Johnston, Dominic Kwiatkowski, Cordelia Langford, John Sillitoe on behalf of the Wellcome Sanger Institute COVID-19 Surveillance Team |
| EPI_ISL_3338323 | Lighthouse Lab in Alderley Park | Wellcome Sanger Institute for the COVID-19 Genomics UK (COG-UK) Consortium | Jacquelyn Wynn, Mairead Hyland, The Lighthouse Lab in Alderley Park and Alex Alderton, Roberto Amato, Jeffrey Barrett, Sonia Goncalves, Ewan Harrison, David K. Jackson, Ian Johnston, Dominic Kwiatkowski, Cordelia Langford, John Sillitoe on behalf of the Wellcome Sanger Institute COVID-19 Surveillance Team |
| EPI_ISL_3338324 | Lighthouse Laboratory Plymouth | Wellcome Sanger Institute for the COVID-19 Genomics UK (COG-UK) Consortium | Lighthouse Laboratory Plymouth and Alex Alderton, Roberto Amato, Jeffrey Barrett, Sonia Goncalves, Ewan Harrison, David K. Jackson, Ian Johnston, Dominic Kwiatkowski, Cordelia Langford, John Sillitoe on behalf of the Wellcome Sanger Institute COVID-19 Surveillance Team |
| EPI_ISL_3338325 | Lighthouse Lab in Glasgow | Wellcome Sanger Institute for the COVID-19 Genomics UK (COG-UK) Consortium | Harper VanSteenhouse, Yumi Kasai, David Gray, Carol Clugston, Anna Dominiczak and Alex Alderton, Roberto Amato, Jeffrey Barrett, Sonia Goncalves, Ewan Harrison, David K. Jackson, Ian Johnston, Dominic Kwiatkowski, Cordelia Langford, John Sillitoe on behalf of the Wellcome Sanger Institute COVID-19 Surveillance Team |
| EPI_ISL_3338326 | Lighthouse Lab in Alderley Park | Wellcome Sanger Institute for the COVID-19 Genomics UK (COG-UK) Consortium | Jacquelyn Wynn, Mairead Hyland, The Lighthouse Lab in Alderley Park and Alex Alderton, Roberto Amato, Jeffrey Barrett, Sonia Goncalves, Ewan Harrison, David K. Jackson, Ian Johnston, Dominic Kwiatkowski, Cordelia Langford, John Sillitoe on behalf of the Wellcome Sanger Institute COVID-19 Surveillance Team |

| | | | |
|---|---|---|---|
| EPI_ISL_3338327, EPI_ISL_3338328 | Lighthouse Lab in Glasgow | Wellcome Sanger Institute for the COVID-19 Genomics UK (COG-UK) Consortium | Harper VanSteenhouse, Yumi Kasai, David Gray, Carol Clugston, Anna Dominiczak and Alex Alderton, Roberto Amato, Jeffrey Barrett, Sonia Goncalves, Ewan Harrison, David K. Jackson, Ian Johnston, Dominic Kwiatkowski, Cordelia Langford, John Sillitoe on behalf of the Wellcome Sanger Institute COVID-19 Surveillance Team |
| EPI_ISL_3338329 | Lighthouse Lab in Alderley Park | Wellcome Sanger Institute for the COVID-19 Genomics UK (COG-UK) Consortium | Jacquelyn Wynn, Mairead Hyland, The Lighthouse Lab in Alderley Park and Alex Alderton, Roberto Amato, Jeffrey Barrett, Sonia Goncalves, Ewan Harrison, David K. Jackson, Ian Johnston, Dominic Kwiatkowski, Cordelia Langford, John Sillitoe on behalf of the Wellcome Sanger Institute COVID-19 Surveillance Team |
| EPI_ISL_3338330, EPI_ISL_3338331 | Lighthouse Lab in Glasgow | Wellcome Sanger Institute for the COVID-19 Genomics UK (COG-UK) Consortium | Harper VanSteenhouse, Yumi Kasai, David Gray, Carol Clugston, Anna Dominiczak and Alex Alderton, Roberto Amato, Jeffrey Barrett, Sonia Goncalves, Ewan Harrison, David K. Jackson, Ian Johnston, Dominic Kwiatkowski, Cordelia Langford, John Sillitoe on behalf of the Wellcome Sanger Institute COVID-19 Surveillance Team |
| EPI_ISL_3338332, EPI_ISL_3338333 | Lighthouse Lab in Alderley Park | Wellcome Sanger Institute for the COVID-19 Genomics UK (COG-UK) Consortium | Jacquelyn Wynn, Mairead Hyland, The Lighthouse Lab in Alderley Park and Alex Alderton, Roberto Amato, Jeffrey Barrett, Sonia Goncalves, Ewan Harrison, David K. Jackson, Ian Johnston, Dominic Kwiatkowski, Cordelia Langford, John Sillitoe on behalf of the Wellcome Sanger Institute COVID-19 Surveillance Team |
| EPI_ISL_3338334 | Lighthouse Laboratory Plymouth | Wellcome Sanger Institute for the COVID-19 Genomics UK (COG-UK) Consortium | Lighthouse Laboratory Plymouth and Alex Alderton, Roberto Amato, Jeffrey Barrett, Sonia Goncalves, Ewan Harrison, David K. Jackson, Ian Johnston, Dominic Kwiatkowski, Cordelia Langford, John Sillitoe on behalf of the Wellcome Sanger Institute COVID-19 Surveillance Team |
| EPI_ISL_3338335 | Lighthouse Lab in Glasgow | Wellcome Sanger Institute for the COVID-19 Genomics UK (COG-UK) Consortium | Harper VanSteenhouse, Yumi Kasai, David Gray, Carol Clugston, Anna Dominiczak and Alex Alderton, Roberto Amato, Jeffrey Barrett, Sonia Goncalves, Ewan Harrison, David K. Jackson, Ian Johnston, Dominic Kwiatkowski, Cordelia Langford, John Sillitoe on behalf of the Wellcome Sanger Institute COVID-19 Surveillance Team |
| EPI_ISL_3338336 | Lighthouse Laboratory Plymouth | Wellcome Sanger Institute for the COVID-19 Genomics UK (COG-UK) Consortium | Lighthouse Laboratory Plymouth and Alex Alderton, Roberto Amato, Jeffrey Barrett, Sonia Goncalves, Ewan Harrison, David K. Jackson, Ian Johnston, Dominic Kwiatkowski, Cordelia Langford, John Sillitoe on behalf of the Wellcome Sanger Institute COVID-19 Surveillance Team |
| EPI_ISL_3338337 | Lighthouse Lab in Glasgow | Wellcome Sanger Institute for the COVID-19 Genomics UK (COG-UK) Consortium | Harper VanSteenhouse, Yumi Kasai, David Gray, Carol Clugston, Anna Dominiczak and Alex Alderton, Roberto Amato, Jeffrey Barrett, Sonia Goncalves, Ewan Harrison, David K. Jackson, Ian Johnston, Dominic Kwiatkowski, Cordelia Langford, John Sillitoe on behalf of the Wellcome Sanger Institute COVID-19 Surveillance Team |
| EPI_ISL_3338338, EPI_ISL_3338339 | Lighthouse Lab in Alderley Park | Wellcome Sanger Institute for the COVID-19 Genomics UK (COG-UK) Consortium | Jacquelyn Wynn, Mairead Hyland, The Lighthouse Lab in Alderley Park and Alex Alderton, Roberto Amato, Jeffrey Barrett, Sonia Goncalves, Ewan Harrison, David K. Jackson, Ian Johnston, Dominic Kwiatkowski, Cordelia Langford, John Sillitoe on behalf of the Wellcome Sanger Institute COVID-19 Surveillance Team |
| EPI_ISL_3338340, EPI_ISL_3338341 | Lighthouse Lab in Glasgow | Wellcome Sanger Institute for the COVID-19 Genomics UK (COG-UK) Consortium | Harper VanSteenhouse, Yumi Kasai, David Gray, Carol Clugston, Anna Dominiczak and Alex Alderton, Roberto Amato, Jeffrey Barrett, Sonia Goncalves, Ewan Harrison, David K. Jackson, Ian Johnston, Dominic Kwiatkowski, Cordelia Langford, John Sillitoe on behalf of the Wellcome Sanger Institute COVID-19 Surveillance Team |
| EPI_ISL_3338342 | Lighthouse Lab in Alderley Park | Wellcome Sanger Institute for the COVID-19 Genomics UK (COG-UK) Consortium | Jacquelyn Wynn, Mairead Hyland, The Lighthouse Lab in Alderley Park and Alex Alderton, Roberto Amato, Jeffrey Barrett, Sonia Goncalves, Ewan Harrison, David K. Jackson, Ian Johnston, Dominic Kwiatkowski, Cordelia Langford, John Sillitoe on behalf of the Wellcome Sanger Institute COVID-19 Surveillance Team |
| EPI_ISL_3338343, EPI_ISL_3338344 | Lighthouse Laboratory Plymouth | Wellcome Sanger Institute for the COVID-19 Genomics UK (COG-UK) Consortium | Lighthouse Laboratory Plymouth and Alex Alderton, Roberto Amato, Jeffrey Barrett, Sonia Goncalves, Ewan Harrison, David K. Jackson, Ian Johnston, Dominic Kwiatkowski, Cordelia Langford, John Sillitoe on behalf of the Wellcome Sanger Institute COVID-19 Surveillance Team |
| EPI_ISL_3338345, EPI_ISL_3338346, EPI_ISL_3338347, EPI_ISL_3338348 | Lighthouse Lab in Alderley Park | Wellcome Sanger Institute for the COVID-19 Genomics UK (COG-UK) Consortium | Jacquelyn Wynn, Mairead Hyland, The Lighthouse Lab in Alderley Park and Alex Alderton, Roberto Amato, Jeffrey Barrett, Sonia Goncalves, Ewan Harrison, David K. Jackson, Ian Johnston, Dominic Kwiatkowski, Cordelia Langford, John Sillitoe on behalf of the Wellcome Sanger Institute COVID-19 Surveillance Team |
| EPI_ISL_3338349 | Lighthouse Lab in Glasgow | Wellcome Sanger Institute for the COVID-19 Genomics UK (COG-UK) Consortium | Harper VanSteenhouse, Yumi Kasai, David Gray, Carol Clugston, Anna Dominiczak and Alex Alderton, Roberto Amato, Jeffrey Barrett, Sonia Goncalves, Ewan Harrison, David K. Jackson, Ian Johnston, Dominic Kwiatkowski, Cordelia Langford, John Sillitoe on behalf of the Wellcome Sanger Institute COVID-19 Surveillance Team |
| EPI_ISL_3338350 | Lighthouse Laboratory Plymouth | Wellcome Sanger Institute for the COVID-19 Genomics UK (COG-UK) Consortium | Lighthouse Laboratory Plymouth and Alex Alderton, Roberto Amato, Jeffrey Barrett, Sonia Goncalves, Ewan Harrison, David K. Jackson, Ian Johnston, Dominic Kwiatkowski, Cordelia Langford, John Sillitoe on behalf of the Wellcome Sanger Institute COVID-19 Surveillance Team |
| EPI_ISL_3338351 | Lighthouse Lab in Alderley Park | Wellcome Sanger Institute for the COVID-19 Genomics UK (COG-UK) Consortium | Jacquelyn Wynn, Mairead Hyland, The Lighthouse Lab in Alderley Park and Alex Alderton, Roberto Amato, Jeffrey Barrett, Sonia Goncalves, Ewan Harrison, David K. Jackson, Ian Johnston, Dominic Kwiatkowski, Cordelia Langford, John Sillitoe on behalf of the Wellcome Sanger Institute COVID-19 Surveillance Team |
| EPI_ISL_3338352 | Lighthouse Lab in Glasgow | Wellcome Sanger Institute for the COVID-19 Genomics UK (COG-UK) Consortium | Harper VanSteenhouse, Yumi Kasai, David Gray, Carol Clugston, Anna Dominiczak and Alex Alderton, Roberto Amato, Jeffrey Barrett, Sonia Goncalves, Ewan Harrison, David K. Jackson, Ian Johnston, Dominic Kwiatkowski, Cordelia Langford, John Sillitoe on behalf of the Wellcome Sanger Institute COVID-19 Surveillance Team |
| EPI_ISL_3338353, EPI_ISL_3338354, EPI_ISL_3338355 | Lighthouse Lab in Alderley Park | Wellcome Sanger Institute for the COVID-19 Genomics UK (COG-UK) Consortium | Jacquelyn Wynn, Mairead Hyland, The Lighthouse Lab in Alderley Park and Alex Alderton, Roberto Amato, Jeffrey Barrett, Sonia Goncalves, Ewan Harrison, David K. Jackson, Ian Johnston, Dominic Kwiatkowski, Cordelia Langford, John Sillitoe on behalf of the Wellcome Sanger Institute COVID-19 Surveillance Team |
| EPI_ISL_3338356 | Lighthouse Lab in Glasgow | Wellcome Sanger Institute for the COVID-19 Genomics UK (COG-UK) Consortium | Harper VanSteenhouse, Yumi Kasai, David Gray, Carol Clugston, Anna Dominiczak and Alex Alderton, Roberto Amato, Jeffrey Barrett, Sonia Goncalves, Ewan Harrison, David K. Jackson, Ian Johnston, Dominic Kwiatkowski, Cordelia Langford, John Sillitoe on behalf of the Wellcome Sanger Institute COVID-19 Surveillance Team |
| EPI_ISL_3338357 | Lighthouse Laboratory Plymouth | Wellcome Sanger Institute for the COVID-19 Genomics UK (COG-UK) Consortium | Lighthouse Laboratory Plymouth and Alex Alderton, Roberto Amato, Jeffrey Barrett, Sonia Goncalves, Ewan Harrison, David K. Jackson, Ian Johnston, Dominic Kwiatkowski, Cordelia Langford, John Sillitoe on behalf of the Wellcome Sanger Institute COVID-19 Surveillance Team |
| EPI_ISL_3338358, EPI_ISL_3338359 | Lighthouse Lab in Alderley Park | Wellcome Sanger Institute for the COVID-19 Genomics UK (COG-UK) Consortium | Jacquelyn Wynn, Mairead Hyland, The Lighthouse Lab in Alderley Park and Alex Alderton, Roberto Amato, Jeffrey Barrett, Sonia Goncalves, Ewan Harrison, David K. Jackson, Ian Johnston, Dominic Kwiatkowski, Cordelia Langford, John Sillitoe on behalf of the Wellcome Sanger Institute COVID-19 Surveillance Team |
| EPI_ISL_3338360 | Lighthouse Lab in Glasgow | Wellcome Sanger Institute for the COVID-19 Genomics UK (COG-UK) Consortium | Harper VanSteenhouse, Yumi Kasai, David Gray, Carol Clugston, Anna Dominiczak and Alex Alderton, Roberto Amato, Jeffrey Barrett, Sonia Goncalves, Ewan Harrison, David K. Jackson, Ian Johnston, Dominic Kwiatkowski, Cordelia Langford, John Sillitoe on behalf of the Wellcome Sanger Institute COVID-19 Surveillance Team |
| EPI_ISL_3338361, EPI_ISL_3338362, EPI_ISL_3338363 | Lighthouse Lab in Alderley Park | Wellcome Sanger Institute for the COVID-19 Genomics UK (COG-UK) Consortium | Jacquelyn Wynn, Mairead Hyland, The Lighthouse Lab in Alderley Park and Alex Alderton, Roberto Amato, Jeffrey Barrett, Sonia Goncalves, Ewan Harrison, David K. Jackson, Ian Johnston, Dominic Kwiatkowski, Cordelia Langford, John Sillitoe on behalf of the Wellcome Sanger Institute COVID-19 Surveillance Team |
| EPI_ISL_3338364 | Lighthouse Lab in Glasgow | Wellcome Sanger Institute for the COVID-19 Genomics UK (COG-UK) Consortium | Harper VanSteenhouse, Yumi Kasai, David Gray, Carol Clugston, Anna Dominiczak and Alex Alderton, Roberto Amato, Jeffrey Barrett, Sonia Goncalves, Ewan Harrison, David K. Jackson, Ian Johnston, Dominic Kwiatkowski, Cordelia Langford, John Sillitoe on behalf of the Wellcome Sanger Institute COVID-19 Surveillance Team |
| EPI_ISL_3338365 | Lighthouse Lab in Alderley Park | Wellcome Sanger Institute for the COVID-19 Genomics UK (COG-UK) Consortium | Jacquelyn Wynn, Mairead Hyland, The Lighthouse Lab in Alderley Park and Alex Alderton, Roberto Amato, Jeffrey Barrett, Sonia Goncalves, Ewan Harrison, David K. Jackson, Ian Johnston, Dominic Kwiatkowski, Cordelia Langford, John Sillitoe on behalf of the Wellcome Sanger Institute COVID-19 Surveillance Team |
| EPI_ISL_3338366 | Lighthouse Laboratory Plymouth | Wellcome Sanger Institute for the COVID-19 Genomics UK (COG-UK) Consortium | Lighthouse Laboratory Plymouth and Alex Alderton, Roberto Amato, Jeffrey Barrett, Sonia Goncalves, Ewan Harrison, David K. Jackson, Ian Johnston, Dominic Kwiatkowski, Cordelia Langford, John Sillitoe on behalf of the Wellcome Sanger Institute COVID-19 Surveillance Team |
| EPI_ISL_3338367, EPI_ISL_3338368 | Lighthouse Lab in Alderley Park | Wellcome Sanger Institute for the COVID-19 Genomics UK (COG-UK) Consortium | Jacquelyn Wynn, Mairead Hyland, The Lighthouse Lab in Alderley Park and Alex Alderton, Roberto Amato, Jeffrey Barrett, Sonia Goncalves, Ewan Harrison, David K. Jackson, Ian Johnston, Dominic Kwiatkowski, Cordelia Langford, John Sillitoe on behalf of the Wellcome Sanger Institute COVID-19 Surveillance Team |
| EPI_ISL_3338369 | Lighthouse Laboratory Plymouth | Wellcome Sanger Institute for the COVID-19 Genomics UK (COG-UK) Consortium | Lighthouse Laboratory Plymouth and Alex Alderton, Roberto Amato, Jeffrey Barrett, Sonia Goncalves, Ewan Harrison, David K. Jackson, Ian Johnston, Dominic Kwiatkowski, Cordelia Langford, John Sillitoe on behalf of the Wellcome Sanger Institute COVID-19 Surveillance Team |
| EPI_ISL_3338370, EPI_ISL_3338371 | Lighthouse Lab in Alderley Park | Wellcome Sanger Institute for the COVID-19 Genomics UK (COG-UK) Consortium | Jacquelyn Wynn, Mairead Hyland, The Lighthouse Lab in Alderley Park and Alex Alderton, Roberto Amato, Jeffrey Barrett, Sonia Goncalves, Ewan Harrison, David K. Jackson, Ian Johnston, Dominic Kwiatkowski, Cordelia Langford, John Sillitoe on behalf of the Wellcome Sanger Institute COVID-19 Surveillance Team |
| EPI_ISL_3338372 | Lighthouse Lab in Glasgow | Wellcome Sanger Institute for the COVID-19 Genomics UK (COG-UK) Consortium | Harper VanSteenhouse, Yumi Kasai, David Gray, Carol Clugston, Anna Dominiczak and Alex Alderton, Roberto Amato, Jeffrey Barrett, Sonia Goncalves, Ewan Harrison, David K. Jackson, Ian Johnston, Dominic Kwiatkowski, Cordelia Langford, John Sillitoe on behalf of the Wellcome Sanger Institute COVID-19 Surveillance Team |
| EPI_ISL_3338373, EPI_ISL_3338374 | Lighthouse Lab in Alderley Park | Wellcome Sanger Institute for the COVID-19 Genomics UK (COG-UK) Consortium | Jacquelyn Wynn, Mairead Hyland, The Lighthouse Lab in Alderley Park and Alex Alderton, Roberto Amato, Jeffrey Barrett, Sonia Goncalves, Ewan Harrison, David K. Jackson, Ian Johnston, Dominic Kwiatkowski, Cordelia Langford, John Sillitoe on behalf of the Wellcome Sanger Institute COVID-19 Surveillance Team |
| EPI_ISL_3338375 | Lighthouse Laboratory Plymouth | Wellcome Sanger Institute for the COVID-19 Genomics UK (COG-UK) Consortium | Lighthouse Laboratory Plymouth and Alex Alderton, Roberto Amato, Jeffrey Barrett, Sonia Goncalves, Ewan Harrison, David K. Jackson, Ian Johnston, Dominic Kwiatkowski, Cordelia Langford, John Sillitoe on behalf of the Wellcome Sanger Institute COVID-19 Surveillance Team |
| EPI_ISL_3338376 | Lighthouse Lab in Glasgow | Wellcome Sanger Institute for the COVID-19 Genomics UK (COG-UK) Consortium | Harper VanSteenhouse, Yumi Kasai, David Gray, Carol Clugston, Anna Dominiczak and Alex Alderton, Roberto Amato, Jeffrey Barrett, Sonia Goncalves, Ewan Harrison, David K. Jackson, Ian Johnston, Dominic Kwiatkowski, Cordelia Langford, John Sillitoe on behalf of the Wellcome Sanger Institute COVID-19 Surveillance Team |
| EPI_ISL_3338377 | Lighthouse Lab in Alderley Park | Wellcome Sanger Institute for the COVID-19 Genomics UK (COG-UK) Consortium | Jacquelyn Wynn, Mairead Hyland, The Lighthouse Lab in Alderley Park and Alex Alderton, Roberto Amato, Jeffrey Barrett, Sonia Goncalves, Ewan Harrison, David K. Jackson, Ian Johnston, Dominic Kwiatkowski, Cordelia Langford, John Sillitoe on behalf of the Wellcome Sanger Institute COVID-19 Surveillance Team |
| EPI_ISL_3338378 | Lighthouse Laboratory Plymouth | Wellcome Sanger Institute for the COVID-19 Genomics UK (COG-UK) Consortium | Lighthouse Laboratory Plymouth and Alex Alderton, Roberto Amato, Jeffrey Barrett, Sonia Goncalves, Ewan Harrison, David K. Jackson, Ian Johnston, Dominic Kwiatkowski, Cordelia Langford, John Sillitoe on behalf of the Wellcome Sanger Institute COVID-19 Surveillance Team |
| EPI_ISL_3338379, EPI_ISL_3338380, EPI_ISL_3338381 | Lighthouse Lab in Alderley Park | Wellcome Sanger Institute for the COVID-19 Genomics UK (COG-UK) Consortium | Jacquelyn Wynn, Mairead Hyland, The Lighthouse Lab in Alderley Park and Alex Alderton, Roberto Amato, Jeffrey Barrett, Sonia Goncalves, Ewan Harrison, David K. Jackson, Ian Johnston, Dominic Kwiatkowski, Cordelia Langford, John Sillitoe on behalf of the Wellcome Sanger Institute COVID-19 Surveillance Team |
| EPI_ISL_3338382 | Lighthouse Laboratory Plymouth | Wellcome Sanger Institute for the COVID-19 Genomics UK (COG-UK) Consortium | Lighthouse Laboratory Plymouth and Alex Alderton, Roberto Amato, Jeffrey Barrett, Sonia Goncalves, Ewan Harrison, David K. Jackson, Ian Johnston, Dominic Kwiatkowski, Cordelia Langford, John Sillitoe on behalf of the Wellcome Sanger Institute COVID-19 Surveillance Team |
| EPI_ISL_3338383 | Lighthouse Lab in Alderley Park | Wellcome Sanger Institute for the COVID-19 Genomics UK (COG-UK) Consortium | Jacquelyn Wynn, Mairead Hyland, The Lighthouse Lab in Alderley Park and Alex Alderton, Roberto Amato, Jeffrey Barrett, Sonia Goncalves, Ewan Harrison, David K. Jackson, Ian Johnston, Dominic Kwiatkowski, Cordelia Langford, John Sillitoe on behalf of the Wellcome Sanger Institute COVID-19 Surveillance Team |
| EPI_ISL_3338384, EPI_ISL_3338385 | Lighthouse Laboratory Plymouth | Wellcome Sanger Institute for the COVID-19 Genomics UK (COG-UK) Consortium | Lighthouse Laboratory Plymouth and Alex Alderton, Roberto Amato, Jeffrey Barrett, Sonia Goncalves, Ewan Harrison, David K. Jackson, Ian Johnston, Dominic Kwiatkowski, Cordelia Langford, John Sillitoe on behalf of the Wellcome Sanger Institute COVID-19 Surveillance Team |
| EPI_ISL_3338386 | Lighthouse Lab in Alderley Park | Wellcome Sanger Institute for the COVID-19 Genomics UK (COG-UK) Consortium | Jacquelyn Wynn, Mairead Hyland, The Lighthouse Lab in Alderley Park and Alex Alderton, Roberto Amato, Jeffrey Barrett, Sonia Goncalves, Ewan Harrison, David K. Jackson, Ian Johnston, Dominic Kwiatkowski, Cordelia Langford, John Sillitoe on behalf of the Wellcome Sanger Institute COVID-19 Surveillance Team |
| EPI_ISL_3338387 | Lighthouse Laboratory Plymouth | Wellcome Sanger Institute for the COVID-19 Genomics UK (COG-UK) Consortium | Lighthouse Laboratory Plymouth and Alex Alderton, Roberto Amato, Jeffrey Barrett, Sonia Goncalves, Ewan Harrison, David K. Jackson, Ian Johnston, Dominic Kwiatkowski, Cordelia Langford, John Sillitoe on behalf of the Wellcome Sanger Institute COVID-19 Surveillance Team |
| EPI_ISL_3338388, EPI_ISL_3338389, EPI_ISL_3338390, EPI_ISL_3338391, EPI_ISL_3338392 | Lighthouse Lab in Alderley Park | Wellcome Sanger Institute for the COVID-19 Genomics UK (COG-UK) Consortium | Jacquelyn Wynn, Mairead Hyland, The Lighthouse Lab in Alderley Park and Alex Alderton, Roberto Amato, Jeffrey Barrett, Sonia Goncalves, Ewan Harrison, David K. Jackson, Ian Johnston, Dominic Kwiatkowski, Cordelia Langford, John Sillitoe on behalf of the Wellcome Sanger Institute COVID-19 Surveillance Team |

| | | | |
|---|---|---|---|
| EPI_ISL_3338393 | Lighthouse Laboratory Plymouth | Wellcome Sanger Institute for the COVID-19 Genomics UK (COG-UK) Consortium | Lighthouse Laboratory Plymouth and Alex Alderton, Roberto Amato, Jeffrey Barrett, Sonia Goncalves, Ewan Harrison, David K. Jackson, Ian Johnston, Dominic Kwiatkowski, Cordelia Langford, John Sillitoe on behalf of the Wellcome Sanger Institute COVID-19 Surveillance Team |
| EPI_ISL_3338394, EPI_ISL_3338395, EPI_ISL_3338396 | Lighthouse Lab in Alderley Park | Wellcome Sanger Institute for the COVID-19 Genomics UK (COG-UK) Consortium | Jacquelyn Wynn, Mairead Hyland, The Lighthouse Lab in Alderley Park and Alex Alderton, Roberto Amato, Jeffrey Barrett, Sonia Goncalves, Ewan Harrison, David K. Jackson, Ian Johnston, Dominic Kwiatkowski, Cordelia Langford, John Sillitoe on behalf of the Wellcome Sanger Institute COVID-19 Surveillance Team |
| EPI_ISL_3338397 | Lighthouse Laboratory Plymouth | Wellcome Sanger Institute for the COVID-19 Genomics UK (COG-UK) Consortium | Lighthouse Laboratory Plymouth and Alex Alderton, Roberto Amato, Jeffrey Barrett, Sonia Goncalves, Ewan Harrison, David K. Jackson, Ian Johnston, Dominic Kwiatkowski, Cordelia Langford, John Sillitoe on behalf of the Wellcome Sanger Institute COVID-19 Surveillance Team |
| EPI_ISL_3338398, EPI_ISL_3338399, EPI_ISL_3338400, EPI_ISL_3338401 | Lighthouse Lab in Alderley Park | Wellcome Sanger Institute for the COVID-19 Genomics UK (COG-UK) Consortium | Jacquelyn Wynn, Mairead Hyland, The Lighthouse Lab in Alderley Park and Alex Alderton, Roberto Amato, Jeffrey Barrett, Sonia Goncalves, Ewan Harrison, David K. Jackson, Ian Johnston, Dominic Kwiatkowski, Cordelia Langford, John Sillitoe on behalf of the Wellcome Sanger Institute COVID-19 Surveillance Team |
| EPI_ISL_3338402 | Lighthouse Laboratory Plymouth | Wellcome Sanger Institute for the COVID-19 Genomics UK (COG-UK) Consortium | Lighthouse Laboratory Plymouth and Alex Alderton, Roberto Amato, Jeffrey Barrett, Sonia Goncalves, Ewan Harrison, David K. Jackson, Ian Johnston, Dominic Kwiatkowski, Cordelia Langford, John Sillitoe on behalf of the Wellcome Sanger Institute COVID-19 Surveillance Team |
| EPI_ISL_3338403, EPI_ISL_3338404 | Lighthouse Lab in Alderley Park | Wellcome Sanger Institute for the COVID-19 Genomics UK (COG-UK) Consortium | Jacquelyn Wynn, Mairead Hyland, The Lighthouse Lab in Alderley Park and Alex Alderton, Roberto Amato, Jeffrey Barrett, Sonia Goncalves, Ewan Harrison, David K. Jackson, Ian Johnston, Dominic Kwiatkowski, Cordelia Langford, John Sillitoe on behalf of the Wellcome Sanger Institute COVID-19 Surveillance Team |
| EPI_ISL_3338405 | Lighthouse Laboratory Plymouth | Wellcome Sanger Institute for the COVID-19 Genomics UK (COG-UK) Consortium | Lighthouse Laboratory Plymouth and Alex Alderton, Roberto Amato, Jeffrey Barrett, Sonia Goncalves, Ewan Harrison, David K. Jackson, Ian Johnston, Dominic Kwiatkowski, Cordelia Langford, John Sillitoe on behalf of the Wellcome Sanger Institute COVID-19 Surveillance Team |
| EPI_ISL_3338406, EPI_ISL_3338407, EPI_ISL_3338408 | Lighthouse Lab in Alderley Park | Wellcome Sanger Institute for the COVID-19 Genomics UK (COG-UK) Consortium | Jacquelyn Wynn, Mairead Hyland, The Lighthouse Lab in Alderley Park and Alex Alderton, Roberto Amato, Jeffrey Barrett, Sonia Goncalves, Ewan Harrison, David K. Jackson, Ian Johnston, Dominic Kwiatkowski, Cordelia Langford, John Sillitoe on behalf of the Wellcome Sanger Institute COVID-19 Surveillance Team |
| EPI_ISL_3338409 | Lighthouse Laboratory Plymouth | Wellcome Sanger Institute for the COVID-19 Genomics UK (COG-UK) Consortium | Lighthouse Laboratory Plymouth and Alex Alderton, Roberto Amato, Jeffrey Barrett, Sonia Goncalves, Ewan Harrison, David K. Jackson, Ian Johnston, Dominic Kwiatkowski, Cordelia Langford, John Sillitoe on behalf of the Wellcome Sanger Institute COVID-19 Surveillance Team |
| EPI_ISL_3338410, EPI_ISL_3338411, EPI_ISL_3338412, EPI_ISL_3338413 | Lighthouse Lab in Glasgow | Wellcome Sanger Institute for the COVID-19 Genomics UK (COG-UK) Consortium | Harper VanSteenhouse, Yumi Kasai, David Gray, Carol Clugston, Anna Dominiczak and Alex Alderton, Roberto Amato, Jeffrey Barrett, Sonia Goncalves, Ewan Harrison, David K. Jackson, Ian Johnston, Dominic Kwiatkowski, Cordelia Langford, John Sillitoe on behalf of the Wellcome Sanger Institute COVID-19 Surveillance Team |
| EPI_ISL_3338414 | Lighthouse Laboratory Plymouth | Wellcome Sanger Institute for the COVID-19 Genomics UK (COG-UK) Consortium | Lighthouse Laboratory Plymouth and Alex Alderton, Roberto Amato, Jeffrey Barrett, Sonia Goncalves, Ewan Harrison, David K. Jackson, Ian Johnston, Dominic Kwiatkowski, Cordelia Langford, John Sillitoe on behalf of the Wellcome Sanger Institute COVID-19 Surveillance Team |
| EPI_ISL_3338415, EPI_ISL_3338416 | Lighthouse Lab in Alderley Park | Wellcome Sanger Institute for the COVID-19 Genomics UK (COG-UK) Consortium | Jacquelyn Wynn, Mairead Hyland, The Lighthouse Lab in Alderley Park and Alex Alderton, Roberto Amato, Jeffrey Barrett, Sonia Goncalves, Ewan Harrison, David K. Jackson, Ian Johnston, Dominic Kwiatkowski, Cordelia Langford, John Sillitoe on behalf of the Wellcome Sanger Institute COVID-19 Surveillance Team |
| EPI_ISL_3338417, EPI_ISL_3338418 | Lighthouse Laboratory Plymouth | Wellcome Sanger Institute for the COVID-19 Genomics UK (COG-UK) Consortium | Lighthouse Laboratory Plymouth and Alex Alderton, Roberto Amato, Jeffrey Barrett, Sonia Goncalves, Ewan Harrison, David K. Jackson, Ian Johnston, Dominic Kwiatkowski, Cordelia Langford, John Sillitoe on behalf of the Wellcome Sanger Institute COVID-19 Surveillance Team |
| EPI_ISL_3338419, EPI_ISL_3338420, EPI_ISL_3338421 | Lighthouse Lab in Alderley Park | Wellcome Sanger Institute for the COVID-19 Genomics UK (COG-UK) Consortium | Jacquelyn Wynn, Mairead Hyland, The Lighthouse Lab in Alderley Park and Alex Alderton, Roberto Amato, Jeffrey Barrett, Sonia Goncalves, Ewan Harrison, David K. Jackson, Ian Johnston, Dominic Kwiatkowski, Cordelia Langford, John Sillitoe on behalf of the Wellcome Sanger Institute COVID-19 Surveillance Team |
| EPI_ISL_3338422 | Lighthouse Lab in Glasgow | Wellcome Sanger Institute for the COVID-19 Genomics UK (COG-UK) Consortium | Harper VanSteenhouse, Yumi Kasai, David Gray, Carol Clugston, Anna Dominiczak and Alex Alderton, Roberto Amato, Jeffrey Barrett, Sonia Goncalves, Ewan Harrison, David K. Jackson, Ian Johnston, Dominic Kwiatkowski, Cordelia Langford, John Sillitoe on behalf of the Wellcome Sanger Institute COVID-19 Surveillance Team |
| EPI_ISL_3338423 | Lighthouse Laboratory Plymouth | Wellcome Sanger Institute for the COVID-19 Genomics UK (COG-UK) Consortium | Lighthouse Laboratory Plymouth and Alex Alderton, Roberto Amato, Jeffrey Barrett, Sonia Goncalves, Ewan Harrison, David K. Jackson, Ian Johnston, Dominic Kwiatkowski, Cordelia Langford, John Sillitoe on behalf of the Wellcome Sanger Institute COVID-19 Surveillance Team |
| EPI_ISL_3338424 | Lighthouse Lab in Alderley Park | Wellcome Sanger Institute for the COVID-19 Genomics UK (COG-UK) Consortium | Jacquelyn Wynn, Mairead Hyland, The Lighthouse Lab in Alderley Park and Alex Alderton, Roberto Amato, Jeffrey Barrett, Sonia Goncalves, Ewan Harrison, David K. Jackson, Ian Johnston, Dominic Kwiatkowski, Cordelia Langford, John Sillitoe on behalf of the Wellcome Sanger Institute COVID-19 Surveillance Team |
| EPI_ISL_3338425, EPI_ISL_3338426 | Lighthouse Laboratory Plymouth | Wellcome Sanger Institute for the COVID-19 Genomics UK (COG-UK) Consortium | Lighthouse Laboratory Plymouth and Alex Alderton, Roberto Amato, Jeffrey Barrett, Sonia Goncalves, Ewan Harrison, David K. Jackson, Ian Johnston, Dominic Kwiatkowski, Cordelia Langford, John Sillitoe on behalf of the Wellcome Sanger Institute COVID-19 Surveillance Team |
| EPI_ISL_3338427 | Lighthouse Lab in Glasgow | Wellcome Sanger Institute for the COVID-19 Genomics UK (COG-UK) Consortium | Harper VanSteenhouse, Yumi Kasai, David Gray, Carol Clugston, Anna Dominiczak and Alex Alderton, Roberto Amato, Jeffrey Barrett, Sonia Goncalves, Ewan Harrison, David K. Jackson, Ian Johnston, Dominic Kwiatkowski, Cordelia Langford, John Sillitoe on behalf of the Wellcome Sanger Institute COVID-19 Surveillance Team |
| EPI_ISL_3338428, EPI_ISL_3338429, EPI_ISL_3338430 | Lighthouse Lab in Alderley Park | Wellcome Sanger Institute for the COVID-19 Genomics UK (COG-UK) Consortium | Jacquelyn Wynn, Mairead Hyland, The Lighthouse Lab in Alderley Park and Alex Alderton, Roberto Amato, Jeffrey Barrett, Sonia Goncalves, Ewan Harrison, David K. Jackson, Ian Johnston, Dominic Kwiatkowski, Cordelia Langford, John Sillitoe on behalf of the Wellcome Sanger Institute COVID-19 Surveillance Team |
| EPI_ISL_3338431 | Lighthouse Laboratory Plymouth | Wellcome Sanger Institute for the COVID-19 Genomics UK (COG-UK) Consortium | Lighthouse Laboratory Plymouth and Alex Alderton, Roberto Amato, Jeffrey Barrett, Sonia Goncalves, Ewan Harrison, David K. Jackson, Ian Johnston, Dominic Kwiatkowski, Cordelia Langford, John Sillitoe on behalf of the Wellcome Sanger Institute COVID-19 Surveillance Team |
| EPI_ISL_3338432 | Lighthouse Lab in Glasgow | Wellcome Sanger Institute for the COVID-19 Genomics UK (COG-UK) Consortium | Harper VanSteenhouse, Yumi Kasai, David Gray, Carol Clugston, Anna Dominiczak and Alex Alderton, Roberto Amato, Jeffrey Barrett, Sonia Goncalves, Ewan Harrison, David K. Jackson, Ian Johnston, Dominic Kwiatkowski, Cordelia Langford, John Sillitoe on behalf of the Wellcome Sanger Institute COVID-19 Surveillance Team |
| EPI_ISL_3338433, EPI_ISL_3338434 | Lighthouse Lab in Alderley Park | Wellcome Sanger Institute for the COVID-19 Genomics UK (COG-UK) Consortium | Jacquelyn Wynn, Mairead Hyland, The Lighthouse Lab in Alderley Park and Alex Alderton, Roberto Amato, Jeffrey Barrett, Sonia Goncalves, Ewan Harrison, David K. Jackson, Ian Johnston, Dominic Kwiatkowski, Cordelia Langford, John Sillitoe on behalf of the Wellcome Sanger Institute COVID-19 Surveillance Team |
| EPI_ISL_3338435 | Lighthouse Lab in Glasgow | Wellcome Sanger Institute for the COVID-19 Genomics UK (COG-UK) Consortium | Harper VanSteenhouse, Yumi Kasai, David Gray, Carol Clugston, Anna Dominiczak and Alex Alderton, Roberto Amato, Jeffrey Barrett, Sonia Goncalves, Ewan Harrison, David K. Jackson, Ian Johnston, Dominic Kwiatkowski, Cordelia Langford, John Sillitoe on behalf of the Wellcome Sanger Institute COVID-19 Surveillance Team |
| EPI_ISL_3338436 | Lighthouse Lab in Alderley Park | Wellcome Sanger Institute for the COVID-19 Genomics UK (COG-UK) Consortium | Jacquelyn Wynn, Mairead Hyland, The Lighthouse Lab in Alderley Park and Alex Alderton, Roberto Amato, Jeffrey Barrett, Sonia Goncalves, Ewan Harrison, David K. Jackson, Ian Johnston, Dominic Kwiatkowski, Cordelia Langford, John Sillitoe on behalf of the Wellcome Sanger Institute COVID-19 Surveillance Team |
| EPI_ISL_3338437, EPI_ISL_3338438 | Lighthouse Laboratory Plymouth | Wellcome Sanger Institute for the COVID-19 Genomics UK (COG-UK) Consortium | Lighthouse Laboratory Plymouth and Alex Alderton, Roberto Amato, Jeffrey Barrett, Sonia Goncalves, Ewan Harrison, David K. Jackson, Ian Johnston, Dominic Kwiatkowski, Cordelia Langford, John Sillitoe on behalf of the Wellcome Sanger Institute COVID-19 Surveillance Team |
| EPI_ISL_3338439 | Lighthouse Lab in Alderley Park | Wellcome Sanger Institute for the COVID-19 Genomics UK (COG-UK) Consortium | Jacquelyn Wynn, Mairead Hyland, The Lighthouse Lab in Alderley Park and Alex Alderton, Roberto Amato, Jeffrey Barrett, Sonia Goncalves, Ewan Harrison, David K. Jackson, Ian Johnston, Dominic Kwiatkowski, Cordelia Langford, John Sillitoe on behalf of the Wellcome Sanger Institute COVID-19 Surveillance Team |
| EPI_ISL_3338440 | Lighthouse Laboratory Plymouth | Wellcome Sanger Institute for the COVID-19 Genomics UK (COG-UK) Consortium | Lighthouse Laboratory Plymouth and Alex Alderton, Roberto Amato, Jeffrey Barrett, Sonia Goncalves, Ewan Harrison, David K. Jackson, Ian Johnston, Dominic Kwiatkowski, Cordelia Langford, John Sillitoe on behalf of the Wellcome Sanger Institute COVID-19 Surveillance Team |
| EPI_ISL_3338441 | Lighthouse Lab in Alderley Park | Wellcome Sanger Institute for the COVID-19 Genomics UK (COG-UK) Consortium | Jacquelyn Wynn, Mairead Hyland, The Lighthouse Lab in Alderley Park and Alex Alderton, Roberto Amato, Jeffrey Barrett, Sonia Goncalves, Ewan Harrison, David K. Jackson, Ian Johnston, Dominic Kwiatkowski, Cordelia Langford, John Sillitoe on behalf of the Wellcome Sanger Institute COVID-19 Surveillance Team |
| EPI_ISL_3338442 | Lighthouse Laboratory Plymouth | Wellcome Sanger Institute for the COVID-19 Genomics UK (COG-UK) Consortium | Lighthouse Laboratory Plymouth and Alex Alderton, Roberto Amato, Jeffrey Barrett, Sonia Goncalves, Ewan Harrison, David K. Jackson, Ian Johnston, Dominic Kwiatkowski, Cordelia Langford, John Sillitoe on behalf of the Wellcome Sanger Institute COVID-19 Surveillance Team |
| EPI_ISL_3338443 | Lighthouse Lab in Alderley Park | Wellcome Sanger Institute for the COVID-19 Genomics UK (COG-UK) Consortium | Jacquelyn Wynn, Mairead Hyland, The Lighthouse Lab in Alderley Park and Alex Alderton, Roberto Amato, Jeffrey Barrett, Sonia Goncalves, Ewan Harrison, David K. Jackson, Ian Johnston, Dominic Kwiatkowski, Cordelia Langford, John Sillitoe on behalf of the Wellcome Sanger Institute COVID-19 Surveillance Team |
| EPI_ISL_3338444, EPI_ISL_3338445 | Lighthouse Laboratory Plymouth | Wellcome Sanger Institute for the COVID-19 Genomics UK (COG-UK) Consortium | Lighthouse Laboratory Plymouth and Alex Alderton, Roberto Amato, Jeffrey Barrett, Sonia Goncalves, Ewan Harrison, David K. Jackson, Ian Johnston, Dominic Kwiatkowski, Cordelia Langford, John Sillitoe on behalf of the Wellcome Sanger Institute COVID-19 Surveillance Team |
| EPI_ISL_3338446 | Lighthouse Lab in Glasgow | Wellcome Sanger Institute for the COVID-19 Genomics UK (COG-UK) Consortium | Harper VanSteenhouse, Yumi Kasai, David Gray, Carol Clugston, Anna Dominiczak and Alex Alderton, Roberto Amato, Jeffrey Barrett, Sonia Goncalves, Ewan Harrison, David K. Jackson, Ian Johnston, Dominic Kwiatkowski, Cordelia Langford, John Sillitoe on behalf of the Wellcome Sanger Institute COVID-19 Surveillance Team |
| EPI_ISL_3338447 | Lighthouse Lab in Alderley Park | Wellcome Sanger Institute for the COVID-19 Genomics UK (COG-UK) Consortium | Jacquelyn Wynn, Mairead Hyland, The Lighthouse Lab in Alderley Park and Alex Alderton, Roberto Amato, Jeffrey Barrett, Sonia Goncalves, Ewan Harrison, David K. Jackson, Ian Johnston, Dominic Kwiatkowski, Cordelia Langford, John Sillitoe on behalf of the Wellcome Sanger Institute COVID-19 Surveillance Team |
| EPI_ISL_3338448 | Lighthouse Laboratory Plymouth | Wellcome Sanger Institute for the COVID-19 Genomics UK (COG-UK) Consortium | Lighthouse Laboratory Plymouth and Alex Alderton, Roberto Amato, Jeffrey Barrett, Sonia Goncalves, Ewan Harrison, David K. Jackson, Ian Johnston, Dominic Kwiatkowski, Cordelia Langford, John Sillitoe on behalf of the Wellcome Sanger Institute COVID-19 Surveillance Team |
| EPI_ISL_3338449, EPI_ISL_3338450, EPI_ISL_3338451 | Lighthouse Lab in Alderley Park | Wellcome Sanger Institute for the COVID-19 Genomics UK (COG-UK) Consortium | Jacquelyn Wynn, Mairead Hyland, The Lighthouse Lab in Alderley Park and Alex Alderton, Roberto Amato, Jeffrey Barrett, Sonia Goncalves, Ewan Harrison, David K. Jackson, Ian Johnston, Dominic Kwiatkowski, Cordelia Langford, John Sillitoe on behalf of the Wellcome Sanger Institute COVID-19 Surveillance Team |
| EPI_ISL_3338452 | Lighthouse Lab in Glasgow | Wellcome Sanger Institute for the COVID-19 Genomics UK (COG-UK) Consortium | Harper VanSteenhouse, Yumi Kasai, David Gray, Carol Clugston, Anna Dominiczak and Alex Alderton, Roberto Amato, Jeffrey Barrett, Sonia Goncalves, Ewan Harrison, David K. Jackson, Ian Johnston, Dominic Kwiatkowski, Cordelia Langford, John Sillitoe on behalf of the Wellcome Sanger Institute COVID-19 Surveillance Team |
| EPI_ISL_3338453 | Lighthouse Lab in Alderley Park | Wellcome Sanger Institute for the COVID-19 Genomics UK (COG-UK) Consortium | Jacquelyn Wynn, Mairead Hyland, The Lighthouse Lab in Alderley Park and Alex Alderton, Roberto Amato, Jeffrey Barrett, Sonia Goncalves, Ewan Harrison, David K. Jackson, Ian Johnston, Dominic Kwiatkowski, Cordelia Langford, John Sillitoe on behalf of the Wellcome Sanger Institute COVID-19 Surveillance Team |
| EPI_ISL_3338454 | Lighthouse Laboratory Plymouth | Wellcome Sanger Institute for the COVID-19 Genomics UK (COG-UK) Consortium | Lighthouse Laboratory Plymouth and Alex Alderton, Roberto Amato, Jeffrey Barrett, Sonia Goncalves, Ewan Harrison, David K. Jackson, Ian Johnston, Dominic Kwiatkowski, Cordelia Langford, John Sillitoe on behalf of the Wellcome Sanger Institute COVID-19 Surveillance Team |
| EPI_ISL_3338455 | Lighthouse Lab in Glasgow | Wellcome Sanger Institute for the COVID-19 Genomics UK (COG-UK) Consortium | Harper VanSteenhouse, Yumi Kasai, David Gray, Carol Clugston, Anna Dominiczak and Alex Alderton, Roberto Amato, Jeffrey Barrett, Sonia Goncalves, Ewan Harrison, David K. Jackson, Ian Johnston, Dominic Kwiatkowski, Cordelia Langford, John Sillitoe on behalf of the Wellcome Sanger Institute COVID-19 Surveillance Team |
| EPI_ISL_3338456, EPI_ISL_3338457 | Lighthouse Laboratory Plymouth | Wellcome Sanger Institute for the COVID-19 Genomics UK (COG-UK) Consortium | Lighthouse Laboratory Plymouth and Alex Alderton, Roberto Amato, Jeffrey Barrett, Sonia Goncalves, Ewan Harrison, David K. Jackson, Ian Johnston, Dominic Kwiatkowski, Cordelia Langford, John Sillitoe on behalf of the Wellcome Sanger Institute COVID-19 Surveillance Team |

| | | | |
|---|---|---|---|
| EPI_ISL_3338458 | Lighthouse Lab in Alderley Park | Wellcome Sanger Institute for the COVID-19 Genomics UK (COG-UK) Consortium | Jacquelyn Wynn, Mairead Hyland, The Lighthouse Lab in Alderley Park and Alex Alderton, Roberto Amato, Jeffrey Barrett, Sonia Goncalves, Ewan Harrison, David K. Jackson, Ian Johnston, Dominic Kwiatkowski, Cordelia Langford, John Sillitoe on behalf of the Wellcome Sanger Institute COVID-19 Surveillance Team |
| EPI_ISL_3338459 | Lighthouse Lab in Glasgow | Wellcome Sanger Institute for the COVID-19 Genomics UK (COG-UK) Consortium | Harper VanSteenhouse, Yumi Kasai, David Gray, Carol Clugston, Anna Dominiczak and Alex Alderton, Roberto Amato, Jeffrey Barrett, Sonia Goncalves, Ewan Harrison, David K. Jackson, Ian Johnston, Dominic Kwiatkowski, Cordelia Langford, John Sillitoe on behalf of the Wellcome Sanger Institute COVID-19 Surveillance Team |
| EPI_ISL_3338460 | Lighthouse Lab in Alderley Park | Wellcome Sanger Institute for the COVID-19 Genomics UK (COG-UK) Consortium | Jacquelyn Wynn, Mairead Hyland, The Lighthouse Lab in Alderley Park and Alex Alderton, Roberto Amato, Jeffrey Barrett, Sonia Goncalves, Ewan Harrison, David K. Jackson, Ian Johnston, Dominic Kwiatkowski, Cordelia Langford, John Sillitoe on behalf of the Wellcome Sanger Institute COVID-19 Surveillance Team |
| EPI_ISL_3338461 | Lighthouse Lab in Glasgow | Wellcome Sanger Institute for the COVID-19 Genomics UK (COG-UK) Consortium | Harper VanSteenhouse, Yumi Kasai, David Gray, Carol Clugston, Anna Dominiczak and Alex Alderton, Roberto Amato, Jeffrey Barrett, Sonia Goncalves, Ewan Harrison, David K. Jackson, Ian Johnston, Dominic Kwiatkowski, Cordelia Langford, John Sillitoe on behalf of the Wellcome Sanger Institute COVID-19 Surveillance Team |
| EPI_ISL_3338462 | Lighthouse Laboratory Plymouth | Wellcome Sanger Institute for the COVID-19 Genomics UK (COG-UK) Consortium | Lighthouse Laboratory Plymouth and Alex Alderton, Roberto Amato, Jeffrey Barrett, Sonia Goncalves, Ewan Harrison, David K. Jackson, Ian Johnston, Dominic Kwiatkowski, Cordelia Langford, John Sillitoe on behalf of the Wellcome Sanger Institute COVID-19 Surveillance Team |
| EPI_ISL_3338463, EPI_ISL_3338464, EPI_ISL_3338465, EPI_ISL_3338466, EPI_ISL_3338467, EPI_ISL_3338468 | Lighthouse Lab in Alderley Park | Wellcome Sanger Institute for the COVID-19 Genomics UK (COG-UK) Consortium | Jacquelyn Wynn, Mairead Hyland, The Lighthouse Lab in Alderley Park and Alex Alderton, Roberto Amato, Jeffrey Barrett, Sonia Goncalves, Ewan Harrison, David K. Jackson, Ian Johnston, Dominic Kwiatkowski, Cordelia Langford, John Sillitoe on behalf of the Wellcome Sanger Institute COVID-19 Surveillance Team |
| EPI_ISL_3338469, EPI_ISL_3338470 | Lighthouse Laboratory Plymouth | Wellcome Sanger Institute for the COVID-19 Genomics UK (COG-UK) Consortium | Lighthouse Laboratory Plymouth and Alex Alderton, Roberto Amato, Jeffrey Barrett, Sonia Goncalves, Ewan Harrison, David K. Jackson, Ian Johnston, Dominic Kwiatkowski, Cordelia Langford, John Sillitoe on behalf of the Wellcome Sanger Institute COVID-19 Surveillance Team |
| EPI_ISL_3338471, EPI_ISL_3338472, EPI_ISL_3338473 | Lighthouse Lab in Alderley Park | Wellcome Sanger Institute for the COVID-19 Genomics UK (COG-UK) Consortium | Jacquelyn Wynn, Mairead Hyland, The Lighthouse Lab in Alderley Park and Alex Alderton, Roberto Amato, Jeffrey Barrett, Sonia Goncalves, Ewan Harrison, David K. Jackson, Ian Johnston, Dominic Kwiatkowski, Cordelia Langford, John Sillitoe on behalf of the Wellcome Sanger Institute COVID-19 Surveillance Team |
| EPI_ISL_3338474, EPI_ISL_3338475, EPI_ISL_3338476 | Lighthouse Laboratory Plymouth | Wellcome Sanger Institute for the COVID-19 Genomics UK (COG-UK) Consortium | Lighthouse Laboratory Plymouth and Alex Alderton, Roberto Amato, Jeffrey Barrett, Sonia Goncalves, Ewan Harrison, David K. Jackson, Ian Johnston, Dominic Kwiatkowski, Cordelia Langford, John Sillitoe on behalf of the Wellcome Sanger Institute COVID-19 Surveillance Team |
| EPI_ISL_3338477 | Lighthouse Lab in Alderley Park | Wellcome Sanger Institute for the COVID-19 Genomics UK (COG-UK) Consortium | Jacquelyn Wynn, Mairead Hyland, The Lighthouse Lab in Alderley Park and Alex Alderton, Roberto Amato, Jeffrey Barrett, Sonia Goncalves, Ewan Harrison, David K. Jackson, Ian Johnston, Dominic Kwiatkowski, Cordelia Langford, John Sillitoe on behalf of the Wellcome Sanger Institute COVID-19 Surveillance Team |
| EPI_ISL_3338478, EPI_ISL_3338479 | Lighthouse Laboratory Plymouth | Wellcome Sanger Institute for the COVID-19 Genomics UK (COG-UK) Consortium | Lighthouse Laboratory Plymouth and Alex Alderton, Roberto Amato, Jeffrey Barrett, Sonia Goncalves, Ewan Harrison, David K. Jackson, Ian Johnston, Dominic Kwiatkowski, Cordelia Langford, John Sillitoe on behalf of the Wellcome Sanger Institute COVID-19 Surveillance Team |
| EPI_ISL_3338480 | Lighthouse Lab in Alderley Park | Wellcome Sanger Institute for the COVID-19 Genomics UK (COG-UK) Consortium | Jacquelyn Wynn, Mairead Hyland, The Lighthouse Lab in Alderley Park and Alex Alderton, Roberto Amato, Jeffrey Barrett, Sonia Goncalves, Ewan Harrison, David K. Jackson, Ian Johnston, Dominic Kwiatkowski, Cordelia Langford, John Sillitoe on behalf of the Wellcome Sanger Institute COVID-19 Surveillance Team |
| EPI_ISL_3338481 | Lighthouse Lab in Glasgow | Wellcome Sanger Institute for the COVID-19 Genomics UK (COG-UK) Consortium | Harper VanSteenhouse, Yumi Kasai, David Gray, Carol Clugston, Anna Dominiczak and Alex Alderton, Roberto Amato, Jeffrey Barrett, Sonia Goncalves, Ewan Harrison, David K. Jackson, Ian Johnston, Dominic Kwiatkowski, Cordelia Langford, John Sillitoe on behalf of the Wellcome Sanger Institute COVID-19 Surveillance Team |
| EPI_ISL_3338482 | Lighthouse Laboratory Plymouth | Wellcome Sanger Institute for the COVID-19 Genomics UK (COG-UK) Consortium | Lighthouse Laboratory Plymouth and Alex Alderton, Roberto Amato, Jeffrey Barrett, Sonia Goncalves, Ewan Harrison, David K. Jackson, Ian Johnston, Dominic Kwiatkowski, Cordelia Langford, John Sillitoe on behalf of the Wellcome Sanger Institute COVID-19 Surveillance Team |
| EPI_ISL_3338483 | Lighthouse Lab in Alderley Park | Wellcome Sanger Institute for the COVID-19 Genomics UK (COG-UK) Consortium | Jacquelyn Wynn, Mairead Hyland, The Lighthouse Lab in Alderley Park and Alex Alderton, Roberto Amato, Jeffrey Barrett, Sonia Goncalves, Ewan Harrison, David K. Jackson, Ian Johnston, Dominic Kwiatkowski, Cordelia Langford, John Sillitoe on behalf of the Wellcome Sanger Institute COVID-19 Surveillance Team |
| EPI_ISL_3338484 | Lighthouse Laboratory Plymouth | Wellcome Sanger Institute for the COVID-19 Genomics UK (COG-UK) Consortium | Lighthouse Laboratory Plymouth and Alex Alderton, Roberto Amato, Jeffrey Barrett, Sonia Goncalves, Ewan Harrison, David K. Jackson, Ian Johnston, Dominic Kwiatkowski, Cordelia Langford, John Sillitoe on behalf of the Wellcome Sanger Institute COVID-19 Surveillance Team |
| EPI_ISL_3338485 | Lighthouse Lab in Alderley Park | Wellcome Sanger Institute for the COVID-19 Genomics UK (COG-UK) Consortium | Jacquelyn Wynn, Mairead Hyland, The Lighthouse Lab in Alderley Park and Alex Alderton, Roberto Amato, Jeffrey Barrett, Sonia Goncalves, Ewan Harrison, David K. Jackson, Ian Johnston, Dominic Kwiatkowski, Cordelia Langford, John Sillitoe on behalf of the Wellcome Sanger Institute COVID-19 Surveillance Team |
| EPI_ISL_3338486 | Lighthouse Laboratory Plymouth | Wellcome Sanger Institute for the COVID-19 Genomics UK (COG-UK) Consortium | Lighthouse Laboratory Plymouth and Alex Alderton, Roberto Amato, Jeffrey Barrett, Sonia Goncalves, Ewan Harrison, David K. Jackson, Ian Johnston, Dominic Kwiatkowski, Cordelia Langford, John Sillitoe on behalf of the Wellcome Sanger Institute COVID-19 Surveillance Team |
| EPI_ISL_3338487 | Lighthouse Lab in Alderley Park | Wellcome Sanger Institute for the COVID-19 Genomics UK (COG-UK) Consortium | Jacquelyn Wynn, Mairead Hyland, The Lighthouse Lab in Alderley Park and Alex Alderton, Roberto Amato, Jeffrey Barrett, Sonia Goncalves, Ewan Harrison, David K. Jackson, Ian Johnston, Dominic Kwiatkowski, Cordelia Langford, John Sillitoe on behalf of the Wellcome Sanger Institute COVID-19 Surveillance Team |
| EPI_ISL_3338488 | Lighthouse Lab in Glasgow | Wellcome Sanger Institute for the COVID-19 Genomics UK (COG-UK) Consortium | Harper VanSteenhouse, Yumi Kasai, David Gray, Carol Clugston, Anna Dominiczak and Alex Alderton, Roberto Amato, Jeffrey Barrett, Sonia Goncalves, Ewan Harrison, David K. Jackson, Ian Johnston, Dominic Kwiatkowski, Cordelia Langford, John Sillitoe on behalf of the Wellcome Sanger Institute COVID-19 Surveillance Team |
| EPI_ISL_3338489 | Lighthouse Laboratory Plymouth | Wellcome Sanger Institute for the COVID-19 Genomics UK (COG-UK) Consortium | Lighthouse Laboratory Plymouth and Alex Alderton, Roberto Amato, Jeffrey Barrett, Sonia Goncalves, Ewan Harrison, David K. Jackson, Ian Johnston, Dominic Kwiatkowski, Cordelia Langford, John Sillitoe on behalf of the Wellcome Sanger Institute COVID-19 Surveillance Team |
| EPI_ISL_3338490 | Lighthouse Lab in Alderley Park | Wellcome Sanger Institute for the COVID-19 Genomics UK (COG-UK) Consortium | Jacquelyn Wynn, Mairead Hyland, The Lighthouse Lab in Alderley Park and Alex Alderton, Roberto Amato, Jeffrey Barrett, Sonia Goncalves, Ewan Harrison, David K. Jackson, Ian Johnston, Dominic Kwiatkowski, Cordelia Langford, John Sillitoe on behalf of the Wellcome Sanger Institute COVID-19 Surveillance Team |
| EPI_ISL_3338491, EPI_ISL_3338492, EPI_ISL_3338493 | Lighthouse Laboratory Plymouth | Wellcome Sanger Institute for the COVID-19 Genomics UK (COG-UK) Consortium | Lighthouse Laboratory Plymouth and Alex Alderton, Roberto Amato, Jeffrey Barrett, Sonia Goncalves, Ewan Harrison, David K. Jackson, Ian Johnston, Dominic Kwiatkowski, Cordelia Langford, John Sillitoe on behalf of the Wellcome Sanger Institute COVID-19 Surveillance Team |
| EPI_ISL_3338494 | Lighthouse Lab in Alderley Park | Wellcome Sanger Institute for the COVID-19 Genomics UK (COG-UK) Consortium | Jacquelyn Wynn, Mairead Hyland, The Lighthouse Lab in Alderley Park and Alex Alderton, Roberto Amato, Jeffrey Barrett, Sonia Goncalves, Ewan Harrison, David K. Jackson, Ian Johnston, Dominic Kwiatkowski, Cordelia Langford, John Sillitoe on behalf of the Wellcome Sanger Institute COVID-19 Surveillance Team |
| EPI_ISL_3338495 | Lighthouse Laboratory Plymouth | Wellcome Sanger Institute for the COVID-19 Genomics UK (COG-UK) Consortium | Lighthouse Laboratory Plymouth and Alex Alderton, Roberto Amato, Jeffrey Barrett, Sonia Goncalves, Ewan Harrison, David K. Jackson, Ian Johnston, Dominic Kwiatkowski, Cordelia Langford, John Sillitoe on behalf of the Wellcome Sanger Institute COVID-19 Surveillance Team |
| EPI_ISL_3338496, EPI_ISL_3338497 | Lighthouse Lab in Alderley Park | Wellcome Sanger Institute for the COVID-19 Genomics UK (COG-UK) Consortium | Jacquelyn Wynn, Mairead Hyland, The Lighthouse Lab in Alderley Park and Alex Alderton, Roberto Amato, Jeffrey Barrett, Sonia Goncalves, Ewan Harrison, David K. Jackson, Ian Johnston, Dominic Kwiatkowski, Cordelia Langford, John Sillitoe on behalf of the Wellcome Sanger Institute COVID-19 Surveillance Team |
| EPI_ISL_3338498, EPI_ISL_3338499, EPI_ISL_3338500 | Lighthouse Laboratory Plymouth | Wellcome Sanger Institute for the COVID-19 Genomics UK (COG-UK) Consortium | Lighthouse Laboratory Plymouth and Alex Alderton, Roberto Amato, Jeffrey Barrett, Sonia Goncalves, Ewan Harrison, David K. Jackson, Ian Johnston, Dominic Kwiatkowski, Cordelia Langford, John Sillitoe on behalf of the Wellcome Sanger Institute COVID-19 Surveillance Team |
| EPI_ISL_3338501, EPI_ISL_3338502, EPI_ISL_3338503, EPI_ISL_3338504, EPI_ISL_3338505 | Lighthouse Lab in Alderley Park | Wellcome Sanger Institute for the COVID-19 Genomics UK (COG-UK) Consortium | Jacquelyn Wynn, Mairead Hyland, The Lighthouse Lab in Alderley Park and Alex Alderton, Roberto Amato, Jeffrey Barrett, Sonia Goncalves, Ewan Harrison, David K. Jackson, Ian Johnston, Dominic Kwiatkowski, Cordelia Langford, John Sillitoe on behalf of the Wellcome Sanger Institute COVID-19 Surveillance Team |
| EPI_ISL_3338506 | Lighthouse Laboratory Plymouth | Wellcome Sanger Institute for the COVID-19 Genomics UK (COG-UK) Consortium | Lighthouse Laboratory Plymouth and Alex Alderton, Roberto Amato, Jeffrey Barrett, Sonia Goncalves, Ewan Harrison, David K. Jackson, Ian Johnston, Dominic Kwiatkowski, Cordelia Langford, John Sillitoe on behalf of the Wellcome Sanger Institute COVID-19 Surveillance Team |
| EPI_ISL_3338507, EPI_ISL_3338508 | Lighthouse Lab in Glasgow | Wellcome Sanger Institute for the COVID-19 Genomics UK (COG-UK) Consortium | Harper VanSteenhouse, Yumi Kasai, David Gray, Carol Clugston, Anna Dominiczak and Alex Alderton, Roberto Amato, Jeffrey Barrett, Sonia Goncalves, Ewan Harrison, David K. Jackson, Ian Johnston, Dominic Kwiatkowski, Cordelia Langford, John Sillitoe on behalf of the Wellcome Sanger Institute COVID-19 Surveillance Team |
| EPI_ISL_3338509 | Lighthouse Laboratory Plymouth | Wellcome Sanger Institute for the COVID-19 Genomics UK (COG-UK) Consortium | Lighthouse Laboratory Plymouth and Alex Alderton, Roberto Amato, Jeffrey Barrett, Sonia Goncalves, Ewan Harrison, David K. Jackson, Ian Johnston, Dominic Kwiatkowski, Cordelia Langford, John Sillitoe on behalf of the Wellcome Sanger Institute COVID-19 Surveillance Team |
| EPI_ISL_3338510 | Lighthouse Lab in Glasgow | Wellcome Sanger Institute for the COVID-19 Genomics UK (COG-UK) Consortium | Harper VanSteenhouse, Yumi Kasai, David Gray, Carol Clugston, Anna Dominiczak and Alex Alderton, Roberto Amato, Jeffrey Barrett, Sonia Goncalves, Ewan Harrison, David K. Jackson, Ian Johnston, Dominic Kwiatkowski, Cordelia Langford, John Sillitoe on behalf of the Wellcome Sanger Institute COVID-19 Surveillance Team |
| EPI_ISL_3338511 | Lighthouse Laboratory Plymouth | Wellcome Sanger Institute for the COVID-19 Genomics UK (COG-UK) Consortium | Lighthouse Laboratory Plymouth and Alex Alderton, Roberto Amato, Jeffrey Barrett, Sonia Goncalves, Ewan Harrison, David K. Jackson, Ian Johnston, Dominic Kwiatkowski, Cordelia Langford, John Sillitoe on behalf of the Wellcome Sanger Institute COVID-19 Surveillance Team |
| EPI_ISL_3338512, EPI_ISL_3338513, EPI_ISL_3338514 | Lighthouse Lab in Alderley Park | Wellcome Sanger Institute for the COVID-19 Genomics UK (COG-UK) Consortium | Jacquelyn Wynn, Mairead Hyland, The Lighthouse Lab in Alderley Park and Alex Alderton, Roberto Amato, Jeffrey Barrett, Sonia Goncalves, Ewan Harrison, David K. Jackson, Ian Johnston, Dominic Kwiatkowski, Cordelia Langford, John Sillitoe on behalf of the Wellcome Sanger Institute COVID-19 Surveillance Team |
| EPI_ISL_3338515 | Lighthouse Lab in Glasgow | Wellcome Sanger Institute for the COVID-19 Genomics UK (COG-UK) Consortium | Harper VanSteenhouse, Yumi Kasai, David Gray, Carol Clugston, Anna Dominiczak and Alex Alderton, Roberto Amato, Jeffrey Barrett, Sonia Goncalves, Ewan Harrison, David K. Jackson, Ian Johnston, Dominic Kwiatkowski, Cordelia Langford, John Sillitoe on behalf of the Wellcome Sanger Institute COVID-19 Surveillance Team |
| EPI_ISL_3338516 | Lighthouse Lab in Alderley Park | Wellcome Sanger Institute for the COVID-19 Genomics UK (COG-UK) Consortium | Jacquelyn Wynn, Mairead Hyland, The Lighthouse Lab in Alderley Park and Alex Alderton, Roberto Amato, Jeffrey Barrett, Sonia Goncalves, Ewan Harrison, David K. Jackson, Ian Johnston, Dominic Kwiatkowski, Cordelia Langford, John Sillitoe on behalf of the Wellcome Sanger Institute COVID-19 Surveillance Team |
| EPI_ISL_3338517 | Lighthouse Laboratory Plymouth | Wellcome Sanger Institute for the COVID-19 Genomics UK (COG-UK) Consortium | Lighthouse Laboratory Plymouth and Alex Alderton, Roberto Amato, Jeffrey Barrett, Sonia Goncalves, Ewan Harrison, David K. Jackson, Ian Johnston, Dominic Kwiatkowski, Cordelia Langford, John Sillitoe on behalf of the Wellcome Sanger Institute COVID-19 Surveillance Team |
| EPI_ISL_3338518, EPI_ISL_3338519, EPI_ISL_3338520, EPI_ISL_3338521, EPI_ISL_3338522 | Lighthouse Lab in Alderley Park | Wellcome Sanger Institute for the COVID-19 Genomics UK (COG-UK) Consortium | Jacquelyn Wynn, Mairead Hyland, The Lighthouse Lab in Alderley Park and Alex Alderton, Roberto Amato, Jeffrey Barrett, Sonia Goncalves, Ewan Harrison, David K. Jackson, Ian Johnston, Dominic Kwiatkowski, Cordelia Langford, John Sillitoe on behalf of the Wellcome Sanger Institute COVID-19 Surveillance Team |
| EPI_ISL_3338523, EPI_ISL_3338524 | Lighthouse Lab in Glasgow | Wellcome Sanger Institute for the COVID-19 Genomics UK (COG-UK) Consortium | Harper VanSteenhouse, Yumi Kasai, David Gray, Carol Clugston, Anna Dominiczak and Alex Alderton, Roberto Amato, Jeffrey Barrett, Sonia Goncalves, Ewan Harrison, David K. Jackson, Ian Johnston, Dominic Kwiatkowski, Cordelia Langford, John Sillitoe on behalf of the Wellcome Sanger Institute COVID-19 Surveillance Team |

| | | | |
|---|---|---|---|
| EPI_ISL_3338525 | Lighthouse Laboratory Plymouth | Wellcome Sanger Institute for the COVID-19 Genomics UK (COG-UK) Consortium | Lighthouse Laboratory Plymouth and Alex Alderton, Roberto Amato, Jeffrey Barrett, Sonia Goncalves, Ewan Harrison, David K. Jackson, Ian Johnston, Dominic Kwiatkowski, Cordelia Langford, John Sillitoe on behalf of the Wellcome Sanger Institute COVID-19 Surveillance Team |
| EPI_ISL_3338526, EPI_ISL_3338527 | Lighthouse Lab in Alderley Park | Wellcome Sanger Institute for the COVID-19 Genomics UK (COG-UK) Consortium | Jacquelyn Wynn, Mairead Hyland, The Lighthouse Lab in Alderley Park and Alex Alderton, Roberto Amato, Jeffrey Barrett, Sonia Goncalves, Ewan Harrison, David K. Jackson, Ian Johnston, Dominic Kwiatkowski, Cordelia Langford, John Sillitoe on behalf of the Wellcome Sanger Institute COVID-19 Surveillance Team |
| EPI_ISL_3338528 | Lighthouse Lab in Glasgow | Wellcome Sanger Institute for the COVID-19 Genomics UK (COG-UK) Consortium | Harper VanSteenhouse, Yumi Kasai, David Gray, Carol Clugston, Anna Dominiczak and Alex Alderton, Roberto Amato, Jeffrey Barrett, Sonia Goncalves, Ewan Harrison, David K. Jackson, Ian Johnston, Dominic Kwiatkowski, Cordelia Langford, John Sillitoe on behalf of the Wellcome Sanger Institute COVID-19 Surveillance Team |
| EPI_ISL_3338529 | Lighthouse Lab in Alderley Park | Wellcome Sanger Institute for the COVID-19 Genomics UK (COG-UK) Consortium | Jacquelyn Wynn, Mairead Hyland, The Lighthouse Lab in Alderley Park and Alex Alderton, Roberto Amato, Jeffrey Barrett, Sonia Goncalves, Ewan Harrison, David K. Jackson, Ian Johnston, Dominic Kwiatkowski, Cordelia Langford, John Sillitoe on behalf of the Wellcome Sanger Institute COVID-19 Surveillance Team |
| EPI_ISL_3338530 | Lighthouse Lab in Glasgow | Wellcome Sanger Institute for the COVID-19 Genomics UK (COG-UK) Consortium | Harper VanSteenhouse, Yumi Kasai, David Gray, Carol Clugston, Anna Dominiczak and Alex Alderton, Roberto Amato, Jeffrey Barrett, Sonia Goncalves, Ewan Harrison, David K. Jackson, Ian Johnston, Dominic Kwiatkowski, Cordelia Langford, John Sillitoe on behalf of the Wellcome Sanger Institute COVID-19 Surveillance Team |
| EPI_ISL_3338531 | Lighthouse Lab in Alderley Park | Wellcome Sanger Institute for the COVID-19 Genomics UK (COG-UK) Consortium | Jacquelyn Wynn, Mairead Hyland, The Lighthouse Lab in Alderley Park and Alex Alderton, Roberto Amato, Jeffrey Barrett, Sonia Goncalves, Ewan Harrison, David K. Jackson, Ian Johnston, Dominic Kwiatkowski, Cordelia Langford, John Sillitoe on behalf of the Wellcome Sanger Institute COVID-19 Surveillance Team |
| EPI_ISL_3338532, EPI_ISL_3338533 | Lighthouse Lab in Glasgow | Wellcome Sanger Institute for the COVID-19 Genomics UK (COG-UK) Consortium | Harper VanSteenhouse, Yumi Kasai, David Gray, Carol Clugston, Anna Dominiczak and Alex Alderton, Roberto Amato, Jeffrey Barrett, Sonia Goncalves, Ewan Harrison, David K. Jackson, Ian Johnston, Dominic Kwiatkowski, Cordelia Langford, John Sillitoe on behalf of the Wellcome Sanger Institute COVID-19 Surveillance Team |
| EPI_ISL_3338534 | Lighthouse Lab in Alderley Park | Wellcome Sanger Institute for the COVID-19 Genomics UK (COG-UK) Consortium | Jacquelyn Wynn, Mairead Hyland, The Lighthouse Lab in Alderley Park and Alex Alderton, Roberto Amato, Jeffrey Barrett, Sonia Goncalves, Ewan Harrison, David K. Jackson, Ian Johnston, Dominic Kwiatkowski, Cordelia Langford, John Sillitoe on behalf of the Wellcome Sanger Institute COVID-19 Surveillance Team |
| EPI_ISL_3338535 | Lighthouse Laboratory Plymouth | Wellcome Sanger Institute for the COVID-19 Genomics UK (COG-UK) Consortium | Lighthouse Laboratory Plymouth and Alex Alderton, Roberto Amato, Jeffrey Barrett, Sonia Goncalves, Ewan Harrison, David K. Jackson, Ian Johnston, Dominic Kwiatkowski, Cordelia Langford, John Sillitoe on behalf of the Wellcome Sanger Institute COVID-19 Surveillance Team |
| EPI_ISL_3338536 | Lighthouse Lab in Alderley Park | Wellcome Sanger Institute for the COVID-19 Genomics UK (COG-UK) Consortium | Jacquelyn Wynn, Mairead Hyland, The Lighthouse Lab in Alderley Park and Alex Alderton, Roberto Amato, Jeffrey Barrett, Sonia Goncalves, Ewan Harrison, David K. Jackson, Ian Johnston, Dominic Kwiatkowski, Cordelia Langford, John Sillitoe on behalf of the Wellcome Sanger Institute COVID-19 Surveillance Team |
| EPI_ISL_3338537, EPI_ISL_3338538 | Lighthouse Laboratory Plymouth | Wellcome Sanger Institute for the COVID-19 Genomics UK (COG-UK) Consortium | Lighthouse Laboratory Plymouth and Alex Alderton, Roberto Amato, Jeffrey Barrett, Sonia Goncalves, Ewan Harrison, David K. Jackson, Ian Johnston, Dominic Kwiatkowski, Cordelia Langford, John Sillitoe on behalf of the Wellcome Sanger Institute COVID-19 Surveillance Team |
| EPI_ISL_3338539, EPI_ISL_3338540 | Lighthouse Lab in Alderley Park | Wellcome Sanger Institute for the COVID-19 Genomics UK (COG-UK) Consortium | Jacquelyn Wynn, Mairead Hyland, The Lighthouse Lab in Alderley Park and Alex Alderton, Roberto Amato, Jeffrey Barrett, Sonia Goncalves, Ewan Harrison, David K. Jackson, Ian Johnston, Dominic Kwiatkowski, Cordelia Langford, John Sillitoe on behalf of the Wellcome Sanger Institute COVID-19 Surveillance Team |
| EPI_ISL_3338541 | Lighthouse Lab in Glasgow | Wellcome Sanger Institute for the COVID-19 Genomics UK (COG-UK) Consortium | Harper VanSteenhouse, Yumi Kasai, David Gray, Carol Clugston, Anna Dominiczak and Alex Alderton, Roberto Amato, Jeffrey Barrett, Sonia Goncalves, Ewan Harrison, David K. Jackson, Ian Johnston, Dominic Kwiatkowski, Cordelia Langford, John Sillitoe on behalf of the Wellcome Sanger Institute COVID-19 Surveillance Team |
| EPI_ISL_3338542 | Lighthouse Laboratory Plymouth | Wellcome Sanger Institute for the COVID-19 Genomics UK (COG-UK) Consortium | Lighthouse Laboratory Plymouth and Alex Alderton, Roberto Amato, Jeffrey Barrett, Sonia Goncalves, Ewan Harrison, David K. Jackson, Ian Johnston, Dominic Kwiatkowski, Cordelia Langford, John Sillitoe on behalf of the Wellcome Sanger Institute COVID-19 Surveillance Team |
| EPI_ISL_3338543 | Lighthouse Lab in Alderley Park | Wellcome Sanger Institute for the COVID-19 Genomics UK (COG-UK) Consortium | Jacquelyn Wynn, Mairead Hyland, The Lighthouse Lab in Alderley Park and Alex Alderton, Roberto Amato, Jeffrey Barrett, Sonia Goncalves, Ewan Harrison, David K. Jackson, Ian Johnston, Dominic Kwiatkowski, Cordelia Langford, John Sillitoe on behalf of the Wellcome Sanger Institute COVID-19 Surveillance Team |
| EPI_ISL_3338544 | Lighthouse Lab in Glasgow | Wellcome Sanger Institute for the COVID-19 Genomics UK (COG-UK) Consortium | Harper VanSteenhouse, Yumi Kasai, David Gray, Carol Clugston, Anna Dominiczak and Alex Alderton, Roberto Amato, Jeffrey Barrett, Sonia Goncalves, Ewan Harrison, David K. Jackson, Ian Johnston, Dominic Kwiatkowski, Cordelia Langford, John Sillitoe on behalf of the Wellcome Sanger Institute COVID-19 Surveillance Team |
| EPI_ISL_3338545 | Lighthouse Lab in Alderley Park | Wellcome Sanger Institute for the COVID-19 Genomics UK (COG-UK) Consortium | Jacquelyn Wynn, Mairead Hyland, The Lighthouse Lab in Alderley Park and Alex Alderton, Roberto Amato, Jeffrey Barrett, Sonia Goncalves, Ewan Harrison, David K. Jackson, Ian Johnston, Dominic Kwiatkowski, Cordelia Langford, John Sillitoe on behalf of the Wellcome Sanger Institute COVID-19 Surveillance Team |
| EPI_ISL_3338546 | Lighthouse Laboratory Plymouth | Wellcome Sanger Institute for the COVID-19 Genomics UK (COG-UK) Consortium | Lighthouse Laboratory Plymouth and Alex Alderton, Roberto Amato, Jeffrey Barrett, Sonia Goncalves, Ewan Harrison, David K. Jackson, Ian Johnston, Dominic Kwiatkowski, Cordelia Langford, John Sillitoe on behalf of the Wellcome Sanger Institute COVID-19 Surveillance Team |
| EPI_ISL_3338547 | Lighthouse Lab in Glasgow | Wellcome Sanger Institute for the COVID-19 Genomics UK (COG-UK) Consortium | Harper VanSteenhouse, Yumi Kasai, David Gray, Carol Clugston, Anna Dominiczak and Alex Alderton, Roberto Amato, Jeffrey Barrett, Sonia Goncalves, Ewan Harrison, David K. Jackson, Ian Johnston, Dominic Kwiatkowski, Cordelia Langford, John Sillitoe on behalf of the Wellcome Sanger Institute COVID-19 Surveillance Team |
| EPI_ISL_3338548, EPI_ISL_3338549, EPI_ISL_3338550, EPI_ISL_3338551, EPI_ISL_3338552 | Lighthouse Lab in Alderley Park | Wellcome Sanger Institute for the COVID-19 Genomics UK (COG-UK) Consortium | Jacquelyn Wynn, Mairead Hyland, The Lighthouse Lab in Alderley Park and Alex Alderton, Roberto Amato, Jeffrey Barrett, Sonia Goncalves, Ewan Harrison, David K. Jackson, Ian Johnston, Dominic Kwiatkowski, Cordelia Langford, John Sillitoe on behalf of the Wellcome Sanger Institute COVID-19 Surveillance Team |
| EPI_ISL_3338553 | Lighthouse Lab in Glasgow | Wellcome Sanger Institute for the COVID-19 Genomics UK (COG-UK) Consortium | Harper VanSteenhouse, Yumi Kasai, David Gray, Carol Clugston, Anna Dominiczak and Alex Alderton, Roberto Amato, Jeffrey Barrett, Sonia Goncalves, Ewan Harrison, David K. Jackson, Ian Johnston, Dominic Kwiatkowski, Cordelia Langford, John Sillitoe on behalf of the Wellcome Sanger Institute COVID-19 Surveillance Team |
| EPI_ISL_3338554, EPI_ISL_3338555 | Lighthouse Laboratory Plymouth | Wellcome Sanger Institute for the COVID-19 Genomics UK (COG-UK) Consortium | Lighthouse Laboratory Plymouth and Alex Alderton, Roberto Amato, Jeffrey Barrett, Sonia Goncalves, Ewan Harrison, David K. Jackson, Ian Johnston, Dominic Kwiatkowski, Cordelia Langford, John Sillitoe on behalf of the Wellcome Sanger Institute COVID-19 Surveillance Team |
| EPI_ISL_3338556 | Lighthouse Lab in Glasgow | Wellcome Sanger Institute for the COVID-19 Genomics UK (COG-UK) Consortium | Harper VanSteenhouse, Yumi Kasai, David Gray, Carol Clugston, Anna Dominiczak and Alex Alderton, Roberto Amato, Jeffrey Barrett, Sonia Goncalves, Ewan Harrison, David K. Jackson, Ian Johnston, Dominic Kwiatkowski, Cordelia Langford, John Sillitoe on behalf of the Wellcome Sanger Institute COVID-19 Surveillance Team |
| EPI_ISL_3338557 | Lighthouse Laboratory Plymouth | Wellcome Sanger Institute for the COVID-19 Genomics UK (COG-UK) Consortium | Lighthouse Laboratory Plymouth and Alex Alderton, Roberto Amato, Jeffrey Barrett, Sonia Goncalves, Ewan Harrison, David K. Jackson, Ian Johnston, Dominic Kwiatkowski, Cordelia Langford, John Sillitoe on behalf of the Wellcome Sanger Institute COVID-19 Surveillance Team |
| EPI_ISL_3338558 | Lighthouse Lab in Alderley Park | Wellcome Sanger Institute for the COVID-19 Genomics UK (COG-UK) Consortium | Jacquelyn Wynn, Mairead Hyland, The Lighthouse Lab in Alderley Park and Alex Alderton, Roberto Amato, Jeffrey Barrett, Sonia Goncalves, Ewan Harrison, David K. Jackson, Ian Johnston, Dominic Kwiatkowski, Cordelia Langford, John Sillitoe on behalf of the Wellcome Sanger Institute COVID-19 Surveillance Team |
| EPI_ISL_3338559 | Lighthouse Laboratory Plymouth | Wellcome Sanger Institute for the COVID-19 Genomics UK (COG-UK) Consortium | Lighthouse Laboratory Plymouth and Alex Alderton, Roberto Amato, Jeffrey Barrett, Sonia Goncalves, Ewan Harrison, David K. Jackson, Ian Johnston, Dominic Kwiatkowski, Cordelia Langford, John Sillitoe on behalf of the Wellcome Sanger Institute COVID-19 Surveillance Team |
| EPI_ISL_3338560 | Lighthouse Lab in Glasgow | Wellcome Sanger Institute for the COVID-19 Genomics UK (COG-UK) Consortium | Harper VanSteenhouse, Yumi Kasai, David Gray, Carol Clugston, Anna Dominiczak and Alex Alderton, Roberto Amato, Jeffrey Barrett, Sonia Goncalves, Ewan Harrison, David K. Jackson, Ian Johnston, Dominic Kwiatkowski, Cordelia Langford, John Sillitoe on behalf of the Wellcome Sanger Institute COVID-19 Surveillance Team |
| EPI_ISL_3338561, EPI_ISL_3338562, EPI_ISL_3338563, EPI_ISL_3338564 | Lighthouse Lab in Alderley Park | Wellcome Sanger Institute for the COVID-19 Genomics UK (COG-UK) Consortium | Jacquelyn Wynn, Mairead Hyland, The Lighthouse Lab in Alderley Park and Alex Alderton, Roberto Amato, Jeffrey Barrett, Sonia Goncalves, Ewan Harrison, David K. Jackson, Ian Johnston, Dominic Kwiatkowski, Cordelia Langford, John Sillitoe on behalf of the Wellcome Sanger Institute COVID-19 Surveillance Team |
| EPI_ISL_3338565 | Lighthouse Laboratory Plymouth | Wellcome Sanger Institute for the COVID-19 Genomics UK (COG-UK) Consortium | Lighthouse Laboratory Plymouth and Alex Alderton, Roberto Amato, Jeffrey Barrett, Sonia Goncalves, Ewan Harrison, David K. Jackson, Ian Johnston, Dominic Kwiatkowski, Cordelia Langford, John Sillitoe on behalf of the Wellcome Sanger Institute COVID-19 Surveillance Team |
| EPI_ISL_3338566 | Lighthouse Lab in Alderley Park | Wellcome Sanger Institute for the COVID-19 Genomics UK (COG-UK) Consortium | Jacquelyn Wynn, Mairead Hyland, The Lighthouse Lab in Alderley Park and Alex Alderton, Roberto Amato, Jeffrey Barrett, Sonia Goncalves, Ewan Harrison, David K. Jackson, Ian Johnston, Dominic Kwiatkowski, Cordelia Langford, John Sillitoe on behalf of the Wellcome Sanger Institute COVID-19 Surveillance Team |
| EPI_ISL_3338567 | Lighthouse Laboratory Plymouth | Wellcome Sanger Institute for the COVID-19 Genomics UK (COG-UK) Consortium | Lighthouse Laboratory Plymouth and Alex Alderton, Roberto Amato, Jeffrey Barrett, Sonia Goncalves, Ewan Harrison, David K. Jackson, Ian Johnston, Dominic Kwiatkowski, Cordelia Langford, John Sillitoe on behalf of the Wellcome Sanger Institute COVID-19 Surveillance Team |
| EPI_ISL_3338568 | Lighthouse Lab in Alderley Park | Wellcome Sanger Institute for the COVID-19 Genomics UK (COG-UK) Consortium | Jacquelyn Wynn, Mairead Hyland, The Lighthouse Lab in Alderley Park and Alex Alderton, Roberto Amato, Jeffrey Barrett, Sonia Goncalves, Ewan Harrison, David K. Jackson, Ian Johnston, Dominic Kwiatkowski, Cordelia Langford, John Sillitoe on behalf of the Wellcome Sanger Institute COVID-19 Surveillance Team |
| EPI_ISL_3338569 | Lighthouse Lab in Glasgow | Wellcome Sanger Institute for the COVID-19 Genomics UK (COG-UK) Consortium | Harper VanSteenhouse, Yumi Kasai, David Gray, Carol Clugston, Anna Dominiczak and Alex Alderton, Roberto Amato, Jeffrey Barrett, Sonia Goncalves, Ewan Harrison, David K. Jackson, Ian Johnston, Dominic Kwiatkowski, Cordelia Langford, John Sillitoe on behalf of the Wellcome Sanger Institute COVID-19 Surveillance Team |
| EPI_ISL_3338570, EPI_ISL_3338571 | Lighthouse Lab in Alderley Park | Wellcome Sanger Institute for the COVID-19 Genomics UK (COG-UK) Consortium | Jacquelyn Wynn, Mairead Hyland, The Lighthouse Lab in Alderley Park and Alex Alderton, Roberto Amato, Jeffrey Barrett, Sonia Goncalves, Ewan Harrison, David K. Jackson, Ian Johnston, Dominic Kwiatkowski, Cordelia Langford, John Sillitoe on behalf of the Wellcome Sanger Institute COVID-19 Surveillance Team |
| EPI_ISL_3338572 | Lighthouse Laboratory Plymouth | Wellcome Sanger Institute for the COVID-19 Genomics UK (COG-UK) Consortium | Lighthouse Laboratory Plymouth and Alex Alderton, Roberto Amato, Jeffrey Barrett, Sonia Goncalves, Ewan Harrison, David K. Jackson, Ian Johnston, Dominic Kwiatkowski, Cordelia Langford, John Sillitoe on behalf of the Wellcome Sanger Institute COVID-19 Surveillance Team |
| EPI_ISL_3338573, EPI_ISL_3338574, EPI_ISL_3338575, EPI_ISL_3338576, EPI_ISL_3338577, EPI_ISL_3338578 | Lighthouse Lab in Alderley Park | Wellcome Sanger Institute for the COVID-19 Genomics UK (COG-UK) Consortium | Jacquelyn Wynn, Mairead Hyland, The Lighthouse Lab in Alderley Park and Alex Alderton, Roberto Amato, Jeffrey Barrett, Sonia Goncalves, Ewan Harrison, David K. Jackson, Ian Johnston, Dominic Kwiatkowski, Cordelia Langford, John Sillitoe on behalf of the Wellcome Sanger Institute COVID-19 Surveillance Team |
| EPI_ISL_3338579, EPI_ISL_3338580, EPI_ISL_3338581, EPI_ISL_3338582 | Lighthouse Lab in Milton Keynes | Wellcome Sanger Institute for the COVID-19 Genomics UK (COG-UK) Consortium | The Lighthouse Lab in Milton Keynes and Alex Alderton, Roberto Amato, Jeffrey Barrett, Sonia Goncalves, Ewan Harrison, David K. Jackson, Ian Johnston, Dominic Kwiatkowski, Cordelia Langford, John Sillitoe on behalf of the Wellcome Sanger Institute COVID-19 Surveillance Team |
| EPI_ISL_3338583 | Lighthouse Lab in Alderley Park | Wellcome Sanger Institute for the COVID-19 Genomics UK (COG-UK) Consortium | Jacquelyn Wynn, Mairead Hyland, The Lighthouse Lab in Alderley Park and Alex Alderton, Roberto Amato, Jeffrey Barrett, Sonia Goncalves, Ewan Harrison, David K. Jackson, Ian Johnston, Dominic Kwiatkowski, Cordelia Langford, John Sillitoe on behalf of the Wellcome Sanger Institute COVID-19 Surveillance Team |
| EPI_ISL_3338584, EPI_ISL_3338585 | Lighthouse Lab in Milton Keynes | Wellcome Sanger Institute for the COVID-19 Genomics UK (COG-UK) Consortium | The Lighthouse Lab in Milton Keynes and Alex Alderton, Roberto Amato, Jeffrey Barrett, Sonia Goncalves, Ewan Harrison, David K. Jackson, Ian Johnston, Dominic Kwiatkowski, Cordelia Langford, John Sillitoe on behalf of the Wellcome Sanger Institute COVID-19 Surveillance Team |
| EPI_ISL_3338586 | Lighthouse Lab in Glasgow | Wellcome Sanger Institute for the COVID-19 Genomics UK (COG-UK) Consortium | Harper VanSteenhouse, Yumi Kasai, David Gray, Carol Clugston, Anna Dominiczak and Alex Alderton, Roberto Amato, Jeffrey Barrett, Sonia Goncalves, Ewan Harrison, David K. Jackson, Ian Johnston, Dominic Kwiatkowski, Cordelia Langford, John Sillitoe on behalf of the Wellcome Sanger Institute COVID-19 Surveillance Team |
| EPI_ISL_3338587, | Lighthouse Lab in Milton Keynes | Wellcome Sanger Institute for the COVID-19 Genomics UK (COG-UK) | The Lighthouse Lab in Milton Keynes and Alex Alderton, Roberto Amato, Jeffrey Barrett, Sonia Goncalves, Ewan Harrison, David K. Jackson, Ian Johnston, Dominic Kwiatkowski, Cordelia Langford, John Sillitoe on behalf of the Wellcome Sanger |

| | | | |
|---|---|---|---|
| EPI_ISL_3338588, EPI_ISL_3338589, EPI_ISL_3338590 | | Consortium | Institute COVID-19 Surveillance Team |
| EPI_ISL_3338591 | Lighthouse Laboratory Plymouth | Wellcome Sanger Institute for the COVID-19 Genomics UK (COG-UK) Consortium | Lighthouse Laboratory Plymouth and Alex Alderton, Roberto Amato, Jeffrey Barrett, Sonia Goncalves, Ewan Harrison, David K. Jackson, Ian Johnston, Dominic Kwiatkowski, Cordelia Langford, John Sillitoe on behalf of the Wellcome Sanger Institute COVID-19 Surveillance Team |
| EPI_ISL_3338592 | Lighthouse Lab in Glasgow | Wellcome Sanger Institute for the COVID-19 Genomics UK (COG-UK) Consortium | Harper VanSteenhouse, Yumi Kasai, David Gray, Carol Clugston, Anna Dominiczak and Alex Alderton, Roberto Amato, Jeffrey Barrett, Sonia Goncalves, Ewan Harrison, David K. Jackson, Ian Johnston, Dominic Kwiatkowski, Cordelia Langford, John Sillitoe on behalf of the Wellcome Sanger Institute COVID-19 Surveillance Team |
| EPI_ISL_3338593 | Lighthouse Lab in Milton Keynes | Wellcome Sanger Institute for the COVID-19 Genomics UK (COG-UK) Consortium | The Lighthouse Lab in Milton Keynes and Alex Alderton, Roberto Amato, Jeffrey Barrett, Sonia Goncalves, Ewan Harrison, David K. Jackson, Ian Johnston, Dominic Kwiatkowski, Cordelia Langford, John Sillitoe on behalf of the Wellcome Sanger Institute COVID-19 Surveillance Team |
| EPI_ISL_3338594, EPI_ISL_3338595 | Lighthouse Lab in Glasgow | Wellcome Sanger Institute for the COVID-19 Genomics UK (COG-UK) Consortium | Harper VanSteenhouse, Yumi Kasai, David Gray, Carol Clugston, Anna Dominiczak and Alex Alderton, Roberto Amato, Jeffrey Barrett, Sonia Goncalves, Ewan Harrison, David K. Jackson, Ian Johnston, Dominic Kwiatkowski, Cordelia Langford, John Sillitoe on behalf of the Wellcome Sanger Institute COVID-19 Surveillance Team |
| EPI_ISL_3338596, EPI_ISL_3338597, EPI_ISL_3338598 | Lighthouse Lab in Milton Keynes | Wellcome Sanger Institute for the COVID-19 Genomics UK (COG-UK) Consortium | The Lighthouse Lab in Milton Keynes and Alex Alderton, Roberto Amato, Jeffrey Barrett, Sonia Goncalves, Ewan Harrison, David K. Jackson, Ian Johnston, Dominic Kwiatkowski, Cordelia Langford, John Sillitoe on behalf of the Wellcome Sanger Institute COVID-19 Surveillance Team |
| EPI_ISL_3338599 | Lighthouse Laboratory Plymouth | Wellcome Sanger Institute for the COVID-19 Genomics UK (COG-UK) Consortium | Lighthouse Laboratory Plymouth and Alex Alderton, Roberto Amato, Jeffrey Barrett, Sonia Goncalves, Ewan Harrison, David K. Jackson, Ian Johnston, Dominic Kwiatkowski, Cordelia Langford, John Sillitoe on behalf of the Wellcome Sanger Institute COVID-19 Surveillance Team |
| EPI_ISL_3338600 | Lighthouse Lab in Glasgow | Wellcome Sanger Institute for the COVID-19 Genomics UK (COG-UK) Consortium | Harper VanSteenhouse, Yumi Kasai, David Gray, Carol Clugston, Anna Dominiczak and Alex Alderton, Roberto Amato, Jeffrey Barrett, Sonia Goncalves, Ewan Harrison, David K. Jackson, Ian Johnston, Dominic Kwiatkowski, Cordelia Langford, John Sillitoe on behalf of the Wellcome Sanger Institute COVID-19 Surveillance Team |
| EPI_ISL_3338601 | Lighthouse Laboratory Plymouth | Wellcome Sanger Institute for the COVID-19 Genomics UK (COG-UK) Consortium | Lighthouse Laboratory Plymouth and Alex Alderton, Roberto Amato, Jeffrey Barrett, Sonia Goncalves, Ewan Harrison, David K. Jackson, Ian Johnston, Dominic Kwiatkowski, Cordelia Langford, John Sillitoe on behalf of the Wellcome Sanger Institute COVID-19 Surveillance Team |
| EPI_ISL_3338602 | Lighthouse Lab in Glasgow | Wellcome Sanger Institute for the COVID-19 Genomics UK (COG-UK) Consortium | Harper VanSteenhouse, Yumi Kasai, David Gray, Carol Clugston, Anna Dominiczak and Alex Alderton, Roberto Amato, Jeffrey Barrett, Sonia Goncalves, Ewan Harrison, David K. Jackson, Ian Johnston, Dominic Kwiatkowski, Cordelia Langford, John Sillitoe on behalf of the Wellcome Sanger Institute COVID-19 Surveillance Team |
| EPI_ISL_3338603 | Lighthouse Lab in Milton Keynes | Wellcome Sanger Institute for the COVID-19 Genomics UK (COG-UK) Consortium | The Lighthouse Lab in Milton Keynes and Alex Alderton, Roberto Amato, Jeffrey Barrett, Sonia Goncalves, Ewan Harrison, David K. Jackson, Ian Johnston, Dominic Kwiatkowski, Cordelia Langford, John Sillitoe on behalf of the Wellcome Sanger Institute COVID-19 Surveillance Team |
| EPI_ISL_3338604 | Lighthouse Lab in Glasgow | Wellcome Sanger Institute for the COVID-19 Genomics UK (COG-UK) Consortium | Harper VanSteenhouse, Yumi Kasai, David Gray, Carol Clugston, Anna Dominiczak and Alex Alderton, Roberto Amato, Jeffrey Barrett, Sonia Goncalves, Ewan Harrison, David K. Jackson, Ian Johnston, Dominic Kwiatkowski, Cordelia Langford, John Sillitoe on behalf of the Wellcome Sanger Institute COVID-19 Surveillance Team |
| EPI_ISL_3338605 | Lighthouse Laboratory Plymouth | Wellcome Sanger Institute for the COVID-19 Genomics UK (COG-UK) Consortium | Lighthouse Laboratory Plymouth and Alex Alderton, Roberto Amato, Jeffrey Barrett, Sonia Goncalves, Ewan Harrison, David K. Jackson, Ian Johnston, Dominic Kwiatkowski, Cordelia Langford, John Sillitoe on behalf of the Wellcome Sanger Institute COVID-19 Surveillance Team |
| EPI_ISL_3338606 | Lighthouse Lab in Milton Keynes | Wellcome Sanger Institute for the COVID-19 Genomics UK (COG-UK) Consortium | The Lighthouse Lab in Milton Keynes and Alex Alderton, Roberto Amato, Jeffrey Barrett, Sonia Goncalves, Ewan Harrison, David K. Jackson, Ian Johnston, Dominic Kwiatkowski, Cordelia Langford, John Sillitoe on behalf of the Wellcome Sanger Institute COVID-19 Surveillance Team |
| EPI_ISL_3338607 | Lighthouse Lab in Glasgow | Wellcome Sanger Institute for the COVID-19 Genomics UK (COG-UK) Consortium | Harper VanSteenhouse, Yumi Kasai, David Gray, Carol Clugston, Anna Dominiczak and Alex Alderton, Roberto Amato, Jeffrey Barrett, Sonia Goncalves, Ewan Harrison, David K. Jackson, Ian Johnston, Dominic Kwiatkowski, Cordelia Langford, John Sillitoe on behalf of the Wellcome Sanger Institute COVID-19 Surveillance Team |
| EPI_ISL_3338608 | Lighthouse Lab in Milton Keynes | Wellcome Sanger Institute for the COVID-19 Genomics UK (COG-UK) Consortium | The Lighthouse Lab in Milton Keynes and Alex Alderton, Roberto Amato, Jeffrey Barrett, Sonia Goncalves, Ewan Harrison, David K. Jackson, Ian Johnston, Dominic Kwiatkowski, Cordelia Langford, John Sillitoe on behalf of the Wellcome Sanger Institute COVID-19 Surveillance Team |
| EPI_ISL_3338609 | Lighthouse Lab in Glasgow | Wellcome Sanger Institute for the COVID-19 Genomics UK (COG-UK) Consortium | Harper VanSteenhouse, Yumi Kasai, David Gray, Carol Clugston, Anna Dominiczak and Alex Alderton, Roberto Amato, Jeffrey Barrett, Sonia Goncalves, Ewan Harrison, David K. Jackson, Ian Johnston, Dominic Kwiatkowski, Cordelia Langford, John Sillitoe on behalf of the Wellcome Sanger Institute COVID-19 Surveillance Team |
| EPI_ISL_3338610 | Lighthouse Lab in Milton Keynes | Wellcome Sanger Institute for the COVID-19 Genomics UK (COG-UK) Consortium | The Lighthouse Lab in Milton Keynes and Alex Alderton, Roberto Amato, Jeffrey Barrett, Sonia Goncalves, Ewan Harrison, David K. Jackson, Ian Johnston, Dominic Kwiatkowski, Cordelia Langford, John Sillitoe on behalf of the Wellcome Sanger Institute COVID-19 Surveillance Team |
| EPI_ISL_3338611 | Lighthouse Laboratory Plymouth | Wellcome Sanger Institute for the COVID-19 Genomics UK (COG-UK) Consortium | Lighthouse Laboratory Plymouth and Alex Alderton, Roberto Amato, Jeffrey Barrett, Sonia Goncalves, Ewan Harrison, David K. Jackson, Ian Johnston, Dominic Kwiatkowski, Cordelia Langford, John Sillitoe on behalf of the Wellcome Sanger Institute COVID-19 Surveillance Team |
| EPI_ISL_3338612, EPI_ISL_3338613 | Lighthouse Lab in Milton Keynes | Wellcome Sanger Institute for the COVID-19 Genomics UK (COG-UK) Consortium | The Lighthouse Lab in Milton Keynes and Alex Alderton, Roberto Amato, Jeffrey Barrett, Sonia Goncalves, Ewan Harrison, David K. Jackson, Ian Johnston, Dominic Kwiatkowski, Cordelia Langford, John Sillitoe on behalf of the Wellcome Sanger Institute COVID-19 Surveillance Team |
| EPI_ISL_3338614, EPI_ISL_3338615 | Lighthouse Laboratory Plymouth | Wellcome Sanger Institute for the COVID-19 Genomics UK (COG-UK) Consortium | Lighthouse Laboratory Plymouth and Alex Alderton, Roberto Amato, Jeffrey Barrett, Sonia Goncalves, Ewan Harrison, David K. Jackson, Ian Johnston, Dominic Kwiatkowski, Cordelia Langford, John Sillitoe on behalf of the Wellcome Sanger Institute COVID-19 Surveillance Team |
| EPI_ISL_3338616 | Lighthouse Lab in Milton Keynes | Wellcome Sanger Institute for the COVID-19 Genomics UK (COG-UK) Consortium | The Lighthouse Lab in Milton Keynes and Alex Alderton, Roberto Amato, Jeffrey Barrett, Sonia Goncalves, Ewan Harrison, David K. Jackson, Ian Johnston, Dominic Kwiatkowski, Cordelia Langford, John Sillitoe on behalf of the Wellcome Sanger Institute COVID-19 Surveillance Team |
| EPI_ISL_3338617, EPI_ISL_3338618 | Lighthouse Laboratory Plymouth | Wellcome Sanger Institute for the COVID-19 Genomics UK (COG-UK) Consortium | Lighthouse Laboratory Plymouth and Alex Alderton, Roberto Amato, Jeffrey Barrett, Sonia Goncalves, Ewan Harrison, David K. Jackson, Ian Johnston, Dominic Kwiatkowski, Cordelia Langford, John Sillitoe on behalf of the Wellcome Sanger Institute COVID-19 Surveillance Team |
| EPI_ISL_3338619 | Lighthouse Lab in Glasgow | Wellcome Sanger Institute for the COVID-19 Genomics UK (COG-UK) Consortium | Harper VanSteenhouse, Yumi Kasai, David Gray, Carol Clugston, Anna Dominiczak and Alex Alderton, Roberto Amato, Jeffrey Barrett, Sonia Goncalves, Ewan Harrison, David K. Jackson, Ian Johnston, Dominic Kwiatkowski, Cordelia Langford, John Sillitoe on behalf of the Wellcome Sanger Institute COVID-19 Surveillance Team |
| EPI_ISL_3338620, EPI_ISL_3338621, EPI_ISL_3338622 | Lighthouse Laboratory Plymouth | Wellcome Sanger Institute for the COVID-19 Genomics UK (COG-UK) Consortium | Lighthouse Laboratory Plymouth and Alex Alderton, Roberto Amato, Jeffrey Barrett, Sonia Goncalves, Ewan Harrison, David K. Jackson, Ian Johnston, Dominic Kwiatkowski, Cordelia Langford, John Sillitoe on behalf of the Wellcome Sanger Institute COVID-19 Surveillance Team |
| EPI_ISL_3338623 | Lighthouse Lab in Glasgow | Wellcome Sanger Institute for the COVID-19 Genomics UK (COG-UK) Consortium | Harper VanSteenhouse, Yumi Kasai, David Gray, Carol Clugston, Anna Dominiczak and Alex Alderton, Roberto Amato, Jeffrey Barrett, Sonia Goncalves, Ewan Harrison, David K. Jackson, Ian Johnston, Dominic Kwiatkowski, Cordelia Langford, John Sillitoe on behalf of the Wellcome Sanger Institute COVID-19 Surveillance Team |
| EPI_ISL_3338624 | Lighthouse Lab in Milton Keynes | Wellcome Sanger Institute for the COVID-19 Genomics UK (COG-UK) Consortium | The Lighthouse Lab in Milton Keynes and Alex Alderton, Roberto Amato, Jeffrey Barrett, Sonia Goncalves, Ewan Harrison, David K. Jackson, Ian Johnston, Dominic Kwiatkowski, Cordelia Langford, John Sillitoe on behalf of the Wellcome Sanger Institute COVID-19 Surveillance Team |
| EPI_ISL_3338625 | Lighthouse Laboratory Plymouth | Wellcome Sanger Institute for the COVID-19 Genomics UK (COG-UK) Consortium | Lighthouse Laboratory Plymouth and Alex Alderton, Roberto Amato, Jeffrey Barrett, Sonia Goncalves, Ewan Harrison, David K. Jackson, Ian Johnston, Dominic Kwiatkowski, Cordelia Langford, John Sillitoe on behalf of the Wellcome Sanger Institute COVID-19 Surveillance Team |
| EPI_ISL_3338626, EPI_ISL_3338627, EPI_ISL_3338628 | Lighthouse Lab in Milton Keynes | Wellcome Sanger Institute for the COVID-19 Genomics UK (COG-UK) Consortium | The Lighthouse Lab in Milton Keynes and Alex Alderton, Roberto Amato, Jeffrey Barrett, Sonia Goncalves, Ewan Harrison, David K. Jackson, Ian Johnston, Dominic Kwiatkowski, Cordelia Langford, John Sillitoe on behalf of the Wellcome Sanger Institute COVID-19 Surveillance Team |
| EPI_ISL_3338629, EPI_ISL_3338630 | Lighthouse Laboratory Plymouth | Wellcome Sanger Institute for the COVID-19 Genomics UK (COG-UK) Consortium | Lighthouse Laboratory Plymouth and Alex Alderton, Roberto Amato, Jeffrey Barrett, Sonia Goncalves, Ewan Harrison, David K. Jackson, Ian Johnston, Dominic Kwiatkowski, Cordelia Langford, John Sillitoe on behalf of the Wellcome Sanger Institute COVID-19 Surveillance Team |
| EPI_ISL_3338631, EPI_ISL_3338632 | Lighthouse Lab in Milton Keynes | Wellcome Sanger Institute for the COVID-19 Genomics UK (COG-UK) Consortium | The Lighthouse Lab in Milton Keynes and Alex Alderton, Roberto Amato, Jeffrey Barrett, Sonia Goncalves, Ewan Harrison, David K. Jackson, Ian Johnston, Dominic Kwiatkowski, Cordelia Langford, John Sillitoe on behalf of the Wellcome Sanger Institute COVID-19 Surveillance Team |
| EPI_ISL_3338633, EPI_ISL_3338634 | Lighthouse Laboratory Plymouth | Wellcome Sanger Institute for the COVID-19 Genomics UK (COG-UK) Consortium | Lighthouse Laboratory Plymouth and Alex Alderton, Roberto Amato, Jeffrey Barrett, Sonia Goncalves, Ewan Harrison, David K. Jackson, Ian Johnston, Dominic Kwiatkowski, Cordelia Langford, John Sillitoe on behalf of the Wellcome Sanger Institute COVID-19 Surveillance Team |
| EPI_ISL_3338635, EPI_ISL_3338636, EPI_ISL_3338637, EPI_ISL_3338638, EPI_ISL_3338639 | Lighthouse Lab in Milton Keynes | Wellcome Sanger Institute for the COVID-19 Genomics UK (COG-UK) Consortium | The Lighthouse Lab in Milton Keynes and Alex Alderton, Roberto Amato, Jeffrey Barrett, Sonia Goncalves, Ewan Harrison, David K. Jackson, Ian Johnston, Dominic Kwiatkowski, Cordelia Langford, John Sillitoe on behalf of the Wellcome Sanger Institute COVID-19 Surveillance Team |
| EPI_ISL_3338640, EPI_ISL_3338641, EPI_ISL_3338642 | Lighthouse Laboratory Plymouth | Wellcome Sanger Institute for the COVID-19 Genomics UK (COG-UK) Consortium | Lighthouse Laboratory Plymouth and Alex Alderton, Roberto Amato, Jeffrey Barrett, Sonia Goncalves, Ewan Harrison, David K. Jackson, Ian Johnston, Dominic Kwiatkowski, Cordelia Langford, John Sillitoe on behalf of the Wellcome Sanger Institute COVID-19 Surveillance Team |
| EPI_ISL_3338643 | Lighthouse Lab in Glasgow | Wellcome Sanger Institute for the COVID-19 Genomics UK (COG-UK) Consortium | Harper VanSteenhouse, Yumi Kasai, David Gray, Carol Clugston, Anna Dominiczak and Alex Alderton, Roberto Amato, Jeffrey Barrett, Sonia Goncalves, Ewan Harrison, David K. Jackson, Ian Johnston, Dominic Kwiatkowski, Cordelia Langford, John Sillitoe on behalf of the Wellcome Sanger Institute COVID-19 Surveillance Team |
| EPI_ISL_3338644 | Lighthouse Lab in Milton Keynes | Wellcome Sanger Institute for the COVID-19 Genomics UK (COG-UK) Consortium | The Lighthouse Lab in Milton Keynes and Alex Alderton, Roberto Amato, Jeffrey Barrett, Sonia Goncalves, Ewan Harrison, David K. Jackson, Ian Johnston, Dominic Kwiatkowski, Cordelia Langford, John Sillitoe on behalf of the Wellcome Sanger Institute COVID-19 Surveillance Team |
| EPI_ISL_3338645 | Lighthouse Laboratory Plymouth | Wellcome Sanger Institute for the COVID-19 Genomics UK (COG-UK) Consortium | Lighthouse Laboratory Plymouth and Alex Alderton, Roberto Amato, Jeffrey Barrett, Sonia Goncalves, Ewan Harrison, David K. Jackson, Ian Johnston, Dominic Kwiatkowski, Cordelia Langford, John Sillitoe on behalf of the Wellcome Sanger Institute COVID-19 Surveillance Team |
| EPI_ISL_3338646, EPI_ISL_3338647 | Lighthouse Lab in Glasgow | Wellcome Sanger Institute for the COVID-19 Genomics UK (COG-UK) Consortium | Harper VanSteenhouse, Yumi Kasai, David Gray, Carol Clugston, Anna Dominiczak and Alex Alderton, Roberto Amato, Jeffrey Barrett, Sonia Goncalves, Ewan Harrison, David K. Jackson, Ian Johnston, Dominic Kwiatkowski, Cordelia Langford, John Sillitoe on behalf of the Wellcome Sanger Institute COVID-19 Surveillance Team |
| EPI_ISL_3338648 | Lighthouse Laboratory Plymouth | Wellcome Sanger Institute for the COVID-19 Genomics UK (COG-UK) Consortium | Lighthouse Laboratory Plymouth and Alex Alderton, Roberto Amato, Jeffrey Barrett, Sonia Goncalves, Ewan Harrison, David K. Jackson, Ian Johnston, Dominic Kwiatkowski, Cordelia Langford, John Sillitoe on behalf of the Wellcome Sanger Institute COVID-19 Surveillance Team |
| EPI_ISL_3338649 | Lighthouse Lab in Milton Keynes | Wellcome Sanger Institute for the COVID-19 Genomics UK (COG-UK) Consortium | The Lighthouse Lab in Milton Keynes and Alex Alderton, Roberto Amato, Jeffrey Barrett, Sonia Goncalves, Ewan Harrison, David K. Jackson, Ian Johnston, Dominic Kwiatkowski, Cordelia Langford, John Sillitoe on behalf of the Wellcome Sanger Institute COVID-19 Surveillance Team |
| EPI_ISL_3338650, EPI_ISL_3338651 | Lighthouse Lab in Glasgow | Wellcome Sanger Institute for the COVID-19 Genomics UK (COG-UK) Consortium | Harper VanSteenhouse, Yumi Kasai, David Gray, Carol Clugston, Anna Dominiczak and Alex Alderton, Roberto Amato, Jeffrey Barrett, Sonia Goncalves, Ewan Harrison, David K. Jackson, Ian Johnston, Dominic Kwiatkowski, Cordelia Langford, John Sillitoe on behalf of the Wellcome Sanger Institute COVID-19 Surveillance Team |
| EPI_ISL_3338652 | Lighthouse Lab in Milton Keynes | Wellcome Sanger Institute for the COVID-19 Genomics UK (COG-UK) | The Lighthouse Lab in Milton Keynes and Alex Alderton, Roberto Amato, Jeffrey Barrett, Sonia Goncalves, Ewan Harrison, David K. Jackson, Ian Johnston, Dominic Kwiatkowski, Cordelia Langford, John Sillitoe on behalf of the Wellcome Sanger |

| | | | |
|---|---|---|---|
| | | Consortium | Institute COVID-19 Surveillance Team |
| EPI_ISL_3338653 | Lighthouse Laboratory Plymouth | Wellcome Sanger Institute for the COVID-19 Genomics UK (COG-UK) Consortium | Lighthouse Laboratory Plymouth and Alex Alderton, Roberto Amato, Jeffrey Barrett, Sonia Goncalves, Ewan Harrison, David K. Jackson, Ian Johnston, Dominic Kwiatkowski, Cordelia Langford, John Sillitoe on behalf of the Wellcome Sanger Institute COVID-19 Surveillance Team |
| EPI_ISL_3338654 | Lighthouse Lab in Glasgow | Wellcome Sanger Institute for the COVID-19 Genomics UK (COG-UK) Consortium | Harper VanSteenhouse, Yumi Kasai, David Gray, Carol Clugston, Anna Dominiczak and Alex Alderton, Roberto Amato, Jeffrey Barrett, Sonia Goncalves, Ewan Harrison, David K. Jackson, Ian Johnston, Dominic Kwiatkowski, Cordelia Langford, John Sillitoe on behalf of the Wellcome Sanger Institute COVID-19 Surveillance Team |
| EPI_ISL_3338655 | Lighthouse Lab in Milton Keynes | Wellcome Sanger Institute for the COVID-19 Genomics UK (COG-UK) Consortium | The Lighthouse Lab in Milton Keynes and Alex Alderton, Roberto Amato, Jeffrey Barrett, Sonia Goncalves, Ewan Harrison, David K. Jackson, Ian Johnston, Dominic Kwiatkowski, Cordelia Langford, John Sillitoe on behalf of the Wellcome Sanger Institute COVID-19 Surveillance Team |
| EPI_ISL_3338656 | Lighthouse Laboratory Plymouth | Wellcome Sanger Institute for the COVID-19 Genomics UK (COG-UK) Consortium | Lighthouse Laboratory Plymouth and Alex Alderton, Roberto Amato, Jeffrey Barrett, Sonia Goncalves, Ewan Harrison, David K. Jackson, Ian Johnston, Dominic Kwiatkowski, Cordelia Langford, John Sillitoe on behalf of the Wellcome Sanger Institute COVID-19 Surveillance Team |
| EPI_ISL_3338657, EPI_ISL_3338658 | Lighthouse Lab in Milton Keynes | Wellcome Sanger Institute for the COVID-19 Genomics UK (COG-UK) Consortium | The Lighthouse Lab in Milton Keynes and Alex Alderton, Roberto Amato, Jeffrey Barrett, Sonia Goncalves, Ewan Harrison, David K. Jackson, Ian Johnston, Dominic Kwiatkowski, Cordelia Langford, John Sillitoe on behalf of the Wellcome Sanger Institute COVID-19 Surveillance Team |
| EPI_ISL_3338659, EPI_ISL_3338660 | Lighthouse Laboratory Plymouth | Wellcome Sanger Institute for the COVID-19 Genomics UK (COG-UK) Consortium | Lighthouse Laboratory Plymouth and Alex Alderton, Roberto Amato, Jeffrey Barrett, Sonia Goncalves, Ewan Harrison, David K. Jackson, Ian Johnston, Dominic Kwiatkowski, Cordelia Langford, John Sillitoe on behalf of the Wellcome Sanger Institute COVID-19 Surveillance Team |
| EPI_ISL_3338661 | Lighthouse Lab in Milton Keynes | Wellcome Sanger Institute for the COVID-19 Genomics UK (COG-UK) Consortium | The Lighthouse Lab in Milton Keynes and Alex Alderton, Roberto Amato, Jeffrey Barrett, Sonia Goncalves, Ewan Harrison, David K. Jackson, Ian Johnston, Dominic Kwiatkowski, Cordelia Langford, John Sillitoe on behalf of the Wellcome Sanger Institute COVID-19 Surveillance Team |
| EPI_ISL_3338662, EPI_ISL_3338663 | Lighthouse Laboratory Plymouth | Wellcome Sanger Institute for the COVID-19 Genomics UK (COG-UK) Consortium | Lighthouse Laboratory Plymouth and Alex Alderton, Roberto Amato, Jeffrey Barrett, Sonia Goncalves, Ewan Harrison, David K. Jackson, Ian Johnston, Dominic Kwiatkowski, Cordelia Langford, John Sillitoe on behalf of the Wellcome Sanger Institute COVID-19 Surveillance Team |
| EPI_ISL_3338664, EPI_ISL_3338665 | Lighthouse Lab in Glasgow | Wellcome Sanger Institute for the COVID-19 Genomics UK (COG-UK) Consortium | Harper VanSteenhouse, Yumi Kasai, David Gray, Carol Clugston, Anna Dominiczak and Alex Alderton, Roberto Amato, Jeffrey Barrett, Sonia Goncalves, Ewan Harrison, David K. Jackson, Ian Johnston, Dominic Kwiatkowski, Cordelia Langford, John Sillitoe on behalf of the Wellcome Sanger Institute COVID-19 Surveillance Team |
| EPI_ISL_3338666 | Lighthouse Laboratory Plymouth | Wellcome Sanger Institute for the COVID-19 Genomics UK (COG-UK) Consortium | Lighthouse Laboratory Plymouth and Alex Alderton, Roberto Amato, Jeffrey Barrett, Sonia Goncalves, Ewan Harrison, David K. Jackson, Ian Johnston, Dominic Kwiatkowski, Cordelia Langford, John Sillitoe on behalf of the Wellcome Sanger Institute COVID-19 Surveillance Team |
| EPI_ISL_3338667 | Lighthouse Lab in Glasgow | Wellcome Sanger Institute for the COVID-19 Genomics UK (COG-UK) Consortium | Harper VanSteenhouse, Yumi Kasai, David Gray, Carol Clugston, Anna Dominiczak and Alex Alderton, Roberto Amato, Jeffrey Barrett, Sonia Goncalves, Ewan Harrison, David K. Jackson, Ian Johnston, Dominic Kwiatkowski, Cordelia Langford, John Sillitoe on behalf of the Wellcome Sanger Institute COVID-19 Surveillance Team |
| EPI_ISL_3338668 | Lighthouse Lab in Milton Keynes | Wellcome Sanger Institute for the COVID-19 Genomics UK (COG-UK) Consortium | The Lighthouse Lab in Milton Keynes and Alex Alderton, Roberto Amato, Jeffrey Barrett, Sonia Goncalves, Ewan Harrison, David K. Jackson, Ian Johnston, Dominic Kwiatkowski, Cordelia Langford, John Sillitoe on behalf of the Wellcome Sanger Institute COVID-19 Surveillance Team |
| EPI_ISL_3338669 | Lighthouse Lab in Glasgow | Wellcome Sanger Institute for the COVID-19 Genomics UK (COG-UK) Consortium | Harper VanSteenhouse, Yumi Kasai, David Gray, Carol Clugston, Anna Dominiczak and Alex Alderton, Roberto Amato, Jeffrey Barrett, Sonia Goncalves, Ewan Harrison, David K. Jackson, Ian Johnston, Dominic Kwiatkowski, Cordelia Langford, John Sillitoe on behalf of the Wellcome Sanger Institute COVID-19 Surveillance Team |
| EPI_ISL_3338670 | Lighthouse Laboratory Plymouth | Wellcome Sanger Institute for the COVID-19 Genomics UK (COG-UK) Consortium | Lighthouse Laboratory Plymouth and Alex Alderton, Roberto Amato, Jeffrey Barrett, Sonia Goncalves, Ewan Harrison, David K. Jackson, Ian Johnston, Dominic Kwiatkowski, Cordelia Langford, John Sillitoe on behalf of the Wellcome Sanger Institute COVID-19 Surveillance Team |
| EPI_ISL_3338671, EPI_ISL_3338672, EPI_ISL_3338673, EPI_ISL_3338674, EPI_ISL_3338675 | Lighthouse Lab in Glasgow | Wellcome Sanger Institute for the COVID-19 Genomics UK (COG-UK) Consortium | Harper VanSteenhouse, Yumi Kasai, David Gray, Carol Clugston, Anna Dominiczak and Alex Alderton, Roberto Amato, Jeffrey Barrett, Sonia Goncalves, Ewan Harrison, David K. Jackson, Ian Johnston, Dominic Kwiatkowski, Cordelia Langford, John Sillitoe on behalf of the Wellcome Sanger Institute COVID-19 Surveillance Team |
| EPI_ISL_3338676 | Lighthouse Laboratory Plymouth | Wellcome Sanger Institute for the COVID-19 Genomics UK (COG-UK) Consortium | Lighthouse Laboratory Plymouth and Alex Alderton, Roberto Amato, Jeffrey Barrett, Sonia Goncalves, Ewan Harrison, David K. Jackson, Ian Johnston, Dominic Kwiatkowski, Cordelia Langford, John Sillitoe on behalf of the Wellcome Sanger Institute COVID-19 Surveillance Team |
| EPI_ISL_3338677, EPI_ISL_3338678, EPI_ISL_3338679, EPI_ISL_3338680 | Lighthouse Lab in Milton Keynes | Wellcome Sanger Institute for the COVID-19 Genomics UK (COG-UK) Consortium | The Lighthouse Lab in Milton Keynes and Alex Alderton, Roberto Amato, Jeffrey Barrett, Sonia Goncalves, Ewan Harrison, David K. Jackson, Ian Johnston, Dominic Kwiatkowski, Cordelia Langford, John Sillitoe on behalf of the Wellcome Sanger Institute COVID-19 Surveillance Team |
| EPI_ISL_3338681 | Lighthouse Laboratory Plymouth | Wellcome Sanger Institute for the COVID-19 Genomics UK (COG-UK) Consortium | Lighthouse Laboratory Plymouth and Alex Alderton, Roberto Amato, Jeffrey Barrett, Sonia Goncalves, Ewan Harrison, David K. Jackson, Ian Johnston, Dominic Kwiatkowski, Cordelia Langford, John Sillitoe on behalf of the Wellcome Sanger Institute COVID-19 Surveillance Team |
| EPI_ISL_3338682, EPI_ISL_3338683 | Lighthouse Lab in Milton Keynes | Wellcome Sanger Institute for the COVID-19 Genomics UK (COG-UK) Consortium | The Lighthouse Lab in Milton Keynes and Alex Alderton, Roberto Amato, Jeffrey Barrett, Sonia Goncalves, Ewan Harrison, David K. Jackson, Ian Johnston, Dominic Kwiatkowski, Cordelia Langford, John Sillitoe on behalf of the Wellcome Sanger Institute COVID-19 Surveillance Team |
| EPI_ISL_3338684 | Lighthouse Laboratory Plymouth | Wellcome Sanger Institute for the COVID-19 Genomics UK (COG-UK) Consortium | Lighthouse Laboratory Plymouth and Alex Alderton, Roberto Amato, Jeffrey Barrett, Sonia Goncalves, Ewan Harrison, David K. Jackson, Ian Johnston, Dominic Kwiatkowski, Cordelia Langford, John Sillitoe on behalf of the Wellcome Sanger Institute COVID-19 Surveillance Team |
| EPI_ISL_3338685 | Lighthouse Lab in Glasgow | Wellcome Sanger Institute for the COVID-19 Genomics UK (COG-UK) Consortium | Harper VanSteenhouse, Yumi Kasai, David Gray, Carol Clugston, Anna Dominiczak and Alex Alderton, Roberto Amato, Jeffrey Barrett, Sonia Goncalves, Ewan Harrison, David K. Jackson, Ian Johnston, Dominic Kwiatkowski, Cordelia Langford, John Sillitoe on behalf of the Wellcome Sanger Institute COVID-19 Surveillance Team |
| EPI_ISL_3338686 | Lighthouse Laboratory Plymouth | Wellcome Sanger Institute for the COVID-19 Genomics UK (COG-UK) Consortium | Lighthouse Laboratory Plymouth and Alex Alderton, Roberto Amato, Jeffrey Barrett, Sonia Goncalves, Ewan Harrison, David K. Jackson, Ian Johnston, Dominic Kwiatkowski, Cordelia Langford, John Sillitoe on behalf of the Wellcome Sanger Institute COVID-19 Surveillance Team |
| EPI_ISL_3338687, EPI_ISL_3338688 | Lighthouse Lab in Milton Keynes | Wellcome Sanger Institute for the COVID-19 Genomics UK (COG-UK) Consortium | The Lighthouse Lab in Milton Keynes and Alex Alderton, Roberto Amato, Jeffrey Barrett, Sonia Goncalves, Ewan Harrison, David K. Jackson, Ian Johnston, Dominic Kwiatkowski, Cordelia Langford, John Sillitoe on behalf of the Wellcome Sanger Institute COVID-19 Surveillance Team |
| EPI_ISL_3338689 | Lighthouse Lab in Glasgow | Wellcome Sanger Institute for the COVID-19 Genomics UK (COG-UK) Consortium | Harper VanSteenhouse, Yumi Kasai, David Gray, Carol Clugston, Anna Dominiczak and Alex Alderton, Roberto Amato, Jeffrey Barrett, Sonia Goncalves, Ewan Harrison, David K. Jackson, Ian Johnston, Dominic Kwiatkowski, Cordelia Langford, John Sillitoe on behalf of the Wellcome Sanger Institute COVID-19 Surveillance Team |
| EPI_ISL_3338690, EPI_ISL_3338691 | Lighthouse Laboratory Plymouth | Wellcome Sanger Institute for the COVID-19 Genomics UK (COG-UK) Consortium | Lighthouse Laboratory Plymouth and Alex Alderton, Roberto Amato, Jeffrey Barrett, Sonia Goncalves, Ewan Harrison, David K. Jackson, Ian Johnston, Dominic Kwiatkowski, Cordelia Langford, John Sillitoe on behalf of the Wellcome Sanger Institute COVID-19 Surveillance Team |
| EPI_ISL_3338692, EPI_ISL_3338693, EPI_ISL_3338694 | Lighthouse Lab in Glasgow | Wellcome Sanger Institute for the COVID-19 Genomics UK (COG-UK) Consortium | Harper VanSteenhouse, Yumi Kasai, David Gray, Carol Clugston, Anna Dominiczak and Alex Alderton, Roberto Amato, Jeffrey Barrett, Sonia Goncalves, Ewan Harrison, David K. Jackson, Ian Johnston, Dominic Kwiatkowski, Cordelia Langford, John Sillitoe on behalf of the Wellcome Sanger Institute COVID-19 Surveillance Team |
| EPI_ISL_3338695 | Lighthouse Laboratory Plymouth | Wellcome Sanger Institute for the COVID-19 Genomics UK (COG-UK) Consortium | Lighthouse Laboratory Plymouth and Alex Alderton, Roberto Amato, Jeffrey Barrett, Sonia Goncalves, Ewan Harrison, David K. Jackson, Ian Johnston, Dominic Kwiatkowski, Cordelia Langford, John Sillitoe on behalf of the Wellcome Sanger Institute COVID-19 Surveillance Team |
| EPI_ISL_3338696, EPI_ISL_3338697, EPI_ISL_3338698, EPI_ISL_3338699, EPI_ISL_3338700 | Lighthouse Lab in Milton Keynes | Wellcome Sanger Institute for the COVID-19 Genomics UK (COG-UK) Consortium | The Lighthouse Lab in Milton Keynes and Alex Alderton, Roberto Amato, Jeffrey Barrett, Sonia Goncalves, Ewan Harrison, David K. Jackson, Ian Johnston, Dominic Kwiatkowski, Cordelia Langford, John Sillitoe on behalf of the Wellcome Sanger Institute COVID-19 Surveillance Team |
| EPI_ISL_3338701 | Lighthouse Laboratory Plymouth | Wellcome Sanger Institute for the COVID-19 Genomics UK (COG-UK) Consortium | Lighthouse Laboratory Plymouth and Alex Alderton, Roberto Amato, Jeffrey Barrett, Sonia Goncalves, Ewan Harrison, David K. Jackson, Ian Johnston, Dominic Kwiatkowski, Cordelia Langford, John Sillitoe on behalf of the Wellcome Sanger Institute COVID-19 Surveillance Team |
| EPI_ISL_3338702, EPI_ISL_3338703 | Lighthouse Lab in Milton Keynes | Wellcome Sanger Institute for the COVID-19 Genomics UK (COG-UK) Consortium | The Lighthouse Lab in Milton Keynes and Alex Alderton, Roberto Amato, Jeffrey Barrett, Sonia Goncalves, Ewan Harrison, David K. Jackson, Ian Johnston, Dominic Kwiatkowski, Cordelia Langford, John Sillitoe on behalf of the Wellcome Sanger Institute COVID-19 Surveillance Team |
| EPI_ISL_3338704 | Lighthouse Laboratory Plymouth | Wellcome Sanger Institute for the COVID-19 Genomics UK (COG-UK) Consortium | Lighthouse Laboratory Plymouth and Alex Alderton, Roberto Amato, Jeffrey Barrett, Sonia Goncalves, Ewan Harrison, David K. Jackson, Ian Johnston, Dominic Kwiatkowski, Cordelia Langford, John Sillitoe on behalf of the Wellcome Sanger Institute COVID-19 Surveillance Team |
| EPI_ISL_3338705 | Lighthouse Lab in Glasgow | Wellcome Sanger Institute for the COVID-19 Genomics UK (COG-UK) Consortium | Harper VanSteenhouse, Yumi Kasai, David Gray, Carol Clugston, Anna Dominiczak and Alex Alderton, Roberto Amato, Jeffrey Barrett, Sonia Goncalves, Ewan Harrison, David K. Jackson, Ian Johnston, Dominic Kwiatkowski, Cordelia Langford, John Sillitoe on behalf of the Wellcome Sanger Institute COVID-19 Surveillance Team |
| EPI_ISL_3338706, EPI_ISL_3338707, EPI_ISL_3338708 | Lighthouse Lab in Milton Keynes | Wellcome Sanger Institute for the COVID-19 Genomics UK (COG-UK) Consortium | The Lighthouse Lab in Milton Keynes and Alex Alderton, Roberto Amato, Jeffrey Barrett, Sonia Goncalves, Ewan Harrison, David K. Jackson, Ian Johnston, Dominic Kwiatkowski, Cordelia Langford, John Sillitoe on behalf of the Wellcome Sanger Institute COVID-19 Surveillance Team |
| EPI_ISL_3338709 | Lighthouse Lab in Glasgow | Wellcome Sanger Institute for the COVID-19 Genomics UK (COG-UK) Consortium | Harper VanSteenhouse, Yumi Kasai, David Gray, Carol Clugston, Anna Dominiczak and Alex Alderton, Roberto Amato, Jeffrey Barrett, Sonia Goncalves, Ewan Harrison, David K. Jackson, Ian Johnston, Dominic Kwiatkowski, Cordelia Langford, John Sillitoe on behalf of the Wellcome Sanger Institute COVID-19 Surveillance Team |
| EPI_ISL_3338710 | Lighthouse Laboratory Plymouth | Wellcome Sanger Institute for the COVID-19 Genomics UK (COG-UK) Consortium | Lighthouse Laboratory Plymouth and Alex Alderton, Roberto Amato, Jeffrey Barrett, Sonia Goncalves, Ewan Harrison, David K. Jackson, Ian Johnston, Dominic Kwiatkowski, Cordelia Langford, John Sillitoe on behalf of the Wellcome Sanger Institute COVID-19 Surveillance Team |
| EPI_ISL_3338711 | Lighthouse Lab in Milton Keynes | Wellcome Sanger Institute for the COVID-19 Genomics UK (COG-UK) Consortium | The Lighthouse Lab in Milton Keynes and Alex Alderton, Roberto Amato, Jeffrey Barrett, Sonia Goncalves, Ewan Harrison, David K. Jackson, Ian Johnston, Dominic Kwiatkowski, Cordelia Langford, John Sillitoe on behalf of the Wellcome Sanger Institute COVID-19 Surveillance Team |
| EPI_ISL_3338712 | Lighthouse Lab in Glasgow | Wellcome Sanger Institute for the COVID-19 Genomics UK (COG-UK) Consortium | Harper VanSteenhouse, Yumi Kasai, David Gray, Carol Clugston, Anna Dominiczak and Alex Alderton, Roberto Amato, Jeffrey Barrett, Sonia Goncalves, Ewan Harrison, David K. Jackson, Ian Johnston, Dominic Kwiatkowski, Cordelia Langford, John Sillitoe on behalf of the Wellcome Sanger Institute COVID-19 Surveillance Team |
| EPI_ISL_3338713, EPI_ISL_3338714 | Lighthouse Lab in Milton Keynes | Wellcome Sanger Institute for the COVID-19 Genomics UK (COG-UK) Consortium | The Lighthouse Lab in Milton Keynes and Alex Alderton, Roberto Amato, Jeffrey Barrett, Sonia Goncalves, Ewan Harrison, David K. Jackson, Ian Johnston, Dominic Kwiatkowski, Cordelia Langford, John Sillitoe on behalf of the Wellcome Sanger Institute COVID-19 Surveillance Team |
| EPI_ISL_3338715, EPI_ISL_3338716 | Lighthouse Lab in Glasgow | Wellcome Sanger Institute for the COVID-19 Genomics UK (COG-UK) Consortium | Harper VanSteenhouse, Yumi Kasai, David Gray, Carol Clugston, Anna Dominiczak and Alex Alderton, Roberto Amato, Jeffrey Barrett, Sonia Goncalves, Ewan Harrison, David K. Jackson, Ian Johnston, Dominic Kwiatkowski, Cordelia Langford, John Sillitoe on behalf of the Wellcome Sanger Institute COVID-19 Surveillance Team |
| EPI_ISL_3338717, EPI_ISL_3338718 | Lighthouse Lab in Milton Keynes | Wellcome Sanger Institute for the COVID-19 Genomics UK (COG-UK) Consortium | The Lighthouse Lab in Milton Keynes and Alex Alderton, Roberto Amato, Jeffrey Barrett, Sonia Goncalves, Ewan Harrison, David K. Jackson, Ian Johnston, Dominic Kwiatkowski, Cordelia Langford, John Sillitoe on behalf of the Wellcome Sanger Institute COVID-19 Surveillance Team |
| EPI_ISL_3338719 | Lighthouse Laboratory Plymouth | Wellcome Sanger Institute for the COVID-19 Genomics UK (COG-UK) | Lighthouse Laboratory Plymouth and Alex Alderton, Roberto Amato, Jeffrey Barrett, Sonia Goncalves, Ewan Harrison, David K. Jackson, Ian Johnston, Dominic Kwiatkowski, Cordelia Langford, John Sillitoe on behalf of the Wellcome Sanger |

| | | | |
|---|---|---|---|
| | | Consortium | Institute COVID-19 Surveillance Team |
| EPI_ISL_3338720 | Lighthouse Lab in Milton Keynes | Wellcome Sanger Institute for the COVID-19 Genomics UK (COG-UK) Consortium | The Lighthouse Lab in Milton Keynes and Alex Alderton, Roberto Amato, Jeffrey Barrett, Sonia Goncalves, Ewan Harrison, David K. Jackson, Ian Johnston, Dominic Kwiatkowski, Cordelia Langford, John Sillitoe on behalf of the Wellcome Sanger Institute COVID-19 Surveillance Team |
| EPI_ISL_3338721 | Lighthouse Lab in Glasgow | Wellcome Sanger Institute for the COVID-19 Genomics UK (COG-UK) Consortium | Harper VanSteenhouse, Yumi Kasai, David Gray, Carol Clugston, Anna Dominiczak and Alex Alderton, Roberto Amato, Jeffrey Barrett, Sonia Goncalves, Ewan Harrison, David K. Jackson, Ian Johnston, Dominic Kwiatkowski, Cordelia Langford, John Sillitoe on behalf of the Wellcome Sanger Institute COVID-19 Surveillance Team |
| EPI_ISL_3338722, EPI_ISL_3338723 | Lighthouse Lab in Milton Keynes | Wellcome Sanger Institute for the COVID-19 Genomics UK (COG-UK) Consortium | The Lighthouse Lab in Milton Keynes and Alex Alderton, Roberto Amato, Jeffrey Barrett, Sonia Goncalves, Ewan Harrison, David K. Jackson, Ian Johnston, Dominic Kwiatkowski, Cordelia Langford, John Sillitoe on behalf of the Wellcome Sanger Institute COVID-19 Surveillance Team |
| EPI_ISL_3338724 | Lighthouse Laboratory Plymouth | Wellcome Sanger Institute for the COVID-19 Genomics UK (COG-UK) Consortium | Lighthouse Laboratory Plymouth and Alex Alderton, Roberto Amato, Jeffrey Barrett, Sonia Goncalves, Ewan Harrison, David K. Jackson, Ian Johnston, Dominic Kwiatkowski, Cordelia Langford, John Sillitoe on behalf of the Wellcome Sanger Institute COVID-19 Surveillance Team |
| EPI_ISL_3338725, EPI_ISL_3338726 | Lighthouse Lab in Milton Keynes | Wellcome Sanger Institute for the COVID-19 Genomics UK (COG-UK) Consortium | The Lighthouse Lab in Milton Keynes and Alex Alderton, Roberto Amato, Jeffrey Barrett, Sonia Goncalves, Ewan Harrison, David K. Jackson, Ian Johnston, Dominic Kwiatkowski, Cordelia Langford, John Sillitoe on behalf of the Wellcome Sanger Institute COVID-19 Surveillance Team |
| EPI_ISL_3338727 | Lighthouse Laboratory Plymouth | Wellcome Sanger Institute for the COVID-19 Genomics UK (COG-UK) Consortium | Lighthouse Laboratory Plymouth and Alex Alderton, Roberto Amato, Jeffrey Barrett, Sonia Goncalves, Ewan Harrison, David K. Jackson, Ian Johnston, Dominic Kwiatkowski, Cordelia Langford, John Sillitoe on behalf of the Wellcome Sanger Institute COVID-19 Surveillance Team |
| EPI_ISL_3338728, EPI_ISL_3338729 | Lighthouse Lab in Milton Keynes | Wellcome Sanger Institute for the COVID-19 Genomics UK (COG-UK) Consortium | The Lighthouse Lab in Milton Keynes and Alex Alderton, Roberto Amato, Jeffrey Barrett, Sonia Goncalves, Ewan Harrison, David K. Jackson, Ian Johnston, Dominic Kwiatkowski, Cordelia Langford, John Sillitoe on behalf of the Wellcome Sanger Institute COVID-19 Surveillance Team |
| EPI_ISL_3338730 | Lighthouse Laboratory Plymouth | Wellcome Sanger Institute for the COVID-19 Genomics UK (COG-UK) Consortium | Lighthouse Laboratory Plymouth and Alex Alderton, Roberto Amato, Jeffrey Barrett, Sonia Goncalves, Ewan Harrison, David K. Jackson, Ian Johnston, Dominic Kwiatkowski, Cordelia Langford, John Sillitoe on behalf of the Wellcome Sanger Institute COVID-19 Surveillance Team |
| EPI_ISL_3338731 | Lighthouse Lab in Glasgow | Wellcome Sanger Institute for the COVID-19 Genomics UK (COG-UK) Consortium | Harper VanSteenhouse, Yumi Kasai, David Gray, Carol Clugston, Anna Dominiczak and Alex Alderton, Roberto Amato, Jeffrey Barrett, Sonia Goncalves, Ewan Harrison, David K. Jackson, Ian Johnston, Dominic Kwiatkowski, Cordelia Langford, John Sillitoe on behalf of the Wellcome Sanger Institute COVID-19 Surveillance Team |
| EPI_ISL_3338732 | Lighthouse Lab in Milton Keynes | Wellcome Sanger Institute for the COVID-19 Genomics UK (COG-UK) Consortium | The Lighthouse Lab in Milton Keynes and Alex Alderton, Roberto Amato, Jeffrey Barrett, Sonia Goncalves, Ewan Harrison, David K. Jackson, Ian Johnston, Dominic Kwiatkowski, Cordelia Langford, John Sillitoe on behalf of the Wellcome Sanger Institute COVID-19 Surveillance Team |
| EPI_ISL_3338733, EPI_ISL_3338734 | Lighthouse Lab in Glasgow | Wellcome Sanger Institute for the COVID-19 Genomics UK (COG-UK) Consortium | Harper VanSteenhouse, Yumi Kasai, David Gray, Carol Clugston, Anna Dominiczak and Alex Alderton, Roberto Amato, Jeffrey Barrett, Sonia Goncalves, Ewan Harrison, David K. Jackson, Ian Johnston, Dominic Kwiatkowski, Cordelia Langford, John Sillitoe on behalf of the Wellcome Sanger Institute COVID-19 Surveillance Team |
| EPI_ISL_3338735 | Lighthouse Laboratory Plymouth | Wellcome Sanger Institute for the COVID-19 Genomics UK (COG-UK) Consortium | Lighthouse Laboratory Plymouth and Alex Alderton, Roberto Amato, Jeffrey Barrett, Sonia Goncalves, Ewan Harrison, David K. Jackson, Ian Johnston, Dominic Kwiatkowski, Cordelia Langford, John Sillitoe on behalf of the Wellcome Sanger Institute COVID-19 Surveillance Team |
| EPI_ISL_3338736 | Lighthouse Lab in Milton Keynes | Wellcome Sanger Institute for the COVID-19 Genomics UK (COG-UK) Consortium | The Lighthouse Lab in Milton Keynes and Alex Alderton, Roberto Amato, Jeffrey Barrett, Sonia Goncalves, Ewan Harrison, David K. Jackson, Ian Johnston, Dominic Kwiatkowski, Cordelia Langford, John Sillitoe on behalf of the Wellcome Sanger Institute COVID-19 Surveillance Team |
| EPI_ISL_3338737 | Lighthouse Lab in Glasgow | Wellcome Sanger Institute for the COVID-19 Genomics UK (COG-UK) Consortium | Harper VanSteenhouse, Yumi Kasai, David Gray, Carol Clugston, Anna Dominiczak and Alex Alderton, Roberto Amato, Jeffrey Barrett, Sonia Goncalves, Ewan Harrison, David K. Jackson, Ian Johnston, Dominic Kwiatkowski, Cordelia Langford, John Sillitoe on behalf of the Wellcome Sanger Institute COVID-19 Surveillance Team |
| EPI_ISL_3338738 | Lighthouse Laboratory Plymouth | Wellcome Sanger Institute for the COVID-19 Genomics UK (COG-UK) Consortium | Lighthouse Laboratory Plymouth and Alex Alderton, Roberto Amato, Jeffrey Barrett, Sonia Goncalves, Ewan Harrison, David K. Jackson, Ian Johnston, Dominic Kwiatkowski, Cordelia Langford, John Sillitoe on behalf of the Wellcome Sanger Institute COVID-19 Surveillance Team |
| EPI_ISL_3338739, EPI_ISL_3338740, EPI_ISL_3338741, EPI_ISL_3338742, EPI_ISL_3338743 | Lighthouse Lab in Milton Keynes | Wellcome Sanger Institute for the COVID-19 Genomics UK (COG-UK) Consortium | The Lighthouse Lab in Milton Keynes and Alex Alderton, Roberto Amato, Jeffrey Barrett, Sonia Goncalves, Ewan Harrison, David K. Jackson, Ian Johnston, Dominic Kwiatkowski, Cordelia Langford, John Sillitoe on behalf of the Wellcome Sanger Institute COVID-19 Surveillance Team |
| EPI_ISL_3338744, EPI_ISL_3338745 | Lighthouse Lab in Glasgow | Wellcome Sanger Institute for the COVID-19 Genomics UK (COG-UK) Consortium | Harper VanSteenhouse, Yumi Kasai, David Gray, Carol Clugston, Anna Dominiczak and Alex Alderton, Roberto Amato, Jeffrey Barrett, Sonia Goncalves, Ewan Harrison, David K. Jackson, Ian Johnston, Dominic Kwiatkowski, Cordelia Langford, John Sillitoe on behalf of the Wellcome Sanger Institute COVID-19 Surveillance Team |
| EPI_ISL_3338746 | Lighthouse Lab in Milton Keynes | Wellcome Sanger Institute for the COVID-19 Genomics UK (COG-UK) Consortium | The Lighthouse Lab in Milton Keynes and Alex Alderton, Roberto Amato, Jeffrey Barrett, Sonia Goncalves, Ewan Harrison, David K. Jackson, Ian Johnston, Dominic Kwiatkowski, Cordelia Langford, John Sillitoe on behalf of the Wellcome Sanger Institute COVID-19 Surveillance Team |
| EPI_ISL_3338747 | Lighthouse Laboratory Plymouth | Wellcome Sanger Institute for the COVID-19 Genomics UK (COG-UK) Consortium | Lighthouse Laboratory Plymouth and Alex Alderton, Roberto Amato, Jeffrey Barrett, Sonia Goncalves, Ewan Harrison, David K. Jackson, Ian Johnston, Dominic Kwiatkowski, Cordelia Langford, John Sillitoe on behalf of the Wellcome Sanger Institute COVID-19 Surveillance Team |
| EPI_ISL_3338748 | Lighthouse Lab in Glasgow | Wellcome Sanger Institute for the COVID-19 Genomics UK (COG-UK) Consortium | Harper VanSteenhouse, Yumi Kasai, David Gray, Carol Clugston, Anna Dominiczak and Alex Alderton, Roberto Amato, Jeffrey Barrett, Sonia Goncalves, Ewan Harrison, David K. Jackson, Ian Johnston, Dominic Kwiatkowski, Cordelia Langford, John Sillitoe on behalf of the Wellcome Sanger Institute COVID-19 Surveillance Team |
| EPI_ISL_3338749 | Lighthouse Laboratory Plymouth | Wellcome Sanger Institute for the COVID-19 Genomics UK (COG-UK) Consortium | Lighthouse Laboratory Plymouth and Alex Alderton, Roberto Amato, Jeffrey Barrett, Sonia Goncalves, Ewan Harrison, David K. Jackson, Ian Johnston, Dominic Kwiatkowski, Cordelia Langford, John Sillitoe on behalf of the Wellcome Sanger Institute COVID-19 Surveillance Team |
| EPI_ISL_3338750, EPI_ISL_3338751, EPI_ISL_3338752 | Lighthouse Lab in Glasgow | Wellcome Sanger Institute for the COVID-19 Genomics UK (COG-UK) Consortium | Harper VanSteenhouse, Yumi Kasai, David Gray, Carol Clugston, Anna Dominiczak and Alex Alderton, Roberto Amato, Jeffrey Barrett, Sonia Goncalves, Ewan Harrison, David K. Jackson, Ian Johnston, Dominic Kwiatkowski, Cordelia Langford, John Sillitoe on behalf of the Wellcome Sanger Institute COVID-19 Surveillance Team |
| EPI_ISL_3338753 | Lighthouse Lab in Milton Keynes | Wellcome Sanger Institute for the COVID-19 Genomics UK (COG-UK) Consortium | The Lighthouse Lab in Milton Keynes and Alex Alderton, Roberto Amato, Jeffrey Barrett, Sonia Goncalves, Ewan Harrison, David K. Jackson, Ian Johnston, Dominic Kwiatkowski, Cordelia Langford, John Sillitoe on behalf of the Wellcome Sanger Institute COVID-19 Surveillance Team |
| EPI_ISL_3338754, EPI_ISL_3338755 | Lighthouse Laboratory Plymouth | Wellcome Sanger Institute for the COVID-19 Genomics UK (COG-UK) Consortium | Lighthouse Laboratory Plymouth and Alex Alderton, Roberto Amato, Jeffrey Barrett, Sonia Goncalves, Ewan Harrison, David K. Jackson, Ian Johnston, Dominic Kwiatkowski, Cordelia Langford, John Sillitoe on behalf of the Wellcome Sanger Institute COVID-19 Surveillance Team |
| EPI_ISL_3338756, EPI_ISL_3338757, EPI_ISL_3338758 | Lighthouse Lab in Milton Keynes | Wellcome Sanger Institute for the COVID-19 Genomics UK (COG-UK) Consortium | The Lighthouse Lab in Milton Keynes and Alex Alderton, Roberto Amato, Jeffrey Barrett, Sonia Goncalves, Ewan Harrison, David K. Jackson, Ian Johnston, Dominic Kwiatkowski, Cordelia Langford, John Sillitoe on behalf of the Wellcome Sanger Institute COVID-19 Surveillance Team |
| EPI_ISL_3338759 | Lighthouse Laboratory Plymouth | Wellcome Sanger Institute for the COVID-19 Genomics UK (COG-UK) Consortium | Lighthouse Laboratory Plymouth and Alex Alderton, Roberto Amato, Jeffrey Barrett, Sonia Goncalves, Ewan Harrison, David K. Jackson, Ian Johnston, Dominic Kwiatkowski, Cordelia Langford, John Sillitoe on behalf of the Wellcome Sanger Institute COVID-19 Surveillance Team |
| EPI_ISL_3338760 | Lighthouse Lab in Milton Keynes | Wellcome Sanger Institute for the COVID-19 Genomics UK (COG-UK) Consortium | The Lighthouse Lab in Milton Keynes and Alex Alderton, Roberto Amato, Jeffrey Barrett, Sonia Goncalves, Ewan Harrison, David K. Jackson, Ian Johnston, Dominic Kwiatkowski, Cordelia Langford, John Sillitoe on behalf of the Wellcome Sanger Institute COVID-19 Surveillance Team |
| EPI_ISL_3338761 | Lighthouse Lab in Glasgow | Wellcome Sanger Institute for the COVID-19 Genomics UK (COG-UK) Consortium | Harper VanSteenhouse, Yumi Kasai, David Gray, Carol Clugston, Anna Dominiczak and Alex Alderton, Roberto Amato, Jeffrey Barrett, Sonia Goncalves, Ewan Harrison, David K. Jackson, Ian Johnston, Dominic Kwiatkowski, Cordelia Langford, John Sillitoe on behalf of the Wellcome Sanger Institute COVID-19 Surveillance Team |
| EPI_ISL_3338762 | Lighthouse Lab in Milton Keynes | Wellcome Sanger Institute for the COVID-19 Genomics UK (COG-UK) Consortium | The Lighthouse Lab in Milton Keynes and Alex Alderton, Roberto Amato, Jeffrey Barrett, Sonia Goncalves, Ewan Harrison, David K. Jackson, Ian Johnston, Dominic Kwiatkowski, Cordelia Langford, John Sillitoe on behalf of the Wellcome Sanger Institute COVID-19 Surveillance Team |
| EPI_ISL_3338763 | Lighthouse Laboratory Plymouth | Wellcome Sanger Institute for the COVID-19 Genomics UK (COG-UK) Consortium | Lighthouse Laboratory Plymouth and Alex Alderton, Roberto Amato, Jeffrey Barrett, Sonia Goncalves, Ewan Harrison, David K. Jackson, Ian Johnston, Dominic Kwiatkowski, Cordelia Langford, John Sillitoe on behalf of the Wellcome Sanger Institute COVID-19 Surveillance Team |
| EPI_ISL_3338764, EPI_ISL_3338765, EPI_ISL_3338766, EPI_ISL_3338767, EPI_ISL_3338768, EPI_ISL_3338769 | Lighthouse Lab in Milton Keynes | Wellcome Sanger Institute for the COVID-19 Genomics UK (COG-UK) Consortium | The Lighthouse Lab in Milton Keynes and Alex Alderton, Roberto Amato, Jeffrey Barrett, Sonia Goncalves, Ewan Harrison, David K. Jackson, Ian Johnston, Dominic Kwiatkowski, Cordelia Langford, John Sillitoe on behalf of the Wellcome Sanger Institute COVID-19 Surveillance Team |
| EPI_ISL_3338770, EPI_ISL_3338771, EPI_ISL_3338772 | Lighthouse Laboratory Plymouth | Wellcome Sanger Institute for the COVID-19 Genomics UK (COG-UK) Consortium | Lighthouse Laboratory Plymouth and Alex Alderton, Roberto Amato, Jeffrey Barrett, Sonia Goncalves, Ewan Harrison, David K. Jackson, Ian Johnston, Dominic Kwiatkowski, Cordelia Langford, John Sillitoe on behalf of the Wellcome Sanger Institute COVID-19 Surveillance Team |
| EPI_ISL_3338773 | Lighthouse Lab in Glasgow | Wellcome Sanger Institute for the COVID-19 Genomics UK (COG-UK) Consortium | Harper VanSteenhouse, Yumi Kasai, David Gray, Carol Clugston, Anna Dominiczak and Alex Alderton, Roberto Amato, Jeffrey Barrett, Sonia Goncalves, Ewan Harrison, David K. Jackson, Ian Johnston, Dominic Kwiatkowski, Cordelia Langford, John Sillitoe on behalf of the Wellcome Sanger Institute COVID-19 Surveillance Team |
| EPI_ISL_3338774 | Lighthouse Laboratory Plymouth | Wellcome Sanger Institute for the COVID-19 Genomics UK (COG-UK) Consortium | Lighthouse Laboratory Plymouth and Alex Alderton, Roberto Amato, Jeffrey Barrett, Sonia Goncalves, Ewan Harrison, David K. Jackson, Ian Johnston, Dominic Kwiatkowski, Cordelia Langford, John Sillitoe on behalf of the Wellcome Sanger Institute COVID-19 Surveillance Team |
| EPI_ISL_3338775 | Lighthouse Lab in Milton Keynes | Wellcome Sanger Institute for the COVID-19 Genomics UK (COG-UK) Consortium | The Lighthouse Lab in Milton Keynes and Alex Alderton, Roberto Amato, Jeffrey Barrett, Sonia Goncalves, Ewan Harrison, David K. Jackson, Ian Johnston, Dominic Kwiatkowski, Cordelia Langford, John Sillitoe on behalf of the Wellcome Sanger Institute COVID-19 Surveillance Team |
| EPI_ISL_3338776 | Lighthouse Lab in Glasgow | Wellcome Sanger Institute for the COVID-19 Genomics UK (COG-UK) Consortium | Harper VanSteenhouse, Yumi Kasai, David Gray, Carol Clugston, Anna Dominiczak and Alex Alderton, Roberto Amato, Jeffrey Barrett, Sonia Goncalves, Ewan Harrison, David K. Jackson, Ian Johnston, Dominic Kwiatkowski, Cordelia Langford, John Sillitoe on behalf of the Wellcome Sanger Institute COVID-19 Surveillance Team |
| EPI_ISL_3338777 | Lighthouse Lab in Milton Keynes | Wellcome Sanger Institute for the COVID-19 Genomics UK (COG-UK) Consortium | The Lighthouse Lab in Milton Keynes and Alex Alderton, Roberto Amato, Jeffrey Barrett, Sonia Goncalves, Ewan Harrison, David K. Jackson, Ian Johnston, Dominic Kwiatkowski, Cordelia Langford, John Sillitoe on behalf of the Wellcome Sanger Institute COVID-19 Surveillance Team |
| EPI_ISL_3338778, EPI_ISL_3338779 | Lighthouse Lab in Glasgow | Wellcome Sanger Institute for the COVID-19 Genomics UK (COG-UK) Consortium | Harper VanSteenhouse, Yumi Kasai, David Gray, Carol Clugston, Anna Dominiczak and Alex Alderton, Roberto Amato, Jeffrey Barrett, Sonia Goncalves, Ewan Harrison, David K. Jackson, Ian Johnston, Dominic Kwiatkowski, Cordelia Langford, John Sillitoe on behalf of the Wellcome Sanger Institute COVID-19 Surveillance Team |
| EPI_ISL_3338780 | Lighthouse Laboratory Plymouth | Wellcome Sanger Institute for the COVID-19 Genomics UK (COG-UK) Consortium | Lighthouse Laboratory Plymouth and Alex Alderton, Roberto Amato, Jeffrey Barrett, Sonia Goncalves, Ewan Harrison, David K. Jackson, Ian Johnston, Dominic Kwiatkowski, Cordelia Langford, John Sillitoe on behalf of the Wellcome Sanger Institute COVID-19 Surveillance Team |
| EPI_ISL_3338781 | Lighthouse Lab in Milton Keynes | Wellcome Sanger Institute for the COVID-19 Genomics UK (COG-UK) Consortium | The Lighthouse Lab in Milton Keynes and Alex Alderton, Roberto Amato, Jeffrey Barrett, Sonia Goncalves, Ewan Harrison, David K. Jackson, Ian Johnston, Dominic Kwiatkowski, Cordelia Langford, John Sillitoe on behalf of the Wellcome Sanger Institute COVID-19 Surveillance Team |
| EPI_ISL_3338782 | Lighthouse Lab in Glasgow | Wellcome Sanger Institute for the COVID-19 Genomics UK (COG-UK) | Harper VanSteenhouse, Yumi Kasai, David Gray, Carol Clugston, Anna Dominiczak and Alex Alderton, Roberto Amato, Jeffrey Barrett, Sonia Goncalves, Ewan Harrison, David K. Jackson, Ian Johnston, Dominic Kwiatkowski, Cordelia Langford, |

| | | | |
|---|---|---|---|
| | | Consortium | John Sillitoe on behalf of the Wellcome Sanger Institute COVID-19 Surveillance Team |
| EPI_ISL_3338783 | Lighthouse Laboratory Plymouth | Wellcome Sanger Institute for the COVID-19 Genomics UK (COG-UK) Consortium | Lighthouse Laboratory Plymouth and Alex Alderton, Roberto Amato, Jeffrey Barrett, Sonia Goncalves, Ewan Harrison, David K. Jackson, Ian Johnston, Dominic Kwiatkowski, Cordelia Langford, John Sillitoe on behalf of the Wellcome Sanger Institute COVID-19 Surveillance Team |
| EPI_ISL_3338784, EPI_ISL_3338785, EPI_ISL_3338786 | Lighthouse Lab in Milton Keynes | Wellcome Sanger Institute for the COVID-19 Genomics UK (COG-UK) Consortium | The Lighthouse Lab in Milton Keynes and Alex Alderton, Roberto Amato, Jeffrey Barrett, Sonia Goncalves, Ewan Harrison, David K. Jackson, Ian Johnston, Dominic Kwiatkowski, Cordelia Langford, John Sillitoe on behalf of the Wellcome Sanger Institute COVID-19 Surveillance Team |
| EPI_ISL_3338787, EPI_ISL_3338788 | Lighthouse Laboratory Plymouth | Wellcome Sanger Institute for the COVID-19 Genomics UK (COG-UK) Consortium | Lighthouse Laboratory Plymouth and Alex Alderton, Roberto Amato, Jeffrey Barrett, Sonia Goncalves, Ewan Harrison, David K. Jackson, Ian Johnston, Dominic Kwiatkowski, Cordelia Langford, John Sillitoe on behalf of the Wellcome Sanger Institute COVID-19 Surveillance Team |
| EPI_ISL_3338789 | Lighthouse Lab in Glasgow | Wellcome Sanger Institute for the COVID-19 Genomics UK (COG-UK) Consortium | Harper VanSteenhouse, Yumi Kasai, David Gray, Carol Clugston, Anna Dominiczak and Alex Alderton, Roberto Amato, Jeffrey Barrett, Sonia Goncalves, Ewan Harrison, David K. Jackson, Ian Johnston, Dominic Kwiatkowski, Cordelia Langford, John Sillitoe on behalf of the Wellcome Sanger Institute COVID-19 Surveillance Team |
| EPI_ISL_3338790, EPI_ISL_3338791, EPI_ISL_3338792 | Lighthouse Lab in Milton Keynes | Wellcome Sanger Institute for the COVID-19 Genomics UK (COG-UK) Consortium | The Lighthouse Lab in Milton Keynes and Alex Alderton, Roberto Amato, Jeffrey Barrett, Sonia Goncalves, Ewan Harrison, David K. Jackson, Ian Johnston, Dominic Kwiatkowski, Cordelia Langford, John Sillitoe on behalf of the Wellcome Sanger Institute COVID-19 Surveillance Team |
| EPI_ISL_3338793 | Lighthouse Laboratory Plymouth | Wellcome Sanger Institute for the COVID-19 Genomics UK (COG-UK) Consortium | Lighthouse Laboratory Plymouth and Alex Alderton, Roberto Amato, Jeffrey Barrett, Sonia Goncalves, Ewan Harrison, David K. Jackson, Ian Johnston, Dominic Kwiatkowski, Cordelia Langford, John Sillitoe on behalf of the Wellcome Sanger Institute COVID-19 Surveillance Team |
| EPI_ISL_3338794 | Lighthouse Lab in Milton Keynes | Wellcome Sanger Institute for the COVID-19 Genomics UK (COG-UK) Consortium | The Lighthouse Lab in Milton Keynes and Alex Alderton, Roberto Amato, Jeffrey Barrett, Sonia Goncalves, Ewan Harrison, David K. Jackson, Ian Johnston, Dominic Kwiatkowski, Cordelia Langford, John Sillitoe on behalf of the Wellcome Sanger Institute COVID-19 Surveillance Team |
| EPI_ISL_3338795, EPI_ISL_3338796 | Lighthouse Lab in Glasgow | Wellcome Sanger Institute for the COVID-19 Genomics UK (COG-UK) Consortium | Harper VanSteenhouse, Yumi Kasai, David Gray, Carol Clugston, Anna Dominiczak and Alex Alderton, Roberto Amato, Jeffrey Barrett, Sonia Goncalves, Ewan Harrison, David K. Jackson, Ian Johnston, Dominic Kwiatkowski, Cordelia Langford, John Sillitoe on behalf of the Wellcome Sanger Institute COVID-19 Surveillance Team |
| EPI_ISL_3338797 | Lighthouse Laboratory Plymouth | Wellcome Sanger Institute for the COVID-19 Genomics UK (COG-UK) Consortium | Lighthouse Laboratory Plymouth and Alex Alderton, Roberto Amato, Jeffrey Barrett, Sonia Goncalves, Ewan Harrison, David K. Jackson, Ian Johnston, Dominic Kwiatkowski, Cordelia Langford, John Sillitoe on behalf of the Wellcome Sanger Institute COVID-19 Surveillance Team |
| EPI_ISL_3338798, EPI_ISL_3338799 | Lighthouse Lab in Glasgow | Wellcome Sanger Institute for the COVID-19 Genomics UK (COG-UK) Consortium | Harper VanSteenhouse, Yumi Kasai, David Gray, Carol Clugston, Anna Dominiczak and Alex Alderton, Roberto Amato, Jeffrey Barrett, Sonia Goncalves, Ewan Harrison, David K. Jackson, Ian Johnston, Dominic Kwiatkowski, Cordelia Langford, John Sillitoe on behalf of the Wellcome Sanger Institute COVID-19 Surveillance Team |
| EPI_ISL_3338800, EPI_ISL_3338801, EPI_ISL_3338802 | Lighthouse Lab in Milton Keynes | Wellcome Sanger Institute for the COVID-19 Genomics UK (COG-UK) Consortium | The Lighthouse Lab in Milton Keynes and Alex Alderton, Roberto Amato, Jeffrey Barrett, Sonia Goncalves, Ewan Harrison, David K. Jackson, Ian Johnston, Dominic Kwiatkowski, Cordelia Langford, John Sillitoe on behalf of the Wellcome Sanger Institute COVID-19 Surveillance Team |
| EPI_ISL_3338803 | Lighthouse Lab in Glasgow | Wellcome Sanger Institute for the COVID-19 Genomics UK (COG-UK) Consortium | Harper VanSteenhouse, Yumi Kasai, David Gray, Carol Clugston, Anna Dominiczak and Alex Alderton, Roberto Amato, Jeffrey Barrett, Sonia Goncalves, Ewan Harrison, David K. Jackson, Ian Johnston, Dominic Kwiatkowski, Cordelia Langford, John Sillitoe on behalf of the Wellcome Sanger Institute COVID-19 Surveillance Team |
| EPI_ISL_3338804 | Lighthouse Laboratory Plymouth | Wellcome Sanger Institute for the COVID-19 Genomics UK (COG-UK) Consortium | Lighthouse Laboratory Plymouth and Alex Alderton, Roberto Amato, Jeffrey Barrett, Sonia Goncalves, Ewan Harrison, David K. Jackson, Ian Johnston, Dominic Kwiatkowski, Cordelia Langford, John Sillitoe on behalf of the Wellcome Sanger Institute COVID-19 Surveillance Team |
| EPI_ISL_3338805, EPI_ISL_3338806, EPI_ISL_3338807, EPI_ISL_3338808, EPI_ISL_3338809 | Lighthouse Lab in Milton Keynes | Wellcome Sanger Institute for the COVID-19 Genomics UK (COG-UK) Consortium | The Lighthouse Lab in Milton Keynes and Alex Alderton, Roberto Amato, Jeffrey Barrett, Sonia Goncalves, Ewan Harrison, David K. Jackson, Ian Johnston, Dominic Kwiatkowski, Cordelia Langford, John Sillitoe on behalf of the Wellcome Sanger Institute COVID-19 Surveillance Team |
| EPI_ISL_3338810 | Lighthouse Lab in Glasgow | Wellcome Sanger Institute for the COVID-19 Genomics UK (COG-UK) Consortium | Harper VanSteenhouse, Yumi Kasai, David Gray, Carol Clugston, Anna Dominiczak and Alex Alderton, Roberto Amato, Jeffrey Barrett, Sonia Goncalves, Ewan Harrison, David K. Jackson, Ian Johnston, Dominic Kwiatkowski, Cordelia Langford, John Sillitoe on behalf of the Wellcome Sanger Institute COVID-19 Surveillance Team |
| EPI_ISL_3338811, EPI_ISL_3338812, EPI_ISL_3338813, EPI_ISL_3338814, EPI_ISL_3338815, EPI_ISL_3338816, EPI_ISL_3338817, EPI_ISL_3338818 | Lighthouse Lab in Milton Keynes | Wellcome Sanger Institute for the COVID-19 Genomics UK (COG-UK) Consortium | The Lighthouse Lab in Milton Keynes and Alex Alderton, Roberto Amato, Jeffrey Barrett, Sonia Goncalves, Ewan Harrison, David K. Jackson, Ian Johnston, Dominic Kwiatkowski, Cordelia Langford, John Sillitoe on behalf of the Wellcome Sanger Institute COVID-19 Surveillance Team |
| EPI_ISL_3338819, EPI_ISL_3338820 | Lighthouse Lab in Glasgow | Wellcome Sanger Institute for the COVID-19 Genomics UK (COG-UK) Consortium | Harper VanSteenhouse, Yumi Kasai, David Gray, Carol Clugston, Anna Dominiczak and Alex Alderton, Roberto Amato, Jeffrey Barrett, Sonia Goncalves, Ewan Harrison, David K. Jackson, Ian Johnston, Dominic Kwiatkowski, Cordelia Langford, John Sillitoe on behalf of the Wellcome Sanger Institute COVID-19 Surveillance Team |
| EPI_ISL_3338821, EPI_ISL_3338822 | Lighthouse Laboratory Plymouth | Wellcome Sanger Institute for the COVID-19 Genomics UK (COG-UK) Consortium | Lighthouse Laboratory Plymouth and Alex Alderton, Roberto Amato, Jeffrey Barrett, Sonia Goncalves, Ewan Harrison, David K. Jackson, Ian Johnston, Dominic Kwiatkowski, Cordelia Langford, John Sillitoe on behalf of the Wellcome Sanger Institute COVID-19 Surveillance Team |
| EPI_ISL_3338823 | Lighthouse Lab in Glasgow | Wellcome Sanger Institute for the COVID-19 Genomics UK (COG-UK) Consortium | Harper VanSteenhouse, Yumi Kasai, David Gray, Carol Clugston, Anna Dominiczak and Alex Alderton, Roberto Amato, Jeffrey Barrett, Sonia Goncalves, Ewan Harrison, David K. Jackson, Ian Johnston, Dominic Kwiatkowski, Cordelia Langford, John Sillitoe on behalf of the Wellcome Sanger Institute COVID-19 Surveillance Team |
| EPI_ISL_3338824, EPI_ISL_3338825 | Lighthouse Laboratory Plymouth | Wellcome Sanger Institute for the COVID-19 Genomics UK (COG-UK) Consortium | Lighthouse Laboratory Plymouth and Alex Alderton, Roberto Amato, Jeffrey Barrett, Sonia Goncalves, Ewan Harrison, David K. Jackson, Ian Johnston, Dominic Kwiatkowski, Cordelia Langford, John Sillitoe on behalf of the Wellcome Sanger Institute COVID-19 Surveillance Team |
| EPI_ISL_3338826 | Lighthouse Lab in Glasgow | Wellcome Sanger Institute for the COVID-19 Genomics UK (COG-UK) Consortium | Harper VanSteenhouse, Yumi Kasai, David Gray, Carol Clugston, Anna Dominiczak and Alex Alderton, Roberto Amato, Jeffrey Barrett, Sonia Goncalves, Ewan Harrison, David K. Jackson, Ian Johnston, Dominic Kwiatkowski, Cordelia Langford, John Sillitoe on behalf of the Wellcome Sanger Institute COVID-19 Surveillance Team |
| EPI_ISL_3338827 | Lighthouse Laboratory Plymouth | Wellcome Sanger Institute for the COVID-19 Genomics UK (COG-UK) Consortium | Lighthouse Laboratory Plymouth and Alex Alderton, Roberto Amato, Jeffrey Barrett, Sonia Goncalves, Ewan Harrison, David K. Jackson, Ian Johnston, Dominic Kwiatkowski, Cordelia Langford, John Sillitoe on behalf of the Wellcome Sanger Institute COVID-19 Surveillance Team |
| EPI_ISL_3338828, EPI_ISL_3338829 | Lighthouse Lab in Milton Keynes | Wellcome Sanger Institute for the COVID-19 Genomics UK (COG-UK) Consortium | The Lighthouse Lab in Milton Keynes and Alex Alderton, Roberto Amato, Jeffrey Barrett, Sonia Goncalves, Ewan Harrison, David K. Jackson, Ian Johnston, Dominic Kwiatkowski, Cordelia Langford, John Sillitoe on behalf of the Wellcome Sanger Institute COVID-19 Surveillance Team |
| EPI_ISL_3338830 | Lighthouse Lab in Glasgow | Wellcome Sanger Institute for the COVID-19 Genomics UK (COG-UK) Consortium | Harper VanSteenhouse, Yumi Kasai, David Gray, Carol Clugston, Anna Dominiczak and Alex Alderton, Roberto Amato, Jeffrey Barrett, Sonia Goncalves, Ewan Harrison, David K. Jackson, Ian Johnston, Dominic Kwiatkowski, Cordelia Langford, John Sillitoe on behalf of the Wellcome Sanger Institute COVID-19 Surveillance Team |
| EPI_ISL_3338831, EPI_ISL_3338832, EPI_ISL_3338833 | Lighthouse Lab in Milton Keynes | Wellcome Sanger Institute for the COVID-19 Genomics UK (COG-UK) Consortium | The Lighthouse Lab in Milton Keynes and Alex Alderton, Roberto Amato, Jeffrey Barrett, Sonia Goncalves, Ewan Harrison, David K. Jackson, Ian Johnston, Dominic Kwiatkowski, Cordelia Langford, John Sillitoe on behalf of the Wellcome Sanger Institute COVID-19 Surveillance Team |
| EPI_ISL_3338834 | Lighthouse Laboratory Plymouth | Wellcome Sanger Institute for the COVID-19 Genomics UK (COG-UK) Consortium | Lighthouse Laboratory Plymouth and Alex Alderton, Roberto Amato, Jeffrey Barrett, Sonia Goncalves, Ewan Harrison, David K. Jackson, Ian Johnston, Dominic Kwiatkowski, Cordelia Langford, John Sillitoe on behalf of the Wellcome Sanger Institute COVID-19 Surveillance Team |
| EPI_ISL_3338835 | Lighthouse Lab in Glasgow | Wellcome Sanger Institute for the COVID-19 Genomics UK (COG-UK) Consortium | Harper VanSteenhouse, Yumi Kasai, David Gray, Carol Clugston, Anna Dominiczak and Alex Alderton, Roberto Amato, Jeffrey Barrett, Sonia Goncalves, Ewan Harrison, David K. Jackson, Ian Johnston, Dominic Kwiatkowski, Cordelia Langford, John Sillitoe on behalf of the Wellcome Sanger Institute COVID-19 Surveillance Team |
| EPI_ISL_3338836 | Lighthouse Lab in Milton Keynes | Wellcome Sanger Institute for the COVID-19 Genomics UK (COG-UK) Consortium | The Lighthouse Lab in Milton Keynes and Alex Alderton, Roberto Amato, Jeffrey Barrett, Sonia Goncalves, Ewan Harrison, David K. Jackson, Ian Johnston, Dominic Kwiatkowski, Cordelia Langford, John Sillitoe on behalf of the Wellcome Sanger Institute COVID-19 Surveillance Team |
| EPI_ISL_3338837 | Lighthouse Lab in Glasgow | Wellcome Sanger Institute for the COVID-19 Genomics UK (COG-UK) Consortium | Harper VanSteenhouse, Yumi Kasai, David Gray, Carol Clugston, Anna Dominiczak and Alex Alderton, Roberto Amato, Jeffrey Barrett, Sonia Goncalves, Ewan Harrison, David K. Jackson, Ian Johnston, Dominic Kwiatkowski, Cordelia Langford, John Sillitoe on behalf of the Wellcome Sanger Institute COVID-19 Surveillance Team |
| EPI_ISL_3338838, EPI_ISL_3338839 | Lighthouse Lab in Milton Keynes | Wellcome Sanger Institute for the COVID-19 Genomics UK (COG-UK) Consortium | The Lighthouse Lab in Milton Keynes and Alex Alderton, Roberto Amato, Jeffrey Barrett, Sonia Goncalves, Ewan Harrison, David K. Jackson, Ian Johnston, Dominic Kwiatkowski, Cordelia Langford, John Sillitoe on behalf of the Wellcome Sanger Institute COVID-19 Surveillance Team |
| EPI_ISL_3338840, EPI_ISL_3338841, EPI_ISL_3338842 | Lighthouse Laboratory Plymouth | Wellcome Sanger Institute for the COVID-19 Genomics UK (COG-UK) Consortium | Lighthouse Laboratory Plymouth and Alex Alderton, Roberto Amato, Jeffrey Barrett, Sonia Goncalves, Ewan Harrison, David K. Jackson, Ian Johnston, Dominic Kwiatkowski, Cordelia Langford, John Sillitoe on behalf of the Wellcome Sanger Institute COVID-19 Surveillance Team |
| EPI_ISL_3338843, EPI_ISL_3338844, EPI_ISL_3338845, EPI_ISL_3338846 | Lighthouse Lab in Milton Keynes | Wellcome Sanger Institute for the COVID-19 Genomics UK (COG-UK) Consortium | The Lighthouse Lab in Milton Keynes and Alex Alderton, Roberto Amato, Jeffrey Barrett, Sonia Goncalves, Ewan Harrison, David K. Jackson, Ian Johnston, Dominic Kwiatkowski, Cordelia Langford, John Sillitoe on behalf of the Wellcome Sanger Institute COVID-19 Surveillance Team |
| EPI_ISL_3338847, EPI_ISL_3338848 | Lighthouse Laboratory Plymouth | Wellcome Sanger Institute for the COVID-19 Genomics UK (COG-UK) Consortium | Lighthouse Laboratory Plymouth and Alex Alderton, Roberto Amato, Jeffrey Barrett, Sonia Goncalves, Ewan Harrison, David K. Jackson, Ian Johnston, Dominic Kwiatkowski, Cordelia Langford, John Sillitoe on behalf of the Wellcome Sanger Institute COVID-19 Surveillance Team |
| EPI_ISL_3338849 | Lighthouse Lab in Glasgow | Wellcome Sanger Institute for the COVID-19 Genomics UK (COG-UK) Consortium | Harper VanSteenhouse, Yumi Kasai, David Gray, Carol Clugston, Anna Dominiczak and Alex Alderton, Roberto Amato, Jeffrey Barrett, Sonia Goncalves, Ewan Harrison, David K. Jackson, Ian Johnston, Dominic Kwiatkowski, Cordelia Langford, John Sillitoe on behalf of the Wellcome Sanger Institute COVID-19 Surveillance Team |
| EPI_ISL_3338850 | Lighthouse Lab in Milton Keynes | Wellcome Sanger Institute for the COVID-19 Genomics UK (COG-UK) Consortium | The Lighthouse Lab in Milton Keynes and Alex Alderton, Roberto Amato, Jeffrey Barrett, Sonia Goncalves, Ewan Harrison, David K. Jackson, Ian Johnston, Dominic Kwiatkowski, Cordelia Langford, John Sillitoe on behalf of the Wellcome Sanger Institute COVID-19 Surveillance Team |
| EPI_ISL_3338851 | Lighthouse Lab in Glasgow | Wellcome Sanger Institute for the COVID-19 Genomics UK (COG-UK) Consortium | Harper VanSteenhouse, Yumi Kasai, David Gray, Carol Clugston, Anna Dominiczak and Alex Alderton, Roberto Amato, Jeffrey Barrett, Sonia Goncalves, Ewan Harrison, David K. Jackson, Ian Johnston, Dominic Kwiatkowski, Cordelia Langford, John Sillitoe on behalf of the Wellcome Sanger Institute COVID-19 Surveillance Team |
| EPI_ISL_3338852 | Lighthouse Lab in Milton Keynes | Wellcome Sanger Institute for the COVID-19 Genomics UK (COG-UK) Consortium | The Lighthouse Lab in Milton Keynes and Alex Alderton, Roberto Amato, Jeffrey Barrett, Sonia Goncalves, Ewan Harrison, David K. Jackson, Ian Johnston, Dominic Kwiatkowski, Cordelia Langford, John Sillitoe on behalf of the Wellcome Sanger Institute COVID-19 Surveillance Team |
| EPI_ISL_3338853, EPI_ISL_3338854, | Lighthouse Laboratory Plymouth | Wellcome Sanger Institute for the COVID-19 Genomics UK (COG-UK) Consortium | Lighthouse Laboratory Plymouth and Alex Alderton, Roberto Amato, Jeffrey Barrett, Sonia Goncalves, Ewan Harrison, David K. Jackson, Ian Johnston, Dominic Kwiatkowski, Cordelia Langford, John Sillitoe on behalf of the Wellcome Sanger Institute COVID-19 Surveillance Team |

| | | | |
|---|---|---|---|
| EPI_ISL_3338855, EPI_ISL_3338856, EPI_ISL_3338857 | | | |
| EPI_ISL_3338858, EPI_ISL_3338859, EPI_ISL_3338860 | Lighthouse Lab in Milton Keynes | Wellcome Sanger Institute for the COVID-19 Genomics UK (COG-UK) Consortium | The Lighthouse Lab in Milton Keynes and Alex Alderton, Roberto Amato, Jeffrey Barrett, Sonia Goncalves, Ewan Harrison, David K. Jackson, Ian Johnston, Dominic Kwiatkowski, Cordelia Langford, John Sillitoe on behalf of the Wellcome Sanger Institute COVID-19 Surveillance Team |
| EPI_ISL_3338861, EPI_ISL_3338862 | Lighthouse Lab in Alderley Park | Wellcome Sanger Institute for the COVID-19 Genomics UK (COG-UK) Consortium | Jacquelyn Wynn, Mairead Hyland, The Lighthouse Lab in Alderley Park and Alex Alderton, Roberto Amato, Jeffrey Barrett, Sonia Goncalves, Ewan Harrison, David K. Jackson, Ian Johnston, Dominic Kwiatkowski, Cordelia Langford, John Sillitoe on behalf of the Wellcome Sanger Institute COVID-19 Surveillance Team |
| EPI_ISL_3338863, EPI_ISL_3338864 | Lighthouse Lab in Milton Keynes | Wellcome Sanger Institute for the COVID-19 Genomics UK (COG-UK) Consortium | The Lighthouse Lab in Milton Keynes and Alex Alderton, Roberto Amato, Jeffrey Barrett, Sonia Goncalves, Ewan Harrison, David K. Jackson, Ian Johnston, Dominic Kwiatkowski, Cordelia Langford, John Sillitoe on behalf of the Wellcome Sanger Institute COVID-19 Surveillance Team |
| EPI_ISL_3338865 | Berkshire and Surrey Pathology Services Lighthouse Laboratory | Wellcome Sanger Institute for the COVID-19 Genomics UK (COG-UK) Consortium | Berkshire and Surrey Pathology Services Lighthouse Laboratory and Alex Alderton, Roberto Amato, Jeffrey Barrett, Sonia Goncalves, Ewan Harrison, David K. Jackson, Ian Johnston, Dominic Kwiatkowski, Cordelia Langford, John Sillitoe on behalf of the Wellcome Sanger Institute COVID-19 Surveillance Team |
| EPI_ISL_3338866 | Lighthouse Lab in Glasgow | Wellcome Sanger Institute for the COVID-19 Genomics UK (COG-UK) Consortium | Harper VanSteenhouse, Yumi Kasai, David Gray, Carol Clugston, Anna Dominiczak and Alex Alderton, Roberto Amato, Jeffrey Barrett, Sonia Goncalves, Ewan Harrison, David K. Jackson, Ian Johnston, Dominic Kwiatkowski, Cordelia Langford, John Sillitoe on behalf of the Wellcome Sanger Institute COVID-19 Surveillance Team |
| EPI_ISL_3338867, EPI_ISL_3338868 | Lighthouse Lab in Alderley Park | Wellcome Sanger Institute for the COVID-19 Genomics UK (COG-UK) Consortium | Jacquelyn Wynn, Mairead Hyland, The Lighthouse Lab in Alderley Park and Alex Alderton, Roberto Amato, Jeffrey Barrett, Sonia Goncalves, Ewan Harrison, David K. Jackson, Ian Johnston, Dominic Kwiatkowski, Cordelia Langford, John Sillitoe on behalf of the Wellcome Sanger Institute COVID-19 Surveillance Team |
| EPI_ISL_3338869 | Randox Laboratories | Wellcome Sanger Institute for the COVID-19 Genomics UK (COG-UK) Consortium | Randox Laboratories and Alex Alderton, Roberto Amato, Jeffrey Barrett, Sonia Goncalves, Ewan Harrison, David K. Jackson, Ian Johnston, Dominic Kwiatkowski, Cordelia Langford, John Sillitoe on behalf of the Wellcome Sanger Institute COVID-19 Surveillance Team |
| EPI_ISL_3338870, EPI_ISL_3338871, EPI_ISL_3338872 | Lighthouse Lab in Alderley Park | Wellcome Sanger Institute for the COVID-19 Genomics UK (COG-UK) Consortium | Jacquelyn Wynn, Mairead Hyland, The Lighthouse Lab in Alderley Park and Alex Alderton, Roberto Amato, Jeffrey Barrett, Sonia Goncalves, Ewan Harrison, David K. Jackson, Ian Johnston, Dominic Kwiatkowski, Cordelia Langford, John Sillitoe on behalf of the Wellcome Sanger Institute COVID-19 Surveillance Team |
| EPI_ISL_3338873 | Lighthouse Lab in Glasgow | Wellcome Sanger Institute for the COVID-19 Genomics UK (COG-UK) Consortium | Harper VanSteenhouse, Yumi Kasai, David Gray, Carol Clugston, Anna Dominiczak and Alex Alderton, Roberto Amato, Jeffrey Barrett, Sonia Goncalves, Ewan Harrison, David K. Jackson, Ian Johnston, Dominic Kwiatkowski, Cordelia Langford, John Sillitoe on behalf of the Wellcome Sanger Institute COVID-19 Surveillance Team |
| EPI_ISL_3338874 | Lighthouse Lab in Alderley Park | Wellcome Sanger Institute for the COVID-19 Genomics UK (COG-UK) Consortium | Jacquelyn Wynn, Mairead Hyland, The Lighthouse Lab in Alderley Park and Alex Alderton, Roberto Amato, Jeffrey Barrett, Sonia Goncalves, Ewan Harrison, David K. Jackson, Ian Johnston, Dominic Kwiatkowski, Cordelia Langford, John Sillitoe on behalf of the Wellcome Sanger Institute COVID-19 Surveillance Team |
| EPI_ISL_3338875 | Lighthouse Lab in Glasgow | Wellcome Sanger Institute for the COVID-19 Genomics UK (COG-UK) Consortium | Harper VanSteenhouse, Yumi Kasai, David Gray, Carol Clugston, Anna Dominiczak and Alex Alderton, Roberto Amato, Jeffrey Barrett, Sonia Goncalves, Ewan Harrison, David K. Jackson, Ian Johnston, Dominic Kwiatkowski, Cordelia Langford, John Sillitoe on behalf of the Wellcome Sanger Institute COVID-19 Surveillance Team |
| EPI_ISL_3338876 | Lighthouse Lab in Alderley Park | Wellcome Sanger Institute for the COVID-19 Genomics UK (COG-UK) Consortium | Jacquelyn Wynn, Mairead Hyland, The Lighthouse Lab in Alderley Park and Alex Alderton, Roberto Amato, Jeffrey Barrett, Sonia Goncalves, Ewan Harrison, David K. Jackson, Ian Johnston, Dominic Kwiatkowski, Cordelia Langford, John Sillitoe on behalf of the Wellcome Sanger Institute COVID-19 Surveillance Team |
| EPI_ISL_3338877 | Lighthouse Lab in Glasgow | Wellcome Sanger Institute for the COVID-19 Genomics UK (COG-UK) Consortium | Harper VanSteenhouse, Yumi Kasai, David Gray, Carol Clugston, Anna Dominiczak and Alex Alderton, Roberto Amato, Jeffrey Barrett, Sonia Goncalves, Ewan Harrison, David K. Jackson, Ian Johnston, Dominic Kwiatkowski, Cordelia Langford, John Sillitoe on behalf of the Wellcome Sanger Institute COVID-19 Surveillance Team |
| EPI_ISL_3338878, EPI_ISL_3338879, EPI_ISL_3338880 | Lighthouse Lab in Alderley Park | Wellcome Sanger Institute for the COVID-19 Genomics UK (COG-UK) Consortium | Jacquelyn Wynn, Mairead Hyland, The Lighthouse Lab in Alderley Park and Alex Alderton, Roberto Amato, Jeffrey Barrett, Sonia Goncalves, Ewan Harrison, David K. Jackson, Ian Johnston, Dominic Kwiatkowski, Cordelia Langford, John Sillitoe on behalf of the Wellcome Sanger Institute COVID-19 Surveillance Team |
| EPI_ISL_3338881 | Lighthouse Lab in Milton Keynes | Wellcome Sanger Institute for the COVID-19 Genomics UK (COG-UK) Consortium | The Lighthouse Lab in Milton Keynes and Alex Alderton, Roberto Amato, Jeffrey Barrett, Sonia Goncalves, Ewan Harrison, David K. Jackson, Ian Johnston, Dominic Kwiatkowski, Cordelia Langford, John Sillitoe on behalf of the Wellcome Sanger Institute COVID-19 Surveillance Team |
| EPI_ISL_3338882, EPI_ISL_3338883 | Lighthouse Lab in Alderley Park | Wellcome Sanger Institute for the COVID-19 Genomics UK (COG-UK) Consortium | Jacquelyn Wynn, Mairead Hyland, The Lighthouse Lab in Alderley Park and Alex Alderton, Roberto Amato, Jeffrey Barrett, Sonia Goncalves, Ewan Harrison, David K. Jackson, Ian Johnston, Dominic Kwiatkowski, Cordelia Langford, John Sillitoe on behalf of the Wellcome Sanger Institute COVID-19 Surveillance Team |
| EPI_ISL_3338884 | Lighthouse Lab in Milton Keynes | Wellcome Sanger Institute for the COVID-19 Genomics UK (COG-UK) Consortium | The Lighthouse Lab in Milton Keynes and Alex Alderton, Roberto Amato, Jeffrey Barrett, Sonia Goncalves, Ewan Harrison, David K. Jackson, Ian Johnston, Dominic Kwiatkowski, Cordelia Langford, John Sillitoe on behalf of the Wellcome Sanger Institute COVID-19 Surveillance Team |
| EPI_ISL_3338885 | Lighthouse Lab in Alderley Park | Wellcome Sanger Institute for the COVID-19 Genomics UK (COG-UK) Consortium | Jacquelyn Wynn, Mairead Hyland, The Lighthouse Lab in Alderley Park and Alex Alderton, Roberto Amato, Jeffrey Barrett, Sonia Goncalves, Ewan Harrison, David K. Jackson, Ian Johnston, Dominic Kwiatkowski, Cordelia Langford, John Sillitoe on behalf of the Wellcome Sanger Institute COVID-19 Surveillance Team |
| EPI_ISL_3338886, EPI_ISL_3338887 | Lighthouse Lab in Milton Keynes | Wellcome Sanger Institute for the COVID-19 Genomics UK (COG-UK) Consortium | The Lighthouse Lab in Milton Keynes and Alex Alderton, Roberto Amato, Jeffrey Barrett, Sonia Goncalves, Ewan Harrison, David K. Jackson, Ian Johnston, Dominic Kwiatkowski, Cordelia Langford, John Sillitoe on behalf of the Wellcome Sanger Institute COVID-19 Surveillance Team |
| EPI_ISL_3338888, EPI_ISL_3338889 | Lighthouse Lab in Alderley Park | Wellcome Sanger Institute for the COVID-19 Genomics UK (COG-UK) Consortium | Jacquelyn Wynn, Mairead Hyland, The Lighthouse Lab in Alderley Park and Alex Alderton, Roberto Amato, Jeffrey Barrett, Sonia Goncalves, Ewan Harrison, David K. Jackson, Ian Johnston, Dominic Kwiatkowski, Cordelia Langford, John Sillitoe on behalf of the Wellcome Sanger Institute COVID-19 Surveillance Team |
| EPI_ISL_3338890, EPI_ISL_3338891, EPI_ISL_3338892 | Lighthouse Lab in Milton Keynes | Wellcome Sanger Institute for the COVID-19 Genomics UK (COG-UK) Consortium | The Lighthouse Lab in Milton Keynes and Alex Alderton, Roberto Amato, Jeffrey Barrett, Sonia Goncalves, Ewan Harrison, David K. Jackson, Ian Johnston, Dominic Kwiatkowski, Cordelia Langford, John Sillitoe on behalf of the Wellcome Sanger Institute COVID-19 Surveillance Team |
| EPI_ISL_3338893, EPI_ISL_3338894, EPI_ISL_3338895 | Lighthouse Lab in Alderley Park | Wellcome Sanger Institute for the COVID-19 Genomics UK (COG-UK) Consortium | Jacquelyn Wynn, Mairead Hyland, The Lighthouse Lab in Alderley Park and Alex Alderton, Roberto Amato, Jeffrey Barrett, Sonia Goncalves, Ewan Harrison, David K. Jackson, Ian Johnston, Dominic Kwiatkowski, Cordelia Langford, John Sillitoe on behalf of the Wellcome Sanger Institute COVID-19 Surveillance Team |
| EPI_ISL_3338896 | Lighthouse Lab in Milton Keynes | Wellcome Sanger Institute for the COVID-19 Genomics UK (COG-UK) Consortium | The Lighthouse Lab in Milton Keynes and Alex Alderton, Roberto Amato, Jeffrey Barrett, Sonia Goncalves, Ewan Harrison, David K. Jackson, Ian Johnston, Dominic Kwiatkowski, Cordelia Langford, John Sillitoe on behalf of the Wellcome Sanger Institute COVID-19 Surveillance Team |
| EPI_ISL_3338897 | Lighthouse Lab in Alderley Park | Wellcome Sanger Institute for the COVID-19 Genomics UK (COG-UK) Consortium | Jacquelyn Wynn, Mairead Hyland, The Lighthouse Lab in Alderley Park and Alex Alderton, Roberto Amato, Jeffrey Barrett, Sonia Goncalves, Ewan Harrison, David K. Jackson, Ian Johnston, Dominic Kwiatkowski, Cordelia Langford, John Sillitoe on behalf of the Wellcome Sanger Institute COVID-19 Surveillance Team |
| EPI_ISL_3338898 | Lighthouse Lab in Glasgow | Wellcome Sanger Institute for the COVID-19 Genomics UK (COG-UK) Consortium | Harper VanSteenhouse, Yumi Kasai, David Gray, Carol Clugston, Anna Dominiczak and Alex Alderton, Roberto Amato, Jeffrey Barrett, Sonia Goncalves, Ewan Harrison, David K. Jackson, Ian Johnston, Dominic Kwiatkowski, Cordelia Langford, John Sillitoe on behalf of the Wellcome Sanger Institute COVID-19 Surveillance Team |
| EPI_ISL_3338899 | Lighthouse Lab in Milton Keynes | Wellcome Sanger Institute for the COVID-19 Genomics UK (COG-UK) Consortium | The Lighthouse Lab in Milton Keynes and Alex Alderton, Roberto Amato, Jeffrey Barrett, Sonia Goncalves, Ewan Harrison, David K. Jackson, Ian Johnston, Dominic Kwiatkowski, Cordelia Langford, John Sillitoe on behalf of the Wellcome Sanger Institute COVID-19 Surveillance Team |
| EPI_ISL_3338900, EPI_ISL_3338901, EPI_ISL_3338902, EPI_ISL_3338903 | Lighthouse Lab in Alderley Park | Wellcome Sanger Institute for the COVID-19 Genomics UK (COG-UK) Consortium | Jacquelyn Wynn, Mairead Hyland, The Lighthouse Lab in Alderley Park and Alex Alderton, Roberto Amato, Jeffrey Barrett, Sonia Goncalves, Ewan Harrison, David K. Jackson, Ian Johnston, Dominic Kwiatkowski, Cordelia Langford, John Sillitoe on behalf of the Wellcome Sanger Institute COVID-19 Surveillance Team |
| EPI_ISL_3338904 | Lighthouse Lab in Milton Keynes | Wellcome Sanger Institute for the COVID-19 Genomics UK (COG-UK) Consortium | The Lighthouse Lab in Milton Keynes and Alex Alderton, Roberto Amato, Jeffrey Barrett, Sonia Goncalves, Ewan Harrison, David K. Jackson, Ian Johnston, Dominic Kwiatkowski, Cordelia Langford, John Sillitoe on behalf of the Wellcome Sanger Institute COVID-19 Surveillance Team |
| EPI_ISL_3338905 | Lighthouse Lab in Alderley Park | Wellcome Sanger Institute for the COVID-19 Genomics UK (COG-UK) Consortium | Jacquelyn Wynn, Mairead Hyland, The Lighthouse Lab in Alderley Park and Alex Alderton, Roberto Amato, Jeffrey Barrett, Sonia Goncalves, Ewan Harrison, David K. Jackson, Ian Johnston, Dominic Kwiatkowski, Cordelia Langford, John Sillitoe on behalf of the Wellcome Sanger Institute COVID-19 Surveillance Team |
| EPI_ISL_3338906, EPI_ISL_3338907 | Lighthouse Lab in Milton Keynes | Wellcome Sanger Institute for the COVID-19 Genomics UK (COG-UK) Consortium | The Lighthouse Lab in Milton Keynes and Alex Alderton, Roberto Amato, Jeffrey Barrett, Sonia Goncalves, Ewan Harrison, David K. Jackson, Ian Johnston, Dominic Kwiatkowski, Cordelia Langford, John Sillitoe on behalf of the Wellcome Sanger Institute COVID-19 Surveillance Team |
| EPI_ISL_3338908, EPI_ISL_3338909 | Lighthouse Lab in Alderley Park | Wellcome Sanger Institute for the COVID-19 Genomics UK (COG-UK) Consortium | Jacquelyn Wynn, Mairead Hyland, The Lighthouse Lab in Alderley Park and Alex Alderton, Roberto Amato, Jeffrey Barrett, Sonia Goncalves, Ewan Harrison, David K. Jackson, Ian Johnston, Dominic Kwiatkowski, Cordelia Langford, John Sillitoe on behalf of the Wellcome Sanger Institute COVID-19 Surveillance Team |
| EPI_ISL_3338910, EPI_ISL_3338911 | Lighthouse Lab in Glasgow | Wellcome Sanger Institute for the COVID-19 Genomics UK (COG-UK) Consortium | Harper VanSteenhouse, Yumi Kasai, David Gray, Carol Clugston, Anna Dominiczak and Alex Alderton, Roberto Amato, Jeffrey Barrett, Sonia Goncalves, Ewan Harrison, David K. Jackson, Ian Johnston, Dominic Kwiatkowski, Cordelia Langford, John Sillitoe on behalf of the Wellcome Sanger Institute COVID-19 Surveillance Team |
| EPI_ISL_3338912 | Lighthouse Lab in Alderley Park | Wellcome Sanger Institute for the COVID-19 Genomics UK (COG-UK) Consortium | Jacquelyn Wynn, Mairead Hyland, The Lighthouse Lab in Alderley Park and Alex Alderton, Roberto Amato, Jeffrey Barrett, Sonia Goncalves, Ewan Harrison, David K. Jackson, Ian Johnston, Dominic Kwiatkowski, Cordelia Langford, John Sillitoe on behalf of the Wellcome Sanger Institute COVID-19 Surveillance Team |
| EPI_ISL_3338913 | Randox Laboratories | Wellcome Sanger Institute for the COVID-19 Genomics UK (COG-UK) Consortium | Randox Laboratories and Alex Alderton, Roberto Amato, Jeffrey Barrett, Sonia Goncalves, Ewan Harrison, David K. Jackson, Ian Johnston, Dominic Kwiatkowski, Cordelia Langford, John Sillitoe on behalf of the Wellcome Sanger Institute COVID-19 Surveillance Team |
| EPI_ISL_3338914, EPI_ISL_3338915 | Lighthouse Lab in Milton Keynes | Wellcome Sanger Institute for the COVID-19 Genomics UK (COG-UK) Consortium | The Lighthouse Lab in Milton Keynes and Alex Alderton, Roberto Amato, Jeffrey Barrett, Sonia Goncalves, Ewan Harrison, David K. Jackson, Ian Johnston, Dominic Kwiatkowski, Cordelia Langford, John Sillitoe on behalf of the Wellcome Sanger Institute COVID-19 Surveillance Team |
| EPI_ISL_3338916 | Lighthouse Lab in Glasgow | Wellcome Sanger Institute for the COVID-19 Genomics UK (COG-UK) Consortium | Harper VanSteenhouse, Yumi Kasai, David Gray, Carol Clugston, Anna Dominiczak and Alex Alderton, Roberto Amato, Jeffrey Barrett, Sonia Goncalves, Ewan Harrison, David K. Jackson, Ian Johnston, Dominic Kwiatkowski, Cordelia Langford, John Sillitoe on behalf of the Wellcome Sanger Institute COVID-19 Surveillance Team |
| EPI_ISL_3338917 | Lighthouse Lab in Alderley Park | Wellcome Sanger Institute for the COVID-19 Genomics UK (COG-UK) Consortium | Jacquelyn Wynn, Mairead Hyland, The Lighthouse Lab in Alderley Park and Alex Alderton, Roberto Amato, Jeffrey Barrett, Sonia Goncalves, Ewan Harrison, David K. Jackson, Ian Johnston, Dominic Kwiatkowski, Cordelia Langford, John Sillitoe on behalf of the Wellcome Sanger Institute COVID-19 Surveillance Team |
| EPI_ISL_3338918 | Randox Laboratories | Wellcome Sanger Institute for the COVID-19 Genomics UK (COG-UK) Consortium | Randox Laboratories and Alex Alderton, Roberto Amato, Jeffrey Barrett, Sonia Goncalves, Ewan Harrison, David K. Jackson, Ian Johnston, Dominic Kwiatkowski, Cordelia Langford, John Sillitoe on behalf of the Wellcome Sanger Institute COVID-19 Surveillance Team |
| EPI_ISL_3338919, EPI_ISL_3338920 | Lighthouse Lab in Alderley Park | Wellcome Sanger Institute for the COVID-19 Genomics UK (COG-UK) Consortium | Jacquelyn Wynn, Mairead Hyland, The Lighthouse Lab in Alderley Park and Alex Alderton, Roberto Amato, Jeffrey Barrett, Sonia Goncalves, Ewan Harrison, David K. Jackson, Ian Johnston, Dominic Kwiatkowski, Cordelia Langford, John Sillitoe on behalf of the Wellcome Sanger Institute COVID-19 Surveillance Team |
| EPI_ISL_3338921 | Lighthouse Lab in Glasgow | Wellcome Sanger Institute for the COVID-19 Genomics UK (COG-UK) Consortium | Harper VanSteenhouse, Yumi Kasai, David Gray, Carol Clugston, Anna Dominiczak and Alex Alderton, Roberto Amato, Jeffrey Barrett, Sonia Goncalves, Ewan Harrison, David K. Jackson, Ian Johnston, Dominic Kwiatkowski, Cordelia Langford, John Sillitoe on behalf of the Wellcome Sanger Institute COVID-19 Surveillance Team |
| EPI_ISL_3338922 | Lighthouse Lab in Alderley Park | Wellcome Sanger Institute for the COVID-19 Genomics UK (COG-UK) | Jacquelyn Wynn, Mairead Hyland, The Lighthouse Lab in Alderley Park and Alex Alderton, Roberto Amato, Jeffrey Barrett, Sonia Goncalves, Ewan Harrison, David K. Jackson, Ian Johnston, Dominic Kwiatkowski, Cordelia Langford, John Sillitoe on |

| | | | |
|---|---|---|---|
| | | Consortium | behalf of the Wellcome Sanger Institute COVID-19 Surveillance Team |
| EPI_ISL_3338923 | Lighthouse Lab in Milton Keynes | Wellcome Sanger Institute for the COVID-19 Genomics UK (COG-UK) Consortium | The Lighthouse Lab in Milton Keynes and Alex Alderton, Roberto Amato, Jeffrey Barrett, Sonia Goncalves, Ewan Harrison, David K. Jackson, Ian Johnston, Dominic Kwiatkowski, Cordelia Langford, John Sillitoe on behalf of the Wellcome Sanger Institute COVID-19 Surveillance Team |
| EPI_ISL_3338924 | Lighthouse Lab in Alderley Park | Wellcome Sanger Institute for the COVID-19 Genomics UK (COG-UK) Consortium | Jacquelyn Wynn, Mairead Hyland, The Lighthouse Lab in Alderley Park and Alex Alderton, Roberto Amato, Jeffrey Barrett, Sonia Goncalves, Ewan Harrison, David K. Jackson, Ian Johnston, Dominic Kwiatkowski, Cordelia Langford, John Sillitoe on behalf of the Wellcome Sanger Institute COVID-19 Surveillance Team |
| EPI_ISL_3338925 | Lighthouse Lab in Glasgow | Wellcome Sanger Institute for the COVID-19 Genomics UK (COG-UK) Consortium | Harper VanSteenhouse, Yumi Kasai, David Gray, Carol Clugston, Anna Dominiczak and Alex Alderton, Roberto Amato, Jeffrey Barrett, Sonia Goncalves, Ewan Harrison, David K. Jackson, Ian Johnston, Dominic Kwiatkowski, Cordelia Langford, John Sillitoe on behalf of the Wellcome Sanger Institute COVID-19 Surveillance Team |
| EPI_ISL_3338926 | Lighthouse Lab in Alderley Park | Wellcome Sanger Institute for the COVID-19 Genomics UK (COG-UK) Consortium | Jacquelyn Wynn, Mairead Hyland, The Lighthouse Lab in Alderley Park and Alex Alderton, Roberto Amato, Jeffrey Barrett, Sonia Goncalves, Ewan Harrison, David K. Jackson, Ian Johnston, Dominic Kwiatkowski, Cordelia Langford, John Sillitoe on behalf of the Wellcome Sanger Institute COVID-19 Surveillance Team |
| EPI_ISL_3338927, EPI_ISL_3338928 | Lighthouse Lab in Milton Keynes | Wellcome Sanger Institute for the COVID-19 Genomics UK (COG-UK) Consortium | The Lighthouse Lab in Milton Keynes and Alex Alderton, Roberto Amato, Jeffrey Barrett, Sonia Goncalves, Ewan Harrison, David K. Jackson, Ian Johnston, Dominic Kwiatkowski, Cordelia Langford, John Sillitoe on behalf of the Wellcome Sanger Institute COVID-19 Surveillance Team |
| EPI_ISL_3338929, EPI_ISL_3338930 | Lighthouse Lab in Alderley Park | Wellcome Sanger Institute for the COVID-19 Genomics UK (COG-UK) Consortium | Jacquelyn Wynn, Mairead Hyland, The Lighthouse Lab in Alderley Park and Alex Alderton, Roberto Amato, Jeffrey Barrett, Sonia Goncalves, Ewan Harrison, David K. Jackson, Ian Johnston, Dominic Kwiatkowski, Cordelia Langford, John Sillitoe on behalf of the Wellcome Sanger Institute COVID-19 Surveillance Team |
| EPI_ISL_3338931 | Lighthouse Lab in Milton Keynes | Wellcome Sanger Institute for the COVID-19 Genomics UK (COG-UK) Consortium | The Lighthouse Lab in Milton Keynes and Alex Alderton, Roberto Amato, Jeffrey Barrett, Sonia Goncalves, Ewan Harrison, David K. Jackson, Ian Johnston, Dominic Kwiatkowski, Cordelia Langford, John Sillitoe on behalf of the Wellcome Sanger Institute COVID-19 Surveillance Team |
| EPI_ISL_3338932 | Lighthouse Lab in Alderley Park | Wellcome Sanger Institute for the COVID-19 Genomics UK (COG-UK) Consortium | Jacquelyn Wynn, Mairead Hyland, The Lighthouse Lab in Alderley Park and Alex Alderton, Roberto Amato, Jeffrey Barrett, Sonia Goncalves, Ewan Harrison, David K. Jackson, Ian Johnston, Dominic Kwiatkowski, Cordelia Langford, John Sillitoe on behalf of the Wellcome Sanger Institute COVID-19 Surveillance Team |
| EPI_ISL_3338933 | Lighthouse Lab in Milton Keynes | Wellcome Sanger Institute for the COVID-19 Genomics UK (COG-UK) Consortium | The Lighthouse Lab in Milton Keynes and Alex Alderton, Roberto Amato, Jeffrey Barrett, Sonia Goncalves, Ewan Harrison, David K. Jackson, Ian Johnston, Dominic Kwiatkowski, Cordelia Langford, John Sillitoe on behalf of the Wellcome Sanger Institute COVID-19 Surveillance Team |
| EPI_ISL_3338934 | Lighthouse Lab in Alderley Park | Wellcome Sanger Institute for the COVID-19 Genomics UK (COG-UK) Consortium | Jacquelyn Wynn, Mairead Hyland, The Lighthouse Lab in Alderley Park and Alex Alderton, Roberto Amato, Jeffrey Barrett, Sonia Goncalves, Ewan Harrison, David K. Jackson, Ian Johnston, Dominic Kwiatkowski, Cordelia Langford, John Sillitoe on behalf of the Wellcome Sanger Institute COVID-19 Surveillance Team |
| EPI_ISL_3338935 | Lighthouse Lab in Milton Keynes | Wellcome Sanger Institute for the COVID-19 Genomics UK (COG-UK) Consortium | The Lighthouse Lab in Milton Keynes and Alex Alderton, Roberto Amato, Jeffrey Barrett, Sonia Goncalves, Ewan Harrison, David K. Jackson, Ian Johnston, Dominic Kwiatkowski, Cordelia Langford, John Sillitoe on behalf of the Wellcome Sanger Institute COVID-19 Surveillance Team |
| EPI_ISL_3338936 | Lighthouse Lab in Alderley Park | Wellcome Sanger Institute for the COVID-19 Genomics UK (COG-UK) Consortium | Jacquelyn Wynn, Mairead Hyland, The Lighthouse Lab in Alderley Park and Alex Alderton, Roberto Amato, Jeffrey Barrett, Sonia Goncalves, Ewan Harrison, David K. Jackson, Ian Johnston, Dominic Kwiatkowski, Cordelia Langford, John Sillitoe on behalf of the Wellcome Sanger Institute COVID-19 Surveillance Team |
| EPI_ISL_3338937 | Lighthouse Lab in Milton Keynes | Wellcome Sanger Institute for the COVID-19 Genomics UK (COG-UK) Consortium | The Lighthouse Lab in Milton Keynes and Alex Alderton, Roberto Amato, Jeffrey Barrett, Sonia Goncalves, Ewan Harrison, David K. Jackson, Ian Johnston, Dominic Kwiatkowski, Cordelia Langford, John Sillitoe on behalf of the Wellcome Sanger Institute COVID-19 Surveillance Team |
| EPI_ISL_3338938, EPI_ISL_3338939 | Lighthouse Lab in Alderley Park | Wellcome Sanger Institute for the COVID-19 Genomics UK (COG-UK) Consortium | Jacquelyn Wynn, Mairead Hyland, The Lighthouse Lab in Alderley Park and Alex Alderton, Roberto Amato, Jeffrey Barrett, Sonia Goncalves, Ewan Harrison, David K. Jackson, Ian Johnston, Dominic Kwiatkowski, Cordelia Langford, John Sillitoe on behalf of the Wellcome Sanger Institute COVID-19 Surveillance Team |
| EPI_ISL_3338940 | Lighthouse Lab in Milton Keynes | Wellcome Sanger Institute for the COVID-19 Genomics UK (COG-UK) Consortium | The Lighthouse Lab in Milton Keynes and Alex Alderton, Roberto Amato, Jeffrey Barrett, Sonia Goncalves, Ewan Harrison, David K. Jackson, Ian Johnston, Dominic Kwiatkowski, Cordelia Langford, John Sillitoe on behalf of the Wellcome Sanger Institute COVID-19 Surveillance Team |
| EPI_ISL_3338941 | Lighthouse Lab in Alderley Park | Wellcome Sanger Institute for the COVID-19 Genomics UK (COG-UK) Consortium | Jacquelyn Wynn, Mairead Hyland, The Lighthouse Lab in Alderley Park and Alex Alderton, Roberto Amato, Jeffrey Barrett, Sonia Goncalves, Ewan Harrison, David K. Jackson, Ian Johnston, Dominic Kwiatkowski, Cordelia Langford, John Sillitoe on behalf of the Wellcome Sanger Institute COVID-19 Surveillance Team |
| EPI_ISL_3338942 | Lighthouse Lab in Glasgow | Wellcome Sanger Institute for the COVID-19 Genomics UK (COG-UK) Consortium | Harper VanSteenhouse, Yumi Kasai, David Gray, Carol Clugston, Anna Dominiczak and Alex Alderton, Roberto Amato, Jeffrey Barrett, Sonia Goncalves, Ewan Harrison, David K. Jackson, Ian Johnston, Dominic Kwiatkowski, Cordelia Langford, John Sillitoe on behalf of the Wellcome Sanger Institute COVID-19 Surveillance Team |
| EPI_ISL_3338943, EPI_ISL_3338944, EPI_ISL_3338945, EPI_ISL_3338946 | Lighthouse Lab in Alderley Park | Wellcome Sanger Institute for the COVID-19 Genomics UK (COG-UK) Consortium | Jacquelyn Wynn, Mairead Hyland, The Lighthouse Lab in Alderley Park and Alex Alderton, Roberto Amato, Jeffrey Barrett, Sonia Goncalves, Ewan Harrison, David K. Jackson, Ian Johnston, Dominic Kwiatkowski, Cordelia Langford, John Sillitoe on behalf of the Wellcome Sanger Institute COVID-19 Surveillance Team |
| EPI_ISL_3338947 | Lighthouse Lab in Milton Keynes | Wellcome Sanger Institute for the COVID-19 Genomics UK (COG-UK) Consortium | The Lighthouse Lab in Milton Keynes and Alex Alderton, Roberto Amato, Jeffrey Barrett, Sonia Goncalves, Ewan Harrison, David K. Jackson, Ian Johnston, Dominic Kwiatkowski, Cordelia Langford, John Sillitoe on behalf of the Wellcome Sanger Institute COVID-19 Surveillance Team |
| EPI_ISL_3338948 | Lighthouse Lab in Alderley Park | Wellcome Sanger Institute for the COVID-19 Genomics UK (COG-UK) Consortium | Jacquelyn Wynn, Mairead Hyland, The Lighthouse Lab in Alderley Park and Alex Alderton, Roberto Amato, Jeffrey Barrett, Sonia Goncalves, Ewan Harrison, David K. Jackson, Ian Johnston, Dominic Kwiatkowski, Cordelia Langford, John Sillitoe on behalf of the Wellcome Sanger Institute COVID-19 Surveillance Team |
| EPI_ISL_3338949, EPI_ISL_3338950, EPI_ISL_3338951 | Lighthouse Lab in Milton Keynes | Wellcome Sanger Institute for the COVID-19 Genomics UK (COG-UK) Consortium | The Lighthouse Lab in Milton Keynes and Alex Alderton, Roberto Amato, Jeffrey Barrett, Sonia Goncalves, Ewan Harrison, David K. Jackson, Ian Johnston, Dominic Kwiatkowski, Cordelia Langford, John Sillitoe on behalf of the Wellcome Sanger Institute COVID-19 Surveillance Team |
| EPI_ISL_3338952, EPI_ISL_3338953, EPI_ISL_3338954 | Lighthouse Lab in Alderley Park | Wellcome Sanger Institute for the COVID-19 Genomics UK (COG-UK) Consortium | Jacquelyn Wynn, Mairead Hyland, The Lighthouse Lab in Alderley Park and Alex Alderton, Roberto Amato, Jeffrey Barrett, Sonia Goncalves, Ewan Harrison, David K. Jackson, Ian Johnston, Dominic Kwiatkowski, Cordelia Langford, John Sillitoe on behalf of the Wellcome Sanger Institute COVID-19 Surveillance Team |
| EPI_ISL_3338955 | Lighthouse Lab in Glasgow | Wellcome Sanger Institute for the COVID-19 Genomics UK (COG-UK) Consortium | Harper VanSteenhouse, Yumi Kasai, David Gray, Carol Clugston, Anna Dominiczak and Alex Alderton, Roberto Amato, Jeffrey Barrett, Sonia Goncalves, Ewan Harrison, David K. Jackson, Ian Johnston, Dominic Kwiatkowski, Cordelia Langford, John Sillitoe on behalf of the Wellcome Sanger Institute COVID-19 Surveillance Team |
| EPI_ISL_3338956, EPI_ISL_3338957, EPI_ISL_3338958, EPI_ISL_3338959 | Lighthouse Lab in Alderley Park | Wellcome Sanger Institute for the COVID-19 Genomics UK (COG-UK) Consortium | Jacquelyn Wynn, Mairead Hyland, The Lighthouse Lab in Alderley Park and Alex Alderton, Roberto Amato, Jeffrey Barrett, Sonia Goncalves, Ewan Harrison, David K. Jackson, Ian Johnston, Dominic Kwiatkowski, Cordelia Langford, John Sillitoe on behalf of the Wellcome Sanger Institute COVID-19 Surveillance Team |
| EPI_ISL_3338960 | Lighthouse Lab in Milton Keynes | Wellcome Sanger Institute for the COVID-19 Genomics UK (COG-UK) Consortium | The Lighthouse Lab in Milton Keynes and Alex Alderton, Roberto Amato, Jeffrey Barrett, Sonia Goncalves, Ewan Harrison, David K. Jackson, Ian Johnston, Dominic Kwiatkowski, Cordelia Langford, John Sillitoe on behalf of the Wellcome Sanger Institute COVID-19 Surveillance Team |
| EPI_ISL_3338961, EPI_ISL_3338962, EPI_ISL_3338963 | Lighthouse Lab in Alderley Park | Wellcome Sanger Institute for the COVID-19 Genomics UK (COG-UK) Consortium | Jacquelyn Wynn, Mairead Hyland, The Lighthouse Lab in Alderley Park and Alex Alderton, Roberto Amato, Jeffrey Barrett, Sonia Goncalves, Ewan Harrison, David K. Jackson, Ian Johnston, Dominic Kwiatkowski, Cordelia Langford, John Sillitoe on behalf of the Wellcome Sanger Institute COVID-19 Surveillance Team |
| EPI_ISL_3338964 | Lighthouse Lab in Glasgow | Wellcome Sanger Institute for the COVID-19 Genomics UK (COG-UK) Consortium | Harper VanSteenhouse, Yumi Kasai, David Gray, Carol Clugston, Anna Dominiczak and Alex Alderton, Roberto Amato, Jeffrey Barrett, Sonia Goncalves, Ewan Harrison, David K. Jackson, Ian Johnston, Dominic Kwiatkowski, Cordelia Langford, John Sillitoe on behalf of the Wellcome Sanger Institute COVID-19 Surveillance Team |
| EPI_ISL_3338965 | Lighthouse Lab in Alderley Park | Wellcome Sanger Institute for the COVID-19 Genomics UK (COG-UK) Consortium | Jacquelyn Wynn, Mairead Hyland, The Lighthouse Lab in Alderley Park and Alex Alderton, Roberto Amato, Jeffrey Barrett, Sonia Goncalves, Ewan Harrison, David K. Jackson, Ian Johnston, Dominic Kwiatkowski, Cordelia Langford, John Sillitoe on behalf of the Wellcome Sanger Institute COVID-19 Surveillance Team |
| EPI_ISL_3338966 | Lighthouse Lab in Glasgow | Wellcome Sanger Institute for the COVID-19 Genomics UK (COG-UK) Consortium | Harper VanSteenhouse, Yumi Kasai, David Gray, Carol Clugston, Anna Dominiczak and Alex Alderton, Roberto Amato, Jeffrey Barrett, Sonia Goncalves, Ewan Harrison, David K. Jackson, Ian Johnston, Dominic Kwiatkowski, Cordelia Langford, John Sillitoe on behalf of the Wellcome Sanger Institute COVID-19 Surveillance Team |
| EPI_ISL_3338967 | Lighthouse Lab in Alderley Park | Wellcome Sanger Institute for the COVID-19 Genomics UK (COG-UK) Consortium | Jacquelyn Wynn, Mairead Hyland, The Lighthouse Lab in Alderley Park and Alex Alderton, Roberto Amato, Jeffrey Barrett, Sonia Goncalves, Ewan Harrison, David K. Jackson, Ian Johnston, Dominic Kwiatkowski, Cordelia Langford, John Sillitoe on behalf of the Wellcome Sanger Institute COVID-19 Surveillance Team |
| EPI_ISL_3338968 | Lighthouse Lab in Milton Keynes | Wellcome Sanger Institute for the COVID-19 Genomics UK (COG-UK) Consortium | The Lighthouse Lab in Milton Keynes and Alex Alderton, Roberto Amato, Jeffrey Barrett, Sonia Goncalves, Ewan Harrison, David K. Jackson, Ian Johnston, Dominic Kwiatkowski, Cordelia Langford, John Sillitoe on behalf of the Wellcome Sanger Institute COVID-19 Surveillance Team |
| EPI_ISL_3338969 | Lighthouse Lab in Glasgow | Wellcome Sanger Institute for the COVID-19 Genomics UK (COG-UK) Consortium | Harper VanSteenhouse, Yumi Kasai, David Gray, Carol Clugston, Anna Dominiczak and Alex Alderton, Roberto Amato, Jeffrey Barrett, Sonia Goncalves, Ewan Harrison, David K. Jackson, Ian Johnston, Dominic Kwiatkowski, Cordelia Langford, John Sillitoe on behalf of the Wellcome Sanger Institute COVID-19 Surveillance Team |
| EPI_ISL_3338970, EPI_ISL_3338971, EPI_ISL_3338972 | Lighthouse Lab in Milton Keynes | Wellcome Sanger Institute for the COVID-19 Genomics UK (COG-UK) Consortium | The Lighthouse Lab in Milton Keynes and Alex Alderton, Roberto Amato, Jeffrey Barrett, Sonia Goncalves, Ewan Harrison, David K. Jackson, Ian Johnston, Dominic Kwiatkowski, Cordelia Langford, John Sillitoe on behalf of the Wellcome Sanger Institute COVID-19 Surveillance Team |
| EPI_ISL_3338973 | Lighthouse Lab in Glasgow | Wellcome Sanger Institute for the COVID-19 Genomics UK (COG-UK) Consortium | Harper VanSteenhouse, Yumi Kasai, David Gray, Carol Clugston, Anna Dominiczak and Alex Alderton, Roberto Amato, Jeffrey Barrett, Sonia Goncalves, Ewan Harrison, David K. Jackson, Ian Johnston, Dominic Kwiatkowski, Cordelia Langford, John Sillitoe on behalf of the Wellcome Sanger Institute COVID-19 Surveillance Team |
| EPI_ISL_3338974 | Lighthouse Lab in Milton Keynes | Wellcome Sanger Institute for the COVID-19 Genomics UK (COG-UK) Consortium | The Lighthouse Lab in Milton Keynes and Alex Alderton, Roberto Amato, Jeffrey Barrett, Sonia Goncalves, Ewan Harrison, David K. Jackson, Ian Johnston, Dominic Kwiatkowski, Cordelia Langford, John Sillitoe on behalf of the Wellcome Sanger Institute COVID-19 Surveillance Team |
| EPI_ISL_3338975 | Lighthouse Lab in Alderley Park | Wellcome Sanger Institute for the COVID-19 Genomics UK (COG-UK) Consortium | Jacquelyn Wynn, Mairead Hyland, The Lighthouse Lab in Alderley Park and Alex Alderton, Roberto Amato, Jeffrey Barrett, Sonia Goncalves, Ewan Harrison, David K. Jackson, Ian Johnston, Dominic Kwiatkowski, Cordelia Langford, John Sillitoe on behalf of the Wellcome Sanger Institute COVID-19 Surveillance Team |
| EPI_ISL_3338976 | Lighthouse Lab in Milton Keynes | Wellcome Sanger Institute for the COVID-19 Genomics UK (COG-UK) Consortium | The Lighthouse Lab in Milton Keynes and Alex Alderton, Roberto Amato, Jeffrey Barrett, Sonia Goncalves, Ewan Harrison, David K. Jackson, Ian Johnston, Dominic Kwiatkowski, Cordelia Langford, John Sillitoe on behalf of the Wellcome Sanger Institute COVID-19 Surveillance Team |
| EPI_ISL_3338977 | Lighthouse Lab in Glasgow | Wellcome Sanger Institute for the COVID-19 Genomics UK (COG-UK) Consortium | Harper VanSteenhouse, Yumi Kasai, David Gray, Carol Clugston, Anna Dominiczak and Alex Alderton, Roberto Amato, Jeffrey Barrett, Sonia Goncalves, Ewan Harrison, David K. Jackson, Ian Johnston, Dominic Kwiatkowski, Cordelia Langford, John Sillitoe on behalf of the Wellcome Sanger Institute COVID-19 Surveillance Team |
| EPI_ISL_3338978 | Lighthouse Lab in Milton Keynes | Wellcome Sanger Institute for the COVID-19 Genomics UK (COG-UK) Consortium | The Lighthouse Lab in Milton Keynes and Alex Alderton, Roberto Amato, Jeffrey Barrett, Sonia Goncalves, Ewan Harrison, David K. Jackson, Ian Johnston, Dominic Kwiatkowski, Cordelia Langford, John Sillitoe on behalf of the Wellcome Sanger Institute COVID-19 Surveillance Team |
| EPI_ISL_3338979 | Lighthouse Lab in Alderley Park | Wellcome Sanger Institute for the COVID-19 Genomics UK (COG-UK) Consortium | Jacquelyn Wynn, Mairead Hyland, The Lighthouse Lab in Alderley Park and Alex Alderton, Roberto Amato, Jeffrey Barrett, Sonia Goncalves, Ewan Harrison, David K. Jackson, Ian Johnston, Dominic Kwiatkowski, Cordelia Langford, John Sillitoe on behalf of the Wellcome Sanger Institute COVID-19 Surveillance Team |
| EPI_ISL_3338980 | Lighthouse Lab in Milton Keynes | Wellcome Sanger Institute for the COVID-19 Genomics UK (COG-UK) Consortium | The Lighthouse Lab in Milton Keynes and Alex Alderton, Roberto Amato, Jeffrey Barrett, Sonia Goncalves, Ewan Harrison, David K. Jackson, Ian Johnston, Dominic Kwiatkowski, Cordelia Langford, John Sillitoe on behalf of the Wellcome Sanger Institute COVID-19 Surveillance Team |

| | | | |
|---|---|---|---|
| EPI_ISL_3338981 | Lighthouse Lab in Alderley Park | Wellcome Sanger Institute for the COVID-19 Genomics UK (COG-UK) Consortium | Jacquelyn Wynn, Mairead Hyland, The Lighthouse Lab in Alderley Park and Alex Alderton, Roberto Amato, Jeffrey Barrett, Sonia Goncalves, Ewan Harrison, David K. Jackson, Ian Johnston, Dominic Kwiatkowski, Cordelia Langford, John Sillitoe on behalf of the Wellcome Sanger Institute COVID-19 Surveillance Team |
| EPI_ISL_3338982 | Lighthouse Lab in Milton Keynes | Wellcome Sanger Institute for the COVID-19 Genomics UK (COG-UK) Consortium | The Lighthouse Lab in Milton Keynes and Alex Alderton, Roberto Amato, Jeffrey Barrett, Sonia Goncalves, Ewan Harrison, David K. Jackson, Ian Johnston, Dominic Kwiatkowski, Cordelia Langford, John Sillitoe on behalf of the Wellcome Sanger Institute COVID-19 Surveillance Team |
| EPI_ISL_3338983 | Lighthouse Lab in Glasgow | Wellcome Sanger Institute for the COVID-19 Genomics UK (COG-UK) Consortium | Harper VanSteenhouse, Yumi Kasai, David Gray, Carol Clugston, Anna Dominiczak and Alex Alderton, Roberto Amato, Jeffrey Barrett, Sonia Goncalves, Ewan Harrison, David K. Jackson, Ian Johnston, Dominic Kwiatkowski, Cordelia Langford, John Sillitoe on behalf of the Wellcome Sanger Institute COVID-19 Surveillance Team |
| EPI_ISL_3338984 | Lighthouse Lab in Milton Keynes | Wellcome Sanger Institute for the COVID-19 Genomics UK (COG-UK) Consortium | The Lighthouse Lab in Milton Keynes and Alex Alderton, Roberto Amato, Jeffrey Barrett, Sonia Goncalves, Ewan Harrison, David K. Jackson, Ian Johnston, Dominic Kwiatkowski, Cordelia Langford, John Sillitoe on behalf of the Wellcome Sanger Institute COVID-19 Surveillance Team |
| EPI_ISL_3338985, EPI_ISL_3338986, EPI_ISL_3338987, EPI_ISL_3338988, EPI_ISL_3338989, EPI_ISL_3338990 | Lighthouse Lab in Alderley Park | Wellcome Sanger Institute for the COVID-19 Genomics UK (COG-UK) Consortium | Jacquelyn Wynn, Mairead Hyland, The Lighthouse Lab in Alderley Park and Alex Alderton, Roberto Amato, Jeffrey Barrett, Sonia Goncalves, Ewan Harrison, David K. Jackson, Ian Johnston, Dominic Kwiatkowski, Cordelia Langford, John Sillitoe on behalf of the Wellcome Sanger Institute COVID-19 Surveillance Team |
| EPI_ISL_3338991 | Lighthouse Lab in Milton Keynes | Wellcome Sanger Institute for the COVID-19 Genomics UK (COG-UK) Consortium | The Lighthouse Lab in Milton Keynes and Alex Alderton, Roberto Amato, Jeffrey Barrett, Sonia Goncalves, Ewan Harrison, David K. Jackson, Ian Johnston, Dominic Kwiatkowski, Cordelia Langford, John Sillitoe on behalf of the Wellcome Sanger Institute COVID-19 Surveillance Team |
| EPI_ISL_3338992 | Lighthouse Lab in Alderley Park | Wellcome Sanger Institute for the COVID-19 Genomics UK (COG-UK) Consortium | Jacquelyn Wynn, Mairead Hyland, The Lighthouse Lab in Alderley Park and Alex Alderton, Roberto Amato, Jeffrey Barrett, Sonia Goncalves, Ewan Harrison, David K. Jackson, Ian Johnston, Dominic Kwiatkowski, Cordelia Langford, John Sillitoe on behalf of the Wellcome Sanger Institute COVID-19 Surveillance Team |
| EPI_ISL_3338993 | Lighthouse Lab in Milton Keynes | Wellcome Sanger Institute for the COVID-19 Genomics UK (COG-UK) Consortium | The Lighthouse Lab in Milton Keynes and Alex Alderton, Roberto Amato, Jeffrey Barrett, Sonia Goncalves, Ewan Harrison, David K. Jackson, Ian Johnston, Dominic Kwiatkowski, Cordelia Langford, John Sillitoe on behalf of the Wellcome Sanger Institute COVID-19 Surveillance Team |
| EPI_ISL_3338994 | Lighthouse Lab in Glasgow | Wellcome Sanger Institute for the COVID-19 Genomics UK (COG-UK) Consortium | Harper VanSteenhouse, Yumi Kasai, David Gray, Carol Clugston, Anna Dominiczak and Alex Alderton, Roberto Amato, Jeffrey Barrett, Sonia Goncalves, Ewan Harrison, David K. Jackson, Ian Johnston, Dominic Kwiatkowski, Cordelia Langford, John Sillitoe on behalf of the Wellcome Sanger Institute COVID-19 Surveillance Team |
| EPI_ISL_3338995 | Lighthouse Lab in Alderley Park | Wellcome Sanger Institute for the COVID-19 Genomics UK (COG-UK) Consortium | Jacquelyn Wynn, Mairead Hyland, The Lighthouse Lab in Alderley Park and Alex Alderton, Roberto Amato, Jeffrey Barrett, Sonia Goncalves, Ewan Harrison, David K. Jackson, Ian Johnston, Dominic Kwiatkowski, Cordelia Langford, John Sillitoe on behalf of the Wellcome Sanger Institute COVID-19 Surveillance Team |
| EPI_ISL_3338996 | Lighthouse Lab in Milton Keynes | Wellcome Sanger Institute for the COVID-19 Genomics UK (COG-UK) Consortium | The Lighthouse Lab in Milton Keynes and Alex Alderton, Roberto Amato, Jeffrey Barrett, Sonia Goncalves, Ewan Harrison, David K. Jackson, Ian Johnston, Dominic Kwiatkowski, Cordelia Langford, John Sillitoe on behalf of the Wellcome Sanger Institute COVID-19 Surveillance Team |
| EPI_ISL_3338997 | Lighthouse Lab in Alderley Park | Wellcome Sanger Institute for the COVID-19 Genomics UK (COG-UK) Consortium | Jacquelyn Wynn, Mairead Hyland, The Lighthouse Lab in Alderley Park and Alex Alderton, Roberto Amato, Jeffrey Barrett, Sonia Goncalves, Ewan Harrison, David K. Jackson, Ian Johnston, Dominic Kwiatkowski, Cordelia Langford, John Sillitoe on behalf of the Wellcome Sanger Institute COVID-19 Surveillance Team |
| EPI_ISL_3338998 | Lighthouse Lab in Milton Keynes | Wellcome Sanger Institute for the COVID-19 Genomics UK (COG-UK) Consortium | The Lighthouse Lab in Milton Keynes and Alex Alderton, Roberto Amato, Jeffrey Barrett, Sonia Goncalves, Ewan Harrison, David K. Jackson, Ian Johnston, Dominic Kwiatkowski, Cordelia Langford, John Sillitoe on behalf of the Wellcome Sanger Institute COVID-19 Surveillance Team |
| EPI_ISL_3338999, EPI_ISL_3339000, EPI_ISL_3339001, EPI_ISL_3339002, EPI_ISL_3339003, EPI_ISL_3339004, EPI_ISL_3339005 | Lighthouse Lab in Alderley Park | Wellcome Sanger Institute for the COVID-19 Genomics UK (COG-UK) Consortium | Jacquelyn Wynn, Mairead Hyland, The Lighthouse Lab in Alderley Park and Alex Alderton, Roberto Amato, Jeffrey Barrett, Sonia Goncalves, Ewan Harrison, David K. Jackson, Ian Johnston, Dominic Kwiatkowski, Cordelia Langford, John Sillitoe on behalf of the Wellcome Sanger Institute COVID-19 Surveillance Team |
| EPI_ISL_3339006 | SYNLAB | GIGA Medical Genomics | Keith Durkin, Maria Artesi, Bouchra Boujemla, Nathalie Renotte, Nadine Cambisano, Sébastien Bontems, Cécile Meex, Claire Gourzonès, Olivier Ek, Laurent Gillet, Marie-Pierre Hayette, Vincent Bours |
| EPI_ISL_3339007, EPI_ISL_3339008 | Lighthouse Lab in Milton Keynes | Wellcome Sanger Institute for the COVID-19 Genomics UK (COG-UK) Consortium | The Lighthouse Lab in Milton Keynes and Alex Alderton, Roberto Amato, Jeffrey Barrett, Sonia Goncalves, Ewan Harrison, David K. Jackson, Ian Johnston, Dominic Kwiatkowski, Cordelia Langford, John Sillitoe on behalf of the Wellcome Sanger Institute COVID-19 Surveillance Team |
| EPI_ISL_3339009 | SYNLAB | GIGA Medical Genomics | Keith Durkin, Maria Artesi, Bouchra Boujemla, Nathalie Renotte, Nadine Cambisano, Sébastien Bontems, Cécile Meex, Claire Gourzonès, Olivier Ek, Laurent Gillet, Marie-Pierre Hayette, Vincent Bours |
| EPI_ISL_3339010 | Lighthouse Lab in Milton Keynes | Wellcome Sanger Institute for the COVID-19 Genomics UK (COG-UK) Consortium | The Lighthouse Lab in Milton Keynes and Alex Alderton, Roberto Amato, Jeffrey Barrett, Sonia Goncalves, Ewan Harrison, David K. Jackson, Ian Johnston, Dominic Kwiatkowski, Cordelia Langford, John Sillitoe on behalf of the Wellcome Sanger Institute COVID-19 Surveillance Team |
| EPI_ISL_3339011 | Lighthouse Lab in Glasgow | Wellcome Sanger Institute for the COVID-19 Genomics UK (COG-UK) Consortium | Harper VanSteenhouse, Yumi Kasai, David Gray, Carol Clugston, Anna Dominiczak and Alex Alderton, Roberto Amato, Jeffrey Barrett, Sonia Goncalves, Ewan Harrison, David K. Jackson, Ian Johnston, Dominic Kwiatkowski, Cordelia Langford, John Sillitoe on behalf of the Wellcome Sanger Institute COVID-19 Surveillance Team |
| EPI_ISL_3339012 | Lighthouse Lab in Milton Keynes | Wellcome Sanger Institute for the COVID-19 Genomics UK (COG-UK) Consortium | The Lighthouse Lab in Milton Keynes and Alex Alderton, Roberto Amato, Jeffrey Barrett, Sonia Goncalves, Ewan Harrison, David K. Jackson, Ian Johnston, Dominic Kwiatkowski, Cordelia Langford, John Sillitoe on behalf of the Wellcome Sanger Institute COVID-19 Surveillance Team |
| EPI_ISL_3339013 | SYNLAB | GIGA Medical Genomics | Keith Durkin, Maria Artesi, Bouchra Boujemla, Nathalie Renotte, Nadine Cambisano, Sébastien Bontems, Cécile Meex, Claire Gourzonès, Olivier Ek, Laurent Gillet, Marie-Pierre Hayette, Vincent Bours |
| EPI_ISL_3339014 | Lighthouse Lab in Alderley Park | Wellcome Sanger Institute for the COVID-19 Genomics UK (COG-UK) Consortium | Jacquelyn Wynn, Mairead Hyland, The Lighthouse Lab in Alderley Park and Alex Alderton, Roberto Amato, Jeffrey Barrett, Sonia Goncalves, Ewan Harrison, David K. Jackson, Ian Johnston, Dominic Kwiatkowski, Cordelia Langford, John Sillitoe on behalf of the Wellcome Sanger Institute COVID-19 Surveillance Team |
| EPI_ISL_3339015 | Lighthouse Lab in Milton Keynes | Wellcome Sanger Institute for the COVID-19 Genomics UK (COG-UK) Consortium | The Lighthouse Lab in Milton Keynes and Alex Alderton, Roberto Amato, Jeffrey Barrett, Sonia Goncalves, Ewan Harrison, David K. Jackson, Ian Johnston, Dominic Kwiatkowski, Cordelia Langford, John Sillitoe on behalf of the Wellcome Sanger Institute COVID-19 Surveillance Team |
| EPI_ISL_3339016 | SYNLAB | GIGA Medical Genomics | Keith Durkin, Maria Artesi, Bouchra Boujemla, Nathalie Renotte, Nadine Cambisano, Sébastien Bontems, Cécile Meex, Claire Gourzonès, Olivier Ek, Laurent Gillet, Marie-Pierre Hayette, Vincent Bours |
| EPI_ISL_3339017 | Lighthouse Lab in Alderley Park | Wellcome Sanger Institute for the COVID-19 Genomics UK (COG-UK) Consortium | Jacquelyn Wynn, Mairead Hyland, The Lighthouse Lab in Alderley Park and Alex Alderton, Roberto Amato, Jeffrey Barrett, Sonia Goncalves, Ewan Harrison, David K. Jackson, Ian Johnston, Dominic Kwiatkowski, Cordelia Langford, John Sillitoe on behalf of the Wellcome Sanger Institute COVID-19 Surveillance Team |
| EPI_ISL_3339018 | Lighthouse Lab in Milton Keynes | Wellcome Sanger Institute for the COVID-19 Genomics UK (COG-UK) Consortium | The Lighthouse Lab in Milton Keynes and Alex Alderton, Roberto Amato, Jeffrey Barrett, Sonia Goncalves, Ewan Harrison, David K. Jackson, Ian Johnston, Dominic Kwiatkowski, Cordelia Langford, John Sillitoe on behalf of the Wellcome Sanger Institute COVID-19 Surveillance Team |
| EPI_ISL_3339020 | Lighthouse Lab in Glasgow | Wellcome Sanger Institute for the COVID-19 Genomics UK (COG-UK) Consortium | Harper VanSteenhouse, Yumi Kasai, David Gray, Carol Clugston, Anna Dominiczak and Alex Alderton, Roberto Amato, Jeffrey Barrett, Sonia Goncalves, Ewan Harrison, David K. Jackson, Ian Johnston, Dominic Kwiatkowski, Cordelia Langford, John Sillitoe on behalf of the Wellcome Sanger Institute COVID-19 Surveillance Team |
| EPI_ISL_3339021 | Lighthouse Lab in Alderley Park | Wellcome Sanger Institute for the COVID-19 Genomics UK (COG-UK) Consortium | Jacquelyn Wynn, Mairead Hyland, The Lighthouse Lab in Alderley Park and Alex Alderton, Roberto Amato, Jeffrey Barrett, Sonia Goncalves, Ewan Harrison, David K. Jackson, Ian Johnston, Dominic Kwiatkowski, Cordelia Langford, John Sillitoe on behalf of the Wellcome Sanger Institute COVID-19 Surveillance Team |
| EPI_ISL_3339022 | SYNLAB | GIGA Medical Genomics | Keith Durkin, Maria Artesi, Bouchra Boujemla, Nathalie Renotte, Nadine Cambisano, Sébastien Bontems, Cécile Meex, Claire Gourzonès, Olivier Ek, Laurent Gillet, Marie-Pierre Hayette, Vincent Bours |
| EPI_ISL_3339023, EPI_ISL_3339024 | Lighthouse Lab in Milton Keynes | Wellcome Sanger Institute for the COVID-19 Genomics UK (COG-UK) Consortium | The Lighthouse Lab in Milton Keynes and Alex Alderton, Roberto Amato, Jeffrey Barrett, Sonia Goncalves, Ewan Harrison, David K. Jackson, Ian Johnston, Dominic Kwiatkowski, Cordelia Langford, John Sillitoe on behalf of the Wellcome Sanger Institute COVID-19 Surveillance Team |
| EPI_ISL_3339025 | Lighthouse Lab in Alderley Park | Wellcome Sanger Institute for the COVID-19 Genomics UK (COG-UK) Consortium | Jacquelyn Wynn, Mairead Hyland, The Lighthouse Lab in Alderley Park and Alex Alderton, Roberto Amato, Jeffrey Barrett, Sonia Goncalves, Ewan Harrison, David K. Jackson, Ian Johnston, Dominic Kwiatkowski, Cordelia Langford, John Sillitoe on behalf of the Wellcome Sanger Institute COVID-19 Surveillance Team |
| EPI_ISL_3339026 | SYNLAB | GIGA Medical Genomics | Keith Durkin, Maria Artesi, Bouchra Boujemla, Nathalie Renotte, Nadine Cambisano, Sébastien Bontems, Cécile Meex, Claire Gourzonès, Olivier Ek, Laurent Gillet, Marie-Pierre Hayette, Vincent Bours |
| EPI_ISL_3339027, EPI_ISL_3339028, EPI_ISL_3339029 | Lighthouse Lab in Alderley Park | Wellcome Sanger Institute for the COVID-19 Genomics UK (COG-UK) Consortium | Jacquelyn Wynn, Mairead Hyland, The Lighthouse Lab in Alderley Park and Alex Alderton, Roberto Amato, Jeffrey Barrett, Sonia Goncalves, Ewan Harrison, David K. Jackson, Ian Johnston, Dominic Kwiatkowski, Cordelia Langford, John Sillitoe on behalf of the Wellcome Sanger Institute COVID-19 Surveillance Team |
| EPI_ISL_3339030 | SYNLAB | GIGA Medical Genomics | Keith Durkin, Maria Artesi, Bouchra Boujemla, Nathalie Renotte, Nadine Cambisano, Sébastien Bontems, Cécile Meex, Claire Gourzonès, Olivier Ek, Laurent Gillet, Marie-Pierre Hayette, Vincent Bours |
| EPI_ISL_3339031 | Lighthouse Lab in Milton Keynes | Wellcome Sanger Institute for the COVID-19 Genomics UK (COG-UK) Consortium | The Lighthouse Lab in Milton Keynes and Alex Alderton, Roberto Amato, Jeffrey Barrett, Sonia Goncalves, Ewan Harrison, David K. Jackson, Ian Johnston, Dominic Kwiatkowski, Cordelia Langford, John Sillitoe on behalf of the Wellcome Sanger Institute COVID-19 Surveillance Team |
| EPI_ISL_3339032 | Lighthouse Lab in Alderley Park | Wellcome Sanger Institute for the COVID-19 Genomics UK (COG-UK) Consortium | Jacquelyn Wynn, Mairead Hyland, The Lighthouse Lab in Alderley Park and Alex Alderton, Roberto Amato, Jeffrey Barrett, Sonia Goncalves, Ewan Harrison, David K. Jackson, Ian Johnston, Dominic Kwiatkowski, Cordelia Langford, John Sillitoe on behalf of the Wellcome Sanger Institute COVID-19 Surveillance Team |
| EPI_ISL_3339033 | SYNLAB | GIGA Medical Genomics | Keith Durkin, Maria Artesi, Bouchra Boujemla, Nathalie Renotte, Nadine Cambisano, Sébastien Bontems, Cécile Meex, Claire Gourzonès, Olivier Ek, Laurent Gillet, Marie-Pierre Hayette, Vincent Bours |
| EPI_ISL_3339034 | Lighthouse Lab in Alderley Park | Wellcome Sanger Institute for the COVID-19 Genomics UK (COG-UK) Consortium | Jacquelyn Wynn, Mairead Hyland, The Lighthouse Lab in Alderley Park and Alex Alderton, Roberto Amato, Jeffrey Barrett, Sonia Goncalves, Ewan Harrison, David K. Jackson, Ian Johnston, Dominic Kwiatkowski, Cordelia Langford, John Sillitoe on behalf of the Wellcome Sanger Institute COVID-19 Surveillance Team |
| EPI_ISL_3339035 | Lighthouse Lab in Milton Keynes | Wellcome Sanger Institute for the COVID-19 Genomics UK (COG-UK) Consortium | The Lighthouse Lab in Milton Keynes and Alex Alderton, Roberto Amato, Jeffrey Barrett, Sonia Goncalves, Ewan Harrison, David K. Jackson, Ian Johnston, Dominic Kwiatkowski, Cordelia Langford, John Sillitoe on behalf of the Wellcome Sanger Institute COVID-19 Surveillance Team |
| EPI_ISL_3339036 | SYNLAB | GIGA Medical Genomics | Keith Durkin, Maria Artesi, Bouchra Boujemla, Nathalie Renotte, Nadine Cambisano, Sébastien Bontems, Cécile Meex, Claire Gourzonès, Olivier Ek, Laurent Gillet, Marie-Pierre Hayette, Vincent Bours |
| EPI_ISL_3339037 | Lighthouse Lab in Milton Keynes | Wellcome Sanger Institute for the COVID-19 Genomics UK (COG-UK) Consortium | The Lighthouse Lab in Milton Keynes and Alex Alderton, Roberto Amato, Jeffrey Barrett, Sonia Goncalves, Ewan Harrison, David K. Jackson, Ian Johnston, Dominic Kwiatkowski, Cordelia Langford, John Sillitoe on behalf of the Wellcome Sanger Institute COVID-19 Surveillance Team |
| EPI_ISL_3339038 | Randox Laboratories | Wellcome Sanger Institute for the COVID-19 Genomics UK (COG-UK) Consortium | Randox Laboratories and Alex Alderton, Roberto Amato, Jeffrey Barrett, Sonia Goncalves, Ewan Harrison, David K. Jackson, Ian Johnston, Dominic Kwiatkowski, Cordelia Langford, John Sillitoe on behalf of the Wellcome Sanger Institute COVID-19 Surveillance Team |
| EPI_ISL_3339039 | SYNLAB | GIGA Medical Genomics | Keith Durkin, Maria Artesi, Bouchra Boujemla, Nathalie Renotte, Nadine Cambisano, Sébastien Bontems, Cécile Meex, Claire Gourzonès, Olivier Ek, Laurent Gillet, Marie-Pierre Hayette, Vincent Bours |
| EPI_ISL_3339040 | Randox Laboratories | Wellcome Sanger Institute for the COVID-19 Genomics UK (COG-UK) Consortium | Randox Laboratories and Alex Alderton, Roberto Amato, Jeffrey Barrett, Sonia Goncalves, Ewan Harrison, David K. Jackson, Ian Johnston, Dominic Kwiatkowski, Cordelia Langford, John Sillitoe on behalf of the Wellcome Sanger Institute COVID-19 Surveillance Team |
| EPI_ISL_3339041, EPI_ISL_3339042 | Lighthouse Lab in Glasgow | Wellcome Sanger Institute for the COVID-19 Genomics UK (COG-UK) Consortium | Harper VanSteenhouse, Yumi Kasai, David Gray, Carol Clugston, Anna Dominiczak and Alex Alderton, Roberto Amato, Jeffrey Barrett, Sonia Goncalves, Ewan Harrison, David K. Jackson, Ian Johnston, Dominic Kwiatkowski, Cordelia Langford, John Sillitoe on behalf of the Wellcome Sanger Institute COVID-19 Surveillance Team |

| | | | |
|---|---|---|---|
| EPI_ISL_3339043 | SYNLAB | GIGA Medical Genomics | Keith Durkin, Maria Artesi, Bouchra Boujemla, Nathalie Renotte, Nadine Cambisano, Sébastien Bontems, Cécile Meex, Claire Gourzonès, Olivier Ek, Laurent Gillet, Marie-Pierre Hayette, Vincent Bours |
| EPI_ISL_3339044 | Randox Laboratories | Wellcome Sanger Institute for the COVID-19 Genomics UK (COG-UK) Consortium | Randox Laboratories and Alex Alderton, Roberto Amato, Jeffrey Barrett, Sonia Goncalves, Ewan Harrison, David K. Jackson, Ian Johnston, Dominic Kwiatkowski, Cordelia Langford, John Sillitoe on behalf of the Wellcome Sanger Institute COVID-19 Surveillance Team |
| EPI_ISL_3339045, EPI_ISL_3339046 | Lighthouse Lab in Alderley Park | Wellcome Sanger Institute for the COVID-19 Genomics UK (COG-UK) Consortium | Jacquelyn Wynn, Mairead Hyland, The Lighthouse Lab in Alderley Park and Alex Alderton, Roberto Amato, Jeffrey Barrett, Sonia Goncalves, Ewan Harrison, David K. Jackson, Ian Johnston, Dominic Kwiatkowski, Cordelia Langford, John Sillitoe on behalf of the Wellcome Sanger Institute COVID-19 Surveillance Team |
| EPI_ISL_3339047 | SYNLAB | GIGA Medical Genomics | Keith Durkin, Maria Artesi, Bouchra Boujemla, Nathalie Renotte, Nadine Cambisano, Sébastien Bontems, Cécile Meex, Claire Gourzonès, Olivier Ek, Laurent Gillet, Marie-Pierre Hayette, Vincent Bours |
| EPI_ISL_3339048 | Lighthouse Lab in Alderley Park | Wellcome Sanger Institute for the COVID-19 Genomics UK (COG-UK) Consortium | Jacquelyn Wynn, Mairead Hyland, The Lighthouse Lab in Alderley Park and Alex Alderton, Roberto Amato, Jeffrey Barrett, Sonia Goncalves, Ewan Harrison, David K. Jackson, Ian Johnston, Dominic Kwiatkowski, Cordelia Langford, John Sillitoe on behalf of the Wellcome Sanger Institute COVID-19 Surveillance Team |
| EPI_ISL_3339049 | Lighthouse Lab in Milton Keynes | Wellcome Sanger Institute for the COVID-19 Genomics UK (COG-UK) Consortium | The Lighthouse Lab in Milton Keynes and Alex Alderton, Roberto Amato, Jeffrey Barrett, Sonia Goncalves, Ewan Harrison, David K. Jackson, Ian Johnston, Dominic Kwiatkowski, Cordelia Langford, John Sillitoe on behalf of the Wellcome Sanger Institute COVID-19 Surveillance Team |
| EPI_ISL_3339050 | SYNLAB | GIGA Medical Genomics | Keith Durkin, Maria Artesi, Bouchra Boujemla, Nathalie Renotte, Nadine Cambisano, Sébastien Bontems, Cécile Meex, Claire Gourzonès, Olivier Ek, Laurent Gillet, Marie-Pierre Hayette, Vincent Bours |
| EPI_ISL_3339051 | Lighthouse Lab in Alderley Park | Wellcome Sanger Institute for the COVID-19 Genomics UK (COG-UK) Consortium | Jacquelyn Wynn, Mairead Hyland, The Lighthouse Lab in Alderley Park and Alex Alderton, Roberto Amato, Jeffrey Barrett, Sonia Goncalves, Ewan Harrison, David K. Jackson, Ian Johnston, Dominic Kwiatkowski, Cordelia Langford, John Sillitoe on behalf of the Wellcome Sanger Institute COVID-19 Surveillance Team |
| EPI_ISL_3339052 | Lighthouse Lab in Milton Keynes | Wellcome Sanger Institute for the COVID-19 Genomics UK (COG-UK) Consortium | The Lighthouse Lab in Milton Keynes and Alex Alderton, Roberto Amato, Jeffrey Barrett, Sonia Goncalves, Ewan Harrison, David K. Jackson, Ian Johnston, Dominic Kwiatkowski, Cordelia Langford, John Sillitoe on behalf of the Wellcome Sanger Institute COVID-19 Surveillance Team |
| EPI_ISL_3339053 | SYNLAB | GIGA Medical Genomics | Keith Durkin, Maria Artesi, Bouchra Boujemla, Nathalie Renotte, Nadine Cambisano, Sébastien Bontems, Cécile Meex, Claire Gourzonès, Olivier Ek, Laurent Gillet, Marie-Pierre Hayette, Vincent Bours |
| EPI_ISL_3339054, EPI_ISL_3339055, EPI_ISL_3339056 | Lighthouse Lab in Milton Keynes | Wellcome Sanger Institute for the COVID-19 Genomics UK (COG-UK) Consortium | The Lighthouse Lab in Milton Keynes and Alex Alderton, Roberto Amato, Jeffrey Barrett, Sonia Goncalves, Ewan Harrison, David K. Jackson, Ian Johnston, Dominic Kwiatkowski, Cordelia Langford, John Sillitoe on behalf of the Wellcome Sanger Institute COVID-19 Surveillance Team |
| EPI_ISL_3339057 | SYNLAB | GIGA Medical Genomics | Keith Durkin, Maria Artesi, Bouchra Boujemla, Nathalie Renotte, Nadine Cambisano, Sébastien Bontems, Cécile Meex, Claire Gourzonès, Olivier Ek, Laurent Gillet, Marie-Pierre Hayette, Vincent Bours |
| EPI_ISL_3339058, EPI_ISL_3339059 | Lighthouse Lab in Milton Keynes | Wellcome Sanger Institute for the COVID-19 Genomics UK (COG-UK) Consortium | The Lighthouse Lab in Milton Keynes and Alex Alderton, Roberto Amato, Jeffrey Barrett, Sonia Goncalves, Ewan Harrison, David K. Jackson, Ian Johnston, Dominic Kwiatkowski, Cordelia Langford, John Sillitoe on behalf of the Wellcome Sanger Institute COVID-19 Surveillance Team |
| EPI_ISL_3339060 | SYNLAB | GIGA Medical Genomics | Keith Durkin, Maria Artesi, Bouchra Boujemla, Nathalie Renotte, Nadine Cambisano, Sébastien Bontems, Cécile Meex, Claire Gourzonès, Olivier Ek, Laurent Gillet, Marie-Pierre Hayette, Vincent Bours |
| EPI_ISL_3339061 | Lighthouse Lab in Milton Keynes | Wellcome Sanger Institute for the COVID-19 Genomics UK (COG-UK) Consortium | The Lighthouse Lab in Milton Keynes and Alex Alderton, Roberto Amato, Jeffrey Barrett, Sonia Goncalves, Ewan Harrison, David K. Jackson, Ian Johnston, Dominic Kwiatkowski, Cordelia Langford, John Sillitoe on behalf of the Wellcome Sanger Institute COVID-19 Surveillance Team |
| EPI_ISL_3339062, EPI_ISL_3339063 | Lighthouse Lab in Alderley Park | Wellcome Sanger Institute for the COVID-19 Genomics UK (COG-UK) Consortium | Jacquelyn Wynn, Mairead Hyland, The Lighthouse Lab in Alderley Park and Alex Alderton, Roberto Amato, Jeffrey Barrett, Sonia Goncalves, Ewan Harrison, David K. Jackson, Ian Johnston, Dominic Kwiatkowski, Cordelia Langford, John Sillitoe on behalf of the Wellcome Sanger Institute COVID-19 Surveillance Team |
| EPI_ISL_3339064 | SYNLAB | GIGA Medical Genomics | Keith Durkin, Maria Artesi, Bouchra Boujemla, Nathalie Renotte, Nadine Cambisano, Sébastien Bontems, Cécile Meex, Claire Gourzonès, Olivier Ek, Laurent Gillet, Marie-Pierre Hayette, Vincent Bours |
| EPI_ISL_3339065, EPI_ISL_3339066 | Lighthouse Lab in Milton Keynes | Wellcome Sanger Institute for the COVID-19 Genomics UK (COG-UK) Consortium | The Lighthouse Lab in Milton Keynes and Alex Alderton, Roberto Amato, Jeffrey Barrett, Sonia Goncalves, Ewan Harrison, David K. Jackson, Ian Johnston, Dominic Kwiatkowski, Cordelia Langford, John Sillitoe on behalf of the Wellcome Sanger Institute COVID-19 Surveillance Team |
| EPI_ISL_3339067 | SYNLAB | GIGA Medical Genomics | Keith Durkin, Maria Artesi, Bouchra Boujemla, Nathalie Renotte, Nadine Cambisano, Sébastien Bontems, Cécile Meex, Claire Gourzonès, Olivier Ek, Laurent Gillet, Marie-Pierre Hayette, Vincent Bours |
| EPI_ISL_3339068 | Lighthouse Lab in Glasgow | Wellcome Sanger Institute for the COVID-19 Genomics UK (COG-UK) Consortium | Harper VanSteenhouse, Yumi Kasai, David Gray, Carol Clugston, Anna Dominiczak and Alex Alderton, Roberto Amato, Jeffrey Barrett, Sonia Goncalves, Ewan Harrison, David K. Jackson, Ian Johnston, Dominic Kwiatkowski, Cordelia Langford, John Sillitoe on behalf of the Wellcome Sanger Institute COVID-19 Surveillance Team |
| EPI_ISL_3339069 | Randox Laboratories | Wellcome Sanger Institute for the COVID-19 Genomics UK (COG-UK) Consortium | Randox Laboratories and Alex Alderton, Roberto Amato, Jeffrey Barrett, Sonia Goncalves, Ewan Harrison, David K. Jackson, Ian Johnston, Dominic Kwiatkowski, Cordelia Langford, John Sillitoe on behalf of the Wellcome Sanger Institute COVID-19 Surveillance Team |
| EPI_ISL_3339070 | Lighthouse Lab in Alderley Park | Wellcome Sanger Institute for the COVID-19 Genomics UK (COG-UK) Consortium | Jacquelyn Wynn, Mairead Hyland, The Lighthouse Lab in Alderley Park and Alex Alderton, Roberto Amato, Jeffrey Barrett, Sonia Goncalves, Ewan Harrison, David K. Jackson, Ian Johnston, Dominic Kwiatkowski, Cordelia Langford, John Sillitoe on behalf of the Wellcome Sanger Institute COVID-19 Surveillance Team |
| EPI_ISL_3339071 | SYNLAB | GIGA Medical Genomics | Keith Durkin, Maria Artesi, Bouchra Boujemla, Nathalie Renotte, Nadine Cambisano, Sébastien Bontems, Cécile Meex, Claire Gourzonès, Olivier Ek, Laurent Gillet, Marie-Pierre Hayette, Vincent Bours |
| EPI_ISL_3339072 | Lighthouse Lab in Glasgow | Wellcome Sanger Institute for the COVID-19 Genomics UK (COG-UK) Consortium | Harper VanSteenhouse, Yumi Kasai, David Gray, Carol Clugston, Anna Dominiczak and Alex Alderton, Roberto Amato, Jeffrey Barrett, Sonia Goncalves, Ewan Harrison, David K. Jackson, Ian Johnston, Dominic Kwiatkowski, Cordelia Langford, John Sillitoe on behalf of the Wellcome Sanger Institute COVID-19 Surveillance Team |
| EPI_ISL_3339073 | Lighthouse Lab in Alderley Park | Wellcome Sanger Institute for the COVID-19 Genomics UK (COG-UK) Consortium | Jacquelyn Wynn, Mairead Hyland, The Lighthouse Lab in Alderley Park and Alex Alderton, Roberto Amato, Jeffrey Barrett, Sonia Goncalves, Ewan Harrison, David K. Jackson, Ian Johnston, Dominic Kwiatkowski, Cordelia Langford, John Sillitoe on behalf of the Wellcome Sanger Institute COVID-19 Surveillance Team |
| EPI_ISL_3339074 | SYNLAB | GIGA Medical Genomics | Keith Durkin, Maria Artesi, Bouchra Boujemla, Nathalie Renotte, Nadine Cambisano, Sébastien Bontems, Cécile Meex, Claire Gourzonès, Olivier Ek, Laurent Gillet, Marie-Pierre Hayette, Vincent Bours |
| EPI_ISL_3339075 | Lighthouse Lab in Alderley Park | Wellcome Sanger Institute for the COVID-19 Genomics UK (COG-UK) Consortium | Jacquelyn Wynn, Mairead Hyland, The Lighthouse Lab in Alderley Park and Alex Alderton, Roberto Amato, Jeffrey Barrett, Sonia Goncalves, Ewan Harrison, David K. Jackson, Ian Johnston, Dominic Kwiatkowski, Cordelia Langford, John Sillitoe on behalf of the Wellcome Sanger Institute COVID-19 Surveillance Team |
| EPI_ISL_3339076 | Lighthouse Lab in Glasgow | Wellcome Sanger Institute for the COVID-19 Genomics UK (COG-UK) Consortium | Harper VanSteenhouse, Yumi Kasai, David Gray, Carol Clugston, Anna Dominiczak and Alex Alderton, Roberto Amato, Jeffrey Barrett, Sonia Goncalves, Ewan Harrison, David K. Jackson, Ian Johnston, Dominic Kwiatkowski, Cordelia Langford, John Sillitoe on behalf of the Wellcome Sanger Institute COVID-19 Surveillance Team |
| EPI_ISL_3339077 | SYNLAB | GIGA Medical Genomics | Keith Durkin, Maria Artesi, Bouchra Boujemla, Nathalie Renotte, Nadine Cambisano, Sébastien Bontems, Cécile Meex, Claire Gourzonès, Olivier Ek, Laurent Gillet, Marie-Pierre Hayette, Vincent Bours |
| EPI_ISL_3339078 | Lighthouse Lab in Milton Keynes | Wellcome Sanger Institute for the COVID-19 Genomics UK (COG-UK) Consortium | The Lighthouse Lab in Milton Keynes and Alex Alderton, Roberto Amato, Jeffrey Barrett, Sonia Goncalves, Ewan Harrison, David K. Jackson, Ian Johnston, Dominic Kwiatkowski, Cordelia Langford, John Sillitoe on behalf of the Wellcome Sanger Institute COVID-19 Surveillance Team |
| EPI_ISL_3339079 | Lighthouse Lab in Alderley Park | Wellcome Sanger Institute for the COVID-19 Genomics UK (COG-UK) Consortium | Jacquelyn Wynn, Mairead Hyland, The Lighthouse Lab in Alderley Park and Alex Alderton, Roberto Amato, Jeffrey Barrett, Sonia Goncalves, Ewan Harrison, David K. Jackson, Ian Johnston, Dominic Kwiatkowski, Cordelia Langford, John Sillitoe on behalf of the Wellcome Sanger Institute COVID-19 Surveillance Team |
| EPI_ISL_3339080 | Lighthouse Lab in Glasgow | Wellcome Sanger Institute for the COVID-19 Genomics UK (COG-UK) Consortium | Harper VanSteenhouse, Yumi Kasai, David Gray, Carol Clugston, Anna Dominiczak and Alex Alderton, Roberto Amato, Jeffrey Barrett, Sonia Goncalves, Ewan Harrison, David K. Jackson, Ian Johnston, Dominic Kwiatkowski, Cordelia Langford, John Sillitoe on behalf of the Wellcome Sanger Institute COVID-19 Surveillance Team |
| EPI_ISL_3339081 | SYNLAB | GIGA Medical Genomics | Keith Durkin, Maria Artesi, Bouchra Boujemla, Nathalie Renotte, Nadine Cambisano, Sébastien Bontems, Cécile Meex, Claire Gourzonès, Olivier Ek, Laurent Gillet, Marie-Pierre Hayette, Vincent Bours |
| EPI_ISL_3339082, EPI_ISL_3339083 | Lighthouse Lab in Alderley Park | Wellcome Sanger Institute for the COVID-19 Genomics UK (COG-UK) Consortium | Jacquelyn Wynn, Mairead Hyland, The Lighthouse Lab in Alderley Park and Alex Alderton, Roberto Amato, Jeffrey Barrett, Sonia Goncalves, Ewan Harrison, David K. Jackson, Ian Johnston, Dominic Kwiatkowski, Cordelia Langford, John Sillitoe on behalf of the Wellcome Sanger Institute COVID-19 Surveillance Team |
| EPI_ISL_3339084 | SYNLAB | GIGA Medical Genomics | Keith Durkin, Maria Artesi, Bouchra Boujemla, Nathalie Renotte, Nadine Cambisano, Sébastien Bontems, Cécile Meex, Claire Gourzonès, Olivier Ek, Laurent Gillet, Marie-Pierre Hayette, Vincent Bours |
| EPI_ISL_3339085 | Lighthouse Lab in Milton Keynes | Wellcome Sanger Institute for the COVID-19 Genomics UK (COG-UK) Consortium | The Lighthouse Lab in Milton Keynes and Alex Alderton, Roberto Amato, Jeffrey Barrett, Sonia Goncalves, Ewan Harrison, David K. Jackson, Ian Johnston, Dominic Kwiatkowski, Cordelia Langford, John Sillitoe on behalf of the Wellcome Sanger Institute COVID-19 Surveillance Team |
| EPI_ISL_3339086, EPI_ISL_3339087 | Lighthouse Lab in Alderley Park | Wellcome Sanger Institute for the COVID-19 Genomics UK (COG-UK) Consortium | Jacquelyn Wynn, Mairead Hyland, The Lighthouse Lab in Alderley Park and Alex Alderton, Roberto Amato, Jeffrey Barrett, Sonia Goncalves, Ewan Harrison, David K. Jackson, Ian Johnston, Dominic Kwiatkowski, Cordelia Langford, John Sillitoe on behalf of the Wellcome Sanger Institute COVID-19 Surveillance Team |
| EPI_ISL_3339088 | SYNLAB | GIGA Medical Genomics | Keith Durkin, Maria Artesi, Bouchra Boujemla, Nathalie Renotte, Nadine Cambisano, Sébastien Bontems, Cécile Meex, Claire Gourzonès, Olivier Ek, Laurent Gillet, Marie-Pierre Hayette, Vincent Bours |
| EPI_ISL_3339089 | Lighthouse Lab in Milton Keynes | Wellcome Sanger Institute for the COVID-19 Genomics UK (COG-UK) Consortium | The Lighthouse Lab in Milton Keynes and Alex Alderton, Roberto Amato, Jeffrey Barrett, Sonia Goncalves, Ewan Harrison, David K. Jackson, Ian Johnston, Dominic Kwiatkowski, Cordelia Langford, John Sillitoe on behalf of the Wellcome Sanger Institute COVID-19 Surveillance Team |
| EPI_ISL_3339090 | Lighthouse Lab in Alderley Park | Wellcome Sanger Institute for the COVID-19 Genomics UK (COG-UK) Consortium | Jacquelyn Wynn, Mairead Hyland, The Lighthouse Lab in Alderley Park and Alex Alderton, Roberto Amato, Jeffrey Barrett, Sonia Goncalves, Ewan Harrison, David K. Jackson, Ian Johnston, Dominic Kwiatkowski, Cordelia Langford, John Sillitoe on behalf of the Wellcome Sanger Institute COVID-19 Surveillance Team |
| EPI_ISL_3339091 | SYNLAB | GIGA Medical Genomics | Keith Durkin, Maria Artesi, Bouchra Boujemla, Nathalie Renotte, Nadine Cambisano, Sébastien Bontems, Cécile Meex, Claire Gourzonès, Olivier Ek, Laurent Gillet, Marie-Pierre Hayette, Vincent Bours |
| EPI_ISL_3339092, EPI_ISL_3339093, EPI_ISL_3339094 | Lighthouse Lab in Alderley Park | Wellcome Sanger Institute for the COVID-19 Genomics UK (COG-UK) Consortium | Jacquelyn Wynn, Mairead Hyland, The Lighthouse Lab in Alderley Park and Alex Alderton, Roberto Amato, Jeffrey Barrett, Sonia Goncalves, Ewan Harrison, David K. Jackson, Ian Johnston, Dominic Kwiatkowski, Cordelia Langford, John Sillitoe on behalf of the Wellcome Sanger Institute COVID-19 Surveillance Team |
| EPI_ISL_3339095 | SYNLAB | GIGA Medical Genomics | Keith Durkin, Maria Artesi, Bouchra Boujemla, Nathalie Renotte, Nadine Cambisano, Sébastien Bontems, Cécile Meex, Claire Gourzonès, Olivier Ek, Laurent Gillet, Marie-Pierre Hayette, Vincent Bours |
| EPI_ISL_3339096 | Lighthouse Lab in Milton Keynes | Wellcome Sanger Institute for the COVID-19 Genomics UK (COG-UK) Consortium | The Lighthouse Lab in Milton Keynes and Alex Alderton, Roberto Amato, Jeffrey Barrett, Sonia Goncalves, Ewan Harrison, David K. Jackson, Ian Johnston, Dominic Kwiatkowski, Cordelia Langford, John Sillitoe on behalf of the Wellcome Sanger Institute COVID-19 Surveillance Team |
| EPI_ISL_3339097 | Lighthouse Lab in Alderley Park | Wellcome Sanger Institute for the COVID-19 Genomics UK (COG-UK) Consortium | Jacquelyn Wynn, Mairead Hyland, The Lighthouse Lab in Alderley Park and Alex Alderton, Roberto Amato, Jeffrey Barrett, Sonia Goncalves, Ewan Harrison, David K. Jackson, Ian Johnston, Dominic Kwiatkowski, Cordelia Langford, John Sillitoe on behalf of the Wellcome Sanger Institute COVID-19 Surveillance Team |
| EPI_ISL_3339098 | SYNLAB | GIGA Medical Genomics | Keith Durkin, Maria Artesi, Bouchra Boujemla, Nathalie Renotte, Nadine Cambisano, Sébastien Bontems, Cécile Meex, Claire Gourzonès, Olivier Ek, Laurent Gillet, Marie-Pierre Hayette, Vincent Bours |
| EPI_ISL_3339099 | Lighthouse Lab in Alderley Park | Wellcome Sanger Institute for the COVID-19 Genomics UK (COG-UK) Consortium | Jacquelyn Wynn, Mairead Hyland, The Lighthouse Lab in Alderley Park and Alex Alderton, Roberto Amato, Jeffrey Barrett, Sonia Goncalves, Ewan Harrison, David K. Jackson, Ian Johnston, Dominic Kwiatkowski, Cordelia Langford, John Sillitoe on behalf of the Wellcome Sanger Institute COVID-19 Surveillance Team |
| EPI_ISL_3339100 | Lighthouse Lab in Milton Keynes | Wellcome Sanger Institute for the COVID-19 Genomics UK (COG-UK) Consortium | The Lighthouse Lab in Milton Keynes and Alex Alderton, Roberto Amato, Jeffrey Barrett, Sonia Goncalves, Ewan Harrison, David K. Jackson, Ian Johnston, Dominic Kwiatkowski, Cordelia Langford, John Sillitoe on behalf of the Wellcome Sanger Institute COVID-19 Surveillance Team |
| EPI_ISL_3339101 | Lighthouse Lab in Alderley Park | Wellcome Sanger Institute for the COVID-19 Genomics UK (COG-UK) Consortium | Jacquelyn Wynn, Mairead Hyland, The Lighthouse Lab in Alderley Park and Alex Alderton, Roberto Amato, Jeffrey Barrett, Sonia Goncalves, Ewan Harrison, David K. Jackson, Ian Johnston, Dominic Kwiatkowski, Cordelia Langford, John Sillitoe on behalf of the Wellcome Sanger Institute COVID-19 Surveillance Team |
| EPI_ISL_3339102 | SYNLAB | GIGA Medical Genomics | Keith Durkin, Maria Artesi, Bouchra Boujemla, Nathalie Renotte, Nadine Cambisano, Sébastien Bontems, Cécile Meex, Claire Gourzonès, Olivier Ek, Laurent Gillet, Marie-Pierre Hayette, Vincent Bours |
| EPI_ISL_3339103, EPI_ISL_3339104 | Lighthouse Lab in Alderley Park | Wellcome Sanger Institute for the COVID-19 Genomics UK (COG-UK) Consortium | Jacquelyn Wynn, Mairead Hyland, The Lighthouse Lab in Alderley Park and Alex Alderton, Roberto Amato, Jeffrey Barrett, Sonia Goncalves, Ewan Harrison, David K. Jackson, Ian Johnston, Dominic Kwiatkowski, Cordelia Langford, John Sillitoe on behalf of the Wellcome Sanger Institute COVID-19 Surveillance Team |
| EPI_ISL_3339105 | Randox Laboratories | Wellcome Sanger Institute for the COVID-19 Genomics UK (COG-UK) | Randox Laboratories and Alex Alderton, Roberto Amato, Jeffrey Barrett, Sonia Goncalves, Ewan Harrison, David K. Jackson, Ian Johnston, Dominic Kwiatkowski, Cordelia Langford, John Sillitoe on behalf of the Wellcome Sanger Institute COVID- |

| | | | |
|---|---|---|---|
| | | Consortium | 19 Surveillance Team |
| EPI_ISL_3339106 | SYNLAB | GIGA Medical Genomics | Keith Durkin, Maria Artesi, Bouchra Boujemla, Nathalie Renotte, Nadine Cambisano, Sébastien Bontems, Cécile Meex, Claire Gourzonès, Olivier Ek, Laurent Gillet, Marie-Pierre Hayette, Vincent Bours |
| EPI_ISL_3339107 | Lighthouse Lab in Alderley Park | Wellcome Sanger Institute for the COVID-19 Genomics UK (COG-UK) Consortium | Jacquelyn Wynn, Mairead Hyland, The Lighthouse Lab in Alderley Park and Alex Alderton, Roberto Amato, Jeffrey Barrett, Sonia Goncalves, Ewan Harrison, David K. Jackson, Ian Johnston, Dominic Kwiatkowski, Cordelia Langford, John Sillitoe on behalf of the Wellcome Sanger Institute COVID-19 Surveillance Team |
| EPI_ISL_3339108 | Lighthouse Lab in Glasgow | Wellcome Sanger Institute for the COVID-19 Genomics UK (COG-UK) Consortium | Harper VanSteenhouse, Yumi Kasai, David Gray, Carol Clugston, Anna Dominiczak and Alex Alderton, Roberto Amato, Jeffrey Barrett, Sonia Goncalves, Ewan Harrison, David K. Jackson, Ian Johnston, Dominic Kwiatkowski, Cordelia Langford, John Sillitoe on behalf of the Wellcome Sanger Institute COVID-19 Surveillance Team |
| EPI_ISL_3339109 | SYNLAB | GIGA Medical Genomics | Keith Durkin, Maria Artesi, Bouchra Boujemla, Nathalie Renotte, Nadine Cambisano, Sébastien Bontems, Cécile Meex, Claire Gourzonès, Olivier Ek, Laurent Gillet, Marie-Pierre Hayette, Vincent Bours |
| EPI_ISL_3339110, EPI_ISL_3339111 | Lighthouse Lab in Alderley Park | Wellcome Sanger Institute for the COVID-19 Genomics UK (COG-UK) Consortium | Jacquelyn Wynn, Mairead Hyland, The Lighthouse Lab in Alderley Park and Alex Alderton, Roberto Amato, Jeffrey Barrett, Sonia Goncalves, Ewan Harrison, David K. Jackson, Ian Johnston, Dominic Kwiatkowski, Cordelia Langford, John Sillitoe on behalf of the Wellcome Sanger Institute COVID-19 Surveillance Team |
| EPI_ISL_3339112 | SYNLAB | GIGA Medical Genomics | Keith Durkin, Maria Artesi, Bouchra Boujemla, Nathalie Renotte, Nadine Cambisano, Sébastien Bontems, Cécile Meex, Claire Gourzonès, Olivier Ek, Laurent Gillet, Marie-Pierre Hayette, Vincent Bours |
| EPI_ISL_3339113 | Lighthouse Lab in Alderley Park | Wellcome Sanger Institute for the COVID-19 Genomics UK (COG-UK) Consortium | Jacquelyn Wynn, Mairead Hyland, The Lighthouse Lab in Alderley Park and Alex Alderton, Roberto Amato, Jeffrey Barrett, Sonia Goncalves, Ewan Harrison, David K. Jackson, Ian Johnston, Dominic Kwiatkowski, Cordelia Langford, John Sillitoe on behalf of the Wellcome Sanger Institute COVID-19 Surveillance Team |
| EPI_ISL_3339114 | Randox Laboratories | Wellcome Sanger Institute for the COVID-19 Genomics UK (COG-UK) Consortium | Randox Laboratories and Alex Alderton, Roberto Amato, Jeffrey Barrett, Sonia Goncalves, Ewan Harrison, David K. Jackson, Ian Johnston, Dominic Kwiatkowski, Cordelia Langford, John Sillitoe on behalf of the Wellcome Sanger Institute COVID-19 Surveillance Team |
| EPI_ISL_3339115 | Lighthouse Lab in Milton Keynes | Wellcome Sanger Institute for the COVID-19 Genomics UK (COG-UK) Consortium | The Lighthouse Lab in Milton Keynes and Alex Alderton, Roberto Amato, Jeffrey Barrett, Sonia Goncalves, Ewan Harrison, David K. Jackson, Ian Johnston, Dominic Kwiatkowski, Cordelia Langford, John Sillitoe on behalf of the Wellcome Sanger Institute COVID-19 Surveillance Team |
| EPI_ISL_3339116 | SYNLAB | GIGA Medical Genomics | Keith Durkin, Maria Artesi, Bouchra Boujemla, Nathalie Renotte, Nadine Cambisano, Sébastien Bontems, Cécile Meex, Claire Gourzonès, Olivier Ek, Laurent Gillet, Marie-Pierre Hayette, Vincent Bours |
| EPI_ISL_3339117 | Lighthouse Lab in Alderley Park | Wellcome Sanger Institute for the COVID-19 Genomics UK (COG-UK) Consortium | Jacquelyn Wynn, Mairead Hyland, The Lighthouse Lab in Alderley Park and Alex Alderton, Roberto Amato, Jeffrey Barrett, Sonia Goncalves, Ewan Harrison, David K. Jackson, Ian Johnston, Dominic Kwiatkowski, Cordelia Langford, John Sillitoe on behalf of the Wellcome Sanger Institute COVID-19 Surveillance Team |
| EPI_ISL_3339118 | Lighthouse Lab in Milton Keynes | Wellcome Sanger Institute for the COVID-19 Genomics UK (COG-UK) Consortium | The Lighthouse Lab in Milton Keynes and Alex Alderton, Roberto Amato, Jeffrey Barrett, Sonia Goncalves, Ewan Harrison, David K. Jackson, Ian Johnston, Dominic Kwiatkowski, Cordelia Langford, John Sillitoe on behalf of the Wellcome Sanger Institute COVID-19 Surveillance Team |
| EPI_ISL_3339119 | SYNLAB | GIGA Medical Genomics | Keith Durkin, Maria Artesi, Bouchra Boujemla, Nathalie Renotte, Nadine Cambisano, Sébastien Bontems, Cécile Meex, Claire Gourzonès, Olivier Ek, Laurent Gillet, Marie-Pierre Hayette, Vincent Bours |
| EPI_ISL_3339120 | Lighthouse Lab in Milton Keynes | Wellcome Sanger Institute for the COVID-19 Genomics UK (COG-UK) Consortium | The Lighthouse Lab in Milton Keynes and Alex Alderton, Roberto Amato, Jeffrey Barrett, Sonia Goncalves, Ewan Harrison, David K. Jackson, Ian Johnston, Dominic Kwiatkowski, Cordelia Langford, John Sillitoe on behalf of the Wellcome Sanger Institute COVID-19 Surveillance Team |
| EPI_ISL_3339121 | Lighthouse Lab in Alderley Park | Wellcome Sanger Institute for the COVID-19 Genomics UK (COG-UK) Consortium | Jacquelyn Wynn, Mairead Hyland, The Lighthouse Lab in Alderley Park and Alex Alderton, Roberto Amato, Jeffrey Barrett, Sonia Goncalves, Ewan Harrison, David K. Jackson, Ian Johnston, Dominic Kwiatkowski, Cordelia Langford, John Sillitoe on behalf of the Wellcome Sanger Institute COVID-19 Surveillance Team |
| EPI_ISL_3339122 | SYNLAB | GIGA Medical Genomics | Keith Durkin, Maria Artesi, Bouchra Boujemla, Nathalie Renotte, Nadine Cambisano, Sébastien Bontems, Cécile Meex, Claire Gourzonès, Olivier Ek, Laurent Gillet, Marie-Pierre Hayette, Vincent Bours |
| EPI_ISL_3339123, EPI_ISL_3339124 | Lighthouse Lab in Milton Keynes | Wellcome Sanger Institute for the COVID-19 Genomics UK (COG-UK) Consortium | The Lighthouse Lab in Milton Keynes and Alex Alderton, Roberto Amato, Jeffrey Barrett, Sonia Goncalves, Ewan Harrison, David K. Jackson, Ian Johnston, Dominic Kwiatkowski, Cordelia Langford, John Sillitoe on behalf of the Wellcome Sanger Institute COVID-19 Surveillance Team |
| EPI_ISL_3339125 | Lighthouse Lab in Alderley Park | Wellcome Sanger Institute for the COVID-19 Genomics UK (COG-UK) Consortium | Jacquelyn Wynn, Mairead Hyland, The Lighthouse Lab in Alderley Park and Alex Alderton, Roberto Amato, Jeffrey Barrett, Sonia Goncalves, Ewan Harrison, David K. Jackson, Ian Johnston, Dominic Kwiatkowski, Cordelia Langford, John Sillitoe on behalf of the Wellcome Sanger Institute COVID-19 Surveillance Team |
| EPI_ISL_3339126 | SYNLAB | GIGA Medical Genomics | Keith Durkin, Maria Artesi, Bouchra Boujemla, Nathalie Renotte, Nadine Cambisano, Sébastien Bontems, Cécile Meex, Claire Gourzonès, Olivier Ek, Laurent Gillet, Marie-Pierre Hayette, Vincent Bours |
| EPI_ISL_3339127 | Lighthouse Lab in Glasgow | Wellcome Sanger Institute for the COVID-19 Genomics UK (COG-UK) Consortium | Harper VanSteenhouse, Yumi Kasai, David Gray, Carol Clugston, Anna Dominiczak and Alex Alderton, Roberto Amato, Jeffrey Barrett, Sonia Goncalves, Ewan Harrison, David K. Jackson, Ian Johnston, Dominic Kwiatkowski, Cordelia Langford, John Sillitoe on behalf of the Wellcome Sanger Institute COVID-19 Surveillance Team |
| EPI_ISL_3339128 | Lighthouse Lab in Alderley Park | Wellcome Sanger Institute for the COVID-19 Genomics UK (COG-UK) Consortium | Jacquelyn Wynn, Mairead Hyland, The Lighthouse Lab in Alderley Park and Alex Alderton, Roberto Amato, Jeffrey Barrett, Sonia Goncalves, Ewan Harrison, David K. Jackson, Ian Johnston, Dominic Kwiatkowski, Cordelia Langford, John Sillitoe on behalf of the Wellcome Sanger Institute COVID-19 Surveillance Team |
| EPI_ISL_3339129 | SYNLAB | GIGA Medical Genomics | Keith Durkin, Maria Artesi, Bouchra Boujemla, Nathalie Renotte, Nadine Cambisano, Sébastien Bontems, Cécile Meex, Claire Gourzonès, Olivier Ek, Laurent Gillet, Marie-Pierre Hayette, Vincent Bours |
| EPI_ISL_3339130 | Lighthouse Lab in Milton Keynes | Wellcome Sanger Institute for the COVID-19 Genomics UK (COG-UK) Consortium | The Lighthouse Lab in Milton Keynes and Alex Alderton, Roberto Amato, Jeffrey Barrett, Sonia Goncalves, Ewan Harrison, David K. Jackson, Ian Johnston, Dominic Kwiatkowski, Cordelia Langford, John Sillitoe on behalf of the Wellcome Sanger Institute COVID-19 Surveillance Team |
| EPI_ISL_3339131 | Lighthouse Lab in Glasgow | Wellcome Sanger Institute for the COVID-19 Genomics UK (COG-UK) Consortium | Harper VanSteenhouse, Yumi Kasai, David Gray, Carol Clugston, Anna Dominiczak and Alex Alderton, Roberto Amato, Jeffrey Barrett, Sonia Goncalves, Ewan Harrison, David K. Jackson, Ian Johnston, Dominic Kwiatkowski, Cordelia Langford, John Sillitoe on behalf of the Wellcome Sanger Institute COVID-19 Surveillance Team |
| EPI_ISL_3339132 | SYNLAB | GIGA Medical Genomics | Keith Durkin, Maria Artesi, Bouchra Boujemla, Nathalie Renotte, Nadine Cambisano, Sébastien Bontems, Cécile Meex, Claire Gourzonès, Olivier Ek, Laurent Gillet, Marie-Pierre Hayette, Vincent Bours |
| EPI_ISL_3339133 | Lighthouse Lab in Alderley Park | Wellcome Sanger Institute for the COVID-19 Genomics UK (COG-UK) Consortium | Jacquelyn Wynn, Mairead Hyland, The Lighthouse Lab in Alderley Park and Alex Alderton, Roberto Amato, Jeffrey Barrett, Sonia Goncalves, Ewan Harrison, David K. Jackson, Ian Johnston, Dominic Kwiatkowski, Cordelia Langford, John Sillitoe on behalf of the Wellcome Sanger Institute COVID-19 Surveillance Team |
| EPI_ISL_3339134 | Lighthouse Lab in Glasgow | Wellcome Sanger Institute for the COVID-19 Genomics UK (COG-UK) Consortium | Harper VanSteenhouse, Yumi Kasai, David Gray, Carol Clugston, Anna Dominiczak and Alex Alderton, Roberto Amato, Jeffrey Barrett, Sonia Goncalves, Ewan Harrison, David K. Jackson, Ian Johnston, Dominic Kwiatkowski, Cordelia Langford, John Sillitoe on behalf of the Wellcome Sanger Institute COVID-19 Surveillance Team |
| EPI_ISL_3339135 | Lighthouse Lab in Alderley Park | Wellcome Sanger Institute for the COVID-19 Genomics UK (COG-UK) Consortium | Jacquelyn Wynn, Mairead Hyland, The Lighthouse Lab in Alderley Park and Alex Alderton, Roberto Amato, Jeffrey Barrett, Sonia Goncalves, Ewan Harrison, David K. Jackson, Ian Johnston, Dominic Kwiatkowski, Cordelia Langford, John Sillitoe on behalf of the Wellcome Sanger Institute COVID-19 Surveillance Team |
| EPI_ISL_3339136 | SYNLAB | GIGA Medical Genomics | Keith Durkin, Maria Artesi, Bouchra Boujemla, Nathalie Renotte, Nadine Cambisano, Sébastien Bontems, Cécile Meex, Claire Gourzonès, Olivier Ek, Laurent Gillet, Marie-Pierre Hayette, Vincent Bours |
| EPI_ISL_3339137, EPI_ISL_3339138 | Lighthouse Lab in Alderley Park | Wellcome Sanger Institute for the COVID-19 Genomics UK (COG-UK) Consortium | Jacquelyn Wynn, Mairead Hyland, The Lighthouse Lab in Alderley Park and Alex Alderton, Roberto Amato, Jeffrey Barrett, Sonia Goncalves, Ewan Harrison, David K. Jackson, Ian Johnston, Dominic Kwiatkowski, Cordelia Langford, John Sillitoe on behalf of the Wellcome Sanger Institute COVID-19 Surveillance Team |
| EPI_ISL_3339139 | SYNLAB | GIGA Medical Genomics | Keith Durkin, Maria Artesi, Bouchra Boujemla, Nathalie Renotte, Nadine Cambisano, Sébastien Bontems, Cécile Meex, Claire Gourzonès, Olivier Ek, Laurent Gillet, Marie-Pierre Hayette, Vincent Bours |
| EPI_ISL_3339140 | Lighthouse Lab in Glasgow | Wellcome Sanger Institute for the COVID-19 Genomics UK (COG-UK) Consortium | Harper VanSteenhouse, Yumi Kasai, David Gray, Carol Clugston, Anna Dominiczak and Alex Alderton, Roberto Amato, Jeffrey Barrett, Sonia Goncalves, Ewan Harrison, David K. Jackson, Ian Johnston, Dominic Kwiatkowski, Cordelia Langford, John Sillitoe on behalf of the Wellcome Sanger Institute COVID-19 Surveillance Team |
| EPI_ISL_3339141 | Lighthouse Lab in Alderley Park | Wellcome Sanger Institute for the COVID-19 Genomics UK (COG-UK) Consortium | Jacquelyn Wynn, Mairead Hyland, The Lighthouse Lab in Alderley Park and Alex Alderton, Roberto Amato, Jeffrey Barrett, Sonia Goncalves, Ewan Harrison, David K. Jackson, Ian Johnston, Dominic Kwiatkowski, Cordelia Langford, John Sillitoe on behalf of the Wellcome Sanger Institute COVID-19 Surveillance Team |
| EPI_ISL_3339142 | SYNLAB | GIGA Medical Genomics | Keith Durkin, Maria Artesi, Bouchra Boujemla, Nathalie Renotte, Nadine Cambisano, Sébastien Bontems, Cécile Meex, Claire Gourzonès, Olivier Ek, Laurent Gillet, Marie-Pierre Hayette, Vincent Bours |
| EPI_ISL_3339143, EPI_ISL_3339144, EPI_ISL_3339145 | Lighthouse Lab in Milton Keynes | Wellcome Sanger Institute for the COVID-19 Genomics UK (COG-UK) Consortium | The Lighthouse Lab in Milton Keynes and Alex Alderton, Roberto Amato, Jeffrey Barrett, Sonia Goncalves, Ewan Harrison, David K. Jackson, Ian Johnston, Dominic Kwiatkowski, Cordelia Langford, John Sillitoe on behalf of the Wellcome Sanger Institute COVID-19 Surveillance Team |
| EPI_ISL_3339146 | SYNLAB | GIGA Medical Genomics | Keith Durkin, Maria Artesi, Bouchra Boujemla, Nathalie Renotte, Nadine Cambisano, Sébastien Bontems, Cécile Meex, Claire Gourzonès, Olivier Ek, Laurent Gillet, Marie-Pierre Hayette, Vincent Bours |
| EPI_ISL_3339147, EPI_ISL_3339148 | Lighthouse Lab in Alderley Park | Wellcome Sanger Institute for the COVID-19 Genomics UK (COG-UK) Consortium | Jacquelyn Wynn, Mairead Hyland, The Lighthouse Lab in Alderley Park and Alex Alderton, Roberto Amato, Jeffrey Barrett, Sonia Goncalves, Ewan Harrison, David K. Jackson, Ian Johnston, Dominic Kwiatkowski, Cordelia Langford, John Sillitoe on behalf of the Wellcome Sanger Institute COVID-19 Surveillance Team |
| EPI_ISL_3339149 | Department of Clinical Microbiology | GIGA Medical Genomics | Keith Durkin, Maria Artesi, Bouchra Boujemla, Nathalie Renotte, Nadine Cambisano, Sébastien Bontems, Cécile Meex, Claire Gourzonès, Olivier Ek, Laurent Gillet, Marie-Pierre Hayette, Vincent Bours |
| EPI_ISL_3339150 | Lighthouse Lab in Milton Keynes | Wellcome Sanger Institute for the COVID-19 Genomics UK (COG-UK) Consortium | The Lighthouse Lab in Milton Keynes and Alex Alderton, Roberto Amato, Jeffrey Barrett, Sonia Goncalves, Ewan Harrison, David K. Jackson, Ian Johnston, Dominic Kwiatkowski, Cordelia Langford, John Sillitoe on behalf of the Wellcome Sanger Institute COVID-19 Surveillance Team |
| EPI_ISL_3339151 | Lighthouse Lab in Alderley Park | Wellcome Sanger Institute for the COVID-19 Genomics UK (COG-UK) Consortium | Jacquelyn Wynn, Mairead Hyland, The Lighthouse Lab in Alderley Park and Alex Alderton, Roberto Amato, Jeffrey Barrett, Sonia Goncalves, Ewan Harrison, David K. Jackson, Ian Johnston, Dominic Kwiatkowski, Cordelia Langford, John Sillitoe on behalf of the Wellcome Sanger Institute COVID-19 Surveillance Team |
| EPI_ISL_3339152 | Department of Clinical Microbiology | GIGA Medical Genomics | Keith Durkin, Maria Artesi, Bouchra Boujemla, Nathalie Renotte, Nadine Cambisano, Sébastien Bontems, Cécile Meex, Claire Gourzonès, Olivier Ek, Laurent Gillet, Marie-Pierre Hayette, Vincent Bours |
| EPI_ISL_3339153 | Lighthouse Lab in Glasgow | Wellcome Sanger Institute for the COVID-19 Genomics UK (COG-UK) Consortium | Harper VanSteenhouse, Yumi Kasai, David Gray, Carol Clugston, Anna Dominiczak and Alex Alderton, Roberto Amato, Jeffrey Barrett, Sonia Goncalves, Ewan Harrison, David K. Jackson, Ian Johnston, Dominic Kwiatkowski, Cordelia Langford, John Sillitoe on behalf of the Wellcome Sanger Institute COVID-19 Surveillance Team |
| EPI_ISL_3339154 | Lighthouse Lab in Alderley Park | Wellcome Sanger Institute for the COVID-19 Genomics UK (COG-UK) Consortium | Jacquelyn Wynn, Mairead Hyland, The Lighthouse Lab in Alderley Park and Alex Alderton, Roberto Amato, Jeffrey Barrett, Sonia Goncalves, Ewan Harrison, David K. Jackson, Ian Johnston, Dominic Kwiatkowski, Cordelia Langford, John Sillitoe on behalf of the Wellcome Sanger Institute COVID-19 Surveillance Team |
| EPI_ISL_3339155 | SYNLAB | GIGA Medical Genomics | Keith Durkin, Maria Artesi, Bouchra Boujemla, Nathalie Renotte, Nadine Cambisano, Sébastien Bontems, Cécile Meex, Claire Gourzonès, Olivier Ek, Laurent Gillet, Marie-Pierre Hayette, Vincent Bours |
| EPI_ISL_3339156, EPI_ISL_3339157, EPI_ISL_3339158 | Lighthouse Lab in Alderley Park | Wellcome Sanger Institute for the COVID-19 Genomics UK (COG-UK) Consortium | Jacquelyn Wynn, Mairead Hyland, The Lighthouse Lab in Alderley Park and Alex Alderton, Roberto Amato, Jeffrey Barrett, Sonia Goncalves, Ewan Harrison, David K. Jackson, Ian Johnston, Dominic Kwiatkowski, Cordelia Langford, John Sillitoe on behalf of the Wellcome Sanger Institute COVID-19 Surveillance Team |
| EPI_ISL_3339159 | SYNLAB | GIGA Medical Genomics | Keith Durkin, Maria Artesi, Bouchra Boujemla, Nathalie Renotte, Nadine Cambisano, Sébastien Bontems, Cécile Meex, Claire Gourzonès, Olivier Ek, Laurent Gillet, Marie-Pierre Hayette, Vincent Bours |
| EPI_ISL_3339160, EPI_ISL_3339161 | Lighthouse Lab in Alderley Park | Wellcome Sanger Institute for the COVID-19 Genomics UK (COG-UK) Consortium | Jacquelyn Wynn, Mairead Hyland, The Lighthouse Lab in Alderley Park and Alex Alderton, Roberto Amato, Jeffrey Barrett, Sonia Goncalves, Ewan Harrison, David K. Jackson, Ian Johnston, Dominic Kwiatkowski, Cordelia Langford, John Sillitoe on behalf of the Wellcome Sanger Institute COVID-19 Surveillance Team |
| EPI_ISL_3339162 | SYNLAB | GIGA Medical Genomics | Keith Durkin, Maria Artesi, Bouchra Boujemla, Nathalie Renotte, Nadine Cambisano, Sébastien Bontems, Cécile Meex, Claire Gourzonès, Olivier Ek, Laurent Gillet, Marie-Pierre Hayette, Vincent Bours |
| EPI_ISL_3339163 | Lighthouse Lab in Glasgow | Wellcome Sanger Institute for the COVID-19 Genomics UK (COG-UK) Consortium | Harper VanSteenhouse, Yumi Kasai, David Gray, Carol Clugston, Anna Dominiczak and Alex Alderton, Roberto Amato, Jeffrey Barrett, Sonia Goncalves, Ewan Harrison, David K. Jackson, Ian Johnston, Dominic Kwiatkowski, Cordelia Langford, John Sillitoe on behalf of the Wellcome Sanger Institute COVID-19 Surveillance Team |
| EPI_ISL_3339164 | Lighthouse Lab in Alderley Park | Wellcome Sanger Institute for the COVID-19 Genomics UK (COG-UK) Consortium | Jacquelyn Wynn, Mairead Hyland, The Lighthouse Lab in Alderley Park and Alex Alderton, Roberto Amato, Jeffrey Barrett, Sonia Goncalves, Ewan Harrison, David K. Jackson, Ian Johnston, Dominic Kwiatkowski, Cordelia Langford, John Sillitoe on behalf of the Wellcome Sanger Institute COVID-19 Surveillance Team |
| EPI_ISL_3339165 | SYNLAB | GIGA Medical Genomics | Keith Durkin, Maria Artesi, Bouchra Boujemla, Nathalie Renotte, Nadine Cambisano, Sébastien Bontems, Cécile Meex, Claire Gourzonès, Olivier Ek, Laurent Gillet, Marie-Pierre Hayette, Vincent Bours |
| EPI_ISL_3339166, | Lighthouse Lab in Alderley Park | Wellcome Sanger Institute for the COVID-19 Genomics UK (COG-UK) | Jacquelyn Wynn, Mairead Hyland, The Lighthouse Lab in Alderley Park and Alex Alderton, Roberto Amato, Jeffrey Barrett, Sonia Goncalves, Ewan Harrison, David K. Jackson, Ian Johnston, Dominic Kwiatkowski, Cordelia Langford, John Sillitoe on |

| | | | |
|---|---|---|---|
| EPI_ISL_3339167 | | Consortium | behalf of the Wellcome Sanger Institute COVID-19 Surveillance Team |
| EPI_ISL_3339168 | Lighthouse Lab in Milton Keynes | Wellcome Sanger Institute for the COVID-19 Genomics UK (COG-UK) Consortium | The Lighthouse Lab in Milton Keynes and Alex Alderton, Roberto Amato, Jeffrey Barrett, Sonia Goncalves, Ewan Harrison, David K. Jackson, Ian Johnston, Dominic Kwiatkowski, Cordelia Langford, John Sillitoe on behalf of the Wellcome Sanger Institute COVID-19 Surveillance Team |
| EPI_ISL_3339169 | SYNLAB | GIGA Medical Genomics | Keith Durkin, Maria Artesi, Bouchra Boujemla, Nathalie Renotte, Nadine Cambisano, Sébastien Bontems, Cécile Meex, Claire Gourzonès, Olivier Ek, Laurent Gillet, Marie-Pierre Hayette, Vincent Bours |
| EPI_ISL_3339170 | Lighthouse Lab in Alderley Park | Wellcome Sanger Institute for the COVID-19 Genomics UK (COG-UK) Consortium | Jacquelyn Wynn, Mairead Hyland, The Lighthouse Lab in Alderley Park and Alex Alderton, Roberto Amato, Jeffrey Barrett, Sonia Goncalves, Ewan Harrison, David K. Jackson, Ian Johnston, Dominic Kwiatkowski, Cordelia Langford, John Sillitoe on behalf of the Wellcome Sanger Institute COVID-19 Surveillance Team |
| EPI_ISL_3339171 | Lighthouse Lab in Milton Keynes | Wellcome Sanger Institute for the COVID-19 Genomics UK (COG-UK) Consortium | The Lighthouse Lab in Milton Keynes and Alex Alderton, Roberto Amato, Jeffrey Barrett, Sonia Goncalves, Ewan Harrison, David K. Jackson, Ian Johnston, Dominic Kwiatkowski, Cordelia Langford, John Sillitoe on behalf of the Wellcome Sanger Institute COVID-19 Surveillance Team |
| EPI_ISL_3339172 | SYNLAB | GIGA Medical Genomics | Keith Durkin, Maria Artesi, Bouchra Boujemla, Nathalie Renotte, Nadine Cambisano, Sébastien Bontems, Cécile Meex, Claire Gourzonès, Olivier Ek, Laurent Gillet, Marie-Pierre Hayette, Vincent Bours |
| EPI_ISL_3339173, EPI_ISL_3339174 | Lighthouse Lab in Alderley Park | Wellcome Sanger Institute for the COVID-19 Genomics UK (COG-UK) Consortium | Jacquelyn Wynn, Mairead Hyland, The Lighthouse Lab in Alderley Park and Alex Alderton, Roberto Amato, Jeffrey Barrett, Sonia Goncalves, Ewan Harrison, David K. Jackson, Ian Johnston, Dominic Kwiatkowski, Cordelia Langford, John Sillitoe on behalf of the Wellcome Sanger Institute COVID-19 Surveillance Team |
| EPI_ISL_3339175 | Lighthouse Lab in Milton Keynes | Wellcome Sanger Institute for the COVID-19 Genomics UK (COG-UK) Consortium | The Lighthouse Lab in Milton Keynes and Alex Alderton, Roberto Amato, Jeffrey Barrett, Sonia Goncalves, Ewan Harrison, David K. Jackson, Ian Johnston, Dominic Kwiatkowski, Cordelia Langford, John Sillitoe on behalf of the Wellcome Sanger Institute COVID-19 Surveillance Team |
| EPI_ISL_3339176 | SYNLAB | GIGA Medical Genomics | Keith Durkin, Maria Artesi, Bouchra Boujemla, Nathalie Renotte, Nadine Cambisano, Sébastien Bontems, Cécile Meex, Claire Gourzonès, Olivier Ek, Laurent Gillet, Marie-Pierre Hayette, Vincent Bours |
| EPI_ISL_3339177 | Lighthouse Lab in Milton Keynes | Wellcome Sanger Institute for the COVID-19 Genomics UK (COG-UK) Consortium | The Lighthouse Lab in Milton Keynes and Alex Alderton, Roberto Amato, Jeffrey Barrett, Sonia Goncalves, Ewan Harrison, David K. Jackson, Ian Johnston, Dominic Kwiatkowski, Cordelia Langford, John Sillitoe on behalf of the Wellcome Sanger Institute COVID-19 Surveillance Team |
| EPI_ISL_3339178 | Lighthouse Lab in Alderley Park | Wellcome Sanger Institute for the COVID-19 Genomics UK (COG-UK) Consortium | Jacquelyn Wynn, Mairead Hyland, The Lighthouse Lab in Alderley Park and Alex Alderton, Roberto Amato, Jeffrey Barrett, Sonia Goncalves, Ewan Harrison, David K. Jackson, Ian Johnston, Dominic Kwiatkowski, Cordelia Langford, John Sillitoe on behalf of the Wellcome Sanger Institute COVID-19 Surveillance Team |
| EPI_ISL_3339179 | SYNLAB | GIGA Medical Genomics | Keith Durkin, Maria Artesi, Bouchra Boujemla, Nathalie Renotte, Nadine Cambisano, Sébastien Bontems, Cécile Meex, Claire Gourzonès, Olivier Ek, Laurent Gillet, Marie-Pierre Hayette, Vincent Bours |
| EPI_ISL_3339180 | Lighthouse Lab in Alderley Park | Wellcome Sanger Institute for the COVID-19 Genomics UK (COG-UK) Consortium | Jacquelyn Wynn, Mairead Hyland, The Lighthouse Lab in Alderley Park and Alex Alderton, Roberto Amato, Jeffrey Barrett, Sonia Goncalves, Ewan Harrison, David K. Jackson, Ian Johnston, Dominic Kwiatkowski, Cordelia Langford, John Sillitoe on behalf of the Wellcome Sanger Institute COVID-19 Surveillance Team |
| EPI_ISL_3339181, EPI_ISL_3339182 | Lighthouse Lab in Milton Keynes | Wellcome Sanger Institute for the COVID-19 Genomics UK (COG-UK) Consortium | The Lighthouse Lab in Milton Keynes and Alex Alderton, Roberto Amato, Jeffrey Barrett, Sonia Goncalves, Ewan Harrison, David K. Jackson, Ian Johnston, Dominic Kwiatkowski, Cordelia Langford, John Sillitoe on behalf of the Wellcome Sanger Institute COVID-19 Surveillance Team |
| EPI_ISL_3339183 | SYNLAB | GIGA Medical Genomics | Keith Durkin, Maria Artesi, Bouchra Boujemla, Nathalie Renotte, Nadine Cambisano, Sébastien Bontems, Cécile Meex, Claire Gourzonès, Olivier Ek, Laurent Gillet, Marie-Pierre Hayette, Vincent Bours |
| EPI_ISL_3339184, EPI_ISL_3339185 | Lighthouse Lab in Milton Keynes | Wellcome Sanger Institute for the COVID-19 Genomics UK (COG-UK) Consortium | The Lighthouse Lab in Milton Keynes and Alex Alderton, Roberto Amato, Jeffrey Barrett, Sonia Goncalves, Ewan Harrison, David K. Jackson, Ian Johnston, Dominic Kwiatkowski, Cordelia Langford, John Sillitoe on behalf of the Wellcome Sanger Institute COVID-19 Surveillance Team |
| EPI_ISL_3339186 | SYNLAB | GIGA Medical Genomics | Keith Durkin, Maria Artesi, Bouchra Boujemla, Nathalie Renotte, Nadine Cambisano, Sébastien Bontems, Cécile Meex, Claire Gourzonès, Olivier Ek, Laurent Gillet, Marie-Pierre Hayette, Vincent Bours |
| EPI_ISL_3339187 | Lighthouse Laboratory Plymouth | Wellcome Sanger Institute for the COVID-19 Genomics UK (COG-UK) Consortium | Lighthouse Laboratory Plymouth and Alex Alderton, Roberto Amato, Jeffrey Barrett, Sonia Goncalves, Ewan Harrison, David K. Jackson, Ian Johnston, Dominic Kwiatkowski, Cordelia Langford, John Sillitoe on behalf of the Wellcome Sanger Institute COVID-19 Surveillance Team |
| EPI_ISL_3339188, EPI_ISL_3339189 | Lighthouse Lab in Alderley Park | Wellcome Sanger Institute for the COVID-19 Genomics UK (COG-UK) Consortium | Jacquelyn Wynn, Mairead Hyland, The Lighthouse Lab in Alderley Park and Alex Alderton, Roberto Amato, Jeffrey Barrett, Sonia Goncalves, Ewan Harrison, David K. Jackson, Ian Johnston, Dominic Kwiatkowski, Cordelia Langford, John Sillitoe on behalf of the Wellcome Sanger Institute COVID-19 Surveillance Team |
| EPI_ISL_3339190 | SYNLAB | GIGA Medical Genomics | Keith Durkin, Maria Artesi, Bouchra Boujemla, Nathalie Renotte, Nadine Cambisano, Sébastien Bontems, Cécile Meex, Claire Gourzonès, Olivier Ek, Laurent Gillet, Marie-Pierre Hayette, Vincent Bours |
| EPI_ISL_3339191, EPI_ISL_3339192 | Lighthouse Lab in Milton Keynes | Wellcome Sanger Institute for the COVID-19 Genomics UK (COG-UK) Consortium | The Lighthouse Lab in Milton Keynes and Alex Alderton, Roberto Amato, Jeffrey Barrett, Sonia Goncalves, Ewan Harrison, David K. Jackson, Ian Johnston, Dominic Kwiatkowski, Cordelia Langford, John Sillitoe on behalf of the Wellcome Sanger Institute COVID-19 Surveillance Team |
| EPI_ISL_3339193 | SYNLAB | GIGA Medical Genomics | Keith Durkin, Maria Artesi, Bouchra Boujemla, Nathalie Renotte, Nadine Cambisano, Sébastien Bontems, Cécile Meex, Claire Gourzonès, Olivier Ek, Laurent Gillet, Marie-Pierre Hayette, Vincent Bours |
| EPI_ISL_3339194, EPI_ISL_3339195 | Lighthouse Lab in Alderley Park | Wellcome Sanger Institute for the COVID-19 Genomics UK (COG-UK) Consortium | Jacquelyn Wynn, Mairead Hyland, The Lighthouse Lab in Alderley Park and Alex Alderton, Roberto Amato, Jeffrey Barrett, Sonia Goncalves, Ewan Harrison, David K. Jackson, Ian Johnston, Dominic Kwiatkowski, Cordelia Langford, John Sillitoe on behalf of the Wellcome Sanger Institute COVID-19 Surveillance Team |
| EPI_ISL_3339196 | SYNLAB | GIGA Medical Genomics | Keith Durkin, Maria Artesi, Bouchra Boujemla, Nathalie Renotte, Nadine Cambisano, Sébastien Bontems, Cécile Meex, Claire Gourzonès, Olivier Ek, Laurent Gillet, Marie-Pierre Hayette, Vincent Bours |
| EPI_ISL_3339197 | Lighthouse Lab in Alderley Park | Wellcome Sanger Institute for the COVID-19 Genomics UK (COG-UK) Consortium | Jacquelyn Wynn, Mairead Hyland, The Lighthouse Lab in Alderley Park and Alex Alderton, Roberto Amato, Jeffrey Barrett, Sonia Goncalves, Ewan Harrison, David K. Jackson, Ian Johnston, Dominic Kwiatkowski, Cordelia Langford, John Sillitoe on behalf of the Wellcome Sanger Institute COVID-19 Surveillance Team |
| EPI_ISL_3339198 | Lighthouse Lab in Milton Keynes | Wellcome Sanger Institute for the COVID-19 Genomics UK (COG-UK) Consortium | The Lighthouse Lab in Milton Keynes and Alex Alderton, Roberto Amato, Jeffrey Barrett, Sonia Goncalves, Ewan Harrison, David K. Jackson, Ian Johnston, Dominic Kwiatkowski, Cordelia Langford, John Sillitoe on behalf of the Wellcome Sanger Institute COVID-19 Surveillance Team |
| EPI_ISL_3339199 | SYNLAB | GIGA Medical Genomics | Keith Durkin, Maria Artesi, Bouchra Boujemla, Nathalie Renotte, Nadine Cambisano, Sébastien Bontems, Cécile Meex, Claire Gourzonès, Olivier Ek, Laurent Gillet, Marie-Pierre Hayette, Vincent Bours |
| EPI_ISL_3339200, EPI_ISL_3339201 | Lighthouse Lab in Alderley Park | Wellcome Sanger Institute for the COVID-19 Genomics UK (COG-UK) Consortium | Jacquelyn Wynn, Mairead Hyland, The Lighthouse Lab in Alderley Park and Alex Alderton, Roberto Amato, Jeffrey Barrett, Sonia Goncalves, Ewan Harrison, David K. Jackson, Ian Johnston, Dominic Kwiatkowski, Cordelia Langford, John Sillitoe on behalf of the Wellcome Sanger Institute COVID-19 Surveillance Team |
| EPI_ISL_3339202 | Lighthouse Laboratory Plymouth | Wellcome Sanger Institute for the COVID-19 Genomics UK (COG-UK) Consortium | Lighthouse Laboratory Plymouth and Alex Alderton, Roberto Amato, Jeffrey Barrett, Sonia Goncalves, Ewan Harrison, David K. Jackson, Ian Johnston, Dominic Kwiatkowski, Cordelia Langford, John Sillitoe on behalf of the Wellcome Sanger Institute COVID-19 Surveillance Team |
| EPI_ISL_3339203 | SYNLAB | GIGA Medical Genomics | Keith Durkin, Maria Artesi, Bouchra Boujemla, Nathalie Renotte, Nadine Cambisano, Sébastien Bontems, Cécile Meex, Claire Gourzonès, Olivier Ek, Laurent Gillet, Marie-Pierre Hayette, Vincent Bours |
| EPI_ISL_3339204 | Lighthouse Laboratory Plymouth | Wellcome Sanger Institute for the COVID-19 Genomics UK (COG-UK) Consortium | Lighthouse Laboratory Plymouth and Alex Alderton, Roberto Amato, Jeffrey Barrett, Sonia Goncalves, Ewan Harrison, David K. Jackson, Ian Johnston, Dominic Kwiatkowski, Cordelia Langford, John Sillitoe on behalf of the Wellcome Sanger Institute COVID-19 Surveillance Team |
| EPI_ISL_3339205 | Lighthouse Lab in Milton Keynes | Wellcome Sanger Institute for the COVID-19 Genomics UK (COG-UK) Consortium | The Lighthouse Lab in Milton Keynes and Alex Alderton, Roberto Amato, Jeffrey Barrett, Sonia Goncalves, Ewan Harrison, David K. Jackson, Ian Johnston, Dominic Kwiatkowski, Cordelia Langford, John Sillitoe on behalf of the Wellcome Sanger Institute COVID-19 Surveillance Team |
| EPI_ISL_3339206 | SYNLAB | GIGA Medical Genomics | Keith Durkin, Maria Artesi, Bouchra Boujemla, Nathalie Renotte, Nadine Cambisano, Sébastien Bontems, Cécile Meex, Claire Gourzonès, Olivier Ek, Laurent Gillet, Marie-Pierre Hayette, Vincent Bours |
| EPI_ISL_3339207, EPI_ISL_3339208 | Lighthouse Lab in Milton Keynes | Wellcome Sanger Institute for the COVID-19 Genomics UK (COG-UK) Consortium | The Lighthouse Lab in Milton Keynes and Alex Alderton, Roberto Amato, Jeffrey Barrett, Sonia Goncalves, Ewan Harrison, David K. Jackson, Ian Johnston, Dominic Kwiatkowski, Cordelia Langford, John Sillitoe on behalf of the Wellcome Sanger Institute COVID-19 Surveillance Team |
| EPI_ISL_3339209 | SYNLAB | GIGA Medical Genomics | Keith Durkin, Maria Artesi, Bouchra Boujemla, Nathalie Renotte, Nadine Cambisano, Sébastien Bontems, Cécile Meex, Claire Gourzonès, Olivier Ek, Laurent Gillet, Marie-Pierre Hayette, Vincent Bours |
| EPI_ISL_3339210, EPI_ISL_3339211 | Lighthouse Laboratory Plymouth | Wellcome Sanger Institute for the COVID-19 Genomics UK (COG-UK) Consortium | Lighthouse Laboratory Plymouth and Alex Alderton, Roberto Amato, Jeffrey Barrett, Sonia Goncalves, Ewan Harrison, David K. Jackson, Ian Johnston, Dominic Kwiatkowski, Cordelia Langford, John Sillitoe on behalf of the Wellcome Sanger Institute COVID-19 Surveillance Team |
| EPI_ISL_3339212 | SYNLAB | GIGA Medical Genomics | Keith Durkin, Maria Artesi, Bouchra Boujemla, Nathalie Renotte, Nadine Cambisano, Sébastien Bontems, Cécile Meex, Claire Gourzonès, Olivier Ek, Laurent Gillet, Marie-Pierre Hayette, Vincent Bours |
| EPI_ISL_3339213 | Lighthouse Laboratory Plymouth | Wellcome Sanger Institute for the COVID-19 Genomics UK (COG-UK) Consortium | Lighthouse Laboratory Plymouth and Alex Alderton, Roberto Amato, Jeffrey Barrett, Sonia Goncalves, Ewan Harrison, David K. Jackson, Ian Johnston, Dominic Kwiatkowski, Cordelia Langford, John Sillitoe on behalf of the Wellcome Sanger Institute COVID-19 Surveillance Team |
| EPI_ISL_3339214 | Lighthouse Lab in Milton Keynes | Wellcome Sanger Institute for the COVID-19 Genomics UK (COG-UK) Consortium | The Lighthouse Lab in Milton Keynes and Alex Alderton, Roberto Amato, Jeffrey Barrett, Sonia Goncalves, Ewan Harrison, David K. Jackson, Ian Johnston, Dominic Kwiatkowski, Cordelia Langford, John Sillitoe on behalf of the Wellcome Sanger Institute COVID-19 Surveillance Team |
| EPI_ISL_3339215 | SYNLAB | GIGA Medical Genomics | Keith Durkin, Maria Artesi, Bouchra Boujemla, Nathalie Renotte, Nadine Cambisano, Sébastien Bontems, Cécile Meex, Claire Gourzonès, Olivier Ek, Laurent Gillet, Marie-Pierre Hayette, Vincent Bours |
| EPI_ISL_3339216, EPI_ISL_3339217, EPI_ISL_3339218 | Lighthouse Lab in Milton Keynes | Wellcome Sanger Institute for the COVID-19 Genomics UK (COG-UK) Consortium | The Lighthouse Lab in Milton Keynes and Alex Alderton, Roberto Amato, Jeffrey Barrett, Sonia Goncalves, Ewan Harrison, David K. Jackson, Ian Johnston, Dominic Kwiatkowski, Cordelia Langford, John Sillitoe on behalf of the Wellcome Sanger Institute COVID-19 Surveillance Team |
| EPI_ISL_3339219 | SYNLAB | GIGA Medical Genomics | Keith Durkin, Maria Artesi, Bouchra Boujemla, Nathalie Renotte, Nadine Cambisano, Sébastien Bontems, Cécile Meex, Claire Gourzonès, Olivier Ek, Laurent Gillet, Marie-Pierre Hayette, Vincent Bours |
| EPI_ISL_3339220, EPI_ISL_3339221 | Lighthouse Lab in Milton Keynes | Wellcome Sanger Institute for the COVID-19 Genomics UK (COG-UK) Consortium | The Lighthouse Lab in Milton Keynes and Alex Alderton, Roberto Amato, Jeffrey Barrett, Sonia Goncalves, Ewan Harrison, David K. Jackson, Ian Johnston, Dominic Kwiatkowski, Cordelia Langford, John Sillitoe on behalf of the Wellcome Sanger Institute COVID-19 Surveillance Team |
| EPI_ISL_3339222 | Department of Clinical Microbiology | GIGA Medical Genomics | Keith Durkin, Maria Artesi, Bouchra Boujemla, Nathalie Renotte, Nadine Cambisano, Sébastien Bontems, Cécile Meex, Claire Gourzonès, Olivier Ek, Laurent Gillet, Marie-Pierre Hayette, Vincent Bours |
| EPI_ISL_3339223 | Lighthouse Lab in Milton Keynes | Wellcome Sanger Institute for the COVID-19 Genomics UK (COG-UK) Consortium | The Lighthouse Lab in Milton Keynes and Alex Alderton, Roberto Amato, Jeffrey Barrett, Sonia Goncalves, Ewan Harrison, David K. Jackson, Ian Johnston, Dominic Kwiatkowski, Cordelia Langford, John Sillitoe on behalf of the Wellcome Sanger Institute COVID-19 Surveillance Team |
| EPI_ISL_3339224 | Lighthouse Laboratory Plymouth | Wellcome Sanger Institute for the COVID-19 Genomics UK (COG-UK) Consortium | Lighthouse Laboratory Plymouth and Alex Alderton, Roberto Amato, Jeffrey Barrett, Sonia Goncalves, Ewan Harrison, David K. Jackson, Ian Johnston, Dominic Kwiatkowski, Cordelia Langford, John Sillitoe on behalf of the Wellcome Sanger Institute COVID-19 Surveillance Team |
| EPI_ISL_3339225 | Lighthouse Lab in Alderley Park | Wellcome Sanger Institute for the COVID-19 Genomics UK (COG-UK) Consortium | Jacquelyn Wynn, Mairead Hyland, The Lighthouse Lab in Alderley Park and Alex Alderton, Roberto Amato, Jeffrey Barrett, Sonia Goncalves, Ewan Harrison, David K. Jackson, Ian Johnston, Dominic Kwiatkowski, Cordelia Langford, John Sillitoe on behalf of the Wellcome Sanger Institute COVID-19 Surveillance Team |
| EPI_ISL_3339226, EPI_ISL_3339227 | Lighthouse Lab in Milton Keynes | Wellcome Sanger Institute for the COVID-19 Genomics UK (COG-UK) Consortium | The Lighthouse Lab in Milton Keynes and Alex Alderton, Roberto Amato, Jeffrey Barrett, Sonia Goncalves, Ewan Harrison, David K. Jackson, Ian Johnston, Dominic Kwiatkowski, Cordelia Langford, John Sillitoe on behalf of the Wellcome Sanger Institute COVID-19 Surveillance Team |
| EPI_ISL_3339228 | Lighthouse Laboratory Plymouth | Wellcome Sanger Institute for the COVID-19 Genomics UK (COG-UK) Consortium | Lighthouse Laboratory Plymouth and Alex Alderton, Roberto Amato, Jeffrey Barrett, Sonia Goncalves, Ewan Harrison, David K. Jackson, Ian Johnston, Dominic Kwiatkowski, Cordelia Langford, John Sillitoe on behalf of the Wellcome Sanger Institute COVID-19 Surveillance Team |
| EPI_ISL_3339229, EPI_ISL_3339230, EPI_ISL_3339231 | Lighthouse Lab in Milton Keynes | Wellcome Sanger Institute for the COVID-19 Genomics UK (COG-UK) Consortium | The Lighthouse Lab in Milton Keynes and Alex Alderton, Roberto Amato, Jeffrey Barrett, Sonia Goncalves, Ewan Harrison, David K. Jackson, Ian Johnston, Dominic Kwiatkowski, Cordelia Langford, John Sillitoe on behalf of the Wellcome Sanger Institute COVID-19 Surveillance Team |
| EPI_ISL_3339232 | Lighthouse Laboratory Plymouth | Wellcome Sanger Institute for the COVID-19 Genomics UK (COG-UK) Consortium | Lighthouse Laboratory Plymouth and Alex Alderton, Roberto Amato, Jeffrey Barrett, Sonia Goncalves, Ewan Harrison, David K. Jackson, Ian Johnston, Dominic Kwiatkowski, Cordelia Langford, John Sillitoe on behalf of the Wellcome Sanger Institute COVID-19 Surveillance Team |
| EPI_ISL_3339233, | Lighthouse Lab in Alderley Park | Wellcome Sanger Institute for the COVID-19 Genomics UK (COG-UK) | Jacquelyn Wynn, Mairead Hyland, The Lighthouse Lab in Alderley Park and Alex Alderton, Roberto Amato, Jeffrey Barrett, Sonia Goncalves, Ewan Harrison, David K. Jackson, Ian Johnston, Dominic Kwiatkowski, Cordelia Langford, John Sillitoe on |

| | | | |
|---|---|---|---|
| EPI_ISL_3339234 | | Consortium | behalf of the Wellcome Sanger Institute COVID-19 Surveillance Team |
| EPI_ISL_3339235 | Lighthouse Laboratory Plymouth | Wellcome Sanger Institute for the COVID-19 Genomics UK (COG-UK) Consortium | Lighthouse Laboratory Plymouth and Alex Alderton, Roberto Amato, Jeffrey Barrett, Sonia Goncalves, Ewan Harrison, David K. Jackson, Ian Johnston, Dominic Kwiatkowski, Cordelia Langford, John Sillitoe on behalf of the Wellcome Sanger Institute COVID-19 Surveillance Team |
| EPI_ISL_3339236 | Lighthouse Lab in Milton Keynes | Wellcome Sanger Institute for the COVID-19 Genomics UK (COG-UK) Consortium | The Lighthouse Lab in Milton Keynes and Alex Alderton, Roberto Amato, Jeffrey Barrett, Sonia Goncalves, Ewan Harrison, David K. Jackson, Ian Johnston, Dominic Kwiatkowski, Cordelia Langford, John Sillitoe on behalf of the Wellcome Sanger Institute COVID-19 Surveillance Team |
| EPI_ISL_3339237, EPI_ISL_3339238 | Lighthouse Laboratory Plymouth | Wellcome Sanger Institute for the COVID-19 Genomics UK (COG-UK) Consortium | Lighthouse Laboratory Plymouth and Alex Alderton, Roberto Amato, Jeffrey Barrett, Sonia Goncalves, Ewan Harrison, David K. Jackson, Ian Johnston, Dominic Kwiatkowski, Cordelia Langford, John Sillitoe on behalf of the Wellcome Sanger Institute COVID-19 Surveillance Team |
| EPI_ISL_3339239, EPI_ISL_3339240, EPI_ISL_3339241, EPI_ISL_3339242 | Lighthouse Lab in Milton Keynes | Wellcome Sanger Institute for the COVID-19 Genomics UK (COG-UK) Consortium | The Lighthouse Lab in Milton Keynes and Alex Alderton, Roberto Amato, Jeffrey Barrett, Sonia Goncalves, Ewan Harrison, David K. Jackson, Ian Johnston, Dominic Kwiatkowski, Cordelia Langford, John Sillitoe on behalf of the Wellcome Sanger Institute COVID-19 Surveillance Team |
| EPI_ISL_3339243 | Lighthouse Laboratory Plymouth | Wellcome Sanger Institute for the COVID-19 Genomics UK (COG-UK) Consortium | Lighthouse Laboratory Plymouth and Alex Alderton, Roberto Amato, Jeffrey Barrett, Sonia Goncalves, Ewan Harrison, David K. Jackson, Ian Johnston, Dominic Kwiatkowski, Cordelia Langford, John Sillitoe on behalf of the Wellcome Sanger Institute COVID-19 Surveillance Team |
| EPI_ISL_3339244 | Lighthouse Lab in Alderley Park | Wellcome Sanger Institute for the COVID-19 Genomics UK (COG-UK) Consortium | Jacquelyn Wynn, Mairead Hyland, The Lighthouse Lab in Alderley Park and Alex Alderton, Roberto Amato, Jeffrey Barrett, Sonia Goncalves, Ewan Harrison, David K. Jackson, Ian Johnston, Dominic Kwiatkowski, Cordelia Langford, John Sillitoe on behalf of the Wellcome Sanger Institute COVID-19 Surveillance Team |
| EPI_ISL_3339245, EPI_ISL_3339246, EPI_ISL_3339247, EPI_ISL_3339248, EPI_ISL_3339249 | Lighthouse Lab in Milton Keynes | Wellcome Sanger Institute for the COVID-19 Genomics UK (COG-UK) Consortium | The Lighthouse Lab in Milton Keynes and Alex Alderton, Roberto Amato, Jeffrey Barrett, Sonia Goncalves, Ewan Harrison, David K. Jackson, Ian Johnston, Dominic Kwiatkowski, Cordelia Langford, John Sillitoe on behalf of the Wellcome Sanger Institute COVID-19 Surveillance Team |
| EPI_ISL_3339250 | Lighthouse Lab in Alderley Park | Wellcome Sanger Institute for the COVID-19 Genomics UK (COG-UK) Consortium | Jacquelyn Wynn, Mairead Hyland, The Lighthouse Lab in Alderley Park and Alex Alderton, Roberto Amato, Jeffrey Barrett, Sonia Goncalves, Ewan Harrison, David K. Jackson, Ian Johnston, Dominic Kwiatkowski, Cordelia Langford, John Sillitoe on behalf of the Wellcome Sanger Institute COVID-19 Surveillance Team |
| EPI_ISL_3339251 | Lighthouse Lab in Milton Keynes | Wellcome Sanger Institute for the COVID-19 Genomics UK (COG-UK) Consortium | The Lighthouse Lab in Milton Keynes and Alex Alderton, Roberto Amato, Jeffrey Barrett, Sonia Goncalves, Ewan Harrison, David K. Jackson, Ian Johnston, Dominic Kwiatkowski, Cordelia Langford, John Sillitoe on behalf of the Wellcome Sanger Institute COVID-19 Surveillance Team |
| EPI_ISL_3339252 | Lighthouse Lab in Alderley Park | Wellcome Sanger Institute for the COVID-19 Genomics UK (COG-UK) Consortium | Jacquelyn Wynn, Mairead Hyland, The Lighthouse Lab in Alderley Park and Alex Alderton, Roberto Amato, Jeffrey Barrett, Sonia Goncalves, Ewan Harrison, David K. Jackson, Ian Johnston, Dominic Kwiatkowski, Cordelia Langford, John Sillitoe on behalf of the Wellcome Sanger Institute COVID-19 Surveillance Team |
| EPI_ISL_3339253, EPI_ISL_3339254 | Lighthouse Lab in Milton Keynes | Wellcome Sanger Institute for the COVID-19 Genomics UK (COG-UK) Consortium | The Lighthouse Lab in Milton Keynes and Alex Alderton, Roberto Amato, Jeffrey Barrett, Sonia Goncalves, Ewan Harrison, David K. Jackson, Ian Johnston, Dominic Kwiatkowski, Cordelia Langford, John Sillitoe on behalf of the Wellcome Sanger Institute COVID-19 Surveillance Team |
| EPI_ISL_3339255, EPI_ISL_3339256, EPI_ISL_3339257 | Lighthouse Lab in Alderley Park | Wellcome Sanger Institute for the COVID-19 Genomics UK (COG-UK) Consortium | Jacquelyn Wynn, Mairead Hyland, The Lighthouse Lab in Alderley Park and Alex Alderton, Roberto Amato, Jeffrey Barrett, Sonia Goncalves, Ewan Harrison, David K. Jackson, Ian Johnston, Dominic Kwiatkowski, Cordelia Langford, John Sillitoe on behalf of the Wellcome Sanger Institute COVID-19 Surveillance Team |
| EPI_ISL_3339258 | Lighthouse Lab in Milton Keynes | Wellcome Sanger Institute for the COVID-19 Genomics UK (COG-UK) Consortium | The Lighthouse Lab in Milton Keynes and Alex Alderton, Roberto Amato, Jeffrey Barrett, Sonia Goncalves, Ewan Harrison, David K. Jackson, Ian Johnston, Dominic Kwiatkowski, Cordelia Langford, John Sillitoe on behalf of the Wellcome Sanger Institute COVID-19 Surveillance Team |
| EPI_ISL_3339259 | Lighthouse Lab in Alderley Park | Wellcome Sanger Institute for the COVID-19 Genomics UK (COG-UK) Consortium | Jacquelyn Wynn, Mairead Hyland, The Lighthouse Lab in Alderley Park and Alex Alderton, Roberto Amato, Jeffrey Barrett, Sonia Goncalves, Ewan Harrison, David K. Jackson, Ian Johnston, Dominic Kwiatkowski, Cordelia Langford, John Sillitoe on behalf of the Wellcome Sanger Institute COVID-19 Surveillance Team |
| EPI_ISL_3339260 | Lighthouse Lab in Milton Keynes | Wellcome Sanger Institute for the COVID-19 Genomics UK (COG-UK) Consortium | The Lighthouse Lab in Milton Keynes and Alex Alderton, Roberto Amato, Jeffrey Barrett, Sonia Goncalves, Ewan Harrison, David K. Jackson, Ian Johnston, Dominic Kwiatkowski, Cordelia Langford, John Sillitoe on behalf of the Wellcome Sanger Institute COVID-19 Surveillance Team |
| EPI_ISL_3339261, EPI_ISL_3339262, EPI_ISL_3339263, EPI_ISL_3339264, EPI_ISL_3339265 | Lighthouse Laboratory Plymouth | Wellcome Sanger Institute for the COVID-19 Genomics UK (COG-UK) Consortium | Lighthouse Laboratory Plymouth and Alex Alderton, Roberto Amato, Jeffrey Barrett, Sonia Goncalves, Ewan Harrison, David K. Jackson, Ian Johnston, Dominic Kwiatkowski, Cordelia Langford, John Sillitoe on behalf of the Wellcome Sanger Institute COVID-19 Surveillance Team |
| EPI_ISL_3339266 | Lighthouse Lab in Alderley Park | Wellcome Sanger Institute for the COVID-19 Genomics UK (COG-UK) Consortium | Jacquelyn Wynn, Mairead Hyland, The Lighthouse Lab in Alderley Park and Alex Alderton, Roberto Amato, Jeffrey Barrett, Sonia Goncalves, Ewan Harrison, David K. Jackson, Ian Johnston, Dominic Kwiatkowski, Cordelia Langford, John Sillitoe on behalf of the Wellcome Sanger Institute COVID-19 Surveillance Team |
| EPI_ISL_3339267, EPI_ISL_3339268 | Lighthouse Lab in Milton Keynes | Wellcome Sanger Institute for the COVID-19 Genomics UK (COG-UK) Consortium | The Lighthouse Lab in Milton Keynes and Alex Alderton, Roberto Amato, Jeffrey Barrett, Sonia Goncalves, Ewan Harrison, David K. Jackson, Ian Johnston, Dominic Kwiatkowski, Cordelia Langford, John Sillitoe on behalf of the Wellcome Sanger Institute COVID-19 Surveillance Team |
| EPI_ISL_3339269 | Lighthouse Laboratory Plymouth | Wellcome Sanger Institute for the COVID-19 Genomics UK (COG-UK) Consortium | Lighthouse Laboratory Plymouth and Alex Alderton, Roberto Amato, Jeffrey Barrett, Sonia Goncalves, Ewan Harrison, David K. Jackson, Ian Johnston, Dominic Kwiatkowski, Cordelia Langford, John Sillitoe on behalf of the Wellcome Sanger Institute COVID-19 Surveillance Team |
| EPI_ISL_3339270, EPI_ISL_3339271, EPI_ISL_3339272, EPI_ISL_3339273 | Lighthouse Lab in Milton Keynes | Wellcome Sanger Institute for the COVID-19 Genomics UK (COG-UK) Consortium | The Lighthouse Lab in Milton Keynes and Alex Alderton, Roberto Amato, Jeffrey Barrett, Sonia Goncalves, Ewan Harrison, David K. Jackson, Ian Johnston, Dominic Kwiatkowski, Cordelia Langford, John Sillitoe on behalf of the Wellcome Sanger Institute COVID-19 Surveillance Team |
| EPI_ISL_3339274 | Lighthouse Lab in Alderley Park | Wellcome Sanger Institute for the COVID-19 Genomics UK (COG-UK) Consortium | Jacquelyn Wynn, Mairead Hyland, The Lighthouse Lab in Alderley Park and Alex Alderton, Roberto Amato, Jeffrey Barrett, Sonia Goncalves, Ewan Harrison, David K. Jackson, Ian Johnston, Dominic Kwiatkowski, Cordelia Langford, John Sillitoe on behalf of the Wellcome Sanger Institute COVID-19 Surveillance Team |
| EPI_ISL_3339275 | Lighthouse Lab in Milton Keynes | Wellcome Sanger Institute for the COVID-19 Genomics UK (COG-UK) Consortium | The Lighthouse Lab in Milton Keynes and Alex Alderton, Roberto Amato, Jeffrey Barrett, Sonia Goncalves, Ewan Harrison, David K. Jackson, Ian Johnston, Dominic Kwiatkowski, Cordelia Langford, John Sillitoe on behalf of the Wellcome Sanger Institute COVID-19 Surveillance Team |
| EPI_ISL_3339276 | Lighthouse Laboratory Plymouth | Wellcome Sanger Institute for the COVID-19 Genomics UK (COG-UK) Consortium | Lighthouse Laboratory Plymouth and Alex Alderton, Roberto Amato, Jeffrey Barrett, Sonia Goncalves, Ewan Harrison, David K. Jackson, Ian Johnston, Dominic Kwiatkowski, Cordelia Langford, John Sillitoe on behalf of the Wellcome Sanger Institute COVID-19 Surveillance Team |
| EPI_ISL_3339277 | Lighthouse Lab in Milton Keynes | Wellcome Sanger Institute for the COVID-19 Genomics UK (COG-UK) Consortium | The Lighthouse Lab in Milton Keynes and Alex Alderton, Roberto Amato, Jeffrey Barrett, Sonia Goncalves, Ewan Harrison, David K. Jackson, Ian Johnston, Dominic Kwiatkowski, Cordelia Langford, John Sillitoe on behalf of the Wellcome Sanger Institute COVID-19 Surveillance Team |
| EPI_ISL_3339278 | Lighthouse Lab in Alderley Park | Wellcome Sanger Institute for the COVID-19 Genomics UK (COG-UK) Consortium | Jacquelyn Wynn, Mairead Hyland, The Lighthouse Lab in Alderley Park and Alex Alderton, Roberto Amato, Jeffrey Barrett, Sonia Goncalves, Ewan Harrison, David K. Jackson, Ian Johnston, Dominic Kwiatkowski, Cordelia Langford, John Sillitoe on behalf of the Wellcome Sanger Institute COVID-19 Surveillance Team |
| EPI_ISL_3339279 | Lighthouse Laboratory Plymouth | Wellcome Sanger Institute for the COVID-19 Genomics UK (COG-UK) Consortium | Lighthouse Laboratory Plymouth and Alex Alderton, Roberto Amato, Jeffrey Barrett, Sonia Goncalves, Ewan Harrison, David K. Jackson, Ian Johnston, Dominic Kwiatkowski, Cordelia Langford, John Sillitoe on behalf of the Wellcome Sanger Institute COVID-19 Surveillance Team |
| EPI_ISL_3339280 | Lighthouse Lab in Milton Keynes | Wellcome Sanger Institute for the COVID-19 Genomics UK (COG-UK) Consortium | The Lighthouse Lab in Milton Keynes and Alex Alderton, Roberto Amato, Jeffrey Barrett, Sonia Goncalves, Ewan Harrison, David K. Jackson, Ian Johnston, Dominic Kwiatkowski, Cordelia Langford, John Sillitoe on behalf of the Wellcome Sanger Institute COVID-19 Surveillance Team |
| EPI_ISL_3339281 | Lighthouse Lab in Alderley Park | Wellcome Sanger Institute for the COVID-19 Genomics UK (COG-UK) Consortium | Jacquelyn Wynn, Mairead Hyland, The Lighthouse Lab in Alderley Park and Alex Alderton, Roberto Amato, Jeffrey Barrett, Sonia Goncalves, Ewan Harrison, David K. Jackson, Ian Johnston, Dominic Kwiatkowski, Cordelia Langford, John Sillitoe on behalf of the Wellcome Sanger Institute COVID-19 Surveillance Team |
| EPI_ISL_3339282, EPI_ISL_3339283, EPI_ISL_3339284 | Lighthouse Lab in Milton Keynes | Wellcome Sanger Institute for the COVID-19 Genomics UK (COG-UK) Consortium | The Lighthouse Lab in Milton Keynes and Alex Alderton, Roberto Amato, Jeffrey Barrett, Sonia Goncalves, Ewan Harrison, David K. Jackson, Ian Johnston, Dominic Kwiatkowski, Cordelia Langford, John Sillitoe on behalf of the Wellcome Sanger Institute COVID-19 Surveillance Team |
| EPI_ISL_3339285 | Lighthouse Lab in Alderley Park | Wellcome Sanger Institute for the COVID-19 Genomics UK (COG-UK) Consortium | Jacquelyn Wynn, Mairead Hyland, The Lighthouse Lab in Alderley Park and Alex Alderton, Roberto Amato, Jeffrey Barrett, Sonia Goncalves, Ewan Harrison, David K. Jackson, Ian Johnston, Dominic Kwiatkowski, Cordelia Langford, John Sillitoe on behalf of the Wellcome Sanger Institute COVID-19 Surveillance Team |
| EPI_ISL_3339286 | Lighthouse Lab in Milton Keynes | Wellcome Sanger Institute for the COVID-19 Genomics UK (COG-UK) Consortium | The Lighthouse Lab in Milton Keynes and Alex Alderton, Roberto Amato, Jeffrey Barrett, Sonia Goncalves, Ewan Harrison, David K. Jackson, Ian Johnston, Dominic Kwiatkowski, Cordelia Langford, John Sillitoe on behalf of the Wellcome Sanger Institute COVID-19 Surveillance Team |
| EPI_ISL_3339287 | Lighthouse Lab in Alderley Park | Wellcome Sanger Institute for the COVID-19 Genomics UK (COG-UK) Consortium | Jacquelyn Wynn, Mairead Hyland, The Lighthouse Lab in Alderley Park and Alex Alderton, Roberto Amato, Jeffrey Barrett, Sonia Goncalves, Ewan Harrison, David K. Jackson, Ian Johnston, Dominic Kwiatkowski, Cordelia Langford, John Sillitoe on behalf of the Wellcome Sanger Institute COVID-19 Surveillance Team |
| EPI_ISL_3339288, EPI_ISL_3339289, EPI_ISL_3339290 | Lighthouse Lab in Milton Keynes | Wellcome Sanger Institute for the COVID-19 Genomics UK (COG-UK) Consortium | The Lighthouse Lab in Milton Keynes and Alex Alderton, Roberto Amato, Jeffrey Barrett, Sonia Goncalves, Ewan Harrison, David K. Jackson, Ian Johnston, Dominic Kwiatkowski, Cordelia Langford, John Sillitoe on behalf of the Wellcome Sanger Institute COVID-19 Surveillance Team |
| EPI_ISL_3339291 | Lighthouse Laboratory Plymouth | Wellcome Sanger Institute for the COVID-19 Genomics UK (COG-UK) Consortium | Lighthouse Laboratory Plymouth and Alex Alderton, Roberto Amato, Jeffrey Barrett, Sonia Goncalves, Ewan Harrison, David K. Jackson, Ian Johnston, Dominic Kwiatkowski, Cordelia Langford, John Sillitoe on behalf of the Wellcome Sanger Institute COVID-19 Surveillance Team |
| EPI_ISL_3339292 | Lighthouse Lab in Milton Keynes | Wellcome Sanger Institute for the COVID-19 Genomics UK (COG-UK) Consortium | The Lighthouse Lab in Milton Keynes and Alex Alderton, Roberto Amato, Jeffrey Barrett, Sonia Goncalves, Ewan Harrison, David K. Jackson, Ian Johnston, Dominic Kwiatkowski, Cordelia Langford, John Sillitoe on behalf of the Wellcome Sanger Institute COVID-19 Surveillance Team |
| EPI_ISL_3339293 | Lighthouse Lab in Alderley Park | Wellcome Sanger Institute for the COVID-19 Genomics UK (COG-UK) Consortium | Jacquelyn Wynn, Mairead Hyland, The Lighthouse Lab in Alderley Park and Alex Alderton, Roberto Amato, Jeffrey Barrett, Sonia Goncalves, Ewan Harrison, David K. Jackson, Ian Johnston, Dominic Kwiatkowski, Cordelia Langford, John Sillitoe on behalf of the Wellcome Sanger Institute COVID-19 Surveillance Team |
| EPI_ISL_3339294 | Lighthouse Lab in Milton Keynes | Wellcome Sanger Institute for the COVID-19 Genomics UK (COG-UK) Consortium | The Lighthouse Lab in Milton Keynes and Alex Alderton, Roberto Amato, Jeffrey Barrett, Sonia Goncalves, Ewan Harrison, David K. Jackson, Ian Johnston, Dominic Kwiatkowski, Cordelia Langford, John Sillitoe on behalf of the Wellcome Sanger Institute COVID-19 Surveillance Team |
| EPI_ISL_3339295 | Lighthouse Laboratory Plymouth | Wellcome Sanger Institute for the COVID-19 Genomics UK (COG-UK) Consortium | Lighthouse Laboratory Plymouth and Alex Alderton, Roberto Amato, Jeffrey Barrett, Sonia Goncalves, Ewan Harrison, David K. Jackson, Ian Johnston, Dominic Kwiatkowski, Cordelia Langford, John Sillitoe on behalf of the Wellcome Sanger Institute COVID-19 Surveillance Team |
| EPI_ISL_3339296 | Lighthouse Lab in Alderley Park | Wellcome Sanger Institute for the COVID-19 Genomics UK (COG-UK) Consortium | Jacquelyn Wynn, Mairead Hyland, The Lighthouse Lab in Alderley Park and Alex Alderton, Roberto Amato, Jeffrey Barrett, Sonia Goncalves, Ewan Harrison, David K. Jackson, Ian Johnston, Dominic Kwiatkowski, Cordelia Langford, John Sillitoe on behalf of the Wellcome Sanger Institute COVID-19 Surveillance Team |

| | | | |
|---|---|---|---|
| EPI_ISL_3339297, EPI_ISL_3339298 | Lighthouse Lab in Milton Keynes | Wellcome Sanger Institute for the COVID-19 Genomics UK (COG-UK) Consortium | The Lighthouse Lab in Milton Keynes and Alex Alderton, Roberto Amato, Jeffrey Barrett, Sonia Goncalves, Ewan Harrison, David K. Jackson, Ian Johnston, Dominic Kwiatkowski, Cordelia Langford, John Sillitoe on behalf of the Wellcome Sanger Institute COVID-19 Surveillance Team |
| EPI_ISL_3339299 | Lighthouse Laboratory Plymouth | Wellcome Sanger Institute for the COVID-19 Genomics UK (COG-UK) Consortium | Lighthouse Laboratory Plymouth and Alex Alderton, Roberto Amato, Jeffrey Barrett, Sonia Goncalves, Ewan Harrison, David K. Jackson, Ian Johnston, Dominic Kwiatkowski, Cordelia Langford, John Sillitoe on behalf of the Wellcome Sanger Institute COVID-19 Surveillance Team |
| EPI_ISL_3339300, EPI_ISL_3339301, EPI_ISL_3339302 | Lighthouse Lab in Milton Keynes | Wellcome Sanger Institute for the COVID-19 Genomics UK (COG-UK) Consortium | The Lighthouse Lab in Milton Keynes and Alex Alderton, Roberto Amato, Jeffrey Barrett, Sonia Goncalves, Ewan Harrison, David K. Jackson, Ian Johnston, Dominic Kwiatkowski, Cordelia Langford, John Sillitoe on behalf of the Wellcome Sanger Institute COVID-19 Surveillance Team |
| EPI_ISL_3339303 | Lighthouse Laboratory Plymouth | Wellcome Sanger Institute for the COVID-19 Genomics UK (COG-UK) Consortium | Lighthouse Laboratory Plymouth and Alex Alderton, Roberto Amato, Jeffrey Barrett, Sonia Goncalves, Ewan Harrison, David K. Jackson, Ian Johnston, Dominic Kwiatkowski, Cordelia Langford, John Sillitoe on behalf of the Wellcome Sanger Institute COVID-19 Surveillance Team |
| EPI_ISL_3339304, EPI_ISL_3339305, EPI_ISL_3339306 | Lighthouse Lab in Milton Keynes | Wellcome Sanger Institute for the COVID-19 Genomics UK (COG-UK) Consortium | The Lighthouse Lab in Milton Keynes and Alex Alderton, Roberto Amato, Jeffrey Barrett, Sonia Goncalves, Ewan Harrison, David K. Jackson, Ian Johnston, Dominic Kwiatkowski, Cordelia Langford, John Sillitoe on behalf of the Wellcome Sanger Institute COVID-19 Surveillance Team |
| EPI_ISL_3339307 | Lighthouse Lab in Alderley Park | Wellcome Sanger Institute for the COVID-19 Genomics UK (COG-UK) Consortium | Jacquelyn Wynn, Mairead Hyland, The Lighthouse Lab in Alderley Park and Alex Alderton, Roberto Amato, Jeffrey Barrett, Sonia Goncalves, Ewan Harrison, David K. Jackson, Ian Johnston, Dominic Kwiatkowski, Cordelia Langford, John Sillitoe on behalf of the Wellcome Sanger Institute COVID-19 Surveillance Team |
| EPI_ISL_3339308 | Lighthouse Laboratory Plymouth | Wellcome Sanger Institute for the COVID-19 Genomics UK (COG-UK) Consortium | Lighthouse Laboratory Plymouth and Alex Alderton, Roberto Amato, Jeffrey Barrett, Sonia Goncalves, Ewan Harrison, David K. Jackson, Ian Johnston, Dominic Kwiatkowski, Cordelia Langford, John Sillitoe on behalf of the Wellcome Sanger Institute COVID-19 Surveillance Team |
| EPI_ISL_3339309 | Lighthouse Lab in Alderley Park | Wellcome Sanger Institute for the COVID-19 Genomics UK (COG-UK) Consortium | Jacquelyn Wynn, Mairead Hyland, The Lighthouse Lab in Alderley Park and Alex Alderton, Roberto Amato, Jeffrey Barrett, Sonia Goncalves, Ewan Harrison, David K. Jackson, Ian Johnston, Dominic Kwiatkowski, Cordelia Langford, John Sillitoe on behalf of the Wellcome Sanger Institute COVID-19 Surveillance Team |
| EPI_ISL_3339310 | Lighthouse Laboratory Plymouth | Wellcome Sanger Institute for the COVID-19 Genomics UK (COG-UK) Consortium | Lighthouse Laboratory Plymouth and Alex Alderton, Roberto Amato, Jeffrey Barrett, Sonia Goncalves, Ewan Harrison, David K. Jackson, Ian Johnston, Dominic Kwiatkowski, Cordelia Langford, John Sillitoe on behalf of the Wellcome Sanger Institute COVID-19 Surveillance Team |
| EPI_ISL_3339311, EPI_ISL_3339312, EPI_ISL_3339313, EPI_ISL_3339314, EPI_ISL_3339315, EPI_ISL_3339316 | Lighthouse Lab in Milton Keynes | Wellcome Sanger Institute for the COVID-19 Genomics UK (COG-UK) Consortium | The Lighthouse Lab in Milton Keynes and Alex Alderton, Roberto Amato, Jeffrey Barrett, Sonia Goncalves, Ewan Harrison, David K. Jackson, Ian Johnston, Dominic Kwiatkowski, Cordelia Langford, John Sillitoe on behalf of the Wellcome Sanger Institute COVID-19 Surveillance Team |
| EPI_ISL_3339317 | Lighthouse Lab in Alderley Park | Wellcome Sanger Institute for the COVID-19 Genomics UK (COG-UK) Consortium | Jacquelyn Wynn, Mairead Hyland, The Lighthouse Lab in Alderley Park and Alex Alderton, Roberto Amato, Jeffrey Barrett, Sonia Goncalves, Ewan Harrison, David K. Jackson, Ian Johnston, Dominic Kwiatkowski, Cordelia Langford, John Sillitoe on behalf of the Wellcome Sanger Institute COVID-19 Surveillance Team |
| EPI_ISL_3339318, EPI_ISL_3339319 | Lighthouse Lab in Milton Keynes | Wellcome Sanger Institute for the COVID-19 Genomics UK (COG-UK) Consortium | The Lighthouse Lab in Milton Keynes and Alex Alderton, Roberto Amato, Jeffrey Barrett, Sonia Goncalves, Ewan Harrison, David K. Jackson, Ian Johnston, Dominic Kwiatkowski, Cordelia Langford, John Sillitoe on behalf of the Wellcome Sanger Institute COVID-19 Surveillance Team |
| EPI_ISL_3339320 | Lighthouse Laboratory Plymouth | Wellcome Sanger Institute for the COVID-19 Genomics UK (COG-UK) Consortium | Lighthouse Laboratory Plymouth and Alex Alderton, Roberto Amato, Jeffrey Barrett, Sonia Goncalves, Ewan Harrison, David K. Jackson, Ian Johnston, Dominic Kwiatkowski, Cordelia Langford, John Sillitoe on behalf of the Wellcome Sanger Institute COVID-19 Surveillance Team |
| EPI_ISL_3339321, EPI_ISL_3339322, EPI_ISL_3339323, EPI_ISL_3339324, EPI_ISL_3339325, EPI_ISL_3339326 | Lighthouse Lab in Milton Keynes | Wellcome Sanger Institute for the COVID-19 Genomics UK (COG-UK) Consortium | The Lighthouse Lab in Milton Keynes and Alex Alderton, Roberto Amato, Jeffrey Barrett, Sonia Goncalves, Ewan Harrison, David K. Jackson, Ian Johnston, Dominic Kwiatkowski, Cordelia Langford, John Sillitoe on behalf of the Wellcome Sanger Institute COVID-19 Surveillance Team |
| EPI_ISL_3339327 | Lighthouse Laboratory Plymouth | Wellcome Sanger Institute for the COVID-19 Genomics UK (COG-UK) Consortium | Lighthouse Laboratory Plymouth and Alex Alderton, Roberto Amato, Jeffrey Barrett, Sonia Goncalves, Ewan Harrison, David K. Jackson, Ian Johnston, Dominic Kwiatkowski, Cordelia Langford, John Sillitoe on behalf of the Wellcome Sanger Institute COVID-19 Surveillance Team |
| EPI_ISL_3339328 | Lighthouse Lab in Alderley Park | Wellcome Sanger Institute for the COVID-19 Genomics UK (COG-UK) Consortium | Jacquelyn Wynn, Mairead Hyland, The Lighthouse Lab in Alderley Park and Alex Alderton, Roberto Amato, Jeffrey Barrett, Sonia Goncalves, Ewan Harrison, David K. Jackson, Ian Johnston, Dominic Kwiatkowski, Cordelia Langford, John Sillitoe on behalf of the Wellcome Sanger Institute COVID-19 Surveillance Team |
| EPI_ISL_3339329 | Lighthouse Laboratory Plymouth | Wellcome Sanger Institute for the COVID-19 Genomics UK (COG-UK) Consortium | Lighthouse Laboratory Plymouth and Alex Alderton, Roberto Amato, Jeffrey Barrett, Sonia Goncalves, Ewan Harrison, David K. Jackson, Ian Johnston, Dominic Kwiatkowski, Cordelia Langford, John Sillitoe on behalf of the Wellcome Sanger Institute COVID-19 Surveillance Team |
| EPI_ISL_3339330, EPI_ISL_3339331 | Lighthouse Lab in Milton Keynes | Wellcome Sanger Institute for the COVID-19 Genomics UK (COG-UK) Consortium | The Lighthouse Lab in Milton Keynes and Alex Alderton, Roberto Amato, Jeffrey Barrett, Sonia Goncalves, Ewan Harrison, David K. Jackson, Ian Johnston, Dominic Kwiatkowski, Cordelia Langford, John Sillitoe on behalf of the Wellcome Sanger Institute COVID-19 Surveillance Team |
| EPI_ISL_3339332 | Lighthouse Lab in Alderley Park | Wellcome Sanger Institute for the COVID-19 Genomics UK (COG-UK) Consortium | Jacquelyn Wynn, Mairead Hyland, The Lighthouse Lab in Alderley Park and Alex Alderton, Roberto Amato, Jeffrey Barrett, Sonia Goncalves, Ewan Harrison, David K. Jackson, Ian Johnston, Dominic Kwiatkowski, Cordelia Langford, John Sillitoe on behalf of the Wellcome Sanger Institute COVID-19 Surveillance Team |
| EPI_ISL_3339333, EPI_ISL_3339334, EPI_ISL_3339335 | Lighthouse Lab in Milton Keynes | Wellcome Sanger Institute for the COVID-19 Genomics UK (COG-UK) Consortium | The Lighthouse Lab in Milton Keynes and Alex Alderton, Roberto Amato, Jeffrey Barrett, Sonia Goncalves, Ewan Harrison, David K. Jackson, Ian Johnston, Dominic Kwiatkowski, Cordelia Langford, John Sillitoe on behalf of the Wellcome Sanger Institute COVID-19 Surveillance Team |
| EPI_ISL_3339336 | Lighthouse Lab in Alderley Park | Wellcome Sanger Institute for the COVID-19 Genomics UK (COG-UK) Consortium | Jacquelyn Wynn, Mairead Hyland, The Lighthouse Lab in Alderley Park and Alex Alderton, Roberto Amato, Jeffrey Barrett, Sonia Goncalves, Ewan Harrison, David K. Jackson, Ian Johnston, Dominic Kwiatkowski, Cordelia Langford, John Sillitoe on behalf of the Wellcome Sanger Institute COVID-19 Surveillance Team |
| EPI_ISL_3339337 | Lighthouse Laboratory Plymouth | Wellcome Sanger Institute for the COVID-19 Genomics UK (COG-UK) Consortium | Lighthouse Laboratory Plymouth and Alex Alderton, Roberto Amato, Jeffrey Barrett, Sonia Goncalves, Ewan Harrison, David K. Jackson, Ian Johnston, Dominic Kwiatkowski, Cordelia Langford, John Sillitoe on behalf of the Wellcome Sanger Institute COVID-19 Surveillance Team |
| EPI_ISL_3339338 | Lighthouse Lab in Milton Keynes | Wellcome Sanger Institute for the COVID-19 Genomics UK (COG-UK) Consortium | The Lighthouse Lab in Milton Keynes and Alex Alderton, Roberto Amato, Jeffrey Barrett, Sonia Goncalves, Ewan Harrison, David K. Jackson, Ian Johnston, Dominic Kwiatkowski, Cordelia Langford, John Sillitoe on behalf of the Wellcome Sanger Institute COVID-19 Surveillance Team |
| EPI_ISL_3339339 | Lighthouse Lab in Alderley Park | Wellcome Sanger Institute for the COVID-19 Genomics UK (COG-UK) Consortium | Jacquelyn Wynn, Mairead Hyland, The Lighthouse Lab in Alderley Park and Alex Alderton, Roberto Amato, Jeffrey Barrett, Sonia Goncalves, Ewan Harrison, David K. Jackson, Ian Johnston, Dominic Kwiatkowski, Cordelia Langford, John Sillitoe on behalf of the Wellcome Sanger Institute COVID-19 Surveillance Team |
| EPI_ISL_3339340, EPI_ISL_3339341 | Lighthouse Lab in Milton Keynes | Wellcome Sanger Institute for the COVID-19 Genomics UK (COG-UK) Consortium | The Lighthouse Lab in Milton Keynes and Alex Alderton, Roberto Amato, Jeffrey Barrett, Sonia Goncalves, Ewan Harrison, David K. Jackson, Ian Johnston, Dominic Kwiatkowski, Cordelia Langford, John Sillitoe on behalf of the Wellcome Sanger Institute COVID-19 Surveillance Team |
| EPI_ISL_3339342 | Lighthouse Laboratory Plymouth | Wellcome Sanger Institute for the COVID-19 Genomics UK (COG-UK) Consortium | Lighthouse Laboratory Plymouth and Alex Alderton, Roberto Amato, Jeffrey Barrett, Sonia Goncalves, Ewan Harrison, David K. Jackson, Ian Johnston, Dominic Kwiatkowski, Cordelia Langford, John Sillitoe on behalf of the Wellcome Sanger Institute COVID-19 Surveillance Team |
| EPI_ISL_3339343 | Lighthouse Lab in Alderley Park | Wellcome Sanger Institute for the COVID-19 Genomics UK (COG-UK) Consortium | Jacquelyn Wynn, Mairead Hyland, The Lighthouse Lab in Alderley Park and Alex Alderton, Roberto Amato, Jeffrey Barrett, Sonia Goncalves, Ewan Harrison, David K. Jackson, Ian Johnston, Dominic Kwiatkowski, Cordelia Langford, John Sillitoe on behalf of the Wellcome Sanger Institute COVID-19 Surveillance Team |
| EPI_ISL_3339344, EPI_ISL_3339345 | Lighthouse Laboratory Plymouth | Wellcome Sanger Institute for the COVID-19 Genomics UK (COG-UK) Consortium | Lighthouse Laboratory Plymouth and Alex Alderton, Roberto Amato, Jeffrey Barrett, Sonia Goncalves, Ewan Harrison, David K. Jackson, Ian Johnston, Dominic Kwiatkowski, Cordelia Langford, John Sillitoe on behalf of the Wellcome Sanger Institute COVID-19 Surveillance Team |
| EPI_ISL_3339346 | Lighthouse Lab in Milton Keynes | Wellcome Sanger Institute for the COVID-19 Genomics UK (COG-UK) Consortium | The Lighthouse Lab in Milton Keynes and Alex Alderton, Roberto Amato, Jeffrey Barrett, Sonia Goncalves, Ewan Harrison, David K. Jackson, Ian Johnston, Dominic Kwiatkowski, Cordelia Langford, John Sillitoe on behalf of the Wellcome Sanger Institute COVID-19 Surveillance Team |
| EPI_ISL_3339347 | Lighthouse Laboratory Plymouth | Wellcome Sanger Institute for the COVID-19 Genomics UK (COG-UK) Consortium | Lighthouse Laboratory Plymouth and Alex Alderton, Roberto Amato, Jeffrey Barrett, Sonia Goncalves, Ewan Harrison, David K. Jackson, Ian Johnston, Dominic Kwiatkowski, Cordelia Langford, John Sillitoe on behalf of the Wellcome Sanger Institute COVID-19 Surveillance Team |
| EPI_ISL_3339348, EPI_ISL_3339349 | Lighthouse Lab in Milton Keynes | Wellcome Sanger Institute for the COVID-19 Genomics UK (COG-UK) Consortium | The Lighthouse Lab in Milton Keynes and Alex Alderton, Roberto Amato, Jeffrey Barrett, Sonia Goncalves, Ewan Harrison, David K. Jackson, Ian Johnston, Dominic Kwiatkowski, Cordelia Langford, John Sillitoe on behalf of the Wellcome Sanger Institute COVID-19 Surveillance Team |
| EPI_ISL_3339350 | Lighthouse Lab in Alderley Park | Wellcome Sanger Institute for the COVID-19 Genomics UK (COG-UK) Consortium | Jacquelyn Wynn, Mairead Hyland, The Lighthouse Lab in Alderley Park and Alex Alderton, Roberto Amato, Jeffrey Barrett, Sonia Goncalves, Ewan Harrison, David K. Jackson, Ian Johnston, Dominic Kwiatkowski, Cordelia Langford, John Sillitoe on behalf of the Wellcome Sanger Institute COVID-19 Surveillance Team |
| EPI_ISL_3339351 | Lighthouse Lab in Milton Keynes | Wellcome Sanger Institute for the COVID-19 Genomics UK (COG-UK) Consortium | The Lighthouse Lab in Milton Keynes and Alex Alderton, Roberto Amato, Jeffrey Barrett, Sonia Goncalves, Ewan Harrison, David K. Jackson, Ian Johnston, Dominic Kwiatkowski, Cordelia Langford, John Sillitoe on behalf of the Wellcome Sanger Institute COVID-19 Surveillance Team |
| EPI_ISL_3339352, EPI_ISL_3339353 | Lighthouse Lab in Alderley Park | Wellcome Sanger Institute for the COVID-19 Genomics UK (COG-UK) Consortium | Jacquelyn Wynn, Mairead Hyland, The Lighthouse Lab in Alderley Park and Alex Alderton, Roberto Amato, Jeffrey Barrett, Sonia Goncalves, Ewan Harrison, David K. Jackson, Ian Johnston, Dominic Kwiatkowski, Cordelia Langford, John Sillitoe on behalf of the Wellcome Sanger Institute COVID-19 Surveillance Team |
| EPI_ISL_3339354 | Lighthouse Laboratory Plymouth | Wellcome Sanger Institute for the COVID-19 Genomics UK (COG-UK) Consortium | Lighthouse Laboratory Plymouth and Alex Alderton, Roberto Amato, Jeffrey Barrett, Sonia Goncalves, Ewan Harrison, David K. Jackson, Ian Johnston, Dominic Kwiatkowski, Cordelia Langford, John Sillitoe on behalf of the Wellcome Sanger Institute COVID-19 Surveillance Team |
| EPI_ISL_3339355 | Lighthouse Lab in Alderley Park | Wellcome Sanger Institute for the COVID-19 Genomics UK (COG-UK) Consortium | Jacquelyn Wynn, Mairead Hyland, The Lighthouse Lab in Alderley Park and Alex Alderton, Roberto Amato, Jeffrey Barrett, Sonia Goncalves, Ewan Harrison, David K. Jackson, Ian Johnston, Dominic Kwiatkowski, Cordelia Langford, John Sillitoe on behalf of the Wellcome Sanger Institute COVID-19 Surveillance Team |
| EPI_ISL_3339356 | Lighthouse Laboratory Plymouth | Wellcome Sanger Institute for the COVID-19 Genomics UK (COG-UK) Consortium | Lighthouse Laboratory Plymouth and Alex Alderton, Roberto Amato, Jeffrey Barrett, Sonia Goncalves, Ewan Harrison, David K. Jackson, Ian Johnston, Dominic Kwiatkowski, Cordelia Langford, John Sillitoe on behalf of the Wellcome Sanger Institute COVID-19 Surveillance Team |
| EPI_ISL_3339357 | Lighthouse Lab in Milton Keynes | Wellcome Sanger Institute for the COVID-19 Genomics UK (COG-UK) Consortium | The Lighthouse Lab in Milton Keynes and Alex Alderton, Roberto Amato, Jeffrey Barrett, Sonia Goncalves, Ewan Harrison, David K. Jackson, Ian Johnston, Dominic Kwiatkowski, Cordelia Langford, John Sillitoe on behalf of the Wellcome Sanger Institute COVID-19 Surveillance Team |
| EPI_ISL_3339358, EPI_ISL_3339359 | Lighthouse Lab in Alderley Park | Wellcome Sanger Institute for the COVID-19 Genomics UK (COG-UK) Consortium | Jacquelyn Wynn, Mairead Hyland, The Lighthouse Lab in Alderley Park and Alex Alderton, Roberto Amato, Jeffrey Barrett, Sonia Goncalves, Ewan Harrison, David K. Jackson, Ian Johnston, Dominic Kwiatkowski, Cordelia Langford, John Sillitoe on behalf of the Wellcome Sanger Institute COVID-19 Surveillance Team |
| EPI_ISL_3339360 | Lighthouse Laboratory Plymouth | Wellcome Sanger Institute for the COVID-19 Genomics UK (COG-UK) Consortium | Lighthouse Laboratory Plymouth and Alex Alderton, Roberto Amato, Jeffrey Barrett, Sonia Goncalves, Ewan Harrison, David K. Jackson, Ian Johnston, Dominic Kwiatkowski, Cordelia Langford, John Sillitoe on behalf of the Wellcome Sanger Institute COVID-19 Surveillance Team |
| EPI_ISL_3339361 | Lighthouse Lab in Alderley Park | Wellcome Sanger Institute for the COVID-19 Genomics UK (COG-UK) Consortium | Jacquelyn Wynn, Mairead Hyland, The Lighthouse Lab in Alderley Park and Alex Alderton, Roberto Amato, Jeffrey Barrett, Sonia Goncalves, Ewan Harrison, David K. Jackson, Ian Johnston, Dominic Kwiatkowski, Cordelia Langford, John Sillitoe on behalf of the Wellcome Sanger Institute COVID-19 Surveillance Team |

| | | | |
|---|---|---|---|
| EPI_ISL_3339362 | Lighthouse Lab in Milton Keynes | Wellcome Sanger Institute for the COVID-19 Genomics UK (COG-UK) Consortium | The Lighthouse Lab in Milton Keynes and Alex Alderton, Roberto Amato, Jeffrey Barrett, Sonia Goncalves, Ewan Harrison, David K. Jackson, Ian Johnston, Dominic Kwiatkowski, Cordelia Langford, John Sillitoe on behalf of the Wellcome Sanger Institute COVID-19 Surveillance Team |
| EPI_ISL_3339363 | Lighthouse Lab in Alderley Park | Wellcome Sanger Institute for the COVID-19 Genomics UK (COG-UK) Consortium | Jacquelyn Wynn, Mairead Hyland, The Lighthouse Lab in Alderley Park and Alex Alderton, Roberto Amato, Jeffrey Barrett, Sonia Goncalves, Ewan Harrison, David K. Jackson, Ian Johnston, Dominic Kwiatkowski, Cordelia Langford, John Sillitoe on behalf of the Wellcome Sanger Institute COVID-19 Surveillance Team |
| EPI_ISL_3339364 | Lighthouse Lab in Milton Keynes | Wellcome Sanger Institute for the COVID-19 Genomics UK (COG-UK) Consortium | The Lighthouse Lab in Milton Keynes and Alex Alderton, Roberto Amato, Jeffrey Barrett, Sonia Goncalves, Ewan Harrison, David K. Jackson, Ian Johnston, Dominic Kwiatkowski, Cordelia Langford, John Sillitoe on behalf of the Wellcome Sanger Institute COVID-19 Surveillance Team |
| EPI_ISL_3339365 | Lighthouse Laboratory Plymouth | Wellcome Sanger Institute for the COVID-19 Genomics UK (COG-UK) Consortium | Lighthouse Laboratory Plymouth and Alex Alderton, Roberto Amato, Jeffrey Barrett, Sonia Goncalves, Ewan Harrison, David K. Jackson, Ian Johnston, Dominic Kwiatkowski, Cordelia Langford, John Sillitoe on behalf of the Wellcome Sanger Institute COVID-19 Surveillance Team |
| EPI_ISL_3339366 | Lighthouse Lab in Milton Keynes | Wellcome Sanger Institute for the COVID-19 Genomics UK (COG-UK) Consortium | The Lighthouse Lab in Milton Keynes and Alex Alderton, Roberto Amato, Jeffrey Barrett, Sonia Goncalves, Ewan Harrison, David K. Jackson, Ian Johnston, Dominic Kwiatkowski, Cordelia Langford, John Sillitoe on behalf of the Wellcome Sanger Institute COVID-19 Surveillance Team |
| EPI_ISL_3339367 | Lighthouse Laboratory Plymouth | Wellcome Sanger Institute for the COVID-19 Genomics UK (COG-UK) Consortium | Lighthouse Laboratory Plymouth and Alex Alderton, Roberto Amato, Jeffrey Barrett, Sonia Goncalves, Ewan Harrison, David K. Jackson, Ian Johnston, Dominic Kwiatkowski, Cordelia Langford, John Sillitoe on behalf of the Wellcome Sanger Institute COVID-19 Surveillance Team |
| EPI_ISL_3339368, EPI_ISL_3339369, EPI_ISL_3339370, EPI_ISL_3339371 | Lighthouse Lab in Milton Keynes | Wellcome Sanger Institute for the COVID-19 Genomics UK (COG-UK) Consortium | The Lighthouse Lab in Milton Keynes and Alex Alderton, Roberto Amato, Jeffrey Barrett, Sonia Goncalves, Ewan Harrison, David K. Jackson, Ian Johnston, Dominic Kwiatkowski, Cordelia Langford, John Sillitoe on behalf of the Wellcome Sanger Institute COVID-19 Surveillance Team |
| EPI_ISL_3339372 | Lighthouse Lab in Alderley Park | Wellcome Sanger Institute for the COVID-19 Genomics UK (COG-UK) Consortium | Jacquelyn Wynn, Mairead Hyland, The Lighthouse Lab in Alderley Park and Alex Alderton, Roberto Amato, Jeffrey Barrett, Sonia Goncalves, Ewan Harrison, David K. Jackson, Ian Johnston, Dominic Kwiatkowski, Cordelia Langford, John Sillitoe on behalf of the Wellcome Sanger Institute COVID-19 Surveillance Team |
| EPI_ISL_3339373, EPI_ISL_3339374, EPI_ISL_3339375, EPI_ISL_3339376, EPI_ISL_3339377, EPI_ISL_3339378, EPI_ISL_3339379, EPI_ISL_3339380, EPI_ISL_3339381 | Lighthouse Lab in Milton Keynes | Wellcome Sanger Institute for the COVID-19 Genomics UK (COG-UK) Consortium | The Lighthouse Lab in Milton Keynes and Alex Alderton, Roberto Amato, Jeffrey Barrett, Sonia Goncalves, Ewan Harrison, David K. Jackson, Ian Johnston, Dominic Kwiatkowski, Cordelia Langford, John Sillitoe on behalf of the Wellcome Sanger Institute COVID-19 Surveillance Team |
| EPI_ISL_3339382, EPI_ISL_3339383 | Lighthouse Laboratory Plymouth | Wellcome Sanger Institute for the COVID-19 Genomics UK (COG-UK) Consortium | Lighthouse Laboratory Plymouth and Alex Alderton, Roberto Amato, Jeffrey Barrett, Sonia Goncalves, Ewan Harrison, David K. Jackson, Ian Johnston, Dominic Kwiatkowski, Cordelia Langford, John Sillitoe on behalf of the Wellcome Sanger Institute COVID-19 Surveillance Team |
| EPI_ISL_3339384 | Lighthouse Lab in Alderley Park | Wellcome Sanger Institute for the COVID-19 Genomics UK (COG-UK) Consortium | Jacquelyn Wynn, Mairead Hyland, The Lighthouse Lab in Alderley Park and Alex Alderton, Roberto Amato, Jeffrey Barrett, Sonia Goncalves, Ewan Harrison, David K. Jackson, Ian Johnston, Dominic Kwiatkowski, Cordelia Langford, John Sillitoe on behalf of the Wellcome Sanger Institute COVID-19 Surveillance Team |
| EPI_ISL_3339385, EPI_ISL_3339386 | Lighthouse Laboratory Plymouth | Wellcome Sanger Institute for the COVID-19 Genomics UK (COG-UK) Consortium | Lighthouse Laboratory Plymouth and Alex Alderton, Roberto Amato, Jeffrey Barrett, Sonia Goncalves, Ewan Harrison, David K. Jackson, Ian Johnston, Dominic Kwiatkowski, Cordelia Langford, John Sillitoe on behalf of the Wellcome Sanger Institute COVID-19 Surveillance Team |
| EPI_ISL_3339387, EPI_ISL_3339388 | Lighthouse Lab in Milton Keynes | Wellcome Sanger Institute for the COVID-19 Genomics UK (COG-UK) Consortium | The Lighthouse Lab in Milton Keynes and Alex Alderton, Roberto Amato, Jeffrey Barrett, Sonia Goncalves, Ewan Harrison, David K. Jackson, Ian Johnston, Dominic Kwiatkowski, Cordelia Langford, John Sillitoe on behalf of the Wellcome Sanger Institute COVID-19 Surveillance Team |
| EPI_ISL_3339389 | Lighthouse Lab in Alderley Park | Wellcome Sanger Institute for the COVID-19 Genomics UK (COG-UK) Consortium | Jacquelyn Wynn, Mairead Hyland, The Lighthouse Lab in Alderley Park and Alex Alderton, Roberto Amato, Jeffrey Barrett, Sonia Goncalves, Ewan Harrison, David K. Jackson, Ian Johnston, Dominic Kwiatkowski, Cordelia Langford, John Sillitoe on behalf of the Wellcome Sanger Institute COVID-19 Surveillance Team |
| EPI_ISL_3339390, EPI_ISL_3339391, EPI_ISL_3339392, EPI_ISL_3339393, EPI_ISL_3339394, EPI_ISL_3339395, EPI_ISL_3339396, EPI_ISL_3339397, EPI_ISL_3339398 | Lighthouse Lab in Milton Keynes | Wellcome Sanger Institute for the COVID-19 Genomics UK (COG-UK) Consortium | The Lighthouse Lab in Milton Keynes and Alex Alderton, Roberto Amato, Jeffrey Barrett, Sonia Goncalves, Ewan Harrison, David K. Jackson, Ian Johnston, Dominic Kwiatkowski, Cordelia Langford, John Sillitoe on behalf of the Wellcome Sanger Institute COVID-19 Surveillance Team |
| EPI_ISL_3339399 | Lighthouse Lab in Alderley Park | Wellcome Sanger Institute for the COVID-19 Genomics UK (COG-UK) Consortium | Jacquelyn Wynn, Mairead Hyland, The Lighthouse Lab in Alderley Park and Alex Alderton, Roberto Amato, Jeffrey Barrett, Sonia Goncalves, Ewan Harrison, David K. Jackson, Ian Johnston, Dominic Kwiatkowski, Cordelia Langford, John Sillitoe on behalf of the Wellcome Sanger Institute COVID-19 Surveillance Team |
| EPI_ISL_3339400 | Lighthouse Lab in Milton Keynes | Wellcome Sanger Institute for the COVID-19 Genomics UK (COG-UK) Consortium | The Lighthouse Lab in Milton Keynes and Alex Alderton, Roberto Amato, Jeffrey Barrett, Sonia Goncalves, Ewan Harrison, David K. Jackson, Ian Johnston, Dominic Kwiatkowski, Cordelia Langford, John Sillitoe on behalf of the Wellcome Sanger Institute COVID-19 Surveillance Team |
| EPI_ISL_3339401 | Lighthouse Lab in Alderley Park | Wellcome Sanger Institute for the COVID-19 Genomics UK (COG-UK) Consortium | Jacquelyn Wynn, Mairead Hyland, The Lighthouse Lab in Alderley Park and Alex Alderton, Roberto Amato, Jeffrey Barrett, Sonia Goncalves, Ewan Harrison, David K. Jackson, Ian Johnston, Dominic Kwiatkowski, Cordelia Langford, John Sillitoe on behalf of the Wellcome Sanger Institute COVID-19 Surveillance Team |
| EPI_ISL_3339402 | Lighthouse Laboratory Plymouth | Wellcome Sanger Institute for the COVID-19 Genomics UK (COG-UK) Consortium | Lighthouse Laboratory Plymouth and Alex Alderton, Roberto Amato, Jeffrey Barrett, Sonia Goncalves, Ewan Harrison, David K. Jackson, Ian Johnston, Dominic Kwiatkowski, Cordelia Langford, John Sillitoe on behalf of the Wellcome Sanger Institute COVID-19 Surveillance Team |
| EPI_ISL_3339403 | Lighthouse Lab in Alderley Park | Wellcome Sanger Institute for the COVID-19 Genomics UK (COG-UK) Consortium | Jacquelyn Wynn, Mairead Hyland, The Lighthouse Lab in Alderley Park and Alex Alderton, Roberto Amato, Jeffrey Barrett, Sonia Goncalves, Ewan Harrison, David K. Jackson, Ian Johnston, Dominic Kwiatkowski, Cordelia Langford, John Sillitoe on behalf of the Wellcome Sanger Institute COVID-19 Surveillance Team |
| EPI_ISL_3339404, EPI_ISL_3339405 | Lighthouse Lab in Milton Keynes | Wellcome Sanger Institute for the COVID-19 Genomics UK (COG-UK) Consortium | The Lighthouse Lab in Milton Keynes and Alex Alderton, Roberto Amato, Jeffrey Barrett, Sonia Goncalves, Ewan Harrison, David K. Jackson, Ian Johnston, Dominic Kwiatkowski, Cordelia Langford, John Sillitoe on behalf of the Wellcome Sanger Institute COVID-19 Surveillance Team |
| EPI_ISL_3339406, EPI_ISL_3339407, EPI_ISL_3339408, EPI_ISL_3339409 | Lighthouse Laboratory Plymouth | Wellcome Sanger Institute for the COVID-19 Genomics UK (COG-UK) Consortium | Lighthouse Laboratory Plymouth and Alex Alderton, Roberto Amato, Jeffrey Barrett, Sonia Goncalves, Ewan Harrison, David K. Jackson, Ian Johnston, Dominic Kwiatkowski, Cordelia Langford, John Sillitoe on behalf of the Wellcome Sanger Institute COVID-19 Surveillance Team |
| EPI_ISL_3339410 | Lighthouse Lab in Milton Keynes | Wellcome Sanger Institute for the COVID-19 Genomics UK (COG-UK) Consortium | The Lighthouse Lab in Milton Keynes and Alex Alderton, Roberto Amato, Jeffrey Barrett, Sonia Goncalves, Ewan Harrison, David K. Jackson, Ian Johnston, Dominic Kwiatkowski, Cordelia Langford, John Sillitoe on behalf of the Wellcome Sanger Institute COVID-19 Surveillance Team |
| EPI_ISL_3339411 | Lighthouse Lab in Alderley Park | Wellcome Sanger Institute for the COVID-19 Genomics UK (COG-UK) Consortium | Jacquelyn Wynn, Mairead Hyland, The Lighthouse Lab in Alderley Park and Alex Alderton, Roberto Amato, Jeffrey Barrett, Sonia Goncalves, Ewan Harrison, David K. Jackson, Ian Johnston, Dominic Kwiatkowski, Cordelia Langford, John Sillitoe on behalf of the Wellcome Sanger Institute COVID-19 Surveillance Team |
| EPI_ISL_3339412, EPI_ISL_3339413, EPI_ISL_3339414 | Lighthouse Lab in Milton Keynes | Wellcome Sanger Institute for the COVID-19 Genomics UK (COG-UK) Consortium | The Lighthouse Lab in Milton Keynes and Alex Alderton, Roberto Amato, Jeffrey Barrett, Sonia Goncalves, Ewan Harrison, David K. Jackson, Ian Johnston, Dominic Kwiatkowski, Cordelia Langford, John Sillitoe on behalf of the Wellcome Sanger Institute COVID-19 Surveillance Team |
| EPI_ISL_3339415 | Lighthouse Laboratory Plymouth | Wellcome Sanger Institute for the COVID-19 Genomics UK (COG-UK) Consortium | Lighthouse Laboratory Plymouth and Alex Alderton, Roberto Amato, Jeffrey Barrett, Sonia Goncalves, Ewan Harrison, David K. Jackson, Ian Johnston, Dominic Kwiatkowski, Cordelia Langford, John Sillitoe on behalf of the Wellcome Sanger Institute COVID-19 Surveillance Team |
| EPI_ISL_3339416 | Lighthouse Lab in Milton Keynes | Wellcome Sanger Institute for the COVID-19 Genomics UK (COG-UK) Consortium | The Lighthouse Lab in Milton Keynes and Alex Alderton, Roberto Amato, Jeffrey Barrett, Sonia Goncalves, Ewan Harrison, David K. Jackson, Ian Johnston, Dominic Kwiatkowski, Cordelia Langford, John Sillitoe on behalf of the Wellcome Sanger Institute COVID-19 Surveillance Team |
| EPI_ISL_3339417, EPI_ISL_3339418 | Lighthouse Laboratory Plymouth | Wellcome Sanger Institute for the COVID-19 Genomics UK (COG-UK) Consortium | Lighthouse Laboratory Plymouth and Alex Alderton, Roberto Amato, Jeffrey Barrett, Sonia Goncalves, Ewan Harrison, David K. Jackson, Ian Johnston, Dominic Kwiatkowski, Cordelia Langford, John Sillitoe on behalf of the Wellcome Sanger Institute COVID-19 Surveillance Team |
| EPI_ISL_3339419, EPI_ISL_3339420, EPI_ISL_3339421 | Lighthouse Lab in Milton Keynes | Wellcome Sanger Institute for the COVID-19 Genomics UK (COG-UK) Consortium | The Lighthouse Lab in Milton Keynes and Alex Alderton, Roberto Amato, Jeffrey Barrett, Sonia Goncalves, Ewan Harrison, David K. Jackson, Ian Johnston, Dominic Kwiatkowski, Cordelia Langford, John Sillitoe on behalf of the Wellcome Sanger Institute COVID-19 Surveillance Team |
| EPI_ISL_3339422 | Lighthouse Lab in Alderley Park | Wellcome Sanger Institute for the COVID-19 Genomics UK (COG-UK) Consortium | Jacquelyn Wynn, Mairead Hyland, The Lighthouse Lab in Alderley Park and Alex Alderton, Roberto Amato, Jeffrey Barrett, Sonia Goncalves, Ewan Harrison, David K. Jackson, Ian Johnston, Dominic Kwiatkowski, Cordelia Langford, John Sillitoe on behalf of the Wellcome Sanger Institute COVID-19 Surveillance Team |
| EPI_ISL_3339423, EPI_ISL_3339424, EPI_ISL_3339425 | Lighthouse Laboratory Plymouth | Wellcome Sanger Institute for the COVID-19 Genomics UK (COG-UK) Consortium | Lighthouse Laboratory Plymouth and Alex Alderton, Roberto Amato, Jeffrey Barrett, Sonia Goncalves, Ewan Harrison, David K. Jackson, Ian Johnston, Dominic Kwiatkowski, Cordelia Langford, John Sillitoe on behalf of the Wellcome Sanger Institute COVID-19 Surveillance Team |
| EPI_ISL_3339426, EPI_ISL_3339427, EPI_ISL_3339428 | Lighthouse Lab in Milton Keynes | Wellcome Sanger Institute for the COVID-19 Genomics UK (COG-UK) Consortium | The Lighthouse Lab in Milton Keynes and Alex Alderton, Roberto Amato, Jeffrey Barrett, Sonia Goncalves, Ewan Harrison, David K. Jackson, Ian Johnston, Dominic Kwiatkowski, Cordelia Langford, John Sillitoe on behalf of the Wellcome Sanger Institute COVID-19 Surveillance Team |
| EPI_ISL_3339429 | Lighthouse Lab in Alderley Park | Wellcome Sanger Institute for the COVID-19 Genomics UK (COG-UK) Consortium | Jacquelyn Wynn, Mairead Hyland, The Lighthouse Lab in Alderley Park and Alex Alderton, Roberto Amato, Jeffrey Barrett, Sonia Goncalves, Ewan Harrison, David K. Jackson, Ian Johnston, Dominic Kwiatkowski, Cordelia Langford, John Sillitoe on behalf of the Wellcome Sanger Institute COVID-19 Surveillance Team |
| EPI_ISL_3339430 | Lighthouse Lab in Milton Keynes | Wellcome Sanger Institute for the COVID-19 Genomics UK (COG-UK) Consortium | The Lighthouse Lab in Milton Keynes and Alex Alderton, Roberto Amato, Jeffrey Barrett, Sonia Goncalves, Ewan Harrison, David K. Jackson, Ian Johnston, Dominic Kwiatkowski, Cordelia Langford, John Sillitoe on behalf of the Wellcome Sanger Institute COVID-19 Surveillance Team |
| EPI_ISL_3339431, EPI_ISL_3339432 | Lighthouse Laboratory Plymouth | Wellcome Sanger Institute for the COVID-19 Genomics UK (COG-UK) Consortium | Lighthouse Laboratory Plymouth and Alex Alderton, Roberto Amato, Jeffrey Barrett, Sonia Goncalves, Ewan Harrison, David K. Jackson, Ian Johnston, Dominic Kwiatkowski, Cordelia Langford, John Sillitoe on behalf of the Wellcome Sanger Institute COVID-19 Surveillance Team |
| EPI_ISL_3339433 | Lighthouse Lab in Milton Keynes | Wellcome Sanger Institute for the COVID-19 Genomics UK (COG-UK) | The Lighthouse Lab in Milton Keynes and Alex Alderton, Roberto Amato, Jeffrey Barrett, Sonia Goncalves, Ewan Harrison, David K. Jackson, Ian Johnston, Dominic Kwiatkowski, Cordelia Langford, John Sillitoe on behalf of the Wellcome Sanger |

| | | | |
|---|---|---|---|
| | | Consortium | Institute COVID-19 Surveillance Team |
| EPI_ISL_3339434 | Lighthouse Laboratory Plymouth | Wellcome Sanger Institute for the COVID-19 Genomics UK (COG-UK) Consortium | Lighthouse Laboratory Plymouth and Alex Alderton, Roberto Amato, Jeffrey Barrett, Sonia Goncalves, Ewan Harrison, David K. Jackson, Ian Johnston, Dominic Kwiatkowski, Cordelia Langford, John Sillitoe on behalf of the Wellcome Sanger Institute COVID-19 Surveillance Team |
| EPI_ISL_3339435 | Lighthouse Lab in Alderley Park | Wellcome Sanger Institute for the COVID-19 Genomics UK (COG-UK) Consortium | Jacquelyn Wynn, Mairead Hyland, The Lighthouse Lab in Alderley Park and Alex Alderton, Roberto Amato, Jeffrey Barrett, Sonia Goncalves, Ewan Harrison, David K. Jackson, Ian Johnston, Dominic Kwiatkowski, Cordelia Langford, John Sillitoe on behalf of the Wellcome Sanger Institute COVID-19 Surveillance Team |
| EPI_ISL_3339436, EPI_ISL_3339437, EPI_ISL_3339438 | Lighthouse Laboratory Plymouth | Wellcome Sanger Institute for the COVID-19 Genomics UK (COG-UK) Consortium | Lighthouse Laboratory Plymouth and Alex Alderton, Roberto Amato, Jeffrey Barrett, Sonia Goncalves, Ewan Harrison, David K. Jackson, Ian Johnston, Dominic Kwiatkowski, Cordelia Langford, John Sillitoe on behalf of the Wellcome Sanger Institute COVID-19 Surveillance Team |
| EPI_ISL_3339439, EPI_ISL_3339440 | Lighthouse Lab in Milton Keynes | Wellcome Sanger Institute for the COVID-19 Genomics UK (COG-UK) Consortium | The Lighthouse Lab in Milton Keynes and Alex Alderton, Roberto Amato, Jeffrey Barrett, Sonia Goncalves, Ewan Harrison, David K. Jackson, Ian Johnston, Dominic Kwiatkowski, Cordelia Langford, John Sillitoe on behalf of the Wellcome Sanger Institute COVID-19 Surveillance Team |
| EPI_ISL_3339441 | Lighthouse Laboratory Plymouth | Wellcome Sanger Institute for the COVID-19 Genomics UK (COG-UK) Consortium | Lighthouse Laboratory Plymouth and Alex Alderton, Roberto Amato, Jeffrey Barrett, Sonia Goncalves, Ewan Harrison, David K. Jackson, Ian Johnston, Dominic Kwiatkowski, Cordelia Langford, John Sillitoe on behalf of the Wellcome Sanger Institute COVID-19 Surveillance Team |
| EPI_ISL_3339442 | Lighthouse Lab in Milton Keynes | Wellcome Sanger Institute for the COVID-19 Genomics UK (COG-UK) Consortium | The Lighthouse Lab in Milton Keynes and Alex Alderton, Roberto Amato, Jeffrey Barrett, Sonia Goncalves, Ewan Harrison, David K. Jackson, Ian Johnston, Dominic Kwiatkowski, Cordelia Langford, John Sillitoe on behalf of the Wellcome Sanger Institute COVID-19 Surveillance Team |
| EPI_ISL_3339443, EPI_ISL_3339444 | Lighthouse Lab in Alderley Park | Wellcome Sanger Institute for the COVID-19 Genomics UK (COG-UK) Consortium | Jacquelyn Wynn, Mairead Hyland, The Lighthouse Lab in Alderley Park and Alex Alderton, Roberto Amato, Jeffrey Barrett, Sonia Goncalves, Ewan Harrison, David K. Jackson, Ian Johnston, Dominic Kwiatkowski, Cordelia Langford, John Sillitoe on behalf of the Wellcome Sanger Institute COVID-19 Surveillance Team |
| EPI_ISL_3339445 | Lighthouse Lab in Milton Keynes | Wellcome Sanger Institute for the COVID-19 Genomics UK (COG-UK) Consortium | The Lighthouse Lab in Milton Keynes and Alex Alderton, Roberto Amato, Jeffrey Barrett, Sonia Goncalves, Ewan Harrison, David K. Jackson, Ian Johnston, Dominic Kwiatkowski, Cordelia Langford, John Sillitoe on behalf of the Wellcome Sanger Institute COVID-19 Surveillance Team |
| EPI_ISL_3339446 | Lighthouse Laboratory Plymouth | Wellcome Sanger Institute for the COVID-19 Genomics UK (COG-UK) Consortium | Lighthouse Laboratory Plymouth and Alex Alderton, Roberto Amato, Jeffrey Barrett, Sonia Goncalves, Ewan Harrison, David K. Jackson, Ian Johnston, Dominic Kwiatkowski, Cordelia Langford, John Sillitoe on behalf of the Wellcome Sanger Institute COVID-19 Surveillance Team |
| EPI_ISL_3339447 | Lighthouse Lab in Milton Keynes | Wellcome Sanger Institute for the COVID-19 Genomics UK (COG-UK) Consortium | The Lighthouse Lab in Milton Keynes and Alex Alderton, Roberto Amato, Jeffrey Barrett, Sonia Goncalves, Ewan Harrison, David K. Jackson, Ian Johnston, Dominic Kwiatkowski, Cordelia Langford, John Sillitoe on behalf of the Wellcome Sanger Institute COVID-19 Surveillance Team |
| EPI_ISL_3339448 | Lighthouse Lab in Alderley Park | Wellcome Sanger Institute for the COVID-19 Genomics UK (COG-UK) Consortium | Jacquelyn Wynn, Mairead Hyland, The Lighthouse Lab in Alderley Park and Alex Alderton, Roberto Amato, Jeffrey Barrett, Sonia Goncalves, Ewan Harrison, David K. Jackson, Ian Johnston, Dominic Kwiatkowski, Cordelia Langford, John Sillitoe on behalf of the Wellcome Sanger Institute COVID-19 Surveillance Team |
| EPI_ISL_3339449 | Lighthouse Laboratory Plymouth | Wellcome Sanger Institute for the COVID-19 Genomics UK (COG-UK) Consortium | Lighthouse Laboratory Plymouth and Alex Alderton, Roberto Amato, Jeffrey Barrett, Sonia Goncalves, Ewan Harrison, David K. Jackson, Ian Johnston, Dominic Kwiatkowski, Cordelia Langford, John Sillitoe on behalf of the Wellcome Sanger Institute COVID-19 Surveillance Team |
| EPI_ISL_3339450 | Lighthouse Lab in Alderley Park | Wellcome Sanger Institute for the COVID-19 Genomics UK (COG-UK) Consortium | Jacquelyn Wynn, Mairead Hyland, The Lighthouse Lab in Alderley Park and Alex Alderton, Roberto Amato, Jeffrey Barrett, Sonia Goncalves, Ewan Harrison, David K. Jackson, Ian Johnston, Dominic Kwiatkowski, Cordelia Langford, John Sillitoe on behalf of the Wellcome Sanger Institute COVID-19 Surveillance Team |
| EPI_ISL_3339451 | Lighthouse Lab in Milton Keynes | Wellcome Sanger Institute for the COVID-19 Genomics UK (COG-UK) Consortium | The Lighthouse Lab in Milton Keynes and Alex Alderton, Roberto Amato, Jeffrey Barrett, Sonia Goncalves, Ewan Harrison, David K. Jackson, Ian Johnston, Dominic Kwiatkowski, Cordelia Langford, John Sillitoe on behalf of the Wellcome Sanger Institute COVID-19 Surveillance Team |
| EPI_ISL_3339452 | Lighthouse Lab in Alderley Park | Wellcome Sanger Institute for the COVID-19 Genomics UK (COG-UK) Consortium | Jacquelyn Wynn, Mairead Hyland, The Lighthouse Lab in Alderley Park and Alex Alderton, Roberto Amato, Jeffrey Barrett, Sonia Goncalves, Ewan Harrison, David K. Jackson, Ian Johnston, Dominic Kwiatkowski, Cordelia Langford, John Sillitoe on behalf of the Wellcome Sanger Institute COVID-19 Surveillance Team |
| EPI_ISL_3339453 | Lighthouse Laboratory Plymouth | Wellcome Sanger Institute for the COVID-19 Genomics UK (COG-UK) Consortium | Lighthouse Laboratory Plymouth and Alex Alderton, Roberto Amato, Jeffrey Barrett, Sonia Goncalves, Ewan Harrison, David K. Jackson, Ian Johnston, Dominic Kwiatkowski, Cordelia Langford, John Sillitoe on behalf of the Wellcome Sanger Institute COVID-19 Surveillance Team |
| EPI_ISL_3339454 | Lighthouse Lab in Milton Keynes | Wellcome Sanger Institute for the COVID-19 Genomics UK (COG-UK) Consortium | The Lighthouse Lab in Milton Keynes and Alex Alderton, Roberto Amato, Jeffrey Barrett, Sonia Goncalves, Ewan Harrison, David K. Jackson, Ian Johnston, Dominic Kwiatkowski, Cordelia Langford, John Sillitoe on behalf of the Wellcome Sanger Institute COVID-19 Surveillance Team |
| EPI_ISL_3339455 | Lighthouse Laboratory Plymouth | Wellcome Sanger Institute for the COVID-19 Genomics UK (COG-UK) Consortium | Lighthouse Laboratory Plymouth and Alex Alderton, Roberto Amato, Jeffrey Barrett, Sonia Goncalves, Ewan Harrison, David K. Jackson, Ian Johnston, Dominic Kwiatkowski, Cordelia Langford, John Sillitoe on behalf of the Wellcome Sanger Institute COVID-19 Surveillance Team |
| EPI_ISL_3339456 | Lighthouse Lab in Milton Keynes | Wellcome Sanger Institute for the COVID-19 Genomics UK (COG-UK) Consortium | The Lighthouse Lab in Milton Keynes and Alex Alderton, Roberto Amato, Jeffrey Barrett, Sonia Goncalves, Ewan Harrison, David K. Jackson, Ian Johnston, Dominic Kwiatkowski, Cordelia Langford, John Sillitoe on behalf of the Wellcome Sanger Institute COVID-19 Surveillance Team |
| EPI_ISL_3339457 | Lighthouse Laboratory Plymouth | Wellcome Sanger Institute for the COVID-19 Genomics UK (COG-UK) Consortium | Lighthouse Laboratory Plymouth and Alex Alderton, Roberto Amato, Jeffrey Barrett, Sonia Goncalves, Ewan Harrison, David K. Jackson, Ian Johnston, Dominic Kwiatkowski, Cordelia Langford, John Sillitoe on behalf of the Wellcome Sanger Institute COVID-19 Surveillance Team |
| EPI_ISL_3339458 | Lighthouse Lab in Alderley Park | Wellcome Sanger Institute for the COVID-19 Genomics UK (COG-UK) Consortium | Jacquelyn Wynn, Mairead Hyland, The Lighthouse Lab in Alderley Park and Alex Alderton, Roberto Amato, Jeffrey Barrett, Sonia Goncalves, Ewan Harrison, David K. Jackson, Ian Johnston, Dominic Kwiatkowski, Cordelia Langford, John Sillitoe on behalf of the Wellcome Sanger Institute COVID-19 Surveillance Team |
| EPI_ISL_3339459, EPI_ISL_3339460 | Lighthouse Laboratory Plymouth | Wellcome Sanger Institute for the COVID-19 Genomics UK (COG-UK) Consortium | Lighthouse Laboratory Plymouth and Alex Alderton, Roberto Amato, Jeffrey Barrett, Sonia Goncalves, Ewan Harrison, David K. Jackson, Ian Johnston, Dominic Kwiatkowski, Cordelia Langford, John Sillitoe on behalf of the Wellcome Sanger Institute COVID-19 Surveillance Team |
| EPI_ISL_3339461 | Lighthouse Lab in Milton Keynes | Wellcome Sanger Institute for the COVID-19 Genomics UK (COG-UK) Consortium | The Lighthouse Lab in Milton Keynes and Alex Alderton, Roberto Amato, Jeffrey Barrett, Sonia Goncalves, Ewan Harrison, David K. Jackson, Ian Johnston, Dominic Kwiatkowski, Cordelia Langford, John Sillitoe on behalf of the Wellcome Sanger Institute COVID-19 Surveillance Team |
| EPI_ISL_3339462 | Lighthouse Lab in Alderley Park | Wellcome Sanger Institute for the COVID-19 Genomics UK (COG-UK) Consortium | Jacquelyn Wynn, Mairead Hyland, The Lighthouse Lab in Alderley Park and Alex Alderton, Roberto Amato, Jeffrey Barrett, Sonia Goncalves, Ewan Harrison, David K. Jackson, Ian Johnston, Dominic Kwiatkowski, Cordelia Langford, John Sillitoe on behalf of the Wellcome Sanger Institute COVID-19 Surveillance Team |
| EPI_ISL_3339463 | Lighthouse Lab in Milton Keynes | Wellcome Sanger Institute for the COVID-19 Genomics UK (COG-UK) Consortium | The Lighthouse Lab in Milton Keynes and Alex Alderton, Roberto Amato, Jeffrey Barrett, Sonia Goncalves, Ewan Harrison, David K. Jackson, Ian Johnston, Dominic Kwiatkowski, Cordelia Langford, John Sillitoe on behalf of the Wellcome Sanger Institute COVID-19 Surveillance Team |
| EPI_ISL_3339464, EPI_ISL_3339465 | Lighthouse Laboratory Plymouth | Wellcome Sanger Institute for the COVID-19 Genomics UK (COG-UK) Consortium | Lighthouse Laboratory Plymouth and Alex Alderton, Roberto Amato, Jeffrey Barrett, Sonia Goncalves, Ewan Harrison, David K. Jackson, Ian Johnston, Dominic Kwiatkowski, Cordelia Langford, John Sillitoe on behalf of the Wellcome Sanger Institute COVID-19 Surveillance Team |
| EPI_ISL_3339466 | Lighthouse Lab in Milton Keynes | Wellcome Sanger Institute for the COVID-19 Genomics UK (COG-UK) Consortium | The Lighthouse Lab in Milton Keynes and Alex Alderton, Roberto Amato, Jeffrey Barrett, Sonia Goncalves, Ewan Harrison, David K. Jackson, Ian Johnston, Dominic Kwiatkowski, Cordelia Langford, John Sillitoe on behalf of the Wellcome Sanger Institute COVID-19 Surveillance Team |
| EPI_ISL_3339467 | Lighthouse Laboratory Plymouth | Wellcome Sanger Institute for the COVID-19 Genomics UK (COG-UK) Consortium | Lighthouse Laboratory Plymouth and Alex Alderton, Roberto Amato, Jeffrey Barrett, Sonia Goncalves, Ewan Harrison, David K. Jackson, Ian Johnston, Dominic Kwiatkowski, Cordelia Langford, John Sillitoe on behalf of the Wellcome Sanger Institute COVID-19 Surveillance Team |
| EPI_ISL_3339468 | Lighthouse Lab in Milton Keynes | Wellcome Sanger Institute for the COVID-19 Genomics UK (COG-UK) Consortium | The Lighthouse Lab in Milton Keynes and Alex Alderton, Roberto Amato, Jeffrey Barrett, Sonia Goncalves, Ewan Harrison, David K. Jackson, Ian Johnston, Dominic Kwiatkowski, Cordelia Langford, John Sillitoe on behalf of the Wellcome Sanger Institute COVID-19 Surveillance Team |
| EPI_ISL_3339469, EPI_ISL_3339470 | Lighthouse Laboratory Plymouth | Wellcome Sanger Institute for the COVID-19 Genomics UK (COG-UK) Consortium | Lighthouse Laboratory Plymouth and Alex Alderton, Roberto Amato, Jeffrey Barrett, Sonia Goncalves, Ewan Harrison, David K. Jackson, Ian Johnston, Dominic Kwiatkowski, Cordelia Langford, John Sillitoe on behalf of the Wellcome Sanger Institute COVID-19 Surveillance Team |
| EPI_ISL_3339471 | Lighthouse Lab in Alderley Park | Wellcome Sanger Institute for the COVID-19 Genomics UK (COG-UK) Consortium | Jacquelyn Wynn, Mairead Hyland, The Lighthouse Lab in Alderley Park and Alex Alderton, Roberto Amato, Jeffrey Barrett, Sonia Goncalves, Ewan Harrison, David K. Jackson, Ian Johnston, Dominic Kwiatkowski, Cordelia Langford, John Sillitoe on behalf of the Wellcome Sanger Institute COVID-19 Surveillance Team |
| EPI_ISL_3339472 | Lighthouse Lab in Milton Keynes | Wellcome Sanger Institute for the COVID-19 Genomics UK (COG-UK) Consortium | The Lighthouse Lab in Milton Keynes and Alex Alderton, Roberto Amato, Jeffrey Barrett, Sonia Goncalves, Ewan Harrison, David K. Jackson, Ian Johnston, Dominic Kwiatkowski, Cordelia Langford, John Sillitoe on behalf of the Wellcome Sanger Institute COVID-19 Surveillance Team |
| EPI_ISL_3339473, EPI_ISL_3339474, EPI_ISL_3339475 | Lighthouse Lab in Alderley Park | Wellcome Sanger Institute for the COVID-19 Genomics UK (COG-UK) Consortium | Jacquelyn Wynn, Mairead Hyland, The Lighthouse Lab in Alderley Park and Alex Alderton, Roberto Amato, Jeffrey Barrett, Sonia Goncalves, Ewan Harrison, David K. Jackson, Ian Johnston, Dominic Kwiatkowski, Cordelia Langford, John Sillitoe on behalf of the Wellcome Sanger Institute COVID-19 Surveillance Team |
| EPI_ISL_3339476 | Lighthouse Lab in Milton Keynes | Wellcome Sanger Institute for the COVID-19 Genomics UK (COG-UK) Consortium | The Lighthouse Lab in Milton Keynes and Alex Alderton, Roberto Amato, Jeffrey Barrett, Sonia Goncalves, Ewan Harrison, David K. Jackson, Ian Johnston, Dominic Kwiatkowski, Cordelia Langford, John Sillitoe on behalf of the Wellcome Sanger Institute COVID-19 Surveillance Team |
| EPI_ISL_3339477 | Lighthouse Laboratory Plymouth | Wellcome Sanger Institute for the COVID-19 Genomics UK (COG-UK) Consortium | Lighthouse Laboratory Plymouth and Alex Alderton, Roberto Amato, Jeffrey Barrett, Sonia Goncalves, Ewan Harrison, David K. Jackson, Ian Johnston, Dominic Kwiatkowski, Cordelia Langford, John Sillitoe on behalf of the Wellcome Sanger Institute COVID-19 Surveillance Team |
| EPI_ISL_3339478 | Lighthouse Lab in Milton Keynes | Wellcome Sanger Institute for the COVID-19 Genomics UK (COG-UK) Consortium | The Lighthouse Lab in Milton Keynes and Alex Alderton, Roberto Amato, Jeffrey Barrett, Sonia Goncalves, Ewan Harrison, David K. Jackson, Ian Johnston, Dominic Kwiatkowski, Cordelia Langford, John Sillitoe on behalf of the Wellcome Sanger Institute COVID-19 Surveillance Team |
| EPI_ISL_3339479 | Lighthouse Lab in Alderley Park | Wellcome Sanger Institute for the COVID-19 Genomics UK (COG-UK) Consortium | Jacquelyn Wynn, Mairead Hyland, The Lighthouse Lab in Alderley Park and Alex Alderton, Roberto Amato, Jeffrey Barrett, Sonia Goncalves, Ewan Harrison, David K. Jackson, Ian Johnston, Dominic Kwiatkowski, Cordelia Langford, John Sillitoe on behalf of the Wellcome Sanger Institute COVID-19 Surveillance Team |
| EPI_ISL_3339480 | Lighthouse Lab in Milton Keynes | Wellcome Sanger Institute for the COVID-19 Genomics UK (COG-UK) Consortium | The Lighthouse Lab in Milton Keynes and Alex Alderton, Roberto Amato, Jeffrey Barrett, Sonia Goncalves, Ewan Harrison, David K. Jackson, Ian Johnston, Dominic Kwiatkowski, Cordelia Langford, John Sillitoe on behalf of the Wellcome Sanger Institute COVID-19 Surveillance Team |
| EPI_ISL_3339481 | Lighthouse Laboratory Plymouth | Wellcome Sanger Institute for the COVID-19 Genomics UK (COG-UK) Consortium | Lighthouse Laboratory Plymouth and Alex Alderton, Roberto Amato, Jeffrey Barrett, Sonia Goncalves, Ewan Harrison, David K. Jackson, Ian Johnston, Dominic Kwiatkowski, Cordelia Langford, John Sillitoe on behalf of the Wellcome Sanger Institute COVID-19 Surveillance Team |
| EPI_ISL_3339482, EPI_ISL_3339483, EPI_ISL_3339484 | Lighthouse Lab in Milton Keynes | Wellcome Sanger Institute for the COVID-19 Genomics UK (COG-UK) Consortium | The Lighthouse Lab in Milton Keynes and Alex Alderton, Roberto Amato, Jeffrey Barrett, Sonia Goncalves, Ewan Harrison, David K. Jackson, Ian Johnston, Dominic Kwiatkowski, Cordelia Langford, John Sillitoe on behalf of the Wellcome Sanger Institute COVID-19 Surveillance Team |
| EPI_ISL_3339485 | Lighthouse Laboratory Plymouth | Wellcome Sanger Institute for the COVID-19 Genomics UK (COG-UK) Consortium | Lighthouse Laboratory Plymouth and Alex Alderton, Roberto Amato, Jeffrey Barrett, Sonia Goncalves, Ewan Harrison, David K. Jackson, Ian Johnston, Dominic Kwiatkowski, Cordelia Langford, John Sillitoe on behalf of the Wellcome Sanger Institute COVID-19 Surveillance Team |
| EPI_ISL_3339486, EPI_ISL_3339487 | Lighthouse Lab in Alderley Park | Wellcome Sanger Institute for the COVID-19 Genomics UK (COG-UK) Consortium | Jacquelyn Wynn, Mairead Hyland, The Lighthouse Lab in Alderley Park and Alex Alderton, Roberto Amato, Jeffrey Barrett, Sonia Goncalves, Ewan Harrison, David K. Jackson, Ian Johnston, Dominic Kwiatkowski, Cordelia Langford, John Sillitoe on behalf of the Wellcome Sanger Institute COVID-19 Surveillance Team |
| EPI_ISL_3339488 | Lighthouse Lab in Milton Keynes | Wellcome Sanger Institute for the COVID-19 Genomics UK (COG-UK) Consortium | The Lighthouse Lab in Milton Keynes and Alex Alderton, Roberto Amato, Jeffrey Barrett, Sonia Goncalves, Ewan Harrison, David K. Jackson, Ian Johnston, Dominic Kwiatkowski, Cordelia Langford, John Sillitoe on behalf of the Wellcome Sanger Institute COVID-19 Surveillance Team |
| EPI_ISL_3339489, | Lighthouse Laboratory Plymouth | Wellcome Sanger Institute for the COVID-19 Genomics UK (COG-UK) | Lighthouse Laboratory Plymouth and Alex Alderton, Roberto Amato, Jeffrey Barrett, Sonia Goncalves, Ewan Harrison, David K. Jackson, Ian Johnston, Dominic Kwiatkowski, Cordelia Langford, John Sillitoe on behalf of the Wellcome Sanger |

| | | | |
|---|---|---|---|
| EPI_ISL_3339490 | | Consortium | Institute COVID-19 Surveillance Team |
| EPI_ISL_3339491 | Lighthouse Lab in Alderley Park | Wellcome Sanger Institute for the COVID-19 Genomics UK (COG-UK) Consortium | Jacquelyn Wynn, Mairead Hyland, The Lighthouse Lab in Alderley Park and Alex Alderton, Roberto Amato, Jeffrey Barrett, Sonia Goncalves, Ewan Harrison, David K. Jackson, Ian Johnston, Dominic Kwiatkowski, Cordelia Langford, John Sillitoe on behalf of the Wellcome Sanger Institute COVID-19 Surveillance Team |
| EPI_ISL_3339492 | Lighthouse Lab in Milton Keynes | Wellcome Sanger Institute for the COVID-19 Genomics UK (COG-UK) Consortium | The Lighthouse Lab in Milton Keynes and Alex Alderton, Roberto Amato, Jeffrey Barrett, Sonia Goncalves, Ewan Harrison, David K. Jackson, Ian Johnston, Dominic Kwiatkowski, Cordelia Langford, John Sillitoe on behalf of the Wellcome Sanger Institute COVID-19 Surveillance Team |
| EPI_ISL_3339493, EPI_ISL_3339494, EPI_ISL_3339495, EPI_ISL_3339496 | Lighthouse Lab in Alderley Park | Wellcome Sanger Institute for the COVID-19 Genomics UK (COG-UK) Consortium | Jacquelyn Wynn, Mairead Hyland, The Lighthouse Lab in Alderley Park and Alex Alderton, Roberto Amato, Jeffrey Barrett, Sonia Goncalves, Ewan Harrison, David K. Jackson, Ian Johnston, Dominic Kwiatkowski, Cordelia Langford, John Sillitoe on behalf of the Wellcome Sanger Institute COVID-19 Surveillance Team |
| EPI_ISL_3339497 | Lighthouse Laboratory Plymouth | Wellcome Sanger Institute for the COVID-19 Genomics UK (COG-UK) Consortium | Lighthouse Laboratory Plymouth and Alex Alderton, Roberto Amato, Jeffrey Barrett, Sonia Goncalves, Ewan Harrison, David K. Jackson, Ian Johnston, Dominic Kwiatkowski, Cordelia Langford, John Sillitoe on behalf of the Wellcome Sanger Institute COVID-19 Surveillance Team |
| EPI_ISL_3339498 | Lighthouse Lab in Milton Keynes | Wellcome Sanger Institute for the COVID-19 Genomics UK (COG-UK) Consortium | The Lighthouse Lab in Milton Keynes and Alex Alderton, Roberto Amato, Jeffrey Barrett, Sonia Goncalves, Ewan Harrison, David K. Jackson, Ian Johnston, Dominic Kwiatkowski, Cordelia Langford, John Sillitoe on behalf of the Wellcome Sanger Institute COVID-19 Surveillance Team |
| EPI_ISL_3339499, EPI_ISL_3339500 | Lighthouse Laboratory Plymouth | Wellcome Sanger Institute for the COVID-19 Genomics UK (COG-UK) Consortium | Lighthouse Laboratory Plymouth and Alex Alderton, Roberto Amato, Jeffrey Barrett, Sonia Goncalves, Ewan Harrison, David K. Jackson, Ian Johnston, Dominic Kwiatkowski, Cordelia Langford, John Sillitoe on behalf of the Wellcome Sanger Institute COVID-19 Surveillance Team |
| EPI_ISL_3339501, EPI_ISL_3339502, EPI_ISL_3339503, EPI_ISL_3339504 | Lighthouse Lab in Milton Keynes | Wellcome Sanger Institute for the COVID-19 Genomics UK (COG-UK) Consortium | The Lighthouse Lab in Milton Keynes and Alex Alderton, Roberto Amato, Jeffrey Barrett, Sonia Goncalves, Ewan Harrison, David K. Jackson, Ian Johnston, Dominic Kwiatkowski, Cordelia Langford, John Sillitoe on behalf of the Wellcome Sanger Institute COVID-19 Surveillance Team |
| EPI_ISL_3339505 | Lighthouse Laboratory Plymouth | Wellcome Sanger Institute for the COVID-19 Genomics UK (COG-UK) Consortium | Lighthouse Laboratory Plymouth and Alex Alderton, Roberto Amato, Jeffrey Barrett, Sonia Goncalves, Ewan Harrison, David K. Jackson, Ian Johnston, Dominic Kwiatkowski, Cordelia Langford, John Sillitoe on behalf of the Wellcome Sanger Institute COVID-19 Surveillance Team |
| EPI_ISL_3339506, EPI_ISL_3339507, EPI_ISL_3339508, EPI_ISL_3339509 | Lighthouse Lab in Milton Keynes | Wellcome Sanger Institute for the COVID-19 Genomics UK (COG-UK) Consortium | The Lighthouse Lab in Milton Keynes and Alex Alderton, Roberto Amato, Jeffrey Barrett, Sonia Goncalves, Ewan Harrison, David K. Jackson, Ian Johnston, Dominic Kwiatkowski, Cordelia Langford, John Sillitoe on behalf of the Wellcome Sanger Institute COVID-19 Surveillance Team |
| EPI_ISL_3339510 | Lighthouse Laboratory Plymouth | Wellcome Sanger Institute for the COVID-19 Genomics UK (COG-UK) Consortium | Lighthouse Laboratory Plymouth and Alex Alderton, Roberto Amato, Jeffrey Barrett, Sonia Goncalves, Ewan Harrison, David K. Jackson, Ian Johnston, Dominic Kwiatkowski, Cordelia Langford, John Sillitoe on behalf of the Wellcome Sanger Institute COVID-19 Surveillance Team |
| EPI_ISL_3339511, EPI_ISL_3339512, EPI_ISL_3339513, EPI_ISL_3339514, EPI_ISL_3339515 | Lighthouse Lab in Milton Keynes | Wellcome Sanger Institute for the COVID-19 Genomics UK (COG-UK) Consortium | The Lighthouse Lab in Milton Keynes and Alex Alderton, Roberto Amato, Jeffrey Barrett, Sonia Goncalves, Ewan Harrison, David K. Jackson, Ian Johnston, Dominic Kwiatkowski, Cordelia Langford, John Sillitoe on behalf of the Wellcome Sanger Institute COVID-19 Surveillance Team |
| EPI_ISL_3339516, EPI_ISL_3339517 | Lighthouse Lab in Alderley Park | Wellcome Sanger Institute for the COVID-19 Genomics UK (COG-UK) Consortium | Jacquelyn Wynn, Mairead Hyland, The Lighthouse Lab in Alderley Park and Alex Alderton, Roberto Amato, Jeffrey Barrett, Sonia Goncalves, Ewan Harrison, David K. Jackson, Ian Johnston, Dominic Kwiatkowski, Cordelia Langford, John Sillitoe on behalf of the Wellcome Sanger Institute COVID-19 Surveillance Team |
| EPI_ISL_3339518 | Lighthouse Lab in Milton Keynes | Wellcome Sanger Institute for the COVID-19 Genomics UK (COG-UK) Consortium | The Lighthouse Lab in Milton Keynes and Alex Alderton, Roberto Amato, Jeffrey Barrett, Sonia Goncalves, Ewan Harrison, David K. Jackson, Ian Johnston, Dominic Kwiatkowski, Cordelia Langford, John Sillitoe on behalf of the Wellcome Sanger Institute COVID-19 Surveillance Team |
| EPI_ISL_3339519, EPI_ISL_3339520 | Lighthouse Lab in Alderley Park | Wellcome Sanger Institute for the COVID-19 Genomics UK (COG-UK) Consortium | Jacquelyn Wynn, Mairead Hyland, The Lighthouse Lab in Alderley Park and Alex Alderton, Roberto Amato, Jeffrey Barrett, Sonia Goncalves, Ewan Harrison, David K. Jackson, Ian Johnston, Dominic Kwiatkowski, Cordelia Langford, John Sillitoe on behalf of the Wellcome Sanger Institute COVID-19 Surveillance Team |
| EPI_ISL_3339521 | Lighthouse Lab in Milton Keynes | Wellcome Sanger Institute for the COVID-19 Genomics UK (COG-UK) Consortium | The Lighthouse Lab in Milton Keynes and Alex Alderton, Roberto Amato, Jeffrey Barrett, Sonia Goncalves, Ewan Harrison, David K. Jackson, Ian Johnston, Dominic Kwiatkowski, Cordelia Langford, John Sillitoe on behalf of the Wellcome Sanger Institute COVID-19 Surveillance Team |
| EPI_ISL_3339522 | Lighthouse Lab in Alderley Park | Wellcome Sanger Institute for the COVID-19 Genomics UK (COG-UK) Consortium | Jacquelyn Wynn, Mairead Hyland, The Lighthouse Lab in Alderley Park and Alex Alderton, Roberto Amato, Jeffrey Barrett, Sonia Goncalves, Ewan Harrison, David K. Jackson, Ian Johnston, Dominic Kwiatkowski, Cordelia Langford, John Sillitoe on behalf of the Wellcome Sanger Institute COVID-19 Surveillance Team |
| EPI_ISL_3339523 | Lighthouse Laboratory Plymouth | Wellcome Sanger Institute for the COVID-19 Genomics UK (COG-UK) Consortium | Lighthouse Laboratory Plymouth and Alex Alderton, Roberto Amato, Jeffrey Barrett, Sonia Goncalves, Ewan Harrison, David K. Jackson, Ian Johnston, Dominic Kwiatkowski, Cordelia Langford, John Sillitoe on behalf of the Wellcome Sanger Institute COVID-19 Surveillance Team |
| EPI_ISL_3339524 | Lighthouse Lab in Milton Keynes | Wellcome Sanger Institute for the COVID-19 Genomics UK (COG-UK) Consortium | The Lighthouse Lab in Milton Keynes and Alex Alderton, Roberto Amato, Jeffrey Barrett, Sonia Goncalves, Ewan Harrison, David K. Jackson, Ian Johnston, Dominic Kwiatkowski, Cordelia Langford, John Sillitoe on behalf of the Wellcome Sanger Institute COVID-19 Surveillance Team |
| EPI_ISL_3339525, EPI_ISL_3339526 | Lighthouse Lab in Glasgow | Wellcome Sanger Institute for the COVID-19 Genomics UK (COG-UK) Consortium | Harper VanSteenhouse, Yumi Kasai, David Gray, Carol Clugston, Anna Dominiczak and Alex Alderton, Roberto Amato, Jeffrey Barrett, Sonia Goncalves, Ewan Harrison, David K. Jackson, Ian Johnston, Dominic Kwiatkowski, Cordelia Langford, John Sillitoe on behalf of the Wellcome Sanger Institute COVID-19 Surveillance Team |
| EPI_ISL_3339527, EPI_ISL_3339528 | Lighthouse Lab in Milton Keynes | Wellcome Sanger Institute for the COVID-19 Genomics UK (COG-UK) Consortium | The Lighthouse Lab in Milton Keynes and Alex Alderton, Roberto Amato, Jeffrey Barrett, Sonia Goncalves, Ewan Harrison, David K. Jackson, Ian Johnston, Dominic Kwiatkowski, Cordelia Langford, John Sillitoe on behalf of the Wellcome Sanger Institute COVID-19 Surveillance Team |
| EPI_ISL_3339529 | Lighthouse Lab in Glasgow | Wellcome Sanger Institute for the COVID-19 Genomics UK (COG-UK) Consortium | Harper VanSteenhouse, Yumi Kasai, David Gray, Carol Clugston, Anna Dominiczak and Alex Alderton, Roberto Amato, Jeffrey Barrett, Sonia Goncalves, Ewan Harrison, David K. Jackson, Ian Johnston, Dominic Kwiatkowski, Cordelia Langford, John Sillitoe on behalf of the Wellcome Sanger Institute COVID-19 Surveillance Team |
| EPI_ISL_3339530, EPI_ISL_3339531, EPI_ISL_3339532, EPI_ISL_3339533, EPI_ISL_3339535, EPI_ISL_3339537, EPI_ISL_3339538, EPI_ISL_3339539, EPI_ISL_3339540, EPI_ISL_3339541, EPI_ISL_3339543, EPI_ISL_3339544, EPI_ISL_3339545, EPI_ISL_3339546, EPI_ISL_3339547, EPI_ISL_3339548, EPI_ISL_3339549, EPI_ISL_3339550, EPI_ISL_3339551, EPI_ISL_3339552, EPI_ISL_3339559, EPI_ISL_3339560, EPI_ISL_3339561, EPI_ISL_3339562, EPI_ISL_3339563 | | | |
| see above | Pandemic Response Lab - NYC | Pandemic Response Lab, R&D | Henry Lee, Michael Hammerling, Melissa Hopkins, Cybill del Castillo, Shinyoung Clair Kang, William Ward, Pradeep Bugga, Sol Rey, Dylan Law, Alex Carpio, Isabel Fernandez Escapa, Haiping Hao, Jon Laurent |
| EPI_ISL_3339565, EPI_ISL_3339566, EPI_ISL_3339567, EPI_ISL_3339568, EPI_ISL_3339569, EPI_ISL_3339570, EPI_ISL_3339571, EPI_ISL_3339572, EPI_ISL_3339573, EPI_ISL_3339574, EPI_ISL_3339575, EPI_ISL_3339576, EPI_ISL_3339577, EPI_ISL_3339578, EPI_ISL_3339579, EPI_ISL_3339580, EPI_ISL_3339581, EPI_ISL_3339582, EPI_ISL_3339583, EPI_ISL_3339584, EPI_ISL_3339585, EPI_ISL_3339586, EPI_ISL_3339587, EPI_ISL_3339588, EPI_ISL_3339589, EPI_ISL_3339590, EPI_ISL_3339591, EPI_ISL_3339592, EPI_ISL_3339593, EPI_ISL_3339594, EPI_ISL_3339595, EPI_ISL_3339596, EPI_ISL_3339597, EPI_ISL_3339598, EPI_ISL_3339599, EPI_ISL_3339600, EPI_ISL_3339601, EPI_ISL_3339602, EPI_ISL_3339603, EPI_ISL_3339604 | | | |
| see above | UAB InMedica | Lithuanian University of Health Sciences | Arnoldas Pautienius, Kamile Tamusauskaite, Gediminas Alzbutas, Dovydas Gecys, Lukas Zemaitis, Vaiva Lesauskaite |
| EPI_ISL_3339605, EPI_ISL_3339606, EPI_ISL_3339607, EPI_ISL_3339608, EPI_ISL_3339609, EPI_ISL_3339610, EPI_ISL_3339611, EPI_ISL_3339612, EPI_ISL_3339613, EPI_ISL_3339614, EPI_ISL_3339615, EPI_ISL_3339616, EPI_ISL_3339617, EPI_ISL_3339618, EPI_ISL_3339619, EPI_ISL_3339620, EPI_ISL_3339621, EPI_ISL_3339622, EPI_ISL_3339623, EPI_ISL_3339624, EPI_ISL_3339625, EPI_ISL_3339628 | | | |
| see above | Department of Bacteria, Parasites and Fungi, Statens Serum Institut, Copenhagen, Denmark | Statens Serum Institut Bioinformatics and Microbial Genomics | Danish Covid-19 Genome Consortium |
| EPI_ISL_3339629 | Molekylær Medicinsk Afdeling, Aarhus University Hospital, Aarhus, Denmark | Statens Serum Institut Bioinformatics and Microbial Genomics | Danish Covid-19 Genome Consortium |
| EPI_ISL_3339630, EPI_ISL_3339631, EPI_ISL_3339632, EPI_ISL_3339633, EPI_ISL_3339634, EPI_ISL_3339635, EPI_ISL_3339636 | Department of Bacteria, Parasites and Fungi, Statens Serum Institut, Copenhagen, Denmark | Statens Serum Institut Bioinformatics and Microbial Genomics | Danish Covid-19 Genome Consortium |
| EPI_ISL_3339637 | Department of Clinical Microbiology, Odense University Hospital, Odense, Denmark | Statens Serum Institut Bioinformatics and Microbial Genomics | Danish Covid-19 Genome Consortium |
| EPI_ISL_3339638, EPI_ISL_3339639, EPI_ISL_3339640, EPI_ISL_3339641, EPI_ISL_3339642, EPI_ISL_3339643, EPI_ISL_3339644, EPI_ISL_3339645, EPI_ISL_3339646, EPI_ISL_3339647, EPI_ISL_3339648, EPI_ISL_3339649, EPI_ISL_3339650, EPI_ISL_3339651, EPI_ISL_3339652, EPI_ISL_3339654, EPI_ISL_3339655, EPI_ISL_3339656, EPI_ISL_3339657, EPI_ISL_3339658, EPI_ISL_3339659, EPI_ISL_3339660, EPI_ISL_3339661, EPI_ISL_3339662, EPI_ISL_3339663, EPI_ISL_3339664, EPI_ISL_3339665, EPI_ISL_3339666, EPI_ISL_3339667, EPI_ISL_3339668, EPI_ISL_3339669, EPI_ISL_3339670, EPI_ISL_3339671, EPI_ISL_3339672, EPI_ISL_3339673, EPI_ISL_3339674, EPI_ISL_3339675, EPI_ISL_3339676, EPI_ISL_3339677, EPI_ISL_3339678, EPI_ISL_3339679, EPI_ISL_3339680, EPI_ISL_3339681, EPI_ISL_3339682, EPI_ISL_3339683, EPI_ISL_3339684, EPI_ISL_3339685, EPI_ISL_3339686, EPI_ISL_3339687, EPI_ISL_3339688, EPI_ISL_3339689, EPI_ISL_3339690, EPI_ISL_3339692, EPI_ISL_3339693, EPI_ISL_3339694, EPI_ISL_3339695, EPI_ISL_3339696, EPI_ISL_3339697, EPI_ISL_3339698, EPI_ISL_3339699, EPI_ISL_3339700, EPI_ISL_3339701, EPI_ISL_3339702, EPI_ISL_3339703, EPI_ISL_3339704, EPI_ISL_3339705, EPI_ISL_3339706, EPI_ISL_3339707, EPI_ISL_3339708, EPI_ISL_3339709, EPI_ISL_3339710, EPI_ISL_3339711, EPI_ISL_3339712, EPI_ISL_3339714, EPI_ISL_3339715, EPI_ISL_3339716, EPI_ISL_3339717, EPI_ISL_3339718, EPI_ISL_3339719, EPI_ISL_3339720, EPI_ISL_3339721, EPI_ISL_3339722, EPI_ISL_3339723, EPI_ISL_3339724, EPI_ISL_3339725, EPI_ISL_3339726, EPI_ISL_3339727, EPI_ISL_3339728, EPI_ISL_3339729, EPI_ISL_3339730, EPI_ISL_3339731, EPI_ISL_3339732, EPI_ISL_3339733, EPI_ISL_3339734, EPI_ISL_3339735, EPI_ISL_3339736, EPI_ISL_3339737, EPI_ISL_3339738, EPI_ISL_3339739, EPI_ISL_3339740, EPI_ISL_3339741, EPI_ISL_3339742, EPI_ISL_3339743, EPI_ISL_3339744, EPI_ISL_3339745, EPI_ISL_3339746, EPI_ISL_3339747, EPI_ISL_3339748, EPI_ISL_3339749, EPI_ISL_3339750, EPI_ISL_3339751, EPI_ISL_3339752, EPI_ISL_3339753, EPI_ISL_3339754, EPI_ISL_3339755, EPI_ISL_3339756, EPI_ISL_3339757, EPI_ISL_3339758, EPI_ISL_3339759, EPI_ISL_3339760, EPI_ISL_3339761, EPI_ISL_3339762, EPI_ISL_3339763, EPI_ISL_3339764, EPI_ISL_3339765, EPI_ISL_3339766, EPI_ISL_3339767, EPI_ISL_3339769, EPI_ISL_3339770, EPI_ISL_3339771, EPI_ISL_3339772, EPI_ISL_3339774, EPI_ISL_3339775, EPI_ISL_3339776, EPI_ISL_3339777, EPI_ISL_3339778, EPI_ISL_3339779, EPI_ISL_3339780, EPI_ISL_3339781, EPI_ISL_3339782, EPI_ISL_3339783, EPI_ISL_3339784, EPI_ISL_3339785, EPI_ISL_3339786, EPI_ISL_3339787, EPI_ISL_3339788, EPI_ISL_3339789, EPI_ISL_3339790, EPI_ISL_3339791, EPI_ISL_3339792, EPI_ISL_3339793, EPI_ISL_3339794, EPI_ISL_3339795, EPI_ISL_3339796, EPI_ISL_3339797, EPI_ISL_3339798, EPI_ISL_3339799, EPI_ISL_3339800, EPI_ISL_3339801, EPI_ISL_3339802, EPI_ISL_3339803, EPI_ISL_3339804, EPI_ISL_3339805, EPI_ISL_3339806, EPI_ISL_3339807, EPI_ISL_3339808, EPI_ISL_3339809, EPI_ISL_3339810, EPI_ISL_3339812, EPI_ISL_3339813, EPI_ISL_3339814, EPI_ISL_3339815, EPI_ISL_3339816, EPI_ISL_3339817, EPI_ISL_3339818, EPI_ISL_3339819, EPI_ISL_3339820, EPI_ISL_3339822, EPI_ISL_3339823, EPI_ISL_3339824, EPI_ISL_3339825, EPI_ISL_3339826, EPI_ISL_3339827, EPI_ISL_3339828, EPI_ISL_3339829, EPI_ISL_3339830, EPI_ISL_3339831, EPI_ISL_3339832, EPI_ISL_3339833, EPI_ISL_3339834, EPI_ISL_3339835, EPI_ISL_3339836, EPI_ISL_3339837, EPI_ISL_3339838, EPI_ISL_3339839, EPI_ISL_3339840 | | | |
| see above | Department of Bacteria, Parasites and Fungi, Statens Serum Institut, Copenhagen, Denmark | Statens Serum Institut Bioinformatics and Microbial Genomics | Danish Covid-19 Genome Consortium |
| EPI_ISL_3339841 | Department of Clinical Microbiology, Odense University Hospital, Odense, Denmark | Statens Serum Institut Bioinformatics and Microbial Genomics | Danish Covid-19 Genome Consortium |
| EPI_ISL_3339842, EPI_ISL_3339843, EPI_ISL_3339845, EPI_ISL_3339846, EPI_ISL_3339847, EPI_ISL_3339848, EPI_ISL_3339849, EPI_ISL_3339850, EPI_ISL_3339851, EPI_ISL_3339853, EPI_ISL_3339854, EPI_ISL_3339855, EPI_ISL_3339856, EPI_ISL_3339857, EPI_ISL_3339858, EPI_ISL_3339859, EPI_ISL_3339860, EPI_ISL_3339861, EPI_ISL_3339862, EPI_ISL_3339863, EPI_ISL_3339864, EPI_ISL_3339865, EPI_ISL_3339866, EPI_ISL_3339867, EPI_ISL_3339868, EPI_ISL_3339869, EPI_ISL_3339870, EPI_ISL_3339871, EPI_ISL_3339872, EPI_ISL_3339873, EPI_ISL_3339874, EPI_ISL_3339875, EPI_ISL_3339876, EPI_ISL_3339877, EPI_ISL_3339878, EPI_ISL_3339879, EPI_ISL_3339880, EPI_ISL_3339881, EPI_ISL_3339882, EPI_ISL_3339883, EPI_ISL_3339884, EPI_ISL_3339885, EPI_ISL_3339886, EPI_ISL_3339887, EPI_ISL_3339888, EPI_ISL_3339889, EPI_ISL_3339890, EPI_ISL_3339891, EPI_ISL_3339892, EPI_ISL_3339893, EPI_ISL_3339894, EPI_ISL_3339895, EPI_ISL_3339896, EPI_ISL_3339897, EPI_ISL_3339898, EPI_ISL_3339899, EPI_ISL_3339900, EPI_ISL_3339901, EPI_ISL_3339902, EPI_ISL_3339903, EPI_ISL_3339904, EPI_ISL_3339905, EPI_ISL_3339906, EPI_ISL_3339907, EPI_ISL_3339908, EPI_ISL_3339909, EPI_ISL_3339910, EPI_ISL_3339911, EPI_ISL_3339912, EPI_ISL_3339913, EPI_ISL_3339915, EPI_ISL_3339916, EPI_ISL_3339917, EPI_ISL_3339918, EPI_ISL_3339919, EPI_ISL_3339920, EPI_ISL_3339921 | | | |
| see above | Department of Bacteria, Parasites and Fungi, Statens Serum Institut, Copenhagen, Denmark | Statens Serum Institut Bioinformatics and Microbial Genomics | Danish Covid-19 Genome Consortium |
| EPI_ISL_3339922 | Molekylær Medicinsk Afdeling, Aarhus University Hospital, Aarhus, Denmark | Statens Serum Institut Bioinformatics and Microbial Genomics | Danish Covid-19 Genome Consortium |
| EPI_ISL_3339923, | Department of Bacteria, Parasites and Fungi, Statens Serum | Statens Serum Institut Bioinformatics and Microbial Genomics | Danish Covid-19 Genome Consortium |

| | | | |
|---|---|---|---|
| EPI_ISL_3339924, EPI_ISL_3339925, EPI_ISL_3339926 | Institut, Copenhagen, Denmark | | |
| EPI_ISL_3339927 | Department of Clinical Microbiology, Odense University Hospital, Odense, Denmark | Statens Serum Institut Bioinformatics and Microbial Genomics | Danish Covid-19 Genome Consortium |
| EPI_ISL_3339928, EPI_ISL_3339929, EPI_ISL_3339930, EPI_ISL_3339931, EPI_ISL_3339932, EPI_ISL_3339933, EPI_ISL_3339934, EPI_ISL_3339935, EPI_ISL_3339936, EPI_ISL_3339937, EPI_ISL_3339938, EPI_ISL_3339939, EPI_ISL_3339940, EPI_ISL_3339941, EPI_ISL_3339942 | | | |
| see above | Department of Bacteria, Parasites and Fungi, Statens Serum Institut, Copenhagen, Denmark | Statens Serum Institut Bioinformatics and Microbial Genomics | Danish Covid-19 Genome Consortium |
| EPI_ISL_3339943 | Molekylær Medicinsk Afdeling, Aarhus University Hospital, Aarhus, Denmark | Statens Serum Institut Bioinformatics and Microbial Genomics | Danish Covid-19 Genome Consortium |
| EPI_ISL_3339944, EPI_ISL_3339945, EPI_ISL_3339946, EPI_ISL_3339947, EPI_ISL_3339948, EPI_ISL_3339949, EPI_ISL_3339950, EPI_ISL_3339951, EPI_ISL_3339952, EPI_ISL_3339953, EPI_ISL_3339954, EPI_ISL_3339955, EPI_ISL_3339956, EPI_ISL_3339957, EPI_ISL_3339958, EPI_ISL_3339959 | | | |
| see above | Department of Bacteria, Parasites and Fungi, Statens Serum Institut, Copenhagen, Denmark | Statens Serum Institut Bioinformatics and Microbial Genomics | Danish Covid-19 Genome Consortium |
| EPI_ISL_3339960 | Molekylær Medicinsk Afdeling, Aarhus University Hospital, Aarhus, Denmark | Statens Serum Institut Bioinformatics and Microbial Genomics | Danish Covid-19 Genome Consortium |
| EPI_ISL_3339961, EPI_ISL_3339962, EPI_ISL_3339963, EPI_ISL_3339964 | Department of Bacteria, Parasites and Fungi, Statens Serum Institut, Copenhagen, Denmark | Statens Serum Institut Bioinformatics and Microbial Genomics | Danish Covid-19 Genome Consortium |
| EPI_ISL_3339965 | Viešoji įstaiga Vilniaus universiteto ligoninė Santaros klinikos | Vilnius University Hospital Santaros Klinikos, Center of Laboratory Medicine | Ingrida Olendraite, Gytis Dudas, Daniel Naumovas, Ligita Raugaite, Monika Katenaite, Rimvydas Norvilas, Dovile Juozapaite |
| EPI_ISL_3339966 | Department of Bacteria, Parasites and Fungi, Statens Serum Institut, Copenhagen, Denmark | Statens Serum Institut Bioinformatics and Microbial Genomics | Danish Covid-19 Genome Consortium |
| EPI_ISL_3339967 | Viešoji įstaiga Vilniaus universiteto ligoninė Santaros klinikos | Vilnius University Hospital Santaros Klinikos, Center of Laboratory Medicine | Ingrida Olendraite, Gytis Dudas, Daniel Naumovas, Ligita Raugaite, Monika Katenaite, Rimvydas Norvilas, Dovile Juozapaite |
| EPI_ISL_3339968 | Department of Bacteria, Parasites and Fungi, Statens Serum Institut, Copenhagen, Denmark | Statens Serum Institut Bioinformatics and Microbial Genomics | Danish Covid-19 Genome Consortium |
| EPI_ISL_3339969 | Viešoji įstaiga Klaipėdos universitetinė ligoninė | Vilnius University Hospital Santaros Klinikos, Center of Laboratory Medicine | Ingrida Olendraite, Gytis Dudas, Daniel Naumovas, Ligita Raugaite, Monika Katenaite, Rimvydas Norvilas, Dovile Juozapaite |
| EPI_ISL_3339970 | Department of Bacteria, Parasites and Fungi, Statens Serum Institut, Copenhagen, Denmark | Statens Serum Institut Bioinformatics and Microbial Genomics | Danish Covid-19 Genome Consortium |
| EPI_ISL_3339971 | Viešoji įstaiga Vilniaus universiteto ligoninė Santaros klinikos | Vilnius University Hospital Santaros Klinikos, Center of Laboratory Medicine | Ingrida Olendraite, Gytis Dudas, Daniel Naumovas, Ligita Raugaite, Monika Katenaite, Rimvydas Norvilas, Dovile Juozapaite |
| EPI_ISL_3339972 | Department of Bacteria, Parasites and Fungi, Statens Serum Institut, Copenhagen, Denmark | Statens Serum Institut Bioinformatics and Microbial Genomics | Danish Covid-19 Genome Consortium |
| EPI_ISL_3339973 | Viešoji įstaiga Vilniaus universiteto ligoninė Santaros klinikos | Vilnius University Hospital Santaros Klinikos, Center of Laboratory Medicine | Ingrida Olendraite, Gytis Dudas, Daniel Naumovas, Ligita Raugaite, Monika Katenaite, Rimvydas Norvilas, Dovile Juozapaite |
| EPI_ISL_3339974 | Department of Bacteria, Parasites and Fungi, Statens Serum Institut, Copenhagen, Denmark | Statens Serum Institut Bioinformatics and Microbial Genomics | Danish Covid-19 Genome Consortium |
| EPI_ISL_3339975 | Viešoji įstaiga Klaipėdos universitetinė ligoninė | Vilnius University Hospital Santaros Klinikos, Center of Laboratory Medicine | Ingrida Olendraite, Gytis Dudas, Daniel Naumovas, Ligita Raugaite, Monika Katenaite, Rimvydas Norvilas, Dovile Juozapaite |
| EPI_ISL_3339976 | Department of Bacteria, Parasites and Fungi, Statens Serum Institut, Copenhagen, Denmark | Statens Serum Institut Bioinformatics and Microbial Genomics | Danish Covid-19 Genome Consortium |
| EPI_ISL_3339977 | Viešoji įstaiga Vilniaus universiteto ligoninė Santaros klinikos | Vilnius University Hospital Santaros Klinikos, Center of Laboratory Medicine | Ingrida Olendraite, Gytis Dudas, Daniel Naumovas, Ligita Raugaite, Monika Katenaite, Rimvydas Norvilas, Dovile Juozapaite |
| EPI_ISL_3339978 | Department of Bacteria, Parasites and Fungi, Statens Serum Institut, Copenhagen, Denmark | Statens Serum Institut Bioinformatics and Microbial Genomics | Danish Covid-19 Genome Consortium |
| EPI_ISL_3339979 | Viešoji įstaiga Klaipėdos universitetinė ligoninė | Vilnius University Hospital Santaros Klinikos, Center of Laboratory Medicine | Ingrida Olendraite, Gytis Dudas, Daniel Naumovas, Ligita Raugaite, Monika Katenaite, Rimvydas Norvilas, Dovile Juozapaite |
| EPI_ISL_3339980 | Department of Bacteria, Parasites and Fungi, Statens Serum Institut, Copenhagen, Denmark | Statens Serum Institut Bioinformatics and Microbial Genomics | Danish Covid-19 Genome Consortium |
| EPI_ISL_3339981 | Viešoji įstaiga Vilniaus universiteto ligoninė Santaros klinikos | Vilnius University Hospital Santaros Klinikos, Center of Laboratory Medicine | Ingrida Olendraite, Gytis Dudas, Daniel Naumovas, Ligita Raugaite, Monika Katenaite, Rimvydas Norvilas, Dovile Juozapaite |
| EPI_ISL_3339982 | Department of Bacteria, Parasites and Fungi, Statens Serum Institut, Copenhagen, Denmark | Statens Serum Institut Bioinformatics and Microbial Genomics | Danish Covid-19 Genome Consortium |
| EPI_ISL_3339983 | Viešoji įstaiga Klaipėdos universitetinė ligoninė | Vilnius University Hospital Santaros Klinikos, Center of Laboratory Medicine | Ingrida Olendraite, Gytis Dudas, Daniel Naumovas, Ligita Raugaite, Monika Katenaite, Rimvydas Norvilas, Dovile Juozapaite |
| EPI_ISL_3339984 | Department of Bacteria, Parasites and Fungi, Statens Serum Institut, Copenhagen, Denmark | Statens Serum Institut Bioinformatics and Microbial Genomics | Danish Covid-19 Genome Consortium |
| EPI_ISL_3339985 | Viešoji įstaiga Klaipėdos universitetinė ligoninė | Vilnius University Hospital Santaros Klinikos, Center of Laboratory Medicine | Ingrida Olendraite, Gytis Dudas, Daniel Naumovas, Ligita Raugaite, Monika Katenaite, Rimvydas Norvilas, Dovile Juozapaite |
| EPI_ISL_3339986 | Department of Bacteria, Parasites and Fungi, Statens Serum Institut, Copenhagen, Denmark | Statens Serum Institut Bioinformatics and Microbial Genomics | Danish Covid-19 Genome Consortium |
| EPI_ISL_3339987 | Viešoji įstaiga Klaipėdos universitetinė ligoninė | Vilnius University Hospital Santaros Klinikos, Center of Laboratory Medicine | Ingrida Olendraite, Gytis Dudas, Daniel Naumovas, Ligita Raugaite, Monika Katenaite, Rimvydas Norvilas, Dovile Juozapaite |
| EPI_ISL_3339988, EPI_ISL_3339990 | Department of Bacteria, Parasites and Fungi, Statens Serum Institut, Copenhagen, Denmark | Statens Serum Institut Bioinformatics and Microbial Genomics | Danish Covid-19 Genome Consortium |
| EPI_ISL_3339991 | Viešoji įstaiga Vilniaus universiteto ligoninė Santaros klinikos | Vilnius University Hospital Santaros Klinikos, Center of Laboratory Medicine | Ingrida Olendraite, Gytis Dudas, Daniel Naumovas, Ligita Raugaite, Monika Katenaite, Rimvydas Norvilas, Dovile Juozapaite |
| EPI_ISL_3339992 | Department of Bacteria, Parasites and Fungi, Statens Serum Institut, Copenhagen, Denmark | Statens Serum Institut Bioinformatics and Microbial Genomics | Danish Covid-19 Genome Consortium |
| EPI_ISL_3339993 | Viešoji įstaiga Klaipėdos universitetinė ligoninė | Vilnius University Hospital Santaros Klinikos, Center of Laboratory Medicine | Ingrida Olendraite, Gytis Dudas, Daniel Naumovas, Ligita Raugaite, Monika Katenaite, Rimvydas Norvilas, Dovile Juozapaite |
| EPI_ISL_3339994 | Department of Bacteria, Parasites and Fungi, Statens Serum Institut, Copenhagen, Denmark | Statens Serum Institut Bioinformatics and Microbial Genomics | Danish Covid-19 Genome Consortium |
| EPI_ISL_3339995 | Viešoji įstaiga Vilniaus universiteto ligoninė Santaros klinikos | Vilnius University Hospital Santaros Klinikos, Center of Laboratory Medicine | Ingrida Olendraite, Gytis Dudas, Daniel Naumovas, Ligita Raugaite, Monika Katenaite, Rimvydas Norvilas, Dovile Juozapaite |
| EPI_ISL_3339996, EPI_ISL_3339997 | Department of Bacteria, Parasites and Fungi, Statens Serum Institut, Copenhagen, Denmark | Statens Serum Institut Bioinformatics and Microbial Genomics | Danish Covid-19 Genome Consortium |
| EPI_ISL_3339998 | Viešoji įstaiga Vilniaus universiteto ligoninė Santaros klinikos | Vilnius University Hospital Santaros Klinikos, Center of Laboratory Medicine | Ingrida Olendraite, Gytis Dudas, Daniel Naumovas, Ligita Raugaite, Monika Katenaite, Rimvydas Norvilas, Dovile Juozapaite |
| EPI_ISL_3339999 | Department of Bacteria, Parasites and Fungi, Statens Serum Institut, Copenhagen, Denmark | Statens Serum Institut Bioinformatics and Microbial Genomics | Danish Covid-19 Genome Consortium |
| EPI_ISL_3340000 | Viešoji įstaiga Vilniaus universiteto ligoninė Santaros klinikos | Vilnius University Hospital Santaros Klinikos, Center of Laboratory Medicine | Ingrida Olendraite, Gytis Dudas, Daniel Naumovas, Ligita Raugaite, Monika Katenaite, Rimvydas Norvilas, Dovile Juozapaite |
| EPI_ISL_3340001 | Department of Bacteria, Parasites and Fungi, Statens Serum Institut, Copenhagen, Denmark | Statens Serum Institut Bioinformatics and Microbial Genomics | Danish Covid-19 Genome Consortium |
| EPI_ISL_3340002 | Viešoji įstaiga Vilniaus universiteto ligoninė Santaros klinikos | Vilnius University Hospital Santaros Klinikos, Center of Laboratory Medicine | Ingrida Olendraite, Gytis Dudas, Daniel Naumovas, Ligita Raugaite, Monika Katenaite, Rimvydas Norvilas, Dovile Juozapaite |
| EPI_ISL_3340003 | Department of Bacteria, Parasites and Fungi, Statens Serum Institut, Copenhagen, Denmark | Statens Serum Institut Bioinformatics and Microbial Genomics | Danish Covid-19 Genome Consortium |

| | | | |
|---|---|---|---|
| EPI_ISL_3340004 | Viešoji įstaiga Vilniaus universiteto ligoninė Santaros klinikos | Vilnius University Hospital Santaros Klinikos, Center of Laboratory Medicine | Ingrida Olendraite, Gytis Dudas, Daniel Naumovas, Ligita Raugaite, Monika Katenaite, Rimvydas Norvilas, Dovile Juozapaite |
| EPI_ISL_3340005 | Department of Bacteria, Parasites and Fungi, Statens Serum Institut, Copenhagen, Denmark | Statens Serum Institut Bioinformatics and Microbial Genomics | Danish Covid-19 Genome Consortium |
| EPI_ISL_3340006 | Viešoji įstaiga Vilniaus universiteto ligoninė Santaros klinikos | Vilnius University Hospital Santaros Klinikos, Center of Laboratory Medicine | Ingrida Olendraite, Gytis Dudas, Daniel Naumovas, Ligita Raugaite, Monika Katenaite, Rimvydas Norvilas, Dovile Juozapaite |
| EPI_ISL_3340007, EPI_ISL_3340009, EPI_ISL_3340010 | Department of Bacteria, Parasites and Fungi, Statens Serum Institut, Copenhagen, Denmark | Statens Serum Institut Bioinformatics and Microbial Genomics | Danish Covid-19 Genome Consortium |
| EPI_ISL_3340011 | Viešoji įstaiga Vilniaus universiteto ligoninė Santaros klinikos | Vilnius University Hospital Santaros Klinikos, Center of Laboratory Medicine | Ingrida Olendraite, Gytis Dudas, Daniel Naumovas, Ligita Raugaite, Monika Katenaite, Rimvydas Norvilas, Dovile Juozapaite |
| EPI_ISL_3340012 | Department of Bacteria, Parasites and Fungi, Statens Serum Institut, Copenhagen, Denmark | Statens Serum Institut Bioinformatics and Microbial Genomics | Danish Covid-19 Genome Consortium |
| EPI_ISL_3340013, EPI_ISL_3340015 | Viešoji įstaiga Vilniaus universiteto ligoninė Santaros klinikos | Vilnius University Hospital Santaros Klinikos, Center of Laboratory Medicine | Ingrida Olendraite, Gytis Dudas, Daniel Naumovas, Ligita Raugaite, Monika Katenaite, Rimvydas Norvilas, Dovile Juozapaite |
| EPI_ISL_3340017 | Viešoji įstaiga Klaipėdos universitetinė ligoninė | Vilnius University Hospital Santaros Klinikos, Center of Laboratory Medicine | Ingrida Olendraite, Gytis Dudas, Daniel Naumovas, Ligita Raugaite, Monika Katenaite, Rimvydas Norvilas, Dovile Juozapaite |
| EPI_ISL_3340018 | Department of Bacteria, Parasites and Fungi, Statens Serum Institut, Copenhagen, Denmark | Statens Serum Institut Bioinformatics and Microbial Genomics | Danish Covid-19 Genome Consortium |
| EPI_ISL_3340019 | Viešoji įstaiga Vilniaus universiteto ligoninė Santaros klinikos | Vilnius University Hospital Santaros Klinikos, Center of Laboratory Medicine | Ingrida Olendraite, Gytis Dudas, Daniel Naumovas, Ligita Raugaite, Monika Katenaite, Rimvydas Norvilas, Dovile Juozapaite |
| EPI_ISL_3340020 | Department of Bacteria, Parasites and Fungi, Statens Serum Institut, Copenhagen, Denmark | Statens Serum Institut Bioinformatics and Microbial Genomics | Danish Covid-19 Genome Consortium |
| EPI_ISL_3340021 | Viešoji įstaiga Vilniaus universiteto ligoninė Santaros klinikos | Vilnius University Hospital Santaros Klinikos, Center of Laboratory Medicine | Ingrida Olendraite, Gytis Dudas, Daniel Naumovas, Ligita Raugaite, Monika Katenaite, Rimvydas Norvilas, Dovile Juozapaite |
| EPI_ISL_3340022 | Department of Bacteria, Parasites and Fungi, Statens Serum Institut, Copenhagen, Denmark | Statens Serum Institut Bioinformatics and Microbial Genomics | Danish Covid-19 Genome Consortium |
| EPI_ISL_3340023 | Viešoji įstaiga Vilniaus universiteto ligoninė Santaros klinikos | Vilnius University Hospital Santaros Klinikos, Center of Laboratory Medicine | Ingrida Olendraite, Gytis Dudas, Daniel Naumovas, Ligita Raugaite, Monika Katenaite, Rimvydas Norvilas, Dovile Juozapaite |
| EPI_ISL_3340024, EPI_ISL_3340026 | Department of Bacteria, Parasites and Fungi, Statens Serum Institut, Copenhagen, Denmark | Statens Serum Institut Bioinformatics and Microbial Genomics | Danish Covid-19 Genome Consortium |
| EPI_ISL_3340027 | Molekylær Medicinsk Afdeling, Aarhus University Hospital, Aarhus, Denmark | Statens Serum Institut Bioinformatics and Microbial Genomics | Danish Covid-19 Genome Consortium |
| EPI_ISL_3340028 | Viešoji įstaiga Klaipėdos universitetinė ligoninė | Vilnius University Hospital Santaros Klinikos, Center of Laboratory Medicine | Ingrida Olendraite, Gytis Dudas, Daniel Naumovas, Ligita Raugaite, Monika Katenaite, Rimvydas Norvilas, Dovile Juozapaite |
| EPI_ISL_3340029 | Department of Bacteria, Parasites and Fungi, Statens Serum Institut, Copenhagen, Denmark | Statens Serum Institut Bioinformatics and Microbial Genomics | Danish Covid-19 Genome Consortium |
| EPI_ISL_3340030 | Viešoji įstaiga Klaipėdos universitetinė ligoninė | Vilnius University Hospital Santaros Klinikos, Center of Laboratory Medicine | Ingrida Olendraite, Gytis Dudas, Daniel Naumovas, Ligita Raugaite, Monika Katenaite, Rimvydas Norvilas, Dovile Juozapaite |
| EPI_ISL_3340031 | Department of Bacteria, Parasites and Fungi, Statens Serum Institut, Copenhagen, Denmark | Statens Serum Institut Bioinformatics and Microbial Genomics | Danish Covid-19 Genome Consortium |
| EPI_ISL_3340032 | Viešoji įstaiga Klaipėdos universitetinė ligoninė | Vilnius University Hospital Santaros Klinikos, Center of Laboratory Medicine | Ingrida Olendraite, Gytis Dudas, Daniel Naumovas, Ligita Raugaite, Monika Katenaite, Rimvydas Norvilas, Dovile Juozapaite |
| EPI_ISL_3340033 | Department of Bacteria, Parasites and Fungi, Statens Serum Institut, Copenhagen, Denmark | Statens Serum Institut Bioinformatics and Microbial Genomics | Danish Covid-19 Genome Consortium |
| EPI_ISL_3340034 | Viešoji įstaiga Klaipėdos universitetinė ligoninė | Vilnius University Hospital Santaros Klinikos, Center of Laboratory Medicine | Ingrida Olendraite, Gytis Dudas, Daniel Naumovas, Ligita Raugaite, Monika Katenaite, Rimvydas Norvilas, Dovile Juozapaite |
| EPI_ISL_3340035 | Department of Bacteria, Parasites and Fungi, Statens Serum Institut, Copenhagen, Denmark | Statens Serum Institut Bioinformatics and Microbial Genomics | Danish Covid-19 Genome Consortium |
| EPI_ISL_3340036 | Viešoji įstaiga Klaipėdos universitetinė ligoninė | Vilnius University Hospital Santaros Klinikos, Center of Laboratory Medicine | Ingrida Olendraite, Gytis Dudas, Daniel Naumovas, Ligita Raugaite, Monika Katenaite, Rimvydas Norvilas, Dovile Juozapaite |
| EPI_ISL_3340037 | Department of Bacteria, Parasites and Fungi, Statens Serum Institut, Copenhagen, Denmark | Statens Serum Institut Bioinformatics and Microbial Genomics | Danish Covid-19 Genome Consortium |
| EPI_ISL_3340038 | Viešoji įstaiga Klaipėdos universitetinė ligoninė | Vilnius University Hospital Santaros Klinikos, Center of Laboratory Medicine | Ingrida Olendraite, Gytis Dudas, Daniel Naumovas, Ligita Raugaite, Monika Katenaite, Rimvydas Norvilas, Dovile Juozapaite |
| EPI_ISL_3340039 | Department of Bacteria, Parasites and Fungi, Statens Serum Institut, Copenhagen, Denmark | Statens Serum Institut Bioinformatics and Microbial Genomics | Danish Covid-19 Genome Consortium |
| EPI_ISL_3340040 | Viešoji įstaiga Vilniaus universiteto ligoninė Santaros klinikos | Vilnius University Hospital Santaros Klinikos, Center of Laboratory Medicine | Ingrida Olendraite, Gytis Dudas, Daniel Naumovas, Ligita Raugaite, Monika Katenaite, Rimvydas Norvilas, Dovile Juozapaite |
| EPI_ISL_3340041 | Department of Bacteria, Parasites and Fungi, Statens Serum Institut, Copenhagen, Denmark | Statens Serum Institut Bioinformatics and Microbial Genomics | Danish Covid-19 Genome Consortium |
| EPI_ISL_3340042 | Viešoji įstaiga Vilniaus universiteto ligoninė Santaros klinikos | Vilnius University Hospital Santaros Klinikos, Center of Laboratory Medicine | Ingrida Olendraite, Gytis Dudas, Daniel Naumovas, Ligita Raugaite, Monika Katenaite, Rimvydas Norvilas, Dovile Juozapaite |
| EPI_ISL_3340044, EPI_ISL_3340046 | Viešoji įstaiga Klaipėdos universitetinė ligoninė | Vilnius University Hospital Santaros Klinikos, Center of Laboratory Medicine | Ingrida Olendraite, Gytis Dudas, Daniel Naumovas, Ligita Raugaite, Monika Katenaite, Rimvydas Norvilas, Dovile Juozapaite |
| EPI_ISL_3340047, EPI_ISL_3340048 | Department of Bacteria, Parasites and Fungi, Statens Serum Institut, Copenhagen, Denmark | Statens Serum Institut Bioinformatics and Microbial Genomics | Danish Covid-19 Genome Consortium |
| EPI_ISL_3340049 | Viešoji įstaiga Klaipėdos universitetinė ligoninė | Vilnius University Hospital Santaros Klinikos, Center of Laboratory Medicine | Ingrida Olendraite, Gytis Dudas, Daniel Naumovas, Ligita Raugaite, Monika Katenaite, Rimvydas Norvilas, Dovile Juozapaite |
| EPI_ISL_3340050 | Department of Bacteria, Parasites and Fungi, Statens Serum Institut, Copenhagen, Denmark | Statens Serum Institut Bioinformatics and Microbial Genomics | Danish Covid-19 Genome Consortium |
| EPI_ISL_3340051 | Viešoji įstaiga Vilniaus universiteto ligoninė Santaros klinikos | Vilnius University Hospital Santaros Klinikos, Center of Laboratory Medicine | Ingrida Olendraite, Gytis Dudas, Daniel Naumovas, Ligita Raugaite, Monika Katenaite, Rimvydas Norvilas, Dovile Juozapaite |
| EPI_ISL_3340052 | Department of Bacteria, Parasites and Fungi, Statens Serum Institut, Copenhagen, Denmark | Statens Serum Institut Bioinformatics and Microbial Genomics | Danish Covid-19 Genome Consortium |
| EPI_ISL_3340053 | Viešoji įstaiga Vilniaus universiteto ligoninė Santaros klinikos | Vilnius University Hospital Santaros Klinikos, Center of Laboratory Medicine | Ingrida Olendraite, Gytis Dudas, Daniel Naumovas, Ligita Raugaite, Monika Katenaite, Rimvydas Norvilas, Dovile Juozapaite |
| EPI_ISL_3340054 | Department of Bacteria, Parasites and Fungi, Statens Serum Institut, Copenhagen, Denmark | Statens Serum Institut Bioinformatics and Microbial Genomics | Danish Covid-19 Genome Consortium |
| EPI_ISL_3340055 | Viešoji įstaiga Vilniaus universiteto ligoninė Santaros klinikos | Vilnius University Hospital Santaros Klinikos, Center of Laboratory Medicine | Ingrida Olendraite, Gytis Dudas, Daniel Naumovas, Ligita Raugaite, Monika Katenaite, Rimvydas Norvilas, Dovile Juozapaite |
| EPI_ISL_3340056, EPI_ISL_3340058 | Department of Bacteria, Parasites and Fungi, Statens Serum Institut, Copenhagen, Denmark | Statens Serum Institut Bioinformatics and Microbial Genomics | Danish Covid-19 Genome Consortium |
| EPI_ISL_3340059 | Viešoji įstaiga Vilniaus universiteto ligoninė Santaros klinikos | Vilnius University Hospital Santaros Klinikos, Center of Laboratory Medicine | Ingrida Olendraite, Gytis Dudas, Daniel Naumovas, Ligita Raugaite, Monika Katenaite, Rimvydas Norvilas, Dovile Juozapaite |
| EPI_ISL_3340060 | Department of Bacteria, Parasites and Fungi, Statens Serum Institut, Copenhagen, Denmark | Statens Serum Institut Bioinformatics and Microbial Genomics | Danish Covid-19 Genome Consortium |
| EPI_ISL_3340061 | Viešoji įstaiga Klaipėdos universitetinė ligoninė | Vilnius University Hospital Santaros Klinikos, Center of Laboratory Medicine | Ingrida Olendraite, Gytis Dudas, Daniel Naumovas, Ligita Raugaite, Monika Katenaite, Rimvydas Norvilas, Dovile Juozapaite |
| EPI_ISL_3340062, | Department of Bacteria, Parasites and Fungi, Statens Serum | Statens Serum Institut Bioinformatics and Microbial Genomics | Danish Covid-19 Genome Consortium |

| | | | |
|---|---|---|---|
| EPI_ISL_3340063 | Institut, Copenhagen, Denmark | | |
| EPI_ISL_3340064 | Viešoji įstaiga Vilniaus universiteto ligoninė Santaros klinikos | Vilnius University Hospital Santaros Klinikos, Center of Laboratory Medicine | Ingrida Olendraite, Gytis Dudas, Daniel Naumovas, Ligita Raugaite, Monika Katenaite, Rimvydas Norvilas, Dovile Juozapaite |
| EPI_ISL_3340065 | Department of Bacteria, Parasites and Fungi, Statens Serum Institut, Copenhagen, Denmark | Statens Serum Institut Bioinformatics and Microbial Genomics | Danish Covid-19 Genome Consortium |
| EPI_ISL_3340066 | Viešoji įstaiga Klaipėdos universitetinė ligoninė | Vilnius University Hospital Santaros Klinikos, Center of Laboratory Medicine | Ingrida Olendraite, Gytis Dudas, Daniel Naumovas, Ligita Raugaite, Monika Katenaite, Rimvydas Norvilas, Dovile Juozapaite |
| EPI_ISL_3340067 | Department of Bacteria, Parasites and Fungi, Statens Serum Institut, Copenhagen, Denmark | Statens Serum Institut Bioinformatics and Microbial Genomics | Danish Covid-19 Genome Consortium |
| EPI_ISL_3340068 | Viešoji įstaiga Klaipėdos universitetinė ligoninė | Vilnius University Hospital Santaros Klinikos, Center of Laboratory Medicine | Ingrida Olendraite, Gytis Dudas, Daniel Naumovas, Ligita Raugaite, Monika Katenaite, Rimvydas Norvilas, Dovile Juozapaite |
| EPI_ISL_3340069 | Department of Bacteria, Parasites and Fungi, Statens Serum Institut, Copenhagen, Denmark | Statens Serum Institut Bioinformatics and Microbial Genomics | Danish Covid-19 Genome Consortium |
| EPI_ISL_3340070 | Viešoji įstaiga Klaipėdos universitetinė ligoninė | Vilnius University Hospital Santaros Klinikos, Center of Laboratory Medicine | Ingrida Olendraite, Gytis Dudas, Daniel Naumovas, Ligita Raugaite, Monika Katenaite, Rimvydas Norvilas, Dovile Juozapaite |
| EPI_ISL_3340071 | Department of Bacteria, Parasites and Fungi, Statens Serum Institut, Copenhagen, Denmark | Statens Serum Institut Bioinformatics and Microbial Genomics | Danish Covid-19 Genome Consortium |
| EPI_ISL_3340072 | Viešoji įstaiga Vilniaus universiteto ligoninė Santaros klinikos | Vilnius University Hospital Santaros Klinikos, Center of Laboratory Medicine | Ingrida Olendraite, Gytis Dudas, Daniel Naumovas, Ligita Raugaite, Monika Katenaite, Rimvydas Norvilas, Dovile Juozapaite |
| EPI_ISL_3340073 | Department of Bacteria, Parasites and Fungi, Statens Serum Institut, Copenhagen, Denmark | Statens Serum Institut Bioinformatics and Microbial Genomics | Danish Covid-19 Genome Consortium |
| EPI_ISL_3340074 | Viešoji įstaiga Klaipėdos universitetinė ligoninė | Vilnius University Hospital Santaros Klinikos, Center of Laboratory Medicine | Ingrida Olendraite, Gytis Dudas, Daniel Naumovas, Ligita Raugaite, Monika Katenaite, Rimvydas Norvilas, Dovile Juozapaite |
| EPI_ISL_3340075 | Department of Bacteria, Parasites and Fungi, Statens Serum Institut, Copenhagen, Denmark | Statens Serum Institut Bioinformatics and Microbial Genomics | Danish Covid-19 Genome Consortium |
| EPI_ISL_3340076 | Viešoji įstaiga Klaipėdos universitetinė ligoninė | Vilnius University Hospital Santaros Klinikos, Center of Laboratory Medicine | Ingrida Olendraite, Gytis Dudas, Daniel Naumovas, Ligita Raugaite, Monika Katenaite, Rimvydas Norvilas, Dovile Juozapaite |
| EPI_ISL_3340077 | Department of Bacteria, Parasites and Fungi, Statens Serum Institut, Copenhagen, Denmark | Statens Serum Institut Bioinformatics and Microbial Genomics | Danish Covid-19 Genome Consortium |
| EPI_ISL_3340078 | Viešoji įstaiga Klaipėdos universitetinė ligoninė | Vilnius University Hospital Santaros Klinikos, Center of Laboratory Medicine | Ingrida Olendraite, Gytis Dudas, Daniel Naumovas, Ligita Raugaite, Monika Katenaite, Rimvydas Norvilas, Dovile Juozapaite |
| EPI_ISL_3340079 | Department of Bacteria, Parasites and Fungi, Statens Serum Institut, Copenhagen, Denmark | Statens Serum Institut Bioinformatics and Microbial Genomics | Danish Covid-19 Genome Consortium |
| EPI_ISL_3340080 | Viešoji įstaiga Klaipėdos universitetinė ligoninė | Vilnius University Hospital Santaros Klinikos, Center of Laboratory Medicine | Ingrida Olendraite, Gytis Dudas, Daniel Naumovas, Ligita Raugaite, Monika Katenaite, Rimvydas Norvilas, Dovile Juozapaite |
| EPI_ISL_3340081 | Department of Bacteria, Parasites and Fungi, Statens Serum Institut, Copenhagen, Denmark | Statens Serum Institut Bioinformatics and Microbial Genomics | Danish Covid-19 Genome Consortium |
| EPI_ISL_3340082 | Viešoji įstaiga Klaipėdos universitetinė ligoninė | Vilnius University Hospital Santaros Klinikos, Center of Laboratory Medicine | Ingrida Olendraite, Gytis Dudas, Daniel Naumovas, Ligita Raugaite, Monika Katenaite, Rimvydas Norvilas, Dovile Juozapaite |
| EPI_ISL_3340083, EPI_ISL_3340084 | Department of Bacteria, Parasites and Fungi, Statens Serum Institut, Copenhagen, Denmark | Statens Serum Institut Bioinformatics and Microbial Genomics | Danish Covid-19 Genome Consortium |
| EPI_ISL_3340085 | Viešoji įstaiga Vilniaus universiteto ligoninė Santaros klinikos | Vilnius University Hospital Santaros Klinikos, Center of Laboratory Medicine | Ingrida Olendraite, Gytis Dudas, Daniel Naumovas, Ligita Raugaite, Monika Katenaite, Rimvydas Norvilas, Dovile Juozapaite |
| EPI_ISL_3340086 | Department of Bacteria, Parasites and Fungi, Statens Serum Institut, Copenhagen, Denmark | Statens Serum Institut Bioinformatics and Microbial Genomics | Danish Covid-19 Genome Consortium |
| EPI_ISL_3340087 | Viešoji įstaiga Klaipėdos universitetinė ligoninė | Vilnius University Hospital Santaros Klinikos, Center of Laboratory Medicine | Ingrida Olendraite, Gytis Dudas, Daniel Naumovas, Ligita Raugaite, Monika Katenaite, Rimvydas Norvilas, Dovile Juozapaite |
| EPI_ISL_3340088 | Department of Bacteria, Parasites and Fungi, Statens Serum Institut, Copenhagen, Denmark | Statens Serum Institut Bioinformatics and Microbial Genomics | Danish Covid-19 Genome Consortium |
| EPI_ISL_3340089 | Viešoji įstaiga Vilniaus universiteto ligoninė Santaros klinikos | Vilnius University Hospital Santaros Klinikos, Center of Laboratory Medicine | Ingrida Olendraite, Gytis Dudas, Daniel Naumovas, Ligita Raugaite, Monika Katenaite, Rimvydas Norvilas, Dovile Juozapaite |
| EPI_ISL_3340090 | Department of Bacteria, Parasites and Fungi, Statens Serum Institut, Copenhagen, Denmark | Statens Serum Institut Bioinformatics and Microbial Genomics | Danish Covid-19 Genome Consortium |
| EPI_ISL_3340091 | Viešoji įstaiga Klaipėdos universitetinė ligoninė | Vilnius University Hospital Santaros Klinikos, Center of Laboratory Medicine | Ingrida Olendraite, Gytis Dudas, Daniel Naumovas, Ligita Raugaite, Monika Katenaite, Rimvydas Norvilas, Dovile Juozapaite |
| EPI_ISL_3340092, EPI_ISL_3340094, EPI_ISL_3340095 | Department of Bacteria, Parasites and Fungi, Statens Serum Institut, Copenhagen, Denmark | Statens Serum Institut Bioinformatics and Microbial Genomics | Danish Covid-19 Genome Consortium |
| EPI_ISL_3340096 | Viešoji įstaiga Vilniaus universiteto ligoninė Santaros klinikos | Vilnius University Hospital Santaros Klinikos, Center of Laboratory Medicine | Ingrida Olendraite, Gytis Dudas, Daniel Naumovas, Ligita Raugaite, Monika Katenaite, Rimvydas Norvilas, Dovile Juozapaite |
| EPI_ISL_3340097 | Department of Bacteria, Parasites and Fungi, Statens Serum Institut, Copenhagen, Denmark | Statens Serum Institut Bioinformatics and Microbial Genomics | Danish Covid-19 Genome Consortium |
| EPI_ISL_3340098 | Viešoji įstaiga Vilniaus universiteto ligoninė Santaros klinikos | Vilnius University Hospital Santaros Klinikos, Center of Laboratory Medicine | Ingrida Olendraite, Gytis Dudas, Daniel Naumovas, Ligita Raugaite, Monika Katenaite, Rimvydas Norvilas, Dovile Juozapaite |
| EPI_ISL_3340099 | Department of Bacteria, Parasites and Fungi, Statens Serum Institut, Copenhagen, Denmark | Statens Serum Institut Bioinformatics and Microbial Genomics | Danish Covid-19 Genome Consortium |
| EPI_ISL_3340100 | Viešoji įstaiga Klaipėdos universitetinė ligoninė | Vilnius University Hospital Santaros Klinikos, Center of Laboratory Medicine | Ingrida Olendraite, Gytis Dudas, Daniel Naumovas, Ligita Raugaite, Monika Katenaite, Rimvydas Norvilas, Dovile Juozapaite |
| EPI_ISL_3340101 | Molekylær Medicinsk Afdeling, Aarhus University Hospital, Aarhus, Denmark | Statens Serum Institut Bioinformatics and Microbial Genomics | Danish Covid-19 Genome Consortium |
| EPI_ISL_3340102 | Viešoji įstaiga Vilniaus universiteto ligoninė Santaros klinikos | Vilnius University Hospital Santaros Klinikos, Center of Laboratory Medicine | Ingrida Olendraite, Gytis Dudas, Daniel Naumovas, Ligita Raugaite, Monika Katenaite, Rimvydas Norvilas, Dovile Juozapaite |
| EPI_ISL_3340103 | Department of Bacteria, Parasites and Fungi, Statens Serum Institut, Copenhagen, Denmark | Statens Serum Institut Bioinformatics and Microbial Genomics | Danish Covid-19 Genome Consortium |
| EPI_ISL_3340104 | Viešoji įstaiga Vilniaus universiteto ligoninė Santaros klinikos | Vilnius University Hospital Santaros Klinikos, Center of Laboratory Medicine | Ingrida Olendraite, Gytis Dudas, Daniel Naumovas, Ligita Raugaite, Monika Katenaite, Rimvydas Norvilas, Dovile Juozapaite |
| EPI_ISL_3340105 | Department of Bacteria, Parasites and Fungi, Statens Serum Institut, Copenhagen, Denmark | Statens Serum Institut Bioinformatics and Microbial Genomics | Danish Covid-19 Genome Consortium |
| EPI_ISL_3340106 | Viešoji įstaiga Vilniaus universiteto ligoninė Santaros klinikos | Vilnius University Hospital Santaros Klinikos, Center of Laboratory Medicine | Ingrida Olendraite, Gytis Dudas, Daniel Naumovas, Ligita Raugaite, Monika Katenaite, Rimvydas Norvilas, Dovile Juozapaite |
| EPI_ISL_3340107 | Department of Bacteria, Parasites and Fungi, Statens Serum Institut, Copenhagen, Denmark | Statens Serum Institut Bioinformatics and Microbial Genomics | Danish Covid-19 Genome Consortium |
| EPI_ISL_3340108 | Viešoji įstaiga Klaipėdos universitetinė ligoninė | Vilnius University Hospital Santaros Klinikos, Center of Laboratory Medicine | Ingrida Olendraite, Gytis Dudas, Daniel Naumovas, Ligita Raugaite, Monika Katenaite, Rimvydas Norvilas, Dovile Juozapaite |
| EPI_ISL_3340109, EPI_ISL_3340110 | Department of Bacteria, Parasites and Fungi, Statens Serum Institut, Copenhagen, Denmark | Statens Serum Institut Bioinformatics and Microbial Genomics | Danish Covid-19 Genome Consortium |
| EPI_ISL_3340111 | Viešoji įstaiga Vilniaus universiteto ligoninė Santaros klinikos | Vilnius University Hospital Santaros Klinikos, Center of Laboratory Medicine | Ingrida Olendraite, Gytis Dudas, Daniel Naumovas, Ligita Raugaite, Monika Katenaite, Rimvydas Norvilas, Dovile Juozapaite |
| EPI_ISL_3340112 | Department of Bacteria, Parasites and Fungi, Statens Serum Institut, Copenhagen, Denmark | Statens Serum Institut Bioinformatics and Microbial Genomics | Danish Covid-19 Genome Consortium |

| | | | |
|---|---|---|---|
| EPI_ISL_3340113 | Viešoji įstaiga Vilniaus universiteto ligoninė Santaros klinikos | Vilnius University Hospital Santaros Klinikos, Center of Laboratory Medicine | Ingrida Olendraite, Gytis Dudas, Daniel Naumovas, Ligita Raugaite, Monika Katenaite, Rimvydas Norvilas, Dovile Juozapaite |
| EPI_ISL_3340114 | Department of Bacteria, Parasites and Fungi, Statens Serum Institut, Copenhagen, Denmark | Statens Serum Institut Bioinformatics and Microbial Genomics | Danish Covid-19 Genome Consortium |
| EPI_ISL_3340115 | Viešoji įstaiga Klaipėdos universitetinė ligoninė | Vilnius University Hospital Santaros Klinikos, Center of Laboratory Medicine | Ingrida Olendraite, Gytis Dudas, Daniel Naumovas, Ligita Raugaite, Monika Katenaite, Rimvydas Norvilas, Dovile Juozapaite |
| EPI_ISL_3340116 | Department of Bacteria, Parasites and Fungi, Statens Serum Institut, Copenhagen, Denmark | Statens Serum Institut Bioinformatics and Microbial Genomics | Danish Covid-19 Genome Consortium |
| EPI_ISL_3340117 | Viešoji įstaiga Klaipėdos universitetinė ligoninė | Vilnius University Hospital Santaros Klinikos, Center of Laboratory Medicine | Ingrida Olendraite, Gytis Dudas, Daniel Naumovas, Ligita Raugaite, Monika Katenaite, Rimvydas Norvilas, Dovile Juozapaite |
| EPI_ISL_3340118 | Department of Bacteria, Parasites and Fungi, Statens Serum Institut, Copenhagen, Denmark | Statens Serum Institut Bioinformatics and Microbial Genomics | Danish Covid-19 Genome Consortium |
| EPI_ISL_3340119 | Viešoji įstaiga Klaipėdos universitetinė ligoninė | Vilnius University Hospital Santaros Klinikos, Center of Laboratory Medicine | Ingrida Olendraite, Gytis Dudas, Daniel Naumovas, Ligita Raugaite, Monika Katenaite, Rimvydas Norvilas, Dovile Juozapaite |
| EPI_ISL_3340120 | Department of Bacteria, Parasites and Fungi, Statens Serum Institut, Copenhagen, Denmark | Statens Serum Institut Bioinformatics and Microbial Genomics | Danish Covid-19 Genome Consortium |
| EPI_ISL_3340121 | Viešoji įstaiga Klaipėdos universitetinė ligoninė | Vilnius University Hospital Santaros Klinikos, Center of Laboratory Medicine | Ingrida Olendraite, Gytis Dudas, Daniel Naumovas, Ligita Raugaite, Monika Katenaite, Rimvydas Norvilas, Dovile Juozapaite |
| EPI_ISL_3340122 | Department of Bacteria, Parasites and Fungi, Statens Serum Institut, Copenhagen, Denmark | Statens Serum Institut Bioinformatics and Microbial Genomics | Danish Covid-19 Genome Consortium |
| EPI_ISL_3340123 | Viešoji įstaiga Klaipėdos universitetinė ligoninė | Vilnius University Hospital Santaros Klinikos, Center of Laboratory Medicine | Ingrida Olendraite, Gytis Dudas, Daniel Naumovas, Ligita Raugaite, Monika Katenaite, Rimvydas Norvilas, Dovile Juozapaite |
| EPI_ISL_3340124 | Department of Bacteria, Parasites and Fungi, Statens Serum Institut, Copenhagen, Denmark | Statens Serum Institut Bioinformatics and Microbial Genomics | Danish Covid-19 Genome Consortium |
| EPI_ISL_3340125 | Viešoji įstaiga Vilniaus universiteto ligoninė Santaros klinikos | Vilnius University Hospital Santaros Klinikos, Center of Laboratory Medicine | Ingrida Olendraite, Gytis Dudas, Daniel Naumovas, Ligita Raugaite, Monika Katenaite, Rimvydas Norvilas, Dovile Juozapaite |
| EPI_ISL_3340127 | Department of Bacteria, Parasites and Fungi, Statens Serum Institut, Copenhagen, Denmark | Statens Serum Institut Bioinformatics and Microbial Genomics | Danish Covid-19 Genome Consortium |
| EPI_ISL_3340128 | Viešoji įstaiga Klaipėdos universitetinė ligoninė | Vilnius University Hospital Santaros Klinikos, Center of Laboratory Medicine | Ingrida Olendraite, Gytis Dudas, Daniel Naumovas, Ligita Raugaite, Monika Katenaite, Rimvydas Norvilas, Dovile Juozapaite |
| EPI_ISL_3340129 | Department of Bacteria, Parasites and Fungi, Statens Serum Institut, Copenhagen, Denmark | Statens Serum Institut Bioinformatics and Microbial Genomics | Danish Covid-19 Genome Consortium |
| EPI_ISL_3340130 | Viešoji įstaiga Vilniaus universiteto ligoninė Santaros klinikos | Vilnius University Hospital Santaros Klinikos, Center of Laboratory Medicine | Ingrida Olendraite, Gytis Dudas, Daniel Naumovas, Ligita Raugaite, Monika Katenaite, Rimvydas Norvilas, Dovile Juozapaite |
| EPI_ISL_3340131 | Department of Bacteria, Parasites and Fungi, Statens Serum Institut, Copenhagen, Denmark | Statens Serum Institut Bioinformatics and Microbial Genomics | Danish Covid-19 Genome Consortium |
| EPI_ISL_3340132 | Viešoji įstaiga Vilniaus universiteto ligoninė Santaros klinikos | Vilnius University Hospital Santaros Klinikos, Center of Laboratory Medicine | Ingrida Olendraite, Gytis Dudas, Daniel Naumovas, Ligita Raugaite, Monika Katenaite, Rimvydas Norvilas, Dovile Juozapaite |
| EPI_ISL_3340133, EPI_ISL_3340136, EPI_ISL_3340138 | Department of Bacteria, Parasites and Fungi, Statens Serum Institut, Copenhagen, Denmark | Statens Serum Institut Bioinformatics and Microbial Genomics | Danish Covid-19 Genome Consortium |
| EPI_ISL_3340139 | Viešoji įstaiga Klaipėdos universitetinė ligoninė | Vilnius University Hospital Santaros Klinikos, Center of Laboratory Medicine | Ingrida Olendraite, Gytis Dudas, Daniel Naumovas, Ligita Raugaite, Monika Katenaite, Rimvydas Norvilas, Dovile Juozapaite |
| EPI_ISL_3340140 | Department of Bacteria, Parasites and Fungi, Statens Serum Institut, Copenhagen, Denmark | Statens Serum Institut Bioinformatics and Microbial Genomics | Danish Covid-19 Genome Consortium |
| EPI_ISL_3340141 | Viešoji įstaiga Vilniaus universiteto ligoninė Santaros klinikos | Vilnius University Hospital Santaros Klinikos, Center of Laboratory Medicine | Ingrida Olendraite, Gytis Dudas, Daniel Naumovas, Ligita Raugaite, Monika Katenaite, Rimvydas Norvilas, Dovile Juozapaite |
| EPI_ISL_3340142 | Department of Bacteria, Parasites and Fungi, Statens Serum Institut, Copenhagen, Denmark | Statens Serum Institut Bioinformatics and Microbial Genomics | Danish Covid-19 Genome Consortium |
| EPI_ISL_3340143 | Viešoji įstaiga Vilniaus universiteto ligoninė Santaros klinikos | Vilnius University Hospital Santaros Klinikos, Center of Laboratory Medicine | Ingrida Olendraite, Gytis Dudas, Daniel Naumovas, Ligita Raugaite, Monika Katenaite, Rimvydas Norvilas, Dovile Juozapaite |
| EPI_ISL_3340144 | Department of Bacteria, Parasites and Fungi, Statens Serum Institut, Copenhagen, Denmark | Statens Serum Institut Bioinformatics and Microbial Genomics | Danish Covid-19 Genome Consortium |
| EPI_ISL_3340145 | Viešoji įstaiga Vilniaus universiteto ligoninė Santaros klinikos | Vilnius University Hospital Santaros Klinikos, Center of Laboratory Medicine | Ingrida Olendraite, Gytis Dudas, Daniel Naumovas, Ligita Raugaite, Monika Katenaite, Rimvydas Norvilas, Dovile Juozapaite |
| EPI_ISL_3340146, EPI_ISL_3340147 | Department of Bacteria, Parasites and Fungi, Statens Serum Institut, Copenhagen, Denmark | Statens Serum Institut Bioinformatics and Microbial Genomics | Danish Covid-19 Genome Consortium |
| EPI_ISL_3340148 | Viešoji įstaiga Klaipėdos universitetinė ligoninė | Vilnius University Hospital Santaros Klinikos, Center of Laboratory Medicine | Ingrida Olendraite, Gytis Dudas, Daniel Naumovas, Ligita Raugaite, Monika Katenaite, Rimvydas Norvilas, Dovile Juozapaite |
| EPI_ISL_3340149, EPI_ISL_3340150, EPI_ISL_3340151, EPI_ISL_3340152, EPI_ISL_3340153, EPI_ISL_3340154, EPI_ISL_3340155, EPI_ISL_3340157, EPI_ISL_3340158, EPI_ISL_3340159, EPI_ISL_3340160, EPI_ISL_3340161, EPI_ISL_3340162, EPI_ISL_3340163, EPI_ISL_3340164, EPI_ISL_3340165, EPI_ISL_3340166, EPI_ISL_3340167, EPI_ISL_3340168, EPI_ISL_3340169, EPI_ISL_3340170, EPI_ISL_3340171, EPI_ISL_3340172, EPI_ISL_3340173, EPI_ISL_3340174, EPI_ISL_3340175, EPI_ISL_3340176, EPI_ISL_3340177, EPI_ISL_3340178, EPI_ISL_3340179, EPI_ISL_3340180, EPI_ISL_3340181, EPI_ISL_3340182, EPI_ISL_3340183, EPI_ISL_3340184, EPI_ISL_3340185, EPI_ISL_3340186, EPI_ISL_3340187, EPI_ISL_3340188, EPI_ISL_3340189, EPI_ISL_3340190, EPI_ISL_3340191, EPI_ISL_3340192 | | | |
| see above | Department of Bacteria, Parasites and Fungi, Statens Serum Institut, Copenhagen, Denmark | Statens Serum Institut Bioinformatics and Microbial Genomics | Danish Covid-19 Genome Consortium |
| EPI_ISL_3340193 | Molekylær Medicinsk Afdeling, Aarhus University Hospital, Aarhus, Denmark | Statens Serum Institut Bioinformatics and Microbial Genomics | Danish Covid-19 Genome Consortium |
| EPI_ISL_3340194, EPI_ISL_3340195, EPI_ISL_3340196, EPI_ISL_3340197, EPI_ISL_3340198, EPI_ISL_3340199, EPI_ISL_3340200, EPI_ISL_3340201, EPI_ISL_3340202, EPI_ISL_3340203, EPI_ISL_3340204, EPI_ISL_3340205, EPI_ISL_3340206, EPI_ISL_3340207, EPI_ISL_3340208, EPI_ISL_3340209, EPI_ISL_3340211, EPI_ISL_3340212, EPI_ISL_3340213, EPI_ISL_3340214, EPI_ISL_3340215 | | | |
| see above | Department of Bacteria, Parasites and Fungi, Statens Serum Institut, Copenhagen, Denmark | Statens Serum Institut Bioinformatics and Microbial Genomics | Danish Covid-19 Genome Consortium |
| EPI_ISL_3340216 | Molekylær Medicinsk Afdeling, Aarhus University Hospital, Aarhus, Denmark | Statens Serum Institut Bioinformatics and Microbial Genomics | Danish Covid-19 Genome Consortium |
| EPI_ISL_3340217, EPI_ISL_3340218, EPI_ISL_3340219, EPI_ISL_3340220, EPI_ISL_3340221, EPI_ISL_3340222, EPI_ISL_3340223, EPI_ISL_3340224, EPI_ISL_3340225, EPI_ISL_3340226, EPI_ISL_3340227, EPI_ISL_3340228, EPI_ISL_3340229, EPI_ISL_3340230, EPI_ISL_3340232, EPI_ISL_3340233, EPI_ISL_3340234, EPI_ISL_3340235, EPI_ISL_3340236, EPI_ISL_3340237, EPI_ISL_3340238, EPI_ISL_3340239, EPI_ISL_3340241, EPI_ISL_3340242, EPI_ISL_3340243, EPI_ISL_3340244, EPI_ISL_3340246, EPI_ISL_3340248, EPI_ISL_3340249, EPI_ISL_3340250, EPI_ISL_3340251, EPI_ISL_3340252, EPI_ISL_3340253, EPI_ISL_3340254, EPI_ISL_3340255, EPI_ISL_3340256, EPI_ISL_3340257, EPI_ISL_3340258, EPI_ISL_3340259, EPI_ISL_3340260, EPI_ISL_3340261, EPI_ISL_3340262, EPI_ISL_3340263, EPI_ISL_3340264, EPI_ISL_3340265, EPI_ISL_3340266, EPI_ISL_3340267, EPI_ISL_3340268, EPI_ISL_3340269, EPI_ISL_3340270, EPI_ISL_3340271, EPI_ISL_3340272, EPI_ISL_3340273, EPI_ISL_3340274, EPI_ISL_3340275, EPI_ISL_3340276, EPI_ISL_3340277, EPI_ISL_3340278, EPI_ISL_3340279, EPI_ISL_3340280, EPI_ISL_3340281, EPI_ISL_3340282, EPI_ISL_3340283, EPI_ISL_3340284, EPI_ISL_3340285, EPI_ISL_3340286, EPI_ISL_3340287, EPI_ISL_3340288, EPI_ISL_3340289, EPI_ISL_3340290, EPI_ISL_3340291, EPI_ISL_3340292, EPI_ISL_3340293, EPI_ISL_3340294, EPI_ISL_3340295, EPI_ISL_3340296, EPI_ISL_3340297, EPI_ISL_3340298, EPI_ISL_3340299, EPI_ISL_3340300, EPI_ISL_3340301 | | | |
| see above | Department of Bacteria, Parasites and Fungi, Statens Serum Institut, Copenhagen, Denmark | Statens Serum Institut Bioinformatics and Microbial Genomics | Danish Covid-19 Genome Consortium |
| EPI_ISL_3340303 | Department of Clinical Microbiology, Odense University Hospital, Odense, Denmark | Statens Serum Institut Bioinformatics and Microbial Genomics | Danish Covid-19 Genome Consortium |
| EPI_ISL_3340304, EPI_ISL_3340305, EPI_ISL_3340306, EPI_ISL_3340307, EPI_ISL_3340308, EPI_ISL_3340309, EPI_ISL_3340310, EPI_ISL_3340311, EPI_ISL_3340312, EPI_ISL_3340313, EPI_ISL_3340314, EPI_ISL_3340315, EPI_ISL_3340316, EPI_ISL_3340317, EPI_ISL_3340318, EPI_ISL_3340319, EPI_ISL_3340320, EPI_ISL_3340322, EPI_ISL_3340323, EPI_ISL_3340324, EPI_ISL_3340325, EPI_ISL_3340326, EPI_ISL_3340327, EPI_ISL_3340328, EPI_ISL_3340329, EPI_ISL_3340330, EPI_ISL_3340331, EPI_ISL_3340332, EPI_ISL_3340333, EPI_ISL_3340334, EPI_ISL_3340335, EPI_ISL_3340336, EPI_ISL_3340337, EPI_ISL_3340338, EPI_ISL_3340339, EPI_ISL_3340340, EPI_ISL_3340341, EPI_ISL_3340342, EPI_ISL_3340343, EPI_ISL_3340344, EPI_ISL_3340345, EPI_ISL_3340346, EPI_ISL_3340348, EPI_ISL_3340349, EPI_ISL_3340350, EPI_ISL_3340351, EPI_ISL_3340352, EPI_ISL_3340353, EPI_ISL_3340354, EPI_ISL_3340355, EPI_ISL_3340356, EPI_ISL_3340357, EPI_ISL_3340358, EPI_ISL_3340359, EPI_ISL_3340360, EPI_ISL_3340361, EPI_ISL_3340362, EPI_ISL_3340363, EPI_ISL_3340364, EPI_ISL_3340365, EPI_ISL_3340366, EPI_ISL_3340367, EPI_ISL_3340368, EPI_ISL_3340369, EPI_ISL_3340370, EPI_ISL_3340371, EPI_ISL_3340373, EPI_ISL_3340374, EPI_ISL_3340375, EPI_ISL_3340376, EPI_ISL_3340377, EPI_ISL_3340378, EPI_ISL_3340379, EPI_ISL_3340380, EPI_ISL_3340381, EPI_ISL_3340382, EPI_ISL_3340383, EPI_ISL_3340384, EPI_ISL_3340386, EPI_ISL_3340387, EPI_ISL_3340388, EPI_ISL_3340389, EPI_ISL_3340390, EPI_ISL_3340391 | | | |
| see above | Department of Bacteria, Parasites and Fungi, Statens Serum Institut, Copenhagen, Denmark | Statens Serum Institut Bioinformatics and Microbial Genomics | Danish Covid-19 Genome Consortium |
| EPI_ISL_3340392 | Molekylær Medicinsk Afdeling, Aarhus University Hospital, Aarhus, Denmark | Statens Serum Institut Bioinformatics and Microbial Genomics | Danish Covid-19 Genome Consortium |
| EPI_ISL_3340393, EPI_ISL_3340394, EPI_ISL_3340395, EPI_ISL_3340396, EPI_ISL_3340397, EPI_ISL_3340398, EPI_ISL_3340399, EPI_ISL_3340400, EPI_ISL_3340402, EPI_ISL_3340403, EPI_ISL_3340404, EPI_ISL_3340405, EPI_ISL_3340406, EPI_ISL_3340407, EPI_ISL_3340409, EPI_ISL_3340410, EPI_ISL_3340411, EPI_ISL_3340412, EPI_ISL_3340413, EPI_ISL_3340414, EPI_ISL_3340415, EPI_ISL_3340416, EPI_ISL_3340417, EPI_ISL_3340418, EPI_ISL_3340419, EPI_ISL_3340420, EPI_ISL_3340421, EPI_ISL_3340422, EPI_ISL_3340423, EPI_ISL_3340424, EPI_ISL_3340425, EPI_ISL_3340426, EPI_ISL_3340427, EPI_ISL_3340428, EPI_ISL_3340429 | | | |
| see above | Department of Bacteria, Parasites and Fungi, Statens Serum Institut, Copenhagen, Denmark | Statens Serum Institut Bioinformatics and Microbial Genomics | Danish Covid-19 Genome Consortium |
| EPI_ISL_3340430 | Department of Clinical Microbiology, Odense University Hospital, | Statens Serum Institut Bioinformatics and Microbial Genomics | Danish Covid-19 Genome Consortium |

| | | | |
|---|---|---|---|
| | Odense, Denmark | | |
| EPI_ISL_3340431, EPI_ISL_3340432, EPI_ISL_3340433, EPI_ISL_3340434, EPI_ISL_3340435, EPI_ISL_3340436, EPI_ISL_3340437, EPI_ISL_3340438, EPI_ISL_3340439, EPI_ISL_3340440, EPI_ISL_3340441, EPI_ISL_3340442, EPI_ISL_3340443, EPI_ISL_3340444, EPI_ISL_3340446, EPI_ISL_3340447, EPI_ISL_3340448, EPI_ISL_3340449, EPI_ISL_3340450, EPI_ISL_3340451, EPI_ISL_3340452, EPI_ISL_3340453, EPI_ISL_3340454 | | | |
| see above | Department of Bacteria, Parasites and Fungi, Statens Serum Institut, Copenhagen, Denmark | Statens Serum Institut Bioinformatics and Microbial Genomics | Danish Covid-19 Genome Consortium |
| EPI_ISL_3340455 | Molekylær Medicinsk Afdeling, Aarhus University Hospital, Aarhus, Denmark | Statens Serum Institut Bioinformatics and Microbial Genomics | Danish Covid-19 Genome Consortium |
| EPI_ISL_3340456, EPI_ISL_3340457, EPI_ISL_3340458, EPI_ISL_3340459, EPI_ISL_3340460, EPI_ISL_3340461, EPI_ISL_3340462, EPI_ISL_3340463, EPI_ISL_3340464, EPI_ISL_3340465, EPI_ISL_3340466, EPI_ISL_3340467, EPI_ISL_3340468, EPI_ISL_3340469, EPI_ISL_3340470, EPI_ISL_3340471, EPI_ISL_3340472, EPI_ISL_3340473, EPI_ISL_3340474, EPI_ISL_3340475, EPI_ISL_3340476, EPI_ISL_3340477, EPI_ISL_3340478, EPI_ISL_3340479, EPI_ISL_3340480, EPI_ISL_3340482, EPI_ISL_3340483, EPI_ISL_3340484, EPI_ISL_3340485, EPI_ISL_3340487, EPI_ISL_3340488, EPI_ISL_3340489, EPI_ISL_3340490, EPI_ISL_3340491, EPI_ISL_3340492, EPI_ISL_3340493, EPI_ISL_3340494, EPI_ISL_3340495, EPI_ISL_3340496, EPI_ISL_3340497, EPI_ISL_3340498, EPI_ISL_3340499, EPI_ISL_3340500, EPI_ISL_3340501, EPI_ISL_3340502, EPI_ISL_3340503, EPI_ISL_3340504, EPI_ISL_3340505, EPI_ISL_3340506, EPI_ISL_3340507, EPI_ISL_3340508, EPI_ISL_3340509, EPI_ISL_3340510, EPI_ISL_3340511, EPI_ISL_3340512, EPI_ISL_3340513, EPI_ISL_3340514, EPI_ISL_3340515, EPI_ISL_3340516, EPI_ISL_3340517, EPI_ISL_3340518, EPI_ISL_3340519, EPI_ISL_3340521, EPI_ISL_3340522, EPI_ISL_3340523, EPI_ISL_3340524, EPI_ISL_3340527, EPI_ISL_3340528, EPI_ISL_3340529, EPI_ISL_3340530, EPI_ISL_3340531, EPI_ISL_3340532, EPI_ISL_3340533, EPI_ISL_3340534, EPI_ISL_3340535, EPI_ISL_3340536, EPI_ISL_3340537, EPI_ISL_3340538, EPI_ISL_3340539, EPI_ISL_3340540, EPI_ISL_3340541, EPI_ISL_3340542, EPI_ISL_3340543, EPI_ISL_3340544, EPI_ISL_3340545, EPI_ISL_3340546, EPI_ISL_3340547, EPI_ISL_3340548, EPI_ISL_3340549, EPI_ISL_3340550, EPI_ISL_3340551, EPI_ISL_3340552, EPI_ISL_3340553, EPI_ISL_3340554, EPI_ISL_3340555, EPI_ISL_3340556, EPI_ISL_3340557, EPI_ISL_3340558, EPI_ISL_3340559, EPI_ISL_3340560, EPI_ISL_3340561, EPI_ISL_3340562, EPI_ISL_3340563, EPI_ISL_3340564, EPI_ISL_3340565, EPI_ISL_3340566, EPI_ISL_3340567, EPI_ISL_3340568, EPI_ISL_3340569, EPI_ISL_3340570, EPI_ISL_3340571, EPI_ISL_3340572, EPI_ISL_3340573, EPI_ISL_3340574, EPI_ISL_3340575, EPI_ISL_3340576, EPI_ISL_3340577, EPI_ISL_3340578, EPI_ISL_3340579, EPI_ISL_3340580, EPI_ISL_3340581, EPI_ISL_3340583, EPI_ISL_3340585, EPI_ISL_3340586, EPI_ISL_3340587, EPI_ISL_3340588, EPI_ISL_3340589, EPI_ISL_3340590, EPI_ISL_3340591, EPI_ISL_3340592, EPI_ISL_3340593, EPI_ISL_3340594 | | | |
| see above | Department of Bacteria, Parasites and Fungi, Statens Serum Institut, Copenhagen, Denmark | Statens Serum Institut Bioinformatics and Microbial Genomics | Danish Covid-19 Genome Consortium |
| EPI_ISL_3340595 | Molekylær Medicinsk Afdeling, Aarhus University Hospital, Aarhus, Denmark | Statens Serum Institut Bioinformatics and Microbial Genomics | Danish Covid-19 Genome Consortium |
| EPI_ISL_3340596, EPI_ISL_3340597, EPI_ISL_3340598, EPI_ISL_3340599, EPI_ISL_3340600, EPI_ISL_3340601, EPI_ISL_3340602, EPI_ISL_3340603, EPI_ISL_3340604, EPI_ISL_3340606, EPI_ISL_3340607, EPI_ISL_3340608, EPI_ISL_3340609, EPI_ISL_3340610, EPI_ISL_3340611, EPI_ISL_3340612, EPI_ISL_3340613, EPI_ISL_3340614, EPI_ISL_3340615, EPI_ISL_3340616, EPI_ISL_3340618, EPI_ISL_3340619, EPI_ISL_3340620, EPI_ISL_3340621, EPI_ISL_3340623, EPI_ISL_3340624, EPI_ISL_3340625, EPI_ISL_3340627 | | | |
| see above | Department of Bacteria, Parasites and Fungi, Statens Serum Institut, Copenhagen, Denmark | Statens Serum Institut Bioinformatics and Microbial Genomics | Danish Covid-19 Genome Consortium |
| EPI_ISL_3340628 | Molekylær Medicinsk Afdeling, Aarhus University Hospital, Aarhus, Denmark | Statens Serum Institut Bioinformatics and Microbial Genomics | Danish Covid-19 Genome Consortium |
| EPI_ISL_3340629, EPI_ISL_3340630, EPI_ISL_3340631, EPI_ISL_3340632, EPI_ISL_3340633, EPI_ISL_3340634, EPI_ISL_3340635, EPI_ISL_3340636, EPI_ISL_3340637, EPI_ISL_3340638, EPI_ISL_3340639, EPI_ISL_3340640, EPI_ISL_3340641, EPI_ISL_3340642, EPI_ISL_3340643, EPI_ISL_3340644, EPI_ISL_3340645, EPI_ISL_3340646, EPI_ISL_3340647, EPI_ISL_3340648, EPI_ISL_3340649, EPI_ISL_3340650, EPI_ISL_3340651, EPI_ISL_3340652, EPI_ISL_3340653, EPI_ISL_3340654, EPI_ISL_3340655, EPI_ISL_3340656, EPI_ISL_3340657, EPI_ISL_3340658, EPI_ISL_3340659, EPI_ISL_3340660, EPI_ISL_3340661, EPI_ISL_3340662, EPI_ISL_3340663, EPI_ISL_3340664, EPI_ISL_3340665, EPI_ISL_3340666, EPI_ISL_3340667, EPI_ISL_3340668, EPI_ISL_3340670, EPI_ISL_3340671, EPI_ISL_3340672, EPI_ISL_3340673 | | | |
| see above | Department of Bacteria, Parasites and Fungi, Statens Serum Institut, Copenhagen, Denmark | Statens Serum Institut Bioinformatics and Microbial Genomics | Danish Covid-19 Genome Consortium |
| EPI_ISL_3340674 | Molekylær Medicinsk Afdeling, Aarhus University Hospital, Aarhus, Denmark | Statens Serum Institut Bioinformatics and Microbial Genomics | Danish Covid-19 Genome Consortium |
| EPI_ISL_3340675, EPI_ISL_3340676, EPI_ISL_3340677, EPI_ISL_3340678, EPI_ISL_3340679 | Department of Bacteria, Parasites and Fungi, Statens Serum Institut, Copenhagen, Denmark | Statens Serum Institut Bioinformatics and Microbial Genomics | Danish Covid-19 Genome Consortium |
| EPI_ISL_3340680 | UAB InMedica | unknown | Rasa Ugenskiene, Darius Cereskevicius, Inga Nasvytiene, Zivile Zemeckiene, Mantas Sarauskas, Marius Sukys, Rima Vainoriene, Astra Vitkauskiene, Renaldas Jurkevicius |
| EPI_ISL_3340681 | UAB Medicina practica laboratorija | unknown | Rasa Ugenskiene, Darius Cereskevicius, Inga Nasvytiene, Zivile Zemeckiene, Mantas Sarauskas, Marius Sukys, Rima Vainoriene, Astra Vitkauskiene, Renaldas Jurkevicius |
| EPI_ISL_3340682, EPI_ISL_3340683 | UAB InMedica | unknown | Rasa Ugenskiene, Darius Cereskevicius, Inga Nasvytiene, Zivile Zemeckiene, Mantas Sarauskas, Marius Sukys, Rima Vainoriene, Astra Vitkauskiene, Renaldas Jurkevicius |
| EPI_ISL_3340684, EPI_ISL_3340685 | UAB Medicina practica laboratorija | unknown | Rasa Ugenskiene, Darius Cereskevicius, Inga Nasvytiene, Zivile Zemeckiene, Mantas Sarauskas, Marius Sukys, Rima Vainoriene, Astra Vitkauskiene, Renaldas Jurkevicius |
| EPI_ISL_3340686 | UAB InMedica | unknown | Rasa Ugenskiene, Darius Cereskevicius, Inga Nasvytiene, Zivile Zemeckiene, Mantas Sarauskas, Marius Sukys, Rima Vainoriene, Astra Vitkauskiene, Renaldas Jurkevicius |
| EPI_ISL_3340687 | UAB Medicina practica laboratorija | unknown | Rasa Ugenskiene, Darius Cereskevicius, Inga Nasvytiene, Zivile Zemeckiene, Mantas Sarauskas, Marius Sukys, Rima Vainoriene, Astra Vitkauskiene, Renaldas Jurkevicius |
| EPI_ISL_3340688 | UAB InMedica | unknown | Rasa Ugenskiene, Darius Cereskevicius, Inga Nasvytiene, Zivile Zemeckiene, Mantas Sarauskas, Marius Sukys, Rima Vainoriene, Astra Vitkauskiene, Renaldas Jurkevicius |
| EPI_ISL_3340689 | UAB Medicina practica laboratorija | unknown | Rasa Ugenskiene, Darius Cereskevicius, Inga Nasvytiene, Zivile Zemeckiene, Mantas Sarauskas, Marius Sukys, Rima Vainoriene, Astra Vitkauskiene, Renaldas Jurkevicius |
| EPI_ISL_3340690 | UAB InMedica | unknown | Rasa Ugenskiene, Darius Cereskevicius, Inga Nasvytiene, Zivile Zemeckiene, Mantas Sarauskas, Marius Sukys, Rima Vainoriene, Astra Vitkauskiene, Renaldas Jurkevicius |
| EPI_ISL_3340691 | UAB Medicina practica laboratorija | unknown | Rasa Ugenskiene, Darius Cereskevicius, Inga Nasvytiene, Zivile Zemeckiene, Mantas Sarauskas, Marius Sukys, Rima Vainoriene, Astra Vitkauskiene, Renaldas Jurkevicius |
| EPI_ISL_3340692 | UAB InMedica | unknown | Rasa Ugenskiene, Darius Cereskevicius, Inga Nasvytiene, Zivile Zemeckiene, Mantas Sarauskas, Marius Sukys, Rima Vainoriene, Astra Vitkauskiene, Renaldas Jurkevicius |
| EPI_ISL_3340693, EPI_ISL_3340694 | UAB Medicina practica laboratorija | unknown | Rasa Ugenskiene, Darius Cereskevicius, Inga Nasvytiene, Zivile Zemeckiene, Mantas Sarauskas, Marius Sukys, Rima Vainoriene, Astra Vitkauskiene, Renaldas Jurkevicius |
| EPI_ISL_3340695, EPI_ISL_3340696 | UAB InMedica | unknown | Rasa Ugenskiene, Darius Cereskevicius, Inga Nasvytiene, Zivile Zemeckiene, Mantas Sarauskas, Marius Sukys, Rima Vainoriene, Astra Vitkauskiene, Renaldas Jurkevicius |
| EPI_ISL_3340697, EPI_ISL_3340698 | UAB Medicina practica laboratorija | unknown | Rasa Ugenskiene, Darius Cereskevicius, Inga Nasvytiene, Zivile Zemeckiene, Mantas Sarauskas, Marius Sukys, Rima Vainoriene, Astra Vitkauskiene, Renaldas Jurkevicius |
| EPI_ISL_3340699 | UAB InMedica | unknown | Rasa Ugenskiene, Darius Cereskevicius, Inga Nasvytiene, Zivile Zemeckiene, Mantas Sarauskas, Marius Sukys, Rima Vainoriene, Astra Vitkauskiene, Renaldas Jurkevicius |
| EPI_ISL_3340700, EPI_ISL_3340701, EPI_ISL_3340702 | UAB Medicina practica laboratorija | unknown | Rasa Ugenskiene, Darius Cereskevicius, Inga Nasvytiene, Zivile Zemeckiene, Mantas Sarauskas, Marius Sukys, Rima Vainoriene, Astra Vitkauskiene, Renaldas Jurkevicius |
| EPI_ISL_3340703, EPI_ISL_3340704 | UAB InMedica | unknown | Rasa Ugenskiene, Darius Cereskevicius, Inga Nasvytiene, Zivile Zemeckiene, Mantas Sarauskas, Marius Sukys, Rima Vainoriene, Astra Vitkauskiene, Renaldas Jurkevicius |
| EPI_ISL_3340705 | UAB Medicina practica laboratorija | unknown | Rasa Ugenskiene, Darius Cereskevicius, Inga Nasvytiene, Zivile Zemeckiene, Mantas Sarauskas, Marius Sukys, Rima Vainoriene, Astra Vitkauskiene, Renaldas Jurkevicius |
| EPI_ISL_3340706, EPI_ISL_3340707 | UAB InMedica | unknown | Rasa Ugenskiene, Darius Cereskevicius, Inga Nasvytiene, Zivile Zemeckiene, Mantas Sarauskas, Marius Sukys, Rima Vainoriene, Astra Vitkauskiene, Renaldas Jurkevicius |
| EPI_ISL_3340708, EPI_ISL_3340709, EPI_ISL_3340710, EPI_ISL_3340711, EPI_ISL_3340712 | UAB Medicina practica laboratorija | unknown | Rasa Ugenskiene, Darius Cereskevicius, Inga Nasvytiene, Zivile Zemeckiene, Mantas Sarauskas, Marius Sukys, Rima Vainoriene, Astra Vitkauskiene, Renaldas Jurkevicius |
| EPI_ISL_3340713, EPI_ISL_3340714 | UAB InMedica | unknown | Rasa Ugenskiene, Darius Cereskevicius, Inga Nasvytiene, Zivile Zemeckiene, Mantas Sarauskas, Marius Sukys, Rima Vainoriene, Astra Vitkauskiene, Renaldas Jurkevicius |
| EPI_ISL_3340715 | UAB Medicina practica laboratorija | unknown | Rasa Ugenskiene, Darius Cereskevicius, Inga Nasvytiene, Zivile Zemeckiene, Mantas Sarauskas, Marius Sukys, Rima Vainoriene, Astra Vitkauskiene, Renaldas Jurkevicius |
| EPI_ISL_3340716, EPI_ISL_3340717, EPI_ISL_3340718, EPI_ISL_3340719 | UAB InMedica | unknown | Rasa Ugenskiene, Darius Cereskevicius, Inga Nasvytiene, Zivile Zemeckiene, Mantas Sarauskas, Marius Sukys, Rima Vainoriene, Astra Vitkauskiene, Renaldas Jurkevicius |
| EPI_ISL_3340720 | UAB Medicina practica laboratorija | unknown | Rasa Ugenskiene, Darius Cereskevicius, Inga Nasvytiene, Zivile Zemeckiene, Mantas Sarauskas, Marius Sukys, Rima Vainoriene, Astra Vitkauskiene, Renaldas Jurkevicius |
| EPI_ISL_3340721, EPI_ISL_3340722 | UAB InMedica | unknown | Rasa Ugenskiene, Darius Cereskevicius, Inga Nasvytiene, Zivile Zemeckiene, Mantas Sarauskas, Marius Sukys, Rima Vainoriene, Astra Vitkauskiene, Renaldas Jurkevicius |
| EPI_ISL_3340723, EPI_ISL_3340724 | UAB Medicina practica laboratorija | unknown | Rasa Ugenskiene, Darius Cereskevicius, Inga Nasvytiene, Zivile Zemeckiene, Mantas Sarauskas, Marius Sukys, Rima Vainoriene, Astra Vitkauskiene, Renaldas Jurkevicius |
| EPI_ISL_3340725 | UAB InMedica | unknown | Rasa Ugenskiene, Darius Cereskevicius, Inga Nasvytiene, Zivile Zemeckiene, Mantas Sarauskas, Marius Sukys, Rima Vainoriene, Astra Vitkauskiene, Renaldas Jurkevicius |
| EPI_ISL_3340726 | UAB Medicina practica laboratorija | unknown | Rasa Ugenskiene, Darius Cereskevicius, Inga Nasvytiene, Zivile Zemeckiene, Mantas Sarauskas, Marius Sukys, Rima Vainoriene, Astra Vitkauskiene, Renaldas Jurkevicius |
| EPI_ISL_3340727 | UAB InMedica | unknown | Rasa Ugenskiene, Darius Cereskevicius, Inga Nasvytiene, Zivile Zemeckiene, Mantas Sarauskas, Marius Sukys, Rima Vainoriene, Astra Vitkauskiene, Renaldas Jurkevicius |
| EPI_ISL_3340728, EPI_ISL_3340729, EPI_ISL_3340730, EPI_ISL_3340731, EPI_ISL_3340732 | UAB Medicina practica laboratorija | unknown | Rasa Ugenskiene, Darius Cereskevicius, Inga Nasvytiene, Zivile Zemeckiene, Mantas Sarauskas, Marius Sukys, Rima Vainoriene, Astra Vitkauskiene, Renaldas Jurkevicius |
| EPI_ISL_3340733, EPI_ISL_3340734, EPI_ISL_3340735 | UAB InMedica | unknown | Rasa Ugenskiene, Darius Cereskevicius, Inga Nasvytiene, Zivile Zemeckiene, Mantas Sarauskas, Marius Sukys, Rima Vainoriene, Astra Vitkauskiene, Renaldas Jurkevicius |

| | | | |
|---|---|---|---|
| EPI_ISL_3340736 | UAB Medicina practica laboratorija | unknown | Rasa Ugenskiene, Darius Cereskevicius, Inga Nasvytiene, Zivile Zemeckiene, Mantas Sarauskas, Marius Sukys, Rima Vainoriene, Astra Vitkauskiene, Renaldas Jurkevicius |
| EPI_ISL_3340737, EPI_ISL_3340738 | UAB InMedica | unknown | Rasa Ugenskiene, Darius Cereskevicius, Inga Nasvytiene, Zivile Zemeckiene, Mantas Sarauskas, Marius Sukys, Rima Vainoriene, Astra Vitkauskiene, Renaldas Jurkevicius |
| EPI_ISL_3340739 | UAB Medicina practica laboratorija | unknown | Rasa Ugenskiene, Darius Cereskevicius, Inga Nasvytiene, Zivile Zemeckiene, Mantas Sarauskas, Marius Sukys, Rima Vainoriene, Astra Vitkauskiene, Renaldas Jurkevicius |
| EPI_ISL_3340740, EPI_ISL_3340741 | UAB InMedica | unknown | Rasa Ugenskiene, Darius Cereskevicius, Inga Nasvytiene, Zivile Zemeckiene, Mantas Sarauskas, Marius Sukys, Rima Vainoriene, Astra Vitkauskiene, Renaldas Jurkevicius |
| EPI_ISL_3340742 | UAB Medicina practica laboratorija | unknown | Rasa Ugenskiene, Darius Cereskevicius, Inga Nasvytiene, Zivile Zemeckiene, Mantas Sarauskas, Marius Sukys, Rima Vainoriene, Astra Vitkauskiene, Renaldas Jurkevicius |
| EPI_ISL_3340743, EPI_ISL_3340744 | UAB InMedica | unknown | Rasa Ugenskiene, Darius Cereskevicius, Inga Nasvytiene, Zivile Zemeckiene, Mantas Sarauskas, Marius Sukys, Rima Vainoriene, Astra Vitkauskiene, Renaldas Jurkevicius |
| EPI_ISL_3340745 | UAB Medicina practica laboratorija | unknown | Rasa Ugenskiene, Darius Cereskevicius, Inga Nasvytiene, Zivile Zemeckiene, Mantas Sarauskas, Marius Sukys, Rima Vainoriene, Astra Vitkauskiene, Renaldas Jurkevicius |
| EPI_ISL_3340746, EPI_ISL_3340747, EPI_ISL_3340748 | UAB InMedica | unknown | Rasa Ugenskiene, Darius Cereskevicius, Inga Nasvytiene, Zivile Zemeckiene, Mantas Sarauskas, Marius Sukys, Rima Vainoriene, Astra Vitkauskiene, Renaldas Jurkevicius |
| EPI_ISL_3340749, EPI_ISL_3340750 | UAB Medicina practica laboratorija | unknown | Rasa Ugenskiene, Darius Cereskevicius, Inga Nasvytiene, Zivile Zemeckiene, Mantas Sarauskas, Marius Sukys, Rima Vainoriene, Astra Vitkauskiene, Renaldas Jurkevicius |
| EPI_ISL_3340751 | UAB InMedica | unknown | Rasa Ugenskiene, Darius Cereskevicius, Inga Nasvytiene, Zivile Zemeckiene, Mantas Sarauskas, Marius Sukys, Rima Vainoriene, Astra Vitkauskiene, Renaldas Jurkevicius |
| EPI_ISL_3340752 | UAB Medicina practica laboratorija | unknown | Rasa Ugenskiene, Darius Cereskevicius, Inga Nasvytiene, Zivile Zemeckiene, Mantas Sarauskas, Marius Sukys, Rima Vainoriene, Astra Vitkauskiene, Renaldas Jurkevicius |
| EPI_ISL_3340753 | UAB InMedica | unknown | Rasa Ugenskiene, Darius Cereskevicius, Inga Nasvytiene, Zivile Zemeckiene, Mantas Sarauskas, Marius Sukys, Rima Vainoriene, Astra Vitkauskiene, Renaldas Jurkevicius |
| EPI_ISL_3340754, EPI_ISL_3340755, EPI_ISL_3340756, EPI_ISL_3340757 | UAB Medicina practica laboratorija | unknown | Rasa Ugenskiene, Darius Cereskevicius, Inga Nasvytiene, Zivile Zemeckiene, Mantas Sarauskas, Marius Sukys, Rima Vainoriene, Astra Vitkauskiene, Renaldas Jurkevicius |
| EPI_ISL_3340758 | UAB InMedica | unknown | Rasa Ugenskiene, Darius Cereskevicius, Inga Nasvytiene, Zivile Zemeckiene, Mantas Sarauskas, Marius Sukys, Rima Vainoriene, Astra Vitkauskiene, Renaldas Jurkevicius |
| EPI_ISL_3340759, EPI_ISL_3340760, EPI_ISL_3340761, EPI_ISL_3340762, EPI_ISL_3340763, EPI_ISL_3340764 | UAB Medicina practica laboratorija | unknown | Rasa Ugenskiene, Darius Cereskevicius, Inga Nasvytiene, Zivile Zemeckiene, Mantas Sarauskas, Marius Sukys, Rima Vainoriene, Astra Vitkauskiene, Renaldas Jurkevicius |
| EPI_ISL_3340765, EPI_ISL_3340766, EPI_ISL_3340767, EPI_ISL_3340768, EPI_ISL_3340769, EPI_ISL_3340770, EPI_ISL_3340771 | UAB InMedica | unknown | Rasa Ugenskiene, Darius Cereskevicius, Inga Nasvytiene, Zivile Zemeckiene, Mantas Sarauskas, Marius Sukys, Rima Vainoriene, Astra Vitkauskiene, Renaldas Jurkevicius |
| EPI_ISL_3340772 | UAB Medicina practica laboratorija | unknown | Rasa Ugenskiene, Darius Cereskevicius, Inga Nasvytiene, Zivile Zemeckiene, Mantas Sarauskas, Marius Sukys, Rima Vainoriene, Astra Vitkauskiene, Renaldas Jurkevicius |
| EPI_ISL_3340773, EPI_ISL_3340774 | UAB InMedica | unknown | Rasa Ugenskiene, Darius Cereskevicius, Inga Nasvytiene, Zivile Zemeckiene, Mantas Sarauskas, Marius Sukys, Rima Vainoriene, Astra Vitkauskiene, Renaldas Jurkevicius |
| EPI_ISL_3340775, EPI_ISL_3340776, EPI_ISL_3340777 | UAB Medicina practica laboratorija | unknown | Rasa Ugenskiene, Darius Cereskevicius, Inga Nasvytiene, Zivile Zemeckiene, Mantas Sarauskas, Marius Sukys, Rima Vainoriene, Astra Vitkauskiene, Renaldas Jurkevicius |
| EPI_ISL_3340778 | UAB InMedica | unknown | Rasa Ugenskiene, Darius Cereskevicius, Inga Nasvytiene, Zivile Zemeckiene, Mantas Sarauskas, Marius Sukys, Rima Vainoriene, Astra Vitkauskiene, Renaldas Jurkevicius |
| EPI_ISL_3340779, EPI_ISL_3340780 | UAB Medicina practica laboratorija | unknown | Rasa Ugenskiene, Darius Cereskevicius, Inga Nasvytiene, Zivile Zemeckiene, Mantas Sarauskas, Marius Sukys, Rima Vainoriene, Astra Vitkauskiene, Renaldas Jurkevicius |
| EPI_ISL_3340781, EPI_ISL_3340782, EPI_ISL_3340783, EPI_ISL_3340784, EPI_ISL_3340785, EPI_ISL_3340786, EPI_ISL_3340787, EPI_ISL_3340788 | UAB InMedica | unknown | Rasa Ugenskiene, Darius Cereskevicius, Inga Nasvytiene, Zivile Zemeckiene, Mantas Sarauskas, Marius Sukys, Rima Vainoriene, Astra Vitkauskiene, Renaldas Jurkevicius |
| EPI_ISL_3340789 | UAB Medicina practica laboratorija | unknown | Rasa Ugenskiene, Darius Cereskevicius, Inga Nasvytiene, Zivile Zemeckiene, Mantas Sarauskas, Marius Sukys, Rima Vainoriene, Astra Vitkauskiene, Renaldas Jurkevicius |
| EPI_ISL_3340790 | UAB InMedica | unknown | Rasa Ugenskiene, Darius Cereskevicius, Inga Nasvytiene, Zivile Zemeckiene, Mantas Sarauskas, Marius Sukys, Rima Vainoriene, Astra Vitkauskiene, Renaldas Jurkevicius |
| EPI_ISL_3340791, EPI_ISL_3340792, EPI_ISL_3340793, EPI_ISL_3340794, EPI_ISL_3340795, EPI_ISL_3340796, EPI_ISL_3340797, EPI_ISL_3340798, EPI_ISL_3340799 | UAB Medicina practica laboratorija | unknown | Rasa Ugenskiene, Darius Cereskevicius, Inga Nasvytiene, Zivile Zemeckiene, Mantas Sarauskas, Marius Sukys, Rima Vainoriene, Astra Vitkauskiene, Renaldas Jurkevicius |
| EPI_ISL_3340800 | UAB InMedica | unknown | Rasa Ugenskiene, Darius Cereskevicius, Inga Nasvytiene, Zivile Zemeckiene, Mantas Sarauskas, Marius Sukys, Rima Vainoriene, Astra Vitkauskiene, Renaldas Jurkevicius |
| EPI_ISL_3340801, EPI_ISL_3340802 | UAB Medicina practica laboratorija | unknown | Rasa Ugenskiene, Darius Cereskevicius, Inga Nasvytiene, Zivile Zemeckiene, Mantas Sarauskas, Marius Sukys, Rima Vainoriene, Astra Vitkauskiene, Renaldas Jurkevicius |
| EPI_ISL_3340803, EPI_ISL_3340804 | UAB InMedica | unknown | Rasa Ugenskiene, Darius Cereskevicius, Inga Nasvytiene, Zivile Zemeckiene, Mantas Sarauskas, Marius Sukys, Rima Vainoriene, Astra Vitkauskiene, Renaldas Jurkevicius |
| EPI_ISL_3340805 | UAB Medicina practica laboratorija | unknown | Rasa Ugenskiene, Darius Cereskevicius, Inga Nasvytiene, Zivile Zemeckiene, Mantas Sarauskas, Marius Sukys, Rima Vainoriene, Astra Vitkauskiene, Renaldas Jurkevicius |
| EPI_ISL_3340806, EPI_ISL_3340807, EPI_ISL_3340808, EPI_ISL_3340809, EPI_ISL_3340810 | UAB InMedica | unknown | Rasa Ugenskiene, Darius Cereskevicius, Inga Nasvytiene, Zivile Zemeckiene, Mantas Sarauskas, Marius Sukys, Rima Vainoriene, Astra Vitkauskiene, Renaldas Jurkevicius |
| EPI_ISL_3340811 | UAB Medicina practica laboratorija | unknown | Rasa Ugenskiene, Darius Cereskevicius, Inga Nasvytiene, Zivile Zemeckiene, Mantas Sarauskas, Marius Sukys, Rima Vainoriene, Astra Vitkauskiene, Renaldas Jurkevicius |
| EPI_ISL_3340812 | UAB InMedica | unknown | Rasa Ugenskiene, Darius Cereskevicius, Inga Nasvytiene, Zivile Zemeckiene, Mantas Sarauskas, Marius Sukys, Rima Vainoriene, Astra Vitkauskiene, Renaldas Jurkevicius |
| EPI_ISL_3340813, EPI_ISL_3340814, EPI_ISL_3340815 | UAB Medicina practica laboratorija | unknown | Rasa Ugenskiene, Darius Cereskevicius, Inga Nasvytiene, Zivile Zemeckiene, Mantas Sarauskas, Marius Sukys, Rima Vainoriene, Astra Vitkauskiene, Renaldas Jurkevicius |
| EPI_ISL_3340816, EPI_ISL_3340817 | UAB InMedica | unknown | Rasa Ugenskiene, Darius Cereskevicius, Inga Nasvytiene, Zivile Zemeckiene, Mantas Sarauskas, Marius Sukys, Rima Vainoriene, Astra Vitkauskiene, Renaldas Jurkevicius |
| EPI_ISL_3340818, EPI_ISL_3340819, EPI_ISL_3340820, EPI_ISL_3340821, EPI_ISL_3340822 | UAB Medicina practica laboratorija | unknown | Rasa Ugenskiene, Darius Cereskevicius, Inga Nasvytiene, Zivile Zemeckiene, Mantas Sarauskas, Marius Sukys, Rima Vainoriene, Astra Vitkauskiene, Renaldas Jurkevicius |
| EPI_ISL_3340823 | UAB InMedica | unknown | Rasa Ugenskiene, Darius Cereskevicius, Inga Nasvytiene, Zivile Zemeckiene, Mantas Sarauskas, Marius Sukys, Rima Vainoriene, Astra Vitkauskiene, Renaldas Jurkevicius |
| EPI_ISL_3340824, EPI_ISL_3340825 | UAB Medicina practica laboratorija | unknown | Rasa Ugenskiene, Darius Cereskevicius, Inga Nasvytiene, Zivile Zemeckiene, Mantas Sarauskas, Marius Sukys, Rima Vainoriene, Astra Vitkauskiene, Renaldas Jurkevicius |
| EPI_ISL_3340826, EPI_ISL_3340827 | UAB InMedica | unknown | Rasa Ugenskiene, Darius Cereskevicius, Inga Nasvytiene, Zivile Zemeckiene, Mantas Sarauskas, Marius Sukys, Rima Vainoriene, Astra Vitkauskiene, Renaldas Jurkevicius |
| EPI_ISL_3340828, | UAB Medicina practica laboratorija | unknown | Rasa Ugenskiene, Darius Cereskevicius, Inga Nasvytiene, Zivile Zemeckiene, Mantas Sarauskas, Marius Sukys, Rima Vainoriene, Astra Vitkauskiene, Renaldas Jurkevicius |

| | | | |
|---|---|---|---|
| EPI_ISL_3340829, EPI_ISL_3340830 | | | |
| EPI_ISL_3340831, EPI_ISL_3340832, EPI_ISL_3340833 | UAB InMedica | unknown | Rasa Ugenskiene, Darius Cereskevicius, Inga Nasvytiene, Zivile Zemeckiene, Mantas Sarauskas, Marius Sukys, Rima Vainoriene, Astra Vitkauskiene, Renaldas Jurkevicius |
| EPI_ISL_3340834, EPI_ISL_3340835 | UAB Medicina practica laboratorija | unknown | Rasa Ugenskiene, Darius Cereskevicius, Inga Nasvytiene, Zivile Zemeckiene, Mantas Sarauskas, Marius Sukys, Rima Vainoriene, Astra Vitkauskiene, Renaldas Jurkevicius |
| EPI_ISL_3340836 | UAB InMedica | unknown | Rasa Ugenskiene, Darius Cereskevicius, Inga Nasvytiene, Zivile Zemeckiene, Mantas Sarauskas, Marius Sukys, Rima Vainoriene, Astra Vitkauskiene, Renaldas Jurkevicius |
| EPI_ISL_3340837, EPI_ISL_3340838, EPI_ISL_3340839, EPI_ISL_3340840 | UAB Medicina practica laboratorija | unknown | Rasa Ugenskiene, Darius Cereskevicius, Inga Nasvytiene, Zivile Zemeckiene, Mantas Sarauskas, Marius Sukys, Rima Vainoriene, Astra Vitkauskiene, Renaldas Jurkevicius |
| EPI_ISL_3340841, EPI_ISL_3340842, EPI_ISL_3340843 | UAB InMedica | unknown | Rasa Ugenskiene, Darius Cereskevicius, Inga Nasvytiene, Zivile Zemeckiene, Mantas Sarauskas, Marius Sukys, Rima Vainoriene, Astra Vitkauskiene, Renaldas Jurkevicius |
| EPI_ISL_3340844, EPI_ISL_3340845 | UAB Medicina practica laboratorija | unknown | Rasa Ugenskiene, Darius Cereskevicius, Inga Nasvytiene, Zivile Zemeckiene, Mantas Sarauskas, Marius Sukys, Rima Vainoriene, Astra Vitkauskiene, Renaldas Jurkevicius |
| EPI_ISL_3340846 | UAB InMedica | unknown | Rasa Ugenskiene, Darius Cereskevicius, Inga Nasvytiene, Zivile Zemeckiene, Mantas Sarauskas, Marius Sukys, Rima Vainoriene, Astra Vitkauskiene, Renaldas Jurkevicius |
| EPI_ISL_3340847, EPI_ISL_3340848 | UAB Medicina practica laboratorija | unknown | Rasa Ugenskiene, Darius Cereskevicius, Inga Nasvytiene, Zivile Zemeckiene, Mantas Sarauskas, Marius Sukys, Rima Vainoriene, Astra Vitkauskiene, Renaldas Jurkevicius |
| EPI_ISL_3340849 | UAB InMedica | unknown | Rasa Ugenskiene, Darius Cereskevicius, Inga Nasvytiene, Zivile Zemeckiene, Mantas Sarauskas, Marius Sukys, Rima Vainoriene, Astra Vitkauskiene, Renaldas Jurkevicius |
| EPI_ISL_3340850 | Viešoji įstaiga Klaipėdos universitetinė ligoninė | National Public Health Surveillance Laboratory | Lukas Zemaitis, Migle Gabrielaite, Jelena Razmuk, Svajune Muralyte, Ana Steponkiene, Lukas Vasionis, Danas Baksa |
| EPI_ISL_3340851 | UAB Diagnostikos laboratorija | National Public Health Surveillance Laboratory | Lukas Zemaitis, Migle Gabrielaite, Jelena Razmuk, Svajune Muralyte, Ana Steponkiene, Lukas Vasionis, Danas Baksa |
| EPI_ISL_3340852 | Viešoji įstaiga Klaipėdos universitetinė ligoninė | National Public Health Surveillance Laboratory | Lukas Zemaitis, Migle Gabrielaite, Jelena Razmuk, Svajune Muralyte, Ana Steponkiene, Lukas Vasionis, Danas Baksa |
| EPI_ISL_3340853, EPI_ISL_3340854 | UAB Diagnostikos laboratorija | National Public Health Surveillance Laboratory | Lukas Zemaitis, Migle Gabrielaite, Jelena Razmuk, Svajune Muralyte, Ana Steponkiene, Lukas Vasionis, Danas Baksa |
| EPI_ISL_3340855 | UAB "Baltic Medics" | National Public Health Surveillance Laboratory | Lukas Zemaitis, Migle Gabrielaite, Jelena Razmuk, Svajune Muralyte, Ana Steponkiene, Lukas Vasionis, Danas Baksa |
| EPI_ISL_3340856 | Viešoji įstaiga Vilniaus universiteto ligoninė Santaros klinikos | National Public Health Surveillance Laboratory | Lukas Zemaitis, Migle Gabrielaite, Jelena Razmuk, Svajune Muralyte, Ana Steponkiene, Lukas Vasionis, Danas Baksa |
| EPI_ISL_3340857, EPI_ISL_3340858, EPI_ISL_3340859, EPI_ISL_3340860 | UAB Diagnostikos laboratorija | National Public Health Surveillance Laboratory | Lukas Zemaitis, Migle Gabrielaite, Jelena Razmuk, Svajune Muralyte, Ana Steponkiene, Lukas Vasionis, Danas Baksa |
| EPI_ISL_3340861, EPI_ISL_3340862 | Viešoji įstaiga Klaipėdos universitetinė ligoninė | National Public Health Surveillance Laboratory | Lukas Zemaitis, Migle Gabrielaite, Jelena Razmuk, Svajune Muralyte, Ana Steponkiene, Lukas Vasionis, Danas Baksa |
| EPI_ISL_3340863 | UAB Diagnostikos laboratorija | National Public Health Surveillance Laboratory | Lukas Zemaitis, Migle Gabrielaite, Jelena Razmuk, Svajune Muralyte, Ana Steponkiene, Lukas Vasionis, Danas Baksa |
| EPI_ISL_3340864, EPI_ISL_3340865, EPI_ISL_3340866 | Viešoji įstaiga Klaipėdos universitetinė ligoninė | National Public Health Surveillance Laboratory | Lukas Zemaitis, Migle Gabrielaite, Jelena Razmuk, Svajune Muralyte, Ana Steponkiene, Lukas Vasionis, Danas Baksa |
| EPI_ISL_3340867 | UAB "Rezus.lt" | National Public Health Surveillance Laboratory | Lukas Zemaitis, Migle Gabrielaite, Jelena Razmuk, Svajune Muralyte, Ana Steponkiene, Lukas Vasionis, Danas Baksa |
| EPI_ISL_3340868, EPI_ISL_3340869 | UAB Diagnostikos laboratorija | National Public Health Surveillance Laboratory | Lukas Zemaitis, Migle Gabrielaite, Jelena Razmuk, Svajune Muralyte, Ana Steponkiene, Lukas Vasionis, Danas Baksa |
| EPI_ISL_3340870 | Viešoji įstaiga Klaipėdos universitetinė ligoninė | National Public Health Surveillance Laboratory | Lukas Zemaitis, Migle Gabrielaite, Jelena Razmuk, Svajune Muralyte, Ana Steponkiene, Lukas Vasionis, Danas Baksa |
| EPI_ISL_3340871 | UAB Diagnostikos laboratorija | National Public Health Surveillance Laboratory | Lukas Zemaitis, Migle Gabrielaite, Jelena Razmuk, Svajune Muralyte, Ana Steponkiene, Lukas Vasionis, Danas Baksa |
| EPI_ISL_3340872, EPI_ISL_3340873, EPI_ISL_3340874 | Viešoji įstaiga Klaipėdos universitetinė ligoninė | National Public Health Surveillance Laboratory | Lukas Zemaitis, Migle Gabrielaite, Jelena Razmuk, Svajune Muralyte, Ana Steponkiene, Lukas Vasionis, Danas Baksa |
| EPI_ISL_3340875 | UAB Diagnostikos laboratorija | National Public Health Surveillance Laboratory | Lukas Zemaitis, Migle Gabrielaite, Jelena Razmuk, Svajune Muralyte, Ana Steponkiene, Lukas Vasionis, Danas Baksa |
| EPI_ISL_3340876 | UAB "Rezus.lt" | National Public Health Surveillance Laboratory | Lukas Zemaitis, Migle Gabrielaite, Jelena Razmuk, Svajune Muralyte, Ana Steponkiene, Lukas Vasionis, Danas Baksa |
| EPI_ISL_3340877 | UAB Diagnostikos laboratorija | National Public Health Surveillance Laboratory | Lukas Zemaitis, Migle Gabrielaite, Jelena Razmuk, Svajune Muralyte, Ana Steponkiene, Lukas Vasionis, Danas Baksa |
| EPI_ISL_3340878 | Viešoji įstaiga Vilniaus universiteto ligoninė Santaros klinikos | National Public Health Surveillance Laboratory | Lukas Zemaitis, Migle Gabrielaite, Jelena Razmuk, Svajune Muralyte, Ana Steponkiene, Lukas Vasionis, Danas Baksa |
| EPI_ISL_3340879 | Viešoji įstaiga Klaipėdos universitetinė ligoninė | National Public Health Surveillance Laboratory | Lukas Zemaitis, Migle Gabrielaite, Jelena Razmuk, Svajune Muralyte, Ana Steponkiene, Lukas Vasionis, Danas Baksa |
| EPI_ISL_3340880 | UAB Diagnostikos laboratorija | National Public Health Surveillance Laboratory | Lukas Zemaitis, Migle Gabrielaite, Jelena Razmuk, Svajune Muralyte, Ana Steponkiene, Lukas Vasionis, Danas Baksa |
| EPI_ISL_3340881, EPI_ISL_3340882 | Viešoji įstaiga Klaipėdos universitetinė ligoninė | National Public Health Surveillance Laboratory | Lukas Zemaitis, Migle Gabrielaite, Jelena Razmuk, Svajune Muralyte, Ana Steponkiene, Lukas Vasionis, Danas Baksa |
| EPI_ISL_3340883 | Viešoji įstaiga Vilniaus universiteto ligoninė Santaros klinikos | National Public Health Surveillance Laboratory | Lukas Zemaitis, Migle Gabrielaite, Jelena Razmuk, Svajune Muralyte, Ana Steponkiene, Lukas Vasionis, Danas Baksa |
| EPI_ISL_3340884 | UAB Diagnostikos laboratorija | National Public Health Surveillance Laboratory | Lukas Zemaitis, Migle Gabrielaite, Jelena Razmuk, Svajune Muralyte, Ana Steponkiene, Lukas Vasionis, Danas Baksa |
| EPI_ISL_3340885 | Viešoji įstaiga Klaipėdos universitetinė ligoninė | National Public Health Surveillance Laboratory | Lukas Zemaitis, Migle Gabrielaite, Jelena Razmuk, Svajune Muralyte, Ana Steponkiene, Lukas Vasionis, Danas Baksa |
| EPI_ISL_3340886 | Viešoji įstaiga Vilniaus universiteto ligoninė Santaros klinikos | National Public Health Surveillance Laboratory | Lukas Zemaitis, Migle Gabrielaite, Jelena Razmuk, Svajune Muralyte, Ana Steponkiene, Lukas Vasionis, Danas Baksa |
| EPI_ISL_3340887 | Viešoji įstaiga Klaipėdos universitetinė ligoninė | National Public Health Surveillance Laboratory | Lukas Zemaitis, Migle Gabrielaite, Jelena Razmuk, Svajune Muralyte, Ana Steponkiene, Lukas Vasionis, Danas Baksa |
| EPI_ISL_3340888 | UAB Diagnostikos laboratorija | National Public Health Surveillance Laboratory | Lukas Zemaitis, Migle Gabrielaite, Jelena Razmuk, Svajune Muralyte, Ana Steponkiene, Lukas Vasionis, Danas Baksa |
| EPI_ISL_3340889, EPI_ISL_3340890, EPI_ISL_3340891 | Viešoji įstaiga Klaipėdos universitetinė ligoninė | National Public Health Surveillance Laboratory | Lukas Zemaitis, Migle Gabrielaite, Jelena Razmuk, Svajune Muralyte, Ana Steponkiene, Lukas Vasionis, Danas Baksa |
| EPI_ISL_3340892, EPI_ISL_3340893 | Viešoji įstaiga Respublikinė Šiaulių ligoninė | National Public Health Surveillance Laboratory | Lukas Zemaitis, Migle Gabrielaite, Jelena Razmuk, Svajune Muralyte, Ana Steponkiene, Lukas Vasionis, Danas Baksa |
| EPI_ISL_3340894 | SYNLAB Lietuva UAB | National Public Health Surveillance Laboratory | Lukas Zemaitis, Migle Gabrielaite, Jelena Razmuk, Svajune Muralyte, Ana Steponkiene, Lukas Vasionis, Danas Baksa |
| EPI_ISL_3340895, EPI_ISL_3340896 | Viešoji įstaiga Klaipėdos universitetinė ligoninė | National Public Health Surveillance Laboratory | Lukas Zemaitis, Migle Gabrielaite, Jelena Razmuk, Svajune Muralyte, Ana Steponkiene, Lukas Vasionis, Danas Baksa |
| EPI_ISL_3340897 | UAB "Rezus.lt" | National Public Health Surveillance Laboratory | Lukas Zemaitis, Migle Gabrielaite, Jelena Razmuk, Svajune Muralyte, Ana Steponkiene, Lukas Vasionis, Danas Baksa |
| EPI_ISL_3340898, EPI_ISL_3340899 | UAB Diagnostikos laboratorija | National Public Health Surveillance Laboratory | Lukas Zemaitis, Migle Gabrielaite, Jelena Razmuk, Svajune Muralyte, Ana Steponkiene, Lukas Vasionis, Danas Baksa |
| EPI_ISL_3340900 | Viešoji įstaiga Vilniaus universiteto ligoninė Santaros klinikos | National Public Health Surveillance Laboratory | Lukas Zemaitis, Migle Gabrielaite, Jelena Razmuk, Svajune Muralyte, Ana Steponkiene, Lukas Vasionis, Danas Baksa |
| EPI_ISL_3340901 | UAB "Rezus.lt" | National Public Health Surveillance Laboratory | Lukas Zemaitis, Migle Gabrielaite, Jelena Razmuk, Svajune Muralyte, Ana Steponkiene, Lukas Vasionis, Danas Baksa |
| EPI_ISL_3340902 | Viešoji įstaiga Klaipėdos universitetinė ligoninė | National Public Health Surveillance Laboratory | Lukas Zemaitis, Migle Gabrielaite, Jelena Razmuk, Svajune Muralyte, Ana Steponkiene, Lukas Vasionis, Danas Baksa |
| EPI_ISL_3340903 | UAB "Rezus.lt" | National Public Health Surveillance Laboratory | Lukas Zemaitis, Migle Gabrielaite, Jelena Razmuk, Svajune Muralyte, Ana Steponkiene, Lukas Vasionis, Danas Baksa |
| EPI_ISL_3340904 | Viešoji įstaiga Klaipėdos universitetinė ligoninė | National Public Health Surveillance Laboratory | Lukas Zemaitis, Migle Gabrielaite, Jelena Razmuk, Svajune Muralyte, Ana Steponkiene, Lukas Vasionis, Danas Baksa |
| EPI_ISL_3340905 | UAB Diagnostikos laboratorija | National Public Health Surveillance Laboratory | Lukas Zemaitis, Migle Gabrielaite, Jelena Razmuk, Svajune Muralyte, Ana Steponkiene, Lukas Vasionis, Danas Baksa |
| EPI_ISL_3340906, EPI_ISL_3340907 | Viešoji įstaiga Klaipėdos universitetinė ligoninė | National Public Health Surveillance Laboratory | Lukas Zemaitis, Migle Gabrielaite, Jelena Razmuk, Svajune Muralyte, Ana Steponkiene, Lukas Vasionis, Danas Baksa |
| EPI_ISL_3340908 | Viešoji įstaiga Vilniaus universiteto ligoninė Santaros klinikos | National Public Health Surveillance Laboratory | Lukas Zemaitis, Migle Gabrielaite, Jelena Razmuk, Svajune Muralyte, Ana Steponkiene, Lukas Vasionis, Danas Baksa |
| EPI_ISL_3340909, EPI_ISL_3340910 | UAB "Rezus.lt" | National Public Health Surveillance Laboratory | Lukas Zemaitis, Migle Gabrielaite, Jelena Razmuk, Svajune Muralyte, Ana Steponkiene, Lukas Vasionis, Danas Baksa |
| EPI_ISL_3340911 | SYNLAB Lietuva UAB | National Public Health Surveillance Laboratory | Lukas Zemaitis, Migle Gabrielaite, Jelena Razmuk, Svajune Muralyte, Ana Steponkiene, Lukas Vasionis, Danas Baksa |
| EPI_ISL_3340912 | Viešoji įstaiga Respublikinė Šiaulių ligoninė | National Public Health Surveillance Laboratory | Lukas Zemaitis, Migle Gabrielaite, Jelena Razmuk, Svajune Muralyte, Ana Steponkiene, Lukas Vasionis, Danas Baksa |
| EPI_ISL_3340913 | Viešoji įstaiga Klaipėdos universitetinė ligoninė | National Public Health Surveillance Laboratory | Lukas Zemaitis, Migle Gabrielaite, Jelena Razmuk, Svajune Muralyte, Ana Steponkiene, Lukas Vasionis, Danas Baksa |

| | | | |
|---|---|---|---|
| EPI_ISL_3340914 | UAB "Rezus.lt" | National Public Health Surveillance Laboratory | Lukas Zemaitis, Migle Gabrielaite, Jelena Razmuk, Svajune Muralyte, Ana Steponkiene, Lukas Vasionis, Danas Baksa |
| EPI_ISL_3340915 | UAB Diagnostikos laboratorija | National Public Health Surveillance Laboratory | Lukas Zemaitis, Migle Gabrielaite, Jelena Razmuk, Svajune Muralyte, Ana Steponkiene, Lukas Vasionis, Danas Baksa |
| EPI_ISL_3340916, EPI_ISL_3340917 | Viešoji įstaiga Klaipėdos universitetinė ligoninė | National Public Health Surveillance Laboratory | Lukas Zemaitis, Migle Gabrielaite, Jelena Razmuk, Svajune Muralyte, Ana Steponkiene, Lukas Vasionis, Danas Baksa |
| EPI_ISL_3340918, EPI_ISL_3340919, EPI_ISL_3340920 | UAB Diagnostikos laboratorija | National Public Health Surveillance Laboratory | Lukas Zemaitis, Migle Gabrielaite, Jelena Razmuk, Svajune Muralyte, Ana Steponkiene, Lukas Vasionis, Danas Baksa |
| EPI_ISL_3340921 | Viešoji įstaiga Klaipėdos universitetinė ligoninė | National Public Health Surveillance Laboratory | Lukas Zemaitis, Migle Gabrielaite, Jelena Razmuk, Svajune Muralyte, Ana Steponkiene, Lukas Vasionis, Danas Baksa |
| EPI_ISL_3340922, EPI_ISL_3340923 | UAB "Rezus.lt" | National Public Health Surveillance Laboratory | Lukas Zemaitis, Migle Gabrielaite, Jelena Razmuk, Svajune Muralyte, Ana Steponkiene, Lukas Vasionis, Danas Baksa |
| EPI_ISL_3340924, EPI_ISL_3340925 | UAB Diagnostikos laboratorija | National Public Health Surveillance Laboratory | Lukas Zemaitis, Migle Gabrielaite, Jelena Razmuk, Svajune Muralyte, Ana Steponkiene, Lukas Vasionis, Danas Baksa |
| EPI_ISL_3340926 | Viešoji įstaiga Klaipėdos universitetinė ligoninė | National Public Health Surveillance Laboratory | Lukas Zemaitis, Migle Gabrielaite, Jelena Razmuk, Svajune Muralyte, Ana Steponkiene, Lukas Vasionis, Danas Baksa |
| EPI_ISL_3340927 | UAB Diagnostikos laboratorija | National Public Health Surveillance Laboratory | Lukas Zemaitis, Migle Gabrielaite, Jelena Razmuk, Svajune Muralyte, Ana Steponkiene, Lukas Vasionis, Danas Baksa |
| EPI_ISL_3340928 | Viešoji įstaiga Klaipėdos universitetinė ligoninė | National Public Health Surveillance Laboratory | Lukas Zemaitis, Migle Gabrielaite, Jelena Razmuk, Svajune Muralyte, Ana Steponkiene, Lukas Vasionis, Danas Baksa |
| EPI_ISL_3340929 | UAB Diagnostikos laboratorija | National Public Health Surveillance Laboratory | Lukas Zemaitis, Migle Gabrielaite, Jelena Razmuk, Svajune Muralyte, Ana Steponkiene, Lukas Vasionis, Danas Baksa |
| EPI_ISL_3340930 | Viešoji įstaiga Klaipėdos universitetinė ligoninė | National Public Health Surveillance Laboratory | Lukas Zemaitis, Migle Gabrielaite, Jelena Razmuk, Svajune Muralyte, Ana Steponkiene, Lukas Vasionis, Danas Baksa |
| EPI_ISL_3340931 | UAB "Rezus.lt" | National Public Health Surveillance Laboratory | Lukas Zemaitis, Migle Gabrielaite, Jelena Razmuk, Svajune Muralyte, Ana Steponkiene, Lukas Vasionis, Danas Baksa |
| EPI_ISL_3340932 | UAB Diagnostikos laboratorija | National Public Health Surveillance Laboratory | Lukas Zemaitis, Migle Gabrielaite, Jelena Razmuk, Svajune Muralyte, Ana Steponkiene, Lukas Vasionis, Danas Baksa |
| EPI_ISL_3340933 | Viešoji įstaiga Klaipėdos universitetinė ligoninė | National Public Health Surveillance Laboratory | Lukas Zemaitis, Migle Gabrielaite, Jelena Razmuk, Svajune Muralyte, Ana Steponkiene, Lukas Vasionis, Danas Baksa |
| EPI_ISL_3340934, EPI_ISL_3340935 | UAB "Rezus.lt" | National Public Health Surveillance Laboratory | Lukas Zemaitis, Migle Gabrielaite, Jelena Razmuk, Svajune Muralyte, Ana Steponkiene, Lukas Vasionis, Danas Baksa |
| EPI_ISL_3340936 | Viešoji įstaiga Klaipėdos universitetinė ligoninė | National Public Health Surveillance Laboratory | Lukas Zemaitis, Migle Gabrielaite, Jelena Razmuk, Svajune Muralyte, Ana Steponkiene, Lukas Vasionis, Danas Baksa |
| EPI_ISL_3340937, EPI_ISL_3340938 | UAB Diagnostikos laboratorija | National Public Health Surveillance Laboratory | Lukas Zemaitis, Migle Gabrielaite, Jelena Razmuk, Svajune Muralyte, Ana Steponkiene, Lukas Vasionis, Danas Baksa |
| EPI_ISL_3340939 | Viešoji įstaiga Klaipėdos universitetinė ligoninė | National Public Health Surveillance Laboratory | Lukas Zemaitis, Migle Gabrielaite, Jelena Razmuk, Svajune Muralyte, Ana Steponkiene, Lukas Vasionis, Danas Baksa |
| EPI_ISL_3340940 | UAB Diagnostikos laboratorija | National Public Health Surveillance Laboratory | Lukas Zemaitis, Migle Gabrielaite, Jelena Razmuk, Svajune Muralyte, Ana Steponkiene, Lukas Vasionis, Danas Baksa |
| EPI_ISL_3340941 | UAB "Baltic Medics" | National Public Health Surveillance Laboratory | Lukas Zemaitis, Migle Gabrielaite, Jelena Razmuk, Svajune Muralyte, Ana Steponkiene, Lukas Vasionis, Danas Baksa |
| EPI_ISL_3340942 | Viešoji įstaiga Klaipėdos universitetinė ligoninė | National Public Health Surveillance Laboratory | Lukas Zemaitis, Migle Gabrielaite, Jelena Razmuk, Svajune Muralyte, Ana Steponkiene, Lukas Vasionis, Danas Baksa |
| EPI_ISL_3340943 | UAB "Rezus.lt" | National Public Health Surveillance Laboratory | Lukas Zemaitis, Migle Gabrielaite, Jelena Razmuk, Svajune Muralyte, Ana Steponkiene, Lukas Vasionis, Danas Baksa |
| EPI_ISL_3340944 | UAB Diagnostikos laboratorija | National Public Health Surveillance Laboratory | Lukas Zemaitis, Migle Gabrielaite, Jelena Razmuk, Svajune Muralyte, Ana Steponkiene, Lukas Vasionis, Danas Baksa |
| EPI_ISL_3340945 | Viešoji įstaiga Klaipėdos universitetinė ligoninė | National Public Health Surveillance Laboratory | Lukas Zemaitis, Migle Gabrielaite, Jelena Razmuk, Svajune Muralyte, Ana Steponkiene, Lukas Vasionis, Danas Baksa |
| EPI_ISL_3340946 | UAB "Baltic Medics" | National Public Health Surveillance Laboratory | Lukas Zemaitis, Migle Gabrielaite, Jelena Razmuk, Svajune Muralyte, Ana Steponkiene, Lukas Vasionis, Danas Baksa |
| EPI_ISL_3340947, EPI_ISL_3340948, EPI_ISL_3340949 | UAB Diagnostikos laboratorija | National Public Health Surveillance Laboratory | Lukas Zemaitis, Migle Gabrielaite, Jelena Razmuk, Svajune Muralyte, Ana Steponkiene, Lukas Vasionis, Danas Baksa |
| EPI_ISL_3340950 | Viešoji įstaiga Klaipėdos universitetinė ligoninė | National Public Health Surveillance Laboratory | Lukas Zemaitis, Migle Gabrielaite, Jelena Razmuk, Svajune Muralyte, Ana Steponkiene, Lukas Vasionis, Danas Baksa |
| EPI_ISL_3340951 | SYNLAB Lietuva UAB | National Public Health Surveillance Laboratory | Lukas Zemaitis, Migle Gabrielaite, Jelena Razmuk, Svajune Muralyte, Ana Steponkiene, Lukas Vasionis, Danas Baksa |
| EPI_ISL_3340952 | UAB "Baltic Medics" | National Public Health Surveillance Laboratory | Lukas Zemaitis, Migle Gabrielaite, Jelena Razmuk, Svajune Muralyte, Ana Steponkiene, Lukas Vasionis, Danas Baksa |
| EPI_ISL_3340953 | UAB "Rezus.lt" | National Public Health Surveillance Laboratory | Lukas Zemaitis, Migle Gabrielaite, Jelena Razmuk, Svajune Muralyte, Ana Steponkiene, Lukas Vasionis, Danas Baksa |
| EPI_ISL_3340954 | Viešoji įstaiga Klaipėdos universitetinė ligoninė | National Public Health Surveillance Laboratory | Lukas Zemaitis, Migle Gabrielaite, Jelena Razmuk, Svajune Muralyte, Ana Steponkiene, Lukas Vasionis, Danas Baksa |
| EPI_ISL_3340955 | UAB Diagnostikos laboratorija | National Public Health Surveillance Laboratory | Lukas Zemaitis, Migle Gabrielaite, Jelena Razmuk, Svajune Muralyte, Ana Steponkiene, Lukas Vasionis, Danas Baksa |
| EPI_ISL_3340956 | UAB "Rezus.lt" | National Public Health Surveillance Laboratory | Lukas Zemaitis, Migle Gabrielaite, Jelena Razmuk, Svajune Muralyte, Ana Steponkiene, Lukas Vasionis, Danas Baksa |
| EPI_ISL_3340957 | Viešoji įstaiga Vilniaus universiteto ligoninė Santaros klinikos | National Public Health Surveillance Laboratory | Lukas Zemaitis, Migle Gabrielaite, Jelena Razmuk, Svajune Muralyte, Ana Steponkiene, Lukas Vasionis, Danas Baksa |
| EPI_ISL_3340958, EPI_ISL_3340959 | UAB Diagnostikos laboratorija | National Public Health Surveillance Laboratory | Lukas Zemaitis, Migle Gabrielaite, Jelena Razmuk, Svajune Muralyte, Ana Steponkiene, Lukas Vasionis, Danas Baksa |
| EPI_ISL_3340960 | Viešoji įstaiga Klaipėdos universitetinė ligoninė | National Public Health Surveillance Laboratory | Lukas Zemaitis, Migle Gabrielaite, Jelena Razmuk, Svajune Muralyte, Ana Steponkiene, Lukas Vasionis, Danas Baksa |
| EPI_ISL_3340961, EPI_ISL_3340962 | UAB Diagnostikos laboratorija | National Public Health Surveillance Laboratory | Lukas Zemaitis, Migle Gabrielaite, Jelena Razmuk, Svajune Muralyte, Ana Steponkiene, Lukas Vasionis, Danas Baksa |
| EPI_ISL_3340963 | UAB "Baltic Medics" | National Public Health Surveillance Laboratory | Lukas Zemaitis, Migle Gabrielaite, Jelena Razmuk, Svajune Muralyte, Ana Steponkiene, Lukas Vasionis, Danas Baksa |
| EPI_ISL_3340964, EPI_ISL_3340965 | Viešoji įstaiga Klaipėdos universitetinė ligoninė | National Public Health Surveillance Laboratory | Lukas Zemaitis, Migle Gabrielaite, Jelena Razmuk, Svajune Muralyte, Ana Steponkiene, Lukas Vasionis, Danas Baksa |
| EPI_ISL_3340966, EPI_ISL_3340967 | UAB "Rezus.lt" | National Public Health Surveillance Laboratory | Lukas Zemaitis, Migle Gabrielaite, Jelena Razmuk, Svajune Muralyte, Ana Steponkiene, Lukas Vasionis, Danas Baksa |
| EPI_ISL_3340968, EPI_ISL_3340969 | UAB Diagnostikos laboratorija | National Public Health Surveillance Laboratory | Lukas Zemaitis, Migle Gabrielaite, Jelena Razmuk, Svajune Muralyte, Ana Steponkiene, Lukas Vasionis, Danas Baksa |
| EPI_ISL_3340970, EPI_ISL_3340971 | Viešoji įstaiga Klaipėdos universitetinė ligoninė | National Public Health Surveillance Laboratory | Lukas Zemaitis, Migle Gabrielaite, Jelena Razmuk, Svajune Muralyte, Ana Steponkiene, Lukas Vasionis, Danas Baksa |
| EPI_ISL_3340972 | UAB Diagnostikos laboratorija | National Public Health Surveillance Laboratory | Lukas Zemaitis, Migle Gabrielaite, Jelena Razmuk, Svajune Muralyte, Ana Steponkiene, Lukas Vasionis, Danas Baksa |
| EPI_ISL_3340973 | Viešoji įstaiga Klaipėdos universitetinė ligoninė | National Public Health Surveillance Laboratory | Lukas Zemaitis, Migle Gabrielaite, Jelena Razmuk, Svajune Muralyte, Ana Steponkiene, Lukas Vasionis, Danas Baksa |
| EPI_ISL_3340974 | SYNLAB Lietuva UAB | National Public Health Surveillance Laboratory | Lukas Zemaitis, Migle Gabrielaite, Jelena Razmuk, Svajune Muralyte, Ana Steponkiene, Lukas Vasionis, Danas Baksa |
| EPI_ISL_3340975 | Viešoji įstaiga Vilniaus universiteto ligoninė Santaros klinikos | National Public Health Surveillance Laboratory | Lukas Zemaitis, Migle Gabrielaite, Jelena Razmuk, Svajune Muralyte, Ana Steponkiene, Lukas Vasionis, Danas Baksa |
| EPI_ISL_3340976, EPI_ISL_3340977 | UAB Diagnostikos laboratorija | National Public Health Surveillance Laboratory | Lukas Zemaitis, Migle Gabrielaite, Jelena Razmuk, Svajune Muralyte, Ana Steponkiene, Lukas Vasionis, Danas Baksa |
| EPI_ISL_3340978 | UAB "Baltic Medics" | National Public Health Surveillance Laboratory | Lukas Zemaitis, Migle Gabrielaite, Jelena Razmuk, Svajune Muralyte, Ana Steponkiene, Lukas Vasionis, Danas Baksa |
| EPI_ISL_3340979, EPI_ISL_3340980, EPI_ISL_3340981 | UAB Diagnostikos laboratorija | National Public Health Surveillance Laboratory | Lukas Zemaitis, Migle Gabrielaite, Jelena Razmuk, Svajune Muralyte, Ana Steponkiene, Lukas Vasionis, Danas Baksa |
| EPI_ISL_3340982 | Viešoji įstaiga Vilniaus universiteto ligoninė Santaros klinikos | National Public Health Surveillance Laboratory | Lukas Zemaitis, Migle Gabrielaite, Jelena Razmuk, Svajune Muralyte, Ana Steponkiene, Lukas Vasionis, Danas Baksa |
| EPI_ISL_3340983 | UAB Diagnostikos laboratorija | National Public Health Surveillance Laboratory | Lukas Zemaitis, Migle Gabrielaite, Jelena Razmuk, Svajune Muralyte, Ana Steponkiene, Lukas Vasionis, Danas Baksa |
| EPI_ISL_3340984 | UAB "Rezus.lt" | National Public Health Surveillance Laboratory | Lukas Zemaitis, Migle Gabrielaite, Jelena Razmuk, Svajune Muralyte, Ana Steponkiene, Lukas Vasionis, Danas Baksa |
| EPI_ISL_3340985, EPI_ISL_3340986, EPI_ISL_3340987 | UAB Diagnostikos laboratorija | National Public Health Surveillance Laboratory | Lukas Zemaitis, Migle Gabrielaite, Jelena Razmuk, Svajune Muralyte, Ana Steponkiene, Lukas Vasionis, Danas Baksa |
| EPI_ISL_3340988, EPI_ISL_3340989 | Viešoji įstaiga Klaipėdos universitetinė ligoninė | National Public Health Surveillance Laboratory | Lukas Zemaitis, Migle Gabrielaite, Jelena Razmuk, Svajune Muralyte, Ana Steponkiene, Lukas Vasionis, Danas Baksa |
| EPI_ISL_3340990 | SYNLAB Lietuva UAB | National Public Health Surveillance Laboratory | Lukas Zemaitis, Migle Gabrielaite, Jelena Razmuk, Svajune Muralyte, Ana Steponkiene, Lukas Vasionis, Danas Baksa |
| EPI_ISL_3340991 | UAB "Rezus.lt" | National Public Health Surveillance Laboratory | Lukas Zemaitis, Migle Gabrielaite, Jelena Razmuk, Svajune Muralyte, Ana Steponkiene, Lukas Vasionis, Danas Baksa |
| EPI_ISL_3340992 | UAB "Baltic Medics" | National Public Health Surveillance Laboratory | Lukas Zemaitis, Migle Gabrielaite, Jelena Razmuk, Svajune Muralyte, Ana Steponkiene, Lukas Vasionis, Danas Baksa |
| EPI_ISL_3340993 | Viešoji įstaiga Klaipėdos universitetinė ligoninė | National Public Health Surveillance Laboratory | Lukas Zemaitis, Migle Gabrielaite, Jelena Razmuk, Svajune Muralyte, Ana Steponkiene, Lukas Vasionis, Danas Baksa |
| EPI_ISL_3340994 | Viešoji įstaiga Respublikinė Šiaulių ligoninė | National Public Health Surveillance Laboratory | Lukas Zemaitis, Migle Gabrielaite, Jelena Razmuk, Svajune Muralyte, Ana Steponkiene, Lukas Vasionis, Danas Baksa |
| EPI_ISL_3340995 | Viešoji įstaiga Vilniaus universiteto ligoninė Santaros klinikos | National Public Health Surveillance Laboratory | Lukas Zemaitis, Migle Gabrielaite, Jelena Razmuk, Svajune Muralyte, Ana Steponkiene, Lukas Vasionis, Danas Baksa |
| EPI_ISL_3340996 | Viešoji įstaiga Klaipėdos universitetinė ligoninė | National Public Health Surveillance Laboratory | Lukas Zemaitis, Migle Gabrielaite, Jelena Razmuk, Svajune Muralyte, Ana Steponkiene, Lukas Vasionis, Danas Baksa |

| | | | |
|---|---|---|---|
| EPI_ISL_3340997, EPI_ISL_3340998 | UAB Diagnostikos laboratorija | National Public Health Surveillance Laboratory | Lukas Zemaitis, Migle Gabrielaite, Jelena Razmuk, Svajune Muralyte, Ana Steponkiene, Lukas Vasionis, Danas Baksa |
| EPI_ISL_3340999, EPI_ISL_3341000 | Viešoji įstaiga Klaipėdos universitetinė ligoninė | National Public Health Surveillance Laboratory | Lukas Zemaitis, Migle Gabrielaite, Jelena Razmuk, Svajune Muralyte, Ana Steponkiene, Lukas Vasionis, Danas Baksa |
| EPI_ISL_3341001 | Viešoji įstaiga Vilniaus universiteto ligoninė Santaros klinikos | National Public Health Surveillance Laboratory | Lukas Zemaitis, Migle Gabrielaite, Jelena Razmuk, Svajune Muralyte, Ana Steponkiene, Lukas Vasionis, Danas Baksa |
| EPI_ISL_3341002, EPI_ISL_3341003, EPI_ISL_3341004 | Viešoji įstaiga Klaipėdos universitetinė ligoninė | National Public Health Surveillance Laboratory | Lukas Zemaitis, Migle Gabrielaite, Jelena Razmuk, Svajune Muralyte, Ana Steponkiene, Lukas Vasionis, Danas Baksa |
| EPI_ISL_3341005 | UAB Diagnostikos laboratorija | National Public Health Surveillance Laboratory | Lukas Zemaitis, Migle Gabrielaite, Jelena Razmuk, Svajune Muralyte, Ana Steponkiene, Lukas Vasionis, Danas Baksa |
| EPI_ISL_3341006 | Viešoji įstaiga Klaipėdos universitetinė ligoninė | National Public Health Surveillance Laboratory | Lukas Zemaitis, Migle Gabrielaite, Jelena Razmuk, Svajune Muralyte, Ana Steponkiene, Lukas Vasionis, Danas Baksa |
| EPI_ISL_3341007, EPI_ISL_3341008, EPI_ISL_3341009 | UAB Diagnostikos laboratorija | National Public Health Surveillance Laboratory | Lukas Zemaitis, Migle Gabrielaite, Jelena Razmuk, Svajune Muralyte, Ana Steponkiene, Lukas Vasionis, Danas Baksa |
| EPI_ISL_3341010 | UAB "Rezus.lt" | National Public Health Surveillance Laboratory | Lukas Zemaitis, Migle Gabrielaite, Jelena Razmuk, Svajune Muralyte, Ana Steponkiene, Lukas Vasionis, Danas Baksa |
| EPI_ISL_3341011, EPI_ISL_3341012 | Viešoji įstaiga Klaipėdos universitetinė ligoninė | National Public Health Surveillance Laboratory | Lukas Zemaitis, Migle Gabrielaite, Jelena Razmuk, Svajune Muralyte, Ana Steponkiene, Lukas Vasionis, Danas Baksa |
| EPI_ISL_3341013 | UAB Diagnostikos laboratorija | National Public Health Surveillance Laboratory | Lukas Zemaitis, Migle Gabrielaite, Jelena Razmuk, Svajune Muralyte, Ana Steponkiene, Lukas Vasionis, Danas Baksa |
| EPI_ISL_3341014 | Viešoji įstaiga Vilniaus universiteto ligoninė Santaros klinikos | National Public Health Surveillance Laboratory | Lukas Zemaitis, Migle Gabrielaite, Jelena Razmuk, Svajune Muralyte, Ana Steponkiene, Lukas Vasionis, Danas Baksa |
| EPI_ISL_3341015 | Viešoji įstaiga Klaipėdos universitetinė ligoninė | National Public Health Surveillance Laboratory | Lukas Zemaitis, Migle Gabrielaite, Jelena Razmuk, Svajune Muralyte, Ana Steponkiene, Lukas Vasionis, Danas Baksa |
| EPI_ISL_3341016 | UAB Diagnostikos laboratorija | National Public Health Surveillance Laboratory | Lukas Zemaitis, Migle Gabrielaite, Jelena Razmuk, Svajune Muralyte, Ana Steponkiene, Lukas Vasionis, Danas Baksa |
| EPI_ISL_3341017 | Viešoji įstaiga Klaipėdos universitetinė ligoninė | National Public Health Surveillance Laboratory | Lukas Zemaitis, Migle Gabrielaite, Jelena Razmuk, Svajune Muralyte, Ana Steponkiene, Lukas Vasionis, Danas Baksa |
| EPI_ISL_3341018 | UAB Diagnostikos laboratorija | National Public Health Surveillance Laboratory | Lukas Zemaitis, Migle Gabrielaite, Jelena Razmuk, Svajune Muralyte, Ana Steponkiene, Lukas Vasionis, Danas Baksa |
| EPI_ISL_3341019, EPI_ISL_3341020, EPI_ISL_3341021, EPI_ISL_3341022 | Viešoji įstaiga Klaipėdos universitetinė ligoninė | National Public Health Surveillance Laboratory | Lukas Zemaitis, Migle Gabrielaite, Jelena Razmuk, Svajune Muralyte, Ana Steponkiene, Lukas Vasionis, Danas Baksa |
| EPI_ISL_3341023 | UAB "Rezus.lt" | National Public Health Surveillance Laboratory | Lukas Zemaitis, Migle Gabrielaite, Jelena Razmuk, Svajune Muralyte, Ana Steponkiene, Lukas Vasionis, Danas Baksa |
| EPI_ISL_3341024 | Viešoji įstaiga Klaipėdos universitetinė ligoninė | National Public Health Surveillance Laboratory | Lukas Zemaitis, Migle Gabrielaite, Jelena Razmuk, Svajune Muralyte, Ana Steponkiene, Lukas Vasionis, Danas Baksa |
| EPI_ISL_3341025 | UAB Diagnostikos laboratorija | National Public Health Surveillance Laboratory | Lukas Zemaitis, Migle Gabrielaite, Jelena Razmuk, Svajune Muralyte, Ana Steponkiene, Lukas Vasionis, Danas Baksa |
| EPI_ISL_3341026 | Viešoji įstaiga Klaipėdos universitetinė ligoninė | National Public Health Surveillance Laboratory | Lukas Zemaitis, Migle Gabrielaite, Jelena Razmuk, Svajune Muralyte, Ana Steponkiene, Lukas Vasionis, Danas Baksa |
| EPI_ISL_3341027 | UAB Diagnostikos laboratorija | National Public Health Surveillance Laboratory | Lukas Zemaitis, Migle Gabrielaite, Jelena Razmuk, Svajune Muralyte, Ana Steponkiene, Lukas Vasionis, Danas Baksa |
| EPI_ISL_3341028 | UAB "Baltic Medics" | National Public Health Surveillance Laboratory | Lukas Zemaitis, Migle Gabrielaite, Jelena Razmuk, Svajune Muralyte, Ana Steponkiene, Lukas Vasionis, Danas Baksa |
| EPI_ISL_3341029, EPI_ISL_3341030 | UAB Diagnostikos laboratorija | National Public Health Surveillance Laboratory | Lukas Zemaitis, Migle Gabrielaite, Jelena Razmuk, Svajune Muralyte, Ana Steponkiene, Lukas Vasionis, Danas Baksa |
| EPI_ISL_3341031 | Viešoji įstaiga Vilniaus universiteto ligoninė Santaros klinikos | National Public Health Surveillance Laboratory | Lukas Zemaitis, Migle Gabrielaite, Jelena Razmuk, Svajune Muralyte, Ana Steponkiene, Lukas Vasionis, Danas Baksa |
| EPI_ISL_3341032, EPI_ISL_3341033 | Viešoji įstaiga Klaipėdos universitetinė ligoninė | National Public Health Surveillance Laboratory | Lukas Zemaitis, Migle Gabrielaite, Jelena Razmuk, Svajune Muralyte, Ana Steponkiene, Lukas Vasionis, Danas Baksa |
| EPI_ISL_3341034, EPI_ISL_3341035 | UAB Diagnostikos laboratorija | National Public Health Surveillance Laboratory | Lukas Zemaitis, Migle Gabrielaite, Jelena Razmuk, Svajune Muralyte, Ana Steponkiene, Lukas Vasionis, Danas Baksa |
| EPI_ISL_3341036 | Viešoji įstaiga Klaipėdos universitetinė ligoninė | National Public Health Surveillance Laboratory | Lukas Zemaitis, Migle Gabrielaite, Jelena Razmuk, Svajune Muralyte, Ana Steponkiene, Lukas Vasionis, Danas Baksa |
| EPI_ISL_3341037 | UAB "Rezus.lt" | National Public Health Surveillance Laboratory | Lukas Zemaitis, Migle Gabrielaite, Jelena Razmuk, Svajune Muralyte, Ana Steponkiene, Lukas Vasionis, Danas Baksa |
| EPI_ISL_3341038 | UAB "Baltic Medics" | National Public Health Surveillance Laboratory | Lukas Zemaitis, Migle Gabrielaite, Jelena Razmuk, Svajune Muralyte, Ana Steponkiene, Lukas Vasionis, Danas Baksa |
| EPI_ISL_3341039, EPI_ISL_3341040, EPI_ISL_3341041, EPI_ISL_3341042 | Viešoji įstaiga Klaipėdos universitetinė ligoninė | National Public Health Surveillance Laboratory | Lukas Zemaitis, Migle Gabrielaite, Jelena Razmuk, Svajune Muralyte, Ana Steponkiene, Lukas Vasionis, Danas Baksa |
| EPI_ISL_3341043 | UAB Diagnostikos laboratorija | National Public Health Surveillance Laboratory | Lukas Zemaitis, Migle Gabrielaite, Jelena Razmuk, Svajune Muralyte, Ana Steponkiene, Lukas Vasionis, Danas Baksa |
| EPI_ISL_3341044 | Viešoji įstaiga Klaipėdos universitetinė ligoninė | National Public Health Surveillance Laboratory | Lukas Zemaitis, Migle Gabrielaite, Jelena Razmuk, Svajune Muralyte, Ana Steponkiene, Lukas Vasionis, Danas Baksa |
| EPI_ISL_3341045 | UAB Diagnostikos laboratorija | National Public Health Surveillance Laboratory | Lukas Zemaitis, Migle Gabrielaite, Jelena Razmuk, Svajune Muralyte, Ana Steponkiene, Lukas Vasionis, Danas Baksa |
| EPI_ISL_3341046, EPI_ISL_3341047 | Viešoji įstaiga Klaipėdos universitetinė ligoninė | National Public Health Surveillance Laboratory | Lukas Zemaitis, Migle Gabrielaite, Jelena Razmuk, Svajune Muralyte, Ana Steponkiene, Lukas Vasionis, Danas Baksa |
| EPI_ISL_3341048 | UAB "Rezus.lt" | National Public Health Surveillance Laboratory | Lukas Zemaitis, Migle Gabrielaite, Jelena Razmuk, Svajune Muralyte, Ana Steponkiene, Lukas Vasionis, Danas Baksa |
| EPI_ISL_3341049 | Viešoji įstaiga Klaipėdos universitetinė ligoninė | National Public Health Surveillance Laboratory | Lukas Zemaitis, Migle Gabrielaite, Jelena Razmuk, Svajune Muralyte, Ana Steponkiene, Lukas Vasionis, Danas Baksa |
| EPI_ISL_3341050 | UAB "Baltic Medics" | National Public Health Surveillance Laboratory | Lukas Zemaitis, Migle Gabrielaite, Jelena Razmuk, Svajune Muralyte, Ana Steponkiene, Lukas Vasionis, Danas Baksa |
| EPI_ISL_3341051, EPI_ISL_3341052 | UAB Diagnostikos laboratorija | National Public Health Surveillance Laboratory | Lukas Zemaitis, Migle Gabrielaite, Jelena Razmuk, Svajune Muralyte, Ana Steponkiene, Lukas Vasionis, Danas Baksa |
| EPI_ISL_3341053 | Viešoji įstaiga Klaipėdos universitetinė ligoninė | National Public Health Surveillance Laboratory | Lukas Zemaitis, Migle Gabrielaite, Jelena Razmuk, Svajune Muralyte, Ana Steponkiene, Lukas Vasionis, Danas Baksa |
| EPI_ISL_3341054 | UAB "Rezus.lt" | National Public Health Surveillance Laboratory | Lukas Zemaitis, Migle Gabrielaite, Jelena Razmuk, Svajune Muralyte, Ana Steponkiene, Lukas Vasionis, Danas Baksa |
| EPI_ISL_3341055 | Viešoji įstaiga Vilniaus universiteto ligoninė Santaros klinikos | National Public Health Surveillance Laboratory | Lukas Zemaitis, Migle Gabrielaite, Jelena Razmuk, Svajune Muralyte, Ana Steponkiene, Lukas Vasionis, Danas Baksa |
| EPI_ISL_3341056 | Viešoji įstaiga Klaipėdos universitetinė ligoninė | National Public Health Surveillance Laboratory | Lukas Zemaitis, Migle Gabrielaite, Jelena Razmuk, Svajune Muralyte, Ana Steponkiene, Lukas Vasionis, Danas Baksa |
| EPI_ISL_3341057 | UAB "Baltic Medics" | National Public Health Surveillance Laboratory | Lukas Zemaitis, Migle Gabrielaite, Jelena Razmuk, Svajune Muralyte, Ana Steponkiene, Lukas Vasionis, Danas Baksa |
| EPI_ISL_3341058, EPI_ISL_3341059 | UAB Diagnostikos laboratorija | National Public Health Surveillance Laboratory | Lukas Zemaitis, Migle Gabrielaite, Jelena Razmuk, Svajune Muralyte, Ana Steponkiene, Lukas Vasionis, Danas Baksa |
| EPI_ISL_3341060 | Viešoji įstaiga Klaipėdos universitetinė ligoninė | National Public Health Surveillance Laboratory | Lukas Zemaitis, Migle Gabrielaite, Jelena Razmuk, Svajune Muralyte, Ana Steponkiene, Lukas Vasionis, Danas Baksa |
| EPI_ISL_3341061 | UAB Diagnostikos laboratorija | National Public Health Surveillance Laboratory | Lukas Zemaitis, Migle Gabrielaite, Jelena Razmuk, Svajune Muralyte, Ana Steponkiene, Lukas Vasionis, Danas Baksa |
| EPI_ISL_3341062, EPI_ISL_3341063 | Viešoji įstaiga Klaipėdos universitetinė ligoninė | National Public Health Surveillance Laboratory | Lukas Zemaitis, Migle Gabrielaite, Jelena Razmuk, Svajune Muralyte, Ana Steponkiene, Lukas Vasionis, Danas Baksa |
| EPI_ISL_3341064 | UAB "SK IMPEKS MEDICINOS DIAGNOSTIKOS CENTRAS" | National Public Health Surveillance Laboratory | Lukas Zemaitis, Migle Gabrielaite, Jelena Razmuk, Svajune Muralyte, Ana Steponkiene, Lukas Vasionis, Danas Baksa |
| EPI_ISL_3341065, EPI_ISL_3341066 | UAB Diagnostikos laboratorija | National Public Health Surveillance Laboratory | Lukas Zemaitis, Migle Gabrielaite, Jelena Razmuk, Svajune Muralyte, Ana Steponkiene, Lukas Vasionis, Danas Baksa |
| EPI_ISL_3341067 | Viešoji įstaiga Klaipėdos universitetinė ligoninė | National Public Health Surveillance Laboratory | Lukas Zemaitis, Migle Gabrielaite, Jelena Razmuk, Svajune Muralyte, Ana Steponkiene, Lukas Vasionis, Danas Baksa |
| EPI_ISL_3341068 | UAB Diagnostikos laboratorija | National Public Health Surveillance Laboratory | Lukas Zemaitis, Migle Gabrielaite, Jelena Razmuk, Svajune Muralyte, Ana Steponkiene, Lukas Vasionis, Danas Baksa |
| EPI_ISL_3341069 | Viešoji įstaiga Klaipėdos universitetinė ligoninė | National Public Health Surveillance Laboratory | Lukas Zemaitis, Migle Gabrielaite, Jelena Razmuk, Svajune Muralyte, Ana Steponkiene, Lukas Vasionis, Danas Baksa |
| EPI_ISL_3341070 | UAB Diagnostikos laboratorija | National Public Health Surveillance Laboratory | Lukas Zemaitis, Migle Gabrielaite, Jelena Razmuk, Svajune Muralyte, Ana Steponkiene, Lukas Vasionis, Danas Baksa |
| EPI_ISL_3341071 | UAB "Rezus.lt" | National Public Health Surveillance Laboratory | Lukas Zemaitis, Migle Gabrielaite, Jelena Razmuk, Svajune Muralyte, Ana Steponkiene, Lukas Vasionis, Danas Baksa |
| EPI_ISL_3341072, EPI_ISL_3341073 | UAB Diagnostikos laboratorija | National Public Health Surveillance Laboratory | Lukas Zemaitis, Migle Gabrielaite, Jelena Razmuk, Svajune Muralyte, Ana Steponkiene, Lukas Vasionis, Danas Baksa |
| EPI_ISL_3341074 | Viešoji įstaiga Klaipėdos universitetinė ligoninė | National Public Health Surveillance Laboratory | Lukas Zemaitis, Migle Gabrielaite, Jelena Razmuk, Svajune Muralyte, Ana Steponkiene, Lukas Vasionis, Danas Baksa |
| EPI_ISL_3341075 | Viešoji įstaiga Respublikinė Šiaulių ligoninė | National Public Health Surveillance Laboratory | Lukas Zemaitis, Migle Gabrielaite, Jelena Razmuk, Svajune Muralyte, Ana Steponkiene, Lukas Vasionis, Danas Baksa |
| EPI_ISL_3341076 | UAB "Rezus.lt" | National Public Health Surveillance Laboratory | Lukas Zemaitis, Migle Gabrielaite, Jelena Razmuk, Svajune Muralyte, Ana Steponkiene, Lukas Vasionis, Danas Baksa |
| EPI_ISL_3341077, EPI_ISL_3341078 | Viešoji įstaiga Klaipėdos universitetinė ligoninė | National Public Health Surveillance Laboratory | Lukas Zemaitis, Migle Gabrielaite, Jelena Razmuk, Svajune Muralyte, Ana Steponkiene, Lukas Vasionis, Danas Baksa |
| EPI_ISL_3341079 | UAB "Rezus.lt" | National Public Health Surveillance Laboratory | Lukas Zemaitis, Migle Gabrielaite, Jelena Razmuk, Svajune Muralyte, Ana Steponkiene, Lukas Vasionis, Danas Baksa |
| EPI_ISL_3341080, | UAB Diagnostikos laboratorija | National Public Health Surveillance Laboratory | Lukas Zemaitis, Migle Gabrielaite, Jelena Razmuk, Svajune Muralyte, Ana Steponkiene, Lukas Vasionis, Danas Baksa |

| | | | |
|---|---|---|---|
| EPI_ISL_3341081 | | | |
| EPI_ISL_3341082 | UAB "Baltic Medics" | National Public Health Surveillance Laboratory | Lukas Zemaitis, Migle Gabrielaite, Jelena Razmuk, Svajune Muralyte, Ana Steponkiene, Lukas Vasionis, Danas Baksa |
| EPI_ISL_3341083, EPI_ISL_3341084 | UAB Diagnostikos laboratorija | National Public Health Surveillance Laboratory | Lukas Zemaitis, Migle Gabrielaite, Jelena Razmuk, Svajune Muralyte, Ana Steponkiene, Lukas Vasionis, Danas Baksa |
| EPI_ISL_3341085 | Viešoji įstaiga Vilniaus universiteto ligoninė Santaros klinikos | National Public Health Surveillance Laboratory | Lukas Zemaitis, Migle Gabrielaite, Jelena Razmuk, Svajune Muralyte, Ana Steponkiene, Lukas Vasionis, Danas Baksa |
| EPI_ISL_3341086 | Viešoji įstaiga Klaipėdos universitetinė ligoninė | National Public Health Surveillance Laboratory | Lukas Zemaitis, Migle Gabrielaite, Jelena Razmuk, Svajune Muralyte, Ana Steponkiene, Lukas Vasionis, Danas Baksa |
| EPI_ISL_3341087 | Viešoji įstaiga Vilniaus universiteto ligoninė Santaros klinikos | National Public Health Surveillance Laboratory | Lukas Zemaitis, Migle Gabrielaite, Jelena Razmuk, Svajune Muralyte, Ana Steponkiene, Lukas Vasionis, Danas Baksa |
| EPI_ISL_3341088 | UAB "Rezus.lt" | National Public Health Surveillance Laboratory | Lukas Zemaitis, Migle Gabrielaite, Jelena Razmuk, Svajune Muralyte, Ana Steponkiene, Lukas Vasionis, Danas Baksa |
| EPI_ISL_3341089 | UAB Diagnostikos laboratorija | National Public Health Surveillance Laboratory | Lukas Zemaitis, Migle Gabrielaite, Jelena Razmuk, Svajune Muralyte, Ana Steponkiene, Lukas Vasionis, Danas Baksa |
| EPI_ISL_3341090, EPI_ISL_3341091, EPI_ISL_3341092 | UAB "Rezus.lt" | National Public Health Surveillance Laboratory | Lukas Zemaitis, Migle Gabrielaite, Jelena Razmuk, Svajune Muralyte, Ana Steponkiene, Lukas Vasionis, Danas Baksa |
| EPI_ISL_3341093 | Viešoji įstaiga Klaipėdos universitetinė ligoninė | National Public Health Surveillance Laboratory | Lukas Zemaitis, Migle Gabrielaite, Jelena Razmuk, Svajune Muralyte, Ana Steponkiene, Lukas Vasionis, Danas Baksa |
| EPI_ISL_3341094 | UAB Diagnostikos laboratorija | National Public Health Surveillance Laboratory | Lukas Zemaitis, Migle Gabrielaite, Jelena Razmuk, Svajune Muralyte, Ana Steponkiene, Lukas Vasionis, Danas Baksa |
| EPI_ISL_3341095 | UAB "Rezus.lt" | National Public Health Surveillance Laboratory | Lukas Zemaitis, Migle Gabrielaite, Jelena Razmuk, Svajune Muralyte, Ana Steponkiene, Lukas Vasionis, Danas Baksa |
| EPI_ISL_3341096, EPI_ISL_3341097, EPI_ISL_3341098 | Viešoji įstaiga Klaipėdos universitetinė ligoninė | National Public Health Surveillance Laboratory | Lukas Zemaitis, Migle Gabrielaite, Jelena Razmuk, Svajune Muralyte, Ana Steponkiene, Lukas Vasionis, Danas Baksa |
| EPI_ISL_3341099 | UAB Diagnostikos laboratorija | National Public Health Surveillance Laboratory | Lukas Zemaitis, Migle Gabrielaite, Jelena Razmuk, Svajune Muralyte, Ana Steponkiene, Lukas Vasionis, Danas Baksa |
| EPI_ISL_3341100, EPI_ISL_3341101 | Viešoji įstaiga Vilniaus universiteto ligoninė Santaros klinikos | National Public Health Surveillance Laboratory | Lukas Zemaitis, Migle Gabrielaite, Jelena Razmuk, Svajune Muralyte, Ana Steponkiene, Lukas Vasionis, Danas Baksa |
| EPI_ISL_3341102 | UAB Diagnostikos laboratorija | National Public Health Surveillance Laboratory | Lukas Zemaitis, Migle Gabrielaite, Jelena Razmuk, Svajune Muralyte, Ana Steponkiene, Lukas Vasionis, Danas Baksa |
| EPI_ISL_3341103, EPI_ISL_3341104 | Viešoji įstaiga Klaipėdos universitetinė ligoninė | National Public Health Surveillance Laboratory | Lukas Zemaitis, Migle Gabrielaite, Jelena Razmuk, Svajune Muralyte, Ana Steponkiene, Lukas Vasionis, Danas Baksa |
| EPI_ISL_3341105, EPI_ISL_3341106, EPI_ISL_3341107 | UAB Diagnostikos laboratorija | National Public Health Surveillance Laboratory | Lukas Zemaitis, Migle Gabrielaite, Jelena Razmuk, Svajune Muralyte, Ana Steponkiene, Lukas Vasionis, Danas Baksa |
| EPI_ISL_3341108 | UAB "Rezus.lt" | National Public Health Surveillance Laboratory | Lukas Zemaitis, Migle Gabrielaite, Jelena Razmuk, Svajune Muralyte, Ana Steponkiene, Lukas Vasionis, Danas Baksa |
| EPI_ISL_3341109 | Viešoji įstaiga Vilniaus universiteto ligoninė Santaros klinikos | National Public Health Surveillance Laboratory | Lukas Zemaitis, Migle Gabrielaite, Jelena Razmuk, Svajune Muralyte, Ana Steponkiene, Lukas Vasionis, Danas Baksa |
| EPI_ISL_3341110, EPI_ISL_3341111, EPI_ISL_3341112 | Viešoji įstaiga Klaipėdos universitetinė ligoninė | National Public Health Surveillance Laboratory | Lukas Zemaitis, Migle Gabrielaite, Jelena Razmuk, Svajune Muralyte, Ana Steponkiene, Lukas Vasionis, Danas Baksa |
| EPI_ISL_3341113, EPI_ISL_3341114 | UAB Diagnostikos laboratorija | National Public Health Surveillance Laboratory | Lukas Zemaitis, Migle Gabrielaite, Jelena Razmuk, Svajune Muralyte, Ana Steponkiene, Lukas Vasionis, Danas Baksa |
| EPI_ISL_3341115 | Viešoji įstaiga Klaipėdos universitetinė ligoninė | National Public Health Surveillance Laboratory | Lukas Zemaitis, Migle Gabrielaite, Jelena Razmuk, Svajune Muralyte, Ana Steponkiene, Lukas Vasionis, Danas Baksa |
| EPI_ISL_3341116 | Viešoji įstaiga Vilniaus universiteto ligoninė Santaros klinikos | National Public Health Surveillance Laboratory | Lukas Zemaitis, Migle Gabrielaite, Jelena Razmuk, Svajune Muralyte, Ana Steponkiene, Lukas Vasionis, Danas Baksa |
| EPI_ISL_3341117 | UAB "Rezus.lt" | National Public Health Surveillance Laboratory | Lukas Zemaitis, Migle Gabrielaite, Jelena Razmuk, Svajune Muralyte, Ana Steponkiene, Lukas Vasionis, Danas Baksa |
| EPI_ISL_3341118 | Viešoji įstaiga Klaipėdos universitetinė ligoninė | National Public Health Surveillance Laboratory | Lukas Zemaitis, Migle Gabrielaite, Jelena Razmuk, Svajune Muralyte, Ana Steponkiene, Lukas Vasionis, Danas Baksa |
| EPI_ISL_3341119 | UAB Diagnostikos laboratorija | National Public Health Surveillance Laboratory | Lukas Zemaitis, Migle Gabrielaite, Jelena Razmuk, Svajune Muralyte, Ana Steponkiene, Lukas Vasionis, Danas Baksa |
| EPI_ISL_3341120, EPI_ISL_3341121 | Viešoji įstaiga Klaipėdos universitetinė ligoninė | National Public Health Surveillance Laboratory | Lukas Zemaitis, Migle Gabrielaite, Jelena Razmuk, Svajune Muralyte, Ana Steponkiene, Lukas Vasionis, Danas Baksa |
| EPI_ISL_3341122, EPI_ISL_3341123, EPI_ISL_3341124 | UAB Diagnostikos laboratorija | National Public Health Surveillance Laboratory | Lukas Zemaitis, Migle Gabrielaite, Jelena Razmuk, Svajune Muralyte, Ana Steponkiene, Lukas Vasionis, Danas Baksa |
| EPI_ISL_3341125, EPI_ISL_3341126 | SYNLAB Lietuva UAB | National Public Health Surveillance Laboratory | Lukas Zemaitis, Migle Gabrielaite, Jelena Razmuk, Svajune Muralyte, Ana Steponkiene, Lukas Vasionis, Danas Baksa |
| EPI_ISL_3341127 | Viešoji įstaiga Klaipėdos universitetinė ligoninė | National Public Health Surveillance Laboratory | Lukas Zemaitis, Migle Gabrielaite, Jelena Razmuk, Svajune Muralyte, Ana Steponkiene, Lukas Vasionis, Danas Baksa |
| EPI_ISL_3341128 | Viešoji įstaiga Vilniaus universiteto ligoninė Santaros klinikos | National Public Health Surveillance Laboratory | Lukas Zemaitis, Migle Gabrielaite, Jelena Razmuk, Svajune Muralyte, Ana Steponkiene, Lukas Vasionis, Danas Baksa |
| EPI_ISL_3341129, EPI_ISL_3341130, EPI_ISL_3341131 | Viešoji įstaiga Klaipėdos universitetinė ligoninė | National Public Health Surveillance Laboratory | Lukas Zemaitis, Migle Gabrielaite, Jelena Razmuk, Svajune Muralyte, Ana Steponkiene, Lukas Vasionis, Danas Baksa |
| EPI_ISL_3341132, EPI_ISL_3341133 | Viešoji įstaiga Vilniaus universiteto ligoninė Santaros klinikos | National Public Health Surveillance Laboratory | Lukas Zemaitis, Migle Gabrielaite, Jelena Razmuk, Svajune Muralyte, Ana Steponkiene, Lukas Vasionis, Danas Baksa |
| EPI_ISL_3341134, EPI_ISL_3341135, EPI_ISL_3341136 | Viešoji įstaiga Klaipėdos universitetinė ligoninė | National Public Health Surveillance Laboratory | Lukas Zemaitis, Migle Gabrielaite, Jelena Razmuk, Svajune Muralyte, Ana Steponkiene, Lukas Vasionis, Danas Baksa |
| EPI_ISL_3341137 | UAB Diagnostikos laboratorija | National Public Health Surveillance Laboratory | Lukas Zemaitis, Migle Gabrielaite, Jelena Razmuk, Svajune Muralyte, Ana Steponkiene, Lukas Vasionis, Danas Baksa |
| EPI_ISL_3341138 | Viešoji įstaiga Klaipėdos universitetinė ligoninė | National Public Health Surveillance Laboratory | Lukas Zemaitis, Migle Gabrielaite, Jelena Razmuk, Svajune Muralyte, Ana Steponkiene, Lukas Vasionis, Danas Baksa |
| EPI_ISL_3341139 | UAB Diagnostikos laboratorija | National Public Health Surveillance Laboratory | Lukas Zemaitis, Migle Gabrielaite, Jelena Razmuk, Svajune Muralyte, Ana Steponkiene, Lukas Vasionis, Danas Baksa |
| EPI_ISL_3341140 | UAB "Rezus.lt" | National Public Health Surveillance Laboratory | Lukas Zemaitis, Migle Gabrielaite, Jelena Razmuk, Svajune Muralyte, Ana Steponkiene, Lukas Vasionis, Danas Baksa |
| EPI_ISL_3341141, EPI_ISL_3341142, EPI_ISL_3341143 | Viešoji įstaiga Klaipėdos universitetinė ligoninė | National Public Health Surveillance Laboratory | Lukas Zemaitis, Migle Gabrielaite, Jelena Razmuk, Svajune Muralyte, Ana Steponkiene, Lukas Vasionis, Danas Baksa |
| EPI_ISL_3341144, EPI_ISL_3341145 | UAB Diagnostikos laboratorija | National Public Health Surveillance Laboratory | Lukas Zemaitis, Migle Gabrielaite, Jelena Razmuk, Svajune Muralyte, Ana Steponkiene, Lukas Vasionis, Danas Baksa |
| EPI_ISL_3341146 | SYNLAB Lietuva UAB | National Public Health Surveillance Laboratory | Lukas Zemaitis, Migle Gabrielaite, Jelena Razmuk, Svajune Muralyte, Ana Steponkiene, Lukas Vasionis, Danas Baksa |
| EPI_ISL_3341147 | Viešoji įstaiga Klaipėdos universitetinė ligoninė | National Public Health Surveillance Laboratory | Lukas Zemaitis, Migle Gabrielaite, Jelena Razmuk, Svajune Muralyte, Ana Steponkiene, Lukas Vasionis, Danas Baksa |
| EPI_ISL_3341148, EPI_ISL_3341149 | UAB Diagnostikos laboratorija | National Public Health Surveillance Laboratory | Lukas Zemaitis, Migle Gabrielaite, Jelena Razmuk, Svajune Muralyte, Ana Steponkiene, Lukas Vasionis, Danas Baksa |
| EPI_ISL_3341150 | UAB "Baltic Medics" | National Public Health Surveillance Laboratory | Lukas Zemaitis, Migle Gabrielaite, Jelena Razmuk, Svajune Muralyte, Ana Steponkiene, Lukas Vasionis, Danas Baksa |
| EPI_ISL_3341151 | UAB "Rezus.lt" | National Public Health Surveillance Laboratory | Lukas Zemaitis, Migle Gabrielaite, Jelena Razmuk, Svajune Muralyte, Ana Steponkiene, Lukas Vasionis, Danas Baksa |
| EPI_ISL_3341152, EPI_ISL_3341153 | UAB Diagnostikos laboratorija | National Public Health Surveillance Laboratory | Lukas Zemaitis, Migle Gabrielaite, Jelena Razmuk, Svajune Muralyte, Ana Steponkiene, Lukas Vasionis, Danas Baksa |
| EPI_ISL_3341154, EPI_ISL_3341155, EPI_ISL_3341156 | UAB "Rezus.lt" | National Public Health Surveillance Laboratory | Lukas Zemaitis, Migle Gabrielaite, Jelena Razmuk, Svajune Muralyte, Ana Steponkiene, Lukas Vasionis, Danas Baksa |
| EPI_ISL_3341157 | UAB Diagnostikos laboratorija | National Public Health Surveillance Laboratory | Lukas Zemaitis, Migle Gabrielaite, Jelena Razmuk, Svajune Muralyte, Ana Steponkiene, Lukas Vasionis, Danas Baksa |
| EPI_ISL_3341158 | Viešoji įstaiga Klaipėdos universitetinė ligoninė | National Public Health Surveillance Laboratory | Lukas Zemaitis, Migle Gabrielaite, Jelena Razmuk, Svajune Muralyte, Ana Steponkiene, Lukas Vasionis, Danas Baksa |
| EPI_ISL_3341159 | Viešoji įstaiga Vilniaus universiteto ligoninė Santaros klinikos | National Public Health Surveillance Laboratory | Lukas Zemaitis, Migle Gabrielaite, Jelena Razmuk, Svajune Muralyte, Ana Steponkiene, Lukas Vasionis, Danas Baksa |
| EPI_ISL_3341161 | UAB Diagnostikos laboratorija | National Public Health Surveillance Laboratory | Lukas Zemaitis, Migle Gabrielaite, Jelena Razmuk, Svajune Muralyte, Ana Steponkiene, Lukas Vasionis, Danas Baksa |
| EPI_ISL_3341162, EPI_ISL_3341163 | Viešoji įstaiga Klaipėdos universitetinė ligoninė | National Public Health Surveillance Laboratory | Lukas Zemaitis, Migle Gabrielaite, Jelena Razmuk, Svajune Muralyte, Ana Steponkiene, Lukas Vasionis, Danas Baksa |
| EPI_ISL_3341164, EPI_ISL_3341165 | UAB "Rezus.lt" | National Public Health Surveillance Laboratory | Lukas Zemaitis, Migle Gabrielaite, Jelena Razmuk, Svajune Muralyte, Ana Steponkiene, Lukas Vasionis, Danas Baksa |
| EPI_ISL_3341166, EPI_ISL_3341167 | Viešoji įstaiga Klaipėdos universitetinė ligoninė | National Public Health Surveillance Laboratory | Lukas Zemaitis, Migle Gabrielaite, Jelena Razmuk, Svajune Muralyte, Ana Steponkiene, Lukas Vasionis, Danas Baksa |

| | | | |
|---|---|---|---|
| EPI_ISL_3341168 | UAB "Rezus.lt" | National Public Health Surveillance Laboratory | Lukas Zemaitis, Migle Gabrielaite, Jelena Razmuk, Svajune Muralyte, Ana Steponkiene, Lukas Vasionis, Danas Baksa |
| EPI_ISL_3341169 | Viešoji įstaiga Vilniaus universiteto ligoninė Santaros klinikos | National Public Health Surveillance Laboratory | Lukas Zemaitis, Migle Gabrielaite, Jelena Razmuk, Svajune Muralyte, Ana Steponkiene, Lukas Vasionis, Danas Baksa |
| EPI_ISL_3341170, EPI_ISL_3341171 | UAB Diagnostikos laboratorija | National Public Health Surveillance Laboratory | Lukas Zemaitis, Migle Gabrielaite, Jelena Razmuk, Svajune Muralyte, Ana Steponkiene, Lukas Vasionis, Danas Baksa |
| EPI_ISL_3341172 | Viešoji įstaiga Vilniaus universiteto ligoninė Santaros klinikos | National Public Health Surveillance Laboratory | Lukas Zemaitis, Migle Gabrielaite, Jelena Razmuk, Svajune Muralyte, Ana Steponkiene, Lukas Vasionis, Danas Baksa |
| EPI_ISL_3341173 | Viešoji įstaiga Klaipėdos universitetinė ligoninė | National Public Health Surveillance Laboratory | Lukas Zemaitis, Migle Gabrielaite, Jelena Razmuk, Svajune Muralyte, Ana Steponkiene, Lukas Vasionis, Danas Baksa |
| EPI_ISL_3341174 | UAB Diagnostikos laboratorija | National Public Health Surveillance Laboratory | Lukas Zemaitis, Migle Gabrielaite, Jelena Razmuk, Svajune Muralyte, Ana Steponkiene, Lukas Vasionis, Danas Baksa |
| EPI_ISL_3341175 | UAB "Baltic Medics" | National Public Health Surveillance Laboratory | Lukas Zemaitis, Migle Gabrielaite, Jelena Razmuk, Svajune Muralyte, Ana Steponkiene, Lukas Vasionis, Danas Baksa |
| EPI_ISL_3341176 | Viešoji įstaiga Klaipėdos universitetinė ligoninė | National Public Health Surveillance Laboratory | Lukas Zemaitis, Migle Gabrielaite, Jelena Razmuk, Svajune Muralyte, Ana Steponkiene, Lukas Vasionis, Danas Baksa |
| EPI_ISL_3341177 | UAB Diagnostikos laboratorija | National Public Health Surveillance Laboratory | Lukas Zemaitis, Migle Gabrielaite, Jelena Razmuk, Svajune Muralyte, Ana Steponkiene, Lukas Vasionis, Danas Baksa |
| EPI_ISL_3341178, EPI_ISL_3341179 | Viešoji įstaiga Klaipėdos universitetinė ligoninė | National Public Health Surveillance Laboratory | Lukas Zemaitis, Migle Gabrielaite, Jelena Razmuk, Svajune Muralyte, Ana Steponkiene, Lukas Vasionis, Danas Baksa |
| EPI_ISL_3341180 | UAB "Rezus.lt" | National Public Health Surveillance Laboratory | Lukas Zemaitis, Migle Gabrielaite, Jelena Razmuk, Svajune Muralyte, Ana Steponkiene, Lukas Vasionis, Danas Baksa |
| EPI_ISL_3341181 | Viešoji įstaiga Klaipėdos universitetinė ligoninė | National Public Health Surveillance Laboratory | Lukas Zemaitis, Migle Gabrielaite, Jelena Razmuk, Svajune Muralyte, Ana Steponkiene, Lukas Vasionis, Danas Baksa |
| EPI_ISL_3341182 | UAB "Baltic Medics" | National Public Health Surveillance Laboratory | Lukas Zemaitis, Migle Gabrielaite, Jelena Razmuk, Svajune Muralyte, Ana Steponkiene, Lukas Vasionis, Danas Baksa |
| EPI_ISL_3341183, EPI_ISL_3341184 | Viešoji įstaiga Klaipėdos universitetinė ligoninė | National Public Health Surveillance Laboratory | Lukas Zemaitis, Migle Gabrielaite, Jelena Razmuk, Svajune Muralyte, Ana Steponkiene, Lukas Vasionis, Danas Baksa |
| EPI_ISL_3341185 | UAB "Rezus.lt" | National Public Health Surveillance Laboratory | Lukas Zemaitis, Migle Gabrielaite, Jelena Razmuk, Svajune Muralyte, Ana Steponkiene, Lukas Vasionis, Danas Baksa |
| EPI_ISL_3341186 | Viešoji įstaiga Respublikinė Šiaulių ligoninė | National Public Health Surveillance Laboratory | Lukas Zemaitis, Migle Gabrielaite, Jelena Razmuk, Svajune Muralyte, Ana Steponkiene, Lukas Vasionis, Danas Baksa |
| EPI_ISL_3341187, EPI_ISL_3341188 | Viešoji įstaiga Klaipėdos universitetinė ligoninė | National Public Health Surveillance Laboratory | Lukas Zemaitis, Migle Gabrielaite, Jelena Razmuk, Svajune Muralyte, Ana Steponkiene, Lukas Vasionis, Danas Baksa |
| EPI_ISL_3341189 | UAB Diagnostikos laboratorija | National Public Health Surveillance Laboratory | Lukas Zemaitis, Migle Gabrielaite, Jelena Razmuk, Svajune Muralyte, Ana Steponkiene, Lukas Vasionis, Danas Baksa |
| EPI_ISL_3341190 | Viešoji įstaiga Klaipėdos universitetinė ligoninė | National Public Health Surveillance Laboratory | Lukas Zemaitis, Migle Gabrielaite, Jelena Razmuk, Svajune Muralyte, Ana Steponkiene, Lukas Vasionis, Danas Baksa |
| EPI_ISL_3341191 | UAB "Rezus.lt" | National Public Health Surveillance Laboratory | Lukas Zemaitis, Migle Gabrielaite, Jelena Razmuk, Svajune Muralyte, Ana Steponkiene, Lukas Vasionis, Danas Baksa |
| EPI_ISL_3341192 | UAB Diagnostikos laboratorija | National Public Health Surveillance Laboratory | Lukas Zemaitis, Migle Gabrielaite, Jelena Razmuk, Svajune Muralyte, Ana Steponkiene, Lukas Vasionis, Danas Baksa |
| EPI_ISL_3341193 | UAB "Rezus.lt" | National Public Health Surveillance Laboratory | Lukas Zemaitis, Migle Gabrielaite, Jelena Razmuk, Svajune Muralyte, Ana Steponkiene, Lukas Vasionis, Danas Baksa |
| EPI_ISL_3341194 | Viešoji įstaiga Vilniaus universiteto ligoninė Santaros klinikos | National Public Health Surveillance Laboratory | Lukas Zemaitis, Migle Gabrielaite, Jelena Razmuk, Svajune Muralyte, Ana Steponkiene, Lukas Vasionis, Danas Baksa |
| EPI_ISL_3341195 | UAB Diagnostikos laboratorija | National Public Health Surveillance Laboratory | Lukas Zemaitis, Migle Gabrielaite, Jelena Razmuk, Svajune Muralyte, Ana Steponkiene, Lukas Vasionis, Danas Baksa |
| EPI_ISL_3341196 | UAB "Rezus.lt" | National Public Health Surveillance Laboratory | Lukas Zemaitis, Migle Gabrielaite, Jelena Razmuk, Svajune Muralyte, Ana Steponkiene, Lukas Vasionis, Danas Baksa |
| EPI_ISL_3341197 | UAB Diagnostikos laboratorija | National Public Health Surveillance Laboratory | Lukas Zemaitis, Migle Gabrielaite, Jelena Razmuk, Svajune Muralyte, Ana Steponkiene, Lukas Vasionis, Danas Baksa |
| EPI_ISL_3341198 | UAB "Rezus.lt" | National Public Health Surveillance Laboratory | Lukas Zemaitis, Migle Gabrielaite, Jelena Razmuk, Svajune Muralyte, Ana Steponkiene, Lukas Vasionis, Danas Baksa |
| EPI_ISL_3341199 | Viešoji įstaiga Vilniaus universiteto ligoninė Santaros klinikos | National Public Health Surveillance Laboratory | Lukas Zemaitis, Migle Gabrielaite, Jelena Razmuk, Svajune Muralyte, Ana Steponkiene, Lukas Vasionis, Danas Baksa |
| EPI_ISL_3341200 | UAB Diagnostikos laboratorija | National Public Health Surveillance Laboratory | Lukas Zemaitis, Migle Gabrielaite, Jelena Razmuk, Svajune Muralyte, Ana Steponkiene, Lukas Vasionis, Danas Baksa |
| EPI_ISL_3341201, EPI_ISL_3341202, EPI_ISL_3341203 | Viešoji įstaiga Klaipėdos universitetinė ligoninė | National Public Health Surveillance Laboratory | Lukas Zemaitis, Migle Gabrielaite, Jelena Razmuk, Svajune Muralyte, Ana Steponkiene, Lukas Vasionis, Danas Baksa |
| EPI_ISL_3341204 | Viešoji įstaiga Respublikinė Šiaulių ligoninė | National Public Health Surveillance Laboratory | Lukas Zemaitis, Migle Gabrielaite, Jelena Razmuk, Svajune Muralyte, Ana Steponkiene, Lukas Vasionis, Danas Baksa |
| EPI_ISL_3341205 | Viešoji įstaiga Klaipėdos universitetinė ligoninė | National Public Health Surveillance Laboratory | Lukas Zemaitis, Migle Gabrielaite, Jelena Razmuk, Svajune Muralyte, Ana Steponkiene, Lukas Vasionis, Danas Baksa |
| EPI_ISL_3341206, EPI_ISL_3341207 | UAB Diagnostikos laboratorija | National Public Health Surveillance Laboratory | Lukas Zemaitis, Migle Gabrielaite, Jelena Razmuk, Svajune Muralyte, Ana Steponkiene, Lukas Vasionis, Danas Baksa |
| EPI_ISL_3341208 | UAB "Rezus.lt" | National Public Health Surveillance Laboratory | Lukas Zemaitis, Migle Gabrielaite, Jelena Razmuk, Svajune Muralyte, Ana Steponkiene, Lukas Vasionis, Danas Baksa |
| EPI_ISL_3341209, EPI_ISL_3341210 | UAB Diagnostikos laboratorija | National Public Health Surveillance Laboratory | Lukas Zemaitis, Migle Gabrielaite, Jelena Razmuk, Svajune Muralyte, Ana Steponkiene, Lukas Vasionis, Danas Baksa |
| EPI_ISL_3341211, EPI_ISL_3341212 | Viešoji įstaiga Klaipėdos universitetinė ligoninė | National Public Health Surveillance Laboratory | Lukas Zemaitis, Migle Gabrielaite, Jelena Razmuk, Svajune Muralyte, Ana Steponkiene, Lukas Vasionis, Danas Baksa |
| EPI_ISL_3341213 | UAB Diagnostikos laboratorija | National Public Health Surveillance Laboratory | Lukas Zemaitis, Migle Gabrielaite, Jelena Razmuk, Svajune Muralyte, Ana Steponkiene, Lukas Vasionis, Danas Baksa |
| EPI_ISL_3341214, EPI_ISL_3341215 | Viešoji įstaiga Klaipėdos universitetinė ligoninė | National Public Health Surveillance Laboratory | Lukas Zemaitis, Migle Gabrielaite, Jelena Razmuk, Svajune Muralyte, Ana Steponkiene, Lukas Vasionis, Danas Baksa |
| EPI_ISL_3341216 | Viešoji įstaiga Vilniaus universiteto ligoninė Santaros klinikos | National Public Health Surveillance Laboratory | Lukas Zemaitis, Migle Gabrielaite, Jelena Razmuk, Svajune Muralyte, Ana Steponkiene, Lukas Vasionis, Danas Baksa |
| EPI_ISL_3341217, EPI_ISL_3341218 | Viešoji įstaiga Klaipėdos universitetinė ligoninė | National Public Health Surveillance Laboratory | Lukas Zemaitis, Migle Gabrielaite, Jelena Razmuk, Svajune Muralyte, Ana Steponkiene, Lukas Vasionis, Danas Baksa |
| EPI_ISL_3341219 | Viešoji įstaiga Vilniaus universiteto ligoninė Santaros klinikos | National Public Health Surveillance Laboratory | Lukas Zemaitis, Migle Gabrielaite, Jelena Razmuk, Svajune Muralyte, Ana Steponkiene, Lukas Vasionis, Danas Baksa |
| EPI_ISL_3341220, EPI_ISL_3341221, EPI_ISL_3341222, EPI_ISL_3341223 | UAB Diagnostikos laboratorija | National Public Health Surveillance Laboratory | Lukas Zemaitis, Migle Gabrielaite, Jelena Razmuk, Svajune Muralyte, Ana Steponkiene, Lukas Vasionis, Danas Baksa |
| EPI_ISL_3341224 | UAB "Baltic Medics" | National Public Health Surveillance Laboratory | Lukas Zemaitis, Migle Gabrielaite, Jelena Razmuk, Svajune Muralyte, Ana Steponkiene, Lukas Vasionis, Danas Baksa |
| EPI_ISL_3341225 | UAB "Rezus.lt" | National Public Health Surveillance Laboratory | Lukas Zemaitis, Migle Gabrielaite, Jelena Razmuk, Svajune Muralyte, Ana Steponkiene, Lukas Vasionis, Danas Baksa |
| EPI_ISL_3341226 | Viešoji įstaiga Klaipėdos universitetinė ligoninė | National Public Health Surveillance Laboratory | Lukas Zemaitis, Migle Gabrielaite, Jelena Razmuk, Svajune Muralyte, Ana Steponkiene, Lukas Vasionis, Danas Baksa |
| EPI_ISL_3341227 | Viešoji įstaiga Vilniaus universiteto ligoninė Santaros klinikos | National Public Health Surveillance Laboratory | Lukas Zemaitis, Migle Gabrielaite, Jelena Razmuk, Svajune Muralyte, Ana Steponkiene, Lukas Vasionis, Danas Baksa |
| EPI_ISL_3341228 | UAB Diagnostikos laboratorija | National Public Health Surveillance Laboratory | Lukas Zemaitis, Migle Gabrielaite, Jelena Razmuk, Svajune Muralyte, Ana Steponkiene, Lukas Vasionis, Danas Baksa |
| EPI_ISL_3341229 | Viešoji įstaiga Respublikinė Šiaulių ligoninė | National Public Health Surveillance Laboratory | Lukas Zemaitis, Migle Gabrielaite, Jelena Razmuk, Svajune Muralyte, Ana Steponkiene, Lukas Vasionis, Danas Baksa |
| EPI_ISL_3341230 | UAB "Rezus.lt" | National Public Health Surveillance Laboratory | Lukas Zemaitis, Migle Gabrielaite, Jelena Razmuk, Svajune Muralyte, Ana Steponkiene, Lukas Vasionis, Danas Baksa |
| EPI_ISL_3341231 | Viešoji įstaiga Vilniaus universiteto ligoninė Santaros klinikos | National Public Health Surveillance Laboratory | Lukas Zemaitis, Migle Gabrielaite, Jelena Razmuk, Svajune Muralyte, Ana Steponkiene, Lukas Vasionis, Danas Baksa |
| EPI_ISL_3341232 | Viešoji įstaiga Klaipėdos universitetinė ligoninė | National Public Health Surveillance Laboratory | Lukas Zemaitis, Migle Gabrielaite, Jelena Razmuk, Svajune Muralyte, Ana Steponkiene, Lukas Vasionis, Danas Baksa |
| EPI_ISL_3341233 | UAB Diagnostikos laboratorija | National Public Health Surveillance Laboratory | Lukas Zemaitis, Migle Gabrielaite, Jelena Razmuk, Svajune Muralyte, Ana Steponkiene, Lukas Vasionis, Danas Baksa |
| EPI_ISL_3341234, EPI_ISL_3341235 | Viešoji įstaiga Klaipėdos universitetinė ligoninė | National Public Health Surveillance Laboratory | Lukas Zemaitis, Migle Gabrielaite, Jelena Razmuk, Svajune Muralyte, Ana Steponkiene, Lukas Vasionis, Danas Baksa |
| EPI_ISL_3341236 | UAB "Rezus.lt" | National Public Health Surveillance Laboratory | Lukas Zemaitis, Migle Gabrielaite, Jelena Razmuk, Svajune Muralyte, Ana Steponkiene, Lukas Vasionis, Danas Baksa |
| EPI_ISL_3341237 | Viešoji įstaiga Klaipėdos universitetinė ligoninė | National Public Health Surveillance Laboratory | Lukas Zemaitis, Migle Gabrielaite, Jelena Razmuk, Svajune Muralyte, Ana Steponkiene, Lukas Vasionis, Danas Baksa |
| EPI_ISL_3341238 | UAB "Rezus.lt" | National Public Health Surveillance Laboratory | Lukas Zemaitis, Migle Gabrielaite, Jelena Razmuk, Svajune Muralyte, Ana Steponkiene, Lukas Vasionis, Danas Baksa |
| EPI_ISL_3341239 | UAB Diagnostikos laboratorija | National Public Health Surveillance Laboratory | Lukas Zemaitis, Migle Gabrielaite, Jelena Razmuk, Svajune Muralyte, Ana Steponkiene, Lukas Vasionis, Danas Baksa |
| EPI_ISL_3341240, EPI_ISL_3341241 | UAB "Rezus.lt" | National Public Health Surveillance Laboratory | Lukas Zemaitis, Migle Gabrielaite, Jelena Razmuk, Svajune Muralyte, Ana Steponkiene, Lukas Vasionis, Danas Baksa |
| EPI_ISL_3341242 | UAB Diagnostikos laboratorija | National Public Health Surveillance Laboratory | Lukas Zemaitis, Migle Gabrielaite, Jelena Razmuk, Svajune Muralyte, Ana Steponkiene, Lukas Vasionis, Danas Baksa |
| EPI_ISL_3341243 | UAB "Rezus.lt" | National Public Health Surveillance Laboratory | Lukas Zemaitis, Migle Gabrielaite, Jelena Razmuk, Svajune Muralyte, Ana Steponkiene, Lukas Vasionis, Danas Baksa |
| EPI_ISL_3341244 | Viešoji įstaiga Klaipėdos universitetinė ligoninė | National Public Health Surveillance Laboratory | Lukas Zemaitis, Migle Gabrielaite, Jelena Razmuk, Svajune Muralyte, Ana Steponkiene, Lukas Vasionis, Danas Baksa |
| EPI_ISL_3341245 | UAB "Rezus.lt" | National Public Health Surveillance Laboratory | Lukas Zemaitis, Migle Gabrielaite, Jelena Razmuk, Svajune Muralyte, Ana Steponkiene, Lukas Vasionis, Danas Baksa |
| EPI_ISL_3341246 | UAB Diagnostikos laboratorija | National Public Health Surveillance Laboratory | Lukas Zemaitis, Migle Gabrielaite, Jelena Razmuk, Svajune Muralyte, Ana Steponkiene, Lukas Vasionis, Danas Baksa |
| EPI_ISL_3341247 | UAB "Rezus.lt" | National Public Health Surveillance Laboratory | Lukas Zemaitis, Migle Gabrielaite, Jelena Razmuk, Svajune Muralyte, Ana Steponkiene, Lukas Vasionis, Danas Baksa |
| EPI_ISL_3341248 | Viešoji įstaiga Vilniaus universiteto ligoninė Santaros klinikos | National Public Health Surveillance Laboratory | Lukas Zemaitis, Migle Gabrielaite, Jelena Razmuk, Svajune Muralyte, Ana Steponkiene, Lukas Vasionis, Danas Baksa |

| | | | |
|---|---|---|---|
| EPI_ISL_3341249, EPI_ISL_3341250 | Viešoji įstaiga Klaipėdos universitetinė ligoninė | National Public Health Surveillance Laboratory | Lukas Zemaitis, Migle Gabrielaite, Jelena Razmuk, Svajune Muralyte, Ana Steponkiene, Lukas Vasionis, Danas Baksa |
| EPI_ISL_3341251, EPI_ISL_3341252 | UAB Diagnostikos laboratorija | National Public Health Surveillance Laboratory | Lukas Zemaitis, Migle Gabrielaite, Jelena Razmuk, Svajune Muralyte, Ana Steponkiene, Lukas Vasionis, Danas Baksa |
| EPI_ISL_3341253 | Viešoji įstaiga Klaipėdos universitetinė ligoninė | National Public Health Surveillance Laboratory | Lukas Zemaitis, Migle Gabrielaite, Jelena Razmuk, Svajune Muralyte, Ana Steponkiene, Lukas Vasionis, Danas Baksa |
| EPI_ISL_3341254, EPI_ISL_3341255 | UAB "Rezus.lt" | National Public Health Surveillance Laboratory | Lukas Zemaitis, Migle Gabrielaite, Jelena Razmuk, Svajune Muralyte, Ana Steponkiene, Lukas Vasionis, Danas Baksa |
| EPI_ISL_3341256 | Viešoji įstaiga Vilniaus universiteto ligoninė Santaros klinikos | National Public Health Surveillance Laboratory | Lukas Zemaitis, Migle Gabrielaite, Jelena Razmuk, Svajune Muralyte, Ana Steponkiene, Lukas Vasionis, Danas Baksa |
| EPI_ISL_3341257 | Viešoji įstaiga Klaipėdos universitetinė ligoninė | National Public Health Surveillance Laboratory | Lukas Zemaitis, Migle Gabrielaite, Jelena Razmuk, Svajune Muralyte, Ana Steponkiene, Lukas Vasionis, Danas Baksa |
| EPI_ISL_3341258 | UAB "Baltic Medics" | National Public Health Surveillance Laboratory | Lukas Zemaitis, Migle Gabrielaite, Jelena Razmuk, Svajune Muralyte, Ana Steponkiene, Lukas Vasionis, Danas Baksa |
| EPI_ISL_3341259 | Viešoji įstaiga Klaipėdos universitetinė ligoninė | National Public Health Surveillance Laboratory | Lukas Zemaitis, Migle Gabrielaite, Jelena Razmuk, Svajune Muralyte, Ana Steponkiene, Lukas Vasionis, Danas Baksa |
| EPI_ISL_3341260, EPI_ISL_3341261 | UAB Diagnostikos laboratorija | National Public Health Surveillance Laboratory | Lukas Zemaitis, Migle Gabrielaite, Jelena Razmuk, Svajune Muralyte, Ana Steponkiene, Lukas Vasionis, Danas Baksa |
| EPI_ISL_3341262, EPI_ISL_3341263 | Viešoji įstaiga Klaipėdos universitetinė ligoninė | National Public Health Surveillance Laboratory | Lukas Zemaitis, Migle Gabrielaite, Jelena Razmuk, Svajune Muralyte, Ana Steponkiene, Lukas Vasionis, Danas Baksa |
| EPI_ISL_3341264 | Viešoji įstaiga Vilniaus universiteto ligoninė Santaros klinikos | National Public Health Surveillance Laboratory | Lukas Zemaitis, Migle Gabrielaite, Jelena Razmuk, Svajune Muralyte, Ana Steponkiene, Lukas Vasionis, Danas Baksa |
| EPI_ISL_3341265, EPI_ISL_3341266, EPI_ISL_3341267, EPI_ISL_3341268 | UAB Diagnostikos laboratorija | National Public Health Surveillance Laboratory | Lukas Zemaitis, Migle Gabrielaite, Jelena Razmuk, Svajune Muralyte, Ana Steponkiene, Lukas Vasionis, Danas Baksa |
| EPI_ISL_3341269 | UAB "Rezus.lt" | National Public Health Surveillance Laboratory | Lukas Zemaitis, Migle Gabrielaite, Jelena Razmuk, Svajune Muralyte, Ana Steponkiene, Lukas Vasionis, Danas Baksa |
| EPI_ISL_3341270 | Viešoji įstaiga Klaipėdos universitetinė ligoninė | National Public Health Surveillance Laboratory | Lukas Zemaitis, Migle Gabrielaite, Jelena Razmuk, Svajune Muralyte, Ana Steponkiene, Lukas Vasionis, Danas Baksa |
| EPI_ISL_3341271 | UAB "Rezus.lt" | National Public Health Surveillance Laboratory | Lukas Zemaitis, Migle Gabrielaite, Jelena Razmuk, Svajune Muralyte, Ana Steponkiene, Lukas Vasionis, Danas Baksa |
| EPI_ISL_3341272 | Viešoji įstaiga Klaipėdos universitetinė ligoninė | National Public Health Surveillance Laboratory | Lukas Zemaitis, Migle Gabrielaite, Jelena Razmuk, Svajune Muralyte, Ana Steponkiene, Lukas Vasionis, Danas Baksa |
| EPI_ISL_3341273 | UAB Diagnostikos laboratorija | National Public Health Surveillance Laboratory | Lukas Zemaitis, Migle Gabrielaite, Jelena Razmuk, Svajune Muralyte, Ana Steponkiene, Lukas Vasionis, Danas Baksa |
| EPI_ISL_3341274 | Viešoji įstaiga Klaipėdos universitetinė ligoninė | National Public Health Surveillance Laboratory | Lukas Zemaitis, Migle Gabrielaite, Jelena Razmuk, Svajune Muralyte, Ana Steponkiene, Lukas Vasionis, Danas Baksa |
| EPI_ISL_3341275 | UAB "SK IMPEKS MEDICINOS DIAGNOSTIKOS CENTRAS" | National Public Health Surveillance Laboratory | Lukas Zemaitis, Migle Gabrielaite, Jelena Razmuk, Svajune Muralyte, Ana Steponkiene, Lukas Vasionis, Danas Baksa |
| EPI_ISL_3341276, EPI_ISL_3341277, EPI_ISL_3341278 | Viešoji įstaiga Klaipėdos universitetinė ligoninė | National Public Health Surveillance Laboratory | Lukas Zemaitis, Migle Gabrielaite, Jelena Razmuk, Svajune Muralyte, Ana Steponkiene, Lukas Vasionis, Danas Baksa |
| EPI_ISL_3341279 | UAB "Baltic Medics" | National Public Health Surveillance Laboratory | Lukas Zemaitis, Migle Gabrielaite, Jelena Razmuk, Svajune Muralyte, Ana Steponkiene, Lukas Vasionis, Danas Baksa |
| EPI_ISL_3341280 | Viešoji įstaiga Klaipėdos universitetinė ligoninė | National Public Health Surveillance Laboratory | Lukas Zemaitis, Migle Gabrielaite, Jelena Razmuk, Svajune Muralyte, Ana Steponkiene, Lukas Vasionis, Danas Baksa |
| EPI_ISL_3341281 | UAB Diagnostikos laboratorija | National Public Health Surveillance Laboratory | Lukas Zemaitis, Migle Gabrielaite, Jelena Razmuk, Svajune Muralyte, Ana Steponkiene, Lukas Vasionis, Danas Baksa |
| EPI_ISL_3341282 | Viešoji įstaiga Klaipėdos universitetinė ligoninė | National Public Health Surveillance Laboratory | Lukas Zemaitis, Migle Gabrielaite, Jelena Razmuk, Svajune Muralyte, Ana Steponkiene, Lukas Vasionis, Danas Baksa |
| EPI_ISL_3341283 | UAB Diagnostikos laboratorija | National Public Health Surveillance Laboratory | Lukas Zemaitis, Migle Gabrielaite, Jelena Razmuk, Svajune Muralyte, Ana Steponkiene, Lukas Vasionis, Danas Baksa |
| EPI_ISL_3341284, EPI_ISL_3341285, EPI_ISL_3341286 | Viešoji įstaiga Klaipėdos universitetinė ligoninė | National Public Health Surveillance Laboratory | Lukas Zemaitis, Migle Gabrielaite, Jelena Razmuk, Svajune Muralyte, Ana Steponkiene, Lukas Vasionis, Danas Baksa |
| EPI_ISL_3341287 | UAB "Rezus.lt" | National Public Health Surveillance Laboratory | Lukas Zemaitis, Migle Gabrielaite, Jelena Razmuk, Svajune Muralyte, Ana Steponkiene, Lukas Vasionis, Danas Baksa |
| EPI_ISL_3341288, EPI_ISL_3341289, EPI_ISL_3341290, EPI_ISL_3341291, EPI_ISL_3341292, EPI_ISL_3341293, EPI_ISL_3341294, EPI_ISL_3341295, EPI_ISL_3341296, EPI_ISL_3341297, EPI_ISL_3341298, EPI_ISL_3341299, EPI_ISL_3341300, EPI_ISL_3341301, EPI_ISL_3341303, EPI_ISL_3341304, EPI_ISL_3341305, EPI_ISL_3341306, EPI_ISL_3341307, EPI_ISL_3341308, EPI_ISL_3341310, EPI_ISL_3341311, EPI_ISL_3341312, EPI_ISL_3341313, EPI_ISL_3341314, EPI_ISL_3341315, EPI_ISL_3341316, EPI_ISL_3341318, EPI_ISL_3341319, EPI_ISL_3341320, EPI_ISL_3341321, EPI_ISL_3341322, EPI_ISL_3341326, EPI_ISL_3341327, EPI_ISL_3341328, EPI_ISL_3341329, EPI_ISL_3341330, EPI_ISL_3341331, EPI_ISL_3341332, EPI_ISL_3341333, EPI_ISL_3341334, EPI_ISL_3341335, EPI_ISL_3341336, EPI_ISL_3341337, EPI_ISL_3341338, EPI_ISL_3341339, EPI_ISL_3341340, EPI_ISL_3341341, EPI_ISL_3341342, EPI_ISL_3341343, EPI_ISL_3341344, EPI_ISL_3341345, EPI_ISL_3341346, EPI_ISL_3341347, EPI_ISL_3341348, EPI_ISL_3341349, EPI_ISL_3341350, EPI_ISL_3341351, EPI_ISL_3341352, EPI_ISL_3341353, EPI_ISL_3341354, EPI_ISL_3341355, EPI_ISL_3341356, EPI_ISL_3341357, EPI_ISL_3341358, EPI_ISL_3341359, EPI_ISL_3341360, EPI_ISL_3341361, EPI_ISL_3341362, EPI_ISL_3341363, EPI_ISL_3341364, EPI_ISL_3341365, EPI_ISL_3341366, EPI_ISL_3341367, EPI_ISL_3341368, EPI_ISL_3341369, EPI_ISL_3341370, EPI_ISL_3341371, EPI_ISL_3341372, EPI_ISL_3341373, EPI_ISL_3341374, EPI_ISL_3341375, EPI_ISL_3341376, EPI_ISL_3341377, EPI_ISL_3341378, EPI_ISL_3341379, EPI_ISL_3341380, EPI_ISL_3341381, EPI_ISL_3341382, EPI_ISL_3341383, EPI_ISL_3341384, EPI_ISL_3341385, EPI_ISL_3341386, EPI_ISL_3341387, EPI_ISL_3341388, EPI_ISL_3341389, EPI_ISL_3341390, EPI_ISL_3341391, EPI_ISL_3341392, EPI_ISL_3341393, EPI_ISL_3341394, EPI_ISL_3341395, EPI_ISL_3341396, EPI_ISL_3341397 | | | |
| see above | Gravity Diagnostics, LLC | Gravity Diagnostics, LLC | Gravity Diagnostics |
| EPI_ISL_3341398, EPI_ISL_3341399 | CDFD | CDFD | Murali Bashyam, Asmita Gupta, Reelina Basu, Vinay Donipadi, Nagamani Kammili, Divya Vashisht, Ashwin Dalal |
| EPI_ISL_3341405, EPI_ISL_3341406, EPI_ISL_3341407, EPI_ISL_3341408, EPI_ISL_3341409, EPI_ISL_3341410, EPI_ISL_3341411, EPI_ISL_3341412, EPI_ISL_3341413, EPI_ISL_3341414, EPI_ISL_3341416, EPI_ISL_3341417, EPI_ISL_3341418, EPI_ISL_3341420, EPI_ISL_3341421, EPI_ISL_3341422, EPI_ISL_3341423, EPI_ISL_3341424, EPI_ISL_3341425, EPI_ISL_3341426, EPI_ISL_3341428, EPI_ISL_3341429, EPI_ISL_3341430, EPI_ISL_3341431, EPI_ISL_3341432, EPI_ISL_3341433, EPI_ISL_3341434, EPI_ISL_3341436, EPI_ISL_3341438, EPI_ISL_3341439, EPI_ISL_3341440, EPI_ISL_3341441, EPI_ISL_3341442, EPI_ISL_3341443, EPI_ISL_3341445, EPI_ISL_3341446, EPI_ISL_3341447, EPI_ISL_3341448, EPI_ISL_3341449, EPI_ISL_3341450, EPI_ISL_3341451, EPI_ISL_3341453, EPI_ISL_3341454, EPI_ISL_3341457, EPI_ISL_3341458, EPI_ISL_3341460, EPI_ISL_3341462, EPI_ISL_3341464, EPI_ISL_3341465, EPI_ISL_3341466, EPI_ISL_3341467, EPI_ISL_3341468, EPI_ISL_3341470, EPI_ISL_3341471, EPI_ISL_3341472, EPI_ISL_3341473, EPI_ISL_3341474, EPI_ISL_3341475, EPI_ISL_3341476, EPI_ISL_3341477, EPI_ISL_3341478, EPI_ISL_3341479, EPI_ISL_3341480, EPI_ISL_3341481, EPI_ISL_3341482, EPI_ISL_3341484, EPI_ISL_3341486, EPI_ISL_3341487, EPI_ISL_3341488, EPI_ISL_3341489, EPI_ISL_3341490, EPI_ISL_3341491, EPI_ISL_3341492, EPI_ISL_3341493, EPI_ISL_3341499, EPI_ISL_3341500, EPI_ISL_3341501, EPI_ISL_3341502, EPI_ISL_3341503, EPI_ISL_3341504, EPI_ISL_3341505, EPI_ISL_3341506, EPI_ISL_3341507, EPI_ISL_3341509, EPI_ISL_3341510, EPI_ISL_3341511, EPI_ISL_3341512, EPI_ISL_3341513, EPI_ISL_3341514, EPI_ISL_3341515, EPI_ISL_3341516, EPI_ISL_3341517, EPI_ISL_3341518, EPI_ISL_3341520, EPI_ISL_3341521, EPI_ISL_3341522, EPI_ISL_3341523, EPI_ISL_3341524, EPI_ISL_3341525, EPI_ISL_3341526, EPI_ISL_3341528, EPI_ISL_3341529, EPI_ISL_3341530, EPI_ISL_3341531, EPI_ISL_3341532, EPI_ISL_3341533, EPI_ISL_3341534, EPI_ISL_3341535, EPI_ISL_3341536, EPI_ISL_3341537, EPI_ISL_3341538, EPI_ISL_3341539, EPI_ISL_3341540, EPI_ISL_3341541, EPI_ISL_3341542, EPI_ISL_3341543, EPI_ISL_3341544, EPI_ISL_3341545, EPI_ISL_3341546, EPI_ISL_3341547, EPI_ISL_3341548, EPI_ISL_3341549, EPI_ISL_3341550, EPI_ISL_3341551, EPI_ISL_3341552, EPI_ISL_3341553, EPI_ISL_3341554, EPI_ISL_3341555, EPI_ISL_3341556, EPI_ISL_3341557, EPI_ISL_3341558, EPI_ISL_3341559, EPI_ISL_3341560, EPI_ISL_3341561, EPI_ISL_3341562, EPI_ISL_3341563, EPI_ISL_3341564, EPI_ISL_3341565, EPI_ISL_3341566, EPI_ISL_3341568, EPI_ISL_3341569, EPI_ISL_3341570, EPI_ISL_3341571, EPI_ISL_3341572, EPI_ISL_3341573, EPI_ISL_3341574, EPI_ISL_3341575, EPI_ISL_3341576, EPI_ISL_3341577, EPI_ISL_3341578, EPI_ISL_3341579, EPI_ISL_3341580, EPI_ISL_3341581, EPI_ISL_3341582, EPI_ISL_3341583, EPI_ISL_3341584, EPI_ISL_3341585, EPI_ISL_3341586, EPI_ISL_3341587, EPI_ISL_3341588, EPI_ISL_3341589, EPI_ISL_3341590, EPI_ISL_3341591, EPI_ISL_3341592, EPI_ISL_3341593, EPI_ISL_3341594, EPI_ISL_3341598, EPI_ISL_3341599, EPI_ISL_3341600, EPI_ISL_3341601, EPI_ISL_3341602, EPI_ISL_3341603, EPI_ISL_3341604, EPI_ISL_3341605, EPI_ISL_3341606, EPI_ISL_3341607, EPI_ISL_3341608, EPI_ISL_3341609, EPI_ISL_3341612, EPI_ISL_3341613, EPI_ISL_3341614, EPI_ISL_3341615, EPI_ISL_3341616, EPI_ISL_3341617, EPI_ISL_3341618, EPI_ISL_3341619, EPI_ISL_3341620, EPI_ISL_3341621, EPI_ISL_3341622, EPI_ISL_3341623, EPI_ISL_3341624, EPI_ISL_3341626, EPI_ISL_3341627, EPI_ISL_3341628, EPI_ISL_3341629, EPI_ISL_3341630, EPI_ISL_3341631, EPI_ISL_3341632, EPI_ISL_3341633, EPI_ISL_3341634, EPI_ISL_3341635, EPI_ISL_3341636, EPI_ISL_3341637, EPI_ISL_3341638, EPI_ISL_3341639, EPI_ISL_3341640, EPI_ISL_3341641, EPI_ISL_3341642, EPI_ISL_3341643, EPI_ISL_3341644, EPI_ISL_3341646, EPI_ISL_3341647, EPI_ISL_3341649, EPI_ISL_3341650, EPI_ISL_3341651, EPI_ISL_3341652, EPI_ISL_3341653, EPI_ISL_3341654, EPI_ISL_3341655, EPI_ISL_3341656, EPI_ISL_3341657, EPI_ISL_3341658, EPI_ISL_3341659, EPI_ISL_3341660, EPI_ISL_3341661, EPI_ISL_3341663, EPI_ISL_3341664, EPI_ISL_3341665, EPI_ISL_3341666, EPI_ISL_3341667, EPI_ISL_3341668, EPI_ISL_3341669, EPI_ISL_3341670, EPI_ISL_3341671, EPI_ISL_3341672, EPI_ISL_3341673, EPI_ISL_3341674, EPI_ISL_3341675, EPI_ISL_3341676, EPI_ISL_3341680, EPI_ISL_3341681, EPI_ISL_3341682, EPI_ISL_3341683, EPI_ISL_3341684, EPI_ISL_3341685, EPI_ISL_3341686, EPI_ISL_3341687, EPI_ISL_3341688, EPI_ISL_3341689, EPI_ISL_3341690, EPI_ISL_3341691, EPI_ISL_3341692, EPI_ISL_3341693, EPI_ISL_3341694, EPI_ISL_3341695, EPI_ISL_3341696, EPI_ISL_3341697, EPI_ISL_3341698, EPI_ISL_3341699, EPI_ISL_3341700, EPI_ISL_3341701, EPI_ISL_3341703, EPI_ISL_3341704, EPI_ISL_3341705, EPI_ISL_3341706, EPI_ISL_3341707, EPI_ISL_3341709, EPI_ISL_3341710, EPI_ISL_3341712, EPI_ISL_3341713, EPI_ISL_3341714, EPI_ISL_3341715, EPI_ISL_3341716, EPI_ISL_3341719, EPI_ISL_3341720, EPI_ISL_3341721, EPI_ISL_3341722, EPI_ISL_3341723, EPI_ISL_3341724, EPI_ISL_3341725, EPI_ISL_3341726, EPI_ISL_3341727, EPI_ISL_3341728, EPI_ISL_3341729, EPI_ISL_3341730, EPI_ISL_3341731, EPI_ISL_3341732, EPI_ISL_3341733, EPI_ISL_3341735, EPI_ISL_3341736, EPI_ISL_3341737, EPI_ISL_3341738, EPI_ISL_3341739, EPI_ISL_3341740, EPI_ISL_3341741, EPI_ISL_3341742, EPI_ISL_3341743, EPI_ISL_3341744, EPI_ISL_3341745, EPI_ISL_3341746, EPI_ISL_3341747, EPI_ISL_3341748, EPI_ISL_3341749, EPI_ISL_3341750, EPI_ISL_3341751, EPI_ISL_3341752, EPI_ISL_3341753, EPI_ISL_3341754, EPI_ISL_3341755, EPI_ISL_3341756, EPI_ISL_3341757, EPI_ISL_3341758, EPI_ISL_3341759, EPI_ISL_3341760, EPI_ISL_3341761, EPI_ISL_3341762, EPI_ISL_3341763, EPI_ISL_3341764, EPI_ISL_3341766, EPI_ISL_3341767, EPI_ISL_3341768, EPI_ISL_3341769, EPI_ISL_3341771, EPI_ISL_3341772, EPI_ISL_3341773, EPI_ISL_3341774, EPI_ISL_3341775, EPI_ISL_3341776, EPI_ISL_3341777, EPI_ISL_3341778, EPI_ISL_3341779, EPI_ISL_3341780, EPI_ISL_3341781, EPI_ISL_3341782, EPI_ISL_3341783, EPI_ISL_3341784, EPI_ISL_3341785, EPI_ISL_3341786, EPI_ISL_3341787, EPI_ISL_3341788, EPI_ISL_3341789, EPI_ISL_3341790, EPI_ISL_3341791, EPI_ISL_3341792, EPI_ISL_3341793, EPI_ISL_3341794, EPI_ISL_3341795, EPI_ISL_3341796, EPI_ISL_3341797, EPI_ISL_3341798, EPI_ISL_3341799, EPI_ISL_3341800, EPI_ISL_3341801, EPI_ISL_3341802, EPI_ISL_3341803, EPI_ISL_3341804, EPI_ISL_3341805, EPI_ISL_3341806, EPI_ISL_3341807, EPI_ISL_3341808, EPI_ISL_3341809, EPI_ISL_3341810, EPI_ISL_3341811, EPI_ISL_3341812, EPI_ISL_3341813, EPI_ISL_3341814, EPI_ISL_3341815, EPI_ISL_3341816, EPI_ISL_3341817, EPI_ISL_3341818, EPI_ISL_3341819, EPI_ISL_3341820, EPI_ISL_3341821, EPI_ISL_3341822, EPI_ISL_3341823, EPI_ISL_3341824, EPI_ISL_3341826, EPI_ISL_3341827, EPI_ISL_3341828, EPI_ISL_3341829, EPI_ISL_3341830, EPI_ISL_3341831, EPI_ISL_3341832, EPI_ISL_3341833, EPI_ISL_3341834, EPI_ISL_3341835, EPI_ISL_3341837, EPI_ISL_3341838, EPI_ISL_3341839, EPI_ISL_3341840, EPI_ISL_3341841, EPI_ISL_3341842, EPI_ISL_3341843, EPI_ISL_3341844, EPI_ISL_3341845, EPI_ISL_3341846, EPI_ISL_3341847, EPI_ISL_3341848, EPI_ISL_3341849, EPI_ISL_3341850, EPI_ISL_3341851, EPI_ISL_3341852, EPI_ISL_3341853, EPI_ISL_3341854, EPI_ISL_3341855, EPI_ISL_3341856 | | | |
| see above | Public Health Ontario Laboratory | Public Health Ontario Laboratory | Vanessa G Allen, Philip Banh, Yao Chen, Richard de Borja, Alireza Eshaghi, Nahuel Fittipaldi, Christine Frantz, Jonathan B Gubbay, Jennifer L Guthrie, Lawrence Heisler, Esha Joshi, Michael Laszloffy, Aimin Li, Michael CY Li, Dean Maxwell, Sandeep Nagra, Samir N Patel, Jared Simpson, Karthikeyan Sivaraman, Ashleigh Sullivan, Yogi Sundaravadanam, Sarah Teatero, Andre Villegas, Matthew Watson, Sandra Zittermann |
| EPI_ISL_3341857, EPI_ISL_3341859, EPI_ISL_3341862, EPI_ISL_3341863, EPI_ISL_3341864, EPI_ISL_3341866 | Genetica Molecular and Subdepartamento de Virologia ISP Chile | Instituto de Salud Publica de Chile | Karen Orostica, Constanza Campano, Barbara Parra, Loredana Arata, Gisselle Barra, Patricia Bustos, Rodrigo Fasce, Javier Tognarelli, Andres Castillo, Soledad Ulloa, Jorge Fernandez |
| EPI_ISL_3341871 | CGB laboratoř, a.s. | Institute of Molecular and Translational Medicine / Laboratory of Experimental Medicine, Faculty of Medicine and Dentistry, Palacky University | Rastislav Slavkovský, Hana Jaworek, Vladimíra Koudeláková, Barbora Blumová, Marián Hajdúch |
| EPI_ISL_3341880, EPI_ISL_3341881 | Nemocnice Šumperk, a.s. | Institute of Molecular and Translational Medicine / Laboratory of Experimental Medicine, Faculty of Medicine and Dentistry, Palacky University | Rastislav Slavkovský, Hana Jaworek, Vladimíra Koudeláková, Barbora Blumová, Marián Hajdúch |
| EPI_ISL_3341882, EPI_ISL_3341883, EPI_ISL_3341884, EPI_ISL_3341885, EPI_ISL_3341887, EPI_ISL_3341888, EPI_ISL_3341889, EPI_ISL_3341890, EPI_ISL_3341891, EPI_ISL_3341892, EPI_ISL_3341893, EPI_ISL_3341894, EPI_ISL_3341895, EPI_ISL_3341896, EPI_ISL_3341897, EPI_ISL_3341898, EPI_ISL_3341899, EPI_ISL_3341900, EPI_ISL_3341901, EPI_ISL_3341902 | | | |
| see above | Institute of Molecular and Translational Medicine / Laboratory of Experimental Medicine, Faculty of Medicine and Dentistry, Palacky University and University Hospital Olomouc | Institute of Molecular and Translational Medicine / Laboratory of Experimental Medicine, Faculty of Medicine and Dentistry, Palacky University | Rastislav Slavkovský, Hana Jaworek, Vladimíra Koudeláková, Barbora Blumová, Marián Hajdúch |
| EPI_ISL_3341903 | UAB "Rezus.lt" | National Public Health Surveillance Laboratory | Lukas Zemaitis, Migle Gabrielaite, Jelena Razmuk, Svajune Muralyte, Ana Steponkiene, Lukas Vasionis, Danas Baksa |
| EPI_ISL_3341904 | Viešoji įstaiga Vilniaus universiteto ligoninė Santaros klinikos | National Public Health Surveillance Laboratory | Lukas Zemaitis, Migle Gabrielaite, Jelena Razmuk, Svajune Muralyte, Ana Steponkiene, Lukas Vasionis, Danas Baksa |
| EPI_ISL_3341905 | UAB Diagnostikos laboratorija | National Public Health Surveillance Laboratory | Lukas Zemaitis, Migle Gabrielaite, Jelena Razmuk, Svajune Muralyte, Ana Steponkiene, Lukas Vasionis, Danas Baksa |
| EPI_ISL_3341906 | Viešoji įstaiga Klaipėdos universitetinė ligoninė | National Public Health Surveillance Laboratory | Lukas Zemaitis, Migle Gabrielaite, Jelena Razmuk, Svajune Muralyte, Ana Steponkiene, Lukas Vasionis, Danas Baksa |
| EPI_ISL_3341917, EPI_ISL_3341918, EPI_ISL_3341919, | Genetica Molecular and Subdepartamento de Virologia ISP Chile | Instituto de Salud Publica de Chile | Karen Orostica, Constanza Campano, Barbara Parra, Loredana Arata, Gisselle Barra, Patricia Bustos, Rodrigo Fasce, Javier Tognarelli, Andres Castillo, Soledad Ulloa, Jorge Fernandez |

| | | | |
|---|---|---|---|
| EPI_ISL_3341920, EPI_ISL_3341921, EPI_ISL_3341922 | | | |
| EPI_ISL_3341927, EPI_ISL_3341928, EPI_ISL_3341929, EPI_ISL_3341930, EPI_ISL_3341931, EPI_ISL_3341932, EPI_ISL_3341933, EPI_ISL_3341934 | UPMC Clinical Microbiology Laboratory | Microbial Genome Sequencing Center; Microbial Genomic Epidemiology Laboratory, University of Pittsburgh | Lee H. Harrison, Jane W. Marsh, Marissa P. Griffith, Vatsala R. Srinivasa, Kady D. Waggle, Daniel J. Snyder, Vaughn S. Cooper |
| EPI_ISL_3341943, EPI_ISL_3341944, EPI_ISL_3341945, EPI_ISL_3341946, EPI_ISL_3341947, EPI_ISL_3341948, EPI_ISL_3341949 | Genetica Molecular and Subdepartamento de Virologia ISP Chile | Instituto de Salud Publica de Chile | Karen Orostica, Constanza Campano, Barbara Parra, Loredana Arata, Gisselle Barra, Patricia Bustos, Rodrigo Fasce, Javier Tognarelli, Andres Castillo, Soledad Ulloa, Jorge Fernandez |
| EPI_ISL_3341951, EPI_ISL_3341952, EPI_ISL_3341953, EPI_ISL_3341955, EPI_ISL_3341956, EPI_ISL_3341957, EPI_ISL_3341958, EPI_ISL_3341959, EPI_ISL_3341962, EPI_ISL_3341963, EPI_ISL_3341964, EPI_ISL_3341965, EPI_ISL_3341967, EPI_ISL_3341968, EPI_ISL_3341970, EPI_ISL_3341974, EPI_ISL_3341977, EPI_ISL_3341978, EPI_ISL_3341979, EPI_ISL_3341980, EPI_ISL_3341981, EPI_ISL_3341982, EPI_ISL_3341983, EPI_ISL_3341984, EPI_ISL_3341985, EPI_ISL_3341988 | | | |
| see above | Child Health Research Foundation | Child Health Research Foundation | CHRF Bangladesh Genomics Team |
| EPI_ISL_3341989, EPI_ISL_3341990, EPI_ISL_3341991, EPI_ISL_3341992, EPI_ISL_3341993, EPI_ISL_3341994, EPI_ISL_3341995, EPI_ISL_3341996 | Victor Babes Hospital | National Institute of Infectious Diseases-Prof. Dr. Matei Bals Molecular Diagnostics Laboratory | Corina Casangiu, Leontina Banica, Marius Surleac, Ovidiu Vlaicu, Petre Milu, Robert Hohan, Simona Paraschiv, Dan Otelea |
| EPI_ISL_3341997, EPI_ISL_3341998 | Department of Public Health Tulcea | National Institute of Infectious Diseases-Prof. Dr. Matei Bals Molecular Diagnostics Laboratory | Corina Casangiu, Leontina Banica, Marius Surleac, Ovidiu Vlaicu, Petre Milu, Robert Hohan, Simona Paraschiv, Dan Otelea |
| EPI_ISL_3341999 | Department of Public Health Constanta | National Institute of Infectious Diseases-Prof. Dr. Matei Bals Molecular Diagnostics Laboratory | Corina Casangiu, Leontina Banica, Marius Surleac, Ovidiu Vlaicu, Petre Milu, Robert Hohan, Simona Paraschiv, Dan Otelea |
| EPI_ISL_3342000, EPI_ISL_3342001, EPI_ISL_3342002, EPI_ISL_3342003, EPI_ISL_3342004, EPI_ISL_3342005 | National Institute of Infectious Diseases-Prof. Dr. Matei Bals Molecular Diagnostics Laboratory | National Institute of Infectious Diseases-Prof. Dr. Matei Bals Molecular Diagnostics Laboratory | Corina Casangiu, Leontina Banica, Marius Surleac, Ovidiu Vlaicu, Petre Milu, Robert Hohan, Simona Paraschiv, Dan Otelea |
| EPI_ISL_3342006, EPI_ISL_3342007, EPI_ISL_3342008, EPI_ISL_3342009, EPI_ISL_3342010 | Victor Babes Hospital | National Institute of Infectious Diseases-Prof. Dr. Matei Bals Molecular Diagnostics Laboratory | Corina Casangiu, Leontina Banica, Marius Surleac, Ovidiu Vlaicu, Petre Milu, Robert Hohan, Simona Paraschiv, Dan Otelea |
| EPI_ISL_3342011, EPI_ISL_3342012, EPI_ISL_3342013, EPI_ISL_3342014 | Department of Public Health Tulcea | National Institute of Infectious Diseases-Prof. Dr. Matei Bals Molecular Diagnostics Laboratory | Corina Casangiu, Leontina Banica, Marius Surleac, Ovidiu Vlaicu, Petre Milu, Robert Hohan, Simona Paraschiv, Dan Otelea |
| EPI_ISL_3342015, EPI_ISL_3342016, EPI_ISL_3342017, EPI_ISL_3342018, EPI_ISL_3342019, EPI_ISL_3342020, EPI_ISL_3342021, EPI_ISL_3342022, EPI_ISL_3342023 | Department of Public Health Constanta | National Institute of Infectious Diseases-Prof. Dr. Matei Bals Molecular Diagnostics Laboratory | Corina Casangiu, Leontina Banica, Marius Surleac, Ovidiu Vlaicu, Petre Milu, Robert Hohan, Simona Paraschiv, Dan Otelea |
| EPI_ISL_3342024, EPI_ISL_3342025, EPI_ISL_3342026 | Department of Public Health Ilfov | National Institute of Infectious Diseases-Prof. Dr. Matei Bals Molecular Diagnostics Laboratory | Corina Casangiu, Leontina Banica, Marius Surleac, Ovidiu Vlaicu, Petre Milu, Robert Hohan, Simona Paraschiv, Dan Otelea |
| EPI_ISL_3342027 | Department of Public Health Vrancea | National Institute of Infectious Diseases-Prof. Dr. Matei Bals Molecular Diagnostics Laboratory | Corina Casangiu, Leontina Banica, Marius Surleac, Ovidiu Vlaicu, Petre Milu, Robert Hohan, Simona Paraschiv, Dan Otelea |
| EPI_ISL_3342028, EPI_ISL_3342029, EPI_ISL_3342030, EPI_ISL_3342031, EPI_ISL_3342032, EPI_ISL_3342033, EPI_ISL_3342034, EPI_ISL_3342035, EPI_ISL_3342036, EPI_ISL_3342037, EPI_ISL_3342038, EPI_ISL_3342039 | | | |
| see above | Department of Public Health Iasi | National Institute of Infectious Diseases-Prof. Dr. Matei Bals Molecular Diagnostics Laboratory | Corina Casangiu, Leontina Banica, Marius Surleac, Ovidiu Vlaicu, Petre Milu, Robert Hohan, Simona Paraschiv, Dan Otelea |
| EPI_ISL_3342046, EPI_ISL_3342047 | Genetica Molecular and Subdepartamento de Virologia ISP Chile | Instituto de Salud Publica de Chile | Karen Orostica, Constanza Campano, Barbara Parra, Loredana Arata, Gisselle Barra, Patricia Bustos, Rodrigo Fasce, Javier Tognarelli, Andres Castillo, Soledad Ulloa, Jorge Fernandez |
| EPI_ISL_3342052, EPI_ISL_3342053, EPI_ISL_3342054, EPI_ISL_3342055, EPI_ISL_3342056, EPI_ISL_3342057, EPI_ISL_3342058, EPI_ISL_3342060 | Indian Council of Medical Research-National Institute of Virology, Microbial Containment Complex | Indian Council of Medical Research-National Institute of Virology, Microbial Containment Complex | Pragya D. Yadav |
| EPI_ISL_3342061 | Regional Medical Sciences Center 5 Samut Songkhram | National Institute of Health, Department of Medical Sciences, Ministry of Public Health, Thailand | Pilailuk Okada; Siripaporn Phuygun; Nuttida Thongpramul; Sittiporn Parnmen; Ratana Tacharoenmuang; Pakorn Piromtong; Natchaya Khiadsang; Thanutsapa Thanadachakul; Warawan Wongboot; sirikanda wimol; Sunthareeya Waicharoen; |
| EPI_ISL_3342062, EPI_ISL_3342063 | Regional Medical Sciences Center 3 Nakhonsawan | National Institute of Health, Department of Medical Sciences, Ministry of Public Health, Thailand | Pilailuk Okada; Siripaporn Phuygun; Nuttida Thongpramul; Sittiporn Parnmen; Ratana Tacharoenmuang; Pakorn Piromtong; Natchaya Khiadsang; Thanutsapa Thanadachakul; Warawan Wongboot; sirikanda wimol; Sunthareeya Waicharoen; |
| EPI_ISL_3342064, EPI_ISL_3342065 | Urban Institute for Disease Prevention and Control | National Institute of Health, Department of Medical Sciences, Ministry of Public Health, Thailand | Pilailuk Okada; Siripaporn Phuygun; Nuttida Thongpramul; Sittiporn Parnmen; Ratana Tacharoenmuang; Pakorn Piromtong; Natchaya Khiadsang; Thanutsapa Thanadachakul; Warawan Wongboot; sirikanda wimol; Sunthareeya Waicharoen; |
| EPI_ISL_3342066 | Regional Medical Sciences Center 9 Nakhon Ratchasima | National Institute of Health, Department of Medical Sciences, Ministry of Public Health, Thailand | Pilailuk Okada; Siripaporn Phuygun; Nuttida Thongpramul; Sittiporn Parnmen; Ratana Tacharoenmuang; Pakorn Piromtong; Natchaya Khiadsang; Thanutsapa Thanadachakul; Warawan Wongboot; sirikanda wimol; Sunthareeya Waicharoen; |
| EPI_ISL_3342069 | Regional Medical Sciences Center 8 Udonthani | National Institute of Health, Department of Medical Sciences, Ministry of Public Health, Thailand | Pilailuk Okada; Siripaporn Phuygun; Nuttida Thongpramul; Sittiporn Parnmen; Ratana Tacharoenmuang; Pakorn Piromtong; Natchaya Khiadsang; Thanutsapa Thanadachakul; Warawan Wongboot; sirikanda wimol; Sunthareeya Waicharoen; |
| EPI_ISL_3342070, EPI_ISL_3342071, EPI_ISL_3342072, EPI_ISL_3342073 | Laboratory of Molecular Biology and Cancer Immunology, Faculty of Sciences, Lebanese University | Microbial Pathogenomics Lab - LAU | Sima tokajian, Georgi Merhi, Jad Koweyes, Bassam Badran, Fadi Abdel Sater, Rawan Makki, Tatiana Tohmeh, Nada Ghosn, Hamad Hassan |
| EPI_ISL_3342075, EPI_ISL_3342076, EPI_ISL_3342077, EPI_ISL_3342078, EPI_ISL_3342079, EPI_ISL_3342080, EPI_ISL_3342081, EPI_ISL_3342082, EPI_ISL_3342083, EPI_ISL_3342084, EPI_ISL_3342085, EPI_ISL_3342086, EPI_ISL_3342087, EPI_ISL_3342088, EPI_ISL_3342089, EPI_ISL_3342090, EPI_ISL_3342091, EPI_ISL_3342092, EPI_ISL_3342093, EPI_ISL_3342094, EPI_ISL_3342095, EPI_ISL_3342096, EPI_ISL_3342097, EPI_ISL_3342098, EPI_ISL_3342099, EPI_ISL_3342100, EPI_ISL_3342101, EPI_ISL_3342102, EPI_ISL_3342103, EPI_ISL_3342104, EPI_ISL_3342105, EPI_ISL_3342106, EPI_ISL_3342107, EPI_ISL_3342108, EPI_ISL_3342109, EPI_ISL_3342110, EPI_ISL_3342111, EPI_ISL_3342112, EPI_ISL_3342113, EPI_ISL_3342114, EPI_ISL_3342115, EPI_ISL_3342116, EPI_ISL_3342117, EPI_ISL_3342118, EPI_ISL_3342119, EPI_ISL_3342120, EPI_ISL_3342121, EPI_ISL_3342122, EPI_ISL_3342123, EPI_ISL_3342124, EPI_ISL_3342125, EPI_ISL_3342126, EPI_ISL_3342127, EPI_ISL_3342128, EPI_ISL_3342129, EPI_ISL_3342130, EPI_ISL_3342131, EPI_ISL_3342132, EPI_ISL_3342133, EPI_ISL_3342134, EPI_ISL_3342135, EPI_ISL_3342136, EPI_ISL_3342137, EPI_ISL_3342138, EPI_ISL_3342139, EPI_ISL_3342140, EPI_ISL_3342141, EPI_ISL_3342142, EPI_ISL_3342143, EPI_ISL_3342144, EPI_ISL_3342145, EPI_ISL_3342146 | | | |
| see above | Baylor Scott & White-Temple | Baylor Scott & White-Temple | Ari Rao, Linden Morales, Kimberly Walker, Marcus Volz, Shelby Johnson |
| EPI_ISL_3342153, EPI_ISL_3342154, EPI_ISL_3342155, EPI_ISL_3342156, EPI_ISL_3342157, EPI_ISL_3342158, EPI_ISL_3342159, EPI_ISL_3342160, EPI_ISL_3342161, EPI_ISL_3342162, EPI_ISL_3342163, EPI_ISL_3342164, EPI_ISL_3342165, EPI_ISL_3342166, EPI_ISL_3342167, EPI_ISL_3342168, EPI_ISL_3342169, EPI_ISL_3342170, EPI_ISL_3342171, EPI_ISL_3342172, EPI_ISL_3342173, EPI_ISL_3342174, EPI_ISL_3342175, EPI_ISL_3342176, EPI_ISL_3342178, EPI_ISL_3342179, EPI_ISL_3342180, EPI_ISL_3342181, EPI_ISL_3342182, EPI_ISL_3342185, EPI_ISL_3342186, EPI_ISL_3342188, EPI_ISL_3342189, EPI_ISL_3342191, EPI_ISL_3342192, EPI_ISL_3342193, EPI_ISL_3342194, EPI_ISL_3342195, EPI_ISL_3342196, EPI_ISL_3342198, EPI_ISL_3342199, EPI_ISL_3342200, EPI_ISL_3342201, EPI_ISL_3342202, EPI_ISL_3342203, EPI_ISL_3342204, EPI_ISL_3342205, EPI_ISL_3342206, EPI_ISL_3342207, EPI_ISL_3342208, EPI_ISL_3342209, EPI_ISL_3342210, EPI_ISL_3342212, EPI_ISL_3342213, EPI_ISL_3342214, EPI_ISL_3342215, EPI_ISL_3342216, EPI_ISL_3342217, EPI_ISL_3342218, EPI_ISL_3342219, EPI_ISL_3342220, EPI_ISL_3342221 | | | |
| see above | TSHEPONG LABORATORY | National Institute for Communicable Diseases of the National Health | Amoako DG, Everatt J, Scheepers C, Mohale T, Ntuli N, Mahlangu B, Mnguni A, Ismail A, Bhiman JN |

| | | | |
|---|---|---|---|
| | | Laboratory Service | |
| EPI_ISL_3342222, EPI_ISL_3342223, EPI_ISL_3342225, EPI_ISL_3342226, EPI_ISL_3342228, EPI_ISL_3342229, EPI_ISL_3342230, EPI_ISL_3342232, EPI_ISL_3342233, EPI_ISL_3342235, EPI_ISL_3342236, EPI_ISL_3342237, EPI_ISL_3342238, EPI_ISL_3342239, EPI_ISL_3342240, EPI_ISL_3342241, EPI_ISL_3342242, EPI_ISL_3342243, EPI_ISL_3342245, EPI_ISL_3342246, EPI_ISL_3342248, EPI_ISL_3342249, EPI_ISL_3342251, EPI_ISL_3342253, EPI_ISL_3342254, EPI_ISL_3342255, EPI_ISL_3342256 | | | |
| see above | KOPANONG LABORATORY | National Institute for Communicable Diseases of the National Health Laboratory Service | Amoako DG, Everatt J, Scheepers C, Mohale T, Ntuli N, Mahlangu B, Mnguni A, Ismail A, Bhiman JN |
| EPI_ISL_3342257, EPI_ISL_3342258, EPI_ISL_3342260, EPI_ISL_3342261, EPI_ISL_3342262, EPI_ISL_3342263, EPI_ISL_3342264, EPI_ISL_3342265 | TAMBO MEMORIAL LABORATORY | National Institute for Communicable Diseases of the National Health Laboratory Service | Amoako DG, Everatt J, Scheepers C, Mohale T, Ntuli N, Mahlangu B, Mnguni A, Ismail A, Bhiman JN |
| EPI_ISL_3342266, EPI_ISL_3342267, EPI_ISL_3342268, EPI_ISL_3342269, EPI_ISL_3342270, EPI_ISL_3342271, EPI_ISL_3342272, EPI_ISL_3342273, EPI_ISL_3342274, EPI_ISL_3342275, EPI_ISL_3342276, EPI_ISL_3342277, EPI_ISL_3342278, EPI_ISL_3342279, EPI_ISL_3342280, EPI_ISL_3342281, EPI_ISL_3342282, EPI_ISL_3342283 | | | |
| see above | POLOKWANE LABORATORY | National Institute for Communicable Diseases of the National Health Laboratory Service | Amoako DG, Everatt J, Scheepers C, Mohale T, Ntuli N, Mahlangu B, Mnguni A, Ismail A, Bhiman JN |
| EPI_ISL_3342284, EPI_ISL_3342285, EPI_ISL_3342286, EPI_ISL_3342287 | LEBOWAKGOMO LABORATORY | National Institute for Communicable Diseases of the National Health Laboratory Service | Amoako DG, Everatt J, Scheepers C, Mohale T, Ntuli N, Mahlangu B, Mnguni A, Ismail A, Bhiman JN |
| EPI_ISL_3342288 | MAFIKENG LABORATORY | National Institute for Communicable Diseases of the National Health Laboratory Service | Amoako DG, Everatt J, Scheepers C, Mohale T, Ntuli N, Mahlangu B, Mnguni A, Ismail A, Bhiman JN |
| EPI_ISL_3342289, EPI_ISL_3342290, EPI_ISL_3342291, EPI_ISL_3342292, EPI_ISL_3342293, EPI_ISL_3342294, EPI_ISL_3342295 | ROB FERREIRA LABORATORY | National Institute for Communicable Diseases of the National Health Laboratory Service | Amoako DG, Everatt J, Scheepers C, Mohale T, Ntuli N, Mahlangu B, Mnguni A, Ismail A, Bhiman JN |
| EPI_ISL_3342296, EPI_ISL_3342297, EPI_ISL_3342298, EPI_ISL_3342299, EPI_ISL_3342300, EPI_ISL_3342302, EPI_ISL_3342303, EPI_ISL_3342304, EPI_ISL_3342305, EPI_ISL_3342306, EPI_ISL_3342307, EPI_ISL_3342308, EPI_ISL_3342309, EPI_ISL_3342310, EPI_ISL_3342311, EPI_ISL_3342312, EPI_ISL_3342313, EPI_ISL_3342314, EPI_ISL_3342315, EPI_ISL_3342316, EPI_ISL_3342317, EPI_ISL_3342318, EPI_ISL_3342319, EPI_ISL_3342320, EPI_ISL_3342321, EPI_ISL_3342322, EPI_ISL_3342323, EPI_ISL_3342324, EPI_ISL_3342325, EPI_ISL_3342326, EPI_ISL_3342327, EPI_ISL_3342328, EPI_ISL_3342329, EPI_ISL_3342330 | | | |
| see above | POLOKWANE LABORATORY | National Institute for Communicable Diseases of the National Health Laboratory Service | Amoako DG, Everatt J, Scheepers C, Mohale T, Ntuli N, Mahlangu B, Mnguni A, Ismail A, Bhiman JN |
| EPI_ISL_3342423, EPI_ISL_3342424, EPI_ISL_3342425, EPI_ISL_3342426, EPI_ISL_3342427, EPI_ISL_3342428, EPI_ISL_3342429, EPI_ISL_3342430, EPI_ISL_3342431, EPI_ISL_3342433, EPI_ISL_3342434, EPI_ISL_3342435, EPI_ISL_3342436, EPI_ISL_3342438, EPI_ISL_3342440, EPI_ISL_3342441, EPI_ISL_3342442, EPI_ISL_3342443, EPI_ISL_3342444, EPI_ISL_3342445 | | | |
| see above | National Institute for Communicable Diseases of the National Health Laboratory Service | National Institute for Communicable Diseases of the National Health Laboratory Service | Amoako DG, Everatt J, Scheepers C, Mohale T, Ntuli N, Mahlangu B, Mnguni A, Ismail A, Bhiman JN |
| EPI_ISL_3342446 | LKO | Jessa | Marijke Raymaekers et al. on behalf of the Jessa_cmdLab |
| EPI_ISL_3342447 | Jessa | Jessa | Marijke Raymaekers et al. on behalf of the Jessa_cmdLab |
| EPI_ISL_3342448, EPI_ISL_3342449, EPI_ISL_3342450, EPI_ISL_3342451, EPI_ISL_3342452, EPI_ISL_3342453, EPI_ISL_3342454, EPI_ISL_3342455, EPI_ISL_3342456, EPI_ISL_3342457, EPI_ISL_3342458, EPI_ISL_3342459, EPI_ISL_3342460, EPI_ISL_3342461, EPI_ISL_3342462, EPI_ISL_3342463, EPI_ISL_3342464, EPI_ISL_3342465, EPI_ISL_3342466, EPI_ISL_3342467, EPI_ISL_3342468, EPI_ISL_3342469, EPI_ISL_3342470, EPI_ISL_3342471, EPI_ISL_3342472, EPI_ISL_3342473, EPI_ISL_3342474 | | | |
| see above | ZOL | Jessa | Marijke Raymaekers et al. on behalf of the Jessa_cmdLab |
| EPI_ISL_3342475, EPI_ISL_3342476, EPI_ISL_3342477, EPI_ISL_3342478, EPI_ISL_3342479, EPI_ISL_3342480, EPI_ISL_3342481, EPI_ISL_3342482, EPI_ISL_3342483, EPI_ISL_3342484, EPI_ISL_3342485 | | | |
| see above | Jessa | Jessa | Marijke Raymaekers et al. on behalf of the Jessa_cmdLab |
| EPI_ISL_3342487, EPI_ISL_3342488, EPI_ISL_3342489, EPI_ISL_3342490, EPI_ISL_3342491, EPI_ISL_3342492, EPI_ISL_3342493, EPI_ISL_3342494, EPI_ISL_3342495, EPI_ISL_3342496, EPI_ISL_3342497, EPI_ISL_3342498, EPI_ISL_3342499, EPI_ISL_3342500, EPI_ISL_3342501 | | | |
| see above | INNO Diagnostics Reference Laboratory | RCMI-Center for Research Resources, Ponce Research Institute | Vanessa Rivera-Amill, Andrea Arias-García, Raphael Sánchez-Torres |
| EPI_ISL_3342502, EPI_ISL_3342503, EPI_ISL_3342505, EPI_ISL_3342506, EPI_ISL_3342507, EPI_ISL_3342508, EPI_ISL_3342509, EPI_ISL_3342510, EPI_ISL_3342511, EPI_ISL_3342512, EPI_ISL_3342513, EPI_ISL_3342514, EPI_ISL_3342515, EPI_ISL_3342516, EPI_ISL_3342517, EPI_ISL_3342518, EPI_ISL_3342519, EPI_ISL_3342520, EPI_ISL_3342521, EPI_ISL_3342522, EPI_ISL_3342523, EPI_ISL_3342525, EPI_ISL_3342526, EPI_ISL_3342527, EPI_ISL_3342528, EPI_ISL_3342529, EPI_ISL_3342530, EPI_ISL_3342531, EPI_ISL_3342533, EPI_ISL_3342534, EPI_ISL_3342535, EPI_ISL_3342536, EPI_ISL_3342537, EPI_ISL_3342538, EPI_ISL_3342539, EPI_ISL_3342540, EPI_ISL_3342541, EPI_ISL_3342542, EPI_ISL_3342543, EPI_ISL_3342544 | | | |
| see above | ILSPPR | RCMI-Center for Research Resources, Ponce Research Institute | Vanessa Rivera-Amill, Andrea Arias-García, Raphael Sánchez-Torres |
| EPI_ISL_3342545, EPI_ISL_3342546, EPI_ISL_3342547, EPI_ISL_3342548, EPI_ISL_3342549, EPI_ISL_3342550, EPI_ISL_3342551, EPI_ISL_3342552 | Laboratorio Clinico Toledo | RCMI-Center for Research Resources, Ponce Research Institute | Vanessa Rivera-Amill, Andrea Arias-García, Raphael Sánchez-Torres |
| EPI_ISL_3342553, EPI_ISL_3342575 | Ohio Department of Health Laboratory | Ohio Department of Health Laboratory | Holmes, Jennifer; Eric Brandt, Keoni Omura, Glen McGillivary, Caitlin McDonnell, Jade Mowery, Stephanie Mccracken, Tyler Payne, Kirtana Ramadugu, Erica Leasure, Brent Lee, Kelsey Florek, Heather Blankenship, Allison Black, Quanta Brown, and Tammy Bannerman |
| EPI_ISL_3342576, EPI_ISL_3342577, EPI_ISL_3342578, EPI_ISL_3342579, EPI_ISL_3342580, EPI_ISL_3342581, EPI_ISL_3342582, EPI_ISL_3342583, EPI_ISL_3342584, EPI_ISL_3342585, EPI_ISL_3342586, EPI_ISL_3342587, EPI_ISL_3342588, EPI_ISL_3342589, EPI_ISL_3342591, EPI_ISL_3342592, EPI_ISL_3342593, EPI_ISL_3342594, EPI_ISL_3342595, EPI_ISL_3342596, EPI_ISL_3342597, EPI_ISL_3342598, EPI_ISL_3342599, EPI_ISL_3342600, EPI_ISL_3342601, EPI_ISL_3342602, EPI_ISL_3342603, EPI_ISL_3342604, EPI_ISL_3342605, EPI_ISL_3342606, EPI_ISL_3342608, EPI_ISL_3342609, EPI_ISL_3342610, EPI_ISL_3342611, EPI_ISL_3342612, EPI_ISL_3342613, EPI_ISL_3342614, EPI_ISL_3342615, EPI_ISL_3342616, EPI_ISL_3342617, EPI_ISL_3342618, EPI_ISL_3342619, EPI_ISL_3342620, EPI_ISL_3342621, EPI_ISL_3342622, EPI_ISL_3342623, EPI_ISL_3342624, EPI_ISL_3342625, EPI_ISL_3342626, EPI_ISL_3342627, EPI_ISL_3342628, EPI_ISL_3342629, EPI_ISL_3342630, EPI_ISL_3342631, EPI_ISL_3342632, EPI_ISL_3342633, EPI_ISL_3342634, EPI_ISL_3342635, EPI_ISL_3342636, EPI_ISL_3342637, EPI_ISL_3342638, EPI_ISL_3342639, EPI_ISL_3342640, EPI_ISL_3342641, EPI_ISL_3342642, EPI_ISL_3342643, EPI_ISL_3342644, EPI_ISL_3342645, EPI_ISL_3342646, EPI_ISL_3342647, EPI_ISL_3342648, EPI_ISL_3342649, EPI_ISL_3342650, EPI_ISL_3342651, EPI_ISL_3342652 | | | |
| see above | Alaska State Virology Laboratory | Alaska State Virology Laboratory | Stephanie DeRonde, Elva House, Jacob Zidek, Lisa Smith, Ph.D., Jack Chen, Ph.D. |
| EPI_ISL_3342653, EPI_ISL_3342654, EPI_ISL_3342655, EPI_ISL_3342657, EPI_ISL_3342658, EPI_ISL_3342659, EPI_ISL_3342660, EPI_ISL_3342661, EPI_ISL_3342662, EPI_ISL_3342663, EPI_ISL_3342664, EPI_ISL_3342665, EPI_ISL_3342666, EPI_ISL_3342667, EPI_ISL_3342668, EPI_ISL_3342669, EPI_ISL_3342670, EPI_ISL_3342671, EPI_ISL_3342672, EPI_ISL_3342673, EPI_ISL_3342674, EPI_ISL_3342675, EPI_ISL_3342676, EPI_ISL_3342677, EPI_ISL_3342678, EPI_ISL_3342679, EPI_ISL_3342680 | | | |
| see above | Laboratorio Central de Epidemiología (LCE) | Instituto de Biotecnología de la UNAM | Consorcio Mexicano de Vigilancia Genómica (CoViGen-Mex). Authors (in alphabetical order): Julio Elias Alvarado-Yaah, Carlos F. Arias, Santiago Ávila-Ríos, Eduardo Becerril-Vargas, Víctor Hugo Borja-Aburto, Celia Boukadida, Cristóbal Cháidez-Quiróz, Juan Bautista Chale-Dzul, Ricardo Ciria Merce, Andreu Comas-García, Célida Duque Molina, Julissa Enciso-Ibarra, José Antonio Enciso-Moreno, Gloria Elena Espinosa-Ayala, Fernando Fontove-Herrera, Daniel Fregoso-Rueda, Victor Eduardo García-Arias, Alejandra García-Gasca, Bruno Gómez-Gil, Jean Pierre González, Irvin González-López, Concepción Grajales-Muñiz, Ricardo Grande, Rosa María Gutiérrez Rios, Jesús Hernández, Alejandra Hernández-Terán, Alfredo Herrera-Estrella, Carla Ivón Herrera-Najera, Pavel Isa, Verónica Mata-Haro, Daniel Lira Morales, Susana Lopez, Antonio Loza Román, Brenda Irasema Maldonado-Meza, Bernardo Martínez-Miguel, José Arturo Martínez-Orozco, Célida Martínez-Rodríguez, Margarita Matías-Florentino, Fidencio Mejía-Nepomuceno, María Guadalupe de Jesús Mireles-Rivera, Gloria María Molina-Salinas, Hector Montoya-Fuentes, , Mario Mújica-Sánchez, José Esteban Muñoz-Medina, José de Jesús Nuñez-Contreras, Alicia Ocaña-Mondragón, Luis Alberto Ochoa-Carrera, Hector Esteban Paz-Juárez, Marissa Perez-Garcia, Francisco Pulido, Helen Haydee Fernanda Ramírez-Plascencia, Jorge Salas-Hernández, Angel Gustavo Salas-Lais, Alejandro Sánchez-Flores, Clara Esperanza Santacruz-Tinoco, María Guadalupe Santiago-Mauricio, Selene Zárate, Nelly Sélem-Mojica, Blanca Taboada, Kathia Elizabeth Tapia-Diaz, Gloria Vazquez, Joel Armando Vázquez-Pérez). |
| EPI_ISL_3342681, EPI_ISL_3342682, EPI_ISL_3342683, EPI_ISL_3342684, EPI_ISL_3342685, EPI_ISL_3342686 | Baylor Scott & White-Temple | Baylor Scott & White-Temple | Ari Rao, Linden Morales, Kimberly Walker, Marcus Volz, Shelby Johnson |
| EPI_ISL_3342687, EPI_ISL_3342688, EPI_ISL_3342720, EPI_ISL_3342722 | Ohio Department of Health Laboratory | Ohio Department of Health Laboratory | Holmes, Jennifer; Eric Brandt, Keoni Omura, Glen McGillivary, Caitlin McDonnell, Jade Mowery, Stephanie Mccracken, Tyler Payne, Kirtana Ramadugu, Erica Leasure, Brent Lee, Kelsey Florek, Heather Blankenship, Allison Black, Quanta Brown, and Tammy Bannerman |
| EPI_ISL_3342724, EPI_ISL_3342725, EPI_ISL_3342726, EPI_ISL_3342727 | The Jackson Laboratory | The Jackson Laboratory | Uvalic J, Long J, Renzette N, Adams M, Omerza G, Kelly K, Li L |
| EPI_ISL_3342728 | Ohio Department of Health Laboratory | Ohio Department of Health Laboratory | Holmes, Jennifer; Eric Brandt, Keoni Omura, Glen McGillivary, Caitlin McDonnell, Jade Mowery, Stephanie Mccracken, Tyler Payne, Kirtana Ramadugu, Erica Leasure, Brent Lee, Kelsey Florek, Heather Blankenship, Allison Black, Quanta Brown, and Tammy Bannerman |
| EPI_ISL_3342739, EPI_ISL_3342740, EPI_ISL_3342741, EPI_ISL_3342742, EPI_ISL_3342743, EPI_ISL_3342744, EPI_ISL_3342745, EPI_ISL_3342746, EPI_ISL_3342747, EPI_ISL_3342748, EPI_ISL_3342749, EPI_ISL_3342750, EPI_ISL_3342751, EPI_ISL_3342752, EPI_ISL_3342753, EPI_ISL_3342754, EPI_ISL_3342755, EPI_ISL_3342756, EPI_ISL_3342757, EPI_ISL_3342758, EPI_ISL_3342759, EPI_ISL_3342760, EPI_ISL_3342761, EPI_ISL_3342762, EPI_ISL_3342763, EPI_ISL_3342764, EPI_ISL_3342765, EPI_ISL_3342766, EPI_ISL_3342767, EPI_ISL_3342768, EPI_ISL_3342769, EPI_ISL_3342770, EPI_ISL_3342771, EPI_ISL_3342772, EPI_ISL_3342773, EPI_ISL_3342774, EPI_ISL_3342775, EPI_ISL_3342776, EPI_ISL_3342777, EPI_ISL_3342778, EPI_ISL_3342779, EPI_ISL_3342780, EPI_ISL_3342781 | | | |
| see above | Saint Louis County Department of Public Health Environmental Health Laboratory | University of Minnesota Genomics Center | Daryl M. Gohl, John Garbe, Jaquelyn Kuriger-Laber, Corbin Dirkx |
| EPI_ISL_3342783, EPI_ISL_3342784, EPI_ISL_3342785, EPI_ISL_3342786, EPI_ISL_3342787, EPI_ISL_3342788, EPI_ISL_3342789, EPI_ISL_3342790, EPI_ISL_3342791, EPI_ISL_3342792, EPI_ISL_3342793, EPI_ISL_3342794, EPI_ISL_3342795, EPI_ISL_3342796, EPI_ISL_3342797, EPI_ISL_3342798, EPI_ISL_3342799, EPI_ISL_3342800, EPI_ISL_3342801, EPI_ISL_3342802, EPI_ISL_3342803, EPI_ISL_3342804, EPI_ISL_3342805, EPI_ISL_3342806, EPI_ISL_3342807, EPI_ISL_3342808, EPI_ISL_3342809, EPI_ISL_3342810, EPI_ISL_3342811, EPI_ISL_3342812, EPI_ISL_3342813, EPI_ISL_3342814, EPI_ISL_3342815, EPI_ISL_3342817, EPI_ISL_3342818, EPI_ISL_3342819, EPI_ISL_3342820, EPI_ISL_3342821, EPI_ISL_3342822, EPI_ISL_3342823, EPI_ISL_3342824, EPI_ISL_3342825, EPI_ISL_3342826, EPI_ISL_3342827, | | | |

| | | | |
|---|---|---|---|
| EPI_ISL_3342828, EPI_ISL_3342829, EPI_ISL_3342830, EPI_ISL_3342831, EPI_ISL_3342832, EPI_ISL_3342833, EPI_ISL_3342834, EPI_ISL_3342835, EPI_ISL_3342836, EPI_ISL_3342837, EPI_ISL_3342838, EPI_ISL_3342839, EPI_ISL_3342840, EPI_ISL_3342841, EPI_ISL_3342842, EPI_ISL_3342843, EPI_ISL_3342844, EPI_ISL_3342845, EPI_ISL_3342846, EPI_ISL_3342847, EPI_ISL_3342848, EPI_ISL_3342849, EPI_ISL_3342850, EPI_ISL_3342851, EPI_ISL_3342852, EPI_ISL_3342853, EPI_ISL_3342854, EPI_ISL_3342855, EPI_ISL_3342856, EPI_ISL_3342857, EPI_ISL_3342858, EPI_ISL_3342859, EPI_ISL_3342860, EPI_ISL_3342861, EPI_ISL_3342862, EPI_ISL_3342863, EPI_ISL_3342864, EPI_ISL_3342865, EPI_ISL_3342866, EPI_ISL_3342867, EPI_ISL_3342868, EPI_ISL_3342869, EPI_ISL_3342870, EPI_ISL_3342871, EPI_ISL_3342872, EPI_ISL_3342873, EPI_ISL_3342874, EPI_ISL_3342875, EPI_ISL_3342876, EPI_ISL_3342877, EPI_ISL_3342878, EPI_ISL_3342879, EPI_ISL_3342880, EPI_ISL_3342881, EPI_ISL_3342882, EPI_ISL_3342883, EPI_ISL_3342885, EPI_ISL_3342886, EPI_ISL_3342887, EPI_ISL_3342888, EPI_ISL_3342889, EPI_ISL_3342890, EPI_ISL_3342891, EPI_ISL_3342892, EPI_ISL_3342893, EPI_ISL_3342894, EPI_ISL_3342895, EPI_ISL_3342896, EPI_ISL_3342897 | | | |
| see above | Michigan Department of Health and Human Services, Bureau of Laboratories | Michigan Department of Health and Human Services, Bureau of Laboratories | Blankenship HM, Riner D, Soehnlen MK |
| EPI_ISL_3342900 | Microbiology Department, Laboratori Clínic Metropolitana Nord. Hospital Universitari Germans Trias i Pujol | IrsiCaixa | Marc Noguera-Julian, Mariona Parera, Maria Casadellà, Pilar Armengol, Francesc Catala-Moll, Roger Paredes, Bonaventura Clotet Marc Noguera-Julian, Pilar Armengol, Ignacio Blanco, Antoni E Bordoy, Francesc Catala-Moll, Pere-Joan Cardona, Maria Casadellà, Cristina Casañ, Gemma Clara, Bonaventura Clotet, Montserrat Giménez, Mercedes Guerrero, Anna Not, David Panisello, Roger Paredes, Mariona Parera, Alexia París, Verónica Saludes, Alba Sánchez, Laia Soler, Lauro Sumoy, and Elisa Martró on behalf of the Can Ruti SARS-CoV-2 Sequencing Hub |
| EPI_ISL_3342901 | UOC Microbiology and Virology Policlinico Umberto I Rome, Italy | University Campus Bio-Medico of Rome (UCBM) | Fogolari M., De Florio L., Riva E., Francesconi M., Lintas C., Veralli R., Angeletti S. |
| EPI_ISL_3344068, EPI_ISL_3344069, EPI_ISL_3344070, EPI_ISL_3344071, EPI_ISL_3344072 | Biolytix AG | Clinical Bacteriology | Tim Roloff, Fanny Wegner, Helena MB Seth-Smith, Alfredo Mari, Karoline Leuzinger, Julia Bielicki, Manuel Battegay, Hans Hirsch, Adrian Egli |
| EPI_ISL_3344074, EPI_ISL_3344075, EPI_ISL_3344076, EPI_ISL_3344077, EPI_ISL_3344079 | Clinical Virology | Clinical Bacteriology | Tim Roloff, Fanny Wegner, Helena MB Seth-Smith, Alfredo Mari, Karoline Leuzinger, Julia Bielicki, Manuel Battegay, Hans Hirsch, Adrian Egli |
| EPI_ISL_3344080, EPI_ISL_3344081, EPI_ISL_3344082 | Synlab Suisse SA | Clinical Bacteriology | Tim Roloff, Fanny Wegner, Helena MB Seth-Smith, Alfredo Mari, Karoline Leuzinger, Julia Bielicki, Manuel Battegay, Hans Hirsch, Adrian Egli |
| EPI_ISL_3344083, EPI_ISL_3344084, EPI_ISL_3344086, EPI_ISL_3344087, EPI_ISL_3344088 | Clinical Virology | Clinical Bacteriology | Tim Roloff, Fanny Wegner, Helena MB Seth-Smith, Alfredo Mari, Karoline Leuzinger, Julia Bielicki, Manuel Battegay, Hans Hirsch, Adrian Egli |
| EPI_ISL_3344089 | Synlab Suisse SA | Clinical Bacteriology | Tim Roloff, Fanny Wegner, Helena MB Seth-Smith, Alfredo Mari, Karoline Leuzinger, Julia Bielicki, Manuel Battegay, Hans Hirsch, Adrian Egli |
| EPI_ISL_3344090, EPI_ISL_3344091, EPI_ISL_3344092, EPI_ISL_3344093, EPI_ISL_3344094 | Rothen Medizinische Laboratorien AG | Clinical Bacteriology | Tim Roloff, Fanny Wegner, Helena MB Seth-Smith, Alfredo Mari, Karoline Leuzinger, Julia Bielicki, Ingrid Steffen, Manuel Battegay, Hans Hirsch, Adrian Egli |
| EPI_ISL_3344095, EPI_ISL_3344096, EPI_ISL_3344097, EPI_ISL_3344098, EPI_ISL_3344099, EPI_ISL_3344100 | Clinical Virology | Clinical Bacteriology | Tim Roloff, Fanny Wegner, Helena MB Seth-Smith, Alfredo Mari, Karoline Leuzinger, Julia Bielicki, Manuel Battegay, Hans Hirsch, Adrian Egli |
| EPI_ISL_3344101 | Rothen Medizinische Laboratorien AG | Clinical Bacteriology | Tim Roloff, Fanny Wegner, Helena MB Seth-Smith, Alfredo Mari, Karoline Leuzinger, Julia Bielicki, Ingrid Steffen, Manuel Battegay, Hans Hirsch, Adrian Egli |
| EPI_ISL_3344102, EPI_ISL_3344103, EPI_ISL_3344104, EPI_ISL_3344105, EPI_ISL_3344106, EPI_ISL_3344107, EPI_ISL_3344108, EPI_ISL_3344109, EPI_ISL_3344110, EPI_ISL_3344111, EPI_ISL_3344112, EPI_ISL_3344113, EPI_ISL_3344114, EPI_ISL_3344115, EPI_ISL_3344116, EPI_ISL_3344117, EPI_ISL_3344118 | | | |
| see above | Clinical Virology | Clinical Bacteriology | Tim Roloff, Fanny Wegner, Helena MB Seth-Smith, Alfredo Mari, Karoline Leuzinger, Julia Bielicki, Manuel Battegay, Hans Hirsch, Adrian Egli |
| EPI_ISL_3344119, EPI_ISL_3344120, EPI_ISL_3344121 | Synlab Suisse SA | Clinical Bacteriology | Tim Roloff, Fanny Wegner, Helena MB Seth-Smith, Alfredo Mari, Karoline Leuzinger, Julia Bielicki, Manuel Battegay, Hans Hirsch, Adrian Egli |
| EPI_ISL_3344122 | Labormedizinisches Zentrum Dr Risch | Clinical Bacteriology | Tim Roloff, Fanny Wegner, Helena MB Seth-Smith, Alfredo Mari, Karoline Leuzinger, Julia Bielicki, Nadia Wohlwend,Martin Risch, Lorenz Risch, Manuel Battegay, Hans Hirsch, Adrian Egli |
| EPI_ISL_3344123, EPI_ISL_3344124, EPI_ISL_3344125 | Clinical Virology | Clinical Bacteriology | Tim Roloff, Fanny Wegner, Helena MB Seth-Smith, Alfredo Mari, Karoline Leuzinger, Julia Bielicki, Manuel Battegay, Hans Hirsch, Adrian Egli |
| EPI_ISL_3344129, EPI_ISL_3344130, EPI_ISL_3344131, EPI_ISL_3344132, EPI_ISL_3344133, EPI_ISL_3344134, EPI_ISL_3344135, EPI_ISL_3344136, EPI_ISL_3344137, EPI_ISL_3344138, EPI_ISL_3344139, EPI_ISL_3344140, EPI_ISL_3344141, EPI_ISL_3344142, EPI_ISL_3344143, EPI_ISL_3344144, EPI_ISL_3344145, EPI_ISL_3344146, EPI_ISL_3344147, EPI_ISL_3344148, EPI_ISL_3344149, EPI_ISL_3344150, EPI_ISL_3344151, EPI_ISL_3344152, EPI_ISL_3344153, EPI_ISL_3344154, EPI_ISL_3344155, EPI_ISL_3344156, EPI_ISL_3344157, EPI_ISL_3344158, EPI_ISL_3344159, EPI_ISL_3344160, EPI_ISL_3344161, EPI_ISL_3344162, EPI_ISL_3344163, EPI_ISL_3344164, EPI_ISL_3344165, EPI_ISL_3344166, EPI_ISL_3344167, EPI_ISL_3344168, EPI_ISL_3344169, EPI_ISL_3344170, EPI_ISL_3344171, EPI_ISL_3344172, EPI_ISL_3344174, EPI_ISL_3344175, EPI_ISL_3344176, EPI_ISL_3344177, EPI_ISL_3344178, EPI_ISL_3344179, EPI_ISL_3344180, EPI_ISL_3344181, EPI_ISL_3344182, EPI_ISL_3344183, EPI_ISL_3344184, EPI_ISL_3344185, EPI_ISL_3344186, EPI_ISL_3344187, EPI_ISL_3344188, EPI_ISL_3344189, EPI_ISL_3344190, EPI_ISL_3344191, EPI_ISL_3344192, EPI_ISL_3344193, EPI_ISL_3344194, EPI_ISL_3344195, EPI_ISL_3344196, EPI_ISL_3344197, EPI_ISL_3344198, EPI_ISL_3344199, EPI_ISL_3344200, EPI_ISL_3344201, EPI_ISL_3344203, EPI_ISL_3344204, EPI_ISL_3344205, EPI_ISL_3344206, EPI_ISL_3344207, EPI_ISL_3344208, EPI_ISL_3344209, EPI_ISL_3344210, EPI_ISL_3344211, EPI_ISL_3344212, EPI_ISL_3344213, EPI_ISL_3344214, EPI_ISL_3344215, EPI_ISL_3344216, EPI_ISL_3344217, EPI_ISL_3344218, EPI_ISL_3344219, EPI_ISL_3344220, EPI_ISL_3344221, EPI_ISL_3344222, EPI_ISL_3344223, EPI_ISL_3344224, EPI_ISL_3344225, EPI_ISL_3344226, EPI_ISL_3344227, EPI_ISL_3344228, EPI_ISL_3344229, EPI_ISL_3344230, EPI_ISL_3344231, EPI_ISL_3344232 | | | |
| see above | Fulgent Genetics | Centers for Disease Control and Prevention Division of Viral Diseases, Pathogen Discovery | Dakota Howard, Dhwani Batra, Peter W. Cook, Kara Moser, Adrian Paskey, Jason Caravas, Benjamin Rambo-Martin, Shatavia Morrison, Christopher Gulvick, Scott Sammons, Yvette Unoarumhi, Darlene Wagner, Matthew Schmerer, Harry Gao, Mickey Li, John Gao, Joseph Fierro, Benafsh Sapra, Becky Tsai, Yan Meng, Doreen Ng, James Xie, Clinton R. Paden, Duncan MacCannell |
| EPI_ISL_3344233 | UOC Microbiology and Virology Policlinico Umberto I Rome, Italy | University Campus Bio-Medico of Rome (UCBM) | Fogolari M., De Florio L., Riva E., Francesconi M., Lintas C., Veralli R., Angeletti S. |
| EPI_ISL_3344234, EPI_ISL_3344235, EPI_ISL_3344236, EPI_ISL_3344237, EPI_ISL_3344238, EPI_ISL_3344239, EPI_ISL_3344240, EPI_ISL_3344241, EPI_ISL_3344242, EPI_ISL_3344243, EPI_ISL_3344244, EPI_ISL_3344245, EPI_ISL_3344246, EPI_ISL_3344247, EPI_ISL_3344248, EPI_ISL_3344249, EPI_ISL_3344250, EPI_ISL_3344251, EPI_ISL_3344252, EPI_ISL_3344253, EPI_ISL_3344254, EPI_ISL_3344255, EPI_ISL_3344256, EPI_ISL_3344257, EPI_ISL_3344258, EPI_ISL_3344259, EPI_ISL_3344260, EPI_ISL_3344261, EPI_ISL_3344262, EPI_ISL_3344263, EPI_ISL_3344264, EPI_ISL_3344265, EPI_ISL_3344266, EPI_ISL_3344267, EPI_ISL_3344268, EPI_ISL_3344269, EPI_ISL_3344270, EPI_ISL_3344271, EPI_ISL_3344272, EPI_ISL_3344273, EPI_ISL_3344274, EPI_ISL_3344275, EPI_ISL_3344276, EPI_ISL_3344277, EPI_ISL_3344278, EPI_ISL_3344279, EPI_ISL_3344280, EPI_ISL_3344281, EPI_ISL_3344282, EPI_ISL_3344283, EPI_ISL_3344284, EPI_ISL_3344285, EPI_ISL_3344286, EPI_ISL_3344287, EPI_ISL_3344288, EPI_ISL_3344289, EPI_ISL_3344290, EPI_ISL_3344291, EPI_ISL_3344292, EPI_ISL_3344293, EPI_ISL_3344294, EPI_ISL_3344296, EPI_ISL_3344297, EPI_ISL_3344298, EPI_ISL_3344299, EPI_ISL_3344300, EPI_ISL_3344301, EPI_ISL_3344302, EPI_ISL_3344303, EPI_ISL_3344304, EPI_ISL_3344306, EPI_ISL_3344307, EPI_ISL_3344308, EPI_ISL_3344309, EPI_ISL_3344310, EPI_ISL_3344311, EPI_ISL_3344312, EPI_ISL_3344313, EPI_ISL_3344314, EPI_ISL_3344315, EPI_ISL_3344316, EPI_ISL_3344317, EPI_ISL_3344318, EPI_ISL_3344319, EPI_ISL_3344320, EPI_ISL_3344321, EPI_ISL_3344322, EPI_ISL_3344323, EPI_ISL_3344324, EPI_ISL_3344325, EPI_ISL_3344326, EPI_ISL_3344327, EPI_ISL_3344328, EPI_ISL_3344329, EPI_ISL_3344330, EPI_ISL_3344331, EPI_ISL_3344332, EPI_ISL_3344333, EPI_ISL_3344334, EPI_ISL_3344335, EPI_ISL_3344336, EPI_ISL_3344337, EPI_ISL_3344338, EPI_ISL_3344339, EPI_ISL_3344340, EPI_ISL_3344341, EPI_ISL_3344342, EPI_ISL_3344343, EPI_ISL_3344344, EPI_ISL_3344345, EPI_ISL_3344346, EPI_ISL_3344347, EPI_ISL_3344348, EPI_ISL_3344349, EPI_ISL_3344350, EPI_ISL_3344351, EPI_ISL_3344352, EPI_ISL_3344353, EPI_ISL_3344354, EPI_ISL_3344355, EPI_ISL_3344356, EPI_ISL_3344357, EPI_ISL_3344358, EPI_ISL_3344359, EPI_ISL_3344360, EPI_ISL_3344361, EPI_ISL_3344362, EPI_ISL_3344363, EPI_ISL_3344364, EPI_ISL_3344365, EPI_ISL_3344366, EPI_ISL_3344367, EPI_ISL_3344368, EPI_ISL_3344369, EPI_ISL_3344370, EPI_ISL_3344371, EPI_ISL_3344372, EPI_ISL_3344373, EPI_ISL_3344374, EPI_ISL_3344375, EPI_ISL_3344376, EPI_ISL_3344377, EPI_ISL_3344378, EPI_ISL_3344379, EPI_ISL_3344380, EPI_ISL_3344381, EPI_ISL_3344382, EPI_ISL_3344383, EPI_ISL_3344384, EPI_ISL_3344385, EPI_ISL_3344386, EPI_ISL_3344387, EPI_ISL_3344388, EPI_ISL_3344389, EPI_ISL_3344390, EPI_ISL_3344391, EPI_ISL_3344392, EPI_ISL_3344393, EPI_ISL_3344394, EPI_ISL_3344395, EPI_ISL_3344396, EPI_ISL_3344397, EPI_ISL_3344398, EPI_ISL_3344399, EPI_ISL_3344400, EPI_ISL_3344401, EPI_ISL_3344402, EPI_ISL_3344403, EPI_ISL_3344404, EPI_ISL_3344405, EPI_ISL_3344406, EPI_ISL_3344407, EPI_ISL_3344408, EPI_ISL_3344409, EPI_ISL_3344410, EPI_ISL_3344411, EPI_ISL_3344412, EPI_ISL_3344413, EPI_ISL_3344414, EPI_ISL_3344415, EPI_ISL_3344416, EPI_ISL_3344417, EPI_ISL_3344418, EPI_ISL_3344419, EPI_ISL_3344420, EPI_ISL_3344421, EPI_ISL_3344422, EPI_ISL_3344423, EPI_ISL_3344424, EPI_ISL_3344426, EPI_ISL_3344427, EPI_ISL_3344428, EPI_ISL_3344429, EPI_ISL_3344430, EPI_ISL_3344431, EPI_ISL_3344432, EPI_ISL_3344433, EPI_ISL_3344435, EPI_ISL_3344436, EPI_ISL_3344437, EPI_ISL_3344438, EPI_ISL_3344439, EPI_ISL_3344440, EPI_ISL_3344441, EPI_ISL_3344442, EPI_ISL_3344443, EPI_ISL_3344444, EPI_ISL_3344445, EPI_ISL_3344446, EPI_ISL_3344447, EPI_ISL_3344448, EPI_ISL_3344449, EPI_ISL_3344450, EPI_ISL_3344451, EPI_ISL_3344452, EPI_ISL_3344453, EPI_ISL_3344454, EPI_ISL_3344455, EPI_ISL_3344456, EPI_ISL_3344457, EPI_ISL_3344458, EPI_ISL_3344459, EPI_ISL_3344460, EPI_ISL_3344461, EPI_ISL_3344462, EPI_ISL_3344463, EPI_ISL_3344464, EPI_ISL_3344465, EPI_ISL_3344466, EPI_ISL_3344467, EPI_ISL_3344468, EPI_ISL_3344469, EPI_ISL_3344470, EPI_ISL_3344471, EPI_ISL_3344472, EPI_ISL_3344473, EPI_ISL_3344474, EPI_ISL_3344475, EPI_ISL_3344476, EPI_ISL_3344477, EPI_ISL_3344478, EPI_ISL_3344479, EPI_ISL_3344480, EPI_ISL_3344481, EPI_ISL_3344482, EPI_ISL_3344483, EPI_ISL_3344484, EPI_ISL_3344485, EPI_ISL_3344486, EPI_ISL_3344487, EPI_ISL_3344488, EPI_ISL_3344489, EPI_ISL_3344490, EPI_ISL_3344491, EPI_ISL_3344492, EPI_ISL_3344493, EPI_ISL_3344494, EPI_ISL_3344495, EPI_ISL_3344496, EPI_ISL_3344497, EPI_ISL_3344498, EPI_ISL_3344499, EPI_ISL_3344500, EPI_ISL_3344501, EPI_ISL_3344502, EPI_ISL_3344503, EPI_ISL_3344504, EPI_ISL_3344505, EPI_ISL_3344506, EPI_ISL_3344507, EPI_ISL_3344508, EPI_ISL_3344509, EPI_ISL_3344510, EPI_ISL_3344511, EPI_ISL_3344512, EPI_ISL_3344513, EPI_ISL_3344514, EPI_ISL_3344515, EPI_ISL_3344516, EPI_ISL_3344517, EPI_ISL_3344518, EPI_ISL_3344519, EPI_ISL_3344520, EPI_ISL_3344521, EPI_ISL_3344522, EPI_ISL_3344523, EPI_ISL_3344524, EPI_ISL_3344525, EPI_ISL_3344526, EPI_ISL_3344527, EPI_ISL_3344528, EPI_ISL_3344529, EPI_ISL_3344530, EPI_ISL_3344531, EPI_ISL_3344532, EPI_ISL_3344533, EPI_ISL_3344534, EPI_ISL_3344535, EPI_ISL_3344536, EPI_ISL_3344537, EPI_ISL_3344538, EPI_ISL_3344539, EPI_ISL_3344540, EPI_ISL_3344541, EPI_ISL_3344542, EPI_ISL_3344543, EPI_ISL_3344544, EPI_ISL_3344545, EPI_ISL_3344546, EPI_ISL_3344547, EPI_ISL_3344548, EPI_ISL_3344549, EPI_ISL_3344550, EPI_ISL_3344551, EPI_ISL_3344552, EPI_ISL_3344553, EPI_ISL_3344554, EPI_ISL_3344555, EPI_ISL_3344556, EPI_ISL_3344557, EPI_ISL_3344558, EPI_ISL_3344559, EPI_ISL_3344560, EPI_ISL_3344561, EPI_ISL_3344562, EPI_ISL_3344563, EPI_ISL_3344564, EPI_ISL_3344565, EPI_ISL_3344566, EPI_ISL_3344567, EPI_ISL_3344568, EPI_ISL_3344569, EPI_ISL_3344570, EPI_ISL_3344571, EPI_ISL_3344572, EPI_ISL_3344573, EPI_ISL_3344574, EPI_ISL_3344575, EPI_ISL_3344576, EPI_ISL_3344577, EPI_ISL_3344578, EPI_ISL_3344579, EPI_ISL_3344580, EPI_ISL_3344581, EPI_ISL_3344582, EPI_ISL_3344583, EPI_ISL_3344584, EPI_ISL_3344585, EPI_ISL_3344586, EPI_ISL_3344587, EPI_ISL_3344588, EPI_ISL_3344589, EPI_ISL_3344591, EPI_ISL_3344592, EPI_ISL_3344593, EPI_ISL_3344594, EPI_ISL_3344595, EPI_ISL_3344596, EPI_ISL_3344597, EPI_ISL_3344598, EPI_ISL_3344599, EPI_ISL_3344600, EPI_ISL_3344601, EPI_ISL_3344602, EPI_ISL_3344603, EPI_ISL_3344604, EPI_ISL_3344605, EPI_ISL_3344606, EPI_ISL_3344607, EPI_ISL_3344608, EPI_ISL_3344609, EPI_ISL_3344610, EPI_ISL_3344611, EPI_ISL_3344612, EPI_ISL_3344613, EPI_ISL_3344614, EPI_ISL_3344615, EPI_ISL_3344616, EPI_ISL_3344617, EPI_ISL_3344618, EPI_ISL_3344619, EPI_ISL_3344620, EPI_ISL_3344621, EPI_ISL_3344622, EPI_ISL_3344623, EPI_ISL_3344624, EPI_ISL_3344625, EPI_ISL_3344626, EPI_ISL_3344627, EPI_ISL_3344628, EPI_ISL_3344629, EPI_ISL_3344630, EPI_ISL_3344631, EPI_ISL_3344632, EPI_ISL_3344633, EPI_ISL_3344634, EPI_ISL_3344635, EPI_ISL_3344636, EPI_ISL_3344637, EPI_ISL_3344638, EPI_ISL_3344639, EPI_ISL_3344640, EPI_ISL_3344641, EPI_ISL_3344642, EPI_ISL_3344643, EPI_ISL_3344644, EPI_ISL_3344645, EPI_ISL_3344646, EPI_ISL_3344647, EPI_ISL_3344648, EPI_ISL_3344649, EPI_ISL_3344650, EPI_ISL_3344651, EPI_ISL_3344652, EPI_ISL_3344653, EPI_ISL_3344654, EPI_ISL_3344656, EPI_ISL_3344657, EPI_ISL_3344658, EPI_ISL_3344659, EPI_ISL_3344660, EPI_ISL_3344661, EPI_ISL_3344662, EPI_ISL_3344663, EPI_ISL_3344664, EPI_ISL_3344666, EPI_ISL_3344667, EPI_ISL_3344668, EPI_ISL_3344669, EPI_ISL_3344670, EPI_ISL_3344671, EPI_ISL_3344672, EPI_ISL_3344673, EPI_ISL_3344674, EPI_ISL_3344675, EPI_ISL_3344676, EPI_ISL_3344677, EPI_ISL_3344678, EPI_ISL_3344679, EPI_ISL_3344680, EPI_ISL_3344681, EPI_ISL_3344682, EPI_ISL_3344683, EPI_ISL_3344684, EPI_ISL_3344685, EPI_ISL_3344686, EPI_ISL_3344687, EPI_ISL_3344688, EPI_ISL_3344689, EPI_ISL_3344690, EPI_ISL_3344691, EPI_ISL_3344693, EPI_ISL_3344696, EPI_ISL_3344697, EPI_ISL_3344698, EPI_ISL_3344699, EPI_ISL_3344700, EPI_ISL_3344701, EPI_ISL_3344702, EPI_ISL_3344703, EPI_ISL_3344704, EPI_ISL_3344705, EPI_ISL_3344706, EPI_ISL_3344707, EPI_ISL_3344708, EPI_ISL_3344709, EPI_ISL_3344710, EPI_ISL_3344711, EPI_ISL_3344712, EPI_ISL_3344713, EPI_ISL_3344714, EPI_ISL_3344716, EPI_ISL_3344717, EPI_ISL_3344718, EPI_ISL_3344719, EPI_ISL_3344720, EPI_ISL_3344721, EPI_ISL_3344722 | | | |
| see above | Fulgent Genetics | Centers for Disease Control and Prevention Division of Viral Diseases, Pathogen Discovery | Dakota Howard, Dhwani Batra, Peter W. Cook, Kara Moser, Adrian Paskey, Jason Caravas, Benjamin Rambo-Martin, Shatavia Morrison, Christopher Gulvick, Scott Sammons, Yvette Unoarumhi, Darlene Wagner, Matthew Schmerer, Harry Gao, Mickey Li, John Gao, Joseph Fierro, Benafsh Sapra, Becky Tsai, Yan Meng, Doreen Ng, James Xie, Clinton R. Paden, Duncan MacCannell |
| EPI_ISL_3344723 | Spital Dornach | Clinical Bacteriology | Tim Roloff, Fanny Wegner, Helena MB Seth-Smith, Alfredo Mari, Karoline Leuzinger, Julia Bielicki, Manuel Battegay, Hans Hirsch, Adrian Egli |
| EPI_ISL_3344724, EPI_ISL_3344725, EPI_ISL_3344726, EPI_ISL_3344727, EPI_ISL_3344728, EPI_ISL_3344729, EPI_ISL_3344730, EPI_ISL_3344731, EPI_ISL_3344732, EPI_ISL_3344733, EPI_ISL_3344734, EPI_ISL_3344735, EPI_ISL_3344736, EPI_ISL_3344737, EPI_ISL_3344738, EPI_ISL_3344739, EPI_ISL_3344740, EPI_ISL_3344741, EPI_ISL_3344742, EPI_ISL_3344743, EPI_ISL_3344744, EPI_ISL_3344745, EPI_ISL_3344746, EPI_ISL_3344747, EPI_ISL_3344748, EPI_ISL_3344749, EPI_ISL_3344750, EPI_ISL_3344751, EPI_ISL_3344752, EPI_ISL_3344753, EPI_ISL_3344754, EPI_ISL_3344755, EPI_ISL_3344756, EPI_ISL_3344757, EPI_ISL_3344758, EPI_ISL_3344759, EPI_ISL_3344760, EPI_ISL_3344761, EPI_ISL_3344762, EPI_ISL_3344763, EPI_ISL_3344764, EPI_ISL_3344765, EPI_ISL_3344766, EPI_ISL_3344767, EPI_ISL_3344768, EPI_ISL_3344769, EPI_ISL_3344770, EPI_ISL_3344771, EPI_ISL_3344772, EPI_ISL_3344773, EPI_ISL_3344774, EPI_ISL_3344775, EPI_ISL_3344776, EPI_ISL_3344777, EPI_ISL_3344778, EPI_ISL_3344779, EPI_ISL_3344780, EPI_ISL_3344781, EPI_ISL_3344782, EPI_ISL_3344783, EPI_ISL_3344784, EPI_ISL_3344785, EPI_ISL_3344786, EPI_ISL_3344787, EPI_ISL_3344788, EPI_ISL_3344789, EPI_ISL_3344790, EPI_ISL_3344791, EPI_ISL_3344792, EPI_ISL_3344793, EPI_ISL_3344794, EPI_ISL_3344795, EPI_ISL_3344796, EPI_ISL_3344797, EPI_ISL_3344798, EPI_ISL_3344799, EPI_ISL_3344800, EPI_ISL_3344801, EPI_ISL_3344802, EPI_ISL_3344803, EPI_ISL_3344804, EPI_ISL_3344805, EPI_ISL_3344806, EPI_ISL_3344807, EPI_ISL_3344808, EPI_ISL_3344809, EPI_ISL_3344810, EPI_ISL_3344811, EPI_ISL_3344812, EPI_ISL_3344813, EPI_ISL_3344814, EPI_ISL_3344815, EPI_ISL_3344816, EPI_ISL_3344817, EPI_ISL_3344818, EPI_ISL_3344819, EPI_ISL_3344820, EPI_ISL_3344821, EPI_ISL_3344822, EPI_ISL_3344823, EPI_ISL_3344824, EPI_ISL_3344825, EPI_ISL_3344826, EPI_ISL_3344827, EPI_ISL_3344828, EPI_ISL_3344829, EPI_ISL_3344830, EPI_ISL_3344831, EPI_ISL_3344832, EPI_ISL_3344833, EPI_ISL_3344834, EPI_ISL_3344835, EPI_ISL_3344836, EPI_ISL_3344837, EPI_ISL_3344838, EPI_ISL_3344839, EPI_ISL_3344840, EPI_ISL_3344841, EPI_ISL_3344842, EPI_ISL_3344843, EPI_ISL_3344844, EPI_ISL_3344845, EPI_ISL_3344846, EPI_ISL_3344847, EPI_ISL_3344848, EPI_ISL_3344849, EPI_ISL_3344850, EPI_ISL_3344851, EPI_ISL_3344852, EPI_ISL_3344853, EPI_ISL_3344854, EPI_ISL_3344855, EPI_ISL_3344856, EPI_ISL_3344857, EPI_ISL_3344858, EPI_ISL_3344859, EPI_ISL_3344860, EPI_ISL_3344861, EPI_ISL_3344862, EPI_ISL_3344863 | | | |
| see above | Fulgent Genetics | Centers for Disease Control and Prevention Division of Viral Diseases, Pathogen Discovery | Dakota Howard, Dhwani Batra, Peter W. Cook, Kara Moser, Adrian Paskey, Jason Caravas, Benjamin Rambo-Martin, Shatavia Morrison, Christopher Gulvick, Scott Sammons, Yvette Unoarumhi, Darlene Wagner, Matthew Schmerer, Harry Gao, Mickey Li, John Gao, Joseph Fierro, Benafsh Sapra, Becky Tsai, Yan Meng, Doreen Ng, James Xie, Clinton R. Paden, Duncan MacCannell |
| EPI_ISL_3344864 | MPHL | MPHL | Beata Karowlewicz |
| EPI_ISL_3344865, EPI_ISL_3344866, EPI_ISL_3344867, EPI_ISL_3344868, EPI_ISL_3344869, EPI_ISL_3344870, EPI_ISL_3344871, EPI_ISL_3344873, EPI_ISL_3344874, EPI_ISL_3344875, EPI_ISL_3344876, EPI_ISL_3344877, EPI_ISL_3344878, EPI_ISL_3344879, EPI_ISL_3344880, EPI_ISL_3344881, EPI_ISL_3344882, EPI_ISL_3344883, EPI_ISL_3344884, EPI_ISL_3344885, EPI_ISL_3344886, EPI_ISL_3344887, EPI_ISL_3344888, EPI_ISL_3344889, EPI_ISL_3344890, EPI_ISL_3344891, EPI_ISL_3344892, EPI_ISL_3344893, EPI_ISL_3344894, EPI_ISL_3344895, EPI_ISL_3344896, EPI_ISL_3344897, EPI_ISL_3344898, EPI_ISL_3344899, EPI_ISL_3344900, EPI_ISL_3344901, EPI_ISL_3344902, EPI_ISL_3344903, EPI_ISL_3344904, EPI_ISL_3344905, EPI_ISL_3344906, EPI_ISL_3344907, EPI_ISL_3344908, EPI_ISL_3344909, EPI_ISL_3344910, EPI_ISL_3344911, EPI_ISL_3344912, EPI_ISL_3344913, EPI_ISL_3344914, EPI_ISL_3344915, EPI_ISL_3344916, EPI_ISL_3344917, EPI_ISL_3344918, EPI_ISL_3344919, EPI_ISL_3344920, EPI_ISL_3344921, EPI_ISL_3344922, EPI_ISL_3344923, EPI_ISL_3344924, EPI_ISL_3344926, EPI_ISL_3344927, EPI_ISL_3344928, EPI_ISL_3344929, EPI_ISL_3344930, EPI_ISL_3344931, EPI_ISL_3344932, EPI_ISL_3344933, EPI_ISL_3344934, EPI_ISL_3344935, EPI_ISL_3344936, EPI_ISL_3344937, EPI_ISL_3344938, EPI_ISL_3344939, EPI_ISL_3344940, EPI_ISL_3344941, EPI_ISL_3344942, EPI_ISL_3344943, EPI_ISL_3344944, EPI_ISL_3344945, EPI_ISL_3344946, EPI_ISL_3344947, EPI_ISL_3344948, EPI_ISL_3344949, EPI_ISL_3344950, EPI_ISL_3344952, EPI_ISL_3344953, EPI_ISL_3344954, EPI_ISL_3344955, EPI_ISL_3344956, EPI_ISL_3344957, EPI_ISL_3344958, EPI_ISL_3344959, EPI_ISL_3344960, EPI_ISL_3344961, EPI_ISL_3344962, EPI_ISL_3344963, EPI_ISL_3344964, EPI_ISL_3344965, EPI_ISL_3344966, EPI_ISL_3344967, EPI_ISL_3344968, EPI_ISL_3344969, EPI_ISL_3344970, EPI_ISL_3344971, EPI_ISL_3344972, EPI_ISL_3344973, EPI_ISL_3344974, EPI_ISL_3344975, EPI_ISL_3344976, EPI_ISL_3344977, EPI_ISL_3344978, EPI_ISL_3344979, EPI_ISL_3344980, EPI_ISL_3344981, EPI_ISL_3344982, EPI_ISL_3344983, EPI_ISL_3344984, EPI_ISL_3344985, EPI_ISL_3344986, EPI_ISL_3344987, EPI_ISL_3344988, EPI_ISL_3344989, EPI_ISL_3344990, EPI_ISL_3344991, EPI_ISL_3344992, EPI_ISL_3344993, EPI_ISL_3344994, EPI_ISL_3344995, EPI_ISL_3344996, EPI_ISL_3344997, EPI_ISL_3344998, EPI_ISL_3344999, | | | |

EPI_ISL_3345000, EPI_ISL_3345001, EPI_ISL_3345002, EPI_ISL_3345003, EPI_ISL_3345004, EPI_ISL_3345005, EPI_ISL_3345006, EPI_ISL_3345007, EPI_ISL_3345008, EPI_ISL_3345009, EPI_ISL_3345010, EPI_ISL_3345011, EPI_ISL_3345012, EPI_ISL_3345014, EPI_ISL_3345015, EPI_ISL_3345016, EPI_ISL_3345017, EPI_ISL_3345018, EPI_ISL_3345019, EPI_ISL_3345020, EPI_ISL_3345021, EPI_ISL_3345022, EPI_ISL_3345023, EPI_ISL_3345024, EPI_ISL_3345025, EPI_ISL_3345026, EPI_ISL_3345027, EPI_ISL_3345028, EPI_ISL_3345029, EPI_ISL_3345030, EPI_ISL_3345031, EPI_ISL_3345032, EPI_ISL_3345033, EPI_ISL_3345034, EPI_ISL_3345035, EPI_ISL_3345036, EPI_ISL_3345037, EPI_ISL_3345038, EPI_ISL_3345039, EPI_ISL_3345040, EPI_ISL_3345041, EPI_ISL_3345042, EPI_ISL_3345043, EPI_ISL_3345044, EPI_ISL_3345045, EPI_ISL_3345046, EPI_ISL_3345047, EPI_ISL_3345048, EPI_ISL_3345049, EPI_ISL_3345050, EPI_ISL_3345051, EPI_ISL_3345052, EPI_ISL_3345053, EPI_ISL_3345054, EPI_ISL_3345055, EPI_ISL_3345056, EPI_ISL_3345057, EPI_ISL_3345058, EPI_ISL_3345059, EPI_ISL_3345060, EPI_ISL_3345061, EPI_ISL_3345062, EPI_ISL_3345063, EPI_ISL_3345064, EPI_ISL_3345065, EPI_ISL_3345066, EPI_ISL_3345067, EPI_ISL_3345068, EPI_ISL_3345069, EPI_ISL_3345070, EPI_ISL_3345071, EPI_ISL_3345072, EPI_ISL_3345073, EPI_ISL_3345074, EPI_ISL_3345075, EPI_ISL_3345076, EPI_ISL_3345077, EPI_ISL_3345078, EPI_ISL_3345079, EPI_ISL_3345080, EPI_ISL_3345081, EPI_ISL_3345082, EPI_ISL_3345083, EPI_ISL_3345084, EPI_ISL_3345085, EPI_ISL_3345086, EPI_ISL_3345087, EPI_ISL_3345088, EPI_ISL_3345089, EPI_ISL_3345090, EPI_ISL_3345091, EPI_ISL_3345092, EPI_ISL_3345093, EPI_ISL_3345094, EPI_ISL_3345095, EPI_ISL_3345096, EPI_ISL_3345097, EPI_ISL_3345098, EPI_ISL_3345099, EPI_ISL_3345100, EPI_ISL_3345101, EPI_ISL_3345102, EPI_ISL_3345103, EPI_ISL_3345104, EPI_ISL_3345105, EPI_ISL_3345106, EPI_ISL_3345107, EPI_ISL_3345108, EPI_ISL_3345109, EPI_ISL_3345110, EPI_ISL_3345111, EPI_ISL_3345112, EPI_ISL_3345113, EPI_ISL_3345114, EPI_ISL_3345115, EPI_ISL_3345117, EPI_ISL_3345118, EPI_ISL_3345119, EPI_ISL_3345120, EPI_ISL_3345121, EPI_ISL_3345122, EPI_ISL_3345123, EPI_ISL_3345124, EPI_ISL_3345125, EPI_ISL_3345126, EPI_ISL_3345127, EPI_ISL_3345128, EPI_ISL_3345129, EPI_ISL_3345130, EPI_ISL_3345131, EPI_ISL_3345132, EPI_ISL_3345133, EPI_ISL_3345134, EPI_ISL_3345135, EPI_ISL_3345136, EPI_ISL_3345137, EPI_ISL_3345138, EPI_ISL_3345139, EPI_ISL_3345140, EPI_ISL_3345141, EPI_ISL_3345142, EPI_ISL_3345143, EPI_ISL_3345144, EPI_ISL_3345145, EPI_ISL_3345146, EPI_ISL_3345147, EPI_ISL_3345148, EPI_ISL_3345149, EPI_ISL_3345150, EPI_ISL_3345151, EPI_ISL_3345152, EPI_ISL_3345154, EPI_ISL_3345155, EPI_ISL_3345156, EPI_ISL_3345157, EPI_ISL_3345158, EPI_ISL_3345159, EPI_ISL_3345160, EPI_ISL_3345161, EPI_ISL_3345162, EPI_ISL_3345163, EPI_ISL_3345164, EPI_ISL_3345165, EPI_ISL_3345166, EPI_ISL_3345167, EPI_ISL_3345168, EPI_ISL_3345169, EPI_ISL_3345170, EPI_ISL_3345171, EPI_ISL_3345172, EPI_ISL_3345173, EPI_ISL_3345175, EPI_ISL_3345176, EPI_ISL_3345177, EPI_ISL_3345178, EPI_ISL_3345179, EPI_ISL_3345180, EPI_ISL_3345181, EPI_ISL_3345182, EPI_ISL_3345183, EPI_ISL_3345184, EPI_ISL_3345185, EPI_ISL_3345186, EPI_ISL_3345187, EPI_ISL_3345188, EPI_ISL_3345189, EPI_ISL_3345190, EPI_ISL_3345191, EPI_ISL_3345192, EPI_ISL_3345193, EPI_ISL_3345194, EPI_ISL_3345195, EPI_ISL_3345196, EPI_ISL_3345197, EPI_ISL_3345198, EPI_ISL_3345199, EPI_ISL_3345200, EPI_ISL_3345201, EPI_ISL_3345202, EPI_ISL_3345203, EPI_ISL_3345204, EPI_ISL_3345205, EPI_ISL_3345206, EPI_ISL_3345207, EPI_ISL_3345208, EPI_ISL_3345209, EPI_ISL_3345210, EPI_ISL_3345211, EPI_ISL_3345212, EPI_ISL_3345213, EPI_ISL_3345214, EPI_ISL_3345215, EPI_ISL_3345216, EPI_ISL_3345217, EPI_ISL_3345218, EPI_ISL_3345219, EPI_ISL_3345220, EPI_ISL_3345221, EPI_ISL_3345222, EPI_ISL_3345223, EPI_ISL_3345224, EPI_ISL_3345225, EPI_ISL_3345226, EPI_ISL_3345227, EPI_ISL_3345228, EPI_ISL_3345229, EPI_ISL_3345230, EPI_ISL_3345231, EPI_ISL_3345232, EPI_ISL_3345233, EPI_ISL_3345234, EPI_ISL_3345235, EPI_ISL_3345236, EPI_ISL_3345237, EPI_ISL_3345238, EPI_ISL_3345239, EPI_ISL_3345240, EPI_ISL_3345241, EPI_ISL_3345242, EPI_ISL_3345243, EPI_ISL_3345244, EPI_ISL_3345245, EPI_ISL_3345246, EPI_ISL_3345247, EPI_ISL_3345248, EPI_ISL_3345249, EPI_ISL_3345250, EPI_ISL_3345251, EPI_ISL_3345252, EPI_ISL_3345253, EPI_ISL_3345254, EPI_ISL_3345255, EPI_ISL_3345256, EPI_ISL_3345257, EPI_ISL_3345258, EPI_ISL_3345259, EPI_ISL_3345260, EPI_ISL_3345261, EPI_ISL_3345262, EPI_ISL_3345263, EPI_ISL_3345264, EPI_ISL_3345265, EPI_ISL_3345266, EPI_ISL_3345267, EPI_ISL_3345268, EPI_ISL_3345269, EPI_ISL_3345270, EPI_ISL_3345271, EPI_ISL_3345272, EPI_ISL_3345273, EPI_ISL_3345274, EPI_ISL_3345275, EPI_ISL_3345276, EPI_ISL_3345277, EPI_ISL_3345278, EPI_ISL_3345279, EPI_ISL_3345280, EPI_ISL_3345281, EPI_ISL_3345282, EPI_ISL_3345283, EPI_ISL_3345284, EPI_ISL_3345285, EPI_ISL_3345286, EPI_ISL_3345287, EPI_ISL_3345288, EPI_ISL_3345289, EPI_ISL_3345290, EPI_ISL_3345291, EPI_ISL_3345292, EPI_ISL_3345293, EPI_ISL_3345294, EPI_ISL_3345295, EPI_ISL_3345296, EPI_ISL_3345297, EPI_ISL_3345298, EPI_ISL_3345299, EPI_ISL_3345300, EPI_ISL_3345301, EPI_ISL_3345302, EPI_ISL_3345303, EPI_ISL_3345304, EPI_ISL_3345305, EPI_ISL_3345306, EPI_ISL_3345307, EPI_ISL_3345308, EPI_ISL_3345309, EPI_ISL_3345310, EPI_ISL_3345311, EPI_ISL_3345312, EPI_ISL_3345313, EPI_ISL_3345314, EPI_ISL_3345315, EPI_ISL_3345316, EPI_ISL_3345317, EPI_ISL_3345318, EPI_ISL_3345319, EPI_ISL_3345320, EPI_ISL_3345321, EPI_ISL_3345322, EPI_ISL_3345323, EPI_ISL_3345324, EPI_ISL_3345325, EPI_ISL_3345326, EPI_ISL_3345327, EPI_ISL_3345328, EPI_ISL_3345329, EPI_ISL_3345330, EPI_ISL_3345331, EPI_ISL_3345332, EPI_ISL_3345333, EPI_ISL_3345334, EPI_ISL_3345335, EPI_ISL_3345336, EPI_ISL_3345337, EPI_ISL_3345338, EPI_ISL_3345339, EPI_ISL_3345340, EPI_ISL_3345341, EPI_ISL_3345342, EPI_ISL_3345343, EPI_ISL_3345344, EPI_ISL_3345345, EPI_ISL_3345346, EPI_ISL_3345347, EPI_ISL_3345348, EPI_ISL_3345349, EPI_ISL_3345350, EPI_ISL_3345351, EPI_ISL_3345352, EPI_ISL_3345353, EPI_ISL_3345354, EPI_ISL_3345355, EPI_ISL_3345356, EPI_ISL_3345357, EPI_ISL_3345358, EPI_ISL_3345359, EPI_ISL_3345360, EPI_ISL_3345361, EPI_ISL_3345362, EPI_ISL_3345363, EPI_ISL_3345364, EPI_ISL_3345365, EPI_ISL_3345366, EPI_ISL_3345367, EPI_ISL_3345368, EPI_ISL_3345369, EPI_ISL_3345370, EPI_ISL_3345371, EPI_ISL_3345372, EPI_ISL_3345373, EPI_ISL_3345374, EPI_ISL_3345375, EPI_ISL_3345376, EPI_ISL_3345377, EPI_ISL_3345378, EPI_ISL_3345379, EPI_ISL_3345380, EPI_ISL_3345381, EPI_ISL_3345383, EPI_ISL_3345384, EPI_ISL_3345385, EPI_ISL_3345386, EPI_ISL_3345387, EPI_ISL_3345388, EPI_ISL_3345389, EPI_ISL_3345390, EPI_ISL_3345391, EPI_ISL_3345392, EPI_ISL_3345393, EPI_ISL_3345394, EPI_ISL_3345395, EPI_ISL_3345396, EPI_ISL_3345397, EPI_ISL_3345398, EPI_ISL_3345400, EPI_ISL_3345401, EPI_ISL_3345402, EPI_ISL_3345403, EPI_ISL_3345404, EPI_ISL_3345405, EPI_ISL_3345406, EPI_ISL_3345407, EPI_ISL_3345408, EPI_ISL_3345409, EPI_ISL_3345410, EPI_ISL_3345411, EPI_ISL_3345412, EPI_ISL_3345413, EPI_ISL_3345414, EPI_ISL_3345415, EPI_ISL_3345416, EPI_ISL_3345417, EPI_ISL_3345418, EPI_ISL_3345419, EPI_ISL_3345420, EPI_ISL_3345421, EPI_ISL_3345422, EPI_ISL_3345423, EPI_ISL_3345424, EPI_ISL_3345425, EPI_ISL_3345426, EPI_ISL_3345427, EPI_ISL_3345428, EPI_ISL_3345429, EPI_ISL_3345430, EPI_ISL_3345431, EPI_ISL_3345432, EPI_ISL_3345433, EPI_ISL_3345434, EPI_ISL_3345435, EPI_ISL_3345436, EPI_ISL_3345437, EPI_ISL_3345438, EPI_ISL_3345439, EPI_ISL_3345440, EPI_ISL_3345441, EPI_ISL_3345442, EPI_ISL_3345443, EPI_ISL_3345444, EPI_ISL_3345445, EPI_ISL_3345446, EPI_ISL_3345447, EPI_ISL_3345448, EPI_ISL_3345449, EPI_ISL_3345450, EPI_ISL_3345451, EPI_ISL_3345452, EPI_ISL_3345453, EPI_ISL_3345454, EPI_ISL_3345455, EPI_ISL_3345456, EPI_ISL_3345457, EPI_ISL_3345458, EPI_ISL_3345459, EPI_ISL_3345460, EPI_ISL_3345461, EPI_ISL_3345463, EPI_ISL_3345464, EPI_ISL_3345466, EPI_ISL_3345467, EPI_ISL_3345468, EPI_ISL_3345469, EPI_ISL_3345470, EPI_ISL_3345471, EPI_ISL_3345473, EPI_ISL_3345474, EPI_ISL_3345475, EPI_ISL_3345476, EPI_ISL_3345477, EPI_ISL_3345478, EPI_ISL_3345479, EPI_ISL_3345480, EPI_ISL_3345481, EPI_ISL_3345482, EPI_ISL_3345483, EPI_ISL_3345484, EPI_ISL_3345485, EPI_ISL_3345486, EPI_ISL_3345487, EPI_ISL_3345488, EPI_ISL_3345489, EPI_ISL_3345491, EPI_ISL_3345492, EPI_ISL_3345493, EPI_ISL_3345494, EPI_ISL_3345495, EPI_ISL_3345496, EPI_ISL_3345497, EPI_ISL_3345498, EPI_ISL_3345499, EPI_ISL_3345500, EPI_ISL_3345501, EPI_ISL_3345502, EPI_ISL_3345503, EPI_ISL_3345504, EPI_ISL_3345505, EPI_ISL_3345506, EPI_ISL_3345507, EPI_ISL_3345508, EPI_ISL_3345509, EPI_ISL_3345510, EPI_ISL_3345511, EPI_ISL_3345512, EPI_ISL_3345513, EPI_ISL_3345514, EPI_ISL_3345515, EPI_ISL_3345516, EPI_ISL_3345517, EPI_ISL_3345518, EPI_ISL_3345519, EPI_ISL_3345520, EPI_ISL_3345521, EPI_ISL_3345522, EPI_ISL_3345523, EPI_ISL_3345524, EPI_ISL_3345525, EPI_ISL_3345526, EPI_ISL_3345527, EPI_ISL_3345528, EPI_ISL_3345529, EPI_ISL_3345530, EPI_ISL_3345531, EPI_ISL_3345532, EPI_ISL_3345533, EPI_ISL_3345534, EPI_ISL_3345535, EPI_ISL_3345536, EPI_ISL_3345537, EPI_ISL_3345538, EPI_ISL_3345539, EPI_ISL_3345540, EPI_ISL_3345541, EPI_ISL_3345542, EPI_ISL_3345543, EPI_ISL_3345544, EPI_ISL_3345545, EPI_ISL_3345546, EPI_ISL_3345547, EPI_ISL_3345548, EPI_ISL_3345549, EPI_ISL_3345550, EPI_ISL_3345551, EPI_ISL_3345552, EPI_ISL_3345553, EPI_ISL_3345554, EPI_ISL_3345555, EPI_ISL_3345556, EPI_ISL_3345557, EPI_ISL_3345558, EPI_ISL_3345559, EPI_ISL_3345560, EPI_ISL_3345561, EPI_ISL_3345562, EPI_ISL_3345563, EPI_ISL_3345564, EPI_ISL_3345565, EPI_ISL_3345566, EPI_ISL_3345567, EPI_ISL_3345568, EPI_ISL_3345569, EPI_ISL_3345570, EPI_ISL_3345571, EPI_ISL_3345572, EPI_ISL_3345573, EPI_ISL_3345574, EPI_ISL_3345575, EPI_ISL_3345577, EPI_ISL_3345578, EPI_ISL_3345579, EPI_ISL_3345580, EPI_ISL_3345581, EPI_ISL_3345582, EPI_ISL_3345583, EPI_ISL_3345584, EPI_ISL_3345585

| | | | |
|---|---|---|---|
| see above | Fulgent Genetics | Centers for Disease Control and Prevention Division of Viral Diseases, Pathogen Discovery | Dakota Howard, Dhwani Batra, Peter W. Cook, Kara Moser, Adrian Paskey, Jason Caravas, Benjamin Rambo-Martin, Shatavia Morrison, Christopher Gulvick, Scott Sammons, Yvette Unoarumhi, Darlene Wagner, Matthew Schmerer, Harry Gao, Mickey Li, John Gao, Joseph Fierro, Benafsh Sapra, Becky Tsai, Yan Meng, Doreen Ng, James Xie, Clinton R. Paden, Duncan MacCannell |
| EPI_ISL_3345586 | Viollier AG | Department of Biosystems Science and Engineering, ETH Zürich | Chaoran Chen, Sarah Nadeau, Louis du Plessis, Catharine Aquino, Ivan Topolsky, Kim Philipp Jablonski, Lara Fuhrmann, Daniel Ehrsam, Isabel Stürmer, Andreia Cabral de Gouvea, Maria Domenica Moccia, Simon Grüter, Timothy Sykes, Lennart Opitz, Griffin White, Laura Neff, Doris Popovic, Andrea Patrignani, Jay Tracy, Ralph Schlapbach, Christiane Beckmann, Maurice Redondo, Olivier Kobel, Christoph Noppen, Niko Beerenwinkel, Tanja Stadler |
| EPI_ISL_3345587 | Viollier AG | Department of Biosystems Science and Engineering, ETH Zürich | Christian Beisel, Sarah Nadeau, Chaoran Chen, Louis du Plessis, Ivan Topolsky, Kim Philipp Jablonski, Lara Fuhrmann, Rebecca Denes, Mirjam Feldkamp, Ina Nissen, Natascha Santacroce, Elodie Burcklen, Christiane Beckmann, Maurice Redondo, Olivier Kobel, Christoph Noppen, Niko Beerenwinkel, Tanja Stadler |
| EPI_ISL_3345588 | Fulgent Genetics | Centers for Disease Control and Prevention Division of Viral Diseases, Pathogen Discovery | Dakota Howard, Dhwani Batra, Peter W. Cook, Kara Moser, Adrian Paskey, Jason Caravas, Benjamin Rambo-Martin, Shatavia Morrison, Christopher Gulvick, Scott Sammons, Yvette Unoarumhi, Darlene Wagner, Matthew Schmerer, Harry Gao, Mickey Li, John Gao, Joseph Fierro, Benafsh Sapra, Becky Tsai, Yan Meng, Doreen Ng, James Xie, Clinton R. Paden, Duncan MacCannell |
| EPI_ISL_3345589 | Viollier AG | Department of Biosystems Science and Engineering, ETH Zürich | Chaoran Chen, Sarah Nadeau, Louis du Plessis, Catharine Aquino, Ivan Topolsky, Kim Philipp Jablonski, Lara Fuhrmann, Daniel Ehrsam, Isabel Stürmer, Andreia Cabral de Gouvea, Maria Domenica Moccia, Simon Grüter, Timothy Sykes, Lennart Opitz, Griffin White, Laura Neff, Doris Popovic, Andrea Patrignani, Jay Tracy, Ralph Schlapbach, Christiane Beckmann, Maurice Redondo, Olivier Kobel, Christoph Noppen, Niko Beerenwinkel, Tanja Stadler |
| EPI_ISL_3345590 | Fulgent Genetics | Centers for Disease Control and Prevention Division of Viral Diseases, Pathogen Discovery | Dakota Howard, Dhwani Batra, Peter W. Cook, Kara Moser, Adrian Paskey, Jason Caravas, Benjamin Rambo-Martin, Shatavia Morrison, Christopher Gulvick, Scott Sammons, Yvette Unoarumhi, Darlene Wagner, Matthew Schmerer, Harry Gao, Mickey Li, John Gao, Joseph Fierro, Benafsh Sapra, Becky Tsai, Yan Meng, Doreen Ng, James Xie, Clinton R. Paden, Duncan MacCannell |
| EPI_ISL_3345591 | Viollier AG | Department of Biosystems Science and Engineering, ETH Zürich | Chaoran Chen, Sarah Nadeau, Louis du Plessis, Catharine Aquino, Ivan Topolsky, Kim Philipp Jablonski, Lara Fuhrmann, Daniel Ehrsam, Isabel Stürmer, Andreia Cabral de Gouvea, Maria Domenica Moccia, Simon Grüter, Timothy Sykes, Lennart Opitz, Griffin White, Laura Neff, Doris Popovic, Andrea Patrignani, Jay Tracy, Ralph Schlapbach, Christiane Beckmann, Maurice Redondo, Olivier Kobel, Christoph Noppen, Niko Beerenwinkel, Tanja Stadler |
| EPI_ISL_3345592 | Viollier AG | Department of Biosystems Science and Engineering, ETH Zürich | Christian Beisel, Sarah Nadeau, Chaoran Chen, Louis du Plessis, Ivan Topolsky, Kim Philipp Jablonski, Lara Fuhrmann, Rebecca Denes, Mirjam Feldkamp, Ina Nissen, Natascha Santacroce, Elodie Burcklen, Christiane Beckmann, Maurice Redondo, Olivier Kobel, Christoph Noppen, Niko Beerenwinkel, Tanja Stadler |
| EPI_ISL_3345593 | Fulgent Genetics | Centers for Disease Control and Prevention Division of Viral Diseases, Pathogen Discovery | Dakota Howard, Dhwani Batra, Peter W. Cook, Kara Moser, Adrian Paskey, Jason Caravas, Benjamin Rambo-Martin, Shatavia Morrison, Christopher Gulvick, Scott Sammons, Yvette Unoarumhi, Darlene Wagner, Matthew Schmerer, Harry Gao, Mickey Li, John Gao, Joseph Fierro, Benafsh Sapra, Becky Tsai, Yan Meng, Doreen Ng, James Xie, Clinton R. Paden, Duncan MacCannell |
| EPI_ISL_3345594 | Viollier AG | Department of Biosystems Science and Engineering, ETH Zürich | Chaoran Chen, Sarah Nadeau, Louis du Plessis, Catharine Aquino, Ivan Topolsky, Kim Philipp Jablonski, Lara Fuhrmann, Daniel Ehrsam, Isabel Stürmer, Andreia Cabral de Gouvea, Maria Domenica Moccia, Simon Grüter, Timothy Sykes, Lennart Opitz, Griffin White, Laura Neff, Doris Popovic, Andrea Patrignani, Jay Tracy, Ralph Schlapbach, Christiane Beckmann, Maurice Redondo, Olivier Kobel, Christoph Noppen, Niko Beerenwinkel, Tanja Stadler |
| EPI_ISL_3345595 | Viollier AG | Department of Biosystems Science and Engineering, ETH Zürich | Christian Beisel, Sarah Nadeau, Chaoran Chen, Louis du Plessis, Ivan Topolsky, Kim Philipp Jablonski, Lara Fuhrmann, Rebecca Denes, Mirjam Feldkamp, Ina Nissen, Natascha Santacroce, Elodie Burcklen, Christiane Beckmann, Maurice Redondo, Olivier Kobel, Christoph Noppen, Niko Beerenwinkel, Tanja Stadler |
| EPI_ISL_3345596 | Fulgent Genetics | Centers for Disease Control and Prevention Division of Viral Diseases, Pathogen Discovery | Dakota Howard, Dhwani Batra, Peter W. Cook, Kara Moser, Adrian Paskey, Jason Caravas, Benjamin Rambo-Martin, Shatavia Morrison, Christopher Gulvick, Scott Sammons, Yvette Unoarumhi, Darlene Wagner, Matthew Schmerer, Harry Gao, Mickey Li, John Gao, Joseph Fierro, Benafsh Sapra, Becky Tsai, Yan Meng, Doreen Ng, James Xie, Clinton R. Paden, Duncan MacCannell |
| EPI_ISL_3345597, EPI_ISL_3345598 | Viollier AG | Department of Biosystems Science and Engineering, ETH Zürich | Chaoran Chen, Sarah Nadeau, Louis du Plessis, Catharine Aquino, Ivan Topolsky, Kim Philipp Jablonski, Lara Fuhrmann, Daniel Ehrsam, Isabel Stürmer, Andreia Cabral de Gouvea, Maria Domenica Moccia, Simon Grüter, Timothy Sykes, Lennart Opitz, Griffin White, Laura Neff, Doris Popovic, Andrea Patrignani, Jay Tracy, Ralph Schlapbach, Christiane Beckmann, Maurice Redondo, Olivier Kobel, Christoph Noppen, Niko Beerenwinkel, Tanja Stadler |
| EPI_ISL_3345599 | Fulgent Genetics | Centers for Disease Control and Prevention Division of Viral Diseases, Pathogen Discovery | Dakota Howard, Dhwani Batra, Peter W. Cook, Kara Moser, Adrian Paskey, Jason Caravas, Benjamin Rambo-Martin, Shatavia Morrison, Christopher Gulvick, Scott Sammons, Yvette Unoarumhi, Darlene Wagner, Matthew Schmerer, Harry Gao, Mickey Li, John Gao, Joseph Fierro, Benafsh Sapra, Becky Tsai, Yan Meng, Doreen Ng, James Xie, Clinton R. Paden, Duncan MacCannell |
| EPI_ISL_3345600 | Viollier AG | Department of Biosystems Science and Engineering, ETH Zürich | Chaoran Chen, Sarah Nadeau, Louis du Plessis, Catharine Aquino, Ivan Topolsky, Kim Philipp Jablonski, Lara Fuhrmann, Daniel Ehrsam, Isabel Stürmer, Andreia Cabral de Gouvea, Maria Domenica Moccia, Simon Grüter, Timothy Sykes, Lennart Opitz, Griffin White, Laura Neff, Doris Popovic, Andrea Patrignani, Jay Tracy, Ralph Schlapbach, Christiane Beckmann, Maurice Redondo, Olivier Kobel, Christoph Noppen, Niko Beerenwinkel, Tanja Stadler |
| EPI_ISL_3345601 | Fulgent Genetics | Centers for Disease Control and Prevention Division of Viral Diseases, Pathogen Discovery | Dakota Howard, Dhwani Batra, Peter W. Cook, Kara Moser, Adrian Paskey, Jason Caravas, Benjamin Rambo-Martin, Shatavia Morrison, Christopher Gulvick, Scott Sammons, Yvette Unoarumhi, Darlene Wagner, Matthew Schmerer, Harry Gao, Mickey Li, John Gao, Joseph Fierro, Benafsh Sapra, Becky Tsai, Yan Meng, Doreen Ng, James Xie, Clinton R. Paden, Duncan MacCannell |
| EPI_ISL_3345602, EPI_ISL_3345603 | Viollier AG | Department of Biosystems Science and Engineering, ETH Zürich | Chaoran Chen, Sarah Nadeau, Louis du Plessis, Catharine Aquino, Ivan Topolsky, Kim Philipp Jablonski, Lara Fuhrmann, Daniel Ehrsam, Isabel Stürmer, Andreia Cabral de Gouvea, Maria Domenica Moccia, Simon Grüter, Timothy Sykes, Lennart Opitz, Griffin White, Laura Neff, Doris Popovic, Andrea Patrignani, Jay Tracy, Ralph Schlapbach, Christiane Beckmann, Maurice Redondo, Olivier Kobel, Christoph Noppen, Niko Beerenwinkel, Tanja Stadler |
| EPI_ISL_3345604 | Fulgent Genetics | Centers for Disease Control and Prevention Division of Viral Diseases, Pathogen Discovery | Dakota Howard, Dhwani Batra, Peter W. Cook, Kara Moser, Adrian Paskey, Jason Caravas, Benjamin Rambo-Martin, Shatavia Morrison, Christopher Gulvick, Scott Sammons, Yvette Unoarumhi, Darlene Wagner, Matthew Schmerer, Harry Gao, Mickey Li, John Gao, Joseph Fierro, Benafsh Sapra, Becky Tsai, Yan Meng, Doreen Ng, James Xie, Clinton R. Paden, Duncan MacCannell |
| EPI_ISL_3345605, EPI_ISL_3345606 | Viollier AG | Department of Biosystems Science and Engineering, ETH Zürich | Chaoran Chen, Sarah Nadeau, Louis du Plessis, Catharine Aquino, Ivan Topolsky, Kim Philipp Jablonski, Lara Fuhrmann, Daniel Ehrsam, Isabel Stürmer, Andreia Cabral de Gouvea, Maria Domenica Moccia, Simon Grüter, Timothy Sykes, Lennart Opitz, Griffin White, Laura Neff, Doris Popovic, Andrea Patrignani, Jay Tracy, Ralph Schlapbach, Christiane Beckmann, Maurice Redondo, Olivier Kobel, Christoph Noppen, Niko Beerenwinkel, Tanja Stadler |
| EPI_ISL_3345607 | Fulgent Genetics | Centers for Disease Control and Prevention Division of Viral Diseases, Pathogen Discovery | Dakota Howard, Dhwani Batra, Peter W. Cook, Kara Moser, Adrian Paskey, Jason Caravas, Benjamin Rambo-Martin, Shatavia Morrison, Christopher Gulvick, Scott Sammons, Yvette Unoarumhi, Darlene Wagner, Matthew Schmerer, Harry Gao, Mickey Li, John Gao, Joseph Fierro, Benafsh Sapra, Becky Tsai, Yan Meng, Doreen Ng, James Xie, Clinton R. Paden, Duncan MacCannell |
| EPI_ISL_3345608 | Viollier AG | Department of Biosystems Science and Engineering, ETH Zürich | Christian Beisel, Sarah Nadeau, Chaoran Chen, Louis du Plessis, Ivan Topolsky, Kim Philipp Jablonski, Lara Fuhrmann, Rebecca Denes, Mirjam Feldkamp, Ina Nissen, Natascha Santacroce, Elodie Burcklen, Christiane Beckmann, Maurice Redondo, Olivier Kobel, Christoph Noppen, Niko Beerenwinkel, Tanja Stadler |
| EPI_ISL_3345609, EPI_ISL_3345611 | Viollier AG | Department of Biosystems Science and Engineering, ETH Zürich | Chaoran Chen, Sarah Nadeau, Louis du Plessis, Catharine Aquino, Ivan Topolsky, Kim Philipp Jablonski, Lara Fuhrmann, Daniel Ehrsam, Isabel Stürmer, Andreia Cabral de Gouvea, Maria Domenica Moccia, Simon Grüter, Timothy Sykes, Lennart Opitz, Griffin White, Laura Neff, Doris Popovic, Andrea Patrignani, Jay Tracy, Ralph Schlapbach, Christiane Beckmann, Maurice Redondo, Olivier Kobel, Christoph Noppen, Niko Beerenwinkel, Tanja Stadler |
| EPI_ISL_3345612 | Fulgent Genetics | Centers for Disease Control and Prevention Division of Viral Diseases, Pathogen Discovery | Dakota Howard, Dhwani Batra, Peter W. Cook, Kara Moser, Adrian Paskey, Jason Caravas, Benjamin Rambo-Martin, Shatavia Morrison, Christopher Gulvick, Scott Sammons, Yvette Unoarumhi, Darlene Wagner, Matthew Schmerer, Harry Gao, Mickey Li, John Gao, Joseph Fierro, Benafsh Sapra, Becky Tsai, Yan Meng, Doreen Ng, James Xie, Clinton R. Paden, Duncan MacCannell |
| EPI_ISL_3345613 | Viollier AG | Department of Biosystems Science and Engineering, ETH Zürich | Chaoran Chen, Sarah Nadeau, Louis du Plessis, Catharine Aquino, Ivan Topolsky, Kim Philipp Jablonski, Lara Fuhrmann, Daniel Ehrsam, Isabel Stürmer, Andreia Cabral de Gouvea, Maria Domenica Moccia, Simon Grüter, Timothy Sykes, Lennart Opitz, Griffin White, Laura Neff, Doris Popovic, Andrea Patrignani, Jay Tracy, Ralph Schlapbach, Christiane Beckmann, Maurice Redondo, Olivier Kobel, Christoph Noppen, Niko Beerenwinkel, Tanja Stadler |
| EPI_ISL_3345614 | Fulgent Genetics | Centers for Disease Control and Prevention Division of Viral Diseases, Pathogen Discovery | Dakota Howard, Dhwani Batra, Peter W. Cook, Kara Moser, Adrian Paskey, Jason Caravas, Benjamin Rambo-Martin, Shatavia Morrison, Christopher Gulvick, Scott Sammons, Yvette Unoarumhi, Darlene Wagner, Matthew Schmerer, Harry Gao, Mickey Li, John Gao, Joseph Fierro, Benafsh Sapra, Becky Tsai, Yan Meng, Doreen Ng, James Xie, Clinton R. Paden, Duncan MacCannell |
| EPI_ISL_3345615, EPI_ISL_3345616 | Viollier AG | Department of Biosystems Science and Engineering, ETH Zürich | Chaoran Chen, Sarah Nadeau, Louis du Plessis, Catharine Aquino, Ivan Topolsky, Kim Philipp Jablonski, Lara Fuhrmann, Daniel Ehrsam, Isabel Stürmer, Andreia Cabral de Gouvea, Maria Domenica Moccia, Simon Grüter, Timothy Sykes, Lennart Opitz, Griffin White, Laura Neff, Doris Popovic, Andrea Patrignani, Jay Tracy, Ralph Schlapbach, Christiane Beckmann, Maurice Redondo, Olivier Kobel, Christoph Noppen, Niko Beerenwinkel, Tanja Stadler |
| EPI_ISL_3345617 | Fulgent Genetics | Centers for Disease Control and Prevention Division of Viral Diseases, Pathogen Discovery | Dakota Howard, Dhwani Batra, Peter W. Cook, Kara Moser, Adrian Paskey, Jason Caravas, Benjamin Rambo-Martin, Shatavia Morrison, Christopher Gulvick, Scott Sammons, Yvette Unoarumhi, Darlene Wagner, Matthew Schmerer, Harry Gao, Mickey Li, John Gao, Joseph Fierro, Benafsh Sapra, Becky Tsai, Yan Meng, Doreen Ng, James Xie, Clinton R. Paden, Duncan MacCannell |
| EPI_ISL_3345618 | Viollier AG | Department of Biosystems Science and Engineering, ETH Zürich | Chaoran Chen, Sarah Nadeau, Louis du Plessis, Catharine Aquino, Ivan Topolsky, Kim Philipp Jablonski, Lara Fuhrmann, Daniel Ehrsam, Isabel Stürmer, Andreia Cabral de Gouvea, Maria Domenica Moccia, Simon Grüter, Timothy Sykes, Lennart Opitz, Griffin White, Laura Neff, Doris Popovic, Andrea Patrignani, Jay Tracy, Ralph Schlapbach, Christiane Beckmann, Maurice Redondo, Olivier Kobel, Christoph Noppen, Niko Beerenwinkel, Tanja Stadler |
| EPI_ISL_3345619 | Viollier AG | Department of Biosystems Science and Engineering, ETH Zürich | Christian Beisel, Sarah Nadeau, Chaoran Chen, Louis du Plessis, Ivan Topolsky, Kim Philipp Jablonski, Lara Fuhrmann, Rebecca Denes, Mirjam Feldkamp, Ina Nissen, Natascha Santacroce, Elodie Burcklen, Christiane Beckmann, Maurice Redondo, Olivier Kobel, Christoph Noppen, Niko Beerenwinkel, Tanja Stadler |
| EPI_ISL_3345621 | Viollier AG | Department of Biosystems Science and Engineering, ETH Zürich | Chaoran Chen, Sarah Nadeau, Louis du Plessis, Catharine Aquino, Ivan Topolsky, Kim Philipp Jablonski, Lara Fuhrmann, Daniel Ehrsam, Isabel Stürmer, Andreia Cabral de Gouvea, Maria Domenica Moccia, Simon Grüter, Timothy Sykes, Lennart Opitz, Griffin White, Laura Neff, Doris Popovic, Andrea Patrignani, Jay Tracy, Ralph Schlapbach, Christiane Beckmann, Maurice Redondo, Olivier Kobel, Christoph Noppen, Niko Beerenwinkel, Tanja Stadler |
| EPI_ISL_3345622 | Fulgent Genetics | Centers for Disease Control and Prevention Division of Viral Diseases, Pathogen Discovery | Dakota Howard, Dhwani Batra, Peter W. Cook, Kara Moser, Adrian Paskey, Jason Caravas, Benjamin Rambo-Martin, Shatavia Morrison, Christopher Gulvick, Scott Sammons, Yvette Unoarumhi, Darlene Wagner, Matthew Schmerer, Harry Gao, Mickey Li, John Gao, Joseph Fierro, Benafsh Sapra, Becky Tsai, Yan Meng, Doreen Ng, James Xie, Clinton R. Paden, Duncan MacCannell |
| EPI_ISL_3345623, EPI_ISL_3345624, EPI_ISL_3345625 | Viollier AG | Department of Biosystems Science and Engineering, ETH Zürich | Chaoran Chen, Sarah Nadeau, Louis du Plessis, Catharine Aquino, Ivan Topolsky, Kim Philipp Jablonski, Lara Fuhrmann, Daniel Ehrsam, Isabel Stürmer, Andreia Cabral de Gouvea, Maria Domenica Moccia, Simon Grüter, Timothy Sykes, Lennart Opitz, Griffin White, Laura Neff, Doris Popovic, Andrea Patrignani, Jay Tracy, Ralph Schlapbach, Christiane Beckmann, Maurice Redondo, Olivier Kobel, Christoph Noppen, Niko Beerenwinkel, Tanja Stadler |
| EPI_ISL_3345626 | Fulgent Genetics | Centers for Disease Control and Prevention Division of Viral Diseases, Pathogen Discovery | Dakota Howard, Dhwani Batra, Peter W. Cook, Kara Moser, Adrian Paskey, Jason Caravas, Benjamin Rambo-Martin, Shatavia Morrison, Christopher Gulvick, Scott Sammons, Yvette Unoarumhi, Darlene Wagner, Matthew Schmerer, Harry Gao, Mickey Li, John Gao, Joseph Fierro, Benafsh Sapra, Becky Tsai, Yan Meng, Doreen Ng, James Xie, Clinton R. Paden, Duncan MacCannell |
| EPI_ISL_3345627 | Viollier AG | Department of Biosystems Science and Engineering, ETH Zürich | Chaoran Chen, Sarah Nadeau, Louis du Plessis, Catharine Aquino, Ivan Topolsky, Kim Philipp Jablonski, Lara Fuhrmann, Daniel Ehrsam, Isabel Stürmer, Andreia Cabral de Gouvea, Maria Domenica Moccia, Simon Grüter, Timothy Sykes, Lennart |

| | | | |
|---|---|---|---|
| | | | Opitz, Griffin White, Laura Neff, Doris Popovic, Andrea Patrignani, Jay Tracy, Ralph Schlapbach, Christiane Beckmann, Maurice Redondo, Olivier Kobel, Christoph Noppen, Niko Beerenwinkel, Tanja Stadler |
| EPI_ISL_3345628 | Fulgent Genetics | Centers for Disease Control and Prevention Division of Viral Diseases, Pathogen Discovery | Dakota Howard, Dhwani Batra, Peter W. Cook, Kara Moser, Adrian Paskey, Jason Caravas, Benjamin Rambo-Martin, Shatavia Morrison, Christopher Gulvick, Scott Sammons, Yvette Unoarumhi, Darlene Wagner, Matthew Schmerer, Harry Gao, Mickey Li, John Gao, Joseph Fierro, Benafsh Sapra, Becky Tsai, Yan Meng, Doreen Ng, James Xie, Clinton R. Paden, Duncan MacCannell |
| EPI_ISL_3345629, EPI_ISL_3345630 | Viollier AG | Department of Biosystems Science and Engineering, ETH Zürich | Chaoran Chen, Sarah Nadeau, Louis du Plessis, Catharine Aquino, Ivan Topolsky, Kim Philipp Jablonski, Lara Fuhrmann, Daniel Ehrsam, Isabel Stürmer, Andreia Cabral de Gouvea, Maria Domenica Moccia, Simon Grüter, Timothy Sykes, Lennart Opitz, Griffin White, Laura Neff, Doris Popovic, Andrea Patrignani, Jay Tracy, Ralph Schlapbach, Christiane Beckmann, Maurice Redondo, Olivier Kobel, Christoph Noppen, Niko Beerenwinkel, Tanja Stadler |
| EPI_ISL_3345631 | Fulgent Genetics | Centers for Disease Control and Prevention Division of Viral Diseases, Pathogen Discovery | Dakota Howard, Dhwani Batra, Peter W. Cook, Kara Moser, Adrian Paskey, Jason Caravas, Benjamin Rambo-Martin, Shatavia Morrison, Christopher Gulvick, Scott Sammons, Yvette Unoarumhi, Darlene Wagner, Matthew Schmerer, Harry Gao, Mickey Li, John Gao, Joseph Fierro, Benafsh Sapra, Becky Tsai, Yan Meng, Doreen Ng, James Xie, Clinton R. Paden, Duncan MacCannell |
| EPI_ISL_3345632 | Viollier AG | Department of Biosystems Science and Engineering, ETH Zürich | Chaoran Chen, Sarah Nadeau, Louis du Plessis, Catharine Aquino, Ivan Topolsky, Kim Philipp Jablonski, Lara Fuhrmann, Daniel Ehrsam, Isabel Stürmer, Andreia Cabral de Gouvea, Maria Domenica Moccia, Simon Grüter, Timothy Sykes, Lennart Opitz, Griffin White, Laura Neff, Doris Popovic, Andrea Patrignani, Jay Tracy, Ralph Schlapbach, Christiane Beckmann, Maurice Redondo, Olivier Kobel, Christoph Noppen, Niko Beerenwinkel, Tanja Stadler |
| EPI_ISL_3345633 | Fulgent Genetics | Centers for Disease Control and Prevention Division of Viral Diseases, Pathogen Discovery | Dakota Howard, Dhwani Batra, Peter W. Cook, Kara Moser, Adrian Paskey, Jason Caravas, Benjamin Rambo-Martin, Shatavia Morrison, Christopher Gulvick, Scott Sammons, Yvette Unoarumhi, Darlene Wagner, Matthew Schmerer, Harry Gao, Mickey Li, John Gao, Joseph Fierro, Benafsh Sapra, Becky Tsai, Yan Meng, Doreen Ng, James Xie, Clinton R. Paden, Duncan MacCannell |
| EPI_ISL_3345634, EPI_ISL_3345635 | Viollier AG | Department of Biosystems Science and Engineering, ETH Zürich | Chaoran Chen, Sarah Nadeau, Louis du Plessis, Catharine Aquino, Ivan Topolsky, Kim Philipp Jablonski, Lara Fuhrmann, Daniel Ehrsam, Isabel Stürmer, Andreia Cabral de Gouvea, Maria Domenica Moccia, Simon Grüter, Timothy Sykes, Lennart Opitz, Griffin White, Laura Neff, Doris Popovic, Andrea Patrignani, Jay Tracy, Ralph Schlapbach, Christiane Beckmann, Maurice Redondo, Olivier Kobel, Christoph Noppen, Niko Beerenwinkel, Tanja Stadler |
| EPI_ISL_3345636 | Fulgent Genetics | Centers for Disease Control and Prevention Division of Viral Diseases, Pathogen Discovery | Dakota Howard, Dhwani Batra, Peter W. Cook, Kara Moser, Adrian Paskey, Jason Caravas, Benjamin Rambo-Martin, Shatavia Morrison, Christopher Gulvick, Scott Sammons, Yvette Unoarumhi, Darlene Wagner, Matthew Schmerer, Harry Gao, Mickey Li, John Gao, Joseph Fierro, Benafsh Sapra, Becky Tsai, Yan Meng, Doreen Ng, James Xie, Clinton R. Paden, Duncan MacCannell |
| EPI_ISL_3345637, EPI_ISL_3345638 | Viollier AG | Department of Biosystems Science and Engineering, ETH Zürich | Chaoran Chen, Sarah Nadeau, Louis du Plessis, Catharine Aquino, Ivan Topolsky, Kim Philipp Jablonski, Lara Fuhrmann, Daniel Ehrsam, Isabel Stürmer, Andreia Cabral de Gouvea, Maria Domenica Moccia, Simon Grüter, Timothy Sykes, Lennart Opitz, Griffin White, Laura Neff, Doris Popovic, Andrea Patrignani, Jay Tracy, Ralph Schlapbach, Christiane Beckmann, Maurice Redondo, Olivier Kobel, Christoph Noppen, Niko Beerenwinkel, Tanja Stadler |
| EPI_ISL_3345639 | Fulgent Genetics | Centers for Disease Control and Prevention Division of Viral Diseases, Pathogen Discovery | Dakota Howard, Dhwani Batra, Peter W. Cook, Kara Moser, Adrian Paskey, Jason Caravas, Benjamin Rambo-Martin, Shatavia Morrison, Christopher Gulvick, Scott Sammons, Yvette Unoarumhi, Darlene Wagner, Matthew Schmerer, Harry Gao, Mickey Li, John Gao, Joseph Fierro, Benafsh Sapra, Becky Tsai, Yan Meng, Doreen Ng, James Xie, Clinton R. Paden, Duncan MacCannell |
| EPI_ISL_3345640 | Viollier AG | Department of Biosystems Science and Engineering, ETH Zürich | Chaoran Chen, Sarah Nadeau, Louis du Plessis, Catharine Aquino, Ivan Topolsky, Kim Philipp Jablonski, Lara Fuhrmann, Daniel Ehrsam, Isabel Stürmer, Andreia Cabral de Gouvea, Maria Domenica Moccia, Simon Grüter, Timothy Sykes, Lennart Opitz, Griffin White, Laura Neff, Doris Popovic, Andrea Patrignani, Jay Tracy, Ralph Schlapbach, Christiane Beckmann, Maurice Redondo, Olivier Kobel, Christoph Noppen, Niko Beerenwinkel, Tanja Stadler |
| EPI_ISL_3345641 | Fulgent Genetics | Centers for Disease Control and Prevention Division of Viral Diseases, Pathogen Discovery | Dakota Howard, Dhwani Batra, Peter W. Cook, Kara Moser, Adrian Paskey, Jason Caravas, Benjamin Rambo-Martin, Shatavia Morrison, Christopher Gulvick, Scott Sammons, Yvette Unoarumhi, Darlene Wagner, Matthew Schmerer, Harry Gao, Mickey Li, John Gao, Joseph Fierro, Benafsh Sapra, Becky Tsai, Yan Meng, Doreen Ng, James Xie, Clinton R. Paden, Duncan MacCannell |
| EPI_ISL_3345642, EPI_ISL_3345643 | Viollier AG | Department of Biosystems Science and Engineering, ETH Zürich | Chaoran Chen, Sarah Nadeau, Louis du Plessis, Catharine Aquino, Ivan Topolsky, Kim Philipp Jablonski, Lara Fuhrmann, Daniel Ehrsam, Isabel Stürmer, Andreia Cabral de Gouvea, Maria Domenica Moccia, Simon Grüter, Timothy Sykes, Lennart Opitz, Griffin White, Laura Neff, Doris Popovic, Andrea Patrignani, Jay Tracy, Ralph Schlapbach, Christiane Beckmann, Maurice Redondo, Olivier Kobel, Christoph Noppen, Niko Beerenwinkel, Tanja Stadler |
| EPI_ISL_3345644 | Fulgent Genetics | Centers for Disease Control and Prevention Division of Viral Diseases, Pathogen Discovery | Dakota Howard, Dhwani Batra, Peter W. Cook, Kara Moser, Adrian Paskey, Jason Caravas, Benjamin Rambo-Martin, Shatavia Morrison, Christopher Gulvick, Scott Sammons, Yvette Unoarumhi, Darlene Wagner, Matthew Schmerer, Harry Gao, Mickey Li, John Gao, Joseph Fierro, Benafsh Sapra, Becky Tsai, Yan Meng, Doreen Ng, James Xie, Clinton R. Paden, Duncan MacCannell |
| EPI_ISL_3345645, EPI_ISL_3345646 | Viollier AG | Department of Biosystems Science and Engineering, ETH Zürich | Chaoran Chen, Sarah Nadeau, Louis du Plessis, Catharine Aquino, Ivan Topolsky, Kim Philipp Jablonski, Lara Fuhrmann, Daniel Ehrsam, Isabel Stürmer, Andreia Cabral de Gouvea, Maria Domenica Moccia, Simon Grüter, Timothy Sykes, Lennart Opitz, Griffin White, Laura Neff, Doris Popovic, Andrea Patrignani, Jay Tracy, Ralph Schlapbach, Christiane Beckmann, Maurice Redondo, Olivier Kobel, Christoph Noppen, Niko Beerenwinkel, Tanja Stadler |
| EPI_ISL_3345647 | Fulgent Genetics | Centers for Disease Control and Prevention Division of Viral Diseases, Pathogen Discovery | Dakota Howard, Dhwani Batra, Peter W. Cook, Kara Moser, Adrian Paskey, Jason Caravas, Benjamin Rambo-Martin, Shatavia Morrison, Christopher Gulvick, Scott Sammons, Yvette Unoarumhi, Darlene Wagner, Matthew Schmerer, Harry Gao, Mickey Li, John Gao, Joseph Fierro, Benafsh Sapra, Becky Tsai, Yan Meng, Doreen Ng, James Xie, Clinton R. Paden, Duncan MacCannell |
| EPI_ISL_3345648 | Viollier AG | Department of Biosystems Science and Engineering, ETH Zürich | Chaoran Chen, Sarah Nadeau, Louis du Plessis, Catharine Aquino, Ivan Topolsky, Kim Philipp Jablonski, Lara Fuhrmann, Daniel Ehrsam, Isabel Stürmer, Andreia Cabral de Gouvea, Maria Domenica Moccia, Simon Grüter, Timothy Sykes, Lennart Opitz, Griffin White, Laura Neff, Doris Popovic, Andrea Patrignani, Jay Tracy, Ralph Schlapbach, Christiane Beckmann, Maurice Redondo, Olivier Kobel, Christoph Noppen, Niko Beerenwinkel, Tanja Stadler |
| EPI_ISL_3345649 | Viollier AG | Department of Biosystems Science and Engineering, ETH Zürich | Christian Beisel, Sarah Nadeau, Chaoran Chen, Louis du Plessis, Ivan Topolsky, Kim Philipp Jablonski, Lara Fuhrmann, Rebecca Denes, Mirjam Feldkamp, Ina Nissen, Natascha Santacroce, Elodie Burcklen, Christiane Beckmann, Maurice Redondo, Olivier Kobel, Christoph Noppen, Niko Beerenwinkel, Tanja Stadler |
| EPI_ISL_3345650 | Fulgent Genetics | Centers for Disease Control and Prevention Division of Viral Diseases, Pathogen Discovery | Dakota Howard, Dhwani Batra, Peter W. Cook, Kara Moser, Adrian Paskey, Jason Caravas, Benjamin Rambo-Martin, Shatavia Morrison, Christopher Gulvick, Scott Sammons, Yvette Unoarumhi, Darlene Wagner, Matthew Schmerer, Harry Gao, Mickey Li, John Gao, Joseph Fierro, Benafsh Sapra, Becky Tsai, Yan Meng, Doreen Ng, James Xie, Clinton R. Paden, Duncan MacCannell |
| EPI_ISL_3345651 | Viollier AG | Department of Biosystems Science and Engineering, ETH Zürich | Chaoran Chen, Sarah Nadeau, Louis du Plessis, Catharine Aquino, Ivan Topolsky, Kim Philipp Jablonski, Lara Fuhrmann, Daniel Ehrsam, Isabel Stürmer, Andreia Cabral de Gouvea, Maria Domenica Moccia, Simon Grüter, Timothy Sykes, Lennart Opitz, Griffin White, Laura Neff, Doris Popovic, Andrea Patrignani, Jay Tracy, Ralph Schlapbach, Christiane Beckmann, Maurice Redondo, Olivier Kobel, Christoph Noppen, Niko Beerenwinkel, Tanja Stadler |
| EPI_ISL_3345652 | Fulgent Genetics | Centers for Disease Control and Prevention Division of Viral Diseases, Pathogen Discovery | Dakota Howard, Dhwani Batra, Peter W. Cook, Kara Moser, Adrian Paskey, Jason Caravas, Benjamin Rambo-Martin, Shatavia Morrison, Christopher Gulvick, Scott Sammons, Yvette Unoarumhi, Darlene Wagner, Matthew Schmerer, Harry Gao, Mickey Li, John Gao, Joseph Fierro, Benafsh Sapra, Becky Tsai, Yan Meng, Doreen Ng, James Xie, Clinton R. Paden, Duncan MacCannell |
| EPI_ISL_3345653 | Viollier AG | Department of Biosystems Science and Engineering, ETH Zürich | Chaoran Chen, Sarah Nadeau, Louis du Plessis, Catharine Aquino, Ivan Topolsky, Kim Philipp Jablonski, Lara Fuhrmann, Daniel Ehrsam, Isabel Stürmer, Andreia Cabral de Gouvea, Maria Domenica Moccia, Simon Grüter, Timothy Sykes, Lennart Opitz, Griffin White, Laura Neff, Doris Popovic, Andrea Patrignani, Jay Tracy, Ralph Schlapbach, Christiane Beckmann, Maurice Redondo, Olivier Kobel, Christoph Noppen, Niko Beerenwinkel, Tanja Stadler |
| EPI_ISL_3345654 | Fulgent Genetics | Centers for Disease Control and Prevention Division of Viral Diseases, Pathogen Discovery | Dakota Howard, Dhwani Batra, Peter W. Cook, Kara Moser, Adrian Paskey, Jason Caravas, Benjamin Rambo-Martin, Shatavia Morrison, Christopher Gulvick, Scott Sammons, Yvette Unoarumhi, Darlene Wagner, Matthew Schmerer, Harry Gao, Mickey Li, John Gao, Joseph Fierro, Benafsh Sapra, Becky Tsai, Yan Meng, Doreen Ng, James Xie, Clinton R. Paden, Duncan MacCannell |
| EPI_ISL_3345655 | Viollier AG | Department of Biosystems Science and Engineering, ETH Zürich | Christian Beisel, Sarah Nadeau, Chaoran Chen, Louis du Plessis, Ivan Topolsky, Kim Philipp Jablonski, Lara Fuhrmann, Rebecca Denes, Mirjam Feldkamp, Ina Nissen, Natascha Santacroce, Elodie Burcklen, Christiane Beckmann, Maurice Redondo, Olivier Kobel, Christoph Noppen, Niko Beerenwinkel, Tanja Stadler |
| EPI_ISL_3345656 | Viollier AG | Department of Biosystems Science and Engineering, ETH Zürich | Chaoran Chen, Sarah Nadeau, Louis du Plessis, Catharine Aquino, Ivan Topolsky, Kim Philipp Jablonski, Lara Fuhrmann, Daniel Ehrsam, Isabel Stürmer, Andreia Cabral de Gouvea, Maria Domenica Moccia, Simon Grüter, Timothy Sykes, Lennart Opitz, Griffin White, Laura Neff, Doris Popovic, Andrea Patrignani, Jay Tracy, Ralph Schlapbach, Christiane Beckmann, Maurice Redondo, Olivier Kobel, Christoph Noppen, Niko Beerenwinkel, Tanja Stadler |
| EPI_ISL_3345657 | Fulgent Genetics | Centers for Disease Control and Prevention Division of Viral Diseases, Pathogen Discovery | Dakota Howard, Dhwani Batra, Peter W. Cook, Kara Moser, Adrian Paskey, Jason Caravas, Benjamin Rambo-Martin, Shatavia Morrison, Christopher Gulvick, Scott Sammons, Yvette Unoarumhi, Darlene Wagner, Matthew Schmerer, Harry Gao, Mickey Li, John Gao, Joseph Fierro, Benafsh Sapra, Becky Tsai, Yan Meng, Doreen Ng, James Xie, Clinton R. Paden, Duncan MacCannell |
| EPI_ISL_3345658, EPI_ISL_3345659 | Viollier AG | Department of Biosystems Science and Engineering, ETH Zürich | Chaoran Chen, Sarah Nadeau, Louis du Plessis, Catharine Aquino, Ivan Topolsky, Kim Philipp Jablonski, Lara Fuhrmann, Daniel Ehrsam, Isabel Stürmer, Andreia Cabral de Gouvea, Maria Domenica Moccia, Simon Grüter, Timothy Sykes, Lennart Opitz, Griffin White, Laura Neff, Doris Popovic, Andrea Patrignani, Jay Tracy, Ralph Schlapbach, Christiane Beckmann, Maurice Redondo, Olivier Kobel, Christoph Noppen, Niko Beerenwinkel, Tanja Stadler |
| EPI_ISL_3345660 | Fulgent Genetics | Centers for Disease Control and Prevention Division of Viral Diseases, Pathogen Discovery | Dakota Howard, Dhwani Batra, Peter W. Cook, Kara Moser, Adrian Paskey, Jason Caravas, Benjamin Rambo-Martin, Shatavia Morrison, Christopher Gulvick, Scott Sammons, Yvette Unoarumhi, Darlene Wagner, Matthew Schmerer, Harry Gao, Mickey Li, John Gao, Joseph Fierro, Benafsh Sapra, Becky Tsai, Yan Meng, Doreen Ng, James Xie, Clinton R. Paden, Duncan MacCannell |
| EPI_ISL_3345661 | Viollier AG | Department of Biosystems Science and Engineering, ETH Zürich | Chaoran Chen, Sarah Nadeau, Louis du Plessis, Catharine Aquino, Ivan Topolsky, Kim Philipp Jablonski, Lara Fuhrmann, Daniel Ehrsam, Isabel Stürmer, Andreia Cabral de Gouvea, Maria Domenica Moccia, Simon Grüter, Timothy Sykes, Lennart Opitz, Griffin White, Laura Neff, Doris Popovic, Andrea Patrignani, Jay Tracy, Ralph Schlapbach, Christiane Beckmann, Maurice Redondo, Olivier Kobel, Christoph Noppen, Niko Beerenwinkel, Tanja Stadler |
| EPI_ISL_3345662 | Fulgent Genetics | Centers for Disease Control and Prevention Division of Viral Diseases, Pathogen Discovery | Dakota Howard, Dhwani Batra, Peter W. Cook, Kara Moser, Adrian Paskey, Jason Caravas, Benjamin Rambo-Martin, Shatavia Morrison, Christopher Gulvick, Scott Sammons, Yvette Unoarumhi, Darlene Wagner, Matthew Schmerer, Harry Gao, Mickey Li, John Gao, Joseph Fierro, Benafsh Sapra, Becky Tsai, Yan Meng, Doreen Ng, James Xie, Clinton R. Paden, Duncan MacCannell |
| EPI_ISL_3345663 | Viollier AG | Department of Biosystems Science and Engineering, ETH Zürich | Chaoran Chen, Sarah Nadeau, Louis du Plessis, Catharine Aquino, Ivan Topolsky, Kim Philipp Jablonski, Lara Fuhrmann, Daniel Ehrsam, Isabel Stürmer, Andreia Cabral de Gouvea, Maria Domenica Moccia, Simon Grüter, Timothy Sykes, Lennart Opitz, Griffin White, Laura Neff, Doris Popovic, Andrea Patrignani, Jay Tracy, Ralph Schlapbach, Christiane Beckmann, Maurice Redondo, Olivier Kobel, Christoph Noppen, Niko Beerenwinkel, Tanja Stadler |
| EPI_ISL_3345664 | Viollier AG | Department of Biosystems Science and Engineering, ETH Zürich | Christian Beisel, Sarah Nadeau, Chaoran Chen, Louis du Plessis, Ivan Topolsky, Kim Philipp Jablonski, Lara Fuhrmann, Rebecca Denes, Mirjam Feldkamp, Ina Nissen, Natascha Santacroce, Elodie Burcklen, Christiane Beckmann, Maurice Redondo, Olivier Kobel, Christoph Noppen, Niko Beerenwinkel, Tanja Stadler |
| EPI_ISL_3345665 | Fulgent Genetics | Centers for Disease Control and Prevention Division of Viral Diseases, Pathogen Discovery | Dakota Howard, Dhwani Batra, Peter W. Cook, Kara Moser, Adrian Paskey, Jason Caravas, Benjamin Rambo-Martin, Shatavia Morrison, Christopher Gulvick, Scott Sammons, Yvette Unoarumhi, Darlene Wagner, Matthew Schmerer, Harry Gao, Mickey Li, John Gao, Joseph Fierro, Benafsh Sapra, Becky Tsai, Yan Meng, Doreen Ng, James Xie, Clinton R. Paden, Duncan MacCannell |
| EPI_ISL_3345666, EPI_ISL_3345667 | Viollier AG | Department of Biosystems Science and Engineering, ETH Zürich | Chaoran Chen, Sarah Nadeau, Louis du Plessis, Catharine Aquino, Ivan Topolsky, Kim Philipp Jablonski, Lara Fuhrmann, Daniel Ehrsam, Isabel Stürmer, Andreia Cabral de Gouvea, Maria Domenica Moccia, Simon Grüter, Timothy Sykes, Lennart Opitz, Griffin White, Laura Neff, Doris Popovic, Andrea Patrignani, Jay Tracy, Ralph Schlapbach, Christiane Beckmann, Maurice Redondo, Olivier Kobel, Christoph Noppen, Niko Beerenwinkel, Tanja Stadler |
| EPI_ISL_3345668 | Fulgent Genetics | Centers for Disease Control and Prevention Division of Viral Diseases, Pathogen Discovery | Dakota Howard, Dhwani Batra, Peter W. Cook, Kara Moser, Adrian Paskey, Jason Caravas, Benjamin Rambo-Martin, Shatavia Morrison, Christopher Gulvick, Scott Sammons, Yvette Unoarumhi, Darlene Wagner, Matthew Schmerer, Harry Gao, Mickey Li, John Gao, Joseph Fierro, Benafsh Sapra, Becky Tsai, Yan Meng, Doreen Ng, James Xie, Clinton R. Paden, Duncan MacCannell |
| EPI_ISL_3345669 | Viollier AG | Department of Biosystems Science and Engineering, ETH Zürich | Chaoran Chen, Sarah Nadeau, Louis du Plessis, Catharine Aquino, Ivan Topolsky, Kim Philipp Jablonski, Lara Fuhrmann, Daniel Ehrsam, Isabel Stürmer, Andreia Cabral de Gouvea, Maria Domenica Moccia, Simon Grüter, Timothy Sykes, Lennart Opitz, Griffin White, Laura Neff, Doris Popovic, Andrea Patrignani, Jay Tracy, Ralph Schlapbach, Christiane Beckmann, Maurice Redondo, Olivier Kobel, Christoph Noppen, Niko Beerenwinkel, Tanja Stadler |
| EPI_ISL_3345670 | Viollier AG | Department of Biosystems Science and Engineering, ETH Zürich | Christian Beisel, Sarah Nadeau, Chaoran Chen, Louis du Plessis, Ivan Topolsky, Kim Philipp Jablonski, Lara Fuhrmann, Rebecca Denes, Mirjam Feldkamp, Ina Nissen, Natascha Santacroce, Elodie Burcklen, Christiane Beckmann, Maurice Redondo, Olivier Kobel, Christoph Noppen, Niko Beerenwinkel, Tanja Stadler |
| EPI_ISL_3345671 | Fulgent Genetics | Centers for Disease Control and Prevention Division of Viral Diseases, Pathogen Discovery | Dakota Howard, Dhwani Batra, Peter W. Cook, Kara Moser, Adrian Paskey, Jason Caravas, Benjamin Rambo-Martin, Shatavia Morrison, Christopher Gulvick, Scott Sammons, Yvette Unoarumhi, Darlene Wagner, Matthew Schmerer, Harry Gao, Mickey Li, John Gao, Joseph Fierro, Benafsh Sapra, Becky Tsai, Yan Meng, Doreen Ng, James Xie, Clinton R. Paden, Duncan MacCannell |
| EPI_ISL_3345672, EPI_ISL_3345673 | Viollier AG | Department of Biosystems Science and Engineering, ETH Zürich | Chaoran Chen, Sarah Nadeau, Louis du Plessis, Catharine Aquino, Ivan Topolsky, Kim Philipp Jablonski, Lara Fuhrmann, Daniel Ehrsam, Isabel Stürmer, Andreia Cabral de Gouvea, Maria Domenica Moccia, Simon Grüter, Timothy Sykes, Lennart Opitz, Griffin White, Laura Neff, Doris Popovic, Andrea Patrignani, Jay Tracy, Ralph Schlapbach, Christiane Beckmann, Maurice Redondo, Olivier Kobel, Christoph Noppen, Niko Beerenwinkel, Tanja Stadler |
| EPI_ISL_3345674 | Fulgent Genetics | Centers for Disease Control and Prevention Division of Viral Diseases, Pathogen Discovery | Dakota Howard, Dhwani Batra, Peter W. Cook, Kara Moser, Adrian Paskey, Jason Caravas, Benjamin Rambo-Martin, Shatavia Morrison, Christopher Gulvick, Scott Sammons, Yvette Unoarumhi, Darlene Wagner, Matthew Schmerer, Harry Gao, Mickey Li, John Gao, Joseph Fierro, Benafsh Sapra, Becky Tsai, Yan Meng, Doreen Ng, James Xie, Clinton R. Paden, Duncan MacCannell |
| EPI_ISL_3345675 | Viollier AG | Department of Biosystems Science and Engineering, ETH Zürich | Chaoran Chen, Sarah Nadeau, Louis du Plessis, Catharine Aquino, Ivan Topolsky, Kim Philipp Jablonski, Lara Fuhrmann, Daniel Ehrsam, Isabel Stürmer, Andreia Cabral de Gouvea, Maria Domenica Moccia, Simon Grüter, Timothy Sykes, Lennart Opitz, Griffin White, Laura Neff, Doris Popovic, Andrea Patrignani, Jay Tracy, Ralph Schlapbach, Christiane Beckmann, Maurice Redondo, Olivier Kobel, Christoph Noppen, Niko Beerenwinkel, Tanja Stadler |
| EPI_ISL_3345676 | Viollier AG | Department of Biosystems Science and Engineering, ETH Zürich | Christian Beisel, Sarah Nadeau, Chaoran Chen, Louis du Plessis, Ivan Topolsky, Kim Philipp Jablonski, Lara Fuhrmann, Rebecca Denes, Mirjam Feldkamp, Ina Nissen, Natascha Santacroce, Elodie Burcklen, Christiane Beckmann, Maurice Redondo, Olivier Kobel, Christoph Noppen, Niko Beerenwinkel, Tanja Stadler |
| EPI_ISL_3345677 | Fulgent Genetics | Centers for Disease Control and Prevention Division of Viral Diseases, Pathogen Discovery | Dakota Howard, Dhwani Batra, Peter W. Cook, Kara Moser, Adrian Paskey, Jason Caravas, Benjamin Rambo-Martin, Shatavia Morrison, Christopher Gulvick, Scott Sammons, Yvette Unoarumhi, Darlene Wagner, Matthew Schmerer, Harry Gao, Mickey Li, John Gao, Joseph Fierro, Benafsh Sapra, Becky Tsai, Yan Meng, Doreen Ng, James Xie, Clinton R. Paden, Duncan MacCannell |
| EPI_ISL_3345678, EPI_ISL_3345680, EPI_ISL_3345681 | Viollier AG | Department of Biosystems Science and Engineering, ETH Zürich | Chaoran Chen, Sarah Nadeau, Louis du Plessis, Catharine Aquino, Ivan Topolsky, Kim Philipp Jablonski, Lara Fuhrmann, Daniel Ehrsam, Isabel Stürmer, Andreia Cabral de Gouvea, Maria Domenica Moccia, Simon Grüter, Timothy Sykes, Lennart Opitz, Griffin White, Laura Neff, Doris Popovic, Andrea Patrignani, Jay Tracy, Ralph Schlapbach, Christiane Beckmann, Maurice Redondo, Olivier Kobel, Christoph Noppen, Niko Beerenwinkel, Tanja Stadler |
| EPI_ISL_3345682 | Fulgent Genetics | Centers for Disease Control and Prevention Division of Viral Diseases, Pathogen Discovery | Dakota Howard, Dhwani Batra, Peter W. Cook, Kara Moser, Adrian Paskey, Jason Caravas, Benjamin Rambo-Martin, Shatavia Morrison, Christopher Gulvick, Scott Sammons, Yvette Unoarumhi, Darlene Wagner, Matthew Schmerer, Harry Gao, Mickey Li, John Gao, Joseph Fierro, Benafsh Sapra, Becky Tsai, Yan Meng, Doreen Ng, James Xie, Clinton R. Paden, Duncan MacCannell |

| | | | |
|---|---|---|---|
| EPI_ISL_3345683 | Viollier AG | Department of Biosystems Science and Engineering, ETH Zürich | Christian Beisel, Sarah Nadeau, Chaoran Chen, Louis du Plessis, Ivan Topolsky, Kim Philipp Jablonski, Lara Fuhrmann, Rebecca Denes, Mirjam Feldkamp, Ina Nissen, Natascha Santacroce, Elodie Burcklen, Christiane Beckmann, Maurice Redondo, Olivier Kobel, Christoph Noppen, Niko Beerenwinkel, Tanja Stadler |
| EPI_ISL_3345684 | Fulgent Genetics | Centers for Disease Control and Prevention Division of Viral Diseases, Pathogen Discovery | Dakota Howard, Dhwani Batra, Peter W. Cook, Kara Moser, Adrian Paskey, Jason Caravas, Benjamin Rambo-Martin, Shatavia Morrison, Christopher Gulvick, Scott Sammons, Yvette Unoarumhi, Darlene Wagner, Matthew Schmerer, Harry Gao, Mickey Li, John Gao, Joseph Fierro, Benafsh Sapra, Becky Tsai, Yan Meng, Doreen Ng, James Xie, Clinton R. Paden, Duncan MacCannell |
| EPI_ISL_3345685, EPI_ISL_3345686 | Viollier AG | Department of Biosystems Science and Engineering, ETH Zürich | Chaoran Chen, Sarah Nadeau, Louis du Plessis, Catharine Aquino, Ivan Topolsky, Kim Philipp Jablonski, Lara Fuhrmann, Daniel Ehrsam, Isabel Stürmer, Andreia Cabral de Gouvea, Maria Domenica Moccia, Simon Grüter, Timothy Sykes, Lennart Opitz, Griffin White, Laura Neff, Doris Popovic, Andrea Patrignani, Jay Tracy, Ralph Schlapbach, Christiane Beckmann, Maurice Redondo, Olivier Kobel, Christoph Noppen, Niko Beerenwinkel, Tanja Stadler |
| EPI_ISL_3345687 | Fulgent Genetics | Centers for Disease Control and Prevention Division of Viral Diseases, Pathogen Discovery | Dakota Howard, Dhwani Batra, Peter W. Cook, Kara Moser, Adrian Paskey, Jason Caravas, Benjamin Rambo-Martin, Shatavia Morrison, Christopher Gulvick, Scott Sammons, Yvette Unoarumhi, Darlene Wagner, Matthew Schmerer, Harry Gao, Mickey Li, John Gao, Joseph Fierro, Benafsh Sapra, Becky Tsai, Yan Meng, Doreen Ng, James Xie, Clinton R. Paden, Duncan MacCannell |
| EPI_ISL_3345688 | Viollier AG | Department of Biosystems Science and Engineering, ETH Zürich | Chaoran Chen, Sarah Nadeau, Louis du Plessis, Catharine Aquino, Ivan Topolsky, Kim Philipp Jablonski, Lara Fuhrmann, Daniel Ehrsam, Isabel Stürmer, Andreia Cabral de Gouvea, Maria Domenica Moccia, Simon Grüter, Timothy Sykes, Lennart Opitz, Griffin White, Laura Neff, Doris Popovic, Andrea Patrignani, Jay Tracy, Ralph Schlapbach, Christiane Beckmann, Maurice Redondo, Olivier Kobel, Christoph Noppen, Niko Beerenwinkel, Tanja Stadler |
| EPI_ISL_3345689 | Fulgent Genetics | Centers for Disease Control and Prevention Division of Viral Diseases, Pathogen Discovery | Dakota Howard, Dhwani Batra, Peter W. Cook, Kara Moser, Adrian Paskey, Jason Caravas, Benjamin Rambo-Martin, Shatavia Morrison, Christopher Gulvick, Scott Sammons, Yvette Unoarumhi, Darlene Wagner, Matthew Schmerer, Harry Gao, Mickey Li, John Gao, Joseph Fierro, Benafsh Sapra, Becky Tsai, Yan Meng, Doreen Ng, James Xie, Clinton R. Paden, Duncan MacCannell |
| EPI_ISL_3345690, EPI_ISL_3345691 | Viollier AG | Department of Biosystems Science and Engineering, ETH Zürich | Chaoran Chen, Sarah Nadeau, Louis du Plessis, Catharine Aquino, Ivan Topolsky, Kim Philipp Jablonski, Lara Fuhrmann, Daniel Ehrsam, Isabel Stürmer, Andreia Cabral de Gouvea, Maria Domenica Moccia, Simon Grüter, Timothy Sykes, Lennart Opitz, Griffin White, Laura Neff, Doris Popovic, Andrea Patrignani, Jay Tracy, Ralph Schlapbach, Christiane Beckmann, Maurice Redondo, Olivier Kobel, Christoph Noppen, Niko Beerenwinkel, Tanja Stadler |
| EPI_ISL_3345692 | Fulgent Genetics | Centers for Disease Control and Prevention Division of Viral Diseases, Pathogen Discovery | Dakota Howard, Dhwani Batra, Peter W. Cook, Kara Moser, Adrian Paskey, Jason Caravas, Benjamin Rambo-Martin, Shatavia Morrison, Christopher Gulvick, Scott Sammons, Yvette Unoarumhi, Darlene Wagner, Matthew Schmerer, Harry Gao, Mickey Li, John Gao, Joseph Fierro, Benafsh Sapra, Becky Tsai, Yan Meng, Doreen Ng, James Xie, Clinton R. Paden, Duncan MacCannell |
| EPI_ISL_3345693, EPI_ISL_3345694 | Viollier AG | Department of Biosystems Science and Engineering, ETH Zürich | Chaoran Chen, Sarah Nadeau, Louis du Plessis, Catharine Aquino, Ivan Topolsky, Kim Philipp Jablonski, Lara Fuhrmann, Daniel Ehrsam, Isabel Stürmer, Andreia Cabral de Gouvea, Maria Domenica Moccia, Simon Grüter, Timothy Sykes, Lennart Opitz, Griffin White, Laura Neff, Doris Popovic, Andrea Patrignani, Jay Tracy, Ralph Schlapbach, Christiane Beckmann, Maurice Redondo, Olivier Kobel, Christoph Noppen, Niko Beerenwinkel, Tanja Stadler |
| EPI_ISL_3345695 | Fulgent Genetics | Centers for Disease Control and Prevention Division of Viral Diseases, Pathogen Discovery | Dakota Howard, Dhwani Batra, Peter W. Cook, Kara Moser, Adrian Paskey, Jason Caravas, Benjamin Rambo-Martin, Shatavia Morrison, Christopher Gulvick, Scott Sammons, Yvette Unoarumhi, Darlene Wagner, Matthew Schmerer, Harry Gao, Mickey Li, John Gao, Joseph Fierro, Benafsh Sapra, Becky Tsai, Yan Meng, Doreen Ng, James Xie, Clinton R. Paden, Duncan MacCannell |
| EPI_ISL_3345697 | Viollier AG | Department of Biosystems Science and Engineering, ETH Zürich | Chaoran Chen, Sarah Nadeau, Louis du Plessis, Catharine Aquino, Ivan Topolsky, Kim Philipp Jablonski, Lara Fuhrmann, Daniel Ehrsam, Isabel Stürmer, Andreia Cabral de Gouvea, Maria Domenica Moccia, Simon Grüter, Timothy Sykes, Lennart Opitz, Griffin White, Laura Neff, Doris Popovic, Andrea Patrignani, Jay Tracy, Ralph Schlapbach, Christiane Beckmann, Maurice Redondo, Olivier Kobel, Christoph Noppen, Niko Beerenwinkel, Tanja Stadler |
| EPI_ISL_3345698 | Fulgent Genetics | Centers for Disease Control and Prevention Division of Viral Diseases, Pathogen Discovery | Dakota Howard, Dhwani Batra, Peter W. Cook, Kara Moser, Adrian Paskey, Jason Caravas, Benjamin Rambo-Martin, Shatavia Morrison, Christopher Gulvick, Scott Sammons, Yvette Unoarumhi, Darlene Wagner, Matthew Schmerer, Harry Gao, Mickey Li, John Gao, Joseph Fierro, Benafsh Sapra, Becky Tsai, Yan Meng, Doreen Ng, James Xie, Clinton R. Paden, Duncan MacCannell |
| EPI_ISL_3345699, EPI_ISL_3345700 | Viollier AG | Department of Biosystems Science and Engineering, ETH Zürich | Chaoran Chen, Sarah Nadeau, Louis du Plessis, Catharine Aquino, Ivan Topolsky, Kim Philipp Jablonski, Lara Fuhrmann, Daniel Ehrsam, Isabel Stürmer, Andreia Cabral de Gouvea, Maria Domenica Moccia, Simon Grüter, Timothy Sykes, Lennart Opitz, Griffin White, Laura Neff, Doris Popovic, Andrea Patrignani, Jay Tracy, Ralph Schlapbach, Christiane Beckmann, Maurice Redondo, Olivier Kobel, Christoph Noppen, Niko Beerenwinkel, Tanja Stadler |
| EPI_ISL_3345701 | Fulgent Genetics | Centers for Disease Control and Prevention Division of Viral Diseases, Pathogen Discovery | Dakota Howard, Dhwani Batra, Peter W. Cook, Kara Moser, Adrian Paskey, Jason Caravas, Benjamin Rambo-Martin, Shatavia Morrison, Christopher Gulvick, Scott Sammons, Yvette Unoarumhi, Darlene Wagner, Matthew Schmerer, Harry Gao, Mickey Li, John Gao, Joseph Fierro, Benafsh Sapra, Becky Tsai, Yan Meng, Doreen Ng, James Xie, Clinton R. Paden, Duncan MacCannell |
| EPI_ISL_3345702 | Viollier AG | Department of Biosystems Science and Engineering, ETH Zürich | Chaoran Chen, Sarah Nadeau, Louis du Plessis, Catharine Aquino, Ivan Topolsky, Kim Philipp Jablonski, Lara Fuhrmann, Daniel Ehrsam, Isabel Stürmer, Andreia Cabral de Gouvea, Maria Domenica Moccia, Simon Grüter, Timothy Sykes, Lennart Opitz, Griffin White, Laura Neff, Doris Popovic, Andrea Patrignani, Jay Tracy, Ralph Schlapbach, Christiane Beckmann, Maurice Redondo, Olivier Kobel, Christoph Noppen, Niko Beerenwinkel, Tanja Stadler |
| EPI_ISL_3345703 | Viollier AG | Department of Biosystems Science and Engineering, ETH Zürich | Christian Beisel, Sarah Nadeau, Chaoran Chen, Louis du Plessis, Ivan Topolsky, Kim Philipp Jablonski, Lara Fuhrmann, Rebecca Denes, Mirjam Feldkamp, Ina Nissen, Natascha Santacroce, Elodie Burcklen, Christiane Beckmann, Maurice Redondo, Olivier Kobel, Christoph Noppen, Niko Beerenwinkel, Tanja Stadler |
| EPI_ISL_3345704, EPI_ISL_3345706 | Fulgent Genetics | Centers for Disease Control and Prevention Division of Viral Diseases, Pathogen Discovery | Dakota Howard, Dhwani Batra, Peter W. Cook, Kara Moser, Adrian Paskey, Jason Caravas, Benjamin Rambo-Martin, Shatavia Morrison, Christopher Gulvick, Scott Sammons, Yvette Unoarumhi, Darlene Wagner, Matthew Schmerer, Harry Gao, Mickey Li, John Gao, Joseph Fierro, Benafsh Sapra, Becky Tsai, Yan Meng, Doreen Ng, James Xie, Clinton R. Paden, Duncan MacCannell |
| EPI_ISL_3345707, EPI_ISL_3345708 | Viollier AG | Department of Biosystems Science and Engineering, ETH Zürich | Chaoran Chen, Sarah Nadeau, Louis du Plessis, Catharine Aquino, Ivan Topolsky, Kim Philipp Jablonski, Lara Fuhrmann, Daniel Ehrsam, Isabel Stürmer, Andreia Cabral de Gouvea, Maria Domenica Moccia, Simon Grüter, Timothy Sykes, Lennart Opitz, Griffin White, Laura Neff, Doris Popovic, Andrea Patrignani, Jay Tracy, Ralph Schlapbach, Christiane Beckmann, Maurice Redondo, Olivier Kobel, Christoph Noppen, Niko Beerenwinkel, Tanja Stadler |
| EPI_ISL_3345709 | Fulgent Genetics | Centers for Disease Control and Prevention Division of Viral Diseases, Pathogen Discovery | Dakota Howard, Dhwani Batra, Peter W. Cook, Kara Moser, Adrian Paskey, Jason Caravas, Benjamin Rambo-Martin, Shatavia Morrison, Christopher Gulvick, Scott Sammons, Yvette Unoarumhi, Darlene Wagner, Matthew Schmerer, Harry Gao, Mickey Li, John Gao, Joseph Fierro, Benafsh Sapra, Becky Tsai, Yan Meng, Doreen Ng, James Xie, Clinton R. Paden, Duncan MacCannell |
| EPI_ISL_3345710 | Viollier AG | Department of Biosystems Science and Engineering, ETH Zürich | Christian Beisel, Sarah Nadeau, Chaoran Chen, Louis du Plessis, Ivan Topolsky, Kim Philipp Jablonski, Lara Fuhrmann, Rebecca Denes, Mirjam Feldkamp, Ina Nissen, Natascha Santacroce, Elodie Burcklen, Christiane Beckmann, Maurice Redondo, Olivier Kobel, Christoph Noppen, Niko Beerenwinkel, Tanja Stadler |
| EPI_ISL_3345711 | Viollier AG | Department of Biosystems Science and Engineering, ETH Zürich | Chaoran Chen, Sarah Nadeau, Louis du Plessis, Catharine Aquino, Ivan Topolsky, Kim Philipp Jablonski, Lara Fuhrmann, Daniel Ehrsam, Isabel Stürmer, Andreia Cabral de Gouvea, Maria Domenica Moccia, Simon Grüter, Timothy Sykes, Lennart Opitz, Griffin White, Laura Neff, Doris Popovic, Andrea Patrignani, Jay Tracy, Ralph Schlapbach, Christiane Beckmann, Maurice Redondo, Olivier Kobel, Christoph Noppen, Niko Beerenwinkel, Tanja Stadler |
| EPI_ISL_3345712 | Fulgent Genetics | Centers for Disease Control and Prevention Division of Viral Diseases, Pathogen Discovery | Dakota Howard, Dhwani Batra, Peter W. Cook, Kara Moser, Adrian Paskey, Jason Caravas, Benjamin Rambo-Martin, Shatavia Morrison, Christopher Gulvick, Scott Sammons, Yvette Unoarumhi, Darlene Wagner, Matthew Schmerer, Harry Gao, Mickey Li, John Gao, Joseph Fierro, Benafsh Sapra, Becky Tsai, Yan Meng, Doreen Ng, James Xie, Clinton R. Paden, Duncan MacCannell |
| EPI_ISL_3345713, EPI_ISL_3345714 | Viollier AG | Department of Biosystems Science and Engineering, ETH Zürich | Chaoran Chen, Sarah Nadeau, Louis du Plessis, Catharine Aquino, Ivan Topolsky, Kim Philipp Jablonski, Lara Fuhrmann, Daniel Ehrsam, Isabel Stürmer, Andreia Cabral de Gouvea, Maria Domenica Moccia, Simon Grüter, Timothy Sykes, Lennart Opitz, Griffin White, Laura Neff, Doris Popovic, Andrea Patrignani, Jay Tracy, Ralph Schlapbach, Christiane Beckmann, Maurice Redondo, Olivier Kobel, Christoph Noppen, Niko Beerenwinkel, Tanja Stadler |
| EPI_ISL_3345715 | Fulgent Genetics | Centers for Disease Control and Prevention Division of Viral Diseases, Pathogen Discovery | Dakota Howard, Dhwani Batra, Peter W. Cook, Kara Moser, Adrian Paskey, Jason Caravas, Benjamin Rambo-Martin, Shatavia Morrison, Christopher Gulvick, Scott Sammons, Yvette Unoarumhi, Darlene Wagner, Matthew Schmerer, Harry Gao, Mickey Li, John Gao, Joseph Fierro, Benafsh Sapra, Becky Tsai, Yan Meng, Doreen Ng, James Xie, Clinton R. Paden, Duncan MacCannell |
| EPI_ISL_3345716 | Viollier AG | Department of Biosystems Science and Engineering, ETH Zürich | Christian Beisel, Sarah Nadeau, Chaoran Chen, Louis du Plessis, Ivan Topolsky, Kim Philipp Jablonski, Lara Fuhrmann, Rebecca Denes, Mirjam Feldkamp, Ina Nissen, Natascha Santacroce, Elodie Burcklen, Christiane Beckmann, Maurice Redondo, Olivier Kobel, Christoph Noppen, Niko Beerenwinkel, Tanja Stadler |
| EPI_ISL_3345717 | Viollier AG | Department of Biosystems Science and Engineering, ETH Zürich | Chaoran Chen, Sarah Nadeau, Louis du Plessis, Catharine Aquino, Ivan Topolsky, Kim Philipp Jablonski, Lara Fuhrmann, Daniel Ehrsam, Isabel Stürmer, Andreia Cabral de Gouvea, Maria Domenica Moccia, Simon Grüter, Timothy Sykes, Lennart Opitz, Griffin White, Laura Neff, Doris Popovic, Andrea Patrignani, Jay Tracy, Ralph Schlapbach, Christiane Beckmann, Maurice Redondo, Olivier Kobel, Christoph Noppen, Niko Beerenwinkel, Tanja Stadler |
| EPI_ISL_3345718 | Fulgent Genetics | Centers for Disease Control and Prevention Division of Viral Diseases, Pathogen Discovery | Dakota Howard, Dhwani Batra, Peter W. Cook, Kara Moser, Adrian Paskey, Jason Caravas, Benjamin Rambo-Martin, Shatavia Morrison, Christopher Gulvick, Scott Sammons, Yvette Unoarumhi, Darlene Wagner, Matthew Schmerer, Harry Gao, Mickey Li, John Gao, Joseph Fierro, Benafsh Sapra, Becky Tsai, Yan Meng, Doreen Ng, James Xie, Clinton R. Paden, Duncan MacCannell |
| EPI_ISL_3345719 | Viollier AG | Department of Biosystems Science and Engineering, ETH Zürich | Chaoran Chen, Sarah Nadeau, Louis du Plessis, Catharine Aquino, Ivan Topolsky, Kim Philipp Jablonski, Lara Fuhrmann, Daniel Ehrsam, Isabel Stürmer, Andreia Cabral de Gouvea, Maria Domenica Moccia, Simon Grüter, Timothy Sykes, Lennart Opitz, Griffin White, Laura Neff, Doris Popovic, Andrea Patrignani, Jay Tracy, Ralph Schlapbach, Christiane Beckmann, Maurice Redondo, Olivier Kobel, Christoph Noppen, Niko Beerenwinkel, Tanja Stadler |
| EPI_ISL_3345720 | Fulgent Genetics | Centers for Disease Control and Prevention Division of Viral Diseases, Pathogen Discovery | Dakota Howard, Dhwani Batra, Peter W. Cook, Kara Moser, Adrian Paskey, Jason Caravas, Benjamin Rambo-Martin, Shatavia Morrison, Christopher Gulvick, Scott Sammons, Yvette Unoarumhi, Darlene Wagner, Matthew Schmerer, Harry Gao, Mickey Li, John Gao, Joseph Fierro, Benafsh Sapra, Becky Tsai, Yan Meng, Doreen Ng, James Xie, Clinton R. Paden, Duncan MacCannell |
| EPI_ISL_3345721 | Viollier AG | Department of Biosystems Science and Engineering, ETH Zürich | Christian Beisel, Sarah Nadeau, Chaoran Chen, Louis du Plessis, Ivan Topolsky, Kim Philipp Jablonski, Lara Fuhrmann, Rebecca Denes, Mirjam Feldkamp, Ina Nissen, Natascha Santacroce, Elodie Burcklen, Christiane Beckmann, Maurice Redondo, Olivier Kobel, Christoph Noppen, Niko Beerenwinkel, Tanja Stadler |
| EPI_ISL_3345722 | Viollier AG | Department of Biosystems Science and Engineering, ETH Zürich | Chaoran Chen, Sarah Nadeau, Louis du Plessis, Catharine Aquino, Ivan Topolsky, Kim Philipp Jablonski, Lara Fuhrmann, Daniel Ehrsam, Isabel Stürmer, Andreia Cabral de Gouvea, Maria Domenica Moccia, Simon Grüter, Timothy Sykes, Lennart Opitz, Griffin White, Laura Neff, Doris Popovic, Andrea Patrignani, Jay Tracy, Ralph Schlapbach, Christiane Beckmann, Maurice Redondo, Olivier Kobel, Christoph Noppen, Niko Beerenwinkel, Tanja Stadler |
| EPI_ISL_3345723 | Fulgent Genetics | Centers for Disease Control and Prevention Division of Viral Diseases, Pathogen Discovery | Dakota Howard, Dhwani Batra, Peter W. Cook, Kara Moser, Adrian Paskey, Jason Caravas, Benjamin Rambo-Martin, Shatavia Morrison, Christopher Gulvick, Scott Sammons, Yvette Unoarumhi, Darlene Wagner, Matthew Schmerer, Harry Gao, Mickey Li, John Gao, Joseph Fierro, Benafsh Sapra, Becky Tsai, Yan Meng, Doreen Ng, James Xie, Clinton R. Paden, Duncan MacCannell |
| EPI_ISL_3345724, EPI_ISL_3345725 | Viollier AG | Department of Biosystems Science and Engineering, ETH Zürich | Chaoran Chen, Sarah Nadeau, Louis du Plessis, Catharine Aquino, Ivan Topolsky, Kim Philipp Jablonski, Lara Fuhrmann, Daniel Ehrsam, Isabel Stürmer, Andreia Cabral de Gouvea, Maria Domenica Moccia, Simon Grüter, Timothy Sykes, Lennart Opitz, Griffin White, Laura Neff, Doris Popovic, Andrea Patrignani, Jay Tracy, Ralph Schlapbach, Christiane Beckmann, Maurice Redondo, Olivier Kobel, Christoph Noppen, Niko Beerenwinkel, Tanja Stadler |
| EPI_ISL_3345726 | Fulgent Genetics | Centers for Disease Control and Prevention Division of Viral Diseases, Pathogen Discovery | Dakota Howard, Dhwani Batra, Peter W. Cook, Kara Moser, Adrian Paskey, Jason Caravas, Benjamin Rambo-Martin, Shatavia Morrison, Christopher Gulvick, Scott Sammons, Yvette Unoarumhi, Darlene Wagner, Matthew Schmerer, Harry Gao, Mickey Li, John Gao, Joseph Fierro, Benafsh Sapra, Becky Tsai, Yan Meng, Doreen Ng, James Xie, Clinton R. Paden, Duncan MacCannell |
| EPI_ISL_3345727, EPI_ISL_3345728 | Viollier AG | Department of Biosystems Science and Engineering, ETH Zürich | Chaoran Chen, Sarah Nadeau, Louis du Plessis, Catharine Aquino, Ivan Topolsky, Kim Philipp Jablonski, Lara Fuhrmann, Daniel Ehrsam, Isabel Stürmer, Andreia Cabral de Gouvea, Maria Domenica Moccia, Simon Grüter, Timothy Sykes, Lennart Opitz, Griffin White, Laura Neff, Doris Popovic, Andrea Patrignani, Jay Tracy, Ralph Schlapbach, Christiane Beckmann, Maurice Redondo, Olivier Kobel, Christoph Noppen, Niko Beerenwinkel, Tanja Stadler |
| EPI_ISL_3345729 | Fulgent Genetics | Centers for Disease Control and Prevention Division of Viral Diseases, Pathogen Discovery | Dakota Howard, Dhwani Batra, Peter W. Cook, Kara Moser, Adrian Paskey, Jason Caravas, Benjamin Rambo-Martin, Shatavia Morrison, Christopher Gulvick, Scott Sammons, Yvette Unoarumhi, Darlene Wagner, Matthew Schmerer, Harry Gao, Mickey Li, John Gao, Joseph Fierro, Benafsh Sapra, Becky Tsai, Yan Meng, Doreen Ng, James Xie, Clinton R. Paden, Duncan MacCannell |
| EPI_ISL_3345730 | Viollier AG | Department of Biosystems Science and Engineering, ETH Zürich | Chaoran Chen, Sarah Nadeau, Louis du Plessis, Catharine Aquino, Ivan Topolsky, Kim Philipp Jablonski, Lara Fuhrmann, Daniel Ehrsam, Isabel Stürmer, Andreia Cabral de Gouvea, Maria Domenica Moccia, Simon Grüter, Timothy Sykes, Lennart Opitz, Griffin White, Laura Neff, Doris Popovic, Andrea Patrignani, Jay Tracy, Ralph Schlapbach, Christiane Beckmann, Maurice Redondo, Olivier Kobel, Christoph Noppen, Niko Beerenwinkel, Tanja Stadler |
| EPI_ISL_3345731 | Fulgent Genetics | Centers for Disease Control and Prevention Division of Viral Diseases, Pathogen Discovery | Dakota Howard, Dhwani Batra, Peter W. Cook, Kara Moser, Adrian Paskey, Jason Caravas, Benjamin Rambo-Martin, Shatavia Morrison, Christopher Gulvick, Scott Sammons, Yvette Unoarumhi, Darlene Wagner, Matthew Schmerer, Harry Gao, Mickey Li, John Gao, Joseph Fierro, Benafsh Sapra, Becky Tsai, Yan Meng, Doreen Ng, James Xie, Clinton R. Paden, Duncan MacCannell |
| EPI_ISL_3345732 | Viollier AG | Department of Biosystems Science and Engineering, ETH Zürich | Christian Beisel, Sarah Nadeau, Chaoran Chen, Louis du Plessis, Ivan Topolsky, Kim Philipp Jablonski, Lara Fuhrmann, Rebecca Denes, Mirjam Feldkamp, Ina Nissen, Natascha Santacroce, Elodie Burcklen, Christiane Beckmann, Maurice Redondo, Olivier Kobel, Christoph Noppen, Niko Beerenwinkel, Tanja Stadler |
| EPI_ISL_3345733 | Viollier AG | Department of Biosystems Science and Engineering, ETH Zürich | Chaoran Chen, Sarah Nadeau, Louis du Plessis, Catharine Aquino, Ivan Topolsky, Kim Philipp Jablonski, Lara Fuhrmann, Daniel Ehrsam, Isabel Stürmer, Andreia Cabral de Gouvea, Maria Domenica Moccia, Simon Grüter, Timothy Sykes, Lennart Opitz, Griffin White, Laura Neff, Doris Popovic, Andrea Patrignani, Jay Tracy, Ralph Schlapbach, Christiane Beckmann, Maurice Redondo, Olivier Kobel, Christoph Noppen, Niko Beerenwinkel, Tanja Stadler |
| EPI_ISL_3345734 | Fulgent Genetics | Centers for Disease Control and Prevention Division of Viral Diseases, Pathogen Discovery | Dakota Howard, Dhwani Batra, Peter W. Cook, Kara Moser, Adrian Paskey, Jason Caravas, Benjamin Rambo-Martin, Shatavia Morrison, Christopher Gulvick, Scott Sammons, Yvette Unoarumhi, Darlene Wagner, Matthew Schmerer, Harry Gao, Mickey Li, John Gao, Joseph Fierro, Benafsh Sapra, Becky Tsai, Yan Meng, Doreen Ng, James Xie, Clinton R. Paden, Duncan MacCannell |
| EPI_ISL_3345735, EPI_ISL_3345736 | Viollier AG | Department of Biosystems Science and Engineering, ETH Zürich | Christian Beisel, Sarah Nadeau, Chaoran Chen, Louis du Plessis, Ivan Topolsky, Kim Philipp Jablonski, Lara Fuhrmann, Rebecca Denes, Mirjam Feldkamp, Ina Nissen, Natascha Santacroce, Elodie Burcklen, Christiane Beckmann, Maurice Redondo, Olivier Kobel, Christoph Noppen, Niko Beerenwinkel, Tanja Stadler |
| EPI_ISL_3345738 | Viollier AG | Department of Biosystems Science and Engineering, ETH Zürich | Chaoran Chen, Sarah Nadeau, Louis du Plessis, Catharine Aquino, Ivan Topolsky, Kim Philipp Jablonski, Lara Fuhrmann, Daniel Ehrsam, Isabel Stürmer, Andreia Cabral de Gouvea, Maria Domenica Moccia, Simon Grüter, Timothy Sykes, Lennart Opitz, Griffin White, Laura Neff, Doris Popovic, Andrea Patrignani, Jay Tracy, Ralph Schlapbach, Christiane Beckmann, Maurice Redondo, Olivier Kobel, Christoph Noppen, Niko Beerenwinkel, Tanja Stadler |
| EPI_ISL_3345739 | Viollier AG | Department of Biosystems Science and Engineering, ETH Zürich | Christian Beisel, Sarah Nadeau, Chaoran Chen, Louis du Plessis, Ivan Topolsky, Kim Philipp Jablonski, Lara Fuhrmann, Rebecca Denes, Mirjam Feldkamp, Ina Nissen, Natascha Santacroce, Elodie Burcklen, Christiane Beckmann, Maurice Redondo, Olivier Kobel, Christoph Noppen, Niko Beerenwinkel, Tanja Stadler |
| EPI_ISL_3345740 | Fulgent Genetics | Centers for Disease Control and Prevention Division of Viral Diseases, Pathogen Discovery | Dakota Howard, Dhwani Batra, Peter W. Cook, Kara Moser, Adrian Paskey, Jason Caravas, Benjamin Rambo-Martin, Shatavia Morrison, Christopher Gulvick, Scott Sammons, Yvette Unoarumhi, Darlene Wagner, Matthew Schmerer, Harry Gao, Mickey Li, John Gao, Joseph Fierro, Benafsh Sapra, Becky Tsai, Yan Meng, Doreen Ng, James Xie, Clinton R. Paden, Duncan MacCannell |

| | | | |
|---|---|---|---|
| EPI_ISL_3345741 | Viollier AG | Department of Biosystems Science and Engineering, ETH Zürich | Christian Beisel, Sarah Nadeau, Chaoran Chen, Louis du Plessis, Ivan Topolsky, Kim Philipp Jablonski, Lara Fuhrmann, Rebecca Denes, Mirjam Feldkamp, Ina Nissen, Natascha Santacroce, Elodie Burcklen, Christiane Beckmann, Maurice Redondo, Olivier Kobel, Christoph Noppen, Niko Beerenwinkel, Tanja Stadler |
| EPI_ISL_3345742 | Fulgent Genetics | Centers for Disease Control and Prevention Division of Viral Diseases, Pathogen Discovery | Dakota Howard, Dhwani Batra, Peter W. Cook, Kara Moser, Adrian Paskey, Jason Caravas, Benjamin Rambo-Martin, Shatavia Morrison, Christopher Gulvick, Scott Sammons, Yvette Unoarumhi, Darlene Wagner, Matthew Schmerer, Harry Gao, Mickey Li, John Gao, Joseph Fierro, Benafsh Sapra, Becky Tsai, Yan Meng, Doreen Ng, James Xie, Clinton R. Paden, Duncan MacCannell |
| EPI_ISL_3345743 | Viollier AG | Department of Biosystems Science and Engineering, ETH Zürich | Chaoran Chen, Sarah Nadeau, Louis du Plessis, Catharine Aquino, Ivan Topolsky, Kim Philipp Jablonski, Lara Fuhrmann, Daniel Ehrsam, Isabel Stürmer, Andreia Cabral de Gouvea, Maria Domenica Moccia, Simon Grüter, Timothy Sykes, Lennart Opitz, Griffin White, Laura Neff, Doris Popovic, Andrea Patrignani, Jay Tracy, Ralph Schlapbach, Christiane Beckmann, Maurice Redondo, Olivier Kobel, Christoph Noppen, Niko Beerenwinkel, Tanja Stadler |
| EPI_ISL_3345744 | Viollier AG | Department of Biosystems Science and Engineering, ETH Zürich | Christian Beisel, Sarah Nadeau, Chaoran Chen, Louis du Plessis, Ivan Topolsky, Kim Philipp Jablonski, Lara Fuhrmann, Rebecca Denes, Mirjam Feldkamp, Ina Nissen, Natascha Santacroce, Elodie Burcklen, Christiane Beckmann, Maurice Redondo, Olivier Kobel, Christoph Noppen, Niko Beerenwinkel, Tanja Stadler |
| EPI_ISL_3345745 | Fulgent Genetics | Centers for Disease Control and Prevention Division of Viral Diseases, Pathogen Discovery | Dakota Howard, Dhwani Batra, Peter W. Cook, Kara Moser, Adrian Paskey, Jason Caravas, Benjamin Rambo-Martin, Shatavia Morrison, Christopher Gulvick, Scott Sammons, Yvette Unoarumhi, Darlene Wagner, Matthew Schmerer, Harry Gao, Mickey Li, John Gao, Joseph Fierro, Benafsh Sapra, Becky Tsai, Yan Meng, Doreen Ng, James Xie, Clinton R. Paden, Duncan MacCannell |
| EPI_ISL_3345747 | Viollier AG | Department of Biosystems Science and Engineering, ETH Zürich | Chaoran Chen, Sarah Nadeau, Louis du Plessis, Catharine Aquino, Ivan Topolsky, Kim Philipp Jablonski, Lara Fuhrmann, Daniel Ehrsam, Isabel Stürmer, Andreia Cabral de Gouvea, Maria Domenica Moccia, Simon Grüter, Timothy Sykes, Lennart Opitz, Griffin White, Laura Neff, Doris Popovic, Andrea Patrignani, Jay Tracy, Ralph Schlapbach, Christiane Beckmann, Maurice Redondo, Olivier Kobel, Christoph Noppen, Niko Beerenwinkel, Tanja Stadler |
| EPI_ISL_3345748 | Fulgent Genetics | Centers for Disease Control and Prevention Division of Viral Diseases, Pathogen Discovery | Dakota Howard, Dhwani Batra, Peter W. Cook, Kara Moser, Adrian Paskey, Jason Caravas, Benjamin Rambo-Martin, Shatavia Morrison, Christopher Gulvick, Scott Sammons, Yvette Unoarumhi, Darlene Wagner, Matthew Schmerer, Harry Gao, Mickey Li, John Gao, Joseph Fierro, Benafsh Sapra, Becky Tsai, Yan Meng, Doreen Ng, James Xie, Clinton R. Paden, Duncan MacCannell |
| EPI_ISL_3345749 | Viollier AG | Department of Biosystems Science and Engineering, ETH Zürich | Chaoran Chen, Sarah Nadeau, Louis du Plessis, Catharine Aquino, Ivan Topolsky, Kim Philipp Jablonski, Lara Fuhrmann, Daniel Ehrsam, Isabel Stürmer, Andreia Cabral de Gouvea, Maria Domenica Moccia, Simon Grüter, Timothy Sykes, Lennart Opitz, Griffin White, Laura Neff, Doris Popovic, Andrea Patrignani, Jay Tracy, Ralph Schlapbach, Christiane Beckmann, Maurice Redondo, Olivier Kobel, Christoph Noppen, Niko Beerenwinkel, Tanja Stadler |
| EPI_ISL_3345750 | Fulgent Genetics | Centers for Disease Control and Prevention Division of Viral Diseases, Pathogen Discovery | Dakota Howard, Dhwani Batra, Peter W. Cook, Kara Moser, Adrian Paskey, Jason Caravas, Benjamin Rambo-Martin, Shatavia Morrison, Christopher Gulvick, Scott Sammons, Yvette Unoarumhi, Darlene Wagner, Matthew Schmerer, Harry Gao, Mickey Li, John Gao, Joseph Fierro, Benafsh Sapra, Becky Tsai, Yan Meng, Doreen Ng, James Xie, Clinton R. Paden, Duncan MacCannell |
| EPI_ISL_3345751 | Viollier AG | Department of Biosystems Science and Engineering, ETH Zürich | Chaoran Chen, Sarah Nadeau, Louis du Plessis, Catharine Aquino, Ivan Topolsky, Kim Philipp Jablonski, Lara Fuhrmann, Daniel Ehrsam, Isabel Stürmer, Andreia Cabral de Gouvea, Maria Domenica Moccia, Simon Grüter, Timothy Sykes, Lennart Opitz, Griffin White, Laura Neff, Doris Popovic, Andrea Patrignani, Jay Tracy, Ralph Schlapbach, Christiane Beckmann, Maurice Redondo, Olivier Kobel, Christoph Noppen, Niko Beerenwinkel, Tanja Stadler |
| EPI_ISL_3345753 | Fulgent Genetics | Centers for Disease Control and Prevention Division of Viral Diseases, Pathogen Discovery | Dakota Howard, Dhwani Batra, Peter W. Cook, Kara Moser, Adrian Paskey, Jason Caravas, Benjamin Rambo-Martin, Shatavia Morrison, Christopher Gulvick, Scott Sammons, Yvette Unoarumhi, Darlene Wagner, Matthew Schmerer, Harry Gao, Mickey Li, John Gao, Joseph Fierro, Benafsh Sapra, Becky Tsai, Yan Meng, Doreen Ng, James Xie, Clinton R. Paden, Duncan MacCannell |
| EPI_ISL_3345754 | Viollier AG | Department of Biosystems Science and Engineering, ETH Zürich | Chaoran Chen, Sarah Nadeau, Louis du Plessis, Catharine Aquino, Ivan Topolsky, Kim Philipp Jablonski, Lara Fuhrmann, Daniel Ehrsam, Isabel Stürmer, Andreia Cabral de Gouvea, Maria Domenica Moccia, Simon Grüter, Timothy Sykes, Lennart Opitz, Griffin White, Laura Neff, Doris Popovic, Andrea Patrignani, Jay Tracy, Ralph Schlapbach, Christiane Beckmann, Maurice Redondo, Olivier Kobel, Christoph Noppen, Niko Beerenwinkel, Tanja Stadler |
| EPI_ISL_3345755 | Viollier AG | Department of Biosystems Science and Engineering, ETH Zürich | Christian Beisel, Sarah Nadeau, Chaoran Chen, Louis du Plessis, Ivan Topolsky, Kim Philipp Jablonski, Lara Fuhrmann, Rebecca Denes, Mirjam Feldkamp, Ina Nissen, Natascha Santacroce, Elodie Burcklen, Christiane Beckmann, Maurice Redondo, Olivier Kobel, Christoph Noppen, Niko Beerenwinkel, Tanja Stadler |
| EPI_ISL_3345756 | Fulgent Genetics | Centers for Disease Control and Prevention Division of Viral Diseases, Pathogen Discovery | Dakota Howard, Dhwani Batra, Peter W. Cook, Kara Moser, Adrian Paskey, Jason Caravas, Benjamin Rambo-Martin, Shatavia Morrison, Christopher Gulvick, Scott Sammons, Yvette Unoarumhi, Darlene Wagner, Matthew Schmerer, Harry Gao, Mickey Li, John Gao, Joseph Fierro, Benafsh Sapra, Becky Tsai, Yan Meng, Doreen Ng, James Xie, Clinton R. Paden, Duncan MacCannell |
| EPI_ISL_3345757 | Viollier AG | Department of Biosystems Science and Engineering, ETH Zürich | Chaoran Chen, Sarah Nadeau, Louis du Plessis, Catharine Aquino, Ivan Topolsky, Kim Philipp Jablonski, Lara Fuhrmann, Daniel Ehrsam, Isabel Stürmer, Andreia Cabral de Gouvea, Maria Domenica Moccia, Simon Grüter, Timothy Sykes, Lennart Opitz, Griffin White, Laura Neff, Doris Popovic, Andrea Patrignani, Jay Tracy, Ralph Schlapbach, Christiane Beckmann, Maurice Redondo, Olivier Kobel, Christoph Noppen, Niko Beerenwinkel, Tanja Stadler |
| EPI_ISL_3345758 | Fulgent Genetics | Centers for Disease Control and Prevention Division of Viral Diseases, Pathogen Discovery | Dakota Howard, Dhwani Batra, Peter W. Cook, Kara Moser, Adrian Paskey, Jason Caravas, Benjamin Rambo-Martin, Shatavia Morrison, Christopher Gulvick, Scott Sammons, Yvette Unoarumhi, Darlene Wagner, Matthew Schmerer, Harry Gao, Mickey Li, John Gao, Joseph Fierro, Benafsh Sapra, Becky Tsai, Yan Meng, Doreen Ng, James Xie, Clinton R. Paden, Duncan MacCannell |
| EPI_ISL_3345759 | Viollier AG | Department of Biosystems Science and Engineering, ETH Zürich | Chaoran Chen, Sarah Nadeau, Louis du Plessis, Catharine Aquino, Ivan Topolsky, Kim Philipp Jablonski, Lara Fuhrmann, Daniel Ehrsam, Isabel Stürmer, Andreia Cabral de Gouvea, Maria Domenica Moccia, Simon Grüter, Timothy Sykes, Lennart Opitz, Griffin White, Laura Neff, Doris Popovic, Andrea Patrignani, Jay Tracy, Ralph Schlapbach, Christiane Beckmann, Maurice Redondo, Olivier Kobel, Christoph Noppen, Niko Beerenwinkel, Tanja Stadler |
| EPI_ISL_3345760 | Viollier AG | Department of Biosystems Science and Engineering, ETH Zürich | Christian Beisel, Sarah Nadeau, Chaoran Chen, Louis du Plessis, Ivan Topolsky, Kim Philipp Jablonski, Lara Fuhrmann, Rebecca Denes, Mirjam Feldkamp, Ina Nissen, Natascha Santacroce, Elodie Burcklen, Christiane Beckmann, Maurice Redondo, Olivier Kobel, Christoph Noppen, Niko Beerenwinkel, Tanja Stadler |
| EPI_ISL_3345761 | Fulgent Genetics | Centers for Disease Control and Prevention Division of Viral Diseases, Pathogen Discovery | Dakota Howard, Dhwani Batra, Peter W. Cook, Kara Moser, Adrian Paskey, Jason Caravas, Benjamin Rambo-Martin, Shatavia Morrison, Christopher Gulvick, Scott Sammons, Yvette Unoarumhi, Darlene Wagner, Matthew Schmerer, Harry Gao, Mickey Li, John Gao, Joseph Fierro, Benafsh Sapra, Becky Tsai, Yan Meng, Doreen Ng, James Xie, Clinton R. Paden, Duncan MacCannell |
| EPI_ISL_3345762, EPI_ISL_3345763 | Viollier AG | Department of Biosystems Science and Engineering, ETH Zürich | Chaoran Chen, Sarah Nadeau, Louis du Plessis, Catharine Aquino, Ivan Topolsky, Kim Philipp Jablonski, Lara Fuhrmann, Daniel Ehrsam, Isabel Stürmer, Andreia Cabral de Gouvea, Maria Domenica Moccia, Simon Grüter, Timothy Sykes, Lennart Opitz, Griffin White, Laura Neff, Doris Popovic, Andrea Patrignani, Jay Tracy, Ralph Schlapbach, Christiane Beckmann, Maurice Redondo, Olivier Kobel, Christoph Noppen, Niko Beerenwinkel, Tanja Stadler |
| EPI_ISL_3345764 | Fulgent Genetics | Centers for Disease Control and Prevention Division of Viral Diseases, Pathogen Discovery | Dakota Howard, Dhwani Batra, Peter W. Cook, Kara Moser, Adrian Paskey, Jason Caravas, Benjamin Rambo-Martin, Shatavia Morrison, Christopher Gulvick, Scott Sammons, Yvette Unoarumhi, Darlene Wagner, Matthew Schmerer, Harry Gao, Mickey Li, John Gao, Joseph Fierro, Benafsh Sapra, Becky Tsai, Yan Meng, Doreen Ng, James Xie, Clinton R. Paden, Duncan MacCannell |
| EPI_ISL_3345765 | Viollier AG | Department of Biosystems Science and Engineering, ETH Zürich | Christian Beisel, Sarah Nadeau, Chaoran Chen, Louis du Plessis, Ivan Topolsky, Kim Philipp Jablonski, Lara Fuhrmann, Rebecca Denes, Mirjam Feldkamp, Ina Nissen, Natascha Santacroce, Elodie Burcklen, Christiane Beckmann, Maurice Redondo, Olivier Kobel, Christoph Noppen, Niko Beerenwinkel, Tanja Stadler |
| EPI_ISL_3345766 | Fulgent Genetics | Centers for Disease Control and Prevention Division of Viral Diseases, Pathogen Discovery | Dakota Howard, Dhwani Batra, Peter W. Cook, Kara Moser, Adrian Paskey, Jason Caravas, Benjamin Rambo-Martin, Shatavia Morrison, Christopher Gulvick, Scott Sammons, Yvette Unoarumhi, Darlene Wagner, Matthew Schmerer, Harry Gao, Mickey Li, John Gao, Joseph Fierro, Benafsh Sapra, Becky Tsai, Yan Meng, Doreen Ng, James Xie, Clinton R. Paden, Duncan MacCannell |
| EPI_ISL_3345767 | Viollier AG | Department of Biosystems Science and Engineering, ETH Zürich | Christian Beisel, Sarah Nadeau, Chaoran Chen, Louis du Plessis, Ivan Topolsky, Kim Philipp Jablonski, Lara Fuhrmann, Rebecca Denes, Mirjam Feldkamp, Ina Nissen, Natascha Santacroce, Elodie Burcklen, Christiane Beckmann, Maurice Redondo, Olivier Kobel, Christoph Noppen, Niko Beerenwinkel, Tanja Stadler |
| EPI_ISL_3345768 | Viollier AG | Department of Biosystems Science and Engineering, ETH Zürich | Chaoran Chen, Sarah Nadeau, Louis du Plessis, Catharine Aquino, Ivan Topolsky, Kim Philipp Jablonski, Lara Fuhrmann, Daniel Ehrsam, Isabel Stürmer, Andreia Cabral de Gouvea, Maria Domenica Moccia, Simon Grüter, Timothy Sykes, Lennart Opitz, Griffin White, Laura Neff, Doris Popovic, Andrea Patrignani, Jay Tracy, Ralph Schlapbach, Christiane Beckmann, Maurice Redondo, Olivier Kobel, Christoph Noppen, Niko Beerenwinkel, Tanja Stadler |
| EPI_ISL_3345769 | Fulgent Genetics | Centers for Disease Control and Prevention Division of Viral Diseases, Pathogen Discovery | Dakota Howard, Dhwani Batra, Peter W. Cook, Kara Moser, Adrian Paskey, Jason Caravas, Benjamin Rambo-Martin, Shatavia Morrison, Christopher Gulvick, Scott Sammons, Yvette Unoarumhi, Darlene Wagner, Matthew Schmerer, Harry Gao, Mickey Li, John Gao, Joseph Fierro, Benafsh Sapra, Becky Tsai, Yan Meng, Doreen Ng, James Xie, Clinton R. Paden, Duncan MacCannell |
| EPI_ISL_3345770, EPI_ISL_3345771 | Viollier AG | Department of Biosystems Science and Engineering, ETH Zürich | Chaoran Chen, Sarah Nadeau, Louis du Plessis, Catharine Aquino, Ivan Topolsky, Kim Philipp Jablonski, Lara Fuhrmann, Daniel Ehrsam, Isabel Stürmer, Andreia Cabral de Gouvea, Maria Domenica Moccia, Simon Grüter, Timothy Sykes, Lennart Opitz, Griffin White, Laura Neff, Doris Popovic, Andrea Patrignani, Jay Tracy, Ralph Schlapbach, Christiane Beckmann, Maurice Redondo, Olivier Kobel, Christoph Noppen, Niko Beerenwinkel, Tanja Stadler |
| EPI_ISL_3345772 | Fulgent Genetics | Centers for Disease Control and Prevention Division of Viral Diseases, Pathogen Discovery | Dakota Howard, Dhwani Batra, Peter W. Cook, Kara Moser, Adrian Paskey, Jason Caravas, Benjamin Rambo-Martin, Shatavia Morrison, Christopher Gulvick, Scott Sammons, Yvette Unoarumhi, Darlene Wagner, Matthew Schmerer, Harry Gao, Mickey Li, John Gao, Joseph Fierro, Benafsh Sapra, Becky Tsai, Yan Meng, Doreen Ng, James Xie, Clinton R. Paden, Duncan MacCannell |
| EPI_ISL_3345773 | Viollier AG | Department of Biosystems Science and Engineering, ETH Zürich | Christian Beisel, Sarah Nadeau, Chaoran Chen, Louis du Plessis, Ivan Topolsky, Kim Philipp Jablonski, Lara Fuhrmann, Rebecca Denes, Mirjam Feldkamp, Ina Nissen, Natascha Santacroce, Elodie Burcklen, Christiane Beckmann, Maurice Redondo, Olivier Kobel, Christoph Noppen, Niko Beerenwinkel, Tanja Stadler |
| EPI_ISL_3345774 | Fulgent Genetics | Centers for Disease Control and Prevention Division of Viral Diseases, Pathogen Discovery | Dakota Howard, Dhwani Batra, Peter W. Cook, Kara Moser, Adrian Paskey, Jason Caravas, Benjamin Rambo-Martin, Shatavia Morrison, Christopher Gulvick, Scott Sammons, Yvette Unoarumhi, Darlene Wagner, Matthew Schmerer, Harry Gao, Mickey Li, John Gao, Joseph Fierro, Benafsh Sapra, Becky Tsai, Yan Meng, Doreen Ng, James Xie, Clinton R. Paden, Duncan MacCannell |
| EPI_ISL_3345775, EPI_ISL_3345776 | Viollier AG | Department of Biosystems Science and Engineering, ETH Zürich | Chaoran Chen, Sarah Nadeau, Louis du Plessis, Catharine Aquino, Ivan Topolsky, Kim Philipp Jablonski, Lara Fuhrmann, Daniel Ehrsam, Isabel Stürmer, Andreia Cabral de Gouvea, Maria Domenica Moccia, Simon Grüter, Timothy Sykes, Lennart Opitz, Griffin White, Laura Neff, Doris Popovic, Andrea Patrignani, Jay Tracy, Ralph Schlapbach, Christiane Beckmann, Maurice Redondo, Olivier Kobel, Christoph Noppen, Niko Beerenwinkel, Tanja Stadler |
| EPI_ISL_3345777 | Fulgent Genetics | Centers for Disease Control and Prevention Division of Viral Diseases, Pathogen Discovery | Dakota Howard, Dhwani Batra, Peter W. Cook, Kara Moser, Adrian Paskey, Jason Caravas, Benjamin Rambo-Martin, Shatavia Morrison, Christopher Gulvick, Scott Sammons, Yvette Unoarumhi, Darlene Wagner, Matthew Schmerer, Harry Gao, Mickey Li, John Gao, Joseph Fierro, Benafsh Sapra, Becky Tsai, Yan Meng, Doreen Ng, James Xie, Clinton R. Paden, Duncan MacCannell |
| EPI_ISL_3345778, EPI_ISL_3345779 | Viollier AG | Department of Biosystems Science and Engineering, ETH Zürich | Chaoran Chen, Sarah Nadeau, Louis du Plessis, Catharine Aquino, Ivan Topolsky, Kim Philipp Jablonski, Lara Fuhrmann, Daniel Ehrsam, Isabel Stürmer, Andreia Cabral de Gouvea, Maria Domenica Moccia, Simon Grüter, Timothy Sykes, Lennart Opitz, Griffin White, Laura Neff, Doris Popovic, Andrea Patrignani, Jay Tracy, Ralph Schlapbach, Christiane Beckmann, Maurice Redondo, Olivier Kobel, Christoph Noppen, Niko Beerenwinkel, Tanja Stadler |
| EPI_ISL_3345780 | Fulgent Genetics | Centers for Disease Control and Prevention Division of Viral Diseases, Pathogen Discovery | Dakota Howard, Dhwani Batra, Peter W. Cook, Kara Moser, Adrian Paskey, Jason Caravas, Benjamin Rambo-Martin, Shatavia Morrison, Christopher Gulvick, Scott Sammons, Yvette Unoarumhi, Darlene Wagner, Matthew Schmerer, Harry Gao, Mickey Li, John Gao, Joseph Fierro, Benafsh Sapra, Becky Tsai, Yan Meng, Doreen Ng, James Xie, Clinton R. Paden, Duncan MacCannell |
| EPI_ISL_3345782 | Viollier AG | Department of Biosystems Science and Engineering, ETH Zürich | Chaoran Chen, Sarah Nadeau, Louis du Plessis, Catharine Aquino, Ivan Topolsky, Kim Philipp Jablonski, Lara Fuhrmann, Daniel Ehrsam, Isabel Stürmer, Andreia Cabral de Gouvea, Maria Domenica Moccia, Simon Grüter, Timothy Sykes, Lennart Opitz, Griffin White, Laura Neff, Doris Popovic, Andrea Patrignani, Jay Tracy, Ralph Schlapbach, Christiane Beckmann, Maurice Redondo, Olivier Kobel, Christoph Noppen, Niko Beerenwinkel, Tanja Stadler |
| EPI_ISL_3345783, EPI_ISL_3345785 | Fulgent Genetics | Centers for Disease Control and Prevention Division of Viral Diseases, Pathogen Discovery | Dakota Howard, Dhwani Batra, Peter W. Cook, Kara Moser, Adrian Paskey, Jason Caravas, Benjamin Rambo-Martin, Shatavia Morrison, Christopher Gulvick, Scott Sammons, Yvette Unoarumhi, Darlene Wagner, Matthew Schmerer, Harry Gao, Mickey Li, John Gao, Joseph Fierro, Benafsh Sapra, Becky Tsai, Yan Meng, Doreen Ng, James Xie, Clinton R. Paden, Duncan MacCannell |
| EPI_ISL_3345786 | Viollier AG | Department of Biosystems Science and Engineering, ETH Zürich | Chaoran Chen, Sarah Nadeau, Louis du Plessis, Catharine Aquino, Ivan Topolsky, Kim Philipp Jablonski, Lara Fuhrmann, Daniel Ehrsam, Isabel Stürmer, Andreia Cabral de Gouvea, Maria Domenica Moccia, Simon Grüter, Timothy Sykes, Lennart Opitz, Griffin White, Laura Neff, Doris Popovic, Andrea Patrignani, Jay Tracy, Ralph Schlapbach, Christiane Beckmann, Maurice Redondo, Olivier Kobel, Christoph Noppen, Niko Beerenwinkel, Tanja Stadler |
| EPI_ISL_3345787 | Viollier AG | Department of Biosystems Science and Engineering, ETH Zürich | Christian Beisel, Sarah Nadeau, Chaoran Chen, Louis du Plessis, Ivan Topolsky, Kim Philipp Jablonski, Lara Fuhrmann, Rebecca Denes, Mirjam Feldkamp, Ina Nissen, Natascha Santacroce, Elodie Burcklen, Christiane Beckmann, Maurice Redondo, Olivier Kobel, Christoph Noppen, Niko Beerenwinkel, Tanja Stadler |
| EPI_ISL_3345788 | Fulgent Genetics | Centers for Disease Control and Prevention Division of Viral Diseases, Pathogen Discovery | Dakota Howard, Dhwani Batra, Peter W. Cook, Kara Moser, Adrian Paskey, Jason Caravas, Benjamin Rambo-Martin, Shatavia Morrison, Christopher Gulvick, Scott Sammons, Yvette Unoarumhi, Darlene Wagner, Matthew Schmerer, Harry Gao, Mickey Li, John Gao, Joseph Fierro, Benafsh Sapra, Becky Tsai, Yan Meng, Doreen Ng, James Xie, Clinton R. Paden, Duncan MacCannell |
| EPI_ISL_3345789, EPI_ISL_3345790 | Viollier AG | Department of Biosystems Science and Engineering, ETH Zürich | Chaoran Chen, Sarah Nadeau, Louis du Plessis, Catharine Aquino, Ivan Topolsky, Kim Philipp Jablonski, Lara Fuhrmann, Daniel Ehrsam, Isabel Stürmer, Andreia Cabral de Gouvea, Maria Domenica Moccia, Simon Grüter, Timothy Sykes, Lennart Opitz, Griffin White, Laura Neff, Doris Popovic, Andrea Patrignani, Jay Tracy, Ralph Schlapbach, Christiane Beckmann, Maurice Redondo, Olivier Kobel, Christoph Noppen, Niko Beerenwinkel, Tanja Stadler |
| EPI_ISL_3345791 | Fulgent Genetics | Centers for Disease Control and Prevention Division of Viral Diseases, Pathogen Discovery | Dakota Howard, Dhwani Batra, Peter W. Cook, Kara Moser, Adrian Paskey, Jason Caravas, Benjamin Rambo-Martin, Shatavia Morrison, Christopher Gulvick, Scott Sammons, Yvette Unoarumhi, Darlene Wagner, Matthew Schmerer, Harry Gao, Mickey Li, John Gao, Joseph Fierro, Benafsh Sapra, Becky Tsai, Yan Meng, Doreen Ng, James Xie, Clinton R. Paden, Duncan MacCannell |
| EPI_ISL_3345792, EPI_ISL_3345793 | Viollier AG | Department of Biosystems Science and Engineering, ETH Zürich | Chaoran Chen, Sarah Nadeau, Louis du Plessis, Catharine Aquino, Ivan Topolsky, Kim Philipp Jablonski, Lara Fuhrmann, Daniel Ehrsam, Isabel Stürmer, Andreia Cabral de Gouvea, Maria Domenica Moccia, Simon Grüter, Timothy Sykes, Lennart Opitz, Griffin White, Laura Neff, Doris Popovic, Andrea Patrignani, Jay Tracy, Ralph Schlapbach, Christiane Beckmann, Maurice Redondo, Olivier Kobel, Christoph Noppen, Niko Beerenwinkel, Tanja Stadler |
| EPI_ISL_3345794 | Fulgent Genetics | Centers for Disease Control and Prevention Division of Viral Diseases, Pathogen Discovery | Dakota Howard, Dhwani Batra, Peter W. Cook, Kara Moser, Adrian Paskey, Jason Caravas, Benjamin Rambo-Martin, Shatavia Morrison, Christopher Gulvick, Scott Sammons, Yvette Unoarumhi, Darlene Wagner, Matthew Schmerer, Harry Gao, Mickey Li, John Gao, Joseph Fierro, Benafsh Sapra, Becky Tsai, Yan Meng, Doreen Ng, James Xie, Clinton R. Paden, Duncan MacCannell |
| EPI_ISL_3345795 | Viollier AG | Department of Biosystems Science and Engineering, ETH Zürich | Chaoran Chen, Sarah Nadeau, Louis du Plessis, Catharine Aquino, Ivan Topolsky, Kim Philipp Jablonski, Lara Fuhrmann, Daniel Ehrsam, Isabel Stürmer, Andreia Cabral de Gouvea, Maria Domenica Moccia, Simon Grüter, Timothy Sykes, Lennart Opitz, Griffin White, Laura Neff, Doris Popovic, Andrea Patrignani, Jay Tracy, Ralph Schlapbach, Christiane Beckmann, Maurice Redondo, Olivier Kobel, Christoph Noppen, Niko Beerenwinkel, Tanja Stadler |
| EPI_ISL_3345796 | Fulgent Genetics | Centers for Disease Control and Prevention Division of Viral Diseases, Pathogen Discovery | Dakota Howard, Dhwani Batra, Peter W. Cook, Kara Moser, Adrian Paskey, Jason Caravas, Benjamin Rambo-Martin, Shatavia Morrison, Christopher Gulvick, Scott Sammons, Yvette Unoarumhi, Darlene Wagner, Matthew Schmerer, Harry Gao, Mickey Li, John Gao, Joseph Fierro, Benafsh Sapra, Becky Tsai, Yan Meng, Doreen Ng, James Xie, Clinton R. Paden, Duncan MacCannell |

| | | | |
|---|---|---|---|
| EPI_ISL_3345797 | Viollier AG | Department of Biosystems Science and Engineering, ETH Zürich | Christian Beisel, Sarah Nadeau, Chaoran Chen, Louis du Plessis, Ivan Topolsky, Kim Philipp Jablonski, Lara Fuhrmann, Rebecca Denes, Mirjam Feldkamp, Ina Nissen, Natascha Santacroce, Elodie Burcklen, Christiane Beckmann, Maurice Redondo, Olivier Kobel, Christoph Noppen, Niko Beerenwinkel, Tanja Stadler |
| EPI_ISL_3345798 | Viollier AG | Department of Biosystems Science and Engineering, ETH Zürich | Chaoran Chen, Sarah Nadeau, Louis du Plessis, Catharine Aquino, Ivan Topolsky, Kim Philipp Jablonski, Lara Fuhrmann, Daniel Ehrsam, Isabel Stürmer, Andreia Cabral de Gouvea, Maria Domenica Moccia, Simon Grüter, Timothy Sykes, Lennart Opitz, Griffin White, Laura Neff, Doris Popovic, Andrea Patrignani, Jay Tracy, Ralph Schlapbach, Christiane Beckmann, Maurice Redondo, Olivier Kobel, Christoph Noppen, Niko Beerenwinkel, Tanja Stadler |
| EPI_ISL_3345799 | Fulgent Genetics | Centers for Disease Control and Prevention Division of Viral Diseases, Pathogen Discovery | Dakota Howard, Dhwani Batra, Peter W. Cook, Kara Moser, Adrian Paskey, Jason Caravas, Benjamin Rambo-Martin, Shatavia Morrison, Christopher Gulvick, Scott Sammons, Yvette Unoarumhi, Darlene Wagner, Matthew Schmerer, Harry Gao, Mickey Li, John Gao, Joseph Fierro, Benafsh Sapra, Becky Tsai, Yan Meng, Doreen Ng, James Xie, Clinton R. Paden, Duncan MacCannell |
| EPI_ISL_3345800, EPI_ISL_3345801 | Viollier AG | Department of Biosystems Science and Engineering, ETH Zürich | Christian Beisel, Sarah Nadeau, Chaoran Chen, Louis du Plessis, Ivan Topolsky, Kim Philipp Jablonski, Lara Fuhrmann, Rebecca Denes, Mirjam Feldkamp, Ina Nissen, Natascha Santacroce, Elodie Burcklen, Christiane Beckmann, Maurice Redondo, Olivier Kobel, Christoph Noppen, Niko Beerenwinkel, Tanja Stadler |
| EPI_ISL_3345802 | Fulgent Genetics | Centers for Disease Control and Prevention Division of Viral Diseases, Pathogen Discovery | Dakota Howard, Dhwani Batra, Peter W. Cook, Kara Moser, Adrian Paskey, Jason Caravas, Benjamin Rambo-Martin, Shatavia Morrison, Christopher Gulvick, Scott Sammons, Yvette Unoarumhi, Darlene Wagner, Matthew Schmerer, Harry Gao, Mickey Li, John Gao, Joseph Fierro, Benafsh Sapra, Becky Tsai, Yan Meng, Doreen Ng, James Xie, Clinton R. Paden, Duncan MacCannell |
| EPI_ISL_3345803 | Viollier AG | Department of Biosystems Science and Engineering, ETH Zürich | Chaoran Chen, Sarah Nadeau, Louis du Plessis, Catharine Aquino, Ivan Topolsky, Kim Philipp Jablonski, Lara Fuhrmann, Daniel Ehrsam, Isabel Stürmer, Andreia Cabral de Gouvea, Maria Domenica Moccia, Simon Grüter, Timothy Sykes, Lennart Opitz, Griffin White, Laura Neff, Doris Popovic, Andrea Patrignani, Jay Tracy, Ralph Schlapbach, Christiane Beckmann, Maurice Redondo, Olivier Kobel, Christoph Noppen, Niko Beerenwinkel, Tanja Stadler |
| EPI_ISL_3345804 | Fulgent Genetics | Centers for Disease Control and Prevention Division of Viral Diseases, Pathogen Discovery | Dakota Howard, Dhwani Batra, Peter W. Cook, Kara Moser, Adrian Paskey, Jason Caravas, Benjamin Rambo-Martin, Shatavia Morrison, Christopher Gulvick, Scott Sammons, Yvette Unoarumhi, Darlene Wagner, Matthew Schmerer, Harry Gao, Mickey Li, John Gao, Joseph Fierro, Benafsh Sapra, Becky Tsai, Yan Meng, Doreen Ng, James Xie, Clinton R. Paden, Duncan MacCannell |
| EPI_ISL_3345805, EPI_ISL_3345806 | Viollier AG | Department of Biosystems Science and Engineering, ETH Zürich | Chaoran Chen, Sarah Nadeau, Louis du Plessis, Catharine Aquino, Ivan Topolsky, Kim Philipp Jablonski, Lara Fuhrmann, Daniel Ehrsam, Isabel Stürmer, Andreia Cabral de Gouvea, Maria Domenica Moccia, Simon Grüter, Timothy Sykes, Lennart Opitz, Griffin White, Laura Neff, Doris Popovic, Andrea Patrignani, Jay Tracy, Ralph Schlapbach, Christiane Beckmann, Maurice Redondo, Olivier Kobel, Christoph Noppen, Niko Beerenwinkel, Tanja Stadler |
| EPI_ISL_3345807 | Fulgent Genetics | Centers for Disease Control and Prevention Division of Viral Diseases, Pathogen Discovery | Dakota Howard, Dhwani Batra, Peter W. Cook, Kara Moser, Adrian Paskey, Jason Caravas, Benjamin Rambo-Martin, Shatavia Morrison, Christopher Gulvick, Scott Sammons, Yvette Unoarumhi, Darlene Wagner, Matthew Schmerer, Harry Gao, Mickey Li, John Gao, Joseph Fierro, Benafsh Sapra, Becky Tsai, Yan Meng, Doreen Ng, James Xie, Clinton R. Paden, Duncan MacCannell |
| EPI_ISL_3345808, EPI_ISL_3345809 | Viollier AG | Department of Biosystems Science and Engineering, ETH Zürich | Chaoran Chen, Sarah Nadeau, Louis du Plessis, Catharine Aquino, Ivan Topolsky, Kim Philipp Jablonski, Lara Fuhrmann, Daniel Ehrsam, Isabel Stürmer, Andreia Cabral de Gouvea, Maria Domenica Moccia, Simon Grüter, Timothy Sykes, Lennart Opitz, Griffin White, Laura Neff, Doris Popovic, Andrea Patrignani, Jay Tracy, Ralph Schlapbach, Christiane Beckmann, Maurice Redondo, Olivier Kobel, Christoph Noppen, Niko Beerenwinkel, Tanja Stadler |
| EPI_ISL_3345810 | Fulgent Genetics | Centers for Disease Control and Prevention Division of Viral Diseases, Pathogen Discovery | Dakota Howard, Dhwani Batra, Peter W. Cook, Kara Moser, Adrian Paskey, Jason Caravas, Benjamin Rambo-Martin, Shatavia Morrison, Christopher Gulvick, Scott Sammons, Yvette Unoarumhi, Darlene Wagner, Matthew Schmerer, Harry Gao, Mickey Li, John Gao, Joseph Fierro, Benafsh Sapra, Becky Tsai, Yan Meng, Doreen Ng, James Xie, Clinton R. Paden, Duncan MacCannell |
| EPI_ISL_3345811 | Viollier AG | Department of Biosystems Science and Engineering, ETH Zürich | Christian Beisel, Sarah Nadeau, Chaoran Chen, Louis du Plessis, Ivan Topolsky, Kim Philipp Jablonski, Lara Fuhrmann, Rebecca Denes, Mirjam Feldkamp, Ina Nissen, Natascha Santacroce, Elodie Burcklen, Christiane Beckmann, Maurice Redondo, Olivier Kobel, Christoph Noppen, Niko Beerenwinkel, Tanja Stadler |
| EPI_ISL_3345812 | Fulgent Genetics | Centers for Disease Control and Prevention Division of Viral Diseases, Pathogen Discovery | Dakota Howard, Dhwani Batra, Peter W. Cook, Kara Moser, Adrian Paskey, Jason Caravas, Benjamin Rambo-Martin, Shatavia Morrison, Christopher Gulvick, Scott Sammons, Yvette Unoarumhi, Darlene Wagner, Matthew Schmerer, Harry Gao, Mickey Li, John Gao, Joseph Fierro, Benafsh Sapra, Becky Tsai, Yan Meng, Doreen Ng, James Xie, Clinton R. Paden, Duncan MacCannell |
| EPI_ISL_3345813, EPI_ISL_3345814 | Viollier AG | Department of Biosystems Science and Engineering, ETH Zürich | Chaoran Chen, Sarah Nadeau, Louis du Plessis, Catharine Aquino, Ivan Topolsky, Kim Philipp Jablonski, Lara Fuhrmann, Daniel Ehrsam, Isabel Stürmer, Andreia Cabral de Gouvea, Maria Domenica Moccia, Simon Grüter, Timothy Sykes, Lennart Opitz, Griffin White, Laura Neff, Doris Popovic, Andrea Patrignani, Jay Tracy, Ralph Schlapbach, Christiane Beckmann, Maurice Redondo, Olivier Kobel, Christoph Noppen, Niko Beerenwinkel, Tanja Stadler |
| EPI_ISL_3345815 | Fulgent Genetics | Centers for Disease Control and Prevention Division of Viral Diseases, Pathogen Discovery | Dakota Howard, Dhwani Batra, Peter W. Cook, Kara Moser, Adrian Paskey, Jason Caravas, Benjamin Rambo-Martin, Shatavia Morrison, Christopher Gulvick, Scott Sammons, Yvette Unoarumhi, Darlene Wagner, Matthew Schmerer, Harry Gao, Mickey Li, John Gao, Joseph Fierro, Benafsh Sapra, Becky Tsai, Yan Meng, Doreen Ng, James Xie, Clinton R. Paden, Duncan MacCannell |
| EPI_ISL_3345816 | Viollier AG | Department of Biosystems Science and Engineering, ETH Zürich | Christian Beisel, Sarah Nadeau, Chaoran Chen, Louis du Plessis, Ivan Topolsky, Kim Philipp Jablonski, Lara Fuhrmann, Rebecca Denes, Mirjam Feldkamp, Ina Nissen, Natascha Santacroce, Elodie Burcklen, Christiane Beckmann, Maurice Redondo, Olivier Kobel, Christoph Noppen, Niko Beerenwinkel, Tanja Stadler |
| EPI_ISL_3345817 | Viollier AG | Department of Biosystems Science and Engineering, ETH Zürich | Chaoran Chen, Sarah Nadeau, Louis du Plessis, Catharine Aquino, Ivan Topolsky, Kim Philipp Jablonski, Lara Fuhrmann, Daniel Ehrsam, Isabel Stürmer, Andreia Cabral de Gouvea, Maria Domenica Moccia, Simon Grüter, Timothy Sykes, Lennart Opitz, Griffin White, Laura Neff, Doris Popovic, Andrea Patrignani, Jay Tracy, Ralph Schlapbach, Christiane Beckmann, Maurice Redondo, Olivier Kobel, Christoph Noppen, Niko Beerenwinkel, Tanja Stadler |
| EPI_ISL_3345818 | Fulgent Genetics | Centers for Disease Control and Prevention Division of Viral Diseases, Pathogen Discovery | Dakota Howard, Dhwani Batra, Peter W. Cook, Kara Moser, Adrian Paskey, Jason Caravas, Benjamin Rambo-Martin, Shatavia Morrison, Christopher Gulvick, Scott Sammons, Yvette Unoarumhi, Darlene Wagner, Matthew Schmerer, Harry Gao, Mickey Li, John Gao, Joseph Fierro, Benafsh Sapra, Becky Tsai, Yan Meng, Doreen Ng, James Xie, Clinton R. Paden, Duncan MacCannell |
| EPI_ISL_3345819 | Viollier AG | Department of Biosystems Science and Engineering, ETH Zürich | Chaoran Chen, Sarah Nadeau, Louis du Plessis, Catharine Aquino, Ivan Topolsky, Kim Philipp Jablonski, Lara Fuhrmann, Daniel Ehrsam, Isabel Stürmer, Andreia Cabral de Gouvea, Maria Domenica Moccia, Simon Grüter, Timothy Sykes, Lennart Opitz, Griffin White, Laura Neff, Doris Popovic, Andrea Patrignani, Jay Tracy, Ralph Schlapbach, Christiane Beckmann, Maurice Redondo, Olivier Kobel, Christoph Noppen, Niko Beerenwinkel, Tanja Stadler |
| EPI_ISL_3345820 | Fulgent Genetics | Centers for Disease Control and Prevention Division of Viral Diseases, Pathogen Discovery | Dakota Howard, Dhwani Batra, Peter W. Cook, Kara Moser, Adrian Paskey, Jason Caravas, Benjamin Rambo-Martin, Shatavia Morrison, Christopher Gulvick, Scott Sammons, Yvette Unoarumhi, Darlene Wagner, Matthew Schmerer, Harry Gao, Mickey Li, John Gao, Joseph Fierro, Benafsh Sapra, Becky Tsai, Yan Meng, Doreen Ng, James Xie, Clinton R. Paden, Duncan MacCannell |
| EPI_ISL_3345821, EPI_ISL_3345822 | Viollier AG | Department of Biosystems Science and Engineering, ETH Zürich | Chaoran Chen, Sarah Nadeau, Louis du Plessis, Catharine Aquino, Ivan Topolsky, Kim Philipp Jablonski, Lara Fuhrmann, Daniel Ehrsam, Isabel Stürmer, Andreia Cabral de Gouvea, Maria Domenica Moccia, Simon Grüter, Timothy Sykes, Lennart Opitz, Griffin White, Laura Neff, Doris Popovic, Andrea Patrignani, Jay Tracy, Ralph Schlapbach, Christiane Beckmann, Maurice Redondo, Olivier Kobel, Christoph Noppen, Niko Beerenwinkel, Tanja Stadler |
| EPI_ISL_3345823 | Fulgent Genetics | Centers for Disease Control and Prevention Division of Viral Diseases, Pathogen Discovery | Dakota Howard, Dhwani Batra, Peter W. Cook, Kara Moser, Adrian Paskey, Jason Caravas, Benjamin Rambo-Martin, Shatavia Morrison, Christopher Gulvick, Scott Sammons, Yvette Unoarumhi, Darlene Wagner, Matthew Schmerer, Harry Gao, Mickey Li, John Gao, Joseph Fierro, Benafsh Sapra, Becky Tsai, Yan Meng, Doreen Ng, James Xie, Clinton R. Paden, Duncan MacCannell |
| EPI_ISL_3345824 | Viollier AG | Department of Biosystems Science and Engineering, ETH Zürich | Christian Beisel, Sarah Nadeau, Chaoran Chen, Louis du Plessis, Ivan Topolsky, Kim Philipp Jablonski, Lara Fuhrmann, Rebecca Denes, Mirjam Feldkamp, Ina Nissen, Natascha Santacroce, Elodie Burcklen, Christiane Beckmann, Maurice Redondo, Olivier Kobel, Christoph Noppen, Niko Beerenwinkel, Tanja Stadler |
| EPI_ISL_3345825 | Viollier AG | Department of Biosystems Science and Engineering, ETH Zürich | Chaoran Chen, Sarah Nadeau, Louis du Plessis, Catharine Aquino, Ivan Topolsky, Kim Philipp Jablonski, Lara Fuhrmann, Daniel Ehrsam, Isabel Stürmer, Andreia Cabral de Gouvea, Maria Domenica Moccia, Simon Grüter, Timothy Sykes, Lennart Opitz, Griffin White, Laura Neff, Doris Popovic, Andrea Patrignani, Jay Tracy, Ralph Schlapbach, Christiane Beckmann, Maurice Redondo, Olivier Kobel, Christoph Noppen, Niko Beerenwinkel, Tanja Stadler |
| EPI_ISL_3345826 | Fulgent Genetics | Centers for Disease Control and Prevention Division of Viral Diseases, Pathogen Discovery | Dakota Howard, Dhwani Batra, Peter W. Cook, Kara Moser, Adrian Paskey, Jason Caravas, Benjamin Rambo-Martin, Shatavia Morrison, Christopher Gulvick, Scott Sammons, Yvette Unoarumhi, Darlene Wagner, Matthew Schmerer, Harry Gao, Mickey Li, John Gao, Joseph Fierro, Benafsh Sapra, Becky Tsai, Yan Meng, Doreen Ng, James Xie, Clinton R. Paden, Duncan MacCannell |
| EPI_ISL_3345827, EPI_ISL_3345828 | Viollier AG | Department of Biosystems Science and Engineering, ETH Zürich | Chaoran Chen, Sarah Nadeau, Louis du Plessis, Catharine Aquino, Ivan Topolsky, Kim Philipp Jablonski, Lara Fuhrmann, Daniel Ehrsam, Isabel Stürmer, Andreia Cabral de Gouvea, Maria Domenica Moccia, Simon Grüter, Timothy Sykes, Lennart Opitz, Griffin White, Laura Neff, Doris Popovic, Andrea Patrignani, Jay Tracy, Ralph Schlapbach, Christiane Beckmann, Maurice Redondo, Olivier Kobel, Christoph Noppen, Niko Beerenwinkel, Tanja Stadler |
| EPI_ISL_3345829 | Fulgent Genetics | Centers for Disease Control and Prevention Division of Viral Diseases, Pathogen Discovery | Dakota Howard, Dhwani Batra, Peter W. Cook, Kara Moser, Adrian Paskey, Jason Caravas, Benjamin Rambo-Martin, Shatavia Morrison, Christopher Gulvick, Scott Sammons, Yvette Unoarumhi, Darlene Wagner, Matthew Schmerer, Harry Gao, Mickey Li, John Gao, Joseph Fierro, Benafsh Sapra, Becky Tsai, Yan Meng, Doreen Ng, James Xie, Clinton R. Paden, Duncan MacCannell |
| EPI_ISL_3345830 | Viollier AG | Department of Biosystems Science and Engineering, ETH Zürich | Christian Beisel, Sarah Nadeau, Chaoran Chen, Louis du Plessis, Ivan Topolsky, Kim Philipp Jablonski, Lara Fuhrmann, Rebecca Denes, Mirjam Feldkamp, Ina Nissen, Natascha Santacroce, Elodie Burcklen, Christiane Beckmann, Maurice Redondo, Olivier Kobel, Christoph Noppen, Niko Beerenwinkel, Tanja Stadler |
| EPI_ISL_3345831 | Viollier AG | Department of Biosystems Science and Engineering, ETH Zürich | Chaoran Chen, Sarah Nadeau, Louis du Plessis, Catharine Aquino, Ivan Topolsky, Kim Philipp Jablonski, Lara Fuhrmann, Daniel Ehrsam, Isabel Stürmer, Andreia Cabral de Gouvea, Maria Domenica Moccia, Simon Grüter, Timothy Sykes, Lennart Opitz, Griffin White, Laura Neff, Doris Popovic, Andrea Patrignani, Jay Tracy, Ralph Schlapbach, Christiane Beckmann, Maurice Redondo, Olivier Kobel, Christoph Noppen, Niko Beerenwinkel, Tanja Stadler |
| EPI_ISL_3345832 | Fulgent Genetics | Centers for Disease Control and Prevention Division of Viral Diseases, Pathogen Discovery | Dakota Howard, Dhwani Batra, Peter W. Cook, Kara Moser, Adrian Paskey, Jason Caravas, Benjamin Rambo-Martin, Shatavia Morrison, Christopher Gulvick, Scott Sammons, Yvette Unoarumhi, Darlene Wagner, Matthew Schmerer, Harry Gao, Mickey Li, John Gao, Joseph Fierro, Benafsh Sapra, Becky Tsai, Yan Meng, Doreen Ng, James Xie, Clinton R. Paden, Duncan MacCannell |
| EPI_ISL_3345833, EPI_ISL_3345834 | Viollier AG | Department of Biosystems Science and Engineering, ETH Zürich | Chaoran Chen, Sarah Nadeau, Louis du Plessis, Catharine Aquino, Ivan Topolsky, Kim Philipp Jablonski, Lara Fuhrmann, Daniel Ehrsam, Isabel Stürmer, Andreia Cabral de Gouvea, Maria Domenica Moccia, Simon Grüter, Timothy Sykes, Lennart Opitz, Griffin White, Laura Neff, Doris Popovic, Andrea Patrignani, Jay Tracy, Ralph Schlapbach, Christiane Beckmann, Maurice Redondo, Olivier Kobel, Christoph Noppen, Niko Beerenwinkel, Tanja Stadler |
| EPI_ISL_3345835 | Fulgent Genetics | Centers for Disease Control and Prevention Division of Viral Diseases, Pathogen Discovery | Dakota Howard, Dhwani Batra, Peter W. Cook, Kara Moser, Adrian Paskey, Jason Caravas, Benjamin Rambo-Martin, Shatavia Morrison, Christopher Gulvick, Scott Sammons, Yvette Unoarumhi, Darlene Wagner, Matthew Schmerer, Harry Gao, Mickey Li, John Gao, Joseph Fierro, Benafsh Sapra, Becky Tsai, Yan Meng, Doreen Ng, James Xie, Clinton R. Paden, Duncan MacCannell |
| EPI_ISL_3345836 | Viollier AG | Department of Biosystems Science and Engineering, ETH Zürich | Chaoran Chen, Sarah Nadeau, Louis du Plessis, Catharine Aquino, Ivan Topolsky, Kim Philipp Jablonski, Lara Fuhrmann, Daniel Ehrsam, Isabel Stürmer, Andreia Cabral de Gouvea, Maria Domenica Moccia, Simon Grüter, Timothy Sykes, Lennart Opitz, Griffin White, Laura Neff, Doris Popovic, Andrea Patrignani, Jay Tracy, Ralph Schlapbach, Christiane Beckmann, Maurice Redondo, Olivier Kobel, Christoph Noppen, Niko Beerenwinkel, Tanja Stadler |
| EPI_ISL_3345837 | Fulgent Genetics | Centers for Disease Control and Prevention Division of Viral Diseases, Pathogen Discovery | Dakota Howard, Dhwani Batra, Peter W. Cook, Kara Moser, Adrian Paskey, Jason Caravas, Benjamin Rambo-Martin, Shatavia Morrison, Christopher Gulvick, Scott Sammons, Yvette Unoarumhi, Darlene Wagner, Matthew Schmerer, Harry Gao, Mickey Li, John Gao, Joseph Fierro, Benafsh Sapra, Becky Tsai, Yan Meng, Doreen Ng, James Xie, Clinton R. Paden, Duncan MacCannell |
| EPI_ISL_3345838, EPI_ISL_3345839 | Viollier AG | Department of Biosystems Science and Engineering, ETH Zürich | Chaoran Chen, Sarah Nadeau, Louis du Plessis, Catharine Aquino, Ivan Topolsky, Kim Philipp Jablonski, Lara Fuhrmann, Daniel Ehrsam, Isabel Stürmer, Andreia Cabral de Gouvea, Maria Domenica Moccia, Simon Grüter, Timothy Sykes, Lennart Opitz, Griffin White, Laura Neff, Doris Popovic, Andrea Patrignani, Jay Tracy, Ralph Schlapbach, Christiane Beckmann, Maurice Redondo, Olivier Kobel, Christoph Noppen, Niko Beerenwinkel, Tanja Stadler |
| EPI_ISL_3345840 | Fulgent Genetics | Centers for Disease Control and Prevention Division of Viral Diseases, Pathogen Discovery | Dakota Howard, Dhwani Batra, Peter W. Cook, Kara Moser, Adrian Paskey, Jason Caravas, Benjamin Rambo-Martin, Shatavia Morrison, Christopher Gulvick, Scott Sammons, Yvette Unoarumhi, Darlene Wagner, Matthew Schmerer, Harry Gao, Mickey Li, John Gao, Joseph Fierro, Benafsh Sapra, Becky Tsai, Yan Meng, Doreen Ng, James Xie, Clinton R. Paden, Duncan MacCannell |
| EPI_ISL_3345841 | Viollier AG | Department of Biosystems Science and Engineering, ETH Zürich | Chaoran Chen, Sarah Nadeau, Louis du Plessis, Catharine Aquino, Ivan Topolsky, Kim Philipp Jablonski, Lara Fuhrmann, Daniel Ehrsam, Isabel Stürmer, Andreia Cabral de Gouvea, Maria Domenica Moccia, Simon Grüter, Timothy Sykes, Lennart Opitz, Griffin White, Laura Neff, Doris Popovic, Andrea Patrignani, Jay Tracy, Ralph Schlapbach, Christiane Beckmann, Maurice Redondo, Olivier Kobel, Christoph Noppen, Niko Beerenwinkel, Tanja Stadler |
| EPI_ISL_3345842 | Viollier AG | Department of Biosystems Science and Engineering, ETH Zürich | Christian Beisel, Sarah Nadeau, Chaoran Chen, Louis du Plessis, Ivan Topolsky, Kim Philipp Jablonski, Lara Fuhrmann, Rebecca Denes, Mirjam Feldkamp, Ina Nissen, Natascha Santacroce, Elodie Burcklen, Christiane Beckmann, Maurice Redondo, Olivier Kobel, Christoph Noppen, Niko Beerenwinkel, Tanja Stadler |
| EPI_ISL_3345843 | Fulgent Genetics | Centers for Disease Control and Prevention Division of Viral Diseases, Pathogen Discovery | Dakota Howard, Dhwani Batra, Peter W. Cook, Kara Moser, Adrian Paskey, Jason Caravas, Benjamin Rambo-Martin, Shatavia Morrison, Christopher Gulvick, Scott Sammons, Yvette Unoarumhi, Darlene Wagner, Matthew Schmerer, Harry Gao, Mickey Li, John Gao, Joseph Fierro, Benafsh Sapra, Becky Tsai, Yan Meng, Doreen Ng, James Xie, Clinton R. Paden, Duncan MacCannell |
| EPI_ISL_3345844 | Viollier AG | Department of Biosystems Science and Engineering, ETH Zürich | Chaoran Chen, Sarah Nadeau, Louis du Plessis, Catharine Aquino, Ivan Topolsky, Kim Philipp Jablonski, Lara Fuhrmann, Daniel Ehrsam, Isabel Stürmer, Andreia Cabral de Gouvea, Maria Domenica Moccia, Simon Grüter, Timothy Sykes, Lennart Opitz, Griffin White, Laura Neff, Doris Popovic, Andrea Patrignani, Jay Tracy, Ralph Schlapbach, Christiane Beckmann, Maurice Redondo, Olivier Kobel, Christoph Noppen, Niko Beerenwinkel, Tanja Stadler |
| EPI_ISL_3345845 | Fulgent Genetics | Centers for Disease Control and Prevention Division of Viral Diseases, Pathogen Discovery | Dakota Howard, Dhwani Batra, Peter W. Cook, Kara Moser, Adrian Paskey, Jason Caravas, Benjamin Rambo-Martin, Shatavia Morrison, Christopher Gulvick, Scott Sammons, Yvette Unoarumhi, Darlene Wagner, Matthew Schmerer, Harry Gao, Mickey Li, John Gao, Joseph Fierro, Benafsh Sapra, Becky Tsai, Yan Meng, Doreen Ng, James Xie, Clinton R. Paden, Duncan MacCannell |
| EPI_ISL_3345846, EPI_ISL_3345847 | Viollier AG | Department of Biosystems Science and Engineering, ETH Zürich | Chaoran Chen, Sarah Nadeau, Louis du Plessis, Catharine Aquino, Ivan Topolsky, Kim Philipp Jablonski, Lara Fuhrmann, Daniel Ehrsam, Isabel Stürmer, Andreia Cabral de Gouvea, Maria Domenica Moccia, Simon Grüter, Timothy Sykes, Lennart Opitz, Griffin White, Laura Neff, Doris Popovic, Andrea Patrignani, Jay Tracy, Ralph Schlapbach, Christiane Beckmann, Maurice Redondo, Olivier Kobel, Christoph Noppen, Niko Beerenwinkel, Tanja Stadler |
| EPI_ISL_3345848 | Fulgent Genetics | Centers for Disease Control and Prevention Division of Viral Diseases, Pathogen Discovery | Dakota Howard, Dhwani Batra, Peter W. Cook, Kara Moser, Adrian Paskey, Jason Caravas, Benjamin Rambo-Martin, Shatavia Morrison, Christopher Gulvick, Scott Sammons, Yvette Unoarumhi, Darlene Wagner, Matthew Schmerer, Harry Gao, Mickey Li, John Gao, Joseph Fierro, Benafsh Sapra, Becky Tsai, Yan Meng, Doreen Ng, James Xie, Clinton R. Paden, Duncan MacCannell |
| EPI_ISL_3345849, EPI_ISL_3345850 | Viollier AG | Department of Biosystems Science and Engineering, ETH Zürich | Chaoran Chen, Sarah Nadeau, Louis du Plessis, Catharine Aquino, Ivan Topolsky, Kim Philipp Jablonski, Lara Fuhrmann, Daniel Ehrsam, Isabel Stürmer, Andreia Cabral de Gouvea, Maria Domenica Moccia, Simon Grüter, Timothy Sykes, Lennart Opitz, Griffin White, Laura Neff, Doris Popovic, Andrea Patrignani, Jay Tracy, Ralph Schlapbach, Christiane Beckmann, Maurice Redondo, Olivier Kobel, Christoph Noppen, Niko Beerenwinkel, Tanja Stadler |
| EPI_ISL_3345851 | Fulgent Genetics | Centers for Disease Control and Prevention Division of Viral Diseases, Pathogen Discovery | Dakota Howard, Dhwani Batra, Peter W. Cook, Kara Moser, Adrian Paskey, Jason Caravas, Benjamin Rambo-Martin, Shatavia Morrison, Christopher Gulvick, Scott Sammons, Yvette Unoarumhi, Darlene Wagner, Matthew Schmerer, Harry Gao, Mickey Li, John Gao, Joseph Fierro, Benafsh Sapra, Becky Tsai, Yan Meng, Doreen Ng, James Xie, Clinton R. Paden, Duncan MacCannell |

| | | | |
|---|---|---|---|
| EPI_ISL_3345852 | Viollier AG | Department of Biosystems Science and Engineering, ETH Zürich | Christian Beisel, Sarah Nadeau, Chaoran Chen, Louis du Plessis, Ivan Topolsky, Kim Philipp Jablonski, Lara Fuhrmann, Rebecca Denes, Mirjam Feldkamp, Ina Nissen, Natascha Santacroce, Elodie Burcklen, Christiane Beckmann, Maurice Redondo, Olivier Kobel, Christoph Noppen, Niko Beerenwinkel, Tanja Stadler |
| EPI_ISL_3345853 | Fulgent Genetics | Centers for Disease Control and Prevention Division of Viral Diseases, Pathogen Discovery | Dakota Howard, Dhwani Batra, Peter W. Cook, Kara Moser, Adrian Paskey, Jason Caravas, Benjamin Rambo-Martin, Shatavia Morrison, Christopher Gulvick, Scott Sammons, Yvette Unoarumhi, Darlene Wagner, Matthew Schmerer, Harry Gao, Mickey Li, John Gao, Joseph Fierro, Benafsh Sapra, Becky Tsai, Yan Meng, Doreen Ng, James Xie, Clinton R. Paden, Duncan MacCannell |
| EPI_ISL_3345854, EPI_ISL_3345855 | Viollier AG | Department of Biosystems Science and Engineering, ETH Zürich | Chaoran Chen, Sarah Nadeau, Louis du Plessis, Catharine Aquino, Ivan Topolsky, Kim Philipp Jablonski, Lara Fuhrmann, Daniel Ehrsam, Isabel Stürmer, Andreia Cabral de Gouvea, Maria Domenica Moccia, Simon Grüter, Timothy Sykes, Lennart Opitz, Griffin White, Laura Neff, Doris Popovic, Andrea Patrignani, Jay Tracy, Ralph Schlapbach, Christiane Beckmann, Maurice Redondo, Olivier Kobel, Christoph Noppen, Niko Beerenwinkel, Tanja Stadler |
| EPI_ISL_3345856 | Fulgent Genetics | Centers for Disease Control and Prevention Division of Viral Diseases, Pathogen Discovery | Dakota Howard, Dhwani Batra, Peter W. Cook, Kara Moser, Adrian Paskey, Jason Caravas, Benjamin Rambo-Martin, Shatavia Morrison, Christopher Gulvick, Scott Sammons, Yvette Unoarumhi, Darlene Wagner, Matthew Schmerer, Harry Gao, Mickey Li, John Gao, Joseph Fierro, Benafsh Sapra, Becky Tsai, Yan Meng, Doreen Ng, James Xie, Clinton R. Paden, Duncan MacCannell |
| EPI_ISL_3345857, EPI_ISL_3345858 | Viollier AG | Department of Biosystems Science and Engineering, ETH Zürich | Chaoran Chen, Sarah Nadeau, Louis du Plessis, Catharine Aquino, Ivan Topolsky, Kim Philipp Jablonski, Lara Fuhrmann, Daniel Ehrsam, Isabel Stürmer, Andreia Cabral de Gouvea, Maria Domenica Moccia, Simon Grüter, Timothy Sykes, Lennart Opitz, Griffin White, Laura Neff, Doris Popovic, Andrea Patrignani, Jay Tracy, Ralph Schlapbach, Christiane Beckmann, Maurice Redondo, Olivier Kobel, Christoph Noppen, Niko Beerenwinkel, Tanja Stadler |
| EPI_ISL_3345859 | Fulgent Genetics | Centers for Disease Control and Prevention Division of Viral Diseases, Pathogen Discovery | Dakota Howard, Dhwani Batra, Peter W. Cook, Kara Moser, Adrian Paskey, Jason Caravas, Benjamin Rambo-Martin, Shatavia Morrison, Christopher Gulvick, Scott Sammons, Yvette Unoarumhi, Darlene Wagner, Matthew Schmerer, Harry Gao, Mickey Li, John Gao, Joseph Fierro, Benafsh Sapra, Becky Tsai, Yan Meng, Doreen Ng, James Xie, Clinton R. Paden, Duncan MacCannell |
| EPI_ISL_3345860 | Viollier AG | Department of Biosystems Science and Engineering, ETH Zürich | Christian Beisel, Sarah Nadeau, Chaoran Chen, Louis du Plessis, Ivan Topolsky, Kim Philipp Jablonski, Lara Fuhrmann, Rebecca Denes, Mirjam Feldkamp, Ina Nissen, Natascha Santacroce, Elodie Burcklen, Christiane Beckmann, Maurice Redondo, Olivier Kobel, Christoph Noppen, Niko Beerenwinkel, Tanja Stadler |
| EPI_ISL_3345861 | Viollier AG | Department of Biosystems Science and Engineering, ETH Zürich | Chaoran Chen, Sarah Nadeau, Louis du Plessis, Catharine Aquino, Ivan Topolsky, Kim Philipp Jablonski, Lara Fuhrmann, Daniel Ehrsam, Isabel Stürmer, Andreia Cabral de Gouvea, Maria Domenica Moccia, Simon Grüter, Timothy Sykes, Lennart Opitz, Griffin White, Laura Neff, Doris Popovic, Andrea Patrignani, Jay Tracy, Ralph Schlapbach, Christiane Beckmann, Maurice Redondo, Olivier Kobel, Christoph Noppen, Niko Beerenwinkel, Tanja Stadler |
| EPI_ISL_3345862 | Fulgent Genetics | Centers for Disease Control and Prevention Division of Viral Diseases, Pathogen Discovery | Dakota Howard, Dhwani Batra, Peter W. Cook, Kara Moser, Adrian Paskey, Jason Caravas, Benjamin Rambo-Martin, Shatavia Morrison, Christopher Gulvick, Scott Sammons, Yvette Unoarumhi, Darlene Wagner, Matthew Schmerer, Harry Gao, Mickey Li, John Gao, Joseph Fierro, Benafsh Sapra, Becky Tsai, Yan Meng, Doreen Ng, James Xie, Clinton R. Paden, Duncan MacCannell |
| EPI_ISL_3345863 | Viollier AG | Department of Biosystems Science and Engineering, ETH Zürich | Chaoran Chen, Sarah Nadeau, Louis du Plessis, Catharine Aquino, Ivan Topolsky, Kim Philipp Jablonski, Lara Fuhrmann, Daniel Ehrsam, Isabel Stürmer, Andreia Cabral de Gouvea, Maria Domenica Moccia, Simon Grüter, Timothy Sykes, Lennart Opitz, Griffin White, Laura Neff, Doris Popovic, Andrea Patrignani, Jay Tracy, Ralph Schlapbach, Christiane Beckmann, Maurice Redondo, Olivier Kobel, Christoph Noppen, Niko Beerenwinkel, Tanja Stadler |
| EPI_ISL_3345864 | Fulgent Genetics | Centers for Disease Control and Prevention Division of Viral Diseases, Pathogen Discovery | Dakota Howard, Dhwani Batra, Peter W. Cook, Kara Moser, Adrian Paskey, Jason Caravas, Benjamin Rambo-Martin, Shatavia Morrison, Christopher Gulvick, Scott Sammons, Yvette Unoarumhi, Darlene Wagner, Matthew Schmerer, Harry Gao, Mickey Li, John Gao, Joseph Fierro, Benafsh Sapra, Becky Tsai, Yan Meng, Doreen Ng, James Xie, Clinton R. Paden, Duncan MacCannell |
| EPI_ISL_3345865, EPI_ISL_3345866 | Viollier AG | Department of Biosystems Science and Engineering, ETH Zürich | Chaoran Chen, Sarah Nadeau, Louis du Plessis, Catharine Aquino, Ivan Topolsky, Kim Philipp Jablonski, Lara Fuhrmann, Daniel Ehrsam, Isabel Stürmer, Andreia Cabral de Gouvea, Maria Domenica Moccia, Simon Grüter, Timothy Sykes, Lennart Opitz, Griffin White, Laura Neff, Doris Popovic, Andrea Patrignani, Jay Tracy, Ralph Schlapbach, Christiane Beckmann, Maurice Redondo, Olivier Kobel, Christoph Noppen, Niko Beerenwinkel, Tanja Stadler |
| EPI_ISL_3345867 | Fulgent Genetics | Centers for Disease Control and Prevention Division of Viral Diseases, Pathogen Discovery | Dakota Howard, Dhwani Batra, Peter W. Cook, Kara Moser, Adrian Paskey, Jason Caravas, Benjamin Rambo-Martin, Shatavia Morrison, Christopher Gulvick, Scott Sammons, Yvette Unoarumhi, Darlene Wagner, Matthew Schmerer, Harry Gao, Mickey Li, John Gao, Joseph Fierro, Benafsh Sapra, Becky Tsai, Yan Meng, Doreen Ng, James Xie, Clinton R. Paden, Duncan MacCannell |
| EPI_ISL_3345868, EPI_ISL_3345869 | Viollier AG | Department of Biosystems Science and Engineering, ETH Zürich | Chaoran Chen, Sarah Nadeau, Louis du Plessis, Catharine Aquino, Ivan Topolsky, Kim Philipp Jablonski, Lara Fuhrmann, Daniel Ehrsam, Isabel Stürmer, Andreia Cabral de Gouvea, Maria Domenica Moccia, Simon Grüter, Timothy Sykes, Lennart Opitz, Griffin White, Laura Neff, Doris Popovic, Andrea Patrignani, Jay Tracy, Ralph Schlapbach, Christiane Beckmann, Maurice Redondo, Olivier Kobel, Christoph Noppen, Niko Beerenwinkel, Tanja Stadler |
| EPI_ISL_3345870 | Fulgent Genetics | Centers for Disease Control and Prevention Division of Viral Diseases, Pathogen Discovery | Dakota Howard, Dhwani Batra, Peter W. Cook, Kara Moser, Adrian Paskey, Jason Caravas, Benjamin Rambo-Martin, Shatavia Morrison, Christopher Gulvick, Scott Sammons, Yvette Unoarumhi, Darlene Wagner, Matthew Schmerer, Harry Gao, Mickey Li, John Gao, Joseph Fierro, Benafsh Sapra, Becky Tsai, Yan Meng, Doreen Ng, James Xie, Clinton R. Paden, Duncan MacCannell |
| EPI_ISL_3345871 | Viollier AG | Department of Biosystems Science and Engineering, ETH Zürich | Chaoran Chen, Sarah Nadeau, Louis du Plessis, Catharine Aquino, Ivan Topolsky, Kim Philipp Jablonski, Lara Fuhrmann, Daniel Ehrsam, Isabel Stürmer, Andreia Cabral de Gouvea, Maria Domenica Moccia, Simon Grüter, Timothy Sykes, Lennart Opitz, Griffin White, Laura Neff, Doris Popovic, Andrea Patrignani, Jay Tracy, Ralph Schlapbach, Christiane Beckmann, Maurice Redondo, Olivier Kobel, Christoph Noppen, Niko Beerenwinkel, Tanja Stadler |
| EPI_ISL_3345872 | Fulgent Genetics | Centers for Disease Control and Prevention Division of Viral Diseases, Pathogen Discovery | Dakota Howard, Dhwani Batra, Peter W. Cook, Kara Moser, Adrian Paskey, Jason Caravas, Benjamin Rambo-Martin, Shatavia Morrison, Christopher Gulvick, Scott Sammons, Yvette Unoarumhi, Darlene Wagner, Matthew Schmerer, Harry Gao, Mickey Li, John Gao, Joseph Fierro, Benafsh Sapra, Becky Tsai, Yan Meng, Doreen Ng, James Xie, Clinton R. Paden, Duncan MacCannell |
| EPI_ISL_3345873, EPI_ISL_3345874 | Viollier AG | Department of Biosystems Science and Engineering, ETH Zürich | Chaoran Chen, Sarah Nadeau, Louis du Plessis, Catharine Aquino, Ivan Topolsky, Kim Philipp Jablonski, Lara Fuhrmann, Daniel Ehrsam, Isabel Stürmer, Andreia Cabral de Gouvea, Maria Domenica Moccia, Simon Grüter, Timothy Sykes, Lennart Opitz, Griffin White, Laura Neff, Doris Popovic, Andrea Patrignani, Jay Tracy, Ralph Schlapbach, Christiane Beckmann, Maurice Redondo, Olivier Kobel, Christoph Noppen, Niko Beerenwinkel, Tanja Stadler |
| EPI_ISL_3345875 | Fulgent Genetics | Centers for Disease Control and Prevention Division of Viral Diseases, Pathogen Discovery | Dakota Howard, Dhwani Batra, Peter W. Cook, Kara Moser, Adrian Paskey, Jason Caravas, Benjamin Rambo-Martin, Shatavia Morrison, Christopher Gulvick, Scott Sammons, Yvette Unoarumhi, Darlene Wagner, Matthew Schmerer, Harry Gao, Mickey Li, John Gao, Joseph Fierro, Benafsh Sapra, Becky Tsai, Yan Meng, Doreen Ng, James Xie, Clinton R. Paden, Duncan MacCannell |
| EPI_ISL_3345876, EPI_ISL_3345877 | Viollier AG | Department of Biosystems Science and Engineering, ETH Zürich | Christian Beisel, Sarah Nadeau, Chaoran Chen, Louis du Plessis, Ivan Topolsky, Kim Philipp Jablonski, Lara Fuhrmann, Rebecca Denes, Mirjam Feldkamp, Ina Nissen, Natascha Santacroce, Elodie Burcklen, Christiane Beckmann, Maurice Redondo, Olivier Kobel, Christoph Noppen, Niko Beerenwinkel, Tanja Stadler |
| EPI_ISL_3345878 | Fulgent Genetics | Centers for Disease Control and Prevention Division of Viral Diseases, Pathogen Discovery | Dakota Howard, Dhwani Batra, Peter W. Cook, Kara Moser, Adrian Paskey, Jason Caravas, Benjamin Rambo-Martin, Shatavia Morrison, Christopher Gulvick, Scott Sammons, Yvette Unoarumhi, Darlene Wagner, Matthew Schmerer, Harry Gao, Mickey Li, John Gao, Joseph Fierro, Benafsh Sapra, Becky Tsai, Yan Meng, Doreen Ng, James Xie, Clinton R. Paden, Duncan MacCannell |
| EPI_ISL_3345879 | Viollier AG | Department of Biosystems Science and Engineering, ETH Zürich | Chaoran Chen, Sarah Nadeau, Louis du Plessis, Catharine Aquino, Ivan Topolsky, Kim Philipp Jablonski, Lara Fuhrmann, Daniel Ehrsam, Isabel Stürmer, Andreia Cabral de Gouvea, Maria Domenica Moccia, Simon Grüter, Timothy Sykes, Lennart Opitz, Griffin White, Laura Neff, Doris Popovic, Andrea Patrignani, Jay Tracy, Ralph Schlapbach, Christiane Beckmann, Maurice Redondo, Olivier Kobel, Christoph Noppen, Niko Beerenwinkel, Tanja Stadler |
| EPI_ISL_3345880 | Fulgent Genetics | Centers for Disease Control and Prevention Division of Viral Diseases, Pathogen Discovery | Dakota Howard, Dhwani Batra, Peter W. Cook, Kara Moser, Adrian Paskey, Jason Caravas, Benjamin Rambo-Martin, Shatavia Morrison, Christopher Gulvick, Scott Sammons, Yvette Unoarumhi, Darlene Wagner, Matthew Schmerer, Harry Gao, Mickey Li, John Gao, Joseph Fierro, Benafsh Sapra, Becky Tsai, Yan Meng, Doreen Ng, James Xie, Clinton R. Paden, Duncan MacCannell |
| EPI_ISL_3345881 | Viollier AG | Department of Biosystems Science and Engineering, ETH Zürich | Chaoran Chen, Sarah Nadeau, Louis du Plessis, Catharine Aquino, Ivan Topolsky, Kim Philipp Jablonski, Lara Fuhrmann, Daniel Ehrsam, Isabel Stürmer, Andreia Cabral de Gouvea, Maria Domenica Moccia, Simon Grüter, Timothy Sykes, Lennart Opitz, Griffin White, Laura Neff, Doris Popovic, Andrea Patrignani, Jay Tracy, Ralph Schlapbach, Christiane Beckmann, Maurice Redondo, Olivier Kobel, Christoph Noppen, Niko Beerenwinkel, Tanja Stadler |
| EPI_ISL_3345882 | Viollier AG | Department of Biosystems Science and Engineering, ETH Zürich | Christian Beisel, Sarah Nadeau, Chaoran Chen, Louis du Plessis, Ivan Topolsky, Kim Philipp Jablonski, Lara Fuhrmann, Rebecca Denes, Mirjam Feldkamp, Ina Nissen, Natascha Santacroce, Elodie Burcklen, Christiane Beckmann, Maurice Redondo, Olivier Kobel, Christoph Noppen, Niko Beerenwinkel, Tanja Stadler |
| EPI_ISL_3345883 | Fulgent Genetics | Centers for Disease Control and Prevention Division of Viral Diseases, Pathogen Discovery | Dakota Howard, Dhwani Batra, Peter W. Cook, Kara Moser, Adrian Paskey, Jason Caravas, Benjamin Rambo-Martin, Shatavia Morrison, Christopher Gulvick, Scott Sammons, Yvette Unoarumhi, Darlene Wagner, Matthew Schmerer, Harry Gao, Mickey Li, John Gao, Joseph Fierro, Benafsh Sapra, Becky Tsai, Yan Meng, Doreen Ng, James Xie, Clinton R. Paden, Duncan MacCannell |
| EPI_ISL_3345885 | Viollier AG | Department of Biosystems Science and Engineering, ETH Zürich | Chaoran Chen, Sarah Nadeau, Louis du Plessis, Catharine Aquino, Ivan Topolsky, Kim Philipp Jablonski, Lara Fuhrmann, Daniel Ehrsam, Isabel Stürmer, Andreia Cabral de Gouvea, Maria Domenica Moccia, Simon Grüter, Timothy Sykes, Lennart Opitz, Griffin White, Laura Neff, Doris Popovic, Andrea Patrignani, Jay Tracy, Ralph Schlapbach, Christiane Beckmann, Maurice Redondo, Olivier Kobel, Christoph Noppen, Niko Beerenwinkel, Tanja Stadler |
| EPI_ISL_3345886 | Fulgent Genetics | Centers for Disease Control and Prevention Division of Viral Diseases, Pathogen Discovery | Dakota Howard, Dhwani Batra, Peter W. Cook, Kara Moser, Adrian Paskey, Jason Caravas, Benjamin Rambo-Martin, Shatavia Morrison, Christopher Gulvick, Scott Sammons, Yvette Unoarumhi, Darlene Wagner, Matthew Schmerer, Harry Gao, Mickey Li, John Gao, Joseph Fierro, Benafsh Sapra, Becky Tsai, Yan Meng, Doreen Ng, James Xie, Clinton R. Paden, Duncan MacCannell |
| EPI_ISL_3345887 | Viollier AG | Department of Biosystems Science and Engineering, ETH Zürich | Chaoran Chen, Sarah Nadeau, Louis du Plessis, Catharine Aquino, Ivan Topolsky, Kim Philipp Jablonski, Lara Fuhrmann, Daniel Ehrsam, Isabel Stürmer, Andreia Cabral de Gouvea, Maria Domenica Moccia, Simon Grüter, Timothy Sykes, Lennart Opitz, Griffin White, Laura Neff, Doris Popovic, Andrea Patrignani, Jay Tracy, Ralph Schlapbach, Christiane Beckmann, Maurice Redondo, Olivier Kobel, Christoph Noppen, Niko Beerenwinkel, Tanja Stadler |
| EPI_ISL_3345888 | Fulgent Genetics | Centers for Disease Control and Prevention Division of Viral Diseases, Pathogen Discovery | Dakota Howard, Dhwani Batra, Peter W. Cook, Kara Moser, Adrian Paskey, Jason Caravas, Benjamin Rambo-Martin, Shatavia Morrison, Christopher Gulvick, Scott Sammons, Yvette Unoarumhi, Darlene Wagner, Matthew Schmerer, Harry Gao, Mickey Li, John Gao, Joseph Fierro, Benafsh Sapra, Becky Tsai, Yan Meng, Doreen Ng, James Xie, Clinton R. Paden, Duncan MacCannell |
| EPI_ISL_3345889 | Viollier AG | Department of Biosystems Science and Engineering, ETH Zürich | Christian Beisel, Sarah Nadeau, Chaoran Chen, Louis du Plessis, Ivan Topolsky, Kim Philipp Jablonski, Lara Fuhrmann, Rebecca Denes, Mirjam Feldkamp, Ina Nissen, Natascha Santacroce, Elodie Burcklen, Christiane Beckmann, Maurice Redondo, Olivier Kobel, Christoph Noppen, Niko Beerenwinkel, Tanja Stadler |
| EPI_ISL_3345890 | Viollier AG | Department of Biosystems Science and Engineering, ETH Zürich | Chaoran Chen, Sarah Nadeau, Louis du Plessis, Catharine Aquino, Ivan Topolsky, Kim Philipp Jablonski, Lara Fuhrmann, Daniel Ehrsam, Isabel Stürmer, Andreia Cabral de Gouvea, Maria Domenica Moccia, Simon Grüter, Timothy Sykes, Lennart Opitz, Griffin White, Laura Neff, Doris Popovic, Andrea Patrignani, Jay Tracy, Ralph Schlapbach, Christiane Beckmann, Maurice Redondo, Olivier Kobel, Christoph Noppen, Niko Beerenwinkel, Tanja Stadler |
| EPI_ISL_3345891 | Fulgent Genetics | Centers for Disease Control and Prevention Division of Viral Diseases, Pathogen Discovery | Dakota Howard, Dhwani Batra, Peter W. Cook, Kara Moser, Adrian Paskey, Jason Caravas, Benjamin Rambo-Martin, Shatavia Morrison, Christopher Gulvick, Scott Sammons, Yvette Unoarumhi, Darlene Wagner, Matthew Schmerer, Harry Gao, Mickey Li, John Gao, Joseph Fierro, Benafsh Sapra, Becky Tsai, Yan Meng, Doreen Ng, James Xie, Clinton R. Paden, Duncan MacCannell |
| EPI_ISL_3345892 | Viollier AG | Department of Biosystems Science and Engineering, ETH Zürich | Christian Beisel, Sarah Nadeau, Chaoran Chen, Louis du Plessis, Ivan Topolsky, Kim Philipp Jablonski, Lara Fuhrmann, Rebecca Denes, Mirjam Feldkamp, Ina Nissen, Natascha Santacroce, Elodie Burcklen, Christiane Beckmann, Maurice Redondo, Olivier Kobel, Christoph Noppen, Niko Beerenwinkel, Tanja Stadler |
| EPI_ISL_3345893 | Viollier AG | Department of Biosystems Science and Engineering, ETH Zürich | Chaoran Chen, Sarah Nadeau, Louis du Plessis, Catharine Aquino, Ivan Topolsky, Kim Philipp Jablonski, Lara Fuhrmann, Daniel Ehrsam, Isabel Stürmer, Andreia Cabral de Gouvea, Maria Domenica Moccia, Simon Grüter, Timothy Sykes, Lennart Opitz, Griffin White, Laura Neff, Doris Popovic, Andrea Patrignani, Jay Tracy, Ralph Schlapbach, Christiane Beckmann, Maurice Redondo, Olivier Kobel, Christoph Noppen, Niko Beerenwinkel, Tanja Stadler |
| EPI_ISL_3345894 | Fulgent Genetics | Centers for Disease Control and Prevention Division of Viral Diseases, Pathogen Discovery | Dakota Howard, Dhwani Batra, Peter W. Cook, Kara Moser, Adrian Paskey, Jason Caravas, Benjamin Rambo-Martin, Shatavia Morrison, Christopher Gulvick, Scott Sammons, Yvette Unoarumhi, Darlene Wagner, Matthew Schmerer, Harry Gao, Mickey Li, John Gao, Joseph Fierro, Benafsh Sapra, Becky Tsai, Yan Meng, Doreen Ng, James Xie, Clinton R. Paden, Duncan MacCannell |
| EPI_ISL_3345895 | Viollier AG | Department of Biosystems Science and Engineering, ETH Zürich | Christian Beisel, Sarah Nadeau, Chaoran Chen, Louis du Plessis, Ivan Topolsky, Kim Philipp Jablonski, Lara Fuhrmann, Rebecca Denes, Mirjam Feldkamp, Ina Nissen, Natascha Santacroce, Elodie Burcklen, Christiane Beckmann, Maurice Redondo, Olivier Kobel, Christoph Noppen, Niko Beerenwinkel, Tanja Stadler |
| EPI_ISL_3345896 | Viollier AG | Department of Biosystems Science and Engineering, ETH Zürich | Chaoran Chen, Sarah Nadeau, Louis du Plessis, Catharine Aquino, Ivan Topolsky, Kim Philipp Jablonski, Lara Fuhrmann, Daniel Ehrsam, Isabel Stürmer, Andreia Cabral de Gouvea, Maria Domenica Moccia, Simon Grüter, Timothy Sykes, Lennart Opitz, Griffin White, Laura Neff, Doris Popovic, Andrea Patrignani, Jay Tracy, Ralph Schlapbach, Christiane Beckmann, Maurice Redondo, Olivier Kobel, Christoph Noppen, Niko Beerenwinkel, Tanja Stadler |
| EPI_ISL_3345897 | Fulgent Genetics | Centers for Disease Control and Prevention Division of Viral Diseases, Pathogen Discovery | Dakota Howard, Dhwani Batra, Peter W. Cook, Kara Moser, Adrian Paskey, Jason Caravas, Benjamin Rambo-Martin, Shatavia Morrison, Christopher Gulvick, Scott Sammons, Yvette Unoarumhi, Darlene Wagner, Matthew Schmerer, Harry Gao, Mickey Li, John Gao, Joseph Fierro, Benafsh Sapra, Becky Tsai, Yan Meng, Doreen Ng, James Xie, Clinton R. Paden, Duncan MacCannell |
| EPI_ISL_3345898 | Viollier AG | Department of Biosystems Science and Engineering, ETH Zürich | Chaoran Chen, Sarah Nadeau, Louis du Plessis, Catharine Aquino, Ivan Topolsky, Kim Philipp Jablonski, Lara Fuhrmann, Daniel Ehrsam, Isabel Stürmer, Andreia Cabral de Gouvea, Maria Domenica Moccia, Simon Grüter, Timothy Sykes, Lennart Opitz, Griffin White, Laura Neff, Doris Popovic, Andrea Patrignani, Jay Tracy, Ralph Schlapbach, Christiane Beckmann, Maurice Redondo, Olivier Kobel, Christoph Noppen, Niko Beerenwinkel, Tanja Stadler |
| EPI_ISL_3345899 | Fulgent Genetics | Centers for Disease Control and Prevention Division of Viral Diseases, Pathogen Discovery | Dakota Howard, Dhwani Batra, Peter W. Cook, Kara Moser, Adrian Paskey, Jason Caravas, Benjamin Rambo-Martin, Shatavia Morrison, Christopher Gulvick, Scott Sammons, Yvette Unoarumhi, Darlene Wagner, Matthew Schmerer, Harry Gao, Mickey Li, John Gao, Joseph Fierro, Benafsh Sapra, Becky Tsai, Yan Meng, Doreen Ng, James Xie, Clinton R. Paden, Duncan MacCannell |
| EPI_ISL_3345900, EPI_ISL_3345901 | Viollier AG | Department of Biosystems Science and Engineering, ETH Zürich | Christian Beisel, Sarah Nadeau, Chaoran Chen, Louis du Plessis, Ivan Topolsky, Kim Philipp Jablonski, Lara Fuhrmann, Rebecca Denes, Mirjam Feldkamp, Ina Nissen, Natascha Santacroce, Elodie Burcklen, Christiane Beckmann, Maurice Redondo, Olivier Kobel, Christoph Noppen, Niko Beerenwinkel, Tanja Stadler |
| EPI_ISL_3345902 | Fulgent Genetics | Centers for Disease Control and Prevention Division of Viral Diseases, Pathogen Discovery | Dakota Howard, Dhwani Batra, Peter W. Cook, Kara Moser, Adrian Paskey, Jason Caravas, Benjamin Rambo-Martin, Shatavia Morrison, Christopher Gulvick, Scott Sammons, Yvette Unoarumhi, Darlene Wagner, Matthew Schmerer, Harry Gao, Mickey Li, John Gao, Joseph Fierro, Benafsh Sapra, Becky Tsai, Yan Meng, Doreen Ng, James Xie, Clinton R. Paden, Duncan MacCannell |
| EPI_ISL_3345903 | Viollier AG | Department of Biosystems Science and Engineering, ETH Zürich | Chaoran Chen, Sarah Nadeau, Louis du Plessis, Catharine Aquino, Ivan Topolsky, Kim Philipp Jablonski, Lara Fuhrmann, Daniel Ehrsam, Isabel Stürmer, Andreia Cabral de Gouvea, Maria Domenica Moccia, Simon Grüter, Timothy Sykes, Lennart Opitz, Griffin White, Laura Neff, Doris Popovic, Andrea Patrignani, Jay Tracy, Ralph Schlapbach, Christiane Beckmann, Maurice Redondo, Olivier Kobel, Christoph Noppen, Niko Beerenwinkel, Tanja Stadler |
| EPI_ISL_3345904 | Viollier AG | Department of Biosystems Science and Engineering, ETH Zürich | Christian Beisel, Sarah Nadeau, Chaoran Chen, Louis du Plessis, Ivan Topolsky, Kim Philipp Jablonski, Lara Fuhrmann, Rebecca Denes, Mirjam Feldkamp, Ina Nissen, Natascha Santacroce, Elodie Burcklen, Christiane Beckmann, Maurice Redondo, Olivier Kobel, Christoph Noppen, Niko Beerenwinkel, Tanja Stadler |

| | | | |
|---|---|---|---|
| EPI_ISL_3345905 | Fulgent Genetics | Centers for Disease Control and Prevention Division of Viral Diseases, Pathogen Discovery | Dakota Howard, Dhwani Batra, Peter W. Cook, Kara Moser, Adrian Paskey, Jason Caravas, Benjamin Rambo-Martin, Shatavia Morrison, Christopher Gulvick, Scott Sammons, Yvette Unoarumhi, Darlene Wagner, Matthew Schmerer, Harry Gao, Mickey Li, John Gao, Joseph Fierro, Benafsh Sapra, Becky Tsai, Yan Meng, Doreen Ng, James Xie, Clinton R. Paden, Duncan MacCannell |
| EPI_ISL_3345906, EPI_ISL_3345907 | Viollier AG | Department of Biosystems Science and Engineering, ETH Zürich | Chaoran Chen, Sarah Nadeau, Louis du Plessis, Catharine Aquino, Ivan Topolsky, Kim Philipp Jablonski, Lara Fuhrmann, Daniel Ehrsam, Isabel Stürmer, Andreia Cabral de Gouvea, Maria Domenica Moccia, Simon Grüter, Timothy Sykes, Lennart Opitz, Griffin White, Laura Neff, Doris Popovic, Andrea Patrignani, Jay Tracy, Ralph Schlapbach, Christiane Beckmann, Maurice Redondo, Olivier Kobel, Christoph Noppen, Niko Beerenwinkel, Tanja Stadler |
| EPI_ISL_3345908 | Fulgent Genetics | Centers for Disease Control and Prevention Division of Viral Diseases, Pathogen Discovery | Dakota Howard, Dhwani Batra, Peter W. Cook, Kara Moser, Adrian Paskey, Jason Caravas, Benjamin Rambo-Martin, Shatavia Morrison, Christopher Gulvick, Scott Sammons, Yvette Unoarumhi, Darlene Wagner, Matthew Schmerer, Harry Gao, Mickey Li, John Gao, Joseph Fierro, Benafsh Sapra, Becky Tsai, Yan Meng, Doreen Ng, James Xie, Clinton R. Paden, Duncan MacCannell |
| EPI_ISL_3345909, EPI_ISL_3345910 | Viollier AG | Department of Biosystems Science and Engineering, ETH Zürich | Christian Beisel, Sarah Nadeau, Chaoran Chen, Louis du Plessis, Ivan Topolsky, Kim Philipp Jablonski, Lara Fuhrmann, Rebecca Denes, Mirjam Feldkamp, Ina Nissen, Natascha Santacroce, Elodie Burcklen, Christiane Beckmann, Maurice Redondo, Olivier Kobel, Christoph Noppen, Niko Beerenwinkel, Tanja Stadler |
| EPI_ISL_3345911 | Fulgent Genetics | Centers for Disease Control and Prevention Division of Viral Diseases, Pathogen Discovery | Dakota Howard, Dhwani Batra, Peter W. Cook, Kara Moser, Adrian Paskey, Jason Caravas, Benjamin Rambo-Martin, Shatavia Morrison, Christopher Gulvick, Scott Sammons, Yvette Unoarumhi, Darlene Wagner, Matthew Schmerer, Harry Gao, Mickey Li, John Gao, Joseph Fierro, Benafsh Sapra, Becky Tsai, Yan Meng, Doreen Ng, James Xie, Clinton R. Paden, Duncan MacCannell |
| EPI_ISL_3345912 | Viollier AG | Department of Biosystems Science and Engineering, ETH Zürich | Chaoran Chen, Sarah Nadeau, Louis du Plessis, Catharine Aquino, Ivan Topolsky, Kim Philipp Jablonski, Lara Fuhrmann, Daniel Ehrsam, Isabel Stürmer, Andreia Cabral de Gouvea, Maria Domenica Moccia, Simon Grüter, Timothy Sykes, Lennart Opitz, Griffin White, Laura Neff, Doris Popovic, Andrea Patrignani, Jay Tracy, Ralph Schlapbach, Christiane Beckmann, Maurice Redondo, Olivier Kobel, Christoph Noppen, Niko Beerenwinkel, Tanja Stadler |
| EPI_ISL_3345913 | Fulgent Genetics | Centers for Disease Control and Prevention Division of Viral Diseases, Pathogen Discovery | Dakota Howard, Dhwani Batra, Peter W. Cook, Kara Moser, Adrian Paskey, Jason Caravas, Benjamin Rambo-Martin, Shatavia Morrison, Christopher Gulvick, Scott Sammons, Yvette Unoarumhi, Darlene Wagner, Matthew Schmerer, Harry Gao, Mickey Li, John Gao, Joseph Fierro, Benafsh Sapra, Becky Tsai, Yan Meng, Doreen Ng, James Xie, Clinton R. Paden, Duncan MacCannell |
| EPI_ISL_3345914 | Viollier AG | Department of Biosystems Science and Engineering, ETH Zürich | Chaoran Chen, Sarah Nadeau, Louis du Plessis, Catharine Aquino, Ivan Topolsky, Kim Philipp Jablonski, Lara Fuhrmann, Daniel Ehrsam, Isabel Stürmer, Andreia Cabral de Gouvea, Maria Domenica Moccia, Simon Grüter, Timothy Sykes, Lennart Opitz, Griffin White, Laura Neff, Doris Popovic, Andrea Patrignani, Jay Tracy, Ralph Schlapbach, Christiane Beckmann, Maurice Redondo, Olivier Kobel, Christoph Noppen, Niko Beerenwinkel, Tanja Stadler |
| EPI_ISL_3345915 | Viollier AG | Department of Biosystems Science and Engineering, ETH Zürich | Christian Beisel, Sarah Nadeau, Chaoran Chen, Louis du Plessis, Ivan Topolsky, Kim Philipp Jablonski, Lara Fuhrmann, Rebecca Denes, Mirjam Feldkamp, Ina Nissen, Natascha Santacroce, Elodie Burcklen, Christiane Beckmann, Maurice Redondo, Olivier Kobel, Christoph Noppen, Niko Beerenwinkel, Tanja Stadler |
| EPI_ISL_3345916 | Fulgent Genetics | Centers for Disease Control and Prevention Division of Viral Diseases, Pathogen Discovery | Dakota Howard, Dhwani Batra, Peter W. Cook, Kara Moser, Adrian Paskey, Jason Caravas, Benjamin Rambo-Martin, Shatavia Morrison, Christopher Gulvick, Scott Sammons, Yvette Unoarumhi, Darlene Wagner, Matthew Schmerer, Harry Gao, Mickey Li, John Gao, Joseph Fierro, Benafsh Sapra, Becky Tsai, Yan Meng, Doreen Ng, James Xie, Clinton R. Paden, Duncan MacCannell |
| EPI_ISL_3345917, EPI_ISL_3345918 | Viollier AG | Department of Biosystems Science and Engineering, ETH Zürich | Chaoran Chen, Sarah Nadeau, Louis du Plessis, Catharine Aquino, Ivan Topolsky, Kim Philipp Jablonski, Lara Fuhrmann, Daniel Ehrsam, Isabel Stürmer, Andreia Cabral de Gouvea, Maria Domenica Moccia, Simon Grüter, Timothy Sykes, Lennart Opitz, Griffin White, Laura Neff, Doris Popovic, Andrea Patrignani, Jay Tracy, Ralph Schlapbach, Christiane Beckmann, Maurice Redondo, Olivier Kobel, Christoph Noppen, Niko Beerenwinkel, Tanja Stadler |
| EPI_ISL_3345919 | Fulgent Genetics | Centers for Disease Control and Prevention Division of Viral Diseases, Pathogen Discovery | Dakota Howard, Dhwani Batra, Peter W. Cook, Kara Moser, Adrian Paskey, Jason Caravas, Benjamin Rambo-Martin, Shatavia Morrison, Christopher Gulvick, Scott Sammons, Yvette Unoarumhi, Darlene Wagner, Matthew Schmerer, Harry Gao, Mickey Li, John Gao, Joseph Fierro, Benafsh Sapra, Becky Tsai, Yan Meng, Doreen Ng, James Xie, Clinton R. Paden, Duncan MacCannell |
| EPI_ISL_3345920 | Viollier AG | Department of Biosystems Science and Engineering, ETH Zürich | Chaoran Chen, Sarah Nadeau, Louis du Plessis, Catharine Aquino, Ivan Topolsky, Kim Philipp Jablonski, Lara Fuhrmann, Daniel Ehrsam, Isabel Stürmer, Andreia Cabral de Gouvea, Maria Domenica Moccia, Simon Grüter, Timothy Sykes, Lennart Opitz, Griffin White, Laura Neff, Doris Popovic, Andrea Patrignani, Jay Tracy, Ralph Schlapbach, Christiane Beckmann, Maurice Redondo, Olivier Kobel, Christoph Noppen, Niko Beerenwinkel, Tanja Stadler |
| EPI_ISL_3345921 | Fulgent Genetics | Centers for Disease Control and Prevention Division of Viral Diseases, Pathogen Discovery | Dakota Howard, Dhwani Batra, Peter W. Cook, Kara Moser, Adrian Paskey, Jason Caravas, Benjamin Rambo-Martin, Shatavia Morrison, Christopher Gulvick, Scott Sammons, Yvette Unoarumhi, Darlene Wagner, Matthew Schmerer, Harry Gao, Mickey Li, John Gao, Joseph Fierro, Benafsh Sapra, Becky Tsai, Yan Meng, Doreen Ng, James Xie, Clinton R. Paden, Duncan MacCannell |
| EPI_ISL_3345922, EPI_ISL_3345923 | Viollier AG | Department of Biosystems Science and Engineering, ETH Zürich | Chaoran Chen, Sarah Nadeau, Louis du Plessis, Catharine Aquino, Ivan Topolsky, Kim Philipp Jablonski, Lara Fuhrmann, Daniel Ehrsam, Isabel Stürmer, Andreia Cabral de Gouvea, Maria Domenica Moccia, Simon Grüter, Timothy Sykes, Lennart Opitz, Griffin White, Laura Neff, Doris Popovic, Andrea Patrignani, Jay Tracy, Ralph Schlapbach, Christiane Beckmann, Maurice Redondo, Olivier Kobel, Christoph Noppen, Niko Beerenwinkel, Tanja Stadler |
| EPI_ISL_3345924 | Fulgent Genetics | Centers for Disease Control and Prevention Division of Viral Diseases, Pathogen Discovery | Dakota Howard, Dhwani Batra, Peter W. Cook, Kara Moser, Adrian Paskey, Jason Caravas, Benjamin Rambo-Martin, Shatavia Morrison, Christopher Gulvick, Scott Sammons, Yvette Unoarumhi, Darlene Wagner, Matthew Schmerer, Harry Gao, Mickey Li, John Gao, Joseph Fierro, Benafsh Sapra, Becky Tsai, Yan Meng, Doreen Ng, James Xie, Clinton R. Paden, Duncan MacCannell |
| EPI_ISL_3345925, EPI_ISL_3345926 | Viollier AG | Department of Biosystems Science and Engineering, ETH Zürich | Chaoran Chen, Sarah Nadeau, Louis du Plessis, Catharine Aquino, Ivan Topolsky, Kim Philipp Jablonski, Lara Fuhrmann, Daniel Ehrsam, Isabel Stürmer, Andreia Cabral de Gouvea, Maria Domenica Moccia, Simon Grüter, Timothy Sykes, Lennart Opitz, Griffin White, Laura Neff, Doris Popovic, Andrea Patrignani, Jay Tracy, Ralph Schlapbach, Christiane Beckmann, Maurice Redondo, Olivier Kobel, Christoph Noppen, Niko Beerenwinkel, Tanja Stadler |
| EPI_ISL_3345927 | Fulgent Genetics | Centers for Disease Control and Prevention Division of Viral Diseases, Pathogen Discovery | Dakota Howard, Dhwani Batra, Peter W. Cook, Kara Moser, Adrian Paskey, Jason Caravas, Benjamin Rambo-Martin, Shatavia Morrison, Christopher Gulvick, Scott Sammons, Yvette Unoarumhi, Darlene Wagner, Matthew Schmerer, Harry Gao, Mickey Li, John Gao, Joseph Fierro, Benafsh Sapra, Becky Tsai, Yan Meng, Doreen Ng, James Xie, Clinton R. Paden, Duncan MacCannell |
| EPI_ISL_3345928, EPI_ISL_3345929 | Viollier AG | Department of Biosystems Science and Engineering, ETH Zürich | Chaoran Chen, Sarah Nadeau, Louis du Plessis, Catharine Aquino, Ivan Topolsky, Kim Philipp Jablonski, Lara Fuhrmann, Daniel Ehrsam, Isabel Stürmer, Andreia Cabral de Gouvea, Maria Domenica Moccia, Simon Grüter, Timothy Sykes, Lennart Opitz, Griffin White, Laura Neff, Doris Popovic, Andrea Patrignani, Jay Tracy, Ralph Schlapbach, Christiane Beckmann, Maurice Redondo, Olivier Kobel, Christoph Noppen, Niko Beerenwinkel, Tanja Stadler |
| EPI_ISL_3345930 | Fulgent Genetics | Centers for Disease Control and Prevention Division of Viral Diseases, Pathogen Discovery | Dakota Howard, Dhwani Batra, Peter W. Cook, Kara Moser, Adrian Paskey, Jason Caravas, Benjamin Rambo-Martin, Shatavia Morrison, Christopher Gulvick, Scott Sammons, Yvette Unoarumhi, Darlene Wagner, Matthew Schmerer, Harry Gao, Mickey Li, John Gao, Joseph Fierro, Benafsh Sapra, Becky Tsai, Yan Meng, Doreen Ng, James Xie, Clinton R. Paden, Duncan MacCannell |
| EPI_ISL_3345931 | Viollier AG | Department of Biosystems Science and Engineering, ETH Zürich | Christian Beisel, Sarah Nadeau, Chaoran Chen, Louis du Plessis, Ivan Topolsky, Kim Philipp Jablonski, Lara Fuhrmann, Rebecca Denes, Mirjam Feldkamp, Ina Nissen, Natascha Santacroce, Elodie Burcklen, Christiane Beckmann, Maurice Redondo, Olivier Kobel, Christoph Noppen, Niko Beerenwinkel, Tanja Stadler |
| EPI_ISL_3345932 | Fulgent Genetics | Centers for Disease Control and Prevention Division of Viral Diseases, Pathogen Discovery | Dakota Howard, Dhwani Batra, Peter W. Cook, Kara Moser, Adrian Paskey, Jason Caravas, Benjamin Rambo-Martin, Shatavia Morrison, Christopher Gulvick, Scott Sammons, Yvette Unoarumhi, Darlene Wagner, Matthew Schmerer, Harry Gao, Mickey Li, John Gao, Joseph Fierro, Benafsh Sapra, Becky Tsai, Yan Meng, Doreen Ng, James Xie, Clinton R. Paden, Duncan MacCannell |
| EPI_ISL_3345933, EPI_ISL_3345934 | Viollier AG | Department of Biosystems Science and Engineering, ETH Zürich | Chaoran Chen, Sarah Nadeau, Louis du Plessis, Catharine Aquino, Ivan Topolsky, Kim Philipp Jablonski, Lara Fuhrmann, Daniel Ehrsam, Isabel Stürmer, Andreia Cabral de Gouvea, Maria Domenica Moccia, Simon Grüter, Timothy Sykes, Lennart Opitz, Griffin White, Laura Neff, Doris Popovic, Andrea Patrignani, Jay Tracy, Ralph Schlapbach, Christiane Beckmann, Maurice Redondo, Olivier Kobel, Christoph Noppen, Niko Beerenwinkel, Tanja Stadler |
| EPI_ISL_3345935 | Fulgent Genetics | Centers for Disease Control and Prevention Division of Viral Diseases, Pathogen Discovery | Dakota Howard, Dhwani Batra, Peter W. Cook, Kara Moser, Adrian Paskey, Jason Caravas, Benjamin Rambo-Martin, Shatavia Morrison, Christopher Gulvick, Scott Sammons, Yvette Unoarumhi, Darlene Wagner, Matthew Schmerer, Harry Gao, Mickey Li, John Gao, Joseph Fierro, Benafsh Sapra, Becky Tsai, Yan Meng, Doreen Ng, James Xie, Clinton R. Paden, Duncan MacCannell |
| EPI_ISL_3345936, EPI_ISL_3345937 | Viollier AG | Department of Biosystems Science and Engineering, ETH Zürich | Chaoran Chen, Sarah Nadeau, Louis du Plessis, Catharine Aquino, Ivan Topolsky, Kim Philipp Jablonski, Lara Fuhrmann, Daniel Ehrsam, Isabel Stürmer, Andreia Cabral de Gouvea, Maria Domenica Moccia, Simon Grüter, Timothy Sykes, Lennart Opitz, Griffin White, Laura Neff, Doris Popovic, Andrea Patrignani, Jay Tracy, Ralph Schlapbach, Christiane Beckmann, Maurice Redondo, Olivier Kobel, Christoph Noppen, Niko Beerenwinkel, Tanja Stadler |
| EPI_ISL_3345938 | Fulgent Genetics | Centers for Disease Control and Prevention Division of Viral Diseases, Pathogen Discovery | Dakota Howard, Dhwani Batra, Peter W. Cook, Kara Moser, Adrian Paskey, Jason Caravas, Benjamin Rambo-Martin, Shatavia Morrison, Christopher Gulvick, Scott Sammons, Yvette Unoarumhi, Darlene Wagner, Matthew Schmerer, Harry Gao, Mickey Li, John Gao, Joseph Fierro, Benafsh Sapra, Becky Tsai, Yan Meng, Doreen Ng, James Xie, Clinton R. Paden, Duncan MacCannell |
| EPI_ISL_3345939, EPI_ISL_3345940 | Viollier AG | Department of Biosystems Science and Engineering, ETH Zürich | Chaoran Chen, Sarah Nadeau, Louis du Plessis, Catharine Aquino, Ivan Topolsky, Kim Philipp Jablonski, Lara Fuhrmann, Daniel Ehrsam, Isabel Stürmer, Andreia Cabral de Gouvea, Maria Domenica Moccia, Simon Grüter, Timothy Sykes, Lennart Opitz, Griffin White, Laura Neff, Doris Popovic, Andrea Patrignani, Jay Tracy, Ralph Schlapbach, Christiane Beckmann, Maurice Redondo, Olivier Kobel, Christoph Noppen, Niko Beerenwinkel, Tanja Stadler |
| EPI_ISL_3345941 | Fulgent Genetics | Centers for Disease Control and Prevention Division of Viral Diseases, Pathogen Discovery | Dakota Howard, Dhwani Batra, Peter W. Cook, Kara Moser, Adrian Paskey, Jason Caravas, Benjamin Rambo-Martin, Shatavia Morrison, Christopher Gulvick, Scott Sammons, Yvette Unoarumhi, Darlene Wagner, Matthew Schmerer, Harry Gao, Mickey Li, John Gao, Joseph Fierro, Benafsh Sapra, Becky Tsai, Yan Meng, Doreen Ng, James Xie, Clinton R. Paden, Duncan MacCannell |
| EPI_ISL_3345942, EPI_ISL_3345943 | Viollier AG | Department of Biosystems Science and Engineering, ETH Zürich | Christian Beisel, Sarah Nadeau, Chaoran Chen, Louis du Plessis, Ivan Topolsky, Kim Philipp Jablonski, Lara Fuhrmann, Rebecca Denes, Mirjam Feldkamp, Ina Nissen, Natascha Santacroce, Elodie Burcklen, Christiane Beckmann, Maurice Redondo, Olivier Kobel, Christoph Noppen, Niko Beerenwinkel, Tanja Stadler |
| EPI_ISL_3345944 | Fulgent Genetics | Centers for Disease Control and Prevention Division of Viral Diseases, Pathogen Discovery | Dakota Howard, Dhwani Batra, Peter W. Cook, Kara Moser, Adrian Paskey, Jason Caravas, Benjamin Rambo-Martin, Shatavia Morrison, Christopher Gulvick, Scott Sammons, Yvette Unoarumhi, Darlene Wagner, Matthew Schmerer, Harry Gao, Mickey Li, John Gao, Joseph Fierro, Benafsh Sapra, Becky Tsai, Yan Meng, Doreen Ng, James Xie, Clinton R. Paden, Duncan MacCannell |
| EPI_ISL_3345945 | Viollier AG | Department of Biosystems Science and Engineering, ETH Zürich | Chaoran Chen, Sarah Nadeau, Louis du Plessis, Catharine Aquino, Ivan Topolsky, Kim Philipp Jablonski, Lara Fuhrmann, Daniel Ehrsam, Isabel Stürmer, Andreia Cabral de Gouvea, Maria Domenica Moccia, Simon Grüter, Timothy Sykes, Lennart Opitz, Griffin White, Laura Neff, Doris Popovic, Andrea Patrignani, Jay Tracy, Ralph Schlapbach, Christiane Beckmann, Maurice Redondo, Olivier Kobel, Christoph Noppen, Niko Beerenwinkel, Tanja Stadler |
| EPI_ISL_3345946 | Fulgent Genetics | Centers for Disease Control and Prevention Division of Viral Diseases, Pathogen Discovery | Dakota Howard, Dhwani Batra, Peter W. Cook, Kara Moser, Adrian Paskey, Jason Caravas, Benjamin Rambo-Martin, Shatavia Morrison, Christopher Gulvick, Scott Sammons, Yvette Unoarumhi, Darlene Wagner, Matthew Schmerer, Harry Gao, Mickey Li, John Gao, Joseph Fierro, Benafsh Sapra, Becky Tsai, Yan Meng, Doreen Ng, James Xie, Clinton R. Paden, Duncan MacCannell |
| EPI_ISL_3345947, EPI_ISL_3345948 | Viollier AG | Department of Biosystems Science and Engineering, ETH Zürich | Chaoran Chen, Sarah Nadeau, Louis du Plessis, Catharine Aquino, Ivan Topolsky, Kim Philipp Jablonski, Lara Fuhrmann, Daniel Ehrsam, Isabel Stürmer, Andreia Cabral de Gouvea, Maria Domenica Moccia, Simon Grüter, Timothy Sykes, Lennart Opitz, Griffin White, Laura Neff, Doris Popovic, Andrea Patrignani, Jay Tracy, Ralph Schlapbach, Christiane Beckmann, Maurice Redondo, Olivier Kobel, Christoph Noppen, Niko Beerenwinkel, Tanja Stadler |
| EPI_ISL_3345949 | Fulgent Genetics | Centers for Disease Control and Prevention Division of Viral Diseases, Pathogen Discovery | Dakota Howard, Dhwani Batra, Peter W. Cook, Kara Moser, Adrian Paskey, Jason Caravas, Benjamin Rambo-Martin, Shatavia Morrison, Christopher Gulvick, Scott Sammons, Yvette Unoarumhi, Darlene Wagner, Matthew Schmerer, Harry Gao, Mickey Li, John Gao, Joseph Fierro, Benafsh Sapra, Becky Tsai, Yan Meng, Doreen Ng, James Xie, Clinton R. Paden, Duncan MacCannell |
| EPI_ISL_3345950 | Viollier AG | Department of Biosystems Science and Engineering, ETH Zürich | Christian Beisel, Sarah Nadeau, Chaoran Chen, Louis du Plessis, Ivan Topolsky, Kim Philipp Jablonski, Lara Fuhrmann, Rebecca Denes, Mirjam Feldkamp, Ina Nissen, Natascha Santacroce, Elodie Burcklen, Christiane Beckmann, Maurice Redondo, Olivier Kobel, Christoph Noppen, Niko Beerenwinkel, Tanja Stadler |
| EPI_ISL_3345951 | Fulgent Genetics | Centers for Disease Control and Prevention Division of Viral Diseases, Pathogen Discovery | Dakota Howard, Dhwani Batra, Peter W. Cook, Kara Moser, Adrian Paskey, Jason Caravas, Benjamin Rambo-Martin, Shatavia Morrison, Christopher Gulvick, Scott Sammons, Yvette Unoarumhi, Darlene Wagner, Matthew Schmerer, Harry Gao, Mickey Li, John Gao, Joseph Fierro, Benafsh Sapra, Becky Tsai, Yan Meng, Doreen Ng, James Xie, Clinton R. Paden, Duncan MacCannell |
| EPI_ISL_3345952, EPI_ISL_3345953, EPI_ISL_3345954, EPI_ISL_3345955, EPI_ISL_3345956, EPI_ISL_3345957, EPI_ISL_3345958 | Viollier AG | Department of Biosystems Science and Engineering, ETH Zürich | Chaoran Chen, Sarah Nadeau, Louis du Plessis, Catharine Aquino, Ivan Topolsky, Kim Philipp Jablonski, Lara Fuhrmann, Daniel Ehrsam, Isabel Stürmer, Andreia Cabral de Gouvea, Maria Domenica Moccia, Simon Grüter, Timothy Sykes, Lennart Opitz, Griffin White, Laura Neff, Doris Popovic, Andrea Patrignani, Jay Tracy, Ralph Schlapbach, Christiane Beckmann, Maurice Redondo, Olivier Kobel, Christoph Noppen, Niko Beerenwinkel, Tanja Stadler |
| EPI_ISL_3345959 | Fulgent Genetics | Centers for Disease Control and Prevention Division of Viral Diseases, Pathogen Discovery | Dakota Howard, Dhwani Batra, Peter W. Cook, Kara Moser, Adrian Paskey, Jason Caravas, Benjamin Rambo-Martin, Shatavia Morrison, Christopher Gulvick, Scott Sammons, Yvette Unoarumhi, Darlene Wagner, Matthew Schmerer, Harry Gao, Mickey Li, John Gao, Joseph Fierro, Benafsh Sapra, Becky Tsai, Yan Meng, Doreen Ng, James Xie, Clinton R. Paden, Duncan MacCannell |
| EPI_ISL_3345960, EPI_ISL_3345961 | Viollier AG | Department of Biosystems Science and Engineering, ETH Zürich | Chaoran Chen, Sarah Nadeau, Louis du Plessis, Catharine Aquino, Ivan Topolsky, Kim Philipp Jablonski, Lara Fuhrmann, Daniel Ehrsam, Isabel Stürmer, Andreia Cabral de Gouvea, Maria Domenica Moccia, Simon Grüter, Timothy Sykes, Lennart Opitz, Griffin White, Laura Neff, Doris Popovic, Andrea Patrignani, Jay Tracy, Ralph Schlapbach, Christiane Beckmann, Maurice Redondo, Olivier Kobel, Christoph Noppen, Niko Beerenwinkel, Tanja Stadler |
| EPI_ISL_3345962 | Fulgent Genetics | Centers for Disease Control and Prevention Division of Viral Diseases, Pathogen Discovery | Dakota Howard, Dhwani Batra, Peter W. Cook, Kara Moser, Adrian Paskey, Jason Caravas, Benjamin Rambo-Martin, Shatavia Morrison, Christopher Gulvick, Scott Sammons, Yvette Unoarumhi, Darlene Wagner, Matthew Schmerer, Harry Gao, Mickey Li, John Gao, Joseph Fierro, Benafsh Sapra, Becky Tsai, Yan Meng, Doreen Ng, James Xie, Clinton R. Paden, Duncan MacCannell |
| EPI_ISL_3345963 | Viollier AG | Department of Biosystems Science and Engineering, ETH Zürich | Christian Beisel, Sarah Nadeau, Chaoran Chen, Louis du Plessis, Ivan Topolsky, Kim Philipp Jablonski, Lara Fuhrmann, Rebecca Denes, Mirjam Feldkamp, Ina Nissen, Natascha Santacroce, Elodie Burcklen, Christiane Beckmann, Maurice Redondo, Olivier Kobel, Christoph Noppen, Niko Beerenwinkel, Tanja Stadler |
| EPI_ISL_3345964 | Viollier AG | Department of Biosystems Science and Engineering, ETH Zürich | Chaoran Chen, Sarah Nadeau, Louis du Plessis, Catharine Aquino, Ivan Topolsky, Kim Philipp Jablonski, Lara Fuhrmann, Daniel Ehrsam, Isabel Stürmer, Andreia Cabral de Gouvea, Maria Domenica Moccia, Simon Grüter, Timothy Sykes, Lennart Opitz, Griffin White, Laura Neff, Doris Popovic, Andrea Patrignani, Jay Tracy, Ralph Schlapbach, Christiane Beckmann, Maurice Redondo, Olivier Kobel, Christoph Noppen, Niko Beerenwinkel, Tanja Stadler |
| EPI_ISL_3345965 | Fulgent Genetics | Centers for Disease Control and Prevention Division of Viral Diseases, Pathogen Discovery | Dakota Howard, Dhwani Batra, Peter W. Cook, Kara Moser, Adrian Paskey, Jason Caravas, Benjamin Rambo-Martin, Shatavia Morrison, Christopher Gulvick, Scott Sammons, Yvette Unoarumhi, Darlene Wagner, Matthew Schmerer, Harry Gao, Mickey Li, John Gao, Joseph Fierro, Benafsh Sapra, Becky Tsai, Yan Meng, Doreen Ng, James Xie, Clinton R. Paden, Duncan MacCannell |

| | | | |
|---|---|---|---|
| EPI_ISL_3345966 | Viollier AG | Department of Biosystems Science and Engineering, ETH Zürich | Chaoran Chen, Sarah Nadeau, Louis du Plessis, Catharine Aquino, Ivan Topolsky, Kim Philipp Jablonski, Lara Fuhrmann, Daniel Ehrsam, Isabel Stürmer, Andreia Cabral de Gouvea, Maria Domenica Moccia, Simon Grüter, Timothy Sykes, Lennart Opitz, Griffin White, Laura Neff, Doris Popovic, Andrea Patrignani, Jay Tracy, Ralph Schlapbach, Christiane Beckmann, Maurice Redondo, Olivier Kobel, Christoph Noppen, Niko Beerenwinkel, Tanja Stadler |
| EPI_ISL_3345967 | Fulgent Genetics | Centers for Disease Control and Prevention Division of Viral Diseases, Pathogen Discovery | Dakota Howard, Dhwani Batra, Peter W. Cook, Kara Moser, Adrian Paskey, Jason Caravas, Benjamin Rambo-Martin, Shatavia Morrison, Christopher Gulvick, Scott Sammons, Yvette Unoarumhi, Darlene Wagner, Matthew Schmerer, Harry Gao, Mickey Li, John Gao, Joseph Fierro, Benafsh Sapra, Becky Tsai, Yan Meng, Doreen Ng, James Xie, Clinton R. Paden, Duncan MacCannell |
| EPI_ISL_3345969 | Viollier AG | Department of Biosystems Science and Engineering, ETH Zürich | Christian Beisel, Sarah Nadeau, Chaoran Chen, Louis du Plessis, Ivan Topolsky, Kim Philipp Jablonski, Lara Fuhrmann, Rebecca Denes, Mirjam Feldkamp, Ina Nissen, Natascha Santacroce, Elodie Burcklen, Christiane Beckmann, Maurice Redondo, Olivier Kobel, Christoph Noppen, Niko Beerenwinkel, Tanja Stadler |
| EPI_ISL_3345970 | Fulgent Genetics | Centers for Disease Control and Prevention Division of Viral Diseases, Pathogen Discovery | Dakota Howard, Dhwani Batra, Peter W. Cook, Kara Moser, Adrian Paskey, Jason Caravas, Benjamin Rambo-Martin, Shatavia Morrison, Christopher Gulvick, Scott Sammons, Yvette Unoarumhi, Darlene Wagner, Matthew Schmerer, Harry Gao, Mickey Li, John Gao, Joseph Fierro, Benafsh Sapra, Becky Tsai, Yan Meng, Doreen Ng, James Xie, Clinton R. Paden, Duncan MacCannell |
| EPI_ISL_3345971 | Viollier AG | Department of Biosystems Science and Engineering, ETH Zürich | Christian Beisel, Sarah Nadeau, Chaoran Chen, Louis du Plessis, Ivan Topolsky, Kim Philipp Jablonski, Lara Fuhrmann, Rebecca Denes, Mirjam Feldkamp, Ina Nissen, Natascha Santacroce, Elodie Burcklen, Christiane Beckmann, Maurice Redondo, Olivier Kobel, Christoph Noppen, Niko Beerenwinkel, Tanja Stadler |
| EPI_ISL_3345972 | Viollier AG | Department of Biosystems Science and Engineering, ETH Zürich | Chaoran Chen, Sarah Nadeau, Louis du Plessis, Catharine Aquino, Ivan Topolsky, Kim Philipp Jablonski, Lara Fuhrmann, Daniel Ehrsam, Isabel Stürmer, Andreia Cabral de Gouvea, Maria Domenica Moccia, Simon Grüter, Timothy Sykes, Lennart Opitz, Griffin White, Laura Neff, Doris Popovic, Andrea Patrignani, Jay Tracy, Ralph Schlapbach, Christiane Beckmann, Maurice Redondo, Olivier Kobel, Christoph Noppen, Niko Beerenwinkel, Tanja Stadler |
| EPI_ISL_3345973 | Fulgent Genetics | Centers for Disease Control and Prevention Division of Viral Diseases, Pathogen Discovery | Dakota Howard, Dhwani Batra, Peter W. Cook, Kara Moser, Adrian Paskey, Jason Caravas, Benjamin Rambo-Martin, Shatavia Morrison, Christopher Gulvick, Scott Sammons, Yvette Unoarumhi, Darlene Wagner, Matthew Schmerer, Harry Gao, Mickey Li, John Gao, Joseph Fierro, Benafsh Sapra, Becky Tsai, Yan Meng, Doreen Ng, James Xie, Clinton R. Paden, Duncan MacCannell |
| EPI_ISL_3345974, EPI_ISL_3345975 | Viollier AG | Department of Biosystems Science and Engineering, ETH Zürich | Chaoran Chen, Sarah Nadeau, Louis du Plessis, Catharine Aquino, Ivan Topolsky, Kim Philipp Jablonski, Lara Fuhrmann, Daniel Ehrsam, Isabel Stürmer, Andreia Cabral de Gouvea, Maria Domenica Moccia, Simon Grüter, Timothy Sykes, Lennart Opitz, Griffin White, Laura Neff, Doris Popovic, Andrea Patrignani, Jay Tracy, Ralph Schlapbach, Christiane Beckmann, Maurice Redondo, Olivier Kobel, Christoph Noppen, Niko Beerenwinkel, Tanja Stadler |
| EPI_ISL_3345976 | Fulgent Genetics | Centers for Disease Control and Prevention Division of Viral Diseases, Pathogen Discovery | Dakota Howard, Dhwani Batra, Peter W. Cook, Kara Moser, Adrian Paskey, Jason Caravas, Benjamin Rambo-Martin, Shatavia Morrison, Christopher Gulvick, Scott Sammons, Yvette Unoarumhi, Darlene Wagner, Matthew Schmerer, Harry Gao, Mickey Li, John Gao, Joseph Fierro, Benafsh Sapra, Becky Tsai, Yan Meng, Doreen Ng, James Xie, Clinton R. Paden, Duncan MacCannell |
| EPI_ISL_3345977 | Viollier AG | Department of Biosystems Science and Engineering, ETH Zürich | Christian Beisel, Sarah Nadeau, Chaoran Chen, Louis du Plessis, Ivan Topolsky, Kim Philipp Jablonski, Lara Fuhrmann, Rebecca Denes, Mirjam Feldkamp, Ina Nissen, Natascha Santacroce, Elodie Burcklen, Christiane Beckmann, Maurice Redondo, Olivier Kobel, Christoph Noppen, Niko Beerenwinkel, Tanja Stadler |
| EPI_ISL_3345978 | Fulgent Genetics | Centers for Disease Control and Prevention Division of Viral Diseases, Pathogen Discovery | Dakota Howard, Dhwani Batra, Peter W. Cook, Kara Moser, Adrian Paskey, Jason Caravas, Benjamin Rambo-Martin, Shatavia Morrison, Christopher Gulvick, Scott Sammons, Yvette Unoarumhi, Darlene Wagner, Matthew Schmerer, Harry Gao, Mickey Li, John Gao, Joseph Fierro, Benafsh Sapra, Becky Tsai, Yan Meng, Doreen Ng, James Xie, Clinton R. Paden, Duncan MacCannell |
| EPI_ISL_3345979 | Viollier AG | Department of Biosystems Science and Engineering, ETH Zürich | Christian Beisel, Sarah Nadeau, Chaoran Chen, Louis du Plessis, Ivan Topolsky, Kim Philipp Jablonski, Lara Fuhrmann, Rebecca Denes, Mirjam Feldkamp, Ina Nissen, Natascha Santacroce, Elodie Burcklen, Christiane Beckmann, Maurice Redondo, Olivier Kobel, Christoph Noppen, Niko Beerenwinkel, Tanja Stadler |
| EPI_ISL_3345980 | Viollier AG | Department of Biosystems Science and Engineering, ETH Zürich | Chaoran Chen, Sarah Nadeau, Louis du Plessis, Catharine Aquino, Ivan Topolsky, Kim Philipp Jablonski, Lara Fuhrmann, Daniel Ehrsam, Isabel Stürmer, Andreia Cabral de Gouvea, Maria Domenica Moccia, Simon Grüter, Timothy Sykes, Lennart Opitz, Griffin White, Laura Neff, Doris Popovic, Andrea Patrignani, Jay Tracy, Ralph Schlapbach, Christiane Beckmann, Maurice Redondo, Olivier Kobel, Christoph Noppen, Niko Beerenwinkel, Tanja Stadler |
| EPI_ISL_3345981 | Fulgent Genetics | Centers for Disease Control and Prevention Division of Viral Diseases, Pathogen Discovery | Dakota Howard, Dhwani Batra, Peter W. Cook, Kara Moser, Adrian Paskey, Jason Caravas, Benjamin Rambo-Martin, Shatavia Morrison, Christopher Gulvick, Scott Sammons, Yvette Unoarumhi, Darlene Wagner, Matthew Schmerer, Harry Gao, Mickey Li, John Gao, Joseph Fierro, Benafsh Sapra, Becky Tsai, Yan Meng, Doreen Ng, James Xie, Clinton R. Paden, Duncan MacCannell |
| EPI_ISL_3345982 | Viollier AG | Department of Biosystems Science and Engineering, ETH Zürich | Christian Beisel, Sarah Nadeau, Chaoran Chen, Louis du Plessis, Ivan Topolsky, Kim Philipp Jablonski, Lara Fuhrmann, Rebecca Denes, Mirjam Feldkamp, Ina Nissen, Natascha Santacroce, Elodie Burcklen, Christiane Beckmann, Maurice Redondo, Olivier Kobel, Christoph Noppen, Niko Beerenwinkel, Tanja Stadler |
| EPI_ISL_3345983 | Fulgent Genetics | Centers for Disease Control and Prevention Division of Viral Diseases, Pathogen Discovery | Dakota Howard, Dhwani Batra, Peter W. Cook, Kara Moser, Adrian Paskey, Jason Caravas, Benjamin Rambo-Martin, Shatavia Morrison, Christopher Gulvick, Scott Sammons, Yvette Unoarumhi, Darlene Wagner, Matthew Schmerer, Harry Gao, Mickey Li, John Gao, Joseph Fierro, Benafsh Sapra, Becky Tsai, Yan Meng, Doreen Ng, James Xie, Clinton R. Paden, Duncan MacCannell |
| EPI_ISL_3345984, EPI_ISL_3345985 | Viollier AG | Department of Biosystems Science and Engineering, ETH Zürich | Chaoran Chen, Sarah Nadeau, Louis du Plessis, Catharine Aquino, Ivan Topolsky, Kim Philipp Jablonski, Lara Fuhrmann, Daniel Ehrsam, Isabel Stürmer, Andreia Cabral de Gouvea, Maria Domenica Moccia, Simon Grüter, Timothy Sykes, Lennart Opitz, Griffin White, Laura Neff, Doris Popovic, Andrea Patrignani, Jay Tracy, Ralph Schlapbach, Christiane Beckmann, Maurice Redondo, Olivier Kobel, Christoph Noppen, Niko Beerenwinkel, Tanja Stadler |
| EPI_ISL_3345986 | Fulgent Genetics | Centers for Disease Control and Prevention Division of Viral Diseases, Pathogen Discovery | Dakota Howard, Dhwani Batra, Peter W. Cook, Kara Moser, Adrian Paskey, Jason Caravas, Benjamin Rambo-Martin, Shatavia Morrison, Christopher Gulvick, Scott Sammons, Yvette Unoarumhi, Darlene Wagner, Matthew Schmerer, Harry Gao, Mickey Li, John Gao, Joseph Fierro, Benafsh Sapra, Becky Tsai, Yan Meng, Doreen Ng, James Xie, Clinton R. Paden, Duncan MacCannell |
| EPI_ISL_3345987 | Viollier AG | Department of Biosystems Science and Engineering, ETH Zürich | Chaoran Chen, Sarah Nadeau, Louis du Plessis, Catharine Aquino, Ivan Topolsky, Kim Philipp Jablonski, Lara Fuhrmann, Daniel Ehrsam, Isabel Stürmer, Andreia Cabral de Gouvea, Maria Domenica Moccia, Simon Grüter, Timothy Sykes, Lennart Opitz, Griffin White, Laura Neff, Doris Popovic, Andrea Patrignani, Jay Tracy, Ralph Schlapbach, Christiane Beckmann, Maurice Redondo, Olivier Kobel, Christoph Noppen, Niko Beerenwinkel, Tanja Stadler |
| EPI_ISL_3345988 | Fulgent Genetics | Centers for Disease Control and Prevention Division of Viral Diseases, Pathogen Discovery | Dakota Howard, Dhwani Batra, Peter W. Cook, Kara Moser, Adrian Paskey, Jason Caravas, Benjamin Rambo-Martin, Shatavia Morrison, Christopher Gulvick, Scott Sammons, Yvette Unoarumhi, Darlene Wagner, Matthew Schmerer, Harry Gao, Mickey Li, John Gao, Joseph Fierro, Benafsh Sapra, Becky Tsai, Yan Meng, Doreen Ng, James Xie, Clinton R. Paden, Duncan MacCannell |
| EPI_ISL_3345989, EPI_ISL_3345990 | Viollier AG | Department of Biosystems Science and Engineering, ETH Zürich | Chaoran Chen, Sarah Nadeau, Louis du Plessis, Catharine Aquino, Ivan Topolsky, Kim Philipp Jablonski, Lara Fuhrmann, Daniel Ehrsam, Isabel Stürmer, Andreia Cabral de Gouvea, Maria Domenica Moccia, Simon Grüter, Timothy Sykes, Lennart Opitz, Griffin White, Laura Neff, Doris Popovic, Andrea Patrignani, Jay Tracy, Ralph Schlapbach, Christiane Beckmann, Maurice Redondo, Olivier Kobel, Christoph Noppen, Niko Beerenwinkel, Tanja Stadler |
| EPI_ISL_3345991 | Fulgent Genetics | Centers for Disease Control and Prevention Division of Viral Diseases, Pathogen Discovery | Dakota Howard, Dhwani Batra, Peter W. Cook, Kara Moser, Adrian Paskey, Jason Caravas, Benjamin Rambo-Martin, Shatavia Morrison, Christopher Gulvick, Scott Sammons, Yvette Unoarumhi, Darlene Wagner, Matthew Schmerer, Harry Gao, Mickey Li, John Gao, Joseph Fierro, Benafsh Sapra, Becky Tsai, Yan Meng, Doreen Ng, James Xie, Clinton R. Paden, Duncan MacCannell |
| EPI_ISL_3345992 | Viollier AG | Department of Biosystems Science and Engineering, ETH Zürich | Chaoran Chen, Sarah Nadeau, Louis du Plessis, Catharine Aquino, Ivan Topolsky, Kim Philipp Jablonski, Lara Fuhrmann, Daniel Ehrsam, Isabel Stürmer, Andreia Cabral de Gouvea, Maria Domenica Moccia, Simon Grüter, Timothy Sykes, Lennart Opitz, Griffin White, Laura Neff, Doris Popovic, Andrea Patrignani, Jay Tracy, Ralph Schlapbach, Christiane Beckmann, Maurice Redondo, Olivier Kobel, Christoph Noppen, Niko Beerenwinkel, Tanja Stadler |
| EPI_ISL_3345994 | Fulgent Genetics | Centers for Disease Control and Prevention Division of Viral Diseases, Pathogen Discovery | Dakota Howard, Dhwani Batra, Peter W. Cook, Kara Moser, Adrian Paskey, Jason Caravas, Benjamin Rambo-Martin, Shatavia Morrison, Christopher Gulvick, Scott Sammons, Yvette Unoarumhi, Darlene Wagner, Matthew Schmerer, Harry Gao, Mickey Li, John Gao, Joseph Fierro, Benafsh Sapra, Becky Tsai, Yan Meng, Doreen Ng, James Xie, Clinton R. Paden, Duncan MacCannell |
| EPI_ISL_3345995, EPI_ISL_3345996 | Viollier AG | Department of Biosystems Science and Engineering, ETH Zürich | Chaoran Chen, Sarah Nadeau, Louis du Plessis, Catharine Aquino, Ivan Topolsky, Kim Philipp Jablonski, Lara Fuhrmann, Daniel Ehrsam, Isabel Stürmer, Andreia Cabral de Gouvea, Maria Domenica Moccia, Simon Grüter, Timothy Sykes, Lennart Opitz, Griffin White, Laura Neff, Doris Popovic, Andrea Patrignani, Jay Tracy, Ralph Schlapbach, Christiane Beckmann, Maurice Redondo, Olivier Kobel, Christoph Noppen, Niko Beerenwinkel, Tanja Stadler |
| EPI_ISL_3345997 | Fulgent Genetics | Centers for Disease Control and Prevention Division of Viral Diseases, Pathogen Discovery | Dakota Howard, Dhwani Batra, Peter W. Cook, Kara Moser, Adrian Paskey, Jason Caravas, Benjamin Rambo-Martin, Shatavia Morrison, Christopher Gulvick, Scott Sammons, Yvette Unoarumhi, Darlene Wagner, Matthew Schmerer, Harry Gao, Mickey Li, John Gao, Joseph Fierro, Benafsh Sapra, Becky Tsai, Yan Meng, Doreen Ng, James Xie, Clinton R. Paden, Duncan MacCannell |
| EPI_ISL_3345998 | Viollier AG | Department of Biosystems Science and Engineering, ETH Zürich | Chaoran Chen, Sarah Nadeau, Louis du Plessis, Catharine Aquino, Ivan Topolsky, Kim Philipp Jablonski, Lara Fuhrmann, Daniel Ehrsam, Isabel Stürmer, Andreia Cabral de Gouvea, Maria Domenica Moccia, Simon Grüter, Timothy Sykes, Lennart Opitz, Griffin White, Laura Neff, Doris Popovic, Andrea Patrignani, Jay Tracy, Ralph Schlapbach, Christiane Beckmann, Maurice Redondo, Olivier Kobel, Christoph Noppen, Niko Beerenwinkel, Tanja Stadler |
| EPI_ISL_3345999 | Viollier AG | Department of Biosystems Science and Engineering, ETH Zürich | Christian Beisel, Sarah Nadeau, Chaoran Chen, Louis du Plessis, Ivan Topolsky, Kim Philipp Jablonski, Lara Fuhrmann, Rebecca Denes, Mirjam Feldkamp, Ina Nissen, Natascha Santacroce, Elodie Burcklen, Christiane Beckmann, Maurice Redondo, Olivier Kobel, Christoph Noppen, Niko Beerenwinkel, Tanja Stadler |
| EPI_ISL_3346000 | Fulgent Genetics | Centers for Disease Control and Prevention Division of Viral Diseases, Pathogen Discovery | Dakota Howard, Dhwani Batra, Peter W. Cook, Kara Moser, Adrian Paskey, Jason Caravas, Benjamin Rambo-Martin, Shatavia Morrison, Christopher Gulvick, Scott Sammons, Yvette Unoarumhi, Darlene Wagner, Matthew Schmerer, Harry Gao, Mickey Li, John Gao, Joseph Fierro, Benafsh Sapra, Becky Tsai, Yan Meng, Doreen Ng, James Xie, Clinton R. Paden, Duncan MacCannell |
| EPI_ISL_3346001, EPI_ISL_3346002 | Viollier AG | Department of Biosystems Science and Engineering, ETH Zürich | Chaoran Chen, Sarah Nadeau, Louis du Plessis, Catharine Aquino, Ivan Topolsky, Kim Philipp Jablonski, Lara Fuhrmann, Daniel Ehrsam, Isabel Stürmer, Andreia Cabral de Gouvea, Maria Domenica Moccia, Simon Grüter, Timothy Sykes, Lennart Opitz, Griffin White, Laura Neff, Doris Popovic, Andrea Patrignani, Jay Tracy, Ralph Schlapbach, Christiane Beckmann, Maurice Redondo, Olivier Kobel, Christoph Noppen, Niko Beerenwinkel, Tanja Stadler |
| EPI_ISL_3346003 | Fulgent Genetics | Centers for Disease Control and Prevention Division of Viral Diseases, Pathogen Discovery | Dakota Howard, Dhwani Batra, Peter W. Cook, Kara Moser, Adrian Paskey, Jason Caravas, Benjamin Rambo-Martin, Shatavia Morrison, Christopher Gulvick, Scott Sammons, Yvette Unoarumhi, Darlene Wagner, Matthew Schmerer, Harry Gao, Mickey Li, John Gao, Joseph Fierro, Benafsh Sapra, Becky Tsai, Yan Meng, Doreen Ng, James Xie, Clinton R. Paden, Duncan MacCannell |
| EPI_ISL_3346004 | Viollier AG | Department of Biosystems Science and Engineering, ETH Zürich | Chaoran Chen, Sarah Nadeau, Louis du Plessis, Catharine Aquino, Ivan Topolsky, Kim Philipp Jablonski, Lara Fuhrmann, Daniel Ehrsam, Isabel Stürmer, Andreia Cabral de Gouvea, Maria Domenica Moccia, Simon Grüter, Timothy Sykes, Lennart Opitz, Griffin White, Laura Neff, Doris Popovic, Andrea Patrignani, Jay Tracy, Ralph Schlapbach, Christiane Beckmann, Maurice Redondo, Olivier Kobel, Christoph Noppen, Niko Beerenwinkel, Tanja Stadler |
| EPI_ISL_3346005 | Fulgent Genetics | Centers for Disease Control and Prevention Division of Viral Diseases, Pathogen Discovery | Dakota Howard, Dhwani Batra, Peter W. Cook, Kara Moser, Adrian Paskey, Jason Caravas, Benjamin Rambo-Martin, Shatavia Morrison, Christopher Gulvick, Scott Sammons, Yvette Unoarumhi, Darlene Wagner, Matthew Schmerer, Harry Gao, Mickey Li, John Gao, Joseph Fierro, Benafsh Sapra, Becky Tsai, Yan Meng, Doreen Ng, James Xie, Clinton R. Paden, Duncan MacCannell |
| EPI_ISL_3346006 | Viollier AG | Department of Biosystems Science and Engineering, ETH Zürich | Christian Beisel, Sarah Nadeau, Chaoran Chen, Louis du Plessis, Ivan Topolsky, Kim Philipp Jablonski, Lara Fuhrmann, Rebecca Denes, Mirjam Feldkamp, Ina Nissen, Natascha Santacroce, Elodie Burcklen, Christiane Beckmann, Maurice Redondo, Olivier Kobel, Christoph Noppen, Niko Beerenwinkel, Tanja Stadler |
| EPI_ISL_3346007 | Viollier AG | Department of Biosystems Science and Engineering, ETH Zürich | Chaoran Chen, Sarah Nadeau, Louis du Plessis, Catharine Aquino, Ivan Topolsky, Kim Philipp Jablonski, Lara Fuhrmann, Daniel Ehrsam, Isabel Stürmer, Andreia Cabral de Gouvea, Maria Domenica Moccia, Simon Grüter, Timothy Sykes, Lennart Opitz, Griffin White, Laura Neff, Doris Popovic, Andrea Patrignani, Jay Tracy, Ralph Schlapbach, Christiane Beckmann, Maurice Redondo, Olivier Kobel, Christoph Noppen, Niko Beerenwinkel, Tanja Stadler |
| EPI_ISL_3346008 | Fulgent Genetics | Centers for Disease Control and Prevention Division of Viral Diseases, Pathogen Discovery | Dakota Howard, Dhwani Batra, Peter W. Cook, Kara Moser, Adrian Paskey, Jason Caravas, Benjamin Rambo-Martin, Shatavia Morrison, Christopher Gulvick, Scott Sammons, Yvette Unoarumhi, Darlene Wagner, Matthew Schmerer, Harry Gao, Mickey Li, John Gao, Joseph Fierro, Benafsh Sapra, Becky Tsai, Yan Meng, Doreen Ng, James Xie, Clinton R. Paden, Duncan MacCannell |
| EPI_ISL_3346009 | Viollier AG | Department of Biosystems Science and Engineering, ETH Zürich | Chaoran Chen, Sarah Nadeau, Louis du Plessis, Catharine Aquino, Ivan Topolsky, Kim Philipp Jablonski, Lara Fuhrmann, Daniel Ehrsam, Isabel Stürmer, Andreia Cabral de Gouvea, Maria Domenica Moccia, Simon Grüter, Timothy Sykes, Lennart Opitz, Griffin White, Laura Neff, Doris Popovic, Andrea Patrignani, Jay Tracy, Ralph Schlapbach, Christiane Beckmann, Maurice Redondo, Olivier Kobel, Christoph Noppen, Niko Beerenwinkel, Tanja Stadler |
| EPI_ISL_3346010 | Fulgent Genetics | Centers for Disease Control and Prevention Division of Viral Diseases, Pathogen Discovery | Dakota Howard, Dhwani Batra, Peter W. Cook, Kara Moser, Adrian Paskey, Jason Caravas, Benjamin Rambo-Martin, Shatavia Morrison, Christopher Gulvick, Scott Sammons, Yvette Unoarumhi, Darlene Wagner, Matthew Schmerer, Harry Gao, Mickey Li, John Gao, Joseph Fierro, Benafsh Sapra, Becky Tsai, Yan Meng, Doreen Ng, James Xie, Clinton R. Paden, Duncan MacCannell |
| EPI_ISL_3346011, EPI_ISL_3346012 | Viollier AG | Department of Biosystems Science and Engineering, ETH Zürich | Chaoran Chen, Sarah Nadeau, Louis du Plessis, Catharine Aquino, Ivan Topolsky, Kim Philipp Jablonski, Lara Fuhrmann, Daniel Ehrsam, Isabel Stürmer, Andreia Cabral de Gouvea, Maria Domenica Moccia, Simon Grüter, Timothy Sykes, Lennart Opitz, Griffin White, Laura Neff, Doris Popovic, Andrea Patrignani, Jay Tracy, Ralph Schlapbach, Christiane Beckmann, Maurice Redondo, Olivier Kobel, Christoph Noppen, Niko Beerenwinkel, Tanja Stadler |
| EPI_ISL_3346013 | Fulgent Genetics | Centers for Disease Control and Prevention Division of Viral Diseases, Pathogen Discovery | Dakota Howard, Dhwani Batra, Peter W. Cook, Kara Moser, Adrian Paskey, Jason Caravas, Benjamin Rambo-Martin, Shatavia Morrison, Christopher Gulvick, Scott Sammons, Yvette Unoarumhi, Darlene Wagner, Matthew Schmerer, Harry Gao, Mickey Li, John Gao, Joseph Fierro, Benafsh Sapra, Becky Tsai, Yan Meng, Doreen Ng, James Xie, Clinton R. Paden, Duncan MacCannell |
| EPI_ISL_3346014, EPI_ISL_3346015 | Viollier AG | Department of Biosystems Science and Engineering, ETH Zürich | Chaoran Chen, Sarah Nadeau, Louis du Plessis, Catharine Aquino, Ivan Topolsky, Kim Philipp Jablonski, Lara Fuhrmann, Daniel Ehrsam, Isabel Stürmer, Andreia Cabral de Gouvea, Maria Domenica Moccia, Simon Grüter, Timothy Sykes, Lennart Opitz, Griffin White, Laura Neff, Doris Popovic, Andrea Patrignani, Jay Tracy, Ralph Schlapbach, Christiane Beckmann, Maurice Redondo, Olivier Kobel, Christoph Noppen, Niko Beerenwinkel, Tanja Stadler |
| EPI_ISL_3346016 | Fulgent Genetics | Centers for Disease Control and Prevention Division of Viral Diseases, Pathogen Discovery | Dakota Howard, Dhwani Batra, Peter W. Cook, Kara Moser, Adrian Paskey, Jason Caravas, Benjamin Rambo-Martin, Shatavia Morrison, Christopher Gulvick, Scott Sammons, Yvette Unoarumhi, Darlene Wagner, Matthew Schmerer, Harry Gao, Mickey Li, John Gao, Joseph Fierro, Benafsh Sapra, Becky Tsai, Yan Meng, Doreen Ng, James Xie, Clinton R. Paden, Duncan MacCannell |
| EPI_ISL_3346017 | Viollier AG | Department of Biosystems Science and Engineering, ETH Zürich | Christian Beisel, Sarah Nadeau, Chaoran Chen, Louis du Plessis, Ivan Topolsky, Kim Philipp Jablonski, Lara Fuhrmann, Rebecca Denes, Mirjam Feldkamp, Ina Nissen, Natascha Santacroce, Elodie Burcklen, Christiane Beckmann, Maurice Redondo, Olivier Kobel, Christoph Noppen, Niko Beerenwinkel, Tanja Stadler |
| EPI_ISL_3346018 | Fulgent Genetics | Centers for Disease Control and Prevention Division of Viral Diseases, Pathogen Discovery | Dakota Howard, Dhwani Batra, Peter W. Cook, Kara Moser, Adrian Paskey, Jason Caravas, Benjamin Rambo-Martin, Shatavia Morrison, Christopher Gulvick, Scott Sammons, Yvette Unoarumhi, Darlene Wagner, Matthew Schmerer, Harry Gao, Mickey Li, John Gao, Joseph Fierro, Benafsh Sapra, Becky Tsai, Yan Meng, Doreen Ng, James Xie, Clinton R. Paden, Duncan MacCannell |
| EPI_ISL_3346019, EPI_ISL_3346020 | Viollier AG | Department of Biosystems Science and Engineering, ETH Zürich | Christian Beisel, Sarah Nadeau, Chaoran Chen, Louis du Plessis, Ivan Topolsky, Kim Philipp Jablonski, Lara Fuhrmann, Rebecca Denes, Mirjam Feldkamp, Ina Nissen, Natascha Santacroce, Elodie Burcklen, Christiane Beckmann, Maurice Redondo, Olivier Kobel, Christoph Noppen, Niko Beerenwinkel, Tanja Stadler |

| | | | |
|---|---|---|---|
| EPI_ISL_3346021 | Fulgent Genetics | Centers for Disease Control and Prevention Division of Viral Diseases, Pathogen Discovery | Dakota Howard, Dhwani Batra, Peter W. Cook, Kara Moser, Adrian Paskey, Jason Caravas, Benjamin Rambo-Martin, Shatavia Morrison, Christopher Gulvick, Scott Sammons, Yvette Unoarumhi, Darlene Wagner, Matthew Schmerer, Harry Gao, Mickey Li, John Gao, Joseph Fierro, Benafsh Sapra, Becky Tsai, Yan Meng, Doreen Ng, James Xie, Clinton R. Paden, Duncan MacCannell |
| EPI_ISL_3346022 | Viollier AG | Department of Biosystems Science and Engineering, ETH Zürich | Christian Beisel, Sarah Nadeau, Chaoran Chen, Louis du Plessis, Ivan Topolsky, Kim Philipp Jablonski, Lara Fuhrmann, Rebecca Denes, Mirjam Feldkamp, Ina Nissen, Natascha Santacroce, Elodie Burcklen, Christiane Beckmann, Maurice Redondo, Olivier Kobel, Christoph Noppen, Niko Beerenwinkel, Tanja Stadler |
| EPI_ISL_3346023 | Fulgent Genetics | Centers for Disease Control and Prevention Division of Viral Diseases, Pathogen Discovery | Dakota Howard, Dhwani Batra, Peter W. Cook, Kara Moser, Adrian Paskey, Jason Caravas, Benjamin Rambo-Martin, Shatavia Morrison, Christopher Gulvick, Scott Sammons, Yvette Unoarumhi, Darlene Wagner, Matthew Schmerer, Harry Gao, Mickey Li, John Gao, Joseph Fierro, Benafsh Sapra, Becky Tsai, Yan Meng, Doreen Ng, James Xie, Clinton R. Paden, Duncan MacCannell |
| EPI_ISL_3346024, EPI_ISL_3346025 | Viollier AG | Department of Biosystems Science and Engineering, ETH Zürich | Chaoran Chen, Sarah Nadeau, Louis du Plessis, Catharine Aquino, Ivan Topolsky, Kim Philipp Jablonski, Lara Fuhrmann, Daniel Ehrsam, Isabel Stürmer, Andreia Cabral de Gouvea, Maria Domenica Moccia, Simon Grüter, Timothy Sykes, Lennart Opitz, Griffin White, Laura Neff, Doris Popovic, Andrea Patrignani, Jay Tracy, Ralph Schlapbach, Christiane Beckmann, Maurice Redondo, Olivier Kobel, Christoph Noppen, Niko Beerenwinkel, Tanja Stadler |
| EPI_ISL_3346026 | Fulgent Genetics | Centers for Disease Control and Prevention Division of Viral Diseases, Pathogen Discovery | Dakota Howard, Dhwani Batra, Peter W. Cook, Kara Moser, Adrian Paskey, Jason Caravas, Benjamin Rambo-Martin, Shatavia Morrison, Christopher Gulvick, Scott Sammons, Yvette Unoarumhi, Darlene Wagner, Matthew Schmerer, Harry Gao, Mickey Li, John Gao, Joseph Fierro, Benafsh Sapra, Becky Tsai, Yan Meng, Doreen Ng, James Xie, Clinton R. Paden, Duncan MacCannell |
| EPI_ISL_3346027, EPI_ISL_3346028 | Viollier AG | Department of Biosystems Science and Engineering, ETH Zürich | Chaoran Chen, Sarah Nadeau, Louis du Plessis, Catharine Aquino, Ivan Topolsky, Kim Philipp Jablonski, Lara Fuhrmann, Daniel Ehrsam, Isabel Stürmer, Andreia Cabral de Gouvea, Maria Domenica Moccia, Simon Grüter, Timothy Sykes, Lennart Opitz, Griffin White, Laura Neff, Doris Popovic, Andrea Patrignani, Jay Tracy, Ralph Schlapbach, Christiane Beckmann, Maurice Redondo, Olivier Kobel, Christoph Noppen, Niko Beerenwinkel, Tanja Stadler |
| EPI_ISL_3346029 | Fulgent Genetics | Centers for Disease Control and Prevention Division of Viral Diseases, Pathogen Discovery | Dakota Howard, Dhwani Batra, Peter W. Cook, Kara Moser, Adrian Paskey, Jason Caravas, Benjamin Rambo-Martin, Shatavia Morrison, Christopher Gulvick, Scott Sammons, Yvette Unoarumhi, Darlene Wagner, Matthew Schmerer, Harry Gao, Mickey Li, John Gao, Joseph Fierro, Benafsh Sapra, Becky Tsai, Yan Meng, Doreen Ng, James Xie, Clinton R. Paden, Duncan MacCannell |
| EPI_ISL_3346030, EPI_ISL_3346031 | Viollier AG | Department of Biosystems Science and Engineering, ETH Zürich | Chaoran Chen, Sarah Nadeau, Louis du Plessis, Catharine Aquino, Ivan Topolsky, Kim Philipp Jablonski, Lara Fuhrmann, Daniel Ehrsam, Isabel Stürmer, Andreia Cabral de Gouvea, Maria Domenica Moccia, Simon Grüter, Timothy Sykes, Lennart Opitz, Griffin White, Laura Neff, Doris Popovic, Andrea Patrignani, Jay Tracy, Ralph Schlapbach, Christiane Beckmann, Maurice Redondo, Olivier Kobel, Christoph Noppen, Niko Beerenwinkel, Tanja Stadler |
| EPI_ISL_3346032 | Fulgent Genetics | Centers for Disease Control and Prevention Division of Viral Diseases, Pathogen Discovery | Dakota Howard, Dhwani Batra, Peter W. Cook, Kara Moser, Adrian Paskey, Jason Caravas, Benjamin Rambo-Martin, Shatavia Morrison, Christopher Gulvick, Scott Sammons, Yvette Unoarumhi, Darlene Wagner, Matthew Schmerer, Harry Gao, Mickey Li, John Gao, Joseph Fierro, Benafsh Sapra, Becky Tsai, Yan Meng, Doreen Ng, James Xie, Clinton R. Paden, Duncan MacCannell |
| EPI_ISL_3346033 | Viollier AG | Department of Biosystems Science and Engineering, ETH Zürich | Christian Beisel, Sarah Nadeau, Chaoran Chen, Louis du Plessis, Ivan Topolsky, Kim Philipp Jablonski, Lara Fuhrmann, Rebecca Denes, Mirjam Feldkamp, Ina Nissen, Natascha Santacroce, Elodie Burcklen, Christiane Beckmann, Maurice Redondo, Olivier Kobel, Christoph Noppen, Niko Beerenwinkel, Tanja Stadler |
| EPI_ISL_3346034 | Fulgent Genetics | Centers for Disease Control and Prevention Division of Viral Diseases, Pathogen Discovery | Dakota Howard, Dhwani Batra, Peter W. Cook, Kara Moser, Adrian Paskey, Jason Caravas, Benjamin Rambo-Martin, Shatavia Morrison, Christopher Gulvick, Scott Sammons, Yvette Unoarumhi, Darlene Wagner, Matthew Schmerer, Harry Gao, Mickey Li, John Gao, Joseph Fierro, Benafsh Sapra, Becky Tsai, Yan Meng, Doreen Ng, James Xie, Clinton R. Paden, Duncan MacCannell |
| EPI_ISL_3346035 | Viollier AG | Department of Biosystems Science and Engineering, ETH Zürich | Chaoran Chen, Sarah Nadeau, Louis du Plessis, Catharine Aquino, Ivan Topolsky, Kim Philipp Jablonski, Lara Fuhrmann, Daniel Ehrsam, Isabel Stürmer, Andreia Cabral de Gouvea, Maria Domenica Moccia, Simon Grüter, Timothy Sykes, Lennart Opitz, Griffin White, Laura Neff, Doris Popovic, Andrea Patrignani, Jay Tracy, Ralph Schlapbach, Christiane Beckmann, Maurice Redondo, Olivier Kobel, Christoph Noppen, Niko Beerenwinkel, Tanja Stadler |
| EPI_ISL_3346036 | Viollier AG | Department of Biosystems Science and Engineering, ETH Zürich | Christian Beisel, Sarah Nadeau, Chaoran Chen, Louis du Plessis, Ivan Topolsky, Kim Philipp Jablonski, Lara Fuhrmann, Rebecca Denes, Mirjam Feldkamp, Ina Nissen, Natascha Santacroce, Elodie Burcklen, Christiane Beckmann, Maurice Redondo, Olivier Kobel, Christoph Noppen, Niko Beerenwinkel, Tanja Stadler |
| EPI_ISL_3346037 | Fulgent Genetics | Centers for Disease Control and Prevention Division of Viral Diseases, Pathogen Discovery | Dakota Howard, Dhwani Batra, Peter W. Cook, Kara Moser, Adrian Paskey, Jason Caravas, Benjamin Rambo-Martin, Shatavia Morrison, Christopher Gulvick, Scott Sammons, Yvette Unoarumhi, Darlene Wagner, Matthew Schmerer, Harry Gao, Mickey Li, John Gao, Joseph Fierro, Benafsh Sapra, Becky Tsai, Yan Meng, Doreen Ng, James Xie, Clinton R. Paden, Duncan MacCannell |
| EPI_ISL_3346038 | Viollier AG | Department of Biosystems Science and Engineering, ETH Zürich | Chaoran Chen, Sarah Nadeau, Louis du Plessis, Catharine Aquino, Ivan Topolsky, Kim Philipp Jablonski, Lara Fuhrmann, Daniel Ehrsam, Isabel Stürmer, Andreia Cabral de Gouvea, Maria Domenica Moccia, Simon Grüter, Timothy Sykes, Lennart Opitz, Griffin White, Laura Neff, Doris Popovic, Andrea Patrignani, Jay Tracy, Ralph Schlapbach, Christiane Beckmann, Maurice Redondo, Olivier Kobel, Christoph Noppen, Niko Beerenwinkel, Tanja Stadler |
| EPI_ISL_3346039, EPI_ISL_3346040 | Viollier AG | Department of Biosystems Science and Engineering, ETH Zürich | Christian Beisel, Sarah Nadeau, Chaoran Chen, Louis du Plessis, Ivan Topolsky, Kim Philipp Jablonski, Lara Fuhrmann, Rebecca Denes, Mirjam Feldkamp, Ina Nissen, Natascha Santacroce, Elodie Burcklen, Christiane Beckmann, Maurice Redondo, Olivier Kobel, Christoph Noppen, Niko Beerenwinkel, Tanja Stadler |
| EPI_ISL_3346041 | Viollier AG | Department of Biosystems Science and Engineering, ETH Zürich | Chaoran Chen, Sarah Nadeau, Louis du Plessis, Catharine Aquino, Ivan Topolsky, Kim Philipp Jablonski, Lara Fuhrmann, Daniel Ehrsam, Isabel Stürmer, Andreia Cabral de Gouvea, Maria Domenica Moccia, Simon Grüter, Timothy Sykes, Lennart Opitz, Griffin White, Laura Neff, Doris Popovic, Andrea Patrignani, Jay Tracy, Ralph Schlapbach, Christiane Beckmann, Maurice Redondo, Olivier Kobel, Christoph Noppen, Niko Beerenwinkel, Tanja Stadler |
| EPI_ISL_3346043 | Viollier AG | Department of Biosystems Science and Engineering, ETH Zürich | Christian Beisel, Sarah Nadeau, Chaoran Chen, Louis du Plessis, Ivan Topolsky, Kim Philipp Jablonski, Lara Fuhrmann, Rebecca Denes, Mirjam Feldkamp, Ina Nissen, Natascha Santacroce, Elodie Burcklen, Christiane Beckmann, Maurice Redondo, Olivier Kobel, Christoph Noppen, Niko Beerenwinkel, Tanja Stadler |
| EPI_ISL_3346044 | Viollier AG | Department of Biosystems Science and Engineering, ETH Zürich | Chaoran Chen, Sarah Nadeau, Louis du Plessis, Catharine Aquino, Ivan Topolsky, Kim Philipp Jablonski, Lara Fuhrmann, Daniel Ehrsam, Isabel Stürmer, Andreia Cabral de Gouvea, Maria Domenica Moccia, Simon Grüter, Timothy Sykes, Lennart Opitz, Griffin White, Laura Neff, Doris Popovic, Andrea Patrignani, Jay Tracy, Ralph Schlapbach, Christiane Beckmann, Maurice Redondo, Olivier Kobel, Christoph Noppen, Niko Beerenwinkel, Tanja Stadler |
| EPI_ISL_3346045 | Viollier AG | Department of Biosystems Science and Engineering, ETH Zürich | Christian Beisel, Sarah Nadeau, Chaoran Chen, Louis du Plessis, Ivan Topolsky, Kim Philipp Jablonski, Lara Fuhrmann, Rebecca Denes, Mirjam Feldkamp, Ina Nissen, Natascha Santacroce, Elodie Burcklen, Christiane Beckmann, Maurice Redondo, Olivier Kobel, Christoph Noppen, Niko Beerenwinkel, Tanja Stadler |
| EPI_ISL_3346046, EPI_ISL_3346047 | Viollier AG | Department of Biosystems Science and Engineering, ETH Zürich | Chaoran Chen, Sarah Nadeau, Louis du Plessis, Catharine Aquino, Ivan Topolsky, Kim Philipp Jablonski, Lara Fuhrmann, Daniel Ehrsam, Isabel Stürmer, Andreia Cabral de Gouvea, Maria Domenica Moccia, Simon Grüter, Timothy Sykes, Lennart Opitz, Griffin White, Laura Neff, Doris Popovic, Andrea Patrignani, Jay Tracy, Ralph Schlapbach, Christiane Beckmann, Maurice Redondo, Olivier Kobel, Christoph Noppen, Niko Beerenwinkel, Tanja Stadler |
| EPI_ISL_3346048 | Viollier AG | Department of Biosystems Science and Engineering, ETH Zürich | Christian Beisel, Sarah Nadeau, Chaoran Chen, Louis du Plessis, Ivan Topolsky, Kim Philipp Jablonski, Lara Fuhrmann, Rebecca Denes, Mirjam Feldkamp, Ina Nissen, Natascha Santacroce, Elodie Burcklen, Christiane Beckmann, Maurice Redondo, Olivier Kobel, Christoph Noppen, Niko Beerenwinkel, Tanja Stadler |
| EPI_ISL_3346049 | Viollier AG | Department of Biosystems Science and Engineering, ETH Zürich | Chaoran Chen, Sarah Nadeau, Louis du Plessis, Catharine Aquino, Ivan Topolsky, Kim Philipp Jablonski, Lara Fuhrmann, Daniel Ehrsam, Isabel Stürmer, Andreia Cabral de Gouvea, Maria Domenica Moccia, Simon Grüter, Timothy Sykes, Lennart Opitz, Griffin White, Laura Neff, Doris Popovic, Andrea Patrignani, Jay Tracy, Ralph Schlapbach, Christiane Beckmann, Maurice Redondo, Olivier Kobel, Christoph Noppen, Niko Beerenwinkel, Tanja Stadler |
| EPI_ISL_3346050 | Viollier AG | Department of Biosystems Science and Engineering, ETH Zürich | Christian Beisel, Sarah Nadeau, Chaoran Chen, Louis du Plessis, Ivan Topolsky, Kim Philipp Jablonski, Lara Fuhrmann, Rebecca Denes, Mirjam Feldkamp, Ina Nissen, Natascha Santacroce, Elodie Burcklen, Christiane Beckmann, Maurice Redondo, Olivier Kobel, Christoph Noppen, Niko Beerenwinkel, Tanja Stadler |
| EPI_ISL_3346051 | Viollier AG | Department of Biosystems Science and Engineering, ETH Zürich | Chaoran Chen, Sarah Nadeau, Louis du Plessis, Catharine Aquino, Ivan Topolsky, Kim Philipp Jablonski, Lara Fuhrmann, Daniel Ehrsam, Isabel Stürmer, Andreia Cabral de Gouvea, Maria Domenica Moccia, Simon Grüter, Timothy Sykes, Lennart Opitz, Griffin White, Laura Neff, Doris Popovic, Andrea Patrignani, Jay Tracy, Ralph Schlapbach, Christiane Beckmann, Maurice Redondo, Olivier Kobel, Christoph Noppen, Niko Beerenwinkel, Tanja Stadler |
| EPI_ISL_3346052, EPI_ISL_3346053 | Viollier AG | Department of Biosystems Science and Engineering, ETH Zürich | Christian Beisel, Sarah Nadeau, Chaoran Chen, Louis du Plessis, Ivan Topolsky, Kim Philipp Jablonski, Lara Fuhrmann, Rebecca Denes, Mirjam Feldkamp, Ina Nissen, Natascha Santacroce, Elodie Burcklen, Christiane Beckmann, Maurice Redondo, Olivier Kobel, Christoph Noppen, Niko Beerenwinkel, Tanja Stadler |
| EPI_ISL_3346054, EPI_ISL_3346055 | Viollier AG | Department of Biosystems Science and Engineering, ETH Zürich | Chaoran Chen, Sarah Nadeau, Louis du Plessis, Catharine Aquino, Ivan Topolsky, Kim Philipp Jablonski, Lara Fuhrmann, Daniel Ehrsam, Isabel Stürmer, Andreia Cabral de Gouvea, Maria Domenica Moccia, Simon Grüter, Timothy Sykes, Lennart Opitz, Griffin White, Laura Neff, Doris Popovic, Andrea Patrignani, Jay Tracy, Ralph Schlapbach, Christiane Beckmann, Maurice Redondo, Olivier Kobel, Christoph Noppen, Niko Beerenwinkel, Tanja Stadler |
| EPI_ISL_3346056 | Viollier AG | Department of Biosystems Science and Engineering, ETH Zürich | Christian Beisel, Sarah Nadeau, Chaoran Chen, Louis du Plessis, Ivan Topolsky, Kim Philipp Jablonski, Lara Fuhrmann, Rebecca Denes, Mirjam Feldkamp, Ina Nissen, Natascha Santacroce, Elodie Burcklen, Christiane Beckmann, Maurice Redondo, Olivier Kobel, Christoph Noppen, Niko Beerenwinkel, Tanja Stadler |
| EPI_ISL_3346057 | Viollier AG | Department of Biosystems Science and Engineering, ETH Zürich | Chaoran Chen, Sarah Nadeau, Louis du Plessis, Catharine Aquino, Ivan Topolsky, Kim Philipp Jablonski, Lara Fuhrmann, Daniel Ehrsam, Isabel Stürmer, Andreia Cabral de Gouvea, Maria Domenica Moccia, Simon Grüter, Timothy Sykes, Lennart Opitz, Griffin White, Laura Neff, Doris Popovic, Andrea Patrignani, Jay Tracy, Ralph Schlapbach, Christiane Beckmann, Maurice Redondo, Olivier Kobel, Christoph Noppen, Niko Beerenwinkel, Tanja Stadler |
| EPI_ISL_3346058 | Viollier AG | Department of Biosystems Science and Engineering, ETH Zürich | Christian Beisel, Sarah Nadeau, Chaoran Chen, Louis du Plessis, Ivan Topolsky, Kim Philipp Jablonski, Lara Fuhrmann, Rebecca Denes, Mirjam Feldkamp, Ina Nissen, Natascha Santacroce, Elodie Burcklen, Christiane Beckmann, Maurice Redondo, Olivier Kobel, Christoph Noppen, Niko Beerenwinkel, Tanja Stadler |
| EPI_ISL_3346059, EPI_ISL_3346060, EPI_ISL_3346061, EPI_ISL_3346062, EPI_ISL_3346063, EPI_ISL_3346064, EPI_ISL_3346065 | Viollier AG | Department of Biosystems Science and Engineering, ETH Zürich | Chaoran Chen, Sarah Nadeau, Louis du Plessis, Catharine Aquino, Ivan Topolsky, Kim Philipp Jablonski, Lara Fuhrmann, Daniel Ehrsam, Isabel Stürmer, Andreia Cabral de Gouvea, Maria Domenica Moccia, Simon Grüter, Timothy Sykes, Lennart Opitz, Griffin White, Laura Neff, Doris Popovic, Andrea Patrignani, Jay Tracy, Ralph Schlapbach, Christiane Beckmann, Maurice Redondo, Olivier Kobel, Christoph Noppen, Niko Beerenwinkel, Tanja Stadler |
| EPI_ISL_3346066, EPI_ISL_3346067 | Viollier AG | Department of Biosystems Science and Engineering, ETH Zürich | Christian Beisel, Sarah Nadeau, Chaoran Chen, Louis du Plessis, Ivan Topolsky, Kim Philipp Jablonski, Lara Fuhrmann, Rebecca Denes, Mirjam Feldkamp, Ina Nissen, Natascha Santacroce, Elodie Burcklen, Christiane Beckmann, Maurice Redondo, Olivier Kobel, Christoph Noppen, Niko Beerenwinkel, Tanja Stadler |
| EPI_ISL_3346068, EPI_ISL_3346069, EPI_ISL_3346070, EPI_ISL_3346071 | Viollier AG | Department of Biosystems Science and Engineering, ETH Zürich | Chaoran Chen, Sarah Nadeau, Louis du Plessis, Catharine Aquino, Ivan Topolsky, Kim Philipp Jablonski, Lara Fuhrmann, Daniel Ehrsam, Isabel Stürmer, Andreia Cabral de Gouvea, Maria Domenica Moccia, Simon Grüter, Timothy Sykes, Lennart Opitz, Griffin White, Laura Neff, Doris Popovic, Andrea Patrignani, Jay Tracy, Ralph Schlapbach, Christiane Beckmann, Maurice Redondo, Olivier Kobel, Christoph Noppen, Niko Beerenwinkel, Tanja Stadler |
| EPI_ISL_3346072, EPI_ISL_3346073 | Viollier AG | Department of Biosystems Science and Engineering, ETH Zürich | Christian Beisel, Sarah Nadeau, Chaoran Chen, Louis du Plessis, Ivan Topolsky, Kim Philipp Jablonski, Lara Fuhrmann, Rebecca Denes, Mirjam Feldkamp, Ina Nissen, Natascha Santacroce, Elodie Burcklen, Christiane Beckmann, Maurice Redondo, Olivier Kobel, Christoph Noppen, Niko Beerenwinkel, Tanja Stadler |
| EPI_ISL_3346074, EPI_ISL_3346075, EPI_ISL_3346076, EPI_ISL_3346077 | Viollier AG | Department of Biosystems Science and Engineering, ETH Zürich | Chaoran Chen, Sarah Nadeau, Louis du Plessis, Catharine Aquino, Ivan Topolsky, Kim Philipp Jablonski, Lara Fuhrmann, Daniel Ehrsam, Isabel Stürmer, Andreia Cabral de Gouvea, Maria Domenica Moccia, Simon Grüter, Timothy Sykes, Lennart Opitz, Griffin White, Laura Neff, Doris Popovic, Andrea Patrignani, Jay Tracy, Ralph Schlapbach, Christiane Beckmann, Maurice Redondo, Olivier Kobel, Christoph Noppen, Niko Beerenwinkel, Tanja Stadler |
| EPI_ISL_3346078 | Viollier AG | Department of Biosystems Science and Engineering, ETH Zürich | Christian Beisel, Sarah Nadeau, Chaoran Chen, Louis du Plessis, Ivan Topolsky, Kim Philipp Jablonski, Lara Fuhrmann, Rebecca Denes, Mirjam Feldkamp, Ina Nissen, Natascha Santacroce, Elodie Burcklen, Christiane Beckmann, Maurice Redondo, Olivier Kobel, Christoph Noppen, Niko Beerenwinkel, Tanja Stadler |
| EPI_ISL_3346079, EPI_ISL_3346080, EPI_ISL_3346081, EPI_ISL_3346082 | Viollier AG | Department of Biosystems Science and Engineering, ETH Zürich | Chaoran Chen, Sarah Nadeau, Louis du Plessis, Catharine Aquino, Ivan Topolsky, Kim Philipp Jablonski, Lara Fuhrmann, Daniel Ehrsam, Isabel Stürmer, Andreia Cabral de Gouvea, Maria Domenica Moccia, Simon Grüter, Timothy Sykes, Lennart Opitz, Griffin White, Laura Neff, Doris Popovic, Andrea Patrignani, Jay Tracy, Ralph Schlapbach, Christiane Beckmann, Maurice Redondo, Olivier Kobel, Christoph Noppen, Niko Beerenwinkel, Tanja Stadler |
| EPI_ISL_3346083, EPI_ISL_3346084, EPI_ISL_3346085, EPI_ISL_3346086 | Viollier AG | Department of Biosystems Science and Engineering, ETH Zürich | Christian Beisel, Sarah Nadeau, Chaoran Chen, Louis du Plessis, Ivan Topolsky, Kim Philipp Jablonski, Lara Fuhrmann, Rebecca Denes, Mirjam Feldkamp, Ina Nissen, Natascha Santacroce, Elodie Burcklen, Christiane Beckmann, Maurice Redondo, Olivier Kobel, Christoph Noppen, Niko Beerenwinkel, Tanja Stadler |
| EPI_ISL_3346087 | Viollier AG | Department of Biosystems Science and Engineering, ETH Zürich | Chaoran Chen, Sarah Nadeau, Louis du Plessis, Catharine Aquino, Ivan Topolsky, Kim Philipp Jablonski, Lara Fuhrmann, Daniel Ehrsam, Isabel Stürmer, Andreia Cabral de Gouvea, Maria Domenica Moccia, Simon Grüter, Timothy Sykes, Lennart Opitz, Griffin White, Laura Neff, Doris Popovic, Andrea Patrignani, Jay Tracy, Ralph Schlapbach, Christiane Beckmann, Maurice Redondo, Olivier Kobel, Christoph Noppen, Niko Beerenwinkel, Tanja Stadler |
| EPI_ISL_3346088 | Viollier AG | Department of Biosystems Science and Engineering, ETH Zürich | Christian Beisel, Sarah Nadeau, Chaoran Chen, Louis du Plessis, Ivan Topolsky, Kim Philipp Jablonski, Lara Fuhrmann, Rebecca Denes, Mirjam Feldkamp, Ina Nissen, Natascha Santacroce, Elodie Burcklen, Christiane Beckmann, Maurice |

| | | | |
|---|---|---|---|
| | | | Redondo, Olivier Kobel, Christoph Noppen, Niko Beerenwinkel, Tanja Stadler |
| EPI_ISL_3346089 | Viollier AG | Department of Biosystems Science and Engineering, ETH Zürich | Chaoran Chen, Sarah Nadeau, Louis du Plessis, Catharine Aquino, Ivan Topolsky, Kim Philipp Jablonski, Lara Fuhrmann, Daniel Ehrsam, Isabel Stürmer, Andreia Cabral de Gouvea, Maria Domenica Moccia, Simon Grüter, Timothy Sykes, Lennart Opitz, Griffin White, Laura Neff, Doris Popovic, Andrea Patrignani, Jay Tracy, Ralph Schlapbach, Christiane Beckmann, Maurice Redondo, Olivier Kobel, Christoph Noppen, Niko Beerenwinkel, Tanja Stadler |
| EPI_ISL_3346090, EPI_ISL_3346091, EPI_ISL_3346092 | Viollier AG | Department of Biosystems Science and Engineering, ETH Zürich | Christian Beisel, Sarah Nadeau, Chaoran Chen, Louis du Plessis, Ivan Topolsky, Kim Philipp Jablonski, Lara Fuhrmann, Rebecca Denes, Mirjam Feldkamp, Ina Nissen, Natascha Santacroce, Elodie Burcklen, Christiane Beckmann, Maurice Redondo, Olivier Kobel, Christoph Noppen, Niko Beerenwinkel, Tanja Stadler |
| EPI_ISL_3346093, EPI_ISL_3346094, EPI_ISL_3346095, EPI_ISL_3346096, EPI_ISL_3346097, EPI_ISL_3346098 | Viollier AG | Department of Biosystems Science and Engineering, ETH Zürich | Chaoran Chen, Sarah Nadeau, Louis du Plessis, Catharine Aquino, Ivan Topolsky, Kim Philipp Jablonski, Lara Fuhrmann, Daniel Ehrsam, Isabel Stürmer, Andreia Cabral de Gouvea, Maria Domenica Moccia, Simon Grüter, Timothy Sykes, Lennart Opitz, Griffin White, Laura Neff, Doris Popovic, Andrea Patrignani, Jay Tracy, Ralph Schlapbach, Christiane Beckmann, Maurice Redondo, Olivier Kobel, Christoph Noppen, Niko Beerenwinkel, Tanja Stadler |
| EPI_ISL_3346099 | Viollier AG | Department of Biosystems Science and Engineering, ETH Zürich | Christian Beisel, Sarah Nadeau, Chaoran Chen, Louis du Plessis, Ivan Topolsky, Kim Philipp Jablonski, Lara Fuhrmann, Rebecca Denes, Mirjam Feldkamp, Ina Nissen, Natascha Santacroce, Elodie Burcklen, Christiane Beckmann, Maurice Redondo, Olivier Kobel, Christoph Noppen, Niko Beerenwinkel, Tanja Stadler |
| EPI_ISL_3346101, EPI_ISL_3346102 | Viollier AG | Department of Biosystems Science and Engineering, ETH Zürich | Chaoran Chen, Sarah Nadeau, Louis du Plessis, Catharine Aquino, Ivan Topolsky, Kim Philipp Jablonski, Lara Fuhrmann, Daniel Ehrsam, Isabel Stürmer, Andreia Cabral de Gouvea, Maria Domenica Moccia, Simon Grüter, Timothy Sykes, Lennart Opitz, Griffin White, Laura Neff, Doris Popovic, Andrea Patrignani, Jay Tracy, Ralph Schlapbach, Christiane Beckmann, Maurice Redondo, Olivier Kobel, Christoph Noppen, Niko Beerenwinkel, Tanja Stadler |
| EPI_ISL_3346103 | Viollier AG | Department of Biosystems Science and Engineering, ETH Zürich | Christian Beisel, Sarah Nadeau, Chaoran Chen, Louis du Plessis, Ivan Topolsky, Kim Philipp Jablonski, Lara Fuhrmann, Rebecca Denes, Mirjam Feldkamp, Ina Nissen, Natascha Santacroce, Elodie Burcklen, Christiane Beckmann, Maurice Redondo, Olivier Kobel, Christoph Noppen, Niko Beerenwinkel, Tanja Stadler |
| EPI_ISL_3346105, EPI_ISL_3346106, EPI_ISL_3346107, EPI_ISL_3346108 | Viollier AG | Department of Biosystems Science and Engineering, ETH Zürich | Chaoran Chen, Sarah Nadeau, Louis du Plessis, Catharine Aquino, Ivan Topolsky, Kim Philipp Jablonski, Lara Fuhrmann, Daniel Ehrsam, Isabel Stürmer, Andreia Cabral de Gouvea, Maria Domenica Moccia, Simon Grüter, Timothy Sykes, Lennart Opitz, Griffin White, Laura Neff, Doris Popovic, Andrea Patrignani, Jay Tracy, Ralph Schlapbach, Christiane Beckmann, Maurice Redondo, Olivier Kobel, Christoph Noppen, Niko Beerenwinkel, Tanja Stadler |
| EPI_ISL_3346109 | Viollier AG | Department of Biosystems Science and Engineering, ETH Zürich | Christian Beisel, Sarah Nadeau, Chaoran Chen, Louis du Plessis, Ivan Topolsky, Kim Philipp Jablonski, Lara Fuhrmann, Rebecca Denes, Mirjam Feldkamp, Ina Nissen, Natascha Santacroce, Elodie Burcklen, Christiane Beckmann, Maurice Redondo, Olivier Kobel, Christoph Noppen, Niko Beerenwinkel, Tanja Stadler |
| EPI_ISL_3346110, EPI_ISL_3346111 | Viollier AG | Department of Biosystems Science and Engineering, ETH Zürich | Chaoran Chen, Sarah Nadeau, Louis du Plessis, Catharine Aquino, Ivan Topolsky, Kim Philipp Jablonski, Lara Fuhrmann, Daniel Ehrsam, Isabel Stürmer, Andreia Cabral de Gouvea, Maria Domenica Moccia, Simon Grüter, Timothy Sykes, Lennart Opitz, Griffin White, Laura Neff, Doris Popovic, Andrea Patrignani, Jay Tracy, Ralph Schlapbach, Christiane Beckmann, Maurice Redondo, Olivier Kobel, Christoph Noppen, Niko Beerenwinkel, Tanja Stadler |
| EPI_ISL_3346112, EPI_ISL_3346113 | Viollier AG | Department of Biosystems Science and Engineering, ETH Zürich | Christian Beisel, Sarah Nadeau, Chaoran Chen, Louis du Plessis, Ivan Topolsky, Kim Philipp Jablonski, Lara Fuhrmann, Rebecca Denes, Mirjam Feldkamp, Ina Nissen, Natascha Santacroce, Elodie Burcklen, Christiane Beckmann, Maurice Redondo, Olivier Kobel, Christoph Noppen, Niko Beerenwinkel, Tanja Stadler |
| EPI_ISL_3346114, EPI_ISL_3346115 | Viollier AG | Department of Biosystems Science and Engineering, ETH Zürich | Chaoran Chen, Sarah Nadeau, Louis du Plessis, Catharine Aquino, Ivan Topolsky, Kim Philipp Jablonski, Lara Fuhrmann, Daniel Ehrsam, Isabel Stürmer, Andreia Cabral de Gouvea, Maria Domenica Moccia, Simon Grüter, Timothy Sykes, Lennart Opitz, Griffin White, Laura Neff, Doris Popovic, Andrea Patrignani, Jay Tracy, Ralph Schlapbach, Christiane Beckmann, Maurice Redondo, Olivier Kobel, Christoph Noppen, Niko Beerenwinkel, Tanja Stadler |
| EPI_ISL_3346116 | Viollier AG | Department of Biosystems Science and Engineering, ETH Zürich | Christian Beisel, Sarah Nadeau, Chaoran Chen, Louis du Plessis, Ivan Topolsky, Kim Philipp Jablonski, Lara Fuhrmann, Rebecca Denes, Mirjam Feldkamp, Ina Nissen, Natascha Santacroce, Elodie Burcklen, Christiane Beckmann, Maurice Redondo, Olivier Kobel, Christoph Noppen, Niko Beerenwinkel, Tanja Stadler |
| EPI_ISL_3346117 | Viollier AG | Department of Biosystems Science and Engineering, ETH Zürich | Chaoran Chen, Sarah Nadeau, Louis du Plessis, Catharine Aquino, Ivan Topolsky, Kim Philipp Jablonski, Lara Fuhrmann, Daniel Ehrsam, Isabel Stürmer, Andreia Cabral de Gouvea, Maria Domenica Moccia, Simon Grüter, Timothy Sykes, Lennart Opitz, Griffin White, Laura Neff, Doris Popovic, Andrea Patrignani, Jay Tracy, Ralph Schlapbach, Christiane Beckmann, Maurice Redondo, Olivier Kobel, Christoph Noppen, Niko Beerenwinkel, Tanja Stadler |
| EPI_ISL_3346118 | Viollier AG | Department of Biosystems Science and Engineering, ETH Zürich | Christian Beisel, Sarah Nadeau, Chaoran Chen, Louis du Plessis, Ivan Topolsky, Kim Philipp Jablonski, Lara Fuhrmann, Rebecca Denes, Mirjam Feldkamp, Ina Nissen, Natascha Santacroce, Elodie Burcklen, Christiane Beckmann, Maurice Redondo, Olivier Kobel, Christoph Noppen, Niko Beerenwinkel, Tanja Stadler |
| EPI_ISL_3346119, EPI_ISL_3346120, EPI_ISL_3346121, EPI_ISL_3346122 | Viollier AG | Department of Biosystems Science and Engineering, ETH Zürich | Chaoran Chen, Sarah Nadeau, Louis du Plessis, Catharine Aquino, Ivan Topolsky, Kim Philipp Jablonski, Lara Fuhrmann, Daniel Ehrsam, Isabel Stürmer, Andreia Cabral de Gouvea, Maria Domenica Moccia, Simon Grüter, Timothy Sykes, Lennart Opitz, Griffin White, Laura Neff, Doris Popovic, Andrea Patrignani, Jay Tracy, Ralph Schlapbach, Christiane Beckmann, Maurice Redondo, Olivier Kobel, Christoph Noppen, Niko Beerenwinkel, Tanja Stadler |
| EPI_ISL_3346123 | Viollier AG | Department of Biosystems Science and Engineering, ETH Zürich | Christian Beisel, Sarah Nadeau, Chaoran Chen, Louis du Plessis, Ivan Topolsky, Kim Philipp Jablonski, Lara Fuhrmann, Rebecca Denes, Mirjam Feldkamp, Ina Nissen, Natascha Santacroce, Elodie Burcklen, Christiane Beckmann, Maurice Redondo, Olivier Kobel, Christoph Noppen, Niko Beerenwinkel, Tanja Stadler |
| EPI_ISL_3346124, EPI_ISL_3346125, EPI_ISL_3346126, EPI_ISL_3346127 | Viollier AG | Department of Biosystems Science and Engineering, ETH Zürich | Chaoran Chen, Sarah Nadeau, Louis du Plessis, Catharine Aquino, Ivan Topolsky, Kim Philipp Jablonski, Lara Fuhrmann, Daniel Ehrsam, Isabel Stürmer, Andreia Cabral de Gouvea, Maria Domenica Moccia, Simon Grüter, Timothy Sykes, Lennart Opitz, Griffin White, Laura Neff, Doris Popovic, Andrea Patrignani, Jay Tracy, Ralph Schlapbach, Christiane Beckmann, Maurice Redondo, Olivier Kobel, Christoph Noppen, Niko Beerenwinkel, Tanja Stadler |
| EPI_ISL_3346128, EPI_ISL_3346129 | Viollier AG | Department of Biosystems Science and Engineering, ETH Zürich | Christian Beisel, Sarah Nadeau, Chaoran Chen, Louis du Plessis, Ivan Topolsky, Kim Philipp Jablonski, Lara Fuhrmann, Rebecca Denes, Mirjam Feldkamp, Ina Nissen, Natascha Santacroce, Elodie Burcklen, Christiane Beckmann, Maurice Redondo, Olivier Kobel, Christoph Noppen, Niko Beerenwinkel, Tanja Stadler |
| EPI_ISL_3346131, EPI_ISL_3346132, EPI_ISL_3346134 | Viollier AG | Department of Biosystems Science and Engineering, ETH Zürich | Chaoran Chen, Sarah Nadeau, Louis du Plessis, Catharine Aquino, Ivan Topolsky, Kim Philipp Jablonski, Lara Fuhrmann, Daniel Ehrsam, Isabel Stürmer, Andreia Cabral de Gouvea, Maria Domenica Moccia, Simon Grüter, Timothy Sykes, Lennart Opitz, Griffin White, Laura Neff, Doris Popovic, Andrea Patrignani, Jay Tracy, Ralph Schlapbach, Christiane Beckmann, Maurice Redondo, Olivier Kobel, Christoph Noppen, Niko Beerenwinkel, Tanja Stadler |
| EPI_ISL_3346135 | Viollier AG | Department of Biosystems Science and Engineering, ETH Zürich | Christian Beisel, Sarah Nadeau, Chaoran Chen, Louis du Plessis, Ivan Topolsky, Kim Philipp Jablonski, Lara Fuhrmann, Rebecca Denes, Mirjam Feldkamp, Ina Nissen, Natascha Santacroce, Elodie Burcklen, Christiane Beckmann, Maurice Redondo, Olivier Kobel, Christoph Noppen, Niko Beerenwinkel, Tanja Stadler |
| EPI_ISL_3346137, EPI_ISL_3346138 | Viollier AG | Department of Biosystems Science and Engineering, ETH Zürich | Chaoran Chen, Sarah Nadeau, Louis du Plessis, Catharine Aquino, Ivan Topolsky, Kim Philipp Jablonski, Lara Fuhrmann, Daniel Ehrsam, Isabel Stürmer, Andreia Cabral de Gouvea, Maria Domenica Moccia, Simon Grüter, Timothy Sykes, Lennart Opitz, Griffin White, Laura Neff, Doris Popovic, Andrea Patrignani, Jay Tracy, Ralph Schlapbach, Christiane Beckmann, Maurice Redondo, Olivier Kobel, Christoph Noppen, Niko Beerenwinkel, Tanja Stadler |
| EPI_ISL_3346139, EPI_ISL_3346140, EPI_ISL_3346141, EPI_ISL_3346142 | Viollier AG | Department of Biosystems Science and Engineering, ETH Zürich | Christian Beisel, Sarah Nadeau, Chaoran Chen, Louis du Plessis, Ivan Topolsky, Kim Philipp Jablonski, Lara Fuhrmann, Rebecca Denes, Mirjam Feldkamp, Ina Nissen, Natascha Santacroce, Elodie Burcklen, Christiane Beckmann, Maurice Redondo, Olivier Kobel, Christoph Noppen, Niko Beerenwinkel, Tanja Stadler |
| EPI_ISL_3346143 | Viollier AG | Department of Biosystems Science and Engineering, ETH Zürich | Chaoran Chen, Sarah Nadeau, Louis du Plessis, Catharine Aquino, Ivan Topolsky, Kim Philipp Jablonski, Lara Fuhrmann, Daniel Ehrsam, Isabel Stürmer, Andreia Cabral de Gouvea, Maria Domenica Moccia, Simon Grüter, Timothy Sykes, Lennart Opitz, Griffin White, Laura Neff, Doris Popovic, Andrea Patrignani, Jay Tracy, Ralph Schlapbach, Christiane Beckmann, Maurice Redondo, Olivier Kobel, Christoph Noppen, Niko Beerenwinkel, Tanja Stadler |
| EPI_ISL_3346144 | Viollier AG | Department of Biosystems Science and Engineering, ETH Zürich | Christian Beisel, Sarah Nadeau, Chaoran Chen, Louis du Plessis, Ivan Topolsky, Kim Philipp Jablonski, Lara Fuhrmann, Rebecca Denes, Mirjam Feldkamp, Ina Nissen, Natascha Santacroce, Elodie Burcklen, Christiane Beckmann, Maurice Redondo, Olivier Kobel, Christoph Noppen, Niko Beerenwinkel, Tanja Stadler |
| EPI_ISL_3346145, EPI_ISL_3346146, EPI_ISL_3346147 | Viollier AG | Department of Biosystems Science and Engineering, ETH Zürich | Chaoran Chen, Sarah Nadeau, Louis du Plessis, Catharine Aquino, Ivan Topolsky, Kim Philipp Jablonski, Lara Fuhrmann, Daniel Ehrsam, Isabel Stürmer, Andreia Cabral de Gouvea, Maria Domenica Moccia, Simon Grüter, Timothy Sykes, Lennart Opitz, Griffin White, Laura Neff, Doris Popovic, Andrea Patrignani, Jay Tracy, Ralph Schlapbach, Christiane Beckmann, Maurice Redondo, Olivier Kobel, Christoph Noppen, Niko Beerenwinkel, Tanja Stadler |
| EPI_ISL_3346148 | Viollier AG | Department of Biosystems Science and Engineering, ETH Zürich | Christian Beisel, Sarah Nadeau, Chaoran Chen, Louis du Plessis, Ivan Topolsky, Kim Philipp Jablonski, Lara Fuhrmann, Rebecca Denes, Mirjam Feldkamp, Ina Nissen, Natascha Santacroce, Elodie Burcklen, Christiane Beckmann, Maurice Redondo, Olivier Kobel, Christoph Noppen, Niko Beerenwinkel, Tanja Stadler |
| EPI_ISL_3346149, EPI_ISL_3346150, EPI_ISL_3346151, EPI_ISL_3346152, EPI_ISL_3346153, EPI_ISL_3346156, EPI_ISL_3346157, EPI_ISL_3346158, EPI_ISL_3346159, EPI_ISL_3346160, EPI_ISL_3346161 | | | |
| see above | Viollier AG | Department of Biosystems Science and Engineering, ETH Zürich | Chaoran Chen, Sarah Nadeau, Louis du Plessis, Catharine Aquino, Ivan Topolsky, Kim Philipp Jablonski, Lara Fuhrmann, Daniel Ehrsam, Isabel Stürmer, Andreia Cabral de Gouvea, Maria Domenica Moccia, Simon Grüter, Timothy Sykes, Lennart Opitz, Griffin White, Laura Neff, Doris Popovic, Andrea Patrignani, Jay Tracy, Ralph Schlapbach, Christiane Beckmann, Maurice Redondo, Olivier Kobel, Christoph Noppen, Niko Beerenwinkel, Tanja Stadler |
| EPI_ISL_3346162 | Viollier AG | Department of Biosystems Science and Engineering, ETH Zürich | Christian Beisel, Sarah Nadeau, Chaoran Chen, Louis du Plessis, Ivan Topolsky, Kim Philipp Jablonski, Lara Fuhrmann, Rebecca Denes, Mirjam Feldkamp, Ina Nissen, Natascha Santacroce, Elodie Burcklen, Christiane Beckmann, Maurice Redondo, Olivier Kobel, Christoph Noppen, Niko Beerenwinkel, Tanja Stadler |
| EPI_ISL_3346163, EPI_ISL_3346164, EPI_ISL_3346165 | Viollier AG | Department of Biosystems Science and Engineering, ETH Zürich | Chaoran Chen, Sarah Nadeau, Louis du Plessis, Catharine Aquino, Ivan Topolsky, Kim Philipp Jablonski, Lara Fuhrmann, Daniel Ehrsam, Isabel Stürmer, Andreia Cabral de Gouvea, Maria Domenica Moccia, Simon Grüter, Timothy Sykes, Lennart Opitz, Griffin White, Laura Neff, Doris Popovic, Andrea Patrignani, Jay Tracy, Ralph Schlapbach, Christiane Beckmann, Maurice Redondo, Olivier Kobel, Christoph Noppen, Niko Beerenwinkel, Tanja Stadler |
| EPI_ISL_3346166, EPI_ISL_3346167 | Viollier AG | Department of Biosystems Science and Engineering, ETH Zürich | Christian Beisel, Sarah Nadeau, Chaoran Chen, Louis du Plessis, Ivan Topolsky, Kim Philipp Jablonski, Lara Fuhrmann, Rebecca Denes, Mirjam Feldkamp, Ina Nissen, Natascha Santacroce, Elodie Burcklen, Christiane Beckmann, Maurice Redondo, Olivier Kobel, Christoph Noppen, Niko Beerenwinkel, Tanja Stadler |
| EPI_ISL_3346168, EPI_ISL_3346169, EPI_ISL_3346170, EPI_ISL_3346171, EPI_ISL_3346172, EPI_ISL_3346173, EPI_ISL_3346174, EPI_ISL_3346175, EPI_ISL_3346176, EPI_ISL_3346177, EPI_ISL_3346178, EPI_ISL_3346179, EPI_ISL_3346180, EPI_ISL_3346181, EPI_ISL_3346182, EPI_ISL_3346183, EPI_ISL_3346184, EPI_ISL_3346185, EPI_ISL_3346186, EPI_ISL_3346187, EPI_ISL_3346188, EPI_ISL_3346189, EPI_ISL_3346190, EPI_ISL_3346191, EPI_ISL_3346192, EPI_ISL_3346193, EPI_ISL_3346194, EPI_ISL_3346195, EPI_ISL_3346197, EPI_ISL_3346198, EPI_ISL_3346200, EPI_ISL_3346201, EPI_ISL_3346203, EPI_ISL_3346204, EPI_ISL_3346206, EPI_ISL_3346208, EPI_ISL_3346209, EPI_ISL_3346211, EPI_ISL_3346212, EPI_ISL_3346214, EPI_ISL_3346215 | | | |
| see above | Viollier AG | Department of Biosystems Science and Engineering, ETH Zürich | Chaoran Chen, Sarah Nadeau, Louis du Plessis, Catharine Aquino, Ivan Topolsky, Kim Philipp Jablonski, Lara Fuhrmann, Daniel Ehrsam, Isabel Stürmer, Andreia Cabral de Gouvea, Maria Domenica Moccia, Simon Grüter, Timothy Sykes, Lennart Opitz, Griffin White, Laura Neff, Doris Popovic, Andrea Patrignani, Jay Tracy, Ralph Schlapbach, Christiane Beckmann, Maurice Redondo, Olivier Kobel, Christoph Noppen, Niko Beerenwinkel, Tanja Stadler |
| EPI_ISL_3346217 | Viollier AG | Department of Biosystems Science and Engineering, ETH Zürich | Christian Beisel, Sarah Nadeau, Chaoran Chen, Louis du Plessis, Ivan Topolsky, Kim Philipp Jablonski, Lara Fuhrmann, Rebecca Denes, Mirjam Feldkamp, Ina Nissen, Natascha Santacroce, Elodie Burcklen, Christiane Beckmann, Maurice Redondo, Olivier Kobel, Christoph Noppen, Niko Beerenwinkel, Tanja Stadler |
| EPI_ISL_3346218, EPI_ISL_3346220, EPI_ISL_3346222, EPI_ISL_3346223, EPI_ISL_3346225 | Viollier AG | Department of Biosystems Science and Engineering, ETH Zürich | Chaoran Chen, Sarah Nadeau, Louis du Plessis, Catharine Aquino, Ivan Topolsky, Kim Philipp Jablonski, Lara Fuhrmann, Daniel Ehrsam, Isabel Stürmer, Andreia Cabral de Gouvea, Maria Domenica Moccia, Simon Grüter, Timothy Sykes, Lennart Opitz, Griffin White, Laura Neff, Doris Popovic, Andrea Patrignani, Jay Tracy, Ralph Schlapbach, Christiane Beckmann, Maurice Redondo, Olivier Kobel, Christoph Noppen, Niko Beerenwinkel, Tanja Stadler |
| EPI_ISL_3346226 | Viollier AG | Department of Biosystems Science and Engineering, ETH Zürich | Christian Beisel, Sarah Nadeau, Chaoran Chen, Louis du Plessis, Ivan Topolsky, Kim Philipp Jablonski, Lara Fuhrmann, Rebecca Denes, Mirjam Feldkamp, Ina Nissen, Natascha Santacroce, Elodie Burcklen, Christiane Beckmann, Maurice Redondo, Olivier Kobel, Christoph Noppen, Niko Beerenwinkel, Tanja Stadler |
| EPI_ISL_3346228, EPI_ISL_3346229, EPI_ISL_3346231, | Viollier AG | Department of Biosystems Science and Engineering, ETH Zürich | Chaoran Chen, Sarah Nadeau, Louis du Plessis, Catharine Aquino, Ivan Topolsky, Kim Philipp Jablonski, Lara Fuhrmann, Daniel Ehrsam, Isabel Stürmer, Andreia Cabral de Gouvea, Maria Domenica Moccia, Simon Grüter, Timothy Sykes, Lennart Opitz, Griffin White, Laura Neff, Doris Popovic, Andrea Patrignani, Jay Tracy, Ralph Schlapbach, Christiane Beckmann, Maurice Redondo, Olivier Kobel, Christoph Noppen, Niko Beerenwinkel, Tanja Stadler |

| | | | |
|---|---|---|---|
| EPI_ISL_3346233 | | | |
| EPI_ISL_3346234 | Viollier AG | Department of Biosystems Science and Engineering, ETH Zürich | Christian Beisel, Sarah Nadeau, Chaoran Chen, Louis du Plessis, Ivan Topolsky, Kim Philipp Jablonski, Lara Fuhrmann, Rebecca Denes, Mirjam Feldkamp, Ina Nissen, Natascha Santacroce, Elodie Burcklen, Christiane Beckmann, Maurice Redondo, Olivier Kobel, Christoph Noppen, Niko Beerenwinkel, Tanja Stadler |
| EPI_ISL_3346236, EPI_ISL_3346238, EPI_ISL_3346239, EPI_ISL_3346241, EPI_ISL_3346242 | Viollier AG | Department of Biosystems Science and Engineering, ETH Zürich | Chaoran Chen, Sarah Nadeau, Louis du Plessis, Catharine Aquino, Ivan Topolsky, Kim Philipp Jablonski, Lara Fuhrmann, Daniel Ehrsam, Isabel Stürmer, Andreia Cabral de Gouvea, Maria Domenica Moccia, Simon Grüter, Timothy Sykes, Lennart Opitz, Griffin White, Laura Neff, Doris Popovic, Andrea Patrignani, Jay Tracy, Ralph Schlapbach, Christiane Beckmann, Maurice Redondo, Olivier Kobel, Christoph Noppen, Niko Beerenwinkel, Tanja Stadler |
| EPI_ISL_3346244, EPI_ISL_3346245, EPI_ISL_3346247 | Viollier AG | Department of Biosystems Science and Engineering, ETH Zürich | Christian Beisel, Sarah Nadeau, Chaoran Chen, Louis du Plessis, Ivan Topolsky, Kim Philipp Jablonski, Lara Fuhrmann, Rebecca Denes, Mirjam Feldkamp, Ina Nissen, Natascha Santacroce, Elodie Burcklen, Christiane Beckmann, Maurice Redondo, Olivier Kobel, Christoph Noppen, Niko Beerenwinkel, Tanja Stadler |
| EPI_ISL_3346248, EPI_ISL_3346250 | Viollier AG | Department of Biosystems Science and Engineering, ETH Zürich | Chaoran Chen, Sarah Nadeau, Louis du Plessis, Catharine Aquino, Ivan Topolsky, Kim Philipp Jablonski, Lara Fuhrmann, Daniel Ehrsam, Isabel Stürmer, Andreia Cabral de Gouvea, Maria Domenica Moccia, Simon Grüter, Timothy Sykes, Lennart Opitz, Griffin White, Laura Neff, Doris Popovic, Andrea Patrignani, Jay Tracy, Ralph Schlapbach, Christiane Beckmann, Maurice Redondo, Olivier Kobel, Christoph Noppen, Niko Beerenwinkel, Tanja Stadler |
| EPI_ISL_3346251 | Viollier AG | Department of Biosystems Science and Engineering, ETH Zürich | Christian Beisel, Sarah Nadeau, Chaoran Chen, Louis du Plessis, Ivan Topolsky, Kim Philipp Jablonski, Lara Fuhrmann, Rebecca Denes, Mirjam Feldkamp, Ina Nissen, Natascha Santacroce, Elodie Burcklen, Christiane Beckmann, Maurice Redondo, Olivier Kobel, Christoph Noppen, Niko Beerenwinkel, Tanja Stadler |
| EPI_ISL_3346252 | Viollier AG | Department of Biosystems Science and Engineering, ETH Zürich | Chaoran Chen, Sarah Nadeau, Louis du Plessis, Catharine Aquino, Ivan Topolsky, Kim Philipp Jablonski, Lara Fuhrmann, Daniel Ehrsam, Isabel Stürmer, Andreia Cabral de Gouvea, Maria Domenica Moccia, Simon Grüter, Timothy Sykes, Lennart Opitz, Griffin White, Laura Neff, Doris Popovic, Andrea Patrignani, Jay Tracy, Ralph Schlapbach, Christiane Beckmann, Maurice Redondo, Olivier Kobel, Christoph Noppen, Niko Beerenwinkel, Tanja Stadler |
| EPI_ISL_3346253, EPI_ISL_3346254 | Viollier AG | Department of Biosystems Science and Engineering, ETH Zürich | Christian Beisel, Sarah Nadeau, Chaoran Chen, Louis du Plessis, Ivan Topolsky, Kim Philipp Jablonski, Lara Fuhrmann, Rebecca Denes, Mirjam Feldkamp, Ina Nissen, Natascha Santacroce, Elodie Burcklen, Christiane Beckmann, Maurice Redondo, Olivier Kobel, Christoph Noppen, Niko Beerenwinkel, Tanja Stadler |
| EPI_ISL_3346255 | Viollier AG | Department of Biosystems Science and Engineering, ETH Zürich | Chaoran Chen, Sarah Nadeau, Louis du Plessis, Catharine Aquino, Ivan Topolsky, Kim Philipp Jablonski, Lara Fuhrmann, Daniel Ehrsam, Isabel Stürmer, Andreia Cabral de Gouvea, Maria Domenica Moccia, Simon Grüter, Timothy Sykes, Lennart Opitz, Griffin White, Laura Neff, Doris Popovic, Andrea Patrignani, Jay Tracy, Ralph Schlapbach, Christiane Beckmann, Maurice Redondo, Olivier Kobel, Christoph Noppen, Niko Beerenwinkel, Tanja Stadler |
| EPI_ISL_3346256 | Viollier AG | Department of Biosystems Science and Engineering, ETH Zürich | Christian Beisel, Sarah Nadeau, Chaoran Chen, Louis du Plessis, Ivan Topolsky, Kim Philipp Jablonski, Lara Fuhrmann, Rebecca Denes, Mirjam Feldkamp, Ina Nissen, Natascha Santacroce, Elodie Burcklen, Christiane Beckmann, Maurice Redondo, Olivier Kobel, Christoph Noppen, Niko Beerenwinkel, Tanja Stadler |
| EPI_ISL_3346257 | Viollier AG | Department of Biosystems Science and Engineering, ETH Zürich | Chaoran Chen, Sarah Nadeau, Louis du Plessis, Catharine Aquino, Ivan Topolsky, Kim Philipp Jablonski, Lara Fuhrmann, Daniel Ehrsam, Isabel Stürmer, Andreia Cabral de Gouvea, Maria Domenica Moccia, Simon Grüter, Timothy Sykes, Lennart Opitz, Griffin White, Laura Neff, Doris Popovic, Andrea Patrignani, Jay Tracy, Ralph Schlapbach, Christiane Beckmann, Maurice Redondo, Olivier Kobel, Christoph Noppen, Niko Beerenwinkel, Tanja Stadler |
| EPI_ISL_3346258 | Viollier AG | Department of Biosystems Science and Engineering, ETH Zürich | Christian Beisel, Sarah Nadeau, Chaoran Chen, Louis du Plessis, Ivan Topolsky, Kim Philipp Jablonski, Lara Fuhrmann, Rebecca Denes, Mirjam Feldkamp, Ina Nissen, Natascha Santacroce, Elodie Burcklen, Christiane Beckmann, Maurice Redondo, Olivier Kobel, Christoph Noppen, Niko Beerenwinkel, Tanja Stadler |
| EPI_ISL_3346259, EPI_ISL_3346260, EPI_ISL_3346261 | Viollier AG | Department of Biosystems Science and Engineering, ETH Zürich | Chaoran Chen, Sarah Nadeau, Louis du Plessis, Catharine Aquino, Ivan Topolsky, Kim Philipp Jablonski, Lara Fuhrmann, Daniel Ehrsam, Isabel Stürmer, Andreia Cabral de Gouvea, Maria Domenica Moccia, Simon Grüter, Timothy Sykes, Lennart Opitz, Griffin White, Laura Neff, Doris Popovic, Andrea Patrignani, Jay Tracy, Ralph Schlapbach, Christiane Beckmann, Maurice Redondo, Olivier Kobel, Christoph Noppen, Niko Beerenwinkel, Tanja Stadler |
| EPI_ISL_3346262 | Viollier AG | Department of Biosystems Science and Engineering, ETH Zürich | Christian Beisel, Sarah Nadeau, Chaoran Chen, Louis du Plessis, Ivan Topolsky, Kim Philipp Jablonski, Lara Fuhrmann, Rebecca Denes, Mirjam Feldkamp, Ina Nissen, Natascha Santacroce, Elodie Burcklen, Christiane Beckmann, Maurice Redondo, Olivier Kobel, Christoph Noppen, Niko Beerenwinkel, Tanja Stadler |
| EPI_ISL_3346263 | Viollier AG | Department of Biosystems Science and Engineering, ETH Zürich | Chaoran Chen, Sarah Nadeau, Louis du Plessis, Catharine Aquino, Ivan Topolsky, Kim Philipp Jablonski, Lara Fuhrmann, Daniel Ehrsam, Isabel Stürmer, Andreia Cabral de Gouvea, Maria Domenica Moccia, Simon Grüter, Timothy Sykes, Lennart Opitz, Griffin White, Laura Neff, Doris Popovic, Andrea Patrignani, Jay Tracy, Ralph Schlapbach, Christiane Beckmann, Maurice Redondo, Olivier Kobel, Christoph Noppen, Niko Beerenwinkel, Tanja Stadler |
| EPI_ISL_3346264 | Viollier AG | Department of Biosystems Science and Engineering, ETH Zürich | Christian Beisel, Sarah Nadeau, Chaoran Chen, Louis du Plessis, Ivan Topolsky, Kim Philipp Jablonski, Lara Fuhrmann, Rebecca Denes, Mirjam Feldkamp, Ina Nissen, Natascha Santacroce, Elodie Burcklen, Christiane Beckmann, Maurice Redondo, Olivier Kobel, Christoph Noppen, Niko Beerenwinkel, Tanja Stadler |
| EPI_ISL_3346265, EPI_ISL_3346266, EPI_ISL_3346267 | Viollier AG | Department of Biosystems Science and Engineering, ETH Zürich | Chaoran Chen, Sarah Nadeau, Louis du Plessis, Catharine Aquino, Ivan Topolsky, Kim Philipp Jablonski, Lara Fuhrmann, Daniel Ehrsam, Isabel Stürmer, Andreia Cabral de Gouvea, Maria Domenica Moccia, Simon Grüter, Timothy Sykes, Lennart Opitz, Griffin White, Laura Neff, Doris Popovic, Andrea Patrignani, Jay Tracy, Ralph Schlapbach, Christiane Beckmann, Maurice Redondo, Olivier Kobel, Christoph Noppen, Niko Beerenwinkel, Tanja Stadler |
| EPI_ISL_3346268, EPI_ISL_3346269 | Viollier AG | Department of Biosystems Science and Engineering, ETH Zürich | Christian Beisel, Sarah Nadeau, Chaoran Chen, Louis du Plessis, Ivan Topolsky, Kim Philipp Jablonski, Lara Fuhrmann, Rebecca Denes, Mirjam Feldkamp, Ina Nissen, Natascha Santacroce, Elodie Burcklen, Christiane Beckmann, Maurice Redondo, Olivier Kobel, Christoph Noppen, Niko Beerenwinkel, Tanja Stadler |
| EPI_ISL_3346270 | Viollier AG | Department of Biosystems Science and Engineering, ETH Zürich | Chaoran Chen, Sarah Nadeau, Louis du Plessis, Catharine Aquino, Ivan Topolsky, Kim Philipp Jablonski, Lara Fuhrmann, Daniel Ehrsam, Isabel Stürmer, Andreia Cabral de Gouvea, Maria Domenica Moccia, Simon Grüter, Timothy Sykes, Lennart Opitz, Griffin White, Laura Neff, Doris Popovic, Andrea Patrignani, Jay Tracy, Ralph Schlapbach, Christiane Beckmann, Maurice Redondo, Olivier Kobel, Christoph Noppen, Niko Beerenwinkel, Tanja Stadler |
| EPI_ISL_3346271 | Viollier AG | Department of Biosystems Science and Engineering, ETH Zürich | Christian Beisel, Sarah Nadeau, Chaoran Chen, Louis du Plessis, Ivan Topolsky, Kim Philipp Jablonski, Lara Fuhrmann, Rebecca Denes, Mirjam Feldkamp, Ina Nissen, Natascha Santacroce, Elodie Burcklen, Christiane Beckmann, Maurice Redondo, Olivier Kobel, Christoph Noppen, Niko Beerenwinkel, Tanja Stadler |
| EPI_ISL_3346272, EPI_ISL_3346273, EPI_ISL_3346274, EPI_ISL_3346275, EPI_ISL_3346276, EPI_ISL_3346277 | Viollier AG | Department of Biosystems Science and Engineering, ETH Zürich | Chaoran Chen, Sarah Nadeau, Louis du Plessis, Catharine Aquino, Ivan Topolsky, Kim Philipp Jablonski, Lara Fuhrmann, Daniel Ehrsam, Isabel Stürmer, Andreia Cabral de Gouvea, Maria Domenica Moccia, Simon Grüter, Timothy Sykes, Lennart Opitz, Griffin White, Laura Neff, Doris Popovic, Andrea Patrignani, Jay Tracy, Ralph Schlapbach, Christiane Beckmann, Maurice Redondo, Olivier Kobel, Christoph Noppen, Niko Beerenwinkel, Tanja Stadler |
| EPI_ISL_3346278, EPI_ISL_3346279, EPI_ISL_3346280 | Viollier AG | Department of Biosystems Science and Engineering, ETH Zürich | Christian Beisel, Sarah Nadeau, Chaoran Chen, Louis du Plessis, Ivan Topolsky, Kim Philipp Jablonski, Lara Fuhrmann, Rebecca Denes, Mirjam Feldkamp, Ina Nissen, Natascha Santacroce, Elodie Burcklen, Christiane Beckmann, Maurice Redondo, Olivier Kobel, Christoph Noppen, Niko Beerenwinkel, Tanja Stadler |
| EPI_ISL_3346281, EPI_ISL_3346282, EPI_ISL_3346283, EPI_ISL_3346284 | Viollier AG | Department of Biosystems Science and Engineering, ETH Zürich | Chaoran Chen, Sarah Nadeau, Louis du Plessis, Catharine Aquino, Ivan Topolsky, Kim Philipp Jablonski, Lara Fuhrmann, Daniel Ehrsam, Isabel Stürmer, Andreia Cabral de Gouvea, Maria Domenica Moccia, Simon Grüter, Timothy Sykes, Lennart Opitz, Griffin White, Laura Neff, Doris Popovic, Andrea Patrignani, Jay Tracy, Ralph Schlapbach, Christiane Beckmann, Maurice Redondo, Olivier Kobel, Christoph Noppen, Niko Beerenwinkel, Tanja Stadler |
| EPI_ISL_3346285 | Viollier AG | Department of Biosystems Science and Engineering, ETH Zürich | Christian Beisel, Sarah Nadeau, Chaoran Chen, Louis du Plessis, Ivan Topolsky, Kim Philipp Jablonski, Lara Fuhrmann, Rebecca Denes, Mirjam Feldkamp, Ina Nissen, Natascha Santacroce, Elodie Burcklen, Christiane Beckmann, Maurice Redondo, Olivier Kobel, Christoph Noppen, Niko Beerenwinkel, Tanja Stadler |
| EPI_ISL_3346286 | Viollier AG | Department of Biosystems Science and Engineering, ETH Zürich | Chaoran Chen, Sarah Nadeau, Louis du Plessis, Catharine Aquino, Ivan Topolsky, Kim Philipp Jablonski, Lara Fuhrmann, Daniel Ehrsam, Isabel Stürmer, Andreia Cabral de Gouvea, Maria Domenica Moccia, Simon Grüter, Timothy Sykes, Lennart Opitz, Griffin White, Laura Neff, Doris Popovic, Andrea Patrignani, Jay Tracy, Ralph Schlapbach, Christiane Beckmann, Maurice Redondo, Olivier Kobel, Christoph Noppen, Niko Beerenwinkel, Tanja Stadler |
| EPI_ISL_3346287 | Viollier AG | Department of Biosystems Science and Engineering, ETH Zürich | Christian Beisel, Sarah Nadeau, Chaoran Chen, Louis du Plessis, Ivan Topolsky, Kim Philipp Jablonski, Lara Fuhrmann, Rebecca Denes, Mirjam Feldkamp, Ina Nissen, Natascha Santacroce, Elodie Burcklen, Christiane Beckmann, Maurice Redondo, Olivier Kobel, Christoph Noppen, Niko Beerenwinkel, Tanja Stadler |
| EPI_ISL_3346289 | Viollier AG | Department of Biosystems Science and Engineering, ETH Zürich | Chaoran Chen, Sarah Nadeau, Louis du Plessis, Catharine Aquino, Ivan Topolsky, Kim Philipp Jablonski, Lara Fuhrmann, Daniel Ehrsam, Isabel Stürmer, Andreia Cabral de Gouvea, Maria Domenica Moccia, Simon Grüter, Timothy Sykes, Lennart Opitz, Griffin White, Laura Neff, Doris Popovic, Andrea Patrignani, Jay Tracy, Ralph Schlapbach, Christiane Beckmann, Maurice Redondo, Olivier Kobel, Christoph Noppen, Niko Beerenwinkel, Tanja Stadler |
| EPI_ISL_3346290, EPI_ISL_3346291, EPI_ISL_3346292, EPI_ISL_3346293 | Viollier AG | Department of Biosystems Science and Engineering, ETH Zürich | Christian Beisel, Sarah Nadeau, Chaoran Chen, Louis du Plessis, Ivan Topolsky, Kim Philipp Jablonski, Lara Fuhrmann, Rebecca Denes, Mirjam Feldkamp, Ina Nissen, Natascha Santacroce, Elodie Burcklen, Christiane Beckmann, Maurice Redondo, Olivier Kobel, Christoph Noppen, Niko Beerenwinkel, Tanja Stadler |
| EPI_ISL_3346294, EPI_ISL_3346295 | Viollier AG | Department of Biosystems Science and Engineering, ETH Zürich | Chaoran Chen, Sarah Nadeau, Louis du Plessis, Catharine Aquino, Ivan Topolsky, Kim Philipp Jablonski, Lara Fuhrmann, Daniel Ehrsam, Isabel Stürmer, Andreia Cabral de Gouvea, Maria Domenica Moccia, Simon Grüter, Timothy Sykes, Lennart Opitz, Griffin White, Laura Neff, Doris Popovic, Andrea Patrignani, Jay Tracy, Ralph Schlapbach, Christiane Beckmann, Maurice Redondo, Olivier Kobel, Christoph Noppen, Niko Beerenwinkel, Tanja Stadler |
| EPI_ISL_3346296 | Viollier AG | Department of Biosystems Science and Engineering, ETH Zürich | Christian Beisel, Sarah Nadeau, Chaoran Chen, Louis du Plessis, Ivan Topolsky, Kim Philipp Jablonski, Lara Fuhrmann, Rebecca Denes, Mirjam Feldkamp, Ina Nissen, Natascha Santacroce, Elodie Burcklen, Christiane Beckmann, Maurice Redondo, Olivier Kobel, Christoph Noppen, Niko Beerenwinkel, Tanja Stadler |
| EPI_ISL_3346297 | Viollier AG | Department of Biosystems Science and Engineering, ETH Zürich | Chaoran Chen, Sarah Nadeau, Louis du Plessis, Catharine Aquino, Ivan Topolsky, Kim Philipp Jablonski, Lara Fuhrmann, Daniel Ehrsam, Isabel Stürmer, Andreia Cabral de Gouvea, Maria Domenica Moccia, Simon Grüter, Timothy Sykes, Lennart Opitz, Griffin White, Laura Neff, Doris Popovic, Andrea Patrignani, Jay Tracy, Ralph Schlapbach, Christiane Beckmann, Maurice Redondo, Olivier Kobel, Christoph Noppen, Niko Beerenwinkel, Tanja Stadler |
| EPI_ISL_3346298 | Viollier AG | Department of Biosystems Science and Engineering, ETH Zürich | Christian Beisel, Sarah Nadeau, Chaoran Chen, Louis du Plessis, Ivan Topolsky, Kim Philipp Jablonski, Lara Fuhrmann, Rebecca Denes, Mirjam Feldkamp, Ina Nissen, Natascha Santacroce, Elodie Burcklen, Christiane Beckmann, Maurice Redondo, Olivier Kobel, Christoph Noppen, Niko Beerenwinkel, Tanja Stadler |
| EPI_ISL_3346299, EPI_ISL_3346300, EPI_ISL_3346301 | Viollier AG | Department of Biosystems Science and Engineering, ETH Zürich | Chaoran Chen, Sarah Nadeau, Louis du Plessis, Catharine Aquino, Ivan Topolsky, Kim Philipp Jablonski, Lara Fuhrmann, Daniel Ehrsam, Isabel Stürmer, Andreia Cabral de Gouvea, Maria Domenica Moccia, Simon Grüter, Timothy Sykes, Lennart Opitz, Griffin White, Laura Neff, Doris Popovic, Andrea Patrignani, Jay Tracy, Ralph Schlapbach, Christiane Beckmann, Maurice Redondo, Olivier Kobel, Christoph Noppen, Niko Beerenwinkel, Tanja Stadler |
| EPI_ISL_3346302 | Viollier AG | Department of Biosystems Science and Engineering, ETH Zürich | Christian Beisel, Sarah Nadeau, Chaoran Chen, Louis du Plessis, Ivan Topolsky, Kim Philipp Jablonski, Lara Fuhrmann, Rebecca Denes, Mirjam Feldkamp, Ina Nissen, Natascha Santacroce, Elodie Burcklen, Christiane Beckmann, Maurice Redondo, Olivier Kobel, Christoph Noppen, Niko Beerenwinkel, Tanja Stadler |
| EPI_ISL_3346303, EPI_ISL_3346304 | Viollier AG | Department of Biosystems Science and Engineering, ETH Zürich | Chaoran Chen, Sarah Nadeau, Louis du Plessis, Catharine Aquino, Ivan Topolsky, Kim Philipp Jablonski, Lara Fuhrmann, Daniel Ehrsam, Isabel Stürmer, Andreia Cabral de Gouvea, Maria Domenica Moccia, Simon Grüter, Timothy Sykes, Lennart Opitz, Griffin White, Laura Neff, Doris Popovic, Andrea Patrignani, Jay Tracy, Ralph Schlapbach, Christiane Beckmann, Maurice Redondo, Olivier Kobel, Christoph Noppen, Niko Beerenwinkel, Tanja Stadler |
| EPI_ISL_3346305, EPI_ISL_3346306, EPI_ISL_3346307, EPI_ISL_3346308 | Viollier AG | Department of Biosystems Science and Engineering, ETH Zürich | Christian Beisel, Sarah Nadeau, Chaoran Chen, Louis du Plessis, Ivan Topolsky, Kim Philipp Jablonski, Lara Fuhrmann, Rebecca Denes, Mirjam Feldkamp, Ina Nissen, Natascha Santacroce, Elodie Burcklen, Christiane Beckmann, Maurice Redondo, Olivier Kobel, Christoph Noppen, Niko Beerenwinkel, Tanja Stadler |
| EPI_ISL_3346309 | Viollier AG | Department of Biosystems Science and Engineering, ETH Zürich | Chaoran Chen, Sarah Nadeau, Louis du Plessis, Catharine Aquino, Ivan Topolsky, Kim Philipp Jablonski, Lara Fuhrmann, Daniel Ehrsam, Isabel Stürmer, Andreia Cabral de Gouvea, Maria Domenica Moccia, Simon Grüter, Timothy Sykes, Lennart Opitz, Griffin White, Laura Neff, Doris Popovic, Andrea Patrignani, Jay Tracy, Ralph Schlapbach, Christiane Beckmann, Maurice Redondo, Olivier Kobel, Christoph Noppen, Niko Beerenwinkel, Tanja Stadler |
| EPI_ISL_3346310 | Viollier AG | Department of Biosystems Science and Engineering, ETH Zürich | Christian Beisel, Sarah Nadeau, Chaoran Chen, Louis du Plessis, Ivan Topolsky, Kim Philipp Jablonski, Lara Fuhrmann, Rebecca Denes, Mirjam Feldkamp, Ina Nissen, Natascha Santacroce, Elodie Burcklen, Christiane Beckmann, Maurice Redondo, Olivier Kobel, Christoph Noppen, Niko Beerenwinkel, Tanja Stadler |

| | | | |
|---|---|---|---|
| EPI_ISL_3346311 | Unilabs | Institute of Medical Virology | Verena Kufner, Gabriela Ziltener, Maryam Zaheri, Stefan Schmutz, Annette Audigé, Maria Grünberg, Kevin Steiner, Jon Huder, Cyril Shah, Riccarda Capaul, Guido Bloemberg, Jürg Böni, Michael Huber, Alexandra Trkola |
| EPI_ISL_3346312, EPI_ISL_3346313 | Kantonsspital Baden AG | Institute of Medical Virology | Verena Kufner, Gabriela Ziltener, Maryam Zaheri, Stefan Schmutz, Annette Audigé, Maria Grünberg, Kevin Steiner, Jon Huder, Cyril Shah, Riccarda Capaul, Guido Bloemberg, Jürg Böni, Michael Huber, Alexandra Trkola |
| EPI_ISL_3346314 | Medica | Institute of Medical Virology | Verena Kufner, Gabriela Ziltener, Maryam Zaheri, Stefan Schmutz, Annette Audigé, Maria Grünberg, Kevin Steiner, Jon Huder, Cyril Shah, Riccarda Capaul, Guido Bloemberg, Jürg Böni, Michael Huber, Alexandra Trkola |
| EPI_ISL_3346315 | Analytica | Institute of Medical Virology | Verena Kufner, Gabriela Ziltener, Maryam Zaheri, Stefan Schmutz, Annette Audigé, Maria Grünberg, Kevin Steiner, Jon Huder, Cyril Shah, Riccarda Capaul, Guido Bloemberg, Jürg Böni, Michael Huber, Alexandra Trkola |
| EPI_ISL_3346316 | Universität Zürich | Institute of Medical Virology | Verena Kufner, Gabriela Ziltener, Maryam Zaheri, Stefan Schmutz, Annette Audigé, Maria Grünberg, Kevin Steiner, Jon Huder, Cyril Shah, Riccarda Capaul, Guido Bloemberg, Jürg Böni, Michael Huber, Alexandra Trkola |
| EPI_ISL_3346317, EPI_ISL_3346318 | UniversitätsSpital Zürich | Institute of Medical Virology | Verena Kufner, Gabriela Ziltener, Maryam Zaheri, Stefan Schmutz, Annette Audigé, Maria Grünberg, Kevin Steiner, Jon Huder, Cyril Shah, Riccarda Capaul, Guido Bloemberg, Jürg Böni, Michael Huber, Alexandra Trkola |
| EPI_ISL_3346319 | Kantonsspital Aarau | Institute of Medical Virology | Verena Kufner, Gabriela Ziltener, Maryam Zaheri, Stefan Schmutz, Annette Audigé, Maria Grünberg, Kevin Steiner, Jon Huder, Cyril Shah, Riccarda Capaul, Guido Bloemberg, Jürg Böni, Michael Huber, Alexandra Trkola |
| EPI_ISL_3346320 | Kantonsspital Baden AG | Institute of Medical Virology | Verena Kufner, Gabriela Ziltener, Maryam Zaheri, Stefan Schmutz, Annette Audigé, Maria Grünberg, Kevin Steiner, Jon Huder, Cyril Shah, Riccarda Capaul, Guido Bloemberg, Jürg Böni, Michael Huber, Alexandra Trkola |
| EPI_ISL_3346321, EPI_ISL_3346322 | Zentrallabor Zürich | Institute of Medical Virology | Verena Kufner, Gabriela Ziltener, Maryam Zaheri, Stefan Schmutz, Annette Audigé, Maria Grünberg, Kevin Steiner, Jon Huder, Cyril Shah, Riccarda Capaul, Guido Bloemberg, Jürg Böni, Michael Huber, Alexandra Trkola |
| EPI_ISL_3346323 | UniversitätsSpital Zürich | Institute of Medical Virology | Verena Kufner, Gabriela Ziltener, Maryam Zaheri, Stefan Schmutz, Annette Audigé, Maria Grünberg, Kevin Steiner, Jon Huder, Cyril Shah, Riccarda Capaul, Guido Bloemberg, Jürg Böni, Michael Huber, Alexandra Trkola |
| EPI_ISL_3346324, EPI_ISL_3346325 | Kantonsspital Aarau | Institute of Medical Virology | Verena Kufner, Gabriela Ziltener, Maryam Zaheri, Stefan Schmutz, Annette Audigé, Maria Grünberg, Kevin Steiner, Jon Huder, Cyril Shah, Riccarda Capaul, Guido Bloemberg, Jürg Böni, Michael Huber, Alexandra Trkola |
| EPI_ISL_3346326 | Kantonsspital Baden AG | Institute of Medical Virology | Verena Kufner, Gabriela Ziltener, Maryam Zaheri, Stefan Schmutz, Annette Audigé, Maria Grünberg, Kevin Steiner, Jon Huder, Cyril Shah, Riccarda Capaul, Guido Bloemberg, Jürg Böni, Michael Huber, Alexandra Trkola |
| EPI_ISL_3346327 | Stadtspital Waid | Institute of Medical Virology | Verena Kufner, Gabriela Ziltener, Maryam Zaheri, Stefan Schmutz, Annette Audigé, Maria Grünberg, Kevin Steiner, Jon Huder, Cyril Shah, Riccarda Capaul, Guido Bloemberg, Jürg Böni, Michael Huber, Alexandra Trkola |
| EPI_ISL_3346328, EPI_ISL_3346329, EPI_ISL_3346330 | Medica | Institute of Medical Virology | Verena Kufner, Gabriela Ziltener, Maryam Zaheri, Stefan Schmutz, Annette Audigé, Maria Grünberg, Kevin Steiner, Jon Huder, Cyril Shah, Riccarda Capaul, Guido Bloemberg, Jürg Böni, Michael Huber, Alexandra Trkola |
| EPI_ISL_3346331, EPI_ISL_3346332 | Analytica | Institute of Medical Virology | Verena Kufner, Gabriela Ziltener, Maryam Zaheri, Stefan Schmutz, Annette Audigé, Maria Grünberg, Kevin Steiner, Jon Huder, Cyril Shah, Riccarda Capaul, Guido Bloemberg, Jürg Böni, Michael Huber, Alexandra Trkola |
| EPI_ISL_3346333, EPI_ISL_3346334, EPI_ISL_3346335 | Zentrum für Labormedizin | Institute of Medical Virology | Verena Kufner, Gabriela Ziltener, Maryam Zaheri, Stefan Schmutz, Annette Audigé, Maria Grünberg, Kevin Steiner, Jon Huder, Cyril Shah, Riccarda Capaul, Guido Bloemberg, Jürg Böni, Michael Huber, Alexandra Trkola |
| EPI_ISL_3346336, EPI_ISL_3346337 | Universität Zürich | Institute of Medical Virology | Verena Kufner, Gabriela Ziltener, Maryam Zaheri, Stefan Schmutz, Annette Audigé, Maria Grünberg, Kevin Steiner, Jon Huder, Cyril Shah, Riccarda Capaul, Guido Bloemberg, Jürg Böni, Michael Huber, Alexandra Trkola |
| EPI_ISL_3346338 | UniversitätsSpital Zürich | Institute of Medical Virology | Verena Kufner, Gabriela Ziltener, Maryam Zaheri, Stefan Schmutz, Annette Audigé, Maria Grünberg, Kevin Steiner, Jon Huder, Cyril Shah, Riccarda Capaul, Guido Bloemberg, Jürg Böni, Michael Huber, Alexandra Trkola |
| EPI_ISL_3346500 | CHC Andrée Rosemon | Institut Pasteur de la Guyane | Anne Lavergne, Dominique Rousset, Antoine Enfissi, Arielle Salmier |
| EPI_ISL_3346501 | CNR Institut Pasteur de la Guyane | Institut Pasteur de la Guyane | Anne Lavergne, Dominique Rousset, Antoine Enfissi, Arielle Salmier |
| EPI_ISL_3346502, EPI_ISL_3346503 | State Hygienic Laboratory at the University of Iowa | State Hygienic Laboratory at the University of Iowa | Valerie Reeb, Erik Twait, Anna Yakos, Mariah Knutson, Kris Eveland, Kristen Zanon, Jeff, Benfer, Davis Rieckenberg, Wes Hottel, Alankar Kampoowale |
| EPI_ISL_3346520 | OHSU-MM Lab | OHSU-MM Lab | Jeannine Lama |
| EPI_ISL_3346523, EPI_ISL_3346524, EPI_ISL_3346525, EPI_ISL_3346526, EPI_ISL_3346527, EPI_ISL_3346528, EPI_ISL_3346529, EPI_ISL_3346530, EPI_ISL_3346531, EPI_ISL_3346532, EPI_ISL_3346533, EPI_ISL_3346534, EPI_ISL_3346535, EPI_ISL_3346536, EPI_ISL_3346537, EPI_ISL_3346538, EPI_ISL_3346539, EPI_ISL_3346540, EPI_ISL_3346541, EPI_ISL_3346542, EPI_ISL_3346543, EPI_ISL_3346544, EPI_ISL_3346545, EPI_ISL_3346546, EPI_ISL_3346547, EPI_ISL_3346548, EPI_ISL_3346549, EPI_ISL_3346550, EPI_ISL_3346551, EPI_ISL_3346552, EPI_ISL_3346553 | | | |
| see above | Nebraska Public Health Laboratory | NPHL COVID-19 Response Team | NPHL COVID-19 Response Team |
| EPI_ISL_3346554 | University Campus Bio-Medico of Rome (UCBM) | University Campus Bio-Medico of Rome (UCBM) | Fogolari M., De Florio L., Riva E., Francesconi M., Lintas C., Veralli R., Angeletti S. |
| EPI_ISL_3346557 | State Hygienic Laboratory at the University of Iowa | State Hygienic Laboratory at the University of Iowa | Valerie Reeb, Erik Twait, Anna Yakos, Mariah Knutson, Kris Eveland, Kristen Zanon, Jeff, Benfer, Davis Rieckenberg, Wes Hottel, Alankar Kampoowale |
| EPI_ISL_3346558, EPI_ISL_3346559, EPI_ISL_3346571 | University Campus Bio-Medico of Rome (UCBM) | University Campus Bio-Medico of Rome (UCBM) | Fogolari M., De Florio L., Riva E., Francesconi M., Lintas C., Veralli R., Angeletti S. |
| EPI_ISL_3346598, EPI_ISL_3346599, EPI_ISL_3346604, EPI_ISL_3346607, EPI_ISL_3346610 | San Diego County Public Health Laboratory | San Diego County Public Health Laboratory | Ashleigh Murphy Schafer, Brett Austin |
| EPI_ISL_3346611, EPI_ISL_3346612, EPI_ISL_3346613, EPI_ISL_3346614, EPI_ISL_3346615, EPI_ISL_3346616, EPI_ISL_3346617, EPI_ISL_3346618, EPI_ISL_3346619, EPI_ISL_3346620, EPI_ISL_3346621, EPI_ISL_3346622, EPI_ISL_3346623, EPI_ISL_3346624, EPI_ISL_3346625, EPI_ISL_3346626, EPI_ISL_3346627, EPI_ISL_3346628, EPI_ISL_3346629, EPI_ISL_3346630, EPI_ISL_3346631, EPI_ISL_3346632, EPI_ISL_3346633, EPI_ISL_3346634, EPI_ISL_3346635, EPI_ISL_3346636, EPI_ISL_3346637, EPI_ISL_3346638, EPI_ISL_3346639, EPI_ISL_3346640, EPI_ISL_3346641, EPI_ISL_3346642, EPI_ISL_3346643, EPI_ISL_3346644, EPI_ISL_3346645, EPI_ISL_3346646, EPI_ISL_3346647, EPI_ISL_3346648, EPI_ISL_3346649, EPI_ISL_3346650, EPI_ISL_3346651, EPI_ISL_3346652, EPI_ISL_3346653, EPI_ISL_3346654, EPI_ISL_3346655, EPI_ISL_3346656, EPI_ISL_3346657, EPI_ISL_3346658, EPI_ISL_3346659, EPI_ISL_3346660, EPI_ISL_3346661, EPI_ISL_3346662, EPI_ISL_3346663, EPI_ISL_3346664, EPI_ISL_3346665, EPI_ISL_3346666, EPI_ISL_3346667, EPI_ISL_3346668, EPI_ISL_3346669, EPI_ISL_3346670, EPI_ISL_3346671, EPI_ISL_3346672, EPI_ISL_3346673, EPI_ISL_3346674, EPI_ISL_3346675, EPI_ISL_3346676, EPI_ISL_3346677, EPI_ISL_3346678, EPI_ISL_3346679, EPI_ISL_3346680, EPI_ISL_3346681, EPI_ISL_3346682, EPI_ISL_3346683, EPI_ISL_3346684, EPI_ISL_3346685, EPI_ISL_3346686, EPI_ISL_3346687, EPI_ISL_3346688, EPI_ISL_3346689, EPI_ISL_3346690, EPI_ISL_3346691, EPI_ISL_3346692, EPI_ISL_3346693, EPI_ISL_3346694, EPI_ISL_3346695, EPI_ISL_3346696, EPI_ISL_3346697, EPI_ISL_3346698, EPI_ISL_3346699, EPI_ISL_3346700, EPI_ISL_3346701, EPI_ISL_3346702, EPI_ISL_3346703, EPI_ISL_3346704, EPI_ISL_3346705, EPI_ISL_3346706, EPI_ISL_3346707, EPI_ISL_3346708, EPI_ISL_3346709, EPI_ISL_3346710, EPI_ISL_3346711, EPI_ISL_3346712, EPI_ISL_3346713, EPI_ISL_3346714, EPI_ISL_3346715, EPI_ISL_3346716, EPI_ISL_3346717, EPI_ISL_3346718, EPI_ISL_3346719, EPI_ISL_3346720, EPI_ISL_3346721, EPI_ISL_3346722, EPI_ISL_3346723, EPI_ISL_3346724, EPI_ISL_3346725, EPI_ISL_3346726, EPI_ISL_3346727, EPI_ISL_3346728, EPI_ISL_3346729, EPI_ISL_3346730, EPI_ISL_3346731, EPI_ISL_3346732, EPI_ISL_3346733, EPI_ISL_3346734, EPI_ISL_3346735, EPI_ISL_3346736, EPI_ISL_3346737, EPI_ISL_3346738, EPI_ISL_3346739, EPI_ISL_3346740, EPI_ISL_3346741, EPI_ISL_3346742, EPI_ISL_3346743, EPI_ISL_3346744, EPI_ISL_3346745, EPI_ISL_3346746, EPI_ISL_3346747, EPI_ISL_3346748, EPI_ISL_3346749, EPI_ISL_3346750, EPI_ISL_3346751, EPI_ISL_3346752, EPI_ISL_3346753, EPI_ISL_3346754, EPI_ISL_3346755, EPI_ISL_3346756, EPI_ISL_3346757, EPI_ISL_3346758, EPI_ISL_3346759, EPI_ISL_3346760, EPI_ISL_3346761, EPI_ISL_3346762, EPI_ISL_3346763, EPI_ISL_3346764, EPI_ISL_3346765, EPI_ISL_3346766, EPI_ISL_3346767, EPI_ISL_3346768, EPI_ISL_3346769, EPI_ISL_3346770, EPI_ISL_3346771, EPI_ISL_3346772, EPI_ISL_3346773, EPI_ISL_3346774, EPI_ISL_3346775, EPI_ISL_3346776, EPI_ISL_3346777, EPI_ISL_3346778, EPI_ISL_3346779, EPI_ISL_3346780, EPI_ISL_3346781, EPI_ISL_3346782, EPI_ISL_3346783, EPI_ISL_3346785, EPI_ISL_3346786, EPI_ISL_3346787, EPI_ISL_3346788, EPI_ISL_3346789, EPI_ISL_3346790, EPI_ISL_3346791, EPI_ISL_3346792, EPI_ISL_3346793, EPI_ISL_3346794, EPI_ISL_3346795, EPI_ISL_3346796, EPI_ISL_3346797, EPI_ISL_3346798, EPI_ISL_3346799, EPI_ISL_3346800, EPI_ISL_3346801, EPI_ISL_3346802, EPI_ISL_3346803, EPI_ISL_3346804, EPI_ISL_3346805, EPI_ISL_3346806, EPI_ISL_3346807, EPI_ISL_3346808, EPI_ISL_3346809, EPI_ISL_3346810, EPI_ISL_3346811, EPI_ISL_3346812, EPI_ISL_3346813, EPI_ISL_3346815, EPI_ISL_3346816, EPI_ISL_3346817, EPI_ISL_3346818, EPI_ISL_3346819, EPI_ISL_3346820, EPI_ISL_3346821, EPI_ISL_3346822, EPI_ISL_3346823, EPI_ISL_3346824, EPI_ISL_3346825, EPI_ISL_3346826, EPI_ISL_3346827, EPI_ISL_3346828, EPI_ISL_3346829, EPI_ISL_3346830, EPI_ISL_3346831, EPI_ISL_3346832, EPI_ISL_3346833, EPI_ISL_3346834, EPI_ISL_3346835, EPI_ISL_3346836, EPI_ISL_3346837, EPI_ISL_3346838, EPI_ISL_3346839, EPI_ISL_3346840, EPI_ISL_3346841, EPI_ISL_3346842, EPI_ISL_3346843, EPI_ISL_3346844, EPI_ISL_3346845, EPI_ISL_3346846, EPI_ISL_3346847, EPI_ISL_3346848, EPI_ISL_3346849, EPI_ISL_3346850, EPI_ISL_3346851, EPI_ISL_3346852, EPI_ISL_3346853, EPI_ISL_3346854, EPI_ISL_3346855, EPI_ISL_3346856 | | | |
| see above | Springfield-Greene County Health Department | University of Minnesota Genomics Center | Daryl M. Gohl, John Garbe, Jaquelyn Kuriger-Laber, Corbin Dirkx |
| EPI_ISL_3346857, EPI_ISL_3346858, EPI_ISL_3346859, EPI_ISL_3346860, EPI_ISL_3346862, EPI_ISL_3346863, EPI_ISL_3346864, EPI_ISL_3346865, EPI_ISL_3346866, EPI_ISL_3346867, EPI_ISL_3346868, EPI_ISL_3346869, EPI_ISL_3346870, EPI_ISL_3346871, EPI_ISL_3346872, EPI_ISL_3346873, EPI_ISL_3346874, EPI_ISL_3346875, EPI_ISL_3346876, EPI_ISL_3346877, EPI_ISL_3346878, EPI_ISL_3346879, EPI_ISL_3346880, EPI_ISL_3346881, EPI_ISL_3346882, EPI_ISL_3346883, EPI_ISL_3346884, EPI_ISL_3346885, EPI_ISL_3346886, EPI_ISL_3346887, EPI_ISL_3346888, EPI_ISL_3346889, EPI_ISL_3346890, EPI_ISL_3346891, EPI_ISL_3346892, EPI_ISL_3346894, EPI_ISL_3346895, EPI_ISL_3346896, EPI_ISL_3346897, EPI_ISL_3346898, EPI_ISL_3346899, EPI_ISL_3346900, EPI_ISL_3346901, EPI_ISL_3346902, EPI_ISL_3346903, EPI_ISL_3346904, EPI_ISL_3346906, EPI_ISL_3346907, EPI_ISL_3346908, EPI_ISL_3346909, EPI_ISL_3346910, EPI_ISL_3346911, EPI_ISL_3346912, EPI_ISL_3346914, EPI_ISL_3346915, EPI_ISL_3346916, EPI_ISL_3346917, EPI_ISL_3346918, EPI_ISL_3346919, EPI_ISL_3346921, EPI_ISL_3346923, EPI_ISL_3346925, EPI_ISL_3346926, EPI_ISL_3346927, EPI_ISL_3346928, EPI_ISL_3346929, EPI_ISL_3346930, EPI_ISL_3346931, EPI_ISL_3346932, EPI_ISL_3346933, EPI_ISL_3346934, EPI_ISL_3346935 | | | |
| see above | Northwest Laboratory | Seattle Flu Study | Deborah A. Nickerson, Chris D. Frazar, Jover Lee, Benjamin Pelle, Erica Ryke, Matthew Richardson, Amanda Adler, Elisabeth Brandstetter, Peter D. Han, Kairsten Fay, Misja Ilcisin, Kirsten Lacombe, Thomas R. Sibley, Melissa Truong, Caitlin R. Wolf, Romesh Gautom, Geoff Melly, Brian Hiatt, Philip Dykema, Scott Lindquist, Michael Boeckh, Janet A. Englund, Michael Famulare, Barry R. Lutz, Mark J. Rieder, Lea M. Starita, Matthew Thompson, Helen Y. Chu, Jay Shendure, Trevor Bedford |
| EPI_ISL_3346936, EPI_ISL_3346937, EPI_ISL_3346938, EPI_ISL_3346939, EPI_ISL_3346940, EPI_ISL_3346941, EPI_ISL_3346942, EPI_ISL_3346943, EPI_ISL_3346944, EPI_ISL_3346945, EPI_ISL_3346946, EPI_ISL_3346947, EPI_ISL_3346948, EPI_ISL_3346949, EPI_ISL_3346950, EPI_ISL_3346951, EPI_ISL_3346952, EPI_ISL_3346953, EPI_ISL_3346954, EPI_ISL_3346955, EPI_ISL_3346956, EPI_ISL_3346957, EPI_ISL_3346958, EPI_ISL_3346959, EPI_ISL_3346960, EPI_ISL_3346961, EPI_ISL_3346962, EPI_ISL_3346963, EPI_ISL_3346964, EPI_ISL_3346965, EPI_ISL_3346966, EPI_ISL_3346967, EPI_ISL_3346968, EPI_ISL_3346969, EPI_ISL_3346970, EPI_ISL_3346971, EPI_ISL_3346972, EPI_ISL_3346973, EPI_ISL_3346974, EPI_ISL_3346975, EPI_ISL_3346976, EPI_ISL_3346977, EPI_ISL_3346978, EPI_ISL_3346979, EPI_ISL_3346981, EPI_ISL_3346982, EPI_ISL_3346983, EPI_ISL_3346984, EPI_ISL_3346985, EPI_ISL_3346986, EPI_ISL_3346987, EPI_ISL_3346988, EPI_ISL_3346989, EPI_ISL_3346990, EPI_ISL_3346991, EPI_ISL_3346992, EPI_ISL_3346993, EPI_ISL_3346994 | | | |
| see above | Atlas Genomics | Seattle Flu Study | Deborah A. Nickerson, Chris D. Frazar, Jover Lee, Benjamin Pelle, Erica Ryke, Matthew Richardson, Amanda Adler, Elisabeth Brandstetter, Peter D. Han, Kairsten Fay, Misja Ilcisin, Kirsten Lacombe, Thomas R. Sibley, Melissa Truong, Caitlin R. Wolf, Romesh Gautom, Geoff Melly, Brian Hiatt, Philip Dykema, Scott Lindquist, Michael Boeckh, Janet A. Englund, Michael Famulare, Barry R. Lutz, Mark J. Rieder, Lea M. Starita, Matthew Thompson, Helen Y. Chu, Jay Shendure, Trevor Bedford |
| EPI_ISL_3346995, EPI_ISL_3346996, EPI_ISL_3346998, EPI_ISL_3347003, EPI_ISL_3347004, EPI_ISL_3347007, EPI_ISL_3347008, EPI_ISL_3347009, EPI_ISL_3347012, EPI_ISL_3347013, EPI_ISL_3347014, EPI_ISL_3347016, EPI_ISL_3347017, EPI_ISL_3347018, EPI_ISL_3347019, EPI_ISL_3347023, EPI_ISL_3347024, EPI_ISL_3347025, EPI_ISL_3347026, EPI_ISL_3347027, EPI_ISL_3347028, EPI_ISL_3347030, EPI_ISL_3347031, EPI_ISL_3347032, EPI_ISL_3347033, EPI_ISL_3347034, EPI_ISL_3347035, EPI_ISL_3347036, EPI_ISL_3347037, EPI_ISL_3347039, EPI_ISL_3347040, EPI_ISL_3347041, EPI_ISL_3347042, EPI_ISL_3347043, EPI_ISL_3347045, EPI_ISL_3347046, EPI_ISL_3347047, EPI_ISL_3347048, EPI_ISL_3347050, EPI_ISL_3347051, EPI_ISL_3347052, EPI_ISL_3347053, EPI_ISL_3347054, EPI_ISL_3347055, EPI_ISL_3347056, EPI_ISL_3347057, EPI_ISL_3347058, EPI_ISL_3347059, EPI_ISL_3347060, EPI_ISL_3347061, EPI_ISL_3347062, EPI_ISL_3347063, EPI_ISL_3347064, EPI_ISL_3347065 | | | |
| see above | Incyte Diagnostics Spokane | Seattle Flu Study | Deborah A. Nickerson, Chris D. Frazar, Jover Lee, Benjamin Pelle, Erica Ryke, Matthew Richardson, Amanda Adler, Elisabeth Brandstetter, Peter D. Han, Kairsten Fay, Misja Ilcisin, Kirsten Lacombe, Thomas R. Sibley, Melissa Truong, Caitlin R. Wolf, Romesh Gautom, Geoff Melly, Brian Hiatt, Philip Dykema, Scott Lindquist, Michael Boeckh, Janet A. Englund, Michael Famulare, Barry R. Lutz, Mark J. Rieder, Lea M. Starita, Matthew Thompson, Helen Y. Chu, Jay Shendure, Trevor Bedford |
| EPI_ISL_3347066, EPI_ISL_3347069, EPI_ISL_3347070, EPI_ISL_3347072, EPI_ISL_3347073, EPI_ISL_3347075, EPI_ISL_3347076, EPI_ISL_3347079, EPI_ISL_3347082, EPI_ISL_3347091, EPI_ISL_3347094, EPI_ISL_3347097, EPI_ISL_3347100, EPI_ISL_3347103, EPI_ISL_3347107, EPI_ISL_3347111, EPI_ISL_3347112, EPI_ISL_3347114, EPI_ISL_3347119, EPI_ISL_3347121, EPI_ISL_3347122 | | | |
| see above | Interpath Laboratory | Seattle Flu Study | Deborah A. Nickerson, Chris D. Frazar, Jover Lee, Benjamin Pelle, Erica Ryke, Matthew Richardson, Amanda Adler, Elisabeth Brandstetter, Peter D. Han, Kairsten Fay, Misja Ilcisin, Kirsten Lacombe, Thomas R. Sibley, Melissa Truong, Caitlin R. Wolf, Romesh Gautom, Geoff Melly, Brian Hiatt, Philip Dykema, Scott Lindquist, Michael Boeckh, Janet A. Englund, Michael Famulare, Barry R. Lutz, Mark J. Rieder, Lea M. Starita, Matthew Thompson, Helen Y. Chu, Jay Shendure, Trevor Bedford |
| EPI_ISL_3347123, EPI_ISL_3347124, EPI_ISL_3347125, EPI_ISL_3347126, EPI_ISL_3347127, EPI_ISL_3347128, EPI_ISL_3347129, EPI_ISL_3347130, EPI_ISL_3347131, EPI_ISL_3347132, EPI_ISL_3347133, EPI_ISL_3347134, EPI_ISL_3347135, EPI_ISL_3347136, EPI_ISL_3347137, EPI_ISL_3347138, EPI_ISL_3347139, EPI_ISL_3347140, EPI_ISL_3347141, EPI_ISL_3347142, EPI_ISL_3347143, EPI_ISL_3347144, EPI_ISL_3347145, EPI_ISL_3347146, EPI_ISL_3347147, EPI_ISL_3347148, EPI_ISL_3347149, EPI_ISL_3347150, EPI_ISL_3347151, EPI_ISL_3347152, EPI_ISL_3347153, EPI_ISL_3347154, EPI_ISL_3347155, EPI_ISL_3347156, EPI_ISL_3347157, EPI_ISL_3347158, EPI_ISL_3347159, EPI_ISL_3347160, EPI_ISL_3347161, EPI_ISL_3347162, EPI_ISL_3347163, EPI_ISL_3347164, EPI_ISL_3347165, EPI_ISL_3347166, EPI_ISL_3347167, EPI_ISL_3347168, EPI_ISL_3347169, EPI_ISL_3347170, EPI_ISL_3347171, EPI_ISL_3347172 | | | |
| see above | Atlas Genomics | Seattle Flu Study | Deborah A. Nickerson, Chris D. Frazar, Jover Lee, Benjamin Pelle, Erica Ryke, Matthew Richardson, Amanda Adler, Elisabeth Brandstetter, Peter D. Han, Kairsten Fay, Misja Ilcisin, Kirsten Lacombe, Thomas R. Sibley, Melissa Truong, Caitlin R. Wolf, Romesh Gautom, Geoff Melly, Brian Hiatt, Philip Dykema, Scott Lindquist, Michael Boeckh, Janet A. Englund, Michael Famulare, Barry R. Lutz, Mark J. Rieder, Lea M. Starita, Matthew Thompson, Helen Y. Chu, Jay Shendure, Trevor Bedford |
| EPI_ISL_3347211, EPI_ISL_3347213, EPI_ISL_3347214 | UCC (Universidad Central del Caribe) | Grubaugh Lab - Yale School of Public Health | Joseph Fauver, Mallery Breban, Isabel Ott, Tara Alpert, Mary Petrone, Anderson Brito, Chantal Vogels, Annie Watkins, Chaney Kalinich, Jessica Rothman, Kendall Billig, Tobias Koch, Fabiola Cruz-Lopez, David Rivera Aponte, Fabiola Fontanet, Natalie Machargo, Andrea Firpo, Vianca Aponte, Ernesto Vazquez, Patricia Serrano, Tania Mitwalli, Karolane Gonzalez, Nathan Grubaugh |
| EPI_ISL_3347215, EPI_ISL_3347216, EPI_ISL_3347217, EPI_ISL_3347218, EPI_ISL_3347220, EPI_ISL_3347221, EPI_ISL_3347222, EPI_ISL_3347223, EPI_ISL_3347224, EPI_ISL_3347226, EPI_ISL_3347228, EPI_ISL_3347231, EPI_ISL_3347232 | | | |
| see above | VIDOH (Virgin Islands Department of Health) | Grubaugh Lab - Yale School of Public Health | Joseph Fauver, Mallery Breban, Isabel Ott, Tara Alpert, Mary Petrone, Anderson Brito, Chantal Vogels, Annie Watkins, Chaney Kalinich, Jessica Rothman, Kendall Billig, Tobias Koch, Jendai Richards, TaLesa Aderohunmu, Esther Ellis, Marlon Lawrence, Brett Ellis, Nathan Grubaugh |
| EPI_ISL_3347233, EPI_ISL_3347234, EPI_ISL_3347235, EPI_ISL_3347236, EPI_ISL_3347237, EPI_ISL_3347238, EPI_ISL_3347239, EPI_ISL_3347240, EPI_ISL_3347241, EPI_ISL_3347242, EPI_ISL_3347243, EPI_ISL_3347244, EPI_ISL_3347245, EPI_ISL_3347246, EPI_ISL_3347247, EPI_ISL_3347248, EPI_ISL_3347249, EPI_ISL_3347250, EPI_ISL_3347251, EPI_ISL_3347252, EPI_ISL_3347253, EPI_ISL_3347254, EPI_ISL_3347255, EPI_ISL_3347256, EPI_ISL_3347257, EPI_ISL_3347258, EPI_ISL_3347259, EPI_ISL_3347260, EPI_ISL_3347261, EPI_ISL_3347262, EPI_ISL_3347263, EPI_ISL_3347264, EPI_ISL_3347265, EPI_ISL_3347266, EPI_ISL_3347267, EPI_ISL_3347268, EPI_ISL_3347269, EPI_ISL_3347270, EPI_ISL_3347271, EPI_ISL_3347272, EPI_ISL_3347273, EPI_ISL_3347274, EPI_ISL_3347275, EPI_ISL_3347276 | | | |

| | | | |
|---|---|---|---|
| see above | Yale Clinical Virology Lab | Grubaugh Lab - Yale School of Public Health | Joseph Fauver, Mallery Breban, Isabel Ott, Tara Alpert, Mary Petrone, Anderson Brito, Chantal Vogels, Annie Watkins, Chaney Kalinich, Jessica Rothman, Kendall Billig, Tobias Koch, Marie L. Landry, Nathan Grubaugh |
| EPI_ISL_3347277, EPI_ISL_3347278, EPI_ISL_3347280, EPI_ISL_3347281, EPI_ISL_3347282, EPI_ISL_3347283, EPI_ISL_3347284, EPI_ISL_3347285, EPI_ISL_3347286, EPI_ISL_3347287, EPI_ISL_3347288, EPI_ISL_3347289, EPI_ISL_3347290, EPI_ISL_3347291, EPI_ISL_3347292, EPI_ISL_3347293, EPI_ISL_3347294, EPI_ISL_3347295, EPI_ISL_3347296, EPI_ISL_3347297, EPI_ISL_3347298, EPI_ISL_3347299, EPI_ISL_3347300, EPI_ISL_3347301, EPI_ISL_3347302, EPI_ISL_3347303, EPI_ISL_3347304, EPI_ISL_3347305, EPI_ISL_3347306, EPI_ISL_3347307, EPI_ISL_3347308, EPI_ISL_3347309, EPI_ISL_3347310, EPI_ISL_3347311, EPI_ISL_3347312, EPI_ISL_3347313, EPI_ISL_3347314, EPI_ISL_3347315, EPI_ISL_3347316, EPI_ISL_3347317, EPI_ISL_3347318, EPI_ISL_3347319, EPI_ISL_3347320, EPI_ISL_3347321, EPI_ISL_3347322, EPI_ISL_3347323, EPI_ISL_3347327, EPI_ISL_3347328, EPI_ISL_3347329, EPI_ISL_3347330 | | | |
| see above | UCC (Universidad Central del Caribe) | Grubaugh Lab - Yale School of Public Health | Joseph Fauver, Mallery Breban, Isabel Ott, Tara Alpert, Mary Petrone, Anderson Brito, Chantal Vogels, Annie Watkins, Chaney Kalinich, Jessica Rothman, Kendall Billig, Tobias Koch, Fabiola Cruz-Lopez, David Rivera Aponte, Fabiola Fontanet, Natalie Machargo, Andrea Firpo, Vianca Aponte, Ernesto Vazquez, Patricia Serrano, Tania Mitwalli, Karolane Gonzalez, Nathan Grubaugh |
| EPI_ISL_3347331, EPI_ISL_3347332, EPI_ISL_3347333, EPI_ISL_3347334, EPI_ISL_3347335, EPI_ISL_3347337, EPI_ISL_3347338, EPI_ISL_3347341, EPI_ISL_3347342, EPI_ISL_3347345, EPI_ISL_3347346, EPI_ISL_3347348, EPI_ISL_3347349, EPI_ISL_3347350, EPI_ISL_3347351, EPI_ISL_3347352 | | | |
| see above | VIDOH (Virgin Islands Department of Health) | Grubaugh Lab - Yale School of Public Health | Joseph Fauver, Mallery Breban, Isabel Ott, Tara Alpert, Mary Petrone, Anderson Brito, Chantal Vogels, Annie Watkins, Chaney Kalinich, Jessica Rothman, Kendall Billig, Tobias Koch, Jendai Richards, TaLesa Aderohunmu, Esther Ellis, Marlon Lawrence, Brett Ellis, Nathan Grubaugh |
| EPI_ISL_3347354, EPI_ISL_3347355, EPI_ISL_3347356, EPI_ISL_3347357, EPI_ISL_3347358, EPI_ISL_3347359, EPI_ISL_3347360, EPI_ISL_3347361, EPI_ISL_3347362, EPI_ISL_3347363, EPI_ISL_3347364, EPI_ISL_3347365, EPI_ISL_3347366 | | | |
| see above | Yale Pathology Lab | Grubaugh Lab - Yale School of Public Health | Joseph Fauver, Mallery Breban, Isabel Ott, Tara Alpert, Mary Petrone, Anderson Brito, Chantal Vogels, Annie Watkins, Chaney Kalinich, Jessica Rothman, Kendall Billig, Tobias Koch, Jianhui Wang, Chen Liu, Pei Hui, Nathan Grubaugh |
| EPI_ISL_3347381, EPI_ISL_3347382, EPI_ISL_3347383, EPI_ISL_3347384, EPI_ISL_3347385, EPI_ISL_3347386, EPI_ISL_3347387, EPI_ISL_3347388, EPI_ISL_3347389, EPI_ISL_3347390, EPI_ISL_3347391, EPI_ISL_3347392, EPI_ISL_3347393, EPI_ISL_3347394, EPI_ISL_3347395, EPI_ISL_3347396, EPI_ISL_3347397, EPI_ISL_3347398, EPI_ISL_3347399, EPI_ISL_3347400, EPI_ISL_3347401, EPI_ISL_3347402, EPI_ISL_3347403, EPI_ISL_3347404, EPI_ISL_3347405, EPI_ISL_3347406 | | | |
| see above | Kansas Health and Environmental Lab | Kansas Health and Environmental Lab | Mike Grose, Jonathan Barnell, Amanda Bradley, and Phil Adam |
| EPI_ISL_3347407, EPI_ISL_3347408, EPI_ISL_3347409, EPI_ISL_3347410, EPI_ISL_3347411, EPI_ISL_3347412, EPI_ISL_3347413, EPI_ISL_3347414, EPI_ISL_3347415, EPI_ISL_3347416, EPI_ISL_3347417, EPI_ISL_3347418, EPI_ISL_3347419, EPI_ISL_3347420, EPI_ISL_3347421, EPI_ISL_3347422, EPI_ISL_3347423, EPI_ISL_3347424, EPI_ISL_3347425, EPI_ISL_3347426, EPI_ISL_3347427, EPI_ISL_3347428, EPI_ISL_3347429, EPI_ISL_3347430, EPI_ISL_3347431, EPI_ISL_3347432, EPI_ISL_3347433, EPI_ISL_3347434, EPI_ISL_3347435, EPI_ISL_3347436, EPI_ISL_3347437, EPI_ISL_3347438, EPI_ISL_3347439, EPI_ISL_3347440, EPI_ISL_3347441, EPI_ISL_3347442, EPI_ISL_3347443, EPI_ISL_3347444, EPI_ISL_3347445, EPI_ISL_3347446, EPI_ISL_3347447, EPI_ISL_3347448, EPI_ISL_3347449, EPI_ISL_3347450, EPI_ISL_3347451, EPI_ISL_3347452, EPI_ISL_3347453, EPI_ISL_3347454, EPI_ISL_3347455, EPI_ISL_3347456, EPI_ISL_3347457, EPI_ISL_3347458, EPI_ISL_3347459, EPI_ISL_3347460, EPI_ISL_3347461, EPI_ISL_3347462, EPI_ISL_3347463, EPI_ISL_3347464, EPI_ISL_3347465, EPI_ISL_3347466, EPI_ISL_3347467, EPI_ISL_3347468, EPI_ISL_3347469, EPI_ISL_3347470, EPI_ISL_3347471, EPI_ISL_3347472, EPI_ISL_3347473, EPI_ISL_3347474, EPI_ISL_3347475, EPI_ISL_3347476 | | | |
| see above | NYU Langone Health | Departments of Pathology and Medicine, New York University School of Medicine | Adriana Heguy, Dacia Dimartino, Emily Guzman, Christian Marier, Peter Meyn, Sitharam Ramaswami, Gael Westby, Paul Zappile, Yutong Zhang, Paolo Cotzia, Guiqing Wang |
| EPI_ISL_3347477, EPI_ISL_3347478, EPI_ISL_3347479, EPI_ISL_3347480, EPI_ISL_3347481, EPI_ISL_3347482, EPI_ISL_3347483, EPI_ISL_3347484, EPI_ISL_3347485, EPI_ISL_3347486, EPI_ISL_3347487, EPI_ISL_3347488, EPI_ISL_3347489, EPI_ISL_3347490, EPI_ISL_3347491, EPI_ISL_3347492, EPI_ISL_3347493, EPI_ISL_3347494, EPI_ISL_3347495, EPI_ISL_3347496, EPI_ISL_3347497, EPI_ISL_3347498, EPI_ISL_3347499, EPI_ISL_3347500, EPI_ISL_3347501, EPI_ISL_3347503, EPI_ISL_3347504, EPI_ISL_3347505, EPI_ISL_3347506 | | | |
| see above | Nebraska Public Health Laboratory | NPHL COVID-19 Response Team | NPHL COVID-19 Response Team |
| EPI_ISL_3347507, EPI_ISL_3347508 | Hospital Herrera Llerandi | Asociación de Salud Integral/Clínica Familiar "Luis Ángel García" | Eduardo Arathoon, Ana S. Gonzalez-Reiche, Claudia Rangel, Luis Rivas, Oscar Bonilla, Narda Medina, Luis Aguirre, Danicela Mercado, Hilda Ruiz, Osmar Gamboa |
| EPI_ISL_3347509 | Laboratorio Central de Epidemiología (LCE) | Instituto de Biotecnología de la UNAM | Consorcio Mexicano de Vigilancia Genómica (CoViGen-Mex). Authors (in alphabetical order): Julio Elias Alvarado-Yaah, Carlos F. Arias, Santiago Ávila-Ríos, Eduardo Becerril-Vargas, Víctor Hugo Borja-Aburto, Celia Boukadida, Cristóbal Cháidez-Quiróz, Juan Bautista Chale-Dzul, Ricardo Ciria Merce, Andreu Comas-García, Célida Duque Molina, Julissa Enciso-Ibarra, José Antonio Enciso-Moreno, Gloria Elena Espinosa-Ayala, Fernando Fontove-Herrera, Daniel Fregoso-Rueda, Victor Eduardo García-Arias, Alejandra García-Gasca, Bruno Gómez-Gil, Jean Pierre González, Irvin González-López, Concepción Grajales-Muñiz, Ricardo Grande, Rosa María Gutiérrez Rios, Jesús Hernández, Alejandra Hernández-Terán, Alfredo Herrera-Estrella, Carla Ivón Herrera-Najera, Pavel Isa, Verónica Mata-Haro, Daniel Lira Morales, Susana Lopez, Antonio Loza Román, Brenda Irasema Maldonado-Meza, Bernardo Martínez-Miguel, José Arturo Martínez-Orozco, Célida Martínez-Rodríguez, Margarita Matías-Florentino, Fidencio Mejía-Nepomuceno, María Guadalupe de Jesús Mireles-Rivera, Gloria María Molina-Salinas, Hector Montoya-Fuentes, , Mario Mújica-Sánchez, José Esteban Muñoz-Medina, José de Jesús Nuñez-Contreras, Alicia Ocaña-Mondragón, Luis Alberto Ochoa-Carrera, Hector Esteban Paz-Juárez, Marissa Perez-Garcia, Francisco Pulido, Helen Haydee Fernanda Ramírez-Plascencia, Jorge Salas-Hernández, Angel Gustavo Salas-Lais, Alejandro Sánchez-Flores, Clara Esperanza Santacruz-Tinoco, María Guadalupe Santiago-Mauricio, Selene Zárate, Nelly Sélem-Mojica, Blanca Taboada, Kathia Elizabeth Tapia-Diaz, Gloria Vazquez, Joel Armando Vázquez-Pérez). |
| EPI_ISL_3347510, EPI_ISL_3347511 | Centro de Investigación Biomédica del Noreste (CIBIN) | Instituto de Biotecnología de la UNAM | Consorcio Mexicano de Vigilancia Genómica (CoViGen-Mex). Authors (in alphabetical order): Julio Elias Alvarado-Yaah, Carlos F. Arias, Santiago Ávila-Ríos, Eduardo Becerril-Vargas, Víctor Hugo Borja-Aburto, Celia Boukadida, Cristóbal Cháidez-Quiróz, Juan Bautista Chale-Dzul, Ricardo Ciria Merce, Andreu Comas-García, Célida Duque Molina, Julissa Enciso-Ibarra, José Antonio Enciso-Moreno, Gloria Elena Espinosa-Ayala, Fernando Fontove-Herrera, Daniel Fregoso-Rueda, Victor Eduardo García-Arias, Alejandra García-Gasca, Bruno Gómez-Gil, Jean Pierre González, Irvin González-López, Concepción Grajales-Muñiz, Ricardo Grande, Rosa María Gutiérrez Rios, Jesús Hernández, Alejandra Hernández-Terán, Alfredo Herrera-Estrella, Carla Ivón Herrera-Najera, Pavel Isa, Verónica Mata-Haro, Daniel Lira Morales, Susana Lopez, Antonio Loza Román, Brenda Irasema Maldonado-Meza, Bernardo Martínez-Miguel, José Arturo Martínez-Orozco, Célida Martínez-Rodríguez, Margarita Matías-Florentino, Fidencio Mejía-Nepomuceno, María Guadalupe de Jesús Mireles-Rivera, Gloria María Molina-Salinas, Hector Montoya-Fuentes, , Mario Mújica-Sánchez, José Esteban Muñoz-Medina, José de Jesús Nuñez-Contreras, Alicia Ocaña-Mondragón, Luis Alberto Ochoa-Carrera, Hector Esteban Paz-Juárez, Marissa Perez-Garcia, Francisco Pulido, Helen Haydee Fernanda Ramírez-Plascencia, Jorge Salas-Hernández, Angel Gustavo Salas-Lais, Alejandro Sánchez-Flores, Clara Esperanza Santacruz-Tinoco, María Guadalupe Santiago-Mauricio, Selene Zárate, Nelly Sélem-Mojica, Blanca Taboada, Kathia Elizabeth Tapia-Diaz, Gloria Vazquez, Joel Armando Vázquez-Pérez). |
| EPI_ISL_3347512, EPI_ISL_3347513 | Laboratorio Central de Epidemiología (LCE) | Instituto de Biotecnología de la UNAM | Consorcio Mexicano de Vigilancia Genómica (CoViGen-Mex). Authors (in alphabetical order): Julio Elias Alvarado-Yaah, Carlos F. Arias, Santiago Ávila-Ríos, Eduardo Becerril-Vargas, Víctor Hugo Borja-Aburto, Celia Boukadida, Cristóbal Cháidez-Quiróz, Juan Bautista Chale-Dzul, Ricardo Ciria Merce, Andreu Comas-García, Célida Duque Molina, Julissa Enciso-Ibarra, José Antonio Enciso-Moreno, Gloria Elena Espinosa-Ayala, Fernando Fontove-Herrera, Daniel Fregoso-Rueda, Victor Eduardo García-Arias, Alejandra García-Gasca, Bruno Gómez-Gil, Jean Pierre González, Irvin González-López, Concepción Grajales-Muñiz, Ricardo Grande, Rosa María Gutiérrez Rios, Jesús Hernández, Alejandra Hernández-Terán, Alfredo Herrera-Estrella, Carla Ivón Herrera-Najera, Pavel Isa, Verónica Mata-Haro, Daniel Lira Morales, Susana Lopez, Antonio Loza Román, Brenda Irasema Maldonado-Meza, Bernardo Martínez-Miguel, José Arturo Martínez-Orozco, Célida Martínez-Rodríguez, Margarita Matías-Florentino, Fidencio Mejía-Nepomuceno, María Guadalupe de Jesús Mireles-Rivera, Gloria María Molina-Salinas, Hector Montoya-Fuentes, , Mario Mújica-Sánchez, José Esteban Muñoz-Medina, José de Jesús Nuñez-Contreras, Alicia Ocaña-Mondragón, Luis Alberto Ochoa-Carrera, Hector Esteban Paz-Juárez, Marissa Perez-Garcia, Francisco Pulido, Helen Haydee Fernanda Ramírez-Plascencia, Jorge Salas-Hernández, Angel Gustavo Salas-Lais, Alejandro Sánchez-Flores, Clara Esperanza Santacruz-Tinoco, María Guadalupe Santiago-Mauricio, Selene Zárate, Nelly Sélem-Mojica, Blanca Taboada, Kathia Elizabeth Tapia-Diaz, Gloria Vazquez, Joel Armando Vázquez-Pérez). |
| EPI_ISL_3347518, EPI_ISL_3347520 | Centro de Investigación Biomédica de Occidente (CIBO) | Instituto de Biotecnología de la UNAM | Consorcio Mexicano de Vigilancia Genómica (CoViGen-Mex). Authors (in alphabetical order): Julio Elias Alvarado-Yaah, Carlos F. Arias, Santiago Ávila-Ríos, Eduardo Becerril-Vargas, Víctor Hugo Borja-Aburto, Celia Boukadida, Cristóbal Cháidez-Quiróz, Juan Bautista Chale-Dzul, Ricardo Ciria Merce, Andreu Comas-García, Célida Duque Molina, Julissa Enciso-Ibarra, José Antonio Enciso-Moreno, Gloria Elena Espinosa-Ayala, Fernando Fontove-Herrera, Daniel Fregoso-Rueda, Victor Eduardo García-Arias, Alejandra García-Gasca, Bruno Gómez-Gil, Jean Pierre González, Irvin González-López, Concepción Grajales-Muñiz, Ricardo Grande, Rosa María Gutiérrez Rios, Jesús Hernández, Alejandra Hernández-Terán, Alfredo Herrera-Estrella, Carla Ivón Herrera-Najera, Pavel Isa, Verónica Mata-Haro, Daniel Lira Morales, Susana Lopez, Antonio Loza Román, Brenda Irasema Maldonado-Meza, Bernardo Martínez-Miguel, José Arturo Martínez-Orozco, Célida Martínez-Rodríguez, Margarita Matías-Florentino, Fidencio Mejía-Nepomuceno, María Guadalupe de Jesús Mireles-Rivera, Gloria María Molina-Salinas, Hector Montoya-Fuentes, , Mario Mújica-Sánchez, José Esteban Muñoz-Medina, José de Jesús Nuñez-Contreras, Alicia Ocaña-Mondragón, Luis Alberto Ochoa-Carrera, Hector Esteban Paz-Juárez, Marissa Perez-Garcia, Francisco Pulido, Helen Haydee Fernanda Ramírez-Plascencia, Jorge Salas-Hernández, Angel Gustavo Salas-Lais, Alejandro Sánchez-Flores, Clara Esperanza Santacruz-Tinoco, María Guadalupe Santiago-Mauricio, Selene Zárate, Nelly Sélem-Mojica, Blanca Taboada, Kathia Elizabeth Tapia-Diaz, Gloria Vazquez, Joel Armando Vázquez-Pérez). |
| EPI_ISL_3347522, EPI_ISL_3347523, EPI_ISL_3347524, EPI_ISL_3347525, EPI_ISL_3347526, EPI_ISL_3347527, EPI_ISL_3347528 | Laboratorio Central de Epidemiología (LCE) | Instituto de Biotecnología de la UNAM | Consorcio Mexicano de Vigilancia Genómica (CoViGen-Mex). Authors (in alphabetical order): Julio Elias Alvarado-Yaah, Carlos F. Arias, Santiago Ávila-Ríos, Eduardo Becerril-Vargas, Víctor Hugo Borja-Aburto, Celia Boukadida, Cristóbal Cháidez-Quiróz, Juan Bautista Chale-Dzul, Ricardo Ciria Merce, Andreu Comas-García, Célida Duque Molina, Julissa Enciso-Ibarra, José Antonio Enciso-Moreno, Gloria Elena Espinosa-Ayala, Fernando Fontove-Herrera, Daniel Fregoso-Rueda, Victor Eduardo García-Arias, Alejandra García-Gasca, Bruno Gómez-Gil, Jean Pierre González, Irvin González-López, Concepción Grajales-Muñiz, Ricardo Grande, Rosa María Gutiérrez Rios, Jesús Hernández, Alejandra Hernández-Terán, Alfredo Herrera-Estrella, Carla Ivón Herrera-Najera, Pavel Isa, Verónica Mata-Haro, Daniel Lira Morales, Susana Lopez, Antonio Loza Román, Brenda Irasema Maldonado-Meza, Bernardo Martínez-Miguel, José Arturo Martínez-Orozco, Célida Martínez-Rodríguez, Margarita Matías-Florentino, Fidencio Mejía-Nepomuceno, María Guadalupe de Jesús Mireles-Rivera, Gloria María Molina-Salinas, Hector Montoya-Fuentes, , Mario Mújica-Sánchez, José Esteban Muñoz-Medina, José de Jesús Nuñez-Contreras, Alicia Ocaña-Mondragón, Luis Alberto Ochoa-Carrera, Hector Esteban Paz-Juárez, Marissa Perez-Garcia, Francisco Pulido, Helen Haydee Fernanda Ramírez-Plascencia, Jorge Salas-Hernández, Angel Gustavo Salas-Lais, Alejandro Sánchez-Flores, Clara Esperanza Santacruz-Tinoco, María Guadalupe Santiago-Mauricio, Selene Zárate, Nelly Sélem-Mojica, Blanca Taboada, Kathia Elizabeth Tapia-Diaz, Gloria Vazquez, Joel Armando Vázquez-Pérez). |
| EPI_ISL_3347565, EPI_ISL_3347566 | Centro de Investigación Biomédica del Noreste (CIBIN) | Instituto de Biotecnología de la UNAM | Consorcio Mexicano de Vigilancia Genómica (CoViGen-Mex). Authors (in alphabetical order): Julio Elias Alvarado-Yaah, Carlos F. Arias, Santiago Ávila-Ríos, Eduardo Becerril-Vargas, Víctor Hugo Borja-Aburto, Celia Boukadida, Cristóbal Cháidez-Quiróz, Juan Bautista Chale-Dzul, Ricardo Ciria Merce, Andreu Comas-García, Célida Duque Molina, Julissa Enciso-Ibarra, José Antonio Enciso-Moreno, Gloria Elena Espinosa-Ayala, Fernando Fontove-Herrera, Daniel Fregoso-Rueda, Victor Eduardo García-Arias, Alejandra García-Gasca, Bruno Gómez-Gil, Jean Pierre González, Irvin González-López, Concepción Grajales-Muñiz, Ricardo Grande, Rosa María Gutiérrez Rios, Jesús Hernández, Alejandra Hernández-Terán, Alfredo Herrera-Estrella, Carla Ivón Herrera-Najera, Pavel Isa, Verónica Mata-Haro, Daniel Lira Morales, Susana Lopez, Antonio Loza Román, Brenda Irasema Maldonado-Meza, Bernardo Martínez-Miguel, José Arturo Martínez-Orozco, Célida Martínez-Rodríguez, Margarita Matías-Florentino, Fidencio Mejía-Nepomuceno, María Guadalupe de Jesús Mireles-Rivera, Gloria María Molina-Salinas, Hector Montoya-Fuentes, , Mario Mújica-Sánchez, José Esteban Muñoz-Medina, José de Jesús Nuñez-Contreras, Alicia Ocaña-Mondragón, Luis Alberto Ochoa-Carrera, Hector Esteban Paz-Juárez, Marissa Perez-Garcia, Francisco Pulido, Helen Haydee Fernanda Ramírez-Plascencia, Jorge Salas-Hernández, Angel Gustavo Salas-Lais, Alejandro Sánchez-Flores, Clara Esperanza Santacruz-Tinoco, María Guadalupe Santiago-Mauricio, Selene Zárate, Nelly Sélem-Mojica, Blanca Taboada, Kathia Elizabeth Tapia-Diaz, Gloria Vazquez, Joel Armando Vázquez-Pérez). |
| EPI_ISL_3347567, EPI_ISL_3347581, EPI_ISL_3347597, EPI_ISL_3347605 | Laboratorio Central de Epidemiología (LCE) | Instituto de Biotecnología de la UNAM | Consorcio Mexicano de Vigilancia Genómica (CoViGen-Mex). Authors (in alphabetical order): Julio Elias Alvarado-Yaah, Carlos F. Arias, Santiago Ávila-Ríos, Eduardo Becerril-Vargas, Víctor Hugo Borja-Aburto, Celia Boukadida, Cristóbal Cháidez-Quiróz, Juan Bautista Chale-Dzul, Ricardo Ciria Merce, Andreu Comas-García, Célida Duque Molina, Julissa Enciso-Ibarra, José Antonio Enciso-Moreno, Gloria Elena Espinosa-Ayala, Fernando Fontove-Herrera, Daniel Fregoso-Rueda, Victor Eduardo García-Arias, Alejandra García-Gasca, Bruno Gómez-Gil, Jean Pierre González, Irvin González-López, Concepción Grajales-Muñiz, Ricardo Grande, Rosa María Gutiérrez Rios, Jesús Hernández, Alejandra Hernández-Terán, Alfredo Herrera-Estrella, Carla Ivón Herrera-Najera, Pavel Isa, Verónica Mata-Haro, Daniel Lira Morales, Susana Lopez, Antonio Loza Román, Brenda Irasema Maldonado-Meza, Bernardo Martínez-Miguel, José Arturo Martínez-Orozco, Célida Martínez-Rodríguez, Margarita Matías-Florentino, Fidencio Mejía-Nepomuceno, María Guadalupe de Jesús Mireles-Rivera, Gloria María Molina-Salinas, Hector Montoya-Fuentes, , Mario Mújica-Sánchez, José Esteban Muñoz-Medina, José de Jesús Nuñez-Contreras, Alicia Ocaña-Mondragón, Luis Alberto Ochoa-Carrera, Hector Esteban Paz-Juárez, Marissa Perez-Garcia, Francisco Pulido, Helen Haydee Fernanda Ramírez-Plascencia, Jorge Salas-Hernández, Angel Gustavo Salas-Lais, Alejandro Sánchez-Flores, Clara Esperanza Santacruz-Tinoco, María Guadalupe Santiago-Mauricio, Selene Zárate, Nelly Sélem-Mojica, Blanca Taboada, Kathia Elizabeth Tapia-Diaz, Gloria Vazquez, Joel Armando Vázquez-Pérez). |
| EPI_ISL_3347607 | Centro de Investigación Biomédica de Occidente (CIBO) | Instituto de Biotecnología de la UNAM | Consorcio Mexicano de Vigilancia Genómica (CoViGen-Mex). Authors (in alphabetical order): Julio Elias Alvarado-Yaah, Carlos F. Arias, Santiago Ávila-Ríos, Eduardo Becerril-Vargas, Víctor Hugo Borja-Aburto, Celia Boukadida, Cristóbal Cháidez-Quiróz, Juan Bautista Chale-Dzul, Ricardo Ciria Merce, Andreu Comas-García, Célida Duque Molina, Julissa Enciso-Ibarra, José Antonio Enciso-Moreno, Gloria Elena Espinosa-Ayala, Fernando Fontove-Herrera, Daniel Fregoso-Rueda, Victor Eduardo García-Arias, Alejandra García-Gasca, Bruno Gómez-Gil, Jean Pierre González, Irvin González-López, Concepción Grajales-Muñiz, Ricardo Grande, Rosa María Gutiérrez Rios, Jesús Hernández, Alejandra Hernández-Terán, Alfredo Herrera-Estrella, Carla Ivón Herrera-Najera, Pavel Isa, Verónica Mata-Haro, Daniel Lira Morales, Susana Lopez, Antonio Loza Román, Brenda Irasema Maldonado-Meza, Bernardo Martínez-Miguel, José Arturo Martínez-Orozco, Célida Martínez-Rodríguez, Margarita Matías-Florentino, Fidencio Mejía-Nepomuceno, María Guadalupe de Jesús Mireles-Rivera, Gloria María Molina-Salinas, Hector Montoya-Fuentes, , Mario Mújica-Sánchez, José Esteban Muñoz-Medina, José de Jesús Nuñez-Contreras, Alicia Ocaña-Mondragón, Luis Alberto Ochoa-Carrera, Hector Esteban Paz-Juárez, Marissa Perez-Garcia, Francisco Pulido, Helen Haydee Fernanda Ramírez-Plascencia, Jorge Salas-Hernández, Angel Gustavo Salas-Lais, Alejandro Sánchez-Flores, Clara Esperanza Santacruz-Tinoco, María Guadalupe Santiago-Mauricio, Selene Zárate, Nelly Sélem-Mojica, Blanca Taboada, Kathia Elizabeth Tapia-Diaz, Gloria Vazquez, Joel Armando Vázquez-Pérez). |
| EPI_ISL_3347609 | Laboratorio Central de Epidemiología (LCE) | Instituto de Biotecnología de la UNAM | Consorcio Mexicano de Vigilancia Genómica (CoViGen-Mex). Authors (in alphabetical order): Julio Elias Alvarado-Yaah, Carlos F. Arias, Santiago Ávila-Ríos, Eduardo Becerril-Vargas, Víctor Hugo Borja-Aburto, Celia Boukadida, Cristóbal Cháidez-Quiróz, Juan Bautista Chale-Dzul, Ricardo Ciria Merce, Andreu Comas-García, Célida Duque Molina, Julissa Enciso-Ibarra, José Antonio Enciso-Moreno, Gloria Elena Espinosa-Ayala, Fernando Fontove-Herrera, Daniel Fregoso-Rueda, Victor Eduardo García-Arias, Alejandra García-Gasca, Bruno Gómez-Gil, Jean Pierre González, Irvin González-López, Concepción Grajales-Muñiz, Ricardo Grande, Rosa María Gutiérrez Rios, Jesús Hernández, Alejandra Hernández-Terán, Alfredo Herrera-Estrella, Carla Ivón Herrera-Najera, Pavel Isa, Verónica Mata-Haro, Daniel Lira Morales, Susana Lopez, Antonio Loza Román, Brenda Irasema Maldonado-Meza, Bernardo Martínez-Miguel, José Arturo Martínez-Orozco, Célida Martínez-Rodríguez, Margarita Matías-Florentino, Fidencio Mejía-Nepomuceno, María Guadalupe de Jesús Mireles-Rivera, Gloria María Molina-Salinas, Hector Montoya-Fuentes, , Mario Mújica-Sánchez, José Esteban Muñoz-Medina, José de Jesús Nuñez-Contreras, Alicia Ocaña-Mondragón, Luis Alberto Ochoa-Carrera, Hector Esteban Paz-Juárez, Marissa Perez-Garcia, Francisco Pulido, Helen Haydee Fernanda Ramírez-Plascencia, Jorge Salas-Hernández, Angel Gustavo Salas-Lais, Alejandro Sánchez-Flores, Clara Esperanza Santacruz-Tinoco, María Guadalupe Santiago-Mauricio, Selene Zárate, Nelly Sélem-Mojica, Blanca Taboada, Kathia Elizabeth Tapia-Diaz, Gloria Vazquez, Joel Armando Vázquez-Pérez). |
| EPI_ISL_3347610, EPI_ISL_3347612, EPI_ISL_3347613, EPI_ISL_3347614, EPI_ISL_3347615, EPI_ISL_3347616, EPI_ISL_3347617, EPI_ISL_3347618, EPI_ISL_3347619, EPI_ISL_3347620, EPI_ISL_3347621, EPI_ISL_3347622, EPI_ISL_3347623, EPI_ISL_3347624, EPI_ISL_3347625, EPI_ISL_3347626, EPI_ISL_3347627, EPI_ISL_3347628, EPI_ISL_3347629, EPI_ISL_3347630, EPI_ISL_3347631, EPI_ISL_3347632, EPI_ISL_3347633, EPI_ISL_3347634, EPI_ISL_3347635, EPI_ISL_3347636, EPI_ISL_3347637, EPI_ISL_3347638, EPI_ISL_3347639, EPI_ISL_3347640, EPI_ISL_3347641, EPI_ISL_3347642, EPI_ISL_3347643, EPI_ISL_3347644, EPI_ISL_3347645, EPI_ISL_3347646, EPI_ISL_3347647, EPI_ISL_3347648, EPI_ISL_3347649, EPI_ISL_3347650, EPI_ISL_3347651, EPI_ISL_3347652, EPI_ISL_3347653, EPI_ISL_3347654, EPI_ISL_3347655, EPI_ISL_3347656, EPI_ISL_3347657, EPI_ISL_3347658, EPI_ISL_3347659, EPI_ISL_3347660, EPI_ISL_3347661, EPI_ISL_3347662, EPI_ISL_3347663, EPI_ISL_3347664, EPI_ISL_3347665, EPI_ISL_3347666, EPI_ISL_3347667, EPI_ISL_3347668, EPI_ISL_3347669, EPI_ISL_3347670, EPI_ISL_3347671, EPI_ISL_3347672, EPI_ISL_3347673, EPI_ISL_3347674, EPI_ISL_3347675 | | | |
| see above | Centro de Investigación Biomédica del Noreste (CIBIN) | Instituto de Biotecnología de la UNAM | Consorcio Mexicano de Vigilancia Genómica (CoViGen-Mex). Authors (in alphabetical order): Julio Elias Alvarado-Yaah, Carlos F. Arias, Santiago Ávila-Ríos, Eduardo Becerril-Vargas, Víctor Hugo Borja-Aburto, Celia Boukadida, Cristóbal Cháidez-Quiróz, Juan Bautista Chale-Dzul, Ricardo Ciria Merce, Andreu Comas-García, Célida Duque Molina, Julissa Enciso-Ibarra, José Antonio Enciso-Moreno, Gloria Elena Espinosa-Ayala, Fernando Fontove-Herrera, Daniel Fregoso-Rueda, Victor Eduardo García-Arias, Alejandra García-Gasca, Bruno Gómez-Gil, Jean Pierre González, Irvin González-López, Concepción Grajales-Muñiz, Ricardo Grande, Rosa María Gutiérrez Rios, Jesús Hernández, Alejandra Hernández-Terán, Alfredo Herrera-Estrella, Carla Ivón Herrera-Najera, Pavel Isa, Verónica Mata-Haro, Daniel Lira Morales, Susana Lopez, Antonio Loza Román, Brenda Irasema Maldonado-Meza, Bernardo Martínez-Miguel, José Arturo Martínez-Orozco, Célida Martínez-Rodríguez, Margarita Matías-Florentino, Fidencio Mejía-Nepomuceno, María Guadalupe de Jesús Mireles-Rivera, Gloria María Molina-Salinas, Hector Montoya-Fuentes, , Mario Mújica-Sánchez, José Esteban Muñoz-Medina, José de Jesús Nuñez-Contreras, Alicia Ocaña-Mondragón, Luis Alberto Ochoa-Carrera, Hector Esteban Paz-Juárez, Marissa Perez-Garcia, Francisco Pulido, Helen Haydee Fernanda Ramírez-Plascencia, Jorge Salas-Hernández, Angel Gustavo Salas-Lais, Alejandro Sánchez-Flores, Clara Esperanza Santacruz-Tinoco, María Guadalupe Santiago-Mauricio, Selene Zárate, Nelly Sélem-Mojica, Blanca Taboada, Kathia Elizabeth Tapia-Diaz, Gloria Vazquez, Joel Armando Vázquez-Pérez). |

| | | | |
|---|---|---|---|
| EPI_ISL_3347676, EPI_ISL_3347677, EPI_ISL_3347678, EPI_ISL_3347679, EPI_ISL_3347680, EPI_ISL_3347681, EPI_ISL_3347682, EPI_ISL_3347683, EPI_ISL_3347684, EPI_ISL_3347685, EPI_ISL_3347686 | | | |
| see above | Unidad de Investigación Médica de Yucatán (UIMY) | Instituto de Biotecnología de la UNAM | Consorcio Mexicano de Vigilancia Genómica (CoViGen-Mex). Authors (in alphabetical order): Julio Elias Alvarado-Yaah, Carlos F. Arias, Santiago Ávila-Ríos, Eduardo Becerril-Vargas, Víctor Hugo Borja-Aburto, Celia Boukadida, Cristóbal Cháidez-Quiróz, Juan Bautista Chale-Dzul, Ricardo Ciria Merce, Andreu Comas-García, Célida Duque Molina, Julissa Enciso-Ibarra, José Antonio Enciso-Moreno, Gloria Elena Espinosa-Ayala, Fernando Fontove-Herrera, Daniel Fregoso-Rueda, Victor Eduardo García-Arias, Alejandra García-Gasca, Bruno Gómez-Gil, Jean Pierre González, Irvin González-López, Concepción Grajales-Muñiz, Ricardo Grande, Rosa María Gutiérrez Rios, Jesús Hernández, Alejandra Hernández-Terán, Alfredo Herrera-Estrella, Carla Ivón Herrera-Najera, Pavel Isa, Verónica Mata-Haro, Daniel Lira Morales, Susana Lopez, Antonio Loza Román, Brenda Irasema Maldonado-Meza, Bernardo Martínez-Miguel, José Arturo Martínez-Orozco, Célida Martínez-Rodríguez, Margarita Matías-Florentino, Fidencio Mejía-Nepomuceno, María Guadalupe de Jesús Mireles-Rivera, Gloria María Molina-Salinas, Hector Montoya-Fuentes, , Mario Mújica-Sánchez, José Esteban Muñoz-Medina, José de Jesús Nuñez-Contreras, Alicia Ocaña-Mondragón, Luis Alberto Ochoa-Carrera, Hector Esteban Paz-Juárez, Marissa Perez-Garcia, Francisco Pulido, Helen Haydee Fernanda Ramírez-Plascencia, Jorge Salas-Hernández, Angel Gustavo Salas-Lais, Alejandro Sánchez-Flores, Clara Esperanza Santacruz-Tinoco, María Guadalupe Santiago-Mauricio, Selene Zárate, Nelly Sélem-Mojica, Blanca Taboada, Kathia Elizabeth Tapia-Diaz, Gloria Vazquez, Joel Armando Vázquez-Pérez). |
| EPI_ISL_3347687, EPI_ISL_3347688, EPI_ISL_3347689, EPI_ISL_3347690, EPI_ISL_3347691, EPI_ISL_3347692, EPI_ISL_3347693, EPI_ISL_3347694, EPI_ISL_3347695, EPI_ISL_3347696, EPI_ISL_3347697, EPI_ISL_3347698, EPI_ISL_3347699, EPI_ISL_3347700, EPI_ISL_3347701, EPI_ISL_3347702, EPI_ISL_3347703, EPI_ISL_3347704, EPI_ISL_3347705, EPI_ISL_3347706, EPI_ISL_3347707, EPI_ISL_3347708, EPI_ISL_3347709, EPI_ISL_3347710, EPI_ISL_3347711, EPI_ISL_3347712, EPI_ISL_3347713, EPI_ISL_3347714, EPI_ISL_3347715, EPI_ISL_3347716, EPI_ISL_3347717, EPI_ISL_3347718, EPI_ISL_3347719, EPI_ISL_3347720, EPI_ISL_3347721, EPI_ISL_3347722, EPI_ISL_3347723, EPI_ISL_3347724, EPI_ISL_3347725, EPI_ISL_3347726, EPI_ISL_3347727, EPI_ISL_3347728, EPI_ISL_3347729, EPI_ISL_3347730, EPI_ISL_3347731, EPI_ISL_3347732, EPI_ISL_3347733, EPI_ISL_3347734, EPI_ISL_3347735, EPI_ISL_3347736, EPI_ISL_3347737, EPI_ISL_3347738, EPI_ISL_3347739, EPI_ISL_3347740, EPI_ISL_3347741, EPI_ISL_3347742, EPI_ISL_3347743, EPI_ISL_3347744, EPI_ISL_3347745, EPI_ISL_3347746, EPI_ISL_3347747, EPI_ISL_3347748, EPI_ISL_3347749, EPI_ISL_3347750, EPI_ISL_3347751, EPI_ISL_3347752, EPI_ISL_3347753, EPI_ISL_3347754, EPI_ISL_3347755, EPI_ISL_3347756, EPI_ISL_3347757, EPI_ISL_3347758, EPI_ISL_3347759, EPI_ISL_3347760, EPI_ISL_3347761, EPI_ISL_3347762, EPI_ISL_3347763, EPI_ISL_3347764, EPI_ISL_3347765, EPI_ISL_3347766, EPI_ISL_3347767, EPI_ISL_3347768, EPI_ISL_3347769, EPI_ISL_3347770, EPI_ISL_3347771, EPI_ISL_3347772, EPI_ISL_3347773, EPI_ISL_3347774, EPI_ISL_3347775, EPI_ISL_3347776, EPI_ISL_3347777, EPI_ISL_3347778, EPI_ISL_3347779, EPI_ISL_3347780, EPI_ISL_3347781, EPI_ISL_3347782, EPI_ISL_3347783, EPI_ISL_3347784, EPI_ISL_3347785, EPI_ISL_3347786, EPI_ISL_3347787, EPI_ISL_3347788, EPI_ISL_3347789, EPI_ISL_3347790, EPI_ISL_3347791, EPI_ISL_3347792, EPI_ISL_3347793, EPI_ISL_3347794, EPI_ISL_3347795, EPI_ISL_3347796, EPI_ISL_3347797, EPI_ISL_3347798, EPI_ISL_3347799, EPI_ISL_3347800, EPI_ISL_3347801, EPI_ISL_3347802, EPI_ISL_3347803, EPI_ISL_3347804, EPI_ISL_3347805, EPI_ISL_3347806, EPI_ISL_3347807, EPI_ISL_3347808 | | | |
| see above | Laboratorio Central de Epidemiología (LCE) | Instituto de Biotecnología de la UNAM | Consorcio Mexicano de Vigilancia Genómica (CoViGen-Mex). Authors (in alphabetical order): Julio Elias Alvarado-Yaah, Carlos F. Arias, Santiago Ávila-Ríos, Eduardo Becerril-Vargas, Víctor Hugo Borja-Aburto, Celia Boukadida, Cristóbal Cháidez-Quiróz, Juan Bautista Chale-Dzul, Ricardo Ciria Merce, Andreu Comas-García, Célida Duque Molina, Julissa Enciso-Ibarra, José Antonio Enciso-Moreno, Gloria Elena Espinosa-Ayala, Fernando Fontove-Herrera, Daniel Fregoso-Rueda, Victor Eduardo García-Arias, Alejandra García-Gasca, Bruno Gómez-Gil, Jean Pierre González, Irvin González-López, Concepción Grajales-Muñiz, Ricardo Grande, Rosa María Gutiérrez Rios, Jesús Hernández, Alejandra Hernández-Terán, Alfredo Herrera-Estrella, Carla Ivón Herrera-Najera, Pavel Isa, Verónica Mata-Haro, Daniel Lira Morales, Susana Lopez, Antonio Loza Román, Brenda Irasema Maldonado-Meza, Bernardo Martínez-Miguel, José Arturo Martínez-Orozco, Célida Martínez-Rodríguez, Margarita Matías-Florentino, Fidencio Mejía-Nepomuceno, María Guadalupe de Jesús Mireles-Rivera, Gloria María Molina-Salinas, Hector Montoya-Fuentes, , Mario Mújica-Sánchez, José Esteban Muñoz-Medina, José de Jesús Nuñez-Contreras, Alicia Ocaña-Mondragón, Luis Alberto Ochoa-Carrera, Hector Esteban Paz-Juárez, Marissa Perez-Garcia, Francisco Pulido, Helen Haydee Fernanda Ramírez-Plascencia, Jorge Salas-Hernández, Angel Gustavo Salas-Lais, Alejandro Sánchez-Flores, Clara Esperanza Santacruz-Tinoco, María Guadalupe Santiago-Mauricio, Selene Zárate, Nelly Sélem-Mojica, Blanca Taboada, Kathia Elizabeth Tapia-Diaz, Gloria Vazquez, Joel Armando Vázquez-Pérez). |
| EPI_ISL_3347809, EPI_ISL_3347810, EPI_ISL_3347811, EPI_ISL_3347812, EPI_ISL_3347813, EPI_ISL_3347814, EPI_ISL_3347815, EPI_ISL_3347816, EPI_ISL_3347817, EPI_ISL_3347818, EPI_ISL_3347819, EPI_ISL_3347820, EPI_ISL_3347821, EPI_ISL_3347822, EPI_ISL_3347823, EPI_ISL_3347824, EPI_ISL_3347825, EPI_ISL_3347826, EPI_ISL_3347827, EPI_ISL_3347828, EPI_ISL_3347829, EPI_ISL_3347830, EPI_ISL_3347831, EPI_ISL_3347832, EPI_ISL_3347833, EPI_ISL_3347834, EPI_ISL_3347835, EPI_ISL_3347836, EPI_ISL_3347837, EPI_ISL_3347838, EPI_ISL_3347839, EPI_ISL_3347840, EPI_ISL_3347841, EPI_ISL_3347842, EPI_ISL_3347843, EPI_ISL_3347844, EPI_ISL_3347845, EPI_ISL_3347846, EPI_ISL_3347847, EPI_ISL_3347848, EPI_ISL_3347849, EPI_ISL_3347850, EPI_ISL_3347851, EPI_ISL_3347852, EPI_ISL_3347853, EPI_ISL_3347854, EPI_ISL_3347855, EPI_ISL_3347856, EPI_ISL_3347857, EPI_ISL_3347858, EPI_ISL_3347859, EPI_ISL_3347860, EPI_ISL_3347861, EPI_ISL_3347862 | | | |
| see above | Centro de Investigación Biomédica de Occidente (CIBO) | Instituto de Biotecnología de la UNAM | Consorcio Mexicano de Vigilancia Genómica (CoViGen-Mex). Authors (in alphabetical order): Julio Elias Alvarado-Yaah, Carlos F. Arias, Santiago Ávila-Ríos, Eduardo Becerril-Vargas, Víctor Hugo Borja-Aburto, Celia Boukadida, Cristóbal Cháidez-Quiróz, Juan Bautista Chale-Dzul, Ricardo Ciria Merce, Andreu Comas-García, Célida Duque Molina, Julissa Enciso-Ibarra, José Antonio Enciso-Moreno, Gloria Elena Espinosa-Ayala, Fernando Fontove-Herrera, Daniel Fregoso-Rueda, Victor Eduardo García-Arias, Alejandra García-Gasca, Bruno Gómez-Gil, Jean Pierre González, Irvin González-López, Concepción Grajales-Muñiz, Ricardo Grande, Rosa María Gutiérrez Rios, Jesús Hernández, Alejandra Hernández-Terán, Alfredo Herrera-Estrella, Carla Ivón Herrera-Najera, Pavel Isa, Verónica Mata-Haro, Daniel Lira Morales, Susana Lopez, Antonio Loza Román, Brenda Irasema Maldonado-Meza, Bernardo Martínez-Miguel, José Arturo Martínez-Orozco, Célida Martínez-Rodríguez, Margarita Matías-Florentino, Fidencio Mejía-Nepomuceno, María Guadalupe de Jesús Mireles-Rivera, Gloria María Molina-Salinas, Hector Montoya-Fuentes, , Mario Mújica-Sánchez, José Esteban Muñoz-Medina, José de Jesús Nuñez-Contreras, Alicia Ocaña-Mondragón, Luis Alberto Ochoa-Carrera, Hector Esteban Paz-Juárez, Marissa Perez-Garcia, Francisco Pulido, Helen Haydee Fernanda Ramírez-Plascencia, Jorge Salas-Hernández, Angel Gustavo Salas-Lais, Alejandro Sánchez-Flores, Clara Esperanza Santacruz-Tinoco, María Guadalupe Santiago-Mauricio, Selene Zárate, Nelly Sélem-Mojica, Blanca Taboada, Kathia Elizabeth Tapia-Diaz, Gloria Vazquez, Joel Armando Vázquez-Pérez). |
| EPI_ISL_3347863 | Hospital Herrera Llerandi | Asociación de Salud Integral/Clínica Familiar "Luis Ángel García" | Eduardo Arathoon, Ana S. Gonzalez-Reiche, Claudia Rangel, Luis Rivas, Oscar Bonilla, Narda Medina, Luis Aguirre, Danicela Mercado, Hilda Ruiz, Osmar Gamboa |
| EPI_ISL_3347864, EPI_ISL_3347865, EPI_ISL_3347866, EPI_ISL_3347867, EPI_ISL_3347868, EPI_ISL_3347869, EPI_ISL_3347870, EPI_ISL_3347871, EPI_ISL_3347872, EPI_ISL_3347873, EPI_ISL_3347874, EPI_ISL_3347875, EPI_ISL_3347876, EPI_ISL_3347877, EPI_ISL_3347878, EPI_ISL_3347879, EPI_ISL_3347880, EPI_ISL_3347881, EPI_ISL_3347882, EPI_ISL_3347883, EPI_ISL_3347884, EPI_ISL_3347885, EPI_ISL_3347886, EPI_ISL_3347887, EPI_ISL_3347888, EPI_ISL_3347889, EPI_ISL_3347890, EPI_ISL_3347891, EPI_ISL_3347892, EPI_ISL_3347893, EPI_ISL_3347894, EPI_ISL_3347895 | | | |
| see above | Centro de Investigación Biomédica de Occidente (CIBO) | Instituto de Biotecnología de la UNAM | Consorcio Mexicano de Vigilancia Genómica (CoViGen-Mex). Authors (in alphabetical order): Julio Elias Alvarado-Yaah, Carlos F. Arias, Santiago Ávila-Ríos, Eduardo Becerril-Vargas, Víctor Hugo Borja-Aburto, Celia Boukadida, Cristóbal Cháidez-Quiróz, Juan Bautista Chale-Dzul, Ricardo Ciria Merce, Andreu Comas-García, Célida Duque Molina, Julissa Enciso-Ibarra, José Antonio Enciso-Moreno, Gloria Elena Espinosa-Ayala, Fernando Fontove-Herrera, Daniel Fregoso-Rueda, Victor Eduardo García-Arias, Alejandra García-Gasca, Bruno Gómez-Gil, Jean Pierre González, Irvin González-López, Concepción Grajales-Muñiz, Ricardo Grande, Rosa María Gutiérrez Rios, Jesús Hernández, Alejandra Hernández-Terán, Alfredo Herrera-Estrella, Carla Ivón Herrera-Najera, Pavel Isa, Verónica Mata-Haro, Daniel Lira Morales, Susana Lopez, Antonio Loza Román, Brenda Irasema Maldonado-Meza, Bernardo Martínez-Miguel, José Arturo Martínez-Orozco, Célida Martínez-Rodríguez, Margarita Matías-Florentino, Fidencio Mejía-Nepomuceno, María Guadalupe de Jesús Mireles-Rivera, Gloria María Molina-Salinas, Hector Montoya-Fuentes, , Mario Mújica-Sánchez, José Esteban Muñoz-Medina, José de Jesús Nuñez-Contreras, Alicia Ocaña-Mondragón, Luis Alberto Ochoa-Carrera, Hector Esteban Paz-Juárez, Marissa Perez-Garcia, Francisco Pulido, Helen Haydee Fernanda Ramírez-Plascencia, Jorge Salas-Hernández, Angel Gustavo Salas-Lais, Alejandro Sánchez-Flores, Clara Esperanza Santacruz-Tinoco, María Guadalupe Santiago-Mauricio, Selene Zárate, Nelly Sélem-Mojica, Blanca Taboada, Kathia Elizabeth Tapia-Diaz, Gloria Vazquez, Joel Armando Vázquez-Pérez). |
| EPI_ISL_3347896, EPI_ISL_3347897, EPI_ISL_3347898, EPI_ISL_3347899, EPI_ISL_3347900, EPI_ISL_3347901, EPI_ISL_3347902, EPI_ISL_3347903, EPI_ISL_3347904, EPI_ISL_3347905, EPI_ISL_3347906, EPI_ISL_3347907, EPI_ISL_3347908, EPI_ISL_3347909, EPI_ISL_3347910, EPI_ISL_3347911, EPI_ISL_3347912 | | | |
| see above | Unidad de Investigación Biomédica de Zacatecas (UIBZ) | Instituto de Biotecnología de la UNAM | Consorcio Mexicano de Vigilancia Genómica (CoViGen-Mex). Authors (in alphabetical order): Julio Elias Alvarado-Yaah, Carlos F. Arias, Santiago Ávila-Ríos, Eduardo Becerril-Vargas, Víctor Hugo Borja-Aburto, Celia Boukadida, Cristóbal Cháidez-Quiróz, Juan Bautista Chale-Dzul, Ricardo Ciria Merce, Andreu Comas-García, Célida Duque Molina, Julissa Enciso-Ibarra, José Antonio Enciso-Moreno, Gloria Elena Espinosa-Ayala, Fernando Fontove-Herrera, Daniel Fregoso-Rueda, Victor Eduardo García-Arias, Alejandra García-Gasca, Bruno Gómez-Gil, Jean Pierre González, Irvin González-López, Concepción Grajales-Muñiz, Ricardo Grande, Rosa María Gutiérrez Rios, Jesús Hernández, Alejandra Hernández-Terán, Alfredo Herrera-Estrella, Carla Ivón Herrera-Najera, Pavel Isa, Verónica Mata-Haro, Daniel Lira Morales, Susana Lopez, Antonio Loza Román, Brenda Irasema Maldonado-Meza, Bernardo Martínez-Miguel, José Arturo Martínez-Orozco, Célida Martínez-Rodríguez, Margarita Matías-Florentino, Fidencio Mejía-Nepomuceno, María Guadalupe de Jesús Mireles-Rivera, Gloria María Molina-Salinas, Hector Montoya-Fuentes, , Mario Mújica-Sánchez, José Esteban Muñoz-Medina, José de Jesús Nuñez-Contreras, Alicia Ocaña-Mondragón, Luis Alberto Ochoa-Carrera, Hector Esteban Paz-Juárez, Marissa Perez-Garcia, Francisco Pulido, Helen Haydee Fernanda Ramírez-Plascencia, Jorge Salas-Hernández, Angel Gustavo Salas-Lais, Alejandro Sánchez-Flores, Clara Esperanza Santacruz-Tinoco, María Guadalupe Santiago-Mauricio, Selene Zárate, Nelly Sélem-Mojica, Blanca Taboada, Kathia Elizabeth Tapia-Diaz, Gloria Vazquez, Joel Armando Vázquez-Pérez). |
| EPI_ISL_3347913, EPI_ISL_3347914, EPI_ISL_3347915, EPI_ISL_3347916, EPI_ISL_3347917 | Centro de Investigación Biomédica del Noreste (CIBIN) | Instituto de Biotecnología de la UNAM | Consorcio Mexicano de Vigilancia Genómica (CoViGen-Mex). Authors (in alphabetical order): Julio Elias Alvarado-Yaah, Carlos F. Arias, Santiago Ávila-Ríos, Eduardo Becerril-Vargas, Víctor Hugo Borja-Aburto, Celia Boukadida, Cristóbal Cháidez-Quiróz, Juan Bautista Chale-Dzul, Ricardo Ciria Merce, Andreu Comas-García, Célida Duque Molina, Julissa Enciso-Ibarra, José Antonio Enciso-Moreno, Gloria Elena Espinosa-Ayala, Fernando Fontove-Herrera, Daniel Fregoso-Rueda, Victor Eduardo García-Arias, Alejandra García-Gasca, Bruno Gómez-Gil, Jean Pierre González, Irvin González-López, Concepción Grajales-Muñiz, Ricardo Grande, Rosa María Gutiérrez Rios, Jesús Hernández, Alejandra Hernández-Terán, Alfredo Herrera-Estrella, Carla Ivón Herrera-Najera, Pavel Isa, Verónica Mata-Haro, Daniel Lira Morales, Susana Lopez, Antonio Loza Román, Brenda Irasema Maldonado-Meza, Bernardo Martínez-Miguel, José Arturo Martínez-Orozco, Célida Martínez-Rodríguez, Margarita Matías-Florentino, Fidencio Mejía-Nepomuceno, María Guadalupe de Jesús Mireles-Rivera, Gloria María Molina-Salinas, Hector Montoya-Fuentes, , Mario Mújica-Sánchez, José Esteban Muñoz-Medina, José de Jesús Nuñez-Contreras, Alicia Ocaña-Mondragón, Luis Alberto Ochoa-Carrera, Hector Esteban Paz-Juárez, Marissa Perez-Garcia, Francisco Pulido, Helen Haydee Fernanda Ramírez-Plascencia, Jorge Salas-Hernández, Angel Gustavo Salas-Lais, Alejandro Sánchez-Flores, Clara Esperanza Santacruz-Tinoco, María Guadalupe Santiago-Mauricio, Selene Zárate, Nelly Sélem-Mojica, Blanca Taboada, Kathia Elizabeth Tapia-Diaz, Gloria Vazquez, Joel Armando Vázquez-Pérez). |
| EPI_ISL_3347918, EPI_ISL_3347919 | Unidad de Investigación Médica de Yucatán (UIMY) | Instituto de Biotecnología de la UNAM | Consorcio Mexicano de Vigilancia Genómica (CoViGen-Mex). Authors (in alphabetical order): Julio Elias Alvarado-Yaah, Carlos F. Arias, Santiago Ávila-Ríos, Eduardo Becerril-Vargas, Víctor Hugo Borja-Aburto, Celia Boukadida, Cristóbal Cháidez-Quiróz, Juan Bautista Chale-Dzul, Ricardo Ciria Merce, Andreu Comas-García, Célida Duque Molina, Julissa Enciso-Ibarra, José Antonio Enciso-Moreno, Gloria Elena Espinosa-Ayala, Fernando Fontove-Herrera, Daniel Fregoso-Rueda, Victor Eduardo García-Arias, Alejandra García-Gasca, Bruno Gómez-Gil, Jean Pierre González, Irvin González-López, Concepción Grajales-Muñiz, Ricardo Grande, Rosa María Gutiérrez Rios, Jesús Hernández, Alejandra Hernández-Terán, Alfredo Herrera-Estrella, Carla Ivón Herrera-Najera, Pavel Isa, Verónica Mata-Haro, Daniel Lira Morales, Susana Lopez, Antonio Loza Román, Brenda Irasema Maldonado-Meza, Bernardo Martínez-Miguel, José Arturo Martínez-Orozco, Célida Martínez-Rodríguez, Margarita Matías-Florentino, Fidencio Mejía-Nepomuceno, María Guadalupe de Jesús Mireles-Rivera, Gloria María Molina-Salinas, Hector Montoya-Fuentes, , Mario Mújica-Sánchez, José Esteban Muñoz-Medina, José de Jesús Nuñez-Contreras, Alicia Ocaña-Mondragón, Luis Alberto Ochoa-Carrera, Hector Esteban Paz-Juárez, Marissa Perez-Garcia, Francisco Pulido, Helen Haydee Fernanda Ramírez-Plascencia, Jorge Salas-Hernández, Angel Gustavo Salas-Lais, Alejandro Sánchez-Flores, Clara Esperanza Santacruz-Tinoco, María Guadalupe Santiago-Mauricio, Selene Zárate, Nelly Sélem-Mojica, Blanca Taboada, Kathia Elizabeth Tapia-Diaz, Gloria Vazquez, Joel Armando Vázquez-Pérez). |
| EPI_ISL_3347920, EPI_ISL_3347921, EPI_ISL_3347922, EPI_ISL_3347923, EPI_ISL_3347924, EPI_ISL_3347925, EPI_ISL_3347926, EPI_ISL_3347927, EPI_ISL_3347928, EPI_ISL_3347929, EPI_ISL_3347930, EPI_ISL_3347931, EPI_ISL_3347932, EPI_ISL_3347933, EPI_ISL_3347934, EPI_ISL_3347935, EPI_ISL_3347936 | | | |
| see above | Laboratorio Central de Epidemiología (LCE) | Instituto de Biotecnología de la UNAM | Consorcio Mexicano de Vigilancia Genómica (CoViGen-Mex). Authors (in alphabetical order): Julio Elias Alvarado-Yaah, Carlos F. Arias, Santiago Ávila-Ríos, Eduardo Becerril-Vargas, Víctor Hugo Borja-Aburto, Celia Boukadida, Cristóbal Cháidez-Quiróz, Juan Bautista Chale-Dzul, Ricardo Ciria Merce, Andreu Comas-García, Célida Duque Molina, Julissa Enciso-Ibarra, José Antonio Enciso-Moreno, Gloria Elena Espinosa-Ayala, Fernando Fontove-Herrera, Daniel Fregoso-Rueda, Victor Eduardo García-Arias, Alejandra García-Gasca, Bruno Gómez-Gil, Jean Pierre González, Irvin González-López, Concepción Grajales-Muñiz, Ricardo Grande, Rosa María Gutiérrez Rios, Jesús Hernández, Alejandra Hernández-Terán, Alfredo Herrera-Estrella, Carla Ivón Herrera-Najera, Pavel Isa, Verónica Mata-Haro, Daniel Lira Morales, Susana Lopez, Antonio Loza Román, Brenda Irasema Maldonado-Meza, Bernardo Martínez-Miguel, José Arturo Martínez-Orozco, Célida Martínez-Rodríguez, Margarita Matías-Florentino, Fidencio Mejía-Nepomuceno, María Guadalupe de Jesús Mireles-Rivera, Gloria María Molina-Salinas, Hector Montoya-Fuentes, , Mario Mújica-Sánchez, José Esteban Muñoz-Medina, José de Jesús Nuñez-Contreras, Alicia Ocaña-Mondragón, Luis Alberto Ochoa-Carrera, Hector Esteban Paz-Juárez, Marissa Perez-Garcia, Francisco Pulido, Helen Haydee Fernanda Ramírez-Plascencia, Jorge Salas-Hernández, Angel Gustavo Salas-Lais, Alejandro Sánchez-Flores, Clara Esperanza Santacruz-Tinoco, María Guadalupe Santiago-Mauricio, Selene Zárate, Nelly Sélem-Mojica, Blanca Taboada, Kathia Elizabeth Tapia-Diaz, Gloria Vazquez, Joel Armando Vázquez-Pérez). |
| EPI_ISL_3347937, EPI_ISL_3347938, EPI_ISL_3347939, EPI_ISL_3347940, EPI_ISL_3347941, EPI_ISL_3347942, EPI_ISL_3347943, EPI_ISL_3347944 | Centro de Investigación Biomédica de Occidente (CIBO) | Instituto de Biotecnología de la UNAM | Consorcio Mexicano de Vigilancia Genómica (CoViGen-Mex). Authors (in alphabetical order): Julio Elias Alvarado-Yaah, Carlos F. Arias, Santiago Ávila-Ríos, Eduardo Becerril-Vargas, Víctor Hugo Borja-Aburto, Celia Boukadida, Cristóbal Cháidez-Quiróz, Juan Bautista Chale-Dzul, Ricardo Ciria Merce, Andreu Comas-García, Célida Duque Molina, Julissa Enciso-Ibarra, José Antonio Enciso-Moreno, Gloria Elena Espinosa-Ayala, Fernando Fontove-Herrera, Daniel Fregoso-Rueda, Victor Eduardo García-Arias, Alejandra García-Gasca, Bruno Gómez-Gil, Jean Pierre González, Irvin González-López, Concepción Grajales-Muñiz, Ricardo Grande, Rosa María Gutiérrez Rios, Jesús Hernández, Alejandra Hernández-Terán, Alfredo Herrera-Estrella, Carla Ivón Herrera-Najera, Pavel Isa, Verónica Mata-Haro, Daniel Lira Morales, Susana Lopez, Antonio Loza Román, Brenda Irasema Maldonado-Meza, Bernardo Martínez-Miguel, José Arturo Martínez-Orozco, Célida Martínez-Rodríguez, Margarita Matías-Florentino, Fidencio Mejía-Nepomuceno, María Guadalupe de Jesús Mireles-Rivera, Gloria María Molina-Salinas, Hector Montoya-Fuentes, , Mario Mújica-Sánchez, José Esteban Muñoz-Medina, José de Jesús Nuñez-Contreras, Alicia Ocaña-Mondragón, Luis Alberto Ochoa-Carrera, Hector Esteban Paz-Juárez, Marissa Perez-Garcia, Francisco Pulido, Helen Haydee Fernanda Ramírez-Plascencia, Jorge Salas-Hernández, Angel Gustavo Salas-Lais, Alejandro Sánchez-Flores, Clara Esperanza Santacruz-Tinoco, María Guadalupe Santiago-Mauricio, Selene Zárate, Nelly Sélem-Mojica, Blanca Taboada, Kathia Elizabeth Tapia-Diaz, Gloria Vazquez, Joel Armando Vázquez-Pérez). |
| EPI_ISL_3347945, EPI_ISL_3347946 | Unidad de Investigación Biomédica de Zacatecas (UIBZ) | Instituto de Biotecnología de la UNAM | Consorcio Mexicano de Vigilancia Genómica (CoViGen-Mex). Authors (in alphabetical order): Julio Elias Alvarado-Yaah, Carlos F. Arias, Santiago Ávila-Ríos, Eduardo Becerril-Vargas, Víctor Hugo Borja-Aburto, Celia Boukadida, Cristóbal Cháidez-Quiróz, Juan Bautista Chale-Dzul, Ricardo Ciria Merce, Andreu Comas-García, Célida Duque Molina, Julissa Enciso-Ibarra, José Antonio Enciso-Moreno, Gloria Elena Espinosa-Ayala, Fernando Fontove-Herrera, Daniel Fregoso-Rueda, Victor Eduardo García-Arias, Alejandra García-Gasca, Bruno Gómez-Gil, Jean Pierre González, Irvin González-López, Concepción Grajales-Muñiz, Ricardo Grande, Rosa María Gutiérrez Rios, Jesús Hernández, Alejandra Hernández-Terán, Alfredo Herrera-Estrella, Carla Ivón Herrera-Najera, Pavel Isa, Verónica Mata-Haro, Daniel Lira Morales, Susana Lopez, Antonio Loza Román, Brenda Irasema Maldonado-Meza, Bernardo Martínez-Miguel, José Arturo Martínez-Orozco, Célida Martínez-Rodríguez, Margarita Matías-Florentino, Fidencio Mejía-Nepomuceno, María Guadalupe de Jesús Mireles-Rivera, Gloria María Molina-Salinas, Hector Montoya-Fuentes, , Mario Mújica-Sánchez, José Esteban Muñoz-Medina, José de Jesús Nuñez-Contreras, Alicia Ocaña-Mondragón, Luis Alberto Ochoa-Carrera, Hector Esteban Paz-Juárez, Marissa Perez-Garcia, Francisco Pulido, Helen Haydee Fernanda Ramírez-Plascencia, Jorge Salas-Hernández, Angel Gustavo Salas-Lais, Alejandro Sánchez-Flores, Clara Esperanza Santacruz-Tinoco, María Guadalupe Santiago-Mauricio, Selene Zárate, Nelly Sélem-Mojica, Blanca Taboada, Kathia Elizabeth Tapia-Diaz, Gloria Vazquez, Joel Armando Vázquez-Pérez). |
| EPI_ISL_3347949 | Centro de Investigación Biomédica del Noreste (CIBIN) | Instituto de Biotecnología de la UNAM | Consorcio Mexicano de Vigilancia Genómica (CoViGen-Mex). Authors (in alphabetical order): Julio Elias Alvarado-Yaah, Carlos F. Arias, Santiago Ávila-Ríos, Eduardo Becerril-Vargas, Víctor Hugo Borja-Aburto, Celia Boukadida, Cristóbal Cháidez-Quiróz, Juan Bautista Chale-Dzul, Ricardo Ciria Merce, Andreu Comas-García, Célida Duque Molina, Julissa Enciso-Ibarra, José Antonio Enciso-Moreno, Gloria Elena Espinosa-Ayala, Fernando Fontove-Herrera, Daniel Fregoso-Rueda, Victor Eduardo García-Arias, Alejandra García-Gasca, Bruno Gómez-Gil, Jean Pierre González, Irvin González-López, Concepción Grajales-Muñiz, Ricardo Grande, Rosa María Gutiérrez Rios, Jesús Hernández, Alejandra Hernández-Terán, Alfredo Herrera-Estrella, Carla Ivón Herrera-Najera, Pavel Isa, Verónica Mata-Haro, Daniel Lira Morales, Susana Lopez, Antonio Loza Román, Brenda Irasema Maldonado-Meza, Bernardo Martínez-Miguel, José Arturo Martínez-Orozco, Célida Martínez-Rodríguez, Margarita Matías-Florentino, Fidencio Mejía-Nepomuceno, María Guadalupe de Jesús Mireles-Rivera, Gloria María Molina-Salinas, Hector Montoya-Fuentes, , Mario Mújica-Sánchez, José Esteban Muñoz-Medina, José de Jesús Nuñez-Contreras, Alicia Ocaña-Mondragón, Luis Alberto Ochoa-Carrera, Hector Esteban Paz-Juárez, Marissa Perez-Garcia, Francisco Pulido, Helen Haydee Fernanda Ramírez-Plascencia, Jorge Salas-Hernández, Angel Gustavo Salas-Lais, Alejandro Sánchez-Flores, Clara Esperanza Santacruz-Tinoco, María Guadalupe Santiago-Mauricio, Selene Zárate, Nelly Sélem-Mojica, Blanca Taboada, Kathia Elizabeth Tapia-Diaz, Gloria Vazquez, Joel Armando Vázquez-Pérez). |
| EPI_ISL_3347954 | Hospital Herrera Llerandi | Asociación de Salud Integral/Clínica Familiar "Luis Ángel García" | Eduardo Arathoon, Ana S. Gonzalez-Reiche, Claudia Rangel, Luis Rivas, Oscar Bonilla, Narda Medina, Luis Aguirre, Danicela Mercado, Hilda Ruiz, Osmar Gamboa |
| EPI_ISL_3347955, EPI_ISL_3347956, EPI_ISL_3347957, EPI_ISL_3347958, EPI_ISL_3347959, EPI_ISL_3347960, EPI_ISL_3347961, EPI_ISL_3347962, EPI_ISL_3347963, EPI_ISL_3347964, EPI_ISL_3347965, EPI_ISL_3347966, EPI_ISL_3347967, EPI_ISL_3347968, EPI_ISL_3347969, EPI_ISL_3347970, EPI_ISL_3347971, EPI_ISL_3347972, EPI_ISL_3347973, EPI_ISL_3347974, EPI_ISL_3347975, EPI_ISL_3347976, EPI_ISL_3347977, EPI_ISL_3347978, EPI_ISL_3347979, EPI_ISL_3347980, EPI_ISL_3347981, EPI_ISL_3347982, EPI_ISL_3347983, EPI_ISL_3347984, EPI_ISL_3347985, EPI_ISL_3347986, EPI_ISL_3347987, EPI_ISL_3347988, EPI_ISL_3347989, EPI_ISL_3347990, EPI_ISL_3347991, EPI_ISL_3347992, EPI_ISL_3347993, EPI_ISL_3347994, EPI_ISL_3347995, EPI_ISL_3347996, EPI_ISL_3347997, EPI_ISL_3347998, EPI_ISL_3347999, EPI_ISL_3348000, EPI_ISL_3348001, EPI_ISL_3348002, EPI_ISL_3348003, EPI_ISL_3348004, EPI_ISL_3348005, EPI_ISL_3348006, EPI_ISL_3348007, EPI_ISL_3348008, EPI_ISL_3348009, EPI_ISL_3348010, EPI_ISL_3348011, EPI_ISL_3348012, EPI_ISL_3348013, EPI_ISL_3348014, EPI_ISL_3348015, EPI_ISL_3348016, EPI_ISL_3348017, EPI_ISL_3348018, EPI_ISL_3348019, EPI_ISL_3348020, EPI_ISL_3348021, EPI_ISL_3348022, EPI_ISL_3348023, EPI_ISL_3348024, EPI_ISL_3348025, EPI_ISL_3348026, EPI_ISL_3348027, EPI_ISL_3348028, EPI_ISL_3348029, EPI_ISL_3348030, EPI_ISL_3348031, EPI_ISL_3348032, EPI_ISL_3348033, EPI_ISL_3348034, EPI_ISL_3348035, EPI_ISL_3348036, EPI_ISL_3348037, EPI_ISL_3348038, EPI_ISL_3348039, EPI_ISL_3348040, EPI_ISL_3348041, EPI_ISL_3348042, EPI_ISL_3348043, EPI_ISL_3348044, EPI_ISL_3348045, EPI_ISL_3348046, EPI_ISL_3348047, EPI_ISL_3348048, EPI_ISL_3348049, EPI_ISL_3348050, EPI_ISL_3348051, EPI_ISL_3348052, EPI_ISL_3348053, EPI_ISL_3348054, EPI_ISL_3348055, EPI_ISL_3348056, EPI_ISL_3348057, EPI_ISL_3348058, EPI_ISL_3348059, EPI_ISL_3348060, EPI_ISL_3348061, EPI_ISL_3348062, EPI_ISL_3348063, EPI_ISL_3348064, EPI_ISL_3348065, EPI_ISL_3348066, EPI_ISL_3348067, EPI_ISL_3348068, EPI_ISL_3348069, EPI_ISL_3348070, EPI_ISL_3348071, EPI_ISL_3348072, EPI_ISL_3348073, EPI_ISL_3348074, EPI_ISL_3348075, EPI_ISL_3348076, EPI_ISL_3348077, EPI_ISL_3348078, EPI_ISL_3348079, EPI_ISL_3348080, EPI_ISL_3348081, EPI_ISL_3348082, EPI_ISL_3348083, EPI_ISL_3348084, EPI_ISL_3348085, EPI_ISL_3348086, EPI_ISL_3348087, EPI_ISL_3348088, EPI_ISL_3348089, EPI_ISL_3348090, EPI_ISL_3348091, EPI_ISL_3348092, EPI_ISL_3348093, EPI_ISL_3348094, EPI_ISL_3348095, EPI_ISL_3348096, EPI_ISL_3348097, EPI_ISL_3348098, EPI_ISL_3348099, EPI_ISL_3348100, EPI_ISL_3348101, EPI_ISL_3348102, EPI_ISL_3348103, EPI_ISL_3348104, EPI_ISL_3348105, EPI_ISL_3348106, EPI_ISL_3348107, EPI_ISL_3348108, EPI_ISL_3348109, | | | |

EPI_ISL_3348110, EPI_ISL_3348111, EPI_ISL_3348112, EPI_ISL_3348113, EPI_ISL_3348114, EPI_ISL_3348115, EPI_ISL_3348116, EPI_ISL_3348117, EPI_ISL_3348118, EPI_ISL_3348119, EPI_ISL_3348120, EPI_ISL_3348121, EPI_ISL_3348122, EPI_ISL_3348123, EPI_ISL_3348124, EPI_ISL_3348125, EPI_ISL_3348126, EPI_ISL_3348127, EPI_ISL_3348128, EPI_ISL_3348129, EPI_ISL_3348130, EPI_ISL_3348131,
EPI_ISL_3348132, EPI_ISL_3348133, EPI_ISL_3348134, EPI_ISL_3348135, EPI_ISL_3348136, EPI_ISL_3348137, EPI_ISL_3348138, EPI_ISL_3348140, EPI_ISL_3348141, EPI_ISL_3348142, EPI_ISL_3348143, EPI_ISL_3348144, EPI_ISL_3348145, EPI_ISL_3348146, EPI_ISL_3348147, EPI_ISL_3348148, EPI_ISL_3348149, EPI_ISL_3348150, EPI_ISL_3348151, EPI_ISL_3348152, EPI_ISL_3348153, EPI_ISL_3348154,
EPI_ISL_3348155, EPI_ISL_3348156, EPI_ISL_3348157, EPI_ISL_3348158, EPI_ISL_3348159, EPI_ISL_3348160, EPI_ISL_3348161, EPI_ISL_3348162, EPI_ISL_3348163, EPI_ISL_3348165, EPI_ISL_3348166, EPI_ISL_3348167, EPI_ISL_3348168, EPI_ISL_3348169, EPI_ISL_3348170, EPI_ISL_3348171, EPI_ISL_3348172, EPI_ISL_3348173, EPI_ISL_3348174, EPI_ISL_3348175, EPI_ISL_3348176, EPI_ISL_3348177,
EPI_ISL_3348178, EPI_ISL_3348179, EPI_ISL_3348180, EPI_ISL_3348181, EPI_ISL_3348182, EPI_ISL_3348183, EPI_ISL_3348184, EPI_ISL_3348185, EPI_ISL_3348186, EPI_ISL_3348187, EPI_ISL_3348188, EPI_ISL_3348189, EPI_ISL_3348190, EPI_ISL_3348191, EPI_ISL_3348192, EPI_ISL_3348193, EPI_ISL_3348194, EPI_ISL_3348195, EPI_ISL_3348196, EPI_ISL_3348197, EPI_ISL_3348198, EPI_ISL_3348199,
EPI_ISL_3348200, EPI_ISL_3348201, EPI_ISL_3348202, EPI_ISL_3348203, EPI_ISL_3348204, EPI_ISL_3348205, EPI_ISL_3348206, EPI_ISL_3348207, EPI_ISL_3348208, EPI_ISL_3348209, EPI_ISL_3348210, EPI_ISL_3348211, EPI_ISL_3348212, EPI_ISL_3348213, EPI_ISL_3348214, EPI_ISL_3348215, EPI_ISL_3348216, EPI_ISL_3348217, EPI_ISL_3348218, EPI_ISL_3348219, EPI_ISL_3348220, EPI_ISL_3348221,
EPI_ISL_3348222, EPI_ISL_3348223, EPI_ISL_3348224, EPI_ISL_3348225, EPI_ISL_3348226, EPI_ISL_3348227, EPI_ISL_3348228, EPI_ISL_3348229, EPI_ISL_3348230, EPI_ISL_3348231, EPI_ISL_3348232, EPI_ISL_3348233, EPI_ISL_3348234, EPI_ISL_3348235, EPI_ISL_3348236, EPI_ISL_3348237, EPI_ISL_3348238, EPI_ISL_3348239, EPI_ISL_3348240, EPI_ISL_3348241, EPI_ISL_3348242, EPI_ISL_3348243,
EPI_ISL_3348244, EPI_ISL_3348245, EPI_ISL_3348246, EPI_ISL_3348247, EPI_ISL_3348248, EPI_ISL_3348249, EPI_ISL_3348250, EPI_ISL_3348251, EPI_ISL_3348252, EPI_ISL_3348253, EPI_ISL_3348254, EPI_ISL_3348255, EPI_ISL_3348256, EPI_ISL_3348257, EPI_ISL_3348258, EPI_ISL_3348259, EPI_ISL_3348260, EPI_ISL_3348261, EPI_ISL_3348262, EPI_ISL_3348263, EPI_ISL_3348264, EPI_ISL_3348265,
EPI_ISL_3348266, EPI_ISL_3348267, EPI_ISL_3348268, EPI_ISL_3348269, EPI_ISL_3348270, EPI_ISL_3348271, EPI_ISL_3348272, EPI_ISL_3348273, EPI_ISL_3348274, EPI_ISL_3348275, EPI_ISL_3348276, EPI_ISL_3348277, EPI_ISL_3348279, EPI_ISL_3348280, EPI_ISL_3348281, EPI_ISL_3348282, EPI_ISL_3348283, EPI_ISL_3348284, EPI_ISL_3348285, EPI_ISL_3348286, EPI_ISL_3348287, EPI_ISL_3348288,
EPI_ISL_3348289, EPI_ISL_3348290, EPI_ISL_3348292, EPI_ISL_3348293, EPI_ISL_3348294, EPI_ISL_3348295, EPI_ISL_3348296, EPI_ISL_3348297, EPI_ISL_3348298, EPI_ISL_3348299, EPI_ISL_3348300, EPI_ISL_3348301, EPI_ISL_3348302, EPI_ISL_3348303, EPI_ISL_3348304, EPI_ISL_3348305, EPI_ISL_3348306, EPI_ISL_3348307, EPI_ISL_3348308, EPI_ISL_3348309, EPI_ISL_3348310, EPI_ISL_3348311,
EPI_ISL_3348312, EPI_ISL_3348313, EPI_ISL_3348314, EPI_ISL_3348315, EPI_ISL_3348316, EPI_ISL_3348317, EPI_ISL_3348318, EPI_ISL_3348319, EPI_ISL_3348320, EPI_ISL_3348321, EPI_ISL_3348322, EPI_ISL_3348323, EPI_ISL_3348324, EPI_ISL_3348325, EPI_ISL_3348326, EPI_ISL_3348327, EPI_ISL_3348328, EPI_ISL_3348329, EPI_ISL_3348330, EPI_ISL_3348331, EPI_ISL_3348332, EPI_ISL_3348333,
EPI_ISL_3348334, EPI_ISL_3348335, EPI_ISL_3348336, EPI_ISL_3348337, EPI_ISL_3348338, EPI_ISL_3348339, EPI_ISL_3348340, EPI_ISL_3348341, EPI_ISL_3348342, EPI_ISL_3348343, EPI_ISL_3348344, EPI_ISL_3348345, EPI_ISL_3348346, EPI_ISL_3348347, EPI_ISL_3348348, EPI_ISL_3348349, EPI_ISL_3348350, EPI_ISL_3348351, EPI_ISL_3348352, EPI_ISL_3348353, EPI_ISL_3348355, EPI_ISL_3348356,
EPI_ISL_3348358, EPI_ISL_3348359, EPI_ISL_3348360, EPI_ISL_3348361, EPI_ISL_3348362, EPI_ISL_3348363, EPI_ISL_3348364, EPI_ISL_3348365, EPI_ISL_3348366, EPI_ISL_3348367, EPI_ISL_3348368, EPI_ISL_3348369, EPI_ISL_3348370, EPI_ISL_3348371, EPI_ISL_3348372, EPI_ISL_3348373, EPI_ISL_3348374, EPI_ISL_3348375, EPI_ISL_3348376, EPI_ISL_3348377, EPI_ISL_3348378, EPI_ISL_3348379,
EPI_ISL_3348380, EPI_ISL_3348381, EPI_ISL_3348382, EPI_ISL_3348383, EPI_ISL_3348384, EPI_ISL_3348385, EPI_ISL_3348386, EPI_ISL_3348387, EPI_ISL_3348388, EPI_ISL_3348389, EPI_ISL_3348390, EPI_ISL_3348391, EPI_ISL_3348392, EPI_ISL_3348393, EPI_ISL_3348394, EPI_ISL_3348395, EPI_ISL_3348396, EPI_ISL_3348397, EPI_ISL_3348398, EPI_ISL_3348399, EPI_ISL_3348400, EPI_ISL_3348401,
EPI_ISL_3348402, EPI_ISL_3348403, EPI_ISL_3348404, EPI_ISL_3348405, EPI_ISL_3348406, EPI_ISL_3348407, EPI_ISL_3348408, EPI_ISL_3348409, EPI_ISL_3348410, EPI_ISL_3348411, EPI_ISL_3348412, EPI_ISL_3348413, EPI_ISL_3348414, EPI_ISL_3348415, EPI_ISL_3348416, EPI_ISL_3348417, EPI_ISL_3348418, EPI_ISL_3348419, EPI_ISL_3348420, EPI_ISL_3348421, EPI_ISL_3348422, EPI_ISL_3348423,
EPI_ISL_3348424, EPI_ISL_3348425, EPI_ISL_3348426, EPI_ISL_3348427, EPI_ISL_3348428, EPI_ISL_3348429, EPI_ISL_3348430, EPI_ISL_3348431, EPI_ISL_3348432, EPI_ISL_3348433, EPI_ISL_3348434, EPI_ISL_3348435, EPI_ISL_3348436, EPI_ISL_3348437, EPI_ISL_3348438, EPI_ISL_3348439, EPI_ISL_3348440, EPI_ISL_3348441, EPI_ISL_3348442, EPI_ISL_3348443, EPI_ISL_3348444, EPI_ISL_3348445,
EPI_ISL_3348446, EPI_ISL_3348447, EPI_ISL_3348448, EPI_ISL_3348449, EPI_ISL_3348450, EPI_ISL_3348451, EPI_ISL_3348452, EPI_ISL_3348453, EPI_ISL_3348454, EPI_ISL_3348455, EPI_ISL_3348456, EPI_ISL_3348457, EPI_ISL_3348458, EPI_ISL_3348459, EPI_ISL_3348460, EPI_ISL_3348461, EPI_ISL_3348462, EPI_ISL_3348463, EPI_ISL_3348464, EPI_ISL_3348465, EPI_ISL_3348466, EPI_ISL_3348467,
EPI_ISL_3348468, EPI_ISL_3348469, EPI_ISL_3348470, EPI_ISL_3348471, EPI_ISL_3348472, EPI_ISL_3348473, EPI_ISL_3348474, EPI_ISL_3348475, EPI_ISL_3348476, EPI_ISL_3348477, EPI_ISL_3348478, EPI_ISL_3348479, EPI_ISL_3348480, EPI_ISL_3348481, EPI_ISL_3348482, EPI_ISL_3348483, EPI_ISL_3348484, EPI_ISL_3348485, EPI_ISL_3348486, EPI_ISL_3348487, EPI_ISL_3348488, EPI_ISL_3348489,
EPI_ISL_3348490, EPI_ISL_3348491, EPI_ISL_3348492, EPI_ISL_3348493, EPI_ISL_3348494, EPI_ISL_3348495, EPI_ISL_3348496, EPI_ISL_3348497, EPI_ISL_3348498, EPI_ISL_3348499, EPI_ISL_3348500, EPI_ISL_3348501, EPI_ISL_3348502, EPI_ISL_3348503, EPI_ISL_3348504, EPI_ISL_3348505, EPI_ISL_3348506, EPI_ISL_3348507, EPI_ISL_3348508, EPI_ISL_3348509, EPI_ISL_3348510, EPI_ISL_3348511,
EPI_ISL_3348512, EPI_ISL_3348513, EPI_ISL_3348514, EPI_ISL_3348515, EPI_ISL_3348516, EPI_ISL_3348517, EPI_ISL_3348518, EPI_ISL_3348519, EPI_ISL_3348521, EPI_ISL_3348522, EPI_ISL_3348523, EPI_ISL_3348524, EPI_ISL_3348525, EPI_ISL_3348526, EPI_ISL_3348527, EPI_ISL_3348528, EPI_ISL_3348529, EPI_ISL_3348530, EPI_ISL_3348531, EPI_ISL_3348532, EPI_ISL_3348533, EPI_ISL_3348534,
EPI_ISL_3348535, EPI_ISL_3348536, EPI_ISL_3348537, EPI_ISL_3348538, EPI_ISL_3348539, EPI_ISL_3348540, EPI_ISL_3348541, EPI_ISL_3348542, EPI_ISL_3348543, EPI_ISL_3348544, EPI_ISL_3348545, EPI_ISL_3348546, EPI_ISL_3348547, EPI_ISL_3348548, EPI_ISL_3348549, EPI_ISL_3348550, EPI_ISL_3348551, EPI_ISL_3348552, EPI_ISL_3348553, EPI_ISL_3348554, EPI_ISL_3348555, EPI_ISL_3348556,
EPI_ISL_3348557, EPI_ISL_3348558, EPI_ISL_3348559, EPI_ISL_3348560, EPI_ISL_3348561, EPI_ISL_3348562, EPI_ISL_3348563, EPI_ISL_3348564, EPI_ISL_3348565, EPI_ISL_3348566, EPI_ISL_3348567, EPI_ISL_3348568, EPI_ISL_3348569, EPI_ISL_3348570, EPI_ISL_3348571, EPI_ISL_3348572, EPI_ISL_3348573, EPI_ISL_3348574, EPI_ISL_3348575, EPI_ISL_3348576, EPI_ISL_3348577, EPI_ISL_3348578,
EPI_ISL_3348579, EPI_ISL_3348580, EPI_ISL_3348581, EPI_ISL_3348582, EPI_ISL_3348583, EPI_ISL_3348584, EPI_ISL_3348585, EPI_ISL_3348586, EPI_ISL_3348587, EPI_ISL_3348588, EPI_ISL_3348589, EPI_ISL_3348590, EPI_ISL_3348591, EPI_ISL_3348592, EPI_ISL_3348593, EPI_ISL_3348594, EPI_ISL_3348595, EPI_ISL_3348596, EPI_ISL_3348597, EPI_ISL_3348598, EPI_ISL_3348599, EPI_ISL_3348600,
EPI_ISL_3348601, EPI_ISL_3348602, EPI_ISL_3348603, EPI_ISL_3348604, EPI_ISL_3348605, EPI_ISL_3348606, EPI_ISL_3348607, EPI_ISL_3348608, EPI_ISL_3348609, EPI_ISL_3348610, EPI_ISL_3348611, EPI_ISL_3348612, EPI_ISL_3348613, EPI_ISL_3348614, EPI_ISL_3348615, EPI_ISL_3348616, EPI_ISL_3348617, EPI_ISL_3348618, EPI_ISL_3348619, EPI_ISL_3348620, EPI_ISL_3348621, EPI_ISL_3348622,
EPI_ISL_3348623, EPI_ISL_3348624, EPI_ISL_3348625, EPI_ISL_3348626, EPI_ISL_3348627, EPI_ISL_3348628, EPI_ISL_3348629, EPI_ISL_3348630, EPI_ISL_3348631, EPI_ISL_3348632, EPI_ISL_3348633, EPI_ISL_3348634, EPI_ISL_3348635, EPI_ISL_3348636, EPI_ISL_3348637, EPI_ISL_3348638, EPI_ISL_3348639, EPI_ISL_3348640, EPI_ISL_3348641, EPI_ISL_3348642, EPI_ISL_3348643, EPI_ISL_3348644,
EPI_ISL_3348645, EPI_ISL_3348646, EPI_ISL_3348647, EPI_ISL_3348648, EPI_ISL_3348649, EPI_ISL_3348650, EPI_ISL_3348651, EPI_ISL_3348652, EPI_ISL_3348653, EPI_ISL_3348654, EPI_ISL_3348655, EPI_ISL_3348656, EPI_ISL_3348657, EPI_ISL_3348658, EPI_ISL_3348659, EPI_ISL_3348660, EPI_ISL_3348661, EPI_ISL_3348662, EPI_ISL_3348663, EPI_ISL_3348664, EPI_ISL_3348665, EPI_ISL_3348666,
EPI_ISL_3348667, EPI_ISL_3348668, EPI_ISL_3348669, EPI_ISL_3348670, EPI_ISL_3348671, EPI_ISL_3348672, EPI_ISL_3348673, EPI_ISL_3348674, EPI_ISL_3348675, EPI_ISL_3348676, EPI_ISL_3348677, EPI_ISL_3348678, EPI_ISL_3348679, EPI_ISL_3348680, EPI_ISL_3348681, EPI_ISL_3348682, EPI_ISL_3348683, EPI_ISL_3348684, EPI_ISL_3348685, EPI_ISL_3348686, EPI_ISL_3348687, EPI_ISL_3348688,
EPI_ISL_3348689, EPI_ISL_3348690, EPI_ISL_3348691, EPI_ISL_3348692, EPI_ISL_3348693, EPI_ISL_3348694, EPI_ISL_3348695, EPI_ISL_3348696, EPI_ISL_3348697, EPI_ISL_3348698, EPI_ISL_3348699, EPI_ISL_3348700, EPI_ISL_3348701, EPI_ISL_3348702, EPI_ISL_3348703, EPI_ISL_3348704, EPI_ISL_3348705, EPI_ISL_3348706, EPI_ISL_3348707, EPI_ISL_3348708, EPI_ISL_3348709, EPI_ISL_3348710,
EPI_ISL_3348711, EPI_ISL_3348712, EPI_ISL_3348713, EPI_ISL_3348714, EPI_ISL_3348715, EPI_ISL_3348716, EPI_ISL_3348717, EPI_ISL_3348718, EPI_ISL_3348719, EPI_ISL_3348720, EPI_ISL_3348721, EPI_ISL_3348722, EPI_ISL_3348723, EPI_ISL_3348724, EPI_ISL_3348725, EPI_ISL_3348726, EPI_ISL_3348727, EPI_ISL_3348728, EPI_ISL_3348729, EPI_ISL_3348730, EPI_ISL_3348731, EPI_ISL_3348732,
EPI_ISL_3348733, EPI_ISL_3348734, EPI_ISL_3348735, EPI_ISL_3348736, EPI_ISL_3348737, EPI_ISL_3348738, EPI_ISL_3348739, EPI_ISL_3348740, EPI_ISL_3348741, EPI_ISL_3348742, EPI_ISL_3348743, EPI_ISL_3348744, EPI_ISL_3348745, EPI_ISL_3348746, EPI_ISL_3348747, EPI_ISL_3348748, EPI_ISL_3348749, EPI_ISL_3348750, EPI_ISL_3348751, EPI_ISL_3348752, EPI_ISL_3348753, EPI_ISL_3348754,
EPI_ISL_3348755, EPI_ISL_3348756, EPI_ISL_3348757, EPI_ISL_3348758, EPI_ISL_3348759, EPI_ISL_3348760, EPI_ISL_3348761, EPI_ISL_3348762, EPI_ISL_3348763, EPI_ISL_3348764, EPI_ISL_3348765, EPI_ISL_3348766, EPI_ISL_3348767, EPI_ISL_3348768, EPI_ISL_3348769, EPI_ISL_3348770, EPI_ISL_3348771, EPI_ISL_3348772, EPI_ISL_3348773, EPI_ISL_3348774, EPI_ISL_3348775, EPI_ISL_3348776,
EPI_ISL_3348777, EPI_ISL_3348778, EPI_ISL_3348779, EPI_ISL_3348780, EPI_ISL_3348781, EPI_ISL_3348782, EPI_ISL_3348783, EPI_ISL_3348784, EPI_ISL_3348785, EPI_ISL_3348786, EPI_ISL_3348787, EPI_ISL_3348788, EPI_ISL_3348789, EPI_ISL_3348790, EPI_ISL_3348791, EPI_ISL_3348792, EPI_ISL_3348793, EPI_ISL_3348794, EPI_ISL_3348795, EPI_ISL_3348796, EPI_ISL_3348797, EPI_ISL_3348798,
EPI_ISL_3348799, EPI_ISL_3348800, EPI_ISL_3348801, EPI_ISL_3348802, EPI_ISL_3348803, EPI_ISL_3348804, EPI_ISL_3348805, EPI_ISL_3348806, EPI_ISL_3348807, EPI_ISL_3348808, EPI_ISL_3348809, EPI_ISL_3348810, EPI_ISL_3348811, EPI_ISL_3348812, EPI_ISL_3348813, EPI_ISL_3348814, EPI_ISL_3348815, EPI_ISL_3348816, EPI_ISL_3348817, EPI_ISL_3348818, EPI_ISL_3348819, EPI_ISL_3348820,
EPI_ISL_3348821, EPI_ISL_3348822, EPI_ISL_3348823, EPI_ISL_3348824, EPI_ISL_3348825, EPI_ISL_3348826, EPI_ISL_3348827, EPI_ISL_3348828, EPI_ISL_3348829, EPI_ISL_3348830, EPI_ISL_3348831, EPI_ISL_3348832, EPI_ISL_3348833, EPI_ISL_3348834, EPI_ISL_3348835, EPI_ISL_3348836, EPI_ISL_3348837, EPI_ISL_3348838, EPI_ISL_3348839, EPI_ISL_3348840, EPI_ISL_3348841, EPI_ISL_3348842,
EPI_ISL_3348843, EPI_ISL_3348844, EPI_ISL_3348845, EPI_ISL_3348846, EPI_ISL_3348847, EPI_ISL_3348848, EPI_ISL_3348849, EPI_ISL_3348850, EPI_ISL_3348851, EPI_ISL_3348852, EPI_ISL_3348853, EPI_ISL_3348854, EPI_ISL_3348855, EPI_ISL_3348856, EPI_ISL_3348857, EPI_ISL_3348858, EPI_ISL_3348859, EPI_ISL_3348860, EPI_ISL_3348861, EPI_ISL_3348862, EPI_ISL_3348863, EPI_ISL_3348864,
EPI_ISL_3348865, EPI_ISL_3348866, EPI_ISL_3348867, EPI_ISL_3348868, EPI_ISL_3348869, EPI_ISL_3348870, EPI_ISL_3348871, EPI_ISL_3348872, EPI_ISL_3348873, EPI_ISL_3348874, EPI_ISL_3348875, EPI_ISL_3348877, EPI_ISL_3348878, EPI_ISL_3348879, EPI_ISL_3348880, EPI_ISL_3348881, EPI_ISL_3348882, EPI_ISL_3348883, EPI_ISL_3348884, EPI_ISL_3348885, EPI_ISL_3348886, EPI_ISL_3348887,
EPI_ISL_3348888, EPI_ISL_3348890, EPI_ISL_3348891, EPI_ISL_3348892, EPI_ISL_3348893, EPI_ISL_3348894, EPI_ISL_3348895, EPI_ISL_3348896, EPI_ISL_3348897, EPI_ISL_3348898, EPI_ISL_3348899, EPI_ISL_3348900, EPI_ISL_3348901, EPI_ISL_3348903, EPI_ISL_3348904, EPI_ISL_3348905, EPI_ISL_3348906, EPI_ISL_3348907, EPI_ISL_3348908, EPI_ISL_3348909, EPI_ISL_3348910, EPI_ISL_3348911,
EPI_ISL_3348912, EPI_ISL_3348913, EPI_ISL_3348914, EPI_ISL_3348915, EPI_ISL_3348916, EPI_ISL_3348917, EPI_ISL_3348919, EPI_ISL_3348920, EPI_ISL_3348921, EPI_ISL_3348922, EPI_ISL_3348923, EPI_ISL_3348924, EPI_ISL_3348925, EPI_ISL_3348926, EPI_ISL_3348927, EPI_ISL_3348928, EPI_ISL_3348929, EPI_ISL_3348930, EPI_ISL_3348931, EPI_ISL_3348932, EPI_ISL_3348933, EPI_ISL_3348934,
EPI_ISL_3348935, EPI_ISL_3348937, EPI_ISL_3348938, EPI_ISL_3348939, EPI_ISL_3348940, EPI_ISL_3348941, EPI_ISL_3348942, EPI_ISL_3348943, EPI_ISL_3348944, EPI_ISL_3348945, EPI_ISL_3348946, EPI_ISL_3348947, EPI_ISL_3348948, EPI_ISL_3348949, EPI_ISL_3348950, EPI_ISL_3348951, EPI_ISL_3348952, EPI_ISL_3348953, EPI_ISL_3348954, EPI_ISL_3348955, EPI_ISL_3348956, EPI_ISL_3348957,
EPI_ISL_3348958, EPI_ISL_3348959, EPI_ISL_3348960, EPI_ISL_3348961, EPI_ISL_3348962, EPI_ISL_3348963, EPI_ISL_3348964, EPI_ISL_3348965, EPI_ISL_3348966, EPI_ISL_3348967, EPI_ISL_3348968, EPI_ISL_3348969, EPI_ISL_3348970, EPI_ISL_3348972, EPI_ISL_3348973, EPI_ISL_3348974, EPI_ISL_3348975, EPI_ISL_3348976, EPI_ISL_3348977, EPI_ISL_3348978, EPI_ISL_3348979, EPI_ISL_3348980,
EPI_ISL_3348981, EPI_ISL_3348982, EPI_ISL_3348983, EPI_ISL_3348984, EPI_ISL_3348985, EPI_ISL_3348986, EPI_ISL_3348987, EPI_ISL_3348988, EPI_ISL_3348989, EPI_ISL_3348990, EPI_ISL_3348991, EPI_ISL_3348992, EPI_ISL_3348993, EPI_ISL_3348994, EPI_ISL_3348995, EPI_ISL_3348996, EPI_ISL_3348997, EPI_ISL_3348998, EPI_ISL_3348999, EPI_ISL_3349000, EPI_ISL_3349001, EPI_ISL_3349002,
EPI_ISL_3349003, EPI_ISL_3349004, EPI_ISL_3349005, EPI_ISL_3349006, EPI_ISL_3349007, EPI_ISL_3349008, EPI_ISL_3349009, EPI_ISL_3349010, EPI_ISL_3349011, EPI_ISL_3349012, EPI_ISL_3349013, EPI_ISL_3349014, EPI_ISL_3349015, EPI_ISL_3349016, EPI_ISL_3349017, EPI_ISL_3349018, EPI_ISL_3349019, EPI_ISL_3349020, EPI_ISL_3349021, EPI_ISL_3349022, EPI_ISL_3349023, EPI_ISL_3349024,
EPI_ISL_3349025, EPI_ISL_3349026, EPI_ISL_3349027, EPI_ISL_3349028, EPI_ISL_3349029, EPI_ISL_3349030, EPI_ISL_3349031, EPI_ISL_3349032, EPI_ISL_3349033, EPI_ISL_3349034, EPI_ISL_3349035, EPI_ISL_3349036, EPI_ISL_3349037, EPI_ISL_3349038, EPI_ISL_3349039, EPI_ISL_3349040, EPI_ISL_3349041, EPI_ISL_3349042, EPI_ISL_3349043, EPI_ISL_3349044, EPI_ISL_3349045, EPI_ISL_3349046,
EPI_ISL_3349047, EPI_ISL_3349048, EPI_ISL_3349049, EPI_ISL_3349050, EPI_ISL_3349051, EPI_ISL_3349052, EPI_ISL_3349053, EPI_ISL_3349054, EPI_ISL_3349055, EPI_ISL_3349056, EPI_ISL_3349057, EPI_ISL_3349058, EPI_ISL_3349059, EPI_ISL_3349060, EPI_ISL_3349061, EPI_ISL_3349062, EPI_ISL_3349063, EPI_ISL_3349064, EPI_ISL_3349065, EPI_ISL_3349066, EPI_ISL_3349067, EPI_ISL_3349068,
EPI_ISL_3349069, EPI_ISL_3349070, EPI_ISL_3349071, EPI_ISL_3349072, EPI_ISL_3349073, EPI_ISL_3349074, EPI_ISL_3349075, EPI_ISL_3349076, EPI_ISL_3349077, EPI_ISL_3349078, EPI_ISL_3349079, EPI_ISL_3349080, EPI_ISL_3349081, EPI_ISL_3349082, EPI_ISL_3349083, EPI_ISL_3349084, EPI_ISL_3349085, EPI_ISL_3349086, EPI_ISL_3349087, EPI_ISL_3349088, EPI_ISL_3349089, EPI_ISL_3349090,
EPI_ISL_3349091, EPI_ISL_3349092, EPI_ISL_3349093, EPI_ISL_3349094, EPI_ISL_3349095, EPI_ISL_3349096, EPI_ISL_3349097, EPI_ISL_3349098, EPI_ISL_3349099, EPI_ISL_3349100, EPI_ISL_3349101, EPI_ISL_3349102, EPI_ISL_3349103, EPI_ISL_3349104, EPI_ISL_3349105, EPI_ISL_3349106, EPI_ISL_3349107, EPI_ISL_3349109, EPI_ISL_3349110, EPI_ISL_3349111, EPI_ISL_3349112, EPI_ISL_3349113,
EPI_ISL_3349114, EPI_ISL_3349115, EPI_ISL_3349116, EPI_ISL_3349117, EPI_ISL_3349118, EPI_ISL_3349119, EPI_ISL_3349120, EPI_ISL_3349121, EPI_ISL_3349122, EPI_ISL_3349123, EPI_ISL_3349124, EPI_ISL_3349125, EPI_ISL_3349126, EPI_ISL_3349127, EPI_ISL_3349128, EPI_ISL_3349129, EPI_ISL_3349130, EPI_ISL_3349131, EPI_ISL_3349132, EPI_ISL_3349133, EPI_ISL_3349134, EPI_ISL_3349135,
EPI_ISL_3349136, EPI_ISL_3349137, EPI_ISL_3349138, EPI_ISL_3349139, EPI_ISL_3349140, EPI_ISL_3349141, EPI_ISL_3349142, EPI_ISL_3349143, EPI_ISL_3349144, EPI_ISL_3349145, EPI_ISL_3349146, EPI_ISL_3349147, EPI_ISL_3349148, EPI_ISL_3349149, EPI_ISL_3349150, EPI_ISL_3349151, EPI_ISL_3349152, EPI_ISL_3349153, EPI_ISL_3349154, EPI_ISL_3349155, EPI_ISL_3349156, EPI_ISL_3349157,
EPI_ISL_3349158, EPI_ISL_3349159, EPI_ISL_3349160, EPI_ISL_3349161, EPI_ISL_3349162, EPI_ISL_3349163, EPI_ISL_3349164, EPI_ISL_3349165, EPI_ISL_3349166, EPI_ISL_3349167, EPI_ISL_3349168, EPI_ISL_3349169, EPI_ISL_3349170, EPI_ISL_3349171, EPI_ISL_3349172, EPI_ISL_3349173, EPI_ISL_3349174, EPI_ISL_3349175, EPI_ISL_3349176, EPI_ISL_3349177, EPI_ISL_3349178, EPI_ISL_3349179,
EPI_ISL_3349180, EPI_ISL_3349181, EPI_ISL_3349182, EPI_ISL_3349183, EPI_ISL_3349184, EPI_ISL_3349185, EPI_ISL_3349186, EPI_ISL_3349187, EPI_ISL_3349188, EPI_ISL_3349189, EPI_ISL_3349190, EPI_ISL_3349191, EPI_ISL_3349192, EPI_ISL_3349193, EPI_ISL_3349194, EPI_ISL_3349195, EPI_ISL_3349196, EPI_ISL_3349197, EPI_ISL_3349198, EPI_ISL_3349199, EPI_ISL_3349200, EPI_ISL_3349201,
EPI_ISL_3349202, EPI_ISL_3349203, EPI_ISL_3349204, EPI_ISL_3349205, EPI_ISL_3349206, EPI_ISL_3349207, EPI_ISL_3349208, EPI_ISL_3349209, EPI_ISL_3349211, EPI_ISL_3349212, EPI_ISL_3349213, EPI_ISL_3349214, EPI_ISL_3349215, EPI_ISL_3349216, EPI_ISL_3349217, EPI_ISL_3349218, EPI_ISL_3349219, EPI_ISL_3349220, EPI_ISL_3349221, EPI_ISL_3349222, EPI_ISL_3349223, EPI_ISL_3349224,
EPI_ISL_3349225, EPI_ISL_3349226, EPI_ISL_3349227, EPI_ISL_3349228, EPI_ISL_3349229, EPI_ISL_3349230, EPI_ISL_3349231, EPI_ISL_3349232, EPI_ISL_3349233, EPI_ISL_3349234, EPI_ISL_3349235, EPI_ISL_3349236, EPI_ISL_3349237, EPI_ISL_3349238, EPI_ISL_3349239, EPI_ISL_3349240, EPI_ISL_3349241, EPI_ISL_3349242, EPI_ISL_3349243, EPI_ISL_3349244, EPI_ISL_3349245, EPI_ISL_3349246,
EPI_ISL_3349247, EPI_ISL_3349248, EPI_ISL_3349249, EPI_ISL_3349250, EPI_ISL_3349251, EPI_ISL_3349252, EPI_ISL_3349253, EPI_ISL_3349254, EPI_ISL_3349255, EPI_ISL_3349256, EPI_ISL_3349257, EPI_ISL_3349258, EPI_ISL_3349259, EPI_ISL_3349260, EPI_ISL_3349261, EPI_ISL_3349262, EPI_ISL_3349263, EPI_ISL_3349264, EPI_ISL_3349265, EPI_ISL_3349266, EPI_ISL_3349267, EPI_ISL_3349268,
EPI_ISL_3349269, EPI_ISL_3349270, EPI_ISL_3349271, EPI_ISL_3349272, EPI_ISL_3349273, EPI_ISL_3349274, EPI_ISL_3349275, EPI_ISL_3349276, EPI_ISL_3349277, EPI_ISL_3349278, EPI_ISL_3349279, EPI_ISL_3349280, EPI_ISL_3349281, EPI_ISL_3349282, EPI_ISL_3349283, EPI_ISL_3349284, EPI_ISL_3349285, EPI_ISL_3349286, EPI_ISL_3349287, EPI_ISL_3349288, EPI_ISL_3349289, EPI_ISL_3349290,
EPI_ISL_3349291, EPI_ISL_3349292, EPI_ISL_3349293, EPI_ISL_3349294, EPI_ISL_3349295, EPI_ISL_3349296, EPI_ISL_3349297, EPI_ISL_3349298, EPI_ISL_3349299, EPI_ISL_3349300, EPI_ISL_3349301, EPI_ISL_3349302, EPI_ISL_3349303, EPI_ISL_3349304, EPI_ISL_3349305, EPI_ISL_3349306, EPI_ISL_3349307, EPI_ISL_3349308, EPI_ISL_3349309, EPI_ISL_3349310, EPI_ISL_3349311, EPI_ISL_3349312,
EPI_ISL_3349313, EPI_ISL_3349314, EPI_ISL_3349315, EPI_ISL_3349316, EPI_ISL_3349317, EPI_ISL_3349318, EPI_ISL_3349319, EPI_ISL_3349320, EPI_ISL_3349321, EPI_ISL_3349322, EPI_ISL_3349323, EPI_ISL_3349324, EPI_ISL_3349325, EPI_ISL_3349326, EPI_ISL_3349327, EPI_ISL_3349328, EPI_ISL_3349329, EPI_ISL_3349330, EPI_ISL_3349331, EPI_ISL_3349332, EPI_ISL_3349333, EPI_ISL_3349335,
EPI_ISL_3349336, EPI_ISL_3349337, EPI_ISL_3349338, EPI_ISL_3349339, EPI_ISL_3349340, EPI_ISL_3349341, EPI_ISL_3349342, EPI_ISL_3349343, EPI_ISL_3349344, EPI_ISL_3349345, EPI_ISL_3349346, EPI_ISL_3349347, EPI_ISL_3349348, EPI_ISL_3349349, EPI_ISL_3349350, EPI_ISL_3349351, EPI_ISL_3349352, EPI_ISL_3349353, EPI_ISL_3349354, EPI_ISL_3349355, EPI_ISL_3349356, EPI_ISL_3349357,
EPI_ISL_3349358, EPI_ISL_3349359, EPI_ISL_3349360, EPI_ISL_3349361, EPI_ISL_3349362, EPI_ISL_3349363, EPI_ISL_3349364, EPI_ISL_3349365, EPI_ISL_3349366, EPI_ISL_3349367, EPI_ISL_3349368, EPI_ISL_3349369, EPI_ISL_3349370, EPI_ISL_3349371, EPI_ISL_3349372, EPI_ISL_3349373, EPI_ISL_3349374, EPI_ISL_3349375, EPI_ISL_3349376, EPI_ISL_3349377, EPI_ISL_3349378, EPI_ISL_3349379,
EPI_ISL_3349380, EPI_ISL_3349381, EPI_ISL_3349382, EPI_ISL_3349383, EPI_ISL_3349384, EPI_ISL_3349385, EPI_ISL_3349386, EPI_ISL_3349387, EPI_ISL_3349388, EPI_ISL_3349389, EPI_ISL_3349390, EPI_ISL_3349391, EPI_ISL_3349392, EPI_ISL_3349393, EPI_ISL_3349394, EPI_ISL_3349395, EPI_ISL_3349396, EPI_ISL_3349397, EPI_ISL_3349398, EPI_ISL_3349399, EPI_ISL_3349400, EPI_ISL_3349401,
EPI_ISL_3349402, EPI_ISL_3349403, EPI_ISL_3349404, EPI_ISL_3349405, EPI_ISL_3349406, EPI_ISL_3349407, EPI_ISL_3349408, EPI_ISL_3349410, EPI_ISL_3349411, EPI_ISL_3349412, EPI_ISL_3349413, EPI_ISL_3349414, EPI_ISL_3349415, EPI_ISL_3349416, EPI_ISL_3349417, EPI_ISL_3349418, EPI_ISL_3349420, EPI_ISL_3349421, EPI_ISL_3349422, EPI_ISL_3349423, EPI_ISL_3349424, EPI_ISL_3349425,
EPI_ISL_3349426, EPI_ISL_3349427, EPI_ISL_3349428, EPI_ISL_3349429, EPI_ISL_3349430, EPI_ISL_3349431, EPI_ISL_3349432, EPI_ISL_3349433, EPI_ISL_3349434, EPI_ISL_3349435, EPI_ISL_3349436, EPI_ISL_3349437, EPI_ISL_3349438, EPI_ISL_3349439, EPI_ISL_3349440, EPI_ISL_3349441, EPI_ISL_3349442, EPI_ISL_3349443, EPI_ISL_3349444, EPI_ISL_3349445, EPI_ISL_3349449, EPI_ISL_3349450,
EPI_ISL_3349451, EPI_ISL_3349452, EPI_ISL_3349453, EPI_ISL_3349454, EPI_ISL_3349455, EPI_ISL_3349456, EPI_ISL_3349457, EPI_ISL_3349459, EPI_ISL_3349461, EPI_ISL_3349462, EPI_ISL_3349463, EPI_ISL_3349464, EPI_ISL_3349465, EPI_ISL_3349466, EPI_ISL_3349467, EPI_ISL_3349468, EPI_ISL_3349469, EPI_ISL_3349470, EPI_ISL_3349471, EPI_ISL_3349472, EPI_ISL_3349473, EPI_ISL_3349474,
EPI_ISL_3349475, EPI_ISL_3349476, EPI_ISL_3349477, EPI_ISL_3349478, EPI_ISL_3349479, EPI_ISL_3349480, EPI_ISL_3349481, EPI_ISL_3349482, EPI_ISL_3349483, EPI_ISL_3349484, EPI_ISL_3349485, EPI_ISL_3349486, EPI_ISL_3349487, EPI_ISL_3349488, EPI_ISL_3349489, EPI_ISL_3349490, EPI_ISL_3349491, EPI_ISL_3349492, EPI_ISL_3349493, EPI_ISL_3349494, EPI_ISL_3349495, EPI_ISL_3349496,
EPI_ISL_3349497, EPI_ISL_3349498, EPI_ISL_3349499, EPI_ISL_3349500, EPI_ISL_3349501, EPI_ISL_3349502, EPI_ISL_3349503, EPI_ISL_3349504, EPI_ISL_3349505, EPI_ISL_3349506, EPI_ISL_3349507, EPI_ISL_3349509, EPI_ISL_3349510, EPI_ISL_3349511, EPI_ISL_3349512, EPI_ISL_3349513, EPI_ISL_3349514, EPI_ISL_3349515, EPI_ISL_3349516, EPI_ISL_3349517, EPI_ISL_3349518, EPI_ISL_3349519,
EPI_ISL_3349520, EPI_ISL_3349521, EPI_ISL_3349522, EPI_ISL_3349523, EPI_ISL_3349524, EPI_ISL_3349525, EPI_ISL_3349526, EPI_ISL_3349527, EPI_ISL_3349528, EPI_ISL_3349529, EPI_ISL_3349530, EPI_ISL_3349531, EPI_ISL_3349532, EPI_ISL_3349533, EPI_ISL_3349534, EPI_ISL_3349535, EPI_ISL_3349536, EPI_ISL_3349537, EPI_ISL_3349538, EPI_ISL_3349539, EPI_ISL_3349540, EPI_ISL_3349541,
EPI_ISL_3349542, EPI_ISL_3349543, EPI_ISL_3349544, EPI_ISL_3349545, EPI_ISL_3349546, EPI_ISL_3349547, EPI_ISL_3349548, EPI_ISL_3349549, EPI_ISL_3349550, EPI_ISL_3349551, EPI_ISL_3349552, EPI_ISL_3349553, EPI_ISL_3349554, EPI_ISL_3349555, EPI_ISL_3349556, EPI_ISL_3349557, EPI_ISL_3349558, EPI_ISL_3349559, EPI_ISL_3349560, EPI_ISL_3349561, EPI_ISL_3349562, EPI_ISL_3349563,
EPI_ISL_3349564, EPI_ISL_3349565, EPI_ISL_3349566, EPI_ISL_3349567, EPI_ISL_3349568, EPI_ISL_3349569, EPI_ISL_3349570, EPI_ISL_3349571, EPI_ISL_3349572, EPI_ISL_3349573, EPI_ISL_3349574, EPI_ISL_3349575, EPI_ISL_3349576, EPI_ISL_3349577, EPI_ISL_3349578, EPI_ISL_3349579, EPI_ISL_3349580, EPI_ISL_3349581, EPI_ISL_3349582, EPI_ISL_3349583, EPI_ISL_3349584, EPI_ISL_3349585,
EPI_ISL_3349586, EPI_ISL_3349587, EPI_ISL_3349588, EPI_ISL_3349589, EPI_ISL_3349591, EPI_ISL_3349592, EPI_ISL_3349593, EPI_ISL_3349594, EPI_ISL_3349595, EPI_ISL_3349596, EPI_ISL_3349597, EPI_ISL_3349598, EPI_ISL_3349599, EPI_ISL_3349600, EPI_ISL_3349601, EPI_ISL_3349602, EPI_ISL_3349603, EPI_ISL_3349604, EPI_ISL_3349605, EPI_ISL_3349606, EPI_ISL_3349607, EPI_ISL_3349608,
EPI_ISL_3349609, EPI_ISL_3349610, EPI_ISL_3349611, EPI_ISL_3349612, EPI_ISL_3349613, EPI_ISL_3349614, EPI_ISL_3349615, EPI_ISL_3349616, EPI_ISL_3349617, EPI_ISL_3349618, EPI_ISL_3349619, EPI_ISL_3349620, EPI_ISL_3349621, EPI_ISL_3349622, EPI_ISL_3349623, EPI_ISL_3349624, EPI_ISL_3349625, EPI_ISL_3349626, EPI_ISL_3349627, EPI_ISL_3349628, EPI_ISL_3349629, EPI_ISL_3349630,
EPI_ISL_3349631, EPI_ISL_3349632, EPI_ISL_3349633, EPI_ISL_3349634, EPI_ISL_3349635, EPI_ISL_3349636, EPI_ISL_3349637, EPI_ISL_3349638, EPI_ISL_3349639, EPI_ISL_3349640, EPI_ISL_3349641, EPI_ISL_3349642, EPI_ISL_3349643, EPI_ISL_3349644, EPI_ISL_3349645, EPI_ISL_3349646, EPI_ISL_3349647, EPI_ISL_3349648, EPI_ISL_3349649, EPI_ISL_3349650, EPI_ISL_3349651, EPI_ISL_3349652,
EPI_ISL_3349653, EPI_ISL_3349654, EPI_ISL_3349655, EPI_ISL_3349656, EPI_ISL_3349657, EPI_ISL_3349658, EPI_ISL_3349659, EPI_ISL_3349660, EPI_ISL_3349661, EPI_ISL_3349662, EPI_ISL_3349663, EPI_ISL_3349664, EPI_ISL_3349665, EPI_ISL_3349666, EPI_ISL_3349667, EPI_ISL_3349668, EPI_ISL_3349669, EPI_ISL_3349670, EPI_ISL_3349671, EPI_ISL_3349672, EPI_ISL_3349673, EPI_ISL_3349674,
EPI_ISL_3349675, EPI_ISL_3349676, EPI_ISL_3349677, EPI_ISL_3349678, EPI_ISL_3349679, EPI_ISL_3349680, EPI_ISL_3349681, EPI_ISL_3349682, EPI_ISL_3349683, EPI_ISL_3349684, EPI_ISL_3349685, EPI_ISL_3349686, EPI_ISL_3349687, EPI_ISL_3349688, EPI_ISL_3349689, EPI_ISL_3349690, EPI_ISL_3349691, EPI_ISL_3349692, EPI_ISL_3349693, EPI_ISL_3349694, EPI_ISL_3349695, EPI_ISL_3349696,
EPI_ISL_3349697, EPI_ISL_3349698, EPI_ISL_3349699, EPI_ISL_3349700, EPI_ISL_3349701, EPI_ISL_3349702, EPI_ISL_3349703, EPI_ISL_3349704, EPI_ISL_3349705, EPI_ISL_3349706, EPI_ISL_3349707, EPI_ISL_3349708, EPI_ISL_3349709, EPI_ISL_3349710, EPI_ISL_3349711, EPI_ISL_3349712, EPI_ISL_3349713, EPI_ISL_3349714, EPI_ISL_3349715, EPI_ISL_3349717, EPI_ISL_3349718, EPI_ISL_3349719,
EPI_ISL_3349720, EPI_ISL_3349721, EPI_ISL_3349722, EPI_ISL_3349723, EPI_ISL_3349724, EPI_ISL_3349725, EPI_ISL_3349726, EPI_ISL_3349727, EPI_ISL_3349728, EPI_ISL_3349729, EPI_ISL_3349730, EPI_ISL_3349731, EPI_ISL_3349732, EPI_ISL_3349733, EPI_ISL_3349734, EPI_ISL_3349735, EPI_ISL_3349736, EPI_ISL_3349737, EPI_ISL_3349738, EPI_ISL_3349739, EPI_ISL_3349740, EPI_ISL_3349741,
EPI_ISL_3349742, EPI_ISL_3349743, EPI_ISL_3349744, EPI_ISL_3349745, EPI_ISL_3349746, EPI_ISL_3349747, EPI_ISL_3349748, EPI_ISL_3349749, EPI_ISL_3349750, EPI_ISL_3349751, EPI_ISL_3349752, EPI_ISL_3349753, EPI_ISL_3349754, EPI_ISL_3349755, EPI_ISL_3349756, EPI_ISL_3349757, EPI_ISL_3349758, EPI_ISL_3349759, EPI_ISL_3349760, EPI_ISL_3349761, EPI_ISL_3349762, EPI_ISL_3349763,
EPI_ISL_3349764, EPI_ISL_3349765, EPI_ISL_3349766, EPI_ISL_3349767, EPI_ISL_3349768, EPI_ISL_3349769, EPI_ISL_3349770, EPI_ISL_3349771, EPI_ISL_3349772, EPI_ISL_3349773, EPI_ISL_3349774, EPI_ISL_3349775, EPI_ISL_3349776, EPI_ISL_3349777, EPI_ISL_3349778, EPI_ISL_3349779, EPI_ISL_3349780, EPI_ISL_3349781, EPI_ISL_3349782, EPI_ISL_3349783, EPI_ISL_3349784, EPI_ISL_3349785,
EPI_ISL_3349786, EPI_ISL_3349787, EPI_ISL_3349788, EPI_ISL_3349789, EPI_ISL_3349790, EPI_ISL_3349791, EPI_ISL_3349792, EPI_ISL_3349793, EPI_ISL_3349794, EPI_ISL_3349795, EPI_ISL_3349796, EPI_ISL_3349797, EPI_ISL_3349798, EPI_ISL_3349799, EPI_ISL_3349800, EPI_ISL_3349801, EPI_ISL_3349802, EPI_ISL_3349803, EPI_ISL_3349804, EPI_ISL_3349805, EPI_ISL_3349806, EPI_ISL_3349807,
EPI_ISL_3349808, EPI_ISL_3349809, EPI_ISL_3349810, EPI_ISL_3349811, EPI_ISL_3349812, EPI_ISL_3349813, EPI_ISL_3349814, EPI_ISL_3349815, EPI_ISL_3349816, EPI_ISL_3349817, EPI_ISL_3349818, EPI_ISL_3349819, EPI_ISL_3349820, EPI_ISL_3349821, EPI_ISL_3349822, EPI_ISL_3349823, EPI_ISL_3349824, EPI_ISL_3349825, EPI_ISL_3349826, EPI_ISL_3349827, EPI_ISL_3349828, EPI_ISL_3349829,
EPI_ISL_3349830, EPI_ISL_3349831, EPI_ISL_3349832, EPI_ISL_3349833, EPI_ISL_3349835, EPI_ISL_3349836, EPI_ISL_3349837, EPI_ISL_3349838, EPI_ISL_3349839, EPI_ISL_3349840, EPI_ISL_3349841, EPI_ISL_3349842, EPI_ISL_3349843, EPI_ISL_3349844, EPI_ISL_3349846, EPI_ISL_3349847, EPI_ISL_3349848, EPI_ISL_3349849, EPI_ISL_3349850, EPI_ISL_3349851, EPI_ISL_3349852, EPI_ISL_3349853,
EPI_ISL_3349854, EPI_ISL_3349855, EPI_ISL_3349856, EPI_ISL_3349857, EPI_ISL_3349858, EPI_ISL_3349859, EPI_ISL_3349860, EPI_ISL_3349861, EPI_ISL_3349862, EPI_ISL_3349863, EPI_ISL_3349864, EPI_ISL_3349865, EPI_ISL_3349866, EPI_ISL_3349867, EPI_ISL_3349868, EPI_ISL_3349869, EPI_ISL_3349870, EPI_ISL_3349871, EPI_ISL_3349872, EPI_ISL_3349873, EPI_ISL_3349874, EPI_ISL_3349875,
EPI_ISL_3349876, EPI_ISL_3349877, EPI_ISL_3349878, EPI_ISL_3349879, EPI_ISL_3349880, EPI_ISL_3349881, EPI_ISL_3349882, EPI_ISL_3349883, EPI_ISL_3349884, EPI_ISL_3349885, EPI_ISL_3349886, EPI_ISL_3349887, EPI_ISL_3349888, EPI_ISL_3349889, EPI_ISL_3349890, EPI_ISL_3349891, EPI_ISL_3349892, EPI_ISL_3349893, EPI_ISL_3349894, EPI_ISL_3349895, EPI_ISL_3349896, EPI_ISL_3349897,
EPI_ISL_3349898, EPI_ISL_3349899, EPI_ISL_3349900, EPI_ISL_3349901, EPI_ISL_3349902, EPI_ISL_3349903, EPI_ISL_3349904, EPI_ISL_3349905, EPI_ISL_3349906, EPI_ISL_3349907, EPI_ISL_3349908, EPI_ISL_3349909, EPI_ISL_3349910, EPI_ISL_3349911, EPI_ISL_3349912, EPI_ISL_3349913, EPI_ISL_3349915, EPI_ISL_3349916, EPI_ISL_3349917, EPI_ISL_3349918, EPI_ISL_3349919, EPI_ISL_3349920,
EPI_ISL_3349921, EPI_ISL_3349922, EPI_ISL_3349923, EPI_ISL_3349924, EPI_ISL_3349925, EPI_ISL_3349926, EPI_ISL_3349927, EPI_ISL_3349928, EPI_ISL_3349929, EPI_ISL_3349930, EPI_ISL_3349931, EPI_ISL_3349932, EPI_ISL_3349933, EPI_ISL_3349934, EPI_ISL_3349935, EPI_ISL_3349936, EPI_ISL_3349937, EPI_ISL_3349938, EPI_ISL_3349939, EPI_ISL_3349940, EPI_ISL_3349941, EPI_ISL_3349942,
EPI_ISL_3349943, EPI_ISL_3349944, EPI_ISL_3349945, EPI_ISL_3349946, EPI_ISL_3349947, EPI_ISL_3349948, EPI_ISL_3349949, EPI_ISL_3349950, EPI_ISL_3349951, EPI_ISL_3349952, EPI_ISL_3349953, EPI_ISL_3349954, EPI_ISL_3349955, EPI_ISL_3349956, EPI_ISL_3349957, EPI_ISL_3349958, EPI_ISL_3349959, EPI_ISL_3349960, EPI_ISL_3349961, EPI_ISL_3349962, EPI_ISL_3349963, EPI_ISL_3349964,
EPI_ISL_3349965, EPI_ISL_3349966, EPI_ISL_3349967, EPI_ISL_3349968, EPI_ISL_3349969, EPI_ISL_3349970, EPI_ISL_3349971, EPI_ISL_3349972, EPI_ISL_3349973, EPI_ISL_3349974, EPI_ISL_3349975, EPI_ISL_3349976, EPI_ISL_3349977, EPI_ISL_3349978, EPI_ISL_3349979, EPI_ISL_3349980, EPI_ISL_3349981, EPI_ISL_3349982, EPI_ISL_3349983, EPI_ISL_3349984, EPI_ISL_3349985, EPI_ISL_3349986,
EPI_ISL_3349987, EPI_ISL_3349988, EPI_ISL_3349989, EPI_ISL_3349990, EPI_ISL_3349991, EPI_ISL_3349992, EPI_ISL_3349993, EPI_ISL_3349994, EPI_ISL_3349995, EPI_ISL_3349996, EPI_ISL_3349997, EPI_ISL_3349998, EPI_ISL_3349999, EPI_ISL_3350000, EPI_ISL_3350001, EPI_ISL_3350002, EPI_ISL_3350003, EPI_ISL_3350004, EPI_ISL_3350005, EPI_ISL_3350006, EPI_ISL_3350007, EPI_ISL_3350008,
EPI_ISL_3350009, EPI_ISL_3350010, EPI_ISL_3350011, EPI_ISL_3350012, EPI_ISL_3350013, EPI_ISL_3350014, EPI_ISL_3350015, EPI_ISL_3350016, EPI_ISL_3350017, EPI_ISL_3350018, EPI_ISL_3350019, EPI_ISL_3350020, EPI_ISL_3350021, EPI_ISL_3350022, EPI_ISL_3350023, EPI_ISL_3350024, EPI_ISL_3350025, EPI_ISL_3350026, EPI_ISL_3350027, EPI_ISL_3350028, EPI_ISL_3350029, EPI_ISL_3350030,
EPI_ISL_3350031, EPI_ISL_3350032, EPI_ISL_3350033, EPI_ISL_3350034, EPI_ISL_3350035, EPI_ISL_3350036, EPI_ISL_3350037, EPI_ISL_3350038, EPI_ISL_3350039, EPI_ISL_3350040, EPI_ISL_3350041, EPI_ISL_3350042, EPI_ISL_3350043, EPI_ISL_3350044, EPI_ISL_3350045, EPI_ISL_3350046, EPI_ISL_3350047, EPI_ISL_3350048, EPI_ISL_3350049, EPI_ISL_3350050, EPI_ISL_3350051, EPI_ISL_3350052,
EPI_ISL_3350053, EPI_ISL_3350054, EPI_ISL_3350055, EPI_ISL_3350056, EPI_ISL_3350057, EPI_ISL_3350058, EPI_ISL_3350059, EPI_ISL_3350060, EPI_ISL_3350061, EPI_ISL_3350062, EPI_ISL_3350063, EPI_ISL_3350064, EPI_ISL_3350065, EPI_ISL_3350066, EPI_ISL_3350067, EPI_ISL_3350068, EPI_ISL_3350069, EPI_ISL_3350070, EPI_ISL_3350071, EPI_ISL_3350072, EPI_ISL_3350073, EPI_ISL_3350074,
EPI_ISL_3350075, EPI_ISL_3350076, EPI_ISL_3350077, EPI_ISL_3350078, EPI_ISL_3350079, EPI_ISL_3350080, EPI_ISL_3350081, EPI_ISL_3350082, EPI_ISL_3350083, EPI_ISL_3350084, EPI_ISL_3350085, EPI_ISL_3350086, EPI_ISL_3350087, EPI_ISL_3350088, EPI_ISL_3350089, EPI_ISL_3350090, EPI_ISL_3350091, EPI_ISL_3350092, EPI_ISL_3350093, EPI_ISL_3350094, EPI_ISL_3350095, EPI_ISL_3350096,
EPI_ISL_3350097, EPI_ISL_3350098, EPI_ISL_3350099, EPI_ISL_3350100, EPI_ISL_3350101, EPI_ISL_3350102, EPI_ISL_3350103, EPI_ISL_3350104, EPI_ISL_3350105, EPI_ISL_3350106, EPI_ISL_3350107, EPI_ISL_3350108, EPI_ISL_3350109, EPI_ISL_3350110, EPI_ISL_3350111, EPI_ISL_3350112, EPI_ISL_3350113, EPI_ISL_3350114, EPI_ISL_3350115, EPI_ISL_3350116, EPI_ISL_3350117, EPI_ISL_3350118,
EPI_ISL_3350119, EPI_ISL_3350120, EPI_ISL_3350121, EPI_ISL_3350122, EPI_ISL_3350123, EPI_ISL_3350124, EPI_ISL_3350125, EPI_ISL_3350126, EPI_ISL_3350127, EPI_ISL_3350128, EPI_ISL_3350129, EPI_ISL_3350130, EPI_ISL_3350131, EPI_ISL_3350132, EPI_ISL_3350133, EPI_ISL_3350134, EPI_ISL_3350135, EPI_ISL_3350136, EPI_ISL_3350137, EPI_ISL_3350138, EPI_ISL_3350139, EPI_ISL_3350140,
EPI_ISL_3350141, EPI_ISL_3350142, EPI_ISL_3350143, EPI_ISL_3350144, EPI_ISL_3350145, EPI_ISL_3350146, EPI_ISL_3350147, EPI_ISL_3350148, EPI_ISL_3350149, EPI_ISL_3350150, EPI_ISL_3350151, EPI_ISL_3350152, EPI_ISL_3350153, EPI_ISL_3350154, EPI_ISL_3350155, EPI_ISL_3350156, EPI_ISL_3350157, EPI_ISL_3350158, EPI_ISL_3350159, EPI_ISL_3350160, EPI_ISL_3350161, EPI_ISL_3350162,
EPI_ISL_3350163, EPI_ISL_3350164, EPI_ISL_3350165, EPI_ISL_3350166, EPI_ISL_3350167, EPI_ISL_3350168, EPI_ISL_3350169, EPI_ISL_3350170, EPI_ISL_3350171, EPI_ISL_3350172, EPI_ISL_3350173, EPI_ISL_3350174, EPI_ISL_3350175, EPI_ISL_3350176, EPI_ISL_3350177, EPI_ISL_3350178, EPI_ISL_3350179, EPI_ISL_3350180, EPI_ISL_3350181, EPI_ISL_3350182, EPI_ISL_3350183, EPI_ISL_3350184,
EPI_ISL_3350185, EPI_ISL_3350186, EPI_ISL_3350187, EPI_ISL_3350188, EPI_ISL_3350189, EPI_ISL_3350190, EPI_ISL_3350191, EPI_ISL_3350192, EPI_ISL_3350193, EPI_ISL_3350194, EPI_ISL_3350195, EPI_ISL_3350196, EPI_ISL_3350197, EPI_ISL_3350198, EPI_ISL_3350199, EPI_ISL_3350200, EPI_ISL_3350201, EPI_ISL_3350202, EPI_ISL_3350203, EPI_ISL_3350204, EPI_ISL_3350205, EPI_ISL_3350206,
EPI_ISL_3350207, EPI_ISL_3350208, EPI_ISL_3350209, EPI_ISL_3350210, EPI_ISL_3350211, EPI_ISL_3350212, EPI_ISL_3350213, EPI_ISL_3350214, EPI_ISL_3350215, EPI_ISL_3350216, EPI_ISL_3350217, EPI_ISL_3350218, EPI_ISL_3350219, EPI_ISL_3350220, EPI_ISL_3350221, EPI_ISL_3350222, EPI_ISL_3350223, EPI_ISL_3350224, EPI_ISL_3350225, EPI_ISL_3350226, EPI_ISL_3350227, EPI_ISL_3350228,
EPI_ISL_3350229, EPI_ISL_3350230, EPI_ISL_3350231, EPI_ISL_3350232, EPI_ISL_3350233, EPI_ISL_3350234, EPI_ISL_3350235, EPI_ISL_3350236, EPI_ISL_3350237, EPI_ISL_3350238, EPI_ISL_3350239, EPI_ISL_3350240, EPI_ISL_3350241, EPI_ISL_3350242, EPI_ISL_3350243, EPI_ISL_3350244, EPI_ISL_3350245, EPI_ISL_3350246, EPI_ISL_3350247, EPI_ISL_3350248, EPI_ISL_3350249, EPI_ISL_3350250,
EPI_ISL_3350251, EPI_ISL_3350252, EPI_ISL_3350254, EPI_ISL_3350255, EPI_ISL_3350256, EPI_ISL_3350257, EPI_ISL_3350258, EPI_ISL_3350259, EPI_ISL_3350260, EPI_ISL_3350261, EPI_ISL_3350262, EPI_ISL_3350263, EPI_ISL_3350264, EPI_ISL_3350265, EPI_ISL_3350266, EPI_ISL_3350267, EPI_ISL_3350268, EPI_ISL_3350269, EPI_ISL_3350270, EPI_ISL_3350271, EPI_ISL_3350272, EPI_ISL_3350273,
EPI_ISL_3350274, EPI_ISL_3350275, EPI_ISL_3350276, EPI_ISL_3350277, EPI_ISL_3350278, EPI_ISL_3350279, EPI_ISL_3350281, EPI_ISL_3350282, EPI_ISL_3350283, EPI_ISL_3350284, EPI_ISL_3350285, EPI_ISL_3350286, EPI_ISL_3350287, EPI_ISL_3350288, EPI_ISL_3350289, EPI_ISL_3350291, EPI_ISL_3350292, EPI_ISL_3350293, EPI_ISL_3350294, EPI_ISL_3350295, EPI_ISL_3350296, EPI_ISL_3350297,
EPI_ISL_3350298, EPI_ISL_3350299, EPI_ISL_3350300, EPI_ISL_3350301, EPI_ISL_3350302, EPI_ISL_3350303, EPI_ISL_3350304, EPI_ISL_3350305, EPI_ISL_3350306, EPI_ISL_3350307, EPI_ISL_3350308, EPI_ISL_3350309, EPI_ISL_3350310, EPI_ISL_3350311, EPI_ISL_3350312, EPI_ISL_3350313, EPI_ISL_3350314, EPI_ISL_3350315, EPI_ISL_3350316, EPI_ISL_3350317, EPI_ISL_3350318, EPI_ISL_3350319,
EPI_ISL_3350320, EPI_ISL_3350321, EPI_ISL_3350322, EPI_ISL_3350323, EPI_ISL_3350324, EPI_ISL_3350325, EPI_ISL_3350326, EPI_ISL_3350327, EPI_ISL_3350328, EPI_ISL_3350329, EPI_ISL_3350330, EPI_ISL_3350331, EPI_ISL_3350332, EPI_ISL_3350333, EPI_ISL_3350334, EPI_ISL_3350335, EPI_ISL_3350336, EPI_ISL_3350337, EPI_ISL_3350338, EPI_ISL_3350339, EPI_ISL_3350340, EPI_ISL_3350341,
EPI_ISL_3350342, EPI_ISL_3350343, EPI_ISL_3350344, EPI_ISL_3350345, EPI_ISL_3350346, EPI_ISL_3350347, EPI_ISL_3350348, EPI_ISL_3350349, EPI_ISL_3350350, EPI_ISL_3350351, EPI_ISL_3350352, EPI_ISL_3350353, EPI_ISL_3350354, EPI_ISL_3350355, EPI_ISL_3350356, EPI_ISL_3350357, EPI_ISL_3350358, EPI_ISL_3350359, EPI_ISL_3350360, EPI_ISL_3350361, EPI_ISL_3350362, EPI_ISL_3350363,
EPI_ISL_3350364, EPI_ISL_3350365, EPI_ISL_3350366, EPI_ISL_3350367, EPI_ISL_3350368, EPI_ISL_3350369, EPI_ISL_3350370, EPI_ISL_3350371, EPI_ISL_3350372, EPI_ISL_3350373, EPI_ISL_3350374, EPI_ISL_3350375, EPI_ISL_3350376, EPI_ISL_3350377, EPI_ISL_3350378, EPI_ISL_3350379, EPI_ISL_3350380, EPI_ISL_3350381, EPI_ISL_3350382, EPI_ISL_3350383, EPI_ISL_3350384, EPI_ISL_3350385,
EPI_ISL_3350386, EPI_ISL_3350387, EPI_ISL_3350388, EPI_ISL_3350389, EPI_ISL_3350390, EPI_ISL_3350391, EPI_ISL_3350392, EPI_ISL_3350393, EPI_ISL_3350394, EPI_ISL_3350395, EPI_ISL_3350396, EPI_ISL_3350397, EPI_ISL_3350398, EPI_ISL_3350399, EPI_ISL_3350400, EPI_ISL_3350401, EPI_ISL_3350402, EPI_ISL_3350403, EPI_ISL_3350404, EPI_ISL_3350405, EPI_ISL_3350406, EPI_ISL_3350407,
EPI_ISL_3350408, EPI_ISL_3350409, EPI_ISL_3350410, EPI_ISL_3350411, EPI_ISL_3350412, EPI_ISL_3350413, EPI_ISL_3350414, EPI_ISL_3350415, EPI_ISL_3350416, EPI_ISL_3350417, EPI_ISL_3350418, EPI_ISL_3350419, EPI_ISL_3350420, EPI_ISL_3350421, EPI_ISL_3350422, EPI_ISL_3350423, EPI_ISL_3350424, EPI_ISL_3350425, EPI_ISL_3350427, EPI_ISL_3350428, EPI_ISL_3350429, EPI_ISL_3350430,
EPI_ISL_3350431, EPI_ISL_3350432, EPI_ISL_3350433, EPI_ISL_3350434, EPI_ISL_3350435, EPI_ISL_3350436, EPI_ISL_3350437, EPI_ISL_3350438, EPI_ISL_3350439, EPI_ISL_3350440, EPI_ISL_3350441, EPI_ISL_3350442, EPI_ISL_3350443, EPI_ISL_3350444, EPI_ISL_3350445, EPI_ISL_3350446, EPI_ISL_3350447, EPI_ISL_3350448, EPI_ISL_3350449, EPI_ISL_3350450, EPI_ISL_3350451, EPI_ISL_3350452,
EPI_ISL_3350453, EPI_ISL_3350455, EPI_ISL_3350456, EPI_ISL_3350457, EPI_ISL_3350458, EPI_ISL_3350459, EPI_ISL_3350460, EPI_ISL_3350461, EPI_ISL_3350462, EPI_ISL_3350463, EPI_ISL_3350464, EPI_ISL_3350465, EPI_ISL_3350466, EPI_ISL_3350467, EPI_ISL_3350468, EPI_ISL_3350469, EPI_ISL_3350470, EPI_ISL_3350471, EPI_ISL_3350472, EPI_ISL_3350473, EPI_ISL_3350474, EPI_ISL_3350475,
EPI_ISL_3350476, EPI_ISL_3350477, EPI_ISL_3350478, EPI_ISL_3350479, EPI_ISL_3350481, EPI_ISL_3350482, EPI_ISL_3350483, EPI_ISL_3350484, EPI_ISL_3350485, EPI_ISL_3350486, EPI_ISL_3350487, EPI_ISL_3350488, EPI_ISL_3350489, EPI_ISL_3350490, EPI_ISL_3350491, EPI_ISL_3350492, EPI_ISL_3350493, EPI_ISL_3350494, EPI_ISL_3350495, EPI_ISL_3350496, EPI_ISL_3350497, EPI_ISL_3350498,
EPI_ISL_3350499, EPI_ISL_3350500, EPI_ISL_3350501, EPI_ISL_3350502, EPI_ISL_3350503, EPI_ISL_3350504, EPI_ISL_3350505, EPI_ISL_3350506, EPI_ISL_3350507, EPI_ISL_3350508, EPI_ISL_3350509, EPI_ISL_3350510, EPI_ISL_3350511, EPI_ISL_3350512, EPI_ISL_3350513, EPI_ISL_3350514, EPI_ISL_3350515, EPI_ISL_3350516, EPI_ISL_3350517, EPI_ISL_3350518, EPI_ISL_3350519, EPI_ISL_3350521,
EPI_ISL_3350522, EPI_ISL_3350523, EPI_ISL_3350524, EPI_ISL_3350525, EPI_ISL_3350526, EPI_ISL_3350527, EPI_ISL_3350528, EPI_ISL_3350529, EPI_ISL_3350530, EPI_ISL_3350531, EPI_ISL_3350532, EPI_ISL_3350533, EPI_ISL_3350534, EPI_ISL_3350535, EPI_ISL_3350536, EPI_ISL_3350537, EPI_ISL_3350538, EPI_ISL_3350539, EPI_ISL_3350540, EPI_ISL_3350541, EPI_ISL_3350542, EPI_ISL_3350543,
EPI_ISL_3350544, EPI_ISL_3350545, EPI_ISL_3350546, EPI_ISL_3350547, EPI_ISL_3350548, EPI_ISL_3350549, EPI_ISL_3350550, EPI_ISL_3350551, EPI_ISL_3350553, EPI_ISL_3350554, EPI_ISL_3350555, EPI_ISL_3350556, EPI_ISL_3350557, EPI_ISL_3350558, EPI_ISL_3350559, EPI_ISL_3350560, EPI_ISL_3350561, EPI_ISL_3350562, EPI_ISL_3350563, EPI_ISL_3350564, EPI_ISL_3350565, EPI_ISL_3350566,

| | | | |
|---|---|---|---|
| EPI_ISL_3350567, EPI_ISL_3350568, EPI_ISL_3350569, EPI_ISL_3350570, EPI_ISL_3350571, EPI_ISL_3350572, EPI_ISL_3350573, EPI_ISL_3350574, EPI_ISL_3350575, EPI_ISL_3350576, EPI_ISL_3350577, EPI_ISL_3350578, EPI_ISL_3350580, EPI_ISL_3350581, EPI_ISL_3350582, EPI_ISL_3350583, EPI_ISL_3350584, EPI_ISL_3350585, EPI_ISL_3350586, EPI_ISL_3350587, EPI_ISL_3350588, EPI_ISL_3350589, EPI_ISL_3350590, EPI_ISL_3350591, EPI_ISL_3350592, EPI_ISL_3350593, EPI_ISL_3350594, EPI_ISL_3350595, EPI_ISL_3350596, EPI_ISL_3350597, EPI_ISL_3350598, EPI_ISL_3350599, EPI_ISL_3350600, EPI_ISL_3350601, EPI_ISL_3350602, EPI_ISL_3350603, EPI_ISL_3350604, EPI_ISL_3350605, EPI_ISL_3350606, EPI_ISL_3350607, EPI_ISL_3350608, EPI_ISL_3350609, EPI_ISL_3350610, EPI_ISL_3350611, EPI_ISL_3350612, EPI_ISL_3350613, EPI_ISL_3350614, EPI_ISL_3350615, EPI_ISL_3350616, EPI_ISL_3350617, EPI_ISL_3350618, EPI_ISL_3350619, EPI_ISL_3350620, EPI_ISL_3350621, EPI_ISL_3350622, EPI_ISL_3350623, EPI_ISL_3350624, EPI_ISL_3350625, EPI_ISL_3350626, EPI_ISL_3350627, EPI_ISL_3350628, EPI_ISL_3350629, EPI_ISL_3350630, EPI_ISL_3350631, EPI_ISL_3350632, EPI_ISL_3350633, EPI_ISL_3350634, EPI_ISL_3350635, EPI_ISL_3350636, EPI_ISL_3350637, EPI_ISL_3350638, EPI_ISL_3350639, EPI_ISL_3350640, EPI_ISL_3350641, EPI_ISL_3350642, EPI_ISL_3350643, EPI_ISL_3350644, EPI_ISL_3350645, EPI_ISL_3350646, EPI_ISL_3350647, EPI_ISL_3350648, EPI_ISL_3350649, EPI_ISL_3350650, EPI_ISL_3350651, EPI_ISL_3350652, EPI_ISL_3350653, EPI_ISL_3350654, EPI_ISL_3350655, EPI_ISL_3350656, EPI_ISL_3350657, EPI_ISL_3350658, EPI_ISL_3350659, EPI_ISL_3350660, EPI_ISL_3350662, EPI_ISL_3350663, EPI_ISL_3350664, EPI_ISL_3350665, EPI_ISL_3350666, EPI_ISL_3350667, EPI_ISL_3350668, EPI_ISL_3350669, EPI_ISL_3350670, EPI_ISL_3350671, EPI_ISL_3350672, EPI_ISL_3350673, EPI_ISL_3350674, EPI_ISL_3350675, EPI_ISL_3350676, EPI_ISL_3350677, EPI_ISL_3350678, EPI_ISL_3350679, EPI_ISL_3350680, EPI_ISL_3350681, EPI_ISL_3350682, EPI_ISL_3350683, EPI_ISL_3350684, EPI_ISL_3350685, EPI_ISL_3350686, EPI_ISL_3350687, EPI_ISL_3350688, EPI_ISL_3350689, EPI_ISL_3350690, EPI_ISL_3350691, EPI_ISL_3350692, EPI_ISL_3350693, EPI_ISL_3350694, EPI_ISL_3350695, EPI_ISL_3350696, EPI_ISL_3350697, EPI_ISL_3350698, EPI_ISL_3350699, EPI_ISL_3350700, EPI_ISL_3350701, EPI_ISL_3350702, EPI_ISL_3350703, EPI_ISL_3350704, EPI_ISL_3350705, EPI_ISL_3350706, EPI_ISL_3350707, EPI_ISL_3350708, EPI_ISL_3350709, EPI_ISL_3350710, EPI_ISL_3350711, EPI_ISL_3350712, EPI_ISL_3350713, EPI_ISL_3350714, EPI_ISL_3350715, EPI_ISL_3350716, EPI_ISL_3350717, EPI_ISL_3350718, EPI_ISL_3350719, EPI_ISL_3350720, EPI_ISL_3350721, EPI_ISL_3350722, EPI_ISL_3350723, EPI_ISL_3350724, EPI_ISL_3350725, EPI_ISL_3350726, EPI_ISL_3350727, EPI_ISL_3350728, EPI_ISL_3350729, EPI_ISL_3350730, EPI_ISL_3350731, EPI_ISL_3350732, EPI_ISL_3350733, EPI_ISL_3350734, EPI_ISL_3350735, EPI_ISL_3350736, EPI_ISL_3350737, EPI_ISL_3350738, EPI_ISL_3350739, EPI_ISL_3350740, EPI_ISL_3350741, EPI_ISL_3350742, EPI_ISL_3350743, EPI_ISL_3350744, EPI_ISL_3350745, EPI_ISL_3350746, EPI_ISL_3350747, EPI_ISL_3350748, EPI_ISL_3350749, EPI_ISL_3350750, EPI_ISL_3350751, EPI_ISL_3350752, EPI_ISL_3350753, EPI_ISL_3350754, EPI_ISL_3350755, EPI_ISL_3350756, EPI_ISL_3350757, EPI_ISL_3350758, EPI_ISL_3350759, EPI_ISL_3350760, EPI_ISL_3350761, EPI_ISL_3350762, EPI_ISL_3350763, EPI_ISL_3350764, EPI_ISL_3350765, EPI_ISL_3350766, EPI_ISL_3350767, EPI_ISL_3350768, EPI_ISL_3350769, EPI_ISL_3350770, EPI_ISL_3350771, EPI_ISL_3350772, EPI_ISL_3350773, EPI_ISL_3350774, EPI_ISL_3350775, EPI_ISL_3350776, EPI_ISL_3350777, EPI_ISL_3350778, EPI_ISL_3350779, EPI_ISL_3350780, EPI_ISL_3350781, EPI_ISL_3350782, EPI_ISL_3350783, EPI_ISL_3350784, EPI_ISL_3350785, EPI_ISL_3350786, EPI_ISL_3350787, EPI_ISL_3350788, EPI_ISL_3350789, EPI_ISL_3350790, EPI_ISL_3350791, EPI_ISL_3350792, EPI_ISL_3350793, EPI_ISL_3350794, EPI_ISL_3350795, EPI_ISL_3350796, EPI_ISL_3350797, EPI_ISL_3350798, EPI_ISL_3350799, EPI_ISL_3350800, EPI_ISL_3350801, EPI_ISL_3350802, EPI_ISL_3350803, EPI_ISL_3350804, EPI_ISL_3350805, EPI_ISL_3350806, EPI_ISL_3350807, EPI_ISL_3350808, EPI_ISL_3350809, EPI_ISL_3350810, EPI_ISL_3350811, EPI_ISL_3350812, EPI_ISL_3350813, EPI_ISL_3350814, EPI_ISL_3350815, EPI_ISL_3350816, EPI_ISL_3350817, EPI_ISL_3350818, EPI_ISL_3350819, EPI_ISL_3350820, EPI_ISL_3350821, EPI_ISL_3350822, EPI_ISL_3350823, EPI_ISL_3350824, EPI_ISL_3350825, EPI_ISL_3350826, EPI_ISL_3350827, EPI_ISL_3350828, EPI_ISL_3350829, EPI_ISL_3350830, EPI_ISL_3350831, EPI_ISL_3350832, EPI_ISL_3350833, EPI_ISL_3350834, EPI_ISL_3350835, EPI_ISL_3350836, EPI_ISL_3350837, EPI_ISL_3350838, EPI_ISL_3350839, EPI_ISL_3350840, EPI_ISL_3350841, EPI_ISL_3350842, EPI_ISL_3350843, EPI_ISL_3350844, EPI_ISL_3350845, EPI_ISL_3350846, EPI_ISL_3350847, EPI_ISL_3350848, EPI_ISL_3350849, EPI_ISL_3350850, EPI_ISL_3350851, EPI_ISL_3350852, EPI_ISL_3350853, EPI_ISL_3350854, EPI_ISL_3350855, EPI_ISL_3350856, EPI_ISL_3350857, EPI_ISL_3350858, EPI_ISL_3350859, EPI_ISL_3350860, EPI_ISL_3350861, EPI_ISL_3350863, EPI_ISL_3350864, EPI_ISL_3350865, EPI_ISL_3350866, EPI_ISL_3350867, EPI_ISL_3350868, EPI_ISL_3350869, EPI_ISL_3350870, EPI_ISL_3350871, EPI_ISL_3350872, EPI_ISL_3350873, EPI_ISL_3350874, EPI_ISL_3350875, EPI_ISL_3350876, EPI_ISL_3350877, EPI_ISL_3350878, EPI_ISL_3350879, EPI_ISL_3350880, EPI_ISL_3350881, EPI_ISL_3350882, EPI_ISL_3350883, EPI_ISL_3350884, EPI_ISL_3350885, EPI_ISL_3350886, EPI_ISL_3350887, EPI_ISL_3350888, EPI_ISL_3350889, EPI_ISL_3350890, EPI_ISL_3350891, EPI_ISL_3350892, EPI_ISL_3350893, EPI_ISL_3350894, EPI_ISL_3350895, EPI_ISL_3350896, EPI_ISL_3350897, EPI_ISL_3350898, EPI_ISL_3350899, EPI_ISL_3350900, EPI_ISL_3350901, EPI_ISL_3350902, EPI_ISL_3350903, EPI_ISL_3350904, EPI_ISL_3350905, EPI_ISL_3350906, EPI_ISL_3350907, EPI_ISL_3350908, EPI_ISL_3350909, EPI_ISL_3350910, EPI_ISL_3350911, EPI_ISL_3350912, EPI_ISL_3350913, EPI_ISL_3350914, EPI_ISL_3350915, EPI_ISL_3350916, EPI_ISL_3350917, EPI_ISL_3350918, EPI_ISL_3350919, EPI_ISL_3350920, EPI_ISL_3350921, EPI_ISL_3350922, EPI_ISL_3350923, EPI_ISL_3350924, EPI_ISL_3350925, EPI_ISL_3350926, EPI_ISL_3350927, EPI_ISL_3350928, EPI_ISL_3350929, EPI_ISL_3350930, EPI_ISL_3350931, EPI_ISL_3350932, EPI_ISL_3350933, EPI_ISL_3350934, EPI_ISL_3350935, EPI_ISL_3350936, EPI_ISL_3350937, EPI_ISL_3350938, EPI_ISL_3350939, EPI_ISL_3350940, EPI_ISL_3350941, EPI_ISL_3350942, EPI_ISL_3350943, EPI_ISL_3350944, EPI_ISL_3350945, EPI_ISL_3350946, EPI_ISL_3350947, EPI_ISL_3350948, EPI_ISL_3350949, EPI_ISL_3350950, EPI_ISL_3350951, EPI_ISL_3350952, EPI_ISL_3350953, EPI_ISL_3350954, EPI_ISL_3350955, EPI_ISL_3350956, EPI_ISL_3350957, EPI_ISL_3350958, EPI_ISL_3350959, EPI_ISL_3350960, EPI_ISL_3350961, EPI_ISL_3350962, EPI_ISL_3350963, EPI_ISL_3350964, EPI_ISL_3350965, EPI_ISL_3350966, EPI_ISL_3350967, EPI_ISL_3350968, EPI_ISL_3350969, EPI_ISL_3350970, EPI_ISL_3350971, EPI_ISL_3350972, EPI_ISL_3350973, EPI_ISL_3350974, EPI_ISL_3350975, EPI_ISL_3350976, EPI_ISL_3350977, EPI_ISL_3350978, EPI_ISL_3350979, EPI_ISL_3350980, EPI_ISL_3350981, EPI_ISL_3350982, EPI_ISL_3350983, EPI_ISL_3350984, EPI_ISL_3350985, EPI_ISL_3350986, EPI_ISL_3350987, EPI_ISL_3350988, EPI_ISL_3350989, EPI_ISL_3350990, EPI_ISL_3350991, EPI_ISL_3350992, EPI_ISL_3350993, EPI_ISL_3350994, EPI_ISL_3350995, EPI_ISL_3350996, EPI_ISL_3350997, EPI_ISL_3350998, EPI_ISL_3350999, EPI_ISL_3351000, EPI_ISL_3351001, EPI_ISL_3351002, EPI_ISL_3351003, EPI_ISL_3351004, EPI_ISL_3351005, EPI_ISL_3351006, EPI_ISL_3351007, EPI_ISL_3351008, EPI_ISL_3351009, EPI_ISL_3351010, EPI_ISL_3351011, EPI_ISL_3351012, EPI_ISL_3351013, EPI_ISL_3351014, EPI_ISL_3351016, EPI_ISL_3351017, EPI_ISL_3351018, EPI_ISL_3351019, EPI_ISL_3351020, EPI_ISL_3351021, EPI_ISL_3351022, EPI_ISL_3351023, EPI_ISL_3351024, EPI_ISL_3351025, EPI_ISL_3351026, EPI_ISL_3351027, EPI_ISL_3351028, EPI_ISL_3351029, EPI_ISL_3351030, EPI_ISL_3351031, EPI_ISL_3351032, EPI_ISL_3351033, EPI_ISL_3351034, EPI_ISL_3351035, EPI_ISL_3351036, EPI_ISL_3351037, EPI_ISL_3351038, EPI_ISL_3351039, EPI_ISL_3351040, EPI_ISL_3351041, EPI_ISL_3351042, EPI_ISL_3351043, EPI_ISL_3351044, EPI_ISL_3351045, EPI_ISL_3351046, EPI_ISL_3351047, EPI_ISL_3351048, EPI_ISL_3351049, EPI_ISL_3351050, EPI_ISL_3351051, EPI_ISL_3351052, EPI_ISL_3351053, EPI_ISL_3351054, EPI_ISL_3351055, EPI_ISL_3351056, EPI_ISL_3351057, EPI_ISL_3351058, EPI_ISL_3351059, EPI_ISL_3351060, EPI_ISL_3351061, EPI_ISL_3351062, EPI_ISL_3351063, EPI_ISL_3351064, EPI_ISL_3351065, EPI_ISL_3351066, EPI_ISL_3351067, EPI_ISL_3351068, EPI_ISL_3351069, EPI_ISL_3351070, EPI_ISL_3351071, EPI_ISL_3351072, EPI_ISL_3351073, EPI_ISL_3351074, EPI_ISL_3351075, EPI_ISL_3351076, EPI_ISL_3351077, EPI_ISL_3351078, EPI_ISL_3351079, EPI_ISL_3351080, EPI_ISL_3351081, EPI_ISL_3351082, EPI_ISL_3351083, EPI_ISL_3351084, EPI_ISL_3351085, EPI_ISL_3351086, EPI_ISL_3351087, EPI_ISL_3351088, EPI_ISL_3351089, EPI_ISL_3351090, EPI_ISL_3351091, EPI_ISL_3351092, EPI_ISL_3351093, EPI_ISL_3351094, EPI_ISL_3351095, EPI_ISL_3351096, EPI_ISL_3351097, EPI_ISL_3351098, EPI_ISL_3351099, EPI_ISL_3351100, EPI_ISL_3351101, EPI_ISL_3351102, EPI_ISL_3351103, EPI_ISL_3351104, EPI_ISL_3351105, EPI_ISL_3351106, EPI_ISL_3351107, EPI_ISL_3351108, EPI_ISL_3351109, EPI_ISL_3351110, EPI_ISL_3351111, EPI_ISL_3351112, EPI_ISL_3351113, EPI_ISL_3351114, EPI_ISL_3351115, EPI_ISL_3351116, EPI_ISL_3351117, EPI_ISL_3351118, EPI_ISL_3351119, EPI_ISL_3351120, EPI_ISL_3351121, EPI_ISL_3351122, EPI_ISL_3351123, EPI_ISL_3351124, EPI_ISL_3351125, EPI_ISL_3351126, EPI_ISL_3351127, EPI_ISL_3351128, EPI_ISL_3351129, EPI_ISL_3351130, EPI_ISL_3351131, EPI_ISL_3351132, EPI_ISL_3351133, EPI_ISL_3351134, EPI_ISL_3351135, EPI_ISL_3351136, EPI_ISL_3351137, EPI_ISL_3351138, EPI_ISL_3351139, EPI_ISL_3351140, EPI_ISL_3351141, EPI_ISL_3351142, EPI_ISL_3351143, EPI_ISL_3351144, EPI_ISL_3351145, EPI_ISL_3351146, EPI_ISL_3351147, EPI_ISL_3351148, EPI_ISL_3351149, EPI_ISL_3351150, EPI_ISL_3351151, EPI_ISL_3351152, EPI_ISL_3351153, EPI_ISL_3351154, EPI_ISL_3351155, EPI_ISL_3351156, EPI_ISL_3351157, EPI_ISL_3351158, EPI_ISL_3351159, EPI_ISL_3351160, EPI_ISL_3351161, EPI_ISL_3351162, EPI_ISL_3351163, EPI_ISL_3351164, EPI_ISL_3351165, EPI_ISL_3351166, EPI_ISL_3351167, EPI_ISL_3351168, EPI_ISL_3351169, EPI_ISL_3351170, EPI_ISL_3351171, EPI_ISL_3351172, EPI_ISL_3351173, EPI_ISL_3351174, EPI_ISL_3351175, EPI_ISL_3351176, EPI_ISL_3351177, EPI_ISL_3351178, EPI_ISL_3351179, EPI_ISL_3351180, EPI_ISL_3351181, EPI_ISL_3351182, EPI_ISL_3351183, EPI_ISL_3351184, EPI_ISL_3351185, EPI_ISL_3351186, EPI_ISL_3351187, EPI_ISL_3351188, EPI_ISL_3351189, EPI_ISL_3351190, EPI_ISL_3351191, EPI_ISL_3351192, EPI_ISL_3351193, EPI_ISL_3351194, EPI_ISL_3351195, EPI_ISL_3351196, EPI_ISL_3351197, EPI_ISL_3351198, EPI_ISL_3351199, EPI_ISL_3351200, EPI_ISL_3351201, EPI_ISL_3351202, EPI_ISL_3351203, EPI_ISL_3351204, EPI_ISL_3351205, EPI_ISL_3351206, EPI_ISL_3351207, EPI_ISL_3351208, EPI_ISL_3351209, EPI_ISL_3351210, EPI_ISL_3351211, EPI_ISL_3351212, EPI_ISL_3351214, EPI_ISL_3351215, EPI_ISL_3351216, EPI_ISL_3351217, EPI_ISL_3351218, EPI_ISL_3351219, EPI_ISL_3351220, EPI_ISL_3351221, EPI_ISL_3351222, EPI_ISL_3351223, EPI_ISL_3351224, EPI_ISL_3351225, EPI_ISL_3351226, EPI_ISL_3351227, EPI_ISL_3351228, EPI_ISL_3351229, EPI_ISL_3351230, EPI_ISL_3351231, EPI_ISL_3351232, EPI_ISL_3351233, EPI_ISL_3351234, EPI_ISL_3351235, EPI_ISL_3351236, EPI_ISL_3351237, EPI_ISL_3351238, EPI_ISL_3351239, EPI_ISL_3351240, EPI_ISL_3351241, EPI_ISL_3351242, EPI_ISL_3351243, EPI_ISL_3351244, EPI_ISL_3351245, EPI_ISL_3351246, EPI_ISL_3351247, EPI_ISL_3351248, EPI_ISL_3351249, EPI_ISL_3351250, EPI_ISL_3351251, EPI_ISL_3351252, EPI_ISL_3351253, EPI_ISL_3351254, EPI_ISL_3351255, EPI_ISL_3351256, EPI_ISL_3351257, EPI_ISL_3351258, EPI_ISL_3351259, EPI_ISL_3351260, EPI_ISL_3351261, EPI_ISL_3351262, EPI_ISL_3351263, EPI_ISL_3351264, EPI_ISL_3351265, EPI_ISL_3351266, EPI_ISL_3351267, EPI_ISL_3351268, EPI_ISL_3351269, EPI_ISL_3351270, EPI_ISL_3351271, EPI_ISL_3351272, EPI_ISL_3351273, EPI_ISL_3351274, EPI_ISL_3351275, EPI_ISL_3351276, EPI_ISL_3351277, EPI_ISL_3351278, EPI_ISL_3351280, EPI_ISL_3351281, EPI_ISL_3351282, EPI_ISL_3351283, EPI_ISL_3351284, EPI_ISL_3351285, EPI_ISL_3351286, EPI_ISL_3351287, EPI_ISL_3351288, EPI_ISL_3351289, EPI_ISL_3351290, EPI_ISL_3351291, EPI_ISL_3351292, EPI_ISL_3351293, EPI_ISL_3351294, EPI_ISL_3351295, EPI_ISL_3351296, EPI_ISL_3351297, EPI_ISL_3351298, EPI_ISL_3351299, EPI_ISL_3351300, EPI_ISL_3351301, EPI_ISL_3351302, EPI_ISL_3351303, EPI_ISL_3351304, EPI_ISL_3351305, EPI_ISL_3351306, EPI_ISL_3351307, EPI_ISL_3351308, EPI_ISL_3351309, EPI_ISL_3351310, EPI_ISL_3351311, EPI_ISL_3351312, EPI_ISL_3351313, EPI_ISL_3351314, EPI_ISL_3351315, EPI_ISL_3351316, EPI_ISL_3351317, EPI_ISL_3351318, EPI_ISL_3351319, EPI_ISL_3351320, EPI_ISL_3351321, EPI_ISL_3351322, EPI_ISL_3351323, EPI_ISL_3351324, EPI_ISL_3351325, EPI_ISL_3351326, EPI_ISL_3351327, EPI_ISL_3351328, EPI_ISL_3351329, EPI_ISL_3351330, EPI_ISL_3351331, EPI_ISL_3351332, EPI_ISL_3351333, EPI_ISL_3351334, EPI_ISL_3351335, EPI_ISL_3351336, EPI_ISL_3351337, EPI_ISL_3351338, EPI_ISL_3351339, EPI_ISL_3351340, EPI_ISL_3351341, EPI_ISL_3351342, EPI_ISL_3351344, EPI_ISL_3351345, EPI_ISL_3351346, EPI_ISL_3351347, EPI_ISL_3351348, EPI_ISL_3351349, EPI_ISL_3351350, EPI_ISL_3351351, EPI_ISL_3351352, EPI_ISL_3351353, EPI_ISL_3351354, EPI_ISL_3351355, EPI_ISL_3351357, EPI_ISL_3351358, EPI_ISL_3351359, EPI_ISL_3351360, EPI_ISL_3351361, EPI_ISL_3351362, EPI_ISL_3351363, EPI_ISL_3351364, EPI_ISL_3351365, EPI_ISL_3351366, EPI_ISL_3351367, EPI_ISL_3351368, EPI_ISL_3351369, EPI_ISL_3351370, EPI_ISL_3351371, EPI_ISL_3351372, EPI_ISL_3351373, EPI_ISL_3351374, EPI_ISL_3351375, EPI_ISL_3351376, EPI_ISL_3351377, EPI_ISL_3351378, EPI_ISL_3351379, EPI_ISL_3351380, EPI_ISL_3351381, EPI_ISL_3351382, EPI_ISL_3351383, EPI_ISL_3351384, EPI_ISL_3351385, EPI_ISL_3351386, EPI_ISL_3351387, EPI_ISL_3351388, EPI_ISL_3351389, EPI_ISL_3351390, EPI_ISL_3351391, EPI_ISL_3351392, EPI_ISL_3351393, EPI_ISL_3351394, EPI_ISL_3351395, EPI_ISL_3351396, EPI_ISL_3351397, EPI_ISL_3351398, EPI_ISL_3351399, EPI_ISL_3351400, EPI_ISL_3351401, EPI_ISL_3351402, EPI_ISL_3351403, EPI_ISL_3351404, EPI_ISL_3351405, EPI_ISL_3351406, EPI_ISL_3351407, EPI_ISL_3351408, EPI_ISL_3351409, EPI_ISL_3351410, EPI_ISL_3351411, EPI_ISL_3351412, EPI_ISL_3351413, EPI_ISL_3351414, EPI_ISL_3351415, EPI_ISL_3351416, EPI_ISL_3351417, EPI_ISL_3351418, EPI_ISL_3351419, EPI_ISL_3351420, EPI_ISL_3351421, EPI_ISL_3351422, EPI_ISL_3351423, EPI_ISL_3351424, EPI_ISL_3351425, EPI_ISL_3351426, EPI_ISL_3351427, EPI_ISL_3351428, EPI_ISL_3351429, EPI_ISL_3351430, EPI_ISL_3351431, EPI_ISL_3351432, EPI_ISL_3351433, EPI_ISL_3351434, EPI_ISL_3351435, EPI_ISL_3351436, EPI_ISL_3351437, EPI_ISL_3351438, EPI_ISL_3351439, EPI_ISL_3351440, EPI_ISL_3351441, EPI_ISL_3351442, EPI_ISL_3351443, EPI_ISL_3351444, EPI_ISL_3351445, EPI_ISL_3351446, EPI_ISL_3351447, EPI_ISL_3351448, EPI_ISL_3351449, EPI_ISL_3351450, EPI_ISL_3351451, EPI_ISL_3351452, EPI_ISL_3351453, EPI_ISL_3351454, EPI_ISL_3351455, EPI_ISL_3351456, EPI_ISL_3351457, EPI_ISL_3351458, EPI_ISL_3351459, EPI_ISL_3351460, EPI_ISL_3351461, EPI_ISL_3351462, EPI_ISL_3351463, EPI_ISL_3351464, EPI_ISL_3351465, EPI_ISL_3351466, EPI_ISL_3351467, EPI_ISL_3351468, EPI_ISL_3351469, EPI_ISL_3351470, EPI_ISL_3351471, EPI_ISL_3351472, EPI_ISL_3351473, EPI_ISL_3351474, EPI_ISL_3351475, EPI_ISL_3351476, EPI_ISL_3351477, EPI_ISL_3351478, EPI_ISL_3351479, EPI_ISL_3351480, EPI_ISL_3351481, EPI_ISL_3351482, EPI_ISL_3351483, EPI_ISL_3351484, EPI_ISL_3351485, EPI_ISL_3351486, EPI_ISL_3351487, EPI_ISL_3351488, EPI_ISL_3351489, EPI_ISL_3351490, EPI_ISL_3351491, EPI_ISL_3351492, EPI_ISL_3351493, EPI_ISL_3351494, EPI_ISL_3351495, EPI_ISL_3351496, EPI_ISL_3351497, EPI_ISL_3351498, EPI_ISL_3351499, EPI_ISL_3351500, EPI_ISL_3351501, EPI_ISL_3351502, EPI_ISL_3351503, EPI_ISL_3351504, EPI_ISL_3351505, EPI_ISL_3351506, EPI_ISL_3351507, EPI_ISL_3351508, EPI_ISL_3351509, EPI_ISL_3351510, EPI_ISL_3351511, EPI_ISL_3351512, EPI_ISL_3351513, EPI_ISL_3351514, EPI_ISL_3351515, EPI_ISL_3351516, EPI_ISL_3351518, EPI_ISL_3351519, EPI_ISL_3351520, EPI_ISL_3351521, EPI_ISL_3351522, EPI_ISL_3351523, EPI_ISL_3351524, EPI_ISL_3351525, EPI_ISL_3351526, EPI_ISL_3351527, EPI_ISL_3351528, EPI_ISL_3351529, EPI_ISL_3351530, EPI_ISL_3351531, EPI_ISL_3351532, EPI_ISL_3351533, EPI_ISL_3351534, EPI_ISL_3351535, EPI_ISL_3351536, EPI_ISL_3351537, EPI_ISL_3351538, EPI_ISL_3351539, EPI_ISL_3351540, EPI_ISL_3351541, EPI_ISL_3351542, EPI_ISL_3351543, EPI_ISL_3351544, EPI_ISL_3351545, EPI_ISL_3351546, EPI_ISL_3351547, EPI_ISL_3351548, EPI_ISL_3351549, EPI_ISL_3351550, EPI_ISL_3351551, EPI_ISL_3351552, EPI_ISL_3351553, EPI_ISL_3351554, EPI_ISL_3351555, EPI_ISL_3351556, EPI_ISL_3351557, EPI_ISL_3351558, EPI_ISL_3351559, EPI_ISL_3351560, EPI_ISL_3351561, EPI_ISL_3351562, EPI_ISL_3351563, EPI_ISL_3351564, EPI_ISL_3351565, EPI_ISL_3351566, EPI_ISL_3351567, EPI_ISL_3351568, EPI_ISL_3351569, EPI_ISL_3351570, EPI_ISL_3351571, EPI_ISL_3351572, EPI_ISL_3351573, EPI_ISL_3351574, EPI_ISL_3351575, EPI_ISL_3351576, EPI_ISL_3351577, EPI_ISL_3351578, EPI_ISL_3351579, EPI_ISL_3351580, EPI_ISL_3351581, EPI_ISL_3351582, EPI_ISL_3351583, EPI_ISL_3351584, EPI_ISL_3351585, EPI_ISL_3351586, EPI_ISL_3351587, EPI_ISL_3351588, EPI_ISL_3351589, EPI_ISL_3351590, EPI_ISL_3351591, EPI_ISL_3351592, EPI_ISL_3351593, EPI_ISL_3351594, EPI_ISL_3351595, EPI_ISL_3351596, EPI_ISL_3351597, EPI_ISL_3351598, EPI_ISL_3351599, EPI_ISL_3351600, EPI_ISL_3351601, EPI_ISL_3351602, EPI_ISL_3351603, EPI_ISL_3351604, EPI_ISL_3351605, EPI_ISL_3351606, EPI_ISL_3351607, EPI_ISL_3351608, EPI_ISL_3351609, EPI_ISL_3351610, EPI_ISL_3351611, EPI_ISL_3351612, EPI_ISL_3351613, EPI_ISL_3351614, EPI_ISL_3351615, EPI_ISL_3351616, EPI_ISL_3351617, EPI_ISL_3351618, EPI_ISL_3351620, EPI_ISL_3351621, EPI_ISL_3351622, EPI_ISL_3351623, EPI_ISL_3351624, EPI_ISL_3351625, EPI_ISL_3351626, EPI_ISL_3351627, EPI_ISL_3351628, EPI_ISL_3351629, EPI_ISL_3351630, EPI_ISL_3351631, EPI_ISL_3351632, EPI_ISL_3351633, EPI_ISL_3351634, EPI_ISL_3351635, EPI_ISL_3351636, EPI_ISL_3351637, EPI_ISL_3351638, EPI_ISL_3351639, EPI_ISL_3351640, EPI_ISL_3351641, EPI_ISL_3351642, EPI_ISL_3351643, EPI_ISL_3351644, EPI_ISL_3351645, EPI_ISL_3351646, EPI_ISL_3351647, EPI_ISL_3351648, EPI_ISL_3351649, EPI_ISL_3351650, EPI_ISL_3351651, EPI_ISL_3351652, EPI_ISL_3351653, EPI_ISL_3351654, EPI_ISL_3351655, EPI_ISL_3351656, EPI_ISL_3351657, EPI_ISL_3351658, EPI_ISL_3351659, EPI_ISL_3351660, EPI_ISL_3351661, EPI_ISL_3351662, EPI_ISL_3351663, EPI_ISL_3351664, EPI_ISL_3351665, EPI_ISL_3351666, EPI_ISL_3351667, EPI_ISL_3351668, EPI_ISL_3351669, EPI_ISL_3351670, EPI_ISL_3351671, EPI_ISL_3351672, EPI_ISL_3351673, EPI_ISL_3351674, EPI_ISL_3351675, EPI_ISL_3351677, EPI_ISL_3351678, EPI_ISL_3351679, EPI_ISL_3351680, EPI_ISL_3351681, EPI_ISL_3351682, EPI_ISL_3351683, EPI_ISL_3351684, EPI_ISL_3351685, EPI_ISL_3351686, EPI_ISL_3351687, EPI_ISL_3351688, EPI_ISL_3351689, EPI_ISL_3351690, EPI_ISL_3351691, EPI_ISL_3351692, EPI_ISL_3351693, EPI_ISL_3351694, EPI_ISL_3351695, EPI_ISL_3351696, EPI_ISL_3351697, EPI_ISL_3351698, EPI_ISL_3351700, EPI_ISL_3351701, EPI_ISL_3351702, EPI_ISL_3351703, EPI_ISL_3351704, EPI_ISL_3351705, EPI_ISL_3351706, EPI_ISL_3351707, EPI_ISL_3351708, EPI_ISL_3351709, EPI_ISL_3351710, EPI_ISL_3351711, EPI_ISL_3351712, EPI_ISL_3351713, EPI_ISL_3351714, EPI_ISL_3351715, EPI_ISL_3351716, EPI_ISL_3351717, EPI_ISL_3351718, EPI_ISL_3351719, EPI_ISL_3351720, EPI_ISL_3351721, EPI_ISL_3351722, EPI_ISL_3351723, EPI_ISL_3351724, EPI_ISL_3351725, EPI_ISL_3351726, EPI_ISL_3351727, EPI_ISL_3351728, EPI_ISL_3351729, EPI_ISL_3351730, EPI_ISL_3351731, EPI_ISL_3351732, EPI_ISL_3351733, EPI_ISL_3351734, EPI_ISL_3351735, EPI_ISL_3351736, EPI_ISL_3351737, EPI_ISL_3351738, EPI_ISL_3351739, EPI_ISL_3351740, EPI_ISL_3351741, EPI_ISL_3351742, EPI_ISL_3351743, EPI_ISL_3351744, EPI_ISL_3351745, EPI_ISL_3351746, EPI_ISL_3351747, EPI_ISL_3351748, EPI_ISL_3351749, EPI_ISL_3351750, EPI_ISL_3351751, EPI_ISL_3351752, EPI_ISL_3351753, EPI_ISL_3351754, EPI_ISL_3351755, EPI_ISL_3351756, EPI_ISL_3351757, EPI_ISL_3351758, EPI_ISL_3351759, EPI_ISL_3351760, EPI_ISL_3351761, EPI_ISL_3351762, EPI_ISL_3351763, EPI_ISL_3351764, EPI_ISL_3351765, EPI_ISL_3351766, EPI_ISL_3351767, EPI_ISL_3351768, EPI_ISL_3351769, EPI_ISL_3351770, EPI_ISL_3351771, EPI_ISL_3351772, EPI_ISL_3351773, EPI_ISL_3351774, EPI_ISL_3351775, EPI_ISL_3351776, EPI_ISL_3351777, EPI_ISL_3351778, EPI_ISL_3351779, EPI_ISL_3351780, EPI_ISL_3351781, EPI_ISL_3351782, EPI_ISL_3351783, EPI_ISL_3351784, EPI_ISL_3351785, EPI_ISL_3351786, EPI_ISL_3351787, EPI_ISL_3351788, EPI_ISL_3351789, EPI_ISL_3351790, EPI_ISL_3351791, EPI_ISL_3351792, EPI_ISL_3351793, EPI_ISL_3351794, EPI_ISL_3351795, EPI_ISL_3351796, EPI_ISL_3351797, EPI_ISL_3351798, EPI_ISL_3351799, EPI_ISL_3351800, EPI_ISL_3351801, EPI_ISL_3351802, EPI_ISL_3351803, EPI_ISL_3351804, EPI_ISL_3351805, EPI_ISL_3351806, EPI_ISL_3351807, EPI_ISL_3351808, EPI_ISL_3351809, EPI_ISL_3351810, EPI_ISL_3351811, EPI_ISL_3351812, EPI_ISL_3351813, EPI_ISL_3351814, EPI_ISL_3351815, EPI_ISL_3351816, EPI_ISL_3351817, EPI_ISL_3351818, EPI_ISL_3351819, EPI_ISL_3351820, EPI_ISL_3351821, EPI_ISL_3351822, EPI_ISL_3351823, EPI_ISL_3351824, EPI_ISL_3351825, EPI_ISL_3351826, EPI_ISL_3351827, EPI_ISL_3351828, EPI_ISL_3351829, EPI_ISL_3351830, EPI_ISL_3351831, EPI_ISL_3351832, EPI_ISL_3351833, EPI_ISL_3351834, EPI_ISL_3351835, EPI_ISL_3351836, EPI_ISL_3351837, EPI_ISL_3351838, EPI_ISL_3351839, EPI_ISL_3351840, EPI_ISL_3351841, EPI_ISL_3351842, EPI_ISL_3351843, EPI_ISL_3351844, EPI_ISL_3351845, EPI_ISL_3351846, EPI_ISL_3351847, EPI_ISL_3351848, EPI_ISL_3351849, EPI_ISL_3351850, EPI_ISL_3351851, EPI_ISL_3351852, EPI_ISL_3351853, EPI_ISL_3351854, EPI_ISL_3351855, EPI_ISL_3351856, EPI_ISL_3351857, EPI_ISL_3351858, EPI_ISL_3351859, EPI_ISL_3351860, EPI_ISL_3351861, EPI_ISL_3351862, EPI_ISL_3351863, EPI_ISL_3351864, EPI_ISL_3351865, EPI_ISL_3351866, EPI_ISL_3351867, EPI_ISL_3351868, EPI_ISL_3351869, EPI_ISL_3351870, EPI_ISL_3351871, EPI_ISL_3351872, EPI_ISL_3351873, EPI_ISL_3351874, EPI_ISL_3351875, EPI_ISL_3351876, EPI_ISL_3351877, EPI_ISL_3351878, EPI_ISL_3351879, EPI_ISL_3351880, EPI_ISL_3351881, EPI_ISL_3351882, EPI_ISL_3351883, EPI_ISL_3351884, EPI_ISL_3351885, EPI_ISL_3351886, EPI_ISL_3351887, EPI_ISL_3351888, EPI_ISL_3351889, EPI_ISL_3351890, EPI_ISL_3351891, EPI_ISL_3351892, EPI_ISL_3351893, EPI_ISL_3351894, EPI_ISL_3351895, EPI_ISL_3351896, EPI_ISL_3351897, EPI_ISL_3351898, EPI_ISL_3351899, EPI_ISL_3351900, EPI_ISL_3351901, EPI_ISL_3351902, EPI_ISL_3351903, EPI_ISL_3351904, EPI_ISL_3351905, EPI_ISL_3351906, EPI_ISL_3351907, EPI_ISL_3351908, EPI_ISL_3351909, EPI_ISL_3351910, EPI_ISL_3351911, EPI_ISL_3351912, EPI_ISL_3351913, EPI_ISL_3351914, EPI_ISL_3351915, EPI_ISL_3351917, EPI_ISL_3351918, EPI_ISL_3351919, EPI_ISL_3351920, EPI_ISL_3351921, EPI_ISL_3351922, EPI_ISL_3351923, EPI_ISL_3351924, EPI_ISL_3351925, EPI_ISL_3351926, EPI_ISL_3351927, EPI_ISL_3351928, EPI_ISL_3351929, EPI_ISL_3351930, EPI_ISL_3351931, EPI_ISL_3351932, EPI_ISL_3351933, EPI_ISL_3351934, EPI_ISL_3351935, EPI_ISL_3351936, EPI_ISL_3351937, EPI_ISL_3351938, EPI_ISL_3351939, EPI_ISL_3351940, EPI_ISL_3351941, EPI_ISL_3351942, EPI_ISL_3351943, EPI_ISL_3351944, EPI_ISL_3351945, EPI_ISL_3351946, EPI_ISL_3351947, EPI_ISL_3351948, EPI_ISL_3351949, EPI_ISL_3351950, EPI_ISL_3351951, EPI_ISL_3351952, EPI_ISL_3351953, EPI_ISL_3351954, EPI_ISL_3351955, EPI_ISL_3351956, EPI_ISL_3351957, EPI_ISL_3351958, EPI_ISL_3351959, EPI_ISL_3351960, EPI_ISL_3351961, EPI_ISL_3351962, EPI_ISL_3351963, EPI_ISL_3351964, EPI_ISL_3351965, EPI_ISL_3351966, EPI_ISL_3351967, EPI_ISL_3351968, EPI_ISL_3351969, EPI_ISL_3351970, EPI_ISL_3351971, EPI_ISL_3351972, EPI_ISL_3351973, EPI_ISL_3351974, EPI_ISL_3351975, EPI_ISL_3351976, EPI_ISL_3351977, EPI_ISL_3351978, EPI_ISL_3351979, EPI_ISL_3351980, EPI_ISL_3351981, EPI_ISL_3351982, EPI_ISL_3351983, EPI_ISL_3351984, EPI_ISL_3351985, EPI_ISL_3351986, EPI_ISL_3351987, EPI_ISL_3351988, EPI_ISL_3351989, EPI_ISL_3351990, EPI_ISL_3351991, EPI_ISL_3351992, EPI_ISL_3351993, EPI_ISL_3351994, EPI_ISL_3351995, EPI_ISL_3351996, EPI_ISL_3351997, EPI_ISL_3351998, EPI_ISL_3351999, EPI_ISL_3352000, EPI_ISL_3352001, EPI_ISL_3352002, EPI_ISL_3352003, EPI_ISL_3352004, EPI_ISL_3352005, EPI_ISL_3352006, EPI_ISL_3352007, EPI_ISL_3352008, EPI_ISL_3352009, EPI_ISL_3352010, EPI_ISL_3352011, EPI_ISL_3352012, EPI_ISL_3352013, EPI_ISL_3352014, EPI_ISL_3352015, EPI_ISL_3352016, EPI_ISL_3352017, EPI_ISL_3352018, EPI_ISL_3352019, EPI_ISL_3352020, EPI_ISL_3352021, EPI_ISL_3352022, EPI_ISL_3352023, EPI_ISL_3352024, EPI_ISL_3352025, EPI_ISL_3352026, EPI_ISL_3352027, EPI_ISL_3352028, EPI_ISL_3352029, EPI_ISL_3352030, EPI_ISL_3352031, EPI_ISL_3352032, EPI_ISL_3352033, EPI_ISL_3352034, EPI_ISL_3352035, EPI_ISL_3352036, EPI_ISL_3352037, EPI_ISL_3352038, EPI_ISL_3352039, EPI_ISL_3352040, EPI_ISL_3352042, EPI_ISL_3352043, EPI_ISL_3352044, EPI_ISL_3352045, EPI_ISL_3352046, EPI_ISL_3352047, EPI_ISL_3352048, EPI_ISL_3352049, EPI_ISL_3352050, EPI_ISL_3352051, EPI_ISL_3352052, EPI_ISL_3352053, EPI_ISL_3352054, EPI_ISL_3352055, EPI_ISL_3352056, EPI_ISL_3352057, EPI_ISL_3352058, EPI_ISL_3352059, EPI_ISL_3352060, EPI_ISL_3352061, EPI_ISL_3352062, EPI_ISL_3352063, EPI_ISL_3352064, EPI_ISL_3352065, EPI_ISL_3352066, EPI_ISL_3352067, EPI_ISL_3352068, EPI_ISL_3352069, EPI_ISL_3352070, EPI_ISL_3352071, EPI_ISL_3352072, EPI_ISL_3352073, EPI_ISL_3352074, EPI_ISL_3352075, EPI_ISL_3352076, EPI_ISL_3352077, EPI_ISL_3352078, EPI_ISL_3352079, EPI_ISL_3352080, EPI_ISL_3352081, EPI_ISL_3352082, EPI_ISL_3352083, EPI_ISL_3352084, EPI_ISL_3352085, EPI_ISL_3352086, EPI_ISL_3352087, EPI_ISL_3352088, EPI_ISL_3352089, EPI_ISL_3352090, EPI_ISL_3352091, EPI_ISL_3352092, EPI_ISL_3352093, EPI_ISL_3352094, EPI_ISL_3352095, EPI_ISL_3352096, EPI_ISL_3352097, EPI_ISL_3352098, EPI_ISL_3352099, EPI_ISL_3352100, EPI_ISL_3352101, EPI_ISL_3352102, EPI_ISL_3352103, EPI_ISL_3352104, EPI_ISL_3352105, EPI_ISL_3352106, EPI_ISL_3352107, EPI_ISL_3352108, EPI_ISL_3352109, EPI_ISL_3352110, EPI_ISL_3352111, EPI_ISL_3352112, EPI_ISL_3352113, EPI_ISL_3352114, EPI_ISL_3352115, EPI_ISL_3352116, EPI_ISL_3352117, EPI_ISL_3352118, EPI_ISL_3352119, EPI_ISL_3352120, EPI_ISL_3352121, EPI_ISL_3352122, EPI_ISL_3352123, EPI_ISL_3352124, EPI_ISL_3352125, EPI_ISL_3352126, EPI_ISL_3352127, EPI_ISL_3352128, EPI_ISL_3352129, EPI_ISL_3352130, EPI_ISL_3352131, EPI_ISL_3352132, EPI_ISL_3352133, EPI_ISL_3352134, EPI_ISL_3352135, EPI_ISL_3352136, EPI_ISL_3352137, EPI_ISL_3352138, EPI_ISL_3352139, EPI_ISL_3352140, EPI_ISL_3352141, EPI_ISL_3352142, EPI_ISL_3352143, EPI_ISL_3352144, EPI_ISL_3352145, EPI_ISL_3352146, EPI_ISL_3352147, EPI_ISL_3352148, EPI_ISL_3352149, EPI_ISL_3352150, EPI_ISL_3352151, EPI_ISL_3352152, EPI_ISL_3352153, EPI_ISL_3352154, EPI_ISL_3352155, EPI_ISL_3352156, EPI_ISL_3352157, EPI_ISL_3352158, EPI_ISL_3352159, EPI_ISL_3352160, EPI_ISL_3352161, EPI_ISL_3352162, EPI_ISL_3352163, EPI_ISL_3352164, EPI_ISL_3352165, EPI_ISL_3352166, EPI_ISL_3352167, EPI_ISL_3352168, EPI_ISL_3352169, EPI_ISL_3352170, EPI_ISL_3352171, EPI_ISL_3352172, EPI_ISL_3352173, EPI_ISL_3352174, EPI_ISL_3352175, EPI_ISL_3352176, EPI_ISL_3352177, EPI_ISL_3352178, EPI_ISL_3352179, EPI_ISL_3352180, EPI_ISL_3352181, EPI_ISL_3352182, EPI_ISL_3352183, EPI_ISL_3352184, EPI_ISL_3352185, EPI_ISL_3352186, EPI_ISL_3352187, EPI_ISL_3352188, EPI_ISL_3352189, EPI_ISL_3352190, EPI_ISL_3352191, EPI_ISL_3352192, EPI_ISL_3352193, EPI_ISL_3352194, EPI_ISL_3352195, EPI_ISL_3352196, EPI_ISL_3352197, EPI_ISL_3352198, EPI_ISL_3352199, EPI_ISL_3352200, EPI_ISL_3352201, EPI_ISL_3352202, EPI_ISL_3352203, EPI_ISL_3352204, EPI_ISL_3352205, EPI_ISL_3352206, EPI_ISL_3352207, EPI_ISL_3352208, EPI_ISL_3352209, EPI_ISL_3352211, EPI_ISL_3352212, EPI_ISL_3352213, EPI_ISL_3352214, EPI_ISL_3352215, EPI_ISL_3352216, EPI_ISL_3352217, EPI_ISL_3352218, EPI_ISL_3352219, EPI_ISL_3352220, EPI_ISL_3352221, EPI_ISL_3352222, EPI_ISL_3352223, EPI_ISL_3352224, EPI_ISL_3352225, EPI_ISL_3352226, EPI_ISL_3352227, EPI_ISL_3352228, EPI_ISL_3352229, EPI_ISL_3352230, EPI_ISL_3352231, EPI_ISL_3352232, EPI_ISL_3352233, EPI_ISL_3352234, EPI_ISL_3352235, EPI_ISL_3352236, EPI_ISL_3352237, EPI_ISL_3352238, EPI_ISL_3352239, EPI_ISL_3352240, EPI_ISL_3352241, EPI_ISL_3352242, EPI_ISL_3352243, EPI_ISL_3352244, EPI_ISL_3352245, EPI_ISL_3352246, EPI_ISL_3352247, EPI_ISL_3352248, EPI_ISL_3352249, EPI_ISL_3352250, EPI_ISL_3352251, EPI_ISL_3352252, EPI_ISL_3352253, EPI_ISL_3352254, EPI_ISL_3352255, EPI_ISL_3352256, EPI_ISL_3352257, EPI_ISL_3352258, EPI_ISL_3352259, EPI_ISL_3352260, EPI_ISL_3352261, EPI_ISL_3352262, EPI_ISL_3352263, EPI_ISL_3352264, EPI_ISL_3352265, EPI_ISL_3352266, EPI_ISL_3352267, EPI_ISL_3352268, EPI_ISL_3352269, EPI_ISL_3352270, EPI_ISL_3352271, EPI_ISL_3352272, EPI_ISL_3352273, EPI_ISL_3352274, EPI_ISL_3352275, EPI_ISL_3352276, EPI_ISL_3352277, EPI_ISL_3352278, EPI_ISL_3352279, EPI_ISL_3352280, EPI_ISL_3352281, EPI_ISL_3352282, EPI_ISL_3352283, EPI_ISL_3352284, EPI_ISL_3352285, EPI_ISL_3352286, EPI_ISL_3352287, EPI_ISL_3352288, EPI_ISL_3352289, EPI_ISL_3352290, EPI_ISL_3352291, EPI_ISL_3352292, EPI_ISL_3352293, EPI_ISL_3352294, EPI_ISL_3352295, EPI_ISL_3352296, EPI_ISL_3352297, EPI_ISL_3352298, EPI_ISL_3352299, EPI_ISL_3352300, EPI_ISL_3352301, EPI_ISL_3352302, EPI_ISL_3352303, EPI_ISL_3352304, EPI_ISL_3352305, EPI_ISL_3352306, EPI_ISL_3352307, EPI_ISL_3352308, EPI_ISL_3352309, EPI_ISL_3352310, EPI_ISL_3352311, EPI_ISL_3352312, EPI_ISL_3352313, EPI_ISL_3352314, EPI_ISL_3352315, EPI_ISL_3352316, EPI_ISL_3352317, EPI_ISL_3352318, EPI_ISL_3352319, EPI_ISL_3352320, EPI_ISL_3352321, EPI_ISL_3352322, EPI_ISL_3352323, EPI_ISL_3352324, EPI_ISL_3352325, EPI_ISL_3352326, EPI_ISL_3352327, EPI_ISL_3352328, EPI_ISL_3352329, EPI_ISL_3352330, EPI_ISL_3352331, EPI_ISL_3352332, EPI_ISL_3352334, EPI_ISL_3352335, EPI_ISL_3352336, EPI_ISL_3352337, EPI_ISL_3352339, EPI_ISL_3352340, EPI_ISL_3352341, EPI_ISL_3352344, EPI_ISL_3352345, EPI_ISL_3352346, EPI_ISL_3352347, EPI_ISL_3352348, EPI_ISL_3352349, EPI_ISL_3352350, EPI_ISL_3352351, EPI_ISL_3352353, EPI_ISL_3352355, EPI_ISL_3352356, EPI_ISL_3352357, EPI_ISL_3352358, EPI_ISL_3352359, EPI_ISL_3352360, EPI_ISL_3352361, EPI_ISL_3352362, EPI_ISL_3352363, EPI_ISL_3352364, EPI_ISL_3352365, EPI_ISL_3352366, EPI_ISL_3352367, EPI_ISL_3352368, EPI_ISL_3352369, EPI_ISL_3352370, EPI_ISL_3352371, EPI_ISL_3352372, EPI_ISL_3352374, EPI_ISL_3352375, EPI_ISL_3352376, EPI_ISL_3352377, EPI_ISL_3352378, EPI_ISL_3352379, EPI_ISL_3352380, EPI_ISL_3352381, EPI_ISL_3352382, EPI_ISL_3352383, EPI_ISL_3352384, EPI_ISL_3352385, EPI_ISL_3352386 | | | |
| see above | Fulgent Genetics | Centers for Disease Control and Prevention Division of Viral Diseases, Pathogen Discovery | Dakota Howard, Dhwani Batra, Peter W. Cook, Kara Moser, Adrian Paskey, Jason Caravas, Benjamin Rambo-Martin, Shatavia Morrison, Christopher Gulvick, Scott Sammons, Yvette Unoarumhi, Darlene Wagner, Matthew Schmerer, Harry Gao, Mickey Li, John Gao, Joseph Fierro, Benafsh Sapra, Becky Tsai, Yan Meng, Doreen Ng, James Xie, Clinton R. Paden, Duncan MacCannell |
| EPI_ISL_3352387 | NYU Langone Health | Departments of Pathology and Medicine, New York University School of Medicine | Adriana Heguy, Dacia Dimartino, Emily Guzman, Christian Marier, Peter Meyn, Sitharam Ramaswami, Gael Westby, Paul Zappile, Yutong Zhang, Paolo Cotzia, Guiqing Wang |
| EPI_ISL_3352388, EPI_ISL_3352389, EPI_ISL_3352390, EPI_ISL_3352391, EPI_ISL_3352392, EPI_ISL_3352393, EPI_ISL_3352394, EPI_ISL_3352395, EPI_ISL_3352396, EPI_ISL_3352397, EPI_ISL_3352398, EPI_ISL_3352399, EPI_ISL_3352400, EPI_ISL_3352401, EPI_ISL_3352402, EPI_ISL_3352403, EPI_ISL_3352404, EPI_ISL_3352405, EPI_ISL_3352406, EPI_ISL_3352407, EPI_ISL_3352408, EPI_ISL_3352409, EPI_ISL_3352410, EPI_ISL_3352411, EPI_ISL_3352412, EPI_ISL_3352413, EPI_ISL_3352414, EPI_ISL_3352415, EPI_ISL_3352416, EPI_ISL_3352417, EPI_ISL_3352418, EPI_ISL_3352419, EPI_ISL_3352420, EPI_ISL_3352421, EPI_ISL_3352422, EPI_ISL_3352423, EPI_ISL_3352424, EPI_ISL_3352425, EPI_ISL_3352426, EPI_ISL_3352427, EPI_ISL_3352428, EPI_ISL_3352429, EPI_ISL_3352430, EPI_ISL_3352431, EPI_ISL_3352432, EPI_ISL_3352433, EPI_ISL_3352434, EPI_ISL_3352435, EPI_ISL_3352436, EPI_ISL_3352437, EPI_ISL_3352438, EPI_ISL_3352439, EPI_ISL_3352440, EPI_ISL_3352441, EPI_ISL_3352442, EPI_ISL_3352443, EPI_ISL_3352444, EPI_ISL_3352445, EPI_ISL_3352446, EPI_ISL_3352447, EPI_ISL_3352448, EPI_ISL_3352449, EPI_ISL_3352450, EPI_ISL_3352451, EPI_ISL_3352453, EPI_ISL_3352454, EPI_ISL_3352455, EPI_ISL_3352456, EPI_ISL_3352457, EPI_ISL_3352458, EPI_ISL_3352459, EPI_ISL_3352460, EPI_ISL_3352461, EPI_ISL_3352462 | | | |
| see above | Fulgent Genetics | Centers for Disease Control and Prevention Division of Viral Diseases, Pathogen Discovery | Dakota Howard, Dhwani Batra, Peter W. Cook, Kara Moser, Adrian Paskey, Jason Caravas, Benjamin Rambo-Martin, Shatavia Morrison, Christopher Gulvick, Scott Sammons, Yvette Unoarumhi, Darlene Wagner, Matthew Schmerer, Harry Gao, Mickey Li, John Gao, Joseph Fierro, Benafsh Sapra, Becky Tsai, Yan Meng, Doreen Ng, James Xie, Clinton R. Paden, Duncan MacCannell |
| EPI_ISL_3352463, EPI_ISL_3352464, EPI_ISL_3352465, EPI_ISL_3352466, EPI_ISL_3352467, EPI_ISL_3352468, EPI_ISL_3352469, EPI_ISL_3352470, EPI_ISL_3352471, EPI_ISL_3352472, EPI_ISL_3352473, EPI_ISL_3352474, EPI_ISL_3352475, EPI_ISL_3352476, EPI_ISL_3352477, EPI_ISL_3352478, EPI_ISL_3352479, EPI_ISL_3352481, EPI_ISL_3352482, EPI_ISL_3352483, EPI_ISL_3352484, EPI_ISL_3352485, EPI_ISL_3352486, EPI_ISL_3352487, EPI_ISL_3352488, EPI_ISL_3352489, EPI_ISL_3352490, EPI_ISL_3352491, EPI_ISL_3352492, EPI_ISL_3352493, EPI_ISL_3352494, EPI_ISL_3352495, EPI_ISL_3352496, EPI_ISL_3352497, EPI_ISL_3352498, EPI_ISL_3352499, EPI_ISL_3352500, EPI_ISL_3352501, EPI_ISL_3352502, EPI_ISL_3352503, EPI_ISL_3352504, EPI_ISL_3352505, EPI_ISL_3352506, EPI_ISL_3352507, EPI_ISL_3352508, EPI_ISL_3352509, EPI_ISL_3352510, EPI_ISL_3352511, EPI_ISL_3352512, EPI_ISL_3352514, EPI_ISL_3352515, EPI_ISL_3352516, EPI_ISL_3352517 | | | |
| see above | Aegis Sciences Corporation | Centers for Disease Control and Prevention Division of Viral Diseases, Pathogen Discovery | Dakota Howard, Dhwani Batra, Peter W. Cook, Kara Moser, Adrian Paskey, Jason Caravas, Benjamin Rambo-Martin, Shatavia Morrison, Christopher Gulvick, Scott Sammons, Yvette Unoarumhi, Darlene Wagner, Matthew Schmerer, Cyndi Clark, Patrick Campbell, Rob Case, Vikramsinha Ghorpade, Holly Houdeshell, Ola Kvalvaag, Dillon Nall, Ethan Sanders, Alec Vest, Shaun Westlund, Matthew Hardison, Clinton R. Paden, Duncan MacCannell |
| EPI_ISL_3352518, EPI_ISL_3352519, EPI_ISL_3352520, EPI_ISL_3352521, EPI_ISL_3352522, EPI_ISL_3352523, EPI_ISL_3352524, EPI_ISL_3352525, EPI_ISL_3352526, EPI_ISL_3352527, EPI_ISL_3352528, EPI_ISL_3352529, EPI_ISL_3352530, EPI_ISL_3352531, EPI_ISL_3352532, EPI_ISL_3352534, EPI_ISL_3352535, EPI_ISL_3352536, EPI_ISL_3352537, EPI_ISL_3352538, EPI_ISL_3352539, EPI_ISL_3352540, EPI_ISL_3352541, EPI_ISL_3352542, EPI_ISL_3352543, EPI_ISL_3352544, EPI_ISL_3352545, EPI_ISL_3352546, EPI_ISL_3352547, EPI_ISL_3352548, EPI_ISL_3352549, EPI_ISL_3352550, EPI_ISL_3352551, EPI_ISL_3352552, EPI_ISL_3352553, EPI_ISL_3352554, EPI_ISL_3352555, EPI_ISL_3352556, EPI_ISL_3352557, EPI_ISL_3352558, EPI_ISL_3352559, EPI_ISL_3352560, EPI_ISL_3352561, EPI_ISL_3352562, EPI_ISL_3352563, EPI_ISL_3352564, EPI_ISL_3352565, EPI_ISL_3352566, EPI_ISL_3352567, EPI_ISL_3352568, EPI_ISL_3352569, EPI_ISL_3352570, EPI_ISL_3352571, EPI_ISL_3352572, EPI_ISL_3352573, EPI_ISL_3352575, EPI_ISL_3352576, EPI_ISL_3352577, EPI_ISL_3352579, EPI_ISL_3352580, EPI_ISL_3352581, EPI_ISL_3352582, EPI_ISL_3352583, EPI_ISL_3352584, EPI_ISL_3352585, EPI_ISL_3352586, EPI_ISL_3352587, EPI_ISL_3352588, EPI_ISL_3352589, EPI_ISL_3352590, EPI_ISL_3352591, EPI_ISL_3352592, EPI_ISL_3352593, EPI_ISL_3352594, EPI_ISL_3352595, EPI_ISL_3352596, EPI_ISL_3352597, EPI_ISL_3352598, EPI_ISL_3352599, EPI_ISL_3352600, EPI_ISL_3352601, EPI_ISL_3352602, EPI_ISL_3352603, EPI_ISL_3352604, EPI_ISL_3352605, EPI_ISL_3352606, EPI_ISL_3352607, EPI_ISL_3352608, EPI_ISL_3352609, EPI_ISL_3352610, EPI_ISL_3352612, EPI_ISL_3352613, EPI_ISL_3352614, EPI_ISL_3352615, EPI_ISL_3352616, EPI_ISL_3352617, EPI_ISL_3352618, EPI_ISL_3352619, EPI_ISL_3352620, EPI_ISL_3352621, EPI_ISL_3352622, EPI_ISL_3352623, EPI_ISL_3352624, EPI_ISL_3352625, EPI_ISL_3352626, EPI_ISL_3352627, EPI_ISL_3352628, EPI_ISL_3352629, EPI_ISL_3352630, EPI_ISL_3352631, EPI_ISL_3352632, EPI_ISL_3352633, EPI_ISL_3352634, EPI_ISL_3352635, EPI_ISL_3352636, EPI_ISL_3352637, EPI_ISL_3352638, EPI_ISL_3352639, EPI_ISL_3352640, EPI_ISL_3352641, EPI_ISL_3352642, EPI_ISL_3352643, EPI_ISL_3352644, EPI_ISL_3352645, EPI_ISL_3352646, EPI_ISL_3352647, EPI_ISL_3352648, EPI_ISL_3352649, EPI_ISL_3352650, EPI_ISL_3352651, EPI_ISL_3352652, EPI_ISL_3352653, EPI_ISL_3352655, EPI_ISL_3352656, EPI_ISL_3352657, EPI_ISL_3352658, EPI_ISL_3352660, EPI_ISL_3352661, EPI_ISL_3352662, EPI_ISL_3352663, EPI_ISL_3352664, EPI_ISL_3352665, EPI_ISL_3352666, EPI_ISL_3352667, EPI_ISL_3352668, EPI_ISL_3352669, EPI_ISL_3352670, EPI_ISL_3352671, EPI_ISL_3352672, EPI_ISL_3352673, EPI_ISL_3352674, EPI_ISL_3352675, EPI_ISL_3352676, EPI_ISL_3352677, EPI_ISL_3352678, EPI_ISL_3352679, EPI_ISL_3352681, EPI_ISL_3352682, EPI_ISL_3352683, EPI_ISL_3352684, EPI_ISL_3352685, EPI_ISL_3352686, EPI_ISL_3352687, EPI_ISL_3352688, EPI_ISL_3352689, EPI_ISL_3352690, EPI_ISL_3352691, EPI_ISL_3352692, EPI_ISL_3352693, EPI_ISL_3352694, EPI_ISL_3352695, EPI_ISL_3352696, EPI_ISL_3352697, EPI_ISL_3352698, EPI_ISL_3352699, EPI_ISL_3352700, EPI_ISL_3352701, EPI_ISL_3352702, EPI_ISL_3352703, EPI_ISL_3352704, EPI_ISL_3352705, EPI_ISL_3352706, EPI_ISL_3352707, EPI_ISL_3352708, EPI_ISL_3352709, EPI_ISL_3352710, EPI_ISL_3352711, EPI_ISL_3352712, EPI_ISL_3352713, EPI_ISL_3352714, EPI_ISL_3352715, EPI_ISL_3352716, EPI_ISL_3352717, EPI_ISL_3352718, EPI_ISL_3352719, EPI_ISL_3352720, EPI_ISL_3352721, EPI_ISL_3352722, EPI_ISL_3352723, EPI_ISL_3352724, EPI_ISL_3352725, EPI_ISL_3352726, EPI_ISL_3352727, EPI_ISL_3352728, EPI_ISL_3352729, EPI_ISL_3352730, EPI_ISL_3352731, EPI_ISL_3352733, EPI_ISL_3352734, EPI_ISL_3352735, EPI_ISL_3352736, EPI_ISL_3352737, EPI_ISL_3352738, EPI_ISL_3352739, EPI_ISL_3352740, EPI_ISL_3352741, EPI_ISL_3352742, EPI_ISL_3352743, EPI_ISL_3352744, EPI_ISL_3352745, | | | |

| | | | |
|---|---|---|---|
| EPI_ISL_3352746, EPI_ISL_3352747, EPI_ISL_3352748, EPI_ISL_3352749, EPI_ISL_3352750, EPI_ISL_3352751, EPI_ISL_3352752, EPI_ISL_3352753, EPI_ISL_3352754, EPI_ISL_3352755, EPI_ISL_3352756, EPI_ISL_3352757, EPI_ISL_3352758, EPI_ISL_3352759, EPI_ISL_3352761, EPI_ISL_3352762, EPI_ISL_3352763, EPI_ISL_3352764, EPI_ISL_3352765, EPI_ISL_3352766, EPI_ISL_3352767, EPI_ISL_3352768, EPI_ISL_3352769, EPI_ISL_3352770, EPI_ISL_3352771, EPI_ISL_3352772, EPI_ISL_3352773, EPI_ISL_3352774, EPI_ISL_3352775, EPI_ISL_3352776, EPI_ISL_3352777, EPI_ISL_3352778, EPI_ISL_3352779, EPI_ISL_3352780, EPI_ISL_3352781, EPI_ISL_3352782, EPI_ISL_3352783, EPI_ISL_3352784, EPI_ISL_3352785, EPI_ISL_3352786, EPI_ISL_3352787, EPI_ISL_3352788, EPI_ISL_3352789, EPI_ISL_3352790, EPI_ISL_3352791, EPI_ISL_3352792, EPI_ISL_3352793, EPI_ISL_3352794, EPI_ISL_3352795, EPI_ISL_3352796, EPI_ISL_3352797, EPI_ISL_3352798, EPI_ISL_3352799, EPI_ISL_3352800, EPI_ISL_3352801, EPI_ISL_3352802, EPI_ISL_3352803, EPI_ISL_3352804, EPI_ISL_3352805, EPI_ISL_3352806, EPI_ISL_3352807, EPI_ISL_3352808, EPI_ISL_3352809, EPI_ISL_3352810, EPI_ISL_3352811, EPI_ISL_3352812, EPI_ISL_3352813, EPI_ISL_3352814, EPI_ISL_3352815, EPI_ISL_3352816, EPI_ISL_3352818, EPI_ISL_3352819, EPI_ISL_3352820, EPI_ISL_3352821, EPI_ISL_3352822, EPI_ISL_3352823, EPI_ISL_3352824, EPI_ISL_3352825, EPI_ISL_3352826, EPI_ISL_3352827, EPI_ISL_3352828, EPI_ISL_3352829, EPI_ISL_3352830, EPI_ISL_3352831, EPI_ISL_3352832, EPI_ISL_3352833, EPI_ISL_3352834, EPI_ISL_3352835, EPI_ISL_3352836, EPI_ISL_3352837, EPI_ISL_3352838, EPI_ISL_3352839, EPI_ISL_3352840, EPI_ISL_3352841, EPI_ISL_3352842, EPI_ISL_3352843, EPI_ISL_3352844, EPI_ISL_3352845, EPI_ISL_3352846, EPI_ISL_3352847, EPI_ISL_3352848, EPI_ISL_3352849, EPI_ISL_3352850, EPI_ISL_3352851, EPI_ISL_3352852, EPI_ISL_3352853, EPI_ISL_3352854, EPI_ISL_3352855, EPI_ISL_3352856, EPI_ISL_3352857, EPI_ISL_3352858, EPI_ISL_3352859, EPI_ISL_3352860, EPI_ISL_3352862, EPI_ISL_3352863, EPI_ISL_3352864, EPI_ISL_3352865, EPI_ISL_3352866, EPI_ISL_3352867, EPI_ISL_3352868, EPI_ISL_3352869, EPI_ISL_3352870, EPI_ISL_3352871, EPI_ISL_3352872, EPI_ISL_3352873, EPI_ISL_3352874, EPI_ISL_3352875, EPI_ISL_3352876, EPI_ISL_3352877, EPI_ISL_3352878, EPI_ISL_3352879, EPI_ISL_3352880, EPI_ISL_3352881, EPI_ISL_3352882, EPI_ISL_3352883, EPI_ISL_3352884, EPI_ISL_3352885, EPI_ISL_3352886, EPI_ISL_3352887, EPI_ISL_3352888, EPI_ISL_3352889, EPI_ISL_3352890, EPI_ISL_3352891, EPI_ISL_3352892, EPI_ISL_3352893, EPI_ISL_3352894, EPI_ISL_3352895, EPI_ISL_3352896, EPI_ISL_3352897, EPI_ISL_3352898, EPI_ISL_3352899, EPI_ISL_3352900, EPI_ISL_3352901, EPI_ISL_3352902, EPI_ISL_3352903, EPI_ISL_3352904, EPI_ISL_3352905, EPI_ISL_3352906, EPI_ISL_3352907, EPI_ISL_3352908, EPI_ISL_3352909, EPI_ISL_3352910, EPI_ISL_3352911, EPI_ISL_3352912, EPI_ISL_3352913, EPI_ISL_3352914, EPI_ISL_3352915, EPI_ISL_3352916, EPI_ISL_3352917, EPI_ISL_3352918, EPI_ISL_3352919, EPI_ISL_3352920, EPI_ISL_3352921, EPI_ISL_3352922, EPI_ISL_3352923, EPI_ISL_3352924, EPI_ISL_3352925, EPI_ISL_3352926, EPI_ISL_3352927, EPI_ISL_3352928, EPI_ISL_3352929, EPI_ISL_3352930, EPI_ISL_3352931, EPI_ISL_3352932, EPI_ISL_3352934, EPI_ISL_3352935, EPI_ISL_3352936, EPI_ISL_3352937, EPI_ISL_3352938, EPI_ISL_3352939, EPI_ISL_3352941, EPI_ISL_3352942, EPI_ISL_3352943, EPI_ISL_3352944, EPI_ISL_3352945, EPI_ISL_3352946, EPI_ISL_3352947, EPI_ISL_3352948, EPI_ISL_3352949, EPI_ISL_3352950, EPI_ISL_3352951, EPI_ISL_3352952, EPI_ISL_3352953, EPI_ISL_3352954, EPI_ISL_3352955, EPI_ISL_3352956, EPI_ISL_3352957, EPI_ISL_3352958, EPI_ISL_3352959, EPI_ISL_3352960, EPI_ISL_3352961, EPI_ISL_3352962, EPI_ISL_3352963, EPI_ISL_3352964, EPI_ISL_3352965, EPI_ISL_3352966, EPI_ISL_3352967, EPI_ISL_3352968, EPI_ISL_3352969, EPI_ISL_3352970, EPI_ISL_3352971, EPI_ISL_3352972, EPI_ISL_3352973, EPI_ISL_3352974, EPI_ISL_3352975, EPI_ISL_3352976, EPI_ISL_3352977, EPI_ISL_3352978, EPI_ISL_3352979, EPI_ISL_3352980, EPI_ISL_3352981, EPI_ISL_3352982, EPI_ISL_3352983, EPI_ISL_3352984, EPI_ISL_3352985, EPI_ISL_3352986, EPI_ISL_3352987, EPI_ISL_3352988, EPI_ISL_3352989, EPI_ISL_3352990, EPI_ISL_3352991, EPI_ISL_3352992, EPI_ISL_3352993, EPI_ISL_3352994, EPI_ISL_3352995, EPI_ISL_3352996, EPI_ISL_3352997, EPI_ISL_3352998, EPI_ISL_3352999, EPI_ISL_3353000, EPI_ISL_3353001, EPI_ISL_3353002, EPI_ISL_3353003, EPI_ISL_3353004, EPI_ISL_3353005, EPI_ISL_3353006, EPI_ISL_3353007, EPI_ISL_3353008, EPI_ISL_3353009, EPI_ISL_3353010, EPI_ISL_3353011, EPI_ISL_3353012, EPI_ISL_3353013, EPI_ISL_3353014, EPI_ISL_3353015, EPI_ISL_3353016, EPI_ISL_3353017, EPI_ISL_3353018, EPI_ISL_3353019, EPI_ISL_3353020, EPI_ISL_3353021, EPI_ISL_3353022, EPI_ISL_3353023, EPI_ISL_3353024, EPI_ISL_3353025, EPI_ISL_3353026, EPI_ISL_3353027, EPI_ISL_3353028, EPI_ISL_3353029, EPI_ISL_3353030, EPI_ISL_3353031, EPI_ISL_3353032, EPI_ISL_3353033, EPI_ISL_3353034, EPI_ISL_3353035, EPI_ISL_3353036, EPI_ISL_3353037, EPI_ISL_3353038, EPI_ISL_3353039, EPI_ISL_3353040, EPI_ISL_3353041, EPI_ISL_3353042, EPI_ISL_3353043, EPI_ISL_3353044, EPI_ISL_3353045, EPI_ISL_3353046, EPI_ISL_3353047, EPI_ISL_3353048, EPI_ISL_3353049, EPI_ISL_3353050, EPI_ISL_3353051, EPI_ISL_3353052, EPI_ISL_3353053, EPI_ISL_3353054, EPI_ISL_3353055, EPI_ISL_3353056, EPI_ISL_3353057, EPI_ISL_3353058, EPI_ISL_3353059, EPI_ISL_3353060, EPI_ISL_3353062, EPI_ISL_3353063, EPI_ISL_3353064, EPI_ISL_3353065, EPI_ISL_3353066, EPI_ISL_3353067, EPI_ISL_3353068, EPI_ISL_3353069, EPI_ISL_3353070, EPI_ISL_3353071, EPI_ISL_3353072, EPI_ISL_3353073, EPI_ISL_3353074, EPI_ISL_3353075, EPI_ISL_3353076, EPI_ISL_3353077, EPI_ISL_3353078, EPI_ISL_3353079, EPI_ISL_3353080, EPI_ISL_3353081, EPI_ISL_3353082, EPI_ISL_3353083, EPI_ISL_3353084, EPI_ISL_3353085, EPI_ISL_3353086, EPI_ISL_3353087, EPI_ISL_3353088, EPI_ISL_3353089, EPI_ISL_3353090, EPI_ISL_3353091, EPI_ISL_3353092, EPI_ISL_3353093, EPI_ISL_3353094, EPI_ISL_3353095, EPI_ISL_3353096, EPI_ISL_3353097, EPI_ISL_3353098, EPI_ISL_3353099, EPI_ISL_3353100, EPI_ISL_3353101, EPI_ISL_3353102, EPI_ISL_3353103, EPI_ISL_3353104, EPI_ISL_3353105, EPI_ISL_3353106, EPI_ISL_3353107, EPI_ISL_3353108, EPI_ISL_3353109, EPI_ISL_3353110, EPI_ISL_3353111, EPI_ISL_3353112, EPI_ISL_3353113, EPI_ISL_3353114, EPI_ISL_3353115, EPI_ISL_3353116, EPI_ISL_3353117, EPI_ISL_3353118, EPI_ISL_3353119, EPI_ISL_3353120, EPI_ISL_3353121, EPI_ISL_3353122, EPI_ISL_3353123, EPI_ISL_3353124, EPI_ISL_3353125, EPI_ISL_3353126, EPI_ISL_3353127, EPI_ISL_3353128, EPI_ISL_3353129, EPI_ISL_3353130, EPI_ISL_3353131, EPI_ISL_3353132, EPI_ISL_3353133, EPI_ISL_3353134, EPI_ISL_3353135, EPI_ISL_3353136, EPI_ISL_3353137, EPI_ISL_3353138, EPI_ISL_3353139, EPI_ISL_3353140, EPI_ISL_3353141, EPI_ISL_3353142, EPI_ISL_3353143, EPI_ISL_3353144, EPI_ISL_3353145, EPI_ISL_3353146, EPI_ISL_3353147, EPI_ISL_3353148, EPI_ISL_3353149, EPI_ISL_3353150, EPI_ISL_3353151, EPI_ISL_3353152, EPI_ISL_3353153, EPI_ISL_3353154, EPI_ISL_3353155, EPI_ISL_3353156, EPI_ISL_3353157, EPI_ISL_3353158, EPI_ISL_3353159, EPI_ISL_3353160, EPI_ISL_3353161, EPI_ISL_3353162, EPI_ISL_3353163, EPI_ISL_3353164, EPI_ISL_3353165, EPI_ISL_3353166, EPI_ISL_3353167, EPI_ISL_3353168, EPI_ISL_3353169, EPI_ISL_3353170, EPI_ISL_3353171, EPI_ISL_3353172, EPI_ISL_3353173, EPI_ISL_3353174, EPI_ISL_3353175, EPI_ISL_3353176, EPI_ISL_3353177, EPI_ISL_3353178, EPI_ISL_3353179, EPI_ISL_3353180, EPI_ISL_3353181, EPI_ISL_3353182, EPI_ISL_3353183, EPI_ISL_3353184, EPI_ISL_3353185, EPI_ISL_3353186, EPI_ISL_3353187, EPI_ISL_3353188, EPI_ISL_3353189, EPI_ISL_3353190, EPI_ISL_3353191, EPI_ISL_3353192, EPI_ISL_3353193, EPI_ISL_3353194, EPI_ISL_3353195, EPI_ISL_3353196, EPI_ISL_3353197, EPI_ISL_3353198, EPI_ISL_3353199, EPI_ISL_3353200, EPI_ISL_3353201, EPI_ISL_3353202, EPI_ISL_3353203, EPI_ISL_3353204, EPI_ISL_3353205, EPI_ISL_3353206, EPI_ISL_3353207, EPI_ISL_3353208, EPI_ISL_3353209, EPI_ISL_3353210, EPI_ISL_3353211, EPI_ISL_3353212, EPI_ISL_3353213, EPI_ISL_3353214, EPI_ISL_3353215, EPI_ISL_3353216, EPI_ISL_3353217, EPI_ISL_3353218, EPI_ISL_3353219, EPI_ISL_3353220, EPI_ISL_3353221, EPI_ISL_3353222, EPI_ISL_3353223, EPI_ISL_3353224, EPI_ISL_3353225, EPI_ISL_3353226, EPI_ISL_3353227, EPI_ISL_3353228, EPI_ISL_3353229, EPI_ISL_3353230, EPI_ISL_3353231, EPI_ISL_3353232, EPI_ISL_3353233, EPI_ISL_3353234, EPI_ISL_3353235, EPI_ISL_3353236, EPI_ISL_3353237, EPI_ISL_3353238, EPI_ISL_3353239, EPI_ISL_3353240, EPI_ISL_3353241, EPI_ISL_3353242, EPI_ISL_3353243, EPI_ISL_3353244, EPI_ISL_3353245, EPI_ISL_3353246, EPI_ISL_3353247, EPI_ISL_3353248, EPI_ISL_3353249, EPI_ISL_3353250, EPI_ISL_3353251, EPI_ISL_3353252, EPI_ISL_3353253, EPI_ISL_3353254, EPI_ISL_3353255, EPI_ISL_3353256, EPI_ISL_3353258, EPI_ISL_3353259, EPI_ISL_3353260, EPI_ISL_3353261, EPI_ISL_3353262, EPI_ISL_3353263, EPI_ISL_3353264, EPI_ISL_3353265, EPI_ISL_3353266, EPI_ISL_3353267, EPI_ISL_3353268, EPI_ISL_3353269, EPI_ISL_3353270, EPI_ISL_3353271, EPI_ISL_3353272, EPI_ISL_3353273, EPI_ISL_3353274, EPI_ISL_3353275, EPI_ISL_3353276, EPI_ISL_3353277, EPI_ISL_3353278, EPI_ISL_3353279, EPI_ISL_3353280, EPI_ISL_3353281, EPI_ISL_3353282, EPI_ISL_3353283, EPI_ISL_3353284, EPI_ISL_3353285, EPI_ISL_3353287, EPI_ISL_3353288, EPI_ISL_3353289, EPI_ISL_3353290, EPI_ISL_3353291, EPI_ISL_3353292, EPI_ISL_3353293, EPI_ISL_3353294, EPI_ISL_3353296, EPI_ISL_3353297, EPI_ISL_3353298, EPI_ISL_3353299, EPI_ISL_3353300, EPI_ISL_3353301, EPI_ISL_3353302, EPI_ISL_3353303, EPI_ISL_3353304, EPI_ISL_3353306, EPI_ISL_3353307, EPI_ISL_3353308, EPI_ISL_3353309, EPI_ISL_3353310, EPI_ISL_3353311, EPI_ISL_3353312, EPI_ISL_3353313, EPI_ISL_3353314, EPI_ISL_3353315, EPI_ISL_3353316, EPI_ISL_3353317, EPI_ISL_3353318, EPI_ISL_3353319, EPI_ISL_3353320, EPI_ISL_3353321, EPI_ISL_3353322, EPI_ISL_3353323, EPI_ISL_3353324, EPI_ISL_3353325, EPI_ISL_3353326, EPI_ISL_3353327, EPI_ISL_3353328, EPI_ISL_3353329, EPI_ISL_3353330, EPI_ISL_3353331, EPI_ISL_3353332, EPI_ISL_3353333, EPI_ISL_3353334, EPI_ISL_3353335, EPI_ISL_3353336, EPI_ISL_3353337, EPI_ISL_3353338, EPI_ISL_3353339, EPI_ISL_3353340, EPI_ISL_3353341, EPI_ISL_3353342, EPI_ISL_3353343, EPI_ISL_3353344, EPI_ISL_3353345, EPI_ISL_3353346, EPI_ISL_3353347, EPI_ISL_3353348, EPI_ISL_3353350, EPI_ISL_3353351, EPI_ISL_3353352, EPI_ISL_3353353, EPI_ISL_3353355, EPI_ISL_3353356, EPI_ISL_3353357, EPI_ISL_3353358, EPI_ISL_3353359, EPI_ISL_3353360, EPI_ISL_3353361, EPI_ISL_3353362, EPI_ISL_3353363, EPI_ISL_3353364, EPI_ISL_3353365, EPI_ISL_3353366, EPI_ISL_3353367, EPI_ISL_3353368, EPI_ISL_3353369 | | | |
| see above | Quest Diagnostics Incorporated | Centers for Disease Control and Prevention Division of Viral Diseases, Pathogen Discovery | Dakota Howard, Dhwani Batra, Peter W. Cook, Kara Moser, Adrian Paskey, Jason Caravas, Benjamin Rambo-Martin, Shatavia Morrison, Christopher Gulvick, Scott Sammons, Yvette Unoarumhi, Darlene Wagner, Matthew Schmerer, S. H. Rosenthal, A. Gerasimova, R. M. Kagan, B. Anderson, M. Hua, Y. Liu, L.E. Bernstein, K.E. Livingston, A. Perez, I. A. Shlyakhter, R. V. Rolando, R. Owen, P. Tanpaiboon, F. Lacbawan, Clinton R. Paden, Duncan MacCannell |
| EPI_ISL_3353370, EPI_ISL_3353371, EPI_ISL_3353372, EPI_ISL_3353373, EPI_ISL_3353374, EPI_ISL_3353375, EPI_ISL_3353376, EPI_ISL_3353377, EPI_ISL_3353378, EPI_ISL_3353379, EPI_ISL_3353380, EPI_ISL_3353381, EPI_ISL_3353382, EPI_ISL_3353383, EPI_ISL_3353384, EPI_ISL_3353385, EPI_ISL_3353386, EPI_ISL_3353387, EPI_ISL_3353388, EPI_ISL_3353389, EPI_ISL_3353390, EPI_ISL_3353391, EPI_ISL_3353392, EPI_ISL_3353393, EPI_ISL_3353394, EPI_ISL_3353395, EPI_ISL_3353396, EPI_ISL_3353397, EPI_ISL_3353398, EPI_ISL_3353399, EPI_ISL_3353400, EPI_ISL_3353401, EPI_ISL_3353402, EPI_ISL_3353403, EPI_ISL_3353404, EPI_ISL_3353405, EPI_ISL_3353406, EPI_ISL_3353407, EPI_ISL_3353408, EPI_ISL_3353409, EPI_ISL_3353410, EPI_ISL_3353411, EPI_ISL_3353412, EPI_ISL_3353413, EPI_ISL_3353414, EPI_ISL_3353415, EPI_ISL_3353416, EPI_ISL_3353417, EPI_ISL_3353418, EPI_ISL_3353419, EPI_ISL_3353420, EPI_ISL_3353421, EPI_ISL_3353422, EPI_ISL_3353423, EPI_ISL_3353424, EPI_ISL_3353425, EPI_ISL_3353426, EPI_ISL_3353427, EPI_ISL_3353428, EPI_ISL_3353429, EPI_ISL_3353430, EPI_ISL_3353431, EPI_ISL_3353432, EPI_ISL_3353433, EPI_ISL_3353434, EPI_ISL_3353435, EPI_ISL_3353436, EPI_ISL_3353437, EPI_ISL_3353438, EPI_ISL_3353439, EPI_ISL_3353440, EPI_ISL_3353441, EPI_ISL_3353442, EPI_ISL_3353443, EPI_ISL_3353444, EPI_ISL_3353445, EPI_ISL_3353446, EPI_ISL_3353447, EPI_ISL_3353448, EPI_ISL_3353449, EPI_ISL_3353450, EPI_ISL_3353451, EPI_ISL_3353452, EPI_ISL_3353453, EPI_ISL_3353454, EPI_ISL_3353455, EPI_ISL_3353456, EPI_ISL_3353457, EPI_ISL_3353458, EPI_ISL_3353459, EPI_ISL_3353460, EPI_ISL_3353461, EPI_ISL_3353462, EPI_ISL_3353463, EPI_ISL_3353464, EPI_ISL_3353465, EPI_ISL_3353466, EPI_ISL_3353467, EPI_ISL_3353468, EPI_ISL_3353469, EPI_ISL_3353470, EPI_ISL_3353471, EPI_ISL_3353472, EPI_ISL_3353473, EPI_ISL_3353474, EPI_ISL_3353475, EPI_ISL_3353476, EPI_ISL_3353477, EPI_ISL_3353478, EPI_ISL_3353479, EPI_ISL_3353480, EPI_ISL_3353481, EPI_ISL_3353482, EPI_ISL_3353483, EPI_ISL_3353484, EPI_ISL_3353485, EPI_ISL_3353486, EPI_ISL_3353487, EPI_ISL_3353488, EPI_ISL_3353489, EPI_ISL_3353490, EPI_ISL_3353491, EPI_ISL_3353492, EPI_ISL_3353493, EPI_ISL_3353494, EPI_ISL_3353495, EPI_ISL_3353496, EPI_ISL_3353497, EPI_ISL_3353499, EPI_ISL_3353500, EPI_ISL_3353501, EPI_ISL_3353502, EPI_ISL_3353503, EPI_ISL_3353504, EPI_ISL_3353505, EPI_ISL_3353506, EPI_ISL_3353507, EPI_ISL_3353508, EPI_ISL_3353509, EPI_ISL_3353510, EPI_ISL_3353511, EPI_ISL_3353512, EPI_ISL_3353513, EPI_ISL_3353514, EPI_ISL_3353515, EPI_ISL_3353516, EPI_ISL_3353518, EPI_ISL_3353519, EPI_ISL_3353520, EPI_ISL_3353521, EPI_ISL_3353522, EPI_ISL_3353523, EPI_ISL_3353524, EPI_ISL_3353525, EPI_ISL_3353526, EPI_ISL_3353527, EPI_ISL_3353528, EPI_ISL_3353529, EPI_ISL_3353530, EPI_ISL_3353531, EPI_ISL_3353532, EPI_ISL_3353533, EPI_ISL_3353534, EPI_ISL_3353535, EPI_ISL_3353536, EPI_ISL_3353537, EPI_ISL_3353538, EPI_ISL_3353539, EPI_ISL_3353540, EPI_ISL_3353541, EPI_ISL_3353542, EPI_ISL_3353543, EPI_ISL_3353544, EPI_ISL_3353545, EPI_ISL_3353546, EPI_ISL_3353547, EPI_ISL_3353548, EPI_ISL_3353549, EPI_ISL_3353550, EPI_ISL_3353551, EPI_ISL_3353552, EPI_ISL_3353553, EPI_ISL_3353554, EPI_ISL_3353555, EPI_ISL_3353556, EPI_ISL_3353557, EPI_ISL_3353558, EPI_ISL_3353559, EPI_ISL_3353560, EPI_ISL_3353561, EPI_ISL_3353562, EPI_ISL_3353563, EPI_ISL_3353564, EPI_ISL_3353565, EPI_ISL_3353566, EPI_ISL_3353567, EPI_ISL_3353568, EPI_ISL_3353569, EPI_ISL_3353570, EPI_ISL_3353571, EPI_ISL_3353572, EPI_ISL_3353573, EPI_ISL_3353574, EPI_ISL_3353575, EPI_ISL_3353576, EPI_ISL_3353577, EPI_ISL_3353578, EPI_ISL_3353579, EPI_ISL_3353580, EPI_ISL_3353581, EPI_ISL_3353582, EPI_ISL_3353583, EPI_ISL_3353584, EPI_ISL_3353585, EPI_ISL_3353586, EPI_ISL_3353587, EPI_ISL_3353588, EPI_ISL_3353589, EPI_ISL_3353590, EPI_ISL_3353591, EPI_ISL_3353592, EPI_ISL_3353593, EPI_ISL_3353594, EPI_ISL_3353595, EPI_ISL_3353596, EPI_ISL_3353597, EPI_ISL_3353598, EPI_ISL_3353599, EPI_ISL_3353600 | | | |
| see above | Infinity Biologix | Centers for Disease Control and Prevention Division of Viral Diseases, Pathogen Discovery | Dakota Howard, Dhwani Batra, Peter W. Cook, Kara Moser, Adrian Paskey, Jason Caravas, Benjamin Rambo-Martin, Shatavia Morrison, Christopher Gulvick, Scott Sammons, Yvette Unoarumhi, Darlene Wagner, Matthew Schmerer, Christian Bixby, Yihe Wang, Jonathan Schultz, Chirayu Goswami, Russ Hager, Robin Grimwood, Clinton R. Paden, Duncan MacCannell |
| EPI_ISL_3353628 | IA State Hygienic Laboratory | Centers for Disease Control and Prevention Division of Viral Diseases, Pathogen Discovery | Mili Sheth,Sarah Nobles,Jasmine Padilla,Alex Burgin,Meghan Bentz,Mark Burroughs,Peter Cook,Clinton Paden,Dhwani Batra,Krista Queen,Kristen Knipe,Dakota Howard,Yvette Unoarumhi,Matthew Schmerer,Ben Rambo-Martin,Kristine Lacek,Sam Shepard,Dave Wentworth,Vivien Dugan,Suxiang Tong,Justin Lee |
| EPI_ISL_3353629 | "AZ SPHL, Arizona Department of Health Services" | Centers for Disease Control and Prevention Division of Viral Diseases, Pathogen Discovery | Mili Sheth,Sarah Nobles,Jasmine Padilla,Alex Burgin,Meghan Bentz,Mark Burroughs,Peter Cook,Clinton Paden,Dhwani Batra,Krista Queen,Kristen Knipe,Dakota Howard,Yvette Unoarumhi,Matthew Schmerer,Ben Rambo-Martin,Kristine Lacek,Sam Shepard,Dave Wentworth,Vivien Dugan,Suxiang Tong,Justin Lee |
| EPI_ISL_3353630 | "AR Dept. of Health-PHL, Molecular Diagnostics" | Centers for Disease Control and Prevention Division of Viral Diseases, Pathogen Discovery | Mili Sheth,Sarah Nobles,Jasmine Padilla,Alex Burgin,Meghan Bentz,Mark Burroughs,Peter Cook,Clinton Paden,Dhwani Batra,Krista Queen,Kristen Knipe,Dakota Howard,Yvette Unoarumhi,Matthew Schmerer,Ben Rambo-Martin,Kristine Lacek,Sam Shepard,Dave Wentworth,Vivien Dugan,Suxiang Tong,Justin Lee |
| EPI_ISL_3353632 | KS Health and Environmental Laboratories | Centers for Disease Control and Prevention Division of Viral Diseases, Pathogen Discovery | Mili Sheth,Sarah Nobles,Jasmine Padilla,Alex Burgin,Meghan Bentz,Mark Burroughs,Peter Cook,Clinton Paden,Dhwani Batra,Krista Queen,Kristen Knipe,Dakota Howard,Yvette Unoarumhi,Matthew Schmerer,Ben Rambo-Martin,Kristine Lacek,Sam Shepard,Dave Wentworth,Vivien Dugan,Suxiang Tong,Justin Lee |
| EPI_ISL_3353633 | "AZ SPHL, Arizona Department of Health Services" | Centers for Disease Control and Prevention Division of Viral Diseases, Pathogen Discovery | Mili Sheth,Sarah Nobles,Jasmine Padilla,Alex Burgin,Meghan Bentz,Mark Burroughs,Peter Cook,Clinton Paden,Dhwani Batra,Krista Queen,Kristen Knipe,Dakota Howard,Yvette Unoarumhi,Matthew Schmerer,Ben Rambo-Martin,Kristine Lacek,Sam Shepard,Dave Wentworth,Vivien Dugan,Suxiang Tong,Justin Lee |
| EPI_ISL_3353635 | ID Bureau of Laboratories | Centers for Disease Control and Prevention Division of Viral Diseases, Pathogen Discovery | Mili Sheth,Sarah Nobles,Jasmine Padilla,Alex Burgin,Meghan Bentz,Mark Burroughs,Peter Cook,Clinton Paden,Dhwani Batra,Krista Queen,Kristen Knipe,Dakota Howard,Yvette Unoarumhi,Matthew Schmerer,Ben Rambo-Martin,Kristine Lacek,Sam Shepard,Dave Wentworth,Vivien Dugan,Suxiang Tong,Justin Lee |
| EPI_ISL_3353636, EPI_ISL_3353637 | FL Bur. of Public Health Laboratories-Jacksonville | Centers for Disease Control and Prevention Division of Viral Diseases, Pathogen Discovery | Mili Sheth,Sarah Nobles,Jasmine Padilla,Alex Burgin,Meghan Bentz,Mark Burroughs,Peter Cook,Clinton Paden,Dhwani Batra,Krista Queen,Kristen Knipe,Dakota Howard,Yvette Unoarumhi,Matthew Schmerer,Ben Rambo-Martin,Kristine Lacek,Sam Shepard,Dave Wentworth,Vivien Dugan,Suxiang Tong,Justin Lee |
| EPI_ISL_3353638 | SD Public Health Laboratory | Centers for Disease Control and Prevention Division of Viral Diseases, Pathogen Discovery | Mili Sheth,Sarah Nobles,Jasmine Padilla,Alex Burgin,Meghan Bentz,Mark Burroughs,Peter Cook,Clinton Paden,Dhwani Batra,Krista Queen,Kristen Knipe,Dakota Howard,Yvette Unoarumhi,Matthew Schmerer,Ben Rambo-Martin,Kristine Lacek,Sam Shepard,Dave Wentworth,Vivien Dugan,Suxiang Tong,Justin Lee |
| EPI_ISL_3354941 | NV State Public Health Laboratory | Centers for Disease Control and Prevention Division of Viral Diseases, Pathogen Discovery | Mili Sheth,Sarah Nobles,Jasmine Padilla,Alex Burgin,Meghan Bentz,Mark Burroughs,Peter Cook,Clinton Paden,Dhwani Batra,Krista Queen,Kristen Knipe,Dakota Howard,Yvette Unoarumhi,Matthew Schmerer,Ben Rambo-Martin,Kristine Lacek,Sam Shepard,Dave Wentworth,Vivien Dugan,Suxiang Tong,Justin Lee |
| EPI_ISL_3355040 | MA State Public Health Laboratory | Centers for Disease Control and Prevention Division of Viral Diseases, Pathogen Discovery | Mili Sheth,Sarah Nobles,Jasmine Padilla,Alex Burgin,Meghan Bentz,Mark Burroughs,Peter Cook,Clinton Paden,Dhwani Batra,Krista Queen,Kristen Knipe,Dakota Howard,Yvette Unoarumhi,Matthew Schmerer,Ben Rambo-Martin,Kristine Lacek,Sam Shepard,Dave Wentworth,Vivien Dugan,Suxiang Tong,Justin Lee |
| EPI_ISL_3355089 | MS Public Health Laboratory | Centers for Disease Control and Prevention Division of Viral Diseases, Pathogen Discovery | Mili Sheth,Sarah Nobles,Jasmine Padilla,Alex Burgin,Meghan Bentz,Mark Burroughs,Peter Cook,Clinton Paden,Dhwani Batra,Krista Queen,Kristen Knipe,Dakota Howard,Yvette Unoarumhi,Matthew Schmerer,Ben Rambo-Martin,Kristine Lacek,Sam Shepard,Dave Wentworth,Vivien Dugan,Suxiang Tong,Justin Lee |
| EPI_ISL_3355092 | AR Dept. of Health-PHL/Chemical Terrorism | Centers for Disease Control and Prevention Division of Viral Diseases, Pathogen Discovery | Mili Sheth,Sarah Nobles,Jasmine Padilla,Alex Burgin,Meghan Bentz,Mark Burroughs,Peter Cook,Clinton Paden,Dhwani Batra,Krista Queen,Kristen Knipe,Dakota Howard,Yvette Unoarumhi,Matthew Schmerer,Ben Rambo-Martin,Kristine Lacek,Sam Shepard,Dave Wentworth,Vivien Dugan,Suxiang Tong,Justin Lee |
| EPI_ISL_3355094 | "AR Dept. of Health-PHL, Molecular Diagnostics" | Centers for Disease Control and Prevention Division of Viral Diseases, Pathogen Discovery | Mili Sheth,Sarah Nobles,Jasmine Padilla,Alex Burgin,Meghan Bentz,Mark Burroughs,Peter Cook,Clinton Paden,Dhwani Batra,Krista Queen,Kristen Knipe,Dakota Howard,Yvette Unoarumhi,Matthew Schmerer,Ben Rambo-Martin,Kristine Lacek,Sam Shepard,Dave Wentworth,Vivien Dugan,Suxiang Tong,Justin Lee |
| EPI_ISL_3355095 | IA State Hygienic Laboratory | Centers for Disease Control and Prevention Division of Viral Diseases, Pathogen Discovery | Mili Sheth,Sarah Nobles,Jasmine Padilla,Alex Burgin,Meghan Bentz,Mark Burroughs,Peter Cook,Clinton Paden,Dhwani Batra,Krista Queen,Kristen Knipe,Dakota Howard,Yvette Unoarumhi,Matthew Schmerer,Ben Rambo-Martin,Kristine Lacek,Sam Shepard,Dave Wentworth,Vivien Dugan,Suxiang Tong,Justin Lee |
| EPI_ISL_3355146, EPI_ISL_3355149, EPI_ISL_3355170, EPI_ISL_3355172, EPI_ISL_3355174, EPI_ISL_3355178 | SD Public Health Laboratory | Centers for Disease Control and Prevention Division of Viral Diseases, Pathogen Discovery | Mili Sheth,Sarah Nobles,Jasmine Padilla,Alex Burgin,Meghan Bentz,Mark Burroughs,Peter Cook,Clinton Paden,Dhwani Batra,Krista Queen,Kristen Knipe,Dakota Howard,Yvette Unoarumhi,Matthew Schmerer,Ben Rambo-Martin,Kristine Lacek,Sam Shepard,Dave Wentworth,Vivien Dugan,Suxiang Tong,Justin Lee |
| EPI_ISL_3355182 | VA-Division of Consolidated Laboratory Services | Centers for Disease Control and Prevention Division of Viral Diseases, Pathogen Discovery | Mili Sheth,Sarah Nobles,Jasmine Padilla,Alex Burgin,Meghan Bentz,Mark Burroughs,Peter Cook,Clinton Paden,Dhwani Batra,Krista Queen,Kristen Knipe,Dakota Howard,Yvette Unoarumhi,Matthew Schmerer,Ben Rambo-Martin,Kristine Lacek,Sam Shepard,Dave Wentworth,Vivien Dugan,Suxiang Tong,Justin Lee |
| EPI_ISL_3355184 | PR Public Health Lab | Centers for Disease Control and Prevention Division of Viral Diseases, Pathogen Discovery | Mili Sheth,Sarah Nobles,Jasmine Padilla,Alex Burgin,Meghan Bentz,Mark Burroughs,Peter Cook,Clinton Paden,Dhwani Batra,Krista Queen,Kristen Knipe,Dakota Howard,Yvette Unoarumhi,Matthew Schmerer,Ben Rambo-Martin,Kristine Lacek,Sam Shepard,Dave Wentworth,Vivien Dugan,Suxiang Tong,Justin Lee |
| EPI_ISL_3355187 | Guam Public Health Laboratory | Centers for Disease Control and Prevention Division of Viral Diseases, Pathogen Discovery | Mili Sheth,Sarah Nobles,Jasmine Padilla,Alex Burgin,Meghan Bentz,Mark Burroughs,Peter Cook,Clinton Paden,Dhwani Batra,Krista Queen,Kristen Knipe,Dakota Howard,Yvette Unoarumhi,Matthew Schmerer,Ben Rambo-Martin,Kristine Lacek,Sam Shepard,Dave Wentworth,Vivien Dugan,Suxiang Tong,Justin Lee |
| EPI_ISL_3355211 | SD Public Health Laboratory | Centers for Disease Control and Prevention Division of Viral Diseases, Pathogen Discovery | Mili Sheth,Sarah Nobles,Jasmine Padilla,Alex Burgin,Meghan Bentz,Mark Burroughs,Peter Cook,Clinton Paden,Dhwani Batra,Krista Queen,Kristen Knipe,Dakota Howard,Yvette Unoarumhi,Matthew Schmerer,Ben Rambo-Martin,Kristine Lacek,Sam Shepard,Dave Wentworth,Vivien Dugan,Suxiang Tong,Justin Lee |
| EPI_ISL_3355216, EPI_ISL_3355226 | "AR Dept. of Health-PHL, Molecular Diagnostics" | Centers for Disease Control and Prevention Division of Viral Diseases, Pathogen Discovery | Mili Sheth,Sarah Nobles,Jasmine Padilla,Alex Burgin,Meghan Bentz,Mark Burroughs,Peter Cook,Clinton Paden,Dhwani Batra,Krista Queen,Kristen Knipe,Dakota Howard,Yvette Unoarumhi,Matthew Schmerer,Ben Rambo-Martin,Kristine Lacek,Sam Shepard,Dave Wentworth,Vivien Dugan,Suxiang Tong,Justin Lee |
| EPI_ISL_3355230 | IA State Hygienic Laboratory | Centers for Disease Control and Prevention Division of Viral Diseases, Pathogen Discovery | Mili Sheth,Sarah Nobles,Jasmine Padilla,Alex Burgin,Meghan Bentz,Mark Burroughs,Peter Cook,Clinton Paden,Dhwani Batra,Krista Queen,Kristen Knipe,Dakota Howard,Yvette Unoarumhi,Matthew Schmerer,Ben Rambo-Martin,Kristine Lacek,Sam Shepard,Dave Wentworth,Vivien Dugan,Suxiang Tong,Justin Lee |
| EPI_ISL_3355237 | KS Health and Environmental Laboratories | Centers for Disease Control and Prevention Division of Viral Diseases, Pathogen Discovery | Mili Sheth,Sarah Nobles,Jasmine Padilla,Alex Burgin,Meghan Bentz,Mark Burroughs,Peter Cook,Clinton Paden,Dhwani Batra,Krista Queen,Kristen Knipe,Dakota Howard,Yvette Unoarumhi,Matthew Schmerer,Ben Rambo-Martin,Kristine Lacek,Sam Shepard,Dave Wentworth,Vivien Dugan,Suxiang Tong,Justin Lee |
| EPI_ISL_3355238 | "AZ SPHL, Arizona Department of Health Services" | Centers for Disease Control and Prevention Division of Viral Diseases, Pathogen Discovery | Mili Sheth,Sarah Nobles,Jasmine Padilla,Alex Burgin,Meghan Bentz,Mark Burroughs,Peter Cook,Clinton Paden,Dhwani Batra,Krista Queen,Kristen Knipe,Dakota Howard,Yvette Unoarumhi,Matthew Schmerer,Ben Rambo-Martin,Kristine Lacek,Sam Shepard,Dave Wentworth,Vivien Dugan,Suxiang Tong,Justin Lee |
| EPI_ISL_3355241 | IA State Hygienic Laboratory | Centers for Disease Control and Prevention Division of Viral Diseases, Pathogen Discovery | Mili Sheth,Sarah Nobles,Jasmine Padilla,Alex Burgin,Meghan Bentz,Mark Burroughs,Peter Cook,Clinton Paden,Dhwani Batra,Krista Queen,Kristen Knipe,Dakota Howard,Yvette Unoarumhi,Matthew Schmerer,Ben Rambo-Martin,Kristine Lacek,Sam Shepard,Dave Wentworth,Vivien Dugan,Suxiang Tong,Justin Lee |
| EPI_ISL_3355245 | "AR Dept. of Health-PHL, Molecular Diagnostics" | Centers for Disease Control and Prevention Division of Viral Diseases, Pathogen Discovery | Mili Sheth,Sarah Nobles,Jasmine Padilla,Alex Burgin,Meghan Bentz,Mark Burroughs,Peter Cook,Clinton Paden,Dhwani Batra,Krista Queen,Kristen Knipe,Dakota Howard,Yvette Unoarumhi,Matthew Schmerer,Ben Rambo-Martin,Kristine Lacek,Sam Shepard,Dave Wentworth,Vivien Dugan,Suxiang Tong,Justin Lee |

| | | | |
|---|---|---|---|
| EPI_ISL_3355246 | IA State Hygienic Laboratory | Centers for Disease Control and Prevention Division of Viral Diseases, Pathogen Discovery | Mili Sheth,Sarah Nobles,Jasmine Padilla,Alex Burgin,Meghan Bentz,Mark Burroughs,Peter Cook,Clinton Paden,Dhwani Batra,Krista Queen,Kristen Knipe,Dakota Howard,Yvette Unoarumhi,Matthew Schmerer,Ben Rambo-Martin,Kristine Lacek,Sam Shepard,Dave Wentworth,Vivien Dugan,Suxiang Tong,Justin Lee |
| EPI_ISL_3355252 | "AR Dept. of Health-PHL, Molecular Diagnostics" | Centers for Disease Control and Prevention Division of Viral Diseases, Pathogen Discovery | Mili Sheth,Sarah Nobles,Jasmine Padilla,Alex Burgin,Meghan Bentz,Mark Burroughs,Peter Cook,Clinton Paden,Dhwani Batra,Krista Queen,Kristen Knipe,Dakota Howard,Yvette Unoarumhi,Matthew Schmerer,Ben Rambo-Martin,Kristine Lacek,Sam Shepard,Dave Wentworth,Vivien Dugan,Suxiang Tong,Justin Lee |
| EPI_ISL_3355253 | IA State Hygienic Laboratory | Centers for Disease Control and Prevention Division of Viral Diseases, Pathogen Discovery | Mili Sheth,Sarah Nobles,Jasmine Padilla,Alex Burgin,Meghan Bentz,Mark Burroughs,Peter Cook,Clinton Paden,Dhwani Batra,Krista Queen,Kristen Knipe,Dakota Howard,Yvette Unoarumhi,Matthew Schmerer,Ben Rambo-Martin,Kristine Lacek,Sam Shepard,Dave Wentworth,Vivien Dugan,Suxiang Tong,Justin Lee |
| EPI_ISL_3355256 | "AR Dept. of Health-PHL, Molecular Diagnostics" | Centers for Disease Control and Prevention Division of Viral Diseases, Pathogen Discovery | Mili Sheth,Sarah Nobles,Jasmine Padilla,Alex Burgin,Meghan Bentz,Mark Burroughs,Peter Cook,Clinton Paden,Dhwani Batra,Krista Queen,Kristen Knipe,Dakota Howard,Yvette Unoarumhi,Matthew Schmerer,Ben Rambo-Martin,Kristine Lacek,Sam Shepard,Dave Wentworth,Vivien Dugan,Suxiang Tong,Justin Lee |
| EPI_ISL_3355257 | IA State Hygienic Laboratory | Centers for Disease Control and Prevention Division of Viral Diseases, Pathogen Discovery | Mili Sheth,Sarah Nobles,Jasmine Padilla,Alex Burgin,Meghan Bentz,Mark Burroughs,Peter Cook,Clinton Paden,Dhwani Batra,Krista Queen,Kristen Knipe,Dakota Howard,Yvette Unoarumhi,Matthew Schmerer,Ben Rambo-Martin,Kristine Lacek,Sam Shepard,Dave Wentworth,Vivien Dugan,Suxiang Tong,Justin Lee |
| EPI_ISL_3355258 | PR Public Health Lab | Centers for Disease Control and Prevention Division of Viral Diseases, Pathogen Discovery | Mili Sheth,Sarah Nobles,Jasmine Padilla,Alex Burgin,Meghan Bentz,Mark Burroughs,Peter Cook,Clinton Paden,Dhwani Batra,Krista Queen,Kristen Knipe,Dakota Howard,Yvette Unoarumhi,Matthew Schmerer,Ben Rambo-Martin,Kristine Lacek,Sam Shepard,Dave Wentworth,Vivien Dugan,Suxiang Tong,Justin Lee |
| EPI_ISL_3355259 | IA State Hygienic Laboratory | Centers for Disease Control and Prevention Division of Viral Diseases, Pathogen Discovery | Mili Sheth,Sarah Nobles,Jasmine Padilla,Alex Burgin,Meghan Bentz,Mark Burroughs,Peter Cook,Clinton Paden,Dhwani Batra,Krista Queen,Kristen Knipe,Dakota Howard,Yvette Unoarumhi,Matthew Schmerer,Ben Rambo-Martin,Kristine Lacek,Sam Shepard,Dave Wentworth,Vivien Dugan,Suxiang Tong,Justin Lee |
| EPI_ISL_3355260 | "AR Dept. of Health-PHL, Molecular Diagnostics" | Centers for Disease Control and Prevention Division of Viral Diseases, Pathogen Discovery | Mili Sheth,Sarah Nobles,Jasmine Padilla,Alex Burgin,Meghan Bentz,Mark Burroughs,Peter Cook,Clinton Paden,Dhwani Batra,Krista Queen,Kristen Knipe,Dakota Howard,Yvette Unoarumhi,Matthew Schmerer,Ben Rambo-Martin,Kristine Lacek,Sam Shepard,Dave Wentworth,Vivien Dugan,Suxiang Tong,Justin Lee |
| EPI_ISL_3355264 | KS Health and Environmental Laboratories | Centers for Disease Control and Prevention Division of Viral Diseases, Pathogen Discovery | Mili Sheth,Sarah Nobles,Jasmine Padilla,Alex Burgin,Meghan Bentz,Mark Burroughs,Peter Cook,Clinton Paden,Dhwani Batra,Krista Queen,Kristen Knipe,Dakota Howard,Yvette Unoarumhi,Matthew Schmerer,Ben Rambo-Martin,Kristine Lacek,Sam Shepard,Dave Wentworth,Vivien Dugan,Suxiang Tong,Justin Lee |
| EPI_ISL_3355290 | "NM Dept. Health, Scientific Laboratory Division " | Centers for Disease Control and Prevention Division of Viral Diseases, Pathogen Discovery | Mili Sheth,Sarah Nobles,Jasmine Padilla,Alex Burgin,Meghan Bentz,Mark Burroughs,Peter Cook,Clinton Paden,Dhwani Batra,Krista Queen,Kristen Knipe,Dakota Howard,Yvette Unoarumhi,Matthew Schmerer,Ben Rambo-Martin,Kristine Lacek,Sam Shepard,Dave Wentworth,Vivien Dugan,Suxiang Tong,Justin Lee |
| EPI_ISL_3355299 | GA Department of Public Health Laboratory | Centers for Disease Control and Prevention Division of Viral Diseases, Pathogen Discovery | Mili Sheth,Sarah Nobles,Jasmine Padilla,Alex Burgin,Meghan Bentz,Mark Burroughs,Peter Cook,Clinton Paden,Dhwani Batra,Krista Queen,Kristen Knipe,Dakota Howard,Yvette Unoarumhi,Matthew Schmerer,Ben Rambo-Martin,Kristine Lacek,Sam Shepard,Dave Wentworth,Vivien Dugan,Suxiang Tong,Justin Lee |
| EPI_ISL_3355318 | "NM Dept. Health, Scientific Laboratory Division " | Centers for Disease Control and Prevention Division of Viral Diseases, Pathogen Discovery | Mili Sheth,Sarah Nobles,Jasmine Padilla,Alex Burgin,Meghan Bentz,Mark Burroughs,Peter Cook,Clinton Paden,Dhwani Batra,Krista Queen,Kristen Knipe,Dakota Howard,Yvette Unoarumhi,Matthew Schmerer,Ben Rambo-Martin,Kristine Lacek,Sam Shepard,Dave Wentworth,Vivien Dugan,Suxiang Tong,Justin Lee |
| EPI_ISL_3355334 | FL Bur. of Public Health Laboratories-Jacksonville | Centers for Disease Control and Prevention Division of Viral Diseases, Pathogen Discovery | Mili Sheth,Sarah Nobles,Jasmine Padilla,Alex Burgin,Meghan Bentz,Mark Burroughs,Peter Cook,Clinton Paden,Dhwani Batra,Krista Queen,Kristen Knipe,Dakota Howard,Yvette Unoarumhi,Matthew Schmerer,Ben Rambo-Martin,Kristine Lacek,Sam Shepard,Dave Wentworth,Vivien Dugan,Suxiang Tong,Justin Lee |
| EPI_ISL_3355335 | FL Bureau of Public Health Laboratories-Tampa | Centers for Disease Control and Prevention Division of Viral Diseases, Pathogen Discovery | Mili Sheth,Sarah Nobles,Jasmine Padilla,Alex Burgin,Meghan Bentz,Mark Burroughs,Peter Cook,Clinton Paden,Dhwani Batra,Krista Queen,Kristen Knipe,Dakota Howard,Yvette Unoarumhi,Matthew Schmerer,Ben Rambo-Martin,Kristine Lacek,Sam Shepard,Dave Wentworth,Vivien Dugan,Suxiang Tong,Justin Lee |
| EPI_ISL_3355336, EPI_ISL_3355337 | "CDPH, Viral and Rickettsial Disease Laboratory" | Centers for Disease Control and Prevention Division of Viral Diseases, Pathogen Discovery | Mili Sheth,Sarah Nobles,Jasmine Padilla,Alex Burgin,Meghan Bentz,Mark Burroughs,Peter Cook,Clinton Paden,Dhwani Batra,Krista Queen,Kristen Knipe,Dakota Howard,Yvette Unoarumhi,Matthew Schmerer,Ben Rambo-Martin,Kristine Lacek,Sam Shepard,Dave Wentworth,Vivien Dugan,Suxiang Tong,Justin Lee |
| EPI_ISL_3355341 | WY Public Health Laboratory | Centers for Disease Control and Prevention Division of Viral Diseases, Pathogen Discovery | Mili Sheth,Sarah Nobles,Jasmine Padilla,Alex Burgin,Meghan Bentz,Mark Burroughs,Peter Cook,Clinton Paden,Dhwani Batra,Krista Queen,Kristen Knipe,Dakota Howard,Yvette Unoarumhi,Matthew Schmerer,Ben Rambo-Martin,Kristine Lacek,Sam Shepard,Dave Wentworth,Vivien Dugan,Suxiang Tong,Justin Lee |
| EPI_ISL_3355342 | "AR Dept. of Health-PHL, Molecular Diagnostics" | Centers for Disease Control and Prevention Division of Viral Diseases, Pathogen Discovery | Mili Sheth,Sarah Nobles,Jasmine Padilla,Alex Burgin,Meghan Bentz,Mark Burroughs,Peter Cook,Clinton Paden,Dhwani Batra,Krista Queen,Kristen Knipe,Dakota Howard,Yvette Unoarumhi,Matthew Schmerer,Ben Rambo-Martin,Kristine Lacek,Sam Shepard,Dave Wentworth,Vivien Dugan,Suxiang Tong,Justin Lee |
| EPI_ISL_3355343, EPI_ISL_3355345 | FL Bur. of Public Health Laboratories-Jacksonville | Centers for Disease Control and Prevention Division of Viral Diseases, Pathogen Discovery | Mili Sheth,Sarah Nobles,Jasmine Padilla,Alex Burgin,Meghan Bentz,Mark Burroughs,Peter Cook,Clinton Paden,Dhwani Batra,Krista Queen,Kristen Knipe,Dakota Howard,Yvette Unoarumhi,Matthew Schmerer,Ben Rambo-Martin,Kristine Lacek,Sam Shepard,Dave Wentworth,Vivien Dugan,Suxiang Tong,Justin Lee |
| EPI_ISL_3355346 | WY Public Health Laboratory | Centers for Disease Control and Prevention Division of Viral Diseases, Pathogen Discovery | Mili Sheth,Sarah Nobles,Jasmine Padilla,Alex Burgin,Meghan Bentz,Mark Burroughs,Peter Cook,Clinton Paden,Dhwani Batra,Krista Queen,Kristen Knipe,Dakota Howard,Yvette Unoarumhi,Matthew Schmerer,Ben Rambo-Martin,Kristine Lacek,Sam Shepard,Dave Wentworth,Vivien Dugan,Suxiang Tong,Justin Lee |
| EPI_ISL_3355347 | "AR Dept. of Health-PHL, Molecular Diagnostics" | Centers for Disease Control and Prevention Division of Viral Diseases, Pathogen Discovery | Mili Sheth,Sarah Nobles,Jasmine Padilla,Alex Burgin,Meghan Bentz,Mark Burroughs,Peter Cook,Clinton Paden,Dhwani Batra,Krista Queen,Kristen Knipe,Dakota Howard,Yvette Unoarumhi,Matthew Schmerer,Ben Rambo-Martin,Kristine Lacek,Sam Shepard,Dave Wentworth,Vivien Dugan,Suxiang Tong,Justin Lee |
| EPI_ISL_3355348 | "CDPH, Viral and Rickettsial Disease Laboratory" | Centers for Disease Control and Prevention Division of Viral Diseases, Pathogen Discovery | Mili Sheth,Sarah Nobles,Jasmine Padilla,Alex Burgin,Meghan Bentz,Mark Burroughs,Peter Cook,Clinton Paden,Dhwani Batra,Krista Queen,Kristen Knipe,Dakota Howard,Yvette Unoarumhi,Matthew Schmerer,Ben Rambo-Martin,Kristine Lacek,Sam Shepard,Dave Wentworth,Vivien Dugan,Suxiang Tong,Justin Lee |
| EPI_ISL_3355350 | FL Bur. of Public Health Laboratories-Jacksonville | Centers for Disease Control and Prevention Division of Viral Diseases, Pathogen Discovery | Mili Sheth,Sarah Nobles,Jasmine Padilla,Alex Burgin,Meghan Bentz,Mark Burroughs,Peter Cook,Clinton Paden,Dhwani Batra,Krista Queen,Kristen Knipe,Dakota Howard,Yvette Unoarumhi,Matthew Schmerer,Ben Rambo-Martin,Kristine Lacek,Sam Shepard,Dave Wentworth,Vivien Dugan,Suxiang Tong,Justin Lee |
| EPI_ISL_3355352, EPI_ISL_3355353 | "CDPH, Viral and Rickettsial Disease Laboratory" | Centers for Disease Control and Prevention Division of Viral Diseases, Pathogen Discovery | Mili Sheth,Sarah Nobles,Jasmine Padilla,Alex Burgin,Meghan Bentz,Mark Burroughs,Peter Cook,Clinton Paden,Dhwani Batra,Krista Queen,Kristen Knipe,Dakota Howard,Yvette Unoarumhi,Matthew Schmerer,Ben Rambo-Martin,Kristine Lacek,Sam Shepard,Dave Wentworth,Vivien Dugan,Suxiang Tong,Justin Lee |
| EPI_ISL_3355357 | "AR Dept. of Health-PHL, Molecular Diagnostics" | Centers for Disease Control and Prevention Division of Viral Diseases, Pathogen Discovery | Mili Sheth,Sarah Nobles,Jasmine Padilla,Alex Burgin,Meghan Bentz,Mark Burroughs,Peter Cook,Clinton Paden,Dhwani Batra,Krista Queen,Kristen Knipe,Dakota Howard,Yvette Unoarumhi,Matthew Schmerer,Ben Rambo-Martin,Kristine Lacek,Sam Shepard,Dave Wentworth,Vivien Dugan,Suxiang Tong,Justin Lee |
| EPI_ISL_3355359 | FL Bur. of Public Health Laboratories-Jacksonville | Centers for Disease Control and Prevention Division of Viral Diseases, Pathogen Discovery | Mili Sheth,Sarah Nobles,Jasmine Padilla,Alex Burgin,Meghan Bentz,Mark Burroughs,Peter Cook,Clinton Paden,Dhwani Batra,Krista Queen,Kristen Knipe,Dakota Howard,Yvette Unoarumhi,Matthew Schmerer,Ben Rambo-Martin,Kristine Lacek,Sam Shepard,Dave Wentworth,Vivien Dugan,Suxiang Tong,Justin Lee |
| EPI_ISL_3355361, EPI_ISL_3355362 | FL Bureau of Public Health Laboratories-Tampa | Centers for Disease Control and Prevention Division of Viral Diseases, Pathogen Discovery | Mili Sheth,Sarah Nobles,Jasmine Padilla,Alex Burgin,Meghan Bentz,Mark Burroughs,Peter Cook,Clinton Paden,Dhwani Batra,Krista Queen,Kristen Knipe,Dakota Howard,Yvette Unoarumhi,Matthew Schmerer,Ben Rambo-Martin,Kristine Lacek,Sam Shepard,Dave Wentworth,Vivien Dugan,Suxiang Tong,Justin Lee |
| EPI_ISL_3355363 | "AR Dept. of Health-PHL, Molecular Diagnostics" | Centers for Disease Control and Prevention Division of Viral Diseases, Pathogen Discovery | Mili Sheth,Sarah Nobles,Jasmine Padilla,Alex Burgin,Meghan Bentz,Mark Burroughs,Peter Cook,Clinton Paden,Dhwani Batra,Krista Queen,Kristen Knipe,Dakota Howard,Yvette Unoarumhi,Matthew Schmerer,Ben Rambo-Martin,Kristine Lacek,Sam Shepard,Dave Wentworth,Vivien Dugan,Suxiang Tong,Justin Lee |
| EPI_ISL_3355364 | WY Public Health Laboratory | Centers for Disease Control and Prevention Division of Viral Diseases, Pathogen Discovery | Mili Sheth,Sarah Nobles,Jasmine Padilla,Alex Burgin,Meghan Bentz,Mark Burroughs,Peter Cook,Clinton Paden,Dhwani Batra,Krista Queen,Kristen Knipe,Dakota Howard,Yvette Unoarumhi,Matthew Schmerer,Ben Rambo-Martin,Kristine Lacek,Sam Shepard,Dave Wentworth,Vivien Dugan,Suxiang Tong,Justin Lee |
| EPI_ISL_3355366 | "AR Dept. of Health-PHL, Molecular Diagnostics" | Centers for Disease Control and Prevention Division of Viral Diseases, Pathogen Discovery | Mili Sheth,Sarah Nobles,Jasmine Padilla,Alex Burgin,Meghan Bentz,Mark Burroughs,Peter Cook,Clinton Paden,Dhwani Batra,Krista Queen,Kristen Knipe,Dakota Howard,Yvette Unoarumhi,Matthew Schmerer,Ben Rambo-Martin,Kristine Lacek,Sam Shepard,Dave Wentworth,Vivien Dugan,Suxiang Tong,Justin Lee |
| EPI_ISL_3355368, EPI_ISL_3355369, EPI_ISL_3355371, EPI_ISL_3355374, EPI_ISL_3355376, EPI_ISL_3355377 | FL Bur. of Public Health Laboratories-Jacksonville | Centers for Disease Control and Prevention Division of Viral Diseases, Pathogen Discovery | Mili Sheth,Sarah Nobles,Jasmine Padilla,Alex Burgin,Meghan Bentz,Mark Burroughs,Peter Cook,Clinton Paden,Dhwani Batra,Krista Queen,Kristen Knipe,Dakota Howard,Yvette Unoarumhi,Matthew Schmerer,Ben Rambo-Martin,Kristine Lacek,Sam Shepard,Dave Wentworth,Vivien Dugan,Suxiang Tong,Justin Lee |
| EPI_ISL_3355378 | WY Public Health Laboratory | Centers for Disease Control and Prevention Division of Viral Diseases, Pathogen Discovery | Mili Sheth,Sarah Nobles,Jasmine Padilla,Alex Burgin,Meghan Bentz,Mark Burroughs,Peter Cook,Clinton Paden,Dhwani Batra,Krista Queen,Kristen Knipe,Dakota Howard,Yvette Unoarumhi,Matthew Schmerer,Ben Rambo-Martin,Kristine Lacek,Sam Shepard,Dave Wentworth,Vivien Dugan,Suxiang Tong,Justin Lee |
| EPI_ISL_3355379, EPI_ISL_3355383 | "AR Dept. of Health-PHL, Molecular Diagnostics" | Centers for Disease Control and Prevention Division of Viral Diseases, Pathogen Discovery | Mili Sheth,Sarah Nobles,Jasmine Padilla,Alex Burgin,Meghan Bentz,Mark Burroughs,Peter Cook,Clinton Paden,Dhwani Batra,Krista Queen,Kristen Knipe,Dakota Howard,Yvette Unoarumhi,Matthew Schmerer,Ben Rambo-Martin,Kristine Lacek,Sam Shepard,Dave Wentworth,Vivien Dugan,Suxiang Tong,Justin Lee |
| EPI_ISL_3355385, EPI_ISL_3355386 | FL Bur. of Public Health Laboratories-Jacksonville | Centers for Disease Control and Prevention Division of Viral Diseases, Pathogen Discovery | Mili Sheth,Sarah Nobles,Jasmine Padilla,Alex Burgin,Meghan Bentz,Mark Burroughs,Peter Cook,Clinton Paden,Dhwani Batra,Krista Queen,Kristen Knipe,Dakota Howard,Yvette Unoarumhi,Matthew Schmerer,Ben Rambo-Martin,Kristine Lacek,Sam Shepard,Dave Wentworth,Vivien Dugan,Suxiang Tong,Justin Lee |
| EPI_ISL_3355387 | RI State Health Laboratories | Centers for Disease Control and Prevention Division of Viral Diseases, Pathogen Discovery | Mili Sheth,Sarah Nobles,Jasmine Padilla,Alex Burgin,Meghan Bentz,Mark Burroughs,Peter Cook,Clinton Paden,Dhwani Batra,Krista Queen,Kristen Knipe,Dakota Howard,Yvette Unoarumhi,Matthew Schmerer,Ben Rambo-Martin,Kristine Lacek,Sam Shepard,Dave Wentworth,Vivien Dugan,Suxiang Tong,Justin Lee |
| EPI_ISL_3355388 | "CDPH, Viral and Rickettsial Disease Laboratory" | Centers for Disease Control and Prevention Division of Viral Diseases, Pathogen Discovery | Mili Sheth,Sarah Nobles,Jasmine Padilla,Alex Burgin,Meghan Bentz,Mark Burroughs,Peter Cook,Clinton Paden,Dhwani Batra,Krista Queen,Kristen Knipe,Dakota Howard,Yvette Unoarumhi,Matthew Schmerer,Ben Rambo-Martin,Kristine Lacek,Sam Shepard,Dave Wentworth,Vivien Dugan,Suxiang Tong,Justin Lee |
| EPI_ISL_3355390 | WY Public Health Laboratory | Centers for Disease Control and Prevention Division of Viral Diseases, Pathogen Discovery | Mili Sheth,Sarah Nobles,Jasmine Padilla,Alex Burgin,Meghan Bentz,Mark Burroughs,Peter Cook,Clinton Paden,Dhwani Batra,Krista Queen,Kristen Knipe,Dakota Howard,Yvette Unoarumhi,Matthew Schmerer,Ben Rambo-Martin,Kristine Lacek,Sam Shepard,Dave Wentworth,Vivien Dugan,Suxiang Tong,Justin Lee |
| EPI_ISL_3355391 | "AR Dept. of Health-PHL, Molecular Diagnostics" | Centers for Disease Control and Prevention Division of Viral Diseases, Pathogen Discovery | Mili Sheth,Sarah Nobles,Jasmine Padilla,Alex Burgin,Meghan Bentz,Mark Burroughs,Peter Cook,Clinton Paden,Dhwani Batra,Krista Queen,Kristen Knipe,Dakota Howard,Yvette Unoarumhi,Matthew Schmerer,Ben Rambo-Martin,Kristine Lacek,Sam Shepard,Dave Wentworth,Vivien Dugan,Suxiang Tong,Justin Lee |
| EPI_ISL_3355394 | FL Bur. of Public Health Laboratories-Jacksonville | Centers for Disease Control and Prevention Division of Viral Diseases, Pathogen Discovery | Mili Sheth,Sarah Nobles,Jasmine Padilla,Alex Burgin,Meghan Bentz,Mark Burroughs,Peter Cook,Clinton Paden,Dhwani Batra,Krista Queen,Kristen Knipe,Dakota Howard,Yvette Unoarumhi,Matthew Schmerer,Ben Rambo-Martin,Kristine Lacek,Sam Shepard,Dave Wentworth,Vivien Dugan,Suxiang Tong,Justin Lee |
| EPI_ISL_3355395 | FL Bureau of Public Health Laboratories-Tampa | Centers for Disease Control and Prevention Division of Viral Diseases, Pathogen Discovery | Mili Sheth,Sarah Nobles,Jasmine Padilla,Alex Burgin,Meghan Bentz,Mark Burroughs,Peter Cook,Clinton Paden,Dhwani Batra,Krista Queen,Kristen Knipe,Dakota Howard,Yvette Unoarumhi,Matthew Schmerer,Ben Rambo-Martin,Kristine Lacek,Sam Shepard,Dave Wentworth,Vivien Dugan,Suxiang Tong,Justin Lee |
| EPI_ISL_3355397 | "CDPH, Viral and Rickettsial Disease Laboratory" | Centers for Disease Control and Prevention Division of Viral Diseases, Pathogen Discovery | Mili Sheth,Sarah Nobles,Jasmine Padilla,Alex Burgin,Meghan Bentz,Mark Burroughs,Peter Cook,Clinton Paden,Dhwani Batra,Krista Queen,Kristen Knipe,Dakota Howard,Yvette Unoarumhi,Matthew Schmerer,Ben Rambo-Martin,Kristine Lacek,Sam Shepard,Dave Wentworth,Vivien Dugan,Suxiang Tong,Justin Lee |
| EPI_ISL_3355398 | "AR Dept. of Health-PHL, Molecular Diagnostics" | Centers for Disease Control and Prevention Division of Viral Diseases, Pathogen Discovery | Mili Sheth,Sarah Nobles,Jasmine Padilla,Alex Burgin,Meghan Bentz,Mark Burroughs,Peter Cook,Clinton Paden,Dhwani Batra,Krista Queen,Kristen Knipe,Dakota Howard,Yvette Unoarumhi,Matthew Schmerer,Ben Rambo-Martin,Kristine Lacek,Sam Shepard,Dave Wentworth,Vivien Dugan,Suxiang Tong,Justin Lee |
| EPI_ISL_3355399, EPI_ISL_3355400 | WY Public Health Laboratory | Centers for Disease Control and Prevention Division of Viral Diseases, Pathogen Discovery | Mili Sheth,Sarah Nobles,Jasmine Padilla,Alex Burgin,Meghan Bentz,Mark Burroughs,Peter Cook,Clinton Paden,Dhwani Batra,Krista Queen,Kristen Knipe,Dakota Howard,Yvette Unoarumhi,Matthew Schmerer,Ben Rambo-Martin,Kristine Lacek,Sam Shepard,Dave Wentworth,Vivien Dugan,Suxiang Tong,Justin Lee |
| EPI_ISL_3355402 | FL Bur. of Public Health Laboratories-Jacksonville | Centers for Disease Control and Prevention Division of Viral Diseases, Pathogen Discovery | Mili Sheth,Sarah Nobles,Jasmine Padilla,Alex Burgin,Meghan Bentz,Mark Burroughs,Peter Cook,Clinton Paden,Dhwani Batra,Krista Queen,Kristen Knipe,Dakota Howard,Yvette Unoarumhi,Matthew Schmerer,Ben Rambo-Martin,Kristine Lacek,Sam Shepard,Dave Wentworth,Vivien Dugan,Suxiang Tong,Justin Lee |
| EPI_ISL_3355403 | "AR Dept. of Health-PHL, Molecular Diagnostics" | Centers for Disease Control and Prevention Division of Viral | Mili Sheth,Sarah Nobles,Jasmine Padilla,Alex Burgin,Meghan Bentz,Mark Burroughs,Peter Cook,Clinton Paden,Dhwani Batra,Krista Queen,Kristen Knipe,Dakota Howard,Yvette Unoarumhi,Matthew Schmerer,Ben Rambo-Martin,Kristine Lacek,Sam |

| | | | |
|---|---|---|---|
| | | Diseases, Pathogen Discovery | Shepard,Dave Wentworth,Vivien Dugan,Suxiang Tong,Justin Lee |
| EPI_ISL_3355404 | WY Public Health Laboratory | Centers for Disease Control and Prevention Division of Viral Diseases, Pathogen Discovery | Mili Sheth,Sarah Nobles,Jasmine Padilla,Alex Burgin,Meghan Bentz,Mark Burroughs,Peter Cook,Clinton Paden,Dhwani Batra,Krista Queen,Kristen Knipe,Dakota Howard,Yvette Unoarumhi,Matthew Schmerer,Ben Rambo-Martin,Kristine Lacek,Sam Shepard,Dave Wentworth,Vivien Dugan,Suxiang Tong,Justin Lee |
| EPI_ISL_3355407 | "CDPH, Viral and Rickettsial Disease Laboratory" | Centers for Disease Control and Prevention Division of Viral Diseases, Pathogen Discovery | Mili Sheth,Sarah Nobles,Jasmine Padilla,Alex Burgin,Meghan Bentz,Mark Burroughs,Peter Cook,Clinton Paden,Dhwani Batra,Krista Queen,Kristen Knipe,Dakota Howard,Yvette Unoarumhi,Matthew Schmerer,Ben Rambo-Martin,Kristine Lacek,Sam Shepard,Dave Wentworth,Vivien Dugan,Suxiang Tong,Justin Lee |
| EPI_ISL_3355410 | RI State Health Laboratories | Centers for Disease Control and Prevention Division of Viral Diseases, Pathogen Discovery | Mili Sheth,Sarah Nobles,Jasmine Padilla,Alex Burgin,Meghan Bentz,Mark Burroughs,Peter Cook,Clinton Paden,Dhwani Batra,Krista Queen,Kristen Knipe,Dakota Howard,Yvette Unoarumhi,Matthew Schmerer,Ben Rambo-Martin,Kristine Lacek,Sam Shepard,Dave Wentworth,Vivien Dugan,Suxiang Tong,Justin Lee |
| EPI_ISL_3355413 | FL Bur. of Public Health Laboratories-Jacksonville | Centers for Disease Control and Prevention Division of Viral Diseases, Pathogen Discovery | Mili Sheth,Sarah Nobles,Jasmine Padilla,Alex Burgin,Meghan Bentz,Mark Burroughs,Peter Cook,Clinton Paden,Dhwani Batra,Krista Queen,Kristen Knipe,Dakota Howard,Yvette Unoarumhi,Matthew Schmerer,Ben Rambo-Martin,Kristine Lacek,Sam Shepard,Dave Wentworth,Vivien Dugan,Suxiang Tong,Justin Lee |
| EPI_ISL_3355415, EPI_ISL_3355416 | MI - Michigan Department of Health and Human Services - Bureau of Laboratories | Centers for Disease Control and Prevention Division of Viral Diseases, Pathogen Discovery | Mili Sheth,Sarah Nobles,Jasmine Padilla,Alex Burgin,Meghan Bentz,Mark Burroughs,Peter Cook,Clinton Paden,Dhwani Batra,Krista Queen,Kristen Knipe,Dakota Howard,Yvette Unoarumhi,Matthew Schmerer,Ben Rambo-Martin,Kristine Lacek,Sam Shepard,Dave Wentworth,Vivien Dugan,Suxiang Tong,Justin Lee |
| EPI_ISL_3355417, EPI_ISL_3355418 | IL Dept. of Public Health Springfield Laboratory | Centers for Disease Control and Prevention Division of Viral Diseases, Pathogen Discovery | Mili Sheth,Sarah Nobles,Jasmine Padilla,Alex Burgin,Meghan Bentz,Mark Burroughs,Peter Cook,Clinton Paden,Dhwani Batra,Krista Queen,Kristen Knipe,Dakota Howard,Yvette Unoarumhi,Matthew Schmerer,Ben Rambo-Martin,Kristine Lacek,Sam Shepard,Dave Wentworth,Vivien Dugan,Suxiang Tong,Justin Lee |
| EPI_ISL_3355420 | ND Dept. of Health Laboratory Services-Microbiology | Centers for Disease Control and Prevention Division of Viral Diseases, Pathogen Discovery | Mili Sheth,Sarah Nobles,Jasmine Padilla,Alex Burgin,Meghan Bentz,Mark Burroughs,Peter Cook,Clinton Paden,Dhwani Batra,Krista Queen,Kristen Knipe,Dakota Howard,Yvette Unoarumhi,Matthew Schmerer,Ben Rambo-Martin,Kristine Lacek,Sam Shepard,Dave Wentworth,Vivien Dugan,Suxiang Tong,Justin Lee |
| EPI_ISL_3355422 | IL Dept. of Public Health Springfield Laboratory | Centers for Disease Control and Prevention Division of Viral Diseases, Pathogen Discovery | Mili Sheth,Sarah Nobles,Jasmine Padilla,Alex Burgin,Meghan Bentz,Mark Burroughs,Peter Cook,Clinton Paden,Dhwani Batra,Krista Queen,Kristen Knipe,Dakota Howard,Yvette Unoarumhi,Matthew Schmerer,Ben Rambo-Martin,Kristine Lacek,Sam Shepard,Dave Wentworth,Vivien Dugan,Suxiang Tong,Justin Lee |
| EPI_ISL_3355423, EPI_ISL_3355424 | SD Public Health Laboratory | Centers for Disease Control and Prevention Division of Viral Diseases, Pathogen Discovery | Mili Sheth,Sarah Nobles,Jasmine Padilla,Alex Burgin,Meghan Bentz,Mark Burroughs,Peter Cook,Clinton Paden,Dhwani Batra,Krista Queen,Kristen Knipe,Dakota Howard,Yvette Unoarumhi,Matthew Schmerer,Ben Rambo-Martin,Kristine Lacek,Sam Shepard,Dave Wentworth,Vivien Dugan,Suxiang Tong,Justin Lee |
| EPI_ISL_3355425 | RI State Health Laboratories | Centers for Disease Control and Prevention Division of Viral Diseases, Pathogen Discovery | Mili Sheth,Sarah Nobles,Jasmine Padilla,Alex Burgin,Meghan Bentz,Mark Burroughs,Peter Cook,Clinton Paden,Dhwani Batra,Krista Queen,Kristen Knipe,Dakota Howard,Yvette Unoarumhi,Matthew Schmerer,Ben Rambo-Martin,Kristine Lacek,Sam Shepard,Dave Wentworth,Vivien Dugan,Suxiang Tong,Justin Lee |
| EPI_ISL_3355427, EPI_ISL_3355428, EPI_ISL_3355431 | IL Dept. of Public Health Springfield Laboratory | Centers for Disease Control and Prevention Division of Viral Diseases, Pathogen Discovery | Mili Sheth,Sarah Nobles,Jasmine Padilla,Alex Burgin,Meghan Bentz,Mark Burroughs,Peter Cook,Clinton Paden,Dhwani Batra,Krista Queen,Kristen Knipe,Dakota Howard,Yvette Unoarumhi,Matthew Schmerer,Ben Rambo-Martin,Kristine Lacek,Sam Shepard,Dave Wentworth,Vivien Dugan,Suxiang Tong,Justin Lee |
| EPI_ISL_3355432 | SD Public Health Laboratory | Centers for Disease Control and Prevention Division of Viral Diseases, Pathogen Discovery | Mili Sheth,Sarah Nobles,Jasmine Padilla,Alex Burgin,Meghan Bentz,Mark Burroughs,Peter Cook,Clinton Paden,Dhwani Batra,Krista Queen,Kristen Knipe,Dakota Howard,Yvette Unoarumhi,Matthew Schmerer,Ben Rambo-Martin,Kristine Lacek,Sam Shepard,Dave Wentworth,Vivien Dugan,Suxiang Tong,Justin Lee |
| EPI_ISL_3355436 | KS Health and Environmental Laboratories | Centers for Disease Control and Prevention Division of Viral Diseases, Pathogen Discovery | Mili Sheth,Sarah Nobles,Jasmine Padilla,Alex Burgin,Meghan Bentz,Mark Burroughs,Peter Cook,Clinton Paden,Dhwani Batra,Krista Queen,Kristen Knipe,Dakota Howard,Yvette Unoarumhi,Matthew Schmerer,Ben Rambo-Martin,Kristine Lacek,Sam Shepard,Dave Wentworth,Vivien Dugan,Suxiang Tong,Justin Lee |
| EPI_ISL_3355437, EPI_ISL_3355441 | ND Dept. of Health Laboratory Services-Microbiology | Centers for Disease Control and Prevention Division of Viral Diseases, Pathogen Discovery | Mili Sheth,Sarah Nobles,Jasmine Padilla,Alex Burgin,Meghan Bentz,Mark Burroughs,Peter Cook,Clinton Paden,Dhwani Batra,Krista Queen,Kristen Knipe,Dakota Howard,Yvette Unoarumhi,Matthew Schmerer,Ben Rambo-Martin,Kristine Lacek,Sam Shepard,Dave Wentworth,Vivien Dugan,Suxiang Tong,Justin Lee |
| EPI_ISL_3355443 | SD Public Health Laboratory | Centers for Disease Control and Prevention Division of Viral Diseases, Pathogen Discovery | Mili Sheth,Sarah Nobles,Jasmine Padilla,Alex Burgin,Meghan Bentz,Mark Burroughs,Peter Cook,Clinton Paden,Dhwani Batra,Krista Queen,Kristen Knipe,Dakota Howard,Yvette Unoarumhi,Matthew Schmerer,Ben Rambo-Martin,Kristine Lacek,Sam Shepard,Dave Wentworth,Vivien Dugan,Suxiang Tong,Justin Lee |
| EPI_ISL_3355444, EPI_ISL_3355447, EPI_ISL_3355448, EPI_ISL_3355450, EPI_ISL_3355451 | IL Dept. of Public Health Springfield Laboratory | Centers for Disease Control and Prevention Division of Viral Diseases, Pathogen Discovery | Mili Sheth,Sarah Nobles,Jasmine Padilla,Alex Burgin,Meghan Bentz,Mark Burroughs,Peter Cook,Clinton Paden,Dhwani Batra,Krista Queen,Kristen Knipe,Dakota Howard,Yvette Unoarumhi,Matthew Schmerer,Ben Rambo-Martin,Kristine Lacek,Sam Shepard,Dave Wentworth,Vivien Dugan,Suxiang Tong,Justin Lee |
| EPI_ISL_3355452 | MI - Michigan Department of Health and Human Services - Bureau of Laboratories | Centers for Disease Control and Prevention Division of Viral Diseases, Pathogen Discovery | Mili Sheth,Sarah Nobles,Jasmine Padilla,Alex Burgin,Meghan Bentz,Mark Burroughs,Peter Cook,Clinton Paden,Dhwani Batra,Krista Queen,Kristen Knipe,Dakota Howard,Yvette Unoarumhi,Matthew Schmerer,Ben Rambo-Martin,Kristine Lacek,Sam Shepard,Dave Wentworth,Vivien Dugan,Suxiang Tong,Justin Lee |
| EPI_ISL_3355455 | KS Health and Environmental Laboratories | Centers for Disease Control and Prevention Division of Viral Diseases, Pathogen Discovery | Mili Sheth,Sarah Nobles,Jasmine Padilla,Alex Burgin,Meghan Bentz,Mark Burroughs,Peter Cook,Clinton Paden,Dhwani Batra,Krista Queen,Kristen Knipe,Dakota Howard,Yvette Unoarumhi,Matthew Schmerer,Ben Rambo-Martin,Kristine Lacek,Sam Shepard,Dave Wentworth,Vivien Dugan,Suxiang Tong,Justin Lee |
| EPI_ISL_3355456 | ND Dept. of Health Laboratory Services-Microbiology | Centers for Disease Control and Prevention Division of Viral Diseases, Pathogen Discovery | Mili Sheth,Sarah Nobles,Jasmine Padilla,Alex Burgin,Meghan Bentz,Mark Burroughs,Peter Cook,Clinton Paden,Dhwani Batra,Krista Queen,Kristen Knipe,Dakota Howard,Yvette Unoarumhi,Matthew Schmerer,Ben Rambo-Martin,Kristine Lacek,Sam Shepard,Dave Wentworth,Vivien Dugan,Suxiang Tong,Justin Lee |
| EPI_ISL_3355464 | Lablink (M) Sdn. Bhd. | Institute for Medical Research, Infectious Disease Research Centre, National Institutes of Health, Ministry of Health Malaysia | Suppiah J, Kamel K, Mohd Zawawi Z, Azizan MA, Ramly N, Robert F, Anasir MI, Thayan R |
| EPI_ISL_3355465 | Lablink (M) Sdn. Bhd | Institute for Medical Research, Infectious Disease Research Centre, National Institutes of Health, Ministry of Health Malaysia | Suppiah J, Kamel K, Mohd Zawawi Z, Azizan MA, Ramly N, Robert F, Anasir MI, Thayan R |
| EPI_ISL_3355519 | Tengku Ampuan Rahimah Hospital | Institute for Medical Research, Infectious Disease Research Centre, National Institutes of Health, Ministry of Health Malaysia | Suppiah J, Kamel K, Mohd Zawawi Z, Azizan MA, Ramly N, Robert F, Anasir MI, Thayan R |
| EPI_ISL_3355522, EPI_ISL_3355523, EPI_ISL_3355524, EPI_ISL_3355525, EPI_ISL_3355526, EPI_ISL_3355528, EPI_ISL_3355530, EPI_ISL_3355531, EPI_ISL_3355532, EPI_ISL_3355533, EPI_ISL_3355534, EPI_ISL_3355535, EPI_ISL_3355536, EPI_ISL_3355537 | | | |
| see above | Department of Microbiology, Yamanashi Institute for Public Health and Environment | Genome Analysis Center, Yamanashi Central Hospital | Yosuke Hirotsu |
| EPI_ISL_3355545, EPI_ISL_3355546, EPI_ISL_3355547, EPI_ISL_3355550, EPI_ISL_3355551, EPI_ISL_3355552, EPI_ISL_3355553 | Genome Analysis Center, Yamanashi Central Hospital | Genome Analysis Center, Yamanashi Central Hospital | Yosuke Hirotsu |
| EPI_ISL_3355556 | Laboratorio Central de Epidemiología (LCE) | Instituto de Biotecnología de la UNAM | Consorcio Mexicano de Vigilancia Genómica (CoViGen-Mex). Authors (in alphabetical order): Julio Elias Alvarado-Yaah, Carlos F. Arias, Santiago Ávila-Ríos, Eduardo Becerril-Vargas, Víctor Hugo Borja-Aburto, Celia Boukadida, Cristóbal Cháidez-Quiróz, Juan Bautista Chale-Dzul, Ricardo Ciria Merce, Andreu Comas-García, Célida Duque Molina, Julissa Enciso-Ibarra, José Antonio Enciso-Moreno, Gloria Elena Espinosa-Ayala, Fernando Fontove-Herrera, Daniel Fregoso-Rueda, Victor Eduardo García-Arias, Alejandra García-Gasca, Bruno Gómez-Gil, Jean Pierre González, Irvin González-López, Concepción Grajales-Muñiz, Ricardo Grande, Rosa María Gutiérrez Rios, Jesús Hernández, Alejandra Hernández-Terán, Alfredo Herrera-Estrella, Carla Ivón Herrera-Najera, Pavel Isa, Verónica Mata-Haro, Daniel Lira Morales, Susana Lopez, Antonio Loza Román, Brenda Irasema Maldonado-Meza, Bernardo Martínez-Miguel, José Arturo Martínez-Orozco, Célida Martínez-Rodríguez, Margarita Matías-Florentino, Fidencio Mejía-Nepomuceno, María Guadalupe de Jesús Mireles-Rivera, Gloria María Molina-Salinas, Hector Montoya-Fuentes, , Mario Mújica-Sánchez, José Esteban Muñoz-Medina, José de Jesús Nuñez-Contreras, Alicia Ocaña-Mondragón, Luis Alberto Ochoa-Carrera, Hector Esteban Paz-Juárez, Marissa Perez-Garcia, Francisco Pulido, Helen Haydee Fernanda Ramírez-Plascencia, Jorge Salas-Hernández, Angel Gustavo Salas-Lais, Alejandro Sánchez-Flores, Clara Esperanza Santacruz-Tinoco, María Guadalupe Santiago-Mauricio, Selene Zárate, Nelly Sélem-Mojica, Blanca Taboada, Gloria Vazquez, Joel Armando Vázquez-Pérez. |
| EPI_ISL_3355605, EPI_ISL_3355606, EPI_ISL_3355607, EPI_ISL_3355608, EPI_ISL_3355609, EPI_ISL_3355610, EPI_ISL_3355611, EPI_ISL_3355612, EPI_ISL_3355613, EPI_ISL_3355614, EPI_ISL_3355615, EPI_ISL_3355616, EPI_ISL_3355617 | | | |
| see above | CLINIQUE N-D DE GRACE | Institut de Pathologie et Genetique (IPG) | Pascale Hilbert, Jérémie Gras |
| EPI_ISL_3355618, EPI_ISL_3355619, EPI_ISL_3355620, EPI_ISL_3355621, EPI_ISL_3355622, EPI_ISL_3355623, EPI_ISL_3355624, EPI_ISL_3355625 | CHWAPI - SITE NOTRE DAME | Institut de Pathologie et Genetique (IPG) | Pascale Hilbert, Jérémie Gras |
| EPI_ISL_3355626, EPI_ISL_3355627 | Labo Analyses Med | National Reference Center for Viruses of Respiratory Infections, Institut Pasteur, Paris | Marion Barbet, Sylvie Behillil, Méline Bizard, Angela Brisebarre, Camille Capel, Vincent Enouf, Louise Lefrançois, Frédéric Lemoine, Christophe Malabat, Corinne Maufrais, Hub de Bioinformatique et Biostatistique, Etienne Simon-Lorière, Maud Vanpeene, Sylvie Van der Werf ,Baptiste Moreira |
| EPI_ISL_3355628, EPI_ISL_3355629, EPI_ISL_3355630, EPI_ISL_3355631, EPI_ISL_3355632, EPI_ISL_3355633, EPI_ISL_3355634, EPI_ISL_3355635, EPI_ISL_3355636, EPI_ISL_3355637, EPI_ISL_3355639, EPI_ISL_3355640, EPI_ISL_3355641, EPI_ISL_3355642, EPI_ISL_3355643 | | | |
| see above | Labo Analyses Med | National Reference Center for Viruses of Respiratory Infections, Institut Pasteur, Paris | Marion Barbet, Sylvie Behillil, Méline Bizard, Angela Brisebarre, Camille Capel, Vincent Enouf, Louise Lefrançois, Frédéric Lemoine, Christophe Malabat, Corinne Maufrais, Hub de Bioinformatique et Biostatistique, Etienne Simon-Lorière, Maud Vanpeene, Sylvie Van der Werf |
| EPI_ISL_3355644, EPI_ISL_3355645, EPI_ISL_3355646, EPI_ISL_3355647, EPI_ISL_3355648, EPI_ISL_3355649, EPI_ISL_3355650, EPI_ISL_3355651, EPI_ISL_3355652, EPI_ISL_3355653, EPI_ISL_3355654, EPI_ISL_3355655, EPI_ISL_3355656, EPI_ISL_3355657, EPI_ISL_3355658, EPI_ISL_3355659, EPI_ISL_3355660, EPI_ISL_3355661, EPI_ISL_3355662 | | | |
| see above | Labo Analyses Med | National Reference Center for Viruses of Respiratory Infections, Institut Pasteur, Paris | Marion Barbet, Sylvie Behillil, Méline Bizard, Angela Brisebarre, Camille Capel, Vincent Enouf, Louise Lefrançois, Frédéric Lemoine, Christophe Malabat, Corinne Maufrais, Hub de Bioinformatique et Biostatistique, Etienne Simon-Lorière, Maud Vanpeene, Sylvie Van der Werf ,C. Lecorre |
| EPI_ISL_3355679, EPI_ISL_3355680 | Hospital | National Reference Center for Viruses of Respiratory Infections, Institut Pasteur, Paris | Marion Barbet, Sylvie Behillil, Méline Bizard, Angela Brisebarre, Camille Capel, Vincent Enouf, Louise Lefrançois, Frédéric Lemoine, Christophe Malabat, Corinne Maufrais, Hub de Bioinformatique et Biostatistique, Etienne Simon-Lorière, Maud Vanpeene, Sylvie Van der Werf ,Mj Dufour |
| EPI_ISL_3355681, EPI_ISL_3355682, EPI_ISL_3355683, EPI_ISL_3355684, EPI_ISL_3355685, EPI_ISL_3355686, EPI_ISL_3355687, EPI_ISL_3355688, EPI_ISL_3355689, EPI_ISL_3355690, EPI_ISL_3355691, EPI_ISL_3355692, EPI_ISL_3355693 | | | |
| see above | Labo Analyses Med | National Reference Center for Viruses of Respiratory Infections, Institut Pasteur, Paris | Marion Barbet, Sylvie Behillil, Méline Bizard, Angela Brisebarre, Camille Capel, Vincent Enouf, Louise Lefrançois, Frédéric Lemoine, Christophe Malabat, Corinne Maufrais, Hub de Bioinformatique et Biostatistique, Etienne Simon-Lorière, Maud Vanpeene, Sylvie Van der Werf ,FrançOis-Xavier Le Guillou |
| EPI_ISL_3355694, EPI_ISL_3355695, EPI_ISL_3355696, EPI_ISL_3355697, EPI_ISL_3355698, EPI_ISL_3355699, EPI_ISL_3355700, EPI_ISL_3355701, EPI_ISL_3355702, EPI_ISL_3355703, EPI_ISL_3355704, EPI_ISL_3355705, EPI_ISL_3355706, EPI_ISL_3355707, EPI_ISL_3355708, EPI_ISL_3355709, EPI_ISL_3355710, EPI_ISL_3355711, EPI_ISL_3355712 | | | |
| see above | Labo Analyses Med | National Reference Center for Viruses of Respiratory Infections, Institut Pasteur, Paris | Marion Barbet, Sylvie Behillil, Méline Bizard, Angela Brisebarre, Camille Capel, Vincent Enouf, Louise Lefrançois, Frédéric Lemoine, Christophe Malabat, Corinne Maufrais, Hub de Bioinformatique et Biostatistique, Etienne Simon-Lorière, Maud Vanpeene, Sylvie Van der Werf ,Sabine Trombert |
| EPI_ISL_3355715 | Hospital | National Reference Center for Viruses of Respiratory Infections, | Marion Barbet, Sylvie Behillil, Méline Bizard, Angela Brisebarre, Camille Capel, Vincent Enouf, Louise Lefrançois, Frédéric Lemoine, Christophe Malabat, Corinne Maufrais, Hub de Bioinformatique et Biostatistique, Etienne Simon-Lorière, Maud |

| | | | |
|---|---|---|---|
| | | Institut Pasteur, Paris | Vanpeene, Sylvie Van der Werf ,Laurence Louvet |
| EPI_ISL_3355717 | Hospital | National Reference Center for Viruses of Respiratory Infections, Institut Pasteur, Paris | Marion Barbet, Sylvie Behillil, Méline Bizard, Angela Brisebarre, Camille Capel, Vincent Enouf, Louise Lefrançois, Frédéric Lemoine, Christophe Malabat, Corinne Maufrais, Hub de Bioinformatique et Biostatistique, Etienne Simon-Lorière, Maud Vanpeene, Sylvie Van der Werf ,José-Manuel Ramirez |
| EPI_ISL_3355718 | Hospital | National Reference Center for Viruses of Respiratory Infections, Institut Pasteur, Paris | Marion Barbet, Sylvie Behillil, Méline Bizard, Angela Brisebarre, Camille Capel, Vincent Enouf, Louise Lefrançois, Frédéric Lemoine, Christophe Malabat, Corinne Maufrais, Hub de Bioinformatique et Biostatistique, Etienne Simon-Lorière, Maud Vanpeene, Sylvie Van der Werf ,Hugues Cordel |
| EPI_ISL_3355719, EPI_ISL_3355720, EPI_ISL_3355721, EPI_ISL_3355722, EPI_ISL_3355723, EPI_ISL_3355724, EPI_ISL_3355725, EPI_ISL_3355726, EPI_ISL_3355727, EPI_ISL_3355728, EPI_ISL_3355729, EPI_ISL_3355730, EPI_ISL_3355731, EPI_ISL_3355732, EPI_ISL_3355733, EPI_ISL_3355734 | | | |
| see above | Labo Analyses Med | National Reference Center for Viruses of Respiratory Infections, Institut Pasteur, Paris | Marion Barbet, Sylvie Behillil, Méline Bizard, Angela Brisebarre, Camille Capel, Vincent Enouf, Louise Lefrançois, Frédéric Lemoine, Christophe Malabat, Corinne Maufrais, Hub de Bioinformatique et Biostatistique, Etienne Simon-Lorière, Maud Vanpeene, Sylvie Van der Werf |
| EPI_ISL_3355735, EPI_ISL_3355736, EPI_ISL_3355737, EPI_ISL_3355739, EPI_ISL_3355741, EPI_ISL_3355743, EPI_ISL_3355744, EPI_ISL_3355745, EPI_ISL_3355746, EPI_ISL_3355747, EPI_ISL_3355748, EPI_ISL_3355749, EPI_ISL_3355750, EPI_ISL_3355755, EPI_ISL_3355756, EPI_ISL_3355757, EPI_ISL_3355758, EPI_ISL_3355760, EPI_ISL_3355761, EPI_ISL_3355762, EPI_ISL_3355763 | | | |
| see above | Outre mer | National Reference Center for Viruses of Respiratory Infections, Institut Pasteur, Paris | Marion Barbet, Sylvie Behillil, Méline Bizard, Angela Brisebarre, Camille Capel, Vincent Enouf, Louise Lefrançois, Frédéric Lemoine, Christophe Malabat, Corinne Maufrais, Hub de Bioinformatique et Biostatistique, Etienne Simon-Lorière, Maud Vanpeene, Sylvie Van der Werf ,Antoine Talarmin |
| EPI_ISL_3355764, EPI_ISL_3355765, EPI_ISL_3355766, EPI_ISL_3355767, EPI_ISL_3355768, EPI_ISL_3355769, EPI_ISL_3355770, EPI_ISL_3355771, EPI_ISL_3355772, EPI_ISL_3355773, EPI_ISL_3355774, EPI_ISL_3355775, EPI_ISL_3355776, EPI_ISL_3355777, EPI_ISL_3355778 | | | |
| see above | Hospital | National Reference Center for Viruses of Respiratory Infections, Institut Pasteur, Paris | Marion Barbet, Sylvie Behillil, Méline Bizard, Angela Brisebarre, Camille Capel, Vincent Enouf, Louise Lefrançois, Frédéric Lemoine, Christophe Malabat, Corinne Maufrais, Hub de Bioinformatique et Biostatistique, Etienne Simon-Lorière, Maud Vanpeene, Sylvie Van der Werf ,Nabil Gastli |
| EPI_ISL_3355779, EPI_ISL_3355780, EPI_ISL_3355781, EPI_ISL_3355782, EPI_ISL_3355783, EPI_ISL_3355784, EPI_ISL_3355785, EPI_ISL_3355786, EPI_ISL_3355787, EPI_ISL_3355788, EPI_ISL_3355789, EPI_ISL_3355790 | | | |
| see above | Hospital | National Reference Center for Viruses of Respiratory Infections, Institut Pasteur, Paris | Marion Barbet, Sylvie Behillil, Méline Bizard, Angela Brisebarre, Camille Capel, Vincent Enouf, Louise Lefrançois, Frédéric Lemoine, Christophe Malabat, Corinne Maufrais, Hub de Bioinformatique et Biostatistique, Etienne Simon-Lorière, Maud Vanpeene, Sylvie Van der Werf ,CéLine Bressollette |
| EPI_ISL_3355791, EPI_ISL_3355792, EPI_ISL_3355793, EPI_ISL_3355794, EPI_ISL_3355795, EPI_ISL_3355796, EPI_ISL_3355797, EPI_ISL_3355798, EPI_ISL_3355799, EPI_ISL_3355800, EPI_ISL_3355801, EPI_ISL_3355802, EPI_ISL_3355803, EPI_ISL_3355804, EPI_ISL_3355805, EPI_ISL_3355806, EPI_ISL_3355807, EPI_ISL_3355808, EPI_ISL_3355809, EPI_ISL_3355810, EPI_ISL_3355811, EPI_ISL_3355812, EPI_ISL_3355813, EPI_ISL_3355814, EPI_ISL_3355815, EPI_ISL_3355816, EPI_ISL_3355817, EPI_ISL_3355818, EPI_ISL_3355819, EPI_ISL_3355820, EPI_ISL_3355821, EPI_ISL_3355822, EPI_ISL_3355823, EPI_ISL_3355824, EPI_ISL_3355825, EPI_ISL_3355826, EPI_ISL_3355828, EPI_ISL_3355829, EPI_ISL_3355830, EPI_ISL_3355831, EPI_ISL_3355832, EPI_ISL_3355833, EPI_ISL_3355834, EPI_ISL_3355835, EPI_ISL_3355836, EPI_ISL_3355837, EPI_ISL_3355838, EPI_ISL_3355839, EPI_ISL_3355840, EPI_ISL_3355841, EPI_ISL_3355842, EPI_ISL_3355843, EPI_ISL_3355844, EPI_ISL_3355845, EPI_ISL_3355846, EPI_ISL_3355847, EPI_ISL_3355848, EPI_ISL_3355849, EPI_ISL_3355850, EPI_ISL_3355851, EPI_ISL_3355852, EPI_ISL_3355853, EPI_ISL_3355854, EPI_ISL_3355855, EPI_ISL_3355856, EPI_ISL_3355857, EPI_ISL_3355858, EPI_ISL_3355859, EPI_ISL_3355860, EPI_ISL_3355861, EPI_ISL_3355862, EPI_ISL_3355863, EPI_ISL_3355864, EPI_ISL_3355865 | | | |
| see above | Labo Analyses Med | National Reference Center for Viruses of Respiratory Infections, Institut Pasteur, Paris | Marion Barbet, Sylvie Behillil, Méline Bizard, Angela Brisebarre, Camille Capel, Vincent Enouf, Louise Lefrançois, Frédéric Lemoine, Christophe Malabat, Corinne Maufrais, Hub de Bioinformatique et Biostatistique, Etienne Simon-Lorière, Maud Vanpeene, Sylvie Van der Werf ,Pierre-Yves Leonard |
| EPI_ISL_3355867, EPI_ISL_3355868 | Labo Analyses Med | National Reference Center for Viruses of Respiratory Infections, Institut Pasteur, Paris | Marion Barbet, Sylvie Behillil, Méline Bizard, Angela Brisebarre, Camille Capel, Vincent Enouf, Louise Lefrançois, Frédéric Lemoine, Christophe Malabat, Corinne Maufrais, Hub de Bioinformatique et Biostatistique, Etienne Simon-Lorière, Maud Vanpeene, Sylvie Van der Werf ,Claire Pottier |
| EPI_ISL_3355869, EPI_ISL_3355870, EPI_ISL_3355871, EPI_ISL_3355873, EPI_ISL_3355874, EPI_ISL_3355875, EPI_ISL_3355876, EPI_ISL_3355877 | Labo Analyses Med | National Reference Center for Viruses of Respiratory Infections, Institut Pasteur, Paris | Marion Barbet, Sylvie Behillil, Méline Bizard, Angela Brisebarre, Camille Capel, Vincent Enouf, Louise Lefrançois, Frédéric Lemoine, Christophe Malabat, Corinne Maufrais, Hub de Bioinformatique et Biostatistique, Etienne Simon-Lorière, Maud Vanpeene, Sylvie Van der Werf ,GwéNolé Prigent |
| EPI_ISL_3355878 | Hospital | National Reference Center for Viruses of Respiratory Infections, Institut Pasteur, Paris | Marion Barbet, Sylvie Behillil, Méline Bizard, Angela Brisebarre, Camille Capel, Vincent Enouf, Louise Lefrançois, Frédéric Lemoine, Christophe Malabat, Corinne Maufrais, Hub de Bioinformatique et Biostatistique, Etienne Simon-Lorière, Maud Vanpeene, Sylvie Van der Werf ,Eric Farfour |
| EPI_ISL_3355879, EPI_ISL_3355880, EPI_ISL_3355881, EPI_ISL_3355882, EPI_ISL_3355883, EPI_ISL_3355884, EPI_ISL_3355885, EPI_ISL_3355886, EPI_ISL_3355887, EPI_ISL_3355888, EPI_ISL_3355889, EPI_ISL_3355890, EPI_ISL_3355891, EPI_ISL_3355892, EPI_ISL_3355893, EPI_ISL_3355894, EPI_ISL_3355895, EPI_ISL_3355896, EPI_ISL_3355897, EPI_ISL_3355898 | | | |
| see above | Labo Analyses Med | National Reference Center for Viruses of Respiratory Infections, Institut Pasteur, Paris | Marion Barbet, Sylvie Behillil, Méline Bizard, Angela Brisebarre, Camille Capel, Vincent Enouf, Louise Lefrançois, Frédéric Lemoine, Christophe Malabat, Corinne Maufrais, Hub de Bioinformatique et Biostatistique, Etienne Simon-Lorière, Maud Vanpeene, Sylvie Van der Werf |
| EPI_ISL_3355901, EPI_ISL_3355902, EPI_ISL_3355904, EPI_ISL_3355905, EPI_ISL_3355906, EPI_ISL_3355907 | Virology Laboratory, Department of Microbiology, King George's Medical University | Virology Laboratory, Department of Microbiology, King George's Medical University | Shantanu Prakash, Hricha Mishra, Suruchi Shukla, Amita Jain |
| EPI_ISL_3355908, EPI_ISL_3355909, EPI_ISL_3355910, EPI_ISL_3355911, EPI_ISL_3355912, EPI_ISL_3355913, EPI_ISL_3355914, EPI_ISL_3355915, EPI_ISL_3355916, EPI_ISL_3355917, EPI_ISL_3355918, EPI_ISL_3355919, EPI_ISL_3355920 | | | |
| see above | Department of Microbiology, SGPGIMS | Virology Laboratory, Department of Microbiology, King George's Medical University | Ujjala Goshal, Shantanu Prakash, Suruchi Shukla, Amita Jain |
| EPI_ISL_3355921, EPI_ISL_3355922, EPI_ISL_3355923 | Sambhavi Diagnostics, Lucknow | Virology Laboratory, Department of Microbiology, King George's Medical University | Shantanu Prakash, Hricha Mishra, Suruchi Shukla, Amita Jain |
| EPI_ISL_3355924, EPI_ISL_3355925 | Department of Microbiology, Mayo Institute of Medical Sciences | Virology Laboratory, Department of Microbiology, King George's Medical University | Monika Rajani, Meenakshi Sharma, Shantanu Prakash, Hricha Mishra, Suruchi Shukla, Amita Jain |
| EPI_ISL_3355926, EPI_ISL_3355927, EPI_ISL_3355928, EPI_ISL_3355929, EPI_ISL_3355930, EPI_ISL_3355931, EPI_ISL_3355932, EPI_ISL_3355933, EPI_ISL_3355934, EPI_ISL_3355935 | Molecular Diagnostic and Research Laboratory, GIMS, Greater Noida | Virology Laboratory, Department of Microbiology, King George's Medical University | Vivek Gupta, Shantanu Prakash, Suruchi Shukla, Amita Jain |
| EPI_ISL_3355936, EPI_ISL_3355937, EPI_ISL_3355938 | Department of Microbiology, LLRM Medical College | Virology Laboratory, Department of Microbiology, King George's Medical University | Amit Garg, Shantanu Prakash, Suruchi Shukla, Amita Jain |
| EPI_ISL_3355939, EPI_ISL_3355940, EPI_ISL_3355941, EPI_ISL_3355942, EPI_ISL_3355943, EPI_ISL_3355944, EPI_ISL_3355945, EPI_ISL_3355946, EPI_ISL_3355947, EPI_ISL_3355948, EPI_ISL_3355949, EPI_ISL_3355950, EPI_ISL_3355951, EPI_ISL_3355952, EPI_ISL_3355953, EPI_ISL_3355954, EPI_ISL_3355955, EPI_ISL_3355956, EPI_ISL_3355957, EPI_ISL_3355958, EPI_ISL_3355959, EPI_ISL_3355960, EPI_ISL_3355961 | | | |
| see above | Department of Microbiology, SMMH, Saharanpur | Virology Laboratory, Department of Microbiology, King George's Medical University | Pragya Shakya, Shantanu Prakash, Suruchi Shukla, Amita Jain |
| EPI_ISL_3355962, EPI_ISL_3355964, EPI_ISL_3355965, EPI_ISL_3355966 | Department of Microbiology, SSPHPGTI | Virology Laboratory, Department of Microbiology, King George's Medical University | Sumi Nandwani, Shantanu Prakash, Suruchi Shukla, Amita Jain |
| EPI_ISL_3355967, EPI_ISL_3355968, EPI_ISL_3355969, EPI_ISL_3355970, EPI_ISL_3355971, EPI_ISL_3355972, EPI_ISL_3355973, EPI_ISL_3355974 | Department of Microbiology, SGPGIMS | Virology Laboratory, Department of Microbiology, King George's Medical University | Ujjala Goshal, Shantanu Prakash, Suruchi Shukla, Amita Jain |
| EPI_ISL_3355975, EPI_ISL_3355976 | Department of Microbiology, Dr. Ram Manohar Lohia Institute of Medical Sciences | Virology Laboratory, Department of Microbiology, King George's Medical University | Jyotsna Agarwal, Shantanu Prakash, Suruchi Shukla, Amita Jain |
| EPI_ISL_3355977 | Department of Microbiology, AIIMS, Raebareli | Virology Laboratory, Department of Microbiology, King George's Medical University | Shefali Gupta, Shantanu Prakash, Suruchi Shukla, Amita Jain |
| EPI_ISL_3355978, EPI_ISL_3355979, EPI_ISL_3355980, EPI_ISL_3355981, EPI_ISL_3355982, EPI_ISL_3355983, EPI_ISL_3355984 | Department of Microbiology, S.N. Medical College | Virology Laboratory, Department of Microbiology, King George's Medical University | Arti Agrawal, Shantanu Prakash, Suruchi Shukla, Amita Jain |
| EPI_ISL_3355985 | Department of Microbiology, Autonomous State Medical College Firozabad | Virology Laboratory, Department of Microbiology, King George's Medical University | Hariom Sharan, Lekha Tuli, Shantanu Prakash, Suruchi Shukla, Amita Jain |

| | | | |
|---|---|---|---|
| EPI_ISL_3355987 | Department of Microbiology, Veterinary College Mathura, DUVASU | Virology Laboratory, Department of Microbiology, King George's Medical University | Rashmi Singh, Shantanu Prakash, Suruchi Shukla, Amita Jain |
| EPI_ISL_3355988 | Virology Laboratory, Department of Microbiology, King George's Medical University | Virology Laboratory, Department of Microbiology, King George's Medical University | Shantanu Prakash, Hricha Mishra, Suruchi Shukla, Amita Jain |
| EPI_ISL_3355990, EPI_ISL_3355991, EPI_ISL_3355992, EPI_ISL_3355993, EPI_ISL_3355994, EPI_ISL_3355995, EPI_ISL_3355996, EPI_ISL_3355997, EPI_ISL_3355998, EPI_ISL_3355999, EPI_ISL_3356000, EPI_ISL_3356001, EPI_ISL_3356002, EPI_ISL_3356003, EPI_ISL_3356004, EPI_ISL_3356005, EPI_ISL_3356006, EPI_ISL_3356007, EPI_ISL_3356008, EPI_ISL_3356009, EPI_ISL_3356010, EPI_ISL_3356011, EPI_ISL_3356012, EPI_ISL_3356014, EPI_ISL_3356016, EPI_ISL_3356017, EPI_ISL_3356018, EPI_ISL_3356019, EPI_ISL_3356020, EPI_ISL_3356021, EPI_ISL_3356022, EPI_ISL_3356023, EPI_ISL_3356025, EPI_ISL_3356026, EPI_ISL_3356027, EPI_ISL_3356029, EPI_ISL_3356030, EPI_ISL_3356031, EPI_ISL_3356032, EPI_ISL_3356033, EPI_ISL_3356034, EPI_ISL_3356035, EPI_ISL_3356036, EPI_ISL_3356037, EPI_ISL_3356038, EPI_ISL_3356039, EPI_ISL_3356040, EPI_ISL_3356041, EPI_ISL_3356042, EPI_ISL_3356043, EPI_ISL_3356044, EPI_ISL_3356045, EPI_ISL_3356046, EPI_ISL_3356047, EPI_ISL_3356048, EPI_ISL_3356049, EPI_ISL_3356050, EPI_ISL_3356051, EPI_ISL_3356052, EPI_ISL_3356053, EPI_ISL_3356054, EPI_ISL_3356055, EPI_ISL_3356056, EPI_ISL_3356057, EPI_ISL_3356058, EPI_ISL_3356059, EPI_ISL_3356060, EPI_ISL_3356061, EPI_ISL_3356062, EPI_ISL_3356063, EPI_ISL_3356064, EPI_ISL_3356065, EPI_ISL_3356066, EPI_ISL_3356067, EPI_ISL_3356068, EPI_ISL_3356069, EPI_ISL_3356070, EPI_ISL_3356071, EPI_ISL_3356072, EPI_ISL_3356074, EPI_ISL_3356075, EPI_ISL_3356076, EPI_ISL_3356077, EPI_ISL_3356079, EPI_ISL_3356080, EPI_ISL_3356081, EPI_ISL_3356082, EPI_ISL_3356083, EPI_ISL_3356084, EPI_ISL_3356085, EPI_ISL_3356086, EPI_ISL_3356087, EPI_ISL_3356088, EPI_ISL_3356089, EPI_ISL_3356090, EPI_ISL_3356091, EPI_ISL_3356092, EPI_ISL_3356093, EPI_ISL_3356094, EPI_ISL_3356095, EPI_ISL_3356096, EPI_ISL_3356097, EPI_ISL_3356098, EPI_ISL_3356099, EPI_ISL_3356100, EPI_ISL_3356101, EPI_ISL_3356102, EPI_ISL_3356103, EPI_ISL_3356104, EPI_ISL_3356105, EPI_ISL_3356106, EPI_ISL_3356107, EPI_ISL_3356108, EPI_ISL_3356109, EPI_ISL_3356110, EPI_ISL_3356111, EPI_ISL_3356112, EPI_ISL_3356113, EPI_ISL_3356114, EPI_ISL_3356115, EPI_ISL_3356116, EPI_ISL_3356117, EPI_ISL_3356118, EPI_ISL_3356119, EPI_ISL_3356120, EPI_ISL_3356121, EPI_ISL_3356122, EPI_ISL_3356123, EPI_ISL_3356124, EPI_ISL_3356125, EPI_ISL_3356126, EPI_ISL_3356127, EPI_ISL_3356128, EPI_ISL_3356129, EPI_ISL_3356130, EPI_ISL_3356131, EPI_ISL_3356132, EPI_ISL_3356133, EPI_ISL_3356134, EPI_ISL_3356135, EPI_ISL_3356136, EPI_ISL_3356137, EPI_ISL_3356138, EPI_ISL_3356139, EPI_ISL_3356140, EPI_ISL_3356141, EPI_ISL_3356142, EPI_ISL_3356143, EPI_ISL_3356144, EPI_ISL_3356145, EPI_ISL_3356146, EPI_ISL_3356147, EPI_ISL_3356148, EPI_ISL_3356149, EPI_ISL_3356150, EPI_ISL_3356151, EPI_ISL_3356152, EPI_ISL_3356153, EPI_ISL_3356154, EPI_ISL_3356155, EPI_ISL_3356156, EPI_ISL_3356157, EPI_ISL_3356158, EPI_ISL_3356159, EPI_ISL_3356160, EPI_ISL_3356161, EPI_ISL_3356162, EPI_ISL_3356163, EPI_ISL_3356164, EPI_ISL_3356165, EPI_ISL_3356166, EPI_ISL_3356167, EPI_ISL_3356168, EPI_ISL_3356169, EPI_ISL_3356170, EPI_ISL_3356171, EPI_ISL_3356172, EPI_ISL_3356173, EPI_ISL_3356174, EPI_ISL_3356175, EPI_ISL_3356176, EPI_ISL_3356178, EPI_ISL_3356179, EPI_ISL_3356180, EPI_ISL_3356181, EPI_ISL_3356182, EPI_ISL_3356183, EPI_ISL_3356184, EPI_ISL_3356185, EPI_ISL_3356186, EPI_ISL_3356187, EPI_ISL_3356188, EPI_ISL_3356189, EPI_ISL_3356190, EPI_ISL_3356192, EPI_ISL_3356193, EPI_ISL_3356194, EPI_ISL_3356196, EPI_ISL_3356197, EPI_ISL_3356198, EPI_ISL_3356199, EPI_ISL_3356200, EPI_ISL_3356201, EPI_ISL_3356202, EPI_ISL_3356203, EPI_ISL_3356204, EPI_ISL_3356205, EPI_ISL_3356206, EPI_ISL_3356207, EPI_ISL_3356208, EPI_ISL_3356209, EPI_ISL_3356210, EPI_ISL_3356211, EPI_ISL_3356212, EPI_ISL_3356213, EPI_ISL_3356214, EPI_ISL_3356215, EPI_ISL_3356216, EPI_ISL_3356217, EPI_ISL_3356219, EPI_ISL_3356220, EPI_ISL_3356221, EPI_ISL_3356222, EPI_ISL_3356223, EPI_ISL_3356224, EPI_ISL_3356225, EPI_ISL_3356226, EPI_ISL_3356227, EPI_ISL_3356228, EPI_ISL_3356229, EPI_ISL_3356230, EPI_ISL_3356231, EPI_ISL_3356232, EPI_ISL_3356233, EPI_ISL_3356234, EPI_ISL_3356235, EPI_ISL_3356236, EPI_ISL_3356237, EPI_ISL_3356238, EPI_ISL_3356239, EPI_ISL_3356240, EPI_ISL_3356241, EPI_ISL_3356242, EPI_ISL_3356243, EPI_ISL_3356244, EPI_ISL_3356245, EPI_ISL_3356246, EPI_ISL_3356247, EPI_ISL_3356248, EPI_ISL_3356249, EPI_ISL_3356250, EPI_ISL_3356251 | | | |
| see above | National Virus Reference Laboratory | National Virus Reference Laboratory | Zoe Yandle, Charlene Bennett, Gabriel Gonzalez, Michael Carr, Jonathan Dean, Cillian F De Gascun |
| EPI_ISL_3356282, EPI_ISL_3356283, EPI_ISL_3356352 | Serdang Hospital | Institute for Medical Research, Infectious Disease Research Centre, National Institutes of Health, Ministry of Health Malaysia | Suppiah J, Kamel K, Mohd Zawawi Z, Azizan MA, Ramly N, Robert F, Anasir MI, Thayan R |
| EPI_ISL_3356378, EPI_ISL_3356379, EPI_ISL_3356380, EPI_ISL_3356381, EPI_ISL_3356382, EPI_ISL_3356383, EPI_ISL_3356384, EPI_ISL_3356385, EPI_ISL_3356386, EPI_ISL_3356387, EPI_ISL_3356388, EPI_ISL_3356389, EPI_ISL_3356390, EPI_ISL_3356391, EPI_ISL_3356392, EPI_ISL_3356393, EPI_ISL_3356394, EPI_ISL_3356395, EPI_ISL_3356398, EPI_ISL_3356399, EPI_ISL_3356402, EPI_ISL_3356403, EPI_ISL_3356404, EPI_ISL_3356405, EPI_ISL_3356406, EPI_ISL_3356407, EPI_ISL_3356408 | | | |
| see above | CDFD | CDFD | Murali Bashyam, Asmita Gupta, Reelina Basu, Vinay Donipadi, Nagamani Kammili, Divya Vashisht, Ashwin Dalal |
| EPI_ISL_3356409, EPI_ISL_3356410, EPI_ISL_3356411, EPI_ISL_3356412, EPI_ISL_3356413, EPI_ISL_3356414, EPI_ISL_3356415, EPI_ISL_3356416, EPI_ISL_3356417, EPI_ISL_3356418, EPI_ISL_3356419, EPI_ISL_3356420, EPI_ISL_3356422, EPI_ISL_3356423, EPI_ISL_3356424, EPI_ISL_3356425, EPI_ISL_3356426, EPI_ISL_3356427, EPI_ISL_3356428, EPI_ISL_3356429, EPI_ISL_3356430, EPI_ISL_3356431, EPI_ISL_3356432, EPI_ISL_3356434, EPI_ISL_3356435, EPI_ISL_3356436, EPI_ISL_3356437, EPI_ISL_3356438, EPI_ISL_3356439, EPI_ISL_3356440, EPI_ISL_3356441, EPI_ISL_3356442, EPI_ISL_3356443, EPI_ISL_3356444, EPI_ISL_3356445, EPI_ISL_3356446, EPI_ISL_3356447, EPI_ISL_3356448, EPI_ISL_3356449, EPI_ISL_3356450, EPI_ISL_3356451, EPI_ISL_3356452, EPI_ISL_3356453, EPI_ISL_3356454, EPI_ISL_3356455, EPI_ISL_3356456, EPI_ISL_3356459, EPI_ISL_3356460, EPI_ISL_3356461, EPI_ISL_3356462, EPI_ISL_3356463, EPI_ISL_3356464, EPI_ISL_3356465, EPI_ISL_3356466, EPI_ISL_3356468, EPI_ISL_3356469, EPI_ISL_3356470, EPI_ISL_3356471, EPI_ISL_3356472, EPI_ISL_3356473, EPI_ISL_3356474, EPI_ISL_3356475, EPI_ISL_3356476, EPI_ISL_3356477, EPI_ISL_3356479, EPI_ISL_3356480, EPI_ISL_3356481, EPI_ISL_3356482, EPI_ISL_3356483, EPI_ISL_3356484, EPI_ISL_3356485, EPI_ISL_3356486, EPI_ISL_3356488, EPI_ISL_3356489, EPI_ISL_3356491, EPI_ISL_3356492, EPI_ISL_3356493, EPI_ISL_3356494, EPI_ISL_3356495, EPI_ISL_3356497, EPI_ISL_3356498, EPI_ISL_3356499, EPI_ISL_3356500 | | | |
| see above | State Public Health Laboratory | CDFD | Murali Bashyam, Asmita Gupta, Reelina Basu, Vinay Donipadi, Divya Vashisht, Ashwin Dalal |
| EPI_ISL_3356501, EPI_ISL_3356502, EPI_ISL_3356504, EPI_ISL_3356505, EPI_ISL_3356506, EPI_ISL_3356507, EPI_ISL_3356508, EPI_ISL_3356510, EPI_ISL_3356511, EPI_ISL_3356512, EPI_ISL_3356515, EPI_ISL_3356516, EPI_ISL_3356517, EPI_ISL_3356518, EPI_ISL_3356519, EPI_ISL_3356520, EPI_ISL_3356521, EPI_ISL_3356522, EPI_ISL_3356523, EPI_ISL_3356525, EPI_ISL_3356526, EPI_ISL_3356527, EPI_ISL_3356528, EPI_ISL_3356529, EPI_ISL_3356530, EPI_ISL_3356532, EPI_ISL_3356533, EPI_ISL_3356534, EPI_ISL_3356535, EPI_ISL_3356536, EPI_ISL_3356537, EPI_ISL_3356539, EPI_ISL_3356540, EPI_ISL_3356541, EPI_ISL_3356542, EPI_ISL_3356543, EPI_ISL_3356544, EPI_ISL_3356546, EPI_ISL_3356547, EPI_ISL_3356548, EPI_ISL_3356549, EPI_ISL_3356551, EPI_ISL_3356552, EPI_ISL_3356553, EPI_ISL_3356554, EPI_ISL_3356555, EPI_ISL_3356556 | | | |
| see above | CDFD | CDFD | Murali Bashyam, Asmita Gupta, Reelina Basu, Vinay Donipadi, Divya Vashisht, Ashwin Dalal |
| EPI_ISL_3356562, EPI_ISL_3356563, EPI_ISL_3356564, EPI_ISL_3356565, EPI_ISL_3356566, EPI_ISL_3356567, EPI_ISL_3356568, EPI_ISL_3356569, EPI_ISL_3356570, EPI_ISL_3356571, EPI_ISL_3356572, EPI_ISL_3356573, EPI_ISL_3356574, EPI_ISL_3356575, EPI_ISL_3356576, EPI_ISL_3356577, EPI_ISL_3356578, EPI_ISL_3356579, EPI_ISL_3356580, EPI_ISL_3356581, EPI_ISL_3356582, EPI_ISL_3356583, EPI_ISL_3356584, EPI_ISL_3356585, EPI_ISL_3356586, EPI_ISL_3356587, EPI_ISL_3356588, EPI_ISL_3356589, EPI_ISL_3356590, EPI_ISL_3356591, EPI_ISL_3356592, EPI_ISL_3356593, EPI_ISL_3356594, EPI_ISL_3356595, EPI_ISL_3356596, EPI_ISL_3356597, EPI_ISL_3356598, EPI_ISL_3356599, EPI_ISL_3356600, EPI_ISL_3356601, EPI_ISL_3356602, EPI_ISL_3356603, EPI_ISL_3356604, EPI_ISL_3356605, EPI_ISL_3356606, EPI_ISL_3356607, EPI_ISL_3356608, EPI_ISL_3356609, EPI_ISL_3356610, EPI_ISL_3356611, EPI_ISL_3356612, EPI_ISL_3356613, EPI_ISL_3356614, EPI_ISL_3356615, EPI_ISL_3356616, EPI_ISL_3356617, EPI_ISL_3356618, EPI_ISL_3356619, EPI_ISL_3356620, EPI_ISL_3356621, EPI_ISL_3356622, EPI_ISL_3356623, EPI_ISL_3356624, EPI_ISL_3356625, EPI_ISL_3356626, EPI_ISL_3356627, EPI_ISL_3356628, EPI_ISL_3356629, EPI_ISL_3356630, EPI_ISL_3356631, EPI_ISL_3356632, EPI_ISL_3356633, EPI_ISL_3356634, EPI_ISL_3356635, EPI_ISL_3356636, EPI_ISL_3356637, EPI_ISL_3356638, EPI_ISL_3356639, EPI_ISL_3356640, EPI_ISL_3356641, EPI_ISL_3356642, EPI_ISL_3356643, EPI_ISL_3356644, EPI_ISL_3356645, EPI_ISL_3356653, EPI_ISL_3356659, EPI_ISL_3356660, EPI_ISL_3356661, EPI_ISL_3356662, EPI_ISL_3356663, EPI_ISL_3356664, EPI_ISL_3356665, EPI_ISL_3356666, EPI_ISL_3356667, EPI_ISL_3356668, EPI_ISL_3356669, EPI_ISL_3356670, EPI_ISL_3356671, EPI_ISL_3356672, EPI_ISL_3356673, EPI_ISL_3356674, EPI_ISL_3356675, EPI_ISL_3356676, EPI_ISL_3356677, EPI_ISL_3356678, EPI_ISL_3356679, EPI_ISL_3356680, EPI_ISL_3356681, EPI_ISL_3356682, EPI_ISL_3356683, EPI_ISL_3356684, EPI_ISL_3356685, EPI_ISL_3356686, EPI_ISL_3356687, EPI_ISL_3356688, EPI_ISL_3356689, EPI_ISL_3356690, EPI_ISL_3356691, EPI_ISL_3356692, EPI_ISL_3356693, EPI_ISL_3356694, EPI_ISL_3356695, EPI_ISL_3356696, EPI_ISL_3356697, EPI_ISL_3356698, EPI_ISL_3356699, EPI_ISL_3356700, EPI_ISL_3356701, EPI_ISL_3356702, EPI_ISL_3356703, EPI_ISL_3356704, EPI_ISL_3356705, EPI_ISL_3356706, EPI_ISL_3356707, EPI_ISL_3356708, EPI_ISL_3356709, EPI_ISL_3356710, EPI_ISL_3356711, EPI_ISL_3356712, EPI_ISL_3356713, EPI_ISL_3356714, EPI_ISL_3356715, EPI_ISL_3356716, EPI_ISL_3356717, EPI_ISL_3356718, EPI_ISL_3356719, EPI_ISL_3356720, EPI_ISL_3356721, EPI_ISL_3356722, EPI_ISL_3356723, EPI_ISL_3356724, EPI_ISL_3356725, EPI_ISL_3356726, EPI_ISL_3356727 | | | |
| see above | Alaska State Virology Laboratory | Alaska State Virology Laboratory | Stephanie DeRonde, Elva House, Jacob Zidek, Lisa Smith, Ph.D., Jack Chen, Ph.D. |
| EPI_ISL_3356819, EPI_ISL_3356821, EPI_ISL_3356822, EPI_ISL_3356833, EPI_ISL_3356834, EPI_ISL_3356835, EPI_ISL_3356836, EPI_ISL_3356837, EPI_ISL_3356838, EPI_ISL_3356839, EPI_ISL_3356840, EPI_ISL_3356841, EPI_ISL_3356845, EPI_ISL_3356846, EPI_ISL_3356847, EPI_ISL_3356848, EPI_ISL_3356849, EPI_ISL_3356850, EPI_ISL_3356851, EPI_ISL_3356852, EPI_ISL_3356853, EPI_ISL_3356859, EPI_ISL_3356860, EPI_ISL_3356861, EPI_ISL_3356862, EPI_ISL_3356864, EPI_ISL_3356865, EPI_ISL_3356868, EPI_ISL_3356869, EPI_ISL_3356870, EPI_ISL_3356872, EPI_ISL_3356873, EPI_ISL_3356874, EPI_ISL_3356876, EPI_ISL_3356877, EPI_ISL_3356878 | | | |
| see above | MB-Cadham Provincial laboratory | National Microbiology Laboratory (NML) | Anna Majer, Shari Tyson, Grace Seo, Philip Mabon, Elsie Grudeski, Rhiannon Huzarewich, Russell Mandes, Anneliese Landgraff, Jennifer Tanner, Natalie Knox, Morag Graham, Gary Van Domselaar, Paul Van Caeseele, Jared Bullard, David Alexander, Kerry Dust, Nathalie Bastien, Yan Li, Timothy Booth, Darian Hole, Madison Chapel, Kirsten Biggar, CanCOGeN's metadata curation team, Public Health Agency of Canada CanCOGeN team |
| EPI_ISL_3356880, EPI_ISL_3356881, EPI_ISL_3356884, EPI_ISL_3356885, EPI_ISL_3356887, EPI_ISL_3356889, EPI_ISL_3356890, EPI_ISL_3356891, EPI_ISL_3356892, EPI_ISL_3356893, EPI_ISL_3356894 | | | |
| see above | MB-Cadham Provincial laboratory | Manitoba Cadham Provincial Laboratory | Anna Majer, Shari Tyson, Grace Seo, Philip Mabon, Elsie Grudeski, Rhiannon Huzarewich, Russell Mandes, Anneliese Landgraff, Jennifer Tanner, Natalie Knox, Morag Graham, Gary Van Domselaar, Paul Van Caeseele, Jared Bullard, David Alexander, Kerry Dust, Nathalie Bastien, Yan Li, Timothy Booth, Darian Hole, Madison Chapel, Kirsten Biggar, CanCOGeN's metadata curation team, Public Health Agency of Canada CanCOGeN team |
| EPI_ISL_3356918, EPI_ISL_3356919, EPI_ISL_3356920, EPI_ISL_3356921, EPI_ISL_3356922, EPI_ISL_3356923 | Hospital General Universitario de Ciudad Real | Hospital General Universitario de Ciudad Real | José Martínez-Alarcón, Soledad Illescas Fernández-Bermejo, Marta Torres-Narbona, Cristina Colmenarejo, Lidia García-Agudo |
| EPI_ISL_3356924, EPI_ISL_3356925 | Hospital General de Valdepeñas | Hospital General Universitario de Ciudad Real | José Martínez-Alarcón, Soledad Illescas Fernández-Bermejo, Marta Torres-Narbona, Cristina Colmenarejo, Lidia García-Agudo |
| EPI_ISL_3356926 | Hospital General Universitario de Ciudad Real | Hospital General Universitario de Ciudad Real | José Martínez-Alarcón, Soledad Illescas Fernández-Bermejo, Marta Torres-Narbona, Cristina Colmenarejo, Lidia García-Agudo |
| EPI_ISL_3356927 | Hospital General de Valdepeñas | Hospital General Universitario de Ciudad Real | José Martínez-Alarcón, Soledad Illescas Fernández-Bermejo, Marta Torres-Narbona, Cristina Colmenarejo, Lidia García-Agudo |
| EPI_ISL_3356929, EPI_ISL_3356930 | Hospital General Universitario de Ciudad Real | Hospital General Universitario de Ciudad Real | José Martínez-Alarcón, Soledad Illescas Fernández-Bermejo, Marta Torres-Narbona, Cristina Colmenarejo, Lidia García-Agudo |
| EPI_ISL_3356931 | Hospital Virgen De Altagracia | Hospital General Universitario de Ciudad Real | José Martínez-Alarcón, Soledad Illescas Fernández-Bermejo, Marta Torres-Narbona, Cristina Colmenarejo, Lidia García-Agudo |
| EPI_ISL_3356932 | Hospital General Universitario de Ciudad Real | Hospital General Universitario de Ciudad Real | José Martínez-Alarcón, Soledad Illescas Fernández-Bermejo, Marta Torres-Narbona, Cristina Colmenarejo, Lidia García-Agudo |
| EPI_ISL_3356933 | Hospital General de Valdepeñas | Hospital General Universitario de Ciudad Real | José Martínez-Alarcón, Soledad Illescas Fernández-Bermejo, Marta Torres-Narbona, Cristina Colmenarejo, Lidia García-Agudo |
| EPI_ISL_3356934, EPI_ISL_3356935, EPI_ISL_3356936, EPI_ISL_3356937, EPI_ISL_3356938, EPI_ISL_3356939, EPI_ISL_3356940, EPI_ISL_3356941, EPI_ISL_3356942, EPI_ISL_3356943, EPI_ISL_3356944, EPI_ISL_3356945, EPI_ISL_3356946, EPI_ISL_3356947, EPI_ISL_3356948, EPI_ISL_3356949, EPI_ISL_3356950, EPI_ISL_3356951, EPI_ISL_3356952, EPI_ISL_3356953, EPI_ISL_3356954, EPI_ISL_3356955, EPI_ISL_3356956, EPI_ISL_3356957, EPI_ISL_3356958, EPI_ISL_3356959, EPI_ISL_3356960, EPI_ISL_3356961, EPI_ISL_3356962, EPI_ISL_3356963, EPI_ISL_3356964, EPI_ISL_3356965, EPI_ISL_3356966, EPI_ISL_3356967, EPI_ISL_3356968, EPI_ISL_3356969, EPI_ISL_3356970, EPI_ISL_3356971, EPI_ISL_3356972, EPI_ISL_3356973, EPI_ISL_3356974, EPI_ISL_3356975, EPI_ISL_3356976, EPI_ISL_3356977, EPI_ISL_3356978, EPI_ISL_3356979, EPI_ISL_3356980, EPI_ISL_3356981, EPI_ISL_3356982, EPI_ISL_3356983, EPI_ISL_3356984, EPI_ISL_3356985, EPI_ISL_3356986, EPI_ISL_3356987, EPI_ISL_3356988, EPI_ISL_3356989, EPI_ISL_3356990, EPI_ISL_3356991, EPI_ISL_3356992, EPI_ISL_3356993, EPI_ISL_3356994, EPI_ISL_3356995, EPI_ISL_3356996, EPI_ISL_3356997, EPI_ISL_3356998, EPI_ISL_3356999, EPI_ISL_3357000, EPI_ISL_3357001, EPI_ISL_3357002, EPI_ISL_3357003, EPI_ISL_3357004, EPI_ISL_3357005, EPI_ISL_3357006, EPI_ISL_3357007, EPI_ISL_3357008, EPI_ISL_3357009, EPI_ISL_3357010, EPI_ISL_3357011, EPI_ISL_3357012, EPI_ISL_3357013, EPI_ISL_3357014, EPI_ISL_3357015, EPI_ISL_3357016, EPI_ISL_3357017, EPI_ISL_3357018, EPI_ISL_3357019, EPI_ISL_3357020, EPI_ISL_3357021, EPI_ISL_3357022, EPI_ISL_3357023, EPI_ISL_3357024, EPI_ISL_3357025, EPI_ISL_3357026, EPI_ISL_3357027, EPI_ISL_3357028, EPI_ISL_3357029, EPI_ISL_3357030, EPI_ISL_3357031, EPI_ISL_3357032, EPI_ISL_3357033, EPI_ISL_3357034, EPI_ISL_3357035, EPI_ISL_3357036, EPI_ISL_3357037, EPI_ISL_3357038, EPI_ISL_3357039, EPI_ISL_3357040, EPI_ISL_3357041, EPI_ISL_3357042, EPI_ISL_3357043, EPI_ISL_3357044, EPI_ISL_3357045, EPI_ISL_3357046, EPI_ISL_3357047, EPI_ISL_3357048, EPI_ISL_3357049, EPI_ISL_3357050, EPI_ISL_3357051, EPI_ISL_3357052, EPI_ISL_3357053, EPI_ISL_3357054, EPI_ISL_3357055, EPI_ISL_3357056, EPI_ISL_3357057, EPI_ISL_3357058, EPI_ISL_3357059, EPI_ISL_3357060, EPI_ISL_3357061, EPI_ISL_3357062, EPI_ISL_3357063, EPI_ISL_3357064, EPI_ISL_3357065, EPI_ISL_3357066, EPI_ISL_3357067, EPI_ISL_3357068, EPI_ISL_3357069, EPI_ISL_3357070, EPI_ISL_3357071, EPI_ISL_3357072, EPI_ISL_3357073, EPI_ISL_3357074, EPI_ISL_3357075, EPI_ISL_3357076, EPI_ISL_3357077, EPI_ISL_3357078, EPI_ISL_3357079, EPI_ISL_3357080, EPI_ISL_3357081, EPI_ISL_3357082, EPI_ISL_3357083, EPI_ISL_3357084, EPI_ISL_3357085, EPI_ISL_3357086, EPI_ISL_3357087, EPI_ISL_3357088, EPI_ISL_3357089, EPI_ISL_3357091, EPI_ISL_3357092, EPI_ISL_3357093, EPI_ISL_3357094, EPI_ISL_3357095, EPI_ISL_3357096, EPI_ISL_3357097, EPI_ISL_3357098, EPI_ISL_3357099, EPI_ISL_3357100, EPI_ISL_3357101, EPI_ISL_3357102, EPI_ISL_3357103, EPI_ISL_3357104, EPI_ISL_3357105, EPI_ISL_3357106, EPI_ISL_3357107, EPI_ISL_3357108, EPI_ISL_3357109, EPI_ISL_3357110, EPI_ISL_3357111, EPI_ISL_3357112, EPI_ISL_3357113, EPI_ISL_3357114, EPI_ISL_3357115, EPI_ISL_3357116, EPI_ISL_3357117, EPI_ISL_3357118, EPI_ISL_3357119, EPI_ISL_3357120, EPI_ISL_3357121, EPI_ISL_3357122, EPI_ISL_3357123, EPI_ISL_3357124, EPI_ISL_3357125, EPI_ISL_3357126, EPI_ISL_3357127, EPI_ISL_3357128, EPI_ISL_3357129, EPI_ISL_3357130, EPI_ISL_3357131, EPI_ISL_3357132, EPI_ISL_3357133, EPI_ISL_3357134, EPI_ISL_3357135, EPI_ISL_3357136, EPI_ISL_3357137, EPI_ISL_3357138, EPI_ISL_3357139, EPI_ISL_3357140, EPI_ISL_3357141, EPI_ISL_3357142, EPI_ISL_3357143, EPI_ISL_3357144, EPI_ISL_3357145, EPI_ISL_3357146, EPI_ISL_3357147, EPI_ISL_3357148, EPI_ISL_3357149, EPI_ISL_3357150, EPI_ISL_3357151, EPI_ISL_3357152, EPI_ISL_3357153, EPI_ISL_3357154, EPI_ISL_3357155, EPI_ISL_3357156, EPI_ISL_3357157, EPI_ISL_3357158, EPI_ISL_3357159, EPI_ISL_3357160, EPI_ISL_3357161, EPI_ISL_3357162, EPI_ISL_3357163, EPI_ISL_3357164, EPI_ISL_3357165, EPI_ISL_3357166, EPI_ISL_3357167, EPI_ISL_3357168, EPI_ISL_3357169, EPI_ISL_3357170, EPI_ISL_3357171, EPI_ISL_3357172, EPI_ISL_3357173, EPI_ISL_3357174, EPI_ISL_3357175, EPI_ISL_3357176, EPI_ISL_3357177, EPI_ISL_3357178, EPI_ISL_3357179, EPI_ISL_3357180, EPI_ISL_3357181, EPI_ISL_3357182, EPI_ISL_3357183, EPI_ISL_3357184, EPI_ISL_3357185, EPI_ISL_3357186, EPI_ISL_3357187, EPI_ISL_3357188, EPI_ISL_3357189, EPI_ISL_3357190, EPI_ISL_3357191, EPI_ISL_3357192, EPI_ISL_3357193, EPI_ISL_3357194, EPI_ISL_3357195, EPI_ISL_3357196, EPI_ISL_3357197, EPI_ISL_3357198, EPI_ISL_3357199, EPI_ISL_3357200, EPI_ISL_3357201, EPI_ISL_3357202, EPI_ISL_3357203, EPI_ISL_3357204, EPI_ISL_3357205, EPI_ISL_3357206, EPI_ISL_3357207, EPI_ISL_3357208, EPI_ISL_3357209, EPI_ISL_3357210, EPI_ISL_3357211, EPI_ISL_3357212, EPI_ISL_3357213, EPI_ISL_3357214, EPI_ISL_3357215, EPI_ISL_3357216, EPI_ISL_3357217, EPI_ISL_3357218, EPI_ISL_3357219, EPI_ISL_3357220, EPI_ISL_3357221, EPI_ISL_3357222, EPI_ISL_3357223, EPI_ISL_3357224 | | | |
| see above | Department of Bacteria, Parasites and Fungi, Statens Serum Institut, Copenhagen, Denmark | Statens Serum Institut Bioinformatics and Microbial Genomics | Danish Covid-19 Genome Consortium |
| EPI_ISL_3357225 | Department of Clinical Microbiology, Odense University Hospital, Odense, Denmark | Statens Serum Institut Bioinformatics and Microbial Genomics | Danish Covid-19 Genome Consortium |
| EPI_ISL_3357226, EPI_ISL_3357227, EPI_ISL_3357228, EPI_ISL_3357230, EPI_ISL_3357231, EPI_ISL_3357232, EPI_ISL_3357233, EPI_ISL_3357234, EPI_ISL_3357235, EPI_ISL_3357236, EPI_ISL_3357237, EPI_ISL_3357238, EPI_ISL_3357239, EPI_ISL_3357240, EPI_ISL_3357241, EPI_ISL_3357242, EPI_ISL_3357243, EPI_ISL_3357244, EPI_ISL_3357245, EPI_ISL_3357246, EPI_ISL_3357247, EPI_ISL_3357248, EPI_ISL_3357249, EPI_ISL_3357250, EPI_ISL_3357251, EPI_ISL_3357252, EPI_ISL_3357253, EPI_ISL_3357255, EPI_ISL_3357256, EPI_ISL_3357257, EPI_ISL_3357258, EPI_ISL_3357259, EPI_ISL_3357260, EPI_ISL_3357261, EPI_ISL_3357262, EPI_ISL_3357263, EPI_ISL_3357264, EPI_ISL_3357265, EPI_ISL_3357266, EPI_ISL_3357267, EPI_ISL_3357268, EPI_ISL_3357269, EPI_ISL_3357270, EPI_ISL_3357271, EPI_ISL_3357272, EPI_ISL_3357273, EPI_ISL_3357274, EPI_ISL_3357275, EPI_ISL_3357277, EPI_ISL_3357278, EPI_ISL_3357279, EPI_ISL_3357280, EPI_ISL_3357281, EPI_ISL_3357282, EPI_ISL_3357283, EPI_ISL_3357284, EPI_ISL_3357285, EPI_ISL_3357286, EPI_ISL_3357287, EPI_ISL_3357288, EPI_ISL_3357289, EPI_ISL_3357290, EPI_ISL_3357291, EPI_ISL_3357292, EPI_ISL_3357293, EPI_ISL_3357294, EPI_ISL_3357295, EPI_ISL_3357296, EPI_ISL_3357297, EPI_ISL_3357298, EPI_ISL_3357299, EPI_ISL_3357300, EPI_ISL_3357301, EPI_ISL_3357302, EPI_ISL_3357303, EPI_ISL_3357304, EPI_ISL_3357305, EPI_ISL_3357306, EPI_ISL_3357307, EPI_ISL_3357308, EPI_ISL_3357309, EPI_ISL_3357310, EPI_ISL_3357311, EPI_ISL_3357312, EPI_ISL_3357313, EPI_ISL_3357314, EPI_ISL_3357315, EPI_ISL_3357316, EPI_ISL_3357317, EPI_ISL_3357318, EPI_ISL_3357319, EPI_ISL_3357320, EPI_ISL_3357321, EPI_ISL_3357322, EPI_ISL_3357323, EPI_ISL_3357324, EPI_ISL_3357325, EPI_ISL_3357326, EPI_ISL_3357327, EPI_ISL_3357329, EPI_ISL_3357330, EPI_ISL_3357331, EPI_ISL_3357332, EPI_ISL_3357333, EPI_ISL_3357334, EPI_ISL_3357335, EPI_ISL_3357336, EPI_ISL_3357337, EPI_ISL_3357338, EPI_ISL_3357339, EPI_ISL_3357340, EPI_ISL_3357341, EPI_ISL_3357342, EPI_ISL_3357343, EPI_ISL_3357344, EPI_ISL_3357345, EPI_ISL_3357346, EPI_ISL_3357347, EPI_ISL_3357348, EPI_ISL_3357349, EPI_ISL_3357350, EPI_ISL_3357352, EPI_ISL_3357353, EPI_ISL_3357354, EPI_ISL_3357355, EPI_ISL_3357356, EPI_ISL_3357357, EPI_ISL_3357358, EPI_ISL_3357359, EPI_ISL_3357360, EPI_ISL_3357361, EPI_ISL_3357362, EPI_ISL_3357363, EPI_ISL_3357364, EPI_ISL_3357365, EPI_ISL_3357366, EPI_ISL_3357367, EPI_ISL_3357368, EPI_ISL_3357369, EPI_ISL_3357370, EPI_ISL_3357371, EPI_ISL_3357372, EPI_ISL_3357373, EPI_ISL_3357374, EPI_ISL_3357375, EPI_ISL_3357376, EPI_ISL_3357377, EPI_ISL_3357378, EPI_ISL_3357379, EPI_ISL_3357380, EPI_ISL_3357381, EPI_ISL_3357382, EPI_ISL_3357383, EPI_ISL_3357384, EPI_ISL_3357385, EPI_ISL_3357386, EPI_ISL_3357387, EPI_ISL_3357388, EPI_ISL_3357389, EPI_ISL_3357390, EPI_ISL_3357391, EPI_ISL_3357392, EPI_ISL_3357393, EPI_ISL_3357394, EPI_ISL_3357395, EPI_ISL_3357396, EPI_ISL_3357397, EPI_ISL_3357398, EPI_ISL_3357399, EPI_ISL_3357400, EPI_ISL_3357401, EPI_ISL_3357402, EPI_ISL_3357403, EPI_ISL_3357404, EPI_ISL_3357405, EPI_ISL_3357406, EPI_ISL_3357407, EPI_ISL_3357408, EPI_ISL_3357409, EPI_ISL_3357410, EPI_ISL_3357411, EPI_ISL_3357412, EPI_ISL_3357413, EPI_ISL_3357414, EPI_ISL_3357415, EPI_ISL_3357416, EPI_ISL_3357417, EPI_ISL_3357418, EPI_ISL_3357419, EPI_ISL_3357420, EPI_ISL_3357421, EPI_ISL_3357422, EPI_ISL_3357423, EPI_ISL_3357424, EPI_ISL_3357425, EPI_ISL_3357426, EPI_ISL_3357427, EPI_ISL_3357428, EPI_ISL_3357429, EPI_ISL_3357430, EPI_ISL_3357431, EPI_ISL_3357432, EPI_ISL_3357433, EPI_ISL_3357434, EPI_ISL_3357435, EPI_ISL_3357436, EPI_ISL_3357437, EPI_ISL_3357438, EPI_ISL_3357439, EPI_ISL_3357440, EPI_ISL_3357441, EPI_ISL_3357442, EPI_ISL_3357443, EPI_ISL_3357444, EPI_ISL_3357445, EPI_ISL_3357446, EPI_ISL_3357447, EPI_ISL_3357448, EPI_ISL_3357449, EPI_ISL_3357450, EPI_ISL_3357451, EPI_ISL_3357452, EPI_ISL_3357453, EPI_ISL_3357454, EPI_ISL_3357455, EPI_ISL_3357456, EPI_ISL_3357457, EPI_ISL_3357458, EPI_ISL_3357460, EPI_ISL_3357461, EPI_ISL_3357462, EPI_ISL_3357463, EPI_ISL_3357464, EPI_ISL_3357465, EPI_ISL_3357466, EPI_ISL_3357468, EPI_ISL_3357469 | | | |
| see above | Department of Bacteria, Parasites and Fungi, Statens Serum Institut, Copenhagen, Denmark | Statens Serum Institut Bioinformatics and Microbial Genomics | Danish Covid-19 Genome Consortium |
| EPI_ISL_3357470 | Department of Clinical Microbiology, Odense University Hospital, Odense, Denmark | Statens Serum Institut Bioinformatics and Microbial Genomics | Danish Covid-19 Genome Consortium |
| EPI_ISL_3357471, EPI_ISL_3357472, EPI_ISL_3357473, EPI_ISL_3357474, EPI_ISL_3357475, EPI_ISL_3357476, EPI_ISL_3357478, EPI_ISL_3357479, EPI_ISL_3357480, EPI_ISL_3357481, EPI_ISL_3357482, EPI_ISL_3357483, EPI_ISL_3357484, EPI_ISL_3357485, EPI_ISL_3357486, EPI_ISL_3357487, EPI_ISL_3357488, EPI_ISL_3357489, EPI_ISL_3357490, EPI_ISL_3357491, EPI_ISL_3357492, EPI_ISL_3357493, EPI_ISL_3357494, EPI_ISL_3357495, EPI_ISL_3357496, EPI_ISL_3357497, EPI_ISL_3357498, EPI_ISL_3357499, EPI_ISL_3357500, EPI_ISL_3357501, EPI_ISL_3357502, EPI_ISL_3357503, EPI_ISL_3357505, EPI_ISL_3357506, EPI_ISL_3357507, EPI_ISL_3357508, EPI_ISL_3357509, EPI_ISL_3357510, EPI_ISL_3357511, EPI_ISL_3357512, EPI_ISL_3357513, EPI_ISL_3357514, EPI_ISL_3357515, EPI_ISL_3357516, | | | |

| | | | |
|---|---|---|---|
| EPI_ISL_3357517, EPI_ISL_3357518, EPI_ISL_3357519, EPI_ISL_3357520, EPI_ISL_3357521, EPI_ISL_3357522, EPI_ISL_3357523, EPI_ISL_3357524, EPI_ISL_3357525, EPI_ISL_3357526, EPI_ISL_3357527, EPI_ISL_3357528, EPI_ISL_3357529, EPI_ISL_3357530, EPI_ISL_3357531, EPI_ISL_3357532, EPI_ISL_3357533, EPI_ISL_3357534, EPI_ISL_3357535, EPI_ISL_3357536, EPI_ISL_3357537, EPI_ISL_3357538, EPI_ISL_3357539, EPI_ISL_3357540, EPI_ISL_3357541, EPI_ISL_3357542, EPI_ISL_3357543, EPI_ISL_3357544, EPI_ISL_3357545, EPI_ISL_3357546, EPI_ISL_3357547, EPI_ISL_3357548, EPI_ISL_3357549, EPI_ISL_3357550, EPI_ISL_3357551, EPI_ISL_3357552, EPI_ISL_3357553, EPI_ISL_3357554, EPI_ISL_3357556, EPI_ISL_3357557, EPI_ISL_3357558, EPI_ISL_3357559, EPI_ISL_3357560, EPI_ISL_3357561, EPI_ISL_3357562, EPI_ISL_3357563, EPI_ISL_3357564, EPI_ISL_3357565, EPI_ISL_3357566, EPI_ISL_3357567, EPI_ISL_3357568, EPI_ISL_3357569, EPI_ISL_3357570, EPI_ISL_3357571, EPI_ISL_3357572, EPI_ISL_3357573, EPI_ISL_3357574, EPI_ISL_3357575, EPI_ISL_3357577, EPI_ISL_3357578, EPI_ISL_3357579, EPI_ISL_3357580, EPI_ISL_3357581, EPI_ISL_3357582, EPI_ISL_3357583, EPI_ISL_3357584, EPI_ISL_3357585, EPI_ISL_3357586, EPI_ISL_3357587, EPI_ISL_3357588, EPI_ISL_3357589, EPI_ISL_3357590, EPI_ISL_3357591, EPI_ISL_3357592, EPI_ISL_3357593, EPI_ISL_3357594, EPI_ISL_3357595, EPI_ISL_3357596, EPI_ISL_3357597, EPI_ISL_3357598, EPI_ISL_3357599, EPI_ISL_3357600, EPI_ISL_3357601, EPI_ISL_3357602, EPI_ISL_3357603, EPI_ISL_3357604, EPI_ISL_3357605, EPI_ISL_3357606, EPI_ISL_3357607, EPI_ISL_3357608, EPI_ISL_3357609, EPI_ISL_3357610, EPI_ISL_3357611, EPI_ISL_3357612, EPI_ISL_3357613, EPI_ISL_3357614, EPI_ISL_3357615, EPI_ISL_3357616, EPI_ISL_3357617, EPI_ISL_3357618, EPI_ISL_3357619, EPI_ISL_3357620, EPI_ISL_3357621, EPI_ISL_3357622, EPI_ISL_3357623, EPI_ISL_3357624, EPI_ISL_3357625, EPI_ISL_3357626, EPI_ISL_3357627, EPI_ISL_3357628, EPI_ISL_3357629, EPI_ISL_3357630, EPI_ISL_3357631, EPI_ISL_3357632, EPI_ISL_3357633, EPI_ISL_3357634, EPI_ISL_3357635, EPI_ISL_3357636, EPI_ISL_3357637, EPI_ISL_3357638, EPI_ISL_3357639, EPI_ISL_3357640, EPI_ISL_3357641, EPI_ISL_3357642, EPI_ISL_3357643, EPI_ISL_3357644, EPI_ISL_3357645, EPI_ISL_3357646, EPI_ISL_3357647, EPI_ISL_3357648, EPI_ISL_3357649, EPI_ISL_3357650, EPI_ISL_3357651, EPI_ISL_3357652, EPI_ISL_3357653, EPI_ISL_3357654, EPI_ISL_3357655, EPI_ISL_3357656, EPI_ISL_3357657, EPI_ISL_3357658, EPI_ISL_3357659, EPI_ISL_3357660, EPI_ISL_3357661, EPI_ISL_3357662, EPI_ISL_3357663, EPI_ISL_3357664, EPI_ISL_3357665, EPI_ISL_3357666, EPI_ISL_3357667, EPI_ISL_3357668, EPI_ISL_3357669, EPI_ISL_3357670, EPI_ISL_3357671, EPI_ISL_3357672, EPI_ISL_3357673, EPI_ISL_3357674, EPI_ISL_3357675, EPI_ISL_3357676, EPI_ISL_3357677, EPI_ISL_3357678, EPI_ISL_3357679, EPI_ISL_3357680, EPI_ISL_3357681, EPI_ISL_3357683, EPI_ISL_3357684, EPI_ISL_3357685, EPI_ISL_3357687, EPI_ISL_3357688, EPI_ISL_3357689, EPI_ISL_3357690, EPI_ISL_3357691, EPI_ISL_3357692, EPI_ISL_3357693, EPI_ISL_3357694, EPI_ISL_3357695, EPI_ISL_3357696, EPI_ISL_3357697, EPI_ISL_3357698, EPI_ISL_3357699, EPI_ISL_3357700, EPI_ISL_3357701, EPI_ISL_3357702, EPI_ISL_3357703, EPI_ISL_3357704, EPI_ISL_3357705, EPI_ISL_3357706, EPI_ISL_3357708, EPI_ISL_3357709, EPI_ISL_3357710, EPI_ISL_3357711, EPI_ISL_3357713, EPI_ISL_3357714, EPI_ISL_3357715, EPI_ISL_3357716, EPI_ISL_3357717, EPI_ISL_3357718, EPI_ISL_3357719, EPI_ISL_3357720, EPI_ISL_3357721, EPI_ISL_3357722, EPI_ISL_3357723, EPI_ISL_3357724, EPI_ISL_3357725, EPI_ISL_3357726, EPI_ISL_3357727, EPI_ISL_3357728, EPI_ISL_3357729, EPI_ISL_3357730, EPI_ISL_3357731, EPI_ISL_3357732, EPI_ISL_3357733, EPI_ISL_3357734, EPI_ISL_3357735, EPI_ISL_3357736, EPI_ISL_3357737, EPI_ISL_3357738, EPI_ISL_3357739, EPI_ISL_3357741, EPI_ISL_3357742, EPI_ISL_3357743, EPI_ISL_3357744, EPI_ISL_3357746, EPI_ISL_3357747, EPI_ISL_3357749, EPI_ISL_3357751, EPI_ISL_3357753, EPI_ISL_3357755, EPI_ISL_3357757, EPI_ISL_3357759, EPI_ISL_3357761, EPI_ISL_3357762, EPI_ISL_3357764, EPI_ISL_3357766, EPI_ISL_3357768, EPI_ISL_3357769, EPI_ISL_3357771, EPI_ISL_3357773, EPI_ISL_3357775, EPI_ISL_3357776, EPI_ISL_3357778, EPI_ISL_3357780, EPI_ISL_3357781, EPI_ISL_3357785, EPI_ISL_3357787, EPI_ISL_3357788, EPI_ISL_3357790, EPI_ISL_3357792, EPI_ISL_3357794, EPI_ISL_3357796, EPI_ISL_3357798, EPI_ISL_3357800, EPI_ISL_3357802, EPI_ISL_3357804, EPI_ISL_3357806, EPI_ISL_3357807, EPI_ISL_3357809, EPI_ISL_3357811, EPI_ISL_3357813, EPI_ISL_3357815, EPI_ISL_3357816, EPI_ISL_3357818, EPI_ISL_3357820, EPI_ISL_3357822, EPI_ISL_3357824, EPI_ISL_3357825, EPI_ISL_3357827, EPI_ISL_3357829, EPI_ISL_3357831, EPI_ISL_3357833, EPI_ISL_3357835, EPI_ISL_3357837, EPI_ISL_3357838, EPI_ISL_3357840, EPI_ISL_3357842, EPI_ISL_3357844, EPI_ISL_3357846, EPI_ISL_3357848, EPI_ISL_3357850, EPI_ISL_3357852, EPI_ISL_3357854, EPI_ISL_3357855, EPI_ISL_3357857, EPI_ISL_3357859, EPI_ISL_3357861, EPI_ISL_3357863, EPI_ISL_3357864, EPI_ISL_3357867, EPI_ISL_3357869, EPI_ISL_3357871, EPI_ISL_3357872, EPI_ISL_3357874, EPI_ISL_3357876, EPI_ISL_3357878, EPI_ISL_3357880, EPI_ISL_3357882, EPI_ISL_3357884, EPI_ISL_3357886, EPI_ISL_3357887, EPI_ISL_3357889, EPI_ISL_3357891, EPI_ISL_3357893, EPI_ISL_3357895, EPI_ISL_3357897, EPI_ISL_3357899, EPI_ISL_3357901, EPI_ISL_3357903, EPI_ISL_3357905, EPI_ISL_3357907, EPI_ISL_3357909, EPI_ISL_3357911, EPI_ISL_3357913, EPI_ISL_3357915, EPI_ISL_3357917, EPI_ISL_3357919, EPI_ISL_3357921, EPI_ISL_3357923, EPI_ISL_3357924, EPI_ISL_3357926, EPI_ISL_3357928, EPI_ISL_3357930, EPI_ISL_3357932, EPI_ISL_3357934, EPI_ISL_3357936, EPI_ISL_3357939, EPI_ISL_3357941, EPI_ISL_3357943, EPI_ISL_3357945, EPI_ISL_3357947, EPI_ISL_3357949, EPI_ISL_3357951, EPI_ISL_3357953, EPI_ISL_3357955 | | | |
| see above | Department of Bacteria, Parasites and Fungi, Statens Serum Institut, Copenhagen, Denmark | Statens Serum Institut Bioinformatics and Microbial Genomics | Danish Covid-19 Genome Consortium |
| EPI_ISL_3357983, EPI_ISL_3357984, EPI_ISL_3357985, EPI_ISL_3357986, EPI_ISL_3357987, EPI_ISL_3357988, EPI_ISL_3357989, EPI_ISL_3357990, EPI_ISL_3357991, EPI_ISL_3357992, EPI_ISL_3357993, EPI_ISL_3357994, EPI_ISL_3357995, EPI_ISL_3357996, EPI_ISL_3357997, EPI_ISL_3357998, EPI_ISL_3357999, EPI_ISL_3358000, EPI_ISL_3358001, EPI_ISL_3358002, EPI_ISL_3358003, EPI_ISL_3358004, EPI_ISL_3358005, EPI_ISL_3358006, EPI_ISL_3358007, EPI_ISL_3358008, EPI_ISL_3358009, EPI_ISL_3358010, EPI_ISL_3358011, EPI_ISL_3358012, EPI_ISL_3358013, EPI_ISL_3358014, EPI_ISL_3358015, EPI_ISL_3358016, EPI_ISL_3358017, EPI_ISL_3358018, EPI_ISL_3358019, EPI_ISL_3358020, EPI_ISL_3358021, EPI_ISL_3358022, EPI_ISL_3358023, EPI_ISL_3358024, EPI_ISL_3358025, EPI_ISL_3358026, EPI_ISL_3358027, EPI_ISL_3358028, EPI_ISL_3358029, EPI_ISL_3358030, EPI_ISL_3358031, EPI_ISL_3358032, EPI_ISL_3358033, EPI_ISL_3358034, EPI_ISL_3358035, EPI_ISL_3358036, EPI_ISL_3358037, EPI_ISL_3358038, EPI_ISL_3358039, EPI_ISL_3358040, EPI_ISL_3358041, EPI_ISL_3358042, EPI_ISL_3358043, EPI_ISL_3358044, EPI_ISL_3358045, EPI_ISL_3358046, EPI_ISL_3358047, EPI_ISL_3358048, EPI_ISL_3358049, EPI_ISL_3358050, EPI_ISL_3358051, EPI_ISL_3358052, EPI_ISL_3358053, EPI_ISL_3358054, EPI_ISL_3358055, EPI_ISL_3358056, EPI_ISL_3358057, EPI_ISL_3358058, EPI_ISL_3358059, EPI_ISL_3358060, EPI_ISL_3358061, EPI_ISL_3358062, EPI_ISL_3358063, EPI_ISL_3358064, EPI_ISL_3358065, EPI_ISL_3358066, EPI_ISL_3358067, EPI_ISL_3358068, EPI_ISL_3358069, EPI_ISL_3358070, EPI_ISL_3358071, EPI_ISL_3358072, EPI_ISL_3358074, EPI_ISL_3358075, EPI_ISL_3358076, EPI_ISL_3358078, EPI_ISL_3358079, EPI_ISL_3358080, EPI_ISL_3358081, EPI_ISL_3358082, EPI_ISL_3358083, EPI_ISL_3358084, EPI_ISL_3358085, EPI_ISL_3358086, EPI_ISL_3358087, EPI_ISL_3358088, EPI_ISL_3358089, EPI_ISL_3358090, EPI_ISL_3358091, EPI_ISL_3358092, EPI_ISL_3358093, EPI_ISL_3358094, EPI_ISL_3358095, EPI_ISL_3358096, EPI_ISL_3358097, EPI_ISL_3358098, EPI_ISL_3358099, EPI_ISL_3358100, EPI_ISL_3358101, EPI_ISL_3358102, EPI_ISL_3358103, EPI_ISL_3358104, EPI_ISL_3358105, EPI_ISL_3358106, EPI_ISL_3358107, EPI_ISL_3358108, EPI_ISL_3358109, EPI_ISL_3358110, EPI_ISL_3358111, EPI_ISL_3358112, EPI_ISL_3358113, EPI_ISL_3358114, EPI_ISL_3358115, EPI_ISL_3358116, EPI_ISL_3358117, EPI_ISL_3358118, EPI_ISL_3358119, EPI_ISL_3358120, EPI_ISL_3358121, EPI_ISL_3358123, EPI_ISL_3358124, EPI_ISL_3358125, EPI_ISL_3358126, EPI_ISL_3358127, EPI_ISL_3358128, EPI_ISL_3358129, EPI_ISL_3358130, EPI_ISL_3358131, EPI_ISL_3358132, EPI_ISL_3358133, EPI_ISL_3358134, EPI_ISL_3358135, EPI_ISL_3358136, EPI_ISL_3358137, EPI_ISL_3358138, EPI_ISL_3358139, EPI_ISL_3358140, EPI_ISL_3358141, EPI_ISL_3358143, EPI_ISL_3358144, EPI_ISL_3358145, EPI_ISL_3358146, EPI_ISL_3358147, EPI_ISL_3358148, EPI_ISL_3358149, EPI_ISL_3358150, EPI_ISL_3358151, EPI_ISL_3358152, EPI_ISL_3358153, EPI_ISL_3358155, EPI_ISL_3358156, EPI_ISL_3358157, EPI_ISL_3358158, EPI_ISL_3358159, EPI_ISL_3358160, EPI_ISL_3358161, EPI_ISL_3358162, EPI_ISL_3358163, EPI_ISL_3358164, EPI_ISL_3358165, EPI_ISL_3358166, EPI_ISL_3358167, EPI_ISL_3358168, EPI_ISL_3358169, EPI_ISL_3358170, EPI_ISL_3358171, EPI_ISL_3358172, EPI_ISL_3358173, EPI_ISL_3358174, EPI_ISL_3358177, EPI_ISL_3358178, EPI_ISL_3358179, EPI_ISL_3358180, EPI_ISL_3358181, EPI_ISL_3358182, EPI_ISL_3358183, EPI_ISL_3358184, EPI_ISL_3358185, EPI_ISL_3358186, EPI_ISL_3358187, EPI_ISL_3358188, EPI_ISL_3358189, EPI_ISL_3358192, EPI_ISL_3358193, EPI_ISL_3358194, EPI_ISL_3358195, EPI_ISL_3358196, EPI_ISL_3358198, EPI_ISL_3358199, EPI_ISL_3358200, EPI_ISL_3358201, EPI_ISL_3358202, EPI_ISL_3358204, EPI_ISL_3358205, EPI_ISL_3358206, EPI_ISL_3358207, EPI_ISL_3358208, EPI_ISL_3358210, EPI_ISL_3358211, EPI_ISL_3358212, EPI_ISL_3358213, EPI_ISL_3358214, EPI_ISL_3358216, EPI_ISL_3358217, EPI_ISL_3358218, EPI_ISL_3358220, EPI_ISL_3358221, EPI_ISL_3358222, EPI_ISL_3358223, EPI_ISL_3358224, EPI_ISL_3358225, EPI_ISL_3358226, EPI_ISL_3358227, EPI_ISL_3358228, EPI_ISL_3358229, EPI_ISL_3358230, EPI_ISL_3358231, EPI_ISL_3358232, EPI_ISL_3358233, EPI_ISL_3358234, EPI_ISL_3358235, EPI_ISL_3358236, EPI_ISL_3358237, EPI_ISL_3358238, EPI_ISL_3358239, EPI_ISL_3358240, EPI_ISL_3358241, EPI_ISL_3358242, EPI_ISL_3358243, EPI_ISL_3358244, EPI_ISL_3358245, EPI_ISL_3358246, EPI_ISL_3358247, EPI_ISL_3358248, EPI_ISL_3358249, EPI_ISL_3358250, EPI_ISL_3358251, EPI_ISL_3358252, EPI_ISL_3358253, EPI_ISL_3358254, EPI_ISL_3358255, EPI_ISL_3358256, EPI_ISL_3358257, EPI_ISL_3358258, EPI_ISL_3358259, EPI_ISL_3358260, EPI_ISL_3358261, EPI_ISL_3358262, EPI_ISL_3358263, EPI_ISL_3358264, EPI_ISL_3358265, EPI_ISL_3358266, EPI_ISL_3358267, EPI_ISL_3358268, EPI_ISL_3358269, EPI_ISL_3358270, EPI_ISL_3358271, EPI_ISL_3358272, EPI_ISL_3358273, EPI_ISL_3358274, EPI_ISL_3358275, EPI_ISL_3358277, EPI_ISL_3358278, EPI_ISL_3358279, EPI_ISL_3358280, EPI_ISL_3358281, EPI_ISL_3358283, EPI_ISL_3358284, EPI_ISL_3358285, EPI_ISL_3358286, EPI_ISL_3358287, EPI_ISL_3358288, EPI_ISL_3358289, EPI_ISL_3358290, EPI_ISL_3358291, EPI_ISL_3358292, EPI_ISL_3358293, EPI_ISL_3358294, EPI_ISL_3358295, EPI_ISL_3358296, EPI_ISL_3358297, EPI_ISL_3358298, EPI_ISL_3358299, EPI_ISL_3358300, EPI_ISL_3358301, EPI_ISL_3358302, EPI_ISL_3358303, EPI_ISL_3358304, EPI_ISL_3358305, EPI_ISL_3358306, EPI_ISL_3358307, EPI_ISL_3358308, EPI_ISL_3358309, EPI_ISL_3358310, EPI_ISL_3358311, EPI_ISL_3358312, EPI_ISL_3358313, EPI_ISL_3358314, EPI_ISL_3358315, EPI_ISL_3358316, EPI_ISL_3358317, EPI_ISL_3358318, EPI_ISL_3358319, EPI_ISL_3358320, EPI_ISL_3358321, EPI_ISL_3358323, EPI_ISL_3358324, EPI_ISL_3358325, EPI_ISL_3358327, EPI_ISL_3358328, EPI_ISL_3358329, EPI_ISL_3358330, EPI_ISL_3358331, EPI_ISL_3358332, EPI_ISL_3358333, EPI_ISL_3358334, EPI_ISL_3358335, EPI_ISL_3358336, EPI_ISL_3358337, EPI_ISL_3358338, EPI_ISL_3358339, EPI_ISL_3358340, EPI_ISL_3358341, EPI_ISL_3358342, EPI_ISL_3358343, EPI_ISL_3358344 | | | |
| see above | Arkansas Public Health Laboratory, Arkansas Department of Health | University of Minnesota Genomics Center | Daryl M. Gohl, John Garbe, Jaquelyn Kuriger-Laber, Corbin Dirkx |
| EPI_ISL_3358345, EPI_ISL_3358346, EPI_ISL_3358347, EPI_ISL_3358348 | Stadtspital Triemli | Institute of Medical Virology | Daniel Ehrsam, Isabel Stürmer, Catharine Aquino, Joel Wirz, Weihong Qi, Hubert Rehrauer, Verena Kufner, Gabriela Ziltener, Maryam Zaheri, Stefan Schmutz, Annette Audigé, Maria Grünberg, Kevin Steiner, Jon Huder, Cyril Shah, Riccarda Capaul, Guido Bloemberg, Jürg Böni, Michael Huber, Alexandra Trkola |
| EPI_ISL_3358349, EPI_ISL_3358350 | UniversitätsSpital Zürich | Institute of Medical Virology | Daniel Ehrsam, Isabel Stürmer, Catharine Aquino, Joel Wirz, Weihong Qi, Hubert Rehrauer, Verena Kufner, Gabriela Ziltener, Maryam Zaheri, Stefan Schmutz, Annette Audigé, Maria Grünberg, Kevin Steiner, Jon Huder, Cyril Shah, Riccarda Capaul, Guido Bloemberg, Jürg Böni, Michael Huber, Alexandra Trkola |
| EPI_ISL_3358351 | Universitäts-Kinderspital Zürich | Institute of Medical Virology | Daniel Ehrsam, Isabel Stürmer, Catharine Aquino, Joel Wirz, Weihong Qi, Hubert Rehrauer, Verena Kufner, Gabriela Ziltener, Maryam Zaheri, Stefan Schmutz, Annette Audigé, Maria Grünberg, Kevin Steiner, Jon Huder, Cyril Shah, Riccarda Capaul, Guido Bloemberg, Jürg Böni, Michael Huber, Alexandra Trkola |
| EPI_ISL_3358352, EPI_ISL_3358353, EPI_ISL_3358354, EPI_ISL_3358355, EPI_ISL_3358356, EPI_ISL_3358357, EPI_ISL_3358358, EPI_ISL_3358359, EPI_ISL_3358360, EPI_ISL_3358361, EPI_ISL_3358362, EPI_ISL_3358363, EPI_ISL_3358364, EPI_ISL_3358365, EPI_ISL_3358366, EPI_ISL_3358367, EPI_ISL_3358368, EPI_ISL_3358369, EPI_ISL_3358370, EPI_ISL_3358371 | | | |
| see above | Kantonsspital Winterthur | Institute of Medical Virology | Daniel Ehrsam, Isabel Stürmer, Catharine Aquino, Joel Wirz, Weihong Qi, Hubert Rehrauer, Verena Kufner, Gabriela Ziltener, Maryam Zaheri, Stefan Schmutz, Annette Audigé, Maria Grünberg, Kevin Steiner, Jon Huder, Cyril Shah, Riccarda Capaul, Guido Bloemberg, Jürg Böni, Michael Huber, Alexandra Trkola |
| EPI_ISL_3358372 | Kantonsspital Baden AG | Institute of Medical Virology | Daniel Ehrsam, Isabel Stürmer, Catharine Aquino, Joel Wirz, Weihong Qi, Hubert Rehrauer, Verena Kufner, Gabriela Ziltener, Maryam Zaheri, Stefan Schmutz, Annette Audigé, Maria Grünberg, Kevin Steiner, Jon Huder, Cyril Shah, Riccarda Capaul, Guido Bloemberg, Jürg Böni, Michael Huber, Alexandra Trkola |
| EPI_ISL_3358373 | Universitäts-Kinderspital Zürich | Institute of Medical Virology | Daniel Ehrsam, Isabel Stürmer, Catharine Aquino, Joel Wirz, Weihong Qi, Hubert Rehrauer, Verena Kufner, Gabriela Ziltener, Maryam Zaheri, Stefan Schmutz, Annette Audigé, Maria Grünberg, Kevin Steiner, Jon Huder, Cyril Shah, Riccarda Capaul, Guido Bloemberg, Jürg Böni, Michael Huber, Alexandra Trkola |
| EPI_ISL_3358374, EPI_ISL_3358376, EPI_ISL_3358377, EPI_ISL_3358378, EPI_ISL_3358379, EPI_ISL_3358380 | Stadtspital Triemli | Institute of Medical Virology | Daniel Ehrsam, Isabel Stürmer, Catharine Aquino, Joel Wirz, Weihong Qi, Hubert Rehrauer, Verena Kufner, Gabriela Ziltener, Maryam Zaheri, Stefan Schmutz, Annette Audigé, Maria Grünberg, Kevin Steiner, Jon Huder, Cyril Shah, Riccarda Capaul, Guido Bloemberg, Jürg Böni, Michael Huber, Alexandra Trkola |
| EPI_ISL_3358381 | Universitäts-Kinderspital Zürich | Institute of Medical Virology | Daniel Ehrsam, Isabel Stürmer, Catharine Aquino, Joel Wirz, Weihong Qi, Hubert Rehrauer, Verena Kufner, Gabriela Ziltener, Maryam Zaheri, Stefan Schmutz, Annette Audigé, Maria Grünberg, Kevin Steiner, Jon Huder, Cyril Shah, Riccarda Capaul, Guido Bloemberg, Jürg Böni, Michael Huber, Alexandra Trkola |
| EPI_ISL_3358382, EPI_ISL_3358383, EPI_ISL_3358384, EPI_ISL_3358385, EPI_ISL_3358386, EPI_ISL_3358387, EPI_ISL_3358388, EPI_ISL_3358389, EPI_ISL_3358390, EPI_ISL_3358391 | Kantonsspital Winterthur | Institute of Medical Virology | Daniel Ehrsam, Isabel Stürmer, Catharine Aquino, Joel Wirz, Weihong Qi, Hubert Rehrauer, Verena Kufner, Gabriela Ziltener, Maryam Zaheri, Stefan Schmutz, Annette Audigé, Maria Grünberg, Kevin Steiner, Jon Huder, Cyril Shah, Riccarda Capaul, Guido Bloemberg, Jürg Böni, Michael Huber, Alexandra Trkola |
| EPI_ISL_3358392, EPI_ISL_3358393 | Universitäts-Kinderspital Zürich | Institute of Medical Virology | Daniel Ehrsam, Isabel Stürmer, Catharine Aquino, Joel Wirz, Weihong Qi, Hubert Rehrauer, Verena Kufner, Gabriela Ziltener, Maryam Zaheri, Stefan Schmutz, Annette Audigé, Maria Grünberg, Kevin Steiner, Jon Huder, Cyril Shah, Riccarda Capaul, Guido Bloemberg, Jürg Böni, Michael Huber, Alexandra Trkola |
| EPI_ISL_3358394, EPI_ISL_3358395, EPI_ISL_3358396, EPI_ISL_3358397 | Zentrallabor Zürich | Institute of Medical Virology | Daniel Ehrsam, Isabel Stürmer, Catharine Aquino, Joel Wirz, Weihong Qi, Hubert Rehrauer, Verena Kufner, Gabriela Ziltener, Maryam Zaheri, Stefan Schmutz, Annette Audigé, Maria Grünberg, Kevin Steiner, Jon Huder, Cyril Shah, Riccarda Capaul, Guido Bloemberg, Jürg Böni, Michael Huber, Alexandra Trkola |
| EPI_ISL_3358398, EPI_ISL_3358399, EPI_ISL_3358400, EPI_ISL_3358401 | UniversitätsSpital Zürich | Institute of Medical Virology | Daniel Ehrsam, Isabel Stürmer, Catharine Aquino, Joel Wirz, Weihong Qi, Hubert Rehrauer, Verena Kufner, Gabriela Ziltener, Maryam Zaheri, Stefan Schmutz, Annette Audigé, Maria Grünberg, Kevin Steiner, Jon Huder, Cyril Shah, Riccarda Capaul, Guido Bloemberg, Jürg Böni, Michael Huber, Alexandra Trkola |
| EPI_ISL_3358402 | Universität Zürich | Institute of Medical Virology | Daniel Ehrsam, Isabel Stürmer, Catharine Aquino, Joel Wirz, Weihong Qi, Hubert Rehrauer, Verena Kufner, Gabriela Ziltener, Maryam Zaheri, Stefan Schmutz, Annette Audigé, Maria Grünberg, Kevin Steiner, Jon Huder, Cyril Shah, Riccarda Capaul, Guido Bloemberg, Jürg Böni, Michael Huber, Alexandra Trkola |
| EPI_ISL_3358403, EPI_ISL_3358404, EPI_ISL_3358405, EPI_ISL_3358406 | Stadtspital Triemli | Institute of Medical Virology | Daniel Ehrsam, Isabel Stürmer, Catharine Aquino, Joel Wirz, Weihong Qi, Hubert Rehrauer, Verena Kufner, Gabriela Ziltener, Maryam Zaheri, Stefan Schmutz, Annette Audigé, Maria Grünberg, Kevin Steiner, Jon Huder, Cyril Shah, Riccarda Capaul, Guido Bloemberg, Jürg Böni, Michael Huber, Alexandra Trkola |
| EPI_ISL_3358407 | Stadtspital Waid | Institute of Medical Virology | Daniel Ehrsam, Isabel Stürmer, Catharine Aquino, Joel Wirz, Weihong Qi, Hubert Rehrauer, Verena Kufner, Gabriela Ziltener, Maryam Zaheri, Stefan Schmutz, Annette Audigé, Maria Grünberg, Kevin Steiner, Jon Huder, Cyril Shah, Riccarda Capaul, Guido Bloemberg, Jürg Böni, Michael Huber, Alexandra Trkola |
| EPI_ISL_3358408 | Unilabs | Institute of Medical Virology | Verena Kufner, Gabriela Ziltener, Maryam Zaheri, Stefan Schmutz, Annette Audigé, Maria Grünberg, Kevin Steiner, Jon Huder, Cyril Shah, Riccarda Capaul, Guido Bloemberg, Jürg Böni, Michael Huber, Alexandra Trkola |
| EPI_ISL_3358409, EPI_ISL_3358410 | Kantonsspital Baden AG | Institute of Medical Virology | Verena Kufner, Gabriela Ziltener, Maryam Zaheri, Stefan Schmutz, Annette Audigé, Maria Grünberg, Kevin Steiner, Jon Huder, Cyril Shah, Riccarda Capaul, Guido Bloemberg, Jürg Böni, Michael Huber, Alexandra Trkola |
| EPI_ISL_3358411 | Kantonsspital Schaffhausen | Institute of Medical Virology | Verena Kufner, Gabriela Ziltener, Maryam Zaheri, Stefan Schmutz, Annette Audigé, Maria Grünberg, Kevin Steiner, Jon Huder, Cyril Shah, Riccarda Capaul, Guido Bloemberg, Jürg Böni, Michael Huber, Alexandra Trkola |
| EPI_ISL_3358412 | Universität Zürich | Institute of Medical Virology | Verena Kufner, Gabriela Ziltener, Maryam Zaheri, Stefan Schmutz, Annette Audigé, Maria Grünberg, Kevin Steiner, Jon Huder, Cyril Shah, Riccarda Capaul, Guido Bloemberg, Jürg Böni, Michael Huber, Alexandra Trkola |
| EPI_ISL_3358413, EPI_ISL_3358414 | Analytica | Institute of Medical Virology | Verena Kufner, Gabriela Ziltener, Maryam Zaheri, Stefan Schmutz, Annette Audigé, Maria Grünberg, Kevin Steiner, Jon Huder, Cyril Shah, Riccarda Capaul, Guido Bloemberg, Jürg Böni, Michael Huber, Alexandra Trkola |
| EPI_ISL_3358415 | Laborgemeinschaft 1 | Institute of Medical Virology | Verena Kufner, Gabriela Ziltener, Maryam Zaheri, Stefan Schmutz, Annette Audigé, Maria Grünberg, Kevin Steiner, Jon Huder, Cyril Shah, Riccarda Capaul, Guido Bloemberg, Jürg Böni, Michael Huber, Alexandra Trkola |
| EPI_ISL_3358416, EPI_ISL_3358417, EPI_ISL_3358418 | Medica | Institute of Medical Virology | Verena Kufner, Gabriela Ziltener, Maryam Zaheri, Stefan Schmutz, Annette Audigé, Maria Grünberg, Kevin Steiner, Jon Huder, Cyril Shah, Riccarda Capaul, Guido Bloemberg, Jürg Böni, Michael Huber, Alexandra Trkola |

| | | | |
|---|---|---|---|
| EPI_ISL_3358419, EPI_ISL_3358420 | UniversitätsSpital Zürich | Institute of Medical Virology | Verena Kufner, Gabriela Ziltener, Maryam Zaheri, Stefan Schmutz, Annette Audigé, Maria Grünberg, Kevin Steiner, Jon Huder, Cyril Shah, Riccarda Capaul, Guido Bloemberg, Jürg Böni, Michael Huber, Alexandra Trkola |
| EPI_ISL_3358421, EPI_ISL_3358422 | Zentrum für Labormedizin | Institute of Medical Virology | Verena Kufner, Gabriela Ziltener, Maryam Zaheri, Stefan Schmutz, Annette Audigé, Maria Grünberg, Kevin Steiner, Jon Huder, Cyril Shah, Riccarda Capaul, Guido Bloemberg, Jürg Böni, Michael Huber, Alexandra Trkola |
| EPI_ISL_3358423, EPI_ISL_3358424 | Kantonsspital Baden AG | Institute of Medical Virology | Verena Kufner, Gabriela Ziltener, Maryam Zaheri, Stefan Schmutz, Annette Audigé, Maria Grünberg, Kevin Steiner, Jon Huder, Cyril Shah, Riccarda Capaul, Guido Bloemberg, Jürg Böni, Michael Huber, Alexandra Trkola |
| EPI_ISL_3358425 | UniversitätsSpital Zürich | Institute of Medical Virology | Verena Kufner, Gabriela Ziltener, Maryam Zaheri, Stefan Schmutz, Annette Audigé, Maria Grünberg, Kevin Steiner, Jon Huder, Cyril Shah, Riccarda Capaul, Guido Bloemberg, Jürg Böni, Michael Huber, Alexandra Trkola |
| EPI_ISL_3358426, EPI_ISL_3358427 | Unilabs | Institute of Medical Virology | Verena Kufner, Gabriela Ziltener, Maryam Zaheri, Stefan Schmutz, Annette Audigé, Maria Grünberg, Kevin Steiner, Jon Huder, Cyril Shah, Riccarda Capaul, Guido Bloemberg, Jürg Böni, Michael Huber, Alexandra Trkola |
| EPI_ISL_3358428, EPI_ISL_3358429 | Analytica | Institute of Medical Virology | Verena Kufner, Gabriela Ziltener, Maryam Zaheri, Stefan Schmutz, Annette Audigé, Maria Grünberg, Kevin Steiner, Jon Huder, Cyril Shah, Riccarda Capaul, Guido Bloemberg, Jürg Böni, Michael Huber, Alexandra Trkola |
| EPI_ISL_3358430 | Medica | Institute of Medical Virology | Verena Kufner, Gabriela Ziltener, Maryam Zaheri, Stefan Schmutz, Annette Audigé, Maria Grünberg, Kevin Steiner, Jon Huder, Cyril Shah, Riccarda Capaul, Guido Bloemberg, Jürg Böni, Michael Huber, Alexandra Trkola |
| EPI_ISL_3358431 | Laborgemeinschaft 1 | Institute of Medical Virology | Verena Kufner, Gabriela Ziltener, Maryam Zaheri, Stefan Schmutz, Annette Audigé, Maria Grünberg, Kevin Steiner, Jon Huder, Cyril Shah, Riccarda Capaul, Guido Bloemberg, Jürg Böni, Michael Huber, Alexandra Trkola |
| EPI_ISL_3358432 | Universität Zürich | Institute of Medical Virology | Verena Kufner, Gabriela Ziltener, Maryam Zaheri, Stefan Schmutz, Annette Audigé, Maria Grünberg, Kevin Steiner, Jon Huder, Cyril Shah, Riccarda Capaul, Guido Bloemberg, Jürg Böni, Michael Huber, Alexandra Trkola |
| EPI_ISL_3358433, EPI_ISL_3358434 | UniversitätsSpital Zürich | Institute of Medical Virology | Verena Kufner, Gabriela Ziltener, Maryam Zaheri, Stefan Schmutz, Annette Audigé, Maria Grünberg, Kevin Steiner, Jon Huder, Cyril Shah, Riccarda Capaul, Guido Bloemberg, Jürg Böni, Michael Huber, Alexandra Trkola |
| EPI_ISL_3358435, EPI_ISL_3358436, EPI_ISL_3358437, EPI_ISL_3358438, EPI_ISL_3358439, EPI_ISL_3358440, EPI_ISL_3358441, EPI_ISL_3358442, EPI_ISL_3358443 | Universität Zürich | Institute of Medical Virology | Verena Kufner, Gabriela Ziltener, Maryam Zaheri, Stefan Schmutz, Annette Audigé, Maria Grünberg, Kevin Steiner, Jon Huder, Cyril Shah, Riccarda Capaul, Guido Bloemberg, Jürg Böni, Michael Huber, Alexandra Trkola |
| EPI_ISL_3358444 | UniversitätsSpital Zürich | Institute of Medical Virology | Verena Kufner, Gabriela Ziltener, Maryam Zaheri, Stefan Schmutz, Annette Audigé, Maria Grünberg, Kevin Steiner, Jon Huder, Cyril Shah, Riccarda Capaul, Guido Bloemberg, Jürg Böni, Michael Huber, Alexandra Trkola |
| EPI_ISL_3358445, EPI_ISL_3358446 | Universitäts-Kinderspital Zürich | Institute of Medical Virology | Verena Kufner, Gabriela Ziltener, Maryam Zaheri, Stefan Schmutz, Annette Audigé, Maria Grünberg, Kevin Steiner, Jon Huder, Cyril Shah, Riccarda Capaul, Guido Bloemberg, Jürg Böni, Michael Huber, Alexandra Trkola |
| EPI_ISL_3358447, EPI_ISL_3358448, EPI_ISL_3358449, EPI_ISL_3358450, EPI_ISL_3358451, EPI_ISL_3358452, EPI_ISL_3358453 | Stadtspital Triemli | Institute of Medical Virology | Verena Kufner, Gabriela Ziltener, Maryam Zaheri, Stefan Schmutz, Annette Audigé, Maria Grünberg, Kevin Steiner, Jon Huder, Cyril Shah, Riccarda Capaul, Guido Bloemberg, Jürg Böni, Michael Huber, Alexandra Trkola |
| EPI_ISL_3358489, EPI_ISL_3358490, EPI_ISL_3358491, EPI_ISL_3358492, EPI_ISL_3358493, EPI_ISL_3358494, EPI_ISL_3358495, EPI_ISL_3358496, EPI_ISL_3358497, EPI_ISL_3358498, EPI_ISL_3358499, EPI_ISL_3358500, EPI_ISL_3358501, EPI_ISL_3358502, EPI_ISL_3358503, EPI_ISL_3358504, EPI_ISL_3358505, EPI_ISL_3358506, EPI_ISL_3358507, EPI_ISL_3358508, EPI_ISL_3358509, EPI_ISL_3358510, EPI_ISL_3358511, EPI_ISL_3358512, EPI_ISL_3358513, EPI_ISL_3358514, EPI_ISL_3358515, EPI_ISL_3358516, EPI_ISL_3358517, EPI_ISL_3358518, EPI_ISL_3358519, EPI_ISL_3358520, EPI_ISL_3358521, EPI_ISL_3358522, EPI_ISL_3358523, EPI_ISL_3358524, EPI_ISL_3358525, EPI_ISL_3358526, EPI_ISL_3358527, EPI_ISL_3358528, EPI_ISL_3358529, EPI_ISL_3358530, EPI_ISL_3358531, EPI_ISL_3358532 | | | |
| see above | Center for Laboratory Medicine | Center for Laboratory Medicine | Yannick Gerth |
| EPI_ISL_3358577 | Lighthouse Lab in Alderley Park | Wellcome Sanger Institute for the COVID-19 Genomics UK (COG-UK) Consortium | Jacquelyn Wynn, Mairead Hyland, The Lighthouse Lab in Alderley Park and Alex Alderton, Roberto Amato, Jeffrey Barrett, Sonia Goncalves, Ewan Harrison, David K. Jackson, Ian Johnston, Dominic Kwiatkowski, Cordelia Langford, John Sillitoe on behalf of the Wellcome Sanger Institute COVID-19 Surveillance Team |
| EPI_ISL_3358578 | Lighthouse Lab in Milton Keynes | Wellcome Sanger Institute for the COVID-19 Genomics UK (COG-UK) Consortium | The Lighthouse Lab in Milton Keynes and Alex Alderton, Roberto Amato, Jeffrey Barrett, Sonia Goncalves, Ewan Harrison, David K. Jackson, Ian Johnston, Dominic Kwiatkowski, Cordelia Langford, John Sillitoe on behalf of the Wellcome Sanger Institute COVID-19 Surveillance Team |
| EPI_ISL_3358579, EPI_ISL_3358580 | Lighthouse Lab in Glasgow | Wellcome Sanger Institute for the COVID-19 Genomics UK (COG-UK) Consortium | Harper VanSteenhouse, Yumi Kasai, David Gray, Carol Clugston, Anna Dominiczak and Alex Alderton, Roberto Amato, Jeffrey Barrett, Sonia Goncalves, Ewan Harrison, David K. Jackson, Ian Johnston, Dominic Kwiatkowski, Cordelia Langford, John Sillitoe on behalf of the Wellcome Sanger Institute COVID-19 Surveillance Team |
| EPI_ISL_3358581, EPI_ISL_3358582, EPI_ISL_3358583 | Lighthouse Lab in Milton Keynes | Wellcome Sanger Institute for the COVID-19 Genomics UK (COG-UK) Consortium | The Lighthouse Lab in Milton Keynes and Alex Alderton, Roberto Amato, Jeffrey Barrett, Sonia Goncalves, Ewan Harrison, David K. Jackson, Ian Johnston, Dominic Kwiatkowski, Cordelia Langford, John Sillitoe on behalf of the Wellcome Sanger Institute COVID-19 Surveillance Team |
| EPI_ISL_3358584, EPI_ISL_3358585 | Lighthouse Lab in Glasgow | Wellcome Sanger Institute for the COVID-19 Genomics UK (COG-UK) Consortium | Harper VanSteenhouse, Yumi Kasai, David Gray, Carol Clugston, Anna Dominiczak and Alex Alderton, Roberto Amato, Jeffrey Barrett, Sonia Goncalves, Ewan Harrison, David K. Jackson, Ian Johnston, Dominic Kwiatkowski, Cordelia Langford, John Sillitoe on behalf of the Wellcome Sanger Institute COVID-19 Surveillance Team |
| EPI_ISL_3358586 | Lighthouse Lab in Milton Keynes | Wellcome Sanger Institute for the COVID-19 Genomics UK (COG-UK) Consortium | The Lighthouse Lab in Milton Keynes and Alex Alderton, Roberto Amato, Jeffrey Barrett, Sonia Goncalves, Ewan Harrison, David K. Jackson, Ian Johnston, Dominic Kwiatkowski, Cordelia Langford, John Sillitoe on behalf of the Wellcome Sanger Institute COVID-19 Surveillance Team |
| EPI_ISL_3358587 | Lighthouse Lab in Glasgow | Wellcome Sanger Institute for the COVID-19 Genomics UK (COG-UK) Consortium | Harper VanSteenhouse, Yumi Kasai, David Gray, Carol Clugston, Anna Dominiczak and Alex Alderton, Roberto Amato, Jeffrey Barrett, Sonia Goncalves, Ewan Harrison, David K. Jackson, Ian Johnston, Dominic Kwiatkowski, Cordelia Langford, John Sillitoe on behalf of the Wellcome Sanger Institute COVID-19 Surveillance Team |
| EPI_ISL_3358588, EPI_ISL_3358589, EPI_ISL_3358590, EPI_ISL_3358591 | Lighthouse Lab in Milton Keynes | Wellcome Sanger Institute for the COVID-19 Genomics UK (COG-UK) Consortium | The Lighthouse Lab in Milton Keynes and Alex Alderton, Roberto Amato, Jeffrey Barrett, Sonia Goncalves, Ewan Harrison, David K. Jackson, Ian Johnston, Dominic Kwiatkowski, Cordelia Langford, John Sillitoe on behalf of the Wellcome Sanger Institute COVID-19 Surveillance Team |
| EPI_ISL_3358592 | Lighthouse Lab in Alderley Park | Wellcome Sanger Institute for the COVID-19 Genomics UK (COG-UK) Consortium | Jacquelyn Wynn, Mairead Hyland, The Lighthouse Lab in Alderley Park and Alex Alderton, Roberto Amato, Jeffrey Barrett, Sonia Goncalves, Ewan Harrison, David K. Jackson, Ian Johnston, Dominic Kwiatkowski, Cordelia Langford, John Sillitoe on behalf of the Wellcome Sanger Institute COVID-19 Surveillance Team |
| EPI_ISL_3358593, EPI_ISL_3358594, EPI_ISL_3358595 | Lighthouse Lab in Milton Keynes | Wellcome Sanger Institute for the COVID-19 Genomics UK (COG-UK) Consortium | The Lighthouse Lab in Milton Keynes and Alex Alderton, Roberto Amato, Jeffrey Barrett, Sonia Goncalves, Ewan Harrison, David K. Jackson, Ian Johnston, Dominic Kwiatkowski, Cordelia Langford, John Sillitoe on behalf of the Wellcome Sanger Institute COVID-19 Surveillance Team |
| EPI_ISL_3358596, EPI_ISL_3358597 | Lighthouse Lab in Glasgow | Wellcome Sanger Institute for the COVID-19 Genomics UK (COG-UK) Consortium | Harper VanSteenhouse, Yumi Kasai, David Gray, Carol Clugston, Anna Dominiczak and Alex Alderton, Roberto Amato, Jeffrey Barrett, Sonia Goncalves, Ewan Harrison, David K. Jackson, Ian Johnston, Dominic Kwiatkowski, Cordelia Langford, John Sillitoe on behalf of the Wellcome Sanger Institute COVID-19 Surveillance Team |
| EPI_ISL_3358598 | Lighthouse Lab in Alderley Park | Wellcome Sanger Institute for the COVID-19 Genomics UK (COG-UK) Consortium | Jacquelyn Wynn, Mairead Hyland, The Lighthouse Lab in Alderley Park and Alex Alderton, Roberto Amato, Jeffrey Barrett, Sonia Goncalves, Ewan Harrison, David K. Jackson, Ian Johnston, Dominic Kwiatkowski, Cordelia Langford, John Sillitoe on behalf of the Wellcome Sanger Institute COVID-19 Surveillance Team |
| EPI_ISL_3358599 | Lighthouse Lab in Milton Keynes | Wellcome Sanger Institute for the COVID-19 Genomics UK (COG-UK) Consortium | The Lighthouse Lab in Milton Keynes and Alex Alderton, Roberto Amato, Jeffrey Barrett, Sonia Goncalves, Ewan Harrison, David K. Jackson, Ian Johnston, Dominic Kwiatkowski, Cordelia Langford, John Sillitoe on behalf of the Wellcome Sanger Institute COVID-19 Surveillance Team |
| EPI_ISL_3358600 | Lighthouse Lab in Alderley Park | Wellcome Sanger Institute for the COVID-19 Genomics UK (COG-UK) Consortium | Jacquelyn Wynn, Mairead Hyland, The Lighthouse Lab in Alderley Park and Alex Alderton, Roberto Amato, Jeffrey Barrett, Sonia Goncalves, Ewan Harrison, David K. Jackson, Ian Johnston, Dominic Kwiatkowski, Cordelia Langford, John Sillitoe on behalf of the Wellcome Sanger Institute COVID-19 Surveillance Team |
| EPI_ISL_3358601 | Lighthouse Lab in Milton Keynes | Wellcome Sanger Institute for the COVID-19 Genomics UK (COG-UK) Consortium | The Lighthouse Lab in Milton Keynes and Alex Alderton, Roberto Amato, Jeffrey Barrett, Sonia Goncalves, Ewan Harrison, David K. Jackson, Ian Johnston, Dominic Kwiatkowski, Cordelia Langford, John Sillitoe on behalf of the Wellcome Sanger Institute COVID-19 Surveillance Team |
| EPI_ISL_3358602, EPI_ISL_3358603 | Lighthouse Lab in Alderley Park | Wellcome Sanger Institute for the COVID-19 Genomics UK (COG-UK) Consortium | Jacquelyn Wynn, Mairead Hyland, The Lighthouse Lab in Alderley Park and Alex Alderton, Roberto Amato, Jeffrey Barrett, Sonia Goncalves, Ewan Harrison, David K. Jackson, Ian Johnston, Dominic Kwiatkowski, Cordelia Langford, John Sillitoe on behalf of the Wellcome Sanger Institute COVID-19 Surveillance Team |
| EPI_ISL_3358604, EPI_ISL_3358605 | Lighthouse Lab in Milton Keynes | Wellcome Sanger Institute for the COVID-19 Genomics UK (COG-UK) Consortium | The Lighthouse Lab in Milton Keynes and Alex Alderton, Roberto Amato, Jeffrey Barrett, Sonia Goncalves, Ewan Harrison, David K. Jackson, Ian Johnston, Dominic Kwiatkowski, Cordelia Langford, John Sillitoe on behalf of the Wellcome Sanger Institute COVID-19 Surveillance Team |
| EPI_ISL_3358606 | Lighthouse Lab in Glasgow | Wellcome Sanger Institute for the COVID-19 Genomics UK (COG-UK) Consortium | Harper VanSteenhouse, Yumi Kasai, David Gray, Carol Clugston, Anna Dominiczak and Alex Alderton, Roberto Amato, Jeffrey Barrett, Sonia Goncalves, Ewan Harrison, David K. Jackson, Ian Johnston, Dominic Kwiatkowski, Cordelia Langford, John Sillitoe on behalf of the Wellcome Sanger Institute COVID-19 Surveillance Team |
| EPI_ISL_3358607, EPI_ISL_3358608 | Lighthouse Lab in Milton Keynes | Wellcome Sanger Institute for the COVID-19 Genomics UK (COG-UK) Consortium | The Lighthouse Lab in Milton Keynes and Alex Alderton, Roberto Amato, Jeffrey Barrett, Sonia Goncalves, Ewan Harrison, David K. Jackson, Ian Johnston, Dominic Kwiatkowski, Cordelia Langford, John Sillitoe on behalf of the Wellcome Sanger Institute COVID-19 Surveillance Team |
| EPI_ISL_3358609, EPI_ISL_3358610 | Lighthouse Lab in Glasgow | Wellcome Sanger Institute for the COVID-19 Genomics UK (COG-UK) Consortium | Harper VanSteenhouse, Yumi Kasai, David Gray, Carol Clugston, Anna Dominiczak and Alex Alderton, Roberto Amato, Jeffrey Barrett, Sonia Goncalves, Ewan Harrison, David K. Jackson, Ian Johnston, Dominic Kwiatkowski, Cordelia Langford, John Sillitoe on behalf of the Wellcome Sanger Institute COVID-19 Surveillance Team |
| EPI_ISL_3358611 | Lighthouse Lab in Milton Keynes | Wellcome Sanger Institute for the COVID-19 Genomics UK (COG-UK) Consortium | The Lighthouse Lab in Milton Keynes and Alex Alderton, Roberto Amato, Jeffrey Barrett, Sonia Goncalves, Ewan Harrison, David K. Jackson, Ian Johnston, Dominic Kwiatkowski, Cordelia Langford, John Sillitoe on behalf of the Wellcome Sanger Institute COVID-19 Surveillance Team |
| EPI_ISL_3358612, EPI_ISL_3358613 | Lighthouse Lab in Alderley Park | Wellcome Sanger Institute for the COVID-19 Genomics UK (COG-UK) Consortium | Jacquelyn Wynn, Mairead Hyland, The Lighthouse Lab in Alderley Park and Alex Alderton, Roberto Amato, Jeffrey Barrett, Sonia Goncalves, Ewan Harrison, David K. Jackson, Ian Johnston, Dominic Kwiatkowski, Cordelia Langford, John Sillitoe on behalf of the Wellcome Sanger Institute COVID-19 Surveillance Team |
| EPI_ISL_3358614, EPI_ISL_3358615, EPI_ISL_3358616, EPI_ISL_3358617 | Lighthouse Lab in Milton Keynes | Wellcome Sanger Institute for the COVID-19 Genomics UK (COG-UK) Consortium | The Lighthouse Lab in Milton Keynes and Alex Alderton, Roberto Amato, Jeffrey Barrett, Sonia Goncalves, Ewan Harrison, David K. Jackson, Ian Johnston, Dominic Kwiatkowski, Cordelia Langford, John Sillitoe on behalf of the Wellcome Sanger Institute COVID-19 Surveillance Team |
| EPI_ISL_3358618 | Lighthouse Lab in Alderley Park | Wellcome Sanger Institute for the COVID-19 Genomics UK (COG-UK) Consortium | Jacquelyn Wynn, Mairead Hyland, The Lighthouse Lab in Alderley Park and Alex Alderton, Roberto Amato, Jeffrey Barrett, Sonia Goncalves, Ewan Harrison, David K. Jackson, Ian Johnston, Dominic Kwiatkowski, Cordelia Langford, John Sillitoe on behalf of the Wellcome Sanger Institute COVID-19 Surveillance Team |
| EPI_ISL_3358619 | Lighthouse Lab in Milton Keynes | Wellcome Sanger Institute for the COVID-19 Genomics UK (COG-UK) | The Lighthouse Lab in Milton Keynes and Alex Alderton, Roberto Amato, Jeffrey Barrett, Sonia Goncalves, Ewan Harrison, David K. Jackson, Ian Johnston, Dominic Kwiatkowski, Cordelia Langford, John Sillitoe on behalf of the Wellcome Sanger |

| | | | |
|---|---|---|---|
| | | Consortium | Institute COVID-19 Surveillance Team |
| EPI_ISL_3358620, EPI_ISL_3358621 | Lighthouse Lab in Alderley Park | Wellcome Sanger Institute for the COVID-19 Genomics UK (COG-UK) Consortium | Jacquelyn Wynn, Mairead Hyland, The Lighthouse Lab in Alderley Park and Alex Alderton, Roberto Amato, Jeffrey Barrett, Sonia Goncalves, Ewan Harrison, David K. Jackson, Ian Johnston, Dominic Kwiatkowski, Cordelia Langford, John Sillitoe on behalf of the Wellcome Sanger Institute COVID-19 Surveillance Team |
| EPI_ISL_3358622 | Lighthouse Lab in Glasgow | Wellcome Sanger Institute for the COVID-19 Genomics UK (COG-UK) Consortium | Harper VanSteenhouse, Yumi Kasai, David Gray, Carol Clugston, Anna Dominiczak and Alex Alderton, Roberto Amato, Jeffrey Barrett, Sonia Goncalves, Ewan Harrison, David K. Jackson, Ian Johnston, Dominic Kwiatkowski, Cordelia Langford, John Sillitoe on behalf of the Wellcome Sanger Institute COVID-19 Surveillance Team |
| EPI_ISL_3358623 | Lighthouse Lab in Alderley Park | Wellcome Sanger Institute for the COVID-19 Genomics UK (COG-UK) Consortium | Jacquelyn Wynn, Mairead Hyland, The Lighthouse Lab in Alderley Park and Alex Alderton, Roberto Amato, Jeffrey Barrett, Sonia Goncalves, Ewan Harrison, David K. Jackson, Ian Johnston, Dominic Kwiatkowski, Cordelia Langford, John Sillitoe on behalf of the Wellcome Sanger Institute COVID-19 Surveillance Team |
| EPI_ISL_3358624, EPI_ISL_3358625, EPI_ISL_3358626, EPI_ISL_3358627 | Lighthouse Lab in Milton Keynes | Wellcome Sanger Institute for the COVID-19 Genomics UK (COG-UK) Consortium | The Lighthouse Lab in Milton Keynes and Alex Alderton, Roberto Amato, Jeffrey Barrett, Sonia Goncalves, Ewan Harrison, David K. Jackson, Ian Johnston, Dominic Kwiatkowski, Cordelia Langford, John Sillitoe on behalf of the Wellcome Sanger Institute COVID-19 Surveillance Team |
| EPI_ISL_3358628 | Lighthouse Lab in Glasgow | Wellcome Sanger Institute for the COVID-19 Genomics UK (COG-UK) Consortium | Harper VanSteenhouse, Yumi Kasai, David Gray, Carol Clugston, Anna Dominiczak and Alex Alderton, Roberto Amato, Jeffrey Barrett, Sonia Goncalves, Ewan Harrison, David K. Jackson, Ian Johnston, Dominic Kwiatkowski, Cordelia Langford, John Sillitoe on behalf of the Wellcome Sanger Institute COVID-19 Surveillance Team |
| EPI_ISL_3358629 | Lighthouse Lab in Milton Keynes | Wellcome Sanger Institute for the COVID-19 Genomics UK (COG-UK) Consortium | The Lighthouse Lab in Milton Keynes and Alex Alderton, Roberto Amato, Jeffrey Barrett, Sonia Goncalves, Ewan Harrison, David K. Jackson, Ian Johnston, Dominic Kwiatkowski, Cordelia Langford, John Sillitoe on behalf of the Wellcome Sanger Institute COVID-19 Surveillance Team |
| EPI_ISL_3358630 | Lighthouse Lab in Alderley Park | Wellcome Sanger Institute for the COVID-19 Genomics UK (COG-UK) Consortium | Jacquelyn Wynn, Mairead Hyland, The Lighthouse Lab in Alderley Park and Alex Alderton, Roberto Amato, Jeffrey Barrett, Sonia Goncalves, Ewan Harrison, David K. Jackson, Ian Johnston, Dominic Kwiatkowski, Cordelia Langford, John Sillitoe on behalf of the Wellcome Sanger Institute COVID-19 Surveillance Team |
| EPI_ISL_3358631 | Lighthouse Lab in Milton Keynes | Wellcome Sanger Institute for the COVID-19 Genomics UK (COG-UK) Consortium | The Lighthouse Lab in Milton Keynes and Alex Alderton, Roberto Amato, Jeffrey Barrett, Sonia Goncalves, Ewan Harrison, David K. Jackson, Ian Johnston, Dominic Kwiatkowski, Cordelia Langford, John Sillitoe on behalf of the Wellcome Sanger Institute COVID-19 Surveillance Team |
| EPI_ISL_3358632 | Lighthouse Lab in Alderley Park | Wellcome Sanger Institute for the COVID-19 Genomics UK (COG-UK) Consortium | Jacquelyn Wynn, Mairead Hyland, The Lighthouse Lab in Alderley Park and Alex Alderton, Roberto Amato, Jeffrey Barrett, Sonia Goncalves, Ewan Harrison, David K. Jackson, Ian Johnston, Dominic Kwiatkowski, Cordelia Langford, John Sillitoe on behalf of the Wellcome Sanger Institute COVID-19 Surveillance Team |
| EPI_ISL_3358633 | Lighthouse Lab in Milton Keynes | Wellcome Sanger Institute for the COVID-19 Genomics UK (COG-UK) Consortium | The Lighthouse Lab in Milton Keynes and Alex Alderton, Roberto Amato, Jeffrey Barrett, Sonia Goncalves, Ewan Harrison, David K. Jackson, Ian Johnston, Dominic Kwiatkowski, Cordelia Langford, John Sillitoe on behalf of the Wellcome Sanger Institute COVID-19 Surveillance Team |
| EPI_ISL_3358634 | Lighthouse Lab in Alderley Park | Wellcome Sanger Institute for the COVID-19 Genomics UK (COG-UK) Consortium | Jacquelyn Wynn, Mairead Hyland, The Lighthouse Lab in Alderley Park and Alex Alderton, Roberto Amato, Jeffrey Barrett, Sonia Goncalves, Ewan Harrison, David K. Jackson, Ian Johnston, Dominic Kwiatkowski, Cordelia Langford, John Sillitoe on behalf of the Wellcome Sanger Institute COVID-19 Surveillance Team |
| EPI_ISL_3358635 | Lighthouse Lab in Glasgow | Wellcome Sanger Institute for the COVID-19 Genomics UK (COG-UK) Consortium | Harper VanSteenhouse, Yumi Kasai, David Gray, Carol Clugston, Anna Dominiczak and Alex Alderton, Roberto Amato, Jeffrey Barrett, Sonia Goncalves, Ewan Harrison, David K. Jackson, Ian Johnston, Dominic Kwiatkowski, Cordelia Langford, John Sillitoe on behalf of the Wellcome Sanger Institute COVID-19 Surveillance Team |
| EPI_ISL_3358636, EPI_ISL_3358637 | Lighthouse Lab in Alderley Park | Wellcome Sanger Institute for the COVID-19 Genomics UK (COG-UK) Consortium | Jacquelyn Wynn, Mairead Hyland, The Lighthouse Lab in Alderley Park and Alex Alderton, Roberto Amato, Jeffrey Barrett, Sonia Goncalves, Ewan Harrison, David K. Jackson, Ian Johnston, Dominic Kwiatkowski, Cordelia Langford, John Sillitoe on behalf of the Wellcome Sanger Institute COVID-19 Surveillance Team |
| EPI_ISL_3358638 | Lighthouse Lab in Glasgow | Wellcome Sanger Institute for the COVID-19 Genomics UK (COG-UK) Consortium | Harper VanSteenhouse, Yumi Kasai, David Gray, Carol Clugston, Anna Dominiczak and Alex Alderton, Roberto Amato, Jeffrey Barrett, Sonia Goncalves, Ewan Harrison, David K. Jackson, Ian Johnston, Dominic Kwiatkowski, Cordelia Langford, John Sillitoe on behalf of the Wellcome Sanger Institute COVID-19 Surveillance Team |
| EPI_ISL_3358639 | Lighthouse Lab in Alderley Park | Wellcome Sanger Institute for the COVID-19 Genomics UK (COG-UK) Consortium | Jacquelyn Wynn, Mairead Hyland, The Lighthouse Lab in Alderley Park and Alex Alderton, Roberto Amato, Jeffrey Barrett, Sonia Goncalves, Ewan Harrison, David K. Jackson, Ian Johnston, Dominic Kwiatkowski, Cordelia Langford, John Sillitoe on behalf of the Wellcome Sanger Institute COVID-19 Surveillance Team |
| EPI_ISL_3358640, EPI_ISL_3358641 | Lighthouse Lab in Milton Keynes | Wellcome Sanger Institute for the COVID-19 Genomics UK (COG-UK) Consortium | The Lighthouse Lab in Milton Keynes and Alex Alderton, Roberto Amato, Jeffrey Barrett, Sonia Goncalves, Ewan Harrison, David K. Jackson, Ian Johnston, Dominic Kwiatkowski, Cordelia Langford, John Sillitoe on behalf of the Wellcome Sanger Institute COVID-19 Surveillance Team |
| EPI_ISL_3358642 | Lighthouse Lab in Glasgow | Wellcome Sanger Institute for the COVID-19 Genomics UK (COG-UK) Consortium | Harper VanSteenhouse, Yumi Kasai, David Gray, Carol Clugston, Anna Dominiczak and Alex Alderton, Roberto Amato, Jeffrey Barrett, Sonia Goncalves, Ewan Harrison, David K. Jackson, Ian Johnston, Dominic Kwiatkowski, Cordelia Langford, John Sillitoe on behalf of the Wellcome Sanger Institute COVID-19 Surveillance Team |
| EPI_ISL_3358643 | Lighthouse Lab in Milton Keynes | Wellcome Sanger Institute for the COVID-19 Genomics UK (COG-UK) Consortium | The Lighthouse Lab in Milton Keynes and Alex Alderton, Roberto Amato, Jeffrey Barrett, Sonia Goncalves, Ewan Harrison, David K. Jackson, Ian Johnston, Dominic Kwiatkowski, Cordelia Langford, John Sillitoe on behalf of the Wellcome Sanger Institute COVID-19 Surveillance Team |
| EPI_ISL_3358644, EPI_ISL_3358645 | Lighthouse Lab in Alderley Park | Wellcome Sanger Institute for the COVID-19 Genomics UK (COG-UK) Consortium | Jacquelyn Wynn, Mairead Hyland, The Lighthouse Lab in Alderley Park and Alex Alderton, Roberto Amato, Jeffrey Barrett, Sonia Goncalves, Ewan Harrison, David K. Jackson, Ian Johnston, Dominic Kwiatkowski, Cordelia Langford, John Sillitoe on behalf of the Wellcome Sanger Institute COVID-19 Surveillance Team |
| EPI_ISL_3358646 | Lighthouse Lab in Milton Keynes | Wellcome Sanger Institute for the COVID-19 Genomics UK (COG-UK) Consortium | The Lighthouse Lab in Milton Keynes and Alex Alderton, Roberto Amato, Jeffrey Barrett, Sonia Goncalves, Ewan Harrison, David K. Jackson, Ian Johnston, Dominic Kwiatkowski, Cordelia Langford, John Sillitoe on behalf of the Wellcome Sanger Institute COVID-19 Surveillance Team |
| EPI_ISL_3358647 | Lighthouse Lab in Glasgow | Wellcome Sanger Institute for the COVID-19 Genomics UK (COG-UK) Consortium | Harper VanSteenhouse, Yumi Kasai, David Gray, Carol Clugston, Anna Dominiczak and Alex Alderton, Roberto Amato, Jeffrey Barrett, Sonia Goncalves, Ewan Harrison, David K. Jackson, Ian Johnston, Dominic Kwiatkowski, Cordelia Langford, John Sillitoe on behalf of the Wellcome Sanger Institute COVID-19 Surveillance Team |
| EPI_ISL_3358648, EPI_ISL_3358649, EPI_ISL_3358650, EPI_ISL_3358651 | Lighthouse Lab in Milton Keynes | Wellcome Sanger Institute for the COVID-19 Genomics UK (COG-UK) Consortium | The Lighthouse Lab in Milton Keynes and Alex Alderton, Roberto Amato, Jeffrey Barrett, Sonia Goncalves, Ewan Harrison, David K. Jackson, Ian Johnston, Dominic Kwiatkowski, Cordelia Langford, John Sillitoe on behalf of the Wellcome Sanger Institute COVID-19 Surveillance Team |
| EPI_ISL_3358652 | Lighthouse Lab in Alderley Park | Wellcome Sanger Institute for the COVID-19 Genomics UK (COG-UK) Consortium | Jacquelyn Wynn, Mairead Hyland, The Lighthouse Lab in Alderley Park and Alex Alderton, Roberto Amato, Jeffrey Barrett, Sonia Goncalves, Ewan Harrison, David K. Jackson, Ian Johnston, Dominic Kwiatkowski, Cordelia Langford, John Sillitoe on behalf of the Wellcome Sanger Institute COVID-19 Surveillance Team |
| EPI_ISL_3358653 | Lighthouse Lab in Glasgow | Wellcome Sanger Institute for the COVID-19 Genomics UK (COG-UK) Consortium | Harper VanSteenhouse, Yumi Kasai, David Gray, Carol Clugston, Anna Dominiczak and Alex Alderton, Roberto Amato, Jeffrey Barrett, Sonia Goncalves, Ewan Harrison, David K. Jackson, Ian Johnston, Dominic Kwiatkowski, Cordelia Langford, John Sillitoe on behalf of the Wellcome Sanger Institute COVID-19 Surveillance Team |
| EPI_ISL_3358654 | Lighthouse Lab in Alderley Park | Wellcome Sanger Institute for the COVID-19 Genomics UK (COG-UK) Consortium | Jacquelyn Wynn, Mairead Hyland, The Lighthouse Lab in Alderley Park and Alex Alderton, Roberto Amato, Jeffrey Barrett, Sonia Goncalves, Ewan Harrison, David K. Jackson, Ian Johnston, Dominic Kwiatkowski, Cordelia Langford, John Sillitoe on behalf of the Wellcome Sanger Institute COVID-19 Surveillance Team |
| EPI_ISL_3358655 | Lighthouse Lab in Milton Keynes | Wellcome Sanger Institute for the COVID-19 Genomics UK (COG-UK) Consortium | The Lighthouse Lab in Milton Keynes and Alex Alderton, Roberto Amato, Jeffrey Barrett, Sonia Goncalves, Ewan Harrison, David K. Jackson, Ian Johnston, Dominic Kwiatkowski, Cordelia Langford, John Sillitoe on behalf of the Wellcome Sanger Institute COVID-19 Surveillance Team |
| EPI_ISL_3358656 | Lighthouse Lab in Alderley Park | Wellcome Sanger Institute for the COVID-19 Genomics UK (COG-UK) Consortium | Jacquelyn Wynn, Mairead Hyland, The Lighthouse Lab in Alderley Park and Alex Alderton, Roberto Amato, Jeffrey Barrett, Sonia Goncalves, Ewan Harrison, David K. Jackson, Ian Johnston, Dominic Kwiatkowski, Cordelia Langford, John Sillitoe on behalf of the Wellcome Sanger Institute COVID-19 Surveillance Team |
| EPI_ISL_3358657, EPI_ISL_3358658 | Lighthouse Lab in Milton Keynes | Wellcome Sanger Institute for the COVID-19 Genomics UK (COG-UK) Consortium | The Lighthouse Lab in Milton Keynes and Alex Alderton, Roberto Amato, Jeffrey Barrett, Sonia Goncalves, Ewan Harrison, David K. Jackson, Ian Johnston, Dominic Kwiatkowski, Cordelia Langford, John Sillitoe on behalf of the Wellcome Sanger Institute COVID-19 Surveillance Team |
| EPI_ISL_3358659 | Lighthouse Lab in Alderley Park | Wellcome Sanger Institute for the COVID-19 Genomics UK (COG-UK) Consortium | Jacquelyn Wynn, Mairead Hyland, The Lighthouse Lab in Alderley Park and Alex Alderton, Roberto Amato, Jeffrey Barrett, Sonia Goncalves, Ewan Harrison, David K. Jackson, Ian Johnston, Dominic Kwiatkowski, Cordelia Langford, John Sillitoe on behalf of the Wellcome Sanger Institute COVID-19 Surveillance Team |
| EPI_ISL_3358660 | Lighthouse Lab in Glasgow | Wellcome Sanger Institute for the COVID-19 Genomics UK (COG-UK) Consortium | Harper VanSteenhouse, Yumi Kasai, David Gray, Carol Clugston, Anna Dominiczak and Alex Alderton, Roberto Amato, Jeffrey Barrett, Sonia Goncalves, Ewan Harrison, David K. Jackson, Ian Johnston, Dominic Kwiatkowski, Cordelia Langford, John Sillitoe on behalf of the Wellcome Sanger Institute COVID-19 Surveillance Team |
| EPI_ISL_3358661 | Lighthouse Lab in Milton Keynes | Wellcome Sanger Institute for the COVID-19 Genomics UK (COG-UK) Consortium | The Lighthouse Lab in Milton Keynes and Alex Alderton, Roberto Amato, Jeffrey Barrett, Sonia Goncalves, Ewan Harrison, David K. Jackson, Ian Johnston, Dominic Kwiatkowski, Cordelia Langford, John Sillitoe on behalf of the Wellcome Sanger Institute COVID-19 Surveillance Team |
| EPI_ISL_3358662 | Lighthouse Lab in Alderley Park | Wellcome Sanger Institute for the COVID-19 Genomics UK (COG-UK) Consortium | Jacquelyn Wynn, Mairead Hyland, The Lighthouse Lab in Alderley Park and Alex Alderton, Roberto Amato, Jeffrey Barrett, Sonia Goncalves, Ewan Harrison, David K. Jackson, Ian Johnston, Dominic Kwiatkowski, Cordelia Langford, John Sillitoe on behalf of the Wellcome Sanger Institute COVID-19 Surveillance Team |
| EPI_ISL_3358663 | Lighthouse Lab in Milton Keynes | Wellcome Sanger Institute for the COVID-19 Genomics UK (COG-UK) Consortium | The Lighthouse Lab in Milton Keynes and Alex Alderton, Roberto Amato, Jeffrey Barrett, Sonia Goncalves, Ewan Harrison, David K. Jackson, Ian Johnston, Dominic Kwiatkowski, Cordelia Langford, John Sillitoe on behalf of the Wellcome Sanger Institute COVID-19 Surveillance Team |
| EPI_ISL_3358664 | Lighthouse Lab in Glasgow | Wellcome Sanger Institute for the COVID-19 Genomics UK (COG-UK) Consortium | Harper VanSteenhouse, Yumi Kasai, David Gray, Carol Clugston, Anna Dominiczak and Alex Alderton, Roberto Amato, Jeffrey Barrett, Sonia Goncalves, Ewan Harrison, David K. Jackson, Ian Johnston, Dominic Kwiatkowski, Cordelia Langford, John Sillitoe on behalf of the Wellcome Sanger Institute COVID-19 Surveillance Team |
| EPI_ISL_3358665 | Lighthouse Lab in Milton Keynes | Wellcome Sanger Institute for the COVID-19 Genomics UK (COG-UK) Consortium | The Lighthouse Lab in Milton Keynes and Alex Alderton, Roberto Amato, Jeffrey Barrett, Sonia Goncalves, Ewan Harrison, David K. Jackson, Ian Johnston, Dominic Kwiatkowski, Cordelia Langford, John Sillitoe on behalf of the Wellcome Sanger Institute COVID-19 Surveillance Team |
| EPI_ISL_3358666 | Lighthouse Lab in Glasgow | Wellcome Sanger Institute for the COVID-19 Genomics UK (COG-UK) Consortium | Harper VanSteenhouse, Yumi Kasai, David Gray, Carol Clugston, Anna Dominiczak and Alex Alderton, Roberto Amato, Jeffrey Barrett, Sonia Goncalves, Ewan Harrison, David K. Jackson, Ian Johnston, Dominic Kwiatkowski, Cordelia Langford, John Sillitoe on behalf of the Wellcome Sanger Institute COVID-19 Surveillance Team |
| EPI_ISL_3358667, EPI_ISL_3358668 | Lighthouse Lab in Milton Keynes | Wellcome Sanger Institute for the COVID-19 Genomics UK (COG-UK) Consortium | The Lighthouse Lab in Milton Keynes and Alex Alderton, Roberto Amato, Jeffrey Barrett, Sonia Goncalves, Ewan Harrison, David K. Jackson, Ian Johnston, Dominic Kwiatkowski, Cordelia Langford, John Sillitoe on behalf of the Wellcome Sanger Institute COVID-19 Surveillance Team |
| EPI_ISL_3358669, EPI_ISL_3358670, EPI_ISL_3358671 | Lighthouse Lab in Alderley Park | Wellcome Sanger Institute for the COVID-19 Genomics UK (COG-UK) Consortium | Jacquelyn Wynn, Mairead Hyland, The Lighthouse Lab in Alderley Park and Alex Alderton, Roberto Amato, Jeffrey Barrett, Sonia Goncalves, Ewan Harrison, David K. Jackson, Ian Johnston, Dominic Kwiatkowski, Cordelia Langford, John Sillitoe on behalf of the Wellcome Sanger Institute COVID-19 Surveillance Team |
| EPI_ISL_3358672, EPI_ISL_3358673, EPI_ISL_3358674, EPI_ISL_3358675, EPI_ISL_3358676 | Lighthouse Lab in Milton Keynes | Wellcome Sanger Institute for the COVID-19 Genomics UK (COG-UK) Consortium | The Lighthouse Lab in Milton Keynes and Alex Alderton, Roberto Amato, Jeffrey Barrett, Sonia Goncalves, Ewan Harrison, David K. Jackson, Ian Johnston, Dominic Kwiatkowski, Cordelia Langford, John Sillitoe on behalf of the Wellcome Sanger Institute COVID-19 Surveillance Team |
| EPI_ISL_3358677 | Lighthouse Lab in Glasgow | Wellcome Sanger Institute for the COVID-19 Genomics UK (COG-UK) Consortium | Harper VanSteenhouse, Yumi Kasai, David Gray, Carol Clugston, Anna Dominiczak and Alex Alderton, Roberto Amato, Jeffrey Barrett, Sonia Goncalves, Ewan Harrison, David K. Jackson, Ian Johnston, Dominic Kwiatkowski, Cordelia Langford, John Sillitoe on behalf of the Wellcome Sanger Institute COVID-19 Surveillance Team |
| EPI_ISL_3358678 | Lighthouse Lab in Alderley Park | Wellcome Sanger Institute for the COVID-19 Genomics UK (COG-UK) Consortium | Jacquelyn Wynn, Mairead Hyland, The Lighthouse Lab in Alderley Park and Alex Alderton, Roberto Amato, Jeffrey Barrett, Sonia Goncalves, Ewan Harrison, David K. Jackson, Ian Johnston, Dominic Kwiatkowski, Cordelia Langford, John Sillitoe on behalf of the Wellcome Sanger Institute COVID-19 Surveillance Team |
| EPI_ISL_3358679, | Lighthouse Lab in Milton Keynes | Wellcome Sanger Institute for the COVID-19 Genomics UK (COG-UK) | The Lighthouse Lab in Milton Keynes and Alex Alderton, Roberto Amato, Jeffrey Barrett, Sonia Goncalves, Ewan Harrison, David K. Jackson, Ian Johnston, Dominic Kwiatkowski, Cordelia Langford, John Sillitoe on behalf of the Wellcome Sanger |

| | | | |
|---|---|---|---|
| EPI_ISL_3358680 | | Consortium | Institute COVID-19 Surveillance Team |
| EPI_ISL_3358681 | Lighthouse Lab in Glasgow | Wellcome Sanger Institute for the COVID-19 Genomics UK (COG-UK) Consortium | Harper VanSteenhouse, Yumi Kasai, David Gray, Carol Clugston, Anna Dominiczak and Alex Alderton, Roberto Amato, Jeffrey Barrett, Sonia Goncalves, Ewan Harrison, David K. Jackson, Ian Johnston, Dominic Kwiatkowski, Cordelia Langford, John Sillitoe on behalf of the Wellcome Sanger Institute COVID-19 Surveillance Team |
| EPI_ISL_3358682, EPI_ISL_3358683, EPI_ISL_3358684 | Lighthouse Lab in Milton Keynes | Wellcome Sanger Institute for the COVID-19 Genomics UK (COG-UK) Consortium | The Lighthouse Lab in Milton Keynes and Alex Alderton, Roberto Amato, Jeffrey Barrett, Sonia Goncalves, Ewan Harrison, David K. Jackson, Ian Johnston, Dominic Kwiatkowski, Cordelia Langford, John Sillitoe on behalf of the Wellcome Sanger Institute COVID-19 Surveillance Team |
| EPI_ISL_3358685 | Lighthouse Lab in Glasgow | Wellcome Sanger Institute for the COVID-19 Genomics UK (COG-UK) Consortium | Harper VanSteenhouse, Yumi Kasai, David Gray, Carol Clugston, Anna Dominiczak and Alex Alderton, Roberto Amato, Jeffrey Barrett, Sonia Goncalves, Ewan Harrison, David K. Jackson, Ian Johnston, Dominic Kwiatkowski, Cordelia Langford, John Sillitoe on behalf of the Wellcome Sanger Institute COVID-19 Surveillance Team |
| EPI_ISL_3358686, EPI_ISL_3358687 | Lighthouse Lab in Alderley Park | Wellcome Sanger Institute for the COVID-19 Genomics UK (COG-UK) Consortium | Jacquelyn Wynn, Mairead Hyland, The Lighthouse Lab in Alderley Park and Alex Alderton, Roberto Amato, Jeffrey Barrett, Sonia Goncalves, Ewan Harrison, David K. Jackson, Ian Johnston, Dominic Kwiatkowski, Cordelia Langford, John Sillitoe on behalf of the Wellcome Sanger Institute COVID-19 Surveillance Team |
| EPI_ISL_3358688, EPI_ISL_3358689 | Lighthouse Lab in Milton Keynes | Wellcome Sanger Institute for the COVID-19 Genomics UK (COG-UK) Consortium | The Lighthouse Lab in Milton Keynes and Alex Alderton, Roberto Amato, Jeffrey Barrett, Sonia Goncalves, Ewan Harrison, David K. Jackson, Ian Johnston, Dominic Kwiatkowski, Cordelia Langford, John Sillitoe on behalf of the Wellcome Sanger Institute COVID-19 Surveillance Team |
| EPI_ISL_3358690 | Lighthouse Lab in Alderley Park | Wellcome Sanger Institute for the COVID-19 Genomics UK (COG-UK) Consortium | Jacquelyn Wynn, Mairead Hyland, The Lighthouse Lab in Alderley Park and Alex Alderton, Roberto Amato, Jeffrey Barrett, Sonia Goncalves, Ewan Harrison, David K. Jackson, Ian Johnston, Dominic Kwiatkowski, Cordelia Langford, John Sillitoe on behalf of the Wellcome Sanger Institute COVID-19 Surveillance Team |
| EPI_ISL_3358691, EPI_ISL_3358692 | Lighthouse Lab in Milton Keynes | Wellcome Sanger Institute for the COVID-19 Genomics UK (COG-UK) Consortium | The Lighthouse Lab in Milton Keynes and Alex Alderton, Roberto Amato, Jeffrey Barrett, Sonia Goncalves, Ewan Harrison, David K. Jackson, Ian Johnston, Dominic Kwiatkowski, Cordelia Langford, John Sillitoe on behalf of the Wellcome Sanger Institute COVID-19 Surveillance Team |
| EPI_ISL_3358693, EPI_ISL_3358694 | Lighthouse Lab in Alderley Park | Wellcome Sanger Institute for the COVID-19 Genomics UK (COG-UK) Consortium | Jacquelyn Wynn, Mairead Hyland, The Lighthouse Lab in Alderley Park and Alex Alderton, Roberto Amato, Jeffrey Barrett, Sonia Goncalves, Ewan Harrison, David K. Jackson, Ian Johnston, Dominic Kwiatkowski, Cordelia Langford, John Sillitoe on behalf of the Wellcome Sanger Institute COVID-19 Surveillance Team |
| EPI_ISL_3358695 | Lighthouse Lab in Milton Keynes | Wellcome Sanger Institute for the COVID-19 Genomics UK (COG-UK) Consortium | The Lighthouse Lab in Milton Keynes and Alex Alderton, Roberto Amato, Jeffrey Barrett, Sonia Goncalves, Ewan Harrison, David K. Jackson, Ian Johnston, Dominic Kwiatkowski, Cordelia Langford, John Sillitoe on behalf of the Wellcome Sanger Institute COVID-19 Surveillance Team |
| EPI_ISL_3358696 | Lighthouse Lab in Glasgow | Wellcome Sanger Institute for the COVID-19 Genomics UK (COG-UK) Consortium | Harper VanSteenhouse, Yumi Kasai, David Gray, Carol Clugston, Anna Dominiczak and Alex Alderton, Roberto Amato, Jeffrey Barrett, Sonia Goncalves, Ewan Harrison, David K. Jackson, Ian Johnston, Dominic Kwiatkowski, Cordelia Langford, John Sillitoe on behalf of the Wellcome Sanger Institute COVID-19 Surveillance Team |
| EPI_ISL_3358697, EPI_ISL_3358698, EPI_ISL_3358699 | Lighthouse Lab in Alderley Park | Wellcome Sanger Institute for the COVID-19 Genomics UK (COG-UK) Consortium | Jacquelyn Wynn, Mairead Hyland, The Lighthouse Lab in Alderley Park and Alex Alderton, Roberto Amato, Jeffrey Barrett, Sonia Goncalves, Ewan Harrison, David K. Jackson, Ian Johnston, Dominic Kwiatkowski, Cordelia Langford, John Sillitoe on behalf of the Wellcome Sanger Institute COVID-19 Surveillance Team |
| EPI_ISL_3358700 | Lighthouse Lab in Milton Keynes | Wellcome Sanger Institute for the COVID-19 Genomics UK (COG-UK) Consortium | The Lighthouse Lab in Milton Keynes and Alex Alderton, Roberto Amato, Jeffrey Barrett, Sonia Goncalves, Ewan Harrison, David K. Jackson, Ian Johnston, Dominic Kwiatkowski, Cordelia Langford, John Sillitoe on behalf of the Wellcome Sanger Institute COVID-19 Surveillance Team |
| EPI_ISL_3358701 | Lighthouse Lab in Alderley Park | Wellcome Sanger Institute for the COVID-19 Genomics UK (COG-UK) Consortium | Jacquelyn Wynn, Mairead Hyland, The Lighthouse Lab in Alderley Park and Alex Alderton, Roberto Amato, Jeffrey Barrett, Sonia Goncalves, Ewan Harrison, David K. Jackson, Ian Johnston, Dominic Kwiatkowski, Cordelia Langford, John Sillitoe on behalf of the Wellcome Sanger Institute COVID-19 Surveillance Team |
| EPI_ISL_3358702, EPI_ISL_3358703, EPI_ISL_3358704 | Lighthouse Lab in Milton Keynes | Wellcome Sanger Institute for the COVID-19 Genomics UK (COG-UK) Consortium | The Lighthouse Lab in Milton Keynes and Alex Alderton, Roberto Amato, Jeffrey Barrett, Sonia Goncalves, Ewan Harrison, David K. Jackson, Ian Johnston, Dominic Kwiatkowski, Cordelia Langford, John Sillitoe on behalf of the Wellcome Sanger Institute COVID-19 Surveillance Team |
| EPI_ISL_3358705 | Lighthouse Lab in Glasgow | Wellcome Sanger Institute for the COVID-19 Genomics UK (COG-UK) Consortium | Harper VanSteenhouse, Yumi Kasai, David Gray, Carol Clugston, Anna Dominiczak and Alex Alderton, Roberto Amato, Jeffrey Barrett, Sonia Goncalves, Ewan Harrison, David K. Jackson, Ian Johnston, Dominic Kwiatkowski, Cordelia Langford, John Sillitoe on behalf of the Wellcome Sanger Institute COVID-19 Surveillance Team |
| EPI_ISL_3358706, EPI_ISL_3358707, EPI_ISL_3358708, EPI_ISL_3358709, EPI_ISL_3358710, EPI_ISL_3358711 | Lighthouse Lab in Milton Keynes | Wellcome Sanger Institute for the COVID-19 Genomics UK (COG-UK) Consortium | The Lighthouse Lab in Milton Keynes and Alex Alderton, Roberto Amato, Jeffrey Barrett, Sonia Goncalves, Ewan Harrison, David K. Jackson, Ian Johnston, Dominic Kwiatkowski, Cordelia Langford, John Sillitoe on behalf of the Wellcome Sanger Institute COVID-19 Surveillance Team |
| EPI_ISL_3358712, EPI_ISL_3358713, EPI_ISL_3358714 | Lighthouse Lab in Alderley Park | Wellcome Sanger Institute for the COVID-19 Genomics UK (COG-UK) Consortium | Jacquelyn Wynn, Mairead Hyland, The Lighthouse Lab in Alderley Park and Alex Alderton, Roberto Amato, Jeffrey Barrett, Sonia Goncalves, Ewan Harrison, David K. Jackson, Ian Johnston, Dominic Kwiatkowski, Cordelia Langford, John Sillitoe on behalf of the Wellcome Sanger Institute COVID-19 Surveillance Team |
| EPI_ISL_3358715 | Lighthouse Lab in Glasgow | Wellcome Sanger Institute for the COVID-19 Genomics UK (COG-UK) Consortium | Harper VanSteenhouse, Yumi Kasai, David Gray, Carol Clugston, Anna Dominiczak and Alex Alderton, Roberto Amato, Jeffrey Barrett, Sonia Goncalves, Ewan Harrison, David K. Jackson, Ian Johnston, Dominic Kwiatkowski, Cordelia Langford, John Sillitoe on behalf of the Wellcome Sanger Institute COVID-19 Surveillance Team |
| EPI_ISL_3358716, EPI_ISL_3358717, EPI_ISL_3358718 | Lighthouse Lab in Milton Keynes | Wellcome Sanger Institute for the COVID-19 Genomics UK (COG-UK) Consortium | The Lighthouse Lab in Milton Keynes and Alex Alderton, Roberto Amato, Jeffrey Barrett, Sonia Goncalves, Ewan Harrison, David K. Jackson, Ian Johnston, Dominic Kwiatkowski, Cordelia Langford, John Sillitoe on behalf of the Wellcome Sanger Institute COVID-19 Surveillance Team |
| EPI_ISL_3358719 | Lighthouse Lab in Alderley Park | Wellcome Sanger Institute for the COVID-19 Genomics UK (COG-UK) Consortium | Jacquelyn Wynn, Mairead Hyland, The Lighthouse Lab in Alderley Park and Alex Alderton, Roberto Amato, Jeffrey Barrett, Sonia Goncalves, Ewan Harrison, David K. Jackson, Ian Johnston, Dominic Kwiatkowski, Cordelia Langford, John Sillitoe on behalf of the Wellcome Sanger Institute COVID-19 Surveillance Team |
| EPI_ISL_3358720 | Lighthouse Lab in Milton Keynes | Wellcome Sanger Institute for the COVID-19 Genomics UK (COG-UK) Consortium | The Lighthouse Lab in Milton Keynes and Alex Alderton, Roberto Amato, Jeffrey Barrett, Sonia Goncalves, Ewan Harrison, David K. Jackson, Ian Johnston, Dominic Kwiatkowski, Cordelia Langford, John Sillitoe on behalf of the Wellcome Sanger Institute COVID-19 Surveillance Team |
| EPI_ISL_3358721 | Lighthouse Lab in Alderley Park | Wellcome Sanger Institute for the COVID-19 Genomics UK (COG-UK) Consortium | Jacquelyn Wynn, Mairead Hyland, The Lighthouse Lab in Alderley Park and Alex Alderton, Roberto Amato, Jeffrey Barrett, Sonia Goncalves, Ewan Harrison, David K. Jackson, Ian Johnston, Dominic Kwiatkowski, Cordelia Langford, John Sillitoe on behalf of the Wellcome Sanger Institute COVID-19 Surveillance Team |
| EPI_ISL_3358722 | Lighthouse Lab in Milton Keynes | Wellcome Sanger Institute for the COVID-19 Genomics UK (COG-UK) Consortium | The Lighthouse Lab in Milton Keynes and Alex Alderton, Roberto Amato, Jeffrey Barrett, Sonia Goncalves, Ewan Harrison, David K. Jackson, Ian Johnston, Dominic Kwiatkowski, Cordelia Langford, John Sillitoe on behalf of the Wellcome Sanger Institute COVID-19 Surveillance Team |
| EPI_ISL_3358723, EPI_ISL_3358724 | Lighthouse Lab in Glasgow | Wellcome Sanger Institute for the COVID-19 Genomics UK (COG-UK) Consortium | Harper VanSteenhouse, Yumi Kasai, David Gray, Carol Clugston, Anna Dominiczak and Alex Alderton, Roberto Amato, Jeffrey Barrett, Sonia Goncalves, Ewan Harrison, David K. Jackson, Ian Johnston, Dominic Kwiatkowski, Cordelia Langford, John Sillitoe on behalf of the Wellcome Sanger Institute COVID-19 Surveillance Team |
| EPI_ISL_3358725, EPI_ISL_3358726 | Lighthouse Lab in Milton Keynes | Wellcome Sanger Institute for the COVID-19 Genomics UK (COG-UK) Consortium | The Lighthouse Lab in Milton Keynes and Alex Alderton, Roberto Amato, Jeffrey Barrett, Sonia Goncalves, Ewan Harrison, David K. Jackson, Ian Johnston, Dominic Kwiatkowski, Cordelia Langford, John Sillitoe on behalf of the Wellcome Sanger Institute COVID-19 Surveillance Team |
| EPI_ISL_3358727 | Lighthouse Lab in Glasgow | Wellcome Sanger Institute for the COVID-19 Genomics UK (COG-UK) Consortium | Harper VanSteenhouse, Yumi Kasai, David Gray, Carol Clugston, Anna Dominiczak and Alex Alderton, Roberto Amato, Jeffrey Barrett, Sonia Goncalves, Ewan Harrison, David K. Jackson, Ian Johnston, Dominic Kwiatkowski, Cordelia Langford, John Sillitoe on behalf of the Wellcome Sanger Institute COVID-19 Surveillance Team |
| EPI_ISL_3358728, EPI_ISL_3358729 | Lighthouse Lab in Milton Keynes | Wellcome Sanger Institute for the COVID-19 Genomics UK (COG-UK) Consortium | The Lighthouse Lab in Milton Keynes and Alex Alderton, Roberto Amato, Jeffrey Barrett, Sonia Goncalves, Ewan Harrison, David K. Jackson, Ian Johnston, Dominic Kwiatkowski, Cordelia Langford, John Sillitoe on behalf of the Wellcome Sanger Institute COVID-19 Surveillance Team |
| EPI_ISL_3358730 | Lighthouse Lab in Alderley Park | Wellcome Sanger Institute for the COVID-19 Genomics UK (COG-UK) Consortium | Jacquelyn Wynn, Mairead Hyland, The Lighthouse Lab in Alderley Park and Alex Alderton, Roberto Amato, Jeffrey Barrett, Sonia Goncalves, Ewan Harrison, David K. Jackson, Ian Johnston, Dominic Kwiatkowski, Cordelia Langford, John Sillitoe on behalf of the Wellcome Sanger Institute COVID-19 Surveillance Team |
| EPI_ISL_3358731, EPI_ISL_3358732 | Lighthouse Lab in Milton Keynes | Wellcome Sanger Institute for the COVID-19 Genomics UK (COG-UK) Consortium | The Lighthouse Lab in Milton Keynes and Alex Alderton, Roberto Amato, Jeffrey Barrett, Sonia Goncalves, Ewan Harrison, David K. Jackson, Ian Johnston, Dominic Kwiatkowski, Cordelia Langford, John Sillitoe on behalf of the Wellcome Sanger Institute COVID-19 Surveillance Team |
| EPI_ISL_3358733 | Lighthouse Lab in Alderley Park | Wellcome Sanger Institute for the COVID-19 Genomics UK (COG-UK) Consortium | Jacquelyn Wynn, Mairead Hyland, The Lighthouse Lab in Alderley Park and Alex Alderton, Roberto Amato, Jeffrey Barrett, Sonia Goncalves, Ewan Harrison, David K. Jackson, Ian Johnston, Dominic Kwiatkowski, Cordelia Langford, John Sillitoe on behalf of the Wellcome Sanger Institute COVID-19 Surveillance Team |
| EPI_ISL_3358734, EPI_ISL_3358735 | Lighthouse Lab in Milton Keynes | Wellcome Sanger Institute for the COVID-19 Genomics UK (COG-UK) Consortium | The Lighthouse Lab in Milton Keynes and Alex Alderton, Roberto Amato, Jeffrey Barrett, Sonia Goncalves, Ewan Harrison, David K. Jackson, Ian Johnston, Dominic Kwiatkowski, Cordelia Langford, John Sillitoe on behalf of the Wellcome Sanger Institute COVID-19 Surveillance Team |
| EPI_ISL_3358736 | Lighthouse Lab in Glasgow | Wellcome Sanger Institute for the COVID-19 Genomics UK (COG-UK) Consortium | Harper VanSteenhouse, Yumi Kasai, David Gray, Carol Clugston, Anna Dominiczak and Alex Alderton, Roberto Amato, Jeffrey Barrett, Sonia Goncalves, Ewan Harrison, David K. Jackson, Ian Johnston, Dominic Kwiatkowski, Cordelia Langford, John Sillitoe on behalf of the Wellcome Sanger Institute COVID-19 Surveillance Team |
| EPI_ISL_3358737 | Lighthouse Lab in Alderley Park | Wellcome Sanger Institute for the COVID-19 Genomics UK (COG-UK) Consortium | Jacquelyn Wynn, Mairead Hyland, The Lighthouse Lab in Alderley Park and Alex Alderton, Roberto Amato, Jeffrey Barrett, Sonia Goncalves, Ewan Harrison, David K. Jackson, Ian Johnston, Dominic Kwiatkowski, Cordelia Langford, John Sillitoe on behalf of the Wellcome Sanger Institute COVID-19 Surveillance Team |
| EPI_ISL_3358738, EPI_ISL_3358739, EPI_ISL_3358740 | Lighthouse Lab in Milton Keynes | Wellcome Sanger Institute for the COVID-19 Genomics UK (COG-UK) Consortium | The Lighthouse Lab in Milton Keynes and Alex Alderton, Roberto Amato, Jeffrey Barrett, Sonia Goncalves, Ewan Harrison, David K. Jackson, Ian Johnston, Dominic Kwiatkowski, Cordelia Langford, John Sillitoe on behalf of the Wellcome Sanger Institute COVID-19 Surveillance Team |
| EPI_ISL_3358741 | Lighthouse Lab in Glasgow | Wellcome Sanger Institute for the COVID-19 Genomics UK (COG-UK) Consortium | Harper VanSteenhouse, Yumi Kasai, David Gray, Carol Clugston, Anna Dominiczak and Alex Alderton, Roberto Amato, Jeffrey Barrett, Sonia Goncalves, Ewan Harrison, David K. Jackson, Ian Johnston, Dominic Kwiatkowski, Cordelia Langford, John Sillitoe on behalf of the Wellcome Sanger Institute COVID-19 Surveillance Team |
| EPI_ISL_3358742 | Lighthouse Lab in Milton Keynes | Wellcome Sanger Institute for the COVID-19 Genomics UK (COG-UK) Consortium | The Lighthouse Lab in Milton Keynes and Alex Alderton, Roberto Amato, Jeffrey Barrett, Sonia Goncalves, Ewan Harrison, David K. Jackson, Ian Johnston, Dominic Kwiatkowski, Cordelia Langford, John Sillitoe on behalf of the Wellcome Sanger Institute COVID-19 Surveillance Team |
| EPI_ISL_3358743 | Lighthouse Lab in Alderley Park | Wellcome Sanger Institute for the COVID-19 Genomics UK (COG-UK) Consortium | Jacquelyn Wynn, Mairead Hyland, The Lighthouse Lab in Alderley Park and Alex Alderton, Roberto Amato, Jeffrey Barrett, Sonia Goncalves, Ewan Harrison, David K. Jackson, Ian Johnston, Dominic Kwiatkowski, Cordelia Langford, John Sillitoe on behalf of the Wellcome Sanger Institute COVID-19 Surveillance Team |
| EPI_ISL_3358744 | Lighthouse Lab in Milton Keynes | Wellcome Sanger Institute for the COVID-19 Genomics UK (COG-UK) Consortium | The Lighthouse Lab in Milton Keynes and Alex Alderton, Roberto Amato, Jeffrey Barrett, Sonia Goncalves, Ewan Harrison, David K. Jackson, Ian Johnston, Dominic Kwiatkowski, Cordelia Langford, John Sillitoe on behalf of the Wellcome Sanger Institute COVID-19 Surveillance Team |
| EPI_ISL_3358745 | Lighthouse Lab in Alderley Park | Wellcome Sanger Institute for the COVID-19 Genomics UK (COG-UK) Consortium | Jacquelyn Wynn, Mairead Hyland, The Lighthouse Lab in Alderley Park and Alex Alderton, Roberto Amato, Jeffrey Barrett, Sonia Goncalves, Ewan Harrison, David K. Jackson, Ian Johnston, Dominic Kwiatkowski, Cordelia Langford, John Sillitoe on behalf of the Wellcome Sanger Institute COVID-19 Surveillance Team |
| EPI_ISL_3358746, EPI_ISL_3358747 | Lighthouse Lab in Milton Keynes | Wellcome Sanger Institute for the COVID-19 Genomics UK (COG-UK) Consortium | The Lighthouse Lab in Milton Keynes and Alex Alderton, Roberto Amato, Jeffrey Barrett, Sonia Goncalves, Ewan Harrison, David K. Jackson, Ian Johnston, Dominic Kwiatkowski, Cordelia Langford, John Sillitoe on behalf of the Wellcome Sanger Institute COVID-19 Surveillance Team |
| EPI_ISL_3358748 | Lighthouse Lab in Alderley Park | Wellcome Sanger Institute for the COVID-19 Genomics UK (COG-UK) | Jacquelyn Wynn, Mairead Hyland, The Lighthouse Lab in Alderley Park and Alex Alderton, Roberto Amato, Jeffrey Barrett, Sonia Goncalves, Ewan Harrison, David K. Jackson, Ian Johnston, Dominic Kwiatkowski, Cordelia Langford, John Sillitoe on |

| | | | |
|---|---|---|---|
| | | Consortium | behalf of the Wellcome Sanger Institute COVID-19 Surveillance Team |
| EPI_ISL_3358749, EPI_ISL_3358750 | Lighthouse Lab in Milton Keynes | Wellcome Sanger Institute for the COVID-19 Genomics UK (COG-UK) Consortium | The Lighthouse Lab in Milton Keynes and Alex Alderton, Roberto Amato, Jeffrey Barrett, Sonia Goncalves, Ewan Harrison, David K. Jackson, Ian Johnston, Dominic Kwiatkowski, Cordelia Langford, John Sillitoe on behalf of the Wellcome Sanger Institute COVID-19 Surveillance Team |
| EPI_ISL_3358751 | Lighthouse Lab in Glasgow | Wellcome Sanger Institute for the COVID-19 Genomics UK (COG-UK) Consortium | Harper VanSteenhouse, Yumi Kasai, David Gray, Carol Clugston, Anna Dominiczak and Alex Alderton, Roberto Amato, Jeffrey Barrett, Sonia Goncalves, Ewan Harrison, David K. Jackson, Ian Johnston, Dominic Kwiatkowski, Cordelia Langford, John Sillitoe on behalf of the Wellcome Sanger Institute COVID-19 Surveillance Team |
| EPI_ISL_3358752 | Lighthouse Lab in Alderley Park | Wellcome Sanger Institute for the COVID-19 Genomics UK (COG-UK) Consortium | Jacquelyn Wynn, Mairead Hyland, The Lighthouse Lab in Alderley Park and Alex Alderton, Roberto Amato, Jeffrey Barrett, Sonia Goncalves, Ewan Harrison, David K. Jackson, Ian Johnston, Dominic Kwiatkowski, Cordelia Langford, John Sillitoe on behalf of the Wellcome Sanger Institute COVID-19 Surveillance Team |
| EPI_ISL_3358753, EPI_ISL_3358754 | Lighthouse Lab in Glasgow | Wellcome Sanger Institute for the COVID-19 Genomics UK (COG-UK) Consortium | Harper VanSteenhouse, Yumi Kasai, David Gray, Carol Clugston, Anna Dominiczak and Alex Alderton, Roberto Amato, Jeffrey Barrett, Sonia Goncalves, Ewan Harrison, David K. Jackson, Ian Johnston, Dominic Kwiatkowski, Cordelia Langford, John Sillitoe on behalf of the Wellcome Sanger Institute COVID-19 Surveillance Team |
| EPI_ISL_3358755, EPI_ISL_3358756, EPI_ISL_3358757 | Lighthouse Lab in Milton Keynes | Wellcome Sanger Institute for the COVID-19 Genomics UK (COG-UK) Consortium | The Lighthouse Lab in Milton Keynes and Alex Alderton, Roberto Amato, Jeffrey Barrett, Sonia Goncalves, Ewan Harrison, David K. Jackson, Ian Johnston, Dominic Kwiatkowski, Cordelia Langford, John Sillitoe on behalf of the Wellcome Sanger Institute COVID-19 Surveillance Team |
| EPI_ISL_3358758, EPI_ISL_3358759 | Lighthouse Lab in Alderley Park | Wellcome Sanger Institute for the COVID-19 Genomics UK (COG-UK) Consortium | Jacquelyn Wynn, Mairead Hyland, The Lighthouse Lab in Alderley Park and Alex Alderton, Roberto Amato, Jeffrey Barrett, Sonia Goncalves, Ewan Harrison, David K. Jackson, Ian Johnston, Dominic Kwiatkowski, Cordelia Langford, John Sillitoe on behalf of the Wellcome Sanger Institute COVID-19 Surveillance Team |
| EPI_ISL_3358760 | Lighthouse Lab in Glasgow | Wellcome Sanger Institute for the COVID-19 Genomics UK (COG-UK) Consortium | Harper VanSteenhouse, Yumi Kasai, David Gray, Carol Clugston, Anna Dominiczak and Alex Alderton, Roberto Amato, Jeffrey Barrett, Sonia Goncalves, Ewan Harrison, David K. Jackson, Ian Johnston, Dominic Kwiatkowski, Cordelia Langford, John Sillitoe on behalf of the Wellcome Sanger Institute COVID-19 Surveillance Team |
| EPI_ISL_3358761 | Lighthouse Lab in Milton Keynes | Wellcome Sanger Institute for the COVID-19 Genomics UK (COG-UK) Consortium | The Lighthouse Lab in Milton Keynes and Alex Alderton, Roberto Amato, Jeffrey Barrett, Sonia Goncalves, Ewan Harrison, David K. Jackson, Ian Johnston, Dominic Kwiatkowski, Cordelia Langford, John Sillitoe on behalf of the Wellcome Sanger Institute COVID-19 Surveillance Team |
| EPI_ISL_3358762, EPI_ISL_3358763, EPI_ISL_3358764, EPI_ISL_3358765 | Lighthouse Lab in Alderley Park | Wellcome Sanger Institute for the COVID-19 Genomics UK (COG-UK) Consortium | Jacquelyn Wynn, Mairead Hyland, The Lighthouse Lab in Alderley Park and Alex Alderton, Roberto Amato, Jeffrey Barrett, Sonia Goncalves, Ewan Harrison, David K. Jackson, Ian Johnston, Dominic Kwiatkowski, Cordelia Langford, John Sillitoe on behalf of the Wellcome Sanger Institute COVID-19 Surveillance Team |
| EPI_ISL_3358766 | Lighthouse Lab in Glasgow | Wellcome Sanger Institute for the COVID-19 Genomics UK (COG-UK) Consortium | Harper VanSteenhouse, Yumi Kasai, David Gray, Carol Clugston, Anna Dominiczak and Alex Alderton, Roberto Amato, Jeffrey Barrett, Sonia Goncalves, Ewan Harrison, David K. Jackson, Ian Johnston, Dominic Kwiatkowski, Cordelia Langford, John Sillitoe on behalf of the Wellcome Sanger Institute COVID-19 Surveillance Team |
| EPI_ISL_3358767, EPI_ISL_3358768 | Lighthouse Lab in Milton Keynes | Wellcome Sanger Institute for the COVID-19 Genomics UK (COG-UK) Consortium | The Lighthouse Lab in Milton Keynes and Alex Alderton, Roberto Amato, Jeffrey Barrett, Sonia Goncalves, Ewan Harrison, David K. Jackson, Ian Johnston, Dominic Kwiatkowski, Cordelia Langford, John Sillitoe on behalf of the Wellcome Sanger Institute COVID-19 Surveillance Team |
| EPI_ISL_3358769 | Lighthouse Lab in Glasgow | Wellcome Sanger Institute for the COVID-19 Genomics UK (COG-UK) Consortium | Harper VanSteenhouse, Yumi Kasai, David Gray, Carol Clugston, Anna Dominiczak and Alex Alderton, Roberto Amato, Jeffrey Barrett, Sonia Goncalves, Ewan Harrison, David K. Jackson, Ian Johnston, Dominic Kwiatkowski, Cordelia Langford, John Sillitoe on behalf of the Wellcome Sanger Institute COVID-19 Surveillance Team |
| EPI_ISL_3358770, EPI_ISL_3358771 | Lighthouse Lab in Milton Keynes | Wellcome Sanger Institute for the COVID-19 Genomics UK (COG-UK) Consortium | The Lighthouse Lab in Milton Keynes and Alex Alderton, Roberto Amato, Jeffrey Barrett, Sonia Goncalves, Ewan Harrison, David K. Jackson, Ian Johnston, Dominic Kwiatkowski, Cordelia Langford, John Sillitoe on behalf of the Wellcome Sanger Institute COVID-19 Surveillance Team |
| EPI_ISL_3358772, EPI_ISL_3358773 | Lighthouse Lab in Glasgow | Wellcome Sanger Institute for the COVID-19 Genomics UK (COG-UK) Consortium | Harper VanSteenhouse, Yumi Kasai, David Gray, Carol Clugston, Anna Dominiczak and Alex Alderton, Roberto Amato, Jeffrey Barrett, Sonia Goncalves, Ewan Harrison, David K. Jackson, Ian Johnston, Dominic Kwiatkowski, Cordelia Langford, John Sillitoe on behalf of the Wellcome Sanger Institute COVID-19 Surveillance Team |
| EPI_ISL_3358774 | Lighthouse Lab in Milton Keynes | Wellcome Sanger Institute for the COVID-19 Genomics UK (COG-UK) Consortium | The Lighthouse Lab in Milton Keynes and Alex Alderton, Roberto Amato, Jeffrey Barrett, Sonia Goncalves, Ewan Harrison, David K. Jackson, Ian Johnston, Dominic Kwiatkowski, Cordelia Langford, John Sillitoe on behalf of the Wellcome Sanger Institute COVID-19 Surveillance Team |
| EPI_ISL_3358775 | Lighthouse Lab in Alderley Park | Wellcome Sanger Institute for the COVID-19 Genomics UK (COG-UK) Consortium | Jacquelyn Wynn, Mairead Hyland, The Lighthouse Lab in Alderley Park and Alex Alderton, Roberto Amato, Jeffrey Barrett, Sonia Goncalves, Ewan Harrison, David K. Jackson, Ian Johnston, Dominic Kwiatkowski, Cordelia Langford, John Sillitoe on behalf of the Wellcome Sanger Institute COVID-19 Surveillance Team |
| EPI_ISL_3358776 | Lighthouse Lab in Milton Keynes | Wellcome Sanger Institute for the COVID-19 Genomics UK (COG-UK) Consortium | The Lighthouse Lab in Milton Keynes and Alex Alderton, Roberto Amato, Jeffrey Barrett, Sonia Goncalves, Ewan Harrison, David K. Jackson, Ian Johnston, Dominic Kwiatkowski, Cordelia Langford, John Sillitoe on behalf of the Wellcome Sanger Institute COVID-19 Surveillance Team |
| EPI_ISL_3358777, EPI_ISL_3358778, EPI_ISL_3358779 | Lighthouse Lab in Alderley Park | Wellcome Sanger Institute for the COVID-19 Genomics UK (COG-UK) Consortium | Jacquelyn Wynn, Mairead Hyland, The Lighthouse Lab in Alderley Park and Alex Alderton, Roberto Amato, Jeffrey Barrett, Sonia Goncalves, Ewan Harrison, David K. Jackson, Ian Johnston, Dominic Kwiatkowski, Cordelia Langford, John Sillitoe on behalf of the Wellcome Sanger Institute COVID-19 Surveillance Team |
| EPI_ISL_3358780 | Lighthouse Lab in Milton Keynes | Wellcome Sanger Institute for the COVID-19 Genomics UK (COG-UK) Consortium | The Lighthouse Lab in Milton Keynes and Alex Alderton, Roberto Amato, Jeffrey Barrett, Sonia Goncalves, Ewan Harrison, David K. Jackson, Ian Johnston, Dominic Kwiatkowski, Cordelia Langford, John Sillitoe on behalf of the Wellcome Sanger Institute COVID-19 Surveillance Team |
| EPI_ISL_3358781 | Lighthouse Lab in Glasgow | Wellcome Sanger Institute for the COVID-19 Genomics UK (COG-UK) Consortium | Harper VanSteenhouse, Yumi Kasai, David Gray, Carol Clugston, Anna Dominiczak and Alex Alderton, Roberto Amato, Jeffrey Barrett, Sonia Goncalves, Ewan Harrison, David K. Jackson, Ian Johnston, Dominic Kwiatkowski, Cordelia Langford, John Sillitoe on behalf of the Wellcome Sanger Institute COVID-19 Surveillance Team |
| EPI_ISL_3358782 | Lighthouse Lab in Alderley Park | Wellcome Sanger Institute for the COVID-19 Genomics UK (COG-UK) Consortium | Jacquelyn Wynn, Mairead Hyland, The Lighthouse Lab in Alderley Park and Alex Alderton, Roberto Amato, Jeffrey Barrett, Sonia Goncalves, Ewan Harrison, David K. Jackson, Ian Johnston, Dominic Kwiatkowski, Cordelia Langford, John Sillitoe on behalf of the Wellcome Sanger Institute COVID-19 Surveillance Team |
| EPI_ISL_3358783 | Lighthouse Lab in Glasgow | Wellcome Sanger Institute for the COVID-19 Genomics UK (COG-UK) Consortium | Harper VanSteenhouse, Yumi Kasai, David Gray, Carol Clugston, Anna Dominiczak and Alex Alderton, Roberto Amato, Jeffrey Barrett, Sonia Goncalves, Ewan Harrison, David K. Jackson, Ian Johnston, Dominic Kwiatkowski, Cordelia Langford, John Sillitoe on behalf of the Wellcome Sanger Institute COVID-19 Surveillance Team |
| EPI_ISL_3358784 | Lighthouse Lab in Milton Keynes | Wellcome Sanger Institute for the COVID-19 Genomics UK (COG-UK) Consortium | The Lighthouse Lab in Milton Keynes and Alex Alderton, Roberto Amato, Jeffrey Barrett, Sonia Goncalves, Ewan Harrison, David K. Jackson, Ian Johnston, Dominic Kwiatkowski, Cordelia Langford, John Sillitoe on behalf of the Wellcome Sanger Institute COVID-19 Surveillance Team |
| EPI_ISL_3358785 | Lighthouse Lab in Alderley Park | Wellcome Sanger Institute for the COVID-19 Genomics UK (COG-UK) Consortium | Jacquelyn Wynn, Mairead Hyland, The Lighthouse Lab in Alderley Park and Alex Alderton, Roberto Amato, Jeffrey Barrett, Sonia Goncalves, Ewan Harrison, David K. Jackson, Ian Johnston, Dominic Kwiatkowski, Cordelia Langford, John Sillitoe on behalf of the Wellcome Sanger Institute COVID-19 Surveillance Team |
| EPI_ISL_3358786 | Lighthouse Lab in Milton Keynes | Wellcome Sanger Institute for the COVID-19 Genomics UK (COG-UK) Consortium | The Lighthouse Lab in Milton Keynes and Alex Alderton, Roberto Amato, Jeffrey Barrett, Sonia Goncalves, Ewan Harrison, David K. Jackson, Ian Johnston, Dominic Kwiatkowski, Cordelia Langford, John Sillitoe on behalf of the Wellcome Sanger Institute COVID-19 Surveillance Team |
| EPI_ISL_3358787 | Lighthouse Lab in Glasgow | Wellcome Sanger Institute for the COVID-19 Genomics UK (COG-UK) Consortium | Harper VanSteenhouse, Yumi Kasai, David Gray, Carol Clugston, Anna Dominiczak and Alex Alderton, Roberto Amato, Jeffrey Barrett, Sonia Goncalves, Ewan Harrison, David K. Jackson, Ian Johnston, Dominic Kwiatkowski, Cordelia Langford, John Sillitoe on behalf of the Wellcome Sanger Institute COVID-19 Surveillance Team |
| EPI_ISL_3358788, EPI_ISL_3358789, EPI_ISL_3358790, EPI_ISL_3358791, EPI_ISL_3358792 | Lighthouse Lab in Milton Keynes | Wellcome Sanger Institute for the COVID-19 Genomics UK (COG-UK) Consortium | The Lighthouse Lab in Milton Keynes and Alex Alderton, Roberto Amato, Jeffrey Barrett, Sonia Goncalves, Ewan Harrison, David K. Jackson, Ian Johnston, Dominic Kwiatkowski, Cordelia Langford, John Sillitoe on behalf of the Wellcome Sanger Institute COVID-19 Surveillance Team |
| EPI_ISL_3358793 | Lighthouse Lab in Alderley Park | Wellcome Sanger Institute for the COVID-19 Genomics UK (COG-UK) Consortium | Jacquelyn Wynn, Mairead Hyland, The Lighthouse Lab in Alderley Park and Alex Alderton, Roberto Amato, Jeffrey Barrett, Sonia Goncalves, Ewan Harrison, David K. Jackson, Ian Johnston, Dominic Kwiatkowski, Cordelia Langford, John Sillitoe on behalf of the Wellcome Sanger Institute COVID-19 Surveillance Team |
| EPI_ISL_3358794, EPI_ISL_3358795 | Lighthouse Lab in Glasgow | Wellcome Sanger Institute for the COVID-19 Genomics UK (COG-UK) Consortium | Harper VanSteenhouse, Yumi Kasai, David Gray, Carol Clugston, Anna Dominiczak and Alex Alderton, Roberto Amato, Jeffrey Barrett, Sonia Goncalves, Ewan Harrison, David K. Jackson, Ian Johnston, Dominic Kwiatkowski, Cordelia Langford, John Sillitoe on behalf of the Wellcome Sanger Institute COVID-19 Surveillance Team |
| EPI_ISL_3358796 | Lighthouse Lab in Alderley Park | Wellcome Sanger Institute for the COVID-19 Genomics UK (COG-UK) Consortium | Jacquelyn Wynn, Mairead Hyland, The Lighthouse Lab in Alderley Park and Alex Alderton, Roberto Amato, Jeffrey Barrett, Sonia Goncalves, Ewan Harrison, David K. Jackson, Ian Johnston, Dominic Kwiatkowski, Cordelia Langford, John Sillitoe on behalf of the Wellcome Sanger Institute COVID-19 Surveillance Team |
| EPI_ISL_3358797 | Lighthouse Lab in Milton Keynes | Wellcome Sanger Institute for the COVID-19 Genomics UK (COG-UK) Consortium | The Lighthouse Lab in Milton Keynes and Alex Alderton, Roberto Amato, Jeffrey Barrett, Sonia Goncalves, Ewan Harrison, David K. Jackson, Ian Johnston, Dominic Kwiatkowski, Cordelia Langford, John Sillitoe on behalf of the Wellcome Sanger Institute COVID-19 Surveillance Team |
| EPI_ISL_3358798 | Lighthouse Lab in Glasgow | Wellcome Sanger Institute for the COVID-19 Genomics UK (COG-UK) Consortium | Harper VanSteenhouse, Yumi Kasai, David Gray, Carol Clugston, Anna Dominiczak and Alex Alderton, Roberto Amato, Jeffrey Barrett, Sonia Goncalves, Ewan Harrison, David K. Jackson, Ian Johnston, Dominic Kwiatkowski, Cordelia Langford, John Sillitoe on behalf of the Wellcome Sanger Institute COVID-19 Surveillance Team |
| EPI_ISL_3358799 | Lighthouse Lab in Milton Keynes | Wellcome Sanger Institute for the COVID-19 Genomics UK (COG-UK) Consortium | The Lighthouse Lab in Milton Keynes and Alex Alderton, Roberto Amato, Jeffrey Barrett, Sonia Goncalves, Ewan Harrison, David K. Jackson, Ian Johnston, Dominic Kwiatkowski, Cordelia Langford, John Sillitoe on behalf of the Wellcome Sanger Institute COVID-19 Surveillance Team |
| EPI_ISL_3358800 | Lighthouse Lab in Glasgow | Wellcome Sanger Institute for the COVID-19 Genomics UK (COG-UK) Consortium | Harper VanSteenhouse, Yumi Kasai, David Gray, Carol Clugston, Anna Dominiczak and Alex Alderton, Roberto Amato, Jeffrey Barrett, Sonia Goncalves, Ewan Harrison, David K. Jackson, Ian Johnston, Dominic Kwiatkowski, Cordelia Langford, John Sillitoe on behalf of the Wellcome Sanger Institute COVID-19 Surveillance Team |
| EPI_ISL_3358801, EPI_ISL_3358802, EPI_ISL_3358803, EPI_ISL_3358804 | Lighthouse Lab in Milton Keynes | Wellcome Sanger Institute for the COVID-19 Genomics UK (COG-UK) Consortium | The Lighthouse Lab in Milton Keynes and Alex Alderton, Roberto Amato, Jeffrey Barrett, Sonia Goncalves, Ewan Harrison, David K. Jackson, Ian Johnston, Dominic Kwiatkowski, Cordelia Langford, John Sillitoe on behalf of the Wellcome Sanger Institute COVID-19 Surveillance Team |
| EPI_ISL_3358805 | Lighthouse Lab in Glasgow | Wellcome Sanger Institute for the COVID-19 Genomics UK (COG-UK) Consortium | Harper VanSteenhouse, Yumi Kasai, David Gray, Carol Clugston, Anna Dominiczak and Alex Alderton, Roberto Amato, Jeffrey Barrett, Sonia Goncalves, Ewan Harrison, David K. Jackson, Ian Johnston, Dominic Kwiatkowski, Cordelia Langford, John Sillitoe on behalf of the Wellcome Sanger Institute COVID-19 Surveillance Team |
| EPI_ISL_3358806, EPI_ISL_3358807, EPI_ISL_3358808, EPI_ISL_3358809 | Lighthouse Lab in Milton Keynes | Wellcome Sanger Institute for the COVID-19 Genomics UK (COG-UK) Consortium | The Lighthouse Lab in Milton Keynes and Alex Alderton, Roberto Amato, Jeffrey Barrett, Sonia Goncalves, Ewan Harrison, David K. Jackson, Ian Johnston, Dominic Kwiatkowski, Cordelia Langford, John Sillitoe on behalf of the Wellcome Sanger Institute COVID-19 Surveillance Team |
| EPI_ISL_3358810 | Lighthouse Lab in Glasgow | Wellcome Sanger Institute for the COVID-19 Genomics UK (COG-UK) Consortium | Harper VanSteenhouse, Yumi Kasai, David Gray, Carol Clugston, Anna Dominiczak and Alex Alderton, Roberto Amato, Jeffrey Barrett, Sonia Goncalves, Ewan Harrison, David K. Jackson, Ian Johnston, Dominic Kwiatkowski, Cordelia Langford, John Sillitoe on behalf of the Wellcome Sanger Institute COVID-19 Surveillance Team |
| EPI_ISL_3358811, EPI_ISL_3358812, EPI_ISL_3358813 | Lighthouse Lab in Milton Keynes | Wellcome Sanger Institute for the COVID-19 Genomics UK (COG-UK) Consortium | The Lighthouse Lab in Milton Keynes and Alex Alderton, Roberto Amato, Jeffrey Barrett, Sonia Goncalves, Ewan Harrison, David K. Jackson, Ian Johnston, Dominic Kwiatkowski, Cordelia Langford, John Sillitoe on behalf of the Wellcome Sanger Institute COVID-19 Surveillance Team |
| EPI_ISL_3358814 | Lighthouse Lab in Alderley Park | Wellcome Sanger Institute for the COVID-19 Genomics UK (COG-UK) | Jacquelyn Wynn, Mairead Hyland, The Lighthouse Lab in Alderley Park and Alex Alderton, Roberto Amato, Jeffrey Barrett, Sonia Goncalves, Ewan Harrison, David K. Jackson, Ian Johnston, Dominic Kwiatkowski, Cordelia Langford, John Sillitoe on |

| | | | |
|---|---|---|---|
| | | Consortium | behalf of the Wellcome Sanger Institute COVID-19 Surveillance Team |
| EPI_ISL_3358815, EPI_ISL_3358816, EPI_ISL_3358817 | Lighthouse Lab in Milton Keynes | Wellcome Sanger Institute for the COVID-19 Genomics UK (COG-UK) Consortium | The Lighthouse Lab in Milton Keynes and Alex Alderton, Roberto Amato, Jeffrey Barrett, Sonia Goncalves, Ewan Harrison, David K. Jackson, Ian Johnston, Dominic Kwiatkowski, Cordelia Langford, John Sillitoe on behalf of the Wellcome Sanger Institute COVID-19 Surveillance Team |
| EPI_ISL_3358818, EPI_ISL_3358819 | Lighthouse Lab in Alderley Park | Wellcome Sanger Institute for the COVID-19 Genomics UK (COG-UK) Consortium | Jacquelyn Wynn, Mairead Hyland, The Lighthouse Lab in Alderley Park and Alex Alderton, Roberto Amato, Jeffrey Barrett, Sonia Goncalves, Ewan Harrison, David K. Jackson, Ian Johnston, Dominic Kwiatkowski, Cordelia Langford, John Sillitoe on behalf of the Wellcome Sanger Institute COVID-19 Surveillance Team |
| EPI_ISL_3358820 | Lighthouse Lab in Milton Keynes | Wellcome Sanger Institute for the COVID-19 Genomics UK (COG-UK) Consortium | The Lighthouse Lab in Milton Keynes and Alex Alderton, Roberto Amato, Jeffrey Barrett, Sonia Goncalves, Ewan Harrison, David K. Jackson, Ian Johnston, Dominic Kwiatkowski, Cordelia Langford, John Sillitoe on behalf of the Wellcome Sanger Institute COVID-19 Surveillance Team |
| EPI_ISL_3358821 | Lighthouse Lab in Alderley Park | Wellcome Sanger Institute for the COVID-19 Genomics UK (COG-UK) Consortium | Jacquelyn Wynn, Mairead Hyland, The Lighthouse Lab in Alderley Park and Alex Alderton, Roberto Amato, Jeffrey Barrett, Sonia Goncalves, Ewan Harrison, David K. Jackson, Ian Johnston, Dominic Kwiatkowski, Cordelia Langford, John Sillitoe on behalf of the Wellcome Sanger Institute COVID-19 Surveillance Team |
| EPI_ISL_3358822, EPI_ISL_3358823 | Lighthouse Lab in Milton Keynes | Wellcome Sanger Institute for the COVID-19 Genomics UK (COG-UK) Consortium | The Lighthouse Lab in Milton Keynes and Alex Alderton, Roberto Amato, Jeffrey Barrett, Sonia Goncalves, Ewan Harrison, David K. Jackson, Ian Johnston, Dominic Kwiatkowski, Cordelia Langford, John Sillitoe on behalf of the Wellcome Sanger Institute COVID-19 Surveillance Team |
| EPI_ISL_3358824, EPI_ISL_3358825, EPI_ISL_3358826, EPI_ISL_3358827 | Lighthouse Lab in Alderley Park | Wellcome Sanger Institute for the COVID-19 Genomics UK (COG-UK) Consortium | Jacquelyn Wynn, Mairead Hyland, The Lighthouse Lab in Alderley Park and Alex Alderton, Roberto Amato, Jeffrey Barrett, Sonia Goncalves, Ewan Harrison, David K. Jackson, Ian Johnston, Dominic Kwiatkowski, Cordelia Langford, John Sillitoe on behalf of the Wellcome Sanger Institute COVID-19 Surveillance Team |
| EPI_ISL_3358828 | Lighthouse Lab in Milton Keynes | Wellcome Sanger Institute for the COVID-19 Genomics UK (COG-UK) Consortium | The Lighthouse Lab in Milton Keynes and Alex Alderton, Roberto Amato, Jeffrey Barrett, Sonia Goncalves, Ewan Harrison, David K. Jackson, Ian Johnston, Dominic Kwiatkowski, Cordelia Langford, John Sillitoe on behalf of the Wellcome Sanger Institute COVID-19 Surveillance Team |
| EPI_ISL_3358829 | Lighthouse Lab in Glasgow | Wellcome Sanger Institute for the COVID-19 Genomics UK (COG-UK) Consortium | Harper VanSteenhouse, Yumi Kasai, David Gray, Carol Clugston, Anna Dominiczak and Alex Alderton, Roberto Amato, Jeffrey Barrett, Sonia Goncalves, Ewan Harrison, David K. Jackson, Ian Johnston, Dominic Kwiatkowski, Cordelia Langford, John Sillitoe on behalf of the Wellcome Sanger Institute COVID-19 Surveillance Team |
| EPI_ISL_3358830 | Lighthouse Lab in Milton Keynes | Wellcome Sanger Institute for the COVID-19 Genomics UK (COG-UK) Consortium | The Lighthouse Lab in Milton Keynes and Alex Alderton, Roberto Amato, Jeffrey Barrett, Sonia Goncalves, Ewan Harrison, David K. Jackson, Ian Johnston, Dominic Kwiatkowski, Cordelia Langford, John Sillitoe on behalf of the Wellcome Sanger Institute COVID-19 Surveillance Team |
| EPI_ISL_3358831 | Lighthouse Lab in Alderley Park | Wellcome Sanger Institute for the COVID-19 Genomics UK (COG-UK) Consortium | Jacquelyn Wynn, Mairead Hyland, The Lighthouse Lab in Alderley Park and Alex Alderton, Roberto Amato, Jeffrey Barrett, Sonia Goncalves, Ewan Harrison, David K. Jackson, Ian Johnston, Dominic Kwiatkowski, Cordelia Langford, John Sillitoe on behalf of the Wellcome Sanger Institute COVID-19 Surveillance Team |
| EPI_ISL_3358832 | Lighthouse Lab in Glasgow | Wellcome Sanger Institute for the COVID-19 Genomics UK (COG-UK) Consortium | Harper VanSteenhouse, Yumi Kasai, David Gray, Carol Clugston, Anna Dominiczak and Alex Alderton, Roberto Amato, Jeffrey Barrett, Sonia Goncalves, Ewan Harrison, David K. Jackson, Ian Johnston, Dominic Kwiatkowski, Cordelia Langford, John Sillitoe on behalf of the Wellcome Sanger Institute COVID-19 Surveillance Team |
| EPI_ISL_3358833, EPI_ISL_3358834 | Lighthouse Lab in Milton Keynes | Wellcome Sanger Institute for the COVID-19 Genomics UK (COG-UK) Consortium | The Lighthouse Lab in Milton Keynes and Alex Alderton, Roberto Amato, Jeffrey Barrett, Sonia Goncalves, Ewan Harrison, David K. Jackson, Ian Johnston, Dominic Kwiatkowski, Cordelia Langford, John Sillitoe on behalf of the Wellcome Sanger Institute COVID-19 Surveillance Team |
| EPI_ISL_3358835 | Lighthouse Lab in Glasgow | Wellcome Sanger Institute for the COVID-19 Genomics UK (COG-UK) Consortium | Harper VanSteenhouse, Yumi Kasai, David Gray, Carol Clugston, Anna Dominiczak and Alex Alderton, Roberto Amato, Jeffrey Barrett, Sonia Goncalves, Ewan Harrison, David K. Jackson, Ian Johnston, Dominic Kwiatkowski, Cordelia Langford, John Sillitoe on behalf of the Wellcome Sanger Institute COVID-19 Surveillance Team |
| EPI_ISL_3358836, EPI_ISL_3358837, EPI_ISL_3358838 | Lighthouse Lab in Milton Keynes | Wellcome Sanger Institute for the COVID-19 Genomics UK (COG-UK) Consortium | The Lighthouse Lab in Milton Keynes and Alex Alderton, Roberto Amato, Jeffrey Barrett, Sonia Goncalves, Ewan Harrison, David K. Jackson, Ian Johnston, Dominic Kwiatkowski, Cordelia Langford, John Sillitoe on behalf of the Wellcome Sanger Institute COVID-19 Surveillance Team |
| EPI_ISL_3358839 | Lighthouse Lab in Alderley Park | Wellcome Sanger Institute for the COVID-19 Genomics UK (COG-UK) Consortium | Jacquelyn Wynn, Mairead Hyland, The Lighthouse Lab in Alderley Park and Alex Alderton, Roberto Amato, Jeffrey Barrett, Sonia Goncalves, Ewan Harrison, David K. Jackson, Ian Johnston, Dominic Kwiatkowski, Cordelia Langford, John Sillitoe on behalf of the Wellcome Sanger Institute COVID-19 Surveillance Team |
| EPI_ISL_3358840 | Lighthouse Lab in Milton Keynes | Wellcome Sanger Institute for the COVID-19 Genomics UK (COG-UK) Consortium | The Lighthouse Lab in Milton Keynes and Alex Alderton, Roberto Amato, Jeffrey Barrett, Sonia Goncalves, Ewan Harrison, David K. Jackson, Ian Johnston, Dominic Kwiatkowski, Cordelia Langford, John Sillitoe on behalf of the Wellcome Sanger Institute COVID-19 Surveillance Team |
| EPI_ISL_3358841 | Lighthouse Lab in Alderley Park | Wellcome Sanger Institute for the COVID-19 Genomics UK (COG-UK) Consortium | Jacquelyn Wynn, Mairead Hyland, The Lighthouse Lab in Alderley Park and Alex Alderton, Roberto Amato, Jeffrey Barrett, Sonia Goncalves, Ewan Harrison, David K. Jackson, Ian Johnston, Dominic Kwiatkowski, Cordelia Langford, John Sillitoe on behalf of the Wellcome Sanger Institute COVID-19 Surveillance Team |
| EPI_ISL_3358842 | Lighthouse Lab in Milton Keynes | Wellcome Sanger Institute for the COVID-19 Genomics UK (COG-UK) Consortium | The Lighthouse Lab in Milton Keynes and Alex Alderton, Roberto Amato, Jeffrey Barrett, Sonia Goncalves, Ewan Harrison, David K. Jackson, Ian Johnston, Dominic Kwiatkowski, Cordelia Langford, John Sillitoe on behalf of the Wellcome Sanger Institute COVID-19 Surveillance Team |
| EPI_ISL_3358843, EPI_ISL_3358844 | Lighthouse Lab in Alderley Park | Wellcome Sanger Institute for the COVID-19 Genomics UK (COG-UK) Consortium | Jacquelyn Wynn, Mairead Hyland, The Lighthouse Lab in Alderley Park and Alex Alderton, Roberto Amato, Jeffrey Barrett, Sonia Goncalves, Ewan Harrison, David K. Jackson, Ian Johnston, Dominic Kwiatkowski, Cordelia Langford, John Sillitoe on behalf of the Wellcome Sanger Institute COVID-19 Surveillance Team |
| EPI_ISL_3358845 | Lighthouse Lab in Glasgow | Wellcome Sanger Institute for the COVID-19 Genomics UK (COG-UK) Consortium | Harper VanSteenhouse, Yumi Kasai, David Gray, Carol Clugston, Anna Dominiczak and Alex Alderton, Roberto Amato, Jeffrey Barrett, Sonia Goncalves, Ewan Harrison, David K. Jackson, Ian Johnston, Dominic Kwiatkowski, Cordelia Langford, John Sillitoe on behalf of the Wellcome Sanger Institute COVID-19 Surveillance Team |
| EPI_ISL_3358846 | Lighthouse Lab in Milton Keynes | Wellcome Sanger Institute for the COVID-19 Genomics UK (COG-UK) Consortium | The Lighthouse Lab in Milton Keynes and Alex Alderton, Roberto Amato, Jeffrey Barrett, Sonia Goncalves, Ewan Harrison, David K. Jackson, Ian Johnston, Dominic Kwiatkowski, Cordelia Langford, John Sillitoe on behalf of the Wellcome Sanger Institute COVID-19 Surveillance Team |
| EPI_ISL_3358847 | Lighthouse Lab in Glasgow | Wellcome Sanger Institute for the COVID-19 Genomics UK (COG-UK) Consortium | Harper VanSteenhouse, Yumi Kasai, David Gray, Carol Clugston, Anna Dominiczak and Alex Alderton, Roberto Amato, Jeffrey Barrett, Sonia Goncalves, Ewan Harrison, David K. Jackson, Ian Johnston, Dominic Kwiatkowski, Cordelia Langford, John Sillitoe on behalf of the Wellcome Sanger Institute COVID-19 Surveillance Team |
| EPI_ISL_3358848 | Lighthouse Lab in Milton Keynes | Wellcome Sanger Institute for the COVID-19 Genomics UK (COG-UK) Consortium | The Lighthouse Lab in Milton Keynes and Alex Alderton, Roberto Amato, Jeffrey Barrett, Sonia Goncalves, Ewan Harrison, David K. Jackson, Ian Johnston, Dominic Kwiatkowski, Cordelia Langford, John Sillitoe on behalf of the Wellcome Sanger Institute COVID-19 Surveillance Team |
| EPI_ISL_3358849 | Lighthouse Lab in Glasgow | Wellcome Sanger Institute for the COVID-19 Genomics UK (COG-UK) Consortium | Harper VanSteenhouse, Yumi Kasai, David Gray, Carol Clugston, Anna Dominiczak and Alex Alderton, Roberto Amato, Jeffrey Barrett, Sonia Goncalves, Ewan Harrison, David K. Jackson, Ian Johnston, Dominic Kwiatkowski, Cordelia Langford, John Sillitoe on behalf of the Wellcome Sanger Institute COVID-19 Surveillance Team |
| EPI_ISL_3358850 | Lighthouse Lab in Milton Keynes | Wellcome Sanger Institute for the COVID-19 Genomics UK (COG-UK) Consortium | The Lighthouse Lab in Milton Keynes and Alex Alderton, Roberto Amato, Jeffrey Barrett, Sonia Goncalves, Ewan Harrison, David K. Jackson, Ian Johnston, Dominic Kwiatkowski, Cordelia Langford, John Sillitoe on behalf of the Wellcome Sanger Institute COVID-19 Surveillance Team |
| EPI_ISL_3358851 | Lighthouse Lab in Alderley Park | Wellcome Sanger Institute for the COVID-19 Genomics UK (COG-UK) Consortium | Jacquelyn Wynn, Mairead Hyland, The Lighthouse Lab in Alderley Park and Alex Alderton, Roberto Amato, Jeffrey Barrett, Sonia Goncalves, Ewan Harrison, David K. Jackson, Ian Johnston, Dominic Kwiatkowski, Cordelia Langford, John Sillitoe on behalf of the Wellcome Sanger Institute COVID-19 Surveillance Team |
| EPI_ISL_3358852, EPI_ISL_3358853 | Lighthouse Lab in Milton Keynes | Wellcome Sanger Institute for the COVID-19 Genomics UK (COG-UK) Consortium | The Lighthouse Lab in Milton Keynes and Alex Alderton, Roberto Amato, Jeffrey Barrett, Sonia Goncalves, Ewan Harrison, David K. Jackson, Ian Johnston, Dominic Kwiatkowski, Cordelia Langford, John Sillitoe on behalf of the Wellcome Sanger Institute COVID-19 Surveillance Team |
| EPI_ISL_3358854 | Lighthouse Lab in Alderley Park | Wellcome Sanger Institute for the COVID-19 Genomics UK (COG-UK) Consortium | Jacquelyn Wynn, Mairead Hyland, The Lighthouse Lab in Alderley Park and Alex Alderton, Roberto Amato, Jeffrey Barrett, Sonia Goncalves, Ewan Harrison, David K. Jackson, Ian Johnston, Dominic Kwiatkowski, Cordelia Langford, John Sillitoe on behalf of the Wellcome Sanger Institute COVID-19 Surveillance Team |
| EPI_ISL_3358855 | Lighthouse Lab in Milton Keynes | Wellcome Sanger Institute for the COVID-19 Genomics UK (COG-UK) Consortium | The Lighthouse Lab in Milton Keynes and Alex Alderton, Roberto Amato, Jeffrey Barrett, Sonia Goncalves, Ewan Harrison, David K. Jackson, Ian Johnston, Dominic Kwiatkowski, Cordelia Langford, John Sillitoe on behalf of the Wellcome Sanger Institute COVID-19 Surveillance Team |
| EPI_ISL_3358856 | Lighthouse Lab in Glasgow | Wellcome Sanger Institute for the COVID-19 Genomics UK (COG-UK) Consortium | Harper VanSteenhouse, Yumi Kasai, David Gray, Carol Clugston, Anna Dominiczak and Alex Alderton, Roberto Amato, Jeffrey Barrett, Sonia Goncalves, Ewan Harrison, David K. Jackson, Ian Johnston, Dominic Kwiatkowski, Cordelia Langford, John Sillitoe on behalf of the Wellcome Sanger Institute COVID-19 Surveillance Team |
| EPI_ISL_3358857 | Lighthouse Lab in Milton Keynes | Wellcome Sanger Institute for the COVID-19 Genomics UK (COG-UK) Consortium | The Lighthouse Lab in Milton Keynes and Alex Alderton, Roberto Amato, Jeffrey Barrett, Sonia Goncalves, Ewan Harrison, David K. Jackson, Ian Johnston, Dominic Kwiatkowski, Cordelia Langford, John Sillitoe on behalf of the Wellcome Sanger Institute COVID-19 Surveillance Team |
| EPI_ISL_3358858 | Lighthouse Lab in Glasgow | Wellcome Sanger Institute for the COVID-19 Genomics UK (COG-UK) Consortium | Harper VanSteenhouse, Yumi Kasai, David Gray, Carol Clugston, Anna Dominiczak and Alex Alderton, Roberto Amato, Jeffrey Barrett, Sonia Goncalves, Ewan Harrison, David K. Jackson, Ian Johnston, Dominic Kwiatkowski, Cordelia Langford, John Sillitoe on behalf of the Wellcome Sanger Institute COVID-19 Surveillance Team |
| EPI_ISL_3358859 | Lighthouse Lab in Alderley Park | Wellcome Sanger Institute for the COVID-19 Genomics UK (COG-UK) Consortium | Jacquelyn Wynn, Mairead Hyland, The Lighthouse Lab in Alderley Park and Alex Alderton, Roberto Amato, Jeffrey Barrett, Sonia Goncalves, Ewan Harrison, David K. Jackson, Ian Johnston, Dominic Kwiatkowski, Cordelia Langford, John Sillitoe on behalf of the Wellcome Sanger Institute COVID-19 Surveillance Team |
| EPI_ISL_3358860, EPI_ISL_3358861, EPI_ISL_3358862 | Lighthouse Lab in Milton Keynes | Wellcome Sanger Institute for the COVID-19 Genomics UK (COG-UK) Consortium | The Lighthouse Lab in Milton Keynes and Alex Alderton, Roberto Amato, Jeffrey Barrett, Sonia Goncalves, Ewan Harrison, David K. Jackson, Ian Johnston, Dominic Kwiatkowski, Cordelia Langford, John Sillitoe on behalf of the Wellcome Sanger Institute COVID-19 Surveillance Team |
| EPI_ISL_3358863 | Lighthouse Lab in Alderley Park | Wellcome Sanger Institute for the COVID-19 Genomics UK (COG-UK) Consortium | Jacquelyn Wynn, Mairead Hyland, The Lighthouse Lab in Alderley Park and Alex Alderton, Roberto Amato, Jeffrey Barrett, Sonia Goncalves, Ewan Harrison, David K. Jackson, Ian Johnston, Dominic Kwiatkowski, Cordelia Langford, John Sillitoe on behalf of the Wellcome Sanger Institute COVID-19 Surveillance Team |
| EPI_ISL_3358864, EPI_ISL_3358865, EPI_ISL_3358866 | Lighthouse Lab in Milton Keynes | Wellcome Sanger Institute for the COVID-19 Genomics UK (COG-UK) Consortium | The Lighthouse Lab in Milton Keynes and Alex Alderton, Roberto Amato, Jeffrey Barrett, Sonia Goncalves, Ewan Harrison, David K. Jackson, Ian Johnston, Dominic Kwiatkowski, Cordelia Langford, John Sillitoe on behalf of the Wellcome Sanger Institute COVID-19 Surveillance Team |
| EPI_ISL_3358867 | Lighthouse Lab in Alderley Park | Wellcome Sanger Institute for the COVID-19 Genomics UK (COG-UK) Consortium | Jacquelyn Wynn, Mairead Hyland, The Lighthouse Lab in Alderley Park and Alex Alderton, Roberto Amato, Jeffrey Barrett, Sonia Goncalves, Ewan Harrison, David K. Jackson, Ian Johnston, Dominic Kwiatkowski, Cordelia Langford, John Sillitoe on behalf of the Wellcome Sanger Institute COVID-19 Surveillance Team |
| EPI_ISL_3358868, EPI_ISL_3358869, EPI_ISL_3358870, EPI_ISL_3358871 | Lighthouse Lab in Milton Keynes | Wellcome Sanger Institute for the COVID-19 Genomics UK (COG-UK) Consortium | The Lighthouse Lab in Milton Keynes and Alex Alderton, Roberto Amato, Jeffrey Barrett, Sonia Goncalves, Ewan Harrison, David K. Jackson, Ian Johnston, Dominic Kwiatkowski, Cordelia Langford, John Sillitoe on behalf of the Wellcome Sanger Institute COVID-19 Surveillance Team |
| EPI_ISL_3358872 | Lighthouse Lab in Alderley Park | Wellcome Sanger Institute for the COVID-19 Genomics UK (COG-UK) Consortium | Jacquelyn Wynn, Mairead Hyland, The Lighthouse Lab in Alderley Park and Alex Alderton, Roberto Amato, Jeffrey Barrett, Sonia Goncalves, Ewan Harrison, David K. Jackson, Ian Johnston, Dominic Kwiatkowski, Cordelia Langford, John Sillitoe on behalf of the Wellcome Sanger Institute COVID-19 Surveillance Team |
| EPI_ISL_3358873 | Lighthouse Lab in Glasgow | Wellcome Sanger Institute for the COVID-19 Genomics UK (COG-UK) Consortium | Harper VanSteenhouse, Yumi Kasai, David Gray, Carol Clugston, Anna Dominiczak and Alex Alderton, Roberto Amato, Jeffrey Barrett, Sonia Goncalves, Ewan Harrison, David K. Jackson, Ian Johnston, Dominic Kwiatkowski, Cordelia Langford, John Sillitoe on behalf of the Wellcome Sanger Institute COVID-19 Surveillance Team |
| EPI_ISL_3358874, EPI_ISL_3358875 | Lighthouse Lab in Milton Keynes | Wellcome Sanger Institute for the COVID-19 Genomics UK (COG-UK) Consortium | The Lighthouse Lab in Milton Keynes and Alex Alderton, Roberto Amato, Jeffrey Barrett, Sonia Goncalves, Ewan Harrison, David K. Jackson, Ian Johnston, Dominic Kwiatkowski, Cordelia Langford, John Sillitoe on behalf of the Wellcome Sanger Institute COVID-19 Surveillance Team |
| EPI_ISL_3358876 | Lighthouse Lab in Alderley Park | Wellcome Sanger Institute for the COVID-19 Genomics UK (COG-UK) | Jacquelyn Wynn, Mairead Hyland, The Lighthouse Lab in Alderley Park and Alex Alderton, Roberto Amato, Jeffrey Barrett, Sonia Goncalves, Ewan Harrison, David K. Jackson, Ian Johnston, Dominic Kwiatkowski, Cordelia Langford, John Sillitoe on |

| | | | |
|---|---|---|---|
| | | Consortium | behalf of the Wellcome Sanger Institute COVID-19 Surveillance Team |
| EPI_ISL_3358877 | Lighthouse Lab in Milton Keynes | Wellcome Sanger Institute for the COVID-19 Genomics UK (COG-UK) Consortium | The Lighthouse Lab in Milton Keynes and Alex Alderton, Roberto Amato, Jeffrey Barrett, Sonia Goncalves, Ewan Harrison, David K. Jackson, Ian Johnston, Dominic Kwiatkowski, Cordelia Langford, John Sillitoe on behalf of the Wellcome Sanger Institute COVID-19 Surveillance Team |
| EPI_ISL_3358878 | Lighthouse Lab in Alderley Park | Wellcome Sanger Institute for the COVID-19 Genomics UK (COG-UK) Consortium | Jacquelyn Wynn, Mairead Hyland, The Lighthouse Lab in Alderley Park and Alex Alderton, Roberto Amato, Jeffrey Barrett, Sonia Goncalves, Ewan Harrison, David K. Jackson, Ian Johnston, Dominic Kwiatkowski, Cordelia Langford, John Sillitoe on behalf of the Wellcome Sanger Institute COVID-19 Surveillance Team |
| EPI_ISL_3358879, EPI_ISL_3358880, EPI_ISL_3358881, EPI_ISL_3358882, EPI_ISL_3358883, EPI_ISL_3358884, EPI_ISL_3358885, EPI_ISL_3358886 | Lighthouse Lab in Milton Keynes | Wellcome Sanger Institute for the COVID-19 Genomics UK (COG-UK) Consortium | The Lighthouse Lab in Milton Keynes and Alex Alderton, Roberto Amato, Jeffrey Barrett, Sonia Goncalves, Ewan Harrison, David K. Jackson, Ian Johnston, Dominic Kwiatkowski, Cordelia Langford, John Sillitoe on behalf of the Wellcome Sanger Institute COVID-19 Surveillance Team |
| EPI_ISL_3358887, EPI_ISL_3358888 | Lighthouse Lab in Glasgow | Wellcome Sanger Institute for the COVID-19 Genomics UK (COG-UK) Consortium | Harper VanSteenhouse, Yumi Kasai, David Gray, Carol Clugston, Anna Dominiczak and Alex Alderton, Roberto Amato, Jeffrey Barrett, Sonia Goncalves, Ewan Harrison, David K. Jackson, Ian Johnston, Dominic Kwiatkowski, Cordelia Langford, John Sillitoe on behalf of the Wellcome Sanger Institute COVID-19 Surveillance Team |
| EPI_ISL_3358889, EPI_ISL_3358890 | Lighthouse Lab in Milton Keynes | Wellcome Sanger Institute for the COVID-19 Genomics UK (COG-UK) Consortium | The Lighthouse Lab in Milton Keynes and Alex Alderton, Roberto Amato, Jeffrey Barrett, Sonia Goncalves, Ewan Harrison, David K. Jackson, Ian Johnston, Dominic Kwiatkowski, Cordelia Langford, John Sillitoe on behalf of the Wellcome Sanger Institute COVID-19 Surveillance Team |
| EPI_ISL_3358891, EPI_ISL_3358892 | Lighthouse Lab in Glasgow | Wellcome Sanger Institute for the COVID-19 Genomics UK (COG-UK) Consortium | Harper VanSteenhouse, Yumi Kasai, David Gray, Carol Clugston, Anna Dominiczak and Alex Alderton, Roberto Amato, Jeffrey Barrett, Sonia Goncalves, Ewan Harrison, David K. Jackson, Ian Johnston, Dominic Kwiatkowski, Cordelia Langford, John Sillitoe on behalf of the Wellcome Sanger Institute COVID-19 Surveillance Team |
| EPI_ISL_3358893 | Lighthouse Lab in Alderley Park | Wellcome Sanger Institute for the COVID-19 Genomics UK (COG-UK) Consortium | Jacquelyn Wynn, Mairead Hyland, The Lighthouse Lab in Alderley Park and Alex Alderton, Roberto Amato, Jeffrey Barrett, Sonia Goncalves, Ewan Harrison, David K. Jackson, Ian Johnston, Dominic Kwiatkowski, Cordelia Langford, John Sillitoe on behalf of the Wellcome Sanger Institute COVID-19 Surveillance Team |
| EPI_ISL_3358894 | Lighthouse Lab in Milton Keynes | Wellcome Sanger Institute for the COVID-19 Genomics UK (COG-UK) Consortium | The Lighthouse Lab in Milton Keynes and Alex Alderton, Roberto Amato, Jeffrey Barrett, Sonia Goncalves, Ewan Harrison, David K. Jackson, Ian Johnston, Dominic Kwiatkowski, Cordelia Langford, John Sillitoe on behalf of the Wellcome Sanger Institute COVID-19 Surveillance Team |
| EPI_ISL_3358895 | Lighthouse Lab in Alderley Park | Wellcome Sanger Institute for the COVID-19 Genomics UK (COG-UK) Consortium | Jacquelyn Wynn, Mairead Hyland, The Lighthouse Lab in Alderley Park and Alex Alderton, Roberto Amato, Jeffrey Barrett, Sonia Goncalves, Ewan Harrison, David K. Jackson, Ian Johnston, Dominic Kwiatkowski, Cordelia Langford, John Sillitoe on behalf of the Wellcome Sanger Institute COVID-19 Surveillance Team |
| EPI_ISL_3358896, EPI_ISL_3358897 | Lighthouse Lab in Milton Keynes | Wellcome Sanger Institute for the COVID-19 Genomics UK (COG-UK) Consortium | The Lighthouse Lab in Milton Keynes and Alex Alderton, Roberto Amato, Jeffrey Barrett, Sonia Goncalves, Ewan Harrison, David K. Jackson, Ian Johnston, Dominic Kwiatkowski, Cordelia Langford, John Sillitoe on behalf of the Wellcome Sanger Institute COVID-19 Surveillance Team |
| EPI_ISL_3358898 | Lighthouse Lab in Alderley Park | Wellcome Sanger Institute for the COVID-19 Genomics UK (COG-UK) Consortium | Jacquelyn Wynn, Mairead Hyland, The Lighthouse Lab in Alderley Park and Alex Alderton, Roberto Amato, Jeffrey Barrett, Sonia Goncalves, Ewan Harrison, David K. Jackson, Ian Johnston, Dominic Kwiatkowski, Cordelia Langford, John Sillitoe on behalf of the Wellcome Sanger Institute COVID-19 Surveillance Team |
| EPI_ISL_3358899, EPI_ISL_3358900 | Lighthouse Lab in Milton Keynes | Wellcome Sanger Institute for the COVID-19 Genomics UK (COG-UK) Consortium | The Lighthouse Lab in Milton Keynes and Alex Alderton, Roberto Amato, Jeffrey Barrett, Sonia Goncalves, Ewan Harrison, David K. Jackson, Ian Johnston, Dominic Kwiatkowski, Cordelia Langford, John Sillitoe on behalf of the Wellcome Sanger Institute COVID-19 Surveillance Team |
| EPI_ISL_3358901, EPI_ISL_3358902, EPI_ISL_3358903, EPI_ISL_3358904, EPI_ISL_3358905 | Lighthouse Lab in Alderley Park | Wellcome Sanger Institute for the COVID-19 Genomics UK (COG-UK) Consortium | Jacquelyn Wynn, Mairead Hyland, The Lighthouse Lab in Alderley Park and Alex Alderton, Roberto Amato, Jeffrey Barrett, Sonia Goncalves, Ewan Harrison, David K. Jackson, Ian Johnston, Dominic Kwiatkowski, Cordelia Langford, John Sillitoe on behalf of the Wellcome Sanger Institute COVID-19 Surveillance Team |
| EPI_ISL_3358906, EPI_ISL_3358907 | Lighthouse Lab in Milton Keynes | Wellcome Sanger Institute for the COVID-19 Genomics UK (COG-UK) Consortium | The Lighthouse Lab in Milton Keynes and Alex Alderton, Roberto Amato, Jeffrey Barrett, Sonia Goncalves, Ewan Harrison, David K. Jackson, Ian Johnston, Dominic Kwiatkowski, Cordelia Langford, John Sillitoe on behalf of the Wellcome Sanger Institute COVID-19 Surveillance Team |
| EPI_ISL_3358908 | Lighthouse Lab in Alderley Park | Wellcome Sanger Institute for the COVID-19 Genomics UK (COG-UK) Consortium | Jacquelyn Wynn, Mairead Hyland, The Lighthouse Lab in Alderley Park and Alex Alderton, Roberto Amato, Jeffrey Barrett, Sonia Goncalves, Ewan Harrison, David K. Jackson, Ian Johnston, Dominic Kwiatkowski, Cordelia Langford, John Sillitoe on behalf of the Wellcome Sanger Institute COVID-19 Surveillance Team |
| EPI_ISL_3358909, EPI_ISL_3358910, EPI_ISL_3358911 | Lighthouse Lab in Milton Keynes | Wellcome Sanger Institute for the COVID-19 Genomics UK (COG-UK) Consortium | The Lighthouse Lab in Milton Keynes and Alex Alderton, Roberto Amato, Jeffrey Barrett, Sonia Goncalves, Ewan Harrison, David K. Jackson, Ian Johnston, Dominic Kwiatkowski, Cordelia Langford, John Sillitoe on behalf of the Wellcome Sanger Institute COVID-19 Surveillance Team |
| EPI_ISL_3358912 | Berkshire and Surrey Pathology Services Lighthouse Laboratory | Wellcome Sanger Institute for the COVID-19 Genomics UK (COG-UK) Consortium | Berkshire and Surrey Pathology Services Lighthouse Laboratory and Alex Alderton, Roberto Amato, Jeffrey Barrett, Sonia Goncalves, Ewan Harrison, David K. Jackson, Ian Johnston, Dominic Kwiatkowski, Cordelia Langford, John Sillitoe on behalf of the Wellcome Sanger Institute COVID-19 Surveillance Team |
| EPI_ISL_3358913 | Lighthouse Lab in Alderley Park | Wellcome Sanger Institute for the COVID-19 Genomics UK (COG-UK) Consortium | Jacquelyn Wynn, Mairead Hyland, The Lighthouse Lab in Alderley Park and Alex Alderton, Roberto Amato, Jeffrey Barrett, Sonia Goncalves, Ewan Harrison, David K. Jackson, Ian Johnston, Dominic Kwiatkowski, Cordelia Langford, John Sillitoe on behalf of the Wellcome Sanger Institute COVID-19 Surveillance Team |
| EPI_ISL_3358914 | Berkshire and Surrey Pathology Services Lighthouse Laboratory | Wellcome Sanger Institute for the COVID-19 Genomics UK (COG-UK) Consortium | Berkshire and Surrey Pathology Services Lighthouse Laboratory and Alex Alderton, Roberto Amato, Jeffrey Barrett, Sonia Goncalves, Ewan Harrison, David K. Jackson, Ian Johnston, Dominic Kwiatkowski, Cordelia Langford, John Sillitoe on behalf of the Wellcome Sanger Institute COVID-19 Surveillance Team |
| EPI_ISL_3358915, EPI_ISL_3358916, EPI_ISL_3358917 | Lighthouse Lab in Alderley Park | Wellcome Sanger Institute for the COVID-19 Genomics UK (COG-UK) Consortium | Jacquelyn Wynn, Mairead Hyland, The Lighthouse Lab in Alderley Park and Alex Alderton, Roberto Amato, Jeffrey Barrett, Sonia Goncalves, Ewan Harrison, David K. Jackson, Ian Johnston, Dominic Kwiatkowski, Cordelia Langford, John Sillitoe on behalf of the Wellcome Sanger Institute COVID-19 Surveillance Team |
| EPI_ISL_3358918 | Lighthouse Lab in Milton Keynes | Wellcome Sanger Institute for the COVID-19 Genomics UK (COG-UK) Consortium | The Lighthouse Lab in Milton Keynes and Alex Alderton, Roberto Amato, Jeffrey Barrett, Sonia Goncalves, Ewan Harrison, David K. Jackson, Ian Johnston, Dominic Kwiatkowski, Cordelia Langford, John Sillitoe on behalf of the Wellcome Sanger Institute COVID-19 Surveillance Team |
| EPI_ISL_3358919, EPI_ISL_3358920, EPI_ISL_3358921 | Lighthouse Lab in Alderley Park | Wellcome Sanger Institute for the COVID-19 Genomics UK (COG-UK) Consortium | Jacquelyn Wynn, Mairead Hyland, The Lighthouse Lab in Alderley Park and Alex Alderton, Roberto Amato, Jeffrey Barrett, Sonia Goncalves, Ewan Harrison, David K. Jackson, Ian Johnston, Dominic Kwiatkowski, Cordelia Langford, John Sillitoe on behalf of the Wellcome Sanger Institute COVID-19 Surveillance Team |
| EPI_ISL_3358922, EPI_ISL_3358923, EPI_ISL_3358924, EPI_ISL_3358925, EPI_ISL_3358926, EPI_ISL_3358927, EPI_ISL_3358928 | Lighthouse Lab in Milton Keynes | Wellcome Sanger Institute for the COVID-19 Genomics UK (COG-UK) Consortium | The Lighthouse Lab in Milton Keynes and Alex Alderton, Roberto Amato, Jeffrey Barrett, Sonia Goncalves, Ewan Harrison, David K. Jackson, Ian Johnston, Dominic Kwiatkowski, Cordelia Langford, John Sillitoe on behalf of the Wellcome Sanger Institute COVID-19 Surveillance Team |
| EPI_ISL_3358929 | Lighthouse Lab in Alderley Park | Wellcome Sanger Institute for the COVID-19 Genomics UK (COG-UK) Consortium | Jacquelyn Wynn, Mairead Hyland, The Lighthouse Lab in Alderley Park and Alex Alderton, Roberto Amato, Jeffrey Barrett, Sonia Goncalves, Ewan Harrison, David K. Jackson, Ian Johnston, Dominic Kwiatkowski, Cordelia Langford, John Sillitoe on behalf of the Wellcome Sanger Institute COVID-19 Surveillance Team |
| EPI_ISL_3358930, EPI_ISL_3358931, EPI_ISL_3358932, EPI_ISL_3358933 | Lighthouse Lab in Milton Keynes | Wellcome Sanger Institute for the COVID-19 Genomics UK (COG-UK) Consortium | The Lighthouse Lab in Milton Keynes and Alex Alderton, Roberto Amato, Jeffrey Barrett, Sonia Goncalves, Ewan Harrison, David K. Jackson, Ian Johnston, Dominic Kwiatkowski, Cordelia Langford, John Sillitoe on behalf of the Wellcome Sanger Institute COVID-19 Surveillance Team |
| EPI_ISL_3358934, EPI_ISL_3358935, EPI_ISL_3358936 | Lighthouse Lab in Alderley Park | Wellcome Sanger Institute for the COVID-19 Genomics UK (COG-UK) Consortium | Jacquelyn Wynn, Mairead Hyland, The Lighthouse Lab in Alderley Park and Alex Alderton, Roberto Amato, Jeffrey Barrett, Sonia Goncalves, Ewan Harrison, David K. Jackson, Ian Johnston, Dominic Kwiatkowski, Cordelia Langford, John Sillitoe on behalf of the Wellcome Sanger Institute COVID-19 Surveillance Team |
| EPI_ISL_3358937 | Lighthouse Lab in Milton Keynes | Wellcome Sanger Institute for the COVID-19 Genomics UK (COG-UK) Consortium | The Lighthouse Lab in Milton Keynes and Alex Alderton, Roberto Amato, Jeffrey Barrett, Sonia Goncalves, Ewan Harrison, David K. Jackson, Ian Johnston, Dominic Kwiatkowski, Cordelia Langford, John Sillitoe on behalf of the Wellcome Sanger Institute COVID-19 Surveillance Team |
| EPI_ISL_3358938 | Lighthouse Lab in Alderley Park | Wellcome Sanger Institute for the COVID-19 Genomics UK (COG-UK) Consortium | Jacquelyn Wynn, Mairead Hyland, The Lighthouse Lab in Alderley Park and Alex Alderton, Roberto Amato, Jeffrey Barrett, Sonia Goncalves, Ewan Harrison, David K. Jackson, Ian Johnston, Dominic Kwiatkowski, Cordelia Langford, John Sillitoe on behalf of the Wellcome Sanger Institute COVID-19 Surveillance Team |
| EPI_ISL_3358939, EPI_ISL_3358940, EPI_ISL_3358941 | Lighthouse Lab in Milton Keynes | Wellcome Sanger Institute for the COVID-19 Genomics UK (COG-UK) Consortium | The Lighthouse Lab in Milton Keynes and Alex Alderton, Roberto Amato, Jeffrey Barrett, Sonia Goncalves, Ewan Harrison, David K. Jackson, Ian Johnston, Dominic Kwiatkowski, Cordelia Langford, John Sillitoe on behalf of the Wellcome Sanger Institute COVID-19 Surveillance Team |
| EPI_ISL_3358942 | Lighthouse Lab in Alderley Park | Wellcome Sanger Institute for the COVID-19 Genomics UK (COG-UK) Consortium | Jacquelyn Wynn, Mairead Hyland, The Lighthouse Lab in Alderley Park and Alex Alderton, Roberto Amato, Jeffrey Barrett, Sonia Goncalves, Ewan Harrison, David K. Jackson, Ian Johnston, Dominic Kwiatkowski, Cordelia Langford, John Sillitoe on behalf of the Wellcome Sanger Institute COVID-19 Surveillance Team |
| EPI_ISL_3358943 | Lighthouse Lab in Milton Keynes | Wellcome Sanger Institute for the COVID-19 Genomics UK (COG-UK) Consortium | The Lighthouse Lab in Milton Keynes and Alex Alderton, Roberto Amato, Jeffrey Barrett, Sonia Goncalves, Ewan Harrison, David K. Jackson, Ian Johnston, Dominic Kwiatkowski, Cordelia Langford, John Sillitoe on behalf of the Wellcome Sanger Institute COVID-19 Surveillance Team |
| EPI_ISL_3358944 | Berkshire and Surrey Pathology Services Lighthouse Laboratory | Wellcome Sanger Institute for the COVID-19 Genomics UK (COG-UK) Consortium | Berkshire and Surrey Pathology Services Lighthouse Laboratory and Alex Alderton, Roberto Amato, Jeffrey Barrett, Sonia Goncalves, Ewan Harrison, David K. Jackson, Ian Johnston, Dominic Kwiatkowski, Cordelia Langford, John Sillitoe on behalf of the Wellcome Sanger Institute COVID-19 Surveillance Team |
| EPI_ISL_3358945 | Lighthouse Lab in Milton Keynes | Wellcome Sanger Institute for the COVID-19 Genomics UK (COG-UK) Consortium | The Lighthouse Lab in Milton Keynes and Alex Alderton, Roberto Amato, Jeffrey Barrett, Sonia Goncalves, Ewan Harrison, David K. Jackson, Ian Johnston, Dominic Kwiatkowski, Cordelia Langford, John Sillitoe on behalf of the Wellcome Sanger Institute COVID-19 Surveillance Team |
| EPI_ISL_3358946 | Lighthouse Lab in Alderley Park | Wellcome Sanger Institute for the COVID-19 Genomics UK (COG-UK) Consortium | Jacquelyn Wynn, Mairead Hyland, The Lighthouse Lab in Alderley Park and Alex Alderton, Roberto Amato, Jeffrey Barrett, Sonia Goncalves, Ewan Harrison, David K. Jackson, Ian Johnston, Dominic Kwiatkowski, Cordelia Langford, John Sillitoe on behalf of the Wellcome Sanger Institute COVID-19 Surveillance Team |
| EPI_ISL_3358947, EPI_ISL_3358948 | Lighthouse Lab in Milton Keynes | Wellcome Sanger Institute for the COVID-19 Genomics UK (COG-UK) Consortium | The Lighthouse Lab in Milton Keynes and Alex Alderton, Roberto Amato, Jeffrey Barrett, Sonia Goncalves, Ewan Harrison, David K. Jackson, Ian Johnston, Dominic Kwiatkowski, Cordelia Langford, John Sillitoe on behalf of the Wellcome Sanger Institute COVID-19 Surveillance Team |
| EPI_ISL_3358949 | Lighthouse Lab in Alderley Park | Wellcome Sanger Institute for the COVID-19 Genomics UK (COG-UK) Consortium | Jacquelyn Wynn, Mairead Hyland, The Lighthouse Lab in Alderley Park and Alex Alderton, Roberto Amato, Jeffrey Barrett, Sonia Goncalves, Ewan Harrison, David K. Jackson, Ian Johnston, Dominic Kwiatkowski, Cordelia Langford, John Sillitoe on behalf of the Wellcome Sanger Institute COVID-19 Surveillance Team |

| | | | |
|---|---|---|---|
| EPI_ISL_3358950 | Lighthouse Lab in Milton Keynes | Wellcome Sanger Institute for the COVID-19 Genomics UK (COG-UK) Consortium | The Lighthouse Lab in Milton Keynes and Alex Alderton, Roberto Amato, Jeffrey Barrett, Sonia Goncalves, Ewan Harrison, David K. Jackson, Ian Johnston, Dominic Kwiatkowski, Cordelia Langford, John Sillitoe on behalf of the Wellcome Sanger Institute COVID-19 Surveillance Team |
| EPI_ISL_3358951, EPI_ISL_3358952 | Lighthouse Lab in Alderley Park | Wellcome Sanger Institute for the COVID-19 Genomics UK (COG-UK) Consortium | Jacquelyn Wynn, Mairead Hyland, The Lighthouse Lab in Alderley Park and Alex Alderton, Roberto Amato, Jeffrey Barrett, Sonia Goncalves, Ewan Harrison, David K. Jackson, Ian Johnston, Dominic Kwiatkowski, Cordelia Langford, John Sillitoe on behalf of the Wellcome Sanger Institute COVID-19 Surveillance Team |
| EPI_ISL_3358953 | Lighthouse Lab in Milton Keynes | Wellcome Sanger Institute for the COVID-19 Genomics UK (COG-UK) Consortium | The Lighthouse Lab in Milton Keynes and Alex Alderton, Roberto Amato, Jeffrey Barrett, Sonia Goncalves, Ewan Harrison, David K. Jackson, Ian Johnston, Dominic Kwiatkowski, Cordelia Langford, John Sillitoe on behalf of the Wellcome Sanger Institute COVID-19 Surveillance Team |
| EPI_ISL_3358954, EPI_ISL_3358955, EPI_ISL_3358956, EPI_ISL_3358957 | Lighthouse Lab in Alderley Park | Wellcome Sanger Institute for the COVID-19 Genomics UK (COG-UK) Consortium | Jacquelyn Wynn, Mairead Hyland, The Lighthouse Lab in Alderley Park and Alex Alderton, Roberto Amato, Jeffrey Barrett, Sonia Goncalves, Ewan Harrison, David K. Jackson, Ian Johnston, Dominic Kwiatkowski, Cordelia Langford, John Sillitoe on behalf of the Wellcome Sanger Institute COVID-19 Surveillance Team |
| EPI_ISL_3358958, EPI_ISL_3358959 | Lighthouse Lab in Milton Keynes | Wellcome Sanger Institute for the COVID-19 Genomics UK (COG-UK) Consortium | The Lighthouse Lab in Milton Keynes and Alex Alderton, Roberto Amato, Jeffrey Barrett, Sonia Goncalves, Ewan Harrison, David K. Jackson, Ian Johnston, Dominic Kwiatkowski, Cordelia Langford, John Sillitoe on behalf of the Wellcome Sanger Institute COVID-19 Surveillance Team |
| EPI_ISL_3358960, EPI_ISL_3358961, EPI_ISL_3358962, EPI_ISL_3358963, EPI_ISL_3358964 | Lighthouse Lab in Alderley Park | Wellcome Sanger Institute for the COVID-19 Genomics UK (COG-UK) Consortium | Jacquelyn Wynn, Mairead Hyland, The Lighthouse Lab in Alderley Park and Alex Alderton, Roberto Amato, Jeffrey Barrett, Sonia Goncalves, Ewan Harrison, David K. Jackson, Ian Johnston, Dominic Kwiatkowski, Cordelia Langford, John Sillitoe on behalf of the Wellcome Sanger Institute COVID-19 Surveillance Team |
| EPI_ISL_3358965, EPI_ISL_3358966, EPI_ISL_3358967, EPI_ISL_3358968, EPI_ISL_3358969 | Lighthouse Lab in Milton Keynes | Wellcome Sanger Institute for the COVID-19 Genomics UK (COG-UK) Consortium | The Lighthouse Lab in Milton Keynes and Alex Alderton, Roberto Amato, Jeffrey Barrett, Sonia Goncalves, Ewan Harrison, David K. Jackson, Ian Johnston, Dominic Kwiatkowski, Cordelia Langford, John Sillitoe on behalf of the Wellcome Sanger Institute COVID-19 Surveillance Team |
| EPI_ISL_3358970, EPI_ISL_3358971 | Lighthouse Lab in Alderley Park | Wellcome Sanger Institute for the COVID-19 Genomics UK (COG-UK) Consortium | Jacquelyn Wynn, Mairead Hyland, The Lighthouse Lab in Alderley Park and Alex Alderton, Roberto Amato, Jeffrey Barrett, Sonia Goncalves, Ewan Harrison, David K. Jackson, Ian Johnston, Dominic Kwiatkowski, Cordelia Langford, John Sillitoe on behalf of the Wellcome Sanger Institute COVID-19 Surveillance Team |
| EPI_ISL_3358973, EPI_ISL_3358974, EPI_ISL_3358975 | Lighthouse Lab in Milton Keynes | Wellcome Sanger Institute for the COVID-19 Genomics UK (COG-UK) Consortium | The Lighthouse Lab in Milton Keynes and Alex Alderton, Roberto Amato, Jeffrey Barrett, Sonia Goncalves, Ewan Harrison, David K. Jackson, Ian Johnston, Dominic Kwiatkowski, Cordelia Langford, John Sillitoe on behalf of the Wellcome Sanger Institute COVID-19 Surveillance Team |
| EPI_ISL_3358976, EPI_ISL_3358978, EPI_ISL_3358979 | Lighthouse Lab in Alderley Park | Wellcome Sanger Institute for the COVID-19 Genomics UK (COG-UK) Consortium | Jacquelyn Wynn, Mairead Hyland, The Lighthouse Lab in Alderley Park and Alex Alderton, Roberto Amato, Jeffrey Barrett, Sonia Goncalves, Ewan Harrison, David K. Jackson, Ian Johnston, Dominic Kwiatkowski, Cordelia Langford, John Sillitoe on behalf of the Wellcome Sanger Institute COVID-19 Surveillance Team |
| EPI_ISL_3358981 | Lighthouse Lab in Milton Keynes | Wellcome Sanger Institute for the COVID-19 Genomics UK (COG-UK) Consortium | The Lighthouse Lab in Milton Keynes and Alex Alderton, Roberto Amato, Jeffrey Barrett, Sonia Goncalves, Ewan Harrison, David K. Jackson, Ian Johnston, Dominic Kwiatkowski, Cordelia Langford, John Sillitoe on behalf of the Wellcome Sanger Institute COVID-19 Surveillance Team |
| EPI_ISL_3358982 | Lighthouse Lab in Alderley Park | Wellcome Sanger Institute for the COVID-19 Genomics UK (COG-UK) Consortium | Jacquelyn Wynn, Mairead Hyland, The Lighthouse Lab in Alderley Park and Alex Alderton, Roberto Amato, Jeffrey Barrett, Sonia Goncalves, Ewan Harrison, David K. Jackson, Ian Johnston, Dominic Kwiatkowski, Cordelia Langford, John Sillitoe on behalf of the Wellcome Sanger Institute COVID-19 Surveillance Team |
| EPI_ISL_3358983 | Lighthouse Lab in Milton Keynes | Wellcome Sanger Institute for the COVID-19 Genomics UK (COG-UK) Consortium | The Lighthouse Lab in Milton Keynes and Alex Alderton, Roberto Amato, Jeffrey Barrett, Sonia Goncalves, Ewan Harrison, David K. Jackson, Ian Johnston, Dominic Kwiatkowski, Cordelia Langford, John Sillitoe on behalf of the Wellcome Sanger Institute COVID-19 Surveillance Team |
| EPI_ISL_3358985 | Berkshire and Surrey Pathology Services Lighthouse Laboratory | Wellcome Sanger Institute for the COVID-19 Genomics UK (COG-UK) Consortium | Berkshire and Surrey Pathology Services Lighthouse Laboratory and Alex Alderton, Roberto Amato, Jeffrey Barrett, Sonia Goncalves, Ewan Harrison, David K. Jackson, Ian Johnston, Dominic Kwiatkowski, Cordelia Langford, John Sillitoe on behalf of the Wellcome Sanger Institute COVID-19 Surveillance Team |
| EPI_ISL_3358986, EPI_ISL_3358988 | Lighthouse Lab in Alderley Park | Wellcome Sanger Institute for the COVID-19 Genomics UK (COG-UK) Consortium | Jacquelyn Wynn, Mairead Hyland, The Lighthouse Lab in Alderley Park and Alex Alderton, Roberto Amato, Jeffrey Barrett, Sonia Goncalves, Ewan Harrison, David K. Jackson, Ian Johnston, Dominic Kwiatkowski, Cordelia Langford, John Sillitoe on behalf of the Wellcome Sanger Institute COVID-19 Surveillance Team |
| EPI_ISL_3358989 | Lighthouse Lab in Milton Keynes | Wellcome Sanger Institute for the COVID-19 Genomics UK (COG-UK) Consortium | The Lighthouse Lab in Milton Keynes and Alex Alderton, Roberto Amato, Jeffrey Barrett, Sonia Goncalves, Ewan Harrison, David K. Jackson, Ian Johnston, Dominic Kwiatkowski, Cordelia Langford, John Sillitoe on behalf of the Wellcome Sanger Institute COVID-19 Surveillance Team |
| EPI_ISL_3358990 | Lighthouse Lab in Alderley Park | Wellcome Sanger Institute for the COVID-19 Genomics UK (COG-UK) Consortium | Jacquelyn Wynn, Mairead Hyland, The Lighthouse Lab in Alderley Park and Alex Alderton, Roberto Amato, Jeffrey Barrett, Sonia Goncalves, Ewan Harrison, David K. Jackson, Ian Johnston, Dominic Kwiatkowski, Cordelia Langford, John Sillitoe on behalf of the Wellcome Sanger Institute COVID-19 Surveillance Team |
| EPI_ISL_3358992 | Lighthouse Lab in Milton Keynes | Wellcome Sanger Institute for the COVID-19 Genomics UK (COG-UK) Consortium | The Lighthouse Lab in Milton Keynes and Alex Alderton, Roberto Amato, Jeffrey Barrett, Sonia Goncalves, Ewan Harrison, David K. Jackson, Ian Johnston, Dominic Kwiatkowski, Cordelia Langford, John Sillitoe on behalf of the Wellcome Sanger Institute COVID-19 Surveillance Team |
| EPI_ISL_3358993 | Lighthouse Lab in Alderley Park | Wellcome Sanger Institute for the COVID-19 Genomics UK (COG-UK) Consortium | Jacquelyn Wynn, Mairead Hyland, The Lighthouse Lab in Alderley Park and Alex Alderton, Roberto Amato, Jeffrey Barrett, Sonia Goncalves, Ewan Harrison, David K. Jackson, Ian Johnston, Dominic Kwiatkowski, Cordelia Langford, John Sillitoe on behalf of the Wellcome Sanger Institute COVID-19 Surveillance Team |
| EPI_ISL_3358995 | Lighthouse Lab in Milton Keynes | Wellcome Sanger Institute for the COVID-19 Genomics UK (COG-UK) Consortium | The Lighthouse Lab in Milton Keynes and Alex Alderton, Roberto Amato, Jeffrey Barrett, Sonia Goncalves, Ewan Harrison, David K. Jackson, Ian Johnston, Dominic Kwiatkowski, Cordelia Langford, John Sillitoe on behalf of the Wellcome Sanger Institute COVID-19 Surveillance Team |
| EPI_ISL_3358996, EPI_ISL_3358998 | Lighthouse Lab in Alderley Park | Wellcome Sanger Institute for the COVID-19 Genomics UK (COG-UK) Consortium | Jacquelyn Wynn, Mairead Hyland, The Lighthouse Lab in Alderley Park and Alex Alderton, Roberto Amato, Jeffrey Barrett, Sonia Goncalves, Ewan Harrison, David K. Jackson, Ian Johnston, Dominic Kwiatkowski, Cordelia Langford, John Sillitoe on behalf of the Wellcome Sanger Institute COVID-19 Surveillance Team |
| EPI_ISL_3358999, EPI_ISL_3359001 | Lighthouse Lab in Milton Keynes | Wellcome Sanger Institute for the COVID-19 Genomics UK (COG-UK) Consortium | The Lighthouse Lab in Milton Keynes and Alex Alderton, Roberto Amato, Jeffrey Barrett, Sonia Goncalves, Ewan Harrison, David K. Jackson, Ian Johnston, Dominic Kwiatkowski, Cordelia Langford, John Sillitoe on behalf of the Wellcome Sanger Institute COVID-19 Surveillance Team |
| EPI_ISL_3359002 | Lighthouse Lab in Alderley Park | Wellcome Sanger Institute for the COVID-19 Genomics UK (COG-UK) Consortium | Jacquelyn Wynn, Mairead Hyland, The Lighthouse Lab in Alderley Park and Alex Alderton, Roberto Amato, Jeffrey Barrett, Sonia Goncalves, Ewan Harrison, David K. Jackson, Ian Johnston, Dominic Kwiatkowski, Cordelia Langford, John Sillitoe on behalf of the Wellcome Sanger Institute COVID-19 Surveillance Team |
| EPI_ISL_3359003 | Lighthouse Lab in Milton Keynes | Wellcome Sanger Institute for the COVID-19 Genomics UK (COG-UK) Consortium | The Lighthouse Lab in Milton Keynes and Alex Alderton, Roberto Amato, Jeffrey Barrett, Sonia Goncalves, Ewan Harrison, David K. Jackson, Ian Johnston, Dominic Kwiatkowski, Cordelia Langford, John Sillitoe on behalf of the Wellcome Sanger Institute COVID-19 Surveillance Team |
| EPI_ISL_3359005 | Lighthouse Lab in Alderley Park | Wellcome Sanger Institute for the COVID-19 Genomics UK (COG-UK) Consortium | Jacquelyn Wynn, Mairead Hyland, The Lighthouse Lab in Alderley Park and Alex Alderton, Roberto Amato, Jeffrey Barrett, Sonia Goncalves, Ewan Harrison, David K. Jackson, Ian Johnston, Dominic Kwiatkowski, Cordelia Langford, John Sillitoe on behalf of the Wellcome Sanger Institute COVID-19 Surveillance Team |
| EPI_ISL_3359006 | Lighthouse Lab in Milton Keynes | Wellcome Sanger Institute for the COVID-19 Genomics UK (COG-UK) Consortium | The Lighthouse Lab in Milton Keynes and Alex Alderton, Roberto Amato, Jeffrey Barrett, Sonia Goncalves, Ewan Harrison, David K. Jackson, Ian Johnston, Dominic Kwiatkowski, Cordelia Langford, John Sillitoe on behalf of the Wellcome Sanger Institute COVID-19 Surveillance Team |
| EPI_ISL_3359007, EPI_ISL_3359009, EPI_ISL_3359010, EPI_ISL_3359011, EPI_ISL_3359013 | Lighthouse Lab in Alderley Park | Wellcome Sanger Institute for the COVID-19 Genomics UK (COG-UK) Consortium | Jacquelyn Wynn, Mairead Hyland, The Lighthouse Lab in Alderley Park and Alex Alderton, Roberto Amato, Jeffrey Barrett, Sonia Goncalves, Ewan Harrison, David K. Jackson, Ian Johnston, Dominic Kwiatkowski, Cordelia Langford, John Sillitoe on behalf of the Wellcome Sanger Institute COVID-19 Surveillance Team |
| EPI_ISL_3359015, EPI_ISL_3359016 | Lighthouse Lab in Milton Keynes | Wellcome Sanger Institute for the COVID-19 Genomics UK (COG-UK) Consortium | The Lighthouse Lab in Milton Keynes and Alex Alderton, Roberto Amato, Jeffrey Barrett, Sonia Goncalves, Ewan Harrison, David K. Jackson, Ian Johnston, Dominic Kwiatkowski, Cordelia Langford, John Sillitoe on behalf of the Wellcome Sanger Institute COVID-19 Surveillance Team |
| EPI_ISL_3359017, EPI_ISL_3359019 | Lighthouse Lab in Alderley Park | Wellcome Sanger Institute for the COVID-19 Genomics UK (COG-UK) Consortium | Jacquelyn Wynn, Mairead Hyland, The Lighthouse Lab in Alderley Park and Alex Alderton, Roberto Amato, Jeffrey Barrett, Sonia Goncalves, Ewan Harrison, David K. Jackson, Ian Johnston, Dominic Kwiatkowski, Cordelia Langford, John Sillitoe on behalf of the Wellcome Sanger Institute COVID-19 Surveillance Team |
| EPI_ISL_3359020 | Lighthouse Lab in Milton Keynes | Wellcome Sanger Institute for the COVID-19 Genomics UK (COG-UK) Consortium | The Lighthouse Lab in Milton Keynes and Alex Alderton, Roberto Amato, Jeffrey Barrett, Sonia Goncalves, Ewan Harrison, David K. Jackson, Ian Johnston, Dominic Kwiatkowski, Cordelia Langford, John Sillitoe on behalf of the Wellcome Sanger Institute COVID-19 Surveillance Team |
| EPI_ISL_3359022 | Lighthouse Lab in Alderley Park | Wellcome Sanger Institute for the COVID-19 Genomics UK (COG-UK) Consortium | Jacquelyn Wynn, Mairead Hyland, The Lighthouse Lab in Alderley Park and Alex Alderton, Roberto Amato, Jeffrey Barrett, Sonia Goncalves, Ewan Harrison, David K. Jackson, Ian Johnston, Dominic Kwiatkowski, Cordelia Langford, John Sillitoe on behalf of the Wellcome Sanger Institute COVID-19 Surveillance Team |
| EPI_ISL_3359023, EPI_ISL_3359024 | Lighthouse Lab in Milton Keynes | Wellcome Sanger Institute for the COVID-19 Genomics UK (COG-UK) Consortium | The Lighthouse Lab in Milton Keynes and Alex Alderton, Roberto Amato, Jeffrey Barrett, Sonia Goncalves, Ewan Harrison, David K. Jackson, Ian Johnston, Dominic Kwiatkowski, Cordelia Langford, John Sillitoe on behalf of the Wellcome Sanger Institute COVID-19 Surveillance Team |
| EPI_ISL_3359026 | Lighthouse Lab in Alderley Park | Wellcome Sanger Institute for the COVID-19 Genomics UK (COG-UK) Consortium | Jacquelyn Wynn, Mairead Hyland, The Lighthouse Lab in Alderley Park and Alex Alderton, Roberto Amato, Jeffrey Barrett, Sonia Goncalves, Ewan Harrison, David K. Jackson, Ian Johnston, Dominic Kwiatkowski, Cordelia Langford, John Sillitoe on behalf of the Wellcome Sanger Institute COVID-19 Surveillance Team |
| EPI_ISL_3359027 | Berkshire and Surrey Pathology Services Lighthouse Laboratory | Wellcome Sanger Institute for the COVID-19 Genomics UK (COG-UK) Consortium | Berkshire and Surrey Pathology Services Lighthouse Laboratory and Alex Alderton, Roberto Amato, Jeffrey Barrett, Sonia Goncalves, Ewan Harrison, David K. Jackson, Ian Johnston, Dominic Kwiatkowski, Cordelia Langford, John Sillitoe on behalf of the Wellcome Sanger Institute COVID-19 Surveillance Team |
| EPI_ISL_3359029 | Lighthouse Lab in Milton Keynes | Wellcome Sanger Institute for the COVID-19 Genomics UK (COG-UK) Consortium | The Lighthouse Lab in Milton Keynes and Alex Alderton, Roberto Amato, Jeffrey Barrett, Sonia Goncalves, Ewan Harrison, David K. Jackson, Ian Johnston, Dominic Kwiatkowski, Cordelia Langford, John Sillitoe on behalf of the Wellcome Sanger Institute COVID-19 Surveillance Team |
| EPI_ISL_3359030 | Lighthouse Lab in Alderley Park | Wellcome Sanger Institute for the COVID-19 Genomics UK (COG-UK) Consortium | Jacquelyn Wynn, Mairead Hyland, The Lighthouse Lab in Alderley Park and Alex Alderton, Roberto Amato, Jeffrey Barrett, Sonia Goncalves, Ewan Harrison, David K. Jackson, Ian Johnston, Dominic Kwiatkowski, Cordelia Langford, John Sillitoe on behalf of the Wellcome Sanger Institute COVID-19 Surveillance Team |
| EPI_ISL_3359031, EPI_ISL_3359033, EPI_ISL_3359035 | Lighthouse Lab in Milton Keynes | Wellcome Sanger Institute for the COVID-19 Genomics UK (COG-UK) Consortium | The Lighthouse Lab in Milton Keynes and Alex Alderton, Roberto Amato, Jeffrey Barrett, Sonia Goncalves, Ewan Harrison, David K. Jackson, Ian Johnston, Dominic Kwiatkowski, Cordelia Langford, John Sillitoe on behalf of the Wellcome Sanger Institute COVID-19 Surveillance Team |
| EPI_ISL_3359036, EPI_ISL_3359038 | Lighthouse Lab in Alderley Park | Wellcome Sanger Institute for the COVID-19 Genomics UK (COG-UK) Consortium | Jacquelyn Wynn, Mairead Hyland, The Lighthouse Lab in Alderley Park and Alex Alderton, Roberto Amato, Jeffrey Barrett, Sonia Goncalves, Ewan Harrison, David K. Jackson, Ian Johnston, Dominic Kwiatkowski, Cordelia Langford, John Sillitoe on behalf of the Wellcome Sanger Institute COVID-19 Surveillance Team |
| EPI_ISL_3359039, EPI_ISL_3359041, EPI_ISL_3359042 | Lighthouse Lab in Milton Keynes | Wellcome Sanger Institute for the COVID-19 Genomics UK (COG-UK) Consortium | The Lighthouse Lab in Milton Keynes and Alex Alderton, Roberto Amato, Jeffrey Barrett, Sonia Goncalves, Ewan Harrison, David K. Jackson, Ian Johnston, Dominic Kwiatkowski, Cordelia Langford, John Sillitoe on behalf of the Wellcome Sanger Institute COVID-19 Surveillance Team |

| | | | |
|---|---|---|---|
| EPI_ISL_3359043, EPI_ISL_3359045 | Lighthouse Lab in Alderley Park | Wellcome Sanger Institute for the COVID-19 Genomics UK (COG-UK) Consortium | Jacquelyn Wynn, Mairead Hyland, The Lighthouse Lab in Alderley Park and Alex Alderton, Roberto Amato, Jeffrey Barrett, Sonia Goncalves, Ewan Harrison, David K. Jackson, Ian Johnston, Dominic Kwiatkowski, Cordelia Langford, John Sillitoe on behalf of the Wellcome Sanger Institute COVID-19 Surveillance Team |
| EPI_ISL_3359046 | Lighthouse Lab in Milton Keynes | Wellcome Sanger Institute for the COVID-19 Genomics UK (COG-UK) Consortium | The Lighthouse Lab in Milton Keynes and Alex Alderton, Roberto Amato, Jeffrey Barrett, Sonia Goncalves, Ewan Harrison, David K. Jackson, Ian Johnston, Dominic Kwiatkowski, Cordelia Langford, John Sillitoe on behalf of the Wellcome Sanger Institute COVID-19 Surveillance Team |
| EPI_ISL_3359048, EPI_ISL_3359049 | Lighthouse Lab in Alderley Park | Wellcome Sanger Institute for the COVID-19 Genomics UK (COG-UK) Consortium | Jacquelyn Wynn, Mairead Hyland, The Lighthouse Lab in Alderley Park and Alex Alderton, Roberto Amato, Jeffrey Barrett, Sonia Goncalves, Ewan Harrison, David K. Jackson, Ian Johnston, Dominic Kwiatkowski, Cordelia Langford, John Sillitoe on behalf of the Wellcome Sanger Institute COVID-19 Surveillance Team |
| EPI_ISL_3359051, EPI_ISL_3359052 | Lighthouse Lab in Milton Keynes | Wellcome Sanger Institute for the COVID-19 Genomics UK (COG-UK) Consortium | The Lighthouse Lab in Milton Keynes and Alex Alderton, Roberto Amato, Jeffrey Barrett, Sonia Goncalves, Ewan Harrison, David K. Jackson, Ian Johnston, Dominic Kwiatkowski, Cordelia Langford, John Sillitoe on behalf of the Wellcome Sanger Institute COVID-19 Surveillance Team |
| EPI_ISL_3359054 | Lighthouse Lab in Alderley Park | Wellcome Sanger Institute for the COVID-19 Genomics UK (COG-UK) Consortium | Jacquelyn Wynn, Mairead Hyland, The Lighthouse Lab in Alderley Park and Alex Alderton, Roberto Amato, Jeffrey Barrett, Sonia Goncalves, Ewan Harrison, David K. Jackson, Ian Johnston, Dominic Kwiatkowski, Cordelia Langford, John Sillitoe on behalf of the Wellcome Sanger Institute COVID-19 Surveillance Team |
| EPI_ISL_3359055, EPI_ISL_3359057 | Lighthouse Lab in Milton Keynes | Wellcome Sanger Institute for the COVID-19 Genomics UK (COG-UK) Consortium | The Lighthouse Lab in Milton Keynes and Alex Alderton, Roberto Amato, Jeffrey Barrett, Sonia Goncalves, Ewan Harrison, David K. Jackson, Ian Johnston, Dominic Kwiatkowski, Cordelia Langford, John Sillitoe on behalf of the Wellcome Sanger Institute COVID-19 Surveillance Team |
| EPI_ISL_3359058, EPI_ISL_3359060, EPI_ISL_3359061, EPI_ISL_3359063 | Lighthouse Lab in Alderley Park | Wellcome Sanger Institute for the COVID-19 Genomics UK (COG-UK) Consortium | Jacquelyn Wynn, Mairead Hyland, The Lighthouse Lab in Alderley Park and Alex Alderton, Roberto Amato, Jeffrey Barrett, Sonia Goncalves, Ewan Harrison, David K. Jackson, Ian Johnston, Dominic Kwiatkowski, Cordelia Langford, John Sillitoe on behalf of the Wellcome Sanger Institute COVID-19 Surveillance Team |
| EPI_ISL_3359064, EPI_ISL_3359065, EPI_ISL_3359067, EPI_ISL_3359068 | Lighthouse Lab in Milton Keynes | Wellcome Sanger Institute for the COVID-19 Genomics UK (COG-UK) Consortium | The Lighthouse Lab in Milton Keynes and Alex Alderton, Roberto Amato, Jeffrey Barrett, Sonia Goncalves, Ewan Harrison, David K. Jackson, Ian Johnston, Dominic Kwiatkowski, Cordelia Langford, John Sillitoe on behalf of the Wellcome Sanger Institute COVID-19 Surveillance Team |
| EPI_ISL_3359070, EPI_ISL_3359071 | Lighthouse Lab in Alderley Park | Wellcome Sanger Institute for the COVID-19 Genomics UK (COG-UK) Consortium | Jacquelyn Wynn, Mairead Hyland, The Lighthouse Lab in Alderley Park and Alex Alderton, Roberto Amato, Jeffrey Barrett, Sonia Goncalves, Ewan Harrison, David K. Jackson, Ian Johnston, Dominic Kwiatkowski, Cordelia Langford, John Sillitoe on behalf of the Wellcome Sanger Institute COVID-19 Surveillance Team |
| EPI_ISL_3359073, EPI_ISL_3359075 | Lighthouse Lab in Milton Keynes | Wellcome Sanger Institute for the COVID-19 Genomics UK (COG-UK) Consortium | The Lighthouse Lab in Milton Keynes and Alex Alderton, Roberto Amato, Jeffrey Barrett, Sonia Goncalves, Ewan Harrison, David K. Jackson, Ian Johnston, Dominic Kwiatkowski, Cordelia Langford, John Sillitoe on behalf of the Wellcome Sanger Institute COVID-19 Surveillance Team |
| EPI_ISL_3359076, EPI_ISL_3359078 | Lighthouse Lab in Alderley Park | Wellcome Sanger Institute for the COVID-19 Genomics UK (COG-UK) Consortium | Jacquelyn Wynn, Mairead Hyland, The Lighthouse Lab in Alderley Park and Alex Alderton, Roberto Amato, Jeffrey Barrett, Sonia Goncalves, Ewan Harrison, David K. Jackson, Ian Johnston, Dominic Kwiatkowski, Cordelia Langford, John Sillitoe on behalf of the Wellcome Sanger Institute COVID-19 Surveillance Team |
| EPI_ISL_3359079 | Lighthouse Lab in Milton Keynes | Wellcome Sanger Institute for the COVID-19 Genomics UK (COG-UK) Consortium | The Lighthouse Lab in Milton Keynes and Alex Alderton, Roberto Amato, Jeffrey Barrett, Sonia Goncalves, Ewan Harrison, David K. Jackson, Ian Johnston, Dominic Kwiatkowski, Cordelia Langford, John Sillitoe on behalf of the Wellcome Sanger Institute COVID-19 Surveillance Team |
| EPI_ISL_3359080 | Lighthouse Lab in Alderley Park | Wellcome Sanger Institute for the COVID-19 Genomics UK (COG-UK) Consortium | Jacquelyn Wynn, Mairead Hyland, The Lighthouse Lab in Alderley Park and Alex Alderton, Roberto Amato, Jeffrey Barrett, Sonia Goncalves, Ewan Harrison, David K. Jackson, Ian Johnston, Dominic Kwiatkowski, Cordelia Langford, John Sillitoe on behalf of the Wellcome Sanger Institute COVID-19 Surveillance Team |
| EPI_ISL_3359082 | Lighthouse Lab in Milton Keynes | Wellcome Sanger Institute for the COVID-19 Genomics UK (COG-UK) Consortium | The Lighthouse Lab in Milton Keynes and Alex Alderton, Roberto Amato, Jeffrey Barrett, Sonia Goncalves, Ewan Harrison, David K. Jackson, Ian Johnston, Dominic Kwiatkowski, Cordelia Langford, John Sillitoe on behalf of the Wellcome Sanger Institute COVID-19 Surveillance Team |
| EPI_ISL_3359083 | Lighthouse Lab in Alderley Park | Wellcome Sanger Institute for the COVID-19 Genomics UK (COG-UK) Consortium | Jacquelyn Wynn, Mairead Hyland, The Lighthouse Lab in Alderley Park and Alex Alderton, Roberto Amato, Jeffrey Barrett, Sonia Goncalves, Ewan Harrison, David K. Jackson, Ian Johnston, Dominic Kwiatkowski, Cordelia Langford, John Sillitoe on behalf of the Wellcome Sanger Institute COVID-19 Surveillance Team |
| EPI_ISL_3359085, EPI_ISL_3359086 | Lighthouse Lab in Milton Keynes | Wellcome Sanger Institute for the COVID-19 Genomics UK (COG-UK) Consortium | The Lighthouse Lab in Milton Keynes and Alex Alderton, Roberto Amato, Jeffrey Barrett, Sonia Goncalves, Ewan Harrison, David K. Jackson, Ian Johnston, Dominic Kwiatkowski, Cordelia Langford, John Sillitoe on behalf of the Wellcome Sanger Institute COVID-19 Surveillance Team |
| EPI_ISL_3359088 | Lighthouse Lab in Alderley Park | Wellcome Sanger Institute for the COVID-19 Genomics UK (COG-UK) Consortium | Jacquelyn Wynn, Mairead Hyland, The Lighthouse Lab in Alderley Park and Alex Alderton, Roberto Amato, Jeffrey Barrett, Sonia Goncalves, Ewan Harrison, David K. Jackson, Ian Johnston, Dominic Kwiatkowski, Cordelia Langford, John Sillitoe on behalf of the Wellcome Sanger Institute COVID-19 Surveillance Team |
| EPI_ISL_3359089 | Lighthouse Lab in Milton Keynes | Wellcome Sanger Institute for the COVID-19 Genomics UK (COG-UK) Consortium | The Lighthouse Lab in Milton Keynes and Alex Alderton, Roberto Amato, Jeffrey Barrett, Sonia Goncalves, Ewan Harrison, David K. Jackson, Ian Johnston, Dominic Kwiatkowski, Cordelia Langford, John Sillitoe on behalf of the Wellcome Sanger Institute COVID-19 Surveillance Team |
| EPI_ISL_3359090 | Lighthouse Lab in Alderley Park | Wellcome Sanger Institute for the COVID-19 Genomics UK (COG-UK) Consortium | Jacquelyn Wynn, Mairead Hyland, The Lighthouse Lab in Alderley Park and Alex Alderton, Roberto Amato, Jeffrey Barrett, Sonia Goncalves, Ewan Harrison, David K. Jackson, Ian Johnston, Dominic Kwiatkowski, Cordelia Langford, John Sillitoe on behalf of the Wellcome Sanger Institute COVID-19 Surveillance Team |
| EPI_ISL_3359092 | Berkshire and Surrey Pathology Services Lighthouse Laboratory | Wellcome Sanger Institute for the COVID-19 Genomics UK (COG-UK) Consortium | Berkshire and Surrey Pathology Services Lighthouse Laboratory and Alex Alderton, Roberto Amato, Jeffrey Barrett, Sonia Goncalves, Ewan Harrison, David K. Jackson, Ian Johnston, Dominic Kwiatkowski, Cordelia Langford, John Sillitoe on behalf of the Wellcome Sanger Institute COVID-19 Surveillance Team |
| EPI_ISL_3359093, EPI_ISL_3359095, EPI_ISL_3359096, EPI_ISL_3359098, EPI_ISL_3359099 | Lighthouse Lab in Alderley Park | Wellcome Sanger Institute for the COVID-19 Genomics UK (COG-UK) Consortium | Jacquelyn Wynn, Mairead Hyland, The Lighthouse Lab in Alderley Park and Alex Alderton, Roberto Amato, Jeffrey Barrett, Sonia Goncalves, Ewan Harrison, David K. Jackson, Ian Johnston, Dominic Kwiatkowski, Cordelia Langford, John Sillitoe on behalf of the Wellcome Sanger Institute COVID-19 Surveillance Team |
| EPI_ISL_3359101 | Lighthouse Lab in Milton Keynes | Wellcome Sanger Institute for the COVID-19 Genomics UK (COG-UK) Consortium | The Lighthouse Lab in Milton Keynes and Alex Alderton, Roberto Amato, Jeffrey Barrett, Sonia Goncalves, Ewan Harrison, David K. Jackson, Ian Johnston, Dominic Kwiatkowski, Cordelia Langford, John Sillitoe on behalf of the Wellcome Sanger Institute COVID-19 Surveillance Team |
| EPI_ISL_3359102, EPI_ISL_3359104 | Lighthouse Lab in Alderley Park | Wellcome Sanger Institute for the COVID-19 Genomics UK (COG-UK) Consortium | Jacquelyn Wynn, Mairead Hyland, The Lighthouse Lab in Alderley Park and Alex Alderton, Roberto Amato, Jeffrey Barrett, Sonia Goncalves, Ewan Harrison, David K. Jackson, Ian Johnston, Dominic Kwiatkowski, Cordelia Langford, John Sillitoe on behalf of the Wellcome Sanger Institute COVID-19 Surveillance Team |
| EPI_ISL_3359105, EPI_ISL_3359107 | Lighthouse Lab in Milton Keynes | Wellcome Sanger Institute for the COVID-19 Genomics UK (COG-UK) Consortium | The Lighthouse Lab in Milton Keynes and Alex Alderton, Roberto Amato, Jeffrey Barrett, Sonia Goncalves, Ewan Harrison, David K. Jackson, Ian Johnston, Dominic Kwiatkowski, Cordelia Langford, John Sillitoe on behalf of the Wellcome Sanger Institute COVID-19 Surveillance Team |
| EPI_ISL_3359108 | Berkshire and Surrey Pathology Services Lighthouse Laboratory | Wellcome Sanger Institute for the COVID-19 Genomics UK (COG-UK) Consortium | Berkshire and Surrey Pathology Services Lighthouse Laboratory and Alex Alderton, Roberto Amato, Jeffrey Barrett, Sonia Goncalves, Ewan Harrison, David K. Jackson, Ian Johnston, Dominic Kwiatkowski, Cordelia Langford, John Sillitoe on behalf of the Wellcome Sanger Institute COVID-19 Surveillance Team |
| EPI_ISL_3359109 | Lighthouse Lab in Milton Keynes | Wellcome Sanger Institute for the COVID-19 Genomics UK (COG-UK) Consortium | The Lighthouse Lab in Milton Keynes and Alex Alderton, Roberto Amato, Jeffrey Barrett, Sonia Goncalves, Ewan Harrison, David K. Jackson, Ian Johnston, Dominic Kwiatkowski, Cordelia Langford, John Sillitoe on behalf of the Wellcome Sanger Institute COVID-19 Surveillance Team |
| EPI_ISL_3359111 | Berkshire and Surrey Pathology Services Lighthouse Laboratory | Wellcome Sanger Institute for the COVID-19 Genomics UK (COG-UK) Consortium | Berkshire and Surrey Pathology Services Lighthouse Laboratory and Alex Alderton, Roberto Amato, Jeffrey Barrett, Sonia Goncalves, Ewan Harrison, David K. Jackson, Ian Johnston, Dominic Kwiatkowski, Cordelia Langford, John Sillitoe on behalf of the Wellcome Sanger Institute COVID-19 Surveillance Team |
| EPI_ISL_3359112, EPI_ISL_3359113 | Lighthouse Laboratory Plymouth | Wellcome Sanger Institute for the COVID-19 Genomics UK (COG-UK) Consortium | Lighthouse Laboratory Plymouth and Alex Alderton, Roberto Amato, Jeffrey Barrett, Sonia Goncalves, Ewan Harrison, David K. Jackson, Ian Johnston, Dominic Kwiatkowski, Cordelia Langford, John Sillitoe on behalf of the Wellcome Sanger Institute COVID-19 Surveillance Team |
| EPI_ISL_3359115 | Lighthouse Lab in Milton Keynes | Wellcome Sanger Institute for the COVID-19 Genomics UK (COG-UK) Consortium | The Lighthouse Lab in Milton Keynes and Alex Alderton, Roberto Amato, Jeffrey Barrett, Sonia Goncalves, Ewan Harrison, David K. Jackson, Ian Johnston, Dominic Kwiatkowski, Cordelia Langford, John Sillitoe on behalf of the Wellcome Sanger Institute COVID-19 Surveillance Team |
| EPI_ISL_3359116, EPI_ISL_3359118, EPI_ISL_3359120 | Lighthouse Laboratory Plymouth | Wellcome Sanger Institute for the COVID-19 Genomics UK (COG-UK) Consortium | Lighthouse Laboratory Plymouth and Alex Alderton, Roberto Amato, Jeffrey Barrett, Sonia Goncalves, Ewan Harrison, David K. Jackson, Ian Johnston, Dominic Kwiatkowski, Cordelia Langford, John Sillitoe on behalf of the Wellcome Sanger Institute COVID-19 Surveillance Team |
| EPI_ISL_3359121 | Berkshire and Surrey Pathology Services Lighthouse Laboratory | Wellcome Sanger Institute for the COVID-19 Genomics UK (COG-UK) Consortium | Berkshire and Surrey Pathology Services Lighthouse Laboratory and Alex Alderton, Roberto Amato, Jeffrey Barrett, Sonia Goncalves, Ewan Harrison, David K. Jackson, Ian Johnston, Dominic Kwiatkowski, Cordelia Langford, John Sillitoe on behalf of the Wellcome Sanger Institute COVID-19 Surveillance Team |
| EPI_ISL_3359122 | Lighthouse Lab in Milton Keynes | Wellcome Sanger Institute for the COVID-19 Genomics UK (COG-UK) Consortium | The Lighthouse Lab in Milton Keynes and Alex Alderton, Roberto Amato, Jeffrey Barrett, Sonia Goncalves, Ewan Harrison, David K. Jackson, Ian Johnston, Dominic Kwiatkowski, Cordelia Langford, John Sillitoe on behalf of the Wellcome Sanger Institute COVID-19 Surveillance Team |
| EPI_ISL_3359174, EPI_ISL_3359176 | Lighthouse Laboratory Plymouth | Wellcome Sanger Institute for the COVID-19 Genomics UK (COG-UK) Consortium | Lighthouse Laboratory Plymouth and Alex Alderton, Roberto Amato, Jeffrey Barrett, Sonia Goncalves, Ewan Harrison, David K. Jackson, Ian Johnston, Dominic Kwiatkowski, Cordelia Langford, John Sillitoe on behalf of the Wellcome Sanger Institute COVID-19 Surveillance Team |
| EPI_ISL_3359177 | Berkshire and Surrey Pathology Services Lighthouse Laboratory | Wellcome Sanger Institute for the COVID-19 Genomics UK (COG-UK) Consortium | Berkshire and Surrey Pathology Services Lighthouse Laboratory and Alex Alderton, Roberto Amato, Jeffrey Barrett, Sonia Goncalves, Ewan Harrison, David K. Jackson, Ian Johnston, Dominic Kwiatkowski, Cordelia Langford, John Sillitoe on behalf of the Wellcome Sanger Institute COVID-19 Surveillance Team |
| EPI_ISL_3359179, EPI_ISL_3359180, EPI_ISL_3359182 | Lighthouse Lab in Milton Keynes | Wellcome Sanger Institute for the COVID-19 Genomics UK (COG-UK) Consortium | The Lighthouse Lab in Milton Keynes and Alex Alderton, Roberto Amato, Jeffrey Barrett, Sonia Goncalves, Ewan Harrison, David K. Jackson, Ian Johnston, Dominic Kwiatkowski, Cordelia Langford, John Sillitoe on behalf of the Wellcome Sanger Institute COVID-19 Surveillance Team |
| EPI_ISL_3359183, EPI_ISL_3359185 | Lighthouse Laboratory Plymouth | Wellcome Sanger Institute for the COVID-19 Genomics UK (COG-UK) Consortium | Lighthouse Laboratory Plymouth and Alex Alderton, Roberto Amato, Jeffrey Barrett, Sonia Goncalves, Ewan Harrison, David K. Jackson, Ian Johnston, Dominic Kwiatkowski, Cordelia Langford, John Sillitoe on behalf of the Wellcome Sanger Institute COVID-19 Surveillance Team |
| EPI_ISL_3359186 | Berkshire and Surrey Pathology Services Lighthouse Laboratory | Wellcome Sanger Institute for the COVID-19 Genomics UK (COG-UK) Consortium | Berkshire and Surrey Pathology Services Lighthouse Laboratory and Alex Alderton, Roberto Amato, Jeffrey Barrett, Sonia Goncalves, Ewan Harrison, David K. Jackson, Ian Johnston, Dominic Kwiatkowski, Cordelia Langford, John Sillitoe on behalf of the Wellcome Sanger Institute COVID-19 Surveillance Team |
| EPI_ISL_3359188 | Lighthouse Lab in Milton Keynes | Wellcome Sanger Institute for the COVID-19 Genomics UK (COG-UK) Consortium | The Lighthouse Lab in Milton Keynes and Alex Alderton, Roberto Amato, Jeffrey Barrett, Sonia Goncalves, Ewan Harrison, David K. Jackson, Ian Johnston, Dominic Kwiatkowski, Cordelia Langford, John Sillitoe on behalf of the Wellcome Sanger Institute COVID-19 Surveillance Team |
| EPI_ISL_3359189 | Berkshire and Surrey Pathology Services Lighthouse Laboratory | Wellcome Sanger Institute for the COVID-19 Genomics UK (COG-UK) Consortium | Berkshire and Surrey Pathology Services Lighthouse Laboratory and Alex Alderton, Roberto Amato, Jeffrey Barrett, Sonia Goncalves, Ewan Harrison, David K. Jackson, Ian Johnston, Dominic Kwiatkowski, Cordelia Langford, John Sillitoe on behalf of the Wellcome Sanger Institute COVID-19 Surveillance Team |
| EPI_ISL_3359191 | Lighthouse Laboratory Plymouth | Wellcome Sanger Institute for the COVID-19 Genomics UK (COG-UK) Consortium | Lighthouse Laboratory Plymouth and Alex Alderton, Roberto Amato, Jeffrey Barrett, Sonia Goncalves, Ewan Harrison, David K. Jackson, Ian Johnston, Dominic Kwiatkowski, Cordelia Langford, John Sillitoe on behalf of the Wellcome Sanger Institute COVID-19 Surveillance Team |
| EPI_ISL_3359192 | Lighthouse Lab in Milton Keynes | Wellcome Sanger Institute for the COVID-19 Genomics UK (COG-UK) Consortium | The Lighthouse Lab in Milton Keynes and Alex Alderton, Roberto Amato, Jeffrey Barrett, Sonia Goncalves, Ewan Harrison, David K. Jackson, Ian Johnston, Dominic Kwiatkowski, Cordelia Langford, John Sillitoe on behalf of the Wellcome Sanger Institute COVID-19 Surveillance Team |
| EPI_ISL_3359194 | Lighthouse Laboratory Plymouth | Wellcome Sanger Institute for the COVID-19 Genomics UK (COG-UK) | Lighthouse Laboratory Plymouth and Alex Alderton, Roberto Amato, Jeffrey Barrett, Sonia Goncalves, Ewan Harrison, David K. Jackson, Ian Johnston, Dominic Kwiatkowski, Cordelia Langford, John Sillitoe on behalf of the Wellcome Sanger |

| | | | |
|---|---|---|---|
| | | Consortium | Institute COVID-19 Surveillance Team |
| EPI_ISL_3359195 | Berkshire and Surrey Pathology Services Lighthouse Laboratory | Wellcome Sanger Institute for the COVID-19 Genomics UK (COG-UK) Consortium | Berkshire and Surrey Pathology Services Lighthouse Laboratory and Alex Alderton, Roberto Amato, Jeffrey Barrett, Sonia Goncalves, Ewan Harrison, David K. Jackson, Ian Johnston, Dominic Kwiatkowski, Cordelia Langford, John Sillitoe on behalf of the Wellcome Sanger Institute COVID-19 Surveillance Team |
| EPI_ISL_3359196 | Lighthouse Lab in Milton Keynes | Wellcome Sanger Institute for the COVID-19 Genomics UK (COG-UK) Consortium | The Lighthouse Lab in Milton Keynes and Alex Alderton, Roberto Amato, Jeffrey Barrett, Sonia Goncalves, Ewan Harrison, David K. Jackson, Ian Johnston, Dominic Kwiatkowski, Cordelia Langford, John Sillitoe on behalf of the Wellcome Sanger Institute COVID-19 Surveillance Team |
| EPI_ISL_3359198, EPI_ISL_3359199 | Lighthouse Laboratory Plymouth | Wellcome Sanger Institute for the COVID-19 Genomics UK (COG-UK) Consortium | Lighthouse Laboratory Plymouth and Alex Alderton, Roberto Amato, Jeffrey Barrett, Sonia Goncalves, Ewan Harrison, David K. Jackson, Ian Johnston, Dominic Kwiatkowski, Cordelia Langford, John Sillitoe on behalf of the Wellcome Sanger Institute COVID-19 Surveillance Team |
| EPI_ISL_3359200 | Berkshire and Surrey Pathology Services Lighthouse Laboratory | Wellcome Sanger Institute for the COVID-19 Genomics UK (COG-UK) Consortium | Berkshire and Surrey Pathology Services Lighthouse Laboratory and Alex Alderton, Roberto Amato, Jeffrey Barrett, Sonia Goncalves, Ewan Harrison, David K. Jackson, Ian Johnston, Dominic Kwiatkowski, Cordelia Langford, John Sillitoe on behalf of the Wellcome Sanger Institute COVID-19 Surveillance Team |
| EPI_ISL_3359202, EPI_ISL_3359203 | Lighthouse Laboratory Plymouth | Wellcome Sanger Institute for the COVID-19 Genomics UK (COG-UK) Consortium | Lighthouse Laboratory Plymouth and Alex Alderton, Roberto Amato, Jeffrey Barrett, Sonia Goncalves, Ewan Harrison, David K. Jackson, Ian Johnston, Dominic Kwiatkowski, Cordelia Langford, John Sillitoe on behalf of the Wellcome Sanger Institute COVID-19 Surveillance Team |
| EPI_ISL_3359205 | Lighthouse Lab in Milton Keynes | Wellcome Sanger Institute for the COVID-19 Genomics UK (COG-UK) Consortium | The Lighthouse Lab in Milton Keynes and Alex Alderton, Roberto Amato, Jeffrey Barrett, Sonia Goncalves, Ewan Harrison, David K. Jackson, Ian Johnston, Dominic Kwiatkowski, Cordelia Langford, John Sillitoe on behalf of the Wellcome Sanger Institute COVID-19 Surveillance Team |
| EPI_ISL_3359206, EPI_ISL_3359207, EPI_ISL_3359209 | Lighthouse Laboratory Plymouth | Wellcome Sanger Institute for the COVID-19 Genomics UK (COG-UK) Consortium | Lighthouse Laboratory Plymouth and Alex Alderton, Roberto Amato, Jeffrey Barrett, Sonia Goncalves, Ewan Harrison, David K. Jackson, Ian Johnston, Dominic Kwiatkowski, Cordelia Langford, John Sillitoe on behalf of the Wellcome Sanger Institute COVID-19 Surveillance Team |
| EPI_ISL_3359210, EPI_ISL_3359212 | Lighthouse Lab in Milton Keynes | Wellcome Sanger Institute for the COVID-19 Genomics UK (COG-UK) Consortium | The Lighthouse Lab in Milton Keynes and Alex Alderton, Roberto Amato, Jeffrey Barrett, Sonia Goncalves, Ewan Harrison, David K. Jackson, Ian Johnston, Dominic Kwiatkowski, Cordelia Langford, John Sillitoe on behalf of the Wellcome Sanger Institute COVID-19 Surveillance Team |
| EPI_ISL_3359213, EPI_ISL_3359215, EPI_ISL_3359216, EPI_ISL_3359217, EPI_ISL_3359219 | Lighthouse Laboratory Plymouth | Wellcome Sanger Institute for the COVID-19 Genomics UK (COG-UK) Consortium | Lighthouse Laboratory Plymouth and Alex Alderton, Roberto Amato, Jeffrey Barrett, Sonia Goncalves, Ewan Harrison, David K. Jackson, Ian Johnston, Dominic Kwiatkowski, Cordelia Langford, John Sillitoe on behalf of the Wellcome Sanger Institute COVID-19 Surveillance Team |
| EPI_ISL_3359221 | Lighthouse Lab in Milton Keynes | Wellcome Sanger Institute for the COVID-19 Genomics UK (COG-UK) Consortium | The Lighthouse Lab in Milton Keynes and Alex Alderton, Roberto Amato, Jeffrey Barrett, Sonia Goncalves, Ewan Harrison, David K. Jackson, Ian Johnston, Dominic Kwiatkowski, Cordelia Langford, John Sillitoe on behalf of the Wellcome Sanger Institute COVID-19 Surveillance Team |
| EPI_ISL_3359223, EPI_ISL_3359224 | Lighthouse Laboratory Plymouth | Wellcome Sanger Institute for the COVID-19 Genomics UK (COG-UK) Consortium | Lighthouse Laboratory Plymouth and Alex Alderton, Roberto Amato, Jeffrey Barrett, Sonia Goncalves, Ewan Harrison, David K. Jackson, Ian Johnston, Dominic Kwiatkowski, Cordelia Langford, John Sillitoe on behalf of the Wellcome Sanger Institute COVID-19 Surveillance Team |
| EPI_ISL_3359225 | Berkshire and Surrey Pathology Services Lighthouse Laboratory | Wellcome Sanger Institute for the COVID-19 Genomics UK (COG-UK) Consortium | Berkshire and Surrey Pathology Services Lighthouse Laboratory and Alex Alderton, Roberto Amato, Jeffrey Barrett, Sonia Goncalves, Ewan Harrison, David K. Jackson, Ian Johnston, Dominic Kwiatkowski, Cordelia Langford, John Sillitoe on behalf of the Wellcome Sanger Institute COVID-19 Surveillance Team |
| EPI_ISL_3359227 | Lighthouse Lab in Milton Keynes | Wellcome Sanger Institute for the COVID-19 Genomics UK (COG-UK) Consortium | The Lighthouse Lab in Milton Keynes and Alex Alderton, Roberto Amato, Jeffrey Barrett, Sonia Goncalves, Ewan Harrison, David K. Jackson, Ian Johnston, Dominic Kwiatkowski, Cordelia Langford, John Sillitoe on behalf of the Wellcome Sanger Institute COVID-19 Surveillance Team |
| EPI_ISL_3359229 | Berkshire and Surrey Pathology Services Lighthouse Laboratory | Wellcome Sanger Institute for the COVID-19 Genomics UK (COG-UK) Consortium | Berkshire and Surrey Pathology Services Lighthouse Laboratory and Alex Alderton, Roberto Amato, Jeffrey Barrett, Sonia Goncalves, Ewan Harrison, David K. Jackson, Ian Johnston, Dominic Kwiatkowski, Cordelia Langford, John Sillitoe on behalf of the Wellcome Sanger Institute COVID-19 Surveillance Team |
| EPI_ISL_3359230 | Lighthouse Laboratory Plymouth | Wellcome Sanger Institute for the COVID-19 Genomics UK (COG-UK) Consortium | Lighthouse Laboratory Plymouth and Alex Alderton, Roberto Amato, Jeffrey Barrett, Sonia Goncalves, Ewan Harrison, David K. Jackson, Ian Johnston, Dominic Kwiatkowski, Cordelia Langford, John Sillitoe on behalf of the Wellcome Sanger Institute COVID-19 Surveillance Team |
| EPI_ISL_3359231 | Lighthouse Lab in Milton Keynes | Wellcome Sanger Institute for the COVID-19 Genomics UK (COG-UK) Consortium | The Lighthouse Lab in Milton Keynes and Alex Alderton, Roberto Amato, Jeffrey Barrett, Sonia Goncalves, Ewan Harrison, David K. Jackson, Ian Johnston, Dominic Kwiatkowski, Cordelia Langford, John Sillitoe on behalf of the Wellcome Sanger Institute COVID-19 Surveillance Team |
| EPI_ISL_3359233, EPI_ISL_3359234 | Lighthouse Laboratory Plymouth | Wellcome Sanger Institute for the COVID-19 Genomics UK (COG-UK) Consortium | Lighthouse Laboratory Plymouth and Alex Alderton, Roberto Amato, Jeffrey Barrett, Sonia Goncalves, Ewan Harrison, David K. Jackson, Ian Johnston, Dominic Kwiatkowski, Cordelia Langford, John Sillitoe on behalf of the Wellcome Sanger Institute COVID-19 Surveillance Team |
| EPI_ISL_3359236 | Berkshire and Surrey Pathology Services Lighthouse Laboratory | Wellcome Sanger Institute for the COVID-19 Genomics UK (COG-UK) Consortium | Berkshire and Surrey Pathology Services Lighthouse Laboratory and Alex Alderton, Roberto Amato, Jeffrey Barrett, Sonia Goncalves, Ewan Harrison, David K. Jackson, Ian Johnston, Dominic Kwiatkowski, Cordelia Langford, John Sillitoe on behalf of the Wellcome Sanger Institute COVID-19 Surveillance Team |
| EPI_ISL_3359237 | Lighthouse Laboratory Plymouth | Wellcome Sanger Institute for the COVID-19 Genomics UK (COG-UK) Consortium | Lighthouse Laboratory Plymouth and Alex Alderton, Roberto Amato, Jeffrey Barrett, Sonia Goncalves, Ewan Harrison, David K. Jackson, Ian Johnston, Dominic Kwiatkowski, Cordelia Langford, John Sillitoe on behalf of the Wellcome Sanger Institute COVID-19 Surveillance Team |
| EPI_ISL_3359238 | Lighthouse Lab in Milton Keynes | Wellcome Sanger Institute for the COVID-19 Genomics UK (COG-UK) Consortium | The Lighthouse Lab in Milton Keynes and Alex Alderton, Roberto Amato, Jeffrey Barrett, Sonia Goncalves, Ewan Harrison, David K. Jackson, Ian Johnston, Dominic Kwiatkowski, Cordelia Langford, John Sillitoe on behalf of the Wellcome Sanger Institute COVID-19 Surveillance Team |
| EPI_ISL_3359240, EPI_ISL_3359241 | Lighthouse Laboratory Plymouth | Wellcome Sanger Institute for the COVID-19 Genomics UK (COG-UK) Consortium | Lighthouse Laboratory Plymouth and Alex Alderton, Roberto Amato, Jeffrey Barrett, Sonia Goncalves, Ewan Harrison, David K. Jackson, Ian Johnston, Dominic Kwiatkowski, Cordelia Langford, John Sillitoe on behalf of the Wellcome Sanger Institute COVID-19 Surveillance Team |
| EPI_ISL_3359243 | Berkshire and Surrey Pathology Services Lighthouse Laboratory | Wellcome Sanger Institute for the COVID-19 Genomics UK (COG-UK) Consortium | Berkshire and Surrey Pathology Services Lighthouse Laboratory and Alex Alderton, Roberto Amato, Jeffrey Barrett, Sonia Goncalves, Ewan Harrison, David K. Jackson, Ian Johnston, Dominic Kwiatkowski, Cordelia Langford, John Sillitoe on behalf of the Wellcome Sanger Institute COVID-19 Surveillance Team |
| EPI_ISL_3359244, EPI_ISL_3359246 | Lighthouse Laboratory Plymouth | Wellcome Sanger Institute for the COVID-19 Genomics UK (COG-UK) Consortium | Lighthouse Laboratory Plymouth and Alex Alderton, Roberto Amato, Jeffrey Barrett, Sonia Goncalves, Ewan Harrison, David K. Jackson, Ian Johnston, Dominic Kwiatkowski, Cordelia Langford, John Sillitoe on behalf of the Wellcome Sanger Institute COVID-19 Surveillance Team |
| EPI_ISL_3359247 | Berkshire and Surrey Pathology Services Lighthouse Laboratory | Wellcome Sanger Institute for the COVID-19 Genomics UK (COG-UK) Consortium | Berkshire and Surrey Pathology Services Lighthouse Laboratory and Alex Alderton, Roberto Amato, Jeffrey Barrett, Sonia Goncalves, Ewan Harrison, David K. Jackson, Ian Johnston, Dominic Kwiatkowski, Cordelia Langford, John Sillitoe on behalf of the Wellcome Sanger Institute COVID-19 Surveillance Team |
| EPI_ISL_3359249, EPI_ISL_3359250, EPI_ISL_3359252 | Lighthouse Lab in Milton Keynes | Wellcome Sanger Institute for the COVID-19 Genomics UK (COG-UK) Consortium | The Lighthouse Lab in Milton Keynes and Alex Alderton, Roberto Amato, Jeffrey Barrett, Sonia Goncalves, Ewan Harrison, David K. Jackson, Ian Johnston, Dominic Kwiatkowski, Cordelia Langford, John Sillitoe on behalf of the Wellcome Sanger Institute COVID-19 Surveillance Team |
| EPI_ISL_3359253 | Lighthouse Laboratory Plymouth | Wellcome Sanger Institute for the COVID-19 Genomics UK (COG-UK) Consortium | Lighthouse Laboratory Plymouth and Alex Alderton, Roberto Amato, Jeffrey Barrett, Sonia Goncalves, Ewan Harrison, David K. Jackson, Ian Johnston, Dominic Kwiatkowski, Cordelia Langford, John Sillitoe on behalf of the Wellcome Sanger Institute COVID-19 Surveillance Team |
| EPI_ISL_3359255 | Berkshire and Surrey Pathology Services Lighthouse Laboratory | Wellcome Sanger Institute for the COVID-19 Genomics UK (COG-UK) Consortium | Berkshire and Surrey Pathology Services Lighthouse Laboratory and Alex Alderton, Roberto Amato, Jeffrey Barrett, Sonia Goncalves, Ewan Harrison, David K. Jackson, Ian Johnston, Dominic Kwiatkowski, Cordelia Langford, John Sillitoe on behalf of the Wellcome Sanger Institute COVID-19 Surveillance Team |
| EPI_ISL_3359256, EPI_ISL_3359257, EPI_ISL_3359259, EPI_ISL_3359260 | Lighthouse Laboratory Plymouth | Wellcome Sanger Institute for the COVID-19 Genomics UK (COG-UK) Consortium | Lighthouse Laboratory Plymouth and Alex Alderton, Roberto Amato, Jeffrey Barrett, Sonia Goncalves, Ewan Harrison, David K. Jackson, Ian Johnston, Dominic Kwiatkowski, Cordelia Langford, John Sillitoe on behalf of the Wellcome Sanger Institute COVID-19 Surveillance Team |
| EPI_ISL_3359262 | Berkshire and Surrey Pathology Services Lighthouse Laboratory | Wellcome Sanger Institute for the COVID-19 Genomics UK (COG-UK) Consortium | Berkshire and Surrey Pathology Services Lighthouse Laboratory and Alex Alderton, Roberto Amato, Jeffrey Barrett, Sonia Goncalves, Ewan Harrison, David K. Jackson, Ian Johnston, Dominic Kwiatkowski, Cordelia Langford, John Sillitoe on behalf of the Wellcome Sanger Institute COVID-19 Surveillance Team |
| EPI_ISL_3359263 | Lighthouse Laboratory Plymouth | Wellcome Sanger Institute for the COVID-19 Genomics UK (COG-UK) Consortium | Lighthouse Laboratory Plymouth and Alex Alderton, Roberto Amato, Jeffrey Barrett, Sonia Goncalves, Ewan Harrison, David K. Jackson, Ian Johnston, Dominic Kwiatkowski, Cordelia Langford, John Sillitoe on behalf of the Wellcome Sanger Institute COVID-19 Surveillance Team |
| EPI_ISL_3359264 | Berkshire and Surrey Pathology Services Lighthouse Laboratory | Wellcome Sanger Institute for the COVID-19 Genomics UK (COG-UK) Consortium | Berkshire and Surrey Pathology Services Lighthouse Laboratory and Alex Alderton, Roberto Amato, Jeffrey Barrett, Sonia Goncalves, Ewan Harrison, David K. Jackson, Ian Johnston, Dominic Kwiatkowski, Cordelia Langford, John Sillitoe on behalf of the Wellcome Sanger Institute COVID-19 Surveillance Team |
| EPI_ISL_3359266 | Lighthouse Laboratory Plymouth | Wellcome Sanger Institute for the COVID-19 Genomics UK (COG-UK) Consortium | Lighthouse Laboratory Plymouth and Alex Alderton, Roberto Amato, Jeffrey Barrett, Sonia Goncalves, Ewan Harrison, David K. Jackson, Ian Johnston, Dominic Kwiatkowski, Cordelia Langford, John Sillitoe on behalf of the Wellcome Sanger Institute COVID-19 Surveillance Team |
| EPI_ISL_3359267 | Lighthouse Lab in Milton Keynes | Wellcome Sanger Institute for the COVID-19 Genomics UK (COG-UK) Consortium | The Lighthouse Lab in Milton Keynes and Alex Alderton, Roberto Amato, Jeffrey Barrett, Sonia Goncalves, Ewan Harrison, David K. Jackson, Ian Johnston, Dominic Kwiatkowski, Cordelia Langford, John Sillitoe on behalf of the Wellcome Sanger Institute COVID-19 Surveillance Team |
| EPI_ISL_3359269, EPI_ISL_3359270 | Lighthouse Laboratory Plymouth | Wellcome Sanger Institute for the COVID-19 Genomics UK (COG-UK) Consortium | Lighthouse Laboratory Plymouth and Alex Alderton, Roberto Amato, Jeffrey Barrett, Sonia Goncalves, Ewan Harrison, David K. Jackson, Ian Johnston, Dominic Kwiatkowski, Cordelia Langford, John Sillitoe on behalf of the Wellcome Sanger Institute COVID-19 Surveillance Team |
| EPI_ISL_3359271, EPI_ISL_3359272 | Lighthouse Lab in Milton Keynes | Wellcome Sanger Institute for the COVID-19 Genomics UK (COG-UK) Consortium | The Lighthouse Lab in Milton Keynes and Alex Alderton, Roberto Amato, Jeffrey Barrett, Sonia Goncalves, Ewan Harrison, David K. Jackson, Ian Johnston, Dominic Kwiatkowski, Cordelia Langford, John Sillitoe on behalf of the Wellcome Sanger Institute COVID-19 Surveillance Team |
| EPI_ISL_3359274 | Lighthouse Laboratory Plymouth | Wellcome Sanger Institute for the COVID-19 Genomics UK (COG-UK) Consortium | Lighthouse Laboratory Plymouth and Alex Alderton, Roberto Amato, Jeffrey Barrett, Sonia Goncalves, Ewan Harrison, David K. Jackson, Ian Johnston, Dominic Kwiatkowski, Cordelia Langford, John Sillitoe on behalf of the Wellcome Sanger Institute COVID-19 Surveillance Team |
| EPI_ISL_3359275 | Lighthouse Lab in Milton Keynes | Wellcome Sanger Institute for the COVID-19 Genomics UK (COG-UK) Consortium | The Lighthouse Lab in Milton Keynes and Alex Alderton, Roberto Amato, Jeffrey Barrett, Sonia Goncalves, Ewan Harrison, David K. Jackson, Ian Johnston, Dominic Kwiatkowski, Cordelia Langford, John Sillitoe on behalf of the Wellcome Sanger Institute COVID-19 Surveillance Team |
| EPI_ISL_3359277, EPI_ISL_3359279 | Berkshire and Surrey Pathology Services Lighthouse Laboratory | Wellcome Sanger Institute for the COVID-19 Genomics UK (COG-UK) Consortium | Berkshire and Surrey Pathology Services Lighthouse Laboratory and Alex Alderton, Roberto Amato, Jeffrey Barrett, Sonia Goncalves, Ewan Harrison, David K. Jackson, Ian Johnston, Dominic Kwiatkowski, Cordelia Langford, John Sillitoe on behalf of the Wellcome Sanger Institute COVID-19 Surveillance Team |
| EPI_ISL_3359280 | Lighthouse Lab in Milton Keynes | Wellcome Sanger Institute for the COVID-19 Genomics UK (COG-UK) Consortium | The Lighthouse Lab in Milton Keynes and Alex Alderton, Roberto Amato, Jeffrey Barrett, Sonia Goncalves, Ewan Harrison, David K. Jackson, Ian Johnston, Dominic Kwiatkowski, Cordelia Langford, John Sillitoe on behalf of the Wellcome Sanger Institute COVID-19 Surveillance Team |
| EPI_ISL_3359282, EPI_ISL_3359283, EPI_ISL_3359284 | Lighthouse Laboratory Plymouth | Wellcome Sanger Institute for the COVID-19 Genomics UK (COG-UK) Consortium | Lighthouse Laboratory Plymouth and Alex Alderton, Roberto Amato, Jeffrey Barrett, Sonia Goncalves, Ewan Harrison, David K. Jackson, Ian Johnston, Dominic Kwiatkowski, Cordelia Langford, John Sillitoe on behalf of the Wellcome Sanger Institute COVID-19 Surveillance Team |
| EPI_ISL_3359286 | Berkshire and Surrey Pathology Services Lighthouse Laboratory | Wellcome Sanger Institute for the COVID-19 Genomics UK (COG-UK) Consortium | Berkshire and Surrey Pathology Services Lighthouse Laboratory and Alex Alderton, Roberto Amato, Jeffrey Barrett, Sonia Goncalves, Ewan Harrison, David K. Jackson, Ian Johnston, Dominic Kwiatkowski, Cordelia Langford, John Sillitoe on behalf of the Wellcome Sanger Institute COVID-19 Surveillance Team |

| | | | |
|---|---|---|---|
| EPI_ISL_3359288 | Lighthouse Laboratory Plymouth | Wellcome Sanger Institute for the COVID-19 Genomics UK (COG-UK) Consortium | Lighthouse Laboratory Plymouth and Alex Alderton, Roberto Amato, Jeffrey Barrett, Sonia Goncalves, Ewan Harrison, David K. Jackson, Ian Johnston, Dominic Kwiatkowski, Cordelia Langford, John Sillitoe on behalf of the Wellcome Sanger Institute COVID-19 Surveillance Team |
| EPI_ISL_3359289 | Lighthouse Lab in Milton Keynes | Wellcome Sanger Institute for the COVID-19 Genomics UK (COG-UK) Consortium | The Lighthouse Lab in Milton Keynes and Alex Alderton, Roberto Amato, Jeffrey Barrett, Sonia Goncalves, Ewan Harrison, David K. Jackson, Ian Johnston, Dominic Kwiatkowski, Cordelia Langford, John Sillitoe on behalf of the Wellcome Sanger Institute COVID-19 Surveillance Team |
| EPI_ISL_3359290, EPI_ISL_3359292, EPI_ISL_3359293 | Lighthouse Laboratory Plymouth | Wellcome Sanger Institute for the COVID-19 Genomics UK (COG-UK) Consortium | Lighthouse Laboratory Plymouth and Alex Alderton, Roberto Amato, Jeffrey Barrett, Sonia Goncalves, Ewan Harrison, David K. Jackson, Ian Johnston, Dominic Kwiatkowski, Cordelia Langford, John Sillitoe on behalf of the Wellcome Sanger Institute COVID-19 Surveillance Team |
| EPI_ISL_3359295 | Berkshire and Surrey Pathology Services Lighthouse Laboratory | Wellcome Sanger Institute for the COVID-19 Genomics UK (COG-UK) Consortium | Berkshire and Surrey Pathology Services Lighthouse Laboratory and Alex Alderton, Roberto Amato, Jeffrey Barrett, Sonia Goncalves, Ewan Harrison, David K. Jackson, Ian Johnston, Dominic Kwiatkowski, Cordelia Langford, John Sillitoe on behalf of the Wellcome Sanger Institute COVID-19 Surveillance Team |
| EPI_ISL_3359296, EPI_ISL_3359297, EPI_ISL_3359299, EPI_ISL_3359300, EPI_ISL_3359301, EPI_ISL_3359303, EPI_ISL_3359304, EPI_ISL_3359306, EPI_ISL_3359307, EPI_ISL_3359308, EPI_ISL_3359310, EPI_ISL_3359311 | | | |
| see above | Lighthouse Laboratory Plymouth | Wellcome Sanger Institute for the COVID-19 Genomics UK (COG-UK) Consortium | Lighthouse Laboratory Plymouth and Alex Alderton, Roberto Amato, Jeffrey Barrett, Sonia Goncalves, Ewan Harrison, David K. Jackson, Ian Johnston, Dominic Kwiatkowski, Cordelia Langford, John Sillitoe on behalf of the Wellcome Sanger Institute COVID-19 Surveillance Team |
| EPI_ISL_3359313 | Berkshire and Surrey Pathology Services Lighthouse Laboratory | Wellcome Sanger Institute for the COVID-19 Genomics UK (COG-UK) Consortium | Berkshire and Surrey Pathology Services Lighthouse Laboratory and Alex Alderton, Roberto Amato, Jeffrey Barrett, Sonia Goncalves, Ewan Harrison, David K. Jackson, Ian Johnston, Dominic Kwiatkowski, Cordelia Langford, John Sillitoe on behalf of the Wellcome Sanger Institute COVID-19 Surveillance Team |
| EPI_ISL_3359314, EPI_ISL_3359316, EPI_ISL_3359317, EPI_ISL_3359318, EPI_ISL_3359320 | Lighthouse Laboratory Plymouth | Wellcome Sanger Institute for the COVID-19 Genomics UK (COG-UK) Consortium | Lighthouse Laboratory Plymouth and Alex Alderton, Roberto Amato, Jeffrey Barrett, Sonia Goncalves, Ewan Harrison, David K. Jackson, Ian Johnston, Dominic Kwiatkowski, Cordelia Langford, John Sillitoe on behalf of the Wellcome Sanger Institute COVID-19 Surveillance Team |
| EPI_ISL_3359321, EPI_ISL_3359322, EPI_ISL_3359324 | Berkshire and Surrey Pathology Services Lighthouse Laboratory | Wellcome Sanger Institute for the COVID-19 Genomics UK (COG-UK) Consortium | Berkshire and Surrey Pathology Services Lighthouse Laboratory and Alex Alderton, Roberto Amato, Jeffrey Barrett, Sonia Goncalves, Ewan Harrison, David K. Jackson, Ian Johnston, Dominic Kwiatkowski, Cordelia Langford, John Sillitoe on behalf of the Wellcome Sanger Institute COVID-19 Surveillance Team |
| EPI_ISL_3359325, EPI_ISL_3359327, EPI_ISL_3359328, EPI_ISL_3359329 | Lighthouse Laboratory Plymouth | Wellcome Sanger Institute for the COVID-19 Genomics UK (COG-UK) Consortium | Lighthouse Laboratory Plymouth and Alex Alderton, Roberto Amato, Jeffrey Barrett, Sonia Goncalves, Ewan Harrison, David K. Jackson, Ian Johnston, Dominic Kwiatkowski, Cordelia Langford, John Sillitoe on behalf of the Wellcome Sanger Institute COVID-19 Surveillance Team |
| EPI_ISL_3359331, EPI_ISL_3359332 | Berkshire and Surrey Pathology Services Lighthouse Laboratory | Wellcome Sanger Institute for the COVID-19 Genomics UK (COG-UK) Consortium | Berkshire and Surrey Pathology Services Lighthouse Laboratory and Alex Alderton, Roberto Amato, Jeffrey Barrett, Sonia Goncalves, Ewan Harrison, David K. Jackson, Ian Johnston, Dominic Kwiatkowski, Cordelia Langford, John Sillitoe on behalf of the Wellcome Sanger Institute COVID-19 Surveillance Team |
| EPI_ISL_3359334, EPI_ISL_3359335, EPI_ISL_3359337, EPI_ISL_3359338 | Lighthouse Laboratory Plymouth | Wellcome Sanger Institute for the COVID-19 Genomics UK (COG-UK) Consortium | Lighthouse Laboratory Plymouth and Alex Alderton, Roberto Amato, Jeffrey Barrett, Sonia Goncalves, Ewan Harrison, David K. Jackson, Ian Johnston, Dominic Kwiatkowski, Cordelia Langford, John Sillitoe on behalf of the Wellcome Sanger Institute COVID-19 Surveillance Team |
| EPI_ISL_3359340 | Lighthouse Lab in Milton Keynes | Wellcome Sanger Institute for the COVID-19 Genomics UK (COG-UK) Consortium | The Lighthouse Lab in Milton Keynes and Alex Alderton, Roberto Amato, Jeffrey Barrett, Sonia Goncalves, Ewan Harrison, David K. Jackson, Ian Johnston, Dominic Kwiatkowski, Cordelia Langford, John Sillitoe on behalf of the Wellcome Sanger Institute COVID-19 Surveillance Team |
| EPI_ISL_3359341, EPI_ISL_3359342 | Berkshire and Surrey Pathology Services Lighthouse Laboratory | Wellcome Sanger Institute for the COVID-19 Genomics UK (COG-UK) Consortium | Berkshire and Surrey Pathology Services Lighthouse Laboratory and Alex Alderton, Roberto Amato, Jeffrey Barrett, Sonia Goncalves, Ewan Harrison, David K. Jackson, Ian Johnston, Dominic Kwiatkowski, Cordelia Langford, John Sillitoe on behalf of the Wellcome Sanger Institute COVID-19 Surveillance Team |
| EPI_ISL_3359344 | Lighthouse Laboratory Plymouth | Wellcome Sanger Institute for the COVID-19 Genomics UK (COG-UK) Consortium | Lighthouse Laboratory Plymouth and Alex Alderton, Roberto Amato, Jeffrey Barrett, Sonia Goncalves, Ewan Harrison, David K. Jackson, Ian Johnston, Dominic Kwiatkowski, Cordelia Langford, John Sillitoe on behalf of the Wellcome Sanger Institute COVID-19 Surveillance Team |
| EPI_ISL_3359345 | Lighthouse Lab in Milton Keynes | Wellcome Sanger Institute for the COVID-19 Genomics UK (COG-UK) Consortium | The Lighthouse Lab in Milton Keynes and Alex Alderton, Roberto Amato, Jeffrey Barrett, Sonia Goncalves, Ewan Harrison, David K. Jackson, Ian Johnston, Dominic Kwiatkowski, Cordelia Langford, John Sillitoe on behalf of the Wellcome Sanger Institute COVID-19 Surveillance Team |
| EPI_ISL_3359346 | Lighthouse Laboratory Plymouth | Wellcome Sanger Institute for the COVID-19 Genomics UK (COG-UK) Consortium | Lighthouse Laboratory Plymouth and Alex Alderton, Roberto Amato, Jeffrey Barrett, Sonia Goncalves, Ewan Harrison, David K. Jackson, Ian Johnston, Dominic Kwiatkowski, Cordelia Langford, John Sillitoe on behalf of the Wellcome Sanger Institute COVID-19 Surveillance Team |
| EPI_ISL_3359348 | Lighthouse Lab in Milton Keynes | Wellcome Sanger Institute for the COVID-19 Genomics UK (COG-UK) Consortium | The Lighthouse Lab in Milton Keynes and Alex Alderton, Roberto Amato, Jeffrey Barrett, Sonia Goncalves, Ewan Harrison, David K. Jackson, Ian Johnston, Dominic Kwiatkowski, Cordelia Langford, John Sillitoe on behalf of the Wellcome Sanger Institute COVID-19 Surveillance Team |
| EPI_ISL_3359349 | Berkshire and Surrey Pathology Services Lighthouse Laboratory | Wellcome Sanger Institute for the COVID-19 Genomics UK (COG-UK) Consortium | Berkshire and Surrey Pathology Services Lighthouse Laboratory and Alex Alderton, Roberto Amato, Jeffrey Barrett, Sonia Goncalves, Ewan Harrison, David K. Jackson, Ian Johnston, Dominic Kwiatkowski, Cordelia Langford, John Sillitoe on behalf of the Wellcome Sanger Institute COVID-19 Surveillance Team |
| EPI_ISL_3359351 | Lighthouse Lab in Milton Keynes | Wellcome Sanger Institute for the COVID-19 Genomics UK (COG-UK) Consortium | The Lighthouse Lab in Milton Keynes and Alex Alderton, Roberto Amato, Jeffrey Barrett, Sonia Goncalves, Ewan Harrison, David K. Jackson, Ian Johnston, Dominic Kwiatkowski, Cordelia Langford, John Sillitoe on behalf of the Wellcome Sanger Institute COVID-19 Surveillance Team |
| EPI_ISL_3359353, EPI_ISL_3359354 | Lighthouse Laboratory Plymouth | Wellcome Sanger Institute for the COVID-19 Genomics UK (COG-UK) Consortium | Lighthouse Laboratory Plymouth and Alex Alderton, Roberto Amato, Jeffrey Barrett, Sonia Goncalves, Ewan Harrison, David K. Jackson, Ian Johnston, Dominic Kwiatkowski, Cordelia Langford, John Sillitoe on behalf of the Wellcome Sanger Institute COVID-19 Surveillance Team |
| EPI_ISL_3359355 | Berkshire and Surrey Pathology Services Lighthouse Laboratory | Wellcome Sanger Institute for the COVID-19 Genomics UK (COG-UK) Consortium | Berkshire and Surrey Pathology Services Lighthouse Laboratory and Alex Alderton, Roberto Amato, Jeffrey Barrett, Sonia Goncalves, Ewan Harrison, David K. Jackson, Ian Johnston, Dominic Kwiatkowski, Cordelia Langford, John Sillitoe on behalf of the Wellcome Sanger Institute COVID-19 Surveillance Team |
| EPI_ISL_3359357, EPI_ISL_3359358 | Lighthouse Laboratory Plymouth | Wellcome Sanger Institute for the COVID-19 Genomics UK (COG-UK) Consortium | Lighthouse Laboratory Plymouth and Alex Alderton, Roberto Amato, Jeffrey Barrett, Sonia Goncalves, Ewan Harrison, David K. Jackson, Ian Johnston, Dominic Kwiatkowski, Cordelia Langford, John Sillitoe on behalf of the Wellcome Sanger Institute COVID-19 Surveillance Team |
| EPI_ISL_3359360 | Berkshire and Surrey Pathology Services Lighthouse Laboratory | Wellcome Sanger Institute for the COVID-19 Genomics UK (COG-UK) Consortium | Berkshire and Surrey Pathology Services Lighthouse Laboratory and Alex Alderton, Roberto Amato, Jeffrey Barrett, Sonia Goncalves, Ewan Harrison, David K. Jackson, Ian Johnston, Dominic Kwiatkowski, Cordelia Langford, John Sillitoe on behalf of the Wellcome Sanger Institute COVID-19 Surveillance Team |
| EPI_ISL_3359361, EPI_ISL_3359362 | Lighthouse Laboratory Plymouth | Wellcome Sanger Institute for the COVID-19 Genomics UK (COG-UK) Consortium | Lighthouse Laboratory Plymouth and Alex Alderton, Roberto Amato, Jeffrey Barrett, Sonia Goncalves, Ewan Harrison, David K. Jackson, Ian Johnston, Dominic Kwiatkowski, Cordelia Langford, John Sillitoe on behalf of the Wellcome Sanger Institute COVID-19 Surveillance Team |
| EPI_ISL_3359364, EPI_ISL_3359365 | Lighthouse Lab in Milton Keynes | Wellcome Sanger Institute for the COVID-19 Genomics UK (COG-UK) Consortium | The Lighthouse Lab in Milton Keynes and Alex Alderton, Roberto Amato, Jeffrey Barrett, Sonia Goncalves, Ewan Harrison, David K. Jackson, Ian Johnston, Dominic Kwiatkowski, Cordelia Langford, John Sillitoe on behalf of the Wellcome Sanger Institute COVID-19 Surveillance Team |
| EPI_ISL_3359367 | Berkshire and Surrey Pathology Services Lighthouse Laboratory | Wellcome Sanger Institute for the COVID-19 Genomics UK (COG-UK) Consortium | Berkshire and Surrey Pathology Services Lighthouse Laboratory and Alex Alderton, Roberto Amato, Jeffrey Barrett, Sonia Goncalves, Ewan Harrison, David K. Jackson, Ian Johnston, Dominic Kwiatkowski, Cordelia Langford, John Sillitoe on behalf of the Wellcome Sanger Institute COVID-19 Surveillance Team |
| EPI_ISL_3359368 | Lighthouse Lab in Milton Keynes | Wellcome Sanger Institute for the COVID-19 Genomics UK (COG-UK) Consortium | The Lighthouse Lab in Milton Keynes and Alex Alderton, Roberto Amato, Jeffrey Barrett, Sonia Goncalves, Ewan Harrison, David K. Jackson, Ian Johnston, Dominic Kwiatkowski, Cordelia Langford, John Sillitoe on behalf of the Wellcome Sanger Institute COVID-19 Surveillance Team |
| EPI_ISL_3359369 | Lighthouse Laboratory Plymouth | Wellcome Sanger Institute for the COVID-19 Genomics UK (COG-UK) Consortium | Lighthouse Laboratory Plymouth and Alex Alderton, Roberto Amato, Jeffrey Barrett, Sonia Goncalves, Ewan Harrison, David K. Jackson, Ian Johnston, Dominic Kwiatkowski, Cordelia Langford, John Sillitoe on behalf of the Wellcome Sanger Institute COVID-19 Surveillance Team |
| EPI_ISL_3359371 | Lighthouse Lab in Milton Keynes | Wellcome Sanger Institute for the COVID-19 Genomics UK (COG-UK) Consortium | The Lighthouse Lab in Milton Keynes and Alex Alderton, Roberto Amato, Jeffrey Barrett, Sonia Goncalves, Ewan Harrison, David K. Jackson, Ian Johnston, Dominic Kwiatkowski, Cordelia Langford, John Sillitoe on behalf of the Wellcome Sanger Institute COVID-19 Surveillance Team |
| EPI_ISL_3359372, EPI_ISL_3359373, EPI_ISL_3359375, EPI_ISL_3359376, EPI_ISL_3359378 | Lighthouse Laboratory Plymouth | Wellcome Sanger Institute for the COVID-19 Genomics UK (COG-UK) Consortium | Lighthouse Laboratory Plymouth and Alex Alderton, Roberto Amato, Jeffrey Barrett, Sonia Goncalves, Ewan Harrison, David K. Jackson, Ian Johnston, Dominic Kwiatkowski, Cordelia Langford, John Sillitoe on behalf of the Wellcome Sanger Institute COVID-19 Surveillance Team |
| EPI_ISL_3359379 | Lighthouse Lab in Milton Keynes | Wellcome Sanger Institute for the COVID-19 Genomics UK (COG-UK) Consortium | The Lighthouse Lab in Milton Keynes and Alex Alderton, Roberto Amato, Jeffrey Barrett, Sonia Goncalves, Ewan Harrison, David K. Jackson, Ian Johnston, Dominic Kwiatkowski, Cordelia Langford, John Sillitoe on behalf of the Wellcome Sanger Institute COVID-19 Surveillance Team |
| EPI_ISL_3359381, EPI_ISL_3359382, EPI_ISL_3359384 | Lighthouse Laboratory Plymouth | Wellcome Sanger Institute for the COVID-19 Genomics UK (COG-UK) Consortium | Lighthouse Laboratory Plymouth and Alex Alderton, Roberto Amato, Jeffrey Barrett, Sonia Goncalves, Ewan Harrison, David K. Jackson, Ian Johnston, Dominic Kwiatkowski, Cordelia Langford, John Sillitoe on behalf of the Wellcome Sanger Institute COVID-19 Surveillance Team |
| EPI_ISL_3359385, EPI_ISL_3359386 | Lighthouse Lab in Milton Keynes | Wellcome Sanger Institute for the COVID-19 Genomics UK (COG-UK) Consortium | The Lighthouse Lab in Milton Keynes and Alex Alderton, Roberto Amato, Jeffrey Barrett, Sonia Goncalves, Ewan Harrison, David K. Jackson, Ian Johnston, Dominic Kwiatkowski, Cordelia Langford, John Sillitoe on behalf of the Wellcome Sanger Institute COVID-19 Surveillance Team |
| EPI_ISL_3359388, EPI_ISL_3359390, EPI_ISL_3359391 | Lighthouse Laboratory Plymouth | Wellcome Sanger Institute for the COVID-19 Genomics UK (COG-UK) Consortium | Lighthouse Laboratory Plymouth and Alex Alderton, Roberto Amato, Jeffrey Barrett, Sonia Goncalves, Ewan Harrison, David K. Jackson, Ian Johnston, Dominic Kwiatkowski, Cordelia Langford, John Sillitoe on behalf of the Wellcome Sanger Institute COVID-19 Surveillance Team |
| EPI_ISL_3359392 | Lighthouse Lab in Milton Keynes | Wellcome Sanger Institute for the COVID-19 Genomics UK (COG-UK) Consortium | The Lighthouse Lab in Milton Keynes and Alex Alderton, Roberto Amato, Jeffrey Barrett, Sonia Goncalves, Ewan Harrison, David K. Jackson, Ian Johnston, Dominic Kwiatkowski, Cordelia Langford, John Sillitoe on behalf of the Wellcome Sanger Institute COVID-19 Surveillance Team |
| EPI_ISL_3359394, EPI_ISL_3359395 | Lighthouse Laboratory Plymouth | Wellcome Sanger Institute for the COVID-19 Genomics UK (COG-UK) Consortium | Lighthouse Laboratory Plymouth and Alex Alderton, Roberto Amato, Jeffrey Barrett, Sonia Goncalves, Ewan Harrison, David K. Jackson, Ian Johnston, Dominic Kwiatkowski, Cordelia Langford, John Sillitoe on behalf of the Wellcome Sanger Institute COVID-19 Surveillance Team |
| EPI_ISL_3359397 | Lighthouse Lab in Milton Keynes | Wellcome Sanger Institute for the COVID-19 Genomics UK (COG-UK) Consortium | The Lighthouse Lab in Milton Keynes and Alex Alderton, Roberto Amato, Jeffrey Barrett, Sonia Goncalves, Ewan Harrison, David K. Jackson, Ian Johnston, Dominic Kwiatkowski, Cordelia Langford, John Sillitoe on behalf of the Wellcome Sanger Institute COVID-19 Surveillance Team |
| EPI_ISL_3359398 | Lighthouse Laboratory Plymouth | Wellcome Sanger Institute for the COVID-19 Genomics UK (COG-UK) Consortium | Lighthouse Laboratory Plymouth and Alex Alderton, Roberto Amato, Jeffrey Barrett, Sonia Goncalves, Ewan Harrison, David K. Jackson, Ian Johnston, Dominic Kwiatkowski, Cordelia Langford, John Sillitoe on behalf of the Wellcome Sanger Institute COVID-19 Surveillance Team |
| EPI_ISL_3359400, EPI_ISL_3359401 | Berkshire and Surrey Pathology Services Lighthouse Laboratory | Wellcome Sanger Institute for the COVID-19 Genomics UK (COG-UK) Consortium | Berkshire and Surrey Pathology Services Lighthouse Laboratory and Alex Alderton, Roberto Amato, Jeffrey Barrett, Sonia Goncalves, Ewan Harrison, David K. Jackson, Ian Johnston, Dominic Kwiatkowski, Cordelia Langford, John Sillitoe on behalf of the Wellcome Sanger Institute COVID-19 Surveillance Team |

| | | | |
|---|---|---|---|
| EPI_ISL_3359403 | Lighthouse Laboratory Plymouth | Wellcome Sanger Institute for the COVID-19 Genomics UK (COG-UK) Consortium | Lighthouse Laboratory Plymouth and Alex Alderton, Roberto Amato, Jeffrey Barrett, Sonia Goncalves, Ewan Harrison, David K. Jackson, Ian Johnston, Dominic Kwiatkowski, Cordelia Langford, John Sillitoe on behalf of the Wellcome Sanger Institute COVID-19 Surveillance Team |
| EPI_ISL_3359404 | Berkshire and Surrey Pathology Services Lighthouse Laboratory | Wellcome Sanger Institute for the COVID-19 Genomics UK (COG-UK) Consortium | Berkshire and Surrey Pathology Services Lighthouse Laboratory and Alex Alderton, Roberto Amato, Jeffrey Barrett, Sonia Goncalves, Ewan Harrison, David K. Jackson, Ian Johnston, Dominic Kwiatkowski, Cordelia Langford, John Sillitoe on behalf of the Wellcome Sanger Institute COVID-19 Surveillance Team |
| EPI_ISL_3359406, EPI_ISL_3359407 | Lighthouse Laboratory Plymouth | Wellcome Sanger Institute for the COVID-19 Genomics UK (COG-UK) Consortium | Lighthouse Laboratory Plymouth and Alex Alderton, Roberto Amato, Jeffrey Barrett, Sonia Goncalves, Ewan Harrison, David K. Jackson, Ian Johnston, Dominic Kwiatkowski, Cordelia Langford, John Sillitoe on behalf of the Wellcome Sanger Institute COVID-19 Surveillance Team |
| EPI_ISL_3359409 | Lighthouse Lab in Milton Keynes | Wellcome Sanger Institute for the COVID-19 Genomics UK (COG-UK) Consortium | The Lighthouse Lab in Milton Keynes and Alex Alderton, Roberto Amato, Jeffrey Barrett, Sonia Goncalves, Ewan Harrison, David K. Jackson, Ian Johnston, Dominic Kwiatkowski, Cordelia Langford, John Sillitoe on behalf of the Wellcome Sanger Institute COVID-19 Surveillance Team |
| EPI_ISL_3359411, EPI_ISL_3359412, EPI_ISL_3359413 | Lighthouse Laboratory Plymouth | Wellcome Sanger Institute for the COVID-19 Genomics UK (COG-UK) Consortium | Lighthouse Laboratory Plymouth and Alex Alderton, Roberto Amato, Jeffrey Barrett, Sonia Goncalves, Ewan Harrison, David K. Jackson, Ian Johnston, Dominic Kwiatkowski, Cordelia Langford, John Sillitoe on behalf of the Wellcome Sanger Institute COVID-19 Surveillance Team |
| EPI_ISL_3359415 | Lighthouse Lab in Milton Keynes | Wellcome Sanger Institute for the COVID-19 Genomics UK (COG-UK) Consortium | The Lighthouse Lab in Milton Keynes and Alex Alderton, Roberto Amato, Jeffrey Barrett, Sonia Goncalves, Ewan Harrison, David K. Jackson, Ian Johnston, Dominic Kwiatkowski, Cordelia Langford, John Sillitoe on behalf of the Wellcome Sanger Institute COVID-19 Surveillance Team |
| EPI_ISL_3359416 | Berkshire and Surrey Pathology Services Lighthouse Laboratory | Wellcome Sanger Institute for the COVID-19 Genomics UK (COG-UK) Consortium | Berkshire and Surrey Pathology Services Lighthouse Laboratory and Alex Alderton, Roberto Amato, Jeffrey Barrett, Sonia Goncalves, Ewan Harrison, David K. Jackson, Ian Johnston, Dominic Kwiatkowski, Cordelia Langford, John Sillitoe on behalf of the Wellcome Sanger Institute COVID-19 Surveillance Team |
| EPI_ISL_3359417, EPI_ISL_3359419 | Lighthouse Lab in Milton Keynes | Wellcome Sanger Institute for the COVID-19 Genomics UK (COG-UK) Consortium | The Lighthouse Lab in Milton Keynes and Alex Alderton, Roberto Amato, Jeffrey Barrett, Sonia Goncalves, Ewan Harrison, David K. Jackson, Ian Johnston, Dominic Kwiatkowski, Cordelia Langford, John Sillitoe on behalf of the Wellcome Sanger Institute COVID-19 Surveillance Team |
| EPI_ISL_3359420 | Berkshire and Surrey Pathology Services Lighthouse Laboratory | Wellcome Sanger Institute for the COVID-19 Genomics UK (COG-UK) Consortium | Berkshire and Surrey Pathology Services Lighthouse Laboratory and Alex Alderton, Roberto Amato, Jeffrey Barrett, Sonia Goncalves, Ewan Harrison, David K. Jackson, Ian Johnston, Dominic Kwiatkowski, Cordelia Langford, John Sillitoe on behalf of the Wellcome Sanger Institute COVID-19 Surveillance Team |
| EPI_ISL_3359422 | Lighthouse Laboratory Plymouth | Wellcome Sanger Institute for the COVID-19 Genomics UK (COG-UK) Consortium | Lighthouse Laboratory Plymouth and Alex Alderton, Roberto Amato, Jeffrey Barrett, Sonia Goncalves, Ewan Harrison, David K. Jackson, Ian Johnston, Dominic Kwiatkowski, Cordelia Langford, John Sillitoe on behalf of the Wellcome Sanger Institute COVID-19 Surveillance Team |
| EPI_ISL_3359424 | Lighthouse Lab in Milton Keynes | Wellcome Sanger Institute for the COVID-19 Genomics UK (COG-UK) Consortium | The Lighthouse Lab in Milton Keynes and Alex Alderton, Roberto Amato, Jeffrey Barrett, Sonia Goncalves, Ewan Harrison, David K. Jackson, Ian Johnston, Dominic Kwiatkowski, Cordelia Langford, John Sillitoe on behalf of the Wellcome Sanger Institute COVID-19 Surveillance Team |
| EPI_ISL_3359425, EPI_ISL_3359426 | Lighthouse Laboratory Plymouth | Wellcome Sanger Institute for the COVID-19 Genomics UK (COG-UK) Consortium | Lighthouse Laboratory Plymouth and Alex Alderton, Roberto Amato, Jeffrey Barrett, Sonia Goncalves, Ewan Harrison, David K. Jackson, Ian Johnston, Dominic Kwiatkowski, Cordelia Langford, John Sillitoe on behalf of the Wellcome Sanger Institute COVID-19 Surveillance Team |
| EPI_ISL_3359428 | Berkshire and Surrey Pathology Services Lighthouse Laboratory | Wellcome Sanger Institute for the COVID-19 Genomics UK (COG-UK) Consortium | Berkshire and Surrey Pathology Services Lighthouse Laboratory and Alex Alderton, Roberto Amato, Jeffrey Barrett, Sonia Goncalves, Ewan Harrison, David K. Jackson, Ian Johnston, Dominic Kwiatkowski, Cordelia Langford, John Sillitoe on behalf of the Wellcome Sanger Institute COVID-19 Surveillance Team |
| EPI_ISL_3359429 | Lighthouse Lab in Milton Keynes | Wellcome Sanger Institute for the COVID-19 Genomics UK (COG-UK) Consortium | The Lighthouse Lab in Milton Keynes and Alex Alderton, Roberto Amato, Jeffrey Barrett, Sonia Goncalves, Ewan Harrison, David K. Jackson, Ian Johnston, Dominic Kwiatkowski, Cordelia Langford, John Sillitoe on behalf of the Wellcome Sanger Institute COVID-19 Surveillance Team |
| EPI_ISL_3359431 | Lighthouse Laboratory Plymouth | Wellcome Sanger Institute for the COVID-19 Genomics UK (COG-UK) Consortium | Lighthouse Laboratory Plymouth and Alex Alderton, Roberto Amato, Jeffrey Barrett, Sonia Goncalves, Ewan Harrison, David K. Jackson, Ian Johnston, Dominic Kwiatkowski, Cordelia Langford, John Sillitoe on behalf of the Wellcome Sanger Institute COVID-19 Surveillance Team |
| EPI_ISL_3359432 | Berkshire and Surrey Pathology Services Lighthouse Laboratory | Wellcome Sanger Institute for the COVID-19 Genomics UK (COG-UK) Consortium | Berkshire and Surrey Pathology Services Lighthouse Laboratory and Alex Alderton, Roberto Amato, Jeffrey Barrett, Sonia Goncalves, Ewan Harrison, David K. Jackson, Ian Johnston, Dominic Kwiatkowski, Cordelia Langford, John Sillitoe on behalf of the Wellcome Sanger Institute COVID-19 Surveillance Team |
| EPI_ISL_3359434 | Lighthouse Lab in Milton Keynes | Wellcome Sanger Institute for the COVID-19 Genomics UK (COG-UK) Consortium | The Lighthouse Lab in Milton Keynes and Alex Alderton, Roberto Amato, Jeffrey Barrett, Sonia Goncalves, Ewan Harrison, David K. Jackson, Ian Johnston, Dominic Kwiatkowski, Cordelia Langford, John Sillitoe on behalf of the Wellcome Sanger Institute COVID-19 Surveillance Team |
| EPI_ISL_3359435 | Lighthouse Laboratory Plymouth | Wellcome Sanger Institute for the COVID-19 Genomics UK (COG-UK) Consortium | Lighthouse Laboratory Plymouth and Alex Alderton, Roberto Amato, Jeffrey Barrett, Sonia Goncalves, Ewan Harrison, David K. Jackson, Ian Johnston, Dominic Kwiatkowski, Cordelia Langford, John Sillitoe on behalf of the Wellcome Sanger Institute COVID-19 Surveillance Team |
| EPI_ISL_3359437, EPI_ISL_3359439 | Lighthouse Lab in Milton Keynes | Wellcome Sanger Institute for the COVID-19 Genomics UK (COG-UK) Consortium | The Lighthouse Lab in Milton Keynes and Alex Alderton, Roberto Amato, Jeffrey Barrett, Sonia Goncalves, Ewan Harrison, David K. Jackson, Ian Johnston, Dominic Kwiatkowski, Cordelia Langford, John Sillitoe on behalf of the Wellcome Sanger Institute COVID-19 Surveillance Team |
| EPI_ISL_3359440 | Lighthouse Laboratory Plymouth | Wellcome Sanger Institute for the COVID-19 Genomics UK (COG-UK) Consortium | Lighthouse Laboratory Plymouth and Alex Alderton, Roberto Amato, Jeffrey Barrett, Sonia Goncalves, Ewan Harrison, David K. Jackson, Ian Johnston, Dominic Kwiatkowski, Cordelia Langford, John Sillitoe on behalf of the Wellcome Sanger Institute COVID-19 Surveillance Team |
| EPI_ISL_3359441 | Lighthouse Lab in Milton Keynes | Wellcome Sanger Institute for the COVID-19 Genomics UK (COG-UK) Consortium | The Lighthouse Lab in Milton Keynes and Alex Alderton, Roberto Amato, Jeffrey Barrett, Sonia Goncalves, Ewan Harrison, David K. Jackson, Ian Johnston, Dominic Kwiatkowski, Cordelia Langford, John Sillitoe on behalf of the Wellcome Sanger Institute COVID-19 Surveillance Team |
| EPI_ISL_3359443 | Lighthouse Laboratory Plymouth | Wellcome Sanger Institute for the COVID-19 Genomics UK (COG-UK) Consortium | Lighthouse Laboratory Plymouth and Alex Alderton, Roberto Amato, Jeffrey Barrett, Sonia Goncalves, Ewan Harrison, David K. Jackson, Ian Johnston, Dominic Kwiatkowski, Cordelia Langford, John Sillitoe on behalf of the Wellcome Sanger Institute COVID-19 Surveillance Team |
| EPI_ISL_3359444 | Berkshire and Surrey Pathology Services Lighthouse Laboratory | Wellcome Sanger Institute for the COVID-19 Genomics UK (COG-UK) Consortium | Berkshire and Surrey Pathology Services Lighthouse Laboratory and Alex Alderton, Roberto Amato, Jeffrey Barrett, Sonia Goncalves, Ewan Harrison, David K. Jackson, Ian Johnston, Dominic Kwiatkowski, Cordelia Langford, John Sillitoe on behalf of the Wellcome Sanger Institute COVID-19 Surveillance Team |
| EPI_ISL_3359446 | Lighthouse Laboratory Plymouth | Wellcome Sanger Institute for the COVID-19 Genomics UK (COG-UK) Consortium | Lighthouse Laboratory Plymouth and Alex Alderton, Roberto Amato, Jeffrey Barrett, Sonia Goncalves, Ewan Harrison, David K. Jackson, Ian Johnston, Dominic Kwiatkowski, Cordelia Langford, John Sillitoe on behalf of the Wellcome Sanger Institute COVID-19 Surveillance Team |
| EPI_ISL_3359447, EPI_ISL_3359449 | Berkshire and Surrey Pathology Services Lighthouse Laboratory | Wellcome Sanger Institute for the COVID-19 Genomics UK (COG-UK) Consortium | Berkshire and Surrey Pathology Services Lighthouse Laboratory and Alex Alderton, Roberto Amato, Jeffrey Barrett, Sonia Goncalves, Ewan Harrison, David K. Jackson, Ian Johnston, Dominic Kwiatkowski, Cordelia Langford, John Sillitoe on behalf of the Wellcome Sanger Institute COVID-19 Surveillance Team |
| EPI_ISL_3359450, EPI_ISL_3359452, EPI_ISL_3359453, EPI_ISL_3359454, EPI_ISL_3359456, EPI_ISL_3359458, EPI_ISL_3359460, EPI_ISL_3359461 | Lighthouse Laboratory Plymouth | Wellcome Sanger Institute for the COVID-19 Genomics UK (COG-UK) Consortium | Lighthouse Laboratory Plymouth and Alex Alderton, Roberto Amato, Jeffrey Barrett, Sonia Goncalves, Ewan Harrison, David K. Jackson, Ian Johnston, Dominic Kwiatkowski, Cordelia Langford, John Sillitoe on behalf of the Wellcome Sanger Institute COVID-19 Surveillance Team |
| EPI_ISL_3359463 | Lighthouse Lab in Milton Keynes | Wellcome Sanger Institute for the COVID-19 Genomics UK (COG-UK) Consortium | The Lighthouse Lab in Milton Keynes and Alex Alderton, Roberto Amato, Jeffrey Barrett, Sonia Goncalves, Ewan Harrison, David K. Jackson, Ian Johnston, Dominic Kwiatkowski, Cordelia Langford, John Sillitoe on behalf of the Wellcome Sanger Institute COVID-19 Surveillance Team |
| EPI_ISL_3359464 | Berkshire and Surrey Pathology Services Lighthouse Laboratory | Wellcome Sanger Institute for the COVID-19 Genomics UK (COG-UK) Consortium | Berkshire and Surrey Pathology Services Lighthouse Laboratory and Alex Alderton, Roberto Amato, Jeffrey Barrett, Sonia Goncalves, Ewan Harrison, David K. Jackson, Ian Johnston, Dominic Kwiatkowski, Cordelia Langford, John Sillitoe on behalf of the Wellcome Sanger Institute COVID-19 Surveillance Team |
| EPI_ISL_3359465 | Lighthouse Laboratory Plymouth | Wellcome Sanger Institute for the COVID-19 Genomics UK (COG-UK) Consortium | Lighthouse Laboratory Plymouth and Alex Alderton, Roberto Amato, Jeffrey Barrett, Sonia Goncalves, Ewan Harrison, David K. Jackson, Ian Johnston, Dominic Kwiatkowski, Cordelia Langford, John Sillitoe on behalf of the Wellcome Sanger Institute COVID-19 Surveillance Team |
| EPI_ISL_3359467 | Berkshire and Surrey Pathology Services Lighthouse Laboratory | Wellcome Sanger Institute for the COVID-19 Genomics UK (COG-UK) Consortium | Berkshire and Surrey Pathology Services Lighthouse Laboratory and Alex Alderton, Roberto Amato, Jeffrey Barrett, Sonia Goncalves, Ewan Harrison, David K. Jackson, Ian Johnston, Dominic Kwiatkowski, Cordelia Langford, John Sillitoe on behalf of the Wellcome Sanger Institute COVID-19 Surveillance Team |
| EPI_ISL_3359468 | Lighthouse Lab in Milton Keynes | Wellcome Sanger Institute for the COVID-19 Genomics UK (COG-UK) Consortium | The Lighthouse Lab in Milton Keynes and Alex Alderton, Roberto Amato, Jeffrey Barrett, Sonia Goncalves, Ewan Harrison, David K. Jackson, Ian Johnston, Dominic Kwiatkowski, Cordelia Langford, John Sillitoe on behalf of the Wellcome Sanger Institute COVID-19 Surveillance Team |
| EPI_ISL_3359470, EPI_ISL_3359471 | Lighthouse Laboratory Plymouth | Wellcome Sanger Institute for the COVID-19 Genomics UK (COG-UK) Consortium | Lighthouse Laboratory Plymouth and Alex Alderton, Roberto Amato, Jeffrey Barrett, Sonia Goncalves, Ewan Harrison, David K. Jackson, Ian Johnston, Dominic Kwiatkowski, Cordelia Langford, John Sillitoe on behalf of the Wellcome Sanger Institute COVID-19 Surveillance Team |
| EPI_ISL_3359473 | Berkshire and Surrey Pathology Services Lighthouse Laboratory | Wellcome Sanger Institute for the COVID-19 Genomics UK (COG-UK) Consortium | Berkshire and Surrey Pathology Services Lighthouse Laboratory and Alex Alderton, Roberto Amato, Jeffrey Barrett, Sonia Goncalves, Ewan Harrison, David K. Jackson, Ian Johnston, Dominic Kwiatkowski, Cordelia Langford, John Sillitoe on behalf of the Wellcome Sanger Institute COVID-19 Surveillance Team |
| EPI_ISL_3359474 | Lighthouse Laboratory Plymouth | Wellcome Sanger Institute for the COVID-19 Genomics UK (COG-UK) Consortium | Lighthouse Laboratory Plymouth and Alex Alderton, Roberto Amato, Jeffrey Barrett, Sonia Goncalves, Ewan Harrison, David K. Jackson, Ian Johnston, Dominic Kwiatkowski, Cordelia Langford, John Sillitoe on behalf of the Wellcome Sanger Institute COVID-19 Surveillance Team |
| EPI_ISL_3359476 | Lighthouse Lab in Milton Keynes | Wellcome Sanger Institute for the COVID-19 Genomics UK (COG-UK) Consortium | The Lighthouse Lab in Milton Keynes and Alex Alderton, Roberto Amato, Jeffrey Barrett, Sonia Goncalves, Ewan Harrison, David K. Jackson, Ian Johnston, Dominic Kwiatkowski, Cordelia Langford, John Sillitoe on behalf of the Wellcome Sanger Institute COVID-19 Surveillance Team |
| EPI_ISL_3359477 | Berkshire and Surrey Pathology Services Lighthouse Laboratory | Wellcome Sanger Institute for the COVID-19 Genomics UK (COG-UK) Consortium | Berkshire and Surrey Pathology Services Lighthouse Laboratory and Alex Alderton, Roberto Amato, Jeffrey Barrett, Sonia Goncalves, Ewan Harrison, David K. Jackson, Ian Johnston, Dominic Kwiatkowski, Cordelia Langford, John Sillitoe on behalf of the Wellcome Sanger Institute COVID-19 Surveillance Team |
| EPI_ISL_3359479 | Lighthouse Laboratory Plymouth | Wellcome Sanger Institute for the COVID-19 Genomics UK (COG-UK) Consortium | Lighthouse Laboratory Plymouth and Alex Alderton, Roberto Amato, Jeffrey Barrett, Sonia Goncalves, Ewan Harrison, David K. Jackson, Ian Johnston, Dominic Kwiatkowski, Cordelia Langford, John Sillitoe on behalf of the Wellcome Sanger Institute COVID-19 Surveillance Team |
| EPI_ISL_3359480 | Berkshire and Surrey Pathology Services Lighthouse Laboratory | Wellcome Sanger Institute for the COVID-19 Genomics UK (COG-UK) Consortium | Berkshire and Surrey Pathology Services Lighthouse Laboratory and Alex Alderton, Roberto Amato, Jeffrey Barrett, Sonia Goncalves, Ewan Harrison, David K. Jackson, Ian Johnston, Dominic Kwiatkowski, Cordelia Langford, John Sillitoe on behalf of the Wellcome Sanger Institute COVID-19 Surveillance Team |
| EPI_ISL_3359481, EPI_ISL_3359483 | Lighthouse Laboratory Plymouth | Wellcome Sanger Institute for the COVID-19 Genomics UK (COG-UK) Consortium | Lighthouse Laboratory Plymouth and Alex Alderton, Roberto Amato, Jeffrey Barrett, Sonia Goncalves, Ewan Harrison, David K. Jackson, Ian Johnston, Dominic Kwiatkowski, Cordelia Langford, John Sillitoe on behalf of the Wellcome Sanger Institute COVID-19 Surveillance Team |
| EPI_ISL_3359484 | Berkshire and Surrey Pathology Services Lighthouse Laboratory | Wellcome Sanger Institute for the COVID-19 Genomics UK (COG-UK) Consortium | Berkshire and Surrey Pathology Services Lighthouse Laboratory and Alex Alderton, Roberto Amato, Jeffrey Barrett, Sonia Goncalves, Ewan Harrison, David K. Jackson, Ian Johnston, Dominic Kwiatkowski, Cordelia Langford, John Sillitoe on behalf of the Wellcome Sanger Institute COVID-19 Surveillance Team |
| EPI_ISL_3359486 | Lighthouse Lab in Milton Keynes | Wellcome Sanger Institute for the COVID-19 Genomics UK (COG-UK) Consortium | The Lighthouse Lab in Milton Keynes and Alex Alderton, Roberto Amato, Jeffrey Barrett, Sonia Goncalves, Ewan Harrison, David K. Jackson, Ian Johnston, Dominic Kwiatkowski, Cordelia Langford, John Sillitoe on behalf of the Wellcome Sanger Institute COVID-19 Surveillance Team |
| EPI_ISL_3359487, EPI_ISL_3359488 | Lighthouse Laboratory Plymouth | Wellcome Sanger Institute for the COVID-19 Genomics UK (COG-UK) Consortium | Lighthouse Laboratory Plymouth and Alex Alderton, Roberto Amato, Jeffrey Barrett, Sonia Goncalves, Ewan Harrison, David K. Jackson, Ian Johnston, Dominic Kwiatkowski, Cordelia Langford, John Sillitoe on behalf of the Wellcome Sanger Institute COVID-19 Surveillance Team |
| EPI_ISL_3359490 | Lighthouse Lab in Milton Keynes | Wellcome Sanger Institute for the COVID-19 Genomics UK (COG-UK) | The Lighthouse Lab in Milton Keynes and Alex Alderton, Roberto Amato, Jeffrey Barrett, Sonia Goncalves, Ewan Harrison, David K. Jackson, Ian Johnston, Dominic Kwiatkowski, Cordelia Langford, John Sillitoe on behalf of the Wellcome Sanger |

| | | | |
|---|---|---|---|
| | | Consortium | Institute COVID-19 Surveillance Team |
| EPI_ISL_3359491, EPI_ISL_3359493, EPI_ISL_3359494, EPI_ISL_3359495, EPI_ISL_3359496, EPI_ISL_3359498, EPI_ISL_3359500 | Lighthouse Laboratory Plymouth | Wellcome Sanger Institute for the COVID-19 Genomics UK (COG-UK) Consortium | Lighthouse Laboratory Plymouth and Alex Alderton, Roberto Amato, Jeffrey Barrett, Sonia Goncalves, Ewan Harrison, David K. Jackson, Ian Johnston, Dominic Kwiatkowski, Cordelia Langford, John Sillitoe on behalf of the Wellcome Sanger Institute COVID-19 Surveillance Team |
| EPI_ISL_3359501 | Berkshire and Surrey Pathology Services Lighthouse Laboratory | Wellcome Sanger Institute for the COVID-19 Genomics UK (COG-UK) Consortium | Berkshire and Surrey Pathology Services Lighthouse Laboratory and Alex Alderton, Roberto Amato, Jeffrey Barrett, Sonia Goncalves, Ewan Harrison, David K. Jackson, Ian Johnston, Dominic Kwiatkowski, Cordelia Langford, John Sillitoe on behalf of the Wellcome Sanger Institute COVID-19 Surveillance Team |
| EPI_ISL_3359503, EPI_ISL_3359504 | Lighthouse Lab in Milton Keynes | Wellcome Sanger Institute for the COVID-19 Genomics UK (COG-UK) Consortium | The Lighthouse Lab in Milton Keynes and Alex Alderton, Roberto Amato, Jeffrey Barrett, Sonia Goncalves, Ewan Harrison, David K. Jackson, Ian Johnston, Dominic Kwiatkowski, Cordelia Langford, John Sillitoe on behalf of the Wellcome Sanger Institute COVID-19 Surveillance Team |
| EPI_ISL_3359506 | Berkshire and Surrey Pathology Services Lighthouse Laboratory | Wellcome Sanger Institute for the COVID-19 Genomics UK (COG-UK) Consortium | Berkshire and Surrey Pathology Services Lighthouse Laboratory and Alex Alderton, Roberto Amato, Jeffrey Barrett, Sonia Goncalves, Ewan Harrison, David K. Jackson, Ian Johnston, Dominic Kwiatkowski, Cordelia Langford, John Sillitoe on behalf of the Wellcome Sanger Institute COVID-19 Surveillance Team |
| EPI_ISL_3359507, EPI_ISL_3359508, EPI_ISL_3359510, EPI_ISL_3359511, EPI_ISL_3359513, EPI_ISL_3359515, EPI_ISL_3359516, EPI_ISL_3359518, EPI_ISL_3359520 | Lighthouse Laboratory Plymouth | Wellcome Sanger Institute for the COVID-19 Genomics UK (COG-UK) Consortium | Lighthouse Laboratory Plymouth and Alex Alderton, Roberto Amato, Jeffrey Barrett, Sonia Goncalves, Ewan Harrison, David K. Jackson, Ian Johnston, Dominic Kwiatkowski, Cordelia Langford, John Sillitoe on behalf of the Wellcome Sanger Institute COVID-19 Surveillance Team |
| EPI_ISL_3359521 | Berkshire and Surrey Pathology Services Lighthouse Laboratory | Wellcome Sanger Institute for the COVID-19 Genomics UK (COG-UK) Consortium | Berkshire and Surrey Pathology Services Lighthouse Laboratory and Alex Alderton, Roberto Amato, Jeffrey Barrett, Sonia Goncalves, Ewan Harrison, David K. Jackson, Ian Johnston, Dominic Kwiatkowski, Cordelia Langford, John Sillitoe on behalf of the Wellcome Sanger Institute COVID-19 Surveillance Team |
| EPI_ISL_3359522 | Lighthouse Lab in Milton Keynes | Wellcome Sanger Institute for the COVID-19 Genomics UK (COG-UK) Consortium | The Lighthouse Lab in Milton Keynes and Alex Alderton, Roberto Amato, Jeffrey Barrett, Sonia Goncalves, Ewan Harrison, David K. Jackson, Ian Johnston, Dominic Kwiatkowski, Cordelia Langford, John Sillitoe on behalf of the Wellcome Sanger Institute COVID-19 Surveillance Team |
| EPI_ISL_3359524, EPI_ISL_3359525, EPI_ISL_3359527 | Berkshire and Surrey Pathology Services Lighthouse Laboratory | Wellcome Sanger Institute for the COVID-19 Genomics UK (COG-UK) Consortium | Berkshire and Surrey Pathology Services Lighthouse Laboratory and Alex Alderton, Roberto Amato, Jeffrey Barrett, Sonia Goncalves, Ewan Harrison, David K. Jackson, Ian Johnston, Dominic Kwiatkowski, Cordelia Langford, John Sillitoe on behalf of the Wellcome Sanger Institute COVID-19 Surveillance Team |
| EPI_ISL_3359528 | Lighthouse Lab in Milton Keynes | Wellcome Sanger Institute for the COVID-19 Genomics UK (COG-UK) Consortium | The Lighthouse Lab in Milton Keynes and Alex Alderton, Roberto Amato, Jeffrey Barrett, Sonia Goncalves, Ewan Harrison, David K. Jackson, Ian Johnston, Dominic Kwiatkowski, Cordelia Langford, John Sillitoe on behalf of the Wellcome Sanger Institute COVID-19 Surveillance Team |
| EPI_ISL_3359530, EPI_ISL_3359531 | Lighthouse Laboratory Plymouth | Wellcome Sanger Institute for the COVID-19 Genomics UK (COG-UK) Consortium | Lighthouse Laboratory Plymouth and Alex Alderton, Roberto Amato, Jeffrey Barrett, Sonia Goncalves, Ewan Harrison, David K. Jackson, Ian Johnston, Dominic Kwiatkowski, Cordelia Langford, John Sillitoe on behalf of the Wellcome Sanger Institute COVID-19 Surveillance Team |
| EPI_ISL_3359533 | Lighthouse Lab in Milton Keynes | Wellcome Sanger Institute for the COVID-19 Genomics UK (COG-UK) Consortium | The Lighthouse Lab in Milton Keynes and Alex Alderton, Roberto Amato, Jeffrey Barrett, Sonia Goncalves, Ewan Harrison, David K. Jackson, Ian Johnston, Dominic Kwiatkowski, Cordelia Langford, John Sillitoe on behalf of the Wellcome Sanger Institute COVID-19 Surveillance Team |
| EPI_ISL_3359534 | Lighthouse Laboratory Plymouth | Wellcome Sanger Institute for the COVID-19 Genomics UK (COG-UK) Consortium | Lighthouse Laboratory Plymouth and Alex Alderton, Roberto Amato, Jeffrey Barrett, Sonia Goncalves, Ewan Harrison, David K. Jackson, Ian Johnston, Dominic Kwiatkowski, Cordelia Langford, John Sillitoe on behalf of the Wellcome Sanger Institute COVID-19 Surveillance Team |
| EPI_ISL_3359535, EPI_ISL_3359537 | Lighthouse Lab in Milton Keynes | Wellcome Sanger Institute for the COVID-19 Genomics UK (COG-UK) Consortium | The Lighthouse Lab in Milton Keynes and Alex Alderton, Roberto Amato, Jeffrey Barrett, Sonia Goncalves, Ewan Harrison, David K. Jackson, Ian Johnston, Dominic Kwiatkowski, Cordelia Langford, John Sillitoe on behalf of the Wellcome Sanger Institute COVID-19 Surveillance Team |
| EPI_ISL_3359538, EPI_ISL_3359540 | Berkshire and Surrey Pathology Services Lighthouse Laboratory | Wellcome Sanger Institute for the COVID-19 Genomics UK (COG-UK) Consortium | Berkshire and Surrey Pathology Services Lighthouse Laboratory and Alex Alderton, Roberto Amato, Jeffrey Barrett, Sonia Goncalves, Ewan Harrison, David K. Jackson, Ian Johnston, Dominic Kwiatkowski, Cordelia Langford, John Sillitoe on behalf of the Wellcome Sanger Institute COVID-19 Surveillance Team |
| EPI_ISL_3359541, EPI_ISL_3359542, EPI_ISL_3359544, EPI_ISL_3359545 | Lighthouse Laboratory Plymouth | Wellcome Sanger Institute for the COVID-19 Genomics UK (COG-UK) Consortium | Lighthouse Laboratory Plymouth and Alex Alderton, Roberto Amato, Jeffrey Barrett, Sonia Goncalves, Ewan Harrison, David K. Jackson, Ian Johnston, Dominic Kwiatkowski, Cordelia Langford, John Sillitoe on behalf of the Wellcome Sanger Institute COVID-19 Surveillance Team |
| EPI_ISL_3359547 | Berkshire and Surrey Pathology Services Lighthouse Laboratory | Wellcome Sanger Institute for the COVID-19 Genomics UK (COG-UK) Consortium | Berkshire and Surrey Pathology Services Lighthouse Laboratory and Alex Alderton, Roberto Amato, Jeffrey Barrett, Sonia Goncalves, Ewan Harrison, David K. Jackson, Ian Johnston, Dominic Kwiatkowski, Cordelia Langford, John Sillitoe on behalf of the Wellcome Sanger Institute COVID-19 Surveillance Team |
| EPI_ISL_3359548, EPI_ISL_3359550 | Lighthouse Laboratory Plymouth | Wellcome Sanger Institute for the COVID-19 Genomics UK (COG-UK) Consortium | Lighthouse Laboratory Plymouth and Alex Alderton, Roberto Amato, Jeffrey Barrett, Sonia Goncalves, Ewan Harrison, David K. Jackson, Ian Johnston, Dominic Kwiatkowski, Cordelia Langford, John Sillitoe on behalf of the Wellcome Sanger Institute COVID-19 Surveillance Team |
| EPI_ISL_3359551 | Lighthouse Lab in Milton Keynes | Wellcome Sanger Institute for the COVID-19 Genomics UK (COG-UK) Consortium | The Lighthouse Lab in Milton Keynes and Alex Alderton, Roberto Amato, Jeffrey Barrett, Sonia Goncalves, Ewan Harrison, David K. Jackson, Ian Johnston, Dominic Kwiatkowski, Cordelia Langford, John Sillitoe on behalf of the Wellcome Sanger Institute COVID-19 Surveillance Team |
| EPI_ISL_3359552 | Berkshire and Surrey Pathology Services Lighthouse Laboratory | Wellcome Sanger Institute for the COVID-19 Genomics UK (COG-UK) Consortium | Berkshire and Surrey Pathology Services Lighthouse Laboratory and Alex Alderton, Roberto Amato, Jeffrey Barrett, Sonia Goncalves, Ewan Harrison, David K. Jackson, Ian Johnston, Dominic Kwiatkowski, Cordelia Langford, John Sillitoe on behalf of the Wellcome Sanger Institute COVID-19 Surveillance Team |
| EPI_ISL_3359554 | Lighthouse Lab in Milton Keynes | Wellcome Sanger Institute for the COVID-19 Genomics UK (COG-UK) Consortium | The Lighthouse Lab in Milton Keynes and Alex Alderton, Roberto Amato, Jeffrey Barrett, Sonia Goncalves, Ewan Harrison, David K. Jackson, Ian Johnston, Dominic Kwiatkowski, Cordelia Langford, John Sillitoe on behalf of the Wellcome Sanger Institute COVID-19 Surveillance Team |
| EPI_ISL_3359555, EPI_ISL_3359557, EPI_ISL_3359558, EPI_ISL_3359560 | Lighthouse Laboratory Plymouth | Wellcome Sanger Institute for the COVID-19 Genomics UK (COG-UK) Consortium | Lighthouse Laboratory Plymouth and Alex Alderton, Roberto Amato, Jeffrey Barrett, Sonia Goncalves, Ewan Harrison, David K. Jackson, Ian Johnston, Dominic Kwiatkowski, Cordelia Langford, John Sillitoe on behalf of the Wellcome Sanger Institute COVID-19 Surveillance Team |
| EPI_ISL_3359561, EPI_ISL_3359563 | Berkshire and Surrey Pathology Services Lighthouse Laboratory | Wellcome Sanger Institute for the COVID-19 Genomics UK (COG-UK) Consortium | Berkshire and Surrey Pathology Services Lighthouse Laboratory and Alex Alderton, Roberto Amato, Jeffrey Barrett, Sonia Goncalves, Ewan Harrison, David K. Jackson, Ian Johnston, Dominic Kwiatkowski, Cordelia Langford, John Sillitoe on behalf of the Wellcome Sanger Institute COVID-19 Surveillance Team |
| EPI_ISL_3359564 | Lighthouse Laboratory Plymouth | Wellcome Sanger Institute for the COVID-19 Genomics UK (COG-UK) Consortium | Lighthouse Laboratory Plymouth and Alex Alderton, Roberto Amato, Jeffrey Barrett, Sonia Goncalves, Ewan Harrison, David K. Jackson, Ian Johnston, Dominic Kwiatkowski, Cordelia Langford, John Sillitoe on behalf of the Wellcome Sanger Institute COVID-19 Surveillance Team |
| EPI_ISL_3359565 | Lighthouse Lab in Milton Keynes | Wellcome Sanger Institute for the COVID-19 Genomics UK (COG-UK) Consortium | The Lighthouse Lab in Milton Keynes and Alex Alderton, Roberto Amato, Jeffrey Barrett, Sonia Goncalves, Ewan Harrison, David K. Jackson, Ian Johnston, Dominic Kwiatkowski, Cordelia Langford, John Sillitoe on behalf of the Wellcome Sanger Institute COVID-19 Surveillance Team |
| EPI_ISL_3359567, EPI_ISL_3359568, EPI_ISL_3359570 | Berkshire and Surrey Pathology Services Lighthouse Laboratory | Wellcome Sanger Institute for the COVID-19 Genomics UK (COG-UK) Consortium | Berkshire and Surrey Pathology Services Lighthouse Laboratory and Alex Alderton, Roberto Amato, Jeffrey Barrett, Sonia Goncalves, Ewan Harrison, David K. Jackson, Ian Johnston, Dominic Kwiatkowski, Cordelia Langford, John Sillitoe on behalf of the Wellcome Sanger Institute COVID-19 Surveillance Team |
| EPI_ISL_3359571, EPI_ISL_3359572, EPI_ISL_3359574, EPI_ISL_3359575, EPI_ISL_3359576, EPI_ISL_3359578, EPI_ISL_3359579, EPI_ISL_3359581, EPI_ISL_3359582 | Lighthouse Laboratory Plymouth | Wellcome Sanger Institute for the COVID-19 Genomics UK (COG-UK) Consortium | Lighthouse Laboratory Plymouth and Alex Alderton, Roberto Amato, Jeffrey Barrett, Sonia Goncalves, Ewan Harrison, David K. Jackson, Ian Johnston, Dominic Kwiatkowski, Cordelia Langford, John Sillitoe on behalf of the Wellcome Sanger Institute COVID-19 Surveillance Team |
| EPI_ISL_3359583 | Lighthouse Lab in Milton Keynes | Wellcome Sanger Institute for the COVID-19 Genomics UK (COG-UK) Consortium | The Lighthouse Lab in Milton Keynes and Alex Alderton, Roberto Amato, Jeffrey Barrett, Sonia Goncalves, Ewan Harrison, David K. Jackson, Ian Johnston, Dominic Kwiatkowski, Cordelia Langford, John Sillitoe on behalf of the Wellcome Sanger Institute COVID-19 Surveillance Team |
| EPI_ISL_3359585, EPI_ISL_3359586 | Lighthouse Lab in Alderley Park | Wellcome Sanger Institute for the COVID-19 Genomics UK (COG-UK) Consortium | Jacquelyn Wynn, Mairead Hyland, The Lighthouse Lab in Alderley Park and Alex Alderton, Roberto Amato, Jeffrey Barrett, Sonia Goncalves, Ewan Harrison, David K. Jackson, Ian Johnston, Dominic Kwiatkowski, Cordelia Langford, John Sillitoe on behalf of the Wellcome Sanger Institute COVID-19 Surveillance Team |
| EPI_ISL_3359588 | Lighthouse Lab in Glasgow | Wellcome Sanger Institute for the COVID-19 Genomics UK (COG-UK) Consortium | Harper VanSteenhouse, Yumi Kasai, David Gray, Carol Clugston, Anna Dominiczak and Alex Alderton, Roberto Amato, Jeffrey Barrett, Sonia Goncalves, Ewan Harrison, David K. Jackson, Ian Johnston, Dominic Kwiatkowski, Cordelia Langford, John Sillitoe on behalf of the Wellcome Sanger Institute COVID-19 Surveillance Team |
| EPI_ISL_3359589, EPI_ISL_3359590, EPI_ISL_3359592, EPI_ISL_3359593 | Lighthouse Laboratory Plymouth | Wellcome Sanger Institute for the COVID-19 Genomics UK (COG-UK) Consortium | Lighthouse Laboratory Plymouth and Alex Alderton, Roberto Amato, Jeffrey Barrett, Sonia Goncalves, Ewan Harrison, David K. Jackson, Ian Johnston, Dominic Kwiatkowski, Cordelia Langford, John Sillitoe on behalf of the Wellcome Sanger Institute COVID-19 Surveillance Team |
| EPI_ISL_3359595, EPI_ISL_3359596, EPI_ISL_3359598 | Lighthouse Lab in Alderley Park | Wellcome Sanger Institute for the COVID-19 Genomics UK (COG-UK) Consortium | Jacquelyn Wynn, Mairead Hyland, The Lighthouse Lab in Alderley Park and Alex Alderton, Roberto Amato, Jeffrey Barrett, Sonia Goncalves, Ewan Harrison, David K. Jackson, Ian Johnston, Dominic Kwiatkowski, Cordelia Langford, John Sillitoe on behalf of the Wellcome Sanger Institute COVID-19 Surveillance Team |
| EPI_ISL_3359599 | Lighthouse Lab in Glasgow | Wellcome Sanger Institute for the COVID-19 Genomics UK (COG-UK) Consortium | Harper VanSteenhouse, Yumi Kasai, David Gray, Carol Clugston, Anna Dominiczak and Alex Alderton, Roberto Amato, Jeffrey Barrett, Sonia Goncalves, Ewan Harrison, David K. Jackson, Ian Johnston, Dominic Kwiatkowski, Cordelia Langford, John Sillitoe on behalf of the Wellcome Sanger Institute COVID-19 Surveillance Team |
| EPI_ISL_3359600 | Lighthouse Lab in Alderley Park | Wellcome Sanger Institute for the COVID-19 Genomics UK (COG-UK) Consortium | Jacquelyn Wynn, Mairead Hyland, The Lighthouse Lab in Alderley Park and Alex Alderton, Roberto Amato, Jeffrey Barrett, Sonia Goncalves, Ewan Harrison, David K. Jackson, Ian Johnston, Dominic Kwiatkowski, Cordelia Langford, John Sillitoe on behalf of the Wellcome Sanger Institute COVID-19 Surveillance Team |

| | | | |
|---|---|---|---|
| EPI_ISL_3359602 | Lighthouse Lab in Glasgow | Wellcome Sanger Institute for the COVID-19 Genomics UK (COG-UK) Consortium | Harper VanSteenhouse, Yumi Kasai, David Gray, Carol Clugston, Anna Dominiczak and Alex Alderton, Roberto Amato, Jeffrey Barrett, Sonia Goncalves, Ewan Harrison, David K. Jackson, Ian Johnston, Dominic Kwiatkowski, Cordelia Langford, John Sillitoe on behalf of the Wellcome Sanger Institute COVID-19 Surveillance Team |
| EPI_ISL_3359603 | Lighthouse Laboratory Plymouth | Wellcome Sanger Institute for the COVID-19 Genomics UK (COG-UK) Consortium | Lighthouse Laboratory Plymouth and Alex Alderton, Roberto Amato, Jeffrey Barrett, Sonia Goncalves, Ewan Harrison, David K. Jackson, Ian Johnston, Dominic Kwiatkowski, Cordelia Langford, John Sillitoe on behalf of the Wellcome Sanger Institute COVID-19 Surveillance Team |
| EPI_ISL_3359604 | Lighthouse Lab in Glasgow | Wellcome Sanger Institute for the COVID-19 Genomics UK (COG-UK) Consortium | Harper VanSteenhouse, Yumi Kasai, David Gray, Carol Clugston, Anna Dominiczak and Alex Alderton, Roberto Amato, Jeffrey Barrett, Sonia Goncalves, Ewan Harrison, David K. Jackson, Ian Johnston, Dominic Kwiatkowski, Cordelia Langford, John Sillitoe on behalf of the Wellcome Sanger Institute COVID-19 Surveillance Team |
| EPI_ISL_3359606, EPI_ISL_3359607, EPI_ISL_3359609 | Lighthouse Lab in Alderley Park | Wellcome Sanger Institute for the COVID-19 Genomics UK (COG-UK) Consortium | Jacquelyn Wynn, Mairead Hyland, The Lighthouse Lab in Alderley Park and Alex Alderton, Roberto Amato, Jeffrey Barrett, Sonia Goncalves, Ewan Harrison, David K. Jackson, Ian Johnston, Dominic Kwiatkowski, Cordelia Langford, John Sillitoe on behalf of the Wellcome Sanger Institute COVID-19 Surveillance Team |
| EPI_ISL_3359610 | Lighthouse Laboratory Plymouth | Wellcome Sanger Institute for the COVID-19 Genomics UK (COG-UK) Consortium | Lighthouse Laboratory Plymouth and Alex Alderton, Roberto Amato, Jeffrey Barrett, Sonia Goncalves, Ewan Harrison, David K. Jackson, Ian Johnston, Dominic Kwiatkowski, Cordelia Langford, John Sillitoe on behalf of the Wellcome Sanger Institute COVID-19 Surveillance Team |
| EPI_ISL_3359612, EPI_ISL_3359613, EPI_ISL_3359614, EPI_ISL_3359616, EPI_ISL_3359617 | Lighthouse Lab in Alderley Park | Wellcome Sanger Institute for the COVID-19 Genomics UK (COG-UK) Consortium | Jacquelyn Wynn, Mairead Hyland, The Lighthouse Lab in Alderley Park and Alex Alderton, Roberto Amato, Jeffrey Barrett, Sonia Goncalves, Ewan Harrison, David K. Jackson, Ian Johnston, Dominic Kwiatkowski, Cordelia Langford, John Sillitoe on behalf of the Wellcome Sanger Institute COVID-19 Surveillance Team |
| EPI_ISL_3359619, EPI_ISL_3359620, EPI_ISL_3359622 | Lighthouse Laboratory Plymouth | Wellcome Sanger Institute for the COVID-19 Genomics UK (COG-UK) Consortium | Lighthouse Laboratory Plymouth and Alex Alderton, Roberto Amato, Jeffrey Barrett, Sonia Goncalves, Ewan Harrison, David K. Jackson, Ian Johnston, Dominic Kwiatkowski, Cordelia Langford, John Sillitoe on behalf of the Wellcome Sanger Institute COVID-19 Surveillance Team |
| EPI_ISL_3359623, EPI_ISL_3359624, EPI_ISL_3359626, EPI_ISL_3359627 | Lighthouse Lab in Alderley Park | Wellcome Sanger Institute for the COVID-19 Genomics UK (COG-UK) Consortium | Jacquelyn Wynn, Mairead Hyland, The Lighthouse Lab in Alderley Park and Alex Alderton, Roberto Amato, Jeffrey Barrett, Sonia Goncalves, Ewan Harrison, David K. Jackson, Ian Johnston, Dominic Kwiatkowski, Cordelia Langford, John Sillitoe on behalf of the Wellcome Sanger Institute COVID-19 Surveillance Team |
| EPI_ISL_3359629 | Lighthouse Laboratory Plymouth | Wellcome Sanger Institute for the COVID-19 Genomics UK (COG-UK) Consortium | Lighthouse Laboratory Plymouth and Alex Alderton, Roberto Amato, Jeffrey Barrett, Sonia Goncalves, Ewan Harrison, David K. Jackson, Ian Johnston, Dominic Kwiatkowski, Cordelia Langford, John Sillitoe on behalf of the Wellcome Sanger Institute COVID-19 Surveillance Team |
| EPI_ISL_3359630, EPI_ISL_3359631 | Lighthouse Lab in Alderley Park | Wellcome Sanger Institute for the COVID-19 Genomics UK (COG-UK) Consortium | Jacquelyn Wynn, Mairead Hyland, The Lighthouse Lab in Alderley Park and Alex Alderton, Roberto Amato, Jeffrey Barrett, Sonia Goncalves, Ewan Harrison, David K. Jackson, Ian Johnston, Dominic Kwiatkowski, Cordelia Langford, John Sillitoe on behalf of the Wellcome Sanger Institute COVID-19 Surveillance Team |
| EPI_ISL_3359633 | Lighthouse Laboratory Plymouth | Wellcome Sanger Institute for the COVID-19 Genomics UK (COG-UK) Consortium | Lighthouse Laboratory Plymouth and Alex Alderton, Roberto Amato, Jeffrey Barrett, Sonia Goncalves, Ewan Harrison, David K. Jackson, Ian Johnston, Dominic Kwiatkowski, Cordelia Langford, John Sillitoe on behalf of the Wellcome Sanger Institute COVID-19 Surveillance Team |
| EPI_ISL_3359634, EPI_ISL_3359636 | Lighthouse Lab in Alderley Park | Wellcome Sanger Institute for the COVID-19 Genomics UK (COG-UK) Consortium | Jacquelyn Wynn, Mairead Hyland, The Lighthouse Lab in Alderley Park and Alex Alderton, Roberto Amato, Jeffrey Barrett, Sonia Goncalves, Ewan Harrison, David K. Jackson, Ian Johnston, Dominic Kwiatkowski, Cordelia Langford, John Sillitoe on behalf of the Wellcome Sanger Institute COVID-19 Surveillance Team |
| EPI_ISL_3359637, EPI_ISL_3359639 | Lighthouse Laboratory Plymouth | Wellcome Sanger Institute for the COVID-19 Genomics UK (COG-UK) Consortium | Lighthouse Laboratory Plymouth and Alex Alderton, Roberto Amato, Jeffrey Barrett, Sonia Goncalves, Ewan Harrison, David K. Jackson, Ian Johnston, Dominic Kwiatkowski, Cordelia Langford, John Sillitoe on behalf of the Wellcome Sanger Institute COVID-19 Surveillance Team |
| EPI_ISL_3359640 | Lighthouse Lab in Glasgow | Wellcome Sanger Institute for the COVID-19 Genomics UK (COG-UK) Consortium | Harper VanSteenhouse, Yumi Kasai, David Gray, Carol Clugston, Anna Dominiczak and Alex Alderton, Roberto Amato, Jeffrey Barrett, Sonia Goncalves, Ewan Harrison, David K. Jackson, Ian Johnston, Dominic Kwiatkowski, Cordelia Langford, John Sillitoe on behalf of the Wellcome Sanger Institute COVID-19 Surveillance Team |
| EPI_ISL_3359641, EPI_ISL_3359643, EPI_ISL_3359644 | Lighthouse Laboratory Plymouth | Wellcome Sanger Institute for the COVID-19 Genomics UK (COG-UK) Consortium | Lighthouse Laboratory Plymouth and Alex Alderton, Roberto Amato, Jeffrey Barrett, Sonia Goncalves, Ewan Harrison, David K. Jackson, Ian Johnston, Dominic Kwiatkowski, Cordelia Langford, John Sillitoe on behalf of the Wellcome Sanger Institute COVID-19 Surveillance Team |
| EPI_ISL_3359646 | Lighthouse Lab in Alderley Park | Wellcome Sanger Institute for the COVID-19 Genomics UK (COG-UK) Consortium | Jacquelyn Wynn, Mairead Hyland, The Lighthouse Lab in Alderley Park and Alex Alderton, Roberto Amato, Jeffrey Barrett, Sonia Goncalves, Ewan Harrison, David K. Jackson, Ian Johnston, Dominic Kwiatkowski, Cordelia Langford, John Sillitoe on behalf of the Wellcome Sanger Institute COVID-19 Surveillance Team |
| EPI_ISL_3359647 | Lighthouse Laboratory Plymouth | Wellcome Sanger Institute for the COVID-19 Genomics UK (COG-UK) Consortium | Lighthouse Laboratory Plymouth and Alex Alderton, Roberto Amato, Jeffrey Barrett, Sonia Goncalves, Ewan Harrison, David K. Jackson, Ian Johnston, Dominic Kwiatkowski, Cordelia Langford, John Sillitoe on behalf of the Wellcome Sanger Institute COVID-19 Surveillance Team |
| EPI_ISL_3359649, EPI_ISL_3359650 | Lighthouse Lab in Alderley Park | Wellcome Sanger Institute for the COVID-19 Genomics UK (COG-UK) Consortium | Jacquelyn Wynn, Mairead Hyland, The Lighthouse Lab in Alderley Park and Alex Alderton, Roberto Amato, Jeffrey Barrett, Sonia Goncalves, Ewan Harrison, David K. Jackson, Ian Johnston, Dominic Kwiatkowski, Cordelia Langford, John Sillitoe on behalf of the Wellcome Sanger Institute COVID-19 Surveillance Team |
| EPI_ISL_3359652 | Lighthouse Laboratory Plymouth | Wellcome Sanger Institute for the COVID-19 Genomics UK (COG-UK) Consortium | Lighthouse Laboratory Plymouth and Alex Alderton, Roberto Amato, Jeffrey Barrett, Sonia Goncalves, Ewan Harrison, David K. Jackson, Ian Johnston, Dominic Kwiatkowski, Cordelia Langford, John Sillitoe on behalf of the Wellcome Sanger Institute COVID-19 Surveillance Team |
| EPI_ISL_3359653, EPI_ISL_3359655 | Lighthouse Lab in Alderley Park | Wellcome Sanger Institute for the COVID-19 Genomics UK (COG-UK) Consortium | Jacquelyn Wynn, Mairead Hyland, The Lighthouse Lab in Alderley Park and Alex Alderton, Roberto Amato, Jeffrey Barrett, Sonia Goncalves, Ewan Harrison, David K. Jackson, Ian Johnston, Dominic Kwiatkowski, Cordelia Langford, John Sillitoe on behalf of the Wellcome Sanger Institute COVID-19 Surveillance Team |
| EPI_ISL_3359656, EPI_ISL_3359657 | Lighthouse Lab in Glasgow | Wellcome Sanger Institute for the COVID-19 Genomics UK (COG-UK) Consortium | Harper VanSteenhouse, Yumi Kasai, David Gray, Carol Clugston, Anna Dominiczak and Alex Alderton, Roberto Amato, Jeffrey Barrett, Sonia Goncalves, Ewan Harrison, David K. Jackson, Ian Johnston, Dominic Kwiatkowski, Cordelia Langford, John Sillitoe on behalf of the Wellcome Sanger Institute COVID-19 Surveillance Team |
| EPI_ISL_3359659 | Lighthouse Laboratory Plymouth | Wellcome Sanger Institute for the COVID-19 Genomics UK (COG-UK) Consortium | Lighthouse Laboratory Plymouth and Alex Alderton, Roberto Amato, Jeffrey Barrett, Sonia Goncalves, Ewan Harrison, David K. Jackson, Ian Johnston, Dominic Kwiatkowski, Cordelia Langford, John Sillitoe on behalf of the Wellcome Sanger Institute COVID-19 Surveillance Team |
| EPI_ISL_3359660, EPI_ISL_3359662 | Lighthouse Lab in Alderley Park | Wellcome Sanger Institute for the COVID-19 Genomics UK (COG-UK) Consortium | Jacquelyn Wynn, Mairead Hyland, The Lighthouse Lab in Alderley Park and Alex Alderton, Roberto Amato, Jeffrey Barrett, Sonia Goncalves, Ewan Harrison, David K. Jackson, Ian Johnston, Dominic Kwiatkowski, Cordelia Langford, John Sillitoe on behalf of the Wellcome Sanger Institute COVID-19 Surveillance Team |
| EPI_ISL_3359663 | Lighthouse Laboratory Plymouth | Wellcome Sanger Institute for the COVID-19 Genomics UK (COG-UK) Consortium | Lighthouse Laboratory Plymouth and Alex Alderton, Roberto Amato, Jeffrey Barrett, Sonia Goncalves, Ewan Harrison, David K. Jackson, Ian Johnston, Dominic Kwiatkowski, Cordelia Langford, John Sillitoe on behalf of the Wellcome Sanger Institute COVID-19 Surveillance Team |
| EPI_ISL_3359665, EPI_ISL_3359666, EPI_ISL_3359668, EPI_ISL_3359669, EPI_ISL_3359671 | Lighthouse Lab in Alderley Park | Wellcome Sanger Institute for the COVID-19 Genomics UK (COG-UK) Consortium | Jacquelyn Wynn, Mairead Hyland, The Lighthouse Lab in Alderley Park and Alex Alderton, Roberto Amato, Jeffrey Barrett, Sonia Goncalves, Ewan Harrison, David K. Jackson, Ian Johnston, Dominic Kwiatkowski, Cordelia Langford, John Sillitoe on behalf of the Wellcome Sanger Institute COVID-19 Surveillance Team |
| EPI_ISL_3359672 | Lighthouse Laboratory Plymouth | Wellcome Sanger Institute for the COVID-19 Genomics UK (COG-UK) Consortium | Lighthouse Laboratory Plymouth and Alex Alderton, Roberto Amato, Jeffrey Barrett, Sonia Goncalves, Ewan Harrison, David K. Jackson, Ian Johnston, Dominic Kwiatkowski, Cordelia Langford, John Sillitoe on behalf of the Wellcome Sanger Institute COVID-19 Surveillance Team |
| EPI_ISL_3359673 | Lighthouse Lab in Alderley Park | Wellcome Sanger Institute for the COVID-19 Genomics UK (COG-UK) Consortium | Jacquelyn Wynn, Mairead Hyland, The Lighthouse Lab in Alderley Park and Alex Alderton, Roberto Amato, Jeffrey Barrett, Sonia Goncalves, Ewan Harrison, David K. Jackson, Ian Johnston, Dominic Kwiatkowski, Cordelia Langford, John Sillitoe on behalf of the Wellcome Sanger Institute COVID-19 Surveillance Team |
| EPI_ISL_3359675 | Lighthouse Laboratory Plymouth | Wellcome Sanger Institute for the COVID-19 Genomics UK (COG-UK) Consortium | Lighthouse Laboratory Plymouth and Alex Alderton, Roberto Amato, Jeffrey Barrett, Sonia Goncalves, Ewan Harrison, David K. Jackson, Ian Johnston, Dominic Kwiatkowski, Cordelia Langford, John Sillitoe on behalf of the Wellcome Sanger Institute COVID-19 Surveillance Team |
| EPI_ISL_3359676, EPI_ISL_3359677 | Lighthouse Lab in Alderley Park | Wellcome Sanger Institute for the COVID-19 Genomics UK (COG-UK) Consortium | Jacquelyn Wynn, Mairead Hyland, The Lighthouse Lab in Alderley Park and Alex Alderton, Roberto Amato, Jeffrey Barrett, Sonia Goncalves, Ewan Harrison, David K. Jackson, Ian Johnston, Dominic Kwiatkowski, Cordelia Langford, John Sillitoe on behalf of the Wellcome Sanger Institute COVID-19 Surveillance Team |
| EPI_ISL_3359679, EPI_ISL_3359680 | Lighthouse Lab in Glasgow | Wellcome Sanger Institute for the COVID-19 Genomics UK (COG-UK) Consortium | Harper VanSteenhouse, Yumi Kasai, David Gray, Carol Clugston, Anna Dominiczak and Alex Alderton, Roberto Amato, Jeffrey Barrett, Sonia Goncalves, Ewan Harrison, David K. Jackson, Ian Johnston, Dominic Kwiatkowski, Cordelia Langford, John Sillitoe on behalf of the Wellcome Sanger Institute COVID-19 Surveillance Team |
| EPI_ISL_3359682 | Lighthouse Laboratory Plymouth | Wellcome Sanger Institute for the COVID-19 Genomics UK (COG-UK) Consortium | Lighthouse Laboratory Plymouth and Alex Alderton, Roberto Amato, Jeffrey Barrett, Sonia Goncalves, Ewan Harrison, David K. Jackson, Ian Johnston, Dominic Kwiatkowski, Cordelia Langford, John Sillitoe on behalf of the Wellcome Sanger Institute COVID-19 Surveillance Team |
| EPI_ISL_3359683, EPI_ISL_3359684, EPI_ISL_3359686 | Lighthouse Lab in Alderley Park | Wellcome Sanger Institute for the COVID-19 Genomics UK (COG-UK) Consortium | Jacquelyn Wynn, Mairead Hyland, The Lighthouse Lab in Alderley Park and Alex Alderton, Roberto Amato, Jeffrey Barrett, Sonia Goncalves, Ewan Harrison, David K. Jackson, Ian Johnston, Dominic Kwiatkowski, Cordelia Langford, John Sillitoe on behalf of the Wellcome Sanger Institute COVID-19 Surveillance Team |
| EPI_ISL_3359687, EPI_ISL_3359689, EPI_ISL_3359690 | Lighthouse Lab in Glasgow | Wellcome Sanger Institute for the COVID-19 Genomics UK (COG-UK) Consortium | Harper VanSteenhouse, Yumi Kasai, David Gray, Carol Clugston, Anna Dominiczak and Alex Alderton, Roberto Amato, Jeffrey Barrett, Sonia Goncalves, Ewan Harrison, David K. Jackson, Ian Johnston, Dominic Kwiatkowski, Cordelia Langford, John Sillitoe on behalf of the Wellcome Sanger Institute COVID-19 Surveillance Team |
| EPI_ISL_3359692 | Lighthouse Laboratory Plymouth | Wellcome Sanger Institute for the COVID-19 Genomics UK (COG-UK) Consortium | Lighthouse Laboratory Plymouth and Alex Alderton, Roberto Amato, Jeffrey Barrett, Sonia Goncalves, Ewan Harrison, David K. Jackson, Ian Johnston, Dominic Kwiatkowski, Cordelia Langford, John Sillitoe on behalf of the Wellcome Sanger Institute COVID-19 Surveillance Team |
| EPI_ISL_3359693, EPI_ISL_3359694 | Lighthouse Lab in Alderley Park | Wellcome Sanger Institute for the COVID-19 Genomics UK (COG-UK) Consortium | Jacquelyn Wynn, Mairead Hyland, The Lighthouse Lab in Alderley Park and Alex Alderton, Roberto Amato, Jeffrey Barrett, Sonia Goncalves, Ewan Harrison, David K. Jackson, Ian Johnston, Dominic Kwiatkowski, Cordelia Langford, John Sillitoe on behalf of the Wellcome Sanger Institute COVID-19 Surveillance Team |
| EPI_ISL_3359696 | Lighthouse Lab in Glasgow | Wellcome Sanger Institute for the COVID-19 Genomics UK (COG-UK) Consortium | Harper VanSteenhouse, Yumi Kasai, David Gray, Carol Clugston, Anna Dominiczak and Alex Alderton, Roberto Amato, Jeffrey Barrett, Sonia Goncalves, Ewan Harrison, David K. Jackson, Ian Johnston, Dominic Kwiatkowski, Cordelia Langford, John Sillitoe on behalf of the Wellcome Sanger Institute COVID-19 Surveillance Team |
| EPI_ISL_3359697, EPI_ISL_3359699 | Lighthouse Laboratory Plymouth | Wellcome Sanger Institute for the COVID-19 Genomics UK (COG-UK) Consortium | Lighthouse Laboratory Plymouth and Alex Alderton, Roberto Amato, Jeffrey Barrett, Sonia Goncalves, Ewan Harrison, David K. Jackson, Ian Johnston, Dominic Kwiatkowski, Cordelia Langford, John Sillitoe on behalf of the Wellcome Sanger Institute COVID-19 Surveillance Team |
| EPI_ISL_3359700, EPI_ISL_3359701 | Lighthouse Lab in Alderley Park | Wellcome Sanger Institute for the COVID-19 Genomics UK (COG-UK) Consortium | Jacquelyn Wynn, Mairead Hyland, The Lighthouse Lab in Alderley Park and Alex Alderton, Roberto Amato, Jeffrey Barrett, Sonia Goncalves, Ewan Harrison, David K. Jackson, Ian Johnston, Dominic Kwiatkowski, Cordelia Langford, John Sillitoe on behalf of the Wellcome Sanger Institute COVID-19 Surveillance Team |
| EPI_ISL_3359703 | Lighthouse Lab in Glasgow | Wellcome Sanger Institute for the COVID-19 Genomics UK (COG-UK) Consortium | Harper VanSteenhouse, Yumi Kasai, David Gray, Carol Clugston, Anna Dominiczak and Alex Alderton, Roberto Amato, Jeffrey Barrett, Sonia Goncalves, Ewan Harrison, David K. Jackson, Ian Johnston, Dominic Kwiatkowski, Cordelia Langford, John Sillitoe on behalf of the Wellcome Sanger Institute COVID-19 Surveillance Team |
| EPI_ISL_3359704, | Lighthouse Laboratory Plymouth | Wellcome Sanger Institute for the COVID-19 Genomics UK (COG-UK) | Lighthouse Laboratory Plymouth and Alex Alderton, Roberto Amato, Jeffrey Barrett, Sonia Goncalves, Ewan Harrison, David K. Jackson, Ian Johnston, Dominic Kwiatkowski, Cordelia Langford, John Sillitoe on behalf of the Wellcome Sanger |

| | | | |
|---|---|---|---|
| EPI_ISL_3359706, EPI_ISL_3359707 | | Consortium | Institute COVID-19 Surveillance Team |
| EPI_ISL_3359708 | Lighthouse Lab in Glasgow | Wellcome Sanger Institute for the COVID-19 Genomics UK (COG-UK) Consortium | Harper VanSteenhouse, Yumi Kasai, David Gray, Carol Clugston, Anna Dominiczak and Alex Alderton, Roberto Amato, Jeffrey Barrett, Sonia Goncalves, Ewan Harrison, David K. Jackson, Ian Johnston, Dominic Kwiatkowski, Cordelia Langford, John Sillitoe on behalf of the Wellcome Sanger Institute COVID-19 Surveillance Team |
| EPI_ISL_3359710 | Lighthouse Lab in Alderley Park | Wellcome Sanger Institute for the COVID-19 Genomics UK (COG-UK) Consortium | Jacquelyn Wynn, Mairead Hyland, The Lighthouse Lab in Alderley Park and Alex Alderton, Roberto Amato, Jeffrey Barrett, Sonia Goncalves, Ewan Harrison, David K. Jackson, Ian Johnston, Dominic Kwiatkowski, Cordelia Langford, John Sillitoe on behalf of the Wellcome Sanger Institute COVID-19 Surveillance Team |
| EPI_ISL_3359711 | Lighthouse Lab in Glasgow | Wellcome Sanger Institute for the COVID-19 Genomics UK (COG-UK) Consortium | Harper VanSteenhouse, Yumi Kasai, David Gray, Carol Clugston, Anna Dominiczak and Alex Alderton, Roberto Amato, Jeffrey Barrett, Sonia Goncalves, Ewan Harrison, David K. Jackson, Ian Johnston, Dominic Kwiatkowski, Cordelia Langford, John Sillitoe on behalf of the Wellcome Sanger Institute COVID-19 Surveillance Team |
| EPI_ISL_3359712 | Lighthouse Lab in Alderley Park | Wellcome Sanger Institute for the COVID-19 Genomics UK (COG-UK) Consortium | Jacquelyn Wynn, Mairead Hyland, The Lighthouse Lab in Alderley Park and Alex Alderton, Roberto Amato, Jeffrey Barrett, Sonia Goncalves, Ewan Harrison, David K. Jackson, Ian Johnston, Dominic Kwiatkowski, Cordelia Langford, John Sillitoe on behalf of the Wellcome Sanger Institute COVID-19 Surveillance Team |
| EPI_ISL_3359714 | Lighthouse Laboratory Plymouth | Wellcome Sanger Institute for the COVID-19 Genomics UK (COG-UK) Consortium | Lighthouse Laboratory Plymouth and Alex Alderton, Roberto Amato, Jeffrey Barrett, Sonia Goncalves, Ewan Harrison, David K. Jackson, Ian Johnston, Dominic Kwiatkowski, Cordelia Langford, John Sillitoe on behalf of the Wellcome Sanger Institute COVID-19 Surveillance Team |
| EPI_ISL_3359715 | Lighthouse Lab in Alderley Park | Wellcome Sanger Institute for the COVID-19 Genomics UK (COG-UK) Consortium | Jacquelyn Wynn, Mairead Hyland, The Lighthouse Lab in Alderley Park and Alex Alderton, Roberto Amato, Jeffrey Barrett, Sonia Goncalves, Ewan Harrison, David K. Jackson, Ian Johnston, Dominic Kwiatkowski, Cordelia Langford, John Sillitoe on behalf of the Wellcome Sanger Institute COVID-19 Surveillance Team |
| EPI_ISL_3359717, EPI_ISL_3359718 | Lighthouse Lab in Glasgow | Wellcome Sanger Institute for the COVID-19 Genomics UK (COG-UK) Consortium | Harper VanSteenhouse, Yumi Kasai, David Gray, Carol Clugston, Anna Dominiczak and Alex Alderton, Roberto Amato, Jeffrey Barrett, Sonia Goncalves, Ewan Harrison, David K. Jackson, Ian Johnston, Dominic Kwiatkowski, Cordelia Langford, John Sillitoe on behalf of the Wellcome Sanger Institute COVID-19 Surveillance Team |
| EPI_ISL_3359719 | Lighthouse Lab in Alderley Park | Wellcome Sanger Institute for the COVID-19 Genomics UK (COG-UK) Consortium | Jacquelyn Wynn, Mairead Hyland, The Lighthouse Lab in Alderley Park and Alex Alderton, Roberto Amato, Jeffrey Barrett, Sonia Goncalves, Ewan Harrison, David K. Jackson, Ian Johnston, Dominic Kwiatkowski, Cordelia Langford, John Sillitoe on behalf of the Wellcome Sanger Institute COVID-19 Surveillance Team |
| EPI_ISL_3359721, EPI_ISL_3359722 | Lighthouse Laboratory Plymouth | Wellcome Sanger Institute for the COVID-19 Genomics UK (COG-UK) Consortium | Lighthouse Laboratory Plymouth and Alex Alderton, Roberto Amato, Jeffrey Barrett, Sonia Goncalves, Ewan Harrison, David K. Jackson, Ian Johnston, Dominic Kwiatkowski, Cordelia Langford, John Sillitoe on behalf of the Wellcome Sanger Institute COVID-19 Surveillance Team |
| EPI_ISL_3359723, EPI_ISL_3359725, EPI_ISL_3359726, EPI_ISL_3359727, EPI_ISL_3359729, EPI_ISL_3359730 | Lighthouse Lab in Alderley Park | Wellcome Sanger Institute for the COVID-19 Genomics UK (COG-UK) Consortium | Jacquelyn Wynn, Mairead Hyland, The Lighthouse Lab in Alderley Park and Alex Alderton, Roberto Amato, Jeffrey Barrett, Sonia Goncalves, Ewan Harrison, David K. Jackson, Ian Johnston, Dominic Kwiatkowski, Cordelia Langford, John Sillitoe on behalf of the Wellcome Sanger Institute COVID-19 Surveillance Team |
| EPI_ISL_3359732, EPI_ISL_3359733 | Lighthouse Laboratory Plymouth | Wellcome Sanger Institute for the COVID-19 Genomics UK (COG-UK) Consortium | Lighthouse Laboratory Plymouth and Alex Alderton, Roberto Amato, Jeffrey Barrett, Sonia Goncalves, Ewan Harrison, David K. Jackson, Ian Johnston, Dominic Kwiatkowski, Cordelia Langford, John Sillitoe on behalf of the Wellcome Sanger Institute COVID-19 Surveillance Team |
| EPI_ISL_3359735, EPI_ISL_3359736, EPI_ISL_3359738, EPI_ISL_3359739, EPI_ISL_3359741 | Lighthouse Lab in Alderley Park | Wellcome Sanger Institute for the COVID-19 Genomics UK (COG-UK) Consortium | Jacquelyn Wynn, Mairead Hyland, The Lighthouse Lab in Alderley Park and Alex Alderton, Roberto Amato, Jeffrey Barrett, Sonia Goncalves, Ewan Harrison, David K. Jackson, Ian Johnston, Dominic Kwiatkowski, Cordelia Langford, John Sillitoe on behalf of the Wellcome Sanger Institute COVID-19 Surveillance Team |
| EPI_ISL_3359742 | Lighthouse Laboratory Plymouth | Wellcome Sanger Institute for the COVID-19 Genomics UK (COG-UK) Consortium | Lighthouse Laboratory Plymouth and Alex Alderton, Roberto Amato, Jeffrey Barrett, Sonia Goncalves, Ewan Harrison, David K. Jackson, Ian Johnston, Dominic Kwiatkowski, Cordelia Langford, John Sillitoe on behalf of the Wellcome Sanger Institute COVID-19 Surveillance Team |
| EPI_ISL_3359743 | Lighthouse Lab in Alderley Park | Wellcome Sanger Institute for the COVID-19 Genomics UK (COG-UK) Consortium | Jacquelyn Wynn, Mairead Hyland, The Lighthouse Lab in Alderley Park and Alex Alderton, Roberto Amato, Jeffrey Barrett, Sonia Goncalves, Ewan Harrison, David K. Jackson, Ian Johnston, Dominic Kwiatkowski, Cordelia Langford, John Sillitoe on behalf of the Wellcome Sanger Institute COVID-19 Surveillance Team |
| EPI_ISL_3359745 | Lighthouse Lab in Glasgow | Wellcome Sanger Institute for the COVID-19 Genomics UK (COG-UK) Consortium | Harper VanSteenhouse, Yumi Kasai, David Gray, Carol Clugston, Anna Dominiczak and Alex Alderton, Roberto Amato, Jeffrey Barrett, Sonia Goncalves, Ewan Harrison, David K. Jackson, Ian Johnston, Dominic Kwiatkowski, Cordelia Langford, John Sillitoe on behalf of the Wellcome Sanger Institute COVID-19 Surveillance Team |
| EPI_ISL_3359746 | Lighthouse Laboratory Plymouth | Wellcome Sanger Institute for the COVID-19 Genomics UK (COG-UK) Consortium | Lighthouse Laboratory Plymouth and Alex Alderton, Roberto Amato, Jeffrey Barrett, Sonia Goncalves, Ewan Harrison, David K. Jackson, Ian Johnston, Dominic Kwiatkowski, Cordelia Langford, John Sillitoe on behalf of the Wellcome Sanger Institute COVID-19 Surveillance Team |
| EPI_ISL_3359748, EPI_ISL_3359749, EPI_ISL_3359750, EPI_ISL_3359752, EPI_ISL_3359753 | Lighthouse Lab in Alderley Park | Wellcome Sanger Institute for the COVID-19 Genomics UK (COG-UK) Consortium | Jacquelyn Wynn, Mairead Hyland, The Lighthouse Lab in Alderley Park and Alex Alderton, Roberto Amato, Jeffrey Barrett, Sonia Goncalves, Ewan Harrison, David K. Jackson, Ian Johnston, Dominic Kwiatkowski, Cordelia Langford, John Sillitoe on behalf of the Wellcome Sanger Institute COVID-19 Surveillance Team |
| EPI_ISL_3359755 | Lighthouse Lab in Glasgow | Wellcome Sanger Institute for the COVID-19 Genomics UK (COG-UK) Consortium | Harper VanSteenhouse, Yumi Kasai, David Gray, Carol Clugston, Anna Dominiczak and Alex Alderton, Roberto Amato, Jeffrey Barrett, Sonia Goncalves, Ewan Harrison, David K. Jackson, Ian Johnston, Dominic Kwiatkowski, Cordelia Langford, John Sillitoe on behalf of the Wellcome Sanger Institute COVID-19 Surveillance Team |
| EPI_ISL_3359756 | Lighthouse Laboratory Plymouth | Wellcome Sanger Institute for the COVID-19 Genomics UK (COG-UK) Consortium | Lighthouse Laboratory Plymouth and Alex Alderton, Roberto Amato, Jeffrey Barrett, Sonia Goncalves, Ewan Harrison, David K. Jackson, Ian Johnston, Dominic Kwiatkowski, Cordelia Langford, John Sillitoe on behalf of the Wellcome Sanger Institute COVID-19 Surveillance Team |
| EPI_ISL_3359757, EPI_ISL_3359758 | Lighthouse Lab in Alderley Park | Wellcome Sanger Institute for the COVID-19 Genomics UK (COG-UK) Consortium | Jacquelyn Wynn, Mairead Hyland, The Lighthouse Lab in Alderley Park and Alex Alderton, Roberto Amato, Jeffrey Barrett, Sonia Goncalves, Ewan Harrison, David K. Jackson, Ian Johnston, Dominic Kwiatkowski, Cordelia Langford, John Sillitoe on behalf of the Wellcome Sanger Institute COVID-19 Surveillance Team |
| EPI_ISL_3359760 | Lighthouse Laboratory Plymouth | Wellcome Sanger Institute for the COVID-19 Genomics UK (COG-UK) Consortium | Lighthouse Laboratory Plymouth and Alex Alderton, Roberto Amato, Jeffrey Barrett, Sonia Goncalves, Ewan Harrison, David K. Jackson, Ian Johnston, Dominic Kwiatkowski, Cordelia Langford, John Sillitoe on behalf of the Wellcome Sanger Institute COVID-19 Surveillance Team |
| EPI_ISL_3359761 | Lighthouse Lab in Alderley Park | Wellcome Sanger Institute for the COVID-19 Genomics UK (COG-UK) Consortium | Jacquelyn Wynn, Mairead Hyland, The Lighthouse Lab in Alderley Park and Alex Alderton, Roberto Amato, Jeffrey Barrett, Sonia Goncalves, Ewan Harrison, David K. Jackson, Ian Johnston, Dominic Kwiatkowski, Cordelia Langford, John Sillitoe on behalf of the Wellcome Sanger Institute COVID-19 Surveillance Team |
| EPI_ISL_3359763 | Lighthouse Lab in Glasgow | Wellcome Sanger Institute for the COVID-19 Genomics UK (COG-UK) Consortium | Harper VanSteenhouse, Yumi Kasai, David Gray, Carol Clugston, Anna Dominiczak and Alex Alderton, Roberto Amato, Jeffrey Barrett, Sonia Goncalves, Ewan Harrison, David K. Jackson, Ian Johnston, Dominic Kwiatkowski, Cordelia Langford, John Sillitoe on behalf of the Wellcome Sanger Institute COVID-19 Surveillance Team |
| EPI_ISL_3359764 | Lighthouse Lab in Alderley Park | Wellcome Sanger Institute for the COVID-19 Genomics UK (COG-UK) Consortium | Jacquelyn Wynn, Mairead Hyland, The Lighthouse Lab in Alderley Park and Alex Alderton, Roberto Amato, Jeffrey Barrett, Sonia Goncalves, Ewan Harrison, David K. Jackson, Ian Johnston, Dominic Kwiatkowski, Cordelia Langford, John Sillitoe on behalf of the Wellcome Sanger Institute COVID-19 Surveillance Team |
| EPI_ISL_3359765 | Lighthouse Laboratory Plymouth | Wellcome Sanger Institute for the COVID-19 Genomics UK (COG-UK) Consortium | Lighthouse Laboratory Plymouth and Alex Alderton, Roberto Amato, Jeffrey Barrett, Sonia Goncalves, Ewan Harrison, David K. Jackson, Ian Johnston, Dominic Kwiatkowski, Cordelia Langford, John Sillitoe on behalf of the Wellcome Sanger Institute COVID-19 Surveillance Team |
| EPI_ISL_3359767, EPI_ISL_3359768 | Lighthouse Lab in Glasgow | Wellcome Sanger Institute for the COVID-19 Genomics UK (COG-UK) Consortium | Harper VanSteenhouse, Yumi Kasai, David Gray, Carol Clugston, Anna Dominiczak and Alex Alderton, Roberto Amato, Jeffrey Barrett, Sonia Goncalves, Ewan Harrison, David K. Jackson, Ian Johnston, Dominic Kwiatkowski, Cordelia Langford, John Sillitoe on behalf of the Wellcome Sanger Institute COVID-19 Surveillance Team |
| EPI_ISL_3359770, EPI_ISL_3359771 | Lighthouse Lab in Alderley Park | Wellcome Sanger Institute for the COVID-19 Genomics UK (COG-UK) Consortium | Jacquelyn Wynn, Mairead Hyland, The Lighthouse Lab in Alderley Park and Alex Alderton, Roberto Amato, Jeffrey Barrett, Sonia Goncalves, Ewan Harrison, David K. Jackson, Ian Johnston, Dominic Kwiatkowski, Cordelia Langford, John Sillitoe on behalf of the Wellcome Sanger Institute COVID-19 Surveillance Team |
| EPI_ISL_3359773, EPI_ISL_3359774, EPI_ISL_3359775 | Lighthouse Laboratory Plymouth | Wellcome Sanger Institute for the COVID-19 Genomics UK (COG-UK) Consortium | Lighthouse Laboratory Plymouth and Alex Alderton, Roberto Amato, Jeffrey Barrett, Sonia Goncalves, Ewan Harrison, David K. Jackson, Ian Johnston, Dominic Kwiatkowski, Cordelia Langford, John Sillitoe on behalf of the Wellcome Sanger Institute COVID-19 Surveillance Team |
| EPI_ISL_3359777, EPI_ISL_3359778, EPI_ISL_3359780, EPI_ISL_3359782, EPI_ISL_3359783 | Lighthouse Lab in Alderley Park | Wellcome Sanger Institute for the COVID-19 Genomics UK (COG-UK) Consortium | Jacquelyn Wynn, Mairead Hyland, The Lighthouse Lab in Alderley Park and Alex Alderton, Roberto Amato, Jeffrey Barrett, Sonia Goncalves, Ewan Harrison, David K. Jackson, Ian Johnston, Dominic Kwiatkowski, Cordelia Langford, John Sillitoe on behalf of the Wellcome Sanger Institute COVID-19 Surveillance Team |
| EPI_ISL_3359784 | Lighthouse Laboratory Plymouth | Wellcome Sanger Institute for the COVID-19 Genomics UK (COG-UK) Consortium | Lighthouse Laboratory Plymouth and Alex Alderton, Roberto Amato, Jeffrey Barrett, Sonia Goncalves, Ewan Harrison, David K. Jackson, Ian Johnston, Dominic Kwiatkowski, Cordelia Langford, John Sillitoe on behalf of the Wellcome Sanger Institute COVID-19 Surveillance Team |
| EPI_ISL_3359786, EPI_ISL_3359787 | Lighthouse Lab in Alderley Park | Wellcome Sanger Institute for the COVID-19 Genomics UK (COG-UK) Consortium | Jacquelyn Wynn, Mairead Hyland, The Lighthouse Lab in Alderley Park and Alex Alderton, Roberto Amato, Jeffrey Barrett, Sonia Goncalves, Ewan Harrison, David K. Jackson, Ian Johnston, Dominic Kwiatkowski, Cordelia Langford, John Sillitoe on behalf of the Wellcome Sanger Institute COVID-19 Surveillance Team |
| EPI_ISL_3359789 | Lighthouse Laboratory Plymouth | Wellcome Sanger Institute for the COVID-19 Genomics UK (COG-UK) Consortium | Lighthouse Laboratory Plymouth and Alex Alderton, Roberto Amato, Jeffrey Barrett, Sonia Goncalves, Ewan Harrison, David K. Jackson, Ian Johnston, Dominic Kwiatkowski, Cordelia Langford, John Sillitoe on behalf of the Wellcome Sanger Institute COVID-19 Surveillance Team |
| EPI_ISL_3359790 | Lighthouse Lab in Alderley Park | Wellcome Sanger Institute for the COVID-19 Genomics UK (COG-UK) Consortium | Jacquelyn Wynn, Mairead Hyland, The Lighthouse Lab in Alderley Park and Alex Alderton, Roberto Amato, Jeffrey Barrett, Sonia Goncalves, Ewan Harrison, David K. Jackson, Ian Johnston, Dominic Kwiatkowski, Cordelia Langford, John Sillitoe on behalf of the Wellcome Sanger Institute COVID-19 Surveillance Team |
| EPI_ISL_3359791 | Lighthouse Laboratory Plymouth | Wellcome Sanger Institute for the COVID-19 Genomics UK (COG-UK) Consortium | Lighthouse Laboratory Plymouth and Alex Alderton, Roberto Amato, Jeffrey Barrett, Sonia Goncalves, Ewan Harrison, David K. Jackson, Ian Johnston, Dominic Kwiatkowski, Cordelia Langford, John Sillitoe on behalf of the Wellcome Sanger Institute COVID-19 Surveillance Team |
| EPI_ISL_3359793 | Lighthouse Lab in Alderley Park | Wellcome Sanger Institute for the COVID-19 Genomics UK (COG-UK) Consortium | Jacquelyn Wynn, Mairead Hyland, The Lighthouse Lab in Alderley Park and Alex Alderton, Roberto Amato, Jeffrey Barrett, Sonia Goncalves, Ewan Harrison, David K. Jackson, Ian Johnston, Dominic Kwiatkowski, Cordelia Langford, John Sillitoe on behalf of the Wellcome Sanger Institute COVID-19 Surveillance Team |
| EPI_ISL_3359794, EPI_ISL_3359796 | Lighthouse Lab in Glasgow | Wellcome Sanger Institute for the COVID-19 Genomics UK (COG-UK) Consortium | Harper VanSteenhouse, Yumi Kasai, David Gray, Carol Clugston, Anna Dominiczak and Alex Alderton, Roberto Amato, Jeffrey Barrett, Sonia Goncalves, Ewan Harrison, David K. Jackson, Ian Johnston, Dominic Kwiatkowski, Cordelia Langford, John Sillitoe on behalf of the Wellcome Sanger Institute COVID-19 Surveillance Team |
| EPI_ISL_3359798 | Lighthouse Lab in Alderley Park | Wellcome Sanger Institute for the COVID-19 Genomics UK (COG-UK) Consortium | Jacquelyn Wynn, Mairead Hyland, The Lighthouse Lab in Alderley Park and Alex Alderton, Roberto Amato, Jeffrey Barrett, Sonia Goncalves, Ewan Harrison, David K. Jackson, Ian Johnston, Dominic Kwiatkowski, Cordelia Langford, John Sillitoe on behalf of the Wellcome Sanger Institute COVID-19 Surveillance Team |
| EPI_ISL_3359799 | Lighthouse Laboratory Plymouth | Wellcome Sanger Institute for the COVID-19 Genomics UK (COG-UK) Consortium | Lighthouse Laboratory Plymouth and Alex Alderton, Roberto Amato, Jeffrey Barrett, Sonia Goncalves, Ewan Harrison, David K. Jackson, Ian Johnston, Dominic Kwiatkowski, Cordelia Langford, John Sillitoe on behalf of the Wellcome Sanger Institute COVID-19 Surveillance Team |

| | | | |
|---|---|---|---|
| EPI_ISL_3359801 | Lighthouse Lab in Glasgow | Wellcome Sanger Institute for the COVID-19 Genomics UK (COG-UK) Consortium | Harper VanSteenhouse, Yumi Kasai, David Gray, Carol Clugston, Anna Dominiczak and Alex Alderton, Roberto Amato, Jeffrey Barrett, Sonia Goncalves, Ewan Harrison, David K. Jackson, Ian Johnston, Dominic Kwiatkowski, Cordelia Langford, John Sillitoe on behalf of the Wellcome Sanger Institute COVID-19 Surveillance Team |
| EPI_ISL_3359802, EPI_ISL_3359803 | Lighthouse Lab in Alderley Park | Wellcome Sanger Institute for the COVID-19 Genomics UK (COG-UK) Consortium | Jacquelyn Wynn, Mairead Hyland, The Lighthouse Lab in Alderley Park and Alex Alderton, Roberto Amato, Jeffrey Barrett, Sonia Goncalves, Ewan Harrison, David K. Jackson, Ian Johnston, Dominic Kwiatkowski, Cordelia Langford, John Sillitoe on behalf of the Wellcome Sanger Institute COVID-19 Surveillance Team |
| EPI_ISL_3359805 | Lighthouse Lab in Glasgow | Wellcome Sanger Institute for the COVID-19 Genomics UK (COG-UK) Consortium | Harper VanSteenhouse, Yumi Kasai, David Gray, Carol Clugston, Anna Dominiczak and Alex Alderton, Roberto Amato, Jeffrey Barrett, Sonia Goncalves, Ewan Harrison, David K. Jackson, Ian Johnston, Dominic Kwiatkowski, Cordelia Langford, John Sillitoe on behalf of the Wellcome Sanger Institute COVID-19 Surveillance Team |
| EPI_ISL_3359806 | Lighthouse Laboratory Plymouth | Wellcome Sanger Institute for the COVID-19 Genomics UK (COG-UK) Consortium | Lighthouse Laboratory Plymouth and Alex Alderton, Roberto Amato, Jeffrey Barrett, Sonia Goncalves, Ewan Harrison, David K. Jackson, Ian Johnston, Dominic Kwiatkowski, Cordelia Langford, John Sillitoe on behalf of the Wellcome Sanger Institute COVID-19 Surveillance Team |
| EPI_ISL_3359808, EPI_ISL_3359809 | Lighthouse Lab in Alderley Park | Wellcome Sanger Institute for the COVID-19 Genomics UK (COG-UK) Consortium | Jacquelyn Wynn, Mairead Hyland, The Lighthouse Lab in Alderley Park and Alex Alderton, Roberto Amato, Jeffrey Barrett, Sonia Goncalves, Ewan Harrison, David K. Jackson, Ian Johnston, Dominic Kwiatkowski, Cordelia Langford, John Sillitoe on behalf of the Wellcome Sanger Institute COVID-19 Surveillance Team |
| EPI_ISL_3359811 | Lighthouse Laboratory Plymouth | Wellcome Sanger Institute for the COVID-19 Genomics UK (COG-UK) Consortium | Lighthouse Laboratory Plymouth and Alex Alderton, Roberto Amato, Jeffrey Barrett, Sonia Goncalves, Ewan Harrison, David K. Jackson, Ian Johnston, Dominic Kwiatkowski, Cordelia Langford, John Sillitoe on behalf of the Wellcome Sanger Institute COVID-19 Surveillance Team |
| EPI_ISL_3359812 | Lighthouse Lab in Alderley Park | Wellcome Sanger Institute for the COVID-19 Genomics UK (COG-UK) Consortium | Jacquelyn Wynn, Mairead Hyland, The Lighthouse Lab in Alderley Park and Alex Alderton, Roberto Amato, Jeffrey Barrett, Sonia Goncalves, Ewan Harrison, David K. Jackson, Ian Johnston, Dominic Kwiatkowski, Cordelia Langford, John Sillitoe on behalf of the Wellcome Sanger Institute COVID-19 Surveillance Team |
| EPI_ISL_3359814, EPI_ISL_3359815 | Lighthouse Laboratory Plymouth | Wellcome Sanger Institute for the COVID-19 Genomics UK (COG-UK) Consortium | Lighthouse Laboratory Plymouth and Alex Alderton, Roberto Amato, Jeffrey Barrett, Sonia Goncalves, Ewan Harrison, David K. Jackson, Ian Johnston, Dominic Kwiatkowski, Cordelia Langford, John Sillitoe on behalf of the Wellcome Sanger Institute COVID-19 Surveillance Team |
| EPI_ISL_3359816 | Lighthouse Lab in Alderley Park | Wellcome Sanger Institute for the COVID-19 Genomics UK (COG-UK) Consortium | Jacquelyn Wynn, Mairead Hyland, The Lighthouse Lab in Alderley Park and Alex Alderton, Roberto Amato, Jeffrey Barrett, Sonia Goncalves, Ewan Harrison, David K. Jackson, Ian Johnston, Dominic Kwiatkowski, Cordelia Langford, John Sillitoe on behalf of the Wellcome Sanger Institute COVID-19 Surveillance Team |
| EPI_ISL_3359818 | Lighthouse Lab in Glasgow | Wellcome Sanger Institute for the COVID-19 Genomics UK (COG-UK) Consortium | Harper VanSteenhouse, Yumi Kasai, David Gray, Carol Clugston, Anna Dominiczak and Alex Alderton, Roberto Amato, Jeffrey Barrett, Sonia Goncalves, Ewan Harrison, David K. Jackson, Ian Johnston, Dominic Kwiatkowski, Cordelia Langford, John Sillitoe on behalf of the Wellcome Sanger Institute COVID-19 Surveillance Team |
| EPI_ISL_3359819, EPI_ISL_3359821, EPI_ISL_3359822, EPI_ISL_3359824 | Lighthouse Lab in Alderley Park | Wellcome Sanger Institute for the COVID-19 Genomics UK (COG-UK) Consortium | Jacquelyn Wynn, Mairead Hyland, The Lighthouse Lab in Alderley Park and Alex Alderton, Roberto Amato, Jeffrey Barrett, Sonia Goncalves, Ewan Harrison, David K. Jackson, Ian Johnston, Dominic Kwiatkowski, Cordelia Langford, John Sillitoe on behalf of the Wellcome Sanger Institute COVID-19 Surveillance Team |
| EPI_ISL_3359825 | Lighthouse Lab in Glasgow | Wellcome Sanger Institute for the COVID-19 Genomics UK (COG-UK) Consortium | Harper VanSteenhouse, Yumi Kasai, David Gray, Carol Clugston, Anna Dominiczak and Alex Alderton, Roberto Amato, Jeffrey Barrett, Sonia Goncalves, Ewan Harrison, David K. Jackson, Ian Johnston, Dominic Kwiatkowski, Cordelia Langford, John Sillitoe on behalf of the Wellcome Sanger Institute COVID-19 Surveillance Team |
| EPI_ISL_3359827 | Lighthouse Laboratory Plymouth | Wellcome Sanger Institute for the COVID-19 Genomics UK (COG-UK) Consortium | Lighthouse Laboratory Plymouth and Alex Alderton, Roberto Amato, Jeffrey Barrett, Sonia Goncalves, Ewan Harrison, David K. Jackson, Ian Johnston, Dominic Kwiatkowski, Cordelia Langford, John Sillitoe on behalf of the Wellcome Sanger Institute COVID-19 Surveillance Team |
| EPI_ISL_3359828 | Lighthouse Lab in Alderley Park | Wellcome Sanger Institute for the COVID-19 Genomics UK (COG-UK) Consortium | Jacquelyn Wynn, Mairead Hyland, The Lighthouse Lab in Alderley Park and Alex Alderton, Roberto Amato, Jeffrey Barrett, Sonia Goncalves, Ewan Harrison, David K. Jackson, Ian Johnston, Dominic Kwiatkowski, Cordelia Langford, John Sillitoe on behalf of the Wellcome Sanger Institute COVID-19 Surveillance Team |
| EPI_ISL_3359830, EPI_ISL_3359831, EPI_ISL_3359833 | Lighthouse Laboratory Plymouth | Wellcome Sanger Institute for the COVID-19 Genomics UK (COG-UK) Consortium | Lighthouse Laboratory Plymouth and Alex Alderton, Roberto Amato, Jeffrey Barrett, Sonia Goncalves, Ewan Harrison, David K. Jackson, Ian Johnston, Dominic Kwiatkowski, Cordelia Langford, John Sillitoe on behalf of the Wellcome Sanger Institute COVID-19 Surveillance Team |
| EPI_ISL_3359834 | Lighthouse Lab in Glasgow | Wellcome Sanger Institute for the COVID-19 Genomics UK (COG-UK) Consortium | Harper VanSteenhouse, Yumi Kasai, David Gray, Carol Clugston, Anna Dominiczak and Alex Alderton, Roberto Amato, Jeffrey Barrett, Sonia Goncalves, Ewan Harrison, David K. Jackson, Ian Johnston, Dominic Kwiatkowski, Cordelia Langford, John Sillitoe on behalf of the Wellcome Sanger Institute COVID-19 Surveillance Team |
| EPI_ISL_3359836 | Lighthouse Laboratory Plymouth | Wellcome Sanger Institute for the COVID-19 Genomics UK (COG-UK) Consortium | Lighthouse Laboratory Plymouth and Alex Alderton, Roberto Amato, Jeffrey Barrett, Sonia Goncalves, Ewan Harrison, David K. Jackson, Ian Johnston, Dominic Kwiatkowski, Cordelia Langford, John Sillitoe on behalf of the Wellcome Sanger Institute COVID-19 Surveillance Team |
| EPI_ISL_3359837, EPI_ISL_3359839, EPI_ISL_3359840 | Lighthouse Lab in Alderley Park | Wellcome Sanger Institute for the COVID-19 Genomics UK (COG-UK) Consortium | Jacquelyn Wynn, Mairead Hyland, The Lighthouse Lab in Alderley Park and Alex Alderton, Roberto Amato, Jeffrey Barrett, Sonia Goncalves, Ewan Harrison, David K. Jackson, Ian Johnston, Dominic Kwiatkowski, Cordelia Langford, John Sillitoe on behalf of the Wellcome Sanger Institute COVID-19 Surveillance Team |
| EPI_ISL_3359842 | Lighthouse Laboratory Plymouth | Wellcome Sanger Institute for the COVID-19 Genomics UK (COG-UK) Consortium | Lighthouse Laboratory Plymouth and Alex Alderton, Roberto Amato, Jeffrey Barrett, Sonia Goncalves, Ewan Harrison, David K. Jackson, Ian Johnston, Dominic Kwiatkowski, Cordelia Langford, John Sillitoe on behalf of the Wellcome Sanger Institute COVID-19 Surveillance Team |
| EPI_ISL_3359843 | Lighthouse Lab in Alderley Park | Wellcome Sanger Institute for the COVID-19 Genomics UK (COG-UK) Consortium | Jacquelyn Wynn, Mairead Hyland, The Lighthouse Lab in Alderley Park and Alex Alderton, Roberto Amato, Jeffrey Barrett, Sonia Goncalves, Ewan Harrison, David K. Jackson, Ian Johnston, Dominic Kwiatkowski, Cordelia Langford, John Sillitoe on behalf of the Wellcome Sanger Institute COVID-19 Surveillance Team |
| EPI_ISL_3359845 | Lighthouse Laboratory Plymouth | Wellcome Sanger Institute for the COVID-19 Genomics UK (COG-UK) Consortium | Lighthouse Laboratory Plymouth and Alex Alderton, Roberto Amato, Jeffrey Barrett, Sonia Goncalves, Ewan Harrison, David K. Jackson, Ian Johnston, Dominic Kwiatkowski, Cordelia Langford, John Sillitoe on behalf of the Wellcome Sanger Institute COVID-19 Surveillance Team |
| EPI_ISL_3359846 | Lighthouse Lab in Glasgow | Wellcome Sanger Institute for the COVID-19 Genomics UK (COG-UK) Consortium | Harper VanSteenhouse, Yumi Kasai, David Gray, Carol Clugston, Anna Dominiczak and Alex Alderton, Roberto Amato, Jeffrey Barrett, Sonia Goncalves, Ewan Harrison, David K. Jackson, Ian Johnston, Dominic Kwiatkowski, Cordelia Langford, John Sillitoe on behalf of the Wellcome Sanger Institute COVID-19 Surveillance Team |
| EPI_ISL_3359847 | Lighthouse Laboratory Plymouth | Wellcome Sanger Institute for the COVID-19 Genomics UK (COG-UK) Consortium | Lighthouse Laboratory Plymouth and Alex Alderton, Roberto Amato, Jeffrey Barrett, Sonia Goncalves, Ewan Harrison, David K. Jackson, Ian Johnston, Dominic Kwiatkowski, Cordelia Langford, John Sillitoe on behalf of the Wellcome Sanger Institute COVID-19 Surveillance Team |
| EPI_ISL_3359848, EPI_ISL_3359850 | Lighthouse Lab in Alderley Park | Wellcome Sanger Institute for the COVID-19 Genomics UK (COG-UK) Consortium | Jacquelyn Wynn, Mairead Hyland, The Lighthouse Lab in Alderley Park and Alex Alderton, Roberto Amato, Jeffrey Barrett, Sonia Goncalves, Ewan Harrison, David K. Jackson, Ian Johnston, Dominic Kwiatkowski, Cordelia Langford, John Sillitoe on behalf of the Wellcome Sanger Institute COVID-19 Surveillance Team |
| EPI_ISL_3359851 | Lighthouse Laboratory Plymouth | Wellcome Sanger Institute for the COVID-19 Genomics UK (COG-UK) Consortium | Lighthouse Laboratory Plymouth and Alex Alderton, Roberto Amato, Jeffrey Barrett, Sonia Goncalves, Ewan Harrison, David K. Jackson, Ian Johnston, Dominic Kwiatkowski, Cordelia Langford, John Sillitoe on behalf of the Wellcome Sanger Institute COVID-19 Surveillance Team |
| EPI_ISL_3359852, EPI_ISL_3359854 | Lighthouse Lab in Alderley Park | Wellcome Sanger Institute for the COVID-19 Genomics UK (COG-UK) Consortium | Jacquelyn Wynn, Mairead Hyland, The Lighthouse Lab in Alderley Park and Alex Alderton, Roberto Amato, Jeffrey Barrett, Sonia Goncalves, Ewan Harrison, David K. Jackson, Ian Johnston, Dominic Kwiatkowski, Cordelia Langford, John Sillitoe on behalf of the Wellcome Sanger Institute COVID-19 Surveillance Team |
| EPI_ISL_3359855, EPI_ISL_3359857, EPI_ISL_3359858 | Lighthouse Lab in Glasgow | Wellcome Sanger Institute for the COVID-19 Genomics UK (COG-UK) Consortium | Harper VanSteenhouse, Yumi Kasai, David Gray, Carol Clugston, Anna Dominiczak and Alex Alderton, Roberto Amato, Jeffrey Barrett, Sonia Goncalves, Ewan Harrison, David K. Jackson, Ian Johnston, Dominic Kwiatkowski, Cordelia Langford, John Sillitoe on behalf of the Wellcome Sanger Institute COVID-19 Surveillance Team |
| EPI_ISL_3359860 | Lighthouse Lab in Alderley Park | Wellcome Sanger Institute for the COVID-19 Genomics UK (COG-UK) Consortium | Jacquelyn Wynn, Mairead Hyland, The Lighthouse Lab in Alderley Park and Alex Alderton, Roberto Amato, Jeffrey Barrett, Sonia Goncalves, Ewan Harrison, David K. Jackson, Ian Johnston, Dominic Kwiatkowski, Cordelia Langford, John Sillitoe on behalf of the Wellcome Sanger Institute COVID-19 Surveillance Team |
| EPI_ISL_3359862 | Lighthouse Laboratory Plymouth | Wellcome Sanger Institute for the COVID-19 Genomics UK (COG-UK) Consortium | Lighthouse Laboratory Plymouth and Alex Alderton, Roberto Amato, Jeffrey Barrett, Sonia Goncalves, Ewan Harrison, David K. Jackson, Ian Johnston, Dominic Kwiatkowski, Cordelia Langford, John Sillitoe on behalf of the Wellcome Sanger Institute COVID-19 Surveillance Team |
| EPI_ISL_3359863 | Lighthouse Lab in Glasgow | Wellcome Sanger Institute for the COVID-19 Genomics UK (COG-UK) Consortium | Harper VanSteenhouse, Yumi Kasai, David Gray, Carol Clugston, Anna Dominiczak and Alex Alderton, Roberto Amato, Jeffrey Barrett, Sonia Goncalves, Ewan Harrison, David K. Jackson, Ian Johnston, Dominic Kwiatkowski, Cordelia Langford, John Sillitoe on behalf of the Wellcome Sanger Institute COVID-19 Surveillance Team |
| EPI_ISL_3359865 | Lighthouse Laboratory Plymouth | Wellcome Sanger Institute for the COVID-19 Genomics UK (COG-UK) Consortium | Lighthouse Laboratory Plymouth and Alex Alderton, Roberto Amato, Jeffrey Barrett, Sonia Goncalves, Ewan Harrison, David K. Jackson, Ian Johnston, Dominic Kwiatkowski, Cordelia Langford, John Sillitoe on behalf of the Wellcome Sanger Institute COVID-19 Surveillance Team |
| EPI_ISL_3359866, EPI_ISL_3359867, EPI_ISL_3359869 | Lighthouse Lab in Alderley Park | Wellcome Sanger Institute for the COVID-19 Genomics UK (COG-UK) Consortium | Jacquelyn Wynn, Mairead Hyland, The Lighthouse Lab in Alderley Park and Alex Alderton, Roberto Amato, Jeffrey Barrett, Sonia Goncalves, Ewan Harrison, David K. Jackson, Ian Johnston, Dominic Kwiatkowski, Cordelia Langford, John Sillitoe on behalf of the Wellcome Sanger Institute COVID-19 Surveillance Team |
| EPI_ISL_3359870 | Lighthouse Lab in Glasgow | Wellcome Sanger Institute for the COVID-19 Genomics UK (COG-UK) Consortium | Harper VanSteenhouse, Yumi Kasai, David Gray, Carol Clugston, Anna Dominiczak and Alex Alderton, Roberto Amato, Jeffrey Barrett, Sonia Goncalves, Ewan Harrison, David K. Jackson, Ian Johnston, Dominic Kwiatkowski, Cordelia Langford, John Sillitoe on behalf of the Wellcome Sanger Institute COVID-19 Surveillance Team |
| EPI_ISL_3359872 | Lighthouse Laboratory Plymouth | Wellcome Sanger Institute for the COVID-19 Genomics UK (COG-UK) Consortium | Lighthouse Laboratory Plymouth and Alex Alderton, Roberto Amato, Jeffrey Barrett, Sonia Goncalves, Ewan Harrison, David K. Jackson, Ian Johnston, Dominic Kwiatkowski, Cordelia Langford, John Sillitoe on behalf of the Wellcome Sanger Institute COVID-19 Surveillance Team |
| EPI_ISL_3359873 | Lighthouse Lab in Alderley Park | Wellcome Sanger Institute for the COVID-19 Genomics UK (COG-UK) Consortium | Jacquelyn Wynn, Mairead Hyland, The Lighthouse Lab in Alderley Park and Alex Alderton, Roberto Amato, Jeffrey Barrett, Sonia Goncalves, Ewan Harrison, David K. Jackson, Ian Johnston, Dominic Kwiatkowski, Cordelia Langford, John Sillitoe on behalf of the Wellcome Sanger Institute COVID-19 Surveillance Team |
| EPI_ISL_3359875 | Lighthouse Laboratory Plymouth | Wellcome Sanger Institute for the COVID-19 Genomics UK (COG-UK) Consortium | Lighthouse Laboratory Plymouth and Alex Alderton, Roberto Amato, Jeffrey Barrett, Sonia Goncalves, Ewan Harrison, David K. Jackson, Ian Johnston, Dominic Kwiatkowski, Cordelia Langford, John Sillitoe on behalf of the Wellcome Sanger Institute COVID-19 Surveillance Team |
| EPI_ISL_3359877, EPI_ISL_3359878, EPI_ISL_3359879 | Lighthouse Lab in Alderley Park | Wellcome Sanger Institute for the COVID-19 Genomics UK (COG-UK) Consortium | Jacquelyn Wynn, Mairead Hyland, The Lighthouse Lab in Alderley Park and Alex Alderton, Roberto Amato, Jeffrey Barrett, Sonia Goncalves, Ewan Harrison, David K. Jackson, Ian Johnston, Dominic Kwiatkowski, Cordelia Langford, John Sillitoe on behalf of the Wellcome Sanger Institute COVID-19 Surveillance Team |
| EPI_ISL_3359881 | Lighthouse Lab in Glasgow | Wellcome Sanger Institute for the COVID-19 Genomics UK (COG-UK) Consortium | Harper VanSteenhouse, Yumi Kasai, David Gray, Carol Clugston, Anna Dominiczak and Alex Alderton, Roberto Amato, Jeffrey Barrett, Sonia Goncalves, Ewan Harrison, David K. Jackson, Ian Johnston, Dominic Kwiatkowski, Cordelia Langford, John Sillitoe on behalf of the Wellcome Sanger Institute COVID-19 Surveillance Team |
| EPI_ISL_3359882, EPI_ISL_3359884, EPI_ISL_3359885 | Lighthouse Lab in Alderley Park | Wellcome Sanger Institute for the COVID-19 Genomics UK (COG-UK) Consortium | Jacquelyn Wynn, Mairead Hyland, The Lighthouse Lab in Alderley Park and Alex Alderton, Roberto Amato, Jeffrey Barrett, Sonia Goncalves, Ewan Harrison, David K. Jackson, Ian Johnston, Dominic Kwiatkowski, Cordelia Langford, John Sillitoe on behalf of the Wellcome Sanger Institute COVID-19 Surveillance Team |
| EPI_ISL_3359887 | Lighthouse Laboratory Plymouth | Wellcome Sanger Institute for the COVID-19 Genomics UK (COG-UK) Consortium | Lighthouse Laboratory Plymouth and Alex Alderton, Roberto Amato, Jeffrey Barrett, Sonia Goncalves, Ewan Harrison, David K. Jackson, Ian Johnston, Dominic Kwiatkowski, Cordelia Langford, John Sillitoe on behalf of the Wellcome Sanger Institute COVID-19 Surveillance Team |
| EPI_ISL_3359888, EPI_ISL_3359890, EPI_ISL_3359891 | Lighthouse Lab in Alderley Park | Wellcome Sanger Institute for the COVID-19 Genomics UK (COG-UK) Consortium | Jacquelyn Wynn, Mairead Hyland, The Lighthouse Lab in Alderley Park and Alex Alderton, Roberto Amato, Jeffrey Barrett, Sonia Goncalves, Ewan Harrison, David K. Jackson, Ian Johnston, Dominic Kwiatkowski, Cordelia Langford, John Sillitoe on behalf of the Wellcome Sanger Institute COVID-19 Surveillance Team |

| | | | |
|---|---|---|---|
| EPI_ISL_3359892 | Lighthouse Lab in Glasgow | Wellcome Sanger Institute for the COVID-19 Genomics UK (COG-UK) Consortium | Harper VanSteenhouse, Yumi Kasai, David Gray, Carol Clugston, Anna Dominiczak and Alex Alderton, Roberto Amato, Jeffrey Barrett, Sonia Goncalves, Ewan Harrison, David K. Jackson, Ian Johnston, Dominic Kwiatkowski, Cordelia Langford, John Sillitoe on behalf of the Wellcome Sanger Institute COVID-19 Surveillance Team |
| EPI_ISL_3359946 | Lighthouse Lab in Alderley Park | Wellcome Sanger Institute for the COVID-19 Genomics UK (COG-UK) Consortium | Jacquelyn Wynn, Mairead Hyland, The Lighthouse Lab in Alderley Park and Alex Alderton, Roberto Amato, Jeffrey Barrett, Sonia Goncalves, Ewan Harrison, David K. Jackson, Ian Johnston, Dominic Kwiatkowski, Cordelia Langford, John Sillitoe on behalf of the Wellcome Sanger Institute COVID-19 Surveillance Team |
| EPI_ISL_3359948 | Lighthouse Laboratory Plymouth | Wellcome Sanger Institute for the COVID-19 Genomics UK (COG-UK) Consortium | Lighthouse Laboratory Plymouth and Alex Alderton, Roberto Amato, Jeffrey Barrett, Sonia Goncalves, Ewan Harrison, David K. Jackson, Ian Johnston, Dominic Kwiatkowski, Cordelia Langford, John Sillitoe on behalf of the Wellcome Sanger Institute COVID-19 Surveillance Team |
| EPI_ISL_3359949, EPI_ISL_3359951 | Lighthouse Lab in Alderley Park | Wellcome Sanger Institute for the COVID-19 Genomics UK (COG-UK) Consortium | Jacquelyn Wynn, Mairead Hyland, The Lighthouse Lab in Alderley Park and Alex Alderton, Roberto Amato, Jeffrey Barrett, Sonia Goncalves, Ewan Harrison, David K. Jackson, Ian Johnston, Dominic Kwiatkowski, Cordelia Langford, John Sillitoe on behalf of the Wellcome Sanger Institute COVID-19 Surveillance Team |
| EPI_ISL_3359952 | Lighthouse Laboratory Plymouth | Wellcome Sanger Institute for the COVID-19 Genomics UK (COG-UK) Consortium | Lighthouse Laboratory Plymouth and Alex Alderton, Roberto Amato, Jeffrey Barrett, Sonia Goncalves, Ewan Harrison, David K. Jackson, Ian Johnston, Dominic Kwiatkowski, Cordelia Langford, John Sillitoe on behalf of the Wellcome Sanger Institute COVID-19 Surveillance Team |
| EPI_ISL_3359954 | Lighthouse Lab in Glasgow | Wellcome Sanger Institute for the COVID-19 Genomics UK (COG-UK) Consortium | Harper VanSteenhouse, Yumi Kasai, David Gray, Carol Clugston, Anna Dominiczak and Alex Alderton, Roberto Amato, Jeffrey Barrett, Sonia Goncalves, Ewan Harrison, David K. Jackson, Ian Johnston, Dominic Kwiatkowski, Cordelia Langford, John Sillitoe on behalf of the Wellcome Sanger Institute COVID-19 Surveillance Team |
| EPI_ISL_3359955 | Lighthouse Lab in Alderley Park | Wellcome Sanger Institute for the COVID-19 Genomics UK (COG-UK) Consortium | Jacquelyn Wynn, Mairead Hyland, The Lighthouse Lab in Alderley Park and Alex Alderton, Roberto Amato, Jeffrey Barrett, Sonia Goncalves, Ewan Harrison, David K. Jackson, Ian Johnston, Dominic Kwiatkowski, Cordelia Langford, John Sillitoe on behalf of the Wellcome Sanger Institute COVID-19 Surveillance Team |
| EPI_ISL_3359956 | Lighthouse Lab in Glasgow | Wellcome Sanger Institute for the COVID-19 Genomics UK (COG-UK) Consortium | Harper VanSteenhouse, Yumi Kasai, David Gray, Carol Clugston, Anna Dominiczak and Alex Alderton, Roberto Amato, Jeffrey Barrett, Sonia Goncalves, Ewan Harrison, David K. Jackson, Ian Johnston, Dominic Kwiatkowski, Cordelia Langford, John Sillitoe on behalf of the Wellcome Sanger Institute COVID-19 Surveillance Team |
| EPI_ISL_3359958 | Lighthouse Laboratory Plymouth | Wellcome Sanger Institute for the COVID-19 Genomics UK (COG-UK) Consortium | Lighthouse Laboratory Plymouth and Alex Alderton, Roberto Amato, Jeffrey Barrett, Sonia Goncalves, Ewan Harrison, David K. Jackson, Ian Johnston, Dominic Kwiatkowski, Cordelia Langford, John Sillitoe on behalf of the Wellcome Sanger Institute COVID-19 Surveillance Team |
| EPI_ISL_3359959, EPI_ISL_3359961, EPI_ISL_3359962, EPI_ISL_3359963 | Lighthouse Lab in Glasgow | Wellcome Sanger Institute for the COVID-19 Genomics UK (COG-UK) Consortium | Harper VanSteenhouse, Yumi Kasai, David Gray, Carol Clugston, Anna Dominiczak and Alex Alderton, Roberto Amato, Jeffrey Barrett, Sonia Goncalves, Ewan Harrison, David K. Jackson, Ian Johnston, Dominic Kwiatkowski, Cordelia Langford, John Sillitoe on behalf of the Wellcome Sanger Institute COVID-19 Surveillance Team |
| EPI_ISL_3359965, EPI_ISL_3359966, EPI_ISL_3359968, EPI_ISL_3359969, EPI_ISL_3359971, EPI_ISL_3359972, EPI_ISL_3359974 | Lighthouse Lab in Alderley Park | Wellcome Sanger Institute for the COVID-19 Genomics UK (COG-UK) Consortium | Jacquelyn Wynn, Mairead Hyland, The Lighthouse Lab in Alderley Park and Alex Alderton, Roberto Amato, Jeffrey Barrett, Sonia Goncalves, Ewan Harrison, David K. Jackson, Ian Johnston, Dominic Kwiatkowski, Cordelia Langford, John Sillitoe on behalf of the Wellcome Sanger Institute COVID-19 Surveillance Team |
| EPI_ISL_3359975 | Lighthouse Lab in Glasgow | Wellcome Sanger Institute for the COVID-19 Genomics UK (COG-UK) Consortium | Harper VanSteenhouse, Yumi Kasai, David Gray, Carol Clugston, Anna Dominiczak and Alex Alderton, Roberto Amato, Jeffrey Barrett, Sonia Goncalves, Ewan Harrison, David K. Jackson, Ian Johnston, Dominic Kwiatkowski, Cordelia Langford, John Sillitoe on behalf of the Wellcome Sanger Institute COVID-19 Surveillance Team |
| EPI_ISL_3359977 | Lighthouse Laboratory Plymouth | Wellcome Sanger Institute for the COVID-19 Genomics UK (COG-UK) Consortium | Lighthouse Laboratory Plymouth and Alex Alderton, Roberto Amato, Jeffrey Barrett, Sonia Goncalves, Ewan Harrison, David K. Jackson, Ian Johnston, Dominic Kwiatkowski, Cordelia Langford, John Sillitoe on behalf of the Wellcome Sanger Institute COVID-19 Surveillance Team |
| EPI_ISL_3359978 | Lighthouse Lab in Alderley Park | Wellcome Sanger Institute for the COVID-19 Genomics UK (COG-UK) Consortium | Jacquelyn Wynn, Mairead Hyland, The Lighthouse Lab in Alderley Park and Alex Alderton, Roberto Amato, Jeffrey Barrett, Sonia Goncalves, Ewan Harrison, David K. Jackson, Ian Johnston, Dominic Kwiatkowski, Cordelia Langford, John Sillitoe on behalf of the Wellcome Sanger Institute COVID-19 Surveillance Team |
| EPI_ISL_3359980, EPI_ISL_3359981 | Lighthouse Lab in Glasgow | Wellcome Sanger Institute for the COVID-19 Genomics UK (COG-UK) Consortium | Harper VanSteenhouse, Yumi Kasai, David Gray, Carol Clugston, Anna Dominiczak and Alex Alderton, Roberto Amato, Jeffrey Barrett, Sonia Goncalves, Ewan Harrison, David K. Jackson, Ian Johnston, Dominic Kwiatkowski, Cordelia Langford, John Sillitoe on behalf of the Wellcome Sanger Institute COVID-19 Surveillance Team |
| EPI_ISL_3359982, EPI_ISL_3359984 | Lighthouse Lab in Alderley Park | Wellcome Sanger Institute for the COVID-19 Genomics UK (COG-UK) Consortium | Jacquelyn Wynn, Mairead Hyland, The Lighthouse Lab in Alderley Park and Alex Alderton, Roberto Amato, Jeffrey Barrett, Sonia Goncalves, Ewan Harrison, David K. Jackson, Ian Johnston, Dominic Kwiatkowski, Cordelia Langford, John Sillitoe on behalf of the Wellcome Sanger Institute COVID-19 Surveillance Team |
| EPI_ISL_3359985, EPI_ISL_3359987 | Lighthouse Laboratory Plymouth | Wellcome Sanger Institute for the COVID-19 Genomics UK (COG-UK) Consortium | Lighthouse Laboratory Plymouth and Alex Alderton, Roberto Amato, Jeffrey Barrett, Sonia Goncalves, Ewan Harrison, David K. Jackson, Ian Johnston, Dominic Kwiatkowski, Cordelia Langford, John Sillitoe on behalf of the Wellcome Sanger Institute COVID-19 Surveillance Team |
| EPI_ISL_3359988 | Lighthouse Lab in Glasgow | Wellcome Sanger Institute for the COVID-19 Genomics UK (COG-UK) Consortium | Harper VanSteenhouse, Yumi Kasai, David Gray, Carol Clugston, Anna Dominiczak and Alex Alderton, Roberto Amato, Jeffrey Barrett, Sonia Goncalves, Ewan Harrison, David K. Jackson, Ian Johnston, Dominic Kwiatkowski, Cordelia Langford, John Sillitoe on behalf of the Wellcome Sanger Institute COVID-19 Surveillance Team |
| EPI_ISL_3359989 | Lighthouse Lab in Alderley Park | Wellcome Sanger Institute for the COVID-19 Genomics UK (COG-UK) Consortium | Jacquelyn Wynn, Mairead Hyland, The Lighthouse Lab in Alderley Park and Alex Alderton, Roberto Amato, Jeffrey Barrett, Sonia Goncalves, Ewan Harrison, David K. Jackson, Ian Johnston, Dominic Kwiatkowski, Cordelia Langford, John Sillitoe on behalf of the Wellcome Sanger Institute COVID-19 Surveillance Team |
| EPI_ISL_3359991 | Lighthouse Lab in Glasgow | Wellcome Sanger Institute for the COVID-19 Genomics UK (COG-UK) Consortium | Harper VanSteenhouse, Yumi Kasai, David Gray, Carol Clugston, Anna Dominiczak and Alex Alderton, Roberto Amato, Jeffrey Barrett, Sonia Goncalves, Ewan Harrison, David K. Jackson, Ian Johnston, Dominic Kwiatkowski, Cordelia Langford, John Sillitoe on behalf of the Wellcome Sanger Institute COVID-19 Surveillance Team |
| EPI_ISL_3359992 | Lighthouse Lab in Alderley Park | Wellcome Sanger Institute for the COVID-19 Genomics UK (COG-UK) Consortium | Jacquelyn Wynn, Mairead Hyland, The Lighthouse Lab in Alderley Park and Alex Alderton, Roberto Amato, Jeffrey Barrett, Sonia Goncalves, Ewan Harrison, David K. Jackson, Ian Johnston, Dominic Kwiatkowski, Cordelia Langford, John Sillitoe on behalf of the Wellcome Sanger Institute COVID-19 Surveillance Team |
| EPI_ISL_3359994, EPI_ISL_3359995 | Lighthouse Laboratory Plymouth | Wellcome Sanger Institute for the COVID-19 Genomics UK (COG-UK) Consortium | Lighthouse Laboratory Plymouth and Alex Alderton, Roberto Amato, Jeffrey Barrett, Sonia Goncalves, Ewan Harrison, David K. Jackson, Ian Johnston, Dominic Kwiatkowski, Cordelia Langford, John Sillitoe on behalf of the Wellcome Sanger Institute COVID-19 Surveillance Team |
| EPI_ISL_3359996 | Lighthouse Lab in Alderley Park | Wellcome Sanger Institute for the COVID-19 Genomics UK (COG-UK) Consortium | Jacquelyn Wynn, Mairead Hyland, The Lighthouse Lab in Alderley Park and Alex Alderton, Roberto Amato, Jeffrey Barrett, Sonia Goncalves, Ewan Harrison, David K. Jackson, Ian Johnston, Dominic Kwiatkowski, Cordelia Langford, John Sillitoe on behalf of the Wellcome Sanger Institute COVID-19 Surveillance Team |
| EPI_ISL_3359998, EPI_ISL_3359999 | Lighthouse Laboratory Plymouth | Wellcome Sanger Institute for the COVID-19 Genomics UK (COG-UK) Consortium | Lighthouse Laboratory Plymouth and Alex Alderton, Roberto Amato, Jeffrey Barrett, Sonia Goncalves, Ewan Harrison, David K. Jackson, Ian Johnston, Dominic Kwiatkowski, Cordelia Langford, John Sillitoe on behalf of the Wellcome Sanger Institute COVID-19 Surveillance Team |
| EPI_ISL_3360000, EPI_ISL_3360002 | Lighthouse Lab in Alderley Park | Wellcome Sanger Institute for the COVID-19 Genomics UK (COG-UK) Consortium | Jacquelyn Wynn, Mairead Hyland, The Lighthouse Lab in Alderley Park and Alex Alderton, Roberto Amato, Jeffrey Barrett, Sonia Goncalves, Ewan Harrison, David K. Jackson, Ian Johnston, Dominic Kwiatkowski, Cordelia Langford, John Sillitoe on behalf of the Wellcome Sanger Institute COVID-19 Surveillance Team |
| EPI_ISL_3360003 | Lighthouse Lab in Glasgow | Wellcome Sanger Institute for the COVID-19 Genomics UK (COG-UK) Consortium | Harper VanSteenhouse, Yumi Kasai, David Gray, Carol Clugston, Anna Dominiczak and Alex Alderton, Roberto Amato, Jeffrey Barrett, Sonia Goncalves, Ewan Harrison, David K. Jackson, Ian Johnston, Dominic Kwiatkowski, Cordelia Langford, John Sillitoe on behalf of the Wellcome Sanger Institute COVID-19 Surveillance Team |
| EPI_ISL_3360004, EPI_ISL_3360006 | Lighthouse Lab in Alderley Park | Wellcome Sanger Institute for the COVID-19 Genomics UK (COG-UK) Consortium | Jacquelyn Wynn, Mairead Hyland, The Lighthouse Lab in Alderley Park and Alex Alderton, Roberto Amato, Jeffrey Barrett, Sonia Goncalves, Ewan Harrison, David K. Jackson, Ian Johnston, Dominic Kwiatkowski, Cordelia Langford, John Sillitoe on behalf of the Wellcome Sanger Institute COVID-19 Surveillance Team |
| EPI_ISL_3360007 | Lighthouse Lab in Glasgow | Wellcome Sanger Institute for the COVID-19 Genomics UK (COG-UK) Consortium | Harper VanSteenhouse, Yumi Kasai, David Gray, Carol Clugston, Anna Dominiczak and Alex Alderton, Roberto Amato, Jeffrey Barrett, Sonia Goncalves, Ewan Harrison, David K. Jackson, Ian Johnston, Dominic Kwiatkowski, Cordelia Langford, John Sillitoe on behalf of the Wellcome Sanger Institute COVID-19 Surveillance Team |
| EPI_ISL_3360008 | Lighthouse Laboratory Plymouth | Wellcome Sanger Institute for the COVID-19 Genomics UK (COG-UK) Consortium | Lighthouse Laboratory Plymouth and Alex Alderton, Roberto Amato, Jeffrey Barrett, Sonia Goncalves, Ewan Harrison, David K. Jackson, Ian Johnston, Dominic Kwiatkowski, Cordelia Langford, John Sillitoe on behalf of the Wellcome Sanger Institute COVID-19 Surveillance Team |
| EPI_ISL_3360010 | Lighthouse Lab in Alderley Park | Wellcome Sanger Institute for the COVID-19 Genomics UK (COG-UK) Consortium | Jacquelyn Wynn, Mairead Hyland, The Lighthouse Lab in Alderley Park and Alex Alderton, Roberto Amato, Jeffrey Barrett, Sonia Goncalves, Ewan Harrison, David K. Jackson, Ian Johnston, Dominic Kwiatkowski, Cordelia Langford, John Sillitoe on behalf of the Wellcome Sanger Institute COVID-19 Surveillance Team |
| EPI_ISL_3360011 | Lighthouse Lab in Glasgow | Wellcome Sanger Institute for the COVID-19 Genomics UK (COG-UK) Consortium | Harper VanSteenhouse, Yumi Kasai, David Gray, Carol Clugston, Anna Dominiczak and Alex Alderton, Roberto Amato, Jeffrey Barrett, Sonia Goncalves, Ewan Harrison, David K. Jackson, Ian Johnston, Dominic Kwiatkowski, Cordelia Langford, John Sillitoe on behalf of the Wellcome Sanger Institute COVID-19 Surveillance Team |
| EPI_ISL_3360013 | Lighthouse Lab in Alderley Park | Wellcome Sanger Institute for the COVID-19 Genomics UK (COG-UK) Consortium | Jacquelyn Wynn, Mairead Hyland, The Lighthouse Lab in Alderley Park and Alex Alderton, Roberto Amato, Jeffrey Barrett, Sonia Goncalves, Ewan Harrison, David K. Jackson, Ian Johnston, Dominic Kwiatkowski, Cordelia Langford, John Sillitoe on behalf of the Wellcome Sanger Institute COVID-19 Surveillance Team |
| EPI_ISL_3360014 | Lighthouse Lab in Glasgow | Wellcome Sanger Institute for the COVID-19 Genomics UK (COG-UK) Consortium | Harper VanSteenhouse, Yumi Kasai, David Gray, Carol Clugston, Anna Dominiczak and Alex Alderton, Roberto Amato, Jeffrey Barrett, Sonia Goncalves, Ewan Harrison, David K. Jackson, Ian Johnston, Dominic Kwiatkowski, Cordelia Langford, John Sillitoe on behalf of the Wellcome Sanger Institute COVID-19 Surveillance Team |
| EPI_ISL_3360016, EPI_ISL_3360017 | Lighthouse Lab in Alderley Park | Wellcome Sanger Institute for the COVID-19 Genomics UK (COG-UK) Consortium | Jacquelyn Wynn, Mairead Hyland, The Lighthouse Lab in Alderley Park and Alex Alderton, Roberto Amato, Jeffrey Barrett, Sonia Goncalves, Ewan Harrison, David K. Jackson, Ian Johnston, Dominic Kwiatkowski, Cordelia Langford, John Sillitoe on behalf of the Wellcome Sanger Institute COVID-19 Surveillance Team |
| EPI_ISL_3360018 | Lighthouse Laboratory Plymouth | Wellcome Sanger Institute for the COVID-19 Genomics UK (COG-UK) Consortium | Lighthouse Laboratory Plymouth and Alex Alderton, Roberto Amato, Jeffrey Barrett, Sonia Goncalves, Ewan Harrison, David K. Jackson, Ian Johnston, Dominic Kwiatkowski, Cordelia Langford, John Sillitoe on behalf of the Wellcome Sanger Institute COVID-19 Surveillance Team |
| EPI_ISL_3360020, EPI_ISL_3360021, EPI_ISL_3360023 | Lighthouse Lab in Alderley Park | Wellcome Sanger Institute for the COVID-19 Genomics UK (COG-UK) Consortium | Jacquelyn Wynn, Mairead Hyland, The Lighthouse Lab in Alderley Park and Alex Alderton, Roberto Amato, Jeffrey Barrett, Sonia Goncalves, Ewan Harrison, David K. Jackson, Ian Johnston, Dominic Kwiatkowski, Cordelia Langford, John Sillitoe on behalf of the Wellcome Sanger Institute COVID-19 Surveillance Team |
| EPI_ISL_3360024, EPI_ISL_3360025, EPI_ISL_3360027, EPI_ISL_3360028, EPI_ISL_3360030 | Lighthouse Laboratory Plymouth | Wellcome Sanger Institute for the COVID-19 Genomics UK (COG-UK) Consortium | Lighthouse Laboratory Plymouth and Alex Alderton, Roberto Amato, Jeffrey Barrett, Sonia Goncalves, Ewan Harrison, David K. Jackson, Ian Johnston, Dominic Kwiatkowski, Cordelia Langford, John Sillitoe on behalf of the Wellcome Sanger Institute COVID-19 Surveillance Team |
| EPI_ISL_3360031, EPI_ISL_3360032, EPI_ISL_3360034 | Lighthouse Lab in Alderley Park | Wellcome Sanger Institute for the COVID-19 Genomics UK (COG-UK) Consortium | Jacquelyn Wynn, Mairead Hyland, The Lighthouse Lab in Alderley Park and Alex Alderton, Roberto Amato, Jeffrey Barrett, Sonia Goncalves, Ewan Harrison, David K. Jackson, Ian Johnston, Dominic Kwiatkowski, Cordelia Langford, John Sillitoe on behalf of the Wellcome Sanger Institute COVID-19 Surveillance Team |
| EPI_ISL_3360035, EPI_ISL_3360036 | Lighthouse Lab in Glasgow | Wellcome Sanger Institute for the COVID-19 Genomics UK (COG-UK) Consortium | Harper VanSteenhouse, Yumi Kasai, David Gray, Carol Clugston, Anna Dominiczak and Alex Alderton, Roberto Amato, Jeffrey Barrett, Sonia Goncalves, Ewan Harrison, David K. Jackson, Ian Johnston, Dominic Kwiatkowski, Cordelia Langford, John Sillitoe on behalf of the Wellcome Sanger Institute COVID-19 Surveillance Team |
| EPI_ISL_3360038 | Lighthouse Lab in Alderley Park | Wellcome Sanger Institute for the COVID-19 Genomics UK (COG-UK) Consortium | Jacquelyn Wynn, Mairead Hyland, The Lighthouse Lab in Alderley Park and Alex Alderton, Roberto Amato, Jeffrey Barrett, Sonia Goncalves, Ewan Harrison, David K. Jackson, Ian Johnston, Dominic Kwiatkowski, Cordelia Langford, John Sillitoe on behalf of the Wellcome Sanger Institute COVID-19 Surveillance Team |

| | | | |
|---|---|---|---|
| EPI_ISL_3360039 | Lighthouse Laboratory Plymouth | Wellcome Sanger Institute for the COVID-19 Genomics UK (COG-UK) Consortium | Lighthouse Laboratory Plymouth and Alex Alderton, Roberto Amato, Jeffrey Barrett, Sonia Goncalves, Ewan Harrison, David K. Jackson, Ian Johnston, Dominic Kwiatkowski, Cordelia Langford, John Sillitoe on behalf of the Wellcome Sanger Institute COVID-19 Surveillance Team |
| EPI_ISL_3360041, EPI_ISL_3360042 | Lighthouse Lab in Glasgow | Wellcome Sanger Institute for the COVID-19 Genomics UK (COG-UK) Consortium | Harper VanSteenhouse, Yumi Kasai, David Gray, Carol Clugston, Anna Dominiczak and Alex Alderton, Roberto Amato, Jeffrey Barrett, Sonia Goncalves, Ewan Harrison, David K. Jackson, Ian Johnston, Dominic Kwiatkowski, Cordelia Langford, John Sillitoe on behalf of the Wellcome Sanger Institute COVID-19 Surveillance Team |
| EPI_ISL_3360044, EPI_ISL_3360045 | Lighthouse Lab in Alderley Park | Wellcome Sanger Institute for the COVID-19 Genomics UK (COG-UK) Consortium | Jacquelyn Wynn, Mairead Hyland, The Lighthouse Lab in Alderley Park and Alex Alderton, Roberto Amato, Jeffrey Barrett, Sonia Goncalves, Ewan Harrison, David K. Jackson, Ian Johnston, Dominic Kwiatkowski, Cordelia Langford, John Sillitoe on behalf of the Wellcome Sanger Institute COVID-19 Surveillance Team |
| EPI_ISL_3360046 | Lighthouse Lab in Glasgow | Wellcome Sanger Institute for the COVID-19 Genomics UK (COG-UK) Consortium | Harper VanSteenhouse, Yumi Kasai, David Gray, Carol Clugston, Anna Dominiczak and Alex Alderton, Roberto Amato, Jeffrey Barrett, Sonia Goncalves, Ewan Harrison, David K. Jackson, Ian Johnston, Dominic Kwiatkowski, Cordelia Langford, John Sillitoe on behalf of the Wellcome Sanger Institute COVID-19 Surveillance Team |
| EPI_ISL_3360048, EPI_ISL_3360050, EPI_ISL_3360051, EPI_ISL_3360052, EPI_ISL_3360055, EPI_ISL_3360056 | Lighthouse Lab in Alderley Park | Wellcome Sanger Institute for the COVID-19 Genomics UK (COG-UK) Consortium | Jacquelyn Wynn, Mairead Hyland, The Lighthouse Lab in Alderley Park and Alex Alderton, Roberto Amato, Jeffrey Barrett, Sonia Goncalves, Ewan Harrison, David K. Jackson, Ian Johnston, Dominic Kwiatkowski, Cordelia Langford, John Sillitoe on behalf of the Wellcome Sanger Institute COVID-19 Surveillance Team |
| EPI_ISL_3360058, EPI_ISL_3360059 | Lighthouse Lab in Glasgow | Wellcome Sanger Institute for the COVID-19 Genomics UK (COG-UK) Consortium | Harper VanSteenhouse, Yumi Kasai, David Gray, Carol Clugston, Anna Dominiczak and Alex Alderton, Roberto Amato, Jeffrey Barrett, Sonia Goncalves, Ewan Harrison, David K. Jackson, Ian Johnston, Dominic Kwiatkowski, Cordelia Langford, John Sillitoe on behalf of the Wellcome Sanger Institute COVID-19 Surveillance Team |
| EPI_ISL_3360061 | Lighthouse Lab in Alderley Park | Wellcome Sanger Institute for the COVID-19 Genomics UK (COG-UK) Consortium | Jacquelyn Wynn, Mairead Hyland, The Lighthouse Lab in Alderley Park and Alex Alderton, Roberto Amato, Jeffrey Barrett, Sonia Goncalves, Ewan Harrison, David K. Jackson, Ian Johnston, Dominic Kwiatkowski, Cordelia Langford, John Sillitoe on behalf of the Wellcome Sanger Institute COVID-19 Surveillance Team |
| EPI_ISL_3360062 | Lighthouse Lab in Glasgow | Wellcome Sanger Institute for the COVID-19 Genomics UK (COG-UK) Consortium | Harper VanSteenhouse, Yumi Kasai, David Gray, Carol Clugston, Anna Dominiczak and Alex Alderton, Roberto Amato, Jeffrey Barrett, Sonia Goncalves, Ewan Harrison, David K. Jackson, Ian Johnston, Dominic Kwiatkowski, Cordelia Langford, John Sillitoe on behalf of the Wellcome Sanger Institute COVID-19 Surveillance Team |
| EPI_ISL_3360064, EPI_ISL_3360065, EPI_ISL_3360067, EPI_ISL_3360068, EPI_ISL_3360070, EPI_ISL_3360071, EPI_ISL_3360072, EPI_ISL_3360074, EPI_ISL_3360075 | Lighthouse Lab in Alderley Park | Wellcome Sanger Institute for the COVID-19 Genomics UK (COG-UK) Consortium | Jacquelyn Wynn, Mairead Hyland, The Lighthouse Lab in Alderley Park and Alex Alderton, Roberto Amato, Jeffrey Barrett, Sonia Goncalves, Ewan Harrison, David K. Jackson, Ian Johnston, Dominic Kwiatkowski, Cordelia Langford, John Sillitoe on behalf of the Wellcome Sanger Institute COVID-19 Surveillance Team |
| EPI_ISL_3360077 | Lighthouse Lab in Glasgow | Wellcome Sanger Institute for the COVID-19 Genomics UK (COG-UK) Consortium | Harper VanSteenhouse, Yumi Kasai, David Gray, Carol Clugston, Anna Dominiczak and Alex Alderton, Roberto Amato, Jeffrey Barrett, Sonia Goncalves, Ewan Harrison, David K. Jackson, Ian Johnston, Dominic Kwiatkowski, Cordelia Langford, John Sillitoe on behalf of the Wellcome Sanger Institute COVID-19 Surveillance Team |
| EPI_ISL_3360078, EPI_ISL_3360080, EPI_ISL_3360081, EPI_ISL_3360083, EPI_ISL_3360084 | Lighthouse Lab in Alderley Park | Wellcome Sanger Institute for the COVID-19 Genomics UK (COG-UK) Consortium | Jacquelyn Wynn, Mairead Hyland, The Lighthouse Lab in Alderley Park and Alex Alderton, Roberto Amato, Jeffrey Barrett, Sonia Goncalves, Ewan Harrison, David K. Jackson, Ian Johnston, Dominic Kwiatkowski, Cordelia Langford, John Sillitoe on behalf of the Wellcome Sanger Institute COVID-19 Surveillance Team |
| EPI_ISL_3360086 | Lighthouse Lab in Glasgow | Wellcome Sanger Institute for the COVID-19 Genomics UK (COG-UK) Consortium | Harper VanSteenhouse, Yumi Kasai, David Gray, Carol Clugston, Anna Dominiczak and Alex Alderton, Roberto Amato, Jeffrey Barrett, Sonia Goncalves, Ewan Harrison, David K. Jackson, Ian Johnston, Dominic Kwiatkowski, Cordelia Langford, John Sillitoe on behalf of the Wellcome Sanger Institute COVID-19 Surveillance Team |
| EPI_ISL_3360087, EPI_ISL_3360088, EPI_ISL_3360090, EPI_ISL_3360091 | Lighthouse Lab in Alderley Park | Wellcome Sanger Institute for the COVID-19 Genomics UK (COG-UK) Consortium | Jacquelyn Wynn, Mairead Hyland, The Lighthouse Lab in Alderley Park and Alex Alderton, Roberto Amato, Jeffrey Barrett, Sonia Goncalves, Ewan Harrison, David K. Jackson, Ian Johnston, Dominic Kwiatkowski, Cordelia Langford, John Sillitoe on behalf of the Wellcome Sanger Institute COVID-19 Surveillance Team |
| EPI_ISL_3360093, EPI_ISL_3360094 | Lighthouse Lab in Glasgow | Wellcome Sanger Institute for the COVID-19 Genomics UK (COG-UK) Consortium | Harper VanSteenhouse, Yumi Kasai, David Gray, Carol Clugston, Anna Dominiczak and Alex Alderton, Roberto Amato, Jeffrey Barrett, Sonia Goncalves, Ewan Harrison, David K. Jackson, Ian Johnston, Dominic Kwiatkowski, Cordelia Langford, John Sillitoe on behalf of the Wellcome Sanger Institute COVID-19 Surveillance Team |
| EPI_ISL_3360096 | Lighthouse Lab in Alderley Park | Wellcome Sanger Institute for the COVID-19 Genomics UK (COG-UK) Consortium | Jacquelyn Wynn, Mairead Hyland, The Lighthouse Lab in Alderley Park and Alex Alderton, Roberto Amato, Jeffrey Barrett, Sonia Goncalves, Ewan Harrison, David K. Jackson, Ian Johnston, Dominic Kwiatkowski, Cordelia Langford, John Sillitoe on behalf of the Wellcome Sanger Institute COVID-19 Surveillance Team |
| EPI_ISL_3360097 | Lighthouse Lab in Glasgow | Wellcome Sanger Institute for the COVID-19 Genomics UK (COG-UK) Consortium | Harper VanSteenhouse, Yumi Kasai, David Gray, Carol Clugston, Anna Dominiczak and Alex Alderton, Roberto Amato, Jeffrey Barrett, Sonia Goncalves, Ewan Harrison, David K. Jackson, Ian Johnston, Dominic Kwiatkowski, Cordelia Langford, John Sillitoe on behalf of the Wellcome Sanger Institute COVID-19 Surveillance Team |
| EPI_ISL_3360098, EPI_ISL_3360100, EPI_ISL_3360101, EPI_ISL_3360103 | Lighthouse Lab in Alderley Park | Wellcome Sanger Institute for the COVID-19 Genomics UK (COG-UK) Consortium | Jacquelyn Wynn, Mairead Hyland, The Lighthouse Lab in Alderley Park and Alex Alderton, Roberto Amato, Jeffrey Barrett, Sonia Goncalves, Ewan Harrison, David K. Jackson, Ian Johnston, Dominic Kwiatkowski, Cordelia Langford, John Sillitoe on behalf of the Wellcome Sanger Institute COVID-19 Surveillance Team |
| EPI_ISL_3360104, EPI_ISL_3360106, EPI_ISL_3360107 | Lighthouse Lab in Glasgow | Wellcome Sanger Institute for the COVID-19 Genomics UK (COG-UK) Consortium | Harper VanSteenhouse, Yumi Kasai, David Gray, Carol Clugston, Anna Dominiczak and Alex Alderton, Roberto Amato, Jeffrey Barrett, Sonia Goncalves, Ewan Harrison, David K. Jackson, Ian Johnston, Dominic Kwiatkowski, Cordelia Langford, John Sillitoe on behalf of the Wellcome Sanger Institute COVID-19 Surveillance Team |
| EPI_ISL_3360109, EPI_ISL_3360110, EPI_ISL_3360111, EPI_ISL_3360113 | Lighthouse Lab in Alderley Park | Wellcome Sanger Institute for the COVID-19 Genomics UK (COG-UK) Consortium | Jacquelyn Wynn, Mairead Hyland, The Lighthouse Lab in Alderley Park and Alex Alderton, Roberto Amato, Jeffrey Barrett, Sonia Goncalves, Ewan Harrison, David K. Jackson, Ian Johnston, Dominic Kwiatkowski, Cordelia Langford, John Sillitoe on behalf of the Wellcome Sanger Institute COVID-19 Surveillance Team |
| EPI_ISL_3360114, EPI_ISL_3360116 | Lighthouse Lab in Glasgow | Wellcome Sanger Institute for the COVID-19 Genomics UK (COG-UK) Consortium | Harper VanSteenhouse, Yumi Kasai, David Gray, Carol Clugston, Anna Dominiczak and Alex Alderton, Roberto Amato, Jeffrey Barrett, Sonia Goncalves, Ewan Harrison, David K. Jackson, Ian Johnston, Dominic Kwiatkowski, Cordelia Langford, John Sillitoe on behalf of the Wellcome Sanger Institute COVID-19 Surveillance Team |
| EPI_ISL_3360117 | Lighthouse Lab in Alderley Park | Wellcome Sanger Institute for the COVID-19 Genomics UK (COG-UK) Consortium | Jacquelyn Wynn, Mairead Hyland, The Lighthouse Lab in Alderley Park and Alex Alderton, Roberto Amato, Jeffrey Barrett, Sonia Goncalves, Ewan Harrison, David K. Jackson, Ian Johnston, Dominic Kwiatkowski, Cordelia Langford, John Sillitoe on behalf of the Wellcome Sanger Institute COVID-19 Surveillance Team |
| EPI_ISL_3360119 | Lighthouse Lab in Glasgow | Wellcome Sanger Institute for the COVID-19 Genomics UK (COG-UK) Consortium | Harper VanSteenhouse, Yumi Kasai, David Gray, Carol Clugston, Anna Dominiczak and Alex Alderton, Roberto Amato, Jeffrey Barrett, Sonia Goncalves, Ewan Harrison, David K. Jackson, Ian Johnston, Dominic Kwiatkowski, Cordelia Langford, John Sillitoe on behalf of the Wellcome Sanger Institute COVID-19 Surveillance Team |
| EPI_ISL_3360120, EPI_ISL_3360122, EPI_ISL_3360123 | Lighthouse Lab in Alderley Park | Wellcome Sanger Institute for the COVID-19 Genomics UK (COG-UK) Consortium | Jacquelyn Wynn, Mairead Hyland, The Lighthouse Lab in Alderley Park and Alex Alderton, Roberto Amato, Jeffrey Barrett, Sonia Goncalves, Ewan Harrison, David K. Jackson, Ian Johnston, Dominic Kwiatkowski, Cordelia Langford, John Sillitoe on behalf of the Wellcome Sanger Institute COVID-19 Surveillance Team |
| EPI_ISL_3360125 | Lighthouse Lab in Glasgow | Wellcome Sanger Institute for the COVID-19 Genomics UK (COG-UK) Consortium | Harper VanSteenhouse, Yumi Kasai, David Gray, Carol Clugston, Anna Dominiczak and Alex Alderton, Roberto Amato, Jeffrey Barrett, Sonia Goncalves, Ewan Harrison, David K. Jackson, Ian Johnston, Dominic Kwiatkowski, Cordelia Langford, John Sillitoe on behalf of the Wellcome Sanger Institute COVID-19 Surveillance Team |
| EPI_ISL_3360126, EPI_ISL_3360127, EPI_ISL_3360129 | Lighthouse Lab in Alderley Park | Wellcome Sanger Institute for the COVID-19 Genomics UK (COG-UK) Consortium | Jacquelyn Wynn, Mairead Hyland, The Lighthouse Lab in Alderley Park and Alex Alderton, Roberto Amato, Jeffrey Barrett, Sonia Goncalves, Ewan Harrison, David K. Jackson, Ian Johnston, Dominic Kwiatkowski, Cordelia Langford, John Sillitoe on behalf of the Wellcome Sanger Institute COVID-19 Surveillance Team |
| EPI_ISL_3360130 | Lighthouse Lab in Glasgow | Wellcome Sanger Institute for the COVID-19 Genomics UK (COG-UK) Consortium | Harper VanSteenhouse, Yumi Kasai, David Gray, Carol Clugston, Anna Dominiczak and Alex Alderton, Roberto Amato, Jeffrey Barrett, Sonia Goncalves, Ewan Harrison, David K. Jackson, Ian Johnston, Dominic Kwiatkowski, Cordelia Langford, John Sillitoe on behalf of the Wellcome Sanger Institute COVID-19 Surveillance Team |
| EPI_ISL_3360132, EPI_ISL_3360133 | Lighthouse Lab in Alderley Park | Wellcome Sanger Institute for the COVID-19 Genomics UK (COG-UK) Consortium | Jacquelyn Wynn, Mairead Hyland, The Lighthouse Lab in Alderley Park and Alex Alderton, Roberto Amato, Jeffrey Barrett, Sonia Goncalves, Ewan Harrison, David K. Jackson, Ian Johnston, Dominic Kwiatkowski, Cordelia Langford, John Sillitoe on behalf of the Wellcome Sanger Institute COVID-19 Surveillance Team |
| EPI_ISL_3360135 | Lighthouse Lab in Glasgow | Wellcome Sanger Institute for the COVID-19 Genomics UK (COG-UK) Consortium | Harper VanSteenhouse, Yumi Kasai, David Gray, Carol Clugston, Anna Dominiczak and Alex Alderton, Roberto Amato, Jeffrey Barrett, Sonia Goncalves, Ewan Harrison, David K. Jackson, Ian Johnston, Dominic Kwiatkowski, Cordelia Langford, John Sillitoe on behalf of the Wellcome Sanger Institute COVID-19 Surveillance Team |
| EPI_ISL_3360136, EPI_ISL_3360138 | Lighthouse Lab in Alderley Park | Wellcome Sanger Institute for the COVID-19 Genomics UK (COG-UK) Consortium | Jacquelyn Wynn, Mairead Hyland, The Lighthouse Lab in Alderley Park and Alex Alderton, Roberto Amato, Jeffrey Barrett, Sonia Goncalves, Ewan Harrison, David K. Jackson, Ian Johnston, Dominic Kwiatkowski, Cordelia Langford, John Sillitoe on behalf of the Wellcome Sanger Institute COVID-19 Surveillance Team |
| EPI_ISL_3360139, EPI_ISL_3360141 | Lighthouse Lab in Glasgow | Wellcome Sanger Institute for the COVID-19 Genomics UK (COG-UK) Consortium | Harper VanSteenhouse, Yumi Kasai, David Gray, Carol Clugston, Anna Dominiczak and Alex Alderton, Roberto Amato, Jeffrey Barrett, Sonia Goncalves, Ewan Harrison, David K. Jackson, Ian Johnston, Dominic Kwiatkowski, Cordelia Langford, John Sillitoe on behalf of the Wellcome Sanger Institute COVID-19 Surveillance Team |
| EPI_ISL_3360142, EPI_ISL_3360144, EPI_ISL_3360145 | Lighthouse Lab in Alderley Park | Wellcome Sanger Institute for the COVID-19 Genomics UK (COG-UK) Consortium | Jacquelyn Wynn, Mairead Hyland, The Lighthouse Lab in Alderley Park and Alex Alderton, Roberto Amato, Jeffrey Barrett, Sonia Goncalves, Ewan Harrison, David K. Jackson, Ian Johnston, Dominic Kwiatkowski, Cordelia Langford, John Sillitoe on behalf of the Wellcome Sanger Institute COVID-19 Surveillance Team |
| EPI_ISL_3360147 | Lighthouse Lab in Glasgow | Wellcome Sanger Institute for the COVID-19 Genomics UK (COG-UK) Consortium | Harper VanSteenhouse, Yumi Kasai, David Gray, Carol Clugston, Anna Dominiczak and Alex Alderton, Roberto Amato, Jeffrey Barrett, Sonia Goncalves, Ewan Harrison, David K. Jackson, Ian Johnston, Dominic Kwiatkowski, Cordelia Langford, John Sillitoe on behalf of the Wellcome Sanger Institute COVID-19 Surveillance Team |
| EPI_ISL_3360148, EPI_ISL_3360150, EPI_ISL_3360151, EPI_ISL_3360152, EPI_ISL_3360154, EPI_ISL_3360155, EPI_ISL_3360157, EPI_ISL_3360158 | Lighthouse Lab in Alderley Park | Wellcome Sanger Institute for the COVID-19 Genomics UK (COG-UK) Consortium | Jacquelyn Wynn, Mairead Hyland, The Lighthouse Lab in Alderley Park and Alex Alderton, Roberto Amato, Jeffrey Barrett, Sonia Goncalves, Ewan Harrison, David K. Jackson, Ian Johnston, Dominic Kwiatkowski, Cordelia Langford, John Sillitoe on behalf of the Wellcome Sanger Institute COVID-19 Surveillance Team |

| | | | |
|---|---|---|---|
| EPI_ISL_3360159 | Lighthouse Lab in Glasgow | Wellcome Sanger Institute for the COVID-19 Genomics UK (COG-UK) Consortium | Harper VanSteenhouse, Yumi Kasai, David Gray, Carol Clugston, Anna Dominiczak and Alex Alderton, Roberto Amato, Jeffrey Barrett, Sonia Goncalves, Ewan Harrison, David K. Jackson, Ian Johnston, Dominic Kwiatkowski, Cordelia Langford, John Sillitoe on behalf of the Wellcome Sanger Institute COVID-19 Surveillance Team |
| EPI_ISL_3360161 | Lighthouse Lab in Alderley Park | Wellcome Sanger Institute for the COVID-19 Genomics UK (COG-UK) Consortium | Jacquelyn Wynn, Mairead Hyland, The Lighthouse Lab in Alderley Park and Alex Alderton, Roberto Amato, Jeffrey Barrett, Sonia Goncalves, Ewan Harrison, David K. Jackson, Ian Johnston, Dominic Kwiatkowski, Cordelia Langford, John Sillitoe on behalf of the Wellcome Sanger Institute COVID-19 Surveillance Team |
| EPI_ISL_3360162, EPI_ISL_3360164 | Lighthouse Lab in Glasgow | Wellcome Sanger Institute for the COVID-19 Genomics UK (COG-UK) Consortium | Harper VanSteenhouse, Yumi Kasai, David Gray, Carol Clugston, Anna Dominiczak and Alex Alderton, Roberto Amato, Jeffrey Barrett, Sonia Goncalves, Ewan Harrison, David K. Jackson, Ian Johnston, Dominic Kwiatkowski, Cordelia Langford, John Sillitoe on behalf of the Wellcome Sanger Institute COVID-19 Surveillance Team |
| EPI_ISL_3360165, EPI_ISL_3360166, EPI_ISL_3360167, EPI_ISL_3360169, EPI_ISL_3360170, EPI_ISL_3360172, EPI_ISL_3360173, EPI_ISL_3360175 | Lighthouse Lab in Alderley Park | Wellcome Sanger Institute for the COVID-19 Genomics UK (COG-UK) Consortium | Jacquelyn Wynn, Mairead Hyland, The Lighthouse Lab in Alderley Park and Alex Alderton, Roberto Amato, Jeffrey Barrett, Sonia Goncalves, Ewan Harrison, David K. Jackson, Ian Johnston, Dominic Kwiatkowski, Cordelia Langford, John Sillitoe on behalf of the Wellcome Sanger Institute COVID-19 Surveillance Team |
| EPI_ISL_3360176 | Lighthouse Lab in Glasgow | Wellcome Sanger Institute for the COVID-19 Genomics UK (COG-UK) Consortium | Harper VanSteenhouse, Yumi Kasai, David Gray, Carol Clugston, Anna Dominiczak and Alex Alderton, Roberto Amato, Jeffrey Barrett, Sonia Goncalves, Ewan Harrison, David K. Jackson, Ian Johnston, Dominic Kwiatkowski, Cordelia Langford, John Sillitoe on behalf of the Wellcome Sanger Institute COVID-19 Surveillance Team |
| EPI_ISL_3360178, EPI_ISL_3360179, EPI_ISL_3360181, EPI_ISL_3360182, EPI_ISL_3360184, EPI_ISL_3360185, EPI_ISL_3360186 | Lighthouse Lab in Alderley Park | Wellcome Sanger Institute for the COVID-19 Genomics UK (COG-UK) Consortium | Jacquelyn Wynn, Mairead Hyland, The Lighthouse Lab in Alderley Park and Alex Alderton, Roberto Amato, Jeffrey Barrett, Sonia Goncalves, Ewan Harrison, David K. Jackson, Ian Johnston, Dominic Kwiatkowski, Cordelia Langford, John Sillitoe on behalf of the Wellcome Sanger Institute COVID-19 Surveillance Team |
| EPI_ISL_3360188 | Lighthouse Lab in Glasgow | Wellcome Sanger Institute for the COVID-19 Genomics UK (COG-UK) Consortium | Harper VanSteenhouse, Yumi Kasai, David Gray, Carol Clugston, Anna Dominiczak and Alex Alderton, Roberto Amato, Jeffrey Barrett, Sonia Goncalves, Ewan Harrison, David K. Jackson, Ian Johnston, Dominic Kwiatkowski, Cordelia Langford, John Sillitoe on behalf of the Wellcome Sanger Institute COVID-19 Surveillance Team |
| EPI_ISL_3360189, EPI_ISL_3360191, EPI_ISL_3360192, EPI_ISL_3360194, EPI_ISL_3360195, EPI_ISL_3360196, EPI_ISL_3360198, EPI_ISL_3360199 | Lighthouse Lab in Alderley Park | Wellcome Sanger Institute for the COVID-19 Genomics UK (COG-UK) Consortium | Jacquelyn Wynn, Mairead Hyland, The Lighthouse Lab in Alderley Park and Alex Alderton, Roberto Amato, Jeffrey Barrett, Sonia Goncalves, Ewan Harrison, David K. Jackson, Ian Johnston, Dominic Kwiatkowski, Cordelia Langford, John Sillitoe on behalf of the Wellcome Sanger Institute COVID-19 Surveillance Team |
| EPI_ISL_3360201 | Lighthouse Lab in Glasgow | Wellcome Sanger Institute for the COVID-19 Genomics UK (COG-UK) Consortium | Harper VanSteenhouse, Yumi Kasai, David Gray, Carol Clugston, Anna Dominiczak and Alex Alderton, Roberto Amato, Jeffrey Barrett, Sonia Goncalves, Ewan Harrison, David K. Jackson, Ian Johnston, Dominic Kwiatkowski, Cordelia Langford, John Sillitoe on behalf of the Wellcome Sanger Institute COVID-19 Surveillance Team |
| EPI_ISL_3360202 | Lighthouse Lab in Alderley Park | Wellcome Sanger Institute for the COVID-19 Genomics UK (COG-UK) Consortium | Jacquelyn Wynn, Mairead Hyland, The Lighthouse Lab in Alderley Park and Alex Alderton, Roberto Amato, Jeffrey Barrett, Sonia Goncalves, Ewan Harrison, David K. Jackson, Ian Johnston, Dominic Kwiatkowski, Cordelia Langford, John Sillitoe on behalf of the Wellcome Sanger Institute COVID-19 Surveillance Team |
| EPI_ISL_3360204, EPI_ISL_3360205 | Lighthouse Lab in Glasgow | Wellcome Sanger Institute for the COVID-19 Genomics UK (COG-UK) Consortium | Harper VanSteenhouse, Yumi Kasai, David Gray, Carol Clugston, Anna Dominiczak and Alex Alderton, Roberto Amato, Jeffrey Barrett, Sonia Goncalves, Ewan Harrison, David K. Jackson, Ian Johnston, Dominic Kwiatkowski, Cordelia Langford, John Sillitoe on behalf of the Wellcome Sanger Institute COVID-19 Surveillance Team |
| EPI_ISL_3360207, EPI_ISL_3360208, EPI_ISL_3360210, EPI_ISL_3360211, EPI_ISL_3360212, EPI_ISL_3360214, EPI_ISL_3360215, EPI_ISL_3360216 | Lighthouse Lab in Alderley Park | Wellcome Sanger Institute for the COVID-19 Genomics UK (COG-UK) Consortium | Jacquelyn Wynn, Mairead Hyland, The Lighthouse Lab in Alderley Park and Alex Alderton, Roberto Amato, Jeffrey Barrett, Sonia Goncalves, Ewan Harrison, David K. Jackson, Ian Johnston, Dominic Kwiatkowski, Cordelia Langford, John Sillitoe on behalf of the Wellcome Sanger Institute COVID-19 Surveillance Team |
| EPI_ISL_3360218 | Lighthouse Lab in Glasgow | Wellcome Sanger Institute for the COVID-19 Genomics UK (COG-UK) Consortium | Harper VanSteenhouse, Yumi Kasai, David Gray, Carol Clugston, Anna Dominiczak and Alex Alderton, Roberto Amato, Jeffrey Barrett, Sonia Goncalves, Ewan Harrison, David K. Jackson, Ian Johnston, Dominic Kwiatkowski, Cordelia Langford, John Sillitoe on behalf of the Wellcome Sanger Institute COVID-19 Surveillance Team |
| EPI_ISL_3360220, EPI_ISL_3360221, EPI_ISL_3360222, EPI_ISL_3360224, EPI_ISL_3360225, EPI_ISL_3360226, EPI_ISL_3360228 | Lighthouse Lab in Alderley Park | Wellcome Sanger Institute for the COVID-19 Genomics UK (COG-UK) Consortium | Jacquelyn Wynn, Mairead Hyland, The Lighthouse Lab in Alderley Park and Alex Alderton, Roberto Amato, Jeffrey Barrett, Sonia Goncalves, Ewan Harrison, David K. Jackson, Ian Johnston, Dominic Kwiatkowski, Cordelia Langford, John Sillitoe on behalf of the Wellcome Sanger Institute COVID-19 Surveillance Team |
| EPI_ISL_3360230, EPI_ISL_3360231 | Lighthouse Lab in Glasgow | Wellcome Sanger Institute for the COVID-19 Genomics UK (COG-UK) Consortium | Harper VanSteenhouse, Yumi Kasai, David Gray, Carol Clugston, Anna Dominiczak and Alex Alderton, Roberto Amato, Jeffrey Barrett, Sonia Goncalves, Ewan Harrison, David K. Jackson, Ian Johnston, Dominic Kwiatkowski, Cordelia Langford, John Sillitoe on behalf of the Wellcome Sanger Institute COVID-19 Surveillance Team |
| EPI_ISL_3360232, EPI_ISL_3360234 | Lighthouse Lab in Alderley Park | Wellcome Sanger Institute for the COVID-19 Genomics UK (COG-UK) Consortium | Jacquelyn Wynn, Mairead Hyland, The Lighthouse Lab in Alderley Park and Alex Alderton, Roberto Amato, Jeffrey Barrett, Sonia Goncalves, Ewan Harrison, David K. Jackson, Ian Johnston, Dominic Kwiatkowski, Cordelia Langford, John Sillitoe on behalf of the Wellcome Sanger Institute COVID-19 Surveillance Team |
| EPI_ISL_3360235 | Lighthouse Lab in Glasgow | Wellcome Sanger Institute for the COVID-19 Genomics UK (COG-UK) Consortium | Harper VanSteenhouse, Yumi Kasai, David Gray, Carol Clugston, Anna Dominiczak and Alex Alderton, Roberto Amato, Jeffrey Barrett, Sonia Goncalves, Ewan Harrison, David K. Jackson, Ian Johnston, Dominic Kwiatkowski, Cordelia Langford, John Sillitoe on behalf of the Wellcome Sanger Institute COVID-19 Surveillance Team |
| EPI_ISL_3360237, EPI_ISL_3360238, EPI_ISL_3360239, EPI_ISL_3360241 | Lighthouse Lab in Alderley Park | Wellcome Sanger Institute for the COVID-19 Genomics UK (COG-UK) Consortium | Jacquelyn Wynn, Mairead Hyland, The Lighthouse Lab in Alderley Park and Alex Alderton, Roberto Amato, Jeffrey Barrett, Sonia Goncalves, Ewan Harrison, David K. Jackson, Ian Johnston, Dominic Kwiatkowski, Cordelia Langford, John Sillitoe on behalf of the Wellcome Sanger Institute COVID-19 Surveillance Team |
| EPI_ISL_3360242 | Lighthouse Lab in Glasgow | Wellcome Sanger Institute for the COVID-19 Genomics UK (COG-UK) Consortium | Harper VanSteenhouse, Yumi Kasai, David Gray, Carol Clugston, Anna Dominiczak and Alex Alderton, Roberto Amato, Jeffrey Barrett, Sonia Goncalves, Ewan Harrison, David K. Jackson, Ian Johnston, Dominic Kwiatkowski, Cordelia Langford, John Sillitoe on behalf of the Wellcome Sanger Institute COVID-19 Surveillance Team |
| EPI_ISL_3360244, EPI_ISL_3360245, EPI_ISL_3360247, EPI_ISL_3360248, EPI_ISL_3360250, EPI_ISL_3360251, EPI_ISL_3360253, EPI_ISL_3360254, EPI_ISL_3360256, EPI_ISL_3360257, EPI_ISL_3360259 | | | |
| see above | Lighthouse Lab in Alderley Park | Wellcome Sanger Institute for the COVID-19 Genomics UK (COG-UK) Consortium | Jacquelyn Wynn, Mairead Hyland, The Lighthouse Lab in Alderley Park and Alex Alderton, Roberto Amato, Jeffrey Barrett, Sonia Goncalves, Ewan Harrison, David K. Jackson, Ian Johnston, Dominic Kwiatkowski, Cordelia Langford, John Sillitoe on behalf of the Wellcome Sanger Institute COVID-19 Surveillance Team |
| EPI_ISL_3360260 | UAB InMedica | Lithuanian University of Health Sciences | Arnoldas Pautienius, Kamile Tamusauskaite, Gediminas Alzbutas, Dovydas Gecys, Lukas Zemaitis, Vaiva Lesauskaite |
| EPI_ISL_3360261, EPI_ISL_3360263, EPI_ISL_3360264, EPI_ISL_3360266, EPI_ISL_3360267, EPI_ISL_3360268 | Lighthouse Lab in Alderley Park | Wellcome Sanger Institute for the COVID-19 Genomics UK (COG-UK) Consortium | Jacquelyn Wynn, Mairead Hyland, The Lighthouse Lab in Alderley Park and Alex Alderton, Roberto Amato, Jeffrey Barrett, Sonia Goncalves, Ewan Harrison, David K. Jackson, Ian Johnston, Dominic Kwiatkowski, Cordelia Langford, John Sillitoe on behalf of the Wellcome Sanger Institute COVID-19 Surveillance Team |
| EPI_ISL_3360270 | Lighthouse Lab in Glasgow | Wellcome Sanger Institute for the COVID-19 Genomics UK (COG-UK) Consortium | Harper VanSteenhouse, Yumi Kasai, David Gray, Carol Clugston, Anna Dominiczak and Alex Alderton, Roberto Amato, Jeffrey Barrett, Sonia Goncalves, Ewan Harrison, David K. Jackson, Ian Johnston, Dominic Kwiatkowski, Cordelia Langford, John Sillitoe on behalf of the Wellcome Sanger Institute COVID-19 Surveillance Team |
| EPI_ISL_3360272, EPI_ISL_3360273, EPI_ISL_3360275, EPI_ISL_3360276, EPI_ISL_3360278, EPI_ISL_3360279, EPI_ISL_3360281, EPI_ISL_3360282, EPI_ISL_3360284, EPI_ISL_3360285 | Lighthouse Lab in Alderley Park | Wellcome Sanger Institute for the COVID-19 Genomics UK (COG-UK) Consortium | Jacquelyn Wynn, Mairead Hyland, The Lighthouse Lab in Alderley Park and Alex Alderton, Roberto Amato, Jeffrey Barrett, Sonia Goncalves, Ewan Harrison, David K. Jackson, Ian Johnston, Dominic Kwiatkowski, Cordelia Langford, John Sillitoe on behalf of the Wellcome Sanger Institute COVID-19 Surveillance Team |
| EPI_ISL_3360287 | Lighthouse Lab in Glasgow | Wellcome Sanger Institute for the COVID-19 Genomics UK (COG-UK) Consortium | Harper VanSteenhouse, Yumi Kasai, David Gray, Carol Clugston, Anna Dominiczak and Alex Alderton, Roberto Amato, Jeffrey Barrett, Sonia Goncalves, Ewan Harrison, David K. Jackson, Ian Johnston, Dominic Kwiatkowski, Cordelia Langford, John Sillitoe on behalf of the Wellcome Sanger Institute COVID-19 Surveillance Team |
| EPI_ISL_3360288, EPI_ISL_3360290, EPI_ISL_3360291, EPI_ISL_3360292, EPI_ISL_3360294, EPI_ISL_3360295, EPI_ISL_3360297, | Lighthouse Lab in Alderley Park | Wellcome Sanger Institute for the COVID-19 Genomics UK (COG-UK) Consortium | Jacquelyn Wynn, Mairead Hyland, The Lighthouse Lab in Alderley Park and Alex Alderton, Roberto Amato, Jeffrey Barrett, Sonia Goncalves, Ewan Harrison, David K. Jackson, Ian Johnston, Dominic Kwiatkowski, Cordelia Langford, John Sillitoe on behalf of the Wellcome Sanger Institute COVID-19 Surveillance Team |

| | | | |
|---|---|---|---|
| EPI_ISL_3360298, EPI_ISL_3360300 | | | |
| EPI_ISL_3360301 | Lighthouse Lab in Glasgow | Wellcome Sanger Institute for the COVID-19 Genomics UK (COG-UK) Consortium | Harper VanSteenhouse, Yumi Kasai, David Gray, Carol Clugston, Anna Dominiczak and Alex Alderton, Roberto Amato, Jeffrey Barrett, Sonia Goncalves, Ewan Harrison, David K. Jackson, Ian Johnston, Dominic Kwiatkowski, Cordelia Langford, John Sillitoe on behalf of the Wellcome Sanger Institute COVID-19 Surveillance Team |
| EPI_ISL_3360303, EPI_ISL_3360304, EPI_ISL_3360306 | Lighthouse Lab in Alderley Park | Wellcome Sanger Institute for the COVID-19 Genomics UK (COG-UK) Consortium | Jacquelyn Wynn, Mairead Hyland, The Lighthouse Lab in Alderley Park and Alex Alderton, Roberto Amato, Jeffrey Barrett, Sonia Goncalves, Ewan Harrison, David K. Jackson, Ian Johnston, Dominic Kwiatkowski, Cordelia Langford, John Sillitoe on behalf of the Wellcome Sanger Institute COVID-19 Surveillance Team |
| EPI_ISL_3360307 | Lighthouse Lab in Glasgow | Wellcome Sanger Institute for the COVID-19 Genomics UK (COG-UK) Consortium | Harper VanSteenhouse, Yumi Kasai, David Gray, Carol Clugston, Anna Dominiczak and Alex Alderton, Roberto Amato, Jeffrey Barrett, Sonia Goncalves, Ewan Harrison, David K. Jackson, Ian Johnston, Dominic Kwiatkowski, Cordelia Langford, John Sillitoe on behalf of the Wellcome Sanger Institute COVID-19 Surveillance Team |
| EPI_ISL_3360309, EPI_ISL_3360311, EPI_ISL_3360312 | Lighthouse Lab in Alderley Park | Wellcome Sanger Institute for the COVID-19 Genomics UK (COG-UK) Consortium | Jacquelyn Wynn, Mairead Hyland, The Lighthouse Lab in Alderley Park and Alex Alderton, Roberto Amato, Jeffrey Barrett, Sonia Goncalves, Ewan Harrison, David K. Jackson, Ian Johnston, Dominic Kwiatkowski, Cordelia Langford, John Sillitoe on behalf of the Wellcome Sanger Institute COVID-19 Surveillance Team |
| EPI_ISL_3360314 | Lighthouse Lab in Glasgow | Wellcome Sanger Institute for the COVID-19 Genomics UK (COG-UK) Consortium | Harper VanSteenhouse, Yumi Kasai, David Gray, Carol Clugston, Anna Dominiczak and Alex Alderton, Roberto Amato, Jeffrey Barrett, Sonia Goncalves, Ewan Harrison, David K. Jackson, Ian Johnston, Dominic Kwiatkowski, Cordelia Langford, John Sillitoe on behalf of the Wellcome Sanger Institute COVID-19 Surveillance Team |
| EPI_ISL_3360315, EPI_ISL_3360316, EPI_ISL_3360318, EPI_ISL_3360319, EPI_ISL_3360321, EPI_ISL_3360323, EPI_ISL_3360324, EPI_ISL_3360326, EPI_ISL_3360327, EPI_ISL_3360329, EPI_ISL_3360330 | | | |
| see above | Lighthouse Lab in Alderley Park | Wellcome Sanger Institute for the COVID-19 Genomics UK (COG-UK) Consortium | Jacquelyn Wynn, Mairead Hyland, The Lighthouse Lab in Alderley Park and Alex Alderton, Roberto Amato, Jeffrey Barrett, Sonia Goncalves, Ewan Harrison, David K. Jackson, Ian Johnston, Dominic Kwiatkowski, Cordelia Langford, John Sillitoe on behalf of the Wellcome Sanger Institute COVID-19 Surveillance Team |
| EPI_ISL_3360332 | Lighthouse Lab in Glasgow | Wellcome Sanger Institute for the COVID-19 Genomics UK (COG-UK) Consortium | Harper VanSteenhouse, Yumi Kasai, David Gray, Carol Clugston, Anna Dominiczak and Alex Alderton, Roberto Amato, Jeffrey Barrett, Sonia Goncalves, Ewan Harrison, David K. Jackson, Ian Johnston, Dominic Kwiatkowski, Cordelia Langford, John Sillitoe on behalf of the Wellcome Sanger Institute COVID-19 Surveillance Team |
| EPI_ISL_3360333, EPI_ISL_3360335 | Lighthouse Lab in Alderley Park | Wellcome Sanger Institute for the COVID-19 Genomics UK (COG-UK) Consortium | Jacquelyn Wynn, Mairead Hyland, The Lighthouse Lab in Alderley Park and Alex Alderton, Roberto Amato, Jeffrey Barrett, Sonia Goncalves, Ewan Harrison, David K. Jackson, Ian Johnston, Dominic Kwiatkowski, Cordelia Langford, John Sillitoe on behalf of the Wellcome Sanger Institute COVID-19 Surveillance Team |
| EPI_ISL_3360336 | Lighthouse Lab in Glasgow | Wellcome Sanger Institute for the COVID-19 Genomics UK (COG-UK) Consortium | Harper VanSteenhouse, Yumi Kasai, David Gray, Carol Clugston, Anna Dominiczak and Alex Alderton, Roberto Amato, Jeffrey Barrett, Sonia Goncalves, Ewan Harrison, David K. Jackson, Ian Johnston, Dominic Kwiatkowski, Cordelia Langford, John Sillitoe on behalf of the Wellcome Sanger Institute COVID-19 Surveillance Team |
| EPI_ISL_3360338, EPI_ISL_3360339, EPI_ISL_3360341, EPI_ISL_3360342, EPI_ISL_3360344 | Lighthouse Lab in Alderley Park | Wellcome Sanger Institute for the COVID-19 Genomics UK (COG-UK) Consortium | Jacquelyn Wynn, Mairead Hyland, The Lighthouse Lab in Alderley Park and Alex Alderton, Roberto Amato, Jeffrey Barrett, Sonia Goncalves, Ewan Harrison, David K. Jackson, Ian Johnston, Dominic Kwiatkowski, Cordelia Langford, John Sillitoe on behalf of the Wellcome Sanger Institute COVID-19 Surveillance Team |
| EPI_ISL_3360345 | Lighthouse Lab in Glasgow | Wellcome Sanger Institute for the COVID-19 Genomics UK (COG-UK) Consortium | Harper VanSteenhouse, Yumi Kasai, David Gray, Carol Clugston, Anna Dominiczak and Alex Alderton, Roberto Amato, Jeffrey Barrett, Sonia Goncalves, Ewan Harrison, David K. Jackson, Ian Johnston, Dominic Kwiatkowski, Cordelia Langford, John Sillitoe on behalf of the Wellcome Sanger Institute COVID-19 Surveillance Team |
| EPI_ISL_3360347, EPI_ISL_3360349, EPI_ISL_3360350, EPI_ISL_3360351 | Lighthouse Lab in Alderley Park | Wellcome Sanger Institute for the COVID-19 Genomics UK (COG-UK) Consortium | Jacquelyn Wynn, Mairead Hyland, The Lighthouse Lab in Alderley Park and Alex Alderton, Roberto Amato, Jeffrey Barrett, Sonia Goncalves, Ewan Harrison, David K. Jackson, Ian Johnston, Dominic Kwiatkowski, Cordelia Langford, John Sillitoe on behalf of the Wellcome Sanger Institute COVID-19 Surveillance Team |
| EPI_ISL_3360353 | Lighthouse Lab in Glasgow | Wellcome Sanger Institute for the COVID-19 Genomics UK (COG-UK) Consortium | Harper VanSteenhouse, Yumi Kasai, David Gray, Carol Clugston, Anna Dominiczak and Alex Alderton, Roberto Amato, Jeffrey Barrett, Sonia Goncalves, Ewan Harrison, David K. Jackson, Ian Johnston, Dominic Kwiatkowski, Cordelia Langford, John Sillitoe on behalf of the Wellcome Sanger Institute COVID-19 Surveillance Team |
| EPI_ISL_3360355, EPI_ISL_3360356 | Lighthouse Lab in Alderley Park | Wellcome Sanger Institute for the COVID-19 Genomics UK (COG-UK) Consortium | Jacquelyn Wynn, Mairead Hyland, The Lighthouse Lab in Alderley Park and Alex Alderton, Roberto Amato, Jeffrey Barrett, Sonia Goncalves, Ewan Harrison, David K. Jackson, Ian Johnston, Dominic Kwiatkowski, Cordelia Langford, John Sillitoe on behalf of the Wellcome Sanger Institute COVID-19 Surveillance Team |
| EPI_ISL_3360358 | Lighthouse Lab in Glasgow | Wellcome Sanger Institute for the COVID-19 Genomics UK (COG-UK) Consortium | Harper VanSteenhouse, Yumi Kasai, David Gray, Carol Clugston, Anna Dominiczak and Alex Alderton, Roberto Amato, Jeffrey Barrett, Sonia Goncalves, Ewan Harrison, David K. Jackson, Ian Johnston, Dominic Kwiatkowski, Cordelia Langford, John Sillitoe on behalf of the Wellcome Sanger Institute COVID-19 Surveillance Team |
| EPI_ISL_3360359, EPI_ISL_3360360, EPI_ISL_3360362, EPI_ISL_3360364, EPI_ISL_3360365, EPI_ISL_3360367 | Lighthouse Lab in Alderley Park | Wellcome Sanger Institute for the COVID-19 Genomics UK (COG-UK) Consortium | Jacquelyn Wynn, Mairead Hyland, The Lighthouse Lab in Alderley Park and Alex Alderton, Roberto Amato, Jeffrey Barrett, Sonia Goncalves, Ewan Harrison, David K. Jackson, Ian Johnston, Dominic Kwiatkowski, Cordelia Langford, John Sillitoe on behalf of the Wellcome Sanger Institute COVID-19 Surveillance Team |
| EPI_ISL_3360368 | Lighthouse Lab in Glasgow | Wellcome Sanger Institute for the COVID-19 Genomics UK (COG-UK) Consortium | Harper VanSteenhouse, Yumi Kasai, David Gray, Carol Clugston, Anna Dominiczak and Alex Alderton, Roberto Amato, Jeffrey Barrett, Sonia Goncalves, Ewan Harrison, David K. Jackson, Ian Johnston, Dominic Kwiatkowski, Cordelia Langford, John Sillitoe on behalf of the Wellcome Sanger Institute COVID-19 Surveillance Team |
| EPI_ISL_3360369 | Lighthouse Lab in Alderley Park | Wellcome Sanger Institute for the COVID-19 Genomics UK (COG-UK) Consortium | Jacquelyn Wynn, Mairead Hyland, The Lighthouse Lab in Alderley Park and Alex Alderton, Roberto Amato, Jeffrey Barrett, Sonia Goncalves, Ewan Harrison, David K. Jackson, Ian Johnston, Dominic Kwiatkowski, Cordelia Langford, John Sillitoe on behalf of the Wellcome Sanger Institute COVID-19 Surveillance Team |
| EPI_ISL_3360370, EPI_ISL_3360371, EPI_ISL_3360372, EPI_ISL_3360373, EPI_ISL_3360374 | Lighthouse Lab in Glasgow | Wellcome Sanger Institute for the COVID-19 Genomics UK (COG-UK) Consortium | Harper VanSteenhouse, Yumi Kasai, David Gray, Carol Clugston, Anna Dominiczak and Alex Alderton, Roberto Amato, Jeffrey Barrett, Sonia Goncalves, Ewan Harrison, David K. Jackson, Ian Johnston, Dominic Kwiatkowski, Cordelia Langford, John Sillitoe on behalf of the Wellcome Sanger Institute COVID-19 Surveillance Team |
| EPI_ISL_3360375, EPI_ISL_3360376, EPI_ISL_3360378 | Lighthouse Lab in Alderley Park | Wellcome Sanger Institute for the COVID-19 Genomics UK (COG-UK) Consortium | Jacquelyn Wynn, Mairead Hyland, The Lighthouse Lab in Alderley Park and Alex Alderton, Roberto Amato, Jeffrey Barrett, Sonia Goncalves, Ewan Harrison, David K. Jackson, Ian Johnston, Dominic Kwiatkowski, Cordelia Langford, John Sillitoe on behalf of the Wellcome Sanger Institute COVID-19 Surveillance Team |
| EPI_ISL_3360379 | Lighthouse Lab in Glasgow | Wellcome Sanger Institute for the COVID-19 Genomics UK (COG-UK) Consortium | Harper VanSteenhouse, Yumi Kasai, David Gray, Carol Clugston, Anna Dominiczak and Alex Alderton, Roberto Amato, Jeffrey Barrett, Sonia Goncalves, Ewan Harrison, David K. Jackson, Ian Johnston, Dominic Kwiatkowski, Cordelia Langford, John Sillitoe on behalf of the Wellcome Sanger Institute COVID-19 Surveillance Team |
| EPI_ISL_3360381, EPI_ISL_3360382 | Lighthouse Lab in Alderley Park | Wellcome Sanger Institute for the COVID-19 Genomics UK (COG-UK) Consortium | Jacquelyn Wynn, Mairead Hyland, The Lighthouse Lab in Alderley Park and Alex Alderton, Roberto Amato, Jeffrey Barrett, Sonia Goncalves, Ewan Harrison, David K. Jackson, Ian Johnston, Dominic Kwiatkowski, Cordelia Langford, John Sillitoe on behalf of the Wellcome Sanger Institute COVID-19 Surveillance Team |
| EPI_ISL_3360384 | Lighthouse Lab in Glasgow | Wellcome Sanger Institute for the COVID-19 Genomics UK (COG-UK) Consortium | Harper VanSteenhouse, Yumi Kasai, David Gray, Carol Clugston, Anna Dominiczak and Alex Alderton, Roberto Amato, Jeffrey Barrett, Sonia Goncalves, Ewan Harrison, David K. Jackson, Ian Johnston, Dominic Kwiatkowski, Cordelia Langford, John Sillitoe on behalf of the Wellcome Sanger Institute COVID-19 Surveillance Team |
| EPI_ISL_3360385, EPI_ISL_3360387, EPI_ISL_3360388, EPI_ISL_3360389, EPI_ISL_3360391, EPI_ISL_3360393, EPI_ISL_3360394, EPI_ISL_3360396, EPI_ISL_3360397 | Lighthouse Lab in Alderley Park | Wellcome Sanger Institute for the COVID-19 Genomics UK (COG-UK) Consortium | Jacquelyn Wynn, Mairead Hyland, The Lighthouse Lab in Alderley Park and Alex Alderton, Roberto Amato, Jeffrey Barrett, Sonia Goncalves, Ewan Harrison, David K. Jackson, Ian Johnston, Dominic Kwiatkowski, Cordelia Langford, John Sillitoe on behalf of the Wellcome Sanger Institute COVID-19 Surveillance Team |
| EPI_ISL_3360399 | Lighthouse Lab in Glasgow | Wellcome Sanger Institute for the COVID-19 Genomics UK (COG-UK) Consortium | Harper VanSteenhouse, Yumi Kasai, David Gray, Carol Clugston, Anna Dominiczak and Alex Alderton, Roberto Amato, Jeffrey Barrett, Sonia Goncalves, Ewan Harrison, David K. Jackson, Ian Johnston, Dominic Kwiatkowski, Cordelia Langford, John Sillitoe on behalf of the Wellcome Sanger Institute COVID-19 Surveillance Team |
| EPI_ISL_3360400, EPI_ISL_3360402 | Lighthouse Lab in Alderley Park | Wellcome Sanger Institute for the COVID-19 Genomics UK (COG-UK) Consortium | Jacquelyn Wynn, Mairead Hyland, The Lighthouse Lab in Alderley Park and Alex Alderton, Roberto Amato, Jeffrey Barrett, Sonia Goncalves, Ewan Harrison, David K. Jackson, Ian Johnston, Dominic Kwiatkowski, Cordelia Langford, John Sillitoe on behalf of the Wellcome Sanger Institute COVID-19 Surveillance Team |
| EPI_ISL_3360403 | Lighthouse Lab in Glasgow | Wellcome Sanger Institute for the COVID-19 Genomics UK (COG-UK) Consortium | Harper VanSteenhouse, Yumi Kasai, David Gray, Carol Clugston, Anna Dominiczak and Alex Alderton, Roberto Amato, Jeffrey Barrett, Sonia Goncalves, Ewan Harrison, David K. Jackson, Ian Johnston, Dominic Kwiatkowski, Cordelia Langford, John Sillitoe on behalf of the Wellcome Sanger Institute COVID-19 Surveillance Team |
| EPI_ISL_3360405, EPI_ISL_3360406 | Lighthouse Lab in Alderley Park | Wellcome Sanger Institute for the COVID-19 Genomics UK (COG-UK) Consortium | Jacquelyn Wynn, Mairead Hyland, The Lighthouse Lab in Alderley Park and Alex Alderton, Roberto Amato, Jeffrey Barrett, Sonia Goncalves, Ewan Harrison, David K. Jackson, Ian Johnston, Dominic Kwiatkowski, Cordelia Langford, John Sillitoe on behalf of the Wellcome Sanger Institute COVID-19 Surveillance Team |
| EPI_ISL_3360408 | Lighthouse Lab in Glasgow | Wellcome Sanger Institute for the COVID-19 Genomics UK (COG-UK) Consortium | Harper VanSteenhouse, Yumi Kasai, David Gray, Carol Clugston, Anna Dominiczak and Alex Alderton, Roberto Amato, Jeffrey Barrett, Sonia Goncalves, Ewan Harrison, David K. Jackson, Ian Johnston, Dominic Kwiatkowski, Cordelia Langford, John Sillitoe on behalf of the Wellcome Sanger Institute COVID-19 Surveillance Team |
| EPI_ISL_3360409, EPI_ISL_3360411, EPI_ISL_3360412, EPI_ISL_3360414, EPI_ISL_3360415, EPI_ISL_3360417, EPI_ISL_3360418, EPI_ISL_3360420 | Lighthouse Lab in Alderley Park | Wellcome Sanger Institute for the COVID-19 Genomics UK (COG-UK) Consortium | Jacquelyn Wynn, Mairead Hyland, The Lighthouse Lab in Alderley Park and Alex Alderton, Roberto Amato, Jeffrey Barrett, Sonia Goncalves, Ewan Harrison, David K. Jackson, Ian Johnston, Dominic Kwiatkowski, Cordelia Langford, John Sillitoe on behalf of the Wellcome Sanger Institute COVID-19 Surveillance Team |
| EPI_ISL_3360421 | Lighthouse Lab in Glasgow | Wellcome Sanger Institute for the COVID-19 Genomics UK (COG-UK) Consortium | Harper VanSteenhouse, Yumi Kasai, David Gray, Carol Clugston, Anna Dominiczak and Alex Alderton, Roberto Amato, Jeffrey Barrett, Sonia Goncalves, Ewan Harrison, David K. Jackson, Ian Johnston, Dominic Kwiatkowski, Cordelia Langford, John Sillitoe on behalf of the Wellcome Sanger Institute COVID-19 Surveillance Team |
| EPI_ISL_3360423, EPI_ISL_3360424 | Lighthouse Lab in Alderley Park | Wellcome Sanger Institute for the COVID-19 Genomics UK (COG-UK) Consortium | Jacquelyn Wynn, Mairead Hyland, The Lighthouse Lab in Alderley Park and Alex Alderton, Roberto Amato, Jeffrey Barrett, Sonia Goncalves, Ewan Harrison, David K. Jackson, Ian Johnston, Dominic Kwiatkowski, Cordelia Langford, John Sillitoe on behalf of the Wellcome Sanger Institute COVID-19 Surveillance Team |

| | | | |
|---|---|---|---|
| EPI_ISL_3360426 | Lighthouse Lab in Glasgow | Wellcome Sanger Institute for the COVID-19 Genomics UK (COG-UK) Consortium | Harper VanSteenhouse, Yumi Kasai, David Gray, Carol Clugston, Anna Dominiczak and Alex Alderton, Roberto Amato, Jeffrey Barrett, Sonia Goncalves, Ewan Harrison, David K. Jackson, Ian Johnston, Dominic Kwiatkowski, Cordelia Langford, John Sillitoe on behalf of the Wellcome Sanger Institute COVID-19 Surveillance Team |
| EPI_ISL_3360427 | Lighthouse Lab in Alderley Park | Wellcome Sanger Institute for the COVID-19 Genomics UK (COG-UK) Consortium | Jacquelyn Wynn, Mairead Hyland, The Lighthouse Lab in Alderley Park and Alex Alderton, Roberto Amato, Jeffrey Barrett, Sonia Goncalves, Ewan Harrison, David K. Jackson, Ian Johnston, Dominic Kwiatkowski, Cordelia Langford, John Sillitoe on behalf of the Wellcome Sanger Institute COVID-19 Surveillance Team |
| EPI_ISL_3360429 | Lighthouse Lab in Glasgow | Wellcome Sanger Institute for the COVID-19 Genomics UK (COG-UK) Consortium | Harper VanSteenhouse, Yumi Kasai, David Gray, Carol Clugston, Anna Dominiczak and Alex Alderton, Roberto Amato, Jeffrey Barrett, Sonia Goncalves, Ewan Harrison, David K. Jackson, Ian Johnston, Dominic Kwiatkowski, Cordelia Langford, John Sillitoe on behalf of the Wellcome Sanger Institute COVID-19 Surveillance Team |
| EPI_ISL_3360430 | Lighthouse Lab in Alderley Park | Wellcome Sanger Institute for the COVID-19 Genomics UK (COG-UK) Consortium | Jacquelyn Wynn, Mairead Hyland, The Lighthouse Lab in Alderley Park and Alex Alderton, Roberto Amato, Jeffrey Barrett, Sonia Goncalves, Ewan Harrison, David K. Jackson, Ian Johnston, Dominic Kwiatkowski, Cordelia Langford, John Sillitoe on behalf of the Wellcome Sanger Institute COVID-19 Surveillance Team |
| EPI_ISL_3360432 | Lighthouse Lab in Glasgow | Wellcome Sanger Institute for the COVID-19 Genomics UK (COG-UK) Consortium | Harper VanSteenhouse, Yumi Kasai, David Gray, Carol Clugston, Anna Dominiczak and Alex Alderton, Roberto Amato, Jeffrey Barrett, Sonia Goncalves, Ewan Harrison, David K. Jackson, Ian Johnston, Dominic Kwiatkowski, Cordelia Langford, John Sillitoe on behalf of the Wellcome Sanger Institute COVID-19 Surveillance Team |
| EPI_ISL_3360433, EPI_ISL_3360435 | Lighthouse Lab in Alderley Park | Wellcome Sanger Institute for the COVID-19 Genomics UK (COG-UK) Consortium | Jacquelyn Wynn, Mairead Hyland, The Lighthouse Lab in Alderley Park and Alex Alderton, Roberto Amato, Jeffrey Barrett, Sonia Goncalves, Ewan Harrison, David K. Jackson, Ian Johnston, Dominic Kwiatkowski, Cordelia Langford, John Sillitoe on behalf of the Wellcome Sanger Institute COVID-19 Surveillance Team |
| EPI_ISL_3360436 | Lighthouse Lab in Glasgow | Wellcome Sanger Institute for the COVID-19 Genomics UK (COG-UK) Consortium | Harper VanSteenhouse, Yumi Kasai, David Gray, Carol Clugston, Anna Dominiczak and Alex Alderton, Roberto Amato, Jeffrey Barrett, Sonia Goncalves, Ewan Harrison, David K. Jackson, Ian Johnston, Dominic Kwiatkowski, Cordelia Langford, John Sillitoe on behalf of the Wellcome Sanger Institute COVID-19 Surveillance Team |
| EPI_ISL_3360438, EPI_ISL_3360440, EPI_ISL_3360441 | Lighthouse Lab in Alderley Park | Wellcome Sanger Institute for the COVID-19 Genomics UK (COG-UK) Consortium | Jacquelyn Wynn, Mairead Hyland, The Lighthouse Lab in Alderley Park and Alex Alderton, Roberto Amato, Jeffrey Barrett, Sonia Goncalves, Ewan Harrison, David K. Jackson, Ian Johnston, Dominic Kwiatkowski, Cordelia Langford, John Sillitoe on behalf of the Wellcome Sanger Institute COVID-19 Surveillance Team |
| EPI_ISL_3360442, EPI_ISL_3360444 | Lighthouse Lab in Glasgow | Wellcome Sanger Institute for the COVID-19 Genomics UK (COG-UK) Consortium | Harper VanSteenhouse, Yumi Kasai, David Gray, Carol Clugston, Anna Dominiczak and Alex Alderton, Roberto Amato, Jeffrey Barrett, Sonia Goncalves, Ewan Harrison, David K. Jackson, Ian Johnston, Dominic Kwiatkowski, Cordelia Langford, John Sillitoe on behalf of the Wellcome Sanger Institute COVID-19 Surveillance Team |
| EPI_ISL_3360445, EPI_ISL_3360447 | Lighthouse Lab in Alderley Park | Wellcome Sanger Institute for the COVID-19 Genomics UK (COG-UK) Consortium | Jacquelyn Wynn, Mairead Hyland, The Lighthouse Lab in Alderley Park and Alex Alderton, Roberto Amato, Jeffrey Barrett, Sonia Goncalves, Ewan Harrison, David K. Jackson, Ian Johnston, Dominic Kwiatkowski, Cordelia Langford, John Sillitoe on behalf of the Wellcome Sanger Institute COVID-19 Surveillance Team |
| EPI_ISL_3360448 | Lighthouse Lab in Glasgow | Wellcome Sanger Institute for the COVID-19 Genomics UK (COG-UK) Consortium | Harper VanSteenhouse, Yumi Kasai, David Gray, Carol Clugston, Anna Dominiczak and Alex Alderton, Roberto Amato, Jeffrey Barrett, Sonia Goncalves, Ewan Harrison, David K. Jackson, Ian Johnston, Dominic Kwiatkowski, Cordelia Langford, John Sillitoe on behalf of the Wellcome Sanger Institute COVID-19 Surveillance Team |
| EPI_ISL_3360450, EPI_ISL_3360451 | Lighthouse Lab in Alderley Park | Wellcome Sanger Institute for the COVID-19 Genomics UK (COG-UK) Consortium | Jacquelyn Wynn, Mairead Hyland, The Lighthouse Lab in Alderley Park and Alex Alderton, Roberto Amato, Jeffrey Barrett, Sonia Goncalves, Ewan Harrison, David K. Jackson, Ian Johnston, Dominic Kwiatkowski, Cordelia Langford, John Sillitoe on behalf of the Wellcome Sanger Institute COVID-19 Surveillance Team |
| EPI_ISL_3360452 | Lighthouse Lab in Glasgow | Wellcome Sanger Institute for the COVID-19 Genomics UK (COG-UK) Consortium | Harper VanSteenhouse, Yumi Kasai, David Gray, Carol Clugston, Anna Dominiczak and Alex Alderton, Roberto Amato, Jeffrey Barrett, Sonia Goncalves, Ewan Harrison, David K. Jackson, Ian Johnston, Dominic Kwiatkowski, Cordelia Langford, John Sillitoe on behalf of the Wellcome Sanger Institute COVID-19 Surveillance Team |
| EPI_ISL_3360454, EPI_ISL_3360455 | Lighthouse Lab in Alderley Park | Wellcome Sanger Institute for the COVID-19 Genomics UK (COG-UK) Consortium | Jacquelyn Wynn, Mairead Hyland, The Lighthouse Lab in Alderley Park and Alex Alderton, Roberto Amato, Jeffrey Barrett, Sonia Goncalves, Ewan Harrison, David K. Jackson, Ian Johnston, Dominic Kwiatkowski, Cordelia Langford, John Sillitoe on behalf of the Wellcome Sanger Institute COVID-19 Surveillance Team |
| EPI_ISL_3360457 | Lighthouse Lab in Glasgow | Wellcome Sanger Institute for the COVID-19 Genomics UK (COG-UK) Consortium | Harper VanSteenhouse, Yumi Kasai, David Gray, Carol Clugston, Anna Dominiczak and Alex Alderton, Roberto Amato, Jeffrey Barrett, Sonia Goncalves, Ewan Harrison, David K. Jackson, Ian Johnston, Dominic Kwiatkowski, Cordelia Langford, John Sillitoe on behalf of the Wellcome Sanger Institute COVID-19 Surveillance Team |
| EPI_ISL_3360459, EPI_ISL_3360460 | Lighthouse Lab in Alderley Park | Wellcome Sanger Institute for the COVID-19 Genomics UK (COG-UK) Consortium | Jacquelyn Wynn, Mairead Hyland, The Lighthouse Lab in Alderley Park and Alex Alderton, Roberto Amato, Jeffrey Barrett, Sonia Goncalves, Ewan Harrison, David K. Jackson, Ian Johnston, Dominic Kwiatkowski, Cordelia Langford, John Sillitoe on behalf of the Wellcome Sanger Institute COVID-19 Surveillance Team |
| EPI_ISL_3360461, EPI_ISL_3360463, EPI_ISL_3360464, EPI_ISL_3360466 | Lighthouse Lab in Glasgow | Wellcome Sanger Institute for the COVID-19 Genomics UK (COG-UK) Consortium | Harper VanSteenhouse, Yumi Kasai, David Gray, Carol Clugston, Anna Dominiczak and Alex Alderton, Roberto Amato, Jeffrey Barrett, Sonia Goncalves, Ewan Harrison, David K. Jackson, Ian Johnston, Dominic Kwiatkowski, Cordelia Langford, John Sillitoe on behalf of the Wellcome Sanger Institute COVID-19 Surveillance Team |
| EPI_ISL_3360467, EPI_ISL_3360469, EPI_ISL_3360470, EPI_ISL_3360472, EPI_ISL_3360474 | Lighthouse Lab in Alderley Park | Wellcome Sanger Institute for the COVID-19 Genomics UK (COG-UK) Consortium | Jacquelyn Wynn, Mairead Hyland, The Lighthouse Lab in Alderley Park and Alex Alderton, Roberto Amato, Jeffrey Barrett, Sonia Goncalves, Ewan Harrison, David K. Jackson, Ian Johnston, Dominic Kwiatkowski, Cordelia Langford, John Sillitoe on behalf of the Wellcome Sanger Institute COVID-19 Surveillance Team |
| EPI_ISL_3360475 | Lighthouse Lab in Glasgow | Wellcome Sanger Institute for the COVID-19 Genomics UK (COG-UK) Consortium | Harper VanSteenhouse, Yumi Kasai, David Gray, Carol Clugston, Anna Dominiczak and Alex Alderton, Roberto Amato, Jeffrey Barrett, Sonia Goncalves, Ewan Harrison, David K. Jackson, Ian Johnston, Dominic Kwiatkowski, Cordelia Langford, John Sillitoe on behalf of the Wellcome Sanger Institute COVID-19 Surveillance Team |
| EPI_ISL_3360476, EPI_ISL_3360478, EPI_ISL_3360479, EPI_ISL_3360481 | Lighthouse Lab in Alderley Park | Wellcome Sanger Institute for the COVID-19 Genomics UK (COG-UK) Consortium | Jacquelyn Wynn, Mairead Hyland, The Lighthouse Lab in Alderley Park and Alex Alderton, Roberto Amato, Jeffrey Barrett, Sonia Goncalves, Ewan Harrison, David K. Jackson, Ian Johnston, Dominic Kwiatkowski, Cordelia Langford, John Sillitoe on behalf of the Wellcome Sanger Institute COVID-19 Surveillance Team |
| EPI_ISL_3360482 | Lighthouse Lab in Glasgow | Wellcome Sanger Institute for the COVID-19 Genomics UK (COG-UK) Consortium | Harper VanSteenhouse, Yumi Kasai, David Gray, Carol Clugston, Anna Dominiczak and Alex Alderton, Roberto Amato, Jeffrey Barrett, Sonia Goncalves, Ewan Harrison, David K. Jackson, Ian Johnston, Dominic Kwiatkowski, Cordelia Langford, John Sillitoe on behalf of the Wellcome Sanger Institute COVID-19 Surveillance Team |
| EPI_ISL_3360484 | Lighthouse Lab in Alderley Park | Wellcome Sanger Institute for the COVID-19 Genomics UK (COG-UK) Consortium | Jacquelyn Wynn, Mairead Hyland, The Lighthouse Lab in Alderley Park and Alex Alderton, Roberto Amato, Jeffrey Barrett, Sonia Goncalves, Ewan Harrison, David K. Jackson, Ian Johnston, Dominic Kwiatkowski, Cordelia Langford, John Sillitoe on behalf of the Wellcome Sanger Institute COVID-19 Surveillance Team |
| EPI_ISL_3360485 | Lighthouse Lab in Glasgow | Wellcome Sanger Institute for the COVID-19 Genomics UK (COG-UK) Consortium | Harper VanSteenhouse, Yumi Kasai, David Gray, Carol Clugston, Anna Dominiczak and Alex Alderton, Roberto Amato, Jeffrey Barrett, Sonia Goncalves, Ewan Harrison, David K. Jackson, Ian Johnston, Dominic Kwiatkowski, Cordelia Langford, John Sillitoe on behalf of the Wellcome Sanger Institute COVID-19 Surveillance Team |
| EPI_ISL_3360487, EPI_ISL_3360488, EPI_ISL_3360490 | Lighthouse Lab in Alderley Park | Wellcome Sanger Institute for the COVID-19 Genomics UK (COG-UK) Consortium | Jacquelyn Wynn, Mairead Hyland, The Lighthouse Lab in Alderley Park and Alex Alderton, Roberto Amato, Jeffrey Barrett, Sonia Goncalves, Ewan Harrison, David K. Jackson, Ian Johnston, Dominic Kwiatkowski, Cordelia Langford, John Sillitoe on behalf of the Wellcome Sanger Institute COVID-19 Surveillance Team |
| EPI_ISL_3360491 | Lighthouse Lab in Glasgow | Wellcome Sanger Institute for the COVID-19 Genomics UK (COG-UK) Consortium | Harper VanSteenhouse, Yumi Kasai, David Gray, Carol Clugston, Anna Dominiczak and Alex Alderton, Roberto Amato, Jeffrey Barrett, Sonia Goncalves, Ewan Harrison, David K. Jackson, Ian Johnston, Dominic Kwiatkowski, Cordelia Langford, John Sillitoe on behalf of the Wellcome Sanger Institute COVID-19 Surveillance Team |
| EPI_ISL_3360493, EPI_ISL_3360495, EPI_ISL_3360496, EPI_ISL_3360497, EPI_ISL_3360499, EPI_ISL_3360500, EPI_ISL_3360502, EPI_ISL_3360503 | Lighthouse Lab in Alderley Park | Wellcome Sanger Institute for the COVID-19 Genomics UK (COG-UK) Consortium | Jacquelyn Wynn, Mairead Hyland, The Lighthouse Lab in Alderley Park and Alex Alderton, Roberto Amato, Jeffrey Barrett, Sonia Goncalves, Ewan Harrison, David K. Jackson, Ian Johnston, Dominic Kwiatkowski, Cordelia Langford, John Sillitoe on behalf of the Wellcome Sanger Institute COVID-19 Surveillance Team |
| EPI_ISL_3360505 | Lighthouse Lab in Glasgow | Wellcome Sanger Institute for the COVID-19 Genomics UK (COG-UK) Consortium | Harper VanSteenhouse, Yumi Kasai, David Gray, Carol Clugston, Anna Dominiczak and Alex Alderton, Roberto Amato, Jeffrey Barrett, Sonia Goncalves, Ewan Harrison, David K. Jackson, Ian Johnston, Dominic Kwiatkowski, Cordelia Langford, John Sillitoe on behalf of the Wellcome Sanger Institute COVID-19 Surveillance Team |
| EPI_ISL_3360506, EPI_ISL_3360508, EPI_ISL_3360509, EPI_ISL_3360511, EPI_ISL_3360513, EPI_ISL_3360514 | Lighthouse Lab in Alderley Park | Wellcome Sanger Institute for the COVID-19 Genomics UK (COG-UK) Consortium | Jacquelyn Wynn, Mairead Hyland, The Lighthouse Lab in Alderley Park and Alex Alderton, Roberto Amato, Jeffrey Barrett, Sonia Goncalves, Ewan Harrison, David K. Jackson, Ian Johnston, Dominic Kwiatkowski, Cordelia Langford, John Sillitoe on behalf of the Wellcome Sanger Institute COVID-19 Surveillance Team |
| EPI_ISL_3360516 | Lighthouse Lab in Glasgow | Wellcome Sanger Institute for the COVID-19 Genomics UK (COG-UK) Consortium | Harper VanSteenhouse, Yumi Kasai, David Gray, Carol Clugston, Anna Dominiczak and Alex Alderton, Roberto Amato, Jeffrey Barrett, Sonia Goncalves, Ewan Harrison, David K. Jackson, Ian Johnston, Dominic Kwiatkowski, Cordelia Langford, John Sillitoe on behalf of the Wellcome Sanger Institute COVID-19 Surveillance Team |
| EPI_ISL_3360517 | Lighthouse Lab in Milton Keynes | Wellcome Sanger Institute for the COVID-19 Genomics UK (COG-UK) Consortium | The Lighthouse Lab in Milton Keynes and Alex Alderton, Roberto Amato, Jeffrey Barrett, Sonia Goncalves, Ewan Harrison, David K. Jackson, Ian Johnston, Dominic Kwiatkowski, Cordelia Langford, John Sillitoe on behalf of the Wellcome Sanger Institute COVID-19 Surveillance Team |
| EPI_ISL_3360518 | Lighthouse Lab in Alderley Park | Wellcome Sanger Institute for the COVID-19 Genomics UK (COG-UK) Consortium | Jacquelyn Wynn, Mairead Hyland, The Lighthouse Lab in Alderley Park and Alex Alderton, Roberto Amato, Jeffrey Barrett, Sonia Goncalves, Ewan Harrison, David K. Jackson, Ian Johnston, Dominic Kwiatkowski, Cordelia Langford, John Sillitoe on behalf of the Wellcome Sanger Institute COVID-19 Surveillance Team |
| EPI_ISL_3360520 | Lighthouse Lab in Glasgow | Wellcome Sanger Institute for the COVID-19 Genomics UK (COG-UK) Consortium | Harper VanSteenhouse, Yumi Kasai, David Gray, Carol Clugston, Anna Dominiczak and Alex Alderton, Roberto Amato, Jeffrey Barrett, Sonia Goncalves, Ewan Harrison, David K. Jackson, Ian Johnston, Dominic Kwiatkowski, Cordelia Langford, John Sillitoe on behalf of the Wellcome Sanger Institute COVID-19 Surveillance Team |
| EPI_ISL_3360521 | Lighthouse Lab in Milton Keynes | Wellcome Sanger Institute for the COVID-19 Genomics UK (COG-UK) Consortium | The Lighthouse Lab in Milton Keynes and Alex Alderton, Roberto Amato, Jeffrey Barrett, Sonia Goncalves, Ewan Harrison, David K. Jackson, Ian Johnston, Dominic Kwiatkowski, Cordelia Langford, John Sillitoe on behalf of the Wellcome Sanger Institute COVID-19 Surveillance Team |
| EPI_ISL_3360523 | Lighthouse Lab in Alderley Park | Wellcome Sanger Institute for the COVID-19 Genomics UK (COG-UK) Consortium | Jacquelyn Wynn, Mairead Hyland, The Lighthouse Lab in Alderley Park and Alex Alderton, Roberto Amato, Jeffrey Barrett, Sonia Goncalves, Ewan Harrison, David K. Jackson, Ian Johnston, Dominic Kwiatkowski, Cordelia Langford, John Sillitoe on behalf of the Wellcome Sanger Institute COVID-19 Surveillance Team |
| EPI_ISL_3360524 | Lighthouse Lab in Glasgow | Wellcome Sanger Institute for the COVID-19 Genomics UK (COG-UK) Consortium | Harper VanSteenhouse, Yumi Kasai, David Gray, Carol Clugston, Anna Dominiczak and Alex Alderton, Roberto Amato, Jeffrey Barrett, Sonia Goncalves, Ewan Harrison, David K. Jackson, Ian Johnston, Dominic Kwiatkowski, Cordelia Langford, John Sillitoe on behalf of the Wellcome Sanger Institute COVID-19 Surveillance Team |
| EPI_ISL_3360525, EPI_ISL_3360527 | Health Services Laboratories | Wellcome Sanger Institute for the COVID-19 Genomics UK (COG-UK) Consortium | Health Services Laboratories and Alex Alderton, Roberto Amato, Jeffrey Barrett, Sonia Goncalves, Ewan Harrison, David K. Jackson, Ian Johnston, Dominic Kwiatkowski, Cordelia Langford, John Sillitoe on behalf of the Wellcome Sanger Institute COVID-19 Surveillance Team |
| EPI_ISL_3360528 | Lighthouse Lab in Milton Keynes | Wellcome Sanger Institute for the COVID-19 Genomics UK (COG-UK) Consortium | The Lighthouse Lab in Milton Keynes and Alex Alderton, Roberto Amato, Jeffrey Barrett, Sonia Goncalves, Ewan Harrison, David K. Jackson, Ian Johnston, Dominic Kwiatkowski, Cordelia Langford, John Sillitoe on behalf of the Wellcome Sanger Institute COVID-19 Surveillance Team |
| EPI_ISL_3360530 | Health Services Laboratories | Wellcome Sanger Institute for the COVID-19 Genomics UK (COG-UK) | Health Services Laboratories and Alex Alderton, Roberto Amato, Jeffrey Barrett, Sonia Goncalves, Ewan Harrison, David K. Jackson, Ian Johnston, Dominic Kwiatkowski, Cordelia Langford, John Sillitoe on behalf of the Wellcome Sanger Institute |

| | | | |
|---|---|---|---|
| | | Consortium | COVID-19 Surveillance Team |
| EPI_ISL_3360531 | Lighthouse Lab in Milton Keynes | Wellcome Sanger Institute for the COVID-19 Genomics UK (COG-UK) Consortium | The Lighthouse Lab in Milton Keynes and Alex Alderton, Roberto Amato, Jeffrey Barrett, Sonia Goncalves, Ewan Harrison, David K. Jackson, Ian Johnston, Dominic Kwiatkowski, Cordelia Langford, John Sillitoe on behalf of the Wellcome Sanger Institute COVID-19 Surveillance Team |
| EPI_ISL_3360533 | Lighthouse Lab in Alderley Park | Wellcome Sanger Institute for the COVID-19 Genomics UK (COG-UK) Consortium | Jacquelyn Wynn, Mairead Hyland, The Lighthouse Lab in Alderley Park and Alex Alderton, Roberto Amato, Jeffrey Barrett, Sonia Goncalves, Ewan Harrison, David K. Jackson, Ian Johnston, Dominic Kwiatkowski, Cordelia Langford, John Sillitoe on behalf of the Wellcome Sanger Institute COVID-19 Surveillance Team |
| EPI_ISL_3360534 | Lighthouse Lab in Milton Keynes | Wellcome Sanger Institute for the COVID-19 Genomics UK (COG-UK) Consortium | The Lighthouse Lab in Milton Keynes and Alex Alderton, Roberto Amato, Jeffrey Barrett, Sonia Goncalves, Ewan Harrison, David K. Jackson, Ian Johnston, Dominic Kwiatkowski, Cordelia Langford, John Sillitoe on behalf of the Wellcome Sanger Institute COVID-19 Surveillance Team |
| EPI_ISL_3360535 | Health Services Laboratories | Wellcome Sanger Institute for the COVID-19 Genomics UK (COG-UK) Consortium | Health Services Laboratories and Alex Alderton, Roberto Amato, Jeffrey Barrett, Sonia Goncalves, Ewan Harrison, David K. Jackson, Ian Johnston, Dominic Kwiatkowski, Cordelia Langford, John Sillitoe on behalf of the Wellcome Sanger Institute COVID-19 Surveillance Team |
| EPI_ISL_3360537, EPI_ISL_3360538 | Lighthouse Lab in Milton Keynes | Wellcome Sanger Institute for the COVID-19 Genomics UK (COG-UK) Consortium | The Lighthouse Lab in Milton Keynes and Alex Alderton, Roberto Amato, Jeffrey Barrett, Sonia Goncalves, Ewan Harrison, David K. Jackson, Ian Johnston, Dominic Kwiatkowski, Cordelia Langford, John Sillitoe on behalf of the Wellcome Sanger Institute COVID-19 Surveillance Team |
| EPI_ISL_3360540 | Lighthouse Lab in Alderley Park | Wellcome Sanger Institute for the COVID-19 Genomics UK (COG-UK) Consortium | Jacquelyn Wynn, Mairead Hyland, The Lighthouse Lab in Alderley Park and Alex Alderton, Roberto Amato, Jeffrey Barrett, Sonia Goncalves, Ewan Harrison, David K. Jackson, Ian Johnston, Dominic Kwiatkowski, Cordelia Langford, John Sillitoe on behalf of the Wellcome Sanger Institute COVID-19 Surveillance Team |
| EPI_ISL_3360541, EPI_ISL_3360543 | Health Services Laboratories | Wellcome Sanger Institute for the COVID-19 Genomics UK (COG-UK) Consortium | Health Services Laboratories and Alex Alderton, Roberto Amato, Jeffrey Barrett, Sonia Goncalves, Ewan Harrison, David K. Jackson, Ian Johnston, Dominic Kwiatkowski, Cordelia Langford, John Sillitoe on behalf of the Wellcome Sanger Institute COVID-19 Surveillance Team |
| EPI_ISL_3360544 | Lighthouse Lab in Milton Keynes | Wellcome Sanger Institute for the COVID-19 Genomics UK (COG-UK) Consortium | The Lighthouse Lab in Milton Keynes and Alex Alderton, Roberto Amato, Jeffrey Barrett, Sonia Goncalves, Ewan Harrison, David K. Jackson, Ian Johnston, Dominic Kwiatkowski, Cordelia Langford, John Sillitoe on behalf of the Wellcome Sanger Institute COVID-19 Surveillance Team |
| EPI_ISL_3360546 | Lighthouse Lab in Alderley Park | Wellcome Sanger Institute for the COVID-19 Genomics UK (COG-UK) Consortium | Jacquelyn Wynn, Mairead Hyland, The Lighthouse Lab in Alderley Park and Alex Alderton, Roberto Amato, Jeffrey Barrett, Sonia Goncalves, Ewan Harrison, David K. Jackson, Ian Johnston, Dominic Kwiatkowski, Cordelia Langford, John Sillitoe on behalf of the Wellcome Sanger Institute COVID-19 Surveillance Team |
| EPI_ISL_3360547, EPI_ISL_3360549 | Lighthouse Lab in Milton Keynes | Wellcome Sanger Institute for the COVID-19 Genomics UK (COG-UK) Consortium | The Lighthouse Lab in Milton Keynes and Alex Alderton, Roberto Amato, Jeffrey Barrett, Sonia Goncalves, Ewan Harrison, David K. Jackson, Ian Johnston, Dominic Kwiatkowski, Cordelia Langford, John Sillitoe on behalf of the Wellcome Sanger Institute COVID-19 Surveillance Team |
| EPI_ISL_3360550 | Lighthouse Lab in Alderley Park | Wellcome Sanger Institute for the COVID-19 Genomics UK (COG-UK) Consortium | Jacquelyn Wynn, Mairead Hyland, The Lighthouse Lab in Alderley Park and Alex Alderton, Roberto Amato, Jeffrey Barrett, Sonia Goncalves, Ewan Harrison, David K. Jackson, Ian Johnston, Dominic Kwiatkowski, Cordelia Langford, John Sillitoe on behalf of the Wellcome Sanger Institute COVID-19 Surveillance Team |
| EPI_ISL_3360552 | Lighthouse Lab in Glasgow | Wellcome Sanger Institute for the COVID-19 Genomics UK (COG-UK) Consortium | Harper VanSteenhouse, Yumi Kasai, David Gray, Carol Clugston, Anna Dominiczak and Alex Alderton, Roberto Amato, Jeffrey Barrett, Sonia Goncalves, Ewan Harrison, David K. Jackson, Ian Johnston, Dominic Kwiatkowski, Cordelia Langford, John Sillitoe on behalf of the Wellcome Sanger Institute COVID-19 Surveillance Team |
| EPI_ISL_3360553 | Health Services Laboratories | Wellcome Sanger Institute for the COVID-19 Genomics UK (COG-UK) Consortium | Health Services Laboratories and Alex Alderton, Roberto Amato, Jeffrey Barrett, Sonia Goncalves, Ewan Harrison, David K. Jackson, Ian Johnston, Dominic Kwiatkowski, Cordelia Langford, John Sillitoe on behalf of the Wellcome Sanger Institute COVID-19 Surveillance Team |
| EPI_ISL_3360555, EPI_ISL_3360556, EPI_ISL_3360558 | Lighthouse Lab in Glasgow | Wellcome Sanger Institute for the COVID-19 Genomics UK (COG-UK) Consortium | Harper VanSteenhouse, Yumi Kasai, David Gray, Carol Clugston, Anna Dominiczak and Alex Alderton, Roberto Amato, Jeffrey Barrett, Sonia Goncalves, Ewan Harrison, David K. Jackson, Ian Johnston, Dominic Kwiatkowski, Cordelia Langford, John Sillitoe on behalf of the Wellcome Sanger Institute COVID-19 Surveillance Team |
| EPI_ISL_3360559 | Health Services Laboratories | Wellcome Sanger Institute for the COVID-19 Genomics UK (COG-UK) Consortium | Health Services Laboratories and Alex Alderton, Roberto Amato, Jeffrey Barrett, Sonia Goncalves, Ewan Harrison, David K. Jackson, Ian Johnston, Dominic Kwiatkowski, Cordelia Langford, John Sillitoe on behalf of the Wellcome Sanger Institute COVID-19 Surveillance Team |
| EPI_ISL_3360561 | Lighthouse Lab in Milton Keynes | Wellcome Sanger Institute for the COVID-19 Genomics UK (COG-UK) Consortium | The Lighthouse Lab in Milton Keynes and Alex Alderton, Roberto Amato, Jeffrey Barrett, Sonia Goncalves, Ewan Harrison, David K. Jackson, Ian Johnston, Dominic Kwiatkowski, Cordelia Langford, John Sillitoe on behalf of the Wellcome Sanger Institute COVID-19 Surveillance Team |
| EPI_ISL_3360562 | Health Services Laboratories | Wellcome Sanger Institute for the COVID-19 Genomics UK (COG-UK) Consortium | Health Services Laboratories and Alex Alderton, Roberto Amato, Jeffrey Barrett, Sonia Goncalves, Ewan Harrison, David K. Jackson, Ian Johnston, Dominic Kwiatkowski, Cordelia Langford, John Sillitoe on behalf of the Wellcome Sanger Institute COVID-19 Surveillance Team |
| EPI_ISL_3360564 | Lighthouse Lab in Alderley Park | Wellcome Sanger Institute for the COVID-19 Genomics UK (COG-UK) Consortium | Jacquelyn Wynn, Mairead Hyland, The Lighthouse Lab in Alderley Park and Alex Alderton, Roberto Amato, Jeffrey Barrett, Sonia Goncalves, Ewan Harrison, David K. Jackson, Ian Johnston, Dominic Kwiatkowski, Cordelia Langford, John Sillitoe on behalf of the Wellcome Sanger Institute COVID-19 Surveillance Team |
| EPI_ISL_3360565, EPI_ISL_3360567 | Lighthouse Lab in Milton Keynes | Wellcome Sanger Institute for the COVID-19 Genomics UK (COG-UK) Consortium | The Lighthouse Lab in Milton Keynes and Alex Alderton, Roberto Amato, Jeffrey Barrett, Sonia Goncalves, Ewan Harrison, David K. Jackson, Ian Johnston, Dominic Kwiatkowski, Cordelia Langford, John Sillitoe on behalf of the Wellcome Sanger Institute COVID-19 Surveillance Team |
| EPI_ISL_3360568, EPI_ISL_3360570 | Health Services Laboratories | Wellcome Sanger Institute for the COVID-19 Genomics UK (COG-UK) Consortium | Health Services Laboratories and Alex Alderton, Roberto Amato, Jeffrey Barrett, Sonia Goncalves, Ewan Harrison, David K. Jackson, Ian Johnston, Dominic Kwiatkowski, Cordelia Langford, John Sillitoe on behalf of the Wellcome Sanger Institute COVID-19 Surveillance Team |
| EPI_ISL_3360572 | Lighthouse Lab in Alderley Park | Wellcome Sanger Institute for the COVID-19 Genomics UK (COG-UK) Consortium | Jacquelyn Wynn, Mairead Hyland, The Lighthouse Lab in Alderley Park and Alex Alderton, Roberto Amato, Jeffrey Barrett, Sonia Goncalves, Ewan Harrison, David K. Jackson, Ian Johnston, Dominic Kwiatkowski, Cordelia Langford, John Sillitoe on behalf of the Wellcome Sanger Institute COVID-19 Surveillance Team |
| EPI_ISL_3360573 | Lighthouse Lab in Glasgow | Wellcome Sanger Institute for the COVID-19 Genomics UK (COG-UK) Consortium | Harper VanSteenhouse, Yumi Kasai, David Gray, Carol Clugston, Anna Dominiczak and Alex Alderton, Roberto Amato, Jeffrey Barrett, Sonia Goncalves, Ewan Harrison, David K. Jackson, Ian Johnston, Dominic Kwiatkowski, Cordelia Langford, John Sillitoe on behalf of the Wellcome Sanger Institute COVID-19 Surveillance Team |
| EPI_ISL_3360575 | Health Services Laboratories | Wellcome Sanger Institute for the COVID-19 Genomics UK (COG-UK) Consortium | Health Services Laboratories and Alex Alderton, Roberto Amato, Jeffrey Barrett, Sonia Goncalves, Ewan Harrison, David K. Jackson, Ian Johnston, Dominic Kwiatkowski, Cordelia Langford, John Sillitoe on behalf of the Wellcome Sanger Institute COVID-19 Surveillance Team |
| EPI_ISL_3360576 | Lighthouse Lab in Milton Keynes | Wellcome Sanger Institute for the COVID-19 Genomics UK (COG-UK) Consortium | The Lighthouse Lab in Milton Keynes and Alex Alderton, Roberto Amato, Jeffrey Barrett, Sonia Goncalves, Ewan Harrison, David K. Jackson, Ian Johnston, Dominic Kwiatkowski, Cordelia Langford, John Sillitoe on behalf of the Wellcome Sanger Institute COVID-19 Surveillance Team |
| EPI_ISL_3360577 | Health Services Laboratories | Wellcome Sanger Institute for the COVID-19 Genomics UK (COG-UK) Consortium | Health Services Laboratories and Alex Alderton, Roberto Amato, Jeffrey Barrett, Sonia Goncalves, Ewan Harrison, David K. Jackson, Ian Johnston, Dominic Kwiatkowski, Cordelia Langford, John Sillitoe on behalf of the Wellcome Sanger Institute COVID-19 Surveillance Team |
| EPI_ISL_3360579 | Lighthouse Lab in Milton Keynes | Wellcome Sanger Institute for the COVID-19 Genomics UK (COG-UK) Consortium | The Lighthouse Lab in Milton Keynes and Alex Alderton, Roberto Amato, Jeffrey Barrett, Sonia Goncalves, Ewan Harrison, David K. Jackson, Ian Johnston, Dominic Kwiatkowski, Cordelia Langford, John Sillitoe on behalf of the Wellcome Sanger Institute COVID-19 Surveillance Team |
| EPI_ISL_3360580 | Lighthouse Lab in Alderley Park | Wellcome Sanger Institute for the COVID-19 Genomics UK (COG-UK) Consortium | Jacquelyn Wynn, Mairead Hyland, The Lighthouse Lab in Alderley Park and Alex Alderton, Roberto Amato, Jeffrey Barrett, Sonia Goncalves, Ewan Harrison, David K. Jackson, Ian Johnston, Dominic Kwiatkowski, Cordelia Langford, John Sillitoe on behalf of the Wellcome Sanger Institute COVID-19 Surveillance Team |
| EPI_ISL_3360582, EPI_ISL_3360583 | Lighthouse Lab in Glasgow | Wellcome Sanger Institute for the COVID-19 Genomics UK (COG-UK) Consortium | Harper VanSteenhouse, Yumi Kasai, David Gray, Carol Clugston, Anna Dominiczak and Alex Alderton, Roberto Amato, Jeffrey Barrett, Sonia Goncalves, Ewan Harrison, David K. Jackson, Ian Johnston, Dominic Kwiatkowski, Cordelia Langford, John Sillitoe on behalf of the Wellcome Sanger Institute COVID-19 Surveillance Team |
| EPI_ISL_3360586 | Lighthouse Lab in Alderley Park | Wellcome Sanger Institute for the COVID-19 Genomics UK (COG-UK) Consortium | Jacquelyn Wynn, Mairead Hyland, The Lighthouse Lab in Alderley Park and Alex Alderton, Roberto Amato, Jeffrey Barrett, Sonia Goncalves, Ewan Harrison, David K. Jackson, Ian Johnston, Dominic Kwiatkowski, Cordelia Langford, John Sillitoe on behalf of the Wellcome Sanger Institute COVID-19 Surveillance Team |
| EPI_ISL_3360588 | Lighthouse Lab in Milton Keynes | Wellcome Sanger Institute for the COVID-19 Genomics UK (COG-UK) Consortium | The Lighthouse Lab in Milton Keynes and Alex Alderton, Roberto Amato, Jeffrey Barrett, Sonia Goncalves, Ewan Harrison, David K. Jackson, Ian Johnston, Dominic Kwiatkowski, Cordelia Langford, John Sillitoe on behalf of the Wellcome Sanger Institute COVID-19 Surveillance Team |
| EPI_ISL_3360589 | Lighthouse Lab in Alderley Park | Wellcome Sanger Institute for the COVID-19 Genomics UK (COG-UK) Consortium | Jacquelyn Wynn, Mairead Hyland, The Lighthouse Lab in Alderley Park and Alex Alderton, Roberto Amato, Jeffrey Barrett, Sonia Goncalves, Ewan Harrison, David K. Jackson, Ian Johnston, Dominic Kwiatkowski, Cordelia Langford, John Sillitoe on behalf of the Wellcome Sanger Institute COVID-19 Surveillance Team |
| EPI_ISL_3360591 | Lighthouse Lab in Glasgow | Wellcome Sanger Institute for the COVID-19 Genomics UK (COG-UK) Consortium | Harper VanSteenhouse, Yumi Kasai, David Gray, Carol Clugston, Anna Dominiczak and Alex Alderton, Roberto Amato, Jeffrey Barrett, Sonia Goncalves, Ewan Harrison, David K. Jackson, Ian Johnston, Dominic Kwiatkowski, Cordelia Langford, John Sillitoe on behalf of the Wellcome Sanger Institute COVID-19 Surveillance Team |
| EPI_ISL_3360592 | Lighthouse Lab in Alderley Park | Wellcome Sanger Institute for the COVID-19 Genomics UK (COG-UK) Consortium | Jacquelyn Wynn, Mairead Hyland, The Lighthouse Lab in Alderley Park and Alex Alderton, Roberto Amato, Jeffrey Barrett, Sonia Goncalves, Ewan Harrison, David K. Jackson, Ian Johnston, Dominic Kwiatkowski, Cordelia Langford, John Sillitoe on behalf of the Wellcome Sanger Institute COVID-19 Surveillance Team |
| EPI_ISL_3360594 | Lighthouse Lab in Milton Keynes | Wellcome Sanger Institute for the COVID-19 Genomics UK (COG-UK) Consortium | The Lighthouse Lab in Milton Keynes and Alex Alderton, Roberto Amato, Jeffrey Barrett, Sonia Goncalves, Ewan Harrison, David K. Jackson, Ian Johnston, Dominic Kwiatkowski, Cordelia Langford, John Sillitoe on behalf of the Wellcome Sanger Institute COVID-19 Surveillance Team |
| EPI_ISL_3360595, EPI_ISL_3360597 | Lighthouse Lab in Alderley Park | Wellcome Sanger Institute for the COVID-19 Genomics UK (COG-UK) Consortium | Jacquelyn Wynn, Mairead Hyland, The Lighthouse Lab in Alderley Park and Alex Alderton, Roberto Amato, Jeffrey Barrett, Sonia Goncalves, Ewan Harrison, David K. Jackson, Ian Johnston, Dominic Kwiatkowski, Cordelia Langford, John Sillitoe on behalf of the Wellcome Sanger Institute COVID-19 Surveillance Team |
| EPI_ISL_3360598 | Health Services Laboratories | Wellcome Sanger Institute for the COVID-19 Genomics UK (COG-UK) Consortium | Health Services Laboratories and Alex Alderton, Roberto Amato, Jeffrey Barrett, Sonia Goncalves, Ewan Harrison, David K. Jackson, Ian Johnston, Dominic Kwiatkowski, Cordelia Langford, John Sillitoe on behalf of the Wellcome Sanger Institute COVID-19 Surveillance Team |
| EPI_ISL_3360600 | Lighthouse Lab in Milton Keynes | Wellcome Sanger Institute for the COVID-19 Genomics UK (COG-UK) Consortium | The Lighthouse Lab in Milton Keynes and Alex Alderton, Roberto Amato, Jeffrey Barrett, Sonia Goncalves, Ewan Harrison, David K. Jackson, Ian Johnston, Dominic Kwiatkowski, Cordelia Langford, John Sillitoe on behalf of the Wellcome Sanger Institute COVID-19 Surveillance Team |
| EPI_ISL_3360602 | Health Services Laboratories | Wellcome Sanger Institute for the COVID-19 Genomics UK (COG-UK) Consortium | Health Services Laboratories and Alex Alderton, Roberto Amato, Jeffrey Barrett, Sonia Goncalves, Ewan Harrison, David K. Jackson, Ian Johnston, Dominic Kwiatkowski, Cordelia Langford, John Sillitoe on behalf of the Wellcome Sanger Institute COVID-19 Surveillance Team |
| EPI_ISL_3360603 | Lighthouse Lab in Milton Keynes | Wellcome Sanger Institute for the COVID-19 Genomics UK (COG-UK) Consortium | The Lighthouse Lab in Milton Keynes and Alex Alderton, Roberto Amato, Jeffrey Barrett, Sonia Goncalves, Ewan Harrison, David K. Jackson, Ian Johnston, Dominic Kwiatkowski, Cordelia Langford, John Sillitoe on behalf of the Wellcome Sanger Institute COVID-19 Surveillance Team |
| EPI_ISL_3360605 | Lighthouse Lab in Alderley Park | Wellcome Sanger Institute for the COVID-19 Genomics UK (COG-UK) Consortium | Jacquelyn Wynn, Mairead Hyland, The Lighthouse Lab in Alderley Park and Alex Alderton, Roberto Amato, Jeffrey Barrett, Sonia Goncalves, Ewan Harrison, David K. Jackson, Ian Johnston, Dominic Kwiatkowski, Cordelia Langford, John Sillitoe on behalf of the Wellcome Sanger Institute COVID-19 Surveillance Team |
| EPI_ISL_3360606 | Lighthouse Lab in Glasgow | Wellcome Sanger Institute for the COVID-19 Genomics UK (COG-UK) Consortium | Harper VanSteenhouse, Yumi Kasai, David Gray, Carol Clugston, Anna Dominiczak and Alex Alderton, Roberto Amato, Jeffrey Barrett, Sonia Goncalves, Ewan Harrison, David K. Jackson, Ian Johnston, Dominic Kwiatkowski, Cordelia Langford, John Sillitoe on behalf of the Wellcome Sanger Institute COVID-19 Surveillance Team |
| EPI_ISL_3360608 | Lighthouse Lab in Milton Keynes | Wellcome Sanger Institute for the COVID-19 Genomics UK (COG-UK) Consortium | The Lighthouse Lab in Milton Keynes and Alex Alderton, Roberto Amato, Jeffrey Barrett, Sonia Goncalves, Ewan Harrison, David K. Jackson, Ian Johnston, Dominic Kwiatkowski, Cordelia Langford, John Sillitoe on behalf of the Wellcome Sanger Institute COVID-19 Surveillance Team |
| EPI_ISL_3360609 | Health Services Laboratories | Wellcome Sanger Institute for the COVID-19 Genomics UK (COG-UK) Consortium | Health Services Laboratories and Alex Alderton, Roberto Amato, Jeffrey Barrett, Sonia Goncalves, Ewan Harrison, David K. Jackson, Ian Johnston, Dominic Kwiatkowski, Cordelia Langford, John Sillitoe on behalf of the Wellcome Sanger Institute COVID-19 Surveillance Team |
| EPI_ISL_3360610 | Lighthouse Lab in Glasgow | Wellcome Sanger Institute for the COVID-19 Genomics UK (COG-UK) Consortium | Harper VanSteenhouse, Yumi Kasai, David Gray, Carol Clugston, Anna Dominiczak and Alex Alderton, Roberto Amato, Jeffrey Barrett, Sonia Goncalves, Ewan Harrison, David K. Jackson, Ian Johnston, Dominic Kwiatkowski, Cordelia Langford, John Sillitoe on behalf of the Wellcome Sanger Institute COVID-19 Surveillance Team |

| | | | |
|---|---|---|---|
| EPI_ISL_3360612, EPI_ISL_3360614 | Lighthouse Lab in Milton Keynes | Wellcome Sanger Institute for the COVID-19 Genomics UK (COG-UK) Consortium | The Lighthouse Lab in Milton Keynes and Alex Alderton, Roberto Amato, Jeffrey Barrett, Sonia Goncalves, Ewan Harrison, David K. Jackson, Ian Johnston, Dominic Kwiatkowski, Cordelia Langford, John Sillitoe on behalf of the Wellcome Sanger Institute COVID-19 Surveillance Team |
| EPI_ISL_3360615 | Health Services Laboratories | Wellcome Sanger Institute for the COVID-19 Genomics UK (COG-UK) Consortium | Health Services Laboratories and Alex Alderton, Roberto Amato, Jeffrey Barrett, Sonia Goncalves, Ewan Harrison, David K. Jackson, Ian Johnston, Dominic Kwiatkowski, Cordelia Langford, John Sillitoe on behalf of the Wellcome Sanger Institute COVID-19 Surveillance Team |
| EPI_ISL_3360616 | Lighthouse Lab in Glasgow | Wellcome Sanger Institute for the COVID-19 Genomics UK (COG-UK) Consortium | Harper VanSteenhouse, Yumi Kasai, David Gray, Carol Clugston, Anna Dominiczak and Alex Alderton, Roberto Amato, Jeffrey Barrett, Sonia Goncalves, Ewan Harrison, David K. Jackson, Ian Johnston, Dominic Kwiatkowski, Cordelia Langford, John Sillitoe on behalf of the Wellcome Sanger Institute COVID-19 Surveillance Team |
| EPI_ISL_3360618 | Health Services Laboratories | Wellcome Sanger Institute for the COVID-19 Genomics UK (COG-UK) Consortium | Health Services Laboratories and Alex Alderton, Roberto Amato, Jeffrey Barrett, Sonia Goncalves, Ewan Harrison, David K. Jackson, Ian Johnston, Dominic Kwiatkowski, Cordelia Langford, John Sillitoe on behalf of the Wellcome Sanger Institute COVID-19 Surveillance Team |
| EPI_ISL_3360620, EPI_ISL_3360621 | Lighthouse Lab in Milton Keynes | Wellcome Sanger Institute for the COVID-19 Genomics UK (COG-UK) Consortium | The Lighthouse Lab in Milton Keynes and Alex Alderton, Roberto Amato, Jeffrey Barrett, Sonia Goncalves, Ewan Harrison, David K. Jackson, Ian Johnston, Dominic Kwiatkowski, Cordelia Langford, John Sillitoe on behalf of the Wellcome Sanger Institute COVID-19 Surveillance Team |
| EPI_ISL_3360623 | Health Services Laboratories | Wellcome Sanger Institute for the COVID-19 Genomics UK (COG-UK) Consortium | Health Services Laboratories and Alex Alderton, Roberto Amato, Jeffrey Barrett, Sonia Goncalves, Ewan Harrison, David K. Jackson, Ian Johnston, Dominic Kwiatkowski, Cordelia Langford, John Sillitoe on behalf of the Wellcome Sanger Institute COVID-19 Surveillance Team |
| EPI_ISL_3360624 | Lighthouse Lab in Milton Keynes | Wellcome Sanger Institute for the COVID-19 Genomics UK (COG-UK) Consortium | The Lighthouse Lab in Milton Keynes and Alex Alderton, Roberto Amato, Jeffrey Barrett, Sonia Goncalves, Ewan Harrison, David K. Jackson, Ian Johnston, Dominic Kwiatkowski, Cordelia Langford, John Sillitoe on behalf of the Wellcome Sanger Institute COVID-19 Surveillance Team |
| EPI_ISL_3360626 | Lighthouse Lab in Alderley Park | Wellcome Sanger Institute for the COVID-19 Genomics UK (COG-UK) Consortium | Jacquelyn Wynn, Mairead Hyland, The Lighthouse Lab in Alderley Park and Alex Alderton, Roberto Amato, Jeffrey Barrett, Sonia Goncalves, Ewan Harrison, David K. Jackson, Ian Johnston, Dominic Kwiatkowski, Cordelia Langford, John Sillitoe on behalf of the Wellcome Sanger Institute COVID-19 Surveillance Team |
| EPI_ISL_3360627 | Health Services Laboratories | Wellcome Sanger Institute for the COVID-19 Genomics UK (COG-UK) Consortium | Health Services Laboratories and Alex Alderton, Roberto Amato, Jeffrey Barrett, Sonia Goncalves, Ewan Harrison, David K. Jackson, Ian Johnston, Dominic Kwiatkowski, Cordelia Langford, John Sillitoe on behalf of the Wellcome Sanger Institute COVID-19 Surveillance Team |
| EPI_ISL_3360629 | Lighthouse Lab in Alderley Park | Wellcome Sanger Institute for the COVID-19 Genomics UK (COG-UK) Consortium | Jacquelyn Wynn, Mairead Hyland, The Lighthouse Lab in Alderley Park and Alex Alderton, Roberto Amato, Jeffrey Barrett, Sonia Goncalves, Ewan Harrison, David K. Jackson, Ian Johnston, Dominic Kwiatkowski, Cordelia Langford, John Sillitoe on behalf of the Wellcome Sanger Institute COVID-19 Surveillance Team |
| EPI_ISL_3360631 | Health Services Laboratories | Wellcome Sanger Institute for the COVID-19 Genomics UK (COG-UK) Consortium | Health Services Laboratories and Alex Alderton, Roberto Amato, Jeffrey Barrett, Sonia Goncalves, Ewan Harrison, David K. Jackson, Ian Johnston, Dominic Kwiatkowski, Cordelia Langford, John Sillitoe on behalf of the Wellcome Sanger Institute COVID-19 Surveillance Team |
| EPI_ISL_3360632 | Lighthouse Lab in Glasgow | Wellcome Sanger Institute for the COVID-19 Genomics UK (COG-UK) Consortium | Harper VanSteenhouse, Yumi Kasai, David Gray, Carol Clugston, Anna Dominiczak and Alex Alderton, Roberto Amato, Jeffrey Barrett, Sonia Goncalves, Ewan Harrison, David K. Jackson, Ian Johnston, Dominic Kwiatkowski, Cordelia Langford, John Sillitoe on behalf of the Wellcome Sanger Institute COVID-19 Surveillance Team |
| EPI_ISL_3360634 | Health Services Laboratories | Wellcome Sanger Institute for the COVID-19 Genomics UK (COG-UK) Consortium | Health Services Laboratories and Alex Alderton, Roberto Amato, Jeffrey Barrett, Sonia Goncalves, Ewan Harrison, David K. Jackson, Ian Johnston, Dominic Kwiatkowski, Cordelia Langford, John Sillitoe on behalf of the Wellcome Sanger Institute COVID-19 Surveillance Team |
| EPI_ISL_3360636 | Lighthouse Lab in Milton Keynes | Wellcome Sanger Institute for the COVID-19 Genomics UK (COG-UK) Consortium | The Lighthouse Lab in Milton Keynes and Alex Alderton, Roberto Amato, Jeffrey Barrett, Sonia Goncalves, Ewan Harrison, David K. Jackson, Ian Johnston, Dominic Kwiatkowski, Cordelia Langford, John Sillitoe on behalf of the Wellcome Sanger Institute COVID-19 Surveillance Team |
| EPI_ISL_3360637, EPI_ISL_3360639 | Lighthouse Lab in Glasgow | Wellcome Sanger Institute for the COVID-19 Genomics UK (COG-UK) Consortium | Harper VanSteenhouse, Yumi Kasai, David Gray, Carol Clugston, Anna Dominiczak and Alex Alderton, Roberto Amato, Jeffrey Barrett, Sonia Goncalves, Ewan Harrison, David K. Jackson, Ian Johnston, Dominic Kwiatkowski, Cordelia Langford, John Sillitoe on behalf of the Wellcome Sanger Institute COVID-19 Surveillance Team |
| EPI_ISL_3360640 | Health Services Laboratories | Wellcome Sanger Institute for the COVID-19 Genomics UK (COG-UK) Consortium | Health Services Laboratories and Alex Alderton, Roberto Amato, Jeffrey Barrett, Sonia Goncalves, Ewan Harrison, David K. Jackson, Ian Johnston, Dominic Kwiatkowski, Cordelia Langford, John Sillitoe on behalf of the Wellcome Sanger Institute COVID-19 Surveillance Team |
| EPI_ISL_3360642, EPI_ISL_3360643, EPI_ISL_3360645 | Lighthouse Lab in Alderley Park | Wellcome Sanger Institute for the COVID-19 Genomics UK (COG-UK) Consortium | Jacquelyn Wynn, Mairead Hyland, The Lighthouse Lab in Alderley Park and Alex Alderton, Roberto Amato, Jeffrey Barrett, Sonia Goncalves, Ewan Harrison, David K. Jackson, Ian Johnston, Dominic Kwiatkowski, Cordelia Langford, John Sillitoe on behalf of the Wellcome Sanger Institute COVID-19 Surveillance Team |
| EPI_ISL_3360646 | Lighthouse Lab in Milton Keynes | Wellcome Sanger Institute for the COVID-19 Genomics UK (COG-UK) Consortium | The Lighthouse Lab in Milton Keynes and Alex Alderton, Roberto Amato, Jeffrey Barrett, Sonia Goncalves, Ewan Harrison, David K. Jackson, Ian Johnston, Dominic Kwiatkowski, Cordelia Langford, John Sillitoe on behalf of the Wellcome Sanger Institute COVID-19 Surveillance Team |
| EPI_ISL_3360648 | Lighthouse Lab in Glasgow | Wellcome Sanger Institute for the COVID-19 Genomics UK (COG-UK) Consortium | Harper VanSteenhouse, Yumi Kasai, David Gray, Carol Clugston, Anna Dominiczak and Alex Alderton, Roberto Amato, Jeffrey Barrett, Sonia Goncalves, Ewan Harrison, David K. Jackson, Ian Johnston, Dominic Kwiatkowski, Cordelia Langford, John Sillitoe on behalf of the Wellcome Sanger Institute COVID-19 Surveillance Team |
| EPI_ISL_3360649, EPI_ISL_3360651 | Lighthouse Lab in Alderley Park | Wellcome Sanger Institute for the COVID-19 Genomics UK (COG-UK) Consortium | Jacquelyn Wynn, Mairead Hyland, The Lighthouse Lab in Alderley Park and Alex Alderton, Roberto Amato, Jeffrey Barrett, Sonia Goncalves, Ewan Harrison, David K. Jackson, Ian Johnston, Dominic Kwiatkowski, Cordelia Langford, John Sillitoe on behalf of the Wellcome Sanger Institute COVID-19 Surveillance Team |
| EPI_ISL_3360653, EPI_ISL_3360654 | Lighthouse Lab in Glasgow | Wellcome Sanger Institute for the COVID-19 Genomics UK (COG-UK) Consortium | Harper VanSteenhouse, Yumi Kasai, David Gray, Carol Clugston, Anna Dominiczak and Alex Alderton, Roberto Amato, Jeffrey Barrett, Sonia Goncalves, Ewan Harrison, David K. Jackson, Ian Johnston, Dominic Kwiatkowski, Cordelia Langford, John Sillitoe on behalf of the Wellcome Sanger Institute COVID-19 Surveillance Team |
| EPI_ISL_3360656, EPI_ISL_3360657, EPI_ISL_3360659, EPI_ISL_3360660, EPI_ISL_3360661 | Health Services Laboratories | Wellcome Sanger Institute for the COVID-19 Genomics UK (COG-UK) Consortium | Health Services Laboratories and Alex Alderton, Roberto Amato, Jeffrey Barrett, Sonia Goncalves, Ewan Harrison, David K. Jackson, Ian Johnston, Dominic Kwiatkowski, Cordelia Langford, John Sillitoe on behalf of the Wellcome Sanger Institute COVID-19 Surveillance Team |
| EPI_ISL_3360663 | Lighthouse Lab in Glasgow | Wellcome Sanger Institute for the COVID-19 Genomics UK (COG-UK) Consortium | Harper VanSteenhouse, Yumi Kasai, David Gray, Carol Clugston, Anna Dominiczak and Alex Alderton, Roberto Amato, Jeffrey Barrett, Sonia Goncalves, Ewan Harrison, David K. Jackson, Ian Johnston, Dominic Kwiatkowski, Cordelia Langford, John Sillitoe on behalf of the Wellcome Sanger Institute COVID-19 Surveillance Team |
| EPI_ISL_3360664 | Lighthouse Lab in Alderley Park | Wellcome Sanger Institute for the COVID-19 Genomics UK (COG-UK) Consortium | Jacquelyn Wynn, Mairead Hyland, The Lighthouse Lab in Alderley Park and Alex Alderton, Roberto Amato, Jeffrey Barrett, Sonia Goncalves, Ewan Harrison, David K. Jackson, Ian Johnston, Dominic Kwiatkowski, Cordelia Langford, John Sillitoe on behalf of the Wellcome Sanger Institute COVID-19 Surveillance Team |
| EPI_ISL_3360666 | Health Services Laboratories | Wellcome Sanger Institute for the COVID-19 Genomics UK (COG-UK) Consortium | Health Services Laboratories and Alex Alderton, Roberto Amato, Jeffrey Barrett, Sonia Goncalves, Ewan Harrison, David K. Jackson, Ian Johnston, Dominic Kwiatkowski, Cordelia Langford, John Sillitoe on behalf of the Wellcome Sanger Institute COVID-19 Surveillance Team |
| EPI_ISL_3360667, EPI_ISL_3360669 | Lighthouse Lab in Alderley Park | Wellcome Sanger Institute for the COVID-19 Genomics UK (COG-UK) Consortium | Jacquelyn Wynn, Mairead Hyland, The Lighthouse Lab in Alderley Park and Alex Alderton, Roberto Amato, Jeffrey Barrett, Sonia Goncalves, Ewan Harrison, David K. Jackson, Ian Johnston, Dominic Kwiatkowski, Cordelia Langford, John Sillitoe on behalf of the Wellcome Sanger Institute COVID-19 Surveillance Team |
| EPI_ISL_3360670, EPI_ISL_3360671 | Lighthouse Lab in Milton Keynes | Wellcome Sanger Institute for the COVID-19 Genomics UK (COG-UK) Consortium | The Lighthouse Lab in Milton Keynes and Alex Alderton, Roberto Amato, Jeffrey Barrett, Sonia Goncalves, Ewan Harrison, David K. Jackson, Ian Johnston, Dominic Kwiatkowski, Cordelia Langford, John Sillitoe on behalf of the Wellcome Sanger Institute COVID-19 Surveillance Team |
| EPI_ISL_3360673 | Lighthouse Lab in Glasgow | Wellcome Sanger Institute for the COVID-19 Genomics UK (COG-UK) Consortium | Harper VanSteenhouse, Yumi Kasai, David Gray, Carol Clugston, Anna Dominiczak and Alex Alderton, Roberto Amato, Jeffrey Barrett, Sonia Goncalves, Ewan Harrison, David K. Jackson, Ian Johnston, Dominic Kwiatkowski, Cordelia Langford, John Sillitoe on behalf of the Wellcome Sanger Institute COVID-19 Surveillance Team |
| EPI_ISL_3360675 | Lighthouse Lab in Milton Keynes | Wellcome Sanger Institute for the COVID-19 Genomics UK (COG-UK) Consortium | The Lighthouse Lab in Milton Keynes and Alex Alderton, Roberto Amato, Jeffrey Barrett, Sonia Goncalves, Ewan Harrison, David K. Jackson, Ian Johnston, Dominic Kwiatkowski, Cordelia Langford, John Sillitoe on behalf of the Wellcome Sanger Institute COVID-19 Surveillance Team |
| EPI_ISL_3360676 | Lighthouse Lab in Glasgow | Wellcome Sanger Institute for the COVID-19 Genomics UK (COG-UK) Consortium | Harper VanSteenhouse, Yumi Kasai, David Gray, Carol Clugston, Anna Dominiczak and Alex Alderton, Roberto Amato, Jeffrey Barrett, Sonia Goncalves, Ewan Harrison, David K. Jackson, Ian Johnston, Dominic Kwiatkowski, Cordelia Langford, John Sillitoe on behalf of the Wellcome Sanger Institute COVID-19 Surveillance Team |
| EPI_ISL_3360678 | Health Services Laboratories | Wellcome Sanger Institute for the COVID-19 Genomics UK (COG-UK) Consortium | Health Services Laboratories and Alex Alderton, Roberto Amato, Jeffrey Barrett, Sonia Goncalves, Ewan Harrison, David K. Jackson, Ian Johnston, Dominic Kwiatkowski, Cordelia Langford, John Sillitoe on behalf of the Wellcome Sanger Institute COVID-19 Surveillance Team |
| EPI_ISL_3360679 | Lighthouse Lab in Glasgow | Wellcome Sanger Institute for the COVID-19 Genomics UK (COG-UK) Consortium | Harper VanSteenhouse, Yumi Kasai, David Gray, Carol Clugston, Anna Dominiczak and Alex Alderton, Roberto Amato, Jeffrey Barrett, Sonia Goncalves, Ewan Harrison, David K. Jackson, Ian Johnston, Dominic Kwiatkowski, Cordelia Langford, John Sillitoe on behalf of the Wellcome Sanger Institute COVID-19 Surveillance Team |
| EPI_ISL_3360681 | Health Services Laboratories | Wellcome Sanger Institute for the COVID-19 Genomics UK (COG-UK) Consortium | Health Services Laboratories and Alex Alderton, Roberto Amato, Jeffrey Barrett, Sonia Goncalves, Ewan Harrison, David K. Jackson, Ian Johnston, Dominic Kwiatkowski, Cordelia Langford, John Sillitoe on behalf of the Wellcome Sanger Institute COVID-19 Surveillance Team |
| EPI_ISL_3360736 | Lighthouse Lab in Glasgow | Wellcome Sanger Institute for the COVID-19 Genomics UK (COG-UK) Consortium | Harper VanSteenhouse, Yumi Kasai, David Gray, Carol Clugston, Anna Dominiczak and Alex Alderton, Roberto Amato, Jeffrey Barrett, Sonia Goncalves, Ewan Harrison, David K. Jackson, Ian Johnston, Dominic Kwiatkowski, Cordelia Langford, John Sillitoe on behalf of the Wellcome Sanger Institute COVID-19 Surveillance Team |
| EPI_ISL_3360737 | Lighthouse Lab in Milton Keynes | Wellcome Sanger Institute for the COVID-19 Genomics UK (COG-UK) Consortium | The Lighthouse Lab in Milton Keynes and Alex Alderton, Roberto Amato, Jeffrey Barrett, Sonia Goncalves, Ewan Harrison, David K. Jackson, Ian Johnston, Dominic Kwiatkowski, Cordelia Langford, John Sillitoe on behalf of the Wellcome Sanger Institute COVID-19 Surveillance Team |
| EPI_ISL_3360739 | Randox Laboratories | Wellcome Sanger Institute for the COVID-19 Genomics UK (COG-UK) Consortium | Randox Laboratories and Alex Alderton, Roberto Amato, Jeffrey Barrett, Sonia Goncalves, Ewan Harrison, David K. Jackson, Ian Johnston, Dominic Kwiatkowski, Cordelia Langford, John Sillitoe on behalf of the Wellcome Sanger Institute COVID-19 Surveillance Team |
| EPI_ISL_3360740 | Lighthouse Lab in Alderley Park | Wellcome Sanger Institute for the COVID-19 Genomics UK (COG-UK) Consortium | Jacquelyn Wynn, Mairead Hyland, The Lighthouse Lab in Alderley Park and Alex Alderton, Roberto Amato, Jeffrey Barrett, Sonia Goncalves, Ewan Harrison, David K. Jackson, Ian Johnston, Dominic Kwiatkowski, Cordelia Langford, John Sillitoe on behalf of the Wellcome Sanger Institute COVID-19 Surveillance Team |
| EPI_ISL_3360742 | Lighthouse Lab in Glasgow | Wellcome Sanger Institute for the COVID-19 Genomics UK (COG-UK) Consortium | Harper VanSteenhouse, Yumi Kasai, David Gray, Carol Clugston, Anna Dominiczak and Alex Alderton, Roberto Amato, Jeffrey Barrett, Sonia Goncalves, Ewan Harrison, David K. Jackson, Ian Johnston, Dominic Kwiatkowski, Cordelia Langford, John Sillitoe on behalf of the Wellcome Sanger Institute COVID-19 Surveillance Team |
| EPI_ISL_3360743, EPI_ISL_3360745 | Health Services Laboratories | Wellcome Sanger Institute for the COVID-19 Genomics UK (COG-UK) Consortium | Health Services Laboratories and Alex Alderton, Roberto Amato, Jeffrey Barrett, Sonia Goncalves, Ewan Harrison, David K. Jackson, Ian Johnston, Dominic Kwiatkowski, Cordelia Langford, John Sillitoe on behalf of the Wellcome Sanger Institute COVID-19 Surveillance Team |
| EPI_ISL_3360746 | Lighthouse Lab in Glasgow | Wellcome Sanger Institute for the COVID-19 Genomics UK (COG-UK) Consortium | Harper VanSteenhouse, Yumi Kasai, David Gray, Carol Clugston, Anna Dominiczak and Alex Alderton, Roberto Amato, Jeffrey Barrett, Sonia Goncalves, Ewan Harrison, David K. Jackson, Ian Johnston, Dominic Kwiatkowski, Cordelia Langford, John Sillitoe on behalf of the Wellcome Sanger Institute COVID-19 Surveillance Team |
| EPI_ISL_3360748, EPI_ISL_3360749, EPI_ISL_3360751 | Lighthouse Lab in Alderley Park | Wellcome Sanger Institute for the COVID-19 Genomics UK (COG-UK) Consortium | Jacquelyn Wynn, Mairead Hyland, The Lighthouse Lab in Alderley Park and Alex Alderton, Roberto Amato, Jeffrey Barrett, Sonia Goncalves, Ewan Harrison, David K. Jackson, Ian Johnston, Dominic Kwiatkowski, Cordelia Langford, John Sillitoe on behalf of the Wellcome Sanger Institute COVID-19 Surveillance Team |
| EPI_ISL_3360752 | Health Services Laboratories | Wellcome Sanger Institute for the COVID-19 Genomics UK (COG-UK) Consortium | Health Services Laboratories and Alex Alderton, Roberto Amato, Jeffrey Barrett, Sonia Goncalves, Ewan Harrison, David K. Jackson, Ian Johnston, Dominic Kwiatkowski, Cordelia Langford, John Sillitoe on behalf of the Wellcome Sanger Institute COVID-19 Surveillance Team |
| EPI_ISL_3360754 | Lighthouse Lab in Milton Keynes | Wellcome Sanger Institute for the COVID-19 Genomics UK (COG-UK) Consortium | The Lighthouse Lab in Milton Keynes and Alex Alderton, Roberto Amato, Jeffrey Barrett, Sonia Goncalves, Ewan Harrison, David K. Jackson, Ian Johnston, Dominic Kwiatkowski, Cordelia Langford, John Sillitoe on behalf of the Wellcome Sanger Institute COVID-19 Surveillance Team |

| | | | |
|---|---|---|---|
| EPI_ISL_3360755 | Lighthouse Lab in Glasgow | Wellcome Sanger Institute for the COVID-19 Genomics UK (COG-UK) Consortium | Harper VanSteenhouse, Yumi Kasai, David Gray, Carol Clugston, Anna Dominiczak and Alex Alderton, Roberto Amato, Jeffrey Barrett, Sonia Goncalves, Ewan Harrison, David K. Jackson, Ian Johnston, Dominic Kwiatkowski, Cordelia Langford, John Sillitoe on behalf of the Wellcome Sanger Institute COVID-19 Surveillance Team |
| EPI_ISL_3360757 | Health Services Laboratories | Wellcome Sanger Institute for the COVID-19 Genomics UK (COG-UK) Consortium | Health Services Laboratories and Alex Alderton, Roberto Amato, Jeffrey Barrett, Sonia Goncalves, Ewan Harrison, David K. Jackson, Ian Johnston, Dominic Kwiatkowski, Cordelia Langford, John Sillitoe on behalf of the Wellcome Sanger Institute COVID-19 Surveillance Team |
| EPI_ISL_3360758, EPI_ISL_3360760 | Lighthouse Lab in Alderley Park | Wellcome Sanger Institute for the COVID-19 Genomics UK (COG-UK) Consortium | Jacquelyn Wynn, Mairead Hyland, The Lighthouse Lab in Alderley Park and Alex Alderton, Roberto Amato, Jeffrey Barrett, Sonia Goncalves, Ewan Harrison, David K. Jackson, Ian Johnston, Dominic Kwiatkowski, Cordelia Langford, John Sillitoe on behalf of the Wellcome Sanger Institute COVID-19 Surveillance Team |
| EPI_ISL_3360761, EPI_ISL_3360763, EPI_ISL_3360764, EPI_ISL_3360766 | Lighthouse Lab in Milton Keynes | Wellcome Sanger Institute for the COVID-19 Genomics UK (COG-UK) Consortium | The Lighthouse Lab in Milton Keynes and Alex Alderton, Roberto Amato, Jeffrey Barrett, Sonia Goncalves, Ewan Harrison, David K. Jackson, Ian Johnston, Dominic Kwiatkowski, Cordelia Langford, John Sillitoe on behalf of the Wellcome Sanger Institute COVID-19 Surveillance Team |
| EPI_ISL_3360767 | Lighthouse Lab in Glasgow | Wellcome Sanger Institute for the COVID-19 Genomics UK (COG-UK) Consortium | Harper VanSteenhouse, Yumi Kasai, David Gray, Carol Clugston, Anna Dominiczak and Alex Alderton, Roberto Amato, Jeffrey Barrett, Sonia Goncalves, Ewan Harrison, David K. Jackson, Ian Johnston, Dominic Kwiatkowski, Cordelia Langford, John Sillitoe on behalf of the Wellcome Sanger Institute COVID-19 Surveillance Team |
| EPI_ISL_3360769 | Lighthouse Lab in Milton Keynes | Wellcome Sanger Institute for the COVID-19 Genomics UK (COG-UK) Consortium | The Lighthouse Lab in Milton Keynes and Alex Alderton, Roberto Amato, Jeffrey Barrett, Sonia Goncalves, Ewan Harrison, David K. Jackson, Ian Johnston, Dominic Kwiatkowski, Cordelia Langford, John Sillitoe on behalf of the Wellcome Sanger Institute COVID-19 Surveillance Team |
| EPI_ISL_3360770, EPI_ISL_3360771 | Lighthouse Lab in Alderley Park | Wellcome Sanger Institute for the COVID-19 Genomics UK (COG-UK) Consortium | Jacquelyn Wynn, Mairead Hyland, The Lighthouse Lab in Alderley Park and Alex Alderton, Roberto Amato, Jeffrey Barrett, Sonia Goncalves, Ewan Harrison, David K. Jackson, Ian Johnston, Dominic Kwiatkowski, Cordelia Langford, John Sillitoe on behalf of the Wellcome Sanger Institute COVID-19 Surveillance Team |
| EPI_ISL_3360773 | Lighthouse Lab in Glasgow | Wellcome Sanger Institute for the COVID-19 Genomics UK (COG-UK) Consortium | Harper VanSteenhouse, Yumi Kasai, David Gray, Carol Clugston, Anna Dominiczak and Alex Alderton, Roberto Amato, Jeffrey Barrett, Sonia Goncalves, Ewan Harrison, David K. Jackson, Ian Johnston, Dominic Kwiatkowski, Cordelia Langford, John Sillitoe on behalf of the Wellcome Sanger Institute COVID-19 Surveillance Team |
| EPI_ISL_3360775 | Lighthouse Lab in Milton Keynes | Wellcome Sanger Institute for the COVID-19 Genomics UK (COG-UK) Consortium | The Lighthouse Lab in Milton Keynes and Alex Alderton, Roberto Amato, Jeffrey Barrett, Sonia Goncalves, Ewan Harrison, David K. Jackson, Ian Johnston, Dominic Kwiatkowski, Cordelia Langford, John Sillitoe on behalf of the Wellcome Sanger Institute COVID-19 Surveillance Team |
| EPI_ISL_3360776 | Lighthouse Lab in Glasgow | Wellcome Sanger Institute for the COVID-19 Genomics UK (COG-UK) Consortium | Harper VanSteenhouse, Yumi Kasai, David Gray, Carol Clugston, Anna Dominiczak and Alex Alderton, Roberto Amato, Jeffrey Barrett, Sonia Goncalves, Ewan Harrison, David K. Jackson, Ian Johnston, Dominic Kwiatkowski, Cordelia Langford, John Sillitoe on behalf of the Wellcome Sanger Institute COVID-19 Surveillance Team |
| EPI_ISL_3360778, EPI_ISL_3360779 | Lighthouse Lab in Milton Keynes | Wellcome Sanger Institute for the COVID-19 Genomics UK (COG-UK) Consortium | The Lighthouse Lab in Milton Keynes and Alex Alderton, Roberto Amato, Jeffrey Barrett, Sonia Goncalves, Ewan Harrison, David K. Jackson, Ian Johnston, Dominic Kwiatkowski, Cordelia Langford, John Sillitoe on behalf of the Wellcome Sanger Institute COVID-19 Surveillance Team |
| EPI_ISL_3360780 | Lighthouse Lab in Alderley Park | Wellcome Sanger Institute for the COVID-19 Genomics UK (COG-UK) Consortium | Jacquelyn Wynn, Mairead Hyland, The Lighthouse Lab in Alderley Park and Alex Alderton, Roberto Amato, Jeffrey Barrett, Sonia Goncalves, Ewan Harrison, David K. Jackson, Ian Johnston, Dominic Kwiatkowski, Cordelia Langford, John Sillitoe on behalf of the Wellcome Sanger Institute COVID-19 Surveillance Team |
| EPI_ISL_3360782, EPI_ISL_3360783 | Lighthouse Lab in Glasgow | Wellcome Sanger Institute for the COVID-19 Genomics UK (COG-UK) Consortium | Harper VanSteenhouse, Yumi Kasai, David Gray, Carol Clugston, Anna Dominiczak and Alex Alderton, Roberto Amato, Jeffrey Barrett, Sonia Goncalves, Ewan Harrison, David K. Jackson, Ian Johnston, Dominic Kwiatkowski, Cordelia Langford, John Sillitoe on behalf of the Wellcome Sanger Institute COVID-19 Surveillance Team |
| EPI_ISL_3360785 | Lighthouse Lab in Alderley Park | Wellcome Sanger Institute for the COVID-19 Genomics UK (COG-UK) Consortium | Jacquelyn Wynn, Mairead Hyland, The Lighthouse Lab in Alderley Park and Alex Alderton, Roberto Amato, Jeffrey Barrett, Sonia Goncalves, Ewan Harrison, David K. Jackson, Ian Johnston, Dominic Kwiatkowski, Cordelia Langford, John Sillitoe on behalf of the Wellcome Sanger Institute COVID-19 Surveillance Team |
| EPI_ISL_3360786 | Lighthouse Lab in Milton Keynes | Wellcome Sanger Institute for the COVID-19 Genomics UK (COG-UK) Consortium | The Lighthouse Lab in Milton Keynes and Alex Alderton, Roberto Amato, Jeffrey Barrett, Sonia Goncalves, Ewan Harrison, David K. Jackson, Ian Johnston, Dominic Kwiatkowski, Cordelia Langford, John Sillitoe on behalf of the Wellcome Sanger Institute COVID-19 Surveillance Team |
| EPI_ISL_3360788, EPI_ISL_3360789, EPI_ISL_3360791 | Lighthouse Lab in Alderley Park | Wellcome Sanger Institute for the COVID-19 Genomics UK (COG-UK) Consortium | Jacquelyn Wynn, Mairead Hyland, The Lighthouse Lab in Alderley Park and Alex Alderton, Roberto Amato, Jeffrey Barrett, Sonia Goncalves, Ewan Harrison, David K. Jackson, Ian Johnston, Dominic Kwiatkowski, Cordelia Langford, John Sillitoe on behalf of the Wellcome Sanger Institute COVID-19 Surveillance Team |
| EPI_ISL_3360792 | Health Services Laboratories | Wellcome Sanger Institute for the COVID-19 Genomics UK (COG-UK) Consortium | Health Services Laboratories and Alex Alderton, Roberto Amato, Jeffrey Barrett, Sonia Goncalves, Ewan Harrison, David K. Jackson, Ian Johnston, Dominic Kwiatkowski, Cordelia Langford, John Sillitoe on behalf of the Wellcome Sanger Institute COVID-19 Surveillance Team |
| EPI_ISL_3360793 | Lighthouse Lab in Alderley Park | Wellcome Sanger Institute for the COVID-19 Genomics UK (COG-UK) Consortium | Jacquelyn Wynn, Mairead Hyland, The Lighthouse Lab in Alderley Park and Alex Alderton, Roberto Amato, Jeffrey Barrett, Sonia Goncalves, Ewan Harrison, David K. Jackson, Ian Johnston, Dominic Kwiatkowski, Cordelia Langford, John Sillitoe on behalf of the Wellcome Sanger Institute COVID-19 Surveillance Team |
| EPI_ISL_3360795 | Lighthouse Lab in Milton Keynes | Wellcome Sanger Institute for the COVID-19 Genomics UK (COG-UK) Consortium | The Lighthouse Lab in Milton Keynes and Alex Alderton, Roberto Amato, Jeffrey Barrett, Sonia Goncalves, Ewan Harrison, David K. Jackson, Ian Johnston, Dominic Kwiatkowski, Cordelia Langford, John Sillitoe on behalf of the Wellcome Sanger Institute COVID-19 Surveillance Team |
| EPI_ISL_3360796 | Lighthouse Lab in Glasgow | Wellcome Sanger Institute for the COVID-19 Genomics UK (COG-UK) Consortium | Harper VanSteenhouse, Yumi Kasai, David Gray, Carol Clugston, Anna Dominiczak and Alex Alderton, Roberto Amato, Jeffrey Barrett, Sonia Goncalves, Ewan Harrison, David K. Jackson, Ian Johnston, Dominic Kwiatkowski, Cordelia Langford, John Sillitoe on behalf of the Wellcome Sanger Institute COVID-19 Surveillance Team |
| EPI_ISL_3360797 | Lighthouse Lab in Milton Keynes | Wellcome Sanger Institute for the COVID-19 Genomics UK (COG-UK) Consortium | The Lighthouse Lab in Milton Keynes and Alex Alderton, Roberto Amato, Jeffrey Barrett, Sonia Goncalves, Ewan Harrison, David K. Jackson, Ian Johnston, Dominic Kwiatkowski, Cordelia Langford, John Sillitoe on behalf of the Wellcome Sanger Institute COVID-19 Surveillance Team |
| EPI_ISL_3360799 | Lighthouse Lab in Glasgow | Wellcome Sanger Institute for the COVID-19 Genomics UK (COG-UK) Consortium | Harper VanSteenhouse, Yumi Kasai, David Gray, Carol Clugston, Anna Dominiczak and Alex Alderton, Roberto Amato, Jeffrey Barrett, Sonia Goncalves, Ewan Harrison, David K. Jackson, Ian Johnston, Dominic Kwiatkowski, Cordelia Langford, John Sillitoe on behalf of the Wellcome Sanger Institute COVID-19 Surveillance Team |
| EPI_ISL_3360801 | Lighthouse Lab in Milton Keynes | Wellcome Sanger Institute for the COVID-19 Genomics UK (COG-UK) Consortium | The Lighthouse Lab in Milton Keynes and Alex Alderton, Roberto Amato, Jeffrey Barrett, Sonia Goncalves, Ewan Harrison, David K. Jackson, Ian Johnston, Dominic Kwiatkowski, Cordelia Langford, John Sillitoe on behalf of the Wellcome Sanger Institute COVID-19 Surveillance Team |
| EPI_ISL_3360802 | Lighthouse Lab in Alderley Park | Wellcome Sanger Institute for the COVID-19 Genomics UK (COG-UK) Consortium | Jacquelyn Wynn, Mairead Hyland, The Lighthouse Lab in Alderley Park and Alex Alderton, Roberto Amato, Jeffrey Barrett, Sonia Goncalves, Ewan Harrison, David K. Jackson, Ian Johnston, Dominic Kwiatkowski, Cordelia Langford, John Sillitoe on behalf of the Wellcome Sanger Institute COVID-19 Surveillance Team |
| EPI_ISL_3360804, EPI_ISL_3360805, EPI_ISL_3360807 | Health Services Laboratories | Wellcome Sanger Institute for the COVID-19 Genomics UK (COG-UK) Consortium | Health Services Laboratories and Alex Alderton, Roberto Amato, Jeffrey Barrett, Sonia Goncalves, Ewan Harrison, David K. Jackson, Ian Johnston, Dominic Kwiatkowski, Cordelia Langford, John Sillitoe on behalf of the Wellcome Sanger Institute COVID-19 Surveillance Team |
| EPI_ISL_3360808 | Lighthouse Lab in Glasgow | Wellcome Sanger Institute for the COVID-19 Genomics UK (COG-UK) Consortium | Harper VanSteenhouse, Yumi Kasai, David Gray, Carol Clugston, Anna Dominiczak and Alex Alderton, Roberto Amato, Jeffrey Barrett, Sonia Goncalves, Ewan Harrison, David K. Jackson, Ian Johnston, Dominic Kwiatkowski, Cordelia Langford, John Sillitoe on behalf of the Wellcome Sanger Institute COVID-19 Surveillance Team |
| EPI_ISL_3360810 | Lighthouse Lab in Alderley Park | Wellcome Sanger Institute for the COVID-19 Genomics UK (COG-UK) Consortium | Jacquelyn Wynn, Mairead Hyland, The Lighthouse Lab in Alderley Park and Alex Alderton, Roberto Amato, Jeffrey Barrett, Sonia Goncalves, Ewan Harrison, David K. Jackson, Ian Johnston, Dominic Kwiatkowski, Cordelia Langford, John Sillitoe on behalf of the Wellcome Sanger Institute COVID-19 Surveillance Team |
| EPI_ISL_3360811 | Health Services Laboratories | Wellcome Sanger Institute for the COVID-19 Genomics UK (COG-UK) Consortium | Health Services Laboratories and Alex Alderton, Roberto Amato, Jeffrey Barrett, Sonia Goncalves, Ewan Harrison, David K. Jackson, Ian Johnston, Dominic Kwiatkowski, Cordelia Langford, John Sillitoe on behalf of the Wellcome Sanger Institute COVID-19 Surveillance Team |
| EPI_ISL_3360813 | Lighthouse Lab in Glasgow | Wellcome Sanger Institute for the COVID-19 Genomics UK (COG-UK) Consortium | Harper VanSteenhouse, Yumi Kasai, David Gray, Carol Clugston, Anna Dominiczak and Alex Alderton, Roberto Amato, Jeffrey Barrett, Sonia Goncalves, Ewan Harrison, David K. Jackson, Ian Johnston, Dominic Kwiatkowski, Cordelia Langford, John Sillitoe on behalf of the Wellcome Sanger Institute COVID-19 Surveillance Team |
| EPI_ISL_3360814 | Lighthouse Lab in Alderley Park | Wellcome Sanger Institute for the COVID-19 Genomics UK (COG-UK) Consortium | Jacquelyn Wynn, Mairead Hyland, The Lighthouse Lab in Alderley Park and Alex Alderton, Roberto Amato, Jeffrey Barrett, Sonia Goncalves, Ewan Harrison, David K. Jackson, Ian Johnston, Dominic Kwiatkowski, Cordelia Langford, John Sillitoe on behalf of the Wellcome Sanger Institute COVID-19 Surveillance Team |
| EPI_ISL_3360815 | Health Services Laboratories | Wellcome Sanger Institute for the COVID-19 Genomics UK (COG-UK) Consortium | Health Services Laboratories and Alex Alderton, Roberto Amato, Jeffrey Barrett, Sonia Goncalves, Ewan Harrison, David K. Jackson, Ian Johnston, Dominic Kwiatkowski, Cordelia Langford, John Sillitoe on behalf of the Wellcome Sanger Institute COVID-19 Surveillance Team |
| EPI_ISL_3360817, EPI_ISL_3360818 | Lighthouse Lab in Milton Keynes | Wellcome Sanger Institute for the COVID-19 Genomics UK (COG-UK) Consortium | The Lighthouse Lab in Milton Keynes and Alex Alderton, Roberto Amato, Jeffrey Barrett, Sonia Goncalves, Ewan Harrison, David K. Jackson, Ian Johnston, Dominic Kwiatkowski, Cordelia Langford, John Sillitoe on behalf of the Wellcome Sanger Institute COVID-19 Surveillance Team |
| EPI_ISL_3360820 | Health Services Laboratories | Wellcome Sanger Institute for the COVID-19 Genomics UK (COG-UK) Consortium | Health Services Laboratories and Alex Alderton, Roberto Amato, Jeffrey Barrett, Sonia Goncalves, Ewan Harrison, David K. Jackson, Ian Johnston, Dominic Kwiatkowski, Cordelia Langford, John Sillitoe on behalf of the Wellcome Sanger Institute COVID-19 Surveillance Team |
| EPI_ISL_3360821 | Lighthouse Lab in Milton Keynes | Wellcome Sanger Institute for the COVID-19 Genomics UK (COG-UK) Consortium | The Lighthouse Lab in Milton Keynes and Alex Alderton, Roberto Amato, Jeffrey Barrett, Sonia Goncalves, Ewan Harrison, David K. Jackson, Ian Johnston, Dominic Kwiatkowski, Cordelia Langford, John Sillitoe on behalf of the Wellcome Sanger Institute COVID-19 Surveillance Team |
| EPI_ISL_3360822 | Lighthouse Lab in Glasgow | Wellcome Sanger Institute for the COVID-19 Genomics UK (COG-UK) Consortium | Harper VanSteenhouse, Yumi Kasai, David Gray, Carol Clugston, Anna Dominiczak and Alex Alderton, Roberto Amato, Jeffrey Barrett, Sonia Goncalves, Ewan Harrison, David K. Jackson, Ian Johnston, Dominic Kwiatkowski, Cordelia Langford, John Sillitoe on behalf of the Wellcome Sanger Institute COVID-19 Surveillance Team |
| EPI_ISL_3360824 | Health Services Laboratories | Wellcome Sanger Institute for the COVID-19 Genomics UK (COG-UK) Consortium | Health Services Laboratories and Alex Alderton, Roberto Amato, Jeffrey Barrett, Sonia Goncalves, Ewan Harrison, David K. Jackson, Ian Johnston, Dominic Kwiatkowski, Cordelia Langford, John Sillitoe on behalf of the Wellcome Sanger Institute COVID-19 Surveillance Team |
| EPI_ISL_3360826, EPI_ISL_3360827, EPI_ISL_3360828 | Lighthouse Lab in Milton Keynes | Wellcome Sanger Institute for the COVID-19 Genomics UK (COG-UK) Consortium | The Lighthouse Lab in Milton Keynes and Alex Alderton, Roberto Amato, Jeffrey Barrett, Sonia Goncalves, Ewan Harrison, David K. Jackson, Ian Johnston, Dominic Kwiatkowski, Cordelia Langford, John Sillitoe on behalf of the Wellcome Sanger Institute COVID-19 Surveillance Team |
| EPI_ISL_3360830, EPI_ISL_3360831 | Health Services Laboratories | Wellcome Sanger Institute for the COVID-19 Genomics UK (COG-UK) Consortium | Health Services Laboratories and Alex Alderton, Roberto Amato, Jeffrey Barrett, Sonia Goncalves, Ewan Harrison, David K. Jackson, Ian Johnston, Dominic Kwiatkowski, Cordelia Langford, John Sillitoe on behalf of the Wellcome Sanger Institute COVID-19 Surveillance Team |
| EPI_ISL_3360833, EPI_ISL_3360834 | Lighthouse Lab in Milton Keynes | Wellcome Sanger Institute for the COVID-19 Genomics UK (COG-UK) Consortium | The Lighthouse Lab in Milton Keynes and Alex Alderton, Roberto Amato, Jeffrey Barrett, Sonia Goncalves, Ewan Harrison, David K. Jackson, Ian Johnston, Dominic Kwiatkowski, Cordelia Langford, John Sillitoe on behalf of the Wellcome Sanger Institute COVID-19 Surveillance Team |
| EPI_ISL_3360836 | Health Services Laboratories | Wellcome Sanger Institute for the COVID-19 Genomics UK (COG-UK) Consortium | Health Services Laboratories and Alex Alderton, Roberto Amato, Jeffrey Barrett, Sonia Goncalves, Ewan Harrison, David K. Jackson, Ian Johnston, Dominic Kwiatkowski, Cordelia Langford, John Sillitoe on behalf of the Wellcome Sanger Institute COVID-19 Surveillance Team |
| EPI_ISL_3360837, EPI_ISL_3360839 | Lighthouse Lab in Milton Keynes | Wellcome Sanger Institute for the COVID-19 Genomics UK (COG-UK) Consortium | The Lighthouse Lab in Milton Keynes and Alex Alderton, Roberto Amato, Jeffrey Barrett, Sonia Goncalves, Ewan Harrison, David K. Jackson, Ian Johnston, Dominic Kwiatkowski, Cordelia Langford, John Sillitoe on behalf of the Wellcome Sanger Institute COVID-19 Surveillance Team |
| EPI_ISL_3360840, EPI_ISL_3360842 | Lighthouse Lab in Alderley Park | Wellcome Sanger Institute for the COVID-19 Genomics UK (COG-UK) Consortium | Jacquelyn Wynn, Mairead Hyland, The Lighthouse Lab in Alderley Park and Alex Alderton, Roberto Amato, Jeffrey Barrett, Sonia Goncalves, Ewan Harrison, David K. Jackson, Ian Johnston, Dominic Kwiatkowski, Cordelia Langford, John Sillitoe on behalf of the Wellcome Sanger Institute COVID-19 Surveillance Team |
| EPI_ISL_3360843 | Lighthouse Lab in Glasgow | Wellcome Sanger Institute for the COVID-19 Genomics UK (COG-UK) Consortium | Harper VanSteenhouse, Yumi Kasai, David Gray, Carol Clugston, Anna Dominiczak and Alex Alderton, Roberto Amato, Jeffrey Barrett, Sonia Goncalves, Ewan Harrison, David K. Jackson, Ian Johnston, Dominic Kwiatkowski, Cordelia Langford, John Sillitoe on behalf of the Wellcome Sanger Institute COVID-19 Surveillance Team |

| | | | |
|---|---|---|---|
| EPI_ISL_3360845 | Health Services Laboratories | Wellcome Sanger Institute for the COVID-19 Genomics UK (COG-UK) Consortium | Health Services Laboratories and Alex Alderton, Roberto Amato, Jeffrey Barrett, Sonia Goncalves, Ewan Harrison, David K. Jackson, Ian Johnston, Dominic Kwiatkowski, Cordelia Langford, John Sillitoe on behalf of the Wellcome Sanger Institute COVID-19 Surveillance Team |
| EPI_ISL_3360846 | Lighthouse Lab in Alderley Park | Wellcome Sanger Institute for the COVID-19 Genomics UK (COG-UK) Consortium | Jacquelyn Wynn, Mairead Hyland, The Lighthouse Lab in Alderley Park and Alex Alderton, Roberto Amato, Jeffrey Barrett, Sonia Goncalves, Ewan Harrison, David K. Jackson, Ian Johnston, Dominic Kwiatkowski, Cordelia Langford, John Sillitoe on behalf of the Wellcome Sanger Institute COVID-19 Surveillance Team |
| EPI_ISL_3360848 | Lighthouse Lab in Milton Keynes | Wellcome Sanger Institute for the COVID-19 Genomics UK (COG-UK) Consortium | The Lighthouse Lab in Milton Keynes and Alex Alderton, Roberto Amato, Jeffrey Barrett, Sonia Goncalves, Ewan Harrison, David K. Jackson, Ian Johnston, Dominic Kwiatkowski, Cordelia Langford, John Sillitoe on behalf of the Wellcome Sanger Institute COVID-19 Surveillance Team |
| EPI_ISL_3360849 | Lighthouse Lab in Alderley Park | Wellcome Sanger Institute for the COVID-19 Genomics UK (COG-UK) Consortium | Jacquelyn Wynn, Mairead Hyland, The Lighthouse Lab in Alderley Park and Alex Alderton, Roberto Amato, Jeffrey Barrett, Sonia Goncalves, Ewan Harrison, David K. Jackson, Ian Johnston, Dominic Kwiatkowski, Cordelia Langford, John Sillitoe on behalf of the Wellcome Sanger Institute COVID-19 Surveillance Team |
| EPI_ISL_3360850 | Health Services Laboratories | Wellcome Sanger Institute for the COVID-19 Genomics UK (COG-UK) Consortium | Health Services Laboratories and Alex Alderton, Roberto Amato, Jeffrey Barrett, Sonia Goncalves, Ewan Harrison, David K. Jackson, Ian Johnston, Dominic Kwiatkowski, Cordelia Langford, John Sillitoe on behalf of the Wellcome Sanger Institute COVID-19 Surveillance Team |
| EPI_ISL_3360852 | Lighthouse Lab in Alderley Park | Wellcome Sanger Institute for the COVID-19 Genomics UK (COG-UK) Consortium | Jacquelyn Wynn, Mairead Hyland, The Lighthouse Lab in Alderley Park and Alex Alderton, Roberto Amato, Jeffrey Barrett, Sonia Goncalves, Ewan Harrison, David K. Jackson, Ian Johnston, Dominic Kwiatkowski, Cordelia Langford, John Sillitoe on behalf of the Wellcome Sanger Institute COVID-19 Surveillance Team |
| EPI_ISL_3360853 | Health Services Laboratories | Wellcome Sanger Institute for the COVID-19 Genomics UK (COG-UK) Consortium | Health Services Laboratories and Alex Alderton, Roberto Amato, Jeffrey Barrett, Sonia Goncalves, Ewan Harrison, David K. Jackson, Ian Johnston, Dominic Kwiatkowski, Cordelia Langford, John Sillitoe on behalf of the Wellcome Sanger Institute COVID-19 Surveillance Team |
| EPI_ISL_3360855 | Lighthouse Lab in Alderley Park | Wellcome Sanger Institute for the COVID-19 Genomics UK (COG-UK) Consortium | Jacquelyn Wynn, Mairead Hyland, The Lighthouse Lab in Alderley Park and Alex Alderton, Roberto Amato, Jeffrey Barrett, Sonia Goncalves, Ewan Harrison, David K. Jackson, Ian Johnston, Dominic Kwiatkowski, Cordelia Langford, John Sillitoe on behalf of the Wellcome Sanger Institute COVID-19 Surveillance Team |
| EPI_ISL_3360856 | Lighthouse Lab in Milton Keynes | Wellcome Sanger Institute for the COVID-19 Genomics UK (COG-UK) Consortium | The Lighthouse Lab in Milton Keynes and Alex Alderton, Roberto Amato, Jeffrey Barrett, Sonia Goncalves, Ewan Harrison, David K. Jackson, Ian Johnston, Dominic Kwiatkowski, Cordelia Langford, John Sillitoe on behalf of the Wellcome Sanger Institute COVID-19 Surveillance Team |
| EPI_ISL_3360857 | Lighthouse Lab in Alderley Park | Wellcome Sanger Institute for the COVID-19 Genomics UK (COG-UK) Consortium | Jacquelyn Wynn, Mairead Hyland, The Lighthouse Lab in Alderley Park and Alex Alderton, Roberto Amato, Jeffrey Barrett, Sonia Goncalves, Ewan Harrison, David K. Jackson, Ian Johnston, Dominic Kwiatkowski, Cordelia Langford, John Sillitoe on behalf of the Wellcome Sanger Institute COVID-19 Surveillance Team |
| EPI_ISL_3360859 | Lighthouse Lab in Glasgow | Wellcome Sanger Institute for the COVID-19 Genomics UK (COG-UK) Consortium | Harper VanSteenhouse, Yumi Kasai, David Gray, Carol Clugston, Anna Dominiczak and Alex Alderton, Roberto Amato, Jeffrey Barrett, Sonia Goncalves, Ewan Harrison, David K. Jackson, Ian Johnston, Dominic Kwiatkowski, Cordelia Langford, John Sillitoe on behalf of the Wellcome Sanger Institute COVID-19 Surveillance Team |
| EPI_ISL_3360860, EPI_ISL_3360863, EPI_ISL_3360864, EPI_ISL_3360866 | Lighthouse Lab in Milton Keynes | Wellcome Sanger Institute for the COVID-19 Genomics UK (COG-UK) Consortium | The Lighthouse Lab in Milton Keynes and Alex Alderton, Roberto Amato, Jeffrey Barrett, Sonia Goncalves, Ewan Harrison, David K. Jackson, Ian Johnston, Dominic Kwiatkowski, Cordelia Langford, John Sillitoe on behalf of the Wellcome Sanger Institute COVID-19 Surveillance Team |
| EPI_ISL_3360867 | Health Services Laboratories | Wellcome Sanger Institute for the COVID-19 Genomics UK (COG-UK) Consortium | Health Services Laboratories and Alex Alderton, Roberto Amato, Jeffrey Barrett, Sonia Goncalves, Ewan Harrison, David K. Jackson, Ian Johnston, Dominic Kwiatkowski, Cordelia Langford, John Sillitoe on behalf of the Wellcome Sanger Institute COVID-19 Surveillance Team |
| EPI_ISL_3360869 | Lighthouse Lab in Milton Keynes | Wellcome Sanger Institute for the COVID-19 Genomics UK (COG-UK) Consortium | The Lighthouse Lab in Milton Keynes and Alex Alderton, Roberto Amato, Jeffrey Barrett, Sonia Goncalves, Ewan Harrison, David K. Jackson, Ian Johnston, Dominic Kwiatkowski, Cordelia Langford, John Sillitoe on behalf of the Wellcome Sanger Institute COVID-19 Surveillance Team |
| EPI_ISL_3360870 | Lighthouse Lab in Alderley Park | Wellcome Sanger Institute for the COVID-19 Genomics UK (COG-UK) Consortium | Jacquelyn Wynn, Mairead Hyland, The Lighthouse Lab in Alderley Park and Alex Alderton, Roberto Amato, Jeffrey Barrett, Sonia Goncalves, Ewan Harrison, David K. Jackson, Ian Johnston, Dominic Kwiatkowski, Cordelia Langford, John Sillitoe on behalf of the Wellcome Sanger Institute COVID-19 Surveillance Team |
| EPI_ISL_3360871 | Health Services Laboratories | Wellcome Sanger Institute for the COVID-19 Genomics UK (COG-UK) Consortium | Health Services Laboratories and Alex Alderton, Roberto Amato, Jeffrey Barrett, Sonia Goncalves, Ewan Harrison, David K. Jackson, Ian Johnston, Dominic Kwiatkowski, Cordelia Langford, John Sillitoe on behalf of the Wellcome Sanger Institute COVID-19 Surveillance Team |
| EPI_ISL_3360873, EPI_ISL_3360874, EPI_ISL_3360876 | Lighthouse Lab in Alderley Park | Wellcome Sanger Institute for the COVID-19 Genomics UK (COG-UK) Consortium | Jacquelyn Wynn, Mairead Hyland, The Lighthouse Lab in Alderley Park and Alex Alderton, Roberto Amato, Jeffrey Barrett, Sonia Goncalves, Ewan Harrison, David K. Jackson, Ian Johnston, Dominic Kwiatkowski, Cordelia Langford, John Sillitoe on behalf of the Wellcome Sanger Institute COVID-19 Surveillance Team |
| EPI_ISL_3360877 | Lighthouse Lab in Glasgow | Wellcome Sanger Institute for the COVID-19 Genomics UK (COG-UK) Consortium | Harper VanSteenhouse, Yumi Kasai, David Gray, Carol Clugston, Anna Dominiczak and Alex Alderton, Roberto Amato, Jeffrey Barrett, Sonia Goncalves, Ewan Harrison, David K. Jackson, Ian Johnston, Dominic Kwiatkowski, Cordelia Langford, John Sillitoe on behalf of the Wellcome Sanger Institute COVID-19 Surveillance Team |
| EPI_ISL_3360879, EPI_ISL_3360880, EPI_ISL_3360882 | Lighthouse Lab in Alderley Park | Wellcome Sanger Institute for the COVID-19 Genomics UK (COG-UK) Consortium | Jacquelyn Wynn, Mairead Hyland, The Lighthouse Lab in Alderley Park and Alex Alderton, Roberto Amato, Jeffrey Barrett, Sonia Goncalves, Ewan Harrison, David K. Jackson, Ian Johnston, Dominic Kwiatkowski, Cordelia Langford, John Sillitoe on behalf of the Wellcome Sanger Institute COVID-19 Surveillance Team |
| EPI_ISL_3360883 | Health Services Laboratories | Wellcome Sanger Institute for the COVID-19 Genomics UK (COG-UK) Consortium | Health Services Laboratories and Alex Alderton, Roberto Amato, Jeffrey Barrett, Sonia Goncalves, Ewan Harrison, David K. Jackson, Ian Johnston, Dominic Kwiatkowski, Cordelia Langford, John Sillitoe on behalf of the Wellcome Sanger Institute COVID-19 Surveillance Team |
| EPI_ISL_3360884, EPI_ISL_3360886, EPI_ISL_3360887 | Lighthouse Lab in Milton Keynes | Wellcome Sanger Institute for the COVID-19 Genomics UK (COG-UK) Consortium | The Lighthouse Lab in Milton Keynes and Alex Alderton, Roberto Amato, Jeffrey Barrett, Sonia Goncalves, Ewan Harrison, David K. Jackson, Ian Johnston, Dominic Kwiatkowski, Cordelia Langford, John Sillitoe on behalf of the Wellcome Sanger Institute COVID-19 Surveillance Team |
| EPI_ISL_3360889 | Lighthouse Lab in Glasgow | Wellcome Sanger Institute for the COVID-19 Genomics UK (COG-UK) Consortium | Harper VanSteenhouse, Yumi Kasai, David Gray, Carol Clugston, Anna Dominiczak and Alex Alderton, Roberto Amato, Jeffrey Barrett, Sonia Goncalves, Ewan Harrison, David K. Jackson, Ian Johnston, Dominic Kwiatkowski, Cordelia Langford, John Sillitoe on behalf of the Wellcome Sanger Institute COVID-19 Surveillance Team |
| EPI_ISL_3360890 | Lighthouse Lab in Alderley Park | Wellcome Sanger Institute for the COVID-19 Genomics UK (COG-UK) Consortium | Jacquelyn Wynn, Mairead Hyland, The Lighthouse Lab in Alderley Park and Alex Alderton, Roberto Amato, Jeffrey Barrett, Sonia Goncalves, Ewan Harrison, David K. Jackson, Ian Johnston, Dominic Kwiatkowski, Cordelia Langford, John Sillitoe on behalf of the Wellcome Sanger Institute COVID-19 Surveillance Team |
| EPI_ISL_3360892 | Health Services Laboratories | Wellcome Sanger Institute for the COVID-19 Genomics UK (COG-UK) Consortium | Health Services Laboratories and Alex Alderton, Roberto Amato, Jeffrey Barrett, Sonia Goncalves, Ewan Harrison, David K. Jackson, Ian Johnston, Dominic Kwiatkowski, Cordelia Langford, John Sillitoe on behalf of the Wellcome Sanger Institute COVID-19 Surveillance Team |
| EPI_ISL_3360893 | Lighthouse Lab in Milton Keynes | Wellcome Sanger Institute for the COVID-19 Genomics UK (COG-UK) Consortium | The Lighthouse Lab in Milton Keynes and Alex Alderton, Roberto Amato, Jeffrey Barrett, Sonia Goncalves, Ewan Harrison, David K. Jackson, Ian Johnston, Dominic Kwiatkowski, Cordelia Langford, John Sillitoe on behalf of the Wellcome Sanger Institute COVID-19 Surveillance Team |
| EPI_ISL_3360895 | Lighthouse Lab in Glasgow | Wellcome Sanger Institute for the COVID-19 Genomics UK (COG-UK) Consortium | Harper VanSteenhouse, Yumi Kasai, David Gray, Carol Clugston, Anna Dominiczak and Alex Alderton, Roberto Amato, Jeffrey Barrett, Sonia Goncalves, Ewan Harrison, David K. Jackson, Ian Johnston, Dominic Kwiatkowski, Cordelia Langford, John Sillitoe on behalf of the Wellcome Sanger Institute COVID-19 Surveillance Team |
| EPI_ISL_3360896 | Lighthouse Lab in Milton Keynes | Wellcome Sanger Institute for the COVID-19 Genomics UK (COG-UK) Consortium | The Lighthouse Lab in Milton Keynes and Alex Alderton, Roberto Amato, Jeffrey Barrett, Sonia Goncalves, Ewan Harrison, David K. Jackson, Ian Johnston, Dominic Kwiatkowski, Cordelia Langford, John Sillitoe on behalf of the Wellcome Sanger Institute COVID-19 Surveillance Team |
| EPI_ISL_3360897 | Health Services Laboratories | Wellcome Sanger Institute for the COVID-19 Genomics UK (COG-UK) Consortium | Health Services Laboratories and Alex Alderton, Roberto Amato, Jeffrey Barrett, Sonia Goncalves, Ewan Harrison, David K. Jackson, Ian Johnston, Dominic Kwiatkowski, Cordelia Langford, John Sillitoe on behalf of the Wellcome Sanger Institute COVID-19 Surveillance Team |
| EPI_ISL_3360899 | Lighthouse Lab in Alderley Park | Wellcome Sanger Institute for the COVID-19 Genomics UK (COG-UK) Consortium | Jacquelyn Wynn, Mairead Hyland, The Lighthouse Lab in Alderley Park and Alex Alderton, Roberto Amato, Jeffrey Barrett, Sonia Goncalves, Ewan Harrison, David K. Jackson, Ian Johnston, Dominic Kwiatkowski, Cordelia Langford, John Sillitoe on behalf of the Wellcome Sanger Institute COVID-19 Surveillance Team |
| EPI_ISL_3360901 | Lighthouse Lab in Glasgow | Wellcome Sanger Institute for the COVID-19 Genomics UK (COG-UK) Consortium | Harper VanSteenhouse, Yumi Kasai, David Gray, Carol Clugston, Anna Dominiczak and Alex Alderton, Roberto Amato, Jeffrey Barrett, Sonia Goncalves, Ewan Harrison, David K. Jackson, Ian Johnston, Dominic Kwiatkowski, Cordelia Langford, John Sillitoe on behalf of the Wellcome Sanger Institute COVID-19 Surveillance Team |
| EPI_ISL_3360902 | Lighthouse Lab in Alderley Park | Wellcome Sanger Institute for the COVID-19 Genomics UK (COG-UK) Consortium | Jacquelyn Wynn, Mairead Hyland, The Lighthouse Lab in Alderley Park and Alex Alderton, Roberto Amato, Jeffrey Barrett, Sonia Goncalves, Ewan Harrison, David K. Jackson, Ian Johnston, Dominic Kwiatkowski, Cordelia Langford, John Sillitoe on behalf of the Wellcome Sanger Institute COVID-19 Surveillance Team |
| EPI_ISL_3360904, EPI_ISL_3360905 | Lighthouse Lab in Milton Keynes | Wellcome Sanger Institute for the COVID-19 Genomics UK (COG-UK) Consortium | The Lighthouse Lab in Milton Keynes and Alex Alderton, Roberto Amato, Jeffrey Barrett, Sonia Goncalves, Ewan Harrison, David K. Jackson, Ian Johnston, Dominic Kwiatkowski, Cordelia Langford, John Sillitoe on behalf of the Wellcome Sanger Institute COVID-19 Surveillance Team |
| EPI_ISL_3360907 | Health Services Laboratories | Wellcome Sanger Institute for the COVID-19 Genomics UK (COG-UK) Consortium | Health Services Laboratories and Alex Alderton, Roberto Amato, Jeffrey Barrett, Sonia Goncalves, Ewan Harrison, David K. Jackson, Ian Johnston, Dominic Kwiatkowski, Cordelia Langford, John Sillitoe on behalf of the Wellcome Sanger Institute COVID-19 Surveillance Team |
| EPI_ISL_3360908, EPI_ISL_3360910 | Lighthouse Lab in Alderley Park | Wellcome Sanger Institute for the COVID-19 Genomics UK (COG-UK) Consortium | Jacquelyn Wynn, Mairead Hyland, The Lighthouse Lab in Alderley Park and Alex Alderton, Roberto Amato, Jeffrey Barrett, Sonia Goncalves, Ewan Harrison, David K. Jackson, Ian Johnston, Dominic Kwiatkowski, Cordelia Langford, John Sillitoe on behalf of the Wellcome Sanger Institute COVID-19 Surveillance Team |
| EPI_ISL_3360911, EPI_ISL_3360913 | Lighthouse Lab in Glasgow | Wellcome Sanger Institute for the COVID-19 Genomics UK (COG-UK) Consortium | Harper VanSteenhouse, Yumi Kasai, David Gray, Carol Clugston, Anna Dominiczak and Alex Alderton, Roberto Amato, Jeffrey Barrett, Sonia Goncalves, Ewan Harrison, David K. Jackson, Ian Johnston, Dominic Kwiatkowski, Cordelia Langford, John Sillitoe on behalf of the Wellcome Sanger Institute COVID-19 Surveillance Team |
| EPI_ISL_3360914 | Lighthouse Lab in Milton Keynes | Wellcome Sanger Institute for the COVID-19 Genomics UK (COG-UK) Consortium | The Lighthouse Lab in Milton Keynes and Alex Alderton, Roberto Amato, Jeffrey Barrett, Sonia Goncalves, Ewan Harrison, David K. Jackson, Ian Johnston, Dominic Kwiatkowski, Cordelia Langford, John Sillitoe on behalf of the Wellcome Sanger Institute COVID-19 Surveillance Team |
| EPI_ISL_3360915, EPI_ISL_3360917 | Health Services Laboratories | Wellcome Sanger Institute for the COVID-19 Genomics UK (COG-UK) Consortium | Health Services Laboratories and Alex Alderton, Roberto Amato, Jeffrey Barrett, Sonia Goncalves, Ewan Harrison, David K. Jackson, Ian Johnston, Dominic Kwiatkowski, Cordelia Langford, John Sillitoe on behalf of the Wellcome Sanger Institute COVID-19 Surveillance Team |
| EPI_ISL_3360918 | Lighthouse Lab in Milton Keynes | Wellcome Sanger Institute for the COVID-19 Genomics UK (COG-UK) Consortium | The Lighthouse Lab in Milton Keynes and Alex Alderton, Roberto Amato, Jeffrey Barrett, Sonia Goncalves, Ewan Harrison, David K. Jackson, Ian Johnston, Dominic Kwiatkowski, Cordelia Langford, John Sillitoe on behalf of the Wellcome Sanger Institute COVID-19 Surveillance Team |
| EPI_ISL_3360920 | Lighthouse Lab in Glasgow | Wellcome Sanger Institute for the COVID-19 Genomics UK (COG-UK) Consortium | Harper VanSteenhouse, Yumi Kasai, David Gray, Carol Clugston, Anna Dominiczak and Alex Alderton, Roberto Amato, Jeffrey Barrett, Sonia Goncalves, Ewan Harrison, David K. Jackson, Ian Johnston, Dominic Kwiatkowski, Cordelia Langford, John Sillitoe on behalf of the Wellcome Sanger Institute COVID-19 Surveillance Team |
| EPI_ISL_3360921, EPI_ISL_3360923 | Lighthouse Lab in Milton Keynes | Wellcome Sanger Institute for the COVID-19 Genomics UK (COG-UK) Consortium | The Lighthouse Lab in Milton Keynes and Alex Alderton, Roberto Amato, Jeffrey Barrett, Sonia Goncalves, Ewan Harrison, David K. Jackson, Ian Johnston, Dominic Kwiatkowski, Cordelia Langford, John Sillitoe on behalf of the Wellcome Sanger Institute COVID-19 Surveillance Team |
| EPI_ISL_3360924 | Lighthouse Lab in Alderley Park | Wellcome Sanger Institute for the COVID-19 Genomics UK (COG-UK) Consortium | Jacquelyn Wynn, Mairead Hyland, The Lighthouse Lab in Alderley Park and Alex Alderton, Roberto Amato, Jeffrey Barrett, Sonia Goncalves, Ewan Harrison, David K. Jackson, Ian Johnston, Dominic Kwiatkowski, Cordelia Langford, John Sillitoe on behalf of the Wellcome Sanger Institute COVID-19 Surveillance Team |
| EPI_ISL_3360925 | Lighthouse Lab in Milton Keynes | Wellcome Sanger Institute for the COVID-19 Genomics UK (COG-UK) Consortium | The Lighthouse Lab in Milton Keynes and Alex Alderton, Roberto Amato, Jeffrey Barrett, Sonia Goncalves, Ewan Harrison, David K. Jackson, Ian Johnston, Dominic Kwiatkowski, Cordelia Langford, John Sillitoe on behalf of the Wellcome Sanger Institute COVID-19 Surveillance Team |
| EPI_ISL_3360927 | Health Services Laboratories | Wellcome Sanger Institute for the COVID-19 Genomics UK (COG-UK) Consortium | Health Services Laboratories and Alex Alderton, Roberto Amato, Jeffrey Barrett, Sonia Goncalves, Ewan Harrison, David K. Jackson, Ian Johnston, Dominic Kwiatkowski, Cordelia Langford, John Sillitoe on behalf of the Wellcome Sanger Institute COVID-19 Surveillance Team |

| | | | |
|---|---|---|---|
| EPI_ISL_3360929 | Lighthouse Lab in Glasgow | Wellcome Sanger Institute for the COVID-19 Genomics UK (COG-UK) Consortium | Harper VanSteenhouse, Yumi Kasai, David Gray, Carol Clugston, Anna Dominiczak and Alex Alderton, Roberto Amato, Jeffrey Barrett, Sonia Goncalves, Ewan Harrison, David K. Jackson, Ian Johnston, Dominic Kwiatkowski, Cordelia Langford, John Sillitoe on behalf of the Wellcome Sanger Institute COVID-19 Surveillance Team |
| EPI_ISL_3360930, EPI_ISL_3360932 | Lighthouse Lab in Alderley Park | Wellcome Sanger Institute for the COVID-19 Genomics UK (COG-UK) Consortium | Jacquelyn Wynn, Mairead Hyland, The Lighthouse Lab in Alderley Park and Alex Alderton, Roberto Amato, Jeffrey Barrett, Sonia Goncalves, Ewan Harrison, David K. Jackson, Ian Johnston, Dominic Kwiatkowski, Cordelia Langford, John Sillitoe on behalf of the Wellcome Sanger Institute COVID-19 Surveillance Team |
| EPI_ISL_3360933, EPI_ISL_3360935, EPI_ISL_3360936 | Lighthouse Lab in Glasgow | Wellcome Sanger Institute for the COVID-19 Genomics UK (COG-UK) Consortium | Harper VanSteenhouse, Yumi Kasai, David Gray, Carol Clugston, Anna Dominiczak and Alex Alderton, Roberto Amato, Jeffrey Barrett, Sonia Goncalves, Ewan Harrison, David K. Jackson, Ian Johnston, Dominic Kwiatkowski, Cordelia Langford, John Sillitoe on behalf of the Wellcome Sanger Institute COVID-19 Surveillance Team |
| EPI_ISL_3360938 | Health Services Laboratories | Wellcome Sanger Institute for the COVID-19 Genomics UK (COG-UK) Consortium | Health Services Laboratories and Alex Alderton, Roberto Amato, Jeffrey Barrett, Sonia Goncalves, Ewan Harrison, David K. Jackson, Ian Johnston, Dominic Kwiatkowski, Cordelia Langford, John Sillitoe on behalf of the Wellcome Sanger Institute COVID-19 Surveillance Team |
| EPI_ISL_3360939 | Lighthouse Lab in Glasgow | Wellcome Sanger Institute for the COVID-19 Genomics UK (COG-UK) Consortium | Harper VanSteenhouse, Yumi Kasai, David Gray, Carol Clugston, Anna Dominiczak and Alex Alderton, Roberto Amato, Jeffrey Barrett, Sonia Goncalves, Ewan Harrison, David K. Jackson, Ian Johnston, Dominic Kwiatkowski, Cordelia Langford, John Sillitoe on behalf of the Wellcome Sanger Institute COVID-19 Surveillance Team |
| EPI_ISL_3360941 | Lighthouse Lab in Milton Keynes | Wellcome Sanger Institute for the COVID-19 Genomics UK (COG-UK) Consortium | The Lighthouse Lab in Milton Keynes and Alex Alderton, Roberto Amato, Jeffrey Barrett, Sonia Goncalves, Ewan Harrison, David K. Jackson, Ian Johnston, Dominic Kwiatkowski, Cordelia Langford, John Sillitoe on behalf of the Wellcome Sanger Institute COVID-19 Surveillance Team |
| EPI_ISL_3360942 | Lighthouse Lab in Alderley Park | Wellcome Sanger Institute for the COVID-19 Genomics UK (COG-UK) Consortium | Jacquelyn Wynn, Mairead Hyland, The Lighthouse Lab in Alderley Park and Alex Alderton, Roberto Amato, Jeffrey Barrett, Sonia Goncalves, Ewan Harrison, David K. Jackson, Ian Johnston, Dominic Kwiatkowski, Cordelia Langford, John Sillitoe on behalf of the Wellcome Sanger Institute COVID-19 Surveillance Team |
| EPI_ISL_3360944, EPI_ISL_3360945, EPI_ISL_3360947 | Lighthouse Lab in Milton Keynes | Wellcome Sanger Institute for the COVID-19 Genomics UK (COG-UK) Consortium | The Lighthouse Lab in Milton Keynes and Alex Alderton, Roberto Amato, Jeffrey Barrett, Sonia Goncalves, Ewan Harrison, David K. Jackson, Ian Johnston, Dominic Kwiatkowski, Cordelia Langford, John Sillitoe on behalf of the Wellcome Sanger Institute COVID-19 Surveillance Team |
| EPI_ISL_3360948 | Health Services Laboratories | Wellcome Sanger Institute for the COVID-19 Genomics UK (COG-UK) Consortium | Health Services Laboratories and Alex Alderton, Roberto Amato, Jeffrey Barrett, Sonia Goncalves, Ewan Harrison, David K. Jackson, Ian Johnston, Dominic Kwiatkowski, Cordelia Langford, John Sillitoe on behalf of the Wellcome Sanger Institute COVID-19 Surveillance Team |
| EPI_ISL_3360950, EPI_ISL_3360951, EPI_ISL_3360953, EPI_ISL_3360954 | Lighthouse Lab in Alderley Park | Wellcome Sanger Institute for the COVID-19 Genomics UK (COG-UK) Consortium | Jacquelyn Wynn, Mairead Hyland, The Lighthouse Lab in Alderley Park and Alex Alderton, Roberto Amato, Jeffrey Barrett, Sonia Goncalves, Ewan Harrison, David K. Jackson, Ian Johnston, Dominic Kwiatkowski, Cordelia Langford, John Sillitoe on behalf of the Wellcome Sanger Institute COVID-19 Surveillance Team |
| EPI_ISL_3360956 | Health Services Laboratories | Wellcome Sanger Institute for the COVID-19 Genomics UK (COG-UK) Consortium | Health Services Laboratories and Alex Alderton, Roberto Amato, Jeffrey Barrett, Sonia Goncalves, Ewan Harrison, David K. Jackson, Ian Johnston, Dominic Kwiatkowski, Cordelia Langford, John Sillitoe on behalf of the Wellcome Sanger Institute COVID-19 Surveillance Team |
| EPI_ISL_3360957 | Lighthouse Lab in Milton Keynes | Wellcome Sanger Institute for the COVID-19 Genomics UK (COG-UK) Consortium | The Lighthouse Lab in Milton Keynes and Alex Alderton, Roberto Amato, Jeffrey Barrett, Sonia Goncalves, Ewan Harrison, David K. Jackson, Ian Johnston, Dominic Kwiatkowski, Cordelia Langford, John Sillitoe on behalf of the Wellcome Sanger Institute COVID-19 Surveillance Team |
| EPI_ISL_3360959, EPI_ISL_3360960 | Lighthouse Lab in Alderley Park | Wellcome Sanger Institute for the COVID-19 Genomics UK (COG-UK) Consortium | Jacquelyn Wynn, Mairead Hyland, The Lighthouse Lab in Alderley Park and Alex Alderton, Roberto Amato, Jeffrey Barrett, Sonia Goncalves, Ewan Harrison, David K. Jackson, Ian Johnston, Dominic Kwiatkowski, Cordelia Langford, John Sillitoe on behalf of the Wellcome Sanger Institute COVID-19 Surveillance Team |
| EPI_ISL_3360962 | Lighthouse Lab in Milton Keynes | Wellcome Sanger Institute for the COVID-19 Genomics UK (COG-UK) Consortium | The Lighthouse Lab in Milton Keynes and Alex Alderton, Roberto Amato, Jeffrey Barrett, Sonia Goncalves, Ewan Harrison, David K. Jackson, Ian Johnston, Dominic Kwiatkowski, Cordelia Langford, John Sillitoe on behalf of the Wellcome Sanger Institute COVID-19 Surveillance Team |
| EPI_ISL_3360963 | Health Services Laboratories | Wellcome Sanger Institute for the COVID-19 Genomics UK (COG-UK) Consortium | Health Services Laboratories and Alex Alderton, Roberto Amato, Jeffrey Barrett, Sonia Goncalves, Ewan Harrison, David K. Jackson, Ian Johnston, Dominic Kwiatkowski, Cordelia Langford, John Sillitoe on behalf of the Wellcome Sanger Institute COVID-19 Surveillance Team |
| EPI_ISL_3360965 | Lighthouse Lab in Glasgow | Wellcome Sanger Institute for the COVID-19 Genomics UK (COG-UK) Consortium | Harper VanSteenhouse, Yumi Kasai, David Gray, Carol Clugston, Anna Dominiczak and Alex Alderton, Roberto Amato, Jeffrey Barrett, Sonia Goncalves, Ewan Harrison, David K. Jackson, Ian Johnston, Dominic Kwiatkowski, Cordelia Langford, John Sillitoe on behalf of the Wellcome Sanger Institute COVID-19 Surveillance Team |
| EPI_ISL_3360966 | Lighthouse Lab in Alderley Park | Wellcome Sanger Institute for the COVID-19 Genomics UK (COG-UK) Consortium | Jacquelyn Wynn, Mairead Hyland, The Lighthouse Lab in Alderley Park and Alex Alderton, Roberto Amato, Jeffrey Barrett, Sonia Goncalves, Ewan Harrison, David K. Jackson, Ian Johnston, Dominic Kwiatkowski, Cordelia Langford, John Sillitoe on behalf of the Wellcome Sanger Institute COVID-19 Surveillance Team |
| EPI_ISL_3360968 | Health Services Laboratories | Wellcome Sanger Institute for the COVID-19 Genomics UK (COG-UK) Consortium | Health Services Laboratories and Alex Alderton, Roberto Amato, Jeffrey Barrett, Sonia Goncalves, Ewan Harrison, David K. Jackson, Ian Johnston, Dominic Kwiatkowski, Cordelia Langford, John Sillitoe on behalf of the Wellcome Sanger Institute COVID-19 Surveillance Team |
| EPI_ISL_3360969, EPI_ISL_3360971 | Lighthouse Lab in Glasgow | Wellcome Sanger Institute for the COVID-19 Genomics UK (COG-UK) Consortium | Harper VanSteenhouse, Yumi Kasai, David Gray, Carol Clugston, Anna Dominiczak and Alex Alderton, Roberto Amato, Jeffrey Barrett, Sonia Goncalves, Ewan Harrison, David K. Jackson, Ian Johnston, Dominic Kwiatkowski, Cordelia Langford, John Sillitoe on behalf of the Wellcome Sanger Institute COVID-19 Surveillance Team |
| EPI_ISL_3360972 | Lighthouse Lab in Alderley Park | Wellcome Sanger Institute for the COVID-19 Genomics UK (COG-UK) Consortium | Jacquelyn Wynn, Mairead Hyland, The Lighthouse Lab in Alderley Park and Alex Alderton, Roberto Amato, Jeffrey Barrett, Sonia Goncalves, Ewan Harrison, David K. Jackson, Ian Johnston, Dominic Kwiatkowski, Cordelia Langford, John Sillitoe on behalf of the Wellcome Sanger Institute COVID-19 Surveillance Team |
| EPI_ISL_3360973 | Health Services Laboratories | Wellcome Sanger Institute for the COVID-19 Genomics UK (COG-UK) Consortium | Health Services Laboratories and Alex Alderton, Roberto Amato, Jeffrey Barrett, Sonia Goncalves, Ewan Harrison, David K. Jackson, Ian Johnston, Dominic Kwiatkowski, Cordelia Langford, John Sillitoe on behalf of the Wellcome Sanger Institute COVID-19 Surveillance Team |
| EPI_ISL_3360975, EPI_ISL_3360976, EPI_ISL_3360978 | Lighthouse Lab in Alderley Park | Wellcome Sanger Institute for the COVID-19 Genomics UK (COG-UK) Consortium | Jacquelyn Wynn, Mairead Hyland, The Lighthouse Lab in Alderley Park and Alex Alderton, Roberto Amato, Jeffrey Barrett, Sonia Goncalves, Ewan Harrison, David K. Jackson, Ian Johnston, Dominic Kwiatkowski, Cordelia Langford, John Sillitoe on behalf of the Wellcome Sanger Institute COVID-19 Surveillance Team |
| EPI_ISL_3360980 | Lighthouse Lab in Glasgow | Wellcome Sanger Institute for the COVID-19 Genomics UK (COG-UK) Consortium | Harper VanSteenhouse, Yumi Kasai, David Gray, Carol Clugston, Anna Dominiczak and Alex Alderton, Roberto Amato, Jeffrey Barrett, Sonia Goncalves, Ewan Harrison, David K. Jackson, Ian Johnston, Dominic Kwiatkowski, Cordelia Langford, John Sillitoe on behalf of the Wellcome Sanger Institute COVID-19 Surveillance Team |
| EPI_ISL_3360981 | Health Services Laboratories | Wellcome Sanger Institute for the COVID-19 Genomics UK (COG-UK) Consortium | Health Services Laboratories and Alex Alderton, Roberto Amato, Jeffrey Barrett, Sonia Goncalves, Ewan Harrison, David K. Jackson, Ian Johnston, Dominic Kwiatkowski, Cordelia Langford, John Sillitoe on behalf of the Wellcome Sanger Institute COVID-19 Surveillance Team |
| EPI_ISL_3360983, EPI_ISL_3360984 | Lighthouse Lab in Milton Keynes | Wellcome Sanger Institute for the COVID-19 Genomics UK (COG-UK) Consortium | The Lighthouse Lab in Milton Keynes and Alex Alderton, Roberto Amato, Jeffrey Barrett, Sonia Goncalves, Ewan Harrison, David K. Jackson, Ian Johnston, Dominic Kwiatkowski, Cordelia Langford, John Sillitoe on behalf of the Wellcome Sanger Institute COVID-19 Surveillance Team |
| EPI_ISL_3360986, EPI_ISL_3360987 | Lighthouse Lab in Alderley Park | Wellcome Sanger Institute for the COVID-19 Genomics UK (COG-UK) Consortium | Jacquelyn Wynn, Mairead Hyland, The Lighthouse Lab in Alderley Park and Alex Alderton, Roberto Amato, Jeffrey Barrett, Sonia Goncalves, Ewan Harrison, David K. Jackson, Ian Johnston, Dominic Kwiatkowski, Cordelia Langford, John Sillitoe on behalf of the Wellcome Sanger Institute COVID-19 Surveillance Team |
| EPI_ISL_3360989 | Lighthouse Lab in Milton Keynes | Wellcome Sanger Institute for the COVID-19 Genomics UK (COG-UK) Consortium | The Lighthouse Lab in Milton Keynes and Alex Alderton, Roberto Amato, Jeffrey Barrett, Sonia Goncalves, Ewan Harrison, David K. Jackson, Ian Johnston, Dominic Kwiatkowski, Cordelia Langford, John Sillitoe on behalf of the Wellcome Sanger Institute COVID-19 Surveillance Team |
| EPI_ISL_3360990 | Lighthouse Lab in Glasgow | Wellcome Sanger Institute for the COVID-19 Genomics UK (COG-UK) Consortium | Harper VanSteenhouse, Yumi Kasai, David Gray, Carol Clugston, Anna Dominiczak and Alex Alderton, Roberto Amato, Jeffrey Barrett, Sonia Goncalves, Ewan Harrison, David K. Jackson, Ian Johnston, Dominic Kwiatkowski, Cordelia Langford, John Sillitoe on behalf of the Wellcome Sanger Institute COVID-19 Surveillance Team |
| EPI_ISL_3360991, EPI_ISL_3360993 | Lighthouse Lab in Alderley Park | Wellcome Sanger Institute for the COVID-19 Genomics UK (COG-UK) Consortium | Jacquelyn Wynn, Mairead Hyland, The Lighthouse Lab in Alderley Park and Alex Alderton, Roberto Amato, Jeffrey Barrett, Sonia Goncalves, Ewan Harrison, David K. Jackson, Ian Johnston, Dominic Kwiatkowski, Cordelia Langford, John Sillitoe on behalf of the Wellcome Sanger Institute COVID-19 Surveillance Team |
| EPI_ISL_3360994 | Lighthouse Lab in Milton Keynes | Wellcome Sanger Institute for the COVID-19 Genomics UK (COG-UK) Consortium | The Lighthouse Lab in Milton Keynes and Alex Alderton, Roberto Amato, Jeffrey Barrett, Sonia Goncalves, Ewan Harrison, David K. Jackson, Ian Johnston, Dominic Kwiatkowski, Cordelia Langford, John Sillitoe on behalf of the Wellcome Sanger Institute COVID-19 Surveillance Team |
| EPI_ISL_3360996 | Lighthouse Lab in Alderley Park | Wellcome Sanger Institute for the COVID-19 Genomics UK (COG-UK) Consortium | Jacquelyn Wynn, Mairead Hyland, The Lighthouse Lab in Alderley Park and Alex Alderton, Roberto Amato, Jeffrey Barrett, Sonia Goncalves, Ewan Harrison, David K. Jackson, Ian Johnston, Dominic Kwiatkowski, Cordelia Langford, John Sillitoe on behalf of the Wellcome Sanger Institute COVID-19 Surveillance Team |
| EPI_ISL_3360998 | Lighthouse Lab in Milton Keynes | Wellcome Sanger Institute for the COVID-19 Genomics UK (COG-UK) Consortium | The Lighthouse Lab in Milton Keynes and Alex Alderton, Roberto Amato, Jeffrey Barrett, Sonia Goncalves, Ewan Harrison, David K. Jackson, Ian Johnston, Dominic Kwiatkowski, Cordelia Langford, John Sillitoe on behalf of the Wellcome Sanger Institute COVID-19 Surveillance Team |
| EPI_ISL_3360999 | Lighthouse Lab in Alderley Park | Wellcome Sanger Institute for the COVID-19 Genomics UK (COG-UK) Consortium | Jacquelyn Wynn, Mairead Hyland, The Lighthouse Lab in Alderley Park and Alex Alderton, Roberto Amato, Jeffrey Barrett, Sonia Goncalves, Ewan Harrison, David K. Jackson, Ian Johnston, Dominic Kwiatkowski, Cordelia Langford, John Sillitoe on behalf of the Wellcome Sanger Institute COVID-19 Surveillance Team |
| EPI_ISL_3361001 | Health Services Laboratories | Wellcome Sanger Institute for the COVID-19 Genomics UK (COG-UK) Consortium | Health Services Laboratories and Alex Alderton, Roberto Amato, Jeffrey Barrett, Sonia Goncalves, Ewan Harrison, David K. Jackson, Ian Johnston, Dominic Kwiatkowski, Cordelia Langford, John Sillitoe on behalf of the Wellcome Sanger Institute COVID-19 Surveillance Team |
| EPI_ISL_3361002 | Lighthouse Lab in Glasgow | Wellcome Sanger Institute for the COVID-19 Genomics UK (COG-UK) Consortium | Harper VanSteenhouse, Yumi Kasai, David Gray, Carol Clugston, Anna Dominiczak and Alex Alderton, Roberto Amato, Jeffrey Barrett, Sonia Goncalves, Ewan Harrison, David K. Jackson, Ian Johnston, Dominic Kwiatkowski, Cordelia Langford, John Sillitoe on behalf of the Wellcome Sanger Institute COVID-19 Surveillance Team |
| EPI_ISL_3361004 | Lighthouse Lab in Milton Keynes | Wellcome Sanger Institute for the COVID-19 Genomics UK (COG-UK) Consortium | The Lighthouse Lab in Milton Keynes and Alex Alderton, Roberto Amato, Jeffrey Barrett, Sonia Goncalves, Ewan Harrison, David K. Jackson, Ian Johnston, Dominic Kwiatkowski, Cordelia Langford, John Sillitoe on behalf of the Wellcome Sanger Institute COVID-19 Surveillance Team |
| EPI_ISL_3361005 | Health Services Laboratories | Wellcome Sanger Institute for the COVID-19 Genomics UK (COG-UK) Consortium | Health Services Laboratories and Alex Alderton, Roberto Amato, Jeffrey Barrett, Sonia Goncalves, Ewan Harrison, David K. Jackson, Ian Johnston, Dominic Kwiatkowski, Cordelia Langford, John Sillitoe on behalf of the Wellcome Sanger Institute COVID-19 Surveillance Team |
| EPI_ISL_3361007 | Lighthouse Lab in Alderley Park | Wellcome Sanger Institute for the COVID-19 Genomics UK (COG-UK) Consortium | Jacquelyn Wynn, Mairead Hyland, The Lighthouse Lab in Alderley Park and Alex Alderton, Roberto Amato, Jeffrey Barrett, Sonia Goncalves, Ewan Harrison, David K. Jackson, Ian Johnston, Dominic Kwiatkowski, Cordelia Langford, John Sillitoe on behalf of the Wellcome Sanger Institute COVID-19 Surveillance Team |
| EPI_ISL_3361008 | Lighthouse Lab in Milton Keynes | Wellcome Sanger Institute for the COVID-19 Genomics UK (COG-UK) Consortium | The Lighthouse Lab in Milton Keynes and Alex Alderton, Roberto Amato, Jeffrey Barrett, Sonia Goncalves, Ewan Harrison, David K. Jackson, Ian Johnston, Dominic Kwiatkowski, Cordelia Langford, John Sillitoe on behalf of the Wellcome Sanger Institute COVID-19 Surveillance Team |
| EPI_ISL_3361010 | Lighthouse Lab in Glasgow | Wellcome Sanger Institute for the COVID-19 Genomics UK (COG-UK) Consortium | Harper VanSteenhouse, Yumi Kasai, David Gray, Carol Clugston, Anna Dominiczak and Alex Alderton, Roberto Amato, Jeffrey Barrett, Sonia Goncalves, Ewan Harrison, David K. Jackson, Ian Johnston, Dominic Kwiatkowski, Cordelia Langford, John Sillitoe on behalf of the Wellcome Sanger Institute COVID-19 Surveillance Team |
| EPI_ISL_3361011 | Lighthouse Lab in Milton Keynes | Wellcome Sanger Institute for the COVID-19 Genomics UK (COG-UK) Consortium | The Lighthouse Lab in Milton Keynes and Alex Alderton, Roberto Amato, Jeffrey Barrett, Sonia Goncalves, Ewan Harrison, David K. Jackson, Ian Johnston, Dominic Kwiatkowski, Cordelia Langford, John Sillitoe on behalf of the Wellcome Sanger Institute COVID-19 Surveillance Team |
| EPI_ISL_3361013 | Lighthouse Lab in Alderley Park | Wellcome Sanger Institute for the COVID-19 Genomics UK (COG-UK) Consortium | Jacquelyn Wynn, Mairead Hyland, The Lighthouse Lab in Alderley Park and Alex Alderton, Roberto Amato, Jeffrey Barrett, Sonia Goncalves, Ewan Harrison, David K. Jackson, Ian Johnston, Dominic Kwiatkowski, Cordelia Langford, John Sillitoe on behalf of the Wellcome Sanger Institute COVID-19 Surveillance Team |
| EPI_ISL_3361014 | Health Services Laboratories | Wellcome Sanger Institute for the COVID-19 Genomics UK (COG-UK) Consortium | Health Services Laboratories and Alex Alderton, Roberto Amato, Jeffrey Barrett, Sonia Goncalves, Ewan Harrison, David K. Jackson, Ian Johnston, Dominic Kwiatkowski, Cordelia Langford, John Sillitoe on behalf of the Wellcome Sanger Institute COVID-19 Surveillance Team |

| | | | |
|---|---|---|---|
| EPI_ISL_3361016 | Lighthouse Lab in Alderley Park | Wellcome Sanger Institute for the COVID-19 Genomics UK (COG-UK) Consortium | Jacquelyn Wynn, Mairead Hyland, The Lighthouse Lab in Alderley Park and Alex Alderton, Roberto Amato, Jeffrey Barrett, Sonia Goncalves, Ewan Harrison, David K. Jackson, Ian Johnston, Dominic Kwiatkowski, Cordelia Langford, John Sillitoe on behalf of the Wellcome Sanger Institute COVID-19 Surveillance Team |
| EPI_ISL_3361017 | Lighthouse Lab in Glasgow | Wellcome Sanger Institute for the COVID-19 Genomics UK (COG-UK) Consortium | Harper VanSteenhouse, Yumi Kasai, David Gray, Carol Clugston, Anna Dominiczak and Alex Alderton, Roberto Amato, Jeffrey Barrett, Sonia Goncalves, Ewan Harrison, David K. Jackson, Ian Johnston, Dominic Kwiatkowski, Cordelia Langford, John Sillitoe on behalf of the Wellcome Sanger Institute COVID-19 Surveillance Team |
| EPI_ISL_3361019 | Lighthouse Lab in Alderley Park | Wellcome Sanger Institute for the COVID-19 Genomics UK (COG-UK) Consortium | Jacquelyn Wynn, Mairead Hyland, The Lighthouse Lab in Alderley Park and Alex Alderton, Roberto Amato, Jeffrey Barrett, Sonia Goncalves, Ewan Harrison, David K. Jackson, Ian Johnston, Dominic Kwiatkowski, Cordelia Langford, John Sillitoe on behalf of the Wellcome Sanger Institute COVID-19 Surveillance Team |
| EPI_ISL_3361020 | Lighthouse Lab in Glasgow | Wellcome Sanger Institute for the COVID-19 Genomics UK (COG-UK) Consortium | Harper VanSteenhouse, Yumi Kasai, David Gray, Carol Clugston, Anna Dominiczak and Alex Alderton, Roberto Amato, Jeffrey Barrett, Sonia Goncalves, Ewan Harrison, David K. Jackson, Ian Johnston, Dominic Kwiatkowski, Cordelia Langford, John Sillitoe on behalf of the Wellcome Sanger Institute COVID-19 Surveillance Team |
| EPI_ISL_3361022, EPI_ISL_3361024 | Lighthouse Lab in Milton Keynes | Wellcome Sanger Institute for the COVID-19 Genomics UK (COG-UK) Consortium | The Lighthouse Lab in Milton Keynes and Alex Alderton, Roberto Amato, Jeffrey Barrett, Sonia Goncalves, Ewan Harrison, David K. Jackson, Ian Johnston, Dominic Kwiatkowski, Cordelia Langford, John Sillitoe on behalf of the Wellcome Sanger Institute COVID-19 Surveillance Team |
| EPI_ISL_3361025 | Lighthouse Lab in Alderley Park | Wellcome Sanger Institute for the COVID-19 Genomics UK (COG-UK) Consortium | Jacquelyn Wynn, Mairead Hyland, The Lighthouse Lab in Alderley Park and Alex Alderton, Roberto Amato, Jeffrey Barrett, Sonia Goncalves, Ewan Harrison, David K. Jackson, Ian Johnston, Dominic Kwiatkowski, Cordelia Langford, John Sillitoe on behalf of the Wellcome Sanger Institute COVID-19 Surveillance Team |
| EPI_ISL_3361027 | Lighthouse Lab in Glasgow | Wellcome Sanger Institute for the COVID-19 Genomics UK (COG-UK) Consortium | Harper VanSteenhouse, Yumi Kasai, David Gray, Carol Clugston, Anna Dominiczak and Alex Alderton, Roberto Amato, Jeffrey Barrett, Sonia Goncalves, Ewan Harrison, David K. Jackson, Ian Johnston, Dominic Kwiatkowski, Cordelia Langford, John Sillitoe on behalf of the Wellcome Sanger Institute COVID-19 Surveillance Team |
| EPI_ISL_3361029 | Lighthouse Lab in Alderley Park | Wellcome Sanger Institute for the COVID-19 Genomics UK (COG-UK) Consortium | Jacquelyn Wynn, Mairead Hyland, The Lighthouse Lab in Alderley Park and Alex Alderton, Roberto Amato, Jeffrey Barrett, Sonia Goncalves, Ewan Harrison, David K. Jackson, Ian Johnston, Dominic Kwiatkowski, Cordelia Langford, John Sillitoe on behalf of the Wellcome Sanger Institute COVID-19 Surveillance Team |
| EPI_ISL_3361031 | Berkshire and Surrey Pathology Services Lighthouse Laboratory | Wellcome Sanger Institute for the COVID-19 Genomics UK (COG-UK) Consortium | Berkshire and Surrey Pathology Services Lighthouse Laboratory and Alex Alderton, Roberto Amato, Jeffrey Barrett, Sonia Goncalves, Ewan Harrison, David K. Jackson, Ian Johnston, Dominic Kwiatkowski, Cordelia Langford, John Sillitoe on behalf of the Wellcome Sanger Institute COVID-19 Surveillance Team |
| EPI_ISL_3361032 | Lighthouse Lab in Alderley Park | Wellcome Sanger Institute for the COVID-19 Genomics UK (COG-UK) Consortium | Jacquelyn Wynn, Mairead Hyland, The Lighthouse Lab in Alderley Park and Alex Alderton, Roberto Amato, Jeffrey Barrett, Sonia Goncalves, Ewan Harrison, David K. Jackson, Ian Johnston, Dominic Kwiatkowski, Cordelia Langford, John Sillitoe on behalf of the Wellcome Sanger Institute COVID-19 Surveillance Team |
| EPI_ISL_3361034 | Berkshire and Surrey Pathology Services Lighthouse Laboratory | Wellcome Sanger Institute for the COVID-19 Genomics UK (COG-UK) Consortium | Berkshire and Surrey Pathology Services Lighthouse Laboratory and Alex Alderton, Roberto Amato, Jeffrey Barrett, Sonia Goncalves, Ewan Harrison, David K. Jackson, Ian Johnston, Dominic Kwiatkowski, Cordelia Langford, John Sillitoe on behalf of the Wellcome Sanger Institute COVID-19 Surveillance Team |
| EPI_ISL_3361035 | Lighthouse Lab in Milton Keynes | Wellcome Sanger Institute for the COVID-19 Genomics UK (COG-UK) Consortium | The Lighthouse Lab in Milton Keynes and Alex Alderton, Roberto Amato, Jeffrey Barrett, Sonia Goncalves, Ewan Harrison, David K. Jackson, Ian Johnston, Dominic Kwiatkowski, Cordelia Langford, John Sillitoe on behalf of the Wellcome Sanger Institute COVID-19 Surveillance Team |
| EPI_ISL_3361037 | Lighthouse Lab in Alderley Park | Wellcome Sanger Institute for the COVID-19 Genomics UK (COG-UK) Consortium | Jacquelyn Wynn, Mairead Hyland, The Lighthouse Lab in Alderley Park and Alex Alderton, Roberto Amato, Jeffrey Barrett, Sonia Goncalves, Ewan Harrison, David K. Jackson, Ian Johnston, Dominic Kwiatkowski, Cordelia Langford, John Sillitoe on behalf of the Wellcome Sanger Institute COVID-19 Surveillance Team |
| EPI_ISL_3361038 | Lighthouse Lab in Milton Keynes | Wellcome Sanger Institute for the COVID-19 Genomics UK (COG-UK) Consortium | The Lighthouse Lab in Milton Keynes and Alex Alderton, Roberto Amato, Jeffrey Barrett, Sonia Goncalves, Ewan Harrison, David K. Jackson, Ian Johnston, Dominic Kwiatkowski, Cordelia Langford, John Sillitoe on behalf of the Wellcome Sanger Institute COVID-19 Surveillance Team |
| EPI_ISL_3361040 | Lighthouse Lab in Alderley Park | Wellcome Sanger Institute for the COVID-19 Genomics UK (COG-UK) Consortium | Jacquelyn Wynn, Mairead Hyland, The Lighthouse Lab in Alderley Park and Alex Alderton, Roberto Amato, Jeffrey Barrett, Sonia Goncalves, Ewan Harrison, David K. Jackson, Ian Johnston, Dominic Kwiatkowski, Cordelia Langford, John Sillitoe on behalf of the Wellcome Sanger Institute COVID-19 Surveillance Team |
| EPI_ISL_3361042, EPI_ISL_3361043 | Lighthouse Lab in Milton Keynes | Wellcome Sanger Institute for the COVID-19 Genomics UK (COG-UK) Consortium | The Lighthouse Lab in Milton Keynes and Alex Alderton, Roberto Amato, Jeffrey Barrett, Sonia Goncalves, Ewan Harrison, David K. Jackson, Ian Johnston, Dominic Kwiatkowski, Cordelia Langford, John Sillitoe on behalf of the Wellcome Sanger Institute COVID-19 Surveillance Team |
| EPI_ISL_3361044, EPI_ISL_3361046 | Lighthouse Lab in Alderley Park | Wellcome Sanger Institute for the COVID-19 Genomics UK (COG-UK) Consortium | Jacquelyn Wynn, Mairead Hyland, The Lighthouse Lab in Alderley Park and Alex Alderton, Roberto Amato, Jeffrey Barrett, Sonia Goncalves, Ewan Harrison, David K. Jackson, Ian Johnston, Dominic Kwiatkowski, Cordelia Langford, John Sillitoe on behalf of the Wellcome Sanger Institute COVID-19 Surveillance Team |
| EPI_ISL_3361048, EPI_ISL_3361049 | Lighthouse Lab in Milton Keynes | Wellcome Sanger Institute for the COVID-19 Genomics UK (COG-UK) Consortium | The Lighthouse Lab in Milton Keynes and Alex Alderton, Roberto Amato, Jeffrey Barrett, Sonia Goncalves, Ewan Harrison, David K. Jackson, Ian Johnston, Dominic Kwiatkowski, Cordelia Langford, John Sillitoe on behalf of the Wellcome Sanger Institute COVID-19 Surveillance Team |
| EPI_ISL_3361051, EPI_ISL_3361052, EPI_ISL_3361054, EPI_ISL_3361056 | Lighthouse Lab in Alderley Park | Wellcome Sanger Institute for the COVID-19 Genomics UK (COG-UK) Consortium | Jacquelyn Wynn, Mairead Hyland, The Lighthouse Lab in Alderley Park and Alex Alderton, Roberto Amato, Jeffrey Barrett, Sonia Goncalves, Ewan Harrison, David K. Jackson, Ian Johnston, Dominic Kwiatkowski, Cordelia Langford, John Sillitoe on behalf of the Wellcome Sanger Institute COVID-19 Surveillance Team |
| EPI_ISL_3361057 | Berkshire and Surrey Pathology Services Lighthouse Laboratory | Wellcome Sanger Institute for the COVID-19 Genomics UK (COG-UK) Consortium | Berkshire and Surrey Pathology Services Lighthouse Laboratory and Alex Alderton, Roberto Amato, Jeffrey Barrett, Sonia Goncalves, Ewan Harrison, David K. Jackson, Ian Johnston, Dominic Kwiatkowski, Cordelia Langford, John Sillitoe on behalf of the Wellcome Sanger Institute COVID-19 Surveillance Team |
| EPI_ISL_3361059, EPI_ISL_3361060, EPI_ISL_3361062, EPI_ISL_3361063, EPI_ISL_3361065 | Lighthouse Lab in Alderley Park | Wellcome Sanger Institute for the COVID-19 Genomics UK (COG-UK) Consortium | Jacquelyn Wynn, Mairead Hyland, The Lighthouse Lab in Alderley Park and Alex Alderton, Roberto Amato, Jeffrey Barrett, Sonia Goncalves, Ewan Harrison, David K. Jackson, Ian Johnston, Dominic Kwiatkowski, Cordelia Langford, John Sillitoe on behalf of the Wellcome Sanger Institute COVID-19 Surveillance Team |
| EPI_ISL_3361066 | Lighthouse Lab in Milton Keynes | Wellcome Sanger Institute for the COVID-19 Genomics UK (COG-UK) Consortium | The Lighthouse Lab in Milton Keynes and Alex Alderton, Roberto Amato, Jeffrey Barrett, Sonia Goncalves, Ewan Harrison, David K. Jackson, Ian Johnston, Dominic Kwiatkowski, Cordelia Langford, John Sillitoe on behalf of the Wellcome Sanger Institute COVID-19 Surveillance Team |
| EPI_ISL_3361068, EPI_ISL_3361069 | Lighthouse Lab in Alderley Park | Wellcome Sanger Institute for the COVID-19 Genomics UK (COG-UK) Consortium | Jacquelyn Wynn, Mairead Hyland, The Lighthouse Lab in Alderley Park and Alex Alderton, Roberto Amato, Jeffrey Barrett, Sonia Goncalves, Ewan Harrison, David K. Jackson, Ian Johnston, Dominic Kwiatkowski, Cordelia Langford, John Sillitoe on behalf of the Wellcome Sanger Institute COVID-19 Surveillance Team |
| EPI_ISL_3361071 | Lighthouse Lab in Milton Keynes | Wellcome Sanger Institute for the COVID-19 Genomics UK (COG-UK) Consortium | The Lighthouse Lab in Milton Keynes and Alex Alderton, Roberto Amato, Jeffrey Barrett, Sonia Goncalves, Ewan Harrison, David K. Jackson, Ian Johnston, Dominic Kwiatkowski, Cordelia Langford, John Sillitoe on behalf of the Wellcome Sanger Institute COVID-19 Surveillance Team |
| EPI_ISL_3361072 | Lighthouse Lab in Alderley Park | Wellcome Sanger Institute for the COVID-19 Genomics UK (COG-UK) Consortium | Jacquelyn Wynn, Mairead Hyland, The Lighthouse Lab in Alderley Park and Alex Alderton, Roberto Amato, Jeffrey Barrett, Sonia Goncalves, Ewan Harrison, David K. Jackson, Ian Johnston, Dominic Kwiatkowski, Cordelia Langford, John Sillitoe on behalf of the Wellcome Sanger Institute COVID-19 Surveillance Team |
| EPI_ISL_3361074 | Lighthouse Lab in Milton Keynes | Wellcome Sanger Institute for the COVID-19 Genomics UK (COG-UK) Consortium | The Lighthouse Lab in Milton Keynes and Alex Alderton, Roberto Amato, Jeffrey Barrett, Sonia Goncalves, Ewan Harrison, David K. Jackson, Ian Johnston, Dominic Kwiatkowski, Cordelia Langford, John Sillitoe on behalf of the Wellcome Sanger Institute COVID-19 Surveillance Team |
| EPI_ISL_3361075, EPI_ISL_3361076 | Lighthouse Lab in Alderley Park | Wellcome Sanger Institute for the COVID-19 Genomics UK (COG-UK) Consortium | Jacquelyn Wynn, Mairead Hyland, The Lighthouse Lab in Alderley Park and Alex Alderton, Roberto Amato, Jeffrey Barrett, Sonia Goncalves, Ewan Harrison, David K. Jackson, Ian Johnston, Dominic Kwiatkowski, Cordelia Langford, John Sillitoe on behalf of the Wellcome Sanger Institute COVID-19 Surveillance Team |
| EPI_ISL_3361078, EPI_ISL_3361080 | Lighthouse Lab in Milton Keynes | Wellcome Sanger Institute for the COVID-19 Genomics UK (COG-UK) Consortium | The Lighthouse Lab in Milton Keynes and Alex Alderton, Roberto Amato, Jeffrey Barrett, Sonia Goncalves, Ewan Harrison, David K. Jackson, Ian Johnston, Dominic Kwiatkowski, Cordelia Langford, John Sillitoe on behalf of the Wellcome Sanger Institute COVID-19 Surveillance Team |
| EPI_ISL_3361081, EPI_ISL_3361083 | Lighthouse Lab in Alderley Park | Wellcome Sanger Institute for the COVID-19 Genomics UK (COG-UK) Consortium | Jacquelyn Wynn, Mairead Hyland, The Lighthouse Lab in Alderley Park and Alex Alderton, Roberto Amato, Jeffrey Barrett, Sonia Goncalves, Ewan Harrison, David K. Jackson, Ian Johnston, Dominic Kwiatkowski, Cordelia Langford, John Sillitoe on behalf of the Wellcome Sanger Institute COVID-19 Surveillance Team |
| EPI_ISL_3361084, EPI_ISL_3361086 | Berkshire and Surrey Pathology Services Lighthouse Laboratory | Wellcome Sanger Institute for the COVID-19 Genomics UK (COG-UK) Consortium | Berkshire and Surrey Pathology Services Lighthouse Laboratory and Alex Alderton, Roberto Amato, Jeffrey Barrett, Sonia Goncalves, Ewan Harrison, David K. Jackson, Ian Johnston, Dominic Kwiatkowski, Cordelia Langford, John Sillitoe on behalf of the Wellcome Sanger Institute COVID-19 Surveillance Team |
| EPI_ISL_3361087 | Lighthouse Lab in Milton Keynes | Wellcome Sanger Institute for the COVID-19 Genomics UK (COG-UK) Consortium | The Lighthouse Lab in Milton Keynes and Alex Alderton, Roberto Amato, Jeffrey Barrett, Sonia Goncalves, Ewan Harrison, David K. Jackson, Ian Johnston, Dominic Kwiatkowski, Cordelia Langford, John Sillitoe on behalf of the Wellcome Sanger Institute COVID-19 Surveillance Team |
| EPI_ISL_3361088, EPI_ISL_3361089, EPI_ISL_3361091 | Lighthouse Lab in Alderley Park | Wellcome Sanger Institute for the COVID-19 Genomics UK (COG-UK) Consortium | Jacquelyn Wynn, Mairead Hyland, The Lighthouse Lab in Alderley Park and Alex Alderton, Roberto Amato, Jeffrey Barrett, Sonia Goncalves, Ewan Harrison, David K. Jackson, Ian Johnston, Dominic Kwiatkowski, Cordelia Langford, John Sillitoe on behalf of the Wellcome Sanger Institute COVID-19 Surveillance Team |
| EPI_ISL_3361092 | Lighthouse Lab in Milton Keynes | Wellcome Sanger Institute for the COVID-19 Genomics UK (COG-UK) Consortium | The Lighthouse Lab in Milton Keynes and Alex Alderton, Roberto Amato, Jeffrey Barrett, Sonia Goncalves, Ewan Harrison, David K. Jackson, Ian Johnston, Dominic Kwiatkowski, Cordelia Langford, John Sillitoe on behalf of the Wellcome Sanger Institute COVID-19 Surveillance Team |
| EPI_ISL_3361094, EPI_ISL_3361095, EPI_ISL_3361098, EPI_ISL_3361099, EPI_ISL_3361101 | Lighthouse Lab in Alderley Park | Wellcome Sanger Institute for the COVID-19 Genomics UK (COG-UK) Consortium | Jacquelyn Wynn, Mairead Hyland, The Lighthouse Lab in Alderley Park and Alex Alderton, Roberto Amato, Jeffrey Barrett, Sonia Goncalves, Ewan Harrison, David K. Jackson, Ian Johnston, Dominic Kwiatkowski, Cordelia Langford, John Sillitoe on behalf of the Wellcome Sanger Institute COVID-19 Surveillance Team |
| EPI_ISL_3361102, EPI_ISL_3361104 | Lighthouse Lab in Milton Keynes | Wellcome Sanger Institute for the COVID-19 Genomics UK (COG-UK) Consortium | The Lighthouse Lab in Milton Keynes and Alex Alderton, Roberto Amato, Jeffrey Barrett, Sonia Goncalves, Ewan Harrison, David K. Jackson, Ian Johnston, Dominic Kwiatkowski, Cordelia Langford, John Sillitoe on behalf of the Wellcome Sanger Institute COVID-19 Surveillance Team |
| EPI_ISL_3361105, EPI_ISL_3361106 | Lighthouse Lab in Alderley Park | Wellcome Sanger Institute for the COVID-19 Genomics UK (COG-UK) Consortium | Jacquelyn Wynn, Mairead Hyland, The Lighthouse Lab in Alderley Park and Alex Alderton, Roberto Amato, Jeffrey Barrett, Sonia Goncalves, Ewan Harrison, David K. Jackson, Ian Johnston, Dominic Kwiatkowski, Cordelia Langford, John Sillitoe on behalf of the Wellcome Sanger Institute COVID-19 Surveillance Team |
| EPI_ISL_3361107, EPI_ISL_3361108 | Lighthouse Lab in Milton Keynes | Wellcome Sanger Institute for the COVID-19 Genomics UK (COG-UK) Consortium | The Lighthouse Lab in Milton Keynes and Alex Alderton, Roberto Amato, Jeffrey Barrett, Sonia Goncalves, Ewan Harrison, David K. Jackson, Ian Johnston, Dominic Kwiatkowski, Cordelia Langford, John Sillitoe on behalf of the Wellcome Sanger Institute COVID-19 Surveillance Team |
| EPI_ISL_3361109, EPI_ISL_3361111 | Lighthouse Lab in Alderley Park | Wellcome Sanger Institute for the COVID-19 Genomics UK (COG-UK) Consortium | Jacquelyn Wynn, Mairead Hyland, The Lighthouse Lab in Alderley Park and Alex Alderton, Roberto Amato, Jeffrey Barrett, Sonia Goncalves, Ewan Harrison, David K. Jackson, Ian Johnston, Dominic Kwiatkowski, Cordelia Langford, John Sillitoe on behalf of the Wellcome Sanger Institute COVID-19 Surveillance Team |
| EPI_ISL_3361112 | Berkshire and Surrey Pathology Services Lighthouse Laboratory | Wellcome Sanger Institute for the COVID-19 Genomics UK (COG-UK) Consortium | Berkshire and Surrey Pathology Services Lighthouse Laboratory and Alex Alderton, Roberto Amato, Jeffrey Barrett, Sonia Goncalves, Ewan Harrison, David K. Jackson, Ian Johnston, Dominic Kwiatkowski, Cordelia Langford, John Sillitoe on behalf of the Wellcome Sanger Institute COVID-19 Surveillance Team |
| EPI_ISL_3361114, EPI_ISL_3361115 | Lighthouse Lab in Alderley Park | Wellcome Sanger Institute for the COVID-19 Genomics UK (COG-UK) Consortium | Jacquelyn Wynn, Mairead Hyland, The Lighthouse Lab in Alderley Park and Alex Alderton, Roberto Amato, Jeffrey Barrett, Sonia Goncalves, Ewan Harrison, David K. Jackson, Ian Johnston, Dominic Kwiatkowski, Cordelia Langford, John Sillitoe on behalf of the Wellcome Sanger Institute COVID-19 Surveillance Team |
| EPI_ISL_3361117, EPI_ISL_3361118 | Lighthouse Lab in Milton Keynes | Wellcome Sanger Institute for the COVID-19 Genomics UK (COG-UK) Consortium | The Lighthouse Lab in Milton Keynes and Alex Alderton, Roberto Amato, Jeffrey Barrett, Sonia Goncalves, Ewan Harrison, David K. Jackson, Ian Johnston, Dominic Kwiatkowski, Cordelia Langford, John Sillitoe on behalf of the Wellcome Sanger Institute COVID-19 Surveillance Team |
| EPI_ISL_3361120 | Lighthouse Lab in Alderley Park | Wellcome Sanger Institute for the COVID-19 Genomics UK (COG-UK) Consortium | Jacquelyn Wynn, Mairead Hyland, The Lighthouse Lab in Alderley Park and Alex Alderton, Roberto Amato, Jeffrey Barrett, Sonia Goncalves, Ewan Harrison, David K. Jackson, Ian Johnston, Dominic Kwiatkowski, Cordelia Langford, John Sillitoe on |

| | | | |
|---|---|---|---|
| | | Consortium | behalf of the Wellcome Sanger Institute COVID-19 Surveillance Team |
| EPI_ISL_3361121 | Lighthouse Lab in Milton Keynes | Wellcome Sanger Institute for the COVID-19 Genomics UK (COG-UK) Consortium | The Lighthouse Lab in Milton Keynes and Alex Alderton, Roberto Amato, Jeffrey Barrett, Sonia Goncalves, Ewan Harrison, David K. Jackson, Ian Johnston, Dominic Kwiatkowski, Cordelia Langford, John Sillitoe on behalf of the Wellcome Sanger Institute COVID-19 Surveillance Team |
| EPI_ISL_3361123, EPI_ISL_3361124, EPI_ISL_3361126 | Lighthouse Lab in Alderley Park | Wellcome Sanger Institute for the COVID-19 Genomics UK (COG-UK) Consortium | Jacquelyn Wynn, Mairead Hyland, The Lighthouse Lab in Alderley Park and Alex Alderton, Roberto Amato, Jeffrey Barrett, Sonia Goncalves, Ewan Harrison, David K. Jackson, Ian Johnston, Dominic Kwiatkowski, Cordelia Langford, John Sillitoe on behalf of the Wellcome Sanger Institute COVID-19 Surveillance Team |
| EPI_ISL_3361127, EPI_ISL_3361129 | Berkshire and Surrey Pathology Services Lighthouse Laboratory | Wellcome Sanger Institute for the COVID-19 Genomics UK (COG-UK) Consortium | Berkshire and Surrey Pathology Services Lighthouse Laboratory and Alex Alderton, Roberto Amato, Jeffrey Barrett, Sonia Goncalves, Ewan Harrison, David K. Jackson, Ian Johnston, Dominic Kwiatkowski, Cordelia Langford, John Sillitoe on behalf of the Wellcome Sanger Institute COVID-19 Surveillance Team |
| EPI_ISL_3361130, EPI_ISL_3361131, EPI_ISL_3361132 | Lighthouse Lab in Alderley Park | Wellcome Sanger Institute for the COVID-19 Genomics UK (COG-UK) Consortium | Jacquelyn Wynn, Mairead Hyland, The Lighthouse Lab in Alderley Park and Alex Alderton, Roberto Amato, Jeffrey Barrett, Sonia Goncalves, Ewan Harrison, David K. Jackson, Ian Johnston, Dominic Kwiatkowski, Cordelia Langford, John Sillitoe on behalf of the Wellcome Sanger Institute COVID-19 Surveillance Team |
| EPI_ISL_3361134 | Lighthouse Lab in Milton Keynes | Wellcome Sanger Institute for the COVID-19 Genomics UK (COG-UK) Consortium | The Lighthouse Lab in Milton Keynes and Alex Alderton, Roberto Amato, Jeffrey Barrett, Sonia Goncalves, Ewan Harrison, David K. Jackson, Ian Johnston, Dominic Kwiatkowski, Cordelia Langford, John Sillitoe on behalf of the Wellcome Sanger Institute COVID-19 Surveillance Team |
| EPI_ISL_3361135, EPI_ISL_3361137, EPI_ISL_3361138, EPI_ISL_3361140 | Lighthouse Lab in Alderley Park | Wellcome Sanger Institute for the COVID-19 Genomics UK (COG-UK) Consortium | Jacquelyn Wynn, Mairead Hyland, The Lighthouse Lab in Alderley Park and Alex Alderton, Roberto Amato, Jeffrey Barrett, Sonia Goncalves, Ewan Harrison, David K. Jackson, Ian Johnston, Dominic Kwiatkowski, Cordelia Langford, John Sillitoe on behalf of the Wellcome Sanger Institute COVID-19 Surveillance Team |
| EPI_ISL_3361141 | Berkshire and Surrey Pathology Services Lighthouse Laboratory | Wellcome Sanger Institute for the COVID-19 Genomics UK (COG-UK) Consortium | Berkshire and Surrey Pathology Services Lighthouse Laboratory and Alex Alderton, Roberto Amato, Jeffrey Barrett, Sonia Goncalves, Ewan Harrison, David K. Jackson, Ian Johnston, Dominic Kwiatkowski, Cordelia Langford, John Sillitoe on behalf of the Wellcome Sanger Institute COVID-19 Surveillance Team |
| EPI_ISL_3361142 | Lighthouse Lab in Alderley Park | Wellcome Sanger Institute for the COVID-19 Genomics UK (COG-UK) Consortium | Jacquelyn Wynn, Mairead Hyland, The Lighthouse Lab in Alderley Park and Alex Alderton, Roberto Amato, Jeffrey Barrett, Sonia Goncalves, Ewan Harrison, David K. Jackson, Ian Johnston, Dominic Kwiatkowski, Cordelia Langford, John Sillitoe on behalf of the Wellcome Sanger Institute COVID-19 Surveillance Team |
| EPI_ISL_3361144 | Lighthouse Lab in Milton Keynes | Wellcome Sanger Institute for the COVID-19 Genomics UK (COG-UK) Consortium | The Lighthouse Lab in Milton Keynes and Alex Alderton, Roberto Amato, Jeffrey Barrett, Sonia Goncalves, Ewan Harrison, David K. Jackson, Ian Johnston, Dominic Kwiatkowski, Cordelia Langford, John Sillitoe on behalf of the Wellcome Sanger Institute COVID-19 Surveillance Team |
| EPI_ISL_3361146, EPI_ISL_3361147 | Lighthouse Lab in Alderley Park | Wellcome Sanger Institute for the COVID-19 Genomics UK (COG-UK) Consortium | Jacquelyn Wynn, Mairead Hyland, The Lighthouse Lab in Alderley Park and Alex Alderton, Roberto Amato, Jeffrey Barrett, Sonia Goncalves, Ewan Harrison, David K. Jackson, Ian Johnston, Dominic Kwiatkowski, Cordelia Langford, John Sillitoe on behalf of the Wellcome Sanger Institute COVID-19 Surveillance Team |
| EPI_ISL_3361148, EPI_ISL_3361150 | Lighthouse Lab in Milton Keynes | Wellcome Sanger Institute for the COVID-19 Genomics UK (COG-UK) Consortium | The Lighthouse Lab in Milton Keynes and Alex Alderton, Roberto Amato, Jeffrey Barrett, Sonia Goncalves, Ewan Harrison, David K. Jackson, Ian Johnston, Dominic Kwiatkowski, Cordelia Langford, John Sillitoe on behalf of the Wellcome Sanger Institute COVID-19 Surveillance Team |
| EPI_ISL_3361151, EPI_ISL_3361152 | Lighthouse Lab in Alderley Park | Wellcome Sanger Institute for the COVID-19 Genomics UK (COG-UK) Consortium | Jacquelyn Wynn, Mairead Hyland, The Lighthouse Lab in Alderley Park and Alex Alderton, Roberto Amato, Jeffrey Barrett, Sonia Goncalves, Ewan Harrison, David K. Jackson, Ian Johnston, Dominic Kwiatkowski, Cordelia Langford, John Sillitoe on behalf of the Wellcome Sanger Institute COVID-19 Surveillance Team |
| EPI_ISL_3361154 | Berkshire and Surrey Pathology Services Lighthouse Laboratory | Wellcome Sanger Institute for the COVID-19 Genomics UK (COG-UK) Consortium | Berkshire and Surrey Pathology Services Lighthouse Laboratory and Alex Alderton, Roberto Amato, Jeffrey Barrett, Sonia Goncalves, Ewan Harrison, David K. Jackson, Ian Johnston, Dominic Kwiatkowski, Cordelia Langford, John Sillitoe on behalf of the Wellcome Sanger Institute COVID-19 Surveillance Team |
| EPI_ISL_3361155, EPI_ISL_3361157, EPI_ISL_3361158, EPI_ISL_3361159, EPI_ISL_3361161, EPI_ISL_3361162 | Lighthouse Lab in Alderley Park | Wellcome Sanger Institute for the COVID-19 Genomics UK (COG-UK) Consortium | Jacquelyn Wynn, Mairead Hyland, The Lighthouse Lab in Alderley Park and Alex Alderton, Roberto Amato, Jeffrey Barrett, Sonia Goncalves, Ewan Harrison, David K. Jackson, Ian Johnston, Dominic Kwiatkowski, Cordelia Langford, John Sillitoe on behalf of the Wellcome Sanger Institute COVID-19 Surveillance Team |
| EPI_ISL_3361164, EPI_ISL_3361165 | Lighthouse Lab in Milton Keynes | Wellcome Sanger Institute for the COVID-19 Genomics UK (COG-UK) Consortium | The Lighthouse Lab in Milton Keynes and Alex Alderton, Roberto Amato, Jeffrey Barrett, Sonia Goncalves, Ewan Harrison, David K. Jackson, Ian Johnston, Dominic Kwiatkowski, Cordelia Langford, John Sillitoe on behalf of the Wellcome Sanger Institute COVID-19 Surveillance Team |
| EPI_ISL_3361167 | Berkshire and Surrey Pathology Services Lighthouse Laboratory | Wellcome Sanger Institute for the COVID-19 Genomics UK (COG-UK) Consortium | Berkshire and Surrey Pathology Services Lighthouse Laboratory and Alex Alderton, Roberto Amato, Jeffrey Barrett, Sonia Goncalves, Ewan Harrison, David K. Jackson, Ian Johnston, Dominic Kwiatkowski, Cordelia Langford, John Sillitoe on behalf of the Wellcome Sanger Institute COVID-19 Surveillance Team |
| EPI_ISL_3361168, EPI_ISL_3361170, EPI_ISL_3361171 | Lighthouse Lab in Alderley Park | Wellcome Sanger Institute for the COVID-19 Genomics UK (COG-UK) Consortium | Jacquelyn Wynn, Mairead Hyland, The Lighthouse Lab in Alderley Park and Alex Alderton, Roberto Amato, Jeffrey Barrett, Sonia Goncalves, Ewan Harrison, David K. Jackson, Ian Johnston, Dominic Kwiatkowski, Cordelia Langford, John Sillitoe on behalf of the Wellcome Sanger Institute COVID-19 Surveillance Team |
| EPI_ISL_3361172 | Lighthouse Lab in Milton Keynes | Wellcome Sanger Institute for the COVID-19 Genomics UK (COG-UK) Consortium | The Lighthouse Lab in Milton Keynes and Alex Alderton, Roberto Amato, Jeffrey Barrett, Sonia Goncalves, Ewan Harrison, David K. Jackson, Ian Johnston, Dominic Kwiatkowski, Cordelia Langford, John Sillitoe on behalf of the Wellcome Sanger Institute COVID-19 Surveillance Team |
| EPI_ISL_3361174 | Lighthouse Lab in Alderley Park | Wellcome Sanger Institute for the COVID-19 Genomics UK (COG-UK) Consortium | Jacquelyn Wynn, Mairead Hyland, The Lighthouse Lab in Alderley Park and Alex Alderton, Roberto Amato, Jeffrey Barrett, Sonia Goncalves, Ewan Harrison, David K. Jackson, Ian Johnston, Dominic Kwiatkowski, Cordelia Langford, John Sillitoe on behalf of the Wellcome Sanger Institute COVID-19 Surveillance Team |
| EPI_ISL_3361175 | Lighthouse Lab in Milton Keynes | Wellcome Sanger Institute for the COVID-19 Genomics UK (COG-UK) Consortium | The Lighthouse Lab in Milton Keynes and Alex Alderton, Roberto Amato, Jeffrey Barrett, Sonia Goncalves, Ewan Harrison, David K. Jackson, Ian Johnston, Dominic Kwiatkowski, Cordelia Langford, John Sillitoe on behalf of the Wellcome Sanger Institute COVID-19 Surveillance Team |
| EPI_ISL_3361177 | Lighthouse Lab in Alderley Park | Wellcome Sanger Institute for the COVID-19 Genomics UK (COG-UK) Consortium | Jacquelyn Wynn, Mairead Hyland, The Lighthouse Lab in Alderley Park and Alex Alderton, Roberto Amato, Jeffrey Barrett, Sonia Goncalves, Ewan Harrison, David K. Jackson, Ian Johnston, Dominic Kwiatkowski, Cordelia Langford, John Sillitoe on behalf of the Wellcome Sanger Institute COVID-19 Surveillance Team |
| EPI_ISL_3361178 | Lighthouse Lab in Milton Keynes | Wellcome Sanger Institute for the COVID-19 Genomics UK (COG-UK) Consortium | The Lighthouse Lab in Milton Keynes and Alex Alderton, Roberto Amato, Jeffrey Barrett, Sonia Goncalves, Ewan Harrison, David K. Jackson, Ian Johnston, Dominic Kwiatkowski, Cordelia Langford, John Sillitoe on behalf of the Wellcome Sanger Institute COVID-19 Surveillance Team |
| EPI_ISL_3361180 | Lighthouse Lab in Alderley Park | Wellcome Sanger Institute for the COVID-19 Genomics UK (COG-UK) Consortium | Jacquelyn Wynn, Mairead Hyland, The Lighthouse Lab in Alderley Park and Alex Alderton, Roberto Amato, Jeffrey Barrett, Sonia Goncalves, Ewan Harrison, David K. Jackson, Ian Johnston, Dominic Kwiatkowski, Cordelia Langford, John Sillitoe on behalf of the Wellcome Sanger Institute COVID-19 Surveillance Team |
| EPI_ISL_3361182 | Berkshire and Surrey Pathology Services Lighthouse Laboratory | Wellcome Sanger Institute for the COVID-19 Genomics UK (COG-UK) Consortium | Berkshire and Surrey Pathology Services Lighthouse Laboratory and Alex Alderton, Roberto Amato, Jeffrey Barrett, Sonia Goncalves, Ewan Harrison, David K. Jackson, Ian Johnston, Dominic Kwiatkowski, Cordelia Langford, John Sillitoe on behalf of the Wellcome Sanger Institute COVID-19 Surveillance Team |
| EPI_ISL_3361183, EPI_ISL_3361184, EPI_ISL_3361186, EPI_ISL_3361187, EPI_ISL_3361189, EPI_ISL_3361190, EPI_ISL_3361192 | Lighthouse Lab in Alderley Park | Wellcome Sanger Institute for the COVID-19 Genomics UK (COG-UK) Consortium | Jacquelyn Wynn, Mairead Hyland, The Lighthouse Lab in Alderley Park and Alex Alderton, Roberto Amato, Jeffrey Barrett, Sonia Goncalves, Ewan Harrison, David K. Jackson, Ian Johnston, Dominic Kwiatkowski, Cordelia Langford, John Sillitoe on behalf of the Wellcome Sanger Institute COVID-19 Surveillance Team |
| EPI_ISL_3361193 | Berkshire and Surrey Pathology Services Lighthouse Laboratory | Wellcome Sanger Institute for the COVID-19 Genomics UK (COG-UK) Consortium | Berkshire and Surrey Pathology Services Lighthouse Laboratory and Alex Alderton, Roberto Amato, Jeffrey Barrett, Sonia Goncalves, Ewan Harrison, David K. Jackson, Ian Johnston, Dominic Kwiatkowski, Cordelia Langford, John Sillitoe on behalf of the Wellcome Sanger Institute COVID-19 Surveillance Team |
| EPI_ISL_3361194, EPI_ISL_3361196 | Lighthouse Lab in Alderley Park | Wellcome Sanger Institute for the COVID-19 Genomics UK (COG-UK) Consortium | Jacquelyn Wynn, Mairead Hyland, The Lighthouse Lab in Alderley Park and Alex Alderton, Roberto Amato, Jeffrey Barrett, Sonia Goncalves, Ewan Harrison, David K. Jackson, Ian Johnston, Dominic Kwiatkowski, Cordelia Langford, John Sillitoe on behalf of the Wellcome Sanger Institute COVID-19 Surveillance Team |
| EPI_ISL_3361197 | Lighthouse Lab in Milton Keynes | Wellcome Sanger Institute for the COVID-19 Genomics UK (COG-UK) Consortium | The Lighthouse Lab in Milton Keynes and Alex Alderton, Roberto Amato, Jeffrey Barrett, Sonia Goncalves, Ewan Harrison, David K. Jackson, Ian Johnston, Dominic Kwiatkowski, Cordelia Langford, John Sillitoe on behalf of the Wellcome Sanger Institute COVID-19 Surveillance Team |
| EPI_ISL_3361199, EPI_ISL_3361200 | Lighthouse Lab in Alderley Park | Wellcome Sanger Institute for the COVID-19 Genomics UK (COG-UK) Consortium | Jacquelyn Wynn, Mairead Hyland, The Lighthouse Lab in Alderley Park and Alex Alderton, Roberto Amato, Jeffrey Barrett, Sonia Goncalves, Ewan Harrison, David K. Jackson, Ian Johnston, Dominic Kwiatkowski, Cordelia Langford, John Sillitoe on behalf of the Wellcome Sanger Institute COVID-19 Surveillance Team |
| EPI_ISL_3361202 | Lighthouse Lab in Milton Keynes | Wellcome Sanger Institute for the COVID-19 Genomics UK (COG-UK) Consortium | The Lighthouse Lab in Milton Keynes and Alex Alderton, Roberto Amato, Jeffrey Barrett, Sonia Goncalves, Ewan Harrison, David K. Jackson, Ian Johnston, Dominic Kwiatkowski, Cordelia Langford, John Sillitoe on behalf of the Wellcome Sanger Institute COVID-19 Surveillance Team |
| EPI_ISL_3361203 | Lighthouse Lab in Alderley Park | Wellcome Sanger Institute for the COVID-19 Genomics UK (COG-UK) Consortium | Jacquelyn Wynn, Mairead Hyland, The Lighthouse Lab in Alderley Park and Alex Alderton, Roberto Amato, Jeffrey Barrett, Sonia Goncalves, Ewan Harrison, David K. Jackson, Ian Johnston, Dominic Kwiatkowski, Cordelia Langford, John Sillitoe on behalf of the Wellcome Sanger Institute COVID-19 Surveillance Team |
| EPI_ISL_3361205 | Lighthouse Lab in Milton Keynes | Wellcome Sanger Institute for the COVID-19 Genomics UK (COG-UK) Consortium | The Lighthouse Lab in Milton Keynes and Alex Alderton, Roberto Amato, Jeffrey Barrett, Sonia Goncalves, Ewan Harrison, David K. Jackson, Ian Johnston, Dominic Kwiatkowski, Cordelia Langford, John Sillitoe on behalf of the Wellcome Sanger Institute COVID-19 Surveillance Team |
| EPI_ISL_3361206 | Lighthouse Lab in Alderley Park | Wellcome Sanger Institute for the COVID-19 Genomics UK (COG-UK) Consortium | Jacquelyn Wynn, Mairead Hyland, The Lighthouse Lab in Alderley Park and Alex Alderton, Roberto Amato, Jeffrey Barrett, Sonia Goncalves, Ewan Harrison, David K. Jackson, Ian Johnston, Dominic Kwiatkowski, Cordelia Langford, John Sillitoe on behalf of the Wellcome Sanger Institute COVID-19 Surveillance Team |
| EPI_ISL_3361208 | Berkshire and Surrey Pathology Services Lighthouse Laboratory | Wellcome Sanger Institute for the COVID-19 Genomics UK (COG-UK) Consortium | Berkshire and Surrey Pathology Services Lighthouse Laboratory and Alex Alderton, Roberto Amato, Jeffrey Barrett, Sonia Goncalves, Ewan Harrison, David K. Jackson, Ian Johnston, Dominic Kwiatkowski, Cordelia Langford, John Sillitoe on behalf of the Wellcome Sanger Institute COVID-19 Surveillance Team |
| EPI_ISL_3361209 | Lighthouse Lab in Milton Keynes | Wellcome Sanger Institute for the COVID-19 Genomics UK (COG-UK) Consortium | The Lighthouse Lab in Milton Keynes and Alex Alderton, Roberto Amato, Jeffrey Barrett, Sonia Goncalves, Ewan Harrison, David K. Jackson, Ian Johnston, Dominic Kwiatkowski, Cordelia Langford, John Sillitoe on behalf of the Wellcome Sanger Institute COVID-19 Surveillance Team |
| EPI_ISL_3361211, EPI_ISL_3361212 | Lighthouse Lab in Alderley Park | Wellcome Sanger Institute for the COVID-19 Genomics UK (COG-UK) Consortium | Jacquelyn Wynn, Mairead Hyland, The Lighthouse Lab in Alderley Park and Alex Alderton, Roberto Amato, Jeffrey Barrett, Sonia Goncalves, Ewan Harrison, David K. Jackson, Ian Johnston, Dominic Kwiatkowski, Cordelia Langford, John Sillitoe on behalf of the Wellcome Sanger Institute COVID-19 Surveillance Team |
| EPI_ISL_3361214 | Berkshire and Surrey Pathology Services Lighthouse Laboratory | Wellcome Sanger Institute for the COVID-19 Genomics UK (COG-UK) Consortium | Berkshire and Surrey Pathology Services Lighthouse Laboratory and Alex Alderton, Roberto Amato, Jeffrey Barrett, Sonia Goncalves, Ewan Harrison, David K. Jackson, Ian Johnston, Dominic Kwiatkowski, Cordelia Langford, John Sillitoe on behalf of the Wellcome Sanger Institute COVID-19 Surveillance Team |
| EPI_ISL_3361215 | Lighthouse Lab in Milton Keynes | Wellcome Sanger Institute for the COVID-19 Genomics UK (COG-UK) Consortium | The Lighthouse Lab in Milton Keynes and Alex Alderton, Roberto Amato, Jeffrey Barrett, Sonia Goncalves, Ewan Harrison, David K. Jackson, Ian Johnston, Dominic Kwiatkowski, Cordelia Langford, John Sillitoe on behalf of the Wellcome Sanger Institute COVID-19 Surveillance Team |
| EPI_ISL_3361217, EPI_ISL_3361218, | Lighthouse Lab in Alderley Park | Wellcome Sanger Institute for the COVID-19 Genomics UK (COG-UK) Consortium | Jacquelyn Wynn, Mairead Hyland, The Lighthouse Lab in Alderley Park and Alex Alderton, Roberto Amato, Jeffrey Barrett, Sonia Goncalves, Ewan Harrison, David K. Jackson, Ian Johnston, Dominic Kwiatkowski, Cordelia Langford, John Sillitoe on behalf of the Wellcome Sanger Institute COVID-19 Surveillance Team |

| | | | |
|---|---|---|---|
| EPI_ISL_3361220, EPI_ISL_3361221, EPI_ISL_3361223 | | | |
| EPI_ISL_3361224 | Berkshire and Surrey Pathology Services Lighthouse Laboratory | Wellcome Sanger Institute for the COVID-19 Genomics UK (COG-UK) Consortium | Berkshire and Surrey Pathology Services Lighthouse Laboratory and Alex Alderton, Roberto Amato, Jeffrey Barrett, Sonia Goncalves, Ewan Harrison, David K. Jackson, Ian Johnston, Dominic Kwiatkowski, Cordelia Langford, John Sillitoe on behalf of the Wellcome Sanger Institute COVID-19 Surveillance Team |
| EPI_ISL_3361226, EPI_ISL_3361227 | Lighthouse Lab in Alderley Park | Wellcome Sanger Institute for the COVID-19 Genomics UK (COG-UK) Consortium | Jacquelyn Wynn, Mairead Hyland, The Lighthouse Lab in Alderley Park and Alex Alderton, Roberto Amato, Jeffrey Barrett, Sonia Goncalves, Ewan Harrison, David K. Jackson, Ian Johnston, Dominic Kwiatkowski, Cordelia Langford, John Sillitoe on behalf of the Wellcome Sanger Institute COVID-19 Surveillance Team |
| EPI_ISL_3361229 | Berkshire and Surrey Pathology Services Lighthouse Laboratory | Wellcome Sanger Institute for the COVID-19 Genomics UK (COG-UK) Consortium | Berkshire and Surrey Pathology Services Lighthouse Laboratory and Alex Alderton, Roberto Amato, Jeffrey Barrett, Sonia Goncalves, Ewan Harrison, David K. Jackson, Ian Johnston, Dominic Kwiatkowski, Cordelia Langford, John Sillitoe on behalf of the Wellcome Sanger Institute COVID-19 Surveillance Team |
| EPI_ISL_3361230, EPI_ISL_3361232 | Lighthouse Lab in Milton Keynes | Wellcome Sanger Institute for the COVID-19 Genomics UK (COG-UK) Consortium | The Lighthouse Lab in Milton Keynes and Alex Alderton, Roberto Amato, Jeffrey Barrett, Sonia Goncalves, Ewan Harrison, David K. Jackson, Ian Johnston, Dominic Kwiatkowski, Cordelia Langford, John Sillitoe on behalf of the Wellcome Sanger Institute COVID-19 Surveillance Team |
| EPI_ISL_3361233, EPI_ISL_3361235 | Lighthouse Lab in Alderley Park | Wellcome Sanger Institute for the COVID-19 Genomics UK (COG-UK) Consortium | Jacquelyn Wynn, Mairead Hyland, The Lighthouse Lab in Alderley Park and Alex Alderton, Roberto Amato, Jeffrey Barrett, Sonia Goncalves, Ewan Harrison, David K. Jackson, Ian Johnston, Dominic Kwiatkowski, Cordelia Langford, John Sillitoe on behalf of the Wellcome Sanger Institute COVID-19 Surveillance Team |
| EPI_ISL_3361236 | Lighthouse Lab in Milton Keynes | Wellcome Sanger Institute for the COVID-19 Genomics UK (COG-UK) Consortium | The Lighthouse Lab in Milton Keynes and Alex Alderton, Roberto Amato, Jeffrey Barrett, Sonia Goncalves, Ewan Harrison, David K. Jackson, Ian Johnston, Dominic Kwiatkowski, Cordelia Langford, John Sillitoe on behalf of the Wellcome Sanger Institute COVID-19 Surveillance Team |
| EPI_ISL_3361238, EPI_ISL_3361239, EPI_ISL_3361240, EPI_ISL_3361242 | Lighthouse Lab in Alderley Park | Wellcome Sanger Institute for the COVID-19 Genomics UK (COG-UK) Consortium | Jacquelyn Wynn, Mairead Hyland, The Lighthouse Lab in Alderley Park and Alex Alderton, Roberto Amato, Jeffrey Barrett, Sonia Goncalves, Ewan Harrison, David K. Jackson, Ian Johnston, Dominic Kwiatkowski, Cordelia Langford, John Sillitoe on behalf of the Wellcome Sanger Institute COVID-19 Surveillance Team |
| EPI_ISL_3361243 | Lighthouse Lab in Milton Keynes | Wellcome Sanger Institute for the COVID-19 Genomics UK (COG-UK) Consortium | The Lighthouse Lab in Milton Keynes and Alex Alderton, Roberto Amato, Jeffrey Barrett, Sonia Goncalves, Ewan Harrison, David K. Jackson, Ian Johnston, Dominic Kwiatkowski, Cordelia Langford, John Sillitoe on behalf of the Wellcome Sanger Institute COVID-19 Surveillance Team |
| EPI_ISL_3361245, EPI_ISL_3361246, EPI_ISL_3361248, EPI_ISL_3361249, EPI_ISL_3361250 | Lighthouse Lab in Alderley Park | Wellcome Sanger Institute for the COVID-19 Genomics UK (COG-UK) Consortium | Jacquelyn Wynn, Mairead Hyland, The Lighthouse Lab in Alderley Park and Alex Alderton, Roberto Amato, Jeffrey Barrett, Sonia Goncalves, Ewan Harrison, David K. Jackson, Ian Johnston, Dominic Kwiatkowski, Cordelia Langford, John Sillitoe on behalf of the Wellcome Sanger Institute COVID-19 Surveillance Team |
| EPI_ISL_3361252 | Berkshire and Surrey Pathology Services Lighthouse Laboratory | Wellcome Sanger Institute for the COVID-19 Genomics UK (COG-UK) Consortium | Berkshire and Surrey Pathology Services Lighthouse Laboratory and Alex Alderton, Roberto Amato, Jeffrey Barrett, Sonia Goncalves, Ewan Harrison, David K. Jackson, Ian Johnston, Dominic Kwiatkowski, Cordelia Langford, John Sillitoe on behalf of the Wellcome Sanger Institute COVID-19 Surveillance Team |
| EPI_ISL_3361253, EPI_ISL_3361255, EPI_ISL_3361256, EPI_ISL_3361258 | Lighthouse Lab in Milton Keynes | Wellcome Sanger Institute for the COVID-19 Genomics UK (COG-UK) Consortium | The Lighthouse Lab in Milton Keynes and Alex Alderton, Roberto Amato, Jeffrey Barrett, Sonia Goncalves, Ewan Harrison, David K. Jackson, Ian Johnston, Dominic Kwiatkowski, Cordelia Langford, John Sillitoe on behalf of the Wellcome Sanger Institute COVID-19 Surveillance Team |
| EPI_ISL_3361259, EPI_ISL_3361261 | Lighthouse Lab in Alderley Park | Wellcome Sanger Institute for the COVID-19 Genomics UK (COG-UK) Consortium | Jacquelyn Wynn, Mairead Hyland, The Lighthouse Lab in Alderley Park and Alex Alderton, Roberto Amato, Jeffrey Barrett, Sonia Goncalves, Ewan Harrison, David K. Jackson, Ian Johnston, Dominic Kwiatkowski, Cordelia Langford, John Sillitoe on behalf of the Wellcome Sanger Institute COVID-19 Surveillance Team |
| EPI_ISL_3361262 | Lighthouse Lab in Milton Keynes | Wellcome Sanger Institute for the COVID-19 Genomics UK (COG-UK) Consortium | The Lighthouse Lab in Milton Keynes and Alex Alderton, Roberto Amato, Jeffrey Barrett, Sonia Goncalves, Ewan Harrison, David K. Jackson, Ian Johnston, Dominic Kwiatkowski, Cordelia Langford, John Sillitoe on behalf of the Wellcome Sanger Institute COVID-19 Surveillance Team |
| EPI_ISL_3361264 | Berkshire and Surrey Pathology Services Lighthouse Laboratory | Wellcome Sanger Institute for the COVID-19 Genomics UK (COG-UK) Consortium | Berkshire and Surrey Pathology Services Lighthouse Laboratory and Alex Alderton, Roberto Amato, Jeffrey Barrett, Sonia Goncalves, Ewan Harrison, David K. Jackson, Ian Johnston, Dominic Kwiatkowski, Cordelia Langford, John Sillitoe on behalf of the Wellcome Sanger Institute COVID-19 Surveillance Team |
| EPI_ISL_3361265 | Lighthouse Lab in Milton Keynes | Wellcome Sanger Institute for the COVID-19 Genomics UK (COG-UK) Consortium | The Lighthouse Lab in Milton Keynes and Alex Alderton, Roberto Amato, Jeffrey Barrett, Sonia Goncalves, Ewan Harrison, David K. Jackson, Ian Johnston, Dominic Kwiatkowski, Cordelia Langford, John Sillitoe on behalf of the Wellcome Sanger Institute COVID-19 Surveillance Team |
| EPI_ISL_3361267 | Berkshire and Surrey Pathology Services Lighthouse Laboratory | Wellcome Sanger Institute for the COVID-19 Genomics UK (COG-UK) Consortium | Berkshire and Surrey Pathology Services Lighthouse Laboratory and Alex Alderton, Roberto Amato, Jeffrey Barrett, Sonia Goncalves, Ewan Harrison, David K. Jackson, Ian Johnston, Dominic Kwiatkowski, Cordelia Langford, John Sillitoe on behalf of the Wellcome Sanger Institute COVID-19 Surveillance Team |
| EPI_ISL_3361268, EPI_ISL_3361269 | Lighthouse Lab in Milton Keynes | Wellcome Sanger Institute for the COVID-19 Genomics UK (COG-UK) Consortium | The Lighthouse Lab in Milton Keynes and Alex Alderton, Roberto Amato, Jeffrey Barrett, Sonia Goncalves, Ewan Harrison, David K. Jackson, Ian Johnston, Dominic Kwiatkowski, Cordelia Langford, John Sillitoe on behalf of the Wellcome Sanger Institute COVID-19 Surveillance Team |
| EPI_ISL_3361271 | Lighthouse Lab in Alderley Park | Wellcome Sanger Institute for the COVID-19 Genomics UK (COG-UK) Consortium | Jacquelyn Wynn, Mairead Hyland, The Lighthouse Lab in Alderley Park and Alex Alderton, Roberto Amato, Jeffrey Barrett, Sonia Goncalves, Ewan Harrison, David K. Jackson, Ian Johnston, Dominic Kwiatkowski, Cordelia Langford, John Sillitoe on behalf of the Wellcome Sanger Institute COVID-19 Surveillance Team |
| EPI_ISL_3361272 | Berkshire and Surrey Pathology Services Lighthouse Laboratory | Wellcome Sanger Institute for the COVID-19 Genomics UK (COG-UK) Consortium | Berkshire and Surrey Pathology Services Lighthouse Laboratory and Alex Alderton, Roberto Amato, Jeffrey Barrett, Sonia Goncalves, Ewan Harrison, David K. Jackson, Ian Johnston, Dominic Kwiatkowski, Cordelia Langford, John Sillitoe on behalf of the Wellcome Sanger Institute COVID-19 Surveillance Team |
| EPI_ISL_3361274 | Lighthouse Lab in Alderley Park | Wellcome Sanger Institute for the COVID-19 Genomics UK (COG-UK) Consortium | Jacquelyn Wynn, Mairead Hyland, The Lighthouse Lab in Alderley Park and Alex Alderton, Roberto Amato, Jeffrey Barrett, Sonia Goncalves, Ewan Harrison, David K. Jackson, Ian Johnston, Dominic Kwiatkowski, Cordelia Langford, John Sillitoe on behalf of the Wellcome Sanger Institute COVID-19 Surveillance Team |
| EPI_ISL_3361275 | Lighthouse Lab in Milton Keynes | Wellcome Sanger Institute for the COVID-19 Genomics UK (COG-UK) Consortium | The Lighthouse Lab in Milton Keynes and Alex Alderton, Roberto Amato, Jeffrey Barrett, Sonia Goncalves, Ewan Harrison, David K. Jackson, Ian Johnston, Dominic Kwiatkowski, Cordelia Langford, John Sillitoe on behalf of the Wellcome Sanger Institute COVID-19 Surveillance Team |
| EPI_ISL_3361277 | Berkshire and Surrey Pathology Services Lighthouse Laboratory | Wellcome Sanger Institute for the COVID-19 Genomics UK (COG-UK) Consortium | Berkshire and Surrey Pathology Services Lighthouse Laboratory and Alex Alderton, Roberto Amato, Jeffrey Barrett, Sonia Goncalves, Ewan Harrison, David K. Jackson, Ian Johnston, Dominic Kwiatkowski, Cordelia Langford, John Sillitoe on behalf of the Wellcome Sanger Institute COVID-19 Surveillance Team |
| EPI_ISL_3361279 | Lighthouse Lab in Alderley Park | Wellcome Sanger Institute for the COVID-19 Genomics UK (COG-UK) Consortium | Jacquelyn Wynn, Mairead Hyland, The Lighthouse Lab in Alderley Park and Alex Alderton, Roberto Amato, Jeffrey Barrett, Sonia Goncalves, Ewan Harrison, David K. Jackson, Ian Johnston, Dominic Kwiatkowski, Cordelia Langford, John Sillitoe on behalf of the Wellcome Sanger Institute COVID-19 Surveillance Team |
| EPI_ISL_3361280 | Berkshire and Surrey Pathology Services Lighthouse Laboratory | Wellcome Sanger Institute for the COVID-19 Genomics UK (COG-UK) Consortium | Berkshire and Surrey Pathology Services Lighthouse Laboratory and Alex Alderton, Roberto Amato, Jeffrey Barrett, Sonia Goncalves, Ewan Harrison, David K. Jackson, Ian Johnston, Dominic Kwiatkowski, Cordelia Langford, John Sillitoe on behalf of the Wellcome Sanger Institute COVID-19 Surveillance Team |
| EPI_ISL_3361282, EPI_ISL_3361283, EPI_ISL_3361285 | Lighthouse Lab in Alderley Park | Wellcome Sanger Institute for the COVID-19 Genomics UK (COG-UK) Consortium | Jacquelyn Wynn, Mairead Hyland, The Lighthouse Lab in Alderley Park and Alex Alderton, Roberto Amato, Jeffrey Barrett, Sonia Goncalves, Ewan Harrison, David K. Jackson, Ian Johnston, Dominic Kwiatkowski, Cordelia Langford, John Sillitoe on behalf of the Wellcome Sanger Institute COVID-19 Surveillance Team |
| EPI_ISL_3361286 | Berkshire and Surrey Pathology Services Lighthouse Laboratory | Wellcome Sanger Institute for the COVID-19 Genomics UK (COG-UK) Consortium | Berkshire and Surrey Pathology Services Lighthouse Laboratory and Alex Alderton, Roberto Amato, Jeffrey Barrett, Sonia Goncalves, Ewan Harrison, David K. Jackson, Ian Johnston, Dominic Kwiatkowski, Cordelia Langford, John Sillitoe on behalf of the Wellcome Sanger Institute COVID-19 Surveillance Team |
| EPI_ISL_3361287 | Lighthouse Lab in Alderley Park | Wellcome Sanger Institute for the COVID-19 Genomics UK (COG-UK) Consortium | Jacquelyn Wynn, Mairead Hyland, The Lighthouse Lab in Alderley Park and Alex Alderton, Roberto Amato, Jeffrey Barrett, Sonia Goncalves, Ewan Harrison, David K. Jackson, Ian Johnston, Dominic Kwiatkowski, Cordelia Langford, John Sillitoe on behalf of the Wellcome Sanger Institute COVID-19 Surveillance Team |
| EPI_ISL_3361289 | Lighthouse Lab in Milton Keynes | Wellcome Sanger Institute for the COVID-19 Genomics UK (COG-UK) Consortium | The Lighthouse Lab in Milton Keynes and Alex Alderton, Roberto Amato, Jeffrey Barrett, Sonia Goncalves, Ewan Harrison, David K. Jackson, Ian Johnston, Dominic Kwiatkowski, Cordelia Langford, John Sillitoe on behalf of the Wellcome Sanger Institute COVID-19 Surveillance Team |
| EPI_ISL_3361290 | Berkshire and Surrey Pathology Services Lighthouse Laboratory | Wellcome Sanger Institute for the COVID-19 Genomics UK (COG-UK) Consortium | Berkshire and Surrey Pathology Services Lighthouse Laboratory and Alex Alderton, Roberto Amato, Jeffrey Barrett, Sonia Goncalves, Ewan Harrison, David K. Jackson, Ian Johnston, Dominic Kwiatkowski, Cordelia Langford, John Sillitoe on behalf of the Wellcome Sanger Institute COVID-19 Surveillance Team |
| EPI_ISL_3361292, EPI_ISL_3361293, EPI_ISL_3361295, EPI_ISL_3361296, EPI_ISL_3361297, EPI_ISL_3361299, EPI_ISL_3361300 | Lighthouse Lab in Alderley Park | Wellcome Sanger Institute for the COVID-19 Genomics UK (COG-UK) Consortium | Jacquelyn Wynn, Mairead Hyland, The Lighthouse Lab in Alderley Park and Alex Alderton, Roberto Amato, Jeffrey Barrett, Sonia Goncalves, Ewan Harrison, David K. Jackson, Ian Johnston, Dominic Kwiatkowski, Cordelia Langford, John Sillitoe on behalf of the Wellcome Sanger Institute COVID-19 Surveillance Team |
| EPI_ISL_3361302 | Berkshire and Surrey Pathology Services Lighthouse Laboratory | Wellcome Sanger Institute for the COVID-19 Genomics UK (COG-UK) Consortium | Berkshire and Surrey Pathology Services Lighthouse Laboratory and Alex Alderton, Roberto Amato, Jeffrey Barrett, Sonia Goncalves, Ewan Harrison, David K. Jackson, Ian Johnston, Dominic Kwiatkowski, Cordelia Langford, John Sillitoe on behalf of the Wellcome Sanger Institute COVID-19 Surveillance Team |
| EPI_ISL_3361303, EPI_ISL_3361304 | Lighthouse Lab in Milton Keynes | Wellcome Sanger Institute for the COVID-19 Genomics UK (COG-UK) Consortium | The Lighthouse Lab in Milton Keynes and Alex Alderton, Roberto Amato, Jeffrey Barrett, Sonia Goncalves, Ewan Harrison, David K. Jackson, Ian Johnston, Dominic Kwiatkowski, Cordelia Langford, John Sillitoe on behalf of the Wellcome Sanger Institute COVID-19 Surveillance Team |
| EPI_ISL_3361306 | Lighthouse Lab in Alderley Park | Wellcome Sanger Institute for the COVID-19 Genomics UK (COG-UK) Consortium | Jacquelyn Wynn, Mairead Hyland, The Lighthouse Lab in Alderley Park and Alex Alderton, Roberto Amato, Jeffrey Barrett, Sonia Goncalves, Ewan Harrison, David K. Jackson, Ian Johnston, Dominic Kwiatkowski, Cordelia Langford, John Sillitoe on behalf of the Wellcome Sanger Institute COVID-19 Surveillance Team |
| EPI_ISL_3361307, EPI_ISL_3361309 | Lighthouse Lab in Milton Keynes | Wellcome Sanger Institute for the COVID-19 Genomics UK (COG-UK) Consortium | The Lighthouse Lab in Milton Keynes and Alex Alderton, Roberto Amato, Jeffrey Barrett, Sonia Goncalves, Ewan Harrison, David K. Jackson, Ian Johnston, Dominic Kwiatkowski, Cordelia Langford, John Sillitoe on behalf of the Wellcome Sanger Institute COVID-19 Surveillance Team |
| EPI_ISL_3361310, EPI_ISL_3361312, EPI_ISL_3361313 | Lighthouse Lab in Alderley Park | Wellcome Sanger Institute for the COVID-19 Genomics UK (COG-UK) Consortium | Jacquelyn Wynn, Mairead Hyland, The Lighthouse Lab in Alderley Park and Alex Alderton, Roberto Amato, Jeffrey Barrett, Sonia Goncalves, Ewan Harrison, David K. Jackson, Ian Johnston, Dominic Kwiatkowski, Cordelia Langford, John Sillitoe on behalf of the Wellcome Sanger Institute COVID-19 Surveillance Team |
| EPI_ISL_3361315 | Lighthouse Lab in Milton Keynes | Wellcome Sanger Institute for the COVID-19 Genomics UK (COG-UK) Consortium | The Lighthouse Lab in Milton Keynes and Alex Alderton, Roberto Amato, Jeffrey Barrett, Sonia Goncalves, Ewan Harrison, David K. Jackson, Ian Johnston, Dominic Kwiatkowski, Cordelia Langford, John Sillitoe on behalf of the Wellcome Sanger Institute COVID-19 Surveillance Team |
| EPI_ISL_3361316, EPI_ISL_3361318 | Lighthouse Lab in Alderley Park | Wellcome Sanger Institute for the COVID-19 Genomics UK (COG-UK) Consortium | Jacquelyn Wynn, Mairead Hyland, The Lighthouse Lab in Alderley Park and Alex Alderton, Roberto Amato, Jeffrey Barrett, Sonia Goncalves, Ewan Harrison, David K. Jackson, Ian Johnston, Dominic Kwiatkowski, Cordelia Langford, John Sillitoe on behalf of the Wellcome Sanger Institute COVID-19 Surveillance Team |
| EPI_ISL_3361319, EPI_ISL_3361320, | Lighthouse Lab in Milton Keynes | Wellcome Sanger Institute for the COVID-19 Genomics UK (COG-UK) Consortium | The Lighthouse Lab in Milton Keynes and Alex Alderton, Roberto Amato, Jeffrey Barrett, Sonia Goncalves, Ewan Harrison, David K. Jackson, Ian Johnston, Dominic Kwiatkowski, Cordelia Langford, John Sillitoe on behalf of the Wellcome Sanger Institute COVID-19 Surveillance Team |

| | | | |
|---|---|---|---|
| EPI_ISL_3361322, EPI_ISL_3361323, EPI_ISL_3361325 | | | |
| EPI_ISL_3361326 | Lighthouse Lab in Alderley Park | Wellcome Sanger Institute for the COVID-19 Genomics UK (COG-UK) Consortium | Jacquelyn Wynn, Mairead Hyland, The Lighthouse Lab in Alderley Park and Alex Alderton, Roberto Amato, Jeffrey Barrett, Sonia Goncalves, Ewan Harrison, David K. Jackson, Ian Johnston, Dominic Kwiatkowski, Cordelia Langford, John Sillitoe on behalf of the Wellcome Sanger Institute COVID-19 Surveillance Team |
| EPI_ISL_3361328 | Lighthouse Lab in Milton Keynes | Wellcome Sanger Institute for the COVID-19 Genomics UK (COG-UK) Consortium | The Lighthouse Lab in Milton Keynes and Alex Alderton, Roberto Amato, Jeffrey Barrett, Sonia Goncalves, Ewan Harrison, David K. Jackson, Ian Johnston, Dominic Kwiatkowski, Cordelia Langford, John Sillitoe on behalf of the Wellcome Sanger Institute COVID-19 Surveillance Team |
| EPI_ISL_3361329 | Lighthouse Lab in Alderley Park | Wellcome Sanger Institute for the COVID-19 Genomics UK (COG-UK) Consortium | Jacquelyn Wynn, Mairead Hyland, The Lighthouse Lab in Alderley Park and Alex Alderton, Roberto Amato, Jeffrey Barrett, Sonia Goncalves, Ewan Harrison, David K. Jackson, Ian Johnston, Dominic Kwiatkowski, Cordelia Langford, John Sillitoe on behalf of the Wellcome Sanger Institute COVID-19 Surveillance Team |
| EPI_ISL_3361330, EPI_ISL_3361332, EPI_ISL_3361333, EPI_ISL_3361335, EPI_ISL_3361337 | Lighthouse Lab in Milton Keynes | Wellcome Sanger Institute for the COVID-19 Genomics UK (COG-UK) Consortium | The Lighthouse Lab in Milton Keynes and Alex Alderton, Roberto Amato, Jeffrey Barrett, Sonia Goncalves, Ewan Harrison, David K. Jackson, Ian Johnston, Dominic Kwiatkowski, Cordelia Langford, John Sillitoe on behalf of the Wellcome Sanger Institute COVID-19 Surveillance Team |
| EPI_ISL_3361338 | Lighthouse Lab in Alderley Park | Wellcome Sanger Institute for the COVID-19 Genomics UK (COG-UK) Consortium | Jacquelyn Wynn, Mairead Hyland, The Lighthouse Lab in Alderley Park and Alex Alderton, Roberto Amato, Jeffrey Barrett, Sonia Goncalves, Ewan Harrison, David K. Jackson, Ian Johnston, Dominic Kwiatkowski, Cordelia Langford, John Sillitoe on behalf of the Wellcome Sanger Institute COVID-19 Surveillance Team |
| EPI_ISL_3361339, EPI_ISL_3361341 | Berkshire and Surrey Pathology Services Lighthouse Laboratory | Wellcome Sanger Institute for the COVID-19 Genomics UK (COG-UK) Consortium | Berkshire and Surrey Pathology Services Lighthouse Laboratory and Alex Alderton, Roberto Amato, Jeffrey Barrett, Sonia Goncalves, Ewan Harrison, David K. Jackson, Ian Johnston, Dominic Kwiatkowski, Cordelia Langford, John Sillitoe on behalf of the Wellcome Sanger Institute COVID-19 Surveillance Team |
| EPI_ISL_3361342 | Lighthouse Lab in Alderley Park | Wellcome Sanger Institute for the COVID-19 Genomics UK (COG-UK) Consortium | Jacquelyn Wynn, Mairead Hyland, The Lighthouse Lab in Alderley Park and Alex Alderton, Roberto Amato, Jeffrey Barrett, Sonia Goncalves, Ewan Harrison, David K. Jackson, Ian Johnston, Dominic Kwiatkowski, Cordelia Langford, John Sillitoe on behalf of the Wellcome Sanger Institute COVID-19 Surveillance Team |
| EPI_ISL_3361344 | Lighthouse Lab in Milton Keynes | Wellcome Sanger Institute for the COVID-19 Genomics UK (COG-UK) Consortium | The Lighthouse Lab in Milton Keynes and Alex Alderton, Roberto Amato, Jeffrey Barrett, Sonia Goncalves, Ewan Harrison, David K. Jackson, Ian Johnston, Dominic Kwiatkowski, Cordelia Langford, John Sillitoe on behalf of the Wellcome Sanger Institute COVID-19 Surveillance Team |
| EPI_ISL_3361345, EPI_ISL_3361347 | Lighthouse Lab in Alderley Park | Wellcome Sanger Institute for the COVID-19 Genomics UK (COG-UK) Consortium | Jacquelyn Wynn, Mairead Hyland, The Lighthouse Lab in Alderley Park and Alex Alderton, Roberto Amato, Jeffrey Barrett, Sonia Goncalves, Ewan Harrison, David K. Jackson, Ian Johnston, Dominic Kwiatkowski, Cordelia Langford, John Sillitoe on behalf of the Wellcome Sanger Institute COVID-19 Surveillance Team |
| EPI_ISL_3361348 | Lighthouse Lab in Milton Keynes | Wellcome Sanger Institute for the COVID-19 Genomics UK (COG-UK) Consortium | The Lighthouse Lab in Milton Keynes and Alex Alderton, Roberto Amato, Jeffrey Barrett, Sonia Goncalves, Ewan Harrison, David K. Jackson, Ian Johnston, Dominic Kwiatkowski, Cordelia Langford, John Sillitoe on behalf of the Wellcome Sanger Institute COVID-19 Surveillance Team |
| EPI_ISL_3361349, EPI_ISL_3361351 | Lighthouse Lab in Alderley Park | Wellcome Sanger Institute for the COVID-19 Genomics UK (COG-UK) Consortium | Jacquelyn Wynn, Mairead Hyland, The Lighthouse Lab in Alderley Park and Alex Alderton, Roberto Amato, Jeffrey Barrett, Sonia Goncalves, Ewan Harrison, David K. Jackson, Ian Johnston, Dominic Kwiatkowski, Cordelia Langford, John Sillitoe on behalf of the Wellcome Sanger Institute COVID-19 Surveillance Team |
| EPI_ISL_3361352 | Lighthouse Lab in Milton Keynes | Wellcome Sanger Institute for the COVID-19 Genomics UK (COG-UK) Consortium | The Lighthouse Lab in Milton Keynes and Alex Alderton, Roberto Amato, Jeffrey Barrett, Sonia Goncalves, Ewan Harrison, David K. Jackson, Ian Johnston, Dominic Kwiatkowski, Cordelia Langford, John Sillitoe on behalf of the Wellcome Sanger Institute COVID-19 Surveillance Team |
| EPI_ISL_3361354 | Lighthouse Lab in Alderley Park | Wellcome Sanger Institute for the COVID-19 Genomics UK (COG-UK) Consortium | Jacquelyn Wynn, Mairead Hyland, The Lighthouse Lab in Alderley Park and Alex Alderton, Roberto Amato, Jeffrey Barrett, Sonia Goncalves, Ewan Harrison, David K. Jackson, Ian Johnston, Dominic Kwiatkowski, Cordelia Langford, John Sillitoe on behalf of the Wellcome Sanger Institute COVID-19 Surveillance Team |
| EPI_ISL_3361355 | Lighthouse Lab in Milton Keynes | Wellcome Sanger Institute for the COVID-19 Genomics UK (COG-UK) Consortium | The Lighthouse Lab in Milton Keynes and Alex Alderton, Roberto Amato, Jeffrey Barrett, Sonia Goncalves, Ewan Harrison, David K. Jackson, Ian Johnston, Dominic Kwiatkowski, Cordelia Langford, John Sillitoe on behalf of the Wellcome Sanger Institute COVID-19 Surveillance Team |
| EPI_ISL_3361357, EPI_ISL_3361358, EPI_ISL_3361360, EPI_ISL_3361361 | Lighthouse Lab in Alderley Park | Wellcome Sanger Institute for the COVID-19 Genomics UK (COG-UK) Consortium | Jacquelyn Wynn, Mairead Hyland, The Lighthouse Lab in Alderley Park and Alex Alderton, Roberto Amato, Jeffrey Barrett, Sonia Goncalves, Ewan Harrison, David K. Jackson, Ian Johnston, Dominic Kwiatkowski, Cordelia Langford, John Sillitoe on behalf of the Wellcome Sanger Institute COVID-19 Surveillance Team |
| EPI_ISL_3361363 | Lighthouse Lab in Milton Keynes | Wellcome Sanger Institute for the COVID-19 Genomics UK (COG-UK) Consortium | The Lighthouse Lab in Milton Keynes and Alex Alderton, Roberto Amato, Jeffrey Barrett, Sonia Goncalves, Ewan Harrison, David K. Jackson, Ian Johnston, Dominic Kwiatkowski, Cordelia Langford, John Sillitoe on behalf of the Wellcome Sanger Institute COVID-19 Surveillance Team |
| EPI_ISL_3361364 | Lighthouse Lab in Alderley Park | Wellcome Sanger Institute for the COVID-19 Genomics UK (COG-UK) Consortium | Jacquelyn Wynn, Mairead Hyland, The Lighthouse Lab in Alderley Park and Alex Alderton, Roberto Amato, Jeffrey Barrett, Sonia Goncalves, Ewan Harrison, David K. Jackson, Ian Johnston, Dominic Kwiatkowski, Cordelia Langford, John Sillitoe on behalf of the Wellcome Sanger Institute COVID-19 Surveillance Team |
| EPI_ISL_3361365 | Berkshire and Surrey Pathology Services Lighthouse Laboratory | Wellcome Sanger Institute for the COVID-19 Genomics UK (COG-UK) Consortium | Berkshire and Surrey Pathology Services Lighthouse Laboratory and Alex Alderton, Roberto Amato, Jeffrey Barrett, Sonia Goncalves, Ewan Harrison, David K. Jackson, Ian Johnston, Dominic Kwiatkowski, Cordelia Langford, John Sillitoe on behalf of the Wellcome Sanger Institute COVID-19 Surveillance Team |
| EPI_ISL_3361367 | Lighthouse Lab in Milton Keynes | Wellcome Sanger Institute for the COVID-19 Genomics UK (COG-UK) Consortium | The Lighthouse Lab in Milton Keynes and Alex Alderton, Roberto Amato, Jeffrey Barrett, Sonia Goncalves, Ewan Harrison, David K. Jackson, Ian Johnston, Dominic Kwiatkowski, Cordelia Langford, John Sillitoe on behalf of the Wellcome Sanger Institute COVID-19 Surveillance Team |
| EPI_ISL_3361368, EPI_ISL_3361370, EPI_ISL_3361371 | Lighthouse Lab in Alderley Park | Wellcome Sanger Institute for the COVID-19 Genomics UK (COG-UK) Consortium | Jacquelyn Wynn, Mairead Hyland, The Lighthouse Lab in Alderley Park and Alex Alderton, Roberto Amato, Jeffrey Barrett, Sonia Goncalves, Ewan Harrison, David K. Jackson, Ian Johnston, Dominic Kwiatkowski, Cordelia Langford, John Sillitoe on behalf of the Wellcome Sanger Institute COVID-19 Surveillance Team |
| EPI_ISL_3361373 | Berkshire and Surrey Pathology Services Lighthouse Laboratory | Wellcome Sanger Institute for the COVID-19 Genomics UK (COG-UK) Consortium | Berkshire and Surrey Pathology Services Lighthouse Laboratory and Alex Alderton, Roberto Amato, Jeffrey Barrett, Sonia Goncalves, Ewan Harrison, David K. Jackson, Ian Johnston, Dominic Kwiatkowski, Cordelia Langford, John Sillitoe on behalf of the Wellcome Sanger Institute COVID-19 Surveillance Team |
| EPI_ISL_3361374 | Lighthouse Lab in Alderley Park | Wellcome Sanger Institute for the COVID-19 Genomics UK (COG-UK) Consortium | Jacquelyn Wynn, Mairead Hyland, The Lighthouse Lab in Alderley Park and Alex Alderton, Roberto Amato, Jeffrey Barrett, Sonia Goncalves, Ewan Harrison, David K. Jackson, Ian Johnston, Dominic Kwiatkowski, Cordelia Langford, John Sillitoe on behalf of the Wellcome Sanger Institute COVID-19 Surveillance Team |
| EPI_ISL_3361376 | Lighthouse Lab in Milton Keynes | Wellcome Sanger Institute for the COVID-19 Genomics UK (COG-UK) Consortium | The Lighthouse Lab in Milton Keynes and Alex Alderton, Roberto Amato, Jeffrey Barrett, Sonia Goncalves, Ewan Harrison, David K. Jackson, Ian Johnston, Dominic Kwiatkowski, Cordelia Langford, John Sillitoe on behalf of the Wellcome Sanger Institute COVID-19 Surveillance Team |
| EPI_ISL_3361377 | Lighthouse Lab in Alderley Park | Wellcome Sanger Institute for the COVID-19 Genomics UK (COG-UK) Consortium | Jacquelyn Wynn, Mairead Hyland, The Lighthouse Lab in Alderley Park and Alex Alderton, Roberto Amato, Jeffrey Barrett, Sonia Goncalves, Ewan Harrison, David K. Jackson, Ian Johnston, Dominic Kwiatkowski, Cordelia Langford, John Sillitoe on behalf of the Wellcome Sanger Institute COVID-19 Surveillance Team |
| EPI_ISL_3361378 | Berkshire and Surrey Pathology Services Lighthouse Laboratory | Wellcome Sanger Institute for the COVID-19 Genomics UK (COG-UK) Consortium | Berkshire and Surrey Pathology Services Lighthouse Laboratory and Alex Alderton, Roberto Amato, Jeffrey Barrett, Sonia Goncalves, Ewan Harrison, David K. Jackson, Ian Johnston, Dominic Kwiatkowski, Cordelia Langford, John Sillitoe on behalf of the Wellcome Sanger Institute COVID-19 Surveillance Team |
| EPI_ISL_3361380, EPI_ISL_3361381 | Lighthouse Lab in Alderley Park | Wellcome Sanger Institute for the COVID-19 Genomics UK (COG-UK) Consortium | Jacquelyn Wynn, Mairead Hyland, The Lighthouse Lab in Alderley Park and Alex Alderton, Roberto Amato, Jeffrey Barrett, Sonia Goncalves, Ewan Harrison, David K. Jackson, Ian Johnston, Dominic Kwiatkowski, Cordelia Langford, John Sillitoe on behalf of the Wellcome Sanger Institute COVID-19 Surveillance Team |
| EPI_ISL_3361382 | Lighthouse Lab in Milton Keynes | Wellcome Sanger Institute for the COVID-19 Genomics UK (COG-UK) Consortium | The Lighthouse Lab in Milton Keynes and Alex Alderton, Roberto Amato, Jeffrey Barrett, Sonia Goncalves, Ewan Harrison, David K. Jackson, Ian Johnston, Dominic Kwiatkowski, Cordelia Langford, John Sillitoe on behalf of the Wellcome Sanger Institute COVID-19 Surveillance Team |
| EPI_ISL_3361384 | Lighthouse Lab in Alderley Park | Wellcome Sanger Institute for the COVID-19 Genomics UK (COG-UK) Consortium | Jacquelyn Wynn, Mairead Hyland, The Lighthouse Lab in Alderley Park and Alex Alderton, Roberto Amato, Jeffrey Barrett, Sonia Goncalves, Ewan Harrison, David K. Jackson, Ian Johnston, Dominic Kwiatkowski, Cordelia Langford, John Sillitoe on behalf of the Wellcome Sanger Institute COVID-19 Surveillance Team |
| EPI_ISL_3361385 | Berkshire and Surrey Pathology Services Lighthouse Laboratory | Wellcome Sanger Institute for the COVID-19 Genomics UK (COG-UK) Consortium | Berkshire and Surrey Pathology Services Lighthouse Laboratory and Alex Alderton, Roberto Amato, Jeffrey Barrett, Sonia Goncalves, Ewan Harrison, David K. Jackson, Ian Johnston, Dominic Kwiatkowski, Cordelia Langford, John Sillitoe on behalf of the Wellcome Sanger Institute COVID-19 Surveillance Team |
| EPI_ISL_3361387 | Lighthouse Lab in Alderley Park | Wellcome Sanger Institute for the COVID-19 Genomics UK (COG-UK) Consortium | Jacquelyn Wynn, Mairead Hyland, The Lighthouse Lab in Alderley Park and Alex Alderton, Roberto Amato, Jeffrey Barrett, Sonia Goncalves, Ewan Harrison, David K. Jackson, Ian Johnston, Dominic Kwiatkowski, Cordelia Langford, John Sillitoe on behalf of the Wellcome Sanger Institute COVID-19 Surveillance Team |
| EPI_ISL_3361388, EPI_ISL_3361390 | Lighthouse Lab in Milton Keynes | Wellcome Sanger Institute for the COVID-19 Genomics UK (COG-UK) Consortium | The Lighthouse Lab in Milton Keynes and Alex Alderton, Roberto Amato, Jeffrey Barrett, Sonia Goncalves, Ewan Harrison, David K. Jackson, Ian Johnston, Dominic Kwiatkowski, Cordelia Langford, John Sillitoe on behalf of the Wellcome Sanger Institute COVID-19 Surveillance Team |
| EPI_ISL_3361391, EPI_ISL_3361392, EPI_ISL_3361394, EPI_ISL_3361396, EPI_ISL_3361397 | Lighthouse Lab in Alderley Park | Wellcome Sanger Institute for the COVID-19 Genomics UK (COG-UK) Consortium | Jacquelyn Wynn, Mairead Hyland, The Lighthouse Lab in Alderley Park and Alex Alderton, Roberto Amato, Jeffrey Barrett, Sonia Goncalves, Ewan Harrison, David K. Jackson, Ian Johnston, Dominic Kwiatkowski, Cordelia Langford, John Sillitoe on behalf of the Wellcome Sanger Institute COVID-19 Surveillance Team |
| EPI_ISL_3361399 | Berkshire and Surrey Pathology Services Lighthouse Laboratory | Wellcome Sanger Institute for the COVID-19 Genomics UK (COG-UK) Consortium | Berkshire and Surrey Pathology Services Lighthouse Laboratory and Alex Alderton, Roberto Amato, Jeffrey Barrett, Sonia Goncalves, Ewan Harrison, David K. Jackson, Ian Johnston, Dominic Kwiatkowski, Cordelia Langford, John Sillitoe on behalf of the Wellcome Sanger Institute COVID-19 Surveillance Team |
| EPI_ISL_3361400 | Lighthouse Lab in Alderley Park | Wellcome Sanger Institute for the COVID-19 Genomics UK (COG-UK) Consortium | Jacquelyn Wynn, Mairead Hyland, The Lighthouse Lab in Alderley Park and Alex Alderton, Roberto Amato, Jeffrey Barrett, Sonia Goncalves, Ewan Harrison, David K. Jackson, Ian Johnston, Dominic Kwiatkowski, Cordelia Langford, John Sillitoe on behalf of the Wellcome Sanger Institute COVID-19 Surveillance Team |
| EPI_ISL_3361401 | Lighthouse Lab in Milton Keynes | Wellcome Sanger Institute for the COVID-19 Genomics UK (COG-UK) Consortium | The Lighthouse Lab in Milton Keynes and Alex Alderton, Roberto Amato, Jeffrey Barrett, Sonia Goncalves, Ewan Harrison, David K. Jackson, Ian Johnston, Dominic Kwiatkowski, Cordelia Langford, John Sillitoe on behalf of the Wellcome Sanger Institute COVID-19 Surveillance Team |
| EPI_ISL_3361403, EPI_ISL_3361405 | Lighthouse Lab in Alderley Park | Wellcome Sanger Institute for the COVID-19 Genomics UK (COG-UK) Consortium | Jacquelyn Wynn, Mairead Hyland, The Lighthouse Lab in Alderley Park and Alex Alderton, Roberto Amato, Jeffrey Barrett, Sonia Goncalves, Ewan Harrison, David K. Jackson, Ian Johnston, Dominic Kwiatkowski, Cordelia Langford, John Sillitoe on behalf of the Wellcome Sanger Institute COVID-19 Surveillance Team |
| EPI_ISL_3361406 | Lighthouse Lab in Milton Keynes | Wellcome Sanger Institute for the COVID-19 Genomics UK (COG-UK) Consortium | The Lighthouse Lab in Milton Keynes and Alex Alderton, Roberto Amato, Jeffrey Barrett, Sonia Goncalves, Ewan Harrison, David K. Jackson, Ian Johnston, Dominic Kwiatkowski, Cordelia Langford, John Sillitoe on behalf of the Wellcome Sanger Institute COVID-19 Surveillance Team |
| EPI_ISL_3361408, EPI_ISL_3361409, EPI_ISL_3361411 | Lighthouse Lab in Alderley Park | Wellcome Sanger Institute for the COVID-19 Genomics UK (COG-UK) Consortium | Jacquelyn Wynn, Mairead Hyland, The Lighthouse Lab in Alderley Park and Alex Alderton, Roberto Amato, Jeffrey Barrett, Sonia Goncalves, Ewan Harrison, David K. Jackson, Ian Johnston, Dominic Kwiatkowski, Cordelia Langford, John Sillitoe on behalf of the Wellcome Sanger Institute COVID-19 Surveillance Team |
| EPI_ISL_3361412 | Berkshire and Surrey Pathology Services Lighthouse Laboratory | Wellcome Sanger Institute for the COVID-19 Genomics UK (COG-UK) Consortium | Berkshire and Surrey Pathology Services Lighthouse Laboratory and Alex Alderton, Roberto Amato, Jeffrey Barrett, Sonia Goncalves, Ewan Harrison, David K. Jackson, Ian Johnston, Dominic Kwiatkowski, Cordelia Langford, John Sillitoe on behalf of the Wellcome Sanger Institute COVID-19 Surveillance Team |
| EPI_ISL_3361414 | Lighthouse Lab in Alderley Park | Wellcome Sanger Institute for the COVID-19 Genomics UK (COG-UK) | Jacquelyn Wynn, Mairead Hyland, The Lighthouse Lab in Alderley Park and Alex Alderton, Roberto Amato, Jeffrey Barrett, Sonia Goncalves, Ewan Harrison, David K. Jackson, Ian Johnston, Dominic Kwiatkowski, Cordelia Langford, John Sillitoe on |

| | | | |
|---|---|---|---|
| | | Consortium | behalf of the Wellcome Sanger Institute COVID-19 Surveillance Team |
| EPI_ISL_3361415, EPI_ISL_3361417 | Lighthouse Lab in Milton Keynes | Wellcome Sanger Institute for the COVID-19 Genomics UK (COG-UK) Consortium | The Lighthouse Lab in Milton Keynes and Alex Alderton, Roberto Amato, Jeffrey Barrett, Sonia Goncalves, Ewan Harrison, David K. Jackson, Ian Johnston, Dominic Kwiatkowski, Cordelia Langford, John Sillitoe on behalf of the Wellcome Sanger Institute COVID-19 Surveillance Team |
| EPI_ISL_3361418 | Berkshire and Surrey Pathology Services Lighthouse Laboratory | Wellcome Sanger Institute for the COVID-19 Genomics UK (COG-UK) Consortium | Berkshire and Surrey Pathology Services Lighthouse Laboratory and Alex Alderton, Roberto Amato, Jeffrey Barrett, Sonia Goncalves, Ewan Harrison, David K. Jackson, Ian Johnston, Dominic Kwiatkowski, Cordelia Langford, John Sillitoe on behalf of the Wellcome Sanger Institute COVID-19 Surveillance Team |
| EPI_ISL_3361420, EPI_ISL_3361421, EPI_ISL_3361423, EPI_ISL_3361424, EPI_ISL_3361426 | Lighthouse Lab in Alderley Park | Wellcome Sanger Institute for the COVID-19 Genomics UK (COG-UK) Consortium | Jacquelyn Wynn, Mairead Hyland, The Lighthouse Lab in Alderley Park and Alex Alderton, Roberto Amato, Jeffrey Barrett, Sonia Goncalves, Ewan Harrison, David K. Jackson, Ian Johnston, Dominic Kwiatkowski, Cordelia Langford, John Sillitoe on behalf of the Wellcome Sanger Institute COVID-19 Surveillance Team |
| EPI_ISL_3361427 | Berkshire and Surrey Pathology Services Lighthouse Laboratory | Wellcome Sanger Institute for the COVID-19 Genomics UK (COG-UK) Consortium | Berkshire and Surrey Pathology Services Lighthouse Laboratory and Alex Alderton, Roberto Amato, Jeffrey Barrett, Sonia Goncalves, Ewan Harrison, David K. Jackson, Ian Johnston, Dominic Kwiatkowski, Cordelia Langford, John Sillitoe on behalf of the Wellcome Sanger Institute COVID-19 Surveillance Team |
| EPI_ISL_3361429 | Lighthouse Lab in Milton Keynes | Wellcome Sanger Institute for the COVID-19 Genomics UK (COG-UK) Consortium | The Lighthouse Lab in Milton Keynes and Alex Alderton, Roberto Amato, Jeffrey Barrett, Sonia Goncalves, Ewan Harrison, David K. Jackson, Ian Johnston, Dominic Kwiatkowski, Cordelia Langford, John Sillitoe on behalf of the Wellcome Sanger Institute COVID-19 Surveillance Team |
| EPI_ISL_3361430, EPI_ISL_3361431, EPI_ISL_3361433, EPI_ISL_3361434, EPI_ISL_3361436, EPI_ISL_3361437, EPI_ISL_3361439, EPI_ISL_3361440, EPI_ISL_3361442 | Lighthouse Lab in Alderley Park | Wellcome Sanger Institute for the COVID-19 Genomics UK (COG-UK) Consortium | Jacquelyn Wynn, Mairead Hyland, The Lighthouse Lab in Alderley Park and Alex Alderton, Roberto Amato, Jeffrey Barrett, Sonia Goncalves, Ewan Harrison, David K. Jackson, Ian Johnston, Dominic Kwiatkowski, Cordelia Langford, John Sillitoe on behalf of the Wellcome Sanger Institute COVID-19 Surveillance Team |
| EPI_ISL_3361443 | Lighthouse Lab in Milton Keynes | Wellcome Sanger Institute for the COVID-19 Genomics UK (COG-UK) Consortium | The Lighthouse Lab in Milton Keynes and Alex Alderton, Roberto Amato, Jeffrey Barrett, Sonia Goncalves, Ewan Harrison, David K. Jackson, Ian Johnston, Dominic Kwiatkowski, Cordelia Langford, John Sillitoe on behalf of the Wellcome Sanger Institute COVID-19 Surveillance Team |
| EPI_ISL_3361445, EPI_ISL_3361447, EPI_ISL_3361448, EPI_ISL_3361450, EPI_ISL_3361451, EPI_ISL_3361453, EPI_ISL_3361454, EPI_ISL_3361455, EPI_ISL_3361456, EPI_ISL_3361458 | Lighthouse Lab in Alderley Park | Wellcome Sanger Institute for the COVID-19 Genomics UK (COG-UK) Consortium | Jacquelyn Wynn, Mairead Hyland, The Lighthouse Lab in Alderley Park and Alex Alderton, Roberto Amato, Jeffrey Barrett, Sonia Goncalves, Ewan Harrison, David K. Jackson, Ian Johnston, Dominic Kwiatkowski, Cordelia Langford, John Sillitoe on behalf of the Wellcome Sanger Institute COVID-19 Surveillance Team |
| EPI_ISL_3361459 | Berkshire and Surrey Pathology Services Lighthouse Laboratory | Wellcome Sanger Institute for the COVID-19 Genomics UK (COG-UK) Consortium | Berkshire and Surrey Pathology Services Lighthouse Laboratory and Alex Alderton, Roberto Amato, Jeffrey Barrett, Sonia Goncalves, Ewan Harrison, David K. Jackson, Ian Johnston, Dominic Kwiatkowski, Cordelia Langford, John Sillitoe on behalf of the Wellcome Sanger Institute COVID-19 Surveillance Team |
| EPI_ISL_3361461 | Lighthouse Lab in Milton Keynes | Wellcome Sanger Institute for the COVID-19 Genomics UK (COG-UK) Consortium | The Lighthouse Lab in Milton Keynes and Alex Alderton, Roberto Amato, Jeffrey Barrett, Sonia Goncalves, Ewan Harrison, David K. Jackson, Ian Johnston, Dominic Kwiatkowski, Cordelia Langford, John Sillitoe on behalf of the Wellcome Sanger Institute COVID-19 Surveillance Team |
| EPI_ISL_3361462, EPI_ISL_3361464, EPI_ISL_3361465 | Lighthouse Lab in Alderley Park | Wellcome Sanger Institute for the COVID-19 Genomics UK (COG-UK) Consortium | Jacquelyn Wynn, Mairead Hyland, The Lighthouse Lab in Alderley Park and Alex Alderton, Roberto Amato, Jeffrey Barrett, Sonia Goncalves, Ewan Harrison, David K. Jackson, Ian Johnston, Dominic Kwiatkowski, Cordelia Langford, John Sillitoe on behalf of the Wellcome Sanger Institute COVID-19 Surveillance Team |
| EPI_ISL_3361467 | Berkshire and Surrey Pathology Services Lighthouse Laboratory | Wellcome Sanger Institute for the COVID-19 Genomics UK (COG-UK) Consortium | Berkshire and Surrey Pathology Services Lighthouse Laboratory and Alex Alderton, Roberto Amato, Jeffrey Barrett, Sonia Goncalves, Ewan Harrison, David K. Jackson, Ian Johnston, Dominic Kwiatkowski, Cordelia Langford, John Sillitoe on behalf of the Wellcome Sanger Institute COVID-19 Surveillance Team |
| EPI_ISL_3361468 | Lighthouse Lab in Milton Keynes | Wellcome Sanger Institute for the COVID-19 Genomics UK (COG-UK) Consortium | The Lighthouse Lab in Milton Keynes and Alex Alderton, Roberto Amato, Jeffrey Barrett, Sonia Goncalves, Ewan Harrison, David K. Jackson, Ian Johnston, Dominic Kwiatkowski, Cordelia Langford, John Sillitoe on behalf of the Wellcome Sanger Institute COVID-19 Surveillance Team |
| EPI_ISL_3361470, EPI_ISL_3361471 | Lighthouse Lab in Alderley Park | Wellcome Sanger Institute for the COVID-19 Genomics UK (COG-UK) Consortium | Jacquelyn Wynn, Mairead Hyland, The Lighthouse Lab in Alderley Park and Alex Alderton, Roberto Amato, Jeffrey Barrett, Sonia Goncalves, Ewan Harrison, David K. Jackson, Ian Johnston, Dominic Kwiatkowski, Cordelia Langford, John Sillitoe on behalf of the Wellcome Sanger Institute COVID-19 Surveillance Team |
| EPI_ISL_3361523, EPI_ISL_3361524 | Lighthouse Lab in Milton Keynes | Wellcome Sanger Institute for the COVID-19 Genomics UK (COG-UK) Consortium | The Lighthouse Lab in Milton Keynes and Alex Alderton, Roberto Amato, Jeffrey Barrett, Sonia Goncalves, Ewan Harrison, David K. Jackson, Ian Johnston, Dominic Kwiatkowski, Cordelia Langford, John Sillitoe on behalf of the Wellcome Sanger Institute COVID-19 Surveillance Team |
| EPI_ISL_3361526, EPI_ISL_3361528, EPI_ISL_3361529, EPI_ISL_3361530, EPI_ISL_3361532, EPI_ISL_3361533 | Lighthouse Lab in Alderley Park | Wellcome Sanger Institute for the COVID-19 Genomics UK (COG-UK) Consortium | Jacquelyn Wynn, Mairead Hyland, The Lighthouse Lab in Alderley Park and Alex Alderton, Roberto Amato, Jeffrey Barrett, Sonia Goncalves, Ewan Harrison, David K. Jackson, Ian Johnston, Dominic Kwiatkowski, Cordelia Langford, John Sillitoe on behalf of the Wellcome Sanger Institute COVID-19 Surveillance Team |
| EPI_ISL_3361535 | Berkshire and Surrey Pathology Services Lighthouse Laboratory | Wellcome Sanger Institute for the COVID-19 Genomics UK (COG-UK) Consortium | Berkshire and Surrey Pathology Services Lighthouse Laboratory and Alex Alderton, Roberto Amato, Jeffrey Barrett, Sonia Goncalves, Ewan Harrison, David K. Jackson, Ian Johnston, Dominic Kwiatkowski, Cordelia Langford, John Sillitoe on behalf of the Wellcome Sanger Institute COVID-19 Surveillance Team |
| EPI_ISL_3361536, EPI_ISL_3361538 | Lighthouse Lab in Milton Keynes | Wellcome Sanger Institute for the COVID-19 Genomics UK (COG-UK) Consortium | The Lighthouse Lab in Milton Keynes and Alex Alderton, Roberto Amato, Jeffrey Barrett, Sonia Goncalves, Ewan Harrison, David K. Jackson, Ian Johnston, Dominic Kwiatkowski, Cordelia Langford, John Sillitoe on behalf of the Wellcome Sanger Institute COVID-19 Surveillance Team |
| EPI_ISL_3361539, EPI_ISL_3361540, EPI_ISL_3361542, EPI_ISL_3361543 | Lighthouse Lab in Alderley Park | Wellcome Sanger Institute for the COVID-19 Genomics UK (COG-UK) Consortium | Jacquelyn Wynn, Mairead Hyland, The Lighthouse Lab in Alderley Park and Alex Alderton, Roberto Amato, Jeffrey Barrett, Sonia Goncalves, Ewan Harrison, David K. Jackson, Ian Johnston, Dominic Kwiatkowski, Cordelia Langford, John Sillitoe on behalf of the Wellcome Sanger Institute COVID-19 Surveillance Team |
| EPI_ISL_3361545 | Lighthouse Lab in Milton Keynes | Wellcome Sanger Institute for the COVID-19 Genomics UK (COG-UK) Consortium | The Lighthouse Lab in Milton Keynes and Alex Alderton, Roberto Amato, Jeffrey Barrett, Sonia Goncalves, Ewan Harrison, David K. Jackson, Ian Johnston, Dominic Kwiatkowski, Cordelia Langford, John Sillitoe on behalf of the Wellcome Sanger Institute COVID-19 Surveillance Team |
| EPI_ISL_3361546 | Lighthouse Laboratory Plymouth | Wellcome Sanger Institute for the COVID-19 Genomics UK (COG-UK) Consortium | Lighthouse Laboratory Plymouth and Alex Alderton, Roberto Amato, Jeffrey Barrett, Sonia Goncalves, Ewan Harrison, David K. Jackson, Ian Johnston, Dominic Kwiatkowski, Cordelia Langford, John Sillitoe on behalf of the Wellcome Sanger Institute COVID-19 Surveillance Team |
| EPI_ISL_3361548 | Lighthouse Lab in Milton Keynes | Wellcome Sanger Institute for the COVID-19 Genomics UK (COG-UK) Consortium | The Lighthouse Lab in Milton Keynes and Alex Alderton, Roberto Amato, Jeffrey Barrett, Sonia Goncalves, Ewan Harrison, David K. Jackson, Ian Johnston, Dominic Kwiatkowski, Cordelia Langford, John Sillitoe on behalf of the Wellcome Sanger Institute COVID-19 Surveillance Team |
| EPI_ISL_3361549 | Health Services Laboratories | Wellcome Sanger Institute for the COVID-19 Genomics UK (COG-UK) Consortium | Health Services Laboratories and Alex Alderton, Roberto Amato, Jeffrey Barrett, Sonia Goncalves, Ewan Harrison, David K. Jackson, Ian Johnston, Dominic Kwiatkowski, Cordelia Langford, John Sillitoe on behalf of the Wellcome Sanger Institute COVID-19 Surveillance Team |
| EPI_ISL_3361550, EPI_ISL_3361552, EPI_ISL_3361553 | Lighthouse Lab in Milton Keynes | Wellcome Sanger Institute for the COVID-19 Genomics UK (COG-UK) Consortium | The Lighthouse Lab in Milton Keynes and Alex Alderton, Roberto Amato, Jeffrey Barrett, Sonia Goncalves, Ewan Harrison, David K. Jackson, Ian Johnston, Dominic Kwiatkowski, Cordelia Langford, John Sillitoe on behalf of the Wellcome Sanger Institute COVID-19 Surveillance Team |
| EPI_ISL_3361555 | Lighthouse Laboratory Plymouth | Wellcome Sanger Institute for the COVID-19 Genomics UK (COG-UK) Consortium | Lighthouse Laboratory Plymouth and Alex Alderton, Roberto Amato, Jeffrey Barrett, Sonia Goncalves, Ewan Harrison, David K. Jackson, Ian Johnston, Dominic Kwiatkowski, Cordelia Langford, John Sillitoe on behalf of the Wellcome Sanger Institute COVID-19 Surveillance Team |
| EPI_ISL_3361557 | Lighthouse Lab in Milton Keynes | Wellcome Sanger Institute for the COVID-19 Genomics UK (COG-UK) Consortium | The Lighthouse Lab in Milton Keynes and Alex Alderton, Roberto Amato, Jeffrey Barrett, Sonia Goncalves, Ewan Harrison, David K. Jackson, Ian Johnston, Dominic Kwiatkowski, Cordelia Langford, John Sillitoe on behalf of the Wellcome Sanger Institute COVID-19 Surveillance Team |
| EPI_ISL_3361559 | Lighthouse Laboratory Plymouth | Wellcome Sanger Institute for the COVID-19 Genomics UK (COG-UK) Consortium | Lighthouse Laboratory Plymouth and Alex Alderton, Roberto Amato, Jeffrey Barrett, Sonia Goncalves, Ewan Harrison, David K. Jackson, Ian Johnston, Dominic Kwiatkowski, Cordelia Langford, John Sillitoe on behalf of the Wellcome Sanger Institute COVID-19 Surveillance Team |
| EPI_ISL_3361560, EPI_ISL_3361562 | Lighthouse Lab in Milton Keynes | Wellcome Sanger Institute for the COVID-19 Genomics UK (COG-UK) Consortium | The Lighthouse Lab in Milton Keynes and Alex Alderton, Roberto Amato, Jeffrey Barrett, Sonia Goncalves, Ewan Harrison, David K. Jackson, Ian Johnston, Dominic Kwiatkowski, Cordelia Langford, John Sillitoe on behalf of the Wellcome Sanger Institute COVID-19 Surveillance Team |
| EPI_ISL_3361564, EPI_ISL_3361565 | Lighthouse Laboratory Plymouth | Wellcome Sanger Institute for the COVID-19 Genomics UK (COG-UK) Consortium | Lighthouse Laboratory Plymouth and Alex Alderton, Roberto Amato, Jeffrey Barrett, Sonia Goncalves, Ewan Harrison, David K. Jackson, Ian Johnston, Dominic Kwiatkowski, Cordelia Langford, John Sillitoe on behalf of the Wellcome Sanger Institute COVID-19 Surveillance Team |
| EPI_ISL_3361567 | Health Services Laboratories | Wellcome Sanger Institute for the COVID-19 Genomics UK (COG-UK) Consortium | Health Services Laboratories and Alex Alderton, Roberto Amato, Jeffrey Barrett, Sonia Goncalves, Ewan Harrison, David K. Jackson, Ian Johnston, Dominic Kwiatkowski, Cordelia Langford, John Sillitoe on behalf of the Wellcome Sanger Institute COVID-19 Surveillance Team |
| EPI_ISL_3361569 | Lighthouse Lab in Glasgow | Wellcome Sanger Institute for the COVID-19 Genomics UK (COG-UK) Consortium | Harper VanSteenhouse, Yumi Kasai, David Gray, Carol Clugston, Anna Dominiczak and Alex Alderton, Roberto Amato, Jeffrey Barrett, Sonia Goncalves, Ewan Harrison, David K. Jackson, Ian Johnston, Dominic Kwiatkowski, Cordelia Langford, John Sillitoe on behalf of the Wellcome Sanger Institute COVID-19 Surveillance Team |
| EPI_ISL_3361570 | Lighthouse Laboratory Plymouth | Wellcome Sanger Institute for the COVID-19 Genomics UK (COG-UK) Consortium | Lighthouse Laboratory Plymouth and Alex Alderton, Roberto Amato, Jeffrey Barrett, Sonia Goncalves, Ewan Harrison, David K. Jackson, Ian Johnston, Dominic Kwiatkowski, Cordelia Langford, John Sillitoe on behalf of the Wellcome Sanger Institute COVID-19 Surveillance Team |
| EPI_ISL_3361572 | Lighthouse Lab in Glasgow | Wellcome Sanger Institute for the COVID-19 Genomics UK (COG-UK) Consortium | Harper VanSteenhouse, Yumi Kasai, David Gray, Carol Clugston, Anna Dominiczak and Alex Alderton, Roberto Amato, Jeffrey Barrett, Sonia Goncalves, Ewan Harrison, David K. Jackson, Ian Johnston, Dominic Kwiatkowski, Cordelia Langford, John Sillitoe on behalf of the Wellcome Sanger Institute COVID-19 Surveillance Team |
| EPI_ISL_3361573, EPI_ISL_3361575 | Lighthouse Lab in Milton Keynes | Wellcome Sanger Institute for the COVID-19 Genomics UK (COG-UK) Consortium | The Lighthouse Lab in Milton Keynes and Alex Alderton, Roberto Amato, Jeffrey Barrett, Sonia Goncalves, Ewan Harrison, David K. Jackson, Ian Johnston, Dominic Kwiatkowski, Cordelia Langford, John Sillitoe on behalf of the Wellcome Sanger Institute COVID-19 Surveillance Team |

| | | | |
|---|---|---|---|
| EPI_ISL_3361576 | Lighthouse Laboratory Plymouth | Wellcome Sanger Institute for the COVID-19 Genomics UK (COG-UK) Consortium | Lighthouse Laboratory Plymouth and Alex Alderton, Roberto Amato, Jeffrey Barrett, Sonia Goncalves, Ewan Harrison, David K. Jackson, Ian Johnston, Dominic Kwiatkowski, Cordelia Langford, John Sillitoe on behalf of the Wellcome Sanger Institute COVID-19 Surveillance Team |
| EPI_ISL_3361578 | Health Services Laboratories | Wellcome Sanger Institute for the COVID-19 Genomics UK (COG-UK) Consortium | Health Services Laboratories and Alex Alderton, Roberto Amato, Jeffrey Barrett, Sonia Goncalves, Ewan Harrison, David K. Jackson, Ian Johnston, Dominic Kwiatkowski, Cordelia Langford, John Sillitoe on behalf of the Wellcome Sanger Institute COVID-19 Surveillance Team |
| EPI_ISL_3361580 | Lighthouse Lab in Milton Keynes | Wellcome Sanger Institute for the COVID-19 Genomics UK (COG-UK) Consortium | The Lighthouse Lab in Milton Keynes and Alex Alderton, Roberto Amato, Jeffrey Barrett, Sonia Goncalves, Ewan Harrison, David K. Jackson, Ian Johnston, Dominic Kwiatkowski, Cordelia Langford, John Sillitoe on behalf of the Wellcome Sanger Institute COVID-19 Surveillance Team |
| EPI_ISL_3361582 | Lighthouse Laboratory Plymouth | Wellcome Sanger Institute for the COVID-19 Genomics UK (COG-UK) Consortium | Lighthouse Laboratory Plymouth and Alex Alderton, Roberto Amato, Jeffrey Barrett, Sonia Goncalves, Ewan Harrison, David K. Jackson, Ian Johnston, Dominic Kwiatkowski, Cordelia Langford, John Sillitoe on behalf of the Wellcome Sanger Institute COVID-19 Surveillance Team |
| EPI_ISL_3361583 | Health Services Laboratories | Wellcome Sanger Institute for the COVID-19 Genomics UK (COG-UK) Consortium | Health Services Laboratories and Alex Alderton, Roberto Amato, Jeffrey Barrett, Sonia Goncalves, Ewan Harrison, David K. Jackson, Ian Johnston, Dominic Kwiatkowski, Cordelia Langford, John Sillitoe on behalf of the Wellcome Sanger Institute COVID-19 Surveillance Team |
| EPI_ISL_3361584 | Lighthouse Laboratory Plymouth | Wellcome Sanger Institute for the COVID-19 Genomics UK (COG-UK) Consortium | Lighthouse Laboratory Plymouth and Alex Alderton, Roberto Amato, Jeffrey Barrett, Sonia Goncalves, Ewan Harrison, David K. Jackson, Ian Johnston, Dominic Kwiatkowski, Cordelia Langford, John Sillitoe on behalf of the Wellcome Sanger Institute COVID-19 Surveillance Team |
| EPI_ISL_3361586 | Lighthouse Lab in Milton Keynes | Wellcome Sanger Institute for the COVID-19 Genomics UK (COG-UK) Consortium | The Lighthouse Lab in Milton Keynes and Alex Alderton, Roberto Amato, Jeffrey Barrett, Sonia Goncalves, Ewan Harrison, David K. Jackson, Ian Johnston, Dominic Kwiatkowski, Cordelia Langford, John Sillitoe on behalf of the Wellcome Sanger Institute COVID-19 Surveillance Team |
| EPI_ISL_3361587 | Lighthouse Lab in Glasgow | Wellcome Sanger Institute for the COVID-19 Genomics UK (COG-UK) Consortium | Harper VanSteenhouse, Yumi Kasai, David Gray, Carol Clugston, Anna Dominiczak and Alex Alderton, Roberto Amato, Jeffrey Barrett, Sonia Goncalves, Ewan Harrison, David K. Jackson, Ian Johnston, Dominic Kwiatkowski, Cordelia Langford, John Sillitoe on behalf of the Wellcome Sanger Institute COVID-19 Surveillance Team |
| EPI_ISL_3361589 | Health Services Laboratories | Wellcome Sanger Institute for the COVID-19 Genomics UK (COG-UK) Consortium | Health Services Laboratories and Alex Alderton, Roberto Amato, Jeffrey Barrett, Sonia Goncalves, Ewan Harrison, David K. Jackson, Ian Johnston, Dominic Kwiatkowski, Cordelia Langford, John Sillitoe on behalf of the Wellcome Sanger Institute COVID-19 Surveillance Team |
| EPI_ISL_3361590 | Lighthouse Lab in Glasgow | Wellcome Sanger Institute for the COVID-19 Genomics UK (COG-UK) Consortium | Harper VanSteenhouse, Yumi Kasai, David Gray, Carol Clugston, Anna Dominiczak and Alex Alderton, Roberto Amato, Jeffrey Barrett, Sonia Goncalves, Ewan Harrison, David K. Jackson, Ian Johnston, Dominic Kwiatkowski, Cordelia Langford, John Sillitoe on behalf of the Wellcome Sanger Institute COVID-19 Surveillance Team |
| EPI_ISL_3361591 | Lighthouse Laboratory Plymouth | Wellcome Sanger Institute for the COVID-19 Genomics UK (COG-UK) Consortium | Lighthouse Laboratory Plymouth and Alex Alderton, Roberto Amato, Jeffrey Barrett, Sonia Goncalves, Ewan Harrison, David K. Jackson, Ian Johnston, Dominic Kwiatkowski, Cordelia Langford, John Sillitoe on behalf of the Wellcome Sanger Institute COVID-19 Surveillance Team |
| EPI_ISL_3361593 | Lighthouse Lab in Glasgow | Wellcome Sanger Institute for the COVID-19 Genomics UK (COG-UK) Consortium | Harper VanSteenhouse, Yumi Kasai, David Gray, Carol Clugston, Anna Dominiczak and Alex Alderton, Roberto Amato, Jeffrey Barrett, Sonia Goncalves, Ewan Harrison, David K. Jackson, Ian Johnston, Dominic Kwiatkowski, Cordelia Langford, John Sillitoe on behalf of the Wellcome Sanger Institute COVID-19 Surveillance Team |
| EPI_ISL_3361594 | Lighthouse Laboratory Plymouth | Wellcome Sanger Institute for the COVID-19 Genomics UK (COG-UK) Consortium | Lighthouse Laboratory Plymouth and Alex Alderton, Roberto Amato, Jeffrey Barrett, Sonia Goncalves, Ewan Harrison, David K. Jackson, Ian Johnston, Dominic Kwiatkowski, Cordelia Langford, John Sillitoe on behalf of the Wellcome Sanger Institute COVID-19 Surveillance Team |
| EPI_ISL_3361596 | Lighthouse Lab in Glasgow | Wellcome Sanger Institute for the COVID-19 Genomics UK (COG-UK) Consortium | Harper VanSteenhouse, Yumi Kasai, David Gray, Carol Clugston, Anna Dominiczak and Alex Alderton, Roberto Amato, Jeffrey Barrett, Sonia Goncalves, Ewan Harrison, David K. Jackson, Ian Johnston, Dominic Kwiatkowski, Cordelia Langford, John Sillitoe on behalf of the Wellcome Sanger Institute COVID-19 Surveillance Team |
| EPI_ISL_3361598 | Lighthouse Lab in Milton Keynes | Wellcome Sanger Institute for the COVID-19 Genomics UK (COG-UK) Consortium | The Lighthouse Lab in Milton Keynes and Alex Alderton, Roberto Amato, Jeffrey Barrett, Sonia Goncalves, Ewan Harrison, David K. Jackson, Ian Johnston, Dominic Kwiatkowski, Cordelia Langford, John Sillitoe on behalf of the Wellcome Sanger Institute COVID-19 Surveillance Team |
| EPI_ISL_3361599 | Lighthouse Lab in Glasgow | Wellcome Sanger Institute for the COVID-19 Genomics UK (COG-UK) Consortium | Harper VanSteenhouse, Yumi Kasai, David Gray, Carol Clugston, Anna Dominiczak and Alex Alderton, Roberto Amato, Jeffrey Barrett, Sonia Goncalves, Ewan Harrison, David K. Jackson, Ian Johnston, Dominic Kwiatkowski, Cordelia Langford, John Sillitoe on behalf of the Wellcome Sanger Institute COVID-19 Surveillance Team |
| EPI_ISL_3361601, EPI_ISL_3361602 | Lighthouse Lab in Milton Keynes | Wellcome Sanger Institute for the COVID-19 Genomics UK (COG-UK) Consortium | The Lighthouse Lab in Milton Keynes and Alex Alderton, Roberto Amato, Jeffrey Barrett, Sonia Goncalves, Ewan Harrison, David K. Jackson, Ian Johnston, Dominic Kwiatkowski, Cordelia Langford, John Sillitoe on behalf of the Wellcome Sanger Institute COVID-19 Surveillance Team |
| EPI_ISL_3361603 | Lighthouse Laboratory Plymouth | Wellcome Sanger Institute for the COVID-19 Genomics UK (COG-UK) Consortium | Lighthouse Laboratory Plymouth and Alex Alderton, Roberto Amato, Jeffrey Barrett, Sonia Goncalves, Ewan Harrison, David K. Jackson, Ian Johnston, Dominic Kwiatkowski, Cordelia Langford, John Sillitoe on behalf of the Wellcome Sanger Institute COVID-19 Surveillance Team |
| EPI_ISL_3361605, EPI_ISL_3361606 | Lighthouse Lab in Milton Keynes | Wellcome Sanger Institute for the COVID-19 Genomics UK (COG-UK) Consortium | The Lighthouse Lab in Milton Keynes and Alex Alderton, Roberto Amato, Jeffrey Barrett, Sonia Goncalves, Ewan Harrison, David K. Jackson, Ian Johnston, Dominic Kwiatkowski, Cordelia Langford, John Sillitoe on behalf of the Wellcome Sanger Institute COVID-19 Surveillance Team |
| EPI_ISL_3361608, EPI_ISL_3361609 | Lighthouse Lab in Glasgow | Wellcome Sanger Institute for the COVID-19 Genomics UK (COG-UK) Consortium | Harper VanSteenhouse, Yumi Kasai, David Gray, Carol Clugston, Anna Dominiczak and Alex Alderton, Roberto Amato, Jeffrey Barrett, Sonia Goncalves, Ewan Harrison, David K. Jackson, Ian Johnston, Dominic Kwiatkowski, Cordelia Langford, John Sillitoe on behalf of the Wellcome Sanger Institute COVID-19 Surveillance Team |
| EPI_ISL_3361611, EPI_ISL_3361612, EPI_ISL_3361613, EPI_ISL_3361615 | Lighthouse Laboratory Plymouth | Wellcome Sanger Institute for the COVID-19 Genomics UK (COG-UK) Consortium | Lighthouse Laboratory Plymouth and Alex Alderton, Roberto Amato, Jeffrey Barrett, Sonia Goncalves, Ewan Harrison, David K. Jackson, Ian Johnston, Dominic Kwiatkowski, Cordelia Langford, John Sillitoe on behalf of the Wellcome Sanger Institute COVID-19 Surveillance Team |
| EPI_ISL_3361617 | Lighthouse Lab in Glasgow | Wellcome Sanger Institute for the COVID-19 Genomics UK (COG-UK) Consortium | Harper VanSteenhouse, Yumi Kasai, David Gray, Carol Clugston, Anna Dominiczak and Alex Alderton, Roberto Amato, Jeffrey Barrett, Sonia Goncalves, Ewan Harrison, David K. Jackson, Ian Johnston, Dominic Kwiatkowski, Cordelia Langford, John Sillitoe on behalf of the Wellcome Sanger Institute COVID-19 Surveillance Team |
| EPI_ISL_3361618 | Lighthouse Laboratory Plymouth | Wellcome Sanger Institute for the COVID-19 Genomics UK (COG-UK) Consortium | Lighthouse Laboratory Plymouth and Alex Alderton, Roberto Amato, Jeffrey Barrett, Sonia Goncalves, Ewan Harrison, David K. Jackson, Ian Johnston, Dominic Kwiatkowski, Cordelia Langford, John Sillitoe on behalf of the Wellcome Sanger Institute COVID-19 Surveillance Team |
| EPI_ISL_3361619 | Lighthouse Lab in Milton Keynes | Wellcome Sanger Institute for the COVID-19 Genomics UK (COG-UK) Consortium | The Lighthouse Lab in Milton Keynes and Alex Alderton, Roberto Amato, Jeffrey Barrett, Sonia Goncalves, Ewan Harrison, David K. Jackson, Ian Johnston, Dominic Kwiatkowski, Cordelia Langford, John Sillitoe on behalf of the Wellcome Sanger Institute COVID-19 Surveillance Team |
| EPI_ISL_3361621 | Lighthouse Lab in Alderley Park | Wellcome Sanger Institute for the COVID-19 Genomics UK (COG-UK) Consortium | Jacquelyn Wynn, Mairead Hyland, The Lighthouse Lab in Alderley Park and Alex Alderton, Roberto Amato, Jeffrey Barrett, Sonia Goncalves, Ewan Harrison, David K. Jackson, Ian Johnston, Dominic Kwiatkowski, Cordelia Langford, John Sillitoe on behalf of the Wellcome Sanger Institute COVID-19 Surveillance Team |
| EPI_ISL_3361622 | Lighthouse Laboratory Plymouth | Wellcome Sanger Institute for the COVID-19 Genomics UK (COG-UK) Consortium | Lighthouse Laboratory Plymouth and Alex Alderton, Roberto Amato, Jeffrey Barrett, Sonia Goncalves, Ewan Harrison, David K. Jackson, Ian Johnston, Dominic Kwiatkowski, Cordelia Langford, John Sillitoe on behalf of the Wellcome Sanger Institute COVID-19 Surveillance Team |
| EPI_ISL_3361624 | Lighthouse Lab in Milton Keynes | Wellcome Sanger Institute for the COVID-19 Genomics UK (COG-UK) Consortium | The Lighthouse Lab in Milton Keynes and Alex Alderton, Roberto Amato, Jeffrey Barrett, Sonia Goncalves, Ewan Harrison, David K. Jackson, Ian Johnston, Dominic Kwiatkowski, Cordelia Langford, John Sillitoe on behalf of the Wellcome Sanger Institute COVID-19 Surveillance Team |
| EPI_ISL_3361625 | Lighthouse Laboratory Plymouth | Wellcome Sanger Institute for the COVID-19 Genomics UK (COG-UK) Consortium | Lighthouse Laboratory Plymouth and Alex Alderton, Roberto Amato, Jeffrey Barrett, Sonia Goncalves, Ewan Harrison, David K. Jackson, Ian Johnston, Dominic Kwiatkowski, Cordelia Langford, John Sillitoe on behalf of the Wellcome Sanger Institute COVID-19 Surveillance Team |
| EPI_ISL_3361627 | Lighthouse Lab in Milton Keynes | Wellcome Sanger Institute for the COVID-19 Genomics UK (COG-UK) Consortium | The Lighthouse Lab in Milton Keynes and Alex Alderton, Roberto Amato, Jeffrey Barrett, Sonia Goncalves, Ewan Harrison, David K. Jackson, Ian Johnston, Dominic Kwiatkowski, Cordelia Langford, John Sillitoe on behalf of the Wellcome Sanger Institute COVID-19 Surveillance Team |
| EPI_ISL_3361629 | Lighthouse Lab in Glasgow | Wellcome Sanger Institute for the COVID-19 Genomics UK (COG-UK) Consortium | Harper VanSteenhouse, Yumi Kasai, David Gray, Carol Clugston, Anna Dominiczak and Alex Alderton, Roberto Amato, Jeffrey Barrett, Sonia Goncalves, Ewan Harrison, David K. Jackson, Ian Johnston, Dominic Kwiatkowski, Cordelia Langford, John Sillitoe on behalf of the Wellcome Sanger Institute COVID-19 Surveillance Team |
| EPI_ISL_3361630, EPI_ISL_3361632, EPI_ISL_3361634, EPI_ISL_3361635, EPI_ISL_3361636 | Lighthouse Laboratory Plymouth | Wellcome Sanger Institute for the COVID-19 Genomics UK (COG-UK) Consortium | Lighthouse Laboratory Plymouth and Alex Alderton, Roberto Amato, Jeffrey Barrett, Sonia Goncalves, Ewan Harrison, David K. Jackson, Ian Johnston, Dominic Kwiatkowski, Cordelia Langford, John Sillitoe on behalf of the Wellcome Sanger Institute COVID-19 Surveillance Team |
| EPI_ISL_3361638, EPI_ISL_3361639 | Lighthouse Lab in Glasgow | Wellcome Sanger Institute for the COVID-19 Genomics UK (COG-UK) Consortium | Harper VanSteenhouse, Yumi Kasai, David Gray, Carol Clugston, Anna Dominiczak and Alex Alderton, Roberto Amato, Jeffrey Barrett, Sonia Goncalves, Ewan Harrison, David K. Jackson, Ian Johnston, Dominic Kwiatkowski, Cordelia Langford, John Sillitoe on behalf of the Wellcome Sanger Institute COVID-19 Surveillance Team |
| EPI_ISL_3361641 | Health Services Laboratories | Wellcome Sanger Institute for the COVID-19 Genomics UK (COG-UK) Consortium | Health Services Laboratories and Alex Alderton, Roberto Amato, Jeffrey Barrett, Sonia Goncalves, Ewan Harrison, David K. Jackson, Ian Johnston, Dominic Kwiatkowski, Cordelia Langford, John Sillitoe on behalf of the Wellcome Sanger Institute COVID-19 Surveillance Team |
| EPI_ISL_3361642 | Lighthouse Laboratory Plymouth | Wellcome Sanger Institute for the COVID-19 Genomics UK (COG-UK) Consortium | Lighthouse Laboratory Plymouth and Alex Alderton, Roberto Amato, Jeffrey Barrett, Sonia Goncalves, Ewan Harrison, David K. Jackson, Ian Johnston, Dominic Kwiatkowski, Cordelia Langford, John Sillitoe on behalf of the Wellcome Sanger Institute COVID-19 Surveillance Team |
| EPI_ISL_3361644 | Health Services Laboratories | Wellcome Sanger Institute for the COVID-19 Genomics UK (COG-UK) Consortium | Health Services Laboratories and Alex Alderton, Roberto Amato, Jeffrey Barrett, Sonia Goncalves, Ewan Harrison, David K. Jackson, Ian Johnston, Dominic Kwiatkowski, Cordelia Langford, John Sillitoe on behalf of the Wellcome Sanger Institute COVID-19 Surveillance Team |
| EPI_ISL_3361645 | Lighthouse Laboratory Plymouth | Wellcome Sanger Institute for the COVID-19 Genomics UK (COG-UK) Consortium | Lighthouse Laboratory Plymouth and Alex Alderton, Roberto Amato, Jeffrey Barrett, Sonia Goncalves, Ewan Harrison, David K. Jackson, Ian Johnston, Dominic Kwiatkowski, Cordelia Langford, John Sillitoe on behalf of the Wellcome Sanger Institute COVID-19 Surveillance Team |
| EPI_ISL_3361647 | Lighthouse Lab in Alderley Park | Wellcome Sanger Institute for the COVID-19 Genomics UK (COG-UK) Consortium | Jacquelyn Wynn, Mairead Hyland, The Lighthouse Lab in Alderley Park and Alex Alderton, Roberto Amato, Jeffrey Barrett, Sonia Goncalves, Ewan Harrison, David K. Jackson, Ian Johnston, Dominic Kwiatkowski, Cordelia Langford, John Sillitoe on behalf of the Wellcome Sanger Institute COVID-19 Surveillance Team |
| EPI_ISL_3361648 | Lighthouse Laboratory Plymouth | Wellcome Sanger Institute for the COVID-19 Genomics UK (COG-UK) Consortium | Lighthouse Laboratory Plymouth and Alex Alderton, Roberto Amato, Jeffrey Barrett, Sonia Goncalves, Ewan Harrison, David K. Jackson, Ian Johnston, Dominic Kwiatkowski, Cordelia Langford, John Sillitoe on behalf of the Wellcome Sanger Institute COVID-19 Surveillance Team |
| EPI_ISL_3361650 | Health Services Laboratories | Wellcome Sanger Institute for the COVID-19 Genomics UK (COG-UK) Consortium | Health Services Laboratories and Alex Alderton, Roberto Amato, Jeffrey Barrett, Sonia Goncalves, Ewan Harrison, David K. Jackson, Ian Johnston, Dominic Kwiatkowski, Cordelia Langford, John Sillitoe on behalf of the Wellcome Sanger Institute COVID-19 Surveillance Team |
| EPI_ISL_3361652 | Lighthouse Laboratory Plymouth | Wellcome Sanger Institute for the COVID-19 Genomics UK (COG-UK) Consortium | Lighthouse Laboratory Plymouth and Alex Alderton, Roberto Amato, Jeffrey Barrett, Sonia Goncalves, Ewan Harrison, David K. Jackson, Ian Johnston, Dominic Kwiatkowski, Cordelia Langford, John Sillitoe on behalf of the Wellcome Sanger Institute COVID-19 Surveillance Team |
| EPI_ISL_3361653 | Lighthouse Lab in Milton Keynes | Wellcome Sanger Institute for the COVID-19 Genomics UK (COG-UK) Consortium | The Lighthouse Lab in Milton Keynes and Alex Alderton, Roberto Amato, Jeffrey Barrett, Sonia Goncalves, Ewan Harrison, David K. Jackson, Ian Johnston, Dominic Kwiatkowski, Cordelia Langford, John Sillitoe on behalf of the Wellcome Sanger Institute COVID-19 Surveillance Team |
| EPI_ISL_3361654, EPI_ISL_3361656, EPI_ISL_3361657 | Lighthouse Laboratory Plymouth | Wellcome Sanger Institute for the COVID-19 Genomics UK (COG-UK) Consortium | Lighthouse Laboratory Plymouth and Alex Alderton, Roberto Amato, Jeffrey Barrett, Sonia Goncalves, Ewan Harrison, David K. Jackson, Ian Johnston, Dominic Kwiatkowski, Cordelia Langford, John Sillitoe on behalf of the Wellcome Sanger Institute COVID-19 Surveillance Team |
| EPI_ISL_3361659 | Health Services Laboratories | Wellcome Sanger Institute for the COVID-19 Genomics UK (COG-UK) | Health Services Laboratories and Alex Alderton, Roberto Amato, Jeffrey Barrett, Sonia Goncalves, Ewan Harrison, David K. Jackson, Ian Johnston, Dominic Kwiatkowski, Cordelia Langford, John Sillitoe on behalf of the Wellcome Sanger Institute |

| | | | |
|---|---|---|---|
| | | Consortium | COVID-19 Surveillance Team |
| EPI_ISL_3361660, EPI_ISL_3361661 | Lighthouse Lab in Milton Keynes | Wellcome Sanger Institute for the COVID-19 Genomics UK (COG-UK) Consortium | The Lighthouse Lab in Milton Keynes and Alex Alderton, Roberto Amato, Jeffrey Barrett, Sonia Goncalves, Ewan Harrison, David K. Jackson, Ian Johnston, Dominic Kwiatkowski, Cordelia Langford, John Sillitoe on behalf of the Wellcome Sanger Institute COVID-19 Surveillance Team |
| EPI_ISL_3361663, EPI_ISL_3361664 | Lighthouse Lab in Glasgow | Wellcome Sanger Institute for the COVID-19 Genomics UK (COG-UK) Consortium | Harper VanSteenhouse, Yumi Kasai, David Gray, Carol Clugston, Anna Dominiczak and Alex Alderton, Roberto Amato, Jeffrey Barrett, Sonia Goncalves, Ewan Harrison, David K. Jackson, Ian Johnston, Dominic Kwiatkowski, Cordelia Langford, John Sillitoe on behalf of the Wellcome Sanger Institute COVID-19 Surveillance Team |
| EPI_ISL_3361666 | Lighthouse Lab in Milton Keynes | Wellcome Sanger Institute for the COVID-19 Genomics UK (COG-UK) Consortium | The Lighthouse Lab in Milton Keynes and Alex Alderton, Roberto Amato, Jeffrey Barrett, Sonia Goncalves, Ewan Harrison, David K. Jackson, Ian Johnston, Dominic Kwiatkowski, Cordelia Langford, John Sillitoe on behalf of the Wellcome Sanger Institute COVID-19 Surveillance Team |
| EPI_ISL_3361668 | Lighthouse Lab in Alderley Park | Wellcome Sanger Institute for the COVID-19 Genomics UK (COG-UK) Consortium | Jacquelyn Wynn, Mairead Hyland, The Lighthouse Lab in Alderley Park and Alex Alderton, Roberto Amato, Jeffrey Barrett, Sonia Goncalves, Ewan Harrison, David K. Jackson, Ian Johnston, Dominic Kwiatkowski, Cordelia Langford, John Sillitoe on behalf of the Wellcome Sanger Institute COVID-19 Surveillance Team |
| EPI_ISL_3361669 | Lighthouse Laboratory Plymouth | Wellcome Sanger Institute for the COVID-19 Genomics UK (COG-UK) Consortium | Lighthouse Laboratory Plymouth and Alex Alderton, Roberto Amato, Jeffrey Barrett, Sonia Goncalves, Ewan Harrison, David K. Jackson, Ian Johnston, Dominic Kwiatkowski, Cordelia Langford, John Sillitoe on behalf of the Wellcome Sanger Institute COVID-19 Surveillance Team |
| EPI_ISL_3361671 | Lighthouse Lab in Glasgow | Wellcome Sanger Institute for the COVID-19 Genomics UK (COG-UK) Consortium | Harper VanSteenhouse, Yumi Kasai, David Gray, Carol Clugston, Anna Dominiczak and Alex Alderton, Roberto Amato, Jeffrey Barrett, Sonia Goncalves, Ewan Harrison, David K. Jackson, Ian Johnston, Dominic Kwiatkowski, Cordelia Langford, John Sillitoe on behalf of the Wellcome Sanger Institute COVID-19 Surveillance Team |
| EPI_ISL_3361672 | Lighthouse Lab in Milton Keynes | Wellcome Sanger Institute for the COVID-19 Genomics UK (COG-UK) Consortium | The Lighthouse Lab in Milton Keynes and Alex Alderton, Roberto Amato, Jeffrey Barrett, Sonia Goncalves, Ewan Harrison, David K. Jackson, Ian Johnston, Dominic Kwiatkowski, Cordelia Langford, John Sillitoe on behalf of the Wellcome Sanger Institute COVID-19 Surveillance Team |
| EPI_ISL_3361673 | Lighthouse Lab in Glasgow | Wellcome Sanger Institute for the COVID-19 Genomics UK (COG-UK) Consortium | Harper VanSteenhouse, Yumi Kasai, David Gray, Carol Clugston, Anna Dominiczak and Alex Alderton, Roberto Amato, Jeffrey Barrett, Sonia Goncalves, Ewan Harrison, David K. Jackson, Ian Johnston, Dominic Kwiatkowski, Cordelia Langford, John Sillitoe on behalf of the Wellcome Sanger Institute COVID-19 Surveillance Team |
| EPI_ISL_3361675 | Lighthouse Lab in Milton Keynes | Wellcome Sanger Institute for the COVID-19 Genomics UK (COG-UK) Consortium | The Lighthouse Lab in Milton Keynes and Alex Alderton, Roberto Amato, Jeffrey Barrett, Sonia Goncalves, Ewan Harrison, David K. Jackson, Ian Johnston, Dominic Kwiatkowski, Cordelia Langford, John Sillitoe on behalf of the Wellcome Sanger Institute COVID-19 Surveillance Team |
| EPI_ISL_3361676 | Lighthouse Laboratory Plymouth | Wellcome Sanger Institute for the COVID-19 Genomics UK (COG-UK) Consortium | Lighthouse Laboratory Plymouth and Alex Alderton, Roberto Amato, Jeffrey Barrett, Sonia Goncalves, Ewan Harrison, David K. Jackson, Ian Johnston, Dominic Kwiatkowski, Cordelia Langford, John Sillitoe on behalf of the Wellcome Sanger Institute COVID-19 Surveillance Team |
| EPI_ISL_3361678 | Lighthouse Lab in Glasgow | Wellcome Sanger Institute for the COVID-19 Genomics UK (COG-UK) Consortium | Harper VanSteenhouse, Yumi Kasai, David Gray, Carol Clugston, Anna Dominiczak and Alex Alderton, Roberto Amato, Jeffrey Barrett, Sonia Goncalves, Ewan Harrison, David K. Jackson, Ian Johnston, Dominic Kwiatkowski, Cordelia Langford, John Sillitoe on behalf of the Wellcome Sanger Institute COVID-19 Surveillance Team |
| EPI_ISL_3361679 | Health Services Laboratories | Wellcome Sanger Institute for the COVID-19 Genomics UK (COG-UK) Consortium | Health Services Laboratories and Alex Alderton, Roberto Amato, Jeffrey Barrett, Sonia Goncalves, Ewan Harrison, David K. Jackson, Ian Johnston, Dominic Kwiatkowski, Cordelia Langford, John Sillitoe on behalf of the Wellcome Sanger Institute COVID-19 Surveillance Team |
| EPI_ISL_3361680, EPI_ISL_3361682, EPI_ISL_3361684 | Lighthouse Lab in Milton Keynes | Wellcome Sanger Institute for the COVID-19 Genomics UK (COG-UK) Consortium | The Lighthouse Lab in Milton Keynes and Alex Alderton, Roberto Amato, Jeffrey Barrett, Sonia Goncalves, Ewan Harrison, David K. Jackson, Ian Johnston, Dominic Kwiatkowski, Cordelia Langford, John Sillitoe on behalf of the Wellcome Sanger Institute COVID-19 Surveillance Team |
| EPI_ISL_3361685, EPI_ISL_3361686 | Lighthouse Laboratory Plymouth | Wellcome Sanger Institute for the COVID-19 Genomics UK (COG-UK) Consortium | Lighthouse Laboratory Plymouth and Alex Alderton, Roberto Amato, Jeffrey Barrett, Sonia Goncalves, Ewan Harrison, David K. Jackson, Ian Johnston, Dominic Kwiatkowski, Cordelia Langford, John Sillitoe on behalf of the Wellcome Sanger Institute COVID-19 Surveillance Team |
| EPI_ISL_3361688 | Lighthouse Lab in Milton Keynes | Wellcome Sanger Institute for the COVID-19 Genomics UK (COG-UK) Consortium | The Lighthouse Lab in Milton Keynes and Alex Alderton, Roberto Amato, Jeffrey Barrett, Sonia Goncalves, Ewan Harrison, David K. Jackson, Ian Johnston, Dominic Kwiatkowski, Cordelia Langford, John Sillitoe on behalf of the Wellcome Sanger Institute COVID-19 Surveillance Team |
| EPI_ISL_3361689 | Lighthouse Lab in Glasgow | Wellcome Sanger Institute for the COVID-19 Genomics UK (COG-UK) Consortium | Harper VanSteenhouse, Yumi Kasai, David Gray, Carol Clugston, Anna Dominiczak and Alex Alderton, Roberto Amato, Jeffrey Barrett, Sonia Goncalves, Ewan Harrison, David K. Jackson, Ian Johnston, Dominic Kwiatkowski, Cordelia Langford, John Sillitoe on behalf of the Wellcome Sanger Institute COVID-19 Surveillance Team |
| EPI_ISL_3361691 | Lighthouse Lab in Milton Keynes | Wellcome Sanger Institute for the COVID-19 Genomics UK (COG-UK) Consortium | The Lighthouse Lab in Milton Keynes and Alex Alderton, Roberto Amato, Jeffrey Barrett, Sonia Goncalves, Ewan Harrison, David K. Jackson, Ian Johnston, Dominic Kwiatkowski, Cordelia Langford, John Sillitoe on behalf of the Wellcome Sanger Institute COVID-19 Surveillance Team |
| EPI_ISL_3361692 | Lighthouse Lab in Alderley Park | Wellcome Sanger Institute for the COVID-19 Genomics UK (COG-UK) Consortium | Jacquelyn Wynn, Mairead Hyland, The Lighthouse Lab in Alderley Park and Alex Alderton, Roberto Amato, Jeffrey Barrett, Sonia Goncalves, Ewan Harrison, David K. Jackson, Ian Johnston, Dominic Kwiatkowski, Cordelia Langford, John Sillitoe on behalf of the Wellcome Sanger Institute COVID-19 Surveillance Team |
| EPI_ISL_3361693 | Lighthouse Laboratory Plymouth | Wellcome Sanger Institute for the COVID-19 Genomics UK (COG-UK) Consortium | Lighthouse Laboratory Plymouth and Alex Alderton, Roberto Amato, Jeffrey Barrett, Sonia Goncalves, Ewan Harrison, David K. Jackson, Ian Johnston, Dominic Kwiatkowski, Cordelia Langford, John Sillitoe on behalf of the Wellcome Sanger Institute COVID-19 Surveillance Team |
| EPI_ISL_3361695 | Lighthouse Lab in Alderley Park | Wellcome Sanger Institute for the COVID-19 Genomics UK (COG-UK) Consortium | Jacquelyn Wynn, Mairead Hyland, The Lighthouse Lab in Alderley Park and Alex Alderton, Roberto Amato, Jeffrey Barrett, Sonia Goncalves, Ewan Harrison, David K. Jackson, Ian Johnston, Dominic Kwiatkowski, Cordelia Langford, John Sillitoe on behalf of the Wellcome Sanger Institute COVID-19 Surveillance Team |
| EPI_ISL_3361696 | Lighthouse Laboratory Plymouth | Wellcome Sanger Institute for the COVID-19 Genomics UK (COG-UK) Consortium | Lighthouse Laboratory Plymouth and Alex Alderton, Roberto Amato, Jeffrey Barrett, Sonia Goncalves, Ewan Harrison, David K. Jackson, Ian Johnston, Dominic Kwiatkowski, Cordelia Langford, John Sillitoe on behalf of the Wellcome Sanger Institute COVID-19 Surveillance Team |
| EPI_ISL_3361697 | Health Services Laboratories | Wellcome Sanger Institute for the COVID-19 Genomics UK (COG-UK) Consortium | Health Services Laboratories and Alex Alderton, Roberto Amato, Jeffrey Barrett, Sonia Goncalves, Ewan Harrison, David K. Jackson, Ian Johnston, Dominic Kwiatkowski, Cordelia Langford, John Sillitoe on behalf of the Wellcome Sanger Institute COVID-19 Surveillance Team |
| EPI_ISL_3361698 | Lighthouse Lab in Glasgow | Wellcome Sanger Institute for the COVID-19 Genomics UK (COG-UK) Consortium | Harper VanSteenhouse, Yumi Kasai, David Gray, Carol Clugston, Anna Dominiczak and Alex Alderton, Roberto Amato, Jeffrey Barrett, Sonia Goncalves, Ewan Harrison, David K. Jackson, Ian Johnston, Dominic Kwiatkowski, Cordelia Langford, John Sillitoe on behalf of the Wellcome Sanger Institute COVID-19 Surveillance Team |
| EPI_ISL_3361699 | Lighthouse Lab in Milton Keynes | Wellcome Sanger Institute for the COVID-19 Genomics UK (COG-UK) Consortium | The Lighthouse Lab in Milton Keynes and Alex Alderton, Roberto Amato, Jeffrey Barrett, Sonia Goncalves, Ewan Harrison, David K. Jackson, Ian Johnston, Dominic Kwiatkowski, Cordelia Langford, John Sillitoe on behalf of the Wellcome Sanger Institute COVID-19 Surveillance Team |
| EPI_ISL_3361700 | Lighthouse Lab in Glasgow | Wellcome Sanger Institute for the COVID-19 Genomics UK (COG-UK) Consortium | Harper VanSteenhouse, Yumi Kasai, David Gray, Carol Clugston, Anna Dominiczak and Alex Alderton, Roberto Amato, Jeffrey Barrett, Sonia Goncalves, Ewan Harrison, David K. Jackson, Ian Johnston, Dominic Kwiatkowski, Cordelia Langford, John Sillitoe on behalf of the Wellcome Sanger Institute COVID-19 Surveillance Team |
| EPI_ISL_3361701 | Lighthouse Laboratory Plymouth | Wellcome Sanger Institute for the COVID-19 Genomics UK (COG-UK) Consortium | Lighthouse Laboratory Plymouth and Alex Alderton, Roberto Amato, Jeffrey Barrett, Sonia Goncalves, Ewan Harrison, David K. Jackson, Ian Johnston, Dominic Kwiatkowski, Cordelia Langford, John Sillitoe on behalf of the Wellcome Sanger Institute COVID-19 Surveillance Team |
| EPI_ISL_3361702, EPI_ISL_3361703 | Health Services Laboratories | Wellcome Sanger Institute for the COVID-19 Genomics UK (COG-UK) Consortium | Health Services Laboratories and Alex Alderton, Roberto Amato, Jeffrey Barrett, Sonia Goncalves, Ewan Harrison, David K. Jackson, Ian Johnston, Dominic Kwiatkowski, Cordelia Langford, John Sillitoe on behalf of the Wellcome Sanger Institute COVID-19 Surveillance Team |
| EPI_ISL_3361704 | Lighthouse Lab in Alderley Park | Wellcome Sanger Institute for the COVID-19 Genomics UK (COG-UK) Consortium | Jacquelyn Wynn, Mairead Hyland, The Lighthouse Lab in Alderley Park and Alex Alderton, Roberto Amato, Jeffrey Barrett, Sonia Goncalves, Ewan Harrison, David K. Jackson, Ian Johnston, Dominic Kwiatkowski, Cordelia Langford, John Sillitoe on behalf of the Wellcome Sanger Institute COVID-19 Surveillance Team |
| EPI_ISL_3361706 | Health Services Laboratories | Wellcome Sanger Institute for the COVID-19 Genomics UK (COG-UK) Consortium | Health Services Laboratories and Alex Alderton, Roberto Amato, Jeffrey Barrett, Sonia Goncalves, Ewan Harrison, David K. Jackson, Ian Johnston, Dominic Kwiatkowski, Cordelia Langford, John Sillitoe on behalf of the Wellcome Sanger Institute COVID-19 Surveillance Team |
| EPI_ISL_3361707, EPI_ISL_3361709, EPI_ISL_3361710, EPI_ISL_3361712, EPI_ISL_3361713 | Lighthouse Lab in Milton Keynes | Wellcome Sanger Institute for the COVID-19 Genomics UK (COG-UK) Consortium | The Lighthouse Lab in Milton Keynes and Alex Alderton, Roberto Amato, Jeffrey Barrett, Sonia Goncalves, Ewan Harrison, David K. Jackson, Ian Johnston, Dominic Kwiatkowski, Cordelia Langford, John Sillitoe on behalf of the Wellcome Sanger Institute COVID-19 Surveillance Team |
| EPI_ISL_3361715 | Lighthouse Laboratory Plymouth | Wellcome Sanger Institute for the COVID-19 Genomics UK (COG-UK) Consortium | Lighthouse Laboratory Plymouth and Alex Alderton, Roberto Amato, Jeffrey Barrett, Sonia Goncalves, Ewan Harrison, David K. Jackson, Ian Johnston, Dominic Kwiatkowski, Cordelia Langford, John Sillitoe on behalf of the Wellcome Sanger Institute COVID-19 Surveillance Team |
| EPI_ISL_3361716, EPI_ISL_3361718 | Health Services Laboratories | Wellcome Sanger Institute for the COVID-19 Genomics UK (COG-UK) Consortium | Health Services Laboratories and Alex Alderton, Roberto Amato, Jeffrey Barrett, Sonia Goncalves, Ewan Harrison, David K. Jackson, Ian Johnston, Dominic Kwiatkowski, Cordelia Langford, John Sillitoe on behalf of the Wellcome Sanger Institute COVID-19 Surveillance Team |
| EPI_ISL_3361719 | Lighthouse Lab in Glasgow | Wellcome Sanger Institute for the COVID-19 Genomics UK (COG-UK) Consortium | Harper VanSteenhouse, Yumi Kasai, David Gray, Carol Clugston, Anna Dominiczak and Alex Alderton, Roberto Amato, Jeffrey Barrett, Sonia Goncalves, Ewan Harrison, David K. Jackson, Ian Johnston, Dominic Kwiatkowski, Cordelia Langford, John Sillitoe on behalf of the Wellcome Sanger Institute COVID-19 Surveillance Team |
| EPI_ISL_3361720, EPI_ISL_3361722, EPI_ISL_3361723 | Lighthouse Lab in Alderley Park | Wellcome Sanger Institute for the COVID-19 Genomics UK (COG-UK) Consortium | Jacquelyn Wynn, Mairead Hyland, The Lighthouse Lab in Alderley Park and Alex Alderton, Roberto Amato, Jeffrey Barrett, Sonia Goncalves, Ewan Harrison, David K. Jackson, Ian Johnston, Dominic Kwiatkowski, Cordelia Langford, John Sillitoe on behalf of the Wellcome Sanger Institute COVID-19 Surveillance Team |
| EPI_ISL_3361725 | Lighthouse Lab in Milton Keynes | Wellcome Sanger Institute for the COVID-19 Genomics UK (COG-UK) Consortium | The Lighthouse Lab in Milton Keynes and Alex Alderton, Roberto Amato, Jeffrey Barrett, Sonia Goncalves, Ewan Harrison, David K. Jackson, Ian Johnston, Dominic Kwiatkowski, Cordelia Langford, John Sillitoe on behalf of the Wellcome Sanger Institute COVID-19 Surveillance Team |
| EPI_ISL_3361727 | Lighthouse Laboratory Plymouth | Wellcome Sanger Institute for the COVID-19 Genomics UK (COG-UK) Consortium | Lighthouse Laboratory Plymouth and Alex Alderton, Roberto Amato, Jeffrey Barrett, Sonia Goncalves, Ewan Harrison, David K. Jackson, Ian Johnston, Dominic Kwiatkowski, Cordelia Langford, John Sillitoe on behalf of the Wellcome Sanger Institute COVID-19 Surveillance Team |
| EPI_ISL_3361728 | Health Services Laboratories | Wellcome Sanger Institute for the COVID-19 Genomics UK (COG-UK) Consortium | Health Services Laboratories and Alex Alderton, Roberto Amato, Jeffrey Barrett, Sonia Goncalves, Ewan Harrison, David K. Jackson, Ian Johnston, Dominic Kwiatkowski, Cordelia Langford, John Sillitoe on behalf of the Wellcome Sanger Institute COVID-19 Surveillance Team |
| EPI_ISL_3361730, EPI_ISL_3361731, EPI_ISL_3361733, EPI_ISL_3361734, EPI_ISL_3361736, EPI_ISL_3361737 | Lighthouse Lab in Milton Keynes | Wellcome Sanger Institute for the COVID-19 Genomics UK (COG-UK) Consortium | The Lighthouse Lab in Milton Keynes and Alex Alderton, Roberto Amato, Jeffrey Barrett, Sonia Goncalves, Ewan Harrison, David K. Jackson, Ian Johnston, Dominic Kwiatkowski, Cordelia Langford, John Sillitoe on behalf of the Wellcome Sanger Institute COVID-19 Surveillance Team |
| EPI_ISL_3361739, EPI_ISL_3361740, EPI_ISL_3361742 | Lighthouse Laboratory Plymouth | Wellcome Sanger Institute for the COVID-19 Genomics UK (COG-UK) Consortium | Lighthouse Laboratory Plymouth and Alex Alderton, Roberto Amato, Jeffrey Barrett, Sonia Goncalves, Ewan Harrison, David K. Jackson, Ian Johnston, Dominic Kwiatkowski, Cordelia Langford, John Sillitoe on behalf of the Wellcome Sanger Institute COVID-19 Surveillance Team |
| EPI_ISL_3361744 | Lighthouse Lab in Milton Keynes | Wellcome Sanger Institute for the COVID-19 Genomics UK (COG-UK) Consortium | The Lighthouse Lab in Milton Keynes and Alex Alderton, Roberto Amato, Jeffrey Barrett, Sonia Goncalves, Ewan Harrison, David K. Jackson, Ian Johnston, Dominic Kwiatkowski, Cordelia Langford, John Sillitoe on behalf of the Wellcome Sanger Institute COVID-19 Surveillance Team |
| EPI_ISL_3361746 | Lighthouse Lab in Alderley Park | Wellcome Sanger Institute for the COVID-19 Genomics UK (COG-UK) | Jacquelyn Wynn, Mairead Hyland, The Lighthouse Lab in Alderley Park and Alex Alderton, Roberto Amato, Jeffrey Barrett, Sonia Goncalves, Ewan Harrison, David K. Jackson, Ian Johnston, Dominic Kwiatkowski, Cordelia Langford, John Sillitoe on |

| | | | |
|---|---|---|---|
| | | Consortium | behalf of the Wellcome Sanger Institute COVID-19 Surveillance Team |
| EPI_ISL_3361747 | Lighthouse Lab in Milton Keynes | Wellcome Sanger Institute for the COVID-19 Genomics UK (COG-UK) Consortium | The Lighthouse Lab in Milton Keynes and Alex Alderton, Roberto Amato, Jeffrey Barrett, Sonia Goncalves, Ewan Harrison, David K. Jackson, Ian Johnston, Dominic Kwiatkowski, Cordelia Langford, John Sillitoe on behalf of the Wellcome Sanger Institute COVID-19 Surveillance Team |
| EPI_ISL_3361749, EPI_ISL_3361750, EPI_ISL_3361752 | Lighthouse Laboratory Plymouth | Wellcome Sanger Institute for the COVID-19 Genomics UK (COG-UK) Consortium | Lighthouse Laboratory Plymouth and Alex Alderton, Roberto Amato, Jeffrey Barrett, Sonia Goncalves, Ewan Harrison, David K. Jackson, Ian Johnston, Dominic Kwiatkowski, Cordelia Langford, John Sillitoe on behalf of the Wellcome Sanger Institute COVID-19 Surveillance Team |
| EPI_ISL_3361753 | Lighthouse Lab in Milton Keynes | Wellcome Sanger Institute for the COVID-19 Genomics UK (COG-UK) Consortium | The Lighthouse Lab in Milton Keynes and Alex Alderton, Roberto Amato, Jeffrey Barrett, Sonia Goncalves, Ewan Harrison, David K. Jackson, Ian Johnston, Dominic Kwiatkowski, Cordelia Langford, John Sillitoe on behalf of the Wellcome Sanger Institute COVID-19 Surveillance Team |
| EPI_ISL_3361755 | Health Services Laboratories | Wellcome Sanger Institute for the COVID-19 Genomics UK (COG-UK) Consortium | Health Services Laboratories and Alex Alderton, Roberto Amato, Jeffrey Barrett, Sonia Goncalves, Ewan Harrison, David K. Jackson, Ian Johnston, Dominic Kwiatkowski, Cordelia Langford, John Sillitoe on behalf of the Wellcome Sanger Institute COVID-19 Surveillance Team |
| EPI_ISL_3361756, EPI_ISL_3361758, EPI_ISL_3361760, EPI_ISL_3361761, EPI_ISL_3361763 | Lighthouse Lab in Glasgow | Wellcome Sanger Institute for the COVID-19 Genomics UK (COG-UK) Consortium | Harper VanSteenhouse, Yumi Kasai, David Gray, Carol Clugston, Anna Dominiczak and Alex Alderton, Roberto Amato, Jeffrey Barrett, Sonia Goncalves, Ewan Harrison, David K. Jackson, Ian Johnston, Dominic Kwiatkowski, Cordelia Langford, John Sillitoe on behalf of the Wellcome Sanger Institute COVID-19 Surveillance Team |
| EPI_ISL_3361765 | Lighthouse Laboratory Plymouth | Wellcome Sanger Institute for the COVID-19 Genomics UK (COG-UK) Consortium | Lighthouse Laboratory Plymouth and Alex Alderton, Roberto Amato, Jeffrey Barrett, Sonia Goncalves, Ewan Harrison, David K. Jackson, Ian Johnston, Dominic Kwiatkowski, Cordelia Langford, John Sillitoe on behalf of the Wellcome Sanger Institute COVID-19 Surveillance Team |
| EPI_ISL_3361766 | Lighthouse Lab in Alderley Park | Wellcome Sanger Institute for the COVID-19 Genomics UK (COG-UK) Consortium | Jacquelyn Wynn, Mairead Hyland, The Lighthouse Lab in Alderley Park and Alex Alderton, Roberto Amato, Jeffrey Barrett, Sonia Goncalves, Ewan Harrison, David K. Jackson, Ian Johnston, Dominic Kwiatkowski, Cordelia Langford, John Sillitoe on behalf of the Wellcome Sanger Institute COVID-19 Surveillance Team |
| EPI_ISL_3361768, EPI_ISL_3361769, EPI_ISL_3361771, EPI_ISL_3361772 | Lighthouse Lab in Milton Keynes | Wellcome Sanger Institute for the COVID-19 Genomics UK (COG-UK) Consortium | The Lighthouse Lab in Milton Keynes and Alex Alderton, Roberto Amato, Jeffrey Barrett, Sonia Goncalves, Ewan Harrison, David K. Jackson, Ian Johnston, Dominic Kwiatkowski, Cordelia Langford, John Sillitoe on behalf of the Wellcome Sanger Institute COVID-19 Surveillance Team |
| EPI_ISL_3361774, EPI_ISL_3361775 | Lighthouse Laboratory Plymouth | Wellcome Sanger Institute for the COVID-19 Genomics UK (COG-UK) Consortium | Lighthouse Laboratory Plymouth and Alex Alderton, Roberto Amato, Jeffrey Barrett, Sonia Goncalves, Ewan Harrison, David K. Jackson, Ian Johnston, Dominic Kwiatkowski, Cordelia Langford, John Sillitoe on behalf of the Wellcome Sanger Institute COVID-19 Surveillance Team |
| EPI_ISL_3361776 | Lighthouse Lab in Glasgow | Wellcome Sanger Institute for the COVID-19 Genomics UK (COG-UK) Consortium | Harper VanSteenhouse, Yumi Kasai, David Gray, Carol Clugston, Anna Dominiczak and Alex Alderton, Roberto Amato, Jeffrey Barrett, Sonia Goncalves, Ewan Harrison, David K. Jackson, Ian Johnston, Dominic Kwiatkowski, Cordelia Langford, John Sillitoe on behalf of the Wellcome Sanger Institute COVID-19 Surveillance Team |
| EPI_ISL_3361778, EPI_ISL_3361780 | Lighthouse Laboratory Plymouth | Wellcome Sanger Institute for the COVID-19 Genomics UK (COG-UK) Consortium | Lighthouse Laboratory Plymouth and Alex Alderton, Roberto Amato, Jeffrey Barrett, Sonia Goncalves, Ewan Harrison, David K. Jackson, Ian Johnston, Dominic Kwiatkowski, Cordelia Langford, John Sillitoe on behalf of the Wellcome Sanger Institute COVID-19 Surveillance Team |
| EPI_ISL_3361781 | Health Services Laboratories | Wellcome Sanger Institute for the COVID-19 Genomics UK (COG-UK) Consortium | Health Services Laboratories and Alex Alderton, Roberto Amato, Jeffrey Barrett, Sonia Goncalves, Ewan Harrison, David K. Jackson, Ian Johnston, Dominic Kwiatkowski, Cordelia Langford, John Sillitoe on behalf of the Wellcome Sanger Institute COVID-19 Surveillance Team |
| EPI_ISL_3361783 | Lighthouse Laboratory Plymouth | Wellcome Sanger Institute for the COVID-19 Genomics UK (COG-UK) Consortium | Lighthouse Laboratory Plymouth and Alex Alderton, Roberto Amato, Jeffrey Barrett, Sonia Goncalves, Ewan Harrison, David K. Jackson, Ian Johnston, Dominic Kwiatkowski, Cordelia Langford, John Sillitoe on behalf of the Wellcome Sanger Institute COVID-19 Surveillance Team |
| EPI_ISL_3361784 | Lighthouse Lab in Glasgow | Wellcome Sanger Institute for the COVID-19 Genomics UK (COG-UK) Consortium | Harper VanSteenhouse, Yumi Kasai, David Gray, Carol Clugston, Anna Dominiczak and Alex Alderton, Roberto Amato, Jeffrey Barrett, Sonia Goncalves, Ewan Harrison, David K. Jackson, Ian Johnston, Dominic Kwiatkowski, Cordelia Langford, John Sillitoe on behalf of the Wellcome Sanger Institute COVID-19 Surveillance Team |
| EPI_ISL_3361786 | Health Services Laboratories | Wellcome Sanger Institute for the COVID-19 Genomics UK (COG-UK) Consortium | Health Services Laboratories and Alex Alderton, Roberto Amato, Jeffrey Barrett, Sonia Goncalves, Ewan Harrison, David K. Jackson, Ian Johnston, Dominic Kwiatkowski, Cordelia Langford, John Sillitoe on behalf of the Wellcome Sanger Institute COVID-19 Surveillance Team |
| EPI_ISL_3361787 | Lighthouse Lab in Milton Keynes | Wellcome Sanger Institute for the COVID-19 Genomics UK (COG-UK) Consortium | The Lighthouse Lab in Milton Keynes and Alex Alderton, Roberto Amato, Jeffrey Barrett, Sonia Goncalves, Ewan Harrison, David K. Jackson, Ian Johnston, Dominic Kwiatkowski, Cordelia Langford, John Sillitoe on behalf of the Wellcome Sanger Institute COVID-19 Surveillance Team |
| EPI_ISL_3361789 | Health Services Laboratories | Wellcome Sanger Institute for the COVID-19 Genomics UK (COG-UK) Consortium | Health Services Laboratories and Alex Alderton, Roberto Amato, Jeffrey Barrett, Sonia Goncalves, Ewan Harrison, David K. Jackson, Ian Johnston, Dominic Kwiatkowski, Cordelia Langford, John Sillitoe on behalf of the Wellcome Sanger Institute COVID-19 Surveillance Team |
| EPI_ISL_3361790 | Lighthouse Laboratory Plymouth | Wellcome Sanger Institute for the COVID-19 Genomics UK (COG-UK) Consortium | Lighthouse Laboratory Plymouth and Alex Alderton, Roberto Amato, Jeffrey Barrett, Sonia Goncalves, Ewan Harrison, David K. Jackson, Ian Johnston, Dominic Kwiatkowski, Cordelia Langford, John Sillitoe on behalf of the Wellcome Sanger Institute COVID-19 Surveillance Team |
| EPI_ISL_3361792 | Lighthouse Lab in Glasgow | Wellcome Sanger Institute for the COVID-19 Genomics UK (COG-UK) Consortium | Harper VanSteenhouse, Yumi Kasai, David Gray, Carol Clugston, Anna Dominiczak and Alex Alderton, Roberto Amato, Jeffrey Barrett, Sonia Goncalves, Ewan Harrison, David K. Jackson, Ian Johnston, Dominic Kwiatkowski, Cordelia Langford, John Sillitoe on behalf of the Wellcome Sanger Institute COVID-19 Surveillance Team |
| EPI_ISL_3361793 | Lighthouse Laboratory Plymouth | Wellcome Sanger Institute for the COVID-19 Genomics UK (COG-UK) Consortium | Lighthouse Laboratory Plymouth and Alex Alderton, Roberto Amato, Jeffrey Barrett, Sonia Goncalves, Ewan Harrison, David K. Jackson, Ian Johnston, Dominic Kwiatkowski, Cordelia Langford, John Sillitoe on behalf of the Wellcome Sanger Institute COVID-19 Surveillance Team |
| EPI_ISL_3361798 | Health Services Laboratories | Wellcome Sanger Institute for the COVID-19 Genomics UK (COG-UK) Consortium | Health Services Laboratories and Alex Alderton, Roberto Amato, Jeffrey Barrett, Sonia Goncalves, Ewan Harrison, David K. Jackson, Ian Johnston, Dominic Kwiatkowski, Cordelia Langford, John Sillitoe on behalf of the Wellcome Sanger Institute COVID-19 Surveillance Team |
| EPI_ISL_3361799 | Lighthouse Laboratory Plymouth | Wellcome Sanger Institute for the COVID-19 Genomics UK (COG-UK) Consortium | Lighthouse Laboratory Plymouth and Alex Alderton, Roberto Amato, Jeffrey Barrett, Sonia Goncalves, Ewan Harrison, David K. Jackson, Ian Johnston, Dominic Kwiatkowski, Cordelia Langford, John Sillitoe on behalf of the Wellcome Sanger Institute COVID-19 Surveillance Team |
| EPI_ISL_3361801 | Lighthouse Lab in Glasgow | Wellcome Sanger Institute for the COVID-19 Genomics UK (COG-UK) Consortium | Harper VanSteenhouse, Yumi Kasai, David Gray, Carol Clugston, Anna Dominiczak and Alex Alderton, Roberto Amato, Jeffrey Barrett, Sonia Goncalves, Ewan Harrison, David K. Jackson, Ian Johnston, Dominic Kwiatkowski, Cordelia Langford, John Sillitoe on behalf of the Wellcome Sanger Institute COVID-19 Surveillance Team |
| EPI_ISL_3361802, EPI_ISL_3361803, EPI_ISL_3361805 | Lighthouse Laboratory Plymouth | Wellcome Sanger Institute for the COVID-19 Genomics UK (COG-UK) Consortium | Lighthouse Laboratory Plymouth and Alex Alderton, Roberto Amato, Jeffrey Barrett, Sonia Goncalves, Ewan Harrison, David K. Jackson, Ian Johnston, Dominic Kwiatkowski, Cordelia Langford, John Sillitoe on behalf of the Wellcome Sanger Institute COVID-19 Surveillance Team |
| EPI_ISL_3361806 | Lighthouse Lab in Milton Keynes | Wellcome Sanger Institute for the COVID-19 Genomics UK (COG-UK) Consortium | The Lighthouse Lab in Milton Keynes and Alex Alderton, Roberto Amato, Jeffrey Barrett, Sonia Goncalves, Ewan Harrison, David K. Jackson, Ian Johnston, Dominic Kwiatkowski, Cordelia Langford, John Sillitoe on behalf of the Wellcome Sanger Institute COVID-19 Surveillance Team |
| EPI_ISL_3361808 | Lighthouse Lab in Glasgow | Wellcome Sanger Institute for the COVID-19 Genomics UK (COG-UK) Consortium | Harper VanSteenhouse, Yumi Kasai, David Gray, Carol Clugston, Anna Dominiczak and Alex Alderton, Roberto Amato, Jeffrey Barrett, Sonia Goncalves, Ewan Harrison, David K. Jackson, Ian Johnston, Dominic Kwiatkowski, Cordelia Langford, John Sillitoe on behalf of the Wellcome Sanger Institute COVID-19 Surveillance Team |
| EPI_ISL_3361809, EPI_ISL_3361811 | Lighthouse Laboratory Plymouth | Wellcome Sanger Institute for the COVID-19 Genomics UK (COG-UK) Consortium | Lighthouse Laboratory Plymouth and Alex Alderton, Roberto Amato, Jeffrey Barrett, Sonia Goncalves, Ewan Harrison, David K. Jackson, Ian Johnston, Dominic Kwiatkowski, Cordelia Langford, John Sillitoe on behalf of the Wellcome Sanger Institute COVID-19 Surveillance Team |
| EPI_ISL_3361812, EPI_ISL_3361814 | Health Services Laboratories | Wellcome Sanger Institute for the COVID-19 Genomics UK (COG-UK) Consortium | Health Services Laboratories and Alex Alderton, Roberto Amato, Jeffrey Barrett, Sonia Goncalves, Ewan Harrison, David K. Jackson, Ian Johnston, Dominic Kwiatkowski, Cordelia Langford, John Sillitoe on behalf of the Wellcome Sanger Institute COVID-19 Surveillance Team |
| EPI_ISL_3361815 | Lighthouse Lab in Milton Keynes | Wellcome Sanger Institute for the COVID-19 Genomics UK (COG-UK) Consortium | The Lighthouse Lab in Milton Keynes and Alex Alderton, Roberto Amato, Jeffrey Barrett, Sonia Goncalves, Ewan Harrison, David K. Jackson, Ian Johnston, Dominic Kwiatkowski, Cordelia Langford, John Sillitoe on behalf of the Wellcome Sanger Institute COVID-19 Surveillance Team |
| EPI_ISL_3361817 | Lighthouse Laboratory Plymouth | Wellcome Sanger Institute for the COVID-19 Genomics UK (COG-UK) Consortium | Lighthouse Laboratory Plymouth and Alex Alderton, Roberto Amato, Jeffrey Barrett, Sonia Goncalves, Ewan Harrison, David K. Jackson, Ian Johnston, Dominic Kwiatkowski, Cordelia Langford, John Sillitoe on behalf of the Wellcome Sanger Institute COVID-19 Surveillance Team |
| EPI_ISL_3361818 | Lighthouse Lab in Milton Keynes | Wellcome Sanger Institute for the COVID-19 Genomics UK (COG-UK) Consortium | The Lighthouse Lab in Milton Keynes and Alex Alderton, Roberto Amato, Jeffrey Barrett, Sonia Goncalves, Ewan Harrison, David K. Jackson, Ian Johnston, Dominic Kwiatkowski, Cordelia Langford, John Sillitoe on behalf of the Wellcome Sanger Institute COVID-19 Surveillance Team |
| EPI_ISL_3361820 | Lighthouse Laboratory Plymouth | Wellcome Sanger Institute for the COVID-19 Genomics UK (COG-UK) Consortium | Lighthouse Laboratory Plymouth and Alex Alderton, Roberto Amato, Jeffrey Barrett, Sonia Goncalves, Ewan Harrison, David K. Jackson, Ian Johnston, Dominic Kwiatkowski, Cordelia Langford, John Sillitoe on behalf of the Wellcome Sanger Institute COVID-19 Surveillance Team |
| EPI_ISL_3361821 | Health Services Laboratories | Wellcome Sanger Institute for the COVID-19 Genomics UK (COG-UK) Consortium | Health Services Laboratories and Alex Alderton, Roberto Amato, Jeffrey Barrett, Sonia Goncalves, Ewan Harrison, David K. Jackson, Ian Johnston, Dominic Kwiatkowski, Cordelia Langford, John Sillitoe on behalf of the Wellcome Sanger Institute COVID-19 Surveillance Team |
| EPI_ISL_3361823, EPI_ISL_3361825 | Lighthouse Lab in Milton Keynes | Wellcome Sanger Institute for the COVID-19 Genomics UK (COG-UK) Consortium | The Lighthouse Lab in Milton Keynes and Alex Alderton, Roberto Amato, Jeffrey Barrett, Sonia Goncalves, Ewan Harrison, David K. Jackson, Ian Johnston, Dominic Kwiatkowski, Cordelia Langford, John Sillitoe on behalf of the Wellcome Sanger Institute COVID-19 Surveillance Team |
| EPI_ISL_3361826 | Lighthouse Lab in Alderley Park | Wellcome Sanger Institute for the COVID-19 Genomics UK (COG-UK) Consortium | Jacquelyn Wynn, Mairead Hyland, The Lighthouse Lab in Alderley Park and Alex Alderton, Roberto Amato, Jeffrey Barrett, Sonia Goncalves, Ewan Harrison, David K. Jackson, Ian Johnston, Dominic Kwiatkowski, Cordelia Langford, John Sillitoe on behalf of the Wellcome Sanger Institute COVID-19 Surveillance Team |
| EPI_ISL_3361828 | Lighthouse Lab in Glasgow | Wellcome Sanger Institute for the COVID-19 Genomics UK (COG-UK) Consortium | Harper VanSteenhouse, Yumi Kasai, David Gray, Carol Clugston, Anna Dominiczak and Alex Alderton, Roberto Amato, Jeffrey Barrett, Sonia Goncalves, Ewan Harrison, David K. Jackson, Ian Johnston, Dominic Kwiatkowski, Cordelia Langford, John Sillitoe on behalf of the Wellcome Sanger Institute COVID-19 Surveillance Team |
| EPI_ISL_3361829 | Health Services Laboratories | Wellcome Sanger Institute for the COVID-19 Genomics UK (COG-UK) Consortium | Health Services Laboratories and Alex Alderton, Roberto Amato, Jeffrey Barrett, Sonia Goncalves, Ewan Harrison, David K. Jackson, Ian Johnston, Dominic Kwiatkowski, Cordelia Langford, John Sillitoe on behalf of the Wellcome Sanger Institute COVID-19 Surveillance Team |
| EPI_ISL_3361830 | Lighthouse Laboratory Plymouth | Wellcome Sanger Institute for the COVID-19 Genomics UK (COG-UK) Consortium | Lighthouse Laboratory Plymouth and Alex Alderton, Roberto Amato, Jeffrey Barrett, Sonia Goncalves, Ewan Harrison, David K. Jackson, Ian Johnston, Dominic Kwiatkowski, Cordelia Langford, John Sillitoe on behalf of the Wellcome Sanger Institute COVID-19 Surveillance Team |
| EPI_ISL_3361832 | Lighthouse Lab in Alderley Park | Wellcome Sanger Institute for the COVID-19 Genomics UK (COG-UK) Consortium | Jacquelyn Wynn, Mairead Hyland, The Lighthouse Lab in Alderley Park and Alex Alderton, Roberto Amato, Jeffrey Barrett, Sonia Goncalves, Ewan Harrison, David K. Jackson, Ian Johnston, Dominic Kwiatkowski, Cordelia Langford, John Sillitoe on behalf of the Wellcome Sanger Institute COVID-19 Surveillance Team |
| EPI_ISL_3361833 | Lighthouse Laboratory Plymouth | Wellcome Sanger Institute for the COVID-19 Genomics UK (COG-UK) Consortium | Lighthouse Laboratory Plymouth and Alex Alderton, Roberto Amato, Jeffrey Barrett, Sonia Goncalves, Ewan Harrison, David K. Jackson, Ian Johnston, Dominic Kwiatkowski, Cordelia Langford, John Sillitoe on behalf of the Wellcome Sanger Institute COVID-19 Surveillance Team |
| EPI_ISL_3361835, EPI_ISL_3361836 | Lighthouse Lab in Glasgow | Wellcome Sanger Institute for the COVID-19 Genomics UK (COG-UK) Consortium | Harper VanSteenhouse, Yumi Kasai, David Gray, Carol Clugston, Anna Dominiczak and Alex Alderton, Roberto Amato, Jeffrey Barrett, Sonia Goncalves, Ewan Harrison, David K. Jackson, Ian Johnston, Dominic Kwiatkowski, Cordelia Langford, John Sillitoe on behalf of the Wellcome Sanger Institute COVID-19 Surveillance Team |
| EPI_ISL_3361837 | Lighthouse Lab in Milton Keynes | Wellcome Sanger Institute for the COVID-19 Genomics UK (COG-UK) Consortium | The Lighthouse Lab in Milton Keynes and Alex Alderton, Roberto Amato, Jeffrey Barrett, Sonia Goncalves, Ewan Harrison, David K. Jackson, Ian Johnston, Dominic Kwiatkowski, Cordelia Langford, John Sillitoe on behalf of the Wellcome Sanger Institute COVID-19 Surveillance Team |

| | | | |
|---|---|---|---|
| EPI_ISL_3361839 | Lighthouse Laboratory Plymouth | Wellcome Sanger Institute for the COVID-19 Genomics UK (COG-UK) Consortium | Lighthouse Laboratory Plymouth and Alex Alderton, Roberto Amato, Jeffrey Barrett, Sonia Goncalves, Ewan Harrison, David K. Jackson, Ian Johnston, Dominic Kwiatkowski, Cordelia Langford, John Sillitoe on behalf of the Wellcome Sanger Institute COVID-19 Surveillance Team |
| EPI_ISL_3361840 | Lighthouse Lab in Milton Keynes | Wellcome Sanger Institute for the COVID-19 Genomics UK (COG-UK) Consortium | The Lighthouse Lab in Milton Keynes and Alex Alderton, Roberto Amato, Jeffrey Barrett, Sonia Goncalves, Ewan Harrison, David K. Jackson, Ian Johnston, Dominic Kwiatkowski, Cordelia Langford, John Sillitoe on behalf of the Wellcome Sanger Institute COVID-19 Surveillance Team |
| EPI_ISL_3361842 | Lighthouse Lab in Glasgow | Wellcome Sanger Institute for the COVID-19 Genomics UK (COG-UK) Consortium | Harper VanSteenhouse, Yumi Kasai, David Gray, Carol Clugston, Anna Dominiczak and Alex Alderton, Roberto Amato, Jeffrey Barrett, Sonia Goncalves, Ewan Harrison, David K. Jackson, Ian Johnston, Dominic Kwiatkowski, Cordelia Langford, John Sillitoe on behalf of the Wellcome Sanger Institute COVID-19 Surveillance Team |
| EPI_ISL_3361843 | Lighthouse Laboratory Plymouth | Wellcome Sanger Institute for the COVID-19 Genomics UK (COG-UK) Consortium | Lighthouse Laboratory Plymouth and Alex Alderton, Roberto Amato, Jeffrey Barrett, Sonia Goncalves, Ewan Harrison, David K. Jackson, Ian Johnston, Dominic Kwiatkowski, Cordelia Langford, John Sillitoe on behalf of the Wellcome Sanger Institute COVID-19 Surveillance Team |
| EPI_ISL_3361845 | Lighthouse Lab in Milton Keynes | Wellcome Sanger Institute for the COVID-19 Genomics UK (COG-UK) Consortium | The Lighthouse Lab in Milton Keynes and Alex Alderton, Roberto Amato, Jeffrey Barrett, Sonia Goncalves, Ewan Harrison, David K. Jackson, Ian Johnston, Dominic Kwiatkowski, Cordelia Langford, John Sillitoe on behalf of the Wellcome Sanger Institute COVID-19 Surveillance Team |
| EPI_ISL_3361846 | Lighthouse Lab in Alderley Park | Wellcome Sanger Institute for the COVID-19 Genomics UK (COG-UK) Consortium | Jacquelyn Wynn, Mairead Hyland, The Lighthouse Lab in Alderley Park and Alex Alderton, Roberto Amato, Jeffrey Barrett, Sonia Goncalves, Ewan Harrison, David K. Jackson, Ian Johnston, Dominic Kwiatkowski, Cordelia Langford, John Sillitoe on behalf of the Wellcome Sanger Institute COVID-19 Surveillance Team |
| EPI_ISL_3361848 | Lighthouse Lab in Milton Keynes | Wellcome Sanger Institute for the COVID-19 Genomics UK (COG-UK) Consortium | The Lighthouse Lab in Milton Keynes and Alex Alderton, Roberto Amato, Jeffrey Barrett, Sonia Goncalves, Ewan Harrison, David K. Jackson, Ian Johnston, Dominic Kwiatkowski, Cordelia Langford, John Sillitoe on behalf of the Wellcome Sanger Institute COVID-19 Surveillance Team |
| EPI_ISL_3361849 | Lighthouse Lab in Glasgow | Wellcome Sanger Institute for the COVID-19 Genomics UK (COG-UK) Consortium | Harper VanSteenhouse, Yumi Kasai, David Gray, Carol Clugston, Anna Dominiczak and Alex Alderton, Roberto Amato, Jeffrey Barrett, Sonia Goncalves, Ewan Harrison, David K. Jackson, Ian Johnston, Dominic Kwiatkowski, Cordelia Langford, John Sillitoe on behalf of the Wellcome Sanger Institute COVID-19 Surveillance Team |
| EPI_ISL_3361850 | Lighthouse Lab in Alderley Park | Wellcome Sanger Institute for the COVID-19 Genomics UK (COG-UK) Consortium | Jacquelyn Wynn, Mairead Hyland, The Lighthouse Lab in Alderley Park and Alex Alderton, Roberto Amato, Jeffrey Barrett, Sonia Goncalves, Ewan Harrison, David K. Jackson, Ian Johnston, Dominic Kwiatkowski, Cordelia Langford, John Sillitoe on behalf of the Wellcome Sanger Institute COVID-19 Surveillance Team |
| EPI_ISL_3361852 | Lighthouse Laboratory Plymouth | Wellcome Sanger Institute for the COVID-19 Genomics UK (COG-UK) Consortium | Lighthouse Laboratory Plymouth and Alex Alderton, Roberto Amato, Jeffrey Barrett, Sonia Goncalves, Ewan Harrison, David K. Jackson, Ian Johnston, Dominic Kwiatkowski, Cordelia Langford, John Sillitoe on behalf of the Wellcome Sanger Institute COVID-19 Surveillance Team |
| EPI_ISL_3361853 | Lighthouse Lab in Glasgow | Wellcome Sanger Institute for the COVID-19 Genomics UK (COG-UK) Consortium | Harper VanSteenhouse, Yumi Kasai, David Gray, Carol Clugston, Anna Dominiczak and Alex Alderton, Roberto Amato, Jeffrey Barrett, Sonia Goncalves, Ewan Harrison, David K. Jackson, Ian Johnston, Dominic Kwiatkowski, Cordelia Langford, John Sillitoe on behalf of the Wellcome Sanger Institute COVID-19 Surveillance Team |
| EPI_ISL_3361855, EPI_ISL_3361856 | Lighthouse Lab in Alderley Park | Wellcome Sanger Institute for the COVID-19 Genomics UK (COG-UK) Consortium | Jacquelyn Wynn, Mairead Hyland, The Lighthouse Lab in Alderley Park and Alex Alderton, Roberto Amato, Jeffrey Barrett, Sonia Goncalves, Ewan Harrison, David K. Jackson, Ian Johnston, Dominic Kwiatkowski, Cordelia Langford, John Sillitoe on behalf of the Wellcome Sanger Institute COVID-19 Surveillance Team |
| EPI_ISL_3361858 | Lighthouse Lab in Glasgow | Wellcome Sanger Institute for the COVID-19 Genomics UK (COG-UK) Consortium | Harper VanSteenhouse, Yumi Kasai, David Gray, Carol Clugston, Anna Dominiczak and Alex Alderton, Roberto Amato, Jeffrey Barrett, Sonia Goncalves, Ewan Harrison, David K. Jackson, Ian Johnston, Dominic Kwiatkowski, Cordelia Langford, John Sillitoe on behalf of the Wellcome Sanger Institute COVID-19 Surveillance Team |
| EPI_ISL_3361859 | Lighthouse Lab in Milton Keynes | Wellcome Sanger Institute for the COVID-19 Genomics UK (COG-UK) Consortium | The Lighthouse Lab in Milton Keynes and Alex Alderton, Roberto Amato, Jeffrey Barrett, Sonia Goncalves, Ewan Harrison, David K. Jackson, Ian Johnston, Dominic Kwiatkowski, Cordelia Langford, John Sillitoe on behalf of the Wellcome Sanger Institute COVID-19 Surveillance Team |
| EPI_ISL_3361860 | Lighthouse Lab in Glasgow | Wellcome Sanger Institute for the COVID-19 Genomics UK (COG-UK) Consortium | Harper VanSteenhouse, Yumi Kasai, David Gray, Carol Clugston, Anna Dominiczak and Alex Alderton, Roberto Amato, Jeffrey Barrett, Sonia Goncalves, Ewan Harrison, David K. Jackson, Ian Johnston, Dominic Kwiatkowski, Cordelia Langford, John Sillitoe on behalf of the Wellcome Sanger Institute COVID-19 Surveillance Team |
| EPI_ISL_3361862 | Lighthouse Laboratory Plymouth | Wellcome Sanger Institute for the COVID-19 Genomics UK (COG-UK) Consortium | Lighthouse Laboratory Plymouth and Alex Alderton, Roberto Amato, Jeffrey Barrett, Sonia Goncalves, Ewan Harrison, David K. Jackson, Ian Johnston, Dominic Kwiatkowski, Cordelia Langford, John Sillitoe on behalf of the Wellcome Sanger Institute COVID-19 Surveillance Team |
| EPI_ISL_3361863, EPI_ISL_3361865 | Lighthouse Lab in Milton Keynes | Wellcome Sanger Institute for the COVID-19 Genomics UK (COG-UK) Consortium | The Lighthouse Lab in Milton Keynes and Alex Alderton, Roberto Amato, Jeffrey Barrett, Sonia Goncalves, Ewan Harrison, David K. Jackson, Ian Johnston, Dominic Kwiatkowski, Cordelia Langford, John Sillitoe on behalf of the Wellcome Sanger Institute COVID-19 Surveillance Team |
| EPI_ISL_3361866, EPI_ISL_3361867 | Lighthouse Lab in Glasgow | Wellcome Sanger Institute for the COVID-19 Genomics UK (COG-UK) Consortium | Harper VanSteenhouse, Yumi Kasai, David Gray, Carol Clugston, Anna Dominiczak and Alex Alderton, Roberto Amato, Jeffrey Barrett, Sonia Goncalves, Ewan Harrison, David K. Jackson, Ian Johnston, Dominic Kwiatkowski, Cordelia Langford, John Sillitoe on behalf of the Wellcome Sanger Institute COVID-19 Surveillance Team |
| EPI_ISL_3361869 | Lighthouse Lab in Alderley Park | Wellcome Sanger Institute for the COVID-19 Genomics UK (COG-UK) Consortium | Jacquelyn Wynn, Mairead Hyland, The Lighthouse Lab in Alderley Park and Alex Alderton, Roberto Amato, Jeffrey Barrett, Sonia Goncalves, Ewan Harrison, David K. Jackson, Ian Johnston, Dominic Kwiatkowski, Cordelia Langford, John Sillitoe on behalf of the Wellcome Sanger Institute COVID-19 Surveillance Team |
| EPI_ISL_3361871 | Lighthouse Laboratory Plymouth | Wellcome Sanger Institute for the COVID-19 Genomics UK (COG-UK) Consortium | Lighthouse Laboratory Plymouth and Alex Alderton, Roberto Amato, Jeffrey Barrett, Sonia Goncalves, Ewan Harrison, David K. Jackson, Ian Johnston, Dominic Kwiatkowski, Cordelia Langford, John Sillitoe on behalf of the Wellcome Sanger Institute COVID-19 Surveillance Team |
| EPI_ISL_3361872, EPI_ISL_3361873 | Lighthouse Lab in Milton Keynes | Wellcome Sanger Institute for the COVID-19 Genomics UK (COG-UK) Consortium | The Lighthouse Lab in Milton Keynes and Alex Alderton, Roberto Amato, Jeffrey Barrett, Sonia Goncalves, Ewan Harrison, David K. Jackson, Ian Johnston, Dominic Kwiatkowski, Cordelia Langford, John Sillitoe on behalf of the Wellcome Sanger Institute COVID-19 Surveillance Team |
| EPI_ISL_3361875 | Lighthouse Laboratory Plymouth | Wellcome Sanger Institute for the COVID-19 Genomics UK (COG-UK) Consortium | Lighthouse Laboratory Plymouth and Alex Alderton, Roberto Amato, Jeffrey Barrett, Sonia Goncalves, Ewan Harrison, David K. Jackson, Ian Johnston, Dominic Kwiatkowski, Cordelia Langford, John Sillitoe on behalf of the Wellcome Sanger Institute COVID-19 Surveillance Team |
| EPI_ISL_3361876 | Lighthouse Lab in Alderley Park | Wellcome Sanger Institute for the COVID-19 Genomics UK (COG-UK) Consortium | Jacquelyn Wynn, Mairead Hyland, The Lighthouse Lab in Alderley Park and Alex Alderton, Roberto Amato, Jeffrey Barrett, Sonia Goncalves, Ewan Harrison, David K. Jackson, Ian Johnston, Dominic Kwiatkowski, Cordelia Langford, John Sillitoe on behalf of the Wellcome Sanger Institute COVID-19 Surveillance Team |
| EPI_ISL_3361878 | Health Services Laboratories | Wellcome Sanger Institute for the COVID-19 Genomics UK (COG-UK) Consortium | Health Services Laboratories and Alex Alderton, Roberto Amato, Jeffrey Barrett, Sonia Goncalves, Ewan Harrison, David K. Jackson, Ian Johnston, Dominic Kwiatkowski, Cordelia Langford, John Sillitoe on behalf of the Wellcome Sanger Institute COVID-19 Surveillance Team |
| EPI_ISL_3361879, EPI_ISL_3361881 | Lighthouse Lab in Glasgow | Wellcome Sanger Institute for the COVID-19 Genomics UK (COG-UK) Consortium | Harper VanSteenhouse, Yumi Kasai, David Gray, Carol Clugston, Anna Dominiczak and Alex Alderton, Roberto Amato, Jeffrey Barrett, Sonia Goncalves, Ewan Harrison, David K. Jackson, Ian Johnston, Dominic Kwiatkowski, Cordelia Langford, John Sillitoe on behalf of the Wellcome Sanger Institute COVID-19 Surveillance Team |
| EPI_ISL_3361882 | Lighthouse Lab in Milton Keynes | Wellcome Sanger Institute for the COVID-19 Genomics UK (COG-UK) Consortium | The Lighthouse Lab in Milton Keynes and Alex Alderton, Roberto Amato, Jeffrey Barrett, Sonia Goncalves, Ewan Harrison, David K. Jackson, Ian Johnston, Dominic Kwiatkowski, Cordelia Langford, John Sillitoe on behalf of the Wellcome Sanger Institute COVID-19 Surveillance Team |
| EPI_ISL_3361883 | Lighthouse Lab in Alderley Park | Wellcome Sanger Institute for the COVID-19 Genomics UK (COG-UK) Consortium | Jacquelyn Wynn, Mairead Hyland, The Lighthouse Lab in Alderley Park and Alex Alderton, Roberto Amato, Jeffrey Barrett, Sonia Goncalves, Ewan Harrison, David K. Jackson, Ian Johnston, Dominic Kwiatkowski, Cordelia Langford, John Sillitoe on behalf of the Wellcome Sanger Institute COVID-19 Surveillance Team |
| EPI_ISL_3361885 | Lighthouse Lab in Glasgow | Wellcome Sanger Institute for the COVID-19 Genomics UK (COG-UK) Consortium | Harper VanSteenhouse, Yumi Kasai, David Gray, Carol Clugston, Anna Dominiczak and Alex Alderton, Roberto Amato, Jeffrey Barrett, Sonia Goncalves, Ewan Harrison, David K. Jackson, Ian Johnston, Dominic Kwiatkowski, Cordelia Langford, John Sillitoe on behalf of the Wellcome Sanger Institute COVID-19 Surveillance Team |
| EPI_ISL_3361887, EPI_ISL_3361888 | Lighthouse Lab in Alderley Park | Wellcome Sanger Institute for the COVID-19 Genomics UK (COG-UK) Consortium | Jacquelyn Wynn, Mairead Hyland, The Lighthouse Lab in Alderley Park and Alex Alderton, Roberto Amato, Jeffrey Barrett, Sonia Goncalves, Ewan Harrison, David K. Jackson, Ian Johnston, Dominic Kwiatkowski, Cordelia Langford, John Sillitoe on behalf of the Wellcome Sanger Institute COVID-19 Surveillance Team |
| EPI_ISL_3361889 | Lighthouse Lab in Milton Keynes | Wellcome Sanger Institute for the COVID-19 Genomics UK (COG-UK) Consortium | The Lighthouse Lab in Milton Keynes and Alex Alderton, Roberto Amato, Jeffrey Barrett, Sonia Goncalves, Ewan Harrison, David K. Jackson, Ian Johnston, Dominic Kwiatkowski, Cordelia Langford, John Sillitoe on behalf of the Wellcome Sanger Institute COVID-19 Surveillance Team |
| EPI_ISL_3361891 | Lighthouse Lab in Alderley Park | Wellcome Sanger Institute for the COVID-19 Genomics UK (COG-UK) Consortium | Jacquelyn Wynn, Mairead Hyland, The Lighthouse Lab in Alderley Park and Alex Alderton, Roberto Amato, Jeffrey Barrett, Sonia Goncalves, Ewan Harrison, David K. Jackson, Ian Johnston, Dominic Kwiatkowski, Cordelia Langford, John Sillitoe on behalf of the Wellcome Sanger Institute COVID-19 Surveillance Team |
| EPI_ISL_3361892 | Lighthouse Lab in Glasgow | Wellcome Sanger Institute for the COVID-19 Genomics UK (COG-UK) Consortium | Harper VanSteenhouse, Yumi Kasai, David Gray, Carol Clugston, Anna Dominiczak and Alex Alderton, Roberto Amato, Jeffrey Barrett, Sonia Goncalves, Ewan Harrison, David K. Jackson, Ian Johnston, Dominic Kwiatkowski, Cordelia Langford, John Sillitoe on behalf of the Wellcome Sanger Institute COVID-19 Surveillance Team |
| EPI_ISL_3361894, EPI_ISL_3361895 | Health Services Laboratories | Wellcome Sanger Institute for the COVID-19 Genomics UK (COG-UK) Consortium | Health Services Laboratories and Alex Alderton, Roberto Amato, Jeffrey Barrett, Sonia Goncalves, Ewan Harrison, David K. Jackson, Ian Johnston, Dominic Kwiatkowski, Cordelia Langford, John Sillitoe on behalf of the Wellcome Sanger Institute COVID-19 Surveillance Team |
| EPI_ISL_3361897 | Lighthouse Laboratory Plymouth | Wellcome Sanger Institute for the COVID-19 Genomics UK (COG-UK) Consortium | Lighthouse Laboratory Plymouth and Alex Alderton, Roberto Amato, Jeffrey Barrett, Sonia Goncalves, Ewan Harrison, David K. Jackson, Ian Johnston, Dominic Kwiatkowski, Cordelia Langford, John Sillitoe on behalf of the Wellcome Sanger Institute COVID-19 Surveillance Team |
| EPI_ISL_3361898 | Health Services Laboratories | Wellcome Sanger Institute for the COVID-19 Genomics UK (COG-UK) Consortium | Health Services Laboratories and Alex Alderton, Roberto Amato, Jeffrey Barrett, Sonia Goncalves, Ewan Harrison, David K. Jackson, Ian Johnston, Dominic Kwiatkowski, Cordelia Langford, John Sillitoe on behalf of the Wellcome Sanger Institute COVID-19 Surveillance Team |
| EPI_ISL_3361900 | Lighthouse Lab in Glasgow | Wellcome Sanger Institute for the COVID-19 Genomics UK (COG-UK) Consortium | Harper VanSteenhouse, Yumi Kasai, David Gray, Carol Clugston, Anna Dominiczak and Alex Alderton, Roberto Amato, Jeffrey Barrett, Sonia Goncalves, Ewan Harrison, David K. Jackson, Ian Johnston, Dominic Kwiatkowski, Cordelia Langford, John Sillitoe on behalf of the Wellcome Sanger Institute COVID-19 Surveillance Team |
| EPI_ISL_3361901 | Lighthouse Laboratory Plymouth | Wellcome Sanger Institute for the COVID-19 Genomics UK (COG-UK) Consortium | Lighthouse Laboratory Plymouth and Alex Alderton, Roberto Amato, Jeffrey Barrett, Sonia Goncalves, Ewan Harrison, David K. Jackson, Ian Johnston, Dominic Kwiatkowski, Cordelia Langford, John Sillitoe on behalf of the Wellcome Sanger Institute COVID-19 Surveillance Team |
| EPI_ISL_3361902 | Lighthouse Lab in Alderley Park | Wellcome Sanger Institute for the COVID-19 Genomics UK (COG-UK) Consortium | Jacquelyn Wynn, Mairead Hyland, The Lighthouse Lab in Alderley Park and Alex Alderton, Roberto Amato, Jeffrey Barrett, Sonia Goncalves, Ewan Harrison, David K. Jackson, Ian Johnston, Dominic Kwiatkowski, Cordelia Langford, John Sillitoe on behalf of the Wellcome Sanger Institute COVID-19 Surveillance Team |
| EPI_ISL_3361903, EPI_ISL_3361905 | Lighthouse Lab in Milton Keynes | Wellcome Sanger Institute for the COVID-19 Genomics UK (COG-UK) Consortium | The Lighthouse Lab in Milton Keynes and Alex Alderton, Roberto Amato, Jeffrey Barrett, Sonia Goncalves, Ewan Harrison, David K. Jackson, Ian Johnston, Dominic Kwiatkowski, Cordelia Langford, John Sillitoe on behalf of the Wellcome Sanger Institute COVID-19 Surveillance Team |
| EPI_ISL_3361906, EPI_ISL_3361908 | Lighthouse Laboratory Plymouth | Wellcome Sanger Institute for the COVID-19 Genomics UK (COG-UK) Consortium | Lighthouse Laboratory Plymouth and Alex Alderton, Roberto Amato, Jeffrey Barrett, Sonia Goncalves, Ewan Harrison, David K. Jackson, Ian Johnston, Dominic Kwiatkowski, Cordelia Langford, John Sillitoe on behalf of the Wellcome Sanger Institute COVID-19 Surveillance Team |
| EPI_ISL_3361909 | Health Services Laboratories | Wellcome Sanger Institute for the COVID-19 Genomics UK (COG-UK) Consortium | Health Services Laboratories and Alex Alderton, Roberto Amato, Jeffrey Barrett, Sonia Goncalves, Ewan Harrison, David K. Jackson, Ian Johnston, Dominic Kwiatkowski, Cordelia Langford, John Sillitoe on behalf of the Wellcome Sanger Institute COVID-19 Surveillance Team |
| EPI_ISL_3361911 | Lighthouse Laboratory Plymouth | Wellcome Sanger Institute for the COVID-19 Genomics UK (COG-UK) Consortium | Lighthouse Laboratory Plymouth and Alex Alderton, Roberto Amato, Jeffrey Barrett, Sonia Goncalves, Ewan Harrison, David K. Jackson, Ian Johnston, Dominic Kwiatkowski, Cordelia Langford, John Sillitoe on behalf of the Wellcome Sanger Institute COVID-19 Surveillance Team |
| EPI_ISL_3361912 | Lighthouse Lab in Milton Keynes | Wellcome Sanger Institute for the COVID-19 Genomics UK (COG-UK) Consortium | The Lighthouse Lab in Milton Keynes and Alex Alderton, Roberto Amato, Jeffrey Barrett, Sonia Goncalves, Ewan Harrison, David K. Jackson, Ian Johnston, Dominic Kwiatkowski, Cordelia Langford, John Sillitoe on behalf of the Wellcome Sanger Institute COVID-19 Surveillance Team |
| EPI_ISL_3361913, EPI_ISL_3361915, EPI_ISL_3361916 | Lighthouse Laboratory Plymouth | Wellcome Sanger Institute for the COVID-19 Genomics UK (COG-UK) Consortium | Lighthouse Laboratory Plymouth and Alex Alderton, Roberto Amato, Jeffrey Barrett, Sonia Goncalves, Ewan Harrison, David K. Jackson, Ian Johnston, Dominic Kwiatkowski, Cordelia Langford, John Sillitoe on behalf of the Wellcome Sanger Institute COVID-19 Surveillance Team |
| EPI_ISL_3361918 | Lighthouse Lab in Alderley Park | Wellcome Sanger Institute for the COVID-19 Genomics UK (COG-UK) | Jacquelyn Wynn, Mairead Hyland, The Lighthouse Lab in Alderley Park and Alex Alderton, Roberto Amato, Jeffrey Barrett, Sonia Goncalves, Ewan Harrison, David K. Jackson, Ian Johnston, Dominic Kwiatkowski, Cordelia Langford, John Sillitoe on |

| | | | |
|---|---|---|---|
| | | Consortium | behalf of the Wellcome Sanger Institute COVID-19 Surveillance Team |
| EPI_ISL_3361919 | Health Services Laboratories | Wellcome Sanger Institute for the COVID-19 Genomics UK (COG-UK) Consortium | Health Services Laboratories and Alex Alderton, Roberto Amato, Jeffrey Barrett, Sonia Goncalves, Ewan Harrison, David K. Jackson, Ian Johnston, Dominic Kwiatkowski, Cordelia Langford, John Sillitoe on behalf of the Wellcome Sanger Institute COVID-19 Surveillance Team |
| EPI_ISL_3361920, EPI_ISL_3361922 | Lighthouse Lab in Milton Keynes | Wellcome Sanger Institute for the COVID-19 Genomics UK (COG-UK) Consortium | The Lighthouse Lab in Milton Keynes and Alex Alderton, Roberto Amato, Jeffrey Barrett, Sonia Goncalves, Ewan Harrison, David K. Jackson, Ian Johnston, Dominic Kwiatkowski, Cordelia Langford, John Sillitoe on behalf of the Wellcome Sanger Institute COVID-19 Surveillance Team |
| EPI_ISL_3361923 | Health Services Laboratories | Wellcome Sanger Institute for the COVID-19 Genomics UK (COG-UK) Consortium | Health Services Laboratories and Alex Alderton, Roberto Amato, Jeffrey Barrett, Sonia Goncalves, Ewan Harrison, David K. Jackson, Ian Johnston, Dominic Kwiatkowski, Cordelia Langford, John Sillitoe on behalf of the Wellcome Sanger Institute COVID-19 Surveillance Team |
| EPI_ISL_3361924, EPI_ISL_3361926 | Lighthouse Lab in Milton Keynes | Wellcome Sanger Institute for the COVID-19 Genomics UK (COG-UK) Consortium | The Lighthouse Lab in Milton Keynes and Alex Alderton, Roberto Amato, Jeffrey Barrett, Sonia Goncalves, Ewan Harrison, David K. Jackson, Ian Johnston, Dominic Kwiatkowski, Cordelia Langford, John Sillitoe on behalf of the Wellcome Sanger Institute COVID-19 Surveillance Team |
| EPI_ISL_3361927 | Lighthouse Laboratory Plymouth | Wellcome Sanger Institute for the COVID-19 Genomics UK (COG-UK) Consortium | Lighthouse Laboratory Plymouth and Alex Alderton, Roberto Amato, Jeffrey Barrett, Sonia Goncalves, Ewan Harrison, David K. Jackson, Ian Johnston, Dominic Kwiatkowski, Cordelia Langford, John Sillitoe on behalf of the Wellcome Sanger Institute COVID-19 Surveillance Team |
| EPI_ISL_3361929 | Lighthouse Lab in Milton Keynes | Wellcome Sanger Institute for the COVID-19 Genomics UK (COG-UK) Consortium | The Lighthouse Lab in Milton Keynes and Alex Alderton, Roberto Amato, Jeffrey Barrett, Sonia Goncalves, Ewan Harrison, David K. Jackson, Ian Johnston, Dominic Kwiatkowski, Cordelia Langford, John Sillitoe on behalf of the Wellcome Sanger Institute COVID-19 Surveillance Team |
| EPI_ISL_3361930 | Lighthouse Lab in Glasgow | Wellcome Sanger Institute for the COVID-19 Genomics UK (COG-UK) Consortium | Harper VanSteenhouse, Yumi Kasai, David Gray, Carol Clugston, Anna Dominiczak and Alex Alderton, Roberto Amato, Jeffrey Barrett, Sonia Goncalves, Ewan Harrison, David K. Jackson, Ian Johnston, Dominic Kwiatkowski, Cordelia Langford, John Sillitoe on behalf of the Wellcome Sanger Institute COVID-19 Surveillance Team |
| EPI_ISL_3361932 | Lighthouse Lab in Alderley Park | Wellcome Sanger Institute for the COVID-19 Genomics UK (COG-UK) Consortium | Jacquelyn Wynn, Mairead Hyland, The Lighthouse Lab in Alderley Park and Alex Alderton, Roberto Amato, Jeffrey Barrett, Sonia Goncalves, Ewan Harrison, David K. Jackson, Ian Johnston, Dominic Kwiatkowski, Cordelia Langford, John Sillitoe on behalf of the Wellcome Sanger Institute COVID-19 Surveillance Team |
| EPI_ISL_3361933 | Health Services Laboratories | Wellcome Sanger Institute for the COVID-19 Genomics UK (COG-UK) Consortium | Health Services Laboratories and Alex Alderton, Roberto Amato, Jeffrey Barrett, Sonia Goncalves, Ewan Harrison, David K. Jackson, Ian Johnston, Dominic Kwiatkowski, Cordelia Langford, John Sillitoe on behalf of the Wellcome Sanger Institute COVID-19 Surveillance Team |
| EPI_ISL_3361935 | Lighthouse Laboratory Plymouth | Wellcome Sanger Institute for the COVID-19 Genomics UK (COG-UK) Consortium | Lighthouse Laboratory Plymouth and Alex Alderton, Roberto Amato, Jeffrey Barrett, Sonia Goncalves, Ewan Harrison, David K. Jackson, Ian Johnston, Dominic Kwiatkowski, Cordelia Langford, John Sillitoe on behalf of the Wellcome Sanger Institute COVID-19 Surveillance Team |
| EPI_ISL_3361936 | Lighthouse Lab in Milton Keynes | Wellcome Sanger Institute for the COVID-19 Genomics UK (COG-UK) Consortium | The Lighthouse Lab in Milton Keynes and Alex Alderton, Roberto Amato, Jeffrey Barrett, Sonia Goncalves, Ewan Harrison, David K. Jackson, Ian Johnston, Dominic Kwiatkowski, Cordelia Langford, John Sillitoe on behalf of the Wellcome Sanger Institute COVID-19 Surveillance Team |
| EPI_ISL_3361938 | Lighthouse Laboratory Plymouth | Wellcome Sanger Institute for the COVID-19 Genomics UK (COG-UK) Consortium | Lighthouse Laboratory Plymouth and Alex Alderton, Roberto Amato, Jeffrey Barrett, Sonia Goncalves, Ewan Harrison, David K. Jackson, Ian Johnston, Dominic Kwiatkowski, Cordelia Langford, John Sillitoe on behalf of the Wellcome Sanger Institute COVID-19 Surveillance Team |
| EPI_ISL_3361939 | Lighthouse Lab in Glasgow | Wellcome Sanger Institute for the COVID-19 Genomics UK (COG-UK) Consortium | Harper VanSteenhouse, Yumi Kasai, David Gray, Carol Clugston, Anna Dominiczak and Alex Alderton, Roberto Amato, Jeffrey Barrett, Sonia Goncalves, Ewan Harrison, David K. Jackson, Ian Johnston, Dominic Kwiatkowski, Cordelia Langford, John Sillitoe on behalf of the Wellcome Sanger Institute COVID-19 Surveillance Team |
| EPI_ISL_3361941, EPI_ISL_3361942 | Lighthouse Lab in Milton Keynes | Wellcome Sanger Institute for the COVID-19 Genomics UK (COG-UK) Consortium | The Lighthouse Lab in Milton Keynes and Alex Alderton, Roberto Amato, Jeffrey Barrett, Sonia Goncalves, Ewan Harrison, David K. Jackson, Ian Johnston, Dominic Kwiatkowski, Cordelia Langford, John Sillitoe on behalf of the Wellcome Sanger Institute COVID-19 Surveillance Team |
| EPI_ISL_3361944, EPI_ISL_3361945 | Lighthouse Lab in Glasgow | Wellcome Sanger Institute for the COVID-19 Genomics UK (COG-UK) Consortium | Harper VanSteenhouse, Yumi Kasai, David Gray, Carol Clugston, Anna Dominiczak and Alex Alderton, Roberto Amato, Jeffrey Barrett, Sonia Goncalves, Ewan Harrison, David K. Jackson, Ian Johnston, Dominic Kwiatkowski, Cordelia Langford, John Sillitoe on behalf of the Wellcome Sanger Institute COVID-19 Surveillance Team |
| EPI_ISL_3361946 | Health Services Laboratories | Wellcome Sanger Institute for the COVID-19 Genomics UK (COG-UK) Consortium | Health Services Laboratories and Alex Alderton, Roberto Amato, Jeffrey Barrett, Sonia Goncalves, Ewan Harrison, David K. Jackson, Ian Johnston, Dominic Kwiatkowski, Cordelia Langford, John Sillitoe on behalf of the Wellcome Sanger Institute COVID-19 Surveillance Team |
| EPI_ISL_3361948 | Lighthouse Laboratory Plymouth | Wellcome Sanger Institute for the COVID-19 Genomics UK (COG-UK) Consortium | Lighthouse Laboratory Plymouth and Alex Alderton, Roberto Amato, Jeffrey Barrett, Sonia Goncalves, Ewan Harrison, David K. Jackson, Ian Johnston, Dominic Kwiatkowski, Cordelia Langford, John Sillitoe on behalf of the Wellcome Sanger Institute COVID-19 Surveillance Team |
| EPI_ISL_3361949, EPI_ISL_3361951 | Lighthouse Lab in Milton Keynes | Wellcome Sanger Institute for the COVID-19 Genomics UK (COG-UK) Consortium | The Lighthouse Lab in Milton Keynes and Alex Alderton, Roberto Amato, Jeffrey Barrett, Sonia Goncalves, Ewan Harrison, David K. Jackson, Ian Johnston, Dominic Kwiatkowski, Cordelia Langford, John Sillitoe on behalf of the Wellcome Sanger Institute COVID-19 Surveillance Team |
| EPI_ISL_3361952 | Lighthouse Lab in Alderley Park | Wellcome Sanger Institute for the COVID-19 Genomics UK (COG-UK) Consortium | Jacquelyn Wynn, Mairead Hyland, The Lighthouse Lab in Alderley Park and Alex Alderton, Roberto Amato, Jeffrey Barrett, Sonia Goncalves, Ewan Harrison, David K. Jackson, Ian Johnston, Dominic Kwiatkowski, Cordelia Langford, John Sillitoe on behalf of the Wellcome Sanger Institute COVID-19 Surveillance Team |
| EPI_ISL_3361954 | Lighthouse Lab in Glasgow | Wellcome Sanger Institute for the COVID-19 Genomics UK (COG-UK) Consortium | Harper VanSteenhouse, Yumi Kasai, David Gray, Carol Clugston, Anna Dominiczak and Alex Alderton, Roberto Amato, Jeffrey Barrett, Sonia Goncalves, Ewan Harrison, David K. Jackson, Ian Johnston, Dominic Kwiatkowski, Cordelia Langford, John Sillitoe on behalf of the Wellcome Sanger Institute COVID-19 Surveillance Team |
| EPI_ISL_3361955, EPI_ISL_3361956 | Lighthouse Lab in Milton Keynes | Wellcome Sanger Institute for the COVID-19 Genomics UK (COG-UK) Consortium | The Lighthouse Lab in Milton Keynes and Alex Alderton, Roberto Amato, Jeffrey Barrett, Sonia Goncalves, Ewan Harrison, David K. Jackson, Ian Johnston, Dominic Kwiatkowski, Cordelia Langford, John Sillitoe on behalf of the Wellcome Sanger Institute COVID-19 Surveillance Team |
| EPI_ISL_3361958 | Lighthouse Lab in Glasgow | Wellcome Sanger Institute for the COVID-19 Genomics UK (COG-UK) Consortium | Harper VanSteenhouse, Yumi Kasai, David Gray, Carol Clugston, Anna Dominiczak and Alex Alderton, Roberto Amato, Jeffrey Barrett, Sonia Goncalves, Ewan Harrison, David K. Jackson, Ian Johnston, Dominic Kwiatkowski, Cordelia Langford, John Sillitoe on behalf of the Wellcome Sanger Institute COVID-19 Surveillance Team |
| EPI_ISL_3361959 | Lighthouse Lab in Alderley Park | Wellcome Sanger Institute for the COVID-19 Genomics UK (COG-UK) Consortium | Jacquelyn Wynn, Mairead Hyland, The Lighthouse Lab in Alderley Park and Alex Alderton, Roberto Amato, Jeffrey Barrett, Sonia Goncalves, Ewan Harrison, David K. Jackson, Ian Johnston, Dominic Kwiatkowski, Cordelia Langford, John Sillitoe on behalf of the Wellcome Sanger Institute COVID-19 Surveillance Team |
| EPI_ISL_3361961, EPI_ISL_3361963, EPI_ISL_3361964 | Lighthouse Laboratory Plymouth | Wellcome Sanger Institute for the COVID-19 Genomics UK (COG-UK) Consortium | Lighthouse Laboratory Plymouth and Alex Alderton, Roberto Amato, Jeffrey Barrett, Sonia Goncalves, Ewan Harrison, David K. Jackson, Ian Johnston, Dominic Kwiatkowski, Cordelia Langford, John Sillitoe on behalf of the Wellcome Sanger Institute COVID-19 Surveillance Team |
| EPI_ISL_3361966 | Lighthouse Lab in Glasgow | Wellcome Sanger Institute for the COVID-19 Genomics UK (COG-UK) Consortium | Harper VanSteenhouse, Yumi Kasai, David Gray, Carol Clugston, Anna Dominiczak and Alex Alderton, Roberto Amato, Jeffrey Barrett, Sonia Goncalves, Ewan Harrison, David K. Jackson, Ian Johnston, Dominic Kwiatkowski, Cordelia Langford, John Sillitoe on behalf of the Wellcome Sanger Institute COVID-19 Surveillance Team |
| EPI_ISL_3361967, EPI_ISL_3361969 | Lighthouse Lab in Milton Keynes | Wellcome Sanger Institute for the COVID-19 Genomics UK (COG-UK) Consortium | The Lighthouse Lab in Milton Keynes and Alex Alderton, Roberto Amato, Jeffrey Barrett, Sonia Goncalves, Ewan Harrison, David K. Jackson, Ian Johnston, Dominic Kwiatkowski, Cordelia Langford, John Sillitoe on behalf of the Wellcome Sanger Institute COVID-19 Surveillance Team |
| EPI_ISL_3361970 | Lighthouse Lab in Glasgow | Wellcome Sanger Institute for the COVID-19 Genomics UK (COG-UK) Consortium | Harper VanSteenhouse, Yumi Kasai, David Gray, Carol Clugston, Anna Dominiczak and Alex Alderton, Roberto Amato, Jeffrey Barrett, Sonia Goncalves, Ewan Harrison, David K. Jackson, Ian Johnston, Dominic Kwiatkowski, Cordelia Langford, John Sillitoe on behalf of the Wellcome Sanger Institute COVID-19 Surveillance Team |
| EPI_ISL_3361972, EPI_ISL_3361974 | Lighthouse Lab in Milton Keynes | Wellcome Sanger Institute for the COVID-19 Genomics UK (COG-UK) Consortium | The Lighthouse Lab in Milton Keynes and Alex Alderton, Roberto Amato, Jeffrey Barrett, Sonia Goncalves, Ewan Harrison, David K. Jackson, Ian Johnston, Dominic Kwiatkowski, Cordelia Langford, John Sillitoe on behalf of the Wellcome Sanger Institute COVID-19 Surveillance Team |
| EPI_ISL_3361976 | Health Services Laboratories | Wellcome Sanger Institute for the COVID-19 Genomics UK (COG-UK) Consortium | Health Services Laboratories and Alex Alderton, Roberto Amato, Jeffrey Barrett, Sonia Goncalves, Ewan Harrison, David K. Jackson, Ian Johnston, Dominic Kwiatkowski, Cordelia Langford, John Sillitoe on behalf of the Wellcome Sanger Institute COVID-19 Surveillance Team |
| EPI_ISL_3361977 | Lighthouse Lab in Glasgow | Wellcome Sanger Institute for the COVID-19 Genomics UK (COG-UK) Consortium | Harper VanSteenhouse, Yumi Kasai, David Gray, Carol Clugston, Anna Dominiczak and Alex Alderton, Roberto Amato, Jeffrey Barrett, Sonia Goncalves, Ewan Harrison, David K. Jackson, Ian Johnston, Dominic Kwiatkowski, Cordelia Langford, John Sillitoe on behalf of the Wellcome Sanger Institute COVID-19 Surveillance Team |
| EPI_ISL_3361978, EPI_ISL_3361980, EPI_ISL_3361981 | Lighthouse Lab in Milton Keynes | Wellcome Sanger Institute for the COVID-19 Genomics UK (COG-UK) Consortium | The Lighthouse Lab in Milton Keynes and Alex Alderton, Roberto Amato, Jeffrey Barrett, Sonia Goncalves, Ewan Harrison, David K. Jackson, Ian Johnston, Dominic Kwiatkowski, Cordelia Langford, John Sillitoe on behalf of the Wellcome Sanger Institute COVID-19 Surveillance Team |
| EPI_ISL_3361983, EPI_ISL_3361984 | Lighthouse Lab in Alderley Park | Wellcome Sanger Institute for the COVID-19 Genomics UK (COG-UK) Consortium | Jacquelyn Wynn, Mairead Hyland, The Lighthouse Lab in Alderley Park and Alex Alderton, Roberto Amato, Jeffrey Barrett, Sonia Goncalves, Ewan Harrison, David K. Jackson, Ian Johnston, Dominic Kwiatkowski, Cordelia Langford, John Sillitoe on behalf of the Wellcome Sanger Institute COVID-19 Surveillance Team |
| EPI_ISL_3361986 | Lighthouse Lab in Milton Keynes | Wellcome Sanger Institute for the COVID-19 Genomics UK (COG-UK) Consortium | The Lighthouse Lab in Milton Keynes and Alex Alderton, Roberto Amato, Jeffrey Barrett, Sonia Goncalves, Ewan Harrison, David K. Jackson, Ian Johnston, Dominic Kwiatkowski, Cordelia Langford, John Sillitoe on behalf of the Wellcome Sanger Institute COVID-19 Surveillance Team |
| EPI_ISL_3361987 | Lighthouse Laboratory Plymouth | Wellcome Sanger Institute for the COVID-19 Genomics UK (COG-UK) Consortium | Lighthouse Laboratory Plymouth and Alex Alderton, Roberto Amato, Jeffrey Barrett, Sonia Goncalves, Ewan Harrison, David K. Jackson, Ian Johnston, Dominic Kwiatkowski, Cordelia Langford, John Sillitoe on behalf of the Wellcome Sanger Institute COVID-19 Surveillance Team |
| EPI_ISL_3361989 | Lighthouse Lab in Alderley Park | Wellcome Sanger Institute for the COVID-19 Genomics UK (COG-UK) Consortium | Jacquelyn Wynn, Mairead Hyland, The Lighthouse Lab in Alderley Park and Alex Alderton, Roberto Amato, Jeffrey Barrett, Sonia Goncalves, Ewan Harrison, David K. Jackson, Ian Johnston, Dominic Kwiatkowski, Cordelia Langford, John Sillitoe on behalf of the Wellcome Sanger Institute COVID-19 Surveillance Team |
| EPI_ISL_3361990, EPI_ISL_3361992, EPI_ISL_3361993, EPI_ISL_3361995 | Lighthouse Laboratory Plymouth | Wellcome Sanger Institute for the COVID-19 Genomics UK (COG-UK) Consortium | Lighthouse Laboratory Plymouth and Alex Alderton, Roberto Amato, Jeffrey Barrett, Sonia Goncalves, Ewan Harrison, David K. Jackson, Ian Johnston, Dominic Kwiatkowski, Cordelia Langford, John Sillitoe on behalf of the Wellcome Sanger Institute COVID-19 Surveillance Team |
| EPI_ISL_3361996, EPI_ISL_3361998, EPI_ISL_3361999, EPI_ISL_3362001 | Lighthouse Lab in Milton Keynes | Wellcome Sanger Institute for the COVID-19 Genomics UK (COG-UK) Consortium | The Lighthouse Lab in Milton Keynes and Alex Alderton, Roberto Amato, Jeffrey Barrett, Sonia Goncalves, Ewan Harrison, David K. Jackson, Ian Johnston, Dominic Kwiatkowski, Cordelia Langford, John Sillitoe on behalf of the Wellcome Sanger Institute COVID-19 Surveillance Team |
| EPI_ISL_3362002, EPI_ISL_3362004 | Lighthouse Laboratory Plymouth | Wellcome Sanger Institute for the COVID-19 Genomics UK (COG-UK) Consortium | Lighthouse Laboratory Plymouth and Alex Alderton, Roberto Amato, Jeffrey Barrett, Sonia Goncalves, Ewan Harrison, David K. Jackson, Ian Johnston, Dominic Kwiatkowski, Cordelia Langford, John Sillitoe on behalf of the Wellcome Sanger Institute COVID-19 Surveillance Team |
| EPI_ISL_3362006 | Lighthouse Lab in Milton Keynes | Wellcome Sanger Institute for the COVID-19 Genomics UK (COG-UK) Consortium | The Lighthouse Lab in Milton Keynes and Alex Alderton, Roberto Amato, Jeffrey Barrett, Sonia Goncalves, Ewan Harrison, David K. Jackson, Ian Johnston, Dominic Kwiatkowski, Cordelia Langford, John Sillitoe on behalf of the Wellcome Sanger Institute COVID-19 Surveillance Team |
| EPI_ISL_3362007 | Lighthouse Laboratory Plymouth | Wellcome Sanger Institute for the COVID-19 Genomics UK (COG-UK) Consortium | Lighthouse Laboratory Plymouth and Alex Alderton, Roberto Amato, Jeffrey Barrett, Sonia Goncalves, Ewan Harrison, David K. Jackson, Ian Johnston, Dominic Kwiatkowski, Cordelia Langford, John Sillitoe on behalf of the Wellcome Sanger Institute COVID-19 Surveillance Team |
| EPI_ISL_3362009, EPI_ISL_3362010, EPI_ISL_3362012, EPI_ISL_3362013, EPI_ISL_3362014, | Lighthouse Lab in Milton Keynes | Wellcome Sanger Institute for the COVID-19 Genomics UK (COG-UK) Consortium | The Lighthouse Lab in Milton Keynes and Alex Alderton, Roberto Amato, Jeffrey Barrett, Sonia Goncalves, Ewan Harrison, David K. Jackson, Ian Johnston, Dominic Kwiatkowski, Cordelia Langford, John Sillitoe on behalf of the Wellcome Sanger Institute COVID-19 Surveillance Team |

| | | | |
|---|---|---|---|
| EPI_ISL_3362016 | | | |
| EPI_ISL_3362017 | Lighthouse Lab in Alderley Park | Wellcome Sanger Institute for the COVID-19 Genomics UK (COG-UK) Consortium | Jacquelyn Wynn, Mairead Hyland, The Lighthouse Lab in Alderley Park and Alex Alderton, Roberto Amato, Jeffrey Barrett, Sonia Goncalves, Ewan Harrison, David K. Jackson, Ian Johnston, Dominic Kwiatkowski, Cordelia Langford, John Sillitoe on behalf of the Wellcome Sanger Institute COVID-19 Surveillance Team |
| EPI_ISL_3362019, EPI_ISL_3362020, EPI_ISL_3362021, EPI_ISL_3362023, EPI_ISL_3362025, EPI_ISL_3362026 | Lighthouse Lab in Milton Keynes | Wellcome Sanger Institute for the COVID-19 Genomics UK (COG-UK) Consortium | The Lighthouse Lab in Milton Keynes and Alex Alderton, Roberto Amato, Jeffrey Barrett, Sonia Goncalves, Ewan Harrison, David K. Jackson, Ian Johnston, Dominic Kwiatkowski, Cordelia Langford, John Sillitoe on behalf of the Wellcome Sanger Institute COVID-19 Surveillance Team |
| EPI_ISL_3362028 | Lighthouse Lab in Alderley Park | Wellcome Sanger Institute for the COVID-19 Genomics UK (COG-UK) Consortium | Jacquelyn Wynn, Mairead Hyland, The Lighthouse Lab in Alderley Park and Alex Alderton, Roberto Amato, Jeffrey Barrett, Sonia Goncalves, Ewan Harrison, David K. Jackson, Ian Johnston, Dominic Kwiatkowski, Cordelia Langford, John Sillitoe on behalf of the Wellcome Sanger Institute COVID-19 Surveillance Team |
| EPI_ISL_3362029, EPI_ISL_3362030, EPI_ISL_3362032, EPI_ISL_3362033, EPI_ISL_3362035 | Lighthouse Lab in Milton Keynes | Wellcome Sanger Institute for the COVID-19 Genomics UK (COG-UK) Consortium | The Lighthouse Lab in Milton Keynes and Alex Alderton, Roberto Amato, Jeffrey Barrett, Sonia Goncalves, Ewan Harrison, David K. Jackson, Ian Johnston, Dominic Kwiatkowski, Cordelia Langford, John Sillitoe on behalf of the Wellcome Sanger Institute COVID-19 Surveillance Team |
| EPI_ISL_3362036 | Lighthouse Laboratory Plymouth | Wellcome Sanger Institute for the COVID-19 Genomics UK (COG-UK) Consortium | Lighthouse Laboratory Plymouth and Alex Alderton, Roberto Amato, Jeffrey Barrett, Sonia Goncalves, Ewan Harrison, David K. Jackson, Ian Johnston, Dominic Kwiatkowski, Cordelia Langford, John Sillitoe on behalf of the Wellcome Sanger Institute COVID-19 Surveillance Team |
| EPI_ISL_3362038 | Lighthouse Lab in Alderley Park | Wellcome Sanger Institute for the COVID-19 Genomics UK (COG-UK) Consortium | Jacquelyn Wynn, Mairead Hyland, The Lighthouse Lab in Alderley Park and Alex Alderton, Roberto Amato, Jeffrey Barrett, Sonia Goncalves, Ewan Harrison, David K. Jackson, Ian Johnston, Dominic Kwiatkowski, Cordelia Langford, John Sillitoe on behalf of the Wellcome Sanger Institute COVID-19 Surveillance Team |
| EPI_ISL_3362040, EPI_ISL_3362041, EPI_ISL_3362043, EPI_ISL_3362044, EPI_ISL_3362045 | Lighthouse Lab in Milton Keynes | Wellcome Sanger Institute for the COVID-19 Genomics UK (COG-UK) Consortium | The Lighthouse Lab in Milton Keynes and Alex Alderton, Roberto Amato, Jeffrey Barrett, Sonia Goncalves, Ewan Harrison, David K. Jackson, Ian Johnston, Dominic Kwiatkowski, Cordelia Langford, John Sillitoe on behalf of the Wellcome Sanger Institute COVID-19 Surveillance Team |
| EPI_ISL_3362047 | Lighthouse Lab in Alderley Park | Wellcome Sanger Institute for the COVID-19 Genomics UK (COG-UK) Consortium | Jacquelyn Wynn, Mairead Hyland, The Lighthouse Lab in Alderley Park and Alex Alderton, Roberto Amato, Jeffrey Barrett, Sonia Goncalves, Ewan Harrison, David K. Jackson, Ian Johnston, Dominic Kwiatkowski, Cordelia Langford, John Sillitoe on behalf of the Wellcome Sanger Institute COVID-19 Surveillance Team |
| EPI_ISL_3362049 | Lighthouse Lab in Milton Keynes | Wellcome Sanger Institute for the COVID-19 Genomics UK (COG-UK) Consortium | The Lighthouse Lab in Milton Keynes and Alex Alderton, Roberto Amato, Jeffrey Barrett, Sonia Goncalves, Ewan Harrison, David K. Jackson, Ian Johnston, Dominic Kwiatkowski, Cordelia Langford, John Sillitoe on behalf of the Wellcome Sanger Institute COVID-19 Surveillance Team |
| EPI_ISL_3362050 | Lighthouse Laboratory Plymouth | Wellcome Sanger Institute for the COVID-19 Genomics UK (COG-UK) Consortium | Lighthouse Laboratory Plymouth and Alex Alderton, Roberto Amato, Jeffrey Barrett, Sonia Goncalves, Ewan Harrison, David K. Jackson, Ian Johnston, Dominic Kwiatkowski, Cordelia Langford, John Sillitoe on behalf of the Wellcome Sanger Institute COVID-19 Surveillance Team |
| EPI_ISL_3362051 | Lighthouse Lab in Milton Keynes | Wellcome Sanger Institute for the COVID-19 Genomics UK (COG-UK) Consortium | The Lighthouse Lab in Milton Keynes and Alex Alderton, Roberto Amato, Jeffrey Barrett, Sonia Goncalves, Ewan Harrison, David K. Jackson, Ian Johnston, Dominic Kwiatkowski, Cordelia Langford, John Sillitoe on behalf of the Wellcome Sanger Institute COVID-19 Surveillance Team |
| EPI_ISL_3362053 | Lighthouse Lab in Alderley Park | Wellcome Sanger Institute for the COVID-19 Genomics UK (COG-UK) Consortium | Jacquelyn Wynn, Mairead Hyland, The Lighthouse Lab in Alderley Park and Alex Alderton, Roberto Amato, Jeffrey Barrett, Sonia Goncalves, Ewan Harrison, David K. Jackson, Ian Johnston, Dominic Kwiatkowski, Cordelia Langford, John Sillitoe on behalf of the Wellcome Sanger Institute COVID-19 Surveillance Team |
| EPI_ISL_3362054, EPI_ISL_3362056 | Lighthouse Lab in Milton Keynes | Wellcome Sanger Institute for the COVID-19 Genomics UK (COG-UK) Consortium | The Lighthouse Lab in Milton Keynes and Alex Alderton, Roberto Amato, Jeffrey Barrett, Sonia Goncalves, Ewan Harrison, David K. Jackson, Ian Johnston, Dominic Kwiatkowski, Cordelia Langford, John Sillitoe on behalf of the Wellcome Sanger Institute COVID-19 Surveillance Team |
| EPI_ISL_3362057, EPI_ISL_3362059 | Lighthouse Laboratory Plymouth | Wellcome Sanger Institute for the COVID-19 Genomics UK (COG-UK) Consortium | Lighthouse Laboratory Plymouth and Alex Alderton, Roberto Amato, Jeffrey Barrett, Sonia Goncalves, Ewan Harrison, David K. Jackson, Ian Johnston, Dominic Kwiatkowski, Cordelia Langford, John Sillitoe on behalf of the Wellcome Sanger Institute COVID-19 Surveillance Team |
| EPI_ISL_3362060, EPI_ISL_3362062 | Lighthouse Lab in Alderley Park | Wellcome Sanger Institute for the COVID-19 Genomics UK (COG-UK) Consortium | Jacquelyn Wynn, Mairead Hyland, The Lighthouse Lab in Alderley Park and Alex Alderton, Roberto Amato, Jeffrey Barrett, Sonia Goncalves, Ewan Harrison, David K. Jackson, Ian Johnston, Dominic Kwiatkowski, Cordelia Langford, John Sillitoe on behalf of the Wellcome Sanger Institute COVID-19 Surveillance Team |
| EPI_ISL_3362063 | Lighthouse Laboratory Plymouth | Wellcome Sanger Institute for the COVID-19 Genomics UK (COG-UK) Consortium | Lighthouse Laboratory Plymouth and Alex Alderton, Roberto Amato, Jeffrey Barrett, Sonia Goncalves, Ewan Harrison, David K. Jackson, Ian Johnston, Dominic Kwiatkowski, Cordelia Langford, John Sillitoe on behalf of the Wellcome Sanger Institute COVID-19 Surveillance Team |
| EPI_ISL_3362065 | Lighthouse Lab in Milton Keynes | Wellcome Sanger Institute for the COVID-19 Genomics UK (COG-UK) Consortium | The Lighthouse Lab in Milton Keynes and Alex Alderton, Roberto Amato, Jeffrey Barrett, Sonia Goncalves, Ewan Harrison, David K. Jackson, Ian Johnston, Dominic Kwiatkowski, Cordelia Langford, John Sillitoe on behalf of the Wellcome Sanger Institute COVID-19 Surveillance Team |
| EPI_ISL_3362066, EPI_ISL_3362067 | Lighthouse Laboratory Plymouth | Wellcome Sanger Institute for the COVID-19 Genomics UK (COG-UK) Consortium | Lighthouse Laboratory Plymouth and Alex Alderton, Roberto Amato, Jeffrey Barrett, Sonia Goncalves, Ewan Harrison, David K. Jackson, Ian Johnston, Dominic Kwiatkowski, Cordelia Langford, John Sillitoe on behalf of the Wellcome Sanger Institute COVID-19 Surveillance Team |
| EPI_ISL_3362069, EPI_ISL_3362070 | Lighthouse Lab in Alderley Park | Wellcome Sanger Institute for the COVID-19 Genomics UK (COG-UK) Consortium | Jacquelyn Wynn, Mairead Hyland, The Lighthouse Lab in Alderley Park and Alex Alderton, Roberto Amato, Jeffrey Barrett, Sonia Goncalves, Ewan Harrison, David K. Jackson, Ian Johnston, Dominic Kwiatkowski, Cordelia Langford, John Sillitoe on behalf of the Wellcome Sanger Institute COVID-19 Surveillance Team |
| EPI_ISL_3362072, EPI_ISL_3362074, EPI_ISL_3362075 | Lighthouse Laboratory Plymouth | Wellcome Sanger Institute for the COVID-19 Genomics UK (COG-UK) Consortium | Lighthouse Laboratory Plymouth and Alex Alderton, Roberto Amato, Jeffrey Barrett, Sonia Goncalves, Ewan Harrison, David K. Jackson, Ian Johnston, Dominic Kwiatkowski, Cordelia Langford, John Sillitoe on behalf of the Wellcome Sanger Institute COVID-19 Surveillance Team |
| EPI_ISL_3362077 | Lighthouse Lab in Milton Keynes | Wellcome Sanger Institute for the COVID-19 Genomics UK (COG-UK) Consortium | The Lighthouse Lab in Milton Keynes and Alex Alderton, Roberto Amato, Jeffrey Barrett, Sonia Goncalves, Ewan Harrison, David K. Jackson, Ian Johnston, Dominic Kwiatkowski, Cordelia Langford, John Sillitoe on behalf of the Wellcome Sanger Institute COVID-19 Surveillance Team |
| EPI_ISL_3362078 | Lighthouse Lab in Alderley Park | Wellcome Sanger Institute for the COVID-19 Genomics UK (COG-UK) Consortium | Jacquelyn Wynn, Mairead Hyland, The Lighthouse Lab in Alderley Park and Alex Alderton, Roberto Amato, Jeffrey Barrett, Sonia Goncalves, Ewan Harrison, David K. Jackson, Ian Johnston, Dominic Kwiatkowski, Cordelia Langford, John Sillitoe on behalf of the Wellcome Sanger Institute COVID-19 Surveillance Team |
| EPI_ISL_3362080, EPI_ISL_3362081 | Lighthouse Lab in Milton Keynes | Wellcome Sanger Institute for the COVID-19 Genomics UK (COG-UK) Consortium | The Lighthouse Lab in Milton Keynes and Alex Alderton, Roberto Amato, Jeffrey Barrett, Sonia Goncalves, Ewan Harrison, David K. Jackson, Ian Johnston, Dominic Kwiatkowski, Cordelia Langford, John Sillitoe on behalf of the Wellcome Sanger Institute COVID-19 Surveillance Team |
| EPI_ISL_3362083 | Lighthouse Laboratory Plymouth | Wellcome Sanger Institute for the COVID-19 Genomics UK (COG-UK) Consortium | Lighthouse Laboratory Plymouth and Alex Alderton, Roberto Amato, Jeffrey Barrett, Sonia Goncalves, Ewan Harrison, David K. Jackson, Ian Johnston, Dominic Kwiatkowski, Cordelia Langford, John Sillitoe on behalf of the Wellcome Sanger Institute COVID-19 Surveillance Team |
| EPI_ISL_3362084, EPI_ISL_3362086 | Lighthouse Lab in Alderley Park | Wellcome Sanger Institute for the COVID-19 Genomics UK (COG-UK) Consortium | Jacquelyn Wynn, Mairead Hyland, The Lighthouse Lab in Alderley Park and Alex Alderton, Roberto Amato, Jeffrey Barrett, Sonia Goncalves, Ewan Harrison, David K. Jackson, Ian Johnston, Dominic Kwiatkowski, Cordelia Langford, John Sillitoe on behalf of the Wellcome Sanger Institute COVID-19 Surveillance Team |
| EPI_ISL_3362087 | Lighthouse Lab in Milton Keynes | Wellcome Sanger Institute for the COVID-19 Genomics UK (COG-UK) Consortium | The Lighthouse Lab in Milton Keynes and Alex Alderton, Roberto Amato, Jeffrey Barrett, Sonia Goncalves, Ewan Harrison, David K. Jackson, Ian Johnston, Dominic Kwiatkowski, Cordelia Langford, John Sillitoe on behalf of the Wellcome Sanger Institute COVID-19 Surveillance Team |
| EPI_ISL_3362090 | Lighthouse Laboratory Plymouth | Wellcome Sanger Institute for the COVID-19 Genomics UK (COG-UK) Consortium | Lighthouse Laboratory Plymouth and Alex Alderton, Roberto Amato, Jeffrey Barrett, Sonia Goncalves, Ewan Harrison, David K. Jackson, Ian Johnston, Dominic Kwiatkowski, Cordelia Langford, John Sillitoe on behalf of the Wellcome Sanger Institute COVID-19 Surveillance Team |
| EPI_ISL_3362091 | Lighthouse Lab in Alderley Park | Wellcome Sanger Institute for the COVID-19 Genomics UK (COG-UK) Consortium | Jacquelyn Wynn, Mairead Hyland, The Lighthouse Lab in Alderley Park and Alex Alderton, Roberto Amato, Jeffrey Barrett, Sonia Goncalves, Ewan Harrison, David K. Jackson, Ian Johnston, Dominic Kwiatkowski, Cordelia Langford, John Sillitoe on behalf of the Wellcome Sanger Institute COVID-19 Surveillance Team |
| EPI_ISL_3362093 | Lighthouse Lab in Milton Keynes | Wellcome Sanger Institute for the COVID-19 Genomics UK (COG-UK) Consortium | The Lighthouse Lab in Milton Keynes and Alex Alderton, Roberto Amato, Jeffrey Barrett, Sonia Goncalves, Ewan Harrison, David K. Jackson, Ian Johnston, Dominic Kwiatkowski, Cordelia Langford, John Sillitoe on behalf of the Wellcome Sanger Institute COVID-19 Surveillance Team |
| EPI_ISL_3362094 | Lighthouse Laboratory Plymouth | Wellcome Sanger Institute for the COVID-19 Genomics UK (COG-UK) Consortium | Lighthouse Laboratory Plymouth and Alex Alderton, Roberto Amato, Jeffrey Barrett, Sonia Goncalves, Ewan Harrison, David K. Jackson, Ian Johnston, Dominic Kwiatkowski, Cordelia Langford, John Sillitoe on behalf of the Wellcome Sanger Institute COVID-19 Surveillance Team |
| EPI_ISL_3362095 | Lighthouse Lab in Alderley Park | Wellcome Sanger Institute for the COVID-19 Genomics UK (COG-UK) Consortium | Jacquelyn Wynn, Mairead Hyland, The Lighthouse Lab in Alderley Park and Alex Alderton, Roberto Amato, Jeffrey Barrett, Sonia Goncalves, Ewan Harrison, David K. Jackson, Ian Johnston, Dominic Kwiatkowski, Cordelia Langford, John Sillitoe on behalf of the Wellcome Sanger Institute COVID-19 Surveillance Team |
| EPI_ISL_3362097 | Lighthouse Lab in Milton Keynes | Wellcome Sanger Institute for the COVID-19 Genomics UK (COG-UK) Consortium | The Lighthouse Lab in Milton Keynes and Alex Alderton, Roberto Amato, Jeffrey Barrett, Sonia Goncalves, Ewan Harrison, David K. Jackson, Ian Johnston, Dominic Kwiatkowski, Cordelia Langford, John Sillitoe on behalf of the Wellcome Sanger Institute COVID-19 Surveillance Team |
| EPI_ISL_3362098 | Lighthouse Laboratory Plymouth | Wellcome Sanger Institute for the COVID-19 Genomics UK (COG-UK) Consortium | Lighthouse Laboratory Plymouth and Alex Alderton, Roberto Amato, Jeffrey Barrett, Sonia Goncalves, Ewan Harrison, David K. Jackson, Ian Johnston, Dominic Kwiatkowski, Cordelia Langford, John Sillitoe on behalf of the Wellcome Sanger Institute COVID-19 Surveillance Team |
| EPI_ISL_3362100 | Lighthouse Lab in Alderley Park | Wellcome Sanger Institute for the COVID-19 Genomics UK (COG-UK) Consortium | Jacquelyn Wynn, Mairead Hyland, The Lighthouse Lab in Alderley Park and Alex Alderton, Roberto Amato, Jeffrey Barrett, Sonia Goncalves, Ewan Harrison, David K. Jackson, Ian Johnston, Dominic Kwiatkowski, Cordelia Langford, John Sillitoe on behalf of the Wellcome Sanger Institute COVID-19 Surveillance Team |
| EPI_ISL_3362101 | Lighthouse Laboratory Plymouth | Wellcome Sanger Institute for the COVID-19 Genomics UK (COG-UK) Consortium | Lighthouse Laboratory Plymouth and Alex Alderton, Roberto Amato, Jeffrey Barrett, Sonia Goncalves, Ewan Harrison, David K. Jackson, Ian Johnston, Dominic Kwiatkowski, Cordelia Langford, John Sillitoe on behalf of the Wellcome Sanger Institute COVID-19 Surveillance Team |
| EPI_ISL_3362103 | Lighthouse Lab in Alderley Park | Wellcome Sanger Institute for the COVID-19 Genomics UK (COG-UK) Consortium | Jacquelyn Wynn, Mairead Hyland, The Lighthouse Lab in Alderley Park and Alex Alderton, Roberto Amato, Jeffrey Barrett, Sonia Goncalves, Ewan Harrison, David K. Jackson, Ian Johnston, Dominic Kwiatkowski, Cordelia Langford, John Sillitoe on behalf of the Wellcome Sanger Institute COVID-19 Surveillance Team |
| EPI_ISL_3362104, EPI_ISL_3362106, EPI_ISL_3362107 | Lighthouse Lab in Milton Keynes | Wellcome Sanger Institute for the COVID-19 Genomics UK (COG-UK) Consortium | The Lighthouse Lab in Milton Keynes and Alex Alderton, Roberto Amato, Jeffrey Barrett, Sonia Goncalves, Ewan Harrison, David K. Jackson, Ian Johnston, Dominic Kwiatkowski, Cordelia Langford, John Sillitoe on behalf of the Wellcome Sanger Institute COVID-19 Surveillance Team |
| EPI_ISL_3362108, EPI_ISL_3362110 | Lighthouse Laboratory Plymouth | Wellcome Sanger Institute for the COVID-19 Genomics UK (COG-UK) Consortium | Lighthouse Laboratory Plymouth and Alex Alderton, Roberto Amato, Jeffrey Barrett, Sonia Goncalves, Ewan Harrison, David K. Jackson, Ian Johnston, Dominic Kwiatkowski, Cordelia Langford, John Sillitoe on behalf of the Wellcome Sanger Institute COVID-19 Surveillance Team |
| EPI_ISL_3362111, EPI_ISL_3362113 | Lighthouse Lab in Milton Keynes | Wellcome Sanger Institute for the COVID-19 Genomics UK (COG-UK) Consortium | The Lighthouse Lab in Milton Keynes and Alex Alderton, Roberto Amato, Jeffrey Barrett, Sonia Goncalves, Ewan Harrison, David K. Jackson, Ian Johnston, Dominic Kwiatkowski, Cordelia Langford, John Sillitoe on behalf of the Wellcome Sanger Institute COVID-19 Surveillance Team |
| EPI_ISL_3362114 | Lighthouse Laboratory Plymouth | Wellcome Sanger Institute for the COVID-19 Genomics UK (COG-UK) Consortium | Lighthouse Laboratory Plymouth and Alex Alderton, Roberto Amato, Jeffrey Barrett, Sonia Goncalves, Ewan Harrison, David K. Jackson, Ian Johnston, Dominic Kwiatkowski, Cordelia Langford, John Sillitoe on behalf of the Wellcome Sanger Institute COVID-19 Surveillance Team |

| | | | |
|---|---|---|---|
| EPI_ISL_3362115 | Lighthouse Lab in Milton Keynes | Wellcome Sanger Institute for the COVID-19 Genomics UK (COG-UK) Consortium | The Lighthouse Lab in Milton Keynes and Alex Alderton, Roberto Amato, Jeffrey Barrett, Sonia Goncalves, Ewan Harrison, David K. Jackson, Ian Johnston, Dominic Kwiatkowski, Cordelia Langford, John Sillitoe on behalf of the Wellcome Sanger Institute COVID-19 Surveillance Team |
| EPI_ISL_3362117 | Lighthouse Lab in Alderley Park | Wellcome Sanger Institute for the COVID-19 Genomics UK (COG-UK) Consortium | Jacquelyn Wynn, Mairead Hyland, The Lighthouse Lab in Alderley Park and Alex Alderton, Roberto Amato, Jeffrey Barrett, Sonia Goncalves, Ewan Harrison, David K. Jackson, Ian Johnston, Dominic Kwiatkowski, Cordelia Langford, John Sillitoe on behalf of the Wellcome Sanger Institute COVID-19 Surveillance Team |
| EPI_ISL_3362118 | Lighthouse Laboratory Plymouth | Wellcome Sanger Institute for the COVID-19 Genomics UK (COG-UK) Consortium | Lighthouse Laboratory Plymouth and Alex Alderton, Roberto Amato, Jeffrey Barrett, Sonia Goncalves, Ewan Harrison, David K. Jackson, Ian Johnston, Dominic Kwiatkowski, Cordelia Langford, John Sillitoe on behalf of the Wellcome Sanger Institute COVID-19 Surveillance Team |
| EPI_ISL_3362119, EPI_ISL_3362121 | Lighthouse Lab in Milton Keynes | Wellcome Sanger Institute for the COVID-19 Genomics UK (COG-UK) Consortium | The Lighthouse Lab in Milton Keynes and Alex Alderton, Roberto Amato, Jeffrey Barrett, Sonia Goncalves, Ewan Harrison, David K. Jackson, Ian Johnston, Dominic Kwiatkowski, Cordelia Langford, John Sillitoe on behalf of the Wellcome Sanger Institute COVID-19 Surveillance Team |
| EPI_ISL_3362122 | Lighthouse Lab in Alderley Park | Wellcome Sanger Institute for the COVID-19 Genomics UK (COG-UK) Consortium | Jacquelyn Wynn, Mairead Hyland, The Lighthouse Lab in Alderley Park and Alex Alderton, Roberto Amato, Jeffrey Barrett, Sonia Goncalves, Ewan Harrison, David K. Jackson, Ian Johnston, Dominic Kwiatkowski, Cordelia Langford, John Sillitoe on behalf of the Wellcome Sanger Institute COVID-19 Surveillance Team |
| EPI_ISL_3362124, EPI_ISL_3362125 | Lighthouse Laboratory Plymouth | Wellcome Sanger Institute for the COVID-19 Genomics UK (COG-UK) Consortium | Lighthouse Laboratory Plymouth and Alex Alderton, Roberto Amato, Jeffrey Barrett, Sonia Goncalves, Ewan Harrison, David K. Jackson, Ian Johnston, Dominic Kwiatkowski, Cordelia Langford, John Sillitoe on behalf of the Wellcome Sanger Institute COVID-19 Surveillance Team |
| EPI_ISL_3362127 | Lighthouse Lab in Alderley Park | Wellcome Sanger Institute for the COVID-19 Genomics UK (COG-UK) Consortium | Jacquelyn Wynn, Mairead Hyland, The Lighthouse Lab in Alderley Park and Alex Alderton, Roberto Amato, Jeffrey Barrett, Sonia Goncalves, Ewan Harrison, David K. Jackson, Ian Johnston, Dominic Kwiatkowski, Cordelia Langford, John Sillitoe on behalf of the Wellcome Sanger Institute COVID-19 Surveillance Team |
| EPI_ISL_3362128 | Lighthouse Lab in Milton Keynes | Wellcome Sanger Institute for the COVID-19 Genomics UK (COG-UK) Consortium | The Lighthouse Lab in Milton Keynes and Alex Alderton, Roberto Amato, Jeffrey Barrett, Sonia Goncalves, Ewan Harrison, David K. Jackson, Ian Johnston, Dominic Kwiatkowski, Cordelia Langford, John Sillitoe on behalf of the Wellcome Sanger Institute COVID-19 Surveillance Team |
| EPI_ISL_3362129 | Lighthouse Laboratory Plymouth | Wellcome Sanger Institute for the COVID-19 Genomics UK (COG-UK) Consortium | Lighthouse Laboratory Plymouth and Alex Alderton, Roberto Amato, Jeffrey Barrett, Sonia Goncalves, Ewan Harrison, David K. Jackson, Ian Johnston, Dominic Kwiatkowski, Cordelia Langford, John Sillitoe on behalf of the Wellcome Sanger Institute COVID-19 Surveillance Team |
| EPI_ISL_3362131 | Lighthouse Lab in Milton Keynes | Wellcome Sanger Institute for the COVID-19 Genomics UK (COG-UK) Consortium | The Lighthouse Lab in Milton Keynes and Alex Alderton, Roberto Amato, Jeffrey Barrett, Sonia Goncalves, Ewan Harrison, David K. Jackson, Ian Johnston, Dominic Kwiatkowski, Cordelia Langford, John Sillitoe on behalf of the Wellcome Sanger Institute COVID-19 Surveillance Team |
| EPI_ISL_3362132 | Lighthouse Lab in Alderley Park | Wellcome Sanger Institute for the COVID-19 Genomics UK (COG-UK) Consortium | Jacquelyn Wynn, Mairead Hyland, The Lighthouse Lab in Alderley Park and Alex Alderton, Roberto Amato, Jeffrey Barrett, Sonia Goncalves, Ewan Harrison, David K. Jackson, Ian Johnston, Dominic Kwiatkowski, Cordelia Langford, John Sillitoe on behalf of the Wellcome Sanger Institute COVID-19 Surveillance Team |
| EPI_ISL_3362133 | Lighthouse Laboratory Plymouth | Wellcome Sanger Institute for the COVID-19 Genomics UK (COG-UK) Consortium | Lighthouse Laboratory Plymouth and Alex Alderton, Roberto Amato, Jeffrey Barrett, Sonia Goncalves, Ewan Harrison, David K. Jackson, Ian Johnston, Dominic Kwiatkowski, Cordelia Langford, John Sillitoe on behalf of the Wellcome Sanger Institute COVID-19 Surveillance Team |
| EPI_ISL_3362135, EPI_ISL_3362136 | Lighthouse Lab in Milton Keynes | Wellcome Sanger Institute for the COVID-19 Genomics UK (COG-UK) Consortium | The Lighthouse Lab in Milton Keynes and Alex Alderton, Roberto Amato, Jeffrey Barrett, Sonia Goncalves, Ewan Harrison, David K. Jackson, Ian Johnston, Dominic Kwiatkowski, Cordelia Langford, John Sillitoe on behalf of the Wellcome Sanger Institute COVID-19 Surveillance Team |
| EPI_ISL_3362138 | Lighthouse Lab in Alderley Park | Wellcome Sanger Institute for the COVID-19 Genomics UK (COG-UK) Consortium | Jacquelyn Wynn, Mairead Hyland, The Lighthouse Lab in Alderley Park and Alex Alderton, Roberto Amato, Jeffrey Barrett, Sonia Goncalves, Ewan Harrison, David K. Jackson, Ian Johnston, Dominic Kwiatkowski, Cordelia Langford, John Sillitoe on behalf of the Wellcome Sanger Institute COVID-19 Surveillance Team |
| EPI_ISL_3362139 | Lighthouse Lab in Milton Keynes | Wellcome Sanger Institute for the COVID-19 Genomics UK (COG-UK) Consortium | The Lighthouse Lab in Milton Keynes and Alex Alderton, Roberto Amato, Jeffrey Barrett, Sonia Goncalves, Ewan Harrison, David K. Jackson, Ian Johnston, Dominic Kwiatkowski, Cordelia Langford, John Sillitoe on behalf of the Wellcome Sanger Institute COVID-19 Surveillance Team |
| EPI_ISL_3362141, EPI_ISL_3362142, EPI_ISL_3362144 | Lighthouse Lab in Alderley Park | Wellcome Sanger Institute for the COVID-19 Genomics UK (COG-UK) Consortium | Jacquelyn Wynn, Mairead Hyland, The Lighthouse Lab in Alderley Park and Alex Alderton, Roberto Amato, Jeffrey Barrett, Sonia Goncalves, Ewan Harrison, David K. Jackson, Ian Johnston, Dominic Kwiatkowski, Cordelia Langford, John Sillitoe on behalf of the Wellcome Sanger Institute COVID-19 Surveillance Team |
| EPI_ISL_3362145, EPI_ISL_3362147 | Lighthouse Laboratory Plymouth | Wellcome Sanger Institute for the COVID-19 Genomics UK (COG-UK) Consortium | Lighthouse Laboratory Plymouth and Alex Alderton, Roberto Amato, Jeffrey Barrett, Sonia Goncalves, Ewan Harrison, David K. Jackson, Ian Johnston, Dominic Kwiatkowski, Cordelia Langford, John Sillitoe on behalf of the Wellcome Sanger Institute COVID-19 Surveillance Team |
| EPI_ISL_3362148, EPI_ISL_3362150 | Lighthouse Lab in Milton Keynes | Wellcome Sanger Institute for the COVID-19 Genomics UK (COG-UK) Consortium | The Lighthouse Lab in Milton Keynes and Alex Alderton, Roberto Amato, Jeffrey Barrett, Sonia Goncalves, Ewan Harrison, David K. Jackson, Ian Johnston, Dominic Kwiatkowski, Cordelia Langford, John Sillitoe on behalf of the Wellcome Sanger Institute COVID-19 Surveillance Team |
| EPI_ISL_3362151 | Lighthouse Laboratory Plymouth | Wellcome Sanger Institute for the COVID-19 Genomics UK (COG-UK) Consortium | Lighthouse Laboratory Plymouth and Alex Alderton, Roberto Amato, Jeffrey Barrett, Sonia Goncalves, Ewan Harrison, David K. Jackson, Ian Johnston, Dominic Kwiatkowski, Cordelia Langford, John Sillitoe on behalf of the Wellcome Sanger Institute COVID-19 Surveillance Team |
| EPI_ISL_3362152 | Lighthouse Lab in Milton Keynes | Wellcome Sanger Institute for the COVID-19 Genomics UK (COG-UK) Consortium | The Lighthouse Lab in Milton Keynes and Alex Alderton, Roberto Amato, Jeffrey Barrett, Sonia Goncalves, Ewan Harrison, David K. Jackson, Ian Johnston, Dominic Kwiatkowski, Cordelia Langford, John Sillitoe on behalf of the Wellcome Sanger Institute COVID-19 Surveillance Team |
| EPI_ISL_3362154 | Lighthouse Laboratory Plymouth | Wellcome Sanger Institute for the COVID-19 Genomics UK (COG-UK) Consortium | Lighthouse Laboratory Plymouth and Alex Alderton, Roberto Amato, Jeffrey Barrett, Sonia Goncalves, Ewan Harrison, David K. Jackson, Ian Johnston, Dominic Kwiatkowski, Cordelia Langford, John Sillitoe on behalf of the Wellcome Sanger Institute COVID-19 Surveillance Team |
| EPI_ISL_3362155 | Lighthouse Lab in Milton Keynes | Wellcome Sanger Institute for the COVID-19 Genomics UK (COG-UK) Consortium | The Lighthouse Lab in Milton Keynes and Alex Alderton, Roberto Amato, Jeffrey Barrett, Sonia Goncalves, Ewan Harrison, David K. Jackson, Ian Johnston, Dominic Kwiatkowski, Cordelia Langford, John Sillitoe on behalf of the Wellcome Sanger Institute COVID-19 Surveillance Team |
| EPI_ISL_3362157 | Lighthouse Laboratory Plymouth | Wellcome Sanger Institute for the COVID-19 Genomics UK (COG-UK) Consortium | Lighthouse Laboratory Plymouth and Alex Alderton, Roberto Amato, Jeffrey Barrett, Sonia Goncalves, Ewan Harrison, David K. Jackson, Ian Johnston, Dominic Kwiatkowski, Cordelia Langford, John Sillitoe on behalf of the Wellcome Sanger Institute COVID-19 Surveillance Team |
| EPI_ISL_3362158, EPI_ISL_3362159, EPI_ISL_3362161 | Lighthouse Lab in Milton Keynes | Wellcome Sanger Institute for the COVID-19 Genomics UK (COG-UK) Consortium | The Lighthouse Lab in Milton Keynes and Alex Alderton, Roberto Amato, Jeffrey Barrett, Sonia Goncalves, Ewan Harrison, David K. Jackson, Ian Johnston, Dominic Kwiatkowski, Cordelia Langford, John Sillitoe on behalf of the Wellcome Sanger Institute COVID-19 Surveillance Team |
| EPI_ISL_3362162 | Lighthouse Lab in Alderley Park | Wellcome Sanger Institute for the COVID-19 Genomics UK (COG-UK) Consortium | Jacquelyn Wynn, Mairead Hyland, The Lighthouse Lab in Alderley Park and Alex Alderton, Roberto Amato, Jeffrey Barrett, Sonia Goncalves, Ewan Harrison, David K. Jackson, Ian Johnston, Dominic Kwiatkowski, Cordelia Langford, John Sillitoe on behalf of the Wellcome Sanger Institute COVID-19 Surveillance Team |
| EPI_ISL_3362164, EPI_ISL_3362165 | Lighthouse Laboratory Plymouth | Wellcome Sanger Institute for the COVID-19 Genomics UK (COG-UK) Consortium | Lighthouse Laboratory Plymouth and Alex Alderton, Roberto Amato, Jeffrey Barrett, Sonia Goncalves, Ewan Harrison, David K. Jackson, Ian Johnston, Dominic Kwiatkowski, Cordelia Langford, John Sillitoe on behalf of the Wellcome Sanger Institute COVID-19 Surveillance Team |
| EPI_ISL_3362167, EPI_ISL_3362168, EPI_ISL_3362169, EPI_ISL_3362171 | Lighthouse Lab in Milton Keynes | Wellcome Sanger Institute for the COVID-19 Genomics UK (COG-UK) Consortium | The Lighthouse Lab in Milton Keynes and Alex Alderton, Roberto Amato, Jeffrey Barrett, Sonia Goncalves, Ewan Harrison, David K. Jackson, Ian Johnston, Dominic Kwiatkowski, Cordelia Langford, John Sillitoe on behalf of the Wellcome Sanger Institute COVID-19 Surveillance Team |
| EPI_ISL_3362172 | Lighthouse Laboratory Plymouth | Wellcome Sanger Institute for the COVID-19 Genomics UK (COG-UK) Consortium | Lighthouse Laboratory Plymouth and Alex Alderton, Roberto Amato, Jeffrey Barrett, Sonia Goncalves, Ewan Harrison, David K. Jackson, Ian Johnston, Dominic Kwiatkowski, Cordelia Langford, John Sillitoe on behalf of the Wellcome Sanger Institute COVID-19 Surveillance Team |
| EPI_ISL_3362174 | Lighthouse Lab in Alderley Park | Wellcome Sanger Institute for the COVID-19 Genomics UK (COG-UK) Consortium | Jacquelyn Wynn, Mairead Hyland, The Lighthouse Lab in Alderley Park and Alex Alderton, Roberto Amato, Jeffrey Barrett, Sonia Goncalves, Ewan Harrison, David K. Jackson, Ian Johnston, Dominic Kwiatkowski, Cordelia Langford, John Sillitoe on behalf of the Wellcome Sanger Institute COVID-19 Surveillance Team |
| EPI_ISL_3362176 | Lighthouse Lab in Milton Keynes | Wellcome Sanger Institute for the COVID-19 Genomics UK (COG-UK) Consortium | The Lighthouse Lab in Milton Keynes and Alex Alderton, Roberto Amato, Jeffrey Barrett, Sonia Goncalves, Ewan Harrison, David K. Jackson, Ian Johnston, Dominic Kwiatkowski, Cordelia Langford, John Sillitoe on behalf of the Wellcome Sanger Institute COVID-19 Surveillance Team |
| EPI_ISL_3362177 | Lighthouse Lab in Alderley Park | Wellcome Sanger Institute for the COVID-19 Genomics UK (COG-UK) Consortium | Jacquelyn Wynn, Mairead Hyland, The Lighthouse Lab in Alderley Park and Alex Alderton, Roberto Amato, Jeffrey Barrett, Sonia Goncalves, Ewan Harrison, David K. Jackson, Ian Johnston, Dominic Kwiatkowski, Cordelia Langford, John Sillitoe on behalf of the Wellcome Sanger Institute COVID-19 Surveillance Team |
| EPI_ISL_3362179, EPI_ISL_3362180, EPI_ISL_3362182 | Lighthouse Lab in Milton Keynes | Wellcome Sanger Institute for the COVID-19 Genomics UK (COG-UK) Consortium | The Lighthouse Lab in Milton Keynes and Alex Alderton, Roberto Amato, Jeffrey Barrett, Sonia Goncalves, Ewan Harrison, David K. Jackson, Ian Johnston, Dominic Kwiatkowski, Cordelia Langford, John Sillitoe on behalf of the Wellcome Sanger Institute COVID-19 Surveillance Team |
| EPI_ISL_3362183, EPI_ISL_3362184, EPI_ISL_3362186 | Lighthouse Lab in Alderley Park | Wellcome Sanger Institute for the COVID-19 Genomics UK (COG-UK) Consortium | Jacquelyn Wynn, Mairead Hyland, The Lighthouse Lab in Alderley Park and Alex Alderton, Roberto Amato, Jeffrey Barrett, Sonia Goncalves, Ewan Harrison, David K. Jackson, Ian Johnston, Dominic Kwiatkowski, Cordelia Langford, John Sillitoe on behalf of the Wellcome Sanger Institute COVID-19 Surveillance Team |
| EPI_ISL_3362187, EPI_ISL_3362189, EPI_ISL_3362190, EPI_ISL_3362191 | Lighthouse Lab in Milton Keynes | Wellcome Sanger Institute for the COVID-19 Genomics UK (COG-UK) Consortium | The Lighthouse Lab in Milton Keynes and Alex Alderton, Roberto Amato, Jeffrey Barrett, Sonia Goncalves, Ewan Harrison, David K. Jackson, Ian Johnston, Dominic Kwiatkowski, Cordelia Langford, John Sillitoe on behalf of the Wellcome Sanger Institute COVID-19 Surveillance Team |
| EPI_ISL_3362193 | Lighthouse Lab in Alderley Park | Wellcome Sanger Institute for the COVID-19 Genomics UK (COG-UK) Consortium | Jacquelyn Wynn, Mairead Hyland, The Lighthouse Lab in Alderley Park and Alex Alderton, Roberto Amato, Jeffrey Barrett, Sonia Goncalves, Ewan Harrison, David K. Jackson, Ian Johnston, Dominic Kwiatkowski, Cordelia Langford, John Sillitoe on behalf of the Wellcome Sanger Institute COVID-19 Surveillance Team |
| EPI_ISL_3362195, EPI_ISL_3362197 | Lighthouse Lab in Milton Keynes | Wellcome Sanger Institute for the COVID-19 Genomics UK (COG-UK) Consortium | The Lighthouse Lab in Milton Keynes and Alex Alderton, Roberto Amato, Jeffrey Barrett, Sonia Goncalves, Ewan Harrison, David K. Jackson, Ian Johnston, Dominic Kwiatkowski, Cordelia Langford, John Sillitoe on behalf of the Wellcome Sanger Institute COVID-19 Surveillance Team |
| EPI_ISL_3362198 | Lighthouse Laboratory Plymouth | Wellcome Sanger Institute for the COVID-19 Genomics UK (COG-UK) Consortium | Lighthouse Laboratory Plymouth and Alex Alderton, Roberto Amato, Jeffrey Barrett, Sonia Goncalves, Ewan Harrison, David K. Jackson, Ian Johnston, Dominic Kwiatkowski, Cordelia Langford, John Sillitoe on behalf of the Wellcome Sanger Institute COVID-19 Surveillance Team |
| EPI_ISL_3362199, EPI_ISL_3362201, EPI_ISL_3362202 | Lighthouse Lab in Alderley Park | Wellcome Sanger Institute for the COVID-19 Genomics UK (COG-UK) Consortium | Jacquelyn Wynn, Mairead Hyland, The Lighthouse Lab in Alderley Park and Alex Alderton, Roberto Amato, Jeffrey Barrett, Sonia Goncalves, Ewan Harrison, David K. Jackson, Ian Johnston, Dominic Kwiatkowski, Cordelia Langford, John Sillitoe on behalf of the Wellcome Sanger Institute COVID-19 Surveillance Team |
| EPI_ISL_3362204 | Lighthouse Lab in Milton Keynes | Wellcome Sanger Institute for the COVID-19 Genomics UK (COG-UK) Consortium | The Lighthouse Lab in Milton Keynes and Alex Alderton, Roberto Amato, Jeffrey Barrett, Sonia Goncalves, Ewan Harrison, David K. Jackson, Ian Johnston, Dominic Kwiatkowski, Cordelia Langford, John Sillitoe on behalf of the Wellcome Sanger Institute COVID-19 Surveillance Team |
| EPI_ISL_3362205 | Lighthouse Lab in Alderley Park | Wellcome Sanger Institute for the COVID-19 Genomics UK (COG-UK) Consortium | Jacquelyn Wynn, Mairead Hyland, The Lighthouse Lab in Alderley Park and Alex Alderton, Roberto Amato, Jeffrey Barrett, Sonia Goncalves, Ewan Harrison, David K. Jackson, Ian Johnston, Dominic Kwiatkowski, Cordelia Langford, John Sillitoe on behalf of the Wellcome Sanger Institute COVID-19 Surveillance Team |
| EPI_ISL_3362206 | Lighthouse Lab in Milton Keynes | Wellcome Sanger Institute for the COVID-19 Genomics UK (COG-UK) Consortium | The Lighthouse Lab in Milton Keynes and Alex Alderton, Roberto Amato, Jeffrey Barrett, Sonia Goncalves, Ewan Harrison, David K. Jackson, Ian Johnston, Dominic Kwiatkowski, Cordelia Langford, John Sillitoe on behalf of the Wellcome Sanger Institute COVID-19 Surveillance Team |

| | | | |
|---|---|---|---|
| EPI_ISL_3362208 | Lighthouse Lab in Alderley Park | Wellcome Sanger Institute for the COVID-19 Genomics UK (COG-UK) Consortium | Jacquelyn Wynn, Mairead Hyland, The Lighthouse Lab in Alderley Park and Alex Alderton, Roberto Amato, Jeffrey Barrett, Sonia Goncalves, Ewan Harrison, David K. Jackson, Ian Johnston, Dominic Kwiatkowski, Cordelia Langford, John Sillitoe on behalf of the Wellcome Sanger Institute COVID-19 Surveillance Team |
| EPI_ISL_3362209 | Lighthouse Lab in Milton Keynes | Wellcome Sanger Institute for the COVID-19 Genomics UK (COG-UK) Consortium | The Lighthouse Lab in Milton Keynes and Alex Alderton, Roberto Amato, Jeffrey Barrett, Sonia Goncalves, Ewan Harrison, David K. Jackson, Ian Johnston, Dominic Kwiatkowski, Cordelia Langford, John Sillitoe on behalf of the Wellcome Sanger Institute COVID-19 Surveillance Team |
| EPI_ISL_3362211 | Lighthouse Lab in Alderley Park | Wellcome Sanger Institute for the COVID-19 Genomics UK (COG-UK) Consortium | Jacquelyn Wynn, Mairead Hyland, The Lighthouse Lab in Alderley Park and Alex Alderton, Roberto Amato, Jeffrey Barrett, Sonia Goncalves, Ewan Harrison, David K. Jackson, Ian Johnston, Dominic Kwiatkowski, Cordelia Langford, John Sillitoe on behalf of the Wellcome Sanger Institute COVID-19 Surveillance Team |
| EPI_ISL_3362212 | Lighthouse Lab in Milton Keynes | Wellcome Sanger Institute for the COVID-19 Genomics UK (COG-UK) Consortium | The Lighthouse Lab in Milton Keynes and Alex Alderton, Roberto Amato, Jeffrey Barrett, Sonia Goncalves, Ewan Harrison, David K. Jackson, Ian Johnston, Dominic Kwiatkowski, Cordelia Langford, John Sillitoe on behalf of the Wellcome Sanger Institute COVID-19 Surveillance Team |
| EPI_ISL_3362214 | Lighthouse Lab in Alderley Park | Wellcome Sanger Institute for the COVID-19 Genomics UK (COG-UK) Consortium | Jacquelyn Wynn, Mairead Hyland, The Lighthouse Lab in Alderley Park and Alex Alderton, Roberto Amato, Jeffrey Barrett, Sonia Goncalves, Ewan Harrison, David K. Jackson, Ian Johnston, Dominic Kwiatkowski, Cordelia Langford, John Sillitoe on behalf of the Wellcome Sanger Institute COVID-19 Surveillance Team |
| EPI_ISL_3362215, EPI_ISL_3362217, EPI_ISL_3362218, EPI_ISL_3362219 | Lighthouse Lab in Milton Keynes | Wellcome Sanger Institute for the COVID-19 Genomics UK (COG-UK) Consortium | The Lighthouse Lab in Milton Keynes and Alex Alderton, Roberto Amato, Jeffrey Barrett, Sonia Goncalves, Ewan Harrison, David K. Jackson, Ian Johnston, Dominic Kwiatkowski, Cordelia Langford, John Sillitoe on behalf of the Wellcome Sanger Institute COVID-19 Surveillance Team |
| EPI_ISL_3362221, EPI_ISL_3362222 | Lighthouse Lab in Alderley Park | Wellcome Sanger Institute for the COVID-19 Genomics UK (COG-UK) Consortium | Jacquelyn Wynn, Mairead Hyland, The Lighthouse Lab in Alderley Park and Alex Alderton, Roberto Amato, Jeffrey Barrett, Sonia Goncalves, Ewan Harrison, David K. Jackson, Ian Johnston, Dominic Kwiatkowski, Cordelia Langford, John Sillitoe on behalf of the Wellcome Sanger Institute COVID-19 Surveillance Team |
| EPI_ISL_3362224, EPI_ISL_3362225 | Lighthouse Lab in Milton Keynes | Wellcome Sanger Institute for the COVID-19 Genomics UK (COG-UK) Consortium | The Lighthouse Lab in Milton Keynes and Alex Alderton, Roberto Amato, Jeffrey Barrett, Sonia Goncalves, Ewan Harrison, David K. Jackson, Ian Johnston, Dominic Kwiatkowski, Cordelia Langford, John Sillitoe on behalf of the Wellcome Sanger Institute COVID-19 Surveillance Team |
| EPI_ISL_3362226 | Lighthouse Lab in Alderley Park | Wellcome Sanger Institute for the COVID-19 Genomics UK (COG-UK) Consortium | Jacquelyn Wynn, Mairead Hyland, The Lighthouse Lab in Alderley Park and Alex Alderton, Roberto Amato, Jeffrey Barrett, Sonia Goncalves, Ewan Harrison, David K. Jackson, Ian Johnston, Dominic Kwiatkowski, Cordelia Langford, John Sillitoe on behalf of the Wellcome Sanger Institute COVID-19 Surveillance Team |
| EPI_ISL_3362228, EPI_ISL_3362229, EPI_ISL_3362231, EPI_ISL_3362232 | Lighthouse Lab in Milton Keynes | Wellcome Sanger Institute for the COVID-19 Genomics UK (COG-UK) Consortium | The Lighthouse Lab in Milton Keynes and Alex Alderton, Roberto Amato, Jeffrey Barrett, Sonia Goncalves, Ewan Harrison, David K. Jackson, Ian Johnston, Dominic Kwiatkowski, Cordelia Langford, John Sillitoe on behalf of the Wellcome Sanger Institute COVID-19 Surveillance Team |
| EPI_ISL_3362234 | Lighthouse Laboratory Plymouth | Wellcome Sanger Institute for the COVID-19 Genomics UK (COG-UK) Consortium | Lighthouse Laboratory Plymouth and Alex Alderton, Roberto Amato, Jeffrey Barrett, Sonia Goncalves, Ewan Harrison, David K. Jackson, Ian Johnston, Dominic Kwiatkowski, Cordelia Langford, John Sillitoe on behalf of the Wellcome Sanger Institute COVID-19 Surveillance Team |
| EPI_ISL_3362235 | Lighthouse Lab in Alderley Park | Wellcome Sanger Institute for the COVID-19 Genomics UK (COG-UK) Consortium | Jacquelyn Wynn, Mairead Hyland, The Lighthouse Lab in Alderley Park and Alex Alderton, Roberto Amato, Jeffrey Barrett, Sonia Goncalves, Ewan Harrison, David K. Jackson, Ian Johnston, Dominic Kwiatkowski, Cordelia Langford, John Sillitoe on behalf of the Wellcome Sanger Institute COVID-19 Surveillance Team |
| EPI_ISL_3362237 | Lighthouse Lab in Milton Keynes | Wellcome Sanger Institute for the COVID-19 Genomics UK (COG-UK) Consortium | The Lighthouse Lab in Milton Keynes and Alex Alderton, Roberto Amato, Jeffrey Barrett, Sonia Goncalves, Ewan Harrison, David K. Jackson, Ian Johnston, Dominic Kwiatkowski, Cordelia Langford, John Sillitoe on behalf of the Wellcome Sanger Institute COVID-19 Surveillance Team |
| EPI_ISL_3362238 | Lighthouse Lab in Alderley Park | Wellcome Sanger Institute for the COVID-19 Genomics UK (COG-UK) Consortium | Jacquelyn Wynn, Mairead Hyland, The Lighthouse Lab in Alderley Park and Alex Alderton, Roberto Amato, Jeffrey Barrett, Sonia Goncalves, Ewan Harrison, David K. Jackson, Ian Johnston, Dominic Kwiatkowski, Cordelia Langford, John Sillitoe on behalf of the Wellcome Sanger Institute COVID-19 Surveillance Team |
| EPI_ISL_3362239 | Lighthouse Lab in Milton Keynes | Wellcome Sanger Institute for the COVID-19 Genomics UK (COG-UK) Consortium | The Lighthouse Lab in Milton Keynes and Alex Alderton, Roberto Amato, Jeffrey Barrett, Sonia Goncalves, Ewan Harrison, David K. Jackson, Ian Johnston, Dominic Kwiatkowski, Cordelia Langford, John Sillitoe on behalf of the Wellcome Sanger Institute COVID-19 Surveillance Team |
| EPI_ISL_3362241, EPI_ISL_3362242, EPI_ISL_3362244 | Lighthouse Laboratory Plymouth | Wellcome Sanger Institute for the COVID-19 Genomics UK (COG-UK) Consortium | Lighthouse Laboratory Plymouth and Alex Alderton, Roberto Amato, Jeffrey Barrett, Sonia Goncalves, Ewan Harrison, David K. Jackson, Ian Johnston, Dominic Kwiatkowski, Cordelia Langford, John Sillitoe on behalf of the Wellcome Sanger Institute COVID-19 Surveillance Team |
| EPI_ISL_3362245, EPI_ISL_3362247, EPI_ISL_3362248, EPI_ISL_3362250 | Lighthouse Lab in Milton Keynes | Wellcome Sanger Institute for the COVID-19 Genomics UK (COG-UK) Consortium | The Lighthouse Lab in Milton Keynes and Alex Alderton, Roberto Amato, Jeffrey Barrett, Sonia Goncalves, Ewan Harrison, David K. Jackson, Ian Johnston, Dominic Kwiatkowski, Cordelia Langford, John Sillitoe on behalf of the Wellcome Sanger Institute COVID-19 Surveillance Team |
| EPI_ISL_3362251, EPI_ISL_3362253 | Lighthouse Laboratory Plymouth | Wellcome Sanger Institute for the COVID-19 Genomics UK (COG-UK) Consortium | Lighthouse Laboratory Plymouth and Alex Alderton, Roberto Amato, Jeffrey Barrett, Sonia Goncalves, Ewan Harrison, David K. Jackson, Ian Johnston, Dominic Kwiatkowski, Cordelia Langford, John Sillitoe on behalf of the Wellcome Sanger Institute COVID-19 Surveillance Team |
| EPI_ISL_3362254 | Lighthouse Lab in Milton Keynes | Wellcome Sanger Institute for the COVID-19 Genomics UK (COG-UK) Consortium | The Lighthouse Lab in Milton Keynes and Alex Alderton, Roberto Amato, Jeffrey Barrett, Sonia Goncalves, Ewan Harrison, David K. Jackson, Ian Johnston, Dominic Kwiatkowski, Cordelia Langford, John Sillitoe on behalf of the Wellcome Sanger Institute COVID-19 Surveillance Team |
| EPI_ISL_3362256, EPI_ISL_3362257 | Lighthouse Lab in Alderley Park | Wellcome Sanger Institute for the COVID-19 Genomics UK (COG-UK) Consortium | Jacquelyn Wynn, Mairead Hyland, The Lighthouse Lab in Alderley Park and Alex Alderton, Roberto Amato, Jeffrey Barrett, Sonia Goncalves, Ewan Harrison, David K. Jackson, Ian Johnston, Dominic Kwiatkowski, Cordelia Langford, John Sillitoe on behalf of the Wellcome Sanger Institute COVID-19 Surveillance Team |
| EPI_ISL_3362258, EPI_ISL_3362260 | Lighthouse Laboratory Plymouth | Wellcome Sanger Institute for the COVID-19 Genomics UK (COG-UK) Consortium | Lighthouse Laboratory Plymouth and Alex Alderton, Roberto Amato, Jeffrey Barrett, Sonia Goncalves, Ewan Harrison, David K. Jackson, Ian Johnston, Dominic Kwiatkowski, Cordelia Langford, John Sillitoe on behalf of the Wellcome Sanger Institute COVID-19 Surveillance Team |
| EPI_ISL_3362311 | Lighthouse Lab in Milton Keynes | Wellcome Sanger Institute for the COVID-19 Genomics UK (COG-UK) Consortium | The Lighthouse Lab in Milton Keynes and Alex Alderton, Roberto Amato, Jeffrey Barrett, Sonia Goncalves, Ewan Harrison, David K. Jackson, Ian Johnston, Dominic Kwiatkowski, Cordelia Langford, John Sillitoe on behalf of the Wellcome Sanger Institute COVID-19 Surveillance Team |
| EPI_ISL_3362313 | Lighthouse Lab in Alderley Park | Wellcome Sanger Institute for the COVID-19 Genomics UK (COG-UK) Consortium | Jacquelyn Wynn, Mairead Hyland, The Lighthouse Lab in Alderley Park and Alex Alderton, Roberto Amato, Jeffrey Barrett, Sonia Goncalves, Ewan Harrison, David K. Jackson, Ian Johnston, Dominic Kwiatkowski, Cordelia Langford, John Sillitoe on behalf of the Wellcome Sanger Institute COVID-19 Surveillance Team |
| EPI_ISL_3362314, EPI_ISL_3362316 | Lighthouse Lab in Milton Keynes | Wellcome Sanger Institute for the COVID-19 Genomics UK (COG-UK) Consortium | The Lighthouse Lab in Milton Keynes and Alex Alderton, Roberto Amato, Jeffrey Barrett, Sonia Goncalves, Ewan Harrison, David K. Jackson, Ian Johnston, Dominic Kwiatkowski, Cordelia Langford, John Sillitoe on behalf of the Wellcome Sanger Institute COVID-19 Surveillance Team |
| EPI_ISL_3362317, EPI_ISL_3362319 | Lighthouse Lab in Alderley Park | Wellcome Sanger Institute for the COVID-19 Genomics UK (COG-UK) Consortium | Jacquelyn Wynn, Mairead Hyland, The Lighthouse Lab in Alderley Park and Alex Alderton, Roberto Amato, Jeffrey Barrett, Sonia Goncalves, Ewan Harrison, David K. Jackson, Ian Johnston, Dominic Kwiatkowski, Cordelia Langford, John Sillitoe on behalf of the Wellcome Sanger Institute COVID-19 Surveillance Team |
| EPI_ISL_3362321, EPI_ISL_3362322 | Lighthouse Lab in Milton Keynes | Wellcome Sanger Institute for the COVID-19 Genomics UK (COG-UK) Consortium | The Lighthouse Lab in Milton Keynes and Alex Alderton, Roberto Amato, Jeffrey Barrett, Sonia Goncalves, Ewan Harrison, David K. Jackson, Ian Johnston, Dominic Kwiatkowski, Cordelia Langford, John Sillitoe on behalf of the Wellcome Sanger Institute COVID-19 Surveillance Team |
| EPI_ISL_3362324, EPI_ISL_3362325, EPI_ISL_3362327 | Lighthouse Laboratory Plymouth | Wellcome Sanger Institute for the COVID-19 Genomics UK (COG-UK) Consortium | Lighthouse Laboratory Plymouth and Alex Alderton, Roberto Amato, Jeffrey Barrett, Sonia Goncalves, Ewan Harrison, David K. Jackson, Ian Johnston, Dominic Kwiatkowski, Cordelia Langford, John Sillitoe on behalf of the Wellcome Sanger Institute COVID-19 Surveillance Team |
| EPI_ISL_3362328, EPI_ISL_3362330, EPI_ISL_3362331, EPI_ISL_3362333, EPI_ISL_3362334, EPI_ISL_3362335, EPI_ISL_3362337, EPI_ISL_3362338, EPI_ISL_3362340, EPI_ISL_3362341 | Lighthouse Lab in Milton Keynes | Wellcome Sanger Institute for the COVID-19 Genomics UK (COG-UK) Consortium | The Lighthouse Lab in Milton Keynes and Alex Alderton, Roberto Amato, Jeffrey Barrett, Sonia Goncalves, Ewan Harrison, David K. Jackson, Ian Johnston, Dominic Kwiatkowski, Cordelia Langford, John Sillitoe on behalf of the Wellcome Sanger Institute COVID-19 Surveillance Team |
| EPI_ISL_3362343, EPI_ISL_3362344 | Lighthouse Laboratory Plymouth | Wellcome Sanger Institute for the COVID-19 Genomics UK (COG-UK) Consortium | Lighthouse Laboratory Plymouth and Alex Alderton, Roberto Amato, Jeffrey Barrett, Sonia Goncalves, Ewan Harrison, David K. Jackson, Ian Johnston, Dominic Kwiatkowski, Cordelia Langford, John Sillitoe on behalf of the Wellcome Sanger Institute COVID-19 Surveillance Team |
| EPI_ISL_3362345, EPI_ISL_3362347 | Lighthouse Lab in Alderley Park | Wellcome Sanger Institute for the COVID-19 Genomics UK (COG-UK) Consortium | Jacquelyn Wynn, Mairead Hyland, The Lighthouse Lab in Alderley Park and Alex Alderton, Roberto Amato, Jeffrey Barrett, Sonia Goncalves, Ewan Harrison, David K. Jackson, Ian Johnston, Dominic Kwiatkowski, Cordelia Langford, John Sillitoe on behalf of the Wellcome Sanger Institute COVID-19 Surveillance Team |
| EPI_ISL_3362348 | Lighthouse Lab in Milton Keynes | Wellcome Sanger Institute for the COVID-19 Genomics UK (COG-UK) Consortium | The Lighthouse Lab in Milton Keynes and Alex Alderton, Roberto Amato, Jeffrey Barrett, Sonia Goncalves, Ewan Harrison, David K. Jackson, Ian Johnston, Dominic Kwiatkowski, Cordelia Langford, John Sillitoe on behalf of the Wellcome Sanger Institute COVID-19 Surveillance Team |
| EPI_ISL_3362350 | Lighthouse Lab in Alderley Park | Wellcome Sanger Institute for the COVID-19 Genomics UK (COG-UK) Consortium | Jacquelyn Wynn, Mairead Hyland, The Lighthouse Lab in Alderley Park and Alex Alderton, Roberto Amato, Jeffrey Barrett, Sonia Goncalves, Ewan Harrison, David K. Jackson, Ian Johnston, Dominic Kwiatkowski, Cordelia Langford, John Sillitoe on behalf of the Wellcome Sanger Institute COVID-19 Surveillance Team |
| EPI_ISL_3362351 | Lighthouse Laboratory Plymouth | Wellcome Sanger Institute for the COVID-19 Genomics UK (COG-UK) Consortium | Lighthouse Laboratory Plymouth and Alex Alderton, Roberto Amato, Jeffrey Barrett, Sonia Goncalves, Ewan Harrison, David K. Jackson, Ian Johnston, Dominic Kwiatkowski, Cordelia Langford, John Sillitoe on behalf of the Wellcome Sanger Institute COVID-19 Surveillance Team |
| EPI_ISL_3362353, EPI_ISL_3362354, EPI_ISL_3362356, EPI_ISL_3362357 | Lighthouse Lab in Milton Keynes | Wellcome Sanger Institute for the COVID-19 Genomics UK (COG-UK) Consortium | The Lighthouse Lab in Milton Keynes and Alex Alderton, Roberto Amato, Jeffrey Barrett, Sonia Goncalves, Ewan Harrison, David K. Jackson, Ian Johnston, Dominic Kwiatkowski, Cordelia Langford, John Sillitoe on behalf of the Wellcome Sanger Institute COVID-19 Surveillance Team |
| EPI_ISL_3362358, EPI_ISL_3362360, EPI_ISL_3362362, EPI_ISL_3362363 | Lighthouse Lab in Alderley Park | Wellcome Sanger Institute for the COVID-19 Genomics UK (COG-UK) Consortium | Jacquelyn Wynn, Mairead Hyland, The Lighthouse Lab in Alderley Park and Alex Alderton, Roberto Amato, Jeffrey Barrett, Sonia Goncalves, Ewan Harrison, David K. Jackson, Ian Johnston, Dominic Kwiatkowski, Cordelia Langford, John Sillitoe on behalf of the Wellcome Sanger Institute COVID-19 Surveillance Team |
| EPI_ISL_3362365 | Lighthouse Laboratory Plymouth | Wellcome Sanger Institute for the COVID-19 Genomics UK (COG-UK) Consortium | Lighthouse Laboratory Plymouth and Alex Alderton, Roberto Amato, Jeffrey Barrett, Sonia Goncalves, Ewan Harrison, David K. Jackson, Ian Johnston, Dominic Kwiatkowski, Cordelia Langford, John Sillitoe on behalf of the Wellcome Sanger Institute COVID-19 Surveillance Team |
| EPI_ISL_3362366 | Lighthouse Lab in Alderley Park | Wellcome Sanger Institute for the COVID-19 Genomics UK (COG-UK) | Jacquelyn Wynn, Mairead Hyland, The Lighthouse Lab in Alderley Park and Alex Alderton, Roberto Amato, Jeffrey Barrett, Sonia Goncalves, Ewan Harrison, David K. Jackson, Ian Johnston, Dominic Kwiatkowski, Cordelia Langford, John Sillitoe on |

| | | | |
|---|---|---|---|
| | | Consortium | behalf of the Wellcome Sanger Institute COVID-19 Surveillance Team |
| EPI_ISL_3362367 | Lighthouse Lab in Milton Keynes | Wellcome Sanger Institute for the COVID-19 Genomics UK (COG-UK) Consortium | The Lighthouse Lab in Milton Keynes and Alex Alderton, Roberto Amato, Jeffrey Barrett, Sonia Goncalves, Ewan Harrison, David K. Jackson, Ian Johnston, Dominic Kwiatkowski, Cordelia Langford, John Sillitoe on behalf of the Wellcome Sanger Institute COVID-19 Surveillance Team |
| EPI_ISL_3362369 | Lighthouse Lab in Alderley Park | Wellcome Sanger Institute for the COVID-19 Genomics UK (COG-UK) Consortium | Jacquelyn Wynn, Mairead Hyland, The Lighthouse Lab in Alderley Park and Alex Alderton, Roberto Amato, Jeffrey Barrett, Sonia Goncalves, Ewan Harrison, David K. Jackson, Ian Johnston, Dominic Kwiatkowski, Cordelia Langford, John Sillitoe on behalf of the Wellcome Sanger Institute COVID-19 Surveillance Team |
| EPI_ISL_3362370, EPI_ISL_3362372 | Lighthouse Lab in Milton Keynes | Wellcome Sanger Institute for the COVID-19 Genomics UK (COG-UK) Consortium | The Lighthouse Lab in Milton Keynes and Alex Alderton, Roberto Amato, Jeffrey Barrett, Sonia Goncalves, Ewan Harrison, David K. Jackson, Ian Johnston, Dominic Kwiatkowski, Cordelia Langford, John Sillitoe on behalf of the Wellcome Sanger Institute COVID-19 Surveillance Team |
| EPI_ISL_3362373 | Lighthouse Laboratory Plymouth | Wellcome Sanger Institute for the COVID-19 Genomics UK (COG-UK) Consortium | Lighthouse Laboratory Plymouth and Alex Alderton, Roberto Amato, Jeffrey Barrett, Sonia Goncalves, Ewan Harrison, David K. Jackson, Ian Johnston, Dominic Kwiatkowski, Cordelia Langford, John Sillitoe on behalf of the Wellcome Sanger Institute COVID-19 Surveillance Team |
| EPI_ISL_3362375 | Lighthouse Lab in Alderley Park | Wellcome Sanger Institute for the COVID-19 Genomics UK (COG-UK) Consortium | Jacquelyn Wynn, Mairead Hyland, The Lighthouse Lab in Alderley Park and Alex Alderton, Roberto Amato, Jeffrey Barrett, Sonia Goncalves, Ewan Harrison, David K. Jackson, Ian Johnston, Dominic Kwiatkowski, Cordelia Langford, John Sillitoe on behalf of the Wellcome Sanger Institute COVID-19 Surveillance Team |
| EPI_ISL_3362376 | Lighthouse Lab in Milton Keynes | Wellcome Sanger Institute for the COVID-19 Genomics UK (COG-UK) Consortium | The Lighthouse Lab in Milton Keynes and Alex Alderton, Roberto Amato, Jeffrey Barrett, Sonia Goncalves, Ewan Harrison, David K. Jackson, Ian Johnston, Dominic Kwiatkowski, Cordelia Langford, John Sillitoe on behalf of the Wellcome Sanger Institute COVID-19 Surveillance Team |
| EPI_ISL_3362378, EPI_ISL_3362379 | Lighthouse Lab in Alderley Park | Wellcome Sanger Institute for the COVID-19 Genomics UK (COG-UK) Consortium | Jacquelyn Wynn, Mairead Hyland, The Lighthouse Lab in Alderley Park and Alex Alderton, Roberto Amato, Jeffrey Barrett, Sonia Goncalves, Ewan Harrison, David K. Jackson, Ian Johnston, Dominic Kwiatkowski, Cordelia Langford, John Sillitoe on behalf of the Wellcome Sanger Institute COVID-19 Surveillance Team |
| EPI_ISL_3362380, EPI_ISL_3362382 | Lighthouse Lab in Milton Keynes | Wellcome Sanger Institute for the COVID-19 Genomics UK (COG-UK) Consortium | The Lighthouse Lab in Milton Keynes and Alex Alderton, Roberto Amato, Jeffrey Barrett, Sonia Goncalves, Ewan Harrison, David K. Jackson, Ian Johnston, Dominic Kwiatkowski, Cordelia Langford, John Sillitoe on behalf of the Wellcome Sanger Institute COVID-19 Surveillance Team |
| EPI_ISL_3362383 | Lighthouse Lab in Alderley Park | Wellcome Sanger Institute for the COVID-19 Genomics UK (COG-UK) Consortium | Jacquelyn Wynn, Mairead Hyland, The Lighthouse Lab in Alderley Park and Alex Alderton, Roberto Amato, Jeffrey Barrett, Sonia Goncalves, Ewan Harrison, David K. Jackson, Ian Johnston, Dominic Kwiatkowski, Cordelia Langford, John Sillitoe on behalf of the Wellcome Sanger Institute COVID-19 Surveillance Team |
| EPI_ISL_3362384 | Lighthouse Laboratory Plymouth | Wellcome Sanger Institute for the COVID-19 Genomics UK (COG-UK) Consortium | Lighthouse Laboratory Plymouth and Alex Alderton, Roberto Amato, Jeffrey Barrett, Sonia Goncalves, Ewan Harrison, David K. Jackson, Ian Johnston, Dominic Kwiatkowski, Cordelia Langford, John Sillitoe on behalf of the Wellcome Sanger Institute COVID-19 Surveillance Team |
| EPI_ISL_3362386 | Lighthouse Lab in Alderley Park | Wellcome Sanger Institute for the COVID-19 Genomics UK (COG-UK) Consortium | Jacquelyn Wynn, Mairead Hyland, The Lighthouse Lab in Alderley Park and Alex Alderton, Roberto Amato, Jeffrey Barrett, Sonia Goncalves, Ewan Harrison, David K. Jackson, Ian Johnston, Dominic Kwiatkowski, Cordelia Langford, John Sillitoe on behalf of the Wellcome Sanger Institute COVID-19 Surveillance Team |
| EPI_ISL_3362387, EPI_ISL_3362389 | Lighthouse Lab in Milton Keynes | Wellcome Sanger Institute for the COVID-19 Genomics UK (COG-UK) Consortium | The Lighthouse Lab in Milton Keynes and Alex Alderton, Roberto Amato, Jeffrey Barrett, Sonia Goncalves, Ewan Harrison, David K. Jackson, Ian Johnston, Dominic Kwiatkowski, Cordelia Langford, John Sillitoe on behalf of the Wellcome Sanger Institute COVID-19 Surveillance Team |
| EPI_ISL_3362390 | Lighthouse Lab in Alderley Park | Wellcome Sanger Institute for the COVID-19 Genomics UK (COG-UK) Consortium | Jacquelyn Wynn, Mairead Hyland, The Lighthouse Lab in Alderley Park and Alex Alderton, Roberto Amato, Jeffrey Barrett, Sonia Goncalves, Ewan Harrison, David K. Jackson, Ian Johnston, Dominic Kwiatkowski, Cordelia Langford, John Sillitoe on behalf of the Wellcome Sanger Institute COVID-19 Surveillance Team |
| EPI_ISL_3362392 | Lighthouse Laboratory Plymouth | Wellcome Sanger Institute for the COVID-19 Genomics UK (COG-UK) Consortium | Lighthouse Laboratory Plymouth and Alex Alderton, Roberto Amato, Jeffrey Barrett, Sonia Goncalves, Ewan Harrison, David K. Jackson, Ian Johnston, Dominic Kwiatkowski, Cordelia Langford, John Sillitoe on behalf of the Wellcome Sanger Institute COVID-19 Surveillance Team |
| EPI_ISL_3362393 | Lighthouse Lab in Milton Keynes | Wellcome Sanger Institute for the COVID-19 Genomics UK (COG-UK) Consortium | The Lighthouse Lab in Milton Keynes and Alex Alderton, Roberto Amato, Jeffrey Barrett, Sonia Goncalves, Ewan Harrison, David K. Jackson, Ian Johnston, Dominic Kwiatkowski, Cordelia Langford, John Sillitoe on behalf of the Wellcome Sanger Institute COVID-19 Surveillance Team |
| EPI_ISL_3362395 | Lighthouse Laboratory Plymouth | Wellcome Sanger Institute for the COVID-19 Genomics UK (COG-UK) Consortium | Lighthouse Laboratory Plymouth and Alex Alderton, Roberto Amato, Jeffrey Barrett, Sonia Goncalves, Ewan Harrison, David K. Jackson, Ian Johnston, Dominic Kwiatkowski, Cordelia Langford, John Sillitoe on behalf of the Wellcome Sanger Institute COVID-19 Surveillance Team |
| EPI_ISL_3362396, EPI_ISL_3362398 | Lighthouse Lab in Milton Keynes | Wellcome Sanger Institute for the COVID-19 Genomics UK (COG-UK) Consortium | The Lighthouse Lab in Milton Keynes and Alex Alderton, Roberto Amato, Jeffrey Barrett, Sonia Goncalves, Ewan Harrison, David K. Jackson, Ian Johnston, Dominic Kwiatkowski, Cordelia Langford, John Sillitoe on behalf of the Wellcome Sanger Institute COVID-19 Surveillance Team |
| EPI_ISL_3362399 | Lighthouse Laboratory Plymouth | Wellcome Sanger Institute for the COVID-19 Genomics UK (COG-UK) Consortium | Lighthouse Laboratory Plymouth and Alex Alderton, Roberto Amato, Jeffrey Barrett, Sonia Goncalves, Ewan Harrison, David K. Jackson, Ian Johnston, Dominic Kwiatkowski, Cordelia Langford, John Sillitoe on behalf of the Wellcome Sanger Institute COVID-19 Surveillance Team |
| EPI_ISL_3362400 | Lighthouse Lab in Alderley Park | Wellcome Sanger Institute for the COVID-19 Genomics UK (COG-UK) Consortium | Jacquelyn Wynn, Mairead Hyland, The Lighthouse Lab in Alderley Park and Alex Alderton, Roberto Amato, Jeffrey Barrett, Sonia Goncalves, Ewan Harrison, David K. Jackson, Ian Johnston, Dominic Kwiatkowski, Cordelia Langford, John Sillitoe on behalf of the Wellcome Sanger Institute COVID-19 Surveillance Team |
| EPI_ISL_3362402 | Lighthouse Lab in Milton Keynes | Wellcome Sanger Institute for the COVID-19 Genomics UK (COG-UK) Consortium | The Lighthouse Lab in Milton Keynes and Alex Alderton, Roberto Amato, Jeffrey Barrett, Sonia Goncalves, Ewan Harrison, David K. Jackson, Ian Johnston, Dominic Kwiatkowski, Cordelia Langford, John Sillitoe on behalf of the Wellcome Sanger Institute COVID-19 Surveillance Team |
| EPI_ISL_3362403, EPI_ISL_3362405, EPI_ISL_3362406 | Lighthouse Lab in Alderley Park | Wellcome Sanger Institute for the COVID-19 Genomics UK (COG-UK) Consortium | Jacquelyn Wynn, Mairead Hyland, The Lighthouse Lab in Alderley Park and Alex Alderton, Roberto Amato, Jeffrey Barrett, Sonia Goncalves, Ewan Harrison, David K. Jackson, Ian Johnston, Dominic Kwiatkowski, Cordelia Langford, John Sillitoe on behalf of the Wellcome Sanger Institute COVID-19 Surveillance Team |
| EPI_ISL_3362408 | Lighthouse Laboratory Plymouth | Wellcome Sanger Institute for the COVID-19 Genomics UK (COG-UK) Consortium | Lighthouse Laboratory Plymouth and Alex Alderton, Roberto Amato, Jeffrey Barrett, Sonia Goncalves, Ewan Harrison, David K. Jackson, Ian Johnston, Dominic Kwiatkowski, Cordelia Langford, John Sillitoe on behalf of the Wellcome Sanger Institute COVID-19 Surveillance Team |
| EPI_ISL_3362409 | Lighthouse Lab in Milton Keynes | Wellcome Sanger Institute for the COVID-19 Genomics UK (COG-UK) Consortium | The Lighthouse Lab in Milton Keynes and Alex Alderton, Roberto Amato, Jeffrey Barrett, Sonia Goncalves, Ewan Harrison, David K. Jackson, Ian Johnston, Dominic Kwiatkowski, Cordelia Langford, John Sillitoe on behalf of the Wellcome Sanger Institute COVID-19 Surveillance Team |
| EPI_ISL_3362411 | Lighthouse Lab in Alderley Park | Wellcome Sanger Institute for the COVID-19 Genomics UK (COG-UK) Consortium | Jacquelyn Wynn, Mairead Hyland, The Lighthouse Lab in Alderley Park and Alex Alderton, Roberto Amato, Jeffrey Barrett, Sonia Goncalves, Ewan Harrison, David K. Jackson, Ian Johnston, Dominic Kwiatkowski, Cordelia Langford, John Sillitoe on behalf of the Wellcome Sanger Institute COVID-19 Surveillance Team |
| EPI_ISL_3362412 | Lighthouse Laboratory Plymouth | Wellcome Sanger Institute for the COVID-19 Genomics UK (COG-UK) Consortium | Lighthouse Laboratory Plymouth and Alex Alderton, Roberto Amato, Jeffrey Barrett, Sonia Goncalves, Ewan Harrison, David K. Jackson, Ian Johnston, Dominic Kwiatkowski, Cordelia Langford, John Sillitoe on behalf of the Wellcome Sanger Institute COVID-19 Surveillance Team |
| EPI_ISL_3362413, EPI_ISL_3362414 | Lighthouse Lab in Milton Keynes | Wellcome Sanger Institute for the COVID-19 Genomics UK (COG-UK) Consortium | The Lighthouse Lab in Milton Keynes and Alex Alderton, Roberto Amato, Jeffrey Barrett, Sonia Goncalves, Ewan Harrison, David K. Jackson, Ian Johnston, Dominic Kwiatkowski, Cordelia Langford, John Sillitoe on behalf of the Wellcome Sanger Institute COVID-19 Surveillance Team |
| EPI_ISL_3362416 | Lighthouse Laboratory Plymouth | Wellcome Sanger Institute for the COVID-19 Genomics UK (COG-UK) Consortium | Lighthouse Laboratory Plymouth and Alex Alderton, Roberto Amato, Jeffrey Barrett, Sonia Goncalves, Ewan Harrison, David K. Jackson, Ian Johnston, Dominic Kwiatkowski, Cordelia Langford, John Sillitoe on behalf of the Wellcome Sanger Institute COVID-19 Surveillance Team |
| EPI_ISL_3362417 | Lighthouse Lab in Milton Keynes | Wellcome Sanger Institute for the COVID-19 Genomics UK (COG-UK) Consortium | The Lighthouse Lab in Milton Keynes and Alex Alderton, Roberto Amato, Jeffrey Barrett, Sonia Goncalves, Ewan Harrison, David K. Jackson, Ian Johnston, Dominic Kwiatkowski, Cordelia Langford, John Sillitoe on behalf of the Wellcome Sanger Institute COVID-19 Surveillance Team |
| EPI_ISL_3362419 | Lighthouse Lab in Alderley Park | Wellcome Sanger Institute for the COVID-19 Genomics UK (COG-UK) Consortium | Jacquelyn Wynn, Mairead Hyland, The Lighthouse Lab in Alderley Park and Alex Alderton, Roberto Amato, Jeffrey Barrett, Sonia Goncalves, Ewan Harrison, David K. Jackson, Ian Johnston, Dominic Kwiatkowski, Cordelia Langford, John Sillitoe on behalf of the Wellcome Sanger Institute COVID-19 Surveillance Team |
| EPI_ISL_3362420 | Lighthouse Laboratory Plymouth | Wellcome Sanger Institute for the COVID-19 Genomics UK (COG-UK) Consortium | Lighthouse Laboratory Plymouth and Alex Alderton, Roberto Amato, Jeffrey Barrett, Sonia Goncalves, Ewan Harrison, David K. Jackson, Ian Johnston, Dominic Kwiatkowski, Cordelia Langford, John Sillitoe on behalf of the Wellcome Sanger Institute COVID-19 Surveillance Team |
| EPI_ISL_3362421, EPI_ISL_3362423, EPI_ISL_3362424 | Lighthouse Lab in Alderley Park | Wellcome Sanger Institute for the COVID-19 Genomics UK (COG-UK) Consortium | Jacquelyn Wynn, Mairead Hyland, The Lighthouse Lab in Alderley Park and Alex Alderton, Roberto Amato, Jeffrey Barrett, Sonia Goncalves, Ewan Harrison, David K. Jackson, Ian Johnston, Dominic Kwiatkowski, Cordelia Langford, John Sillitoe on behalf of the Wellcome Sanger Institute COVID-19 Surveillance Team |
| EPI_ISL_3362426 | Lighthouse Laboratory Plymouth | Wellcome Sanger Institute for the COVID-19 Genomics UK (COG-UK) Consortium | Lighthouse Laboratory Plymouth and Alex Alderton, Roberto Amato, Jeffrey Barrett, Sonia Goncalves, Ewan Harrison, David K. Jackson, Ian Johnston, Dominic Kwiatkowski, Cordelia Langford, John Sillitoe on behalf of the Wellcome Sanger Institute COVID-19 Surveillance Team |
| EPI_ISL_3362427 | Lighthouse Lab in Milton Keynes | Wellcome Sanger Institute for the COVID-19 Genomics UK (COG-UK) Consortium | The Lighthouse Lab in Milton Keynes and Alex Alderton, Roberto Amato, Jeffrey Barrett, Sonia Goncalves, Ewan Harrison, David K. Jackson, Ian Johnston, Dominic Kwiatkowski, Cordelia Langford, John Sillitoe on behalf of the Wellcome Sanger Institute COVID-19 Surveillance Team |
| EPI_ISL_3362429 | Lighthouse Lab in Alderley Park | Wellcome Sanger Institute for the COVID-19 Genomics UK (COG-UK) Consortium | Jacquelyn Wynn, Mairead Hyland, The Lighthouse Lab in Alderley Park and Alex Alderton, Roberto Amato, Jeffrey Barrett, Sonia Goncalves, Ewan Harrison, David K. Jackson, Ian Johnston, Dominic Kwiatkowski, Cordelia Langford, John Sillitoe on behalf of the Wellcome Sanger Institute COVID-19 Surveillance Team |
| EPI_ISL_3362430, EPI_ISL_3362432, EPI_ISL_3362433 | Lighthouse Lab in Milton Keynes | Wellcome Sanger Institute for the COVID-19 Genomics UK (COG-UK) Consortium | The Lighthouse Lab in Milton Keynes and Alex Alderton, Roberto Amato, Jeffrey Barrett, Sonia Goncalves, Ewan Harrison, David K. Jackson, Ian Johnston, Dominic Kwiatkowski, Cordelia Langford, John Sillitoe on behalf of the Wellcome Sanger Institute COVID-19 Surveillance Team |
| EPI_ISL_3362435 | Lighthouse Lab in Alderley Park | Wellcome Sanger Institute for the COVID-19 Genomics UK (COG-UK) Consortium | Jacquelyn Wynn, Mairead Hyland, The Lighthouse Lab in Alderley Park and Alex Alderton, Roberto Amato, Jeffrey Barrett, Sonia Goncalves, Ewan Harrison, David K. Jackson, Ian Johnston, Dominic Kwiatkowski, Cordelia Langford, John Sillitoe on behalf of the Wellcome Sanger Institute COVID-19 Surveillance Team |
| EPI_ISL_3362436 | Lighthouse Lab in Milton Keynes | Wellcome Sanger Institute for the COVID-19 Genomics UK (COG-UK) Consortium | The Lighthouse Lab in Milton Keynes and Alex Alderton, Roberto Amato, Jeffrey Barrett, Sonia Goncalves, Ewan Harrison, David K. Jackson, Ian Johnston, Dominic Kwiatkowski, Cordelia Langford, John Sillitoe on behalf of the Wellcome Sanger Institute COVID-19 Surveillance Team |
| EPI_ISL_3362438 | Lighthouse Lab in Alderley Park | Wellcome Sanger Institute for the COVID-19 Genomics UK (COG-UK) Consortium | Jacquelyn Wynn, Mairead Hyland, The Lighthouse Lab in Alderley Park and Alex Alderton, Roberto Amato, Jeffrey Barrett, Sonia Goncalves, Ewan Harrison, David K. Jackson, Ian Johnston, Dominic Kwiatkowski, Cordelia Langford, John Sillitoe on behalf of the Wellcome Sanger Institute COVID-19 Surveillance Team |
| EPI_ISL_3362439, EPI_ISL_3362441 | Lighthouse Laboratory Plymouth | Wellcome Sanger Institute for the COVID-19 Genomics UK (COG-UK) Consortium | Lighthouse Laboratory Plymouth and Alex Alderton, Roberto Amato, Jeffrey Barrett, Sonia Goncalves, Ewan Harrison, David K. Jackson, Ian Johnston, Dominic Kwiatkowski, Cordelia Langford, John Sillitoe on behalf of the Wellcome Sanger Institute COVID-19 Surveillance Team |
| EPI_ISL_3362442, EPI_ISL_3362444, EPI_ISL_3362445 | Lighthouse Lab in Milton Keynes | Wellcome Sanger Institute for the COVID-19 Genomics UK (COG-UK) Consortium | The Lighthouse Lab in Milton Keynes and Alex Alderton, Roberto Amato, Jeffrey Barrett, Sonia Goncalves, Ewan Harrison, David K. Jackson, Ian Johnston, Dominic Kwiatkowski, Cordelia Langford, John Sillitoe on behalf of the Wellcome Sanger Institute COVID-19 Surveillance Team |
| EPI_ISL_3362447 | Lighthouse Laboratory Plymouth | Wellcome Sanger Institute for the COVID-19 Genomics UK (COG-UK) Consortium | Lighthouse Laboratory Plymouth and Alex Alderton, Roberto Amato, Jeffrey Barrett, Sonia Goncalves, Ewan Harrison, David K. Jackson, Ian Johnston, Dominic Kwiatkowski, Cordelia Langford, John Sillitoe on behalf of the Wellcome Sanger Institute COVID-19 Surveillance Team |
| EPI_ISL_3362448, EPI_ISL_3362449 | Lighthouse Lab in Milton Keynes | Wellcome Sanger Institute for the COVID-19 Genomics UK (COG-UK) Consortium | The Lighthouse Lab in Milton Keynes and Alex Alderton, Roberto Amato, Jeffrey Barrett, Sonia Goncalves, Ewan Harrison, David K. Jackson, Ian Johnston, Dominic Kwiatkowski, Cordelia Langford, John Sillitoe on behalf of the Wellcome Sanger Institute COVID-19 Surveillance Team |
| EPI_ISL_3362451 | Lighthouse Laboratory Plymouth | Wellcome Sanger Institute for the COVID-19 Genomics UK (COG-UK) Consortium | Lighthouse Laboratory Plymouth and Alex Alderton, Roberto Amato, Jeffrey Barrett, Sonia Goncalves, Ewan Harrison, David K. Jackson, Ian Johnston, Dominic Kwiatkowski, Cordelia Langford, John Sillitoe on behalf of the Wellcome Sanger Institute COVID-19 Surveillance Team |

| | | | |
|---|---|---|---|
| EPI_ISL_3362452 | Lighthouse Lab in Alderley Park | Wellcome Sanger Institute for the COVID-19 Genomics UK (COG-UK) Consortium | Jacquelyn Wynn, Mairead Hyland, The Lighthouse Lab in Alderley Park and Alex Alderton, Roberto Amato, Jeffrey Barrett, Sonia Goncalves, Ewan Harrison, David K. Jackson, Ian Johnston, Dominic Kwiatkowski, Cordelia Langford, John Sillitoe on behalf of the Wellcome Sanger Institute COVID-19 Surveillance Team |
| EPI_ISL_3362454 | Lighthouse Laboratory Plymouth | Wellcome Sanger Institute for the COVID-19 Genomics UK (COG-UK) Consortium | Lighthouse Laboratory Plymouth and Alex Alderton, Roberto Amato, Jeffrey Barrett, Sonia Goncalves, Ewan Harrison, David K. Jackson, Ian Johnston, Dominic Kwiatkowski, Cordelia Langford, John Sillitoe on behalf of the Wellcome Sanger Institute COVID-19 Surveillance Team |
| EPI_ISL_3362455, EPI_ISL_3362457, EPI_ISL_3362458, EPI_ISL_3362459, EPI_ISL_3362461 | Lighthouse Lab in Milton Keynes | Wellcome Sanger Institute for the COVID-19 Genomics UK (COG-UK) Consortium | The Lighthouse Lab in Milton Keynes and Alex Alderton, Roberto Amato, Jeffrey Barrett, Sonia Goncalves, Ewan Harrison, David K. Jackson, Ian Johnston, Dominic Kwiatkowski, Cordelia Langford, John Sillitoe on behalf of the Wellcome Sanger Institute COVID-19 Surveillance Team |
| EPI_ISL_3362463 | Lighthouse Laboratory Plymouth | Wellcome Sanger Institute for the COVID-19 Genomics UK (COG-UK) Consortium | Lighthouse Laboratory Plymouth and Alex Alderton, Roberto Amato, Jeffrey Barrett, Sonia Goncalves, Ewan Harrison, David K. Jackson, Ian Johnston, Dominic Kwiatkowski, Cordelia Langford, John Sillitoe on behalf of the Wellcome Sanger Institute COVID-19 Surveillance Team |
| EPI_ISL_3362464 | Lighthouse Lab in Milton Keynes | Wellcome Sanger Institute for the COVID-19 Genomics UK (COG-UK) Consortium | The Lighthouse Lab in Milton Keynes and Alex Alderton, Roberto Amato, Jeffrey Barrett, Sonia Goncalves, Ewan Harrison, David K. Jackson, Ian Johnston, Dominic Kwiatkowski, Cordelia Langford, John Sillitoe on behalf of the Wellcome Sanger Institute COVID-19 Surveillance Team |
| EPI_ISL_3362465 | Lighthouse Lab in Alderley Park | Wellcome Sanger Institute for the COVID-19 Genomics UK (COG-UK) Consortium | Jacquelyn Wynn, Mairead Hyland, The Lighthouse Lab in Alderley Park and Alex Alderton, Roberto Amato, Jeffrey Barrett, Sonia Goncalves, Ewan Harrison, David K. Jackson, Ian Johnston, Dominic Kwiatkowski, Cordelia Langford, John Sillitoe on behalf of the Wellcome Sanger Institute COVID-19 Surveillance Team |
| EPI_ISL_3362467 | Lighthouse Laboratory Plymouth | Wellcome Sanger Institute for the COVID-19 Genomics UK (COG-UK) Consortium | Lighthouse Laboratory Plymouth and Alex Alderton, Roberto Amato, Jeffrey Barrett, Sonia Goncalves, Ewan Harrison, David K. Jackson, Ian Johnston, Dominic Kwiatkowski, Cordelia Langford, John Sillitoe on behalf of the Wellcome Sanger Institute COVID-19 Surveillance Team |
| EPI_ISL_3362468, EPI_ISL_3362470, EPI_ISL_3362471 | Lighthouse Lab in Milton Keynes | Wellcome Sanger Institute for the COVID-19 Genomics UK (COG-UK) Consortium | The Lighthouse Lab in Milton Keynes and Alex Alderton, Roberto Amato, Jeffrey Barrett, Sonia Goncalves, Ewan Harrison, David K. Jackson, Ian Johnston, Dominic Kwiatkowski, Cordelia Langford, John Sillitoe on behalf of the Wellcome Sanger Institute COVID-19 Surveillance Team |
| EPI_ISL_3362472 | Lighthouse Laboratory Plymouth | Wellcome Sanger Institute for the COVID-19 Genomics UK (COG-UK) Consortium | Lighthouse Laboratory Plymouth and Alex Alderton, Roberto Amato, Jeffrey Barrett, Sonia Goncalves, Ewan Harrison, David K. Jackson, Ian Johnston, Dominic Kwiatkowski, Cordelia Langford, John Sillitoe on behalf of the Wellcome Sanger Institute COVID-19 Surveillance Team |
| EPI_ISL_3362474 | Lighthouse Lab in Milton Keynes | Wellcome Sanger Institute for the COVID-19 Genomics UK (COG-UK) Consortium | The Lighthouse Lab in Milton Keynes and Alex Alderton, Roberto Amato, Jeffrey Barrett, Sonia Goncalves, Ewan Harrison, David K. Jackson, Ian Johnston, Dominic Kwiatkowski, Cordelia Langford, John Sillitoe on behalf of the Wellcome Sanger Institute COVID-19 Surveillance Team |
| EPI_ISL_3362475 | Lighthouse Lab in Alderley Park | Wellcome Sanger Institute for the COVID-19 Genomics UK (COG-UK) Consortium | Jacquelyn Wynn, Mairead Hyland, The Lighthouse Lab in Alderley Park and Alex Alderton, Roberto Amato, Jeffrey Barrett, Sonia Goncalves, Ewan Harrison, David K. Jackson, Ian Johnston, Dominic Kwiatkowski, Cordelia Langford, John Sillitoe on behalf of the Wellcome Sanger Institute COVID-19 Surveillance Team |
| EPI_ISL_3362477 | Lighthouse Laboratory Plymouth | Wellcome Sanger Institute for the COVID-19 Genomics UK (COG-UK) Consortium | Lighthouse Laboratory Plymouth and Alex Alderton, Roberto Amato, Jeffrey Barrett, Sonia Goncalves, Ewan Harrison, David K. Jackson, Ian Johnston, Dominic Kwiatkowski, Cordelia Langford, John Sillitoe on behalf of the Wellcome Sanger Institute COVID-19 Surveillance Team |
| EPI_ISL_3362478, EPI_ISL_3362479 | Lighthouse Lab in Milton Keynes | Wellcome Sanger Institute for the COVID-19 Genomics UK (COG-UK) Consortium | The Lighthouse Lab in Milton Keynes and Alex Alderton, Roberto Amato, Jeffrey Barrett, Sonia Goncalves, Ewan Harrison, David K. Jackson, Ian Johnston, Dominic Kwiatkowski, Cordelia Langford, John Sillitoe on behalf of the Wellcome Sanger Institute COVID-19 Surveillance Team |
| EPI_ISL_3362481, EPI_ISL_3362482 | Lighthouse Laboratory Plymouth | Wellcome Sanger Institute for the COVID-19 Genomics UK (COG-UK) Consortium | Lighthouse Laboratory Plymouth and Alex Alderton, Roberto Amato, Jeffrey Barrett, Sonia Goncalves, Ewan Harrison, David K. Jackson, Ian Johnston, Dominic Kwiatkowski, Cordelia Langford, John Sillitoe on behalf of the Wellcome Sanger Institute COVID-19 Surveillance Team |
| EPI_ISL_3362484 | Lighthouse Lab in Milton Keynes | Wellcome Sanger Institute for the COVID-19 Genomics UK (COG-UK) Consortium | The Lighthouse Lab in Milton Keynes and Alex Alderton, Roberto Amato, Jeffrey Barrett, Sonia Goncalves, Ewan Harrison, David K. Jackson, Ian Johnston, Dominic Kwiatkowski, Cordelia Langford, John Sillitoe on behalf of the Wellcome Sanger Institute COVID-19 Surveillance Team |
| EPI_ISL_3362485 | Lighthouse Lab in Alderley Park | Wellcome Sanger Institute for the COVID-19 Genomics UK (COG-UK) Consortium | Jacquelyn Wynn, Mairead Hyland, The Lighthouse Lab in Alderley Park and Alex Alderton, Roberto Amato, Jeffrey Barrett, Sonia Goncalves, Ewan Harrison, David K. Jackson, Ian Johnston, Dominic Kwiatkowski, Cordelia Langford, John Sillitoe on behalf of the Wellcome Sanger Institute COVID-19 Surveillance Team |
| EPI_ISL_3362487 | Lighthouse Laboratory Plymouth | Wellcome Sanger Institute for the COVID-19 Genomics UK (COG-UK) Consortium | Lighthouse Laboratory Plymouth and Alex Alderton, Roberto Amato, Jeffrey Barrett, Sonia Goncalves, Ewan Harrison, David K. Jackson, Ian Johnston, Dominic Kwiatkowski, Cordelia Langford, John Sillitoe on behalf of the Wellcome Sanger Institute COVID-19 Surveillance Team |
| EPI_ISL_3362488 | Lighthouse Lab in Milton Keynes | Wellcome Sanger Institute for the COVID-19 Genomics UK (COG-UK) Consortium | The Lighthouse Lab in Milton Keynes and Alex Alderton, Roberto Amato, Jeffrey Barrett, Sonia Goncalves, Ewan Harrison, David K. Jackson, Ian Johnston, Dominic Kwiatkowski, Cordelia Langford, John Sillitoe on behalf of the Wellcome Sanger Institute COVID-19 Surveillance Team |
| EPI_ISL_3362489, EPI_ISL_3362491 | Lighthouse Laboratory Plymouth | Wellcome Sanger Institute for the COVID-19 Genomics UK (COG-UK) Consortium | Lighthouse Laboratory Plymouth and Alex Alderton, Roberto Amato, Jeffrey Barrett, Sonia Goncalves, Ewan Harrison, David K. Jackson, Ian Johnston, Dominic Kwiatkowski, Cordelia Langford, John Sillitoe on behalf of the Wellcome Sanger Institute COVID-19 Surveillance Team |
| EPI_ISL_3362492, EPI_ISL_3362494, EPI_ISL_3362495 | Lighthouse Lab in Milton Keynes | Wellcome Sanger Institute for the COVID-19 Genomics UK (COG-UK) Consortium | The Lighthouse Lab in Milton Keynes and Alex Alderton, Roberto Amato, Jeffrey Barrett, Sonia Goncalves, Ewan Harrison, David K. Jackson, Ian Johnston, Dominic Kwiatkowski, Cordelia Langford, John Sillitoe on behalf of the Wellcome Sanger Institute COVID-19 Surveillance Team |
| EPI_ISL_3362497 | Lighthouse Lab in Alderley Park | Wellcome Sanger Institute for the COVID-19 Genomics UK (COG-UK) Consortium | Jacquelyn Wynn, Mairead Hyland, The Lighthouse Lab in Alderley Park and Alex Alderton, Roberto Amato, Jeffrey Barrett, Sonia Goncalves, Ewan Harrison, David K. Jackson, Ian Johnston, Dominic Kwiatkowski, Cordelia Langford, John Sillitoe on behalf of the Wellcome Sanger Institute COVID-19 Surveillance Team |
| EPI_ISL_3362498, EPI_ISL_3362500, EPI_ISL_3362501, EPI_ISL_3362502 | Lighthouse Lab in Milton Keynes | Wellcome Sanger Institute for the COVID-19 Genomics UK (COG-UK) Consortium | The Lighthouse Lab in Milton Keynes and Alex Alderton, Roberto Amato, Jeffrey Barrett, Sonia Goncalves, Ewan Harrison, David K. Jackson, Ian Johnston, Dominic Kwiatkowski, Cordelia Langford, John Sillitoe on behalf of the Wellcome Sanger Institute COVID-19 Surveillance Team |
| EPI_ISL_3362503, EPI_ISL_3362505 | Lighthouse Lab in Glasgow | Wellcome Sanger Institute for the COVID-19 Genomics UK (COG-UK) Consortium | Harper VanSteenhouse, Yumi Kasai, David Gray, Carol Clugston, Anna Dominiczak and Alex Alderton, Roberto Amato, Jeffrey Barrett, Sonia Goncalves, Ewan Harrison, David K. Jackson, Ian Johnston, Dominic Kwiatkowski, Cordelia Langford, John Sillitoe on behalf of the Wellcome Sanger Institute COVID-19 Surveillance Team |
| EPI_ISL_3362506 | Lighthouse Lab in Milton Keynes | Wellcome Sanger Institute for the COVID-19 Genomics UK (COG-UK) Consortium | The Lighthouse Lab in Milton Keynes and Alex Alderton, Roberto Amato, Jeffrey Barrett, Sonia Goncalves, Ewan Harrison, David K. Jackson, Ian Johnston, Dominic Kwiatkowski, Cordelia Langford, John Sillitoe on behalf of the Wellcome Sanger Institute COVID-19 Surveillance Team |
| EPI_ISL_3362508 | Lighthouse Lab in Glasgow | Wellcome Sanger Institute for the COVID-19 Genomics UK (COG-UK) Consortium | Harper VanSteenhouse, Yumi Kasai, David Gray, Carol Clugston, Anna Dominiczak and Alex Alderton, Roberto Amato, Jeffrey Barrett, Sonia Goncalves, Ewan Harrison, David K. Jackson, Ian Johnston, Dominic Kwiatkowski, Cordelia Langford, John Sillitoe on behalf of the Wellcome Sanger Institute COVID-19 Surveillance Team |
| EPI_ISL_3362509 | Lighthouse Lab in Milton Keynes | Wellcome Sanger Institute for the COVID-19 Genomics UK (COG-UK) Consortium | The Lighthouse Lab in Milton Keynes and Alex Alderton, Roberto Amato, Jeffrey Barrett, Sonia Goncalves, Ewan Harrison, David K. Jackson, Ian Johnston, Dominic Kwiatkowski, Cordelia Langford, John Sillitoe on behalf of the Wellcome Sanger Institute COVID-19 Surveillance Team |
| EPI_ISL_3362511 | Lighthouse Laboratory Plymouth | Wellcome Sanger Institute for the COVID-19 Genomics UK (COG-UK) Consortium | Lighthouse Laboratory Plymouth and Alex Alderton, Roberto Amato, Jeffrey Barrett, Sonia Goncalves, Ewan Harrison, David K. Jackson, Ian Johnston, Dominic Kwiatkowski, Cordelia Langford, John Sillitoe on behalf of the Wellcome Sanger Institute COVID-19 Surveillance Team |
| EPI_ISL_3362512, EPI_ISL_3362514 | Lighthouse Lab in Milton Keynes | Wellcome Sanger Institute for the COVID-19 Genomics UK (COG-UK) Consortium | The Lighthouse Lab in Milton Keynes and Alex Alderton, Roberto Amato, Jeffrey Barrett, Sonia Goncalves, Ewan Harrison, David K. Jackson, Ian Johnston, Dominic Kwiatkowski, Cordelia Langford, John Sillitoe on behalf of the Wellcome Sanger Institute COVID-19 Surveillance Team |
| EPI_ISL_3362515, EPI_ISL_3362516, EPI_ISL_3362518, EPI_ISL_3362519, EPI_ISL_3362521 | Lighthouse Laboratory Plymouth | Wellcome Sanger Institute for the COVID-19 Genomics UK (COG-UK) Consortium | Lighthouse Laboratory Plymouth and Alex Alderton, Roberto Amato, Jeffrey Barrett, Sonia Goncalves, Ewan Harrison, David K. Jackson, Ian Johnston, Dominic Kwiatkowski, Cordelia Langford, John Sillitoe on behalf of the Wellcome Sanger Institute COVID-19 Surveillance Team |
| EPI_ISL_3362522 | Lighthouse Lab in Milton Keynes | Wellcome Sanger Institute for the COVID-19 Genomics UK (COG-UK) Consortium | The Lighthouse Lab in Milton Keynes and Alex Alderton, Roberto Amato, Jeffrey Barrett, Sonia Goncalves, Ewan Harrison, David K. Jackson, Ian Johnston, Dominic Kwiatkowski, Cordelia Langford, John Sillitoe on behalf of the Wellcome Sanger Institute COVID-19 Surveillance Team |
| EPI_ISL_3362524 | Lighthouse Laboratory Plymouth | Wellcome Sanger Institute for the COVID-19 Genomics UK (COG-UK) Consortium | Lighthouse Laboratory Plymouth and Alex Alderton, Roberto Amato, Jeffrey Barrett, Sonia Goncalves, Ewan Harrison, David K. Jackson, Ian Johnston, Dominic Kwiatkowski, Cordelia Langford, John Sillitoe on behalf of the Wellcome Sanger Institute COVID-19 Surveillance Team |
| EPI_ISL_3362525, EPI_ISL_3362527 | Lighthouse Lab in Milton Keynes | Wellcome Sanger Institute for the COVID-19 Genomics UK (COG-UK) Consortium | The Lighthouse Lab in Milton Keynes and Alex Alderton, Roberto Amato, Jeffrey Barrett, Sonia Goncalves, Ewan Harrison, David K. Jackson, Ian Johnston, Dominic Kwiatkowski, Cordelia Langford, John Sillitoe on behalf of the Wellcome Sanger Institute COVID-19 Surveillance Team |
| EPI_ISL_3362528 | Lighthouse Laboratory Plymouth | Wellcome Sanger Institute for the COVID-19 Genomics UK (COG-UK) Consortium | Lighthouse Laboratory Plymouth and Alex Alderton, Roberto Amato, Jeffrey Barrett, Sonia Goncalves, Ewan Harrison, David K. Jackson, Ian Johnston, Dominic Kwiatkowski, Cordelia Langford, John Sillitoe on behalf of the Wellcome Sanger Institute COVID-19 Surveillance Team |
| EPI_ISL_3362529 | Lighthouse Lab in Glasgow | Wellcome Sanger Institute for the COVID-19 Genomics UK (COG-UK) Consortium | Harper VanSteenhouse, Yumi Kasai, David Gray, Carol Clugston, Anna Dominiczak and Alex Alderton, Roberto Amato, Jeffrey Barrett, Sonia Goncalves, Ewan Harrison, David K. Jackson, Ian Johnston, Dominic Kwiatkowski, Cordelia Langford, John Sillitoe on behalf of the Wellcome Sanger Institute COVID-19 Surveillance Team |
| EPI_ISL_3362531 | Lighthouse Laboratory Plymouth | Wellcome Sanger Institute for the COVID-19 Genomics UK (COG-UK) Consortium | Lighthouse Laboratory Plymouth and Alex Alderton, Roberto Amato, Jeffrey Barrett, Sonia Goncalves, Ewan Harrison, David K. Jackson, Ian Johnston, Dominic Kwiatkowski, Cordelia Langford, John Sillitoe on behalf of the Wellcome Sanger Institute COVID-19 Surveillance Team |
| EPI_ISL_3362532 | Lighthouse Lab in Milton Keynes | Wellcome Sanger Institute for the COVID-19 Genomics UK (COG-UK) Consortium | The Lighthouse Lab in Milton Keynes and Alex Alderton, Roberto Amato, Jeffrey Barrett, Sonia Goncalves, Ewan Harrison, David K. Jackson, Ian Johnston, Dominic Kwiatkowski, Cordelia Langford, John Sillitoe on behalf of the Wellcome Sanger Institute COVID-19 Surveillance Team |
| EPI_ISL_3362534 | Lighthouse Laboratory Plymouth | Wellcome Sanger Institute for the COVID-19 Genomics UK (COG-UK) Consortium | Lighthouse Laboratory Plymouth and Alex Alderton, Roberto Amato, Jeffrey Barrett, Sonia Goncalves, Ewan Harrison, David K. Jackson, Ian Johnston, Dominic Kwiatkowski, Cordelia Langford, John Sillitoe on behalf of the Wellcome Sanger Institute COVID-19 Surveillance Team |
| EPI_ISL_3362535 | Lighthouse Lab in Milton Keynes | Wellcome Sanger Institute for the COVID-19 Genomics UK (COG-UK) Consortium | The Lighthouse Lab in Milton Keynes and Alex Alderton, Roberto Amato, Jeffrey Barrett, Sonia Goncalves, Ewan Harrison, David K. Jackson, Ian Johnston, Dominic Kwiatkowski, Cordelia Langford, John Sillitoe on behalf of the Wellcome Sanger Institute COVID-19 Surveillance Team |
| EPI_ISL_3362536 | Lighthouse Laboratory Plymouth | Wellcome Sanger Institute for the COVID-19 Genomics UK (COG-UK) Consortium | Lighthouse Laboratory Plymouth and Alex Alderton, Roberto Amato, Jeffrey Barrett, Sonia Goncalves, Ewan Harrison, David K. Jackson, Ian Johnston, Dominic Kwiatkowski, Cordelia Langford, John Sillitoe on behalf of the Wellcome Sanger Institute COVID-19 Surveillance Team |
| EPI_ISL_3362538 | Lighthouse Lab in Glasgow | Wellcome Sanger Institute for the COVID-19 Genomics UK (COG-UK) Consortium | Harper VanSteenhouse, Yumi Kasai, David Gray, Carol Clugston, Anna Dominiczak and Alex Alderton, Roberto Amato, Jeffrey Barrett, Sonia Goncalves, Ewan Harrison, David K. Jackson, Ian Johnston, Dominic Kwiatkowski, Cordelia Langford, John Sillitoe on behalf of the Wellcome Sanger Institute COVID-19 Surveillance Team |
| EPI_ISL_3362539, EPI_ISL_3362541, EPI_ISL_3362542, | Lighthouse Lab in Milton Keynes | Wellcome Sanger Institute for the COVID-19 Genomics UK (COG-UK) Consortium | The Lighthouse Lab in Milton Keynes and Alex Alderton, Roberto Amato, Jeffrey Barrett, Sonia Goncalves, Ewan Harrison, David K. Jackson, Ian Johnston, Dominic Kwiatkowski, Cordelia Langford, John Sillitoe on behalf of the Wellcome Sanger Institute COVID-19 Surveillance Team |

| | | | |
|---|---|---|---|
| EPI_ISL_3362544, EPI_ISL_3362545 | | | |
| EPI_ISL_3362546, EPI_ISL_3362548 | Lighthouse Laboratory Plymouth | Wellcome Sanger Institute for the COVID-19 Genomics UK (COG-UK) Consortium | Lighthouse Laboratory Plymouth and Alex Alderton, Roberto Amato, Jeffrey Barrett, Sonia Goncalves, Ewan Harrison, David K. Jackson, Ian Johnston, Dominic Kwiatkowski, Cordelia Langford, John Sillitoe on behalf of the Wellcome Sanger Institute COVID-19 Surveillance Team |
| EPI_ISL_3362550, EPI_ISL_3362551, EPI_ISL_3362553 | Lighthouse Lab in Milton Keynes | Wellcome Sanger Institute for the COVID-19 Genomics UK (COG-UK) Consortium | The Lighthouse Lab in Milton Keynes and Alex Alderton, Roberto Amato, Jeffrey Barrett, Sonia Goncalves, Ewan Harrison, David K. Jackson, Ian Johnston, Dominic Kwiatkowski, Cordelia Langford, John Sillitoe on behalf of the Wellcome Sanger Institute COVID-19 Surveillance Team |
| EPI_ISL_3362554 | Lighthouse Laboratory Plymouth | Wellcome Sanger Institute for the COVID-19 Genomics UK (COG-UK) Consortium | Lighthouse Laboratory Plymouth and Alex Alderton, Roberto Amato, Jeffrey Barrett, Sonia Goncalves, Ewan Harrison, David K. Jackson, Ian Johnston, Dominic Kwiatkowski, Cordelia Langford, John Sillitoe on behalf of the Wellcome Sanger Institute COVID-19 Surveillance Team |
| EPI_ISL_3362556 | Lighthouse Lab in Milton Keynes | Wellcome Sanger Institute for the COVID-19 Genomics UK (COG-UK) Consortium | The Lighthouse Lab in Milton Keynes and Alex Alderton, Roberto Amato, Jeffrey Barrett, Sonia Goncalves, Ewan Harrison, David K. Jackson, Ian Johnston, Dominic Kwiatkowski, Cordelia Langford, John Sillitoe on behalf of the Wellcome Sanger Institute COVID-19 Surveillance Team |
| EPI_ISL_3362557, EPI_ISL_3362559 | Lighthouse Laboratory Plymouth | Wellcome Sanger Institute for the COVID-19 Genomics UK (COG-UK) Consortium | Lighthouse Laboratory Plymouth and Alex Alderton, Roberto Amato, Jeffrey Barrett, Sonia Goncalves, Ewan Harrison, David K. Jackson, Ian Johnston, Dominic Kwiatkowski, Cordelia Langford, John Sillitoe on behalf of the Wellcome Sanger Institute COVID-19 Surveillance Team |
| EPI_ISL_3362560 | Lighthouse Lab in Glasgow | Wellcome Sanger Institute for the COVID-19 Genomics UK (COG-UK) Consortium | Harper VanSteenhouse, Yumi Kasai, David Gray, Carol Clugston, Anna Dominiczak and Alex Alderton, Roberto Amato, Jeffrey Barrett, Sonia Goncalves, Ewan Harrison, David K. Jackson, Ian Johnston, Dominic Kwiatkowski, Cordelia Langford, John Sillitoe on behalf of the Wellcome Sanger Institute COVID-19 Surveillance Team |
| EPI_ISL_3362561 | Lighthouse Laboratory Plymouth | Wellcome Sanger Institute for the COVID-19 Genomics UK (COG-UK) Consortium | Lighthouse Laboratory Plymouth and Alex Alderton, Roberto Amato, Jeffrey Barrett, Sonia Goncalves, Ewan Harrison, David K. Jackson, Ian Johnston, Dominic Kwiatkowski, Cordelia Langford, John Sillitoe on behalf of the Wellcome Sanger Institute COVID-19 Surveillance Team |
| EPI_ISL_3362563 | Lighthouse Lab in Glasgow | Wellcome Sanger Institute for the COVID-19 Genomics UK (COG-UK) Consortium | Harper VanSteenhouse, Yumi Kasai, David Gray, Carol Clugston, Anna Dominiczak and Alex Alderton, Roberto Amato, Jeffrey Barrett, Sonia Goncalves, Ewan Harrison, David K. Jackson, Ian Johnston, Dominic Kwiatkowski, Cordelia Langford, John Sillitoe on behalf of the Wellcome Sanger Institute COVID-19 Surveillance Team |
| EPI_ISL_3362564 | Lighthouse Laboratory Plymouth | Wellcome Sanger Institute for the COVID-19 Genomics UK (COG-UK) Consortium | Lighthouse Laboratory Plymouth and Alex Alderton, Roberto Amato, Jeffrey Barrett, Sonia Goncalves, Ewan Harrison, David K. Jackson, Ian Johnston, Dominic Kwiatkowski, Cordelia Langford, John Sillitoe on behalf of the Wellcome Sanger Institute COVID-19 Surveillance Team |
| EPI_ISL_3362566 | Lighthouse Lab in Milton Keynes | Wellcome Sanger Institute for the COVID-19 Genomics UK (COG-UK) Consortium | The Lighthouse Lab in Milton Keynes and Alex Alderton, Roberto Amato, Jeffrey Barrett, Sonia Goncalves, Ewan Harrison, David K. Jackson, Ian Johnston, Dominic Kwiatkowski, Cordelia Langford, John Sillitoe on behalf of the Wellcome Sanger Institute COVID-19 Surveillance Team |
| EPI_ISL_3362567 | Lighthouse Laboratory Plymouth | Wellcome Sanger Institute for the COVID-19 Genomics UK (COG-UK) Consortium | Lighthouse Laboratory Plymouth and Alex Alderton, Roberto Amato, Jeffrey Barrett, Sonia Goncalves, Ewan Harrison, David K. Jackson, Ian Johnston, Dominic Kwiatkowski, Cordelia Langford, John Sillitoe on behalf of the Wellcome Sanger Institute COVID-19 Surveillance Team |
| EPI_ISL_3362569, EPI_ISL_3362570, EPI_ISL_3362572, EPI_ISL_3362573 | Lighthouse Lab in Milton Keynes | Wellcome Sanger Institute for the COVID-19 Genomics UK (COG-UK) Consortium | The Lighthouse Lab in Milton Keynes and Alex Alderton, Roberto Amato, Jeffrey Barrett, Sonia Goncalves, Ewan Harrison, David K. Jackson, Ian Johnston, Dominic Kwiatkowski, Cordelia Langford, John Sillitoe on behalf of the Wellcome Sanger Institute COVID-19 Surveillance Team |
| EPI_ISL_3362575, EPI_ISL_3362576 | Lighthouse Laboratory Plymouth | Wellcome Sanger Institute for the COVID-19 Genomics UK (COG-UK) Consortium | Lighthouse Laboratory Plymouth and Alex Alderton, Roberto Amato, Jeffrey Barrett, Sonia Goncalves, Ewan Harrison, David K. Jackson, Ian Johnston, Dominic Kwiatkowski, Cordelia Langford, John Sillitoe on behalf of the Wellcome Sanger Institute COVID-19 Surveillance Team |
| EPI_ISL_3362578, EPI_ISL_3362579, EPI_ISL_3362580 | Lighthouse Lab in Milton Keynes | Wellcome Sanger Institute for the COVID-19 Genomics UK (COG-UK) Consortium | The Lighthouse Lab in Milton Keynes and Alex Alderton, Roberto Amato, Jeffrey Barrett, Sonia Goncalves, Ewan Harrison, David K. Jackson, Ian Johnston, Dominic Kwiatkowski, Cordelia Langford, John Sillitoe on behalf of the Wellcome Sanger Institute COVID-19 Surveillance Team |
| EPI_ISL_3362582 | Lighthouse Laboratory Plymouth | Wellcome Sanger Institute for the COVID-19 Genomics UK (COG-UK) Consortium | Lighthouse Laboratory Plymouth and Alex Alderton, Roberto Amato, Jeffrey Barrett, Sonia Goncalves, Ewan Harrison, David K. Jackson, Ian Johnston, Dominic Kwiatkowski, Cordelia Langford, John Sillitoe on behalf of the Wellcome Sanger Institute COVID-19 Surveillance Team |
| EPI_ISL_3362584 | Lighthouse Lab in Glasgow | Wellcome Sanger Institute for the COVID-19 Genomics UK (COG-UK) Consortium | Harper VanSteenhouse, Yumi Kasai, David Gray, Carol Clugston, Anna Dominiczak and Alex Alderton, Roberto Amato, Jeffrey Barrett, Sonia Goncalves, Ewan Harrison, David K. Jackson, Ian Johnston, Dominic Kwiatkowski, Cordelia Langford, John Sillitoe on behalf of the Wellcome Sanger Institute COVID-19 Surveillance Team |
| EPI_ISL_3362585, EPI_ISL_3362586, EPI_ISL_3362588, EPI_ISL_3362589, EPI_ISL_3362591 | Lighthouse Lab in Milton Keynes | Wellcome Sanger Institute for the COVID-19 Genomics UK (COG-UK) Consortium | The Lighthouse Lab in Milton Keynes and Alex Alderton, Roberto Amato, Jeffrey Barrett, Sonia Goncalves, Ewan Harrison, David K. Jackson, Ian Johnston, Dominic Kwiatkowski, Cordelia Langford, John Sillitoe on behalf of the Wellcome Sanger Institute COVID-19 Surveillance Team |
| EPI_ISL_3362592, EPI_ISL_3362594 | Lighthouse Laboratory Plymouth | Wellcome Sanger Institute for the COVID-19 Genomics UK (COG-UK) Consortium | Lighthouse Laboratory Plymouth and Alex Alderton, Roberto Amato, Jeffrey Barrett, Sonia Goncalves, Ewan Harrison, David K. Jackson, Ian Johnston, Dominic Kwiatkowski, Cordelia Langford, John Sillitoe on behalf of the Wellcome Sanger Institute COVID-19 Surveillance Team |
| EPI_ISL_3362595, EPI_ISL_3362596, EPI_ISL_3362598 | Lighthouse Lab in Milton Keynes | Wellcome Sanger Institute for the COVID-19 Genomics UK (COG-UK) Consortium | The Lighthouse Lab in Milton Keynes and Alex Alderton, Roberto Amato, Jeffrey Barrett, Sonia Goncalves, Ewan Harrison, David K. Jackson, Ian Johnston, Dominic Kwiatkowski, Cordelia Langford, John Sillitoe on behalf of the Wellcome Sanger Institute COVID-19 Surveillance Team |
| EPI_ISL_3362599 | Lighthouse Lab in Glasgow | Wellcome Sanger Institute for the COVID-19 Genomics UK (COG-UK) Consortium | Harper VanSteenhouse, Yumi Kasai, David Gray, Carol Clugston, Anna Dominiczak and Alex Alderton, Roberto Amato, Jeffrey Barrett, Sonia Goncalves, Ewan Harrison, David K. Jackson, Ian Johnston, Dominic Kwiatkowski, Cordelia Langford, John Sillitoe on behalf of the Wellcome Sanger Institute COVID-19 Surveillance Team |
| EPI_ISL_3362601, EPI_ISL_3362602 | Lighthouse Laboratory Plymouth | Wellcome Sanger Institute for the COVID-19 Genomics UK (COG-UK) Consortium | Lighthouse Laboratory Plymouth and Alex Alderton, Roberto Amato, Jeffrey Barrett, Sonia Goncalves, Ewan Harrison, David K. Jackson, Ian Johnston, Dominic Kwiatkowski, Cordelia Langford, John Sillitoe on behalf of the Wellcome Sanger Institute COVID-19 Surveillance Team |
| EPI_ISL_3362604 | Lighthouse Lab in Milton Keynes | Wellcome Sanger Institute for the COVID-19 Genomics UK (COG-UK) Consortium | The Lighthouse Lab in Milton Keynes and Alex Alderton, Roberto Amato, Jeffrey Barrett, Sonia Goncalves, Ewan Harrison, David K. Jackson, Ian Johnston, Dominic Kwiatkowski, Cordelia Langford, John Sillitoe on behalf of the Wellcome Sanger Institute COVID-19 Surveillance Team |
| EPI_ISL_3362605 | Lighthouse Laboratory Plymouth | Wellcome Sanger Institute for the COVID-19 Genomics UK (COG-UK) Consortium | Lighthouse Laboratory Plymouth and Alex Alderton, Roberto Amato, Jeffrey Barrett, Sonia Goncalves, Ewan Harrison, David K. Jackson, Ian Johnston, Dominic Kwiatkowski, Cordelia Langford, John Sillitoe on behalf of the Wellcome Sanger Institute COVID-19 Surveillance Team |
| EPI_ISL_3362607 | Lighthouse Lab in Milton Keynes | Wellcome Sanger Institute for the COVID-19 Genomics UK (COG-UK) Consortium | The Lighthouse Lab in Milton Keynes and Alex Alderton, Roberto Amato, Jeffrey Barrett, Sonia Goncalves, Ewan Harrison, David K. Jackson, Ian Johnston, Dominic Kwiatkowski, Cordelia Langford, John Sillitoe on behalf of the Wellcome Sanger Institute COVID-19 Surveillance Team |
| EPI_ISL_3362608 | Lighthouse Lab in Glasgow | Wellcome Sanger Institute for the COVID-19 Genomics UK (COG-UK) Consortium | Harper VanSteenhouse, Yumi Kasai, David Gray, Carol Clugston, Anna Dominiczak and Alex Alderton, Roberto Amato, Jeffrey Barrett, Sonia Goncalves, Ewan Harrison, David K. Jackson, Ian Johnston, Dominic Kwiatkowski, Cordelia Langford, John Sillitoe on behalf of the Wellcome Sanger Institute COVID-19 Surveillance Team |
| EPI_ISL_3362609 | Lighthouse Laboratory Plymouth | Wellcome Sanger Institute for the COVID-19 Genomics UK (COG-UK) Consortium | Lighthouse Laboratory Plymouth and Alex Alderton, Roberto Amato, Jeffrey Barrett, Sonia Goncalves, Ewan Harrison, David K. Jackson, Ian Johnston, Dominic Kwiatkowski, Cordelia Langford, John Sillitoe on behalf of the Wellcome Sanger Institute COVID-19 Surveillance Team |
| EPI_ISL_3362611, EPI_ISL_3362612, EPI_ISL_3362614 | Lighthouse Lab in Milton Keynes | Wellcome Sanger Institute for the COVID-19 Genomics UK (COG-UK) Consortium | The Lighthouse Lab in Milton Keynes and Alex Alderton, Roberto Amato, Jeffrey Barrett, Sonia Goncalves, Ewan Harrison, David K. Jackson, Ian Johnston, Dominic Kwiatkowski, Cordelia Langford, John Sillitoe on behalf of the Wellcome Sanger Institute COVID-19 Surveillance Team |
| EPI_ISL_3362615 | Lighthouse Laboratory Plymouth | Wellcome Sanger Institute for the COVID-19 Genomics UK (COG-UK) Consortium | Lighthouse Laboratory Plymouth and Alex Alderton, Roberto Amato, Jeffrey Barrett, Sonia Goncalves, Ewan Harrison, David K. Jackson, Ian Johnston, Dominic Kwiatkowski, Cordelia Langford, John Sillitoe on behalf of the Wellcome Sanger Institute COVID-19 Surveillance Team |
| EPI_ISL_3362617, EPI_ISL_3362618 | Lighthouse Lab in Milton Keynes | Wellcome Sanger Institute for the COVID-19 Genomics UK (COG-UK) Consortium | The Lighthouse Lab in Milton Keynes and Alex Alderton, Roberto Amato, Jeffrey Barrett, Sonia Goncalves, Ewan Harrison, David K. Jackson, Ian Johnston, Dominic Kwiatkowski, Cordelia Langford, John Sillitoe on behalf of the Wellcome Sanger Institute COVID-19 Surveillance Team |
| EPI_ISL_3362619 | Lighthouse Lab in Glasgow | Wellcome Sanger Institute for the COVID-19 Genomics UK (COG-UK) Consortium | Harper VanSteenhouse, Yumi Kasai, David Gray, Carol Clugston, Anna Dominiczak and Alex Alderton, Roberto Amato, Jeffrey Barrett, Sonia Goncalves, Ewan Harrison, David K. Jackson, Ian Johnston, Dominic Kwiatkowski, Cordelia Langford, John Sillitoe on behalf of the Wellcome Sanger Institute COVID-19 Surveillance Team |
| EPI_ISL_3362621 | Lighthouse Laboratory Plymouth | Wellcome Sanger Institute for the COVID-19 Genomics UK (COG-UK) Consortium | Lighthouse Laboratory Plymouth and Alex Alderton, Roberto Amato, Jeffrey Barrett, Sonia Goncalves, Ewan Harrison, David K. Jackson, Ian Johnston, Dominic Kwiatkowski, Cordelia Langford, John Sillitoe on behalf of the Wellcome Sanger Institute COVID-19 Surveillance Team |
| EPI_ISL_3362622 | Lighthouse Lab in Milton Keynes | Wellcome Sanger Institute for the COVID-19 Genomics UK (COG-UK) Consortium | The Lighthouse Lab in Milton Keynes and Alex Alderton, Roberto Amato, Jeffrey Barrett, Sonia Goncalves, Ewan Harrison, David K. Jackson, Ian Johnston, Dominic Kwiatkowski, Cordelia Langford, John Sillitoe on behalf of the Wellcome Sanger Institute COVID-19 Surveillance Team |
| EPI_ISL_3362624 | Lighthouse Laboratory Plymouth | Wellcome Sanger Institute for the COVID-19 Genomics UK (COG-UK) Consortium | Lighthouse Laboratory Plymouth and Alex Alderton, Roberto Amato, Jeffrey Barrett, Sonia Goncalves, Ewan Harrison, David K. Jackson, Ian Johnston, Dominic Kwiatkowski, Cordelia Langford, John Sillitoe on behalf of the Wellcome Sanger Institute COVID-19 Surveillance Team |
| EPI_ISL_3362625 | Lighthouse Lab in Glasgow | Wellcome Sanger Institute for the COVID-19 Genomics UK (COG-UK) Consortium | Harper VanSteenhouse, Yumi Kasai, David Gray, Carol Clugston, Anna Dominiczak and Alex Alderton, Roberto Amato, Jeffrey Barrett, Sonia Goncalves, Ewan Harrison, David K. Jackson, Ian Johnston, Dominic Kwiatkowski, Cordelia Langford, John Sillitoe on behalf of the Wellcome Sanger Institute COVID-19 Surveillance Team |
| EPI_ISL_3362626, EPI_ISL_3362628 | Lighthouse Laboratory Plymouth | Wellcome Sanger Institute for the COVID-19 Genomics UK (COG-UK) Consortium | Lighthouse Laboratory Plymouth and Alex Alderton, Roberto Amato, Jeffrey Barrett, Sonia Goncalves, Ewan Harrison, David K. Jackson, Ian Johnston, Dominic Kwiatkowski, Cordelia Langford, John Sillitoe on behalf of the Wellcome Sanger Institute COVID-19 Surveillance Team |
| EPI_ISL_3362629, EPI_ISL_3362631, EPI_ISL_3362632 | Lighthouse Lab in Glasgow | Wellcome Sanger Institute for the COVID-19 Genomics UK (COG-UK) Consortium | Harper VanSteenhouse, Yumi Kasai, David Gray, Carol Clugston, Anna Dominiczak and Alex Alderton, Roberto Amato, Jeffrey Barrett, Sonia Goncalves, Ewan Harrison, David K. Jackson, Ian Johnston, Dominic Kwiatkowski, Cordelia Langford, John Sillitoe on behalf of the Wellcome Sanger Institute COVID-19 Surveillance Team |
| EPI_ISL_3362634 | Lighthouse Lab in Milton Keynes | Wellcome Sanger Institute for the COVID-19 Genomics UK (COG-UK) Consortium | The Lighthouse Lab in Milton Keynes and Alex Alderton, Roberto Amato, Jeffrey Barrett, Sonia Goncalves, Ewan Harrison, David K. Jackson, Ian Johnston, Dominic Kwiatkowski, Cordelia Langford, John Sillitoe on behalf of the Wellcome Sanger Institute COVID-19 Surveillance Team |
| EPI_ISL_3362635 | Lighthouse Lab in Glasgow | Wellcome Sanger Institute for the COVID-19 Genomics UK (COG-UK) Consortium | Harper VanSteenhouse, Yumi Kasai, David Gray, Carol Clugston, Anna Dominiczak and Alex Alderton, Roberto Amato, Jeffrey Barrett, Sonia Goncalves, Ewan Harrison, David K. Jackson, Ian Johnston, Dominic Kwiatkowski, Cordelia Langford, John Sillitoe on behalf of the Wellcome Sanger Institute COVID-19 Surveillance Team |
| EPI_ISL_3362637, EPI_ISL_3362639 | Lighthouse Lab in Milton Keynes | Wellcome Sanger Institute for the COVID-19 Genomics UK (COG-UK) Consortium | The Lighthouse Lab in Milton Keynes and Alex Alderton, Roberto Amato, Jeffrey Barrett, Sonia Goncalves, Ewan Harrison, David K. Jackson, Ian Johnston, Dominic Kwiatkowski, Cordelia Langford, John Sillitoe on behalf of the Wellcome Sanger Institute COVID-19 Surveillance Team |
| EPI_ISL_3362640 | Lighthouse Lab in Glasgow | Wellcome Sanger Institute for the COVID-19 Genomics UK (COG-UK) Consortium | Harper VanSteenhouse, Yumi Kasai, David Gray, Carol Clugston, Anna Dominiczak and Alex Alderton, Roberto Amato, Jeffrey Barrett, Sonia Goncalves, Ewan Harrison, David K. Jackson, Ian Johnston, Dominic Kwiatkowski, Cordelia Langford, John Sillitoe on behalf of the Wellcome Sanger Institute COVID-19 Surveillance Team |

| | | | |
|---|---|---|---|
| EPI_ISL_3362641 | Lighthouse Laboratory Plymouth | Wellcome Sanger Institute for the COVID-19 Genomics UK (COG-UK) Consortium | Lighthouse Laboratory Plymouth and Alex Alderton, Roberto Amato, Jeffrey Barrett, Sonia Goncalves, Ewan Harrison, David K. Jackson, Ian Johnston, Dominic Kwiatkowski, Cordelia Langford, John Sillitoe on behalf of the Wellcome Sanger Institute COVID-19 Surveillance Team |
| EPI_ISL_3362643 | Lighthouse Lab in Milton Keynes | Wellcome Sanger Institute for the COVID-19 Genomics UK (COG-UK) Consortium | The Lighthouse Lab in Milton Keynes and Alex Alderton, Roberto Amato, Jeffrey Barrett, Sonia Goncalves, Ewan Harrison, David K. Jackson, Ian Johnston, Dominic Kwiatkowski, Cordelia Langford, John Sillitoe on behalf of the Wellcome Sanger Institute COVID-19 Surveillance Team |
| EPI_ISL_3362644 | Lighthouse Laboratory Plymouth | Wellcome Sanger Institute for the COVID-19 Genomics UK (COG-UK) Consortium | Lighthouse Laboratory Plymouth and Alex Alderton, Roberto Amato, Jeffrey Barrett, Sonia Goncalves, Ewan Harrison, David K. Jackson, Ian Johnston, Dominic Kwiatkowski, Cordelia Langford, John Sillitoe on behalf of the Wellcome Sanger Institute COVID-19 Surveillance Team |
| EPI_ISL_3362646 | Lighthouse Lab in Milton Keynes | Wellcome Sanger Institute for the COVID-19 Genomics UK (COG-UK) Consortium | The Lighthouse Lab in Milton Keynes and Alex Alderton, Roberto Amato, Jeffrey Barrett, Sonia Goncalves, Ewan Harrison, David K. Jackson, Ian Johnston, Dominic Kwiatkowski, Cordelia Langford, John Sillitoe on behalf of the Wellcome Sanger Institute COVID-19 Surveillance Team |
| EPI_ISL_3362647 | Lighthouse Laboratory Plymouth | Wellcome Sanger Institute for the COVID-19 Genomics UK (COG-UK) Consortium | Lighthouse Laboratory Plymouth and Alex Alderton, Roberto Amato, Jeffrey Barrett, Sonia Goncalves, Ewan Harrison, David K. Jackson, Ian Johnston, Dominic Kwiatkowski, Cordelia Langford, John Sillitoe on behalf of the Wellcome Sanger Institute COVID-19 Surveillance Team |
| EPI_ISL_3362649 | Lighthouse Lab in Glasgow | Wellcome Sanger Institute for the COVID-19 Genomics UK (COG-UK) Consortium | Harper VanSteenhouse, Yumi Kasai, David Gray, Carol Clugston, Anna Dominiczak and Alex Alderton, Roberto Amato, Jeffrey Barrett, Sonia Goncalves, Ewan Harrison, David K. Jackson, Ian Johnston, Dominic Kwiatkowski, Cordelia Langford, John Sillitoe on behalf of the Wellcome Sanger Institute COVID-19 Surveillance Team |
| EPI_ISL_3362650 | Lighthouse Lab in Milton Keynes | Wellcome Sanger Institute for the COVID-19 Genomics UK (COG-UK) Consortium | The Lighthouse Lab in Milton Keynes and Alex Alderton, Roberto Amato, Jeffrey Barrett, Sonia Goncalves, Ewan Harrison, David K. Jackson, Ian Johnston, Dominic Kwiatkowski, Cordelia Langford, John Sillitoe on behalf of the Wellcome Sanger Institute COVID-19 Surveillance Team |
| EPI_ISL_3362652, EPI_ISL_3362653, EPI_ISL_3362655, EPI_ISL_3362656, EPI_ISL_3362657 | Lighthouse Laboratory Plymouth | Wellcome Sanger Institute for the COVID-19 Genomics UK (COG-UK) Consortium | Lighthouse Laboratory Plymouth and Alex Alderton, Roberto Amato, Jeffrey Barrett, Sonia Goncalves, Ewan Harrison, David K. Jackson, Ian Johnston, Dominic Kwiatkowski, Cordelia Langford, John Sillitoe on behalf of the Wellcome Sanger Institute COVID-19 Surveillance Team |
| EPI_ISL_3362659 | Lighthouse Lab in Glasgow | Wellcome Sanger Institute for the COVID-19 Genomics UK (COG-UK) Consortium | Harper VanSteenhouse, Yumi Kasai, David Gray, Carol Clugston, Anna Dominiczak and Alex Alderton, Roberto Amato, Jeffrey Barrett, Sonia Goncalves, Ewan Harrison, David K. Jackson, Ian Johnston, Dominic Kwiatkowski, Cordelia Langford, John Sillitoe on behalf of the Wellcome Sanger Institute COVID-19 Surveillance Team |
| EPI_ISL_3362660 | Lighthouse Laboratory Plymouth | Wellcome Sanger Institute for the COVID-19 Genomics UK (COG-UK) Consortium | Lighthouse Laboratory Plymouth and Alex Alderton, Roberto Amato, Jeffrey Barrett, Sonia Goncalves, Ewan Harrison, David K. Jackson, Ian Johnston, Dominic Kwiatkowski, Cordelia Langford, John Sillitoe on behalf of the Wellcome Sanger Institute COVID-19 Surveillance Team |
| EPI_ISL_3362662 | Lighthouse Lab in Milton Keynes | Wellcome Sanger Institute for the COVID-19 Genomics UK (COG-UK) Consortium | The Lighthouse Lab in Milton Keynes and Alex Alderton, Roberto Amato, Jeffrey Barrett, Sonia Goncalves, Ewan Harrison, David K. Jackson, Ian Johnston, Dominic Kwiatkowski, Cordelia Langford, John Sillitoe on behalf of the Wellcome Sanger Institute COVID-19 Surveillance Team |
| EPI_ISL_3362663, EPI_ISL_3362665, EPI_ISL_3362666 | Lighthouse Lab in Glasgow | Wellcome Sanger Institute for the COVID-19 Genomics UK (COG-UK) Consortium | Harper VanSteenhouse, Yumi Kasai, David Gray, Carol Clugston, Anna Dominiczak and Alex Alderton, Roberto Amato, Jeffrey Barrett, Sonia Goncalves, Ewan Harrison, David K. Jackson, Ian Johnston, Dominic Kwiatkowski, Cordelia Langford, John Sillitoe on behalf of the Wellcome Sanger Institute COVID-19 Surveillance Team |
| EPI_ISL_3362667 | Lighthouse Lab in Milton Keynes | Wellcome Sanger Institute for the COVID-19 Genomics UK (COG-UK) Consortium | The Lighthouse Lab in Milton Keynes and Alex Alderton, Roberto Amato, Jeffrey Barrett, Sonia Goncalves, Ewan Harrison, David K. Jackson, Ian Johnston, Dominic Kwiatkowski, Cordelia Langford, John Sillitoe on behalf of the Wellcome Sanger Institute COVID-19 Surveillance Team |
| EPI_ISL_3362669, EPI_ISL_3362671 | Lighthouse Lab in Glasgow | Wellcome Sanger Institute for the COVID-19 Genomics UK (COG-UK) Consortium | Harper VanSteenhouse, Yumi Kasai, David Gray, Carol Clugston, Anna Dominiczak and Alex Alderton, Roberto Amato, Jeffrey Barrett, Sonia Goncalves, Ewan Harrison, David K. Jackson, Ian Johnston, Dominic Kwiatkowski, Cordelia Langford, John Sillitoe on behalf of the Wellcome Sanger Institute COVID-19 Surveillance Team |
| EPI_ISL_3362672, EPI_ISL_3362673 | Lighthouse Lab in Milton Keynes | Wellcome Sanger Institute for the COVID-19 Genomics UK (COG-UK) Consortium | The Lighthouse Lab in Milton Keynes and Alex Alderton, Roberto Amato, Jeffrey Barrett, Sonia Goncalves, Ewan Harrison, David K. Jackson, Ian Johnston, Dominic Kwiatkowski, Cordelia Langford, John Sillitoe on behalf of the Wellcome Sanger Institute COVID-19 Surveillance Team |
| EPI_ISL_3362675 | Lighthouse Lab in Glasgow | Wellcome Sanger Institute for the COVID-19 Genomics UK (COG-UK) Consortium | Harper VanSteenhouse, Yumi Kasai, David Gray, Carol Clugston, Anna Dominiczak and Alex Alderton, Roberto Amato, Jeffrey Barrett, Sonia Goncalves, Ewan Harrison, David K. Jackson, Ian Johnston, Dominic Kwiatkowski, Cordelia Langford, John Sillitoe on behalf of the Wellcome Sanger Institute COVID-19 Surveillance Team |
| EPI_ISL_3362676 | Lighthouse Lab in Milton Keynes | Wellcome Sanger Institute for the COVID-19 Genomics UK (COG-UK) Consortium | The Lighthouse Lab in Milton Keynes and Alex Alderton, Roberto Amato, Jeffrey Barrett, Sonia Goncalves, Ewan Harrison, David K. Jackson, Ian Johnston, Dominic Kwiatkowski, Cordelia Langford, John Sillitoe on behalf of the Wellcome Sanger Institute COVID-19 Surveillance Team |
| EPI_ISL_3362678 | Lighthouse Laboratory Plymouth | Wellcome Sanger Institute for the COVID-19 Genomics UK (COG-UK) Consortium | Lighthouse Laboratory Plymouth and Alex Alderton, Roberto Amato, Jeffrey Barrett, Sonia Goncalves, Ewan Harrison, David K. Jackson, Ian Johnston, Dominic Kwiatkowski, Cordelia Langford, John Sillitoe on behalf of the Wellcome Sanger Institute COVID-19 Surveillance Team |
| EPI_ISL_3362679 | Lighthouse Lab in Milton Keynes | Wellcome Sanger Institute for the COVID-19 Genomics UK (COG-UK) Consortium | The Lighthouse Lab in Milton Keynes and Alex Alderton, Roberto Amato, Jeffrey Barrett, Sonia Goncalves, Ewan Harrison, David K. Jackson, Ian Johnston, Dominic Kwiatkowski, Cordelia Langford, John Sillitoe on behalf of the Wellcome Sanger Institute COVID-19 Surveillance Team |
| EPI_ISL_3362681, EPI_ISL_3362682 | Lighthouse Lab in Glasgow | Wellcome Sanger Institute for the COVID-19 Genomics UK (COG-UK) Consortium | Harper VanSteenhouse, Yumi Kasai, David Gray, Carol Clugston, Anna Dominiczak and Alex Alderton, Roberto Amato, Jeffrey Barrett, Sonia Goncalves, Ewan Harrison, David K. Jackson, Ian Johnston, Dominic Kwiatkowski, Cordelia Langford, John Sillitoe on behalf of the Wellcome Sanger Institute COVID-19 Surveillance Team |
| EPI_ISL_3362684 | Lighthouse Lab in Milton Keynes | Wellcome Sanger Institute for the COVID-19 Genomics UK (COG-UK) Consortium | The Lighthouse Lab in Milton Keynes and Alex Alderton, Roberto Amato, Jeffrey Barrett, Sonia Goncalves, Ewan Harrison, David K. Jackson, Ian Johnston, Dominic Kwiatkowski, Cordelia Langford, John Sillitoe on behalf of the Wellcome Sanger Institute COVID-19 Surveillance Team |
| EPI_ISL_3362685 | Lighthouse Laboratory Plymouth | Wellcome Sanger Institute for the COVID-19 Genomics UK (COG-UK) Consortium | Lighthouse Laboratory Plymouth and Alex Alderton, Roberto Amato, Jeffrey Barrett, Sonia Goncalves, Ewan Harrison, David K. Jackson, Ian Johnston, Dominic Kwiatkowski, Cordelia Langford, John Sillitoe on behalf of the Wellcome Sanger Institute COVID-19 Surveillance Team |
| EPI_ISL_3362686 | Lighthouse Lab in Milton Keynes | Wellcome Sanger Institute for the COVID-19 Genomics UK (COG-UK) Consortium | The Lighthouse Lab in Milton Keynes and Alex Alderton, Roberto Amato, Jeffrey Barrett, Sonia Goncalves, Ewan Harrison, David K. Jackson, Ian Johnston, Dominic Kwiatkowski, Cordelia Langford, John Sillitoe on behalf of the Wellcome Sanger Institute COVID-19 Surveillance Team |
| EPI_ISL_3362688, EPI_ISL_3362689, EPI_ISL_3362691 | Lighthouse Laboratory Plymouth | Wellcome Sanger Institute for the COVID-19 Genomics UK (COG-UK) Consortium | Lighthouse Laboratory Plymouth and Alex Alderton, Roberto Amato, Jeffrey Barrett, Sonia Goncalves, Ewan Harrison, David K. Jackson, Ian Johnston, Dominic Kwiatkowski, Cordelia Langford, John Sillitoe on behalf of the Wellcome Sanger Institute COVID-19 Surveillance Team |
| EPI_ISL_3362692 | Lighthouse Lab in Milton Keynes | Wellcome Sanger Institute for the COVID-19 Genomics UK (COG-UK) Consortium | The Lighthouse Lab in Milton Keynes and Alex Alderton, Roberto Amato, Jeffrey Barrett, Sonia Goncalves, Ewan Harrison, David K. Jackson, Ian Johnston, Dominic Kwiatkowski, Cordelia Langford, John Sillitoe on behalf of the Wellcome Sanger Institute COVID-19 Surveillance Team |
| EPI_ISL_3362694 | Lighthouse Laboratory Plymouth | Wellcome Sanger Institute for the COVID-19 Genomics UK (COG-UK) Consortium | Lighthouse Laboratory Plymouth and Alex Alderton, Roberto Amato, Jeffrey Barrett, Sonia Goncalves, Ewan Harrison, David K. Jackson, Ian Johnston, Dominic Kwiatkowski, Cordelia Langford, John Sillitoe on behalf of the Wellcome Sanger Institute COVID-19 Surveillance Team |
| EPI_ISL_3362695, EPI_ISL_3362696 | Lighthouse Lab in Milton Keynes | Wellcome Sanger Institute for the COVID-19 Genomics UK (COG-UK) Consortium | The Lighthouse Lab in Milton Keynes and Alex Alderton, Roberto Amato, Jeffrey Barrett, Sonia Goncalves, Ewan Harrison, David K. Jackson, Ian Johnston, Dominic Kwiatkowski, Cordelia Langford, John Sillitoe on behalf of the Wellcome Sanger Institute COVID-19 Surveillance Team |
| EPI_ISL_3362698 | Lighthouse Laboratory Plymouth | Wellcome Sanger Institute for the COVID-19 Genomics UK (COG-UK) Consortium | Lighthouse Laboratory Plymouth and Alex Alderton, Roberto Amato, Jeffrey Barrett, Sonia Goncalves, Ewan Harrison, David K. Jackson, Ian Johnston, Dominic Kwiatkowski, Cordelia Langford, John Sillitoe on behalf of the Wellcome Sanger Institute COVID-19 Surveillance Team |
| EPI_ISL_3362699 | Lighthouse Lab in Milton Keynes | Wellcome Sanger Institute for the COVID-19 Genomics UK (COG-UK) Consortium | The Lighthouse Lab in Milton Keynes and Alex Alderton, Roberto Amato, Jeffrey Barrett, Sonia Goncalves, Ewan Harrison, David K. Jackson, Ian Johnston, Dominic Kwiatkowski, Cordelia Langford, John Sillitoe on behalf of the Wellcome Sanger Institute COVID-19 Surveillance Team |
| EPI_ISL_3362701 | Lighthouse Lab in Glasgow | Wellcome Sanger Institute for the COVID-19 Genomics UK (COG-UK) Consortium | Harper VanSteenhouse, Yumi Kasai, David Gray, Carol Clugston, Anna Dominiczak and Alex Alderton, Roberto Amato, Jeffrey Barrett, Sonia Goncalves, Ewan Harrison, David K. Jackson, Ian Johnston, Dominic Kwiatkowski, Cordelia Langford, John Sillitoe on behalf of the Wellcome Sanger Institute COVID-19 Surveillance Team |
| EPI_ISL_3362702 | Lighthouse Lab in Milton Keynes | Wellcome Sanger Institute for the COVID-19 Genomics UK (COG-UK) Consortium | The Lighthouse Lab in Milton Keynes and Alex Alderton, Roberto Amato, Jeffrey Barrett, Sonia Goncalves, Ewan Harrison, David K. Jackson, Ian Johnston, Dominic Kwiatkowski, Cordelia Langford, John Sillitoe on behalf of the Wellcome Sanger Institute COVID-19 Surveillance Team |
| EPI_ISL_3362703, EPI_ISL_3362705 | Lighthouse Lab in Glasgow | Wellcome Sanger Institute for the COVID-19 Genomics UK (COG-UK) Consortium | Harper VanSteenhouse, Yumi Kasai, David Gray, Carol Clugston, Anna Dominiczak and Alex Alderton, Roberto Amato, Jeffrey Barrett, Sonia Goncalves, Ewan Harrison, David K. Jackson, Ian Johnston, Dominic Kwiatkowski, Cordelia Langford, John Sillitoe on behalf of the Wellcome Sanger Institute COVID-19 Surveillance Team |
| EPI_ISL_3362706 | Lighthouse Lab in Milton Keynes | Wellcome Sanger Institute for the COVID-19 Genomics UK (COG-UK) Consortium | The Lighthouse Lab in Milton Keynes and Alex Alderton, Roberto Amato, Jeffrey Barrett, Sonia Goncalves, Ewan Harrison, David K. Jackson, Ian Johnston, Dominic Kwiatkowski, Cordelia Langford, John Sillitoe on behalf of the Wellcome Sanger Institute COVID-19 Surveillance Team |
| EPI_ISL_3362708 | Lighthouse Laboratory Plymouth | Wellcome Sanger Institute for the COVID-19 Genomics UK (COG-UK) Consortium | Lighthouse Laboratory Plymouth and Alex Alderton, Roberto Amato, Jeffrey Barrett, Sonia Goncalves, Ewan Harrison, David K. Jackson, Ian Johnston, Dominic Kwiatkowski, Cordelia Langford, John Sillitoe on behalf of the Wellcome Sanger Institute COVID-19 Surveillance Team |
| EPI_ISL_3362709, EPI_ISL_3362711, EPI_ISL_3362712 | Lighthouse Lab in Milton Keynes | Wellcome Sanger Institute for the COVID-19 Genomics UK (COG-UK) Consortium | The Lighthouse Lab in Milton Keynes and Alex Alderton, Roberto Amato, Jeffrey Barrett, Sonia Goncalves, Ewan Harrison, David K. Jackson, Ian Johnston, Dominic Kwiatkowski, Cordelia Langford, John Sillitoe on behalf of the Wellcome Sanger Institute COVID-19 Surveillance Team |
| EPI_ISL_3362714 | Lighthouse Laboratory Plymouth | Wellcome Sanger Institute for the COVID-19 Genomics UK (COG-UK) Consortium | Lighthouse Laboratory Plymouth and Alex Alderton, Roberto Amato, Jeffrey Barrett, Sonia Goncalves, Ewan Harrison, David K. Jackson, Ian Johnston, Dominic Kwiatkowski, Cordelia Langford, John Sillitoe on behalf of the Wellcome Sanger Institute COVID-19 Surveillance Team |
| EPI_ISL_3362715 | Lighthouse Lab in Milton Keynes | Wellcome Sanger Institute for the COVID-19 Genomics UK (COG-UK) Consortium | The Lighthouse Lab in Milton Keynes and Alex Alderton, Roberto Amato, Jeffrey Barrett, Sonia Goncalves, Ewan Harrison, David K. Jackson, Ian Johnston, Dominic Kwiatkowski, Cordelia Langford, John Sillitoe on behalf of the Wellcome Sanger Institute COVID-19 Surveillance Team |
| EPI_ISL_3362716, EPI_ISL_3362718 | Lighthouse Laboratory Plymouth | Wellcome Sanger Institute for the COVID-19 Genomics UK (COG-UK) Consortium | Lighthouse Laboratory Plymouth and Alex Alderton, Roberto Amato, Jeffrey Barrett, Sonia Goncalves, Ewan Harrison, David K. Jackson, Ian Johnston, Dominic Kwiatkowski, Cordelia Langford, John Sillitoe on behalf of the Wellcome Sanger Institute COVID-19 Surveillance Team |
| EPI_ISL_3362719 | Lighthouse Lab in Glasgow | Wellcome Sanger Institute for the COVID-19 Genomics UK (COG-UK) Consortium | Harper VanSteenhouse, Yumi Kasai, David Gray, Carol Clugston, Anna Dominiczak and Alex Alderton, Roberto Amato, Jeffrey Barrett, Sonia Goncalves, Ewan Harrison, David K. Jackson, Ian Johnston, Dominic Kwiatkowski, Cordelia Langford, John Sillitoe on behalf of the Wellcome Sanger Institute COVID-19 Surveillance Team |
| EPI_ISL_3362721 | Lighthouse Laboratory Plymouth | Wellcome Sanger Institute for the COVID-19 Genomics UK (COG-UK) Consortium | Lighthouse Laboratory Plymouth and Alex Alderton, Roberto Amato, Jeffrey Barrett, Sonia Goncalves, Ewan Harrison, David K. Jackson, Ian Johnston, Dominic Kwiatkowski, Cordelia Langford, John Sillitoe on behalf of the Wellcome Sanger Institute COVID-19 Surveillance Team |
| EPI_ISL_3362722, EPI_ISL_3362724 | Lighthouse Lab in Milton Keynes | Wellcome Sanger Institute for the COVID-19 Genomics UK (COG-UK) Consortium | The Lighthouse Lab in Milton Keynes and Alex Alderton, Roberto Amato, Jeffrey Barrett, Sonia Goncalves, Ewan Harrison, David K. Jackson, Ian Johnston, Dominic Kwiatkowski, Cordelia Langford, John Sillitoe on behalf of the Wellcome Sanger Institute COVID-19 Surveillance Team |
| EPI_ISL_3362725, EPI_ISL_3362727 | Lighthouse Laboratory Plymouth | Wellcome Sanger Institute for the COVID-19 Genomics UK (COG-UK) Consortium | Lighthouse Laboratory Plymouth and Alex Alderton, Roberto Amato, Jeffrey Barrett, Sonia Goncalves, Ewan Harrison, David K. Jackson, Ian Johnston, Dominic Kwiatkowski, Cordelia Langford, John Sillitoe on behalf of the Wellcome Sanger Institute COVID-19 Surveillance Team |
| EPI_ISL_3362728 | Lighthouse Lab in Glasgow | Wellcome Sanger Institute for the COVID-19 Genomics UK (COG-UK) | Harper VanSteenhouse, Yumi Kasai, David Gray, Carol Clugston, Anna Dominiczak and Alex Alderton, Roberto Amato, Jeffrey Barrett, Sonia Goncalves, Ewan Harrison, David K. Jackson, Ian Johnston, Dominic Kwiatkowski, Cordelia Langford, |

| | | | |
|---|---|---|---|
| | | Consortium | John Sillitoe on behalf of the Wellcome Sanger Institute COVID-19 Surveillance Team |
| EPI_ISL_3362730 | Lighthouse Laboratory Plymouth | Wellcome Sanger Institute for the COVID-19 Genomics UK (COG-UK) Consortium | Lighthouse Laboratory Plymouth and Alex Alderton, Roberto Amato, Jeffrey Barrett, Sonia Goncalves, Ewan Harrison, David K. Jackson, Ian Johnston, Dominic Kwiatkowski, Cordelia Langford, John Sillitoe on behalf of the Wellcome Sanger Institute COVID-19 Surveillance Team |
| EPI_ISL_3362731, EPI_ISL_3362733, EPI_ISL_3362734 | Lighthouse Lab in Glasgow | Wellcome Sanger Institute for the COVID-19 Genomics UK (COG-UK) Consortium | Harper VanSteenhouse, Yumi Kasai, David Gray, Carol Clugston, Anna Dominiczak and Alex Alderton, Roberto Amato, Jeffrey Barrett, Sonia Goncalves, Ewan Harrison, David K. Jackson, Ian Johnston, Dominic Kwiatkowski, Cordelia Langford, John Sillitoe on behalf of the Wellcome Sanger Institute COVID-19 Surveillance Team |
| EPI_ISL_3362735, EPI_ISL_3362737, EPI_ISL_3362738, EPI_ISL_3362740, EPI_ISL_3362741 | Lighthouse Laboratory Plymouth | Wellcome Sanger Institute for the COVID-19 Genomics UK (COG-UK) Consortium | Lighthouse Laboratory Plymouth and Alex Alderton, Roberto Amato, Jeffrey Barrett, Sonia Goncalves, Ewan Harrison, David K. Jackson, Ian Johnston, Dominic Kwiatkowski, Cordelia Langford, John Sillitoe on behalf of the Wellcome Sanger Institute COVID-19 Surveillance Team |
| EPI_ISL_3362743 | Lighthouse Lab in Milton Keynes | Wellcome Sanger Institute for the COVID-19 Genomics UK (COG-UK) Consortium | The Lighthouse Lab in Milton Keynes and Alex Alderton, Roberto Amato, Jeffrey Barrett, Sonia Goncalves, Ewan Harrison, David K. Jackson, Ian Johnston, Dominic Kwiatkowski, Cordelia Langford, John Sillitoe on behalf of the Wellcome Sanger Institute COVID-19 Surveillance Team |
| EPI_ISL_3362744, EPI_ISL_3362746, EPI_ISL_3362747 | Lighthouse Lab in Glasgow | Wellcome Sanger Institute for the COVID-19 Genomics UK (COG-UK) Consortium | Harper VanSteenhouse, Yumi Kasai, David Gray, Carol Clugston, Anna Dominiczak and Alex Alderton, Roberto Amato, Jeffrey Barrett, Sonia Goncalves, Ewan Harrison, David K. Jackson, Ian Johnston, Dominic Kwiatkowski, Cordelia Langford, John Sillitoe on behalf of the Wellcome Sanger Institute COVID-19 Surveillance Team |
| EPI_ISL_3362748 | Lighthouse Lab in Milton Keynes | Wellcome Sanger Institute for the COVID-19 Genomics UK (COG-UK) Consortium | The Lighthouse Lab in Milton Keynes and Alex Alderton, Roberto Amato, Jeffrey Barrett, Sonia Goncalves, Ewan Harrison, David K. Jackson, Ian Johnston, Dominic Kwiatkowski, Cordelia Langford, John Sillitoe on behalf of the Wellcome Sanger Institute COVID-19 Surveillance Team |
| EPI_ISL_3362750 | Lighthouse Laboratory Plymouth | Wellcome Sanger Institute for the COVID-19 Genomics UK (COG-UK) Consortium | Lighthouse Laboratory Plymouth and Alex Alderton, Roberto Amato, Jeffrey Barrett, Sonia Goncalves, Ewan Harrison, David K. Jackson, Ian Johnston, Dominic Kwiatkowski, Cordelia Langford, John Sillitoe on behalf of the Wellcome Sanger Institute COVID-19 Surveillance Team |
| EPI_ISL_3362751 | Lighthouse Lab in Glasgow | Wellcome Sanger Institute for the COVID-19 Genomics UK (COG-UK) Consortium | Harper VanSteenhouse, Yumi Kasai, David Gray, Carol Clugston, Anna Dominiczak and Alex Alderton, Roberto Amato, Jeffrey Barrett, Sonia Goncalves, Ewan Harrison, David K. Jackson, Ian Johnston, Dominic Kwiatkowski, Cordelia Langford, John Sillitoe on behalf of the Wellcome Sanger Institute COVID-19 Surveillance Team |
| EPI_ISL_3362753 | Lighthouse Laboratory Plymouth | Wellcome Sanger Institute for the COVID-19 Genomics UK (COG-UK) Consortium | Lighthouse Laboratory Plymouth and Alex Alderton, Roberto Amato, Jeffrey Barrett, Sonia Goncalves, Ewan Harrison, David K. Jackson, Ian Johnston, Dominic Kwiatkowski, Cordelia Langford, John Sillitoe on behalf of the Wellcome Sanger Institute COVID-19 Surveillance Team |
| EPI_ISL_3362754, EPI_ISL_3362755, EPI_ISL_3362757 | Lighthouse Lab in Milton Keynes | Wellcome Sanger Institute for the COVID-19 Genomics UK (COG-UK) Consortium | The Lighthouse Lab in Milton Keynes and Alex Alderton, Roberto Amato, Jeffrey Barrett, Sonia Goncalves, Ewan Harrison, David K. Jackson, Ian Johnston, Dominic Kwiatkowski, Cordelia Langford, John Sillitoe on behalf of the Wellcome Sanger Institute COVID-19 Surveillance Team |
| EPI_ISL_3362758, EPI_ISL_3362760 | Lighthouse Laboratory Plymouth | Wellcome Sanger Institute for the COVID-19 Genomics UK (COG-UK) Consortium | Lighthouse Laboratory Plymouth and Alex Alderton, Roberto Amato, Jeffrey Barrett, Sonia Goncalves, Ewan Harrison, David K. Jackson, Ian Johnston, Dominic Kwiatkowski, Cordelia Langford, John Sillitoe on behalf of the Wellcome Sanger Institute COVID-19 Surveillance Team |
| EPI_ISL_3362761 | Lighthouse Lab in Milton Keynes | Wellcome Sanger Institute for the COVID-19 Genomics UK (COG-UK) Consortium | The Lighthouse Lab in Milton Keynes and Alex Alderton, Roberto Amato, Jeffrey Barrett, Sonia Goncalves, Ewan Harrison, David K. Jackson, Ian Johnston, Dominic Kwiatkowski, Cordelia Langford, John Sillitoe on behalf of the Wellcome Sanger Institute COVID-19 Surveillance Team |
| EPI_ISL_3362763 | Lighthouse Lab in Glasgow | Wellcome Sanger Institute for the COVID-19 Genomics UK (COG-UK) Consortium | Harper VanSteenhouse, Yumi Kasai, David Gray, Carol Clugston, Anna Dominiczak and Alex Alderton, Roberto Amato, Jeffrey Barrett, Sonia Goncalves, Ewan Harrison, David K. Jackson, Ian Johnston, Dominic Kwiatkowski, Cordelia Langford, John Sillitoe on behalf of the Wellcome Sanger Institute COVID-19 Surveillance Team |
| EPI_ISL_3362764 | Lighthouse Laboratory Plymouth | Wellcome Sanger Institute for the COVID-19 Genomics UK (COG-UK) Consortium | Lighthouse Laboratory Plymouth and Alex Alderton, Roberto Amato, Jeffrey Barrett, Sonia Goncalves, Ewan Harrison, David K. Jackson, Ian Johnston, Dominic Kwiatkowski, Cordelia Langford, John Sillitoe on behalf of the Wellcome Sanger Institute COVID-19 Surveillance Team |
| EPI_ISL_3362766, EPI_ISL_3362767 | Lighthouse Lab in Milton Keynes | Wellcome Sanger Institute for the COVID-19 Genomics UK (COG-UK) Consortium | The Lighthouse Lab in Milton Keynes and Alex Alderton, Roberto Amato, Jeffrey Barrett, Sonia Goncalves, Ewan Harrison, David K. Jackson, Ian Johnston, Dominic Kwiatkowski, Cordelia Langford, John Sillitoe on behalf of the Wellcome Sanger Institute COVID-19 Surveillance Team |
| EPI_ISL_3362768, EPI_ISL_3362770 | Lighthouse Laboratory Plymouth | Wellcome Sanger Institute for the COVID-19 Genomics UK (COG-UK) Consortium | Lighthouse Laboratory Plymouth and Alex Alderton, Roberto Amato, Jeffrey Barrett, Sonia Goncalves, Ewan Harrison, David K. Jackson, Ian Johnston, Dominic Kwiatkowski, Cordelia Langford, John Sillitoe on behalf of the Wellcome Sanger Institute COVID-19 Surveillance Team |
| EPI_ISL_3362771 | Lighthouse Lab in Glasgow | Wellcome Sanger Institute for the COVID-19 Genomics UK (COG-UK) Consortium | Harper VanSteenhouse, Yumi Kasai, David Gray, Carol Clugston, Anna Dominiczak and Alex Alderton, Roberto Amato, Jeffrey Barrett, Sonia Goncalves, Ewan Harrison, David K. Jackson, Ian Johnston, Dominic Kwiatkowski, Cordelia Langford, John Sillitoe on behalf of the Wellcome Sanger Institute COVID-19 Surveillance Team |
| EPI_ISL_3362773, EPI_ISL_3362774, EPI_ISL_3362776 | Lighthouse Laboratory Plymouth | Wellcome Sanger Institute for the COVID-19 Genomics UK (COG-UK) Consortium | Lighthouse Laboratory Plymouth and Alex Alderton, Roberto Amato, Jeffrey Barrett, Sonia Goncalves, Ewan Harrison, David K. Jackson, Ian Johnston, Dominic Kwiatkowski, Cordelia Langford, John Sillitoe on behalf of the Wellcome Sanger Institute COVID-19 Surveillance Team |
| EPI_ISL_3362777 | Lighthouse Lab in Glasgow | Wellcome Sanger Institute for the COVID-19 Genomics UK (COG-UK) Consortium | Harper VanSteenhouse, Yumi Kasai, David Gray, Carol Clugston, Anna Dominiczak and Alex Alderton, Roberto Amato, Jeffrey Barrett, Sonia Goncalves, Ewan Harrison, David K. Jackson, Ian Johnston, Dominic Kwiatkowski, Cordelia Langford, John Sillitoe on behalf of the Wellcome Sanger Institute COVID-19 Surveillance Team |
| EPI_ISL_3362779 | Lighthouse Lab in Milton Keynes | Wellcome Sanger Institute for the COVID-19 Genomics UK (COG-UK) Consortium | The Lighthouse Lab in Milton Keynes and Alex Alderton, Roberto Amato, Jeffrey Barrett, Sonia Goncalves, Ewan Harrison, David K. Jackson, Ian Johnston, Dominic Kwiatkowski, Cordelia Langford, John Sillitoe on behalf of the Wellcome Sanger Institute COVID-19 Surveillance Team |
| EPI_ISL_3362780, EPI_ISL_3362781, EPI_ISL_3362783, EPI_ISL_3362785 | Lighthouse Lab in Glasgow | Wellcome Sanger Institute for the COVID-19 Genomics UK (COG-UK) Consortium | Harper VanSteenhouse, Yumi Kasai, David Gray, Carol Clugston, Anna Dominiczak and Alex Alderton, Roberto Amato, Jeffrey Barrett, Sonia Goncalves, Ewan Harrison, David K. Jackson, Ian Johnston, Dominic Kwiatkowski, Cordelia Langford, John Sillitoe on behalf of the Wellcome Sanger Institute COVID-19 Surveillance Team |
| EPI_ISL_3362786 | Lighthouse Laboratory Plymouth | Wellcome Sanger Institute for the COVID-19 Genomics UK (COG-UK) Consortium | Lighthouse Laboratory Plymouth and Alex Alderton, Roberto Amato, Jeffrey Barrett, Sonia Goncalves, Ewan Harrison, David K. Jackson, Ian Johnston, Dominic Kwiatkowski, Cordelia Langford, John Sillitoe on behalf of the Wellcome Sanger Institute COVID-19 Surveillance Team |
| EPI_ISL_3362787 | Lighthouse Lab in Milton Keynes | Wellcome Sanger Institute for the COVID-19 Genomics UK (COG-UK) Consortium | The Lighthouse Lab in Milton Keynes and Alex Alderton, Roberto Amato, Jeffrey Barrett, Sonia Goncalves, Ewan Harrison, David K. Jackson, Ian Johnston, Dominic Kwiatkowski, Cordelia Langford, John Sillitoe on behalf of the Wellcome Sanger Institute COVID-19 Surveillance Team |
| EPI_ISL_3362789 | Lighthouse Laboratory Plymouth | Wellcome Sanger Institute for the COVID-19 Genomics UK (COG-UK) Consortium | Lighthouse Laboratory Plymouth and Alex Alderton, Roberto Amato, Jeffrey Barrett, Sonia Goncalves, Ewan Harrison, David K. Jackson, Ian Johnston, Dominic Kwiatkowski, Cordelia Langford, John Sillitoe on behalf of the Wellcome Sanger Institute COVID-19 Surveillance Team |
| EPI_ISL_3362790 | Lighthouse Lab in Milton Keynes | Wellcome Sanger Institute for the COVID-19 Genomics UK (COG-UK) Consortium | The Lighthouse Lab in Milton Keynes and Alex Alderton, Roberto Amato, Jeffrey Barrett, Sonia Goncalves, Ewan Harrison, David K. Jackson, Ian Johnston, Dominic Kwiatkowski, Cordelia Langford, John Sillitoe on behalf of the Wellcome Sanger Institute COVID-19 Surveillance Team |
| EPI_ISL_3362792, EPI_ISL_3362793, EPI_ISL_3362794 | Lighthouse Laboratory Plymouth | Wellcome Sanger Institute for the COVID-19 Genomics UK (COG-UK) Consortium | Lighthouse Laboratory Plymouth and Alex Alderton, Roberto Amato, Jeffrey Barrett, Sonia Goncalves, Ewan Harrison, David K. Jackson, Ian Johnston, Dominic Kwiatkowski, Cordelia Langford, John Sillitoe on behalf of the Wellcome Sanger Institute COVID-19 Surveillance Team |
| EPI_ISL_3362796 | Lighthouse Lab in Glasgow | Wellcome Sanger Institute for the COVID-19 Genomics UK (COG-UK) Consortium | Harper VanSteenhouse, Yumi Kasai, David Gray, Carol Clugston, Anna Dominiczak and Alex Alderton, Roberto Amato, Jeffrey Barrett, Sonia Goncalves, Ewan Harrison, David K. Jackson, Ian Johnston, Dominic Kwiatkowski, Cordelia Langford, John Sillitoe on behalf of the Wellcome Sanger Institute COVID-19 Surveillance Team |
| EPI_ISL_3362798, EPI_ISL_3362799 | Lighthouse Lab in Milton Keynes | Wellcome Sanger Institute for the COVID-19 Genomics UK (COG-UK) Consortium | The Lighthouse Lab in Milton Keynes and Alex Alderton, Roberto Amato, Jeffrey Barrett, Sonia Goncalves, Ewan Harrison, David K. Jackson, Ian Johnston, Dominic Kwiatkowski, Cordelia Langford, John Sillitoe on behalf of the Wellcome Sanger Institute COVID-19 Surveillance Team |
| EPI_ISL_3362800 | Lighthouse Lab in Glasgow | Wellcome Sanger Institute for the COVID-19 Genomics UK (COG-UK) Consortium | Harper VanSteenhouse, Yumi Kasai, David Gray, Carol Clugston, Anna Dominiczak and Alex Alderton, Roberto Amato, Jeffrey Barrett, Sonia Goncalves, Ewan Harrison, David K. Jackson, Ian Johnston, Dominic Kwiatkowski, Cordelia Langford, John Sillitoe on behalf of the Wellcome Sanger Institute COVID-19 Surveillance Team |
| EPI_ISL_3362802 | Lighthouse Lab in Milton Keynes | Wellcome Sanger Institute for the COVID-19 Genomics UK (COG-UK) Consortium | The Lighthouse Lab in Milton Keynes and Alex Alderton, Roberto Amato, Jeffrey Barrett, Sonia Goncalves, Ewan Harrison, David K. Jackson, Ian Johnston, Dominic Kwiatkowski, Cordelia Langford, John Sillitoe on behalf of the Wellcome Sanger Institute COVID-19 Surveillance Team |
| EPI_ISL_3362803 | Lighthouse Lab in Alderley Park | Wellcome Sanger Institute for the COVID-19 Genomics UK (COG-UK) Consortium | Jacquelyn Wynn, Mairead Hyland, The Lighthouse Lab in Alderley Park and Alex Alderton, Roberto Amato, Jeffrey Barrett, Sonia Goncalves, Ewan Harrison, David K. Jackson, Ian Johnston, Dominic Kwiatkowski, Cordelia Langford, John Sillitoe on behalf of the Wellcome Sanger Institute COVID-19 Surveillance Team |
| EPI_ISL_3362805, EPI_ISL_3362806, EPI_ISL_3362808 | Lighthouse Lab in Glasgow | Wellcome Sanger Institute for the COVID-19 Genomics UK (COG-UK) Consortium | Harper VanSteenhouse, Yumi Kasai, David Gray, Carol Clugston, Anna Dominiczak and Alex Alderton, Roberto Amato, Jeffrey Barrett, Sonia Goncalves, Ewan Harrison, David K. Jackson, Ian Johnston, Dominic Kwiatkowski, Cordelia Langford, John Sillitoe on behalf of the Wellcome Sanger Institute COVID-19 Surveillance Team |
| EPI_ISL_3362809 | Lighthouse Lab in Milton Keynes | Wellcome Sanger Institute for the COVID-19 Genomics UK (COG-UK) Consortium | The Lighthouse Lab in Milton Keynes and Alex Alderton, Roberto Amato, Jeffrey Barrett, Sonia Goncalves, Ewan Harrison, David K. Jackson, Ian Johnston, Dominic Kwiatkowski, Cordelia Langford, John Sillitoe on behalf of the Wellcome Sanger Institute COVID-19 Surveillance Team |
| EPI_ISL_3362811, EPI_ISL_3362812, EPI_ISL_3362814, EPI_ISL_3362815 | Lighthouse Lab in Glasgow | Wellcome Sanger Institute for the COVID-19 Genomics UK (COG-UK) Consortium | Harper VanSteenhouse, Yumi Kasai, David Gray, Carol Clugston, Anna Dominiczak and Alex Alderton, Roberto Amato, Jeffrey Barrett, Sonia Goncalves, Ewan Harrison, David K. Jackson, Ian Johnston, Dominic Kwiatkowski, Cordelia Langford, John Sillitoe on behalf of the Wellcome Sanger Institute COVID-19 Surveillance Team |
| EPI_ISL_3362817 | Lighthouse Lab in Alderley Park | Wellcome Sanger Institute for the COVID-19 Genomics UK (COG-UK) Consortium | Jacquelyn Wynn, Mairead Hyland, The Lighthouse Lab in Alderley Park and Alex Alderton, Roberto Amato, Jeffrey Barrett, Sonia Goncalves, Ewan Harrison, David K. Jackson, Ian Johnston, Dominic Kwiatkowski, Cordelia Langford, John Sillitoe on behalf of the Wellcome Sanger Institute COVID-19 Surveillance Team |
| EPI_ISL_3362818, EPI_ISL_3362820 | Lighthouse Lab in Glasgow | Wellcome Sanger Institute for the COVID-19 Genomics UK (COG-UK) Consortium | Harper VanSteenhouse, Yumi Kasai, David Gray, Carol Clugston, Anna Dominiczak and Alex Alderton, Roberto Amato, Jeffrey Barrett, Sonia Goncalves, Ewan Harrison, David K. Jackson, Ian Johnston, Dominic Kwiatkowski, Cordelia Langford, John Sillitoe on behalf of the Wellcome Sanger Institute COVID-19 Surveillance Team |
| EPI_ISL_3362821 | Lighthouse Lab in Milton Keynes | Wellcome Sanger Institute for the COVID-19 Genomics UK (COG-UK) Consortium | The Lighthouse Lab in Milton Keynes and Alex Alderton, Roberto Amato, Jeffrey Barrett, Sonia Goncalves, Ewan Harrison, David K. Jackson, Ian Johnston, Dominic Kwiatkowski, Cordelia Langford, John Sillitoe on behalf of the Wellcome Sanger Institute COVID-19 Surveillance Team |
| EPI_ISL_3362823, EPI_ISL_3362824 | Lighthouse Lab in Alderley Park | Wellcome Sanger Institute for the COVID-19 Genomics UK (COG-UK) Consortium | Jacquelyn Wynn, Mairead Hyland, The Lighthouse Lab in Alderley Park and Alex Alderton, Roberto Amato, Jeffrey Barrett, Sonia Goncalves, Ewan Harrison, David K. Jackson, Ian Johnston, Dominic Kwiatkowski, Cordelia Langford, John Sillitoe on behalf of the Wellcome Sanger Institute COVID-19 Surveillance Team |
| EPI_ISL_3362825, EPI_ISL_3362827 | Lighthouse Lab in Glasgow | Wellcome Sanger Institute for the COVID-19 Genomics UK (COG-UK) Consortium | Harper VanSteenhouse, Yumi Kasai, David Gray, Carol Clugston, Anna Dominiczak and Alex Alderton, Roberto Amato, Jeffrey Barrett, Sonia Goncalves, Ewan Harrison, David K. Jackson, Ian Johnston, Dominic Kwiatkowski, Cordelia Langford, John Sillitoe on behalf of the Wellcome Sanger Institute COVID-19 Surveillance Team |
| EPI_ISL_3362828 | Lighthouse Lab in Milton Keynes | Wellcome Sanger Institute for the COVID-19 Genomics UK (COG-UK) | The Lighthouse Lab in Milton Keynes and Alex Alderton, Roberto Amato, Jeffrey Barrett, Sonia Goncalves, Ewan Harrison, David K. Jackson, Ian Johnston, Dominic Kwiatkowski, Cordelia Langford, John Sillitoe on behalf of the Wellcome Sanger |

| | | | |
|---|---|---|---|
| | | Consortium | Institute COVID-19 Surveillance Team |
| EPI_ISL_3362830, EPI_ISL_3362831 | Lighthouse Lab in Glasgow | Wellcome Sanger Institute for the COVID-19 Genomics UK (COG-UK) Consortium | Harper VanSteenhouse, Yumi Kasai, David Gray, Carol Clugston, Anna Dominiczak and Alex Alderton, Roberto Amato, Jeffrey Barrett, Sonia Goncalves, Ewan Harrison, David K. Jackson, Ian Johnston, Dominic Kwiatkowski, Cordelia Langford, John Sillitoe on behalf of the Wellcome Sanger Institute COVID-19 Surveillance Team |
| EPI_ISL_3362833 | Lighthouse Lab in Milton Keynes | Wellcome Sanger Institute for the COVID-19 Genomics UK (COG-UK) Consortium | The Lighthouse Lab in Milton Keynes and Alex Alderton, Roberto Amato, Jeffrey Barrett, Sonia Goncalves, Ewan Harrison, David K. Jackson, Ian Johnston, Dominic Kwiatkowski, Cordelia Langford, John Sillitoe on behalf of the Wellcome Sanger Institute COVID-19 Surveillance Team |
| EPI_ISL_3362834 | Lighthouse Lab in Alderley Park | Wellcome Sanger Institute for the COVID-19 Genomics UK (COG-UK) Consortium | Jacquelyn Wynn, Mairead Hyland, The Lighthouse Lab in Alderley Park and Alex Alderton, Roberto Amato, Jeffrey Barrett, Sonia Goncalves, Ewan Harrison, David K. Jackson, Ian Johnston, Dominic Kwiatkowski, Cordelia Langford, John Sillitoe on behalf of the Wellcome Sanger Institute COVID-19 Surveillance Team |
| EPI_ISL_3362836, EPI_ISL_3362837, EPI_ISL_3362839 | Lighthouse Lab in Milton Keynes | Wellcome Sanger Institute for the COVID-19 Genomics UK (COG-UK) Consortium | The Lighthouse Lab in Milton Keynes and Alex Alderton, Roberto Amato, Jeffrey Barrett, Sonia Goncalves, Ewan Harrison, David K. Jackson, Ian Johnston, Dominic Kwiatkowski, Cordelia Langford, John Sillitoe on behalf of the Wellcome Sanger Institute COVID-19 Surveillance Team |
| EPI_ISL_3362840 | Lighthouse Lab in Glasgow | Wellcome Sanger Institute for the COVID-19 Genomics UK (COG-UK) Consortium | Harper VanSteenhouse, Yumi Kasai, David Gray, Carol Clugston, Anna Dominiczak and Alex Alderton, Roberto Amato, Jeffrey Barrett, Sonia Goncalves, Ewan Harrison, David K. Jackson, Ian Johnston, Dominic Kwiatkowski, Cordelia Langford, John Sillitoe on behalf of the Wellcome Sanger Institute COVID-19 Surveillance Team |
| EPI_ISL_3362842 | Lighthouse Lab in Milton Keynes | Wellcome Sanger Institute for the COVID-19 Genomics UK (COG-UK) Consortium | The Lighthouse Lab in Milton Keynes and Alex Alderton, Roberto Amato, Jeffrey Barrett, Sonia Goncalves, Ewan Harrison, David K. Jackson, Ian Johnston, Dominic Kwiatkowski, Cordelia Langford, John Sillitoe on behalf of the Wellcome Sanger Institute COVID-19 Surveillance Team |
| EPI_ISL_3362844, EPI_ISL_3362845 | Lighthouse Lab in Glasgow | Wellcome Sanger Institute for the COVID-19 Genomics UK (COG-UK) Consortium | Harper VanSteenhouse, Yumi Kasai, David Gray, Carol Clugston, Anna Dominiczak and Alex Alderton, Roberto Amato, Jeffrey Barrett, Sonia Goncalves, Ewan Harrison, David K. Jackson, Ian Johnston, Dominic Kwiatkowski, Cordelia Langford, John Sillitoe on behalf of the Wellcome Sanger Institute COVID-19 Surveillance Team |
| EPI_ISL_3362846 | Lighthouse Lab in Alderley Park | Wellcome Sanger Institute for the COVID-19 Genomics UK (COG-UK) Consortium | Jacquelyn Wynn, Mairead Hyland, The Lighthouse Lab in Alderley Park and Alex Alderton, Roberto Amato, Jeffrey Barrett, Sonia Goncalves, Ewan Harrison, David K. Jackson, Ian Johnston, Dominic Kwiatkowski, Cordelia Langford, John Sillitoe on behalf of the Wellcome Sanger Institute COVID-19 Surveillance Team |
| EPI_ISL_3362848 | Lighthouse Lab in Milton Keynes | Wellcome Sanger Institute for the COVID-19 Genomics UK (COG-UK) Consortium | The Lighthouse Lab in Milton Keynes and Alex Alderton, Roberto Amato, Jeffrey Barrett, Sonia Goncalves, Ewan Harrison, David K. Jackson, Ian Johnston, Dominic Kwiatkowski, Cordelia Langford, John Sillitoe on behalf of the Wellcome Sanger Institute COVID-19 Surveillance Team |
| EPI_ISL_3362849 | Lighthouse Lab in Glasgow | Wellcome Sanger Institute for the COVID-19 Genomics UK (COG-UK) Consortium | Harper VanSteenhouse, Yumi Kasai, David Gray, Carol Clugston, Anna Dominiczak and Alex Alderton, Roberto Amato, Jeffrey Barrett, Sonia Goncalves, Ewan Harrison, David K. Jackson, Ian Johnston, Dominic Kwiatkowski, Cordelia Langford, John Sillitoe on behalf of the Wellcome Sanger Institute COVID-19 Surveillance Team |
| EPI_ISL_3362851, EPI_ISL_3362852, EPI_ISL_3362853, EPI_ISL_3362855 | Lighthouse Lab in Milton Keynes | Wellcome Sanger Institute for the COVID-19 Genomics UK (COG-UK) Consortium | The Lighthouse Lab in Milton Keynes and Alex Alderton, Roberto Amato, Jeffrey Barrett, Sonia Goncalves, Ewan Harrison, David K. Jackson, Ian Johnston, Dominic Kwiatkowski, Cordelia Langford, John Sillitoe on behalf of the Wellcome Sanger Institute COVID-19 Surveillance Team |
| EPI_ISL_3362856 | Lighthouse Lab in Glasgow | Wellcome Sanger Institute for the COVID-19 Genomics UK (COG-UK) Consortium | Harper VanSteenhouse, Yumi Kasai, David Gray, Carol Clugston, Anna Dominiczak and Alex Alderton, Roberto Amato, Jeffrey Barrett, Sonia Goncalves, Ewan Harrison, David K. Jackson, Ian Johnston, Dominic Kwiatkowski, Cordelia Langford, John Sillitoe on behalf of the Wellcome Sanger Institute COVID-19 Surveillance Team |
| EPI_ISL_3362858 | Lighthouse Lab in Milton Keynes | Wellcome Sanger Institute for the COVID-19 Genomics UK (COG-UK) Consortium | The Lighthouse Lab in Milton Keynes and Alex Alderton, Roberto Amato, Jeffrey Barrett, Sonia Goncalves, Ewan Harrison, David K. Jackson, Ian Johnston, Dominic Kwiatkowski, Cordelia Langford, John Sillitoe on behalf of the Wellcome Sanger Institute COVID-19 Surveillance Team |
| EPI_ISL_3362859 | Lighthouse Lab in Glasgow | Wellcome Sanger Institute for the COVID-19 Genomics UK (COG-UK) Consortium | Harper VanSteenhouse, Yumi Kasai, David Gray, Carol Clugston, Anna Dominiczak and Alex Alderton, Roberto Amato, Jeffrey Barrett, Sonia Goncalves, Ewan Harrison, David K. Jackson, Ian Johnston, Dominic Kwiatkowski, Cordelia Langford, John Sillitoe on behalf of the Wellcome Sanger Institute COVID-19 Surveillance Team |
| EPI_ISL_3362861 | Lighthouse Lab in Alderley Park | Wellcome Sanger Institute for the COVID-19 Genomics UK (COG-UK) Consortium | Jacquelyn Wynn, Mairead Hyland, The Lighthouse Lab in Alderley Park and Alex Alderton, Roberto Amato, Jeffrey Barrett, Sonia Goncalves, Ewan Harrison, David K. Jackson, Ian Johnston, Dominic Kwiatkowski, Cordelia Langford, John Sillitoe on behalf of the Wellcome Sanger Institute COVID-19 Surveillance Team |
| EPI_ISL_3362862, EPI_ISL_3362864 | Lighthouse Lab in Glasgow | Wellcome Sanger Institute for the COVID-19 Genomics UK (COG-UK) Consortium | Harper VanSteenhouse, Yumi Kasai, David Gray, Carol Clugston, Anna Dominiczak and Alex Alderton, Roberto Amato, Jeffrey Barrett, Sonia Goncalves, Ewan Harrison, David K. Jackson, Ian Johnston, Dominic Kwiatkowski, Cordelia Langford, John Sillitoe on behalf of the Wellcome Sanger Institute COVID-19 Surveillance Team |
| EPI_ISL_3362865, EPI_ISL_3362866 | Lighthouse Lab in Milton Keynes | Wellcome Sanger Institute for the COVID-19 Genomics UK (COG-UK) Consortium | The Lighthouse Lab in Milton Keynes and Alex Alderton, Roberto Amato, Jeffrey Barrett, Sonia Goncalves, Ewan Harrison, David K. Jackson, Ian Johnston, Dominic Kwiatkowski, Cordelia Langford, John Sillitoe on behalf of the Wellcome Sanger Institute COVID-19 Surveillance Team |
| EPI_ISL_3362868 | Lighthouse Lab in Alderley Park | Wellcome Sanger Institute for the COVID-19 Genomics UK (COG-UK) Consortium | Jacquelyn Wynn, Mairead Hyland, The Lighthouse Lab in Alderley Park and Alex Alderton, Roberto Amato, Jeffrey Barrett, Sonia Goncalves, Ewan Harrison, David K. Jackson, Ian Johnston, Dominic Kwiatkowski, Cordelia Langford, John Sillitoe on behalf of the Wellcome Sanger Institute COVID-19 Surveillance Team |
| EPI_ISL_3362869 | Lighthouse Lab in Milton Keynes | Wellcome Sanger Institute for the COVID-19 Genomics UK (COG-UK) Consortium | The Lighthouse Lab in Milton Keynes and Alex Alderton, Roberto Amato, Jeffrey Barrett, Sonia Goncalves, Ewan Harrison, David K. Jackson, Ian Johnston, Dominic Kwiatkowski, Cordelia Langford, John Sillitoe on behalf of the Wellcome Sanger Institute COVID-19 Surveillance Team |
| EPI_ISL_3362871, EPI_ISL_3362872, EPI_ISL_3362874 | Lighthouse Lab in Alderley Park | Wellcome Sanger Institute for the COVID-19 Genomics UK (COG-UK) Consortium | Jacquelyn Wynn, Mairead Hyland, The Lighthouse Lab in Alderley Park and Alex Alderton, Roberto Amato, Jeffrey Barrett, Sonia Goncalves, Ewan Harrison, David K. Jackson, Ian Johnston, Dominic Kwiatkowski, Cordelia Langford, John Sillitoe on behalf of the Wellcome Sanger Institute COVID-19 Surveillance Team |
| EPI_ISL_3362875, EPI_ISL_3362876, EPI_ISL_3362878, EPI_ISL_3362879 | Lighthouse Lab in Glasgow | Wellcome Sanger Institute for the COVID-19 Genomics UK (COG-UK) Consortium | Harper VanSteenhouse, Yumi Kasai, David Gray, Carol Clugston, Anna Dominiczak and Alex Alderton, Roberto Amato, Jeffrey Barrett, Sonia Goncalves, Ewan Harrison, David K. Jackson, Ian Johnston, Dominic Kwiatkowski, Cordelia Langford, John Sillitoe on behalf of the Wellcome Sanger Institute COVID-19 Surveillance Team |
| EPI_ISL_3362881 | Lighthouse Lab in Alderley Park | Wellcome Sanger Institute for the COVID-19 Genomics UK (COG-UK) Consortium | Jacquelyn Wynn, Mairead Hyland, The Lighthouse Lab in Alderley Park and Alex Alderton, Roberto Amato, Jeffrey Barrett, Sonia Goncalves, Ewan Harrison, David K. Jackson, Ian Johnston, Dominic Kwiatkowski, Cordelia Langford, John Sillitoe on behalf of the Wellcome Sanger Institute COVID-19 Surveillance Team |
| EPI_ISL_3362882 | Lighthouse Lab in Milton Keynes | Wellcome Sanger Institute for the COVID-19 Genomics UK (COG-UK) Consortium | The Lighthouse Lab in Milton Keynes and Alex Alderton, Roberto Amato, Jeffrey Barrett, Sonia Goncalves, Ewan Harrison, David K. Jackson, Ian Johnston, Dominic Kwiatkowski, Cordelia Langford, John Sillitoe on behalf of the Wellcome Sanger Institute COVID-19 Surveillance Team |
| EPI_ISL_3362884 | Lighthouse Lab in Glasgow | Wellcome Sanger Institute for the COVID-19 Genomics UK (COG-UK) Consortium | Harper VanSteenhouse, Yumi Kasai, David Gray, Carol Clugston, Anna Dominiczak and Alex Alderton, Roberto Amato, Jeffrey Barrett, Sonia Goncalves, Ewan Harrison, David K. Jackson, Ian Johnston, Dominic Kwiatkowski, Cordelia Langford, John Sillitoe on behalf of the Wellcome Sanger Institute COVID-19 Surveillance Team |
| EPI_ISL_3362885 | Lighthouse Lab in Alderley Park | Wellcome Sanger Institute for the COVID-19 Genomics UK (COG-UK) Consortium | Jacquelyn Wynn, Mairead Hyland, The Lighthouse Lab in Alderley Park and Alex Alderton, Roberto Amato, Jeffrey Barrett, Sonia Goncalves, Ewan Harrison, David K. Jackson, Ian Johnston, Dominic Kwiatkowski, Cordelia Langford, John Sillitoe on behalf of the Wellcome Sanger Institute COVID-19 Surveillance Team |
| EPI_ISL_3362887 | Lighthouse Lab in Milton Keynes | Wellcome Sanger Institute for the COVID-19 Genomics UK (COG-UK) Consortium | The Lighthouse Lab in Milton Keynes and Alex Alderton, Roberto Amato, Jeffrey Barrett, Sonia Goncalves, Ewan Harrison, David K. Jackson, Ian Johnston, Dominic Kwiatkowski, Cordelia Langford, John Sillitoe on behalf of the Wellcome Sanger Institute COVID-19 Surveillance Team |
| EPI_ISL_3362888 | Lighthouse Lab in Alderley Park | Wellcome Sanger Institute for the COVID-19 Genomics UK (COG-UK) Consortium | Jacquelyn Wynn, Mairead Hyland, The Lighthouse Lab in Alderley Park and Alex Alderton, Roberto Amato, Jeffrey Barrett, Sonia Goncalves, Ewan Harrison, David K. Jackson, Ian Johnston, Dominic Kwiatkowski, Cordelia Langford, John Sillitoe on behalf of the Wellcome Sanger Institute COVID-19 Surveillance Team |
| EPI_ISL_3362890, EPI_ISL_3362891, EPI_ISL_3362893, EPI_ISL_3362894 | Lighthouse Lab in Milton Keynes | Wellcome Sanger Institute for the COVID-19 Genomics UK (COG-UK) Consortium | The Lighthouse Lab in Milton Keynes and Alex Alderton, Roberto Amato, Jeffrey Barrett, Sonia Goncalves, Ewan Harrison, David K. Jackson, Ian Johnston, Dominic Kwiatkowski, Cordelia Langford, John Sillitoe on behalf of the Wellcome Sanger Institute COVID-19 Surveillance Team |
| EPI_ISL_3362895 | Lighthouse Lab in Glasgow | Wellcome Sanger Institute for the COVID-19 Genomics UK (COG-UK) Consortium | Harper VanSteenhouse, Yumi Kasai, David Gray, Carol Clugston, Anna Dominiczak and Alex Alderton, Roberto Amato, Jeffrey Barrett, Sonia Goncalves, Ewan Harrison, David K. Jackson, Ian Johnston, Dominic Kwiatkowski, Cordelia Langford, John Sillitoe on behalf of the Wellcome Sanger Institute COVID-19 Surveillance Team |
| EPI_ISL_3362896 | Lighthouse Lab in Milton Keynes | Wellcome Sanger Institute for the COVID-19 Genomics UK (COG-UK) Consortium | The Lighthouse Lab in Milton Keynes and Alex Alderton, Roberto Amato, Jeffrey Barrett, Sonia Goncalves, Ewan Harrison, David K. Jackson, Ian Johnston, Dominic Kwiatkowski, Cordelia Langford, John Sillitoe on behalf of the Wellcome Sanger Institute COVID-19 Surveillance Team |
| EPI_ISL_3362898, EPI_ISL_3362899 | Lighthouse Lab in Glasgow | Wellcome Sanger Institute for the COVID-19 Genomics UK (COG-UK) Consortium | Harper VanSteenhouse, Yumi Kasai, David Gray, Carol Clugston, Anna Dominiczak and Alex Alderton, Roberto Amato, Jeffrey Barrett, Sonia Goncalves, Ewan Harrison, David K. Jackson, Ian Johnston, Dominic Kwiatkowski, Cordelia Langford, John Sillitoe on behalf of the Wellcome Sanger Institute COVID-19 Surveillance Team |
| EPI_ISL_3362901 | Lighthouse Lab in Milton Keynes | Wellcome Sanger Institute for the COVID-19 Genomics UK (COG-UK) Consortium | The Lighthouse Lab in Milton Keynes and Alex Alderton, Roberto Amato, Jeffrey Barrett, Sonia Goncalves, Ewan Harrison, David K. Jackson, Ian Johnston, Dominic Kwiatkowski, Cordelia Langford, John Sillitoe on behalf of the Wellcome Sanger Institute COVID-19 Surveillance Team |
| EPI_ISL_3362902 | Lighthouse Lab in Glasgow | Wellcome Sanger Institute for the COVID-19 Genomics UK (COG-UK) Consortium | Harper VanSteenhouse, Yumi Kasai, David Gray, Carol Clugston, Anna Dominiczak and Alex Alderton, Roberto Amato, Jeffrey Barrett, Sonia Goncalves, Ewan Harrison, David K. Jackson, Ian Johnston, Dominic Kwiatkowski, Cordelia Langford, John Sillitoe on behalf of the Wellcome Sanger Institute COVID-19 Surveillance Team |
| EPI_ISL_3362904, EPI_ISL_3362905 | Lighthouse Lab in Alderley Park | Wellcome Sanger Institute for the COVID-19 Genomics UK (COG-UK) Consortium | Jacquelyn Wynn, Mairead Hyland, The Lighthouse Lab in Alderley Park and Alex Alderton, Roberto Amato, Jeffrey Barrett, Sonia Goncalves, Ewan Harrison, David K. Jackson, Ian Johnston, Dominic Kwiatkowski, Cordelia Langford, John Sillitoe on behalf of the Wellcome Sanger Institute COVID-19 Surveillance Team |
| EPI_ISL_3362906, EPI_ISL_3362908 | Lighthouse Lab in Glasgow | Wellcome Sanger Institute for the COVID-19 Genomics UK (COG-UK) Consortium | Harper VanSteenhouse, Yumi Kasai, David Gray, Carol Clugston, Anna Dominiczak and Alex Alderton, Roberto Amato, Jeffrey Barrett, Sonia Goncalves, Ewan Harrison, David K. Jackson, Ian Johnston, Dominic Kwiatkowski, Cordelia Langford, John Sillitoe on behalf of the Wellcome Sanger Institute COVID-19 Surveillance Team |
| EPI_ISL_3362909 | Lighthouse Lab in Milton Keynes | Wellcome Sanger Institute for the COVID-19 Genomics UK (COG-UK) Consortium | The Lighthouse Lab in Milton Keynes and Alex Alderton, Roberto Amato, Jeffrey Barrett, Sonia Goncalves, Ewan Harrison, David K. Jackson, Ian Johnston, Dominic Kwiatkowski, Cordelia Langford, John Sillitoe on behalf of the Wellcome Sanger Institute COVID-19 Surveillance Team |
| EPI_ISL_3362911, EPI_ISL_3362912, EPI_ISL_3362914 | Lighthouse Lab in Glasgow | Wellcome Sanger Institute for the COVID-19 Genomics UK (COG-UK) Consortium | Harper VanSteenhouse, Yumi Kasai, David Gray, Carol Clugston, Anna Dominiczak and Alex Alderton, Roberto Amato, Jeffrey Barrett, Sonia Goncalves, Ewan Harrison, David K. Jackson, Ian Johnston, Dominic Kwiatkowski, Cordelia Langford, John Sillitoe on behalf of the Wellcome Sanger Institute COVID-19 Surveillance Team |
| EPI_ISL_3362915, EPI_ISL_3362917 | Lighthouse Lab in Alderley Park | Wellcome Sanger Institute for the COVID-19 Genomics UK (COG-UK) Consortium | Jacquelyn Wynn, Mairead Hyland, The Lighthouse Lab in Alderley Park and Alex Alderton, Roberto Amato, Jeffrey Barrett, Sonia Goncalves, Ewan Harrison, David K. Jackson, Ian Johnston, Dominic Kwiatkowski, Cordelia Langford, John Sillitoe on behalf of the Wellcome Sanger Institute COVID-19 Surveillance Team |
| EPI_ISL_3362918, EPI_ISL_3362920 | Lighthouse Lab in Glasgow | Wellcome Sanger Institute for the COVID-19 Genomics UK (COG-UK) Consortium | Harper VanSteenhouse, Yumi Kasai, David Gray, Carol Clugston, Anna Dominiczak and Alex Alderton, Roberto Amato, Jeffrey Barrett, Sonia Goncalves, Ewan Harrison, David K. Jackson, Ian Johnston, Dominic Kwiatkowski, Cordelia Langford, John Sillitoe on behalf of the Wellcome Sanger Institute COVID-19 Surveillance Team |
| EPI_ISL_3362921 | Lighthouse Lab in Milton Keynes | Wellcome Sanger Institute for the COVID-19 Genomics UK (COG-UK) Consortium | The Lighthouse Lab in Milton Keynes and Alex Alderton, Roberto Amato, Jeffrey Barrett, Sonia Goncalves, Ewan Harrison, David K. Jackson, Ian Johnston, Dominic Kwiatkowski, Cordelia Langford, John Sillitoe on behalf of the Wellcome Sanger Institute COVID-19 Surveillance Team |
| EPI_ISL_3362923 | Lighthouse Lab in Alderley Park | Wellcome Sanger Institute for the COVID-19 Genomics UK (COG-UK) Consortium | Jacquelyn Wynn, Mairead Hyland, The Lighthouse Lab in Alderley Park and Alex Alderton, Roberto Amato, Jeffrey Barrett, Sonia Goncalves, Ewan Harrison, David K. Jackson, Ian Johnston, Dominic Kwiatkowski, Cordelia Langford, John Sillitoe on behalf of the Wellcome Sanger Institute COVID-19 Surveillance Team |

| | | | |
|---|---|---|---|
| EPI_ISL_3362924, EPI_ISL_3362925, EPI_ISL_3362927 | Lighthouse Lab in Glasgow | Wellcome Sanger Institute for the COVID-19 Genomics UK (COG-UK) Consortium | Harper VanSteenhouse, Yumi Kasai, David Gray, Carol Clugston, Anna Dominiczak and Alex Alderton, Roberto Amato, Jeffrey Barrett, Sonia Goncalves, Ewan Harrison, David K. Jackson, Ian Johnston, Dominic Kwiatkowski, Cordelia Langford, John Sillitoe on behalf of the Wellcome Sanger Institute COVID-19 Surveillance Team |
| EPI_ISL_3362928 | Lighthouse Lab in Alderley Park | Wellcome Sanger Institute for the COVID-19 Genomics UK (COG-UK) Consortium | Jacquelyn Wynn, Mairead Hyland, The Lighthouse Lab in Alderley Park and Alex Alderton, Roberto Amato, Jeffrey Barrett, Sonia Goncalves, Ewan Harrison, David K. Jackson, Ian Johnston, Dominic Kwiatkowski, Cordelia Langford, John Sillitoe on behalf of the Wellcome Sanger Institute COVID-19 Surveillance Team |
| EPI_ISL_3362930 | Lighthouse Lab in Glasgow | Wellcome Sanger Institute for the COVID-19 Genomics UK (COG-UK) Consortium | Harper VanSteenhouse, Yumi Kasai, David Gray, Carol Clugston, Anna Dominiczak and Alex Alderton, Roberto Amato, Jeffrey Barrett, Sonia Goncalves, Ewan Harrison, David K. Jackson, Ian Johnston, Dominic Kwiatkowski, Cordelia Langford, John Sillitoe on behalf of the Wellcome Sanger Institute COVID-19 Surveillance Team |
| EPI_ISL_3362931 | Lighthouse Lab in Alderley Park | Wellcome Sanger Institute for the COVID-19 Genomics UK (COG-UK) Consortium | Jacquelyn Wynn, Mairead Hyland, The Lighthouse Lab in Alderley Park and Alex Alderton, Roberto Amato, Jeffrey Barrett, Sonia Goncalves, Ewan Harrison, David K. Jackson, Ian Johnston, Dominic Kwiatkowski, Cordelia Langford, John Sillitoe on behalf of the Wellcome Sanger Institute COVID-19 Surveillance Team |
| EPI_ISL_3362932, EPI_ISL_3362934, EPI_ISL_3362936 | Lighthouse Lab in Glasgow | Wellcome Sanger Institute for the COVID-19 Genomics UK (COG-UK) Consortium | Harper VanSteenhouse, Yumi Kasai, David Gray, Carol Clugston, Anna Dominiczak and Alex Alderton, Roberto Amato, Jeffrey Barrett, Sonia Goncalves, Ewan Harrison, David K. Jackson, Ian Johnston, Dominic Kwiatkowski, Cordelia Langford, John Sillitoe on behalf of the Wellcome Sanger Institute COVID-19 Surveillance Team |
| EPI_ISL_3362937 | Lighthouse Lab in Milton Keynes | Wellcome Sanger Institute for the COVID-19 Genomics UK (COG-UK) Consortium | The Lighthouse Lab in Milton Keynes and Alex Alderton, Roberto Amato, Jeffrey Barrett, Sonia Goncalves, Ewan Harrison, David K. Jackson, Ian Johnston, Dominic Kwiatkowski, Cordelia Langford, John Sillitoe on behalf of the Wellcome Sanger Institute COVID-19 Surveillance Team |
| EPI_ISL_3362939, EPI_ISL_3362940 | Lighthouse Lab in Alderley Park | Wellcome Sanger Institute for the COVID-19 Genomics UK (COG-UK) Consortium | Jacquelyn Wynn, Mairead Hyland, The Lighthouse Lab in Alderley Park and Alex Alderton, Roberto Amato, Jeffrey Barrett, Sonia Goncalves, Ewan Harrison, David K. Jackson, Ian Johnston, Dominic Kwiatkowski, Cordelia Langford, John Sillitoe on behalf of the Wellcome Sanger Institute COVID-19 Surveillance Team |
| EPI_ISL_3362941, EPI_ISL_3362943, EPI_ISL_3362944 | Lighthouse Lab in Glasgow | Wellcome Sanger Institute for the COVID-19 Genomics UK (COG-UK) Consortium | Harper VanSteenhouse, Yumi Kasai, David Gray, Carol Clugston, Anna Dominiczak and Alex Alderton, Roberto Amato, Jeffrey Barrett, Sonia Goncalves, Ewan Harrison, David K. Jackson, Ian Johnston, Dominic Kwiatkowski, Cordelia Langford, John Sillitoe on behalf of the Wellcome Sanger Institute COVID-19 Surveillance Team |
| EPI_ISL_3362946, EPI_ISL_3362947 | Lighthouse Lab in Alderley Park | Wellcome Sanger Institute for the COVID-19 Genomics UK (COG-UK) Consortium | Jacquelyn Wynn, Mairead Hyland, The Lighthouse Lab in Alderley Park and Alex Alderton, Roberto Amato, Jeffrey Barrett, Sonia Goncalves, Ewan Harrison, David K. Jackson, Ian Johnston, Dominic Kwiatkowski, Cordelia Langford, John Sillitoe on behalf of the Wellcome Sanger Institute COVID-19 Surveillance Team |
| EPI_ISL_3362948 | Lighthouse Lab in Glasgow | Wellcome Sanger Institute for the COVID-19 Genomics UK (COG-UK) Consortium | Harper VanSteenhouse, Yumi Kasai, David Gray, Carol Clugston, Anna Dominiczak and Alex Alderton, Roberto Amato, Jeffrey Barrett, Sonia Goncalves, Ewan Harrison, David K. Jackson, Ian Johnston, Dominic Kwiatkowski, Cordelia Langford, John Sillitoe on behalf of the Wellcome Sanger Institute COVID-19 Surveillance Team |
| EPI_ISL_3362950 | Lighthouse Lab in Milton Keynes | Wellcome Sanger Institute for the COVID-19 Genomics UK (COG-UK) Consortium | The Lighthouse Lab in Milton Keynes and Alex Alderton, Roberto Amato, Jeffrey Barrett, Sonia Goncalves, Ewan Harrison, David K. Jackson, Ian Johnston, Dominic Kwiatkowski, Cordelia Langford, John Sillitoe on behalf of the Wellcome Sanger Institute COVID-19 Surveillance Team |
| EPI_ISL_3362951 | Lighthouse Lab in Alderley Park | Wellcome Sanger Institute for the COVID-19 Genomics UK (COG-UK) Consortium | Jacquelyn Wynn, Mairead Hyland, The Lighthouse Lab in Alderley Park and Alex Alderton, Roberto Amato, Jeffrey Barrett, Sonia Goncalves, Ewan Harrison, David K. Jackson, Ian Johnston, Dominic Kwiatkowski, Cordelia Langford, John Sillitoe on behalf of the Wellcome Sanger Institute COVID-19 Surveillance Team |
| EPI_ISL_3362953, EPI_ISL_3362954 | Lighthouse Lab in Milton Keynes | Wellcome Sanger Institute for the COVID-19 Genomics UK (COG-UK) Consortium | The Lighthouse Lab in Milton Keynes and Alex Alderton, Roberto Amato, Jeffrey Barrett, Sonia Goncalves, Ewan Harrison, David K. Jackson, Ian Johnston, Dominic Kwiatkowski, Cordelia Langford, John Sillitoe on behalf of the Wellcome Sanger Institute COVID-19 Surveillance Team |
| EPI_ISL_3362956, EPI_ISL_3362957 | Lighthouse Lab in Alderley Park | Wellcome Sanger Institute for the COVID-19 Genomics UK (COG-UK) Consortium | Jacquelyn Wynn, Mairead Hyland, The Lighthouse Lab in Alderley Park and Alex Alderton, Roberto Amato, Jeffrey Barrett, Sonia Goncalves, Ewan Harrison, David K. Jackson, Ian Johnston, Dominic Kwiatkowski, Cordelia Langford, John Sillitoe on behalf of the Wellcome Sanger Institute COVID-19 Surveillance Team |
| EPI_ISL_3362959 | Lighthouse Lab in Glasgow | Wellcome Sanger Institute for the COVID-19 Genomics UK (COG-UK) Consortium | Harper VanSteenhouse, Yumi Kasai, David Gray, Carol Clugston, Anna Dominiczak and Alex Alderton, Roberto Amato, Jeffrey Barrett, Sonia Goncalves, Ewan Harrison, David K. Jackson, Ian Johnston, Dominic Kwiatkowski, Cordelia Langford, John Sillitoe on behalf of the Wellcome Sanger Institute COVID-19 Surveillance Team |
| EPI_ISL_3362960 | Lighthouse Lab in Milton Keynes | Wellcome Sanger Institute for the COVID-19 Genomics UK (COG-UK) Consortium | The Lighthouse Lab in Milton Keynes and Alex Alderton, Roberto Amato, Jeffrey Barrett, Sonia Goncalves, Ewan Harrison, David K. Jackson, Ian Johnston, Dominic Kwiatkowski, Cordelia Langford, John Sillitoe on behalf of the Wellcome Sanger Institute COVID-19 Surveillance Team |
| EPI_ISL_3362961, EPI_ISL_3362963, EPI_ISL_3362964 | Lighthouse Lab in Glasgow | Wellcome Sanger Institute for the COVID-19 Genomics UK (COG-UK) Consortium | Harper VanSteenhouse, Yumi Kasai, David Gray, Carol Clugston, Anna Dominiczak and Alex Alderton, Roberto Amato, Jeffrey Barrett, Sonia Goncalves, Ewan Harrison, David K. Jackson, Ian Johnston, Dominic Kwiatkowski, Cordelia Langford, John Sillitoe on behalf of the Wellcome Sanger Institute COVID-19 Surveillance Team |
| EPI_ISL_3362966 | Lighthouse Lab in Alderley Park | Wellcome Sanger Institute for the COVID-19 Genomics UK (COG-UK) Consortium | Jacquelyn Wynn, Mairead Hyland, The Lighthouse Lab in Alderley Park and Alex Alderton, Roberto Amato, Jeffrey Barrett, Sonia Goncalves, Ewan Harrison, David K. Jackson, Ian Johnston, Dominic Kwiatkowski, Cordelia Langford, John Sillitoe on behalf of the Wellcome Sanger Institute COVID-19 Surveillance Team |
| EPI_ISL_3362967, EPI_ISL_3362969 | Lighthouse Lab in Milton Keynes | Wellcome Sanger Institute for the COVID-19 Genomics UK (COG-UK) Consortium | The Lighthouse Lab in Milton Keynes and Alex Alderton, Roberto Amato, Jeffrey Barrett, Sonia Goncalves, Ewan Harrison, David K. Jackson, Ian Johnston, Dominic Kwiatkowski, Cordelia Langford, John Sillitoe on behalf of the Wellcome Sanger Institute COVID-19 Surveillance Team |
| EPI_ISL_3362970, EPI_ISL_3362972 | Lighthouse Lab in Glasgow | Wellcome Sanger Institute for the COVID-19 Genomics UK (COG-UK) Consortium | Harper VanSteenhouse, Yumi Kasai, David Gray, Carol Clugston, Anna Dominiczak and Alex Alderton, Roberto Amato, Jeffrey Barrett, Sonia Goncalves, Ewan Harrison, David K. Jackson, Ian Johnston, Dominic Kwiatkowski, Cordelia Langford, John Sillitoe on behalf of the Wellcome Sanger Institute COVID-19 Surveillance Team |
| EPI_ISL_3362973, EPI_ISL_3362975, EPI_ISL_3362976 | Lighthouse Lab in Milton Keynes | Wellcome Sanger Institute for the COVID-19 Genomics UK (COG-UK) Consortium | The Lighthouse Lab in Milton Keynes and Alex Alderton, Roberto Amato, Jeffrey Barrett, Sonia Goncalves, Ewan Harrison, David K. Jackson, Ian Johnston, Dominic Kwiatkowski, Cordelia Langford, John Sillitoe on behalf of the Wellcome Sanger Institute COVID-19 Surveillance Team |
| EPI_ISL_3362978, EPI_ISL_3362979 | Lighthouse Lab in Alderley Park | Wellcome Sanger Institute for the COVID-19 Genomics UK (COG-UK) Consortium | Jacquelyn Wynn, Mairead Hyland, The Lighthouse Lab in Alderley Park and Alex Alderton, Roberto Amato, Jeffrey Barrett, Sonia Goncalves, Ewan Harrison, David K. Jackson, Ian Johnston, Dominic Kwiatkowski, Cordelia Langford, John Sillitoe on behalf of the Wellcome Sanger Institute COVID-19 Surveillance Team |
| EPI_ISL_3362980, EPI_ISL_3362982 | Lighthouse Lab in Glasgow | Wellcome Sanger Institute for the COVID-19 Genomics UK (COG-UK) Consortium | Harper VanSteenhouse, Yumi Kasai, David Gray, Carol Clugston, Anna Dominiczak and Alex Alderton, Roberto Amato, Jeffrey Barrett, Sonia Goncalves, Ewan Harrison, David K. Jackson, Ian Johnston, Dominic Kwiatkowski, Cordelia Langford, John Sillitoe on behalf of the Wellcome Sanger Institute COVID-19 Surveillance Team |
| EPI_ISL_3362983 | Lighthouse Lab in Milton Keynes | Wellcome Sanger Institute for the COVID-19 Genomics UK (COG-UK) Consortium | The Lighthouse Lab in Milton Keynes and Alex Alderton, Roberto Amato, Jeffrey Barrett, Sonia Goncalves, Ewan Harrison, David K. Jackson, Ian Johnston, Dominic Kwiatkowski, Cordelia Langford, John Sillitoe on behalf of the Wellcome Sanger Institute COVID-19 Surveillance Team |
| EPI_ISL_3362985 | Lighthouse Lab in Alderley Park | Wellcome Sanger Institute for the COVID-19 Genomics UK (COG-UK) Consortium | Jacquelyn Wynn, Mairead Hyland, The Lighthouse Lab in Alderley Park and Alex Alderton, Roberto Amato, Jeffrey Barrett, Sonia Goncalves, Ewan Harrison, David K. Jackson, Ian Johnston, Dominic Kwiatkowski, Cordelia Langford, John Sillitoe on behalf of the Wellcome Sanger Institute COVID-19 Surveillance Team |
| EPI_ISL_3362986 | Lighthouse Lab in Glasgow | Wellcome Sanger Institute for the COVID-19 Genomics UK (COG-UK) Consortium | Harper VanSteenhouse, Yumi Kasai, David Gray, Carol Clugston, Anna Dominiczak and Alex Alderton, Roberto Amato, Jeffrey Barrett, Sonia Goncalves, Ewan Harrison, David K. Jackson, Ian Johnston, Dominic Kwiatkowski, Cordelia Langford, John Sillitoe on behalf of the Wellcome Sanger Institute COVID-19 Surveillance Team |
| EPI_ISL_3362988, EPI_ISL_3362989 | Lighthouse Lab in Alderley Park | Wellcome Sanger Institute for the COVID-19 Genomics UK (COG-UK) Consortium | Jacquelyn Wynn, Mairead Hyland, The Lighthouse Lab in Alderley Park and Alex Alderton, Roberto Amato, Jeffrey Barrett, Sonia Goncalves, Ewan Harrison, David K. Jackson, Ian Johnston, Dominic Kwiatkowski, Cordelia Langford, John Sillitoe on behalf of the Wellcome Sanger Institute COVID-19 Surveillance Team |
| EPI_ISL_3362991, EPI_ISL_3362992 | Lighthouse Lab in Glasgow | Wellcome Sanger Institute for the COVID-19 Genomics UK (COG-UK) Consortium | Harper VanSteenhouse, Yumi Kasai, David Gray, Carol Clugston, Anna Dominiczak and Alex Alderton, Roberto Amato, Jeffrey Barrett, Sonia Goncalves, Ewan Harrison, David K. Jackson, Ian Johnston, Dominic Kwiatkowski, Cordelia Langford, John Sillitoe on behalf of the Wellcome Sanger Institute COVID-19 Surveillance Team |
| EPI_ISL_3362993 | Lighthouse Lab in Milton Keynes | Wellcome Sanger Institute for the COVID-19 Genomics UK (COG-UK) Consortium | The Lighthouse Lab in Milton Keynes and Alex Alderton, Roberto Amato, Jeffrey Barrett, Sonia Goncalves, Ewan Harrison, David K. Jackson, Ian Johnston, Dominic Kwiatkowski, Cordelia Langford, John Sillitoe on behalf of the Wellcome Sanger Institute COVID-19 Surveillance Team |
| EPI_ISL_3362995 | Lighthouse Lab in Glasgow | Wellcome Sanger Institute for the COVID-19 Genomics UK (COG-UK) Consortium | Harper VanSteenhouse, Yumi Kasai, David Gray, Carol Clugston, Anna Dominiczak and Alex Alderton, Roberto Amato, Jeffrey Barrett, Sonia Goncalves, Ewan Harrison, David K. Jackson, Ian Johnston, Dominic Kwiatkowski, Cordelia Langford, John Sillitoe on behalf of the Wellcome Sanger Institute COVID-19 Surveillance Team |
| EPI_ISL_3362996 | Lighthouse Lab in Alderley Park | Wellcome Sanger Institute for the COVID-19 Genomics UK (COG-UK) Consortium | Jacquelyn Wynn, Mairead Hyland, The Lighthouse Lab in Alderley Park and Alex Alderton, Roberto Amato, Jeffrey Barrett, Sonia Goncalves, Ewan Harrison, David K. Jackson, Ian Johnston, Dominic Kwiatkowski, Cordelia Langford, John Sillitoe on behalf of the Wellcome Sanger Institute COVID-19 Surveillance Team |
| EPI_ISL_3362998 | Lighthouse Lab in Glasgow | Wellcome Sanger Institute for the COVID-19 Genomics UK (COG-UK) Consortium | Harper VanSteenhouse, Yumi Kasai, David Gray, Carol Clugston, Anna Dominiczak and Alex Alderton, Roberto Amato, Jeffrey Barrett, Sonia Goncalves, Ewan Harrison, David K. Jackson, Ian Johnston, Dominic Kwiatkowski, Cordelia Langford, John Sillitoe on behalf of the Wellcome Sanger Institute COVID-19 Surveillance Team |
| EPI_ISL_3362999 | Lighthouse Lab in Alderley Park | Wellcome Sanger Institute for the COVID-19 Genomics UK (COG-UK) Consortium | Jacquelyn Wynn, Mairead Hyland, The Lighthouse Lab in Alderley Park and Alex Alderton, Roberto Amato, Jeffrey Barrett, Sonia Goncalves, Ewan Harrison, David K. Jackson, Ian Johnston, Dominic Kwiatkowski, Cordelia Langford, John Sillitoe on behalf of the Wellcome Sanger Institute COVID-19 Surveillance Team |
| EPI_ISL_3363001 | Lighthouse Lab in Milton Keynes | Wellcome Sanger Institute for the COVID-19 Genomics UK (COG-UK) Consortium | The Lighthouse Lab in Milton Keynes and Alex Alderton, Roberto Amato, Jeffrey Barrett, Sonia Goncalves, Ewan Harrison, David K. Jackson, Ian Johnston, Dominic Kwiatkowski, Cordelia Langford, John Sillitoe on behalf of the Wellcome Sanger Institute COVID-19 Surveillance Team |
| EPI_ISL_3363002 | Lighthouse Lab in Alderley Park | Wellcome Sanger Institute for the COVID-19 Genomics UK (COG-UK) Consortium | Jacquelyn Wynn, Mairead Hyland, The Lighthouse Lab in Alderley Park and Alex Alderton, Roberto Amato, Jeffrey Barrett, Sonia Goncalves, Ewan Harrison, David K. Jackson, Ian Johnston, Dominic Kwiatkowski, Cordelia Langford, John Sillitoe on behalf of the Wellcome Sanger Institute COVID-19 Surveillance Team |
| EPI_ISL_3363004, EPI_ISL_3363005 | Lighthouse Lab in Milton Keynes | Wellcome Sanger Institute for the COVID-19 Genomics UK (COG-UK) Consortium | The Lighthouse Lab in Milton Keynes and Alex Alderton, Roberto Amato, Jeffrey Barrett, Sonia Goncalves, Ewan Harrison, David K. Jackson, Ian Johnston, Dominic Kwiatkowski, Cordelia Langford, John Sillitoe on behalf of the Wellcome Sanger Institute COVID-19 Surveillance Team |
| EPI_ISL_3363006 | Lighthouse Lab in Glasgow | Wellcome Sanger Institute for the COVID-19 Genomics UK (COG-UK) Consortium | Harper VanSteenhouse, Yumi Kasai, David Gray, Carol Clugston, Anna Dominiczak and Alex Alderton, Roberto Amato, Jeffrey Barrett, Sonia Goncalves, Ewan Harrison, David K. Jackson, Ian Johnston, Dominic Kwiatkowski, Cordelia Langford, John Sillitoe on behalf of the Wellcome Sanger Institute COVID-19 Surveillance Team |
| EPI_ISL_3363008 | Lighthouse Lab in Alderley Park | Wellcome Sanger Institute for the COVID-19 Genomics UK (COG-UK) Consortium | Jacquelyn Wynn, Mairead Hyland, The Lighthouse Lab in Alderley Park and Alex Alderton, Roberto Amato, Jeffrey Barrett, Sonia Goncalves, Ewan Harrison, David K. Jackson, Ian Johnston, Dominic Kwiatkowski, Cordelia Langford, John Sillitoe on behalf of the Wellcome Sanger Institute COVID-19 Surveillance Team |
| EPI_ISL_3363009, EPI_ISL_3363011 | Lighthouse Lab in Glasgow | Wellcome Sanger Institute for the COVID-19 Genomics UK (COG-UK) Consortium | Harper VanSteenhouse, Yumi Kasai, David Gray, Carol Clugston, Anna Dominiczak and Alex Alderton, Roberto Amato, Jeffrey Barrett, Sonia Goncalves, Ewan Harrison, David K. Jackson, Ian Johnston, Dominic Kwiatkowski, Cordelia Langford, John Sillitoe on behalf of the Wellcome Sanger Institute COVID-19 Surveillance Team |
| EPI_ISL_3363013, EPI_ISL_3363014 | Lighthouse Lab in Alderley Park | Wellcome Sanger Institute for the COVID-19 Genomics UK (COG-UK) Consortium | Jacquelyn Wynn, Mairead Hyland, The Lighthouse Lab in Alderley Park and Alex Alderton, Roberto Amato, Jeffrey Barrett, Sonia Goncalves, Ewan Harrison, David K. Jackson, Ian Johnston, Dominic Kwiatkowski, Cordelia Langford, John Sillitoe on behalf of the Wellcome Sanger Institute COVID-19 Surveillance Team |
| EPI_ISL_3363015, EPI_ISL_3363017, EPI_ISL_3363019 | Lighthouse Lab in Milton Keynes | Wellcome Sanger Institute for the COVID-19 Genomics UK (COG-UK) Consortium | The Lighthouse Lab in Milton Keynes and Alex Alderton, Roberto Amato, Jeffrey Barrett, Sonia Goncalves, Ewan Harrison, David K. Jackson, Ian Johnston, Dominic Kwiatkowski, Cordelia Langford, John Sillitoe on behalf of the Wellcome Sanger Institute COVID-19 Surveillance Team |
| EPI_ISL_3363020 | Lighthouse Lab in Alderley Park | Wellcome Sanger Institute for the COVID-19 Genomics UK (COG-UK) Consortium | Jacquelyn Wynn, Mairead Hyland, The Lighthouse Lab in Alderley Park and Alex Alderton, Roberto Amato, Jeffrey Barrett, Sonia Goncalves, Ewan Harrison, David K. Jackson, Ian Johnston, Dominic Kwiatkowski, Cordelia Langford, John Sillitoe on behalf of the Wellcome Sanger Institute COVID-19 Surveillance Team |
| EPI_ISL_3363022 | Lighthouse Lab in Glasgow | Wellcome Sanger Institute for the COVID-19 Genomics UK (COG-UK) Consortium | Harper VanSteenhouse, Yumi Kasai, David Gray, Carol Clugston, Anna Dominiczak and Alex Alderton, Roberto Amato, Jeffrey Barrett, Sonia Goncalves, Ewan Harrison, David K. Jackson, Ian Johnston, Dominic Kwiatkowski, Cordelia Langford, John Sillitoe on behalf of the Wellcome Sanger Institute COVID-19 Surveillance Team |

| | | | |
|---|---|---|---|
| EPI_ISL_3363023 | Lighthouse Lab in Milton Keynes | Wellcome Sanger Institute for the COVID-19 Genomics UK (COG-UK) Consortium | The Lighthouse Lab in Milton Keynes and Alex Alderton, Roberto Amato, Jeffrey Barrett, Sonia Goncalves, Ewan Harrison, David K. Jackson, Ian Johnston, Dominic Kwiatkowski, Cordelia Langford, John Sillitoe on behalf of the Wellcome Sanger Institute COVID-19 Surveillance Team |
| EPI_ISL_3363025, EPI_ISL_3363027, EPI_ISL_3363028 | Lighthouse Lab in Glasgow | Wellcome Sanger Institute for the COVID-19 Genomics UK (COG-UK) Consortium | Harper VanSteenhouse, Yumi Kasai, David Gray, Carol Clugston, Anna Dominiczak and Alex Alderton, Roberto Amato, Jeffrey Barrett, Sonia Goncalves, Ewan Harrison, David K. Jackson, Ian Johnston, Dominic Kwiatkowski, Cordelia Langford, John Sillitoe on behalf of the Wellcome Sanger Institute COVID-19 Surveillance Team |
| EPI_ISL_3363029 | Lighthouse Lab in Alderley Park | Wellcome Sanger Institute for the COVID-19 Genomics UK (COG-UK) Consortium | Jacquelyn Wynn, Mairead Hyland, The Lighthouse Lab in Alderley Park and Alex Alderton, Roberto Amato, Jeffrey Barrett, Sonia Goncalves, Ewan Harrison, David K. Jackson, Ian Johnston, Dominic Kwiatkowski, Cordelia Langford, John Sillitoe on behalf of the Wellcome Sanger Institute COVID-19 Surveillance Team |
| EPI_ISL_3363031 | Lighthouse Lab in Milton Keynes | Wellcome Sanger Institute for the COVID-19 Genomics UK (COG-UK) Consortium | The Lighthouse Lab in Milton Keynes and Alex Alderton, Roberto Amato, Jeffrey Barrett, Sonia Goncalves, Ewan Harrison, David K. Jackson, Ian Johnston, Dominic Kwiatkowski, Cordelia Langford, John Sillitoe on behalf of the Wellcome Sanger Institute COVID-19 Surveillance Team |
| EPI_ISL_3363032 | Lighthouse Lab in Glasgow | Wellcome Sanger Institute for the COVID-19 Genomics UK (COG-UK) Consortium | Harper VanSteenhouse, Yumi Kasai, David Gray, Carol Clugston, Anna Dominiczak and Alex Alderton, Roberto Amato, Jeffrey Barrett, Sonia Goncalves, Ewan Harrison, David K. Jackson, Ian Johnston, Dominic Kwiatkowski, Cordelia Langford, John Sillitoe on behalf of the Wellcome Sanger Institute COVID-19 Surveillance Team |
| EPI_ISL_3363034, EPI_ISL_3363035 | Lighthouse Lab in Milton Keynes | Wellcome Sanger Institute for the COVID-19 Genomics UK (COG-UK) Consortium | The Lighthouse Lab in Milton Keynes and Alex Alderton, Roberto Amato, Jeffrey Barrett, Sonia Goncalves, Ewan Harrison, David K. Jackson, Ian Johnston, Dominic Kwiatkowski, Cordelia Langford, John Sillitoe on behalf of the Wellcome Sanger Institute COVID-19 Surveillance Team |
| EPI_ISL_3363085 | Lighthouse Lab in Glasgow | Wellcome Sanger Institute for the COVID-19 Genomics UK (COG-UK) Consortium | Harper VanSteenhouse, Yumi Kasai, David Gray, Carol Clugston, Anna Dominiczak and Alex Alderton, Roberto Amato, Jeffrey Barrett, Sonia Goncalves, Ewan Harrison, David K. Jackson, Ian Johnston, Dominic Kwiatkowski, Cordelia Langford, John Sillitoe on behalf of the Wellcome Sanger Institute COVID-19 Surveillance Team |
| EPI_ISL_3363087 | Lighthouse Lab in Milton Keynes | Wellcome Sanger Institute for the COVID-19 Genomics UK (COG-UK) Consortium | The Lighthouse Lab in Milton Keynes and Alex Alderton, Roberto Amato, Jeffrey Barrett, Sonia Goncalves, Ewan Harrison, David K. Jackson, Ian Johnston, Dominic Kwiatkowski, Cordelia Langford, John Sillitoe on behalf of the Wellcome Sanger Institute COVID-19 Surveillance Team |
| EPI_ISL_3363088 | Lighthouse Lab in Alderley Park | Wellcome Sanger Institute for the COVID-19 Genomics UK (COG-UK) Consortium | Jacquelyn Wynn, Mairead Hyland, The Lighthouse Lab in Alderley Park and Alex Alderton, Roberto Amato, Jeffrey Barrett, Sonia Goncalves, Ewan Harrison, David K. Jackson, Ian Johnston, Dominic Kwiatkowski, Cordelia Langford, John Sillitoe on behalf of the Wellcome Sanger Institute COVID-19 Surveillance Team |
| EPI_ISL_3363090 | Lighthouse Lab in Milton Keynes | Wellcome Sanger Institute for the COVID-19 Genomics UK (COG-UK) Consortium | The Lighthouse Lab in Milton Keynes and Alex Alderton, Roberto Amato, Jeffrey Barrett, Sonia Goncalves, Ewan Harrison, David K. Jackson, Ian Johnston, Dominic Kwiatkowski, Cordelia Langford, John Sillitoe on behalf of the Wellcome Sanger Institute COVID-19 Surveillance Team |
| EPI_ISL_3363091 | Lighthouse Lab in Glasgow | Wellcome Sanger Institute for the COVID-19 Genomics UK (COG-UK) Consortium | Harper VanSteenhouse, Yumi Kasai, David Gray, Carol Clugston, Anna Dominiczak and Alex Alderton, Roberto Amato, Jeffrey Barrett, Sonia Goncalves, Ewan Harrison, David K. Jackson, Ian Johnston, Dominic Kwiatkowski, Cordelia Langford, John Sillitoe on behalf of the Wellcome Sanger Institute COVID-19 Surveillance Team |
| EPI_ISL_3363094, EPI_ISL_3363095 | Lighthouse Lab in Alderley Park | Wellcome Sanger Institute for the COVID-19 Genomics UK (COG-UK) Consortium | Jacquelyn Wynn, Mairead Hyland, The Lighthouse Lab in Alderley Park and Alex Alderton, Roberto Amato, Jeffrey Barrett, Sonia Goncalves, Ewan Harrison, David K. Jackson, Ian Johnston, Dominic Kwiatkowski, Cordelia Langford, John Sillitoe on behalf of the Wellcome Sanger Institute COVID-19 Surveillance Team |
| EPI_ISL_3363097 | Lighthouse Lab in Glasgow | Wellcome Sanger Institute for the COVID-19 Genomics UK (COG-UK) Consortium | Harper VanSteenhouse, Yumi Kasai, David Gray, Carol Clugston, Anna Dominiczak and Alex Alderton, Roberto Amato, Jeffrey Barrett, Sonia Goncalves, Ewan Harrison, David K. Jackson, Ian Johnston, Dominic Kwiatkowski, Cordelia Langford, John Sillitoe on behalf of the Wellcome Sanger Institute COVID-19 Surveillance Team |
| EPI_ISL_3363099, EPI_ISL_3363100, EPI_ISL_3363101, EPI_ISL_3363103 | Lighthouse Lab in Alderley Park | Wellcome Sanger Institute for the COVID-19 Genomics UK (COG-UK) Consortium | Jacquelyn Wynn, Mairead Hyland, The Lighthouse Lab in Alderley Park and Alex Alderton, Roberto Amato, Jeffrey Barrett, Sonia Goncalves, Ewan Harrison, David K. Jackson, Ian Johnston, Dominic Kwiatkowski, Cordelia Langford, John Sillitoe on behalf of the Wellcome Sanger Institute COVID-19 Surveillance Team |
| EPI_ISL_3363104 | Lighthouse Lab in Milton Keynes | Wellcome Sanger Institute for the COVID-19 Genomics UK (COG-UK) Consortium | The Lighthouse Lab in Milton Keynes and Alex Alderton, Roberto Amato, Jeffrey Barrett, Sonia Goncalves, Ewan Harrison, David K. Jackson, Ian Johnston, Dominic Kwiatkowski, Cordelia Langford, John Sillitoe on behalf of the Wellcome Sanger Institute COVID-19 Surveillance Team |
| EPI_ISL_3363106 | Lighthouse Lab in Alderley Park | Wellcome Sanger Institute for the COVID-19 Genomics UK (COG-UK) Consortium | Jacquelyn Wynn, Mairead Hyland, The Lighthouse Lab in Alderley Park and Alex Alderton, Roberto Amato, Jeffrey Barrett, Sonia Goncalves, Ewan Harrison, David K. Jackson, Ian Johnston, Dominic Kwiatkowski, Cordelia Langford, John Sillitoe on behalf of the Wellcome Sanger Institute COVID-19 Surveillance Team |
| EPI_ISL_3363107, EPI_ISL_3363108 | Lighthouse Lab in Milton Keynes | Wellcome Sanger Institute for the COVID-19 Genomics UK (COG-UK) Consortium | The Lighthouse Lab in Milton Keynes and Alex Alderton, Roberto Amato, Jeffrey Barrett, Sonia Goncalves, Ewan Harrison, David K. Jackson, Ian Johnston, Dominic Kwiatkowski, Cordelia Langford, John Sillitoe on behalf of the Wellcome Sanger Institute COVID-19 Surveillance Team |
| EPI_ISL_3363110 | Lighthouse Lab in Alderley Park | Wellcome Sanger Institute for the COVID-19 Genomics UK (COG-UK) Consortium | Jacquelyn Wynn, Mairead Hyland, The Lighthouse Lab in Alderley Park and Alex Alderton, Roberto Amato, Jeffrey Barrett, Sonia Goncalves, Ewan Harrison, David K. Jackson, Ian Johnston, Dominic Kwiatkowski, Cordelia Langford, John Sillitoe on behalf of the Wellcome Sanger Institute COVID-19 Surveillance Team |
| EPI_ISL_3363111 | Lighthouse Lab in Glasgow | Wellcome Sanger Institute for the COVID-19 Genomics UK (COG-UK) Consortium | Harper VanSteenhouse, Yumi Kasai, David Gray, Carol Clugston, Anna Dominiczak and Alex Alderton, Roberto Amato, Jeffrey Barrett, Sonia Goncalves, Ewan Harrison, David K. Jackson, Ian Johnston, Dominic Kwiatkowski, Cordelia Langford, John Sillitoe on behalf of the Wellcome Sanger Institute COVID-19 Surveillance Team |
| EPI_ISL_3363113, EPI_ISL_3363114, EPI_ISL_3363116 | Lighthouse Lab in Alderley Park | Wellcome Sanger Institute for the COVID-19 Genomics UK (COG-UK) Consortium | Jacquelyn Wynn, Mairead Hyland, The Lighthouse Lab in Alderley Park and Alex Alderton, Roberto Amato, Jeffrey Barrett, Sonia Goncalves, Ewan Harrison, David K. Jackson, Ian Johnston, Dominic Kwiatkowski, Cordelia Langford, John Sillitoe on behalf of the Wellcome Sanger Institute COVID-19 Surveillance Team |
| EPI_ISL_3363117 | Lighthouse Lab in Milton Keynes | Wellcome Sanger Institute for the COVID-19 Genomics UK (COG-UK) Consortium | The Lighthouse Lab in Milton Keynes and Alex Alderton, Roberto Amato, Jeffrey Barrett, Sonia Goncalves, Ewan Harrison, David K. Jackson, Ian Johnston, Dominic Kwiatkowski, Cordelia Langford, John Sillitoe on behalf of the Wellcome Sanger Institute COVID-19 Surveillance Team |
| EPI_ISL_3363118, EPI_ISL_3363120 | Lighthouse Lab in Alderley Park | Wellcome Sanger Institute for the COVID-19 Genomics UK (COG-UK) Consortium | Jacquelyn Wynn, Mairead Hyland, The Lighthouse Lab in Alderley Park and Alex Alderton, Roberto Amato, Jeffrey Barrett, Sonia Goncalves, Ewan Harrison, David K. Jackson, Ian Johnston, Dominic Kwiatkowski, Cordelia Langford, John Sillitoe on behalf of the Wellcome Sanger Institute COVID-19 Surveillance Team |
| EPI_ISL_3363121 | Lighthouse Lab in Milton Keynes | Wellcome Sanger Institute for the COVID-19 Genomics UK (COG-UK) Consortium | The Lighthouse Lab in Milton Keynes and Alex Alderton, Roberto Amato, Jeffrey Barrett, Sonia Goncalves, Ewan Harrison, David K. Jackson, Ian Johnston, Dominic Kwiatkowski, Cordelia Langford, John Sillitoe on behalf of the Wellcome Sanger Institute COVID-19 Surveillance Team |
| EPI_ISL_3363123, EPI_ISL_3363124 | Lighthouse Lab in Glasgow | Wellcome Sanger Institute for the COVID-19 Genomics UK (COG-UK) Consortium | Harper VanSteenhouse, Yumi Kasai, David Gray, Carol Clugston, Anna Dominiczak and Alex Alderton, Roberto Amato, Jeffrey Barrett, Sonia Goncalves, Ewan Harrison, David K. Jackson, Ian Johnston, Dominic Kwiatkowski, Cordelia Langford, John Sillitoe on behalf of the Wellcome Sanger Institute COVID-19 Surveillance Team |
| EPI_ISL_3363125 | Lighthouse Lab in Milton Keynes | Wellcome Sanger Institute for the COVID-19 Genomics UK (COG-UK) Consortium | The Lighthouse Lab in Milton Keynes and Alex Alderton, Roberto Amato, Jeffrey Barrett, Sonia Goncalves, Ewan Harrison, David K. Jackson, Ian Johnston, Dominic Kwiatkowski, Cordelia Langford, John Sillitoe on behalf of the Wellcome Sanger Institute COVID-19 Surveillance Team |
| EPI_ISL_3363127 | Lighthouse Lab in Alderley Park | Wellcome Sanger Institute for the COVID-19 Genomics UK (COG-UK) Consortium | Jacquelyn Wynn, Mairead Hyland, The Lighthouse Lab in Alderley Park and Alex Alderton, Roberto Amato, Jeffrey Barrett, Sonia Goncalves, Ewan Harrison, David K. Jackson, Ian Johnston, Dominic Kwiatkowski, Cordelia Langford, John Sillitoe on behalf of the Wellcome Sanger Institute COVID-19 Surveillance Team |
| EPI_ISL_3363128, EPI_ISL_3363130 | Lighthouse Lab in Milton Keynes | Wellcome Sanger Institute for the COVID-19 Genomics UK (COG-UK) Consortium | The Lighthouse Lab in Milton Keynes and Alex Alderton, Roberto Amato, Jeffrey Barrett, Sonia Goncalves, Ewan Harrison, David K. Jackson, Ian Johnston, Dominic Kwiatkowski, Cordelia Langford, John Sillitoe on behalf of the Wellcome Sanger Institute COVID-19 Surveillance Team |
| EPI_ISL_3363131 | Lighthouse Lab in Glasgow | Wellcome Sanger Institute for the COVID-19 Genomics UK (COG-UK) Consortium | Harper VanSteenhouse, Yumi Kasai, David Gray, Carol Clugston, Anna Dominiczak and Alex Alderton, Roberto Amato, Jeffrey Barrett, Sonia Goncalves, Ewan Harrison, David K. Jackson, Ian Johnston, Dominic Kwiatkowski, Cordelia Langford, John Sillitoe on behalf of the Wellcome Sanger Institute COVID-19 Surveillance Team |
| EPI_ISL_3363133 | Lighthouse Lab in Alderley Park | Wellcome Sanger Institute for the COVID-19 Genomics UK (COG-UK) Consortium | Jacquelyn Wynn, Mairead Hyland, The Lighthouse Lab in Alderley Park and Alex Alderton, Roberto Amato, Jeffrey Barrett, Sonia Goncalves, Ewan Harrison, David K. Jackson, Ian Johnston, Dominic Kwiatkowski, Cordelia Langford, John Sillitoe on behalf of the Wellcome Sanger Institute COVID-19 Surveillance Team |
| EPI_ISL_3363134, EPI_ISL_3363136 | Lighthouse Lab in Glasgow | Wellcome Sanger Institute for the COVID-19 Genomics UK (COG-UK) Consortium | Harper VanSteenhouse, Yumi Kasai, David Gray, Carol Clugston, Anna Dominiczak and Alex Alderton, Roberto Amato, Jeffrey Barrett, Sonia Goncalves, Ewan Harrison, David K. Jackson, Ian Johnston, Dominic Kwiatkowski, Cordelia Langford, John Sillitoe on behalf of the Wellcome Sanger Institute COVID-19 Surveillance Team |
| EPI_ISL_3363137 | Lighthouse Lab in Milton Keynes | Wellcome Sanger Institute for the COVID-19 Genomics UK (COG-UK) Consortium | The Lighthouse Lab in Milton Keynes and Alex Alderton, Roberto Amato, Jeffrey Barrett, Sonia Goncalves, Ewan Harrison, David K. Jackson, Ian Johnston, Dominic Kwiatkowski, Cordelia Langford, John Sillitoe on behalf of the Wellcome Sanger Institute COVID-19 Surveillance Team |
| EPI_ISL_3363139, EPI_ISL_3363140 | Lighthouse Lab in Glasgow | Wellcome Sanger Institute for the COVID-19 Genomics UK (COG-UK) Consortium | Harper VanSteenhouse, Yumi Kasai, David Gray, Carol Clugston, Anna Dominiczak and Alex Alderton, Roberto Amato, Jeffrey Barrett, Sonia Goncalves, Ewan Harrison, David K. Jackson, Ian Johnston, Dominic Kwiatkowski, Cordelia Langford, John Sillitoe on behalf of the Wellcome Sanger Institute COVID-19 Surveillance Team |
| EPI_ISL_3363142 | Lighthouse Lab in Alderley Park | Wellcome Sanger Institute for the COVID-19 Genomics UK (COG-UK) Consortium | Jacquelyn Wynn, Mairead Hyland, The Lighthouse Lab in Alderley Park and Alex Alderton, Roberto Amato, Jeffrey Barrett, Sonia Goncalves, Ewan Harrison, David K. Jackson, Ian Johnston, Dominic Kwiatkowski, Cordelia Langford, John Sillitoe on behalf of the Wellcome Sanger Institute COVID-19 Surveillance Team |
| EPI_ISL_3363143 | Lighthouse Lab in Glasgow | Wellcome Sanger Institute for the COVID-19 Genomics UK (COG-UK) Consortium | Harper VanSteenhouse, Yumi Kasai, David Gray, Carol Clugston, Anna Dominiczak and Alex Alderton, Roberto Amato, Jeffrey Barrett, Sonia Goncalves, Ewan Harrison, David K. Jackson, Ian Johnston, Dominic Kwiatkowski, Cordelia Langford, John Sillitoe on behalf of the Wellcome Sanger Institute COVID-19 Surveillance Team |
| EPI_ISL_3363144, EPI_ISL_3363145 | Lighthouse Lab in Milton Keynes | Wellcome Sanger Institute for the COVID-19 Genomics UK (COG-UK) Consortium | The Lighthouse Lab in Milton Keynes and Alex Alderton, Roberto Amato, Jeffrey Barrett, Sonia Goncalves, Ewan Harrison, David K. Jackson, Ian Johnston, Dominic Kwiatkowski, Cordelia Langford, John Sillitoe on behalf of the Wellcome Sanger Institute COVID-19 Surveillance Team |
| EPI_ISL_3363147, EPI_ISL_3363148, EPI_ISL_3363149, EPI_ISL_3363151 | Lighthouse Lab in Glasgow | Wellcome Sanger Institute for the COVID-19 Genomics UK (COG-UK) Consortium | Harper VanSteenhouse, Yumi Kasai, David Gray, Carol Clugston, Anna Dominiczak and Alex Alderton, Roberto Amato, Jeffrey Barrett, Sonia Goncalves, Ewan Harrison, David K. Jackson, Ian Johnston, Dominic Kwiatkowski, Cordelia Langford, John Sillitoe on behalf of the Wellcome Sanger Institute COVID-19 Surveillance Team |
| EPI_ISL_3363152 | Lighthouse Lab in Alderley Park | Wellcome Sanger Institute for the COVID-19 Genomics UK (COG-UK) Consortium | Jacquelyn Wynn, Mairead Hyland, The Lighthouse Lab in Alderley Park and Alex Alderton, Roberto Amato, Jeffrey Barrett, Sonia Goncalves, Ewan Harrison, David K. Jackson, Ian Johnston, Dominic Kwiatkowski, Cordelia Langford, John Sillitoe on behalf of the Wellcome Sanger Institute COVID-19 Surveillance Team |
| EPI_ISL_3363154 | Lighthouse Lab in Milton Keynes | Wellcome Sanger Institute for the COVID-19 Genomics UK (COG-UK) Consortium | The Lighthouse Lab in Milton Keynes and Alex Alderton, Roberto Amato, Jeffrey Barrett, Sonia Goncalves, Ewan Harrison, David K. Jackson, Ian Johnston, Dominic Kwiatkowski, Cordelia Langford, John Sillitoe on behalf of the Wellcome Sanger Institute COVID-19 Surveillance Team |
| EPI_ISL_3363155 | Lighthouse Lab in Glasgow | Wellcome Sanger Institute for the COVID-19 Genomics UK (COG-UK) Consortium | Harper VanSteenhouse, Yumi Kasai, David Gray, Carol Clugston, Anna Dominiczak and Alex Alderton, Roberto Amato, Jeffrey Barrett, Sonia Goncalves, Ewan Harrison, David K. Jackson, Ian Johnston, Dominic Kwiatkowski, Cordelia Langford, John Sillitoe on behalf of the Wellcome Sanger Institute COVID-19 Surveillance Team |
| EPI_ISL_3363157 | Lighthouse Lab in Alderley Park | Wellcome Sanger Institute for the COVID-19 Genomics UK (COG-UK) Consortium | Jacquelyn Wynn, Mairead Hyland, The Lighthouse Lab in Alderley Park and Alex Alderton, Roberto Amato, Jeffrey Barrett, Sonia Goncalves, Ewan Harrison, David K. Jackson, Ian Johnston, Dominic Kwiatkowski, Cordelia Langford, John Sillitoe on behalf of the Wellcome Sanger Institute COVID-19 Surveillance Team |
| EPI_ISL_3363158 | Lighthouse Lab in Milton Keynes | Wellcome Sanger Institute for the COVID-19 Genomics UK (COG-UK) Consortium | The Lighthouse Lab in Milton Keynes and Alex Alderton, Roberto Amato, Jeffrey Barrett, Sonia Goncalves, Ewan Harrison, David K. Jackson, Ian Johnston, Dominic Kwiatkowski, Cordelia Langford, John Sillitoe on behalf of the Wellcome Sanger Institute COVID-19 Surveillance Team |
| EPI_ISL_3363160, EPI_ISL_3363161 | Lighthouse Lab in Glasgow | Wellcome Sanger Institute for the COVID-19 Genomics UK (COG-UK) Consortium | Harper VanSteenhouse, Yumi Kasai, David Gray, Carol Clugston, Anna Dominiczak and Alex Alderton, Roberto Amato, Jeffrey Barrett, Sonia Goncalves, Ewan Harrison, David K. Jackson, Ian Johnston, Dominic Kwiatkowski, Cordelia Langford, John Sillitoe on behalf of the Wellcome Sanger Institute COVID-19 Surveillance Team |
| EPI_ISL_3363162, | Lighthouse Lab in Alderley Park | Wellcome Sanger Institute for the COVID-19 Genomics UK (COG-UK) | Jacquelyn Wynn, Mairead Hyland, The Lighthouse Lab in Alderley Park and Alex Alderton, Roberto Amato, Jeffrey Barrett, Sonia Goncalves, Ewan Harrison, David K. Jackson, Ian Johnston, Dominic Kwiatkowski, Cordelia Langford, John Sillitoe on |

| | | | |
|---|---|---|---|
| EPI_ISL_3363164 | | Consortium | behalf of the Wellcome Sanger Institute COVID-19 Surveillance Team |
| EPI_ISL_3363165 | Lighthouse Lab in Milton Keynes | Wellcome Sanger Institute for the COVID-19 Genomics UK (COG-UK) Consortium | The Lighthouse Lab in Milton Keynes and Alex Alderton, Roberto Amato, Jeffrey Barrett, Sonia Goncalves, Ewan Harrison, David K. Jackson, Ian Johnston, Dominic Kwiatkowski, Cordelia Langford, John Sillitoe on behalf of the Wellcome Sanger Institute COVID-19 Surveillance Team |
| EPI_ISL_3363167 | Lighthouse Lab in Glasgow | Wellcome Sanger Institute for the COVID-19 Genomics UK (COG-UK) Consortium | Harper VanSteenhouse, Yumi Kasai, David Gray, Carol Clugston, Anna Dominiczak and Alex Alderton, Roberto Amato, Jeffrey Barrett, Sonia Goncalves, Ewan Harrison, David K. Jackson, Ian Johnston, Dominic Kwiatkowski, Cordelia Langford, John Sillitoe on behalf of the Wellcome Sanger Institute COVID-19 Surveillance Team |
| EPI_ISL_3363168 | Lighthouse Lab in Milton Keynes | Wellcome Sanger Institute for the COVID-19 Genomics UK (COG-UK) Consortium | The Lighthouse Lab in Milton Keynes and Alex Alderton, Roberto Amato, Jeffrey Barrett, Sonia Goncalves, Ewan Harrison, David K. Jackson, Ian Johnston, Dominic Kwiatkowski, Cordelia Langford, John Sillitoe on behalf of the Wellcome Sanger Institute COVID-19 Surveillance Team |
| EPI_ISL_3363169 | Lighthouse Lab in Glasgow | Wellcome Sanger Institute for the COVID-19 Genomics UK (COG-UK) Consortium | Harper VanSteenhouse, Yumi Kasai, David Gray, Carol Clugston, Anna Dominiczak and Alex Alderton, Roberto Amato, Jeffrey Barrett, Sonia Goncalves, Ewan Harrison, David K. Jackson, Ian Johnston, Dominic Kwiatkowski, Cordelia Langford, John Sillitoe on behalf of the Wellcome Sanger Institute COVID-19 Surveillance Team |
| EPI_ISL_3363171, EPI_ISL_3363172, EPI_ISL_3363174 | Lighthouse Lab in Milton Keynes | Wellcome Sanger Institute for the COVID-19 Genomics UK (COG-UK) Consortium | The Lighthouse Lab in Milton Keynes and Alex Alderton, Roberto Amato, Jeffrey Barrett, Sonia Goncalves, Ewan Harrison, David K. Jackson, Ian Johnston, Dominic Kwiatkowski, Cordelia Langford, John Sillitoe on behalf of the Wellcome Sanger Institute COVID-19 Surveillance Team |
| EPI_ISL_3363175, EPI_ISL_3363177 | Lighthouse Lab in Glasgow | Wellcome Sanger Institute for the COVID-19 Genomics UK (COG-UK) Consortium | Harper VanSteenhouse, Yumi Kasai, David Gray, Carol Clugston, Anna Dominiczak and Alex Alderton, Roberto Amato, Jeffrey Barrett, Sonia Goncalves, Ewan Harrison, David K. Jackson, Ian Johnston, Dominic Kwiatkowski, Cordelia Langford, John Sillitoe on behalf of the Wellcome Sanger Institute COVID-19 Surveillance Team |
| EPI_ISL_3363178 | Lighthouse Lab in Alderley Park | Wellcome Sanger Institute for the COVID-19 Genomics UK (COG-UK) Consortium | Jacquelyn Wynn, Mairead Hyland, The Lighthouse Lab in Alderley Park and Alex Alderton, Roberto Amato, Jeffrey Barrett, Sonia Goncalves, Ewan Harrison, David K. Jackson, Ian Johnston, Dominic Kwiatkowski, Cordelia Langford, John Sillitoe on behalf of the Wellcome Sanger Institute COVID-19 Surveillance Team |
| EPI_ISL_3363179, EPI_ISL_3363181, EPI_ISL_3363182 | Lighthouse Lab in Glasgow | Wellcome Sanger Institute for the COVID-19 Genomics UK (COG-UK) Consortium | Harper VanSteenhouse, Yumi Kasai, David Gray, Carol Clugston, Anna Dominiczak and Alex Alderton, Roberto Amato, Jeffrey Barrett, Sonia Goncalves, Ewan Harrison, David K. Jackson, Ian Johnston, Dominic Kwiatkowski, Cordelia Langford, John Sillitoe on behalf of the Wellcome Sanger Institute COVID-19 Surveillance Team |
| EPI_ISL_3363184 | Lighthouse Lab in Alderley Park | Wellcome Sanger Institute for the COVID-19 Genomics UK (COG-UK) Consortium | Jacquelyn Wynn, Mairead Hyland, The Lighthouse Lab in Alderley Park and Alex Alderton, Roberto Amato, Jeffrey Barrett, Sonia Goncalves, Ewan Harrison, David K. Jackson, Ian Johnston, Dominic Kwiatkowski, Cordelia Langford, John Sillitoe on behalf of the Wellcome Sanger Institute COVID-19 Surveillance Team |
| EPI_ISL_3363185 | Lighthouse Lab in Glasgow | Wellcome Sanger Institute for the COVID-19 Genomics UK (COG-UK) Consortium | Harper VanSteenhouse, Yumi Kasai, David Gray, Carol Clugston, Anna Dominiczak and Alex Alderton, Roberto Amato, Jeffrey Barrett, Sonia Goncalves, Ewan Harrison, David K. Jackson, Ian Johnston, Dominic Kwiatkowski, Cordelia Langford, John Sillitoe on behalf of the Wellcome Sanger Institute COVID-19 Surveillance Team |
| EPI_ISL_3363187, EPI_ISL_3363188 | Lighthouse Lab in Milton Keynes | Wellcome Sanger Institute for the COVID-19 Genomics UK (COG-UK) Consortium | The Lighthouse Lab in Milton Keynes and Alex Alderton, Roberto Amato, Jeffrey Barrett, Sonia Goncalves, Ewan Harrison, David K. Jackson, Ian Johnston, Dominic Kwiatkowski, Cordelia Langford, John Sillitoe on behalf of the Wellcome Sanger Institute COVID-19 Surveillance Team |
| EPI_ISL_3363189 | Lighthouse Lab in Glasgow | Wellcome Sanger Institute for the COVID-19 Genomics UK (COG-UK) Consortium | Harper VanSteenhouse, Yumi Kasai, David Gray, Carol Clugston, Anna Dominiczak and Alex Alderton, Roberto Amato, Jeffrey Barrett, Sonia Goncalves, Ewan Harrison, David K. Jackson, Ian Johnston, Dominic Kwiatkowski, Cordelia Langford, John Sillitoe on behalf of the Wellcome Sanger Institute COVID-19 Surveillance Team |
| EPI_ISL_3363191 | Lighthouse Lab in Milton Keynes | Wellcome Sanger Institute for the COVID-19 Genomics UK (COG-UK) Consortium | The Lighthouse Lab in Milton Keynes and Alex Alderton, Roberto Amato, Jeffrey Barrett, Sonia Goncalves, Ewan Harrison, David K. Jackson, Ian Johnston, Dominic Kwiatkowski, Cordelia Langford, John Sillitoe on behalf of the Wellcome Sanger Institute COVID-19 Surveillance Team |
| EPI_ISL_3363192 | Lighthouse Lab in Glasgow | Wellcome Sanger Institute for the COVID-19 Genomics UK (COG-UK) Consortium | Harper VanSteenhouse, Yumi Kasai, David Gray, Carol Clugston, Anna Dominiczak and Alex Alderton, Roberto Amato, Jeffrey Barrett, Sonia Goncalves, Ewan Harrison, David K. Jackson, Ian Johnston, Dominic Kwiatkowski, Cordelia Langford, John Sillitoe on behalf of the Wellcome Sanger Institute COVID-19 Surveillance Team |
| EPI_ISL_3363194 | Lighthouse Lab in Milton Keynes | Wellcome Sanger Institute for the COVID-19 Genomics UK (COG-UK) Consortium | The Lighthouse Lab in Milton Keynes and Alex Alderton, Roberto Amato, Jeffrey Barrett, Sonia Goncalves, Ewan Harrison, David K. Jackson, Ian Johnston, Dominic Kwiatkowski, Cordelia Langford, John Sillitoe on behalf of the Wellcome Sanger Institute COVID-19 Surveillance Team |
| EPI_ISL_3363195 | Lighthouse Lab in Alderley Park | Wellcome Sanger Institute for the COVID-19 Genomics UK (COG-UK) Consortium | Jacquelyn Wynn, Mairead Hyland, The Lighthouse Lab in Alderley Park and Alex Alderton, Roberto Amato, Jeffrey Barrett, Sonia Goncalves, Ewan Harrison, David K. Jackson, Ian Johnston, Dominic Kwiatkowski, Cordelia Langford, John Sillitoe on behalf of the Wellcome Sanger Institute COVID-19 Surveillance Team |
| EPI_ISL_3363197 | Lighthouse Lab in Glasgow | Wellcome Sanger Institute for the COVID-19 Genomics UK (COG-UK) Consortium | Harper VanSteenhouse, Yumi Kasai, David Gray, Carol Clugston, Anna Dominiczak and Alex Alderton, Roberto Amato, Jeffrey Barrett, Sonia Goncalves, Ewan Harrison, David K. Jackson, Ian Johnston, Dominic Kwiatkowski, Cordelia Langford, John Sillitoe on behalf of the Wellcome Sanger Institute COVID-19 Surveillance Team |
| EPI_ISL_3363198, EPI_ISL_3363200, EPI_ISL_3363201 | Lighthouse Lab in Milton Keynes | Wellcome Sanger Institute for the COVID-19 Genomics UK (COG-UK) Consortium | The Lighthouse Lab in Milton Keynes and Alex Alderton, Roberto Amato, Jeffrey Barrett, Sonia Goncalves, Ewan Harrison, David K. Jackson, Ian Johnston, Dominic Kwiatkowski, Cordelia Langford, John Sillitoe on behalf of the Wellcome Sanger Institute COVID-19 Surveillance Team |
| EPI_ISL_3363202 | Lighthouse Lab in Alderley Park | Wellcome Sanger Institute for the COVID-19 Genomics UK (COG-UK) Consortium | Jacquelyn Wynn, Mairead Hyland, The Lighthouse Lab in Alderley Park and Alex Alderton, Roberto Amato, Jeffrey Barrett, Sonia Goncalves, Ewan Harrison, David K. Jackson, Ian Johnston, Dominic Kwiatkowski, Cordelia Langford, John Sillitoe on behalf of the Wellcome Sanger Institute COVID-19 Surveillance Team |
| EPI_ISL_3363204 | Lighthouse Lab in Glasgow | Wellcome Sanger Institute for the COVID-19 Genomics UK (COG-UK) Consortium | Harper VanSteenhouse, Yumi Kasai, David Gray, Carol Clugston, Anna Dominiczak and Alex Alderton, Roberto Amato, Jeffrey Barrett, Sonia Goncalves, Ewan Harrison, David K. Jackson, Ian Johnston, Dominic Kwiatkowski, Cordelia Langford, John Sillitoe on behalf of the Wellcome Sanger Institute COVID-19 Surveillance Team |
| EPI_ISL_3363205, EPI_ISL_3363207, EPI_ISL_3363208, EPI_ISL_3363210 | Lighthouse Lab in Alderley Park | Wellcome Sanger Institute for the COVID-19 Genomics UK (COG-UK) Consortium | Jacquelyn Wynn, Mairead Hyland, The Lighthouse Lab in Alderley Park and Alex Alderton, Roberto Amato, Jeffrey Barrett, Sonia Goncalves, Ewan Harrison, David K. Jackson, Ian Johnston, Dominic Kwiatkowski, Cordelia Langford, John Sillitoe on behalf of the Wellcome Sanger Institute COVID-19 Surveillance Team |
| EPI_ISL_3363211 | Lighthouse Lab in Milton Keynes | Wellcome Sanger Institute for the COVID-19 Genomics UK (COG-UK) Consortium | The Lighthouse Lab in Milton Keynes and Alex Alderton, Roberto Amato, Jeffrey Barrett, Sonia Goncalves, Ewan Harrison, David K. Jackson, Ian Johnston, Dominic Kwiatkowski, Cordelia Langford, John Sillitoe on behalf of the Wellcome Sanger Institute COVID-19 Surveillance Team |
| EPI_ISL_3363212, EPI_ISL_3363214 | Lighthouse Lab in Alderley Park | Wellcome Sanger Institute for the COVID-19 Genomics UK (COG-UK) Consortium | Jacquelyn Wynn, Mairead Hyland, The Lighthouse Lab in Alderley Park and Alex Alderton, Roberto Amato, Jeffrey Barrett, Sonia Goncalves, Ewan Harrison, David K. Jackson, Ian Johnston, Dominic Kwiatkowski, Cordelia Langford, John Sillitoe on behalf of the Wellcome Sanger Institute COVID-19 Surveillance Team |
| EPI_ISL_3363215, EPI_ISL_3363217, EPI_ISL_3363218 | Lighthouse Lab in Glasgow | Wellcome Sanger Institute for the COVID-19 Genomics UK (COG-UK) Consortium | Harper VanSteenhouse, Yumi Kasai, David Gray, Carol Clugston, Anna Dominiczak and Alex Alderton, Roberto Amato, Jeffrey Barrett, Sonia Goncalves, Ewan Harrison, David K. Jackson, Ian Johnston, Dominic Kwiatkowski, Cordelia Langford, John Sillitoe on behalf of the Wellcome Sanger Institute COVID-19 Surveillance Team |
| EPI_ISL_3363220, EPI_ISL_3363221 | Lighthouse Lab in Milton Keynes | Wellcome Sanger Institute for the COVID-19 Genomics UK (COG-UK) Consortium | The Lighthouse Lab in Milton Keynes and Alex Alderton, Roberto Amato, Jeffrey Barrett, Sonia Goncalves, Ewan Harrison, David K. Jackson, Ian Johnston, Dominic Kwiatkowski, Cordelia Langford, John Sillitoe on behalf of the Wellcome Sanger Institute COVID-19 Surveillance Team |
| EPI_ISL_3363222, EPI_ISL_3363224 | Lighthouse Lab in Alderley Park | Wellcome Sanger Institute for the COVID-19 Genomics UK (COG-UK) Consortium | Jacquelyn Wynn, Mairead Hyland, The Lighthouse Lab in Alderley Park and Alex Alderton, Roberto Amato, Jeffrey Barrett, Sonia Goncalves, Ewan Harrison, David K. Jackson, Ian Johnston, Dominic Kwiatkowski, Cordelia Langford, John Sillitoe on behalf of the Wellcome Sanger Institute COVID-19 Surveillance Team |
| EPI_ISL_3363225 | Lighthouse Lab in Milton Keynes | Wellcome Sanger Institute for the COVID-19 Genomics UK (COG-UK) Consortium | The Lighthouse Lab in Milton Keynes and Alex Alderton, Roberto Amato, Jeffrey Barrett, Sonia Goncalves, Ewan Harrison, David K. Jackson, Ian Johnston, Dominic Kwiatkowski, Cordelia Langford, John Sillitoe on behalf of the Wellcome Sanger Institute COVID-19 Surveillance Team |
| EPI_ISL_3363227 | Lighthouse Lab in Alderley Park | Wellcome Sanger Institute for the COVID-19 Genomics UK (COG-UK) Consortium | Jacquelyn Wynn, Mairead Hyland, The Lighthouse Lab in Alderley Park and Alex Alderton, Roberto Amato, Jeffrey Barrett, Sonia Goncalves, Ewan Harrison, David K. Jackson, Ian Johnston, Dominic Kwiatkowski, Cordelia Langford, John Sillitoe on behalf of the Wellcome Sanger Institute COVID-19 Surveillance Team |
| EPI_ISL_3363228 | Lighthouse Lab in Glasgow | Wellcome Sanger Institute for the COVID-19 Genomics UK (COG-UK) Consortium | Harper VanSteenhouse, Yumi Kasai, David Gray, Carol Clugston, Anna Dominiczak and Alex Alderton, Roberto Amato, Jeffrey Barrett, Sonia Goncalves, Ewan Harrison, David K. Jackson, Ian Johnston, Dominic Kwiatkowski, Cordelia Langford, John Sillitoe on behalf of the Wellcome Sanger Institute COVID-19 Surveillance Team |
| EPI_ISL_3363229, EPI_ISL_3363231 | Lighthouse Lab in Alderley Park | Wellcome Sanger Institute for the COVID-19 Genomics UK (COG-UK) Consortium | Jacquelyn Wynn, Mairead Hyland, The Lighthouse Lab in Alderley Park and Alex Alderton, Roberto Amato, Jeffrey Barrett, Sonia Goncalves, Ewan Harrison, David K. Jackson, Ian Johnston, Dominic Kwiatkowski, Cordelia Langford, John Sillitoe on behalf of the Wellcome Sanger Institute COVID-19 Surveillance Team |
| EPI_ISL_3363232 | Lighthouse Lab in Milton Keynes | Wellcome Sanger Institute for the COVID-19 Genomics UK (COG-UK) Consortium | The Lighthouse Lab in Milton Keynes and Alex Alderton, Roberto Amato, Jeffrey Barrett, Sonia Goncalves, Ewan Harrison, David K. Jackson, Ian Johnston, Dominic Kwiatkowski, Cordelia Langford, John Sillitoe on behalf of the Wellcome Sanger Institute COVID-19 Surveillance Team |
| EPI_ISL_3363234 | Lighthouse Lab in Alderley Park | Wellcome Sanger Institute for the COVID-19 Genomics UK (COG-UK) Consortium | Jacquelyn Wynn, Mairead Hyland, The Lighthouse Lab in Alderley Park and Alex Alderton, Roberto Amato, Jeffrey Barrett, Sonia Goncalves, Ewan Harrison, David K. Jackson, Ian Johnston, Dominic Kwiatkowski, Cordelia Langford, John Sillitoe on behalf of the Wellcome Sanger Institute COVID-19 Surveillance Team |
| EPI_ISL_3363235, EPI_ISL_3363237 | Lighthouse Lab in Glasgow | Wellcome Sanger Institute for the COVID-19 Genomics UK (COG-UK) Consortium | Harper VanSteenhouse, Yumi Kasai, David Gray, Carol Clugston, Anna Dominiczak and Alex Alderton, Roberto Amato, Jeffrey Barrett, Sonia Goncalves, Ewan Harrison, David K. Jackson, Ian Johnston, Dominic Kwiatkowski, Cordelia Langford, John Sillitoe on behalf of the Wellcome Sanger Institute COVID-19 Surveillance Team |
| EPI_ISL_3363238, EPI_ISL_3363239 | Lighthouse Lab in Alderley Park | Wellcome Sanger Institute for the COVID-19 Genomics UK (COG-UK) Consortium | Jacquelyn Wynn, Mairead Hyland, The Lighthouse Lab in Alderley Park and Alex Alderton, Roberto Amato, Jeffrey Barrett, Sonia Goncalves, Ewan Harrison, David K. Jackson, Ian Johnston, Dominic Kwiatkowski, Cordelia Langford, John Sillitoe on behalf of the Wellcome Sanger Institute COVID-19 Surveillance Team |
| EPI_ISL_3363241 | Lighthouse Lab in Milton Keynes | Wellcome Sanger Institute for the COVID-19 Genomics UK (COG-UK) Consortium | The Lighthouse Lab in Milton Keynes and Alex Alderton, Roberto Amato, Jeffrey Barrett, Sonia Goncalves, Ewan Harrison, David K. Jackson, Ian Johnston, Dominic Kwiatkowski, Cordelia Langford, John Sillitoe on behalf of the Wellcome Sanger Institute COVID-19 Surveillance Team |
| EPI_ISL_3363242 | Lighthouse Lab in Glasgow | Wellcome Sanger Institute for the COVID-19 Genomics UK (COG-UK) Consortium | Harper VanSteenhouse, Yumi Kasai, David Gray, Carol Clugston, Anna Dominiczak and Alex Alderton, Roberto Amato, Jeffrey Barrett, Sonia Goncalves, Ewan Harrison, David K. Jackson, Ian Johnston, Dominic Kwiatkowski, Cordelia Langford, John Sillitoe on behalf of the Wellcome Sanger Institute COVID-19 Surveillance Team |
| EPI_ISL_3363243, EPI_ISL_3363244, EPI_ISL_3363245, EPI_ISL_3363246, EPI_ISL_3363247 | Lighthouse Laboratory Plymouth | Wellcome Sanger Institute for the COVID-19 Genomics UK (COG-UK) Consortium | Lighthouse Laboratory Plymouth and Alex Alderton, Roberto Amato, Jeffrey Barrett, Sonia Goncalves, Ewan Harrison, David K. Jackson, Ian Johnston, Dominic Kwiatkowski, Cordelia Langford, John Sillitoe on behalf of the Wellcome Sanger Institute COVID-19 Surveillance Team |
| EPI_ISL_3363248 | Randox Laboratories | Wellcome Sanger Institute for the COVID-19 Genomics UK (COG-UK) Consortium | Randox Laboratories and Alex Alderton, Roberto Amato, Jeffrey Barrett, Sonia Goncalves, Ewan Harrison, David K. Jackson, Ian Johnston, Dominic Kwiatkowski, Cordelia Langford, John Sillitoe on behalf of the Wellcome Sanger Institute COVID-19 Surveillance Team |
| EPI_ISL_3363249, EPI_ISL_3363250, EPI_ISL_3363251 | Lighthouse Laboratory Plymouth | Wellcome Sanger Institute for the COVID-19 Genomics UK (COG-UK) Consortium | Lighthouse Laboratory Plymouth and Alex Alderton, Roberto Amato, Jeffrey Barrett, Sonia Goncalves, Ewan Harrison, David K. Jackson, Ian Johnston, Dominic Kwiatkowski, Cordelia Langford, John Sillitoe on behalf of the Wellcome Sanger Institute COVID-19 Surveillance Team |
| EPI_ISL_3363252 | Randox Laboratories | Wellcome Sanger Institute for the COVID-19 Genomics UK (COG-UK) Consortium | Randox Laboratories and Alex Alderton, Roberto Amato, Jeffrey Barrett, Sonia Goncalves, Ewan Harrison, David K. Jackson, Ian Johnston, Dominic Kwiatkowski, Cordelia Langford, John Sillitoe on behalf of the Wellcome Sanger Institute COVID-19 Surveillance Team |
| EPI_ISL_3363253, | Lighthouse Laboratory Plymouth | Wellcome Sanger Institute for the COVID-19 Genomics UK (COG-UK) | Lighthouse Laboratory Plymouth and Alex Alderton, Roberto Amato, Jeffrey Barrett, Sonia Goncalves, Ewan Harrison, David K. Jackson, Ian Johnston, Dominic Kwiatkowski, Cordelia Langford, John Sillitoe on behalf of the Wellcome Sanger |

| | | | |
|---|---|---|---|
| EPI_ISL_3363254, EPI_ISL_3363255, EPI_ISL_3363256, EPI_ISL_3363257 | | Consortium | Institute COVID-19 Surveillance Team |
| EPI_ISL_3363258, EPI_ISL_3363259, EPI_ISL_3363260 | Randox Laboratories | Wellcome Sanger Institute for the COVID-19 Genomics UK (COG-UK) Consortium | Randox Laboratories and Alex Alderton, Roberto Amato, Jeffrey Barrett, Sonia Goncalves, Ewan Harrison, David K. Jackson, Ian Johnston, Dominic Kwiatkowski, Cordelia Langford, John Sillitoe on behalf of the Wellcome Sanger Institute COVID-19 Surveillance Team |
| EPI_ISL_3363261, EPI_ISL_3363262 | Lighthouse Laboratory Plymouth | Wellcome Sanger Institute for the COVID-19 Genomics UK (COG-UK) Consortium | Lighthouse Laboratory Plymouth and Alex Alderton, Roberto Amato, Jeffrey Barrett, Sonia Goncalves, Ewan Harrison, David K. Jackson, Ian Johnston, Dominic Kwiatkowski, Cordelia Langford, John Sillitoe on behalf of the Wellcome Sanger Institute COVID-19 Surveillance Team |
| EPI_ISL_3363263 | Randox Laboratories | Wellcome Sanger Institute for the COVID-19 Genomics UK (COG-UK) Consortium | Randox Laboratories and Alex Alderton, Roberto Amato, Jeffrey Barrett, Sonia Goncalves, Ewan Harrison, David K. Jackson, Ian Johnston, Dominic Kwiatkowski, Cordelia Langford, John Sillitoe on behalf of the Wellcome Sanger Institute COVID-19 Surveillance Team |
| EPI_ISL_3363264, EPI_ISL_3363265, EPI_ISL_3363266, EPI_ISL_3363267 | Lighthouse Laboratory Plymouth | Wellcome Sanger Institute for the COVID-19 Genomics UK (COG-UK) Consortium | Lighthouse Laboratory Plymouth and Alex Alderton, Roberto Amato, Jeffrey Barrett, Sonia Goncalves, Ewan Harrison, David K. Jackson, Ian Johnston, Dominic Kwiatkowski, Cordelia Langford, John Sillitoe on behalf of the Wellcome Sanger Institute COVID-19 Surveillance Team |
| EPI_ISL_3363268, EPI_ISL_3363269 | Randox Laboratories | Wellcome Sanger Institute for the COVID-19 Genomics UK (COG-UK) Consortium | Randox Laboratories and Alex Alderton, Roberto Amato, Jeffrey Barrett, Sonia Goncalves, Ewan Harrison, David K. Jackson, Ian Johnston, Dominic Kwiatkowski, Cordelia Langford, John Sillitoe on behalf of the Wellcome Sanger Institute COVID-19 Surveillance Team |
| EPI_ISL_3363270, EPI_ISL_3363271, EPI_ISL_3363272 | Lighthouse Laboratory Plymouth | Wellcome Sanger Institute for the COVID-19 Genomics UK (COG-UK) Consortium | Lighthouse Laboratory Plymouth and Alex Alderton, Roberto Amato, Jeffrey Barrett, Sonia Goncalves, Ewan Harrison, David K. Jackson, Ian Johnston, Dominic Kwiatkowski, Cordelia Langford, John Sillitoe on behalf of the Wellcome Sanger Institute COVID-19 Surveillance Team |
| EPI_ISL_3363273 | Randox Laboratories | Wellcome Sanger Institute for the COVID-19 Genomics UK (COG-UK) Consortium | Randox Laboratories and Alex Alderton, Roberto Amato, Jeffrey Barrett, Sonia Goncalves, Ewan Harrison, David K. Jackson, Ian Johnston, Dominic Kwiatkowski, Cordelia Langford, John Sillitoe on behalf of the Wellcome Sanger Institute COVID-19 Surveillance Team |
| EPI_ISL_3363274, EPI_ISL_3363275, EPI_ISL_3363276, EPI_ISL_3363277, EPI_ISL_3363278, EPI_ISL_3363279, EPI_ISL_3363280 | Lighthouse Laboratory Plymouth | Wellcome Sanger Institute for the COVID-19 Genomics UK (COG-UK) Consortium | Lighthouse Laboratory Plymouth and Alex Alderton, Roberto Amato, Jeffrey Barrett, Sonia Goncalves, Ewan Harrison, David K. Jackson, Ian Johnston, Dominic Kwiatkowski, Cordelia Langford, John Sillitoe on behalf of the Wellcome Sanger Institute COVID-19 Surveillance Team |
| EPI_ISL_3363281, EPI_ISL_3363282 | Randox Laboratories | Wellcome Sanger Institute for the COVID-19 Genomics UK (COG-UK) Consortium | Randox Laboratories and Alex Alderton, Roberto Amato, Jeffrey Barrett, Sonia Goncalves, Ewan Harrison, David K. Jackson, Ian Johnston, Dominic Kwiatkowski, Cordelia Langford, John Sillitoe on behalf of the Wellcome Sanger Institute COVID-19 Surveillance Team |
| EPI_ISL_3363283, EPI_ISL_3363284, EPI_ISL_3363285, EPI_ISL_3363286 | Lighthouse Laboratory Plymouth | Wellcome Sanger Institute for the COVID-19 Genomics UK (COG-UK) Consortium | Lighthouse Laboratory Plymouth and Alex Alderton, Roberto Amato, Jeffrey Barrett, Sonia Goncalves, Ewan Harrison, David K. Jackson, Ian Johnston, Dominic Kwiatkowski, Cordelia Langford, John Sillitoe on behalf of the Wellcome Sanger Institute COVID-19 Surveillance Team |
| EPI_ISL_3363287 | Randox Laboratories | Wellcome Sanger Institute for the COVID-19 Genomics UK (COG-UK) Consortium | Randox Laboratories and Alex Alderton, Roberto Amato, Jeffrey Barrett, Sonia Goncalves, Ewan Harrison, David K. Jackson, Ian Johnston, Dominic Kwiatkowski, Cordelia Langford, John Sillitoe on behalf of the Wellcome Sanger Institute COVID-19 Surveillance Team |
| EPI_ISL_3363288, EPI_ISL_3363289 | Lighthouse Laboratory Plymouth | Wellcome Sanger Institute for the COVID-19 Genomics UK (COG-UK) Consortium | Lighthouse Laboratory Plymouth and Alex Alderton, Roberto Amato, Jeffrey Barrett, Sonia Goncalves, Ewan Harrison, David K. Jackson, Ian Johnston, Dominic Kwiatkowski, Cordelia Langford, John Sillitoe on behalf of the Wellcome Sanger Institute COVID-19 Surveillance Team |
| EPI_ISL_3363290 | Randox Laboratories | Wellcome Sanger Institute for the COVID-19 Genomics UK (COG-UK) Consortium | Randox Laboratories and Alex Alderton, Roberto Amato, Jeffrey Barrett, Sonia Goncalves, Ewan Harrison, David K. Jackson, Ian Johnston, Dominic Kwiatkowski, Cordelia Langford, John Sillitoe on behalf of the Wellcome Sanger Institute COVID-19 Surveillance Team |
| EPI_ISL_3363291, EPI_ISL_3363292, EPI_ISL_3363293 | Lighthouse Laboratory Plymouth | Wellcome Sanger Institute for the COVID-19 Genomics UK (COG-UK) Consortium | Lighthouse Laboratory Plymouth and Alex Alderton, Roberto Amato, Jeffrey Barrett, Sonia Goncalves, Ewan Harrison, David K. Jackson, Ian Johnston, Dominic Kwiatkowski, Cordelia Langford, John Sillitoe on behalf of the Wellcome Sanger Institute COVID-19 Surveillance Team |
| EPI_ISL_3363294 | Randox Laboratories | Wellcome Sanger Institute for the COVID-19 Genomics UK (COG-UK) Consortium | Randox Laboratories and Alex Alderton, Roberto Amato, Jeffrey Barrett, Sonia Goncalves, Ewan Harrison, David K. Jackson, Ian Johnston, Dominic Kwiatkowski, Cordelia Langford, John Sillitoe on behalf of the Wellcome Sanger Institute COVID-19 Surveillance Team |
| EPI_ISL_3363295, EPI_ISL_3363296, EPI_ISL_3363297, EPI_ISL_3363298, EPI_ISL_3363299, EPI_ISL_3363300, EPI_ISL_3363301, EPI_ISL_3363302 | Lighthouse Laboratory Plymouth | Wellcome Sanger Institute for the COVID-19 Genomics UK (COG-UK) Consortium | Lighthouse Laboratory Plymouth and Alex Alderton, Roberto Amato, Jeffrey Barrett, Sonia Goncalves, Ewan Harrison, David K. Jackson, Ian Johnston, Dominic Kwiatkowski, Cordelia Langford, John Sillitoe on behalf of the Wellcome Sanger Institute COVID-19 Surveillance Team |
| EPI_ISL_3363303, EPI_ISL_3363304, EPI_ISL_3363305 | Randox Laboratories | Wellcome Sanger Institute for the COVID-19 Genomics UK (COG-UK) Consortium | Randox Laboratories and Alex Alderton, Roberto Amato, Jeffrey Barrett, Sonia Goncalves, Ewan Harrison, David K. Jackson, Ian Johnston, Dominic Kwiatkowski, Cordelia Langford, John Sillitoe on behalf of the Wellcome Sanger Institute COVID-19 Surveillance Team |
| EPI_ISL_3363306, EPI_ISL_3363307, EPI_ISL_3363308, EPI_ISL_3363309, EPI_ISL_3363310 | Lighthouse Laboratory Plymouth | Wellcome Sanger Institute for the COVID-19 Genomics UK (COG-UK) Consortium | Lighthouse Laboratory Plymouth and Alex Alderton, Roberto Amato, Jeffrey Barrett, Sonia Goncalves, Ewan Harrison, David K. Jackson, Ian Johnston, Dominic Kwiatkowski, Cordelia Langford, John Sillitoe on behalf of the Wellcome Sanger Institute COVID-19 Surveillance Team |
| EPI_ISL_3363311 | Randox Laboratories | Wellcome Sanger Institute for the COVID-19 Genomics UK (COG-UK) Consortium | Randox Laboratories and Alex Alderton, Roberto Amato, Jeffrey Barrett, Sonia Goncalves, Ewan Harrison, David K. Jackson, Ian Johnston, Dominic Kwiatkowski, Cordelia Langford, John Sillitoe on behalf of the Wellcome Sanger Institute COVID-19 Surveillance Team |
| EPI_ISL_3363312, EPI_ISL_3363313, EPI_ISL_3363314, EPI_ISL_3363315, EPI_ISL_3363316, EPI_ISL_3363317, EPI_ISL_3363318 | Lighthouse Laboratory Plymouth | Wellcome Sanger Institute for the COVID-19 Genomics UK (COG-UK) Consortium | Lighthouse Laboratory Plymouth and Alex Alderton, Roberto Amato, Jeffrey Barrett, Sonia Goncalves, Ewan Harrison, David K. Jackson, Ian Johnston, Dominic Kwiatkowski, Cordelia Langford, John Sillitoe on behalf of the Wellcome Sanger Institute COVID-19 Surveillance Team |
| EPI_ISL_3363319 | Randox Laboratories | Wellcome Sanger Institute for the COVID-19 Genomics UK (COG-UK) Consortium | Randox Laboratories and Alex Alderton, Roberto Amato, Jeffrey Barrett, Sonia Goncalves, Ewan Harrison, David K. Jackson, Ian Johnston, Dominic Kwiatkowski, Cordelia Langford, John Sillitoe on behalf of the Wellcome Sanger Institute COVID-19 Surveillance Team |
| EPI_ISL_3363320 | Lighthouse Laboratory Plymouth | Wellcome Sanger Institute for the COVID-19 Genomics UK (COG-UK) Consortium | Lighthouse Laboratory Plymouth and Alex Alderton, Roberto Amato, Jeffrey Barrett, Sonia Goncalves, Ewan Harrison, David K. Jackson, Ian Johnston, Dominic Kwiatkowski, Cordelia Langford, John Sillitoe on behalf of the Wellcome Sanger Institute COVID-19 Surveillance Team |
| EPI_ISL_3363321, EPI_ISL_3363322, EPI_ISL_3363323 | Randox Laboratories | Wellcome Sanger Institute for the COVID-19 Genomics UK (COG-UK) Consortium | Randox Laboratories and Alex Alderton, Roberto Amato, Jeffrey Barrett, Sonia Goncalves, Ewan Harrison, David K. Jackson, Ian Johnston, Dominic Kwiatkowski, Cordelia Langford, John Sillitoe on behalf of the Wellcome Sanger Institute COVID-19 Surveillance Team |
| EPI_ISL_3363324 | Lighthouse Laboratory Plymouth | Wellcome Sanger Institute for the COVID-19 Genomics UK (COG-UK) Consortium | Lighthouse Laboratory Plymouth and Alex Alderton, Roberto Amato, Jeffrey Barrett, Sonia Goncalves, Ewan Harrison, David K. Jackson, Ian Johnston, Dominic Kwiatkowski, Cordelia Langford, John Sillitoe on behalf of the Wellcome Sanger Institute COVID-19 Surveillance Team |
| EPI_ISL_3363325 | Randox Laboratories | Wellcome Sanger Institute for the COVID-19 Genomics UK (COG-UK) Consortium | Randox Laboratories and Alex Alderton, Roberto Amato, Jeffrey Barrett, Sonia Goncalves, Ewan Harrison, David K. Jackson, Ian Johnston, Dominic Kwiatkowski, Cordelia Langford, John Sillitoe on behalf of the Wellcome Sanger Institute COVID-19 Surveillance Team |
| EPI_ISL_3363326, EPI_ISL_3363327, EPI_ISL_3363328 | Lighthouse Laboratory Plymouth | Wellcome Sanger Institute for the COVID-19 Genomics UK (COG-UK) Consortium | Lighthouse Laboratory Plymouth and Alex Alderton, Roberto Amato, Jeffrey Barrett, Sonia Goncalves, Ewan Harrison, David K. Jackson, Ian Johnston, Dominic Kwiatkowski, Cordelia Langford, John Sillitoe on behalf of the Wellcome Sanger Institute COVID-19 Surveillance Team |
| EPI_ISL_3363329 | Randox Laboratories | Wellcome Sanger Institute for the COVID-19 Genomics UK (COG-UK) Consortium | Randox Laboratories and Alex Alderton, Roberto Amato, Jeffrey Barrett, Sonia Goncalves, Ewan Harrison, David K. Jackson, Ian Johnston, Dominic Kwiatkowski, Cordelia Langford, John Sillitoe on behalf of the Wellcome Sanger Institute COVID-19 Surveillance Team |
| EPI_ISL_3363330 | Lighthouse Laboratory Plymouth | Wellcome Sanger Institute for the COVID-19 Genomics UK (COG-UK) Consortium | Lighthouse Laboratory Plymouth and Alex Alderton, Roberto Amato, Jeffrey Barrett, Sonia Goncalves, Ewan Harrison, David K. Jackson, Ian Johnston, Dominic Kwiatkowski, Cordelia Langford, John Sillitoe on behalf of the Wellcome Sanger Institute COVID-19 Surveillance Team |
| EPI_ISL_3363331 | Randox Laboratories | Wellcome Sanger Institute for the COVID-19 Genomics UK (COG-UK) Consortium | Randox Laboratories and Alex Alderton, Roberto Amato, Jeffrey Barrett, Sonia Goncalves, Ewan Harrison, David K. Jackson, Ian Johnston, Dominic Kwiatkowski, Cordelia Langford, John Sillitoe on behalf of the Wellcome Sanger Institute COVID-19 Surveillance Team |
| EPI_ISL_3363332 | Lighthouse Laboratory Plymouth | Wellcome Sanger Institute for the COVID-19 Genomics UK (COG-UK) Consortium | Lighthouse Laboratory Plymouth and Alex Alderton, Roberto Amato, Jeffrey Barrett, Sonia Goncalves, Ewan Harrison, David K. Jackson, Ian Johnston, Dominic Kwiatkowski, Cordelia Langford, John Sillitoe on behalf of the Wellcome Sanger Institute COVID-19 Surveillance Team |
| EPI_ISL_3363333, EPI_ISL_3363334 | Randox Laboratories | Wellcome Sanger Institute for the COVID-19 Genomics UK (COG-UK) Consortium | Randox Laboratories and Alex Alderton, Roberto Amato, Jeffrey Barrett, Sonia Goncalves, Ewan Harrison, David K. Jackson, Ian Johnston, Dominic Kwiatkowski, Cordelia Langford, John Sillitoe on behalf of the Wellcome Sanger Institute COVID-19 Surveillance Team |
| EPI_ISL_3363335, | Lighthouse Laboratory Plymouth | Wellcome Sanger Institute for the COVID-19 Genomics UK (COG-UK) | Lighthouse Laboratory Plymouth and Alex Alderton, Roberto Amato, Jeffrey Barrett, Sonia Goncalves, Ewan Harrison, David K. Jackson, Ian Johnston, Dominic Kwiatkowski, Cordelia Langford, John Sillitoe on behalf of the Wellcome Sanger |

| | | | |
|---|---|---|---|
| EPI_ISL_3363336 | | Consortium | Institute COVID-19 Surveillance Team |
| EPI_ISL_3363337, EPI_ISL_3363338 | Randox Laboratories | Wellcome Sanger Institute for the COVID-19 Genomics UK (COG-UK) Consortium | Randox Laboratories and Alex Alderton, Roberto Amato, Jeffrey Barrett, Sonia Goncalves, Ewan Harrison, David K. Jackson, Ian Johnston, Dominic Kwiatkowski, Cordelia Langford, John Sillitoe on behalf of the Wellcome Sanger Institute COVID-19 Surveillance Team |
| EPI_ISL_3363339, EPI_ISL_3363340, EPI_ISL_3363341, EPI_ISL_3363342, EPI_ISL_3363343, EPI_ISL_3363344, EPI_ISL_3363345 | Lighthouse Laboratory Plymouth | Wellcome Sanger Institute for the COVID-19 Genomics UK (COG-UK) Consortium | Lighthouse Laboratory Plymouth and Alex Alderton, Roberto Amato, Jeffrey Barrett, Sonia Goncalves, Ewan Harrison, David K. Jackson, Ian Johnston, Dominic Kwiatkowski, Cordelia Langford, John Sillitoe on behalf of the Wellcome Sanger Institute COVID-19 Surveillance Team |
| EPI_ISL_3363346, EPI_ISL_3363347 | Randox Laboratories | Wellcome Sanger Institute for the COVID-19 Genomics UK (COG-UK) Consortium | Randox Laboratories and Alex Alderton, Roberto Amato, Jeffrey Barrett, Sonia Goncalves, Ewan Harrison, David K. Jackson, Ian Johnston, Dominic Kwiatkowski, Cordelia Langford, John Sillitoe on behalf of the Wellcome Sanger Institute COVID-19 Surveillance Team |
| EPI_ISL_3363348, EPI_ISL_3363349, EPI_ISL_3363350, EPI_ISL_3363351 | Lighthouse Laboratory Plymouth | Wellcome Sanger Institute for the COVID-19 Genomics UK (COG-UK) Consortium | Lighthouse Laboratory Plymouth and Alex Alderton, Roberto Amato, Jeffrey Barrett, Sonia Goncalves, Ewan Harrison, David K. Jackson, Ian Johnston, Dominic Kwiatkowski, Cordelia Langford, John Sillitoe on behalf of the Wellcome Sanger Institute COVID-19 Surveillance Team |
| EPI_ISL_3363352 | Randox Laboratories | Wellcome Sanger Institute for the COVID-19 Genomics UK (COG-UK) Consortium | Randox Laboratories and Alex Alderton, Roberto Amato, Jeffrey Barrett, Sonia Goncalves, Ewan Harrison, David K. Jackson, Ian Johnston, Dominic Kwiatkowski, Cordelia Langford, John Sillitoe on behalf of the Wellcome Sanger Institute COVID-19 Surveillance Team |
| EPI_ISL_3363353 | Lighthouse Laboratory Plymouth | Wellcome Sanger Institute for the COVID-19 Genomics UK (COG-UK) Consortium | Lighthouse Laboratory Plymouth and Alex Alderton, Roberto Amato, Jeffrey Barrett, Sonia Goncalves, Ewan Harrison, David K. Jackson, Ian Johnston, Dominic Kwiatkowski, Cordelia Langford, John Sillitoe on behalf of the Wellcome Sanger Institute COVID-19 Surveillance Team |
| EPI_ISL_3363354 | Randox Laboratories | Wellcome Sanger Institute for the COVID-19 Genomics UK (COG-UK) Consortium | Randox Laboratories and Alex Alderton, Roberto Amato, Jeffrey Barrett, Sonia Goncalves, Ewan Harrison, David K. Jackson, Ian Johnston, Dominic Kwiatkowski, Cordelia Langford, John Sillitoe on behalf of the Wellcome Sanger Institute COVID-19 Surveillance Team |
| EPI_ISL_3363355, EPI_ISL_3363356, EPI_ISL_3363357, EPI_ISL_3363358 | Lighthouse Laboratory Plymouth | Wellcome Sanger Institute for the COVID-19 Genomics UK (COG-UK) Consortium | Lighthouse Laboratory Plymouth and Alex Alderton, Roberto Amato, Jeffrey Barrett, Sonia Goncalves, Ewan Harrison, David K. Jackson, Ian Johnston, Dominic Kwiatkowski, Cordelia Langford, John Sillitoe on behalf of the Wellcome Sanger Institute COVID-19 Surveillance Team |
| EPI_ISL_3363359, EPI_ISL_3363360, EPI_ISL_3363361 | Randox Laboratories | Wellcome Sanger Institute for the COVID-19 Genomics UK (COG-UK) Consortium | Randox Laboratories and Alex Alderton, Roberto Amato, Jeffrey Barrett, Sonia Goncalves, Ewan Harrison, David K. Jackson, Ian Johnston, Dominic Kwiatkowski, Cordelia Langford, John Sillitoe on behalf of the Wellcome Sanger Institute COVID-19 Surveillance Team |
| EPI_ISL_3363362 | Lighthouse Laboratory Plymouth | Wellcome Sanger Institute for the COVID-19 Genomics UK (COG-UK) Consortium | Lighthouse Laboratory Plymouth and Alex Alderton, Roberto Amato, Jeffrey Barrett, Sonia Goncalves, Ewan Harrison, David K. Jackson, Ian Johnston, Dominic Kwiatkowski, Cordelia Langford, John Sillitoe on behalf of the Wellcome Sanger Institute COVID-19 Surveillance Team |
| EPI_ISL_3363363 | Randox Laboratories | Wellcome Sanger Institute for the COVID-19 Genomics UK (COG-UK) Consortium | Randox Laboratories and Alex Alderton, Roberto Amato, Jeffrey Barrett, Sonia Goncalves, Ewan Harrison, David K. Jackson, Ian Johnston, Dominic Kwiatkowski, Cordelia Langford, John Sillitoe on behalf of the Wellcome Sanger Institute COVID-19 Surveillance Team |
| EPI_ISL_3363364, EPI_ISL_3363365, EPI_ISL_3363366, EPI_ISL_3363367 | Lighthouse Laboratory Plymouth | Wellcome Sanger Institute for the COVID-19 Genomics UK (COG-UK) Consortium | Lighthouse Laboratory Plymouth and Alex Alderton, Roberto Amato, Jeffrey Barrett, Sonia Goncalves, Ewan Harrison, David K. Jackson, Ian Johnston, Dominic Kwiatkowski, Cordelia Langford, John Sillitoe on behalf of the Wellcome Sanger Institute COVID-19 Surveillance Team |
| EPI_ISL_3363368 | Randox Laboratories | Wellcome Sanger Institute for the COVID-19 Genomics UK (COG-UK) Consortium | Randox Laboratories and Alex Alderton, Roberto Amato, Jeffrey Barrett, Sonia Goncalves, Ewan Harrison, David K. Jackson, Ian Johnston, Dominic Kwiatkowski, Cordelia Langford, John Sillitoe on behalf of the Wellcome Sanger Institute COVID-19 Surveillance Team |
| EPI_ISL_3363369 | Lighthouse Laboratory Plymouth | Wellcome Sanger Institute for the COVID-19 Genomics UK (COG-UK) Consortium | Lighthouse Laboratory Plymouth and Alex Alderton, Roberto Amato, Jeffrey Barrett, Sonia Goncalves, Ewan Harrison, David K. Jackson, Ian Johnston, Dominic Kwiatkowski, Cordelia Langford, John Sillitoe on behalf of the Wellcome Sanger Institute COVID-19 Surveillance Team |
| EPI_ISL_3363370, EPI_ISL_3363371 | Randox Laboratories | Wellcome Sanger Institute for the COVID-19 Genomics UK (COG-UK) Consortium | Randox Laboratories and Alex Alderton, Roberto Amato, Jeffrey Barrett, Sonia Goncalves, Ewan Harrison, David K. Jackson, Ian Johnston, Dominic Kwiatkowski, Cordelia Langford, John Sillitoe on behalf of the Wellcome Sanger Institute COVID-19 Surveillance Team |
| EPI_ISL_3363372 | Lighthouse Laboratory Plymouth | Wellcome Sanger Institute for the COVID-19 Genomics UK (COG-UK) Consortium | Lighthouse Laboratory Plymouth and Alex Alderton, Roberto Amato, Jeffrey Barrett, Sonia Goncalves, Ewan Harrison, David K. Jackson, Ian Johnston, Dominic Kwiatkowski, Cordelia Langford, John Sillitoe on behalf of the Wellcome Sanger Institute COVID-19 Surveillance Team |
| EPI_ISL_3363373 | Randox Laboratories | Wellcome Sanger Institute for the COVID-19 Genomics UK (COG-UK) Consortium | Randox Laboratories and Alex Alderton, Roberto Amato, Jeffrey Barrett, Sonia Goncalves, Ewan Harrison, David K. Jackson, Ian Johnston, Dominic Kwiatkowski, Cordelia Langford, John Sillitoe on behalf of the Wellcome Sanger Institute COVID-19 Surveillance Team |
| EPI_ISL_3363374, EPI_ISL_3363375, EPI_ISL_3363376 | Lighthouse Laboratory Plymouth | Wellcome Sanger Institute for the COVID-19 Genomics UK (COG-UK) Consortium | Lighthouse Laboratory Plymouth and Alex Alderton, Roberto Amato, Jeffrey Barrett, Sonia Goncalves, Ewan Harrison, David K. Jackson, Ian Johnston, Dominic Kwiatkowski, Cordelia Langford, John Sillitoe on behalf of the Wellcome Sanger Institute COVID-19 Surveillance Team |
| EPI_ISL_3363377, EPI_ISL_3363378 | Randox Laboratories | Wellcome Sanger Institute for the COVID-19 Genomics UK (COG-UK) Consortium | Randox Laboratories and Alex Alderton, Roberto Amato, Jeffrey Barrett, Sonia Goncalves, Ewan Harrison, David K. Jackson, Ian Johnston, Dominic Kwiatkowski, Cordelia Langford, John Sillitoe on behalf of the Wellcome Sanger Institute COVID-19 Surveillance Team |
| EPI_ISL_3363379, EPI_ISL_3363380 | Lighthouse Laboratory Plymouth | Wellcome Sanger Institute for the COVID-19 Genomics UK (COG-UK) Consortium | Lighthouse Laboratory Plymouth and Alex Alderton, Roberto Amato, Jeffrey Barrett, Sonia Goncalves, Ewan Harrison, David K. Jackson, Ian Johnston, Dominic Kwiatkowski, Cordelia Langford, John Sillitoe on behalf of the Wellcome Sanger Institute COVID-19 Surveillance Team |
| EPI_ISL_3363381 | Randox Laboratories | Wellcome Sanger Institute for the COVID-19 Genomics UK (COG-UK) Consortium | Randox Laboratories and Alex Alderton, Roberto Amato, Jeffrey Barrett, Sonia Goncalves, Ewan Harrison, David K. Jackson, Ian Johnston, Dominic Kwiatkowski, Cordelia Langford, John Sillitoe on behalf of the Wellcome Sanger Institute COVID-19 Surveillance Team |
| EPI_ISL_3363382, EPI_ISL_3363383, EPI_ISL_3363384, EPI_ISL_3363385, EPI_ISL_3363386, EPI_ISL_3363387, EPI_ISL_3363388 | Lighthouse Laboratory Plymouth | Wellcome Sanger Institute for the COVID-19 Genomics UK (COG-UK) Consortium | Lighthouse Laboratory Plymouth and Alex Alderton, Roberto Amato, Jeffrey Barrett, Sonia Goncalves, Ewan Harrison, David K. Jackson, Ian Johnston, Dominic Kwiatkowski, Cordelia Langford, John Sillitoe on behalf of the Wellcome Sanger Institute COVID-19 Surveillance Team |
| EPI_ISL_3363389 | Randox Laboratories | Wellcome Sanger Institute for the COVID-19 Genomics UK (COG-UK) Consortium | Randox Laboratories and Alex Alderton, Roberto Amato, Jeffrey Barrett, Sonia Goncalves, Ewan Harrison, David K. Jackson, Ian Johnston, Dominic Kwiatkowski, Cordelia Langford, John Sillitoe on behalf of the Wellcome Sanger Institute COVID-19 Surveillance Team |
| EPI_ISL_3363390, EPI_ISL_3363391, EPI_ISL_3363392 | Lighthouse Laboratory Plymouth | Wellcome Sanger Institute for the COVID-19 Genomics UK (COG-UK) Consortium | Lighthouse Laboratory Plymouth and Alex Alderton, Roberto Amato, Jeffrey Barrett, Sonia Goncalves, Ewan Harrison, David K. Jackson, Ian Johnston, Dominic Kwiatkowski, Cordelia Langford, John Sillitoe on behalf of the Wellcome Sanger Institute COVID-19 Surveillance Team |
| EPI_ISL_3363393 | Randox Laboratories | Wellcome Sanger Institute for the COVID-19 Genomics UK (COG-UK) Consortium | Randox Laboratories and Alex Alderton, Roberto Amato, Jeffrey Barrett, Sonia Goncalves, Ewan Harrison, David K. Jackson, Ian Johnston, Dominic Kwiatkowski, Cordelia Langford, John Sillitoe on behalf of the Wellcome Sanger Institute COVID-19 Surveillance Team |
| EPI_ISL_3363394, EPI_ISL_3363395, EPI_ISL_3363396, EPI_ISL_3363397, EPI_ISL_3363398, EPI_ISL_3363399, EPI_ISL_3363400 | Lighthouse Laboratory Plymouth | Wellcome Sanger Institute for the COVID-19 Genomics UK (COG-UK) Consortium | Lighthouse Laboratory Plymouth and Alex Alderton, Roberto Amato, Jeffrey Barrett, Sonia Goncalves, Ewan Harrison, David K. Jackson, Ian Johnston, Dominic Kwiatkowski, Cordelia Langford, John Sillitoe on behalf of the Wellcome Sanger Institute COVID-19 Surveillance Team |
| EPI_ISL_3363401 | Randox Laboratories | Wellcome Sanger Institute for the COVID-19 Genomics UK (COG-UK) Consortium | Randox Laboratories and Alex Alderton, Roberto Amato, Jeffrey Barrett, Sonia Goncalves, Ewan Harrison, David K. Jackson, Ian Johnston, Dominic Kwiatkowski, Cordelia Langford, John Sillitoe on behalf of the Wellcome Sanger Institute COVID-19 Surveillance Team |
| EPI_ISL_3363402, EPI_ISL_3363403, EPI_ISL_3363404, EPI_ISL_3363405, EPI_ISL_3363406, EPI_ISL_3363407, EPI_ISL_3363408, EPI_ISL_3363409 | Lighthouse Laboratory Plymouth | Wellcome Sanger Institute for the COVID-19 Genomics UK (COG-UK) Consortium | Lighthouse Laboratory Plymouth and Alex Alderton, Roberto Amato, Jeffrey Barrett, Sonia Goncalves, Ewan Harrison, David K. Jackson, Ian Johnston, Dominic Kwiatkowski, Cordelia Langford, John Sillitoe on behalf of the Wellcome Sanger Institute COVID-19 Surveillance Team |
| EPI_ISL_3363410, EPI_ISL_3363411 | Randox Laboratories | Wellcome Sanger Institute for the COVID-19 Genomics UK (COG-UK) Consortium | Randox Laboratories and Alex Alderton, Roberto Amato, Jeffrey Barrett, Sonia Goncalves, Ewan Harrison, David K. Jackson, Ian Johnston, Dominic Kwiatkowski, Cordelia Langford, John Sillitoe on behalf of the Wellcome Sanger Institute COVID-19 Surveillance Team |
| EPI_ISL_3363412, EPI_ISL_3363413 | Lighthouse Laboratory Plymouth | Wellcome Sanger Institute for the COVID-19 Genomics UK (COG-UK) Consortium | Lighthouse Laboratory Plymouth and Alex Alderton, Roberto Amato, Jeffrey Barrett, Sonia Goncalves, Ewan Harrison, David K. Jackson, Ian Johnston, Dominic Kwiatkowski, Cordelia Langford, John Sillitoe on behalf of the Wellcome Sanger Institute COVID-19 Surveillance Team |
| EPI_ISL_3363414 | Randox Laboratories | Wellcome Sanger Institute for the COVID-19 Genomics UK (COG-UK) Consortium | Randox Laboratories and Alex Alderton, Roberto Amato, Jeffrey Barrett, Sonia Goncalves, Ewan Harrison, David K. Jackson, Ian Johnston, Dominic Kwiatkowski, Cordelia Langford, John Sillitoe on behalf of the Wellcome Sanger Institute COVID-19 Surveillance Team |
| EPI_ISL_3363415, EPI_ISL_3363416 | Lighthouse Laboratory Plymouth | Wellcome Sanger Institute for the COVID-19 Genomics UK (COG-UK) Consortium | Lighthouse Laboratory Plymouth and Alex Alderton, Roberto Amato, Jeffrey Barrett, Sonia Goncalves, Ewan Harrison, David K. Jackson, Ian Johnston, Dominic Kwiatkowski, Cordelia Langford, John Sillitoe on behalf of the Wellcome Sanger Institute COVID-19 Surveillance Team |
| EPI_ISL_3363417 | Randox Laboratories | Wellcome Sanger Institute for the COVID-19 Genomics UK (COG-UK) | Randox Laboratories and Alex Alderton, Roberto Amato, Jeffrey Barrett, Sonia Goncalves, Ewan Harrison, David K. Jackson, Ian Johnston, Dominic Kwiatkowski, Cordelia Langford, John Sillitoe on behalf of the Wellcome Sanger Institute COVID- |

| | | | |
|---|---|---|---|
| | | Consortium | 19 Surveillance Team |
| EPI_ISL_3363418, EPI_ISL_3363419, EPI_ISL_3363420, EPI_ISL_3363421 | Lighthouse Laboratory Plymouth | Wellcome Sanger Institute for the COVID-19 Genomics UK (COG-UK) Consortium | Lighthouse Laboratory Plymouth and Alex Alderton, Roberto Amato, Jeffrey Barrett, Sonia Goncalves, Ewan Harrison, David K. Jackson, Ian Johnston, Dominic Kwiatkowski, Cordelia Langford, John Sillitoe on behalf of the Wellcome Sanger Institute COVID-19 Surveillance Team |
| EPI_ISL_3363422, EPI_ISL_3363423 | Randox Laboratories | Wellcome Sanger Institute for the COVID-19 Genomics UK (COG-UK) Consortium | Randox Laboratories and Alex Alderton, Roberto Amato, Jeffrey Barrett, Sonia Goncalves, Ewan Harrison, David K. Jackson, Ian Johnston, Dominic Kwiatkowski, Cordelia Langford, John Sillitoe on behalf of the Wellcome Sanger Institute COVID-19 Surveillance Team |
| EPI_ISL_3363424, EPI_ISL_3363425, EPI_ISL_3363426, EPI_ISL_3363427 | Lighthouse Laboratory Plymouth | Wellcome Sanger Institute for the COVID-19 Genomics UK (COG-UK) Consortium | Lighthouse Laboratory Plymouth and Alex Alderton, Roberto Amato, Jeffrey Barrett, Sonia Goncalves, Ewan Harrison, David K. Jackson, Ian Johnston, Dominic Kwiatkowski, Cordelia Langford, John Sillitoe on behalf of the Wellcome Sanger Institute COVID-19 Surveillance Team |
| EPI_ISL_3363428 | Randox Laboratories | Wellcome Sanger Institute for the COVID-19 Genomics UK (COG-UK) Consortium | Randox Laboratories and Alex Alderton, Roberto Amato, Jeffrey Barrett, Sonia Goncalves, Ewan Harrison, David K. Jackson, Ian Johnston, Dominic Kwiatkowski, Cordelia Langford, John Sillitoe on behalf of the Wellcome Sanger Institute COVID-19 Surveillance Team |
| EPI_ISL_3363429, EPI_ISL_3363430 | Lighthouse Laboratory Plymouth | Wellcome Sanger Institute for the COVID-19 Genomics UK (COG-UK) Consortium | Lighthouse Laboratory Plymouth and Alex Alderton, Roberto Amato, Jeffrey Barrett, Sonia Goncalves, Ewan Harrison, David K. Jackson, Ian Johnston, Dominic Kwiatkowski, Cordelia Langford, John Sillitoe on behalf of the Wellcome Sanger Institute COVID-19 Surveillance Team |
| EPI_ISL_3363431 | Randox Laboratories | Wellcome Sanger Institute for the COVID-19 Genomics UK (COG-UK) Consortium | Randox Laboratories and Alex Alderton, Roberto Amato, Jeffrey Barrett, Sonia Goncalves, Ewan Harrison, David K. Jackson, Ian Johnston, Dominic Kwiatkowski, Cordelia Langford, John Sillitoe on behalf of the Wellcome Sanger Institute COVID-19 Surveillance Team |
| EPI_ISL_3363432, EPI_ISL_3363433 | Lighthouse Laboratory Plymouth | Wellcome Sanger Institute for the COVID-19 Genomics UK (COG-UK) Consortium | Lighthouse Laboratory Plymouth and Alex Alderton, Roberto Amato, Jeffrey Barrett, Sonia Goncalves, Ewan Harrison, David K. Jackson, Ian Johnston, Dominic Kwiatkowski, Cordelia Langford, John Sillitoe on behalf of the Wellcome Sanger Institute COVID-19 Surveillance Team |
| EPI_ISL_3363434, EPI_ISL_3363435 | Randox Laboratories | Wellcome Sanger Institute for the COVID-19 Genomics UK (COG-UK) Consortium | Randox Laboratories and Alex Alderton, Roberto Amato, Jeffrey Barrett, Sonia Goncalves, Ewan Harrison, David K. Jackson, Ian Johnston, Dominic Kwiatkowski, Cordelia Langford, John Sillitoe on behalf of the Wellcome Sanger Institute COVID-19 Surveillance Team |
| EPI_ISL_3363436, EPI_ISL_3363437, EPI_ISL_3363438, EPI_ISL_3363439, EPI_ISL_3363440, EPI_ISL_3363441, EPI_ISL_3363442 | Lighthouse Laboratory Plymouth | Wellcome Sanger Institute for the COVID-19 Genomics UK (COG-UK) Consortium | Lighthouse Laboratory Plymouth and Alex Alderton, Roberto Amato, Jeffrey Barrett, Sonia Goncalves, Ewan Harrison, David K. Jackson, Ian Johnston, Dominic Kwiatkowski, Cordelia Langford, John Sillitoe on behalf of the Wellcome Sanger Institute COVID-19 Surveillance Team |
| EPI_ISL_3363443, EPI_ISL_3363444 | Randox Laboratories | Wellcome Sanger Institute for the COVID-19 Genomics UK (COG-UK) Consortium | Randox Laboratories and Alex Alderton, Roberto Amato, Jeffrey Barrett, Sonia Goncalves, Ewan Harrison, David K. Jackson, Ian Johnston, Dominic Kwiatkowski, Cordelia Langford, John Sillitoe on behalf of the Wellcome Sanger Institute COVID-19 Surveillance Team |
| EPI_ISL_3363445, EPI_ISL_3363446, EPI_ISL_3363447 | Lighthouse Laboratory Plymouth | Wellcome Sanger Institute for the COVID-19 Genomics UK (COG-UK) Consortium | Lighthouse Laboratory Plymouth and Alex Alderton, Roberto Amato, Jeffrey Barrett, Sonia Goncalves, Ewan Harrison, David K. Jackson, Ian Johnston, Dominic Kwiatkowski, Cordelia Langford, John Sillitoe on behalf of the Wellcome Sanger Institute COVID-19 Surveillance Team |
| EPI_ISL_3363448 | Randox Laboratories | Wellcome Sanger Institute for the COVID-19 Genomics UK (COG-UK) Consortium | Randox Laboratories and Alex Alderton, Roberto Amato, Jeffrey Barrett, Sonia Goncalves, Ewan Harrison, David K. Jackson, Ian Johnston, Dominic Kwiatkowski, Cordelia Langford, John Sillitoe on behalf of the Wellcome Sanger Institute COVID-19 Surveillance Team |
| EPI_ISL_3363449, EPI_ISL_3363450, EPI_ISL_3363451, EPI_ISL_3363452, EPI_ISL_3363453, EPI_ISL_3363454, EPI_ISL_3363455 | Lighthouse Laboratory Plymouth | Wellcome Sanger Institute for the COVID-19 Genomics UK (COG-UK) Consortium | Lighthouse Laboratory Plymouth and Alex Alderton, Roberto Amato, Jeffrey Barrett, Sonia Goncalves, Ewan Harrison, David K. Jackson, Ian Johnston, Dominic Kwiatkowski, Cordelia Langford, John Sillitoe on behalf of the Wellcome Sanger Institute COVID-19 Surveillance Team |
| EPI_ISL_3363456 | Randox Laboratories | Wellcome Sanger Institute for the COVID-19 Genomics UK (COG-UK) Consortium | Randox Laboratories and Alex Alderton, Roberto Amato, Jeffrey Barrett, Sonia Goncalves, Ewan Harrison, David K. Jackson, Ian Johnston, Dominic Kwiatkowski, Cordelia Langford, John Sillitoe on behalf of the Wellcome Sanger Institute COVID-19 Surveillance Team |
| EPI_ISL_3363457, EPI_ISL_3363458, EPI_ISL_3363459 | Lighthouse Laboratory Plymouth | Wellcome Sanger Institute for the COVID-19 Genomics UK (COG-UK) Consortium | Lighthouse Laboratory Plymouth and Alex Alderton, Roberto Amato, Jeffrey Barrett, Sonia Goncalves, Ewan Harrison, David K. Jackson, Ian Johnston, Dominic Kwiatkowski, Cordelia Langford, John Sillitoe on behalf of the Wellcome Sanger Institute COVID-19 Surveillance Team |
| EPI_ISL_3363460, EPI_ISL_3363461 | Randox Laboratories | Wellcome Sanger Institute for the COVID-19 Genomics UK (COG-UK) Consortium | Randox Laboratories and Alex Alderton, Roberto Amato, Jeffrey Barrett, Sonia Goncalves, Ewan Harrison, David K. Jackson, Ian Johnston, Dominic Kwiatkowski, Cordelia Langford, John Sillitoe on behalf of the Wellcome Sanger Institute COVID-19 Surveillance Team |
| EPI_ISL_3363462, EPI_ISL_3363463 | Lighthouse Laboratory Plymouth | Wellcome Sanger Institute for the COVID-19 Genomics UK (COG-UK) Consortium | Lighthouse Laboratory Plymouth and Alex Alderton, Roberto Amato, Jeffrey Barrett, Sonia Goncalves, Ewan Harrison, David K. Jackson, Ian Johnston, Dominic Kwiatkowski, Cordelia Langford, John Sillitoe on behalf of the Wellcome Sanger Institute COVID-19 Surveillance Team |
| EPI_ISL_3363464 | Randox Laboratories | Wellcome Sanger Institute for the COVID-19 Genomics UK (COG-UK) Consortium | Randox Laboratories and Alex Alderton, Roberto Amato, Jeffrey Barrett, Sonia Goncalves, Ewan Harrison, David K. Jackson, Ian Johnston, Dominic Kwiatkowski, Cordelia Langford, John Sillitoe on behalf of the Wellcome Sanger Institute COVID-19 Surveillance Team |
| EPI_ISL_3363465 | Lighthouse Laboratory Plymouth | Wellcome Sanger Institute for the COVID-19 Genomics UK (COG-UK) Consortium | Lighthouse Laboratory Plymouth and Alex Alderton, Roberto Amato, Jeffrey Barrett, Sonia Goncalves, Ewan Harrison, David K. Jackson, Ian Johnston, Dominic Kwiatkowski, Cordelia Langford, John Sillitoe on behalf of the Wellcome Sanger Institute COVID-19 Surveillance Team |
| EPI_ISL_3363466 | Randox Laboratories | Wellcome Sanger Institute for the COVID-19 Genomics UK (COG-UK) Consortium | Randox Laboratories and Alex Alderton, Roberto Amato, Jeffrey Barrett, Sonia Goncalves, Ewan Harrison, David K. Jackson, Ian Johnston, Dominic Kwiatkowski, Cordelia Langford, John Sillitoe on behalf of the Wellcome Sanger Institute COVID-19 Surveillance Team |
| EPI_ISL_3363467 | Lighthouse Laboratory Plymouth | Wellcome Sanger Institute for the COVID-19 Genomics UK (COG-UK) Consortium | Lighthouse Laboratory Plymouth and Alex Alderton, Roberto Amato, Jeffrey Barrett, Sonia Goncalves, Ewan Harrison, David K. Jackson, Ian Johnston, Dominic Kwiatkowski, Cordelia Langford, John Sillitoe on behalf of the Wellcome Sanger Institute COVID-19 Surveillance Team |
| EPI_ISL_3363468, EPI_ISL_3363469 | Randox Laboratories | Wellcome Sanger Institute for the COVID-19 Genomics UK (COG-UK) Consortium | Randox Laboratories and Alex Alderton, Roberto Amato, Jeffrey Barrett, Sonia Goncalves, Ewan Harrison, David K. Jackson, Ian Johnston, Dominic Kwiatkowski, Cordelia Langford, John Sillitoe on behalf of the Wellcome Sanger Institute COVID-19 Surveillance Team |
| EPI_ISL_3363470, EPI_ISL_3363471 | Lighthouse Laboratory Plymouth | Wellcome Sanger Institute for the COVID-19 Genomics UK (COG-UK) Consortium | Lighthouse Laboratory Plymouth and Alex Alderton, Roberto Amato, Jeffrey Barrett, Sonia Goncalves, Ewan Harrison, David K. Jackson, Ian Johnston, Dominic Kwiatkowski, Cordelia Langford, John Sillitoe on behalf of the Wellcome Sanger Institute COVID-19 Surveillance Team |
| EPI_ISL_3363472, EPI_ISL_3363473 | Randox Laboratories | Wellcome Sanger Institute for the COVID-19 Genomics UK (COG-UK) Consortium | Randox Laboratories and Alex Alderton, Roberto Amato, Jeffrey Barrett, Sonia Goncalves, Ewan Harrison, David K. Jackson, Ian Johnston, Dominic Kwiatkowski, Cordelia Langford, John Sillitoe on behalf of the Wellcome Sanger Institute COVID-19 Surveillance Team |
| EPI_ISL_3363474 | Lighthouse Laboratory Plymouth | Wellcome Sanger Institute for the COVID-19 Genomics UK (COG-UK) Consortium | Lighthouse Laboratory Plymouth and Alex Alderton, Roberto Amato, Jeffrey Barrett, Sonia Goncalves, Ewan Harrison, David K. Jackson, Ian Johnston, Dominic Kwiatkowski, Cordelia Langford, John Sillitoe on behalf of the Wellcome Sanger Institute COVID-19 Surveillance Team |
| EPI_ISL_3363475 | Randox Laboratories | Wellcome Sanger Institute for the COVID-19 Genomics UK (COG-UK) Consortium | Randox Laboratories and Alex Alderton, Roberto Amato, Jeffrey Barrett, Sonia Goncalves, Ewan Harrison, David K. Jackson, Ian Johnston, Dominic Kwiatkowski, Cordelia Langford, John Sillitoe on behalf of the Wellcome Sanger Institute COVID-19 Surveillance Team |
| EPI_ISL_3363476 | Lighthouse Laboratory Plymouth | Wellcome Sanger Institute for the COVID-19 Genomics UK (COG-UK) Consortium | Lighthouse Laboratory Plymouth and Alex Alderton, Roberto Amato, Jeffrey Barrett, Sonia Goncalves, Ewan Harrison, David K. Jackson, Ian Johnston, Dominic Kwiatkowski, Cordelia Langford, John Sillitoe on behalf of the Wellcome Sanger Institute COVID-19 Surveillance Team |
| EPI_ISL_3363477, EPI_ISL_3363478 | Randox Laboratories | Wellcome Sanger Institute for the COVID-19 Genomics UK (COG-UK) Consortium | Randox Laboratories and Alex Alderton, Roberto Amato, Jeffrey Barrett, Sonia Goncalves, Ewan Harrison, David K. Jackson, Ian Johnston, Dominic Kwiatkowski, Cordelia Langford, John Sillitoe on behalf of the Wellcome Sanger Institute COVID-19 Surveillance Team |
| EPI_ISL_3363479 | Lighthouse Laboratory Plymouth | Wellcome Sanger Institute for the COVID-19 Genomics UK (COG-UK) Consortium | Lighthouse Laboratory Plymouth and Alex Alderton, Roberto Amato, Jeffrey Barrett, Sonia Goncalves, Ewan Harrison, David K. Jackson, Ian Johnston, Dominic Kwiatkowski, Cordelia Langford, John Sillitoe on behalf of the Wellcome Sanger Institute COVID-19 Surveillance Team |
| EPI_ISL_3363480 | Randox Laboratories | Wellcome Sanger Institute for the COVID-19 Genomics UK (COG-UK) Consortium | Randox Laboratories and Alex Alderton, Roberto Amato, Jeffrey Barrett, Sonia Goncalves, Ewan Harrison, David K. Jackson, Ian Johnston, Dominic Kwiatkowski, Cordelia Langford, John Sillitoe on behalf of the Wellcome Sanger Institute COVID-19 Surveillance Team |
| EPI_ISL_3363481 | Lighthouse Laboratory Plymouth | Wellcome Sanger Institute for the COVID-19 Genomics UK (COG-UK) Consortium | Lighthouse Laboratory Plymouth and Alex Alderton, Roberto Amato, Jeffrey Barrett, Sonia Goncalves, Ewan Harrison, David K. Jackson, Ian Johnston, Dominic Kwiatkowski, Cordelia Langford, John Sillitoe on behalf of the Wellcome Sanger Institute COVID-19 Surveillance Team |
| EPI_ISL_3363483, EPI_ISL_3363484 | Lighthouse Lab in Milton Keynes | Wellcome Sanger Institute for the COVID-19 Genomics UK (COG-UK) Consortium | The Lighthouse Lab in Milton Keynes and Alex Alderton, Roberto Amato, Jeffrey Barrett, Sonia Goncalves, Ewan Harrison, David K. Jackson, Ian Johnston, Dominic Kwiatkowski, Cordelia Langford, John Sillitoe on behalf of the Wellcome Sanger Institute COVID-19 Surveillance Team |
| EPI_ISL_3363485 | Berkshire and Surrey Pathology Services Lighthouse Laboratory | Wellcome Sanger Institute for the COVID-19 Genomics UK (COG-UK) Consortium | Berkshire and Surrey Pathology Services Lighthouse Laboratory and Alex Alderton, Roberto Amato, Jeffrey Barrett, Sonia Goncalves, Ewan Harrison, David K. Jackson, Ian Johnston, Dominic Kwiatkowski, Cordelia Langford, John Sillitoe on behalf of the Wellcome Sanger Institute COVID-19 Surveillance Team |
| EPI_ISL_3363486 | Lighthouse Lab in Glasgow | Wellcome Sanger Institute for the COVID-19 Genomics UK (COG-UK) Consortium | Harper VanSteenhouse, Yumi Kasai, David Gray, Carol Clugston, Anna Dominiczak and Alex Alderton, Roberto Amato, Jeffrey Barrett, Sonia Goncalves, Ewan Harrison, David K. Jackson, Ian Johnston, Dominic Kwiatkowski, Cordelia Langford, John Sillitoe on behalf of the Wellcome Sanger Institute COVID-19 Surveillance Team |
| EPI_ISL_3363487 | Lighthouse Lab in Alderley Park | Wellcome Sanger Institute for the COVID-19 Genomics UK (COG-UK) Consortium | Jacquelyn Wynn, Mairead Hyland, The Lighthouse Lab in Alderley Park and Alex Alderton, Roberto Amato, Jeffrey Barrett, Sonia Goncalves, Ewan Harrison, David K. Jackson, Ian Johnston, Dominic Kwiatkowski, Cordelia Langford, John Sillitoe on behalf of the Wellcome Sanger Institute COVID-19 Surveillance Team |
| EPI_ISL_3363488 | Berkshire and Surrey Pathology Services Lighthouse Laboratory | Wellcome Sanger Institute for the COVID-19 Genomics UK (COG-UK) Consortium | Berkshire and Surrey Pathology Services Lighthouse Laboratory and Alex Alderton, Roberto Amato, Jeffrey Barrett, Sonia Goncalves, Ewan Harrison, David K. Jackson, Ian Johnston, Dominic Kwiatkowski, Cordelia Langford, John Sillitoe on behalf of the Wellcome Sanger Institute COVID-19 Surveillance Team |
| EPI_ISL_3363489 | Lighthouse Lab in Milton Keynes | Wellcome Sanger Institute for the COVID-19 Genomics UK (COG-UK) Consortium | The Lighthouse Lab in Milton Keynes and Alex Alderton, Roberto Amato, Jeffrey Barrett, Sonia Goncalves, Ewan Harrison, David K. Jackson, Ian Johnston, Dominic Kwiatkowski, Cordelia Langford, John Sillitoe on behalf of the Wellcome Sanger Institute COVID-19 Surveillance Team |
| EPI_ISL_3363491, EPI_ISL_3363492 | Berkshire and Surrey Pathology Services Lighthouse Laboratory | Wellcome Sanger Institute for the COVID-19 Genomics UK (COG-UK) Consortium | Berkshire and Surrey Pathology Services Lighthouse Laboratory and Alex Alderton, Roberto Amato, Jeffrey Barrett, Sonia Goncalves, Ewan Harrison, David K. Jackson, Ian Johnston, Dominic Kwiatkowski, Cordelia Langford, John Sillitoe on behalf of the Wellcome Sanger Institute COVID-19 Surveillance Team |

| | | | |
|---|---|---|---|
| EPI_ISL_3363493 | Lighthouse Lab in Milton Keynes | Wellcome Sanger Institute for the COVID-19 Genomics UK (COG-UK) Consortium | The Lighthouse Lab in Milton Keynes and Alex Alderton, Roberto Amato, Jeffrey Barrett, Sonia Goncalves, Ewan Harrison, David K. Jackson, Ian Johnston, Dominic Kwiatkowski, Cordelia Langford, John Sillitoe on behalf of the Wellcome Sanger Institute COVID-19 Surveillance Team |
| EPI_ISL_3363495 | Lighthouse Lab in Alderley Park | Wellcome Sanger Institute for the COVID-19 Genomics UK (COG-UK) Consortium | Jacquelyn Wynn, Mairead Hyland, The Lighthouse Lab in Alderley Park and Alex Alderton, Roberto Amato, Jeffrey Barrett, Sonia Goncalves, Ewan Harrison, David K. Jackson, Ian Johnston, Dominic Kwiatkowski, Cordelia Langford, John Sillitoe on behalf of the Wellcome Sanger Institute COVID-19 Surveillance Team |
| EPI_ISL_3363496 | Lighthouse Lab in Milton Keynes | Wellcome Sanger Institute for the COVID-19 Genomics UK (COG-UK) Consortium | The Lighthouse Lab in Milton Keynes and Alex Alderton, Roberto Amato, Jeffrey Barrett, Sonia Goncalves, Ewan Harrison, David K. Jackson, Ian Johnston, Dominic Kwiatkowski, Cordelia Langford, John Sillitoe on behalf of the Wellcome Sanger Institute COVID-19 Surveillance Team |
| EPI_ISL_3363497 | Berkshire and Surrey Pathology Services Lighthouse Laboratory | Wellcome Sanger Institute for the COVID-19 Genomics UK (COG-UK) Consortium | Berkshire and Surrey Pathology Services Lighthouse Laboratory and Alex Alderton, Roberto Amato, Jeffrey Barrett, Sonia Goncalves, Ewan Harrison, David K. Jackson, Ian Johnston, Dominic Kwiatkowski, Cordelia Langford, John Sillitoe on behalf of the Wellcome Sanger Institute COVID-19 Surveillance Team |
| EPI_ISL_3363499, EPI_ISL_3363500 | Lighthouse Lab in Milton Keynes | Wellcome Sanger Institute for the COVID-19 Genomics UK (COG-UK) Consortium | The Lighthouse Lab in Milton Keynes and Alex Alderton, Roberto Amato, Jeffrey Barrett, Sonia Goncalves, Ewan Harrison, David K. Jackson, Ian Johnston, Dominic Kwiatkowski, Cordelia Langford, John Sillitoe on behalf of the Wellcome Sanger Institute COVID-19 Surveillance Team |
| EPI_ISL_3363502, EPI_ISL_3363503 | Lighthouse Lab in Alderley Park | Wellcome Sanger Institute for the COVID-19 Genomics UK (COG-UK) Consortium | Jacquelyn Wynn, Mairead Hyland, The Lighthouse Lab in Alderley Park and Alex Alderton, Roberto Amato, Jeffrey Barrett, Sonia Goncalves, Ewan Harrison, David K. Jackson, Ian Johnston, Dominic Kwiatkowski, Cordelia Langford, John Sillitoe on behalf of the Wellcome Sanger Institute COVID-19 Surveillance Team |
| EPI_ISL_3363505 | Lighthouse Lab in Milton Keynes | Wellcome Sanger Institute for the COVID-19 Genomics UK (COG-UK) Consortium | The Lighthouse Lab in Milton Keynes and Alex Alderton, Roberto Amato, Jeffrey Barrett, Sonia Goncalves, Ewan Harrison, David K. Jackson, Ian Johnston, Dominic Kwiatkowski, Cordelia Langford, John Sillitoe on behalf of the Wellcome Sanger Institute COVID-19 Surveillance Team |
| EPI_ISL_3363506 | Lighthouse Lab in Alderley Park | Wellcome Sanger Institute for the COVID-19 Genomics UK (COG-UK) Consortium | Jacquelyn Wynn, Mairead Hyland, The Lighthouse Lab in Alderley Park and Alex Alderton, Roberto Amato, Jeffrey Barrett, Sonia Goncalves, Ewan Harrison, David K. Jackson, Ian Johnston, Dominic Kwiatkowski, Cordelia Langford, John Sillitoe on behalf of the Wellcome Sanger Institute COVID-19 Surveillance Team |
| EPI_ISL_3363507, EPI_ISL_3363508 | Berkshire and Surrey Pathology Services Lighthouse Laboratory | Wellcome Sanger Institute for the COVID-19 Genomics UK (COG-UK) Consortium | Berkshire and Surrey Pathology Services Lighthouse Laboratory and Alex Alderton, Roberto Amato, Jeffrey Barrett, Sonia Goncalves, Ewan Harrison, David K. Jackson, Ian Johnston, Dominic Kwiatkowski, Cordelia Langford, John Sillitoe on behalf of the Wellcome Sanger Institute COVID-19 Surveillance Team |
| EPI_ISL_3363510 | Lighthouse Lab in Glasgow | Wellcome Sanger Institute for the COVID-19 Genomics UK (COG-UK) Consortium | Harper VanSteenhouse, Yumi Kasai, David Gray, Carol Clugston, Anna Dominiczak and Alex Alderton, Roberto Amato, Jeffrey Barrett, Sonia Goncalves, Ewan Harrison, David K. Jackson, Ian Johnston, Dominic Kwiatkowski, Cordelia Langford, John Sillitoe on behalf of the Wellcome Sanger Institute COVID-19 Surveillance Team |
| EPI_ISL_3363511 | Berkshire and Surrey Pathology Services Lighthouse Laboratory | Wellcome Sanger Institute for the COVID-19 Genomics UK (COG-UK) Consortium | Berkshire and Surrey Pathology Services Lighthouse Laboratory and Alex Alderton, Roberto Amato, Jeffrey Barrett, Sonia Goncalves, Ewan Harrison, David K. Jackson, Ian Johnston, Dominic Kwiatkowski, Cordelia Langford, John Sillitoe on behalf of the Wellcome Sanger Institute COVID-19 Surveillance Team |
| EPI_ISL_3363512 | Lighthouse Lab in Glasgow | Wellcome Sanger Institute for the COVID-19 Genomics UK (COG-UK) Consortium | Harper VanSteenhouse, Yumi Kasai, David Gray, Carol Clugston, Anna Dominiczak and Alex Alderton, Roberto Amato, Jeffrey Barrett, Sonia Goncalves, Ewan Harrison, David K. Jackson, Ian Johnston, Dominic Kwiatkowski, Cordelia Langford, John Sillitoe on behalf of the Wellcome Sanger Institute COVID-19 Surveillance Team |
| EPI_ISL_3363513 | Lighthouse Lab in Milton Keynes | Wellcome Sanger Institute for the COVID-19 Genomics UK (COG-UK) Consortium | The Lighthouse Lab in Milton Keynes and Alex Alderton, Roberto Amato, Jeffrey Barrett, Sonia Goncalves, Ewan Harrison, David K. Jackson, Ian Johnston, Dominic Kwiatkowski, Cordelia Langford, John Sillitoe on behalf of the Wellcome Sanger Institute COVID-19 Surveillance Team |
| EPI_ISL_3363514 | Lighthouse Lab in Glasgow | Wellcome Sanger Institute for the COVID-19 Genomics UK (COG-UK) Consortium | Harper VanSteenhouse, Yumi Kasai, David Gray, Carol Clugston, Anna Dominiczak and Alex Alderton, Roberto Amato, Jeffrey Barrett, Sonia Goncalves, Ewan Harrison, David K. Jackson, Ian Johnston, Dominic Kwiatkowski, Cordelia Langford, John Sillitoe on behalf of the Wellcome Sanger Institute COVID-19 Surveillance Team |
| EPI_ISL_3363515 | Lighthouse Lab in Alderley Park | Wellcome Sanger Institute for the COVID-19 Genomics UK (COG-UK) Consortium | Jacquelyn Wynn, Mairead Hyland, The Lighthouse Lab in Alderley Park and Alex Alderton, Roberto Amato, Jeffrey Barrett, Sonia Goncalves, Ewan Harrison, David K. Jackson, Ian Johnston, Dominic Kwiatkowski, Cordelia Langford, John Sillitoe on behalf of the Wellcome Sanger Institute COVID-19 Surveillance Team |
| EPI_ISL_3363516 | Berkshire and Surrey Pathology Services Lighthouse Laboratory | Wellcome Sanger Institute for the COVID-19 Genomics UK (COG-UK) Consortium | Berkshire and Surrey Pathology Services Lighthouse Laboratory and Alex Alderton, Roberto Amato, Jeffrey Barrett, Sonia Goncalves, Ewan Harrison, David K. Jackson, Ian Johnston, Dominic Kwiatkowski, Cordelia Langford, John Sillitoe on behalf of the Wellcome Sanger Institute COVID-19 Surveillance Team |
| EPI_ISL_3363518, EPI_ISL_3363519 | Lighthouse Lab in Milton Keynes | Wellcome Sanger Institute for the COVID-19 Genomics UK (COG-UK) Consortium | The Lighthouse Lab in Milton Keynes and Alex Alderton, Roberto Amato, Jeffrey Barrett, Sonia Goncalves, Ewan Harrison, David K. Jackson, Ian Johnston, Dominic Kwiatkowski, Cordelia Langford, John Sillitoe on behalf of the Wellcome Sanger Institute COVID-19 Surveillance Team |
| EPI_ISL_3363521, EPI_ISL_3363522, EPI_ISL_3363523 | Lighthouse Lab in Glasgow | Wellcome Sanger Institute for the COVID-19 Genomics UK (COG-UK) Consortium | Harper VanSteenhouse, Yumi Kasai, David Gray, Carol Clugston, Anna Dominiczak and Alex Alderton, Roberto Amato, Jeffrey Barrett, Sonia Goncalves, Ewan Harrison, David K. Jackson, Ian Johnston, Dominic Kwiatkowski, Cordelia Langford, John Sillitoe on behalf of the Wellcome Sanger Institute COVID-19 Surveillance Team |
| EPI_ISL_3363524 | Lighthouse Lab in Alderley Park | Wellcome Sanger Institute for the COVID-19 Genomics UK (COG-UK) Consortium | Jacquelyn Wynn, Mairead Hyland, The Lighthouse Lab in Alderley Park and Alex Alderton, Roberto Amato, Jeffrey Barrett, Sonia Goncalves, Ewan Harrison, David K. Jackson, Ian Johnston, Dominic Kwiatkowski, Cordelia Langford, John Sillitoe on behalf of the Wellcome Sanger Institute COVID-19 Surveillance Team |
| EPI_ISL_3363526 | Lighthouse Lab in Milton Keynes | Wellcome Sanger Institute for the COVID-19 Genomics UK (COG-UK) Consortium | The Lighthouse Lab in Milton Keynes and Alex Alderton, Roberto Amato, Jeffrey Barrett, Sonia Goncalves, Ewan Harrison, David K. Jackson, Ian Johnston, Dominic Kwiatkowski, Cordelia Langford, John Sillitoe on behalf of the Wellcome Sanger Institute COVID-19 Surveillance Team |
| EPI_ISL_3363527 | Berkshire and Surrey Pathology Services Lighthouse Laboratory | Wellcome Sanger Institute for the COVID-19 Genomics UK (COG-UK) Consortium | Berkshire and Surrey Pathology Services Lighthouse Laboratory and Alex Alderton, Roberto Amato, Jeffrey Barrett, Sonia Goncalves, Ewan Harrison, David K. Jackson, Ian Johnston, Dominic Kwiatkowski, Cordelia Langford, John Sillitoe on behalf of the Wellcome Sanger Institute COVID-19 Surveillance Team |
| EPI_ISL_3363529 | Lighthouse Lab in Alderley Park | Wellcome Sanger Institute for the COVID-19 Genomics UK (COG-UK) Consortium | Jacquelyn Wynn, Mairead Hyland, The Lighthouse Lab in Alderley Park and Alex Alderton, Roberto Amato, Jeffrey Barrett, Sonia Goncalves, Ewan Harrison, David K. Jackson, Ian Johnston, Dominic Kwiatkowski, Cordelia Langford, John Sillitoe on behalf of the Wellcome Sanger Institute COVID-19 Surveillance Team |
| EPI_ISL_3363530, EPI_ISL_3363531 | Berkshire and Surrey Pathology Services Lighthouse Laboratory | Wellcome Sanger Institute for the COVID-19 Genomics UK (COG-UK) Consortium | Berkshire and Surrey Pathology Services Lighthouse Laboratory and Alex Alderton, Roberto Amato, Jeffrey Barrett, Sonia Goncalves, Ewan Harrison, David K. Jackson, Ian Johnston, Dominic Kwiatkowski, Cordelia Langford, John Sillitoe on behalf of the Wellcome Sanger Institute COVID-19 Surveillance Team |
| EPI_ISL_3363532 | Lighthouse Lab in Milton Keynes | Wellcome Sanger Institute for the COVID-19 Genomics UK (COG-UK) Consortium | The Lighthouse Lab in Milton Keynes and Alex Alderton, Roberto Amato, Jeffrey Barrett, Sonia Goncalves, Ewan Harrison, David K. Jackson, Ian Johnston, Dominic Kwiatkowski, Cordelia Langford, John Sillitoe on behalf of the Wellcome Sanger Institute COVID-19 Surveillance Team |
| EPI_ISL_3363534, EPI_ISL_3363535 | Lighthouse Lab in Alderley Park | Wellcome Sanger Institute for the COVID-19 Genomics UK (COG-UK) Consortium | Jacquelyn Wynn, Mairead Hyland, The Lighthouse Lab in Alderley Park and Alex Alderton, Roberto Amato, Jeffrey Barrett, Sonia Goncalves, Ewan Harrison, David K. Jackson, Ian Johnston, Dominic Kwiatkowski, Cordelia Langford, John Sillitoe on behalf of the Wellcome Sanger Institute COVID-19 Surveillance Team |
| EPI_ISL_3363536, EPI_ISL_3363538 | Lighthouse Lab in Glasgow | Wellcome Sanger Institute for the COVID-19 Genomics UK (COG-UK) Consortium | Harper VanSteenhouse, Yumi Kasai, David Gray, Carol Clugston, Anna Dominiczak and Alex Alderton, Roberto Amato, Jeffrey Barrett, Sonia Goncalves, Ewan Harrison, David K. Jackson, Ian Johnston, Dominic Kwiatkowski, Cordelia Langford, John Sillitoe on behalf of the Wellcome Sanger Institute COVID-19 Surveillance Team |
| EPI_ISL_3363539, EPI_ISL_3363541 | Berkshire and Surrey Pathology Services Lighthouse Laboratory | Wellcome Sanger Institute for the COVID-19 Genomics UK (COG-UK) Consortium | Berkshire and Surrey Pathology Services Lighthouse Laboratory and Alex Alderton, Roberto Amato, Jeffrey Barrett, Sonia Goncalves, Ewan Harrison, David K. Jackson, Ian Johnston, Dominic Kwiatkowski, Cordelia Langford, John Sillitoe on behalf of the Wellcome Sanger Institute COVID-19 Surveillance Team |
| EPI_ISL_3363542 | Lighthouse Lab in Alderley Park | Wellcome Sanger Institute for the COVID-19 Genomics UK (COG-UK) Consortium | Jacquelyn Wynn, Mairead Hyland, The Lighthouse Lab in Alderley Park and Alex Alderton, Roberto Amato, Jeffrey Barrett, Sonia Goncalves, Ewan Harrison, David K. Jackson, Ian Johnston, Dominic Kwiatkowski, Cordelia Langford, John Sillitoe on behalf of the Wellcome Sanger Institute COVID-19 Surveillance Team |
| EPI_ISL_3363543, EPI_ISL_3363545 | Berkshire and Surrey Pathology Services Lighthouse Laboratory | Wellcome Sanger Institute for the COVID-19 Genomics UK (COG-UK) Consortium | Berkshire and Surrey Pathology Services Lighthouse Laboratory and Alex Alderton, Roberto Amato, Jeffrey Barrett, Sonia Goncalves, Ewan Harrison, David K. Jackson, Ian Johnston, Dominic Kwiatkowski, Cordelia Langford, John Sillitoe on behalf of the Wellcome Sanger Institute COVID-19 Surveillance Team |
| EPI_ISL_3363546, EPI_ISL_3363547 | Lighthouse Lab in Alderley Park | Wellcome Sanger Institute for the COVID-19 Genomics UK (COG-UK) Consortium | Jacquelyn Wynn, Mairead Hyland, The Lighthouse Lab in Alderley Park and Alex Alderton, Roberto Amato, Jeffrey Barrett, Sonia Goncalves, Ewan Harrison, David K. Jackson, Ian Johnston, Dominic Kwiatkowski, Cordelia Langford, John Sillitoe on behalf of the Wellcome Sanger Institute COVID-19 Surveillance Team |
| EPI_ISL_3363549, EPI_ISL_3363550 | Berkshire and Surrey Pathology Services Lighthouse Laboratory | Wellcome Sanger Institute for the COVID-19 Genomics UK (COG-UK) Consortium | Berkshire and Surrey Pathology Services Lighthouse Laboratory and Alex Alderton, Roberto Amato, Jeffrey Barrett, Sonia Goncalves, Ewan Harrison, David K. Jackson, Ian Johnston, Dominic Kwiatkowski, Cordelia Langford, John Sillitoe on behalf of the Wellcome Sanger Institute COVID-19 Surveillance Team |
| EPI_ISL_3363552 | Lighthouse Lab in Glasgow | Wellcome Sanger Institute for the COVID-19 Genomics UK (COG-UK) Consortium | Harper VanSteenhouse, Yumi Kasai, David Gray, Carol Clugston, Anna Dominiczak and Alex Alderton, Roberto Amato, Jeffrey Barrett, Sonia Goncalves, Ewan Harrison, David K. Jackson, Ian Johnston, Dominic Kwiatkowski, Cordelia Langford, John Sillitoe on behalf of the Wellcome Sanger Institute COVID-19 Surveillance Team |
| EPI_ISL_3363553 | Berkshire and Surrey Pathology Services Lighthouse Laboratory | Wellcome Sanger Institute for the COVID-19 Genomics UK (COG-UK) Consortium | Berkshire and Surrey Pathology Services Lighthouse Laboratory and Alex Alderton, Roberto Amato, Jeffrey Barrett, Sonia Goncalves, Ewan Harrison, David K. Jackson, Ian Johnston, Dominic Kwiatkowski, Cordelia Langford, John Sillitoe on behalf of the Wellcome Sanger Institute COVID-19 Surveillance Team |
| EPI_ISL_3363554 | Lighthouse Lab in Milton Keynes | Wellcome Sanger Institute for the COVID-19 Genomics UK (COG-UK) Consortium | The Lighthouse Lab in Milton Keynes and Alex Alderton, Roberto Amato, Jeffrey Barrett, Sonia Goncalves, Ewan Harrison, David K. Jackson, Ian Johnston, Dominic Kwiatkowski, Cordelia Langford, John Sillitoe on behalf of the Wellcome Sanger Institute COVID-19 Surveillance Team |
| EPI_ISL_3363556, EPI_ISL_3363557 | Lighthouse Lab in Alderley Park | Wellcome Sanger Institute for the COVID-19 Genomics UK (COG-UK) Consortium | Jacquelyn Wynn, Mairead Hyland, The Lighthouse Lab in Alderley Park and Alex Alderton, Roberto Amato, Jeffrey Barrett, Sonia Goncalves, Ewan Harrison, David K. Jackson, Ian Johnston, Dominic Kwiatkowski, Cordelia Langford, John Sillitoe on behalf of the Wellcome Sanger Institute COVID-19 Surveillance Team |
| EPI_ISL_3363558 | Lighthouse Lab in Glasgow | Wellcome Sanger Institute for the COVID-19 Genomics UK (COG-UK) Consortium | Harper VanSteenhouse, Yumi Kasai, David Gray, Carol Clugston, Anna Dominiczak and Alex Alderton, Roberto Amato, Jeffrey Barrett, Sonia Goncalves, Ewan Harrison, David K. Jackson, Ian Johnston, Dominic Kwiatkowski, Cordelia Langford, John Sillitoe on behalf of the Wellcome Sanger Institute COVID-19 Surveillance Team |
| EPI_ISL_3363560 | Lighthouse Lab in Milton Keynes | Wellcome Sanger Institute for the COVID-19 Genomics UK (COG-UK) Consortium | The Lighthouse Lab in Milton Keynes and Alex Alderton, Roberto Amato, Jeffrey Barrett, Sonia Goncalves, Ewan Harrison, David K. Jackson, Ian Johnston, Dominic Kwiatkowski, Cordelia Langford, John Sillitoe on behalf of the Wellcome Sanger Institute COVID-19 Surveillance Team |
| EPI_ISL_3363561 | Lighthouse Lab in Alderley Park | Wellcome Sanger Institute for the COVID-19 Genomics UK (COG-UK) Consortium | Jacquelyn Wynn, Mairead Hyland, The Lighthouse Lab in Alderley Park and Alex Alderton, Roberto Amato, Jeffrey Barrett, Sonia Goncalves, Ewan Harrison, David K. Jackson, Ian Johnston, Dominic Kwiatkowski, Cordelia Langford, John Sillitoe on behalf of the Wellcome Sanger Institute COVID-19 Surveillance Team |
| EPI_ISL_3363562 | Lighthouse Lab in Glasgow | Wellcome Sanger Institute for the COVID-19 Genomics UK (COG-UK) Consortium | Harper VanSteenhouse, Yumi Kasai, David Gray, Carol Clugston, Anna Dominiczak and Alex Alderton, Roberto Amato, Jeffrey Barrett, Sonia Goncalves, Ewan Harrison, David K. Jackson, Ian Johnston, Dominic Kwiatkowski, Cordelia Langford, John Sillitoe on behalf of the Wellcome Sanger Institute COVID-19 Surveillance Team |
| EPI_ISL_3363563, EPI_ISL_3363564 | Lighthouse Lab in Alderley Park | Wellcome Sanger Institute for the COVID-19 Genomics UK (COG-UK) Consortium | Jacquelyn Wynn, Mairead Hyland, The Lighthouse Lab in Alderley Park and Alex Alderton, Roberto Amato, Jeffrey Barrett, Sonia Goncalves, Ewan Harrison, David K. Jackson, Ian Johnston, Dominic Kwiatkowski, Cordelia Langford, John Sillitoe on behalf of the Wellcome Sanger Institute COVID-19 Surveillance Team |
| EPI_ISL_3363565, EPI_ISL_3363566 | Lighthouse Lab in Milton Keynes | Wellcome Sanger Institute for the COVID-19 Genomics UK (COG-UK) Consortium | The Lighthouse Lab in Milton Keynes and Alex Alderton, Roberto Amato, Jeffrey Barrett, Sonia Goncalves, Ewan Harrison, David K. Jackson, Ian Johnston, Dominic Kwiatkowski, Cordelia Langford, John Sillitoe on behalf of the Wellcome Sanger Institute COVID-19 Surveillance Team |
| EPI_ISL_3363567, EPI_ISL_3363568 | Lighthouse Lab in Glasgow | Wellcome Sanger Institute for the COVID-19 Genomics UK (COG-UK) Consortium | Harper VanSteenhouse, Yumi Kasai, David Gray, Carol Clugston, Anna Dominiczak and Alex Alderton, Roberto Amato, Jeffrey Barrett, Sonia Goncalves, Ewan Harrison, David K. Jackson, Ian Johnston, Dominic Kwiatkowski, Cordelia Langford, John Sillitoe on behalf of the Wellcome Sanger Institute COVID-19 Surveillance Team |
| EPI_ISL_3363570 | Lighthouse Lab in Alderley Park | Wellcome Sanger Institute for the COVID-19 Genomics UK (COG-UK) Consortium | Jacquelyn Wynn, Mairead Hyland, The Lighthouse Lab in Alderley Park and Alex Alderton, Roberto Amato, Jeffrey Barrett, Sonia Goncalves, Ewan Harrison, David K. Jackson, Ian Johnston, Dominic Kwiatkowski, Cordelia Langford, John Sillitoe on behalf of the Wellcome Sanger Institute COVID-19 Surveillance Team |
| EPI_ISL_3363571 | Lighthouse Lab in Milton Keynes | Wellcome Sanger Institute for the COVID-19 Genomics UK (COG-UK) Consortium | The Lighthouse Lab in Milton Keynes and Alex Alderton, Roberto Amato, Jeffrey Barrett, Sonia Goncalves, Ewan Harrison, David K. Jackson, Ian Johnston, Dominic Kwiatkowski, Cordelia Langford, John Sillitoe on behalf of the Wellcome Sanger Institute COVID-19 Surveillance Team |
| EPI_ISL_3363572 | Berkshire and Surrey Pathology Services Lighthouse Laboratory | Wellcome Sanger Institute for the COVID-19 Genomics UK (COG-UK) | Berkshire and Surrey Pathology Services Lighthouse Laboratory and Alex Alderton, Roberto Amato, Jeffrey Barrett, Sonia Goncalves, Ewan Harrison, David K. Jackson, Ian Johnston, Dominic Kwiatkowski, Cordelia Langford, John Sillitoe on behalf |

| | | | |
|---|---|---|---|
| | | Consortium | of the Wellcome Sanger Institute COVID-19 Surveillance Team |
| EPI_ISL_3363573 | Lighthouse Lab in Alderley Park | Wellcome Sanger Institute for the COVID-19 Genomics UK (COG-UK) Consortium | Jacquelyn Wynn, Mairead Hyland, The Lighthouse Lab in Alderley Park and Alex Alderton, Roberto Amato, Jeffrey Barrett, Sonia Goncalves, Ewan Harrison, David K. Jackson, Ian Johnston, Dominic Kwiatkowski, Cordelia Langford, John Sillitoe on behalf of the Wellcome Sanger Institute COVID-19 Surveillance Team |
| EPI_ISL_3363574 | Berkshire and Surrey Pathology Services Lighthouse Laboratory | Wellcome Sanger Institute for the COVID-19 Genomics UK (COG-UK) Consortium | Berkshire and Surrey Pathology Services Lighthouse Laboratory and Alex Alderton, Roberto Amato, Jeffrey Barrett, Sonia Goncalves, Ewan Harrison, David K. Jackson, Ian Johnston, Dominic Kwiatkowski, Cordelia Langford, John Sillitoe on behalf of the Wellcome Sanger Institute COVID-19 Surveillance Team |
| EPI_ISL_3363575 | Lighthouse Lab in Milton Keynes | Wellcome Sanger Institute for the COVID-19 Genomics UK (COG-UK) Consortium | The Lighthouse Lab in Milton Keynes and Alex Alderton, Roberto Amato, Jeffrey Barrett, Sonia Goncalves, Ewan Harrison, David K. Jackson, Ian Johnston, Dominic Kwiatkowski, Cordelia Langford, John Sillitoe on behalf of the Wellcome Sanger Institute COVID-19 Surveillance Team |
| EPI_ISL_3363576, EPI_ISL_3363577 | Lighthouse Lab in Alderley Park | Wellcome Sanger Institute for the COVID-19 Genomics UK (COG-UK) Consortium | Jacquelyn Wynn, Mairead Hyland, The Lighthouse Lab in Alderley Park and Alex Alderton, Roberto Amato, Jeffrey Barrett, Sonia Goncalves, Ewan Harrison, David K. Jackson, Ian Johnston, Dominic Kwiatkowski, Cordelia Langford, John Sillitoe on behalf of the Wellcome Sanger Institute COVID-19 Surveillance Team |
| EPI_ISL_3363579, EPI_ISL_3363580 | Lighthouse Lab in Glasgow | Wellcome Sanger Institute for the COVID-19 Genomics UK (COG-UK) Consortium | Harper VanSteenhouse, Yumi Kasai, David Gray, Carol Clugston, Anna Dominiczak and Alex Alderton, Roberto Amato, Jeffrey Barrett, Sonia Goncalves, Ewan Harrison, David K. Jackson, Ian Johnston, Dominic Kwiatkowski, Cordelia Langford, John Sillitoe on behalf of the Wellcome Sanger Institute COVID-19 Surveillance Team |
| EPI_ISL_3363581 | Berkshire and Surrey Pathology Services Lighthouse Laboratory | Wellcome Sanger Institute for the COVID-19 Genomics UK (COG-UK) Consortium | Berkshire and Surrey Pathology Services Lighthouse Laboratory and Alex Alderton, Roberto Amato, Jeffrey Barrett, Sonia Goncalves, Ewan Harrison, David K. Jackson, Ian Johnston, Dominic Kwiatkowski, Cordelia Langford, John Sillitoe on behalf of the Wellcome Sanger Institute COVID-19 Surveillance Team |
| EPI_ISL_3363583 | Lighthouse Lab in Milton Keynes | Wellcome Sanger Institute for the COVID-19 Genomics UK (COG-UK) Consortium | The Lighthouse Lab in Milton Keynes and Alex Alderton, Roberto Amato, Jeffrey Barrett, Sonia Goncalves, Ewan Harrison, David K. Jackson, Ian Johnston, Dominic Kwiatkowski, Cordelia Langford, John Sillitoe on behalf of the Wellcome Sanger Institute COVID-19 Surveillance Team |
| EPI_ISL_3363584 | Berkshire and Surrey Pathology Services Lighthouse Laboratory | Wellcome Sanger Institute for the COVID-19 Genomics UK (COG-UK) Consortium | Berkshire and Surrey Pathology Services Lighthouse Laboratory and Alex Alderton, Roberto Amato, Jeffrey Barrett, Sonia Goncalves, Ewan Harrison, David K. Jackson, Ian Johnston, Dominic Kwiatkowski, Cordelia Langford, John Sillitoe on behalf of the Wellcome Sanger Institute COVID-19 Surveillance Team |
| EPI_ISL_3363586 | Lighthouse Lab in Alderley Park | Wellcome Sanger Institute for the COVID-19 Genomics UK (COG-UK) Consortium | Jacquelyn Wynn, Mairead Hyland, The Lighthouse Lab in Alderley Park and Alex Alderton, Roberto Amato, Jeffrey Barrett, Sonia Goncalves, Ewan Harrison, David K. Jackson, Ian Johnston, Dominic Kwiatkowski, Cordelia Langford, John Sillitoe on behalf of the Wellcome Sanger Institute COVID-19 Surveillance Team |
| EPI_ISL_3363587, EPI_ISL_3363588 | Lighthouse Lab in Milton Keynes | Wellcome Sanger Institute for the COVID-19 Genomics UK (COG-UK) Consortium | The Lighthouse Lab in Milton Keynes and Alex Alderton, Roberto Amato, Jeffrey Barrett, Sonia Goncalves, Ewan Harrison, David K. Jackson, Ian Johnston, Dominic Kwiatkowski, Cordelia Langford, John Sillitoe on behalf of the Wellcome Sanger Institute COVID-19 Surveillance Team |
| EPI_ISL_3363590, EPI_ISL_3363591 | Lighthouse Lab in Glasgow | Wellcome Sanger Institute for the COVID-19 Genomics UK (COG-UK) Consortium | Harper VanSteenhouse, Yumi Kasai, David Gray, Carol Clugston, Anna Dominiczak and Alex Alderton, Roberto Amato, Jeffrey Barrett, Sonia Goncalves, Ewan Harrison, David K. Jackson, Ian Johnston, Dominic Kwiatkowski, Cordelia Langford, John Sillitoe on behalf of the Wellcome Sanger Institute COVID-19 Surveillance Team |
| EPI_ISL_3363592 | Lighthouse Lab in Alderley Park | Wellcome Sanger Institute for the COVID-19 Genomics UK (COG-UK) Consortium | Jacquelyn Wynn, Mairead Hyland, The Lighthouse Lab in Alderley Park and Alex Alderton, Roberto Amato, Jeffrey Barrett, Sonia Goncalves, Ewan Harrison, David K. Jackson, Ian Johnston, Dominic Kwiatkowski, Cordelia Langford, John Sillitoe on behalf of the Wellcome Sanger Institute COVID-19 Surveillance Team |
| EPI_ISL_3363594 | Berkshire and Surrey Pathology Services Lighthouse Laboratory | Wellcome Sanger Institute for the COVID-19 Genomics UK (COG-UK) Consortium | Berkshire and Surrey Pathology Services Lighthouse Laboratory and Alex Alderton, Roberto Amato, Jeffrey Barrett, Sonia Goncalves, Ewan Harrison, David K. Jackson, Ian Johnston, Dominic Kwiatkowski, Cordelia Langford, John Sillitoe on behalf of the Wellcome Sanger Institute COVID-19 Surveillance Team |
| EPI_ISL_3363595 | Lighthouse Lab in Alderley Park | Wellcome Sanger Institute for the COVID-19 Genomics UK (COG-UK) Consortium | Jacquelyn Wynn, Mairead Hyland, The Lighthouse Lab in Alderley Park and Alex Alderton, Roberto Amato, Jeffrey Barrett, Sonia Goncalves, Ewan Harrison, David K. Jackson, Ian Johnston, Dominic Kwiatkowski, Cordelia Langford, John Sillitoe on behalf of the Wellcome Sanger Institute COVID-19 Surveillance Team |
| EPI_ISL_3363596, EPI_ISL_3363598 | Lighthouse Lab in Milton Keynes | Wellcome Sanger Institute for the COVID-19 Genomics UK (COG-UK) Consortium | The Lighthouse Lab in Milton Keynes and Alex Alderton, Roberto Amato, Jeffrey Barrett, Sonia Goncalves, Ewan Harrison, David K. Jackson, Ian Johnston, Dominic Kwiatkowski, Cordelia Langford, John Sillitoe on behalf of the Wellcome Sanger Institute COVID-19 Surveillance Team |
| EPI_ISL_3363599 | Lighthouse Lab in Alderley Park | Wellcome Sanger Institute for the COVID-19 Genomics UK (COG-UK) Consortium | Jacquelyn Wynn, Mairead Hyland, The Lighthouse Lab in Alderley Park and Alex Alderton, Roberto Amato, Jeffrey Barrett, Sonia Goncalves, Ewan Harrison, David K. Jackson, Ian Johnston, Dominic Kwiatkowski, Cordelia Langford, John Sillitoe on behalf of the Wellcome Sanger Institute COVID-19 Surveillance Team |
| EPI_ISL_3363600 | Lighthouse Lab in Glasgow | Wellcome Sanger Institute for the COVID-19 Genomics UK (COG-UK) Consortium | Harper VanSteenhouse, Yumi Kasai, David Gray, Carol Clugston, Anna Dominiczak and Alex Alderton, Roberto Amato, Jeffrey Barrett, Sonia Goncalves, Ewan Harrison, David K. Jackson, Ian Johnston, Dominic Kwiatkowski, Cordelia Langford, John Sillitoe on behalf of the Wellcome Sanger Institute COVID-19 Surveillance Team |
| EPI_ISL_3363602 | Berkshire and Surrey Pathology Services Lighthouse Laboratory | Wellcome Sanger Institute for the COVID-19 Genomics UK (COG-UK) Consortium | Berkshire and Surrey Pathology Services Lighthouse Laboratory and Alex Alderton, Roberto Amato, Jeffrey Barrett, Sonia Goncalves, Ewan Harrison, David K. Jackson, Ian Johnston, Dominic Kwiatkowski, Cordelia Langford, John Sillitoe on behalf of the Wellcome Sanger Institute COVID-19 Surveillance Team |
| EPI_ISL_3363603 | Lighthouse Lab in Alderley Park | Wellcome Sanger Institute for the COVID-19 Genomics UK (COG-UK) Consortium | Jacquelyn Wynn, Mairead Hyland, The Lighthouse Lab in Alderley Park and Alex Alderton, Roberto Amato, Jeffrey Barrett, Sonia Goncalves, Ewan Harrison, David K. Jackson, Ian Johnston, Dominic Kwiatkowski, Cordelia Langford, John Sillitoe on behalf of the Wellcome Sanger Institute COVID-19 Surveillance Team |
| EPI_ISL_3363604 | Lighthouse Lab in Milton Keynes | Wellcome Sanger Institute for the COVID-19 Genomics UK (COG-UK) Consortium | The Lighthouse Lab in Milton Keynes and Alex Alderton, Roberto Amato, Jeffrey Barrett, Sonia Goncalves, Ewan Harrison, David K. Jackson, Ian Johnston, Dominic Kwiatkowski, Cordelia Langford, John Sillitoe on behalf of the Wellcome Sanger Institute COVID-19 Surveillance Team |
| EPI_ISL_3363605, EPI_ISL_3363606 | Berkshire and Surrey Pathology Services Lighthouse Laboratory | Wellcome Sanger Institute for the COVID-19 Genomics UK (COG-UK) Consortium | Berkshire and Surrey Pathology Services Lighthouse Laboratory and Alex Alderton, Roberto Amato, Jeffrey Barrett, Sonia Goncalves, Ewan Harrison, David K. Jackson, Ian Johnston, Dominic Kwiatkowski, Cordelia Langford, John Sillitoe on behalf of the Wellcome Sanger Institute COVID-19 Surveillance Team |
| EPI_ISL_3363607, EPI_ISL_3363608 | Lighthouse Lab in Milton Keynes | Wellcome Sanger Institute for the COVID-19 Genomics UK (COG-UK) Consortium | The Lighthouse Lab in Milton Keynes and Alex Alderton, Roberto Amato, Jeffrey Barrett, Sonia Goncalves, Ewan Harrison, David K. Jackson, Ian Johnston, Dominic Kwiatkowski, Cordelia Langford, John Sillitoe on behalf of the Wellcome Sanger Institute COVID-19 Surveillance Team |
| EPI_ISL_3363610 | Lighthouse Lab in Alderley Park | Wellcome Sanger Institute for the COVID-19 Genomics UK (COG-UK) Consortium | Jacquelyn Wynn, Mairead Hyland, The Lighthouse Lab in Alderley Park and Alex Alderton, Roberto Amato, Jeffrey Barrett, Sonia Goncalves, Ewan Harrison, David K. Jackson, Ian Johnston, Dominic Kwiatkowski, Cordelia Langford, John Sillitoe on behalf of the Wellcome Sanger Institute COVID-19 Surveillance Team |
| EPI_ISL_3363611 | Berkshire and Surrey Pathology Services Lighthouse Laboratory | Wellcome Sanger Institute for the COVID-19 Genomics UK (COG-UK) Consortium | Berkshire and Surrey Pathology Services Lighthouse Laboratory and Alex Alderton, Roberto Amato, Jeffrey Barrett, Sonia Goncalves, Ewan Harrison, David K. Jackson, Ian Johnston, Dominic Kwiatkowski, Cordelia Langford, John Sillitoe on behalf of the Wellcome Sanger Institute COVID-19 Surveillance Team |
| EPI_ISL_3363612, EPI_ISL_3363613 | Lighthouse Lab in Alderley Park | Wellcome Sanger Institute for the COVID-19 Genomics UK (COG-UK) Consortium | Jacquelyn Wynn, Mairead Hyland, The Lighthouse Lab in Alderley Park and Alex Alderton, Roberto Amato, Jeffrey Barrett, Sonia Goncalves, Ewan Harrison, David K. Jackson, Ian Johnston, Dominic Kwiatkowski, Cordelia Langford, John Sillitoe on behalf of the Wellcome Sanger Institute COVID-19 Surveillance Team |
| EPI_ISL_3363614 | Berkshire and Surrey Pathology Services Lighthouse Laboratory | Wellcome Sanger Institute for the COVID-19 Genomics UK (COG-UK) Consortium | Berkshire and Surrey Pathology Services Lighthouse Laboratory and Alex Alderton, Roberto Amato, Jeffrey Barrett, Sonia Goncalves, Ewan Harrison, David K. Jackson, Ian Johnston, Dominic Kwiatkowski, Cordelia Langford, John Sillitoe on behalf of the Wellcome Sanger Institute COVID-19 Surveillance Team |
| EPI_ISL_3363615, EPI_ISL_3363616 | Lighthouse Lab in Milton Keynes | Wellcome Sanger Institute for the COVID-19 Genomics UK (COG-UK) Consortium | The Lighthouse Lab in Milton Keynes and Alex Alderton, Roberto Amato, Jeffrey Barrett, Sonia Goncalves, Ewan Harrison, David K. Jackson, Ian Johnston, Dominic Kwiatkowski, Cordelia Langford, John Sillitoe on behalf of the Wellcome Sanger Institute COVID-19 Surveillance Team |
| EPI_ISL_3363617, EPI_ISL_3363619 | Lighthouse Lab in Glasgow | Wellcome Sanger Institute for the COVID-19 Genomics UK (COG-UK) Consortium | Harper VanSteenhouse, Yumi Kasai, David Gray, Carol Clugston, Anna Dominiczak and Alex Alderton, Roberto Amato, Jeffrey Barrett, Sonia Goncalves, Ewan Harrison, David K. Jackson, Ian Johnston, Dominic Kwiatkowski, Cordelia Langford, John Sillitoe on behalf of the Wellcome Sanger Institute COVID-19 Surveillance Team |
| EPI_ISL_3363620 | Lighthouse Lab in Alderley Park | Wellcome Sanger Institute for the COVID-19 Genomics UK (COG-UK) Consortium | Jacquelyn Wynn, Mairead Hyland, The Lighthouse Lab in Alderley Park and Alex Alderton, Roberto Amato, Jeffrey Barrett, Sonia Goncalves, Ewan Harrison, David K. Jackson, Ian Johnston, Dominic Kwiatkowski, Cordelia Langford, John Sillitoe on behalf of the Wellcome Sanger Institute COVID-19 Surveillance Team |
| EPI_ISL_3363621, EPI_ISL_3363622 | Lighthouse Lab in Milton Keynes | Wellcome Sanger Institute for the COVID-19 Genomics UK (COG-UK) Consortium | The Lighthouse Lab in Milton Keynes and Alex Alderton, Roberto Amato, Jeffrey Barrett, Sonia Goncalves, Ewan Harrison, David K. Jackson, Ian Johnston, Dominic Kwiatkowski, Cordelia Langford, John Sillitoe on behalf of the Wellcome Sanger Institute COVID-19 Surveillance Team |
| EPI_ISL_3363623, EPI_ISL_3363624 | Lighthouse Lab in Glasgow | Wellcome Sanger Institute for the COVID-19 Genomics UK (COG-UK) Consortium | Harper VanSteenhouse, Yumi Kasai, David Gray, Carol Clugston, Anna Dominiczak and Alex Alderton, Roberto Amato, Jeffrey Barrett, Sonia Goncalves, Ewan Harrison, David K. Jackson, Ian Johnston, Dominic Kwiatkowski, Cordelia Langford, John Sillitoe on behalf of the Wellcome Sanger Institute COVID-19 Surveillance Team |
| EPI_ISL_3363625, EPI_ISL_3363626, EPI_ISL_3363628, EPI_ISL_3363629 | Berkshire and Surrey Pathology Services Lighthouse Laboratory | Wellcome Sanger Institute for the COVID-19 Genomics UK (COG-UK) Consortium | Berkshire and Surrey Pathology Services Lighthouse Laboratory and Alex Alderton, Roberto Amato, Jeffrey Barrett, Sonia Goncalves, Ewan Harrison, David K. Jackson, Ian Johnston, Dominic Kwiatkowski, Cordelia Langford, John Sillitoe on behalf of the Wellcome Sanger Institute COVID-19 Surveillance Team |
| EPI_ISL_3363630 | Lighthouse Lab in Alderley Park | Wellcome Sanger Institute for the COVID-19 Genomics UK (COG-UK) Consortium | Jacquelyn Wynn, Mairead Hyland, The Lighthouse Lab in Alderley Park and Alex Alderton, Roberto Amato, Jeffrey Barrett, Sonia Goncalves, Ewan Harrison, David K. Jackson, Ian Johnston, Dominic Kwiatkowski, Cordelia Langford, John Sillitoe on behalf of the Wellcome Sanger Institute COVID-19 Surveillance Team |
| EPI_ISL_3363632 | Berkshire and Surrey Pathology Services Lighthouse Laboratory | Wellcome Sanger Institute for the COVID-19 Genomics UK (COG-UK) Consortium | Berkshire and Surrey Pathology Services Lighthouse Laboratory and Alex Alderton, Roberto Amato, Jeffrey Barrett, Sonia Goncalves, Ewan Harrison, David K. Jackson, Ian Johnston, Dominic Kwiatkowski, Cordelia Langford, John Sillitoe on behalf of the Wellcome Sanger Institute COVID-19 Surveillance Team |
| EPI_ISL_3363633 | Lighthouse Lab in Glasgow | Wellcome Sanger Institute for the COVID-19 Genomics UK (COG-UK) Consortium | Harper VanSteenhouse, Yumi Kasai, David Gray, Carol Clugston, Anna Dominiczak and Alex Alderton, Roberto Amato, Jeffrey Barrett, Sonia Goncalves, Ewan Harrison, David K. Jackson, Ian Johnston, Dominic Kwiatkowski, Cordelia Langford, John Sillitoe on behalf of the Wellcome Sanger Institute COVID-19 Surveillance Team |
| EPI_ISL_3363634 | Berkshire and Surrey Pathology Services Lighthouse Laboratory | Wellcome Sanger Institute for the COVID-19 Genomics UK (COG-UK) Consortium | Berkshire and Surrey Pathology Services Lighthouse Laboratory and Alex Alderton, Roberto Amato, Jeffrey Barrett, Sonia Goncalves, Ewan Harrison, David K. Jackson, Ian Johnston, Dominic Kwiatkowski, Cordelia Langford, John Sillitoe on behalf of the Wellcome Sanger Institute COVID-19 Surveillance Team |
| EPI_ISL_3363636 | Lighthouse Lab in Glasgow | Wellcome Sanger Institute for the COVID-19 Genomics UK (COG-UK) Consortium | Harper VanSteenhouse, Yumi Kasai, David Gray, Carol Clugston, Anna Dominiczak and Alex Alderton, Roberto Amato, Jeffrey Barrett, Sonia Goncalves, Ewan Harrison, David K. Jackson, Ian Johnston, Dominic Kwiatkowski, Cordelia Langford, John Sillitoe on behalf of the Wellcome Sanger Institute COVID-19 Surveillance Team |
| EPI_ISL_3363637, EPI_ISL_3363638 | Berkshire and Surrey Pathology Services Lighthouse Laboratory | Wellcome Sanger Institute for the COVID-19 Genomics UK (COG-UK) Consortium | Berkshire and Surrey Pathology Services Lighthouse Laboratory and Alex Alderton, Roberto Amato, Jeffrey Barrett, Sonia Goncalves, Ewan Harrison, David K. Jackson, Ian Johnston, Dominic Kwiatkowski, Cordelia Langford, John Sillitoe on behalf of the Wellcome Sanger Institute COVID-19 Surveillance Team |
| EPI_ISL_3363639 | Lighthouse Lab in Glasgow | Wellcome Sanger Institute for the COVID-19 Genomics UK (COG-UK) Consortium | Harper VanSteenhouse, Yumi Kasai, David Gray, Carol Clugston, Anna Dominiczak and Alex Alderton, Roberto Amato, Jeffrey Barrett, Sonia Goncalves, Ewan Harrison, David K. Jackson, Ian Johnston, Dominic Kwiatkowski, Cordelia Langford, John Sillitoe on behalf of the Wellcome Sanger Institute COVID-19 Surveillance Team |
| EPI_ISL_3363640, EPI_ISL_3363641, EPI_ISL_3363642, EPI_ISL_3363643 | Lighthouse Lab in Milton Keynes | Wellcome Sanger Institute for the COVID-19 Genomics UK (COG-UK) Consortium | The Lighthouse Lab in Milton Keynes and Alex Alderton, Roberto Amato, Jeffrey Barrett, Sonia Goncalves, Ewan Harrison, David K. Jackson, Ian Johnston, Dominic Kwiatkowski, Cordelia Langford, John Sillitoe on behalf of the Wellcome Sanger Institute COVID-19 Surveillance Team |
| EPI_ISL_3363644, EPI_ISL_3363645, EPI_ISL_3363647 | Lighthouse Lab in Glasgow | Wellcome Sanger Institute for the COVID-19 Genomics UK (COG-UK) Consortium | Harper VanSteenhouse, Yumi Kasai, David Gray, Carol Clugston, Anna Dominiczak and Alex Alderton, Roberto Amato, Jeffrey Barrett, Sonia Goncalves, Ewan Harrison, David K. Jackson, Ian Johnston, Dominic Kwiatkowski, Cordelia Langford, John Sillitoe on behalf of the Wellcome Sanger Institute COVID-19 Surveillance Team |
| EPI_ISL_3363648, EPI_ISL_3363650 | Berkshire and Surrey Pathology Services Lighthouse Laboratory | Wellcome Sanger Institute for the COVID-19 Genomics UK (COG-UK) Consortium | Berkshire and Surrey Pathology Services Lighthouse Laboratory and Alex Alderton, Roberto Amato, Jeffrey Barrett, Sonia Goncalves, Ewan Harrison, David K. Jackson, Ian Johnston, Dominic Kwiatkowski, Cordelia Langford, John Sillitoe on behalf of the Wellcome Sanger Institute COVID-19 Surveillance Team |

| | | | |
|---|---|---|---|
| EPI_ISL_3363651, EPI_ISL_3363652 | Lighthouse Lab in Glasgow | Wellcome Sanger Institute for the COVID-19 Genomics UK (COG-UK) Consortium | Harper VanSteenhouse, Yumi Kasai, David Gray, Carol Clugston, Anna Dominiczak and Alex Alderton, Roberto Amato, Jeffrey Barrett, Sonia Goncalves, Ewan Harrison, David K. Jackson, Ian Johnston, Dominic Kwiatkowski, Cordelia Langford, John Sillitoe on behalf of the Wellcome Sanger Institute COVID-19 Surveillance Team |
| EPI_ISL_3363654 | Lighthouse Lab in Alderley Park | Wellcome Sanger Institute for the COVID-19 Genomics UK (COG-UK) Consortium | Jacquelyn Wynn, Mairead Hyland, The Lighthouse Lab in Alderley Park and Alex Alderton, Roberto Amato, Jeffrey Barrett, Sonia Goncalves, Ewan Harrison, David K. Jackson, Ian Johnston, Dominic Kwiatkowski, Cordelia Langford, John Sillitoe on behalf of the Wellcome Sanger Institute COVID-19 Surveillance Team |
| EPI_ISL_3363655 | Lighthouse Lab in Glasgow | Wellcome Sanger Institute for the COVID-19 Genomics UK (COG-UK) Consortium | Harper VanSteenhouse, Yumi Kasai, David Gray, Carol Clugston, Anna Dominiczak and Alex Alderton, Roberto Amato, Jeffrey Barrett, Sonia Goncalves, Ewan Harrison, David K. Jackson, Ian Johnston, Dominic Kwiatkowski, Cordelia Langford, John Sillitoe on behalf of the Wellcome Sanger Institute COVID-19 Surveillance Team |
| EPI_ISL_3363657 | Lighthouse Lab in Milton Keynes | Wellcome Sanger Institute for the COVID-19 Genomics UK (COG-UK) Consortium | The Lighthouse Lab in Milton Keynes and Alex Alderton, Roberto Amato, Jeffrey Barrett, Sonia Goncalves, Ewan Harrison, David K. Jackson, Ian Johnston, Dominic Kwiatkowski, Cordelia Langford, John Sillitoe on behalf of the Wellcome Sanger Institute COVID-19 Surveillance Team |
| EPI_ISL_3363658 | Lighthouse Lab in Glasgow | Wellcome Sanger Institute for the COVID-19 Genomics UK (COG-UK) Consortium | Harper VanSteenhouse, Yumi Kasai, David Gray, Carol Clugston, Anna Dominiczak and Alex Alderton, Roberto Amato, Jeffrey Barrett, Sonia Goncalves, Ewan Harrison, David K. Jackson, Ian Johnston, Dominic Kwiatkowski, Cordelia Langford, John Sillitoe on behalf of the Wellcome Sanger Institute COVID-19 Surveillance Team |
| EPI_ISL_3363660 | Lighthouse Lab in Milton Keynes | Wellcome Sanger Institute for the COVID-19 Genomics UK (COG-UK) Consortium | The Lighthouse Lab in Milton Keynes and Alex Alderton, Roberto Amato, Jeffrey Barrett, Sonia Goncalves, Ewan Harrison, David K. Jackson, Ian Johnston, Dominic Kwiatkowski, Cordelia Langford, John Sillitoe on behalf of the Wellcome Sanger Institute COVID-19 Surveillance Team |
| EPI_ISL_3363661 | Lighthouse Lab in Alderley Park | Wellcome Sanger Institute for the COVID-19 Genomics UK (COG-UK) Consortium | Jacquelyn Wynn, Mairead Hyland, The Lighthouse Lab in Alderley Park and Alex Alderton, Roberto Amato, Jeffrey Barrett, Sonia Goncalves, Ewan Harrison, David K. Jackson, Ian Johnston, Dominic Kwiatkowski, Cordelia Langford, John Sillitoe on behalf of the Wellcome Sanger Institute COVID-19 Surveillance Team |
| EPI_ISL_3363663 | Lighthouse Lab in Milton Keynes | Wellcome Sanger Institute for the COVID-19 Genomics UK (COG-UK) Consortium | The Lighthouse Lab in Milton Keynes and Alex Alderton, Roberto Amato, Jeffrey Barrett, Sonia Goncalves, Ewan Harrison, David K. Jackson, Ian Johnston, Dominic Kwiatkowski, Cordelia Langford, John Sillitoe on behalf of the Wellcome Sanger Institute COVID-19 Surveillance Team |
| EPI_ISL_3363664 | Berkshire and Surrey Pathology Services Lighthouse Laboratory | Wellcome Sanger Institute for the COVID-19 Genomics UK (COG-UK) Consortium | Berkshire and Surrey Pathology Services Lighthouse Laboratory and Alex Alderton, Roberto Amato, Jeffrey Barrett, Sonia Goncalves, Ewan Harrison, David K. Jackson, Ian Johnston, Dominic Kwiatkowski, Cordelia Langford, John Sillitoe on behalf of the Wellcome Sanger Institute COVID-19 Surveillance Team |
| EPI_ISL_3363665 | Lighthouse Lab in Milton Keynes | Wellcome Sanger Institute for the COVID-19 Genomics UK (COG-UK) Consortium | The Lighthouse Lab in Milton Keynes and Alex Alderton, Roberto Amato, Jeffrey Barrett, Sonia Goncalves, Ewan Harrison, David K. Jackson, Ian Johnston, Dominic Kwiatkowski, Cordelia Langford, John Sillitoe on behalf of the Wellcome Sanger Institute COVID-19 Surveillance Team |
| EPI_ISL_3363667 | Lighthouse Lab in Alderley Park | Wellcome Sanger Institute for the COVID-19 Genomics UK (COG-UK) Consortium | Jacquelyn Wynn, Mairead Hyland, The Lighthouse Lab in Alderley Park and Alex Alderton, Roberto Amato, Jeffrey Barrett, Sonia Goncalves, Ewan Harrison, David K. Jackson, Ian Johnston, Dominic Kwiatkowski, Cordelia Langford, John Sillitoe on behalf of the Wellcome Sanger Institute COVID-19 Surveillance Team |
| EPI_ISL_3363668 | Lighthouse Lab in Glasgow | Wellcome Sanger Institute for the COVID-19 Genomics UK (COG-UK) Consortium | Harper VanSteenhouse, Yumi Kasai, David Gray, Carol Clugston, Anna Dominiczak and Alex Alderton, Roberto Amato, Jeffrey Barrett, Sonia Goncalves, Ewan Harrison, David K. Jackson, Ian Johnston, Dominic Kwiatkowski, Cordelia Langford, John Sillitoe on behalf of the Wellcome Sanger Institute COVID-19 Surveillance Team |
| EPI_ISL_3363670 | Lighthouse Lab in Alderley Park | Wellcome Sanger Institute for the COVID-19 Genomics UK (COG-UK) Consortium | Jacquelyn Wynn, Mairead Hyland, The Lighthouse Lab in Alderley Park and Alex Alderton, Roberto Amato, Jeffrey Barrett, Sonia Goncalves, Ewan Harrison, David K. Jackson, Ian Johnston, Dominic Kwiatkowski, Cordelia Langford, John Sillitoe on behalf of the Wellcome Sanger Institute COVID-19 Surveillance Team |
| EPI_ISL_3363671 | Lighthouse Lab in Milton Keynes | Wellcome Sanger Institute for the COVID-19 Genomics UK (COG-UK) Consortium | The Lighthouse Lab in Milton Keynes and Alex Alderton, Roberto Amato, Jeffrey Barrett, Sonia Goncalves, Ewan Harrison, David K. Jackson, Ian Johnston, Dominic Kwiatkowski, Cordelia Langford, John Sillitoe on behalf of the Wellcome Sanger Institute COVID-19 Surveillance Team |
| EPI_ISL_3363672 | Lighthouse Lab in Glasgow | Wellcome Sanger Institute for the COVID-19 Genomics UK (COG-UK) Consortium | Harper VanSteenhouse, Yumi Kasai, David Gray, Carol Clugston, Anna Dominiczak and Alex Alderton, Roberto Amato, Jeffrey Barrett, Sonia Goncalves, Ewan Harrison, David K. Jackson, Ian Johnston, Dominic Kwiatkowski, Cordelia Langford, John Sillitoe on behalf of the Wellcome Sanger Institute COVID-19 Surveillance Team |
| EPI_ISL_3363688, EPI_ISL_3363689 | Genetica Molecular and Subdepartamento de Virologia ISP Chile | Instituto de Salud Publica de Chile | Karen Orostica, Constanza Campano, Barbara Parra, Loredana Arata, Gisselle Barra, Patricia Bustos, Rodrigo Fasce, Javier Tognarelli, Andres Castillo, Soledad Ulloa, Jorge Fernandez |
| EPI_ISL_3363690 | Lighthouse Lab in Milton Keynes | Wellcome Sanger Institute for the COVID-19 Genomics UK (COG-UK) Consortium | The Lighthouse Lab in Milton Keynes and Alex Alderton, Roberto Amato, Jeffrey Barrett, Sonia Goncalves, Ewan Harrison, David K. Jackson, Ian Johnston, Dominic Kwiatkowski, Cordelia Langford, John Sillitoe on behalf of the Wellcome Sanger Institute COVID-19 Surveillance Team |
| EPI_ISL_3363691 | Lighthouse Lab in Alderley Park | Wellcome Sanger Institute for the COVID-19 Genomics UK (COG-UK) Consortium | Jacquelyn Wynn, Mairead Hyland, The Lighthouse Lab in Alderley Park and Alex Alderton, Roberto Amato, Jeffrey Barrett, Sonia Goncalves, Ewan Harrison, David K. Jackson, Ian Johnston, Dominic Kwiatkowski, Cordelia Langford, John Sillitoe on behalf of the Wellcome Sanger Institute COVID-19 Surveillance Team |
| EPI_ISL_3363692 | Lighthouse Lab in Milton Keynes | Wellcome Sanger Institute for the COVID-19 Genomics UK (COG-UK) Consortium | The Lighthouse Lab in Milton Keynes and Alex Alderton, Roberto Amato, Jeffrey Barrett, Sonia Goncalves, Ewan Harrison, David K. Jackson, Ian Johnston, Dominic Kwiatkowski, Cordelia Langford, John Sillitoe on behalf of the Wellcome Sanger Institute COVID-19 Surveillance Team |
| EPI_ISL_3363693 | Lighthouse Lab in Alderley Park | Wellcome Sanger Institute for the COVID-19 Genomics UK (COG-UK) Consortium | Jacquelyn Wynn, Mairead Hyland, The Lighthouse Lab in Alderley Park and Alex Alderton, Roberto Amato, Jeffrey Barrett, Sonia Goncalves, Ewan Harrison, David K. Jackson, Ian Johnston, Dominic Kwiatkowski, Cordelia Langford, John Sillitoe on behalf of the Wellcome Sanger Institute COVID-19 Surveillance Team |
| EPI_ISL_3363694, EPI_ISL_3363695 | Lighthouse Lab in Milton Keynes | Wellcome Sanger Institute for the COVID-19 Genomics UK (COG-UK) Consortium | The Lighthouse Lab in Milton Keynes and Alex Alderton, Roberto Amato, Jeffrey Barrett, Sonia Goncalves, Ewan Harrison, David K. Jackson, Ian Johnston, Dominic Kwiatkowski, Cordelia Langford, John Sillitoe on behalf of the Wellcome Sanger Institute COVID-19 Surveillance Team |
| EPI_ISL_3363696, EPI_ISL_3363697, EPI_ISL_3363698 | Lighthouse Lab in Alderley Park | Wellcome Sanger Institute for the COVID-19 Genomics UK (COG-UK) Consortium | Jacquelyn Wynn, Mairead Hyland, The Lighthouse Lab in Alderley Park and Alex Alderton, Roberto Amato, Jeffrey Barrett, Sonia Goncalves, Ewan Harrison, David K. Jackson, Ian Johnston, Dominic Kwiatkowski, Cordelia Langford, John Sillitoe on behalf of the Wellcome Sanger Institute COVID-19 Surveillance Team |
| EPI_ISL_3363699 | Lighthouse Lab in Milton Keynes | Wellcome Sanger Institute for the COVID-19 Genomics UK (COG-UK) Consortium | The Lighthouse Lab in Milton Keynes and Alex Alderton, Roberto Amato, Jeffrey Barrett, Sonia Goncalves, Ewan Harrison, David K. Jackson, Ian Johnston, Dominic Kwiatkowski, Cordelia Langford, John Sillitoe on behalf of the Wellcome Sanger Institute COVID-19 Surveillance Team |
| EPI_ISL_3363700 | Berkshire and Surrey Pathology Services Lighthouse Laboratory | Wellcome Sanger Institute for the COVID-19 Genomics UK (COG-UK) Consortium | Berkshire and Surrey Pathology Services Lighthouse Laboratory and Alex Alderton, Roberto Amato, Jeffrey Barrett, Sonia Goncalves, Ewan Harrison, David K. Jackson, Ian Johnston, Dominic Kwiatkowski, Cordelia Langford, John Sillitoe on behalf of the Wellcome Sanger Institute COVID-19 Surveillance Team |
| EPI_ISL_3363701, EPI_ISL_3363702 | Lighthouse Lab in Glasgow | Wellcome Sanger Institute for the COVID-19 Genomics UK (COG-UK) Consortium | Harper VanSteenhouse, Yumi Kasai, David Gray, Carol Clugston, Anna Dominiczak and Alex Alderton, Roberto Amato, Jeffrey Barrett, Sonia Goncalves, Ewan Harrison, David K. Jackson, Ian Johnston, Dominic Kwiatkowski, Cordelia Langford, John Sillitoe on behalf of the Wellcome Sanger Institute COVID-19 Surveillance Team |
| EPI_ISL_3363703, EPI_ISL_3363704 | Berkshire and Surrey Pathology Services Lighthouse Laboratory | Wellcome Sanger Institute for the COVID-19 Genomics UK (COG-UK) Consortium | Berkshire and Surrey Pathology Services Lighthouse Laboratory and Alex Alderton, Roberto Amato, Jeffrey Barrett, Sonia Goncalves, Ewan Harrison, David K. Jackson, Ian Johnston, Dominic Kwiatkowski, Cordelia Langford, John Sillitoe on behalf of the Wellcome Sanger Institute COVID-19 Surveillance Team |
| EPI_ISL_3363705 | Lighthouse Lab in Milton Keynes | Wellcome Sanger Institute for the COVID-19 Genomics UK (COG-UK) Consortium | The Lighthouse Lab in Milton Keynes and Alex Alderton, Roberto Amato, Jeffrey Barrett, Sonia Goncalves, Ewan Harrison, David K. Jackson, Ian Johnston, Dominic Kwiatkowski, Cordelia Langford, John Sillitoe on behalf of the Wellcome Sanger Institute COVID-19 Surveillance Team |
| EPI_ISL_3363706, EPI_ISL_3363707, EPI_ISL_3363708 | Lighthouse Lab in Alderley Park | Wellcome Sanger Institute for the COVID-19 Genomics UK (COG-UK) Consortium | Jacquelyn Wynn, Mairead Hyland, The Lighthouse Lab in Alderley Park and Alex Alderton, Roberto Amato, Jeffrey Barrett, Sonia Goncalves, Ewan Harrison, David K. Jackson, Ian Johnston, Dominic Kwiatkowski, Cordelia Langford, John Sillitoe on behalf of the Wellcome Sanger Institute COVID-19 Surveillance Team |
| EPI_ISL_3363709 | Lighthouse Lab in Milton Keynes | Wellcome Sanger Institute for the COVID-19 Genomics UK (COG-UK) Consortium | The Lighthouse Lab in Milton Keynes and Alex Alderton, Roberto Amato, Jeffrey Barrett, Sonia Goncalves, Ewan Harrison, David K. Jackson, Ian Johnston, Dominic Kwiatkowski, Cordelia Langford, John Sillitoe on behalf of the Wellcome Sanger Institute COVID-19 Surveillance Team |
| EPI_ISL_3363710 | Berkshire and Surrey Pathology Services Lighthouse Laboratory | Wellcome Sanger Institute for the COVID-19 Genomics UK (COG-UK) Consortium | Berkshire and Surrey Pathology Services Lighthouse Laboratory and Alex Alderton, Roberto Amato, Jeffrey Barrett, Sonia Goncalves, Ewan Harrison, David K. Jackson, Ian Johnston, Dominic Kwiatkowski, Cordelia Langford, John Sillitoe on behalf of the Wellcome Sanger Institute COVID-19 Surveillance Team |
| EPI_ISL_3363711 | Lighthouse Lab in Glasgow | Wellcome Sanger Institute for the COVID-19 Genomics UK (COG-UK) Consortium | Harper VanSteenhouse, Yumi Kasai, David Gray, Carol Clugston, Anna Dominiczak and Alex Alderton, Roberto Amato, Jeffrey Barrett, Sonia Goncalves, Ewan Harrison, David K. Jackson, Ian Johnston, Dominic Kwiatkowski, Cordelia Langford, John Sillitoe on behalf of the Wellcome Sanger Institute COVID-19 Surveillance Team |
| EPI_ISL_3363712 | Lighthouse Lab in Milton Keynes | Wellcome Sanger Institute for the COVID-19 Genomics UK (COG-UK) Consortium | The Lighthouse Lab in Milton Keynes and Alex Alderton, Roberto Amato, Jeffrey Barrett, Sonia Goncalves, Ewan Harrison, David K. Jackson, Ian Johnston, Dominic Kwiatkowski, Cordelia Langford, John Sillitoe on behalf of the Wellcome Sanger Institute COVID-19 Surveillance Team |
| EPI_ISL_3363713 | Lighthouse Lab in Glasgow | Wellcome Sanger Institute for the COVID-19 Genomics UK (COG-UK) Consortium | Harper VanSteenhouse, Yumi Kasai, David Gray, Carol Clugston, Anna Dominiczak and Alex Alderton, Roberto Amato, Jeffrey Barrett, Sonia Goncalves, Ewan Harrison, David K. Jackson, Ian Johnston, Dominic Kwiatkowski, Cordelia Langford, John Sillitoe on behalf of the Wellcome Sanger Institute COVID-19 Surveillance Team |
| EPI_ISL_3363714 | Lighthouse Lab in Milton Keynes | Wellcome Sanger Institute for the COVID-19 Genomics UK (COG-UK) Consortium | The Lighthouse Lab in Milton Keynes and Alex Alderton, Roberto Amato, Jeffrey Barrett, Sonia Goncalves, Ewan Harrison, David K. Jackson, Ian Johnston, Dominic Kwiatkowski, Cordelia Langford, John Sillitoe on behalf of the Wellcome Sanger Institute COVID-19 Surveillance Team |
| EPI_ISL_3363715 | Lighthouse Lab in Alderley Park | Wellcome Sanger Institute for the COVID-19 Genomics UK (COG-UK) Consortium | Jacquelyn Wynn, Mairead Hyland, The Lighthouse Lab in Alderley Park and Alex Alderton, Roberto Amato, Jeffrey Barrett, Sonia Goncalves, Ewan Harrison, David K. Jackson, Ian Johnston, Dominic Kwiatkowski, Cordelia Langford, John Sillitoe on behalf of the Wellcome Sanger Institute COVID-19 Surveillance Team |
| EPI_ISL_3363716, EPI_ISL_3363717 | Lighthouse Lab in Milton Keynes | Wellcome Sanger Institute for the COVID-19 Genomics UK (COG-UK) Consortium | The Lighthouse Lab in Milton Keynes and Alex Alderton, Roberto Amato, Jeffrey Barrett, Sonia Goncalves, Ewan Harrison, David K. Jackson, Ian Johnston, Dominic Kwiatkowski, Cordelia Langford, John Sillitoe on behalf of the Wellcome Sanger Institute COVID-19 Surveillance Team |
| EPI_ISL_3363718 | Lighthouse Lab in Glasgow | Wellcome Sanger Institute for the COVID-19 Genomics UK (COG-UK) Consortium | Harper VanSteenhouse, Yumi Kasai, David Gray, Carol Clugston, Anna Dominiczak and Alex Alderton, Roberto Amato, Jeffrey Barrett, Sonia Goncalves, Ewan Harrison, David K. Jackson, Ian Johnston, Dominic Kwiatkowski, Cordelia Langford, John Sillitoe on behalf of the Wellcome Sanger Institute COVID-19 Surveillance Team |
| EPI_ISL_3363719 | Lighthouse Lab in Alderley Park | Wellcome Sanger Institute for the COVID-19 Genomics UK (COG-UK) Consortium | Jacquelyn Wynn, Mairead Hyland, The Lighthouse Lab in Alderley Park and Alex Alderton, Roberto Amato, Jeffrey Barrett, Sonia Goncalves, Ewan Harrison, David K. Jackson, Ian Johnston, Dominic Kwiatkowski, Cordelia Langford, John Sillitoe on behalf of the Wellcome Sanger Institute COVID-19 Surveillance Team |
| EPI_ISL_3363720 | Lighthouse Lab in Milton Keynes | Wellcome Sanger Institute for the COVID-19 Genomics UK (COG-UK) Consortium | The Lighthouse Lab in Milton Keynes and Alex Alderton, Roberto Amato, Jeffrey Barrett, Sonia Goncalves, Ewan Harrison, David K. Jackson, Ian Johnston, Dominic Kwiatkowski, Cordelia Langford, John Sillitoe on behalf of the Wellcome Sanger Institute COVID-19 Surveillance Team |
| EPI_ISL_3363721, EPI_ISL_3363722 | Lighthouse Lab in Glasgow | Wellcome Sanger Institute for the COVID-19 Genomics UK (COG-UK) Consortium | Harper VanSteenhouse, Yumi Kasai, David Gray, Carol Clugston, Anna Dominiczak and Alex Alderton, Roberto Amato, Jeffrey Barrett, Sonia Goncalves, Ewan Harrison, David K. Jackson, Ian Johnston, Dominic Kwiatkowski, Cordelia Langford, John Sillitoe on behalf of the Wellcome Sanger Institute COVID-19 Surveillance Team |
| EPI_ISL_3363723 | Lighthouse Lab in Alderley Park | Wellcome Sanger Institute for the COVID-19 Genomics UK (COG-UK) Consortium | Jacquelyn Wynn, Mairead Hyland, The Lighthouse Lab in Alderley Park and Alex Alderton, Roberto Amato, Jeffrey Barrett, Sonia Goncalves, Ewan Harrison, David K. Jackson, Ian Johnston, Dominic Kwiatkowski, Cordelia Langford, John Sillitoe on behalf of the Wellcome Sanger Institute COVID-19 Surveillance Team |
| EPI_ISL_3363724, EPI_ISL_3363725, EPI_ISL_3363726 | Berkshire and Surrey Pathology Services Lighthouse Laboratory | Wellcome Sanger Institute for the COVID-19 Genomics UK (COG-UK) Consortium | Berkshire and Surrey Pathology Services Lighthouse Laboratory and Alex Alderton, Roberto Amato, Jeffrey Barrett, Sonia Goncalves, Ewan Harrison, David K. Jackson, Ian Johnston, Dominic Kwiatkowski, Cordelia Langford, John Sillitoe on behalf of the Wellcome Sanger Institute COVID-19 Surveillance Team |
| EPI_ISL_3363727, EPI_ISL_3363728 | Lighthouse Lab in Glasgow | Wellcome Sanger Institute for the COVID-19 Genomics UK (COG-UK) Consortium | Harper VanSteenhouse, Yumi Kasai, David Gray, Carol Clugston, Anna Dominiczak and Alex Alderton, Roberto Amato, Jeffrey Barrett, Sonia Goncalves, Ewan Harrison, David K. Jackson, Ian Johnston, Dominic Kwiatkowski, Cordelia Langford, John Sillitoe on behalf of the Wellcome Sanger Institute COVID-19 Surveillance Team |
| EPI_ISL_3363729 | Lighthouse Lab in Alderley Park | Wellcome Sanger Institute for the COVID-19 Genomics UK (COG-UK) Consortium | Jacquelyn Wynn, Mairead Hyland, The Lighthouse Lab in Alderley Park and Alex Alderton, Roberto Amato, Jeffrey Barrett, Sonia Goncalves, Ewan Harrison, David K. Jackson, Ian Johnston, Dominic Kwiatkowski, Cordelia Langford, John Sillitoe on behalf of the Wellcome Sanger Institute COVID-19 Surveillance Team |

| | | | |
|---|---|---|---|
| EPI_ISL_3363730 | Berkshire and Surrey Pathology Services Lighthouse Laboratory | Wellcome Sanger Institute for the COVID-19 Genomics UK (COG-UK) Consortium | Berkshire and Surrey Pathology Services Lighthouse Laboratory and Alex Alderton, Roberto Amato, Jeffrey Barrett, Sonia Goncalves, Ewan Harrison, David K. Jackson, Ian Johnston, Dominic Kwiatkowski, Cordelia Langford, John Sillitoe on behalf of the Wellcome Sanger Institute COVID-19 Surveillance Team |
| EPI_ISL_3363731, EPI_ISL_3363732 | Lighthouse Lab in Glasgow | Wellcome Sanger Institute for the COVID-19 Genomics UK (COG-UK) Consortium | Harper VanSteenhouse, Yumi Kasai, David Gray, Carol Clugston, Anna Dominiczak and Alex Alderton, Roberto Amato, Jeffrey Barrett, Sonia Goncalves, Ewan Harrison, David K. Jackson, Ian Johnston, Dominic Kwiatkowski, Cordelia Langford, John Sillitoe on behalf of the Wellcome Sanger Institute COVID-19 Surveillance Team |
| EPI_ISL_3363733 | Lighthouse Lab in Alderley Park | Wellcome Sanger Institute for the COVID-19 Genomics UK (COG-UK) Consortium | Jacquelyn Wynn, Mairead Hyland, The Lighthouse Lab in Alderley Park and Alex Alderton, Roberto Amato, Jeffrey Barrett, Sonia Goncalves, Ewan Harrison, David K. Jackson, Ian Johnston, Dominic Kwiatkowski, Cordelia Langford, John Sillitoe on behalf of the Wellcome Sanger Institute COVID-19 Surveillance Team |
| EPI_ISL_3363734 | Berkshire and Surrey Pathology Services Lighthouse Laboratory | Wellcome Sanger Institute for the COVID-19 Genomics UK (COG-UK) Consortium | Berkshire and Surrey Pathology Services Lighthouse Laboratory and Alex Alderton, Roberto Amato, Jeffrey Barrett, Sonia Goncalves, Ewan Harrison, David K. Jackson, Ian Johnston, Dominic Kwiatkowski, Cordelia Langford, John Sillitoe on behalf of the Wellcome Sanger Institute COVID-19 Surveillance Team |
| EPI_ISL_3363735, EPI_ISL_3363736 | Lighthouse Lab in Glasgow | Wellcome Sanger Institute for the COVID-19 Genomics UK (COG-UK) Consortium | Harper VanSteenhouse, Yumi Kasai, David Gray, Carol Clugston, Anna Dominiczak and Alex Alderton, Roberto Amato, Jeffrey Barrett, Sonia Goncalves, Ewan Harrison, David K. Jackson, Ian Johnston, Dominic Kwiatkowski, Cordelia Langford, John Sillitoe on behalf of the Wellcome Sanger Institute COVID-19 Surveillance Team |
| EPI_ISL_3363737 | Lighthouse Lab in Milton Keynes | Wellcome Sanger Institute for the COVID-19 Genomics UK (COG-UK) Consortium | The Lighthouse Lab in Milton Keynes and Alex Alderton, Roberto Amato, Jeffrey Barrett, Sonia Goncalves, Ewan Harrison, David K. Jackson, Ian Johnston, Dominic Kwiatkowski, Cordelia Langford, John Sillitoe on behalf of the Wellcome Sanger Institute COVID-19 Surveillance Team |
| EPI_ISL_3363738 | Berkshire and Surrey Pathology Services Lighthouse Laboratory | Wellcome Sanger Institute for the COVID-19 Genomics UK (COG-UK) Consortium | Berkshire and Surrey Pathology Services Lighthouse Laboratory and Alex Alderton, Roberto Amato, Jeffrey Barrett, Sonia Goncalves, Ewan Harrison, David K. Jackson, Ian Johnston, Dominic Kwiatkowski, Cordelia Langford, John Sillitoe on behalf of the Wellcome Sanger Institute COVID-19 Surveillance Team |
| EPI_ISL_3363739 | Lighthouse Lab in Alderley Park | Wellcome Sanger Institute for the COVID-19 Genomics UK (COG-UK) Consortium | Jacquelyn Wynn, Mairead Hyland, The Lighthouse Lab in Alderley Park and Alex Alderton, Roberto Amato, Jeffrey Barrett, Sonia Goncalves, Ewan Harrison, David K. Jackson, Ian Johnston, Dominic Kwiatkowski, Cordelia Langford, John Sillitoe on behalf of the Wellcome Sanger Institute COVID-19 Surveillance Team |
| EPI_ISL_3363740 | Lighthouse Lab in Milton Keynes | Wellcome Sanger Institute for the COVID-19 Genomics UK (COG-UK) Consortium | The Lighthouse Lab in Milton Keynes and Alex Alderton, Roberto Amato, Jeffrey Barrett, Sonia Goncalves, Ewan Harrison, David K. Jackson, Ian Johnston, Dominic Kwiatkowski, Cordelia Langford, John Sillitoe on behalf of the Wellcome Sanger Institute COVID-19 Surveillance Team |
| EPI_ISL_3363741 | Lighthouse Lab in Alderley Park | Wellcome Sanger Institute for the COVID-19 Genomics UK (COG-UK) Consortium | Jacquelyn Wynn, Mairead Hyland, The Lighthouse Lab in Alderley Park and Alex Alderton, Roberto Amato, Jeffrey Barrett, Sonia Goncalves, Ewan Harrison, David K. Jackson, Ian Johnston, Dominic Kwiatkowski, Cordelia Langford, John Sillitoe on behalf of the Wellcome Sanger Institute COVID-19 Surveillance Team |
| EPI_ISL_3363742 | Berkshire and Surrey Pathology Services Lighthouse Laboratory | Wellcome Sanger Institute for the COVID-19 Genomics UK (COG-UK) Consortium | Berkshire and Surrey Pathology Services Lighthouse Laboratory and Alex Alderton, Roberto Amato, Jeffrey Barrett, Sonia Goncalves, Ewan Harrison, David K. Jackson, Ian Johnston, Dominic Kwiatkowski, Cordelia Langford, John Sillitoe on behalf of the Wellcome Sanger Institute COVID-19 Surveillance Team |
| EPI_ISL_3363743, EPI_ISL_3363744 | Lighthouse Lab in Glasgow | Wellcome Sanger Institute for the COVID-19 Genomics UK (COG-UK) Consortium | Harper VanSteenhouse, Yumi Kasai, David Gray, Carol Clugston, Anna Dominiczak and Alex Alderton, Roberto Amato, Jeffrey Barrett, Sonia Goncalves, Ewan Harrison, David K. Jackson, Ian Johnston, Dominic Kwiatkowski, Cordelia Langford, John Sillitoe on behalf of the Wellcome Sanger Institute COVID-19 Surveillance Team |
| EPI_ISL_3363745 | Lighthouse Lab in Milton Keynes | Wellcome Sanger Institute for the COVID-19 Genomics UK (COG-UK) Consortium | The Lighthouse Lab in Milton Keynes and Alex Alderton, Roberto Amato, Jeffrey Barrett, Sonia Goncalves, Ewan Harrison, David K. Jackson, Ian Johnston, Dominic Kwiatkowski, Cordelia Langford, John Sillitoe on behalf of the Wellcome Sanger Institute COVID-19 Surveillance Team |
| EPI_ISL_3363746, EPI_ISL_3363747 | Berkshire and Surrey Pathology Services Lighthouse Laboratory | Wellcome Sanger Institute for the COVID-19 Genomics UK (COG-UK) Consortium | Berkshire and Surrey Pathology Services Lighthouse Laboratory and Alex Alderton, Roberto Amato, Jeffrey Barrett, Sonia Goncalves, Ewan Harrison, David K. Jackson, Ian Johnston, Dominic Kwiatkowski, Cordelia Langford, John Sillitoe on behalf of the Wellcome Sanger Institute COVID-19 Surveillance Team |
| EPI_ISL_3363748 | Lighthouse Lab in Milton Keynes | Wellcome Sanger Institute for the COVID-19 Genomics UK (COG-UK) Consortium | The Lighthouse Lab in Milton Keynes and Alex Alderton, Roberto Amato, Jeffrey Barrett, Sonia Goncalves, Ewan Harrison, David K. Jackson, Ian Johnston, Dominic Kwiatkowski, Cordelia Langford, John Sillitoe on behalf of the Wellcome Sanger Institute COVID-19 Surveillance Team |
| EPI_ISL_3363749 | Lighthouse Lab in Glasgow | Wellcome Sanger Institute for the COVID-19 Genomics UK (COG-UK) Consortium | Harper VanSteenhouse, Yumi Kasai, David Gray, Carol Clugston, Anna Dominiczak and Alex Alderton, Roberto Amato, Jeffrey Barrett, Sonia Goncalves, Ewan Harrison, David K. Jackson, Ian Johnston, Dominic Kwiatkowski, Cordelia Langford, John Sillitoe on behalf of the Wellcome Sanger Institute COVID-19 Surveillance Team |
| EPI_ISL_3363750 | Lighthouse Lab in Milton Keynes | Wellcome Sanger Institute for the COVID-19 Genomics UK (COG-UK) Consortium | The Lighthouse Lab in Milton Keynes and Alex Alderton, Roberto Amato, Jeffrey Barrett, Sonia Goncalves, Ewan Harrison, David K. Jackson, Ian Johnston, Dominic Kwiatkowski, Cordelia Langford, John Sillitoe on behalf of the Wellcome Sanger Institute COVID-19 Surveillance Team |
| EPI_ISL_3363751 | Lighthouse Lab in Glasgow | Wellcome Sanger Institute for the COVID-19 Genomics UK (COG-UK) Consortium | Harper VanSteenhouse, Yumi Kasai, David Gray, Carol Clugston, Anna Dominiczak and Alex Alderton, Roberto Amato, Jeffrey Barrett, Sonia Goncalves, Ewan Harrison, David K. Jackson, Ian Johnston, Dominic Kwiatkowski, Cordelia Langford, John Sillitoe on behalf of the Wellcome Sanger Institute COVID-19 Surveillance Team |
| EPI_ISL_3363752, EPI_ISL_3363753 | Berkshire and Surrey Pathology Services Lighthouse Laboratory | Wellcome Sanger Institute for the COVID-19 Genomics UK (COG-UK) Consortium | Berkshire and Surrey Pathology Services Lighthouse Laboratory and Alex Alderton, Roberto Amato, Jeffrey Barrett, Sonia Goncalves, Ewan Harrison, David K. Jackson, Ian Johnston, Dominic Kwiatkowski, Cordelia Langford, John Sillitoe on behalf of the Wellcome Sanger Institute COVID-19 Surveillance Team |
| EPI_ISL_3363754 | Lighthouse Lab in Milton Keynes | Wellcome Sanger Institute for the COVID-19 Genomics UK (COG-UK) Consortium | The Lighthouse Lab in Milton Keynes and Alex Alderton, Roberto Amato, Jeffrey Barrett, Sonia Goncalves, Ewan Harrison, David K. Jackson, Ian Johnston, Dominic Kwiatkowski, Cordelia Langford, John Sillitoe on behalf of the Wellcome Sanger Institute COVID-19 Surveillance Team |
| EPI_ISL_3363755 | Lighthouse Lab in Glasgow | Wellcome Sanger Institute for the COVID-19 Genomics UK (COG-UK) Consortium | Harper VanSteenhouse, Yumi Kasai, David Gray, Carol Clugston, Anna Dominiczak and Alex Alderton, Roberto Amato, Jeffrey Barrett, Sonia Goncalves, Ewan Harrison, David K. Jackson, Ian Johnston, Dominic Kwiatkowski, Cordelia Langford, John Sillitoe on behalf of the Wellcome Sanger Institute COVID-19 Surveillance Team |
| EPI_ISL_3363756, EPI_ISL_3363757, EPI_ISL_3363758 | Lighthouse Lab in Alderley Park | Wellcome Sanger Institute for the COVID-19 Genomics UK (COG-UK) Consortium | Jacquelyn Wynn, Mairead Hyland, The Lighthouse Lab in Alderley Park and Alex Alderton, Roberto Amato, Jeffrey Barrett, Sonia Goncalves, Ewan Harrison, David K. Jackson, Ian Johnston, Dominic Kwiatkowski, Cordelia Langford, John Sillitoe on behalf of the Wellcome Sanger Institute COVID-19 Surveillance Team |
| EPI_ISL_3363759, EPI_ISL_3363760 | Lighthouse Lab in Glasgow | Wellcome Sanger Institute for the COVID-19 Genomics UK (COG-UK) Consortium | Harper VanSteenhouse, Yumi Kasai, David Gray, Carol Clugston, Anna Dominiczak and Alex Alderton, Roberto Amato, Jeffrey Barrett, Sonia Goncalves, Ewan Harrison, David K. Jackson, Ian Johnston, Dominic Kwiatkowski, Cordelia Langford, John Sillitoe on behalf of the Wellcome Sanger Institute COVID-19 Surveillance Team |
| EPI_ISL_3363761 | Lighthouse Lab in Alderley Park | Wellcome Sanger Institute for the COVID-19 Genomics UK (COG-UK) Consortium | Jacquelyn Wynn, Mairead Hyland, The Lighthouse Lab in Alderley Park and Alex Alderton, Roberto Amato, Jeffrey Barrett, Sonia Goncalves, Ewan Harrison, David K. Jackson, Ian Johnston, Dominic Kwiatkowski, Cordelia Langford, John Sillitoe on behalf of the Wellcome Sanger Institute COVID-19 Surveillance Team |
| EPI_ISL_3363762 | Lighthouse Lab in Milton Keynes | Wellcome Sanger Institute for the COVID-19 Genomics UK (COG-UK) Consortium | The Lighthouse Lab in Milton Keynes and Alex Alderton, Roberto Amato, Jeffrey Barrett, Sonia Goncalves, Ewan Harrison, David K. Jackson, Ian Johnston, Dominic Kwiatkowski, Cordelia Langford, John Sillitoe on behalf of the Wellcome Sanger Institute COVID-19 Surveillance Team |
| EPI_ISL_3363763 | Lighthouse Lab in Alderley Park | Wellcome Sanger Institute for the COVID-19 Genomics UK (COG-UK) Consortium | Jacquelyn Wynn, Mairead Hyland, The Lighthouse Lab in Alderley Park and Alex Alderton, Roberto Amato, Jeffrey Barrett, Sonia Goncalves, Ewan Harrison, David K. Jackson, Ian Johnston, Dominic Kwiatkowski, Cordelia Langford, John Sillitoe on behalf of the Wellcome Sanger Institute COVID-19 Surveillance Team |
| EPI_ISL_3363764 | Berkshire and Surrey Pathology Services Lighthouse Laboratory | Wellcome Sanger Institute for the COVID-19 Genomics UK (COG-UK) Consortium | Berkshire and Surrey Pathology Services Lighthouse Laboratory and Alex Alderton, Roberto Amato, Jeffrey Barrett, Sonia Goncalves, Ewan Harrison, David K. Jackson, Ian Johnston, Dominic Kwiatkowski, Cordelia Langford, John Sillitoe on behalf of the Wellcome Sanger Institute COVID-19 Surveillance Team |
| EPI_ISL_3363765 | Lighthouse Lab in Milton Keynes | Wellcome Sanger Institute for the COVID-19 Genomics UK (COG-UK) Consortium | The Lighthouse Lab in Milton Keynes and Alex Alderton, Roberto Amato, Jeffrey Barrett, Sonia Goncalves, Ewan Harrison, David K. Jackson, Ian Johnston, Dominic Kwiatkowski, Cordelia Langford, John Sillitoe on behalf of the Wellcome Sanger Institute COVID-19 Surveillance Team |
| EPI_ISL_3363766 | Lighthouse Lab in Alderley Park | Wellcome Sanger Institute for the COVID-19 Genomics UK (COG-UK) Consortium | Jacquelyn Wynn, Mairead Hyland, The Lighthouse Lab in Alderley Park and Alex Alderton, Roberto Amato, Jeffrey Barrett, Sonia Goncalves, Ewan Harrison, David K. Jackson, Ian Johnston, Dominic Kwiatkowski, Cordelia Langford, John Sillitoe on behalf of the Wellcome Sanger Institute COVID-19 Surveillance Team |
| EPI_ISL_3363767, EPI_ISL_3363768 | Lighthouse Lab in Glasgow | Wellcome Sanger Institute for the COVID-19 Genomics UK (COG-UK) Consortium | Harper VanSteenhouse, Yumi Kasai, David Gray, Carol Clugston, Anna Dominiczak and Alex Alderton, Roberto Amato, Jeffrey Barrett, Sonia Goncalves, Ewan Harrison, David K. Jackson, Ian Johnston, Dominic Kwiatkowski, Cordelia Langford, John Sillitoe on behalf of the Wellcome Sanger Institute COVID-19 Surveillance Team |
| EPI_ISL_3363769, EPI_ISL_3363770, EPI_ISL_3363771 | Lighthouse Lab in Milton Keynes | Wellcome Sanger Institute for the COVID-19 Genomics UK (COG-UK) Consortium | The Lighthouse Lab in Milton Keynes and Alex Alderton, Roberto Amato, Jeffrey Barrett, Sonia Goncalves, Ewan Harrison, David K. Jackson, Ian Johnston, Dominic Kwiatkowski, Cordelia Langford, John Sillitoe on behalf of the Wellcome Sanger Institute COVID-19 Surveillance Team |
| EPI_ISL_3363772, EPI_ISL_3363773 | Lighthouse Lab in Glasgow | Wellcome Sanger Institute for the COVID-19 Genomics UK (COG-UK) Consortium | Harper VanSteenhouse, Yumi Kasai, David Gray, Carol Clugston, Anna Dominiczak and Alex Alderton, Roberto Amato, Jeffrey Barrett, Sonia Goncalves, Ewan Harrison, David K. Jackson, Ian Johnston, Dominic Kwiatkowski, Cordelia Langford, John Sillitoe on behalf of the Wellcome Sanger Institute COVID-19 Surveillance Team |
| EPI_ISL_3363774 | Lighthouse Lab in Alderley Park | Wellcome Sanger Institute for the COVID-19 Genomics UK (COG-UK) Consortium | Jacquelyn Wynn, Mairead Hyland, The Lighthouse Lab in Alderley Park and Alex Alderton, Roberto Amato, Jeffrey Barrett, Sonia Goncalves, Ewan Harrison, David K. Jackson, Ian Johnston, Dominic Kwiatkowski, Cordelia Langford, John Sillitoe on behalf of the Wellcome Sanger Institute COVID-19 Surveillance Team |
| EPI_ISL_3363775, EPI_ISL_3363776 | Lighthouse Lab in Milton Keynes | Wellcome Sanger Institute for the COVID-19 Genomics UK (COG-UK) Consortium | The Lighthouse Lab in Milton Keynes and Alex Alderton, Roberto Amato, Jeffrey Barrett, Sonia Goncalves, Ewan Harrison, David K. Jackson, Ian Johnston, Dominic Kwiatkowski, Cordelia Langford, John Sillitoe on behalf of the Wellcome Sanger Institute COVID-19 Surveillance Team |
| EPI_ISL_3363777 | Lighthouse Lab in Alderley Park | Wellcome Sanger Institute for the COVID-19 Genomics UK (COG-UK) Consortium | Jacquelyn Wynn, Mairead Hyland, The Lighthouse Lab in Alderley Park and Alex Alderton, Roberto Amato, Jeffrey Barrett, Sonia Goncalves, Ewan Harrison, David K. Jackson, Ian Johnston, Dominic Kwiatkowski, Cordelia Langford, John Sillitoe on behalf of the Wellcome Sanger Institute COVID-19 Surveillance Team |
| EPI_ISL_3363778, EPI_ISL_3363779 | Lighthouse Lab in Glasgow | Wellcome Sanger Institute for the COVID-19 Genomics UK (COG-UK) Consortium | Harper VanSteenhouse, Yumi Kasai, David Gray, Carol Clugston, Anna Dominiczak and Alex Alderton, Roberto Amato, Jeffrey Barrett, Sonia Goncalves, Ewan Harrison, David K. Jackson, Ian Johnston, Dominic Kwiatkowski, Cordelia Langford, John Sillitoe on behalf of the Wellcome Sanger Institute COVID-19 Surveillance Team |
| EPI_ISL_3363780 | Lighthouse Lab in Alderley Park | Wellcome Sanger Institute for the COVID-19 Genomics UK (COG-UK) Consortium | Jacquelyn Wynn, Mairead Hyland, The Lighthouse Lab in Alderley Park and Alex Alderton, Roberto Amato, Jeffrey Barrett, Sonia Goncalves, Ewan Harrison, David K. Jackson, Ian Johnston, Dominic Kwiatkowski, Cordelia Langford, John Sillitoe on behalf of the Wellcome Sanger Institute COVID-19 Surveillance Team |
| EPI_ISL_3363781, EPI_ISL_3363782 | Lighthouse Lab in Glasgow | Wellcome Sanger Institute for the COVID-19 Genomics UK (COG-UK) Consortium | Harper VanSteenhouse, Yumi Kasai, David Gray, Carol Clugston, Anna Dominiczak and Alex Alderton, Roberto Amato, Jeffrey Barrett, Sonia Goncalves, Ewan Harrison, David K. Jackson, Ian Johnston, Dominic Kwiatkowski, Cordelia Langford, John Sillitoe on behalf of the Wellcome Sanger Institute COVID-19 Surveillance Team |
| EPI_ISL_3363783, EPI_ISL_3363784 | Berkshire and Surrey Pathology Services Lighthouse Laboratory | Wellcome Sanger Institute for the COVID-19 Genomics UK (COG-UK) Consortium | Berkshire and Surrey Pathology Services Lighthouse Laboratory and Alex Alderton, Roberto Amato, Jeffrey Barrett, Sonia Goncalves, Ewan Harrison, David K. Jackson, Ian Johnston, Dominic Kwiatkowski, Cordelia Langford, John Sillitoe on behalf of the Wellcome Sanger Institute COVID-19 Surveillance Team |
| EPI_ISL_3363785 | Lighthouse Lab in Milton Keynes | Wellcome Sanger Institute for the COVID-19 Genomics UK (COG-UK) Consortium | The Lighthouse Lab in Milton Keynes and Alex Alderton, Roberto Amato, Jeffrey Barrett, Sonia Goncalves, Ewan Harrison, David K. Jackson, Ian Johnston, Dominic Kwiatkowski, Cordelia Langford, John Sillitoe on behalf of the Wellcome Sanger Institute COVID-19 Surveillance Team |
| EPI_ISL_3363786 | Lighthouse Lab in Alderley Park | Wellcome Sanger Institute for the COVID-19 Genomics UK (COG-UK) Consortium | Jacquelyn Wynn, Mairead Hyland, The Lighthouse Lab in Alderley Park and Alex Alderton, Roberto Amato, Jeffrey Barrett, Sonia Goncalves, Ewan Harrison, David K. Jackson, Ian Johnston, Dominic Kwiatkowski, Cordelia Langford, John Sillitoe on behalf of the Wellcome Sanger Institute COVID-19 Surveillance Team |
| EPI_ISL_3363787 | Lighthouse Lab in Glasgow | Wellcome Sanger Institute for the COVID-19 Genomics UK (COG-UK) Consortium | Harper VanSteenhouse, Yumi Kasai, David Gray, Carol Clugston, Anna Dominiczak and Alex Alderton, Roberto Amato, Jeffrey Barrett, Sonia Goncalves, Ewan Harrison, David K. Jackson, Ian Johnston, Dominic Kwiatkowski, Cordelia Langford, John Sillitoe on behalf of the Wellcome Sanger Institute COVID-19 Surveillance Team |
| EPI_ISL_3363788 | Berkshire and Surrey Pathology Services Lighthouse Laboratory | Wellcome Sanger Institute for the COVID-19 Genomics UK (COG-UK) Consortium | Berkshire and Surrey Pathology Services Lighthouse Laboratory and Alex Alderton, Roberto Amato, Jeffrey Barrett, Sonia Goncalves, Ewan Harrison, David K. Jackson, Ian Johnston, Dominic Kwiatkowski, Cordelia Langford, John Sillitoe on behalf of the Wellcome Sanger Institute COVID-19 Surveillance Team |
| EPI_ISL_3363789 | Lighthouse Lab in Milton Keynes | Wellcome Sanger Institute for the COVID-19 Genomics UK (COG-UK) Consortium | The Lighthouse Lab in Milton Keynes and Alex Alderton, Roberto Amato, Jeffrey Barrett, Sonia Goncalves, Ewan Harrison, David K. Jackson, Ian Johnston, Dominic Kwiatkowski, Cordelia Langford, John Sillitoe on behalf of the Wellcome Sanger Institute COVID-19 Surveillance Team |

| | | | |
|---|---|---|---|
| EPI_ISL_3363790 | Lighthouse Lab in Glasgow | Wellcome Sanger Institute for the COVID-19 Genomics UK (COG-UK) Consortium | Harper VanSteenhouse, Yumi Kasai, David Gray, Carol Clugston, Anna Dominiczak and Alex Alderton, Roberto Amato, Jeffrey Barrett, Sonia Goncalves, Ewan Harrison, David K. Jackson, Ian Johnston, Dominic Kwiatkowski, Cordelia Langford, John Sillitoe on behalf of the Wellcome Sanger Institute COVID-19 Surveillance Team |
| EPI_ISL_3363791 | Lighthouse Lab in Milton Keynes | Wellcome Sanger Institute for the COVID-19 Genomics UK (COG-UK) Consortium | The Lighthouse Lab in Milton Keynes and Alex Alderton, Roberto Amato, Jeffrey Barrett, Sonia Goncalves, Ewan Harrison, David K. Jackson, Ian Johnston, Dominic Kwiatkowski, Cordelia Langford, John Sillitoe on behalf of the Wellcome Sanger Institute COVID-19 Surveillance Team |
| EPI_ISL_3363792 | Lighthouse Lab in Glasgow | Wellcome Sanger Institute for the COVID-19 Genomics UK (COG-UK) Consortium | Harper VanSteenhouse, Yumi Kasai, David Gray, Carol Clugston, Anna Dominiczak and Alex Alderton, Roberto Amato, Jeffrey Barrett, Sonia Goncalves, Ewan Harrison, David K. Jackson, Ian Johnston, Dominic Kwiatkowski, Cordelia Langford, John Sillitoe on behalf of the Wellcome Sanger Institute COVID-19 Surveillance Team |
| EPI_ISL_3363793 | Lighthouse Lab in Alderley Park | Wellcome Sanger Institute for the COVID-19 Genomics UK (COG-UK) Consortium | Jacquelyn Wynn, Mairead Hyland, The Lighthouse Lab in Alderley Park and Alex Alderton, Roberto Amato, Jeffrey Barrett, Sonia Goncalves, Ewan Harrison, David K. Jackson, Ian Johnston, Dominic Kwiatkowski, Cordelia Langford, John Sillitoe on behalf of the Wellcome Sanger Institute COVID-19 Surveillance Team |
| EPI_ISL_3363794 | Lighthouse Lab in Glasgow | Wellcome Sanger Institute for the COVID-19 Genomics UK (COG-UK) Consortium | Harper VanSteenhouse, Yumi Kasai, David Gray, Carol Clugston, Anna Dominiczak and Alex Alderton, Roberto Amato, Jeffrey Barrett, Sonia Goncalves, Ewan Harrison, David K. Jackson, Ian Johnston, Dominic Kwiatkowski, Cordelia Langford, John Sillitoe on behalf of the Wellcome Sanger Institute COVID-19 Surveillance Team |
| EPI_ISL_3363795 | Lighthouse Lab in Alderley Park | Wellcome Sanger Institute for the COVID-19 Genomics UK (COG-UK) Consortium | Jacquelyn Wynn, Mairead Hyland, The Lighthouse Lab in Alderley Park and Alex Alderton, Roberto Amato, Jeffrey Barrett, Sonia Goncalves, Ewan Harrison, David K. Jackson, Ian Johnston, Dominic Kwiatkowski, Cordelia Langford, John Sillitoe on behalf of the Wellcome Sanger Institute COVID-19 Surveillance Team |
| EPI_ISL_3363796 | Lighthouse Lab in Milton Keynes | Wellcome Sanger Institute for the COVID-19 Genomics UK (COG-UK) Consortium | The Lighthouse Lab in Milton Keynes and Alex Alderton, Roberto Amato, Jeffrey Barrett, Sonia Goncalves, Ewan Harrison, David K. Jackson, Ian Johnston, Dominic Kwiatkowski, Cordelia Langford, John Sillitoe on behalf of the Wellcome Sanger Institute COVID-19 Surveillance Team |
| EPI_ISL_3363797, EPI_ISL_3363798, EPI_ISL_3363799 | Berkshire and Surrey Pathology Services Lighthouse Laboratory | Wellcome Sanger Institute for the COVID-19 Genomics UK (COG-UK) Consortium | Berkshire and Surrey Pathology Services Lighthouse Laboratory and Alex Alderton, Roberto Amato, Jeffrey Barrett, Sonia Goncalves, Ewan Harrison, David K. Jackson, Ian Johnston, Dominic Kwiatkowski, Cordelia Langford, John Sillitoe on behalf of the Wellcome Sanger Institute COVID-19 Surveillance Team |
| EPI_ISL_3363800 | Lighthouse Lab in Glasgow | Wellcome Sanger Institute for the COVID-19 Genomics UK (COG-UK) Consortium | Harper VanSteenhouse, Yumi Kasai, David Gray, Carol Clugston, Anna Dominiczak and Alex Alderton, Roberto Amato, Jeffrey Barrett, Sonia Goncalves, Ewan Harrison, David K. Jackson, Ian Johnston, Dominic Kwiatkowski, Cordelia Langford, John Sillitoe on behalf of the Wellcome Sanger Institute COVID-19 Surveillance Team |
| EPI_ISL_3363801, EPI_ISL_3363802, EPI_ISL_3363803 | Lighthouse Lab in Alderley Park | Wellcome Sanger Institute for the COVID-19 Genomics UK (COG-UK) Consortium | Jacquelyn Wynn, Mairead Hyland, The Lighthouse Lab in Alderley Park and Alex Alderton, Roberto Amato, Jeffrey Barrett, Sonia Goncalves, Ewan Harrison, David K. Jackson, Ian Johnston, Dominic Kwiatkowski, Cordelia Langford, John Sillitoe on behalf of the Wellcome Sanger Institute COVID-19 Surveillance Team |
| EPI_ISL_3363804 | Lighthouse Lab in Milton Keynes | Wellcome Sanger Institute for the COVID-19 Genomics UK (COG-UK) Consortium | The Lighthouse Lab in Milton Keynes and Alex Alderton, Roberto Amato, Jeffrey Barrett, Sonia Goncalves, Ewan Harrison, David K. Jackson, Ian Johnston, Dominic Kwiatkowski, Cordelia Langford, John Sillitoe on behalf of the Wellcome Sanger Institute COVID-19 Surveillance Team |
| EPI_ISL_3363805 | Lighthouse Lab in Alderley Park | Wellcome Sanger Institute for the COVID-19 Genomics UK (COG-UK) Consortium | Jacquelyn Wynn, Mairead Hyland, The Lighthouse Lab in Alderley Park and Alex Alderton, Roberto Amato, Jeffrey Barrett, Sonia Goncalves, Ewan Harrison, David K. Jackson, Ian Johnston, Dominic Kwiatkowski, Cordelia Langford, John Sillitoe on behalf of the Wellcome Sanger Institute COVID-19 Surveillance Team |
| EPI_ISL_3363806, EPI_ISL_3363807, EPI_ISL_3363808 | Lighthouse Lab in Milton Keynes | Wellcome Sanger Institute for the COVID-19 Genomics UK (COG-UK) Consortium | The Lighthouse Lab in Milton Keynes and Alex Alderton, Roberto Amato, Jeffrey Barrett, Sonia Goncalves, Ewan Harrison, David K. Jackson, Ian Johnston, Dominic Kwiatkowski, Cordelia Langford, John Sillitoe on behalf of the Wellcome Sanger Institute COVID-19 Surveillance Team |
| EPI_ISL_3363809 | Berkshire and Surrey Pathology Services Lighthouse Laboratory | Wellcome Sanger Institute for the COVID-19 Genomics UK (COG-UK) Consortium | Berkshire and Surrey Pathology Services Lighthouse Laboratory and Alex Alderton, Roberto Amato, Jeffrey Barrett, Sonia Goncalves, Ewan Harrison, David K. Jackson, Ian Johnston, Dominic Kwiatkowski, Cordelia Langford, John Sillitoe on behalf of the Wellcome Sanger Institute COVID-19 Surveillance Team |
| EPI_ISL_3363810, EPI_ISL_3363811, EPI_ISL_3363812, EPI_ISL_3363813 | Lighthouse Lab in Milton Keynes | Wellcome Sanger Institute for the COVID-19 Genomics UK (COG-UK) Consortium | The Lighthouse Lab in Milton Keynes and Alex Alderton, Roberto Amato, Jeffrey Barrett, Sonia Goncalves, Ewan Harrison, David K. Jackson, Ian Johnston, Dominic Kwiatkowski, Cordelia Langford, John Sillitoe on behalf of the Wellcome Sanger Institute COVID-19 Surveillance Team |
| EPI_ISL_3363814 | Lighthouse Lab in Alderley Park | Wellcome Sanger Institute for the COVID-19 Genomics UK (COG-UK) Consortium | Jacquelyn Wynn, Mairead Hyland, The Lighthouse Lab in Alderley Park and Alex Alderton, Roberto Amato, Jeffrey Barrett, Sonia Goncalves, Ewan Harrison, David K. Jackson, Ian Johnston, Dominic Kwiatkowski, Cordelia Langford, John Sillitoe on behalf of the Wellcome Sanger Institute COVID-19 Surveillance Team |
| EPI_ISL_3363815 | Lighthouse Lab in Milton Keynes | Wellcome Sanger Institute for the COVID-19 Genomics UK (COG-UK) Consortium | The Lighthouse Lab in Milton Keynes and Alex Alderton, Roberto Amato, Jeffrey Barrett, Sonia Goncalves, Ewan Harrison, David K. Jackson, Ian Johnston, Dominic Kwiatkowski, Cordelia Langford, John Sillitoe on behalf of the Wellcome Sanger Institute COVID-19 Surveillance Team |
| EPI_ISL_3363816 | Lighthouse Lab in Glasgow | Wellcome Sanger Institute for the COVID-19 Genomics UK (COG-UK) Consortium | Harper VanSteenhouse, Yumi Kasai, David Gray, Carol Clugston, Anna Dominiczak and Alex Alderton, Roberto Amato, Jeffrey Barrett, Sonia Goncalves, Ewan Harrison, David K. Jackson, Ian Johnston, Dominic Kwiatkowski, Cordelia Langford, John Sillitoe on behalf of the Wellcome Sanger Institute COVID-19 Surveillance Team |
| EPI_ISL_3363817 | Lighthouse Lab in Milton Keynes | Wellcome Sanger Institute for the COVID-19 Genomics UK (COG-UK) Consortium | The Lighthouse Lab in Milton Keynes and Alex Alderton, Roberto Amato, Jeffrey Barrett, Sonia Goncalves, Ewan Harrison, David K. Jackson, Ian Johnston, Dominic Kwiatkowski, Cordelia Langford, John Sillitoe on behalf of the Wellcome Sanger Institute COVID-19 Surveillance Team |
| EPI_ISL_3363818 | Lighthouse Lab in Alderley Park | Wellcome Sanger Institute for the COVID-19 Genomics UK (COG-UK) Consortium | Jacquelyn Wynn, Mairead Hyland, The Lighthouse Lab in Alderley Park and Alex Alderton, Roberto Amato, Jeffrey Barrett, Sonia Goncalves, Ewan Harrison, David K. Jackson, Ian Johnston, Dominic Kwiatkowski, Cordelia Langford, John Sillitoe on behalf of the Wellcome Sanger Institute COVID-19 Surveillance Team |
| EPI_ISL_3363819 | Lighthouse Lab in Milton Keynes | Wellcome Sanger Institute for the COVID-19 Genomics UK (COG-UK) Consortium | The Lighthouse Lab in Milton Keynes and Alex Alderton, Roberto Amato, Jeffrey Barrett, Sonia Goncalves, Ewan Harrison, David K. Jackson, Ian Johnston, Dominic Kwiatkowski, Cordelia Langford, John Sillitoe on behalf of the Wellcome Sanger Institute COVID-19 Surveillance Team |
| EPI_ISL_3363820 | Berkshire and Surrey Pathology Services Lighthouse Laboratory | Wellcome Sanger Institute for the COVID-19 Genomics UK (COG-UK) Consortium | Berkshire and Surrey Pathology Services Lighthouse Laboratory and Alex Alderton, Roberto Amato, Jeffrey Barrett, Sonia Goncalves, Ewan Harrison, David K. Jackson, Ian Johnston, Dominic Kwiatkowski, Cordelia Langford, John Sillitoe on behalf of the Wellcome Sanger Institute COVID-19 Surveillance Team |
| EPI_ISL_3363821 | Lighthouse Lab in Milton Keynes | Wellcome Sanger Institute for the COVID-19 Genomics UK (COG-UK) Consortium | The Lighthouse Lab in Milton Keynes and Alex Alderton, Roberto Amato, Jeffrey Barrett, Sonia Goncalves, Ewan Harrison, David K. Jackson, Ian Johnston, Dominic Kwiatkowski, Cordelia Langford, John Sillitoe on behalf of the Wellcome Sanger Institute COVID-19 Surveillance Team |
| EPI_ISL_3363822 | Lighthouse Lab in Glasgow | Wellcome Sanger Institute for the COVID-19 Genomics UK (COG-UK) Consortium | Harper VanSteenhouse, Yumi Kasai, David Gray, Carol Clugston, Anna Dominiczak and Alex Alderton, Roberto Amato, Jeffrey Barrett, Sonia Goncalves, Ewan Harrison, David K. Jackson, Ian Johnston, Dominic Kwiatkowski, Cordelia Langford, John Sillitoe on behalf of the Wellcome Sanger Institute COVID-19 Surveillance Team |
| EPI_ISL_3363823 | Berkshire and Surrey Pathology Services Lighthouse Laboratory | Wellcome Sanger Institute for the COVID-19 Genomics UK (COG-UK) Consortium | Berkshire and Surrey Pathology Services Lighthouse Laboratory and Alex Alderton, Roberto Amato, Jeffrey Barrett, Sonia Goncalves, Ewan Harrison, David K. Jackson, Ian Johnston, Dominic Kwiatkowski, Cordelia Langford, John Sillitoe on behalf of the Wellcome Sanger Institute COVID-19 Surveillance Team |
| EPI_ISL_3363824 | Lighthouse Lab in Glasgow | Wellcome Sanger Institute for the COVID-19 Genomics UK (COG-UK) Consortium | Harper VanSteenhouse, Yumi Kasai, David Gray, Carol Clugston, Anna Dominiczak and Alex Alderton, Roberto Amato, Jeffrey Barrett, Sonia Goncalves, Ewan Harrison, David K. Jackson, Ian Johnston, Dominic Kwiatkowski, Cordelia Langford, John Sillitoe on behalf of the Wellcome Sanger Institute COVID-19 Surveillance Team |
| EPI_ISL_3363825, EPI_ISL_3363826 | Berkshire and Surrey Pathology Services Lighthouse Laboratory | Wellcome Sanger Institute for the COVID-19 Genomics UK (COG-UK) Consortium | Berkshire and Surrey Pathology Services Lighthouse Laboratory and Alex Alderton, Roberto Amato, Jeffrey Barrett, Sonia Goncalves, Ewan Harrison, David K. Jackson, Ian Johnston, Dominic Kwiatkowski, Cordelia Langford, John Sillitoe on behalf of the Wellcome Sanger Institute COVID-19 Surveillance Team |
| EPI_ISL_3363827 | Lighthouse Lab in Milton Keynes | Wellcome Sanger Institute for the COVID-19 Genomics UK (COG-UK) Consortium | The Lighthouse Lab in Milton Keynes and Alex Alderton, Roberto Amato, Jeffrey Barrett, Sonia Goncalves, Ewan Harrison, David K. Jackson, Ian Johnston, Dominic Kwiatkowski, Cordelia Langford, John Sillitoe on behalf of the Wellcome Sanger Institute COVID-19 Surveillance Team |
| EPI_ISL_3363828 | Berkshire and Surrey Pathology Services Lighthouse Laboratory | Wellcome Sanger Institute for the COVID-19 Genomics UK (COG-UK) Consortium | Berkshire and Surrey Pathology Services Lighthouse Laboratory and Alex Alderton, Roberto Amato, Jeffrey Barrett, Sonia Goncalves, Ewan Harrison, David K. Jackson, Ian Johnston, Dominic Kwiatkowski, Cordelia Langford, John Sillitoe on behalf of the Wellcome Sanger Institute COVID-19 Surveillance Team |
| EPI_ISL_3363829 | Lighthouse Lab in Milton Keynes | Wellcome Sanger Institute for the COVID-19 Genomics UK (COG-UK) Consortium | The Lighthouse Lab in Milton Keynes and Alex Alderton, Roberto Amato, Jeffrey Barrett, Sonia Goncalves, Ewan Harrison, David K. Jackson, Ian Johnston, Dominic Kwiatkowski, Cordelia Langford, John Sillitoe on behalf of the Wellcome Sanger Institute COVID-19 Surveillance Team |
| EPI_ISL_3363830 | Berkshire and Surrey Pathology Services Lighthouse Laboratory | Wellcome Sanger Institute for the COVID-19 Genomics UK (COG-UK) Consortium | Berkshire and Surrey Pathology Services Lighthouse Laboratory and Alex Alderton, Roberto Amato, Jeffrey Barrett, Sonia Goncalves, Ewan Harrison, David K. Jackson, Ian Johnston, Dominic Kwiatkowski, Cordelia Langford, John Sillitoe on behalf of the Wellcome Sanger Institute COVID-19 Surveillance Team |
| EPI_ISL_3363831, EPI_ISL_3363832, EPI_ISL_3363833 | Lighthouse Lab in Glasgow | Wellcome Sanger Institute for the COVID-19 Genomics UK (COG-UK) Consortium | Harper VanSteenhouse, Yumi Kasai, David Gray, Carol Clugston, Anna Dominiczak and Alex Alderton, Roberto Amato, Jeffrey Barrett, Sonia Goncalves, Ewan Harrison, David K. Jackson, Ian Johnston, Dominic Kwiatkowski, Cordelia Langford, John Sillitoe on behalf of the Wellcome Sanger Institute COVID-19 Surveillance Team |
| EPI_ISL_3363834, EPI_ISL_3363835 | Lighthouse Lab in Alderley Park | Wellcome Sanger Institute for the COVID-19 Genomics UK (COG-UK) Consortium | Jacquelyn Wynn, Mairead Hyland, The Lighthouse Lab in Alderley Park and Alex Alderton, Roberto Amato, Jeffrey Barrett, Sonia Goncalves, Ewan Harrison, David K. Jackson, Ian Johnston, Dominic Kwiatkowski, Cordelia Langford, John Sillitoe on behalf of the Wellcome Sanger Institute COVID-19 Surveillance Team |
| EPI_ISL_3363836 | Lighthouse Lab in Milton Keynes | Wellcome Sanger Institute for the COVID-19 Genomics UK (COG-UK) Consortium | The Lighthouse Lab in Milton Keynes and Alex Alderton, Roberto Amato, Jeffrey Barrett, Sonia Goncalves, Ewan Harrison, David K. Jackson, Ian Johnston, Dominic Kwiatkowski, Cordelia Langford, John Sillitoe on behalf of the Wellcome Sanger Institute COVID-19 Surveillance Team |
| EPI_ISL_3363837 | Lighthouse Lab in Glasgow | Wellcome Sanger Institute for the COVID-19 Genomics UK (COG-UK) Consortium | Harper VanSteenhouse, Yumi Kasai, David Gray, Carol Clugston, Anna Dominiczak and Alex Alderton, Roberto Amato, Jeffrey Barrett, Sonia Goncalves, Ewan Harrison, David K. Jackson, Ian Johnston, Dominic Kwiatkowski, Cordelia Langford, John Sillitoe on behalf of the Wellcome Sanger Institute COVID-19 Surveillance Team |
| EPI_ISL_3363838, EPI_ISL_3363839 | Berkshire and Surrey Pathology Services Lighthouse Laboratory | Wellcome Sanger Institute for the COVID-19 Genomics UK (COG-UK) Consortium | Berkshire and Surrey Pathology Services Lighthouse Laboratory and Alex Alderton, Roberto Amato, Jeffrey Barrett, Sonia Goncalves, Ewan Harrison, David K. Jackson, Ian Johnston, Dominic Kwiatkowski, Cordelia Langford, John Sillitoe on behalf of the Wellcome Sanger Institute COVID-19 Surveillance Team |
| EPI_ISL_3363840 | Lighthouse Lab in Alderley Park | Wellcome Sanger Institute for the COVID-19 Genomics UK (COG-UK) Consortium | Jacquelyn Wynn, Mairead Hyland, The Lighthouse Lab in Alderley Park and Alex Alderton, Roberto Amato, Jeffrey Barrett, Sonia Goncalves, Ewan Harrison, David K. Jackson, Ian Johnston, Dominic Kwiatkowski, Cordelia Langford, John Sillitoe on behalf of the Wellcome Sanger Institute COVID-19 Surveillance Team |
| EPI_ISL_3363841 | Lighthouse Lab in Milton Keynes | Wellcome Sanger Institute for the COVID-19 Genomics UK (COG-UK) Consortium | The Lighthouse Lab in Milton Keynes and Alex Alderton, Roberto Amato, Jeffrey Barrett, Sonia Goncalves, Ewan Harrison, David K. Jackson, Ian Johnston, Dominic Kwiatkowski, Cordelia Langford, John Sillitoe on behalf of the Wellcome Sanger Institute COVID-19 Surveillance Team |
| EPI_ISL_3363842 | Lighthouse Lab in Alderley Park | Wellcome Sanger Institute for the COVID-19 Genomics UK (COG-UK) Consortium | Jacquelyn Wynn, Mairead Hyland, The Lighthouse Lab in Alderley Park and Alex Alderton, Roberto Amato, Jeffrey Barrett, Sonia Goncalves, Ewan Harrison, David K. Jackson, Ian Johnston, Dominic Kwiatkowski, Cordelia Langford, John Sillitoe on behalf of the Wellcome Sanger Institute COVID-19 Surveillance Team |
| EPI_ISL_3363843 | Lighthouse Lab in Glasgow | Wellcome Sanger Institute for the COVID-19 Genomics UK (COG-UK) Consortium | Harper VanSteenhouse, Yumi Kasai, David Gray, Carol Clugston, Anna Dominiczak and Alex Alderton, Roberto Amato, Jeffrey Barrett, Sonia Goncalves, Ewan Harrison, David K. Jackson, Ian Johnston, Dominic Kwiatkowski, Cordelia Langford, John Sillitoe on behalf of the Wellcome Sanger Institute COVID-19 Surveillance Team |
| EPI_ISL_3363844 | Lighthouse Lab in Milton Keynes | Wellcome Sanger Institute for the COVID-19 Genomics UK (COG-UK) Consortium | The Lighthouse Lab in Milton Keynes and Alex Alderton, Roberto Amato, Jeffrey Barrett, Sonia Goncalves, Ewan Harrison, David K. Jackson, Ian Johnston, Dominic Kwiatkowski, Cordelia Langford, John Sillitoe on behalf of the Wellcome Sanger Institute COVID-19 Surveillance Team |
| EPI_ISL_3363845 | Lighthouse Lab in Alderley Park | Wellcome Sanger Institute for the COVID-19 Genomics UK (COG-UK) Consortium | Jacquelyn Wynn, Mairead Hyland, The Lighthouse Lab in Alderley Park and Alex Alderton, Roberto Amato, Jeffrey Barrett, Sonia Goncalves, Ewan Harrison, David K. Jackson, Ian Johnston, Dominic Kwiatkowski, Cordelia Langford, John Sillitoe on behalf of the Wellcome Sanger Institute COVID-19 Surveillance Team |
| EPI_ISL_3363846 | Berkshire and Surrey Pathology Services Lighthouse Laboratory | Wellcome Sanger Institute for the COVID-19 Genomics UK (COG-UK) | Berkshire and Surrey Pathology Services Lighthouse Laboratory and Alex Alderton, Roberto Amato, Jeffrey Barrett, Sonia Goncalves, Ewan Harrison, David K. Jackson, Ian Johnston, Dominic Kwiatkowski, Cordelia Langford, John Sillitoe on behalf |

| | | | |
|---|---|---|---|
| | | Consortium | of the Wellcome Sanger Institute COVID-19 Surveillance Team |
| EPI_ISL_3363847, EPI_ISL_3363848 | Lighthouse Lab in Glasgow | Wellcome Sanger Institute for the COVID-19 Genomics UK (COG-UK) Consortium | Harper VanSteenhouse, Yumi Kasai, David Gray, Carol Clugston, Anna Dominiczak and Alex Alderton, Roberto Amato, Jeffrey Barrett, Sonia Goncalves, Ewan Harrison, David K. Jackson, Ian Johnston, Dominic Kwiatkowski, Cordelia Langford, John Sillitoe on behalf of the Wellcome Sanger Institute COVID-19 Surveillance Team |
| EPI_ISL_3363849, EPI_ISL_3363850 | Lighthouse Lab in Alderley Park | Wellcome Sanger Institute for the COVID-19 Genomics UK (COG-UK) Consortium | Jacquelyn Wynn, Mairead Hyland, The Lighthouse Lab in Alderley Park and Alex Alderton, Roberto Amato, Jeffrey Barrett, Sonia Goncalves, Ewan Harrison, David K. Jackson, Ian Johnston, Dominic Kwiatkowski, Cordelia Langford, John Sillitoe on behalf of the Wellcome Sanger Institute COVID-19 Surveillance Team |
| EPI_ISL_3363851 | Lighthouse Lab in Milton Keynes | Wellcome Sanger Institute for the COVID-19 Genomics UK (COG-UK) Consortium | The Lighthouse Lab in Milton Keynes and Alex Alderton, Roberto Amato, Jeffrey Barrett, Sonia Goncalves, Ewan Harrison, David K. Jackson, Ian Johnston, Dominic Kwiatkowski, Cordelia Langford, John Sillitoe on behalf of the Wellcome Sanger Institute COVID-19 Surveillance Team |
| EPI_ISL_3363852 | Lighthouse Lab in Alderley Park | Wellcome Sanger Institute for the COVID-19 Genomics UK (COG-UK) Consortium | Jacquelyn Wynn, Mairead Hyland, The Lighthouse Lab in Alderley Park and Alex Alderton, Roberto Amato, Jeffrey Barrett, Sonia Goncalves, Ewan Harrison, David K. Jackson, Ian Johnston, Dominic Kwiatkowski, Cordelia Langford, John Sillitoe on behalf of the Wellcome Sanger Institute COVID-19 Surveillance Team |
| EPI_ISL_3363853 | Lighthouse Lab in Milton Keynes | Wellcome Sanger Institute for the COVID-19 Genomics UK (COG-UK) Consortium | The Lighthouse Lab in Milton Keynes and Alex Alderton, Roberto Amato, Jeffrey Barrett, Sonia Goncalves, Ewan Harrison, David K. Jackson, Ian Johnston, Dominic Kwiatkowski, Cordelia Langford, John Sillitoe on behalf of the Wellcome Sanger Institute COVID-19 Surveillance Team |
| EPI_ISL_3363854 | Lighthouse Lab in Glasgow | Wellcome Sanger Institute for the COVID-19 Genomics UK (COG-UK) Consortium | Harper VanSteenhouse, Yumi Kasai, David Gray, Carol Clugston, Anna Dominiczak and Alex Alderton, Roberto Amato, Jeffrey Barrett, Sonia Goncalves, Ewan Harrison, David K. Jackson, Ian Johnston, Dominic Kwiatkowski, Cordelia Langford, John Sillitoe on behalf of the Wellcome Sanger Institute COVID-19 Surveillance Team |
| EPI_ISL_3363855, EPI_ISL_3363856 | Lighthouse Lab in Alderley Park | Wellcome Sanger Institute for the COVID-19 Genomics UK (COG-UK) Consortium | Jacquelyn Wynn, Mairead Hyland, The Lighthouse Lab in Alderley Park and Alex Alderton, Roberto Amato, Jeffrey Barrett, Sonia Goncalves, Ewan Harrison, David K. Jackson, Ian Johnston, Dominic Kwiatkowski, Cordelia Langford, John Sillitoe on behalf of the Wellcome Sanger Institute COVID-19 Surveillance Team |
| EPI_ISL_3363857 | Lighthouse Lab in Glasgow | Wellcome Sanger Institute for the COVID-19 Genomics UK (COG-UK) Consortium | Harper VanSteenhouse, Yumi Kasai, David Gray, Carol Clugston, Anna Dominiczak and Alex Alderton, Roberto Amato, Jeffrey Barrett, Sonia Goncalves, Ewan Harrison, David K. Jackson, Ian Johnston, Dominic Kwiatkowski, Cordelia Langford, John Sillitoe on behalf of the Wellcome Sanger Institute COVID-19 Surveillance Team |
| EPI_ISL_3363858, EPI_ISL_3363859 | Lighthouse Lab in Alderley Park | Wellcome Sanger Institute for the COVID-19 Genomics UK (COG-UK) Consortium | Jacquelyn Wynn, Mairead Hyland, The Lighthouse Lab in Alderley Park and Alex Alderton, Roberto Amato, Jeffrey Barrett, Sonia Goncalves, Ewan Harrison, David K. Jackson, Ian Johnston, Dominic Kwiatkowski, Cordelia Langford, John Sillitoe on behalf of the Wellcome Sanger Institute COVID-19 Surveillance Team |
| EPI_ISL_3363860 | Berkshire and Surrey Pathology Services Lighthouse Laboratory | Wellcome Sanger Institute for the COVID-19 Genomics UK (COG-UK) Consortium | Berkshire and Surrey Pathology Services Lighthouse Laboratory and Alex Alderton, Roberto Amato, Jeffrey Barrett, Sonia Goncalves, Ewan Harrison, David K. Jackson, Ian Johnston, Dominic Kwiatkowski, Cordelia Langford, John Sillitoe on behalf of the Wellcome Sanger Institute COVID-19 Surveillance Team |
| EPI_ISL_3363861 | Lighthouse Lab in Alderley Park | Wellcome Sanger Institute for the COVID-19 Genomics UK (COG-UK) Consortium | Jacquelyn Wynn, Mairead Hyland, The Lighthouse Lab in Alderley Park and Alex Alderton, Roberto Amato, Jeffrey Barrett, Sonia Goncalves, Ewan Harrison, David K. Jackson, Ian Johnston, Dominic Kwiatkowski, Cordelia Langford, John Sillitoe on behalf of the Wellcome Sanger Institute COVID-19 Surveillance Team |
| EPI_ISL_3363862 | Berkshire and Surrey Pathology Services Lighthouse Laboratory | Wellcome Sanger Institute for the COVID-19 Genomics UK (COG-UK) Consortium | Berkshire and Surrey Pathology Services Lighthouse Laboratory and Alex Alderton, Roberto Amato, Jeffrey Barrett, Sonia Goncalves, Ewan Harrison, David K. Jackson, Ian Johnston, Dominic Kwiatkowski, Cordelia Langford, John Sillitoe on behalf of the Wellcome Sanger Institute COVID-19 Surveillance Team |
| EPI_ISL_3363863, EPI_ISL_3363864 | Lighthouse Lab in Alderley Park | Wellcome Sanger Institute for the COVID-19 Genomics UK (COG-UK) Consortium | Jacquelyn Wynn, Mairead Hyland, The Lighthouse Lab in Alderley Park and Alex Alderton, Roberto Amato, Jeffrey Barrett, Sonia Goncalves, Ewan Harrison, David K. Jackson, Ian Johnston, Dominic Kwiatkowski, Cordelia Langford, John Sillitoe on behalf of the Wellcome Sanger Institute COVID-19 Surveillance Team |
| EPI_ISL_3363865 | Berkshire and Surrey Pathology Services Lighthouse Laboratory | Wellcome Sanger Institute for the COVID-19 Genomics UK (COG-UK) Consortium | Berkshire and Surrey Pathology Services Lighthouse Laboratory and Alex Alderton, Roberto Amato, Jeffrey Barrett, Sonia Goncalves, Ewan Harrison, David K. Jackson, Ian Johnston, Dominic Kwiatkowski, Cordelia Langford, John Sillitoe on behalf of the Wellcome Sanger Institute COVID-19 Surveillance Team |
| EPI_ISL_3363866, EPI_ISL_3363867, EPI_ISL_3363868, EPI_ISL_3363869 | Lighthouse Lab in Milton Keynes | Wellcome Sanger Institute for the COVID-19 Genomics UK (COG-UK) Consortium | The Lighthouse Lab in Milton Keynes and Alex Alderton, Roberto Amato, Jeffrey Barrett, Sonia Goncalves, Ewan Harrison, David K. Jackson, Ian Johnston, Dominic Kwiatkowski, Cordelia Langford, John Sillitoe on behalf of the Wellcome Sanger Institute COVID-19 Surveillance Team |
| EPI_ISL_3363870 | Lighthouse Lab in Alderley Park | Wellcome Sanger Institute for the COVID-19 Genomics UK (COG-UK) Consortium | Jacquelyn Wynn, Mairead Hyland, The Lighthouse Lab in Alderley Park and Alex Alderton, Roberto Amato, Jeffrey Barrett, Sonia Goncalves, Ewan Harrison, David K. Jackson, Ian Johnston, Dominic Kwiatkowski, Cordelia Langford, John Sillitoe on behalf of the Wellcome Sanger Institute COVID-19 Surveillance Team |
| EPI_ISL_3363871, EPI_ISL_3363872, EPI_ISL_3363873 | Lighthouse Lab in Milton Keynes | Wellcome Sanger Institute for the COVID-19 Genomics UK (COG-UK) Consortium | The Lighthouse Lab in Milton Keynes and Alex Alderton, Roberto Amato, Jeffrey Barrett, Sonia Goncalves, Ewan Harrison, David K. Jackson, Ian Johnston, Dominic Kwiatkowski, Cordelia Langford, John Sillitoe on behalf of the Wellcome Sanger Institute COVID-19 Surveillance Team |
| EPI_ISL_3363874 | Lighthouse Lab in Alderley Park | Wellcome Sanger Institute for the COVID-19 Genomics UK (COG-UK) Consortium | Jacquelyn Wynn, Mairead Hyland, The Lighthouse Lab in Alderley Park and Alex Alderton, Roberto Amato, Jeffrey Barrett, Sonia Goncalves, Ewan Harrison, David K. Jackson, Ian Johnston, Dominic Kwiatkowski, Cordelia Langford, John Sillitoe on behalf of the Wellcome Sanger Institute COVID-19 Surveillance Team |
| EPI_ISL_3363875, EPI_ISL_3363876, EPI_ISL_3363877 | Lighthouse Lab in Milton Keynes | Wellcome Sanger Institute for the COVID-19 Genomics UK (COG-UK) Consortium | The Lighthouse Lab in Milton Keynes and Alex Alderton, Roberto Amato, Jeffrey Barrett, Sonia Goncalves, Ewan Harrison, David K. Jackson, Ian Johnston, Dominic Kwiatkowski, Cordelia Langford, John Sillitoe on behalf of the Wellcome Sanger Institute COVID-19 Surveillance Team |
| EPI_ISL_3363878, EPI_ISL_3363879, EPI_ISL_3363880 | Lighthouse Lab in Alderley Park | Wellcome Sanger Institute for the COVID-19 Genomics UK (COG-UK) Consortium | Jacquelyn Wynn, Mairead Hyland, The Lighthouse Lab in Alderley Park and Alex Alderton, Roberto Amato, Jeffrey Barrett, Sonia Goncalves, Ewan Harrison, David K. Jackson, Ian Johnston, Dominic Kwiatkowski, Cordelia Langford, John Sillitoe on behalf of the Wellcome Sanger Institute COVID-19 Surveillance Team |
| EPI_ISL_3363881 | Berkshire and Surrey Pathology Services Lighthouse Laboratory | Wellcome Sanger Institute for the COVID-19 Genomics UK (COG-UK) Consortium | Berkshire and Surrey Pathology Services Lighthouse Laboratory and Alex Alderton, Roberto Amato, Jeffrey Barrett, Sonia Goncalves, Ewan Harrison, David K. Jackson, Ian Johnston, Dominic Kwiatkowski, Cordelia Langford, John Sillitoe on behalf of the Wellcome Sanger Institute COVID-19 Surveillance Team |
| EPI_ISL_3363882 | Lighthouse Lab in Milton Keynes | Wellcome Sanger Institute for the COVID-19 Genomics UK (COG-UK) Consortium | The Lighthouse Lab in Milton Keynes and Alex Alderton, Roberto Amato, Jeffrey Barrett, Sonia Goncalves, Ewan Harrison, David K. Jackson, Ian Johnston, Dominic Kwiatkowski, Cordelia Langford, John Sillitoe on behalf of the Wellcome Sanger Institute COVID-19 Surveillance Team |
| EPI_ISL_3363883, EPI_ISL_3363884 | Berkshire and Surrey Pathology Services Lighthouse Laboratory | Wellcome Sanger Institute for the COVID-19 Genomics UK (COG-UK) Consortium | Berkshire and Surrey Pathology Services Lighthouse Laboratory and Alex Alderton, Roberto Amato, Jeffrey Barrett, Sonia Goncalves, Ewan Harrison, David K. Jackson, Ian Johnston, Dominic Kwiatkowski, Cordelia Langford, John Sillitoe on behalf of the Wellcome Sanger Institute COVID-19 Surveillance Team |
| EPI_ISL_3363885 | Lighthouse Lab in Alderley Park | Wellcome Sanger Institute for the COVID-19 Genomics UK (COG-UK) Consortium | Jacquelyn Wynn, Mairead Hyland, The Lighthouse Lab in Alderley Park and Alex Alderton, Roberto Amato, Jeffrey Barrett, Sonia Goncalves, Ewan Harrison, David K. Jackson, Ian Johnston, Dominic Kwiatkowski, Cordelia Langford, John Sillitoe on behalf of the Wellcome Sanger Institute COVID-19 Surveillance Team |
| EPI_ISL_3363886 | Berkshire and Surrey Pathology Services Lighthouse Laboratory | Wellcome Sanger Institute for the COVID-19 Genomics UK (COG-UK) Consortium | Berkshire and Surrey Pathology Services Lighthouse Laboratory and Alex Alderton, Roberto Amato, Jeffrey Barrett, Sonia Goncalves, Ewan Harrison, David K. Jackson, Ian Johnston, Dominic Kwiatkowski, Cordelia Langford, John Sillitoe on behalf of the Wellcome Sanger Institute COVID-19 Surveillance Team |
| EPI_ISL_3363887, EPI_ISL_3363888 | Lighthouse Lab in Alderley Park | Wellcome Sanger Institute for the COVID-19 Genomics UK (COG-UK) Consortium | Jacquelyn Wynn, Mairead Hyland, The Lighthouse Lab in Alderley Park and Alex Alderton, Roberto Amato, Jeffrey Barrett, Sonia Goncalves, Ewan Harrison, David K. Jackson, Ian Johnston, Dominic Kwiatkowski, Cordelia Langford, John Sillitoe on behalf of the Wellcome Sanger Institute COVID-19 Surveillance Team |
| EPI_ISL_3363889 | Berkshire and Surrey Pathology Services Lighthouse Laboratory | Wellcome Sanger Institute for the COVID-19 Genomics UK (COG-UK) Consortium | Berkshire and Surrey Pathology Services Lighthouse Laboratory and Alex Alderton, Roberto Amato, Jeffrey Barrett, Sonia Goncalves, Ewan Harrison, David K. Jackson, Ian Johnston, Dominic Kwiatkowski, Cordelia Langford, John Sillitoe on behalf of the Wellcome Sanger Institute COVID-19 Surveillance Team |
| EPI_ISL_3363890 | Lighthouse Lab in Alderley Park | Wellcome Sanger Institute for the COVID-19 Genomics UK (COG-UK) Consortium | Jacquelyn Wynn, Mairead Hyland, The Lighthouse Lab in Alderley Park and Alex Alderton, Roberto Amato, Jeffrey Barrett, Sonia Goncalves, Ewan Harrison, David K. Jackson, Ian Johnston, Dominic Kwiatkowski, Cordelia Langford, John Sillitoe on behalf of the Wellcome Sanger Institute COVID-19 Surveillance Team |
| EPI_ISL_3363891 | Lighthouse Lab in Milton Keynes | Wellcome Sanger Institute for the COVID-19 Genomics UK (COG-UK) Consortium | The Lighthouse Lab in Milton Keynes and Alex Alderton, Roberto Amato, Jeffrey Barrett, Sonia Goncalves, Ewan Harrison, David K. Jackson, Ian Johnston, Dominic Kwiatkowski, Cordelia Langford, John Sillitoe on behalf of the Wellcome Sanger Institute COVID-19 Surveillance Team |
| EPI_ISL_3363892 | Lighthouse Lab in Alderley Park | Wellcome Sanger Institute for the COVID-19 Genomics UK (COG-UK) Consortium | Jacquelyn Wynn, Mairead Hyland, The Lighthouse Lab in Alderley Park and Alex Alderton, Roberto Amato, Jeffrey Barrett, Sonia Goncalves, Ewan Harrison, David K. Jackson, Ian Johnston, Dominic Kwiatkowski, Cordelia Langford, John Sillitoe on behalf of the Wellcome Sanger Institute COVID-19 Surveillance Team |
| EPI_ISL_3363893 | Berkshire and Surrey Pathology Services Lighthouse Laboratory | Wellcome Sanger Institute for the COVID-19 Genomics UK (COG-UK) Consortium | Berkshire and Surrey Pathology Services Lighthouse Laboratory and Alex Alderton, Roberto Amato, Jeffrey Barrett, Sonia Goncalves, Ewan Harrison, David K. Jackson, Ian Johnston, Dominic Kwiatkowski, Cordelia Langford, John Sillitoe on behalf of the Wellcome Sanger Institute COVID-19 Surveillance Team |
| EPI_ISL_3363894 | Lighthouse Lab in Milton Keynes | Wellcome Sanger Institute for the COVID-19 Genomics UK (COG-UK) Consortium | The Lighthouse Lab in Milton Keynes and Alex Alderton, Roberto Amato, Jeffrey Barrett, Sonia Goncalves, Ewan Harrison, David K. Jackson, Ian Johnston, Dominic Kwiatkowski, Cordelia Langford, John Sillitoe on behalf of the Wellcome Sanger Institute COVID-19 Surveillance Team |
| EPI_ISL_3363895, EPI_ISL_3363896 | Berkshire and Surrey Pathology Services Lighthouse Laboratory | Wellcome Sanger Institute for the COVID-19 Genomics UK (COG-UK) Consortium | Berkshire and Surrey Pathology Services Lighthouse Laboratory and Alex Alderton, Roberto Amato, Jeffrey Barrett, Sonia Goncalves, Ewan Harrison, David K. Jackson, Ian Johnston, Dominic Kwiatkowski, Cordelia Langford, John Sillitoe on behalf of the Wellcome Sanger Institute COVID-19 Surveillance Team |
| EPI_ISL_3363897 | Lighthouse Lab in Milton Keynes | Wellcome Sanger Institute for the COVID-19 Genomics UK (COG-UK) Consortium | The Lighthouse Lab in Milton Keynes and Alex Alderton, Roberto Amato, Jeffrey Barrett, Sonia Goncalves, Ewan Harrison, David K. Jackson, Ian Johnston, Dominic Kwiatkowski, Cordelia Langford, John Sillitoe on behalf of the Wellcome Sanger Institute COVID-19 Surveillance Team |
| EPI_ISL_3363898, EPI_ISL_3363899, EPI_ISL_3363900, EPI_ISL_3363901 | Lighthouse Lab in Alderley Park | Wellcome Sanger Institute for the COVID-19 Genomics UK (COG-UK) Consortium | Jacquelyn Wynn, Mairead Hyland, The Lighthouse Lab in Alderley Park and Alex Alderton, Roberto Amato, Jeffrey Barrett, Sonia Goncalves, Ewan Harrison, David K. Jackson, Ian Johnston, Dominic Kwiatkowski, Cordelia Langford, John Sillitoe on behalf of the Wellcome Sanger Institute COVID-19 Surveillance Team |
| EPI_ISL_3363902 | Berkshire and Surrey Pathology Services Lighthouse Laboratory | Wellcome Sanger Institute for the COVID-19 Genomics UK (COG-UK) Consortium | Berkshire and Surrey Pathology Services Lighthouse Laboratory and Alex Alderton, Roberto Amato, Jeffrey Barrett, Sonia Goncalves, Ewan Harrison, David K. Jackson, Ian Johnston, Dominic Kwiatkowski, Cordelia Langford, John Sillitoe on behalf of the Wellcome Sanger Institute COVID-19 Surveillance Team |
| EPI_ISL_3363903 | Lighthouse Lab in Alderley Park | Wellcome Sanger Institute for the COVID-19 Genomics UK (COG-UK) Consortium | Jacquelyn Wynn, Mairead Hyland, The Lighthouse Lab in Alderley Park and Alex Alderton, Roberto Amato, Jeffrey Barrett, Sonia Goncalves, Ewan Harrison, David K. Jackson, Ian Johnston, Dominic Kwiatkowski, Cordelia Langford, John Sillitoe on behalf of the Wellcome Sanger Institute COVID-19 Surveillance Team |
| EPI_ISL_3363904 | Lighthouse Lab in Milton Keynes | Wellcome Sanger Institute for the COVID-19 Genomics UK (COG-UK) Consortium | The Lighthouse Lab in Milton Keynes and Alex Alderton, Roberto Amato, Jeffrey Barrett, Sonia Goncalves, Ewan Harrison, David K. Jackson, Ian Johnston, Dominic Kwiatkowski, Cordelia Langford, John Sillitoe on behalf of the Wellcome Sanger Institute COVID-19 Surveillance Team |
| EPI_ISL_3363905 | Lighthouse Lab in Alderley Park | Wellcome Sanger Institute for the COVID-19 Genomics UK (COG-UK) Consortium | Jacquelyn Wynn, Mairead Hyland, The Lighthouse Lab in Alderley Park and Alex Alderton, Roberto Amato, Jeffrey Barrett, Sonia Goncalves, Ewan Harrison, David K. Jackson, Ian Johnston, Dominic Kwiatkowski, Cordelia Langford, John Sillitoe on behalf of the Wellcome Sanger Institute COVID-19 Surveillance Team |
| EPI_ISL_3363906 | Lighthouse Lab in Milton Keynes | Wellcome Sanger Institute for the COVID-19 Genomics UK (COG-UK) Consortium | The Lighthouse Lab in Milton Keynes and Alex Alderton, Roberto Amato, Jeffrey Barrett, Sonia Goncalves, Ewan Harrison, David K. Jackson, Ian Johnston, Dominic Kwiatkowski, Cordelia Langford, John Sillitoe on behalf of the Wellcome Sanger Institute COVID-19 Surveillance Team |
| EPI_ISL_3363907 | Lighthouse Lab in Alderley Park | Wellcome Sanger Institute for the COVID-19 Genomics UK (COG-UK) Consortium | Jacquelyn Wynn, Mairead Hyland, The Lighthouse Lab in Alderley Park and Alex Alderton, Roberto Amato, Jeffrey Barrett, Sonia Goncalves, Ewan Harrison, David K. Jackson, Ian Johnston, Dominic Kwiatkowski, Cordelia Langford, John Sillitoe on behalf of the Wellcome Sanger Institute COVID-19 Surveillance Team |
| EPI_ISL_3363908 | Lighthouse Lab in Milton Keynes | Wellcome Sanger Institute for the COVID-19 Genomics UK (COG-UK) | The Lighthouse Lab in Milton Keynes and Alex Alderton, Roberto Amato, Jeffrey Barrett, Sonia Goncalves, Ewan Harrison, David K. Jackson, Ian Johnston, Dominic Kwiatkowski, Cordelia Langford, John Sillitoe on behalf of the Wellcome Sanger |

| | | | |
|---|---|---|---|
| | | Consortium | Institute COVID-19 Surveillance Team |
| EPI_ISL_3363909 | Lighthouse Lab in Alderley Park | Wellcome Sanger Institute for the COVID-19 Genomics UK (COG-UK) Consortium | Jacquelyn Wynn, Mairead Hyland, The Lighthouse Lab in Alderley Park and Alex Alderton, Roberto Amato, Jeffrey Barrett, Sonia Goncalves, Ewan Harrison, David K. Jackson, Ian Johnston, Dominic Kwiatkowski, Cordelia Langford, John Sillitoe on behalf of the Wellcome Sanger Institute COVID-19 Surveillance Team |
| EPI_ISL_3363910 | Lighthouse Lab in Milton Keynes | Wellcome Sanger Institute for the COVID-19 Genomics UK (COG-UK) Consortium | The Lighthouse Lab in Milton Keynes and Alex Alderton, Roberto Amato, Jeffrey Barrett, Sonia Goncalves, Ewan Harrison, David K. Jackson, Ian Johnston, Dominic Kwiatkowski, Cordelia Langford, John Sillitoe on behalf of the Wellcome Sanger Institute COVID-19 Surveillance Team |
| EPI_ISL_3363911, EPI_ISL_3363912, EPI_ISL_3363913 | Lighthouse Lab in Alderley Park | Wellcome Sanger Institute for the COVID-19 Genomics UK (COG-UK) Consortium | Jacquelyn Wynn, Mairead Hyland, The Lighthouse Lab in Alderley Park and Alex Alderton, Roberto Amato, Jeffrey Barrett, Sonia Goncalves, Ewan Harrison, David K. Jackson, Ian Johnston, Dominic Kwiatkowski, Cordelia Langford, John Sillitoe on behalf of the Wellcome Sanger Institute COVID-19 Surveillance Team |
| EPI_ISL_3363914 | Berkshire and Surrey Pathology Services Lighthouse Laboratory | Wellcome Sanger Institute for the COVID-19 Genomics UK (COG-UK) Consortium | Berkshire and Surrey Pathology Services Lighthouse Laboratory and Alex Alderton, Roberto Amato, Jeffrey Barrett, Sonia Goncalves, Ewan Harrison, David K. Jackson, Ian Johnston, Dominic Kwiatkowski, Cordelia Langford, John Sillitoe on behalf of the Wellcome Sanger Institute COVID-19 Surveillance Team |
| EPI_ISL_3363915, EPI_ISL_3363916 | Lighthouse Lab in Milton Keynes | Wellcome Sanger Institute for the COVID-19 Genomics UK (COG-UK) Consortium | The Lighthouse Lab in Milton Keynes and Alex Alderton, Roberto Amato, Jeffrey Barrett, Sonia Goncalves, Ewan Harrison, David K. Jackson, Ian Johnston, Dominic Kwiatkowski, Cordelia Langford, John Sillitoe on behalf of the Wellcome Sanger Institute COVID-19 Surveillance Team |
| EPI_ISL_3363917, EPI_ISL_3363918 | Lighthouse Lab in Alderley Park | Wellcome Sanger Institute for the COVID-19 Genomics UK (COG-UK) Consortium | Jacquelyn Wynn, Mairead Hyland, The Lighthouse Lab in Alderley Park and Alex Alderton, Roberto Amato, Jeffrey Barrett, Sonia Goncalves, Ewan Harrison, David K. Jackson, Ian Johnston, Dominic Kwiatkowski, Cordelia Langford, John Sillitoe on behalf of the Wellcome Sanger Institute COVID-19 Surveillance Team |
| EPI_ISL_3363919 | Lighthouse Lab in Milton Keynes | Wellcome Sanger Institute for the COVID-19 Genomics UK (COG-UK) Consortium | The Lighthouse Lab in Milton Keynes and Alex Alderton, Roberto Amato, Jeffrey Barrett, Sonia Goncalves, Ewan Harrison, David K. Jackson, Ian Johnston, Dominic Kwiatkowski, Cordelia Langford, John Sillitoe on behalf of the Wellcome Sanger Institute COVID-19 Surveillance Team |
| EPI_ISL_3363920 | Lighthouse Lab in Alderley Park | Wellcome Sanger Institute for the COVID-19 Genomics UK (COG-UK) Consortium | Jacquelyn Wynn, Mairead Hyland, The Lighthouse Lab in Alderley Park and Alex Alderton, Roberto Amato, Jeffrey Barrett, Sonia Goncalves, Ewan Harrison, David K. Jackson, Ian Johnston, Dominic Kwiatkowski, Cordelia Langford, John Sillitoe on behalf of the Wellcome Sanger Institute COVID-19 Surveillance Team |
| EPI_ISL_3363921 | Lighthouse Lab in Milton Keynes | Wellcome Sanger Institute for the COVID-19 Genomics UK (COG-UK) Consortium | The Lighthouse Lab in Milton Keynes and Alex Alderton, Roberto Amato, Jeffrey Barrett, Sonia Goncalves, Ewan Harrison, David K. Jackson, Ian Johnston, Dominic Kwiatkowski, Cordelia Langford, John Sillitoe on behalf of the Wellcome Sanger Institute COVID-19 Surveillance Team |
| EPI_ISL_3363922, EPI_ISL_3363923 | Lighthouse Lab in Alderley Park | Wellcome Sanger Institute for the COVID-19 Genomics UK (COG-UK) Consortium | Jacquelyn Wynn, Mairead Hyland, The Lighthouse Lab in Alderley Park and Alex Alderton, Roberto Amato, Jeffrey Barrett, Sonia Goncalves, Ewan Harrison, David K. Jackson, Ian Johnston, Dominic Kwiatkowski, Cordelia Langford, John Sillitoe on behalf of the Wellcome Sanger Institute COVID-19 Surveillance Team |
| EPI_ISL_3363924, EPI_ISL_3363925 | Berkshire and Surrey Pathology Services Lighthouse Laboratory | Wellcome Sanger Institute for the COVID-19 Genomics UK (COG-UK) Consortium | Berkshire and Surrey Pathology Services Lighthouse Laboratory and Alex Alderton, Roberto Amato, Jeffrey Barrett, Sonia Goncalves, Ewan Harrison, David K. Jackson, Ian Johnston, Dominic Kwiatkowski, Cordelia Langford, John Sillitoe on behalf of the Wellcome Sanger Institute COVID-19 Surveillance Team |
| EPI_ISL_3363926 | Lighthouse Lab in Milton Keynes | Wellcome Sanger Institute for the COVID-19 Genomics UK (COG-UK) Consortium | The Lighthouse Lab in Milton Keynes and Alex Alderton, Roberto Amato, Jeffrey Barrett, Sonia Goncalves, Ewan Harrison, David K. Jackson, Ian Johnston, Dominic Kwiatkowski, Cordelia Langford, John Sillitoe on behalf of the Wellcome Sanger Institute COVID-19 Surveillance Team |
| EPI_ISL_3363927 | Berkshire and Surrey Pathology Services Lighthouse Laboratory | Wellcome Sanger Institute for the COVID-19 Genomics UK (COG-UK) Consortium | Berkshire and Surrey Pathology Services Lighthouse Laboratory and Alex Alderton, Roberto Amato, Jeffrey Barrett, Sonia Goncalves, Ewan Harrison, David K. Jackson, Ian Johnston, Dominic Kwiatkowski, Cordelia Langford, John Sillitoe on behalf of the Wellcome Sanger Institute COVID-19 Surveillance Team |
| EPI_ISL_3363928, EPI_ISL_3363929, EPI_ISL_3363930, EPI_ISL_3363931, EPI_ISL_3363932, EPI_ISL_3363933 | Lighthouse Lab in Alderley Park | Wellcome Sanger Institute for the COVID-19 Genomics UK (COG-UK) Consortium | Jacquelyn Wynn, Mairead Hyland, The Lighthouse Lab in Alderley Park and Alex Alderton, Roberto Amato, Jeffrey Barrett, Sonia Goncalves, Ewan Harrison, David K. Jackson, Ian Johnston, Dominic Kwiatkowski, Cordelia Langford, John Sillitoe on behalf of the Wellcome Sanger Institute COVID-19 Surveillance Team |
| EPI_ISL_3363934, EPI_ISL_3363935, EPI_ISL_3363936 | Lighthouse Lab in Milton Keynes | Wellcome Sanger Institute for the COVID-19 Genomics UK (COG-UK) Consortium | The Lighthouse Lab in Milton Keynes and Alex Alderton, Roberto Amato, Jeffrey Barrett, Sonia Goncalves, Ewan Harrison, David K. Jackson, Ian Johnston, Dominic Kwiatkowski, Cordelia Langford, John Sillitoe on behalf of the Wellcome Sanger Institute COVID-19 Surveillance Team |
| EPI_ISL_3363937 | Berkshire and Surrey Pathology Services Lighthouse Laboratory | Wellcome Sanger Institute for the COVID-19 Genomics UK (COG-UK) Consortium | Berkshire and Surrey Pathology Services Lighthouse Laboratory and Alex Alderton, Roberto Amato, Jeffrey Barrett, Sonia Goncalves, Ewan Harrison, David K. Jackson, Ian Johnston, Dominic Kwiatkowski, Cordelia Langford, John Sillitoe on behalf of the Wellcome Sanger Institute COVID-19 Surveillance Team |
| EPI_ISL_3363938, EPI_ISL_3363939 | Lighthouse Lab in Milton Keynes | Wellcome Sanger Institute for the COVID-19 Genomics UK (COG-UK) Consortium | The Lighthouse Lab in Milton Keynes and Alex Alderton, Roberto Amato, Jeffrey Barrett, Sonia Goncalves, Ewan Harrison, David K. Jackson, Ian Johnston, Dominic Kwiatkowski, Cordelia Langford, John Sillitoe on behalf of the Wellcome Sanger Institute COVID-19 Surveillance Team |
| EPI_ISL_3363940, EPI_ISL_3363941 | Lighthouse Lab in Alderley Park | Wellcome Sanger Institute for the COVID-19 Genomics UK (COG-UK) Consortium | Jacquelyn Wynn, Mairead Hyland, The Lighthouse Lab in Alderley Park and Alex Alderton, Roberto Amato, Jeffrey Barrett, Sonia Goncalves, Ewan Harrison, David K. Jackson, Ian Johnston, Dominic Kwiatkowski, Cordelia Langford, John Sillitoe on behalf of the Wellcome Sanger Institute COVID-19 Surveillance Team |
| EPI_ISL_3363942, EPI_ISL_3363943 | Lighthouse Lab in Milton Keynes | Wellcome Sanger Institute for the COVID-19 Genomics UK (COG-UK) Consortium | The Lighthouse Lab in Milton Keynes and Alex Alderton, Roberto Amato, Jeffrey Barrett, Sonia Goncalves, Ewan Harrison, David K. Jackson, Ian Johnston, Dominic Kwiatkowski, Cordelia Langford, John Sillitoe on behalf of the Wellcome Sanger Institute COVID-19 Surveillance Team |
| EPI_ISL_3363944 | Berkshire and Surrey Pathology Services Lighthouse Laboratory | Wellcome Sanger Institute for the COVID-19 Genomics UK (COG-UK) Consortium | Berkshire and Surrey Pathology Services Lighthouse Laboratory and Alex Alderton, Roberto Amato, Jeffrey Barrett, Sonia Goncalves, Ewan Harrison, David K. Jackson, Ian Johnston, Dominic Kwiatkowski, Cordelia Langford, John Sillitoe on behalf of the Wellcome Sanger Institute COVID-19 Surveillance Team |
| EPI_ISL_3363945 | Lighthouse Lab in Alderley Park | Wellcome Sanger Institute for the COVID-19 Genomics UK (COG-UK) Consortium | Jacquelyn Wynn, Mairead Hyland, The Lighthouse Lab in Alderley Park and Alex Alderton, Roberto Amato, Jeffrey Barrett, Sonia Goncalves, Ewan Harrison, David K. Jackson, Ian Johnston, Dominic Kwiatkowski, Cordelia Langford, John Sillitoe on behalf of the Wellcome Sanger Institute COVID-19 Surveillance Team |
| EPI_ISL_3363946 | Lighthouse Lab in Milton Keynes | Wellcome Sanger Institute for the COVID-19 Genomics UK (COG-UK) Consortium | The Lighthouse Lab in Milton Keynes and Alex Alderton, Roberto Amato, Jeffrey Barrett, Sonia Goncalves, Ewan Harrison, David K. Jackson, Ian Johnston, Dominic Kwiatkowski, Cordelia Langford, John Sillitoe on behalf of the Wellcome Sanger Institute COVID-19 Surveillance Team |
| EPI_ISL_3363947, EPI_ISL_3363948 | Lighthouse Lab in Alderley Park | Wellcome Sanger Institute for the COVID-19 Genomics UK (COG-UK) Consortium | Jacquelyn Wynn, Mairead Hyland, The Lighthouse Lab in Alderley Park and Alex Alderton, Roberto Amato, Jeffrey Barrett, Sonia Goncalves, Ewan Harrison, David K. Jackson, Ian Johnston, Dominic Kwiatkowski, Cordelia Langford, John Sillitoe on behalf of the Wellcome Sanger Institute COVID-19 Surveillance Team |
| EPI_ISL_3363949, EPI_ISL_3363950, EPI_ISL_3363951 | Berkshire and Surrey Pathology Services Lighthouse Laboratory | Wellcome Sanger Institute for the COVID-19 Genomics UK (COG-UK) Consortium | Berkshire and Surrey Pathology Services Lighthouse Laboratory and Alex Alderton, Roberto Amato, Jeffrey Barrett, Sonia Goncalves, Ewan Harrison, David K. Jackson, Ian Johnston, Dominic Kwiatkowski, Cordelia Langford, John Sillitoe on behalf of the Wellcome Sanger Institute COVID-19 Surveillance Team |
| EPI_ISL_3363952 | Lighthouse Lab in Alderley Park | Wellcome Sanger Institute for the COVID-19 Genomics UK (COG-UK) Consortium | Jacquelyn Wynn, Mairead Hyland, The Lighthouse Lab in Alderley Park and Alex Alderton, Roberto Amato, Jeffrey Barrett, Sonia Goncalves, Ewan Harrison, David K. Jackson, Ian Johnston, Dominic Kwiatkowski, Cordelia Langford, John Sillitoe on behalf of the Wellcome Sanger Institute COVID-19 Surveillance Team |
| EPI_ISL_3363953 | Berkshire and Surrey Pathology Services Lighthouse Laboratory | Wellcome Sanger Institute for the COVID-19 Genomics UK (COG-UK) Consortium | Berkshire and Surrey Pathology Services Lighthouse Laboratory and Alex Alderton, Roberto Amato, Jeffrey Barrett, Sonia Goncalves, Ewan Harrison, David K. Jackson, Ian Johnston, Dominic Kwiatkowski, Cordelia Langford, John Sillitoe on behalf of the Wellcome Sanger Institute COVID-19 Surveillance Team |
| EPI_ISL_3363954 | Lighthouse Lab in Milton Keynes | Wellcome Sanger Institute for the COVID-19 Genomics UK (COG-UK) Consortium | The Lighthouse Lab in Milton Keynes and Alex Alderton, Roberto Amato, Jeffrey Barrett, Sonia Goncalves, Ewan Harrison, David K. Jackson, Ian Johnston, Dominic Kwiatkowski, Cordelia Langford, John Sillitoe on behalf of the Wellcome Sanger Institute COVID-19 Surveillance Team |
| EPI_ISL_3363955 | Berkshire and Surrey Pathology Services Lighthouse Laboratory | Wellcome Sanger Institute for the COVID-19 Genomics UK (COG-UK) Consortium | Berkshire and Surrey Pathology Services Lighthouse Laboratory and Alex Alderton, Roberto Amato, Jeffrey Barrett, Sonia Goncalves, Ewan Harrison, David K. Jackson, Ian Johnston, Dominic Kwiatkowski, Cordelia Langford, John Sillitoe on behalf of the Wellcome Sanger Institute COVID-19 Surveillance Team |
| EPI_ISL_3363956, EPI_ISL_3363957 | Lighthouse Lab in Alderley Park | Wellcome Sanger Institute for the COVID-19 Genomics UK (COG-UK) Consortium | Jacquelyn Wynn, Mairead Hyland, The Lighthouse Lab in Alderley Park and Alex Alderton, Roberto Amato, Jeffrey Barrett, Sonia Goncalves, Ewan Harrison, David K. Jackson, Ian Johnston, Dominic Kwiatkowski, Cordelia Langford, John Sillitoe on behalf of the Wellcome Sanger Institute COVID-19 Surveillance Team |
| EPI_ISL_3363958 | Lighthouse Lab in Milton Keynes | Wellcome Sanger Institute for the COVID-19 Genomics UK (COG-UK) Consortium | The Lighthouse Lab in Milton Keynes and Alex Alderton, Roberto Amato, Jeffrey Barrett, Sonia Goncalves, Ewan Harrison, David K. Jackson, Ian Johnston, Dominic Kwiatkowski, Cordelia Langford, John Sillitoe on behalf of the Wellcome Sanger Institute COVID-19 Surveillance Team |
| EPI_ISL_3363959 | Lighthouse Lab in Alderley Park | Wellcome Sanger Institute for the COVID-19 Genomics UK (COG-UK) Consortium | Jacquelyn Wynn, Mairead Hyland, The Lighthouse Lab in Alderley Park and Alex Alderton, Roberto Amato, Jeffrey Barrett, Sonia Goncalves, Ewan Harrison, David K. Jackson, Ian Johnston, Dominic Kwiatkowski, Cordelia Langford, John Sillitoe on behalf of the Wellcome Sanger Institute COVID-19 Surveillance Team |
| EPI_ISL_3363960, EPI_ISL_3363961 | Berkshire and Surrey Pathology Services Lighthouse Laboratory | Wellcome Sanger Institute for the COVID-19 Genomics UK (COG-UK) Consortium | Berkshire and Surrey Pathology Services Lighthouse Laboratory and Alex Alderton, Roberto Amato, Jeffrey Barrett, Sonia Goncalves, Ewan Harrison, David K. Jackson, Ian Johnston, Dominic Kwiatkowski, Cordelia Langford, John Sillitoe on behalf of the Wellcome Sanger Institute COVID-19 Surveillance Team |
| EPI_ISL_3363962 | Lighthouse Lab in Milton Keynes | Wellcome Sanger Institute for the COVID-19 Genomics UK (COG-UK) Consortium | The Lighthouse Lab in Milton Keynes and Alex Alderton, Roberto Amato, Jeffrey Barrett, Sonia Goncalves, Ewan Harrison, David K. Jackson, Ian Johnston, Dominic Kwiatkowski, Cordelia Langford, John Sillitoe on behalf of the Wellcome Sanger Institute COVID-19 Surveillance Team |
| EPI_ISL_3363963 | Berkshire and Surrey Pathology Services Lighthouse Laboratory | Wellcome Sanger Institute for the COVID-19 Genomics UK (COG-UK) Consortium | Berkshire and Surrey Pathology Services Lighthouse Laboratory and Alex Alderton, Roberto Amato, Jeffrey Barrett, Sonia Goncalves, Ewan Harrison, David K. Jackson, Ian Johnston, Dominic Kwiatkowski, Cordelia Langford, John Sillitoe on behalf of the Wellcome Sanger Institute COVID-19 Surveillance Team |
| EPI_ISL_3363964, EPI_ISL_3363965, EPI_ISL_3363966, EPI_ISL_3363967, EPI_ISL_3363968, EPI_ISL_3363969, EPI_ISL_3363970, EPI_ISL_3363971 | Lighthouse Lab in Alderley Park | Wellcome Sanger Institute for the COVID-19 Genomics UK (COG-UK) Consortium | Jacquelyn Wynn, Mairead Hyland, The Lighthouse Lab in Alderley Park and Alex Alderton, Roberto Amato, Jeffrey Barrett, Sonia Goncalves, Ewan Harrison, David K. Jackson, Ian Johnston, Dominic Kwiatkowski, Cordelia Langford, John Sillitoe on behalf of the Wellcome Sanger Institute COVID-19 Surveillance Team |
| EPI_ISL_3363972 | Lighthouse Lab in Milton Keynes | Wellcome Sanger Institute for the COVID-19 Genomics UK (COG-UK) Consortium | The Lighthouse Lab in Milton Keynes and Alex Alderton, Roberto Amato, Jeffrey Barrett, Sonia Goncalves, Ewan Harrison, David K. Jackson, Ian Johnston, Dominic Kwiatkowski, Cordelia Langford, John Sillitoe on behalf of the Wellcome Sanger Institute COVID-19 Surveillance Team |
| EPI_ISL_3363973 | Lighthouse Lab in Alderley Park | Wellcome Sanger Institute for the COVID-19 Genomics UK (COG-UK) Consortium | Jacquelyn Wynn, Mairead Hyland, The Lighthouse Lab in Alderley Park and Alex Alderton, Roberto Amato, Jeffrey Barrett, Sonia Goncalves, Ewan Harrison, David K. Jackson, Ian Johnston, Dominic Kwiatkowski, Cordelia Langford, John Sillitoe on behalf of the Wellcome Sanger Institute COVID-19 Surveillance Team |
| EPI_ISL_3363974 | Lighthouse Lab in Milton Keynes | Wellcome Sanger Institute for the COVID-19 Genomics UK (COG-UK) Consortium | The Lighthouse Lab in Milton Keynes and Alex Alderton, Roberto Amato, Jeffrey Barrett, Sonia Goncalves, Ewan Harrison, David K. Jackson, Ian Johnston, Dominic Kwiatkowski, Cordelia Langford, John Sillitoe on behalf of the Wellcome Sanger Institute COVID-19 Surveillance Team |
| EPI_ISL_3363975, EPI_ISL_3363976 | Lighthouse Lab in Alderley Park | Wellcome Sanger Institute for the COVID-19 Genomics UK (COG-UK) Consortium | Jacquelyn Wynn, Mairead Hyland, The Lighthouse Lab in Alderley Park and Alex Alderton, Roberto Amato, Jeffrey Barrett, Sonia Goncalves, Ewan Harrison, David K. Jackson, Ian Johnston, Dominic Kwiatkowski, Cordelia Langford, John Sillitoe on behalf of the Wellcome Sanger Institute COVID-19 Surveillance Team |

| | | | |
|---|---|---|---|
| EPI_ISL_3363977 | Lighthouse Lab in Milton Keynes | Wellcome Sanger Institute for the COVID-19 Genomics UK (COG-UK) Consortium | The Lighthouse Lab in Milton Keynes and Alex Alderton, Roberto Amato, Jeffrey Barrett, Sonia Goncalves, Ewan Harrison, David K. Jackson, Ian Johnston, Dominic Kwiatkowski, Cordelia Langford, John Sillitoe on behalf of the Wellcome Sanger Institute COVID-19 Surveillance Team |
| EPI_ISL_3363978 | Berkshire and Surrey Pathology Services Lighthouse Laboratory | Wellcome Sanger Institute for the COVID-19 Genomics UK (COG-UK) Consortium | Berkshire and Surrey Pathology Services Lighthouse Laboratory and Alex Alderton, Roberto Amato, Jeffrey Barrett, Sonia Goncalves, Ewan Harrison, David K. Jackson, Ian Johnston, Dominic Kwiatkowski, Cordelia Langford, John Sillitoe on behalf of the Wellcome Sanger Institute COVID-19 Surveillance Team |
| EPI_ISL_3363979 | Lighthouse Lab in Milton Keynes | Wellcome Sanger Institute for the COVID-19 Genomics UK (COG-UK) Consortium | The Lighthouse Lab in Milton Keynes and Alex Alderton, Roberto Amato, Jeffrey Barrett, Sonia Goncalves, Ewan Harrison, David K. Jackson, Ian Johnston, Dominic Kwiatkowski, Cordelia Langford, John Sillitoe on behalf of the Wellcome Sanger Institute COVID-19 Surveillance Team |
| EPI_ISL_3363980 | Berkshire and Surrey Pathology Services Lighthouse Laboratory | Wellcome Sanger Institute for the COVID-19 Genomics UK (COG-UK) Consortium | Berkshire and Surrey Pathology Services Lighthouse Laboratory and Alex Alderton, Roberto Amato, Jeffrey Barrett, Sonia Goncalves, Ewan Harrison, David K. Jackson, Ian Johnston, Dominic Kwiatkowski, Cordelia Langford, John Sillitoe on behalf of the Wellcome Sanger Institute COVID-19 Surveillance Team |
| EPI_ISL_3363981, EPI_ISL_3363982, EPI_ISL_3363983, EPI_ISL_3363984 | Lighthouse Lab in Alderley Park | Wellcome Sanger Institute for the COVID-19 Genomics UK (COG-UK) Consortium | Jacquelyn Wynn, Mairead Hyland, The Lighthouse Lab in Alderley Park and Alex Alderton, Roberto Amato, Jeffrey Barrett, Sonia Goncalves, Ewan Harrison, David K. Jackson, Ian Johnston, Dominic Kwiatkowski, Cordelia Langford, John Sillitoe on behalf of the Wellcome Sanger Institute COVID-19 Surveillance Team |
| EPI_ISL_3363985 | Berkshire and Surrey Pathology Services Lighthouse Laboratory | Wellcome Sanger Institute for the COVID-19 Genomics UK (COG-UK) Consortium | Berkshire and Surrey Pathology Services Lighthouse Laboratory and Alex Alderton, Roberto Amato, Jeffrey Barrett, Sonia Goncalves, Ewan Harrison, David K. Jackson, Ian Johnston, Dominic Kwiatkowski, Cordelia Langford, John Sillitoe on behalf of the Wellcome Sanger Institute COVID-19 Surveillance Team |
| EPI_ISL_3363986 | Lighthouse Lab in Alderley Park | Wellcome Sanger Institute for the COVID-19 Genomics UK (COG-UK) Consortium | Jacquelyn Wynn, Mairead Hyland, The Lighthouse Lab in Alderley Park and Alex Alderton, Roberto Amato, Jeffrey Barrett, Sonia Goncalves, Ewan Harrison, David K. Jackson, Ian Johnston, Dominic Kwiatkowski, Cordelia Langford, John Sillitoe on behalf of the Wellcome Sanger Institute COVID-19 Surveillance Team |
| EPI_ISL_3363987, EPI_ISL_3363988 | Lighthouse Lab in Milton Keynes | Wellcome Sanger Institute for the COVID-19 Genomics UK (COG-UK) Consortium | The Lighthouse Lab in Milton Keynes and Alex Alderton, Roberto Amato, Jeffrey Barrett, Sonia Goncalves, Ewan Harrison, David K. Jackson, Ian Johnston, Dominic Kwiatkowski, Cordelia Langford, John Sillitoe on behalf of the Wellcome Sanger Institute COVID-19 Surveillance Team |
| EPI_ISL_3363989, EPI_ISL_3363990 | Lighthouse Lab in Alderley Park | Wellcome Sanger Institute for the COVID-19 Genomics UK (COG-UK) Consortium | Jacquelyn Wynn, Mairead Hyland, The Lighthouse Lab in Alderley Park and Alex Alderton, Roberto Amato, Jeffrey Barrett, Sonia Goncalves, Ewan Harrison, David K. Jackson, Ian Johnston, Dominic Kwiatkowski, Cordelia Langford, John Sillitoe on behalf of the Wellcome Sanger Institute COVID-19 Surveillance Team |
| EPI_ISL_3363991, EPI_ISL_3363992 | Berkshire and Surrey Pathology Services Lighthouse Laboratory | Wellcome Sanger Institute for the COVID-19 Genomics UK (COG-UK) Consortium | Berkshire and Surrey Pathology Services Lighthouse Laboratory and Alex Alderton, Roberto Amato, Jeffrey Barrett, Sonia Goncalves, Ewan Harrison, David K. Jackson, Ian Johnston, Dominic Kwiatkowski, Cordelia Langford, John Sillitoe on behalf of the Wellcome Sanger Institute COVID-19 Surveillance Team |
| EPI_ISL_3363993 | Lighthouse Lab in Alderley Park | Wellcome Sanger Institute for the COVID-19 Genomics UK (COG-UK) Consortium | Jacquelyn Wynn, Mairead Hyland, The Lighthouse Lab in Alderley Park and Alex Alderton, Roberto Amato, Jeffrey Barrett, Sonia Goncalves, Ewan Harrison, David K. Jackson, Ian Johnston, Dominic Kwiatkowski, Cordelia Langford, John Sillitoe on behalf of the Wellcome Sanger Institute COVID-19 Surveillance Team |
| EPI_ISL_3363994 | Lighthouse Lab in Milton Keynes | Wellcome Sanger Institute for the COVID-19 Genomics UK (COG-UK) Consortium | The Lighthouse Lab in Milton Keynes and Alex Alderton, Roberto Amato, Jeffrey Barrett, Sonia Goncalves, Ewan Harrison, David K. Jackson, Ian Johnston, Dominic Kwiatkowski, Cordelia Langford, John Sillitoe on behalf of the Wellcome Sanger Institute COVID-19 Surveillance Team |
| EPI_ISL_3363995 | Lighthouse Lab in Alderley Park | Wellcome Sanger Institute for the COVID-19 Genomics UK (COG-UK) Consortium | Jacquelyn Wynn, Mairead Hyland, The Lighthouse Lab in Alderley Park and Alex Alderton, Roberto Amato, Jeffrey Barrett, Sonia Goncalves, Ewan Harrison, David K. Jackson, Ian Johnston, Dominic Kwiatkowski, Cordelia Langford, John Sillitoe on behalf of the Wellcome Sanger Institute COVID-19 Surveillance Team |
| EPI_ISL_3363996, EPI_ISL_3363997, EPI_ISL_3363998, EPI_ISL_3363999, EPI_ISL_3364000 | Berkshire and Surrey Pathology Services Lighthouse Laboratory | Wellcome Sanger Institute for the COVID-19 Genomics UK (COG-UK) Consortium | Berkshire and Surrey Pathology Services Lighthouse Laboratory and Alex Alderton, Roberto Amato, Jeffrey Barrett, Sonia Goncalves, Ewan Harrison, David K. Jackson, Ian Johnston, Dominic Kwiatkowski, Cordelia Langford, John Sillitoe on behalf of the Wellcome Sanger Institute COVID-19 Surveillance Team |
| EPI_ISL_3364001 | Lighthouse Lab in Milton Keynes | Wellcome Sanger Institute for the COVID-19 Genomics UK (COG-UK) Consortium | The Lighthouse Lab in Milton Keynes and Alex Alderton, Roberto Amato, Jeffrey Barrett, Sonia Goncalves, Ewan Harrison, David K. Jackson, Ian Johnston, Dominic Kwiatkowski, Cordelia Langford, John Sillitoe on behalf of the Wellcome Sanger Institute COVID-19 Surveillance Team |
| EPI_ISL_3364002 | Lighthouse Lab in Alderley Park | Wellcome Sanger Institute for the COVID-19 Genomics UK (COG-UK) Consortium | Jacquelyn Wynn, Mairead Hyland, The Lighthouse Lab in Alderley Park and Alex Alderton, Roberto Amato, Jeffrey Barrett, Sonia Goncalves, Ewan Harrison, David K. Jackson, Ian Johnston, Dominic Kwiatkowski, Cordelia Langford, John Sillitoe on behalf of the Wellcome Sanger Institute COVID-19 Surveillance Team |
| EPI_ISL_3364003 | Lighthouse Lab in Milton Keynes | Wellcome Sanger Institute for the COVID-19 Genomics UK (COG-UK) Consortium | The Lighthouse Lab in Milton Keynes and Alex Alderton, Roberto Amato, Jeffrey Barrett, Sonia Goncalves, Ewan Harrison, David K. Jackson, Ian Johnston, Dominic Kwiatkowski, Cordelia Langford, John Sillitoe on behalf of the Wellcome Sanger Institute COVID-19 Surveillance Team |
| EPI_ISL_3364004 | Berkshire and Surrey Pathology Services Lighthouse Laboratory | Wellcome Sanger Institute for the COVID-19 Genomics UK (COG-UK) Consortium | Berkshire and Surrey Pathology Services Lighthouse Laboratory and Alex Alderton, Roberto Amato, Jeffrey Barrett, Sonia Goncalves, Ewan Harrison, David K. Jackson, Ian Johnston, Dominic Kwiatkowski, Cordelia Langford, John Sillitoe on behalf of the Wellcome Sanger Institute COVID-19 Surveillance Team |
| EPI_ISL_3364005, EPI_ISL_3364006, EPI_ISL_3364008, EPI_ISL_3364009, EPI_ISL_3364010 | Lighthouse Lab in Alderley Park | Wellcome Sanger Institute for the COVID-19 Genomics UK (COG-UK) Consortium | Jacquelyn Wynn, Mairead Hyland, The Lighthouse Lab in Alderley Park and Alex Alderton, Roberto Amato, Jeffrey Barrett, Sonia Goncalves, Ewan Harrison, David K. Jackson, Ian Johnston, Dominic Kwiatkowski, Cordelia Langford, John Sillitoe on behalf of the Wellcome Sanger Institute COVID-19 Surveillance Team |
| EPI_ISL_3364012 | Lighthouse Lab in Milton Keynes | Wellcome Sanger Institute for the COVID-19 Genomics UK (COG-UK) Consortium | The Lighthouse Lab in Milton Keynes and Alex Alderton, Roberto Amato, Jeffrey Barrett, Sonia Goncalves, Ewan Harrison, David K. Jackson, Ian Johnston, Dominic Kwiatkowski, Cordelia Langford, John Sillitoe on behalf of the Wellcome Sanger Institute COVID-19 Surveillance Team |
| EPI_ISL_3364013 | Berkshire and Surrey Pathology Services Lighthouse Laboratory | Wellcome Sanger Institute for the COVID-19 Genomics UK (COG-UK) Consortium | Berkshire and Surrey Pathology Services Lighthouse Laboratory and Alex Alderton, Roberto Amato, Jeffrey Barrett, Sonia Goncalves, Ewan Harrison, David K. Jackson, Ian Johnston, Dominic Kwiatkowski, Cordelia Langford, John Sillitoe on behalf of the Wellcome Sanger Institute COVID-19 Surveillance Team |
| EPI_ISL_3364014 | Lighthouse Lab in Milton Keynes | Wellcome Sanger Institute for the COVID-19 Genomics UK (COG-UK) Consortium | The Lighthouse Lab in Milton Keynes and Alex Alderton, Roberto Amato, Jeffrey Barrett, Sonia Goncalves, Ewan Harrison, David K. Jackson, Ian Johnston, Dominic Kwiatkowski, Cordelia Langford, John Sillitoe on behalf of the Wellcome Sanger Institute COVID-19 Surveillance Team |
| EPI_ISL_3364016, EPI_ISL_3364017 | Lighthouse Lab in Alderley Park | Wellcome Sanger Institute for the COVID-19 Genomics UK (COG-UK) Consortium | Jacquelyn Wynn, Mairead Hyland, The Lighthouse Lab in Alderley Park and Alex Alderton, Roberto Amato, Jeffrey Barrett, Sonia Goncalves, Ewan Harrison, David K. Jackson, Ian Johnston, Dominic Kwiatkowski, Cordelia Langford, John Sillitoe on behalf of the Wellcome Sanger Institute COVID-19 Surveillance Team |
| EPI_ISL_3364019 | Lighthouse Lab in Milton Keynes | Wellcome Sanger Institute for the COVID-19 Genomics UK (COG-UK) Consortium | The Lighthouse Lab in Milton Keynes and Alex Alderton, Roberto Amato, Jeffrey Barrett, Sonia Goncalves, Ewan Harrison, David K. Jackson, Ian Johnston, Dominic Kwiatkowski, Cordelia Langford, John Sillitoe on behalf of the Wellcome Sanger Institute COVID-19 Surveillance Team |
| EPI_ISL_3364020, EPI_ISL_3364021 | Berkshire and Surrey Pathology Services Lighthouse Laboratory | Wellcome Sanger Institute for the COVID-19 Genomics UK (COG-UK) Consortium | Berkshire and Surrey Pathology Services Lighthouse Laboratory and Alex Alderton, Roberto Amato, Jeffrey Barrett, Sonia Goncalves, Ewan Harrison, David K. Jackson, Ian Johnston, Dominic Kwiatkowski, Cordelia Langford, John Sillitoe on behalf of the Wellcome Sanger Institute COVID-19 Surveillance Team |
| EPI_ISL_3364023, EPI_ISL_3364024 | Lighthouse Lab in Milton Keynes | Wellcome Sanger Institute for the COVID-19 Genomics UK (COG-UK) Consortium | The Lighthouse Lab in Milton Keynes and Alex Alderton, Roberto Amato, Jeffrey Barrett, Sonia Goncalves, Ewan Harrison, David K. Jackson, Ian Johnston, Dominic Kwiatkowski, Cordelia Langford, John Sillitoe on behalf of the Wellcome Sanger Institute COVID-19 Surveillance Team |
| EPI_ISL_3364026 | Berkshire and Surrey Pathology Services Lighthouse Laboratory | Wellcome Sanger Institute for the COVID-19 Genomics UK (COG-UK) Consortium | Berkshire and Surrey Pathology Services Lighthouse Laboratory and Alex Alderton, Roberto Amato, Jeffrey Barrett, Sonia Goncalves, Ewan Harrison, David K. Jackson, Ian Johnston, Dominic Kwiatkowski, Cordelia Langford, John Sillitoe on behalf of the Wellcome Sanger Institute COVID-19 Surveillance Team |
| EPI_ISL_3364027 | Lighthouse Lab in Milton Keynes | Wellcome Sanger Institute for the COVID-19 Genomics UK (COG-UK) Consortium | The Lighthouse Lab in Milton Keynes and Alex Alderton, Roberto Amato, Jeffrey Barrett, Sonia Goncalves, Ewan Harrison, David K. Jackson, Ian Johnston, Dominic Kwiatkowski, Cordelia Langford, John Sillitoe on behalf of the Wellcome Sanger Institute COVID-19 Surveillance Team |
| EPI_ISL_3364028, EPI_ISL_3364030 | Lighthouse Lab in Alderley Park | Wellcome Sanger Institute for the COVID-19 Genomics UK (COG-UK) Consortium | Jacquelyn Wynn, Mairead Hyland, The Lighthouse Lab in Alderley Park and Alex Alderton, Roberto Amato, Jeffrey Barrett, Sonia Goncalves, Ewan Harrison, David K. Jackson, Ian Johnston, Dominic Kwiatkowski, Cordelia Langford, John Sillitoe on behalf of the Wellcome Sanger Institute COVID-19 Surveillance Team |
| EPI_ISL_3364031 | Berkshire and Surrey Pathology Services Lighthouse Laboratory | Wellcome Sanger Institute for the COVID-19 Genomics UK (COG-UK) Consortium | Berkshire and Surrey Pathology Services Lighthouse Laboratory and Alex Alderton, Roberto Amato, Jeffrey Barrett, Sonia Goncalves, Ewan Harrison, David K. Jackson, Ian Johnston, Dominic Kwiatkowski, Cordelia Langford, John Sillitoe on behalf of the Wellcome Sanger Institute COVID-19 Surveillance Team |
| EPI_ISL_3364033, EPI_ISL_3364034, EPI_ISL_3364035, EPI_ISL_3364037, EPI_ISL_3364038 | Lighthouse Lab in Alderley Park | Wellcome Sanger Institute for the COVID-19 Genomics UK (COG-UK) Consortium | Jacquelyn Wynn, Mairead Hyland, The Lighthouse Lab in Alderley Park and Alex Alderton, Roberto Amato, Jeffrey Barrett, Sonia Goncalves, Ewan Harrison, David K. Jackson, Ian Johnston, Dominic Kwiatkowski, Cordelia Langford, John Sillitoe on behalf of the Wellcome Sanger Institute COVID-19 Surveillance Team |
| EPI_ISL_3364040 | Berkshire and Surrey Pathology Services Lighthouse Laboratory | Wellcome Sanger Institute for the COVID-19 Genomics UK (COG-UK) Consortium | Berkshire and Surrey Pathology Services Lighthouse Laboratory and Alex Alderton, Roberto Amato, Jeffrey Barrett, Sonia Goncalves, Ewan Harrison, David K. Jackson, Ian Johnston, Dominic Kwiatkowski, Cordelia Langford, John Sillitoe on behalf of the Wellcome Sanger Institute COVID-19 Surveillance Team |
| EPI_ISL_3364041, EPI_ISL_3364042, EPI_ISL_3364044, EPI_ISL_3364045, EPI_ISL_3364046 | Lighthouse Lab in Alderley Park | Wellcome Sanger Institute for the COVID-19 Genomics UK (COG-UK) Consortium | Jacquelyn Wynn, Mairead Hyland, The Lighthouse Lab in Alderley Park and Alex Alderton, Roberto Amato, Jeffrey Barrett, Sonia Goncalves, Ewan Harrison, David K. Jackson, Ian Johnston, Dominic Kwiatkowski, Cordelia Langford, John Sillitoe on behalf of the Wellcome Sanger Institute COVID-19 Surveillance Team |
| EPI_ISL_3364048 | Berkshire and Surrey Pathology Services Lighthouse Laboratory | Wellcome Sanger Institute for the COVID-19 Genomics UK (COG-UK) Consortium | Berkshire and Surrey Pathology Services Lighthouse Laboratory and Alex Alderton, Roberto Amato, Jeffrey Barrett, Sonia Goncalves, Ewan Harrison, David K. Jackson, Ian Johnston, Dominic Kwiatkowski, Cordelia Langford, John Sillitoe on behalf of the Wellcome Sanger Institute COVID-19 Surveillance Team |
| EPI_ISL_3364049 | Lighthouse Lab in Alderley Park | Wellcome Sanger Institute for the COVID-19 Genomics UK (COG-UK) Consortium | Jacquelyn Wynn, Mairead Hyland, The Lighthouse Lab in Alderley Park and Alex Alderton, Roberto Amato, Jeffrey Barrett, Sonia Goncalves, Ewan Harrison, David K. Jackson, Ian Johnston, Dominic Kwiatkowski, Cordelia Langford, John Sillitoe on behalf of the Wellcome Sanger Institute COVID-19 Surveillance Team |
| EPI_ISL_3364051 | Lighthouse Lab in Milton Keynes | Wellcome Sanger Institute for the COVID-19 Genomics UK (COG-UK) Consortium | The Lighthouse Lab in Milton Keynes and Alex Alderton, Roberto Amato, Jeffrey Barrett, Sonia Goncalves, Ewan Harrison, David K. Jackson, Ian Johnston, Dominic Kwiatkowski, Cordelia Langford, John Sillitoe on behalf of the Wellcome Sanger Institute COVID-19 Surveillance Team |
| EPI_ISL_3364052 | Lighthouse Lab in Alderley Park | Wellcome Sanger Institute for the COVID-19 Genomics UK (COG-UK) Consortium | Jacquelyn Wynn, Mairead Hyland, The Lighthouse Lab in Alderley Park and Alex Alderton, Roberto Amato, Jeffrey Barrett, Sonia Goncalves, Ewan Harrison, David K. Jackson, Ian Johnston, Dominic Kwiatkowski, Cordelia Langford, John Sillitoe on behalf of the Wellcome Sanger Institute COVID-19 Surveillance Team |
| EPI_ISL_3364053 | Lighthouse Lab in Milton Keynes | Wellcome Sanger Institute for the COVID-19 Genomics UK (COG-UK) Consortium | The Lighthouse Lab in Milton Keynes and Alex Alderton, Roberto Amato, Jeffrey Barrett, Sonia Goncalves, Ewan Harrison, David K. Jackson, Ian Johnston, Dominic Kwiatkowski, Cordelia Langford, John Sillitoe on behalf of the Wellcome Sanger Institute COVID-19 Surveillance Team |
| EPI_ISL_3364055, EPI_ISL_3364056 | Lighthouse Lab in Alderley Park | Wellcome Sanger Institute for the COVID-19 Genomics UK (COG-UK) Consortium | Jacquelyn Wynn, Mairead Hyland, The Lighthouse Lab in Alderley Park and Alex Alderton, Roberto Amato, Jeffrey Barrett, Sonia Goncalves, Ewan Harrison, David K. Jackson, Ian Johnston, Dominic Kwiatkowski, Cordelia Langford, John Sillitoe on behalf of the Wellcome Sanger Institute COVID-19 Surveillance Team |
| EPI_ISL_3364058 | Berkshire and Surrey Pathology Services Lighthouse Laboratory | Wellcome Sanger Institute for the COVID-19 Genomics UK (COG-UK) | Berkshire and Surrey Pathology Services Lighthouse Laboratory and Alex Alderton, Roberto Amato, Jeffrey Barrett, Sonia Goncalves, Ewan Harrison, David K. Jackson, Ian Johnston, Dominic Kwiatkowski, Cordelia Langford, John Sillitoe on behalf |

| | | | |
|---|---|---|---|
| | | Consortium | of the Wellcome Sanger Institute COVID-19 Surveillance Team |
| EPI_ISL_3364059 | Lighthouse Lab in Milton Keynes | Wellcome Sanger Institute for the COVID-19 Genomics UK (COG-UK) Consortium | The Lighthouse Lab in Milton Keynes and Alex Alderton, Roberto Amato, Jeffrey Barrett, Sonia Goncalves, Ewan Harrison, David K. Jackson, Ian Johnston, Dominic Kwiatkowski, Cordelia Langford, John Sillitoe on behalf of the Wellcome Sanger Institute COVID-19 Surveillance Team |
| EPI_ISL_3364060 | Berkshire and Surrey Pathology Services Lighthouse Laboratory | Wellcome Sanger Institute for the COVID-19 Genomics UK (COG-UK) Consortium | Berkshire and Surrey Pathology Services Lighthouse Laboratory and Alex Alderton, Roberto Amato, Jeffrey Barrett, Sonia Goncalves, Ewan Harrison, David K. Jackson, Ian Johnston, Dominic Kwiatkowski, Cordelia Langford, John Sillitoe on behalf of the Wellcome Sanger Institute COVID-19 Surveillance Team |
| EPI_ISL_3364062 | Lighthouse Lab in Alderley Park | Wellcome Sanger Institute for the COVID-19 Genomics UK (COG-UK) Consortium | Jacquelyn Wynn, Mairead Hyland, The Lighthouse Lab in Alderley Park and Alex Alderton, Roberto Amato, Jeffrey Barrett, Sonia Goncalves, Ewan Harrison, David K. Jackson, Ian Johnston, Dominic Kwiatkowski, Cordelia Langford, John Sillitoe on behalf of the Wellcome Sanger Institute COVID-19 Surveillance Team |
| EPI_ISL_3364063 | Lighthouse Lab in Milton Keynes | Wellcome Sanger Institute for the COVID-19 Genomics UK (COG-UK) Consortium | The Lighthouse Lab in Milton Keynes and Alex Alderton, Roberto Amato, Jeffrey Barrett, Sonia Goncalves, Ewan Harrison, David K. Jackson, Ian Johnston, Dominic Kwiatkowski, Cordelia Langford, John Sillitoe on behalf of the Wellcome Sanger Institute COVID-19 Surveillance Team |
| EPI_ISL_3364064, EPI_ISL_3364065, EPI_ISL_3364067 | Lighthouse Lab in Alderley Park | Wellcome Sanger Institute for the COVID-19 Genomics UK (COG-UK) Consortium | Jacquelyn Wynn, Mairead Hyland, The Lighthouse Lab in Alderley Park and Alex Alderton, Roberto Amato, Jeffrey Barrett, Sonia Goncalves, Ewan Harrison, David K. Jackson, Ian Johnston, Dominic Kwiatkowski, Cordelia Langford, John Sillitoe on behalf of the Wellcome Sanger Institute COVID-19 Surveillance Team |
| EPI_ISL_3364068 | Lighthouse Lab in Milton Keynes | Wellcome Sanger Institute for the COVID-19 Genomics UK (COG-UK) Consortium | The Lighthouse Lab in Milton Keynes and Alex Alderton, Roberto Amato, Jeffrey Barrett, Sonia Goncalves, Ewan Harrison, David K. Jackson, Ian Johnston, Dominic Kwiatkowski, Cordelia Langford, John Sillitoe on behalf of the Wellcome Sanger Institute COVID-19 Surveillance Team |
| EPI_ISL_3364069, EPI_ISL_3364070 | Lighthouse Lab in Alderley Park | Wellcome Sanger Institute for the COVID-19 Genomics UK (COG-UK) Consortium | Jacquelyn Wynn, Mairead Hyland, The Lighthouse Lab in Alderley Park and Alex Alderton, Roberto Amato, Jeffrey Barrett, Sonia Goncalves, Ewan Harrison, David K. Jackson, Ian Johnston, Dominic Kwiatkowski, Cordelia Langford, John Sillitoe on behalf of the Wellcome Sanger Institute COVID-19 Surveillance Team |
| EPI_ISL_3364072, EPI_ISL_3364073 | Lighthouse Lab in Milton Keynes | Wellcome Sanger Institute for the COVID-19 Genomics UK (COG-UK) Consortium | The Lighthouse Lab in Milton Keynes and Alex Alderton, Roberto Amato, Jeffrey Barrett, Sonia Goncalves, Ewan Harrison, David K. Jackson, Ian Johnston, Dominic Kwiatkowski, Cordelia Langford, John Sillitoe on behalf of the Wellcome Sanger Institute COVID-19 Surveillance Team |
| EPI_ISL_3364074 | Berkshire and Surrey Pathology Services Lighthouse Laboratory | Wellcome Sanger Institute for the COVID-19 Genomics UK (COG-UK) Consortium | Berkshire and Surrey Pathology Services Lighthouse Laboratory and Alex Alderton, Roberto Amato, Jeffrey Barrett, Sonia Goncalves, Ewan Harrison, David K. Jackson, Ian Johnston, Dominic Kwiatkowski, Cordelia Langford, John Sillitoe on behalf of the Wellcome Sanger Institute COVID-19 Surveillance Team |
| EPI_ISL_3364075, EPI_ISL_3364077, EPI_ISL_3364078, EPI_ISL_3364079 | Lighthouse Lab in Alderley Park | Wellcome Sanger Institute for the COVID-19 Genomics UK (COG-UK) Consortium | Jacquelyn Wynn, Mairead Hyland, The Lighthouse Lab in Alderley Park and Alex Alderton, Roberto Amato, Jeffrey Barrett, Sonia Goncalves, Ewan Harrison, David K. Jackson, Ian Johnston, Dominic Kwiatkowski, Cordelia Langford, John Sillitoe on behalf of the Wellcome Sanger Institute COVID-19 Surveillance Team |
| EPI_ISL_3364081 | Lighthouse Lab in Milton Keynes | Wellcome Sanger Institute for the COVID-19 Genomics UK (COG-UK) Consortium | The Lighthouse Lab in Milton Keynes and Alex Alderton, Roberto Amato, Jeffrey Barrett, Sonia Goncalves, Ewan Harrison, David K. Jackson, Ian Johnston, Dominic Kwiatkowski, Cordelia Langford, John Sillitoe on behalf of the Wellcome Sanger Institute COVID-19 Surveillance Team |
| EPI_ISL_3364082 | Lighthouse Lab in Alderley Park | Wellcome Sanger Institute for the COVID-19 Genomics UK (COG-UK) Consortium | Jacquelyn Wynn, Mairead Hyland, The Lighthouse Lab in Alderley Park and Alex Alderton, Roberto Amato, Jeffrey Barrett, Sonia Goncalves, Ewan Harrison, David K. Jackson, Ian Johnston, Dominic Kwiatkowski, Cordelia Langford, John Sillitoe on behalf of the Wellcome Sanger Institute COVID-19 Surveillance Team |
| EPI_ISL_3364083 | Lighthouse Lab in Milton Keynes | Wellcome Sanger Institute for the COVID-19 Genomics UK (COG-UK) Consortium | The Lighthouse Lab in Milton Keynes and Alex Alderton, Roberto Amato, Jeffrey Barrett, Sonia Goncalves, Ewan Harrison, David K. Jackson, Ian Johnston, Dominic Kwiatkowski, Cordelia Langford, John Sillitoe on behalf of the Wellcome Sanger Institute COVID-19 Surveillance Team |
| EPI_ISL_3364084 | Berkshire and Surrey Pathology Services Lighthouse Laboratory | Wellcome Sanger Institute for the COVID-19 Genomics UK (COG-UK) Consortium | Berkshire and Surrey Pathology Services Lighthouse Laboratory and Alex Alderton, Roberto Amato, Jeffrey Barrett, Sonia Goncalves, Ewan Harrison, David K. Jackson, Ian Johnston, Dominic Kwiatkowski, Cordelia Langford, John Sillitoe on behalf of the Wellcome Sanger Institute COVID-19 Surveillance Team |
| EPI_ISL_3364085 | Lighthouse Lab in Alderley Park | Wellcome Sanger Institute for the COVID-19 Genomics UK (COG-UK) Consortium | Jacquelyn Wynn, Mairead Hyland, The Lighthouse Lab in Alderley Park and Alex Alderton, Roberto Amato, Jeffrey Barrett, Sonia Goncalves, Ewan Harrison, David K. Jackson, Ian Johnston, Dominic Kwiatkowski, Cordelia Langford, John Sillitoe on behalf of the Wellcome Sanger Institute COVID-19 Surveillance Team |
| EPI_ISL_3364086, EPI_ISL_3364087 | Berkshire and Surrey Pathology Services Lighthouse Laboratory | Wellcome Sanger Institute for the COVID-19 Genomics UK (COG-UK) Consortium | Berkshire and Surrey Pathology Services Lighthouse Laboratory and Alex Alderton, Roberto Amato, Jeffrey Barrett, Sonia Goncalves, Ewan Harrison, David K. Jackson, Ian Johnston, Dominic Kwiatkowski, Cordelia Langford, John Sillitoe on behalf of the Wellcome Sanger Institute COVID-19 Surveillance Team |
| EPI_ISL_3364088 | Lighthouse Lab in Alderley Park | Wellcome Sanger Institute for the COVID-19 Genomics UK (COG-UK) Consortium | Jacquelyn Wynn, Mairead Hyland, The Lighthouse Lab in Alderley Park and Alex Alderton, Roberto Amato, Jeffrey Barrett, Sonia Goncalves, Ewan Harrison, David K. Jackson, Ian Johnston, Dominic Kwiatkowski, Cordelia Langford, John Sillitoe on behalf of the Wellcome Sanger Institute COVID-19 Surveillance Team |
| EPI_ISL_3364090 | Berkshire and Surrey Pathology Services Lighthouse Laboratory | Wellcome Sanger Institute for the COVID-19 Genomics UK (COG-UK) Consortium | Berkshire and Surrey Pathology Services Lighthouse Laboratory and Alex Alderton, Roberto Amato, Jeffrey Barrett, Sonia Goncalves, Ewan Harrison, David K. Jackson, Ian Johnston, Dominic Kwiatkowski, Cordelia Langford, John Sillitoe on behalf of the Wellcome Sanger Institute COVID-19 Surveillance Team |
| EPI_ISL_3364091 | Lighthouse Lab in Milton Keynes | Wellcome Sanger Institute for the COVID-19 Genomics UK (COG-UK) Consortium | The Lighthouse Lab in Milton Keynes and Alex Alderton, Roberto Amato, Jeffrey Barrett, Sonia Goncalves, Ewan Harrison, David K. Jackson, Ian Johnston, Dominic Kwiatkowski, Cordelia Langford, John Sillitoe on behalf of the Wellcome Sanger Institute COVID-19 Surveillance Team |
| EPI_ISL_3364092 | Lighthouse Lab in Alderley Park | Wellcome Sanger Institute for the COVID-19 Genomics UK (COG-UK) Consortium | Jacquelyn Wynn, Mairead Hyland, The Lighthouse Lab in Alderley Park and Alex Alderton, Roberto Amato, Jeffrey Barrett, Sonia Goncalves, Ewan Harrison, David K. Jackson, Ian Johnston, Dominic Kwiatkowski, Cordelia Langford, John Sillitoe on behalf of the Wellcome Sanger Institute COVID-19 Surveillance Team |
| EPI_ISL_3364093 | Lighthouse Lab in Milton Keynes | Wellcome Sanger Institute for the COVID-19 Genomics UK (COG-UK) Consortium | The Lighthouse Lab in Milton Keynes and Alex Alderton, Roberto Amato, Jeffrey Barrett, Sonia Goncalves, Ewan Harrison, David K. Jackson, Ian Johnston, Dominic Kwiatkowski, Cordelia Langford, John Sillitoe on behalf of the Wellcome Sanger Institute COVID-19 Surveillance Team |
| EPI_ISL_3364094 | Lighthouse Lab in Alderley Park | Wellcome Sanger Institute for the COVID-19 Genomics UK (COG-UK) Consortium | Jacquelyn Wynn, Mairead Hyland, The Lighthouse Lab in Alderley Park and Alex Alderton, Roberto Amato, Jeffrey Barrett, Sonia Goncalves, Ewan Harrison, David K. Jackson, Ian Johnston, Dominic Kwiatkowski, Cordelia Langford, John Sillitoe on behalf of the Wellcome Sanger Institute COVID-19 Surveillance Team |
| EPI_ISL_3364095, EPI_ISL_3364096 | Berkshire and Surrey Pathology Services Lighthouse Laboratory | Wellcome Sanger Institute for the COVID-19 Genomics UK (COG-UK) Consortium | Berkshire and Surrey Pathology Services Lighthouse Laboratory and Alex Alderton, Roberto Amato, Jeffrey Barrett, Sonia Goncalves, Ewan Harrison, David K. Jackson, Ian Johnston, Dominic Kwiatkowski, Cordelia Langford, John Sillitoe on behalf of the Wellcome Sanger Institute COVID-19 Surveillance Team |
| EPI_ISL_3364098 | Lighthouse Lab in Milton Keynes | Wellcome Sanger Institute for the COVID-19 Genomics UK (COG-UK) Consortium | The Lighthouse Lab in Milton Keynes and Alex Alderton, Roberto Amato, Jeffrey Barrett, Sonia Goncalves, Ewan Harrison, David K. Jackson, Ian Johnston, Dominic Kwiatkowski, Cordelia Langford, John Sillitoe on behalf of the Wellcome Sanger Institute COVID-19 Surveillance Team |
| EPI_ISL_3364099, EPI_ISL_3364100, EPI_ISL_3364101, EPI_ISL_3364102 | Lighthouse Lab in Alderley Park | Wellcome Sanger Institute for the COVID-19 Genomics UK (COG-UK) Consortium | Jacquelyn Wynn, Mairead Hyland, The Lighthouse Lab in Alderley Park and Alex Alderton, Roberto Amato, Jeffrey Barrett, Sonia Goncalves, Ewan Harrison, David K. Jackson, Ian Johnston, Dominic Kwiatkowski, Cordelia Langford, John Sillitoe on behalf of the Wellcome Sanger Institute COVID-19 Surveillance Team |
| EPI_ISL_3364103 | Lighthouse Lab in Milton Keynes | Wellcome Sanger Institute for the COVID-19 Genomics UK (COG-UK) Consortium | The Lighthouse Lab in Milton Keynes and Alex Alderton, Roberto Amato, Jeffrey Barrett, Sonia Goncalves, Ewan Harrison, David K. Jackson, Ian Johnston, Dominic Kwiatkowski, Cordelia Langford, John Sillitoe on behalf of the Wellcome Sanger Institute COVID-19 Surveillance Team |
| EPI_ISL_3364104, EPI_ISL_3364105 | Lighthouse Lab in Alderley Park | Wellcome Sanger Institute for the COVID-19 Genomics UK (COG-UK) Consortium | Jacquelyn Wynn, Mairead Hyland, The Lighthouse Lab in Alderley Park and Alex Alderton, Roberto Amato, Jeffrey Barrett, Sonia Goncalves, Ewan Harrison, David K. Jackson, Ian Johnston, Dominic Kwiatkowski, Cordelia Langford, John Sillitoe on behalf of the Wellcome Sanger Institute COVID-19 Surveillance Team |
| EPI_ISL_3364107 | Berkshire and Surrey Pathology Services Lighthouse Laboratory | Wellcome Sanger Institute for the COVID-19 Genomics UK (COG-UK) Consortium | Berkshire and Surrey Pathology Services Lighthouse Laboratory and Alex Alderton, Roberto Amato, Jeffrey Barrett, Sonia Goncalves, Ewan Harrison, David K. Jackson, Ian Johnston, Dominic Kwiatkowski, Cordelia Langford, John Sillitoe on behalf of the Wellcome Sanger Institute COVID-19 Surveillance Team |
| EPI_ISL_3364108, EPI_ISL_3364109, EPI_ISL_3364111, EPI_ISL_3364112 | Lighthouse Lab in Alderley Park | Wellcome Sanger Institute for the COVID-19 Genomics UK (COG-UK) Consortium | Jacquelyn Wynn, Mairead Hyland, The Lighthouse Lab in Alderley Park and Alex Alderton, Roberto Amato, Jeffrey Barrett, Sonia Goncalves, Ewan Harrison, David K. Jackson, Ian Johnston, Dominic Kwiatkowski, Cordelia Langford, John Sillitoe on behalf of the Wellcome Sanger Institute COVID-19 Surveillance Team |
| EPI_ISL_3364114 | Lighthouse Lab in Milton Keynes | Wellcome Sanger Institute for the COVID-19 Genomics UK (COG-UK) Consortium | The Lighthouse Lab in Milton Keynes and Alex Alderton, Roberto Amato, Jeffrey Barrett, Sonia Goncalves, Ewan Harrison, David K. Jackson, Ian Johnston, Dominic Kwiatkowski, Cordelia Langford, John Sillitoe on behalf of the Wellcome Sanger Institute COVID-19 Surveillance Team |
| EPI_ISL_3364115, EPI_ISL_3364116 | Lighthouse Lab in Alderley Park | Wellcome Sanger Institute for the COVID-19 Genomics UK (COG-UK) Consortium | Jacquelyn Wynn, Mairead Hyland, The Lighthouse Lab in Alderley Park and Alex Alderton, Roberto Amato, Jeffrey Barrett, Sonia Goncalves, Ewan Harrison, David K. Jackson, Ian Johnston, Dominic Kwiatkowski, Cordelia Langford, John Sillitoe on behalf of the Wellcome Sanger Institute COVID-19 Surveillance Team |
| EPI_ISL_3364118 | Berkshire and Surrey Pathology Services Lighthouse Laboratory | Wellcome Sanger Institute for the COVID-19 Genomics UK (COG-UK) Consortium | Berkshire and Surrey Pathology Services Lighthouse Laboratory and Alex Alderton, Roberto Amato, Jeffrey Barrett, Sonia Goncalves, Ewan Harrison, David K. Jackson, Ian Johnston, Dominic Kwiatkowski, Cordelia Langford, John Sillitoe on behalf of the Wellcome Sanger Institute COVID-19 Surveillance Team |
| EPI_ISL_3364119, EPI_ISL_3364120, EPI_ISL_3364122, EPI_ISL_3364123, EPI_ISL_3364124 | Lighthouse Lab in Alderley Park | Wellcome Sanger Institute for the COVID-19 Genomics UK (COG-UK) Consortium | Jacquelyn Wynn, Mairead Hyland, The Lighthouse Lab in Alderley Park and Alex Alderton, Roberto Amato, Jeffrey Barrett, Sonia Goncalves, Ewan Harrison, David K. Jackson, Ian Johnston, Dominic Kwiatkowski, Cordelia Langford, John Sillitoe on behalf of the Wellcome Sanger Institute COVID-19 Surveillance Team |
| EPI_ISL_3364126 | Lighthouse Lab in Milton Keynes | Wellcome Sanger Institute for the COVID-19 Genomics UK (COG-UK) Consortium | The Lighthouse Lab in Milton Keynes and Alex Alderton, Roberto Amato, Jeffrey Barrett, Sonia Goncalves, Ewan Harrison, David K. Jackson, Ian Johnston, Dominic Kwiatkowski, Cordelia Langford, John Sillitoe on behalf of the Wellcome Sanger Institute COVID-19 Surveillance Team |
| EPI_ISL_3364127 | Lighthouse Lab in Alderley Park | Wellcome Sanger Institute for the COVID-19 Genomics UK (COG-UK) Consortium | Jacquelyn Wynn, Mairead Hyland, The Lighthouse Lab in Alderley Park and Alex Alderton, Roberto Amato, Jeffrey Barrett, Sonia Goncalves, Ewan Harrison, David K. Jackson, Ian Johnston, Dominic Kwiatkowski, Cordelia Langford, John Sillitoe on behalf of the Wellcome Sanger Institute COVID-19 Surveillance Team |
| EPI_ISL_3364128 | Lighthouse Lab in Milton Keynes | Wellcome Sanger Institute for the COVID-19 Genomics UK (COG-UK) Consortium | The Lighthouse Lab in Milton Keynes and Alex Alderton, Roberto Amato, Jeffrey Barrett, Sonia Goncalves, Ewan Harrison, David K. Jackson, Ian Johnston, Dominic Kwiatkowski, Cordelia Langford, John Sillitoe on behalf of the Wellcome Sanger Institute COVID-19 Surveillance Team |
| EPI_ISL_3364129, EPI_ISL_3364130 | Lighthouse Lab in Alderley Park | Wellcome Sanger Institute for the COVID-19 Genomics UK (COG-UK) Consortium | Jacquelyn Wynn, Mairead Hyland, The Lighthouse Lab in Alderley Park and Alex Alderton, Roberto Amato, Jeffrey Barrett, Sonia Goncalves, Ewan Harrison, David K. Jackson, Ian Johnston, Dominic Kwiatkowski, Cordelia Langford, John Sillitoe on behalf of the Wellcome Sanger Institute COVID-19 Surveillance Team |
| EPI_ISL_3364131 | Lighthouse Lab in Milton Keynes | Wellcome Sanger Institute for the COVID-19 Genomics UK (COG-UK) Consortium | The Lighthouse Lab in Milton Keynes and Alex Alderton, Roberto Amato, Jeffrey Barrett, Sonia Goncalves, Ewan Harrison, David K. Jackson, Ian Johnston, Dominic Kwiatkowski, Cordelia Langford, John Sillitoe on behalf of the Wellcome Sanger Institute COVID-19 Surveillance Team |
| EPI_ISL_3364132, EPI_ISL_3364133, EPI_ISL_3364134 | Lighthouse Lab in Alderley Park | Wellcome Sanger Institute for the COVID-19 Genomics UK (COG-UK) Consortium | Jacquelyn Wynn, Mairead Hyland, The Lighthouse Lab in Alderley Park and Alex Alderton, Roberto Amato, Jeffrey Barrett, Sonia Goncalves, Ewan Harrison, David K. Jackson, Ian Johnston, Dominic Kwiatkowski, Cordelia Langford, John Sillitoe on behalf of the Wellcome Sanger Institute COVID-19 Surveillance Team |
| EPI_ISL_3364136 | Lighthouse Lab in Milton Keynes | Wellcome Sanger Institute for the COVID-19 Genomics UK (COG-UK) Consortium | The Lighthouse Lab in Milton Keynes and Alex Alderton, Roberto Amato, Jeffrey Barrett, Sonia Goncalves, Ewan Harrison, David K. Jackson, Ian Johnston, Dominic Kwiatkowski, Cordelia Langford, John Sillitoe on behalf of the Wellcome Sanger Institute COVID-19 Surveillance Team |

| | | | |
|---|---|---|---|
| EPI_ISL_3364137, EPI_ISL_3364138, EPI_ISL_3364140 | Lighthouse Lab in Alderley Park | Wellcome Sanger Institute for the COVID-19 Genomics UK (COG-UK) Consortium | Jacquelyn Wynn, Mairead Hyland, The Lighthouse Lab in Alderley Park and Alex Alderton, Roberto Amato, Jeffrey Barrett, Sonia Goncalves, Ewan Harrison, David K. Jackson, Ian Johnston, Dominic Kwiatkowski, Cordelia Langford, John Sillitoe on behalf of the Wellcome Sanger Institute COVID-19 Surveillance Team |
| EPI_ISL_3364141 | Berkshire and Surrey Pathology Services Lighthouse Laboratory | Wellcome Sanger Institute for the COVID-19 Genomics UK (COG-UK) Consortium | Berkshire and Surrey Pathology Services Lighthouse Laboratory and Alex Alderton, Roberto Amato, Jeffrey Barrett, Sonia Goncalves, Ewan Harrison, David K. Jackson, Ian Johnston, Dominic Kwiatkowski, Cordelia Langford, John Sillitoe on behalf of the Wellcome Sanger Institute COVID-19 Surveillance Team |
| EPI_ISL_3364142 | Lighthouse Lab in Milton Keynes | Wellcome Sanger Institute for the COVID-19 Genomics UK (COG-UK) Consortium | The Lighthouse Lab in Milton Keynes and Alex Alderton, Roberto Amato, Jeffrey Barrett, Sonia Goncalves, Ewan Harrison, David K. Jackson, Ian Johnston, Dominic Kwiatkowski, Cordelia Langford, John Sillitoe on behalf of the Wellcome Sanger Institute COVID-19 Surveillance Team |
| EPI_ISL_3364144 | Berkshire and Surrey Pathology Services Lighthouse Laboratory | Wellcome Sanger Institute for the COVID-19 Genomics UK (COG-UK) Consortium | Berkshire and Surrey Pathology Services Lighthouse Laboratory and Alex Alderton, Roberto Amato, Jeffrey Barrett, Sonia Goncalves, Ewan Harrison, David K. Jackson, Ian Johnston, Dominic Kwiatkowski, Cordelia Langford, John Sillitoe on behalf of the Wellcome Sanger Institute COVID-19 Surveillance Team |
| EPI_ISL_3364145, EPI_ISL_3364147, EPI_ISL_3364148 | Lighthouse Lab in Alderley Park | Wellcome Sanger Institute for the COVID-19 Genomics UK (COG-UK) Consortium | Jacquelyn Wynn, Mairead Hyland, The Lighthouse Lab in Alderley Park and Alex Alderton, Roberto Amato, Jeffrey Barrett, Sonia Goncalves, Ewan Harrison, David K. Jackson, Ian Johnston, Dominic Kwiatkowski, Cordelia Langford, John Sillitoe on behalf of the Wellcome Sanger Institute COVID-19 Surveillance Team |
| EPI_ISL_3364149 | Lighthouse Lab in Milton Keynes | Wellcome Sanger Institute for the COVID-19 Genomics UK (COG-UK) Consortium | The Lighthouse Lab in Milton Keynes and Alex Alderton, Roberto Amato, Jeffrey Barrett, Sonia Goncalves, Ewan Harrison, David K. Jackson, Ian Johnston, Dominic Kwiatkowski, Cordelia Langford, John Sillitoe on behalf of the Wellcome Sanger Institute COVID-19 Surveillance Team |
| EPI_ISL_3364151, EPI_ISL_3364152 | Lighthouse Lab in Alderley Park | Wellcome Sanger Institute for the COVID-19 Genomics UK (COG-UK) Consortium | Jacquelyn Wynn, Mairead Hyland, The Lighthouse Lab in Alderley Park and Alex Alderton, Roberto Amato, Jeffrey Barrett, Sonia Goncalves, Ewan Harrison, David K. Jackson, Ian Johnston, Dominic Kwiatkowski, Cordelia Langford, John Sillitoe on behalf of the Wellcome Sanger Institute COVID-19 Surveillance Team |
| EPI_ISL_3364154 | Berkshire and Surrey Pathology Services Lighthouse Laboratory | Wellcome Sanger Institute for the COVID-19 Genomics UK (COG-UK) Consortium | Berkshire and Surrey Pathology Services Lighthouse Laboratory and Alex Alderton, Roberto Amato, Jeffrey Barrett, Sonia Goncalves, Ewan Harrison, David K. Jackson, Ian Johnston, Dominic Kwiatkowski, Cordelia Langford, John Sillitoe on behalf of the Wellcome Sanger Institute COVID-19 Surveillance Team |
| EPI_ISL_3364155, EPI_ISL_3364157 | Lighthouse Lab in Milton Keynes | Wellcome Sanger Institute for the COVID-19 Genomics UK (COG-UK) Consortium | The Lighthouse Lab in Milton Keynes and Alex Alderton, Roberto Amato, Jeffrey Barrett, Sonia Goncalves, Ewan Harrison, David K. Jackson, Ian Johnston, Dominic Kwiatkowski, Cordelia Langford, John Sillitoe on behalf of the Wellcome Sanger Institute COVID-19 Surveillance Team |
| EPI_ISL_3364158, EPI_ISL_3364159, EPI_ISL_3364161, EPI_ISL_3364162, EPI_ISL_3364164, EPI_ISL_3364165, EPI_ISL_3364166, EPI_ISL_3364168 | Lighthouse Lab in Alderley Park | Wellcome Sanger Institute for the COVID-19 Genomics UK (COG-UK) Consortium | Jacquelyn Wynn, Mairead Hyland, The Lighthouse Lab in Alderley Park and Alex Alderton, Roberto Amato, Jeffrey Barrett, Sonia Goncalves, Ewan Harrison, David K. Jackson, Ian Johnston, Dominic Kwiatkowski, Cordelia Langford, John Sillitoe on behalf of the Wellcome Sanger Institute COVID-19 Surveillance Team |
| EPI_ISL_3364169 | Berkshire and Surrey Pathology Services Lighthouse Laboratory | Wellcome Sanger Institute for the COVID-19 Genomics UK (COG-UK) Consortium | Berkshire and Surrey Pathology Services Lighthouse Laboratory and Alex Alderton, Roberto Amato, Jeffrey Barrett, Sonia Goncalves, Ewan Harrison, David K. Jackson, Ian Johnston, Dominic Kwiatkowski, Cordelia Langford, John Sillitoe on behalf of the Wellcome Sanger Institute COVID-19 Surveillance Team |
| EPI_ISL_3364170 | Lighthouse Lab in Alderley Park | Wellcome Sanger Institute for the COVID-19 Genomics UK (COG-UK) Consortium | Jacquelyn Wynn, Mairead Hyland, The Lighthouse Lab in Alderley Park and Alex Alderton, Roberto Amato, Jeffrey Barrett, Sonia Goncalves, Ewan Harrison, David K. Jackson, Ian Johnston, Dominic Kwiatkowski, Cordelia Langford, John Sillitoe on behalf of the Wellcome Sanger Institute COVID-19 Surveillance Team |
| EPI_ISL_3364172, EPI_ISL_3364173 | Berkshire and Surrey Pathology Services Lighthouse Laboratory | Wellcome Sanger Institute for the COVID-19 Genomics UK (COG-UK) Consortium | Berkshire and Surrey Pathology Services Lighthouse Laboratory and Alex Alderton, Roberto Amato, Jeffrey Barrett, Sonia Goncalves, Ewan Harrison, David K. Jackson, Ian Johnston, Dominic Kwiatkowski, Cordelia Langford, John Sillitoe on behalf of the Wellcome Sanger Institute COVID-19 Surveillance Team |
| EPI_ISL_3364175, EPI_ISL_3364176 | Lighthouse Lab in Milton Keynes | Wellcome Sanger Institute for the COVID-19 Genomics UK (COG-UK) Consortium | The Lighthouse Lab in Milton Keynes and Alex Alderton, Roberto Amato, Jeffrey Barrett, Sonia Goncalves, Ewan Harrison, David K. Jackson, Ian Johnston, Dominic Kwiatkowski, Cordelia Langford, John Sillitoe on behalf of the Wellcome Sanger Institute COVID-19 Surveillance Team |
| EPI_ISL_3364177, EPI_ISL_3364179, EPI_ISL_3364180, EPI_ISL_3364181, EPI_ISL_3364183 | Lighthouse Lab in Alderley Park | Wellcome Sanger Institute for the COVID-19 Genomics UK (COG-UK) Consortium | Jacquelyn Wynn, Mairead Hyland, The Lighthouse Lab in Alderley Park and Alex Alderton, Roberto Amato, Jeffrey Barrett, Sonia Goncalves, Ewan Harrison, David K. Jackson, Ian Johnston, Dominic Kwiatkowski, Cordelia Langford, John Sillitoe on behalf of the Wellcome Sanger Institute COVID-19 Surveillance Team |
| EPI_ISL_3364184 | Lighthouse Lab in Milton Keynes | Wellcome Sanger Institute for the COVID-19 Genomics UK (COG-UK) Consortium | The Lighthouse Lab in Milton Keynes and Alex Alderton, Roberto Amato, Jeffrey Barrett, Sonia Goncalves, Ewan Harrison, David K. Jackson, Ian Johnston, Dominic Kwiatkowski, Cordelia Langford, John Sillitoe on behalf of the Wellcome Sanger Institute COVID-19 Surveillance Team |
| EPI_ISL_3364186 | Berkshire and Surrey Pathology Services Lighthouse Laboratory | Wellcome Sanger Institute for the COVID-19 Genomics UK (COG-UK) Consortium | Berkshire and Surrey Pathology Services Lighthouse Laboratory and Alex Alderton, Roberto Amato, Jeffrey Barrett, Sonia Goncalves, Ewan Harrison, David K. Jackson, Ian Johnston, Dominic Kwiatkowski, Cordelia Langford, John Sillitoe on behalf of the Wellcome Sanger Institute COVID-19 Surveillance Team |
| EPI_ISL_3364187, EPI_ISL_3364189 | Lighthouse Lab in Alderley Park | Wellcome Sanger Institute for the COVID-19 Genomics UK (COG-UK) Consortium | Jacquelyn Wynn, Mairead Hyland, The Lighthouse Lab in Alderley Park and Alex Alderton, Roberto Amato, Jeffrey Barrett, Sonia Goncalves, Ewan Harrison, David K. Jackson, Ian Johnston, Dominic Kwiatkowski, Cordelia Langford, John Sillitoe on behalf of the Wellcome Sanger Institute COVID-19 Surveillance Team |
| EPI_ISL_3364190 | Berkshire and Surrey Pathology Services Lighthouse Laboratory | Wellcome Sanger Institute for the COVID-19 Genomics UK (COG-UK) Consortium | Berkshire and Surrey Pathology Services Lighthouse Laboratory and Alex Alderton, Roberto Amato, Jeffrey Barrett, Sonia Goncalves, Ewan Harrison, David K. Jackson, Ian Johnston, Dominic Kwiatkowski, Cordelia Langford, John Sillitoe on behalf of the Wellcome Sanger Institute COVID-19 Surveillance Team |
| EPI_ISL_3364191 | Lighthouse Lab in Alderley Park | Wellcome Sanger Institute for the COVID-19 Genomics UK (COG-UK) Consortium | Jacquelyn Wynn, Mairead Hyland, The Lighthouse Lab in Alderley Park and Alex Alderton, Roberto Amato, Jeffrey Barrett, Sonia Goncalves, Ewan Harrison, David K. Jackson, Ian Johnston, Dominic Kwiatkowski, Cordelia Langford, John Sillitoe on behalf of the Wellcome Sanger Institute COVID-19 Surveillance Team |
| EPI_ISL_3364192 | Berkshire and Surrey Pathology Services Lighthouse Laboratory | Wellcome Sanger Institute for the COVID-19 Genomics UK (COG-UK) Consortium | Berkshire and Surrey Pathology Services Lighthouse Laboratory and Alex Alderton, Roberto Amato, Jeffrey Barrett, Sonia Goncalves, Ewan Harrison, David K. Jackson, Ian Johnston, Dominic Kwiatkowski, Cordelia Langford, John Sillitoe on behalf of the Wellcome Sanger Institute COVID-19 Surveillance Team |
| EPI_ISL_3364194 | Lighthouse Lab in Alderley Park | Wellcome Sanger Institute for the COVID-19 Genomics UK (COG-UK) Consortium | Jacquelyn Wynn, Mairead Hyland, The Lighthouse Lab in Alderley Park and Alex Alderton, Roberto Amato, Jeffrey Barrett, Sonia Goncalves, Ewan Harrison, David K. Jackson, Ian Johnston, Dominic Kwiatkowski, Cordelia Langford, John Sillitoe on behalf of the Wellcome Sanger Institute COVID-19 Surveillance Team |
| EPI_ISL_3364195 | Lighthouse Lab in Milton Keynes | Wellcome Sanger Institute for the COVID-19 Genomics UK (COG-UK) Consortium | The Lighthouse Lab in Milton Keynes and Alex Alderton, Roberto Amato, Jeffrey Barrett, Sonia Goncalves, Ewan Harrison, David K. Jackson, Ian Johnston, Dominic Kwiatkowski, Cordelia Langford, John Sillitoe on behalf of the Wellcome Sanger Institute COVID-19 Surveillance Team |
| EPI_ISL_3364197 | Berkshire and Surrey Pathology Services Lighthouse Laboratory | Wellcome Sanger Institute for the COVID-19 Genomics UK (COG-UK) Consortium | Berkshire and Surrey Pathology Services Lighthouse Laboratory and Alex Alderton, Roberto Amato, Jeffrey Barrett, Sonia Goncalves, Ewan Harrison, David K. Jackson, Ian Johnston, Dominic Kwiatkowski, Cordelia Langford, John Sillitoe on behalf of the Wellcome Sanger Institute COVID-19 Surveillance Team |
| EPI_ISL_3364198, EPI_ISL_3364199, EPI_ISL_3364200 | Lighthouse Lab in Alderley Park | Wellcome Sanger Institute for the COVID-19 Genomics UK (COG-UK) Consortium | Jacquelyn Wynn, Mairead Hyland, The Lighthouse Lab in Alderley Park and Alex Alderton, Roberto Amato, Jeffrey Barrett, Sonia Goncalves, Ewan Harrison, David K. Jackson, Ian Johnston, Dominic Kwiatkowski, Cordelia Langford, John Sillitoe on behalf of the Wellcome Sanger Institute COVID-19 Surveillance Team |
| EPI_ISL_3364201 | Lighthouse Lab in Milton Keynes | Wellcome Sanger Institute for the COVID-19 Genomics UK (COG-UK) Consortium | The Lighthouse Lab in Milton Keynes and Alex Alderton, Roberto Amato, Jeffrey Barrett, Sonia Goncalves, Ewan Harrison, David K. Jackson, Ian Johnston, Dominic Kwiatkowski, Cordelia Langford, John Sillitoe on behalf of the Wellcome Sanger Institute COVID-19 Surveillance Team |
| EPI_ISL_3364202 | Lighthouse Lab in Alderley Park | Wellcome Sanger Institute for the COVID-19 Genomics UK (COG-UK) Consortium | Jacquelyn Wynn, Mairead Hyland, The Lighthouse Lab in Alderley Park and Alex Alderton, Roberto Amato, Jeffrey Barrett, Sonia Goncalves, Ewan Harrison, David K. Jackson, Ian Johnston, Dominic Kwiatkowski, Cordelia Langford, John Sillitoe on behalf of the Wellcome Sanger Institute COVID-19 Surveillance Team |
| EPI_ISL_3364203 | Berkshire and Surrey Pathology Services Lighthouse Laboratory | Wellcome Sanger Institute for the COVID-19 Genomics UK (COG-UK) Consortium | Berkshire and Surrey Pathology Services Lighthouse Laboratory and Alex Alderton, Roberto Amato, Jeffrey Barrett, Sonia Goncalves, Ewan Harrison, David K. Jackson, Ian Johnston, Dominic Kwiatkowski, Cordelia Langford, John Sillitoe on behalf of the Wellcome Sanger Institute COVID-19 Surveillance Team |
| EPI_ISL_3364204 | Lighthouse Lab in Milton Keynes | Wellcome Sanger Institute for the COVID-19 Genomics UK (COG-UK) Consortium | The Lighthouse Lab in Milton Keynes and Alex Alderton, Roberto Amato, Jeffrey Barrett, Sonia Goncalves, Ewan Harrison, David K. Jackson, Ian Johnston, Dominic Kwiatkowski, Cordelia Langford, John Sillitoe on behalf of the Wellcome Sanger Institute COVID-19 Surveillance Team |
| EPI_ISL_3364206, EPI_ISL_3364207, EPI_ISL_3364208, EPI_ISL_3364210 | Lighthouse Lab in Alderley Park | Wellcome Sanger Institute for the COVID-19 Genomics UK (COG-UK) Consortium | Jacquelyn Wynn, Mairead Hyland, The Lighthouse Lab in Alderley Park and Alex Alderton, Roberto Amato, Jeffrey Barrett, Sonia Goncalves, Ewan Harrison, David K. Jackson, Ian Johnston, Dominic Kwiatkowski, Cordelia Langford, John Sillitoe on behalf of the Wellcome Sanger Institute COVID-19 Surveillance Team |
| EPI_ISL_3364211 | Lighthouse Lab in Milton Keynes | Wellcome Sanger Institute for the COVID-19 Genomics UK (COG-UK) Consortium | The Lighthouse Lab in Milton Keynes and Alex Alderton, Roberto Amato, Jeffrey Barrett, Sonia Goncalves, Ewan Harrison, David K. Jackson, Ian Johnston, Dominic Kwiatkowski, Cordelia Langford, John Sillitoe on behalf of the Wellcome Sanger Institute COVID-19 Surveillance Team |
| EPI_ISL_3364213 | Lighthouse Lab in Alderley Park | Wellcome Sanger Institute for the COVID-19 Genomics UK (COG-UK) Consortium | Jacquelyn Wynn, Mairead Hyland, The Lighthouse Lab in Alderley Park and Alex Alderton, Roberto Amato, Jeffrey Barrett, Sonia Goncalves, Ewan Harrison, David K. Jackson, Ian Johnston, Dominic Kwiatkowski, Cordelia Langford, John Sillitoe on behalf of the Wellcome Sanger Institute COVID-19 Surveillance Team |
| EPI_ISL_3364214 | Lighthouse Lab in Milton Keynes | Wellcome Sanger Institute for the COVID-19 Genomics UK (COG-UK) Consortium | The Lighthouse Lab in Milton Keynes and Alex Alderton, Roberto Amato, Jeffrey Barrett, Sonia Goncalves, Ewan Harrison, David K. Jackson, Ian Johnston, Dominic Kwiatkowski, Cordelia Langford, John Sillitoe on behalf of the Wellcome Sanger Institute COVID-19 Surveillance Team |
| EPI_ISL_3364216 | Lighthouse Lab in Alderley Park | Wellcome Sanger Institute for the COVID-19 Genomics UK (COG-UK) Consortium | Jacquelyn Wynn, Mairead Hyland, The Lighthouse Lab in Alderley Park and Alex Alderton, Roberto Amato, Jeffrey Barrett, Sonia Goncalves, Ewan Harrison, David K. Jackson, Ian Johnston, Dominic Kwiatkowski, Cordelia Langford, John Sillitoe on behalf of the Wellcome Sanger Institute COVID-19 Surveillance Team |
| EPI_ISL_3364217 | Lighthouse Lab in Milton Keynes | Wellcome Sanger Institute for the COVID-19 Genomics UK (COG-UK) Consortium | The Lighthouse Lab in Milton Keynes and Alex Alderton, Roberto Amato, Jeffrey Barrett, Sonia Goncalves, Ewan Harrison, David K. Jackson, Ian Johnston, Dominic Kwiatkowski, Cordelia Langford, John Sillitoe on behalf of the Wellcome Sanger Institute COVID-19 Surveillance Team |
| EPI_ISL_3364218, EPI_ISL_3364220 | Lighthouse Lab in Alderley Park | Wellcome Sanger Institute for the COVID-19 Genomics UK (COG-UK) Consortium | Jacquelyn Wynn, Mairead Hyland, The Lighthouse Lab in Alderley Park and Alex Alderton, Roberto Amato, Jeffrey Barrett, Sonia Goncalves, Ewan Harrison, David K. Jackson, Ian Johnston, Dominic Kwiatkowski, Cordelia Langford, John Sillitoe on behalf of the Wellcome Sanger Institute COVID-19 Surveillance Team |
| EPI_ISL_3364221 | Berkshire and Surrey Pathology Services Lighthouse Laboratory | Wellcome Sanger Institute for the COVID-19 Genomics UK (COG-UK) Consortium | Berkshire and Surrey Pathology Services Lighthouse Laboratory and Alex Alderton, Roberto Amato, Jeffrey Barrett, Sonia Goncalves, Ewan Harrison, David K. Jackson, Ian Johnston, Dominic Kwiatkowski, Cordelia Langford, John Sillitoe on behalf of the Wellcome Sanger Institute COVID-19 Surveillance Team |
| EPI_ISL_3364222 | Lighthouse Lab in Milton Keynes | Wellcome Sanger Institute for the COVID-19 Genomics UK (COG-UK) Consortium | The Lighthouse Lab in Milton Keynes and Alex Alderton, Roberto Amato, Jeffrey Barrett, Sonia Goncalves, Ewan Harrison, David K. Jackson, Ian Johnston, Dominic Kwiatkowski, Cordelia Langford, John Sillitoe on behalf of the Wellcome Sanger Institute COVID-19 Surveillance Team |
| EPI_ISL_3364224, EPI_ISL_3364225 | Lighthouse Lab in Alderley Park | Wellcome Sanger Institute for the COVID-19 Genomics UK (COG-UK) Consortium | Jacquelyn Wynn, Mairead Hyland, The Lighthouse Lab in Alderley Park and Alex Alderton, Roberto Amato, Jeffrey Barrett, Sonia Goncalves, Ewan Harrison, David K. Jackson, Ian Johnston, Dominic Kwiatkowski, Cordelia Langford, John Sillitoe on behalf of the Wellcome Sanger Institute COVID-19 Surveillance Team |
| EPI_ISL_3364227 | Berkshire and Surrey Pathology Services Lighthouse Laboratory | Wellcome Sanger Institute for the COVID-19 Genomics UK (COG-UK) | Berkshire and Surrey Pathology Services Lighthouse Laboratory and Alex Alderton, Roberto Amato, Jeffrey Barrett, Sonia Goncalves, Ewan Harrison, David K. Jackson, Ian Johnston, Dominic Kwiatkowski, Cordelia Langford, John Sillitoe on behalf |

| | | | |
|---|---|---|---|
| | | Consortium | of the Wellcome Sanger Institute COVID-19 Surveillance Team |
| EPI_ISL_3364228, EPI_ISL_3364229, EPI_ISL_3364231 | Lighthouse Lab in Alderley Park | Wellcome Sanger Institute for the COVID-19 Genomics UK (COG-UK) Consortium | Jacquelyn Wynn, Mairead Hyland, The Lighthouse Lab in Alderley Park and Alex Alderton, Roberto Amato, Jeffrey Barrett, Sonia Goncalves, Ewan Harrison, David K. Jackson, Ian Johnston, Dominic Kwiatkowski, Cordelia Langford, John Sillitoe on behalf of the Wellcome Sanger Institute COVID-19 Surveillance Team |
| EPI_ISL_3364232, EPI_ISL_3364234 | Lighthouse Lab in Milton Keynes | Wellcome Sanger Institute for the COVID-19 Genomics UK (COG-UK) Consortium | The Lighthouse Lab in Milton Keynes and Alex Alderton, Roberto Amato, Jeffrey Barrett, Sonia Goncalves, Ewan Harrison, David K. Jackson, Ian Johnston, Dominic Kwiatkowski, Cordelia Langford, John Sillitoe on behalf of the Wellcome Sanger Institute COVID-19 Surveillance Team |
| EPI_ISL_3364235 | Lighthouse Lab in Alderley Park | Wellcome Sanger Institute for the COVID-19 Genomics UK (COG-UK) Consortium | Jacquelyn Wynn, Mairead Hyland, The Lighthouse Lab in Alderley Park and Alex Alderton, Roberto Amato, Jeffrey Barrett, Sonia Goncalves, Ewan Harrison, David K. Jackson, Ian Johnston, Dominic Kwiatkowski, Cordelia Langford, John Sillitoe on behalf of the Wellcome Sanger Institute COVID-19 Surveillance Team |
| EPI_ISL_3364236, EPI_ISL_3364238 | Lighthouse Lab in Milton Keynes | Wellcome Sanger Institute for the COVID-19 Genomics UK (COG-UK) Consortium | The Lighthouse Lab in Milton Keynes and Alex Alderton, Roberto Amato, Jeffrey Barrett, Sonia Goncalves, Ewan Harrison, David K. Jackson, Ian Johnston, Dominic Kwiatkowski, Cordelia Langford, John Sillitoe on behalf of the Wellcome Sanger Institute COVID-19 Surveillance Team |
| EPI_ISL_3364239 | Lighthouse Lab in Alderley Park | Wellcome Sanger Institute for the COVID-19 Genomics UK (COG-UK) Consortium | Jacquelyn Wynn, Mairead Hyland, The Lighthouse Lab in Alderley Park and Alex Alderton, Roberto Amato, Jeffrey Barrett, Sonia Goncalves, Ewan Harrison, David K. Jackson, Ian Johnston, Dominic Kwiatkowski, Cordelia Langford, John Sillitoe on behalf of the Wellcome Sanger Institute COVID-19 Surveillance Team |
| EPI_ISL_3364241 | Lighthouse Lab in Milton Keynes | Wellcome Sanger Institute for the COVID-19 Genomics UK (COG-UK) Consortium | The Lighthouse Lab in Milton Keynes and Alex Alderton, Roberto Amato, Jeffrey Barrett, Sonia Goncalves, Ewan Harrison, David K. Jackson, Ian Johnston, Dominic Kwiatkowski, Cordelia Langford, John Sillitoe on behalf of the Wellcome Sanger Institute COVID-19 Surveillance Team |
| EPI_ISL_3364242 | Berkshire and Surrey Pathology Services Lighthouse Laboratory | Wellcome Sanger Institute for the COVID-19 Genomics UK (COG-UK) Consortium | Berkshire and Surrey Pathology Services Lighthouse Laboratory and Alex Alderton, Roberto Amato, Jeffrey Barrett, Sonia Goncalves, Ewan Harrison, David K. Jackson, Ian Johnston, Dominic Kwiatkowski, Cordelia Langford, John Sillitoe on behalf of the Wellcome Sanger Institute COVID-19 Surveillance Team |
| EPI_ISL_3364243 | Lighthouse Lab in Alderley Park | Wellcome Sanger Institute for the COVID-19 Genomics UK (COG-UK) Consortium | Jacquelyn Wynn, Mairead Hyland, The Lighthouse Lab in Alderley Park and Alex Alderton, Roberto Amato, Jeffrey Barrett, Sonia Goncalves, Ewan Harrison, David K. Jackson, Ian Johnston, Dominic Kwiatkowski, Cordelia Langford, John Sillitoe on behalf of the Wellcome Sanger Institute COVID-19 Surveillance Team |
| EPI_ISL_3364245 | Lighthouse Lab in Milton Keynes | Wellcome Sanger Institute for the COVID-19 Genomics UK (COG-UK) Consortium | The Lighthouse Lab in Milton Keynes and Alex Alderton, Roberto Amato, Jeffrey Barrett, Sonia Goncalves, Ewan Harrison, David K. Jackson, Ian Johnston, Dominic Kwiatkowski, Cordelia Langford, John Sillitoe on behalf of the Wellcome Sanger Institute COVID-19 Surveillance Team |
| EPI_ISL_3364246, EPI_ISL_3364248 | Lighthouse Lab in Alderley Park | Wellcome Sanger Institute for the COVID-19 Genomics UK (COG-UK) Consortium | Jacquelyn Wynn, Mairead Hyland, The Lighthouse Lab in Alderley Park and Alex Alderton, Roberto Amato, Jeffrey Barrett, Sonia Goncalves, Ewan Harrison, David K. Jackson, Ian Johnston, Dominic Kwiatkowski, Cordelia Langford, John Sillitoe on behalf of the Wellcome Sanger Institute COVID-19 Surveillance Team |
| EPI_ISL_3364296, EPI_ISL_3364298 | Lighthouse Lab in Milton Keynes | Wellcome Sanger Institute for the COVID-19 Genomics UK (COG-UK) Consortium | The Lighthouse Lab in Milton Keynes and Alex Alderton, Roberto Amato, Jeffrey Barrett, Sonia Goncalves, Ewan Harrison, David K. Jackson, Ian Johnston, Dominic Kwiatkowski, Cordelia Langford, John Sillitoe on behalf of the Wellcome Sanger Institute COVID-19 Surveillance Team |
| EPI_ISL_3364299 | Lighthouse Lab in Alderley Park | Wellcome Sanger Institute for the COVID-19 Genomics UK (COG-UK) Consortium | Jacquelyn Wynn, Mairead Hyland, The Lighthouse Lab in Alderley Park and Alex Alderton, Roberto Amato, Jeffrey Barrett, Sonia Goncalves, Ewan Harrison, David K. Jackson, Ian Johnston, Dominic Kwiatkowski, Cordelia Langford, John Sillitoe on behalf of the Wellcome Sanger Institute COVID-19 Surveillance Team |
| EPI_ISL_3364301, EPI_ISL_3364302, EPI_ISL_3364304 | Lighthouse Lab in Milton Keynes | Wellcome Sanger Institute for the COVID-19 Genomics UK (COG-UK) Consortium | The Lighthouse Lab in Milton Keynes and Alex Alderton, Roberto Amato, Jeffrey Barrett, Sonia Goncalves, Ewan Harrison, David K. Jackson, Ian Johnston, Dominic Kwiatkowski, Cordelia Langford, John Sillitoe on behalf of the Wellcome Sanger Institute COVID-19 Surveillance Team |
| EPI_ISL_3364305 | Lighthouse Lab in Alderley Park | Wellcome Sanger Institute for the COVID-19 Genomics UK (COG-UK) Consortium | Jacquelyn Wynn, Mairead Hyland, The Lighthouse Lab in Alderley Park and Alex Alderton, Roberto Amato, Jeffrey Barrett, Sonia Goncalves, Ewan Harrison, David K. Jackson, Ian Johnston, Dominic Kwiatkowski, Cordelia Langford, John Sillitoe on behalf of the Wellcome Sanger Institute COVID-19 Surveillance Team |
| EPI_ISL_3364306 | Berkshire and Surrey Pathology Services Lighthouse Laboratory | Wellcome Sanger Institute for the COVID-19 Genomics UK (COG-UK) Consortium | Berkshire and Surrey Pathology Services Lighthouse Laboratory and Alex Alderton, Roberto Amato, Jeffrey Barrett, Sonia Goncalves, Ewan Harrison, David K. Jackson, Ian Johnston, Dominic Kwiatkowski, Cordelia Langford, John Sillitoe on behalf of the Wellcome Sanger Institute COVID-19 Surveillance Team |
| EPI_ISL_3364308, EPI_ISL_3364309, EPI_ISL_3364311 | Lighthouse Lab in Alderley Park | Wellcome Sanger Institute for the COVID-19 Genomics UK (COG-UK) Consortium | Jacquelyn Wynn, Mairead Hyland, The Lighthouse Lab in Alderley Park and Alex Alderton, Roberto Amato, Jeffrey Barrett, Sonia Goncalves, Ewan Harrison, David K. Jackson, Ian Johnston, Dominic Kwiatkowski, Cordelia Langford, John Sillitoe on behalf of the Wellcome Sanger Institute COVID-19 Surveillance Team |
| EPI_ISL_3364312 | Lighthouse Lab in Milton Keynes | Wellcome Sanger Institute for the COVID-19 Genomics UK (COG-UK) Consortium | The Lighthouse Lab in Milton Keynes and Alex Alderton, Roberto Amato, Jeffrey Barrett, Sonia Goncalves, Ewan Harrison, David K. Jackson, Ian Johnston, Dominic Kwiatkowski, Cordelia Langford, John Sillitoe on behalf of the Wellcome Sanger Institute COVID-19 Surveillance Team |
| EPI_ISL_3364313 | Berkshire and Surrey Pathology Services Lighthouse Laboratory | Wellcome Sanger Institute for the COVID-19 Genomics UK (COG-UK) Consortium | Berkshire and Surrey Pathology Services Lighthouse Laboratory and Alex Alderton, Roberto Amato, Jeffrey Barrett, Sonia Goncalves, Ewan Harrison, David K. Jackson, Ian Johnston, Dominic Kwiatkowski, Cordelia Langford, John Sillitoe on behalf of the Wellcome Sanger Institute COVID-19 Surveillance Team |
| EPI_ISL_3364315 | Lighthouse Lab in Milton Keynes | Wellcome Sanger Institute for the COVID-19 Genomics UK (COG-UK) Consortium | The Lighthouse Lab in Milton Keynes and Alex Alderton, Roberto Amato, Jeffrey Barrett, Sonia Goncalves, Ewan Harrison, David K. Jackson, Ian Johnston, Dominic Kwiatkowski, Cordelia Langford, John Sillitoe on behalf of the Wellcome Sanger Institute COVID-19 Surveillance Team |
| EPI_ISL_3364316 | Berkshire and Surrey Pathology Services Lighthouse Laboratory | Wellcome Sanger Institute for the COVID-19 Genomics UK (COG-UK) Consortium | Berkshire and Surrey Pathology Services Lighthouse Laboratory and Alex Alderton, Roberto Amato, Jeffrey Barrett, Sonia Goncalves, Ewan Harrison, David K. Jackson, Ian Johnston, Dominic Kwiatkowski, Cordelia Langford, John Sillitoe on behalf of the Wellcome Sanger Institute COVID-19 Surveillance Team |
| EPI_ISL_3364318, EPI_ISL_3364319, EPI_ISL_3364320 | Lighthouse Lab in Milton Keynes | Wellcome Sanger Institute for the COVID-19 Genomics UK (COG-UK) Consortium | The Lighthouse Lab in Milton Keynes and Alex Alderton, Roberto Amato, Jeffrey Barrett, Sonia Goncalves, Ewan Harrison, David K. Jackson, Ian Johnston, Dominic Kwiatkowski, Cordelia Langford, John Sillitoe on behalf of the Wellcome Sanger Institute COVID-19 Surveillance Team |
| EPI_ISL_3364322 | Berkshire and Surrey Pathology Services Lighthouse Laboratory | Wellcome Sanger Institute for the COVID-19 Genomics UK (COG-UK) Consortium | Berkshire and Surrey Pathology Services Lighthouse Laboratory and Alex Alderton, Roberto Amato, Jeffrey Barrett, Sonia Goncalves, Ewan Harrison, David K. Jackson, Ian Johnston, Dominic Kwiatkowski, Cordelia Langford, John Sillitoe on behalf of the Wellcome Sanger Institute COVID-19 Surveillance Team |
| EPI_ISL_3364323 | Lighthouse Lab in Milton Keynes | Wellcome Sanger Institute for the COVID-19 Genomics UK (COG-UK) Consortium | The Lighthouse Lab in Milton Keynes and Alex Alderton, Roberto Amato, Jeffrey Barrett, Sonia Goncalves, Ewan Harrison, David K. Jackson, Ian Johnston, Dominic Kwiatkowski, Cordelia Langford, John Sillitoe on behalf of the Wellcome Sanger Institute COVID-19 Surveillance Team |
| EPI_ISL_3364325, EPI_ISL_3364326 | Berkshire and Surrey Pathology Services Lighthouse Laboratory | Wellcome Sanger Institute for the COVID-19 Genomics UK (COG-UK) Consortium | Berkshire and Surrey Pathology Services Lighthouse Laboratory and Alex Alderton, Roberto Amato, Jeffrey Barrett, Sonia Goncalves, Ewan Harrison, David K. Jackson, Ian Johnston, Dominic Kwiatkowski, Cordelia Langford, John Sillitoe on behalf of the Wellcome Sanger Institute COVID-19 Surveillance Team |
| EPI_ISL_3364327, EPI_ISL_3364329 | Lighthouse Lab in Milton Keynes | Wellcome Sanger Institute for the COVID-19 Genomics UK (COG-UK) Consortium | The Lighthouse Lab in Milton Keynes and Alex Alderton, Roberto Amato, Jeffrey Barrett, Sonia Goncalves, Ewan Harrison, David K. Jackson, Ian Johnston, Dominic Kwiatkowski, Cordelia Langford, John Sillitoe on behalf of the Wellcome Sanger Institute COVID-19 Surveillance Team |
| EPI_ISL_3364330 | Berkshire and Surrey Pathology Services Lighthouse Laboratory | Wellcome Sanger Institute for the COVID-19 Genomics UK (COG-UK) Consortium | Berkshire and Surrey Pathology Services Lighthouse Laboratory and Alex Alderton, Roberto Amato, Jeffrey Barrett, Sonia Goncalves, Ewan Harrison, David K. Jackson, Ian Johnston, Dominic Kwiatkowski, Cordelia Langford, John Sillitoe on behalf of the Wellcome Sanger Institute COVID-19 Surveillance Team |
| EPI_ISL_3364331, EPI_ISL_3364333, EPI_ISL_3364334 | Lighthouse Lab in Milton Keynes | Wellcome Sanger Institute for the COVID-19 Genomics UK (COG-UK) Consortium | The Lighthouse Lab in Milton Keynes and Alex Alderton, Roberto Amato, Jeffrey Barrett, Sonia Goncalves, Ewan Harrison, David K. Jackson, Ian Johnston, Dominic Kwiatkowski, Cordelia Langford, John Sillitoe on behalf of the Wellcome Sanger Institute COVID-19 Surveillance Team |
| EPI_ISL_3364336 | Berkshire and Surrey Pathology Services Lighthouse Laboratory | Wellcome Sanger Institute for the COVID-19 Genomics UK (COG-UK) Consortium | Berkshire and Surrey Pathology Services Lighthouse Laboratory and Alex Alderton, Roberto Amato, Jeffrey Barrett, Sonia Goncalves, Ewan Harrison, David K. Jackson, Ian Johnston, Dominic Kwiatkowski, Cordelia Langford, John Sillitoe on behalf of the Wellcome Sanger Institute COVID-19 Surveillance Team |
| EPI_ISL_3364337, EPI_ISL_3364338, EPI_ISL_3364340 | Lighthouse Lab in Milton Keynes | Wellcome Sanger Institute for the COVID-19 Genomics UK (COG-UK) Consortium | The Lighthouse Lab in Milton Keynes and Alex Alderton, Roberto Amato, Jeffrey Barrett, Sonia Goncalves, Ewan Harrison, David K. Jackson, Ian Johnston, Dominic Kwiatkowski, Cordelia Langford, John Sillitoe on behalf of the Wellcome Sanger Institute COVID-19 Surveillance Team |
| EPI_ISL_3364341 | Berkshire and Surrey Pathology Services Lighthouse Laboratory | Wellcome Sanger Institute for the COVID-19 Genomics UK (COG-UK) Consortium | Berkshire and Surrey Pathology Services Lighthouse Laboratory and Alex Alderton, Roberto Amato, Jeffrey Barrett, Sonia Goncalves, Ewan Harrison, David K. Jackson, Ian Johnston, Dominic Kwiatkowski, Cordelia Langford, John Sillitoe on behalf of the Wellcome Sanger Institute COVID-19 Surveillance Team |
| EPI_ISL_3364342 | Lighthouse Lab in Milton Keynes | Wellcome Sanger Institute for the COVID-19 Genomics UK (COG-UK) Consortium | The Lighthouse Lab in Milton Keynes and Alex Alderton, Roberto Amato, Jeffrey Barrett, Sonia Goncalves, Ewan Harrison, David K. Jackson, Ian Johnston, Dominic Kwiatkowski, Cordelia Langford, John Sillitoe on behalf of the Wellcome Sanger Institute COVID-19 Surveillance Team |
| EPI_ISL_3364344 | Berkshire and Surrey Pathology Services Lighthouse Laboratory | Wellcome Sanger Institute for the COVID-19 Genomics UK (COG-UK) Consortium | Berkshire and Surrey Pathology Services Lighthouse Laboratory and Alex Alderton, Roberto Amato, Jeffrey Barrett, Sonia Goncalves, Ewan Harrison, David K. Jackson, Ian Johnston, Dominic Kwiatkowski, Cordelia Langford, John Sillitoe on behalf of the Wellcome Sanger Institute COVID-19 Surveillance Team |
| EPI_ISL_3364345, EPI_ISL_3364347, EPI_ISL_3364348 | Lighthouse Lab in Milton Keynes | Wellcome Sanger Institute for the COVID-19 Genomics UK (COG-UK) Consortium | The Lighthouse Lab in Milton Keynes and Alex Alderton, Roberto Amato, Jeffrey Barrett, Sonia Goncalves, Ewan Harrison, David K. Jackson, Ian Johnston, Dominic Kwiatkowski, Cordelia Langford, John Sillitoe on behalf of the Wellcome Sanger Institute COVID-19 Surveillance Team |
| EPI_ISL_3364350, EPI_ISL_3364351, EPI_ISL_3364352, EPI_ISL_3364354 | Berkshire and Surrey Pathology Services Lighthouse Laboratory | Wellcome Sanger Institute for the COVID-19 Genomics UK (COG-UK) Consortium | Berkshire and Surrey Pathology Services Lighthouse Laboratory and Alex Alderton, Roberto Amato, Jeffrey Barrett, Sonia Goncalves, Ewan Harrison, David K. Jackson, Ian Johnston, Dominic Kwiatkowski, Cordelia Langford, John Sillitoe on behalf of the Wellcome Sanger Institute COVID-19 Surveillance Team |
| EPI_ISL_3364355 | Lighthouse Lab in Milton Keynes | Wellcome Sanger Institute for the COVID-19 Genomics UK (COG-UK) Consortium | The Lighthouse Lab in Milton Keynes and Alex Alderton, Roberto Amato, Jeffrey Barrett, Sonia Goncalves, Ewan Harrison, David K. Jackson, Ian Johnston, Dominic Kwiatkowski, Cordelia Langford, John Sillitoe on behalf of the Wellcome Sanger Institute COVID-19 Surveillance Team |
| EPI_ISL_3364356 | Berkshire and Surrey Pathology Services Lighthouse Laboratory | Wellcome Sanger Institute for the COVID-19 Genomics UK (COG-UK) Consortium | Berkshire and Surrey Pathology Services Lighthouse Laboratory and Alex Alderton, Roberto Amato, Jeffrey Barrett, Sonia Goncalves, Ewan Harrison, David K. Jackson, Ian Johnston, Dominic Kwiatkowski, Cordelia Langford, John Sillitoe on behalf of the Wellcome Sanger Institute COVID-19 Surveillance Team |
| EPI_ISL_3364358, EPI_ISL_3364359, EPI_ISL_3364361 | Lighthouse Lab in Milton Keynes | Wellcome Sanger Institute for the COVID-19 Genomics UK (COG-UK) Consortium | The Lighthouse Lab in Milton Keynes and Alex Alderton, Roberto Amato, Jeffrey Barrett, Sonia Goncalves, Ewan Harrison, David K. Jackson, Ian Johnston, Dominic Kwiatkowski, Cordelia Langford, John Sillitoe on behalf of the Wellcome Sanger Institute COVID-19 Surveillance Team |
| EPI_ISL_3364362 | Berkshire and Surrey Pathology Services Lighthouse Laboratory | Wellcome Sanger Institute for the COVID-19 Genomics UK (COG-UK) Consortium | Berkshire and Surrey Pathology Services Lighthouse Laboratory and Alex Alderton, Roberto Amato, Jeffrey Barrett, Sonia Goncalves, Ewan Harrison, David K. Jackson, Ian Johnston, Dominic Kwiatkowski, Cordelia Langford, John Sillitoe on behalf of the Wellcome Sanger Institute COVID-19 Surveillance Team |
| EPI_ISL_3364363, EPI_ISL_3364365, EPI_ISL_3364366, EPI_ISL_3364368, EPI_ISL_3364369, EPI_ISL_3364370 | Lighthouse Lab in Milton Keynes | Wellcome Sanger Institute for the COVID-19 Genomics UK (COG-UK) Consortium | The Lighthouse Lab in Milton Keynes and Alex Alderton, Roberto Amato, Jeffrey Barrett, Sonia Goncalves, Ewan Harrison, David K. Jackson, Ian Johnston, Dominic Kwiatkowski, Cordelia Langford, John Sillitoe on behalf of the Wellcome Sanger Institute COVID-19 Surveillance Team |

| | | | |
|---|---|---|---|
| EPI_ISL_3364372 | Berkshire and Surrey Pathology Services Lighthouse Laboratory | Wellcome Sanger Institute for the COVID-19 Genomics UK (COG-UK) Consortium | Berkshire and Surrey Pathology Services Lighthouse Laboratory and Alex Alderton, Roberto Amato, Jeffrey Barrett, Sonia Goncalves, Ewan Harrison, David K. Jackson, Ian Johnston, Dominic Kwiatkowski, Cordelia Langford, John Sillitoe on behalf of the Wellcome Sanger Institute COVID-19 Surveillance Team |
| EPI_ISL_3364373 | Lighthouse Lab in Milton Keynes | Wellcome Sanger Institute for the COVID-19 Genomics UK (COG-UK) Consortium | The Lighthouse Lab in Milton Keynes and Alex Alderton, Roberto Amato, Jeffrey Barrett, Sonia Goncalves, Ewan Harrison, David K. Jackson, Ian Johnston, Dominic Kwiatkowski, Cordelia Langford, John Sillitoe on behalf of the Wellcome Sanger Institute COVID-19 Surveillance Team |
| EPI_ISL_3364374, EPI_ISL_3364376, EPI_ISL_3364377, EPI_ISL_3364379, EPI_ISL_3364380, EPI_ISL_3364381 | Berkshire and Surrey Pathology Services Lighthouse Laboratory | Wellcome Sanger Institute for the COVID-19 Genomics UK (COG-UK) Consortium | Berkshire and Surrey Pathology Services Lighthouse Laboratory and Alex Alderton, Roberto Amato, Jeffrey Barrett, Sonia Goncalves, Ewan Harrison, David K. Jackson, Ian Johnston, Dominic Kwiatkowski, Cordelia Langford, John Sillitoe on behalf of the Wellcome Sanger Institute COVID-19 Surveillance Team |
| EPI_ISL_3364383 | Lighthouse Lab in Milton Keynes | Wellcome Sanger Institute for the COVID-19 Genomics UK (COG-UK) Consortium | The Lighthouse Lab in Milton Keynes and Alex Alderton, Roberto Amato, Jeffrey Barrett, Sonia Goncalves, Ewan Harrison, David K. Jackson, Ian Johnston, Dominic Kwiatkowski, Cordelia Langford, John Sillitoe on behalf of the Wellcome Sanger Institute COVID-19 Surveillance Team |
| EPI_ISL_3364384 | Berkshire and Surrey Pathology Services Lighthouse Laboratory | Wellcome Sanger Institute for the COVID-19 Genomics UK (COG-UK) Consortium | Berkshire and Surrey Pathology Services Lighthouse Laboratory and Alex Alderton, Roberto Amato, Jeffrey Barrett, Sonia Goncalves, Ewan Harrison, David K. Jackson, Ian Johnston, Dominic Kwiatkowski, Cordelia Langford, John Sillitoe on behalf of the Wellcome Sanger Institute COVID-19 Surveillance Team |
| EPI_ISL_3364386, EPI_ISL_3364387, EPI_ISL_3364388 | Lighthouse Lab in Milton Keynes | Wellcome Sanger Institute for the COVID-19 Genomics UK (COG-UK) Consortium | The Lighthouse Lab in Milton Keynes and Alex Alderton, Roberto Amato, Jeffrey Barrett, Sonia Goncalves, Ewan Harrison, David K. Jackson, Ian Johnston, Dominic Kwiatkowski, Cordelia Langford, John Sillitoe on behalf of the Wellcome Sanger Institute COVID-19 Surveillance Team |
| EPI_ISL_3364390, EPI_ISL_3364391 | Berkshire and Surrey Pathology Services Lighthouse Laboratory | Wellcome Sanger Institute for the COVID-19 Genomics UK (COG-UK) Consortium | Berkshire and Surrey Pathology Services Lighthouse Laboratory and Alex Alderton, Roberto Amato, Jeffrey Barrett, Sonia Goncalves, Ewan Harrison, David K. Jackson, Ian Johnston, Dominic Kwiatkowski, Cordelia Langford, John Sillitoe on behalf of the Wellcome Sanger Institute COVID-19 Surveillance Team |
| EPI_ISL_3364392, EPI_ISL_3364394, EPI_ISL_3364395 | Lighthouse Lab in Milton Keynes | Wellcome Sanger Institute for the COVID-19 Genomics UK (COG-UK) Consortium | The Lighthouse Lab in Milton Keynes and Alex Alderton, Roberto Amato, Jeffrey Barrett, Sonia Goncalves, Ewan Harrison, David K. Jackson, Ian Johnston, Dominic Kwiatkowski, Cordelia Langford, John Sillitoe on behalf of the Wellcome Sanger Institute COVID-19 Surveillance Team |
| EPI_ISL_3364397 | Berkshire and Surrey Pathology Services Lighthouse Laboratory | Wellcome Sanger Institute for the COVID-19 Genomics UK (COG-UK) Consortium | Berkshire and Surrey Pathology Services Lighthouse Laboratory and Alex Alderton, Roberto Amato, Jeffrey Barrett, Sonia Goncalves, Ewan Harrison, David K. Jackson, Ian Johnston, Dominic Kwiatkowski, Cordelia Langford, John Sillitoe on behalf of the Wellcome Sanger Institute COVID-19 Surveillance Team |
| EPI_ISL_3364398 | Lighthouse Lab in Milton Keynes | Wellcome Sanger Institute for the COVID-19 Genomics UK (COG-UK) Consortium | The Lighthouse Lab in Milton Keynes and Alex Alderton, Roberto Amato, Jeffrey Barrett, Sonia Goncalves, Ewan Harrison, David K. Jackson, Ian Johnston, Dominic Kwiatkowski, Cordelia Langford, John Sillitoe on behalf of the Wellcome Sanger Institute COVID-19 Surveillance Team |
| EPI_ISL_3364399, EPI_ISL_3364401 | Berkshire and Surrey Pathology Services Lighthouse Laboratory | Wellcome Sanger Institute for the COVID-19 Genomics UK (COG-UK) Consortium | Berkshire and Surrey Pathology Services Lighthouse Laboratory and Alex Alderton, Roberto Amato, Jeffrey Barrett, Sonia Goncalves, Ewan Harrison, David K. Jackson, Ian Johnston, Dominic Kwiatkowski, Cordelia Langford, John Sillitoe on behalf of the Wellcome Sanger Institute COVID-19 Surveillance Team |
| EPI_ISL_3364402 | Lighthouse Lab in Milton Keynes | Wellcome Sanger Institute for the COVID-19 Genomics UK (COG-UK) Consortium | The Lighthouse Lab in Milton Keynes and Alex Alderton, Roberto Amato, Jeffrey Barrett, Sonia Goncalves, Ewan Harrison, David K. Jackson, Ian Johnston, Dominic Kwiatkowski, Cordelia Langford, John Sillitoe on behalf of the Wellcome Sanger Institute COVID-19 Surveillance Team |
| EPI_ISL_3364404 | Berkshire and Surrey Pathology Services Lighthouse Laboratory | Wellcome Sanger Institute for the COVID-19 Genomics UK (COG-UK) Consortium | Berkshire and Surrey Pathology Services Lighthouse Laboratory and Alex Alderton, Roberto Amato, Jeffrey Barrett, Sonia Goncalves, Ewan Harrison, David K. Jackson, Ian Johnston, Dominic Kwiatkowski, Cordelia Langford, John Sillitoe on behalf of the Wellcome Sanger Institute COVID-19 Surveillance Team |
| EPI_ISL_3364405, EPI_ISL_3364407, EPI_ISL_3364408 | Lighthouse Lab in Milton Keynes | Wellcome Sanger Institute for the COVID-19 Genomics UK (COG-UK) Consortium | The Lighthouse Lab in Milton Keynes and Alex Alderton, Roberto Amato, Jeffrey Barrett, Sonia Goncalves, Ewan Harrison, David K. Jackson, Ian Johnston, Dominic Kwiatkowski, Cordelia Langford, John Sillitoe on behalf of the Wellcome Sanger Institute COVID-19 Surveillance Team |
| EPI_ISL_3364409 | Berkshire and Surrey Pathology Services Lighthouse Laboratory | Wellcome Sanger Institute for the COVID-19 Genomics UK (COG-UK) Consortium | Berkshire and Surrey Pathology Services Lighthouse Laboratory and Alex Alderton, Roberto Amato, Jeffrey Barrett, Sonia Goncalves, Ewan Harrison, David K. Jackson, Ian Johnston, Dominic Kwiatkowski, Cordelia Langford, John Sillitoe on behalf of the Wellcome Sanger Institute COVID-19 Surveillance Team |
| EPI_ISL_3364411, EPI_ISL_3364412, EPI_ISL_3364414, EPI_ISL_3364415 | Lighthouse Lab in Milton Keynes | Wellcome Sanger Institute for the COVID-19 Genomics UK (COG-UK) Consortium | The Lighthouse Lab in Milton Keynes and Alex Alderton, Roberto Amato, Jeffrey Barrett, Sonia Goncalves, Ewan Harrison, David K. Jackson, Ian Johnston, Dominic Kwiatkowski, Cordelia Langford, John Sillitoe on behalf of the Wellcome Sanger Institute COVID-19 Surveillance Team |
| EPI_ISL_3364417 | Berkshire and Surrey Pathology Services Lighthouse Laboratory | Wellcome Sanger Institute for the COVID-19 Genomics UK (COG-UK) Consortium | Berkshire and Surrey Pathology Services Lighthouse Laboratory and Alex Alderton, Roberto Amato, Jeffrey Barrett, Sonia Goncalves, Ewan Harrison, David K. Jackson, Ian Johnston, Dominic Kwiatkowski, Cordelia Langford, John Sillitoe on behalf of the Wellcome Sanger Institute COVID-19 Surveillance Team |
| EPI_ISL_3364418, EPI_ISL_3364419 | Lighthouse Lab in Milton Keynes | Wellcome Sanger Institute for the COVID-19 Genomics UK (COG-UK) Consortium | The Lighthouse Lab in Milton Keynes and Alex Alderton, Roberto Amato, Jeffrey Barrett, Sonia Goncalves, Ewan Harrison, David K. Jackson, Ian Johnston, Dominic Kwiatkowski, Cordelia Langford, John Sillitoe on behalf of the Wellcome Sanger Institute COVID-19 Surveillance Team |
| EPI_ISL_3364421 | Berkshire and Surrey Pathology Services Lighthouse Laboratory | Wellcome Sanger Institute for the COVID-19 Genomics UK (COG-UK) Consortium | Berkshire and Surrey Pathology Services Lighthouse Laboratory and Alex Alderton, Roberto Amato, Jeffrey Barrett, Sonia Goncalves, Ewan Harrison, David K. Jackson, Ian Johnston, Dominic Kwiatkowski, Cordelia Langford, John Sillitoe on behalf of the Wellcome Sanger Institute COVID-19 Surveillance Team |
| EPI_ISL_3364422 | Lighthouse Lab in Milton Keynes | Wellcome Sanger Institute for the COVID-19 Genomics UK (COG-UK) Consortium | The Lighthouse Lab in Milton Keynes and Alex Alderton, Roberto Amato, Jeffrey Barrett, Sonia Goncalves, Ewan Harrison, David K. Jackson, Ian Johnston, Dominic Kwiatkowski, Cordelia Langford, John Sillitoe on behalf of the Wellcome Sanger Institute COVID-19 Surveillance Team |
| EPI_ISL_3364423, EPI_ISL_3364425, EPI_ISL_3364426, EPI_ISL_3364427 | Berkshire and Surrey Pathology Services Lighthouse Laboratory | Wellcome Sanger Institute for the COVID-19 Genomics UK (COG-UK) Consortium | Berkshire and Surrey Pathology Services Lighthouse Laboratory and Alex Alderton, Roberto Amato, Jeffrey Barrett, Sonia Goncalves, Ewan Harrison, David K. Jackson, Ian Johnston, Dominic Kwiatkowski, Cordelia Langford, John Sillitoe on behalf of the Wellcome Sanger Institute COVID-19 Surveillance Team |
| EPI_ISL_3364429 | Lighthouse Lab in Milton Keynes | Wellcome Sanger Institute for the COVID-19 Genomics UK (COG-UK) Consortium | The Lighthouse Lab in Milton Keynes and Alex Alderton, Roberto Amato, Jeffrey Barrett, Sonia Goncalves, Ewan Harrison, David K. Jackson, Ian Johnston, Dominic Kwiatkowski, Cordelia Langford, John Sillitoe on behalf of the Wellcome Sanger Institute COVID-19 Surveillance Team |
| EPI_ISL_3364430, EPI_ISL_3364434 | Berkshire and Surrey Pathology Services Lighthouse Laboratory | Wellcome Sanger Institute for the COVID-19 Genomics UK (COG-UK) Consortium | Berkshire and Surrey Pathology Services Lighthouse Laboratory and Alex Alderton, Roberto Amato, Jeffrey Barrett, Sonia Goncalves, Ewan Harrison, David K. Jackson, Ian Johnston, Dominic Kwiatkowski, Cordelia Langford, John Sillitoe on behalf of the Wellcome Sanger Institute COVID-19 Surveillance Team |
| EPI_ISL_3364435 | Lighthouse Lab in Milton Keynes | Wellcome Sanger Institute for the COVID-19 Genomics UK (COG-UK) Consortium | The Lighthouse Lab in Milton Keynes and Alex Alderton, Roberto Amato, Jeffrey Barrett, Sonia Goncalves, Ewan Harrison, David K. Jackson, Ian Johnston, Dominic Kwiatkowski, Cordelia Langford, John Sillitoe on behalf of the Wellcome Sanger Institute COVID-19 Surveillance Team |
| EPI_ISL_3364436, EPI_ISL_3364438, EPI_ISL_3364439 | Berkshire and Surrey Pathology Services Lighthouse Laboratory | Wellcome Sanger Institute for the COVID-19 Genomics UK (COG-UK) Consortium | Berkshire and Surrey Pathology Services Lighthouse Laboratory and Alex Alderton, Roberto Amato, Jeffrey Barrett, Sonia Goncalves, Ewan Harrison, David K. Jackson, Ian Johnston, Dominic Kwiatkowski, Cordelia Langford, John Sillitoe on behalf of the Wellcome Sanger Institute COVID-19 Surveillance Team |
| EPI_ISL_3364440, EPI_ISL_3364442 | Lighthouse Lab in Milton Keynes | Wellcome Sanger Institute for the COVID-19 Genomics UK (COG-UK) Consortium | The Lighthouse Lab in Milton Keynes and Alex Alderton, Roberto Amato, Jeffrey Barrett, Sonia Goncalves, Ewan Harrison, David K. Jackson, Ian Johnston, Dominic Kwiatkowski, Cordelia Langford, John Sillitoe on behalf of the Wellcome Sanger Institute COVID-19 Surveillance Team |
| EPI_ISL_3364443 | Berkshire and Surrey Pathology Services Lighthouse Laboratory | Wellcome Sanger Institute for the COVID-19 Genomics UK (COG-UK) Consortium | Berkshire and Surrey Pathology Services Lighthouse Laboratory and Alex Alderton, Roberto Amato, Jeffrey Barrett, Sonia Goncalves, Ewan Harrison, David K. Jackson, Ian Johnston, Dominic Kwiatkowski, Cordelia Langford, John Sillitoe on behalf of the Wellcome Sanger Institute COVID-19 Surveillance Team |
| EPI_ISL_3364445 | Lighthouse Lab in Milton Keynes | Wellcome Sanger Institute for the COVID-19 Genomics UK (COG-UK) Consortium | The Lighthouse Lab in Milton Keynes and Alex Alderton, Roberto Amato, Jeffrey Barrett, Sonia Goncalves, Ewan Harrison, David K. Jackson, Ian Johnston, Dominic Kwiatkowski, Cordelia Langford, John Sillitoe on behalf of the Wellcome Sanger Institute COVID-19 Surveillance Team |
| EPI_ISL_3364446, EPI_ISL_3364448, EPI_ISL_3364449, EPI_ISL_3364450, EPI_ISL_3364452 | Berkshire and Surrey Pathology Services Lighthouse Laboratory | Wellcome Sanger Institute for the COVID-19 Genomics UK (COG-UK) Consortium | Berkshire and Surrey Pathology Services Lighthouse Laboratory and Alex Alderton, Roberto Amato, Jeffrey Barrett, Sonia Goncalves, Ewan Harrison, David K. Jackson, Ian Johnston, Dominic Kwiatkowski, Cordelia Langford, John Sillitoe on behalf of the Wellcome Sanger Institute COVID-19 Surveillance Team |
| EPI_ISL_3364453, EPI_ISL_3364454 | Lighthouse Lab in Milton Keynes | Wellcome Sanger Institute for the COVID-19 Genomics UK (COG-UK) Consortium | The Lighthouse Lab in Milton Keynes and Alex Alderton, Roberto Amato, Jeffrey Barrett, Sonia Goncalves, Ewan Harrison, David K. Jackson, Ian Johnston, Dominic Kwiatkowski, Cordelia Langford, John Sillitoe on behalf of the Wellcome Sanger Institute COVID-19 Surveillance Team |
| EPI_ISL_3364456 | Berkshire and Surrey Pathology Services Lighthouse Laboratory | Wellcome Sanger Institute for the COVID-19 Genomics UK (COG-UK) Consortium | Berkshire and Surrey Pathology Services Lighthouse Laboratory and Alex Alderton, Roberto Amato, Jeffrey Barrett, Sonia Goncalves, Ewan Harrison, David K. Jackson, Ian Johnston, Dominic Kwiatkowski, Cordelia Langford, John Sillitoe on behalf of the Wellcome Sanger Institute COVID-19 Surveillance Team |
| EPI_ISL_3364457 | Lighthouse Lab in Milton Keynes | Wellcome Sanger Institute for the COVID-19 Genomics UK (COG-UK) Consortium | The Lighthouse Lab in Milton Keynes and Alex Alderton, Roberto Amato, Jeffrey Barrett, Sonia Goncalves, Ewan Harrison, David K. Jackson, Ian Johnston, Dominic Kwiatkowski, Cordelia Langford, John Sillitoe on behalf of the Wellcome Sanger Institute COVID-19 Surveillance Team |
| EPI_ISL_3364459 | Berkshire and Surrey Pathology Services Lighthouse Laboratory | Wellcome Sanger Institute for the COVID-19 Genomics UK (COG-UK) Consortium | Berkshire and Surrey Pathology Services Lighthouse Laboratory and Alex Alderton, Roberto Amato, Jeffrey Barrett, Sonia Goncalves, Ewan Harrison, David K. Jackson, Ian Johnston, Dominic Kwiatkowski, Cordelia Langford, John Sillitoe on behalf of the Wellcome Sanger Institute COVID-19 Surveillance Team |
| EPI_ISL_3364460, EPI_ISL_3364461 | Lighthouse Lab in Milton Keynes | Wellcome Sanger Institute for the COVID-19 Genomics UK (COG-UK) Consortium | The Lighthouse Lab in Milton Keynes and Alex Alderton, Roberto Amato, Jeffrey Barrett, Sonia Goncalves, Ewan Harrison, David K. Jackson, Ian Johnston, Dominic Kwiatkowski, Cordelia Langford, John Sillitoe on behalf of the Wellcome Sanger Institute COVID-19 Surveillance Team |
| EPI_ISL_3364463, EPI_ISL_3364464, EPI_ISL_3364466, EPI_ISL_3364467 | Berkshire and Surrey Pathology Services Lighthouse Laboratory | Wellcome Sanger Institute for the COVID-19 Genomics UK (COG-UK) Consortium | Berkshire and Surrey Pathology Services Lighthouse Laboratory and Alex Alderton, Roberto Amato, Jeffrey Barrett, Sonia Goncalves, Ewan Harrison, David K. Jackson, Ian Johnston, Dominic Kwiatkowski, Cordelia Langford, John Sillitoe on behalf of the Wellcome Sanger Institute COVID-19 Surveillance Team |
| EPI_ISL_3364468, EPI_ISL_3364470, EPI_ISL_3364471, EPI_ISL_3364473 | Lighthouse Lab in Milton Keynes | Wellcome Sanger Institute for the COVID-19 Genomics UK (COG-UK) Consortium | The Lighthouse Lab in Milton Keynes and Alex Alderton, Roberto Amato, Jeffrey Barrett, Sonia Goncalves, Ewan Harrison, David K. Jackson, Ian Johnston, Dominic Kwiatkowski, Cordelia Langford, John Sillitoe on behalf of the Wellcome Sanger Institute COVID-19 Surveillance Team |
| EPI_ISL_3364474, EPI_ISL_3364476 | Berkshire and Surrey Pathology Services Lighthouse Laboratory | Wellcome Sanger Institute for the COVID-19 Genomics UK (COG-UK) Consortium | Berkshire and Surrey Pathology Services Lighthouse Laboratory and Alex Alderton, Roberto Amato, Jeffrey Barrett, Sonia Goncalves, Ewan Harrison, David K. Jackson, Ian Johnston, Dominic Kwiatkowski, Cordelia Langford, John Sillitoe on behalf of the Wellcome Sanger Institute COVID-19 Surveillance Team |
| EPI_ISL_3364477 | Lighthouse Lab in Milton Keynes | Wellcome Sanger Institute for the COVID-19 Genomics UK (COG-UK) | The Lighthouse Lab in Milton Keynes and Alex Alderton, Roberto Amato, Jeffrey Barrett, Sonia Goncalves, Ewan Harrison, David K. Jackson, Ian Johnston, Dominic Kwiatkowski, Cordelia Langford, John Sillitoe on behalf of the Wellcome Sanger |

| | | | |
|---|---|---|---|
| | | Consortium | Institute COVID-19 Surveillance Team |
| EPI_ISL_3364478, EPI_ISL_3364480 | Berkshire and Surrey Pathology Services Lighthouse Laboratory | Wellcome Sanger Institute for the COVID-19 Genomics UK (COG-UK) Consortium | Berkshire and Surrey Pathology Services Lighthouse Laboratory and Alex Alderton, Roberto Amato, Jeffrey Barrett, Sonia Goncalves, Ewan Harrison, David K. Jackson, Ian Johnston, Dominic Kwiatkowski, Cordelia Langford, John Sillitoe on behalf of the Wellcome Sanger Institute COVID-19 Surveillance Team |
| EPI_ISL_3364481 | Lighthouse Lab in Milton Keynes | Wellcome Sanger Institute for the COVID-19 Genomics UK (COG-UK) Consortium | The Lighthouse Lab in Milton Keynes and Alex Alderton, Roberto Amato, Jeffrey Barrett, Sonia Goncalves, Ewan Harrison, David K. Jackson, Ian Johnston, Dominic Kwiatkowski, Cordelia Langford, John Sillitoe on behalf of the Wellcome Sanger Institute COVID-19 Surveillance Team |
| EPI_ISL_3364483 | Berkshire and Surrey Pathology Services Lighthouse Laboratory | Wellcome Sanger Institute for the COVID-19 Genomics UK (COG-UK) Consortium | Berkshire and Surrey Pathology Services Lighthouse Laboratory and Alex Alderton, Roberto Amato, Jeffrey Barrett, Sonia Goncalves, Ewan Harrison, David K. Jackson, Ian Johnston, Dominic Kwiatkowski, Cordelia Langford, John Sillitoe on behalf of the Wellcome Sanger Institute COVID-19 Surveillance Team |
| EPI_ISL_3364484 | Lighthouse Lab in Milton Keynes | Wellcome Sanger Institute for the COVID-19 Genomics UK (COG-UK) Consortium | The Lighthouse Lab in Milton Keynes and Alex Alderton, Roberto Amato, Jeffrey Barrett, Sonia Goncalves, Ewan Harrison, David K. Jackson, Ian Johnston, Dominic Kwiatkowski, Cordelia Langford, John Sillitoe on behalf of the Wellcome Sanger Institute COVID-19 Surveillance Team |
| EPI_ISL_3364485 | Berkshire and Surrey Pathology Services Lighthouse Laboratory | Wellcome Sanger Institute for the COVID-19 Genomics UK (COG-UK) Consortium | Berkshire and Surrey Pathology Services Lighthouse Laboratory and Alex Alderton, Roberto Amato, Jeffrey Barrett, Sonia Goncalves, Ewan Harrison, David K. Jackson, Ian Johnston, Dominic Kwiatkowski, Cordelia Langford, John Sillitoe on behalf of the Wellcome Sanger Institute COVID-19 Surveillance Team |
| EPI_ISL_3364487, EPI_ISL_3364488 | Lighthouse Lab in Milton Keynes | Wellcome Sanger Institute for the COVID-19 Genomics UK (COG-UK) Consortium | The Lighthouse Lab in Milton Keynes and Alex Alderton, Roberto Amato, Jeffrey Barrett, Sonia Goncalves, Ewan Harrison, David K. Jackson, Ian Johnston, Dominic Kwiatkowski, Cordelia Langford, John Sillitoe on behalf of the Wellcome Sanger Institute COVID-19 Surveillance Team |
| EPI_ISL_3364490 | Berkshire and Surrey Pathology Services Lighthouse Laboratory | Wellcome Sanger Institute for the COVID-19 Genomics UK (COG-UK) Consortium | Berkshire and Surrey Pathology Services Lighthouse Laboratory and Alex Alderton, Roberto Amato, Jeffrey Barrett, Sonia Goncalves, Ewan Harrison, David K. Jackson, Ian Johnston, Dominic Kwiatkowski, Cordelia Langford, John Sillitoe on behalf of the Wellcome Sanger Institute COVID-19 Surveillance Team |
| EPI_ISL_3364491 | Lighthouse Lab in Milton Keynes | Wellcome Sanger Institute for the COVID-19 Genomics UK (COG-UK) Consortium | The Lighthouse Lab in Milton Keynes and Alex Alderton, Roberto Amato, Jeffrey Barrett, Sonia Goncalves, Ewan Harrison, David K. Jackson, Ian Johnston, Dominic Kwiatkowski, Cordelia Langford, John Sillitoe on behalf of the Wellcome Sanger Institute COVID-19 Surveillance Team |
| EPI_ISL_3364493 | Berkshire and Surrey Pathology Services Lighthouse Laboratory | Wellcome Sanger Institute for the COVID-19 Genomics UK (COG-UK) Consortium | Berkshire and Surrey Pathology Services Lighthouse Laboratory and Alex Alderton, Roberto Amato, Jeffrey Barrett, Sonia Goncalves, Ewan Harrison, David K. Jackson, Ian Johnston, Dominic Kwiatkowski, Cordelia Langford, John Sillitoe on behalf of the Wellcome Sanger Institute COVID-19 Surveillance Team |
| EPI_ISL_3364494 | Lighthouse Lab in Milton Keynes | Wellcome Sanger Institute for the COVID-19 Genomics UK (COG-UK) Consortium | The Lighthouse Lab in Milton Keynes and Alex Alderton, Roberto Amato, Jeffrey Barrett, Sonia Goncalves, Ewan Harrison, David K. Jackson, Ian Johnston, Dominic Kwiatkowski, Cordelia Langford, John Sillitoe on behalf of the Wellcome Sanger Institute COVID-19 Surveillance Team |
| EPI_ISL_3364495 | Berkshire and Surrey Pathology Services Lighthouse Laboratory | Wellcome Sanger Institute for the COVID-19 Genomics UK (COG-UK) Consortium | Berkshire and Surrey Pathology Services Lighthouse Laboratory and Alex Alderton, Roberto Amato, Jeffrey Barrett, Sonia Goncalves, Ewan Harrison, David K. Jackson, Ian Johnston, Dominic Kwiatkowski, Cordelia Langford, John Sillitoe on behalf of the Wellcome Sanger Institute COVID-19 Surveillance Team |
| EPI_ISL_3364497, EPI_ISL_3364498 | Lighthouse Lab in Milton Keynes | Wellcome Sanger Institute for the COVID-19 Genomics UK (COG-UK) Consortium | The Lighthouse Lab in Milton Keynes and Alex Alderton, Roberto Amato, Jeffrey Barrett, Sonia Goncalves, Ewan Harrison, David K. Jackson, Ian Johnston, Dominic Kwiatkowski, Cordelia Langford, John Sillitoe on behalf of the Wellcome Sanger Institute COVID-19 Surveillance Team |
| EPI_ISL_3364499 | Berkshire and Surrey Pathology Services Lighthouse Laboratory | Wellcome Sanger Institute for the COVID-19 Genomics UK (COG-UK) Consortium | Berkshire and Surrey Pathology Services Lighthouse Laboratory and Alex Alderton, Roberto Amato, Jeffrey Barrett, Sonia Goncalves, Ewan Harrison, David K. Jackson, Ian Johnston, Dominic Kwiatkowski, Cordelia Langford, John Sillitoe on behalf of the Wellcome Sanger Institute COVID-19 Surveillance Team |
| EPI_ISL_3364501 | Lighthouse Lab in Milton Keynes | Wellcome Sanger Institute for the COVID-19 Genomics UK (COG-UK) Consortium | The Lighthouse Lab in Milton Keynes and Alex Alderton, Roberto Amato, Jeffrey Barrett, Sonia Goncalves, Ewan Harrison, David K. Jackson, Ian Johnston, Dominic Kwiatkowski, Cordelia Langford, John Sillitoe on behalf of the Wellcome Sanger Institute COVID-19 Surveillance Team |
| EPI_ISL_3364502 | Berkshire and Surrey Pathology Services Lighthouse Laboratory | Wellcome Sanger Institute for the COVID-19 Genomics UK (COG-UK) Consortium | Berkshire and Surrey Pathology Services Lighthouse Laboratory and Alex Alderton, Roberto Amato, Jeffrey Barrett, Sonia Goncalves, Ewan Harrison, David K. Jackson, Ian Johnston, Dominic Kwiatkowski, Cordelia Langford, John Sillitoe on behalf of the Wellcome Sanger Institute COVID-19 Surveillance Team |
| EPI_ISL_3364503 | Lighthouse Lab in Milton Keynes | Wellcome Sanger Institute for the COVID-19 Genomics UK (COG-UK) Consortium | The Lighthouse Lab in Milton Keynes and Alex Alderton, Roberto Amato, Jeffrey Barrett, Sonia Goncalves, Ewan Harrison, David K. Jackson, Ian Johnston, Dominic Kwiatkowski, Cordelia Langford, John Sillitoe on behalf of the Wellcome Sanger Institute COVID-19 Surveillance Team |
| EPI_ISL_3364505 | Berkshire and Surrey Pathology Services Lighthouse Laboratory | Wellcome Sanger Institute for the COVID-19 Genomics UK (COG-UK) Consortium | Berkshire and Surrey Pathology Services Lighthouse Laboratory and Alex Alderton, Roberto Amato, Jeffrey Barrett, Sonia Goncalves, Ewan Harrison, David K. Jackson, Ian Johnston, Dominic Kwiatkowski, Cordelia Langford, John Sillitoe on behalf of the Wellcome Sanger Institute COVID-19 Surveillance Team |
| EPI_ISL_3364506 | Lighthouse Lab in Milton Keynes | Wellcome Sanger Institute for the COVID-19 Genomics UK (COG-UK) Consortium | The Lighthouse Lab in Milton Keynes and Alex Alderton, Roberto Amato, Jeffrey Barrett, Sonia Goncalves, Ewan Harrison, David K. Jackson, Ian Johnston, Dominic Kwiatkowski, Cordelia Langford, John Sillitoe on behalf of the Wellcome Sanger Institute COVID-19 Surveillance Team |
| EPI_ISL_3364507 | Berkshire and Surrey Pathology Services Lighthouse Laboratory | Wellcome Sanger Institute for the COVID-19 Genomics UK (COG-UK) Consortium | Berkshire and Surrey Pathology Services Lighthouse Laboratory and Alex Alderton, Roberto Amato, Jeffrey Barrett, Sonia Goncalves, Ewan Harrison, David K. Jackson, Ian Johnston, Dominic Kwiatkowski, Cordelia Langford, John Sillitoe on behalf of the Wellcome Sanger Institute COVID-19 Surveillance Team |
| EPI_ISL_3364509 | Lighthouse Lab in Milton Keynes | Wellcome Sanger Institute for the COVID-19 Genomics UK (COG-UK) Consortium | The Lighthouse Lab in Milton Keynes and Alex Alderton, Roberto Amato, Jeffrey Barrett, Sonia Goncalves, Ewan Harrison, David K. Jackson, Ian Johnston, Dominic Kwiatkowski, Cordelia Langford, John Sillitoe on behalf of the Wellcome Sanger Institute COVID-19 Surveillance Team |
| EPI_ISL_3364510, EPI_ISL_3364512, EPI_ISL_3364513, EPI_ISL_3364514, EPI_ISL_3364516, EPI_ISL_3364517, EPI_ISL_3364519 | Berkshire and Surrey Pathology Services Lighthouse Laboratory | Wellcome Sanger Institute for the COVID-19 Genomics UK (COG-UK) Consortium | Berkshire and Surrey Pathology Services Lighthouse Laboratory and Alex Alderton, Roberto Amato, Jeffrey Barrett, Sonia Goncalves, Ewan Harrison, David K. Jackson, Ian Johnston, Dominic Kwiatkowski, Cordelia Langford, John Sillitoe on behalf of the Wellcome Sanger Institute COVID-19 Surveillance Team |
| EPI_ISL_3364520, EPI_ISL_3364521 | Lighthouse Lab in Milton Keynes | Wellcome Sanger Institute for the COVID-19 Genomics UK (COG-UK) Consortium | The Lighthouse Lab in Milton Keynes and Alex Alderton, Roberto Amato, Jeffrey Barrett, Sonia Goncalves, Ewan Harrison, David K. Jackson, Ian Johnston, Dominic Kwiatkowski, Cordelia Langford, John Sillitoe on behalf of the Wellcome Sanger Institute COVID-19 Surveillance Team |
| EPI_ISL_3364523, EPI_ISL_3364524 | Berkshire and Surrey Pathology Services Lighthouse Laboratory | Wellcome Sanger Institute for the COVID-19 Genomics UK (COG-UK) Consortium | Berkshire and Surrey Pathology Services Lighthouse Laboratory and Alex Alderton, Roberto Amato, Jeffrey Barrett, Sonia Goncalves, Ewan Harrison, David K. Jackson, Ian Johnston, Dominic Kwiatkowski, Cordelia Langford, John Sillitoe on behalf of the Wellcome Sanger Institute COVID-19 Surveillance Team |
| EPI_ISL_3364525 | Lighthouse Lab in Milton Keynes | Wellcome Sanger Institute for the COVID-19 Genomics UK (COG-UK) Consortium | The Lighthouse Lab in Milton Keynes and Alex Alderton, Roberto Amato, Jeffrey Barrett, Sonia Goncalves, Ewan Harrison, David K. Jackson, Ian Johnston, Dominic Kwiatkowski, Cordelia Langford, John Sillitoe on behalf of the Wellcome Sanger Institute COVID-19 Surveillance Team |
| EPI_ISL_3364527, EPI_ISL_3364528, EPI_ISL_3364529 | Berkshire and Surrey Pathology Services Lighthouse Laboratory | Wellcome Sanger Institute for the COVID-19 Genomics UK (COG-UK) Consortium | Berkshire and Surrey Pathology Services Lighthouse Laboratory and Alex Alderton, Roberto Amato, Jeffrey Barrett, Sonia Goncalves, Ewan Harrison, David K. Jackson, Ian Johnston, Dominic Kwiatkowski, Cordelia Langford, John Sillitoe on behalf of the Wellcome Sanger Institute COVID-19 Surveillance Team |
| EPI_ISL_3364530, EPI_ISL_3364531, EPI_ISL_3364532, EPI_ISL_3364533 | Lighthouse Lab in Milton Keynes | Wellcome Sanger Institute for the COVID-19 Genomics UK (COG-UK) Consortium | The Lighthouse Lab in Milton Keynes and Alex Alderton, Roberto Amato, Jeffrey Barrett, Sonia Goncalves, Ewan Harrison, David K. Jackson, Ian Johnston, Dominic Kwiatkowski, Cordelia Langford, John Sillitoe on behalf of the Wellcome Sanger Institute COVID-19 Surveillance Team |
| EPI_ISL_3364534 | Berkshire and Surrey Pathology Services Lighthouse Laboratory | Wellcome Sanger Institute for the COVID-19 Genomics UK (COG-UK) Consortium | Berkshire and Surrey Pathology Services Lighthouse Laboratory and Alex Alderton, Roberto Amato, Jeffrey Barrett, Sonia Goncalves, Ewan Harrison, David K. Jackson, Ian Johnston, Dominic Kwiatkowski, Cordelia Langford, John Sillitoe on behalf of the Wellcome Sanger Institute COVID-19 Surveillance Team |
| EPI_ISL_3364535, EPI_ISL_3364537, EPI_ISL_3364538 | Lighthouse Lab in Milton Keynes | Wellcome Sanger Institute for the COVID-19 Genomics UK (COG-UK) Consortium | The Lighthouse Lab in Milton Keynes and Alex Alderton, Roberto Amato, Jeffrey Barrett, Sonia Goncalves, Ewan Harrison, David K. Jackson, Ian Johnston, Dominic Kwiatkowski, Cordelia Langford, John Sillitoe on behalf of the Wellcome Sanger Institute COVID-19 Surveillance Team |
| EPI_ISL_3364540 | Berkshire and Surrey Pathology Services Lighthouse Laboratory | Wellcome Sanger Institute for the COVID-19 Genomics UK (COG-UK) Consortium | Berkshire and Surrey Pathology Services Lighthouse Laboratory and Alex Alderton, Roberto Amato, Jeffrey Barrett, Sonia Goncalves, Ewan Harrison, David K. Jackson, Ian Johnston, Dominic Kwiatkowski, Cordelia Langford, John Sillitoe on behalf of the Wellcome Sanger Institute COVID-19 Surveillance Team |
| EPI_ISL_3364541, EPI_ISL_3364543 | Lighthouse Lab in Milton Keynes | Wellcome Sanger Institute for the COVID-19 Genomics UK (COG-UK) Consortium | The Lighthouse Lab in Milton Keynes and Alex Alderton, Roberto Amato, Jeffrey Barrett, Sonia Goncalves, Ewan Harrison, David K. Jackson, Ian Johnston, Dominic Kwiatkowski, Cordelia Langford, John Sillitoe on behalf of the Wellcome Sanger Institute COVID-19 Surveillance Team |
| EPI_ISL_3364544, EPI_ISL_3364546 | Berkshire and Surrey Pathology Services Lighthouse Laboratory | Wellcome Sanger Institute for the COVID-19 Genomics UK (COG-UK) Consortium | Berkshire and Surrey Pathology Services Lighthouse Laboratory and Alex Alderton, Roberto Amato, Jeffrey Barrett, Sonia Goncalves, Ewan Harrison, David K. Jackson, Ian Johnston, Dominic Kwiatkowski, Cordelia Langford, John Sillitoe on behalf of the Wellcome Sanger Institute COVID-19 Surveillance Team |
| EPI_ISL_3364547 | Lighthouse Lab in Milton Keynes | Wellcome Sanger Institute for the COVID-19 Genomics UK (COG-UK) Consortium | The Lighthouse Lab in Milton Keynes and Alex Alderton, Roberto Amato, Jeffrey Barrett, Sonia Goncalves, Ewan Harrison, David K. Jackson, Ian Johnston, Dominic Kwiatkowski, Cordelia Langford, John Sillitoe on behalf of the Wellcome Sanger Institute COVID-19 Surveillance Team |
| EPI_ISL_3364549 | Berkshire and Surrey Pathology Services Lighthouse Laboratory | Wellcome Sanger Institute for the COVID-19 Genomics UK (COG-UK) Consortium | Berkshire and Surrey Pathology Services Lighthouse Laboratory and Alex Alderton, Roberto Amato, Jeffrey Barrett, Sonia Goncalves, Ewan Harrison, David K. Jackson, Ian Johnston, Dominic Kwiatkowski, Cordelia Langford, John Sillitoe on behalf of the Wellcome Sanger Institute COVID-19 Surveillance Team |
| EPI_ISL_3364550, EPI_ISL_3364551, EPI_ISL_3364553 | Lighthouse Laboratory Plymouth | Wellcome Sanger Institute for the COVID-19 Genomics UK (COG-UK) Consortium | Lighthouse Laboratory Plymouth and Alex Alderton, Roberto Amato, Jeffrey Barrett, Sonia Goncalves, Ewan Harrison, David K. Jackson, Ian Johnston, Dominic Kwiatkowski, Cordelia Langford, John Sillitoe on behalf of the Wellcome Sanger Institute COVID-19 Surveillance Team |
| EPI_ISL_3364554, EPI_ISL_3364555, EPI_ISL_3364557, EPI_ISL_3364558, EPI_ISL_3364560, EPI_ISL_3364561, EPI_ISL_3364563, EPI_ISL_3364564 | Lighthouse Lab in Milton Keynes | Wellcome Sanger Institute for the COVID-19 Genomics UK (COG-UK) Consortium | The Lighthouse Lab in Milton Keynes and Alex Alderton, Roberto Amato, Jeffrey Barrett, Sonia Goncalves, Ewan Harrison, David K. Jackson, Ian Johnston, Dominic Kwiatkowski, Cordelia Langford, John Sillitoe on behalf of the Wellcome Sanger Institute COVID-19 Surveillance Team |
| EPI_ISL_3364566, EPI_ISL_3364567, EPI_ISL_3364568 | Lighthouse Laboratory Plymouth | Wellcome Sanger Institute for the COVID-19 Genomics UK (COG-UK) Consortium | Lighthouse Laboratory Plymouth and Alex Alderton, Roberto Amato, Jeffrey Barrett, Sonia Goncalves, Ewan Harrison, David K. Jackson, Ian Johnston, Dominic Kwiatkowski, Cordelia Langford, John Sillitoe on behalf of the Wellcome Sanger Institute COVID-19 Surveillance Team |
| EPI_ISL_3364570, EPI_ISL_3364571 | Lighthouse Lab in Milton Keynes | Wellcome Sanger Institute for the COVID-19 Genomics UK (COG-UK) Consortium | The Lighthouse Lab in Milton Keynes and Alex Alderton, Roberto Amato, Jeffrey Barrett, Sonia Goncalves, Ewan Harrison, David K. Jackson, Ian Johnston, Dominic Kwiatkowski, Cordelia Langford, John Sillitoe on behalf of the Wellcome Sanger Institute COVID-19 Surveillance Team |
| EPI_ISL_3364573, | Lighthouse Laboratory Plymouth | Wellcome Sanger Institute for the COVID-19 Genomics UK (COG-UK) | Lighthouse Laboratory Plymouth and Alex Alderton, Roberto Amato, Jeffrey Barrett, Sonia Goncalves, Ewan Harrison, David K. Jackson, Ian Johnston, Dominic Kwiatkowski, Cordelia Langford, John Sillitoe on behalf of the Wellcome Sanger |

| | | | |
|---|---|---|---|
| EPI_ISL_3364574, EPI_ISL_3364575 | | Consortium | Institute COVID-19 Surveillance Team |
| EPI_ISL_3364576 | Biotia | Biotia | Dorottya Nagy-Szakal, Mara Couto-Rodriguez, Xavier Jirau Serrano, Marilyne Debieu, David Danko, Christopher Mason, Niamh O'Hara |
| EPI_ISL_3364577, EPI_ISL_3364578, EPI_ISL_3364579, EPI_ISL_3364580 | Lighthouse Laboratory Plymouth | Wellcome Sanger Institute for the COVID-19 Genomics UK (COG-UK) Consortium | Lighthouse Laboratory Plymouth and Alex Alderton, Roberto Amato, Jeffrey Barrett, Sonia Goncalves, Ewan Harrison, David K. Jackson, Ian Johnston, Dominic Kwiatkowski, Cordelia Langford, John Sillitoe on behalf of the Wellcome Sanger Institute COVID-19 Surveillance Team |
| EPI_ISL_3364581, EPI_ISL_3364582 | Lighthouse Lab in Milton Keynes | Wellcome Sanger Institute for the COVID-19 Genomics UK (COG-UK) Consortium | The Lighthouse Lab in Milton Keynes and Alex Alderton, Roberto Amato, Jeffrey Barrett, Sonia Goncalves, Ewan Harrison, David K. Jackson, Ian Johnston, Dominic Kwiatkowski, Cordelia Langford, John Sillitoe on behalf of the Wellcome Sanger Institute COVID-19 Surveillance Team |
| EPI_ISL_3364583 | Lighthouse Laboratory Plymouth | Wellcome Sanger Institute for the COVID-19 Genomics UK (COG-UK) Consortium | Lighthouse Laboratory Plymouth and Alex Alderton, Roberto Amato, Jeffrey Barrett, Sonia Goncalves, Ewan Harrison, David K. Jackson, Ian Johnston, Dominic Kwiatkowski, Cordelia Langford, John Sillitoe on behalf of the Wellcome Sanger Institute COVID-19 Surveillance Team |
| EPI_ISL_3364584 | Biotia | Biotia | Dorottya Nagy-Szakal, Mara Couto-Rodriguez, Xavier Jirau Serrano, Marilyne Debieu, David Danko, Christopher Mason, Niamh O'Hara |
| EPI_ISL_3364585 | Lighthouse Laboratory Plymouth | Wellcome Sanger Institute for the COVID-19 Genomics UK (COG-UK) Consortium | Lighthouse Laboratory Plymouth and Alex Alderton, Roberto Amato, Jeffrey Barrett, Sonia Goncalves, Ewan Harrison, David K. Jackson, Ian Johnston, Dominic Kwiatkowski, Cordelia Langford, John Sillitoe on behalf of the Wellcome Sanger Institute COVID-19 Surveillance Team |
| EPI_ISL_3364586, EPI_ISL_3364587 | Lighthouse Lab in Milton Keynes | Wellcome Sanger Institute for the COVID-19 Genomics UK (COG-UK) Consortium | The Lighthouse Lab in Milton Keynes and Alex Alderton, Roberto Amato, Jeffrey Barrett, Sonia Goncalves, Ewan Harrison, David K. Jackson, Ian Johnston, Dominic Kwiatkowski, Cordelia Langford, John Sillitoe on behalf of the Wellcome Sanger Institute COVID-19 Surveillance Team |
| EPI_ISL_3364589, EPI_ISL_3364590, EPI_ISL_3364591 | Lighthouse Laboratory Plymouth | Wellcome Sanger Institute for the COVID-19 Genomics UK (COG-UK) Consortium | Lighthouse Laboratory Plymouth and Alex Alderton, Roberto Amato, Jeffrey Barrett, Sonia Goncalves, Ewan Harrison, David K. Jackson, Ian Johnston, Dominic Kwiatkowski, Cordelia Langford, John Sillitoe on behalf of the Wellcome Sanger Institute COVID-19 Surveillance Team |
| EPI_ISL_3364592 | Biotia | Biotia | Dorottya Nagy-Szakal, Mara Couto-Rodriguez, Xavier Jirau Serrano, Marilyne Debieu, David Danko, Christopher Mason, Niamh O'Hara |
| EPI_ISL_3364593 | Lighthouse Lab in Milton Keynes | Wellcome Sanger Institute for the COVID-19 Genomics UK (COG-UK) Consortium | The Lighthouse Lab in Milton Keynes and Alex Alderton, Roberto Amato, Jeffrey Barrett, Sonia Goncalves, Ewan Harrison, David K. Jackson, Ian Johnston, Dominic Kwiatkowski, Cordelia Langford, John Sillitoe on behalf of the Wellcome Sanger Institute COVID-19 Surveillance Team |
| EPI_ISL_3364594 | Lighthouse Laboratory Plymouth | Wellcome Sanger Institute for the COVID-19 Genomics UK (COG-UK) Consortium | Lighthouse Laboratory Plymouth and Alex Alderton, Roberto Amato, Jeffrey Barrett, Sonia Goncalves, Ewan Harrison, David K. Jackson, Ian Johnston, Dominic Kwiatkowski, Cordelia Langford, John Sillitoe on behalf of the Wellcome Sanger Institute COVID-19 Surveillance Team |
| EPI_ISL_3364595 | Biotia | Biotia | Dorottya Nagy-Szakal, Mara Couto-Rodriguez, Xavier Jirau Serrano, Marilyne Debieu, David Danko, Christopher Mason, Niamh O'Hara |
| EPI_ISL_3364596 | Lighthouse Laboratory Plymouth | Wellcome Sanger Institute for the COVID-19 Genomics UK (COG-UK) Consortium | Lighthouse Laboratory Plymouth and Alex Alderton, Roberto Amato, Jeffrey Barrett, Sonia Goncalves, Ewan Harrison, David K. Jackson, Ian Johnston, Dominic Kwiatkowski, Cordelia Langford, John Sillitoe on behalf of the Wellcome Sanger Institute COVID-19 Surveillance Team |
| EPI_ISL_3364597 | Lighthouse Lab in Milton Keynes | Wellcome Sanger Institute for the COVID-19 Genomics UK (COG-UK) Consortium | The Lighthouse Lab in Milton Keynes and Alex Alderton, Roberto Amato, Jeffrey Barrett, Sonia Goncalves, Ewan Harrison, David K. Jackson, Ian Johnston, Dominic Kwiatkowski, Cordelia Langford, John Sillitoe on behalf of the Wellcome Sanger Institute COVID-19 Surveillance Team |
| EPI_ISL_3364598 | Lighthouse Laboratory Plymouth | Wellcome Sanger Institute for the COVID-19 Genomics UK (COG-UK) Consortium | Lighthouse Laboratory Plymouth and Alex Alderton, Roberto Amato, Jeffrey Barrett, Sonia Goncalves, Ewan Harrison, David K. Jackson, Ian Johnston, Dominic Kwiatkowski, Cordelia Langford, John Sillitoe on behalf of the Wellcome Sanger Institute COVID-19 Surveillance Team |
| EPI_ISL_3364599 | Biotia | Biotia | Dorottya Nagy-Szakal, Mara Couto-Rodriguez, Xavier Jirau Serrano, Marilyne Debieu, David Danko, Christopher Mason, Niamh O'Hara |
| EPI_ISL_3364600, EPI_ISL_3364601, EPI_ISL_3364602 | Lighthouse Laboratory Plymouth | Wellcome Sanger Institute for the COVID-19 Genomics UK (COG-UK) Consortium | Lighthouse Laboratory Plymouth and Alex Alderton, Roberto Amato, Jeffrey Barrett, Sonia Goncalves, Ewan Harrison, David K. Jackson, Ian Johnston, Dominic Kwiatkowski, Cordelia Langford, John Sillitoe on behalf of the Wellcome Sanger Institute COVID-19 Surveillance Team |
| EPI_ISL_3364603, EPI_ISL_3364604 | Lighthouse Lab in Milton Keynes | Wellcome Sanger Institute for the COVID-19 Genomics UK (COG-UK) Consortium | The Lighthouse Lab in Milton Keynes and Alex Alderton, Roberto Amato, Jeffrey Barrett, Sonia Goncalves, Ewan Harrison, David K. Jackson, Ian Johnston, Dominic Kwiatkowski, Cordelia Langford, John Sillitoe on behalf of the Wellcome Sanger Institute COVID-19 Surveillance Team |
| EPI_ISL_3364605 | Lighthouse Laboratory Plymouth | Wellcome Sanger Institute for the COVID-19 Genomics UK (COG-UK) Consortium | Lighthouse Laboratory Plymouth and Alex Alderton, Roberto Amato, Jeffrey Barrett, Sonia Goncalves, Ewan Harrison, David K. Jackson, Ian Johnston, Dominic Kwiatkowski, Cordelia Langford, John Sillitoe on behalf of the Wellcome Sanger Institute COVID-19 Surveillance Team |
| EPI_ISL_3364606 | Lighthouse Lab in Milton Keynes | Wellcome Sanger Institute for the COVID-19 Genomics UK (COG-UK) Consortium | The Lighthouse Lab in Milton Keynes and Alex Alderton, Roberto Amato, Jeffrey Barrett, Sonia Goncalves, Ewan Harrison, David K. Jackson, Ian Johnston, Dominic Kwiatkowski, Cordelia Langford, John Sillitoe on behalf of the Wellcome Sanger Institute COVID-19 Surveillance Team |
| EPI_ISL_3364607 | Lighthouse Laboratory Plymouth | Wellcome Sanger Institute for the COVID-19 Genomics UK (COG-UK) Consortium | Lighthouse Laboratory Plymouth and Alex Alderton, Roberto Amato, Jeffrey Barrett, Sonia Goncalves, Ewan Harrison, David K. Jackson, Ian Johnston, Dominic Kwiatkowski, Cordelia Langford, John Sillitoe on behalf of the Wellcome Sanger Institute COVID-19 Surveillance Team |
| EPI_ISL_3364608, EPI_ISL_3364609 | Lighthouse Lab in Milton Keynes | Wellcome Sanger Institute for the COVID-19 Genomics UK (COG-UK) Consortium | The Lighthouse Lab in Milton Keynes and Alex Alderton, Roberto Amato, Jeffrey Barrett, Sonia Goncalves, Ewan Harrison, David K. Jackson, Ian Johnston, Dominic Kwiatkowski, Cordelia Langford, John Sillitoe on behalf of the Wellcome Sanger Institute COVID-19 Surveillance Team |
| EPI_ISL_3364610 | Lighthouse Laboratory Plymouth | Wellcome Sanger Institute for the COVID-19 Genomics UK (COG-UK) Consortium | Lighthouse Laboratory Plymouth and Alex Alderton, Roberto Amato, Jeffrey Barrett, Sonia Goncalves, Ewan Harrison, David K. Jackson, Ian Johnston, Dominic Kwiatkowski, Cordelia Langford, John Sillitoe on behalf of the Wellcome Sanger Institute COVID-19 Surveillance Team |
| EPI_ISL_3364611, EPI_ISL_3364612, EPI_ISL_3364613 | Lighthouse Lab in Milton Keynes | Wellcome Sanger Institute for the COVID-19 Genomics UK (COG-UK) Consortium | The Lighthouse Lab in Milton Keynes and Alex Alderton, Roberto Amato, Jeffrey Barrett, Sonia Goncalves, Ewan Harrison, David K. Jackson, Ian Johnston, Dominic Kwiatkowski, Cordelia Langford, John Sillitoe on behalf of the Wellcome Sanger Institute COVID-19 Surveillance Team |
| EPI_ISL_3364614, EPI_ISL_3364615 | Lighthouse Laboratory Plymouth | Wellcome Sanger Institute for the COVID-19 Genomics UK (COG-UK) Consortium | Lighthouse Laboratory Plymouth and Alex Alderton, Roberto Amato, Jeffrey Barrett, Sonia Goncalves, Ewan Harrison, David K. Jackson, Ian Johnston, Dominic Kwiatkowski, Cordelia Langford, John Sillitoe on behalf of the Wellcome Sanger Institute COVID-19 Surveillance Team |
| EPI_ISL_3364616 | Lighthouse Lab in Milton Keynes | Wellcome Sanger Institute for the COVID-19 Genomics UK (COG-UK) Consortium | The Lighthouse Lab in Milton Keynes and Alex Alderton, Roberto Amato, Jeffrey Barrett, Sonia Goncalves, Ewan Harrison, David K. Jackson, Ian Johnston, Dominic Kwiatkowski, Cordelia Langford, John Sillitoe on behalf of the Wellcome Sanger Institute COVID-19 Surveillance Team |
| EPI_ISL_3364617, EPI_ISL_3364618, EPI_ISL_3364619, EPI_ISL_3364620 | Lighthouse Laboratory Plymouth | Wellcome Sanger Institute for the COVID-19 Genomics UK (COG-UK) Consortium | Lighthouse Laboratory Plymouth and Alex Alderton, Roberto Amato, Jeffrey Barrett, Sonia Goncalves, Ewan Harrison, David K. Jackson, Ian Johnston, Dominic Kwiatkowski, Cordelia Langford, John Sillitoe on behalf of the Wellcome Sanger Institute COVID-19 Surveillance Team |
| EPI_ISL_3364621, EPI_ISL_3364622 | Lighthouse Lab in Milton Keynes | Wellcome Sanger Institute for the COVID-19 Genomics UK (COG-UK) Consortium | The Lighthouse Lab in Milton Keynes and Alex Alderton, Roberto Amato, Jeffrey Barrett, Sonia Goncalves, Ewan Harrison, David K. Jackson, Ian Johnston, Dominic Kwiatkowski, Cordelia Langford, John Sillitoe on behalf of the Wellcome Sanger Institute COVID-19 Surveillance Team |
| EPI_ISL_3364623, EPI_ISL_3364624 | Lighthouse Laboratory Plymouth | Wellcome Sanger Institute for the COVID-19 Genomics UK (COG-UK) Consortium | Lighthouse Laboratory Plymouth and Alex Alderton, Roberto Amato, Jeffrey Barrett, Sonia Goncalves, Ewan Harrison, David K. Jackson, Ian Johnston, Dominic Kwiatkowski, Cordelia Langford, John Sillitoe on behalf of the Wellcome Sanger Institute COVID-19 Surveillance Team |
| EPI_ISL_3364625, EPI_ISL_3364626 | Lighthouse Lab in Milton Keynes | Wellcome Sanger Institute for the COVID-19 Genomics UK (COG-UK) Consortium | The Lighthouse Lab in Milton Keynes and Alex Alderton, Roberto Amato, Jeffrey Barrett, Sonia Goncalves, Ewan Harrison, David K. Jackson, Ian Johnston, Dominic Kwiatkowski, Cordelia Langford, John Sillitoe on behalf of the Wellcome Sanger Institute COVID-19 Surveillance Team |
| EPI_ISL_3364627 | Lighthouse Laboratory Plymouth | Wellcome Sanger Institute for the COVID-19 Genomics UK (COG-UK) Consortium | Lighthouse Laboratory Plymouth and Alex Alderton, Roberto Amato, Jeffrey Barrett, Sonia Goncalves, Ewan Harrison, David K. Jackson, Ian Johnston, Dominic Kwiatkowski, Cordelia Langford, John Sillitoe on behalf of the Wellcome Sanger Institute COVID-19 Surveillance Team |
| EPI_ISL_3364628, EPI_ISL_3364629, EPI_ISL_3364630, EPI_ISL_3364631 | Lighthouse Lab in Milton Keynes | Wellcome Sanger Institute for the COVID-19 Genomics UK (COG-UK) Consortium | The Lighthouse Lab in Milton Keynes and Alex Alderton, Roberto Amato, Jeffrey Barrett, Sonia Goncalves, Ewan Harrison, David K. Jackson, Ian Johnston, Dominic Kwiatkowski, Cordelia Langford, John Sillitoe on behalf of the Wellcome Sanger Institute COVID-19 Surveillance Team |
| EPI_ISL_3364632 | Lighthouse Laboratory Plymouth | Wellcome Sanger Institute for the COVID-19 Genomics UK (COG-UK) Consortium | Lighthouse Laboratory Plymouth and Alex Alderton, Roberto Amato, Jeffrey Barrett, Sonia Goncalves, Ewan Harrison, David K. Jackson, Ian Johnston, Dominic Kwiatkowski, Cordelia Langford, John Sillitoe on behalf of the Wellcome Sanger Institute COVID-19 Surveillance Team |
| EPI_ISL_3364633 | Lighthouse Lab in Milton Keynes | Wellcome Sanger Institute for the COVID-19 Genomics UK (COG-UK) Consortium | The Lighthouse Lab in Milton Keynes and Alex Alderton, Roberto Amato, Jeffrey Barrett, Sonia Goncalves, Ewan Harrison, David K. Jackson, Ian Johnston, Dominic Kwiatkowski, Cordelia Langford, John Sillitoe on behalf of the Wellcome Sanger Institute COVID-19 Surveillance Team |
| EPI_ISL_3364634, EPI_ISL_3364635, EPI_ISL_3364636 | Lighthouse Laboratory Plymouth | Wellcome Sanger Institute for the COVID-19 Genomics UK (COG-UK) Consortium | Lighthouse Laboratory Plymouth and Alex Alderton, Roberto Amato, Jeffrey Barrett, Sonia Goncalves, Ewan Harrison, David K. Jackson, Ian Johnston, Dominic Kwiatkowski, Cordelia Langford, John Sillitoe on behalf of the Wellcome Sanger Institute COVID-19 Surveillance Team |
| EPI_ISL_3364637, EPI_ISL_3364638 | Lighthouse Lab in Milton Keynes | Wellcome Sanger Institute for the COVID-19 Genomics UK (COG-UK) Consortium | The Lighthouse Lab in Milton Keynes and Alex Alderton, Roberto Amato, Jeffrey Barrett, Sonia Goncalves, Ewan Harrison, David K. Jackson, Ian Johnston, Dominic Kwiatkowski, Cordelia Langford, John Sillitoe on behalf of the Wellcome Sanger Institute COVID-19 Surveillance Team |
| EPI_ISL_3364639, EPI_ISL_3364640 | Lighthouse Laboratory Plymouth | Wellcome Sanger Institute for the COVID-19 Genomics UK (COG-UK) Consortium | Lighthouse Laboratory Plymouth and Alex Alderton, Roberto Amato, Jeffrey Barrett, Sonia Goncalves, Ewan Harrison, David K. Jackson, Ian Johnston, Dominic Kwiatkowski, Cordelia Langford, John Sillitoe on behalf of the Wellcome Sanger Institute COVID-19 Surveillance Team |
| EPI_ISL_3364641 | Lighthouse Lab in Milton Keynes | Wellcome Sanger Institute for the COVID-19 Genomics UK (COG-UK) Consortium | The Lighthouse Lab in Milton Keynes and Alex Alderton, Roberto Amato, Jeffrey Barrett, Sonia Goncalves, Ewan Harrison, David K. Jackson, Ian Johnston, Dominic Kwiatkowski, Cordelia Langford, John Sillitoe on behalf of the Wellcome Sanger Institute COVID-19 Surveillance Team |
| EPI_ISL_3364642 | Lighthouse Laboratory Plymouth | Wellcome Sanger Institute for the COVID-19 Genomics UK (COG-UK) Consortium | Lighthouse Laboratory Plymouth and Alex Alderton, Roberto Amato, Jeffrey Barrett, Sonia Goncalves, Ewan Harrison, David K. Jackson, Ian Johnston, Dominic Kwiatkowski, Cordelia Langford, John Sillitoe on behalf of the Wellcome Sanger Institute COVID-19 Surveillance Team |
| EPI_ISL_3364643, EPI_ISL_3364644 | Lighthouse Lab in Milton Keynes | Wellcome Sanger Institute for the COVID-19 Genomics UK (COG-UK) Consortium | The Lighthouse Lab in Milton Keynes and Alex Alderton, Roberto Amato, Jeffrey Barrett, Sonia Goncalves, Ewan Harrison, David K. Jackson, Ian Johnston, Dominic Kwiatkowski, Cordelia Langford, John Sillitoe on behalf of the Wellcome Sanger Institute COVID-19 Surveillance Team |
| EPI_ISL_3364645 | Lighthouse Laboratory Plymouth | Wellcome Sanger Institute for the COVID-19 Genomics UK (COG-UK) Consortium | Lighthouse Laboratory Plymouth and Alex Alderton, Roberto Amato, Jeffrey Barrett, Sonia Goncalves, Ewan Harrison, David K. Jackson, Ian Johnston, Dominic Kwiatkowski, Cordelia Langford, John Sillitoe on behalf of the Wellcome Sanger Institute COVID-19 Surveillance Team |
| EPI_ISL_3364646 | Lighthouse Lab in Milton Keynes | Wellcome Sanger Institute for the COVID-19 Genomics UK (COG-UK) | The Lighthouse Lab in Milton Keynes and Alex Alderton, Roberto Amato, Jeffrey Barrett, Sonia Goncalves, Ewan Harrison, David K. Jackson, Ian Johnston, Dominic Kwiatkowski, Cordelia Langford, John Sillitoe on behalf of the Wellcome Sanger |

| | | | |
|---|---|---|---|
| | | Consortium | Institute COVID-19 Surveillance Team |
| EPI_ISL_3364647, EPI_ISL_3364648, EPI_ISL_3364649, EPI_ISL_3364650 | Lighthouse Laboratory Plymouth | Wellcome Sanger Institute for the COVID-19 Genomics UK (COG-UK) Consortium | Lighthouse Laboratory Plymouth and Alex Alderton, Roberto Amato, Jeffrey Barrett, Sonia Goncalves, Ewan Harrison, David K. Jackson, Ian Johnston, Dominic Kwiatkowski, Cordelia Langford, John Sillitoe on behalf of the Wellcome Sanger Institute COVID-19 Surveillance Team |
| EPI_ISL_3364651 | Lighthouse Lab in Milton Keynes | Wellcome Sanger Institute for the COVID-19 Genomics UK (COG-UK) Consortium | The Lighthouse Lab in Milton Keynes and Alex Alderton, Roberto Amato, Jeffrey Barrett, Sonia Goncalves, Ewan Harrison, David K. Jackson, Ian Johnston, Dominic Kwiatkowski, Cordelia Langford, John Sillitoe on behalf of the Wellcome Sanger Institute COVID-19 Surveillance Team |
| EPI_ISL_3364652, EPI_ISL_3364653 | Lighthouse Laboratory Plymouth | Wellcome Sanger Institute for the COVID-19 Genomics UK (COG-UK) Consortium | Lighthouse Laboratory Plymouth and Alex Alderton, Roberto Amato, Jeffrey Barrett, Sonia Goncalves, Ewan Harrison, David K. Jackson, Ian Johnston, Dominic Kwiatkowski, Cordelia Langford, John Sillitoe on behalf of the Wellcome Sanger Institute COVID-19 Surveillance Team |
| EPI_ISL_3364654 | Lighthouse Lab in Milton Keynes | Wellcome Sanger Institute for the COVID-19 Genomics UK (COG-UK) Consortium | The Lighthouse Lab in Milton Keynes and Alex Alderton, Roberto Amato, Jeffrey Barrett, Sonia Goncalves, Ewan Harrison, David K. Jackson, Ian Johnston, Dominic Kwiatkowski, Cordelia Langford, John Sillitoe on behalf of the Wellcome Sanger Institute COVID-19 Surveillance Team |
| EPI_ISL_3364655, EPI_ISL_3364656 | Lighthouse Laboratory Plymouth | Wellcome Sanger Institute for the COVID-19 Genomics UK (COG-UK) Consortium | Lighthouse Laboratory Plymouth and Alex Alderton, Roberto Amato, Jeffrey Barrett, Sonia Goncalves, Ewan Harrison, David K. Jackson, Ian Johnston, Dominic Kwiatkowski, Cordelia Langford, John Sillitoe on behalf of the Wellcome Sanger Institute COVID-19 Surveillance Team |
| EPI_ISL_3364657 | Lighthouse Lab in Milton Keynes | Wellcome Sanger Institute for the COVID-19 Genomics UK (COG-UK) Consortium | The Lighthouse Lab in Milton Keynes and Alex Alderton, Roberto Amato, Jeffrey Barrett, Sonia Goncalves, Ewan Harrison, David K. Jackson, Ian Johnston, Dominic Kwiatkowski, Cordelia Langford, John Sillitoe on behalf of the Wellcome Sanger Institute COVID-19 Surveillance Team |
| EPI_ISL_3364658, EPI_ISL_3364659 | Lighthouse Laboratory Plymouth | Wellcome Sanger Institute for the COVID-19 Genomics UK (COG-UK) Consortium | Lighthouse Laboratory Plymouth and Alex Alderton, Roberto Amato, Jeffrey Barrett, Sonia Goncalves, Ewan Harrison, David K. Jackson, Ian Johnston, Dominic Kwiatkowski, Cordelia Langford, John Sillitoe on behalf of the Wellcome Sanger Institute COVID-19 Surveillance Team |
| EPI_ISL_3364660, EPI_ISL_3364661, EPI_ISL_3364662 | Lighthouse Lab in Milton Keynes | Wellcome Sanger Institute for the COVID-19 Genomics UK (COG-UK) Consortium | The Lighthouse Lab in Milton Keynes and Alex Alderton, Roberto Amato, Jeffrey Barrett, Sonia Goncalves, Ewan Harrison, David K. Jackson, Ian Johnston, Dominic Kwiatkowski, Cordelia Langford, John Sillitoe on behalf of the Wellcome Sanger Institute COVID-19 Surveillance Team |
| EPI_ISL_3364663 | Lighthouse Laboratory Plymouth | Wellcome Sanger Institute for the COVID-19 Genomics UK (COG-UK) Consortium | Lighthouse Laboratory Plymouth and Alex Alderton, Roberto Amato, Jeffrey Barrett, Sonia Goncalves, Ewan Harrison, David K. Jackson, Ian Johnston, Dominic Kwiatkowski, Cordelia Langford, John Sillitoe on behalf of the Wellcome Sanger Institute COVID-19 Surveillance Team |
| EPI_ISL_3364664, EPI_ISL_3364665, EPI_ISL_3364666 | Lighthouse Lab in Milton Keynes | Wellcome Sanger Institute for the COVID-19 Genomics UK (COG-UK) Consortium | The Lighthouse Lab in Milton Keynes and Alex Alderton, Roberto Amato, Jeffrey Barrett, Sonia Goncalves, Ewan Harrison, David K. Jackson, Ian Johnston, Dominic Kwiatkowski, Cordelia Langford, John Sillitoe on behalf of the Wellcome Sanger Institute COVID-19 Surveillance Team |
| EPI_ISL_3364667 | Lighthouse Laboratory Plymouth | Wellcome Sanger Institute for the COVID-19 Genomics UK (COG-UK) Consortium | Lighthouse Laboratory Plymouth and Alex Alderton, Roberto Amato, Jeffrey Barrett, Sonia Goncalves, Ewan Harrison, David K. Jackson, Ian Johnston, Dominic Kwiatkowski, Cordelia Langford, John Sillitoe on behalf of the Wellcome Sanger Institute COVID-19 Surveillance Team |
| EPI_ISL_3364668 | Lighthouse Lab in Milton Keynes | Wellcome Sanger Institute for the COVID-19 Genomics UK (COG-UK) Consortium | The Lighthouse Lab in Milton Keynes and Alex Alderton, Roberto Amato, Jeffrey Barrett, Sonia Goncalves, Ewan Harrison, David K. Jackson, Ian Johnston, Dominic Kwiatkowski, Cordelia Langford, John Sillitoe on behalf of the Wellcome Sanger Institute COVID-19 Surveillance Team |
| EPI_ISL_3364669, EPI_ISL_3364670, EPI_ISL_3364671, EPI_ISL_3364672 | Lighthouse Laboratory Plymouth | Wellcome Sanger Institute for the COVID-19 Genomics UK (COG-UK) Consortium | Lighthouse Laboratory Plymouth and Alex Alderton, Roberto Amato, Jeffrey Barrett, Sonia Goncalves, Ewan Harrison, David K. Jackson, Ian Johnston, Dominic Kwiatkowski, Cordelia Langford, John Sillitoe on behalf of the Wellcome Sanger Institute COVID-19 Surveillance Team |
| EPI_ISL_3364673 | Lighthouse Lab in Milton Keynes | Wellcome Sanger Institute for the COVID-19 Genomics UK (COG-UK) Consortium | The Lighthouse Lab in Milton Keynes and Alex Alderton, Roberto Amato, Jeffrey Barrett, Sonia Goncalves, Ewan Harrison, David K. Jackson, Ian Johnston, Dominic Kwiatkowski, Cordelia Langford, John Sillitoe on behalf of the Wellcome Sanger Institute COVID-19 Surveillance Team |
| EPI_ISL_3364674 | Lighthouse Laboratory Plymouth | Wellcome Sanger Institute for the COVID-19 Genomics UK (COG-UK) Consortium | Lighthouse Laboratory Plymouth and Alex Alderton, Roberto Amato, Jeffrey Barrett, Sonia Goncalves, Ewan Harrison, David K. Jackson, Ian Johnston, Dominic Kwiatkowski, Cordelia Langford, John Sillitoe on behalf of the Wellcome Sanger Institute COVID-19 Surveillance Team |
| EPI_ISL_3364675, EPI_ISL_3364676, EPI_ISL_3364677 | Lighthouse Lab in Milton Keynes | Wellcome Sanger Institute for the COVID-19 Genomics UK (COG-UK) Consortium | The Lighthouse Lab in Milton Keynes and Alex Alderton, Roberto Amato, Jeffrey Barrett, Sonia Goncalves, Ewan Harrison, David K. Jackson, Ian Johnston, Dominic Kwiatkowski, Cordelia Langford, John Sillitoe on behalf of the Wellcome Sanger Institute COVID-19 Surveillance Team |
| EPI_ISL_3364678 | Lighthouse Laboratory Plymouth | Wellcome Sanger Institute for the COVID-19 Genomics UK (COG-UK) Consortium | Lighthouse Laboratory Plymouth and Alex Alderton, Roberto Amato, Jeffrey Barrett, Sonia Goncalves, Ewan Harrison, David K. Jackson, Ian Johnston, Dominic Kwiatkowski, Cordelia Langford, John Sillitoe on behalf of the Wellcome Sanger Institute COVID-19 Surveillance Team |
| EPI_ISL_3364679, EPI_ISL_3364680, EPI_ISL_3364681, EPI_ISL_3364682, EPI_ISL_3364683, EPI_ISL_3364684 | Lighthouse Lab in Milton Keynes | Wellcome Sanger Institute for the COVID-19 Genomics UK (COG-UK) Consortium | The Lighthouse Lab in Milton Keynes and Alex Alderton, Roberto Amato, Jeffrey Barrett, Sonia Goncalves, Ewan Harrison, David K. Jackson, Ian Johnston, Dominic Kwiatkowski, Cordelia Langford, John Sillitoe on behalf of the Wellcome Sanger Institute COVID-19 Surveillance Team |
| EPI_ISL_3364685, EPI_ISL_3364686 | Lighthouse Laboratory Plymouth | Wellcome Sanger Institute for the COVID-19 Genomics UK (COG-UK) Consortium | Lighthouse Laboratory Plymouth and Alex Alderton, Roberto Amato, Jeffrey Barrett, Sonia Goncalves, Ewan Harrison, David K. Jackson, Ian Johnston, Dominic Kwiatkowski, Cordelia Langford, John Sillitoe on behalf of the Wellcome Sanger Institute COVID-19 Surveillance Team |
| EPI_ISL_3364687, EPI_ISL_3364688 | Lighthouse Lab in Milton Keynes | Wellcome Sanger Institute for the COVID-19 Genomics UK (COG-UK) Consortium | The Lighthouse Lab in Milton Keynes and Alex Alderton, Roberto Amato, Jeffrey Barrett, Sonia Goncalves, Ewan Harrison, David K. Jackson, Ian Johnston, Dominic Kwiatkowski, Cordelia Langford, John Sillitoe on behalf of the Wellcome Sanger Institute COVID-19 Surveillance Team |
| EPI_ISL_3364689 | Lighthouse Laboratory Plymouth | Wellcome Sanger Institute for the COVID-19 Genomics UK (COG-UK) Consortium | Lighthouse Laboratory Plymouth and Alex Alderton, Roberto Amato, Jeffrey Barrett, Sonia Goncalves, Ewan Harrison, David K. Jackson, Ian Johnston, Dominic Kwiatkowski, Cordelia Langford, John Sillitoe on behalf of the Wellcome Sanger Institute COVID-19 Surveillance Team |
| EPI_ISL_3364690 | Lighthouse Lab in Milton Keynes | Wellcome Sanger Institute for the COVID-19 Genomics UK (COG-UK) Consortium | The Lighthouse Lab in Milton Keynes and Alex Alderton, Roberto Amato, Jeffrey Barrett, Sonia Goncalves, Ewan Harrison, David K. Jackson, Ian Johnston, Dominic Kwiatkowski, Cordelia Langford, John Sillitoe on behalf of the Wellcome Sanger Institute COVID-19 Surveillance Team |
| EPI_ISL_3364691, EPI_ISL_3364692 | Lighthouse Laboratory Plymouth | Wellcome Sanger Institute for the COVID-19 Genomics UK (COG-UK) Consortium | Lighthouse Laboratory Plymouth and Alex Alderton, Roberto Amato, Jeffrey Barrett, Sonia Goncalves, Ewan Harrison, David K. Jackson, Ian Johnston, Dominic Kwiatkowski, Cordelia Langford, John Sillitoe on behalf of the Wellcome Sanger Institute COVID-19 Surveillance Team |
| EPI_ISL_3364693, EPI_ISL_3364694, EPI_ISL_3364695 | Lighthouse Lab in Milton Keynes | Wellcome Sanger Institute for the COVID-19 Genomics UK (COG-UK) Consortium | The Lighthouse Lab in Milton Keynes and Alex Alderton, Roberto Amato, Jeffrey Barrett, Sonia Goncalves, Ewan Harrison, David K. Jackson, Ian Johnston, Dominic Kwiatkowski, Cordelia Langford, John Sillitoe on behalf of the Wellcome Sanger Institute COVID-19 Surveillance Team |
| EPI_ISL_3364696 | Lighthouse Laboratory Plymouth | Wellcome Sanger Institute for the COVID-19 Genomics UK (COG-UK) Consortium | Lighthouse Laboratory Plymouth and Alex Alderton, Roberto Amato, Jeffrey Barrett, Sonia Goncalves, Ewan Harrison, David K. Jackson, Ian Johnston, Dominic Kwiatkowski, Cordelia Langford, John Sillitoe on behalf of the Wellcome Sanger Institute COVID-19 Surveillance Team |
| EPI_ISL_3364697 | Lighthouse Lab in Milton Keynes | Wellcome Sanger Institute for the COVID-19 Genomics UK (COG-UK) Consortium | The Lighthouse Lab in Milton Keynes and Alex Alderton, Roberto Amato, Jeffrey Barrett, Sonia Goncalves, Ewan Harrison, David K. Jackson, Ian Johnston, Dominic Kwiatkowski, Cordelia Langford, John Sillitoe on behalf of the Wellcome Sanger Institute COVID-19 Surveillance Team |
| EPI_ISL_3364698 | Lighthouse Laboratory Plymouth | Wellcome Sanger Institute for the COVID-19 Genomics UK (COG-UK) Consortium | Lighthouse Laboratory Plymouth and Alex Alderton, Roberto Amato, Jeffrey Barrett, Sonia Goncalves, Ewan Harrison, David K. Jackson, Ian Johnston, Dominic Kwiatkowski, Cordelia Langford, John Sillitoe on behalf of the Wellcome Sanger Institute COVID-19 Surveillance Team |
| EPI_ISL_3364699, EPI_ISL_3364700 | Lighthouse Lab in Milton Keynes | Wellcome Sanger Institute for the COVID-19 Genomics UK (COG-UK) Consortium | The Lighthouse Lab in Milton Keynes and Alex Alderton, Roberto Amato, Jeffrey Barrett, Sonia Goncalves, Ewan Harrison, David K. Jackson, Ian Johnston, Dominic Kwiatkowski, Cordelia Langford, John Sillitoe on behalf of the Wellcome Sanger Institute COVID-19 Surveillance Team |
| EPI_ISL_3364701 | Lighthouse Laboratory Plymouth | Wellcome Sanger Institute for the COVID-19 Genomics UK (COG-UK) Consortium | Lighthouse Laboratory Plymouth and Alex Alderton, Roberto Amato, Jeffrey Barrett, Sonia Goncalves, Ewan Harrison, David K. Jackson, Ian Johnston, Dominic Kwiatkowski, Cordelia Langford, John Sillitoe on behalf of the Wellcome Sanger Institute COVID-19 Surveillance Team |
| EPI_ISL_3364702 | Lighthouse Lab in Milton Keynes | Wellcome Sanger Institute for the COVID-19 Genomics UK (COG-UK) Consortium | The Lighthouse Lab in Milton Keynes and Alex Alderton, Roberto Amato, Jeffrey Barrett, Sonia Goncalves, Ewan Harrison, David K. Jackson, Ian Johnston, Dominic Kwiatkowski, Cordelia Langford, John Sillitoe on behalf of the Wellcome Sanger Institute COVID-19 Surveillance Team |
| EPI_ISL_3364703 | Lighthouse Laboratory Plymouth | Wellcome Sanger Institute for the COVID-19 Genomics UK (COG-UK) Consortium | Lighthouse Laboratory Plymouth and Alex Alderton, Roberto Amato, Jeffrey Barrett, Sonia Goncalves, Ewan Harrison, David K. Jackson, Ian Johnston, Dominic Kwiatkowski, Cordelia Langford, John Sillitoe on behalf of the Wellcome Sanger Institute COVID-19 Surveillance Team |
| EPI_ISL_3364704, EPI_ISL_3364705 | Lighthouse Lab in Milton Keynes | Wellcome Sanger Institute for the COVID-19 Genomics UK (COG-UK) Consortium | The Lighthouse Lab in Milton Keynes and Alex Alderton, Roberto Amato, Jeffrey Barrett, Sonia Goncalves, Ewan Harrison, David K. Jackson, Ian Johnston, Dominic Kwiatkowski, Cordelia Langford, John Sillitoe on behalf of the Wellcome Sanger Institute COVID-19 Surveillance Team |
| EPI_ISL_3364706, EPI_ISL_3364707, EPI_ISL_3364708, EPI_ISL_3364709 | Lighthouse Laboratory Plymouth | Wellcome Sanger Institute for the COVID-19 Genomics UK (COG-UK) Consortium | Lighthouse Laboratory Plymouth and Alex Alderton, Roberto Amato, Jeffrey Barrett, Sonia Goncalves, Ewan Harrison, David K. Jackson, Ian Johnston, Dominic Kwiatkowski, Cordelia Langford, John Sillitoe on behalf of the Wellcome Sanger Institute COVID-19 Surveillance Team |
| EPI_ISL_3364710 | Lighthouse Lab in Milton Keynes | Wellcome Sanger Institute for the COVID-19 Genomics UK (COG-UK) Consortium | The Lighthouse Lab in Milton Keynes and Alex Alderton, Roberto Amato, Jeffrey Barrett, Sonia Goncalves, Ewan Harrison, David K. Jackson, Ian Johnston, Dominic Kwiatkowski, Cordelia Langford, John Sillitoe on behalf of the Wellcome Sanger Institute COVID-19 Surveillance Team |
| EPI_ISL_3364711, EPI_ISL_3364712, EPI_ISL_3364713 | Lighthouse Laboratory Plymouth | Wellcome Sanger Institute for the COVID-19 Genomics UK (COG-UK) Consortium | Lighthouse Laboratory Plymouth and Alex Alderton, Roberto Amato, Jeffrey Barrett, Sonia Goncalves, Ewan Harrison, David K. Jackson, Ian Johnston, Dominic Kwiatkowski, Cordelia Langford, John Sillitoe on behalf of the Wellcome Sanger Institute COVID-19 Surveillance Team |
| EPI_ISL_3364714 | Lighthouse Lab in Milton Keynes | Wellcome Sanger Institute for the COVID-19 Genomics UK (COG-UK) Consortium | The Lighthouse Lab in Milton Keynes and Alex Alderton, Roberto Amato, Jeffrey Barrett, Sonia Goncalves, Ewan Harrison, David K. Jackson, Ian Johnston, Dominic Kwiatkowski, Cordelia Langford, John Sillitoe on behalf of the Wellcome Sanger Institute COVID-19 Surveillance Team |
| EPI_ISL_3364715 | Lighthouse Laboratory Plymouth | Wellcome Sanger Institute for the COVID-19 Genomics UK (COG-UK) Consortium | Lighthouse Laboratory Plymouth and Alex Alderton, Roberto Amato, Jeffrey Barrett, Sonia Goncalves, Ewan Harrison, David K. Jackson, Ian Johnston, Dominic Kwiatkowski, Cordelia Langford, John Sillitoe on behalf of the Wellcome Sanger Institute COVID-19 Surveillance Team |
| EPI_ISL_3364716 | Lighthouse Lab in Milton Keynes | Wellcome Sanger Institute for the COVID-19 Genomics UK (COG-UK) Consortium | The Lighthouse Lab in Milton Keynes and Alex Alderton, Roberto Amato, Jeffrey Barrett, Sonia Goncalves, Ewan Harrison, David K. Jackson, Ian Johnston, Dominic Kwiatkowski, Cordelia Langford, John Sillitoe on behalf of the Wellcome Sanger Institute COVID-19 Surveillance Team |
| EPI_ISL_3364717 | Lighthouse Laboratory Plymouth | Wellcome Sanger Institute for the COVID-19 Genomics UK (COG-UK) | Lighthouse Laboratory Plymouth and Alex Alderton, Roberto Amato, Jeffrey Barrett, Sonia Goncalves, Ewan Harrison, David K. Jackson, Ian Johnston, Dominic Kwiatkowski, Cordelia Langford, John Sillitoe on behalf of the Wellcome Sanger |

| | | | |
|---|---|---|---|
| | | Consortium | Institute COVID-19 Surveillance Team |
| EPI_ISL_3364718, EPI_ISL_3364719 | Lighthouse Lab in Milton Keynes | Wellcome Sanger Institute for the COVID-19 Genomics UK (COG-UK) Consortium | The Lighthouse Lab in Milton Keynes and Alex Alderton, Roberto Amato, Jeffrey Barrett, Sonia Goncalves, Ewan Harrison, David K. Jackson, Ian Johnston, Dominic Kwiatkowski, Cordelia Langford, John Sillitoe on behalf of the Wellcome Sanger Institute COVID-19 Surveillance Team |
| EPI_ISL_3364720, EPI_ISL_3364721 | Lighthouse Laboratory Plymouth | Wellcome Sanger Institute for the COVID-19 Genomics UK (COG-UK) Consortium | Lighthouse Laboratory Plymouth and Alex Alderton, Roberto Amato, Jeffrey Barrett, Sonia Goncalves, Ewan Harrison, David K. Jackson, Ian Johnston, Dominic Kwiatkowski, Cordelia Langford, John Sillitoe on behalf of the Wellcome Sanger Institute COVID-19 Surveillance Team |
| EPI_ISL_3364722 | Lighthouse Lab in Milton Keynes | Wellcome Sanger Institute for the COVID-19 Genomics UK (COG-UK) Consortium | The Lighthouse Lab in Milton Keynes and Alex Alderton, Roberto Amato, Jeffrey Barrett, Sonia Goncalves, Ewan Harrison, David K. Jackson, Ian Johnston, Dominic Kwiatkowski, Cordelia Langford, John Sillitoe on behalf of the Wellcome Sanger Institute COVID-19 Surveillance Team |
| EPI_ISL_3364723 | Lighthouse Laboratory Plymouth | Wellcome Sanger Institute for the COVID-19 Genomics UK (COG-UK) Consortium | Lighthouse Laboratory Plymouth and Alex Alderton, Roberto Amato, Jeffrey Barrett, Sonia Goncalves, Ewan Harrison, David K. Jackson, Ian Johnston, Dominic Kwiatkowski, Cordelia Langford, John Sillitoe on behalf of the Wellcome Sanger Institute COVID-19 Surveillance Team |
| EPI_ISL_3364724 | Lighthouse Lab in Milton Keynes | Wellcome Sanger Institute for the COVID-19 Genomics UK (COG-UK) Consortium | The Lighthouse Lab in Milton Keynes and Alex Alderton, Roberto Amato, Jeffrey Barrett, Sonia Goncalves, Ewan Harrison, David K. Jackson, Ian Johnston, Dominic Kwiatkowski, Cordelia Langford, John Sillitoe on behalf of the Wellcome Sanger Institute COVID-19 Surveillance Team |
| EPI_ISL_3364725, EPI_ISL_3364726 | Lighthouse Laboratory Plymouth | Wellcome Sanger Institute for the COVID-19 Genomics UK (COG-UK) Consortium | Lighthouse Laboratory Plymouth and Alex Alderton, Roberto Amato, Jeffrey Barrett, Sonia Goncalves, Ewan Harrison, David K. Jackson, Ian Johnston, Dominic Kwiatkowski, Cordelia Langford, John Sillitoe on behalf of the Wellcome Sanger Institute COVID-19 Surveillance Team |
| EPI_ISL_3364727, EPI_ISL_3364728, EPI_ISL_3364729, EPI_ISL_3364730, EPI_ISL_3364731, EPI_ISL_3364732 | Lighthouse Lab in Milton Keynes | Wellcome Sanger Institute for the COVID-19 Genomics UK (COG-UK) Consortium | The Lighthouse Lab in Milton Keynes and Alex Alderton, Roberto Amato, Jeffrey Barrett, Sonia Goncalves, Ewan Harrison, David K. Jackson, Ian Johnston, Dominic Kwiatkowski, Cordelia Langford, John Sillitoe on behalf of the Wellcome Sanger Institute COVID-19 Surveillance Team |
| EPI_ISL_3364733, EPI_ISL_3364734, EPI_ISL_3364735 | Lighthouse Laboratory Plymouth | Wellcome Sanger Institute for the COVID-19 Genomics UK (COG-UK) Consortium | Lighthouse Laboratory Plymouth and Alex Alderton, Roberto Amato, Jeffrey Barrett, Sonia Goncalves, Ewan Harrison, David K. Jackson, Ian Johnston, Dominic Kwiatkowski, Cordelia Langford, John Sillitoe on behalf of the Wellcome Sanger Institute COVID-19 Surveillance Team |
| EPI_ISL_3364736, EPI_ISL_3364737, EPI_ISL_3364738 | Lighthouse Lab in Milton Keynes | Wellcome Sanger Institute for the COVID-19 Genomics UK (COG-UK) Consortium | The Lighthouse Lab in Milton Keynes and Alex Alderton, Roberto Amato, Jeffrey Barrett, Sonia Goncalves, Ewan Harrison, David K. Jackson, Ian Johnston, Dominic Kwiatkowski, Cordelia Langford, John Sillitoe on behalf of the Wellcome Sanger Institute COVID-19 Surveillance Team |
| EPI_ISL_3364739 | Lighthouse Lab in Alderley Park | Wellcome Sanger Institute for the COVID-19 Genomics UK (COG-UK) Consortium | Jacquelyn Wynn, Mairead Hyland, The Lighthouse Lab in Alderley Park and Alex Alderton, Roberto Amato, Jeffrey Barrett, Sonia Goncalves, Ewan Harrison, David K. Jackson, Ian Johnston, Dominic Kwiatkowski, Cordelia Langford, John Sillitoe on behalf of the Wellcome Sanger Institute COVID-19 Surveillance Team |
| EPI_ISL_3364740, EPI_ISL_3364741, EPI_ISL_3364742, EPI_ISL_3364743, EPI_ISL_3364744, EPI_ISL_3364745, EPI_ISL_3364746, EPI_ISL_3364747, EPI_ISL_3364748, EPI_ISL_3364749, EPI_ISL_3364750, EPI_ISL_3364751, EPI_ISL_3364752, EPI_ISL_3364753, EPI_ISL_3364754, EPI_ISL_3364755, EPI_ISL_3364756 | | | |
| see above | Lighthouse Lab in Milton Keynes | Wellcome Sanger Institute for the COVID-19 Genomics UK (COG-UK) Consortium | The Lighthouse Lab in Milton Keynes and Alex Alderton, Roberto Amato, Jeffrey Barrett, Sonia Goncalves, Ewan Harrison, David K. Jackson, Ian Johnston, Dominic Kwiatkowski, Cordelia Langford, John Sillitoe on behalf of the Wellcome Sanger Institute COVID-19 Surveillance Team |
| EPI_ISL_3364757 | Lighthouse Lab in Alderley Park | Wellcome Sanger Institute for the COVID-19 Genomics UK (COG-UK) Consortium | Jacquelyn Wynn, Mairead Hyland, The Lighthouse Lab in Alderley Park and Alex Alderton, Roberto Amato, Jeffrey Barrett, Sonia Goncalves, Ewan Harrison, David K. Jackson, Ian Johnston, Dominic Kwiatkowski, Cordelia Langford, John Sillitoe on behalf of the Wellcome Sanger Institute COVID-19 Surveillance Team |
| EPI_ISL_3364758, EPI_ISL_3364759 | Lighthouse Lab in Milton Keynes | Wellcome Sanger Institute for the COVID-19 Genomics UK (COG-UK) Consortium | The Lighthouse Lab in Milton Keynes and Alex Alderton, Roberto Amato, Jeffrey Barrett, Sonia Goncalves, Ewan Harrison, David K. Jackson, Ian Johnston, Dominic Kwiatkowski, Cordelia Langford, John Sillitoe on behalf of the Wellcome Sanger Institute COVID-19 Surveillance Team |
| EPI_ISL_3364760 | Lighthouse Lab in Alderley Park | Wellcome Sanger Institute for the COVID-19 Genomics UK (COG-UK) Consortium | Jacquelyn Wynn, Mairead Hyland, The Lighthouse Lab in Alderley Park and Alex Alderton, Roberto Amato, Jeffrey Barrett, Sonia Goncalves, Ewan Harrison, David K. Jackson, Ian Johnston, Dominic Kwiatkowski, Cordelia Langford, John Sillitoe on behalf of the Wellcome Sanger Institute COVID-19 Surveillance Team |
| EPI_ISL_3364761, EPI_ISL_3364762, EPI_ISL_3364763, EPI_ISL_3364764, EPI_ISL_3364765, EPI_ISL_3364766, EPI_ISL_3364767, EPI_ISL_3364768, EPI_ISL_3364769, EPI_ISL_3364770, EPI_ISL_3364771, EPI_ISL_3364772, EPI_ISL_3364773, EPI_ISL_3364774 | | | |
| see above | Lighthouse Lab in Milton Keynes | Wellcome Sanger Institute for the COVID-19 Genomics UK (COG-UK) Consortium | The Lighthouse Lab in Milton Keynes and Alex Alderton, Roberto Amato, Jeffrey Barrett, Sonia Goncalves, Ewan Harrison, David K. Jackson, Ian Johnston, Dominic Kwiatkowski, Cordelia Langford, John Sillitoe on behalf of the Wellcome Sanger Institute COVID-19 Surveillance Team |
| EPI_ISL_3364775 | Lighthouse Lab in Alderley Park | Wellcome Sanger Institute for the COVID-19 Genomics UK (COG-UK) Consortium | Jacquelyn Wynn, Mairead Hyland, The Lighthouse Lab in Alderley Park and Alex Alderton, Roberto Amato, Jeffrey Barrett, Sonia Goncalves, Ewan Harrison, David K. Jackson, Ian Johnston, Dominic Kwiatkowski, Cordelia Langford, John Sillitoe on behalf of the Wellcome Sanger Institute COVID-19 Surveillance Team |
| EPI_ISL_3364776, EPI_ISL_3364777 | Lighthouse Lab in Milton Keynes | Wellcome Sanger Institute for the COVID-19 Genomics UK (COG-UK) Consortium | The Lighthouse Lab in Milton Keynes and Alex Alderton, Roberto Amato, Jeffrey Barrett, Sonia Goncalves, Ewan Harrison, David K. Jackson, Ian Johnston, Dominic Kwiatkowski, Cordelia Langford, John Sillitoe on behalf of the Wellcome Sanger Institute COVID-19 Surveillance Team |
| EPI_ISL_3364778 | Lighthouse Lab in Alderley Park | Wellcome Sanger Institute for the COVID-19 Genomics UK (COG-UK) Consortium | Jacquelyn Wynn, Mairead Hyland, The Lighthouse Lab in Alderley Park and Alex Alderton, Roberto Amato, Jeffrey Barrett, Sonia Goncalves, Ewan Harrison, David K. Jackson, Ian Johnston, Dominic Kwiatkowski, Cordelia Langford, John Sillitoe on behalf of the Wellcome Sanger Institute COVID-19 Surveillance Team |
| EPI_ISL_3364779, EPI_ISL_3364780, EPI_ISL_3364781, EPI_ISL_3364782, EPI_ISL_3364783, EPI_ISL_3364784, EPI_ISL_3364785, EPI_ISL_3364786, EPI_ISL_3364787, EPI_ISL_3364788, EPI_ISL_3364789, EPI_ISL_3364790, EPI_ISL_3364791, EPI_ISL_3364792, EPI_ISL_3364793, EPI_ISL_3364794, EPI_ISL_3364795, EPI_ISL_3364796, EPI_ISL_3364797, EPI_ISL_3364798, EPI_ISL_3364799, EPI_ISL_3364800, EPI_ISL_3364801, EPI_ISL_3364802, EPI_ISL_3364803, EPI_ISL_3364804, EPI_ISL_3364805, EPI_ISL_3364806, EPI_ISL_3364807, EPI_ISL_3364808, EPI_ISL_3364809, EPI_ISL_3364810, EPI_ISL_3364811, EPI_ISL_3364812, EPI_ISL_3364813, EPI_ISL_3364814, EPI_ISL_3364815, EPI_ISL_3364816, EPI_ISL_3364817, EPI_ISL_3364818 | | | |
| see above | Lighthouse Lab in Milton Keynes | Wellcome Sanger Institute for the COVID-19 Genomics UK (COG-UK) Consortium | The Lighthouse Lab in Milton Keynes and Alex Alderton, Roberto Amato, Jeffrey Barrett, Sonia Goncalves, Ewan Harrison, David K. Jackson, Ian Johnston, Dominic Kwiatkowski, Cordelia Langford, John Sillitoe on behalf of the Wellcome Sanger Institute COVID-19 Surveillance Team |
| EPI_ISL_3364819 | Lighthouse Lab in Alderley Park | Wellcome Sanger Institute for the COVID-19 Genomics UK (COG-UK) Consortium | Jacquelyn Wynn, Mairead Hyland, The Lighthouse Lab in Alderley Park and Alex Alderton, Roberto Amato, Jeffrey Barrett, Sonia Goncalves, Ewan Harrison, David K. Jackson, Ian Johnston, Dominic Kwiatkowski, Cordelia Langford, John Sillitoe on behalf of the Wellcome Sanger Institute COVID-19 Surveillance Team |
| EPI_ISL_3364820, EPI_ISL_3364821, EPI_ISL_3364822, EPI_ISL_3364823, EPI_ISL_3364824, EPI_ISL_3364825, EPI_ISL_3364826, EPI_ISL_3364827, EPI_ISL_3364828, EPI_ISL_3364829, EPI_ISL_3364830, EPI_ISL_3364831, EPI_ISL_3364832, EPI_ISL_3364833, EPI_ISL_3364834, EPI_ISL_3364835 | | | |
| see above | Lighthouse Lab in Milton Keynes | Wellcome Sanger Institute for the COVID-19 Genomics UK (COG-UK) Consortium | The Lighthouse Lab in Milton Keynes and Alex Alderton, Roberto Amato, Jeffrey Barrett, Sonia Goncalves, Ewan Harrison, David K. Jackson, Ian Johnston, Dominic Kwiatkowski, Cordelia Langford, John Sillitoe on behalf of the Wellcome Sanger Institute COVID-19 Surveillance Team |
| EPI_ISL_3364836 | Lighthouse Lab in Alderley Park | Wellcome Sanger Institute for the COVID-19 Genomics UK (COG-UK) Consortium | Jacquelyn Wynn, Mairead Hyland, The Lighthouse Lab in Alderley Park and Alex Alderton, Roberto Amato, Jeffrey Barrett, Sonia Goncalves, Ewan Harrison, David K. Jackson, Ian Johnston, Dominic Kwiatkowski, Cordelia Langford, John Sillitoe on behalf of the Wellcome Sanger Institute COVID-19 Surveillance Team |
| EPI_ISL_3364837, EPI_ISL_3364838, EPI_ISL_3364839, EPI_ISL_3364840 | Lighthouse Lab in Milton Keynes | Wellcome Sanger Institute for the COVID-19 Genomics UK (COG-UK) Consortium | The Lighthouse Lab in Milton Keynes and Alex Alderton, Roberto Amato, Jeffrey Barrett, Sonia Goncalves, Ewan Harrison, David K. Jackson, Ian Johnston, Dominic Kwiatkowski, Cordelia Langford, John Sillitoe on behalf of the Wellcome Sanger Institute COVID-19 Surveillance Team |
| EPI_ISL_3364841 | Lighthouse Lab in Alderley Park | Wellcome Sanger Institute for the COVID-19 Genomics UK (COG-UK) Consortium | Jacquelyn Wynn, Mairead Hyland, The Lighthouse Lab in Alderley Park and Alex Alderton, Roberto Amato, Jeffrey Barrett, Sonia Goncalves, Ewan Harrison, David K. Jackson, Ian Johnston, Dominic Kwiatkowski, Cordelia Langford, John Sillitoe on behalf of the Wellcome Sanger Institute COVID-19 Surveillance Team |
| EPI_ISL_3364842, EPI_ISL_3364843, EPI_ISL_3364844 | Lighthouse Lab in Milton Keynes | Wellcome Sanger Institute for the COVID-19 Genomics UK (COG-UK) Consortium | The Lighthouse Lab in Milton Keynes and Alex Alderton, Roberto Amato, Jeffrey Barrett, Sonia Goncalves, Ewan Harrison, David K. Jackson, Ian Johnston, Dominic Kwiatkowski, Cordelia Langford, John Sillitoe on behalf of the Wellcome Sanger Institute COVID-19 Surveillance Team |
| EPI_ISL_3364845 | Lighthouse Lab in Alderley Park | Wellcome Sanger Institute for the COVID-19 Genomics UK (COG-UK) Consortium | Jacquelyn Wynn, Mairead Hyland, The Lighthouse Lab in Alderley Park and Alex Alderton, Roberto Amato, Jeffrey Barrett, Sonia Goncalves, Ewan Harrison, David K. Jackson, Ian Johnston, Dominic Kwiatkowski, Cordelia Langford, John Sillitoe on behalf of the Wellcome Sanger Institute COVID-19 Surveillance Team |
| EPI_ISL_3364846, EPI_ISL_3364847, EPI_ISL_3364848, EPI_ISL_3364849, EPI_ISL_3364850, EPI_ISL_3364851, EPI_ISL_3364852, EPI_ISL_3364853, EPI_ISL_3364854, EPI_ISL_3364855, EPI_ISL_3364856, EPI_ISL_3364857, EPI_ISL_3364858, EPI_ISL_3364859, EPI_ISL_3364860, EPI_ISL_3364861 | | | |
| see above | Lighthouse Lab in Milton Keynes | Wellcome Sanger Institute for the COVID-19 Genomics UK (COG-UK) Consortium | The Lighthouse Lab in Milton Keynes and Alex Alderton, Roberto Amato, Jeffrey Barrett, Sonia Goncalves, Ewan Harrison, David K. Jackson, Ian Johnston, Dominic Kwiatkowski, Cordelia Langford, John Sillitoe on behalf of the Wellcome Sanger Institute COVID-19 Surveillance Team |
| EPI_ISL_3364862, EPI_ISL_3364863 | Lighthouse Lab in Alderley Park | Wellcome Sanger Institute for the COVID-19 Genomics UK (COG-UK) Consortium | Jacquelyn Wynn, Mairead Hyland, The Lighthouse Lab in Alderley Park and Alex Alderton, Roberto Amato, Jeffrey Barrett, Sonia Goncalves, Ewan Harrison, David K. Jackson, Ian Johnston, Dominic Kwiatkowski, Cordelia Langford, John Sillitoe on behalf of the Wellcome Sanger Institute COVID-19 Surveillance Team |
| EPI_ISL_3364864, EPI_ISL_3364865 | Lighthouse Lab in Milton Keynes | Wellcome Sanger Institute for the COVID-19 Genomics UK (COG-UK) Consortium | The Lighthouse Lab in Milton Keynes and Alex Alderton, Roberto Amato, Jeffrey Barrett, Sonia Goncalves, Ewan Harrison, David K. Jackson, Ian Johnston, Dominic Kwiatkowski, Cordelia Langford, John Sillitoe on behalf of the Wellcome Sanger Institute COVID-19 Surveillance Team |
| EPI_ISL_3364866 | Lighthouse Lab in Alderley Park | Wellcome Sanger Institute for the COVID-19 Genomics UK (COG-UK) Consortium | Jacquelyn Wynn, Mairead Hyland, The Lighthouse Lab in Alderley Park and Alex Alderton, Roberto Amato, Jeffrey Barrett, Sonia Goncalves, Ewan Harrison, David K. Jackson, Ian Johnston, Dominic Kwiatkowski, Cordelia Langford, John Sillitoe on behalf of the Wellcome Sanger Institute COVID-19 Surveillance Team |
| EPI_ISL_3364867, EPI_ISL_3364868, EPI_ISL_3364869, EPI_ISL_3364870, EPI_ISL_3364871, EPI_ISL_3364872, EPI_ISL_3364873, EPI_ISL_3364874, EPI_ISL_3364875, EPI_ISL_3364876, EPI_ISL_3364877, EPI_ISL_3364878, EPI_ISL_3364879, EPI_ISL_3364880, EPI_ISL_3364883, EPI_ISL_3364884 | | | |
| see above | Lighthouse Lab in Milton Keynes | Wellcome Sanger Institute for the COVID-19 Genomics UK (COG-UK) Consortium | The Lighthouse Lab in Milton Keynes and Alex Alderton, Roberto Amato, Jeffrey Barrett, Sonia Goncalves, Ewan Harrison, David K. Jackson, Ian Johnston, Dominic Kwiatkowski, Cordelia Langford, John Sillitoe on behalf of the Wellcome Sanger Institute COVID-19 Surveillance Team |
| EPI_ISL_3364885 | Genetica Molecular and Subdepartamento de Virologia ISP Chile | Instituto de Salud Publica de Chile | Karen Orostica, Constanza Campano, Barbara Parra, Loredana Arata, Gisselle Barra, Patricia Bustos, Rodrigo Fasce, Javier Tognarelli, Andres Castillo, Soledad Ulloa, Jorge Fernandez |
| EPI_ISL_3364886, EPI_ISL_3364887, EPI_ISL_3364888, EPI_ISL_3364890, EPI_ISL_3364891, EPI_ISL_3364893, EPI_ISL_3364894, EPI_ISL_3364895 | Lighthouse Lab in Milton Keynes | Wellcome Sanger Institute for the COVID-19 Genomics UK (COG-UK) Consortium | The Lighthouse Lab in Milton Keynes and Alex Alderton, Roberto Amato, Jeffrey Barrett, Sonia Goncalves, Ewan Harrison, David K. Jackson, Ian Johnston, Dominic Kwiatkowski, Cordelia Langford, John Sillitoe on behalf of the Wellcome Sanger Institute COVID-19 Surveillance Team |
| EPI_ISL_3364897 | Lighthouse Lab in Alderley Park | Wellcome Sanger Institute for the COVID-19 Genomics UK (COG-UK) Consortium | Jacquelyn Wynn, Mairead Hyland, The Lighthouse Lab in Alderley Park and Alex Alderton, Roberto Amato, Jeffrey Barrett, Sonia Goncalves, Ewan Harrison, David K. Jackson, Ian Johnston, Dominic Kwiatkowski, Cordelia Langford, John Sillitoe on behalf of the Wellcome Sanger Institute COVID-19 Surveillance Team |
| EPI_ISL_3364898, EPI_ISL_3364899, | Lighthouse Lab in Milton Keynes | Wellcome Sanger Institute for the COVID-19 Genomics UK (COG-UK) Consortium | The Lighthouse Lab in Milton Keynes and Alex Alderton, Roberto Amato, Jeffrey Barrett, Sonia Goncalves, Ewan Harrison, David K. Jackson, Ian Johnston, Dominic Kwiatkowski, Cordelia Langford, John Sillitoe on behalf of the Wellcome Sanger Institute COVID-19 Surveillance Team |

| | | | |
|---|---|---|---|
| EPI_ISL_3364901, EPI_ISL_3364902, EPI_ISL_3364903, EPI_ISL_3364905, EPI_ISL_3364906, EPI_ISL_3364908, EPI_ISL_3364909 | | | |
| EPI_ISL_3364910 | Lighthouse Lab in Alderley Park | Wellcome Sanger Institute for the COVID-19 Genomics UK (COG-UK) Consortium | Jacquelyn Wynn, Mairead Hyland, The Lighthouse Lab in Alderley Park and Alex Alderton, Roberto Amato, Jeffrey Barrett, Sonia Goncalves, Ewan Harrison, David K. Jackson, Ian Johnston, Dominic Kwiatkowski, Cordelia Langford, John Sillitoe on behalf of the Wellcome Sanger Institute COVID-19 Surveillance Team |
| EPI_ISL_3364912, EPI_ISL_3364913, EPI_ISL_3364914, EPI_ISL_3364916, EPI_ISL_3364917, EPI_ISL_3364918, EPI_ISL_3364920, EPI_ISL_3364921, EPI_ISL_3364922, EPI_ISL_3364924, EPI_ISL_3364925, EPI_ISL_3364927, EPI_ISL_3364928, EPI_ISL_3364930, EPI_ISL_3364931, EPI_ISL_3364932, EPI_ISL_3364934, EPI_ISL_3364935, EPI_ISL_3364937, EPI_ISL_3364938, EPI_ISL_3364939, EPI_ISL_3364941 | | | |
| see above | Lighthouse Lab in Milton Keynes | Wellcome Sanger Institute for the COVID-19 Genomics UK (COG-UK) Consortium | The Lighthouse Lab in Milton Keynes and Alex Alderton, Roberto Amato, Jeffrey Barrett, Sonia Goncalves, Ewan Harrison, David K. Jackson, Ian Johnston, Dominic Kwiatkowski, Cordelia Langford, John Sillitoe on behalf of the Wellcome Sanger Institute COVID-19 Surveillance Team |
| EPI_ISL_3364942 | Lighthouse Lab in Alderley Park | Wellcome Sanger Institute for the COVID-19 Genomics UK (COG-UK) Consortium | Jacquelyn Wynn, Mairead Hyland, The Lighthouse Lab in Alderley Park and Alex Alderton, Roberto Amato, Jeffrey Barrett, Sonia Goncalves, Ewan Harrison, David K. Jackson, Ian Johnston, Dominic Kwiatkowski, Cordelia Langford, John Sillitoe on behalf of the Wellcome Sanger Institute COVID-19 Surveillance Team |
| EPI_ISL_3364944 | Berkshire and Surrey Pathology Services Lighthouse Laboratory | Wellcome Sanger Institute for the COVID-19 Genomics UK (COG-UK) Consortium | Berkshire and Surrey Pathology Services Lighthouse Laboratory and Alex Alderton, Roberto Amato, Jeffrey Barrett, Sonia Goncalves, Ewan Harrison, David K. Jackson, Ian Johnston, Dominic Kwiatkowski, Cordelia Langford, John Sillitoe on behalf of the Wellcome Sanger Institute COVID-19 Surveillance Team |
| EPI_ISL_3364945, EPI_ISL_3364946 | Lighthouse Lab in Alderley Park | Wellcome Sanger Institute for the COVID-19 Genomics UK (COG-UK) Consortium | Jacquelyn Wynn, Mairead Hyland, The Lighthouse Lab in Alderley Park and Alex Alderton, Roberto Amato, Jeffrey Barrett, Sonia Goncalves, Ewan Harrison, David K. Jackson, Ian Johnston, Dominic Kwiatkowski, Cordelia Langford, John Sillitoe on behalf of the Wellcome Sanger Institute COVID-19 Surveillance Team |
| EPI_ISL_3364948 | Berkshire and Surrey Pathology Services Lighthouse Laboratory | Wellcome Sanger Institute for the COVID-19 Genomics UK (COG-UK) Consortium | Berkshire and Surrey Pathology Services Lighthouse Laboratory and Alex Alderton, Roberto Amato, Jeffrey Barrett, Sonia Goncalves, Ewan Harrison, David K. Jackson, Ian Johnston, Dominic Kwiatkowski, Cordelia Langford, John Sillitoe on behalf of the Wellcome Sanger Institute COVID-19 Surveillance Team |
| EPI_ISL_3364949, EPI_ISL_3364950 | Lighthouse Lab in Milton Keynes | Wellcome Sanger Institute for the COVID-19 Genomics UK (COG-UK) Consortium | The Lighthouse Lab in Milton Keynes and Alex Alderton, Roberto Amato, Jeffrey Barrett, Sonia Goncalves, Ewan Harrison, David K. Jackson, Ian Johnston, Dominic Kwiatkowski, Cordelia Langford, John Sillitoe on behalf of the Wellcome Sanger Institute COVID-19 Surveillance Team |
| EPI_ISL_3364952, EPI_ISL_3364953, EPI_ISL_3364955 | Lighthouse Lab in Alderley Park | Wellcome Sanger Institute for the COVID-19 Genomics UK (COG-UK) Consortium | Jacquelyn Wynn, Mairead Hyland, The Lighthouse Lab in Alderley Park and Alex Alderton, Roberto Amato, Jeffrey Barrett, Sonia Goncalves, Ewan Harrison, David K. Jackson, Ian Johnston, Dominic Kwiatkowski, Cordelia Langford, John Sillitoe on behalf of the Wellcome Sanger Institute COVID-19 Surveillance Team |
| EPI_ISL_3364956 | Berkshire and Surrey Pathology Services Lighthouse Laboratory | Wellcome Sanger Institute for the COVID-19 Genomics UK (COG-UK) Consortium | Berkshire and Surrey Pathology Services Lighthouse Laboratory and Alex Alderton, Roberto Amato, Jeffrey Barrett, Sonia Goncalves, Ewan Harrison, David K. Jackson, Ian Johnston, Dominic Kwiatkowski, Cordelia Langford, John Sillitoe on behalf of the Wellcome Sanger Institute COVID-19 Surveillance Team |
| EPI_ISL_3364957 | Lighthouse Lab in Milton Keynes | Wellcome Sanger Institute for the COVID-19 Genomics UK (COG-UK) Consortium | The Lighthouse Lab in Milton Keynes and Alex Alderton, Roberto Amato, Jeffrey Barrett, Sonia Goncalves, Ewan Harrison, David K. Jackson, Ian Johnston, Dominic Kwiatkowski, Cordelia Langford, John Sillitoe on behalf of the Wellcome Sanger Institute COVID-19 Surveillance Team |
| EPI_ISL_3364959, EPI_ISL_3364960, EPI_ISL_3364961 | Lighthouse Lab in Alderley Park | Wellcome Sanger Institute for the COVID-19 Genomics UK (COG-UK) Consortium | Jacquelyn Wynn, Mairead Hyland, The Lighthouse Lab in Alderley Park and Alex Alderton, Roberto Amato, Jeffrey Barrett, Sonia Goncalves, Ewan Harrison, David K. Jackson, Ian Johnston, Dominic Kwiatkowski, Cordelia Langford, John Sillitoe on behalf of the Wellcome Sanger Institute COVID-19 Surveillance Team |
| EPI_ISL_3364962 | Berkshire and Surrey Pathology Services Lighthouse Laboratory | Wellcome Sanger Institute for the COVID-19 Genomics UK (COG-UK) Consortium | Berkshire and Surrey Pathology Services Lighthouse Laboratory and Alex Alderton, Roberto Amato, Jeffrey Barrett, Sonia Goncalves, Ewan Harrison, David K. Jackson, Ian Johnston, Dominic Kwiatkowski, Cordelia Langford, John Sillitoe on behalf of the Wellcome Sanger Institute COVID-19 Surveillance Team |
| EPI_ISL_3364964 | Lighthouse Lab in Milton Keynes | Wellcome Sanger Institute for the COVID-19 Genomics UK (COG-UK) Consortium | The Lighthouse Lab in Milton Keynes and Alex Alderton, Roberto Amato, Jeffrey Barrett, Sonia Goncalves, Ewan Harrison, David K. Jackson, Ian Johnston, Dominic Kwiatkowski, Cordelia Langford, John Sillitoe on behalf of the Wellcome Sanger Institute COVID-19 Surveillance Team |
| EPI_ISL_3364965 | Lighthouse Lab in Alderley Park | Wellcome Sanger Institute for the COVID-19 Genomics UK (COG-UK) Consortium | Jacquelyn Wynn, Mairead Hyland, The Lighthouse Lab in Alderley Park and Alex Alderton, Roberto Amato, Jeffrey Barrett, Sonia Goncalves, Ewan Harrison, David K. Jackson, Ian Johnston, Dominic Kwiatkowski, Cordelia Langford, John Sillitoe on behalf of the Wellcome Sanger Institute COVID-19 Surveillance Team |
| EPI_ISL_3364966 | Lighthouse Lab in Milton Keynes | Wellcome Sanger Institute for the COVID-19 Genomics UK (COG-UK) Consortium | The Lighthouse Lab in Milton Keynes and Alex Alderton, Roberto Amato, Jeffrey Barrett, Sonia Goncalves, Ewan Harrison, David K. Jackson, Ian Johnston, Dominic Kwiatkowski, Cordelia Langford, John Sillitoe on behalf of the Wellcome Sanger Institute COVID-19 Surveillance Team |
| EPI_ISL_3364968, EPI_ISL_3364969 | Lighthouse Lab in Alderley Park | Wellcome Sanger Institute for the COVID-19 Genomics UK (COG-UK) Consortium | Jacquelyn Wynn, Mairead Hyland, The Lighthouse Lab in Alderley Park and Alex Alderton, Roberto Amato, Jeffrey Barrett, Sonia Goncalves, Ewan Harrison, David K. Jackson, Ian Johnston, Dominic Kwiatkowski, Cordelia Langford, John Sillitoe on behalf of the Wellcome Sanger Institute COVID-19 Surveillance Team |
| EPI_ISL_3364970 | Berkshire and Surrey Pathology Services Lighthouse Laboratory | Wellcome Sanger Institute for the COVID-19 Genomics UK (COG-UK) Consortium | Berkshire and Surrey Pathology Services Lighthouse Laboratory and Alex Alderton, Roberto Amato, Jeffrey Barrett, Sonia Goncalves, Ewan Harrison, David K. Jackson, Ian Johnston, Dominic Kwiatkowski, Cordelia Langford, John Sillitoe on behalf of the Wellcome Sanger Institute COVID-19 Surveillance Team |
| EPI_ISL_3364972, EPI_ISL_3364973, EPI_ISL_3364974, EPI_ISL_3364976, EPI_ISL_3364977 | Lighthouse Lab in Alderley Park | Wellcome Sanger Institute for the COVID-19 Genomics UK (COG-UK) Consortium | Jacquelyn Wynn, Mairead Hyland, The Lighthouse Lab in Alderley Park and Alex Alderton, Roberto Amato, Jeffrey Barrett, Sonia Goncalves, Ewan Harrison, David K. Jackson, Ian Johnston, Dominic Kwiatkowski, Cordelia Langford, John Sillitoe on behalf of the Wellcome Sanger Institute COVID-19 Surveillance Team |
| EPI_ISL_3364978 | Berkshire and Surrey Pathology Services Lighthouse Laboratory | Wellcome Sanger Institute for the COVID-19 Genomics UK (COG-UK) Consortium | Berkshire and Surrey Pathology Services Lighthouse Laboratory and Alex Alderton, Roberto Amato, Jeffrey Barrett, Sonia Goncalves, Ewan Harrison, David K. Jackson, Ian Johnston, Dominic Kwiatkowski, Cordelia Langford, John Sillitoe on behalf of the Wellcome Sanger Institute COVID-19 Surveillance Team |
| EPI_ISL_3364980 | Lighthouse Lab in Alderley Park | Wellcome Sanger Institute for the COVID-19 Genomics UK (COG-UK) Consortium | Jacquelyn Wynn, Mairead Hyland, The Lighthouse Lab in Alderley Park and Alex Alderton, Roberto Amato, Jeffrey Barrett, Sonia Goncalves, Ewan Harrison, David K. Jackson, Ian Johnston, Dominic Kwiatkowski, Cordelia Langford, John Sillitoe on behalf of the Wellcome Sanger Institute COVID-19 Surveillance Team |
| EPI_ISL_3364981, EPI_ISL_3364982 | Berkshire and Surrey Pathology Services Lighthouse Laboratory | Wellcome Sanger Institute for the COVID-19 Genomics UK (COG-UK) Consortium | Berkshire and Surrey Pathology Services Lighthouse Laboratory and Alex Alderton, Roberto Amato, Jeffrey Barrett, Sonia Goncalves, Ewan Harrison, David K. Jackson, Ian Johnston, Dominic Kwiatkowski, Cordelia Langford, John Sillitoe on behalf of the Wellcome Sanger Institute COVID-19 Surveillance Team |
| EPI_ISL_3364984, EPI_ISL_3364985, EPI_ISL_3364986 | Lighthouse Lab in Alderley Park | Wellcome Sanger Institute for the COVID-19 Genomics UK (COG-UK) Consortium | Jacquelyn Wynn, Mairead Hyland, The Lighthouse Lab in Alderley Park and Alex Alderton, Roberto Amato, Jeffrey Barrett, Sonia Goncalves, Ewan Harrison, David K. Jackson, Ian Johnston, Dominic Kwiatkowski, Cordelia Langford, John Sillitoe on behalf of the Wellcome Sanger Institute COVID-19 Surveillance Team |
| EPI_ISL_3364988, EPI_ISL_3364989 | Berkshire and Surrey Pathology Services Lighthouse Laboratory | Wellcome Sanger Institute for the COVID-19 Genomics UK (COG-UK) Consortium | Berkshire and Surrey Pathology Services Lighthouse Laboratory and Alex Alderton, Roberto Amato, Jeffrey Barrett, Sonia Goncalves, Ewan Harrison, David K. Jackson, Ian Johnston, Dominic Kwiatkowski, Cordelia Langford, John Sillitoe on behalf of the Wellcome Sanger Institute COVID-19 Surveillance Team |
| EPI_ISL_3364990, EPI_ISL_3364992, EPI_ISL_3364993 | Lighthouse Lab in Alderley Park | Wellcome Sanger Institute for the COVID-19 Genomics UK (COG-UK) Consortium | Jacquelyn Wynn, Mairead Hyland, The Lighthouse Lab in Alderley Park and Alex Alderton, Roberto Amato, Jeffrey Barrett, Sonia Goncalves, Ewan Harrison, David K. Jackson, Ian Johnston, Dominic Kwiatkowski, Cordelia Langford, John Sillitoe on behalf of the Wellcome Sanger Institute COVID-19 Surveillance Team |
| EPI_ISL_3364994, EPI_ISL_3364996 | Berkshire and Surrey Pathology Services Lighthouse Laboratory | Wellcome Sanger Institute for the COVID-19 Genomics UK (COG-UK) Consortium | Berkshire and Surrey Pathology Services Lighthouse Laboratory and Alex Alderton, Roberto Amato, Jeffrey Barrett, Sonia Goncalves, Ewan Harrison, David K. Jackson, Ian Johnston, Dominic Kwiatkowski, Cordelia Langford, John Sillitoe on behalf of the Wellcome Sanger Institute COVID-19 Surveillance Team |
| EPI_ISL_3364997 | Lighthouse Lab in Alderley Park | Wellcome Sanger Institute for the COVID-19 Genomics UK (COG-UK) Consortium | Jacquelyn Wynn, Mairead Hyland, The Lighthouse Lab in Alderley Park and Alex Alderton, Roberto Amato, Jeffrey Barrett, Sonia Goncalves, Ewan Harrison, David K. Jackson, Ian Johnston, Dominic Kwiatkowski, Cordelia Langford, John Sillitoe on behalf of the Wellcome Sanger Institute COVID-19 Surveillance Team |
| EPI_ISL_3364998, EPI_ISL_3365000, EPI_ISL_3365001 | Berkshire and Surrey Pathology Services Lighthouse Laboratory | Wellcome Sanger Institute for the COVID-19 Genomics UK (COG-UK) Consortium | Berkshire and Surrey Pathology Services Lighthouse Laboratory and Alex Alderton, Roberto Amato, Jeffrey Barrett, Sonia Goncalves, Ewan Harrison, David K. Jackson, Ian Johnston, Dominic Kwiatkowski, Cordelia Langford, John Sillitoe on behalf of the Wellcome Sanger Institute COVID-19 Surveillance Team |
| EPI_ISL_3365002 | Lighthouse Lab in Alderley Park | Wellcome Sanger Institute for the COVID-19 Genomics UK (COG-UK) Consortium | Jacquelyn Wynn, Mairead Hyland, The Lighthouse Lab in Alderley Park and Alex Alderton, Roberto Amato, Jeffrey Barrett, Sonia Goncalves, Ewan Harrison, David K. Jackson, Ian Johnston, Dominic Kwiatkowski, Cordelia Langford, John Sillitoe on behalf of the Wellcome Sanger Institute COVID-19 Surveillance Team |
| EPI_ISL_3365004 | Berkshire and Surrey Pathology Services Lighthouse Laboratory | Wellcome Sanger Institute for the COVID-19 Genomics UK (COG-UK) Consortium | Berkshire and Surrey Pathology Services Lighthouse Laboratory and Alex Alderton, Roberto Amato, Jeffrey Barrett, Sonia Goncalves, Ewan Harrison, David K. Jackson, Ian Johnston, Dominic Kwiatkowski, Cordelia Langford, John Sillitoe on behalf of the Wellcome Sanger Institute COVID-19 Surveillance Team |
| EPI_ISL_3365005 | Lighthouse Lab in Alderley Park | Wellcome Sanger Institute for the COVID-19 Genomics UK (COG-UK) Consortium | Jacquelyn Wynn, Mairead Hyland, The Lighthouse Lab in Alderley Park and Alex Alderton, Roberto Amato, Jeffrey Barrett, Sonia Goncalves, Ewan Harrison, David K. Jackson, Ian Johnston, Dominic Kwiatkowski, Cordelia Langford, John Sillitoe on behalf of the Wellcome Sanger Institute COVID-19 Surveillance Team |
| EPI_ISL_3365006 | Berkshire and Surrey Pathology Services Lighthouse Laboratory | Wellcome Sanger Institute for the COVID-19 Genomics UK (COG-UK) Consortium | Berkshire and Surrey Pathology Services Lighthouse Laboratory and Alex Alderton, Roberto Amato, Jeffrey Barrett, Sonia Goncalves, Ewan Harrison, David K. Jackson, Ian Johnston, Dominic Kwiatkowski, Cordelia Langford, John Sillitoe on behalf of the Wellcome Sanger Institute COVID-19 Surveillance Team |
| EPI_ISL_3365008, EPI_ISL_3365009, EPI_ISL_3365010 | Lighthouse Lab in Alderley Park | Wellcome Sanger Institute for the COVID-19 Genomics UK (COG-UK) Consortium | Jacquelyn Wynn, Mairead Hyland, The Lighthouse Lab in Alderley Park and Alex Alderton, Roberto Amato, Jeffrey Barrett, Sonia Goncalves, Ewan Harrison, David K. Jackson, Ian Johnston, Dominic Kwiatkowski, Cordelia Langford, John Sillitoe on behalf of the Wellcome Sanger Institute COVID-19 Surveillance Team |
| EPI_ISL_3365012, EPI_ISL_3365013, EPI_ISL_3365015 | Berkshire and Surrey Pathology Services Lighthouse Laboratory | Wellcome Sanger Institute for the COVID-19 Genomics UK (COG-UK) Consortium | Berkshire and Surrey Pathology Services Lighthouse Laboratory and Alex Alderton, Roberto Amato, Jeffrey Barrett, Sonia Goncalves, Ewan Harrison, David K. Jackson, Ian Johnston, Dominic Kwiatkowski, Cordelia Langford, John Sillitoe on behalf of the Wellcome Sanger Institute COVID-19 Surveillance Team |
| EPI_ISL_3365016 | Lighthouse Lab in Alderley Park | Wellcome Sanger Institute for the COVID-19 Genomics UK (COG-UK) Consortium | Jacquelyn Wynn, Mairead Hyland, The Lighthouse Lab in Alderley Park and Alex Alderton, Roberto Amato, Jeffrey Barrett, Sonia Goncalves, Ewan Harrison, David K. Jackson, Ian Johnston, Dominic Kwiatkowski, Cordelia Langford, John Sillitoe on behalf of the Wellcome Sanger Institute COVID-19 Surveillance Team |
| EPI_ISL_3365017, EPI_ISL_3365019 | Lighthouse Lab in Milton Keynes | Wellcome Sanger Institute for the COVID-19 Genomics UK (COG-UK) Consortium | The Lighthouse Lab in Milton Keynes and Alex Alderton, Roberto Amato, Jeffrey Barrett, Sonia Goncalves, Ewan Harrison, David K. Jackson, Ian Johnston, Dominic Kwiatkowski, Cordelia Langford, John Sillitoe on behalf of the Wellcome Sanger Institute COVID-19 Surveillance Team |
| EPI_ISL_3365020 | Berkshire and Surrey Pathology Services Lighthouse Laboratory | Wellcome Sanger Institute for the COVID-19 Genomics UK (COG-UK) Consortium | Berkshire and Surrey Pathology Services Lighthouse Laboratory and Alex Alderton, Roberto Amato, Jeffrey Barrett, Sonia Goncalves, Ewan Harrison, David K. Jackson, Ian Johnston, Dominic Kwiatkowski, Cordelia Langford, John Sillitoe on behalf of the Wellcome Sanger Institute COVID-19 Surveillance Team |
| EPI_ISL_3365021 | Lighthouse Lab in Alderley Park | Wellcome Sanger Institute for the COVID-19 Genomics UK (COG-UK) Consortium | Jacquelyn Wynn, Mairead Hyland, The Lighthouse Lab in Alderley Park and Alex Alderton, Roberto Amato, Jeffrey Barrett, Sonia Goncalves, Ewan Harrison, David K. Jackson, Ian Johnston, Dominic Kwiatkowski, Cordelia Langford, John Sillitoe on behalf of the Wellcome Sanger Institute COVID-19 Surveillance Team |

| | | | |
|---|---|---|---|
| EPI_ISL_3365023 | Lighthouse Lab in Milton Keynes | Wellcome Sanger Institute for the COVID-19 Genomics UK (COG-UK) Consortium | The Lighthouse Lab in Milton Keynes and Alex Alderton, Roberto Amato, Jeffrey Barrett, Sonia Goncalves, Ewan Harrison, David K. Jackson, Ian Johnston, Dominic Kwiatkowski, Cordelia Langford, John Sillitoe on behalf of the Wellcome Sanger Institute COVID-19 Surveillance Team |
| EPI_ISL_3365024 | Lighthouse Lab in Alderley Park | Wellcome Sanger Institute for the COVID-19 Genomics UK (COG-UK) Consortium | Jacquelyn Wynn, Mairead Hyland, The Lighthouse Lab in Alderley Park and Alex Alderton, Roberto Amato, Jeffrey Barrett, Sonia Goncalves, Ewan Harrison, David K. Jackson, Ian Johnston, Dominic Kwiatkowski, Cordelia Langford, John Sillitoe on behalf of the Wellcome Sanger Institute COVID-19 Surveillance Team |
| EPI_ISL_3365025 | Lighthouse Lab in Milton Keynes | Wellcome Sanger Institute for the COVID-19 Genomics UK (COG-UK) Consortium | The Lighthouse Lab in Milton Keynes and Alex Alderton, Roberto Amato, Jeffrey Barrett, Sonia Goncalves, Ewan Harrison, David K. Jackson, Ian Johnston, Dominic Kwiatkowski, Cordelia Langford, John Sillitoe on behalf of the Wellcome Sanger Institute COVID-19 Surveillance Team |
| EPI_ISL_3365027 | Berkshire and Surrey Pathology Services Lighthouse Laboratory | Wellcome Sanger Institute for the COVID-19 Genomics UK (COG-UK) Consortium | Berkshire and Surrey Pathology Services Lighthouse Laboratory and Alex Alderton, Roberto Amato, Jeffrey Barrett, Sonia Goncalves, Ewan Harrison, David K. Jackson, Ian Johnston, Dominic Kwiatkowski, Cordelia Langford, John Sillitoe on behalf of the Wellcome Sanger Institute COVID-19 Surveillance Team |
| EPI_ISL_3365028, EPI_ISL_3365029 | Lighthouse Lab in Alderley Park | Wellcome Sanger Institute for the COVID-19 Genomics UK (COG-UK) Consortium | Jacquelyn Wynn, Mairead Hyland, The Lighthouse Lab in Alderley Park and Alex Alderton, Roberto Amato, Jeffrey Barrett, Sonia Goncalves, Ewan Harrison, David K. Jackson, Ian Johnston, Dominic Kwiatkowski, Cordelia Langford, John Sillitoe on behalf of the Wellcome Sanger Institute COVID-19 Surveillance Team |
| EPI_ISL_3365031 | Lighthouse Lab in Milton Keynes | Wellcome Sanger Institute for the COVID-19 Genomics UK (COG-UK) Consortium | The Lighthouse Lab in Milton Keynes and Alex Alderton, Roberto Amato, Jeffrey Barrett, Sonia Goncalves, Ewan Harrison, David K. Jackson, Ian Johnston, Dominic Kwiatkowski, Cordelia Langford, John Sillitoe on behalf of the Wellcome Sanger Institute COVID-19 Surveillance Team |
| EPI_ISL_3365032 | Berkshire and Surrey Pathology Services Lighthouse Laboratory | Wellcome Sanger Institute for the COVID-19 Genomics UK (COG-UK) Consortium | Berkshire and Surrey Pathology Services Lighthouse Laboratory and Alex Alderton, Roberto Amato, Jeffrey Barrett, Sonia Goncalves, Ewan Harrison, David K. Jackson, Ian Johnston, Dominic Kwiatkowski, Cordelia Langford, John Sillitoe on behalf of the Wellcome Sanger Institute COVID-19 Surveillance Team |
| EPI_ISL_3365033 | Lighthouse Lab in Milton Keynes | Wellcome Sanger Institute for the COVID-19 Genomics UK (COG-UK) Consortium | The Lighthouse Lab in Milton Keynes and Alex Alderton, Roberto Amato, Jeffrey Barrett, Sonia Goncalves, Ewan Harrison, David K. Jackson, Ian Johnston, Dominic Kwiatkowski, Cordelia Langford, John Sillitoe on behalf of the Wellcome Sanger Institute COVID-19 Surveillance Team |
| EPI_ISL_3365034, EPI_ISL_3365036 | Lighthouse Lab in Alderley Park | Wellcome Sanger Institute for the COVID-19 Genomics UK (COG-UK) Consortium | Jacquelyn Wynn, Mairead Hyland, The Lighthouse Lab in Alderley Park and Alex Alderton, Roberto Amato, Jeffrey Barrett, Sonia Goncalves, Ewan Harrison, David K. Jackson, Ian Johnston, Dominic Kwiatkowski, Cordelia Langford, John Sillitoe on behalf of the Wellcome Sanger Institute COVID-19 Surveillance Team |
| EPI_ISL_3365037 | Berkshire and Surrey Pathology Services Lighthouse Laboratory | Wellcome Sanger Institute for the COVID-19 Genomics UK (COG-UK) Consortium | Berkshire and Surrey Pathology Services Lighthouse Laboratory and Alex Alderton, Roberto Amato, Jeffrey Barrett, Sonia Goncalves, Ewan Harrison, David K. Jackson, Ian Johnston, Dominic Kwiatkowski, Cordelia Langford, John Sillitoe on behalf of the Wellcome Sanger Institute COVID-19 Surveillance Team |
| EPI_ISL_3365039, EPI_ISL_3365040 | Lighthouse Lab in Milton Keynes | Wellcome Sanger Institute for the COVID-19 Genomics UK (COG-UK) Consortium | The Lighthouse Lab in Milton Keynes and Alex Alderton, Roberto Amato, Jeffrey Barrett, Sonia Goncalves, Ewan Harrison, David K. Jackson, Ian Johnston, Dominic Kwiatkowski, Cordelia Langford, John Sillitoe on behalf of the Wellcome Sanger Institute COVID-19 Surveillance Team |
| EPI_ISL_3365041, EPI_ISL_3365042, EPI_ISL_3365043, EPI_ISL_3365044, EPI_ISL_3365046 | Lighthouse Lab in Alderley Park | Wellcome Sanger Institute for the COVID-19 Genomics UK (COG-UK) Consortium | Jacquelyn Wynn, Mairead Hyland, The Lighthouse Lab in Alderley Park and Alex Alderton, Roberto Amato, Jeffrey Barrett, Sonia Goncalves, Ewan Harrison, David K. Jackson, Ian Johnston, Dominic Kwiatkowski, Cordelia Langford, John Sillitoe on behalf of the Wellcome Sanger Institute COVID-19 Surveillance Team |
| EPI_ISL_3365047 | Berkshire and Surrey Pathology Services Lighthouse Laboratory | Wellcome Sanger Institute for the COVID-19 Genomics UK (COG-UK) Consortium | Berkshire and Surrey Pathology Services Lighthouse Laboratory and Alex Alderton, Roberto Amato, Jeffrey Barrett, Sonia Goncalves, Ewan Harrison, David K. Jackson, Ian Johnston, Dominic Kwiatkowski, Cordelia Langford, John Sillitoe on behalf of the Wellcome Sanger Institute COVID-19 Surveillance Team |
| EPI_ISL_3365048, EPI_ISL_3365050 | Lighthouse Lab in Alderley Park | Wellcome Sanger Institute for the COVID-19 Genomics UK (COG-UK) Consortium | Jacquelyn Wynn, Mairead Hyland, The Lighthouse Lab in Alderley Park and Alex Alderton, Roberto Amato, Jeffrey Barrett, Sonia Goncalves, Ewan Harrison, David K. Jackson, Ian Johnston, Dominic Kwiatkowski, Cordelia Langford, John Sillitoe on behalf of the Wellcome Sanger Institute COVID-19 Surveillance Team |
| EPI_ISL_3365051 | Lighthouse Lab in Milton Keynes | Wellcome Sanger Institute for the COVID-19 Genomics UK (COG-UK) Consortium | The Lighthouse Lab in Milton Keynes and Alex Alderton, Roberto Amato, Jeffrey Barrett, Sonia Goncalves, Ewan Harrison, David K. Jackson, Ian Johnston, Dominic Kwiatkowski, Cordelia Langford, John Sillitoe on behalf of the Wellcome Sanger Institute COVID-19 Surveillance Team |
| EPI_ISL_3365053, EPI_ISL_3365054, EPI_ISL_3365055, EPI_ISL_3365057 | Lighthouse Lab in Alderley Park | Wellcome Sanger Institute for the COVID-19 Genomics UK (COG-UK) Consortium | Jacquelyn Wynn, Mairead Hyland, The Lighthouse Lab in Alderley Park and Alex Alderton, Roberto Amato, Jeffrey Barrett, Sonia Goncalves, Ewan Harrison, David K. Jackson, Ian Johnston, Dominic Kwiatkowski, Cordelia Langford, John Sillitoe on behalf of the Wellcome Sanger Institute COVID-19 Surveillance Team |
| EPI_ISL_3365058 | Berkshire and Surrey Pathology Services Lighthouse Laboratory | Wellcome Sanger Institute for the COVID-19 Genomics UK (COG-UK) Consortium | Berkshire and Surrey Pathology Services Lighthouse Laboratory and Alex Alderton, Roberto Amato, Jeffrey Barrett, Sonia Goncalves, Ewan Harrison, David K. Jackson, Ian Johnston, Dominic Kwiatkowski, Cordelia Langford, John Sillitoe on behalf of the Wellcome Sanger Institute COVID-19 Surveillance Team |
| EPI_ISL_3365059, EPI_ISL_3365061 | Lighthouse Lab in Alderley Park | Wellcome Sanger Institute for the COVID-19 Genomics UK (COG-UK) Consortium | Jacquelyn Wynn, Mairead Hyland, The Lighthouse Lab in Alderley Park and Alex Alderton, Roberto Amato, Jeffrey Barrett, Sonia Goncalves, Ewan Harrison, David K. Jackson, Ian Johnston, Dominic Kwiatkowski, Cordelia Langford, John Sillitoe on behalf of the Wellcome Sanger Institute COVID-19 Surveillance Team |
| EPI_ISL_3365062 | Lighthouse Lab in Milton Keynes | Wellcome Sanger Institute for the COVID-19 Genomics UK (COG-UK) Consortium | The Lighthouse Lab in Milton Keynes and Alex Alderton, Roberto Amato, Jeffrey Barrett, Sonia Goncalves, Ewan Harrison, David K. Jackson, Ian Johnston, Dominic Kwiatkowski, Cordelia Langford, John Sillitoe on behalf of the Wellcome Sanger Institute COVID-19 Surveillance Team |
| EPI_ISL_3365063 | Berkshire and Surrey Pathology Services Lighthouse Laboratory | Wellcome Sanger Institute for the COVID-19 Genomics UK (COG-UK) Consortium | Berkshire and Surrey Pathology Services Lighthouse Laboratory and Alex Alderton, Roberto Amato, Jeffrey Barrett, Sonia Goncalves, Ewan Harrison, David K. Jackson, Ian Johnston, Dominic Kwiatkowski, Cordelia Langford, John Sillitoe on behalf of the Wellcome Sanger Institute COVID-19 Surveillance Team |
| EPI_ISL_3365065, EPI_ISL_3365066, EPI_ISL_3365068 | Lighthouse Lab in Alderley Park | Wellcome Sanger Institute for the COVID-19 Genomics UK (COG-UK) Consortium | Jacquelyn Wynn, Mairead Hyland, The Lighthouse Lab in Alderley Park and Alex Alderton, Roberto Amato, Jeffrey Barrett, Sonia Goncalves, Ewan Harrison, David K. Jackson, Ian Johnston, Dominic Kwiatkowski, Cordelia Langford, John Sillitoe on behalf of the Wellcome Sanger Institute COVID-19 Surveillance Team |
| EPI_ISL_3365069, EPI_ISL_3365070, EPI_ISL_3365072, EPI_ISL_3365073 | Berkshire and Surrey Pathology Services Lighthouse Laboratory | Wellcome Sanger Institute for the COVID-19 Genomics UK (COG-UK) Consortium | Berkshire and Surrey Pathology Services Lighthouse Laboratory and Alex Alderton, Roberto Amato, Jeffrey Barrett, Sonia Goncalves, Ewan Harrison, David K. Jackson, Ian Johnston, Dominic Kwiatkowski, Cordelia Langford, John Sillitoe on behalf of the Wellcome Sanger Institute COVID-19 Surveillance Team |
| EPI_ISL_3365075 | Lighthouse Lab in Alderley Park | Wellcome Sanger Institute for the COVID-19 Genomics UK (COG-UK) Consortium | Jacquelyn Wynn, Mairead Hyland, The Lighthouse Lab in Alderley Park and Alex Alderton, Roberto Amato, Jeffrey Barrett, Sonia Goncalves, Ewan Harrison, David K. Jackson, Ian Johnston, Dominic Kwiatkowski, Cordelia Langford, John Sillitoe on behalf of the Wellcome Sanger Institute COVID-19 Surveillance Team |
| EPI_ISL_3365076, EPI_ISL_3365077 | Lighthouse Lab in Milton Keynes | Wellcome Sanger Institute for the COVID-19 Genomics UK (COG-UK) Consortium | The Lighthouse Lab in Milton Keynes and Alex Alderton, Roberto Amato, Jeffrey Barrett, Sonia Goncalves, Ewan Harrison, David K. Jackson, Ian Johnston, Dominic Kwiatkowski, Cordelia Langford, John Sillitoe on behalf of the Wellcome Sanger Institute COVID-19 Surveillance Team |
| EPI_ISL_3365079 | Lighthouse Lab in Alderley Park | Wellcome Sanger Institute for the COVID-19 Genomics UK (COG-UK) Consortium | Jacquelyn Wynn, Mairead Hyland, The Lighthouse Lab in Alderley Park and Alex Alderton, Roberto Amato, Jeffrey Barrett, Sonia Goncalves, Ewan Harrison, David K. Jackson, Ian Johnston, Dominic Kwiatkowski, Cordelia Langford, John Sillitoe on behalf of the Wellcome Sanger Institute COVID-19 Surveillance Team |
| EPI_ISL_3365080 | Lighthouse Lab in Milton Keynes | Wellcome Sanger Institute for the COVID-19 Genomics UK (COG-UK) Consortium | The Lighthouse Lab in Milton Keynes and Alex Alderton, Roberto Amato, Jeffrey Barrett, Sonia Goncalves, Ewan Harrison, David K. Jackson, Ian Johnston, Dominic Kwiatkowski, Cordelia Langford, John Sillitoe on behalf of the Wellcome Sanger Institute COVID-19 Surveillance Team |
| EPI_ISL_3365081 | Lighthouse Lab in Alderley Park | Wellcome Sanger Institute for the COVID-19 Genomics UK (COG-UK) Consortium | Jacquelyn Wynn, Mairead Hyland, The Lighthouse Lab in Alderley Park and Alex Alderton, Roberto Amato, Jeffrey Barrett, Sonia Goncalves, Ewan Harrison, David K. Jackson, Ian Johnston, Dominic Kwiatkowski, Cordelia Langford, John Sillitoe on behalf of the Wellcome Sanger Institute COVID-19 Surveillance Team |
| EPI_ISL_3365083, EPI_ISL_3365084 | Lighthouse Lab in Milton Keynes | Wellcome Sanger Institute for the COVID-19 Genomics UK (COG-UK) Consortium | The Lighthouse Lab in Milton Keynes and Alex Alderton, Roberto Amato, Jeffrey Barrett, Sonia Goncalves, Ewan Harrison, David K. Jackson, Ian Johnston, Dominic Kwiatkowski, Cordelia Langford, John Sillitoe on behalf of the Wellcome Sanger Institute COVID-19 Surveillance Team |
| EPI_ISL_3365086 | Berkshire and Surrey Pathology Services Lighthouse Laboratory | Wellcome Sanger Institute for the COVID-19 Genomics UK (COG-UK) Consortium | Berkshire and Surrey Pathology Services Lighthouse Laboratory and Alex Alderton, Roberto Amato, Jeffrey Barrett, Sonia Goncalves, Ewan Harrison, David K. Jackson, Ian Johnston, Dominic Kwiatkowski, Cordelia Langford, John Sillitoe on behalf of the Wellcome Sanger Institute COVID-19 Surveillance Team |
| EPI_ISL_3365087 | Lighthouse Lab in Alderley Park | Wellcome Sanger Institute for the COVID-19 Genomics UK (COG-UK) Consortium | Jacquelyn Wynn, Mairead Hyland, The Lighthouse Lab in Alderley Park and Alex Alderton, Roberto Amato, Jeffrey Barrett, Sonia Goncalves, Ewan Harrison, David K. Jackson, Ian Johnston, Dominic Kwiatkowski, Cordelia Langford, John Sillitoe on behalf of the Wellcome Sanger Institute COVID-19 Surveillance Team |
| EPI_ISL_3365088 | Lighthouse Lab in Milton Keynes | Wellcome Sanger Institute for the COVID-19 Genomics UK (COG-UK) Consortium | The Lighthouse Lab in Milton Keynes and Alex Alderton, Roberto Amato, Jeffrey Barrett, Sonia Goncalves, Ewan Harrison, David K. Jackson, Ian Johnston, Dominic Kwiatkowski, Cordelia Langford, John Sillitoe on behalf of the Wellcome Sanger Institute COVID-19 Surveillance Team |
| EPI_ISL_3365090, EPI_ISL_3365091, EPI_ISL_3365092 | Lighthouse Lab in Alderley Park | Wellcome Sanger Institute for the COVID-19 Genomics UK (COG-UK) Consortium | Jacquelyn Wynn, Mairead Hyland, The Lighthouse Lab in Alderley Park and Alex Alderton, Roberto Amato, Jeffrey Barrett, Sonia Goncalves, Ewan Harrison, David K. Jackson, Ian Johnston, Dominic Kwiatkowski, Cordelia Langford, John Sillitoe on behalf of the Wellcome Sanger Institute COVID-19 Surveillance Team |
| EPI_ISL_3365094 | Berkshire and Surrey Pathology Services Lighthouse Laboratory | Wellcome Sanger Institute for the COVID-19 Genomics UK (COG-UK) Consortium | Berkshire and Surrey Pathology Services Lighthouse Laboratory and Alex Alderton, Roberto Amato, Jeffrey Barrett, Sonia Goncalves, Ewan Harrison, David K. Jackson, Ian Johnston, Dominic Kwiatkowski, Cordelia Langford, John Sillitoe on behalf of the Wellcome Sanger Institute COVID-19 Surveillance Team |
| EPI_ISL_3365095 | Lighthouse Lab in Milton Keynes | Wellcome Sanger Institute for the COVID-19 Genomics UK (COG-UK) Consortium | The Lighthouse Lab in Milton Keynes and Alex Alderton, Roberto Amato, Jeffrey Barrett, Sonia Goncalves, Ewan Harrison, David K. Jackson, Ian Johnston, Dominic Kwiatkowski, Cordelia Langford, John Sillitoe on behalf of the Wellcome Sanger Institute COVID-19 Surveillance Team |
| EPI_ISL_3365097, EPI_ISL_3365098, EPI_ISL_3365099 | Lighthouse Lab in Alderley Park | Wellcome Sanger Institute for the COVID-19 Genomics UK (COG-UK) Consortium | Jacquelyn Wynn, Mairead Hyland, The Lighthouse Lab in Alderley Park and Alex Alderton, Roberto Amato, Jeffrey Barrett, Sonia Goncalves, Ewan Harrison, David K. Jackson, Ian Johnston, Dominic Kwiatkowski, Cordelia Langford, John Sillitoe on behalf of the Wellcome Sanger Institute COVID-19 Surveillance Team |
| EPI_ISL_3365101 | Berkshire and Surrey Pathology Services Lighthouse Laboratory | Wellcome Sanger Institute for the COVID-19 Genomics UK (COG-UK) Consortium | Berkshire and Surrey Pathology Services Lighthouse Laboratory and Alex Alderton, Roberto Amato, Jeffrey Barrett, Sonia Goncalves, Ewan Harrison, David K. Jackson, Ian Johnston, Dominic Kwiatkowski, Cordelia Langford, John Sillitoe on behalf of the Wellcome Sanger Institute COVID-19 Surveillance Team |
| EPI_ISL_3365102 | Lighthouse Lab in Alderley Park | Wellcome Sanger Institute for the COVID-19 Genomics UK (COG-UK) Consortium | Jacquelyn Wynn, Mairead Hyland, The Lighthouse Lab in Alderley Park and Alex Alderton, Roberto Amato, Jeffrey Barrett, Sonia Goncalves, Ewan Harrison, David K. Jackson, Ian Johnston, Dominic Kwiatkowski, Cordelia Langford, John Sillitoe on behalf of the Wellcome Sanger Institute COVID-19 Surveillance Team |
| EPI_ISL_3365103 | Lighthouse Lab in Milton Keynes | Wellcome Sanger Institute for the COVID-19 Genomics UK (COG-UK) Consortium | The Lighthouse Lab in Milton Keynes and Alex Alderton, Roberto Amato, Jeffrey Barrett, Sonia Goncalves, Ewan Harrison, David K. Jackson, Ian Johnston, Dominic Kwiatkowski, Cordelia Langford, John Sillitoe on behalf of the Wellcome Sanger Institute COVID-19 Surveillance Team |
| EPI_ISL_3365104 | Lighthouse Lab in Alderley Park | Wellcome Sanger Institute for the COVID-19 Genomics UK (COG-UK) Consortium | Jacquelyn Wynn, Mairead Hyland, The Lighthouse Lab in Alderley Park and Alex Alderton, Roberto Amato, Jeffrey Barrett, Sonia Goncalves, Ewan Harrison, David K. Jackson, Ian Johnston, Dominic Kwiatkowski, Cordelia Langford, John Sillitoe on behalf of the Wellcome Sanger Institute COVID-19 Surveillance Team |
| EPI_ISL_3365105 | Genetica Molecular and Subdepartamento de Virologia ISP Chile | Instituto de Salud Publica de Chile | Karen Orostica, Constanza Campano, Barbara Parra, Loredana Arata, Gisselle Barra, Patricia Bustos, Rodrigo Fasce, Javier Tognarelli, Andres Castillo, Soledad Ulloa, Jorge Fernandez |
| EPI_ISL_3365106, EPI_ISL_3365107 | Lighthouse Lab in Alderley Park | Wellcome Sanger Institute for the COVID-19 Genomics UK (COG-UK) Consortium | Jacquelyn Wynn, Mairead Hyland, The Lighthouse Lab in Alderley Park and Alex Alderton, Roberto Amato, Jeffrey Barrett, Sonia Goncalves, Ewan Harrison, David K. Jackson, Ian Johnston, Dominic Kwiatkowski, Cordelia Langford, John Sillitoe on behalf of the Wellcome Sanger Institute COVID-19 Surveillance Team |
| EPI_ISL_3365108 | Berkshire and Surrey Pathology Services Lighthouse Laboratory | Wellcome Sanger Institute for the COVID-19 Genomics UK (COG-UK) | Berkshire and Surrey Pathology Services Lighthouse Laboratory and Alex Alderton, Roberto Amato, Jeffrey Barrett, Sonia Goncalves, Ewan Harrison, David K. Jackson, Ian Johnston, Dominic Kwiatkowski, Cordelia Langford, John Sillitoe on behalf |

| | | | |
|---|---|---|---|
| | | Consortium | of the Wellcome Sanger Institute COVID-19 Surveillance Team |
| EPI_ISL_3365109 | Genetica Molecular and Subdepartamento de Virologia ISP Chile | Instituto de Salud Publica de Chile | Karen Orostica, Constanza Campano, Barbara Parra, Loredana Arata, Gisselle Barra, Patricia Bustos, Rodrigo Fasce, Javier Tognarelli, Andres Castillo, Soledad Ulloa, Jorge Fernandez |
| EPI_ISL_3365110 | Lighthouse Lab in Alderley Park | Wellcome Sanger Institute for the COVID-19 Genomics UK (COG-UK) Consortium | Jacquelyn Wynn, Mairead Hyland, The Lighthouse Lab in Alderley Park and Alex Alderton, Roberto Amato, Jeffrey Barrett, Sonia Goncalves, Ewan Harrison, David K. Jackson, Ian Johnston, Dominic Kwiatkowski, Cordelia Langford, John Sillitoe on behalf of the Wellcome Sanger Institute COVID-19 Surveillance Team |
| EPI_ISL_3365111 | Lighthouse Lab in Milton Keynes | Wellcome Sanger Institute for the COVID-19 Genomics UK (COG-UK) Consortium | The Lighthouse Lab in Milton Keynes and Alex Alderton, Roberto Amato, Jeffrey Barrett, Sonia Goncalves, Ewan Harrison, David K. Jackson, Ian Johnston, Dominic Kwiatkowski, Cordelia Langford, John Sillitoe on behalf of the Wellcome Sanger Institute COVID-19 Surveillance Team |
| EPI_ISL_3365112 | Lighthouse Lab in Alderley Park | Wellcome Sanger Institute for the COVID-19 Genomics UK (COG-UK) Consortium | Jacquelyn Wynn, Mairead Hyland, The Lighthouse Lab in Alderley Park and Alex Alderton, Roberto Amato, Jeffrey Barrett, Sonia Goncalves, Ewan Harrison, David K. Jackson, Ian Johnston, Dominic Kwiatkowski, Cordelia Langford, John Sillitoe on behalf of the Wellcome Sanger Institute COVID-19 Surveillance Team |
| EPI_ISL_3365113 | Genetica Molecular and Subdepartamento de Virologia ISP Chile | Instituto de Salud Publica de Chile | Karen Orostica, Constanza Campano, Barbara Parra, Loredana Arata, Gisselle Barra, Patricia Bustos, Rodrigo Fasce, Javier Tognarelli, Andres Castillo, Soledad Ulloa, Jorge Fernandez |
| EPI_ISL_3365114 | Lighthouse Lab in Milton Keynes | Wellcome Sanger Institute for the COVID-19 Genomics UK (COG-UK) Consortium | The Lighthouse Lab in Milton Keynes and Alex Alderton, Roberto Amato, Jeffrey Barrett, Sonia Goncalves, Ewan Harrison, David K. Jackson, Ian Johnston, Dominic Kwiatkowski, Cordelia Langford, John Sillitoe on behalf of the Wellcome Sanger Institute COVID-19 Surveillance Team |
| EPI_ISL_3365115 | Berkshire and Surrey Pathology Services Lighthouse Laboratory | Wellcome Sanger Institute for the COVID-19 Genomics UK (COG-UK) Consortium | Berkshire and Surrey Pathology Services Lighthouse Laboratory and Alex Alderton, Roberto Amato, Jeffrey Barrett, Sonia Goncalves, Ewan Harrison, David K. Jackson, Ian Johnston, Dominic Kwiatkowski, Cordelia Langford, John Sillitoe on behalf of the Wellcome Sanger Institute COVID-19 Surveillance Team |
| EPI_ISL_3365116 | Lighthouse Lab in Milton Keynes | Wellcome Sanger Institute for the COVID-19 Genomics UK (COG-UK) Consortium | The Lighthouse Lab in Milton Keynes and Alex Alderton, Roberto Amato, Jeffrey Barrett, Sonia Goncalves, Ewan Harrison, David K. Jackson, Ian Johnston, Dominic Kwiatkowski, Cordelia Langford, John Sillitoe on behalf of the Wellcome Sanger Institute COVID-19 Surveillance Team |
| EPI_ISL_3365117 | Genetica Molecular and Subdepartamento de Virologia ISP Chile | Instituto de Salud Publica de Chile | Karen Orostica, Constanza Campano, Barbara Parra, Loredana Arata, Gisselle Barra, Patricia Bustos, Rodrigo Fasce, Javier Tognarelli, Andres Castillo, Soledad Ulloa, Jorge Fernandez |
| EPI_ISL_3365118 | Lighthouse Lab in Milton Keynes | Wellcome Sanger Institute for the COVID-19 Genomics UK (COG-UK) Consortium | The Lighthouse Lab in Milton Keynes and Alex Alderton, Roberto Amato, Jeffrey Barrett, Sonia Goncalves, Ewan Harrison, David K. Jackson, Ian Johnston, Dominic Kwiatkowski, Cordelia Langford, John Sillitoe on behalf of the Wellcome Sanger Institute COVID-19 Surveillance Team |
| EPI_ISL_3365119, EPI_ISL_3365120 | Lighthouse Lab in Alderley Park | Wellcome Sanger Institute for the COVID-19 Genomics UK (COG-UK) Consortium | Jacquelyn Wynn, Mairead Hyland, The Lighthouse Lab in Alderley Park and Alex Alderton, Roberto Amato, Jeffrey Barrett, Sonia Goncalves, Ewan Harrison, David K. Jackson, Ian Johnston, Dominic Kwiatkowski, Cordelia Langford, John Sillitoe on behalf of the Wellcome Sanger Institute COVID-19 Surveillance Team |
| EPI_ISL_3365121, EPI_ISL_3365122 | Lighthouse Lab in Milton Keynes | Wellcome Sanger Institute for the COVID-19 Genomics UK (COG-UK) Consortium | The Lighthouse Lab in Milton Keynes and Alex Alderton, Roberto Amato, Jeffrey Barrett, Sonia Goncalves, Ewan Harrison, David K. Jackson, Ian Johnston, Dominic Kwiatkowski, Cordelia Langford, John Sillitoe on behalf of the Wellcome Sanger Institute COVID-19 Surveillance Team |
| EPI_ISL_3365123, EPI_ISL_3365124 | Berkshire and Surrey Pathology Services Lighthouse Laboratory | Wellcome Sanger Institute for the COVID-19 Genomics UK (COG-UK) Consortium | Berkshire and Surrey Pathology Services Lighthouse Laboratory and Alex Alderton, Roberto Amato, Jeffrey Barrett, Sonia Goncalves, Ewan Harrison, David K. Jackson, Ian Johnston, Dominic Kwiatkowski, Cordelia Langford, John Sillitoe on behalf of the Wellcome Sanger Institute COVID-19 Surveillance Team |
| EPI_ISL_3365125, EPI_ISL_3365126 | Lighthouse Lab in Milton Keynes | Wellcome Sanger Institute for the COVID-19 Genomics UK (COG-UK) Consortium | The Lighthouse Lab in Milton Keynes and Alex Alderton, Roberto Amato, Jeffrey Barrett, Sonia Goncalves, Ewan Harrison, David K. Jackson, Ian Johnston, Dominic Kwiatkowski, Cordelia Langford, John Sillitoe on behalf of the Wellcome Sanger Institute COVID-19 Surveillance Team |
| EPI_ISL_3365127, EPI_ISL_3365128 | Lighthouse Lab in Alderley Park | Wellcome Sanger Institute for the COVID-19 Genomics UK (COG-UK) Consortium | Jacquelyn Wynn, Mairead Hyland, The Lighthouse Lab in Alderley Park and Alex Alderton, Roberto Amato, Jeffrey Barrett, Sonia Goncalves, Ewan Harrison, David K. Jackson, Ian Johnston, Dominic Kwiatkowski, Cordelia Langford, John Sillitoe on behalf of the Wellcome Sanger Institute COVID-19 Surveillance Team |
| EPI_ISL_3365129 | Berkshire and Surrey Pathology Services Lighthouse Laboratory | Wellcome Sanger Institute for the COVID-19 Genomics UK (COG-UK) Consortium | Berkshire and Surrey Pathology Services Lighthouse Laboratory and Alex Alderton, Roberto Amato, Jeffrey Barrett, Sonia Goncalves, Ewan Harrison, David K. Jackson, Ian Johnston, Dominic Kwiatkowski, Cordelia Langford, John Sillitoe on behalf of the Wellcome Sanger Institute COVID-19 Surveillance Team |
| EPI_ISL_3365130, EPI_ISL_3365131 | Lighthouse Lab in Milton Keynes | Wellcome Sanger Institute for the COVID-19 Genomics UK (COG-UK) Consortium | The Lighthouse Lab in Milton Keynes and Alex Alderton, Roberto Amato, Jeffrey Barrett, Sonia Goncalves, Ewan Harrison, David K. Jackson, Ian Johnston, Dominic Kwiatkowski, Cordelia Langford, John Sillitoe on behalf of the Wellcome Sanger Institute COVID-19 Surveillance Team |
| EPI_ISL_3365132, EPI_ISL_3365133 | Lighthouse Lab in Alderley Park | Wellcome Sanger Institute for the COVID-19 Genomics UK (COG-UK) Consortium | Jacquelyn Wynn, Mairead Hyland, The Lighthouse Lab in Alderley Park and Alex Alderton, Roberto Amato, Jeffrey Barrett, Sonia Goncalves, Ewan Harrison, David K. Jackson, Ian Johnston, Dominic Kwiatkowski, Cordelia Langford, John Sillitoe on behalf of the Wellcome Sanger Institute COVID-19 Surveillance Team |
| EPI_ISL_3365134 | Lighthouse Lab in Milton Keynes | Wellcome Sanger Institute for the COVID-19 Genomics UK (COG-UK) Consortium | The Lighthouse Lab in Milton Keynes and Alex Alderton, Roberto Amato, Jeffrey Barrett, Sonia Goncalves, Ewan Harrison, David K. Jackson, Ian Johnston, Dominic Kwiatkowski, Cordelia Langford, John Sillitoe on behalf of the Wellcome Sanger Institute COVID-19 Surveillance Team |
| EPI_ISL_3365135 | Berkshire and Surrey Pathology Services Lighthouse Laboratory | Wellcome Sanger Institute for the COVID-19 Genomics UK (COG-UK) Consortium | Berkshire and Surrey Pathology Services Lighthouse Laboratory and Alex Alderton, Roberto Amato, Jeffrey Barrett, Sonia Goncalves, Ewan Harrison, David K. Jackson, Ian Johnston, Dominic Kwiatkowski, Cordelia Langford, John Sillitoe on behalf of the Wellcome Sanger Institute COVID-19 Surveillance Team |
| EPI_ISL_3365136, EPI_ISL_3365137, EPI_ISL_3365138 | Lighthouse Lab in Milton Keynes | Wellcome Sanger Institute for the COVID-19 Genomics UK (COG-UK) Consortium | The Lighthouse Lab in Milton Keynes and Alex Alderton, Roberto Amato, Jeffrey Barrett, Sonia Goncalves, Ewan Harrison, David K. Jackson, Ian Johnston, Dominic Kwiatkowski, Cordelia Langford, John Sillitoe on behalf of the Wellcome Sanger Institute COVID-19 Surveillance Team |
| EPI_ISL_3365139, EPI_ISL_3365140, EPI_ISL_3365141, EPI_ISL_3365142 | Lighthouse Lab in Alderley Park | Wellcome Sanger Institute for the COVID-19 Genomics UK (COG-UK) Consortium | Jacquelyn Wynn, Mairead Hyland, The Lighthouse Lab in Alderley Park and Alex Alderton, Roberto Amato, Jeffrey Barrett, Sonia Goncalves, Ewan Harrison, David K. Jackson, Ian Johnston, Dominic Kwiatkowski, Cordelia Langford, John Sillitoe on behalf of the Wellcome Sanger Institute COVID-19 Surveillance Team |
| EPI_ISL_3365143 | Berkshire and Surrey Pathology Services Lighthouse Laboratory | Wellcome Sanger Institute for the COVID-19 Genomics UK (COG-UK) Consortium | Berkshire and Surrey Pathology Services Lighthouse Laboratory and Alex Alderton, Roberto Amato, Jeffrey Barrett, Sonia Goncalves, Ewan Harrison, David K. Jackson, Ian Johnston, Dominic Kwiatkowski, Cordelia Langford, John Sillitoe on behalf of the Wellcome Sanger Institute COVID-19 Surveillance Team |
| EPI_ISL_3365144, EPI_ISL_3365145, EPI_ISL_3365146, EPI_ISL_3365147 | Lighthouse Lab in Alderley Park | Wellcome Sanger Institute for the COVID-19 Genomics UK (COG-UK) Consortium | Jacquelyn Wynn, Mairead Hyland, The Lighthouse Lab in Alderley Park and Alex Alderton, Roberto Amato, Jeffrey Barrett, Sonia Goncalves, Ewan Harrison, David K. Jackson, Ian Johnston, Dominic Kwiatkowski, Cordelia Langford, John Sillitoe on behalf of the Wellcome Sanger Institute COVID-19 Surveillance Team |
| EPI_ISL_3365148 | Berkshire and Surrey Pathology Services Lighthouse Laboratory | Wellcome Sanger Institute for the COVID-19 Genomics UK (COG-UK) Consortium | Berkshire and Surrey Pathology Services Lighthouse Laboratory and Alex Alderton, Roberto Amato, Jeffrey Barrett, Sonia Goncalves, Ewan Harrison, David K. Jackson, Ian Johnston, Dominic Kwiatkowski, Cordelia Langford, John Sillitoe on behalf of the Wellcome Sanger Institute COVID-19 Surveillance Team |
| EPI_ISL_3365149 | Lighthouse Lab in Milton Keynes | Wellcome Sanger Institute for the COVID-19 Genomics UK (COG-UK) Consortium | The Lighthouse Lab in Milton Keynes and Alex Alderton, Roberto Amato, Jeffrey Barrett, Sonia Goncalves, Ewan Harrison, David K. Jackson, Ian Johnston, Dominic Kwiatkowski, Cordelia Langford, John Sillitoe on behalf of the Wellcome Sanger Institute COVID-19 Surveillance Team |
| EPI_ISL_3365150, EPI_ISL_3365151, EPI_ISL_3365152 | Lighthouse Lab in Alderley Park | Wellcome Sanger Institute for the COVID-19 Genomics UK (COG-UK) Consortium | Jacquelyn Wynn, Mairead Hyland, The Lighthouse Lab in Alderley Park and Alex Alderton, Roberto Amato, Jeffrey Barrett, Sonia Goncalves, Ewan Harrison, David K. Jackson, Ian Johnston, Dominic Kwiatkowski, Cordelia Langford, John Sillitoe on behalf of the Wellcome Sanger Institute COVID-19 Surveillance Team |
| EPI_ISL_3365153, EPI_ISL_3365154 | Lighthouse Lab in Milton Keynes | Wellcome Sanger Institute for the COVID-19 Genomics UK (COG-UK) Consortium | The Lighthouse Lab in Milton Keynes and Alex Alderton, Roberto Amato, Jeffrey Barrett, Sonia Goncalves, Ewan Harrison, David K. Jackson, Ian Johnston, Dominic Kwiatkowski, Cordelia Langford, John Sillitoe on behalf of the Wellcome Sanger Institute COVID-19 Surveillance Team |
| EPI_ISL_3365155, EPI_ISL_3365156, EPI_ISL_3365157, EPI_ISL_3365158, EPI_ISL_3365159 | Lighthouse Lab in Alderley Park | Wellcome Sanger Institute for the COVID-19 Genomics UK (COG-UK) Consortium | Jacquelyn Wynn, Mairead Hyland, The Lighthouse Lab in Alderley Park and Alex Alderton, Roberto Amato, Jeffrey Barrett, Sonia Goncalves, Ewan Harrison, David K. Jackson, Ian Johnston, Dominic Kwiatkowski, Cordelia Langford, John Sillitoe on behalf of the Wellcome Sanger Institute COVID-19 Surveillance Team |
| EPI_ISL_3365160, EPI_ISL_3365161 | Lighthouse Lab in Milton Keynes | Wellcome Sanger Institute for the COVID-19 Genomics UK (COG-UK) Consortium | The Lighthouse Lab in Milton Keynes and Alex Alderton, Roberto Amato, Jeffrey Barrett, Sonia Goncalves, Ewan Harrison, David K. Jackson, Ian Johnston, Dominic Kwiatkowski, Cordelia Langford, John Sillitoe on behalf of the Wellcome Sanger Institute COVID-19 Surveillance Team |
| EPI_ISL_3365162 | Lighthouse Lab in Alderley Park | Wellcome Sanger Institute for the COVID-19 Genomics UK (COG-UK) Consortium | Jacquelyn Wynn, Mairead Hyland, The Lighthouse Lab in Alderley Park and Alex Alderton, Roberto Amato, Jeffrey Barrett, Sonia Goncalves, Ewan Harrison, David K. Jackson, Ian Johnston, Dominic Kwiatkowski, Cordelia Langford, John Sillitoe on behalf of the Wellcome Sanger Institute COVID-19 Surveillance Team |
| EPI_ISL_3365163 | Berkshire and Surrey Pathology Services Lighthouse Laboratory | Wellcome Sanger Institute for the COVID-19 Genomics UK (COG-UK) Consortium | Berkshire and Surrey Pathology Services Lighthouse Laboratory and Alex Alderton, Roberto Amato, Jeffrey Barrett, Sonia Goncalves, Ewan Harrison, David K. Jackson, Ian Johnston, Dominic Kwiatkowski, Cordelia Langford, John Sillitoe on behalf of the Wellcome Sanger Institute COVID-19 Surveillance Team |
| EPI_ISL_3365164, EPI_ISL_3365165 | Lighthouse Lab in Alderley Park | Wellcome Sanger Institute for the COVID-19 Genomics UK (COG-UK) Consortium | Jacquelyn Wynn, Mairead Hyland, The Lighthouse Lab in Alderley Park and Alex Alderton, Roberto Amato, Jeffrey Barrett, Sonia Goncalves, Ewan Harrison, David K. Jackson, Ian Johnston, Dominic Kwiatkowski, Cordelia Langford, John Sillitoe on behalf of the Wellcome Sanger Institute COVID-19 Surveillance Team |
| EPI_ISL_3365166, EPI_ISL_3365167 | Lighthouse Lab in Milton Keynes | Wellcome Sanger Institute for the COVID-19 Genomics UK (COG-UK) Consortium | The Lighthouse Lab in Milton Keynes and Alex Alderton, Roberto Amato, Jeffrey Barrett, Sonia Goncalves, Ewan Harrison, David K. Jackson, Ian Johnston, Dominic Kwiatkowski, Cordelia Langford, John Sillitoe on behalf of the Wellcome Sanger Institute COVID-19 Surveillance Team |
| EPI_ISL_3365168 | Berkshire and Surrey Pathology Services Lighthouse Laboratory | Wellcome Sanger Institute for the COVID-19 Genomics UK (COG-UK) Consortium | Berkshire and Surrey Pathology Services Lighthouse Laboratory and Alex Alderton, Roberto Amato, Jeffrey Barrett, Sonia Goncalves, Ewan Harrison, David K. Jackson, Ian Johnston, Dominic Kwiatkowski, Cordelia Langford, John Sillitoe on behalf of the Wellcome Sanger Institute COVID-19 Surveillance Team |
| EPI_ISL_3365169 | Lighthouse Lab in Milton Keynes | Wellcome Sanger Institute for the COVID-19 Genomics UK (COG-UK) Consortium | The Lighthouse Lab in Milton Keynes and Alex Alderton, Roberto Amato, Jeffrey Barrett, Sonia Goncalves, Ewan Harrison, David K. Jackson, Ian Johnston, Dominic Kwiatkowski, Cordelia Langford, John Sillitoe on behalf of the Wellcome Sanger Institute COVID-19 Surveillance Team |
| EPI_ISL_3365170 | Berkshire and Surrey Pathology Services Lighthouse Laboratory | Wellcome Sanger Institute for the COVID-19 Genomics UK (COG-UK) Consortium | Berkshire and Surrey Pathology Services Lighthouse Laboratory and Alex Alderton, Roberto Amato, Jeffrey Barrett, Sonia Goncalves, Ewan Harrison, David K. Jackson, Ian Johnston, Dominic Kwiatkowski, Cordelia Langford, John Sillitoe on behalf of the Wellcome Sanger Institute COVID-19 Surveillance Team |
| EPI_ISL_3365171 | Lighthouse Lab in Alderley Park | Wellcome Sanger Institute for the COVID-19 Genomics UK (COG-UK) Consortium | Jacquelyn Wynn, Mairead Hyland, The Lighthouse Lab in Alderley Park and Alex Alderton, Roberto Amato, Jeffrey Barrett, Sonia Goncalves, Ewan Harrison, David K. Jackson, Ian Johnston, Dominic Kwiatkowski, Cordelia Langford, John Sillitoe on behalf of the Wellcome Sanger Institute COVID-19 Surveillance Team |
| EPI_ISL_3365172, EPI_ISL_3365173 | Lighthouse Lab in Milton Keynes | Wellcome Sanger Institute for the COVID-19 Genomics UK (COG-UK) Consortium | The Lighthouse Lab in Milton Keynes and Alex Alderton, Roberto Amato, Jeffrey Barrett, Sonia Goncalves, Ewan Harrison, David K. Jackson, Ian Johnston, Dominic Kwiatkowski, Cordelia Langford, John Sillitoe on behalf of the Wellcome Sanger Institute COVID-19 Surveillance Team |
| EPI_ISL_3365174 | Berkshire and Surrey Pathology Services Lighthouse Laboratory | Wellcome Sanger Institute for the COVID-19 Genomics UK (COG-UK) Consortium | Berkshire and Surrey Pathology Services Lighthouse Laboratory and Alex Alderton, Roberto Amato, Jeffrey Barrett, Sonia Goncalves, Ewan Harrison, David K. Jackson, Ian Johnston, Dominic Kwiatkowski, Cordelia Langford, John Sillitoe on behalf of the Wellcome Sanger Institute COVID-19 Surveillance Team |
| EPI_ISL_3365175 | Lighthouse Lab in Alderley Park | Wellcome Sanger Institute for the COVID-19 Genomics UK (COG-UK) Consortium | Jacquelyn Wynn, Mairead Hyland, The Lighthouse Lab in Alderley Park and Alex Alderton, Roberto Amato, Jeffrey Barrett, Sonia Goncalves, Ewan Harrison, David K. Jackson, Ian Johnston, Dominic Kwiatkowski, Cordelia Langford, John Sillitoe on behalf of the Wellcome Sanger Institute COVID-19 Surveillance Team |
| EPI_ISL_3365176, EPI_ISL_3365177 | Berkshire and Surrey Pathology Services Lighthouse Laboratory | Wellcome Sanger Institute for the COVID-19 Genomics UK (COG-UK) Consortium | Berkshire and Surrey Pathology Services Lighthouse Laboratory and Alex Alderton, Roberto Amato, Jeffrey Barrett, Sonia Goncalves, Ewan Harrison, David K. Jackson, Ian Johnston, Dominic Kwiatkowski, Cordelia Langford, John Sillitoe on behalf of the Wellcome Sanger Institute COVID-19 Surveillance Team |

| | | | |
|---|---|---|---|
| EPI_ISL_3365178 | Lighthouse Lab in Alderley Park | Wellcome Sanger Institute for the COVID-19 Genomics UK (COG-UK) Consortium | Jacquelyn Wynn, Mairead Hyland, The Lighthouse Lab in Alderley Park and Alex Alderton, Roberto Amato, Jeffrey Barrett, Sonia Goncalves, Ewan Harrison, David K. Jackson, Ian Johnston, Dominic Kwiatkowski, Cordelia Langford, John Sillitoe on behalf of the Wellcome Sanger Institute COVID-19 Surveillance Team |
| EPI_ISL_3365179, EPI_ISL_3365180 | Berkshire and Surrey Pathology Services Lighthouse Laboratory | Wellcome Sanger Institute for the COVID-19 Genomics UK (COG-UK) Consortium | Berkshire and Surrey Pathology Services Lighthouse Laboratory and Alex Alderton, Roberto Amato, Jeffrey Barrett, Sonia Goncalves, Ewan Harrison, David K. Jackson, Ian Johnston, Dominic Kwiatkowski, Cordelia Langford, John Sillitoe on behalf of the Wellcome Sanger Institute COVID-19 Surveillance Team |
| EPI_ISL_3365181 | Lighthouse Lab in Milton Keynes | Wellcome Sanger Institute for the COVID-19 Genomics UK (COG-UK) Consortium | The Lighthouse Lab in Milton Keynes and Alex Alderton, Roberto Amato, Jeffrey Barrett, Sonia Goncalves, Ewan Harrison, David K. Jackson, Ian Johnston, Dominic Kwiatkowski, Cordelia Langford, John Sillitoe on behalf of the Wellcome Sanger Institute COVID-19 Surveillance Team |
| EPI_ISL_3365182, EPI_ISL_3365183 | Berkshire and Surrey Pathology Services Lighthouse Laboratory | Wellcome Sanger Institute for the COVID-19 Genomics UK (COG-UK) Consortium | Berkshire and Surrey Pathology Services Lighthouse Laboratory and Alex Alderton, Roberto Amato, Jeffrey Barrett, Sonia Goncalves, Ewan Harrison, David K. Jackson, Ian Johnston, Dominic Kwiatkowski, Cordelia Langford, John Sillitoe on behalf of the Wellcome Sanger Institute COVID-19 Surveillance Team |
| EPI_ISL_3365184, EPI_ISL_3365185, EPI_ISL_3365186 | Lighthouse Lab in Alderley Park | Wellcome Sanger Institute for the COVID-19 Genomics UK (COG-UK) Consortium | Jacquelyn Wynn, Mairead Hyland, The Lighthouse Lab in Alderley Park and Alex Alderton, Roberto Amato, Jeffrey Barrett, Sonia Goncalves, Ewan Harrison, David K. Jackson, Ian Johnston, Dominic Kwiatkowski, Cordelia Langford, John Sillitoe on behalf of the Wellcome Sanger Institute COVID-19 Surveillance Team |
| EPI_ISL_3365187, EPI_ISL_3365188 | Berkshire and Surrey Pathology Services Lighthouse Laboratory | Wellcome Sanger Institute for the COVID-19 Genomics UK (COG-UK) Consortium | Berkshire and Surrey Pathology Services Lighthouse Laboratory and Alex Alderton, Roberto Amato, Jeffrey Barrett, Sonia Goncalves, Ewan Harrison, David K. Jackson, Ian Johnston, Dominic Kwiatkowski, Cordelia Langford, John Sillitoe on behalf of the Wellcome Sanger Institute COVID-19 Surveillance Team |
| EPI_ISL_3365189 | Lighthouse Lab in Milton Keynes | Wellcome Sanger Institute for the COVID-19 Genomics UK (COG-UK) Consortium | The Lighthouse Lab in Milton Keynes and Alex Alderton, Roberto Amato, Jeffrey Barrett, Sonia Goncalves, Ewan Harrison, David K. Jackson, Ian Johnston, Dominic Kwiatkowski, Cordelia Langford, John Sillitoe on behalf of the Wellcome Sanger Institute COVID-19 Surveillance Team |
| EPI_ISL_3365190, EPI_ISL_3365191 | Lighthouse Lab in Alderley Park | Wellcome Sanger Institute for the COVID-19 Genomics UK (COG-UK) Consortium | Jacquelyn Wynn, Mairead Hyland, The Lighthouse Lab in Alderley Park and Alex Alderton, Roberto Amato, Jeffrey Barrett, Sonia Goncalves, Ewan Harrison, David K. Jackson, Ian Johnston, Dominic Kwiatkowski, Cordelia Langford, John Sillitoe on behalf of the Wellcome Sanger Institute COVID-19 Surveillance Team |
| EPI_ISL_3365192 | Lighthouse Lab in Milton Keynes | Wellcome Sanger Institute for the COVID-19 Genomics UK (COG-UK) Consortium | The Lighthouse Lab in Milton Keynes and Alex Alderton, Roberto Amato, Jeffrey Barrett, Sonia Goncalves, Ewan Harrison, David K. Jackson, Ian Johnston, Dominic Kwiatkowski, Cordelia Langford, John Sillitoe on behalf of the Wellcome Sanger Institute COVID-19 Surveillance Team |
| EPI_ISL_3365193, EPI_ISL_3365194, EPI_ISL_3365195, EPI_ISL_3365196, EPI_ISL_3365197, EPI_ISL_3365198 | Lighthouse Lab in Alderley Park | Wellcome Sanger Institute for the COVID-19 Genomics UK (COG-UK) Consortium | Jacquelyn Wynn, Mairead Hyland, The Lighthouse Lab in Alderley Park and Alex Alderton, Roberto Amato, Jeffrey Barrett, Sonia Goncalves, Ewan Harrison, David K. Jackson, Ian Johnston, Dominic Kwiatkowski, Cordelia Langford, John Sillitoe on behalf of the Wellcome Sanger Institute COVID-19 Surveillance Team |
| EPI_ISL_3365199 | Lighthouse Lab in Milton Keynes | Wellcome Sanger Institute for the COVID-19 Genomics UK (COG-UK) Consortium | The Lighthouse Lab in Milton Keynes and Alex Alderton, Roberto Amato, Jeffrey Barrett, Sonia Goncalves, Ewan Harrison, David K. Jackson, Ian Johnston, Dominic Kwiatkowski, Cordelia Langford, John Sillitoe on behalf of the Wellcome Sanger Institute COVID-19 Surveillance Team |
| EPI_ISL_3365201, EPI_ISL_3365202, EPI_ISL_3365203, EPI_ISL_3365204, EPI_ISL_3365205 | Lighthouse Lab in Alderley Park | Wellcome Sanger Institute for the COVID-19 Genomics UK (COG-UK) Consortium | Jacquelyn Wynn, Mairead Hyland, The Lighthouse Lab in Alderley Park and Alex Alderton, Roberto Amato, Jeffrey Barrett, Sonia Goncalves, Ewan Harrison, David K. Jackson, Ian Johnston, Dominic Kwiatkowski, Cordelia Langford, John Sillitoe on behalf of the Wellcome Sanger Institute COVID-19 Surveillance Team |
| EPI_ISL_3365206 | Lighthouse Lab in Milton Keynes | Wellcome Sanger Institute for the COVID-19 Genomics UK (COG-UK) Consortium | The Lighthouse Lab in Milton Keynes and Alex Alderton, Roberto Amato, Jeffrey Barrett, Sonia Goncalves, Ewan Harrison, David K. Jackson, Ian Johnston, Dominic Kwiatkowski, Cordelia Langford, John Sillitoe on behalf of the Wellcome Sanger Institute COVID-19 Surveillance Team |
| EPI_ISL_3365207 | Berkshire and Surrey Pathology Services Lighthouse Laboratory | Wellcome Sanger Institute for the COVID-19 Genomics UK (COG-UK) Consortium | Berkshire and Surrey Pathology Services Lighthouse Laboratory and Alex Alderton, Roberto Amato, Jeffrey Barrett, Sonia Goncalves, Ewan Harrison, David K. Jackson, Ian Johnston, Dominic Kwiatkowski, Cordelia Langford, John Sillitoe on behalf of the Wellcome Sanger Institute COVID-19 Surveillance Team |
| EPI_ISL_3365208 | Lighthouse Lab in Milton Keynes | Wellcome Sanger Institute for the COVID-19 Genomics UK (COG-UK) Consortium | The Lighthouse Lab in Milton Keynes and Alex Alderton, Roberto Amato, Jeffrey Barrett, Sonia Goncalves, Ewan Harrison, David K. Jackson, Ian Johnston, Dominic Kwiatkowski, Cordelia Langford, John Sillitoe on behalf of the Wellcome Sanger Institute COVID-19 Surveillance Team |
| EPI_ISL_3365209, EPI_ISL_3365210 | Berkshire and Surrey Pathology Services Lighthouse Laboratory | Wellcome Sanger Institute for the COVID-19 Genomics UK (COG-UK) Consortium | Berkshire and Surrey Pathology Services Lighthouse Laboratory and Alex Alderton, Roberto Amato, Jeffrey Barrett, Sonia Goncalves, Ewan Harrison, David K. Jackson, Ian Johnston, Dominic Kwiatkowski, Cordelia Langford, John Sillitoe on behalf of the Wellcome Sanger Institute COVID-19 Surveillance Team |
| EPI_ISL_3365211 | Lighthouse Lab in Alderley Park | Wellcome Sanger Institute for the COVID-19 Genomics UK (COG-UK) Consortium | Jacquelyn Wynn, Mairead Hyland, The Lighthouse Lab in Alderley Park and Alex Alderton, Roberto Amato, Jeffrey Barrett, Sonia Goncalves, Ewan Harrison, David K. Jackson, Ian Johnston, Dominic Kwiatkowski, Cordelia Langford, John Sillitoe on behalf of the Wellcome Sanger Institute COVID-19 Surveillance Team |
| EPI_ISL_3365212 | Berkshire and Surrey Pathology Services Lighthouse Laboratory | Wellcome Sanger Institute for the COVID-19 Genomics UK (COG-UK) Consortium | Berkshire and Surrey Pathology Services Lighthouse Laboratory and Alex Alderton, Roberto Amato, Jeffrey Barrett, Sonia Goncalves, Ewan Harrison, David K. Jackson, Ian Johnston, Dominic Kwiatkowski, Cordelia Langford, John Sillitoe on behalf of the Wellcome Sanger Institute COVID-19 Surveillance Team |
| EPI_ISL_3365213, EPI_ISL_3365214 | Lighthouse Lab in Alderley Park | Wellcome Sanger Institute for the COVID-19 Genomics UK (COG-UK) Consortium | Jacquelyn Wynn, Mairead Hyland, The Lighthouse Lab in Alderley Park and Alex Alderton, Roberto Amato, Jeffrey Barrett, Sonia Goncalves, Ewan Harrison, David K. Jackson, Ian Johnston, Dominic Kwiatkowski, Cordelia Langford, John Sillitoe on behalf of the Wellcome Sanger Institute COVID-19 Surveillance Team |
| EPI_ISL_3365215, EPI_ISL_3365216 | Lighthouse Lab in Milton Keynes | Wellcome Sanger Institute for the COVID-19 Genomics UK (COG-UK) Consortium | The Lighthouse Lab in Milton Keynes and Alex Alderton, Roberto Amato, Jeffrey Barrett, Sonia Goncalves, Ewan Harrison, David K. Jackson, Ian Johnston, Dominic Kwiatkowski, Cordelia Langford, John Sillitoe on behalf of the Wellcome Sanger Institute COVID-19 Surveillance Team |
| EPI_ISL_3365217 | Lighthouse Lab in Alderley Park | Wellcome Sanger Institute for the COVID-19 Genomics UK (COG-UK) Consortium | Jacquelyn Wynn, Mairead Hyland, The Lighthouse Lab in Alderley Park and Alex Alderton, Roberto Amato, Jeffrey Barrett, Sonia Goncalves, Ewan Harrison, David K. Jackson, Ian Johnston, Dominic Kwiatkowski, Cordelia Langford, John Sillitoe on behalf of the Wellcome Sanger Institute COVID-19 Surveillance Team |
| EPI_ISL_3365218 | Lighthouse Lab in Milton Keynes | Wellcome Sanger Institute for the COVID-19 Genomics UK (COG-UK) Consortium | The Lighthouse Lab in Milton Keynes and Alex Alderton, Roberto Amato, Jeffrey Barrett, Sonia Goncalves, Ewan Harrison, David K. Jackson, Ian Johnston, Dominic Kwiatkowski, Cordelia Langford, John Sillitoe on behalf of the Wellcome Sanger Institute COVID-19 Surveillance Team |
| EPI_ISL_3365219, EPI_ISL_3365220, EPI_ISL_3365221, EPI_ISL_3365222, EPI_ISL_3365223 | Lighthouse Lab in Alderley Park | Wellcome Sanger Institute for the COVID-19 Genomics UK (COG-UK) Consortium | Jacquelyn Wynn, Mairead Hyland, The Lighthouse Lab in Alderley Park and Alex Alderton, Roberto Amato, Jeffrey Barrett, Sonia Goncalves, Ewan Harrison, David K. Jackson, Ian Johnston, Dominic Kwiatkowski, Cordelia Langford, John Sillitoe on behalf of the Wellcome Sanger Institute COVID-19 Surveillance Team |
| EPI_ISL_3365224 | Berkshire and Surrey Pathology Services Lighthouse Laboratory | Wellcome Sanger Institute for the COVID-19 Genomics UK (COG-UK) Consortium | Berkshire and Surrey Pathology Services Lighthouse Laboratory and Alex Alderton, Roberto Amato, Jeffrey Barrett, Sonia Goncalves, Ewan Harrison, David K. Jackson, Ian Johnston, Dominic Kwiatkowski, Cordelia Langford, John Sillitoe on behalf of the Wellcome Sanger Institute COVID-19 Surveillance Team |
| EPI_ISL_3365225, EPI_ISL_3365226, EPI_ISL_3365227 | Lighthouse Lab in Milton Keynes | Wellcome Sanger Institute for the COVID-19 Genomics UK (COG-UK) Consortium | The Lighthouse Lab in Milton Keynes and Alex Alderton, Roberto Amato, Jeffrey Barrett, Sonia Goncalves, Ewan Harrison, David K. Jackson, Ian Johnston, Dominic Kwiatkowski, Cordelia Langford, John Sillitoe on behalf of the Wellcome Sanger Institute COVID-19 Surveillance Team |
| EPI_ISL_3365228 | Lighthouse Lab in Alderley Park | Wellcome Sanger Institute for the COVID-19 Genomics UK (COG-UK) Consortium | Jacquelyn Wynn, Mairead Hyland, The Lighthouse Lab in Alderley Park and Alex Alderton, Roberto Amato, Jeffrey Barrett, Sonia Goncalves, Ewan Harrison, David K. Jackson, Ian Johnston, Dominic Kwiatkowski, Cordelia Langford, John Sillitoe on behalf of the Wellcome Sanger Institute COVID-19 Surveillance Team |
| EPI_ISL_3365229 | Berkshire and Surrey Pathology Services Lighthouse Laboratory | Wellcome Sanger Institute for the COVID-19 Genomics UK (COG-UK) Consortium | Berkshire and Surrey Pathology Services Lighthouse Laboratory and Alex Alderton, Roberto Amato, Jeffrey Barrett, Sonia Goncalves, Ewan Harrison, David K. Jackson, Ian Johnston, Dominic Kwiatkowski, Cordelia Langford, John Sillitoe on behalf of the Wellcome Sanger Institute COVID-19 Surveillance Team |
| EPI_ISL_3365230, EPI_ISL_3365231 | Lighthouse Lab in Alderley Park | Wellcome Sanger Institute for the COVID-19 Genomics UK (COG-UK) Consortium | Jacquelyn Wynn, Mairead Hyland, The Lighthouse Lab in Alderley Park and Alex Alderton, Roberto Amato, Jeffrey Barrett, Sonia Goncalves, Ewan Harrison, David K. Jackson, Ian Johnston, Dominic Kwiatkowski, Cordelia Langford, John Sillitoe on behalf of the Wellcome Sanger Institute COVID-19 Surveillance Team |
| EPI_ISL_3365232 | Lighthouse Lab in Milton Keynes | Wellcome Sanger Institute for the COVID-19 Genomics UK (COG-UK) Consortium | The Lighthouse Lab in Milton Keynes and Alex Alderton, Roberto Amato, Jeffrey Barrett, Sonia Goncalves, Ewan Harrison, David K. Jackson, Ian Johnston, Dominic Kwiatkowski, Cordelia Langford, John Sillitoe on behalf of the Wellcome Sanger Institute COVID-19 Surveillance Team |
| EPI_ISL_3365233, EPI_ISL_3365234, EPI_ISL_3365235, EPI_ISL_3365236 | Lighthouse Lab in Alderley Park | Wellcome Sanger Institute for the COVID-19 Genomics UK (COG-UK) Consortium | Jacquelyn Wynn, Mairead Hyland, The Lighthouse Lab in Alderley Park and Alex Alderton, Roberto Amato, Jeffrey Barrett, Sonia Goncalves, Ewan Harrison, David K. Jackson, Ian Johnston, Dominic Kwiatkowski, Cordelia Langford, John Sillitoe on behalf of the Wellcome Sanger Institute COVID-19 Surveillance Team |
| EPI_ISL_3365237 | Lighthouse Lab in Milton Keynes | Wellcome Sanger Institute for the COVID-19 Genomics UK (COG-UK) Consortium | The Lighthouse Lab in Milton Keynes and Alex Alderton, Roberto Amato, Jeffrey Barrett, Sonia Goncalves, Ewan Harrison, David K. Jackson, Ian Johnston, Dominic Kwiatkowski, Cordelia Langford, John Sillitoe on behalf of the Wellcome Sanger Institute COVID-19 Surveillance Team |
| EPI_ISL_3365238 | Berkshire and Surrey Pathology Services Lighthouse Laboratory | Wellcome Sanger Institute for the COVID-19 Genomics UK (COG-UK) Consortium | Berkshire and Surrey Pathology Services Lighthouse Laboratory and Alex Alderton, Roberto Amato, Jeffrey Barrett, Sonia Goncalves, Ewan Harrison, David K. Jackson, Ian Johnston, Dominic Kwiatkowski, Cordelia Langford, John Sillitoe on behalf of the Wellcome Sanger Institute COVID-19 Surveillance Team |
| EPI_ISL_3365239 | Lighthouse Lab in Milton Keynes | Wellcome Sanger Institute for the COVID-19 Genomics UK (COG-UK) Consortium | The Lighthouse Lab in Milton Keynes and Alex Alderton, Roberto Amato, Jeffrey Barrett, Sonia Goncalves, Ewan Harrison, David K. Jackson, Ian Johnston, Dominic Kwiatkowski, Cordelia Langford, John Sillitoe on behalf of the Wellcome Sanger Institute COVID-19 Surveillance Team |
| EPI_ISL_3365240 | Lighthouse Lab in Alderley Park | Wellcome Sanger Institute for the COVID-19 Genomics UK (COG-UK) Consortium | Jacquelyn Wynn, Mairead Hyland, The Lighthouse Lab in Alderley Park and Alex Alderton, Roberto Amato, Jeffrey Barrett, Sonia Goncalves, Ewan Harrison, David K. Jackson, Ian Johnston, Dominic Kwiatkowski, Cordelia Langford, John Sillitoe on behalf of the Wellcome Sanger Institute COVID-19 Surveillance Team |
| EPI_ISL_3365241, EPI_ISL_3365242 | Berkshire and Surrey Pathology Services Lighthouse Laboratory | Wellcome Sanger Institute for the COVID-19 Genomics UK (COG-UK) Consortium | Berkshire and Surrey Pathology Services Lighthouse Laboratory and Alex Alderton, Roberto Amato, Jeffrey Barrett, Sonia Goncalves, Ewan Harrison, David K. Jackson, Ian Johnston, Dominic Kwiatkowski, Cordelia Langford, John Sillitoe on behalf of the Wellcome Sanger Institute COVID-19 Surveillance Team |
| EPI_ISL_3365243 | Lighthouse Lab in Alderley Park | Wellcome Sanger Institute for the COVID-19 Genomics UK (COG-UK) Consortium | Jacquelyn Wynn, Mairead Hyland, The Lighthouse Lab in Alderley Park and Alex Alderton, Roberto Amato, Jeffrey Barrett, Sonia Goncalves, Ewan Harrison, David K. Jackson, Ian Johnston, Dominic Kwiatkowski, Cordelia Langford, John Sillitoe on behalf of the Wellcome Sanger Institute COVID-19 Surveillance Team |
| EPI_ISL_3365244 | Berkshire and Surrey Pathology Services Lighthouse Laboratory | Wellcome Sanger Institute for the COVID-19 Genomics UK (COG-UK) Consortium | Berkshire and Surrey Pathology Services Lighthouse Laboratory and Alex Alderton, Roberto Amato, Jeffrey Barrett, Sonia Goncalves, Ewan Harrison, David K. Jackson, Ian Johnston, Dominic Kwiatkowski, Cordelia Langford, John Sillitoe on behalf of the Wellcome Sanger Institute COVID-19 Surveillance Team |
| EPI_ISL_3365245, EPI_ISL_3365246 | Lighthouse Lab in Milton Keynes | Wellcome Sanger Institute for the COVID-19 Genomics UK (COG-UK) Consortium | The Lighthouse Lab in Milton Keynes and Alex Alderton, Roberto Amato, Jeffrey Barrett, Sonia Goncalves, Ewan Harrison, David K. Jackson, Ian Johnston, Dominic Kwiatkowski, Cordelia Langford, John Sillitoe on behalf of the Wellcome Sanger Institute COVID-19 Surveillance Team |
| EPI_ISL_3365247, EPI_ISL_3365248 | Lighthouse Lab in Alderley Park | Wellcome Sanger Institute for the COVID-19 Genomics UK (COG-UK) Consortium | Jacquelyn Wynn, Mairead Hyland, The Lighthouse Lab in Alderley Park and Alex Alderton, Roberto Amato, Jeffrey Barrett, Sonia Goncalves, Ewan Harrison, David K. Jackson, Ian Johnston, Dominic Kwiatkowski, Cordelia Langford, John Sillitoe on behalf of the Wellcome Sanger Institute COVID-19 Surveillance Team |
| EPI_ISL_3365249 | Lighthouse Lab in Milton Keynes | Wellcome Sanger Institute for the COVID-19 Genomics UK (COG-UK) | The Lighthouse Lab in Milton Keynes and Alex Alderton, Roberto Amato, Jeffrey Barrett, Sonia Goncalves, Ewan Harrison, David K. Jackson, Ian Johnston, Dominic Kwiatkowski, Cordelia Langford, John Sillitoe on behalf of the Wellcome Sanger |

| | | | |
|---|---|---|---|
| | | Consortium | Institute COVID-19 Surveillance Team |
| EPI_ISL_3365250 | Berkshire and Surrey Pathology Services Lighthouse Laboratory | Wellcome Sanger Institute for the COVID-19 Genomics UK (COG-UK) Consortium | Berkshire and Surrey Pathology Services Lighthouse Laboratory and Alex Alderton, Roberto Amato, Jeffrey Barrett, Sonia Goncalves, Ewan Harrison, David K. Jackson, Ian Johnston, Dominic Kwiatkowski, Cordelia Langford, John Sillitoe on behalf of the Wellcome Sanger Institute COVID-19 Surveillance Team |
| EPI_ISL_3365251, EPI_ISL_3365252, EPI_ISL_3365253, EPI_ISL_3365254, EPI_ISL_3365255, EPI_ISL_3365256 | Lighthouse Lab in Alderley Park | Wellcome Sanger Institute for the COVID-19 Genomics UK (COG-UK) Consortium | Jacquelyn Wynn, Mairead Hyland, The Lighthouse Lab in Alderley Park and Alex Alderton, Roberto Amato, Jeffrey Barrett, Sonia Goncalves, Ewan Harrison, David K. Jackson, Ian Johnston, Dominic Kwiatkowski, Cordelia Langford, John Sillitoe on behalf of the Wellcome Sanger Institute COVID-19 Surveillance Team |
| EPI_ISL_3365257 | Lighthouse Lab in Milton Keynes | Wellcome Sanger Institute for the COVID-19 Genomics UK (COG-UK) Consortium | The Lighthouse Lab in Milton Keynes and Alex Alderton, Roberto Amato, Jeffrey Barrett, Sonia Goncalves, Ewan Harrison, David K. Jackson, Ian Johnston, Dominic Kwiatkowski, Cordelia Langford, John Sillitoe on behalf of the Wellcome Sanger Institute COVID-19 Surveillance Team |
| EPI_ISL_3365258, EPI_ISL_3365259 | Berkshire and Surrey Pathology Services Lighthouse Laboratory | Wellcome Sanger Institute for the COVID-19 Genomics UK (COG-UK) Consortium | Berkshire and Surrey Pathology Services Lighthouse Laboratory and Alex Alderton, Roberto Amato, Jeffrey Barrett, Sonia Goncalves, Ewan Harrison, David K. Jackson, Ian Johnston, Dominic Kwiatkowski, Cordelia Langford, John Sillitoe on behalf of the Wellcome Sanger Institute COVID-19 Surveillance Team |
| EPI_ISL_3365260 | Lighthouse Lab in Alderley Park | Wellcome Sanger Institute for the COVID-19 Genomics UK (COG-UK) Consortium | Jacquelyn Wynn, Mairead Hyland, The Lighthouse Lab in Alderley Park and Alex Alderton, Roberto Amato, Jeffrey Barrett, Sonia Goncalves, Ewan Harrison, David K. Jackson, Ian Johnston, Dominic Kwiatkowski, Cordelia Langford, John Sillitoe on behalf of the Wellcome Sanger Institute COVID-19 Surveillance Team |
| EPI_ISL_3365261 | Lighthouse Lab in Milton Keynes | Wellcome Sanger Institute for the COVID-19 Genomics UK (COG-UK) Consortium | The Lighthouse Lab in Milton Keynes and Alex Alderton, Roberto Amato, Jeffrey Barrett, Sonia Goncalves, Ewan Harrison, David K. Jackson, Ian Johnston, Dominic Kwiatkowski, Cordelia Langford, John Sillitoe on behalf of the Wellcome Sanger Institute COVID-19 Surveillance Team |
| EPI_ISL_3365262 | Lighthouse Lab in Alderley Park | Wellcome Sanger Institute for the COVID-19 Genomics UK (COG-UK) Consortium | Jacquelyn Wynn, Mairead Hyland, The Lighthouse Lab in Alderley Park and Alex Alderton, Roberto Amato, Jeffrey Barrett, Sonia Goncalves, Ewan Harrison, David K. Jackson, Ian Johnston, Dominic Kwiatkowski, Cordelia Langford, John Sillitoe on behalf of the Wellcome Sanger Institute COVID-19 Surveillance Team |
| EPI_ISL_3365263, EPI_ISL_3365264, EPI_ISL_3365265 | Berkshire and Surrey Pathology Services Lighthouse Laboratory | Wellcome Sanger Institute for the COVID-19 Genomics UK (COG-UK) Consortium | Berkshire and Surrey Pathology Services Lighthouse Laboratory and Alex Alderton, Roberto Amato, Jeffrey Barrett, Sonia Goncalves, Ewan Harrison, David K. Jackson, Ian Johnston, Dominic Kwiatkowski, Cordelia Langford, John Sillitoe on behalf of the Wellcome Sanger Institute COVID-19 Surveillance Team |
| EPI_ISL_3365266, EPI_ISL_3365267 | Lighthouse Lab in Alderley Park | Wellcome Sanger Institute for the COVID-19 Genomics UK (COG-UK) Consortium | Jacquelyn Wynn, Mairead Hyland, The Lighthouse Lab in Alderley Park and Alex Alderton, Roberto Amato, Jeffrey Barrett, Sonia Goncalves, Ewan Harrison, David K. Jackson, Ian Johnston, Dominic Kwiatkowski, Cordelia Langford, John Sillitoe on behalf of the Wellcome Sanger Institute COVID-19 Surveillance Team |
| EPI_ISL_3365268 | Lighthouse Lab in Milton Keynes | Wellcome Sanger Institute for the COVID-19 Genomics UK (COG-UK) Consortium | The Lighthouse Lab in Milton Keynes and Alex Alderton, Roberto Amato, Jeffrey Barrett, Sonia Goncalves, Ewan Harrison, David K. Jackson, Ian Johnston, Dominic Kwiatkowski, Cordelia Langford, John Sillitoe on behalf of the Wellcome Sanger Institute COVID-19 Surveillance Team |
| EPI_ISL_3365269 | Lighthouse Lab in Alderley Park | Wellcome Sanger Institute for the COVID-19 Genomics UK (COG-UK) Consortium | Jacquelyn Wynn, Mairead Hyland, The Lighthouse Lab in Alderley Park and Alex Alderton, Roberto Amato, Jeffrey Barrett, Sonia Goncalves, Ewan Harrison, David K. Jackson, Ian Johnston, Dominic Kwiatkowski, Cordelia Langford, John Sillitoe on behalf of the Wellcome Sanger Institute COVID-19 Surveillance Team |
| EPI_ISL_3365270, EPI_ISL_3365271 | Lighthouse Lab in Milton Keynes | Wellcome Sanger Institute for the COVID-19 Genomics UK (COG-UK) Consortium | The Lighthouse Lab in Milton Keynes and Alex Alderton, Roberto Amato, Jeffrey Barrett, Sonia Goncalves, Ewan Harrison, David K. Jackson, Ian Johnston, Dominic Kwiatkowski, Cordelia Langford, John Sillitoe on behalf of the Wellcome Sanger Institute COVID-19 Surveillance Team |
| EPI_ISL_3365272 | Lighthouse Lab in Alderley Park | Wellcome Sanger Institute for the COVID-19 Genomics UK (COG-UK) Consortium | Jacquelyn Wynn, Mairead Hyland, The Lighthouse Lab in Alderley Park and Alex Alderton, Roberto Amato, Jeffrey Barrett, Sonia Goncalves, Ewan Harrison, David K. Jackson, Ian Johnston, Dominic Kwiatkowski, Cordelia Langford, John Sillitoe on behalf of the Wellcome Sanger Institute COVID-19 Surveillance Team |
| EPI_ISL_3365273, EPI_ISL_3365274 | Lighthouse Lab in Milton Keynes | Wellcome Sanger Institute for the COVID-19 Genomics UK (COG-UK) Consortium | The Lighthouse Lab in Milton Keynes and Alex Alderton, Roberto Amato, Jeffrey Barrett, Sonia Goncalves, Ewan Harrison, David K. Jackson, Ian Johnston, Dominic Kwiatkowski, Cordelia Langford, John Sillitoe on behalf of the Wellcome Sanger Institute COVID-19 Surveillance Team |
| EPI_ISL_3365275 | Lighthouse Lab in Alderley Park | Wellcome Sanger Institute for the COVID-19 Genomics UK (COG-UK) Consortium | Jacquelyn Wynn, Mairead Hyland, The Lighthouse Lab in Alderley Park and Alex Alderton, Roberto Amato, Jeffrey Barrett, Sonia Goncalves, Ewan Harrison, David K. Jackson, Ian Johnston, Dominic Kwiatkowski, Cordelia Langford, John Sillitoe on behalf of the Wellcome Sanger Institute COVID-19 Surveillance Team |
| EPI_ISL_3365276 | Lighthouse Lab in Milton Keynes | Wellcome Sanger Institute for the COVID-19 Genomics UK (COG-UK) Consortium | The Lighthouse Lab in Milton Keynes and Alex Alderton, Roberto Amato, Jeffrey Barrett, Sonia Goncalves, Ewan Harrison, David K. Jackson, Ian Johnston, Dominic Kwiatkowski, Cordelia Langford, John Sillitoe on behalf of the Wellcome Sanger Institute COVID-19 Surveillance Team |
| EPI_ISL_3365277, EPI_ISL_3365278, EPI_ISL_3365279, EPI_ISL_3365280, EPI_ISL_3365281, EPI_ISL_3365282 | Lighthouse Lab in Alderley Park | Wellcome Sanger Institute for the COVID-19 Genomics UK (COG-UK) Consortium | Jacquelyn Wynn, Mairead Hyland, The Lighthouse Lab in Alderley Park and Alex Alderton, Roberto Amato, Jeffrey Barrett, Sonia Goncalves, Ewan Harrison, David K. Jackson, Ian Johnston, Dominic Kwiatkowski, Cordelia Langford, John Sillitoe on behalf of the Wellcome Sanger Institute COVID-19 Surveillance Team |
| EPI_ISL_3365283 | Berkshire and Surrey Pathology Services Lighthouse Laboratory | Wellcome Sanger Institute for the COVID-19 Genomics UK (COG-UK) Consortium | Berkshire and Surrey Pathology Services Lighthouse Laboratory and Alex Alderton, Roberto Amato, Jeffrey Barrett, Sonia Goncalves, Ewan Harrison, David K. Jackson, Ian Johnston, Dominic Kwiatkowski, Cordelia Langford, John Sillitoe on behalf of the Wellcome Sanger Institute COVID-19 Surveillance Team |
| EPI_ISL_3365284, EPI_ISL_3365285 | Lighthouse Lab in Alderley Park | Wellcome Sanger Institute for the COVID-19 Genomics UK (COG-UK) Consortium | Jacquelyn Wynn, Mairead Hyland, The Lighthouse Lab in Alderley Park and Alex Alderton, Roberto Amato, Jeffrey Barrett, Sonia Goncalves, Ewan Harrison, David K. Jackson, Ian Johnston, Dominic Kwiatkowski, Cordelia Langford, John Sillitoe on behalf of the Wellcome Sanger Institute COVID-19 Surveillance Team |
| EPI_ISL_3365286 | Berkshire and Surrey Pathology Services Lighthouse Laboratory | Wellcome Sanger Institute for the COVID-19 Genomics UK (COG-UK) Consortium | Berkshire and Surrey Pathology Services Lighthouse Laboratory and Alex Alderton, Roberto Amato, Jeffrey Barrett, Sonia Goncalves, Ewan Harrison, David K. Jackson, Ian Johnston, Dominic Kwiatkowski, Cordelia Langford, John Sillitoe on behalf of the Wellcome Sanger Institute COVID-19 Surveillance Team |
| EPI_ISL_3365287, EPI_ISL_3365288, EPI_ISL_3365289 | Lighthouse Lab in Alderley Park | Wellcome Sanger Institute for the COVID-19 Genomics UK (COG-UK) Consortium | Jacquelyn Wynn, Mairead Hyland, The Lighthouse Lab in Alderley Park and Alex Alderton, Roberto Amato, Jeffrey Barrett, Sonia Goncalves, Ewan Harrison, David K. Jackson, Ian Johnston, Dominic Kwiatkowski, Cordelia Langford, John Sillitoe on behalf of the Wellcome Sanger Institute COVID-19 Surveillance Team |
| EPI_ISL_3365290 | Berkshire and Surrey Pathology Services Lighthouse Laboratory | Wellcome Sanger Institute for the COVID-19 Genomics UK (COG-UK) Consortium | Berkshire and Surrey Pathology Services Lighthouse Laboratory and Alex Alderton, Roberto Amato, Jeffrey Barrett, Sonia Goncalves, Ewan Harrison, David K. Jackson, Ian Johnston, Dominic Kwiatkowski, Cordelia Langford, John Sillitoe on behalf of the Wellcome Sanger Institute COVID-19 Surveillance Team |
| EPI_ISL_3365291, EPI_ISL_3365292 | Lighthouse Lab in Alderley Park | Wellcome Sanger Institute for the COVID-19 Genomics UK (COG-UK) Consortium | Jacquelyn Wynn, Mairead Hyland, The Lighthouse Lab in Alderley Park and Alex Alderton, Roberto Amato, Jeffrey Barrett, Sonia Goncalves, Ewan Harrison, David K. Jackson, Ian Johnston, Dominic Kwiatkowski, Cordelia Langford, John Sillitoe on behalf of the Wellcome Sanger Institute COVID-19 Surveillance Team |
| EPI_ISL_3365293 | Berkshire and Surrey Pathology Services Lighthouse Laboratory | Wellcome Sanger Institute for the COVID-19 Genomics UK (COG-UK) Consortium | Berkshire and Surrey Pathology Services Lighthouse Laboratory and Alex Alderton, Roberto Amato, Jeffrey Barrett, Sonia Goncalves, Ewan Harrison, David K. Jackson, Ian Johnston, Dominic Kwiatkowski, Cordelia Langford, John Sillitoe on behalf of the Wellcome Sanger Institute COVID-19 Surveillance Team |
| EPI_ISL_3365294, EPI_ISL_3365295, EPI_ISL_3365296, EPI_ISL_3365297, EPI_ISL_3365298, EPI_ISL_3365299 | Lighthouse Lab in Alderley Park | Wellcome Sanger Institute for the COVID-19 Genomics UK (COG-UK) Consortium | Jacquelyn Wynn, Mairead Hyland, The Lighthouse Lab in Alderley Park and Alex Alderton, Roberto Amato, Jeffrey Barrett, Sonia Goncalves, Ewan Harrison, David K. Jackson, Ian Johnston, Dominic Kwiatkowski, Cordelia Langford, John Sillitoe on behalf of the Wellcome Sanger Institute COVID-19 Surveillance Team |
| EPI_ISL_3365300 | Berkshire and Surrey Pathology Services Lighthouse Laboratory | Wellcome Sanger Institute for the COVID-19 Genomics UK (COG-UK) Consortium | Berkshire and Surrey Pathology Services Lighthouse Laboratory and Alex Alderton, Roberto Amato, Jeffrey Barrett, Sonia Goncalves, Ewan Harrison, David K. Jackson, Ian Johnston, Dominic Kwiatkowski, Cordelia Langford, John Sillitoe on behalf of the Wellcome Sanger Institute COVID-19 Surveillance Team |
| EPI_ISL_3365301 | Lighthouse Lab in Alderley Park | Wellcome Sanger Institute for the COVID-19 Genomics UK (COG-UK) Consortium | Jacquelyn Wynn, Mairead Hyland, The Lighthouse Lab in Alderley Park and Alex Alderton, Roberto Amato, Jeffrey Barrett, Sonia Goncalves, Ewan Harrison, David K. Jackson, Ian Johnston, Dominic Kwiatkowski, Cordelia Langford, John Sillitoe on behalf of the Wellcome Sanger Institute COVID-19 Surveillance Team |
| EPI_ISL_3365302, EPI_ISL_3365303 | Berkshire and Surrey Pathology Services Lighthouse Laboratory | Wellcome Sanger Institute for the COVID-19 Genomics UK (COG-UK) Consortium | Berkshire and Surrey Pathology Services Lighthouse Laboratory and Alex Alderton, Roberto Amato, Jeffrey Barrett, Sonia Goncalves, Ewan Harrison, David K. Jackson, Ian Johnston, Dominic Kwiatkowski, Cordelia Langford, John Sillitoe on behalf of the Wellcome Sanger Institute COVID-19 Surveillance Team |
| EPI_ISL_3365304, EPI_ISL_3365305 | Lighthouse Lab in Milton Keynes | Wellcome Sanger Institute for the COVID-19 Genomics UK (COG-UK) Consortium | The Lighthouse Lab in Milton Keynes and Alex Alderton, Roberto Amato, Jeffrey Barrett, Sonia Goncalves, Ewan Harrison, David K. Jackson, Ian Johnston, Dominic Kwiatkowski, Cordelia Langford, John Sillitoe on behalf of the Wellcome Sanger Institute COVID-19 Surveillance Team |
| EPI_ISL_3365306 | Lighthouse Lab in Alderley Park | Wellcome Sanger Institute for the COVID-19 Genomics UK (COG-UK) Consortium | Jacquelyn Wynn, Mairead Hyland, The Lighthouse Lab in Alderley Park and Alex Alderton, Roberto Amato, Jeffrey Barrett, Sonia Goncalves, Ewan Harrison, David K. Jackson, Ian Johnston, Dominic Kwiatkowski, Cordelia Langford, John Sillitoe on behalf of the Wellcome Sanger Institute COVID-19 Surveillance Team |
| EPI_ISL_3365307, EPI_ISL_3365308 | Lighthouse Lab in Milton Keynes | Wellcome Sanger Institute for the COVID-19 Genomics UK (COG-UK) Consortium | The Lighthouse Lab in Milton Keynes and Alex Alderton, Roberto Amato, Jeffrey Barrett, Sonia Goncalves, Ewan Harrison, David K. Jackson, Ian Johnston, Dominic Kwiatkowski, Cordelia Langford, John Sillitoe on behalf of the Wellcome Sanger Institute COVID-19 Surveillance Team |
| EPI_ISL_3365309 | Berkshire and Surrey Pathology Services Lighthouse Laboratory | Wellcome Sanger Institute for the COVID-19 Genomics UK (COG-UK) Consortium | Berkshire and Surrey Pathology Services Lighthouse Laboratory and Alex Alderton, Roberto Amato, Jeffrey Barrett, Sonia Goncalves, Ewan Harrison, David K. Jackson, Ian Johnston, Dominic Kwiatkowski, Cordelia Langford, John Sillitoe on behalf of the Wellcome Sanger Institute COVID-19 Surveillance Team |
| EPI_ISL_3365310 | Lighthouse Lab in Alderley Park | Wellcome Sanger Institute for the COVID-19 Genomics UK (COG-UK) Consortium | Jacquelyn Wynn, Mairead Hyland, The Lighthouse Lab in Alderley Park and Alex Alderton, Roberto Amato, Jeffrey Barrett, Sonia Goncalves, Ewan Harrison, David K. Jackson, Ian Johnston, Dominic Kwiatkowski, Cordelia Langford, John Sillitoe on behalf of the Wellcome Sanger Institute COVID-19 Surveillance Team |
| EPI_ISL_3365311 | Lighthouse Lab in Milton Keynes | Wellcome Sanger Institute for the COVID-19 Genomics UK (COG-UK) Consortium | The Lighthouse Lab in Milton Keynes and Alex Alderton, Roberto Amato, Jeffrey Barrett, Sonia Goncalves, Ewan Harrison, David K. Jackson, Ian Johnston, Dominic Kwiatkowski, Cordelia Langford, John Sillitoe on behalf of the Wellcome Sanger Institute COVID-19 Surveillance Team |
| EPI_ISL_3365312 | Berkshire and Surrey Pathology Services Lighthouse Laboratory | Wellcome Sanger Institute for the COVID-19 Genomics UK (COG-UK) Consortium | Berkshire and Surrey Pathology Services Lighthouse Laboratory and Alex Alderton, Roberto Amato, Jeffrey Barrett, Sonia Goncalves, Ewan Harrison, David K. Jackson, Ian Johnston, Dominic Kwiatkowski, Cordelia Langford, John Sillitoe on behalf of the Wellcome Sanger Institute COVID-19 Surveillance Team |
| EPI_ISL_3365313, EPI_ISL_3365314 | Lighthouse Lab in Alderley Park | Wellcome Sanger Institute for the COVID-19 Genomics UK (COG-UK) Consortium | Jacquelyn Wynn, Mairead Hyland, The Lighthouse Lab in Alderley Park and Alex Alderton, Roberto Amato, Jeffrey Barrett, Sonia Goncalves, Ewan Harrison, David K. Jackson, Ian Johnston, Dominic Kwiatkowski, Cordelia Langford, John Sillitoe on behalf of the Wellcome Sanger Institute COVID-19 Surveillance Team |
| EPI_ISL_3365315 | Lighthouse Lab in Milton Keynes | Wellcome Sanger Institute for the COVID-19 Genomics UK (COG-UK) Consortium | The Lighthouse Lab in Milton Keynes and Alex Alderton, Roberto Amato, Jeffrey Barrett, Sonia Goncalves, Ewan Harrison, David K. Jackson, Ian Johnston, Dominic Kwiatkowski, Cordelia Langford, John Sillitoe on behalf of the Wellcome Sanger Institute COVID-19 Surveillance Team |
| EPI_ISL_3365316, EPI_ISL_3365317 | Lighthouse Lab in Alderley Park | Wellcome Sanger Institute for the COVID-19 Genomics UK (COG-UK) Consortium | Jacquelyn Wynn, Mairead Hyland, The Lighthouse Lab in Alderley Park and Alex Alderton, Roberto Amato, Jeffrey Barrett, Sonia Goncalves, Ewan Harrison, David K. Jackson, Ian Johnston, Dominic Kwiatkowski, Cordelia Langford, John Sillitoe on behalf of the Wellcome Sanger Institute COVID-19 Surveillance Team |
| EPI_ISL_3365318 | Lighthouse Lab in Milton Keynes | Wellcome Sanger Institute for the COVID-19 Genomics UK (COG-UK) Consortium | The Lighthouse Lab in Milton Keynes and Alex Alderton, Roberto Amato, Jeffrey Barrett, Sonia Goncalves, Ewan Harrison, David K. Jackson, Ian Johnston, Dominic Kwiatkowski, Cordelia Langford, John Sillitoe on behalf of the Wellcome Sanger Institute COVID-19 Surveillance Team |

| | | | |
|---|---|---|---|
| EPI_ISL_3365319 | Berkshire and Surrey Pathology Services Lighthouse Laboratory | Wellcome Sanger Institute for the COVID-19 Genomics UK (COG-UK) Consortium | Berkshire and Surrey Pathology Services Lighthouse Laboratory and Alex Alderton, Roberto Amato, Jeffrey Barrett, Sonia Goncalves, Ewan Harrison, David K. Jackson, Ian Johnston, Dominic Kwiatkowski, Cordelia Langford, John Sillitoe on behalf of the Wellcome Sanger Institute COVID-19 Surveillance Team |
| EPI_ISL_3365320, EPI_ISL_3365321, EPI_ISL_3365322 | Lighthouse Lab in Alderley Park | Wellcome Sanger Institute for the COVID-19 Genomics UK (COG-UK) Consortium | Jacquelyn Wynn, Mairead Hyland, The Lighthouse Lab in Alderley Park and Alex Alderton, Roberto Amato, Jeffrey Barrett, Sonia Goncalves, Ewan Harrison, David K. Jackson, Ian Johnston, Dominic Kwiatkowski, Cordelia Langford, John Sillitoe on behalf of the Wellcome Sanger Institute COVID-19 Surveillance Team |
| EPI_ISL_3365323 | Lighthouse Lab in Milton Keynes | Wellcome Sanger Institute for the COVID-19 Genomics UK (COG-UK) Consortium | The Lighthouse Lab in Milton Keynes and Alex Alderton, Roberto Amato, Jeffrey Barrett, Sonia Goncalves, Ewan Harrison, David K. Jackson, Ian Johnston, Dominic Kwiatkowski, Cordelia Langford, John Sillitoe on behalf of the Wellcome Sanger Institute COVID-19 Surveillance Team |
| EPI_ISL_3365324 | Lighthouse Lab in Alderley Park | Wellcome Sanger Institute for the COVID-19 Genomics UK (COG-UK) Consortium | Jacquelyn Wynn, Mairead Hyland, The Lighthouse Lab in Alderley Park and Alex Alderton, Roberto Amato, Jeffrey Barrett, Sonia Goncalves, Ewan Harrison, David K. Jackson, Ian Johnston, Dominic Kwiatkowski, Cordelia Langford, John Sillitoe on behalf of the Wellcome Sanger Institute COVID-19 Surveillance Team |
| EPI_ISL_3365325 | Lighthouse Lab in Milton Keynes | Wellcome Sanger Institute for the COVID-19 Genomics UK (COG-UK) Consortium | The Lighthouse Lab in Milton Keynes and Alex Alderton, Roberto Amato, Jeffrey Barrett, Sonia Goncalves, Ewan Harrison, David K. Jackson, Ian Johnston, Dominic Kwiatkowski, Cordelia Langford, John Sillitoe on behalf of the Wellcome Sanger Institute COVID-19 Surveillance Team |
| EPI_ISL_3365326 | Lighthouse Lab in Alderley Park | Wellcome Sanger Institute for the COVID-19 Genomics UK (COG-UK) Consortium | Jacquelyn Wynn, Mairead Hyland, The Lighthouse Lab in Alderley Park and Alex Alderton, Roberto Amato, Jeffrey Barrett, Sonia Goncalves, Ewan Harrison, David K. Jackson, Ian Johnston, Dominic Kwiatkowski, Cordelia Langford, John Sillitoe on behalf of the Wellcome Sanger Institute COVID-19 Surveillance Team |
| EPI_ISL_3365327 | Berkshire and Surrey Pathology Services Lighthouse Laboratory | Wellcome Sanger Institute for the COVID-19 Genomics UK (COG-UK) Consortium | Berkshire and Surrey Pathology Services Lighthouse Laboratory and Alex Alderton, Roberto Amato, Jeffrey Barrett, Sonia Goncalves, Ewan Harrison, David K. Jackson, Ian Johnston, Dominic Kwiatkowski, Cordelia Langford, John Sillitoe on behalf of the Wellcome Sanger Institute COVID-19 Surveillance Team |
| EPI_ISL_3365328 | Lighthouse Lab in Milton Keynes | Wellcome Sanger Institute for the COVID-19 Genomics UK (COG-UK) Consortium | The Lighthouse Lab in Milton Keynes and Alex Alderton, Roberto Amato, Jeffrey Barrett, Sonia Goncalves, Ewan Harrison, David K. Jackson, Ian Johnston, Dominic Kwiatkowski, Cordelia Langford, John Sillitoe on behalf of the Wellcome Sanger Institute COVID-19 Surveillance Team |
| EPI_ISL_3365329 | Lighthouse Lab in Alderley Park | Wellcome Sanger Institute for the COVID-19 Genomics UK (COG-UK) Consortium | Jacquelyn Wynn, Mairead Hyland, The Lighthouse Lab in Alderley Park and Alex Alderton, Roberto Amato, Jeffrey Barrett, Sonia Goncalves, Ewan Harrison, David K. Jackson, Ian Johnston, Dominic Kwiatkowski, Cordelia Langford, John Sillitoe on behalf of the Wellcome Sanger Institute COVID-19 Surveillance Team |
| EPI_ISL_3365330 | Lighthouse Lab in Milton Keynes | Wellcome Sanger Institute for the COVID-19 Genomics UK (COG-UK) Consortium | The Lighthouse Lab in Milton Keynes and Alex Alderton, Roberto Amato, Jeffrey Barrett, Sonia Goncalves, Ewan Harrison, David K. Jackson, Ian Johnston, Dominic Kwiatkowski, Cordelia Langford, John Sillitoe on behalf of the Wellcome Sanger Institute COVID-19 Surveillance Team |
| EPI_ISL_3365331, EPI_ISL_3365332 | Lighthouse Lab in Alderley Park | Wellcome Sanger Institute for the COVID-19 Genomics UK (COG-UK) Consortium | Jacquelyn Wynn, Mairead Hyland, The Lighthouse Lab in Alderley Park and Alex Alderton, Roberto Amato, Jeffrey Barrett, Sonia Goncalves, Ewan Harrison, David K. Jackson, Ian Johnston, Dominic Kwiatkowski, Cordelia Langford, John Sillitoe on behalf of the Wellcome Sanger Institute COVID-19 Surveillance Team |
| EPI_ISL_3365333, EPI_ISL_3365334, EPI_ISL_3365335, EPI_ISL_3365336, EPI_ISL_3365337 | Berkshire and Surrey Pathology Services Lighthouse Laboratory | Wellcome Sanger Institute for the COVID-19 Genomics UK (COG-UK) Consortium | Berkshire and Surrey Pathology Services Lighthouse Laboratory and Alex Alderton, Roberto Amato, Jeffrey Barrett, Sonia Goncalves, Ewan Harrison, David K. Jackson, Ian Johnston, Dominic Kwiatkowski, Cordelia Langford, John Sillitoe on behalf of the Wellcome Sanger Institute COVID-19 Surveillance Team |
| EPI_ISL_3365338, EPI_ISL_3365339 | Lighthouse Lab in Milton Keynes | Wellcome Sanger Institute for the COVID-19 Genomics UK (COG-UK) Consortium | The Lighthouse Lab in Milton Keynes and Alex Alderton, Roberto Amato, Jeffrey Barrett, Sonia Goncalves, Ewan Harrison, David K. Jackson, Ian Johnston, Dominic Kwiatkowski, Cordelia Langford, John Sillitoe on behalf of the Wellcome Sanger Institute COVID-19 Surveillance Team |
| EPI_ISL_3365340, EPI_ISL_3365341 | Lighthouse Lab in Alderley Park | Wellcome Sanger Institute for the COVID-19 Genomics UK (COG-UK) Consortium | Jacquelyn Wynn, Mairead Hyland, The Lighthouse Lab in Alderley Park and Alex Alderton, Roberto Amato, Jeffrey Barrett, Sonia Goncalves, Ewan Harrison, David K. Jackson, Ian Johnston, Dominic Kwiatkowski, Cordelia Langford, John Sillitoe on behalf of the Wellcome Sanger Institute COVID-19 Surveillance Team |
| EPI_ISL_3365342 | Berkshire and Surrey Pathology Services Lighthouse Laboratory | Wellcome Sanger Institute for the COVID-19 Genomics UK (COG-UK) Consortium | Berkshire and Surrey Pathology Services Lighthouse Laboratory and Alex Alderton, Roberto Amato, Jeffrey Barrett, Sonia Goncalves, Ewan Harrison, David K. Jackson, Ian Johnston, Dominic Kwiatkowski, Cordelia Langford, John Sillitoe on behalf of the Wellcome Sanger Institute COVID-19 Surveillance Team |
| EPI_ISL_3365343 | Lighthouse Lab in Alderley Park | Wellcome Sanger Institute for the COVID-19 Genomics UK (COG-UK) Consortium | Jacquelyn Wynn, Mairead Hyland, The Lighthouse Lab in Alderley Park and Alex Alderton, Roberto Amato, Jeffrey Barrett, Sonia Goncalves, Ewan Harrison, David K. Jackson, Ian Johnston, Dominic Kwiatkowski, Cordelia Langford, John Sillitoe on behalf of the Wellcome Sanger Institute COVID-19 Surveillance Team |
| EPI_ISL_3365344 | Berkshire and Surrey Pathology Services Lighthouse Laboratory | Wellcome Sanger Institute for the COVID-19 Genomics UK (COG-UK) Consortium | Berkshire and Surrey Pathology Services Lighthouse Laboratory and Alex Alderton, Roberto Amato, Jeffrey Barrett, Sonia Goncalves, Ewan Harrison, David K. Jackson, Ian Johnston, Dominic Kwiatkowski, Cordelia Langford, John Sillitoe on behalf of the Wellcome Sanger Institute COVID-19 Surveillance Team |
| EPI_ISL_3365345 | Lighthouse Lab in Milton Keynes | Wellcome Sanger Institute for the COVID-19 Genomics UK (COG-UK) Consortium | The Lighthouse Lab in Milton Keynes and Alex Alderton, Roberto Amato, Jeffrey Barrett, Sonia Goncalves, Ewan Harrison, David K. Jackson, Ian Johnston, Dominic Kwiatkowski, Cordelia Langford, John Sillitoe on behalf of the Wellcome Sanger Institute COVID-19 Surveillance Team |
| EPI_ISL_3365346 | Lighthouse Lab in Alderley Park | Wellcome Sanger Institute for the COVID-19 Genomics UK (COG-UK) Consortium | Jacquelyn Wynn, Mairead Hyland, The Lighthouse Lab in Alderley Park and Alex Alderton, Roberto Amato, Jeffrey Barrett, Sonia Goncalves, Ewan Harrison, David K. Jackson, Ian Johnston, Dominic Kwiatkowski, Cordelia Langford, John Sillitoe on behalf of the Wellcome Sanger Institute COVID-19 Surveillance Team |
| EPI_ISL_3365347 | Berkshire and Surrey Pathology Services Lighthouse Laboratory | Wellcome Sanger Institute for the COVID-19 Genomics UK (COG-UK) Consortium | Berkshire and Surrey Pathology Services Lighthouse Laboratory and Alex Alderton, Roberto Amato, Jeffrey Barrett, Sonia Goncalves, Ewan Harrison, David K. Jackson, Ian Johnston, Dominic Kwiatkowski, Cordelia Langford, John Sillitoe on behalf of the Wellcome Sanger Institute COVID-19 Surveillance Team |
| EPI_ISL_3365348 | Lighthouse Lab in Alderley Park | Wellcome Sanger Institute for the COVID-19 Genomics UK (COG-UK) Consortium | Jacquelyn Wynn, Mairead Hyland, The Lighthouse Lab in Alderley Park and Alex Alderton, Roberto Amato, Jeffrey Barrett, Sonia Goncalves, Ewan Harrison, David K. Jackson, Ian Johnston, Dominic Kwiatkowski, Cordelia Langford, John Sillitoe on behalf of the Wellcome Sanger Institute COVID-19 Surveillance Team |
| EPI_ISL_3365349, EPI_ISL_3365350 | Lighthouse Lab in Milton Keynes | Wellcome Sanger Institute for the COVID-19 Genomics UK (COG-UK) Consortium | The Lighthouse Lab in Milton Keynes and Alex Alderton, Roberto Amato, Jeffrey Barrett, Sonia Goncalves, Ewan Harrison, David K. Jackson, Ian Johnston, Dominic Kwiatkowski, Cordelia Langford, John Sillitoe on behalf of the Wellcome Sanger Institute COVID-19 Surveillance Team |
| EPI_ISL_3365351, EPI_ISL_3365352 | Berkshire and Surrey Pathology Services Lighthouse Laboratory | Wellcome Sanger Institute for the COVID-19 Genomics UK (COG-UK) Consortium | Berkshire and Surrey Pathology Services Lighthouse Laboratory and Alex Alderton, Roberto Amato, Jeffrey Barrett, Sonia Goncalves, Ewan Harrison, David K. Jackson, Ian Johnston, Dominic Kwiatkowski, Cordelia Langford, John Sillitoe on behalf of the Wellcome Sanger Institute COVID-19 Surveillance Team |
| EPI_ISL_3365353, EPI_ISL_3365354, EPI_ISL_3365355 | Lighthouse Lab in Milton Keynes | Wellcome Sanger Institute for the COVID-19 Genomics UK (COG-UK) Consortium | The Lighthouse Lab in Milton Keynes and Alex Alderton, Roberto Amato, Jeffrey Barrett, Sonia Goncalves, Ewan Harrison, David K. Jackson, Ian Johnston, Dominic Kwiatkowski, Cordelia Langford, John Sillitoe on behalf of the Wellcome Sanger Institute COVID-19 Surveillance Team |
| EPI_ISL_3365356, EPI_ISL_3365357, EPI_ISL_3365358 | Berkshire and Surrey Pathology Services Lighthouse Laboratory | Wellcome Sanger Institute for the COVID-19 Genomics UK (COG-UK) Consortium | Berkshire and Surrey Pathology Services Lighthouse Laboratory and Alex Alderton, Roberto Amato, Jeffrey Barrett, Sonia Goncalves, Ewan Harrison, David K. Jackson, Ian Johnston, Dominic Kwiatkowski, Cordelia Langford, John Sillitoe on behalf of the Wellcome Sanger Institute COVID-19 Surveillance Team |
| EPI_ISL_3365359 | Lighthouse Lab in Alderley Park | Wellcome Sanger Institute for the COVID-19 Genomics UK (COG-UK) Consortium | Jacquelyn Wynn, Mairead Hyland, The Lighthouse Lab in Alderley Park and Alex Alderton, Roberto Amato, Jeffrey Barrett, Sonia Goncalves, Ewan Harrison, David K. Jackson, Ian Johnston, Dominic Kwiatkowski, Cordelia Langford, John Sillitoe on behalf of the Wellcome Sanger Institute COVID-19 Surveillance Team |
| EPI_ISL_3365360 | Lighthouse Lab in Milton Keynes | Wellcome Sanger Institute for the COVID-19 Genomics UK (COG-UK) Consortium | The Lighthouse Lab in Milton Keynes and Alex Alderton, Roberto Amato, Jeffrey Barrett, Sonia Goncalves, Ewan Harrison, David K. Jackson, Ian Johnston, Dominic Kwiatkowski, Cordelia Langford, John Sillitoe on behalf of the Wellcome Sanger Institute COVID-19 Surveillance Team |
| EPI_ISL_3365361 | Berkshire and Surrey Pathology Services Lighthouse Laboratory | Wellcome Sanger Institute for the COVID-19 Genomics UK (COG-UK) Consortium | Berkshire and Surrey Pathology Services Lighthouse Laboratory and Alex Alderton, Roberto Amato, Jeffrey Barrett, Sonia Goncalves, Ewan Harrison, David K. Jackson, Ian Johnston, Dominic Kwiatkowski, Cordelia Langford, John Sillitoe on behalf of the Wellcome Sanger Institute COVID-19 Surveillance Team |
| EPI_ISL_3365362 | Lighthouse Lab in Milton Keynes | Wellcome Sanger Institute for the COVID-19 Genomics UK (COG-UK) Consortium | The Lighthouse Lab in Milton Keynes and Alex Alderton, Roberto Amato, Jeffrey Barrett, Sonia Goncalves, Ewan Harrison, David K. Jackson, Ian Johnston, Dominic Kwiatkowski, Cordelia Langford, John Sillitoe on behalf of the Wellcome Sanger Institute COVID-19 Surveillance Team |
| EPI_ISL_3365363 | Berkshire and Surrey Pathology Services Lighthouse Laboratory | Wellcome Sanger Institute for the COVID-19 Genomics UK (COG-UK) Consortium | Berkshire and Surrey Pathology Services Lighthouse Laboratory and Alex Alderton, Roberto Amato, Jeffrey Barrett, Sonia Goncalves, Ewan Harrison, David K. Jackson, Ian Johnston, Dominic Kwiatkowski, Cordelia Langford, John Sillitoe on behalf of the Wellcome Sanger Institute COVID-19 Surveillance Team |
| EPI_ISL_3365364, EPI_ISL_3365365 | Lighthouse Lab in Alderley Park | Wellcome Sanger Institute for the COVID-19 Genomics UK (COG-UK) Consortium | Jacquelyn Wynn, Mairead Hyland, The Lighthouse Lab in Alderley Park and Alex Alderton, Roberto Amato, Jeffrey Barrett, Sonia Goncalves, Ewan Harrison, David K. Jackson, Ian Johnston, Dominic Kwiatkowski, Cordelia Langford, John Sillitoe on behalf of the Wellcome Sanger Institute COVID-19 Surveillance Team |
| EPI_ISL_3365366, EPI_ISL_3365367, EPI_ISL_3365368, EPI_ISL_3365369 | Berkshire and Surrey Pathology Services Lighthouse Laboratory | Wellcome Sanger Institute for the COVID-19 Genomics UK (COG-UK) Consortium | Berkshire and Surrey Pathology Services Lighthouse Laboratory and Alex Alderton, Roberto Amato, Jeffrey Barrett, Sonia Goncalves, Ewan Harrison, David K. Jackson, Ian Johnston, Dominic Kwiatkowski, Cordelia Langford, John Sillitoe on behalf of the Wellcome Sanger Institute COVID-19 Surveillance Team |
| EPI_ISL_3365370 | Lighthouse Lab in Alderley Park | Wellcome Sanger Institute for the COVID-19 Genomics UK (COG-UK) Consortium | Jacquelyn Wynn, Mairead Hyland, The Lighthouse Lab in Alderley Park and Alex Alderton, Roberto Amato, Jeffrey Barrett, Sonia Goncalves, Ewan Harrison, David K. Jackson, Ian Johnston, Dominic Kwiatkowski, Cordelia Langford, John Sillitoe on behalf of the Wellcome Sanger Institute COVID-19 Surveillance Team |
| EPI_ISL_3365371, EPI_ISL_3365372, EPI_ISL_3365373 | Berkshire and Surrey Pathology Services Lighthouse Laboratory | Wellcome Sanger Institute for the COVID-19 Genomics UK (COG-UK) Consortium | Berkshire and Surrey Pathology Services Lighthouse Laboratory and Alex Alderton, Roberto Amato, Jeffrey Barrett, Sonia Goncalves, Ewan Harrison, David K. Jackson, Ian Johnston, Dominic Kwiatkowski, Cordelia Langford, John Sillitoe on behalf of the Wellcome Sanger Institute COVID-19 Surveillance Team |
| EPI_ISL_3365374 | Lighthouse Lab in Milton Keynes | Wellcome Sanger Institute for the COVID-19 Genomics UK (COG-UK) Consortium | The Lighthouse Lab in Milton Keynes and Alex Alderton, Roberto Amato, Jeffrey Barrett, Sonia Goncalves, Ewan Harrison, David K. Jackson, Ian Johnston, Dominic Kwiatkowski, Cordelia Langford, John Sillitoe on behalf of the Wellcome Sanger Institute COVID-19 Surveillance Team |
| EPI_ISL_3365375, EPI_ISL_3365376 | Lighthouse Lab in Alderley Park | Wellcome Sanger Institute for the COVID-19 Genomics UK (COG-UK) Consortium | Jacquelyn Wynn, Mairead Hyland, The Lighthouse Lab in Alderley Park and Alex Alderton, Roberto Amato, Jeffrey Barrett, Sonia Goncalves, Ewan Harrison, David K. Jackson, Ian Johnston, Dominic Kwiatkowski, Cordelia Langford, John Sillitoe on behalf of the Wellcome Sanger Institute COVID-19 Surveillance Team |
| EPI_ISL_3365377 | Berkshire and Surrey Pathology Services Lighthouse Laboratory | Wellcome Sanger Institute for the COVID-19 Genomics UK (COG-UK) Consortium | Berkshire and Surrey Pathology Services Lighthouse Laboratory and Alex Alderton, Roberto Amato, Jeffrey Barrett, Sonia Goncalves, Ewan Harrison, David K. Jackson, Ian Johnston, Dominic Kwiatkowski, Cordelia Langford, John Sillitoe on behalf of the Wellcome Sanger Institute COVID-19 Surveillance Team |
| EPI_ISL_3365378, EPI_ISL_3365379 | Lighthouse Lab in Alderley Park | Wellcome Sanger Institute for the COVID-19 Genomics UK (COG-UK) Consortium | Jacquelyn Wynn, Mairead Hyland, The Lighthouse Lab in Alderley Park and Alex Alderton, Roberto Amato, Jeffrey Barrett, Sonia Goncalves, Ewan Harrison, David K. Jackson, Ian Johnston, Dominic Kwiatkowski, Cordelia Langford, John Sillitoe on behalf of the Wellcome Sanger Institute COVID-19 Surveillance Team |
| EPI_ISL_3365380 | Berkshire and Surrey Pathology Services Lighthouse Laboratory | Wellcome Sanger Institute for the COVID-19 Genomics UK (COG-UK) Consortium | Berkshire and Surrey Pathology Services Lighthouse Laboratory and Alex Alderton, Roberto Amato, Jeffrey Barrett, Sonia Goncalves, Ewan Harrison, David K. Jackson, Ian Johnston, Dominic Kwiatkowski, Cordelia Langford, John Sillitoe on behalf of the Wellcome Sanger Institute COVID-19 Surveillance Team |
| EPI_ISL_3365381 | Lighthouse Lab in Alderley Park | Wellcome Sanger Institute for the COVID-19 Genomics UK (COG-UK) Consortium | Jacquelyn Wynn, Mairead Hyland, The Lighthouse Lab in Alderley Park and Alex Alderton, Roberto Amato, Jeffrey Barrett, Sonia Goncalves, Ewan Harrison, David K. Jackson, Ian Johnston, Dominic Kwiatkowski, Cordelia Langford, John Sillitoe on behalf of the Wellcome Sanger Institute COVID-19 Surveillance Team |
| EPI_ISL_3365382 | Lighthouse Lab in Milton Keynes | Wellcome Sanger Institute for the COVID-19 Genomics UK (COG-UK) Consortium | The Lighthouse Lab in Milton Keynes and Alex Alderton, Roberto Amato, Jeffrey Barrett, Sonia Goncalves, Ewan Harrison, David K. Jackson, Ian Johnston, Dominic Kwiatkowski, Cordelia Langford, John Sillitoe on behalf of the Wellcome Sanger Institute COVID-19 Surveillance Team |

| | | | |
|---|---|---|---|
| EPI_ISL_3365383, EPI_ISL_3365384, EPI_ISL_3365385, EPI_ISL_3365386 | Berkshire and Surrey Pathology Services Lighthouse Laboratory | Wellcome Sanger Institute for the COVID-19 Genomics UK (COG-UK) Consortium | Berkshire and Surrey Pathology Services Lighthouse Laboratory and Alex Alderton, Roberto Amato, Jeffrey Barrett, Sonia Goncalves, Ewan Harrison, David K. Jackson, Ian Johnston, Dominic Kwiatkowski, Cordelia Langford, John Sillitoe on behalf of the Wellcome Sanger Institute COVID-19 Surveillance Team |
| EPI_ISL_3365387, EPI_ISL_3365388 | Lighthouse Lab in Milton Keynes | Wellcome Sanger Institute for the COVID-19 Genomics UK (COG-UK) Consortium | The Lighthouse Lab in Milton Keynes and Alex Alderton, Roberto Amato, Jeffrey Barrett, Sonia Goncalves, Ewan Harrison, David K. Jackson, Ian Johnston, Dominic Kwiatkowski, Cordelia Langford, John Sillitoe on behalf of the Wellcome Sanger Institute COVID-19 Surveillance Team |
| EPI_ISL_3365389, EPI_ISL_3365390, EPI_ISL_3365391 | Berkshire and Surrey Pathology Services Lighthouse Laboratory | Wellcome Sanger Institute for the COVID-19 Genomics UK (COG-UK) Consortium | Berkshire and Surrey Pathology Services Lighthouse Laboratory and Alex Alderton, Roberto Amato, Jeffrey Barrett, Sonia Goncalves, Ewan Harrison, David K. Jackson, Ian Johnston, Dominic Kwiatkowski, Cordelia Langford, John Sillitoe on behalf of the Wellcome Sanger Institute COVID-19 Surveillance Team |
| EPI_ISL_3365392 | Lighthouse Lab in Alderley Park | Wellcome Sanger Institute for the COVID-19 Genomics UK (COG-UK) Consortium | Jacquelyn Wynn, Mairead Hyland, The Lighthouse Lab in Alderley Park and Alex Alderton, Roberto Amato, Jeffrey Barrett, Sonia Goncalves, Ewan Harrison, David K. Jackson, Ian Johnston, Dominic Kwiatkowski, Cordelia Langford, John Sillitoe on behalf of the Wellcome Sanger Institute COVID-19 Surveillance Team |
| EPI_ISL_3365393, EPI_ISL_3365394 | Lighthouse Lab in Milton Keynes | Wellcome Sanger Institute for the COVID-19 Genomics UK (COG-UK) Consortium | The Lighthouse Lab in Milton Keynes and Alex Alderton, Roberto Amato, Jeffrey Barrett, Sonia Goncalves, Ewan Harrison, David K. Jackson, Ian Johnston, Dominic Kwiatkowski, Cordelia Langford, John Sillitoe on behalf of the Wellcome Sanger Institute COVID-19 Surveillance Team |
| EPI_ISL_3365395 | Berkshire and Surrey Pathology Services Lighthouse Laboratory | Wellcome Sanger Institute for the COVID-19 Genomics UK (COG-UK) Consortium | Berkshire and Surrey Pathology Services Lighthouse Laboratory and Alex Alderton, Roberto Amato, Jeffrey Barrett, Sonia Goncalves, Ewan Harrison, David K. Jackson, Ian Johnston, Dominic Kwiatkowski, Cordelia Langford, John Sillitoe on behalf of the Wellcome Sanger Institute COVID-19 Surveillance Team |
| EPI_ISL_3365396 | Lighthouse Lab in Milton Keynes | Wellcome Sanger Institute for the COVID-19 Genomics UK (COG-UK) Consortium | The Lighthouse Lab in Milton Keynes and Alex Alderton, Roberto Amato, Jeffrey Barrett, Sonia Goncalves, Ewan Harrison, David K. Jackson, Ian Johnston, Dominic Kwiatkowski, Cordelia Langford, John Sillitoe on behalf of the Wellcome Sanger Institute COVID-19 Surveillance Team |
| EPI_ISL_3365397, EPI_ISL_3365398 | Lighthouse Lab in Alderley Park | Wellcome Sanger Institute for the COVID-19 Genomics UK (COG-UK) Consortium | Jacquelyn Wynn, Mairead Hyland, The Lighthouse Lab in Alderley Park and Alex Alderton, Roberto Amato, Jeffrey Barrett, Sonia Goncalves, Ewan Harrison, David K. Jackson, Ian Johnston, Dominic Kwiatkowski, Cordelia Langford, John Sillitoe on behalf of the Wellcome Sanger Institute COVID-19 Surveillance Team |
| EPI_ISL_3365399, EPI_ISL_3365400 | Berkshire and Surrey Pathology Services Lighthouse Laboratory | Wellcome Sanger Institute for the COVID-19 Genomics UK (COG-UK) Consortium | Berkshire and Surrey Pathology Services Lighthouse Laboratory and Alex Alderton, Roberto Amato, Jeffrey Barrett, Sonia Goncalves, Ewan Harrison, David K. Jackson, Ian Johnston, Dominic Kwiatkowski, Cordelia Langford, John Sillitoe on behalf of the Wellcome Sanger Institute COVID-19 Surveillance Team |
| EPI_ISL_3365401 | Lighthouse Lab in Alderley Park | Wellcome Sanger Institute for the COVID-19 Genomics UK (COG-UK) Consortium | Jacquelyn Wynn, Mairead Hyland, The Lighthouse Lab in Alderley Park and Alex Alderton, Roberto Amato, Jeffrey Barrett, Sonia Goncalves, Ewan Harrison, David K. Jackson, Ian Johnston, Dominic Kwiatkowski, Cordelia Langford, John Sillitoe on behalf of the Wellcome Sanger Institute COVID-19 Surveillance Team |
| EPI_ISL_3365402, EPI_ISL_3365403 | Berkshire and Surrey Pathology Services Lighthouse Laboratory | Wellcome Sanger Institute for the COVID-19 Genomics UK (COG-UK) Consortium | Berkshire and Surrey Pathology Services Lighthouse Laboratory and Alex Alderton, Roberto Amato, Jeffrey Barrett, Sonia Goncalves, Ewan Harrison, David K. Jackson, Ian Johnston, Dominic Kwiatkowski, Cordelia Langford, John Sillitoe on behalf of the Wellcome Sanger Institute COVID-19 Surveillance Team |
| EPI_ISL_3365404 | Lighthouse Lab in Milton Keynes | Wellcome Sanger Institute for the COVID-19 Genomics UK (COG-UK) Consortium | The Lighthouse Lab in Milton Keynes and Alex Alderton, Roberto Amato, Jeffrey Barrett, Sonia Goncalves, Ewan Harrison, David K. Jackson, Ian Johnston, Dominic Kwiatkowski, Cordelia Langford, John Sillitoe on behalf of the Wellcome Sanger Institute COVID-19 Surveillance Team |
| EPI_ISL_3365405, EPI_ISL_3365406 | Lighthouse Lab in Alderley Park | Wellcome Sanger Institute for the COVID-19 Genomics UK (COG-UK) Consortium | Jacquelyn Wynn, Mairead Hyland, The Lighthouse Lab in Alderley Park and Alex Alderton, Roberto Amato, Jeffrey Barrett, Sonia Goncalves, Ewan Harrison, David K. Jackson, Ian Johnston, Dominic Kwiatkowski, Cordelia Langford, John Sillitoe on behalf of the Wellcome Sanger Institute COVID-19 Surveillance Team |
| EPI_ISL_3365407 | Lighthouse Lab in Milton Keynes | Wellcome Sanger Institute for the COVID-19 Genomics UK (COG-UK) Consortium | The Lighthouse Lab in Milton Keynes and Alex Alderton, Roberto Amato, Jeffrey Barrett, Sonia Goncalves, Ewan Harrison, David K. Jackson, Ian Johnston, Dominic Kwiatkowski, Cordelia Langford, John Sillitoe on behalf of the Wellcome Sanger Institute COVID-19 Surveillance Team |
| EPI_ISL_3365408 | Berkshire and Surrey Pathology Services Lighthouse Laboratory | Wellcome Sanger Institute for the COVID-19 Genomics UK (COG-UK) Consortium | Berkshire and Surrey Pathology Services Lighthouse Laboratory and Alex Alderton, Roberto Amato, Jeffrey Barrett, Sonia Goncalves, Ewan Harrison, David K. Jackson, Ian Johnston, Dominic Kwiatkowski, Cordelia Langford, John Sillitoe on behalf of the Wellcome Sanger Institute COVID-19 Surveillance Team |
| EPI_ISL_3365409 | Lighthouse Lab in Milton Keynes | Wellcome Sanger Institute for the COVID-19 Genomics UK (COG-UK) Consortium | The Lighthouse Lab in Milton Keynes and Alex Alderton, Roberto Amato, Jeffrey Barrett, Sonia Goncalves, Ewan Harrison, David K. Jackson, Ian Johnston, Dominic Kwiatkowski, Cordelia Langford, John Sillitoe on behalf of the Wellcome Sanger Institute COVID-19 Surveillance Team |
| EPI_ISL_3365410 | Berkshire and Surrey Pathology Services Lighthouse Laboratory | Wellcome Sanger Institute for the COVID-19 Genomics UK (COG-UK) Consortium | Berkshire and Surrey Pathology Services Lighthouse Laboratory and Alex Alderton, Roberto Amato, Jeffrey Barrett, Sonia Goncalves, Ewan Harrison, David K. Jackson, Ian Johnston, Dominic Kwiatkowski, Cordelia Langford, John Sillitoe on behalf of the Wellcome Sanger Institute COVID-19 Surveillance Team |
| EPI_ISL_3365411 | Lighthouse Lab in Alderley Park | Wellcome Sanger Institute for the COVID-19 Genomics UK (COG-UK) Consortium | Jacquelyn Wynn, Mairead Hyland, The Lighthouse Lab in Alderley Park and Alex Alderton, Roberto Amato, Jeffrey Barrett, Sonia Goncalves, Ewan Harrison, David K. Jackson, Ian Johnston, Dominic Kwiatkowski, Cordelia Langford, John Sillitoe on behalf of the Wellcome Sanger Institute COVID-19 Surveillance Team |
| EPI_ISL_3365412 | Lighthouse Lab in Milton Keynes | Wellcome Sanger Institute for the COVID-19 Genomics UK (COG-UK) Consortium | The Lighthouse Lab in Milton Keynes and Alex Alderton, Roberto Amato, Jeffrey Barrett, Sonia Goncalves, Ewan Harrison, David K. Jackson, Ian Johnston, Dominic Kwiatkowski, Cordelia Langford, John Sillitoe on behalf of the Wellcome Sanger Institute COVID-19 Surveillance Team |
| EPI_ISL_3365413 | Berkshire and Surrey Pathology Services Lighthouse Laboratory | Wellcome Sanger Institute for the COVID-19 Genomics UK (COG-UK) Consortium | Berkshire and Surrey Pathology Services Lighthouse Laboratory and Alex Alderton, Roberto Amato, Jeffrey Barrett, Sonia Goncalves, Ewan Harrison, David K. Jackson, Ian Johnston, Dominic Kwiatkowski, Cordelia Langford, John Sillitoe on behalf of the Wellcome Sanger Institute COVID-19 Surveillance Team |
| EPI_ISL_3365414 | Lighthouse Lab in Alderley Park | Wellcome Sanger Institute for the COVID-19 Genomics UK (COG-UK) Consortium | Jacquelyn Wynn, Mairead Hyland, The Lighthouse Lab in Alderley Park and Alex Alderton, Roberto Amato, Jeffrey Barrett, Sonia Goncalves, Ewan Harrison, David K. Jackson, Ian Johnston, Dominic Kwiatkowski, Cordelia Langford, John Sillitoe on behalf of the Wellcome Sanger Institute COVID-19 Surveillance Team |
| EPI_ISL_3365415, EPI_ISL_3365416, EPI_ISL_3365417, EPI_ISL_3365418, EPI_ISL_3365419 | Berkshire and Surrey Pathology Services Lighthouse Laboratory | Wellcome Sanger Institute for the COVID-19 Genomics UK (COG-UK) Consortium | Berkshire and Surrey Pathology Services Lighthouse Laboratory and Alex Alderton, Roberto Amato, Jeffrey Barrett, Sonia Goncalves, Ewan Harrison, David K. Jackson, Ian Johnston, Dominic Kwiatkowski, Cordelia Langford, John Sillitoe on behalf of the Wellcome Sanger Institute COVID-19 Surveillance Team |
| EPI_ISL_3365420 | Lighthouse Lab in Milton Keynes | Wellcome Sanger Institute for the COVID-19 Genomics UK (COG-UK) Consortium | The Lighthouse Lab in Milton Keynes and Alex Alderton, Roberto Amato, Jeffrey Barrett, Sonia Goncalves, Ewan Harrison, David K. Jackson, Ian Johnston, Dominic Kwiatkowski, Cordelia Langford, John Sillitoe on behalf of the Wellcome Sanger Institute COVID-19 Surveillance Team |
| EPI_ISL_3365421 | Lighthouse Lab in Alderley Park | Wellcome Sanger Institute for the COVID-19 Genomics UK (COG-UK) Consortium | Jacquelyn Wynn, Mairead Hyland, The Lighthouse Lab in Alderley Park and Alex Alderton, Roberto Amato, Jeffrey Barrett, Sonia Goncalves, Ewan Harrison, David K. Jackson, Ian Johnston, Dominic Kwiatkowski, Cordelia Langford, John Sillitoe on behalf of the Wellcome Sanger Institute COVID-19 Surveillance Team |
| EPI_ISL_3365422 | Lighthouse Lab in Milton Keynes | Wellcome Sanger Institute for the COVID-19 Genomics UK (COG-UK) Consortium | The Lighthouse Lab in Milton Keynes and Alex Alderton, Roberto Amato, Jeffrey Barrett, Sonia Goncalves, Ewan Harrison, David K. Jackson, Ian Johnston, Dominic Kwiatkowski, Cordelia Langford, John Sillitoe on behalf of the Wellcome Sanger Institute COVID-19 Surveillance Team |
| EPI_ISL_3365423, EPI_ISL_3365424 | Berkshire and Surrey Pathology Services Lighthouse Laboratory | Wellcome Sanger Institute for the COVID-19 Genomics UK (COG-UK) Consortium | Berkshire and Surrey Pathology Services Lighthouse Laboratory and Alex Alderton, Roberto Amato, Jeffrey Barrett, Sonia Goncalves, Ewan Harrison, David K. Jackson, Ian Johnston, Dominic Kwiatkowski, Cordelia Langford, John Sillitoe on behalf of the Wellcome Sanger Institute COVID-19 Surveillance Team |
| EPI_ISL_3365425, EPI_ISL_3365426, EPI_ISL_3365427 | Lighthouse Lab in Alderley Park | Wellcome Sanger Institute for the COVID-19 Genomics UK (COG-UK) Consortium | Jacquelyn Wynn, Mairead Hyland, The Lighthouse Lab in Alderley Park and Alex Alderton, Roberto Amato, Jeffrey Barrett, Sonia Goncalves, Ewan Harrison, David K. Jackson, Ian Johnston, Dominic Kwiatkowski, Cordelia Langford, John Sillitoe on behalf of the Wellcome Sanger Institute COVID-19 Surveillance Team |
| EPI_ISL_3365428, EPI_ISL_3365429, EPI_ISL_3365430 | Berkshire and Surrey Pathology Services Lighthouse Laboratory | Wellcome Sanger Institute for the COVID-19 Genomics UK (COG-UK) Consortium | Berkshire and Surrey Pathology Services Lighthouse Laboratory and Alex Alderton, Roberto Amato, Jeffrey Barrett, Sonia Goncalves, Ewan Harrison, David K. Jackson, Ian Johnston, Dominic Kwiatkowski, Cordelia Langford, John Sillitoe on behalf of the Wellcome Sanger Institute COVID-19 Surveillance Team |
| EPI_ISL_3365431 | Lighthouse Lab in Milton Keynes | Wellcome Sanger Institute for the COVID-19 Genomics UK (COG-UK) Consortium | The Lighthouse Lab in Milton Keynes and Alex Alderton, Roberto Amato, Jeffrey Barrett, Sonia Goncalves, Ewan Harrison, David K. Jackson, Ian Johnston, Dominic Kwiatkowski, Cordelia Langford, John Sillitoe on behalf of the Wellcome Sanger Institute COVID-19 Surveillance Team |
| EPI_ISL_3365432 | Lighthouse Lab in Alderley Park | Wellcome Sanger Institute for the COVID-19 Genomics UK (COG-UK) Consortium | Jacquelyn Wynn, Mairead Hyland, The Lighthouse Lab in Alderley Park and Alex Alderton, Roberto Amato, Jeffrey Barrett, Sonia Goncalves, Ewan Harrison, David K. Jackson, Ian Johnston, Dominic Kwiatkowski, Cordelia Langford, John Sillitoe on behalf of the Wellcome Sanger Institute COVID-19 Surveillance Team |
| EPI_ISL_3365433 | Berkshire and Surrey Pathology Services Lighthouse Laboratory | Wellcome Sanger Institute for the COVID-19 Genomics UK (COG-UK) Consortium | Berkshire and Surrey Pathology Services Lighthouse Laboratory and Alex Alderton, Roberto Amato, Jeffrey Barrett, Sonia Goncalves, Ewan Harrison, David K. Jackson, Ian Johnston, Dominic Kwiatkowski, Cordelia Langford, John Sillitoe on behalf of the Wellcome Sanger Institute COVID-19 Surveillance Team |
| EPI_ISL_3365434, EPI_ISL_3365435 | Lighthouse Lab in Milton Keynes | Wellcome Sanger Institute for the COVID-19 Genomics UK (COG-UK) Consortium | The Lighthouse Lab in Milton Keynes and Alex Alderton, Roberto Amato, Jeffrey Barrett, Sonia Goncalves, Ewan Harrison, David K. Jackson, Ian Johnston, Dominic Kwiatkowski, Cordelia Langford, John Sillitoe on behalf of the Wellcome Sanger Institute COVID-19 Surveillance Team |
| EPI_ISL_3365436 | Berkshire and Surrey Pathology Services Lighthouse Laboratory | Wellcome Sanger Institute for the COVID-19 Genomics UK (COG-UK) Consortium | Berkshire and Surrey Pathology Services Lighthouse Laboratory and Alex Alderton, Roberto Amato, Jeffrey Barrett, Sonia Goncalves, Ewan Harrison, David K. Jackson, Ian Johnston, Dominic Kwiatkowski, Cordelia Langford, John Sillitoe on behalf of the Wellcome Sanger Institute COVID-19 Surveillance Team |
| EPI_ISL_3365437 | Lighthouse Lab in Milton Keynes | Wellcome Sanger Institute for the COVID-19 Genomics UK (COG-UK) Consortium | The Lighthouse Lab in Milton Keynes and Alex Alderton, Roberto Amato, Jeffrey Barrett, Sonia Goncalves, Ewan Harrison, David K. Jackson, Ian Johnston, Dominic Kwiatkowski, Cordelia Langford, John Sillitoe on behalf of the Wellcome Sanger Institute COVID-19 Surveillance Team |
| EPI_ISL_3365438, EPI_ISL_3365439 | Lighthouse Lab in Alderley Park | Wellcome Sanger Institute for the COVID-19 Genomics UK (COG-UK) Consortium | Jacquelyn Wynn, Mairead Hyland, The Lighthouse Lab in Alderley Park and Alex Alderton, Roberto Amato, Jeffrey Barrett, Sonia Goncalves, Ewan Harrison, David K. Jackson, Ian Johnston, Dominic Kwiatkowski, Cordelia Langford, John Sillitoe on behalf of the Wellcome Sanger Institute COVID-19 Surveillance Team |
| EPI_ISL_3365440 | Berkshire and Surrey Pathology Services Lighthouse Laboratory | Wellcome Sanger Institute for the COVID-19 Genomics UK (COG-UK) Consortium | Berkshire and Surrey Pathology Services Lighthouse Laboratory and Alex Alderton, Roberto Amato, Jeffrey Barrett, Sonia Goncalves, Ewan Harrison, David K. Jackson, Ian Johnston, Dominic Kwiatkowski, Cordelia Langford, John Sillitoe on behalf of the Wellcome Sanger Institute COVID-19 Surveillance Team |
| EPI_ISL_3365441, EPI_ISL_3365442, EPI_ISL_3365443 | Lighthouse Lab in Milton Keynes | Wellcome Sanger Institute for the COVID-19 Genomics UK (COG-UK) Consortium | The Lighthouse Lab in Milton Keynes and Alex Alderton, Roberto Amato, Jeffrey Barrett, Sonia Goncalves, Ewan Harrison, David K. Jackson, Ian Johnston, Dominic Kwiatkowski, Cordelia Langford, John Sillitoe on behalf of the Wellcome Sanger Institute COVID-19 Surveillance Team |
| EPI_ISL_3365444 | Lighthouse Lab in Alderley Park | Wellcome Sanger Institute for the COVID-19 Genomics UK (COG-UK) Consortium | Jacquelyn Wynn, Mairead Hyland, The Lighthouse Lab in Alderley Park and Alex Alderton, Roberto Amato, Jeffrey Barrett, Sonia Goncalves, Ewan Harrison, David K. Jackson, Ian Johnston, Dominic Kwiatkowski, Cordelia Langford, John Sillitoe on behalf of the Wellcome Sanger Institute COVID-19 Surveillance Team |
| EPI_ISL_3365445, EPI_ISL_3365446 | Berkshire and Surrey Pathology Services Lighthouse Laboratory | Wellcome Sanger Institute for the COVID-19 Genomics UK (COG-UK) Consortium | Berkshire and Surrey Pathology Services Lighthouse Laboratory and Alex Alderton, Roberto Amato, Jeffrey Barrett, Sonia Goncalves, Ewan Harrison, David K. Jackson, Ian Johnston, Dominic Kwiatkowski, Cordelia Langford, John Sillitoe on behalf of the Wellcome Sanger Institute COVID-19 Surveillance Team |
| EPI_ISL_3365447, EPI_ISL_3365448, EPI_ISL_3365449 | Lighthouse Lab in Milton Keynes | Wellcome Sanger Institute for the COVID-19 Genomics UK (COG-UK) Consortium | The Lighthouse Lab in Milton Keynes and Alex Alderton, Roberto Amato, Jeffrey Barrett, Sonia Goncalves, Ewan Harrison, David K. Jackson, Ian Johnston, Dominic Kwiatkowski, Cordelia Langford, John Sillitoe on behalf of the Wellcome Sanger Institute COVID-19 Surveillance Team |
| EPI_ISL_3365450, EPI_ISL_3365451, | Berkshire and Surrey Pathology Services Lighthouse Laboratory | Wellcome Sanger Institute for the COVID-19 Genomics UK (COG-UK) Consortium | Berkshire and Surrey Pathology Services Lighthouse Laboratory and Alex Alderton, Roberto Amato, Jeffrey Barrett, Sonia Goncalves, Ewan Harrison, David K. Jackson, Ian Johnston, Dominic Kwiatkowski, Cordelia Langford, John Sillitoe on behalf of the Wellcome Sanger Institute COVID-19 Surveillance Team |

| | | | |
|---|---|---|---|
| EPI_ISL_3365452, EPI_ISL_3365453, EPI_ISL_3365454 | | | |
| EPI_ISL_3365455 | Lighthouse Lab in Milton Keynes | Wellcome Sanger Institute for the COVID-19 Genomics UK (COG-UK) Consortium | The Lighthouse Lab in Milton Keynes and Alex Alderton, Roberto Amato, Jeffrey Barrett, Sonia Goncalves, Ewan Harrison, David K. Jackson, Ian Johnston, Dominic Kwiatkowski, Cordelia Langford, John Sillitoe on behalf of the Wellcome Sanger Institute COVID-19 Surveillance Team |
| EPI_ISL_3365456 | Berkshire and Surrey Pathology Services Lighthouse Laboratory | Wellcome Sanger Institute for the COVID-19 Genomics UK (COG-UK) Consortium | Berkshire and Surrey Pathology Services Lighthouse Laboratory and Alex Alderton, Roberto Amato, Jeffrey Barrett, Sonia Goncalves, Ewan Harrison, David K. Jackson, Ian Johnston, Dominic Kwiatkowski, Cordelia Langford, John Sillitoe on behalf of the Wellcome Sanger Institute COVID-19 Surveillance Team |
| EPI_ISL_3365457, EPI_ISL_3365458, EPI_ISL_3365459 | Lighthouse Lab in Milton Keynes | Wellcome Sanger Institute for the COVID-19 Genomics UK (COG-UK) Consortium | The Lighthouse Lab in Milton Keynes and Alex Alderton, Roberto Amato, Jeffrey Barrett, Sonia Goncalves, Ewan Harrison, David K. Jackson, Ian Johnston, Dominic Kwiatkowski, Cordelia Langford, John Sillitoe on behalf of the Wellcome Sanger Institute COVID-19 Surveillance Team |
| EPI_ISL_3365460 | Lighthouse Lab in Alderley Park | Wellcome Sanger Institute for the COVID-19 Genomics UK (COG-UK) Consortium | Jacquelyn Wynn, Mairead Hyland, The Lighthouse Lab in Alderley Park and Alex Alderton, Roberto Amato, Jeffrey Barrett, Sonia Goncalves, Ewan Harrison, David K. Jackson, Ian Johnston, Dominic Kwiatkowski, Cordelia Langford, John Sillitoe on behalf of the Wellcome Sanger Institute COVID-19 Surveillance Team |
| EPI_ISL_3365461 | Berkshire and Surrey Pathology Services Lighthouse Laboratory | Wellcome Sanger Institute for the COVID-19 Genomics UK (COG-UK) Consortium | Berkshire and Surrey Pathology Services Lighthouse Laboratory and Alex Alderton, Roberto Amato, Jeffrey Barrett, Sonia Goncalves, Ewan Harrison, David K. Jackson, Ian Johnston, Dominic Kwiatkowski, Cordelia Langford, John Sillitoe on behalf of the Wellcome Sanger Institute COVID-19 Surveillance Team |
| EPI_ISL_3365462 | Lighthouse Lab in Milton Keynes | Wellcome Sanger Institute for the COVID-19 Genomics UK (COG-UK) Consortium | The Lighthouse Lab in Milton Keynes and Alex Alderton, Roberto Amato, Jeffrey Barrett, Sonia Goncalves, Ewan Harrison, David K. Jackson, Ian Johnston, Dominic Kwiatkowski, Cordelia Langford, John Sillitoe on behalf of the Wellcome Sanger Institute COVID-19 Surveillance Team |
| EPI_ISL_3365463 | Lighthouse Lab in Alderley Park | Wellcome Sanger Institute for the COVID-19 Genomics UK (COG-UK) Consortium | Jacquelyn Wynn, Mairead Hyland, The Lighthouse Lab in Alderley Park and Alex Alderton, Roberto Amato, Jeffrey Barrett, Sonia Goncalves, Ewan Harrison, David K. Jackson, Ian Johnston, Dominic Kwiatkowski, Cordelia Langford, John Sillitoe on behalf of the Wellcome Sanger Institute COVID-19 Surveillance Team |
| EPI_ISL_3365464, EPI_ISL_3365465, EPI_ISL_3365466, EPI_ISL_3365467, EPI_ISL_3365468 | Berkshire and Surrey Pathology Services Lighthouse Laboratory | Wellcome Sanger Institute for the COVID-19 Genomics UK (COG-UK) Consortium | Berkshire and Surrey Pathology Services Lighthouse Laboratory and Alex Alderton, Roberto Amato, Jeffrey Barrett, Sonia Goncalves, Ewan Harrison, David K. Jackson, Ian Johnston, Dominic Kwiatkowski, Cordelia Langford, John Sillitoe on behalf of the Wellcome Sanger Institute COVID-19 Surveillance Team |
| EPI_ISL_3365469, EPI_ISL_3365470 | Lighthouse Lab in Milton Keynes | Wellcome Sanger Institute for the COVID-19 Genomics UK (COG-UK) Consortium | The Lighthouse Lab in Milton Keynes and Alex Alderton, Roberto Amato, Jeffrey Barrett, Sonia Goncalves, Ewan Harrison, David K. Jackson, Ian Johnston, Dominic Kwiatkowski, Cordelia Langford, John Sillitoe on behalf of the Wellcome Sanger Institute COVID-19 Surveillance Team |
| EPI_ISL_3365471, EPI_ISL_3365472, EPI_ISL_3365473 | Berkshire and Surrey Pathology Services Lighthouse Laboratory | Wellcome Sanger Institute for the COVID-19 Genomics UK (COG-UK) Consortium | Berkshire and Surrey Pathology Services Lighthouse Laboratory and Alex Alderton, Roberto Amato, Jeffrey Barrett, Sonia Goncalves, Ewan Harrison, David K. Jackson, Ian Johnston, Dominic Kwiatkowski, Cordelia Langford, John Sillitoe on behalf of the Wellcome Sanger Institute COVID-19 Surveillance Team |
| EPI_ISL_3365474 | Lighthouse Lab in Milton Keynes | Wellcome Sanger Institute for the COVID-19 Genomics UK (COG-UK) Consortium | The Lighthouse Lab in Milton Keynes and Alex Alderton, Roberto Amato, Jeffrey Barrett, Sonia Goncalves, Ewan Harrison, David K. Jackson, Ian Johnston, Dominic Kwiatkowski, Cordelia Langford, John Sillitoe on behalf of the Wellcome Sanger Institute COVID-19 Surveillance Team |
| EPI_ISL_3365475 | Lighthouse Lab in Alderley Park | Wellcome Sanger Institute for the COVID-19 Genomics UK (COG-UK) Consortium | Jacquelyn Wynn, Mairead Hyland, The Lighthouse Lab in Alderley Park and Alex Alderton, Roberto Amato, Jeffrey Barrett, Sonia Goncalves, Ewan Harrison, David K. Jackson, Ian Johnston, Dominic Kwiatkowski, Cordelia Langford, John Sillitoe on behalf of the Wellcome Sanger Institute COVID-19 Surveillance Team |
| EPI_ISL_3365476, EPI_ISL_3365477, EPI_ISL_3365478 | Berkshire and Surrey Pathology Services Lighthouse Laboratory | Wellcome Sanger Institute for the COVID-19 Genomics UK (COG-UK) Consortium | Berkshire and Surrey Pathology Services Lighthouse Laboratory and Alex Alderton, Roberto Amato, Jeffrey Barrett, Sonia Goncalves, Ewan Harrison, David K. Jackson, Ian Johnston, Dominic Kwiatkowski, Cordelia Langford, John Sillitoe on behalf of the Wellcome Sanger Institute COVID-19 Surveillance Team |
| EPI_ISL_3365479, EPI_ISL_3365480 | Lighthouse Lab in Alderley Park | Wellcome Sanger Institute for the COVID-19 Genomics UK (COG-UK) Consortium | Jacquelyn Wynn, Mairead Hyland, The Lighthouse Lab in Alderley Park and Alex Alderton, Roberto Amato, Jeffrey Barrett, Sonia Goncalves, Ewan Harrison, David K. Jackson, Ian Johnston, Dominic Kwiatkowski, Cordelia Langford, John Sillitoe on behalf of the Wellcome Sanger Institute COVID-19 Surveillance Team |
| EPI_ISL_3365481 | Lighthouse Lab in Milton Keynes | Wellcome Sanger Institute for the COVID-19 Genomics UK (COG-UK) Consortium | The Lighthouse Lab in Milton Keynes and Alex Alderton, Roberto Amato, Jeffrey Barrett, Sonia Goncalves, Ewan Harrison, David K. Jackson, Ian Johnston, Dominic Kwiatkowski, Cordelia Langford, John Sillitoe on behalf of the Wellcome Sanger Institute COVID-19 Surveillance Team |
| EPI_ISL_3365482, EPI_ISL_3365483 | Lighthouse Lab in Alderley Park | Wellcome Sanger Institute for the COVID-19 Genomics UK (COG-UK) Consortium | Jacquelyn Wynn, Mairead Hyland, The Lighthouse Lab in Alderley Park and Alex Alderton, Roberto Amato, Jeffrey Barrett, Sonia Goncalves, Ewan Harrison, David K. Jackson, Ian Johnston, Dominic Kwiatkowski, Cordelia Langford, John Sillitoe on behalf of the Wellcome Sanger Institute COVID-19 Surveillance Team |
| EPI_ISL_3365484 | Lighthouse Lab in Milton Keynes | Wellcome Sanger Institute for the COVID-19 Genomics UK (COG-UK) Consortium | The Lighthouse Lab in Milton Keynes and Alex Alderton, Roberto Amato, Jeffrey Barrett, Sonia Goncalves, Ewan Harrison, David K. Jackson, Ian Johnston, Dominic Kwiatkowski, Cordelia Langford, John Sillitoe on behalf of the Wellcome Sanger Institute COVID-19 Surveillance Team |
| EPI_ISL_3365485, EPI_ISL_3365486, EPI_ISL_3365487 | Berkshire and Surrey Pathology Services Lighthouse Laboratory | Wellcome Sanger Institute for the COVID-19 Genomics UK (COG-UK) Consortium | Berkshire and Surrey Pathology Services Lighthouse Laboratory and Alex Alderton, Roberto Amato, Jeffrey Barrett, Sonia Goncalves, Ewan Harrison, David K. Jackson, Ian Johnston, Dominic Kwiatkowski, Cordelia Langford, John Sillitoe on behalf of the Wellcome Sanger Institute COVID-19 Surveillance Team |
| EPI_ISL_3365488, EPI_ISL_3365489 | Lighthouse Lab in Alderley Park | Wellcome Sanger Institute for the COVID-19 Genomics UK (COG-UK) Consortium | Jacquelyn Wynn, Mairead Hyland, The Lighthouse Lab in Alderley Park and Alex Alderton, Roberto Amato, Jeffrey Barrett, Sonia Goncalves, Ewan Harrison, David K. Jackson, Ian Johnston, Dominic Kwiatkowski, Cordelia Langford, John Sillitoe on behalf of the Wellcome Sanger Institute COVID-19 Surveillance Team |
| EPI_ISL_3365490, EPI_ISL_3365491, EPI_ISL_3365492, EPI_ISL_3365493, EPI_ISL_3365494, EPI_ISL_3365495 | Berkshire and Surrey Pathology Services Lighthouse Laboratory | Wellcome Sanger Institute for the COVID-19 Genomics UK (COG-UK) Consortium | Berkshire and Surrey Pathology Services Lighthouse Laboratory and Alex Alderton, Roberto Amato, Jeffrey Barrett, Sonia Goncalves, Ewan Harrison, David K. Jackson, Ian Johnston, Dominic Kwiatkowski, Cordelia Langford, John Sillitoe on behalf of the Wellcome Sanger Institute COVID-19 Surveillance Team |
| EPI_ISL_3365496, EPI_ISL_3365497 | Lighthouse Lab in Milton Keynes | Wellcome Sanger Institute for the COVID-19 Genomics UK (COG-UK) Consortium | The Lighthouse Lab in Milton Keynes and Alex Alderton, Roberto Amato, Jeffrey Barrett, Sonia Goncalves, Ewan Harrison, David K. Jackson, Ian Johnston, Dominic Kwiatkowski, Cordelia Langford, John Sillitoe on behalf of the Wellcome Sanger Institute COVID-19 Surveillance Team |
| EPI_ISL_3365498, EPI_ISL_3365499, EPI_ISL_3365500 | Berkshire and Surrey Pathology Services Lighthouse Laboratory | Wellcome Sanger Institute for the COVID-19 Genomics UK (COG-UK) Consortium | Berkshire and Surrey Pathology Services Lighthouse Laboratory and Alex Alderton, Roberto Amato, Jeffrey Barrett, Sonia Goncalves, Ewan Harrison, David K. Jackson, Ian Johnston, Dominic Kwiatkowski, Cordelia Langford, John Sillitoe on behalf of the Wellcome Sanger Institute COVID-19 Surveillance Team |
| EPI_ISL_3365501 | Lighthouse Lab in Alderley Park | Wellcome Sanger Institute for the COVID-19 Genomics UK (COG-UK) Consortium | Jacquelyn Wynn, Mairead Hyland, The Lighthouse Lab in Alderley Park and Alex Alderton, Roberto Amato, Jeffrey Barrett, Sonia Goncalves, Ewan Harrison, David K. Jackson, Ian Johnston, Dominic Kwiatkowski, Cordelia Langford, John Sillitoe on behalf of the Wellcome Sanger Institute COVID-19 Surveillance Team |
| EPI_ISL_3365502 | Lighthouse Lab in Milton Keynes | Wellcome Sanger Institute for the COVID-19 Genomics UK (COG-UK) Consortium | The Lighthouse Lab in Milton Keynes and Alex Alderton, Roberto Amato, Jeffrey Barrett, Sonia Goncalves, Ewan Harrison, David K. Jackson, Ian Johnston, Dominic Kwiatkowski, Cordelia Langford, John Sillitoe on behalf of the Wellcome Sanger Institute COVID-19 Surveillance Team |
| EPI_ISL_3365503 | Lighthouse Lab in Alderley Park | Wellcome Sanger Institute for the COVID-19 Genomics UK (COG-UK) Consortium | Jacquelyn Wynn, Mairead Hyland, The Lighthouse Lab in Alderley Park and Alex Alderton, Roberto Amato, Jeffrey Barrett, Sonia Goncalves, Ewan Harrison, David K. Jackson, Ian Johnston, Dominic Kwiatkowski, Cordelia Langford, John Sillitoe on behalf of the Wellcome Sanger Institute COVID-19 Surveillance Team |
| EPI_ISL_3365504 | Lighthouse Lab in Milton Keynes | Wellcome Sanger Institute for the COVID-19 Genomics UK (COG-UK) Consortium | The Lighthouse Lab in Milton Keynes and Alex Alderton, Roberto Amato, Jeffrey Barrett, Sonia Goncalves, Ewan Harrison, David K. Jackson, Ian Johnston, Dominic Kwiatkowski, Cordelia Langford, John Sillitoe on behalf of the Wellcome Sanger Institute COVID-19 Surveillance Team |
| EPI_ISL_3365505 | Berkshire and Surrey Pathology Services Lighthouse Laboratory | Wellcome Sanger Institute for the COVID-19 Genomics UK (COG-UK) Consortium | Berkshire and Surrey Pathology Services Lighthouse Laboratory and Alex Alderton, Roberto Amato, Jeffrey Barrett, Sonia Goncalves, Ewan Harrison, David K. Jackson, Ian Johnston, Dominic Kwiatkowski, Cordelia Langford, John Sillitoe on behalf of the Wellcome Sanger Institute COVID-19 Surveillance Team |
| EPI_ISL_3365506 | Lighthouse Lab in Milton Keynes | Wellcome Sanger Institute for the COVID-19 Genomics UK (COG-UK) Consortium | The Lighthouse Lab in Milton Keynes and Alex Alderton, Roberto Amato, Jeffrey Barrett, Sonia Goncalves, Ewan Harrison, David K. Jackson, Ian Johnston, Dominic Kwiatkowski, Cordelia Langford, John Sillitoe on behalf of the Wellcome Sanger Institute COVID-19 Surveillance Team |
| EPI_ISL_3365507 | Lighthouse Lab in Alderley Park | Wellcome Sanger Institute for the COVID-19 Genomics UK (COG-UK) Consortium | Jacquelyn Wynn, Mairead Hyland, The Lighthouse Lab in Alderley Park and Alex Alderton, Roberto Amato, Jeffrey Barrett, Sonia Goncalves, Ewan Harrison, David K. Jackson, Ian Johnston, Dominic Kwiatkowski, Cordelia Langford, John Sillitoe on behalf of the Wellcome Sanger Institute COVID-19 Surveillance Team |
| EPI_ISL_3365508 | Lighthouse Lab in Milton Keynes | Wellcome Sanger Institute for the COVID-19 Genomics UK (COG-UK) Consortium | The Lighthouse Lab in Milton Keynes and Alex Alderton, Roberto Amato, Jeffrey Barrett, Sonia Goncalves, Ewan Harrison, David K. Jackson, Ian Johnston, Dominic Kwiatkowski, Cordelia Langford, John Sillitoe on behalf of the Wellcome Sanger Institute COVID-19 Surveillance Team |
| EPI_ISL_3365509, EPI_ISL_3365510, EPI_ISL_3365511 | Berkshire and Surrey Pathology Services Lighthouse Laboratory | Wellcome Sanger Institute for the COVID-19 Genomics UK (COG-UK) Consortium | Berkshire and Surrey Pathology Services Lighthouse Laboratory and Alex Alderton, Roberto Amato, Jeffrey Barrett, Sonia Goncalves, Ewan Harrison, David K. Jackson, Ian Johnston, Dominic Kwiatkowski, Cordelia Langford, John Sillitoe on behalf of the Wellcome Sanger Institute COVID-19 Surveillance Team |
| EPI_ISL_3365512 | Lighthouse Lab in Milton Keynes | Wellcome Sanger Institute for the COVID-19 Genomics UK (COG-UK) Consortium | The Lighthouse Lab in Milton Keynes and Alex Alderton, Roberto Amato, Jeffrey Barrett, Sonia Goncalves, Ewan Harrison, David K. Jackson, Ian Johnston, Dominic Kwiatkowski, Cordelia Langford, John Sillitoe on behalf of the Wellcome Sanger Institute COVID-19 Surveillance Team |
| EPI_ISL_3365513, EPI_ISL_3365514, EPI_ISL_3365515 | Lighthouse Lab in Alderley Park | Wellcome Sanger Institute for the COVID-19 Genomics UK (COG-UK) Consortium | Jacquelyn Wynn, Mairead Hyland, The Lighthouse Lab in Alderley Park and Alex Alderton, Roberto Amato, Jeffrey Barrett, Sonia Goncalves, Ewan Harrison, David K. Jackson, Ian Johnston, Dominic Kwiatkowski, Cordelia Langford, John Sillitoe on behalf of the Wellcome Sanger Institute COVID-19 Surveillance Team |
| EPI_ISL_3365516 | Berkshire and Surrey Pathology Services Lighthouse Laboratory | Wellcome Sanger Institute for the COVID-19 Genomics UK (COG-UK) Consortium | Berkshire and Surrey Pathology Services Lighthouse Laboratory and Alex Alderton, Roberto Amato, Jeffrey Barrett, Sonia Goncalves, Ewan Harrison, David K. Jackson, Ian Johnston, Dominic Kwiatkowski, Cordelia Langford, John Sillitoe on behalf of the Wellcome Sanger Institute COVID-19 Surveillance Team |
| EPI_ISL_3365517, EPI_ISL_3365518 | Lighthouse Lab in Alderley Park | Wellcome Sanger Institute for the COVID-19 Genomics UK (COG-UK) Consortium | Jacquelyn Wynn, Mairead Hyland, The Lighthouse Lab in Alderley Park and Alex Alderton, Roberto Amato, Jeffrey Barrett, Sonia Goncalves, Ewan Harrison, David K. Jackson, Ian Johnston, Dominic Kwiatkowski, Cordelia Langford, John Sillitoe on behalf of the Wellcome Sanger Institute COVID-19 Surveillance Team |
| EPI_ISL_3365519, EPI_ISL_3365520 | Berkshire and Surrey Pathology Services Lighthouse Laboratory | Wellcome Sanger Institute for the COVID-19 Genomics UK (COG-UK) Consortium | Berkshire and Surrey Pathology Services Lighthouse Laboratory and Alex Alderton, Roberto Amato, Jeffrey Barrett, Sonia Goncalves, Ewan Harrison, David K. Jackson, Ian Johnston, Dominic Kwiatkowski, Cordelia Langford, John Sillitoe on behalf of the Wellcome Sanger Institute COVID-19 Surveillance Team |
| EPI_ISL_3365521, EPI_ISL_3365522 | Lighthouse Lab in Alderley Park | Wellcome Sanger Institute for the COVID-19 Genomics UK (COG-UK) Consortium | Jacquelyn Wynn, Mairead Hyland, The Lighthouse Lab in Alderley Park and Alex Alderton, Roberto Amato, Jeffrey Barrett, Sonia Goncalves, Ewan Harrison, David K. Jackson, Ian Johnston, Dominic Kwiatkowski, Cordelia Langford, John Sillitoe on behalf of the Wellcome Sanger Institute COVID-19 Surveillance Team |
| EPI_ISL_3365523 | Lighthouse Lab in Milton Keynes | Wellcome Sanger Institute for the COVID-19 Genomics UK (COG-UK) Consortium | The Lighthouse Lab in Milton Keynes and Alex Alderton, Roberto Amato, Jeffrey Barrett, Sonia Goncalves, Ewan Harrison, David K. Jackson, Ian Johnston, Dominic Kwiatkowski, Cordelia Langford, John Sillitoe on behalf of the Wellcome Sanger Institute COVID-19 Surveillance Team |

| | | | |
|---|---|---|---|
| EPI_ISL_3365524 | Berkshire and Surrey Pathology Services Lighthouse Laboratory | Wellcome Sanger Institute for the COVID-19 Genomics UK (COG-UK) Consortium | Berkshire and Surrey Pathology Services Lighthouse Laboratory and Alex Alderton, Roberto Amato, Jeffrey Barrett, Sonia Goncalves, Ewan Harrison, David K. Jackson, Ian Johnston, Dominic Kwiatkowski, Cordelia Langford, John Sillitoe on behalf of the Wellcome Sanger Institute COVID-19 Surveillance Team |
| EPI_ISL_3365525 | Lighthouse Lab in Alderley Park | Wellcome Sanger Institute for the COVID-19 Genomics UK (COG-UK) Consortium | Jacquelyn Wynn, Mairead Hyland, The Lighthouse Lab in Alderley Park and Alex Alderton, Roberto Amato, Jeffrey Barrett, Sonia Goncalves, Ewan Harrison, David K. Jackson, Ian Johnston, Dominic Kwiatkowski, Cordelia Langford, John Sillitoe on behalf of the Wellcome Sanger Institute COVID-19 Surveillance Team |
| EPI_ISL_3365526 | Berkshire and Surrey Pathology Services Lighthouse Laboratory | Wellcome Sanger Institute for the COVID-19 Genomics UK (COG-UK) Consortium | Berkshire and Surrey Pathology Services Lighthouse Laboratory and Alex Alderton, Roberto Amato, Jeffrey Barrett, Sonia Goncalves, Ewan Harrison, David K. Jackson, Ian Johnston, Dominic Kwiatkowski, Cordelia Langford, John Sillitoe on behalf of the Wellcome Sanger Institute COVID-19 Surveillance Team |
| EPI_ISL_3365527 | Lighthouse Lab in Milton Keynes | Wellcome Sanger Institute for the COVID-19 Genomics UK (COG-UK) Consortium | The Lighthouse Lab in Milton Keynes and Alex Alderton, Roberto Amato, Jeffrey Barrett, Sonia Goncalves, Ewan Harrison, David K. Jackson, Ian Johnston, Dominic Kwiatkowski, Cordelia Langford, John Sillitoe on behalf of the Wellcome Sanger Institute COVID-19 Surveillance Team |
| EPI_ISL_3365528 | Berkshire and Surrey Pathology Services Lighthouse Laboratory | Wellcome Sanger Institute for the COVID-19 Genomics UK (COG-UK) Consortium | Berkshire and Surrey Pathology Services Lighthouse Laboratory and Alex Alderton, Roberto Amato, Jeffrey Barrett, Sonia Goncalves, Ewan Harrison, David K. Jackson, Ian Johnston, Dominic Kwiatkowski, Cordelia Langford, John Sillitoe on behalf of the Wellcome Sanger Institute COVID-19 Surveillance Team |
| EPI_ISL_3365529 | Lighthouse Lab in Alderley Park | Wellcome Sanger Institute for the COVID-19 Genomics UK (COG-UK) Consortium | Jacquelyn Wynn, Mairead Hyland, The Lighthouse Lab in Alderley Park and Alex Alderton, Roberto Amato, Jeffrey Barrett, Sonia Goncalves, Ewan Harrison, David K. Jackson, Ian Johnston, Dominic Kwiatkowski, Cordelia Langford, John Sillitoe on behalf of the Wellcome Sanger Institute COVID-19 Surveillance Team |
| EPI_ISL_3365530 | Berkshire and Surrey Pathology Services Lighthouse Laboratory | Wellcome Sanger Institute for the COVID-19 Genomics UK (COG-UK) Consortium | Berkshire and Surrey Pathology Services Lighthouse Laboratory and Alex Alderton, Roberto Amato, Jeffrey Barrett, Sonia Goncalves, Ewan Harrison, David K. Jackson, Ian Johnston, Dominic Kwiatkowski, Cordelia Langford, John Sillitoe on behalf of the Wellcome Sanger Institute COVID-19 Surveillance Team |
| EPI_ISL_3365531 | Lighthouse Lab in Alderley Park | Wellcome Sanger Institute for the COVID-19 Genomics UK (COG-UK) Consortium | Jacquelyn Wynn, Mairead Hyland, The Lighthouse Lab in Alderley Park and Alex Alderton, Roberto Amato, Jeffrey Barrett, Sonia Goncalves, Ewan Harrison, David K. Jackson, Ian Johnston, Dominic Kwiatkowski, Cordelia Langford, John Sillitoe on behalf of the Wellcome Sanger Institute COVID-19 Surveillance Team |
| EPI_ISL_3365532 | Berkshire and Surrey Pathology Services Lighthouse Laboratory | Wellcome Sanger Institute for the COVID-19 Genomics UK (COG-UK) Consortium | Berkshire and Surrey Pathology Services Lighthouse Laboratory and Alex Alderton, Roberto Amato, Jeffrey Barrett, Sonia Goncalves, Ewan Harrison, David K. Jackson, Ian Johnston, Dominic Kwiatkowski, Cordelia Langford, John Sillitoe on behalf of the Wellcome Sanger Institute COVID-19 Surveillance Team |
| EPI_ISL_3365533 | Lighthouse Lab in Alderley Park | Wellcome Sanger Institute for the COVID-19 Genomics UK (COG-UK) Consortium | Jacquelyn Wynn, Mairead Hyland, The Lighthouse Lab in Alderley Park and Alex Alderton, Roberto Amato, Jeffrey Barrett, Sonia Goncalves, Ewan Harrison, David K. Jackson, Ian Johnston, Dominic Kwiatkowski, Cordelia Langford, John Sillitoe on behalf of the Wellcome Sanger Institute COVID-19 Surveillance Team |
| EPI_ISL_3365534 | Lighthouse Lab in Milton Keynes | Wellcome Sanger Institute for the COVID-19 Genomics UK (COG-UK) Consortium | The Lighthouse Lab in Milton Keynes and Alex Alderton, Roberto Amato, Jeffrey Barrett, Sonia Goncalves, Ewan Harrison, David K. Jackson, Ian Johnston, Dominic Kwiatkowski, Cordelia Langford, John Sillitoe on behalf of the Wellcome Sanger Institute COVID-19 Surveillance Team |
| EPI_ISL_3365535 | Berkshire and Surrey Pathology Services Lighthouse Laboratory | Wellcome Sanger Institute for the COVID-19 Genomics UK (COG-UK) Consortium | Berkshire and Surrey Pathology Services Lighthouse Laboratory and Alex Alderton, Roberto Amato, Jeffrey Barrett, Sonia Goncalves, Ewan Harrison, David K. Jackson, Ian Johnston, Dominic Kwiatkowski, Cordelia Langford, John Sillitoe on behalf of the Wellcome Sanger Institute COVID-19 Surveillance Team |
| EPI_ISL_3365536 | Lighthouse Lab in Alderley Park | Wellcome Sanger Institute for the COVID-19 Genomics UK (COG-UK) Consortium | Jacquelyn Wynn, Mairead Hyland, The Lighthouse Lab in Alderley Park and Alex Alderton, Roberto Amato, Jeffrey Barrett, Sonia Goncalves, Ewan Harrison, David K. Jackson, Ian Johnston, Dominic Kwiatkowski, Cordelia Langford, John Sillitoe on behalf of the Wellcome Sanger Institute COVID-19 Surveillance Team |
| EPI_ISL_3365537, EPI_ISL_3365538, EPI_ISL_3365539, EPI_ISL_3365540 | Berkshire and Surrey Pathology Services Lighthouse Laboratory | Wellcome Sanger Institute for the COVID-19 Genomics UK (COG-UK) Consortium | Berkshire and Surrey Pathology Services Lighthouse Laboratory and Alex Alderton, Roberto Amato, Jeffrey Barrett, Sonia Goncalves, Ewan Harrison, David K. Jackson, Ian Johnston, Dominic Kwiatkowski, Cordelia Langford, John Sillitoe on behalf of the Wellcome Sanger Institute COVID-19 Surveillance Team |
| EPI_ISL_3365541, EPI_ISL_3365542 | Lighthouse Lab in Milton Keynes | Wellcome Sanger Institute for the COVID-19 Genomics UK (COG-UK) Consortium | The Lighthouse Lab in Milton Keynes and Alex Alderton, Roberto Amato, Jeffrey Barrett, Sonia Goncalves, Ewan Harrison, David K. Jackson, Ian Johnston, Dominic Kwiatkowski, Cordelia Langford, John Sillitoe on behalf of the Wellcome Sanger Institute COVID-19 Surveillance Team |
| EPI_ISL_3365543 | Berkshire and Surrey Pathology Services Lighthouse Laboratory | Wellcome Sanger Institute for the COVID-19 Genomics UK (COG-UK) Consortium | Berkshire and Surrey Pathology Services Lighthouse Laboratory and Alex Alderton, Roberto Amato, Jeffrey Barrett, Sonia Goncalves, Ewan Harrison, David K. Jackson, Ian Johnston, Dominic Kwiatkowski, Cordelia Langford, John Sillitoe on behalf of the Wellcome Sanger Institute COVID-19 Surveillance Team |
| EPI_ISL_3365544 | Lighthouse Lab in Alderley Park | Wellcome Sanger Institute for the COVID-19 Genomics UK (COG-UK) Consortium | Jacquelyn Wynn, Mairead Hyland, The Lighthouse Lab in Alderley Park and Alex Alderton, Roberto Amato, Jeffrey Barrett, Sonia Goncalves, Ewan Harrison, David K. Jackson, Ian Johnston, Dominic Kwiatkowski, Cordelia Langford, John Sillitoe on behalf of the Wellcome Sanger Institute COVID-19 Surveillance Team |
| EPI_ISL_3365545 | Berkshire and Surrey Pathology Services Lighthouse Laboratory | Wellcome Sanger Institute for the COVID-19 Genomics UK (COG-UK) Consortium | Berkshire and Surrey Pathology Services Lighthouse Laboratory and Alex Alderton, Roberto Amato, Jeffrey Barrett, Sonia Goncalves, Ewan Harrison, David K. Jackson, Ian Johnston, Dominic Kwiatkowski, Cordelia Langford, John Sillitoe on behalf of the Wellcome Sanger Institute COVID-19 Surveillance Team |
| EPI_ISL_3365546 | Lighthouse Lab in Milton Keynes | Wellcome Sanger Institute for the COVID-19 Genomics UK (COG-UK) Consortium | The Lighthouse Lab in Milton Keynes and Alex Alderton, Roberto Amato, Jeffrey Barrett, Sonia Goncalves, Ewan Harrison, David K. Jackson, Ian Johnston, Dominic Kwiatkowski, Cordelia Langford, John Sillitoe on behalf of the Wellcome Sanger Institute COVID-19 Surveillance Team |
| EPI_ISL_3365547, EPI_ISL_3365548 | Lighthouse Lab in Alderley Park | Wellcome Sanger Institute for the COVID-19 Genomics UK (COG-UK) Consortium | Jacquelyn Wynn, Mairead Hyland, The Lighthouse Lab in Alderley Park and Alex Alderton, Roberto Amato, Jeffrey Barrett, Sonia Goncalves, Ewan Harrison, David K. Jackson, Ian Johnston, Dominic Kwiatkowski, Cordelia Langford, John Sillitoe on behalf of the Wellcome Sanger Institute COVID-19 Surveillance Team |
| EPI_ISL_3365549, EPI_ISL_3365550 | Lighthouse Lab in Milton Keynes | Wellcome Sanger Institute for the COVID-19 Genomics UK (COG-UK) Consortium | The Lighthouse Lab in Milton Keynes and Alex Alderton, Roberto Amato, Jeffrey Barrett, Sonia Goncalves, Ewan Harrison, David K. Jackson, Ian Johnston, Dominic Kwiatkowski, Cordelia Langford, John Sillitoe on behalf of the Wellcome Sanger Institute COVID-19 Surveillance Team |
| EPI_ISL_3365551 | Berkshire and Surrey Pathology Services Lighthouse Laboratory | Wellcome Sanger Institute for the COVID-19 Genomics UK (COG-UK) Consortium | Berkshire and Surrey Pathology Services Lighthouse Laboratory and Alex Alderton, Roberto Amato, Jeffrey Barrett, Sonia Goncalves, Ewan Harrison, David K. Jackson, Ian Johnston, Dominic Kwiatkowski, Cordelia Langford, John Sillitoe on behalf of the Wellcome Sanger Institute COVID-19 Surveillance Team |
| EPI_ISL_3365552 | Lighthouse Lab in Milton Keynes | Wellcome Sanger Institute for the COVID-19 Genomics UK (COG-UK) Consortium | The Lighthouse Lab in Milton Keynes and Alex Alderton, Roberto Amato, Jeffrey Barrett, Sonia Goncalves, Ewan Harrison, David K. Jackson, Ian Johnston, Dominic Kwiatkowski, Cordelia Langford, John Sillitoe on behalf of the Wellcome Sanger Institute COVID-19 Surveillance Team |
| EPI_ISL_3365553 | Lighthouse Lab in Alderley Park | Wellcome Sanger Institute for the COVID-19 Genomics UK (COG-UK) Consortium | Jacquelyn Wynn, Mairead Hyland, The Lighthouse Lab in Alderley Park and Alex Alderton, Roberto Amato, Jeffrey Barrett, Sonia Goncalves, Ewan Harrison, David K. Jackson, Ian Johnston, Dominic Kwiatkowski, Cordelia Langford, John Sillitoe on behalf of the Wellcome Sanger Institute COVID-19 Surveillance Team |
| EPI_ISL_3365554, EPI_ISL_3365555, EPI_ISL_3365556 | Berkshire and Surrey Pathology Services Lighthouse Laboratory | Wellcome Sanger Institute for the COVID-19 Genomics UK (COG-UK) Consortium | Berkshire and Surrey Pathology Services Lighthouse Laboratory and Alex Alderton, Roberto Amato, Jeffrey Barrett, Sonia Goncalves, Ewan Harrison, David K. Jackson, Ian Johnston, Dominic Kwiatkowski, Cordelia Langford, John Sillitoe on behalf of the Wellcome Sanger Institute COVID-19 Surveillance Team |
| EPI_ISL_3365557 | Lighthouse Lab in Alderley Park | Wellcome Sanger Institute for the COVID-19 Genomics UK (COG-UK) Consortium | Jacquelyn Wynn, Mairead Hyland, The Lighthouse Lab in Alderley Park and Alex Alderton, Roberto Amato, Jeffrey Barrett, Sonia Goncalves, Ewan Harrison, David K. Jackson, Ian Johnston, Dominic Kwiatkowski, Cordelia Langford, John Sillitoe on behalf of the Wellcome Sanger Institute COVID-19 Surveillance Team |
| EPI_ISL_3365558 | Berkshire and Surrey Pathology Services Lighthouse Laboratory | Wellcome Sanger Institute for the COVID-19 Genomics UK (COG-UK) Consortium | Berkshire and Surrey Pathology Services Lighthouse Laboratory and Alex Alderton, Roberto Amato, Jeffrey Barrett, Sonia Goncalves, Ewan Harrison, David K. Jackson, Ian Johnston, Dominic Kwiatkowski, Cordelia Langford, John Sillitoe on behalf of the Wellcome Sanger Institute COVID-19 Surveillance Team |
| EPI_ISL_3365559 | Lighthouse Lab in Milton Keynes | Wellcome Sanger Institute for the COVID-19 Genomics UK (COG-UK) Consortium | The Lighthouse Lab in Milton Keynes and Alex Alderton, Roberto Amato, Jeffrey Barrett, Sonia Goncalves, Ewan Harrison, David K. Jackson, Ian Johnston, Dominic Kwiatkowski, Cordelia Langford, John Sillitoe on behalf of the Wellcome Sanger Institute COVID-19 Surveillance Team |
| EPI_ISL_3365560 | Berkshire and Surrey Pathology Services Lighthouse Laboratory | Wellcome Sanger Institute for the COVID-19 Genomics UK (COG-UK) Consortium | Berkshire and Surrey Pathology Services Lighthouse Laboratory and Alex Alderton, Roberto Amato, Jeffrey Barrett, Sonia Goncalves, Ewan Harrison, David K. Jackson, Ian Johnston, Dominic Kwiatkowski, Cordelia Langford, John Sillitoe on behalf of the Wellcome Sanger Institute COVID-19 Surveillance Team |
| EPI_ISL_3365561 | Lighthouse Lab in Alderley Park | Wellcome Sanger Institute for the COVID-19 Genomics UK (COG-UK) Consortium | Jacquelyn Wynn, Mairead Hyland, The Lighthouse Lab in Alderley Park and Alex Alderton, Roberto Amato, Jeffrey Barrett, Sonia Goncalves, Ewan Harrison, David K. Jackson, Ian Johnston, Dominic Kwiatkowski, Cordelia Langford, John Sillitoe on behalf of the Wellcome Sanger Institute COVID-19 Surveillance Team |
| EPI_ISL_3365562 | Berkshire and Surrey Pathology Services Lighthouse Laboratory | Wellcome Sanger Institute for the COVID-19 Genomics UK (COG-UK) Consortium | Berkshire and Surrey Pathology Services Lighthouse Laboratory and Alex Alderton, Roberto Amato, Jeffrey Barrett, Sonia Goncalves, Ewan Harrison, David K. Jackson, Ian Johnston, Dominic Kwiatkowski, Cordelia Langford, John Sillitoe on behalf of the Wellcome Sanger Institute COVID-19 Surveillance Team |
| EPI_ISL_3365563 | Lighthouse Lab in Alderley Park | Wellcome Sanger Institute for the COVID-19 Genomics UK (COG-UK) Consortium | Jacquelyn Wynn, Mairead Hyland, The Lighthouse Lab in Alderley Park and Alex Alderton, Roberto Amato, Jeffrey Barrett, Sonia Goncalves, Ewan Harrison, David K. Jackson, Ian Johnston, Dominic Kwiatkowski, Cordelia Langford, John Sillitoe on behalf of the Wellcome Sanger Institute COVID-19 Surveillance Team |
| EPI_ISL_3365564, EPI_ISL_3365565 | Berkshire and Surrey Pathology Services Lighthouse Laboratory | Wellcome Sanger Institute for the COVID-19 Genomics UK (COG-UK) Consortium | Berkshire and Surrey Pathology Services Lighthouse Laboratory and Alex Alderton, Roberto Amato, Jeffrey Barrett, Sonia Goncalves, Ewan Harrison, David K. Jackson, Ian Johnston, Dominic Kwiatkowski, Cordelia Langford, John Sillitoe on behalf of the Wellcome Sanger Institute COVID-19 Surveillance Team |
| EPI_ISL_3365566, EPI_ISL_3365567 | Lighthouse Lab in Milton Keynes | Wellcome Sanger Institute for the COVID-19 Genomics UK (COG-UK) Consortium | The Lighthouse Lab in Milton Keynes and Alex Alderton, Roberto Amato, Jeffrey Barrett, Sonia Goncalves, Ewan Harrison, David K. Jackson, Ian Johnston, Dominic Kwiatkowski, Cordelia Langford, John Sillitoe on behalf of the Wellcome Sanger Institute COVID-19 Surveillance Team |
| EPI_ISL_3365568, EPI_ISL_3365569, EPI_ISL_3365570, EPI_ISL_3365571 | Berkshire and Surrey Pathology Services Lighthouse Laboratory | Wellcome Sanger Institute for the COVID-19 Genomics UK (COG-UK) Consortium | Berkshire and Surrey Pathology Services Lighthouse Laboratory and Alex Alderton, Roberto Amato, Jeffrey Barrett, Sonia Goncalves, Ewan Harrison, David K. Jackson, Ian Johnston, Dominic Kwiatkowski, Cordelia Langford, John Sillitoe on behalf of the Wellcome Sanger Institute COVID-19 Surveillance Team |
| EPI_ISL_3365572 | Lighthouse Lab in Alderley Park | Wellcome Sanger Institute for the COVID-19 Genomics UK (COG-UK) Consortium | Jacquelyn Wynn, Mairead Hyland, The Lighthouse Lab in Alderley Park and Alex Alderton, Roberto Amato, Jeffrey Barrett, Sonia Goncalves, Ewan Harrison, David K. Jackson, Ian Johnston, Dominic Kwiatkowski, Cordelia Langford, John Sillitoe on behalf of the Wellcome Sanger Institute COVID-19 Surveillance Team |
| EPI_ISL_3365573, EPI_ISL_3365574 | Berkshire and Surrey Pathology Services Lighthouse Laboratory | Wellcome Sanger Institute for the COVID-19 Genomics UK (COG-UK) Consortium | Berkshire and Surrey Pathology Services Lighthouse Laboratory and Alex Alderton, Roberto Amato, Jeffrey Barrett, Sonia Goncalves, Ewan Harrison, David K. Jackson, Ian Johnston, Dominic Kwiatkowski, Cordelia Langford, John Sillitoe on behalf of the Wellcome Sanger Institute COVID-19 Surveillance Team |
| EPI_ISL_3365575, EPI_ISL_3365576 | Lighthouse Lab in Alderley Park | Wellcome Sanger Institute for the COVID-19 Genomics UK (COG-UK) Consortium | Jacquelyn Wynn, Mairead Hyland, The Lighthouse Lab in Alderley Park and Alex Alderton, Roberto Amato, Jeffrey Barrett, Sonia Goncalves, Ewan Harrison, David K. Jackson, Ian Johnston, Dominic Kwiatkowski, Cordelia Langford, John Sillitoe on behalf of the Wellcome Sanger Institute COVID-19 Surveillance Team |
| EPI_ISL_3365577 | Berkshire and Surrey Pathology Services Lighthouse Laboratory | Wellcome Sanger Institute for the COVID-19 Genomics UK (COG-UK) Consortium | Berkshire and Surrey Pathology Services Lighthouse Laboratory and Alex Alderton, Roberto Amato, Jeffrey Barrett, Sonia Goncalves, Ewan Harrison, David K. Jackson, Ian Johnston, Dominic Kwiatkowski, Cordelia Langford, John Sillitoe on behalf of the Wellcome Sanger Institute COVID-19 Surveillance Team |
| EPI_ISL_3365578 | Lighthouse Lab in Milton Keynes | Wellcome Sanger Institute for the COVID-19 Genomics UK (COG-UK) Consortium | The Lighthouse Lab in Milton Keynes and Alex Alderton, Roberto Amato, Jeffrey Barrett, Sonia Goncalves, Ewan Harrison, David K. Jackson, Ian Johnston, Dominic Kwiatkowski, Cordelia Langford, John Sillitoe on behalf of the Wellcome Sanger Institute COVID-19 Surveillance Team |
| EPI_ISL_3365579 | Berkshire and Surrey Pathology Services Lighthouse Laboratory | Wellcome Sanger Institute for the COVID-19 Genomics UK (COG-UK) Consortium | Berkshire and Surrey Pathology Services Lighthouse Laboratory and Alex Alderton, Roberto Amato, Jeffrey Barrett, Sonia Goncalves, Ewan Harrison, David K. Jackson, Ian Johnston, Dominic Kwiatkowski, Cordelia Langford, John Sillitoe on behalf of the Wellcome Sanger Institute COVID-19 Surveillance Team |
| EPI_ISL_3365580, EPI_ISL_3365581 | Lighthouse Lab in Milton Keynes | Wellcome Sanger Institute for the COVID-19 Genomics UK (COG-UK) Consortium | The Lighthouse Lab in Milton Keynes and Alex Alderton, Roberto Amato, Jeffrey Barrett, Sonia Goncalves, Ewan Harrison, David K. Jackson, Ian Johnston, Dominic Kwiatkowski, Cordelia Langford, John Sillitoe on behalf of the Wellcome Sanger Institute COVID-19 Surveillance Team |
| EPI_ISL_3365582, EPI_ISL_3365583 | Berkshire and Surrey Pathology Services Lighthouse Laboratory | Wellcome Sanger Institute for the COVID-19 Genomics UK (COG-UK) Consortium | Berkshire and Surrey Pathology Services Lighthouse Laboratory and Alex Alderton, Roberto Amato, Jeffrey Barrett, Sonia Goncalves, Ewan Harrison, David K. Jackson, Ian Johnston, Dominic Kwiatkowski, Cordelia Langford, John Sillitoe on behalf of the Wellcome Sanger Institute COVID-19 Surveillance Team |

| | | | |
|---|---|---|---|
| EPI_ISL_3365584 | Lighthouse Lab in Milton Keynes | Wellcome Sanger Institute for the COVID-19 Genomics UK (COG-UK) Consortium | The Lighthouse Lab in Milton Keynes and Alex Alderton, Roberto Amato, Jeffrey Barrett, Sonia Goncalves, Ewan Harrison, David K. Jackson, Ian Johnston, Dominic Kwiatkowski, Cordelia Langford, John Sillitoe on behalf of the Wellcome Sanger Institute COVID-19 Surveillance Team |
| EPI_ISL_3365585, EPI_ISL_3365586 | Berkshire and Surrey Pathology Services Lighthouse Laboratory | Wellcome Sanger Institute for the COVID-19 Genomics UK (COG-UK) Consortium | Berkshire and Surrey Pathology Services Lighthouse Laboratory and Alex Alderton, Roberto Amato, Jeffrey Barrett, Sonia Goncalves, Ewan Harrison, David K. Jackson, Ian Johnston, Dominic Kwiatkowski, Cordelia Langford, John Sillitoe on behalf of the Wellcome Sanger Institute COVID-19 Surveillance Team |
| EPI_ISL_3365587 | Lighthouse Lab in Milton Keynes | Wellcome Sanger Institute for the COVID-19 Genomics UK (COG-UK) Consortium | The Lighthouse Lab in Milton Keynes and Alex Alderton, Roberto Amato, Jeffrey Barrett, Sonia Goncalves, Ewan Harrison, David K. Jackson, Ian Johnston, Dominic Kwiatkowski, Cordelia Langford, John Sillitoe on behalf of the Wellcome Sanger Institute COVID-19 Surveillance Team |
| EPI_ISL_3365588 | Berkshire and Surrey Pathology Services Lighthouse Laboratory | Wellcome Sanger Institute for the COVID-19 Genomics UK (COG-UK) Consortium | Berkshire and Surrey Pathology Services Lighthouse Laboratory and Alex Alderton, Roberto Amato, Jeffrey Barrett, Sonia Goncalves, Ewan Harrison, David K. Jackson, Ian Johnston, Dominic Kwiatkowski, Cordelia Langford, John Sillitoe on behalf of the Wellcome Sanger Institute COVID-19 Surveillance Team |
| EPI_ISL_3365589 | Lighthouse Lab in Alderley Park | Wellcome Sanger Institute for the COVID-19 Genomics UK (COG-UK) Consortium | Jacquelyn Wynn, Mairead Hyland, The Lighthouse Lab in Alderley Park and Alex Alderton, Roberto Amato, Jeffrey Barrett, Sonia Goncalves, Ewan Harrison, David K. Jackson, Ian Johnston, Dominic Kwiatkowski, Cordelia Langford, John Sillitoe on behalf of the Wellcome Sanger Institute COVID-19 Surveillance Team |
| EPI_ISL_3365590, EPI_ISL_3365591 | Berkshire and Surrey Pathology Services Lighthouse Laboratory | Wellcome Sanger Institute for the COVID-19 Genomics UK (COG-UK) Consortium | Berkshire and Surrey Pathology Services Lighthouse Laboratory and Alex Alderton, Roberto Amato, Jeffrey Barrett, Sonia Goncalves, Ewan Harrison, David K. Jackson, Ian Johnston, Dominic Kwiatkowski, Cordelia Langford, John Sillitoe on behalf of the Wellcome Sanger Institute COVID-19 Surveillance Team |
| EPI_ISL_3365592 | Lighthouse Lab in Milton Keynes | Wellcome Sanger Institute for the COVID-19 Genomics UK (COG-UK) Consortium | The Lighthouse Lab in Milton Keynes and Alex Alderton, Roberto Amato, Jeffrey Barrett, Sonia Goncalves, Ewan Harrison, David K. Jackson, Ian Johnston, Dominic Kwiatkowski, Cordelia Langford, John Sillitoe on behalf of the Wellcome Sanger Institute COVID-19 Surveillance Team |
| EPI_ISL_3365593 | Lighthouse Lab in Alderley Park | Wellcome Sanger Institute for the COVID-19 Genomics UK (COG-UK) Consortium | Jacquelyn Wynn, Mairead Hyland, The Lighthouse Lab in Alderley Park and Alex Alderton, Roberto Amato, Jeffrey Barrett, Sonia Goncalves, Ewan Harrison, David K. Jackson, Ian Johnston, Dominic Kwiatkowski, Cordelia Langford, John Sillitoe on behalf of the Wellcome Sanger Institute COVID-19 Surveillance Team |
| EPI_ISL_3365594, EPI_ISL_3365595, EPI_ISL_3365596 | Lighthouse Lab in Milton Keynes | Wellcome Sanger Institute for the COVID-19 Genomics UK (COG-UK) Consortium | The Lighthouse Lab in Milton Keynes and Alex Alderton, Roberto Amato, Jeffrey Barrett, Sonia Goncalves, Ewan Harrison, David K. Jackson, Ian Johnston, Dominic Kwiatkowski, Cordelia Langford, John Sillitoe on behalf of the Wellcome Sanger Institute COVID-19 Surveillance Team |
| EPI_ISL_3365597 | Berkshire and Surrey Pathology Services Lighthouse Laboratory | Wellcome Sanger Institute for the COVID-19 Genomics UK (COG-UK) Consortium | Berkshire and Surrey Pathology Services Lighthouse Laboratory and Alex Alderton, Roberto Amato, Jeffrey Barrett, Sonia Goncalves, Ewan Harrison, David K. Jackson, Ian Johnston, Dominic Kwiatkowski, Cordelia Langford, John Sillitoe on behalf of the Wellcome Sanger Institute COVID-19 Surveillance Team |
| EPI_ISL_3365598, EPI_ISL_3365599 | Lighthouse Lab in Alderley Park | Wellcome Sanger Institute for the COVID-19 Genomics UK (COG-UK) Consortium | Jacquelyn Wynn, Mairead Hyland, The Lighthouse Lab in Alderley Park and Alex Alderton, Roberto Amato, Jeffrey Barrett, Sonia Goncalves, Ewan Harrison, David K. Jackson, Ian Johnston, Dominic Kwiatkowski, Cordelia Langford, John Sillitoe on behalf of the Wellcome Sanger Institute COVID-19 Surveillance Team |
| EPI_ISL_3365600 | Berkshire and Surrey Pathology Services Lighthouse Laboratory | Wellcome Sanger Institute for the COVID-19 Genomics UK (COG-UK) Consortium | Berkshire and Surrey Pathology Services Lighthouse Laboratory and Alex Alderton, Roberto Amato, Jeffrey Barrett, Sonia Goncalves, Ewan Harrison, David K. Jackson, Ian Johnston, Dominic Kwiatkowski, Cordelia Langford, John Sillitoe on behalf of the Wellcome Sanger Institute COVID-19 Surveillance Team |
| EPI_ISL_3365601 | Lighthouse Lab in Milton Keynes | Wellcome Sanger Institute for the COVID-19 Genomics UK (COG-UK) Consortium | The Lighthouse Lab in Milton Keynes and Alex Alderton, Roberto Amato, Jeffrey Barrett, Sonia Goncalves, Ewan Harrison, David K. Jackson, Ian Johnston, Dominic Kwiatkowski, Cordelia Langford, John Sillitoe on behalf of the Wellcome Sanger Institute COVID-19 Surveillance Team |
| EPI_ISL_3365602 | Berkshire and Surrey Pathology Services Lighthouse Laboratory | Wellcome Sanger Institute for the COVID-19 Genomics UK (COG-UK) Consortium | Berkshire and Surrey Pathology Services Lighthouse Laboratory and Alex Alderton, Roberto Amato, Jeffrey Barrett, Sonia Goncalves, Ewan Harrison, David K. Jackson, Ian Johnston, Dominic Kwiatkowski, Cordelia Langford, John Sillitoe on behalf of the Wellcome Sanger Institute COVID-19 Surveillance Team |
| EPI_ISL_3365603 | Lighthouse Lab in Milton Keynes | Wellcome Sanger Institute for the COVID-19 Genomics UK (COG-UK) Consortium | The Lighthouse Lab in Milton Keynes and Alex Alderton, Roberto Amato, Jeffrey Barrett, Sonia Goncalves, Ewan Harrison, David K. Jackson, Ian Johnston, Dominic Kwiatkowski, Cordelia Langford, John Sillitoe on behalf of the Wellcome Sanger Institute COVID-19 Surveillance Team |
| EPI_ISL_3365604 | Berkshire and Surrey Pathology Services Lighthouse Laboratory | Wellcome Sanger Institute for the COVID-19 Genomics UK (COG-UK) Consortium | Berkshire and Surrey Pathology Services Lighthouse Laboratory and Alex Alderton, Roberto Amato, Jeffrey Barrett, Sonia Goncalves, Ewan Harrison, David K. Jackson, Ian Johnston, Dominic Kwiatkowski, Cordelia Langford, John Sillitoe on behalf of the Wellcome Sanger Institute COVID-19 Surveillance Team |
| EPI_ISL_3365605, EPI_ISL_3365606 | Lighthouse Lab in Milton Keynes | Wellcome Sanger Institute for the COVID-19 Genomics UK (COG-UK) Consortium | The Lighthouse Lab in Milton Keynes and Alex Alderton, Roberto Amato, Jeffrey Barrett, Sonia Goncalves, Ewan Harrison, David K. Jackson, Ian Johnston, Dominic Kwiatkowski, Cordelia Langford, John Sillitoe on behalf of the Wellcome Sanger Institute COVID-19 Surveillance Team |
| EPI_ISL_3365607, EPI_ISL_3365608 | Berkshire and Surrey Pathology Services Lighthouse Laboratory | Wellcome Sanger Institute for the COVID-19 Genomics UK (COG-UK) Consortium | Berkshire and Surrey Pathology Services Lighthouse Laboratory and Alex Alderton, Roberto Amato, Jeffrey Barrett, Sonia Goncalves, Ewan Harrison, David K. Jackson, Ian Johnston, Dominic Kwiatkowski, Cordelia Langford, John Sillitoe on behalf of the Wellcome Sanger Institute COVID-19 Surveillance Team |
| EPI_ISL_3365609 | Lighthouse Lab in Alderley Park | Wellcome Sanger Institute for the COVID-19 Genomics UK (COG-UK) Consortium | Jacquelyn Wynn, Mairead Hyland, The Lighthouse Lab in Alderley Park and Alex Alderton, Roberto Amato, Jeffrey Barrett, Sonia Goncalves, Ewan Harrison, David K. Jackson, Ian Johnston, Dominic Kwiatkowski, Cordelia Langford, John Sillitoe on behalf of the Wellcome Sanger Institute COVID-19 Surveillance Team |
| EPI_ISL_3365610, EPI_ISL_3365611 | Lighthouse Lab in Milton Keynes | Wellcome Sanger Institute for the COVID-19 Genomics UK (COG-UK) Consortium | The Lighthouse Lab in Milton Keynes and Alex Alderton, Roberto Amato, Jeffrey Barrett, Sonia Goncalves, Ewan Harrison, David K. Jackson, Ian Johnston, Dominic Kwiatkowski, Cordelia Langford, John Sillitoe on behalf of the Wellcome Sanger Institute COVID-19 Surveillance Team |
| EPI_ISL_3365612 | Lighthouse Lab in Alderley Park | Wellcome Sanger Institute for the COVID-19 Genomics UK (COG-UK) Consortium | Jacquelyn Wynn, Mairead Hyland, The Lighthouse Lab in Alderley Park and Alex Alderton, Roberto Amato, Jeffrey Barrett, Sonia Goncalves, Ewan Harrison, David K. Jackson, Ian Johnston, Dominic Kwiatkowski, Cordelia Langford, John Sillitoe on behalf of the Wellcome Sanger Institute COVID-19 Surveillance Team |
| EPI_ISL_3365613, EPI_ISL_3365614 | Berkshire and Surrey Pathology Services Lighthouse Laboratory | Wellcome Sanger Institute for the COVID-19 Genomics UK (COG-UK) Consortium | Berkshire and Surrey Pathology Services Lighthouse Laboratory and Alex Alderton, Roberto Amato, Jeffrey Barrett, Sonia Goncalves, Ewan Harrison, David K. Jackson, Ian Johnston, Dominic Kwiatkowski, Cordelia Langford, John Sillitoe on behalf of the Wellcome Sanger Institute COVID-19 Surveillance Team |
| EPI_ISL_3365615 | Lighthouse Lab in Alderley Park | Wellcome Sanger Institute for the COVID-19 Genomics UK (COG-UK) Consortium | Jacquelyn Wynn, Mairead Hyland, The Lighthouse Lab in Alderley Park and Alex Alderton, Roberto Amato, Jeffrey Barrett, Sonia Goncalves, Ewan Harrison, David K. Jackson, Ian Johnston, Dominic Kwiatkowski, Cordelia Langford, John Sillitoe on behalf of the Wellcome Sanger Institute COVID-19 Surveillance Team |
| EPI_ISL_3365616 | Berkshire and Surrey Pathology Services Lighthouse Laboratory | Wellcome Sanger Institute for the COVID-19 Genomics UK (COG-UK) Consortium | Berkshire and Surrey Pathology Services Lighthouse Laboratory and Alex Alderton, Roberto Amato, Jeffrey Barrett, Sonia Goncalves, Ewan Harrison, David K. Jackson, Ian Johnston, Dominic Kwiatkowski, Cordelia Langford, John Sillitoe on behalf of the Wellcome Sanger Institute COVID-19 Surveillance Team |
| EPI_ISL_3365617, EPI_ISL_3365618 | Lighthouse Lab in Alderley Park | Wellcome Sanger Institute for the COVID-19 Genomics UK (COG-UK) Consortium | Jacquelyn Wynn, Mairead Hyland, The Lighthouse Lab in Alderley Park and Alex Alderton, Roberto Amato, Jeffrey Barrett, Sonia Goncalves, Ewan Harrison, David K. Jackson, Ian Johnston, Dominic Kwiatkowski, Cordelia Langford, John Sillitoe on behalf of the Wellcome Sanger Institute COVID-19 Surveillance Team |
| EPI_ISL_3365619 | Lighthouse Lab in Milton Keynes | Wellcome Sanger Institute for the COVID-19 Genomics UK (COG-UK) Consortium | The Lighthouse Lab in Milton Keynes and Alex Alderton, Roberto Amato, Jeffrey Barrett, Sonia Goncalves, Ewan Harrison, David K. Jackson, Ian Johnston, Dominic Kwiatkowski, Cordelia Langford, John Sillitoe on behalf of the Wellcome Sanger Institute COVID-19 Surveillance Team |
| EPI_ISL_3365620 | Berkshire and Surrey Pathology Services Lighthouse Laboratory | Wellcome Sanger Institute for the COVID-19 Genomics UK (COG-UK) Consortium | Berkshire and Surrey Pathology Services Lighthouse Laboratory and Alex Alderton, Roberto Amato, Jeffrey Barrett, Sonia Goncalves, Ewan Harrison, David K. Jackson, Ian Johnston, Dominic Kwiatkowski, Cordelia Langford, John Sillitoe on behalf of the Wellcome Sanger Institute COVID-19 Surveillance Team |
| EPI_ISL_3365621 | Lighthouse Lab in Milton Keynes | Wellcome Sanger Institute for the COVID-19 Genomics UK (COG-UK) Consortium | The Lighthouse Lab in Milton Keynes and Alex Alderton, Roberto Amato, Jeffrey Barrett, Sonia Goncalves, Ewan Harrison, David K. Jackson, Ian Johnston, Dominic Kwiatkowski, Cordelia Langford, John Sillitoe on behalf of the Wellcome Sanger Institute COVID-19 Surveillance Team |
| EPI_ISL_3365622, EPI_ISL_3365623, EPI_ISL_3365624, EPI_ISL_3365625, EPI_ISL_3365626, EPI_ISL_3365627 | Berkshire and Surrey Pathology Services Lighthouse Laboratory | Wellcome Sanger Institute for the COVID-19 Genomics UK (COG-UK) Consortium | Berkshire and Surrey Pathology Services Lighthouse Laboratory and Alex Alderton, Roberto Amato, Jeffrey Barrett, Sonia Goncalves, Ewan Harrison, David K. Jackson, Ian Johnston, Dominic Kwiatkowski, Cordelia Langford, John Sillitoe on behalf of the Wellcome Sanger Institute COVID-19 Surveillance Team |
| EPI_ISL_3365628 | Lighthouse Lab in Alderley Park | Wellcome Sanger Institute for the COVID-19 Genomics UK (COG-UK) Consortium | Jacquelyn Wynn, Mairead Hyland, The Lighthouse Lab in Alderley Park and Alex Alderton, Roberto Amato, Jeffrey Barrett, Sonia Goncalves, Ewan Harrison, David K. Jackson, Ian Johnston, Dominic Kwiatkowski, Cordelia Langford, John Sillitoe on behalf of the Wellcome Sanger Institute COVID-19 Surveillance Team |
| EPI_ISL_3365629 | Lighthouse Lab in Milton Keynes | Wellcome Sanger Institute for the COVID-19 Genomics UK (COG-UK) Consortium | The Lighthouse Lab in Milton Keynes and Alex Alderton, Roberto Amato, Jeffrey Barrett, Sonia Goncalves, Ewan Harrison, David K. Jackson, Ian Johnston, Dominic Kwiatkowski, Cordelia Langford, John Sillitoe on behalf of the Wellcome Sanger Institute COVID-19 Surveillance Team |
| EPI_ISL_3365630, EPI_ISL_3365631 | Lighthouse Lab in Alderley Park | Wellcome Sanger Institute for the COVID-19 Genomics UK (COG-UK) Consortium | Jacquelyn Wynn, Mairead Hyland, The Lighthouse Lab in Alderley Park and Alex Alderton, Roberto Amato, Jeffrey Barrett, Sonia Goncalves, Ewan Harrison, David K. Jackson, Ian Johnston, Dominic Kwiatkowski, Cordelia Langford, John Sillitoe on behalf of the Wellcome Sanger Institute COVID-19 Surveillance Team |
| EPI_ISL_3365632 | Berkshire and Surrey Pathology Services Lighthouse Laboratory | Wellcome Sanger Institute for the COVID-19 Genomics UK (COG-UK) Consortium | Berkshire and Surrey Pathology Services Lighthouse Laboratory and Alex Alderton, Roberto Amato, Jeffrey Barrett, Sonia Goncalves, Ewan Harrison, David K. Jackson, Ian Johnston, Dominic Kwiatkowski, Cordelia Langford, John Sillitoe on behalf of the Wellcome Sanger Institute COVID-19 Surveillance Team |
| EPI_ISL_3365633 | Lighthouse Lab in Milton Keynes | Wellcome Sanger Institute for the COVID-19 Genomics UK (COG-UK) Consortium | The Lighthouse Lab in Milton Keynes and Alex Alderton, Roberto Amato, Jeffrey Barrett, Sonia Goncalves, Ewan Harrison, David K. Jackson, Ian Johnston, Dominic Kwiatkowski, Cordelia Langford, John Sillitoe on behalf of the Wellcome Sanger Institute COVID-19 Surveillance Team |
| EPI_ISL_3365634 | Lighthouse Lab in Alderley Park | Wellcome Sanger Institute for the COVID-19 Genomics UK (COG-UK) Consortium | Jacquelyn Wynn, Mairead Hyland, The Lighthouse Lab in Alderley Park and Alex Alderton, Roberto Amato, Jeffrey Barrett, Sonia Goncalves, Ewan Harrison, David K. Jackson, Ian Johnston, Dominic Kwiatkowski, Cordelia Langford, John Sillitoe on behalf of the Wellcome Sanger Institute COVID-19 Surveillance Team |
| EPI_ISL_3365635 | Lighthouse Lab in Milton Keynes | Wellcome Sanger Institute for the COVID-19 Genomics UK (COG-UK) Consortium | The Lighthouse Lab in Milton Keynes and Alex Alderton, Roberto Amato, Jeffrey Barrett, Sonia Goncalves, Ewan Harrison, David K. Jackson, Ian Johnston, Dominic Kwiatkowski, Cordelia Langford, John Sillitoe on behalf of the Wellcome Sanger Institute COVID-19 Surveillance Team |
| EPI_ISL_3365636, EPI_ISL_3365637 | Berkshire and Surrey Pathology Services Lighthouse Laboratory | Wellcome Sanger Institute for the COVID-19 Genomics UK (COG-UK) Consortium | Berkshire and Surrey Pathology Services Lighthouse Laboratory and Alex Alderton, Roberto Amato, Jeffrey Barrett, Sonia Goncalves, Ewan Harrison, David K. Jackson, Ian Johnston, Dominic Kwiatkowski, Cordelia Langford, John Sillitoe on behalf of the Wellcome Sanger Institute COVID-19 Surveillance Team |
| EPI_ISL_3365638, EPI_ISL_3365639, EPI_ISL_3365640, EPI_ISL_3365641 | Lighthouse Lab in Alderley Park | Wellcome Sanger Institute for the COVID-19 Genomics UK (COG-UK) Consortium | Jacquelyn Wynn, Mairead Hyland, The Lighthouse Lab in Alderley Park and Alex Alderton, Roberto Amato, Jeffrey Barrett, Sonia Goncalves, Ewan Harrison, David K. Jackson, Ian Johnston, Dominic Kwiatkowski, Cordelia Langford, John Sillitoe on behalf of the Wellcome Sanger Institute COVID-19 Surveillance Team |
| EPI_ISL_3365642, EPI_ISL_3365643, EPI_ISL_3365644 | Berkshire and Surrey Pathology Services Lighthouse Laboratory | Wellcome Sanger Institute for the COVID-19 Genomics UK (COG-UK) Consortium | Berkshire and Surrey Pathology Services Lighthouse Laboratory and Alex Alderton, Roberto Amato, Jeffrey Barrett, Sonia Goncalves, Ewan Harrison, David K. Jackson, Ian Johnston, Dominic Kwiatkowski, Cordelia Langford, John Sillitoe on behalf of the Wellcome Sanger Institute COVID-19 Surveillance Team |
| EPI_ISL_3365645, EPI_ISL_3365646 | Lighthouse Lab in Milton Keynes | Wellcome Sanger Institute for the COVID-19 Genomics UK (COG-UK) Consortium | The Lighthouse Lab in Milton Keynes and Alex Alderton, Roberto Amato, Jeffrey Barrett, Sonia Goncalves, Ewan Harrison, David K. Jackson, Ian Johnston, Dominic Kwiatkowski, Cordelia Langford, John Sillitoe on behalf of the Wellcome Sanger Institute COVID-19 Surveillance Team |
| EPI_ISL_3365647, EPI_ISL_3365648 | Berkshire and Surrey Pathology Services Lighthouse Laboratory | Wellcome Sanger Institute for the COVID-19 Genomics UK (COG-UK) Consortium | Berkshire and Surrey Pathology Services Lighthouse Laboratory and Alex Alderton, Roberto Amato, Jeffrey Barrett, Sonia Goncalves, Ewan Harrison, David K. Jackson, Ian Johnston, Dominic Kwiatkowski, Cordelia Langford, John Sillitoe on behalf of the Wellcome Sanger Institute COVID-19 Surveillance Team |
| EPI_ISL_3365649, EPI_ISL_3365650 | Lighthouse Lab in Milton Keynes | Wellcome Sanger Institute for the COVID-19 Genomics UK (COG-UK) Consortium | The Lighthouse Lab in Milton Keynes and Alex Alderton, Roberto Amato, Jeffrey Barrett, Sonia Goncalves, Ewan Harrison, David K. Jackson, Ian Johnston, Dominic Kwiatkowski, Cordelia Langford, John Sillitoe on behalf of the Wellcome Sanger Institute COVID-19 Surveillance Team |

| | | | |
|---|---|---|---|
| EPI_ISL_3365651 | Lighthouse Lab in Alderley Park | Wellcome Sanger Institute for the COVID-19 Genomics UK (COG-UK) Consortium | Jacquelyn Wynn, Mairead Hyland, The Lighthouse Lab in Alderley Park and Alex Alderton, Roberto Amato, Jeffrey Barrett, Sonia Goncalves, Ewan Harrison, David K. Jackson, Ian Johnston, Dominic Kwiatkowski, Cordelia Langford, John Sillitoe on behalf of the Wellcome Sanger Institute COVID-19 Surveillance Team |
| EPI_ISL_3365652 | Berkshire and Surrey Pathology Services Lighthouse Laboratory | Wellcome Sanger Institute for the COVID-19 Genomics UK (COG-UK) Consortium | Berkshire and Surrey Pathology Services Lighthouse Laboratory and Alex Alderton, Roberto Amato, Jeffrey Barrett, Sonia Goncalves, Ewan Harrison, David K. Jackson, Ian Johnston, Dominic Kwiatkowski, Cordelia Langford, John Sillitoe on behalf of the Wellcome Sanger Institute COVID-19 Surveillance Team |
| EPI_ISL_3365653 | Lighthouse Lab in Alderley Park | Wellcome Sanger Institute for the COVID-19 Genomics UK (COG-UK) Consortium | Jacquelyn Wynn, Mairead Hyland, The Lighthouse Lab in Alderley Park and Alex Alderton, Roberto Amato, Jeffrey Barrett, Sonia Goncalves, Ewan Harrison, David K. Jackson, Ian Johnston, Dominic Kwiatkowski, Cordelia Langford, John Sillitoe on behalf of the Wellcome Sanger Institute COVID-19 Surveillance Team |
| EPI_ISL_3365654 | Berkshire and Surrey Pathology Services Lighthouse Laboratory | Wellcome Sanger Institute for the COVID-19 Genomics UK (COG-UK) Consortium | Berkshire and Surrey Pathology Services Lighthouse Laboratory and Alex Alderton, Roberto Amato, Jeffrey Barrett, Sonia Goncalves, Ewan Harrison, David K. Jackson, Ian Johnston, Dominic Kwiatkowski, Cordelia Langford, John Sillitoe on behalf of the Wellcome Sanger Institute COVID-19 Surveillance Team |
| EPI_ISL_3365655 | Lighthouse Lab in Alderley Park | Wellcome Sanger Institute for the COVID-19 Genomics UK (COG-UK) Consortium | Jacquelyn Wynn, Mairead Hyland, The Lighthouse Lab in Alderley Park and Alex Alderton, Roberto Amato, Jeffrey Barrett, Sonia Goncalves, Ewan Harrison, David K. Jackson, Ian Johnston, Dominic Kwiatkowski, Cordelia Langford, John Sillitoe on behalf of the Wellcome Sanger Institute COVID-19 Surveillance Team |
| EPI_ISL_3365656 | Lighthouse Lab in Milton Keynes | Wellcome Sanger Institute for the COVID-19 Genomics UK (COG-UK) Consortium | The Lighthouse Lab in Milton Keynes and Alex Alderton, Roberto Amato, Jeffrey Barrett, Sonia Goncalves, Ewan Harrison, David K. Jackson, Ian Johnston, Dominic Kwiatkowski, Cordelia Langford, John Sillitoe on behalf of the Wellcome Sanger Institute COVID-19 Surveillance Team |
| EPI_ISL_3365657 | Lighthouse Lab in Alderley Park | Wellcome Sanger Institute for the COVID-19 Genomics UK (COG-UK) Consortium | Jacquelyn Wynn, Mairead Hyland, The Lighthouse Lab in Alderley Park and Alex Alderton, Roberto Amato, Jeffrey Barrett, Sonia Goncalves, Ewan Harrison, David K. Jackson, Ian Johnston, Dominic Kwiatkowski, Cordelia Langford, John Sillitoe on behalf of the Wellcome Sanger Institute COVID-19 Surveillance Team |
| EPI_ISL_3365658 | Lighthouse Lab in Milton Keynes | Wellcome Sanger Institute for the COVID-19 Genomics UK (COG-UK) Consortium | The Lighthouse Lab in Milton Keynes and Alex Alderton, Roberto Amato, Jeffrey Barrett, Sonia Goncalves, Ewan Harrison, David K. Jackson, Ian Johnston, Dominic Kwiatkowski, Cordelia Langford, John Sillitoe on behalf of the Wellcome Sanger Institute COVID-19 Surveillance Team |
| EPI_ISL_3365659, EPI_ISL_3365660, EPI_ISL_3365661 | Lighthouse Laboratory Plymouth | Wellcome Sanger Institute for the COVID-19 Genomics UK (COG-UK) Consortium | Lighthouse Laboratory Plymouth and Alex Alderton, Roberto Amato, Jeffrey Barrett, Sonia Goncalves, Ewan Harrison, David K. Jackson, Ian Johnston, Dominic Kwiatkowski, Cordelia Langford, John Sillitoe on behalf of the Wellcome Sanger Institute COVID-19 Surveillance Team |
| EPI_ISL_3365662 | Lighthouse Lab in Alderley Park | Wellcome Sanger Institute for the COVID-19 Genomics UK (COG-UK) Consortium | Jacquelyn Wynn, Mairead Hyland, The Lighthouse Lab in Alderley Park and Alex Alderton, Roberto Amato, Jeffrey Barrett, Sonia Goncalves, Ewan Harrison, David K. Jackson, Ian Johnston, Dominic Kwiatkowski, Cordelia Langford, John Sillitoe on behalf of the Wellcome Sanger Institute COVID-19 Surveillance Team |
| EPI_ISL_3365663 | Lighthouse Laboratory Plymouth | Wellcome Sanger Institute for the COVID-19 Genomics UK (COG-UK) Consortium | Lighthouse Laboratory Plymouth and Alex Alderton, Roberto Amato, Jeffrey Barrett, Sonia Goncalves, Ewan Harrison, David K. Jackson, Ian Johnston, Dominic Kwiatkowski, Cordelia Langford, John Sillitoe on behalf of the Wellcome Sanger Institute COVID-19 Surveillance Team |
| EPI_ISL_3365664, EPI_ISL_3365665 | Lighthouse Lab in Milton Keynes | Wellcome Sanger Institute for the COVID-19 Genomics UK (COG-UK) Consortium | The Lighthouse Lab in Milton Keynes and Alex Alderton, Roberto Amato, Jeffrey Barrett, Sonia Goncalves, Ewan Harrison, David K. Jackson, Ian Johnston, Dominic Kwiatkowski, Cordelia Langford, John Sillitoe on behalf of the Wellcome Sanger Institute COVID-19 Surveillance Team |
| EPI_ISL_3365666, EPI_ISL_3365667 | Lighthouse Laboratory Plymouth | Wellcome Sanger Institute for the COVID-19 Genomics UK (COG-UK) Consortium | Lighthouse Laboratory Plymouth and Alex Alderton, Roberto Amato, Jeffrey Barrett, Sonia Goncalves, Ewan Harrison, David K. Jackson, Ian Johnston, Dominic Kwiatkowski, Cordelia Langford, John Sillitoe on behalf of the Wellcome Sanger Institute COVID-19 Surveillance Team |
| EPI_ISL_3365668 | Lighthouse Lab in Milton Keynes | Wellcome Sanger Institute for the COVID-19 Genomics UK (COG-UK) Consortium | The Lighthouse Lab in Milton Keynes and Alex Alderton, Roberto Amato, Jeffrey Barrett, Sonia Goncalves, Ewan Harrison, David K. Jackson, Ian Johnston, Dominic Kwiatkowski, Cordelia Langford, John Sillitoe on behalf of the Wellcome Sanger Institute COVID-19 Surveillance Team |
| EPI_ISL_3365669, EPI_ISL_3365670, EPI_ISL_3365671 | Lighthouse Laboratory Plymouth | Wellcome Sanger Institute for the COVID-19 Genomics UK (COG-UK) Consortium | Lighthouse Laboratory Plymouth and Alex Alderton, Roberto Amato, Jeffrey Barrett, Sonia Goncalves, Ewan Harrison, David K. Jackson, Ian Johnston, Dominic Kwiatkowski, Cordelia Langford, John Sillitoe on behalf of the Wellcome Sanger Institute COVID-19 Surveillance Team |
| EPI_ISL_3365672, EPI_ISL_3365673, EPI_ISL_3365674 | Lighthouse Lab in Alderley Park | Wellcome Sanger Institute for the COVID-19 Genomics UK (COG-UK) Consortium | Jacquelyn Wynn, Mairead Hyland, The Lighthouse Lab in Alderley Park and Alex Alderton, Roberto Amato, Jeffrey Barrett, Sonia Goncalves, Ewan Harrison, David K. Jackson, Ian Johnston, Dominic Kwiatkowski, Cordelia Langford, John Sillitoe on behalf of the Wellcome Sanger Institute COVID-19 Surveillance Team |
| EPI_ISL_3365675, EPI_ISL_3365676 | Lighthouse Lab in Milton Keynes | Wellcome Sanger Institute for the COVID-19 Genomics UK (COG-UK) Consortium | The Lighthouse Lab in Milton Keynes and Alex Alderton, Roberto Amato, Jeffrey Barrett, Sonia Goncalves, Ewan Harrison, David K. Jackson, Ian Johnston, Dominic Kwiatkowski, Cordelia Langford, John Sillitoe on behalf of the Wellcome Sanger Institute COVID-19 Surveillance Team |
| EPI_ISL_3365677 | Lighthouse Lab in Alderley Park | Wellcome Sanger Institute for the COVID-19 Genomics UK (COG-UK) Consortium | Jacquelyn Wynn, Mairead Hyland, The Lighthouse Lab in Alderley Park and Alex Alderton, Roberto Amato, Jeffrey Barrett, Sonia Goncalves, Ewan Harrison, David K. Jackson, Ian Johnston, Dominic Kwiatkowski, Cordelia Langford, John Sillitoe on behalf of the Wellcome Sanger Institute COVID-19 Surveillance Team |
| EPI_ISL_3365678, EPI_ISL_3365679 | Lighthouse Laboratory Plymouth | Wellcome Sanger Institute for the COVID-19 Genomics UK (COG-UK) Consortium | Lighthouse Laboratory Plymouth and Alex Alderton, Roberto Amato, Jeffrey Barrett, Sonia Goncalves, Ewan Harrison, David K. Jackson, Ian Johnston, Dominic Kwiatkowski, Cordelia Langford, John Sillitoe on behalf of the Wellcome Sanger Institute COVID-19 Surveillance Team |
| EPI_ISL_3365680 | Lighthouse Lab in Alderley Park | Wellcome Sanger Institute for the COVID-19 Genomics UK (COG-UK) Consortium | Jacquelyn Wynn, Mairead Hyland, The Lighthouse Lab in Alderley Park and Alex Alderton, Roberto Amato, Jeffrey Barrett, Sonia Goncalves, Ewan Harrison, David K. Jackson, Ian Johnston, Dominic Kwiatkowski, Cordelia Langford, John Sillitoe on behalf of the Wellcome Sanger Institute COVID-19 Surveillance Team |
| EPI_ISL_3365681 | Lighthouse Lab in Milton Keynes | Wellcome Sanger Institute for the COVID-19 Genomics UK (COG-UK) Consortium | The Lighthouse Lab in Milton Keynes and Alex Alderton, Roberto Amato, Jeffrey Barrett, Sonia Goncalves, Ewan Harrison, David K. Jackson, Ian Johnston, Dominic Kwiatkowski, Cordelia Langford, John Sillitoe on behalf of the Wellcome Sanger Institute COVID-19 Surveillance Team |
| EPI_ISL_3365682 | Lighthouse Laboratory Plymouth | Wellcome Sanger Institute for the COVID-19 Genomics UK (COG-UK) Consortium | Lighthouse Laboratory Plymouth and Alex Alderton, Roberto Amato, Jeffrey Barrett, Sonia Goncalves, Ewan Harrison, David K. Jackson, Ian Johnston, Dominic Kwiatkowski, Cordelia Langford, John Sillitoe on behalf of the Wellcome Sanger Institute COVID-19 Surveillance Team |
| EPI_ISL_3365683 | Lighthouse Lab in Alderley Park | Wellcome Sanger Institute for the COVID-19 Genomics UK (COG-UK) Consortium | Jacquelyn Wynn, Mairead Hyland, The Lighthouse Lab in Alderley Park and Alex Alderton, Roberto Amato, Jeffrey Barrett, Sonia Goncalves, Ewan Harrison, David K. Jackson, Ian Johnston, Dominic Kwiatkowski, Cordelia Langford, John Sillitoe on behalf of the Wellcome Sanger Institute COVID-19 Surveillance Team |
| EPI_ISL_3365684 | Lighthouse Laboratory Plymouth | Wellcome Sanger Institute for the COVID-19 Genomics UK (COG-UK) Consortium | Lighthouse Laboratory Plymouth and Alex Alderton, Roberto Amato, Jeffrey Barrett, Sonia Goncalves, Ewan Harrison, David K. Jackson, Ian Johnston, Dominic Kwiatkowski, Cordelia Langford, John Sillitoe on behalf of the Wellcome Sanger Institute COVID-19 Surveillance Team |
| EPI_ISL_3365685 | Lighthouse Lab in Milton Keynes | Wellcome Sanger Institute for the COVID-19 Genomics UK (COG-UK) Consortium | The Lighthouse Lab in Milton Keynes and Alex Alderton, Roberto Amato, Jeffrey Barrett, Sonia Goncalves, Ewan Harrison, David K. Jackson, Ian Johnston, Dominic Kwiatkowski, Cordelia Langford, John Sillitoe on behalf of the Wellcome Sanger Institute COVID-19 Surveillance Team |
| EPI_ISL_3365686, EPI_ISL_3365687, EPI_ISL_3365688 | Lighthouse Laboratory Plymouth | Wellcome Sanger Institute for the COVID-19 Genomics UK (COG-UK) Consortium | Lighthouse Laboratory Plymouth and Alex Alderton, Roberto Amato, Jeffrey Barrett, Sonia Goncalves, Ewan Harrison, David K. Jackson, Ian Johnston, Dominic Kwiatkowski, Cordelia Langford, John Sillitoe on behalf of the Wellcome Sanger Institute COVID-19 Surveillance Team |
| EPI_ISL_3365689 | Lighthouse Lab in Alderley Park | Wellcome Sanger Institute for the COVID-19 Genomics UK (COG-UK) Consortium | Jacquelyn Wynn, Mairead Hyland, The Lighthouse Lab in Alderley Park and Alex Alderton, Roberto Amato, Jeffrey Barrett, Sonia Goncalves, Ewan Harrison, David K. Jackson, Ian Johnston, Dominic Kwiatkowski, Cordelia Langford, John Sillitoe on behalf of the Wellcome Sanger Institute COVID-19 Surveillance Team |
| EPI_ISL_3365690, EPI_ISL_3365691 | Lighthouse Lab in Milton Keynes | Wellcome Sanger Institute for the COVID-19 Genomics UK (COG-UK) Consortium | The Lighthouse Lab in Milton Keynes and Alex Alderton, Roberto Amato, Jeffrey Barrett, Sonia Goncalves, Ewan Harrison, David K. Jackson, Ian Johnston, Dominic Kwiatkowski, Cordelia Langford, John Sillitoe on behalf of the Wellcome Sanger Institute COVID-19 Surveillance Team |
| EPI_ISL_3365692, EPI_ISL_3365693 | Lighthouse Laboratory Plymouth | Wellcome Sanger Institute for the COVID-19 Genomics UK (COG-UK) Consortium | Lighthouse Laboratory Plymouth and Alex Alderton, Roberto Amato, Jeffrey Barrett, Sonia Goncalves, Ewan Harrison, David K. Jackson, Ian Johnston, Dominic Kwiatkowski, Cordelia Langford, John Sillitoe on behalf of the Wellcome Sanger Institute COVID-19 Surveillance Team |
| EPI_ISL_3365694, EPI_ISL_3365695, EPI_ISL_3365696, EPI_ISL_3365697 | Lighthouse Lab in Milton Keynes | Wellcome Sanger Institute for the COVID-19 Genomics UK (COG-UK) Consortium | The Lighthouse Lab in Milton Keynes and Alex Alderton, Roberto Amato, Jeffrey Barrett, Sonia Goncalves, Ewan Harrison, David K. Jackson, Ian Johnston, Dominic Kwiatkowski, Cordelia Langford, John Sillitoe on behalf of the Wellcome Sanger Institute COVID-19 Surveillance Team |
| EPI_ISL_3365698 | Lighthouse Lab in Alderley Park | Wellcome Sanger Institute for the COVID-19 Genomics UK (COG-UK) Consortium | Jacquelyn Wynn, Mairead Hyland, The Lighthouse Lab in Alderley Park and Alex Alderton, Roberto Amato, Jeffrey Barrett, Sonia Goncalves, Ewan Harrison, David K. Jackson, Ian Johnston, Dominic Kwiatkowski, Cordelia Langford, John Sillitoe on behalf of the Wellcome Sanger Institute COVID-19 Surveillance Team |
| EPI_ISL_3365699, EPI_ISL_3365700 | Lighthouse Lab in Milton Keynes | Wellcome Sanger Institute for the COVID-19 Genomics UK (COG-UK) Consortium | The Lighthouse Lab in Milton Keynes and Alex Alderton, Roberto Amato, Jeffrey Barrett, Sonia Goncalves, Ewan Harrison, David K. Jackson, Ian Johnston, Dominic Kwiatkowski, Cordelia Langford, John Sillitoe on behalf of the Wellcome Sanger Institute COVID-19 Surveillance Team |
| EPI_ISL_3365701, EPI_ISL_3365702 | Lighthouse Lab in Alderley Park | Wellcome Sanger Institute for the COVID-19 Genomics UK (COG-UK) Consortium | Jacquelyn Wynn, Mairead Hyland, The Lighthouse Lab in Alderley Park and Alex Alderton, Roberto Amato, Jeffrey Barrett, Sonia Goncalves, Ewan Harrison, David K. Jackson, Ian Johnston, Dominic Kwiatkowski, Cordelia Langford, John Sillitoe on behalf of the Wellcome Sanger Institute COVID-19 Surveillance Team |
| EPI_ISL_3365703, EPI_ISL_3365704 | Lighthouse Laboratory Plymouth | Wellcome Sanger Institute for the COVID-19 Genomics UK (COG-UK) Consortium | Lighthouse Laboratory Plymouth and Alex Alderton, Roberto Amato, Jeffrey Barrett, Sonia Goncalves, Ewan Harrison, David K. Jackson, Ian Johnston, Dominic Kwiatkowski, Cordelia Langford, John Sillitoe on behalf of the Wellcome Sanger Institute COVID-19 Surveillance Team |
| EPI_ISL_3365705, EPI_ISL_3365706 | Lighthouse Lab in Milton Keynes | Wellcome Sanger Institute for the COVID-19 Genomics UK (COG-UK) Consortium | The Lighthouse Lab in Milton Keynes and Alex Alderton, Roberto Amato, Jeffrey Barrett, Sonia Goncalves, Ewan Harrison, David K. Jackson, Ian Johnston, Dominic Kwiatkowski, Cordelia Langford, John Sillitoe on behalf of the Wellcome Sanger Institute COVID-19 Surveillance Team |
| EPI_ISL_3365707 | Lighthouse Lab in Alderley Park | Wellcome Sanger Institute for the COVID-19 Genomics UK (COG-UK) Consortium | Jacquelyn Wynn, Mairead Hyland, The Lighthouse Lab in Alderley Park and Alex Alderton, Roberto Amato, Jeffrey Barrett, Sonia Goncalves, Ewan Harrison, David K. Jackson, Ian Johnston, Dominic Kwiatkowski, Cordelia Langford, John Sillitoe on behalf of the Wellcome Sanger Institute COVID-19 Surveillance Team |
| EPI_ISL_3365708 | Lighthouse Lab in Milton Keynes | Wellcome Sanger Institute for the COVID-19 Genomics UK (COG-UK) Consortium | The Lighthouse Lab in Milton Keynes and Alex Alderton, Roberto Amato, Jeffrey Barrett, Sonia Goncalves, Ewan Harrison, David K. Jackson, Ian Johnston, Dominic Kwiatkowski, Cordelia Langford, John Sillitoe on behalf of the Wellcome Sanger Institute COVID-19 Surveillance Team |
| EPI_ISL_3365709 | Lighthouse Laboratory Plymouth | Wellcome Sanger Institute for the COVID-19 Genomics UK (COG-UK) Consortium | Lighthouse Laboratory Plymouth and Alex Alderton, Roberto Amato, Jeffrey Barrett, Sonia Goncalves, Ewan Harrison, David K. Jackson, Ian Johnston, Dominic Kwiatkowski, Cordelia Langford, John Sillitoe on behalf of the Wellcome Sanger Institute COVID-19 Surveillance Team |
| EPI_ISL_3365710, EPI_ISL_3365711, EPI_ISL_3365712, EPI_ISL_3365713 | Lighthouse Lab in Alderley Park | Wellcome Sanger Institute for the COVID-19 Genomics UK (COG-UK) Consortium | Jacquelyn Wynn, Mairead Hyland, The Lighthouse Lab in Alderley Park and Alex Alderton, Roberto Amato, Jeffrey Barrett, Sonia Goncalves, Ewan Harrison, David K. Jackson, Ian Johnston, Dominic Kwiatkowski, Cordelia Langford, John Sillitoe on behalf of the Wellcome Sanger Institute COVID-19 Surveillance Team |
| EPI_ISL_3365714 | Lighthouse Laboratory Plymouth | Wellcome Sanger Institute for the COVID-19 Genomics UK (COG-UK) Consortium | Lighthouse Laboratory Plymouth and Alex Alderton, Roberto Amato, Jeffrey Barrett, Sonia Goncalves, Ewan Harrison, David K. Jackson, Ian Johnston, Dominic Kwiatkowski, Cordelia Langford, John Sillitoe on behalf of the Wellcome Sanger Institute COVID-19 Surveillance Team |
| EPI_ISL_3365715 | Lighthouse Lab in Alderley Park | Wellcome Sanger Institute for the COVID-19 Genomics UK (COG-UK) Consortium | Jacquelyn Wynn, Mairead Hyland, The Lighthouse Lab in Alderley Park and Alex Alderton, Roberto Amato, Jeffrey Barrett, Sonia Goncalves, Ewan Harrison, David K. Jackson, Ian Johnston, Dominic Kwiatkowski, Cordelia Langford, John Sillitoe on behalf of the Wellcome Sanger Institute COVID-19 Surveillance Team |
| EPI_ISL_3365716 | Lighthouse Lab in Milton Keynes | Wellcome Sanger Institute for the COVID-19 Genomics UK (COG-UK) Consortium | The Lighthouse Lab in Milton Keynes and Alex Alderton, Roberto Amato, Jeffrey Barrett, Sonia Goncalves, Ewan Harrison, David K. Jackson, Ian Johnston, Dominic Kwiatkowski, Cordelia Langford, John Sillitoe on behalf of the Wellcome Sanger Institute COVID-19 Surveillance Team |

| | | | |
|---|---|---|---|
| EPI_ISL_3365717 | Lighthouse Laboratory Plymouth | Wellcome Sanger Institute for the COVID-19 Genomics UK (COG-UK) Consortium | Lighthouse Laboratory Plymouth and Alex Alderton, Roberto Amato, Jeffrey Barrett, Sonia Goncalves, Ewan Harrison, David K. Jackson, Ian Johnston, Dominic Kwiatkowski, Cordelia Langford, John Sillitoe on behalf of the Wellcome Sanger Institute COVID-19 Surveillance Team |
| EPI_ISL_3365718 | Lighthouse Lab in Alderley Park | Wellcome Sanger Institute for the COVID-19 Genomics UK (COG-UK) Consortium | Jacquelyn Wynn, Mairead Hyland, The Lighthouse Lab in Alderley Park and Alex Alderton, Roberto Amato, Jeffrey Barrett, Sonia Goncalves, Ewan Harrison, David K. Jackson, Ian Johnston, Dominic Kwiatkowski, Cordelia Langford, John Sillitoe on behalf of the Wellcome Sanger Institute COVID-19 Surveillance Team |
| EPI_ISL_3365719, EPI_ISL_3365720 | Lighthouse Laboratory Plymouth | Wellcome Sanger Institute for the COVID-19 Genomics UK (COG-UK) Consortium | Lighthouse Laboratory Plymouth and Alex Alderton, Roberto Amato, Jeffrey Barrett, Sonia Goncalves, Ewan Harrison, David K. Jackson, Ian Johnston, Dominic Kwiatkowski, Cordelia Langford, John Sillitoe on behalf of the Wellcome Sanger Institute COVID-19 Surveillance Team |
| EPI_ISL_3365721, EPI_ISL_3365722, EPI_ISL_3365723 | Lighthouse Lab in Alderley Park | Wellcome Sanger Institute for the COVID-19 Genomics UK (COG-UK) Consortium | Jacquelyn Wynn, Mairead Hyland, The Lighthouse Lab in Alderley Park and Alex Alderton, Roberto Amato, Jeffrey Barrett, Sonia Goncalves, Ewan Harrison, David K. Jackson, Ian Johnston, Dominic Kwiatkowski, Cordelia Langford, John Sillitoe on behalf of the Wellcome Sanger Institute COVID-19 Surveillance Team |
| EPI_ISL_3365724 | Lighthouse Laboratory Plymouth | Wellcome Sanger Institute for the COVID-19 Genomics UK (COG-UK) Consortium | Lighthouse Laboratory Plymouth and Alex Alderton, Roberto Amato, Jeffrey Barrett, Sonia Goncalves, Ewan Harrison, David K. Jackson, Ian Johnston, Dominic Kwiatkowski, Cordelia Langford, John Sillitoe on behalf of the Wellcome Sanger Institute COVID-19 Surveillance Team |
| EPI_ISL_3365725 | Lighthouse Lab in Alderley Park | Wellcome Sanger Institute for the COVID-19 Genomics UK (COG-UK) Consortium | Jacquelyn Wynn, Mairead Hyland, The Lighthouse Lab in Alderley Park and Alex Alderton, Roberto Amato, Jeffrey Barrett, Sonia Goncalves, Ewan Harrison, David K. Jackson, Ian Johnston, Dominic Kwiatkowski, Cordelia Langford, John Sillitoe on behalf of the Wellcome Sanger Institute COVID-19 Surveillance Team |
| EPI_ISL_3365726 | Lighthouse Laboratory Plymouth | Wellcome Sanger Institute for the COVID-19 Genomics UK (COG-UK) Consortium | Lighthouse Laboratory Plymouth and Alex Alderton, Roberto Amato, Jeffrey Barrett, Sonia Goncalves, Ewan Harrison, David K. Jackson, Ian Johnston, Dominic Kwiatkowski, Cordelia Langford, John Sillitoe on behalf of the Wellcome Sanger Institute COVID-19 Surveillance Team |
| EPI_ISL_3365727 | Lighthouse Lab in Alderley Park | Wellcome Sanger Institute for the COVID-19 Genomics UK (COG-UK) Consortium | Jacquelyn Wynn, Mairead Hyland, The Lighthouse Lab in Alderley Park and Alex Alderton, Roberto Amato, Jeffrey Barrett, Sonia Goncalves, Ewan Harrison, David K. Jackson, Ian Johnston, Dominic Kwiatkowski, Cordelia Langford, John Sillitoe on behalf of the Wellcome Sanger Institute COVID-19 Surveillance Team |
| EPI_ISL_3365728, EPI_ISL_3365729 | Lighthouse Lab in Milton Keynes | Wellcome Sanger Institute for the COVID-19 Genomics UK (COG-UK) Consortium | The Lighthouse Lab in Milton Keynes and Alex Alderton, Roberto Amato, Jeffrey Barrett, Sonia Goncalves, Ewan Harrison, David K. Jackson, Ian Johnston, Dominic Kwiatkowski, Cordelia Langford, John Sillitoe on behalf of the Wellcome Sanger Institute COVID-19 Surveillance Team |
| EPI_ISL_3365730, EPI_ISL_3365731 | Lighthouse Laboratory Plymouth | Wellcome Sanger Institute for the COVID-19 Genomics UK (COG-UK) Consortium | Lighthouse Laboratory Plymouth and Alex Alderton, Roberto Amato, Jeffrey Barrett, Sonia Goncalves, Ewan Harrison, David K. Jackson, Ian Johnston, Dominic Kwiatkowski, Cordelia Langford, John Sillitoe on behalf of the Wellcome Sanger Institute COVID-19 Surveillance Team |
| EPI_ISL_3365732, EPI_ISL_3365733, EPI_ISL_3365734, EPI_ISL_3365735 | Lighthouse Lab in Milton Keynes | Wellcome Sanger Institute for the COVID-19 Genomics UK (COG-UK) Consortium | The Lighthouse Lab in Milton Keynes and Alex Alderton, Roberto Amato, Jeffrey Barrett, Sonia Goncalves, Ewan Harrison, David K. Jackson, Ian Johnston, Dominic Kwiatkowski, Cordelia Langford, John Sillitoe on behalf of the Wellcome Sanger Institute COVID-19 Surveillance Team |
| EPI_ISL_3365736, EPI_ISL_3365737, EPI_ISL_3365738 | Lighthouse Laboratory Plymouth | Wellcome Sanger Institute for the COVID-19 Genomics UK (COG-UK) Consortium | Lighthouse Laboratory Plymouth and Alex Alderton, Roberto Amato, Jeffrey Barrett, Sonia Goncalves, Ewan Harrison, David K. Jackson, Ian Johnston, Dominic Kwiatkowski, Cordelia Langford, John Sillitoe on behalf of the Wellcome Sanger Institute COVID-19 Surveillance Team |
| EPI_ISL_3365739 | Lighthouse Lab in Milton Keynes | Wellcome Sanger Institute for the COVID-19 Genomics UK (COG-UK) Consortium | The Lighthouse Lab in Milton Keynes and Alex Alderton, Roberto Amato, Jeffrey Barrett, Sonia Goncalves, Ewan Harrison, David K. Jackson, Ian Johnston, Dominic Kwiatkowski, Cordelia Langford, John Sillitoe on behalf of the Wellcome Sanger Institute COVID-19 Surveillance Team |
| EPI_ISL_3365740 | Lighthouse Lab in Alderley Park | Wellcome Sanger Institute for the COVID-19 Genomics UK (COG-UK) Consortium | Jacquelyn Wynn, Mairead Hyland, The Lighthouse Lab in Alderley Park and Alex Alderton, Roberto Amato, Jeffrey Barrett, Sonia Goncalves, Ewan Harrison, David K. Jackson, Ian Johnston, Dominic Kwiatkowski, Cordelia Langford, John Sillitoe on behalf of the Wellcome Sanger Institute COVID-19 Surveillance Team |
| EPI_ISL_3365741 | Lighthouse Laboratory Plymouth | Wellcome Sanger Institute for the COVID-19 Genomics UK (COG-UK) Consortium | Lighthouse Laboratory Plymouth and Alex Alderton, Roberto Amato, Jeffrey Barrett, Sonia Goncalves, Ewan Harrison, David K. Jackson, Ian Johnston, Dominic Kwiatkowski, Cordelia Langford, John Sillitoe on behalf of the Wellcome Sanger Institute COVID-19 Surveillance Team |
| EPI_ISL_3365742 | Lighthouse Lab in Alderley Park | Wellcome Sanger Institute for the COVID-19 Genomics UK (COG-UK) Consortium | Jacquelyn Wynn, Mairead Hyland, The Lighthouse Lab in Alderley Park and Alex Alderton, Roberto Amato, Jeffrey Barrett, Sonia Goncalves, Ewan Harrison, David K. Jackson, Ian Johnston, Dominic Kwiatkowski, Cordelia Langford, John Sillitoe on behalf of the Wellcome Sanger Institute COVID-19 Surveillance Team |
| EPI_ISL_3365743 | Lighthouse Lab in Milton Keynes | Wellcome Sanger Institute for the COVID-19 Genomics UK (COG-UK) Consortium | The Lighthouse Lab in Milton Keynes and Alex Alderton, Roberto Amato, Jeffrey Barrett, Sonia Goncalves, Ewan Harrison, David K. Jackson, Ian Johnston, Dominic Kwiatkowski, Cordelia Langford, John Sillitoe on behalf of the Wellcome Sanger Institute COVID-19 Surveillance Team |
| EPI_ISL_3365744 | Lighthouse Lab in Alderley Park | Wellcome Sanger Institute for the COVID-19 Genomics UK (COG-UK) Consortium | Jacquelyn Wynn, Mairead Hyland, The Lighthouse Lab in Alderley Park and Alex Alderton, Roberto Amato, Jeffrey Barrett, Sonia Goncalves, Ewan Harrison, David K. Jackson, Ian Johnston, Dominic Kwiatkowski, Cordelia Langford, John Sillitoe on behalf of the Wellcome Sanger Institute COVID-19 Surveillance Team |
| EPI_ISL_3365745, EPI_ISL_3365746 | Lighthouse Lab in Milton Keynes | Wellcome Sanger Institute for the COVID-19 Genomics UK (COG-UK) Consortium | The Lighthouse Lab in Milton Keynes and Alex Alderton, Roberto Amato, Jeffrey Barrett, Sonia Goncalves, Ewan Harrison, David K. Jackson, Ian Johnston, Dominic Kwiatkowski, Cordelia Langford, John Sillitoe on behalf of the Wellcome Sanger Institute COVID-19 Surveillance Team |
| EPI_ISL_3365747 | Lighthouse Lab in Alderley Park | Wellcome Sanger Institute for the COVID-19 Genomics UK (COG-UK) Consortium | Jacquelyn Wynn, Mairead Hyland, The Lighthouse Lab in Alderley Park and Alex Alderton, Roberto Amato, Jeffrey Barrett, Sonia Goncalves, Ewan Harrison, David K. Jackson, Ian Johnston, Dominic Kwiatkowski, Cordelia Langford, John Sillitoe on behalf of the Wellcome Sanger Institute COVID-19 Surveillance Team |
| EPI_ISL_3365748 | Lighthouse Lab in Milton Keynes | Wellcome Sanger Institute for the COVID-19 Genomics UK (COG-UK) Consortium | The Lighthouse Lab in Milton Keynes and Alex Alderton, Roberto Amato, Jeffrey Barrett, Sonia Goncalves, Ewan Harrison, David K. Jackson, Ian Johnston, Dominic Kwiatkowski, Cordelia Langford, John Sillitoe on behalf of the Wellcome Sanger Institute COVID-19 Surveillance Team |
| EPI_ISL_3365749, EPI_ISL_3365750, EPI_ISL_3365751 | Lighthouse Lab in Alderley Park | Wellcome Sanger Institute for the COVID-19 Genomics UK (COG-UK) Consortium | Jacquelyn Wynn, Mairead Hyland, The Lighthouse Lab in Alderley Park and Alex Alderton, Roberto Amato, Jeffrey Barrett, Sonia Goncalves, Ewan Harrison, David K. Jackson, Ian Johnston, Dominic Kwiatkowski, Cordelia Langford, John Sillitoe on behalf of the Wellcome Sanger Institute COVID-19 Surveillance Team |
| EPI_ISL_3365752 | Berkshire and Surrey Pathology Services Lighthouse Laboratory | Wellcome Sanger Institute for the COVID-19 Genomics UK (COG-UK) Consortium | Berkshire and Surrey Pathology Services Lighthouse Laboratory and Alex Alderton, Roberto Amato, Jeffrey Barrett, Sonia Goncalves, Ewan Harrison, David K. Jackson, Ian Johnston, Dominic Kwiatkowski, Cordelia Langford, John Sillitoe on behalf of the Wellcome Sanger Institute COVID-19 Surveillance Team |
| EPI_ISL_3365753, EPI_ISL_3365754 | Lighthouse Lab in Milton Keynes | Wellcome Sanger Institute for the COVID-19 Genomics UK (COG-UK) Consortium | The Lighthouse Lab in Milton Keynes and Alex Alderton, Roberto Amato, Jeffrey Barrett, Sonia Goncalves, Ewan Harrison, David K. Jackson, Ian Johnston, Dominic Kwiatkowski, Cordelia Langford, John Sillitoe on behalf of the Wellcome Sanger Institute COVID-19 Surveillance Team |
| EPI_ISL_3365755 | Lighthouse Laboratory Plymouth | Wellcome Sanger Institute for the COVID-19 Genomics UK (COG-UK) Consortium | Lighthouse Laboratory Plymouth and Alex Alderton, Roberto Amato, Jeffrey Barrett, Sonia Goncalves, Ewan Harrison, David K. Jackson, Ian Johnston, Dominic Kwiatkowski, Cordelia Langford, John Sillitoe on behalf of the Wellcome Sanger Institute COVID-19 Surveillance Team |
| EPI_ISL_3365756 | Randox Laboratories | Wellcome Sanger Institute for the COVID-19 Genomics UK (COG-UK) Consortium | Randox Laboratories and Alex Alderton, Roberto Amato, Jeffrey Barrett, Sonia Goncalves, Ewan Harrison, David K. Jackson, Ian Johnston, Dominic Kwiatkowski, Cordelia Langford, John Sillitoe on behalf of the Wellcome Sanger Institute COVID-19 Surveillance Team |
| EPI_ISL_3365757 | Lighthouse Laboratory Plymouth | Wellcome Sanger Institute for the COVID-19 Genomics UK (COG-UK) Consortium | Lighthouse Laboratory Plymouth and Alex Alderton, Roberto Amato, Jeffrey Barrett, Sonia Goncalves, Ewan Harrison, David K. Jackson, Ian Johnston, Dominic Kwiatkowski, Cordelia Langford, John Sillitoe on behalf of the Wellcome Sanger Institute COVID-19 Surveillance Team |
| EPI_ISL_3365758, EPI_ISL_3365759 | Lighthouse Lab in Milton Keynes | Wellcome Sanger Institute for the COVID-19 Genomics UK (COG-UK) Consortium | The Lighthouse Lab in Milton Keynes and Alex Alderton, Roberto Amato, Jeffrey Barrett, Sonia Goncalves, Ewan Harrison, David K. Jackson, Ian Johnston, Dominic Kwiatkowski, Cordelia Langford, John Sillitoe on behalf of the Wellcome Sanger Institute COVID-19 Surveillance Team |
| EPI_ISL_3365760 | Lighthouse Lab in Alderley Park | Wellcome Sanger Institute for the COVID-19 Genomics UK (COG-UK) Consortium | Jacquelyn Wynn, Mairead Hyland, The Lighthouse Lab in Alderley Park and Alex Alderton, Roberto Amato, Jeffrey Barrett, Sonia Goncalves, Ewan Harrison, David K. Jackson, Ian Johnston, Dominic Kwiatkowski, Cordelia Langford, John Sillitoe on behalf of the Wellcome Sanger Institute COVID-19 Surveillance Team |
| EPI_ISL_3365761, EPI_ISL_3365762 | Lighthouse Laboratory Plymouth | Wellcome Sanger Institute for the COVID-19 Genomics UK (COG-UK) Consortium | Lighthouse Laboratory Plymouth and Alex Alderton, Roberto Amato, Jeffrey Barrett, Sonia Goncalves, Ewan Harrison, David K. Jackson, Ian Johnston, Dominic Kwiatkowski, Cordelia Langford, John Sillitoe on behalf of the Wellcome Sanger Institute COVID-19 Surveillance Team |
| EPI_ISL_3365763 | Lighthouse Lab in Alderley Park | Wellcome Sanger Institute for the COVID-19 Genomics UK (COG-UK) Consortium | Jacquelyn Wynn, Mairead Hyland, The Lighthouse Lab in Alderley Park and Alex Alderton, Roberto Amato, Jeffrey Barrett, Sonia Goncalves, Ewan Harrison, David K. Jackson, Ian Johnston, Dominic Kwiatkowski, Cordelia Langford, John Sillitoe on behalf of the Wellcome Sanger Institute COVID-19 Surveillance Team |
| EPI_ISL_3365764, EPI_ISL_3365765, EPI_ISL_3365766 | Lighthouse Laboratory Plymouth | Wellcome Sanger Institute for the COVID-19 Genomics UK (COG-UK) Consortium | Lighthouse Laboratory Plymouth and Alex Alderton, Roberto Amato, Jeffrey Barrett, Sonia Goncalves, Ewan Harrison, David K. Jackson, Ian Johnston, Dominic Kwiatkowski, Cordelia Langford, John Sillitoe on behalf of the Wellcome Sanger Institute COVID-19 Surveillance Team |
| EPI_ISL_3365767 | Lighthouse Lab in Milton Keynes | Wellcome Sanger Institute for the COVID-19 Genomics UK (COG-UK) Consortium | The Lighthouse Lab in Milton Keynes and Alex Alderton, Roberto Amato, Jeffrey Barrett, Sonia Goncalves, Ewan Harrison, David K. Jackson, Ian Johnston, Dominic Kwiatkowski, Cordelia Langford, John Sillitoe on behalf of the Wellcome Sanger Institute COVID-19 Surveillance Team |
| EPI_ISL_3365768 | Lighthouse Laboratory Plymouth | Wellcome Sanger Institute for the COVID-19 Genomics UK (COG-UK) Consortium | Lighthouse Laboratory Plymouth and Alex Alderton, Roberto Amato, Jeffrey Barrett, Sonia Goncalves, Ewan Harrison, David K. Jackson, Ian Johnston, Dominic Kwiatkowski, Cordelia Langford, John Sillitoe on behalf of the Wellcome Sanger Institute COVID-19 Surveillance Team |
| EPI_ISL_3365769 | Lighthouse Lab in Alderley Park | Wellcome Sanger Institute for the COVID-19 Genomics UK (COG-UK) Consortium | Jacquelyn Wynn, Mairead Hyland, The Lighthouse Lab in Alderley Park and Alex Alderton, Roberto Amato, Jeffrey Barrett, Sonia Goncalves, Ewan Harrison, David K. Jackson, Ian Johnston, Dominic Kwiatkowski, Cordelia Langford, John Sillitoe on behalf of the Wellcome Sanger Institute COVID-19 Surveillance Team |
| EPI_ISL_3365770 | Lighthouse Laboratory Plymouth | Wellcome Sanger Institute for the COVID-19 Genomics UK (COG-UK) Consortium | Lighthouse Laboratory Plymouth and Alex Alderton, Roberto Amato, Jeffrey Barrett, Sonia Goncalves, Ewan Harrison, David K. Jackson, Ian Johnston, Dominic Kwiatkowski, Cordelia Langford, John Sillitoe on behalf of the Wellcome Sanger Institute COVID-19 Surveillance Team |
| EPI_ISL_3365771 | Lighthouse Lab in Alderley Park | Wellcome Sanger Institute for the COVID-19 Genomics UK (COG-UK) Consortium | Jacquelyn Wynn, Mairead Hyland, The Lighthouse Lab in Alderley Park and Alex Alderton, Roberto Amato, Jeffrey Barrett, Sonia Goncalves, Ewan Harrison, David K. Jackson, Ian Johnston, Dominic Kwiatkowski, Cordelia Langford, John Sillitoe on behalf of the Wellcome Sanger Institute COVID-19 Surveillance Team |
| EPI_ISL_3365772 | Lighthouse Laboratory Plymouth | Wellcome Sanger Institute for the COVID-19 Genomics UK (COG-UK) Consortium | Lighthouse Laboratory Plymouth and Alex Alderton, Roberto Amato, Jeffrey Barrett, Sonia Goncalves, Ewan Harrison, David K. Jackson, Ian Johnston, Dominic Kwiatkowski, Cordelia Langford, John Sillitoe on behalf of the Wellcome Sanger Institute COVID-19 Surveillance Team |
| EPI_ISL_3365773, EPI_ISL_3365774 | Lighthouse Lab in Milton Keynes | Wellcome Sanger Institute for the COVID-19 Genomics UK (COG-UK) Consortium | The Lighthouse Lab in Milton Keynes and Alex Alderton, Roberto Amato, Jeffrey Barrett, Sonia Goncalves, Ewan Harrison, David K. Jackson, Ian Johnston, Dominic Kwiatkowski, Cordelia Langford, John Sillitoe on behalf of the Wellcome Sanger Institute COVID-19 Surveillance Team |
| EPI_ISL_3365775 | Lighthouse Laboratory Plymouth | Wellcome Sanger Institute for the COVID-19 Genomics UK (COG-UK) Consortium | Lighthouse Laboratory Plymouth and Alex Alderton, Roberto Amato, Jeffrey Barrett, Sonia Goncalves, Ewan Harrison, David K. Jackson, Ian Johnston, Dominic Kwiatkowski, Cordelia Langford, John Sillitoe on behalf of the Wellcome Sanger Institute COVID-19 Surveillance Team |
| EPI_ISL_3365776 | Lighthouse Lab in Milton Keynes | Wellcome Sanger Institute for the COVID-19 Genomics UK (COG-UK) Consortium | The Lighthouse Lab in Milton Keynes and Alex Alderton, Roberto Amato, Jeffrey Barrett, Sonia Goncalves, Ewan Harrison, David K. Jackson, Ian Johnston, Dominic Kwiatkowski, Cordelia Langford, John Sillitoe on behalf of the Wellcome Sanger Institute COVID-19 Surveillance Team |
| EPI_ISL_3365777 | Lighthouse Lab in Alderley Park | Wellcome Sanger Institute for the COVID-19 Genomics UK (COG-UK) Consortium | Jacquelyn Wynn, Mairead Hyland, The Lighthouse Lab in Alderley Park and Alex Alderton, Roberto Amato, Jeffrey Barrett, Sonia Goncalves, Ewan Harrison, David K. Jackson, Ian Johnston, Dominic Kwiatkowski, Cordelia Langford, John Sillitoe on behalf of the Wellcome Sanger Institute COVID-19 Surveillance Team |
| EPI_ISL_3365778 | Lighthouse Lab in Milton Keynes | Wellcome Sanger Institute for the COVID-19 Genomics UK (COG-UK) | The Lighthouse Lab in Milton Keynes and Alex Alderton, Roberto Amato, Jeffrey Barrett, Sonia Goncalves, Ewan Harrison, David K. Jackson, Ian Johnston, Dominic Kwiatkowski, Cordelia Langford, John Sillitoe on behalf of the Wellcome Sanger |

| | | | |
|---|---|---|---|
| | | Consortium | Institute COVID-19 Surveillance Team |
| EPI_ISL_3365779 | Lighthouse Laboratory Plymouth | Wellcome Sanger Institute for the COVID-19 Genomics UK (COG-UK) Consortium | Lighthouse Laboratory Plymouth and Alex Alderton, Roberto Amato, Jeffrey Barrett, Sonia Goncalves, Ewan Harrison, David K. Jackson, Ian Johnston, Dominic Kwiatkowski, Cordelia Langford, John Sillitoe on behalf of the Wellcome Sanger Institute COVID-19 Surveillance Team |
| EPI_ISL_3365780 | Lighthouse Lab in Milton Keynes | Wellcome Sanger Institute for the COVID-19 Genomics UK (COG-UK) Consortium | The Lighthouse Lab in Milton Keynes and Alex Alderton, Roberto Amato, Jeffrey Barrett, Sonia Goncalves, Ewan Harrison, David K. Jackson, Ian Johnston, Dominic Kwiatkowski, Cordelia Langford, John Sillitoe on behalf of the Wellcome Sanger Institute COVID-19 Surveillance Team |
| EPI_ISL_3365781 | Lighthouse Laboratory Plymouth | Wellcome Sanger Institute for the COVID-19 Genomics UK (COG-UK) Consortium | Lighthouse Laboratory Plymouth and Alex Alderton, Roberto Amato, Jeffrey Barrett, Sonia Goncalves, Ewan Harrison, David K. Jackson, Ian Johnston, Dominic Kwiatkowski, Cordelia Langford, John Sillitoe on behalf of the Wellcome Sanger Institute COVID-19 Surveillance Team |
| EPI_ISL_3365782, EPI_ISL_3365783, EPI_ISL_3365784, EPI_ISL_3365785 | Lighthouse Lab in Alderley Park | Wellcome Sanger Institute for the COVID-19 Genomics UK (COG-UK) Consortium | Jacquelyn Wynn, Mairead Hyland, The Lighthouse Lab in Alderley Park and Alex Alderton, Roberto Amato, Jeffrey Barrett, Sonia Goncalves, Ewan Harrison, David K. Jackson, Ian Johnston, Dominic Kwiatkowski, Cordelia Langford, John Sillitoe on behalf of the Wellcome Sanger Institute COVID-19 Surveillance Team |
| EPI_ISL_3365786 | Lighthouse Laboratory Plymouth | Wellcome Sanger Institute for the COVID-19 Genomics UK (COG-UK) Consortium | Lighthouse Laboratory Plymouth and Alex Alderton, Roberto Amato, Jeffrey Barrett, Sonia Goncalves, Ewan Harrison, David K. Jackson, Ian Johnston, Dominic Kwiatkowski, Cordelia Langford, John Sillitoe on behalf of the Wellcome Sanger Institute COVID-19 Surveillance Team |
| EPI_ISL_3365787 | Lighthouse Lab in Milton Keynes | Wellcome Sanger Institute for the COVID-19 Genomics UK (COG-UK) Consortium | The Lighthouse Lab in Milton Keynes and Alex Alderton, Roberto Amato, Jeffrey Barrett, Sonia Goncalves, Ewan Harrison, David K. Jackson, Ian Johnston, Dominic Kwiatkowski, Cordelia Langford, John Sillitoe on behalf of the Wellcome Sanger Institute COVID-19 Surveillance Team |
| EPI_ISL_3365788 | Lighthouse Laboratory Plymouth | Wellcome Sanger Institute for the COVID-19 Genomics UK (COG-UK) Consortium | Lighthouse Laboratory Plymouth and Alex Alderton, Roberto Amato, Jeffrey Barrett, Sonia Goncalves, Ewan Harrison, David K. Jackson, Ian Johnston, Dominic Kwiatkowski, Cordelia Langford, John Sillitoe on behalf of the Wellcome Sanger Institute COVID-19 Surveillance Team |
| EPI_ISL_3365789 | Lighthouse Lab in Milton Keynes | Wellcome Sanger Institute for the COVID-19 Genomics UK (COG-UK) Consortium | The Lighthouse Lab in Milton Keynes and Alex Alderton, Roberto Amato, Jeffrey Barrett, Sonia Goncalves, Ewan Harrison, David K. Jackson, Ian Johnston, Dominic Kwiatkowski, Cordelia Langford, John Sillitoe on behalf of the Wellcome Sanger Institute COVID-19 Surveillance Team |
| EPI_ISL_3365790, EPI_ISL_3365791 | Lighthouse Lab in Alderley Park | Wellcome Sanger Institute for the COVID-19 Genomics UK (COG-UK) Consortium | Jacquelyn Wynn, Mairead Hyland, The Lighthouse Lab in Alderley Park and Alex Alderton, Roberto Amato, Jeffrey Barrett, Sonia Goncalves, Ewan Harrison, David K. Jackson, Ian Johnston, Dominic Kwiatkowski, Cordelia Langford, John Sillitoe on behalf of the Wellcome Sanger Institute COVID-19 Surveillance Team |
| EPI_ISL_3365792 | Lighthouse Lab in Milton Keynes | Wellcome Sanger Institute for the COVID-19 Genomics UK (COG-UK) Consortium | The Lighthouse Lab in Milton Keynes and Alex Alderton, Roberto Amato, Jeffrey Barrett, Sonia Goncalves, Ewan Harrison, David K. Jackson, Ian Johnston, Dominic Kwiatkowski, Cordelia Langford, John Sillitoe on behalf of the Wellcome Sanger Institute COVID-19 Surveillance Team |
| EPI_ISL_3365793, EPI_ISL_3365794, EPI_ISL_3365795 | Lighthouse Lab in Alderley Park | Wellcome Sanger Institute for the COVID-19 Genomics UK (COG-UK) Consortium | Jacquelyn Wynn, Mairead Hyland, The Lighthouse Lab in Alderley Park and Alex Alderton, Roberto Amato, Jeffrey Barrett, Sonia Goncalves, Ewan Harrison, David K. Jackson, Ian Johnston, Dominic Kwiatkowski, Cordelia Langford, John Sillitoe on behalf of the Wellcome Sanger Institute COVID-19 Surveillance Team |
| EPI_ISL_3365796 | Lighthouse Lab in Milton Keynes | Wellcome Sanger Institute for the COVID-19 Genomics UK (COG-UK) Consortium | The Lighthouse Lab in Milton Keynes and Alex Alderton, Roberto Amato, Jeffrey Barrett, Sonia Goncalves, Ewan Harrison, David K. Jackson, Ian Johnston, Dominic Kwiatkowski, Cordelia Langford, John Sillitoe on behalf of the Wellcome Sanger Institute COVID-19 Surveillance Team |
| EPI_ISL_3365797 | Lighthouse Lab in Alderley Park | Wellcome Sanger Institute for the COVID-19 Genomics UK (COG-UK) Consortium | Jacquelyn Wynn, Mairead Hyland, The Lighthouse Lab in Alderley Park and Alex Alderton, Roberto Amato, Jeffrey Barrett, Sonia Goncalves, Ewan Harrison, David K. Jackson, Ian Johnston, Dominic Kwiatkowski, Cordelia Langford, John Sillitoe on behalf of the Wellcome Sanger Institute COVID-19 Surveillance Team |
| EPI_ISL_3365798 | Lighthouse Laboratory Plymouth | Wellcome Sanger Institute for the COVID-19 Genomics UK (COG-UK) Consortium | Lighthouse Laboratory Plymouth and Alex Alderton, Roberto Amato, Jeffrey Barrett, Sonia Goncalves, Ewan Harrison, David K. Jackson, Ian Johnston, Dominic Kwiatkowski, Cordelia Langford, John Sillitoe on behalf of the Wellcome Sanger Institute COVID-19 Surveillance Team |
| EPI_ISL_3365799 | Lighthouse Lab in Alderley Park | Wellcome Sanger Institute for the COVID-19 Genomics UK (COG-UK) Consortium | Jacquelyn Wynn, Mairead Hyland, The Lighthouse Lab in Alderley Park and Alex Alderton, Roberto Amato, Jeffrey Barrett, Sonia Goncalves, Ewan Harrison, David K. Jackson, Ian Johnston, Dominic Kwiatkowski, Cordelia Langford, John Sillitoe on behalf of the Wellcome Sanger Institute COVID-19 Surveillance Team |
| EPI_ISL_3365800, EPI_ISL_3365801, EPI_ISL_3365802 | Lighthouse Lab in Milton Keynes | Wellcome Sanger Institute for the COVID-19 Genomics UK (COG-UK) Consortium | The Lighthouse Lab in Milton Keynes and Alex Alderton, Roberto Amato, Jeffrey Barrett, Sonia Goncalves, Ewan Harrison, David K. Jackson, Ian Johnston, Dominic Kwiatkowski, Cordelia Langford, John Sillitoe on behalf of the Wellcome Sanger Institute COVID-19 Surveillance Team |
| EPI_ISL_3365803, EPI_ISL_3365804 | Lighthouse Lab in Alderley Park | Wellcome Sanger Institute for the COVID-19 Genomics UK (COG-UK) Consortium | Jacquelyn Wynn, Mairead Hyland, The Lighthouse Lab in Alderley Park and Alex Alderton, Roberto Amato, Jeffrey Barrett, Sonia Goncalves, Ewan Harrison, David K. Jackson, Ian Johnston, Dominic Kwiatkowski, Cordelia Langford, John Sillitoe on behalf of the Wellcome Sanger Institute COVID-19 Surveillance Team |
| EPI_ISL_3365805 | Lighthouse Lab in Milton Keynes | Wellcome Sanger Institute for the COVID-19 Genomics UK (COG-UK) Consortium | The Lighthouse Lab in Milton Keynes and Alex Alderton, Roberto Amato, Jeffrey Barrett, Sonia Goncalves, Ewan Harrison, David K. Jackson, Ian Johnston, Dominic Kwiatkowski, Cordelia Langford, John Sillitoe on behalf of the Wellcome Sanger Institute COVID-19 Surveillance Team |
| EPI_ISL_3365806, EPI_ISL_3365807 | Lighthouse Laboratory Plymouth | Wellcome Sanger Institute for the COVID-19 Genomics UK (COG-UK) Consortium | Lighthouse Laboratory Plymouth and Alex Alderton, Roberto Amato, Jeffrey Barrett, Sonia Goncalves, Ewan Harrison, David K. Jackson, Ian Johnston, Dominic Kwiatkowski, Cordelia Langford, John Sillitoe on behalf of the Wellcome Sanger Institute COVID-19 Surveillance Team |
| EPI_ISL_3365808, EPI_ISL_3365809, EPI_ISL_3365810 | Lighthouse Lab in Milton Keynes | Wellcome Sanger Institute for the COVID-19 Genomics UK (COG-UK) Consortium | The Lighthouse Lab in Milton Keynes and Alex Alderton, Roberto Amato, Jeffrey Barrett, Sonia Goncalves, Ewan Harrison, David K. Jackson, Ian Johnston, Dominic Kwiatkowski, Cordelia Langford, John Sillitoe on behalf of the Wellcome Sanger Institute COVID-19 Surveillance Team |
| EPI_ISL_3365811, EPI_ISL_3365812, EPI_ISL_3365813 | Lighthouse Laboratory Plymouth | Wellcome Sanger Institute for the COVID-19 Genomics UK (COG-UK) Consortium | Lighthouse Laboratory Plymouth and Alex Alderton, Roberto Amato, Jeffrey Barrett, Sonia Goncalves, Ewan Harrison, David K. Jackson, Ian Johnston, Dominic Kwiatkowski, Cordelia Langford, John Sillitoe on behalf of the Wellcome Sanger Institute COVID-19 Surveillance Team |
| EPI_ISL_3365814, EPI_ISL_3365815 | Lighthouse Lab in Alderley Park | Wellcome Sanger Institute for the COVID-19 Genomics UK (COG-UK) Consortium | Jacquelyn Wynn, Mairead Hyland, The Lighthouse Lab in Alderley Park and Alex Alderton, Roberto Amato, Jeffrey Barrett, Sonia Goncalves, Ewan Harrison, David K. Jackson, Ian Johnston, Dominic Kwiatkowski, Cordelia Langford, John Sillitoe on behalf of the Wellcome Sanger Institute COVID-19 Surveillance Team |
| EPI_ISL_3365816 | Lighthouse Lab in Milton Keynes | Wellcome Sanger Institute for the COVID-19 Genomics UK (COG-UK) Consortium | The Lighthouse Lab in Milton Keynes and Alex Alderton, Roberto Amato, Jeffrey Barrett, Sonia Goncalves, Ewan Harrison, David K. Jackson, Ian Johnston, Dominic Kwiatkowski, Cordelia Langford, John Sillitoe on behalf of the Wellcome Sanger Institute COVID-19 Surveillance Team |
| EPI_ISL_3365817 | Lighthouse Lab in Alderley Park | Wellcome Sanger Institute for the COVID-19 Genomics UK (COG-UK) Consortium | Jacquelyn Wynn, Mairead Hyland, The Lighthouse Lab in Alderley Park and Alex Alderton, Roberto Amato, Jeffrey Barrett, Sonia Goncalves, Ewan Harrison, David K. Jackson, Ian Johnston, Dominic Kwiatkowski, Cordelia Langford, John Sillitoe on behalf of the Wellcome Sanger Institute COVID-19 Surveillance Team |
| EPI_ISL_3365818, EPI_ISL_3365819 | Lighthouse Laboratory Plymouth | Wellcome Sanger Institute for the COVID-19 Genomics UK (COG-UK) Consortium | Lighthouse Laboratory Plymouth and Alex Alderton, Roberto Amato, Jeffrey Barrett, Sonia Goncalves, Ewan Harrison, David K. Jackson, Ian Johnston, Dominic Kwiatkowski, Cordelia Langford, John Sillitoe on behalf of the Wellcome Sanger Institute COVID-19 Surveillance Team |
| EPI_ISL_3365820 | Lighthouse Lab in Milton Keynes | Wellcome Sanger Institute for the COVID-19 Genomics UK (COG-UK) Consortium | The Lighthouse Lab in Milton Keynes and Alex Alderton, Roberto Amato, Jeffrey Barrett, Sonia Goncalves, Ewan Harrison, David K. Jackson, Ian Johnston, Dominic Kwiatkowski, Cordelia Langford, John Sillitoe on behalf of the Wellcome Sanger Institute COVID-19 Surveillance Team |
| EPI_ISL_3365821 | Lighthouse Lab in Alderley Park | Wellcome Sanger Institute for the COVID-19 Genomics UK (COG-UK) Consortium | Jacquelyn Wynn, Mairead Hyland, The Lighthouse Lab in Alderley Park and Alex Alderton, Roberto Amato, Jeffrey Barrett, Sonia Goncalves, Ewan Harrison, David K. Jackson, Ian Johnston, Dominic Kwiatkowski, Cordelia Langford, John Sillitoe on behalf of the Wellcome Sanger Institute COVID-19 Surveillance Team |
| EPI_ISL_3365822, EPI_ISL_3365823, EPI_ISL_3365824, EPI_ISL_3365825 | Lighthouse Laboratory Plymouth | Wellcome Sanger Institute for the COVID-19 Genomics UK (COG-UK) Consortium | Lighthouse Laboratory Plymouth and Alex Alderton, Roberto Amato, Jeffrey Barrett, Sonia Goncalves, Ewan Harrison, David K. Jackson, Ian Johnston, Dominic Kwiatkowski, Cordelia Langford, John Sillitoe on behalf of the Wellcome Sanger Institute COVID-19 Surveillance Team |
| EPI_ISL_3365826, EPI_ISL_3365827 | Lighthouse Lab in Alderley Park | Wellcome Sanger Institute for the COVID-19 Genomics UK (COG-UK) Consortium | Jacquelyn Wynn, Mairead Hyland, The Lighthouse Lab in Alderley Park and Alex Alderton, Roberto Amato, Jeffrey Barrett, Sonia Goncalves, Ewan Harrison, David K. Jackson, Ian Johnston, Dominic Kwiatkowski, Cordelia Langford, John Sillitoe on behalf of the Wellcome Sanger Institute COVID-19 Surveillance Team |
| EPI_ISL_3365828 | Lighthouse Laboratory Plymouth | Wellcome Sanger Institute for the COVID-19 Genomics UK (COG-UK) Consortium | Lighthouse Laboratory Plymouth and Alex Alderton, Roberto Amato, Jeffrey Barrett, Sonia Goncalves, Ewan Harrison, David K. Jackson, Ian Johnston, Dominic Kwiatkowski, Cordelia Langford, John Sillitoe on behalf of the Wellcome Sanger Institute COVID-19 Surveillance Team |
| EPI_ISL_3365829, EPI_ISL_3365830, EPI_ISL_3365831, EPI_ISL_3365832, EPI_ISL_3365833, EPI_ISL_3365834, EPI_ISL_3365835 | Lighthouse Lab in Milton Keynes | Wellcome Sanger Institute for the COVID-19 Genomics UK (COG-UK) Consortium | The Lighthouse Lab in Milton Keynes and Alex Alderton, Roberto Amato, Jeffrey Barrett, Sonia Goncalves, Ewan Harrison, David K. Jackson, Ian Johnston, Dominic Kwiatkowski, Cordelia Langford, John Sillitoe on behalf of the Wellcome Sanger Institute COVID-19 Surveillance Team |
| EPI_ISL_3365836 | Lighthouse Lab in Alderley Park | Wellcome Sanger Institute for the COVID-19 Genomics UK (COG-UK) Consortium | Jacquelyn Wynn, Mairead Hyland, The Lighthouse Lab in Alderley Park and Alex Alderton, Roberto Amato, Jeffrey Barrett, Sonia Goncalves, Ewan Harrison, David K. Jackson, Ian Johnston, Dominic Kwiatkowski, Cordelia Langford, John Sillitoe on behalf of the Wellcome Sanger Institute COVID-19 Surveillance Team |
| EPI_ISL_3365837 | Lighthouse Laboratory Plymouth | Wellcome Sanger Institute for the COVID-19 Genomics UK (COG-UK) Consortium | Lighthouse Laboratory Plymouth and Alex Alderton, Roberto Amato, Jeffrey Barrett, Sonia Goncalves, Ewan Harrison, David K. Jackson, Ian Johnston, Dominic Kwiatkowski, Cordelia Langford, John Sillitoe on behalf of the Wellcome Sanger Institute COVID-19 Surveillance Team |
| EPI_ISL_3365838 | Lighthouse Lab in Alderley Park | Wellcome Sanger Institute for the COVID-19 Genomics UK (COG-UK) Consortium | Jacquelyn Wynn, Mairead Hyland, The Lighthouse Lab in Alderley Park and Alex Alderton, Roberto Amato, Jeffrey Barrett, Sonia Goncalves, Ewan Harrison, David K. Jackson, Ian Johnston, Dominic Kwiatkowski, Cordelia Langford, John Sillitoe on behalf of the Wellcome Sanger Institute COVID-19 Surveillance Team |
| EPI_ISL_3365839 | Lighthouse Lab in Milton Keynes | Wellcome Sanger Institute for the COVID-19 Genomics UK (COG-UK) Consortium | The Lighthouse Lab in Milton Keynes and Alex Alderton, Roberto Amato, Jeffrey Barrett, Sonia Goncalves, Ewan Harrison, David K. Jackson, Ian Johnston, Dominic Kwiatkowski, Cordelia Langford, John Sillitoe on behalf of the Wellcome Sanger Institute COVID-19 Surveillance Team |
| EPI_ISL_3365840 | Lighthouse Laboratory Plymouth | Wellcome Sanger Institute for the COVID-19 Genomics UK (COG-UK) Consortium | Lighthouse Laboratory Plymouth and Alex Alderton, Roberto Amato, Jeffrey Barrett, Sonia Goncalves, Ewan Harrison, David K. Jackson, Ian Johnston, Dominic Kwiatkowski, Cordelia Langford, John Sillitoe on behalf of the Wellcome Sanger Institute COVID-19 Surveillance Team |
| EPI_ISL_3365841 | Lighthouse Lab in Alderley Park | Wellcome Sanger Institute for the COVID-19 Genomics UK (COG-UK) Consortium | Jacquelyn Wynn, Mairead Hyland, The Lighthouse Lab in Alderley Park and Alex Alderton, Roberto Amato, Jeffrey Barrett, Sonia Goncalves, Ewan Harrison, David K. Jackson, Ian Johnston, Dominic Kwiatkowski, Cordelia Langford, John Sillitoe on behalf of the Wellcome Sanger Institute COVID-19 Surveillance Team |
| EPI_ISL_3365842 | Lighthouse Laboratory Plymouth | Wellcome Sanger Institute for the COVID-19 Genomics UK (COG-UK) Consortium | Lighthouse Laboratory Plymouth and Alex Alderton, Roberto Amato, Jeffrey Barrett, Sonia Goncalves, Ewan Harrison, David K. Jackson, Ian Johnston, Dominic Kwiatkowski, Cordelia Langford, John Sillitoe on behalf of the Wellcome Sanger Institute COVID-19 Surveillance Team |
| EPI_ISL_3365843 | Lighthouse Lab in Alderley Park | Wellcome Sanger Institute for the COVID-19 Genomics UK (COG-UK) Consortium | Jacquelyn Wynn, Mairead Hyland, The Lighthouse Lab in Alderley Park and Alex Alderton, Roberto Amato, Jeffrey Barrett, Sonia Goncalves, Ewan Harrison, David K. Jackson, Ian Johnston, Dominic Kwiatkowski, Cordelia Langford, John Sillitoe on behalf of the Wellcome Sanger Institute COVID-19 Surveillance Team |

| | | | |
|---|---|---|---|
| EPI_ISL_3365844 | Lighthouse Lab in Milton Keynes | Wellcome Sanger Institute for the COVID-19 Genomics UK (COG-UK) Consortium | The Lighthouse Lab in Milton Keynes and Alex Alderton, Roberto Amato, Jeffrey Barrett, Sonia Goncalves, Ewan Harrison, David K. Jackson, Ian Johnston, Dominic Kwiatkowski, Cordelia Langford, John Sillitoe on behalf of the Wellcome Sanger Institute COVID-19 Surveillance Team |
| EPI_ISL_3365846 | Lighthouse Laboratory Plymouth | Wellcome Sanger Institute for the COVID-19 Genomics UK (COG-UK) Consortium | Lighthouse Laboratory Plymouth and Alex Alderton, Roberto Amato, Jeffrey Barrett, Sonia Goncalves, Ewan Harrison, David K. Jackson, Ian Johnston, Dominic Kwiatkowski, Cordelia Langford, John Sillitoe on behalf of the Wellcome Sanger Institute COVID-19 Surveillance Team |
| EPI_ISL_3365847 | Lighthouse Lab in Milton Keynes | Wellcome Sanger Institute for the COVID-19 Genomics UK (COG-UK) Consortium | The Lighthouse Lab in Milton Keynes and Alex Alderton, Roberto Amato, Jeffrey Barrett, Sonia Goncalves, Ewan Harrison, David K. Jackson, Ian Johnston, Dominic Kwiatkowski, Cordelia Langford, John Sillitoe on behalf of the Wellcome Sanger Institute COVID-19 Surveillance Team |
| EPI_ISL_3365848, EPI_ISL_3365849 | Lighthouse Lab in Alderley Park | Wellcome Sanger Institute for the COVID-19 Genomics UK (COG-UK) Consortium | Jacquelyn Wynn, Mairead Hyland, The Lighthouse Lab in Alderley Park and Alex Alderton, Roberto Amato, Jeffrey Barrett, Sonia Goncalves, Ewan Harrison, David K. Jackson, Ian Johnston, Dominic Kwiatkowski, Cordelia Langford, John Sillitoe on behalf of the Wellcome Sanger Institute COVID-19 Surveillance Team |
| EPI_ISL_3365850, EPI_ISL_3365851, EPI_ISL_3365852 | Lighthouse Laboratory Plymouth | Wellcome Sanger Institute for the COVID-19 Genomics UK (COG-UK) Consortium | Lighthouse Laboratory Plymouth and Alex Alderton, Roberto Amato, Jeffrey Barrett, Sonia Goncalves, Ewan Harrison, David K. Jackson, Ian Johnston, Dominic Kwiatkowski, Cordelia Langford, John Sillitoe on behalf of the Wellcome Sanger Institute COVID-19 Surveillance Team |
| EPI_ISL_3365853 | Lighthouse Lab in Milton Keynes | Wellcome Sanger Institute for the COVID-19 Genomics UK (COG-UK) Consortium | The Lighthouse Lab in Milton Keynes and Alex Alderton, Roberto Amato, Jeffrey Barrett, Sonia Goncalves, Ewan Harrison, David K. Jackson, Ian Johnston, Dominic Kwiatkowski, Cordelia Langford, John Sillitoe on behalf of the Wellcome Sanger Institute COVID-19 Surveillance Team |
| EPI_ISL_3365854 | Lighthouse Lab in Alderley Park | Wellcome Sanger Institute for the COVID-19 Genomics UK (COG-UK) Consortium | Jacquelyn Wynn, Mairead Hyland, The Lighthouse Lab in Alderley Park and Alex Alderton, Roberto Amato, Jeffrey Barrett, Sonia Goncalves, Ewan Harrison, David K. Jackson, Ian Johnston, Dominic Kwiatkowski, Cordelia Langford, John Sillitoe on behalf of the Wellcome Sanger Institute COVID-19 Surveillance Team |
| EPI_ISL_3365855 | Lighthouse Laboratory Plymouth | Wellcome Sanger Institute for the COVID-19 Genomics UK (COG-UK) Consortium | Lighthouse Laboratory Plymouth and Alex Alderton, Roberto Amato, Jeffrey Barrett, Sonia Goncalves, Ewan Harrison, David K. Jackson, Ian Johnston, Dominic Kwiatkowski, Cordelia Langford, John Sillitoe on behalf of the Wellcome Sanger Institute COVID-19 Surveillance Team |
| EPI_ISL_3365856 | Lighthouse Lab in Alderley Park | Wellcome Sanger Institute for the COVID-19 Genomics UK (COG-UK) Consortium | Jacquelyn Wynn, Mairead Hyland, The Lighthouse Lab in Alderley Park and Alex Alderton, Roberto Amato, Jeffrey Barrett, Sonia Goncalves, Ewan Harrison, David K. Jackson, Ian Johnston, Dominic Kwiatkowski, Cordelia Langford, John Sillitoe on behalf of the Wellcome Sanger Institute COVID-19 Surveillance Team |
| EPI_ISL_3365857 | Lighthouse Lab in Milton Keynes | Wellcome Sanger Institute for the COVID-19 Genomics UK (COG-UK) Consortium | The Lighthouse Lab in Milton Keynes and Alex Alderton, Roberto Amato, Jeffrey Barrett, Sonia Goncalves, Ewan Harrison, David K. Jackson, Ian Johnston, Dominic Kwiatkowski, Cordelia Langford, John Sillitoe on behalf of the Wellcome Sanger Institute COVID-19 Surveillance Team |
| EPI_ISL_3365858 | Lighthouse Lab in Alderley Park | Wellcome Sanger Institute for the COVID-19 Genomics UK (COG-UK) Consortium | Jacquelyn Wynn, Mairead Hyland, The Lighthouse Lab in Alderley Park and Alex Alderton, Roberto Amato, Jeffrey Barrett, Sonia Goncalves, Ewan Harrison, David K. Jackson, Ian Johnston, Dominic Kwiatkowski, Cordelia Langford, John Sillitoe on behalf of the Wellcome Sanger Institute COVID-19 Surveillance Team |
| EPI_ISL_3365859, EPI_ISL_3365860 | Lighthouse Lab in Milton Keynes | Wellcome Sanger Institute for the COVID-19 Genomics UK (COG-UK) Consortium | The Lighthouse Lab in Milton Keynes and Alex Alderton, Roberto Amato, Jeffrey Barrett, Sonia Goncalves, Ewan Harrison, David K. Jackson, Ian Johnston, Dominic Kwiatkowski, Cordelia Langford, John Sillitoe on behalf of the Wellcome Sanger Institute COVID-19 Surveillance Team |
| EPI_ISL_3365861 | Lighthouse Laboratory Plymouth | Wellcome Sanger Institute for the COVID-19 Genomics UK (COG-UK) Consortium | Lighthouse Laboratory Plymouth and Alex Alderton, Roberto Amato, Jeffrey Barrett, Sonia Goncalves, Ewan Harrison, David K. Jackson, Ian Johnston, Dominic Kwiatkowski, Cordelia Langford, John Sillitoe on behalf of the Wellcome Sanger Institute COVID-19 Surveillance Team |
| EPI_ISL_3365862, EPI_ISL_3365863, EPI_ISL_3365864, EPI_ISL_3365865, EPI_ISL_3365866 | Lighthouse Lab in Alderley Park | Wellcome Sanger Institute for the COVID-19 Genomics UK (COG-UK) Consortium | Jacquelyn Wynn, Mairead Hyland, The Lighthouse Lab in Alderley Park and Alex Alderton, Roberto Amato, Jeffrey Barrett, Sonia Goncalves, Ewan Harrison, David K. Jackson, Ian Johnston, Dominic Kwiatkowski, Cordelia Langford, John Sillitoe on behalf of the Wellcome Sanger Institute COVID-19 Surveillance Team |
| EPI_ISL_3365867, EPI_ISL_3365868 | Lighthouse Laboratory Plymouth | Wellcome Sanger Institute for the COVID-19 Genomics UK (COG-UK) Consortium | Lighthouse Laboratory Plymouth and Alex Alderton, Roberto Amato, Jeffrey Barrett, Sonia Goncalves, Ewan Harrison, David K. Jackson, Ian Johnston, Dominic Kwiatkowski, Cordelia Langford, John Sillitoe on behalf of the Wellcome Sanger Institute COVID-19 Surveillance Team |
| EPI_ISL_3365869 | Lighthouse Lab in Milton Keynes | Wellcome Sanger Institute for the COVID-19 Genomics UK (COG-UK) Consortium | The Lighthouse Lab in Milton Keynes and Alex Alderton, Roberto Amato, Jeffrey Barrett, Sonia Goncalves, Ewan Harrison, David K. Jackson, Ian Johnston, Dominic Kwiatkowski, Cordelia Langford, John Sillitoe on behalf of the Wellcome Sanger Institute COVID-19 Surveillance Team |
| EPI_ISL_3365870 | Lighthouse Laboratory Plymouth | Wellcome Sanger Institute for the COVID-19 Genomics UK (COG-UK) Consortium | Lighthouse Laboratory Plymouth and Alex Alderton, Roberto Amato, Jeffrey Barrett, Sonia Goncalves, Ewan Harrison, David K. Jackson, Ian Johnston, Dominic Kwiatkowski, Cordelia Langford, John Sillitoe on behalf of the Wellcome Sanger Institute COVID-19 Surveillance Team |
| EPI_ISL_3365871 | Lighthouse Lab in Milton Keynes | Wellcome Sanger Institute for the COVID-19 Genomics UK (COG-UK) Consortium | The Lighthouse Lab in Milton Keynes and Alex Alderton, Roberto Amato, Jeffrey Barrett, Sonia Goncalves, Ewan Harrison, David K. Jackson, Ian Johnston, Dominic Kwiatkowski, Cordelia Langford, John Sillitoe on behalf of the Wellcome Sanger Institute COVID-19 Surveillance Team |
| EPI_ISL_3365872 | Berkshire and Surrey Pathology Services Lighthouse Laboratory | Wellcome Sanger Institute for the COVID-19 Genomics UK (COG-UK) Consortium | Berkshire and Surrey Pathology Services Lighthouse Laboratory and Alex Alderton, Roberto Amato, Jeffrey Barrett, Sonia Goncalves, Ewan Harrison, David K. Jackson, Ian Johnston, Dominic Kwiatkowski, Cordelia Langford, John Sillitoe on behalf of the Wellcome Sanger Institute COVID-19 Surveillance Team |
| EPI_ISL_3365873 | Lighthouse Laboratory Plymouth | Wellcome Sanger Institute for the COVID-19 Genomics UK (COG-UK) Consortium | Lighthouse Laboratory Plymouth and Alex Alderton, Roberto Amato, Jeffrey Barrett, Sonia Goncalves, Ewan Harrison, David K. Jackson, Ian Johnston, Dominic Kwiatkowski, Cordelia Langford, John Sillitoe on behalf of the Wellcome Sanger Institute COVID-19 Surveillance Team |
| EPI_ISL_3365874 | Lighthouse Lab in Alderley Park | Wellcome Sanger Institute for the COVID-19 Genomics UK (COG-UK) Consortium | Jacquelyn Wynn, Mairead Hyland, The Lighthouse Lab in Alderley Park and Alex Alderton, Roberto Amato, Jeffrey Barrett, Sonia Goncalves, Ewan Harrison, David K. Jackson, Ian Johnston, Dominic Kwiatkowski, Cordelia Langford, John Sillitoe on behalf of the Wellcome Sanger Institute COVID-19 Surveillance Team |
| EPI_ISL_3365875 | Lighthouse Lab in Milton Keynes | Wellcome Sanger Institute for the COVID-19 Genomics UK (COG-UK) Consortium | The Lighthouse Lab in Milton Keynes and Alex Alderton, Roberto Amato, Jeffrey Barrett, Sonia Goncalves, Ewan Harrison, David K. Jackson, Ian Johnston, Dominic Kwiatkowski, Cordelia Langford, John Sillitoe on behalf of the Wellcome Sanger Institute COVID-19 Surveillance Team |
| EPI_ISL_3365876 | Lighthouse Lab in Alderley Park | Wellcome Sanger Institute for the COVID-19 Genomics UK (COG-UK) Consortium | Jacquelyn Wynn, Mairead Hyland, The Lighthouse Lab in Alderley Park and Alex Alderton, Roberto Amato, Jeffrey Barrett, Sonia Goncalves, Ewan Harrison, David K. Jackson, Ian Johnston, Dominic Kwiatkowski, Cordelia Langford, John Sillitoe on behalf of the Wellcome Sanger Institute COVID-19 Surveillance Team |
| EPI_ISL_3365877 | Lighthouse Laboratory Plymouth | Wellcome Sanger Institute for the COVID-19 Genomics UK (COG-UK) Consortium | Lighthouse Laboratory Plymouth and Alex Alderton, Roberto Amato, Jeffrey Barrett, Sonia Goncalves, Ewan Harrison, David K. Jackson, Ian Johnston, Dominic Kwiatkowski, Cordelia Langford, John Sillitoe on behalf of the Wellcome Sanger Institute COVID-19 Surveillance Team |
| EPI_ISL_3365878, EPI_ISL_3365879 | Lighthouse Lab in Alderley Park | Wellcome Sanger Institute for the COVID-19 Genomics UK (COG-UK) Consortium | Jacquelyn Wynn, Mairead Hyland, The Lighthouse Lab in Alderley Park and Alex Alderton, Roberto Amato, Jeffrey Barrett, Sonia Goncalves, Ewan Harrison, David K. Jackson, Ian Johnston, Dominic Kwiatkowski, Cordelia Langford, John Sillitoe on behalf of the Wellcome Sanger Institute COVID-19 Surveillance Team |
| EPI_ISL_3365880 | Lighthouse Laboratory Plymouth | Wellcome Sanger Institute for the COVID-19 Genomics UK (COG-UK) Consortium | Lighthouse Laboratory Plymouth and Alex Alderton, Roberto Amato, Jeffrey Barrett, Sonia Goncalves, Ewan Harrison, David K. Jackson, Ian Johnston, Dominic Kwiatkowski, Cordelia Langford, John Sillitoe on behalf of the Wellcome Sanger Institute COVID-19 Surveillance Team |
| EPI_ISL_3365881, EPI_ISL_3365882, EPI_ISL_3365883, EPI_ISL_3365884 | Lighthouse Lab in Milton Keynes | Wellcome Sanger Institute for the COVID-19 Genomics UK (COG-UK) Consortium | The Lighthouse Lab in Milton Keynes and Alex Alderton, Roberto Amato, Jeffrey Barrett, Sonia Goncalves, Ewan Harrison, David K. Jackson, Ian Johnston, Dominic Kwiatkowski, Cordelia Langford, John Sillitoe on behalf of the Wellcome Sanger Institute COVID-19 Surveillance Team |
| EPI_ISL_3365885 | Lighthouse Lab in Alderley Park | Wellcome Sanger Institute for the COVID-19 Genomics UK (COG-UK) Consortium | Jacquelyn Wynn, Mairead Hyland, The Lighthouse Lab in Alderley Park and Alex Alderton, Roberto Amato, Jeffrey Barrett, Sonia Goncalves, Ewan Harrison, David K. Jackson, Ian Johnston, Dominic Kwiatkowski, Cordelia Langford, John Sillitoe on behalf of the Wellcome Sanger Institute COVID-19 Surveillance Team |
| EPI_ISL_3365886, EPI_ISL_3365887 | Lighthouse Laboratory Plymouth | Wellcome Sanger Institute for the COVID-19 Genomics UK (COG-UK) Consortium | Lighthouse Laboratory Plymouth and Alex Alderton, Roberto Amato, Jeffrey Barrett, Sonia Goncalves, Ewan Harrison, David K. Jackson, Ian Johnston, Dominic Kwiatkowski, Cordelia Langford, John Sillitoe on behalf of the Wellcome Sanger Institute COVID-19 Surveillance Team |
| EPI_ISL_3365888, EPI_ISL_3365889 | Lighthouse Lab in Alderley Park | Wellcome Sanger Institute for the COVID-19 Genomics UK (COG-UK) Consortium | Jacquelyn Wynn, Mairead Hyland, The Lighthouse Lab in Alderley Park and Alex Alderton, Roberto Amato, Jeffrey Barrett, Sonia Goncalves, Ewan Harrison, David K. Jackson, Ian Johnston, Dominic Kwiatkowski, Cordelia Langford, John Sillitoe on behalf of the Wellcome Sanger Institute COVID-19 Surveillance Team |
| EPI_ISL_3365890 | Lighthouse Lab in Milton Keynes | Wellcome Sanger Institute for the COVID-19 Genomics UK (COG-UK) Consortium | The Lighthouse Lab in Milton Keynes and Alex Alderton, Roberto Amato, Jeffrey Barrett, Sonia Goncalves, Ewan Harrison, David K. Jackson, Ian Johnston, Dominic Kwiatkowski, Cordelia Langford, John Sillitoe on behalf of the Wellcome Sanger Institute COVID-19 Surveillance Team |
| EPI_ISL_3365891 | Lighthouse Lab in Alderley Park | Wellcome Sanger Institute for the COVID-19 Genomics UK (COG-UK) Consortium | Jacquelyn Wynn, Mairead Hyland, The Lighthouse Lab in Alderley Park and Alex Alderton, Roberto Amato, Jeffrey Barrett, Sonia Goncalves, Ewan Harrison, David K. Jackson, Ian Johnston, Dominic Kwiatkowski, Cordelia Langford, John Sillitoe on behalf of the Wellcome Sanger Institute COVID-19 Surveillance Team |
| EPI_ISL_3365892 | Lighthouse Lab in Milton Keynes | Wellcome Sanger Institute for the COVID-19 Genomics UK (COG-UK) Consortium | The Lighthouse Lab in Milton Keynes and Alex Alderton, Roberto Amato, Jeffrey Barrett, Sonia Goncalves, Ewan Harrison, David K. Jackson, Ian Johnston, Dominic Kwiatkowski, Cordelia Langford, John Sillitoe on behalf of the Wellcome Sanger Institute COVID-19 Surveillance Team |
| EPI_ISL_3365893 | Lighthouse Lab in Alderley Park | Wellcome Sanger Institute for the COVID-19 Genomics UK (COG-UK) Consortium | Jacquelyn Wynn, Mairead Hyland, The Lighthouse Lab in Alderley Park and Alex Alderton, Roberto Amato, Jeffrey Barrett, Sonia Goncalves, Ewan Harrison, David K. Jackson, Ian Johnston, Dominic Kwiatkowski, Cordelia Langford, John Sillitoe on behalf of the Wellcome Sanger Institute COVID-19 Surveillance Team |
| EPI_ISL_3365894, EPI_ISL_3365895 | Lighthouse Laboratory Plymouth | Wellcome Sanger Institute for the COVID-19 Genomics UK (COG-UK) Consortium | Lighthouse Laboratory Plymouth and Alex Alderton, Roberto Amato, Jeffrey Barrett, Sonia Goncalves, Ewan Harrison, David K. Jackson, Ian Johnston, Dominic Kwiatkowski, Cordelia Langford, John Sillitoe on behalf of the Wellcome Sanger Institute COVID-19 Surveillance Team |
| EPI_ISL_3365896 | Lighthouse Lab in Alderley Park | Wellcome Sanger Institute for the COVID-19 Genomics UK (COG-UK) Consortium | Jacquelyn Wynn, Mairead Hyland, The Lighthouse Lab in Alderley Park and Alex Alderton, Roberto Amato, Jeffrey Barrett, Sonia Goncalves, Ewan Harrison, David K. Jackson, Ian Johnston, Dominic Kwiatkowski, Cordelia Langford, John Sillitoe on behalf of the Wellcome Sanger Institute COVID-19 Surveillance Team |
| EPI_ISL_3365897, EPI_ISL_3365898, EPI_ISL_3365899 | Lighthouse Lab in Milton Keynes | Wellcome Sanger Institute for the COVID-19 Genomics UK (COG-UK) Consortium | The Lighthouse Lab in Milton Keynes and Alex Alderton, Roberto Amato, Jeffrey Barrett, Sonia Goncalves, Ewan Harrison, David K. Jackson, Ian Johnston, Dominic Kwiatkowski, Cordelia Langford, John Sillitoe on behalf of the Wellcome Sanger Institute COVID-19 Surveillance Team |
| EPI_ISL_3365900, EPI_ISL_3365901 | Lighthouse Laboratory Plymouth | Wellcome Sanger Institute for the COVID-19 Genomics UK (COG-UK) Consortium | Lighthouse Laboratory Plymouth and Alex Alderton, Roberto Amato, Jeffrey Barrett, Sonia Goncalves, Ewan Harrison, David K. Jackson, Ian Johnston, Dominic Kwiatkowski, Cordelia Langford, John Sillitoe on behalf of the Wellcome Sanger Institute COVID-19 Surveillance Team |
| EPI_ISL_3365902 | Lighthouse Lab in Milton Keynes | Wellcome Sanger Institute for the COVID-19 Genomics UK (COG-UK) Consortium | The Lighthouse Lab in Milton Keynes and Alex Alderton, Roberto Amato, Jeffrey Barrett, Sonia Goncalves, Ewan Harrison, David K. Jackson, Ian Johnston, Dominic Kwiatkowski, Cordelia Langford, John Sillitoe on behalf of the Wellcome Sanger Institute COVID-19 Surveillance Team |
| EPI_ISL_3365903 | Lighthouse Laboratory Plymouth | Wellcome Sanger Institute for the COVID-19 Genomics UK (COG-UK) Consortium | Lighthouse Laboratory Plymouth and Alex Alderton, Roberto Amato, Jeffrey Barrett, Sonia Goncalves, Ewan Harrison, David K. Jackson, Ian Johnston, Dominic Kwiatkowski, Cordelia Langford, John Sillitoe on behalf of the Wellcome Sanger Institute COVID-19 Surveillance Team |
| EPI_ISL_3365904, EPI_ISL_3365905 | Lighthouse Lab in Milton Keynes | Wellcome Sanger Institute for the COVID-19 Genomics UK (COG-UK) Consortium | The Lighthouse Lab in Milton Keynes and Alex Alderton, Roberto Amato, Jeffrey Barrett, Sonia Goncalves, Ewan Harrison, David K. Jackson, Ian Johnston, Dominic Kwiatkowski, Cordelia Langford, John Sillitoe on behalf of the Wellcome Sanger Institute COVID-19 Surveillance Team |
| EPI_ISL_3365906, EPI_ISL_3365907 | Lighthouse Laboratory Plymouth | Wellcome Sanger Institute for the COVID-19 Genomics UK (COG-UK) Consortium | Lighthouse Laboratory Plymouth and Alex Alderton, Roberto Amato, Jeffrey Barrett, Sonia Goncalves, Ewan Harrison, David K. Jackson, Ian Johnston, Dominic Kwiatkowski, Cordelia Langford, John Sillitoe on behalf of the Wellcome Sanger Institute COVID-19 Surveillance Team |
| EPI_ISL_3365908 | Lighthouse Lab in Milton Keynes | Wellcome Sanger Institute for the COVID-19 Genomics UK (COG-UK) | The Lighthouse Lab in Milton Keynes and Alex Alderton, Roberto Amato, Jeffrey Barrett, Sonia Goncalves, Ewan Harrison, David K. Jackson, Ian Johnston, Dominic Kwiatkowski, Cordelia Langford, John Sillitoe on behalf of the Wellcome Sanger |

| | | | |
|---|---|---|---|
| | | Consortium | Institute COVID-19 Surveillance Team |
| EPI_ISL_3365909 | Lighthouse Laboratory Plymouth | Wellcome Sanger Institute for the COVID-19 Genomics UK (COG-UK) Consortium | Lighthouse Laboratory Plymouth and Alex Alderton, Roberto Amato, Jeffrey Barrett, Sonia Goncalves, Ewan Harrison, David K. Jackson, Ian Johnston, Dominic Kwiatkowski, Cordelia Langford, John Sillitoe on behalf of the Wellcome Sanger Institute COVID-19 Surveillance Team |
| EPI_ISL_3365910, EPI_ISL_3365911, EPI_ISL_3365912, EPI_ISL_3365913, EPI_ISL_3365914, EPI_ISL_3365915, EPI_ISL_3365916, EPI_ISL_3365917, EPI_ISL_3365918, EPI_ISL_3365919, EPI_ISL_3365920, EPI_ISL_3365921, EPI_ISL_3365922, EPI_ISL_3365923, EPI_ISL_3365924, EPI_ISL_3365925, EPI_ISL_3365926, EPI_ISL_3365927, EPI_ISL_3365928 | | | |
| see above | Lighthouse Lab in Milton Keynes | Wellcome Sanger Institute for the COVID-19 Genomics UK (COG-UK) Consortium | The Lighthouse Lab in Milton Keynes and Alex Alderton, Roberto Amato, Jeffrey Barrett, Sonia Goncalves, Ewan Harrison, David K. Jackson, Ian Johnston, Dominic Kwiatkowski, Cordelia Langford, John Sillitoe on behalf of the Wellcome Sanger Institute COVID-19 Surveillance Team |
| EPI_ISL_3365929 | Lighthouse Lab in Alderley Park | Wellcome Sanger Institute for the COVID-19 Genomics UK (COG-UK) Consortium | Jacquelyn Wynn, Mairead Hyland, The Lighthouse Lab in Alderley Park and Alex Alderton, Roberto Amato, Jeffrey Barrett, Sonia Goncalves, Ewan Harrison, David K. Jackson, Ian Johnston, Dominic Kwiatkowski, Cordelia Langford, John Sillitoe on behalf of the Wellcome Sanger Institute COVID-19 Surveillance Team |
| EPI_ISL_3365930, EPI_ISL_3365931 | Health Services Laboratories | Wellcome Sanger Institute for the COVID-19 Genomics UK (COG-UK) Consortium | Health Services Laboratories and Alex Alderton, Roberto Amato, Jeffrey Barrett, Sonia Goncalves, Ewan Harrison, David K. Jackson, Ian Johnston, Dominic Kwiatkowski, Cordelia Langford, John Sillitoe on behalf of the Wellcome Sanger Institute COVID-19 Surveillance Team |
| EPI_ISL_3365932 | Lighthouse Lab in Alderley Park | Wellcome Sanger Institute for the COVID-19 Genomics UK (COG-UK) Consortium | Jacquelyn Wynn, Mairead Hyland, The Lighthouse Lab in Alderley Park and Alex Alderton, Roberto Amato, Jeffrey Barrett, Sonia Goncalves, Ewan Harrison, David K. Jackson, Ian Johnston, Dominic Kwiatkowski, Cordelia Langford, John Sillitoe on behalf of the Wellcome Sanger Institute COVID-19 Surveillance Team |
| EPI_ISL_3365933, EPI_ISL_3365934 | Health Services Laboratories | Wellcome Sanger Institute for the COVID-19 Genomics UK (COG-UK) Consortium | Health Services Laboratories and Alex Alderton, Roberto Amato, Jeffrey Barrett, Sonia Goncalves, Ewan Harrison, David K. Jackson, Ian Johnston, Dominic Kwiatkowski, Cordelia Langford, John Sillitoe on behalf of the Wellcome Sanger Institute COVID-19 Surveillance Team |
| EPI_ISL_3365935 | Lighthouse Lab in Milton Keynes | Wellcome Sanger Institute for the COVID-19 Genomics UK (COG-UK) Consortium | The Lighthouse Lab in Milton Keynes and Alex Alderton, Roberto Amato, Jeffrey Barrett, Sonia Goncalves, Ewan Harrison, David K. Jackson, Ian Johnston, Dominic Kwiatkowski, Cordelia Langford, John Sillitoe on behalf of the Wellcome Sanger Institute COVID-19 Surveillance Team |
| EPI_ISL_3365936, EPI_ISL_3365937, EPI_ISL_3365938, EPI_ISL_3365939, EPI_ISL_3365940, EPI_ISL_3365941, EPI_ISL_3365942, EPI_ISL_3365943, EPI_ISL_3365944, EPI_ISL_3365945, EPI_ISL_3365946, EPI_ISL_3365947, EPI_ISL_3365948, EPI_ISL_3365949, EPI_ISL_3365950, EPI_ISL_3365951, EPI_ISL_3365952 | | | |
| see above | Health Services Laboratories | Wellcome Sanger Institute for the COVID-19 Genomics UK (COG-UK) Consortium | Health Services Laboratories and Alex Alderton, Roberto Amato, Jeffrey Barrett, Sonia Goncalves, Ewan Harrison, David K. Jackson, Ian Johnston, Dominic Kwiatkowski, Cordelia Langford, John Sillitoe on behalf of the Wellcome Sanger Institute COVID-19 Surveillance Team |
| EPI_ISL_3365953 | Lighthouse Lab in Alderley Park | Wellcome Sanger Institute for the COVID-19 Genomics UK (COG-UK) Consortium | Jacquelyn Wynn, Mairead Hyland, The Lighthouse Lab in Alderley Park and Alex Alderton, Roberto Amato, Jeffrey Barrett, Sonia Goncalves, Ewan Harrison, David K. Jackson, Ian Johnston, Dominic Kwiatkowski, Cordelia Langford, John Sillitoe on behalf of the Wellcome Sanger Institute COVID-19 Surveillance Team |
| EPI_ISL_3365954, EPI_ISL_3365955 | Health Services Laboratories | Wellcome Sanger Institute for the COVID-19 Genomics UK (COG-UK) Consortium | Health Services Laboratories and Alex Alderton, Roberto Amato, Jeffrey Barrett, Sonia Goncalves, Ewan Harrison, David K. Jackson, Ian Johnston, Dominic Kwiatkowski, Cordelia Langford, John Sillitoe on behalf of the Wellcome Sanger Institute COVID-19 Surveillance Team |
| EPI_ISL_3365956 | Lighthouse Lab in Milton Keynes | Wellcome Sanger Institute for the COVID-19 Genomics UK (COG-UK) Consortium | The Lighthouse Lab in Milton Keynes and Alex Alderton, Roberto Amato, Jeffrey Barrett, Sonia Goncalves, Ewan Harrison, David K. Jackson, Ian Johnston, Dominic Kwiatkowski, Cordelia Langford, John Sillitoe on behalf of the Wellcome Sanger Institute COVID-19 Surveillance Team |
| EPI_ISL_3365957, EPI_ISL_3365958, EPI_ISL_3365959, EPI_ISL_3365960, EPI_ISL_3365961 | Health Services Laboratories | Wellcome Sanger Institute for the COVID-19 Genomics UK (COG-UK) Consortium | Health Services Laboratories and Alex Alderton, Roberto Amato, Jeffrey Barrett, Sonia Goncalves, Ewan Harrison, David K. Jackson, Ian Johnston, Dominic Kwiatkowski, Cordelia Langford, John Sillitoe on behalf of the Wellcome Sanger Institute COVID-19 Surveillance Team |
| EPI_ISL_3365962, EPI_ISL_3365963, EPI_ISL_3365964, EPI_ISL_3365965 | Lighthouse Lab in Milton Keynes | Wellcome Sanger Institute for the COVID-19 Genomics UK (COG-UK) Consortium | The Lighthouse Lab in Milton Keynes and Alex Alderton, Roberto Amato, Jeffrey Barrett, Sonia Goncalves, Ewan Harrison, David K. Jackson, Ian Johnston, Dominic Kwiatkowski, Cordelia Langford, John Sillitoe on behalf of the Wellcome Sanger Institute COVID-19 Surveillance Team |
| EPI_ISL_3365966 | Lighthouse Lab in Alderley Park | Wellcome Sanger Institute for the COVID-19 Genomics UK (COG-UK) Consortium | Jacquelyn Wynn, Mairead Hyland, The Lighthouse Lab in Alderley Park and Alex Alderton, Roberto Amato, Jeffrey Barrett, Sonia Goncalves, Ewan Harrison, David K. Jackson, Ian Johnston, Dominic Kwiatkowski, Cordelia Langford, John Sillitoe on behalf of the Wellcome Sanger Institute COVID-19 Surveillance Team |
| EPI_ISL_3365967, EPI_ISL_3365968 | Lighthouse Lab in Milton Keynes | Wellcome Sanger Institute for the COVID-19 Genomics UK (COG-UK) Consortium | The Lighthouse Lab in Milton Keynes and Alex Alderton, Roberto Amato, Jeffrey Barrett, Sonia Goncalves, Ewan Harrison, David K. Jackson, Ian Johnston, Dominic Kwiatkowski, Cordelia Langford, John Sillitoe on behalf of the Wellcome Sanger Institute COVID-19 Surveillance Team |
| EPI_ISL_3365969 | Lighthouse Lab in Alderley Park | Wellcome Sanger Institute for the COVID-19 Genomics UK (COG-UK) Consortium | Jacquelyn Wynn, Mairead Hyland, The Lighthouse Lab in Alderley Park and Alex Alderton, Roberto Amato, Jeffrey Barrett, Sonia Goncalves, Ewan Harrison, David K. Jackson, Ian Johnston, Dominic Kwiatkowski, Cordelia Langford, John Sillitoe on behalf of the Wellcome Sanger Institute COVID-19 Surveillance Team |
| EPI_ISL_3365970 | Lighthouse Lab in Milton Keynes | Wellcome Sanger Institute for the COVID-19 Genomics UK (COG-UK) Consortium | The Lighthouse Lab in Milton Keynes and Alex Alderton, Roberto Amato, Jeffrey Barrett, Sonia Goncalves, Ewan Harrison, David K. Jackson, Ian Johnston, Dominic Kwiatkowski, Cordelia Langford, John Sillitoe on behalf of the Wellcome Sanger Institute COVID-19 Surveillance Team |
| EPI_ISL_3365971 | Randox Laboratories | Wellcome Sanger Institute for the COVID-19 Genomics UK (COG-UK) Consortium | Randox Laboratories and Alex Alderton, Roberto Amato, Jeffrey Barrett, Sonia Goncalves, Ewan Harrison, David K. Jackson, Ian Johnston, Dominic Kwiatkowski, Cordelia Langford, John Sillitoe on behalf of the Wellcome Sanger Institute COVID-19 Surveillance Team |
| EPI_ISL_3365972, EPI_ISL_3365973 | Lighthouse Lab in Milton Keynes | Wellcome Sanger Institute for the COVID-19 Genomics UK (COG-UK) Consortium | The Lighthouse Lab in Milton Keynes and Alex Alderton, Roberto Amato, Jeffrey Barrett, Sonia Goncalves, Ewan Harrison, David K. Jackson, Ian Johnston, Dominic Kwiatkowski, Cordelia Langford, John Sillitoe on behalf of the Wellcome Sanger Institute COVID-19 Surveillance Team |
| EPI_ISL_3365974 | Lighthouse Lab in Alderley Park | Wellcome Sanger Institute for the COVID-19 Genomics UK (COG-UK) Consortium | Jacquelyn Wynn, Mairead Hyland, The Lighthouse Lab in Alderley Park and Alex Alderton, Roberto Amato, Jeffrey Barrett, Sonia Goncalves, Ewan Harrison, David K. Jackson, Ian Johnston, Dominic Kwiatkowski, Cordelia Langford, John Sillitoe on behalf of the Wellcome Sanger Institute COVID-19 Surveillance Team |
| EPI_ISL_3365975 | Lighthouse Lab in Milton Keynes | Wellcome Sanger Institute for the COVID-19 Genomics UK (COG-UK) Consortium | The Lighthouse Lab in Milton Keynes and Alex Alderton, Roberto Amato, Jeffrey Barrett, Sonia Goncalves, Ewan Harrison, David K. Jackson, Ian Johnston, Dominic Kwiatkowski, Cordelia Langford, John Sillitoe on behalf of the Wellcome Sanger Institute COVID-19 Surveillance Team |
| EPI_ISL_3365976 | Randox Laboratories | Wellcome Sanger Institute for the COVID-19 Genomics UK (COG-UK) Consortium | Randox Laboratories and Alex Alderton, Roberto Amato, Jeffrey Barrett, Sonia Goncalves, Ewan Harrison, David K. Jackson, Ian Johnston, Dominic Kwiatkowski, Cordelia Langford, John Sillitoe on behalf of the Wellcome Sanger Institute COVID-19 Surveillance Team |
| EPI_ISL_3365977, EPI_ISL_3365978, EPI_ISL_3365979 | Lighthouse Lab in Milton Keynes | Wellcome Sanger Institute for the COVID-19 Genomics UK (COG-UK) Consortium | The Lighthouse Lab in Milton Keynes and Alex Alderton, Roberto Amato, Jeffrey Barrett, Sonia Goncalves, Ewan Harrison, David K. Jackson, Ian Johnston, Dominic Kwiatkowski, Cordelia Langford, John Sillitoe on behalf of the Wellcome Sanger Institute COVID-19 Surveillance Team |
| EPI_ISL_3365980 | Randox Laboratories | Wellcome Sanger Institute for the COVID-19 Genomics UK (COG-UK) Consortium | Randox Laboratories and Alex Alderton, Roberto Amato, Jeffrey Barrett, Sonia Goncalves, Ewan Harrison, David K. Jackson, Ian Johnston, Dominic Kwiatkowski, Cordelia Langford, John Sillitoe on behalf of the Wellcome Sanger Institute COVID-19 Surveillance Team |
| EPI_ISL_3365981, EPI_ISL_3365982, EPI_ISL_3365983, EPI_ISL_3365984, EPI_ISL_3365985, EPI_ISL_3365986, EPI_ISL_3365987, EPI_ISL_3365988, EPI_ISL_3365989 | Lighthouse Lab in Milton Keynes | Wellcome Sanger Institute for the COVID-19 Genomics UK (COG-UK) Consortium | The Lighthouse Lab in Milton Keynes and Alex Alderton, Roberto Amato, Jeffrey Barrett, Sonia Goncalves, Ewan Harrison, David K. Jackson, Ian Johnston, Dominic Kwiatkowski, Cordelia Langford, John Sillitoe on behalf of the Wellcome Sanger Institute COVID-19 Surveillance Team |
| EPI_ISL_3365990, EPI_ISL_3365991 | Lighthouse Lab in Alderley Park | Wellcome Sanger Institute for the COVID-19 Genomics UK (COG-UK) Consortium | Jacquelyn Wynn, Mairead Hyland, The Lighthouse Lab in Alderley Park and Alex Alderton, Roberto Amato, Jeffrey Barrett, Sonia Goncalves, Ewan Harrison, David K. Jackson, Ian Johnston, Dominic Kwiatkowski, Cordelia Langford, John Sillitoe on behalf of the Wellcome Sanger Institute COVID-19 Surveillance Team |
| EPI_ISL_3365992, EPI_ISL_3365993, EPI_ISL_3365994 | Lighthouse Lab in Milton Keynes | Wellcome Sanger Institute for the COVID-19 Genomics UK (COG-UK) Consortium | The Lighthouse Lab in Milton Keynes and Alex Alderton, Roberto Amato, Jeffrey Barrett, Sonia Goncalves, Ewan Harrison, David K. Jackson, Ian Johnston, Dominic Kwiatkowski, Cordelia Langford, John Sillitoe on behalf of the Wellcome Sanger Institute COVID-19 Surveillance Team |
| EPI_ISL_3365995 | Randox Laboratories | Wellcome Sanger Institute for the COVID-19 Genomics UK (COG-UK) Consortium | Randox Laboratories and Alex Alderton, Roberto Amato, Jeffrey Barrett, Sonia Goncalves, Ewan Harrison, David K. Jackson, Ian Johnston, Dominic Kwiatkowski, Cordelia Langford, John Sillitoe on behalf of the Wellcome Sanger Institute COVID-19 Surveillance Team |
| EPI_ISL_3365996 | Lighthouse Lab in Alderley Park | Wellcome Sanger Institute for the COVID-19 Genomics UK (COG-UK) Consortium | Jacquelyn Wynn, Mairead Hyland, The Lighthouse Lab in Alderley Park and Alex Alderton, Roberto Amato, Jeffrey Barrett, Sonia Goncalves, Ewan Harrison, David K. Jackson, Ian Johnston, Dominic Kwiatkowski, Cordelia Langford, John Sillitoe on behalf of the Wellcome Sanger Institute COVID-19 Surveillance Team |
| EPI_ISL_3365997, EPI_ISL_3365998, EPI_ISL_3365999, EPI_ISL_3366000 | Lighthouse Lab in Milton Keynes | Wellcome Sanger Institute for the COVID-19 Genomics UK (COG-UK) Consortium | The Lighthouse Lab in Milton Keynes and Alex Alderton, Roberto Amato, Jeffrey Barrett, Sonia Goncalves, Ewan Harrison, David K. Jackson, Ian Johnston, Dominic Kwiatkowski, Cordelia Langford, John Sillitoe on behalf of the Wellcome Sanger Institute COVID-19 Surveillance Team |
| EPI_ISL_3366001 | Lighthouse Lab in Alderley Park | Wellcome Sanger Institute for the COVID-19 Genomics UK (COG-UK) Consortium | Jacquelyn Wynn, Mairead Hyland, The Lighthouse Lab in Alderley Park and Alex Alderton, Roberto Amato, Jeffrey Barrett, Sonia Goncalves, Ewan Harrison, David K. Jackson, Ian Johnston, Dominic Kwiatkowski, Cordelia Langford, John Sillitoe on behalf of the Wellcome Sanger Institute COVID-19 Surveillance Team |
| EPI_ISL_3366002 | Lighthouse Lab in Milton Keynes | Wellcome Sanger Institute for the COVID-19 Genomics UK (COG-UK) Consortium | The Lighthouse Lab in Milton Keynes and Alex Alderton, Roberto Amato, Jeffrey Barrett, Sonia Goncalves, Ewan Harrison, David K. Jackson, Ian Johnston, Dominic Kwiatkowski, Cordelia Langford, John Sillitoe on behalf of the Wellcome Sanger Institute COVID-19 Surveillance Team |
| EPI_ISL_3366003, EPI_ISL_3366004 | Lighthouse Lab in Alderley Park | Wellcome Sanger Institute for the COVID-19 Genomics UK (COG-UK) Consortium | Jacquelyn Wynn, Mairead Hyland, The Lighthouse Lab in Alderley Park and Alex Alderton, Roberto Amato, Jeffrey Barrett, Sonia Goncalves, Ewan Harrison, David K. Jackson, Ian Johnston, Dominic Kwiatkowski, Cordelia Langford, John Sillitoe on behalf of the Wellcome Sanger Institute COVID-19 Surveillance Team |
| EPI_ISL_3366005, EPI_ISL_3366006, EPI_ISL_3366007 | Lighthouse Lab in Milton Keynes | Wellcome Sanger Institute for the COVID-19 Genomics UK (COG-UK) Consortium | The Lighthouse Lab in Milton Keynes and Alex Alderton, Roberto Amato, Jeffrey Barrett, Sonia Goncalves, Ewan Harrison, David K. Jackson, Ian Johnston, Dominic Kwiatkowski, Cordelia Langford, John Sillitoe on behalf of the Wellcome Sanger Institute COVID-19 Surveillance Team |
| EPI_ISL_3366008 | Lighthouse Lab in Alderley Park | Wellcome Sanger Institute for the COVID-19 Genomics UK (COG-UK) Consortium | Jacquelyn Wynn, Mairead Hyland, The Lighthouse Lab in Alderley Park and Alex Alderton, Roberto Amato, Jeffrey Barrett, Sonia Goncalves, Ewan Harrison, David K. Jackson, Ian Johnston, Dominic Kwiatkowski, Cordelia Langford, John Sillitoe on behalf of the Wellcome Sanger Institute COVID-19 Surveillance Team |
| EPI_ISL_3366009, EPI_ISL_3366010, EPI_ISL_3366011, | Lighthouse Lab in Milton Keynes | Wellcome Sanger Institute for the COVID-19 Genomics UK (COG-UK) Consortium | The Lighthouse Lab in Milton Keynes and Alex Alderton, Roberto Amato, Jeffrey Barrett, Sonia Goncalves, Ewan Harrison, David K. Jackson, Ian Johnston, Dominic Kwiatkowski, Cordelia Langford, John Sillitoe on behalf of the Wellcome Sanger Institute COVID-19 Surveillance Team |

| | | | |
|---|---|---|---|
| EPI_ISL_3366012 | | | |
| EPI_ISL_3366013 | Lighthouse Lab in Alderley Park | Wellcome Sanger Institute for the COVID-19 Genomics UK (COG-UK) Consortium | Jacquelyn Wynn, Mairead Hyland, The Lighthouse Lab in Alderley Park and Alex Alderton, Roberto Amato, Jeffrey Barrett, Sonia Goncalves, Ewan Harrison, David K. Jackson, Ian Johnston, Dominic Kwiatkowski, Cordelia Langford, John Sillitoe on behalf of the Wellcome Sanger Institute COVID-19 Surveillance Team |
| EPI_ISL_3366014, EPI_ISL_3366015 | Lighthouse Lab in Milton Keynes | Wellcome Sanger Institute for the COVID-19 Genomics UK (COG-UK) Consortium | The Lighthouse Lab in Milton Keynes and Alex Alderton, Roberto Amato, Jeffrey Barrett, Sonia Goncalves, Ewan Harrison, David K. Jackson, Ian Johnston, Dominic Kwiatkowski, Cordelia Langford, John Sillitoe on behalf of the Wellcome Sanger Institute COVID-19 Surveillance Team |
| EPI_ISL_3366016 | Lighthouse Lab in Alderley Park | Wellcome Sanger Institute for the COVID-19 Genomics UK (COG-UK) Consortium | Jacquelyn Wynn, Mairead Hyland, The Lighthouse Lab in Alderley Park and Alex Alderton, Roberto Amato, Jeffrey Barrett, Sonia Goncalves, Ewan Harrison, David K. Jackson, Ian Johnston, Dominic Kwiatkowski, Cordelia Langford, John Sillitoe on behalf of the Wellcome Sanger Institute COVID-19 Surveillance Team |
| EPI_ISL_3366017, EPI_ISL_3366018, EPI_ISL_3366019, EPI_ISL_3366020 | Lighthouse Lab in Milton Keynes | Wellcome Sanger Institute for the COVID-19 Genomics UK (COG-UK) Consortium | The Lighthouse Lab in Milton Keynes and Alex Alderton, Roberto Amato, Jeffrey Barrett, Sonia Goncalves, Ewan Harrison, David K. Jackson, Ian Johnston, Dominic Kwiatkowski, Cordelia Langford, John Sillitoe on behalf of the Wellcome Sanger Institute COVID-19 Surveillance Team |
| EPI_ISL_3366021 | Lighthouse Lab in Alderley Park | Wellcome Sanger Institute for the COVID-19 Genomics UK (COG-UK) Consortium | Jacquelyn Wynn, Mairead Hyland, The Lighthouse Lab in Alderley Park and Alex Alderton, Roberto Amato, Jeffrey Barrett, Sonia Goncalves, Ewan Harrison, David K. Jackson, Ian Johnston, Dominic Kwiatkowski, Cordelia Langford, John Sillitoe on behalf of the Wellcome Sanger Institute COVID-19 Surveillance Team |
| EPI_ISL_3366022, EPI_ISL_3366023 | Randox Laboratories | Wellcome Sanger Institute for the COVID-19 Genomics UK (COG-UK) Consortium | Randox Laboratories and Alex Alderton, Roberto Amato, Jeffrey Barrett, Sonia Goncalves, Ewan Harrison, David K. Jackson, Ian Johnston, Dominic Kwiatkowski, Cordelia Langford, John Sillitoe on behalf of the Wellcome Sanger Institute COVID-19 Surveillance Team |
| EPI_ISL_3366024 | Lighthouse Lab in Alderley Park | Wellcome Sanger Institute for the COVID-19 Genomics UK (COG-UK) Consortium | Jacquelyn Wynn, Mairead Hyland, The Lighthouse Lab in Alderley Park and Alex Alderton, Roberto Amato, Jeffrey Barrett, Sonia Goncalves, Ewan Harrison, David K. Jackson, Ian Johnston, Dominic Kwiatkowski, Cordelia Langford, John Sillitoe on behalf of the Wellcome Sanger Institute COVID-19 Surveillance Team |
| EPI_ISL_3366025, EPI_ISL_3366026, EPI_ISL_3366027 | Lighthouse Lab in Milton Keynes | Wellcome Sanger Institute for the COVID-19 Genomics UK (COG-UK) Consortium | The Lighthouse Lab in Milton Keynes and Alex Alderton, Roberto Amato, Jeffrey Barrett, Sonia Goncalves, Ewan Harrison, David K. Jackson, Ian Johnston, Dominic Kwiatkowski, Cordelia Langford, John Sillitoe on behalf of the Wellcome Sanger Institute COVID-19 Surveillance Team |
| EPI_ISL_3366028 | Randox Laboratories | Wellcome Sanger Institute for the COVID-19 Genomics UK (COG-UK) Consortium | Randox Laboratories and Alex Alderton, Roberto Amato, Jeffrey Barrett, Sonia Goncalves, Ewan Harrison, David K. Jackson, Ian Johnston, Dominic Kwiatkowski, Cordelia Langford, John Sillitoe on behalf of the Wellcome Sanger Institute COVID-19 Surveillance Team |
| EPI_ISL_3366029, EPI_ISL_3366030 | Lighthouse Lab in Milton Keynes | Wellcome Sanger Institute for the COVID-19 Genomics UK (COG-UK) Consortium | The Lighthouse Lab in Milton Keynes and Alex Alderton, Roberto Amato, Jeffrey Barrett, Sonia Goncalves, Ewan Harrison, David K. Jackson, Ian Johnston, Dominic Kwiatkowski, Cordelia Langford, John Sillitoe on behalf of the Wellcome Sanger Institute COVID-19 Surveillance Team |
| EPI_ISL_3366031 | Randox Laboratories | Wellcome Sanger Institute for the COVID-19 Genomics UK (COG-UK) Consortium | Randox Laboratories and Alex Alderton, Roberto Amato, Jeffrey Barrett, Sonia Goncalves, Ewan Harrison, David K. Jackson, Ian Johnston, Dominic Kwiatkowski, Cordelia Langford, John Sillitoe on behalf of the Wellcome Sanger Institute COVID-19 Surveillance Team |
| EPI_ISL_3366032, EPI_ISL_3366033 | Lighthouse Lab in Milton Keynes | Wellcome Sanger Institute for the COVID-19 Genomics UK (COG-UK) Consortium | The Lighthouse Lab in Milton Keynes and Alex Alderton, Roberto Amato, Jeffrey Barrett, Sonia Goncalves, Ewan Harrison, David K. Jackson, Ian Johnston, Dominic Kwiatkowski, Cordelia Langford, John Sillitoe on behalf of the Wellcome Sanger Institute COVID-19 Surveillance Team |
| EPI_ISL_3366034 | Randox Laboratories | Wellcome Sanger Institute for the COVID-19 Genomics UK (COG-UK) Consortium | Randox Laboratories and Alex Alderton, Roberto Amato, Jeffrey Barrett, Sonia Goncalves, Ewan Harrison, David K. Jackson, Ian Johnston, Dominic Kwiatkowski, Cordelia Langford, John Sillitoe on behalf of the Wellcome Sanger Institute COVID-19 Surveillance Team |
| EPI_ISL_3366035 | Lighthouse Lab in Alderley Park | Wellcome Sanger Institute for the COVID-19 Genomics UK (COG-UK) Consortium | Jacquelyn Wynn, Mairead Hyland, The Lighthouse Lab in Alderley Park and Alex Alderton, Roberto Amato, Jeffrey Barrett, Sonia Goncalves, Ewan Harrison, David K. Jackson, Ian Johnston, Dominic Kwiatkowski, Cordelia Langford, John Sillitoe on behalf of the Wellcome Sanger Institute COVID-19 Surveillance Team |
| EPI_ISL_3366036, EPI_ISL_3366037, EPI_ISL_3366038 | Lighthouse Lab in Milton Keynes | Wellcome Sanger Institute for the COVID-19 Genomics UK (COG-UK) Consortium | The Lighthouse Lab in Milton Keynes and Alex Alderton, Roberto Amato, Jeffrey Barrett, Sonia Goncalves, Ewan Harrison, David K. Jackson, Ian Johnston, Dominic Kwiatkowski, Cordelia Langford, John Sillitoe on behalf of the Wellcome Sanger Institute COVID-19 Surveillance Team |
| EPI_ISL_3366039, EPI_ISL_3366040 | Randox Laboratories | Wellcome Sanger Institute for the COVID-19 Genomics UK (COG-UK) Consortium | Randox Laboratories and Alex Alderton, Roberto Amato, Jeffrey Barrett, Sonia Goncalves, Ewan Harrison, David K. Jackson, Ian Johnston, Dominic Kwiatkowski, Cordelia Langford, John Sillitoe on behalf of the Wellcome Sanger Institute COVID-19 Surveillance Team |
| EPI_ISL_3366041 | Lighthouse Lab in Alderley Park | Wellcome Sanger Institute for the COVID-19 Genomics UK (COG-UK) Consortium | Jacquelyn Wynn, Mairead Hyland, The Lighthouse Lab in Alderley Park and Alex Alderton, Roberto Amato, Jeffrey Barrett, Sonia Goncalves, Ewan Harrison, David K. Jackson, Ian Johnston, Dominic Kwiatkowski, Cordelia Langford, John Sillitoe on behalf of the Wellcome Sanger Institute COVID-19 Surveillance Team |
| EPI_ISL_3366042, EPI_ISL_3366043, EPI_ISL_3366044, EPI_ISL_3366045, EPI_ISL_3366046 | Lighthouse Lab in Milton Keynes | Wellcome Sanger Institute for the COVID-19 Genomics UK (COG-UK) Consortium | The Lighthouse Lab in Milton Keynes and Alex Alderton, Roberto Amato, Jeffrey Barrett, Sonia Goncalves, Ewan Harrison, David K. Jackson, Ian Johnston, Dominic Kwiatkowski, Cordelia Langford, John Sillitoe on behalf of the Wellcome Sanger Institute COVID-19 Surveillance Team |
| EPI_ISL_3366047 | Randox Laboratories | Wellcome Sanger Institute for the COVID-19 Genomics UK (COG-UK) Consortium | Randox Laboratories and Alex Alderton, Roberto Amato, Jeffrey Barrett, Sonia Goncalves, Ewan Harrison, David K. Jackson, Ian Johnston, Dominic Kwiatkowski, Cordelia Langford, John Sillitoe on behalf of the Wellcome Sanger Institute COVID-19 Surveillance Team |
| EPI_ISL_3366048 | Lighthouse Lab in Alderley Park | Wellcome Sanger Institute for the COVID-19 Genomics UK (COG-UK) Consortium | Jacquelyn Wynn, Mairead Hyland, The Lighthouse Lab in Alderley Park and Alex Alderton, Roberto Amato, Jeffrey Barrett, Sonia Goncalves, Ewan Harrison, David K. Jackson, Ian Johnston, Dominic Kwiatkowski, Cordelia Langford, John Sillitoe on behalf of the Wellcome Sanger Institute COVID-19 Surveillance Team |
| EPI_ISL_3366049 | Randox Laboratories | Wellcome Sanger Institute for the COVID-19 Genomics UK (COG-UK) Consortium | Randox Laboratories and Alex Alderton, Roberto Amato, Jeffrey Barrett, Sonia Goncalves, Ewan Harrison, David K. Jackson, Ian Johnston, Dominic Kwiatkowski, Cordelia Langford, John Sillitoe on behalf of the Wellcome Sanger Institute COVID-19 Surveillance Team |
| EPI_ISL_3366050, EPI_ISL_3366051 | Lighthouse Lab in Alderley Park | Wellcome Sanger Institute for the COVID-19 Genomics UK (COG-UK) Consortium | Jacquelyn Wynn, Mairead Hyland, The Lighthouse Lab in Alderley Park and Alex Alderton, Roberto Amato, Jeffrey Barrett, Sonia Goncalves, Ewan Harrison, David K. Jackson, Ian Johnston, Dominic Kwiatkowski, Cordelia Langford, John Sillitoe on behalf of the Wellcome Sanger Institute COVID-19 Surveillance Team |
| EPI_ISL_3366052 | Randox Laboratories | Wellcome Sanger Institute for the COVID-19 Genomics UK (COG-UK) Consortium | Randox Laboratories and Alex Alderton, Roberto Amato, Jeffrey Barrett, Sonia Goncalves, Ewan Harrison, David K. Jackson, Ian Johnston, Dominic Kwiatkowski, Cordelia Langford, John Sillitoe on behalf of the Wellcome Sanger Institute COVID-19 Surveillance Team |
| EPI_ISL_3366053, EPI_ISL_3366054 | Lighthouse Lab in Milton Keynes | Wellcome Sanger Institute for the COVID-19 Genomics UK (COG-UK) Consortium | The Lighthouse Lab in Milton Keynes and Alex Alderton, Roberto Amato, Jeffrey Barrett, Sonia Goncalves, Ewan Harrison, David K. Jackson, Ian Johnston, Dominic Kwiatkowski, Cordelia Langford, John Sillitoe on behalf of the Wellcome Sanger Institute COVID-19 Surveillance Team |
| EPI_ISL_3366055, EPI_ISL_3366056, EPI_ISL_3366057 | Lighthouse Lab in Alderley Park | Wellcome Sanger Institute for the COVID-19 Genomics UK (COG-UK) Consortium | Jacquelyn Wynn, Mairead Hyland, The Lighthouse Lab in Alderley Park and Alex Alderton, Roberto Amato, Jeffrey Barrett, Sonia Goncalves, Ewan Harrison, David K. Jackson, Ian Johnston, Dominic Kwiatkowski, Cordelia Langford, John Sillitoe on behalf of the Wellcome Sanger Institute COVID-19 Surveillance Team |
| EPI_ISL_3366058, EPI_ISL_3366059, EPI_ISL_3366060 | Lighthouse Lab in Milton Keynes | Wellcome Sanger Institute for the COVID-19 Genomics UK (COG-UK) Consortium | The Lighthouse Lab in Milton Keynes and Alex Alderton, Roberto Amato, Jeffrey Barrett, Sonia Goncalves, Ewan Harrison, David K. Jackson, Ian Johnston, Dominic Kwiatkowski, Cordelia Langford, John Sillitoe on behalf of the Wellcome Sanger Institute COVID-19 Surveillance Team |
| EPI_ISL_3366061 | Lighthouse Lab in Alderley Park | Wellcome Sanger Institute for the COVID-19 Genomics UK (COG-UK) Consortium | Jacquelyn Wynn, Mairead Hyland, The Lighthouse Lab in Alderley Park and Alex Alderton, Roberto Amato, Jeffrey Barrett, Sonia Goncalves, Ewan Harrison, David K. Jackson, Ian Johnston, Dominic Kwiatkowski, Cordelia Langford, John Sillitoe on behalf of the Wellcome Sanger Institute COVID-19 Surveillance Team |
| EPI_ISL_3366062, EPI_ISL_3366063, EPI_ISL_3366064 | Lighthouse Lab in Milton Keynes | Wellcome Sanger Institute for the COVID-19 Genomics UK (COG-UK) Consortium | The Lighthouse Lab in Milton Keynes and Alex Alderton, Roberto Amato, Jeffrey Barrett, Sonia Goncalves, Ewan Harrison, David K. Jackson, Ian Johnston, Dominic Kwiatkowski, Cordelia Langford, John Sillitoe on behalf of the Wellcome Sanger Institute COVID-19 Surveillance Team |
| EPI_ISL_3366065 | Lighthouse Lab in Alderley Park | Wellcome Sanger Institute for the COVID-19 Genomics UK (COG-UK) Consortium | Jacquelyn Wynn, Mairead Hyland, The Lighthouse Lab in Alderley Park and Alex Alderton, Roberto Amato, Jeffrey Barrett, Sonia Goncalves, Ewan Harrison, David K. Jackson, Ian Johnston, Dominic Kwiatkowski, Cordelia Langford, John Sillitoe on behalf of the Wellcome Sanger Institute COVID-19 Surveillance Team |
| EPI_ISL_3366066 | Lighthouse Lab in Milton Keynes | Wellcome Sanger Institute for the COVID-19 Genomics UK (COG-UK) Consortium | The Lighthouse Lab in Milton Keynes and Alex Alderton, Roberto Amato, Jeffrey Barrett, Sonia Goncalves, Ewan Harrison, David K. Jackson, Ian Johnston, Dominic Kwiatkowski, Cordelia Langford, John Sillitoe on behalf of the Wellcome Sanger Institute COVID-19 Surveillance Team |
| EPI_ISL_3366067 | Lighthouse Lab in Alderley Park | Wellcome Sanger Institute for the COVID-19 Genomics UK (COG-UK) Consortium | Jacquelyn Wynn, Mairead Hyland, The Lighthouse Lab in Alderley Park and Alex Alderton, Roberto Amato, Jeffrey Barrett, Sonia Goncalves, Ewan Harrison, David K. Jackson, Ian Johnston, Dominic Kwiatkowski, Cordelia Langford, John Sillitoe on behalf of the Wellcome Sanger Institute COVID-19 Surveillance Team |
| EPI_ISL_3366068 | Lighthouse Lab in Milton Keynes | Wellcome Sanger Institute for the COVID-19 Genomics UK (COG-UK) Consortium | The Lighthouse Lab in Milton Keynes and Alex Alderton, Roberto Amato, Jeffrey Barrett, Sonia Goncalves, Ewan Harrison, David K. Jackson, Ian Johnston, Dominic Kwiatkowski, Cordelia Langford, John Sillitoe on behalf of the Wellcome Sanger Institute COVID-19 Surveillance Team |
| EPI_ISL_3366069 | Randox Laboratories | Wellcome Sanger Institute for the COVID-19 Genomics UK (COG-UK) Consortium | Randox Laboratories and Alex Alderton, Roberto Amato, Jeffrey Barrett, Sonia Goncalves, Ewan Harrison, David K. Jackson, Ian Johnston, Dominic Kwiatkowski, Cordelia Langford, John Sillitoe on behalf of the Wellcome Sanger Institute COVID-19 Surveillance Team |
| EPI_ISL_3366070, EPI_ISL_3366071 | Lighthouse Lab in Milton Keynes | Wellcome Sanger Institute for the COVID-19 Genomics UK (COG-UK) Consortium | The Lighthouse Lab in Milton Keynes and Alex Alderton, Roberto Amato, Jeffrey Barrett, Sonia Goncalves, Ewan Harrison, David K. Jackson, Ian Johnston, Dominic Kwiatkowski, Cordelia Langford, John Sillitoe on behalf of the Wellcome Sanger Institute COVID-19 Surveillance Team |
| EPI_ISL_3366072 | Lighthouse Lab in Alderley Park | Wellcome Sanger Institute for the COVID-19 Genomics UK (COG-UK) Consortium | Jacquelyn Wynn, Mairead Hyland, The Lighthouse Lab in Alderley Park and Alex Alderton, Roberto Amato, Jeffrey Barrett, Sonia Goncalves, Ewan Harrison, David K. Jackson, Ian Johnston, Dominic Kwiatkowski, Cordelia Langford, John Sillitoe on behalf of the Wellcome Sanger Institute COVID-19 Surveillance Team |
| EPI_ISL_3366073, EPI_ISL_3366074 | Randox Laboratories | Wellcome Sanger Institute for the COVID-19 Genomics UK (COG-UK) Consortium | Randox Laboratories and Alex Alderton, Roberto Amato, Jeffrey Barrett, Sonia Goncalves, Ewan Harrison, David K. Jackson, Ian Johnston, Dominic Kwiatkowski, Cordelia Langford, John Sillitoe on behalf of the Wellcome Sanger Institute COVID-19 Surveillance Team |
| EPI_ISL_3366075 | Lighthouse Lab in Milton Keynes | Wellcome Sanger Institute for the COVID-19 Genomics UK (COG-UK) Consortium | The Lighthouse Lab in Milton Keynes and Alex Alderton, Roberto Amato, Jeffrey Barrett, Sonia Goncalves, Ewan Harrison, David K. Jackson, Ian Johnston, Dominic Kwiatkowski, Cordelia Langford, John Sillitoe on behalf of the Wellcome Sanger Institute COVID-19 Surveillance Team |
| EPI_ISL_3366076 | Randox Laboratories | Wellcome Sanger Institute for the COVID-19 Genomics UK (COG-UK) Consortium | Randox Laboratories and Alex Alderton, Roberto Amato, Jeffrey Barrett, Sonia Goncalves, Ewan Harrison, David K. Jackson, Ian Johnston, Dominic Kwiatkowski, Cordelia Langford, John Sillitoe on behalf of the Wellcome Sanger Institute COVID-19 Surveillance Team |
| EPI_ISL_3366077 | Lighthouse Lab in Milton Keynes | Wellcome Sanger Institute for the COVID-19 Genomics UK (COG-UK) Consortium | The Lighthouse Lab in Milton Keynes and Alex Alderton, Roberto Amato, Jeffrey Barrett, Sonia Goncalves, Ewan Harrison, David K. Jackson, Ian Johnston, Dominic Kwiatkowski, Cordelia Langford, John Sillitoe on behalf of the Wellcome Sanger Institute COVID-19 Surveillance Team |
| EPI_ISL_3366078, EPI_ISL_3366079 | Randox Laboratories | Wellcome Sanger Institute for the COVID-19 Genomics UK (COG-UK) Consortium | Randox Laboratories and Alex Alderton, Roberto Amato, Jeffrey Barrett, Sonia Goncalves, Ewan Harrison, David K. Jackson, Ian Johnston, Dominic Kwiatkowski, Cordelia Langford, John Sillitoe on behalf of the Wellcome Sanger Institute COVID-19 Surveillance Team |
| EPI_ISL_3366080 | Lighthouse Lab in Alderley Park | Wellcome Sanger Institute for the COVID-19 Genomics UK (COG-UK) | Jacquelyn Wynn, Mairead Hyland, The Lighthouse Lab in Alderley Park and Alex Alderton, Roberto Amato, Jeffrey Barrett, Sonia Goncalves, Ewan Harrison, David K. Jackson, Ian Johnston, Dominic Kwiatkowski, Cordelia Langford, John Sillitoe on |

| | | | |
|---|---|---|---|
| | | Consortium | behalf of the Wellcome Sanger Institute COVID-19 Surveillance Team |
| EPI_ISL_3366081, EPI_ISL_3366082 | Lighthouse Lab in Milton Keynes | Wellcome Sanger Institute for the COVID-19 Genomics UK (COG-UK) Consortium | The Lighthouse Lab in Milton Keynes and Alex Alderton, Roberto Amato, Jeffrey Barrett, Sonia Goncalves, Ewan Harrison, David K. Jackson, Ian Johnston, Dominic Kwiatkowski, Cordelia Langford, John Sillitoe on behalf of the Wellcome Sanger Institute COVID-19 Surveillance Team |
| EPI_ISL_3366083 | Randox Laboratories | Wellcome Sanger Institute for the COVID-19 Genomics UK (COG-UK) Consortium | Randox Laboratories and Alex Alderton, Roberto Amato, Jeffrey Barrett, Sonia Goncalves, Ewan Harrison, David K. Jackson, Ian Johnston, Dominic Kwiatkowski, Cordelia Langford, John Sillitoe on behalf of the Wellcome Sanger Institute COVID-19 Surveillance Team |
| EPI_ISL_3366084, EPI_ISL_3366085, EPI_ISL_3366086, EPI_ISL_3366087 | Lighthouse Lab in Milton Keynes | Wellcome Sanger Institute for the COVID-19 Genomics UK (COG-UK) Consortium | The Lighthouse Lab in Milton Keynes and Alex Alderton, Roberto Amato, Jeffrey Barrett, Sonia Goncalves, Ewan Harrison, David K. Jackson, Ian Johnston, Dominic Kwiatkowski, Cordelia Langford, John Sillitoe on behalf of the Wellcome Sanger Institute COVID-19 Surveillance Team |
| EPI_ISL_3366088 | Lighthouse Lab in Alderley Park | Wellcome Sanger Institute for the COVID-19 Genomics UK (COG-UK) Consortium | Jacquelyn Wynn, Mairead Hyland, The Lighthouse Lab in Alderley Park and Alex Alderton, Roberto Amato, Jeffrey Barrett, Sonia Goncalves, Ewan Harrison, David K. Jackson, Ian Johnston, Dominic Kwiatkowski, Cordelia Langford, John Sillitoe on behalf of the Wellcome Sanger Institute COVID-19 Surveillance Team |
| EPI_ISL_3366089, EPI_ISL_3366090, EPI_ISL_3366091 | Lighthouse Lab in Milton Keynes | Wellcome Sanger Institute for the COVID-19 Genomics UK (COG-UK) Consortium | The Lighthouse Lab in Milton Keynes and Alex Alderton, Roberto Amato, Jeffrey Barrett, Sonia Goncalves, Ewan Harrison, David K. Jackson, Ian Johnston, Dominic Kwiatkowski, Cordelia Langford, John Sillitoe on behalf of the Wellcome Sanger Institute COVID-19 Surveillance Team |
| EPI_ISL_3366092, EPI_ISL_3366093, EPI_ISL_3366094, EPI_ISL_3366095 | Lighthouse Lab in Alderley Park | Wellcome Sanger Institute for the COVID-19 Genomics UK (COG-UK) Consortium | Jacquelyn Wynn, Mairead Hyland, The Lighthouse Lab in Alderley Park and Alex Alderton, Roberto Amato, Jeffrey Barrett, Sonia Goncalves, Ewan Harrison, David K. Jackson, Ian Johnston, Dominic Kwiatkowski, Cordelia Langford, John Sillitoe on behalf of the Wellcome Sanger Institute COVID-19 Surveillance Team |
| EPI_ISL_3366096, EPI_ISL_3366097, EPI_ISL_3366098 | Lighthouse Lab in Milton Keynes | Wellcome Sanger Institute for the COVID-19 Genomics UK (COG-UK) Consortium | The Lighthouse Lab in Milton Keynes and Alex Alderton, Roberto Amato, Jeffrey Barrett, Sonia Goncalves, Ewan Harrison, David K. Jackson, Ian Johnston, Dominic Kwiatkowski, Cordelia Langford, John Sillitoe on behalf of the Wellcome Sanger Institute COVID-19 Surveillance Team |
| EPI_ISL_3366099 | Lighthouse Lab in Alderley Park | Wellcome Sanger Institute for the COVID-19 Genomics UK (COG-UK) Consortium | Jacquelyn Wynn, Mairead Hyland, The Lighthouse Lab in Alderley Park and Alex Alderton, Roberto Amato, Jeffrey Barrett, Sonia Goncalves, Ewan Harrison, David K. Jackson, Ian Johnston, Dominic Kwiatkowski, Cordelia Langford, John Sillitoe on behalf of the Wellcome Sanger Institute COVID-19 Surveillance Team |
| EPI_ISL_3366100 | Randox Laboratories | Wellcome Sanger Institute for the COVID-19 Genomics UK (COG-UK) Consortium | Randox Laboratories and Alex Alderton, Roberto Amato, Jeffrey Barrett, Sonia Goncalves, Ewan Harrison, David K. Jackson, Ian Johnston, Dominic Kwiatkowski, Cordelia Langford, John Sillitoe on behalf of the Wellcome Sanger Institute COVID-19 Surveillance Team |
| EPI_ISL_3366101, EPI_ISL_3366102, EPI_ISL_3366103 | Lighthouse Lab in Milton Keynes | Wellcome Sanger Institute for the COVID-19 Genomics UK (COG-UK) Consortium | The Lighthouse Lab in Milton Keynes and Alex Alderton, Roberto Amato, Jeffrey Barrett, Sonia Goncalves, Ewan Harrison, David K. Jackson, Ian Johnston, Dominic Kwiatkowski, Cordelia Langford, John Sillitoe on behalf of the Wellcome Sanger Institute COVID-19 Surveillance Team |
| EPI_ISL_3366104, EPI_ISL_3366105, EPI_ISL_3366106 | Lighthouse Lab in Alderley Park | Wellcome Sanger Institute for the COVID-19 Genomics UK (COG-UK) Consortium | Jacquelyn Wynn, Mairead Hyland, The Lighthouse Lab in Alderley Park and Alex Alderton, Roberto Amato, Jeffrey Barrett, Sonia Goncalves, Ewan Harrison, David K. Jackson, Ian Johnston, Dominic Kwiatkowski, Cordelia Langford, John Sillitoe on behalf of the Wellcome Sanger Institute COVID-19 Surveillance Team |
| EPI_ISL_3366107, EPI_ISL_3366108, EPI_ISL_3366109, EPI_ISL_3366110, EPI_ISL_3366111, EPI_ISL_3366112 | Lighthouse Lab in Milton Keynes | Wellcome Sanger Institute for the COVID-19 Genomics UK (COG-UK) Consortium | The Lighthouse Lab in Milton Keynes and Alex Alderton, Roberto Amato, Jeffrey Barrett, Sonia Goncalves, Ewan Harrison, David K. Jackson, Ian Johnston, Dominic Kwiatkowski, Cordelia Langford, John Sillitoe on behalf of the Wellcome Sanger Institute COVID-19 Surveillance Team |
| EPI_ISL_3366113 | Lighthouse Lab in Alderley Park | Wellcome Sanger Institute for the COVID-19 Genomics UK (COG-UK) Consortium | Jacquelyn Wynn, Mairead Hyland, The Lighthouse Lab in Alderley Park and Alex Alderton, Roberto Amato, Jeffrey Barrett, Sonia Goncalves, Ewan Harrison, David K. Jackson, Ian Johnston, Dominic Kwiatkowski, Cordelia Langford, John Sillitoe on behalf of the Wellcome Sanger Institute COVID-19 Surveillance Team |
| EPI_ISL_3366114, EPI_ISL_3366115, EPI_ISL_3366116 | Lighthouse Lab in Milton Keynes | Wellcome Sanger Institute for the COVID-19 Genomics UK (COG-UK) Consortium | The Lighthouse Lab in Milton Keynes and Alex Alderton, Roberto Amato, Jeffrey Barrett, Sonia Goncalves, Ewan Harrison, David K. Jackson, Ian Johnston, Dominic Kwiatkowski, Cordelia Langford, John Sillitoe on behalf of the Wellcome Sanger Institute COVID-19 Surveillance Team |
| EPI_ISL_3366117, EPI_ISL_3366118, EPI_ISL_3366119 | Lighthouse Lab in Alderley Park | Wellcome Sanger Institute for the COVID-19 Genomics UK (COG-UK) Consortium | Jacquelyn Wynn, Mairead Hyland, The Lighthouse Lab in Alderley Park and Alex Alderton, Roberto Amato, Jeffrey Barrett, Sonia Goncalves, Ewan Harrison, David K. Jackson, Ian Johnston, Dominic Kwiatkowski, Cordelia Langford, John Sillitoe on behalf of the Wellcome Sanger Institute COVID-19 Surveillance Team |
| EPI_ISL_3366120, EPI_ISL_3366121 | Lighthouse Lab in Milton Keynes | Wellcome Sanger Institute for the COVID-19 Genomics UK (COG-UK) Consortium | The Lighthouse Lab in Milton Keynes and Alex Alderton, Roberto Amato, Jeffrey Barrett, Sonia Goncalves, Ewan Harrison, David K. Jackson, Ian Johnston, Dominic Kwiatkowski, Cordelia Langford, John Sillitoe on behalf of the Wellcome Sanger Institute COVID-19 Surveillance Team |
| EPI_ISL_3366122 | Lighthouse Lab in Alderley Park | Wellcome Sanger Institute for the COVID-19 Genomics UK (COG-UK) Consortium | Jacquelyn Wynn, Mairead Hyland, The Lighthouse Lab in Alderley Park and Alex Alderton, Roberto Amato, Jeffrey Barrett, Sonia Goncalves, Ewan Harrison, David K. Jackson, Ian Johnston, Dominic Kwiatkowski, Cordelia Langford, John Sillitoe on behalf of the Wellcome Sanger Institute COVID-19 Surveillance Team |
| EPI_ISL_3366123 | Randox Laboratories | Wellcome Sanger Institute for the COVID-19 Genomics UK (COG-UK) Consortium | Randox Laboratories and Alex Alderton, Roberto Amato, Jeffrey Barrett, Sonia Goncalves, Ewan Harrison, David K. Jackson, Ian Johnston, Dominic Kwiatkowski, Cordelia Langford, John Sillitoe on behalf of the Wellcome Sanger Institute COVID-19 Surveillance Team |
| EPI_ISL_3366124 | Lighthouse Lab in Milton Keynes | Wellcome Sanger Institute for the COVID-19 Genomics UK (COG-UK) Consortium | The Lighthouse Lab in Milton Keynes and Alex Alderton, Roberto Amato, Jeffrey Barrett, Sonia Goncalves, Ewan Harrison, David K. Jackson, Ian Johnston, Dominic Kwiatkowski, Cordelia Langford, John Sillitoe on behalf of the Wellcome Sanger Institute COVID-19 Surveillance Team |
| EPI_ISL_3366125 | Lighthouse Lab in Alderley Park | Wellcome Sanger Institute for the COVID-19 Genomics UK (COG-UK) Consortium | Jacquelyn Wynn, Mairead Hyland, The Lighthouse Lab in Alderley Park and Alex Alderton, Roberto Amato, Jeffrey Barrett, Sonia Goncalves, Ewan Harrison, David K. Jackson, Ian Johnston, Dominic Kwiatkowski, Cordelia Langford, John Sillitoe on behalf of the Wellcome Sanger Institute COVID-19 Surveillance Team |
| EPI_ISL_3366126, EPI_ISL_3366127 | Randox Laboratories | Wellcome Sanger Institute for the COVID-19 Genomics UK (COG-UK) Consortium | Randox Laboratories and Alex Alderton, Roberto Amato, Jeffrey Barrett, Sonia Goncalves, Ewan Harrison, David K. Jackson, Ian Johnston, Dominic Kwiatkowski, Cordelia Langford, John Sillitoe on behalf of the Wellcome Sanger Institute COVID-19 Surveillance Team |
| EPI_ISL_3366128, EPI_ISL_3366129 | Lighthouse Lab in Milton Keynes | Wellcome Sanger Institute for the COVID-19 Genomics UK (COG-UK) Consortium | The Lighthouse Lab in Milton Keynes and Alex Alderton, Roberto Amato, Jeffrey Barrett, Sonia Goncalves, Ewan Harrison, David K. Jackson, Ian Johnston, Dominic Kwiatkowski, Cordelia Langford, John Sillitoe on behalf of the Wellcome Sanger Institute COVID-19 Surveillance Team |
| EPI_ISL_3366130 | Randox Laboratories | Wellcome Sanger Institute for the COVID-19 Genomics UK (COG-UK) Consortium | Randox Laboratories and Alex Alderton, Roberto Amato, Jeffrey Barrett, Sonia Goncalves, Ewan Harrison, David K. Jackson, Ian Johnston, Dominic Kwiatkowski, Cordelia Langford, John Sillitoe on behalf of the Wellcome Sanger Institute COVID-19 Surveillance Team |
| EPI_ISL_3366131, EPI_ISL_3366132, EPI_ISL_3366133 | Lighthouse Lab in Milton Keynes | Wellcome Sanger Institute for the COVID-19 Genomics UK (COG-UK) Consortium | The Lighthouse Lab in Milton Keynes and Alex Alderton, Roberto Amato, Jeffrey Barrett, Sonia Goncalves, Ewan Harrison, David K. Jackson, Ian Johnston, Dominic Kwiatkowski, Cordelia Langford, John Sillitoe on behalf of the Wellcome Sanger Institute COVID-19 Surveillance Team |
| EPI_ISL_3366134, EPI_ISL_3366135 | Lighthouse Lab in Alderley Park | Wellcome Sanger Institute for the COVID-19 Genomics UK (COG-UK) Consortium | Jacquelyn Wynn, Mairead Hyland, The Lighthouse Lab in Alderley Park and Alex Alderton, Roberto Amato, Jeffrey Barrett, Sonia Goncalves, Ewan Harrison, David K. Jackson, Ian Johnston, Dominic Kwiatkowski, Cordelia Langford, John Sillitoe on behalf of the Wellcome Sanger Institute COVID-19 Surveillance Team |
| EPI_ISL_3366136 | Lighthouse Lab in Milton Keynes | Wellcome Sanger Institute for the COVID-19 Genomics UK (COG-UK) Consortium | The Lighthouse Lab in Milton Keynes and Alex Alderton, Roberto Amato, Jeffrey Barrett, Sonia Goncalves, Ewan Harrison, David K. Jackson, Ian Johnston, Dominic Kwiatkowski, Cordelia Langford, John Sillitoe on behalf of the Wellcome Sanger Institute COVID-19 Surveillance Team |
| EPI_ISL_3366137 | Lighthouse Lab in Alderley Park | Wellcome Sanger Institute for the COVID-19 Genomics UK (COG-UK) Consortium | Jacquelyn Wynn, Mairead Hyland, The Lighthouse Lab in Alderley Park and Alex Alderton, Roberto Amato, Jeffrey Barrett, Sonia Goncalves, Ewan Harrison, David K. Jackson, Ian Johnston, Dominic Kwiatkowski, Cordelia Langford, John Sillitoe on behalf of the Wellcome Sanger Institute COVID-19 Surveillance Team |
| EPI_ISL_3366138, EPI_ISL_3366139, EPI_ISL_3366140, EPI_ISL_3366141, EPI_ISL_3366142, EPI_ISL_3366143 | Lighthouse Lab in Milton Keynes | Wellcome Sanger Institute for the COVID-19 Genomics UK (COG-UK) Consortium | The Lighthouse Lab in Milton Keynes and Alex Alderton, Roberto Amato, Jeffrey Barrett, Sonia Goncalves, Ewan Harrison, David K. Jackson, Ian Johnston, Dominic Kwiatkowski, Cordelia Langford, John Sillitoe on behalf of the Wellcome Sanger Institute COVID-19 Surveillance Team |
| EPI_ISL_3366144, EPI_ISL_3366145 | Lighthouse Lab in Alderley Park | Wellcome Sanger Institute for the COVID-19 Genomics UK (COG-UK) Consortium | Jacquelyn Wynn, Mairead Hyland, The Lighthouse Lab in Alderley Park and Alex Alderton, Roberto Amato, Jeffrey Barrett, Sonia Goncalves, Ewan Harrison, David K. Jackson, Ian Johnston, Dominic Kwiatkowski, Cordelia Langford, John Sillitoe on behalf of the Wellcome Sanger Institute COVID-19 Surveillance Team |
| EPI_ISL_3366146 | Randox Laboratories | Wellcome Sanger Institute for the COVID-19 Genomics UK (COG-UK) Consortium | Randox Laboratories and Alex Alderton, Roberto Amato, Jeffrey Barrett, Sonia Goncalves, Ewan Harrison, David K. Jackson, Ian Johnston, Dominic Kwiatkowski, Cordelia Langford, John Sillitoe on behalf of the Wellcome Sanger Institute COVID-19 Surveillance Team |
| EPI_ISL_3366147 | Lighthouse Lab in Milton Keynes | Wellcome Sanger Institute for the COVID-19 Genomics UK (COG-UK) Consortium | The Lighthouse Lab in Milton Keynes and Alex Alderton, Roberto Amato, Jeffrey Barrett, Sonia Goncalves, Ewan Harrison, David K. Jackson, Ian Johnston, Dominic Kwiatkowski, Cordelia Langford, John Sillitoe on behalf of the Wellcome Sanger Institute COVID-19 Surveillance Team |
| EPI_ISL_3366148, EPI_ISL_3366149 | Randox Laboratories | Wellcome Sanger Institute for the COVID-19 Genomics UK (COG-UK) Consortium | Randox Laboratories and Alex Alderton, Roberto Amato, Jeffrey Barrett, Sonia Goncalves, Ewan Harrison, David K. Jackson, Ian Johnston, Dominic Kwiatkowski, Cordelia Langford, John Sillitoe on behalf of the Wellcome Sanger Institute COVID-19 Surveillance Team |
| EPI_ISL_3366150, EPI_ISL_3366151, EPI_ISL_3366152, EPI_ISL_3366153 | Lighthouse Lab in Milton Keynes | Wellcome Sanger Institute for the COVID-19 Genomics UK (COG-UK) Consortium | The Lighthouse Lab in Milton Keynes and Alex Alderton, Roberto Amato, Jeffrey Barrett, Sonia Goncalves, Ewan Harrison, David K. Jackson, Ian Johnston, Dominic Kwiatkowski, Cordelia Langford, John Sillitoe on behalf of the Wellcome Sanger Institute COVID-19 Surveillance Team |
| EPI_ISL_3366154, EPI_ISL_3366155 | Lighthouse Lab in Alderley Park | Wellcome Sanger Institute for the COVID-19 Genomics UK (COG-UK) Consortium | Jacquelyn Wynn, Mairead Hyland, The Lighthouse Lab in Alderley Park and Alex Alderton, Roberto Amato, Jeffrey Barrett, Sonia Goncalves, Ewan Harrison, David K. Jackson, Ian Johnston, Dominic Kwiatkowski, Cordelia Langford, John Sillitoe on behalf of the Wellcome Sanger Institute COVID-19 Surveillance Team |
| EPI_ISL_3366156 | Lighthouse Lab in Milton Keynes | Wellcome Sanger Institute for the COVID-19 Genomics UK (COG-UK) Consortium | The Lighthouse Lab in Milton Keynes and Alex Alderton, Roberto Amato, Jeffrey Barrett, Sonia Goncalves, Ewan Harrison, David K. Jackson, Ian Johnston, Dominic Kwiatkowski, Cordelia Langford, John Sillitoe on behalf of the Wellcome Sanger Institute COVID-19 Surveillance Team |
| EPI_ISL_3366157 | Randox Laboratories | Wellcome Sanger Institute for the COVID-19 Genomics UK (COG-UK) | Randox Laboratories and Alex Alderton, Roberto Amato, Jeffrey Barrett, Sonia Goncalves, Ewan Harrison, David K. Jackson, Ian Johnston, Dominic Kwiatkowski, Cordelia Langford, John Sillitoe on behalf of the Wellcome Sanger Institute COVID- |

| | | | |
|---|---|---|---|
| | | Consortium | 19 Surveillance Team |
| EPI_ISL_3366158, EPI_ISL_3366159 | Lighthouse Lab in Alderley Park | Wellcome Sanger Institute for the COVID-19 Genomics UK (COG-UK) Consortium | Jacquelyn Wynn, Mairead Hyland, The Lighthouse Lab in Alderley Park and Alex Alderton, Roberto Amato, Jeffrey Barrett, Sonia Goncalves, Ewan Harrison, David K. Jackson, Ian Johnston, Dominic Kwiatkowski, Cordelia Langford, John Sillitoe on behalf of the Wellcome Sanger Institute COVID-19 Surveillance Team |
| EPI_ISL_3366160, EPI_ISL_3366161, EPI_ISL_3366162 | Lighthouse Lab in Milton Keynes | Wellcome Sanger Institute for the COVID-19 Genomics UK (COG-UK) Consortium | The Lighthouse Lab in Milton Keynes and Alex Alderton, Roberto Amato, Jeffrey Barrett, Sonia Goncalves, Ewan Harrison, David K. Jackson, Ian Johnston, Dominic Kwiatkowski, Cordelia Langford, John Sillitoe on behalf of the Wellcome Sanger Institute COVID-19 Surveillance Team |
| EPI_ISL_3366163 | Lighthouse Lab in Alderley Park | Wellcome Sanger Institute for the COVID-19 Genomics UK (COG-UK) Consortium | Jacquelyn Wynn, Mairead Hyland, The Lighthouse Lab in Alderley Park and Alex Alderton, Roberto Amato, Jeffrey Barrett, Sonia Goncalves, Ewan Harrison, David K. Jackson, Ian Johnston, Dominic Kwiatkowski, Cordelia Langford, John Sillitoe on behalf of the Wellcome Sanger Institute COVID-19 Surveillance Team |
| EPI_ISL_3366164, EPI_ISL_3366165, EPI_ISL_3366166, EPI_ISL_3366167 | Lighthouse Lab in Milton Keynes | Wellcome Sanger Institute for the COVID-19 Genomics UK (COG-UK) Consortium | The Lighthouse Lab in Milton Keynes and Alex Alderton, Roberto Amato, Jeffrey Barrett, Sonia Goncalves, Ewan Harrison, David K. Jackson, Ian Johnston, Dominic Kwiatkowski, Cordelia Langford, John Sillitoe on behalf of the Wellcome Sanger Institute COVID-19 Surveillance Team |
| EPI_ISL_3366168, EPI_ISL_3366169 | Lighthouse Lab in Alderley Park | Wellcome Sanger Institute for the COVID-19 Genomics UK (COG-UK) Consortium | Jacquelyn Wynn, Mairead Hyland, The Lighthouse Lab in Alderley Park and Alex Alderton, Roberto Amato, Jeffrey Barrett, Sonia Goncalves, Ewan Harrison, David K. Jackson, Ian Johnston, Dominic Kwiatkowski, Cordelia Langford, John Sillitoe on behalf of the Wellcome Sanger Institute COVID-19 Surveillance Team |
| EPI_ISL_3366170 | Randox Laboratories | Wellcome Sanger Institute for the COVID-19 Genomics UK (COG-UK) Consortium | Randox Laboratories and Alex Alderton, Roberto Amato, Jeffrey Barrett, Sonia Goncalves, Ewan Harrison, David K. Jackson, Ian Johnston, Dominic Kwiatkowski, Cordelia Langford, John Sillitoe on behalf of the Wellcome Sanger Institute COVID-19 Surveillance Team |
| EPI_ISL_3366171, EPI_ISL_3366172, EPI_ISL_3366173 | Lighthouse Lab in Milton Keynes | Wellcome Sanger Institute for the COVID-19 Genomics UK (COG-UK) Consortium | The Lighthouse Lab in Milton Keynes and Alex Alderton, Roberto Amato, Jeffrey Barrett, Sonia Goncalves, Ewan Harrison, David K. Jackson, Ian Johnston, Dominic Kwiatkowski, Cordelia Langford, John Sillitoe on behalf of the Wellcome Sanger Institute COVID-19 Surveillance Team |
| EPI_ISL_3366174 | Randox Laboratories | Wellcome Sanger Institute for the COVID-19 Genomics UK (COG-UK) Consortium | Randox Laboratories and Alex Alderton, Roberto Amato, Jeffrey Barrett, Sonia Goncalves, Ewan Harrison, David K. Jackson, Ian Johnston, Dominic Kwiatkowski, Cordelia Langford, John Sillitoe on behalf of the Wellcome Sanger Institute COVID-19 Surveillance Team |
| EPI_ISL_3366175 | Lighthouse Lab in Milton Keynes | Wellcome Sanger Institute for the COVID-19 Genomics UK (COG-UK) Consortium | The Lighthouse Lab in Milton Keynes and Alex Alderton, Roberto Amato, Jeffrey Barrett, Sonia Goncalves, Ewan Harrison, David K. Jackson, Ian Johnston, Dominic Kwiatkowski, Cordelia Langford, John Sillitoe on behalf of the Wellcome Sanger Institute COVID-19 Surveillance Team |
| EPI_ISL_3366176 | Lighthouse Lab in Alderley Park | Wellcome Sanger Institute for the COVID-19 Genomics UK (COG-UK) Consortium | Jacquelyn Wynn, Mairead Hyland, The Lighthouse Lab in Alderley Park and Alex Alderton, Roberto Amato, Jeffrey Barrett, Sonia Goncalves, Ewan Harrison, David K. Jackson, Ian Johnston, Dominic Kwiatkowski, Cordelia Langford, John Sillitoe on behalf of the Wellcome Sanger Institute COVID-19 Surveillance Team |
| EPI_ISL_3366177, EPI_ISL_3366178 | Lighthouse Lab in Milton Keynes | Wellcome Sanger Institute for the COVID-19 Genomics UK (COG-UK) Consortium | The Lighthouse Lab in Milton Keynes and Alex Alderton, Roberto Amato, Jeffrey Barrett, Sonia Goncalves, Ewan Harrison, David K. Jackson, Ian Johnston, Dominic Kwiatkowski, Cordelia Langford, John Sillitoe on behalf of the Wellcome Sanger Institute COVID-19 Surveillance Team |
| EPI_ISL_3366179 | Randox Laboratories | Wellcome Sanger Institute for the COVID-19 Genomics UK (COG-UK) Consortium | Randox Laboratories and Alex Alderton, Roberto Amato, Jeffrey Barrett, Sonia Goncalves, Ewan Harrison, David K. Jackson, Ian Johnston, Dominic Kwiatkowski, Cordelia Langford, John Sillitoe on behalf of the Wellcome Sanger Institute COVID-19 Surveillance Team |
| EPI_ISL_3366180, EPI_ISL_3366181 | Lighthouse Lab in Alderley Park | Wellcome Sanger Institute for the COVID-19 Genomics UK (COG-UK) Consortium | Jacquelyn Wynn, Mairead Hyland, The Lighthouse Lab in Alderley Park and Alex Alderton, Roberto Amato, Jeffrey Barrett, Sonia Goncalves, Ewan Harrison, David K. Jackson, Ian Johnston, Dominic Kwiatkowski, Cordelia Langford, John Sillitoe on behalf of the Wellcome Sanger Institute COVID-19 Surveillance Team |
| EPI_ISL_3366182 | Lighthouse Lab in Milton Keynes | Wellcome Sanger Institute for the COVID-19 Genomics UK (COG-UK) Consortium | The Lighthouse Lab in Milton Keynes and Alex Alderton, Roberto Amato, Jeffrey Barrett, Sonia Goncalves, Ewan Harrison, David K. Jackson, Ian Johnston, Dominic Kwiatkowski, Cordelia Langford, John Sillitoe on behalf of the Wellcome Sanger Institute COVID-19 Surveillance Team |
| EPI_ISL_3366183, EPI_ISL_3366184 | Lighthouse Lab in Alderley Park | Wellcome Sanger Institute for the COVID-19 Genomics UK (COG-UK) Consortium | Jacquelyn Wynn, Mairead Hyland, The Lighthouse Lab in Alderley Park and Alex Alderton, Roberto Amato, Jeffrey Barrett, Sonia Goncalves, Ewan Harrison, David K. Jackson, Ian Johnston, Dominic Kwiatkowski, Cordelia Langford, John Sillitoe on behalf of the Wellcome Sanger Institute COVID-19 Surveillance Team |
| EPI_ISL_3366185 | Randox Laboratories | Wellcome Sanger Institute for the COVID-19 Genomics UK (COG-UK) Consortium | Randox Laboratories and Alex Alderton, Roberto Amato, Jeffrey Barrett, Sonia Goncalves, Ewan Harrison, David K. Jackson, Ian Johnston, Dominic Kwiatkowski, Cordelia Langford, John Sillitoe on behalf of the Wellcome Sanger Institute COVID-19 Surveillance Team |
| EPI_ISL_3366186 | Lighthouse Lab in Alderley Park | Wellcome Sanger Institute for the COVID-19 Genomics UK (COG-UK) Consortium | Jacquelyn Wynn, Mairead Hyland, The Lighthouse Lab in Alderley Park and Alex Alderton, Roberto Amato, Jeffrey Barrett, Sonia Goncalves, Ewan Harrison, David K. Jackson, Ian Johnston, Dominic Kwiatkowski, Cordelia Langford, John Sillitoe on behalf of the Wellcome Sanger Institute COVID-19 Surveillance Team |
| EPI_ISL_3366187, EPI_ISL_3366188, EPI_ISL_3366189, EPI_ISL_3366190 | Lighthouse Lab in Milton Keynes | Wellcome Sanger Institute for the COVID-19 Genomics UK (COG-UK) Consortium | The Lighthouse Lab in Milton Keynes and Alex Alderton, Roberto Amato, Jeffrey Barrett, Sonia Goncalves, Ewan Harrison, David K. Jackson, Ian Johnston, Dominic Kwiatkowski, Cordelia Langford, John Sillitoe on behalf of the Wellcome Sanger Institute COVID-19 Surveillance Team |
| EPI_ISL_3366191 | Randox Laboratories | Wellcome Sanger Institute for the COVID-19 Genomics UK (COG-UK) Consortium | Randox Laboratories and Alex Alderton, Roberto Amato, Jeffrey Barrett, Sonia Goncalves, Ewan Harrison, David K. Jackson, Ian Johnston, Dominic Kwiatkowski, Cordelia Langford, John Sillitoe on behalf of the Wellcome Sanger Institute COVID-19 Surveillance Team |
| EPI_ISL_3366192, EPI_ISL_3366193 | Lighthouse Lab in Milton Keynes | Wellcome Sanger Institute for the COVID-19 Genomics UK (COG-UK) Consortium | The Lighthouse Lab in Milton Keynes and Alex Alderton, Roberto Amato, Jeffrey Barrett, Sonia Goncalves, Ewan Harrison, David K. Jackson, Ian Johnston, Dominic Kwiatkowski, Cordelia Langford, John Sillitoe on behalf of the Wellcome Sanger Institute COVID-19 Surveillance Team |
| EPI_ISL_3366194, EPI_ISL_3366195 | Lighthouse Lab in Alderley Park | Wellcome Sanger Institute for the COVID-19 Genomics UK (COG-UK) Consortium | Jacquelyn Wynn, Mairead Hyland, The Lighthouse Lab in Alderley Park and Alex Alderton, Roberto Amato, Jeffrey Barrett, Sonia Goncalves, Ewan Harrison, David K. Jackson, Ian Johnston, Dominic Kwiatkowski, Cordelia Langford, John Sillitoe on behalf of the Wellcome Sanger Institute COVID-19 Surveillance Team |
| EPI_ISL_3366196, EPI_ISL_3366197 | Lighthouse Lab in Milton Keynes | Wellcome Sanger Institute for the COVID-19 Genomics UK (COG-UK) Consortium | The Lighthouse Lab in Milton Keynes and Alex Alderton, Roberto Amato, Jeffrey Barrett, Sonia Goncalves, Ewan Harrison, David K. Jackson, Ian Johnston, Dominic Kwiatkowski, Cordelia Langford, John Sillitoe on behalf of the Wellcome Sanger Institute COVID-19 Surveillance Team |
| EPI_ISL_3366198, EPI_ISL_3366199, EPI_ISL_3366200 | Lighthouse Lab in Alderley Park | Wellcome Sanger Institute for the COVID-19 Genomics UK (COG-UK) Consortium | Jacquelyn Wynn, Mairead Hyland, The Lighthouse Lab in Alderley Park and Alex Alderton, Roberto Amato, Jeffrey Barrett, Sonia Goncalves, Ewan Harrison, David K. Jackson, Ian Johnston, Dominic Kwiatkowski, Cordelia Langford, John Sillitoe on behalf of the Wellcome Sanger Institute COVID-19 Surveillance Team |
| EPI_ISL_3366201 | Randox Laboratories | Wellcome Sanger Institute for the COVID-19 Genomics UK (COG-UK) Consortium | Randox Laboratories and Alex Alderton, Roberto Amato, Jeffrey Barrett, Sonia Goncalves, Ewan Harrison, David K. Jackson, Ian Johnston, Dominic Kwiatkowski, Cordelia Langford, John Sillitoe on behalf of the Wellcome Sanger Institute COVID-19 Surveillance Team |
| EPI_ISL_3366202, EPI_ISL_3366203 | Lighthouse Lab in Milton Keynes | Wellcome Sanger Institute for the COVID-19 Genomics UK (COG-UK) Consortium | The Lighthouse Lab in Milton Keynes and Alex Alderton, Roberto Amato, Jeffrey Barrett, Sonia Goncalves, Ewan Harrison, David K. Jackson, Ian Johnston, Dominic Kwiatkowski, Cordelia Langford, John Sillitoe on behalf of the Wellcome Sanger Institute COVID-19 Surveillance Team |
| EPI_ISL_3366204 | Lighthouse Lab in Alderley Park | Wellcome Sanger Institute for the COVID-19 Genomics UK (COG-UK) Consortium | Jacquelyn Wynn, Mairead Hyland, The Lighthouse Lab in Alderley Park and Alex Alderton, Roberto Amato, Jeffrey Barrett, Sonia Goncalves, Ewan Harrison, David K. Jackson, Ian Johnston, Dominic Kwiatkowski, Cordelia Langford, John Sillitoe on behalf of the Wellcome Sanger Institute COVID-19 Surveillance Team |
| EPI_ISL_3366205 | Lighthouse Lab in Milton Keynes | Wellcome Sanger Institute for the COVID-19 Genomics UK (COG-UK) Consortium | The Lighthouse Lab in Milton Keynes and Alex Alderton, Roberto Amato, Jeffrey Barrett, Sonia Goncalves, Ewan Harrison, David K. Jackson, Ian Johnston, Dominic Kwiatkowski, Cordelia Langford, John Sillitoe on behalf of the Wellcome Sanger Institute COVID-19 Surveillance Team |
| EPI_ISL_3366206 | Lighthouse Lab in Alderley Park | Wellcome Sanger Institute for the COVID-19 Genomics UK (COG-UK) Consortium | Jacquelyn Wynn, Mairead Hyland, The Lighthouse Lab in Alderley Park and Alex Alderton, Roberto Amato, Jeffrey Barrett, Sonia Goncalves, Ewan Harrison, David K. Jackson, Ian Johnston, Dominic Kwiatkowski, Cordelia Langford, John Sillitoe on behalf of the Wellcome Sanger Institute COVID-19 Surveillance Team |
| EPI_ISL_3366207, EPI_ISL_3366208 | Lighthouse Lab in Milton Keynes | Wellcome Sanger Institute for the COVID-19 Genomics UK (COG-UK) Consortium | The Lighthouse Lab in Milton Keynes and Alex Alderton, Roberto Amato, Jeffrey Barrett, Sonia Goncalves, Ewan Harrison, David K. Jackson, Ian Johnston, Dominic Kwiatkowski, Cordelia Langford, John Sillitoe on behalf of the Wellcome Sanger Institute COVID-19 Surveillance Team |
| EPI_ISL_3366209 | Lighthouse Lab in Alderley Park | Wellcome Sanger Institute for the COVID-19 Genomics UK (COG-UK) Consortium | Jacquelyn Wynn, Mairead Hyland, The Lighthouse Lab in Alderley Park and Alex Alderton, Roberto Amato, Jeffrey Barrett, Sonia Goncalves, Ewan Harrison, David K. Jackson, Ian Johnston, Dominic Kwiatkowski, Cordelia Langford, John Sillitoe on behalf of the Wellcome Sanger Institute COVID-19 Surveillance Team |
| EPI_ISL_3366210, EPI_ISL_3366211, EPI_ISL_3366212 | Lighthouse Lab in Milton Keynes | Wellcome Sanger Institute for the COVID-19 Genomics UK (COG-UK) Consortium | The Lighthouse Lab in Milton Keynes and Alex Alderton, Roberto Amato, Jeffrey Barrett, Sonia Goncalves, Ewan Harrison, David K. Jackson, Ian Johnston, Dominic Kwiatkowski, Cordelia Langford, John Sillitoe on behalf of the Wellcome Sanger Institute COVID-19 Surveillance Team |
| EPI_ISL_3366213 | Lighthouse Lab in Alderley Park | Wellcome Sanger Institute for the COVID-19 Genomics UK (COG-UK) Consortium | Jacquelyn Wynn, Mairead Hyland, The Lighthouse Lab in Alderley Park and Alex Alderton, Roberto Amato, Jeffrey Barrett, Sonia Goncalves, Ewan Harrison, David K. Jackson, Ian Johnston, Dominic Kwiatkowski, Cordelia Langford, John Sillitoe on behalf of the Wellcome Sanger Institute COVID-19 Surveillance Team |
| EPI_ISL_3366214, EPI_ISL_3366215 | Lighthouse Lab in Milton Keynes | Wellcome Sanger Institute for the COVID-19 Genomics UK (COG-UK) Consortium | The Lighthouse Lab in Milton Keynes and Alex Alderton, Roberto Amato, Jeffrey Barrett, Sonia Goncalves, Ewan Harrison, David K. Jackson, Ian Johnston, Dominic Kwiatkowski, Cordelia Langford, John Sillitoe on behalf of the Wellcome Sanger Institute COVID-19 Surveillance Team |
| EPI_ISL_3366216, EPI_ISL_3366217 | Lighthouse Lab in Alderley Park | Wellcome Sanger Institute for the COVID-19 Genomics UK (COG-UK) Consortium | Jacquelyn Wynn, Mairead Hyland, The Lighthouse Lab in Alderley Park and Alex Alderton, Roberto Amato, Jeffrey Barrett, Sonia Goncalves, Ewan Harrison, David K. Jackson, Ian Johnston, Dominic Kwiatkowski, Cordelia Langford, John Sillitoe on behalf of the Wellcome Sanger Institute COVID-19 Surveillance Team |
| EPI_ISL_3366218 | Lighthouse Lab in Milton Keynes | Wellcome Sanger Institute for the COVID-19 Genomics UK (COG-UK) Consortium | The Lighthouse Lab in Milton Keynes and Alex Alderton, Roberto Amato, Jeffrey Barrett, Sonia Goncalves, Ewan Harrison, David K. Jackson, Ian Johnston, Dominic Kwiatkowski, Cordelia Langford, John Sillitoe on behalf of the Wellcome Sanger Institute COVID-19 Surveillance Team |
| EPI_ISL_3366219, EPI_ISL_3366220 | Randox Laboratories | Wellcome Sanger Institute for the COVID-19 Genomics UK (COG-UK) Consortium | Randox Laboratories and Alex Alderton, Roberto Amato, Jeffrey Barrett, Sonia Goncalves, Ewan Harrison, David K. Jackson, Ian Johnston, Dominic Kwiatkowski, Cordelia Langford, John Sillitoe on behalf of the Wellcome Sanger Institute COVID-19 Surveillance Team |
| EPI_ISL_3366221, EPI_ISL_3366222 | Lighthouse Lab in Milton Keynes | Wellcome Sanger Institute for the COVID-19 Genomics UK (COG-UK) Consortium | The Lighthouse Lab in Milton Keynes and Alex Alderton, Roberto Amato, Jeffrey Barrett, Sonia Goncalves, Ewan Harrison, David K. Jackson, Ian Johnston, Dominic Kwiatkowski, Cordelia Langford, John Sillitoe on behalf of the Wellcome Sanger Institute COVID-19 Surveillance Team |
| EPI_ISL_3366223 | Randox Laboratories | Wellcome Sanger Institute for the COVID-19 Genomics UK (COG-UK) Consortium | Randox Laboratories and Alex Alderton, Roberto Amato, Jeffrey Barrett, Sonia Goncalves, Ewan Harrison, David K. Jackson, Ian Johnston, Dominic Kwiatkowski, Cordelia Langford, John Sillitoe on behalf of the Wellcome Sanger Institute COVID-19 Surveillance Team |
| EPI_ISL_3366224, EPI_ISL_3366225, EPI_ISL_3366226 | Lighthouse Lab in Milton Keynes | Wellcome Sanger Institute for the COVID-19 Genomics UK (COG-UK) Consortium | The Lighthouse Lab in Milton Keynes and Alex Alderton, Roberto Amato, Jeffrey Barrett, Sonia Goncalves, Ewan Harrison, David K. Jackson, Ian Johnston, Dominic Kwiatkowski, Cordelia Langford, John Sillitoe on behalf of the Wellcome Sanger Institute COVID-19 Surveillance Team |
| EPI_ISL_3366227 | Lighthouse Lab in Alderley Park | Wellcome Sanger Institute for the COVID-19 Genomics UK (COG-UK) Consortium | Jacquelyn Wynn, Mairead Hyland, The Lighthouse Lab in Alderley Park and Alex Alderton, Roberto Amato, Jeffrey Barrett, Sonia Goncalves, Ewan Harrison, David K. Jackson, Ian Johnston, Dominic Kwiatkowski, Cordelia Langford, John Sillitoe on behalf of the Wellcome Sanger Institute COVID-19 Surveillance Team |
| EPI_ISL_3366228 | Randox Laboratories | Wellcome Sanger Institute for the COVID-19 Genomics UK (COG-UK) Consortium | Randox Laboratories and Alex Alderton, Roberto Amato, Jeffrey Barrett, Sonia Goncalves, Ewan Harrison, David K. Jackson, Ian Johnston, Dominic Kwiatkowski, Cordelia Langford, John Sillitoe on behalf of the Wellcome Sanger Institute COVID-19 Surveillance Team |

| | | | |
|---|---|---|---|
| EPI_ISL_3366229, EPI_ISL_3366230 | Lighthouse Lab in Alderley Park | Wellcome Sanger Institute for the COVID-19 Genomics UK (COG-UK) Consortium | Jacquelyn Wynn, Mairead Hyland, The Lighthouse Lab in Alderley Park and Alex Alderton, Roberto Amato, Jeffrey Barrett, Sonia Goncalves, Ewan Harrison, David K. Jackson, Ian Johnston, Dominic Kwiatkowski, Cordelia Langford, John Sillitoe on behalf of the Wellcome Sanger Institute COVID-19 Surveillance Team |
| EPI_ISL_3366231, EPI_ISL_3366232, EPI_ISL_3366233 | Lighthouse Lab in Milton Keynes | Wellcome Sanger Institute for the COVID-19 Genomics UK (COG-UK) Consortium | The Lighthouse Lab in Milton Keynes and Alex Alderton, Roberto Amato, Jeffrey Barrett, Sonia Goncalves, Ewan Harrison, David K. Jackson, Ian Johnston, Dominic Kwiatkowski, Cordelia Langford, John Sillitoe on behalf of the Wellcome Sanger Institute COVID-19 Surveillance Team |
| EPI_ISL_3366234 | Lighthouse Lab in Alderley Park | Wellcome Sanger Institute for the COVID-19 Genomics UK (COG-UK) Consortium | Jacquelyn Wynn, Mairead Hyland, The Lighthouse Lab in Alderley Park and Alex Alderton, Roberto Amato, Jeffrey Barrett, Sonia Goncalves, Ewan Harrison, David K. Jackson, Ian Johnston, Dominic Kwiatkowski, Cordelia Langford, John Sillitoe on behalf of the Wellcome Sanger Institute COVID-19 Surveillance Team |
| EPI_ISL_3366235, EPI_ISL_3366236, EPI_ISL_3366237, EPI_ISL_3366238 | Lighthouse Lab in Milton Keynes | Wellcome Sanger Institute for the COVID-19 Genomics UK (COG-UK) Consortium | The Lighthouse Lab in Milton Keynes and Alex Alderton, Roberto Amato, Jeffrey Barrett, Sonia Goncalves, Ewan Harrison, David K. Jackson, Ian Johnston, Dominic Kwiatkowski, Cordelia Langford, John Sillitoe on behalf of the Wellcome Sanger Institute COVID-19 Surveillance Team |
| EPI_ISL_3366239 | Randox Laboratories | Wellcome Sanger Institute for the COVID-19 Genomics UK (COG-UK) Consortium | Randox Laboratories and Alex Alderton, Roberto Amato, Jeffrey Barrett, Sonia Goncalves, Ewan Harrison, David K. Jackson, Ian Johnston, Dominic Kwiatkowski, Cordelia Langford, John Sillitoe on behalf of the Wellcome Sanger Institute COVID-19 Surveillance Team |
| EPI_ISL_3366240, EPI_ISL_3366241, EPI_ISL_3366242 | Lighthouse Lab in Milton Keynes | Wellcome Sanger Institute for the COVID-19 Genomics UK (COG-UK) Consortium | The Lighthouse Lab in Milton Keynes and Alex Alderton, Roberto Amato, Jeffrey Barrett, Sonia Goncalves, Ewan Harrison, David K. Jackson, Ian Johnston, Dominic Kwiatkowski, Cordelia Langford, John Sillitoe on behalf of the Wellcome Sanger Institute COVID-19 Surveillance Team |
| EPI_ISL_3366243 | Lighthouse Lab in Alderley Park | Wellcome Sanger Institute for the COVID-19 Genomics UK (COG-UK) Consortium | Jacquelyn Wynn, Mairead Hyland, The Lighthouse Lab in Alderley Park and Alex Alderton, Roberto Amato, Jeffrey Barrett, Sonia Goncalves, Ewan Harrison, David K. Jackson, Ian Johnston, Dominic Kwiatkowski, Cordelia Langford, John Sillitoe on behalf of the Wellcome Sanger Institute COVID-19 Surveillance Team |
| EPI_ISL_3366244, EPI_ISL_3366245, EPI_ISL_3366246 | Lighthouse Lab in Milton Keynes | Wellcome Sanger Institute for the COVID-19 Genomics UK (COG-UK) Consortium | The Lighthouse Lab in Milton Keynes and Alex Alderton, Roberto Amato, Jeffrey Barrett, Sonia Goncalves, Ewan Harrison, David K. Jackson, Ian Johnston, Dominic Kwiatkowski, Cordelia Langford, John Sillitoe on behalf of the Wellcome Sanger Institute COVID-19 Surveillance Team |
| EPI_ISL_3366247 | Lighthouse Lab in Alderley Park | Wellcome Sanger Institute for the COVID-19 Genomics UK (COG-UK) Consortium | Jacquelyn Wynn, Mairead Hyland, The Lighthouse Lab in Alderley Park and Alex Alderton, Roberto Amato, Jeffrey Barrett, Sonia Goncalves, Ewan Harrison, David K. Jackson, Ian Johnston, Dominic Kwiatkowski, Cordelia Langford, John Sillitoe on behalf of the Wellcome Sanger Institute COVID-19 Surveillance Team |
| EPI_ISL_3366248 | Randox Laboratories | Wellcome Sanger Institute for the COVID-19 Genomics UK (COG-UK) Consortium | Randox Laboratories and Alex Alderton, Roberto Amato, Jeffrey Barrett, Sonia Goncalves, Ewan Harrison, David K. Jackson, Ian Johnston, Dominic Kwiatkowski, Cordelia Langford, John Sillitoe on behalf of the Wellcome Sanger Institute COVID-19 Surveillance Team |
| EPI_ISL_3366249, EPI_ISL_3366250 | Lighthouse Lab in Alderley Park | Wellcome Sanger Institute for the COVID-19 Genomics UK (COG-UK) Consortium | Jacquelyn Wynn, Mairead Hyland, The Lighthouse Lab in Alderley Park and Alex Alderton, Roberto Amato, Jeffrey Barrett, Sonia Goncalves, Ewan Harrison, David K. Jackson, Ian Johnston, Dominic Kwiatkowski, Cordelia Langford, John Sillitoe on behalf of the Wellcome Sanger Institute COVID-19 Surveillance Team |
| EPI_ISL_3366251 | Lighthouse Lab in Milton Keynes | Wellcome Sanger Institute for the COVID-19 Genomics UK (COG-UK) Consortium | The Lighthouse Lab in Milton Keynes and Alex Alderton, Roberto Amato, Jeffrey Barrett, Sonia Goncalves, Ewan Harrison, David K. Jackson, Ian Johnston, Dominic Kwiatkowski, Cordelia Langford, John Sillitoe on behalf of the Wellcome Sanger Institute COVID-19 Surveillance Team |
| EPI_ISL_3366252, EPI_ISL_3366253, EPI_ISL_3366254 | Randox Laboratories | Wellcome Sanger Institute for the COVID-19 Genomics UK (COG-UK) Consortium | Randox Laboratories and Alex Alderton, Roberto Amato, Jeffrey Barrett, Sonia Goncalves, Ewan Harrison, David K. Jackson, Ian Johnston, Dominic Kwiatkowski, Cordelia Langford, John Sillitoe on behalf of the Wellcome Sanger Institute COVID-19 Surveillance Team |
| EPI_ISL_3366255, EPI_ISL_3366256, EPI_ISL_3366257, EPI_ISL_3366258, EPI_ISL_3366259, EPI_ISL_3366260, EPI_ISL_3366261, EPI_ISL_3366262 | Lighthouse Lab in Milton Keynes | Wellcome Sanger Institute for the COVID-19 Genomics UK (COG-UK) Consortium | The Lighthouse Lab in Milton Keynes and Alex Alderton, Roberto Amato, Jeffrey Barrett, Sonia Goncalves, Ewan Harrison, David K. Jackson, Ian Johnston, Dominic Kwiatkowski, Cordelia Langford, John Sillitoe on behalf of the Wellcome Sanger Institute COVID-19 Surveillance Team |
| EPI_ISL_3366263 | Randox Laboratories | Wellcome Sanger Institute for the COVID-19 Genomics UK (COG-UK) Consortium | Randox Laboratories and Alex Alderton, Roberto Amato, Jeffrey Barrett, Sonia Goncalves, Ewan Harrison, David K. Jackson, Ian Johnston, Dominic Kwiatkowski, Cordelia Langford, John Sillitoe on behalf of the Wellcome Sanger Institute COVID-19 Surveillance Team |
| EPI_ISL_3366264, EPI_ISL_3366265 | Lighthouse Lab in Milton Keynes | Wellcome Sanger Institute for the COVID-19 Genomics UK (COG-UK) Consortium | The Lighthouse Lab in Milton Keynes and Alex Alderton, Roberto Amato, Jeffrey Barrett, Sonia Goncalves, Ewan Harrison, David K. Jackson, Ian Johnston, Dominic Kwiatkowski, Cordelia Langford, John Sillitoe on behalf of the Wellcome Sanger Institute COVID-19 Surveillance Team |
| EPI_ISL_3366266 | Randox Laboratories | Wellcome Sanger Institute for the COVID-19 Genomics UK (COG-UK) Consortium | Randox Laboratories and Alex Alderton, Roberto Amato, Jeffrey Barrett, Sonia Goncalves, Ewan Harrison, David K. Jackson, Ian Johnston, Dominic Kwiatkowski, Cordelia Langford, John Sillitoe on behalf of the Wellcome Sanger Institute COVID-19 Surveillance Team |
| EPI_ISL_3366267 | Lighthouse Lab in Alderley Park | Wellcome Sanger Institute for the COVID-19 Genomics UK (COG-UK) Consortium | Jacquelyn Wynn, Mairead Hyland, The Lighthouse Lab in Alderley Park and Alex Alderton, Roberto Amato, Jeffrey Barrett, Sonia Goncalves, Ewan Harrison, David K. Jackson, Ian Johnston, Dominic Kwiatkowski, Cordelia Langford, John Sillitoe on behalf of the Wellcome Sanger Institute COVID-19 Surveillance Team |
| EPI_ISL_3366268 | Lighthouse Lab in Milton Keynes | Wellcome Sanger Institute for the COVID-19 Genomics UK (COG-UK) Consortium | The Lighthouse Lab in Milton Keynes and Alex Alderton, Roberto Amato, Jeffrey Barrett, Sonia Goncalves, Ewan Harrison, David K. Jackson, Ian Johnston, Dominic Kwiatkowski, Cordelia Langford, John Sillitoe on behalf of the Wellcome Sanger Institute COVID-19 Surveillance Team |
| EPI_ISL_3366269 | Berkshire and Surrey Pathology Services Lighthouse Laboratory | Wellcome Sanger Institute for the COVID-19 Genomics UK (COG-UK) Consortium | Berkshire and Surrey Pathology Services Lighthouse Laboratory and Alex Alderton, Roberto Amato, Jeffrey Barrett, Sonia Goncalves, Ewan Harrison, David K. Jackson, Ian Johnston, Dominic Kwiatkowski, Cordelia Langford, John Sillitoe on behalf of the Wellcome Sanger Institute COVID-19 Surveillance Team |
| EPI_ISL_3366270 | Lighthouse Lab in Alderley Park | Wellcome Sanger Institute for the COVID-19 Genomics UK (COG-UK) Consortium | Jacquelyn Wynn, Mairead Hyland, The Lighthouse Lab in Alderley Park and Alex Alderton, Roberto Amato, Jeffrey Barrett, Sonia Goncalves, Ewan Harrison, David K. Jackson, Ian Johnston, Dominic Kwiatkowski, Cordelia Langford, John Sillitoe on behalf of the Wellcome Sanger Institute COVID-19 Surveillance Team |
| EPI_ISL_3366271 | Berkshire and Surrey Pathology Services Lighthouse Laboratory | Wellcome Sanger Institute for the COVID-19 Genomics UK (COG-UK) Consortium | Berkshire and Surrey Pathology Services Lighthouse Laboratory and Alex Alderton, Roberto Amato, Jeffrey Barrett, Sonia Goncalves, Ewan Harrison, David K. Jackson, Ian Johnston, Dominic Kwiatkowski, Cordelia Langford, John Sillitoe on behalf of the Wellcome Sanger Institute COVID-19 Surveillance Team |
| EPI_ISL_3366272 | Lighthouse Lab in Alderley Park | Wellcome Sanger Institute for the COVID-19 Genomics UK (COG-UK) Consortium | Jacquelyn Wynn, Mairead Hyland, The Lighthouse Lab in Alderley Park and Alex Alderton, Roberto Amato, Jeffrey Barrett, Sonia Goncalves, Ewan Harrison, David K. Jackson, Ian Johnston, Dominic Kwiatkowski, Cordelia Langford, John Sillitoe on behalf of the Wellcome Sanger Institute COVID-19 Surveillance Team |
| EPI_ISL_3366273 | Berkshire and Surrey Pathology Services Lighthouse Laboratory | Wellcome Sanger Institute for the COVID-19 Genomics UK (COG-UK) Consortium | Berkshire and Surrey Pathology Services Lighthouse Laboratory and Alex Alderton, Roberto Amato, Jeffrey Barrett, Sonia Goncalves, Ewan Harrison, David K. Jackson, Ian Johnston, Dominic Kwiatkowski, Cordelia Langford, John Sillitoe on behalf of the Wellcome Sanger Institute COVID-19 Surveillance Team |
| EPI_ISL_3366274, EPI_ISL_3366275, EPI_ISL_3366276, EPI_ISL_3366277, EPI_ISL_3366278 | Lighthouse Lab in Alderley Park | Wellcome Sanger Institute for the COVID-19 Genomics UK (COG-UK) Consortium | Jacquelyn Wynn, Mairead Hyland, The Lighthouse Lab in Alderley Park and Alex Alderton, Roberto Amato, Jeffrey Barrett, Sonia Goncalves, Ewan Harrison, David K. Jackson, Ian Johnston, Dominic Kwiatkowski, Cordelia Langford, John Sillitoe on behalf of the Wellcome Sanger Institute COVID-19 Surveillance Team |
| EPI_ISL_3366279, EPI_ISL_3366280 | Berkshire and Surrey Pathology Services Lighthouse Laboratory | Wellcome Sanger Institute for the COVID-19 Genomics UK (COG-UK) Consortium | Berkshire and Surrey Pathology Services Lighthouse Laboratory and Alex Alderton, Roberto Amato, Jeffrey Barrett, Sonia Goncalves, Ewan Harrison, David K. Jackson, Ian Johnston, Dominic Kwiatkowski, Cordelia Langford, John Sillitoe on behalf of the Wellcome Sanger Institute COVID-19 Surveillance Team |
| EPI_ISL_3366281 | Lighthouse Lab in Milton Keynes | Wellcome Sanger Institute for the COVID-19 Genomics UK (COG-UK) Consortium | The Lighthouse Lab in Milton Keynes and Alex Alderton, Roberto Amato, Jeffrey Barrett, Sonia Goncalves, Ewan Harrison, David K. Jackson, Ian Johnston, Dominic Kwiatkowski, Cordelia Langford, John Sillitoe on behalf of the Wellcome Sanger Institute COVID-19 Surveillance Team |
| EPI_ISL_3366282, EPI_ISL_3366283, EPI_ISL_3366284 | Berkshire and Surrey Pathology Services Lighthouse Laboratory | Wellcome Sanger Institute for the COVID-19 Genomics UK (COG-UK) Consortium | Berkshire and Surrey Pathology Services Lighthouse Laboratory and Alex Alderton, Roberto Amato, Jeffrey Barrett, Sonia Goncalves, Ewan Harrison, David K. Jackson, Ian Johnston, Dominic Kwiatkowski, Cordelia Langford, John Sillitoe on behalf of the Wellcome Sanger Institute COVID-19 Surveillance Team |
| EPI_ISL_3366285 | Lighthouse Lab in Milton Keynes | Wellcome Sanger Institute for the COVID-19 Genomics UK (COG-UK) Consortium | The Lighthouse Lab in Milton Keynes and Alex Alderton, Roberto Amato, Jeffrey Barrett, Sonia Goncalves, Ewan Harrison, David K. Jackson, Ian Johnston, Dominic Kwiatkowski, Cordelia Langford, John Sillitoe on behalf of the Wellcome Sanger Institute COVID-19 Surveillance Team |
| EPI_ISL_3366286, EPI_ISL_3366287, EPI_ISL_3366288 | Berkshire and Surrey Pathology Services Lighthouse Laboratory | Wellcome Sanger Institute for the COVID-19 Genomics UK (COG-UK) Consortium | Berkshire and Surrey Pathology Services Lighthouse Laboratory and Alex Alderton, Roberto Amato, Jeffrey Barrett, Sonia Goncalves, Ewan Harrison, David K. Jackson, Ian Johnston, Dominic Kwiatkowski, Cordelia Langford, John Sillitoe on behalf of the Wellcome Sanger Institute COVID-19 Surveillance Team |
| EPI_ISL_3366289 | Lighthouse Lab in Milton Keynes | Wellcome Sanger Institute for the COVID-19 Genomics UK (COG-UK) Consortium | The Lighthouse Lab in Milton Keynes and Alex Alderton, Roberto Amato, Jeffrey Barrett, Sonia Goncalves, Ewan Harrison, David K. Jackson, Ian Johnston, Dominic Kwiatkowski, Cordelia Langford, John Sillitoe on behalf of the Wellcome Sanger Institute COVID-19 Surveillance Team |
| EPI_ISL_3366290, EPI_ISL_3366291 | Berkshire and Surrey Pathology Services Lighthouse Laboratory | Wellcome Sanger Institute for the COVID-19 Genomics UK (COG-UK) Consortium | Berkshire and Surrey Pathology Services Lighthouse Laboratory and Alex Alderton, Roberto Amato, Jeffrey Barrett, Sonia Goncalves, Ewan Harrison, David K. Jackson, Ian Johnston, Dominic Kwiatkowski, Cordelia Langford, John Sillitoe on behalf of the Wellcome Sanger Institute COVID-19 Surveillance Team |
| EPI_ISL_3366292, EPI_ISL_3366293, EPI_ISL_3366294, EPI_ISL_3366295 | Health Services Laboratories | Wellcome Sanger Institute for the COVID-19 Genomics UK (COG-UK) Consortium | Health Services Laboratories and Alex Alderton, Roberto Amato, Jeffrey Barrett, Sonia Goncalves, Ewan Harrison, David K. Jackson, Ian Johnston, Dominic Kwiatkowski, Cordelia Langford, John Sillitoe on behalf of the Wellcome Sanger Institute COVID-19 Surveillance Team |
| EPI_ISL_3366296, EPI_ISL_3366297, EPI_ISL_3366298 | Lighthouse Lab in Milton Keynes | Wellcome Sanger Institute for the COVID-19 Genomics UK (COG-UK) Consortium | The Lighthouse Lab in Milton Keynes and Alex Alderton, Roberto Amato, Jeffrey Barrett, Sonia Goncalves, Ewan Harrison, David K. Jackson, Ian Johnston, Dominic Kwiatkowski, Cordelia Langford, John Sillitoe on behalf of the Wellcome Sanger Institute COVID-19 Surveillance Team |
| EPI_ISL_3366299, EPI_ISL_3366300, EPI_ISL_3366301, EPI_ISL_3366302 | Health Services Laboratories | Wellcome Sanger Institute for the COVID-19 Genomics UK (COG-UK) Consortium | Health Services Laboratories and Alex Alderton, Roberto Amato, Jeffrey Barrett, Sonia Goncalves, Ewan Harrison, David K. Jackson, Ian Johnston, Dominic Kwiatkowski, Cordelia Langford, John Sillitoe on behalf of the Wellcome Sanger Institute COVID-19 Surveillance Team |
| EPI_ISL_3366303, EPI_ISL_3366304 | Lighthouse Lab in Milton Keynes | Wellcome Sanger Institute for the COVID-19 Genomics UK (COG-UK) Consortium | The Lighthouse Lab in Milton Keynes and Alex Alderton, Roberto Amato, Jeffrey Barrett, Sonia Goncalves, Ewan Harrison, David K. Jackson, Ian Johnston, Dominic Kwiatkowski, Cordelia Langford, John Sillitoe on behalf of the Wellcome Sanger Institute COVID-19 Surveillance Team |

| | | | |
|---|---|---|---|
| EPI_ISL_3366305 | Berkshire and Surrey Pathology Services Lighthouse Laboratory | Wellcome Sanger Institute for the COVID-19 Genomics UK (COG-UK) Consortium | Berkshire and Surrey Pathology Services Lighthouse Laboratory and Alex Alderton, Roberto Amato, Jeffrey Barrett, Sonia Goncalves, Ewan Harrison, David K. Jackson, Ian Johnston, Dominic Kwiatkowski, Cordelia Langford, John Sillitoe on behalf of the Wellcome Sanger Institute COVID-19 Surveillance Team |
| EPI_ISL_3366306 | Health Services Laboratories | Wellcome Sanger Institute for the COVID-19 Genomics UK (COG-UK) Consortium | Health Services Laboratories and Alex Alderton, Roberto Amato, Jeffrey Barrett, Sonia Goncalves, Ewan Harrison, David K. Jackson, Ian Johnston, Dominic Kwiatkowski, Cordelia Langford, John Sillitoe on behalf of the Wellcome Sanger Institute COVID-19 Surveillance Team |
| EPI_ISL_3366307 | Lighthouse Lab in Milton Keynes | Wellcome Sanger Institute for the COVID-19 Genomics UK (COG-UK) Consortium | The Lighthouse Lab in Milton Keynes and Alex Alderton, Roberto Amato, Jeffrey Barrett, Sonia Goncalves, Ewan Harrison, David K. Jackson, Ian Johnston, Dominic Kwiatkowski, Cordelia Langford, John Sillitoe on behalf of the Wellcome Sanger Institute COVID-19 Surveillance Team |
| EPI_ISL_3366308, EPI_ISL_3366309 | Health Services Laboratories | Wellcome Sanger Institute for the COVID-19 Genomics UK (COG-UK) Consortium | Health Services Laboratories and Alex Alderton, Roberto Amato, Jeffrey Barrett, Sonia Goncalves, Ewan Harrison, David K. Jackson, Ian Johnston, Dominic Kwiatkowski, Cordelia Langford, John Sillitoe on behalf of the Wellcome Sanger Institute COVID-19 Surveillance Team |
| EPI_ISL_3366310 | Berkshire and Surrey Pathology Services Lighthouse Laboratory | Wellcome Sanger Institute for the COVID-19 Genomics UK (COG-UK) Consortium | Berkshire and Surrey Pathology Services Lighthouse Laboratory and Alex Alderton, Roberto Amato, Jeffrey Barrett, Sonia Goncalves, Ewan Harrison, David K. Jackson, Ian Johnston, Dominic Kwiatkowski, Cordelia Langford, John Sillitoe on behalf of the Wellcome Sanger Institute COVID-19 Surveillance Team |
| EPI_ISL_3366311 | Lighthouse Lab in Milton Keynes | Wellcome Sanger Institute for the COVID-19 Genomics UK (COG-UK) Consortium | The Lighthouse Lab in Milton Keynes and Alex Alderton, Roberto Amato, Jeffrey Barrett, Sonia Goncalves, Ewan Harrison, David K. Jackson, Ian Johnston, Dominic Kwiatkowski, Cordelia Langford, John Sillitoe on behalf of the Wellcome Sanger Institute COVID-19 Surveillance Team |
| EPI_ISL_3366312, EPI_ISL_3366313, EPI_ISL_3366314, EPI_ISL_3366315, EPI_ISL_3366316 | Health Services Laboratories | Wellcome Sanger Institute for the COVID-19 Genomics UK (COG-UK) Consortium | Health Services Laboratories and Alex Alderton, Roberto Amato, Jeffrey Barrett, Sonia Goncalves, Ewan Harrison, David K. Jackson, Ian Johnston, Dominic Kwiatkowski, Cordelia Langford, John Sillitoe on behalf of the Wellcome Sanger Institute COVID-19 Surveillance Team |
| EPI_ISL_3366317 | Berkshire and Surrey Pathology Services Lighthouse Laboratory | Wellcome Sanger Institute for the COVID-19 Genomics UK (COG-UK) Consortium | Berkshire and Surrey Pathology Services Lighthouse Laboratory and Alex Alderton, Roberto Amato, Jeffrey Barrett, Sonia Goncalves, Ewan Harrison, David K. Jackson, Ian Johnston, Dominic Kwiatkowski, Cordelia Langford, John Sillitoe on behalf of the Wellcome Sanger Institute COVID-19 Surveillance Team |
| EPI_ISL_3366318 | Health Services Laboratories | Wellcome Sanger Institute for the COVID-19 Genomics UK (COG-UK) Consortium | Health Services Laboratories and Alex Alderton, Roberto Amato, Jeffrey Barrett, Sonia Goncalves, Ewan Harrison, David K. Jackson, Ian Johnston, Dominic Kwiatkowski, Cordelia Langford, John Sillitoe on behalf of the Wellcome Sanger Institute COVID-19 Surveillance Team |
| EPI_ISL_3366319, EPI_ISL_3366320 | Lighthouse Lab in Milton Keynes | Wellcome Sanger Institute for the COVID-19 Genomics UK (COG-UK) Consortium | The Lighthouse Lab in Milton Keynes and Alex Alderton, Roberto Amato, Jeffrey Barrett, Sonia Goncalves, Ewan Harrison, David K. Jackson, Ian Johnston, Dominic Kwiatkowski, Cordelia Langford, John Sillitoe on behalf of the Wellcome Sanger Institute COVID-19 Surveillance Team |
| EPI_ISL_3366321, EPI_ISL_3366322, EPI_ISL_3366323, EPI_ISL_3366324 | Health Services Laboratories | Wellcome Sanger Institute for the COVID-19 Genomics UK (COG-UK) Consortium | Health Services Laboratories and Alex Alderton, Roberto Amato, Jeffrey Barrett, Sonia Goncalves, Ewan Harrison, David K. Jackson, Ian Johnston, Dominic Kwiatkowski, Cordelia Langford, John Sillitoe on behalf of the Wellcome Sanger Institute COVID-19 Surveillance Team |
| EPI_ISL_3366325 | Berkshire and Surrey Pathology Services Lighthouse Laboratory | Wellcome Sanger Institute for the COVID-19 Genomics UK (COG-UK) Consortium | Berkshire and Surrey Pathology Services Lighthouse Laboratory and Alex Alderton, Roberto Amato, Jeffrey Barrett, Sonia Goncalves, Ewan Harrison, David K. Jackson, Ian Johnston, Dominic Kwiatkowski, Cordelia Langford, John Sillitoe on behalf of the Wellcome Sanger Institute COVID-19 Surveillance Team |
| EPI_ISL_3366326 | Health Services Laboratories | Wellcome Sanger Institute for the COVID-19 Genomics UK (COG-UK) Consortium | Health Services Laboratories and Alex Alderton, Roberto Amato, Jeffrey Barrett, Sonia Goncalves, Ewan Harrison, David K. Jackson, Ian Johnston, Dominic Kwiatkowski, Cordelia Langford, John Sillitoe on behalf of the Wellcome Sanger Institute COVID-19 Surveillance Team |
| EPI_ISL_3366327 | Lighthouse Lab in Milton Keynes | Wellcome Sanger Institute for the COVID-19 Genomics UK (COG-UK) Consortium | The Lighthouse Lab in Milton Keynes and Alex Alderton, Roberto Amato, Jeffrey Barrett, Sonia Goncalves, Ewan Harrison, David K. Jackson, Ian Johnston, Dominic Kwiatkowski, Cordelia Langford, John Sillitoe on behalf of the Wellcome Sanger Institute COVID-19 Surveillance Team |
| EPI_ISL_3366328 | Berkshire and Surrey Pathology Services Lighthouse Laboratory | Wellcome Sanger Institute for the COVID-19 Genomics UK (COG-UK) Consortium | Berkshire and Surrey Pathology Services Lighthouse Laboratory and Alex Alderton, Roberto Amato, Jeffrey Barrett, Sonia Goncalves, Ewan Harrison, David K. Jackson, Ian Johnston, Dominic Kwiatkowski, Cordelia Langford, John Sillitoe on behalf of the Wellcome Sanger Institute COVID-19 Surveillance Team |
| EPI_ISL_3366329 | Lighthouse Lab in Milton Keynes | Wellcome Sanger Institute for the COVID-19 Genomics UK (COG-UK) Consortium | The Lighthouse Lab in Milton Keynes and Alex Alderton, Roberto Amato, Jeffrey Barrett, Sonia Goncalves, Ewan Harrison, David K. Jackson, Ian Johnston, Dominic Kwiatkowski, Cordelia Langford, John Sillitoe on behalf of the Wellcome Sanger Institute COVID-19 Surveillance Team |
| EPI_ISL_3366330 | Health Services Laboratories | Wellcome Sanger Institute for the COVID-19 Genomics UK (COG-UK) Consortium | Health Services Laboratories and Alex Alderton, Roberto Amato, Jeffrey Barrett, Sonia Goncalves, Ewan Harrison, David K. Jackson, Ian Johnston, Dominic Kwiatkowski, Cordelia Langford, John Sillitoe on behalf of the Wellcome Sanger Institute COVID-19 Surveillance Team |
| EPI_ISL_3366331 | Lighthouse Lab in Milton Keynes | Wellcome Sanger Institute for the COVID-19 Genomics UK (COG-UK) Consortium | The Lighthouse Lab in Milton Keynes and Alex Alderton, Roberto Amato, Jeffrey Barrett, Sonia Goncalves, Ewan Harrison, David K. Jackson, Ian Johnston, Dominic Kwiatkowski, Cordelia Langford, John Sillitoe on behalf of the Wellcome Sanger Institute COVID-19 Surveillance Team |
| EPI_ISL_3366332, EPI_ISL_3366333, EPI_ISL_3366334, EPI_ISL_3366335, EPI_ISL_3366336 | Health Services Laboratories | Wellcome Sanger Institute for the COVID-19 Genomics UK (COG-UK) Consortium | Health Services Laboratories and Alex Alderton, Roberto Amato, Jeffrey Barrett, Sonia Goncalves, Ewan Harrison, David K. Jackson, Ian Johnston, Dominic Kwiatkowski, Cordelia Langford, John Sillitoe on behalf of the Wellcome Sanger Institute COVID-19 Surveillance Team |
| EPI_ISL_3366337 | Berkshire and Surrey Pathology Services Lighthouse Laboratory | Wellcome Sanger Institute for the COVID-19 Genomics UK (COG-UK) Consortium | Berkshire and Surrey Pathology Services Lighthouse Laboratory and Alex Alderton, Roberto Amato, Jeffrey Barrett, Sonia Goncalves, Ewan Harrison, David K. Jackson, Ian Johnston, Dominic Kwiatkowski, Cordelia Langford, John Sillitoe on behalf of the Wellcome Sanger Institute COVID-19 Surveillance Team |
| EPI_ISL_3366338 | Lighthouse Lab in Milton Keynes | Wellcome Sanger Institute for the COVID-19 Genomics UK (COG-UK) Consortium | The Lighthouse Lab in Milton Keynes and Alex Alderton, Roberto Amato, Jeffrey Barrett, Sonia Goncalves, Ewan Harrison, David K. Jackson, Ian Johnston, Dominic Kwiatkowski, Cordelia Langford, John Sillitoe on behalf of the Wellcome Sanger Institute COVID-19 Surveillance Team |
| EPI_ISL_3366339, EPI_ISL_3366340, EPI_ISL_3366341, EPI_ISL_3366342, EPI_ISL_3366343, EPI_ISL_3366344, EPI_ISL_3366345 | Health Services Laboratories | Wellcome Sanger Institute for the COVID-19 Genomics UK (COG-UK) Consortium | Health Services Laboratories and Alex Alderton, Roberto Amato, Jeffrey Barrett, Sonia Goncalves, Ewan Harrison, David K. Jackson, Ian Johnston, Dominic Kwiatkowski, Cordelia Langford, John Sillitoe on behalf of the Wellcome Sanger Institute COVID-19 Surveillance Team |
| EPI_ISL_3366346, EPI_ISL_3366347, EPI_ISL_3366348 | Lighthouse Lab in Milton Keynes | Wellcome Sanger Institute for the COVID-19 Genomics UK (COG-UK) Consortium | The Lighthouse Lab in Milton Keynes and Alex Alderton, Roberto Amato, Jeffrey Barrett, Sonia Goncalves, Ewan Harrison, David K. Jackson, Ian Johnston, Dominic Kwiatkowski, Cordelia Langford, John Sillitoe on behalf of the Wellcome Sanger Institute COVID-19 Surveillance Team |
| EPI_ISL_3366349, EPI_ISL_3366350 | Lighthouse Lab in Alderley Park | Wellcome Sanger Institute for the COVID-19 Genomics UK (COG-UK) Consortium | Jacquelyn Wynn, Mairead Hyland, The Lighthouse Lab in Alderley Park and Alex Alderton, Roberto Amato, Jeffrey Barrett, Sonia Goncalves, Ewan Harrison, David K. Jackson, Ian Johnston, Dominic Kwiatkowski, Cordelia Langford, John Sillitoe on behalf of the Wellcome Sanger Institute COVID-19 Surveillance Team |
| EPI_ISL_3366351 | Lighthouse Lab in Milton Keynes | Wellcome Sanger Institute for the COVID-19 Genomics UK (COG-UK) Consortium | The Lighthouse Lab in Milton Keynes and Alex Alderton, Roberto Amato, Jeffrey Barrett, Sonia Goncalves, Ewan Harrison, David K. Jackson, Ian Johnston, Dominic Kwiatkowski, Cordelia Langford, John Sillitoe on behalf of the Wellcome Sanger Institute COVID-19 Surveillance Team |
| EPI_ISL_3366352 | Lighthouse Laboratory Plymouth | Wellcome Sanger Institute for the COVID-19 Genomics UK (COG-UK) Consortium | Lighthouse Laboratory Plymouth and Alex Alderton, Roberto Amato, Jeffrey Barrett, Sonia Goncalves, Ewan Harrison, David K. Jackson, Ian Johnston, Dominic Kwiatkowski, Cordelia Langford, John Sillitoe on behalf of the Wellcome Sanger Institute COVID-19 Surveillance Team |
| EPI_ISL_3366353, EPI_ISL_3366354, EPI_ISL_3366355 | Lighthouse Lab in Milton Keynes | Wellcome Sanger Institute for the COVID-19 Genomics UK (COG-UK) Consortium | The Lighthouse Lab in Milton Keynes and Alex Alderton, Roberto Amato, Jeffrey Barrett, Sonia Goncalves, Ewan Harrison, David K. Jackson, Ian Johnston, Dominic Kwiatkowski, Cordelia Langford, John Sillitoe on behalf of the Wellcome Sanger Institute COVID-19 Surveillance Team |
| EPI_ISL_3366356 | Lighthouse Lab in Alderley Park | Wellcome Sanger Institute for the COVID-19 Genomics UK (COG-UK) Consortium | Jacquelyn Wynn, Mairead Hyland, The Lighthouse Lab in Alderley Park and Alex Alderton, Roberto Amato, Jeffrey Barrett, Sonia Goncalves, Ewan Harrison, David K. Jackson, Ian Johnston, Dominic Kwiatkowski, Cordelia Langford, John Sillitoe on behalf of the Wellcome Sanger Institute COVID-19 Surveillance Team |
| EPI_ISL_3366357 | Lighthouse Lab in Milton Keynes | Wellcome Sanger Institute for the COVID-19 Genomics UK (COG-UK) Consortium | The Lighthouse Lab in Milton Keynes and Alex Alderton, Roberto Amato, Jeffrey Barrett, Sonia Goncalves, Ewan Harrison, David K. Jackson, Ian Johnston, Dominic Kwiatkowski, Cordelia Langford, John Sillitoe on behalf of the Wellcome Sanger Institute COVID-19 Surveillance Team |
| EPI_ISL_3366358 | Lighthouse Lab in Alderley Park | Wellcome Sanger Institute for the COVID-19 Genomics UK (COG-UK) Consortium | Jacquelyn Wynn, Mairead Hyland, The Lighthouse Lab in Alderley Park and Alex Alderton, Roberto Amato, Jeffrey Barrett, Sonia Goncalves, Ewan Harrison, David K. Jackson, Ian Johnston, Dominic Kwiatkowski, Cordelia Langford, John Sillitoe on behalf of the Wellcome Sanger Institute COVID-19 Surveillance Team |
| EPI_ISL_3366359, EPI_ISL_3366360, EPI_ISL_3366361, EPI_ISL_3366362 | Lighthouse Lab in Milton Keynes | Wellcome Sanger Institute for the COVID-19 Genomics UK (COG-UK) Consortium | The Lighthouse Lab in Milton Keynes and Alex Alderton, Roberto Amato, Jeffrey Barrett, Sonia Goncalves, Ewan Harrison, David K. Jackson, Ian Johnston, Dominic Kwiatkowski, Cordelia Langford, John Sillitoe on behalf of the Wellcome Sanger Institute COVID-19 Surveillance Team |
| EPI_ISL_3366363 | Berkshire and Surrey Pathology Services Lighthouse Laboratory | Wellcome Sanger Institute for the COVID-19 Genomics UK (COG-UK) Consortium | Berkshire and Surrey Pathology Services Lighthouse Laboratory and Alex Alderton, Roberto Amato, Jeffrey Barrett, Sonia Goncalves, Ewan Harrison, David K. Jackson, Ian Johnston, Dominic Kwiatkowski, Cordelia Langford, John Sillitoe on behalf of the Wellcome Sanger Institute COVID-19 Surveillance Team |
| EPI_ISL_3366364, EPI_ISL_3366365, EPI_ISL_3366366, EPI_ISL_3366367, EPI_ISL_3366368, EPI_ISL_3366369, EPI_ISL_3366370 | Health Services Laboratories | Wellcome Sanger Institute for the COVID-19 Genomics UK (COG-UK) Consortium | Health Services Laboratories and Alex Alderton, Roberto Amato, Jeffrey Barrett, Sonia Goncalves, Ewan Harrison, David K. Jackson, Ian Johnston, Dominic Kwiatkowski, Cordelia Langford, John Sillitoe on behalf of the Wellcome Sanger Institute COVID-19 Surveillance Team |
| EPI_ISL_3366371 | Lighthouse Lab in Milton Keynes | Wellcome Sanger Institute for the COVID-19 Genomics UK (COG-UK) Consortium | The Lighthouse Lab in Milton Keynes and Alex Alderton, Roberto Amato, Jeffrey Barrett, Sonia Goncalves, Ewan Harrison, David K. Jackson, Ian Johnston, Dominic Kwiatkowski, Cordelia Langford, John Sillitoe on behalf of the Wellcome Sanger Institute COVID-19 Surveillance Team |
| EPI_ISL_3366372 | Berkshire and Surrey Pathology Services Lighthouse Laboratory | Wellcome Sanger Institute for the COVID-19 Genomics UK (COG-UK) Consortium | Berkshire and Surrey Pathology Services Lighthouse Laboratory and Alex Alderton, Roberto Amato, Jeffrey Barrett, Sonia Goncalves, Ewan Harrison, David K. Jackson, Ian Johnston, Dominic Kwiatkowski, Cordelia Langford, John Sillitoe on behalf of the Wellcome Sanger Institute COVID-19 Surveillance Team |
| EPI_ISL_3366373 | Health Services Laboratories | Wellcome Sanger Institute for the COVID-19 Genomics UK (COG-UK) Consortium | Health Services Laboratories and Alex Alderton, Roberto Amato, Jeffrey Barrett, Sonia Goncalves, Ewan Harrison, David K. Jackson, Ian Johnston, Dominic Kwiatkowski, Cordelia Langford, John Sillitoe on behalf of the Wellcome Sanger Institute COVID-19 Surveillance Team |

| | | | |
|---|---|---|---|
| EPI_ISL_3366374 | Berkshire and Surrey Pathology Services Lighthouse Laboratory | Wellcome Sanger Institute for the COVID-19 Genomics UK (COG-UK) Consortium | Berkshire and Surrey Pathology Services Lighthouse Laboratory and Alex Alderton, Roberto Amato, Jeffrey Barrett, Sonia Goncalves, Ewan Harrison, David K. Jackson, Ian Johnston, Dominic Kwiatkowski, Cordelia Langford, John Sillitoe on behalf of the Wellcome Sanger Institute COVID-19 Surveillance Team |
| EPI_ISL_3366375, EPI_ISL_3366376, EPI_ISL_3366377 | Health Services Laboratories | Wellcome Sanger Institute for the COVID-19 Genomics UK (COG-UK) Consortium | Health Services Laboratories and Alex Alderton, Roberto Amato, Jeffrey Barrett, Sonia Goncalves, Ewan Harrison, David K. Jackson, Ian Johnston, Dominic Kwiatkowski, Cordelia Langford, John Sillitoe on behalf of the Wellcome Sanger Institute COVID-19 Surveillance Team |
| EPI_ISL_3366378, EPI_ISL_3366379 | Berkshire and Surrey Pathology Services Lighthouse Laboratory | Wellcome Sanger Institute for the COVID-19 Genomics UK (COG-UK) Consortium | Berkshire and Surrey Pathology Services Lighthouse Laboratory and Alex Alderton, Roberto Amato, Jeffrey Barrett, Sonia Goncalves, Ewan Harrison, David K. Jackson, Ian Johnston, Dominic Kwiatkowski, Cordelia Langford, John Sillitoe on behalf of the Wellcome Sanger Institute COVID-19 Surveillance Team |
| EPI_ISL_3366380 | Health Services Laboratories | Wellcome Sanger Institute for the COVID-19 Genomics UK (COG-UK) Consortium | Health Services Laboratories and Alex Alderton, Roberto Amato, Jeffrey Barrett, Sonia Goncalves, Ewan Harrison, David K. Jackson, Ian Johnston, Dominic Kwiatkowski, Cordelia Langford, John Sillitoe on behalf of the Wellcome Sanger Institute COVID-19 Surveillance Team |
| EPI_ISL_3366381, EPI_ISL_3366382 | Berkshire and Surrey Pathology Services Lighthouse Laboratory | Wellcome Sanger Institute for the COVID-19 Genomics UK (COG-UK) Consortium | Berkshire and Surrey Pathology Services Lighthouse Laboratory and Alex Alderton, Roberto Amato, Jeffrey Barrett, Sonia Goncalves, Ewan Harrison, David K. Jackson, Ian Johnston, Dominic Kwiatkowski, Cordelia Langford, John Sillitoe on behalf of the Wellcome Sanger Institute COVID-19 Surveillance Team |
| EPI_ISL_3366383 | Health Services Laboratories | Wellcome Sanger Institute for the COVID-19 Genomics UK (COG-UK) Consortium | Health Services Laboratories and Alex Alderton, Roberto Amato, Jeffrey Barrett, Sonia Goncalves, Ewan Harrison, David K. Jackson, Ian Johnston, Dominic Kwiatkowski, Cordelia Langford, John Sillitoe on behalf of the Wellcome Sanger Institute COVID-19 Surveillance Team |
| EPI_ISL_3366384, EPI_ISL_3366385 | Berkshire and Surrey Pathology Services Lighthouse Laboratory | Wellcome Sanger Institute for the COVID-19 Genomics UK (COG-UK) Consortium | Berkshire and Surrey Pathology Services Lighthouse Laboratory and Alex Alderton, Roberto Amato, Jeffrey Barrett, Sonia Goncalves, Ewan Harrison, David K. Jackson, Ian Johnston, Dominic Kwiatkowski, Cordelia Langford, John Sillitoe on behalf of the Wellcome Sanger Institute COVID-19 Surveillance Team |
| EPI_ISL_3366386, EPI_ISL_3366387 | Health Services Laboratories | Wellcome Sanger Institute for the COVID-19 Genomics UK (COG-UK) Consortium | Health Services Laboratories and Alex Alderton, Roberto Amato, Jeffrey Barrett, Sonia Goncalves, Ewan Harrison, David K. Jackson, Ian Johnston, Dominic Kwiatkowski, Cordelia Langford, John Sillitoe on behalf of the Wellcome Sanger Institute COVID-19 Surveillance Team |
| EPI_ISL_3366388, EPI_ISL_3366389, EPI_ISL_3366390 | Lighthouse Lab in Alderley Park | Wellcome Sanger Institute for the COVID-19 Genomics UK (COG-UK) Consortium | Jacquelyn Wynn, Mairead Hyland, The Lighthouse Lab in Alderley Park and Alex Alderton, Roberto Amato, Jeffrey Barrett, Sonia Goncalves, Ewan Harrison, David K. Jackson, Ian Johnston, Dominic Kwiatkowski, Cordelia Langford, John Sillitoe on behalf of the Wellcome Sanger Institute COVID-19 Surveillance Team |
| EPI_ISL_3366391 | Health Services Laboratories | Wellcome Sanger Institute for the COVID-19 Genomics UK (COG-UK) Consortium | Health Services Laboratories and Alex Alderton, Roberto Amato, Jeffrey Barrett, Sonia Goncalves, Ewan Harrison, David K. Jackson, Ian Johnston, Dominic Kwiatkowski, Cordelia Langford, John Sillitoe on behalf of the Wellcome Sanger Institute COVID-19 Surveillance Team |
| EPI_ISL_3366392 | Lighthouse Lab in Milton Keynes | Wellcome Sanger Institute for the COVID-19 Genomics UK (COG-UK) Consortium | The Lighthouse Lab in Milton Keynes and Alex Alderton, Roberto Amato, Jeffrey Barrett, Sonia Goncalves, Ewan Harrison, David K. Jackson, Ian Johnston, Dominic Kwiatkowski, Cordelia Langford, John Sillitoe on behalf of the Wellcome Sanger Institute COVID-19 Surveillance Team |
| EPI_ISL_3366393 | Health Services Laboratories | Wellcome Sanger Institute for the COVID-19 Genomics UK (COG-UK) Consortium | Health Services Laboratories and Alex Alderton, Roberto Amato, Jeffrey Barrett, Sonia Goncalves, Ewan Harrison, David K. Jackson, Ian Johnston, Dominic Kwiatkowski, Cordelia Langford, John Sillitoe on behalf of the Wellcome Sanger Institute COVID-19 Surveillance Team |
| EPI_ISL_3366394, EPI_ISL_3366395 | Lighthouse Lab in Milton Keynes | Wellcome Sanger Institute for the COVID-19 Genomics UK (COG-UK) Consortium | The Lighthouse Lab in Milton Keynes and Alex Alderton, Roberto Amato, Jeffrey Barrett, Sonia Goncalves, Ewan Harrison, David K. Jackson, Ian Johnston, Dominic Kwiatkowski, Cordelia Langford, John Sillitoe on behalf of the Wellcome Sanger Institute COVID-19 Surveillance Team |
| EPI_ISL_3366396, EPI_ISL_3366397, EPI_ISL_3366398 | Health Services Laboratories | Wellcome Sanger Institute for the COVID-19 Genomics UK (COG-UK) Consortium | Health Services Laboratories and Alex Alderton, Roberto Amato, Jeffrey Barrett, Sonia Goncalves, Ewan Harrison, David K. Jackson, Ian Johnston, Dominic Kwiatkowski, Cordelia Langford, John Sillitoe on behalf of the Wellcome Sanger Institute COVID-19 Surveillance Team |
| EPI_ISL_3366399 | Lighthouse Lab in Milton Keynes | Wellcome Sanger Institute for the COVID-19 Genomics UK (COG-UK) Consortium | The Lighthouse Lab in Milton Keynes and Alex Alderton, Roberto Amato, Jeffrey Barrett, Sonia Goncalves, Ewan Harrison, David K. Jackson, Ian Johnston, Dominic Kwiatkowski, Cordelia Langford, John Sillitoe on behalf of the Wellcome Sanger Institute COVID-19 Surveillance Team |
| EPI_ISL_3366400, EPI_ISL_3366401, EPI_ISL_3366402, EPI_ISL_3366403, EPI_ISL_3366404, EPI_ISL_3366405, EPI_ISL_3366406, EPI_ISL_3366407, EPI_ISL_3366408, EPI_ISL_3366409, EPI_ISL_3366410, EPI_ISL_3366411, EPI_ISL_3366412, EPI_ISL_3366413, EPI_ISL_3366414 | | | |
| see above | Health Services Laboratories | Wellcome Sanger Institute for the COVID-19 Genomics UK (COG-UK) Consortium | Health Services Laboratories and Alex Alderton, Roberto Amato, Jeffrey Barrett, Sonia Goncalves, Ewan Harrison, David K. Jackson, Ian Johnston, Dominic Kwiatkowski, Cordelia Langford, John Sillitoe on behalf of the Wellcome Sanger Institute COVID-19 Surveillance Team |
| EPI_ISL_3366415, EPI_ISL_3366416 | Lighthouse Lab in Milton Keynes | Wellcome Sanger Institute for the COVID-19 Genomics UK (COG-UK) Consortium | The Lighthouse Lab in Milton Keynes and Alex Alderton, Roberto Amato, Jeffrey Barrett, Sonia Goncalves, Ewan Harrison, David K. Jackson, Ian Johnston, Dominic Kwiatkowski, Cordelia Langford, John Sillitoe on behalf of the Wellcome Sanger Institute COVID-19 Surveillance Team |
| EPI_ISL_3366417, EPI_ISL_3366418, EPI_ISL_3366419, EPI_ISL_3366420, EPI_ISL_3366421, EPI_ISL_3366422, EPI_ISL_3366423, EPI_ISL_3366424 | Health Services Laboratories | Wellcome Sanger Institute for the COVID-19 Genomics UK (COG-UK) Consortium | Health Services Laboratories and Alex Alderton, Roberto Amato, Jeffrey Barrett, Sonia Goncalves, Ewan Harrison, David K. Jackson, Ian Johnston, Dominic Kwiatkowski, Cordelia Langford, John Sillitoe on behalf of the Wellcome Sanger Institute COVID-19 Surveillance Team |
| EPI_ISL_3366425, EPI_ISL_3366426, EPI_ISL_3366427, EPI_ISL_3366428, EPI_ISL_3366429, EPI_ISL_3366430, EPI_ISL_3366431 | Lighthouse Lab in Milton Keynes | Wellcome Sanger Institute for the COVID-19 Genomics UK (COG-UK) Consortium | The Lighthouse Lab in Milton Keynes and Alex Alderton, Roberto Amato, Jeffrey Barrett, Sonia Goncalves, Ewan Harrison, David K. Jackson, Ian Johnston, Dominic Kwiatkowski, Cordelia Langford, John Sillitoe on behalf of the Wellcome Sanger Institute COVID-19 Surveillance Team |
| EPI_ISL_3366432, EPI_ISL_3366433, EPI_ISL_3366434, EPI_ISL_3366435 | Lighthouse Lab in Glasgow | Wellcome Sanger Institute for the COVID-19 Genomics UK (COG-UK) Consortium | Harper VanSteenhouse, Yumi Kasai, David Gray, Carol Clugston, Anna Dominiczak and Alex Alderton, Roberto Amato, Jeffrey Barrett, Sonia Goncalves, Ewan Harrison, David K. Jackson, Ian Johnston, Dominic Kwiatkowski, Cordelia Langford, John Sillitoe on behalf of the Wellcome Sanger Institute COVID-19 Surveillance Team |
| EPI_ISL_3366436 | Berkshire and Surrey Pathology Services Lighthouse Laboratory | Wellcome Sanger Institute for the COVID-19 Genomics UK (COG-UK) Consortium | Berkshire and Surrey Pathology Services Lighthouse Laboratory and Alex Alderton, Roberto Amato, Jeffrey Barrett, Sonia Goncalves, Ewan Harrison, David K. Jackson, Ian Johnston, Dominic Kwiatkowski, Cordelia Langford, John Sillitoe on behalf of the Wellcome Sanger Institute COVID-19 Surveillance Team |
| EPI_ISL_3366437, EPI_ISL_3366438, EPI_ISL_3366439, EPI_ISL_3366440, EPI_ISL_3366441, EPI_ISL_3366442, EPI_ISL_3366443, EPI_ISL_3366444 | Lighthouse Lab in Glasgow | Wellcome Sanger Institute for the COVID-19 Genomics UK (COG-UK) Consortium | Harper VanSteenhouse, Yumi Kasai, David Gray, Carol Clugston, Anna Dominiczak and Alex Alderton, Roberto Amato, Jeffrey Barrett, Sonia Goncalves, Ewan Harrison, David K. Jackson, Ian Johnston, Dominic Kwiatkowski, Cordelia Langford, John Sillitoe on behalf of the Wellcome Sanger Institute COVID-19 Surveillance Team |
| EPI_ISL_3366445, EPI_ISL_3366446, EPI_ISL_3366447, EPI_ISL_3366448, EPI_ISL_3366449, EPI_ISL_3366450, EPI_ISL_3366451, EPI_ISL_3366452, EPI_ISL_3366453, EPI_ISL_3366455, EPI_ISL_3366456, EPI_ISL_3366457, EPI_ISL_3366466, EPI_ISL_3366468, EPI_ISL_3366470, EPI_ISL_3366471, EPI_ISL_3366472, EPI_ISL_3366473, EPI_ISL_3366474, EPI_ISL_3366477, EPI_ISL_3366479 | | | |
| see above | Servizo de Microbioloxía. Complexo Hospitalario de Santiago de Compostela | Servizo de Microbioloxía. Complexo Hospitalario de Santiago de Compostela | Xosé Costa, Antonio Aguilera, Gema Barbeito, Amparo Coira, Carlos García_Riestra, Manuela Hernández, José Llovo, Teresa Lopez_Valiño, Laura Millán, Daniel Navarro, Rocío trastoy, Mercedes Treviño, María Luisa Pérez_del_Molino |
| EPI_ISL_3366480 | Lighthouse Lab in Milton Keynes | Wellcome Sanger Institute for the COVID-19 Genomics UK (COG-UK) Consortium | The Lighthouse Lab in Milton Keynes and Alex Alderton, Roberto Amato, Jeffrey Barrett, Sonia Goncalves, Ewan Harrison, David K. Jackson, Ian Johnston, Dominic Kwiatkowski, Cordelia Langford, John Sillitoe on behalf of the Wellcome Sanger Institute COVID-19 Surveillance Team |
| EPI_ISL_3366481, EPI_ISL_3366482, EPI_ISL_3366483, EPI_ISL_3366484, EPI_ISL_3366485 | Lighthouse Lab in Glasgow | Wellcome Sanger Institute for the COVID-19 Genomics UK (COG-UK) Consortium | Harper VanSteenhouse, Yumi Kasai, David Gray, Carol Clugston, Anna Dominiczak and Alex Alderton, Roberto Amato, Jeffrey Barrett, Sonia Goncalves, Ewan Harrison, David K. Jackson, Ian Johnston, Dominic Kwiatkowski, Cordelia Langford, John Sillitoe on behalf of the Wellcome Sanger Institute COVID-19 Surveillance Team |
| EPI_ISL_3366486 | Berkshire and Surrey Pathology Services Lighthouse Laboratory | Wellcome Sanger Institute for the COVID-19 Genomics UK (COG-UK) Consortium | Berkshire and Surrey Pathology Services Lighthouse Laboratory and Alex Alderton, Roberto Amato, Jeffrey Barrett, Sonia Goncalves, Ewan Harrison, David K. Jackson, Ian Johnston, Dominic Kwiatkowski, Cordelia Langford, John Sillitoe on behalf of the Wellcome Sanger Institute COVID-19 Surveillance Team |
| EPI_ISL_3366487, EPI_ISL_3366488, EPI_ISL_3366489, EPI_ISL_3366490, EPI_ISL_3366491, EPI_ISL_3366492, EPI_ISL_3366493, EPI_ISL_3366494, EPI_ISL_3366495, EPI_ISL_3366496, EPI_ISL_3366497, EPI_ISL_3366498, EPI_ISL_3366499 | | | |
| see above | Lighthouse Lab in Glasgow | Wellcome Sanger Institute for the COVID-19 Genomics UK (COG-UK) Consortium | Harper VanSteenhouse, Yumi Kasai, David Gray, Carol Clugston, Anna Dominiczak and Alex Alderton, Roberto Amato, Jeffrey Barrett, Sonia Goncalves, Ewan Harrison, David K. Jackson, Ian Johnston, Dominic Kwiatkowski, Cordelia Langford, John Sillitoe on behalf of the Wellcome Sanger Institute COVID-19 Surveillance Team |
| EPI_ISL_3366500, EPI_ISL_3366501, EPI_ISL_3366502, EPI_ISL_3366503, EPI_ISL_3366504, EPI_ISL_3366505, EPI_ISL_3366506, EPI_ISL_3366507, EPI_ISL_3366508, EPI_ISL_3366509, EPI_ISL_3366510, EPI_ISL_3366511 | | | |
| see above | Health Services Laboratories | Wellcome Sanger Institute for the COVID-19 Genomics UK (COG-UK) Consortium | Health Services Laboratories and Alex Alderton, Roberto Amato, Jeffrey Barrett, Sonia Goncalves, Ewan Harrison, David K. Jackson, Ian Johnston, Dominic Kwiatkowski, Cordelia Langford, John Sillitoe on behalf of the Wellcome Sanger Institute COVID-19 Surveillance Team |
| EPI_ISL_3366512 | Lighthouse Lab in Glasgow | Wellcome Sanger Institute for the COVID-19 Genomics UK (COG-UK) Consortium | Harper VanSteenhouse, Yumi Kasai, David Gray, Carol Clugston, Anna Dominiczak and Alex Alderton, Roberto Amato, Jeffrey Barrett, Sonia Goncalves, Ewan Harrison, David K. Jackson, Ian Johnston, Dominic Kwiatkowski, Cordelia Langford, John Sillitoe on behalf of the Wellcome Sanger Institute COVID-19 Surveillance Team |
| EPI_ISL_3366513, EPI_ISL_3366514, EPI_ISL_3366515, EPI_ISL_3366516, EPI_ISL_3366517, EPI_ISL_3366518, EPI_ISL_3366519, EPI_ISL_3366520, EPI_ISL_3366521, EPI_ISL_3366522, EPI_ISL_3366523, EPI_ISL_3366524, EPI_ISL_3366525, EPI_ISL_3366526, EPI_ISL_3366527, EPI_ISL_3366528, EPI_ISL_3366529, EPI_ISL_3366530, EPI_ISL_3366531, EPI_ISL_3366532, EPI_ISL_3366533, EPI_ISL_3366534, EPI_ISL_3366535, EPI_ISL_3366536, EPI_ISL_3366537, EPI_ISL_3366538, EPI_ISL_3366539 | | | |
| see above | Health Services Laboratories | Wellcome Sanger Institute for the COVID-19 Genomics UK (COG-UK) Consortium | Health Services Laboratories and Alex Alderton, Roberto Amato, Jeffrey Barrett, Sonia Goncalves, Ewan Harrison, David K. Jackson, Ian Johnston, Dominic Kwiatkowski, Cordelia Langford, John Sillitoe on behalf of the Wellcome Sanger Institute COVID-19 Surveillance Team |
| EPI_ISL_3366540 | Lighthouse Lab in Glasgow | Wellcome Sanger Institute for the COVID-19 Genomics UK (COG-UK) Consortium | Harper VanSteenhouse, Yumi Kasai, David Gray, Carol Clugston, Anna Dominiczak and Alex Alderton, Roberto Amato, Jeffrey Barrett, Sonia Goncalves, Ewan Harrison, David K. Jackson, Ian Johnston, Dominic Kwiatkowski, Cordelia Langford, John Sillitoe on behalf of the Wellcome Sanger Institute COVID-19 Surveillance Team |
| EPI_ISL_3366541, EPI_ISL_3366542, EPI_ISL_3366543, EPI_ISL_3366544, EPI_ISL_3366545, EPI_ISL_3366546, EPI_ISL_3366547, EPI_ISL_3366548, EPI_ISL_3366549, EPI_ISL_3366550, EPI_ISL_3366551, EPI_ISL_3366552, EPI_ISL_3366553, EPI_ISL_3366554, EPI_ISL_3366555, EPI_ISL_3366556 | | | |

| | | | |
|---|---|---|---|
| see above | Health Services Laboratories | Wellcome Sanger Institute for the COVID-19 Genomics UK (COG-UK) Consortium | Health Services Laboratories and Alex Alderton, Roberto Amato, Jeffrey Barrett, Sonia Goncalves, Ewan Harrison, David K. Jackson, Ian Johnston, Dominic Kwiatkowski, Cordelia Langford, John Sillitoe on behalf of the Wellcome Sanger Institute COVID-19 Surveillance Team |
| EPI_ISL_3366557, EPI_ISL_3366558 | Lighthouse Lab in Glasgow | Wellcome Sanger Institute for the COVID-19 Genomics UK (COG-UK) Consortium | Harper VanSteenhouse, Yumi Kasai, David Gray, Carol Clugston, Anna Dominiczak and Alex Alderton, Roberto Amato, Jeffrey Barrett, Sonia Goncalves, Ewan Harrison, David K. Jackson, Ian Johnston, Dominic Kwiatkowski, Cordelia Langford, John Sillitoe on behalf of the Wellcome Sanger Institute COVID-19 Surveillance Team |
| EPI_ISL_3366559 | Lighthouse Lab in Milton Keynes | Wellcome Sanger Institute for the COVID-19 Genomics UK (COG-UK) Consortium | The Lighthouse Lab in Milton Keynes and Alex Alderton, Roberto Amato, Jeffrey Barrett, Sonia Goncalves, Ewan Harrison, David K. Jackson, Ian Johnston, Dominic Kwiatkowski, Cordelia Langford, John Sillitoe on behalf of the Wellcome Sanger Institute COVID-19 Surveillance Team |
| EPI_ISL_3366560 | Health Services Laboratories | Wellcome Sanger Institute for the COVID-19 Genomics UK (COG-UK) Consortium | Health Services Laboratories and Alex Alderton, Roberto Amato, Jeffrey Barrett, Sonia Goncalves, Ewan Harrison, David K. Jackson, Ian Johnston, Dominic Kwiatkowski, Cordelia Langford, John Sillitoe on behalf of the Wellcome Sanger Institute COVID-19 Surveillance Team |
| EPI_ISL_3366561, EPI_ISL_3366562 | Lighthouse Lab in Alderley Park | Wellcome Sanger Institute for the COVID-19 Genomics UK (COG-UK) Consortium | Jacquelyn Wynn, Mairead Hyland, The Lighthouse Lab in Alderley Park and Alex Alderton, Roberto Amato, Jeffrey Barrett, Sonia Goncalves, Ewan Harrison, David K. Jackson, Ian Johnston, Dominic Kwiatkowski, Cordelia Langford, John Sillitoe on behalf of the Wellcome Sanger Institute COVID-19 Surveillance Team |
| EPI_ISL_3366563, EPI_ISL_3366564, EPI_ISL_3366565, EPI_ISL_3366566 | Health Services Laboratories | Wellcome Sanger Institute for the COVID-19 Genomics UK (COG-UK) Consortium | Health Services Laboratories and Alex Alderton, Roberto Amato, Jeffrey Barrett, Sonia Goncalves, Ewan Harrison, David K. Jackson, Ian Johnston, Dominic Kwiatkowski, Cordelia Langford, John Sillitoe on behalf of the Wellcome Sanger Institute COVID-19 Surveillance Team |
| EPI_ISL_3366567, EPI_ISL_3366568 | Lighthouse Lab in Alderley Park | Wellcome Sanger Institute for the COVID-19 Genomics UK (COG-UK) Consortium | Jacquelyn Wynn, Mairead Hyland, The Lighthouse Lab in Alderley Park and Alex Alderton, Roberto Amato, Jeffrey Barrett, Sonia Goncalves, Ewan Harrison, David K. Jackson, Ian Johnston, Dominic Kwiatkowski, Cordelia Langford, John Sillitoe on behalf of the Wellcome Sanger Institute COVID-19 Surveillance Team |
| EPI_ISL_3366569 | Health Services Laboratories | Wellcome Sanger Institute for the COVID-19 Genomics UK (COG-UK) Consortium | Health Services Laboratories and Alex Alderton, Roberto Amato, Jeffrey Barrett, Sonia Goncalves, Ewan Harrison, David K. Jackson, Ian Johnston, Dominic Kwiatkowski, Cordelia Langford, John Sillitoe on behalf of the Wellcome Sanger Institute COVID-19 Surveillance Team |
| EPI_ISL_3366570 | Lighthouse Lab in Alderley Park | Wellcome Sanger Institute for the COVID-19 Genomics UK (COG-UK) Consortium | Jacquelyn Wynn, Mairead Hyland, The Lighthouse Lab in Alderley Park and Alex Alderton, Roberto Amato, Jeffrey Barrett, Sonia Goncalves, Ewan Harrison, David K. Jackson, Ian Johnston, Dominic Kwiatkowski, Cordelia Langford, John Sillitoe on behalf of the Wellcome Sanger Institute COVID-19 Surveillance Team |
| EPI_ISL_3366571, EPI_ISL_3366572 | Health Services Laboratories | Wellcome Sanger Institute for the COVID-19 Genomics UK (COG-UK) Consortium | Health Services Laboratories and Alex Alderton, Roberto Amato, Jeffrey Barrett, Sonia Goncalves, Ewan Harrison, David K. Jackson, Ian Johnston, Dominic Kwiatkowski, Cordelia Langford, John Sillitoe on behalf of the Wellcome Sanger Institute COVID-19 Surveillance Team |
| EPI_ISL_3366573, EPI_ISL_3366574 | Lighthouse Lab in Alderley Park | Wellcome Sanger Institute for the COVID-19 Genomics UK (COG-UK) Consortium | Jacquelyn Wynn, Mairead Hyland, The Lighthouse Lab in Alderley Park and Alex Alderton, Roberto Amato, Jeffrey Barrett, Sonia Goncalves, Ewan Harrison, David K. Jackson, Ian Johnston, Dominic Kwiatkowski, Cordelia Langford, John Sillitoe on behalf of the Wellcome Sanger Institute COVID-19 Surveillance Team |
| EPI_ISL_3366575, EPI_ISL_3366576 | Health Services Laboratories | Wellcome Sanger Institute for the COVID-19 Genomics UK (COG-UK) Consortium | Health Services Laboratories and Alex Alderton, Roberto Amato, Jeffrey Barrett, Sonia Goncalves, Ewan Harrison, David K. Jackson, Ian Johnston, Dominic Kwiatkowski, Cordelia Langford, John Sillitoe on behalf of the Wellcome Sanger Institute COVID-19 Surveillance Team |
| EPI_ISL_3366577 | Lighthouse Lab in Alderley Park | Wellcome Sanger Institute for the COVID-19 Genomics UK (COG-UK) Consortium | Jacquelyn Wynn, Mairead Hyland, The Lighthouse Lab in Alderley Park and Alex Alderton, Roberto Amato, Jeffrey Barrett, Sonia Goncalves, Ewan Harrison, David K. Jackson, Ian Johnston, Dominic Kwiatkowski, Cordelia Langford, John Sillitoe on behalf of the Wellcome Sanger Institute COVID-19 Surveillance Team |
| EPI_ISL_3366578 | Lighthouse Laboratory Plymouth | Wellcome Sanger Institute for the COVID-19 Genomics UK (COG-UK) Consortium | Lighthouse Laboratory Plymouth and Alex Alderton, Roberto Amato, Jeffrey Barrett, Sonia Goncalves, Ewan Harrison, David K. Jackson, Ian Johnston, Dominic Kwiatkowski, Cordelia Langford, John Sillitoe on behalf of the Wellcome Sanger Institute COVID-19 Surveillance Team |
| EPI_ISL_3366579 | Health Services Laboratories | Wellcome Sanger Institute for the COVID-19 Genomics UK (COG-UK) Consortium | Health Services Laboratories and Alex Alderton, Roberto Amato, Jeffrey Barrett, Sonia Goncalves, Ewan Harrison, David K. Jackson, Ian Johnston, Dominic Kwiatkowski, Cordelia Langford, John Sillitoe on behalf of the Wellcome Sanger Institute COVID-19 Surveillance Team |
| EPI_ISL_3366580 | Lighthouse Lab in Alderley Park | Wellcome Sanger Institute for the COVID-19 Genomics UK (COG-UK) Consortium | Jacquelyn Wynn, Mairead Hyland, The Lighthouse Lab in Alderley Park and Alex Alderton, Roberto Amato, Jeffrey Barrett, Sonia Goncalves, Ewan Harrison, David K. Jackson, Ian Johnston, Dominic Kwiatkowski, Cordelia Langford, John Sillitoe on behalf of the Wellcome Sanger Institute COVID-19 Surveillance Team |
| EPI_ISL_3366581, EPI_ISL_3366582, EPI_ISL_3366583 | Health Services Laboratories | Wellcome Sanger Institute for the COVID-19 Genomics UK (COG-UK) Consortium | Health Services Laboratories and Alex Alderton, Roberto Amato, Jeffrey Barrett, Sonia Goncalves, Ewan Harrison, David K. Jackson, Ian Johnston, Dominic Kwiatkowski, Cordelia Langford, John Sillitoe on behalf of the Wellcome Sanger Institute COVID-19 Surveillance Team |
| EPI_ISL_3366584 | Lighthouse Lab in Alderley Park | Wellcome Sanger Institute for the COVID-19 Genomics UK (COG-UK) Consortium | Jacquelyn Wynn, Mairead Hyland, The Lighthouse Lab in Alderley Park and Alex Alderton, Roberto Amato, Jeffrey Barrett, Sonia Goncalves, Ewan Harrison, David K. Jackson, Ian Johnston, Dominic Kwiatkowski, Cordelia Langford, John Sillitoe on behalf of the Wellcome Sanger Institute COVID-19 Surveillance Team |
| EPI_ISL_3366585 | Health Services Laboratories | Wellcome Sanger Institute for the COVID-19 Genomics UK (COG-UK) Consortium | Health Services Laboratories and Alex Alderton, Roberto Amato, Jeffrey Barrett, Sonia Goncalves, Ewan Harrison, David K. Jackson, Ian Johnston, Dominic Kwiatkowski, Cordelia Langford, John Sillitoe on behalf of the Wellcome Sanger Institute COVID-19 Surveillance Team |
| EPI_ISL_3366586 | Lighthouse Lab in Alderley Park | Wellcome Sanger Institute for the COVID-19 Genomics UK (COG-UK) Consortium | Jacquelyn Wynn, Mairead Hyland, The Lighthouse Lab in Alderley Park and Alex Alderton, Roberto Amato, Jeffrey Barrett, Sonia Goncalves, Ewan Harrison, David K. Jackson, Ian Johnston, Dominic Kwiatkowski, Cordelia Langford, John Sillitoe on behalf of the Wellcome Sanger Institute COVID-19 Surveillance Team |
| EPI_ISL_3366587, EPI_ISL_3366588 | Health Services Laboratories | Wellcome Sanger Institute for the COVID-19 Genomics UK (COG-UK) Consortium | Health Services Laboratories and Alex Alderton, Roberto Amato, Jeffrey Barrett, Sonia Goncalves, Ewan Harrison, David K. Jackson, Ian Johnston, Dominic Kwiatkowski, Cordelia Langford, John Sillitoe on behalf of the Wellcome Sanger Institute COVID-19 Surveillance Team |
| EPI_ISL_3366589 | Lighthouse Lab in Alderley Park | Wellcome Sanger Institute for the COVID-19 Genomics UK (COG-UK) Consortium | Jacquelyn Wynn, Mairead Hyland, The Lighthouse Lab in Alderley Park and Alex Alderton, Roberto Amato, Jeffrey Barrett, Sonia Goncalves, Ewan Harrison, David K. Jackson, Ian Johnston, Dominic Kwiatkowski, Cordelia Langford, John Sillitoe on behalf of the Wellcome Sanger Institute COVID-19 Surveillance Team |
| EPI_ISL_3366590, EPI_ISL_3366591 | Health Services Laboratories | Wellcome Sanger Institute for the COVID-19 Genomics UK (COG-UK) Consortium | Health Services Laboratories and Alex Alderton, Roberto Amato, Jeffrey Barrett, Sonia Goncalves, Ewan Harrison, David K. Jackson, Ian Johnston, Dominic Kwiatkowski, Cordelia Langford, John Sillitoe on behalf of the Wellcome Sanger Institute COVID-19 Surveillance Team |
| EPI_ISL_3366592, EPI_ISL_3366593, EPI_ISL_3366594 | Lighthouse Lab in Alderley Park | Wellcome Sanger Institute for the COVID-19 Genomics UK (COG-UK) Consortium | Jacquelyn Wynn, Mairead Hyland, The Lighthouse Lab in Alderley Park and Alex Alderton, Roberto Amato, Jeffrey Barrett, Sonia Goncalves, Ewan Harrison, David K. Jackson, Ian Johnston, Dominic Kwiatkowski, Cordelia Langford, John Sillitoe on behalf of the Wellcome Sanger Institute COVID-19 Surveillance Team |
| EPI_ISL_3366595, EPI_ISL_3366596 | Health Services Laboratories | Wellcome Sanger Institute for the COVID-19 Genomics UK (COG-UK) Consortium | Health Services Laboratories and Alex Alderton, Roberto Amato, Jeffrey Barrett, Sonia Goncalves, Ewan Harrison, David K. Jackson, Ian Johnston, Dominic Kwiatkowski, Cordelia Langford, John Sillitoe on behalf of the Wellcome Sanger Institute COVID-19 Surveillance Team |
| EPI_ISL_3366597, EPI_ISL_3366598, EPI_ISL_3366599, EPI_ISL_3366600, EPI_ISL_3366601, EPI_ISL_3366602, EPI_ISL_3366603, EPI_ISL_3366604, EPI_ISL_3366605, EPI_ISL_3366606, EPI_ISL_3366607, EPI_ISL_3366608, EPI_ISL_3366609, EPI_ISL_3366610, EPI_ISL_3366611, EPI_ISL_3366612, EPI_ISL_3366613, EPI_ISL_3366614, EPI_ISL_3366615, EPI_ISL_3366616, EPI_ISL_3366617, EPI_ISL_3366618, EPI_ISL_3366619, EPI_ISL_3366620, EPI_ISL_3366621, EPI_ISL_3366622, EPI_ISL_3366623, EPI_ISL_3366624, EPI_ISL_3366625, EPI_ISL_3366626, EPI_ISL_3366627, EPI_ISL_3366628, EPI_ISL_3366629, EPI_ISL_3366630, EPI_ISL_3366631, EPI_ISL_3366632, EPI_ISL_3366633, EPI_ISL_3366634, EPI_ISL_3366635, EPI_ISL_3366636, EPI_ISL_3366637, EPI_ISL_3366638, EPI_ISL_3366639, EPI_ISL_3366640, EPI_ISL_3366641, EPI_ISL_3366642, EPI_ISL_3366643, EPI_ISL_3366644, EPI_ISL_3366645, EPI_ISL_3366646, EPI_ISL_3366647, EPI_ISL_3366648, EPI_ISL_3366649, EPI_ISL_3366650, EPI_ISL_3366651, EPI_ISL_3366652 | | | |
| see above | Lighthouse Lab in Glasgow | Wellcome Sanger Institute for the COVID-19 Genomics UK (COG-UK) Consortium | Harper VanSteenhouse, Yumi Kasai, David Gray, Carol Clugston, Anna Dominiczak and Alex Alderton, Roberto Amato, Jeffrey Barrett, Sonia Goncalves, Ewan Harrison, David K. Jackson, Ian Johnston, Dominic Kwiatkowski, Cordelia Langford, John Sillitoe on behalf of the Wellcome Sanger Institute COVID-19 Surveillance Team |
| EPI_ISL_3366653, EPI_ISL_3366654 | Lighthouse Lab in Milton Keynes | Wellcome Sanger Institute for the COVID-19 Genomics UK (COG-UK) Consortium | The Lighthouse Lab in Milton Keynes and Alex Alderton, Roberto Amato, Jeffrey Barrett, Sonia Goncalves, Ewan Harrison, David K. Jackson, Ian Johnston, Dominic Kwiatkowski, Cordelia Langford, John Sillitoe on behalf of the Wellcome Sanger Institute COVID-19 Surveillance Team |
| EPI_ISL_3366655, EPI_ISL_3366656 | Lighthouse Laboratory Plymouth | Wellcome Sanger Institute for the COVID-19 Genomics UK (COG-UK) Consortium | Lighthouse Laboratory Plymouth and Alex Alderton, Roberto Amato, Jeffrey Barrett, Sonia Goncalves, Ewan Harrison, David K. Jackson, Ian Johnston, Dominic Kwiatkowski, Cordelia Langford, John Sillitoe on behalf of the Wellcome Sanger Institute COVID-19 Surveillance Team |
| EPI_ISL_3366657, EPI_ISL_3366658, EPI_ISL_3366659 | Lighthouse Lab in Glasgow | Wellcome Sanger Institute for the COVID-19 Genomics UK (COG-UK) Consortium | Harper VanSteenhouse, Yumi Kasai, David Gray, Carol Clugston, Anna Dominiczak and Alex Alderton, Roberto Amato, Jeffrey Barrett, Sonia Goncalves, Ewan Harrison, David K. Jackson, Ian Johnston, Dominic Kwiatkowski, Cordelia Langford, John Sillitoe on behalf of the Wellcome Sanger Institute COVID-19 Surveillance Team |
| EPI_ISL_3366660, EPI_ISL_3366661 | Lighthouse Lab in Alderley Park | Wellcome Sanger Institute for the COVID-19 Genomics UK (COG-UK) Consortium | Jacquelyn Wynn, Mairead Hyland, The Lighthouse Lab in Alderley Park and Alex Alderton, Roberto Amato, Jeffrey Barrett, Sonia Goncalves, Ewan Harrison, David K. Jackson, Ian Johnston, Dominic Kwiatkowski, Cordelia Langford, John Sillitoe on behalf of the Wellcome Sanger Institute COVID-19 Surveillance Team |
| EPI_ISL_3366662 | Lighthouse Lab in Glasgow | Wellcome Sanger Institute for the COVID-19 Genomics UK (COG-UK) Consortium | Harper VanSteenhouse, Yumi Kasai, David Gray, Carol Clugston, Anna Dominiczak and Alex Alderton, Roberto Amato, Jeffrey Barrett, Sonia Goncalves, Ewan Harrison, David K. Jackson, Ian Johnston, Dominic Kwiatkowski, Cordelia Langford, John Sillitoe on behalf of the Wellcome Sanger Institute COVID-19 Surveillance Team |
| EPI_ISL_3366663 | Lighthouse Lab in Alderley Park | Wellcome Sanger Institute for the COVID-19 Genomics UK (COG-UK) Consortium | Jacquelyn Wynn, Mairead Hyland, The Lighthouse Lab in Alderley Park and Alex Alderton, Roberto Amato, Jeffrey Barrett, Sonia Goncalves, Ewan Harrison, David K. Jackson, Ian Johnston, Dominic Kwiatkowski, Cordelia Langford, John Sillitoe on behalf of the Wellcome Sanger Institute COVID-19 Surveillance Team |
| EPI_ISL_3366664 | Lighthouse Lab in Glasgow | Wellcome Sanger Institute for the COVID-19 Genomics UK (COG-UK) Consortium | Harper VanSteenhouse, Yumi Kasai, David Gray, Carol Clugston, Anna Dominiczak and Alex Alderton, Roberto Amato, Jeffrey Barrett, Sonia Goncalves, Ewan Harrison, David K. Jackson, Ian Johnston, Dominic Kwiatkowski, Cordelia Langford, John Sillitoe on behalf of the Wellcome Sanger Institute COVID-19 Surveillance Team |
| EPI_ISL_3366665, EPI_ISL_3366666 | Lighthouse Lab in Alderley Park | Wellcome Sanger Institute for the COVID-19 Genomics UK (COG-UK) Consortium | Jacquelyn Wynn, Mairead Hyland, The Lighthouse Lab in Alderley Park and Alex Alderton, Roberto Amato, Jeffrey Barrett, Sonia Goncalves, Ewan Harrison, David K. Jackson, Ian Johnston, Dominic Kwiatkowski, Cordelia Langford, John Sillitoe on behalf of the Wellcome Sanger Institute COVID-19 Surveillance Team |
| EPI_ISL_3366667 | Lighthouse Lab in Glasgow | Wellcome Sanger Institute for the COVID-19 Genomics UK (COG-UK) Consortium | Harper VanSteenhouse, Yumi Kasai, David Gray, Carol Clugston, Anna Dominiczak and Alex Alderton, Roberto Amato, Jeffrey Barrett, Sonia Goncalves, Ewan Harrison, David K. Jackson, Ian Johnston, Dominic Kwiatkowski, Cordelia Langford, John Sillitoe on behalf of the Wellcome Sanger Institute COVID-19 Surveillance Team |
| EPI_ISL_3366668, EPI_ISL_3366669, EPI_ISL_3366670, EPI_ISL_3366671, EPI_ISL_3366672 | Lighthouse Lab in Alderley Park | Wellcome Sanger Institute for the COVID-19 Genomics UK (COG-UK) Consortium | Jacquelyn Wynn, Mairead Hyland, The Lighthouse Lab in Alderley Park and Alex Alderton, Roberto Amato, Jeffrey Barrett, Sonia Goncalves, Ewan Harrison, David K. Jackson, Ian Johnston, Dominic Kwiatkowski, Cordelia Langford, John Sillitoe on behalf of the Wellcome Sanger Institute COVID-19 Surveillance Team |
| EPI_ISL_3366673 | Lighthouse Lab in Glasgow | Wellcome Sanger Institute for the COVID-19 Genomics UK (COG-UK) Consortium | Harper VanSteenhouse, Yumi Kasai, David Gray, Carol Clugston, Anna Dominiczak and Alex Alderton, Roberto Amato, Jeffrey Barrett, Sonia Goncalves, Ewan Harrison, David K. Jackson, Ian Johnston, Dominic Kwiatkowski, Cordelia Langford, John Sillitoe on behalf of the Wellcome Sanger Institute COVID-19 Surveillance Team |
| EPI_ISL_3366674 | Lighthouse Lab in Alderley Park | Wellcome Sanger Institute for the COVID-19 Genomics UK (COG-UK) Consortium | Jacquelyn Wynn, Mairead Hyland, The Lighthouse Lab in Alderley Park and Alex Alderton, Roberto Amato, Jeffrey Barrett, Sonia Goncalves, Ewan Harrison, David K. Jackson, Ian Johnston, Dominic Kwiatkowski, Cordelia Langford, John Sillitoe on behalf of the Wellcome Sanger Institute COVID-19 Surveillance Team |
| EPI_ISL_3366675, EPI_ISL_3366676, EPI_ISL_3366677 | Lighthouse Lab in Glasgow | Wellcome Sanger Institute for the COVID-19 Genomics UK (COG-UK) Consortium | Harper VanSteenhouse, Yumi Kasai, David Gray, Carol Clugston, Anna Dominiczak and Alex Alderton, Roberto Amato, Jeffrey Barrett, Sonia Goncalves, Ewan Harrison, David K. Jackson, Ian Johnston, Dominic Kwiatkowski, Cordelia Langford, John Sillitoe on behalf of the Wellcome Sanger Institute COVID-19 Surveillance Team |
| EPI_ISL_3366678, | Lighthouse Lab in Alderley Park | Wellcome Sanger Institute for the COVID-19 Genomics UK (COG-UK) Consortium | Jacquelyn Wynn, Mairead Hyland, The Lighthouse Lab in Alderley Park and Alex Alderton, Roberto Amato, Jeffrey Barrett, Sonia Goncalves, Ewan Harrison, David K. Jackson, Ian Johnston, Dominic Kwiatkowski, Cordelia Langford, John Sillitoe on |

| | | | |
|---|---|---|---|
| EPI_ISL_3366679, EPI_ISL_3366680 | | Consortium | behalf of the Wellcome Sanger Institute COVID-19 Surveillance Team |
| EPI_ISL_3366681, EPI_ISL_3366682, EPI_ISL_3366683 | Lighthouse Lab in Glasgow | Wellcome Sanger Institute for the COVID-19 Genomics UK (COG-UK) Consortium | Harper VanSteenhouse, Yumi Kasai, David Gray, Carol Clugston, Anna Dominiczak and Alex Alderton, Roberto Amato, Jeffrey Barrett, Sonia Goncalves, Ewan Harrison, David K. Jackson, Ian Johnston, Dominic Kwiatkowski, Cordelia Langford, John Sillitoe on behalf of the Wellcome Sanger Institute COVID-19 Surveillance Team |
| EPI_ISL_3366684 | Lighthouse Lab in Alderley Park | Wellcome Sanger Institute for the COVID-19 Genomics UK (COG-UK) Consortium | Jacquelyn Wynn, Mairead Hyland, The Lighthouse Lab in Alderley Park and Alex Alderton, Roberto Amato, Jeffrey Barrett, Sonia Goncalves, Ewan Harrison, David K. Jackson, Ian Johnston, Dominic Kwiatkowski, Cordelia Langford, John Sillitoe on behalf of the Wellcome Sanger Institute COVID-19 Surveillance Team |
| EPI_ISL_3366685 | Lighthouse Lab in Glasgow | Wellcome Sanger Institute for the COVID-19 Genomics UK (COG-UK) Consortium | Harper VanSteenhouse, Yumi Kasai, David Gray, Carol Clugston, Anna Dominiczak and Alex Alderton, Roberto Amato, Jeffrey Barrett, Sonia Goncalves, Ewan Harrison, David K. Jackson, Ian Johnston, Dominic Kwiatkowski, Cordelia Langford, John Sillitoe on behalf of the Wellcome Sanger Institute COVID-19 Surveillance Team |
| EPI_ISL_3366686 | Lighthouse Lab in Alderley Park | Wellcome Sanger Institute for the COVID-19 Genomics UK (COG-UK) Consortium | Jacquelyn Wynn, Mairead Hyland, The Lighthouse Lab in Alderley Park and Alex Alderton, Roberto Amato, Jeffrey Barrett, Sonia Goncalves, Ewan Harrison, David K. Jackson, Ian Johnston, Dominic Kwiatkowski, Cordelia Langford, John Sillitoe on behalf of the Wellcome Sanger Institute COVID-19 Surveillance Team |
| EPI_ISL_3366687, EPI_ISL_3366688, EPI_ISL_3366689 | Lighthouse Lab in Glasgow | Wellcome Sanger Institute for the COVID-19 Genomics UK (COG-UK) Consortium | Harper VanSteenhouse, Yumi Kasai, David Gray, Carol Clugston, Anna Dominiczak and Alex Alderton, Roberto Amato, Jeffrey Barrett, Sonia Goncalves, Ewan Harrison, David K. Jackson, Ian Johnston, Dominic Kwiatkowski, Cordelia Langford, John Sillitoe on behalf of the Wellcome Sanger Institute COVID-19 Surveillance Team |
| EPI_ISL_3366690, EPI_ISL_3366691, EPI_ISL_3366692 | Lighthouse Lab in Alderley Park | Wellcome Sanger Institute for the COVID-19 Genomics UK (COG-UK) Consortium | Jacquelyn Wynn, Mairead Hyland, The Lighthouse Lab in Alderley Park and Alex Alderton, Roberto Amato, Jeffrey Barrett, Sonia Goncalves, Ewan Harrison, David K. Jackson, Ian Johnston, Dominic Kwiatkowski, Cordelia Langford, John Sillitoe on behalf of the Wellcome Sanger Institute COVID-19 Surveillance Team |
| EPI_ISL_3366693, EPI_ISL_3366694, EPI_ISL_3366695 | Lighthouse Lab in Milton Keynes | Wellcome Sanger Institute for the COVID-19 Genomics UK (COG-UK) Consortium | The Lighthouse Lab in Milton Keynes and Alex Alderton, Roberto Amato, Jeffrey Barrett, Sonia Goncalves, Ewan Harrison, David K. Jackson, Ian Johnston, Dominic Kwiatkowski, Cordelia Langford, John Sillitoe on behalf of the Wellcome Sanger Institute COVID-19 Surveillance Team |
| EPI_ISL_3366696, EPI_ISL_3366697 | Lighthouse Lab in Alderley Park | Wellcome Sanger Institute for the COVID-19 Genomics UK (COG-UK) Consortium | Jacquelyn Wynn, Mairead Hyland, The Lighthouse Lab in Alderley Park and Alex Alderton, Roberto Amato, Jeffrey Barrett, Sonia Goncalves, Ewan Harrison, David K. Jackson, Ian Johnston, Dominic Kwiatkowski, Cordelia Langford, John Sillitoe on behalf of the Wellcome Sanger Institute COVID-19 Surveillance Team |
| EPI_ISL_3366698 | Lighthouse Lab in Milton Keynes | Wellcome Sanger Institute for the COVID-19 Genomics UK (COG-UK) Consortium | The Lighthouse Lab in Milton Keynes and Alex Alderton, Roberto Amato, Jeffrey Barrett, Sonia Goncalves, Ewan Harrison, David K. Jackson, Ian Johnston, Dominic Kwiatkowski, Cordelia Langford, John Sillitoe on behalf of the Wellcome Sanger Institute COVID-19 Surveillance Team |
| EPI_ISL_3366699, EPI_ISL_3366700 | Lighthouse Lab in Alderley Park | Wellcome Sanger Institute for the COVID-19 Genomics UK (COG-UK) Consortium | Jacquelyn Wynn, Mairead Hyland, The Lighthouse Lab in Alderley Park and Alex Alderton, Roberto Amato, Jeffrey Barrett, Sonia Goncalves, Ewan Harrison, David K. Jackson, Ian Johnston, Dominic Kwiatkowski, Cordelia Langford, John Sillitoe on behalf of the Wellcome Sanger Institute COVID-19 Surveillance Team |
| EPI_ISL_3366701 | Lighthouse Lab in Milton Keynes | Wellcome Sanger Institute for the COVID-19 Genomics UK (COG-UK) Consortium | The Lighthouse Lab in Milton Keynes and Alex Alderton, Roberto Amato, Jeffrey Barrett, Sonia Goncalves, Ewan Harrison, David K. Jackson, Ian Johnston, Dominic Kwiatkowski, Cordelia Langford, John Sillitoe on behalf of the Wellcome Sanger Institute COVID-19 Surveillance Team |
| EPI_ISL_3366702 | Lighthouse Lab in Alderley Park | Wellcome Sanger Institute for the COVID-19 Genomics UK (COG-UK) Consortium | Jacquelyn Wynn, Mairead Hyland, The Lighthouse Lab in Alderley Park and Alex Alderton, Roberto Amato, Jeffrey Barrett, Sonia Goncalves, Ewan Harrison, David K. Jackson, Ian Johnston, Dominic Kwiatkowski, Cordelia Langford, John Sillitoe on behalf of the Wellcome Sanger Institute COVID-19 Surveillance Team |
| EPI_ISL_3366703, EPI_ISL_3366704 | Lighthouse Lab in Milton Keynes | Wellcome Sanger Institute for the COVID-19 Genomics UK (COG-UK) Consortium | The Lighthouse Lab in Milton Keynes and Alex Alderton, Roberto Amato, Jeffrey Barrett, Sonia Goncalves, Ewan Harrison, David K. Jackson, Ian Johnston, Dominic Kwiatkowski, Cordelia Langford, John Sillitoe on behalf of the Wellcome Sanger Institute COVID-19 Surveillance Team |
| EPI_ISL_3366705 | Lighthouse Lab in Alderley Park | Wellcome Sanger Institute for the COVID-19 Genomics UK (COG-UK) Consortium | Jacquelyn Wynn, Mairead Hyland, The Lighthouse Lab in Alderley Park and Alex Alderton, Roberto Amato, Jeffrey Barrett, Sonia Goncalves, Ewan Harrison, David K. Jackson, Ian Johnston, Dominic Kwiatkowski, Cordelia Langford, John Sillitoe on behalf of the Wellcome Sanger Institute COVID-19 Surveillance Team |
| EPI_ISL_3366706, EPI_ISL_3366707 | Lighthouse Lab in Glasgow | Wellcome Sanger Institute for the COVID-19 Genomics UK (COG-UK) Consortium | Harper VanSteenhouse, Yumi Kasai, David Gray, Carol Clugston, Anna Dominiczak and Alex Alderton, Roberto Amato, Jeffrey Barrett, Sonia Goncalves, Ewan Harrison, David K. Jackson, Ian Johnston, Dominic Kwiatkowski, Cordelia Langford, John Sillitoe on behalf of the Wellcome Sanger Institute COVID-19 Surveillance Team |
| EPI_ISL_3366708 | Lighthouse Lab in Alderley Park | Wellcome Sanger Institute for the COVID-19 Genomics UK (COG-UK) Consortium | Jacquelyn Wynn, Mairead Hyland, The Lighthouse Lab in Alderley Park and Alex Alderton, Roberto Amato, Jeffrey Barrett, Sonia Goncalves, Ewan Harrison, David K. Jackson, Ian Johnston, Dominic Kwiatkowski, Cordelia Langford, John Sillitoe on behalf of the Wellcome Sanger Institute COVID-19 Surveillance Team |
| EPI_ISL_3366709, EPI_ISL_3366710 | Lighthouse Lab in Glasgow | Wellcome Sanger Institute for the COVID-19 Genomics UK (COG-UK) Consortium | Harper VanSteenhouse, Yumi Kasai, David Gray, Carol Clugston, Anna Dominiczak and Alex Alderton, Roberto Amato, Jeffrey Barrett, Sonia Goncalves, Ewan Harrison, David K. Jackson, Ian Johnston, Dominic Kwiatkowski, Cordelia Langford, John Sillitoe on behalf of the Wellcome Sanger Institute COVID-19 Surveillance Team |
| EPI_ISL_3366711 | Lighthouse Lab in Alderley Park | Wellcome Sanger Institute for the COVID-19 Genomics UK (COG-UK) Consortium | Jacquelyn Wynn, Mairead Hyland, The Lighthouse Lab in Alderley Park and Alex Alderton, Roberto Amato, Jeffrey Barrett, Sonia Goncalves, Ewan Harrison, David K. Jackson, Ian Johnston, Dominic Kwiatkowski, Cordelia Langford, John Sillitoe on behalf of the Wellcome Sanger Institute COVID-19 Surveillance Team |
| EPI_ISL_3366712, EPI_ISL_3366713 | Lighthouse Lab in Glasgow | Wellcome Sanger Institute for the COVID-19 Genomics UK (COG-UK) Consortium | Harper VanSteenhouse, Yumi Kasai, David Gray, Carol Clugston, Anna Dominiczak and Alex Alderton, Roberto Amato, Jeffrey Barrett, Sonia Goncalves, Ewan Harrison, David K. Jackson, Ian Johnston, Dominic Kwiatkowski, Cordelia Langford, John Sillitoe on behalf of the Wellcome Sanger Institute COVID-19 Surveillance Team |
| EPI_ISL_3366714 | Lighthouse Laboratory Plymouth | Wellcome Sanger Institute for the COVID-19 Genomics UK (COG-UK) Consortium | Lighthouse Laboratory Plymouth and Alex Alderton, Roberto Amato, Jeffrey Barrett, Sonia Goncalves, Ewan Harrison, David K. Jackson, Ian Johnston, Dominic Kwiatkowski, Cordelia Langford, John Sillitoe on behalf of the Wellcome Sanger Institute COVID-19 Surveillance Team |
| EPI_ISL_3366715 | Lighthouse Lab in Alderley Park | Wellcome Sanger Institute for the COVID-19 Genomics UK (COG-UK) Consortium | Jacquelyn Wynn, Mairead Hyland, The Lighthouse Lab in Alderley Park and Alex Alderton, Roberto Amato, Jeffrey Barrett, Sonia Goncalves, Ewan Harrison, David K. Jackson, Ian Johnston, Dominic Kwiatkowski, Cordelia Langford, John Sillitoe on behalf of the Wellcome Sanger Institute COVID-19 Surveillance Team |
| EPI_ISL_3366716 | Lighthouse Laboratory Plymouth | Wellcome Sanger Institute for the COVID-19 Genomics UK (COG-UK) Consortium | Lighthouse Laboratory Plymouth and Alex Alderton, Roberto Amato, Jeffrey Barrett, Sonia Goncalves, Ewan Harrison, David K. Jackson, Ian Johnston, Dominic Kwiatkowski, Cordelia Langford, John Sillitoe on behalf of the Wellcome Sanger Institute COVID-19 Surveillance Team |
| EPI_ISL_3366717, EPI_ISL_3366718, EPI_ISL_3366719, EPI_ISL_3366720, EPI_ISL_3366721, EPI_ISL_3366722 | Lighthouse Lab in Glasgow | Wellcome Sanger Institute for the COVID-19 Genomics UK (COG-UK) Consortium | Harper VanSteenhouse, Yumi Kasai, David Gray, Carol Clugston, Anna Dominiczak and Alex Alderton, Roberto Amato, Jeffrey Barrett, Sonia Goncalves, Ewan Harrison, David K. Jackson, Ian Johnston, Dominic Kwiatkowski, Cordelia Langford, John Sillitoe on behalf of the Wellcome Sanger Institute COVID-19 Surveillance Team |
| EPI_ISL_3366723, EPI_ISL_3366724 | Lighthouse Lab in Alderley Park | Wellcome Sanger Institute for the COVID-19 Genomics UK (COG-UK) Consortium | Jacquelyn Wynn, Mairead Hyland, The Lighthouse Lab in Alderley Park and Alex Alderton, Roberto Amato, Jeffrey Barrett, Sonia Goncalves, Ewan Harrison, David K. Jackson, Ian Johnston, Dominic Kwiatkowski, Cordelia Langford, John Sillitoe on behalf of the Wellcome Sanger Institute COVID-19 Surveillance Team |
| EPI_ISL_3366725, EPI_ISL_3366726, EPI_ISL_3366727 | Lighthouse Lab in Glasgow | Wellcome Sanger Institute for the COVID-19 Genomics UK (COG-UK) Consortium | Harper VanSteenhouse, Yumi Kasai, David Gray, Carol Clugston, Anna Dominiczak and Alex Alderton, Roberto Amato, Jeffrey Barrett, Sonia Goncalves, Ewan Harrison, David K. Jackson, Ian Johnston, Dominic Kwiatkowski, Cordelia Langford, John Sillitoe on behalf of the Wellcome Sanger Institute COVID-19 Surveillance Team |
| EPI_ISL_3366728 | Lighthouse Lab in Alderley Park | Wellcome Sanger Institute for the COVID-19 Genomics UK (COG-UK) Consortium | Jacquelyn Wynn, Mairead Hyland, The Lighthouse Lab in Alderley Park and Alex Alderton, Roberto Amato, Jeffrey Barrett, Sonia Goncalves, Ewan Harrison, David K. Jackson, Ian Johnston, Dominic Kwiatkowski, Cordelia Langford, John Sillitoe on behalf of the Wellcome Sanger Institute COVID-19 Surveillance Team |
| EPI_ISL_3366729, EPI_ISL_3366730 | Lighthouse Lab in Glasgow | Wellcome Sanger Institute for the COVID-19 Genomics UK (COG-UK) Consortium | Harper VanSteenhouse, Yumi Kasai, David Gray, Carol Clugston, Anna Dominiczak and Alex Alderton, Roberto Amato, Jeffrey Barrett, Sonia Goncalves, Ewan Harrison, David K. Jackson, Ian Johnston, Dominic Kwiatkowski, Cordelia Langford, John Sillitoe on behalf of the Wellcome Sanger Institute COVID-19 Surveillance Team |
| EPI_ISL_3366731 | Lighthouse Lab in Alderley Park | Wellcome Sanger Institute for the COVID-19 Genomics UK (COG-UK) Consortium | Jacquelyn Wynn, Mairead Hyland, The Lighthouse Lab in Alderley Park and Alex Alderton, Roberto Amato, Jeffrey Barrett, Sonia Goncalves, Ewan Harrison, David K. Jackson, Ian Johnston, Dominic Kwiatkowski, Cordelia Langford, John Sillitoe on behalf of the Wellcome Sanger Institute COVID-19 Surveillance Team |
| EPI_ISL_3366732, EPI_ISL_3366733 | Lighthouse Lab in Glasgow | Wellcome Sanger Institute for the COVID-19 Genomics UK (COG-UK) Consortium | Harper VanSteenhouse, Yumi Kasai, David Gray, Carol Clugston, Anna Dominiczak and Alex Alderton, Roberto Amato, Jeffrey Barrett, Sonia Goncalves, Ewan Harrison, David K. Jackson, Ian Johnston, Dominic Kwiatkowski, Cordelia Langford, John Sillitoe on behalf of the Wellcome Sanger Institute COVID-19 Surveillance Team |
| EPI_ISL_3366734, EPI_ISL_3366735 | Lighthouse Lab in Alderley Park | Wellcome Sanger Institute for the COVID-19 Genomics UK (COG-UK) Consortium | Jacquelyn Wynn, Mairead Hyland, The Lighthouse Lab in Alderley Park and Alex Alderton, Roberto Amato, Jeffrey Barrett, Sonia Goncalves, Ewan Harrison, David K. Jackson, Ian Johnston, Dominic Kwiatkowski, Cordelia Langford, John Sillitoe on behalf of the Wellcome Sanger Institute COVID-19 Surveillance Team |
| EPI_ISL_3366736 | Lighthouse Lab in Glasgow | Wellcome Sanger Institute for the COVID-19 Genomics UK (COG-UK) Consortium | Harper VanSteenhouse, Yumi Kasai, David Gray, Carol Clugston, Anna Dominiczak and Alex Alderton, Roberto Amato, Jeffrey Barrett, Sonia Goncalves, Ewan Harrison, David K. Jackson, Ian Johnston, Dominic Kwiatkowski, Cordelia Langford, John Sillitoe on behalf of the Wellcome Sanger Institute COVID-19 Surveillance Team |
| EPI_ISL_3366737, EPI_ISL_3366738 | Lighthouse Lab in Alderley Park | Wellcome Sanger Institute for the COVID-19 Genomics UK (COG-UK) Consortium | Jacquelyn Wynn, Mairead Hyland, The Lighthouse Lab in Alderley Park and Alex Alderton, Roberto Amato, Jeffrey Barrett, Sonia Goncalves, Ewan Harrison, David K. Jackson, Ian Johnston, Dominic Kwiatkowski, Cordelia Langford, John Sillitoe on behalf of the Wellcome Sanger Institute COVID-19 Surveillance Team |
| EPI_ISL_3366739 | Lighthouse Lab in Glasgow | Wellcome Sanger Institute for the COVID-19 Genomics UK (COG-UK) Consortium | Harper VanSteenhouse, Yumi Kasai, David Gray, Carol Clugston, Anna Dominiczak and Alex Alderton, Roberto Amato, Jeffrey Barrett, Sonia Goncalves, Ewan Harrison, David K. Jackson, Ian Johnston, Dominic Kwiatkowski, Cordelia Langford, John Sillitoe on behalf of the Wellcome Sanger Institute COVID-19 Surveillance Team |
| EPI_ISL_3366740, EPI_ISL_3366741, EPI_ISL_3366742 | Lighthouse Lab in Alderley Park | Wellcome Sanger Institute for the COVID-19 Genomics UK (COG-UK) Consortium | Jacquelyn Wynn, Mairead Hyland, The Lighthouse Lab in Alderley Park and Alex Alderton, Roberto Amato, Jeffrey Barrett, Sonia Goncalves, Ewan Harrison, David K. Jackson, Ian Johnston, Dominic Kwiatkowski, Cordelia Langford, John Sillitoe on behalf of the Wellcome Sanger Institute COVID-19 Surveillance Team |
| EPI_ISL_3366743, EPI_ISL_3366744 | Lighthouse Lab in Glasgow | Wellcome Sanger Institute for the COVID-19 Genomics UK (COG-UK) Consortium | Harper VanSteenhouse, Yumi Kasai, David Gray, Carol Clugston, Anna Dominiczak and Alex Alderton, Roberto Amato, Jeffrey Barrett, Sonia Goncalves, Ewan Harrison, David K. Jackson, Ian Johnston, Dominic Kwiatkowski, Cordelia Langford, John Sillitoe on behalf of the Wellcome Sanger Institute COVID-19 Surveillance Team |
| EPI_ISL_3366745 | Lighthouse Lab in Alderley Park | Wellcome Sanger Institute for the COVID-19 Genomics UK (COG-UK) Consortium | Jacquelyn Wynn, Mairead Hyland, The Lighthouse Lab in Alderley Park and Alex Alderton, Roberto Amato, Jeffrey Barrett, Sonia Goncalves, Ewan Harrison, David K. Jackson, Ian Johnston, Dominic Kwiatkowski, Cordelia Langford, John Sillitoe on behalf of the Wellcome Sanger Institute COVID-19 Surveillance Team |
| EPI_ISL_3366746, EPI_ISL_3366747, EPI_ISL_3366748 | Lighthouse Lab in Glasgow | Wellcome Sanger Institute for the COVID-19 Genomics UK (COG-UK) Consortium | Harper VanSteenhouse, Yumi Kasai, David Gray, Carol Clugston, Anna Dominiczak and Alex Alderton, Roberto Amato, Jeffrey Barrett, Sonia Goncalves, Ewan Harrison, David K. Jackson, Ian Johnston, Dominic Kwiatkowski, Cordelia Langford, John Sillitoe on behalf of the Wellcome Sanger Institute COVID-19 Surveillance Team |
| EPI_ISL_3366749, EPI_ISL_3366750, EPI_ISL_3366751, EPI_ISL_3366752, EPI_ISL_3366753 | Lighthouse Lab in Alderley Park | Wellcome Sanger Institute for the COVID-19 Genomics UK (COG-UK) Consortium | Jacquelyn Wynn, Mairead Hyland, The Lighthouse Lab in Alderley Park and Alex Alderton, Roberto Amato, Jeffrey Barrett, Sonia Goncalves, Ewan Harrison, David K. Jackson, Ian Johnston, Dominic Kwiatkowski, Cordelia Langford, John Sillitoe on behalf of the Wellcome Sanger Institute COVID-19 Surveillance Team |
| EPI_ISL_3366754, | Lighthouse Lab in Glasgow | Wellcome Sanger Institute for the COVID-19 Genomics UK (COG-UK) | Harper VanSteenhouse, Yumi Kasai, David Gray, Carol Clugston, Anna Dominiczak and Alex Alderton, Roberto Amato, Jeffrey Barrett, Sonia Goncalves, Ewan Harrison, David K. Jackson, Ian Johnston, Dominic Kwiatkowski, Cordelia Langford, |

| | | | |
|---|---|---|---|
| EPI_ISL_3366755 | | Consortium | John Sillitoe on behalf of the Wellcome Sanger Institute COVID-19 Surveillance Team |
| EPI_ISL_3366756 | Lighthouse Lab in Alderley Park | Wellcome Sanger Institute for the COVID-19 Genomics UK (COG-UK) Consortium | Jacquelyn Wynn, Mairead Hyland, The Lighthouse Lab in Alderley Park and Alex Alderton, Roberto Amato, Jeffrey Barrett, Sonia Goncalves, Ewan Harrison, David K. Jackson, Ian Johnston, Dominic Kwiatkowski, Cordelia Langford, John Sillitoe on behalf of the Wellcome Sanger Institute COVID-19 Surveillance Team |
| EPI_ISL_3366757, EPI_ISL_3366758, EPI_ISL_3366759, EPI_ISL_3366760 | Lighthouse Lab in Glasgow | Wellcome Sanger Institute for the COVID-19 Genomics UK (COG-UK) Consortium | Harper VanSteenhouse, Yumi Kasai, David Gray, Carol Clugston, Anna Dominiczak and Alex Alderton, Roberto Amato, Jeffrey Barrett, Sonia Goncalves, Ewan Harrison, David K. Jackson, Ian Johnston, Dominic Kwiatkowski, Cordelia Langford, John Sillitoe on behalf of the Wellcome Sanger Institute COVID-19 Surveillance Team |
| EPI_ISL_3366761 | Lighthouse Lab in Alderley Park | Wellcome Sanger Institute for the COVID-19 Genomics UK (COG-UK) Consortium | Jacquelyn Wynn, Mairead Hyland, The Lighthouse Lab in Alderley Park and Alex Alderton, Roberto Amato, Jeffrey Barrett, Sonia Goncalves, Ewan Harrison, David K. Jackson, Ian Johnston, Dominic Kwiatkowski, Cordelia Langford, John Sillitoe on behalf of the Wellcome Sanger Institute COVID-19 Surveillance Team |
| EPI_ISL_3366762, EPI_ISL_3366763, EPI_ISL_3366764, EPI_ISL_3366765, EPI_ISL_3366766, EPI_ISL_3366767, EPI_ISL_3366768, EPI_ISL_3366769, EPI_ISL_3366770, EPI_ISL_3366771, EPI_ISL_3366772 | | | |
| see above | Lighthouse Lab in Glasgow | Wellcome Sanger Institute for the COVID-19 Genomics UK (COG-UK) Consortium | Harper VanSteenhouse, Yumi Kasai, David Gray, Carol Clugston, Anna Dominiczak and Alex Alderton, Roberto Amato, Jeffrey Barrett, Sonia Goncalves, Ewan Harrison, David K. Jackson, Ian Johnston, Dominic Kwiatkowski, Cordelia Langford, John Sillitoe on behalf of the Wellcome Sanger Institute COVID-19 Surveillance Team |
| EPI_ISL_3366773 | Lighthouse Lab in Milton Keynes | Wellcome Sanger Institute for the COVID-19 Genomics UK (COG-UK) Consortium | The Lighthouse Lab in Milton Keynes and Alex Alderton, Roberto Amato, Jeffrey Barrett, Sonia Goncalves, Ewan Harrison, David K. Jackson, Ian Johnston, Dominic Kwiatkowski, Cordelia Langford, John Sillitoe on behalf of the Wellcome Sanger Institute COVID-19 Surveillance Team |
| EPI_ISL_3366774, EPI_ISL_3366775, EPI_ISL_3366776, EPI_ISL_3366777, EPI_ISL_3366778, EPI_ISL_3366779, EPI_ISL_3366780, EPI_ISL_3366781, EPI_ISL_3366782, EPI_ISL_3366783, EPI_ISL_3366784, EPI_ISL_3366785, EPI_ISL_3366786, EPI_ISL_3366787, EPI_ISL_3366788 | | | |
| see above | Lighthouse Lab in Glasgow | Wellcome Sanger Institute for the COVID-19 Genomics UK (COG-UK) Consortium | Harper VanSteenhouse, Yumi Kasai, David Gray, Carol Clugston, Anna Dominiczak and Alex Alderton, Roberto Amato, Jeffrey Barrett, Sonia Goncalves, Ewan Harrison, David K. Jackson, Ian Johnston, Dominic Kwiatkowski, Cordelia Langford, John Sillitoe on behalf of the Wellcome Sanger Institute COVID-19 Surveillance Team |
| EPI_ISL_3366789, EPI_ISL_3366790, EPI_ISL_3366791, EPI_ISL_3366792, EPI_ISL_3366793, EPI_ISL_3366794, EPI_ISL_3366795, EPI_ISL_3366796, EPI_ISL_3366797 | Lighthouse Lab in Alderley Park | Wellcome Sanger Institute for the COVID-19 Genomics UK (COG-UK) Consortium | Jacquelyn Wynn, Mairead Hyland, The Lighthouse Lab in Alderley Park and Alex Alderton, Roberto Amato, Jeffrey Barrett, Sonia Goncalves, Ewan Harrison, David K. Jackson, Ian Johnston, Dominic Kwiatkowski, Cordelia Langford, John Sillitoe on behalf of the Wellcome Sanger Institute COVID-19 Surveillance Team |
| EPI_ISL_3366798 | Berkshire and Surrey Pathology Services Lighthouse Laboratory | Wellcome Sanger Institute for the COVID-19 Genomics UK (COG-UK) Consortium | Berkshire and Surrey Pathology Services Lighthouse Laboratory and Alex Alderton, Roberto Amato, Jeffrey Barrett, Sonia Goncalves, Ewan Harrison, David K. Jackson, Ian Johnston, Dominic Kwiatkowski, Cordelia Langford, John Sillitoe on behalf of the Wellcome Sanger Institute COVID-19 Surveillance Team |
| EPI_ISL_3366799, EPI_ISL_3366800, EPI_ISL_3366801 | Lighthouse Lab in Alderley Park | Wellcome Sanger Institute for the COVID-19 Genomics UK (COG-UK) Consortium | Jacquelyn Wynn, Mairead Hyland, The Lighthouse Lab in Alderley Park and Alex Alderton, Roberto Amato, Jeffrey Barrett, Sonia Goncalves, Ewan Harrison, David K. Jackson, Ian Johnston, Dominic Kwiatkowski, Cordelia Langford, John Sillitoe on behalf of the Wellcome Sanger Institute COVID-19 Surveillance Team |
| EPI_ISL_3366802 | Berkshire and Surrey Pathology Services Lighthouse Laboratory | Wellcome Sanger Institute for the COVID-19 Genomics UK (COG-UK) Consortium | Berkshire and Surrey Pathology Services Lighthouse Laboratory and Alex Alderton, Roberto Amato, Jeffrey Barrett, Sonia Goncalves, Ewan Harrison, David K. Jackson, Ian Johnston, Dominic Kwiatkowski, Cordelia Langford, John Sillitoe on behalf of the Wellcome Sanger Institute COVID-19 Surveillance Team |
| EPI_ISL_3366803, EPI_ISL_3366804 | Lighthouse Lab in Alderley Park | Wellcome Sanger Institute for the COVID-19 Genomics UK (COG-UK) Consortium | Jacquelyn Wynn, Mairead Hyland, The Lighthouse Lab in Alderley Park and Alex Alderton, Roberto Amato, Jeffrey Barrett, Sonia Goncalves, Ewan Harrison, David K. Jackson, Ian Johnston, Dominic Kwiatkowski, Cordelia Langford, John Sillitoe on behalf of the Wellcome Sanger Institute COVID-19 Surveillance Team |
| EPI_ISL_3366805 | Lighthouse Lab in Milton Keynes | Wellcome Sanger Institute for the COVID-19 Genomics UK (COG-UK) Consortium | The Lighthouse Lab in Milton Keynes and Alex Alderton, Roberto Amato, Jeffrey Barrett, Sonia Goncalves, Ewan Harrison, David K. Jackson, Ian Johnston, Dominic Kwiatkowski, Cordelia Langford, John Sillitoe on behalf of the Wellcome Sanger Institute COVID-19 Surveillance Team |
| EPI_ISL_3366806 | Lighthouse Lab in Alderley Park | Wellcome Sanger Institute for the COVID-19 Genomics UK (COG-UK) Consortium | Jacquelyn Wynn, Mairead Hyland, The Lighthouse Lab in Alderley Park and Alex Alderton, Roberto Amato, Jeffrey Barrett, Sonia Goncalves, Ewan Harrison, David K. Jackson, Ian Johnston, Dominic Kwiatkowski, Cordelia Langford, John Sillitoe on behalf of the Wellcome Sanger Institute COVID-19 Surveillance Team |
| EPI_ISL_3366807, EPI_ISL_3366808 | Lighthouse Lab in Milton Keynes | Wellcome Sanger Institute for the COVID-19 Genomics UK (COG-UK) Consortium | The Lighthouse Lab in Milton Keynes and Alex Alderton, Roberto Amato, Jeffrey Barrett, Sonia Goncalves, Ewan Harrison, David K. Jackson, Ian Johnston, Dominic Kwiatkowski, Cordelia Langford, John Sillitoe on behalf of the Wellcome Sanger Institute COVID-19 Surveillance Team |
| EPI_ISL_3366809 | Lighthouse Lab in Alderley Park | Wellcome Sanger Institute for the COVID-19 Genomics UK (COG-UK) Consortium | Jacquelyn Wynn, Mairead Hyland, The Lighthouse Lab in Alderley Park and Alex Alderton, Roberto Amato, Jeffrey Barrett, Sonia Goncalves, Ewan Harrison, David K. Jackson, Ian Johnston, Dominic Kwiatkowski, Cordelia Langford, John Sillitoe on behalf of the Wellcome Sanger Institute COVID-19 Surveillance Team |
| EPI_ISL_3366810 | Lighthouse Lab in Milton Keynes | Wellcome Sanger Institute for the COVID-19 Genomics UK (COG-UK) Consortium | The Lighthouse Lab in Milton Keynes and Alex Alderton, Roberto Amato, Jeffrey Barrett, Sonia Goncalves, Ewan Harrison, David K. Jackson, Ian Johnston, Dominic Kwiatkowski, Cordelia Langford, John Sillitoe on behalf of the Wellcome Sanger Institute COVID-19 Surveillance Team |
| EPI_ISL_3366811 | Lighthouse Lab in Alderley Park | Wellcome Sanger Institute for the COVID-19 Genomics UK (COG-UK) Consortium | Jacquelyn Wynn, Mairead Hyland, The Lighthouse Lab in Alderley Park and Alex Alderton, Roberto Amato, Jeffrey Barrett, Sonia Goncalves, Ewan Harrison, David K. Jackson, Ian Johnston, Dominic Kwiatkowski, Cordelia Langford, John Sillitoe on behalf of the Wellcome Sanger Institute COVID-19 Surveillance Team |
| EPI_ISL_3366812, EPI_ISL_3366813, EPI_ISL_3366814, EPI_ISL_3366815 | Lighthouse Lab in Milton Keynes | Wellcome Sanger Institute for the COVID-19 Genomics UK (COG-UK) Consortium | The Lighthouse Lab in Milton Keynes and Alex Alderton, Roberto Amato, Jeffrey Barrett, Sonia Goncalves, Ewan Harrison, David K. Jackson, Ian Johnston, Dominic Kwiatkowski, Cordelia Langford, John Sillitoe on behalf of the Wellcome Sanger Institute COVID-19 Surveillance Team |
| EPI_ISL_3366816 | Lighthouse Lab in Alderley Park | Wellcome Sanger Institute for the COVID-19 Genomics UK (COG-UK) Consortium | Jacquelyn Wynn, Mairead Hyland, The Lighthouse Lab in Alderley Park and Alex Alderton, Roberto Amato, Jeffrey Barrett, Sonia Goncalves, Ewan Harrison, David K. Jackson, Ian Johnston, Dominic Kwiatkowski, Cordelia Langford, John Sillitoe on behalf of the Wellcome Sanger Institute COVID-19 Surveillance Team |
| EPI_ISL_3366817, EPI_ISL_3366818, EPI_ISL_3366819, EPI_ISL_3366820 | Lighthouse Lab in Glasgow | Wellcome Sanger Institute for the COVID-19 Genomics UK (COG-UK) Consortium | Harper VanSteenhouse, Yumi Kasai, David Gray, Carol Clugston, Anna Dominiczak and Alex Alderton, Roberto Amato, Jeffrey Barrett, Sonia Goncalves, Ewan Harrison, David K. Jackson, Ian Johnston, Dominic Kwiatkowski, Cordelia Langford, John Sillitoe on behalf of the Wellcome Sanger Institute COVID-19 Surveillance Team |
| EPI_ISL_3366821 | Berkshire and Surrey Pathology Services Lighthouse Laboratory | Wellcome Sanger Institute for the COVID-19 Genomics UK (COG-UK) Consortium | Berkshire and Surrey Pathology Services Lighthouse Laboratory and Alex Alderton, Roberto Amato, Jeffrey Barrett, Sonia Goncalves, Ewan Harrison, David K. Jackson, Ian Johnston, Dominic Kwiatkowski, Cordelia Langford, John Sillitoe on behalf of the Wellcome Sanger Institute COVID-19 Surveillance Team |
| EPI_ISL_3366822, EPI_ISL_3366823 | Lighthouse Lab in Glasgow | Wellcome Sanger Institute for the COVID-19 Genomics UK (COG-UK) Consortium | Harper VanSteenhouse, Yumi Kasai, David Gray, Carol Clugston, Anna Dominiczak and Alex Alderton, Roberto Amato, Jeffrey Barrett, Sonia Goncalves, Ewan Harrison, David K. Jackson, Ian Johnston, Dominic Kwiatkowski, Cordelia Langford, John Sillitoe on behalf of the Wellcome Sanger Institute COVID-19 Surveillance Team |
| EPI_ISL_3366824 | Berkshire and Surrey Pathology Services Lighthouse Laboratory | Wellcome Sanger Institute for the COVID-19 Genomics UK (COG-UK) Consortium | Berkshire and Surrey Pathology Services Lighthouse Laboratory and Alex Alderton, Roberto Amato, Jeffrey Barrett, Sonia Goncalves, Ewan Harrison, David K. Jackson, Ian Johnston, Dominic Kwiatkowski, Cordelia Langford, John Sillitoe on behalf of the Wellcome Sanger Institute COVID-19 Surveillance Team |
| EPI_ISL_3366825, EPI_ISL_3366826, EPI_ISL_3366827, EPI_ISL_3366828, EPI_ISL_3366829, EPI_ISL_3366830, EPI_ISL_3366831, EPI_ISL_3366832, EPI_ISL_3366833, EPI_ISL_3366834, EPI_ISL_3366835, EPI_ISL_3366836, EPI_ISL_3366837, EPI_ISL_3366838, EPI_ISL_3366839, EPI_ISL_3366840, EPI_ISL_3366841, EPI_ISL_3366842, EPI_ISL_3366843, EPI_ISL_3366844, EPI_ISL_3366845, EPI_ISL_3366846, EPI_ISL_3366847, EPI_ISL_3366848, EPI_ISL_3366849, EPI_ISL_3366850, EPI_ISL_3366851, EPI_ISL_3366852, EPI_ISL_3366853, EPI_ISL_3366854, EPI_ISL_3366855, EPI_ISL_3366856, EPI_ISL_3366857, EPI_ISL_3366858, EPI_ISL_3366859, EPI_ISL_3366860, EPI_ISL_3366861, EPI_ISL_3366862, EPI_ISL_3366863, EPI_ISL_3366864, EPI_ISL_3366865, EPI_ISL_3366866, EPI_ISL_3366867, EPI_ISL_3366868, EPI_ISL_3366869, EPI_ISL_3366870, EPI_ISL_3366871, EPI_ISL_3366872, EPI_ISL_3366873, EPI_ISL_3366874, EPI_ISL_3366875, EPI_ISL_3366876, EPI_ISL_3366877, EPI_ISL_3366878, EPI_ISL_3366879, EPI_ISL_3366880, EPI_ISL_3366881, EPI_ISL_3366882, EPI_ISL_3366883, EPI_ISL_3366884, EPI_ISL_3366885, EPI_ISL_3366886, EPI_ISL_3366887, EPI_ISL_3366888, EPI_ISL_3366889, EPI_ISL_3366890, EPI_ISL_3366891, EPI_ISL_3366892, EPI_ISL_3366893, EPI_ISL_3366894, EPI_ISL_3366895, EPI_ISL_3366896, EPI_ISL_3366897, EPI_ISL_3366898, EPI_ISL_3366899, EPI_ISL_3366900, EPI_ISL_3366901, EPI_ISL_3366902, EPI_ISL_3366903, EPI_ISL_3366904, EPI_ISL_3366905 | | | |
| see above | Lighthouse Lab in Glasgow | Wellcome Sanger Institute for the COVID-19 Genomics UK (COG-UK) Consortium | Harper VanSteenhouse, Yumi Kasai, David Gray, Carol Clugston, Anna Dominiczak and Alex Alderton, Roberto Amato, Jeffrey Barrett, Sonia Goncalves, Ewan Harrison, David K. Jackson, Ian Johnston, Dominic Kwiatkowski, Cordelia Langford, John Sillitoe on behalf of the Wellcome Sanger Institute COVID-19 Surveillance Team |
| EPI_ISL_3366906, EPI_ISL_3366907, EPI_ISL_3366908, EPI_ISL_3366909 | Lighthouse Lab in Alderley Park | Wellcome Sanger Institute for the COVID-19 Genomics UK (COG-UK) Consortium | Jacquelyn Wynn, Mairead Hyland, The Lighthouse Lab in Alderley Park and Alex Alderton, Roberto Amato, Jeffrey Barrett, Sonia Goncalves, Ewan Harrison, David K. Jackson, Ian Johnston, Dominic Kwiatkowski, Cordelia Langford, John Sillitoe on behalf of the Wellcome Sanger Institute COVID-19 Surveillance Team |
| EPI_ISL_3366910, EPI_ISL_3366911, EPI_ISL_3366912 | Lighthouse Lab in Milton Keynes | Wellcome Sanger Institute for the COVID-19 Genomics UK (COG-UK) Consortium | The Lighthouse Lab in Milton Keynes and Alex Alderton, Roberto Amato, Jeffrey Barrett, Sonia Goncalves, Ewan Harrison, David K. Jackson, Ian Johnston, Dominic Kwiatkowski, Cordelia Langford, John Sillitoe on behalf of the Wellcome Sanger Institute COVID-19 Surveillance Team |
| EPI_ISL_3366913 | Lighthouse Lab in Alderley Park | Wellcome Sanger Institute for the COVID-19 Genomics UK (COG-UK) Consortium | Jacquelyn Wynn, Mairead Hyland, The Lighthouse Lab in Alderley Park and Alex Alderton, Roberto Amato, Jeffrey Barrett, Sonia Goncalves, Ewan Harrison, David K. Jackson, Ian Johnston, Dominic Kwiatkowski, Cordelia Langford, John Sillitoe on behalf of the Wellcome Sanger Institute COVID-19 Surveillance Team |
| EPI_ISL_3366914 | Lighthouse Lab in Glasgow | Wellcome Sanger Institute for the COVID-19 Genomics UK (COG-UK) Consortium | Harper VanSteenhouse, Yumi Kasai, David Gray, Carol Clugston, Anna Dominiczak and Alex Alderton, Roberto Amato, Jeffrey Barrett, Sonia Goncalves, Ewan Harrison, David K. Jackson, Ian Johnston, Dominic Kwiatkowski, Cordelia Langford, John Sillitoe on behalf of the Wellcome Sanger Institute COVID-19 Surveillance Team |
| EPI_ISL_3366915 | Lighthouse Lab in Milton Keynes | Wellcome Sanger Institute for the COVID-19 Genomics UK (COG-UK) Consortium | The Lighthouse Lab in Milton Keynes and Alex Alderton, Roberto Amato, Jeffrey Barrett, Sonia Goncalves, Ewan Harrison, David K. Jackson, Ian Johnston, Dominic Kwiatkowski, Cordelia Langford, John Sillitoe on behalf of the Wellcome Sanger Institute COVID-19 Surveillance Team |
| EPI_ISL_3366916 | Lighthouse Lab in Glasgow | Wellcome Sanger Institute for the COVID-19 Genomics UK (COG-UK) Consortium | Harper VanSteenhouse, Yumi Kasai, David Gray, Carol Clugston, Anna Dominiczak and Alex Alderton, Roberto Amato, Jeffrey Barrett, Sonia Goncalves, Ewan Harrison, David K. Jackson, Ian Johnston, Dominic Kwiatkowski, Cordelia Langford, John Sillitoe on behalf of the Wellcome Sanger Institute COVID-19 Surveillance Team |
| EPI_ISL_3366917, EPI_ISL_3366918, EPI_ISL_3366919 | Lighthouse Lab in Milton Keynes | Wellcome Sanger Institute for the COVID-19 Genomics UK (COG-UK) Consortium | The Lighthouse Lab in Milton Keynes and Alex Alderton, Roberto Amato, Jeffrey Barrett, Sonia Goncalves, Ewan Harrison, David K. Jackson, Ian Johnston, Dominic Kwiatkowski, Cordelia Langford, John Sillitoe on behalf of the Wellcome Sanger Institute COVID-19 Surveillance Team |
| EPI_ISL_3366920 | Lighthouse Lab in Glasgow | Wellcome Sanger Institute for the COVID-19 Genomics UK (COG-UK) Consortium | Harper VanSteenhouse, Yumi Kasai, David Gray, Carol Clugston, Anna Dominiczak and Alex Alderton, Roberto Amato, Jeffrey Barrett, Sonia Goncalves, Ewan Harrison, David K. Jackson, Ian Johnston, Dominic Kwiatkowski, Cordelia Langford, John Sillitoe on behalf of the Wellcome Sanger Institute COVID-19 Surveillance Team |
| EPI_ISL_3366921, EPI_ISL_3366922 | Lighthouse Lab in Milton Keynes | Wellcome Sanger Institute for the COVID-19 Genomics UK (COG-UK) Consortium | The Lighthouse Lab in Milton Keynes and Alex Alderton, Roberto Amato, Jeffrey Barrett, Sonia Goncalves, Ewan Harrison, David K. Jackson, Ian Johnston, Dominic Kwiatkowski, Cordelia Langford, John Sillitoe on behalf of the Wellcome Sanger Institute COVID-19 Surveillance Team |
| EPI_ISL_3366923, EPI_ISL_3366924 | Lighthouse Lab in Glasgow | Wellcome Sanger Institute for the COVID-19 Genomics UK (COG-UK) Consortium | Harper VanSteenhouse, Yumi Kasai, David Gray, Carol Clugston, Anna Dominiczak and Alex Alderton, Roberto Amato, Jeffrey Barrett, Sonia Goncalves, Ewan Harrison, David K. Jackson, Ian Johnston, Dominic Kwiatkowski, Cordelia Langford, John Sillitoe on behalf of the Wellcome Sanger Institute COVID-19 Surveillance Team |

| | | | |
|---|---|---|---|
| EPI_ISL_3366925, EPI_ISL_3366926 | Lighthouse Lab in Milton Keynes | Wellcome Sanger Institute for the COVID-19 Genomics UK (COG-UK) Consortium | The Lighthouse Lab in Milton Keynes and Alex Alderton, Roberto Amato, Jeffrey Barrett, Sonia Goncalves, Ewan Harrison, David K. Jackson, Ian Johnston, Dominic Kwiatkowski, Cordelia Langford, John Sillitoe on behalf of the Wellcome Sanger Institute COVID-19 Surveillance Team |
| EPI_ISL_3366927, EPI_ISL_3366928, EPI_ISL_3366929, EPI_ISL_3366930, EPI_ISL_3366931, EPI_ISL_3366932, EPI_ISL_3366933 | Lighthouse Lab in Glasgow | Wellcome Sanger Institute for the COVID-19 Genomics UK (COG-UK) Consortium | Harper VanSteenhouse, Yumi Kasai, David Gray, Carol Clugston, Anna Dominiczak and Alex Alderton, Roberto Amato, Jeffrey Barrett, Sonia Goncalves, Ewan Harrison, David K. Jackson, Ian Johnston, Dominic Kwiatkowski, Cordelia Langford, John Sillitoe on behalf of the Wellcome Sanger Institute COVID-19 Surveillance Team |
| EPI_ISL_3366934, EPI_ISL_3366935, EPI_ISL_3366936, EPI_ISL_3366937, EPI_ISL_3366938, EPI_ISL_3366939, EPI_ISL_3366940, EPI_ISL_3366941, EPI_ISL_3366942, EPI_ISL_3366943, EPI_ISL_3366944, EPI_ISL_3366945 | | | |
| see above | Lighthouse Lab in Milton Keynes | Wellcome Sanger Institute for the COVID-19 Genomics UK (COG-UK) Consortium | The Lighthouse Lab in Milton Keynes and Alex Alderton, Roberto Amato, Jeffrey Barrett, Sonia Goncalves, Ewan Harrison, David K. Jackson, Ian Johnston, Dominic Kwiatkowski, Cordelia Langford, John Sillitoe on behalf of the Wellcome Sanger Institute COVID-19 Surveillance Team |
| EPI_ISL_3366946, EPI_ISL_3366947, EPI_ISL_3366948, EPI_ISL_3366949, EPI_ISL_3366950, EPI_ISL_3366951, EPI_ISL_3366952, EPI_ISL_3366953, EPI_ISL_3366954, EPI_ISL_3366955 | Lighthouse Lab in Glasgow | Wellcome Sanger Institute for the COVID-19 Genomics UK (COG-UK) Consortium | Harper VanSteenhouse, Yumi Kasai, David Gray, Carol Clugston, Anna Dominiczak and Alex Alderton, Roberto Amato, Jeffrey Barrett, Sonia Goncalves, Ewan Harrison, David K. Jackson, Ian Johnston, Dominic Kwiatkowski, Cordelia Langford, John Sillitoe on behalf of the Wellcome Sanger Institute COVID-19 Surveillance Team |
| EPI_ISL_3366956 | Berkshire and Surrey Pathology Services Lighthouse Laboratory | Wellcome Sanger Institute for the COVID-19 Genomics UK (COG-UK) Consortium | Berkshire and Surrey Pathology Services Lighthouse Laboratory and Alex Alderton, Roberto Amato, Jeffrey Barrett, Sonia Goncalves, Ewan Harrison, David K. Jackson, Ian Johnston, Dominic Kwiatkowski, Cordelia Langford, John Sillitoe on behalf of the Wellcome Sanger Institute COVID-19 Surveillance Team |
| EPI_ISL_3366957, EPI_ISL_3366958, EPI_ISL_3366959, EPI_ISL_3366960, EPI_ISL_3366961, EPI_ISL_3366962, EPI_ISL_3366963, EPI_ISL_3366964, EPI_ISL_3366965, EPI_ISL_3366966, EPI_ISL_3366967, EPI_ISL_3366968, EPI_ISL_3366969, EPI_ISL_3366970, EPI_ISL_3366971, EPI_ISL_3366972, EPI_ISL_3366973, EPI_ISL_3366974, EPI_ISL_3366975, EPI_ISL_3366976, EPI_ISL_3366977, EPI_ISL_3366978, EPI_ISL_3366979, EPI_ISL_3366980, EPI_ISL_3366981, EPI_ISL_3366982, EPI_ISL_3366983, EPI_ISL_3366984, EPI_ISL_3366985 | | | |
| see above | Lighthouse Lab in Glasgow | Wellcome Sanger Institute for the COVID-19 Genomics UK (COG-UK) Consortium | Harper VanSteenhouse, Yumi Kasai, David Gray, Carol Clugston, Anna Dominiczak and Alex Alderton, Roberto Amato, Jeffrey Barrett, Sonia Goncalves, Ewan Harrison, David K. Jackson, Ian Johnston, Dominic Kwiatkowski, Cordelia Langford, John Sillitoe on behalf of the Wellcome Sanger Institute COVID-19 Surveillance Team |
| EPI_ISL_3366986 | Lighthouse Lab in Alderley Park | Wellcome Sanger Institute for the COVID-19 Genomics UK (COG-UK) Consortium | Jacquelyn Wynn, Mairead Hyland, The Lighthouse Lab in Alderley Park and Alex Alderton, Roberto Amato, Jeffrey Barrett, Sonia Goncalves, Ewan Harrison, David K. Jackson, Ian Johnston, Dominic Kwiatkowski, Cordelia Langford, John Sillitoe on behalf of the Wellcome Sanger Institute COVID-19 Surveillance Team |
| EPI_ISL_3366987, EPI_ISL_3366988, EPI_ISL_3366989, EPI_ISL_3366990, EPI_ISL_3366991, EPI_ISL_3366992, EPI_ISL_3366993, EPI_ISL_3366994, EPI_ISL_3366995, EPI_ISL_3366996, EPI_ISL_3366997, EPI_ISL_3366998, EPI_ISL_3366999, EPI_ISL_3367000 | | | |
| see above | Lighthouse Lab in Glasgow | Wellcome Sanger Institute for the COVID-19 Genomics UK (COG-UK) Consortium | Harper VanSteenhouse, Yumi Kasai, David Gray, Carol Clugston, Anna Dominiczak and Alex Alderton, Roberto Amato, Jeffrey Barrett, Sonia Goncalves, Ewan Harrison, David K. Jackson, Ian Johnston, Dominic Kwiatkowski, Cordelia Langford, John Sillitoe on behalf of the Wellcome Sanger Institute COVID-19 Surveillance Team |
| EPI_ISL_3367001, EPI_ISL_3367002, EPI_ISL_3367003, EPI_ISL_3367004, EPI_ISL_3367005, EPI_ISL_3367006 | Lighthouse Lab in Alderley Park | Wellcome Sanger Institute for the COVID-19 Genomics UK (COG-UK) Consortium | Jacquelyn Wynn, Mairead Hyland, The Lighthouse Lab in Alderley Park and Alex Alderton, Roberto Amato, Jeffrey Barrett, Sonia Goncalves, Ewan Harrison, David K. Jackson, Ian Johnston, Dominic Kwiatkowski, Cordelia Langford, John Sillitoe on behalf of the Wellcome Sanger Institute COVID-19 Surveillance Team |
| EPI_ISL_3367007, EPI_ISL_3367008, EPI_ISL_3367009, EPI_ISL_3367010, EPI_ISL_3367011, EPI_ISL_3367012, EPI_ISL_3367013, EPI_ISL_3367014, EPI_ISL_3367015, EPI_ISL_3367016, EPI_ISL_3367017, EPI_ISL_3367018, EPI_ISL_3367019, EPI_ISL_3367020 | | | |
| see above | Lighthouse Lab in Glasgow | Wellcome Sanger Institute for the COVID-19 Genomics UK (COG-UK) Consortium | Harper VanSteenhouse, Yumi Kasai, David Gray, Carol Clugston, Anna Dominiczak and Alex Alderton, Roberto Amato, Jeffrey Barrett, Sonia Goncalves, Ewan Harrison, David K. Jackson, Ian Johnston, Dominic Kwiatkowski, Cordelia Langford, John Sillitoe on behalf of the Wellcome Sanger Institute COVID-19 Surveillance Team |
| EPI_ISL_3367021 | Lighthouse Lab in Alderley Park | Wellcome Sanger Institute for the COVID-19 Genomics UK (COG-UK) Consortium | Jacquelyn Wynn, Mairead Hyland, The Lighthouse Lab in Alderley Park and Alex Alderton, Roberto Amato, Jeffrey Barrett, Sonia Goncalves, Ewan Harrison, David K. Jackson, Ian Johnston, Dominic Kwiatkowski, Cordelia Langford, John Sillitoe on behalf of the Wellcome Sanger Institute COVID-19 Surveillance Team |
| EPI_ISL_3367022, EPI_ISL_3367023, EPI_ISL_3367024, EPI_ISL_3367025, EPI_ISL_3367026 | Lighthouse Lab in Glasgow | Wellcome Sanger Institute for the COVID-19 Genomics UK (COG-UK) Consortium | Harper VanSteenhouse, Yumi Kasai, David Gray, Carol Clugston, Anna Dominiczak and Alex Alderton, Roberto Amato, Jeffrey Barrett, Sonia Goncalves, Ewan Harrison, David K. Jackson, Ian Johnston, Dominic Kwiatkowski, Cordelia Langford, John Sillitoe on behalf of the Wellcome Sanger Institute COVID-19 Surveillance Team |
| EPI_ISL_3367027 | Lighthouse Lab in Alderley Park | Wellcome Sanger Institute for the COVID-19 Genomics UK (COG-UK) Consortium | Jacquelyn Wynn, Mairead Hyland, The Lighthouse Lab in Alderley Park and Alex Alderton, Roberto Amato, Jeffrey Barrett, Sonia Goncalves, Ewan Harrison, David K. Jackson, Ian Johnston, Dominic Kwiatkowski, Cordelia Langford, John Sillitoe on behalf of the Wellcome Sanger Institute COVID-19 Surveillance Team |
| EPI_ISL_3367028, EPI_ISL_3367029, EPI_ISL_3367030, EPI_ISL_3367031, EPI_ISL_3367032, EPI_ISL_3367033 | Lighthouse Lab in Glasgow | Wellcome Sanger Institute for the COVID-19 Genomics UK (COG-UK) Consortium | Harper VanSteenhouse, Yumi Kasai, David Gray, Carol Clugston, Anna Dominiczak and Alex Alderton, Roberto Amato, Jeffrey Barrett, Sonia Goncalves, Ewan Harrison, David K. Jackson, Ian Johnston, Dominic Kwiatkowski, Cordelia Langford, John Sillitoe on behalf of the Wellcome Sanger Institute COVID-19 Surveillance Team |
| EPI_ISL_3367034 | Lighthouse Lab in Alderley Park | Wellcome Sanger Institute for the COVID-19 Genomics UK (COG-UK) Consortium | Jacquelyn Wynn, Mairead Hyland, The Lighthouse Lab in Alderley Park and Alex Alderton, Roberto Amato, Jeffrey Barrett, Sonia Goncalves, Ewan Harrison, David K. Jackson, Ian Johnston, Dominic Kwiatkowski, Cordelia Langford, John Sillitoe on behalf of the Wellcome Sanger Institute COVID-19 Surveillance Team |
| EPI_ISL_3367035 | Lighthouse Lab in Glasgow | Wellcome Sanger Institute for the COVID-19 Genomics UK (COG-UK) Consortium | Harper VanSteenhouse, Yumi Kasai, David Gray, Carol Clugston, Anna Dominiczak and Alex Alderton, Roberto Amato, Jeffrey Barrett, Sonia Goncalves, Ewan Harrison, David K. Jackson, Ian Johnston, Dominic Kwiatkowski, Cordelia Langford, John Sillitoe on behalf of the Wellcome Sanger Institute COVID-19 Surveillance Team |
| EPI_ISL_3367036 | Lighthouse Lab in Alderley Park | Wellcome Sanger Institute for the COVID-19 Genomics UK (COG-UK) Consortium | Jacquelyn Wynn, Mairead Hyland, The Lighthouse Lab in Alderley Park and Alex Alderton, Roberto Amato, Jeffrey Barrett, Sonia Goncalves, Ewan Harrison, David K. Jackson, Ian Johnston, Dominic Kwiatkowski, Cordelia Langford, John Sillitoe on behalf of the Wellcome Sanger Institute COVID-19 Surveillance Team |
| EPI_ISL_3367037, EPI_ISL_3367038 | Lighthouse Lab in Glasgow | Wellcome Sanger Institute for the COVID-19 Genomics UK (COG-UK) Consortium | Harper VanSteenhouse, Yumi Kasai, David Gray, Carol Clugston, Anna Dominiczak and Alex Alderton, Roberto Amato, Jeffrey Barrett, Sonia Goncalves, Ewan Harrison, David K. Jackson, Ian Johnston, Dominic Kwiatkowski, Cordelia Langford, John Sillitoe on behalf of the Wellcome Sanger Institute COVID-19 Surveillance Team |
| EPI_ISL_3367039 | Lighthouse Lab in Alderley Park | Wellcome Sanger Institute for the COVID-19 Genomics UK (COG-UK) Consortium | Jacquelyn Wynn, Mairead Hyland, The Lighthouse Lab in Alderley Park and Alex Alderton, Roberto Amato, Jeffrey Barrett, Sonia Goncalves, Ewan Harrison, David K. Jackson, Ian Johnston, Dominic Kwiatkowski, Cordelia Langford, John Sillitoe on behalf of the Wellcome Sanger Institute COVID-19 Surveillance Team |
| EPI_ISL_3367040, EPI_ISL_3367041, EPI_ISL_3367042, EPI_ISL_3367043, EPI_ISL_3367044, EPI_ISL_3367045, EPI_ISL_3367046, EPI_ISL_3367047, EPI_ISL_3367048 | Lighthouse Lab in Glasgow | Wellcome Sanger Institute for the COVID-19 Genomics UK (COG-UK) Consortium | Harper VanSteenhouse, Yumi Kasai, David Gray, Carol Clugston, Anna Dominiczak and Alex Alderton, Roberto Amato, Jeffrey Barrett, Sonia Goncalves, Ewan Harrison, David K. Jackson, Ian Johnston, Dominic Kwiatkowski, Cordelia Langford, John Sillitoe on behalf of the Wellcome Sanger Institute COVID-19 Surveillance Team |
| EPI_ISL_3367049 | Lighthouse Lab in Alderley Park | Wellcome Sanger Institute for the COVID-19 Genomics UK (COG-UK) Consortium | Jacquelyn Wynn, Mairead Hyland, The Lighthouse Lab in Alderley Park and Alex Alderton, Roberto Amato, Jeffrey Barrett, Sonia Goncalves, Ewan Harrison, David K. Jackson, Ian Johnston, Dominic Kwiatkowski, Cordelia Langford, John Sillitoe on behalf of the Wellcome Sanger Institute COVID-19 Surveillance Team |
| EPI_ISL_3367050, EPI_ISL_3367051, EPI_ISL_3367052, EPI_ISL_3367053, EPI_ISL_3367054, EPI_ISL_3367055 | Lighthouse Lab in Glasgow | Wellcome Sanger Institute for the COVID-19 Genomics UK (COG-UK) Consortium | Harper VanSteenhouse, Yumi Kasai, David Gray, Carol Clugston, Anna Dominiczak and Alex Alderton, Roberto Amato, Jeffrey Barrett, Sonia Goncalves, Ewan Harrison, David K. Jackson, Ian Johnston, Dominic Kwiatkowski, Cordelia Langford, John Sillitoe on behalf of the Wellcome Sanger Institute COVID-19 Surveillance Team |
| EPI_ISL_3367056 | Lighthouse Lab in Alderley Park | Wellcome Sanger Institute for the COVID-19 Genomics UK (COG-UK) Consortium | Jacquelyn Wynn, Mairead Hyland, The Lighthouse Lab in Alderley Park and Alex Alderton, Roberto Amato, Jeffrey Barrett, Sonia Goncalves, Ewan Harrison, David K. Jackson, Ian Johnston, Dominic Kwiatkowski, Cordelia Langford, John Sillitoe on behalf of the Wellcome Sanger Institute COVID-19 Surveillance Team |
| EPI_ISL_3367057, EPI_ISL_3367058, EPI_ISL_3367059, EPI_ISL_3367060, EPI_ISL_3367061 | Lighthouse Lab in Glasgow | Wellcome Sanger Institute for the COVID-19 Genomics UK (COG-UK) Consortium | Harper VanSteenhouse, Yumi Kasai, David Gray, Carol Clugston, Anna Dominiczak and Alex Alderton, Roberto Amato, Jeffrey Barrett, Sonia Goncalves, Ewan Harrison, David K. Jackson, Ian Johnston, Dominic Kwiatkowski, Cordelia Langford, John Sillitoe on behalf of the Wellcome Sanger Institute COVID-19 Surveillance Team |
| EPI_ISL_3367062, EPI_ISL_3367063, EPI_ISL_3367064, EPI_ISL_3367065, EPI_ISL_3367066, EPI_ISL_3367067, EPI_ISL_3367068, EPI_ISL_3367069, EPI_ISL_3367070, EPI_ISL_3367071, EPI_ISL_3367072, EPI_ISL_3367073, EPI_ISL_3367074, EPI_ISL_3367075, EPI_ISL_3367076, EPI_ISL_3367077, EPI_ISL_3367078, EPI_ISL_3367079, EPI_ISL_3367080, EPI_ISL_3367081, EPI_ISL_3367082, EPI_ISL_3367083, EPI_ISL_3367084, EPI_ISL_3367085, EPI_ISL_3367086, EPI_ISL_3367087, EPI_ISL_3367088, EPI_ISL_3367089, EPI_ISL_3367090, EPI_ISL_3367091, EPI_ISL_3367092, EPI_ISL_3367093, EPI_ISL_3367094, EPI_ISL_3367095, EPI_ISL_3367096, EPI_ISL_3367097, EPI_ISL_3367098 | | | |
| see above | Lighthouse Lab in Milton Keynes | Wellcome Sanger Institute for the COVID-19 Genomics UK (COG-UK) Consortium | The Lighthouse Lab in Milton Keynes and Alex Alderton, Roberto Amato, Jeffrey Barrett, Sonia Goncalves, Ewan Harrison, David K. Jackson, Ian Johnston, Dominic Kwiatkowski, Cordelia Langford, John Sillitoe on behalf of the Wellcome Sanger Institute COVID-19 Surveillance Team |
| EPI_ISL_3367099, EPI_ISL_3367100, EPI_ISL_3367101, EPI_ISL_3367102, EPI_ISL_3367103, EPI_ISL_3367104, EPI_ISL_3367105, EPI_ISL_3367106, EPI_ISL_3367107, EPI_ISL_3367108, EPI_ISL_3367109, EPI_ISL_3367110, EPI_ISL_3367111 | | | |
| see above | Lighthouse Lab in Glasgow | Wellcome Sanger Institute for the COVID-19 Genomics UK (COG-UK) | Harper VanSteenhouse, Yumi Kasai, David Gray, Carol Clugston, Anna Dominiczak and Alex Alderton, Roberto Amato, Jeffrey Barrett, Sonia Goncalves, Ewan Harrison, David K. Jackson, Ian Johnston, Dominic Kwiatkowski, Cordelia Langford, |

| | | | |
|---|---|---|---|
| | | Consortium | John Sillitoe on behalf of the Wellcome Sanger Institute COVID-19 Surveillance Team |
| EPI_ISL_3367112 | Berkshire and Surrey Pathology Services Lighthouse Laboratory | Wellcome Sanger Institute for the COVID-19 Genomics UK (COG-UK) Consortium | Berkshire and Surrey Pathology Services Lighthouse Laboratory and Alex Alderton, Roberto Amato, Jeffrey Barrett, Sonia Goncalves, Ewan Harrison, David K. Jackson, Ian Johnston, Dominic Kwiatkowski, Cordelia Langford, John Sillitoe on behalf of the Wellcome Sanger Institute COVID-19 Surveillance Team |
| EPI_ISL_3367113, EPI_ISL_3367114, EPI_ISL_3367115, EPI_ISL_3367116, EPI_ISL_3367117, EPI_ISL_3367118, EPI_ISL_3367119, EPI_ISL_3367120, EPI_ISL_3367121, EPI_ISL_3367122, EPI_ISL_3367123, EPI_ISL_3367124, EPI_ISL_3367125, EPI_ISL_3367126, EPI_ISL_3367127, EPI_ISL_3367128, EPI_ISL_3367129, EPI_ISL_3367130, EPI_ISL_3367131, EPI_ISL_3367132, EPI_ISL_3367133, EPI_ISL_3367134, EPI_ISL_3367135, EPI_ISL_3367136, EPI_ISL_3367137, EPI_ISL_3367138, EPI_ISL_3367139, EPI_ISL_3367140, EPI_ISL_3367141, EPI_ISL_3367142, EPI_ISL_3367143, EPI_ISL_3367144, EPI_ISL_3367145, EPI_ISL_3367146, EPI_ISL_3367147, EPI_ISL_3367148, EPI_ISL_3367149, EPI_ISL_3367150, EPI_ISL_3367151, EPI_ISL_3367152, EPI_ISL_3367153, EPI_ISL_3367154, EPI_ISL_3367155, EPI_ISL_3367156, EPI_ISL_3367157, EPI_ISL_3367158, EPI_ISL_3367159, EPI_ISL_3367160, EPI_ISL_3367161, EPI_ISL_3367162, EPI_ISL_3367163, EPI_ISL_3367164, EPI_ISL_3367165, EPI_ISL_3367166, EPI_ISL_3367167, EPI_ISL_3367168, EPI_ISL_3367169, EPI_ISL_3367170, EPI_ISL_3367171, EPI_ISL_3367172, EPI_ISL_3367173, EPI_ISL_3367174 | | | |
| see above | Lighthouse Lab in Glasgow | Wellcome Sanger Institute for the COVID-19 Genomics UK (COG-UK) Consortium | Harper VanSteenhouse, Yumi Kasai, David Gray, Carol Clugston, Anna Dominiczak and Alex Alderton, Roberto Amato, Jeffrey Barrett, Sonia Goncalves, Ewan Harrison, David K. Jackson, Ian Johnston, Dominic Kwiatkowski, Cordelia Langford, John Sillitoe on behalf of the Wellcome Sanger Institute COVID-19 Surveillance Team |
| EPI_ISL_3367175, EPI_ISL_3367176 | Berkshire and Surrey Pathology Services Lighthouse Laboratory | Wellcome Sanger Institute for the COVID-19 Genomics UK (COG-UK) Consortium | Berkshire and Surrey Pathology Services Lighthouse Laboratory and Alex Alderton, Roberto Amato, Jeffrey Barrett, Sonia Goncalves, Ewan Harrison, David K. Jackson, Ian Johnston, Dominic Kwiatkowski, Cordelia Langford, John Sillitoe on behalf of the Wellcome Sanger Institute COVID-19 Surveillance Team |
| EPI_ISL_3367177 | Lighthouse Laboratory Plymouth | Wellcome Sanger Institute for the COVID-19 Genomics UK (COG-UK) Consortium | Lighthouse Laboratory Plymouth and Alex Alderton, Roberto Amato, Jeffrey Barrett, Sonia Goncalves, Ewan Harrison, David K. Jackson, Ian Johnston, Dominic Kwiatkowski, Cordelia Langford, John Sillitoe on behalf of the Wellcome Sanger Institute COVID-19 Surveillance Team |
| EPI_ISL_3367178, EPI_ISL_3367179 | Berkshire and Surrey Pathology Services Lighthouse Laboratory | Wellcome Sanger Institute for the COVID-19 Genomics UK (COG-UK) Consortium | Berkshire and Surrey Pathology Services Lighthouse Laboratory and Alex Alderton, Roberto Amato, Jeffrey Barrett, Sonia Goncalves, Ewan Harrison, David K. Jackson, Ian Johnston, Dominic Kwiatkowski, Cordelia Langford, John Sillitoe on behalf of the Wellcome Sanger Institute COVID-19 Surveillance Team |
| EPI_ISL_3367180 | Lighthouse Laboratory Plymouth | Wellcome Sanger Institute for the COVID-19 Genomics UK (COG-UK) Consortium | Lighthouse Laboratory Plymouth and Alex Alderton, Roberto Amato, Jeffrey Barrett, Sonia Goncalves, Ewan Harrison, David K. Jackson, Ian Johnston, Dominic Kwiatkowski, Cordelia Langford, John Sillitoe on behalf of the Wellcome Sanger Institute COVID-19 Surveillance Team |
| EPI_ISL_3367181 | Berkshire and Surrey Pathology Services Lighthouse Laboratory | Wellcome Sanger Institute for the COVID-19 Genomics UK (COG-UK) Consortium | Berkshire and Surrey Pathology Services Lighthouse Laboratory and Alex Alderton, Roberto Amato, Jeffrey Barrett, Sonia Goncalves, Ewan Harrison, David K. Jackson, Ian Johnston, Dominic Kwiatkowski, Cordelia Langford, John Sillitoe on behalf of the Wellcome Sanger Institute COVID-19 Surveillance Team |
| EPI_ISL_3367182 | Lighthouse Lab in Glasgow | Wellcome Sanger Institute for the COVID-19 Genomics UK (COG-UK) Consortium | Harper VanSteenhouse, Yumi Kasai, David Gray, Carol Clugston, Anna Dominiczak and Alex Alderton, Roberto Amato, Jeffrey Barrett, Sonia Goncalves, Ewan Harrison, David K. Jackson, Ian Johnston, Dominic Kwiatkowski, Cordelia Langford, John Sillitoe on behalf of the Wellcome Sanger Institute COVID-19 Surveillance Team |
| EPI_ISL_3367183 | Lighthouse Laboratory Plymouth | Wellcome Sanger Institute for the COVID-19 Genomics UK (COG-UK) Consortium | Lighthouse Laboratory Plymouth and Alex Alderton, Roberto Amato, Jeffrey Barrett, Sonia Goncalves, Ewan Harrison, David K. Jackson, Ian Johnston, Dominic Kwiatkowski, Cordelia Langford, John Sillitoe on behalf of the Wellcome Sanger Institute COVID-19 Surveillance Team |
| EPI_ISL_3367184, EPI_ISL_3367185, EPI_ISL_3367186, EPI_ISL_3367187, EPI_ISL_3367188, EPI_ISL_3367189, EPI_ISL_3367190 | Berkshire and Surrey Pathology Services Lighthouse Laboratory | Wellcome Sanger Institute for the COVID-19 Genomics UK (COG-UK) Consortium | Berkshire and Surrey Pathology Services Lighthouse Laboratory and Alex Alderton, Roberto Amato, Jeffrey Barrett, Sonia Goncalves, Ewan Harrison, David K. Jackson, Ian Johnston, Dominic Kwiatkowski, Cordelia Langford, John Sillitoe on behalf of the Wellcome Sanger Institute COVID-19 Surveillance Team |
| EPI_ISL_3367191 | Lighthouse Laboratory Plymouth | Wellcome Sanger Institute for the COVID-19 Genomics UK (COG-UK) Consortium | Lighthouse Laboratory Plymouth and Alex Alderton, Roberto Amato, Jeffrey Barrett, Sonia Goncalves, Ewan Harrison, David K. Jackson, Ian Johnston, Dominic Kwiatkowski, Cordelia Langford, John Sillitoe on behalf of the Wellcome Sanger Institute COVID-19 Surveillance Team |
| EPI_ISL_3367192 | Berkshire and Surrey Pathology Services Lighthouse Laboratory | Wellcome Sanger Institute for the COVID-19 Genomics UK (COG-UK) Consortium | Berkshire and Surrey Pathology Services Lighthouse Laboratory and Alex Alderton, Roberto Amato, Jeffrey Barrett, Sonia Goncalves, Ewan Harrison, David K. Jackson, Ian Johnston, Dominic Kwiatkowski, Cordelia Langford, John Sillitoe on behalf of the Wellcome Sanger Institute COVID-19 Surveillance Team |
| EPI_ISL_3367193 | Lighthouse Lab in Glasgow | Wellcome Sanger Institute for the COVID-19 Genomics UK (COG-UK) Consortium | Harper VanSteenhouse, Yumi Kasai, David Gray, Carol Clugston, Anna Dominiczak and Alex Alderton, Roberto Amato, Jeffrey Barrett, Sonia Goncalves, Ewan Harrison, David K. Jackson, Ian Johnston, Dominic Kwiatkowski, Cordelia Langford, John Sillitoe on behalf of the Wellcome Sanger Institute COVID-19 Surveillance Team |
| EPI_ISL_3367194, EPI_ISL_3367195, EPI_ISL_3367196 | Berkshire and Surrey Pathology Services Lighthouse Laboratory | Wellcome Sanger Institute for the COVID-19 Genomics UK (COG-UK) Consortium | Berkshire and Surrey Pathology Services Lighthouse Laboratory and Alex Alderton, Roberto Amato, Jeffrey Barrett, Sonia Goncalves, Ewan Harrison, David K. Jackson, Ian Johnston, Dominic Kwiatkowski, Cordelia Langford, John Sillitoe on behalf of the Wellcome Sanger Institute COVID-19 Surveillance Team |
| EPI_ISL_3367197 | Lighthouse Laboratory Plymouth | Wellcome Sanger Institute for the COVID-19 Genomics UK (COG-UK) Consortium | Lighthouse Laboratory Plymouth and Alex Alderton, Roberto Amato, Jeffrey Barrett, Sonia Goncalves, Ewan Harrison, David K. Jackson, Ian Johnston, Dominic Kwiatkowski, Cordelia Langford, John Sillitoe on behalf of the Wellcome Sanger Institute COVID-19 Surveillance Team |
| EPI_ISL_3367198 | Lighthouse Lab in Glasgow | Wellcome Sanger Institute for the COVID-19 Genomics UK (COG-UK) Consortium | Harper VanSteenhouse, Yumi Kasai, David Gray, Carol Clugston, Anna Dominiczak and Alex Alderton, Roberto Amato, Jeffrey Barrett, Sonia Goncalves, Ewan Harrison, David K. Jackson, Ian Johnston, Dominic Kwiatkowski, Cordelia Langford, John Sillitoe on behalf of the Wellcome Sanger Institute COVID-19 Surveillance Team |
| EPI_ISL_3367199 | Lighthouse Laboratory Plymouth | Wellcome Sanger Institute for the COVID-19 Genomics UK (COG-UK) Consortium | Lighthouse Laboratory Plymouth and Alex Alderton, Roberto Amato, Jeffrey Barrett, Sonia Goncalves, Ewan Harrison, David K. Jackson, Ian Johnston, Dominic Kwiatkowski, Cordelia Langford, John Sillitoe on behalf of the Wellcome Sanger Institute COVID-19 Surveillance Team |
| EPI_ISL_3367200, EPI_ISL_3367201 | Berkshire and Surrey Pathology Services Lighthouse Laboratory | Wellcome Sanger Institute for the COVID-19 Genomics UK (COG-UK) Consortium | Berkshire and Surrey Pathology Services Lighthouse Laboratory and Alex Alderton, Roberto Amato, Jeffrey Barrett, Sonia Goncalves, Ewan Harrison, David K. Jackson, Ian Johnston, Dominic Kwiatkowski, Cordelia Langford, John Sillitoe on behalf of the Wellcome Sanger Institute COVID-19 Surveillance Team |
| EPI_ISL_3367202, EPI_ISL_3367203 | Lighthouse Laboratory Plymouth | Wellcome Sanger Institute for the COVID-19 Genomics UK (COG-UK) Consortium | Lighthouse Laboratory Plymouth and Alex Alderton, Roberto Amato, Jeffrey Barrett, Sonia Goncalves, Ewan Harrison, David K. Jackson, Ian Johnston, Dominic Kwiatkowski, Cordelia Langford, John Sillitoe on behalf of the Wellcome Sanger Institute COVID-19 Surveillance Team |
| EPI_ISL_3367204 | Lighthouse Lab in Glasgow | Wellcome Sanger Institute for the COVID-19 Genomics UK (COG-UK) Consortium | Harper VanSteenhouse, Yumi Kasai, David Gray, Carol Clugston, Anna Dominiczak and Alex Alderton, Roberto Amato, Jeffrey Barrett, Sonia Goncalves, Ewan Harrison, David K. Jackson, Ian Johnston, Dominic Kwiatkowski, Cordelia Langford, John Sillitoe on behalf of the Wellcome Sanger Institute COVID-19 Surveillance Team |
| EPI_ISL_3367205 | Lighthouse Laboratory Plymouth | Wellcome Sanger Institute for the COVID-19 Genomics UK (COG-UK) Consortium | Lighthouse Laboratory Plymouth and Alex Alderton, Roberto Amato, Jeffrey Barrett, Sonia Goncalves, Ewan Harrison, David K. Jackson, Ian Johnston, Dominic Kwiatkowski, Cordelia Langford, John Sillitoe on behalf of the Wellcome Sanger Institute COVID-19 Surveillance Team |
| EPI_ISL_3367206, EPI_ISL_3367207 | Lighthouse Lab in Glasgow | Wellcome Sanger Institute for the COVID-19 Genomics UK (COG-UK) Consortium | Harper VanSteenhouse, Yumi Kasai, David Gray, Carol Clugston, Anna Dominiczak and Alex Alderton, Roberto Amato, Jeffrey Barrett, Sonia Goncalves, Ewan Harrison, David K. Jackson, Ian Johnston, Dominic Kwiatkowski, Cordelia Langford, John Sillitoe on behalf of the Wellcome Sanger Institute COVID-19 Surveillance Team |
| EPI_ISL_3367208 | Lighthouse Laboratory Plymouth | Wellcome Sanger Institute for the COVID-19 Genomics UK (COG-UK) Consortium | Lighthouse Laboratory Plymouth and Alex Alderton, Roberto Amato, Jeffrey Barrett, Sonia Goncalves, Ewan Harrison, David K. Jackson, Ian Johnston, Dominic Kwiatkowski, Cordelia Langford, John Sillitoe on behalf of the Wellcome Sanger Institute COVID-19 Surveillance Team |
| EPI_ISL_3367209 | Lighthouse Lab in Glasgow | Wellcome Sanger Institute for the COVID-19 Genomics UK (COG-UK) Consortium | Harper VanSteenhouse, Yumi Kasai, David Gray, Carol Clugston, Anna Dominiczak and Alex Alderton, Roberto Amato, Jeffrey Barrett, Sonia Goncalves, Ewan Harrison, David K. Jackson, Ian Johnston, Dominic Kwiatkowski, Cordelia Langford, John Sillitoe on behalf of the Wellcome Sanger Institute COVID-19 Surveillance Team |
| EPI_ISL_3367210, EPI_ISL_3367211 | Lighthouse Laboratory Plymouth | Wellcome Sanger Institute for the COVID-19 Genomics UK (COG-UK) Consortium | Lighthouse Laboratory Plymouth and Alex Alderton, Roberto Amato, Jeffrey Barrett, Sonia Goncalves, Ewan Harrison, David K. Jackson, Ian Johnston, Dominic Kwiatkowski, Cordelia Langford, John Sillitoe on behalf of the Wellcome Sanger Institute COVID-19 Surveillance Team |
| EPI_ISL_3367212 | Berkshire and Surrey Pathology Services Lighthouse Laboratory | Wellcome Sanger Institute for the COVID-19 Genomics UK (COG-UK) Consortium | Berkshire and Surrey Pathology Services Lighthouse Laboratory and Alex Alderton, Roberto Amato, Jeffrey Barrett, Sonia Goncalves, Ewan Harrison, David K. Jackson, Ian Johnston, Dominic Kwiatkowski, Cordelia Langford, John Sillitoe on behalf of the Wellcome Sanger Institute COVID-19 Surveillance Team |
| EPI_ISL_3367213, EPI_ISL_3367214, EPI_ISL_3367215, EPI_ISL_3367216, EPI_ISL_3367217 | Lighthouse Laboratory Plymouth | Wellcome Sanger Institute for the COVID-19 Genomics UK (COG-UK) Consortium | Lighthouse Laboratory Plymouth and Alex Alderton, Roberto Amato, Jeffrey Barrett, Sonia Goncalves, Ewan Harrison, David K. Jackson, Ian Johnston, Dominic Kwiatkowski, Cordelia Langford, John Sillitoe on behalf of the Wellcome Sanger Institute COVID-19 Surveillance Team |
| EPI_ISL_3367218, EPI_ISL_3367219, EPI_ISL_3367220, EPI_ISL_3367221 | Berkshire and Surrey Pathology Services Lighthouse Laboratory | Wellcome Sanger Institute for the COVID-19 Genomics UK (COG-UK) Consortium | Berkshire and Surrey Pathology Services Lighthouse Laboratory and Alex Alderton, Roberto Amato, Jeffrey Barrett, Sonia Goncalves, Ewan Harrison, David K. Jackson, Ian Johnston, Dominic Kwiatkowski, Cordelia Langford, John Sillitoe on behalf of the Wellcome Sanger Institute COVID-19 Surveillance Team |
| EPI_ISL_3367222 | Lighthouse Lab in Glasgow | Wellcome Sanger Institute for the COVID-19 Genomics UK (COG-UK) Consortium | Harper VanSteenhouse, Yumi Kasai, David Gray, Carol Clugston, Anna Dominiczak and Alex Alderton, Roberto Amato, Jeffrey Barrett, Sonia Goncalves, Ewan Harrison, David K. Jackson, Ian Johnston, Dominic Kwiatkowski, Cordelia Langford, John Sillitoe on behalf of the Wellcome Sanger Institute COVID-19 Surveillance Team |
| EPI_ISL_3367223 | Berkshire and Surrey Pathology Services Lighthouse Laboratory | Wellcome Sanger Institute for the COVID-19 Genomics UK (COG-UK) Consortium | Berkshire and Surrey Pathology Services Lighthouse Laboratory and Alex Alderton, Roberto Amato, Jeffrey Barrett, Sonia Goncalves, Ewan Harrison, David K. Jackson, Ian Johnston, Dominic Kwiatkowski, Cordelia Langford, John Sillitoe on behalf of the Wellcome Sanger Institute COVID-19 Surveillance Team |
| EPI_ISL_3367224 | Lighthouse Lab in Glasgow | Wellcome Sanger Institute for the COVID-19 Genomics UK (COG-UK) Consortium | Harper VanSteenhouse, Yumi Kasai, David Gray, Carol Clugston, Anna Dominiczak and Alex Alderton, Roberto Amato, Jeffrey Barrett, Sonia Goncalves, Ewan Harrison, David K. Jackson, Ian Johnston, Dominic Kwiatkowski, Cordelia Langford, John Sillitoe on behalf of the Wellcome Sanger Institute COVID-19 Surveillance Team |
| EPI_ISL_3367225, EPI_ISL_3367226 | Berkshire and Surrey Pathology Services Lighthouse Laboratory | Wellcome Sanger Institute for the COVID-19 Genomics UK (COG-UK) Consortium | Berkshire and Surrey Pathology Services Lighthouse Laboratory and Alex Alderton, Roberto Amato, Jeffrey Barrett, Sonia Goncalves, Ewan Harrison, David K. Jackson, Ian Johnston, Dominic Kwiatkowski, Cordelia Langford, John Sillitoe on behalf of the Wellcome Sanger Institute COVID-19 Surveillance Team |
| EPI_ISL_3367227 | Lighthouse Laboratory Plymouth | Wellcome Sanger Institute for the COVID-19 Genomics UK (COG-UK) Consortium | Lighthouse Laboratory Plymouth and Alex Alderton, Roberto Amato, Jeffrey Barrett, Sonia Goncalves, Ewan Harrison, David K. Jackson, Ian Johnston, Dominic Kwiatkowski, Cordelia Langford, John Sillitoe on behalf of the Wellcome Sanger Institute COVID-19 Surveillance Team |
| EPI_ISL_3367228, EPI_ISL_3367229 | Berkshire and Surrey Pathology Services Lighthouse Laboratory | Wellcome Sanger Institute for the COVID-19 Genomics UK (COG-UK) Consortium | Berkshire and Surrey Pathology Services Lighthouse Laboratory and Alex Alderton, Roberto Amato, Jeffrey Barrett, Sonia Goncalves, Ewan Harrison, David K. Jackson, Ian Johnston, Dominic Kwiatkowski, Cordelia Langford, John Sillitoe on behalf of the Wellcome Sanger Institute COVID-19 Surveillance Team |
| EPI_ISL_3367230 | Lighthouse Laboratory Plymouth | Wellcome Sanger Institute for the COVID-19 Genomics UK (COG-UK) Consortium | Lighthouse Laboratory Plymouth and Alex Alderton, Roberto Amato, Jeffrey Barrett, Sonia Goncalves, Ewan Harrison, David K. Jackson, Ian Johnston, Dominic Kwiatkowski, Cordelia Langford, John Sillitoe on behalf of the Wellcome Sanger Institute COVID-19 Surveillance Team |
| EPI_ISL_3367231, EPI_ISL_3367232 | Berkshire and Surrey Pathology Services Lighthouse Laboratory | Wellcome Sanger Institute for the COVID-19 Genomics UK (COG-UK) Consortium | Berkshire and Surrey Pathology Services Lighthouse Laboratory and Alex Alderton, Roberto Amato, Jeffrey Barrett, Sonia Goncalves, Ewan Harrison, David K. Jackson, Ian Johnston, Dominic Kwiatkowski, Cordelia Langford, John Sillitoe on behalf of the Wellcome Sanger Institute COVID-19 Surveillance Team |
| EPI_ISL_3367233, EPI_ISL_3367234, EPI_ISL_3367235 | Lighthouse Lab in Glasgow | Wellcome Sanger Institute for the COVID-19 Genomics UK (COG-UK) Consortium | Harper VanSteenhouse, Yumi Kasai, David Gray, Carol Clugston, Anna Dominiczak and Alex Alderton, Roberto Amato, Jeffrey Barrett, Sonia Goncalves, Ewan Harrison, David K. Jackson, Ian Johnston, Dominic Kwiatkowski, Cordelia Langford, John Sillitoe on behalf of the Wellcome Sanger Institute COVID-19 Surveillance Team |
| EPI_ISL_3367236, EPI_ISL_3367237, EPI_ISL_3367238 | Berkshire and Surrey Pathology Services Lighthouse Laboratory | Wellcome Sanger Institute for the COVID-19 Genomics UK (COG-UK) Consortium | Berkshire and Surrey Pathology Services Lighthouse Laboratory and Alex Alderton, Roberto Amato, Jeffrey Barrett, Sonia Goncalves, Ewan Harrison, David K. Jackson, Ian Johnston, Dominic Kwiatkowski, Cordelia Langford, John Sillitoe on behalf of the Wellcome Sanger Institute COVID-19 Surveillance Team |

| | | | |
|---|---|---|---|
| EPI_ISL_3367239 | Lighthouse Laboratory Plymouth | Wellcome Sanger Institute for the COVID-19 Genomics UK (COG-UK) Consortium | Lighthouse Laboratory Plymouth and Alex Alderton, Roberto Amato, Jeffrey Barrett, Sonia Goncalves, Ewan Harrison, David K. Jackson, Ian Johnston, Dominic Kwiatkowski, Cordelia Langford, John Sillitoe on behalf of the Wellcome Sanger Institute COVID-19 Surveillance Team |
| EPI_ISL_3367240 | Berkshire and Surrey Pathology Services Lighthouse Laboratory | Wellcome Sanger Institute for the COVID-19 Genomics UK (COG-UK) Consortium | Berkshire and Surrey Pathology Services Lighthouse Laboratory and Alex Alderton, Roberto Amato, Jeffrey Barrett, Sonia Goncalves, Ewan Harrison, David K. Jackson, Ian Johnston, Dominic Kwiatkowski, Cordelia Langford, John Sillitoe on behalf of the Wellcome Sanger Institute COVID-19 Surveillance Team |
| EPI_ISL_3367241 | Lighthouse Lab in Glasgow | Wellcome Sanger Institute for the COVID-19 Genomics UK (COG-UK) Consortium | Harper VanSteenhouse, Yumi Kasai, David Gray, Carol Clugston, Anna Dominiczak and Alex Alderton, Roberto Amato, Jeffrey Barrett, Sonia Goncalves, Ewan Harrison, David K. Jackson, Ian Johnston, Dominic Kwiatkowski, Cordelia Langford, John Sillitoe on behalf of the Wellcome Sanger Institute COVID-19 Surveillance Team |
| EPI_ISL_3367242 | Lighthouse Laboratory Plymouth | Wellcome Sanger Institute for the COVID-19 Genomics UK (COG-UK) Consortium | Lighthouse Laboratory Plymouth and Alex Alderton, Roberto Amato, Jeffrey Barrett, Sonia Goncalves, Ewan Harrison, David K. Jackson, Ian Johnston, Dominic Kwiatkowski, Cordelia Langford, John Sillitoe on behalf of the Wellcome Sanger Institute COVID-19 Surveillance Team |
| EPI_ISL_3367243, EPI_ISL_3367244, EPI_ISL_3367245 | Lighthouse Lab in Glasgow | Wellcome Sanger Institute for the COVID-19 Genomics UK (COG-UK) Consortium | Harper VanSteenhouse, Yumi Kasai, David Gray, Carol Clugston, Anna Dominiczak and Alex Alderton, Roberto Amato, Jeffrey Barrett, Sonia Goncalves, Ewan Harrison, David K. Jackson, Ian Johnston, Dominic Kwiatkowski, Cordelia Langford, John Sillitoe on behalf of the Wellcome Sanger Institute COVID-19 Surveillance Team |
| EPI_ISL_3367246, EPI_ISL_3367247 | Lighthouse Laboratory Plymouth | Wellcome Sanger Institute for the COVID-19 Genomics UK (COG-UK) Consortium | Lighthouse Laboratory Plymouth and Alex Alderton, Roberto Amato, Jeffrey Barrett, Sonia Goncalves, Ewan Harrison, David K. Jackson, Ian Johnston, Dominic Kwiatkowski, Cordelia Langford, John Sillitoe on behalf of the Wellcome Sanger Institute COVID-19 Surveillance Team |
| EPI_ISL_3367248 | Berkshire and Surrey Pathology Services Lighthouse Laboratory | Wellcome Sanger Institute for the COVID-19 Genomics UK (COG-UK) Consortium | Berkshire and Surrey Pathology Services Lighthouse Laboratory and Alex Alderton, Roberto Amato, Jeffrey Barrett, Sonia Goncalves, Ewan Harrison, David K. Jackson, Ian Johnston, Dominic Kwiatkowski, Cordelia Langford, John Sillitoe on behalf of the Wellcome Sanger Institute COVID-19 Surveillance Team |
| EPI_ISL_3367249, EPI_ISL_3367250 | Lighthouse Laboratory Plymouth | Wellcome Sanger Institute for the COVID-19 Genomics UK (COG-UK) Consortium | Lighthouse Laboratory Plymouth and Alex Alderton, Roberto Amato, Jeffrey Barrett, Sonia Goncalves, Ewan Harrison, David K. Jackson, Ian Johnston, Dominic Kwiatkowski, Cordelia Langford, John Sillitoe on behalf of the Wellcome Sanger Institute COVID-19 Surveillance Team |
| EPI_ISL_3367251, EPI_ISL_3367252, EPI_ISL_3367253 | Berkshire and Surrey Pathology Services Lighthouse Laboratory | Wellcome Sanger Institute for the COVID-19 Genomics UK (COG-UK) Consortium | Berkshire and Surrey Pathology Services Lighthouse Laboratory and Alex Alderton, Roberto Amato, Jeffrey Barrett, Sonia Goncalves, Ewan Harrison, David K. Jackson, Ian Johnston, Dominic Kwiatkowski, Cordelia Langford, John Sillitoe on behalf of the Wellcome Sanger Institute COVID-19 Surveillance Team |
| EPI_ISL_3367254 | Lighthouse Laboratory Plymouth | Wellcome Sanger Institute for the COVID-19 Genomics UK (COG-UK) Consortium | Lighthouse Laboratory Plymouth and Alex Alderton, Roberto Amato, Jeffrey Barrett, Sonia Goncalves, Ewan Harrison, David K. Jackson, Ian Johnston, Dominic Kwiatkowski, Cordelia Langford, John Sillitoe on behalf of the Wellcome Sanger Institute COVID-19 Surveillance Team |
| EPI_ISL_3367255 | Lighthouse Lab in Glasgow | Wellcome Sanger Institute for the COVID-19 Genomics UK (COG-UK) Consortium | Harper VanSteenhouse, Yumi Kasai, David Gray, Carol Clugston, Anna Dominiczak and Alex Alderton, Roberto Amato, Jeffrey Barrett, Sonia Goncalves, Ewan Harrison, David K. Jackson, Ian Johnston, Dominic Kwiatkowski, Cordelia Langford, John Sillitoe on behalf of the Wellcome Sanger Institute COVID-19 Surveillance Team |
| EPI_ISL_3367256, EPI_ISL_3367257, EPI_ISL_3367258 | Berkshire and Surrey Pathology Services Lighthouse Laboratory | Wellcome Sanger Institute for the COVID-19 Genomics UK (COG-UK) Consortium | Berkshire and Surrey Pathology Services Lighthouse Laboratory and Alex Alderton, Roberto Amato, Jeffrey Barrett, Sonia Goncalves, Ewan Harrison, David K. Jackson, Ian Johnston, Dominic Kwiatkowski, Cordelia Langford, John Sillitoe on behalf of the Wellcome Sanger Institute COVID-19 Surveillance Team |
| EPI_ISL_3367259 | Lighthouse Lab in Glasgow | Wellcome Sanger Institute for the COVID-19 Genomics UK (COG-UK) Consortium | Harper VanSteenhouse, Yumi Kasai, David Gray, Carol Clugston, Anna Dominiczak and Alex Alderton, Roberto Amato, Jeffrey Barrett, Sonia Goncalves, Ewan Harrison, David K. Jackson, Ian Johnston, Dominic Kwiatkowski, Cordelia Langford, John Sillitoe on behalf of the Wellcome Sanger Institute COVID-19 Surveillance Team |
| EPI_ISL_3367260 | Berkshire and Surrey Pathology Services Lighthouse Laboratory | Wellcome Sanger Institute for the COVID-19 Genomics UK (COG-UK) Consortium | Berkshire and Surrey Pathology Services Lighthouse Laboratory and Alex Alderton, Roberto Amato, Jeffrey Barrett, Sonia Goncalves, Ewan Harrison, David K. Jackson, Ian Johnston, Dominic Kwiatkowski, Cordelia Langford, John Sillitoe on behalf of the Wellcome Sanger Institute COVID-19 Surveillance Team |
| EPI_ISL_3367261 | Lighthouse Lab in Glasgow | Wellcome Sanger Institute for the COVID-19 Genomics UK (COG-UK) Consortium | Harper VanSteenhouse, Yumi Kasai, David Gray, Carol Clugston, Anna Dominiczak and Alex Alderton, Roberto Amato, Jeffrey Barrett, Sonia Goncalves, Ewan Harrison, David K. Jackson, Ian Johnston, Dominic Kwiatkowski, Cordelia Langford, John Sillitoe on behalf of the Wellcome Sanger Institute COVID-19 Surveillance Team |
| EPI_ISL_3367262 | Lighthouse Laboratory Plymouth | Wellcome Sanger Institute for the COVID-19 Genomics UK (COG-UK) Consortium | Lighthouse Laboratory Plymouth and Alex Alderton, Roberto Amato, Jeffrey Barrett, Sonia Goncalves, Ewan Harrison, David K. Jackson, Ian Johnston, Dominic Kwiatkowski, Cordelia Langford, John Sillitoe on behalf of the Wellcome Sanger Institute COVID-19 Surveillance Team |
| EPI_ISL_3367263 | Lighthouse Lab in Glasgow | Wellcome Sanger Institute for the COVID-19 Genomics UK (COG-UK) Consortium | Harper VanSteenhouse, Yumi Kasai, David Gray, Carol Clugston, Anna Dominiczak and Alex Alderton, Roberto Amato, Jeffrey Barrett, Sonia Goncalves, Ewan Harrison, David K. Jackson, Ian Johnston, Dominic Kwiatkowski, Cordelia Langford, John Sillitoe on behalf of the Wellcome Sanger Institute COVID-19 Surveillance Team |
| EPI_ISL_3367264 | Lighthouse Laboratory Plymouth | Wellcome Sanger Institute for the COVID-19 Genomics UK (COG-UK) Consortium | Lighthouse Laboratory Plymouth and Alex Alderton, Roberto Amato, Jeffrey Barrett, Sonia Goncalves, Ewan Harrison, David K. Jackson, Ian Johnston, Dominic Kwiatkowski, Cordelia Langford, John Sillitoe on behalf of the Wellcome Sanger Institute COVID-19 Surveillance Team |
| EPI_ISL_3367265, EPI_ISL_3367266 | Lighthouse Lab in Glasgow | Wellcome Sanger Institute for the COVID-19 Genomics UK (COG-UK) Consortium | Harper VanSteenhouse, Yumi Kasai, David Gray, Carol Clugston, Anna Dominiczak and Alex Alderton, Roberto Amato, Jeffrey Barrett, Sonia Goncalves, Ewan Harrison, David K. Jackson, Ian Johnston, Dominic Kwiatkowski, Cordelia Langford, John Sillitoe on behalf of the Wellcome Sanger Institute COVID-19 Surveillance Team |
| EPI_ISL_3367267, EPI_ISL_3367268 | Berkshire and Surrey Pathology Services Lighthouse Laboratory | Wellcome Sanger Institute for the COVID-19 Genomics UK (COG-UK) Consortium | Berkshire and Surrey Pathology Services Lighthouse Laboratory and Alex Alderton, Roberto Amato, Jeffrey Barrett, Sonia Goncalves, Ewan Harrison, David K. Jackson, Ian Johnston, Dominic Kwiatkowski, Cordelia Langford, John Sillitoe on behalf of the Wellcome Sanger Institute COVID-19 Surveillance Team |
| EPI_ISL_3367269 | Lighthouse Laboratory Plymouth | Wellcome Sanger Institute for the COVID-19 Genomics UK (COG-UK) Consortium | Lighthouse Laboratory Plymouth and Alex Alderton, Roberto Amato, Jeffrey Barrett, Sonia Goncalves, Ewan Harrison, David K. Jackson, Ian Johnston, Dominic Kwiatkowski, Cordelia Langford, John Sillitoe on behalf of the Wellcome Sanger Institute COVID-19 Surveillance Team |
| EPI_ISL_3367270 | Berkshire and Surrey Pathology Services Lighthouse Laboratory | Wellcome Sanger Institute for the COVID-19 Genomics UK (COG-UK) Consortium | Berkshire and Surrey Pathology Services Lighthouse Laboratory and Alex Alderton, Roberto Amato, Jeffrey Barrett, Sonia Goncalves, Ewan Harrison, David K. Jackson, Ian Johnston, Dominic Kwiatkowski, Cordelia Langford, John Sillitoe on behalf of the Wellcome Sanger Institute COVID-19 Surveillance Team |
| EPI_ISL_3367271 | Lighthouse Lab in Glasgow | Wellcome Sanger Institute for the COVID-19 Genomics UK (COG-UK) Consortium | Harper VanSteenhouse, Yumi Kasai, David Gray, Carol Clugston, Anna Dominiczak and Alex Alderton, Roberto Amato, Jeffrey Barrett, Sonia Goncalves, Ewan Harrison, David K. Jackson, Ian Johnston, Dominic Kwiatkowski, Cordelia Langford, John Sillitoe on behalf of the Wellcome Sanger Institute COVID-19 Surveillance Team |
| EPI_ISL_3367272, EPI_ISL_3367273 | Lighthouse Laboratory Plymouth | Wellcome Sanger Institute for the COVID-19 Genomics UK (COG-UK) Consortium | Lighthouse Laboratory Plymouth and Alex Alderton, Roberto Amato, Jeffrey Barrett, Sonia Goncalves, Ewan Harrison, David K. Jackson, Ian Johnston, Dominic Kwiatkowski, Cordelia Langford, John Sillitoe on behalf of the Wellcome Sanger Institute COVID-19 Surveillance Team |
| EPI_ISL_3367274 | Berkshire and Surrey Pathology Services Lighthouse Laboratory | Wellcome Sanger Institute for the COVID-19 Genomics UK (COG-UK) Consortium | Berkshire and Surrey Pathology Services Lighthouse Laboratory and Alex Alderton, Roberto Amato, Jeffrey Barrett, Sonia Goncalves, Ewan Harrison, David K. Jackson, Ian Johnston, Dominic Kwiatkowski, Cordelia Langford, John Sillitoe on behalf of the Wellcome Sanger Institute COVID-19 Surveillance Team |
| EPI_ISL_3367275 | Lighthouse Laboratory Plymouth | Wellcome Sanger Institute for the COVID-19 Genomics UK (COG-UK) Consortium | Lighthouse Laboratory Plymouth and Alex Alderton, Roberto Amato, Jeffrey Barrett, Sonia Goncalves, Ewan Harrison, David K. Jackson, Ian Johnston, Dominic Kwiatkowski, Cordelia Langford, John Sillitoe on behalf of the Wellcome Sanger Institute COVID-19 Surveillance Team |
| EPI_ISL_3367276 | Berkshire and Surrey Pathology Services Lighthouse Laboratory | Wellcome Sanger Institute for the COVID-19 Genomics UK (COG-UK) Consortium | Berkshire and Surrey Pathology Services Lighthouse Laboratory and Alex Alderton, Roberto Amato, Jeffrey Barrett, Sonia Goncalves, Ewan Harrison, David K. Jackson, Ian Johnston, Dominic Kwiatkowski, Cordelia Langford, John Sillitoe on behalf of the Wellcome Sanger Institute COVID-19 Surveillance Team |
| EPI_ISL_3367277 | Lighthouse Laboratory Plymouth | Wellcome Sanger Institute for the COVID-19 Genomics UK (COG-UK) Consortium | Lighthouse Laboratory Plymouth and Alex Alderton, Roberto Amato, Jeffrey Barrett, Sonia Goncalves, Ewan Harrison, David K. Jackson, Ian Johnston, Dominic Kwiatkowski, Cordelia Langford, John Sillitoe on behalf of the Wellcome Sanger Institute COVID-19 Surveillance Team |
| EPI_ISL_3367278, EPI_ISL_3367279, EPI_ISL_3367280 | Berkshire and Surrey Pathology Services Lighthouse Laboratory | Wellcome Sanger Institute for the COVID-19 Genomics UK (COG-UK) Consortium | Berkshire and Surrey Pathology Services Lighthouse Laboratory and Alex Alderton, Roberto Amato, Jeffrey Barrett, Sonia Goncalves, Ewan Harrison, David K. Jackson, Ian Johnston, Dominic Kwiatkowski, Cordelia Langford, John Sillitoe on behalf of the Wellcome Sanger Institute COVID-19 Surveillance Team |
| EPI_ISL_3367281 | Lighthouse Laboratory Plymouth | Wellcome Sanger Institute for the COVID-19 Genomics UK (COG-UK) Consortium | Lighthouse Laboratory Plymouth and Alex Alderton, Roberto Amato, Jeffrey Barrett, Sonia Goncalves, Ewan Harrison, David K. Jackson, Ian Johnston, Dominic Kwiatkowski, Cordelia Langford, John Sillitoe on behalf of the Wellcome Sanger Institute COVID-19 Surveillance Team |
| EPI_ISL_3367282 | Berkshire and Surrey Pathology Services Lighthouse Laboratory | Wellcome Sanger Institute for the COVID-19 Genomics UK (COG-UK) Consortium | Berkshire and Surrey Pathology Services Lighthouse Laboratory and Alex Alderton, Roberto Amato, Jeffrey Barrett, Sonia Goncalves, Ewan Harrison, David K. Jackson, Ian Johnston, Dominic Kwiatkowski, Cordelia Langford, John Sillitoe on behalf of the Wellcome Sanger Institute COVID-19 Surveillance Team |
| EPI_ISL_3367283 | Lighthouse Laboratory Plymouth | Wellcome Sanger Institute for the COVID-19 Genomics UK (COG-UK) Consortium | Lighthouse Laboratory Plymouth and Alex Alderton, Roberto Amato, Jeffrey Barrett, Sonia Goncalves, Ewan Harrison, David K. Jackson, Ian Johnston, Dominic Kwiatkowski, Cordelia Langford, John Sillitoe on behalf of the Wellcome Sanger Institute COVID-19 Surveillance Team |
| EPI_ISL_3367284, EPI_ISL_3367285, EPI_ISL_3367286, EPI_ISL_3367287 | Berkshire and Surrey Pathology Services Lighthouse Laboratory | Wellcome Sanger Institute for the COVID-19 Genomics UK (COG-UK) Consortium | Berkshire and Surrey Pathology Services Lighthouse Laboratory and Alex Alderton, Roberto Amato, Jeffrey Barrett, Sonia Goncalves, Ewan Harrison, David K. Jackson, Ian Johnston, Dominic Kwiatkowski, Cordelia Langford, John Sillitoe on behalf of the Wellcome Sanger Institute COVID-19 Surveillance Team |
| EPI_ISL_3367288 | Lighthouse Lab in Glasgow | Wellcome Sanger Institute for the COVID-19 Genomics UK (COG-UK) Consortium | Harper VanSteenhouse, Yumi Kasai, David Gray, Carol Clugston, Anna Dominiczak and Alex Alderton, Roberto Amato, Jeffrey Barrett, Sonia Goncalves, Ewan Harrison, David K. Jackson, Ian Johnston, Dominic Kwiatkowski, Cordelia Langford, John Sillitoe on behalf of the Wellcome Sanger Institute COVID-19 Surveillance Team |
| EPI_ISL_3367289 | Berkshire and Surrey Pathology Services Lighthouse Laboratory | Wellcome Sanger Institute for the COVID-19 Genomics UK (COG-UK) Consortium | Berkshire and Surrey Pathology Services Lighthouse Laboratory and Alex Alderton, Roberto Amato, Jeffrey Barrett, Sonia Goncalves, Ewan Harrison, David K. Jackson, Ian Johnston, Dominic Kwiatkowski, Cordelia Langford, John Sillitoe on behalf of the Wellcome Sanger Institute COVID-19 Surveillance Team |
| EPI_ISL_3367290, EPI_ISL_3367291 | Lighthouse Laboratory Plymouth | Wellcome Sanger Institute for the COVID-19 Genomics UK (COG-UK) Consortium | Lighthouse Laboratory Plymouth and Alex Alderton, Roberto Amato, Jeffrey Barrett, Sonia Goncalves, Ewan Harrison, David K. Jackson, Ian Johnston, Dominic Kwiatkowski, Cordelia Langford, John Sillitoe on behalf of the Wellcome Sanger Institute COVID-19 Surveillance Team |
| EPI_ISL_3367292 | Lighthouse Lab in Glasgow | Wellcome Sanger Institute for the COVID-19 Genomics UK (COG-UK) Consortium | Harper VanSteenhouse, Yumi Kasai, David Gray, Carol Clugston, Anna Dominiczak and Alex Alderton, Roberto Amato, Jeffrey Barrett, Sonia Goncalves, Ewan Harrison, David K. Jackson, Ian Johnston, Dominic Kwiatkowski, Cordelia Langford, John Sillitoe on behalf of the Wellcome Sanger Institute COVID-19 Surveillance Team |
| EPI_ISL_3367293, EPI_ISL_3367294 | Berkshire and Surrey Pathology Services Lighthouse Laboratory | Wellcome Sanger Institute for the COVID-19 Genomics UK (COG-UK) Consortium | Berkshire and Surrey Pathology Services Lighthouse Laboratory and Alex Alderton, Roberto Amato, Jeffrey Barrett, Sonia Goncalves, Ewan Harrison, David K. Jackson, Ian Johnston, Dominic Kwiatkowski, Cordelia Langford, John Sillitoe on behalf of the Wellcome Sanger Institute COVID-19 Surveillance Team |
| EPI_ISL_3367295 | Lighthouse Lab in Glasgow | Wellcome Sanger Institute for the COVID-19 Genomics UK (COG-UK) Consortium | Harper VanSteenhouse, Yumi Kasai, David Gray, Carol Clugston, Anna Dominiczak and Alex Alderton, Roberto Amato, Jeffrey Barrett, Sonia Goncalves, Ewan Harrison, David K. Jackson, Ian Johnston, Dominic Kwiatkowski, Cordelia Langford, John Sillitoe on behalf of the Wellcome Sanger Institute COVID-19 Surveillance Team |
| EPI_ISL_3367296, EPI_ISL_3367297, EPI_ISL_3367298, EPI_ISL_3367299 | Berkshire and Surrey Pathology Services Lighthouse Laboratory | Wellcome Sanger Institute for the COVID-19 Genomics UK (COG-UK) Consortium | Berkshire and Surrey Pathology Services Lighthouse Laboratory and Alex Alderton, Roberto Amato, Jeffrey Barrett, Sonia Goncalves, Ewan Harrison, David K. Jackson, Ian Johnston, Dominic Kwiatkowski, Cordelia Langford, John Sillitoe on behalf of the Wellcome Sanger Institute COVID-19 Surveillance Team |
| EPI_ISL_3367300, EPI_ISL_3367301 | Lighthouse Laboratory Plymouth | Wellcome Sanger Institute for the COVID-19 Genomics UK (COG-UK) Consortium | Lighthouse Laboratory Plymouth and Alex Alderton, Roberto Amato, Jeffrey Barrett, Sonia Goncalves, Ewan Harrison, David K. Jackson, Ian Johnston, Dominic Kwiatkowski, Cordelia Langford, John Sillitoe on behalf of the Wellcome Sanger Institute COVID-19 Surveillance Team |
| EPI_ISL_3367302, | Berkshire and Surrey Pathology Services Lighthouse Laboratory | Wellcome Sanger Institute for the COVID-19 Genomics UK (COG-UK) | Berkshire and Surrey Pathology Services Lighthouse Laboratory and Alex Alderton, Roberto Amato, Jeffrey Barrett, Sonia Goncalves, Ewan Harrison, David K. Jackson, Ian Johnston, Dominic Kwiatkowski, Cordelia Langford, John Sillitoe on behalf |

| | | | |
|---|---|---|---|
| EPI_ISL_3367303 | | Consortium | of the Wellcome Sanger Institute COVID-19 Surveillance Team |
| EPI_ISL_3367304 | Lighthouse Laboratory Plymouth | Wellcome Sanger Institute for the COVID-19 Genomics UK (COG-UK) Consortium | Lighthouse Laboratory Plymouth and Alex Alderton, Roberto Amato, Jeffrey Barrett, Sonia Goncalves, Ewan Harrison, David K. Jackson, Ian Johnston, Dominic Kwiatkowski, Cordelia Langford, John Sillitoe on behalf of the Wellcome Sanger Institute COVID-19 Surveillance Team |
| EPI_ISL_3367305, EPI_ISL_3367306 | Berkshire and Surrey Pathology Services Lighthouse Laboratory | Wellcome Sanger Institute for the COVID-19 Genomics UK (COG-UK) Consortium | Berkshire and Surrey Pathology Services Lighthouse Laboratory and Alex Alderton, Roberto Amato, Jeffrey Barrett, Sonia Goncalves, Ewan Harrison, David K. Jackson, Ian Johnston, Dominic Kwiatkowski, Cordelia Langford, John Sillitoe on behalf of the Wellcome Sanger Institute COVID-19 Surveillance Team |
| EPI_ISL_3367307, EPI_ISL_3367308 | Lighthouse Lab in Glasgow | Wellcome Sanger Institute for the COVID-19 Genomics UK (COG-UK) Consortium | Harper VanSteenhouse, Yumi Kasai, David Gray, Carol Clugston, Anna Dominiczak and Alex Alderton, Roberto Amato, Jeffrey Barrett, Sonia Goncalves, Ewan Harrison, David K. Jackson, Ian Johnston, Dominic Kwiatkowski, Cordelia Langford, John Sillitoe on behalf of the Wellcome Sanger Institute COVID-19 Surveillance Team |
| EPI_ISL_3367309, EPI_ISL_3367310, EPI_ISL_3367311 | Berkshire and Surrey Pathology Services Lighthouse Laboratory | Wellcome Sanger Institute for the COVID-19 Genomics UK (COG-UK) Consortium | Berkshire and Surrey Pathology Services Lighthouse Laboratory and Alex Alderton, Roberto Amato, Jeffrey Barrett, Sonia Goncalves, Ewan Harrison, David K. Jackson, Ian Johnston, Dominic Kwiatkowski, Cordelia Langford, John Sillitoe on behalf of the Wellcome Sanger Institute COVID-19 Surveillance Team |
| EPI_ISL_3367312 | Lighthouse Lab in Glasgow | Wellcome Sanger Institute for the COVID-19 Genomics UK (COG-UK) Consortium | Harper VanSteenhouse, Yumi Kasai, David Gray, Carol Clugston, Anna Dominiczak and Alex Alderton, Roberto Amato, Jeffrey Barrett, Sonia Goncalves, Ewan Harrison, David K. Jackson, Ian Johnston, Dominic Kwiatkowski, Cordelia Langford, John Sillitoe on behalf of the Wellcome Sanger Institute COVID-19 Surveillance Team |
| EPI_ISL_3367313 | Berkshire and Surrey Pathology Services Lighthouse Laboratory | Wellcome Sanger Institute for the COVID-19 Genomics UK (COG-UK) Consortium | Berkshire and Surrey Pathology Services Lighthouse Laboratory and Alex Alderton, Roberto Amato, Jeffrey Barrett, Sonia Goncalves, Ewan Harrison, David K. Jackson, Ian Johnston, Dominic Kwiatkowski, Cordelia Langford, John Sillitoe on behalf of the Wellcome Sanger Institute COVID-19 Surveillance Team |
| EPI_ISL_3367314 | Lighthouse Laboratory Plymouth | Wellcome Sanger Institute for the COVID-19 Genomics UK (COG-UK) Consortium | Lighthouse Laboratory Plymouth and Alex Alderton, Roberto Amato, Jeffrey Barrett, Sonia Goncalves, Ewan Harrison, David K. Jackson, Ian Johnston, Dominic Kwiatkowski, Cordelia Langford, John Sillitoe on behalf of the Wellcome Sanger Institute COVID-19 Surveillance Team |
| EPI_ISL_3367315 | Lighthouse Lab in Glasgow | Wellcome Sanger Institute for the COVID-19 Genomics UK (COG-UK) Consortium | Harper VanSteenhouse, Yumi Kasai, David Gray, Carol Clugston, Anna Dominiczak and Alex Alderton, Roberto Amato, Jeffrey Barrett, Sonia Goncalves, Ewan Harrison, David K. Jackson, Ian Johnston, Dominic Kwiatkowski, Cordelia Langford, John Sillitoe on behalf of the Wellcome Sanger Institute COVID-19 Surveillance Team |
| EPI_ISL_3367316 | Berkshire and Surrey Pathology Services Lighthouse Laboratory | Wellcome Sanger Institute for the COVID-19 Genomics UK (COG-UK) Consortium | Berkshire and Surrey Pathology Services Lighthouse Laboratory and Alex Alderton, Roberto Amato, Jeffrey Barrett, Sonia Goncalves, Ewan Harrison, David K. Jackson, Ian Johnston, Dominic Kwiatkowski, Cordelia Langford, John Sillitoe on behalf of the Wellcome Sanger Institute COVID-19 Surveillance Team |
| EPI_ISL_3367317, EPI_ISL_3367318, EPI_ISL_3367319, EPI_ISL_3367320 | Lighthouse Laboratory Plymouth | Wellcome Sanger Institute for the COVID-19 Genomics UK (COG-UK) Consortium | Lighthouse Laboratory Plymouth and Alex Alderton, Roberto Amato, Jeffrey Barrett, Sonia Goncalves, Ewan Harrison, David K. Jackson, Ian Johnston, Dominic Kwiatkowski, Cordelia Langford, John Sillitoe on behalf of the Wellcome Sanger Institute COVID-19 Surveillance Team |
| EPI_ISL_3367321 | Lighthouse Lab in Glasgow | Wellcome Sanger Institute for the COVID-19 Genomics UK (COG-UK) Consortium | Harper VanSteenhouse, Yumi Kasai, David Gray, Carol Clugston, Anna Dominiczak and Alex Alderton, Roberto Amato, Jeffrey Barrett, Sonia Goncalves, Ewan Harrison, David K. Jackson, Ian Johnston, Dominic Kwiatkowski, Cordelia Langford, John Sillitoe on behalf of the Wellcome Sanger Institute COVID-19 Surveillance Team |
| EPI_ISL_3367322 | Lighthouse Laboratory Plymouth | Wellcome Sanger Institute for the COVID-19 Genomics UK (COG-UK) Consortium | Lighthouse Laboratory Plymouth and Alex Alderton, Roberto Amato, Jeffrey Barrett, Sonia Goncalves, Ewan Harrison, David K. Jackson, Ian Johnston, Dominic Kwiatkowski, Cordelia Langford, John Sillitoe on behalf of the Wellcome Sanger Institute COVID-19 Surveillance Team |
| EPI_ISL_3367323 | Berkshire and Surrey Pathology Services Lighthouse Laboratory | Wellcome Sanger Institute for the COVID-19 Genomics UK (COG-UK) Consortium | Berkshire and Surrey Pathology Services Lighthouse Laboratory and Alex Alderton, Roberto Amato, Jeffrey Barrett, Sonia Goncalves, Ewan Harrison, David K. Jackson, Ian Johnston, Dominic Kwiatkowski, Cordelia Langford, John Sillitoe on behalf of the Wellcome Sanger Institute COVID-19 Surveillance Team |
| EPI_ISL_3367324 | Lighthouse Laboratory Plymouth | Wellcome Sanger Institute for the COVID-19 Genomics UK (COG-UK) Consortium | Lighthouse Laboratory Plymouth and Alex Alderton, Roberto Amato, Jeffrey Barrett, Sonia Goncalves, Ewan Harrison, David K. Jackson, Ian Johnston, Dominic Kwiatkowski, Cordelia Langford, John Sillitoe on behalf of the Wellcome Sanger Institute COVID-19 Surveillance Team |
| EPI_ISL_3367325 | Lighthouse Lab in Glasgow | Wellcome Sanger Institute for the COVID-19 Genomics UK (COG-UK) Consortium | Harper VanSteenhouse, Yumi Kasai, David Gray, Carol Clugston, Anna Dominiczak and Alex Alderton, Roberto Amato, Jeffrey Barrett, Sonia Goncalves, Ewan Harrison, David K. Jackson, Ian Johnston, Dominic Kwiatkowski, Cordelia Langford, John Sillitoe on behalf of the Wellcome Sanger Institute COVID-19 Surveillance Team |
| EPI_ISL_3367326 | Lighthouse Laboratory Plymouth | Wellcome Sanger Institute for the COVID-19 Genomics UK (COG-UK) Consortium | Lighthouse Laboratory Plymouth and Alex Alderton, Roberto Amato, Jeffrey Barrett, Sonia Goncalves, Ewan Harrison, David K. Jackson, Ian Johnston, Dominic Kwiatkowski, Cordelia Langford, John Sillitoe on behalf of the Wellcome Sanger Institute COVID-19 Surveillance Team |
| EPI_ISL_3367327, EPI_ISL_3367328, EPI_ISL_3367329, EPI_ISL_3367330 | Berkshire and Surrey Pathology Services Lighthouse Laboratory | Wellcome Sanger Institute for the COVID-19 Genomics UK (COG-UK) Consortium | Berkshire and Surrey Pathology Services Lighthouse Laboratory and Alex Alderton, Roberto Amato, Jeffrey Barrett, Sonia Goncalves, Ewan Harrison, David K. Jackson, Ian Johnston, Dominic Kwiatkowski, Cordelia Langford, John Sillitoe on behalf of the Wellcome Sanger Institute COVID-19 Surveillance Team |
| EPI_ISL_3367331, EPI_ISL_3367332 | Lighthouse Lab in Glasgow | Wellcome Sanger Institute for the COVID-19 Genomics UK (COG-UK) Consortium | Harper VanSteenhouse, Yumi Kasai, David Gray, Carol Clugston, Anna Dominiczak and Alex Alderton, Roberto Amato, Jeffrey Barrett, Sonia Goncalves, Ewan Harrison, David K. Jackson, Ian Johnston, Dominic Kwiatkowski, Cordelia Langford, John Sillitoe on behalf of the Wellcome Sanger Institute COVID-19 Surveillance Team |
| EPI_ISL_3367333, EPI_ISL_3367334 | Berkshire and Surrey Pathology Services Lighthouse Laboratory | Wellcome Sanger Institute for the COVID-19 Genomics UK (COG-UK) Consortium | Berkshire and Surrey Pathology Services Lighthouse Laboratory and Alex Alderton, Roberto Amato, Jeffrey Barrett, Sonia Goncalves, Ewan Harrison, David K. Jackson, Ian Johnston, Dominic Kwiatkowski, Cordelia Langford, John Sillitoe on behalf of the Wellcome Sanger Institute COVID-19 Surveillance Team |
| EPI_ISL_3367335 | Lighthouse Laboratory Plymouth | Wellcome Sanger Institute for the COVID-19 Genomics UK (COG-UK) Consortium | Lighthouse Laboratory Plymouth and Alex Alderton, Roberto Amato, Jeffrey Barrett, Sonia Goncalves, Ewan Harrison, David K. Jackson, Ian Johnston, Dominic Kwiatkowski, Cordelia Langford, John Sillitoe on behalf of the Wellcome Sanger Institute COVID-19 Surveillance Team |
| EPI_ISL_3367336, EPI_ISL_3367337 | Lighthouse Lab in Glasgow | Wellcome Sanger Institute for the COVID-19 Genomics UK (COG-UK) Consortium | Harper VanSteenhouse, Yumi Kasai, David Gray, Carol Clugston, Anna Dominiczak and Alex Alderton, Roberto Amato, Jeffrey Barrett, Sonia Goncalves, Ewan Harrison, David K. Jackson, Ian Johnston, Dominic Kwiatkowski, Cordelia Langford, John Sillitoe on behalf of the Wellcome Sanger Institute COVID-19 Surveillance Team |
| EPI_ISL_3367338 | Berkshire and Surrey Pathology Services Lighthouse Laboratory | Wellcome Sanger Institute for the COVID-19 Genomics UK (COG-UK) Consortium | Berkshire and Surrey Pathology Services Lighthouse Laboratory and Alex Alderton, Roberto Amato, Jeffrey Barrett, Sonia Goncalves, Ewan Harrison, David K. Jackson, Ian Johnston, Dominic Kwiatkowski, Cordelia Langford, John Sillitoe on behalf of the Wellcome Sanger Institute COVID-19 Surveillance Team |
| EPI_ISL_3367339, EPI_ISL_3367340 | Lighthouse Lab in Glasgow | Wellcome Sanger Institute for the COVID-19 Genomics UK (COG-UK) Consortium | Harper VanSteenhouse, Yumi Kasai, David Gray, Carol Clugston, Anna Dominiczak and Alex Alderton, Roberto Amato, Jeffrey Barrett, Sonia Goncalves, Ewan Harrison, David K. Jackson, Ian Johnston, Dominic Kwiatkowski, Cordelia Langford, John Sillitoe on behalf of the Wellcome Sanger Institute COVID-19 Surveillance Team |
| EPI_ISL_3367341 | Lighthouse Laboratory Plymouth | Wellcome Sanger Institute for the COVID-19 Genomics UK (COG-UK) Consortium | Lighthouse Laboratory Plymouth and Alex Alderton, Roberto Amato, Jeffrey Barrett, Sonia Goncalves, Ewan Harrison, David K. Jackson, Ian Johnston, Dominic Kwiatkowski, Cordelia Langford, John Sillitoe on behalf of the Wellcome Sanger Institute COVID-19 Surveillance Team |
| EPI_ISL_3367342 | Lighthouse Lab in Glasgow | Wellcome Sanger Institute for the COVID-19 Genomics UK (COG-UK) Consortium | Harper VanSteenhouse, Yumi Kasai, David Gray, Carol Clugston, Anna Dominiczak and Alex Alderton, Roberto Amato, Jeffrey Barrett, Sonia Goncalves, Ewan Harrison, David K. Jackson, Ian Johnston, Dominic Kwiatkowski, Cordelia Langford, John Sillitoe on behalf of the Wellcome Sanger Institute COVID-19 Surveillance Team |
| EPI_ISL_3367343 | Berkshire and Surrey Pathology Services Lighthouse Laboratory | Wellcome Sanger Institute for the COVID-19 Genomics UK (COG-UK) Consortium | Berkshire and Surrey Pathology Services Lighthouse Laboratory and Alex Alderton, Roberto Amato, Jeffrey Barrett, Sonia Goncalves, Ewan Harrison, David K. Jackson, Ian Johnston, Dominic Kwiatkowski, Cordelia Langford, John Sillitoe on behalf of the Wellcome Sanger Institute COVID-19 Surveillance Team |
| EPI_ISL_3367344 | Lighthouse Laboratory Plymouth | Wellcome Sanger Institute for the COVID-19 Genomics UK (COG-UK) Consortium | Lighthouse Laboratory Plymouth and Alex Alderton, Roberto Amato, Jeffrey Barrett, Sonia Goncalves, Ewan Harrison, David K. Jackson, Ian Johnston, Dominic Kwiatkowski, Cordelia Langford, John Sillitoe on behalf of the Wellcome Sanger Institute COVID-19 Surveillance Team |
| EPI_ISL_3367345, EPI_ISL_3367346, EPI_ISL_3367347 | Berkshire and Surrey Pathology Services Lighthouse Laboratory | Wellcome Sanger Institute for the COVID-19 Genomics UK (COG-UK) Consortium | Berkshire and Surrey Pathology Services Lighthouse Laboratory and Alex Alderton, Roberto Amato, Jeffrey Barrett, Sonia Goncalves, Ewan Harrison, David K. Jackson, Ian Johnston, Dominic Kwiatkowski, Cordelia Langford, John Sillitoe on behalf of the Wellcome Sanger Institute COVID-19 Surveillance Team |
| EPI_ISL_3367348 | Lighthouse Lab in Glasgow | Wellcome Sanger Institute for the COVID-19 Genomics UK (COG-UK) Consortium | Harper VanSteenhouse, Yumi Kasai, David Gray, Carol Clugston, Anna Dominiczak and Alex Alderton, Roberto Amato, Jeffrey Barrett, Sonia Goncalves, Ewan Harrison, David K. Jackson, Ian Johnston, Dominic Kwiatkowski, Cordelia Langford, John Sillitoe on behalf of the Wellcome Sanger Institute COVID-19 Surveillance Team |
| EPI_ISL_3367349 | Lighthouse Laboratory Plymouth | Wellcome Sanger Institute for the COVID-19 Genomics UK (COG-UK) Consortium | Lighthouse Laboratory Plymouth and Alex Alderton, Roberto Amato, Jeffrey Barrett, Sonia Goncalves, Ewan Harrison, David K. Jackson, Ian Johnston, Dominic Kwiatkowski, Cordelia Langford, John Sillitoe on behalf of the Wellcome Sanger Institute COVID-19 Surveillance Team |
| EPI_ISL_3367350, EPI_ISL_3367351 | Berkshire and Surrey Pathology Services Lighthouse Laboratory | Wellcome Sanger Institute for the COVID-19 Genomics UK (COG-UK) Consortium | Berkshire and Surrey Pathology Services Lighthouse Laboratory and Alex Alderton, Roberto Amato, Jeffrey Barrett, Sonia Goncalves, Ewan Harrison, David K. Jackson, Ian Johnston, Dominic Kwiatkowski, Cordelia Langford, John Sillitoe on behalf of the Wellcome Sanger Institute COVID-19 Surveillance Team |
| EPI_ISL_3367352 | Lighthouse Lab in Glasgow | Wellcome Sanger Institute for the COVID-19 Genomics UK (COG-UK) Consortium | Harper VanSteenhouse, Yumi Kasai, David Gray, Carol Clugston, Anna Dominiczak and Alex Alderton, Roberto Amato, Jeffrey Barrett, Sonia Goncalves, Ewan Harrison, David K. Jackson, Ian Johnston, Dominic Kwiatkowski, Cordelia Langford, John Sillitoe on behalf of the Wellcome Sanger Institute COVID-19 Surveillance Team |
| EPI_ISL_3367353 | Berkshire and Surrey Pathology Services Lighthouse Laboratory | Wellcome Sanger Institute for the COVID-19 Genomics UK (COG-UK) Consortium | Berkshire and Surrey Pathology Services Lighthouse Laboratory and Alex Alderton, Roberto Amato, Jeffrey Barrett, Sonia Goncalves, Ewan Harrison, David K. Jackson, Ian Johnston, Dominic Kwiatkowski, Cordelia Langford, John Sillitoe on behalf of the Wellcome Sanger Institute COVID-19 Surveillance Team |
| EPI_ISL_3367354 | Lighthouse Laboratory Plymouth | Wellcome Sanger Institute for the COVID-19 Genomics UK (COG-UK) Consortium | Lighthouse Laboratory Plymouth and Alex Alderton, Roberto Amato, Jeffrey Barrett, Sonia Goncalves, Ewan Harrison, David K. Jackson, Ian Johnston, Dominic Kwiatkowski, Cordelia Langford, John Sillitoe on behalf of the Wellcome Sanger Institute COVID-19 Surveillance Team |
| EPI_ISL_3367355 | Berkshire and Surrey Pathology Services Lighthouse Laboratory | Wellcome Sanger Institute for the COVID-19 Genomics UK (COG-UK) Consortium | Berkshire and Surrey Pathology Services Lighthouse Laboratory and Alex Alderton, Roberto Amato, Jeffrey Barrett, Sonia Goncalves, Ewan Harrison, David K. Jackson, Ian Johnston, Dominic Kwiatkowski, Cordelia Langford, John Sillitoe on behalf of the Wellcome Sanger Institute COVID-19 Surveillance Team |
| EPI_ISL_3367356, EPI_ISL_3367357 | Lighthouse Laboratory Plymouth | Wellcome Sanger Institute for the COVID-19 Genomics UK (COG-UK) Consortium | Lighthouse Laboratory Plymouth and Alex Alderton, Roberto Amato, Jeffrey Barrett, Sonia Goncalves, Ewan Harrison, David K. Jackson, Ian Johnston, Dominic Kwiatkowski, Cordelia Langford, John Sillitoe on behalf of the Wellcome Sanger Institute COVID-19 Surveillance Team |
| EPI_ISL_3367358 | Lighthouse Lab in Glasgow | Wellcome Sanger Institute for the COVID-19 Genomics UK (COG-UK) Consortium | Harper VanSteenhouse, Yumi Kasai, David Gray, Carol Clugston, Anna Dominiczak and Alex Alderton, Roberto Amato, Jeffrey Barrett, Sonia Goncalves, Ewan Harrison, David K. Jackson, Ian Johnston, Dominic Kwiatkowski, Cordelia Langford, John Sillitoe on behalf of the Wellcome Sanger Institute COVID-19 Surveillance Team |
| EPI_ISL_3367359, EPI_ISL_3367360, EPI_ISL_3367361 | Berkshire and Surrey Pathology Services Lighthouse Laboratory | Wellcome Sanger Institute for the COVID-19 Genomics UK (COG-UK) Consortium | Berkshire and Surrey Pathology Services Lighthouse Laboratory and Alex Alderton, Roberto Amato, Jeffrey Barrett, Sonia Goncalves, Ewan Harrison, David K. Jackson, Ian Johnston, Dominic Kwiatkowski, Cordelia Langford, John Sillitoe on behalf of the Wellcome Sanger Institute COVID-19 Surveillance Team |
| EPI_ISL_3367362 | Lighthouse Laboratory Plymouth | Wellcome Sanger Institute for the COVID-19 Genomics UK (COG-UK) Consortium | Lighthouse Laboratory Plymouth and Alex Alderton, Roberto Amato, Jeffrey Barrett, Sonia Goncalves, Ewan Harrison, David K. Jackson, Ian Johnston, Dominic Kwiatkowski, Cordelia Langford, John Sillitoe on behalf of the Wellcome Sanger Institute COVID-19 Surveillance Team |
| EPI_ISL_3367363, EPI_ISL_3367364 | Berkshire and Surrey Pathology Services Lighthouse Laboratory | Wellcome Sanger Institute for the COVID-19 Genomics UK (COG-UK) Consortium | Berkshire and Surrey Pathology Services Lighthouse Laboratory and Alex Alderton, Roberto Amato, Jeffrey Barrett, Sonia Goncalves, Ewan Harrison, David K. Jackson, Ian Johnston, Dominic Kwiatkowski, Cordelia Langford, John Sillitoe on behalf of the Wellcome Sanger Institute COVID-19 Surveillance Team |
| EPI_ISL_3367365 | Lighthouse Laboratory Plymouth | Wellcome Sanger Institute for the COVID-19 Genomics UK (COG-UK) Consortium | Lighthouse Laboratory Plymouth and Alex Alderton, Roberto Amato, Jeffrey Barrett, Sonia Goncalves, Ewan Harrison, David K. Jackson, Ian Johnston, Dominic Kwiatkowski, Cordelia Langford, John Sillitoe on behalf of the Wellcome Sanger Institute COVID-19 Surveillance Team |
| EPI_ISL_3367366, EPI_ISL_3367367 | Berkshire and Surrey Pathology Services Lighthouse Laboratory | Wellcome Sanger Institute for the COVID-19 Genomics UK (COG-UK) Consortium | Berkshire and Surrey Pathology Services Lighthouse Laboratory and Alex Alderton, Roberto Amato, Jeffrey Barrett, Sonia Goncalves, Ewan Harrison, David K. Jackson, Ian Johnston, Dominic Kwiatkowski, Cordelia Langford, John Sillitoe on behalf of the Wellcome Sanger Institute COVID-19 Surveillance Team |

| | | | |
|---|---|---|---|
| EPI_ISL_3367368 | Lighthouse Lab in Glasgow | Wellcome Sanger Institute for the COVID-19 Genomics UK (COG-UK) Consortium | Harper VanSteenhouse, Yumi Kasai, David Gray, Carol Clugston, Anna Dominiczak and Alex Alderton, Roberto Amato, Jeffrey Barrett, Sonia Goncalves, Ewan Harrison, David K. Jackson, Ian Johnston, Dominic Kwiatkowski, Cordelia Langford, John Sillitoe on behalf of the Wellcome Sanger Institute COVID-19 Surveillance Team |
| EPI_ISL_3367369 | Berkshire and Surrey Pathology Services Lighthouse Laboratory | Wellcome Sanger Institute for the COVID-19 Genomics UK (COG-UK) Consortium | Berkshire and Surrey Pathology Services Lighthouse Laboratory and Alex Alderton, Roberto Amato, Jeffrey Barrett, Sonia Goncalves, Ewan Harrison, David K. Jackson, Ian Johnston, Dominic Kwiatkowski, Cordelia Langford, John Sillitoe on behalf of the Wellcome Sanger Institute COVID-19 Surveillance Team |
| EPI_ISL_3367370, EPI_ISL_3367371 | Lighthouse Laboratory Plymouth | Wellcome Sanger Institute for the COVID-19 Genomics UK (COG-UK) Consortium | Lighthouse Laboratory Plymouth and Alex Alderton, Roberto Amato, Jeffrey Barrett, Sonia Goncalves, Ewan Harrison, David K. Jackson, Ian Johnston, Dominic Kwiatkowski, Cordelia Langford, John Sillitoe on behalf of the Wellcome Sanger Institute COVID-19 Surveillance Team |
| EPI_ISL_3367372 | Lighthouse Lab in Glasgow | Wellcome Sanger Institute for the COVID-19 Genomics UK (COG-UK) Consortium | Harper VanSteenhouse, Yumi Kasai, David Gray, Carol Clugston, Anna Dominiczak and Alex Alderton, Roberto Amato, Jeffrey Barrett, Sonia Goncalves, Ewan Harrison, David K. Jackson, Ian Johnston, Dominic Kwiatkowski, Cordelia Langford, John Sillitoe on behalf of the Wellcome Sanger Institute COVID-19 Surveillance Team |
| EPI_ISL_3367373, EPI_ISL_3367374, EPI_ISL_3367375 | Berkshire and Surrey Pathology Services Lighthouse Laboratory | Wellcome Sanger Institute for the COVID-19 Genomics UK (COG-UK) Consortium | Berkshire and Surrey Pathology Services Lighthouse Laboratory and Alex Alderton, Roberto Amato, Jeffrey Barrett, Sonia Goncalves, Ewan Harrison, David K. Jackson, Ian Johnston, Dominic Kwiatkowski, Cordelia Langford, John Sillitoe on behalf of the Wellcome Sanger Institute COVID-19 Surveillance Team |
| EPI_ISL_3367376 | Lighthouse Lab in Glasgow | Wellcome Sanger Institute for the COVID-19 Genomics UK (COG-UK) Consortium | Harper VanSteenhouse, Yumi Kasai, David Gray, Carol Clugston, Anna Dominiczak and Alex Alderton, Roberto Amato, Jeffrey Barrett, Sonia Goncalves, Ewan Harrison, David K. Jackson, Ian Johnston, Dominic Kwiatkowski, Cordelia Langford, John Sillitoe on behalf of the Wellcome Sanger Institute COVID-19 Surveillance Team |
| EPI_ISL_3367377, EPI_ISL_3367378 | Berkshire and Surrey Pathology Services Lighthouse Laboratory | Wellcome Sanger Institute for the COVID-19 Genomics UK (COG-UK) Consortium | Berkshire and Surrey Pathology Services Lighthouse Laboratory and Alex Alderton, Roberto Amato, Jeffrey Barrett, Sonia Goncalves, Ewan Harrison, David K. Jackson, Ian Johnston, Dominic Kwiatkowski, Cordelia Langford, John Sillitoe on behalf of the Wellcome Sanger Institute COVID-19 Surveillance Team |
| EPI_ISL_3367379 | Lighthouse Lab in Glasgow | Wellcome Sanger Institute for the COVID-19 Genomics UK (COG-UK) Consortium | Harper VanSteenhouse, Yumi Kasai, David Gray, Carol Clugston, Anna Dominiczak and Alex Alderton, Roberto Amato, Jeffrey Barrett, Sonia Goncalves, Ewan Harrison, David K. Jackson, Ian Johnston, Dominic Kwiatkowski, Cordelia Langford, John Sillitoe on behalf of the Wellcome Sanger Institute COVID-19 Surveillance Team |
| EPI_ISL_3367380 | Berkshire and Surrey Pathology Services Lighthouse Laboratory | Wellcome Sanger Institute for the COVID-19 Genomics UK (COG-UK) Consortium | Berkshire and Surrey Pathology Services Lighthouse Laboratory and Alex Alderton, Roberto Amato, Jeffrey Barrett, Sonia Goncalves, Ewan Harrison, David K. Jackson, Ian Johnston, Dominic Kwiatkowski, Cordelia Langford, John Sillitoe on behalf of the Wellcome Sanger Institute COVID-19 Surveillance Team |
| EPI_ISL_3367381 | Lighthouse Lab in Glasgow | Wellcome Sanger Institute for the COVID-19 Genomics UK (COG-UK) Consortium | Harper VanSteenhouse, Yumi Kasai, David Gray, Carol Clugston, Anna Dominiczak and Alex Alderton, Roberto Amato, Jeffrey Barrett, Sonia Goncalves, Ewan Harrison, David K. Jackson, Ian Johnston, Dominic Kwiatkowski, Cordelia Langford, John Sillitoe on behalf of the Wellcome Sanger Institute COVID-19 Surveillance Team |
| EPI_ISL_3367382 | Berkshire and Surrey Pathology Services Lighthouse Laboratory | Wellcome Sanger Institute for the COVID-19 Genomics UK (COG-UK) Consortium | Berkshire and Surrey Pathology Services Lighthouse Laboratory and Alex Alderton, Roberto Amato, Jeffrey Barrett, Sonia Goncalves, Ewan Harrison, David K. Jackson, Ian Johnston, Dominic Kwiatkowski, Cordelia Langford, John Sillitoe on behalf of the Wellcome Sanger Institute COVID-19 Surveillance Team |
| EPI_ISL_3367383 | Lighthouse Lab in Glasgow | Wellcome Sanger Institute for the COVID-19 Genomics UK (COG-UK) Consortium | Harper VanSteenhouse, Yumi Kasai, David Gray, Carol Clugston, Anna Dominiczak and Alex Alderton, Roberto Amato, Jeffrey Barrett, Sonia Goncalves, Ewan Harrison, David K. Jackson, Ian Johnston, Dominic Kwiatkowski, Cordelia Langford, John Sillitoe on behalf of the Wellcome Sanger Institute COVID-19 Surveillance Team |
| EPI_ISL_3367384 | Berkshire and Surrey Pathology Services Lighthouse Laboratory | Wellcome Sanger Institute for the COVID-19 Genomics UK (COG-UK) Consortium | Berkshire and Surrey Pathology Services Lighthouse Laboratory and Alex Alderton, Roberto Amato, Jeffrey Barrett, Sonia Goncalves, Ewan Harrison, David K. Jackson, Ian Johnston, Dominic Kwiatkowski, Cordelia Langford, John Sillitoe on behalf of the Wellcome Sanger Institute COVID-19 Surveillance Team |
| EPI_ISL_3367385 | Lighthouse Lab in Glasgow | Wellcome Sanger Institute for the COVID-19 Genomics UK (COG-UK) Consortium | Harper VanSteenhouse, Yumi Kasai, David Gray, Carol Clugston, Anna Dominiczak and Alex Alderton, Roberto Amato, Jeffrey Barrett, Sonia Goncalves, Ewan Harrison, David K. Jackson, Ian Johnston, Dominic Kwiatkowski, Cordelia Langford, John Sillitoe on behalf of the Wellcome Sanger Institute COVID-19 Surveillance Team |
| EPI_ISL_3367386 | Berkshire and Surrey Pathology Services Lighthouse Laboratory | Wellcome Sanger Institute for the COVID-19 Genomics UK (COG-UK) Consortium | Berkshire and Surrey Pathology Services Lighthouse Laboratory and Alex Alderton, Roberto Amato, Jeffrey Barrett, Sonia Goncalves, Ewan Harrison, David K. Jackson, Ian Johnston, Dominic Kwiatkowski, Cordelia Langford, John Sillitoe on behalf of the Wellcome Sanger Institute COVID-19 Surveillance Team |
| EPI_ISL_3367387 | Lighthouse Laboratory Plymouth | Wellcome Sanger Institute for the COVID-19 Genomics UK (COG-UK) Consortium | Lighthouse Laboratory Plymouth and Alex Alderton, Roberto Amato, Jeffrey Barrett, Sonia Goncalves, Ewan Harrison, David K. Jackson, Ian Johnston, Dominic Kwiatkowski, Cordelia Langford, John Sillitoe on behalf of the Wellcome Sanger Institute COVID-19 Surveillance Team |
| EPI_ISL_3367388 | Berkshire and Surrey Pathology Services Lighthouse Laboratory | Wellcome Sanger Institute for the COVID-19 Genomics UK (COG-UK) Consortium | Berkshire and Surrey Pathology Services Lighthouse Laboratory and Alex Alderton, Roberto Amato, Jeffrey Barrett, Sonia Goncalves, Ewan Harrison, David K. Jackson, Ian Johnston, Dominic Kwiatkowski, Cordelia Langford, John Sillitoe on behalf of the Wellcome Sanger Institute COVID-19 Surveillance Team |
| EPI_ISL_3367389 | Lighthouse Lab in Glasgow | Wellcome Sanger Institute for the COVID-19 Genomics UK (COG-UK) Consortium | Harper VanSteenhouse, Yumi Kasai, David Gray, Carol Clugston, Anna Dominiczak and Alex Alderton, Roberto Amato, Jeffrey Barrett, Sonia Goncalves, Ewan Harrison, David K. Jackson, Ian Johnston, Dominic Kwiatkowski, Cordelia Langford, John Sillitoe on behalf of the Wellcome Sanger Institute COVID-19 Surveillance Team |
| EPI_ISL_3367390 | Berkshire and Surrey Pathology Services Lighthouse Laboratory | Wellcome Sanger Institute for the COVID-19 Genomics UK (COG-UK) Consortium | Berkshire and Surrey Pathology Services Lighthouse Laboratory and Alex Alderton, Roberto Amato, Jeffrey Barrett, Sonia Goncalves, Ewan Harrison, David K. Jackson, Ian Johnston, Dominic Kwiatkowski, Cordelia Langford, John Sillitoe on behalf of the Wellcome Sanger Institute COVID-19 Surveillance Team |
| EPI_ISL_3367391 | Lighthouse Laboratory Plymouth | Wellcome Sanger Institute for the COVID-19 Genomics UK (COG-UK) Consortium | Lighthouse Laboratory Plymouth and Alex Alderton, Roberto Amato, Jeffrey Barrett, Sonia Goncalves, Ewan Harrison, David K. Jackson, Ian Johnston, Dominic Kwiatkowski, Cordelia Langford, John Sillitoe on behalf of the Wellcome Sanger Institute COVID-19 Surveillance Team |
| EPI_ISL_3367392 | Lighthouse Lab in Glasgow | Wellcome Sanger Institute for the COVID-19 Genomics UK (COG-UK) Consortium | Harper VanSteenhouse, Yumi Kasai, David Gray, Carol Clugston, Anna Dominiczak and Alex Alderton, Roberto Amato, Jeffrey Barrett, Sonia Goncalves, Ewan Harrison, David K. Jackson, Ian Johnston, Dominic Kwiatkowski, Cordelia Langford, John Sillitoe on behalf of the Wellcome Sanger Institute COVID-19 Surveillance Team |
| EPI_ISL_3367393 | Lighthouse Laboratory Plymouth | Wellcome Sanger Institute for the COVID-19 Genomics UK (COG-UK) Consortium | Lighthouse Laboratory Plymouth and Alex Alderton, Roberto Amato, Jeffrey Barrett, Sonia Goncalves, Ewan Harrison, David K. Jackson, Ian Johnston, Dominic Kwiatkowski, Cordelia Langford, John Sillitoe on behalf of the Wellcome Sanger Institute COVID-19 Surveillance Team |
| EPI_ISL_3367394, EPI_ISL_3367395 | Berkshire and Surrey Pathology Services Lighthouse Laboratory | Wellcome Sanger Institute for the COVID-19 Genomics UK (COG-UK) Consortium | Berkshire and Surrey Pathology Services Lighthouse Laboratory and Alex Alderton, Roberto Amato, Jeffrey Barrett, Sonia Goncalves, Ewan Harrison, David K. Jackson, Ian Johnston, Dominic Kwiatkowski, Cordelia Langford, John Sillitoe on behalf of the Wellcome Sanger Institute COVID-19 Surveillance Team |
| EPI_ISL_3367396 | Lighthouse Lab in Glasgow | Wellcome Sanger Institute for the COVID-19 Genomics UK (COG-UK) Consortium | Harper VanSteenhouse, Yumi Kasai, David Gray, Carol Clugston, Anna Dominiczak and Alex Alderton, Roberto Amato, Jeffrey Barrett, Sonia Goncalves, Ewan Harrison, David K. Jackson, Ian Johnston, Dominic Kwiatkowski, Cordelia Langford, John Sillitoe on behalf of the Wellcome Sanger Institute COVID-19 Surveillance Team |
| EPI_ISL_3367397, EPI_ISL_3367398 | Berkshire and Surrey Pathology Services Lighthouse Laboratory | Wellcome Sanger Institute for the COVID-19 Genomics UK (COG-UK) Consortium | Berkshire and Surrey Pathology Services Lighthouse Laboratory and Alex Alderton, Roberto Amato, Jeffrey Barrett, Sonia Goncalves, Ewan Harrison, David K. Jackson, Ian Johnston, Dominic Kwiatkowski, Cordelia Langford, John Sillitoe on behalf of the Wellcome Sanger Institute COVID-19 Surveillance Team |
| EPI_ISL_3367399 | Lighthouse Lab in Glasgow | Wellcome Sanger Institute for the COVID-19 Genomics UK (COG-UK) Consortium | Harper VanSteenhouse, Yumi Kasai, David Gray, Carol Clugston, Anna Dominiczak and Alex Alderton, Roberto Amato, Jeffrey Barrett, Sonia Goncalves, Ewan Harrison, David K. Jackson, Ian Johnston, Dominic Kwiatkowski, Cordelia Langford, John Sillitoe on behalf of the Wellcome Sanger Institute COVID-19 Surveillance Team |
| EPI_ISL_3367400 | Berkshire and Surrey Pathology Services Lighthouse Laboratory | Wellcome Sanger Institute for the COVID-19 Genomics UK (COG-UK) Consortium | Berkshire and Surrey Pathology Services Lighthouse Laboratory and Alex Alderton, Roberto Amato, Jeffrey Barrett, Sonia Goncalves, Ewan Harrison, David K. Jackson, Ian Johnston, Dominic Kwiatkowski, Cordelia Langford, John Sillitoe on behalf of the Wellcome Sanger Institute COVID-19 Surveillance Team |
| EPI_ISL_3367401 | Lighthouse Laboratory Plymouth | Wellcome Sanger Institute for the COVID-19 Genomics UK (COG-UK) Consortium | Lighthouse Laboratory Plymouth and Alex Alderton, Roberto Amato, Jeffrey Barrett, Sonia Goncalves, Ewan Harrison, David K. Jackson, Ian Johnston, Dominic Kwiatkowski, Cordelia Langford, John Sillitoe on behalf of the Wellcome Sanger Institute COVID-19 Surveillance Team |
| EPI_ISL_3367402 | Berkshire and Surrey Pathology Services Lighthouse Laboratory | Wellcome Sanger Institute for the COVID-19 Genomics UK (COG-UK) Consortium | Berkshire and Surrey Pathology Services Lighthouse Laboratory and Alex Alderton, Roberto Amato, Jeffrey Barrett, Sonia Goncalves, Ewan Harrison, David K. Jackson, Ian Johnston, Dominic Kwiatkowski, Cordelia Langford, John Sillitoe on behalf of the Wellcome Sanger Institute COVID-19 Surveillance Team |
| EPI_ISL_3367403 | Lighthouse Laboratory Plymouth | Wellcome Sanger Institute for the COVID-19 Genomics UK (COG-UK) Consortium | Lighthouse Laboratory Plymouth and Alex Alderton, Roberto Amato, Jeffrey Barrett, Sonia Goncalves, Ewan Harrison, David K. Jackson, Ian Johnston, Dominic Kwiatkowski, Cordelia Langford, John Sillitoe on behalf of the Wellcome Sanger Institute COVID-19 Surveillance Team |
| EPI_ISL_3367404 | Berkshire and Surrey Pathology Services Lighthouse Laboratory | Wellcome Sanger Institute for the COVID-19 Genomics UK (COG-UK) Consortium | Berkshire and Surrey Pathology Services Lighthouse Laboratory and Alex Alderton, Roberto Amato, Jeffrey Barrett, Sonia Goncalves, Ewan Harrison, David K. Jackson, Ian Johnston, Dominic Kwiatkowski, Cordelia Langford, John Sillitoe on behalf of the Wellcome Sanger Institute COVID-19 Surveillance Team |
| EPI_ISL_3367405 | Lighthouse Lab in Glasgow | Wellcome Sanger Institute for the COVID-19 Genomics UK (COG-UK) Consortium | Harper VanSteenhouse, Yumi Kasai, David Gray, Carol Clugston, Anna Dominiczak and Alex Alderton, Roberto Amato, Jeffrey Barrett, Sonia Goncalves, Ewan Harrison, David K. Jackson, Ian Johnston, Dominic Kwiatkowski, Cordelia Langford, John Sillitoe on behalf of the Wellcome Sanger Institute COVID-19 Surveillance Team |
| EPI_ISL_3367406 | Lighthouse Laboratory Plymouth | Wellcome Sanger Institute for the COVID-19 Genomics UK (COG-UK) Consortium | Lighthouse Laboratory Plymouth and Alex Alderton, Roberto Amato, Jeffrey Barrett, Sonia Goncalves, Ewan Harrison, David K. Jackson, Ian Johnston, Dominic Kwiatkowski, Cordelia Langford, John Sillitoe on behalf of the Wellcome Sanger Institute COVID-19 Surveillance Team |
| EPI_ISL_3367407, EPI_ISL_3367408 | Lighthouse Lab in Glasgow | Wellcome Sanger Institute for the COVID-19 Genomics UK (COG-UK) Consortium | Harper VanSteenhouse, Yumi Kasai, David Gray, Carol Clugston, Anna Dominiczak and Alex Alderton, Roberto Amato, Jeffrey Barrett, Sonia Goncalves, Ewan Harrison, David K. Jackson, Ian Johnston, Dominic Kwiatkowski, Cordelia Langford, John Sillitoe on behalf of the Wellcome Sanger Institute COVID-19 Surveillance Team |
| EPI_ISL_3367409 | Berkshire and Surrey Pathology Services Lighthouse Laboratory | Wellcome Sanger Institute for the COVID-19 Genomics UK (COG-UK) Consortium | Berkshire and Surrey Pathology Services Lighthouse Laboratory and Alex Alderton, Roberto Amato, Jeffrey Barrett, Sonia Goncalves, Ewan Harrison, David K. Jackson, Ian Johnston, Dominic Kwiatkowski, Cordelia Langford, John Sillitoe on behalf of the Wellcome Sanger Institute COVID-19 Surveillance Team |
| EPI_ISL_3367410, EPI_ISL_3367411 | Lighthouse Laboratory Plymouth | Wellcome Sanger Institute for the COVID-19 Genomics UK (COG-UK) Consortium | Lighthouse Laboratory Plymouth and Alex Alderton, Roberto Amato, Jeffrey Barrett, Sonia Goncalves, Ewan Harrison, David K. Jackson, Ian Johnston, Dominic Kwiatkowski, Cordelia Langford, John Sillitoe on behalf of the Wellcome Sanger Institute COVID-19 Surveillance Team |
| EPI_ISL_3367412 | Lighthouse Lab in Glasgow | Wellcome Sanger Institute for the COVID-19 Genomics UK (COG-UK) Consortium | Harper VanSteenhouse, Yumi Kasai, David Gray, Carol Clugston, Anna Dominiczak and Alex Alderton, Roberto Amato, Jeffrey Barrett, Sonia Goncalves, Ewan Harrison, David K. Jackson, Ian Johnston, Dominic Kwiatkowski, Cordelia Langford, John Sillitoe on behalf of the Wellcome Sanger Institute COVID-19 Surveillance Team |
| EPI_ISL_3367413, EPI_ISL_3367414, EPI_ISL_3367415 | Berkshire and Surrey Pathology Services Lighthouse Laboratory | Wellcome Sanger Institute for the COVID-19 Genomics UK (COG-UK) Consortium | Berkshire and Surrey Pathology Services Lighthouse Laboratory and Alex Alderton, Roberto Amato, Jeffrey Barrett, Sonia Goncalves, Ewan Harrison, David K. Jackson, Ian Johnston, Dominic Kwiatkowski, Cordelia Langford, John Sillitoe on behalf of the Wellcome Sanger Institute COVID-19 Surveillance Team |
| EPI_ISL_3367416 | Lighthouse Laboratory Plymouth | Wellcome Sanger Institute for the COVID-19 Genomics UK (COG-UK) Consortium | Lighthouse Laboratory Plymouth and Alex Alderton, Roberto Amato, Jeffrey Barrett, Sonia Goncalves, Ewan Harrison, David K. Jackson, Ian Johnston, Dominic Kwiatkowski, Cordelia Langford, John Sillitoe on behalf of the Wellcome Sanger Institute COVID-19 Surveillance Team |
| EPI_ISL_3367417 | Lighthouse Lab in Glasgow | Wellcome Sanger Institute for the COVID-19 Genomics UK (COG-UK) Consortium | Harper VanSteenhouse, Yumi Kasai, David Gray, Carol Clugston, Anna Dominiczak and Alex Alderton, Roberto Amato, Jeffrey Barrett, Sonia Goncalves, Ewan Harrison, David K. Jackson, Ian Johnston, Dominic Kwiatkowski, Cordelia Langford, John Sillitoe on behalf of the Wellcome Sanger Institute COVID-19 Surveillance Team |
| EPI_ISL_3367418 | Berkshire and Surrey Pathology Services Lighthouse Laboratory | Wellcome Sanger Institute for the COVID-19 Genomics UK (COG-UK) Consortium | Berkshire and Surrey Pathology Services Lighthouse Laboratory and Alex Alderton, Roberto Amato, Jeffrey Barrett, Sonia Goncalves, Ewan Harrison, David K. Jackson, Ian Johnston, Dominic Kwiatkowski, Cordelia Langford, John Sillitoe on behalf of the Wellcome Sanger Institute COVID-19 Surveillance Team |
| EPI_ISL_3367419 | Lighthouse Laboratory Plymouth | Wellcome Sanger Institute for the COVID-19 Genomics UK (COG-UK) Consortium | Lighthouse Laboratory Plymouth and Alex Alderton, Roberto Amato, Jeffrey Barrett, Sonia Goncalves, Ewan Harrison, David K. Jackson, Ian Johnston, Dominic Kwiatkowski, Cordelia Langford, John Sillitoe on behalf of the Wellcome Sanger Institute COVID-19 Surveillance Team |
| EPI_ISL_3367420 | Lighthouse Lab in Glasgow | Wellcome Sanger Institute for the COVID-19 Genomics UK (COG-UK) Consortium | Harper VanSteenhouse, Yumi Kasai, David Gray, Carol Clugston, Anna Dominiczak and Alex Alderton, Roberto Amato, Jeffrey Barrett, Sonia Goncalves, Ewan Harrison, David K. Jackson, Ian Johnston, Dominic Kwiatkowski, Cordelia Langford, John Sillitoe on behalf of the Wellcome Sanger Institute COVID-19 Surveillance Team |
| EPI_ISL_3367421 | Lighthouse Laboratory Plymouth | Wellcome Sanger Institute for the COVID-19 Genomics UK (COG-UK) Consortium | Lighthouse Laboratory Plymouth and Alex Alderton, Roberto Amato, Jeffrey Barrett, Sonia Goncalves, Ewan Harrison, David K. Jackson, Ian Johnston, Dominic Kwiatkowski, Cordelia Langford, John Sillitoe on behalf of the Wellcome Sanger Institute COVID-19 Surveillance Team |

| | | | |
|---|---|---|---|
| EPI_ISL_3367422 | Berkshire and Surrey Pathology Services Lighthouse Laboratory | Wellcome Sanger Institute for the COVID-19 Genomics UK (COG-UK) Consortium | Berkshire and Surrey Pathology Services Lighthouse Laboratory and Alex Alderton, Roberto Amato, Jeffrey Barrett, Sonia Goncalves, Ewan Harrison, David K. Jackson, Ian Johnston, Dominic Kwiatkowski, Cordelia Langford, John Sillitoe on behalf of the Wellcome Sanger Institute COVID-19 Surveillance Team |
| EPI_ISL_3367423 | Lighthouse Laboratory Plymouth | Wellcome Sanger Institute for the COVID-19 Genomics UK (COG-UK) Consortium | Lighthouse Laboratory Plymouth and Alex Alderton, Roberto Amato, Jeffrey Barrett, Sonia Goncalves, Ewan Harrison, David K. Jackson, Ian Johnston, Dominic Kwiatkowski, Cordelia Langford, John Sillitoe on behalf of the Wellcome Sanger Institute COVID-19 Surveillance Team |
| EPI_ISL_3367424 | Lighthouse Lab in Glasgow | Wellcome Sanger Institute for the COVID-19 Genomics UK (COG-UK) Consortium | Harper VanSteenhouse, Yumi Kasai, David Gray, Carol Clugston, Anna Dominiczak and Alex Alderton, Roberto Amato, Jeffrey Barrett, Sonia Goncalves, Ewan Harrison, David K. Jackson, Ian Johnston, Dominic Kwiatkowski, Cordelia Langford, John Sillitoe on behalf of the Wellcome Sanger Institute COVID-19 Surveillance Team |
| EPI_ISL_3367425, EPI_ISL_3367426, EPI_ISL_3367427, EPI_ISL_3367428 | Berkshire and Surrey Pathology Services Lighthouse Laboratory | Wellcome Sanger Institute for the COVID-19 Genomics UK (COG-UK) Consortium | Berkshire and Surrey Pathology Services Lighthouse Laboratory and Alex Alderton, Roberto Amato, Jeffrey Barrett, Sonia Goncalves, Ewan Harrison, David K. Jackson, Ian Johnston, Dominic Kwiatkowski, Cordelia Langford, John Sillitoe on behalf of the Wellcome Sanger Institute COVID-19 Surveillance Team |
| EPI_ISL_3367429 | Lighthouse Laboratory Plymouth | Wellcome Sanger Institute for the COVID-19 Genomics UK (COG-UK) Consortium | Lighthouse Laboratory Plymouth and Alex Alderton, Roberto Amato, Jeffrey Barrett, Sonia Goncalves, Ewan Harrison, David K. Jackson, Ian Johnston, Dominic Kwiatkowski, Cordelia Langford, John Sillitoe on behalf of the Wellcome Sanger Institute COVID-19 Surveillance Team |
| EPI_ISL_3367430 | Lighthouse Lab in Glasgow | Wellcome Sanger Institute for the COVID-19 Genomics UK (COG-UK) Consortium | Harper VanSteenhouse, Yumi Kasai, David Gray, Carol Clugston, Anna Dominiczak and Alex Alderton, Roberto Amato, Jeffrey Barrett, Sonia Goncalves, Ewan Harrison, David K. Jackson, Ian Johnston, Dominic Kwiatkowski, Cordelia Langford, John Sillitoe on behalf of the Wellcome Sanger Institute COVID-19 Surveillance Team |
| EPI_ISL_3367431 | Lighthouse Laboratory Plymouth | Wellcome Sanger Institute for the COVID-19 Genomics UK (COG-UK) Consortium | Lighthouse Laboratory Plymouth and Alex Alderton, Roberto Amato, Jeffrey Barrett, Sonia Goncalves, Ewan Harrison, David K. Jackson, Ian Johnston, Dominic Kwiatkowski, Cordelia Langford, John Sillitoe on behalf of the Wellcome Sanger Institute COVID-19 Surveillance Team |
| EPI_ISL_3367432 | Berkshire and Surrey Pathology Services Lighthouse Laboratory | Wellcome Sanger Institute for the COVID-19 Genomics UK (COG-UK) Consortium | Berkshire and Surrey Pathology Services Lighthouse Laboratory and Alex Alderton, Roberto Amato, Jeffrey Barrett, Sonia Goncalves, Ewan Harrison, David K. Jackson, Ian Johnston, Dominic Kwiatkowski, Cordelia Langford, John Sillitoe on behalf of the Wellcome Sanger Institute COVID-19 Surveillance Team |
| EPI_ISL_3367433 | Lighthouse Lab in Glasgow | Wellcome Sanger Institute for the COVID-19 Genomics UK (COG-UK) Consortium | Harper VanSteenhouse, Yumi Kasai, David Gray, Carol Clugston, Anna Dominiczak and Alex Alderton, Roberto Amato, Jeffrey Barrett, Sonia Goncalves, Ewan Harrison, David K. Jackson, Ian Johnston, Dominic Kwiatkowski, Cordelia Langford, John Sillitoe on behalf of the Wellcome Sanger Institute COVID-19 Surveillance Team |
| EPI_ISL_3367434 | Lighthouse Laboratory Plymouth | Wellcome Sanger Institute for the COVID-19 Genomics UK (COG-UK) Consortium | Lighthouse Laboratory Plymouth and Alex Alderton, Roberto Amato, Jeffrey Barrett, Sonia Goncalves, Ewan Harrison, David K. Jackson, Ian Johnston, Dominic Kwiatkowski, Cordelia Langford, John Sillitoe on behalf of the Wellcome Sanger Institute COVID-19 Surveillance Team |
| EPI_ISL_3367435 | Berkshire and Surrey Pathology Services Lighthouse Laboratory | Wellcome Sanger Institute for the COVID-19 Genomics UK (COG-UK) Consortium | Berkshire and Surrey Pathology Services Lighthouse Laboratory and Alex Alderton, Roberto Amato, Jeffrey Barrett, Sonia Goncalves, Ewan Harrison, David K. Jackson, Ian Johnston, Dominic Kwiatkowski, Cordelia Langford, John Sillitoe on behalf of the Wellcome Sanger Institute COVID-19 Surveillance Team |
| EPI_ISL_3367436 | Lighthouse Laboratory Plymouth | Wellcome Sanger Institute for the COVID-19 Genomics UK (COG-UK) Consortium | Lighthouse Laboratory Plymouth and Alex Alderton, Roberto Amato, Jeffrey Barrett, Sonia Goncalves, Ewan Harrison, David K. Jackson, Ian Johnston, Dominic Kwiatkowski, Cordelia Langford, John Sillitoe on behalf of the Wellcome Sanger Institute COVID-19 Surveillance Team |
| EPI_ISL_3367437 | Berkshire and Surrey Pathology Services Lighthouse Laboratory | Wellcome Sanger Institute for the COVID-19 Genomics UK (COG-UK) Consortium | Berkshire and Surrey Pathology Services Lighthouse Laboratory and Alex Alderton, Roberto Amato, Jeffrey Barrett, Sonia Goncalves, Ewan Harrison, David K. Jackson, Ian Johnston, Dominic Kwiatkowski, Cordelia Langford, John Sillitoe on behalf of the Wellcome Sanger Institute COVID-19 Surveillance Team |
| EPI_ISL_3367438 | Lighthouse Lab in Glasgow | Wellcome Sanger Institute for the COVID-19 Genomics UK (COG-UK) Consortium | Harper VanSteenhouse, Yumi Kasai, David Gray, Carol Clugston, Anna Dominiczak and Alex Alderton, Roberto Amato, Jeffrey Barrett, Sonia Goncalves, Ewan Harrison, David K. Jackson, Ian Johnston, Dominic Kwiatkowski, Cordelia Langford, John Sillitoe on behalf of the Wellcome Sanger Institute COVID-19 Surveillance Team |
| EPI_ISL_3367439 | Berkshire and Surrey Pathology Services Lighthouse Laboratory | Wellcome Sanger Institute for the COVID-19 Genomics UK (COG-UK) Consortium | Berkshire and Surrey Pathology Services Lighthouse Laboratory and Alex Alderton, Roberto Amato, Jeffrey Barrett, Sonia Goncalves, Ewan Harrison, David K. Jackson, Ian Johnston, Dominic Kwiatkowski, Cordelia Langford, John Sillitoe on behalf of the Wellcome Sanger Institute COVID-19 Surveillance Team |
| EPI_ISL_3367440, EPI_ISL_3367441 | Lighthouse Laboratory Plymouth | Wellcome Sanger Institute for the COVID-19 Genomics UK (COG-UK) Consortium | Lighthouse Laboratory Plymouth and Alex Alderton, Roberto Amato, Jeffrey Barrett, Sonia Goncalves, Ewan Harrison, David K. Jackson, Ian Johnston, Dominic Kwiatkowski, Cordelia Langford, John Sillitoe on behalf of the Wellcome Sanger Institute COVID-19 Surveillance Team |
| EPI_ISL_3367442 | Lighthouse Lab in Glasgow | Wellcome Sanger Institute for the COVID-19 Genomics UK (COG-UK) Consortium | Harper VanSteenhouse, Yumi Kasai, David Gray, Carol Clugston, Anna Dominiczak and Alex Alderton, Roberto Amato, Jeffrey Barrett, Sonia Goncalves, Ewan Harrison, David K. Jackson, Ian Johnston, Dominic Kwiatkowski, Cordelia Langford, John Sillitoe on behalf of the Wellcome Sanger Institute COVID-19 Surveillance Team |
| EPI_ISL_3367443, EPI_ISL_3367444, EPI_ISL_3367445, EPI_ISL_3367446 | Berkshire and Surrey Pathology Services Lighthouse Laboratory | Wellcome Sanger Institute for the COVID-19 Genomics UK (COG-UK) Consortium | Berkshire and Surrey Pathology Services Lighthouse Laboratory and Alex Alderton, Roberto Amato, Jeffrey Barrett, Sonia Goncalves, Ewan Harrison, David K. Jackson, Ian Johnston, Dominic Kwiatkowski, Cordelia Langford, John Sillitoe on behalf of the Wellcome Sanger Institute COVID-19 Surveillance Team |
| EPI_ISL_3367447 | Lighthouse Laboratory Plymouth | Wellcome Sanger Institute for the COVID-19 Genomics UK (COG-UK) Consortium | Lighthouse Laboratory Plymouth and Alex Alderton, Roberto Amato, Jeffrey Barrett, Sonia Goncalves, Ewan Harrison, David K. Jackson, Ian Johnston, Dominic Kwiatkowski, Cordelia Langford, John Sillitoe on behalf of the Wellcome Sanger Institute COVID-19 Surveillance Team |
| EPI_ISL_3367448 | Berkshire and Surrey Pathology Services Lighthouse Laboratory | Wellcome Sanger Institute for the COVID-19 Genomics UK (COG-UK) Consortium | Berkshire and Surrey Pathology Services Lighthouse Laboratory and Alex Alderton, Roberto Amato, Jeffrey Barrett, Sonia Goncalves, Ewan Harrison, David K. Jackson, Ian Johnston, Dominic Kwiatkowski, Cordelia Langford, John Sillitoe on behalf of the Wellcome Sanger Institute COVID-19 Surveillance Team |
| EPI_ISL_3367449 | Lighthouse Laboratory Plymouth | Wellcome Sanger Institute for the COVID-19 Genomics UK (COG-UK) Consortium | Lighthouse Laboratory Plymouth and Alex Alderton, Roberto Amato, Jeffrey Barrett, Sonia Goncalves, Ewan Harrison, David K. Jackson, Ian Johnston, Dominic Kwiatkowski, Cordelia Langford, John Sillitoe on behalf of the Wellcome Sanger Institute COVID-19 Surveillance Team |
| EPI_ISL_3367450, EPI_ISL_3367451 | Berkshire and Surrey Pathology Services Lighthouse Laboratory | Wellcome Sanger Institute for the COVID-19 Genomics UK (COG-UK) Consortium | Berkshire and Surrey Pathology Services Lighthouse Laboratory and Alex Alderton, Roberto Amato, Jeffrey Barrett, Sonia Goncalves, Ewan Harrison, David K. Jackson, Ian Johnston, Dominic Kwiatkowski, Cordelia Langford, John Sillitoe on behalf of the Wellcome Sanger Institute COVID-19 Surveillance Team |
| EPI_ISL_3367452 | Lighthouse Lab in Glasgow | Wellcome Sanger Institute for the COVID-19 Genomics UK (COG-UK) Consortium | Harper VanSteenhouse, Yumi Kasai, David Gray, Carol Clugston, Anna Dominiczak and Alex Alderton, Roberto Amato, Jeffrey Barrett, Sonia Goncalves, Ewan Harrison, David K. Jackson, Ian Johnston, Dominic Kwiatkowski, Cordelia Langford, John Sillitoe on behalf of the Wellcome Sanger Institute COVID-19 Surveillance Team |
| EPI_ISL_3367453 | Lighthouse Laboratory Plymouth | Wellcome Sanger Institute for the COVID-19 Genomics UK (COG-UK) Consortium | Lighthouse Laboratory Plymouth and Alex Alderton, Roberto Amato, Jeffrey Barrett, Sonia Goncalves, Ewan Harrison, David K. Jackson, Ian Johnston, Dominic Kwiatkowski, Cordelia Langford, John Sillitoe on behalf of the Wellcome Sanger Institute COVID-19 Surveillance Team |
| EPI_ISL_3367454 | Berkshire and Surrey Pathology Services Lighthouse Laboratory | Wellcome Sanger Institute for the COVID-19 Genomics UK (COG-UK) Consortium | Berkshire and Surrey Pathology Services Lighthouse Laboratory and Alex Alderton, Roberto Amato, Jeffrey Barrett, Sonia Goncalves, Ewan Harrison, David K. Jackson, Ian Johnston, Dominic Kwiatkowski, Cordelia Langford, John Sillitoe on behalf of the Wellcome Sanger Institute COVID-19 Surveillance Team |
| EPI_ISL_3367455 | Lighthouse Laboratory Plymouth | Wellcome Sanger Institute for the COVID-19 Genomics UK (COG-UK) Consortium | Lighthouse Laboratory Plymouth and Alex Alderton, Roberto Amato, Jeffrey Barrett, Sonia Goncalves, Ewan Harrison, David K. Jackson, Ian Johnston, Dominic Kwiatkowski, Cordelia Langford, John Sillitoe on behalf of the Wellcome Sanger Institute COVID-19 Surveillance Team |
| EPI_ISL_3367456 | Berkshire and Surrey Pathology Services Lighthouse Laboratory | Wellcome Sanger Institute for the COVID-19 Genomics UK (COG-UK) Consortium | Berkshire and Surrey Pathology Services Lighthouse Laboratory and Alex Alderton, Roberto Amato, Jeffrey Barrett, Sonia Goncalves, Ewan Harrison, David K. Jackson, Ian Johnston, Dominic Kwiatkowski, Cordelia Langford, John Sillitoe on behalf of the Wellcome Sanger Institute COVID-19 Surveillance Team |
| EPI_ISL_3367457 | Lighthouse Laboratory Plymouth | Wellcome Sanger Institute for the COVID-19 Genomics UK (COG-UK) Consortium | Lighthouse Laboratory Plymouth and Alex Alderton, Roberto Amato, Jeffrey Barrett, Sonia Goncalves, Ewan Harrison, David K. Jackson, Ian Johnston, Dominic Kwiatkowski, Cordelia Langford, John Sillitoe on behalf of the Wellcome Sanger Institute COVID-19 Surveillance Team |
| EPI_ISL_3367458, EPI_ISL_3367459, EPI_ISL_3367460 | Berkshire and Surrey Pathology Services Lighthouse Laboratory | Wellcome Sanger Institute for the COVID-19 Genomics UK (COG-UK) Consortium | Berkshire and Surrey Pathology Services Lighthouse Laboratory and Alex Alderton, Roberto Amato, Jeffrey Barrett, Sonia Goncalves, Ewan Harrison, David K. Jackson, Ian Johnston, Dominic Kwiatkowski, Cordelia Langford, John Sillitoe on behalf of the Wellcome Sanger Institute COVID-19 Surveillance Team |
| EPI_ISL_3367461 | Lighthouse Lab in Glasgow | Wellcome Sanger Institute for the COVID-19 Genomics UK (COG-UK) Consortium | Harper VanSteenhouse, Yumi Kasai, David Gray, Carol Clugston, Anna Dominiczak and Alex Alderton, Roberto Amato, Jeffrey Barrett, Sonia Goncalves, Ewan Harrison, David K. Jackson, Ian Johnston, Dominic Kwiatkowski, Cordelia Langford, John Sillitoe on behalf of the Wellcome Sanger Institute COVID-19 Surveillance Team |
| EPI_ISL_3367462, EPI_ISL_3367463, EPI_ISL_3367464, EPI_ISL_3367465 | Berkshire and Surrey Pathology Services Lighthouse Laboratory | Wellcome Sanger Institute for the COVID-19 Genomics UK (COG-UK) Consortium | Berkshire and Surrey Pathology Services Lighthouse Laboratory and Alex Alderton, Roberto Amato, Jeffrey Barrett, Sonia Goncalves, Ewan Harrison, David K. Jackson, Ian Johnston, Dominic Kwiatkowski, Cordelia Langford, John Sillitoe on behalf of the Wellcome Sanger Institute COVID-19 Surveillance Team |
| EPI_ISL_3367466 | Lighthouse Laboratory Plymouth | Wellcome Sanger Institute for the COVID-19 Genomics UK (COG-UK) Consortium | Lighthouse Laboratory Plymouth and Alex Alderton, Roberto Amato, Jeffrey Barrett, Sonia Goncalves, Ewan Harrison, David K. Jackson, Ian Johnston, Dominic Kwiatkowski, Cordelia Langford, John Sillitoe on behalf of the Wellcome Sanger Institute COVID-19 Surveillance Team |
| EPI_ISL_3367467 | Berkshire and Surrey Pathology Services Lighthouse Laboratory | Wellcome Sanger Institute for the COVID-19 Genomics UK (COG-UK) Consortium | Berkshire and Surrey Pathology Services Lighthouse Laboratory and Alex Alderton, Roberto Amato, Jeffrey Barrett, Sonia Goncalves, Ewan Harrison, David K. Jackson, Ian Johnston, Dominic Kwiatkowski, Cordelia Langford, John Sillitoe on behalf of the Wellcome Sanger Institute COVID-19 Surveillance Team |
| EPI_ISL_3367468 | Lighthouse Lab in Glasgow | Wellcome Sanger Institute for the COVID-19 Genomics UK (COG-UK) Consortium | Harper VanSteenhouse, Yumi Kasai, David Gray, Carol Clugston, Anna Dominiczak and Alex Alderton, Roberto Amato, Jeffrey Barrett, Sonia Goncalves, Ewan Harrison, David K. Jackson, Ian Johnston, Dominic Kwiatkowski, Cordelia Langford, John Sillitoe on behalf of the Wellcome Sanger Institute COVID-19 Surveillance Team |
| EPI_ISL_3367469 | Lighthouse Laboratory Plymouth | Wellcome Sanger Institute for the COVID-19 Genomics UK (COG-UK) Consortium | Lighthouse Laboratory Plymouth and Alex Alderton, Roberto Amato, Jeffrey Barrett, Sonia Goncalves, Ewan Harrison, David K. Jackson, Ian Johnston, Dominic Kwiatkowski, Cordelia Langford, John Sillitoe on behalf of the Wellcome Sanger Institute COVID-19 Surveillance Team |
| EPI_ISL_3367470, EPI_ISL_3367471, EPI_ISL_3367472, EPI_ISL_3367473, EPI_ISL_3367474 | Lighthouse Lab in Glasgow | Wellcome Sanger Institute for the COVID-19 Genomics UK (COG-UK) Consortium | Harper VanSteenhouse, Yumi Kasai, David Gray, Carol Clugston, Anna Dominiczak and Alex Alderton, Roberto Amato, Jeffrey Barrett, Sonia Goncalves, Ewan Harrison, David K. Jackson, Ian Johnston, Dominic Kwiatkowski, Cordelia Langford, John Sillitoe on behalf of the Wellcome Sanger Institute COVID-19 Surveillance Team |
| EPI_ISL_3367475, EPI_ISL_3367476 | Lighthouse Laboratory Plymouth | Wellcome Sanger Institute for the COVID-19 Genomics UK (COG-UK) Consortium | Lighthouse Laboratory Plymouth and Alex Alderton, Roberto Amato, Jeffrey Barrett, Sonia Goncalves, Ewan Harrison, David K. Jackson, Ian Johnston, Dominic Kwiatkowski, Cordelia Langford, John Sillitoe on behalf of the Wellcome Sanger Institute COVID-19 Surveillance Team |
| EPI_ISL_3367477 | Lighthouse Lab in Glasgow | Wellcome Sanger Institute for the COVID-19 Genomics UK (COG-UK) Consortium | Harper VanSteenhouse, Yumi Kasai, David Gray, Carol Clugston, Anna Dominiczak and Alex Alderton, Roberto Amato, Jeffrey Barrett, Sonia Goncalves, Ewan Harrison, David K. Jackson, Ian Johnston, Dominic Kwiatkowski, Cordelia Langford, John Sillitoe on behalf of the Wellcome Sanger Institute COVID-19 Surveillance Team |
| EPI_ISL_3367478 | Lighthouse Laboratory Plymouth | Wellcome Sanger Institute for the COVID-19 Genomics UK (COG-UK) Consortium | Lighthouse Laboratory Plymouth and Alex Alderton, Roberto Amato, Jeffrey Barrett, Sonia Goncalves, Ewan Harrison, David K. Jackson, Ian Johnston, Dominic Kwiatkowski, Cordelia Langford, John Sillitoe on behalf of the Wellcome Sanger Institute COVID-19 Surveillance Team |
| EPI_ISL_3367479 | Lighthouse Lab in Glasgow | Wellcome Sanger Institute for the COVID-19 Genomics UK (COG-UK) Consortium | Harper VanSteenhouse, Yumi Kasai, David Gray, Carol Clugston, Anna Dominiczak and Alex Alderton, Roberto Amato, Jeffrey Barrett, Sonia Goncalves, Ewan Harrison, David K. Jackson, Ian Johnston, Dominic Kwiatkowski, Cordelia Langford, John Sillitoe on behalf of the Wellcome Sanger Institute COVID-19 Surveillance Team |
| EPI_ISL_3367480 | Lighthouse Laboratory Plymouth | Wellcome Sanger Institute for the COVID-19 Genomics UK (COG-UK) Consortium | Lighthouse Laboratory Plymouth and Alex Alderton, Roberto Amato, Jeffrey Barrett, Sonia Goncalves, Ewan Harrison, David K. Jackson, Ian Johnston, Dominic Kwiatkowski, Cordelia Langford, John Sillitoe on behalf of the Wellcome Sanger Institute COVID-19 Surveillance Team |

| | | | |
|---|---|---|---|
| EPI_ISL_3367481, EPI_ISL_3367482, EPI_ISL_3367483, EPI_ISL_3367484 | Lighthouse Lab in Glasgow | Wellcome Sanger Institute for the COVID-19 Genomics UK (COG-UK) Consortium | Harper VanSteenhouse, Yumi Kasai, David Gray, Carol Clugston, Anna Dominiczak and Alex Alderton, Roberto Amato, Jeffrey Barrett, Sonia Goncalves, Ewan Harrison, David K. Jackson, Ian Johnston, Dominic Kwiatkowski, Cordelia Langford, John Sillitoe on behalf of the Wellcome Sanger Institute COVID-19 Surveillance Team |
| EPI_ISL_3367485, EPI_ISL_3367486 | Lighthouse Laboratory Plymouth | Wellcome Sanger Institute for the COVID-19 Genomics UK (COG-UK) Consortium | Lighthouse Laboratory Plymouth and Alex Alderton, Roberto Amato, Jeffrey Barrett, Sonia Goncalves, Ewan Harrison, David K. Jackson, Ian Johnston, Dominic Kwiatkowski, Cordelia Langford, John Sillitoe on behalf of the Wellcome Sanger Institute COVID-19 Surveillance Team |
| EPI_ISL_3367487 | Lighthouse Lab in Glasgow | Wellcome Sanger Institute for the COVID-19 Genomics UK (COG-UK) Consortium | Harper VanSteenhouse, Yumi Kasai, David Gray, Carol Clugston, Anna Dominiczak and Alex Alderton, Roberto Amato, Jeffrey Barrett, Sonia Goncalves, Ewan Harrison, David K. Jackson, Ian Johnston, Dominic Kwiatkowski, Cordelia Langford, John Sillitoe on behalf of the Wellcome Sanger Institute COVID-19 Surveillance Team |
| EPI_ISL_3367488 | Lighthouse Laboratory Plymouth | Wellcome Sanger Institute for the COVID-19 Genomics UK (COG-UK) Consortium | Lighthouse Laboratory Plymouth and Alex Alderton, Roberto Amato, Jeffrey Barrett, Sonia Goncalves, Ewan Harrison, David K. Jackson, Ian Johnston, Dominic Kwiatkowski, Cordelia Langford, John Sillitoe on behalf of the Wellcome Sanger Institute COVID-19 Surveillance Team |
| EPI_ISL_3367489, EPI_ISL_3367490, EPI_ISL_3367491 | Lighthouse Lab in Glasgow | Wellcome Sanger Institute for the COVID-19 Genomics UK (COG-UK) Consortium | Harper VanSteenhouse, Yumi Kasai, David Gray, Carol Clugston, Anna Dominiczak and Alex Alderton, Roberto Amato, Jeffrey Barrett, Sonia Goncalves, Ewan Harrison, David K. Jackson, Ian Johnston, Dominic Kwiatkowski, Cordelia Langford, John Sillitoe on behalf of the Wellcome Sanger Institute COVID-19 Surveillance Team |
| EPI_ISL_3367492 | Lighthouse Laboratory Plymouth | Wellcome Sanger Institute for the COVID-19 Genomics UK (COG-UK) Consortium | Lighthouse Laboratory Plymouth and Alex Alderton, Roberto Amato, Jeffrey Barrett, Sonia Goncalves, Ewan Harrison, David K. Jackson, Ian Johnston, Dominic Kwiatkowski, Cordelia Langford, John Sillitoe on behalf of the Wellcome Sanger Institute COVID-19 Surveillance Team |
| EPI_ISL_3367493, EPI_ISL_3367494, EPI_ISL_3367495, EPI_ISL_3367496, EPI_ISL_3367497, EPI_ISL_3367498, EPI_ISL_3367499 | Lighthouse Lab in Glasgow | Wellcome Sanger Institute for the COVID-19 Genomics UK (COG-UK) Consortium | Harper VanSteenhouse, Yumi Kasai, David Gray, Carol Clugston, Anna Dominiczak and Alex Alderton, Roberto Amato, Jeffrey Barrett, Sonia Goncalves, Ewan Harrison, David K. Jackson, Ian Johnston, Dominic Kwiatkowski, Cordelia Langford, John Sillitoe on behalf of the Wellcome Sanger Institute COVID-19 Surveillance Team |
| EPI_ISL_3367500 | Lighthouse Laboratory Plymouth | Wellcome Sanger Institute for the COVID-19 Genomics UK (COG-UK) Consortium | Lighthouse Laboratory Plymouth and Alex Alderton, Roberto Amato, Jeffrey Barrett, Sonia Goncalves, Ewan Harrison, David K. Jackson, Ian Johnston, Dominic Kwiatkowski, Cordelia Langford, John Sillitoe on behalf of the Wellcome Sanger Institute COVID-19 Surveillance Team |
| EPI_ISL_3367501, EPI_ISL_3367502, EPI_ISL_3367503, EPI_ISL_3367504 | Lighthouse Lab in Glasgow | Wellcome Sanger Institute for the COVID-19 Genomics UK (COG-UK) Consortium | Harper VanSteenhouse, Yumi Kasai, David Gray, Carol Clugston, Anna Dominiczak and Alex Alderton, Roberto Amato, Jeffrey Barrett, Sonia Goncalves, Ewan Harrison, David K. Jackson, Ian Johnston, Dominic Kwiatkowski, Cordelia Langford, John Sillitoe on behalf of the Wellcome Sanger Institute COVID-19 Surveillance Team |
| EPI_ISL_3367505, EPI_ISL_3367506, EPI_ISL_3367507, EPI_ISL_3367508, EPI_ISL_3367509 | Lighthouse Laboratory Plymouth | Wellcome Sanger Institute for the COVID-19 Genomics UK (COG-UK) Consortium | Lighthouse Laboratory Plymouth and Alex Alderton, Roberto Amato, Jeffrey Barrett, Sonia Goncalves, Ewan Harrison, David K. Jackson, Ian Johnston, Dominic Kwiatkowski, Cordelia Langford, John Sillitoe on behalf of the Wellcome Sanger Institute COVID-19 Surveillance Team |
| EPI_ISL_3367510 | Lighthouse Lab in Glasgow | Wellcome Sanger Institute for the COVID-19 Genomics UK (COG-UK) Consortium | Harper VanSteenhouse, Yumi Kasai, David Gray, Carol Clugston, Anna Dominiczak and Alex Alderton, Roberto Amato, Jeffrey Barrett, Sonia Goncalves, Ewan Harrison, David K. Jackson, Ian Johnston, Dominic Kwiatkowski, Cordelia Langford, John Sillitoe on behalf of the Wellcome Sanger Institute COVID-19 Surveillance Team |
| EPI_ISL_3367511 | Lighthouse Laboratory Plymouth | Wellcome Sanger Institute for the COVID-19 Genomics UK (COG-UK) Consortium | Lighthouse Laboratory Plymouth and Alex Alderton, Roberto Amato, Jeffrey Barrett, Sonia Goncalves, Ewan Harrison, David K. Jackson, Ian Johnston, Dominic Kwiatkowski, Cordelia Langford, John Sillitoe on behalf of the Wellcome Sanger Institute COVID-19 Surveillance Team |
| EPI_ISL_3367512, EPI_ISL_3367513, EPI_ISL_3367514, EPI_ISL_3367515 | Lighthouse Lab in Glasgow | Wellcome Sanger Institute for the COVID-19 Genomics UK (COG-UK) Consortium | Harper VanSteenhouse, Yumi Kasai, David Gray, Carol Clugston, Anna Dominiczak and Alex Alderton, Roberto Amato, Jeffrey Barrett, Sonia Goncalves, Ewan Harrison, David K. Jackson, Ian Johnston, Dominic Kwiatkowski, Cordelia Langford, John Sillitoe on behalf of the Wellcome Sanger Institute COVID-19 Surveillance Team |
| EPI_ISL_3367516, EPI_ISL_3367517 | Lighthouse Laboratory Plymouth | Wellcome Sanger Institute for the COVID-19 Genomics UK (COG-UK) Consortium | Lighthouse Laboratory Plymouth and Alex Alderton, Roberto Amato, Jeffrey Barrett, Sonia Goncalves, Ewan Harrison, David K. Jackson, Ian Johnston, Dominic Kwiatkowski, Cordelia Langford, John Sillitoe on behalf of the Wellcome Sanger Institute COVID-19 Surveillance Team |
| EPI_ISL_3367518, EPI_ISL_3367519, EPI_ISL_3367520, EPI_ISL_3367521, EPI_ISL_3367522, EPI_ISL_3367523, EPI_ISL_3367524 | Lighthouse Lab in Glasgow | Wellcome Sanger Institute for the COVID-19 Genomics UK (COG-UK) Consortium | Harper VanSteenhouse, Yumi Kasai, David Gray, Carol Clugston, Anna Dominiczak and Alex Alderton, Roberto Amato, Jeffrey Barrett, Sonia Goncalves, Ewan Harrison, David K. Jackson, Ian Johnston, Dominic Kwiatkowski, Cordelia Langford, John Sillitoe on behalf of the Wellcome Sanger Institute COVID-19 Surveillance Team |
| EPI_ISL_3367525 | Lighthouse Laboratory Plymouth | Wellcome Sanger Institute for the COVID-19 Genomics UK (COG-UK) Consortium | Lighthouse Laboratory Plymouth and Alex Alderton, Roberto Amato, Jeffrey Barrett, Sonia Goncalves, Ewan Harrison, David K. Jackson, Ian Johnston, Dominic Kwiatkowski, Cordelia Langford, John Sillitoe on behalf of the Wellcome Sanger Institute COVID-19 Surveillance Team |
| EPI_ISL_3367526, EPI_ISL_3367527, EPI_ISL_3367528, EPI_ISL_3367529, EPI_ISL_3367530, EPI_ISL_3367531, EPI_ISL_3367532, EPI_ISL_3367533, EPI_ISL_3367534, EPI_ISL_3367535, EPI_ISL_3367536, EPI_ISL_3367537, EPI_ISL_3367538 | | | |
| see above | Lighthouse Lab in Glasgow | Wellcome Sanger Institute for the COVID-19 Genomics UK (COG-UK) Consortium | Harper VanSteenhouse, Yumi Kasai, David Gray, Carol Clugston, Anna Dominiczak and Alex Alderton, Roberto Amato, Jeffrey Barrett, Sonia Goncalves, Ewan Harrison, David K. Jackson, Ian Johnston, Dominic Kwiatkowski, Cordelia Langford, John Sillitoe on behalf of the Wellcome Sanger Institute COVID-19 Surveillance Team |
| EPI_ISL_3367539 | Lighthouse Laboratory Plymouth | Wellcome Sanger Institute for the COVID-19 Genomics UK (COG-UK) Consortium | Lighthouse Laboratory Plymouth and Alex Alderton, Roberto Amato, Jeffrey Barrett, Sonia Goncalves, Ewan Harrison, David K. Jackson, Ian Johnston, Dominic Kwiatkowski, Cordelia Langford, John Sillitoe on behalf of the Wellcome Sanger Institute COVID-19 Surveillance Team |
| EPI_ISL_3367540, EPI_ISL_3367541 | Lighthouse Lab in Glasgow | Wellcome Sanger Institute for the COVID-19 Genomics UK (COG-UK) Consortium | Harper VanSteenhouse, Yumi Kasai, David Gray, Carol Clugston, Anna Dominiczak and Alex Alderton, Roberto Amato, Jeffrey Barrett, Sonia Goncalves, Ewan Harrison, David K. Jackson, Ian Johnston, Dominic Kwiatkowski, Cordelia Langford, John Sillitoe on behalf of the Wellcome Sanger Institute COVID-19 Surveillance Team |
| EPI_ISL_3367542 | Lighthouse Laboratory Plymouth | Wellcome Sanger Institute for the COVID-19 Genomics UK (COG-UK) Consortium | Lighthouse Laboratory Plymouth and Alex Alderton, Roberto Amato, Jeffrey Barrett, Sonia Goncalves, Ewan Harrison, David K. Jackson, Ian Johnston, Dominic Kwiatkowski, Cordelia Langford, John Sillitoe on behalf of the Wellcome Sanger Institute COVID-19 Surveillance Team |
| EPI_ISL_3367543, EPI_ISL_3367544, EPI_ISL_3367545, EPI_ISL_3367546, EPI_ISL_3367547, EPI_ISL_3367548, EPI_ISL_3367549, EPI_ISL_3367550, EPI_ISL_3367551, EPI_ISL_3367552, EPI_ISL_3367553, EPI_ISL_3367554 | | | |
| see above | Lighthouse Lab in Glasgow | Wellcome Sanger Institute for the COVID-19 Genomics UK (COG-UK) Consortium | Harper VanSteenhouse, Yumi Kasai, David Gray, Carol Clugston, Anna Dominiczak and Alex Alderton, Roberto Amato, Jeffrey Barrett, Sonia Goncalves, Ewan Harrison, David K. Jackson, Ian Johnston, Dominic Kwiatkowski, Cordelia Langford, John Sillitoe on behalf of the Wellcome Sanger Institute COVID-19 Surveillance Team |
| EPI_ISL_3367555 | Lighthouse Laboratory Plymouth | Wellcome Sanger Institute for the COVID-19 Genomics UK (COG-UK) Consortium | Lighthouse Laboratory Plymouth and Alex Alderton, Roberto Amato, Jeffrey Barrett, Sonia Goncalves, Ewan Harrison, David K. Jackson, Ian Johnston, Dominic Kwiatkowski, Cordelia Langford, John Sillitoe on behalf of the Wellcome Sanger Institute COVID-19 Surveillance Team |
| EPI_ISL_3367556 | Lighthouse Lab in Glasgow | Wellcome Sanger Institute for the COVID-19 Genomics UK (COG-UK) Consortium | Harper VanSteenhouse, Yumi Kasai, David Gray, Carol Clugston, Anna Dominiczak and Alex Alderton, Roberto Amato, Jeffrey Barrett, Sonia Goncalves, Ewan Harrison, David K. Jackson, Ian Johnston, Dominic Kwiatkowski, Cordelia Langford, John Sillitoe on behalf of the Wellcome Sanger Institute COVID-19 Surveillance Team |
| EPI_ISL_3367557, EPI_ISL_3367558 | Lighthouse Laboratory Plymouth | Wellcome Sanger Institute for the COVID-19 Genomics UK (COG-UK) Consortium | Lighthouse Laboratory Plymouth and Alex Alderton, Roberto Amato, Jeffrey Barrett, Sonia Goncalves, Ewan Harrison, David K. Jackson, Ian Johnston, Dominic Kwiatkowski, Cordelia Langford, John Sillitoe on behalf of the Wellcome Sanger Institute COVID-19 Surveillance Team |
| EPI_ISL_3367559, EPI_ISL_3367560, EPI_ISL_3367561, EPI_ISL_3367562, EPI_ISL_3367563 | Lighthouse Lab in Glasgow | Wellcome Sanger Institute for the COVID-19 Genomics UK (COG-UK) Consortium | Harper VanSteenhouse, Yumi Kasai, David Gray, Carol Clugston, Anna Dominiczak and Alex Alderton, Roberto Amato, Jeffrey Barrett, Sonia Goncalves, Ewan Harrison, David K. Jackson, Ian Johnston, Dominic Kwiatkowski, Cordelia Langford, John Sillitoe on behalf of the Wellcome Sanger Institute COVID-19 Surveillance Team |
| EPI_ISL_3367564 | Lighthouse Laboratory Plymouth | Wellcome Sanger Institute for the COVID-19 Genomics UK (COG-UK) Consortium | Lighthouse Laboratory Plymouth and Alex Alderton, Roberto Amato, Jeffrey Barrett, Sonia Goncalves, Ewan Harrison, David K. Jackson, Ian Johnston, Dominic Kwiatkowski, Cordelia Langford, John Sillitoe on behalf of the Wellcome Sanger Institute COVID-19 Surveillance Team |
| EPI_ISL_3367565, EPI_ISL_3367566, EPI_ISL_3367567, EPI_ISL_3367568, EPI_ISL_3367569 | Lighthouse Lab in Glasgow | Wellcome Sanger Institute for the COVID-19 Genomics UK (COG-UK) Consortium | Harper VanSteenhouse, Yumi Kasai, David Gray, Carol Clugston, Anna Dominiczak and Alex Alderton, Roberto Amato, Jeffrey Barrett, Sonia Goncalves, Ewan Harrison, David K. Jackson, Ian Johnston, Dominic Kwiatkowski, Cordelia Langford, John Sillitoe on behalf of the Wellcome Sanger Institute COVID-19 Surveillance Team |
| EPI_ISL_3367570 | Lighthouse Laboratory Plymouth | Wellcome Sanger Institute for the COVID-19 Genomics UK (COG-UK) Consortium | Lighthouse Laboratory Plymouth and Alex Alderton, Roberto Amato, Jeffrey Barrett, Sonia Goncalves, Ewan Harrison, David K. Jackson, Ian Johnston, Dominic Kwiatkowski, Cordelia Langford, John Sillitoe on behalf of the Wellcome Sanger Institute COVID-19 Surveillance Team |
| EPI_ISL_3367571 | Lighthouse Lab in Glasgow | Wellcome Sanger Institute for the COVID-19 Genomics UK (COG-UK) Consortium | Harper VanSteenhouse, Yumi Kasai, David Gray, Carol Clugston, Anna Dominiczak and Alex Alderton, Roberto Amato, Jeffrey Barrett, Sonia Goncalves, Ewan Harrison, David K. Jackson, Ian Johnston, Dominic Kwiatkowski, Cordelia Langford, John Sillitoe on behalf of the Wellcome Sanger Institute COVID-19 Surveillance Team |
| EPI_ISL_3367572, EPI_ISL_3367573, EPI_ISL_3367574, EPI_ISL_3367575 | Lighthouse Laboratory Plymouth | Wellcome Sanger Institute for the COVID-19 Genomics UK (COG-UK) Consortium | Lighthouse Laboratory Plymouth and Alex Alderton, Roberto Amato, Jeffrey Barrett, Sonia Goncalves, Ewan Harrison, David K. Jackson, Ian Johnston, Dominic Kwiatkowski, Cordelia Langford, John Sillitoe on behalf of the Wellcome Sanger Institute COVID-19 Surveillance Team |
| EPI_ISL_3367576, EPI_ISL_3367577 | Lighthouse Lab in Glasgow | Wellcome Sanger Institute for the COVID-19 Genomics UK (COG-UK) Consortium | Harper VanSteenhouse, Yumi Kasai, David Gray, Carol Clugston, Anna Dominiczak and Alex Alderton, Roberto Amato, Jeffrey Barrett, Sonia Goncalves, Ewan Harrison, David K. Jackson, Ian Johnston, Dominic Kwiatkowski, Cordelia Langford, John Sillitoe on behalf of the Wellcome Sanger Institute COVID-19 Surveillance Team |
| EPI_ISL_3367578 | Lighthouse Laboratory Plymouth | Wellcome Sanger Institute for the COVID-19 Genomics UK (COG-UK) Consortium | Lighthouse Laboratory Plymouth and Alex Alderton, Roberto Amato, Jeffrey Barrett, Sonia Goncalves, Ewan Harrison, David K. Jackson, Ian Johnston, Dominic Kwiatkowski, Cordelia Langford, John Sillitoe on behalf of the Wellcome Sanger Institute COVID-19 Surveillance Team |
| EPI_ISL_3367579, | Lighthouse Lab in Glasgow | Wellcome Sanger Institute for the COVID-19 Genomics UK (COG-UK) | Harper VanSteenhouse, Yumi Kasai, David Gray, Carol Clugston, Anna Dominiczak and Alex Alderton, Roberto Amato, Jeffrey Barrett, Sonia Goncalves, Ewan Harrison, David K. Jackson, Ian Johnston, Dominic Kwiatkowski, Cordelia Langford, |

| | | | |
|---|---|---|---|
| EPI_ISL_3367580 | | Consortium | John Sillitoe on behalf of the Wellcome Sanger Institute COVID-19 Surveillance Team |
| EPI_ISL_3367581 | Lighthouse Laboratory Plymouth | Wellcome Sanger Institute for the COVID-19 Genomics UK (COG-UK) Consortium | Lighthouse Laboratory Plymouth and Alex Alderton, Roberto Amato, Jeffrey Barrett, Sonia Goncalves, Ewan Harrison, David K. Jackson, Ian Johnston, Dominic Kwiatkowski, Cordelia Langford, John Sillitoe on behalf of the Wellcome Sanger Institute COVID-19 Surveillance Team |
| EPI_ISL_3367582, EPI_ISL_3367583, EPI_ISL_3367584, EPI_ISL_3367585, EPI_ISL_3367586, EPI_ISL_3367587, EPI_ISL_3367588, EPI_ISL_3367589 | Lighthouse Lab in Glasgow | Wellcome Sanger Institute for the COVID-19 Genomics UK (COG-UK) Consortium | Harper VanSteenhouse, Yumi Kasai, David Gray, Carol Clugston, Anna Dominiczak and Alex Alderton, Roberto Amato, Jeffrey Barrett, Sonia Goncalves, Ewan Harrison, David K. Jackson, Ian Johnston, Dominic Kwiatkowski, Cordelia Langford, John Sillitoe on behalf of the Wellcome Sanger Institute COVID-19 Surveillance Team |
| EPI_ISL_3367590 | Lighthouse Laboratory Plymouth | Wellcome Sanger Institute for the COVID-19 Genomics UK (COG-UK) Consortium | Lighthouse Laboratory Plymouth and Alex Alderton, Roberto Amato, Jeffrey Barrett, Sonia Goncalves, Ewan Harrison, David K. Jackson, Ian Johnston, Dominic Kwiatkowski, Cordelia Langford, John Sillitoe on behalf of the Wellcome Sanger Institute COVID-19 Surveillance Team |
| EPI_ISL_3367591 | Lighthouse Lab in Glasgow | Wellcome Sanger Institute for the COVID-19 Genomics UK (COG-UK) Consortium | Harper VanSteenhouse, Yumi Kasai, David Gray, Carol Clugston, Anna Dominiczak and Alex Alderton, Roberto Amato, Jeffrey Barrett, Sonia Goncalves, Ewan Harrison, David K. Jackson, Ian Johnston, Dominic Kwiatkowski, Cordelia Langford, John Sillitoe on behalf of the Wellcome Sanger Institute COVID-19 Surveillance Team |
| EPI_ISL_3367592 | Lighthouse Laboratory Plymouth | Wellcome Sanger Institute for the COVID-19 Genomics UK (COG-UK) Consortium | Lighthouse Laboratory Plymouth and Alex Alderton, Roberto Amato, Jeffrey Barrett, Sonia Goncalves, Ewan Harrison, David K. Jackson, Ian Johnston, Dominic Kwiatkowski, Cordelia Langford, John Sillitoe on behalf of the Wellcome Sanger Institute COVID-19 Surveillance Team |
| EPI_ISL_3367593, EPI_ISL_3367594 | Lighthouse Lab in Glasgow | Wellcome Sanger Institute for the COVID-19 Genomics UK (COG-UK) Consortium | Harper VanSteenhouse, Yumi Kasai, David Gray, Carol Clugston, Anna Dominiczak and Alex Alderton, Roberto Amato, Jeffrey Barrett, Sonia Goncalves, Ewan Harrison, David K. Jackson, Ian Johnston, Dominic Kwiatkowski, Cordelia Langford, John Sillitoe on behalf of the Wellcome Sanger Institute COVID-19 Surveillance Team |
| EPI_ISL_3367595 | Lighthouse Laboratory Plymouth | Wellcome Sanger Institute for the COVID-19 Genomics UK (COG-UK) Consortium | Lighthouse Laboratory Plymouth and Alex Alderton, Roberto Amato, Jeffrey Barrett, Sonia Goncalves, Ewan Harrison, David K. Jackson, Ian Johnston, Dominic Kwiatkowski, Cordelia Langford, John Sillitoe on behalf of the Wellcome Sanger Institute COVID-19 Surveillance Team |
| EPI_ISL_3367596, EPI_ISL_3367597, EPI_ISL_3367598 | Lighthouse Lab in Glasgow | Wellcome Sanger Institute for the COVID-19 Genomics UK (COG-UK) Consortium | Harper VanSteenhouse, Yumi Kasai, David Gray, Carol Clugston, Anna Dominiczak and Alex Alderton, Roberto Amato, Jeffrey Barrett, Sonia Goncalves, Ewan Harrison, David K. Jackson, Ian Johnston, Dominic Kwiatkowski, Cordelia Langford, John Sillitoe on behalf of the Wellcome Sanger Institute COVID-19 Surveillance Team |
| EPI_ISL_3367599 | Lighthouse Laboratory Plymouth | Wellcome Sanger Institute for the COVID-19 Genomics UK (COG-UK) Consortium | Lighthouse Laboratory Plymouth and Alex Alderton, Roberto Amato, Jeffrey Barrett, Sonia Goncalves, Ewan Harrison, David K. Jackson, Ian Johnston, Dominic Kwiatkowski, Cordelia Langford, John Sillitoe on behalf of the Wellcome Sanger Institute COVID-19 Surveillance Team |
| EPI_ISL_3367600, EPI_ISL_3367601 | Lighthouse Lab in Glasgow | Wellcome Sanger Institute for the COVID-19 Genomics UK (COG-UK) Consortium | Harper VanSteenhouse, Yumi Kasai, David Gray, Carol Clugston, Anna Dominiczak and Alex Alderton, Roberto Amato, Jeffrey Barrett, Sonia Goncalves, Ewan Harrison, David K. Jackson, Ian Johnston, Dominic Kwiatkowski, Cordelia Langford, John Sillitoe on behalf of the Wellcome Sanger Institute COVID-19 Surveillance Team |
| EPI_ISL_3367602 | Lighthouse Laboratory Plymouth | Wellcome Sanger Institute for the COVID-19 Genomics UK (COG-UK) Consortium | Lighthouse Laboratory Plymouth and Alex Alderton, Roberto Amato, Jeffrey Barrett, Sonia Goncalves, Ewan Harrison, David K. Jackson, Ian Johnston, Dominic Kwiatkowski, Cordelia Langford, John Sillitoe on behalf of the Wellcome Sanger Institute COVID-19 Surveillance Team |
| EPI_ISL_3367603 | Lighthouse Lab in Glasgow | Wellcome Sanger Institute for the COVID-19 Genomics UK (COG-UK) Consortium | Harper VanSteenhouse, Yumi Kasai, David Gray, Carol Clugston, Anna Dominiczak and Alex Alderton, Roberto Amato, Jeffrey Barrett, Sonia Goncalves, Ewan Harrison, David K. Jackson, Ian Johnston, Dominic Kwiatkowski, Cordelia Langford, John Sillitoe on behalf of the Wellcome Sanger Institute COVID-19 Surveillance Team |
| EPI_ISL_3367604 | Lighthouse Laboratory Plymouth | Wellcome Sanger Institute for the COVID-19 Genomics UK (COG-UK) Consortium | Lighthouse Laboratory Plymouth and Alex Alderton, Roberto Amato, Jeffrey Barrett, Sonia Goncalves, Ewan Harrison, David K. Jackson, Ian Johnston, Dominic Kwiatkowski, Cordelia Langford, John Sillitoe on behalf of the Wellcome Sanger Institute COVID-19 Surveillance Team |
| EPI_ISL_3367605 | Lighthouse Lab in Glasgow | Wellcome Sanger Institute for the COVID-19 Genomics UK (COG-UK) Consortium | Harper VanSteenhouse, Yumi Kasai, David Gray, Carol Clugston, Anna Dominiczak and Alex Alderton, Roberto Amato, Jeffrey Barrett, Sonia Goncalves, Ewan Harrison, David K. Jackson, Ian Johnston, Dominic Kwiatkowski, Cordelia Langford, John Sillitoe on behalf of the Wellcome Sanger Institute COVID-19 Surveillance Team |
| EPI_ISL_3367606, EPI_ISL_3367607 | Lighthouse Laboratory Plymouth | Wellcome Sanger Institute for the COVID-19 Genomics UK (COG-UK) Consortium | Lighthouse Laboratory Plymouth and Alex Alderton, Roberto Amato, Jeffrey Barrett, Sonia Goncalves, Ewan Harrison, David K. Jackson, Ian Johnston, Dominic Kwiatkowski, Cordelia Langford, John Sillitoe on behalf of the Wellcome Sanger Institute COVID-19 Surveillance Team |
| EPI_ISL_3367608, EPI_ISL_3367609 | Lighthouse Lab in Glasgow | Wellcome Sanger Institute for the COVID-19 Genomics UK (COG-UK) Consortium | Harper VanSteenhouse, Yumi Kasai, David Gray, Carol Clugston, Anna Dominiczak and Alex Alderton, Roberto Amato, Jeffrey Barrett, Sonia Goncalves, Ewan Harrison, David K. Jackson, Ian Johnston, Dominic Kwiatkowski, Cordelia Langford, John Sillitoe on behalf of the Wellcome Sanger Institute COVID-19 Surveillance Team |
| EPI_ISL_3367610 | Lighthouse Laboratory Plymouth | Wellcome Sanger Institute for the COVID-19 Genomics UK (COG-UK) Consortium | Lighthouse Laboratory Plymouth and Alex Alderton, Roberto Amato, Jeffrey Barrett, Sonia Goncalves, Ewan Harrison, David K. Jackson, Ian Johnston, Dominic Kwiatkowski, Cordelia Langford, John Sillitoe on behalf of the Wellcome Sanger Institute COVID-19 Surveillance Team |
| EPI_ISL_3367611 | Lighthouse Lab in Glasgow | Wellcome Sanger Institute for the COVID-19 Genomics UK (COG-UK) Consortium | Harper VanSteenhouse, Yumi Kasai, David Gray, Carol Clugston, Anna Dominiczak and Alex Alderton, Roberto Amato, Jeffrey Barrett, Sonia Goncalves, Ewan Harrison, David K. Jackson, Ian Johnston, Dominic Kwiatkowski, Cordelia Langford, John Sillitoe on behalf of the Wellcome Sanger Institute COVID-19 Surveillance Team |
| EPI_ISL_3367612 | Lighthouse Laboratory Plymouth | Wellcome Sanger Institute for the COVID-19 Genomics UK (COG-UK) Consortium | Lighthouse Laboratory Plymouth and Alex Alderton, Roberto Amato, Jeffrey Barrett, Sonia Goncalves, Ewan Harrison, David K. Jackson, Ian Johnston, Dominic Kwiatkowski, Cordelia Langford, John Sillitoe on behalf of the Wellcome Sanger Institute COVID-19 Surveillance Team |
| EPI_ISL_3367613, EPI_ISL_3367614, EPI_ISL_3367615, EPI_ISL_3367616 | Lighthouse Lab in Glasgow | Wellcome Sanger Institute for the COVID-19 Genomics UK (COG-UK) Consortium | Harper VanSteenhouse, Yumi Kasai, David Gray, Carol Clugston, Anna Dominiczak and Alex Alderton, Roberto Amato, Jeffrey Barrett, Sonia Goncalves, Ewan Harrison, David K. Jackson, Ian Johnston, Dominic Kwiatkowski, Cordelia Langford, John Sillitoe on behalf of the Wellcome Sanger Institute COVID-19 Surveillance Team |
| EPI_ISL_3367617 | Lighthouse Laboratory Plymouth | Wellcome Sanger Institute for the COVID-19 Genomics UK (COG-UK) Consortium | Lighthouse Laboratory Plymouth and Alex Alderton, Roberto Amato, Jeffrey Barrett, Sonia Goncalves, Ewan Harrison, David K. Jackson, Ian Johnston, Dominic Kwiatkowski, Cordelia Langford, John Sillitoe on behalf of the Wellcome Sanger Institute COVID-19 Surveillance Team |
| EPI_ISL_3367618, EPI_ISL_3367619 | Lighthouse Lab in Glasgow | Wellcome Sanger Institute for the COVID-19 Genomics UK (COG-UK) Consortium | Harper VanSteenhouse, Yumi Kasai, David Gray, Carol Clugston, Anna Dominiczak and Alex Alderton, Roberto Amato, Jeffrey Barrett, Sonia Goncalves, Ewan Harrison, David K. Jackson, Ian Johnston, Dominic Kwiatkowski, Cordelia Langford, John Sillitoe on behalf of the Wellcome Sanger Institute COVID-19 Surveillance Team |
| EPI_ISL_3367620 | Lighthouse Laboratory Plymouth | Wellcome Sanger Institute for the COVID-19 Genomics UK (COG-UK) Consortium | Lighthouse Laboratory Plymouth and Alex Alderton, Roberto Amato, Jeffrey Barrett, Sonia Goncalves, Ewan Harrison, David K. Jackson, Ian Johnston, Dominic Kwiatkowski, Cordelia Langford, John Sillitoe on behalf of the Wellcome Sanger Institute COVID-19 Surveillance Team |
| EPI_ISL_3367621 | Lighthouse Lab in Glasgow | Wellcome Sanger Institute for the COVID-19 Genomics UK (COG-UK) Consortium | Harper VanSteenhouse, Yumi Kasai, David Gray, Carol Clugston, Anna Dominiczak and Alex Alderton, Roberto Amato, Jeffrey Barrett, Sonia Goncalves, Ewan Harrison, David K. Jackson, Ian Johnston, Dominic Kwiatkowski, Cordelia Langford, John Sillitoe on behalf of the Wellcome Sanger Institute COVID-19 Surveillance Team |
| EPI_ISL_3367622 | Lighthouse Laboratory Plymouth | Wellcome Sanger Institute for the COVID-19 Genomics UK (COG-UK) Consortium | Lighthouse Laboratory Plymouth and Alex Alderton, Roberto Amato, Jeffrey Barrett, Sonia Goncalves, Ewan Harrison, David K. Jackson, Ian Johnston, Dominic Kwiatkowski, Cordelia Langford, John Sillitoe on behalf of the Wellcome Sanger Institute COVID-19 Surveillance Team |
| EPI_ISL_3367623 | Lighthouse Lab in Glasgow | Wellcome Sanger Institute for the COVID-19 Genomics UK (COG-UK) Consortium | Harper VanSteenhouse, Yumi Kasai, David Gray, Carol Clugston, Anna Dominiczak and Alex Alderton, Roberto Amato, Jeffrey Barrett, Sonia Goncalves, Ewan Harrison, David K. Jackson, Ian Johnston, Dominic Kwiatkowski, Cordelia Langford, John Sillitoe on behalf of the Wellcome Sanger Institute COVID-19 Surveillance Team |
| EPI_ISL_3367624 | Lighthouse Laboratory Plymouth | Wellcome Sanger Institute for the COVID-19 Genomics UK (COG-UK) Consortium | Lighthouse Laboratory Plymouth and Alex Alderton, Roberto Amato, Jeffrey Barrett, Sonia Goncalves, Ewan Harrison, David K. Jackson, Ian Johnston, Dominic Kwiatkowski, Cordelia Langford, John Sillitoe on behalf of the Wellcome Sanger Institute COVID-19 Surveillance Team |
| EPI_ISL_3367625, EPI_ISL_3367626, EPI_ISL_3367627, EPI_ISL_3367628 | Lighthouse Lab in Glasgow | Wellcome Sanger Institute for the COVID-19 Genomics UK (COG-UK) Consortium | Harper VanSteenhouse, Yumi Kasai, David Gray, Carol Clugston, Anna Dominiczak and Alex Alderton, Roberto Amato, Jeffrey Barrett, Sonia Goncalves, Ewan Harrison, David K. Jackson, Ian Johnston, Dominic Kwiatkowski, Cordelia Langford, John Sillitoe on behalf of the Wellcome Sanger Institute COVID-19 Surveillance Team |
| EPI_ISL_3367629 | Lighthouse Laboratory Plymouth | Wellcome Sanger Institute for the COVID-19 Genomics UK (COG-UK) Consortium | Lighthouse Laboratory Plymouth and Alex Alderton, Roberto Amato, Jeffrey Barrett, Sonia Goncalves, Ewan Harrison, David K. Jackson, Ian Johnston, Dominic Kwiatkowski, Cordelia Langford, John Sillitoe on behalf of the Wellcome Sanger Institute COVID-19 Surveillance Team |
| EPI_ISL_3367630, EPI_ISL_3367631, EPI_ISL_3367632, EPI_ISL_3367633, EPI_ISL_3367634 | Lighthouse Lab in Glasgow | Wellcome Sanger Institute for the COVID-19 Genomics UK (COG-UK) Consortium | Harper VanSteenhouse, Yumi Kasai, David Gray, Carol Clugston, Anna Dominiczak and Alex Alderton, Roberto Amato, Jeffrey Barrett, Sonia Goncalves, Ewan Harrison, David K. Jackson, Ian Johnston, Dominic Kwiatkowski, Cordelia Langford, John Sillitoe on behalf of the Wellcome Sanger Institute COVID-19 Surveillance Team |
| EPI_ISL_3367635 | Lighthouse Laboratory Plymouth | Wellcome Sanger Institute for the COVID-19 Genomics UK (COG-UK) Consortium | Lighthouse Laboratory Plymouth and Alex Alderton, Roberto Amato, Jeffrey Barrett, Sonia Goncalves, Ewan Harrison, David K. Jackson, Ian Johnston, Dominic Kwiatkowski, Cordelia Langford, John Sillitoe on behalf of the Wellcome Sanger Institute COVID-19 Surveillance Team |
| EPI_ISL_3367636 | Lighthouse Lab in Glasgow | Wellcome Sanger Institute for the COVID-19 Genomics UK (COG-UK) Consortium | Harper VanSteenhouse, Yumi Kasai, David Gray, Carol Clugston, Anna Dominiczak and Alex Alderton, Roberto Amato, Jeffrey Barrett, Sonia Goncalves, Ewan Harrison, David K. Jackson, Ian Johnston, Dominic Kwiatkowski, Cordelia Langford, John Sillitoe on behalf of the Wellcome Sanger Institute COVID-19 Surveillance Team |
| EPI_ISL_3367637 | Lighthouse Laboratory Plymouth | Wellcome Sanger Institute for the COVID-19 Genomics UK (COG-UK) Consortium | Lighthouse Laboratory Plymouth and Alex Alderton, Roberto Amato, Jeffrey Barrett, Sonia Goncalves, Ewan Harrison, David K. Jackson, Ian Johnston, Dominic Kwiatkowski, Cordelia Langford, John Sillitoe on behalf of the Wellcome Sanger Institute COVID-19 Surveillance Team |
| EPI_ISL_3367638, EPI_ISL_3367639, EPI_ISL_3367640, EPI_ISL_3367641, EPI_ISL_3367642 | Lighthouse Lab in Glasgow | Wellcome Sanger Institute for the COVID-19 Genomics UK (COG-UK) Consortium | Harper VanSteenhouse, Yumi Kasai, David Gray, Carol Clugston, Anna Dominiczak and Alex Alderton, Roberto Amato, Jeffrey Barrett, Sonia Goncalves, Ewan Harrison, David K. Jackson, Ian Johnston, Dominic Kwiatkowski, Cordelia Langford, John Sillitoe on behalf of the Wellcome Sanger Institute COVID-19 Surveillance Team |
| EPI_ISL_3367643 | Lighthouse Laboratory Plymouth | Wellcome Sanger Institute for the COVID-19 Genomics UK (COG-UK) Consortium | Lighthouse Laboratory Plymouth and Alex Alderton, Roberto Amato, Jeffrey Barrett, Sonia Goncalves, Ewan Harrison, David K. Jackson, Ian Johnston, Dominic Kwiatkowski, Cordelia Langford, John Sillitoe on behalf of the Wellcome Sanger Institute COVID-19 Surveillance Team |
| EPI_ISL_3367644, EPI_ISL_3367645, EPI_ISL_3367646 | Lighthouse Lab in Glasgow | Wellcome Sanger Institute for the COVID-19 Genomics UK (COG-UK) Consortium | Harper VanSteenhouse, Yumi Kasai, David Gray, Carol Clugston, Anna Dominiczak and Alex Alderton, Roberto Amato, Jeffrey Barrett, Sonia Goncalves, Ewan Harrison, David K. Jackson, Ian Johnston, Dominic Kwiatkowski, Cordelia Langford, John Sillitoe on behalf of the Wellcome Sanger Institute COVID-19 Surveillance Team |
| EPI_ISL_3367647 | Lighthouse Laboratory Plymouth | Wellcome Sanger Institute for the COVID-19 Genomics UK (COG-UK) Consortium | Lighthouse Laboratory Plymouth and Alex Alderton, Roberto Amato, Jeffrey Barrett, Sonia Goncalves, Ewan Harrison, David K. Jackson, Ian Johnston, Dominic Kwiatkowski, Cordelia Langford, John Sillitoe on behalf of the Wellcome Sanger Institute COVID-19 Surveillance Team |

| | | | |
|---|---|---|---|
| EPI_ISL_3367648, EPI_ISL_3367649, EPI_ISL_3367650, EPI_ISL_3367651, EPI_ISL_3367652, EPI_ISL_3367653, EPI_ISL_3367654, EPI_ISL_3367655 | Lighthouse Lab in Glasgow | Wellcome Sanger Institute for the COVID-19 Genomics UK (COG-UK) Consortium | Harper VanSteenhouse, Yumi Kasai, David Gray, Carol Clugston, Anna Dominiczak and Alex Alderton, Roberto Amato, Jeffrey Barrett, Sonia Goncalves, Ewan Harrison, David K. Jackson, Ian Johnston, Dominic Kwiatkowski, Cordelia Langford, John Sillitoe on behalf of the Wellcome Sanger Institute COVID-19 Surveillance Team |
| EPI_ISL_3367656 | Lighthouse Laboratory Plymouth | Wellcome Sanger Institute for the COVID-19 Genomics UK (COG-UK) Consortium | Lighthouse Laboratory Plymouth and Alex Alderton, Roberto Amato, Jeffrey Barrett, Sonia Goncalves, Ewan Harrison, David K. Jackson, Ian Johnston, Dominic Kwiatkowski, Cordelia Langford, John Sillitoe on behalf of the Wellcome Sanger Institute COVID-19 Surveillance Team |
| EPI_ISL_3367657 | Lighthouse Lab in Glasgow | Wellcome Sanger Institute for the COVID-19 Genomics UK (COG-UK) Consortium | Harper VanSteenhouse, Yumi Kasai, David Gray, Carol Clugston, Anna Dominiczak and Alex Alderton, Roberto Amato, Jeffrey Barrett, Sonia Goncalves, Ewan Harrison, David K. Jackson, Ian Johnston, Dominic Kwiatkowski, Cordelia Langford, John Sillitoe on behalf of the Wellcome Sanger Institute COVID-19 Surveillance Team |
| EPI_ISL_3367658, EPI_ISL_3367659 | Lighthouse Laboratory Plymouth | Wellcome Sanger Institute for the COVID-19 Genomics UK (COG-UK) Consortium | Lighthouse Laboratory Plymouth and Alex Alderton, Roberto Amato, Jeffrey Barrett, Sonia Goncalves, Ewan Harrison, David K. Jackson, Ian Johnston, Dominic Kwiatkowski, Cordelia Langford, John Sillitoe on behalf of the Wellcome Sanger Institute COVID-19 Surveillance Team |
| EPI_ISL_3367660 | Lighthouse Lab in Glasgow | Wellcome Sanger Institute for the COVID-19 Genomics UK (COG-UK) Consortium | Harper VanSteenhouse, Yumi Kasai, David Gray, Carol Clugston, Anna Dominiczak and Alex Alderton, Roberto Amato, Jeffrey Barrett, Sonia Goncalves, Ewan Harrison, David K. Jackson, Ian Johnston, Dominic Kwiatkowski, Cordelia Langford, John Sillitoe on behalf of the Wellcome Sanger Institute COVID-19 Surveillance Team |
| EPI_ISL_3367661 | Lighthouse Laboratory Plymouth | Wellcome Sanger Institute for the COVID-19 Genomics UK (COG-UK) Consortium | Lighthouse Laboratory Plymouth and Alex Alderton, Roberto Amato, Jeffrey Barrett, Sonia Goncalves, Ewan Harrison, David K. Jackson, Ian Johnston, Dominic Kwiatkowski, Cordelia Langford, John Sillitoe on behalf of the Wellcome Sanger Institute COVID-19 Surveillance Team |
| EPI_ISL_3367662, EPI_ISL_3367663, EPI_ISL_3367664, EPI_ISL_3367665, EPI_ISL_3367666, EPI_ISL_3367667, EPI_ISL_3367668 | Lighthouse Lab in Glasgow | Wellcome Sanger Institute for the COVID-19 Genomics UK (COG-UK) Consortium | Harper VanSteenhouse, Yumi Kasai, David Gray, Carol Clugston, Anna Dominiczak and Alex Alderton, Roberto Amato, Jeffrey Barrett, Sonia Goncalves, Ewan Harrison, David K. Jackson, Ian Johnston, Dominic Kwiatkowski, Cordelia Langford, John Sillitoe on behalf of the Wellcome Sanger Institute COVID-19 Surveillance Team |
| EPI_ISL_3367669 | Lighthouse Laboratory Plymouth | Wellcome Sanger Institute for the COVID-19 Genomics UK (COG-UK) Consortium | Lighthouse Laboratory Plymouth and Alex Alderton, Roberto Amato, Jeffrey Barrett, Sonia Goncalves, Ewan Harrison, David K. Jackson, Ian Johnston, Dominic Kwiatkowski, Cordelia Langford, John Sillitoe on behalf of the Wellcome Sanger Institute COVID-19 Surveillance Team |
| EPI_ISL_3367670, EPI_ISL_3367671 | Lighthouse Lab in Glasgow | Wellcome Sanger Institute for the COVID-19 Genomics UK (COG-UK) Consortium | Harper VanSteenhouse, Yumi Kasai, David Gray, Carol Clugston, Anna Dominiczak and Alex Alderton, Roberto Amato, Jeffrey Barrett, Sonia Goncalves, Ewan Harrison, David K. Jackson, Ian Johnston, Dominic Kwiatkowski, Cordelia Langford, John Sillitoe on behalf of the Wellcome Sanger Institute COVID-19 Surveillance Team |
| EPI_ISL_3367672 | Lighthouse Laboratory Plymouth | Wellcome Sanger Institute for the COVID-19 Genomics UK (COG-UK) Consortium | Lighthouse Laboratory Plymouth and Alex Alderton, Roberto Amato, Jeffrey Barrett, Sonia Goncalves, Ewan Harrison, David K. Jackson, Ian Johnston, Dominic Kwiatkowski, Cordelia Langford, John Sillitoe on behalf of the Wellcome Sanger Institute COVID-19 Surveillance Team |
| EPI_ISL_3367673, EPI_ISL_3367674, EPI_ISL_3367675, EPI_ISL_3367676, EPI_ISL_3367677, EPI_ISL_3367678 | Lighthouse Lab in Glasgow | Wellcome Sanger Institute for the COVID-19 Genomics UK (COG-UK) Consortium | Harper VanSteenhouse, Yumi Kasai, David Gray, Carol Clugston, Anna Dominiczak and Alex Alderton, Roberto Amato, Jeffrey Barrett, Sonia Goncalves, Ewan Harrison, David K. Jackson, Ian Johnston, Dominic Kwiatkowski, Cordelia Langford, John Sillitoe on behalf of the Wellcome Sanger Institute COVID-19 Surveillance Team |
| EPI_ISL_3367679, EPI_ISL_3367680 | Lighthouse Laboratory Plymouth | Wellcome Sanger Institute for the COVID-19 Genomics UK (COG-UK) Consortium | Lighthouse Laboratory Plymouth and Alex Alderton, Roberto Amato, Jeffrey Barrett, Sonia Goncalves, Ewan Harrison, David K. Jackson, Ian Johnston, Dominic Kwiatkowski, Cordelia Langford, John Sillitoe on behalf of the Wellcome Sanger Institute COVID-19 Surveillance Team |
| EPI_ISL_3367681, EPI_ISL_3367682, EPI_ISL_3367683, EPI_ISL_3367684, EPI_ISL_3367685 | Lighthouse Lab in Glasgow | Wellcome Sanger Institute for the COVID-19 Genomics UK (COG-UK) Consortium | Harper VanSteenhouse, Yumi Kasai, David Gray, Carol Clugston, Anna Dominiczak and Alex Alderton, Roberto Amato, Jeffrey Barrett, Sonia Goncalves, Ewan Harrison, David K. Jackson, Ian Johnston, Dominic Kwiatkowski, Cordelia Langford, John Sillitoe on behalf of the Wellcome Sanger Institute COVID-19 Surveillance Team |
| EPI_ISL_3367686 | Lighthouse Laboratory Plymouth | Wellcome Sanger Institute for the COVID-19 Genomics UK (COG-UK) Consortium | Lighthouse Laboratory Plymouth and Alex Alderton, Roberto Amato, Jeffrey Barrett, Sonia Goncalves, Ewan Harrison, David K. Jackson, Ian Johnston, Dominic Kwiatkowski, Cordelia Langford, John Sillitoe on behalf of the Wellcome Sanger Institute COVID-19 Surveillance Team |
| EPI_ISL_3367687, EPI_ISL_3367688 | Lighthouse Lab in Glasgow | Wellcome Sanger Institute for the COVID-19 Genomics UK (COG-UK) Consortium | Harper VanSteenhouse, Yumi Kasai, David Gray, Carol Clugston, Anna Dominiczak and Alex Alderton, Roberto Amato, Jeffrey Barrett, Sonia Goncalves, Ewan Harrison, David K. Jackson, Ian Johnston, Dominic Kwiatkowski, Cordelia Langford, John Sillitoe on behalf of the Wellcome Sanger Institute COVID-19 Surveillance Team |
| EPI_ISL_3367689 | Lighthouse Laboratory Plymouth | Wellcome Sanger Institute for the COVID-19 Genomics UK (COG-UK) Consortium | Lighthouse Laboratory Plymouth and Alex Alderton, Roberto Amato, Jeffrey Barrett, Sonia Goncalves, Ewan Harrison, David K. Jackson, Ian Johnston, Dominic Kwiatkowski, Cordelia Langford, John Sillitoe on behalf of the Wellcome Sanger Institute COVID-19 Surveillance Team |
| EPI_ISL_3367690, EPI_ISL_3367691 | Lighthouse Lab in Glasgow | Wellcome Sanger Institute for the COVID-19 Genomics UK (COG-UK) Consortium | Harper VanSteenhouse, Yumi Kasai, David Gray, Carol Clugston, Anna Dominiczak and Alex Alderton, Roberto Amato, Jeffrey Barrett, Sonia Goncalves, Ewan Harrison, David K. Jackson, Ian Johnston, Dominic Kwiatkowski, Cordelia Langford, John Sillitoe on behalf of the Wellcome Sanger Institute COVID-19 Surveillance Team |
| EPI_ISL_3367692, EPI_ISL_3367693, EPI_ISL_3367694 | Lighthouse Laboratory Plymouth | Wellcome Sanger Institute for the COVID-19 Genomics UK (COG-UK) Consortium | Lighthouse Laboratory Plymouth and Alex Alderton, Roberto Amato, Jeffrey Barrett, Sonia Goncalves, Ewan Harrison, David K. Jackson, Ian Johnston, Dominic Kwiatkowski, Cordelia Langford, John Sillitoe on behalf of the Wellcome Sanger Institute COVID-19 Surveillance Team |
| EPI_ISL_3367695 | Lighthouse Lab in Glasgow | Wellcome Sanger Institute for the COVID-19 Genomics UK (COG-UK) Consortium | Harper VanSteenhouse, Yumi Kasai, David Gray, Carol Clugston, Anna Dominiczak and Alex Alderton, Roberto Amato, Jeffrey Barrett, Sonia Goncalves, Ewan Harrison, David K. Jackson, Ian Johnston, Dominic Kwiatkowski, Cordelia Langford, John Sillitoe on behalf of the Wellcome Sanger Institute COVID-19 Surveillance Team |
| EPI_ISL_3367696, EPI_ISL_3367697 | Lighthouse Laboratory Plymouth | Wellcome Sanger Institute for the COVID-19 Genomics UK (COG-UK) Consortium | Lighthouse Laboratory Plymouth and Alex Alderton, Roberto Amato, Jeffrey Barrett, Sonia Goncalves, Ewan Harrison, David K. Jackson, Ian Johnston, Dominic Kwiatkowski, Cordelia Langford, John Sillitoe on behalf of the Wellcome Sanger Institute COVID-19 Surveillance Team |
| EPI_ISL_3367698, EPI_ISL_3367699, EPI_ISL_3367700, EPI_ISL_3367701, EPI_ISL_3367702, EPI_ISL_3367703, EPI_ISL_3367704 | Lighthouse Lab in Glasgow | Wellcome Sanger Institute for the COVID-19 Genomics UK (COG-UK) Consortium | Harper VanSteenhouse, Yumi Kasai, David Gray, Carol Clugston, Anna Dominiczak and Alex Alderton, Roberto Amato, Jeffrey Barrett, Sonia Goncalves, Ewan Harrison, David K. Jackson, Ian Johnston, Dominic Kwiatkowski, Cordelia Langford, John Sillitoe on behalf of the Wellcome Sanger Institute COVID-19 Surveillance Team |
| EPI_ISL_3367705 | Lighthouse Laboratory Plymouth | Wellcome Sanger Institute for the COVID-19 Genomics UK (COG-UK) Consortium | Lighthouse Laboratory Plymouth and Alex Alderton, Roberto Amato, Jeffrey Barrett, Sonia Goncalves, Ewan Harrison, David K. Jackson, Ian Johnston, Dominic Kwiatkowski, Cordelia Langford, John Sillitoe on behalf of the Wellcome Sanger Institute COVID-19 Surveillance Team |
| EPI_ISL_3367706 | Lighthouse Lab in Glasgow | Wellcome Sanger Institute for the COVID-19 Genomics UK (COG-UK) Consortium | Harper VanSteenhouse, Yumi Kasai, David Gray, Carol Clugston, Anna Dominiczak and Alex Alderton, Roberto Amato, Jeffrey Barrett, Sonia Goncalves, Ewan Harrison, David K. Jackson, Ian Johnston, Dominic Kwiatkowski, Cordelia Langford, John Sillitoe on behalf of the Wellcome Sanger Institute COVID-19 Surveillance Team |
| EPI_ISL_3367707, EPI_ISL_3367708 | Lighthouse Laboratory Plymouth | Wellcome Sanger Institute for the COVID-19 Genomics UK (COG-UK) Consortium | Lighthouse Laboratory Plymouth and Alex Alderton, Roberto Amato, Jeffrey Barrett, Sonia Goncalves, Ewan Harrison, David K. Jackson, Ian Johnston, Dominic Kwiatkowski, Cordelia Langford, John Sillitoe on behalf of the Wellcome Sanger Institute COVID-19 Surveillance Team |
| EPI_ISL_3367709, EPI_ISL_3367710, EPI_ISL_3367711, EPI_ISL_3367712 | Lighthouse Lab in Glasgow | Wellcome Sanger Institute for the COVID-19 Genomics UK (COG-UK) Consortium | Harper VanSteenhouse, Yumi Kasai, David Gray, Carol Clugston, Anna Dominiczak and Alex Alderton, Roberto Amato, Jeffrey Barrett, Sonia Goncalves, Ewan Harrison, David K. Jackson, Ian Johnston, Dominic Kwiatkowski, Cordelia Langford, John Sillitoe on behalf of the Wellcome Sanger Institute COVID-19 Surveillance Team |
| EPI_ISL_3367713, EPI_ISL_3367714, EPI_ISL_3367715, EPI_ISL_3367716 | Lighthouse Laboratory Plymouth | Wellcome Sanger Institute for the COVID-19 Genomics UK (COG-UK) Consortium | Lighthouse Laboratory Plymouth and Alex Alderton, Roberto Amato, Jeffrey Barrett, Sonia Goncalves, Ewan Harrison, David K. Jackson, Ian Johnston, Dominic Kwiatkowski, Cordelia Langford, John Sillitoe on behalf of the Wellcome Sanger Institute COVID-19 Surveillance Team |
| EPI_ISL_3367717, EPI_ISL_3367718, EPI_ISL_3367719, EPI_ISL_3367720, EPI_ISL_3367721, EPI_ISL_3367722, EPI_ISL_3367723, EPI_ISL_3367724, EPI_ISL_3367725, EPI_ISL_3367726, EPI_ISL_3367727, EPI_ISL_3367728 | | | |
| see above | Lighthouse Lab in Glasgow | Wellcome Sanger Institute for the COVID-19 Genomics UK (COG-UK) Consortium | Harper VanSteenhouse, Yumi Kasai, David Gray, Carol Clugston, Anna Dominiczak and Alex Alderton, Roberto Amato, Jeffrey Barrett, Sonia Goncalves, Ewan Harrison, David K. Jackson, Ian Johnston, Dominic Kwiatkowski, Cordelia Langford, John Sillitoe on behalf of the Wellcome Sanger Institute COVID-19 Surveillance Team |
| EPI_ISL_3367729 | Lighthouse Laboratory Plymouth | Wellcome Sanger Institute for the COVID-19 Genomics UK (COG-UK) Consortium | Lighthouse Laboratory Plymouth and Alex Alderton, Roberto Amato, Jeffrey Barrett, Sonia Goncalves, Ewan Harrison, David K. Jackson, Ian Johnston, Dominic Kwiatkowski, Cordelia Langford, John Sillitoe on behalf of the Wellcome Sanger Institute COVID-19 Surveillance Team |
| EPI_ISL_3367730 | Lighthouse Lab in Glasgow | Wellcome Sanger Institute for the COVID-19 Genomics UK (COG-UK) Consortium | Harper VanSteenhouse, Yumi Kasai, David Gray, Carol Clugston, Anna Dominiczak and Alex Alderton, Roberto Amato, Jeffrey Barrett, Sonia Goncalves, Ewan Harrison, David K. Jackson, Ian Johnston, Dominic Kwiatkowski, Cordelia Langford, John Sillitoe on behalf of the Wellcome Sanger Institute COVID-19 Surveillance Team |
| EPI_ISL_3367731 | Lighthouse Laboratory Plymouth | Wellcome Sanger Institute for the COVID-19 Genomics UK (COG-UK) Consortium | Lighthouse Laboratory Plymouth and Alex Alderton, Roberto Amato, Jeffrey Barrett, Sonia Goncalves, Ewan Harrison, David K. Jackson, Ian Johnston, Dominic Kwiatkowski, Cordelia Langford, John Sillitoe on behalf of the Wellcome Sanger Institute COVID-19 Surveillance Team |
| EPI_ISL_3367732, EPI_ISL_3367733, EPI_ISL_3367734, EPI_ISL_3367735, EPI_ISL_3367736, EPI_ISL_3367737, EPI_ISL_3367738, EPI_ISL_3367739, EPI_ISL_3367740, EPI_ISL_3367741, EPI_ISL_3367742, EPI_ISL_3367743, EPI_ISL_3367744, EPI_ISL_3367745, EPI_ISL_3367746, EPI_ISL_3367747, EPI_ISL_3367748, EPI_ISL_3367749, EPI_ISL_3367750, EPI_ISL_3367751, EPI_ISL_3367752, EPI_ISL_3367753, EPI_ISL_3367754, EPI_ISL_3367755, EPI_ISL_3367756, EPI_ISL_3367757, EPI_ISL_3367758, EPI_ISL_3367759, EPI_ISL_3367760, EPI_ISL_3367761, EPI_ISL_3367762, EPI_ISL_3367763, EPI_ISL_3367764, EPI_ISL_3367765, EPI_ISL_3367766, EPI_ISL_3367768, EPI_ISL_3367769, EPI_ISL_3367770, EPI_ISL_3367771, EPI_ISL_3367772, EPI_ISL_3367773, EPI_ISL_3367774, EPI_ISL_3367775, EPI_ISL_3367776, EPI_ISL_3367777, EPI_ISL_3367778, EPI_ISL_3367779, EPI_ISL_3367780, EPI_ISL_3367781, EPI_ISL_3367782, EPI_ISL_3367783, EPI_ISL_3367784 | | | |
| see above | Lighthouse Lab in Glasgow | Wellcome Sanger Institute for the COVID-19 Genomics UK (COG-UK) Consortium | Harper VanSteenhouse, Yumi Kasai, David Gray, Carol Clugston, Anna Dominiczak and Alex Alderton, Roberto Amato, Jeffrey Barrett, Sonia Goncalves, Ewan Harrison, David K. Jackson, Ian Johnston, Dominic Kwiatkowski, Cordelia Langford, John Sillitoe on behalf of the Wellcome Sanger Institute COVID-19 Surveillance Team |
| EPI_ISL_3367785, EPI_ISL_3367786 | Lighthouse Lab in Alderley Park | Wellcome Sanger Institute for the COVID-19 Genomics UK (COG-UK) Consortium | Jacquelyn Wynn, Mairead Hyland, The Lighthouse Lab in Alderley Park and Alex Alderton, Roberto Amato, Jeffrey Barrett, Sonia Goncalves, Ewan Harrison, David K. Jackson, Ian Johnston, Dominic Kwiatkowski, Cordelia Langford, John Sillitoe on behalf of the Wellcome Sanger Institute COVID-19 Surveillance Team |

| | | | |
|---|---|---|---|
| EPI_ISL_3367787 | Lighthouse Lab in Glasgow | Wellcome Sanger Institute for the COVID-19 Genomics UK (COG-UK) Consortium | Harper VanSteenhouse, Yumi Kasai, David Gray, Carol Clugston, Anna Dominiczak and Alex Alderton, Roberto Amato, Jeffrey Barrett, Sonia Goncalves, Ewan Harrison, David K. Jackson, Ian Johnston, Dominic Kwiatkowski, Cordelia Langford, John Sillitoe on behalf of the Wellcome Sanger Institute COVID-19 Surveillance Team |
| EPI_ISL_3367788, EPI_ISL_3367789 | Berkshire and Surrey Pathology Services Lighthouse Laboratory | Wellcome Sanger Institute for the COVID-19 Genomics UK (COG-UK) Consortium | Berkshire and Surrey Pathology Services Lighthouse Laboratory and Alex Alderton, Roberto Amato, Jeffrey Barrett, Sonia Goncalves, Ewan Harrison, David K. Jackson, Ian Johnston, Dominic Kwiatkowski, Cordelia Langford, John Sillitoe on behalf of the Wellcome Sanger Institute COVID-19 Surveillance Team |
| EPI_ISL_3367790, EPI_ISL_3367791 | Lighthouse Lab in Milton Keynes | Wellcome Sanger Institute for the COVID-19 Genomics UK (COG-UK) Consortium | The Lighthouse Lab in Milton Keynes and Alex Alderton, Roberto Amato, Jeffrey Barrett, Sonia Goncalves, Ewan Harrison, David K. Jackson, Ian Johnston, Dominic Kwiatkowski, Cordelia Langford, John Sillitoe on behalf of the Wellcome Sanger Institute COVID-19 Surveillance Team |
| EPI_ISL_3367792 | Berkshire and Surrey Pathology Services Lighthouse Laboratory | Wellcome Sanger Institute for the COVID-19 Genomics UK (COG-UK) Consortium | Berkshire and Surrey Pathology Services Lighthouse Laboratory and Alex Alderton, Roberto Amato, Jeffrey Barrett, Sonia Goncalves, Ewan Harrison, David K. Jackson, Ian Johnston, Dominic Kwiatkowski, Cordelia Langford, John Sillitoe on behalf of the Wellcome Sanger Institute COVID-19 Surveillance Team |
| EPI_ISL_3367793 | Lighthouse Laboratory Plymouth | Wellcome Sanger Institute for the COVID-19 Genomics UK (COG-UK) Consortium | Lighthouse Laboratory Plymouth and Alex Alderton, Roberto Amato, Jeffrey Barrett, Sonia Goncalves, Ewan Harrison, David K. Jackson, Ian Johnston, Dominic Kwiatkowski, Cordelia Langford, John Sillitoe on behalf of the Wellcome Sanger Institute COVID-19 Surveillance Team |
| EPI_ISL_3367794 | Berkshire and Surrey Pathology Services Lighthouse Laboratory | Wellcome Sanger Institute for the COVID-19 Genomics UK (COG-UK) Consortium | Berkshire and Surrey Pathology Services Lighthouse Laboratory and Alex Alderton, Roberto Amato, Jeffrey Barrett, Sonia Goncalves, Ewan Harrison, David K. Jackson, Ian Johnston, Dominic Kwiatkowski, Cordelia Langford, John Sillitoe on behalf of the Wellcome Sanger Institute COVID-19 Surveillance Team |
| EPI_ISL_3367795 | Lighthouse Laboratory Plymouth | Wellcome Sanger Institute for the COVID-19 Genomics UK (COG-UK) Consortium | Lighthouse Laboratory Plymouth and Alex Alderton, Roberto Amato, Jeffrey Barrett, Sonia Goncalves, Ewan Harrison, David K. Jackson, Ian Johnston, Dominic Kwiatkowski, Cordelia Langford, John Sillitoe on behalf of the Wellcome Sanger Institute COVID-19 Surveillance Team |
| EPI_ISL_3367796 | Berkshire and Surrey Pathology Services Lighthouse Laboratory | Wellcome Sanger Institute for the COVID-19 Genomics UK (COG-UK) Consortium | Berkshire and Surrey Pathology Services Lighthouse Laboratory and Alex Alderton, Roberto Amato, Jeffrey Barrett, Sonia Goncalves, Ewan Harrison, David K. Jackson, Ian Johnston, Dominic Kwiatkowski, Cordelia Langford, John Sillitoe on behalf of the Wellcome Sanger Institute COVID-19 Surveillance Team |
| EPI_ISL_3367797 | Lighthouse Lab in Milton Keynes | Wellcome Sanger Institute for the COVID-19 Genomics UK (COG-UK) Consortium | The Lighthouse Lab in Milton Keynes and Alex Alderton, Roberto Amato, Jeffrey Barrett, Sonia Goncalves, Ewan Harrison, David K. Jackson, Ian Johnston, Dominic Kwiatkowski, Cordelia Langford, John Sillitoe on behalf of the Wellcome Sanger Institute COVID-19 Surveillance Team |
| EPI_ISL_3367798 | Berkshire and Surrey Pathology Services Lighthouse Laboratory | Wellcome Sanger Institute for the COVID-19 Genomics UK (COG-UK) Consortium | Berkshire and Surrey Pathology Services Lighthouse Laboratory and Alex Alderton, Roberto Amato, Jeffrey Barrett, Sonia Goncalves, Ewan Harrison, David K. Jackson, Ian Johnston, Dominic Kwiatkowski, Cordelia Langford, John Sillitoe on behalf of the Wellcome Sanger Institute COVID-19 Surveillance Team |
| EPI_ISL_3367799 | Lighthouse Lab in Glasgow | Wellcome Sanger Institute for the COVID-19 Genomics UK (COG-UK) Consortium | Harper VanSteenhouse, Yumi Kasai, David Gray, Carol Clugston, Anna Dominiczak and Alex Alderton, Roberto Amato, Jeffrey Barrett, Sonia Goncalves, Ewan Harrison, David K. Jackson, Ian Johnston, Dominic Kwiatkowski, Cordelia Langford, John Sillitoe on behalf of the Wellcome Sanger Institute COVID-19 Surveillance Team |
| EPI_ISL_3367800, EPI_ISL_3367802 | Lighthouse Lab in Milton Keynes | Wellcome Sanger Institute for the COVID-19 Genomics UK (COG-UK) Consortium | The Lighthouse Lab in Milton Keynes and Alex Alderton, Roberto Amato, Jeffrey Barrett, Sonia Goncalves, Ewan Harrison, David K. Jackson, Ian Johnston, Dominic Kwiatkowski, Cordelia Langford, John Sillitoe on behalf of the Wellcome Sanger Institute COVID-19 Surveillance Team |
| EPI_ISL_3367803 | Lighthouse Lab in Glasgow | Wellcome Sanger Institute for the COVID-19 Genomics UK (COG-UK) Consortium | Harper VanSteenhouse, Yumi Kasai, David Gray, Carol Clugston, Anna Dominiczak and Alex Alderton, Roberto Amato, Jeffrey Barrett, Sonia Goncalves, Ewan Harrison, David K. Jackson, Ian Johnston, Dominic Kwiatkowski, Cordelia Langford, John Sillitoe on behalf of the Wellcome Sanger Institute COVID-19 Surveillance Team |
| EPI_ISL_3367804 | Lighthouse Lab in Milton Keynes | Wellcome Sanger Institute for the COVID-19 Genomics UK (COG-UK) Consortium | The Lighthouse Lab in Milton Keynes and Alex Alderton, Roberto Amato, Jeffrey Barrett, Sonia Goncalves, Ewan Harrison, David K. Jackson, Ian Johnston, Dominic Kwiatkowski, Cordelia Langford, John Sillitoe on behalf of the Wellcome Sanger Institute COVID-19 Surveillance Team |
| EPI_ISL_3367805 | Berkshire and Surrey Pathology Services Lighthouse Laboratory | Wellcome Sanger Institute for the COVID-19 Genomics UK (COG-UK) Consortium | Berkshire and Surrey Pathology Services Lighthouse Laboratory and Alex Alderton, Roberto Amato, Jeffrey Barrett, Sonia Goncalves, Ewan Harrison, David K. Jackson, Ian Johnston, Dominic Kwiatkowski, Cordelia Langford, John Sillitoe on behalf of the Wellcome Sanger Institute COVID-19 Surveillance Team |
| EPI_ISL_3367806 | Lighthouse Laboratory Plymouth | Wellcome Sanger Institute for the COVID-19 Genomics UK (COG-UK) Consortium | Lighthouse Laboratory Plymouth and Alex Alderton, Roberto Amato, Jeffrey Barrett, Sonia Goncalves, Ewan Harrison, David K. Jackson, Ian Johnston, Dominic Kwiatkowski, Cordelia Langford, John Sillitoe on behalf of the Wellcome Sanger Institute COVID-19 Surveillance Team |
| EPI_ISL_3367807 | Berkshire and Surrey Pathology Services Lighthouse Laboratory | Wellcome Sanger Institute for the COVID-19 Genomics UK (COG-UK) Consortium | Berkshire and Surrey Pathology Services Lighthouse Laboratory and Alex Alderton, Roberto Amato, Jeffrey Barrett, Sonia Goncalves, Ewan Harrison, David K. Jackson, Ian Johnston, Dominic Kwiatkowski, Cordelia Langford, John Sillitoe on behalf of the Wellcome Sanger Institute COVID-19 Surveillance Team |
| EPI_ISL_3367808 | Lighthouse Lab in Milton Keynes | Wellcome Sanger Institute for the COVID-19 Genomics UK (COG-UK) Consortium | The Lighthouse Lab in Milton Keynes and Alex Alderton, Roberto Amato, Jeffrey Barrett, Sonia Goncalves, Ewan Harrison, David K. Jackson, Ian Johnston, Dominic Kwiatkowski, Cordelia Langford, John Sillitoe on behalf of the Wellcome Sanger Institute COVID-19 Surveillance Team |
| EPI_ISL_3367809 | Berkshire and Surrey Pathology Services Lighthouse Laboratory | Wellcome Sanger Institute for the COVID-19 Genomics UK (COG-UK) Consortium | Berkshire and Surrey Pathology Services Lighthouse Laboratory and Alex Alderton, Roberto Amato, Jeffrey Barrett, Sonia Goncalves, Ewan Harrison, David K. Jackson, Ian Johnston, Dominic Kwiatkowski, Cordelia Langford, John Sillitoe on behalf of the Wellcome Sanger Institute COVID-19 Surveillance Team |
| EPI_ISL_3367810 | Lighthouse Lab in Glasgow | Wellcome Sanger Institute for the COVID-19 Genomics UK (COG-UK) Consortium | Harper VanSteenhouse, Yumi Kasai, David Gray, Carol Clugston, Anna Dominiczak and Alex Alderton, Roberto Amato, Jeffrey Barrett, Sonia Goncalves, Ewan Harrison, David K. Jackson, Ian Johnston, Dominic Kwiatkowski, Cordelia Langford, John Sillitoe on behalf of the Wellcome Sanger Institute COVID-19 Surveillance Team |
| EPI_ISL_3367811 | Lighthouse Laboratory Plymouth | Wellcome Sanger Institute for the COVID-19 Genomics UK (COG-UK) Consortium | Lighthouse Laboratory Plymouth and Alex Alderton, Roberto Amato, Jeffrey Barrett, Sonia Goncalves, Ewan Harrison, David K. Jackson, Ian Johnston, Dominic Kwiatkowski, Cordelia Langford, John Sillitoe on behalf of the Wellcome Sanger Institute COVID-19 Surveillance Team |
| EPI_ISL_3367812 | Lighthouse Lab in Milton Keynes | Wellcome Sanger Institute for the COVID-19 Genomics UK (COG-UK) Consortium | The Lighthouse Lab in Milton Keynes and Alex Alderton, Roberto Amato, Jeffrey Barrett, Sonia Goncalves, Ewan Harrison, David K. Jackson, Ian Johnston, Dominic Kwiatkowski, Cordelia Langford, John Sillitoe on behalf of the Wellcome Sanger Institute COVID-19 Surveillance Team |
| EPI_ISL_3367813, EPI_ISL_3367814, EPI_ISL_3367815, EPI_ISL_3367816 | Lighthouse Laboratory Plymouth | Wellcome Sanger Institute for the COVID-19 Genomics UK (COG-UK) Consortium | Lighthouse Laboratory Plymouth and Alex Alderton, Roberto Amato, Jeffrey Barrett, Sonia Goncalves, Ewan Harrison, David K. Jackson, Ian Johnston, Dominic Kwiatkowski, Cordelia Langford, John Sillitoe on behalf of the Wellcome Sanger Institute COVID-19 Surveillance Team |
| EPI_ISL_3367817, EPI_ISL_3367818 | Lighthouse Lab in Milton Keynes | Wellcome Sanger Institute for the COVID-19 Genomics UK (COG-UK) Consortium | The Lighthouse Lab in Milton Keynes and Alex Alderton, Roberto Amato, Jeffrey Barrett, Sonia Goncalves, Ewan Harrison, David K. Jackson, Ian Johnston, Dominic Kwiatkowski, Cordelia Langford, John Sillitoe on behalf of the Wellcome Sanger Institute COVID-19 Surveillance Team |
| EPI_ISL_3367819, EPI_ISL_3367820, EPI_ISL_3367821 | Lighthouse Lab in Glasgow | Wellcome Sanger Institute for the COVID-19 Genomics UK (COG-UK) Consortium | Harper VanSteenhouse, Yumi Kasai, David Gray, Carol Clugston, Anna Dominiczak and Alex Alderton, Roberto Amato, Jeffrey Barrett, Sonia Goncalves, Ewan Harrison, David K. Jackson, Ian Johnston, Dominic Kwiatkowski, Cordelia Langford, John Sillitoe on behalf of the Wellcome Sanger Institute COVID-19 Surveillance Team |
| EPI_ISL_3367822, EPI_ISL_3367823 | Lighthouse Laboratory Plymouth | Wellcome Sanger Institute for the COVID-19 Genomics UK (COG-UK) Consortium | Lighthouse Laboratory Plymouth and Alex Alderton, Roberto Amato, Jeffrey Barrett, Sonia Goncalves, Ewan Harrison, David K. Jackson, Ian Johnston, Dominic Kwiatkowski, Cordelia Langford, John Sillitoe on behalf of the Wellcome Sanger Institute COVID-19 Surveillance Team |
| EPI_ISL_3367824 | Lighthouse Lab in Milton Keynes | Wellcome Sanger Institute for the COVID-19 Genomics UK (COG-UK) Consortium | The Lighthouse Lab in Milton Keynes and Alex Alderton, Roberto Amato, Jeffrey Barrett, Sonia Goncalves, Ewan Harrison, David K. Jackson, Ian Johnston, Dominic Kwiatkowski, Cordelia Langford, John Sillitoe on behalf of the Wellcome Sanger Institute COVID-19 Surveillance Team |
| EPI_ISL_3367825 | Berkshire and Surrey Pathology Services Lighthouse Laboratory | Wellcome Sanger Institute for the COVID-19 Genomics UK (COG-UK) Consortium | Berkshire and Surrey Pathology Services Lighthouse Laboratory and Alex Alderton, Roberto Amato, Jeffrey Barrett, Sonia Goncalves, Ewan Harrison, David K. Jackson, Ian Johnston, Dominic Kwiatkowski, Cordelia Langford, John Sillitoe on behalf of the Wellcome Sanger Institute COVID-19 Surveillance Team |
| EPI_ISL_3367826, EPI_ISL_3367827 | Lighthouse Laboratory Plymouth | Wellcome Sanger Institute for the COVID-19 Genomics UK (COG-UK) Consortium | Lighthouse Laboratory Plymouth and Alex Alderton, Roberto Amato, Jeffrey Barrett, Sonia Goncalves, Ewan Harrison, David K. Jackson, Ian Johnston, Dominic Kwiatkowski, Cordelia Langford, John Sillitoe on behalf of the Wellcome Sanger Institute COVID-19 Surveillance Team |
| EPI_ISL_3367828, EPI_ISL_3367829 | Lighthouse Lab in Milton Keynes | Wellcome Sanger Institute for the COVID-19 Genomics UK (COG-UK) Consortium | The Lighthouse Lab in Milton Keynes and Alex Alderton, Roberto Amato, Jeffrey Barrett, Sonia Goncalves, Ewan Harrison, David K. Jackson, Ian Johnston, Dominic Kwiatkowski, Cordelia Langford, John Sillitoe on behalf of the Wellcome Sanger Institute COVID-19 Surveillance Team |
| EPI_ISL_3367830 | Lighthouse Laboratory Plymouth | Wellcome Sanger Institute for the COVID-19 Genomics UK (COG-UK) Consortium | Lighthouse Laboratory Plymouth and Alex Alderton, Roberto Amato, Jeffrey Barrett, Sonia Goncalves, Ewan Harrison, David K. Jackson, Ian Johnston, Dominic Kwiatkowski, Cordelia Langford, John Sillitoe on behalf of the Wellcome Sanger Institute COVID-19 Surveillance Team |
| EPI_ISL_3367831 | Lighthouse Lab in Glasgow | Wellcome Sanger Institute for the COVID-19 Genomics UK (COG-UK) Consortium | Harper VanSteenhouse, Yumi Kasai, David Gray, Carol Clugston, Anna Dominiczak and Alex Alderton, Roberto Amato, Jeffrey Barrett, Sonia Goncalves, Ewan Harrison, David K. Jackson, Ian Johnston, Dominic Kwiatkowski, Cordelia Langford, John Sillitoe on behalf of the Wellcome Sanger Institute COVID-19 Surveillance Team |
| EPI_ISL_3367832 | Lighthouse Laboratory Plymouth | Wellcome Sanger Institute for the COVID-19 Genomics UK (COG-UK) Consortium | Lighthouse Laboratory Plymouth and Alex Alderton, Roberto Amato, Jeffrey Barrett, Sonia Goncalves, Ewan Harrison, David K. Jackson, Ian Johnston, Dominic Kwiatkowski, Cordelia Langford, John Sillitoe on behalf of the Wellcome Sanger Institute COVID-19 Surveillance Team |
| EPI_ISL_3367833 | Lighthouse Lab in Glasgow | Wellcome Sanger Institute for the COVID-19 Genomics UK (COG-UK) Consortium | Harper VanSteenhouse, Yumi Kasai, David Gray, Carol Clugston, Anna Dominiczak and Alex Alderton, Roberto Amato, Jeffrey Barrett, Sonia Goncalves, Ewan Harrison, David K. Jackson, Ian Johnston, Dominic Kwiatkowski, Cordelia Langford, John Sillitoe on behalf of the Wellcome Sanger Institute COVID-19 Surveillance Team |
| EPI_ISL_3367834 | Berkshire and Surrey Pathology Services Lighthouse Laboratory | Wellcome Sanger Institute for the COVID-19 Genomics UK (COG-UK) Consortium | Berkshire and Surrey Pathology Services Lighthouse Laboratory and Alex Alderton, Roberto Amato, Jeffrey Barrett, Sonia Goncalves, Ewan Harrison, David K. Jackson, Ian Johnston, Dominic Kwiatkowski, Cordelia Langford, John Sillitoe on behalf of the Wellcome Sanger Institute COVID-19 Surveillance Team |
| EPI_ISL_3367835 | Lighthouse Laboratory Plymouth | Wellcome Sanger Institute for the COVID-19 Genomics UK (COG-UK) Consortium | Lighthouse Laboratory Plymouth and Alex Alderton, Roberto Amato, Jeffrey Barrett, Sonia Goncalves, Ewan Harrison, David K. Jackson, Ian Johnston, Dominic Kwiatkowski, Cordelia Langford, John Sillitoe on behalf of the Wellcome Sanger Institute COVID-19 Surveillance Team |
| EPI_ISL_3367836 | Berkshire and Surrey Pathology Services Lighthouse Laboratory | Wellcome Sanger Institute for the COVID-19 Genomics UK (COG-UK) Consortium | Berkshire and Surrey Pathology Services Lighthouse Laboratory and Alex Alderton, Roberto Amato, Jeffrey Barrett, Sonia Goncalves, Ewan Harrison, David K. Jackson, Ian Johnston, Dominic Kwiatkowski, Cordelia Langford, John Sillitoe on behalf of the Wellcome Sanger Institute COVID-19 Surveillance Team |
| EPI_ISL_3367837 | Lighthouse Laboratory Plymouth | Wellcome Sanger Institute for the COVID-19 Genomics UK (COG-UK) Consortium | Lighthouse Laboratory Plymouth and Alex Alderton, Roberto Amato, Jeffrey Barrett, Sonia Goncalves, Ewan Harrison, David K. Jackson, Ian Johnston, Dominic Kwiatkowski, Cordelia Langford, John Sillitoe on behalf of the Wellcome Sanger Institute COVID-19 Surveillance Team |
| EPI_ISL_3367838 | Lighthouse Lab in Glasgow | Wellcome Sanger Institute for the COVID-19 Genomics UK (COG-UK) Consortium | Harper VanSteenhouse, Yumi Kasai, David Gray, Carol Clugston, Anna Dominiczak and Alex Alderton, Roberto Amato, Jeffrey Barrett, Sonia Goncalves, Ewan Harrison, David K. Jackson, Ian Johnston, Dominic Kwiatkowski, Cordelia Langford, John Sillitoe on behalf of the Wellcome Sanger Institute COVID-19 Surveillance Team |
| EPI_ISL_3367839 | Lighthouse Lab in Milton Keynes | Wellcome Sanger Institute for the COVID-19 Genomics UK (COG-UK) Consortium | The Lighthouse Lab in Milton Keynes and Alex Alderton, Roberto Amato, Jeffrey Barrett, Sonia Goncalves, Ewan Harrison, David K. Jackson, Ian Johnston, Dominic Kwiatkowski, Cordelia Langford, John Sillitoe on behalf of the Wellcome Sanger Institute COVID-19 Surveillance Team |
| EPI_ISL_3367840 | Lighthouse Lab in Glasgow | Wellcome Sanger Institute for the COVID-19 Genomics UK (COG-UK) Consortium | Harper VanSteenhouse, Yumi Kasai, David Gray, Carol Clugston, Anna Dominiczak and Alex Alderton, Roberto Amato, Jeffrey Barrett, Sonia Goncalves, Ewan Harrison, David K. Jackson, Ian Johnston, Dominic Kwiatkowski, Cordelia Langford, John Sillitoe on behalf of the Wellcome Sanger Institute COVID-19 Surveillance Team |
| EPI_ISL_3367841 | Lighthouse Lab in Milton Keynes | Wellcome Sanger Institute for the COVID-19 Genomics UK (COG-UK) Consortium | The Lighthouse Lab in Milton Keynes and Alex Alderton, Roberto Amato, Jeffrey Barrett, Sonia Goncalves, Ewan Harrison, David K. Jackson, Ian Johnston, Dominic Kwiatkowski, Cordelia Langford, John Sillitoe on behalf of the Wellcome Sanger Institute COVID-19 Surveillance Team |
| EPI_ISL_3367842 | Lighthouse Lab in Glasgow | Wellcome Sanger Institute for the COVID-19 Genomics UK (COG-UK) Consortium | Harper VanSteenhouse, Yumi Kasai, David Gray, Carol Clugston, Anna Dominiczak and Alex Alderton, Roberto Amato, Jeffrey Barrett, Sonia Goncalves, Ewan Harrison, David K. Jackson, Ian Johnston, Dominic Kwiatkowski, Cordelia Langford, John Sillitoe on behalf of the Wellcome Sanger Institute COVID-19 Surveillance Team |
| EPI_ISL_3367843 | Berkshire and Surrey Pathology Services Lighthouse Laboratory | Wellcome Sanger Institute for the COVID-19 Genomics UK (COG-UK) | Berkshire and Surrey Pathology Services Lighthouse Laboratory and Alex Alderton, Roberto Amato, Jeffrey Barrett, Sonia Goncalves, Ewan Harrison, David K. Jackson, Ian Johnston, Dominic Kwiatkowski, Cordelia Langford, John Sillitoe on behalf |

| | | | |
|---|---|---|---|
| | | Consortium | of the Wellcome Sanger Institute COVID-19 Surveillance Team |
| EPI_ISL_3367844, EPI_ISL_3367845 | Lighthouse Laboratory Plymouth | Wellcome Sanger Institute for the COVID-19 Genomics UK (COG-UK) Consortium | Lighthouse Laboratory Plymouth and Alex Alderton, Roberto Amato, Jeffrey Barrett, Sonia Goncalves, Ewan Harrison, David K. Jackson, Ian Johnston, Dominic Kwiatkowski, Cordelia Langford, John Sillitoe on behalf of the Wellcome Sanger Institute COVID-19 Surveillance Team |
| EPI_ISL_3367846 | Lighthouse Lab in Milton Keynes | Wellcome Sanger Institute for the COVID-19 Genomics UK (COG-UK) Consortium | The Lighthouse Lab in Milton Keynes and Alex Alderton, Roberto Amato, Jeffrey Barrett, Sonia Goncalves, Ewan Harrison, David K. Jackson, Ian Johnston, Dominic Kwiatkowski, Cordelia Langford, John Sillitoe on behalf of the Wellcome Sanger Institute COVID-19 Surveillance Team |
| EPI_ISL_3367847 | Lighthouse Lab in Glasgow | Wellcome Sanger Institute for the COVID-19 Genomics UK (COG-UK) Consortium | Harper VanSteenhouse, Yumi Kasai, David Gray, Carol Clugston, Anna Dominiczak and Alex Alderton, Roberto Amato, Jeffrey Barrett, Sonia Goncalves, Ewan Harrison, David K. Jackson, Ian Johnston, Dominic Kwiatkowski, Cordelia Langford, John Sillitoe on behalf of the Wellcome Sanger Institute COVID-19 Surveillance Team |
| EPI_ISL_3367848 | Lighthouse Laboratory Plymouth | Wellcome Sanger Institute for the COVID-19 Genomics UK (COG-UK) Consortium | Lighthouse Laboratory Plymouth and Alex Alderton, Roberto Amato, Jeffrey Barrett, Sonia Goncalves, Ewan Harrison, David K. Jackson, Ian Johnston, Dominic Kwiatkowski, Cordelia Langford, John Sillitoe on behalf of the Wellcome Sanger Institute COVID-19 Surveillance Team |
| EPI_ISL_3367849 | Lighthouse Lab in Milton Keynes | Wellcome Sanger Institute for the COVID-19 Genomics UK (COG-UK) Consortium | The Lighthouse Lab in Milton Keynes and Alex Alderton, Roberto Amato, Jeffrey Barrett, Sonia Goncalves, Ewan Harrison, David K. Jackson, Ian Johnston, Dominic Kwiatkowski, Cordelia Langford, John Sillitoe on behalf of the Wellcome Sanger Institute COVID-19 Surveillance Team |
| EPI_ISL_3367850 | Lighthouse Laboratory Plymouth | Wellcome Sanger Institute for the COVID-19 Genomics UK (COG-UK) Consortium | Lighthouse Laboratory Plymouth and Alex Alderton, Roberto Amato, Jeffrey Barrett, Sonia Goncalves, Ewan Harrison, David K. Jackson, Ian Johnston, Dominic Kwiatkowski, Cordelia Langford, John Sillitoe on behalf of the Wellcome Sanger Institute COVID-19 Surveillance Team |
| EPI_ISL_3367851 | Lighthouse Lab in Milton Keynes | Wellcome Sanger Institute for the COVID-19 Genomics UK (COG-UK) Consortium | The Lighthouse Lab in Milton Keynes and Alex Alderton, Roberto Amato, Jeffrey Barrett, Sonia Goncalves, Ewan Harrison, David K. Jackson, Ian Johnston, Dominic Kwiatkowski, Cordelia Langford, John Sillitoe on behalf of the Wellcome Sanger Institute COVID-19 Surveillance Team |
| EPI_ISL_3367852 | Berkshire and Surrey Pathology Services Lighthouse Laboratory | Wellcome Sanger Institute for the COVID-19 Genomics UK (COG-UK) Consortium | Berkshire and Surrey Pathology Services Lighthouse Laboratory and Alex Alderton, Roberto Amato, Jeffrey Barrett, Sonia Goncalves, Ewan Harrison, David K. Jackson, Ian Johnston, Dominic Kwiatkowski, Cordelia Langford, John Sillitoe on behalf of the Wellcome Sanger Institute COVID-19 Surveillance Team |
| EPI_ISL_3367853 | Lighthouse Lab in Glasgow | Wellcome Sanger Institute for the COVID-19 Genomics UK (COG-UK) Consortium | Harper VanSteenhouse, Yumi Kasai, David Gray, Carol Clugston, Anna Dominiczak and Alex Alderton, Roberto Amato, Jeffrey Barrett, Sonia Goncalves, Ewan Harrison, David K. Jackson, Ian Johnston, Dominic Kwiatkowski, Cordelia Langford, John Sillitoe on behalf of the Wellcome Sanger Institute COVID-19 Surveillance Team |
| EPI_ISL_3367854 | Lighthouse Lab in Milton Keynes | Wellcome Sanger Institute for the COVID-19 Genomics UK (COG-UK) Consortium | The Lighthouse Lab in Milton Keynes and Alex Alderton, Roberto Amato, Jeffrey Barrett, Sonia Goncalves, Ewan Harrison, David K. Jackson, Ian Johnston, Dominic Kwiatkowski, Cordelia Langford, John Sillitoe on behalf of the Wellcome Sanger Institute COVID-19 Surveillance Team |
| EPI_ISL_3367855 | Berkshire and Surrey Pathology Services Lighthouse Laboratory | Wellcome Sanger Institute for the COVID-19 Genomics UK (COG-UK) Consortium | Berkshire and Surrey Pathology Services Lighthouse Laboratory and Alex Alderton, Roberto Amato, Jeffrey Barrett, Sonia Goncalves, Ewan Harrison, David K. Jackson, Ian Johnston, Dominic Kwiatkowski, Cordelia Langford, John Sillitoe on behalf of the Wellcome Sanger Institute COVID-19 Surveillance Team |
| EPI_ISL_3367856 | Lighthouse Laboratory Plymouth | Wellcome Sanger Institute for the COVID-19 Genomics UK (COG-UK) Consortium | Lighthouse Laboratory Plymouth and Alex Alderton, Roberto Amato, Jeffrey Barrett, Sonia Goncalves, Ewan Harrison, David K. Jackson, Ian Johnston, Dominic Kwiatkowski, Cordelia Langford, John Sillitoe on behalf of the Wellcome Sanger Institute COVID-19 Surveillance Team |
| EPI_ISL_3367857 | Berkshire and Surrey Pathology Services Lighthouse Laboratory | Wellcome Sanger Institute for the COVID-19 Genomics UK (COG-UK) Consortium | Berkshire and Surrey Pathology Services Lighthouse Laboratory and Alex Alderton, Roberto Amato, Jeffrey Barrett, Sonia Goncalves, Ewan Harrison, David K. Jackson, Ian Johnston, Dominic Kwiatkowski, Cordelia Langford, John Sillitoe on behalf of the Wellcome Sanger Institute COVID-19 Surveillance Team |
| EPI_ISL_3367858, EPI_ISL_3367859 | Lighthouse Laboratory Plymouth | Wellcome Sanger Institute for the COVID-19 Genomics UK (COG-UK) Consortium | Lighthouse Laboratory Plymouth and Alex Alderton, Roberto Amato, Jeffrey Barrett, Sonia Goncalves, Ewan Harrison, David K. Jackson, Ian Johnston, Dominic Kwiatkowski, Cordelia Langford, John Sillitoe on behalf of the Wellcome Sanger Institute COVID-19 Surveillance Team |
| EPI_ISL_3367860, EPI_ISL_3367861 | Lighthouse Lab in Glasgow | Wellcome Sanger Institute for the COVID-19 Genomics UK (COG-UK) Consortium | Harper VanSteenhouse, Yumi Kasai, David Gray, Carol Clugston, Anna Dominiczak and Alex Alderton, Roberto Amato, Jeffrey Barrett, Sonia Goncalves, Ewan Harrison, David K. Jackson, Ian Johnston, Dominic Kwiatkowski, Cordelia Langford, John Sillitoe on behalf of the Wellcome Sanger Institute COVID-19 Surveillance Team |
| EPI_ISL_3367862 | Berkshire and Surrey Pathology Services Lighthouse Laboratory | Wellcome Sanger Institute for the COVID-19 Genomics UK (COG-UK) Consortium | Berkshire and Surrey Pathology Services Lighthouse Laboratory and Alex Alderton, Roberto Amato, Jeffrey Barrett, Sonia Goncalves, Ewan Harrison, David K. Jackson, Ian Johnston, Dominic Kwiatkowski, Cordelia Langford, John Sillitoe on behalf of the Wellcome Sanger Institute COVID-19 Surveillance Team |
| EPI_ISL_3367863 | Lighthouse Lab in Milton Keynes | Wellcome Sanger Institute for the COVID-19 Genomics UK (COG-UK) Consortium | The Lighthouse Lab in Milton Keynes and Alex Alderton, Roberto Amato, Jeffrey Barrett, Sonia Goncalves, Ewan Harrison, David K. Jackson, Ian Johnston, Dominic Kwiatkowski, Cordelia Langford, John Sillitoe on behalf of the Wellcome Sanger Institute COVID-19 Surveillance Team |
| EPI_ISL_3367864, EPI_ISL_3367865 | Berkshire and Surrey Pathology Services Lighthouse Laboratory | Wellcome Sanger Institute for the COVID-19 Genomics UK (COG-UK) Consortium | Berkshire and Surrey Pathology Services Lighthouse Laboratory and Alex Alderton, Roberto Amato, Jeffrey Barrett, Sonia Goncalves, Ewan Harrison, David K. Jackson, Ian Johnston, Dominic Kwiatkowski, Cordelia Langford, John Sillitoe on behalf of the Wellcome Sanger Institute COVID-19 Surveillance Team |
| EPI_ISL_3367866 | Lighthouse Laboratory Plymouth | Wellcome Sanger Institute for the COVID-19 Genomics UK (COG-UK) Consortium | Lighthouse Laboratory Plymouth and Alex Alderton, Roberto Amato, Jeffrey Barrett, Sonia Goncalves, Ewan Harrison, David K. Jackson, Ian Johnston, Dominic Kwiatkowski, Cordelia Langford, John Sillitoe on behalf of the Wellcome Sanger Institute COVID-19 Surveillance Team |
| EPI_ISL_3367867, EPI_ISL_3367869, EPI_ISL_3367870 | Lighthouse Lab in Milton Keynes | Wellcome Sanger Institute for the COVID-19 Genomics UK (COG-UK) Consortium | The Lighthouse Lab in Milton Keynes and Alex Alderton, Roberto Amato, Jeffrey Barrett, Sonia Goncalves, Ewan Harrison, David K. Jackson, Ian Johnston, Dominic Kwiatkowski, Cordelia Langford, John Sillitoe on behalf of the Wellcome Sanger Institute COVID-19 Surveillance Team |
| EPI_ISL_3367871 | Berkshire and Surrey Pathology Services Lighthouse Laboratory | Wellcome Sanger Institute for the COVID-19 Genomics UK (COG-UK) Consortium | Berkshire and Surrey Pathology Services Lighthouse Laboratory and Alex Alderton, Roberto Amato, Jeffrey Barrett, Sonia Goncalves, Ewan Harrison, David K. Jackson, Ian Johnston, Dominic Kwiatkowski, Cordelia Langford, John Sillitoe on behalf of the Wellcome Sanger Institute COVID-19 Surveillance Team |
| EPI_ISL_3367872, EPI_ISL_3367873 | Lighthouse Laboratory Plymouth | Wellcome Sanger Institute for the COVID-19 Genomics UK (COG-UK) Consortium | Lighthouse Laboratory Plymouth and Alex Alderton, Roberto Amato, Jeffrey Barrett, Sonia Goncalves, Ewan Harrison, David K. Jackson, Ian Johnston, Dominic Kwiatkowski, Cordelia Langford, John Sillitoe on behalf of the Wellcome Sanger Institute COVID-19 Surveillance Team |
| EPI_ISL_3367874 | Lighthouse Lab in Milton Keynes | Wellcome Sanger Institute for the COVID-19 Genomics UK (COG-UK) Consortium | The Lighthouse Lab in Milton Keynes and Alex Alderton, Roberto Amato, Jeffrey Barrett, Sonia Goncalves, Ewan Harrison, David K. Jackson, Ian Johnston, Dominic Kwiatkowski, Cordelia Langford, John Sillitoe on behalf of the Wellcome Sanger Institute COVID-19 Surveillance Team |
| EPI_ISL_3367875 | Berkshire and Surrey Pathology Services Lighthouse Laboratory | Wellcome Sanger Institute for the COVID-19 Genomics UK (COG-UK) Consortium | Berkshire and Surrey Pathology Services Lighthouse Laboratory and Alex Alderton, Roberto Amato, Jeffrey Barrett, Sonia Goncalves, Ewan Harrison, David K. Jackson, Ian Johnston, Dominic Kwiatkowski, Cordelia Langford, John Sillitoe on behalf of the Wellcome Sanger Institute COVID-19 Surveillance Team |
| EPI_ISL_3367876 | Lighthouse Laboratory Plymouth | Wellcome Sanger Institute for the COVID-19 Genomics UK (COG-UK) Consortium | Lighthouse Laboratory Plymouth and Alex Alderton, Roberto Amato, Jeffrey Barrett, Sonia Goncalves, Ewan Harrison, David K. Jackson, Ian Johnston, Dominic Kwiatkowski, Cordelia Langford, John Sillitoe on behalf of the Wellcome Sanger Institute COVID-19 Surveillance Team |
| EPI_ISL_3367877 | Lighthouse Lab in Milton Keynes | Wellcome Sanger Institute for the COVID-19 Genomics UK (COG-UK) Consortium | The Lighthouse Lab in Milton Keynes and Alex Alderton, Roberto Amato, Jeffrey Barrett, Sonia Goncalves, Ewan Harrison, David K. Jackson, Ian Johnston, Dominic Kwiatkowski, Cordelia Langford, John Sillitoe on behalf of the Wellcome Sanger Institute COVID-19 Surveillance Team |
| EPI_ISL_3367878 | Berkshire and Surrey Pathology Services Lighthouse Laboratory | Wellcome Sanger Institute for the COVID-19 Genomics UK (COG-UK) Consortium | Berkshire and Surrey Pathology Services Lighthouse Laboratory and Alex Alderton, Roberto Amato, Jeffrey Barrett, Sonia Goncalves, Ewan Harrison, David K. Jackson, Ian Johnston, Dominic Kwiatkowski, Cordelia Langford, John Sillitoe on behalf of the Wellcome Sanger Institute COVID-19 Surveillance Team |
| EPI_ISL_3367879 | Lighthouse Laboratory Plymouth | Wellcome Sanger Institute for the COVID-19 Genomics UK (COG-UK) Consortium | Lighthouse Laboratory Plymouth and Alex Alderton, Roberto Amato, Jeffrey Barrett, Sonia Goncalves, Ewan Harrison, David K. Jackson, Ian Johnston, Dominic Kwiatkowski, Cordelia Langford, John Sillitoe on behalf of the Wellcome Sanger Institute COVID-19 Surveillance Team |
| EPI_ISL_3367880 | Lighthouse Lab in Glasgow | Wellcome Sanger Institute for the COVID-19 Genomics UK (COG-UK) Consortium | Harper VanSteenhouse, Yumi Kasai, David Gray, Carol Clugston, Anna Dominiczak and Alex Alderton, Roberto Amato, Jeffrey Barrett, Sonia Goncalves, Ewan Harrison, David K. Jackson, Ian Johnston, Dominic Kwiatkowski, Cordelia Langford, John Sillitoe on behalf of the Wellcome Sanger Institute COVID-19 Surveillance Team |
| EPI_ISL_3367881 | Lighthouse Laboratory Plymouth | Wellcome Sanger Institute for the COVID-19 Genomics UK (COG-UK) Consortium | Lighthouse Laboratory Plymouth and Alex Alderton, Roberto Amato, Jeffrey Barrett, Sonia Goncalves, Ewan Harrison, David K. Jackson, Ian Johnston, Dominic Kwiatkowski, Cordelia Langford, John Sillitoe on behalf of the Wellcome Sanger Institute COVID-19 Surveillance Team |
| EPI_ISL_3367882 | Berkshire and Surrey Pathology Services Lighthouse Laboratory | Wellcome Sanger Institute for the COVID-19 Genomics UK (COG-UK) Consortium | Berkshire and Surrey Pathology Services Lighthouse Laboratory and Alex Alderton, Roberto Amato, Jeffrey Barrett, Sonia Goncalves, Ewan Harrison, David K. Jackson, Ian Johnston, Dominic Kwiatkowski, Cordelia Langford, John Sillitoe on behalf of the Wellcome Sanger Institute COVID-19 Surveillance Team |
| EPI_ISL_3367883, EPI_ISL_3367884 | Lighthouse Lab in Milton Keynes | Wellcome Sanger Institute for the COVID-19 Genomics UK (COG-UK) Consortium | The Lighthouse Lab in Milton Keynes and Alex Alderton, Roberto Amato, Jeffrey Barrett, Sonia Goncalves, Ewan Harrison, David K. Jackson, Ian Johnston, Dominic Kwiatkowski, Cordelia Langford, John Sillitoe on behalf of the Wellcome Sanger Institute COVID-19 Surveillance Team |
| EPI_ISL_3367885 | Berkshire and Surrey Pathology Services Lighthouse Laboratory | Wellcome Sanger Institute for the COVID-19 Genomics UK (COG-UK) Consortium | Berkshire and Surrey Pathology Services Lighthouse Laboratory and Alex Alderton, Roberto Amato, Jeffrey Barrett, Sonia Goncalves, Ewan Harrison, David K. Jackson, Ian Johnston, Dominic Kwiatkowski, Cordelia Langford, John Sillitoe on behalf of the Wellcome Sanger Institute COVID-19 Surveillance Team |
| EPI_ISL_3367886 | Lighthouse Lab in Milton Keynes | Wellcome Sanger Institute for the COVID-19 Genomics UK (COG-UK) Consortium | The Lighthouse Lab in Milton Keynes and Alex Alderton, Roberto Amato, Jeffrey Barrett, Sonia Goncalves, Ewan Harrison, David K. Jackson, Ian Johnston, Dominic Kwiatkowski, Cordelia Langford, John Sillitoe on behalf of the Wellcome Sanger Institute COVID-19 Surveillance Team |
| EPI_ISL_3367887 | Berkshire and Surrey Pathology Services Lighthouse Laboratory | Wellcome Sanger Institute for the COVID-19 Genomics UK (COG-UK) Consortium | Berkshire and Surrey Pathology Services Lighthouse Laboratory and Alex Alderton, Roberto Amato, Jeffrey Barrett, Sonia Goncalves, Ewan Harrison, David K. Jackson, Ian Johnston, Dominic Kwiatkowski, Cordelia Langford, John Sillitoe on behalf of the Wellcome Sanger Institute COVID-19 Surveillance Team |
| EPI_ISL_3367888 | Lighthouse Lab in Glasgow | Wellcome Sanger Institute for the COVID-19 Genomics UK (COG-UK) Consortium | Harper VanSteenhouse, Yumi Kasai, David Gray, Carol Clugston, Anna Dominiczak and Alex Alderton, Roberto Amato, Jeffrey Barrett, Sonia Goncalves, Ewan Harrison, David K. Jackson, Ian Johnston, Dominic Kwiatkowski, Cordelia Langford, John Sillitoe on behalf of the Wellcome Sanger Institute COVID-19 Surveillance Team |
| EPI_ISL_3367889 | Lighthouse Laboratory Plymouth | Wellcome Sanger Institute for the COVID-19 Genomics UK (COG-UK) Consortium | Lighthouse Laboratory Plymouth and Alex Alderton, Roberto Amato, Jeffrey Barrett, Sonia Goncalves, Ewan Harrison, David K. Jackson, Ian Johnston, Dominic Kwiatkowski, Cordelia Langford, John Sillitoe on behalf of the Wellcome Sanger Institute COVID-19 Surveillance Team |
| EPI_ISL_3367890 | Lighthouse Lab in Milton Keynes | Wellcome Sanger Institute for the COVID-19 Genomics UK (COG-UK) Consortium | The Lighthouse Lab in Milton Keynes and Alex Alderton, Roberto Amato, Jeffrey Barrett, Sonia Goncalves, Ewan Harrison, David K. Jackson, Ian Johnston, Dominic Kwiatkowski, Cordelia Langford, John Sillitoe on behalf of the Wellcome Sanger Institute COVID-19 Surveillance Team |
| EPI_ISL_3367891 | Berkshire and Surrey Pathology Services Lighthouse Laboratory | Wellcome Sanger Institute for the COVID-19 Genomics UK (COG-UK) Consortium | Berkshire and Surrey Pathology Services Lighthouse Laboratory and Alex Alderton, Roberto Amato, Jeffrey Barrett, Sonia Goncalves, Ewan Harrison, David K. Jackson, Ian Johnston, Dominic Kwiatkowski, Cordelia Langford, John Sillitoe on behalf of the Wellcome Sanger Institute COVID-19 Surveillance Team |
| EPI_ISL_3367892 | Lighthouse Lab in Glasgow | Wellcome Sanger Institute for the COVID-19 Genomics UK (COG-UK) Consortium | Harper VanSteenhouse, Yumi Kasai, David Gray, Carol Clugston, Anna Dominiczak and Alex Alderton, Roberto Amato, Jeffrey Barrett, Sonia Goncalves, Ewan Harrison, David K. Jackson, Ian Johnston, Dominic Kwiatkowski, Cordelia Langford, John Sillitoe on behalf of the Wellcome Sanger Institute COVID-19 Surveillance Team |
| EPI_ISL_3367893 | Berkshire and Surrey Pathology Services Lighthouse Laboratory | Wellcome Sanger Institute for the COVID-19 Genomics UK (COG-UK) Consortium | Berkshire and Surrey Pathology Services Lighthouse Laboratory and Alex Alderton, Roberto Amato, Jeffrey Barrett, Sonia Goncalves, Ewan Harrison, David K. Jackson, Ian Johnston, Dominic Kwiatkowski, Cordelia Langford, John Sillitoe on behalf of the Wellcome Sanger Institute COVID-19 Surveillance Team |
| EPI_ISL_3367894 | Lighthouse Laboratory Plymouth | Wellcome Sanger Institute for the COVID-19 Genomics UK (COG-UK) Consortium | Lighthouse Laboratory Plymouth and Alex Alderton, Roberto Amato, Jeffrey Barrett, Sonia Goncalves, Ewan Harrison, David K. Jackson, Ian Johnston, Dominic Kwiatkowski, Cordelia Langford, John Sillitoe on behalf of the Wellcome Sanger Institute COVID-19 Surveillance Team |
| EPI_ISL_3367895 | Berkshire and Surrey Pathology Services Lighthouse Laboratory | Wellcome Sanger Institute for the COVID-19 Genomics UK (COG-UK) Consortium | Berkshire and Surrey Pathology Services Lighthouse Laboratory and Alex Alderton, Roberto Amato, Jeffrey Barrett, Sonia Goncalves, Ewan Harrison, David K. Jackson, Ian Johnston, Dominic Kwiatkowski, Cordelia Langford, John Sillitoe on behalf of the Wellcome Sanger Institute COVID-19 Surveillance Team |
| EPI_ISL_3367896 | Lighthouse Laboratory Plymouth | Wellcome Sanger Institute for the COVID-19 Genomics UK (COG-UK) Consortium | Lighthouse Laboratory Plymouth and Alex Alderton, Roberto Amato, Jeffrey Barrett, Sonia Goncalves, Ewan Harrison, David K. Jackson, Ian Johnston, Dominic Kwiatkowski, Cordelia Langford, John Sillitoe on behalf of the Wellcome Sanger Institute COVID-19 Surveillance Team |

| | | | |
|---|---|---|---|
| EPI_ISL_3367897 | Berkshire and Surrey Pathology Services Lighthouse Laboratory | Wellcome Sanger Institute for the COVID-19 Genomics UK (COG-UK) Consortium | Berkshire and Surrey Pathology Services Lighthouse Laboratory and Alex Alderton, Roberto Amato, Jeffrey Barrett, Sonia Goncalves, Ewan Harrison, David K. Jackson, Ian Johnston, Dominic Kwiatkowski, Cordelia Langford, John Sillitoe on behalf of the Wellcome Sanger Institute COVID-19 Surveillance Team |
| EPI_ISL_3367898, EPI_ISL_3367899 | Lighthouse Lab in Glasgow | Wellcome Sanger Institute for the COVID-19 Genomics UK (COG-UK) Consortium | Harper VanSteenhouse, Yumi Kasai, David Gray, Carol Clugston, Anna Dominiczak and Alex Alderton, Roberto Amato, Jeffrey Barrett, Sonia Goncalves, Ewan Harrison, David K. Jackson, Ian Johnston, Dominic Kwiatkowski, Cordelia Langford, John Sillitoe on behalf of the Wellcome Sanger Institute COVID-19 Surveillance Team |
| EPI_ISL_3367900 | Berkshire and Surrey Pathology Services Lighthouse Laboratory | Wellcome Sanger Institute for the COVID-19 Genomics UK (COG-UK) Consortium | Berkshire and Surrey Pathology Services Lighthouse Laboratory and Alex Alderton, Roberto Amato, Jeffrey Barrett, Sonia Goncalves, Ewan Harrison, David K. Jackson, Ian Johnston, Dominic Kwiatkowski, Cordelia Langford, John Sillitoe on behalf of the Wellcome Sanger Institute COVID-19 Surveillance Team |
| EPI_ISL_3367901 | Lighthouse Lab in Milton Keynes | Wellcome Sanger Institute for the COVID-19 Genomics UK (COG-UK) Consortium | The Lighthouse Lab in Milton Keynes and Alex Alderton, Roberto Amato, Jeffrey Barrett, Sonia Goncalves, Ewan Harrison, David K. Jackson, Ian Johnston, Dominic Kwiatkowski, Cordelia Langford, John Sillitoe on behalf of the Wellcome Sanger Institute COVID-19 Surveillance Team |
| EPI_ISL_3367902 | Berkshire and Surrey Pathology Services Lighthouse Laboratory | Wellcome Sanger Institute for the COVID-19 Genomics UK (COG-UK) Consortium | Berkshire and Surrey Pathology Services Lighthouse Laboratory and Alex Alderton, Roberto Amato, Jeffrey Barrett, Sonia Goncalves, Ewan Harrison, David K. Jackson, Ian Johnston, Dominic Kwiatkowski, Cordelia Langford, John Sillitoe on behalf of the Wellcome Sanger Institute COVID-19 Surveillance Team |
| EPI_ISL_3367903, EPI_ISL_3367904 | Lighthouse Lab in Glasgow | Wellcome Sanger Institute for the COVID-19 Genomics UK (COG-UK) Consortium | Harper VanSteenhouse, Yumi Kasai, David Gray, Carol Clugston, Anna Dominiczak and Alex Alderton, Roberto Amato, Jeffrey Barrett, Sonia Goncalves, Ewan Harrison, David K. Jackson, Ian Johnston, Dominic Kwiatkowski, Cordelia Langford, John Sillitoe on behalf of the Wellcome Sanger Institute COVID-19 Surveillance Team |
| EPI_ISL_3367905 | Lighthouse Laboratory Plymouth | Wellcome Sanger Institute for the COVID-19 Genomics UK (COG-UK) Consortium | Lighthouse Laboratory Plymouth and Alex Alderton, Roberto Amato, Jeffrey Barrett, Sonia Goncalves, Ewan Harrison, David K. Jackson, Ian Johnston, Dominic Kwiatkowski, Cordelia Langford, John Sillitoe on behalf of the Wellcome Sanger Institute COVID-19 Surveillance Team |
| EPI_ISL_3367906 | Lighthouse Lab in Milton Keynes | Wellcome Sanger Institute for the COVID-19 Genomics UK (COG-UK) Consortium | The Lighthouse Lab in Milton Keynes and Alex Alderton, Roberto Amato, Jeffrey Barrett, Sonia Goncalves, Ewan Harrison, David K. Jackson, Ian Johnston, Dominic Kwiatkowski, Cordelia Langford, John Sillitoe on behalf of the Wellcome Sanger Institute COVID-19 Surveillance Team |
| EPI_ISL_3367907 | Lighthouse Lab in Glasgow | Wellcome Sanger Institute for the COVID-19 Genomics UK (COG-UK) Consortium | Harper VanSteenhouse, Yumi Kasai, David Gray, Carol Clugston, Anna Dominiczak and Alex Alderton, Roberto Amato, Jeffrey Barrett, Sonia Goncalves, Ewan Harrison, David K. Jackson, Ian Johnston, Dominic Kwiatkowski, Cordelia Langford, John Sillitoe on behalf of the Wellcome Sanger Institute COVID-19 Surveillance Team |
| EPI_ISL_3367908 | Berkshire and Surrey Pathology Services Lighthouse Laboratory | Wellcome Sanger Institute for the COVID-19 Genomics UK (COG-UK) Consortium | Berkshire and Surrey Pathology Services Lighthouse Laboratory and Alex Alderton, Roberto Amato, Jeffrey Barrett, Sonia Goncalves, Ewan Harrison, David K. Jackson, Ian Johnston, Dominic Kwiatkowski, Cordelia Langford, John Sillitoe on behalf of the Wellcome Sanger Institute COVID-19 Surveillance Team |
| EPI_ISL_3367909, EPI_ISL_3367910 | Lighthouse Lab in Milton Keynes | Wellcome Sanger Institute for the COVID-19 Genomics UK (COG-UK) Consortium | The Lighthouse Lab in Milton Keynes and Alex Alderton, Roberto Amato, Jeffrey Barrett, Sonia Goncalves, Ewan Harrison, David K. Jackson, Ian Johnston, Dominic Kwiatkowski, Cordelia Langford, John Sillitoe on behalf of the Wellcome Sanger Institute COVID-19 Surveillance Team |
| EPI_ISL_3367911, EPI_ISL_3367912 | Lighthouse Laboratory Plymouth | Wellcome Sanger Institute for the COVID-19 Genomics UK (COG-UK) Consortium | Lighthouse Laboratory Plymouth and Alex Alderton, Roberto Amato, Jeffrey Barrett, Sonia Goncalves, Ewan Harrison, David K. Jackson, Ian Johnston, Dominic Kwiatkowski, Cordelia Langford, John Sillitoe on behalf of the Wellcome Sanger Institute COVID-19 Surveillance Team |
| EPI_ISL_3367913 | Lighthouse Lab in Glasgow | Wellcome Sanger Institute for the COVID-19 Genomics UK (COG-UK) Consortium | Harper VanSteenhouse, Yumi Kasai, David Gray, Carol Clugston, Anna Dominiczak and Alex Alderton, Roberto Amato, Jeffrey Barrett, Sonia Goncalves, Ewan Harrison, David K. Jackson, Ian Johnston, Dominic Kwiatkowski, Cordelia Langford, John Sillitoe on behalf of the Wellcome Sanger Institute COVID-19 Surveillance Team |
| EPI_ISL_3367914 | Lighthouse Laboratory Plymouth | Wellcome Sanger Institute for the COVID-19 Genomics UK (COG-UK) Consortium | Lighthouse Laboratory Plymouth and Alex Alderton, Roberto Amato, Jeffrey Barrett, Sonia Goncalves, Ewan Harrison, David K. Jackson, Ian Johnston, Dominic Kwiatkowski, Cordelia Langford, John Sillitoe on behalf of the Wellcome Sanger Institute COVID-19 Surveillance Team |
| EPI_ISL_3367915 | Lighthouse Lab in Glasgow | Wellcome Sanger Institute for the COVID-19 Genomics UK (COG-UK) Consortium | Harper VanSteenhouse, Yumi Kasai, David Gray, Carol Clugston, Anna Dominiczak and Alex Alderton, Roberto Amato, Jeffrey Barrett, Sonia Goncalves, Ewan Harrison, David K. Jackson, Ian Johnston, Dominic Kwiatkowski, Cordelia Langford, John Sillitoe on behalf of the Wellcome Sanger Institute COVID-19 Surveillance Team |
| EPI_ISL_3367916 | Lighthouse Laboratory Plymouth | Wellcome Sanger Institute for the COVID-19 Genomics UK (COG-UK) Consortium | Lighthouse Laboratory Plymouth and Alex Alderton, Roberto Amato, Jeffrey Barrett, Sonia Goncalves, Ewan Harrison, David K. Jackson, Ian Johnston, Dominic Kwiatkowski, Cordelia Langford, John Sillitoe on behalf of the Wellcome Sanger Institute COVID-19 Surveillance Team |
| EPI_ISL_3367917 | Lighthouse Lab in Glasgow | Wellcome Sanger Institute for the COVID-19 Genomics UK (COG-UK) Consortium | Harper VanSteenhouse, Yumi Kasai, David Gray, Carol Clugston, Anna Dominiczak and Alex Alderton, Roberto Amato, Jeffrey Barrett, Sonia Goncalves, Ewan Harrison, David K. Jackson, Ian Johnston, Dominic Kwiatkowski, Cordelia Langford, John Sillitoe on behalf of the Wellcome Sanger Institute COVID-19 Surveillance Team |
| EPI_ISL_3367918 | Berkshire and Surrey Pathology Services Lighthouse Laboratory | Wellcome Sanger Institute for the COVID-19 Genomics UK (COG-UK) Consortium | Berkshire and Surrey Pathology Services Lighthouse Laboratory and Alex Alderton, Roberto Amato, Jeffrey Barrett, Sonia Goncalves, Ewan Harrison, David K. Jackson, Ian Johnston, Dominic Kwiatkowski, Cordelia Langford, John Sillitoe on behalf of the Wellcome Sanger Institute COVID-19 Surveillance Team |
| EPI_ISL_3367919 | Lighthouse Lab in Glasgow | Wellcome Sanger Institute for the COVID-19 Genomics UK (COG-UK) Consortium | Harper VanSteenhouse, Yumi Kasai, David Gray, Carol Clugston, Anna Dominiczak and Alex Alderton, Roberto Amato, Jeffrey Barrett, Sonia Goncalves, Ewan Harrison, David K. Jackson, Ian Johnston, Dominic Kwiatkowski, Cordelia Langford, John Sillitoe on behalf of the Wellcome Sanger Institute COVID-19 Surveillance Team |
| EPI_ISL_3367920, EPI_ISL_3367921, EPI_ISL_3367922 | Lighthouse Lab in Milton Keynes | Wellcome Sanger Institute for the COVID-19 Genomics UK (COG-UK) Consortium | The Lighthouse Lab in Milton Keynes and Alex Alderton, Roberto Amato, Jeffrey Barrett, Sonia Goncalves, Ewan Harrison, David K. Jackson, Ian Johnston, Dominic Kwiatkowski, Cordelia Langford, John Sillitoe on behalf of the Wellcome Sanger Institute COVID-19 Surveillance Team |
| EPI_ISL_3367923 | Berkshire and Surrey Pathology Services Lighthouse Laboratory | Wellcome Sanger Institute for the COVID-19 Genomics UK (COG-UK) Consortium | Berkshire and Surrey Pathology Services Lighthouse Laboratory and Alex Alderton, Roberto Amato, Jeffrey Barrett, Sonia Goncalves, Ewan Harrison, David K. Jackson, Ian Johnston, Dominic Kwiatkowski, Cordelia Langford, John Sillitoe on behalf of the Wellcome Sanger Institute COVID-19 Surveillance Team |
| EPI_ISL_3367924 | Lighthouse Lab in Milton Keynes | Wellcome Sanger Institute for the COVID-19 Genomics UK (COG-UK) Consortium | The Lighthouse Lab in Milton Keynes and Alex Alderton, Roberto Amato, Jeffrey Barrett, Sonia Goncalves, Ewan Harrison, David K. Jackson, Ian Johnston, Dominic Kwiatkowski, Cordelia Langford, John Sillitoe on behalf of the Wellcome Sanger Institute COVID-19 Surveillance Team |
| EPI_ISL_3367925, EPI_ISL_3367926 | Berkshire and Surrey Pathology Services Lighthouse Laboratory | Wellcome Sanger Institute for the COVID-19 Genomics UK (COG-UK) Consortium | Berkshire and Surrey Pathology Services Lighthouse Laboratory and Alex Alderton, Roberto Amato, Jeffrey Barrett, Sonia Goncalves, Ewan Harrison, David K. Jackson, Ian Johnston, Dominic Kwiatkowski, Cordelia Langford, John Sillitoe on behalf of the Wellcome Sanger Institute COVID-19 Surveillance Team |
| EPI_ISL_3367927, EPI_ISL_3367928 | Lighthouse Lab in Glasgow | Wellcome Sanger Institute for the COVID-19 Genomics UK (COG-UK) Consortium | Harper VanSteenhouse, Yumi Kasai, David Gray, Carol Clugston, Anna Dominiczak and Alex Alderton, Roberto Amato, Jeffrey Barrett, Sonia Goncalves, Ewan Harrison, David K. Jackson, Ian Johnston, Dominic Kwiatkowski, Cordelia Langford, John Sillitoe on behalf of the Wellcome Sanger Institute COVID-19 Surveillance Team |
| EPI_ISL_3367929 | Lighthouse Lab in Milton Keynes | Wellcome Sanger Institute for the COVID-19 Genomics UK (COG-UK) Consortium | The Lighthouse Lab in Milton Keynes and Alex Alderton, Roberto Amato, Jeffrey Barrett, Sonia Goncalves, Ewan Harrison, David K. Jackson, Ian Johnston, Dominic Kwiatkowski, Cordelia Langford, John Sillitoe on behalf of the Wellcome Sanger Institute COVID-19 Surveillance Team |
| EPI_ISL_3367930, EPI_ISL_3367931 | Lighthouse Lab in Glasgow | Wellcome Sanger Institute for the COVID-19 Genomics UK (COG-UK) Consortium | Harper VanSteenhouse, Yumi Kasai, David Gray, Carol Clugston, Anna Dominiczak and Alex Alderton, Roberto Amato, Jeffrey Barrett, Sonia Goncalves, Ewan Harrison, David K. Jackson, Ian Johnston, Dominic Kwiatkowski, Cordelia Langford, John Sillitoe on behalf of the Wellcome Sanger Institute COVID-19 Surveillance Team |
| EPI_ISL_3367932 | Lighthouse Laboratory Plymouth | Wellcome Sanger Institute for the COVID-19 Genomics UK (COG-UK) Consortium | Lighthouse Laboratory Plymouth and Alex Alderton, Roberto Amato, Jeffrey Barrett, Sonia Goncalves, Ewan Harrison, David K. Jackson, Ian Johnston, Dominic Kwiatkowski, Cordelia Langford, John Sillitoe on behalf of the Wellcome Sanger Institute COVID-19 Surveillance Team |
| EPI_ISL_3367933, EPI_ISL_3367934, EPI_ISL_3367935, EPI_ISL_3367936, EPI_ISL_3367937 | Lighthouse Lab in Glasgow | Wellcome Sanger Institute for the COVID-19 Genomics UK (COG-UK) Consortium | Harper VanSteenhouse, Yumi Kasai, David Gray, Carol Clugston, Anna Dominiczak and Alex Alderton, Roberto Amato, Jeffrey Barrett, Sonia Goncalves, Ewan Harrison, David K. Jackson, Ian Johnston, Dominic Kwiatkowski, Cordelia Langford, John Sillitoe on behalf of the Wellcome Sanger Institute COVID-19 Surveillance Team |
| EPI_ISL_3367938 | Lighthouse Laboratory Plymouth | Wellcome Sanger Institute for the COVID-19 Genomics UK (COG-UK) Consortium | Lighthouse Laboratory Plymouth and Alex Alderton, Roberto Amato, Jeffrey Barrett, Sonia Goncalves, Ewan Harrison, David K. Jackson, Ian Johnston, Dominic Kwiatkowski, Cordelia Langford, John Sillitoe on behalf of the Wellcome Sanger Institute COVID-19 Surveillance Team |
| EPI_ISL_3367939 | Lighthouse Lab in Glasgow | Wellcome Sanger Institute for the COVID-19 Genomics UK (COG-UK) Consortium | Harper VanSteenhouse, Yumi Kasai, David Gray, Carol Clugston, Anna Dominiczak and Alex Alderton, Roberto Amato, Jeffrey Barrett, Sonia Goncalves, Ewan Harrison, David K. Jackson, Ian Johnston, Dominic Kwiatkowski, Cordelia Langford, John Sillitoe on behalf of the Wellcome Sanger Institute COVID-19 Surveillance Team |
| EPI_ISL_3367940 | Lighthouse Lab in Milton Keynes | Wellcome Sanger Institute for the COVID-19 Genomics UK (COG-UK) Consortium | The Lighthouse Lab in Milton Keynes and Alex Alderton, Roberto Amato, Jeffrey Barrett, Sonia Goncalves, Ewan Harrison, David K. Jackson, Ian Johnston, Dominic Kwiatkowski, Cordelia Langford, John Sillitoe on behalf of the Wellcome Sanger Institute COVID-19 Surveillance Team |
| EPI_ISL_3367941 | Lighthouse Laboratory Plymouth | Wellcome Sanger Institute for the COVID-19 Genomics UK (COG-UK) Consortium | Lighthouse Laboratory Plymouth and Alex Alderton, Roberto Amato, Jeffrey Barrett, Sonia Goncalves, Ewan Harrison, David K. Jackson, Ian Johnston, Dominic Kwiatkowski, Cordelia Langford, John Sillitoe on behalf of the Wellcome Sanger Institute COVID-19 Surveillance Team |
| EPI_ISL_3367942 | Berkshire and Surrey Pathology Services Lighthouse Laboratory | Wellcome Sanger Institute for the COVID-19 Genomics UK (COG-UK) Consortium | Berkshire and Surrey Pathology Services Lighthouse Laboratory and Alex Alderton, Roberto Amato, Jeffrey Barrett, Sonia Goncalves, Ewan Harrison, David K. Jackson, Ian Johnston, Dominic Kwiatkowski, Cordelia Langford, John Sillitoe on behalf of the Wellcome Sanger Institute COVID-19 Surveillance Team |
| EPI_ISL_3367943 | Lighthouse Lab in Glasgow | Wellcome Sanger Institute for the COVID-19 Genomics UK (COG-UK) Consortium | Harper VanSteenhouse, Yumi Kasai, David Gray, Carol Clugston, Anna Dominiczak and Alex Alderton, Roberto Amato, Jeffrey Barrett, Sonia Goncalves, Ewan Harrison, David K. Jackson, Ian Johnston, Dominic Kwiatkowski, Cordelia Langford, John Sillitoe on behalf of the Wellcome Sanger Institute COVID-19 Surveillance Team |
| EPI_ISL_3367944 | Lighthouse Laboratory Plymouth | Wellcome Sanger Institute for the COVID-19 Genomics UK (COG-UK) Consortium | Lighthouse Laboratory Plymouth and Alex Alderton, Roberto Amato, Jeffrey Barrett, Sonia Goncalves, Ewan Harrison, David K. Jackson, Ian Johnston, Dominic Kwiatkowski, Cordelia Langford, John Sillitoe on behalf of the Wellcome Sanger Institute COVID-19 Surveillance Team |
| EPI_ISL_3367945 | Lighthouse Lab in Milton Keynes | Wellcome Sanger Institute for the COVID-19 Genomics UK (COG-UK) Consortium | The Lighthouse Lab in Milton Keynes and Alex Alderton, Roberto Amato, Jeffrey Barrett, Sonia Goncalves, Ewan Harrison, David K. Jackson, Ian Johnston, Dominic Kwiatkowski, Cordelia Langford, John Sillitoe on behalf of the Wellcome Sanger Institute COVID-19 Surveillance Team |
| EPI_ISL_3367946 | Lighthouse Lab in Glasgow | Wellcome Sanger Institute for the COVID-19 Genomics UK (COG-UK) Consortium | Harper VanSteenhouse, Yumi Kasai, David Gray, Carol Clugston, Anna Dominiczak and Alex Alderton, Roberto Amato, Jeffrey Barrett, Sonia Goncalves, Ewan Harrison, David K. Jackson, Ian Johnston, Dominic Kwiatkowski, Cordelia Langford, John Sillitoe on behalf of the Wellcome Sanger Institute COVID-19 Surveillance Team |
| EPI_ISL_3367947 | Berkshire and Surrey Pathology Services Lighthouse Laboratory | Wellcome Sanger Institute for the COVID-19 Genomics UK (COG-UK) Consortium | Berkshire and Surrey Pathology Services Lighthouse Laboratory and Alex Alderton, Roberto Amato, Jeffrey Barrett, Sonia Goncalves, Ewan Harrison, David K. Jackson, Ian Johnston, Dominic Kwiatkowski, Cordelia Langford, John Sillitoe on behalf of the Wellcome Sanger Institute COVID-19 Surveillance Team |
| EPI_ISL_3367948 | Lighthouse Laboratory Plymouth | Wellcome Sanger Institute for the COVID-19 Genomics UK (COG-UK) Consortium | Lighthouse Laboratory Plymouth and Alex Alderton, Roberto Amato, Jeffrey Barrett, Sonia Goncalves, Ewan Harrison, David K. Jackson, Ian Johnston, Dominic Kwiatkowski, Cordelia Langford, John Sillitoe on behalf of the Wellcome Sanger Institute COVID-19 Surveillance Team |
| EPI_ISL_3367949 | Berkshire and Surrey Pathology Services Lighthouse Laboratory | Wellcome Sanger Institute for the COVID-19 Genomics UK (COG-UK) Consortium | Berkshire and Surrey Pathology Services Lighthouse Laboratory and Alex Alderton, Roberto Amato, Jeffrey Barrett, Sonia Goncalves, Ewan Harrison, David K. Jackson, Ian Johnston, Dominic Kwiatkowski, Cordelia Langford, John Sillitoe on behalf of the Wellcome Sanger Institute COVID-19 Surveillance Team |
| EPI_ISL_3367950, EPI_ISL_3367951 | Lighthouse Lab in Milton Keynes | Wellcome Sanger Institute for the COVID-19 Genomics UK (COG-UK) Consortium | The Lighthouse Lab in Milton Keynes and Alex Alderton, Roberto Amato, Jeffrey Barrett, Sonia Goncalves, Ewan Harrison, David K. Jackson, Ian Johnston, Dominic Kwiatkowski, Cordelia Langford, John Sillitoe on behalf of the Wellcome Sanger Institute COVID-19 Surveillance Team |
| EPI_ISL_3367952 | Lighthouse Lab in Glasgow | Wellcome Sanger Institute for the COVID-19 Genomics UK (COG-UK) Consortium | Harper VanSteenhouse, Yumi Kasai, David Gray, Carol Clugston, Anna Dominiczak and Alex Alderton, Roberto Amato, Jeffrey Barrett, Sonia Goncalves, Ewan Harrison, David K. Jackson, Ian Johnston, Dominic Kwiatkowski, Cordelia Langford, John Sillitoe on behalf of the Wellcome Sanger Institute COVID-19 Surveillance Team |
| EPI_ISL_3367953 | Berkshire and Surrey Pathology Services Lighthouse Laboratory | Wellcome Sanger Institute for the COVID-19 Genomics UK (COG-UK) Consortium | Berkshire and Surrey Pathology Services Lighthouse Laboratory and Alex Alderton, Roberto Amato, Jeffrey Barrett, Sonia Goncalves, Ewan Harrison, David K. Jackson, Ian Johnston, Dominic Kwiatkowski, Cordelia Langford, John Sillitoe on behalf of the Wellcome Sanger Institute COVID-19 Surveillance Team |
| EPI_ISL_3367954, | Lighthouse Lab in Milton Keynes | Wellcome Sanger Institute for the COVID-19 Genomics UK (COG-UK) | The Lighthouse Lab in Milton Keynes and Alex Alderton, Roberto Amato, Jeffrey Barrett, Sonia Goncalves, Ewan Harrison, David K. Jackson, Ian Johnston, Dominic Kwiatkowski, Cordelia Langford, John Sillitoe on behalf of the Wellcome Sanger |

| | | | |
|---|---|---|---|
| EPI_ISL_3367955 | | Consortium | Institute COVID-19 Surveillance Team |
| EPI_ISL_3367956 | Lighthouse Laboratory Plymouth | Wellcome Sanger Institute for the COVID-19 Genomics UK (COG-UK) Consortium | Lighthouse Laboratory Plymouth and Alex Alderton, Roberto Amato, Jeffrey Barrett, Sonia Goncalves, Ewan Harrison, David K. Jackson, Ian Johnston, Dominic Kwiatkowski, Cordelia Langford, John Sillitoe on behalf of the Wellcome Sanger Institute COVID-19 Surveillance Team |
| EPI_ISL_3367957, EPI_ISL_3367958 | Lighthouse Lab in Milton Keynes | Wellcome Sanger Institute for the COVID-19 Genomics UK (COG-UK) Consortium | The Lighthouse Lab in Milton Keynes and Alex Alderton, Roberto Amato, Jeffrey Barrett, Sonia Goncalves, Ewan Harrison, David K. Jackson, Ian Johnston, Dominic Kwiatkowski, Cordelia Langford, John Sillitoe on behalf of the Wellcome Sanger Institute COVID-19 Surveillance Team |
| EPI_ISL_3367959 | Lighthouse Laboratory Plymouth | Wellcome Sanger Institute for the COVID-19 Genomics UK (COG-UK) Consortium | Lighthouse Laboratory Plymouth and Alex Alderton, Roberto Amato, Jeffrey Barrett, Sonia Goncalves, Ewan Harrison, David K. Jackson, Ian Johnston, Dominic Kwiatkowski, Cordelia Langford, John Sillitoe on behalf of the Wellcome Sanger Institute COVID-19 Surveillance Team |
| EPI_ISL_3367960 | Lighthouse Lab in Glasgow | Wellcome Sanger Institute for the COVID-19 Genomics UK (COG-UK) Consortium | Harper VanSteenhouse, Yumi Kasai, David Gray, Carol Clugston, Anna Dominiczak and Alex Alderton, Roberto Amato, Jeffrey Barrett, Sonia Goncalves, Ewan Harrison, David K. Jackson, Ian Johnston, Dominic Kwiatkowski, Cordelia Langford, John Sillitoe on behalf of the Wellcome Sanger Institute COVID-19 Surveillance Team |
| EPI_ISL_3367961 | Lighthouse Lab in Milton Keynes | Wellcome Sanger Institute for the COVID-19 Genomics UK (COG-UK) Consortium | The Lighthouse Lab in Milton Keynes and Alex Alderton, Roberto Amato, Jeffrey Barrett, Sonia Goncalves, Ewan Harrison, David K. Jackson, Ian Johnston, Dominic Kwiatkowski, Cordelia Langford, John Sillitoe on behalf of the Wellcome Sanger Institute COVID-19 Surveillance Team |
| EPI_ISL_3367962 | Lighthouse Laboratory Plymouth | Wellcome Sanger Institute for the COVID-19 Genomics UK (COG-UK) Consortium | Lighthouse Laboratory Plymouth and Alex Alderton, Roberto Amato, Jeffrey Barrett, Sonia Goncalves, Ewan Harrison, David K. Jackson, Ian Johnston, Dominic Kwiatkowski, Cordelia Langford, John Sillitoe on behalf of the Wellcome Sanger Institute COVID-19 Surveillance Team |
| EPI_ISL_3367963 | Lighthouse Lab in Glasgow | Wellcome Sanger Institute for the COVID-19 Genomics UK (COG-UK) Consortium | Harper VanSteenhouse, Yumi Kasai, David Gray, Carol Clugston, Anna Dominiczak and Alex Alderton, Roberto Amato, Jeffrey Barrett, Sonia Goncalves, Ewan Harrison, David K. Jackson, Ian Johnston, Dominic Kwiatkowski, Cordelia Langford, John Sillitoe on behalf of the Wellcome Sanger Institute COVID-19 Surveillance Team |
| EPI_ISL_3367964 | Lighthouse Laboratory Plymouth | Wellcome Sanger Institute for the COVID-19 Genomics UK (COG-UK) Consortium | Lighthouse Laboratory Plymouth and Alex Alderton, Roberto Amato, Jeffrey Barrett, Sonia Goncalves, Ewan Harrison, David K. Jackson, Ian Johnston, Dominic Kwiatkowski, Cordelia Langford, John Sillitoe on behalf of the Wellcome Sanger Institute COVID-19 Surveillance Team |
| EPI_ISL_3367965 | Berkshire and Surrey Pathology Services Lighthouse Laboratory | Wellcome Sanger Institute for the COVID-19 Genomics UK (COG-UK) Consortium | Berkshire and Surrey Pathology Services Lighthouse Laboratory and Alex Alderton, Roberto Amato, Jeffrey Barrett, Sonia Goncalves, Ewan Harrison, David K. Jackson, Ian Johnston, Dominic Kwiatkowski, Cordelia Langford, John Sillitoe on behalf of the Wellcome Sanger Institute COVID-19 Surveillance Team |
| EPI_ISL_3367966 | Lighthouse Lab in Glasgow | Wellcome Sanger Institute for the COVID-19 Genomics UK (COG-UK) Consortium | Harper VanSteenhouse, Yumi Kasai, David Gray, Carol Clugston, Anna Dominiczak and Alex Alderton, Roberto Amato, Jeffrey Barrett, Sonia Goncalves, Ewan Harrison, David K. Jackson, Ian Johnston, Dominic Kwiatkowski, Cordelia Langford, John Sillitoe on behalf of the Wellcome Sanger Institute COVID-19 Surveillance Team |
| EPI_ISL_3367967, EPI_ISL_3367968 | Berkshire and Surrey Pathology Services Lighthouse Laboratory | Wellcome Sanger Institute for the COVID-19 Genomics UK (COG-UK) Consortium | Berkshire and Surrey Pathology Services Lighthouse Laboratory and Alex Alderton, Roberto Amato, Jeffrey Barrett, Sonia Goncalves, Ewan Harrison, David K. Jackson, Ian Johnston, Dominic Kwiatkowski, Cordelia Langford, John Sillitoe on behalf of the Wellcome Sanger Institute COVID-19 Surveillance Team |
| EPI_ISL_3367969 | Lighthouse Lab in Milton Keynes | Wellcome Sanger Institute for the COVID-19 Genomics UK (COG-UK) Consortium | The Lighthouse Lab in Milton Keynes and Alex Alderton, Roberto Amato, Jeffrey Barrett, Sonia Goncalves, Ewan Harrison, David K. Jackson, Ian Johnston, Dominic Kwiatkowski, Cordelia Langford, John Sillitoe on behalf of the Wellcome Sanger Institute COVID-19 Surveillance Team |
| EPI_ISL_3367970 | Berkshire and Surrey Pathology Services Lighthouse Laboratory | Wellcome Sanger Institute for the COVID-19 Genomics UK (COG-UK) Consortium | Berkshire and Surrey Pathology Services Lighthouse Laboratory and Alex Alderton, Roberto Amato, Jeffrey Barrett, Sonia Goncalves, Ewan Harrison, David K. Jackson, Ian Johnston, Dominic Kwiatkowski, Cordelia Langford, John Sillitoe on behalf of the Wellcome Sanger Institute COVID-19 Surveillance Team |
| EPI_ISL_3367971 | Lighthouse Lab in Glasgow | Wellcome Sanger Institute for the COVID-19 Genomics UK (COG-UK) Consortium | Harper VanSteenhouse, Yumi Kasai, David Gray, Carol Clugston, Anna Dominiczak and Alex Alderton, Roberto Amato, Jeffrey Barrett, Sonia Goncalves, Ewan Harrison, David K. Jackson, Ian Johnston, Dominic Kwiatkowski, Cordelia Langford, John Sillitoe on behalf of the Wellcome Sanger Institute COVID-19 Surveillance Team |
| EPI_ISL_3367972 | Lighthouse Laboratory Plymouth | Wellcome Sanger Institute for the COVID-19 Genomics UK (COG-UK) Consortium | Lighthouse Laboratory Plymouth and Alex Alderton, Roberto Amato, Jeffrey Barrett, Sonia Goncalves, Ewan Harrison, David K. Jackson, Ian Johnston, Dominic Kwiatkowski, Cordelia Langford, John Sillitoe on behalf of the Wellcome Sanger Institute COVID-19 Surveillance Team |
| EPI_ISL_3367973 | Berkshire and Surrey Pathology Services Lighthouse Laboratory | Wellcome Sanger Institute for the COVID-19 Genomics UK (COG-UK) Consortium | Berkshire and Surrey Pathology Services Lighthouse Laboratory and Alex Alderton, Roberto Amato, Jeffrey Barrett, Sonia Goncalves, Ewan Harrison, David K. Jackson, Ian Johnston, Dominic Kwiatkowski, Cordelia Langford, John Sillitoe on behalf of the Wellcome Sanger Institute COVID-19 Surveillance Team |
| EPI_ISL_3367974 | Lighthouse Laboratory Plymouth | Wellcome Sanger Institute for the COVID-19 Genomics UK (COG-UK) Consortium | Lighthouse Laboratory Plymouth and Alex Alderton, Roberto Amato, Jeffrey Barrett, Sonia Goncalves, Ewan Harrison, David K. Jackson, Ian Johnston, Dominic Kwiatkowski, Cordelia Langford, John Sillitoe on behalf of the Wellcome Sanger Institute COVID-19 Surveillance Team |
| EPI_ISL_3367975 | Lighthouse Lab in Milton Keynes | Wellcome Sanger Institute for the COVID-19 Genomics UK (COG-UK) Consortium | The Lighthouse Lab in Milton Keynes and Alex Alderton, Roberto Amato, Jeffrey Barrett, Sonia Goncalves, Ewan Harrison, David K. Jackson, Ian Johnston, Dominic Kwiatkowski, Cordelia Langford, John Sillitoe on behalf of the Wellcome Sanger Institute COVID-19 Surveillance Team |
| EPI_ISL_3367976 | Berkshire and Surrey Pathology Services Lighthouse Laboratory | Wellcome Sanger Institute for the COVID-19 Genomics UK (COG-UK) Consortium | Berkshire and Surrey Pathology Services Lighthouse Laboratory and Alex Alderton, Roberto Amato, Jeffrey Barrett, Sonia Goncalves, Ewan Harrison, David K. Jackson, Ian Johnston, Dominic Kwiatkowski, Cordelia Langford, John Sillitoe on behalf of the Wellcome Sanger Institute COVID-19 Surveillance Team |
| EPI_ISL_3367977 | Lighthouse Lab in Glasgow | Wellcome Sanger Institute for the COVID-19 Genomics UK (COG-UK) Consortium | Harper VanSteenhouse, Yumi Kasai, David Gray, Carol Clugston, Anna Dominiczak and Alex Alderton, Roberto Amato, Jeffrey Barrett, Sonia Goncalves, Ewan Harrison, David K. Jackson, Ian Johnston, Dominic Kwiatkowski, Cordelia Langford, John Sillitoe on behalf of the Wellcome Sanger Institute COVID-19 Surveillance Team |
| EPI_ISL_3367978 | Berkshire and Surrey Pathology Services Lighthouse Laboratory | Wellcome Sanger Institute for the COVID-19 Genomics UK (COG-UK) Consortium | Berkshire and Surrey Pathology Services Lighthouse Laboratory and Alex Alderton, Roberto Amato, Jeffrey Barrett, Sonia Goncalves, Ewan Harrison, David K. Jackson, Ian Johnston, Dominic Kwiatkowski, Cordelia Langford, John Sillitoe on behalf of the Wellcome Sanger Institute COVID-19 Surveillance Team |
| EPI_ISL_3367979 | Lighthouse Lab in Glasgow | Wellcome Sanger Institute for the COVID-19 Genomics UK (COG-UK) Consortium | Harper VanSteenhouse, Yumi Kasai, David Gray, Carol Clugston, Anna Dominiczak and Alex Alderton, Roberto Amato, Jeffrey Barrett, Sonia Goncalves, Ewan Harrison, David K. Jackson, Ian Johnston, Dominic Kwiatkowski, Cordelia Langford, John Sillitoe on behalf of the Wellcome Sanger Institute COVID-19 Surveillance Team |
| EPI_ISL_3367980 | Lighthouse Lab in Milton Keynes | Wellcome Sanger Institute for the COVID-19 Genomics UK (COG-UK) Consortium | The Lighthouse Lab in Milton Keynes and Alex Alderton, Roberto Amato, Jeffrey Barrett, Sonia Goncalves, Ewan Harrison, David K. Jackson, Ian Johnston, Dominic Kwiatkowski, Cordelia Langford, John Sillitoe on behalf of the Wellcome Sanger Institute COVID-19 Surveillance Team |
| EPI_ISL_3367981 | Lighthouse Lab in Glasgow | Wellcome Sanger Institute for the COVID-19 Genomics UK (COG-UK) Consortium | Harper VanSteenhouse, Yumi Kasai, David Gray, Carol Clugston, Anna Dominiczak and Alex Alderton, Roberto Amato, Jeffrey Barrett, Sonia Goncalves, Ewan Harrison, David K. Jackson, Ian Johnston, Dominic Kwiatkowski, Cordelia Langford, John Sillitoe on behalf of the Wellcome Sanger Institute COVID-19 Surveillance Team |
| EPI_ISL_3367982 | Berkshire and Surrey Pathology Services Lighthouse Laboratory | Wellcome Sanger Institute for the COVID-19 Genomics UK (COG-UK) Consortium | Berkshire and Surrey Pathology Services Lighthouse Laboratory and Alex Alderton, Roberto Amato, Jeffrey Barrett, Sonia Goncalves, Ewan Harrison, David K. Jackson, Ian Johnston, Dominic Kwiatkowski, Cordelia Langford, John Sillitoe on behalf of the Wellcome Sanger Institute COVID-19 Surveillance Team |
| EPI_ISL_3367983 | Lighthouse Lab in Milton Keynes | Wellcome Sanger Institute for the COVID-19 Genomics UK (COG-UK) Consortium | The Lighthouse Lab in Milton Keynes and Alex Alderton, Roberto Amato, Jeffrey Barrett, Sonia Goncalves, Ewan Harrison, David K. Jackson, Ian Johnston, Dominic Kwiatkowski, Cordelia Langford, John Sillitoe on behalf of the Wellcome Sanger Institute COVID-19 Surveillance Team |
| EPI_ISL_3367984, EPI_ISL_3367985 | Lighthouse Lab in Glasgow | Wellcome Sanger Institute for the COVID-19 Genomics UK (COG-UK) Consortium | Harper VanSteenhouse, Yumi Kasai, David Gray, Carol Clugston, Anna Dominiczak and Alex Alderton, Roberto Amato, Jeffrey Barrett, Sonia Goncalves, Ewan Harrison, David K. Jackson, Ian Johnston, Dominic Kwiatkowski, Cordelia Langford, John Sillitoe on behalf of the Wellcome Sanger Institute COVID-19 Surveillance Team |
| EPI_ISL_3367986 | Lighthouse Laboratory Plymouth | Wellcome Sanger Institute for the COVID-19 Genomics UK (COG-UK) Consortium | Lighthouse Laboratory Plymouth and Alex Alderton, Roberto Amato, Jeffrey Barrett, Sonia Goncalves, Ewan Harrison, David K. Jackson, Ian Johnston, Dominic Kwiatkowski, Cordelia Langford, John Sillitoe on behalf of the Wellcome Sanger Institute COVID-19 Surveillance Team |
| EPI_ISL_3367987 | Lighthouse Lab in Glasgow | Wellcome Sanger Institute for the COVID-19 Genomics UK (COG-UK) Consortium | Harper VanSteenhouse, Yumi Kasai, David Gray, Carol Clugston, Anna Dominiczak and Alex Alderton, Roberto Amato, Jeffrey Barrett, Sonia Goncalves, Ewan Harrison, David K. Jackson, Ian Johnston, Dominic Kwiatkowski, Cordelia Langford, John Sillitoe on behalf of the Wellcome Sanger Institute COVID-19 Surveillance Team |
| EPI_ISL_3367988 | Berkshire and Surrey Pathology Services Lighthouse Laboratory | Wellcome Sanger Institute for the COVID-19 Genomics UK (COG-UK) Consortium | Berkshire and Surrey Pathology Services Lighthouse Laboratory and Alex Alderton, Roberto Amato, Jeffrey Barrett, Sonia Goncalves, Ewan Harrison, David K. Jackson, Ian Johnston, Dominic Kwiatkowski, Cordelia Langford, John Sillitoe on behalf of the Wellcome Sanger Institute COVID-19 Surveillance Team |
| EPI_ISL_3367989 | Lighthouse Lab in Milton Keynes | Wellcome Sanger Institute for the COVID-19 Genomics UK (COG-UK) Consortium | The Lighthouse Lab in Milton Keynes and Alex Alderton, Roberto Amato, Jeffrey Barrett, Sonia Goncalves, Ewan Harrison, David K. Jackson, Ian Johnston, Dominic Kwiatkowski, Cordelia Langford, John Sillitoe on behalf of the Wellcome Sanger Institute COVID-19 Surveillance Team |
| EPI_ISL_3367990 | Lighthouse Lab in Glasgow | Wellcome Sanger Institute for the COVID-19 Genomics UK (COG-UK) Consortium | Harper VanSteenhouse, Yumi Kasai, David Gray, Carol Clugston, Anna Dominiczak and Alex Alderton, Roberto Amato, Jeffrey Barrett, Sonia Goncalves, Ewan Harrison, David K. Jackson, Ian Johnston, Dominic Kwiatkowski, Cordelia Langford, John Sillitoe on behalf of the Wellcome Sanger Institute COVID-19 Surveillance Team |
| EPI_ISL_3367991 | Lighthouse Lab in Milton Keynes | Wellcome Sanger Institute for the COVID-19 Genomics UK (COG-UK) Consortium | The Lighthouse Lab in Milton Keynes and Alex Alderton, Roberto Amato, Jeffrey Barrett, Sonia Goncalves, Ewan Harrison, David K. Jackson, Ian Johnston, Dominic Kwiatkowski, Cordelia Langford, John Sillitoe on behalf of the Wellcome Sanger Institute COVID-19 Surveillance Team |
| EPI_ISL_3367992 | Lighthouse Laboratory Plymouth | Wellcome Sanger Institute for the COVID-19 Genomics UK (COG-UK) Consortium | Lighthouse Laboratory Plymouth and Alex Alderton, Roberto Amato, Jeffrey Barrett, Sonia Goncalves, Ewan Harrison, David K. Jackson, Ian Johnston, Dominic Kwiatkowski, Cordelia Langford, John Sillitoe on behalf of the Wellcome Sanger Institute COVID-19 Surveillance Team |
| EPI_ISL_3367993 | Lighthouse Lab in Glasgow | Wellcome Sanger Institute for the COVID-19 Genomics UK (COG-UK) Consortium | Harper VanSteenhouse, Yumi Kasai, David Gray, Carol Clugston, Anna Dominiczak and Alex Alderton, Roberto Amato, Jeffrey Barrett, Sonia Goncalves, Ewan Harrison, David K. Jackson, Ian Johnston, Dominic Kwiatkowski, Cordelia Langford, John Sillitoe on behalf of the Wellcome Sanger Institute COVID-19 Surveillance Team |
| EPI_ISL_3367994 | Lighthouse Lab in Milton Keynes | Wellcome Sanger Institute for the COVID-19 Genomics UK (COG-UK) Consortium | The Lighthouse Lab in Milton Keynes and Alex Alderton, Roberto Amato, Jeffrey Barrett, Sonia Goncalves, Ewan Harrison, David K. Jackson, Ian Johnston, Dominic Kwiatkowski, Cordelia Langford, John Sillitoe on behalf of the Wellcome Sanger Institute COVID-19 Surveillance Team |
| EPI_ISL_3367995, EPI_ISL_3367996 | Lighthouse Laboratory Plymouth | Wellcome Sanger Institute for the COVID-19 Genomics UK (COG-UK) Consortium | Lighthouse Laboratory Plymouth and Alex Alderton, Roberto Amato, Jeffrey Barrett, Sonia Goncalves, Ewan Harrison, David K. Jackson, Ian Johnston, Dominic Kwiatkowski, Cordelia Langford, John Sillitoe on behalf of the Wellcome Sanger Institute COVID-19 Surveillance Team |
| EPI_ISL_3367997, EPI_ISL_3367998, EPI_ISL_3367999, EPI_ISL_3368000 | Lighthouse Lab in Glasgow | Wellcome Sanger Institute for the COVID-19 Genomics UK (COG-UK) Consortium | Harper VanSteenhouse, Yumi Kasai, David Gray, Carol Clugston, Anna Dominiczak and Alex Alderton, Roberto Amato, Jeffrey Barrett, Sonia Goncalves, Ewan Harrison, David K. Jackson, Ian Johnston, Dominic Kwiatkowski, Cordelia Langford, John Sillitoe on behalf of the Wellcome Sanger Institute COVID-19 Surveillance Team |
| EPI_ISL_3368001 | Berkshire and Surrey Pathology Services Lighthouse Laboratory | Wellcome Sanger Institute for the COVID-19 Genomics UK (COG-UK) Consortium | Berkshire and Surrey Pathology Services Lighthouse Laboratory and Alex Alderton, Roberto Amato, Jeffrey Barrett, Sonia Goncalves, Ewan Harrison, David K. Jackson, Ian Johnston, Dominic Kwiatkowski, Cordelia Langford, John Sillitoe on behalf of the Wellcome Sanger Institute COVID-19 Surveillance Team |
| EPI_ISL_3368002 | Lighthouse Lab in Milton Keynes | Wellcome Sanger Institute for the COVID-19 Genomics UK (COG-UK) Consortium | The Lighthouse Lab in Milton Keynes and Alex Alderton, Roberto Amato, Jeffrey Barrett, Sonia Goncalves, Ewan Harrison, David K. Jackson, Ian Johnston, Dominic Kwiatkowski, Cordelia Langford, John Sillitoe on behalf of the Wellcome Sanger Institute COVID-19 Surveillance Team |
| EPI_ISL_3368003 | Berkshire and Surrey Pathology Services Lighthouse Laboratory | Wellcome Sanger Institute for the COVID-19 Genomics UK (COG-UK) Consortium | Berkshire and Surrey Pathology Services Lighthouse Laboratory and Alex Alderton, Roberto Amato, Jeffrey Barrett, Sonia Goncalves, Ewan Harrison, David K. Jackson, Ian Johnston, Dominic Kwiatkowski, Cordelia Langford, John Sillitoe on behalf of the Wellcome Sanger Institute COVID-19 Surveillance Team |
| EPI_ISL_3368004 | Lighthouse Lab in Milton Keynes | Wellcome Sanger Institute for the COVID-19 Genomics UK (COG-UK) Consortium | The Lighthouse Lab in Milton Keynes and Alex Alderton, Roberto Amato, Jeffrey Barrett, Sonia Goncalves, Ewan Harrison, David K. Jackson, Ian Johnston, Dominic Kwiatkowski, Cordelia Langford, John Sillitoe on behalf of the Wellcome Sanger Institute COVID-19 Surveillance Team |
| EPI_ISL_3368005 | Lighthouse Laboratory Plymouth | Wellcome Sanger Institute for the COVID-19 Genomics UK (COG-UK) Consortium | Lighthouse Laboratory Plymouth and Alex Alderton, Roberto Amato, Jeffrey Barrett, Sonia Goncalves, Ewan Harrison, David K. Jackson, Ian Johnston, Dominic Kwiatkowski, Cordelia Langford, John Sillitoe on behalf of the Wellcome Sanger Institute COVID-19 Surveillance Team |
| EPI_ISL_3368006, EPI_ISL_3368007 | Lighthouse Lab in Milton Keynes | Wellcome Sanger Institute for the COVID-19 Genomics UK (COG-UK) Consortium | The Lighthouse Lab in Milton Keynes and Alex Alderton, Roberto Amato, Jeffrey Barrett, Sonia Goncalves, Ewan Harrison, David K. Jackson, Ian Johnston, Dominic Kwiatkowski, Cordelia Langford, John Sillitoe on behalf of the Wellcome Sanger Institute COVID-19 Surveillance Team |

| | | | |
|---|---|---|---|
| EPI_ISL_3368008 | Lighthouse Laboratory Plymouth | Wellcome Sanger Institute for the COVID-19 Genomics UK (COG-UK) Consortium | Lighthouse Laboratory Plymouth and Alex Alderton, Roberto Amato, Jeffrey Barrett, Sonia Goncalves, Ewan Harrison, David K. Jackson, Ian Johnston, Dominic Kwiatkowski, Cordelia Langford, John Sillitoe on behalf of the Wellcome Sanger Institute COVID-19 Surveillance Team |
| EPI_ISL_3368009 | Lighthouse Lab in Glasgow | Wellcome Sanger Institute for the COVID-19 Genomics UK (COG-UK) Consortium | Harper VanSteenhouse, Yumi Kasai, David Gray, Carol Clugston, Anna Dominiczak and Alex Alderton, Roberto Amato, Jeffrey Barrett, Sonia Goncalves, Ewan Harrison, David K. Jackson, Ian Johnston, Dominic Kwiatkowski, Cordelia Langford, John Sillitoe on behalf of the Wellcome Sanger Institute COVID-19 Surveillance Team |
| EPI_ISL_3368010 | Lighthouse Lab in Milton Keynes | Wellcome Sanger Institute for the COVID-19 Genomics UK (COG-UK) Consortium | The Lighthouse Lab in Milton Keynes and Alex Alderton, Roberto Amato, Jeffrey Barrett, Sonia Goncalves, Ewan Harrison, David K. Jackson, Ian Johnston, Dominic Kwiatkowski, Cordelia Langford, John Sillitoe on behalf of the Wellcome Sanger Institute COVID-19 Surveillance Team |
| EPI_ISL_3368011 | Lighthouse Laboratory Plymouth | Wellcome Sanger Institute for the COVID-19 Genomics UK (COG-UK) Consortium | Lighthouse Laboratory Plymouth and Alex Alderton, Roberto Amato, Jeffrey Barrett, Sonia Goncalves, Ewan Harrison, David K. Jackson, Ian Johnston, Dominic Kwiatkowski, Cordelia Langford, John Sillitoe on behalf of the Wellcome Sanger Institute COVID-19 Surveillance Team |
| EPI_ISL_3368012 | Lighthouse Lab in Milton Keynes | Wellcome Sanger Institute for the COVID-19 Genomics UK (COG-UK) Consortium | The Lighthouse Lab in Milton Keynes and Alex Alderton, Roberto Amato, Jeffrey Barrett, Sonia Goncalves, Ewan Harrison, David K. Jackson, Ian Johnston, Dominic Kwiatkowski, Cordelia Langford, John Sillitoe on behalf of the Wellcome Sanger Institute COVID-19 Surveillance Team |
| EPI_ISL_3368013 | Lighthouse Lab in Glasgow | Wellcome Sanger Institute for the COVID-19 Genomics UK (COG-UK) Consortium | Harper VanSteenhouse, Yumi Kasai, David Gray, Carol Clugston, Anna Dominiczak and Alex Alderton, Roberto Amato, Jeffrey Barrett, Sonia Goncalves, Ewan Harrison, David K. Jackson, Ian Johnston, Dominic Kwiatkowski, Cordelia Langford, John Sillitoe on behalf of the Wellcome Sanger Institute COVID-19 Surveillance Team |
| EPI_ISL_3368014 | Lighthouse Laboratory Plymouth | Wellcome Sanger Institute for the COVID-19 Genomics UK (COG-UK) Consortium | Lighthouse Laboratory Plymouth and Alex Alderton, Roberto Amato, Jeffrey Barrett, Sonia Goncalves, Ewan Harrison, David K. Jackson, Ian Johnston, Dominic Kwiatkowski, Cordelia Langford, John Sillitoe on behalf of the Wellcome Sanger Institute COVID-19 Surveillance Team |
| EPI_ISL_3368015 | Lighthouse Lab in Milton Keynes | Wellcome Sanger Institute for the COVID-19 Genomics UK (COG-UK) Consortium | The Lighthouse Lab in Milton Keynes and Alex Alderton, Roberto Amato, Jeffrey Barrett, Sonia Goncalves, Ewan Harrison, David K. Jackson, Ian Johnston, Dominic Kwiatkowski, Cordelia Langford, John Sillitoe on behalf of the Wellcome Sanger Institute COVID-19 Surveillance Team |
| EPI_ISL_3368016, EPI_ISL_3368017 | Berkshire and Surrey Pathology Services Lighthouse Laboratory | Wellcome Sanger Institute for the COVID-19 Genomics UK (COG-UK) Consortium | Berkshire and Surrey Pathology Services Lighthouse Laboratory and Alex Alderton, Roberto Amato, Jeffrey Barrett, Sonia Goncalves, Ewan Harrison, David K. Jackson, Ian Johnston, Dominic Kwiatkowski, Cordelia Langford, John Sillitoe on behalf of the Wellcome Sanger Institute COVID-19 Surveillance Team |
| EPI_ISL_3368018, EPI_ISL_3368019 | Lighthouse Laboratory Plymouth | Wellcome Sanger Institute for the COVID-19 Genomics UK (COG-UK) Consortium | Lighthouse Laboratory Plymouth and Alex Alderton, Roberto Amato, Jeffrey Barrett, Sonia Goncalves, Ewan Harrison, David K. Jackson, Ian Johnston, Dominic Kwiatkowski, Cordelia Langford, John Sillitoe on behalf of the Wellcome Sanger Institute COVID-19 Surveillance Team |
| EPI_ISL_3368020 | Lighthouse Lab in Glasgow | Wellcome Sanger Institute for the COVID-19 Genomics UK (COG-UK) Consortium | Harper VanSteenhouse, Yumi Kasai, David Gray, Carol Clugston, Anna Dominiczak and Alex Alderton, Roberto Amato, Jeffrey Barrett, Sonia Goncalves, Ewan Harrison, David K. Jackson, Ian Johnston, Dominic Kwiatkowski, Cordelia Langford, John Sillitoe on behalf of the Wellcome Sanger Institute COVID-19 Surveillance Team |
| EPI_ISL_3368021 | Lighthouse Lab in Milton Keynes | Wellcome Sanger Institute for the COVID-19 Genomics UK (COG-UK) Consortium | The Lighthouse Lab in Milton Keynes and Alex Alderton, Roberto Amato, Jeffrey Barrett, Sonia Goncalves, Ewan Harrison, David K. Jackson, Ian Johnston, Dominic Kwiatkowski, Cordelia Langford, John Sillitoe on behalf of the Wellcome Sanger Institute COVID-19 Surveillance Team |
| EPI_ISL_3368022 | Berkshire and Surrey Pathology Services Lighthouse Laboratory | Wellcome Sanger Institute for the COVID-19 Genomics UK (COG-UK) Consortium | Berkshire and Surrey Pathology Services Lighthouse Laboratory and Alex Alderton, Roberto Amato, Jeffrey Barrett, Sonia Goncalves, Ewan Harrison, David K. Jackson, Ian Johnston, Dominic Kwiatkowski, Cordelia Langford, John Sillitoe on behalf of the Wellcome Sanger Institute COVID-19 Surveillance Team |
| EPI_ISL_3368023 | Lighthouse Lab in Milton Keynes | Wellcome Sanger Institute for the COVID-19 Genomics UK (COG-UK) Consortium | The Lighthouse Lab in Milton Keynes and Alex Alderton, Roberto Amato, Jeffrey Barrett, Sonia Goncalves, Ewan Harrison, David K. Jackson, Ian Johnston, Dominic Kwiatkowski, Cordelia Langford, John Sillitoe on behalf of the Wellcome Sanger Institute COVID-19 Surveillance Team |
| EPI_ISL_3368024 | Lighthouse Laboratory Plymouth | Wellcome Sanger Institute for the COVID-19 Genomics UK (COG-UK) Consortium | Lighthouse Laboratory Plymouth and Alex Alderton, Roberto Amato, Jeffrey Barrett, Sonia Goncalves, Ewan Harrison, David K. Jackson, Ian Johnston, Dominic Kwiatkowski, Cordelia Langford, John Sillitoe on behalf of the Wellcome Sanger Institute COVID-19 Surveillance Team |
| EPI_ISL_3368025 | Lighthouse Lab in Glasgow | Wellcome Sanger Institute for the COVID-19 Genomics UK (COG-UK) Consortium | Harper VanSteenhouse, Yumi Kasai, David Gray, Carol Clugston, Anna Dominiczak and Alex Alderton, Roberto Amato, Jeffrey Barrett, Sonia Goncalves, Ewan Harrison, David K. Jackson, Ian Johnston, Dominic Kwiatkowski, Cordelia Langford, John Sillitoe on behalf of the Wellcome Sanger Institute COVID-19 Surveillance Team |
| EPI_ISL_3368026 | Lighthouse Laboratory Plymouth | Wellcome Sanger Institute for the COVID-19 Genomics UK (COG-UK) Consortium | Lighthouse Laboratory Plymouth and Alex Alderton, Roberto Amato, Jeffrey Barrett, Sonia Goncalves, Ewan Harrison, David K. Jackson, Ian Johnston, Dominic Kwiatkowski, Cordelia Langford, John Sillitoe on behalf of the Wellcome Sanger Institute COVID-19 Surveillance Team |
| EPI_ISL_3368027, EPI_ISL_3368028 | Lighthouse Lab in Milton Keynes | Wellcome Sanger Institute for the COVID-19 Genomics UK (COG-UK) Consortium | The Lighthouse Lab in Milton Keynes and Alex Alderton, Roberto Amato, Jeffrey Barrett, Sonia Goncalves, Ewan Harrison, David K. Jackson, Ian Johnston, Dominic Kwiatkowski, Cordelia Langford, John Sillitoe on behalf of the Wellcome Sanger Institute COVID-19 Surveillance Team |
| EPI_ISL_3368029 | Lighthouse Laboratory Plymouth | Wellcome Sanger Institute for the COVID-19 Genomics UK (COG-UK) Consortium | Lighthouse Laboratory Plymouth and Alex Alderton, Roberto Amato, Jeffrey Barrett, Sonia Goncalves, Ewan Harrison, David K. Jackson, Ian Johnston, Dominic Kwiatkowski, Cordelia Langford, John Sillitoe on behalf of the Wellcome Sanger Institute COVID-19 Surveillance Team |
| EPI_ISL_3368030, EPI_ISL_3368031 | Berkshire and Surrey Pathology Services Lighthouse Laboratory | Wellcome Sanger Institute for the COVID-19 Genomics UK (COG-UK) Consortium | Berkshire and Surrey Pathology Services Lighthouse Laboratory and Alex Alderton, Roberto Amato, Jeffrey Barrett, Sonia Goncalves, Ewan Harrison, David K. Jackson, Ian Johnston, Dominic Kwiatkowski, Cordelia Langford, John Sillitoe on behalf of the Wellcome Sanger Institute COVID-19 Surveillance Team |
| EPI_ISL_3368032 | Lighthouse Lab in Milton Keynes | Wellcome Sanger Institute for the COVID-19 Genomics UK (COG-UK) Consortium | The Lighthouse Lab in Milton Keynes and Alex Alderton, Roberto Amato, Jeffrey Barrett, Sonia Goncalves, Ewan Harrison, David K. Jackson, Ian Johnston, Dominic Kwiatkowski, Cordelia Langford, John Sillitoe on behalf of the Wellcome Sanger Institute COVID-19 Surveillance Team |
| EPI_ISL_3368033, EPI_ISL_3368034, EPI_ISL_3368035 | Lighthouse Laboratory Plymouth | Wellcome Sanger Institute for the COVID-19 Genomics UK (COG-UK) Consortium | Lighthouse Laboratory Plymouth and Alex Alderton, Roberto Amato, Jeffrey Barrett, Sonia Goncalves, Ewan Harrison, David K. Jackson, Ian Johnston, Dominic Kwiatkowski, Cordelia Langford, John Sillitoe on behalf of the Wellcome Sanger Institute COVID-19 Surveillance Team |
| EPI_ISL_3368036 | Lighthouse Lab in Glasgow | Wellcome Sanger Institute for the COVID-19 Genomics UK (COG-UK) Consortium | Harper VanSteenhouse, Yumi Kasai, David Gray, Carol Clugston, Anna Dominiczak and Alex Alderton, Roberto Amato, Jeffrey Barrett, Sonia Goncalves, Ewan Harrison, David K. Jackson, Ian Johnston, Dominic Kwiatkowski, Cordelia Langford, John Sillitoe on behalf of the Wellcome Sanger Institute COVID-19 Surveillance Team |
| EPI_ISL_3368037, EPI_ISL_3368038 | Berkshire and Surrey Pathology Services Lighthouse Laboratory | Wellcome Sanger Institute for the COVID-19 Genomics UK (COG-UK) Consortium | Berkshire and Surrey Pathology Services Lighthouse Laboratory and Alex Alderton, Roberto Amato, Jeffrey Barrett, Sonia Goncalves, Ewan Harrison, David K. Jackson, Ian Johnston, Dominic Kwiatkowski, Cordelia Langford, John Sillitoe on behalf of the Wellcome Sanger Institute COVID-19 Surveillance Team |
| EPI_ISL_3368039 | Lighthouse Laboratory Plymouth | Wellcome Sanger Institute for the COVID-19 Genomics UK (COG-UK) Consortium | Lighthouse Laboratory Plymouth and Alex Alderton, Roberto Amato, Jeffrey Barrett, Sonia Goncalves, Ewan Harrison, David K. Jackson, Ian Johnston, Dominic Kwiatkowski, Cordelia Langford, John Sillitoe on behalf of the Wellcome Sanger Institute COVID-19 Surveillance Team |
| EPI_ISL_3368040, EPI_ISL_3368041, EPI_ISL_3368042 | Lighthouse Lab in Glasgow | Wellcome Sanger Institute for the COVID-19 Genomics UK (COG-UK) Consortium | Harper VanSteenhouse, Yumi Kasai, David Gray, Carol Clugston, Anna Dominiczak and Alex Alderton, Roberto Amato, Jeffrey Barrett, Sonia Goncalves, Ewan Harrison, David K. Jackson, Ian Johnston, Dominic Kwiatkowski, Cordelia Langford, John Sillitoe on behalf of the Wellcome Sanger Institute COVID-19 Surveillance Team |
| EPI_ISL_3368043 | Lighthouse Lab in Milton Keynes | Wellcome Sanger Institute for the COVID-19 Genomics UK (COG-UK) Consortium | The Lighthouse Lab in Milton Keynes and Alex Alderton, Roberto Amato, Jeffrey Barrett, Sonia Goncalves, Ewan Harrison, David K. Jackson, Ian Johnston, Dominic Kwiatkowski, Cordelia Langford, John Sillitoe on behalf of the Wellcome Sanger Institute COVID-19 Surveillance Team |
| EPI_ISL_3368044 | Berkshire and Surrey Pathology Services Lighthouse Laboratory | Wellcome Sanger Institute for the COVID-19 Genomics UK (COG-UK) Consortium | Berkshire and Surrey Pathology Services Lighthouse Laboratory and Alex Alderton, Roberto Amato, Jeffrey Barrett, Sonia Goncalves, Ewan Harrison, David K. Jackson, Ian Johnston, Dominic Kwiatkowski, Cordelia Langford, John Sillitoe on behalf of the Wellcome Sanger Institute COVID-19 Surveillance Team |
| EPI_ISL_3368045 | Lighthouse Lab in Milton Keynes | Wellcome Sanger Institute for the COVID-19 Genomics UK (COG-UK) Consortium | The Lighthouse Lab in Milton Keynes and Alex Alderton, Roberto Amato, Jeffrey Barrett, Sonia Goncalves, Ewan Harrison, David K. Jackson, Ian Johnston, Dominic Kwiatkowski, Cordelia Langford, John Sillitoe on behalf of the Wellcome Sanger Institute COVID-19 Surveillance Team |
| EPI_ISL_3368046 | Berkshire and Surrey Pathology Services Lighthouse Laboratory | Wellcome Sanger Institute for the COVID-19 Genomics UK (COG-UK) Consortium | Berkshire and Surrey Pathology Services Lighthouse Laboratory and Alex Alderton, Roberto Amato, Jeffrey Barrett, Sonia Goncalves, Ewan Harrison, David K. Jackson, Ian Johnston, Dominic Kwiatkowski, Cordelia Langford, John Sillitoe on behalf of the Wellcome Sanger Institute COVID-19 Surveillance Team |
| EPI_ISL_3368047 | Lighthouse Laboratory Plymouth | Wellcome Sanger Institute for the COVID-19 Genomics UK (COG-UK) Consortium | Lighthouse Laboratory Plymouth and Alex Alderton, Roberto Amato, Jeffrey Barrett, Sonia Goncalves, Ewan Harrison, David K. Jackson, Ian Johnston, Dominic Kwiatkowski, Cordelia Langford, John Sillitoe on behalf of the Wellcome Sanger Institute COVID-19 Surveillance Team |
| EPI_ISL_3368048 | Lighthouse Lab in Glasgow | Wellcome Sanger Institute for the COVID-19 Genomics UK (COG-UK) Consortium | Harper VanSteenhouse, Yumi Kasai, David Gray, Carol Clugston, Anna Dominiczak and Alex Alderton, Roberto Amato, Jeffrey Barrett, Sonia Goncalves, Ewan Harrison, David K. Jackson, Ian Johnston, Dominic Kwiatkowski, Cordelia Langford, John Sillitoe on behalf of the Wellcome Sanger Institute COVID-19 Surveillance Team |
| EPI_ISL_3368049 | Lighthouse Lab in Milton Keynes | Wellcome Sanger Institute for the COVID-19 Genomics UK (COG-UK) Consortium | The Lighthouse Lab in Milton Keynes and Alex Alderton, Roberto Amato, Jeffrey Barrett, Sonia Goncalves, Ewan Harrison, David K. Jackson, Ian Johnston, Dominic Kwiatkowski, Cordelia Langford, John Sillitoe on behalf of the Wellcome Sanger Institute COVID-19 Surveillance Team |
| EPI_ISL_3368050 | Lighthouse Laboratory Plymouth | Wellcome Sanger Institute for the COVID-19 Genomics UK (COG-UK) Consortium | Lighthouse Laboratory Plymouth and Alex Alderton, Roberto Amato, Jeffrey Barrett, Sonia Goncalves, Ewan Harrison, David K. Jackson, Ian Johnston, Dominic Kwiatkowski, Cordelia Langford, John Sillitoe on behalf of the Wellcome Sanger Institute COVID-19 Surveillance Team |
| EPI_ISL_3368051 | Lighthouse Lab in Milton Keynes | Wellcome Sanger Institute for the COVID-19 Genomics UK (COG-UK) Consortium | The Lighthouse Lab in Milton Keynes and Alex Alderton, Roberto Amato, Jeffrey Barrett, Sonia Goncalves, Ewan Harrison, David K. Jackson, Ian Johnston, Dominic Kwiatkowski, Cordelia Langford, John Sillitoe on behalf of the Wellcome Sanger Institute COVID-19 Surveillance Team |
| EPI_ISL_3368052, EPI_ISL_3368053 | Berkshire and Surrey Pathology Services Lighthouse Laboratory | Wellcome Sanger Institute for the COVID-19 Genomics UK (COG-UK) Consortium | Berkshire and Surrey Pathology Services Lighthouse Laboratory and Alex Alderton, Roberto Amato, Jeffrey Barrett, Sonia Goncalves, Ewan Harrison, David K. Jackson, Ian Johnston, Dominic Kwiatkowski, Cordelia Langford, John Sillitoe on behalf of the Wellcome Sanger Institute COVID-19 Surveillance Team |
| EPI_ISL_3368054 | Lighthouse Laboratory Plymouth | Wellcome Sanger Institute for the COVID-19 Genomics UK (COG-UK) Consortium | Lighthouse Laboratory Plymouth and Alex Alderton, Roberto Amato, Jeffrey Barrett, Sonia Goncalves, Ewan Harrison, David K. Jackson, Ian Johnston, Dominic Kwiatkowski, Cordelia Langford, John Sillitoe on behalf of the Wellcome Sanger Institute COVID-19 Surveillance Team |
| EPI_ISL_3368055, EPI_ISL_3368056 | Berkshire and Surrey Pathology Services Lighthouse Laboratory | Wellcome Sanger Institute for the COVID-19 Genomics UK (COG-UK) Consortium | Berkshire and Surrey Pathology Services Lighthouse Laboratory and Alex Alderton, Roberto Amato, Jeffrey Barrett, Sonia Goncalves, Ewan Harrison, David K. Jackson, Ian Johnston, Dominic Kwiatkowski, Cordelia Langford, John Sillitoe on behalf of the Wellcome Sanger Institute COVID-19 Surveillance Team |
| EPI_ISL_3368057 | Lighthouse Lab in Glasgow | Wellcome Sanger Institute for the COVID-19 Genomics UK (COG-UK) Consortium | Harper VanSteenhouse, Yumi Kasai, David Gray, Carol Clugston, Anna Dominiczak and Alex Alderton, Roberto Amato, Jeffrey Barrett, Sonia Goncalves, Ewan Harrison, David K. Jackson, Ian Johnston, Dominic Kwiatkowski, Cordelia Langford, John Sillitoe on behalf of the Wellcome Sanger Institute COVID-19 Surveillance Team |
| EPI_ISL_3368058, EPI_ISL_3368059 | Lighthouse Lab in Milton Keynes | Wellcome Sanger Institute for the COVID-19 Genomics UK (COG-UK) Consortium | The Lighthouse Lab in Milton Keynes and Alex Alderton, Roberto Amato, Jeffrey Barrett, Sonia Goncalves, Ewan Harrison, David K. Jackson, Ian Johnston, Dominic Kwiatkowski, Cordelia Langford, John Sillitoe on behalf of the Wellcome Sanger Institute COVID-19 Surveillance Team |
| EPI_ISL_3368060 | Lighthouse Lab in Glasgow | Wellcome Sanger Institute for the COVID-19 Genomics UK (COG-UK) Consortium | Harper VanSteenhouse, Yumi Kasai, David Gray, Carol Clugston, Anna Dominiczak and Alex Alderton, Roberto Amato, Jeffrey Barrett, Sonia Goncalves, Ewan Harrison, David K. Jackson, Ian Johnston, Dominic Kwiatkowski, Cordelia Langford, John Sillitoe on behalf of the Wellcome Sanger Institute COVID-19 Surveillance Team |
| EPI_ISL_3368061 | Lighthouse Laboratory Plymouth | Wellcome Sanger Institute for the COVID-19 Genomics UK (COG-UK) Consortium | Lighthouse Laboratory Plymouth and Alex Alderton, Roberto Amato, Jeffrey Barrett, Sonia Goncalves, Ewan Harrison, David K. Jackson, Ian Johnston, Dominic Kwiatkowski, Cordelia Langford, John Sillitoe on behalf of the Wellcome Sanger Institute COVID-19 Surveillance Team |
| EPI_ISL_3368062, EPI_ISL_3368063 | Lighthouse Lab in Glasgow | Wellcome Sanger Institute for the COVID-19 Genomics UK (COG-UK) Consortium | Harper VanSteenhouse, Yumi Kasai, David Gray, Carol Clugston, Anna Dominiczak and Alex Alderton, Roberto Amato, Jeffrey Barrett, Sonia Goncalves, Ewan Harrison, David K. Jackson, Ian Johnston, Dominic Kwiatkowski, Cordelia Langford, John Sillitoe on behalf of the Wellcome Sanger Institute COVID-19 Surveillance Team |
| EPI_ISL_3368064, EPI_ISL_3368065, EPI_ISL_3368066 | Lighthouse Laboratory Plymouth | Wellcome Sanger Institute for the COVID-19 Genomics UK (COG-UK) Consortium | Lighthouse Laboratory Plymouth and Alex Alderton, Roberto Amato, Jeffrey Barrett, Sonia Goncalves, Ewan Harrison, David K. Jackson, Ian Johnston, Dominic Kwiatkowski, Cordelia Langford, John Sillitoe on behalf of the Wellcome Sanger Institute COVID-19 Surveillance Team |

| | | | |
|---|---|---|---|
| EPI_ISL_3368067 | Berkshire and Surrey Pathology Services Lighthouse Laboratory | Wellcome Sanger Institute for the COVID-19 Genomics UK (COG-UK) Consortium | Berkshire and Surrey Pathology Services Lighthouse Laboratory and Alex Alderton, Roberto Amato, Jeffrey Barrett, Sonia Goncalves, Ewan Harrison, David K. Jackson, Ian Johnston, Dominic Kwiatkowski, Cordelia Langford, John Sillitoe on behalf of the Wellcome Sanger Institute COVID-19 Surveillance Team |
| EPI_ISL_3368068 | Lighthouse Laboratory Plymouth | Wellcome Sanger Institute for the COVID-19 Genomics UK (COG-UK) Consortium | Lighthouse Laboratory Plymouth and Alex Alderton, Roberto Amato, Jeffrey Barrett, Sonia Goncalves, Ewan Harrison, David K. Jackson, Ian Johnston, Dominic Kwiatkowski, Cordelia Langford, John Sillitoe on behalf of the Wellcome Sanger Institute COVID-19 Surveillance Team |
| EPI_ISL_3368069 | Lighthouse Lab in Glasgow | Wellcome Sanger Institute for the COVID-19 Genomics UK (COG-UK) Consortium | Harper VanSteenhouse, Yumi Kasai, David Gray, Carol Clugston, Anna Dominiczak and Alex Alderton, Roberto Amato, Jeffrey Barrett, Sonia Goncalves, Ewan Harrison, David K. Jackson, Ian Johnston, Dominic Kwiatkowski, Cordelia Langford, John Sillitoe on behalf of the Wellcome Sanger Institute COVID-19 Surveillance Team |
| EPI_ISL_3368070 | Berkshire and Surrey Pathology Services Lighthouse Laboratory | Wellcome Sanger Institute for the COVID-19 Genomics UK (COG-UK) Consortium | Berkshire and Surrey Pathology Services Lighthouse Laboratory and Alex Alderton, Roberto Amato, Jeffrey Barrett, Sonia Goncalves, Ewan Harrison, David K. Jackson, Ian Johnston, Dominic Kwiatkowski, Cordelia Langford, John Sillitoe on behalf of the Wellcome Sanger Institute COVID-19 Surveillance Team |
| EPI_ISL_3368071 | Lighthouse Lab in Milton Keynes | Wellcome Sanger Institute for the COVID-19 Genomics UK (COG-UK) Consortium | The Lighthouse Lab in Milton Keynes and Alex Alderton, Roberto Amato, Jeffrey Barrett, Sonia Goncalves, Ewan Harrison, David K. Jackson, Ian Johnston, Dominic Kwiatkowski, Cordelia Langford, John Sillitoe on behalf of the Wellcome Sanger Institute COVID-19 Surveillance Team |
| EPI_ISL_3368072 | Lighthouse Lab in Glasgow | Wellcome Sanger Institute for the COVID-19 Genomics UK (COG-UK) Consortium | Harper VanSteenhouse, Yumi Kasai, David Gray, Carol Clugston, Anna Dominiczak and Alex Alderton, Roberto Amato, Jeffrey Barrett, Sonia Goncalves, Ewan Harrison, David K. Jackson, Ian Johnston, Dominic Kwiatkowski, Cordelia Langford, John Sillitoe on behalf of the Wellcome Sanger Institute COVID-19 Surveillance Team |
| EPI_ISL_3368073, EPI_ISL_3368074 | Lighthouse Lab in Milton Keynes | Wellcome Sanger Institute for the COVID-19 Genomics UK (COG-UK) Consortium | The Lighthouse Lab in Milton Keynes and Alex Alderton, Roberto Amato, Jeffrey Barrett, Sonia Goncalves, Ewan Harrison, David K. Jackson, Ian Johnston, Dominic Kwiatkowski, Cordelia Langford, John Sillitoe on behalf of the Wellcome Sanger Institute COVID-19 Surveillance Team |
| EPI_ISL_3368075 | Lighthouse Laboratory Plymouth | Wellcome Sanger Institute for the COVID-19 Genomics UK (COG-UK) Consortium | Lighthouse Laboratory Plymouth and Alex Alderton, Roberto Amato, Jeffrey Barrett, Sonia Goncalves, Ewan Harrison, David K. Jackson, Ian Johnston, Dominic Kwiatkowski, Cordelia Langford, John Sillitoe on behalf of the Wellcome Sanger Institute COVID-19 Surveillance Team |
| EPI_ISL_3368076 | Lighthouse Lab in Glasgow | Wellcome Sanger Institute for the COVID-19 Genomics UK (COG-UK) Consortium | Harper VanSteenhouse, Yumi Kasai, David Gray, Carol Clugston, Anna Dominiczak and Alex Alderton, Roberto Amato, Jeffrey Barrett, Sonia Goncalves, Ewan Harrison, David K. Jackson, Ian Johnston, Dominic Kwiatkowski, Cordelia Langford, John Sillitoe on behalf of the Wellcome Sanger Institute COVID-19 Surveillance Team |
| EPI_ISL_3368077 | Lighthouse Laboratory Plymouth | Wellcome Sanger Institute for the COVID-19 Genomics UK (COG-UK) Consortium | Lighthouse Laboratory Plymouth and Alex Alderton, Roberto Amato, Jeffrey Barrett, Sonia Goncalves, Ewan Harrison, David K. Jackson, Ian Johnston, Dominic Kwiatkowski, Cordelia Langford, John Sillitoe on behalf of the Wellcome Sanger Institute COVID-19 Surveillance Team |
| EPI_ISL_3368078 | Lighthouse Lab in Milton Keynes | Wellcome Sanger Institute for the COVID-19 Genomics UK (COG-UK) Consortium | The Lighthouse Lab in Milton Keynes and Alex Alderton, Roberto Amato, Jeffrey Barrett, Sonia Goncalves, Ewan Harrison, David K. Jackson, Ian Johnston, Dominic Kwiatkowski, Cordelia Langford, John Sillitoe on behalf of the Wellcome Sanger Institute COVID-19 Surveillance Team |
| EPI_ISL_3368079 | Lighthouse Laboratory Plymouth | Wellcome Sanger Institute for the COVID-19 Genomics UK (COG-UK) Consortium | Lighthouse Laboratory Plymouth and Alex Alderton, Roberto Amato, Jeffrey Barrett, Sonia Goncalves, Ewan Harrison, David K. Jackson, Ian Johnston, Dominic Kwiatkowski, Cordelia Langford, John Sillitoe on behalf of the Wellcome Sanger Institute COVID-19 Surveillance Team |
| EPI_ISL_3368080 | Lighthouse Lab in Glasgow | Wellcome Sanger Institute for the COVID-19 Genomics UK (COG-UK) Consortium | Harper VanSteenhouse, Yumi Kasai, David Gray, Carol Clugston, Anna Dominiczak and Alex Alderton, Roberto Amato, Jeffrey Barrett, Sonia Goncalves, Ewan Harrison, David K. Jackson, Ian Johnston, Dominic Kwiatkowski, Cordelia Langford, John Sillitoe on behalf of the Wellcome Sanger Institute COVID-19 Surveillance Team |
| EPI_ISL_3368081 | Berkshire and Surrey Pathology Services Lighthouse Laboratory | Wellcome Sanger Institute for the COVID-19 Genomics UK (COG-UK) Consortium | Berkshire and Surrey Pathology Services Lighthouse Laboratory and Alex Alderton, Roberto Amato, Jeffrey Barrett, Sonia Goncalves, Ewan Harrison, David K. Jackson, Ian Johnston, Dominic Kwiatkowski, Cordelia Langford, John Sillitoe on behalf of the Wellcome Sanger Institute COVID-19 Surveillance Team |
| EPI_ISL_3368082 | Lighthouse Lab in Glasgow | Wellcome Sanger Institute for the COVID-19 Genomics UK (COG-UK) Consortium | Harper VanSteenhouse, Yumi Kasai, David Gray, Carol Clugston, Anna Dominiczak and Alex Alderton, Roberto Amato, Jeffrey Barrett, Sonia Goncalves, Ewan Harrison, David K. Jackson, Ian Johnston, Dominic Kwiatkowski, Cordelia Langford, John Sillitoe on behalf of the Wellcome Sanger Institute COVID-19 Surveillance Team |
| EPI_ISL_3368083 | Berkshire and Surrey Pathology Services Lighthouse Laboratory | Wellcome Sanger Institute for the COVID-19 Genomics UK (COG-UK) Consortium | Berkshire and Surrey Pathology Services Lighthouse Laboratory and Alex Alderton, Roberto Amato, Jeffrey Barrett, Sonia Goncalves, Ewan Harrison, David K. Jackson, Ian Johnston, Dominic Kwiatkowski, Cordelia Langford, John Sillitoe on behalf of the Wellcome Sanger Institute COVID-19 Surveillance Team |
| EPI_ISL_3368084, EPI_ISL_3368085, EPI_ISL_3368086 | Lighthouse Lab in Milton Keynes | Wellcome Sanger Institute for the COVID-19 Genomics UK (COG-UK) Consortium | The Lighthouse Lab in Milton Keynes and Alex Alderton, Roberto Amato, Jeffrey Barrett, Sonia Goncalves, Ewan Harrison, David K. Jackson, Ian Johnston, Dominic Kwiatkowski, Cordelia Langford, John Sillitoe on behalf of the Wellcome Sanger Institute COVID-19 Surveillance Team |
| EPI_ISL_3368087 | Berkshire and Surrey Pathology Services Lighthouse Laboratory | Wellcome Sanger Institute for the COVID-19 Genomics UK (COG-UK) Consortium | Berkshire and Surrey Pathology Services Lighthouse Laboratory and Alex Alderton, Roberto Amato, Jeffrey Barrett, Sonia Goncalves, Ewan Harrison, David K. Jackson, Ian Johnston, Dominic Kwiatkowski, Cordelia Langford, John Sillitoe on behalf of the Wellcome Sanger Institute COVID-19 Surveillance Team |
| EPI_ISL_3368088 | Lighthouse Lab in Milton Keynes | Wellcome Sanger Institute for the COVID-19 Genomics UK (COG-UK) Consortium | The Lighthouse Lab in Milton Keynes and Alex Alderton, Roberto Amato, Jeffrey Barrett, Sonia Goncalves, Ewan Harrison, David K. Jackson, Ian Johnston, Dominic Kwiatkowski, Cordelia Langford, John Sillitoe on behalf of the Wellcome Sanger Institute COVID-19 Surveillance Team |
| EPI_ISL_3368089, EPI_ISL_3368090 | Berkshire and Surrey Pathology Services Lighthouse Laboratory | Wellcome Sanger Institute for the COVID-19 Genomics UK (COG-UK) Consortium | Berkshire and Surrey Pathology Services Lighthouse Laboratory and Alex Alderton, Roberto Amato, Jeffrey Barrett, Sonia Goncalves, Ewan Harrison, David K. Jackson, Ian Johnston, Dominic Kwiatkowski, Cordelia Langford, John Sillitoe on behalf of the Wellcome Sanger Institute COVID-19 Surveillance Team |
| EPI_ISL_3368091 | Lighthouse Laboratory Plymouth | Wellcome Sanger Institute for the COVID-19 Genomics UK (COG-UK) Consortium | Lighthouse Laboratory Plymouth and Alex Alderton, Roberto Amato, Jeffrey Barrett, Sonia Goncalves, Ewan Harrison, David K. Jackson, Ian Johnston, Dominic Kwiatkowski, Cordelia Langford, John Sillitoe on behalf of the Wellcome Sanger Institute COVID-19 Surveillance Team |
| EPI_ISL_3368092, EPI_ISL_3368093, EPI_ISL_3368094 | Lighthouse Lab in Milton Keynes | Wellcome Sanger Institute for the COVID-19 Genomics UK (COG-UK) Consortium | The Lighthouse Lab in Milton Keynes and Alex Alderton, Roberto Amato, Jeffrey Barrett, Sonia Goncalves, Ewan Harrison, David K. Jackson, Ian Johnston, Dominic Kwiatkowski, Cordelia Langford, John Sillitoe on behalf of the Wellcome Sanger Institute COVID-19 Surveillance Team |
| EPI_ISL_3368095, EPI_ISL_3368096 | Lighthouse Laboratory Plymouth | Wellcome Sanger Institute for the COVID-19 Genomics UK (COG-UK) Consortium | Lighthouse Laboratory Plymouth and Alex Alderton, Roberto Amato, Jeffrey Barrett, Sonia Goncalves, Ewan Harrison, David K. Jackson, Ian Johnston, Dominic Kwiatkowski, Cordelia Langford, John Sillitoe on behalf of the Wellcome Sanger Institute COVID-19 Surveillance Team |
| EPI_ISL_3368097 | Lighthouse Lab in Glasgow | Wellcome Sanger Institute for the COVID-19 Genomics UK (COG-UK) Consortium | Harper VanSteenhouse, Yumi Kasai, David Gray, Carol Clugston, Anna Dominiczak and Alex Alderton, Roberto Amato, Jeffrey Barrett, Sonia Goncalves, Ewan Harrison, David K. Jackson, Ian Johnston, Dominic Kwiatkowski, Cordelia Langford, John Sillitoe on behalf of the Wellcome Sanger Institute COVID-19 Surveillance Team |
| EPI_ISL_3368098 | Berkshire and Surrey Pathology Services Lighthouse Laboratory | Wellcome Sanger Institute for the COVID-19 Genomics UK (COG-UK) Consortium | Berkshire and Surrey Pathology Services Lighthouse Laboratory and Alex Alderton, Roberto Amato, Jeffrey Barrett, Sonia Goncalves, Ewan Harrison, David K. Jackson, Ian Johnston, Dominic Kwiatkowski, Cordelia Langford, John Sillitoe on behalf of the Wellcome Sanger Institute COVID-19 Surveillance Team |
| EPI_ISL_3368099 | Lighthouse Lab in Glasgow | Wellcome Sanger Institute for the COVID-19 Genomics UK (COG-UK) Consortium | Harper VanSteenhouse, Yumi Kasai, David Gray, Carol Clugston, Anna Dominiczak and Alex Alderton, Roberto Amato, Jeffrey Barrett, Sonia Goncalves, Ewan Harrison, David K. Jackson, Ian Johnston, Dominic Kwiatkowski, Cordelia Langford, John Sillitoe on behalf of the Wellcome Sanger Institute COVID-19 Surveillance Team |
| EPI_ISL_3368100 | Lighthouse Lab in Milton Keynes | Wellcome Sanger Institute for the COVID-19 Genomics UK (COG-UK) Consortium | The Lighthouse Lab in Milton Keynes and Alex Alderton, Roberto Amato, Jeffrey Barrett, Sonia Goncalves, Ewan Harrison, David K. Jackson, Ian Johnston, Dominic Kwiatkowski, Cordelia Langford, John Sillitoe on behalf of the Wellcome Sanger Institute COVID-19 Surveillance Team |
| EPI_ISL_3368101 | Berkshire and Surrey Pathology Services Lighthouse Laboratory | Wellcome Sanger Institute for the COVID-19 Genomics UK (COG-UK) Consortium | Berkshire and Surrey Pathology Services Lighthouse Laboratory and Alex Alderton, Roberto Amato, Jeffrey Barrett, Sonia Goncalves, Ewan Harrison, David K. Jackson, Ian Johnston, Dominic Kwiatkowski, Cordelia Langford, John Sillitoe on behalf of the Wellcome Sanger Institute COVID-19 Surveillance Team |
| EPI_ISL_3368102, EPI_ISL_3368103 | Lighthouse Laboratory Plymouth | Wellcome Sanger Institute for the COVID-19 Genomics UK (COG-UK) Consortium | Lighthouse Laboratory Plymouth and Alex Alderton, Roberto Amato, Jeffrey Barrett, Sonia Goncalves, Ewan Harrison, David K. Jackson, Ian Johnston, Dominic Kwiatkowski, Cordelia Langford, John Sillitoe on behalf of the Wellcome Sanger Institute COVID-19 Surveillance Team |
| EPI_ISL_3368104, EPI_ISL_3368105 | Berkshire and Surrey Pathology Services Lighthouse Laboratory | Wellcome Sanger Institute for the COVID-19 Genomics UK (COG-UK) Consortium | Berkshire and Surrey Pathology Services Lighthouse Laboratory and Alex Alderton, Roberto Amato, Jeffrey Barrett, Sonia Goncalves, Ewan Harrison, David K. Jackson, Ian Johnston, Dominic Kwiatkowski, Cordelia Langford, John Sillitoe on behalf of the Wellcome Sanger Institute COVID-19 Surveillance Team |
| EPI_ISL_3368106 | Lighthouse Laboratory Plymouth | Wellcome Sanger Institute for the COVID-19 Genomics UK (COG-UK) Consortium | Lighthouse Laboratory Plymouth and Alex Alderton, Roberto Amato, Jeffrey Barrett, Sonia Goncalves, Ewan Harrison, David K. Jackson, Ian Johnston, Dominic Kwiatkowski, Cordelia Langford, John Sillitoe on behalf of the Wellcome Sanger Institute COVID-19 Surveillance Team |
| EPI_ISL_3368107 | Berkshire and Surrey Pathology Services Lighthouse Laboratory | Wellcome Sanger Institute for the COVID-19 Genomics UK (COG-UK) Consortium | Berkshire and Surrey Pathology Services Lighthouse Laboratory and Alex Alderton, Roberto Amato, Jeffrey Barrett, Sonia Goncalves, Ewan Harrison, David K. Jackson, Ian Johnston, Dominic Kwiatkowski, Cordelia Langford, John Sillitoe on behalf of the Wellcome Sanger Institute COVID-19 Surveillance Team |
| EPI_ISL_3368108 | Lighthouse Lab in Milton Keynes | Wellcome Sanger Institute for the COVID-19 Genomics UK (COG-UK) Consortium | The Lighthouse Lab in Milton Keynes and Alex Alderton, Roberto Amato, Jeffrey Barrett, Sonia Goncalves, Ewan Harrison, David K. Jackson, Ian Johnston, Dominic Kwiatkowski, Cordelia Langford, John Sillitoe on behalf of the Wellcome Sanger Institute COVID-19 Surveillance Team |
| EPI_ISL_3368109 | Lighthouse Lab in Glasgow | Wellcome Sanger Institute for the COVID-19 Genomics UK (COG-UK) Consortium | Harper VanSteenhouse, Yumi Kasai, David Gray, Carol Clugston, Anna Dominiczak and Alex Alderton, Roberto Amato, Jeffrey Barrett, Sonia Goncalves, Ewan Harrison, David K. Jackson, Ian Johnston, Dominic Kwiatkowski, Cordelia Langford, John Sillitoe on behalf of the Wellcome Sanger Institute COVID-19 Surveillance Team |
| EPI_ISL_3368110 | Lighthouse Lab in Milton Keynes | Wellcome Sanger Institute for the COVID-19 Genomics UK (COG-UK) Consortium | The Lighthouse Lab in Milton Keynes and Alex Alderton, Roberto Amato, Jeffrey Barrett, Sonia Goncalves, Ewan Harrison, David K. Jackson, Ian Johnston, Dominic Kwiatkowski, Cordelia Langford, John Sillitoe on behalf of the Wellcome Sanger Institute COVID-19 Surveillance Team |
| EPI_ISL_3368111 | Berkshire and Surrey Pathology Services Lighthouse Laboratory | Wellcome Sanger Institute for the COVID-19 Genomics UK (COG-UK) Consortium | Berkshire and Surrey Pathology Services Lighthouse Laboratory and Alex Alderton, Roberto Amato, Jeffrey Barrett, Sonia Goncalves, Ewan Harrison, David K. Jackson, Ian Johnston, Dominic Kwiatkowski, Cordelia Langford, John Sillitoe on behalf of the Wellcome Sanger Institute COVID-19 Surveillance Team |
| EPI_ISL_3368112 | Lighthouse Lab in Milton Keynes | Wellcome Sanger Institute for the COVID-19 Genomics UK (COG-UK) Consortium | The Lighthouse Lab in Milton Keynes and Alex Alderton, Roberto Amato, Jeffrey Barrett, Sonia Goncalves, Ewan Harrison, David K. Jackson, Ian Johnston, Dominic Kwiatkowski, Cordelia Langford, John Sillitoe on behalf of the Wellcome Sanger Institute COVID-19 Surveillance Team |
| EPI_ISL_3368113 | Lighthouse Laboratory Plymouth | Wellcome Sanger Institute for the COVID-19 Genomics UK (COG-UK) Consortium | Lighthouse Laboratory Plymouth and Alex Alderton, Roberto Amato, Jeffrey Barrett, Sonia Goncalves, Ewan Harrison, David K. Jackson, Ian Johnston, Dominic Kwiatkowski, Cordelia Langford, John Sillitoe on behalf of the Wellcome Sanger Institute COVID-19 Surveillance Team |
| EPI_ISL_3368114 | Lighthouse Lab in Glasgow | Wellcome Sanger Institute for the COVID-19 Genomics UK (COG-UK) Consortium | Harper VanSteenhouse, Yumi Kasai, David Gray, Carol Clugston, Anna Dominiczak and Alex Alderton, Roberto Amato, Jeffrey Barrett, Sonia Goncalves, Ewan Harrison, David K. Jackson, Ian Johnston, Dominic Kwiatkowski, Cordelia Langford, John Sillitoe on behalf of the Wellcome Sanger Institute COVID-19 Surveillance Team |
| EPI_ISL_3368115 | Lighthouse Laboratory Plymouth | Wellcome Sanger Institute for the COVID-19 Genomics UK (COG-UK) Consortium | Lighthouse Laboratory Plymouth and Alex Alderton, Roberto Amato, Jeffrey Barrett, Sonia Goncalves, Ewan Harrison, David K. Jackson, Ian Johnston, Dominic Kwiatkowski, Cordelia Langford, John Sillitoe on behalf of the Wellcome Sanger Institute COVID-19 Surveillance Team |
| EPI_ISL_3368116, EPI_ISL_3368117, EPI_ISL_3368118, EPI_ISL_3368119, EPI_ISL_3368120, EPI_ISL_3368121, EPI_ISL_3368122, EPI_ISL_3368123, EPI_ISL_3368124, EPI_ISL_3368125, EPI_ISL_3368126, EPI_ISL_3368127, EPI_ISL_3368128, EPI_ISL_3368129, EPI_ISL_3368130, EPI_ISL_3368131, EPI_ISL_3368132, EPI_ISL_3368133, EPI_ISL_3368134, EPI_ISL_3368135, EPI_ISL_3368136, EPI_ISL_3368137, EPI_ISL_3368138, EPI_ISL_3368139, EPI_ISL_3368140, EPI_ISL_3368141, EPI_ISL_3368142, EPI_ISL_3368143, EPI_ISL_3368144, EPI_ISL_3368145, EPI_ISL_3368146, EPI_ISL_3368147, EPI_ISL_3368148, EPI_ISL_3368149, EPI_ISL_3368150, EPI_ISL_3368151, EPI_ISL_3368152, EPI_ISL_3368153, EPI_ISL_3368154, EPI_ISL_3368155, EPI_ISL_3368156, EPI_ISL_3368157, EPI_ISL_3368158, EPI_ISL_3368159, EPI_ISL_3368160, EPI_ISL_3368161, EPI_ISL_3368162, EPI_ISL_3368163, EPI_ISL_3368164, EPI_ISL_3368165, EPI_ISL_3368166, EPI_ISL_3368167, EPI_ISL_3368168, EPI_ISL_3368169, EPI_ISL_3368170, EPI_ISL_3368171, EPI_ISL_3368172, EPI_ISL_3368173, EPI_ISL_3368174, EPI_ISL_3368175, EPI_ISL_3368176, EPI_ISL_3368177, EPI_ISL_3368178, EPI_ISL_3368179, EPI_ISL_3368180, EPI_ISL_3368181, EPI_ISL_3368182, EPI_ISL_3368183, EPI_ISL_3368184, EPI_ISL_3368185, EPI_ISL_3368186, EPI_ISL_3368187, EPI_ISL_3368188, EPI_ISL_3368189, EPI_ISL_3368190, EPI_ISL_3368191, EPI_ISL_3368192, EPI_ISL_3368193, EPI_ISL_3368194, EPI_ISL_3368195, EPI_ISL_3368196, EPI_ISL_3368197, EPI_ISL_3368198, EPI_ISL_3368199, EPI_ISL_3368200, EPI_ISL_3368201, EPI_ISL_3368202, EPI_ISL_3368203, EPI_ISL_3368204, EPI_ISL_3368205, EPI_ISL_3368206, EPI_ISL_3368207, EPI_ISL_3368208, EPI_ISL_3368209, EPI_ISL_3368210, EPI_ISL_3368211, EPI_ISL_3368212, EPI_ISL_3368213, EPI_ISL_3368214, EPI_ISL_3368215, EPI_ISL_3368216, EPI_ISL_3368217, EPI_ISL_3368218, EPI_ISL_3368219, EPI_ISL_3368220, EPI_ISL_3368221, EPI_ISL_3368222, EPI_ISL_3368223, EPI_ISL_3368224, EPI_ISL_3368225, EPI_ISL_3368226, EPI_ISL_3368227, EPI_ISL_3368228, EPI_ISL_3368229, EPI_ISL_3368230, EPI_ISL_3368231, EPI_ISL_3368232, EPI_ISL_3368233, EPI_ISL_3368234, EPI_ISL_3368235, EPI_ISL_3368236, EPI_ISL_3368237, EPI_ISL_3368238, EPI_ISL_3368239, EPI_ISL_3368240, EPI_ISL_3368241, EPI_ISL_3368242, EPI_ISL_3368243, EPI_ISL_3368244, EPI_ISL_3368245, EPI_ISL_3368246, EPI_ISL_3368247, EPI_ISL_3368248, EPI_ISL_3368249, EPI_ISL_3368250, EPI_ISL_3368251, EPI_ISL_3368252, EPI_ISL_3368253, EPI_ISL_3368254, EPI_ISL_3368255, EPI_ISL_3368256, EPI_ISL_3368257, EPI_ISL_3368258, EPI_ISL_3368259, EPI_ISL_3368260, EPI_ISL_3368261, EPI_ISL_3368262, EPI_ISL_3368263, EPI_ISL_3368264, EPI_ISL_3368265 | | | |
| see above | Lighthouse Lab in Glasgow | Wellcome Sanger Institute for the COVID-19 Genomics UK (COG-UK) Consortium | Harper VanSteenhouse, Yumi Kasai, David Gray, Carol Clugston, Anna Dominiczak and Alex Alderton, Roberto Amato, Jeffrey Barrett, Sonia Goncalves, Ewan Harrison, David K. Jackson, Ian Johnston, Dominic Kwiatkowski, Cordelia Langford, John Sillitoe on behalf of the Wellcome Sanger Institute COVID-19 Surveillance Team |

| | | | |
|---|---|---|---|
| EPI_ISL_3368266 | Lighthouse Lab in Alderley Park | Wellcome Sanger Institute for the COVID-19 Genomics UK (COG-UK) Consortium | Jacquelyn Wynn, Mairead Hyland, The Lighthouse Lab in Alderley Park and Alex Alderton, Roberto Amato, Jeffrey Barrett, Sonia Goncalves, Ewan Harrison, David K. Jackson, Ian Johnston, Dominic Kwiatkowski, Cordelia Langford, John Sillitoe on behalf of the Wellcome Sanger Institute COVID-19 Surveillance Team |
| EPI_ISL_3368267 | Lighthouse Lab in Glasgow | Wellcome Sanger Institute for the COVID-19 Genomics UK (COG-UK) Consortium | Harper VanSteenhouse, Yumi Kasai, David Gray, Carol Clugston, Anna Dominiczak and Alex Alderton, Roberto Amato, Jeffrey Barrett, Sonia Goncalves, Ewan Harrison, David K. Jackson, Ian Johnston, Dominic Kwiatkowski, Cordelia Langford, John Sillitoe on behalf of the Wellcome Sanger Institute COVID-19 Surveillance Team |
| EPI_ISL_3368268 | Lighthouse Lab in Alderley Park | Wellcome Sanger Institute for the COVID-19 Genomics UK (COG-UK) Consortium | Jacquelyn Wynn, Mairead Hyland, The Lighthouse Lab in Alderley Park and Alex Alderton, Roberto Amato, Jeffrey Barrett, Sonia Goncalves, Ewan Harrison, David K. Jackson, Ian Johnston, Dominic Kwiatkowski, Cordelia Langford, John Sillitoe on behalf of the Wellcome Sanger Institute COVID-19 Surveillance Team |
| EPI_ISL_3368269 | Lighthouse Lab in Glasgow | Wellcome Sanger Institute for the COVID-19 Genomics UK (COG-UK) Consortium | Harper VanSteenhouse, Yumi Kasai, David Gray, Carol Clugston, Anna Dominiczak and Alex Alderton, Roberto Amato, Jeffrey Barrett, Sonia Goncalves, Ewan Harrison, David K. Jackson, Ian Johnston, Dominic Kwiatkowski, Cordelia Langford, John Sillitoe on behalf of the Wellcome Sanger Institute COVID-19 Surveillance Team |
| EPI_ISL_3368270 | Lighthouse Lab in Alderley Park | Wellcome Sanger Institute for the COVID-19 Genomics UK (COG-UK) Consortium | Jacquelyn Wynn, Mairead Hyland, The Lighthouse Lab in Alderley Park and Alex Alderton, Roberto Amato, Jeffrey Barrett, Sonia Goncalves, Ewan Harrison, David K. Jackson, Ian Johnston, Dominic Kwiatkowski, Cordelia Langford, John Sillitoe on behalf of the Wellcome Sanger Institute COVID-19 Surveillance Team |
| EPI_ISL_3368271 | Lighthouse Lab in Glasgow | Wellcome Sanger Institute for the COVID-19 Genomics UK (COG-UK) Consortium | Harper VanSteenhouse, Yumi Kasai, David Gray, Carol Clugston, Anna Dominiczak and Alex Alderton, Roberto Amato, Jeffrey Barrett, Sonia Goncalves, Ewan Harrison, David K. Jackson, Ian Johnston, Dominic Kwiatkowski, Cordelia Langford, John Sillitoe on behalf of the Wellcome Sanger Institute COVID-19 Surveillance Team |
| EPI_ISL_3368272 | Lighthouse Lab in Milton Keynes | Wellcome Sanger Institute for the COVID-19 Genomics UK (COG-UK) Consortium | The Lighthouse Lab in Milton Keynes and Alex Alderton, Roberto Amato, Jeffrey Barrett, Sonia Goncalves, Ewan Harrison, David K. Jackson, Ian Johnston, Dominic Kwiatkowski, Cordelia Langford, John Sillitoe on behalf of the Wellcome Sanger Institute COVID-19 Surveillance Team |
| EPI_ISL_3368273, EPI_ISL_3368274, EPI_ISL_3368275 | Lighthouse Lab in Glasgow | Wellcome Sanger Institute for the COVID-19 Genomics UK (COG-UK) Consortium | Harper VanSteenhouse, Yumi Kasai, David Gray, Carol Clugston, Anna Dominiczak and Alex Alderton, Roberto Amato, Jeffrey Barrett, Sonia Goncalves, Ewan Harrison, David K. Jackson, Ian Johnston, Dominic Kwiatkowski, Cordelia Langford, John Sillitoe on behalf of the Wellcome Sanger Institute COVID-19 Surveillance Team |
| EPI_ISL_3368276 | Lighthouse Lab in Alderley Park | Wellcome Sanger Institute for the COVID-19 Genomics UK (COG-UK) Consortium | Jacquelyn Wynn, Mairead Hyland, The Lighthouse Lab in Alderley Park and Alex Alderton, Roberto Amato, Jeffrey Barrett, Sonia Goncalves, Ewan Harrison, David K. Jackson, Ian Johnston, Dominic Kwiatkowski, Cordelia Langford, John Sillitoe on behalf of the Wellcome Sanger Institute COVID-19 Surveillance Team |
| EPI_ISL_3368277, EPI_ISL_3368278, EPI_ISL_3368279, EPI_ISL_3368280, EPI_ISL_3368281 | Lighthouse Lab in Glasgow | Wellcome Sanger Institute for the COVID-19 Genomics UK (COG-UK) Consortium | Harper VanSteenhouse, Yumi Kasai, David Gray, Carol Clugston, Anna Dominiczak and Alex Alderton, Roberto Amato, Jeffrey Barrett, Sonia Goncalves, Ewan Harrison, David K. Jackson, Ian Johnston, Dominic Kwiatkowski, Cordelia Langford, John Sillitoe on behalf of the Wellcome Sanger Institute COVID-19 Surveillance Team |
| EPI_ISL_3368282 | Lighthouse Lab in Alderley Park | Wellcome Sanger Institute for the COVID-19 Genomics UK (COG-UK) Consortium | Jacquelyn Wynn, Mairead Hyland, The Lighthouse Lab in Alderley Park and Alex Alderton, Roberto Amato, Jeffrey Barrett, Sonia Goncalves, Ewan Harrison, David K. Jackson, Ian Johnston, Dominic Kwiatkowski, Cordelia Langford, John Sillitoe on behalf of the Wellcome Sanger Institute COVID-19 Surveillance Team |
| EPI_ISL_3368283, EPI_ISL_3368284 | Lighthouse Lab in Glasgow | Wellcome Sanger Institute for the COVID-19 Genomics UK (COG-UK) Consortium | Harper VanSteenhouse, Yumi Kasai, David Gray, Carol Clugston, Anna Dominiczak and Alex Alderton, Roberto Amato, Jeffrey Barrett, Sonia Goncalves, Ewan Harrison, David K. Jackson, Ian Johnston, Dominic Kwiatkowski, Cordelia Langford, John Sillitoe on behalf of the Wellcome Sanger Institute COVID-19 Surveillance Team |
| EPI_ISL_3368285 | Lighthouse Lab in Alderley Park | Wellcome Sanger Institute for the COVID-19 Genomics UK (COG-UK) Consortium | Jacquelyn Wynn, Mairead Hyland, The Lighthouse Lab in Alderley Park and Alex Alderton, Roberto Amato, Jeffrey Barrett, Sonia Goncalves, Ewan Harrison, David K. Jackson, Ian Johnston, Dominic Kwiatkowski, Cordelia Langford, John Sillitoe on behalf of the Wellcome Sanger Institute COVID-19 Surveillance Team |
| EPI_ISL_3368286, EPI_ISL_3368287, EPI_ISL_3368288, EPI_ISL_3368289, EPI_ISL_3368290 | Lighthouse Lab in Glasgow | Wellcome Sanger Institute for the COVID-19 Genomics UK (COG-UK) Consortium | Harper VanSteenhouse, Yumi Kasai, David Gray, Carol Clugston, Anna Dominiczak and Alex Alderton, Roberto Amato, Jeffrey Barrett, Sonia Goncalves, Ewan Harrison, David K. Jackson, Ian Johnston, Dominic Kwiatkowski, Cordelia Langford, John Sillitoe on behalf of the Wellcome Sanger Institute COVID-19 Surveillance Team |
| EPI_ISL_3368291 | Lighthouse Lab in Alderley Park | Wellcome Sanger Institute for the COVID-19 Genomics UK (COG-UK) Consortium | Jacquelyn Wynn, Mairead Hyland, The Lighthouse Lab in Alderley Park and Alex Alderton, Roberto Amato, Jeffrey Barrett, Sonia Goncalves, Ewan Harrison, David K. Jackson, Ian Johnston, Dominic Kwiatkowski, Cordelia Langford, John Sillitoe on behalf of the Wellcome Sanger Institute COVID-19 Surveillance Team |
| EPI_ISL_3368292, EPI_ISL_3368293 | Lighthouse Lab in Glasgow | Wellcome Sanger Institute for the COVID-19 Genomics UK (COG-UK) Consortium | Harper VanSteenhouse, Yumi Kasai, David Gray, Carol Clugston, Anna Dominiczak and Alex Alderton, Roberto Amato, Jeffrey Barrett, Sonia Goncalves, Ewan Harrison, David K. Jackson, Ian Johnston, Dominic Kwiatkowski, Cordelia Langford, John Sillitoe on behalf of the Wellcome Sanger Institute COVID-19 Surveillance Team |
| EPI_ISL_3368294, EPI_ISL_3368295 | Lighthouse Lab in Alderley Park | Wellcome Sanger Institute for the COVID-19 Genomics UK (COG-UK) Consortium | Jacquelyn Wynn, Mairead Hyland, The Lighthouse Lab in Alderley Park and Alex Alderton, Roberto Amato, Jeffrey Barrett, Sonia Goncalves, Ewan Harrison, David K. Jackson, Ian Johnston, Dominic Kwiatkowski, Cordelia Langford, John Sillitoe on behalf of the Wellcome Sanger Institute COVID-19 Surveillance Team |
| EPI_ISL_3368296, EPI_ISL_3368297, EPI_ISL_3368298, EPI_ISL_3368299, EPI_ISL_3368300, EPI_ISL_3368301 | Lighthouse Lab in Glasgow | Wellcome Sanger Institute for the COVID-19 Genomics UK (COG-UK) Consortium | Harper VanSteenhouse, Yumi Kasai, David Gray, Carol Clugston, Anna Dominiczak and Alex Alderton, Roberto Amato, Jeffrey Barrett, Sonia Goncalves, Ewan Harrison, David K. Jackson, Ian Johnston, Dominic Kwiatkowski, Cordelia Langford, John Sillitoe on behalf of the Wellcome Sanger Institute COVID-19 Surveillance Team |
| EPI_ISL_3368302 | Lighthouse Lab in Alderley Park | Wellcome Sanger Institute for the COVID-19 Genomics UK (COG-UK) Consortium | Jacquelyn Wynn, Mairead Hyland, The Lighthouse Lab in Alderley Park and Alex Alderton, Roberto Amato, Jeffrey Barrett, Sonia Goncalves, Ewan Harrison, David K. Jackson, Ian Johnston, Dominic Kwiatkowski, Cordelia Langford, John Sillitoe on behalf of the Wellcome Sanger Institute COVID-19 Surveillance Team |
| EPI_ISL_3368303 | Lighthouse Lab in Glasgow | Wellcome Sanger Institute for the COVID-19 Genomics UK (COG-UK) Consortium | Harper VanSteenhouse, Yumi Kasai, David Gray, Carol Clugston, Anna Dominiczak and Alex Alderton, Roberto Amato, Jeffrey Barrett, Sonia Goncalves, Ewan Harrison, David K. Jackson, Ian Johnston, Dominic Kwiatkowski, Cordelia Langford, John Sillitoe on behalf of the Wellcome Sanger Institute COVID-19 Surveillance Team |
| EPI_ISL_3368304 | Lighthouse Lab in Alderley Park | Wellcome Sanger Institute for the COVID-19 Genomics UK (COG-UK) Consortium | Jacquelyn Wynn, Mairead Hyland, The Lighthouse Lab in Alderley Park and Alex Alderton, Roberto Amato, Jeffrey Barrett, Sonia Goncalves, Ewan Harrison, David K. Jackson, Ian Johnston, Dominic Kwiatkowski, Cordelia Langford, John Sillitoe on behalf of the Wellcome Sanger Institute COVID-19 Surveillance Team |
| EPI_ISL_3368305, EPI_ISL_3368306, EPI_ISL_3368307, EPI_ISL_3368308 | Lighthouse Lab in Glasgow | Wellcome Sanger Institute for the COVID-19 Genomics UK (COG-UK) Consortium | Harper VanSteenhouse, Yumi Kasai, David Gray, Carol Clugston, Anna Dominiczak and Alex Alderton, Roberto Amato, Jeffrey Barrett, Sonia Goncalves, Ewan Harrison, David K. Jackson, Ian Johnston, Dominic Kwiatkowski, Cordelia Langford, John Sillitoe on behalf of the Wellcome Sanger Institute COVID-19 Surveillance Team |
| EPI_ISL_3368309 | Lighthouse Lab in Milton Keynes | Wellcome Sanger Institute for the COVID-19 Genomics UK (COG-UK) Consortium | The Lighthouse Lab in Milton Keynes and Alex Alderton, Roberto Amato, Jeffrey Barrett, Sonia Goncalves, Ewan Harrison, David K. Jackson, Ian Johnston, Dominic Kwiatkowski, Cordelia Langford, John Sillitoe on behalf of the Wellcome Sanger Institute COVID-19 Surveillance Team |
| EPI_ISL_3368310 | Lighthouse Lab in Alderley Park | Wellcome Sanger Institute for the COVID-19 Genomics UK (COG-UK) Consortium | Jacquelyn Wynn, Mairead Hyland, The Lighthouse Lab in Alderley Park and Alex Alderton, Roberto Amato, Jeffrey Barrett, Sonia Goncalves, Ewan Harrison, David K. Jackson, Ian Johnston, Dominic Kwiatkowski, Cordelia Langford, John Sillitoe on behalf of the Wellcome Sanger Institute COVID-19 Surveillance Team |
| EPI_ISL_3368311 | Lighthouse Lab in Glasgow | Wellcome Sanger Institute for the COVID-19 Genomics UK (COG-UK) Consortium | Harper VanSteenhouse, Yumi Kasai, David Gray, Carol Clugston, Anna Dominiczak and Alex Alderton, Roberto Amato, Jeffrey Barrett, Sonia Goncalves, Ewan Harrison, David K. Jackson, Ian Johnston, Dominic Kwiatkowski, Cordelia Langford, John Sillitoe on behalf of the Wellcome Sanger Institute COVID-19 Surveillance Team |
| EPI_ISL_3368312 | Lighthouse Lab in Alderley Park | Wellcome Sanger Institute for the COVID-19 Genomics UK (COG-UK) Consortium | Jacquelyn Wynn, Mairead Hyland, The Lighthouse Lab in Alderley Park and Alex Alderton, Roberto Amato, Jeffrey Barrett, Sonia Goncalves, Ewan Harrison, David K. Jackson, Ian Johnston, Dominic Kwiatkowski, Cordelia Langford, John Sillitoe on behalf of the Wellcome Sanger Institute COVID-19 Surveillance Team |
| EPI_ISL_3368313, EPI_ISL_3368314 | Lighthouse Lab in Glasgow | Wellcome Sanger Institute for the COVID-19 Genomics UK (COG-UK) Consortium | Harper VanSteenhouse, Yumi Kasai, David Gray, Carol Clugston, Anna Dominiczak and Alex Alderton, Roberto Amato, Jeffrey Barrett, Sonia Goncalves, Ewan Harrison, David K. Jackson, Ian Johnston, Dominic Kwiatkowski, Cordelia Langford, John Sillitoe on behalf of the Wellcome Sanger Institute COVID-19 Surveillance Team |
| EPI_ISL_3368315 | Lighthouse Lab in Alderley Park | Wellcome Sanger Institute for the COVID-19 Genomics UK (COG-UK) Consortium | Jacquelyn Wynn, Mairead Hyland, The Lighthouse Lab in Alderley Park and Alex Alderton, Roberto Amato, Jeffrey Barrett, Sonia Goncalves, Ewan Harrison, David K. Jackson, Ian Johnston, Dominic Kwiatkowski, Cordelia Langford, John Sillitoe on behalf of the Wellcome Sanger Institute COVID-19 Surveillance Team |
| EPI_ISL_3368316, EPI_ISL_3368317 | Lighthouse Lab in Glasgow | Wellcome Sanger Institute for the COVID-19 Genomics UK (COG-UK) Consortium | Harper VanSteenhouse, Yumi Kasai, David Gray, Carol Clugston, Anna Dominiczak and Alex Alderton, Roberto Amato, Jeffrey Barrett, Sonia Goncalves, Ewan Harrison, David K. Jackson, Ian Johnston, Dominic Kwiatkowski, Cordelia Langford, John Sillitoe on behalf of the Wellcome Sanger Institute COVID-19 Surveillance Team |
| EPI_ISL_3368318 | Lighthouse Lab in Alderley Park | Wellcome Sanger Institute for the COVID-19 Genomics UK (COG-UK) Consortium | Jacquelyn Wynn, Mairead Hyland, The Lighthouse Lab in Alderley Park and Alex Alderton, Roberto Amato, Jeffrey Barrett, Sonia Goncalves, Ewan Harrison, David K. Jackson, Ian Johnston, Dominic Kwiatkowski, Cordelia Langford, John Sillitoe on behalf of the Wellcome Sanger Institute COVID-19 Surveillance Team |
| EPI_ISL_3368319 | Lighthouse Lab in Glasgow | Wellcome Sanger Institute for the COVID-19 Genomics UK (COG-UK) Consortium | Harper VanSteenhouse, Yumi Kasai, David Gray, Carol Clugston, Anna Dominiczak and Alex Alderton, Roberto Amato, Jeffrey Barrett, Sonia Goncalves, Ewan Harrison, David K. Jackson, Ian Johnston, Dominic Kwiatkowski, Cordelia Langford, John Sillitoe on behalf of the Wellcome Sanger Institute COVID-19 Surveillance Team |
| EPI_ISL_3368320 | Lighthouse Lab in Alderley Park | Wellcome Sanger Institute for the COVID-19 Genomics UK (COG-UK) Consortium | Jacquelyn Wynn, Mairead Hyland, The Lighthouse Lab in Alderley Park and Alex Alderton, Roberto Amato, Jeffrey Barrett, Sonia Goncalves, Ewan Harrison, David K. Jackson, Ian Johnston, Dominic Kwiatkowski, Cordelia Langford, John Sillitoe on behalf of the Wellcome Sanger Institute COVID-19 Surveillance Team |
| EPI_ISL_3368321, EPI_ISL_3368322, EPI_ISL_3368323 | Lighthouse Lab in Glasgow | Wellcome Sanger Institute for the COVID-19 Genomics UK (COG-UK) Consortium | Harper VanSteenhouse, Yumi Kasai, David Gray, Carol Clugston, Anna Dominiczak and Alex Alderton, Roberto Amato, Jeffrey Barrett, Sonia Goncalves, Ewan Harrison, David K. Jackson, Ian Johnston, Dominic Kwiatkowski, Cordelia Langford, John Sillitoe on behalf of the Wellcome Sanger Institute COVID-19 Surveillance Team |
| EPI_ISL_3368324 | Lighthouse Lab in Alderley Park | Wellcome Sanger Institute for the COVID-19 Genomics UK (COG-UK) Consortium | Jacquelyn Wynn, Mairead Hyland, The Lighthouse Lab in Alderley Park and Alex Alderton, Roberto Amato, Jeffrey Barrett, Sonia Goncalves, Ewan Harrison, David K. Jackson, Ian Johnston, Dominic Kwiatkowski, Cordelia Langford, John Sillitoe on behalf of the Wellcome Sanger Institute COVID-19 Surveillance Team |
| EPI_ISL_3368325, EPI_ISL_3368326 | Lighthouse Lab in Glasgow | Wellcome Sanger Institute for the COVID-19 Genomics UK (COG-UK) Consortium | Harper VanSteenhouse, Yumi Kasai, David Gray, Carol Clugston, Anna Dominiczak and Alex Alderton, Roberto Amato, Jeffrey Barrett, Sonia Goncalves, Ewan Harrison, David K. Jackson, Ian Johnston, Dominic Kwiatkowski, Cordelia Langford, John Sillitoe on behalf of the Wellcome Sanger Institute COVID-19 Surveillance Team |
| EPI_ISL_3368327 | Lighthouse Lab in Alderley Park | Wellcome Sanger Institute for the COVID-19 Genomics UK (COG-UK) Consortium | Jacquelyn Wynn, Mairead Hyland, The Lighthouse Lab in Alderley Park and Alex Alderton, Roberto Amato, Jeffrey Barrett, Sonia Goncalves, Ewan Harrison, David K. Jackson, Ian Johnston, Dominic Kwiatkowski, Cordelia Langford, John Sillitoe on behalf of the Wellcome Sanger Institute COVID-19 Surveillance Team |
| EPI_ISL_3368328 | Lighthouse Lab in Milton Keynes | Wellcome Sanger Institute for the COVID-19 Genomics UK (COG-UK) Consortium | The Lighthouse Lab in Milton Keynes and Alex Alderton, Roberto Amato, Jeffrey Barrett, Sonia Goncalves, Ewan Harrison, David K. Jackson, Ian Johnston, Dominic Kwiatkowski, Cordelia Langford, John Sillitoe on behalf of the Wellcome Sanger Institute COVID-19 Surveillance Team |
| EPI_ISL_3368329 | Lighthouse Lab in Alderley Park | Wellcome Sanger Institute for the COVID-19 Genomics UK (COG-UK) Consortium | Jacquelyn Wynn, Mairead Hyland, The Lighthouse Lab in Alderley Park and Alex Alderton, Roberto Amato, Jeffrey Barrett, Sonia Goncalves, Ewan Harrison, David K. Jackson, Ian Johnston, Dominic Kwiatkowski, Cordelia Langford, John Sillitoe on behalf of the Wellcome Sanger Institute COVID-19 Surveillance Team |
| EPI_ISL_3368330, EPI_ISL_3368331, EPI_ISL_3368332 | Lighthouse Lab in Milton Keynes | Wellcome Sanger Institute for the COVID-19 Genomics UK (COG-UK) Consortium | The Lighthouse Lab in Milton Keynes and Alex Alderton, Roberto Amato, Jeffrey Barrett, Sonia Goncalves, Ewan Harrison, David K. Jackson, Ian Johnston, Dominic Kwiatkowski, Cordelia Langford, John Sillitoe on behalf of the Wellcome Sanger Institute COVID-19 Surveillance Team |

| | | | |
|---|---|---|---|
| EPI_ISL_3368333, EPI_ISL_3368334, EPI_ISL_3368335, EPI_ISL_3368336, EPI_ISL_3368337, EPI_ISL_3368338, EPI_ISL_3368339, EPI_ISL_3368340, EPI_ISL_3368341, EPI_ISL_3368342, EPI_ISL_3368343, EPI_ISL_3368344, EPI_ISL_3368345, EPI_ISL_3368346, EPI_ISL_3368347, EPI_ISL_3368348, EPI_ISL_3368349, EPI_ISL_3368350, EPI_ISL_3368351, EPI_ISL_3368352, EPI_ISL_3368353, EPI_ISL_3368354, EPI_ISL_3368355, EPI_ISL_3368356, EPI_ISL_3368357, EPI_ISL_3368358, EPI_ISL_3368359, EPI_ISL_3368360, EPI_ISL_3368361, EPI_ISL_3368362, EPI_ISL_3368363, EPI_ISL_3368364, EPI_ISL_3368365, EPI_ISL_3368366, EPI_ISL_3368367, EPI_ISL_3368368, EPI_ISL_3368369, EPI_ISL_3368370, EPI_ISL_3368371, EPI_ISL_3368372, EPI_ISL_3368373, EPI_ISL_3368374, EPI_ISL_3368375, EPI_ISL_3368376, EPI_ISL_3368377, EPI_ISL_3368378, EPI_ISL_3368379, EPI_ISL_3368380, EPI_ISL_3368381, EPI_ISL_3368382, EPI_ISL_3368383, EPI_ISL_3368384, EPI_ISL_3368385, EPI_ISL_3368386, EPI_ISL_3368387, EPI_ISL_3368388, EPI_ISL_3368389, EPI_ISL_3368390, EPI_ISL_3368391, EPI_ISL_3368392, EPI_ISL_3368393, EPI_ISL_3368394, EPI_ISL_3368395, EPI_ISL_3368396, EPI_ISL_3368397, EPI_ISL_3368398, EPI_ISL_3368399, EPI_ISL_3368400, EPI_ISL_3368401, EPI_ISL_3368402, EPI_ISL_3368403 | | | |
| see above | Lighthouse Lab in Glasgow | Wellcome Sanger Institute for the COVID-19 Genomics UK (COG-UK) Consortium | Harper VanSteenhouse, Yumi Kasai, David Gray, Carol Clugston, Anna Dominiczak and Alex Alderton, Roberto Amato, Jeffrey Barrett, Sonia Goncalves, Ewan Harrison, David K. Jackson, Ian Johnston, Dominic Kwiatkowski, Cordelia Langford, John Sillitoe on behalf of the Wellcome Sanger Institute COVID-19 Surveillance Team |
| EPI_ISL_3368478, EPI_ISL_3368479, EPI_ISL_3368480, EPI_ISL_3368481, EPI_ISL_3368482, EPI_ISL_3368483, EPI_ISL_3368484, EPI_ISL_3368485, EPI_ISL_3368486, EPI_ISL_3368487, EPI_ISL_3368488, EPI_ISL_3368489, EPI_ISL_3368490, EPI_ISL_3368491, EPI_ISL_3368492, EPI_ISL_3368493, EPI_ISL_3368494, EPI_ISL_3368495, EPI_ISL_3368496, EPI_ISL_3368497, EPI_ISL_3368498, EPI_ISL_3368499, EPI_ISL_3368500, EPI_ISL_3368501, EPI_ISL_3368502, EPI_ISL_3368503, EPI_ISL_3368504, EPI_ISL_3368505, EPI_ISL_3368506, EPI_ISL_3368507, EPI_ISL_3368508, EPI_ISL_3368509, EPI_ISL_3368510, EPI_ISL_3368511, EPI_ISL_3368512, EPI_ISL_3368513, EPI_ISL_3368514, EPI_ISL_3368515, EPI_ISL_3368516, EPI_ISL_3368517, EPI_ISL_3368518, EPI_ISL_3368519, EPI_ISL_3368520, EPI_ISL_3368521, EPI_ISL_3368522, EPI_ISL_3368523, EPI_ISL_3368524, EPI_ISL_3368525, EPI_ISL_3368526, EPI_ISL_3368527, EPI_ISL_3368528, EPI_ISL_3368529, EPI_ISL_3368530, EPI_ISL_3368531, EPI_ISL_3368532, EPI_ISL_3368533, EPI_ISL_3368534, EPI_ISL_3368535, EPI_ISL_3368536, EPI_ISL_3368537, EPI_ISL_3368538, EPI_ISL_3368539, EPI_ISL_3368540, EPI_ISL_3368541, EPI_ISL_3368542, EPI_ISL_3368543, EPI_ISL_3368544, EPI_ISL_3368545, EPI_ISL_3368546, EPI_ISL_3368547, EPI_ISL_3368548, EPI_ISL_3368549, EPI_ISL_3368550, EPI_ISL_3368551, EPI_ISL_3368552, EPI_ISL_3368553, EPI_ISL_3368554, EPI_ISL_3368555, EPI_ISL_3368556, EPI_ISL_3368557, EPI_ISL_3368558, EPI_ISL_3368559, EPI_ISL_3368560 | | | |
| see above | Lifebrain Covid Labor GmbH | Dept. of Laboratory Medicine | Harald Esterbauer, Anna Gschaider, Petra Jurkowitsch, Fabian Konig, Sabina Plumer, Robert Strassl, Claudia Weber, Oswald Wagner |
| EPI_ISL_3368608, EPI_ISL_3368609, EPI_ISL_3368614 | Labor Dr. Fenner und Kollegen | Institut für Hygiene und Umwelt Hamburg | Christian Schäfers, Anne Reichstein, Philip Busch, Judith Overhoff, Jens Wild, Andreas Wille |
| EPI_ISL_3368615 | Labor Dr. von Froreich GmbH | Institut für Hygiene und Umwelt Hamburg | Christian Schäfers, Anne Reichstein, Philip Busch, Judith Overhoff, Jens Wild, Andreas Wille |
| EPI_ISL_3368616 | Aesculabor - Hamburg GmbH | Institut für Hygiene und Umwelt Hamburg | Christian Schäfers, Anne Reichstein, Philip Busch, Judith Overhoff, Jens Wild, Andreas Wille |
| EPI_ISL_3368617 | Labor Dr. Heidrich & Kollegen MVZ GmbH | Institut für Hygiene und Umwelt Hamburg | Christian Schäfers, Anne Reichstein, Philip Busch, Judith Overhoff, Jens Wild, Andreas Wille |
| EPI_ISL_3368618, EPI_ISL_3368620 | Labor Dr. Fenner und Kollegen | Institut für Hygiene und Umwelt Hamburg | Christian Schäfers, Anne Reichstein, Philip Busch, Judith Overhoff, Jens Wild, Andreas Wille |
| EPI_ISL_3368621 | Labor Dr. Heidrich & Kollegen MVZ GmbH | Institut für Hygiene und Umwelt Hamburg | Christian Schäfers, Anne Reichstein, Philip Busch, Judith Overhoff, Jens Wild, Andreas Wille |
| EPI_ISL_3368622 | Labor Dr. Fenner und Kollegen | Institut für Hygiene und Umwelt Hamburg | Christian Schäfers, Anne Reichstein, Philip Busch, Judith Overhoff, Jens Wild, Andreas Wille |
| EPI_ISL_3368637, EPI_ISL_3368638, EPI_ISL_3368639, EPI_ISL_3368640, EPI_ISL_3368641, EPI_ISL_3368642, EPI_ISL_3368643, EPI_ISL_3368644, EPI_ISL_3368645, EPI_ISL_3368646, EPI_ISL_3368647, EPI_ISL_3368648, EPI_ISL_3368649, EPI_ISL_3368650, EPI_ISL_3368651, EPI_ISL_3368652, EPI_ISL_3368653, EPI_ISL_3368654, EPI_ISL_3368655, EPI_ISL_3368656, EPI_ISL_3368657, EPI_ISL_3368658, EPI_ISL_3368659, EPI_ISL_3368660, EPI_ISL_3368661, EPI_ISL_3368662, EPI_ISL_3368663, EPI_ISL_3368664, EPI_ISL_3368665, EPI_ISL_3368666, EPI_ISL_3368667, EPI_ISL_3368668, EPI_ISL_3368669, EPI_ISL_3368670, EPI_ISL_3368671, EPI_ISL_3368672, EPI_ISL_3368673, EPI_ISL_3368674, EPI_ISL_3368675, EPI_ISL_3368676, EPI_ISL_3368677, EPI_ISL_3368678, EPI_ISL_3368679, EPI_ISL_3368680, EPI_ISL_3368681, EPI_ISL_3368682, EPI_ISL_3368683, EPI_ISL_3368684, EPI_ISL_3368685, EPI_ISL_3368686, EPI_ISL_3368687, EPI_ISL_3368688, EPI_ISL_3368689, EPI_ISL_3368690, EPI_ISL_3368691, EPI_ISL_3368692, EPI_ISL_3368693, EPI_ISL_3368694, EPI_ISL_3368695, EPI_ISL_3368696, EPI_ISL_3368697, EPI_ISL_3368698, EPI_ISL_3368699, EPI_ISL_3368700, EPI_ISL_3368701, EPI_ISL_3368702, EPI_ISL_3368703, EPI_ISL_3368704, EPI_ISL_3368705, EPI_ISL_3368706, EPI_ISL_3368707, EPI_ISL_3368708, EPI_ISL_3368709, EPI_ISL_3368710, EPI_ISL_3368711, EPI_ISL_3368712, EPI_ISL_3368713, EPI_ISL_3368714, EPI_ISL_3368715, EPI_ISL_3368716, EPI_ISL_3368717, EPI_ISL_3368718, EPI_ISL_3368719, EPI_ISL_3368720, EPI_ISL_3368721, EPI_ISL_3368722, EPI_ISL_3368723, EPI_ISL_3368724, EPI_ISL_3368725, EPI_ISL_3368726, EPI_ISL_3368727, EPI_ISL_3368728, EPI_ISL_3368729, EPI_ISL_3368730, EPI_ISL_3368731, EPI_ISL_3368732, EPI_ISL_3368733, EPI_ISL_3368734, EPI_ISL_3368735, EPI_ISL_3368736, EPI_ISL_3368737, EPI_ISL_3368738, EPI_ISL_3368739, EPI_ISL_3368740, EPI_ISL_3368741, EPI_ISL_3368742, EPI_ISL_3368743, EPI_ISL_3368744, EPI_ISL_3368745, EPI_ISL_3368746, EPI_ISL_3368747, EPI_ISL_3368748, EPI_ISL_3368749, EPI_ISL_3368750, EPI_ISL_3368751, EPI_ISL_3368752, EPI_ISL_3368753, EPI_ISL_3368754, EPI_ISL_3368755, EPI_ISL_3368756 | | | |
| see above | Springfield-Greene County Health Department | University of Minnesota Genomics Center | Daryl M. Gohl, John Garbe, Jaquelyn Kuriger-Laber, Corbin Dirkx |
| EPI_ISL_3368770, EPI_ISL_3368771, EPI_ISL_3368908 | Genetica Molecular and Subdepartamento de Virologia ISP Chile | Instituto de Salud Publica de Chile | Karen Orostica, Constanza Campano, Barbara Parra, Loredana Arata, Gisselle Barra, Patricia Bustos, Rodrigo Fasce, Javier Tognarelli, Andres Castillo, Soledad Ulloa, Jorge Fernandez |
| EPI_ISL_3368914, EPI_ISL_3368916, EPI_ISL_3368918, EPI_ISL_3368924, EPI_ISL_3368932, EPI_ISL_3368935, EPI_ISL_3368939, EPI_ISL_3368962, EPI_ISL_3368984, EPI_ISL_3369001, EPI_ISL_3369007, EPI_ISL_3369008, EPI_ISL_3369023, EPI_ISL_3369027, EPI_ISL_3369028, EPI_ISL_3369029, EPI_ISL_3369030, EPI_ISL_3369031, EPI_ISL_3369032, EPI_ISL_3369033, EPI_ISL_3369034, EPI_ISL_3369035, EPI_ISL_3369036, EPI_ISL_3369037, EPI_ISL_3369038, EPI_ISL_3369039, EPI_ISL_3369040, EPI_ISL_3369041, EPI_ISL_3369042, EPI_ISL_3369043, EPI_ISL_3369044, EPI_ISL_3369045, EPI_ISL_3369046, EPI_ISL_3369047, EPI_ISL_3369048, EPI_ISL_3369049, EPI_ISL_3369050, EPI_ISL_3369051, EPI_ISL_3369052, EPI_ISL_3369053, EPI_ISL_3369054, EPI_ISL_3369055, EPI_ISL_3369056, EPI_ISL_3369057, EPI_ISL_3369058, EPI_ISL_3369059, EPI_ISL_3369060, EPI_ISL_3369061, EPI_ISL_3369062, EPI_ISL_3369063, EPI_ISL_3369064, EPI_ISL_3369065, EPI_ISL_3369066, EPI_ISL_3369067, EPI_ISL_3369068, EPI_ISL_3369069, EPI_ISL_3369070, EPI_ISL_3369071, EPI_ISL_3369072, EPI_ISL_3369073, EPI_ISL_3369074, EPI_ISL_3369075, EPI_ISL_3369076, EPI_ISL_3369077, EPI_ISL_3369078, EPI_ISL_3369079, EPI_ISL_3369080, EPI_ISL_3369081, EPI_ISL_3369082, EPI_ISL_3369083, EPI_ISL_3369084, EPI_ISL_3369085, EPI_ISL_3369086, EPI_ISL_3369087, EPI_ISL_3369088, EPI_ISL_3369089, EPI_ISL_3369090, EPI_ISL_3369091, EPI_ISL_3369092, EPI_ISL_3369093, EPI_ISL_3369094, EPI_ISL_3369095, EPI_ISL_3369096, EPI_ISL_3369097, EPI_ISL_3369098, EPI_ISL_3369099, EPI_ISL_3369100, EPI_ISL_3369101, EPI_ISL_3369102, EPI_ISL_3369103, EPI_ISL_3369104, EPI_ISL_3369105, EPI_ISL_3369106, EPI_ISL_3369107, EPI_ISL_3369108, EPI_ISL_3369109, EPI_ISL_3369110, EPI_ISL_3369111, EPI_ISL_3369112, EPI_ISL_3369113, EPI_ISL_3369114, EPI_ISL_3369115, EPI_ISL_3369116, EPI_ISL_3369117, EPI_ISL_3369118, EPI_ISL_3369119, EPI_ISL_3369120, EPI_ISL_3369121, EPI_ISL_3369122, EPI_ISL_3369123, EPI_ISL_3369124, EPI_ISL_3369125, EPI_ISL_3369126, EPI_ISL_3369127, EPI_ISL_3369128, EPI_ISL_3369129, EPI_ISL_3369130, EPI_ISL_3369131, EPI_ISL_3369132, EPI_ISL_3369133, EPI_ISL_3369134, EPI_ISL_3369135, EPI_ISL_3369136, EPI_ISL_3369137, EPI_ISL_3369138, EPI_ISL_3369139, EPI_ISL_3369140, EPI_ISL_3369141, EPI_ISL_3369142, EPI_ISL_3369143, EPI_ISL_3369144, EPI_ISL_3369145, EPI_ISL_3369146, EPI_ISL_3369147, EPI_ISL_3369148, EPI_ISL_3369149, EPI_ISL_3369150, EPI_ISL_3369151, EPI_ISL_3369152, EPI_ISL_3369153, EPI_ISL_3369154, EPI_ISL_3369155, EPI_ISL_3369156, EPI_ISL_3369157, EPI_ISL_3369158, EPI_ISL_3369159, EPI_ISL_3369160, EPI_ISL_3369161, EPI_ISL_3369162, EPI_ISL_3369163, EPI_ISL_3369164, EPI_ISL_3369165, EPI_ISL_3369166, EPI_ISL_3369167, EPI_ISL_3369168, EPI_ISL_3369169, EPI_ISL_3369170, EPI_ISL_3369171, EPI_ISL_3369172, EPI_ISL_3369173, EPI_ISL_3369174, EPI_ISL_3369175, EPI_ISL_3369176, EPI_ISL_3369177, EPI_ISL_3369178, EPI_ISL_3369179, EPI_ISL_3369180, EPI_ISL_3369181, EPI_ISL_3369182, EPI_ISL_3369183, EPI_ISL_3369184, EPI_ISL_3369185, EPI_ISL_3369199, EPI_ISL_3369200, EPI_ISL_3369201, EPI_ISL_3369202, EPI_ISL_3369203, EPI_ISL_3369204, EPI_ISL_3369205, EPI_ISL_3369206, EPI_ISL_3369207, EPI_ISL_3369208, EPI_ISL_3369209, EPI_ISL_3369210, EPI_ISL_3369211, EPI_ISL_3369212, EPI_ISL_3369213, EPI_ISL_3369214, EPI_ISL_3369215, EPI_ISL_3369216, EPI_ISL_3369217, EPI_ISL_3369218, EPI_ISL_3369219, EPI_ISL_3369220, EPI_ISL_3369221, EPI_ISL_3369222, EPI_ISL_3369228, EPI_ISL_3369230, EPI_ISL_3369231, EPI_ISL_3369232, EPI_ISL_3369233, EPI_ISL_3369234, EPI_ISL_3369235, EPI_ISL_3369236, EPI_ISL_3369237, EPI_ISL_3369238, EPI_ISL_3369239, EPI_ISL_3369240, EPI_ISL_3369241, EPI_ISL_3369242, EPI_ISL_3369243, EPI_ISL_3369244, EPI_ISL_3369245, EPI_ISL_3369246, EPI_ISL_3369247, EPI_ISL_3369248, EPI_ISL_3369268, EPI_ISL_3369269, EPI_ISL_3369270, EPI_ISL_3369272, EPI_ISL_3369274 | | | |
| see above | Division of Emerging Infectious Diseases, Bureau of Infectious Diseases Diagnosis Control, Korea Disease Control and Prevention Agency | Division of Emerging Infectious Diseases, Bureau of Infectious Diseases Diagnosis Control, Korea Disease Control and Prevention Agency | Ae Kyung Park, Il-Hwan Kim, Heui Man Kim, Jeong-Ah Kim, Chae Young Lee, Eun-Jin Kim |
| EPI_ISL_3369277, EPI_ISL_3369285, EPI_ISL_3369330, EPI_ISL_3369331, EPI_ISL_3369332 | Genetica Molecular and Subdepartamento de Virologia ISP Chile | Instituto de Salud Publica de Chile | Karen Orostica, Constanza Campano, Barbara Parra, Loredana Arata, Gisselle Barra, Patricia Bustos, Rodrigo Fasce, Javier Tognarelli, Andres Castillo, Soledad Ulloa, Jorge Fernandez |
| EPI_ISL_3369458, EPI_ISL_3369459, EPI_ISL_3369460, EPI_ISL_3369461, EPI_ISL_3369462, EPI_ISL_3369463, EPI_ISL_3369464, EPI_ISL_3369465, EPI_ISL_3369466, EPI_ISL_3369467, EPI_ISL_3369468, EPI_ISL_3369469, EPI_ISL_3369470, EPI_ISL_3369471, EPI_ISL_3369472, EPI_ISL_3369473, EPI_ISL_3369474, EPI_ISL_3369476, EPI_ISL_3369477, EPI_ISL_3369478, EPI_ISL_3369479, EPI_ISL_3369480, EPI_ISL_3369481, EPI_ISL_3369482, EPI_ISL_3369483, EPI_ISL_3369484, EPI_ISL_3369485, EPI_ISL_3369486 | | | |
| see above | IN State Department of Health Laboratory Services | IN State Department of Health Laboratory Services | Cassandra Campion, Jamie Yeadon, Brian Pope, Lixia Liu, Kyle Brownlee, Melissa Hindenlang, Mark Glazier |
| EPI_ISL_3369491, EPI_ISL_3369495, EPI_ISL_3369496, EPI_ISL_3369497, EPI_ISL_3369498, EPI_ISL_3369499, EPI_ISL_3369500, EPI_ISL_3369501, EPI_ISL_3369502, EPI_ISL_3369503, EPI_ISL_3369504, EPI_ISL_3369505, EPI_ISL_3369506, EPI_ISL_3369507, EPI_ISL_3369508, EPI_ISL_3369509, EPI_ISL_3369510, EPI_ISL_3369511, EPI_ISL_3369512, EPI_ISL_3369513, EPI_ISL_3369514, EPI_ISL_3369515, EPI_ISL_3369516, EPI_ISL_3369517, EPI_ISL_3369518, EPI_ISL_3369519, EPI_ISL_3369520, EPI_ISL_3369521, EPI_ISL_3369522, EPI_ISL_3369523, EPI_ISL_3369524, EPI_ISL_3369526, EPI_ISL_3369527, EPI_ISL_3369528, EPI_ISL_3369529, EPI_ISL_3369530, EPI_ISL_3369531, EPI_ISL_3369532, EPI_ISL_3369533, EPI_ISL_3369534, EPI_ISL_3369535, EPI_ISL_3369536, EPI_ISL_3369537, EPI_ISL_3369538, EPI_ISL_3369539, EPI_ISL_3369540, EPI_ISL_3369541, EPI_ISL_3369542, EPI_ISL_3369543, EPI_ISL_3369544, EPI_ISL_3369545, EPI_ISL_3369546, EPI_ISL_3369547, EPI_ISL_3369548, EPI_ISL_3369549, EPI_ISL_3369550, EPI_ISL_3369551, EPI_ISL_3369552, EPI_ISL_3369553, EPI_ISL_3369554, EPI_ISL_3369555, EPI_ISL_3369556, EPI_ISL_3369557, EPI_ISL_3369558, EPI_ISL_3369559, EPI_ISL_3369560, EPI_ISL_3369561, EPI_ISL_3369562, EPI_ISL_3369563, EPI_ISL_3369564, EPI_ISL_3369565, EPI_ISL_3369566, EPI_ISL_3369567, EPI_ISL_3369568, EPI_ISL_3369569, EPI_ISL_3369572, EPI_ISL_3369573, EPI_ISL_3369574, EPI_ISL_3369575, EPI_ISL_3369576, EPI_ISL_3369577, EPI_ISL_3369578, EPI_ISL_3369579, EPI_ISL_3369580, EPI_ISL_3369581, EPI_ISL_3369582, EPI_ISL_3369583, EPI_ISL_3369584, EPI_ISL_3369585, EPI_ISL_3369586, EPI_ISL_3369589, EPI_ISL_3369590, EPI_ISL_3369591, EPI_ISL_3369592, EPI_ISL_3369593, EPI_ISL_3369594, EPI_ISL_3369595, EPI_ISL_3369596, EPI_ISL_3369597, EPI_ISL_3369598, EPI_ISL_3369599, EPI_ISL_3369600, EPI_ISL_3369601, EPI_ISL_3369602, EPI_ISL_3369603, EPI_ISL_3369604, EPI_ISL_3369605, EPI_ISL_3369606, EPI_ISL_3369607, EPI_ISL_3369608, EPI_ISL_3369609, EPI_ISL_3369610, EPI_ISL_3369611, EPI_ISL_3369612, EPI_ISL_3369613, EPI_ISL_3369614, EPI_ISL_3369615, EPI_ISL_3369616, EPI_ISL_3369617, EPI_ISL_3369618, EPI_ISL_3369619, EPI_ISL_3369620, EPI_ISL_3369621, EPI_ISL_3369622, EPI_ISL_3369623, EPI_ISL_3369624, EPI_ISL_3369625, EPI_ISL_3369626, EPI_ISL_3369627, EPI_ISL_3369628, EPI_ISL_3369629, EPI_ISL_3369630, EPI_ISL_3369631, EPI_ISL_3369632, EPI_ISL_3369633, EPI_ISL_3369634, EPI_ISL_3369635, EPI_ISL_3369636, EPI_ISL_3369637, EPI_ISL_3369638, EPI_ISL_3369639, EPI_ISL_3369640, EPI_ISL_3369641, EPI_ISL_3369642, EPI_ISL_3369643, EPI_ISL_3369644, EPI_ISL_3369645, EPI_ISL_3369646, EPI_ISL_3369647, EPI_ISL_3369648, EPI_ISL_3369649, EPI_ISL_3369650, EPI_ISL_3369651, EPI_ISL_3369652, EPI_ISL_3369653, EPI_ISL_3369654, EPI_ISL_3369655, EPI_ISL_3369656, EPI_ISL_3369657, EPI_ISL_3369658, EPI_ISL_3369659, EPI_ISL_3369660, EPI_ISL_3369661, EPI_ISL_3369662, EPI_ISL_3369663, EPI_ISL_3369664, EPI_ISL_3369665, EPI_ISL_3369666, EPI_ISL_3369667, EPI_ISL_3369668, EPI_ISL_3369669, EPI_ISL_3369670, EPI_ISL_3369671, EPI_ISL_3369672, EPI_ISL_3369673, EPI_ISL_3369674, EPI_ISL_3369675, EPI_ISL_3369676, EPI_ISL_3369677, EPI_ISL_3369678, EPI_ISL_3369679, EPI_ISL_3369680, EPI_ISL_3369681, EPI_ISL_3369682, EPI_ISL_3369683, EPI_ISL_3369684, EPI_ISL_3369685, EPI_ISL_3369686, EPI_ISL_3369687, EPI_ISL_3369688, EPI_ISL_3369689, EPI_ISL_3369690, EPI_ISL_3369691, EPI_ISL_3369692, EPI_ISL_3369693, EPI_ISL_3369694, EPI_ISL_3369695, EPI_ISL_3369696, EPI_ISL_3369697, EPI_ISL_3369698, EPI_ISL_3369699, EPI_ISL_3369700, EPI_ISL_3369701, EPI_ISL_3369702, EPI_ISL_3369703, EPI_ISL_3369704, EPI_ISL_3369705, EPI_ISL_3369706, EPI_ISL_3369707, EPI_ISL_3369708, EPI_ISL_3369709, EPI_ISL_3369710, EPI_ISL_3369711, EPI_ISL_3369712, EPI_ISL_3369713, EPI_ISL_3369714, EPI_ISL_3369715, EPI_ISL_3369716, EPI_ISL_3369717, EPI_ISL_3369718, EPI_ISL_3369719, EPI_ISL_3369720, EPI_ISL_3369721, EPI_ISL_3369722, EPI_ISL_3369723, EPI_ISL_3369724, EPI_ISL_3369725, EPI_ISL_3369726, EPI_ISL_3369727, EPI_ISL_3369728, EPI_ISL_3369729, EPI_ISL_3369730, EPI_ISL_3369731, EPI_ISL_3369732, EPI_ISL_3369733, EPI_ISL_3369734, EPI_ISL_3369735, EPI_ISL_3369736, EPI_ISL_3369737, EPI_ISL_3369738, EPI_ISL_3369739, EPI_ISL_3369740, EPI_ISL_3369741, EPI_ISL_3369742, EPI_ISL_3369743, EPI_ISL_3369744, EPI_ISL_3369745, EPI_ISL_3369746, EPI_ISL_3369747, EPI_ISL_3369748, EPI_ISL_3369749 | | | |
| see above | UW Virology Lab | UW Virology Lab | Pavitra Roychoudhury, Shah Mohamed Bakhash, Ricardo Perez, Hong Xie, Tien V. Nguyen, Maria Lukes, Sean Ellis, Nathan Breit, Robert J. Livingston, Meei-Li Huang, Keith R Jerome, Patrick Mathias, Alexander Greninger |
| EPI_ISL_3369776 | VIDOH (Virgin Islands Department of Health) | Grubaugh Lab - Yale School of Public Health | Joseph Fauver, Mallery Breban, Isabel Ott, Tara Alpert, Mary Petrone, Anderson Brito, Chantal Vogels, Annie Watkins, Chaney Kalinich, Jessica Rothman, Kendall Billig, Tobias Koch, Jendai Richards, TaLesa Aderohunmu, Esther Ellis, Marlon Lawrence, Brett Ellis, Nathan Grubaugh |
| EPI_ISL_3369789, EPI_ISL_3369790 | UCC (Universidad Central del Caribe) | Grubaugh Lab - Yale School of Public Health | Joseph Fauver, Mallery Breban, Isabel Ott, Tara Alpert, Mary Petrone, Anderson Brito, Chantal Vogels, Annie Watkins, Chaney Kalinich, Jessica Rothman, Kendall Billig, Tobias Koch, Fabiola Cruz-Lopez, David Rivera Aponte, Fabiola Fontanet, Natalie Machargo, Andrea Firpo, Vianca Aponte, Ernesto Vazquez, Patricia Serrano, Tania Mitwalli, Karolane Gonzalez, Nathan Grubaugh |
| EPI_ISL_3369800 | Pathology and Laboratory Medicine Institute, Cleveland Clinic, Ohio, USA | Pathology and Laboratory Medicine Institute, Cleveland Clinic, Ohio, USA | Jessica Spildener, Concetta Peck, Jennifer Starbuck, Joy Nakitandwe, Kristen McDonnell, David Plunkett, Zheng Jin Tu, Jay Brock, Yu-Wei Cheng, Gary Procop, Daniel Rhoads, Daniel H. Farkas, David Bosler |
| EPI_ISL_3369801 | University Hospital Brno, OKMI | University Hospital Brno, CMBG | Matej Bezdicek, Jan Svaton, Martina Lengerova, Kristyna Dufkova, Pavlina Volfova |
| EPI_ISL_3369804 | Nemocnice Breclav | University Hospital Brno, CMBG | Matej Bezdicek, Jan Svaton, Martina Lengerova, Kristyna Dufkova, Pavlina Volfova |
| EPI_ISL_3369805 | University Hospital Brno, OKMI | University Hospital Brno, CMBG | Matej Bezdicek, Jan Svaton, Martina Lengerova, Kristyna Dufkova, Pavlina Volfova |
| EPI_ISL_3369806 | Nemocnice Jihlava | University Hospital Brno, CMBG | Matej Bezdicek, Jan Svaton, Martina Lengerova, Kristyna Dufkova, Pavlina Volfova |
| EPI_ISL_3369843 | CEDIBIO-UNILABS OUSTALOUS | CHU Purpan - Laboratoire de Virologie - Institut Fédératif de Biologie | Latour J., Jeanne N., Ranger N., Carcenac R., Harter A., Boyer P., Ferrer V., Tremeaux R., Izopet J. |
| EPI_ISL_3369844 | CEDIBIO UNILABS LESPINET | CHU Purpan - Laboratoire de Virologie - Institut Fédératif de Biologie | Latour J., Jeanne N., Ranger N., Carcenac R., Harter A., Boyer P., Ferrer V., Tremeaux R., Izopet J. |
| EPI_ISL_3369845 | CHU Purpan - Laboratoire de Virologie - Institut Fédératif de Biologie | CHU Purpan - Laboratoire de Virologie - Institut Fédératif de Biologie | Latour J., Jeanne N., Ranger N., Carcenac R., Harter A., Boyer P., Ferrer V., Tremeaux R., Izopet J. |
| EPI_ISL_3369846 | CEDIBIO UNILABS | CHU Purpan - Laboratoire de Virologie - Institut Fédératif de Biologie | Latour J., Jeanne N., Ranger N., Carcenac R., Harter A., Boyer P., Ferrer V., Tremeaux R., Izopet J. |
| EPI_ISL_3369847 | CEDIBIO-UNILABS CASTANET DELHERM | CHU Purpan - Laboratoire de Virologie - Institut Fédératif de Biologie | Latour J., Jeanne N., Ranger N., Carcenac R., Harter A., Boyer P., Ferrer V., Tremeaux R., Izopet J. |
| EPI_ISL_3369848, EPI_ISL_3369849 | CHU Purpan - Laboratoire de Virologie - Institut Fédératif de Biologie | CHU Purpan - Laboratoire de Virologie - Institut Fédératif de Biologie | Latour J., Jeanne N., Ranger N., Carcenac R., Harter A., Boyer P., Ferrer V., Tremeaux R., Izopet J. |
| EPI_ISL_3369850 | CH MONTAUBAN | CHU Purpan - Laboratoire de Virologie - Institut Fédératif de Biologie | Latour J., Jeanne N., Ranger N., Carcenac R., Harter A., Boyer P., Ferrer V., Tremeaux R., Izopet J. |
| EPI_ISL_3369852 | CEDIBIO-UNILABS ST ORENS ARCADES | CHU Purpan - Laboratoire de Virologie - Institut Fédératif de Biologie | Latour J., Jeanne N., Ranger N., Carcenac R., Harter A., Boyer P., Ferrer V., Tremeaux R., Izopet J. |
| EPI_ISL_3369854, EPI_ISL_3369855, EPI_ISL_3369856 | CH ALBI | CHU Purpan - Laboratoire de Virologie - Institut Fédératif de Biologie | Latour J., Jeanne N., Ranger N., Carcenac R., Harter A., Boyer P., Ferrer V., Tremeaux R., Izopet J. |
| EPI_ISL_3369857 | GCS DE BIOLOGIE DU TERRITOIRE DE L'ARIEGE | CHU Purpan - Laboratoire de Virologie - Institut Fédératif de Biologie | Latour J., Jeanne N., Ranger N., Carcenac R., Harter A., Boyer P., Ferrer V., Tremeaux R., Izopet J. |
| EPI_ISL_3369858 | CH CARCASSONE | CHU Purpan - Laboratoire de Virologie - Institut Fédératif de Biologie | Latour J., Jeanne N., Ranger N., Carcenac R., Harter A., Boyer P., Ferrer V., Tremeaux R., Izopet J. |
| EPI_ISL_3369859 | CH MONTAUBAN | CHU Purpan - Laboratoire de Virologie - Institut Fédératif de Biologie | Latour J., Jeanne N., Ranger N., Carcenac R., Harter A., Boyer P., Ferrer V., Tremeaux R., Izopet J. |

| | | | |
|---|---|---|---|
| EPI_ISL_3369860 | LABORATOIRE GCS INTER HOSPITALIER | CHU Purpan - Laboratoire de Virologie - Institut Fédératif de Biologie | Latour J., Jeanne N., Ranger N., Carcenac R., Harter A., Boyer P., Ferrer V., Tremeaux R., Izopet J. |
| EPI_ISL_3369861 | CH MONTAUBAN | CHU Purpan - Laboratoire de Virologie - Institut Fédératif de Biologie | Latour J., Jeanne N., Ranger N., Carcenac R., Harter A., Boyer P., Ferrer V., Tremeaux R., Izopet J. |
| EPI_ISL_3369862 | LABORATOIRE GCS INTER HOSPITALIER | CHU Purpan - Laboratoire de Virologie - Institut Fédératif de Biologie | Latour J., Jeanne N., Ranger N., Carcenac R., Harter A., Boyer P., Ferrer V., Tremeaux R., Izopet J. |
| EPI_ISL_3369863 | CEDIBIO UNILABS BALMA | CHU Purpan - Laboratoire de Virologie - Institut Fédératif de Biologie | Latour J., Jeanne N., Ranger N., Carcenac R., Harter A., Boyer P., Ferrer V., Tremeaux R., Izopet J. |
| EPI_ISL_3369864 | CEDIBIO-UNILABS OUSTALOUS | CHU Purpan - Laboratoire de Virologie - Institut Fédératif de Biologie | Latour J., Jeanne N., Ranger N., Carcenac R., Harter A., Boyer P., Ferrer V., Tremeaux R., Izopet J. |
| EPI_ISL_3369866 | LABORATOIRE GCS INTER HOSPITALIER | CHU Purpan - Laboratoire de Virologie - Institut Fédératif de Biologie | Latour J., Jeanne N., Ranger N., Carcenac R., Harter A., Boyer P., Ferrer V., Tremeaux R., Izopet J. |
| EPI_ISL_3369867 | HOSPILAB 47-VILLENEUVE LOT | CHU Purpan - Laboratoire de Virologie - Institut Fédératif de Biologie | Latour J., Jeanne N., Ranger N., Carcenac R., Harter A., Boyer P., Ferrer V., Tremeaux R., Izopet J. |
| EPI_ISL_3369868 | LABORATOIRE GCS INTER HOSPITALIER | CHU Purpan - Laboratoire de Virologie - Institut Fédératif de Biologie | Latour J., Jeanne N., Ranger N., Carcenac R., Harter A., Boyer P., Ferrer V., Tremeaux R., Izopet J. |
| EPI_ISL_3369869, EPI_ISL_3369870, EPI_ISL_3369871, EPI_ISL_3369872, EPI_ISL_3369873, EPI_ISL_3369874, EPI_ISL_3369875, EPI_ISL_3369880, EPI_ISL_3369881, EPI_ISL_3369882, EPI_ISL_3369883, EPI_ISL_3369884, EPI_ISL_3369885, EPI_ISL_3369886, EPI_ISL_3369887, EPI_ISL_3369888, EPI_ISL_3369889, EPI_ISL_3369890, EPI_ISL_3369891, EPI_ISL_3369892, EPI_ISL_3369894, EPI_ISL_3369895, EPI_ISL_3369896, EPI_ISL_3369897, EPI_ISL_3369899, EPI_ISL_3369900 | | | |
| see above | CHU Purpan - Laboratoire de Virologie - Institut Fédératif de Biologie | CHU Purpan - Laboratoire de Virologie - Institut Fédératif de Biologie | Latour J., Jeanne N., Ranger N., Carcenac R., Harter A., Boyer P., Ferrer V., Tremeaux R., Izopet J. |
| EPI_ISL_3369901, EPI_ISL_3369902, EPI_ISL_3369904 | CH AUCH | CHU Purpan - Laboratoire de Virologie - Institut Fédératif de Biologie | Latour J., Jeanne N., Ranger N., Carcenac R., Harter A., Boyer P., Ferrer V., Tremeaux R., Izopet J. |
| EPI_ISL_3369905, EPI_ISL_3369906, EPI_ISL_3369907, EPI_ISL_3369910, EPI_ISL_3369913, EPI_ISL_3369914, EPI_ISL_3369916, EPI_ISL_3369917, EPI_ISL_3369918 | CHU Purpan - Laboratoire de Virologie - Institut Fédératif de Biologie | CHU Purpan - Laboratoire de Virologie - Institut Fédératif de Biologie | Latour J., Jeanne N., Ranger N., Carcenac R., Harter A., Boyer P., Ferrer V., Tremeaux R., Izopet J. |
| EPI_ISL_3369976, EPI_ISL_3369977, EPI_ISL_3369978, EPI_ISL_3369979, EPI_ISL_3369980, EPI_ISL_3369981, EPI_ISL_3369982, EPI_ISL_3369984, EPI_ISL_3369985, EPI_ISL_3369986, EPI_ISL_3369987, EPI_ISL_3369988, EPI_ISL_3369989, EPI_ISL_3369990 | | | |
| see above | DC Public Health Lab/ Dept. of Forensic Sciences | DC Public Health Lab/ Dept. of Forensic Sciences | Janis Doss, Scott Nguyen, Elizabeth Zelaya, Sarah Scott, Connie Maza, Monica Mann, Brittany Hamilton, David Payne, Jocelyn Hauser |
| EPI_ISL_3370052, EPI_ISL_3370053, EPI_ISL_3370054, EPI_ISL_3370055, EPI_ISL_3370056, EPI_ISL_3370057, EPI_ISL_3370058, EPI_ISL_3370059, EPI_ISL_3370060, EPI_ISL_3370061, EPI_ISL_3370062, EPI_ISL_3370063, EPI_ISL_3370064, EPI_ISL_3370065, EPI_ISL_3370066, EPI_ISL_3370067, EPI_ISL_3370068, EPI_ISL_3370081 | | | |
| see above | Ministry of Health Turkey | Ministry of Health Turkey | Fatma Bayrakdar, Yasemin Coşgun, Süleyman Yalcin, Gülay Korukluoğlu |
| EPI_ISL_3370113, EPI_ISL_3370114, EPI_ISL_3370115, EPI_ISL_3370116, EPI_ISL_3370117, EPI_ISL_3370118, EPI_ISL_3370119, EPI_ISL_3370120, EPI_ISL_3370121, EPI_ISL_3370122, EPI_ISL_3370123, EPI_ISL_3370124, EPI_ISL_3370125, EPI_ISL_3370126, EPI_ISL_3370127, EPI_ISL_3370128, EPI_ISL_3370129, EPI_ISL_3370130, EPI_ISL_3370131, EPI_ISL_3370132, EPI_ISL_3370133, EPI_ISL_3370134, EPI_ISL_3370135, EPI_ISL_3370136, EPI_ISL_3370137, EPI_ISL_3370138, EPI_ISL_3370139, EPI_ISL_3370140, EPI_ISL_3370141, EPI_ISL_3370142, EPI_ISL_3370143, EPI_ISL_3370144, EPI_ISL_3370145, EPI_ISL_3370146, EPI_ISL_3370147, EPI_ISL_3370148, EPI_ISL_3370149, EPI_ISL_3370150, EPI_ISL_3370151, EPI_ISL_3370152, EPI_ISL_3370153, EPI_ISL_3370154, EPI_ISL_3370155, EPI_ISL_3370156, EPI_ISL_3370157, EPI_ISL_3370158, EPI_ISL_3370159, EPI_ISL_3370160, EPI_ISL_3370161, EPI_ISL_3370162, EPI_ISL_3370163, EPI_ISL_3370164, EPI_ISL_3370165, EPI_ISL_3370166, EPI_ISL_3370167, EPI_ISL_3370168, EPI_ISL_3370169, EPI_ISL_3370170, EPI_ISL_3370171, EPI_ISL_3370172, EPI_ISL_3370173, EPI_ISL_3370174, EPI_ISL_3370175, EPI_ISL_3370177, EPI_ISL_3370178, EPI_ISL_3370180, EPI_ISL_3370181, EPI_ISL_3370182, EPI_ISL_3370183, EPI_ISL_3370184, EPI_ISL_3370185, EPI_ISL_3370186, EPI_ISL_3370187, EPI_ISL_3370188, EPI_ISL_3370189, EPI_ISL_3370190, EPI_ISL_3370191, EPI_ISL_3370192, EPI_ISL_3370193, EPI_ISL_3370194, EPI_ISL_3370195, EPI_ISL_3370196, EPI_ISL_3370197, EPI_ISL_3370198, EPI_ISL_3370199, EPI_ISL_3370200, EPI_ISL_3370201, EPI_ISL_3370202, EPI_ISL_3370203, EPI_ISL_3370204, EPI_ISL_3370205, EPI_ISL_3370206, EPI_ISL_3370207, EPI_ISL_3370208, EPI_ISL_3370209, EPI_ISL_3370210, EPI_ISL_3370211, EPI_ISL_3370212, EPI_ISL_3370213, EPI_ISL_3370214, EPI_ISL_3370215, EPI_ISL_3370216, EPI_ISL_3370217, EPI_ISL_3370219, EPI_ISL_3370220, EPI_ISL_3370221, EPI_ISL_3370222, EPI_ISL_3370223, EPI_ISL_3370224, EPI_ISL_3370225, EPI_ISL_3370226, EPI_ISL_3370227, EPI_ISL_3370228, EPI_ISL_3370229, EPI_ISL_3370230, EPI_ISL_3370231, EPI_ISL_3370232, EPI_ISL_3370233, EPI_ISL_3370234, EPI_ISL_3370235, EPI_ISL_3370236, EPI_ISL_3370237, EPI_ISL_3370238, EPI_ISL_3370239, EPI_ISL_3370240, EPI_ISL_3370241, EPI_ISL_3370242, EPI_ISL_3370243, EPI_ISL_3370244, EPI_ISL_3370245, EPI_ISL_3370246, EPI_ISL_3370247, EPI_ISL_3370248, EPI_ISL_3370249, EPI_ISL_3370250, EPI_ISL_3370251, EPI_ISL_3370252, EPI_ISL_3370253, EPI_ISL_3370254, EPI_ISL_3370255, EPI_ISL_3370256, EPI_ISL_3370257, EPI_ISL_3370258, EPI_ISL_3370259, EPI_ISL_3370260, EPI_ISL_3370261, EPI_ISL_3370262, EPI_ISL_3370263, EPI_ISL_3370264, EPI_ISL_3370265, EPI_ISL_3370266, EPI_ISL_3370267, EPI_ISL_3370268 | | | |
| see above | Yale Clinical Virology Lab | Grubaugh Lab - Yale School of Public Health | Joseph Fauver, Mallery Breban, Isabel Ott, Tara Alpert, Mary Petrone, Anderson Brito, Chantal Vogels, Annie Watkins, Chaney Kalinich, Jessica Rothman, Kendall Billig, Tobias Koch, Marie L. Landry, Nathan Grubaugh |
| EPI_ISL_3370271, EPI_ISL_3370272 | Kansas Health and Environmental Lab | Kansas Health and Environmental Lab | Mike Grose, Jonathan Barnell, Amanda Bradley, and Phil Adam |
| EPI_ISL_3370273 | WSU Molecular Diagnostics Laboratory | Kansas Health and Environmental Lab | Mike Grose, Jonathan Barnell, Amanda Bradley, and Phil Adam |
| EPI_ISL_3370274, EPI_ISL_3370275, EPI_ISL_3370276, EPI_ISL_3370277, EPI_ISL_3370278, EPI_ISL_3370280, EPI_ISL_3370281, EPI_ISL_3370282, EPI_ISL_3370283, EPI_ISL_3370284, EPI_ISL_3370285, EPI_ISL_3370286, EPI_ISL_3370287, EPI_ISL_3370288, EPI_ISL_3370289, EPI_ISL_3370290, EPI_ISL_3370291, EPI_ISL_3370292, EPI_ISL_3370293, EPI_ISL_3370294, EPI_ISL_3370295, EPI_ISL_3370296, EPI_ISL_3370297, EPI_ISL_3370298, EPI_ISL_3370299, EPI_ISL_3370300 | | | |
| see above | Kansas Health and Environmental Lab | Kansas Health and Environmental Lab | Mike Grose, Jonathan Barnell, Amanda Bradley, and Phil Adam |
| EPI_ISL_3370302 | Wyoming Public Health Laboratory | Wyoming Public Health Laboratory | Jim Mildenberger, Taylor Fearing, Channing Weber, Ashley Norberg, Chayse Rowley, Marley Goetz, Brian Dominguez, Elliot Thomasson, Sam Britz, Cari Sloma, Robert Petit, and Rob Christensen |
| EPI_ISL_3370324, EPI_ISL_3370325, EPI_ISL_3370326, EPI_ISL_3370327, EPI_ISL_3370328, EPI_ISL_3370329, EPI_ISL_3370330, EPI_ISL_3370331, EPI_ISL_3370332, EPI_ISL_3370333, EPI_ISL_3370334, EPI_ISL_3370335, EPI_ISL_3370336, EPI_ISL_3370337, EPI_ISL_3370338, EPI_ISL_3370339, EPI_ISL_3370340, EPI_ISL_3370341, EPI_ISL_3370342, EPI_ISL_3370343, EPI_ISL_3370344, EPI_ISL_3370345, EPI_ISL_3370346, EPI_ISL_3370347, EPI_ISL_3370348, EPI_ISL_3370349, EPI_ISL_3370350, EPI_ISL_3370351 | | | |
| see above | Kansas Health and Environmental Lab | Kansas Health and Environmental Lab | Mike Grose, Jonathan Barnell, Amanda Bradley, and Phil Adam |
| EPI_ISL_3370352 | Ohio Department of Health Laboratory | Ohio Department of Health Laboratory | Holmes, Jennifer; Eric Brandt, Keoni Omura, Glen McGillivary, Caitlin McDonnell, Jade Mowery, Stephanie Mccracken, Tyler Payne, Kirtana Ramadugu, Erica Leasure, Brent Lee, Kelsey Florek, Heather Blankenship, Allison Black, Quanta Brown, and Tammy Bannerman |
| EPI_ISL_3370394, EPI_ISL_3370395, EPI_ISL_3370396, EPI_ISL_3370397, EPI_ISL_3370398, EPI_ISL_3370399, EPI_ISL_3370400, EPI_ISL_3370401, EPI_ISL_3370402, EPI_ISL_3370403, EPI_ISL_3370404, EPI_ISL_3370405, EPI_ISL_3370406, EPI_ISL_3370407, EPI_ISL_3370408, EPI_ISL_3370409, EPI_ISL_3370410, EPI_ISL_3370411, EPI_ISL_3370412, EPI_ISL_3370413, EPI_ISL_3370414, EPI_ISL_3370415, EPI_ISL_3370416, EPI_ISL_3370417, EPI_ISL_3370418, EPI_ISL_3370419, EPI_ISL_3370421 | | | |
| see above | North Dakota Department of Health, Public Health Laboratory | North Dakota Department of Health, Public Health Laboratory | Lisa Wingerter |
| EPI_ISL_3370463, EPI_ISL_3370464, EPI_ISL_3370465 | Centracare Laboratory Services | Minnesota Department of Health, Public Health Laboratory | Alexandra Lorentz, Jacob Garfin, Matt Plumb, and Xiong Wang |
| EPI_ISL_3370466, EPI_ISL_3370467, EPI_ISL_3370468 | Minnesota Department of Health, Public Health Laboratory | Minnesota Department of Health, Public Health Laboratory | Alexandra Lorentz, Jacob Garfin, Matt Plumb, and Xiong Wang |
| EPI_ISL_3370469, EPI_ISL_3370470, EPI_ISL_3370471, EPI_ISL_3370472, EPI_ISL_3370473, EPI_ISL_3370474 | Centracare Laboratory Services | Minnesota Department of Health, Public Health Laboratory | Alexandra Lorentz, Jacob Garfin, Matt Plumb, and Xiong Wang |
| EPI_ISL_3370475, EPI_ISL_3370476, EPI_ISL_3370477, EPI_ISL_3370478, EPI_ISL_3370479 | Minnesota Department of Health, Public Health Laboratory | Minnesota Department of Health, Public Health Laboratory | Alexandra Lorentz, Jacob Garfin, Matt Plumb, and Xiong Wang |
| EPI_ISL_3370480 | Centracare Laboratory Services | Minnesota Department of Health, Public Health Laboratory | Alexandra Lorentz, Jacob Garfin, Matt Plumb, and Xiong Wang |
| EPI_ISL_3370481, EPI_ISL_3370482 | Minnesota Department of Health, Public Health Laboratory | Minnesota Department of Health, Public Health Laboratory | Alexandra Lorentz, Jacob Garfin, Matt Plumb, and Xiong Wang |
| EPI_ISL_3370483, EPI_ISL_3370484 | Centracare Laboratory Services | Minnesota Department of Health, Public Health Laboratory | Alexandra Lorentz, Jacob Garfin, Matt Plumb, and Xiong Wang |
| EPI_ISL_3370485, EPI_ISL_3370486, EPI_ISL_3370487, EPI_ISL_3370488, EPI_ISL_3370489, EPI_ISL_3370490 | Minnesota Department of Health, Public Health Laboratory | Minnesota Department of Health, Public Health Laboratory | Alexandra Lorentz, Jacob Garfin, Matt Plumb, and Xiong Wang |
| EPI_ISL_3370491, EPI_ISL_3370492, EPI_ISL_3370493, EPI_ISL_3370494, EPI_ISL_3370495 | Centracare Laboratory Services | Minnesota Department of Health, Public Health Laboratory | Alexandra Lorentz, Jacob Garfin, Matt Plumb, and Xiong Wang |
| EPI_ISL_3370496 | Hennepin County Medical Center | Minnesota Department of Health, Public Health Laboratory | Alexandra Lorentz, Jacob Garfin, Matt Plumb, and Xiong Wang |
| EPI_ISL_3370497, EPI_ISL_3370498 | Centracare Laboratory Services | Minnesota Department of Health, Public Health Laboratory | Alexandra Lorentz, Jacob Garfin, Matt Plumb, and Xiong Wang |
| EPI_ISL_3370499 | Minnesota Department of Health, Public Health Laboratory | Minnesota Department of Health, Public Health Laboratory | Alexandra Lorentz, Jacob Garfin, Matt Plumb, and Xiong Wang |
| EPI_ISL_3370500, EPI_ISL_3370501, EPI_ISL_3370502 | Centracare Laboratory Services | Minnesota Department of Health, Public Health Laboratory | Alexandra Lorentz, Jacob Garfin, Matt Plumb, and Xiong Wang |
| EPI_ISL_3370503 | Minnesota Department of Health, Public Health Laboratory | Minnesota Department of Health, Public Health Laboratory | Alexandra Lorentz, Jacob Garfin, Matt Plumb, and Xiong Wang |

| | | | |
|---|---|---|---|
| EPI_ISL_3370504 | Centracare Laboratory Services | Minnesota Department of Health, Public Health Laboratory | Alexandra Lorentz, Jacob Garfin, Matt Plumb, and Xiong Wang |
| EPI_ISL_3370505, EPI_ISL_3370506 | Minnesota Department of Health, Public Health Laboratory | Minnesota Department of Health, Public Health Laboratory | Alexandra Lorentz, Jacob Garfin, Matt Plumb, and Xiong Wang |
| EPI_ISL_3370507 | Centracare Laboratory Services | Minnesota Department of Health, Public Health Laboratory | Alexandra Lorentz, Jacob Garfin, Matt Plumb, and Xiong Wang |
| EPI_ISL_3370509 | Minnesota Department of Health, Public Health Laboratory | Minnesota Department of Health, Public Health Laboratory | Alexandra Lorentz, Jacob Garfin, Matt Plumb, and Xiong Wang |
| EPI_ISL_3370510, EPI_ISL_3370511 | Hennepin County Medical Center | Minnesota Department of Health, Public Health Laboratory | Alexandra Lorentz, Jacob Garfin, Matt Plumb, and Xiong Wang |
| EPI_ISL_3370512 | Centracare Laboratory Services | Minnesota Department of Health, Public Health Laboratory | Alexandra Lorentz, Jacob Garfin, Matt Plumb, and Xiong Wang |
| EPI_ISL_3370513 | Minnesota Department of Health, Public Health Laboratory | Minnesota Department of Health, Public Health Laboratory | Alexandra Lorentz, Jacob Garfin, Matt Plumb, and Xiong Wang |
| EPI_ISL_3370514 | Hennepin County Medical Center | Minnesota Department of Health, Public Health Laboratory | Alexandra Lorentz, Jacob Garfin, Matt Plumb, and Xiong Wang |
| EPI_ISL_3370515 | Minnesota Department of Health, Public Health Laboratory | Minnesota Department of Health, Public Health Laboratory | Alexandra Lorentz, Jacob Garfin, Matt Plumb, and Xiong Wang |
| EPI_ISL_3370516 | Centracare Laboratory Services | Minnesota Department of Health, Public Health Laboratory | Alexandra Lorentz, Jacob Garfin, Matt Plumb, and Xiong Wang |
| EPI_ISL_3370517 | Minnesota Department of Health, Public Health Laboratory | Minnesota Department of Health, Public Health Laboratory | Alexandra Lorentz, Jacob Garfin, Matt Plumb, and Xiong Wang |
| EPI_ISL_3370518, EPI_ISL_3370519, EPI_ISL_3370520 | Centracare Laboratory Services | Minnesota Department of Health, Public Health Laboratory | Alexandra Lorentz, Jacob Garfin, Matt Plumb, and Xiong Wang |
| EPI_ISL_3370521, EPI_ISL_3370522 | Hennepin County Medical Center | Minnesota Department of Health, Public Health Laboratory | Alexandra Lorentz, Jacob Garfin, Matt Plumb, and Xiong Wang |
| EPI_ISL_3370523 | Minnesota Department of Health, Public Health Laboratory | Minnesota Department of Health, Public Health Laboratory | Alexandra Lorentz, Jacob Garfin, Matt Plumb, and Xiong Wang |
| EPI_ISL_3370524 | Centracare Laboratory Services | Minnesota Department of Health, Public Health Laboratory | Alexandra Lorentz, Jacob Garfin, Matt Plumb, and Xiong Wang |
| EPI_ISL_3370525 | Minnesota Department of Health, Public Health Laboratory | Minnesota Department of Health, Public Health Laboratory | Alexandra Lorentz, Jacob Garfin, Matt Plumb, and Xiong Wang |
| EPI_ISL_3370526, EPI_ISL_3370527, EPI_ISL_3370528, EPI_ISL_3370529 | Centracare Laboratory Services | Minnesota Department of Health, Public Health Laboratory | Alexandra Lorentz, Jacob Garfin, Matt Plumb, and Xiong Wang |
| EPI_ISL_3370530, EPI_ISL_3370531, EPI_ISL_3370532, EPI_ISL_3370533, EPI_ISL_3370534, EPI_ISL_3370535, EPI_ISL_3370536, EPI_ISL_3370537, EPI_ISL_3370538, EPI_ISL_3370539 | Minnesota Department of Health, Public Health Laboratory | Minnesota Department of Health, Public Health Laboratory | Alexandra Lorentz, Jacob Garfin, Matt Plumb, and Xiong Wang |
| EPI_ISL_3370540 | Centracare Laboratory Services | Minnesota Department of Health, Public Health Laboratory | Alexandra Lorentz, Jacob Garfin, Matt Plumb, and Xiong Wang |
| EPI_ISL_3370541, EPI_ISL_3370542 | Minnesota Department of Health, Public Health Laboratory | Minnesota Department of Health, Public Health Laboratory | Alexandra Lorentz, Jacob Garfin, Matt Plumb, and Xiong Wang |
| EPI_ISL_3370543, EPI_ISL_3370544, EPI_ISL_3370545, EPI_ISL_3370546, EPI_ISL_3370547 | Centracare Laboratory Services | Minnesota Department of Health, Public Health Laboratory | Alexandra Lorentz, Jacob Garfin, Matt Plumb, and Xiong Wang |
| EPI_ISL_3370548, EPI_ISL_3370549 | Minnesota Department of Health, Public Health Laboratory | Minnesota Department of Health, Public Health Laboratory | Alexandra Lorentz, Jacob Garfin, Matt Plumb, and Xiong Wang |
| EPI_ISL_3370550, EPI_ISL_3370551, EPI_ISL_3370552 | Centracare Laboratory Services | Minnesota Department of Health, Public Health Laboratory | Alexandra Lorentz, Jacob Garfin, Matt Plumb, and Xiong Wang |
| EPI_ISL_3370553, EPI_ISL_3370554, EPI_ISL_3370555 | Minnesota Department of Health, Public Health Laboratory | Minnesota Department of Health, Public Health Laboratory | Alexandra Lorentz, Jacob Garfin, Matt Plumb, and Xiong Wang |
| EPI_ISL_3370556, EPI_ISL_3370557 | Centracare Laboratory Services | Minnesota Department of Health, Public Health Laboratory | Alexandra Lorentz, Jacob Garfin, Matt Plumb, and Xiong Wang |
| EPI_ISL_3370558, EPI_ISL_3370559 | Minnesota Department of Health, Public Health Laboratory | Minnesota Department of Health, Public Health Laboratory | Alexandra Lorentz, Jacob Garfin, Matt Plumb, and Xiong Wang |
| EPI_ISL_3370560 | Centracare Laboratory Services | Minnesota Department of Health, Public Health Laboratory | Alexandra Lorentz, Jacob Garfin, Matt Plumb, and Xiong Wang |
| EPI_ISL_3370561, EPI_ISL_3370562, EPI_ISL_3370563 | Minnesota Department of Health, Public Health Laboratory | Minnesota Department of Health, Public Health Laboratory | Alexandra Lorentz, Jacob Garfin, Matt Plumb, and Xiong Wang |
| EPI_ISL_3370564 | Centracare Laboratory Services | Minnesota Department of Health, Public Health Laboratory | Alexandra Lorentz, Jacob Garfin, Matt Plumb, and Xiong Wang |
| EPI_ISL_3370565, EPI_ISL_3370566, EPI_ISL_3370567, EPI_ISL_3370568 | Minnesota Department of Health, Public Health Laboratory | Minnesota Department of Health, Public Health Laboratory | Alexandra Lorentz, Jacob Garfin, Matt Plumb, and Xiong Wang |
| EPI_ISL_3370569, EPI_ISL_3370570, EPI_ISL_3370571 | Centracare Laboratory Services | Minnesota Department of Health, Public Health Laboratory | Alexandra Lorentz, Jacob Garfin, Matt Plumb, and Xiong Wang |
| EPI_ISL_3370572, EPI_ISL_3370573, EPI_ISL_3370574, EPI_ISL_3370575, EPI_ISL_3370576, EPI_ISL_3370577, EPI_ISL_3370578 | Minnesota Department of Health, Public Health Laboratory | Minnesota Department of Health, Public Health Laboratory | Alexandra Lorentz, Jacob Garfin, Matt Plumb, and Xiong Wang |
| EPI_ISL_3370579 | Centracare Laboratory Services | Minnesota Department of Health, Public Health Laboratory | Alexandra Lorentz, Jacob Garfin, Matt Plumb, and Xiong Wang |
| EPI_ISL_3370580, EPI_ISL_3370581 | Minnesota Department of Health, Public Health Laboratory | Minnesota Department of Health, Public Health Laboratory | Alexandra Lorentz, Jacob Garfin, Matt Plumb, and Xiong Wang |
| EPI_ISL_3370582 | Centracare Laboratory Services | Minnesota Department of Health, Public Health Laboratory | Alexandra Lorentz, Jacob Garfin, Matt Plumb, and Xiong Wang |
| EPI_ISL_3370583, EPI_ISL_3370584, EPI_ISL_3370585, EPI_ISL_3370586, EPI_ISL_3370587, EPI_ISL_3370588, EPI_ISL_3370589, EPI_ISL_3370590, EPI_ISL_3370591, EPI_ISL_3370592, EPI_ISL_3370593, EPI_ISL_3370594, EPI_ISL_3370595, EPI_ISL_3370596, EPI_ISL_3370598 | | | |
| see above | Minnesota Department of Health, Public Health Laboratory | Minnesota Department of Health, Public Health Laboratory | Alexandra Lorentz, Jacob Garfin, Matt Plumb, and Xiong Wang |
| EPI_ISL_3370599, EPI_ISL_3370600, EPI_ISL_3370601, EPI_ISL_3370602 | Centracare Laboratory Services | Minnesota Department of Health, Public Health Laboratory | Alexandra Lorentz, Jacob Garfin, Matt Plumb, and Xiong Wang |
| EPI_ISL_3370603 | Hennepin County Medical Center | Minnesota Department of Health, Public Health Laboratory | Alexandra Lorentz, Jacob Garfin, Matt Plumb, and Xiong Wang |
| EPI_ISL_3370604, EPI_ISL_3370605 | Minnesota Department of Health, Public Health Laboratory | Minnesota Department of Health, Public Health Laboratory | Alexandra Lorentz, Jacob Garfin, Matt Plumb, and Xiong Wang |
| EPI_ISL_3370606, EPI_ISL_3370607, EPI_ISL_3370608, EPI_ISL_3370609 | Centracare Laboratory Services | Minnesota Department of Health, Public Health Laboratory | Alexandra Lorentz, Jacob Garfin, Matt Plumb, and Xiong Wang |

| | | | |
|---|---|---|---|
| EPI_ISL_3370610, EPI_ISL_3370611, EPI_ISL_3370612, EPI_ISL_3370613, EPI_ISL_3370614, EPI_ISL_3370615, EPI_ISL_3370616, EPI_ISL_3370617, EPI_ISL_3370618, EPI_ISL_3370619, EPI_ISL_3370620, EPI_ISL_3370621, EPI_ISL_3370622 | | | |
| see above | Minnesota Department of Health, Public Health Laboratory | Minnesota Department of Health, Public Health Laboratory | Alexandra Lorentz, Jacob Garfin, Matt Plumb, and Xiong Wang |
| EPI_ISL_3370623, EPI_ISL_3370624, EPI_ISL_3370626, EPI_ISL_3370627, EPI_ISL_3370628, EPI_ISL_3370629, EPI_ISL_3370630, EPI_ISL_3370631, EPI_ISL_3370632, EPI_ISL_3370633, EPI_ISL_3370634, EPI_ISL_3370635, EPI_ISL_3370636, EPI_ISL_3370637, EPI_ISL_3370638, EPI_ISL_3370639, EPI_ISL_3370640, EPI_ISL_3370641, EPI_ISL_3370642, EPI_ISL_3370643, EPI_ISL_3370644, EPI_ISL_3370645, EPI_ISL_3370646, EPI_ISL_3370647, EPI_ISL_3370648, EPI_ISL_3370649, EPI_ISL_3370650, EPI_ISL_3370651, EPI_ISL_3370652, EPI_ISL_3370653, EPI_ISL_3370654, EPI_ISL_3370655, EPI_ISL_3370656, EPI_ISL_3370657, EPI_ISL_3370658, EPI_ISL_3370659, EPI_ISL_3370660, EPI_ISL_3370661, EPI_ISL_3370662, EPI_ISL_3370663, EPI_ISL_3370664, EPI_ISL_3370665, EPI_ISL_3370666, EPI_ISL_3370667, EPI_ISL_3370668, EPI_ISL_3370669, EPI_ISL_3370670 | | | |
| see above | NCSLPH | NCSLPH | Chase K, Miller MC, Greene S, Glover W |
| EPI_ISL_3370671 | CH ARMENTIERES | CHU Lille - Laboratoire de Virologie | AIT YAHYA Emilie, ALIDJINOU Enagnon Kazali, BOCKET Laurence, CREPIN Michel, DEMAY Christophe, ENGELMANN Ilka, GEFFROY Sandrine, GUIGON Aurélie, LAMBERT Valérie, LAZREK Mouna, NOBILLIAUX Florian, PREVOST Brigitte, THUILLIER Caroline, TINEZ Claire |
| EPI_ISL_3370672, EPI_ISL_3370673, EPI_ISL_3370674, EPI_ISL_3370675, EPI_ISL_3370676 | CHU LILLE | CHU Lille - Laboratoire de Virologie | AIT YAHYA Emilie, ALIDJINOU Enagnon Kazali, BOCKET Laurence, CREPIN Michel, DEMAY Christophe, ENGELMANN Ilka, GEFFROY Sandrine, GUIGON Aurélie, LAMBERT Valérie, LAZREK Mouna, NOBILLIAUX Florian, PREVOST Brigitte, THUILLIER Caroline, TINEZ Claire |
| EPI_ISL_3370678 | CH TOURCOING | CHU Lille - Laboratoire de Virologie | AIT YAHYA Emilie, ALIDJINOU Enagnon Kazali, BOCKET Laurence, CREPIN Michel, DEMAY Christophe, ENGELMANN Ilka, GEFFROY Sandrine, GUIGON Aurélie, LAMBERT Valérie, LAZREK Mouna, NOBILLIAUX Florian, PREVOST Brigitte, THUILLIER Caroline, TINEZ Claire |
| EPI_ISL_3370679 | CHU LILLE | CHU Lille - Laboratoire de Virologie | AIT YAHYA Emilie, ALIDJINOU Enagnon Kazali, BOCKET Laurence, CREPIN Michel, DEMAY Christophe, ENGELMANN Ilka, GEFFROY Sandrine, GUIGON Aurélie, LAMBERT Valérie, LAZREK Mouna, NOBILLIAUX Florian, PREVOST Brigitte, THUILLIER Caroline, TINEZ Claire |
| EPI_ISL_3370680, EPI_ISL_3370681 | CH VALENCIENNES | CHU Lille - Laboratoire de Virologie | AIT YAHYA Emilie, ALIDJINOU Enagnon Kazali, BOCKET Laurence, CREPIN Michel, DEMAY Christophe, ENGELMANN Ilka, GEFFROY Sandrine, GUIGON Aurélie, LAMBERT Valérie, LAZREK Mouna, NOBILLIAUX Florian, PREVOST Brigitte, THUILLIER Caroline, TINEZ Claire |
| EPI_ISL_3370682, EPI_ISL_3370683, EPI_ISL_3370690, EPI_ISL_3370691, EPI_ISL_3370692, EPI_ISL_3370693, EPI_ISL_3370694 | CHU LILLE | CHU Lille - Laboratoire de Virologie | AIT YAHYA Emilie, ALIDJINOU Enagnon Kazali, BOCKET Laurence, CREPIN Michel, DEMAY Christophe, ENGELMANN Ilka, GEFFROY Sandrine, GUIGON Aurélie, LAMBERT Valérie, LAZREK Mouna, NOBILLIAUX Florian, PREVOST Brigitte, THUILLIER Caroline, TINEZ Claire |
| EPI_ISL_3370695 | CH SAINT-PHILIBERT | CHU Lille - Laboratoire de Virologie | AIT YAHYA Emilie, ALIDJINOU Enagnon Kazali, BOCKET Laurence, CREPIN Michel, DEMAY Christophe, ENGELMANN Ilka, GEFFROY Sandrine, GUIGON Aurélie, LAMBERT Valérie, LAZREK Mouna, NOBILLIAUX Florian, PREVOST Brigitte, THUILLIER Caroline, TINEZ Claire |
| EPI_ISL_3370696, EPI_ISL_3370697, EPI_ISL_3370698 | CHU LILLE | CHU Lille - Laboratoire de Virologie | AIT YAHYA Emilie, ALIDJINOU Enagnon Kazali, BOCKET Laurence, CREPIN Michel, DEMAY Christophe, ENGELMANN Ilka, GEFFROY Sandrine, GUIGON Aurélie, LAMBERT Valérie, LAZREK Mouna, NOBILLIAUX Florian, PREVOST Brigitte, THUILLIER Caroline, TINEZ Claire |
| EPI_ISL_3370699 | CH ROUBAIX | CHU Lille - Laboratoire de Virologie | AIT YAHYA Emilie, ALIDJINOU Enagnon Kazali, BOCKET Laurence, CREPIN Michel, DEMAY Christophe, ENGELMANN Ilka, GEFFROY Sandrine, GUIGON Aurélie, LAMBERT Valérie, LAZREK Mouna, NOBILLIAUX Florian, PREVOST Brigitte, THUILLIER Caroline, TINEZ Claire |
| EPI_ISL_3370700 | CH BOULOGNE-SUR-MER | CHU Lille - Laboratoire de Virologie | AIT YAHYA Emilie, ALIDJINOU Enagnon Kazali, BOCKET Laurence, CREPIN Michel, DEMAY Christophe, ENGELMANN Ilka, GEFFROY Sandrine, GUIGON Aurélie, LAMBERT Valérie, LAZREK Mouna, NOBILLIAUX Florian, PREVOST Brigitte, THUILLIER Caroline, TINEZ Claire |
| EPI_ISL_3370701, EPI_ISL_3370702, EPI_ISL_3370703, EPI_ISL_3370704, EPI_ISL_3370705 | CHU LILLE | CHU Lille - Laboratoire de Virologie | AIT YAHYA Emilie, ALIDJINOU Enagnon Kazali, BOCKET Laurence, CREPIN Michel, DEMAY Christophe, ENGELMANN Ilka, GEFFROY Sandrine, GUIGON Aurélie, LAMBERT Valérie, LAZREK Mouna, NOBILLIAUX Florian, PREVOST Brigitte, THUILLIER Caroline, TINEZ Claire |
| EPI_ISL_3370706 | CH ARRAS | CHU Lille - Laboratoire de Virologie | AIT YAHYA Emilie, ALIDJINOU Enagnon Kazali, BOCKET Laurence, CREPIN Michel, DEMAY Christophe, ENGELMANN Ilka, GEFFROY Sandrine, GUIGON Aurélie, LAMBERT Valérie, LAZREK Mouna, NOBILLIAUX Florian, PREVOST Brigitte, THUILLIER Caroline, TINEZ Claire |
| EPI_ISL_3370707 | CH SAINT-PHILIBERT | CHU Lille - Laboratoire de Virologie | AIT YAHYA Emilie, ALIDJINOU Enagnon Kazali, BOCKET Laurence, CREPIN Michel, DEMAY Christophe, ENGELMANN Ilka, GEFFROY Sandrine, GUIGON Aurélie, LAMBERT Valérie, LAZREK Mouna, NOBILLIAUX Florian, PREVOST Brigitte, THUILLIER Caroline, TINEZ Claire |
| EPI_ISL_3370708 | CHU LILLE | CHU Lille - Laboratoire de Virologie | AIT YAHYA Emilie, ALIDJINOU Enagnon Kazali, BOCKET Laurence, CREPIN Michel, DEMAY Christophe, ENGELMANN Ilka, GEFFROY Sandrine, GUIGON Aurélie, LAMBERT Valérie, LAZREK Mouna, NOBILLIAUX Florian, PREVOST Brigitte, THUILLIER Caroline, TINEZ Claire |
| EPI_ISL_3370709 | CH CALAIS | CHU Lille - Laboratoire de Virologie | AIT YAHYA Emilie, ALIDJINOU Enagnon Kazali, BOCKET Laurence, CREPIN Michel, DEMAY Christophe, ENGELMANN Ilka, GEFFROY Sandrine, GUIGON Aurélie, LAMBERT Valérie, LAZREK Mouna, NOBILLIAUX Florian, PREVOST Brigitte, THUILLIER Caroline, TINEZ Claire |
| EPI_ISL_3370710, EPI_ISL_3370711, EPI_ISL_3370712, EPI_ISL_3370713, EPI_ISL_3370714, EPI_ISL_3370715, EPI_ISL_3370716, EPI_ISL_3370717, EPI_ISL_3370718 | CHU LILLE | CHU Lille - Laboratoire de Virologie | AIT YAHYA Emilie, ALIDJINOU Enagnon Kazali, BOCKET Laurence, CREPIN Michel, DEMAY Christophe, ENGELMANN Ilka, GEFFROY Sandrine, GUIGON Aurélie, LAMBERT Valérie, LAZREK Mouna, NOBILLIAUX Florian, PREVOST Brigitte, THUILLIER Caroline, TINEZ Claire |
| EPI_ISL_3370719, EPI_ISL_3370720 | CH TOURCOING | CHU Lille - Laboratoire de Virologie | AIT YAHYA Emilie, ALIDJINOU Enagnon Kazali, BOCKET Laurence, CREPIN Michel, DEMAY Christophe, ENGELMANN Ilka, GEFFROY Sandrine, GUIGON Aurélie, LAMBERT Valérie, LAZREK Mouna, NOBILLIAUX Florian, PREVOST Brigitte, THUILLIER Caroline, TINEZ Claire |
| EPI_ISL_3370721 | CH ROUBAIX | CHU Lille - Laboratoire de Virologie | AIT YAHYA Emilie, ALIDJINOU Enagnon Kazali, BOCKET Laurence, CREPIN Michel, DEMAY Christophe, ENGELMANN Ilka, GEFFROY Sandrine, GUIGON Aurélie, LAMBERT Valérie, LAZREK Mouna, NOBILLIAUX Florian, PREVOST Brigitte, THUILLIER Caroline, TINEZ Claire |
| EPI_ISL_3370722, EPI_ISL_3370723, EPI_ISL_3370724, EPI_ISL_3370725, EPI_ISL_3370726 | CHU LILLE | CHU Lille - Laboratoire de Virologie | AIT YAHYA Emilie, ALIDJINOU Enagnon Kazali, BOCKET Laurence, CREPIN Michel, DEMAY Christophe, ENGELMANN Ilka, GEFFROY Sandrine, GUIGON Aurélie, LAMBERT Valérie, LAZREK Mouna, NOBILLIAUX Florian, PREVOST Brigitte, THUILLIER Caroline, TINEZ Claire |
| EPI_ISL_3370727, EPI_ISL_3370728, EPI_ISL_3370729, EPI_ISL_3370730, EPI_ISL_3370731, EPI_ISL_3370732, EPI_ISL_3370733, EPI_ISL_3370734, EPI_ISL_3370735 | CH VALENCIENNES | CHU Lille - Laboratoire de Virologie | AIT YAHYA Emilie, ALIDJINOU Enagnon Kazali, BOCKET Laurence, CREPIN Michel, DEMAY Christophe, ENGELMANN Ilka, GEFFROY Sandrine, GUIGON Aurélie, LAMBERT Valérie, LAZREK Mouna, NOBILLIAUX Florian, PREVOST Brigitte, THUILLIER Caroline, TINEZ Claire |
| EPI_ISL_3370736 | CH ROUBAIX | CHU Lille - Laboratoire de Virologie | AIT YAHYA Emilie, ALIDJINOU Enagnon Kazali, BOCKET Laurence, CREPIN Michel, DEMAY Christophe, ENGELMANN Ilka, GEFFROY Sandrine, GUIGON Aurélie, LAMBERT Valérie, LAZREK Mouna, NOBILLIAUX Florian, PREVOST Brigitte, THUILLIER Caroline, TINEZ Claire |
| EPI_ISL_3370737, EPI_ISL_3370738 | CH BETHUNE | CHU Lille - Laboratoire de Virologie | AIT YAHYA Emilie, ALIDJINOU Enagnon Kazali, BOCKET Laurence, CREPIN Michel, DEMAY Christophe, ENGELMANN Ilka, GEFFROY Sandrine, GUIGON Aurélie, LAMBERT Valérie, LAZREK Mouna, NOBILLIAUX Florian, PREVOST Brigitte, THUILLIER Caroline, TINEZ Claire |
| EPI_ISL_3370739 | CH ARMENTIERES | CHU Lille - Laboratoire de Virologie | AIT YAHYA Emilie, ALIDJINOU Enagnon Kazali, BOCKET Laurence, CREPIN Michel, DEMAY Christophe, ENGELMANN Ilka, GEFFROY Sandrine, GUIGON Aurélie, LAMBERT Valérie, LAZREK Mouna, NOBILLIAUX Florian, PREVOST Brigitte, THUILLIER Caroline, TINEZ Claire |
| EPI_ISL_3370740 | CHU LILLE | CHU Lille - Laboratoire de Virologie | AIT YAHYA Emilie, ALIDJINOU Enagnon Kazali, BOCKET Laurence, CREPIN Michel, DEMAY Christophe, ENGELMANN Ilka, GEFFROY Sandrine, GUIGON Aurélie, LAMBERT Valérie, LAZREK Mouna, NOBILLIAUX Florian, PREVOST Brigitte, THUILLIER Caroline, TINEZ Claire |
| EPI_ISL_3370741, EPI_ISL_3370742, EPI_ISL_3370743 | CH TOURCOING | CHU Lille - Laboratoire de Virologie | AIT YAHYA Emilie, ALIDJINOU Enagnon Kazali, BOCKET Laurence, CREPIN Michel, DEMAY Christophe, ENGELMANN Ilka, GEFFROY Sandrine, GUIGON Aurélie, LAMBERT Valérie, LAZREK Mouna, NOBILLIAUX Florian, PREVOST Brigitte, THUILLIER Caroline, TINEZ Claire |
| EPI_ISL_3370744 | CHU LILLE | CHU Lille - Laboratoire de Virologie | AIT YAHYA Emilie, ALIDJINOU Enagnon Kazali, BOCKET Laurence, CREPIN Michel, DEMAY Christophe, ENGELMANN Ilka, GEFFROY Sandrine, GUIGON Aurélie, LAMBERT Valérie, LAZREK Mouna, NOBILLIAUX Florian, PREVOST Brigitte, THUILLIER Caroline, TINEZ Claire |
| EPI_ISL_3370745, EPI_ISL_3370746 | CH VALENCIENNES | CHU Lille - Laboratoire de Virologie | AIT YAHYA Emilie, ALIDJINOU Enagnon Kazali, BOCKET Laurence, CREPIN Michel, DEMAY Christophe, ENGELMANN Ilka, GEFFROY Sandrine, GUIGON Aurélie, LAMBERT Valérie, LAZREK Mouna, NOBILLIAUX Florian, PREVOST Brigitte, THUILLIER Caroline, TINEZ Claire |
| EPI_ISL_3370747, EPI_ISL_3370748, EPI_ISL_3370749, EPI_ISL_3370750 | CHU LILLE | CHU Lille - Laboratoire de Virologie | AIT YAHYA Emilie, ALIDJINOU Enagnon Kazali, BOCKET Laurence, CREPIN Michel, DEMAY Christophe, ENGELMANN Ilka, GEFFROY Sandrine, GUIGON Aurélie, LAMBERT Valérie, LAZREK Mouna, NOBILLIAUX Florian, PREVOST Brigitte, THUILLIER Caroline, TINEZ Claire |
| EPI_ISL_3370752, EPI_ISL_3370753, EPI_ISL_3370754, EPI_ISL_3370755, EPI_ISL_3370756, EPI_ISL_3370757, EPI_ISL_3370758, EPI_ISL_3370759, EPI_ISL_3370760, EPI_ISL_3370761, EPI_ISL_3370762, EPI_ISL_3370763, EPI_ISL_3370764, EPI_ISL_3370765, EPI_ISL_3370766, EPI_ISL_3370767, EPI_ISL_3370768, EPI_ISL_3370769, EPI_ISL_3370770, EPI_ISL_3370771, EPI_ISL_3370772, EPI_ISL_3370773, EPI_ISL_3370774, EPI_ISL_3370775, EPI_ISL_3370776, EPI_ISL_3370777 | | | |

| | | | |
|---|---|---|---|
| see above | Wyoming Public Health Laboratory | Wyoming Public Health Laboratory | Jim Mildenberger, Taylor Fearing, Channing Weber, Ashley Norberg, Chayse Rowley, Marley Goetz, Brian Dominguez, Elliot Thomasson, Sam Britz, Cari Sloma, Robert Petit, and Rob Christensen |
| EPI_ISL_3370778, EPI_ISL_3370779, EPI_ISL_3370780, EPI_ISL_3370782, EPI_ISL_3370783, EPI_ISL_3370784, EPI_ISL_3370785, EPI_ISL_3370786, EPI_ISL_3370787, EPI_ISL_3370788, EPI_ISL_3370789, EPI_ISL_3370790, EPI_ISL_3370791, EPI_ISL_3370792, EPI_ISL_3370794, EPI_ISL_3370795, EPI_ISL_3370796, EPI_ISL_3370797, EPI_ISL_3370798, EPI_ISL_3370799, EPI_ISL_3370800, EPI_ISL_3370801, EPI_ISL_3370802, EPI_ISL_3370803, EPI_ISL_3370804, EPI_ISL_3370805, EPI_ISL_3370806, EPI_ISL_3370807, EPI_ISL_3370808, EPI_ISL_3370809, EPI_ISL_3370810, EPI_ISL_3370811, EPI_ISL_3370812, EPI_ISL_3370813, EPI_ISL_3370814, EPI_ISL_3370815, EPI_ISL_3370816, EPI_ISL_3370817, EPI_ISL_3370818, EPI_ISL_3370819, EPI_ISL_3370820, EPI_ISL_3370821, EPI_ISL_3370823, EPI_ISL_3370824, EPI_ISL_3370825, EPI_ISL_3370826, EPI_ISL_3370827, EPI_ISL_3370828 | | | |
| see above | UW Virology Lab | UW Virology Lab | Pavitra Roychoudhury, Shah Mohamed Bakhash, Ricardo Perez, Hong Xie, Tien V. Nguyen, Maria Lukes, Sean Ellis, Nathan Breit, Robert J. Livingston, Meei-Li Huang, Keith R Jerome, Patrick Mathias, Alexander Greninger |
| EPI_ISL_3370829, EPI_ISL_3370830, EPI_ISL_3370831, EPI_ISL_3370832, EPI_ISL_3370833 | UOC Microbiology and Virology Policlinico Umberto I Rome, Italy | University Campus Bio-Medico of Rome (UCBM) | Fogolari M., De Florio L., Riva E., Francesconi M., Lintas C., Veralli R., Angeletti S. |
| EPI_ISL_3370835, EPI_ISL_3370838, EPI_ISL_3370839, EPI_ISL_3370840, EPI_ISL_3370842, EPI_ISL_3370845, EPI_ISL_3370846, EPI_ISL_3370850 | URMC LABS | Wadsworth Center, New York State Department of Health | Kirsten St. George, Daryl M. Lamson, Alexis Russell, Matthew Shudt, Melissa A Leisner, Jonathan Plitnick, Catharine Prussing, Navjot Singh, John Kelly, Erasmus Schneider, Erica Lasek-Nesselquist |
| EPI_ISL_3370852 | ENZO CLINICAL LABS INC | Wadsworth Center, New York State Department of Health | Kirsten St. George, Daryl M. Lamson, Alexis Russell, Matthew Shudt, Melissa A Leisner, Jonathan Plitnick, Catharine Prussing, Navjot Singh, John Kelly, Erasmus Schneider, Erica Lasek-Nesselquist |
| EPI_ISL_3370853, EPI_ISL_3370854 | STONY BROOK UNIVERSITY HOSPITAL | Wadsworth Center, New York State Department of Health | Kirsten St. George, Daryl M. Lamson, Alexis Russell, Matthew Shudt, Melissa A Leisner, Jonathan Plitnick, Catharine Prussing, Navjot Singh, John Kelly, Erasmus Schneider, Erica Lasek-Nesselquist |
| EPI_ISL_3370859, EPI_ISL_3370860, EPI_ISL_3370861 | URMC LABS | Wadsworth Center, New York State Department of Health | Kirsten St. George, Daryl M. Lamson, Alexis Russell, Matthew Shudt, Melissa A Leisner, Jonathan Plitnick, Catharine Prussing, Navjot Singh, John Kelly, Erasmus Schneider, Erica Lasek-Nesselquist |
| EPI_ISL_3370863, EPI_ISL_3370864 | NORTHWELL HEALTH LABORATORIES | Wadsworth Center, New York State Department of Health | Kirsten St. George, Daryl M. Lamson, Alexis Russell, Matthew Shudt, Melissa A Leisner, Jonathan Plitnick, Catharine Prussing, Navjot Singh, John Kelly, Erasmus Schneider, Erica Lasek-Nesselquist |
| EPI_ISL_3370865, EPI_ISL_3370866, EPI_ISL_3370867, EPI_ISL_3370868, EPI_ISL_3370869, EPI_ISL_3370870, EPI_ISL_3370871 | STONY BROOK UNIVERSITY HOSPITAL | Wadsworth Center, New York State Department of Health | Kirsten St. George, Daryl M. Lamson, Alexis Russell, Matthew Shudt, Melissa A Leisner, Jonathan Plitnick, Catharine Prussing, Navjot Singh, John Kelly, Erasmus Schneider, Erica Lasek-Nesselquist |
| EPI_ISL_3370872, EPI_ISL_3370873, EPI_ISL_3370874, EPI_ISL_3370875 | ALBANY MEDICAL CENTER | Wadsworth Center, New York State Department of Health | Kirsten St. George, Daryl M. Lamson, Alexis Russell, Matthew Shudt, Melissa A Leisner, Jonathan Plitnick, Catharine Prussing, Navjot Singh, John Kelly, Erasmus Schneider, Erica Lasek-Nesselquist |
| EPI_ISL_3370876, EPI_ISL_3370877, EPI_ISL_3370878, EPI_ISL_3370879, EPI_ISL_3370880, EPI_ISL_3370881, EPI_ISL_3370882, EPI_ISL_3370883, EPI_ISL_3370884, EPI_ISL_3370885, EPI_ISL_3370886, EPI_ISL_3370887, EPI_ISL_3370888, EPI_ISL_3370889, EPI_ISL_3370890, EPI_ISL_3370891, EPI_ISL_3370892, EPI_ISL_3370893, EPI_ISL_3370894, EPI_ISL_3370895, EPI_ISL_3370896, EPI_ISL_3370897, EPI_ISL_3370898, EPI_ISL_3370899 | | | |
| see above | URMC LABS | Wadsworth Center, New York State Department of Health | Kirsten St. George, Daryl M. Lamson, Alexis Russell, Matthew Shudt, Melissa A Leisner, Jonathan Plitnick, Catharine Prussing, Navjot Singh, John Kelly, Erasmus Schneider, Erica Lasek-Nesselquist |
| EPI_ISL_3370900, EPI_ISL_3370901, EPI_ISL_3370902, EPI_ISL_3370903 | Saratoga Hospital Laboratory | Wadsworth Center, New York State Department of Health | Kirsten St. George, Daryl M. Lamson, Alexis Russell, Matthew Shudt, Melissa A Leisner, Jonathan Plitnick, Catharine Prussing, Navjot Singh, John Kelly, Erasmus Schneider, Erica Lasek-Nesselquist |
| EPI_ISL_3370905, EPI_ISL_3370906, EPI_ISL_3370907, EPI_ISL_3370908, EPI_ISL_3370909, EPI_ISL_3370911, EPI_ISL_3370912, EPI_ISL_3370913, EPI_ISL_3370914, EPI_ISL_3370915, EPI_ISL_3370916, EPI_ISL_3370917, EPI_ISL_3370918, EPI_ISL_3370919, EPI_ISL_3370920 | | | |
| see above | UCSC Genomics Institute | UCSC Genomics Institute | Mark Akeson, Ikenna Anigbogu, Eric Beraut, Isabel Bjork, Molly Cassatt-Johnstone, Terren Chang, Russell Corbett-Detig, Namrita Dhillon, Maximilian Haeussler, David Haussler, Angie Hinrichs, Miten Jain, Joshua Kapp, A. Marm Kilpatrick, Jakob McBroome, Hugh Olsen, Jeremy Sanford, Beth Shapiro, Michael Stone, Bryan Thornlow, Yatish Turakhia, Ciara Wanket |
| EPI_ISL_3370921, EPI_ISL_3370922, EPI_ISL_3370923, EPI_ISL_3370924, EPI_ISL_3370925, EPI_ISL_3370926, EPI_ISL_3370927, EPI_ISL_3370928, EPI_ISL_3370929, EPI_ISL_3370930, EPI_ISL_3370931, EPI_ISL_3370932, EPI_ISL_3370933, EPI_ISL_3370934, EPI_ISL_3370935, EPI_ISL_3370936, EPI_ISL_3370937, EPI_ISL_3370938, EPI_ISL_3370939, EPI_ISL_3370940, EPI_ISL_3370941, EPI_ISL_3370942, EPI_ISL_3370943, EPI_ISL_3370944, EPI_ISL_3370945, EPI_ISL_3370946, EPI_ISL_3370947, EPI_ISL_3370948, EPI_ISL_3370949, EPI_ISL_3370950, EPI_ISL_3370951, EPI_ISL_3370952, EPI_ISL_3370953, EPI_ISL_3370954, EPI_ISL_3370955 | | | |
| see above | Oregon State Public Health Laboratory | Oregon State Public Health Laboratory | Rafia Razzaque, Eugene Yeboah, Vanda Makris, Laura Tsaknaridis, John Fontana and Shane Sevey |
| EPI_ISL_3370959, EPI_ISL_3370960, EPI_ISL_3370961 | CHC Andrée Rosemon | Institut Pasteur de la Guyane | Anne Lavergne, Dominique Rousset, Antoine Enfissi, Arielle Salmier |
| EPI_ISL_3370963, EPI_ISL_3370964, EPI_ISL_3370965, EPI_ISL_3370966, EPI_ISL_3370967 | CNR Institut Pasteur de la Guyane | Institut Pasteur de la Guyane | Anne Lavergne, Dominique Rousset, Antoine Enfissi, Arielle Salmier |
| EPI_ISL_3370968, EPI_ISL_3370969, EPI_ISL_3370972 | CHC Andrée Rosemon | Institut Pasteur de la Guyane | Anne Lavergne, Dominique Rousset, Antoine Enfissi, Arielle Salmier |
| EPI_ISL_3370974, EPI_ISL_3370975 | CNR Institut Pasteur de la Guyane | Institut Pasteur de la Guyane | Anne Lavergne, Dominique Rousset, Antoine Enfissi, Arielle Salmier |
| EPI_ISL_3370976, EPI_ISL_3370977, EPI_ISL_3370978 | Wyoming Public Health Laboratory | Wyoming Public Health Laboratory | Jim Mildenberger, Taylor Fearing, Channing Weber, Ashley Norberg, Chayse Rowley, Marley Goetz, Brian Dominguez, Elliot Thomasson, Sam Britz, Cari Sloma, Robert Petit, and Rob Christensen |
| EPI_ISL_3371100, EPI_ISL_3371104, EPI_ISL_3371108, EPI_ISL_3371110, EPI_ISL_3371112, EPI_ISL_3371114, EPI_ISL_3371116, EPI_ISL_3371117, EPI_ISL_3371119, EPI_ISL_3371121 | Pennsylvania Department of Health Bureau of Laboratories | Pennsylvania Department of Health Bureau of Laboratories | Dongxiang Xia |
| EPI_ISL_3371415 | Instituto Nacional de Medicina Genomica | Instituto Nacional de Medicina Genomica | Cedro-Tanda A, Cisneros-Villanueva M, Blancas S, Escobar-Arrazola MA, Ramirez-Vega O, Rangel-DeLeon D, Munguia-Garza P, Hidalgo-Miranda A, Mendoza-Vargas A, Reyes-Grajeda JP, Herrera-Montalvo LA. |
| EPI_ISL_3371436, EPI_ISL_3371437, EPI_ISL_3371438, EPI_ISL_3371439, EPI_ISL_3371440, EPI_ISL_3371441, EPI_ISL_3371442, EPI_ISL_3371443, EPI_ISL_3371444, EPI_ISL_3371445, EPI_ISL_3371446, EPI_ISL_3371447, EPI_ISL_3371448, EPI_ISL_3371449, EPI_ISL_3371450, EPI_ISL_3371451, EPI_ISL_3371452, EPI_ISL_3371453, EPI_ISL_3371454, EPI_ISL_3371455, EPI_ISL_3371456, EPI_ISL_3371457, EPI_ISL_3371458, EPI_ISL_3371459, EPI_ISL_3371460, EPI_ISL_3371461, EPI_ISL_3371462, EPI_ISL_3371463, EPI_ISL_3371464, EPI_ISL_3371465, EPI_ISL_3371466, EPI_ISL_3371467, EPI_ISL_3371468, EPI_ISL_3371469, EPI_ISL_3371470, EPI_ISL_3371471, EPI_ISL_3371472, EPI_ISL_3371473, EPI_ISL_3371474, EPI_ISL_3371475, EPI_ISL_3371476, EPI_ISL_3371477, EPI_ISL_3371478, EPI_ISL_3371479, EPI_ISL_3371480, EPI_ISL_3371481, EPI_ISL_3371482, EPI_ISL_3371483, EPI_ISL_3371484, EPI_ISL_3371485, EPI_ISL_3371486, EPI_ISL_3371487, EPI_ISL_3371488, EPI_ISL_3371489, EPI_ISL_3371490, EPI_ISL_3371491, EPI_ISL_3371492, EPI_ISL_3371493, EPI_ISL_3371494, EPI_ISL_3371495, EPI_ISL_3371496, EPI_ISL_3371497, EPI_ISL_3371498, EPI_ISL_3371499, EPI_ISL_3371500, EPI_ISL_3371501, EPI_ISL_3371502, EPI_ISL_3371503, EPI_ISL_3371504, EPI_ISL_3371505, EPI_ISL_3371506, EPI_ISL_3371507, EPI_ISL_3371508, EPI_ISL_3371509, EPI_ISL_3371510, EPI_ISL_3371511, EPI_ISL_3371512, EPI_ISL_3371513, EPI_ISL_3371514, EPI_ISL_3371515, EPI_ISL_3371516, EPI_ISL_3371517, EPI_ISL_3371518, EPI_ISL_3371519, EPI_ISL_3371520, EPI_ISL_3371521, EPI_ISL_3371522, EPI_ISL_3371523, EPI_ISL_3371524, EPI_ISL_3371525, EPI_ISL_3371526, EPI_ISL_3371527, EPI_ISL_3371528, EPI_ISL_3371529, EPI_ISL_3371530, EPI_ISL_3371531, EPI_ISL_3371532, EPI_ISL_3371533, EPI_ISL_3371534, EPI_ISL_3371535, EPI_ISL_3371536, EPI_ISL_3371537, EPI_ISL_3371538, EPI_ISL_3371539, EPI_ISL_3371540, EPI_ISL_3371541, EPI_ISL_3371542, EPI_ISL_3371543, EPI_ISL_3371544, EPI_ISL_3371545, EPI_ISL_3371546, EPI_ISL_3371547, EPI_ISL_3371548, EPI_ISL_3371549, EPI_ISL_3371550, EPI_ISL_3371551, EPI_ISL_3371552, EPI_ISL_3371553, EPI_ISL_3371554, EPI_ISL_3371555, EPI_ISL_3371556, EPI_ISL_3371557, EPI_ISL_3371558, EPI_ISL_3371559, EPI_ISL_3371560, EPI_ISL_3371561, EPI_ISL_3371562, EPI_ISL_3371563, EPI_ISL_3371564, EPI_ISL_3371565, EPI_ISL_3371566, EPI_ISL_3371567, EPI_ISL_3371570, EPI_ISL_3371571, EPI_ISL_3371573, EPI_ISL_3371574, EPI_ISL_3371575, EPI_ISL_3371576, EPI_ISL_3371577, EPI_ISL_3371578, EPI_ISL_3371579, EPI_ISL_3371581, EPI_ISL_3371582, EPI_ISL_3371583, EPI_ISL_3371584, EPI_ISL_3371585, EPI_ISL_3371586, EPI_ISL_3371587, EPI_ISL_3371588, EPI_ISL_3371589, EPI_ISL_3371590, EPI_ISL_3371591, EPI_ISL_3371592, EPI_ISL_3371593, EPI_ISL_3371594, EPI_ISL_3371595, EPI_ISL_3371596, EPI_ISL_3371597, EPI_ISL_3371598, EPI_ISL_3371599, EPI_ISL_3371600, EPI_ISL_3371601, EPI_ISL_3371602, EPI_ISL_3371603, EPI_ISL_3371604, EPI_ISL_3371605, EPI_ISL_3371606, EPI_ISL_3371607, EPI_ISL_3371608, EPI_ISL_3371609, EPI_ISL_3371610, EPI_ISL_3371611, EPI_ISL_3371612, EPI_ISL_3371613, EPI_ISL_3371614 | | | |
| see above | Illinois Department of Public Health - Springfield Lab | Illinois Department of Public Health - Springfield Lab | Bryan Sim, Gordon McCall, Marshall Smock |
| EPI_ISL_3371817, EPI_ISL_3371818 | NCSLPH | NCSLPH | Chase K, Miller MC, Greene S, Glover W |
| EPI_ISL_3371819, EPI_ISL_3371820, EPI_ISL_3371821, EPI_ISL_3371822, EPI_ISL_3371823, EPI_ISL_3371824, EPI_ISL_3371825, EPI_ISL_3371826, EPI_ISL_3371827, EPI_ISL_3371828, EPI_ISL_3371829, EPI_ISL_3371830, EPI_ISL_3371831, EPI_ISL_3371832, EPI_ISL_3371833, EPI_ISL_3371834, EPI_ISL_3371835, EPI_ISL_3371836, EPI_ISL_3371837, EPI_ISL_3371838, EPI_ISL_3371839, EPI_ISL_3371840, EPI_ISL_3371841, EPI_ISL_3371842, EPI_ISL_3371843, EPI_ISL_3371844, EPI_ISL_3371845, EPI_ISL_3371846 | | | |
| see above | IN State Department of Health Laboratory Services | IN State Department of Health Laboratory Services | Cassandra Campion, Jamie Yeadon, Brian Pope, Lixia Liu, Kyle Brownlee, Melissa Hindenlang, Mark Glazier |
| EPI_ISL_3371848 | Provincial Public Health Reference Laboratory (PPHRL) | Provincial Public Health Reference Laboratory (PPHRL) | Andleeb Hanif, Mariam Javed, Hassan Nawaz, Asim Altaf |
| EPI_ISL_3371849 | Provincial Public Health Reference Laboratory (PPHRL) | Provincial Public Health Reference Laboratory (PPHRL) | Mariam Javed, Hassan Nawaz, Andleeb Hanif , Asim Altaf |
| EPI_ISL_3371850, EPI_ISL_3371851, EPI_ISL_3371852, EPI_ISL_3371853, EPI_ISL_3371854, EPI_ISL_3371855, EPI_ISL_3371856, EPI_ISL_3371857, EPI_ISL_3371858, EPI_ISL_3371859, EPI_ISL_3371860, EPI_ISL_3371861, EPI_ISL_3371862, EPI_ISL_3371863, EPI_ISL_3371864, EPI_ISL_3371865, EPI_ISL_3371866, EPI_ISL_3371868, EPI_ISL_3371869, EPI_ISL_3371870, EPI_ISL_3371871, EPI_ISL_3371872, EPI_ISL_3371873, EPI_ISL_3371874, EPI_ISL_3371875, EPI_ISL_3371876, EPI_ISL_3371877, EPI_ISL_3371878, EPI_ISL_3371879, EPI_ISL_3371880, EPI_ISL_3371881, EPI_ISL_3371882, EPI_ISL_3371883, EPI_ISL_3371884, EPI_ISL_3371885, EPI_ISL_3371886, EPI_ISL_3371887, EPI_ISL_3371888, EPI_ISL_3371889, EPI_ISL_3371890, EPI_ISL_3371891, EPI_ISL_3371892, EPI_ISL_3371893, EPI_ISL_3371894, EPI_ISL_3371895, EPI_ISL_3371896, EPI_ISL_3371897, EPI_ISL_3371898, EPI_ISL_3371899, EPI_ISL_3371900, EPI_ISL_3371901, EPI_ISL_3371902, EPI_ISL_3371903, EPI_ISL_3371904, EPI_ISL_3371905, EPI_ISL_3371906, EPI_ISL_3371907, EPI_ISL_3371908, EPI_ISL_3371909, EPI_ISL_3371910, EPI_ISL_3371911, EPI_ISL_3371912, EPI_ISL_3371913, EPI_ISL_3371914, EPI_ISL_3371915, EPI_ISL_3371916, | | | |

| | | | |
|---|---|---|---|
| EPI_ISL_3371917, EPI_ISL_3371918, EPI_ISL_3371919, EPI_ISL_3371920, EPI_ISL_3371921, EPI_ISL_3371922, EPI_ISL_3371924, EPI_ISL_3371925, EPI_ISL_3371926, EPI_ISL_3371927, EPI_ISL_3371928, EPI_ISL_3371929, EPI_ISL_3371930, EPI_ISL_3371931, EPI_ISL_3371932, EPI_ISL_3371933, EPI_ISL_3371934, EPI_ISL_3371935, EPI_ISL_3371936, EPI_ISL_3371937, EPI_ISL_3371938, EPI_ISL_3371939, EPI_ISL_3371940, EPI_ISL_3371941, EPI_ISL_3371942, EPI_ISL_3371943, EPI_ISL_3371944, EPI_ISL_3371945, EPI_ISL_3371946, EPI_ISL_3371947, EPI_ISL_3371948, EPI_ISL_3371949, EPI_ISL_3371950, EPI_ISL_3371951, EPI_ISL_3371952, EPI_ISL_3371953, EPI_ISL_3371954, EPI_ISL_3371955, EPI_ISL_3371956, EPI_ISL_3371957, EPI_ISL_3371958, EPI_ISL_3371959, EPI_ISL_3371960, EPI_ISL_3371961, EPI_ISL_3371962, EPI_ISL_3371963, EPI_ISL_3371964, EPI_ISL_3371965, EPI_ISL_3371966, EPI_ISL_3371967, EPI_ISL_3371968, EPI_ISL_3371969, EPI_ISL_3371970, EPI_ISL_3371971, EPI_ISL_3371972, EPI_ISL_3371973, EPI_ISL_3371974, EPI_ISL_3371977, EPI_ISL_3371978, EPI_ISL_3371979, EPI_ISL_3371980, EPI_ISL_3371981, EPI_ISL_3371982, EPI_ISL_3371983, EPI_ISL_3371984, EPI_ISL_3371985, EPI_ISL_3371986, EPI_ISL_3371987, EPI_ISL_3371988, EPI_ISL_3371989, EPI_ISL_3371990, EPI_ISL_3371991, EPI_ISL_3371992, EPI_ISL_3371993, EPI_ISL_3371994, EPI_ISL_3371995, EPI_ISL_3371996, EPI_ISL_3371997, EPI_ISL_3371998, EPI_ISL_3371999, EPI_ISL_3372000, EPI_ISL_3372001, EPI_ISL_3372002, EPI_ISL_3372003, EPI_ISL_3372004, EPI_ISL_3372005, EPI_ISL_3372006, EPI_ISL_3372007, EPI_ISL_3372008, EPI_ISL_3372009, EPI_ISL_3372010, EPI_ISL_3372011, EPI_ISL_3372012, EPI_ISL_3372013, EPI_ISL_3372014, EPI_ISL_3372015, EPI_ISL_3372016, EPI_ISL_3372017, EPI_ISL_3372018, EPI_ISL_3372019, EPI_ISL_3372020, EPI_ISL_3372021, EPI_ISL_3372022, EPI_ISL_3372023, EPI_ISL_3372024, EPI_ISL_3372025, EPI_ISL_3372026, EPI_ISL_3372027, EPI_ISL_3372028, EPI_ISL_3372029, EPI_ISL_3372030, EPI_ISL_3372031, EPI_ISL_3372032, EPI_ISL_3372033, EPI_ISL_3372034, EPI_ISL_3372035, EPI_ISL_3372036, EPI_ISL_3372037, EPI_ISL_3372038, EPI_ISL_3372041, EPI_ISL_3372042, EPI_ISL_3372044, EPI_ISL_3372046, EPI_ISL_3372047, EPI_ISL_3372048, EPI_ISL_3372049, EPI_ISL_3372050, EPI_ISL_3372051, EPI_ISL_3372052, EPI_ISL_3372053, EPI_ISL_3372054, EPI_ISL_3372055, EPI_ISL_3372056, EPI_ISL_3372057, EPI_ISL_3372058, EPI_ISL_3372059, EPI_ISL_3372060, EPI_ISL_3372061, EPI_ISL_3372062, EPI_ISL_3372063, EPI_ISL_3372064, EPI_ISL_3372065, EPI_ISL_3372066, EPI_ISL_3372067, EPI_ISL_3372068, EPI_ISL_3372069, EPI_ISL_3372070, EPI_ISL_3372071, EPI_ISL_3372072, EPI_ISL_3372073, EPI_ISL_3372075, EPI_ISL_3372076, EPI_ISL_3372077, EPI_ISL_3372078, EPI_ISL_3372079, EPI_ISL_3372080, EPI_ISL_3372081, EPI_ISL_3372082, EPI_ISL_3372083, EPI_ISL_3372084, EPI_ISL_3372085, EPI_ISL_3372086, EPI_ISL_3372087, EPI_ISL_3372088, EPI_ISL_3372089, EPI_ISL_3372090, EPI_ISL_3372091, EPI_ISL_3372093, EPI_ISL_3372094, EPI_ISL_3372095, EPI_ISL_3372096, EPI_ISL_3372097, EPI_ISL_3372098, EPI_ISL_3372101, EPI_ISL_3372102, EPI_ISL_3372103, EPI_ISL_3372104, EPI_ISL_3372105, EPI_ISL_3372106, EPI_ISL_3372107, EPI_ISL_3372108, EPI_ISL_3372109, EPI_ISL_3372110, EPI_ISL_3372111, EPI_ISL_3372112, EPI_ISL_3372113, EPI_ISL_3372114, EPI_ISL_3372115, EPI_ISL_3372116, EPI_ISL_3372117, EPI_ISL_3372118, EPI_ISL_3372119, EPI_ISL_3372120, EPI_ISL_3372121, EPI_ISL_3372122, EPI_ISL_3372123, EPI_ISL_3372124, EPI_ISL_3372125, EPI_ISL_3372126, EPI_ISL_3372127, EPI_ISL_3372128, EPI_ISL_3372129, EPI_ISL_3372130, EPI_ISL_3372131, EPI_ISL_3372132, EPI_ISL_3372133, EPI_ISL_3372134, EPI_ISL_3372135, EPI_ISL_3372136, EPI_ISL_3372137, EPI_ISL_3372138, EPI_ISL_3372139, EPI_ISL_3372140, EPI_ISL_3372141, EPI_ISL_3372142, EPI_ISL_3372143, EPI_ISL_3372144, EPI_ISL_3372145, EPI_ISL_3372146, EPI_ISL_3372147, EPI_ISL_3372148, EPI_ISL_3372149, EPI_ISL_3372150, EPI_ISL_3372151, EPI_ISL_3372152, EPI_ISL_3372153, EPI_ISL_3372154, EPI_ISL_3372155, EPI_ISL_3372156, EPI_ISL_3372157, EPI_ISL_3372158, EPI_ISL_3372159, EPI_ISL_3372160, EPI_ISL_3372161, EPI_ISL_3372163, EPI_ISL_3372164, EPI_ISL_3372165, EPI_ISL_3372166, EPI_ISL_3372167, EPI_ISL_3372168, EPI_ISL_3372169, EPI_ISL_3372170, EPI_ISL_3372171, EPI_ISL_3372172, EPI_ISL_3372173, EPI_ISL_3372174, EPI_ISL_3372175, EPI_ISL_3372176, EPI_ISL_3372177, EPI_ISL_3372178, EPI_ISL_3372179, EPI_ISL_3372180, EPI_ISL_3372181, EPI_ISL_3372182, EPI_ISL_3372183, EPI_ISL_3372184, EPI_ISL_3372185, EPI_ISL_3372186, EPI_ISL_3372187, EPI_ISL_3372188, EPI_ISL_3372189, EPI_ISL_3372190, EPI_ISL_3372191, EPI_ISL_3372192, EPI_ISL_3372193, EPI_ISL_3372194, EPI_ISL_3372195, EPI_ISL_3372196, EPI_ISL_3372197, EPI_ISL_3372198, EPI_ISL_3372199, EPI_ISL_3372200, EPI_ISL_3372201, EPI_ISL_3372203, EPI_ISL_3372204, EPI_ISL_3372205, EPI_ISL_3372206, EPI_ISL_3372207, EPI_ISL_3372208, EPI_ISL_3372209, EPI_ISL_3372210, EPI_ISL_3372211, EPI_ISL_3372212, EPI_ISL_3372213, EPI_ISL_3372215, EPI_ISL_3372216, EPI_ISL_3372217, EPI_ISL_3372218, EPI_ISL_3372219, EPI_ISL_3372220, EPI_ISL_3372221, EPI_ISL_3372222, EPI_ISL_3372223, EPI_ISL_3372224, EPI_ISL_3372225, EPI_ISL_3372226, EPI_ISL_3372227, EPI_ISL_3372228, EPI_ISL_3372229, EPI_ISL_3372230, EPI_ISL_3372231, EPI_ISL_3372232, EPI_ISL_3372233, EPI_ISL_3372234, EPI_ISL_3372236, EPI_ISL_3372237, EPI_ISL_3372238, EPI_ISL_3372239, EPI_ISL_3372240, EPI_ISL_3372241, EPI_ISL_3372242, EPI_ISL_3372243, EPI_ISL_3372244, EPI_ISL_3372245, EPI_ISL_3372246, EPI_ISL_3372247, EPI_ISL_3372248, EPI_ISL_3372249, EPI_ISL_3372250, EPI_ISL_3372251, EPI_ISL_3372252, EPI_ISL_3372253, EPI_ISL_3372254, EPI_ISL_3372256, EPI_ISL_3372257, EPI_ISL_3372258, EPI_ISL_3372259, EPI_ISL_3372260, EPI_ISL_3372261, EPI_ISL_3372262, EPI_ISL_3372263, EPI_ISL_3372264, EPI_ISL_3372265, EPI_ISL_3372266, EPI_ISL_3372267, EPI_ISL_3372268, EPI_ISL_3372269, EPI_ISL_3372270, EPI_ISL_3372274, EPI_ISL_3372275, EPI_ISL_3372276, EPI_ISL_3372277, EPI_ISL_3372278, EPI_ISL_3372279, EPI_ISL_3372280, EPI_ISL_3372281, EPI_ISL_3372282, EPI_ISL_3372283, EPI_ISL_3372284, EPI_ISL_3372285, EPI_ISL_3372286, EPI_ISL_3372287, EPI_ISL_3372288, EPI_ISL_3372289, EPI_ISL_3372290, EPI_ISL_3372291, EPI_ISL_3372292, EPI_ISL_3372293, EPI_ISL_3372294, EPI_ISL_3372295, EPI_ISL_3372296, EPI_ISL_3372297, EPI_ISL_3372298, EPI_ISL_3372299, EPI_ISL_3372300, EPI_ISL_3372301, EPI_ISL_3372302, EPI_ISL_3372303, EPI_ISL_3372304, EPI_ISL_3372305, EPI_ISL_3372306, EPI_ISL_3372307, EPI_ISL_3372308, EPI_ISL_3372309, EPI_ISL_3372310, EPI_ISL_3372311, EPI_ISL_3372312, EPI_ISL_3372313, EPI_ISL_3372314, EPI_ISL_3372315, EPI_ISL_3372316, EPI_ISL_3372317, EPI_ISL_3372318, EPI_ISL_3372319, EPI_ISL_3372320, EPI_ISL_3372321, EPI_ISL_3372322, EPI_ISL_3372323, EPI_ISL_3372324, EPI_ISL_3372325, EPI_ISL_3372326, EPI_ISL_3372327, EPI_ISL_3372328, EPI_ISL_3372329, EPI_ISL_3372330, EPI_ISL_3372331, EPI_ISL_3372332, EPI_ISL_3372333, EPI_ISL_3372334, EPI_ISL_3372335, EPI_ISL_3372336, EPI_ISL_3372337, EPI_ISL_3372338, EPI_ISL_3372339, EPI_ISL_3372340, EPI_ISL_3372341, EPI_ISL_3372342, EPI_ISL_3372343, EPI_ISL_3372344, EPI_ISL_3372345, EPI_ISL_3372346, EPI_ISL_3372347, EPI_ISL_3372348, EPI_ISL_3372349, EPI_ISL_3372350, EPI_ISL_3372351, EPI_ISL_3372352, EPI_ISL_3372353, EPI_ISL_3372354, EPI_ISL_3372355, EPI_ISL_3372356, EPI_ISL_3372357, EPI_ISL_3372358, EPI_ISL_3372359, EPI_ISL_3372360, EPI_ISL_3372361, EPI_ISL_3372362, EPI_ISL_3372363, EPI_ISL_3372364, EPI_ISL_3372365, EPI_ISL_3372366, EPI_ISL_3372367, EPI_ISL_3372368, EPI_ISL_3372369, EPI_ISL_3372370, EPI_ISL_3372371, EPI_ISL_3372372, EPI_ISL_3372373, EPI_ISL_3372374, EPI_ISL_3372375, EPI_ISL_3372376, EPI_ISL_3372377, EPI_ISL_3372378, EPI_ISL_3372379, EPI_ISL_3372380, EPI_ISL_3372381, EPI_ISL_3372382, EPI_ISL_3372383, EPI_ISL_3372384, EPI_ISL_3372385, EPI_ISL_3372386, EPI_ISL_3372388, EPI_ISL_3372389, EPI_ISL_3372390, EPI_ISL_3372391, EPI_ISL_3372392, EPI_ISL_3372393, EPI_ISL_3372394, EPI_ISL_3372395, EPI_ISL_3372396, EPI_ISL_3372397, EPI_ISL_3372398, EPI_ISL_3372399, EPI_ISL_3372400, EPI_ISL_3372401, EPI_ISL_3372402, EPI_ISL_3372403, EPI_ISL_3372404, EPI_ISL_3372405, EPI_ISL_3372406, EPI_ISL_3372407, EPI_ISL_3372408, EPI_ISL_3372409, EPI_ISL_3372410, EPI_ISL_3372411, EPI_ISL_3372412, EPI_ISL_3372413, EPI_ISL_3372414, EPI_ISL_3372415, EPI_ISL_3372416, EPI_ISL_3372417, EPI_ISL_3372418, EPI_ISL_3372419, EPI_ISL_3372420, EPI_ISL_3372421, EPI_ISL_3372422, EPI_ISL_3372423, EPI_ISL_3372424, EPI_ISL_3372425, EPI_ISL_3372426, EPI_ISL_3372427, EPI_ISL_3372428, EPI_ISL_3372429, EPI_ISL_3372430, EPI_ISL_3372431, EPI_ISL_3372432, EPI_ISL_3372433, EPI_ISL_3372434, EPI_ISL_3372435, EPI_ISL_3372436, EPI_ISL_3372437, EPI_ISL_3372438, EPI_ISL_3372439, EPI_ISL_3372440, EPI_ISL_3372441, EPI_ISL_3372442, EPI_ISL_3372443, EPI_ISL_3372444, EPI_ISL_3372445, EPI_ISL_3372446, EPI_ISL_3372447, EPI_ISL_3372448, EPI_ISL_3372449, EPI_ISL_3372450, EPI_ISL_3372451, EPI_ISL_3372452, EPI_ISL_3372453, EPI_ISL_3372454, EPI_ISL_3372455, EPI_ISL_3372456, EPI_ISL_3372457, EPI_ISL_3372458, EPI_ISL_3372459, EPI_ISL_3372460, EPI_ISL_3372461, EPI_ISL_3372462, EPI_ISL_3372463, EPI_ISL_3372464, EPI_ISL_3372465, EPI_ISL_3372466, EPI_ISL_3372467, EPI_ISL_3372468, EPI_ISL_3372469 | | | |
| see above | Nevada State Public Health Laboratory | Nevada State Public Health Laboratory | Andrew Gorzalski, Mark Pandori |
| EPI_ISL_3372471, EPI_ISL_3372472 | Provincial Public Health Reference Laboratory (PPHRL) | Provincial Public Health Reference Laboratory (PPHRL) | Mariam Javed, Hassan Nawaz, Andleeb Hanif, Asim Altaf |
| EPI_ISL_3372473, EPI_ISL_3372474, EPI_ISL_3372475, EPI_ISL_3372476, EPI_ISL_3372477 | Minnesota Department of Health, Public Health Laboratory | Minnesota Department of Health, Public Health Laboratory | Alexandra Lorentz, Jacob Garfin, Matt Plumb, and Xiong Wang |
| EPI_ISL_3372478 | Allina Health Laboratory | Minnesota Department of Health, Public Health Laboratory | Alexandra Lorentz, Jacob Garfin, Matt Plumb, and Xiong Wang |
| EPI_ISL_3372479 | Minnesota Department of Health, Public Health Laboratory | Minnesota Department of Health, Public Health Laboratory | Alexandra Lorentz, Jacob Garfin, Matt Plumb, and Xiong Wang |
| EPI_ISL_3372480 | Allina Health Laboratory | Minnesota Department of Health, Public Health Laboratory | Alexandra Lorentz, Jacob Garfin, Matt Plumb, and Xiong Wang |
| EPI_ISL_3372481, EPI_ISL_3372482, EPI_ISL_3372483, EPI_ISL_3372484 | Minnesota Department of Health, Public Health Laboratory | Minnesota Department of Health, Public Health Laboratory | Alexandra Lorentz, Jacob Garfin, Matt Plumb, and Xiong Wang |
| EPI_ISL_3372485, EPI_ISL_3372486 | Children's Hospitals and Clinics of Minnesota | Minnesota Department of Health, Public Health Laboratory | Alexandra Lorentz, Jacob Garfin, Matt Plumb, and Xiong Wang |
| EPI_ISL_3372487, EPI_ISL_3372488, EPI_ISL_3372489, EPI_ISL_3372490, EPI_ISL_3372491, EPI_ISL_3372492, EPI_ISL_3372493, EPI_ISL_3372494, EPI_ISL_3372495 | Allina Health Laboratory | Minnesota Department of Health, Public Health Laboratory | Alexandra Lorentz, Jacob Garfin, Matt Plumb, and Xiong Wang |
| EPI_ISL_3372496, EPI_ISL_3372497, EPI_ISL_3372498, EPI_ISL_3372499, EPI_ISL_3372500, EPI_ISL_3372501, EPI_ISL_3372502, EPI_ISL_3372503, EPI_ISL_3372504, EPI_ISL_3372505, EPI_ISL_3372506, EPI_ISL_3372507, EPI_ISL_3372508, EPI_ISL_3372509, EPI_ISL_3372510, EPI_ISL_3372511, EPI_ISL_3372512, EPI_ISL_3372513, EPI_ISL_3372514, EPI_ISL_3372515 | | | |
| see above | Minnesota Department of Health, Public Health Laboratory | Minnesota Department of Health, Public Health Laboratory | Alexandra Lorentz, Jacob Garfin, Matt Plumb, and Xiong Wang |
| EPI_ISL_3372516 | Quest Diagnostics | Minnesota Department of Health, Public Health Laboratory | Alexandra Lorentz, Jacob Garfin, Matt Plumb, and Xiong Wang |
| EPI_ISL_3372517, EPI_ISL_3372518, EPI_ISL_3372519, EPI_ISL_3372520, EPI_ISL_3372521 | Minnesota Department of Health, Public Health Laboratory | Minnesota Department of Health, Public Health Laboratory | Alexandra Lorentz, Jacob Garfin, Matt Plumb, and Xiong Wang |
| EPI_ISL_3372522, EPI_ISL_3372523, EPI_ISL_3372524, EPI_ISL_3372525, EPI_ISL_3372526, EPI_ISL_3372527, EPI_ISL_3372528, EPI_ISL_3372529, EPI_ISL_3372530, EPI_ISL_3372531, EPI_ISL_3372532, EPI_ISL_3372533, EPI_ISL_3372534, EPI_ISL_3372535, EPI_ISL_3372536, EPI_ISL_3372537, EPI_ISL_3372538, EPI_ISL_3372539, EPI_ISL_3372540 | | | |
| see above | Centracare Laboratory Services | Minnesota Department of Health, Public Health Laboratory | Alexandra Lorentz, Jacob Garfin, Matt Plumb, and Xiong Wang |
| EPI_ISL_3372541, EPI_ISL_3372542, EPI_ISL_3372543, EPI_ISL_3372544, EPI_ISL_3372545, EPI_ISL_3372546, EPI_ISL_3372547, EPI_ISL_3372548, EPI_ISL_3372549, EPI_ISL_3372550, EPI_ISL_3372551, EPI_ISL_3372552, EPI_ISL_3372553, EPI_ISL_3372554 | | | |
| see above | Minnesota Department of Health, Public Health Laboratory | Minnesota Department of Health, Public Health Laboratory | Alexandra Lorentz, Jacob Garfin, Matt Plumb, and Xiong Wang |
| EPI_ISL_3372555 | Hennepin County Medical Center | Minnesota Department of Health, Public Health Laboratory | Alexandra Lorentz, Jacob Garfin, Matt Plumb, and Xiong Wang |
| EPI_ISL_3372556, EPI_ISL_3372557, EPI_ISL_3372562 | Provincial Public Health Reference Laboratory (PPHRL) | Provincial Public Health Reference Laboratory (PPHRL) | Andleeb Hanif, Mariam Javed, Hassan Nawaz, , Asim Altaf |
| EPI_ISL_3372576, EPI_ISL_3372601, EPI_ISL_3372603 | Dirección regional de salud del Callao (DIRESA-CALLAO) | Centro de Investigaciones Tecnológicas, Biomédicas y Medioambientales (CITBM) | M. Sovero, M. Ramirez, B, Huaman, M. Cuellar, J. Alarcon |
| EPI_ISL_3372609 | Provincial Public Health Reference Laboratory (PPHRL) | Provincial Public Health Reference Laboratory (PPHRL) | Mariam Javed, Hassan Nawaz, Andleeb Hanif , Asim Altaf |
| EPI_ISL_3372614, EPI_ISL_3372615 | DOHMH Central Harlem | New York City Public Health Laboratory | Jade Wang, et al. |
| EPI_ISL_3372622 | Staten Island University Hospital | New York City Public Health Laboratory | Jade Wang, et al. |
| EPI_ISL_3372623 | Richmond University Medical Center | New York City Public Health Laboratory | Jade Wang, et al. |
| EPI_ISL_3372624 | Quest Diagnostics | New York City Public Health Laboratory | Jade Wang, et al. |
| EPI_ISL_3372625 | DOHMH PHL | New York City Public Health Laboratory | Jade Wang, et al. |
| EPI_ISL_3372626 | DOHMH Central Harlem | New York City Public Health Laboratory | Jade Wang, et al. |
| EPI_ISL_3372627 | DOHMH Crown Heights | New York City Public Health Laboratory | Jade Wang, et al. |
| EPI_ISL_3372628 | DOHMH Chelsea | New York City Public Health Laboratory | Jade Wang, et al. |
| EPI_ISL_3372629 | DOHMH Fort Greene | New York City Public Health Laboratory | Jade Wang, et al. |
| EPI_ISL_3372630 | DOHMH Central Harlem | New York City Public Health Laboratory | Jade Wang, et al. |
| EPI_ISL_3372632 | Richmond University Medical Center | New York City Public Health Laboratory | Jade Wang, et al. |
| EPI_ISL_3372633 | Provincial Public Health Reference Laboratory (PPHRL) | Provincial Public Health Reference Laboratory (PPHRL) | Andleeb Hanif, Mariam Javed, Hassan Nawaz, , Asim Altaf |
| EPI_ISL_3372634, EPI_ISL_3372635 | DOHMH PHL | New York City Public Health Laboratory | Jade Wang, et al. |
| EPI_ISL_3372636 | DOHMH Riverside | New York City Public Health Laboratory | Jade Wang, et al. |
| EPI_ISL_3372637 | DOHMH Chelsea | New York City Public Health Laboratory | Jade Wang, et al. |
| EPI_ISL_3372638, EPI_ISL_3372639, EPI_ISL_3372640 | DOHMH Jamaica | New York City Public Health Laboratory | Jade Wang, et al. |
| EPI_ISL_3372641, EPI_ISL_3372642 | DOHMH PHL | New York City Public Health Laboratory | Jade Wang, et al. |

| | | | |
|---|---|---|---|
| EPI_ISL_3372644 | Quest Diagnostics | New York City Public Health Laboratory | Jade Wang, et al. |
| EPI_ISL_3372645, EPI_ISL_3372646, EPI_ISL_3372647 | DOHMH PHL | New York City Public Health Laboratory | Jade Wang, et al. |
| EPI_ISL_3372648, EPI_ISL_3372650 | Provincial Public Health Reference Laboratory (PPHRL) | Provincial Public Health Reference Laboratory (PPHRL) | Andleeb Hanif, Mariam Javed, Hassan Nawaz, , Asim Altaf |
| EPI_ISL_3372651, EPI_ISL_3372652, EPI_ISL_3372653, EPI_ISL_3372654, EPI_ISL_3372655, EPI_ISL_3372656, EPI_ISL_3372657, EPI_ISL_3372658, EPI_ISL_3372659, EPI_ISL_3372660, EPI_ISL_3372661, EPI_ISL_3372662, EPI_ISL_3372663, EPI_ISL_3372664, EPI_ISL_3372665 | | | |
| see above | Institute for Infectious Diseases | Institute for Infectious Diseases | Stefan Neuenschwander, Christian Baumann, Miguel A Terrazos Miani, Cora SÃ¤gesser, Pascal Bittel, Peter Keller, Franziska Suter-Riniker, Stephen L Leib, Alban Ramette |
| EPI_ISL_3372674 | Seattle Flu Study | Seattle Flu Study | Deborah A. Nickerson, Chris D. Frazar, Jover Lee, Benjamin Pelle, Erica Ryke, Matthew Richardson, Amanda Adler, Elisabeth Brandstetter, Peter D. Han, Kairsten Fay, Misja Ilcisin, Kirsten Lacombe, Thomas R. Sibley, Melissa Truong, Caitlin R. Wolf, Michael Boeckh, Janet A. Englund, Michael Famulare, Barry R. Lutz, Mark J. Rieder, Lea M. Starita, Matthew Thompson, Jay Shendure, Trevor Bedford, Helen Y. Chu |
| EPI_ISL_3372675 | Northwest Laboratory | Seattle Flu Study | Deborah A. Nickerson, Chris D. Frazar, Jover Lee, Benjamin Pelle, Erica Ryke, Matthew Richardson, Amanda Adler, Elisabeth Brandstetter, Peter D. Han, Kairsten Fay, Misja Ilcisin, Kirsten Lacombe, Thomas R. Sibley, Melissa Truong, Caitlin R. Wolf, Romesh Gautom, Geoff Melly, Brian Hiatt, Philip Dykema, Scott Lindquist, Michael Boeckh, Janet A. Englund, Michael Famulare, Barry R. Lutz, Mark J. Rieder, Lea M. Starita, Matthew Thompson, Helen Y. Chu, Jay Shendure, Trevor Bedford |
| EPI_ISL_3372676 | Seattle Flu Study | Seattle Flu Study | Deborah A. Nickerson, Chris D. Frazar, Jover Lee, Benjamin Pelle, Erica Ryke, Matthew Richardson, Amanda Adler, Elisabeth Brandstetter, Peter D. Han, Kairsten Fay, Misja Ilcisin, Kirsten Lacombe, Thomas R. Sibley, Melissa Truong, Caitlin R. Wolf, Michael Boeckh, Janet A. Englund, Michael Famulare, Barry R. Lutz, Mark J. Rieder, Lea M. Starita, Matthew Thompson, Jay Shendure, Trevor Bedford, Helen Y. Chu |
| EPI_ISL_3372678 | Atlas Genomics | Seattle Flu Study | Deborah A. Nickerson, Chris D. Frazar, Jover Lee, Benjamin Pelle, Erica Ryke, Matthew Richardson, Amanda Adler, Elisabeth Brandstetter, Peter D. Han, Kairsten Fay, Misja Ilcisin, Kirsten Lacombe, Thomas R. Sibley, Melissa Truong, Caitlin R. Wolf, Romesh Gautom, Geoff Melly, Brian Hiatt, Philip Dykema, Scott Lindquist, Michael Boeckh, Janet A. Englund, Michael Famulare, Barry R. Lutz, Mark J. Rieder, Lea M. Starita, Matthew Thompson, Helen Y. Chu, Jay Shendure, Trevor Bedford |
| EPI_ISL_3372688, EPI_ISL_3372689 | Seattle Flu Study | Seattle Flu Study | Deborah A. Nickerson, Chris D. Frazar, Jover Lee, Benjamin Pelle, Erica Ryke, Matthew Richardson, Amanda Adler, Elisabeth Brandstetter, Peter D. Han, Kairsten Fay, Misja Ilcisin, Kirsten Lacombe, Thomas R. Sibley, Melissa Truong, Caitlin R. Wolf, Karen Cowgill, Stephanie Schrag, Jeff Duchin, Michael Boeckh, Janet A. Englund, Michael Famulare, Barry R. Lutz, Mark J. Rieder, Lea M. Starita, Matthew Thompson, Helen Y. Chu, Trevor Bedford, Jay Shendure |
| EPI_ISL_3372690, EPI_ISL_3372691, EPI_ISL_3372692, EPI_ISL_3372693, EPI_ISL_3372694, EPI_ISL_3372695, EPI_ISL_3372696, EPI_ISL_3372697, EPI_ISL_3372698, EPI_ISL_3372699, EPI_ISL_3372700, EPI_ISL_3372701, EPI_ISL_3372702, EPI_ISL_3372703, EPI_ISL_3372704, EPI_ISL_3372705, EPI_ISL_3372706, EPI_ISL_3372707, EPI_ISL_3372708, EPI_ISL_3372709, EPI_ISL_3372710, EPI_ISL_3372711 | | | |
| see above | Northwest Laboratory | Seattle Flu Study | Deborah A. Nickerson, Chris D. Frazar, Jover Lee, Benjamin Pelle, Erica Ryke, Matthew Richardson, Amanda Adler, Elisabeth Brandstetter, Peter D. Han, Kairsten Fay, Misja Ilcisin, Kirsten Lacombe, Thomas R. Sibley, Melissa Truong, Caitlin R. Wolf, Romesh Gautom, Geoff Melly, Brian Hiatt, Philip Dykema, Scott Lindquist, Michael Boeckh, Janet A. Englund, Michael Famulare, Barry R. Lutz, Mark J. Rieder, Lea M. Starita, Matthew Thompson, Helen Y. Chu, Jay Shendure, Trevor Bedford |
| EPI_ISL_3372712, EPI_ISL_3372713, EPI_ISL_3372714 | Seattle Flu Study | Seattle Flu Study | Deborah A. Nickerson, Chris D. Frazar, Jover Lee, Benjamin Pelle, Erica Ryke, Matthew Richardson, Amanda Adler, Elisabeth Brandstetter, Peter D. Han, Kairsten Fay, Misja Ilcisin, Kirsten Lacombe, Thomas R. Sibley, Melissa Truong, Caitlin R. Wolf, Karen Cowgill, Stephanie Schrag, Jeff Duchin, Michael Boeckh, Janet A. Englund, Michael Famulare, Barry R. Lutz, Mark J. Rieder, Lea M. Starita, Matthew Thompson, Helen Y. Chu, Trevor Bedford, Jay Shendure |
| EPI_ISL_3372715, EPI_ISL_3372716, EPI_ISL_3372717, EPI_ISL_3372718, EPI_ISL_3372719, EPI_ISL_3372720, EPI_ISL_3372721, EPI_ISL_3372722, EPI_ISL_3372723, EPI_ISL_3372724, EPI_ISL_3372725, EPI_ISL_3372726, EPI_ISL_3372727 | | | |
| see above | Northwest Laboratory | Seattle Flu Study | Deborah A. Nickerson, Chris D. Frazar, Jover Lee, Benjamin Pelle, Erica Ryke, Matthew Richardson, Amanda Adler, Elisabeth Brandstetter, Peter D. Han, Kairsten Fay, Misja Ilcisin, Kirsten Lacombe, Thomas R. Sibley, Melissa Truong, Caitlin R. Wolf, Romesh Gautom, Geoff Melly, Brian Hiatt, Philip Dykema, Scott Lindquist, Michael Boeckh, Janet A. Englund, Michael Famulare, Barry R. Lutz, Mark J. Rieder, Lea M. Starita, Matthew Thompson, Helen Y. Chu, Jay Shendure, Trevor Bedford |
| EPI_ISL_3372728, EPI_ISL_3372729, EPI_ISL_3372730, EPI_ISL_3372731 | Seattle Flu Study | Seattle Flu Study | Deborah A. Nickerson, Chris D. Frazar, Jover Lee, Benjamin Pelle, Erica Ryke, Matthew Richardson, Amanda Adler, Elisabeth Brandstetter, Peter D. Han, Kairsten Fay, Misja Ilcisin, Kirsten Lacombe, Thomas R. Sibley, Melissa Truong, Caitlin R. Wolf, Karen Cowgill, Stephanie Schrag, Jeff Duchin, Michael Boeckh, Janet A. Englund, Michael Famulare, Barry R. Lutz, Mark J. Rieder, Lea M. Starita, Matthew Thompson, Helen Y. Chu, Trevor Bedford, Jay Shendure |
| EPI_ISL_3372732, EPI_ISL_3372733 | Seattle Flu Study | Seattle Flu Study | Deborah A. Nickerson, Chris D. Frazar, Jover Lee, Benjamin Pelle, Erica Ryke, Matthew Richardson, Amanda Adler, Elisabeth Brandstetter, Peter D. Han, Kairsten Fay, Misja Ilcisin, Kirsten Lacombe, Thomas R. Sibley, Melissa Truong, Caitlin R. Wolf, Michael Boeckh, Janet A. Englund, Michael Famulare, Barry R. Lutz, Mark J. Rieder, Lea M. Starita, Matthew Thompson, Jay Shendure, Trevor Bedford, Helen Y. Chu |
| EPI_ISL_3372734, EPI_ISL_3372735, EPI_ISL_3372736, EPI_ISL_3372737, EPI_ISL_3372738, EPI_ISL_3372739, EPI_ISL_3372740, EPI_ISL_3372741, EPI_ISL_3372742, EPI_ISL_3372743, EPI_ISL_3372744, EPI_ISL_3372745, EPI_ISL_3372746, EPI_ISL_3372747, EPI_ISL_3372748, EPI_ISL_3372749, EPI_ISL_3372750, EPI_ISL_3372751, EPI_ISL_3372752, EPI_ISL_3372753, EPI_ISL_3372754, EPI_ISL_3372755, EPI_ISL_3372756, EPI_ISL_3372757, EPI_ISL_3372758, EPI_ISL_3372759, EPI_ISL_3372760, EPI_ISL_3372761, EPI_ISL_3372762, EPI_ISL_3372763, EPI_ISL_3372764, EPI_ISL_3372765, EPI_ISL_3372766, EPI_ISL_3372767, EPI_ISL_3372768, EPI_ISL_3372769, EPI_ISL_3372770 | | | |
| see above | Atlas Genomics | Seattle Flu Study | Deborah A. Nickerson, Chris D. Frazar, Jover Lee, Benjamin Pelle, Erica Ryke, Matthew Richardson, Amanda Adler, Elisabeth Brandstetter, Peter D. Han, Kairsten Fay, Misja Ilcisin, Kirsten Lacombe, Thomas R. Sibley, Melissa Truong, Caitlin R. Wolf, Romesh Gautom, Geoff Melly, Brian Hiatt, Philip Dykema, Scott Lindquist, Michael Boeckh, Janet A. Englund, Michael Famulare, Barry R. Lutz, Mark J. Rieder, Lea M. Starita, Matthew Thompson, Helen Y. Chu, Jay Shendure, Trevor Bedford |
| EPI_ISL_3372771 | Seattle Flu Study | Seattle Flu Study | Deborah A. Nickerson, Chris D. Frazar, Jover Lee, Benjamin Pelle, Erica Ryke, Matthew Richardson, Amanda Adler, Elisabeth Brandstetter, Peter D. Han, Kairsten Fay, Misja Ilcisin, Kirsten Lacombe, Thomas R. Sibley, Melissa Truong, Caitlin R. Wolf, Karen Cowgill, Stephanie Schrag, Jeff Duchin, Michael Boeckh, Janet A. Englund, Michael Famulare, Barry R. Lutz, Mark J. Rieder, Lea M. Starita, Matthew Thompson, Helen Y. Chu, Trevor Bedford, Jay Shendure |
| EPI_ISL_3372772 | Atlas Genomics | Seattle Flu Study | Deborah A. Nickerson, Chris D. Frazar, Jover Lee, Benjamin Pelle, Erica Ryke, Matthew Richardson, Amanda Adler, Elisabeth Brandstetter, Peter D. Han, Kairsten Fay, Misja Ilcisin, Kirsten Lacombe, Thomas R. Sibley, Melissa Truong, Caitlin R. Wolf, Romesh Gautom, Geoff Melly, Brian Hiatt, Philip Dykema, Scott Lindquist, Michael Boeckh, Janet A. Englund, Michael Famulare, Barry R. Lutz, Mark J. Rieder, Lea M. Starita, Matthew Thompson, Helen Y. Chu, Jay Shendure, Trevor Bedford |
| EPI_ISL_3372773 | Northwest Laboratory | Seattle Flu Study | Deborah A. Nickerson, Chris D. Frazar, Jover Lee, Benjamin Pelle, Erica Ryke, Matthew Richardson, Amanda Adler, Elisabeth Brandstetter, Peter D. Han, Kairsten Fay, Misja Ilcisin, Kirsten Lacombe, Thomas R. Sibley, Melissa Truong, Caitlin R. Wolf, Romesh Gautom, Geoff Melly, Brian Hiatt, Philip Dykema, Scott Lindquist, Michael Boeckh, Janet A. Englund, Michael Famulare, Barry R. Lutz, Mark J. Rieder, Lea M. Starita, Matthew Thompson, Helen Y. Chu, Jay Shendure, Trevor Bedford |
| EPI_ISL_3372774 | Seattle Flu Study | Seattle Flu Study | Deborah A. Nickerson, Chris D. Frazar, Jover Lee, Benjamin Pelle, Erica Ryke, Matthew Richardson, Amanda Adler, Elisabeth Brandstetter, Peter D. Han, Kairsten Fay, Misja Ilcisin, Kirsten Lacombe, Thomas R. Sibley, Melissa Truong, Caitlin R. Wolf, Karen Cowgill, Stephanie Schrag, Jeff Duchin, Michael Boeckh, Janet A. Englund, Michael Famulare, Barry R. Lutz, Mark J. Rieder, Lea M. Starita, Matthew Thompson, Helen Y. Chu, Trevor Bedford, Jay Shendure |
| EPI_ISL_3372784 | Northwest Laboratory | Seattle Flu Study | Deborah A. Nickerson, Chris D. Frazar, Jover Lee, Benjamin Pelle, Erica Ryke, Matthew Richardson, Amanda Adler, Elisabeth Brandstetter, Peter D. Han, Kairsten Fay, Misja Ilcisin, Kirsten Lacombe, Thomas R. Sibley, Melissa Truong, Caitlin R. Wolf, Romesh Gautom, Geoff Melly, Brian Hiatt, Philip Dykema, Scott Lindquist, Michael Boeckh, Janet A. Englund, Michael Famulare, Barry R. Lutz, Mark J. Rieder, Lea M. Starita, Matthew Thompson, Helen Y. Chu, Jay Shendure, Trevor Bedford |
| EPI_ISL_3372785 | Atlas Genomics | Seattle Flu Study | Deborah A. Nickerson, Chris D. Frazar, Jover Lee, Benjamin Pelle, Erica Ryke, Matthew Richardson, Amanda Adler, Elisabeth Brandstetter, Peter D. Han, Kairsten Fay, Misja Ilcisin, Kirsten Lacombe, Thomas R. Sibley, Melissa Truong, Caitlin R. Wolf, Romesh Gautom, Geoff Melly, Brian Hiatt, Philip Dykema, Scott Lindquist, Michael Boeckh, Janet A. Englund, Michael Famulare, Barry R. Lutz, Mark J. Rieder, Lea M. Starita, Matthew Thompson, Helen Y. Chu, Jay Shendure, Trevor Bedford |
| EPI_ISL_3372787 | Northwest Laboratory | Seattle Flu Study | Deborah A. Nickerson, Chris D. Frazar, Jover Lee, Benjamin Pelle, Erica Ryke, Matthew Richardson, Amanda Adler, Elisabeth Brandstetter, Peter D. Han, Kairsten Fay, Misja Ilcisin, Kirsten Lacombe, Thomas R. Sibley, Melissa Truong, Caitlin R. Wolf, Romesh Gautom, Geoff Melly, Brian Hiatt, Philip Dykema, Scott Lindquist, Michael Boeckh, Janet A. Englund, Michael Famulare, Barry R. Lutz, Mark J. Rieder, Lea M. Starita, Matthew Thompson, Helen Y. Chu, Jay Shendure, Trevor Bedford |
| EPI_ISL_3372788, EPI_ISL_3372789 | Atlas Genomics | Seattle Flu Study | Deborah A. Nickerson, Chris D. Frazar, Jover Lee, Benjamin Pelle, Erica Ryke, Matthew Richardson, Amanda Adler, Elisabeth Brandstetter, Peter D. Han, Kairsten Fay, Misja Ilcisin, Kirsten Lacombe, Thomas R. Sibley, Melissa Truong, Caitlin R. Wolf, Romesh Gautom, Geoff Melly, Brian Hiatt, Philip Dykema, Scott Lindquist, Michael Boeckh, Janet A. Englund, Michael Famulare, Barry R. Lutz, Mark J. Rieder, Lea M. Starita, Matthew Thompson, Helen Y. Chu, Jay Shendure, Trevor Bedford |
| EPI_ISL_3372790, EPI_ISL_3372793, EPI_ISL_3372795, EPI_ISL_3372796, EPI_ISL_3372798 | Northwest Laboratory | Seattle Flu Study | Deborah A. Nickerson, Chris D. Frazar, Jover Lee, Benjamin Pelle, Erica Ryke, Matthew Richardson, Amanda Adler, Elisabeth Brandstetter, Peter D. Han, Kairsten Fay, Misja Ilcisin, Kirsten Lacombe, Thomas R. Sibley, Melissa Truong, Caitlin R. Wolf, Romesh Gautom, Geoff Melly, Brian Hiatt, Philip Dykema, Scott Lindquist, Michael Boeckh, Janet A. Englund, Michael Famulare, Barry R. Lutz, Mark J. Rieder, Lea M. Starita, Matthew Thompson, Helen Y. Chu, Jay Shendure, Trevor Bedford |
| EPI_ISL_3372799 | Seattle Flu Study | Seattle Flu Study | Deborah A. Nickerson, Chris D. Frazar, Jover Lee, Benjamin Pelle, Erica Ryke, Matthew Richardson, Amanda Adler, Elisabeth Brandstetter, Peter D. Han, Kairsten Fay, Misja Ilcisin, Kirsten Lacombe, Thomas R. Sibley, Melissa Truong, Caitlin R. Wolf, Michael Boeckh, Janet A. Englund, Michael Famulare, Barry R. Lutz, Mark J. Rieder, Lea M. Starita, Matthew Thompson, Jay Shendure, Trevor Bedford, Helen Y. Chu |
| EPI_ISL_3372801, EPI_ISL_3372804 | Northwest Laboratory | Seattle Flu Study | Deborah A. Nickerson, Chris D. Frazar, Jover Lee, Benjamin Pelle, Erica Ryke, Matthew Richardson, Amanda Adler, Elisabeth Brandstetter, Peter D. Han, Kairsten Fay, Misja Ilcisin, Kirsten Lacombe, Thomas R. Sibley, Melissa Truong, Caitlin R. Wolf, Romesh Gautom, Geoff Melly, Brian Hiatt, Philip Dykema, Scott Lindquist, Michael Boeckh, Janet A. Englund, Michael Famulare, Barry R. Lutz, Mark J. Rieder, Lea M. Starita, Matthew Thompson, Helen Y. Chu, Jay Shendure, Trevor Bedford |
| EPI_ISL_3372807 | Seattle Flu Study | Seattle Flu Study | Deborah A. Nickerson, Chris D. Frazar, Jover Lee, Benjamin Pelle, Erica Ryke, Matthew Richardson, Amanda Adler, Elisabeth Brandstetter, Peter D. Han, Kairsten Fay, Misja Ilcisin, Kirsten Lacombe, Thomas R. Sibley, Melissa Truong, Caitlin R. Wolf, Karen Cowgill, Stephanie Schrag, Jeff Duchin, Michael Boeckh, Janet A. Englund, Michael Famulare, Barry R. Lutz, Mark J. Rieder, Lea M. Starita, Matthew Thompson, Helen Y. Chu, Trevor Bedford, Jay Shendure |
| EPI_ISL_3372808, EPI_ISL_3372809 | Northwest Laboratory | Seattle Flu Study | Deborah A. Nickerson, Chris D. Frazar, Jover Lee, Benjamin Pelle, Erica Ryke, Matthew Richardson, Amanda Adler, Elisabeth Brandstetter, Peter D. Han, Kairsten Fay, Misja Ilcisin, Kirsten Lacombe, Thomas R. Sibley, Melissa Truong, Caitlin R. Wolf, Romesh Gautom, Geoff Melly, Brian Hiatt, Philip Dykema, Scott Lindquist, Michael Boeckh, Janet A. Englund, Michael Famulare, Barry R. Lutz, Mark J. Rieder, Lea M. Starita, Matthew Thompson, Helen Y. Chu, Jay Shendure, Trevor Bedford |
| EPI_ISL_3372810, EPI_ISL_3372811, EPI_ISL_3372812, EPI_ISL_3372813, EPI_ISL_3372814, EPI_ISL_3372815, EPI_ISL_3372816 | Atlas Genomics | Seattle Flu Study | Deborah A. Nickerson, Chris D. Frazar, Jover Lee, Benjamin Pelle, Erica Ryke, Matthew Richardson, Amanda Adler, Elisabeth Brandstetter, Peter D. Han, Kairsten Fay, Misja Ilcisin, Kirsten Lacombe, Thomas R. Sibley, Melissa Truong, Caitlin R. Wolf, Romesh Gautom, Geoff Melly, Brian Hiatt, Philip Dykema, Scott Lindquist, Michael Boeckh, Janet A. Englund, Michael Famulare, Barry R. Lutz, Mark J. Rieder, Lea M. Starita, Matthew Thompson, Helen Y. Chu, Jay Shendure, Trevor Bedford |
| EPI_ISL_3372817, EPI_ISL_3372818, EPI_ISL_3372819, EPI_ISL_3372820, EPI_ISL_3372821, EPI_ISL_3372822, EPI_ISL_3372823, EPI_ISL_3372824, EPI_ISL_3372825, EPI_ISL_3372831, EPI_ISL_3372832 | | | |
| see above | Northwest Laboratory | Seattle Flu Study | Deborah A. Nickerson, Chris D. Frazar, Jover Lee, Benjamin Pelle, Erica Ryke, Matthew Richardson, Amanda Adler, Elisabeth Brandstetter, Peter D. Han, Kairsten Fay, Misja Ilcisin, Kirsten Lacombe, Thomas R. Sibley, Melissa Truong, Caitlin R. Wolf, Romesh Gautom, Geoff Melly, Brian Hiatt, Philip Dykema, Scott Lindquist, Michael Boeckh, Janet A. Englund, Michael Famulare, Barry R. Lutz, Mark J. Rieder, Lea M. Starita, Matthew Thompson, Helen Y. Chu, Jay Shendure, Trevor Bedford |
| EPI_ISL_3372833 | Seattle Flu Study | Seattle Flu Study | Deborah A. Nickerson, Chris D. Frazar, Jover Lee, Benjamin Pelle, Erica Ryke, Matthew Richardson, Amanda Adler, Elisabeth Brandstetter, Peter D. Han, Kairsten Fay, Misja Ilcisin, Kirsten Lacombe, Thomas R. Sibley, Melissa Truong, Caitlin R. Wolf, Michael Boeckh, Janet A. Englund, Michael Famulare, Barry R. Lutz, Mark J. Rieder, Lea M. Starita, Matthew Thompson, Jay Shendure, Trevor Bedford, Helen Y. Chu |
| EPI_ISL_3372834, EPI_ISL_3372835 | Northwest Laboratory | Seattle Flu Study | Deborah A. Nickerson, Chris D. Frazar, Jover Lee, Benjamin Pelle, Erica Ryke, Matthew Richardson, Amanda Adler, Elisabeth Brandstetter, Peter D. Han, Kairsten Fay, Misja Ilcisin, Kirsten Lacombe, Thomas R. Sibley, Melissa Truong, Caitlin R. Wolf, Romesh Gautom, Geoff Melly, Brian Hiatt, Philip Dykema, Scott Lindquist, Michael Boeckh, Janet A. Englund, Michael Famulare, Barry R. Lutz, Mark J. Rieder, Lea M. Starita, Matthew Thompson, Helen Y. Chu, Jay Shendure, Trevor Bedford |
| EPI_ISL_3372836 | Atlas Genomics | Seattle Flu Study | Deborah A. Nickerson, Chris D. Frazar, Jover Lee, Benjamin Pelle, Erica Ryke, Matthew Richardson, Amanda Adler, Elisabeth Brandstetter, Peter D. Han, Kairsten Fay, Misja Ilcisin, Kirsten Lacombe, Thomas R. Sibley, Melissa Truong, Caitlin R. Wolf, Romesh Gautom, Geoff Melly, Brian Hiatt, Philip Dykema, Scott Lindquist, Michael Boeckh, Janet A. Englund, Michael Famulare, Barry R. Lutz, Mark J. Rieder, Lea M. Starita, Matthew Thompson, Helen Y. Chu, Jay Shendure, Trevor Bedford |
| EPI_ISL_3372838, EPI_ISL_3372839 | Northwest Laboratory | Seattle Flu Study | Deborah A. Nickerson, Chris D. Frazar, Jover Lee, Benjamin Pelle, Erica Ryke, Matthew Richardson, Amanda Adler, Elisabeth Brandstetter, Peter D. Han, Kairsten Fay, Misja Ilcisin, Kirsten Lacombe, Thomas R. Sibley, Melissa Truong, Caitlin R. Wolf, Romesh Gautom, Geoff Melly, Brian Hiatt, Philip Dykema, Scott Lindquist, Michael Boeckh, Janet A. Englund, Michael Famulare, Barry R. Lutz, Mark J. Rieder, Lea M. Starita, Matthew Thompson, Helen Y. Chu, Jay Shendure, Trevor Bedford |
| EPI_ISL_3372840 | Atlas Genomics | Seattle Flu Study | Deborah A. Nickerson, Chris D. Frazar, Jover Lee, Benjamin Pelle, Erica Ryke, Matthew Richardson, Amanda Adler, Elisabeth Brandstetter, Peter D. Han, Kairsten Fay, Misja Ilcisin, Kirsten Lacombe, Thomas R. Sibley, Melissa Truong, Caitlin R. Wolf, Romesh Gautom, Geoff Melly, Brian Hiatt, Philip Dykema, Scott Lindquist, Michael Boeckh, Janet A. Englund, Michael Famulare, Barry R. Lutz, Mark J. Rieder, Lea M. Starita, Matthew Thompson, Helen Y. Chu, Jay Shendure, Trevor Bedford |
| EPI_ISL_3372841 | Seattle Flu Study | Seattle Flu Study | Deborah A. Nickerson, Chris D. Frazar, Jover Lee, Benjamin Pelle, Erica Ryke, Matthew Richardson, Amanda Adler, Elisabeth Brandstetter, Peter D. Han, Kairsten Fay, Misja Ilcisin, Kirsten Lacombe, Thomas R. Sibley, Melissa Truong, Caitlin R. Wolf, Karen Cowgill, Stephanie Schrag, Jeff Duchin, Michael Boeckh, Janet A. Englund, Michael Famulare, Barry R. Lutz, Mark J. Rieder, Lea M. Starita, Matthew Thompson, Helen Y. Chu, Trevor Bedford, Jay Shendure |
| EPI_ISL_3372842 | Northwest Laboratory | Seattle Flu Study | Deborah A. Nickerson, Chris D. Frazar, Jover Lee, Benjamin Pelle, Erica Ryke, Matthew Richardson, Amanda Adler, Elisabeth Brandstetter, Peter D. Han, Kairsten Fay, Misja Ilcisin, Kirsten Lacombe, Thomas R. Sibley, Melissa Truong, Caitlin R. Wolf, Romesh Gautom, Geoff Melly, Brian Hiatt, Philip Dykema, Scott Lindquist, Michael Boeckh, Janet A. Englund, Michael Famulare, Barry R. Lutz, Mark J. Rieder, Lea M. Starita, Matthew Thompson, Helen Y. Chu, Jay Shendure, Trevor Bedford |
| EPI_ISL_3372843 | Atlas Genomics | Seattle Flu Study | Deborah A. Nickerson, Chris D. Frazar, Jover Lee, Benjamin Pelle, Erica Ryke, Matthew Richardson, Amanda Adler, Elisabeth Brandstetter, Peter D. Han, Kairsten Fay, Misja Ilcisin, Kirsten Lacombe, Thomas R. Sibley, Melissa Truong, Caitlin R. |

| | | | |
|---|---|---|---|
| | | | Wolf, Romesh Gautom, Geoff Melly, Brian Hiatt, Philip Dykema, Scott Lindquist, Michael Boeckh, Janet A. Englund, Michael Famulare, Barry R. Lutz, Mark J. Rieder, Lea M. Starita, Matthew Thompson, Helen Y. Chu, Jay Shendure, Trevor Bedford |
| EPI_ISL_3372844, EPI_ISL_3372845, EPI_ISL_3372846, EPI_ISL_3372848, EPI_ISL_3372849 | Northwest Laboratory | Seattle Flu Study | Deborah A. Nickerson, Chris D. Frazar, Jover Lee, Benjamin Pelle, Erica Ryke, Matthew Richardson, Amanda Adler, Elisabeth Brandstetter, Peter D. Han, Kairsten Fay, Misja Ilcisin, Kirsten Lacombe, Thomas R. Sibley, Melissa Truong, Caitlin R. Wolf, Romesh Gautom, Geoff Melly, Brian Hiatt, Philip Dykema, Scott Lindquist, Michael Boeckh, Janet A. Englund, Michael Famulare, Barry R. Lutz, Mark J. Rieder, Lea M. Starita, Matthew Thompson, Helen Y. Chu, Jay Shendure, Trevor Bedford |
| EPI_ISL_3372850 | Seattle Flu Study | Seattle Flu Study | Deborah A. Nickerson, Chris D. Frazar, Jover Lee, Benjamin Pelle, Erica Ryke, Matthew Richardson, Amanda Adler, Elisabeth Brandstetter, Peter D. Han, Kairsten Fay, Misja Ilcisin, Kirsten Lacombe, Thomas R. Sibley, Melissa Truong, Caitlin R. Wolf, Karen Cowgill, Stephanie Schrag, Jeff Duchin, Michael Boeckh, Janet A. Englund, Michael Famulare, Barry R. Lutz, Mark J. Rieder, Lea M. Starita, Matthew Thompson, Helen Y. Chu, Trevor Bedford, Jay Shendure |
| EPI_ISL_3372851 | Northwest Laboratory | Seattle Flu Study | Deborah A. Nickerson, Chris D. Frazar, Jover Lee, Benjamin Pelle, Erica Ryke, Matthew Richardson, Amanda Adler, Elisabeth Brandstetter, Peter D. Han, Kairsten Fay, Misja Ilcisin, Kirsten Lacombe, Thomas R. Sibley, Melissa Truong, Caitlin R. Wolf, Romesh Gautom, Geoff Melly, Brian Hiatt, Philip Dykema, Scott Lindquist, Michael Boeckh, Janet A. Englund, Michael Famulare, Barry R. Lutz, Mark J. Rieder, Lea M. Starita, Matthew Thompson, Helen Y. Chu, Jay Shendure, Trevor Bedford |
| EPI_ISL_3372852 | Atlas Genomics | Seattle Flu Study | Deborah A. Nickerson, Chris D. Frazar, Jover Lee, Benjamin Pelle, Erica Ryke, Matthew Richardson, Amanda Adler, Elisabeth Brandstetter, Peter D. Han, Kairsten Fay, Misja Ilcisin, Kirsten Lacombe, Thomas R. Sibley, Melissa Truong, Caitlin R. Wolf, Romesh Gautom, Geoff Melly, Brian Hiatt, Philip Dykema, Scott Lindquist, Michael Boeckh, Janet A. Englund, Michael Famulare, Barry R. Lutz, Mark J. Rieder, Lea M. Starita, Matthew Thompson, Helen Y. Chu, Jay Shendure, Trevor Bedford |
| EPI_ISL_3372853 | Provincial Public Health Reference Laboratory (PPHRL) | Provincial Public Health Reference Laboratory (PPHRL) | Andleeb Hanif, Mariam Javed, Hassan Nawaz, , Asim Altaf |
| EPI_ISL_3372856 | Seattle Flu Study | Seattle Flu Study | Deborah A. Nickerson, Chris D. Frazar, Jover Lee, Benjamin Pelle, Erica Ryke, Matthew Richardson, Amanda Adler, Elisabeth Brandstetter, Peter D. Han, Kairsten Fay, Misja Ilcisin, Kirsten Lacombe, Thomas R. Sibley, Melissa Truong, Caitlin R. Wolf, Karen Cowgill, Stephanie Schrag, Jeff Duchin, Michael Boeckh, Janet A. Englund, Michael Famulare, Barry R. Lutz, Mark J. Rieder, Lea M. Starita, Matthew Thompson, Helen Y. Chu, Trevor Bedford, Jay Shendure |
| EPI_ISL_3372857, EPI_ISL_3372858, EPI_ISL_3372859, EPI_ISL_3372860 | Northwest Laboratory | Seattle Flu Study | Deborah A. Nickerson, Chris D. Frazar, Jover Lee, Benjamin Pelle, Erica Ryke, Matthew Richardson, Amanda Adler, Elisabeth Brandstetter, Peter D. Han, Kairsten Fay, Misja Ilcisin, Kirsten Lacombe, Thomas R. Sibley, Melissa Truong, Caitlin R. Wolf, Romesh Gautom, Geoff Melly, Brian Hiatt, Philip Dykema, Scott Lindquist, Michael Boeckh, Janet A. Englund, Michael Famulare, Barry R. Lutz, Mark J. Rieder, Lea M. Starita, Matthew Thompson, Helen Y. Chu, Jay Shendure, Trevor Bedford |
| EPI_ISL_3372861 | Seattle Flu Study | Seattle Flu Study | Deborah A. Nickerson, Chris D. Frazar, Jover Lee, Benjamin Pelle, Erica Ryke, Matthew Richardson, Amanda Adler, Elisabeth Brandstetter, Peter D. Han, Kairsten Fay, Misja Ilcisin, Kirsten Lacombe, Thomas R. Sibley, Melissa Truong, Caitlin R. Wolf, Karen Cowgill, Stephanie Schrag, Jeff Duchin, Michael Boeckh, Janet A. Englund, Michael Famulare, Barry R. Lutz, Mark J. Rieder, Lea M. Starita, Matthew Thompson, Helen Y. Chu, Trevor Bedford, Jay Shendure |
| EPI_ISL_3372862 | Atlas Genomics | Seattle Flu Study | Deborah A. Nickerson, Chris D. Frazar, Jover Lee, Benjamin Pelle, Erica Ryke, Matthew Richardson, Amanda Adler, Elisabeth Brandstetter, Peter D. Han, Kairsten Fay, Misja Ilcisin, Kirsten Lacombe, Thomas R. Sibley, Melissa Truong, Caitlin R. Wolf, Romesh Gautom, Geoff Melly, Brian Hiatt, Philip Dykema, Scott Lindquist, Michael Boeckh, Janet A. Englund, Michael Famulare, Barry R. Lutz, Mark J. Rieder, Lea M. Starita, Matthew Thompson, Helen Y. Chu, Jay Shendure, Trevor Bedford |
| EPI_ISL_3372863, EPI_ISL_3372864, EPI_ISL_3372865, EPI_ISL_3372866, EPI_ISL_3372867, EPI_ISL_3372868, EPI_ISL_3372869, EPI_ISL_3372870, EPI_ISL_3372871, EPI_ISL_3372872, EPI_ISL_3372873 | | | |
| see above | Northwest Laboratory | Seattle Flu Study | Deborah A. Nickerson, Chris D. Frazar, Jover Lee, Benjamin Pelle, Erica Ryke, Matthew Richardson, Amanda Adler, Elisabeth Brandstetter, Peter D. Han, Kairsten Fay, Misja Ilcisin, Kirsten Lacombe, Thomas R. Sibley, Melissa Truong, Caitlin R. Wolf, Romesh Gautom, Geoff Melly, Brian Hiatt, Philip Dykema, Scott Lindquist, Michael Boeckh, Janet A. Englund, Michael Famulare, Barry R. Lutz, Mark J. Rieder, Lea M. Starita, Matthew Thompson, Helen Y. Chu, Jay Shendure, Trevor Bedford |
| EPI_ISL_3372874, EPI_ISL_3372875 | Atlas Genomics | Seattle Flu Study | Deborah A. Nickerson, Chris D. Frazar, Jover Lee, Benjamin Pelle, Erica Ryke, Matthew Richardson, Amanda Adler, Elisabeth Brandstetter, Peter D. Han, Kairsten Fay, Misja Ilcisin, Kirsten Lacombe, Thomas R. Sibley, Melissa Truong, Caitlin R. Wolf, Romesh Gautom, Geoff Melly, Brian Hiatt, Philip Dykema, Scott Lindquist, Michael Boeckh, Janet A. Englund, Michael Famulare, Barry R. Lutz, Mark J. Rieder, Lea M. Starita, Matthew Thompson, Helen Y. Chu, Jay Shendure, Trevor Bedford |
| EPI_ISL_3372877, EPI_ISL_3372878, EPI_ISL_3372879, EPI_ISL_3372881, EPI_ISL_3372882, EPI_ISL_3372883, EPI_ISL_3372885, EPI_ISL_3372887, EPI_ISL_3372888 | Northwest Laboratory | Seattle Flu Study | Deborah A. Nickerson, Chris D. Frazar, Jover Lee, Benjamin Pelle, Erica Ryke, Matthew Richardson, Amanda Adler, Elisabeth Brandstetter, Peter D. Han, Kairsten Fay, Misja Ilcisin, Kirsten Lacombe, Thomas R. Sibley, Melissa Truong, Caitlin R. Wolf, Romesh Gautom, Geoff Melly, Brian Hiatt, Philip Dykema, Scott Lindquist, Michael Boeckh, Janet A. Englund, Michael Famulare, Barry R. Lutz, Mark J. Rieder, Lea M. Starita, Matthew Thompson, Helen Y. Chu, Jay Shendure, Trevor Bedford |
| EPI_ISL_3372889 | Atlas Genomics | Seattle Flu Study | Deborah A. Nickerson, Chris D. Frazar, Jover Lee, Benjamin Pelle, Erica Ryke, Matthew Richardson, Amanda Adler, Elisabeth Brandstetter, Peter D. Han, Kairsten Fay, Misja Ilcisin, Kirsten Lacombe, Thomas R. Sibley, Melissa Truong, Caitlin R. Wolf, Romesh Gautom, Geoff Melly, Brian Hiatt, Philip Dykema, Scott Lindquist, Michael Boeckh, Janet A. Englund, Michael Famulare, Barry R. Lutz, Mark J. Rieder, Lea M. Starita, Matthew Thompson, Helen Y. Chu, Jay Shendure, Trevor Bedford |
| EPI_ISL_3372890, EPI_ISL_3372891, EPI_ISL_3372892, EPI_ISL_3372893, EPI_ISL_3372894, EPI_ISL_3372895 | Northwest Laboratory | Seattle Flu Study | Deborah A. Nickerson, Chris D. Frazar, Jover Lee, Benjamin Pelle, Erica Ryke, Matthew Richardson, Amanda Adler, Elisabeth Brandstetter, Peter D. Han, Kairsten Fay, Misja Ilcisin, Kirsten Lacombe, Thomas R. Sibley, Melissa Truong, Caitlin R. Wolf, Romesh Gautom, Geoff Melly, Brian Hiatt, Philip Dykema, Scott Lindquist, Michael Boeckh, Janet A. Englund, Michael Famulare, Barry R. Lutz, Mark J. Rieder, Lea M. Starita, Matthew Thompson, Helen Y. Chu, Jay Shendure, Trevor Bedford |
| EPI_ISL_3372897 | Atlas Genomics | Seattle Flu Study | Deborah A. Nickerson, Chris D. Frazar, Jover Lee, Benjamin Pelle, Erica Ryke, Matthew Richardson, Amanda Adler, Elisabeth Brandstetter, Peter D. Han, Kairsten Fay, Misja Ilcisin, Kirsten Lacombe, Thomas R. Sibley, Melissa Truong, Caitlin R. Wolf, Romesh Gautom, Geoff Melly, Brian Hiatt, Philip Dykema, Scott Lindquist, Michael Boeckh, Janet A. Englund, Michael Famulare, Barry R. Lutz, Mark J. Rieder, Lea M. Starita, Matthew Thompson, Helen Y. Chu, Jay Shendure, Trevor Bedford |
| EPI_ISL_3372899, EPI_ISL_3372900 | Northwest Laboratory | Seattle Flu Study | Deborah A. Nickerson, Chris D. Frazar, Jover Lee, Benjamin Pelle, Erica Ryke, Matthew Richardson, Amanda Adler, Elisabeth Brandstetter, Peter D. Han, Kairsten Fay, Misja Ilcisin, Kirsten Lacombe, Thomas R. Sibley, Melissa Truong, Caitlin R. Wolf, Romesh Gautom, Geoff Melly, Brian Hiatt, Philip Dykema, Scott Lindquist, Michael Boeckh, Janet A. Englund, Michael Famulare, Barry R. Lutz, Mark J. Rieder, Lea M. Starita, Matthew Thompson, Helen Y. Chu, Jay Shendure, Trevor Bedford |
| EPI_ISL_3372901 | Atlas Genomics | Seattle Flu Study | Deborah A. Nickerson, Chris D. Frazar, Jover Lee, Benjamin Pelle, Erica Ryke, Matthew Richardson, Amanda Adler, Elisabeth Brandstetter, Peter D. Han, Kairsten Fay, Misja Ilcisin, Kirsten Lacombe, Thomas R. Sibley, Melissa Truong, Caitlin R. Wolf, Romesh Gautom, Geoff Melly, Brian Hiatt, Philip Dykema, Scott Lindquist, Michael Boeckh, Janet A. Englund, Michael Famulare, Barry R. Lutz, Mark J. Rieder, Lea M. Starita, Matthew Thompson, Helen Y. Chu, Jay Shendure, Trevor Bedford |
| EPI_ISL_3372902 | Northwest Laboratory | Seattle Flu Study | Deborah A. Nickerson, Chris D. Frazar, Jover Lee, Benjamin Pelle, Erica Ryke, Matthew Richardson, Amanda Adler, Elisabeth Brandstetter, Peter D. Han, Kairsten Fay, Misja Ilcisin, Kirsten Lacombe, Thomas R. Sibley, Melissa Truong, Caitlin R. Wolf, Romesh Gautom, Geoff Melly, Brian Hiatt, Philip Dykema, Scott Lindquist, Michael Boeckh, Janet A. Englund, Michael Famulare, Barry R. Lutz, Mark J. Rieder, Lea M. Starita, Matthew Thompson, Helen Y. Chu, Jay Shendure, Trevor Bedford |
| EPI_ISL_3372903 | Atlas Genomics | Seattle Flu Study | Deborah A. Nickerson, Chris D. Frazar, Jover Lee, Benjamin Pelle, Erica Ryke, Matthew Richardson, Amanda Adler, Elisabeth Brandstetter, Peter D. Han, Kairsten Fay, Misja Ilcisin, Kirsten Lacombe, Thomas R. Sibley, Melissa Truong, Caitlin R. Wolf, Romesh Gautom, Geoff Melly, Brian Hiatt, Philip Dykema, Scott Lindquist, Michael Boeckh, Janet A. Englund, Michael Famulare, Barry R. Lutz, Mark J. Rieder, Lea M. Starita, Matthew Thompson, Helen Y. Chu, Jay Shendure, Trevor Bedford |
| EPI_ISL_3372904 | Northwest Laboratory | Seattle Flu Study | Deborah A. Nickerson, Chris D. Frazar, Jover Lee, Benjamin Pelle, Erica Ryke, Matthew Richardson, Amanda Adler, Elisabeth Brandstetter, Peter D. Han, Kairsten Fay, Misja Ilcisin, Kirsten Lacombe, Thomas R. Sibley, Melissa Truong, Caitlin R. Wolf, Romesh Gautom, Geoff Melly, Brian Hiatt, Philip Dykema, Scott Lindquist, Michael Boeckh, Janet A. Englund, Michael Famulare, Barry R. Lutz, Mark J. Rieder, Lea M. Starita, Matthew Thompson, Helen Y. Chu, Jay Shendure, Trevor Bedford |
| EPI_ISL_3372906 | Seattle Flu Study | Seattle Flu Study | Deborah A. Nickerson, Chris D. Frazar, Jover Lee, Benjamin Pelle, Erica Ryke, Matthew Richardson, Amanda Adler, Elisabeth Brandstetter, Peter D. Han, Kairsten Fay, Misja Ilcisin, Kirsten Lacombe, Thomas R. Sibley, Melissa Truong, Caitlin R. Wolf, Michael Boeckh, Janet A. Englund, Michael Famulare, Barry R. Lutz, Mark J. Rieder, Lea M. Starita, Matthew Thompson, Jay Shendure, Trevor Bedford, Helen Y. Chu |
| EPI_ISL_3372907 | Northwest Laboratory | Seattle Flu Study | Deborah A. Nickerson, Chris D. Frazar, Jover Lee, Benjamin Pelle, Erica Ryke, Matthew Richardson, Amanda Adler, Elisabeth Brandstetter, Peter D. Han, Kairsten Fay, Misja Ilcisin, Kirsten Lacombe, Thomas R. Sibley, Melissa Truong, Caitlin R. Wolf, Romesh Gautom, Geoff Melly, Brian Hiatt, Philip Dykema, Scott Lindquist, Michael Boeckh, Janet A. Englund, Michael Famulare, Barry R. Lutz, Mark J. Rieder, Lea M. Starita, Matthew Thompson, Helen Y. Chu, Jay Shendure, Trevor Bedford |
| EPI_ISL_3372909 | Atlas Genomics | Seattle Flu Study | Deborah A. Nickerson, Chris D. Frazar, Jover Lee, Benjamin Pelle, Erica Ryke, Matthew Richardson, Amanda Adler, Elisabeth Brandstetter, Peter D. Han, Kairsten Fay, Misja Ilcisin, Kirsten Lacombe, Thomas R. Sibley, Melissa Truong, Caitlin R. Wolf, Romesh Gautom, Geoff Melly, Brian Hiatt, Philip Dykema, Scott Lindquist, Michael Boeckh, Janet A. Englund, Michael Famulare, Barry R. Lutz, Mark J. Rieder, Lea M. Starita, Matthew Thompson, Helen Y. Chu, Jay Shendure, Trevor Bedford |
| EPI_ISL_3372911, EPI_ISL_3372914 | Northwest Laboratory | Seattle Flu Study | Deborah A. Nickerson, Chris D. Frazar, Jover Lee, Benjamin Pelle, Erica Ryke, Matthew Richardson, Amanda Adler, Elisabeth Brandstetter, Peter D. Han, Kairsten Fay, Misja Ilcisin, Kirsten Lacombe, Thomas R. Sibley, Melissa Truong, Caitlin R. Wolf, Romesh Gautom, Geoff Melly, Brian Hiatt, Philip Dykema, Scott Lindquist, Michael Boeckh, Janet A. Englund, Michael Famulare, Barry R. Lutz, Mark J. Rieder, Lea M. Starita, Matthew Thompson, Helen Y. Chu, Jay Shendure, Trevor Bedford |
| EPI_ISL_3372915 | Atlas Genomics | Seattle Flu Study | Deborah A. Nickerson, Chris D. Frazar, Jover Lee, Benjamin Pelle, Erica Ryke, Matthew Richardson, Amanda Adler, Elisabeth Brandstetter, Peter D. Han, Kairsten Fay, Misja Ilcisin, Kirsten Lacombe, Thomas R. Sibley, Melissa Truong, Caitlin R. Wolf, Romesh Gautom, Geoff Melly, Brian Hiatt, Philip Dykema, Scott Lindquist, Michael Boeckh, Janet A. Englund, Michael Famulare, Barry R. Lutz, Mark J. Rieder, Lea M. Starita, Matthew Thompson, Helen Y. Chu, Jay Shendure, Trevor Bedford |
| EPI_ISL_3372917, EPI_ISL_3372918, EPI_ISL_3372919, EPI_ISL_3372920, EPI_ISL_3372921, EPI_ISL_3372923, EPI_ISL_3372924, EPI_ISL_3372925, EPI_ISL_3372927, EPI_ISL_3372928 | Northwest Laboratory | Seattle Flu Study | Deborah A. Nickerson, Chris D. Frazar, Jover Lee, Benjamin Pelle, Erica Ryke, Matthew Richardson, Amanda Adler, Elisabeth Brandstetter, Peter D. Han, Kairsten Fay, Misja Ilcisin, Kirsten Lacombe, Thomas R. Sibley, Melissa Truong, Caitlin R. Wolf, Romesh Gautom, Geoff Melly, Brian Hiatt, Philip Dykema, Scott Lindquist, Michael Boeckh, Janet A. Englund, Michael Famulare, Barry R. Lutz, Mark J. Rieder, Lea M. Starita, Matthew Thompson, Helen Y. Chu, Jay Shendure, Trevor Bedford |
| EPI_ISL_3372929 | Atlas Genomics | Seattle Flu Study | Deborah A. Nickerson, Chris D. Frazar, Jover Lee, Benjamin Pelle, Erica Ryke, Matthew Richardson, Amanda Adler, Elisabeth Brandstetter, Peter D. Han, Kairsten Fay, Misja Ilcisin, Kirsten Lacombe, Thomas R. Sibley, Melissa Truong, Caitlin R. Wolf, Romesh Gautom, Geoff Melly, Brian Hiatt, Philip Dykema, Scott Lindquist, Michael Boeckh, Janet A. Englund, Michael Famulare, Barry R. Lutz, Mark J. Rieder, Lea M. Starita, Matthew Thompson, Helen Y. Chu, Jay Shendure, Trevor Bedford |
| EPI_ISL_3372931, EPI_ISL_3372933, EPI_ISL_3372934, EPI_ISL_3372936, EPI_ISL_3372937, EPI_ISL_3372938, EPI_ISL_3372939 | Northwest Laboratory | Seattle Flu Study | Deborah A. Nickerson, Chris D. Frazar, Jover Lee, Benjamin Pelle, Erica Ryke, Matthew Richardson, Amanda Adler, Elisabeth Brandstetter, Peter D. Han, Kairsten Fay, Misja Ilcisin, Kirsten Lacombe, Thomas R. Sibley, Melissa Truong, Caitlin R. Wolf, Romesh Gautom, Geoff Melly, Brian Hiatt, Philip Dykema, Scott Lindquist, Michael Boeckh, Janet A. Englund, Michael Famulare, Barry R. Lutz, Mark J. Rieder, Lea M. Starita, Matthew Thompson, Helen Y. Chu, Jay Shendure, Trevor Bedford |
| EPI_ISL_3372940, EPI_ISL_3372941 | Atlas Genomics | Seattle Flu Study | Deborah A. Nickerson, Chris D. Frazar, Jover Lee, Benjamin Pelle, Erica Ryke, Matthew Richardson, Amanda Adler, Elisabeth Brandstetter, Peter D. Han, Kairsten Fay, Misja Ilcisin, Kirsten Lacombe, Thomas R. Sibley, Melissa Truong, Caitlin R. Wolf, Romesh Gautom, Geoff Melly, Brian Hiatt, Philip Dykema, Scott Lindquist, Michael Boeckh, Janet A. Englund, Michael Famulare, Barry R. Lutz, Mark J. Rieder, Lea M. Starita, Matthew Thompson, Helen Y. Chu, Jay Shendure, Trevor Bedford |
| EPI_ISL_3372942, EPI_ISL_3372945 | Northwest Laboratory | Seattle Flu Study | Deborah A. Nickerson, Chris D. Frazar, Jover Lee, Benjamin Pelle, Erica Ryke, Matthew Richardson, Amanda Adler, Elisabeth Brandstetter, Peter D. Han, Kairsten Fay, Misja Ilcisin, Kirsten Lacombe, Thomas R. Sibley, Melissa Truong, Caitlin R. Wolf, Romesh Gautom, Geoff Melly, Brian Hiatt, Philip Dykema, Scott Lindquist, Michael Boeckh, Janet A. Englund, Michael Famulare, Barry R. Lutz, Mark J. Rieder, Lea M. Starita, Matthew Thompson, Helen Y. Chu, Jay Shendure, Trevor Bedford |
| EPI_ISL_3372946 | Seattle Flu Study | Seattle Flu Study | Deborah A. Nickerson, Chris D. Frazar, Jover Lee, Benjamin Pelle, Erica Ryke, Matthew Richardson, Amanda Adler, Elisabeth Brandstetter, Peter D. Han, Kairsten Fay, Misja Ilcisin, Kirsten Lacombe, Thomas R. Sibley, Melissa Truong, Caitlin R. Wolf, Karen Cowgill, Stephanie Schrag, Jeff Duchin, Michael Boeckh, Janet A. Englund, Michael Famulare, Barry R. Lutz, Mark J. Rieder, Lea M. Starita, Matthew Thompson, Helen Y. Chu, Trevor Bedford, Jay Shendure |
| EPI_ISL_3372947, EPI_ISL_3372948, EPI_ISL_3372949, EPI_ISL_3372950, | Northwest Laboratory | Seattle Flu Study | Deborah A. Nickerson, Chris D. Frazar, Jover Lee, Benjamin Pelle, Erica Ryke, Matthew Richardson, Amanda Adler, Elisabeth Brandstetter, Peter D. Han, Kairsten Fay, Misja Ilcisin, Kirsten Lacombe, Thomas R. Sibley, Melissa Truong, Caitlin R. Wolf, Romesh Gautom, Geoff Melly, Brian Hiatt, Philip Dykema, Scott Lindquist, Michael Boeckh, Janet A. Englund, Michael Famulare, Barry R. Lutz, Mark J. Rieder, Lea M. Starita, Matthew Thompson, Helen Y. Chu, Jay Shendure, Trevor Bedford |

| | | | |
|---|---|---|---|
| EPI_ISL_3372951, EPI_ISL_3372952 | | | |
| EPI_ISL_3372953 | Atlas Genomics | Seattle Flu Study | Deborah A. Nickerson, Chris D. Frazar, Jover Lee, Benjamin Pelle, Erica Ryke, Matthew Richardson, Amanda Adler, Elisabeth Brandstetter, Peter D. Han, Kairsten Fay, Misja Ilcisin, Kirsten Lacombe, Thomas R. Sibley, Melissa Truong, Caitlin R. Wolf, Romesh Gautom, Geoff Melly, Brian Hiatt, Philip Dykema, Scott Lindquist, Michael Boeckh, Janet A. Englund, Michael Famulare, Barry R. Lutz, Mark J. Rieder, Lea M. Starita, Matthew Thompson, Helen Y. Chu, Jay Shendure, Trevor Bedford |
| EPI_ISL_3372955 | Northwest Laboratory | Seattle Flu Study | Deborah A. Nickerson, Chris D. Frazar, Jover Lee, Benjamin Pelle, Erica Ryke, Matthew Richardson, Amanda Adler, Elisabeth Brandstetter, Peter D. Han, Kairsten Fay, Misja Ilcisin, Kirsten Lacombe, Thomas R. Sibley, Melissa Truong, Caitlin R. Wolf, Romesh Gautom, Geoff Melly, Brian Hiatt, Philip Dykema, Scott Lindquist, Michael Boeckh, Janet A. Englund, Michael Famulare, Barry R. Lutz, Mark J. Rieder, Lea M. Starita, Matthew Thompson, Helen Y. Chu, Jay Shendure, Trevor Bedford |
| EPI_ISL_3372958 | Seattle Flu Study | Seattle Flu Study | Deborah A. Nickerson, Chris D. Frazar, Jover Lee, Benjamin Pelle, Erica Ryke, Matthew Richardson, Amanda Adler, Elisabeth Brandstetter, Peter D. Han, Kairsten Fay, Misja Ilcisin, Kirsten Lacombe, Thomas R. Sibley, Melissa Truong, Caitlin R. Wolf, Karen Cowgill, Stephanie Schrag, Jeff Duchin, Michael Boeckh, Janet A. Englund, Michael Famulare, Barry R. Lutz, Mark J. Rieder, Lea M. Starita, Matthew Thompson, Helen Y. Chu, Trevor Bedford, Jay Shendure |
| EPI_ISL_3372959 | Atlas Genomics | Seattle Flu Study | Deborah A. Nickerson, Chris D. Frazar, Jover Lee, Benjamin Pelle, Erica Ryke, Matthew Richardson, Amanda Adler, Elisabeth Brandstetter, Peter D. Han, Kairsten Fay, Misja Ilcisin, Kirsten Lacombe, Thomas R. Sibley, Melissa Truong, Caitlin R. Wolf, Romesh Gautom, Geoff Melly, Brian Hiatt, Philip Dykema, Scott Lindquist, Michael Boeckh, Janet A. Englund, Michael Famulare, Barry R. Lutz, Mark J. Rieder, Lea M. Starita, Matthew Thompson, Helen Y. Chu, Jay Shendure, Trevor Bedford |
| EPI_ISL_3372960, EPI_ISL_3372961, EPI_ISL_3372962, EPI_ISL_3372963 | Northwest Laboratory | Seattle Flu Study | Deborah A. Nickerson, Chris D. Frazar, Jover Lee, Benjamin Pelle, Erica Ryke, Matthew Richardson, Amanda Adler, Elisabeth Brandstetter, Peter D. Han, Kairsten Fay, Misja Ilcisin, Kirsten Lacombe, Thomas R. Sibley, Melissa Truong, Caitlin R. Wolf, Romesh Gautom, Geoff Melly, Brian Hiatt, Philip Dykema, Scott Lindquist, Michael Boeckh, Janet A. Englund, Michael Famulare, Barry R. Lutz, Mark J. Rieder, Lea M. Starita, Matthew Thompson, Helen Y. Chu, Jay Shendure, Trevor Bedford |
| EPI_ISL_3372964 | Seattle Flu Study | Seattle Flu Study | Deborah A. Nickerson, Chris D. Frazar, Jover Lee, Benjamin Pelle, Erica Ryke, Matthew Richardson, Amanda Adler, Elisabeth Brandstetter, Peter D. Han, Kairsten Fay, Misja Ilcisin, Kirsten Lacombe, Thomas R. Sibley, Melissa Truong, Caitlin R. Wolf, Karen Cowgill, Stephanie Schrag, Jeff Duchin, Michael Boeckh, Janet A. Englund, Michael Famulare, Barry R. Lutz, Mark J. Rieder, Lea M. Starita, Matthew Thompson, Helen Y. Chu, Trevor Bedford, Jay Shendure |
| EPI_ISL_3372965, EPI_ISL_3372966 | Northwest Laboratory | Seattle Flu Study | Deborah A. Nickerson, Chris D. Frazar, Jover Lee, Benjamin Pelle, Erica Ryke, Matthew Richardson, Amanda Adler, Elisabeth Brandstetter, Peter D. Han, Kairsten Fay, Misja Ilcisin, Kirsten Lacombe, Thomas R. Sibley, Melissa Truong, Caitlin R. Wolf, Romesh Gautom, Geoff Melly, Brian Hiatt, Philip Dykema, Scott Lindquist, Michael Boeckh, Janet A. Englund, Michael Famulare, Barry R. Lutz, Mark J. Rieder, Lea M. Starita, Matthew Thompson, Helen Y. Chu, Jay Shendure, Trevor Bedford |
| EPI_ISL_3372967 | Seattle Flu Study | Seattle Flu Study | Deborah A. Nickerson, Chris D. Frazar, Jover Lee, Benjamin Pelle, Erica Ryke, Matthew Richardson, Amanda Adler, Elisabeth Brandstetter, Peter D. Han, Kairsten Fay, Misja Ilcisin, Kirsten Lacombe, Thomas R. Sibley, Melissa Truong, Caitlin R. Wolf, Michael Boeckh, Janet A. Englund, Michael Famulare, Barry R. Lutz, Mark J. Rieder, Lea M. Starita, Matthew Thompson, Jay Shendure, Trevor Bedford, Helen Y. Chu |
| EPI_ISL_3372968 | Atlas Genomics | Seattle Flu Study | Deborah A. Nickerson, Chris D. Frazar, Jover Lee, Benjamin Pelle, Erica Ryke, Matthew Richardson, Amanda Adler, Elisabeth Brandstetter, Peter D. Han, Kairsten Fay, Misja Ilcisin, Kirsten Lacombe, Thomas R. Sibley, Melissa Truong, Caitlin R. Wolf, Romesh Gautom, Geoff Melly, Brian Hiatt, Philip Dykema, Scott Lindquist, Michael Boeckh, Janet A. Englund, Michael Famulare, Barry R. Lutz, Mark J. Rieder, Lea M. Starita, Matthew Thompson, Helen Y. Chu, Jay Shendure, Trevor Bedford |
| EPI_ISL_3372969, EPI_ISL_3372970, EPI_ISL_3372971 | Northwest Laboratory | Seattle Flu Study | Deborah A. Nickerson, Chris D. Frazar, Jover Lee, Benjamin Pelle, Erica Ryke, Matthew Richardson, Amanda Adler, Elisabeth Brandstetter, Peter D. Han, Kairsten Fay, Misja Ilcisin, Kirsten Lacombe, Thomas R. Sibley, Melissa Truong, Caitlin R. Wolf, Romesh Gautom, Geoff Melly, Brian Hiatt, Philip Dykema, Scott Lindquist, Michael Boeckh, Janet A. Englund, Michael Famulare, Barry R. Lutz, Mark J. Rieder, Lea M. Starita, Matthew Thompson, Helen Y. Chu, Jay Shendure, Trevor Bedford |
| EPI_ISL_3372973, EPI_ISL_3372974 | Atlas Genomics | Seattle Flu Study | Deborah A. Nickerson, Chris D. Frazar, Jover Lee, Benjamin Pelle, Erica Ryke, Matthew Richardson, Amanda Adler, Elisabeth Brandstetter, Peter D. Han, Kairsten Fay, Misja Ilcisin, Kirsten Lacombe, Thomas R. Sibley, Melissa Truong, Caitlin R. Wolf, Romesh Gautom, Geoff Melly, Brian Hiatt, Philip Dykema, Scott Lindquist, Michael Boeckh, Janet A. Englund, Michael Famulare, Barry R. Lutz, Mark J. Rieder, Lea M. Starita, Matthew Thompson, Helen Y. Chu, Jay Shendure, Trevor Bedford |
| EPI_ISL_3372975 | Northwest Laboratory | Seattle Flu Study | Deborah A. Nickerson, Chris D. Frazar, Jover Lee, Benjamin Pelle, Erica Ryke, Matthew Richardson, Amanda Adler, Elisabeth Brandstetter, Peter D. Han, Kairsten Fay, Misja Ilcisin, Kirsten Lacombe, Thomas R. Sibley, Melissa Truong, Caitlin R. Wolf, Romesh Gautom, Geoff Melly, Brian Hiatt, Philip Dykema, Scott Lindquist, Michael Boeckh, Janet A. Englund, Michael Famulare, Barry R. Lutz, Mark J. Rieder, Lea M. Starita, Matthew Thompson, Helen Y. Chu, Jay Shendure, Trevor Bedford |
| EPI_ISL_3372976 | Seattle Flu Study | Seattle Flu Study | Deborah A. Nickerson, Chris D. Frazar, Jover Lee, Benjamin Pelle, Erica Ryke, Matthew Richardson, Amanda Adler, Elisabeth Brandstetter, Peter D. Han, Kairsten Fay, Misja Ilcisin, Kirsten Lacombe, Thomas R. Sibley, Melissa Truong, Caitlin R. Wolf, Karen Cowgill, Stephanie Schrag, Jeff Duchin, Michael Boeckh, Janet A. Englund, Michael Famulare, Barry R. Lutz, Mark J. Rieder, Lea M. Starita, Matthew Thompson, Helen Y. Chu, Trevor Bedford, Jay Shendure |
| EPI_ISL_3372980, EPI_ISL_3372982 | Atlas Genomics | Seattle Flu Study | Deborah A. Nickerson, Chris D. Frazar, Jover Lee, Benjamin Pelle, Erica Ryke, Matthew Richardson, Amanda Adler, Elisabeth Brandstetter, Peter D. Han, Kairsten Fay, Misja Ilcisin, Kirsten Lacombe, Thomas R. Sibley, Melissa Truong, Caitlin R. Wolf, Romesh Gautom, Geoff Melly, Brian Hiatt, Philip Dykema, Scott Lindquist, Michael Boeckh, Janet A. Englund, Michael Famulare, Barry R. Lutz, Mark J. Rieder, Lea M. Starita, Matthew Thompson, Helen Y. Chu, Jay Shendure, Trevor Bedford |
| EPI_ISL_3372985 | Northwest Laboratory | Seattle Flu Study | Deborah A. Nickerson, Chris D. Frazar, Jover Lee, Benjamin Pelle, Erica Ryke, Matthew Richardson, Amanda Adler, Elisabeth Brandstetter, Peter D. Han, Kairsten Fay, Misja Ilcisin, Kirsten Lacombe, Thomas R. Sibley, Melissa Truong, Caitlin R. Wolf, Romesh Gautom, Geoff Melly, Brian Hiatt, Philip Dykema, Scott Lindquist, Michael Boeckh, Janet A. Englund, Michael Famulare, Barry R. Lutz, Mark J. Rieder, Lea M. Starita, Matthew Thompson, Helen Y. Chu, Jay Shendure, Trevor Bedford |
| EPI_ISL_3372986 | Seattle Flu Study | Seattle Flu Study | Deborah A. Nickerson, Chris D. Frazar, Jover Lee, Benjamin Pelle, Erica Ryke, Matthew Richardson, Amanda Adler, Elisabeth Brandstetter, Peter D. Han, Kairsten Fay, Misja Ilcisin, Kirsten Lacombe, Thomas R. Sibley, Melissa Truong, Caitlin R. Wolf, Karen Cowgill, Stephanie Schrag, Jeff Duchin, Michael Boeckh, Janet A. Englund, Michael Famulare, Barry R. Lutz, Mark J. Rieder, Lea M. Starita, Matthew Thompson, Helen Y. Chu, Trevor Bedford, Jay Shendure |
| EPI_ISL_3372988, EPI_ISL_3372989, EPI_ISL_3372990, EPI_ISL_3372991, EPI_ISL_3372992, EPI_ISL_3372993, EPI_ISL_3372994, EPI_ISL_3372995, EPI_ISL_3372996, EPI_ISL_3372997, EPI_ISL_3372998, EPI_ISL_3372999, EPI_ISL_3373000, EPI_ISL_3373001, EPI_ISL_3373002, EPI_ISL_3373003, EPI_ISL_3373004, EPI_ISL_3373005, EPI_ISL_3373006, EPI_ISL_3373007, EPI_ISL_3373008, EPI_ISL_3373009, EPI_ISL_3373010, EPI_ISL_3373011, EPI_ISL_3373012, EPI_ISL_3373013, EPI_ISL_3373014, EPI_ISL_3373015, EPI_ISL_3373016, EPI_ISL_3373017, EPI_ISL_3373018, EPI_ISL_3373019, EPI_ISL_3373020, EPI_ISL_3373021, EPI_ISL_3373022, EPI_ISL_3373023, EPI_ISL_3373024, EPI_ISL_3373025, EPI_ISL_3373026, EPI_ISL_3373027, EPI_ISL_3373028, EPI_ISL_3373029, EPI_ISL_3373030, EPI_ISL_3373031, EPI_ISL_3373032, EPI_ISL_3373033, EPI_ISL_3373034, EPI_ISL_3373035, EPI_ISL_3373036, EPI_ISL_3373037, EPI_ISL_3373038, EPI_ISL_3373039, EPI_ISL_3373040, EPI_ISL_3373041, EPI_ISL_3373042, EPI_ISL_3373043, EPI_ISL_3373044, EPI_ISL_3373045, EPI_ISL_3373046, EPI_ISL_3373047, EPI_ISL_3373048, EPI_ISL_3373049, EPI_ISL_3373050, EPI_ISL_3373051, EPI_ISL_3373052, EPI_ISL_3373053, EPI_ISL_3373054, EPI_ISL_3373055, EPI_ISL_3373056, EPI_ISL_3373057, EPI_ISL_3373058, EPI_ISL_3373059, EPI_ISL_3373060, EPI_ISL_3373061, EPI_ISL_3373062, EPI_ISL_3373063, EPI_ISL_3373064, EPI_ISL_3373065, EPI_ISL_3373066, EPI_ISL_3373067, EPI_ISL_3373069, EPI_ISL_3373070, EPI_ISL_3373071, EPI_ISL_3373072, EPI_ISL_3373073, EPI_ISL_3373074, EPI_ISL_3373075, EPI_ISL_3373076, EPI_ISL_3373077, EPI_ISL_3373078, EPI_ISL_3373079, EPI_ISL_3373080, EPI_ISL_3373081, EPI_ISL_3373082, EPI_ISL_3373083, EPI_ISL_3373084, EPI_ISL_3373085, EPI_ISL_3373086, EPI_ISL_3373087, EPI_ISL_3373089, EPI_ISL_3373090, EPI_ISL_3373091, EPI_ISL_3373092, EPI_ISL_3373093, EPI_ISL_3373094, EPI_ISL_3373095, EPI_ISL_3373096, EPI_ISL_3373098, EPI_ISL_3373099, EPI_ISL_3373100, EPI_ISL_3373101, EPI_ISL_3373102, EPI_ISL_3373103, EPI_ISL_3373104, EPI_ISL_3373105, EPI_ISL_3373106, EPI_ISL_3373108, EPI_ISL_3373109, EPI_ISL_3373110, EPI_ISL_3373111, EPI_ISL_3373112, EPI_ISL_3373113, EPI_ISL_3373115, EPI_ISL_3373116 | | | |
| see above | Utah Public Health Laboratory | Utah Public Health Laboratory | Erin L. Young, Kelly F. Oakeson, Olinto Linares-Perdomo |
| EPI_ISL_3373119, EPI_ISL_3373120, EPI_ISL_3373129, EPI_ISL_3373130, EPI_ISL_3373131, EPI_ISL_3373132, EPI_ISL_3373133, EPI_ISL_3373134, EPI_ISL_3373135, EPI_ISL_3373136, EPI_ISL_3373137, EPI_ISL_3373138, EPI_ISL_3373139, EPI_ISL_3373140, EPI_ISL_3373142, EPI_ISL_3373143, EPI_ISL_3373145, EPI_ISL_3373146, EPI_ISL_3373147, EPI_ISL_3373148, EPI_ISL_3373149, EPI_ISL_3373150, EPI_ISL_3373151, EPI_ISL_3373152, EPI_ISL_3373153, EPI_ISL_3373154, EPI_ISL_3373155, EPI_ISL_3373156, EPI_ISL_3373157, EPI_ISL_3373158, EPI_ISL_3373159, EPI_ISL_3373160, EPI_ISL_3373161, EPI_ISL_3373162, EPI_ISL_3373163, EPI_ISL_3373164, EPI_ISL_3373165, EPI_ISL_3373166, EPI_ISL_3373167, EPI_ISL_3373168, EPI_ISL_3373169, EPI_ISL_3373170, EPI_ISL_3373171, EPI_ISL_3373172, EPI_ISL_3373173, EPI_ISL_3373174, EPI_ISL_3373175, EPI_ISL_3373176, EPI_ISL_3373177, EPI_ISL_3373178, EPI_ISL_3373179, EPI_ISL_3373180, EPI_ISL_3373181, EPI_ISL_3373182, EPI_ISL_3373183, EPI_ISL_3373185, EPI_ISL_3373186, EPI_ISL_3373188, EPI_ISL_3373189, EPI_ISL_3373190, EPI_ISL_3373194, EPI_ISL_3373196, EPI_ISL_3373197, EPI_ISL_3373198, EPI_ISL_3373209, EPI_ISL_3373210, EPI_ISL_3373211, EPI_ISL_3373216, EPI_ISL_3373217, EPI_ISL_3373218, EPI_ISL_3373219, EPI_ISL_3373220, EPI_ISL_3373221, EPI_ISL_3373222, EPI_ISL_3373223, EPI_ISL_3373224, EPI_ISL_3373225, EPI_ISL_3373226, EPI_ISL_3373227, EPI_ISL_3373228, EPI_ISL_3373229, EPI_ISL_3373230, EPI_ISL_3373231, EPI_ISL_3373232, EPI_ISL_3373233, EPI_ISL_3373234, EPI_ISL_3373235, EPI_ISL_3373236, EPI_ISL_3373237, EPI_ISL_3373238, EPI_ISL_3373239, EPI_ISL_3373240, EPI_ISL_3373241, EPI_ISL_3373242, EPI_ISL_3373243, EPI_ISL_3373244, EPI_ISL_3373245, EPI_ISL_3373246, EPI_ISL_3373247, EPI_ISL_3373248, EPI_ISL_3373249, EPI_ISL_3373250, EPI_ISL_3373251, EPI_ISL_3373252, EPI_ISL_3373253, EPI_ISL_3373254, EPI_ISL_3373255 | | | |
| see above | Johns Hopkins Hospital Department of Pathology | Johns Hopkins Hospital Department of Pathology | C. Paul Morris, Chun Huai Luo, David Gaston, Julie M. Norton, Matthew Schwartz, Heba H. Mostafa |
| EPI_ISL_3373333, EPI_ISL_3373337, EPI_ISL_3373407 | Provincial Public Health Reference Laboratory (PPHRL) | Provincial Public Health Reference Laboratory (PPHRL) | Andleeb Hanif, Mariam Javed, Hassan Nawaz, , Asim Altaf |
| EPI_ISL_3373445, EPI_ISL_3373446, EPI_ISL_3373447, EPI_ISL_3373448, EPI_ISL_3373449, EPI_ISL_3373450, EPI_ISL_3373451, EPI_ISL_3373452, EPI_ISL_3373453, EPI_ISL_3373454, EPI_ISL_3373455, EPI_ISL_3373456, EPI_ISL_3373457, EPI_ISL_3373458, EPI_ISL_3373459, EPI_ISL_3373460, EPI_ISL_3373461, EPI_ISL_3373462, EPI_ISL_3373463, EPI_ISL_3373464, EPI_ISL_3373465, EPI_ISL_3373466, EPI_ISL_3373467, EPI_ISL_3373468, EPI_ISL_3373469, EPI_ISL_3373470, EPI_ISL_3373471, EPI_ISL_3373472, EPI_ISL_3373473, EPI_ISL_3373474, EPI_ISL_3373475, EPI_ISL_3373476, EPI_ISL_3373477, EPI_ISL_3373478, EPI_ISL_3373479, EPI_ISL_3373480, EPI_ISL_3373481, EPI_ISL_3373482, EPI_ISL_3373483, EPI_ISL_3373484, EPI_ISL_3373485, EPI_ISL_3373486, EPI_ISL_3373487, EPI_ISL_3373488, EPI_ISL_3373489, EPI_ISL_3373490, EPI_ISL_3373491, EPI_ISL_3373492, EPI_ISL_3373493, EPI_ISL_3373494, EPI_ISL_3373495, EPI_ISL_3373496, EPI_ISL_3373497, EPI_ISL_3373498, EPI_ISL_3373499, EPI_ISL_3373500, EPI_ISL_3373501, EPI_ISL_3373502, EPI_ISL_3373503, EPI_ISL_3373504, EPI_ISL_3373505, EPI_ISL_3373506, EPI_ISL_3373507, EPI_ISL_3373508, EPI_ISL_3373509, EPI_ISL_3373510, EPI_ISL_3373511, EPI_ISL_3373512, EPI_ISL_3373513, EPI_ISL_3373514, EPI_ISL_3373515, EPI_ISL_3373516, EPI_ISL_3373517, EPI_ISL_3373518, EPI_ISL_3373519, EPI_ISL_3373520, EPI_ISL_3373521, EPI_ISL_3373522, EPI_ISL_3373523, EPI_ISL_3373524, EPI_ISL_3373525, EPI_ISL_3373526, EPI_ISL_3373527, EPI_ISL_3373528, EPI_ISL_3373529, EPI_ISL_3373530, EPI_ISL_3373531, EPI_ISL_3373532, EPI_ISL_3373533, EPI_ISL_3373534, EPI_ISL_3373535, EPI_ISL_3373536, EPI_ISL_3373537, EPI_ISL_3373538, EPI_ISL_3373539, EPI_ISL_3373540, EPI_ISL_3373541, EPI_ISL_3373542, EPI_ISL_3373543, EPI_ISL_3373544, EPI_ISL_3373545, EPI_ISL_3373546, EPI_ISL_3373547, EPI_ISL_3373548, EPI_ISL_3373549, EPI_ISL_3373550, EPI_ISL_3373551, EPI_ISL_3373552, EPI_ISL_3373553, EPI_ISL_3373554, EPI_ISL_3373555, EPI_ISL_3373556, EPI_ISL_3373557, EPI_ISL_3373558, EPI_ISL_3373559, EPI_ISL_3373560, EPI_ISL_3373561, EPI_ISL_3373562, EPI_ISL_3373563, EPI_ISL_3373564, EPI_ISL_3373565, EPI_ISL_3373566, EPI_ISL_3373567, EPI_ISL_3373568, EPI_ISL_3373569, EPI_ISL_3373570, EPI_ISL_3373571, EPI_ISL_3373572, EPI_ISL_3373573, EPI_ISL_3373574, EPI_ISL_3373575, EPI_ISL_3373576, EPI_ISL_3373577, EPI_ISL_3373578, EPI_ISL_3373579, EPI_ISL_3373580, EPI_ISL_3373581, EPI_ISL_3373582, EPI_ISL_3373583, EPI_ISL_3373584, EPI_ISL_3373585, EPI_ISL_3373586, EPI_ISL_3373587, EPI_ISL_3373588, EPI_ISL_3373589, EPI_ISL_3373590, EPI_ISL_3373591, EPI_ISL_3373592, EPI_ISL_3373593, EPI_ISL_3373594, EPI_ISL_3373595, EPI_ISL_3373596, EPI_ISL_3373597, EPI_ISL_3373598, EPI_ISL_3373599, EPI_ISL_3373600, EPI_ISL_3373601, EPI_ISL_3373602, EPI_ISL_3373603, EPI_ISL_3373604, EPI_ISL_3373605, EPI_ISL_3373606, EPI_ISL_3373607, EPI_ISL_3373608, EPI_ISL_3373609, EPI_ISL_3373610, EPI_ISL_3373611, EPI_ISL_3373612, EPI_ISL_3373613, EPI_ISL_3373614, EPI_ISL_3373615, EPI_ISL_3373616, EPI_ISL_3373617, EPI_ISL_3373618, EPI_ISL_3373619, EPI_ISL_3373620, EPI_ISL_3373621, EPI_ISL_3373622, EPI_ISL_3373623, EPI_ISL_3373624, EPI_ISL_3373625, EPI_ISL_3373626, EPI_ISL_3373627, EPI_ISL_3373628, EPI_ISL_3373629, EPI_ISL_3373630, EPI_ISL_3373631, EPI_ISL_3373632, EPI_ISL_3373633, EPI_ISL_3373634, EPI_ISL_3373635, EPI_ISL_3373636, EPI_ISL_3373637, EPI_ISL_3373638, EPI_ISL_3373639, EPI_ISL_3373640, EPI_ISL_3373641, EPI_ISL_3373642, EPI_ISL_3373643, EPI_ISL_3373644, EPI_ISL_3373645, EPI_ISL_3373646, EPI_ISL_3373647, EPI_ISL_3373648, EPI_ISL_3373649, EPI_ISL_3373650, EPI_ISL_3373651, EPI_ISL_3373652, EPI_ISL_3373653, EPI_ISL_3373654, EPI_ISL_3373655, EPI_ISL_3373656, EPI_ISL_3373657, EPI_ISL_3373658, EPI_ISL_3373659, EPI_ISL_3373660, EPI_ISL_3373661, EPI_ISL_3373662, EPI_ISL_3373663, EPI_ISL_3373664, EPI_ISL_3373665, EPI_ISL_3373666, EPI_ISL_3373667, EPI_ISL_3373668, EPI_ISL_3373669, EPI_ISL_3373670, EPI_ISL_3373671, EPI_ISL_3373672 | | | |
| see above | Nevada State Public Health Laboratory | Nevada State Public Health Laboratory | Andrew Gorzalski, Mark Pandori |
| EPI_ISL_3373673, EPI_ISL_3373674, EPI_ISL_3373678, EPI_ISL_3373680, EPI_ISL_3373682, EPI_ISL_3373683, EPI_ISL_3373684, EPI_ISL_3373685 | Provincial Public Health Reference Laboratory (PPHRL) | Provincial Public Health Reference Laboratory (PPHRL) | Andleeb Hanif, Mariam Javed, Hassan Nawaz, , Asim Altaf |
| EPI_ISL_3373687, EPI_ISL_3373688, EPI_ISL_3373689 | University Campus Bio-Medico of Rome (UCBM) | University Campus Bio-Medico of Rome (UCBM) | Fogolari M., Riva E., De Florio L., Francesconi M., Lintas C., Veralli R., Angeletti S. |
| EPI_ISL_3373690, EPI_ISL_3373691, EPI_ISL_3373692, EPI_ISL_3373693, EPI_ISL_3373694, EPI_ISL_3373695 | Sharp HealthCare Laboratory | Andersen lab at Scripps Research | SEARCH Alliance San Diego with Aaron Harding, Jacquelyn Berumen, Cathy Woerle, Liam McGinnis, Art Mendoza, Omid Bakhtar |
| EPI_ISL_3373696, EPI_ISL_3373697, EPI_ISL_3373698, EPI_ISL_3373699, EPI_ISL_3373700, EPI_ISL_3373701, EPI_ISL_3373702, EPI_ISL_3373703, EPI_ISL_3373704, EPI_ISL_3373705, EPI_ISL_3373706, EPI_ISL_3373707, EPI_ISL_3373708, EPI_ISL_3373709, EPI_ISL_3373710, EPI_ISL_3373711, EPI_ISL_3373714, EPI_ISL_3373715, EPI_ISL_3373716, EPI_ISL_3373717, EPI_ISL_3373719, EPI_ISL_3373721, EPI_ISL_3373722, EPI_ISL_3373723, EPI_ISL_3373725, EPI_ISL_3373726, EPI_ISL_3373728, EPI_ISL_3373745 | | | |
| see above | Scripps Medical Laboratory | Andersen lab at Scripps Research | SEARCH Alliance San Diego with Michael Quigley, Ellen Stefanski, Ian Mchardy |
| EPI_ISL_3373755, EPI_ISL_3373756, EPI_ISL_3373757, EPI_ISL_3373758, EPI_ISL_3373759, EPI_ISL_3373760, EPI_ISL_3373761, EPI_ISL_3373762, EPI_ISL_3373763, EPI_ISL_3373764, EPI_ISL_3373765, EPI_ISL_3373766, EPI_ISL_3373767, EPI_ISL_3373768, EPI_ISL_3373769, EPI_ISL_3373770, EPI_ISL_3373771, EPI_ISL_3373772, EPI_ISL_3373773, EPI_ISL_3373774, EPI_ISL_3373775, EPI_ISL_3373776, EPI_ISL_3373777, EPI_ISL_3373778, EPI_ISL_3373779, EPI_ISL_3373780, EPI_ISL_3373782, EPI_ISL_3373783, EPI_ISL_3373785, EPI_ISL_3373787, EPI_ISL_3373788, EPI_ISL_3373789, EPI_ISL_3373790, EPI_ISL_3373791, EPI_ISL_3373792, EPI_ISL_3373793, EPI_ISL_3373794, EPI_ISL_3373795, EPI_ISL_3373798, EPI_ISL_3373799, EPI_ISL_3373800, EPI_ISL_3373801, EPI_ISL_3373802, EPI_ISL_3373803, EPI_ISL_3373804, EPI_ISL_3373806, EPI_ISL_3373807, EPI_ISL_3373808, EPI_ISL_3373809, EPI_ISL_3373811, EPI_ISL_3373812, EPI_ISL_3373814, EPI_ISL_3373815, EPI_ISL_3373817, EPI_ISL_3373818, EPI_ISL_3373820, EPI_ISL_3373822, EPI_ISL_3373823, EPI_ISL_3373824, EPI_ISL_3373826, EPI_ISL_3373827, EPI_ISL_3373828, EPI_ISL_3373829, EPI_ISL_3373830, EPI_ISL_3373831, EPI_ISL_3373832, EPI_ISL_3373833, EPI_ISL_3373834, EPI_ISL_3373835, EPI_ISL_3373836, EPI_ISL_3373837, EPI_ISL_3373838, EPI_ISL_3373839, EPI_ISL_3373840, EPI_ISL_3373841, EPI_ISL_3373842, EPI_ISL_3373843, EPI_ISL_3373844, EPI_ISL_3373845, EPI_ISL_3373846, EPI_ISL_3373847, EPI_ISL_3373848, EPI_ISL_3373849, EPI_ISL_3373850, EPI_ISL_3373852, EPI_ISL_3373853, EPI_ISL_3373854, EPI_ISL_3373855, EPI_ISL_3373856, EPI_ISL_3373857, EPI_ISL_3373858, EPI_ISL_3373859, EPI_ISL_3373860, EPI_ISL_3373861, EPI_ISL_3373862, EPI_ISL_3373863, EPI_ISL_3373864, EPI_ISL_3373865, EPI_ISL_3373866, EPI_ISL_3373867, EPI_ISL_3373869, EPI_ISL_3373870, EPI_ISL_3373871, EPI_ISL_3373872, EPI_ISL_3373873, EPI_ISL_3373874, EPI_ISL_3373875, EPI_ISL_3373876, EPI_ISL_3373877, EPI_ISL_3373878, EPI_ISL_3373879, EPI_ISL_3373880, EPI_ISL_3373881, EPI_ISL_3373882, EPI_ISL_3373883, EPI_ISL_3373884, EPI_ISL_3373885, EPI_ISL_3373886, EPI_ISL_3373887, EPI_ISL_3373888, EPI_ISL_3373889, EPI_ISL_3373890, EPI_ISL_3373891, EPI_ISL_3373892, EPI_ISL_3373893, EPI_ISL_3373894, EPI_ISL_3373895, EPI_ISL_3373896, EPI_ISL_3373897, EPI_ISL_3373898, EPI_ISL_3373899, EPI_ISL_3373900, EPI_ISL_3373901, EPI_ISL_3373902, EPI_ISL_3373903, EPI_ISL_3373904, EPI_ISL_3373905, EPI_ISL_3373906, EPI_ISL_3373907, EPI_ISL_3373908, EPI_ISL_3373909, EPI_ISL_3373910, EPI_ISL_3373911, EPI_ISL_3373912, EPI_ISL_3373913, EPI_ISL_3373914, EPI_ISL_3373915, EPI_ISL_3373916, EPI_ISL_3373917, EPI_ISL_3373918, EPI_ISL_3373919, EPI_ISL_3373920, EPI_ISL_3373921, EPI_ISL_3373922, EPI_ISL_3373923, EPI_ISL_3373924, EPI_ISL_3373925, EPI_ISL_3373927, EPI_ISL_3373928, EPI_ISL_3373929, EPI_ISL_3373930, EPI_ISL_3373931, EPI_ISL_3373932, EPI_ISL_3373933, EPI_ISL_3373934, EPI_ISL_3373935, EPI_ISL_3373936, EPI_ISL_3373937, EPI_ISL_3373938, EPI_ISL_3373939, EPI_ISL_3373940, EPI_ISL_3373941, EPI_ISL_3373944, EPI_ISL_3373945, EPI_ISL_3373947, EPI_ISL_3373948, EPI_ISL_3373949, EPI_ISL_3373950, EPI_ISL_3373951, EPI_ISL_3373952, EPI_ISL_3373953, EPI_ISL_3373954, EPI_ISL_3373955, EPI_ISL_3373956, EPI_ISL_3373957, EPI_ISL_3373958, EPI_ISL_3373959, EPI_ISL_3373960, EPI_ISL_3373962, EPI_ISL_3373963, EPI_ISL_3373964, EPI_ISL_3373965, EPI_ISL_3373966, EPI_ISL_3373967, EPI_ISL_3373968, EPI_ISL_3373969, EPI_ISL_3373970, EPI_ISL_3373971, EPI_ISL_3373972, EPI_ISL_3373974 | | | |
| see above | Sharp HealthCare Laboratory | Andersen lab at Scripps Research | SEARCH Alliance San Diego with Aaron Harding, Jacquelyn Berumen, Cathy Woerle, Liam McGinnis, Art Mendoza, Omid Bakhtar |
| EPI_ISL_3373978, EPI_ISL_3373980, EPI_ISL_3373981, EPI_ISL_3373983, EPI_ISL_3373985, EPI_ISL_3373988, EPI_ISL_3373989, EPI_ISL_3373990, EPI_ISL_3373991, EPI_ISL_3373992, EPI_ISL_3373993, EPI_ISL_3373995, EPI_ISL_3373997, EPI_ISL_3373998, EPI_ISL_3373999, EPI_ISL_3374002, EPI_ISL_3374004, EPI_ISL_3374007, EPI_ISL_3374008, EPI_ISL_3374009, EPI_ISL_3374010, EPI_ISL_3374011, | | | |

| | | | |
|---|---|---|---|
| EPI_ISL_3374013, EPI_ISL_3374017, EPI_ISL_3374021, EPI_ISL_3374022, EPI_ISL_3374027, EPI_ISL_3374028, EPI_ISL_3374034, EPI_ISL_3374040, EPI_ISL_3374041 | | | |
| see above | Salud Digna, A.C | Andersen lab at Scripps Research | SEARCH Alliance with Abraham Campos Romero, Miguel A. Fernandez Rojas, Marco Antonio Luna Ruiz-Esparza, Jose Luis Moreno Camacho, Abraham Garcia Gil |
| EPI_ISL_3374046, EPI_ISL_3374047, EPI_ISL_3374064, EPI_ISL_3374065, EPI_ISL_3374067, EPI_ISL_3374070, EPI_ISL_3374072, EPI_ISL_3374073, EPI_ISL_3374074, EPI_ISL_3374075, EPI_ISL_3374076, EPI_ISL_3374081, EPI_ISL_3374082, EPI_ISL_3374084, EPI_ISL_3374085, EPI_ISL_3374086, EPI_ISL_3374088, EPI_ISL_3374090, EPI_ISL_3374091, EPI_ISL_3374093, EPI_ISL_3374095, EPI_ISL_3374097, EPI_ISL_3374098, EPI_ISL_3374099, EPI_ISL_3374100, EPI_ISL_3374101, EPI_ISL_3374102, EPI_ISL_3374103, EPI_ISL_3374104, EPI_ISL_3374105, EPI_ISL_3374107, EPI_ISL_3374108, EPI_ISL_3374111, EPI_ISL_3374115, EPI_ISL_3374116, EPI_ISL_3374117, EPI_ISL_3374119 | | | |
| see above | San Diego County Public Health Laboratory | Andersen lab at Scripps Research | SEARCH Alliance San Diego with Ashleigh Murphy, Jovan Shephard, Brett Austin |
| EPI_ISL_3374151 | Cameron Highlands District Health Office | Institute for Medical Research, Infectious Disease Research Centre, National Institutes of Health, Ministry of Health Malaysia | Suppiah J, Kamel K, Mohd Zawawi Z, Azizan MA, Ramly N, Robert F, Anasir MI, Thayan R |
| EPI_ISL_3374152 | Johor Bahru Public Health Laboratory | Institute for Medical Research, Infectious Disease Research Centre, National Institutes of Health, Ministry of Health Malaysia | Suppiah J, Kamel K, Mohd Zawawi Z, Azizan MA, Ramly N, Robert F, Anasir MI, Thayan R |
| EPI_ISL_3374154 | Putrajaya Health Office | Institute for Medical Research, Infectious Disease Research Centre, National Institutes of Health, Ministry of Health Malaysia | Suppiah J, Kamel K, Mohd Zawawi Z, Azizan MA, Ramly N, Robert F, Anasir MI, Thayan R |
| EPI_ISL_3374155, EPI_ISL_3374156, EPI_ISL_3374157, EPI_ISL_3374158, EPI_ISL_3374159, EPI_ISL_3374160, EPI_ISL_3374161, EPI_ISL_3374162, EPI_ISL_3374163, EPI_ISL_3374164, EPI_ISL_3374165, EPI_ISL_3374166, EPI_ISL_3374167, EPI_ISL_3374168, EPI_ISL_3374169, EPI_ISL_3374170, EPI_ISL_3374171, EPI_ISL_3374172, EPI_ISL_3374173, EPI_ISL_3374174, EPI_ISL_3374175, EPI_ISL_3374176, EPI_ISL_3374177, EPI_ISL_3374178, EPI_ISL_3374179, EPI_ISL_3374180, EPI_ISL_3374181, EPI_ISL_3374182, EPI_ISL_3374183, EPI_ISL_3374184, EPI_ISL_3374185, EPI_ISL_3374186, EPI_ISL_3374187, EPI_ISL_3374188, EPI_ISL_3374189, EPI_ISL_3374190, EPI_ISL_3374191, EPI_ISL_3374192, EPI_ISL_3374193, EPI_ISL_3374194, EPI_ISL_3374195, EPI_ISL_3374196, EPI_ISL_3374197, EPI_ISL_3374198, EPI_ISL_3374199, EPI_ISL_3374200, EPI_ISL_3374201, EPI_ISL_3374202, EPI_ISL_3374203, EPI_ISL_3374205, EPI_ISL_3374206, EPI_ISL_3374207, EPI_ISL_3374208, EPI_ISL_3374209, EPI_ISL_3374210, EPI_ISL_3374211, EPI_ISL_3374212, EPI_ISL_3374213, EPI_ISL_3374214, EPI_ISL_3374215, EPI_ISL_3374216, EPI_ISL_3374217, EPI_ISL_3374218, EPI_ISL_3374219, EPI_ISL_3374220, EPI_ISL_3374221, EPI_ISL_3374222, EPI_ISL_3374223, EPI_ISL_3374224, EPI_ISL_3374225, EPI_ISL_3374226, EPI_ISL_3374227, EPI_ISL_3374228, EPI_ISL_3374230, EPI_ISL_3374231, EPI_ISL_3374232 | | | |
| see above | New South Wales Health Pathology Royal Prince Alfred Hospital | Microbiology RPAH | Foster, C.; Au, J.; Ruiz Silva, M.; Deveson, I.; Bull, R.; Van Hal, S.; Rawlinson, W. |
| EPI_ISL_3374233 | Cheras District Health Office | Institute for Medical Research, Infectious Disease Research Centre, National Institutes of Health, Ministry of Health Malaysia | Suppiah J, Kamel K, Mohd Zawawi Z, Azizan MA, Ramly N, Robert F, Anasir MI, Thayan R |
| EPI_ISL_3374234 | University of Iowa Hospitals and Clinics | State Hygienic Laboratory at the University of Iowa | Valerie Reeb, Erik Twait, Anna Yakos, Mariah Knutson, Kris Eveland, Kristen Zanon, Jeff Benfer, Davis Rieckenberg, Wes Hottel, Alankar Kampoowale |
| EPI_ISL_3374235, EPI_ISL_3374236, EPI_ISL_3374237, EPI_ISL_3374238, EPI_ISL_3374239, EPI_ISL_3374241, EPI_ISL_3374242, EPI_ISL_3374244, EPI_ISL_3374245, EPI_ISL_3374246, EPI_ISL_3374247, EPI_ISL_3374248, EPI_ISL_3374249, EPI_ISL_3374250, EPI_ISL_3374251, EPI_ISL_3374252, EPI_ISL_3374253, EPI_ISL_3374254, EPI_ISL_3374255, EPI_ISL_3374256, EPI_ISL_3374257, EPI_ISL_3374258, EPI_ISL_3374259, EPI_ISL_3374260, EPI_ISL_3374261, EPI_ISL_3374262, EPI_ISL_3374263 | | | |
| see above | State Hygienic Laboratory at the University of Iowa | State Hygienic Laboratory at the University of Iowa | Valerie Reeb, Erik Twait, Anna Yakos, Mariah Knutson, Kris Eveland, Kristen Zanon, Jeff Benfer, Davis Rieckenberg, Wes Hottel, Alankar Kampoowale |
| EPI_ISL_3374264, EPI_ISL_3374265, EPI_ISL_3374266, EPI_ISL_3374267, EPI_ISL_3374268, EPI_ISL_3374270, EPI_ISL_3374271, EPI_ISL_3374272, EPI_ISL_3374273, EPI_ISL_3374274, EPI_ISL_3374275, EPI_ISL_3374276, EPI_ISL_3374277, EPI_ISL_3374278, EPI_ISL_3374279, EPI_ISL_3374280, EPI_ISL_3374281, EPI_ISL_3374282, EPI_ISL_3374283, EPI_ISL_3374284, EPI_ISL_3374285, EPI_ISL_3374286, EPI_ISL_3374287, EPI_ISL_3374288, EPI_ISL_3374289, EPI_ISL_3374290, EPI_ISL_3374291, EPI_ISL_3374292 | | | |
| see above | State Hygienic Laboratory at the University of Iowa | State Hygienic Laboratory at the University of Iowa | Valerie Reeb, Erik Twait, Anna Yakos, Mariah Knutson, Kris Eveland, Kristen Zanon, Jeff, Benfer, Wes Hottel, Alankar Kampoowale |
| EPI_ISL_3374293 | POLICLINICA HORTOLANDIA | Instituto Butantan | Dimas Tadeu Covas, Antonio Jorge Martins, Claudia Renata dos Santos Barros, David Schlesinger, Debora Botequio Moretti, Elaine Cristina Marqueze, Elaine Vieira Santos, Evandra Strazza Rodrigues, Heidge Fukumasu, Jayme Augusto de Souza-Neto, José Salvatore Leister Patané, Luiz Alcantara, Luiz Lehmann Coutinho, Maria Carolina Elias, Maurício Lacerda Nogueira, Rafael dos Santos Bezerra, Raul Machado Neto, Rejane Maria Tommasini Grotto, Ricardo Haddad, Sandra Coccuzzo Sampaio Vessoni, Simone Kashima, Svetoslav Nanev Slavov, Vincent Louis Viala |
| EPI_ISL_3374294 | HOSPITAL DE CAMPANHA COVID 19 MUNICIPIO DE TAUBATE | Instituto Butantan | Dimas Tadeu Covas, Antonio Jorge Martins, Claudia Renata dos Santos Barros, David Schlesinger, Debora Botequio Moretti, Elaine Cristina Marqueze, Elaine Vieira Santos, Evandra Strazza Rodrigues, Heidge Fukumasu, Jayme Augusto de Souza-Neto, José Salvatore Leister Patané, Luiz Alcantara, Luiz Lehmann Coutinho, Maria Carolina Elias, Maurício Lacerda Nogueira, Rafael dos Santos Bezerra, Raul Machado Neto, Rejane Maria Tommasini Grotto, Ricardo Haddad, Sandra Coccuzzo Sampaio Vessoni, Simone Kashima, Svetoslav Nanev Slavov, Vincent Louis Viala |
| EPI_ISL_3374295 | CENTRO DE APOIO EPIDEMIOLOGICO | Instituto Butantan | Dimas Tadeu Covas, Antonio Jorge Martins, Claudia Renata dos Santos Barros, David Schlesinger, Debora Botequio Moretti, Elaine Cristina Marqueze, Elaine Vieira Santos, Evandra Strazza Rodrigues, Heidge Fukumasu, Jayme Augusto de Souza-Neto, José Salvatore Leister Patané, Luiz Alcantara, Luiz Lehmann Coutinho, Maria Carolina Elias, Maurício Lacerda Nogueira, Rafael dos Santos Bezerra, Raul Machado Neto, Rejane Maria Tommasini Grotto, Ricardo Haddad, Sandra Coccuzzo Sampaio Vessoni, Simone Kashima, Svetoslav Nanev Slavov, Vincent Louis Viala |
| EPI_ISL_3374296 | PRONTO SOCORRO VEREADOR LUIZ CARLOS MARQUES | Instituto Butantan | Dimas Tadeu Covas, Antonio Jorge Martins, Claudia Renata dos Santos Barros, David Schlesinger, Debora Botequio Moretti, Elaine Cristina Marqueze, Elaine Vieira Santos, Evandra Strazza Rodrigues, Heidge Fukumasu, Jayme Augusto de Souza-Neto, José Salvatore Leister Patané, Luiz Alcantara, Luiz Lehmann Coutinho, Maria Carolina Elias, Maurício Lacerda Nogueira, Rafael dos Santos Bezerra, Raul Machado Neto, Rejane Maria Tommasini Grotto, Ricardo Haddad, Sandra Coccuzzo Sampaio Vessoni, Simone Kashima, Svetoslav Nanev Slavov, Vincent Louis Viala |
| EPI_ISL_3374297 | HOSPITAL DE CAMPANHA COVID 19 MUNICIPIO DE TAUBATE | Instituto Butantan | Dimas Tadeu Covas, Antonio Jorge Martins, Claudia Renata dos Santos Barros, David Schlesinger, Debora Botequio Moretti, Elaine Cristina Marqueze, Elaine Vieira Santos, Evandra Strazza Rodrigues, Heidge Fukumasu, Jayme Augusto de Souza-Neto, José Salvatore Leister Patané, Luiz Alcantara, Luiz Lehmann Coutinho, Maria Carolina Elias, Maurício Lacerda Nogueira, Rafael dos Santos Bezerra, Raul Machado Neto, Rejane Maria Tommasini Grotto, Ricardo Haddad, Sandra Coccuzzo Sampaio Vessoni, Simone Kashima, Svetoslav Nanev Slavov, Vincent Louis Viala |
| EPI_ISL_3374298 | HOSPITAL DE CAMPANHA COVID 19 CAIEIRAS | Instituto Butantan | Dimas Tadeu Covas, Antonio Jorge Martins, Claudia Renata dos Santos Barros, David Schlesinger, Debora Botequio Moretti, Elaine Cristina Marqueze, Elaine Vieira Santos, Evandra Strazza Rodrigues, Heidge Fukumasu, Jayme Augusto de Souza-Neto, José Salvatore Leister Patané, Luiz Alcantara, Luiz Lehmann Coutinho, Maria Carolina Elias, Maurício Lacerda Nogueira, Rafael dos Santos Bezerra, Raul Machado Neto, Rejane Maria Tommasini Grotto, Ricardo Haddad, Sandra Coccuzzo Sampaio Vessoni, Simone Kashima, Svetoslav Nanev Slavov, Vincent Louis Viala |
| EPI_ISL_3374299 | HOSPITAL REGIONAL DR VIVALDO MARTINS SIMOES OSASCO | Instituto Butantan | Dimas Tadeu Covas, Antonio Jorge Martins, Claudia Renata dos Santos Barros, David Schlesinger, Debora Botequio Moretti, Elaine Cristina Marqueze, Elaine Vieira Santos, Evandra Strazza Rodrigues, Heidge Fukumasu, Jayme Augusto de Souza-Neto, José Salvatore Leister Patané, Luiz Alcantara, Luiz Lehmann Coutinho, Maria Carolina Elias, Maurício Lacerda Nogueira, Rafael dos Santos Bezerra, Raul Machado Neto, Rejane Maria Tommasini Grotto, Ricardo Haddad, Sandra Coccuzzo Sampaio Vessoni, Simone Kashima, Svetoslav Nanev Slavov, Vincent Louis Viala |
| EPI_ISL_3374300 | UBS LAVRAS | Instituto Butantan | Dimas Tadeu Covas, Antonio Jorge Martins, Claudia Renata dos Santos Barros, David Schlesinger, Debora Botequio Moretti, Elaine Cristina Marqueze, Elaine Vieira Santos, Evandra Strazza Rodrigues, Heidge Fukumasu, Jayme Augusto de Souza-Neto, José Salvatore Leister Patané, Luiz Alcantara, Luiz Lehmann Coutinho, Maria Carolina Elias, Maurício Lacerda Nogueira, Rafael dos Santos Bezerra, Raul Machado Neto, Rejane Maria Tommasini Grotto, Ricardo Haddad, Sandra Coccuzzo Sampaio Vessoni, Simone Kashima, Svetoslav Nanev Slavov, Vincent Louis Viala |
| EPI_ISL_3374301 | UNIDADE BASICA DE SAUDE DE PIQUETE | Instituto Butantan | Dimas Tadeu Covas, Antonio Jorge Martins, Claudia Renata dos Santos Barros, David Schlesinger, Debora Botequio Moretti, Elaine Cristina Marqueze, Elaine Vieira Santos, Evandra Strazza Rodrigues, Heidge Fukumasu, Jayme Augusto de Souza-Neto, José Salvatore Leister Patané, Luiz Alcantara, Luiz Lehmann Coutinho, Maria Carolina Elias, Maurício Lacerda Nogueira, Rafael dos Santos Bezerra, Raul Machado Neto, Rejane Maria Tommasini Grotto, Ricardo Haddad, Sandra Coccuzzo Sampaio Vessoni, Simone Kashima, Svetoslav Nanev Slavov, Vincent Louis Viala |
| EPI_ISL_3374302, EPI_ISL_3374303 | LABORATORIO LOCAL DE ITAPECERICA DA SERRA | Instituto Butantan | Dimas Tadeu Covas, Antonio Jorge Martins, Claudia Renata dos Santos Barros, David Schlesinger, Debora Botequio Moretti, Elaine Cristina Marqueze, Elaine Vieira Santos, Evandra Strazza Rodrigues, Heidge Fukumasu, Jayme Augusto de Souza-Neto, José Salvatore Leister Patané, Luiz Alcantara, Luiz Lehmann Coutinho, Maria Carolina Elias, Maurício Lacerda Nogueira, Rafael dos Santos Bezerra, Raul Machado Neto, Rejane Maria Tommasini Grotto, Ricardo Haddad, Sandra Coccuzzo Sampaio Vessoni, Simone Kashima, Svetoslav Nanev Slavov, Vincent Louis Viala |
| EPI_ISL_3374304 | VIGILANCIA EPIDEMIOLOGICA E CONTROLE DE VETORES PIRASSUNUNGA | Instituto Butantan | Dimas Tadeu Covas, Antonio Jorge Martins, Claudia Renata dos Santos Barros, David Schlesinger, Debora Botequio Moretti, Elaine Cristina Marqueze, Elaine Vieira Santos, Evandra Strazza Rodrigues, Heidge Fukumasu, Jayme Augusto de Souza-Neto, José Salvatore Leister Patané, Luiz Alcantara, Luiz Lehmann Coutinho, Maria Carolina Elias, Maurício Lacerda Nogueira, Rafael dos Santos Bezerra, Raul Machado Neto, Rejane Maria Tommasini Grotto, Ricardo Haddad, Sandra Coccuzzo Sampaio Vessoni, Simone Kashima, Svetoslav Nanev Slavov, Vincent Louis Viala |
| EPI_ISL_3374305 | UBS DR HELIO MIGLIARI | Instituto Butantan | Dimas Tadeu Covas, Antonio Jorge Martins, Claudia Renata dos Santos Barros, David Schlesinger, Debora Botequio Moretti, Elaine Cristina Marqueze, Elaine Vieira Santos, Evandra Strazza Rodrigues, Heidge Fukumasu, Jayme Augusto de Souza-Neto, José Salvatore Leister Patané, Luiz Alcantara, Luiz Lehmann Coutinho, Maria Carolina Elias, Maurício Lacerda Nogueira, Rafael dos Santos Bezerra, Raul Machado Neto, Rejane Maria Tommasini Grotto, Ricardo Haddad, Sandra Coccuzzo Sampaio Vessoni, Simone Kashima, Svetoslav Nanev Slavov, Vincent Louis Viala |
| EPI_ISL_3374306 | HOSPITAL MUN DR GUIDO GUIDA | Instituto Butantan | Dimas Tadeu Covas, Antonio Jorge Martins, Claudia Renata dos Santos Barros, David Schlesinger, Debora Botequio Moretti, Elaine Cristina Marqueze, Elaine Vieira Santos, Evandra Strazza Rodrigues, Heidge Fukumasu, Jayme Augusto de Souza-Neto, José Salvatore Leister Patané, Luiz Alcantara, Luiz Lehmann Coutinho, Maria Carolina Elias, Maurício Lacerda Nogueira, Rafael dos Santos Bezerra, Raul Machado Neto, Rejane Maria Tommasini Grotto, Ricardo Haddad, Sandra Coccuzzo Sampaio Vessoni, Simone Kashima, Svetoslav Nanev Slavov, Vincent Louis Viala |
| EPI_ISL_3374307 | CENTRO DE ATENDIMENTO PARA ENFRENTAMENTO A COVID19 DE APARECIDA | Instituto Butantan | Dimas Tadeu Covas, Antonio Jorge Martins, Claudia Renata dos Santos Barros, David Schlesinger, Debora Botequio Moretti, Elaine Cristina Marqueze, Elaine Vieira Santos, Evandra Strazza Rodrigues, Heidge Fukumasu, Jayme Augusto de Souza-Neto, José Salvatore Leister Patané, Luiz Alcantara, Luiz Lehmann Coutinho, Maria Carolina Elias, Maurício Lacerda Nogueira, Rafael dos Santos Bezerra, Raul Machado Neto, Rejane Maria Tommasini Grotto, Ricardo Haddad, Sandra Coccuzzo Sampaio Vessoni, Simone Kashima, Svetoslav Nanev Slavov, Vincent Louis Viala |
| EPI_ISL_3374308, EPI_ISL_3374309, EPI_ISL_3374310 | CENTRO DE APOIO EPIDEMIOLOGICO | Instituto Butantan | Dimas Tadeu Covas, Antonio Jorge Martins, Claudia Renata dos Santos Barros, David Schlesinger, Debora Botequio Moretti, Elaine Cristina Marqueze, Elaine Vieira Santos, Evandra Strazza Rodrigues, Heidge Fukumasu, Jayme Augusto de Souza-Neto, José Salvatore Leister Patané, Luiz Alcantara, Luiz Lehmann Coutinho, Maria Carolina Elias, Maurício Lacerda Nogueira, Rafael dos Santos Bezerra, Raul Machado Neto, Rejane Maria Tommasini Grotto, Ricardo Haddad, Sandra Coccuzzo Sampaio Vessoni, Simone Kashima, Svetoslav Nanev Slavov, Vincent Louis Viala |
| EPI_ISL_3374311 | COMPLEXO HOSPITALAR PADRE BENTO DE GUARULHOS | Instituto Butantan | Dimas Tadeu Covas, Antonio Jorge Martins, Claudia Renata dos Santos Barros, David Schlesinger, Debora Botequio Moretti, Elaine Cristina Marqueze, Elaine Vieira Santos, Evandra Strazza Rodrigues, Heidge Fukumasu, Jayme Augusto de Souza-Neto, José Salvatore Leister Patané, Luiz Alcantara, Luiz Lehmann Coutinho, Maria Carolina Elias, Maurício Lacerda Nogueira, Rafael dos Santos Bezerra, Raul Machado Neto, Rejane Maria Tommasini Grotto, Ricardo Haddad, Sandra Coccuzzo Sampaio Vessoni, Simone Kashima, Svetoslav Nanev Slavov, Vincent Louis Viala |
| EPI_ISL_3374312, EPI_ISL_3374313 | CENTRO DE APOIO EPIDEMIOLOGICO | Instituto Butantan | Dimas Tadeu Covas, Antonio Jorge Martins, Claudia Renata dos Santos Barros, David Schlesinger, Debora Botequio Moretti, Elaine Cristina Marqueze, Elaine Vieira Santos, Evandra Strazza Rodrigues, Heidge Fukumasu, Jayme Augusto de Souza-Neto, José Salvatore Leister Patané, Luiz Alcantara, Luiz Lehmann Coutinho, Maria Carolina Elias, Maurício Lacerda Nogueira, Rafael dos Santos Bezerra, Raul Machado Neto, Rejane Maria Tommasini Grotto, Ricardo Haddad, Sandra Coccuzzo Sampaio Vessoni, Simone Kashima, Svetoslav Nanev Slavov, Vincent Louis Viala |
| EPI_ISL_3374314, EPI_ISL_3374315, EPI_ISL_3374316 | VIGILANCIA SANITARIA E VIG EPIDEMIOLOGICA DE ITAPEVI | Instituto Butantan | Dimas Tadeu Covas, Antonio Jorge Martins, Claudia Renata dos Santos Barros, David Schlesinger, Debora Botequio Moretti, Elaine Cristina Marqueze, Elaine Vieira Santos, Evandra Strazza Rodrigues, Heidge Fukumasu, Jayme Augusto de Souza-Neto, José Salvatore Leister Patané, Luiz Alcantara, Luiz Lehmann Coutinho, Maria Carolina Elias, Maurício Lacerda Nogueira, Rafael dos Santos Bezerra, Raul Machado Neto, Rejane Maria Tommasini Grotto, Ricardo Haddad, Sandra Coccuzzo Sampaio Vessoni, Simone Kashima, Svetoslav Nanev Slavov, Vincent Louis Viala |
| EPI_ISL_3374317 | AMBULATORIO DE ESPECIALIDADE V E MOGI MIRIM | Instituto Butantan | Dimas Tadeu Covas, Antonio Jorge Martins, Claudia Renata dos Santos Barros, David Schlesinger, Debora Botequio Moretti, Elaine Cristina Marqueze, Elaine Vieira Santos, Evandra Strazza Rodrigues, Heidge Fukumasu, Jayme Augusto de Souza-Neto, José Salvatore Leister Patané, Luiz Alcantara, Luiz Lehmann Coutinho, Maria Carolina Elias, Maurício Lacerda Nogueira, Rafael dos Santos Bezerra, Raul Machado Neto, Rejane Maria Tommasini Grotto, Ricardo Haddad, Sandra Coccuzzo Sampaio Vessoni, Simone Kashima, Svetoslav Nanev Slavov, Vincent Louis Viala |
| EPI_ISL_3374318 | USF SANTA ANTONIETA II | Instituto Butantan | Dimas Tadeu Covas, Antonio Jorge Martins, Claudia Renata dos Santos Barros, David Schlesinger, Debora Botequio Moretti, Elaine Cristina Marqueze, Elaine Vieira Santos, Evandra Strazza Rodrigues, Heidge Fukumasu, Jayme Augusto de Souza-Neto, José Salvatore Leister Patané, Luiz Alcantara, Luiz Lehmann Coutinho, Maria Carolina Elias, Maurício Lacerda Nogueira, Rafael dos Santos Bezerra, Raul Machado Neto, Rejane Maria Tommasini Grotto, Ricardo Haddad, Sandra Coccuzzo Sampaio Vessoni, Simone Kashima, Svetoslav Nanev Slavov, Vincent Louis Viala |
| EPI_ISL_3374319, EPI_ISL_3374320, EPI_ISL_3374321 | VIGILANCIA SANITARIA E VIG EPIDEMIOLOGICA DE ITAPEVI | Instituto Butantan | Dimas Tadeu Covas, Antonio Jorge Martins, Claudia Renata dos Santos Barros, David Schlesinger, Debora Botequio Moretti, Elaine Cristina Marqueze, Elaine Vieira Santos, Evandra Strazza Rodrigues, Heidge Fukumasu, Jayme Augusto de Souza-Neto, José Salvatore Leister Patané, Luiz Alcantara, Luiz Lehmann Coutinho, Maria Carolina Elias, Maurício Lacerda Nogueira, Rafael dos Santos Bezerra, Raul Machado Neto, Rejane Maria Tommasini Grotto, Ricardo Haddad, Sandra Coccuzzo Sampaio Vessoni, Simone Kashima, Svetoslav Nanev Slavov, Vincent Louis Viala |
| EPI_ISL_3374322, EPI_ISL_3374323, EPI_ISL_3374324 | SECAO CENTRO DE DIAGNOSTICO SECEDI | Instituto Butantan | Dimas Tadeu Covas, Antonio Jorge Martins, Claudia Renata dos Santos Barros, David Schlesinger, Debora Botequio Moretti, Elaine Cristina Marqueze, Elaine Vieira Santos, Evandra Strazza Rodrigues, Heidge Fukumasu, Jayme Augusto de Souza-Neto, José Salvatore Leister Patané, Luiz Alcantara, Luiz Lehmann Coutinho, Maria Carolina Elias, Maurício Lacerda Nogueira, Rafael dos Santos Bezerra, Raul Machado Neto, Rejane Maria Tommasini Grotto, Ricardo Haddad, Sandra Coccuzzo Sampaio Vessoni, Simone Kashima, Svetoslav Nanev Slavov, Vincent Louis Viala |
| EPI_ISL_3374325 | UNIDADE DE PRONTO ATENDIMENTO UPA 24 HORAS | Instituto Butantan | Dimas Tadeu Covas, Antonio Jorge Martins, Claudia Renata dos Santos Barros, David Schlesinger, Debora Botequio Moretti, Elaine Cristina Marqueze, Elaine Vieira Santos, Evandra Strazza Rodrigues, Heidge Fukumasu, Jayme Augusto de Souza-Neto, José Salvatore Leister Patané, Luiz Alcantara, Luiz Lehmann Coutinho, Maria Carolina Elias, Maurício Lacerda Nogueira, Rafael dos Santos Bezerra, Raul Machado Neto, Rejane Maria Tommasini Grotto, Ricardo Haddad, Sandra Coccuzzo Sampaio Vessoni, Simone Kashima, Svetoslav Nanev Slavov, Vincent Louis Viala |

| | | | |
|---|---|---|---|
| EPI_ISL_3374326 | HOSPITAL DE CAMPANHA COVID 19 MUNICIPIO DE TAUBATE | Instituto Butantan | Dimas Tadeu Covas, Antonio Jorge Martins, Claudia Renata dos Santos Barros, David Schlesinger, Debora Botequio Moretti, Elaine Cristina Marqueze, Elaine Vieira Santos, Evandra Strazza Rodrigues, Heidge Fukumasu, Jayme Augusto de Souza-Neto, José Salvatore Leister Patané, Luiz Alcantara, Luiz Lehmann Coutinho, Maria Carolina Elias, Maurício Lacerda Nogueira, Rafael dos Santos Bezerra, Raul Machado Neto, Rejane Maria Tommasini Grotto, Ricardo Haddad, Sandra Coccuzzo Sampaio Vessoni, Simone Kashima, Svetoslav Nanev Slavov, Vincent Louis Viala |
| EPI_ISL_3374327 | CENTRO DE APOIO EPIDEMIOLOGICO | Instituto Butantan | Dimas Tadeu Covas, Antonio Jorge Martins, Claudia Renata dos Santos Barros, David Schlesinger, Debora Botequio Moretti, Elaine Cristina Marqueze, Elaine Vieira Santos, Evandra Strazza Rodrigues, Heidge Fukumasu, Jayme Augusto de Souza-Neto, José Salvatore Leister Patané, Luiz Alcantara, Luiz Lehmann Coutinho, Maria Carolina Elias, Maurício Lacerda Nogueira, Rafael dos Santos Bezerra, Raul Machado Neto, Rejane Maria Tommasini Grotto, Ricardo Haddad, Sandra Coccuzzo Sampaio Vessoni, Simone Kashima, Svetoslav Nanev Slavov, Vincent Louis Viala |
| EPI_ISL_3374328, EPI_ISL_3374329, EPI_ISL_3374330, EPI_ISL_3374331, EPI_ISL_3374332, EPI_ISL_3374333, EPI_ISL_3374334, EPI_ISL_3374335, EPI_ISL_3374336, EPI_ISL_3374337, EPI_ISL_3374338, EPI_ISL_3374339 | | | |
| see above | AFIP | Instituto Butantan | Dimas Tadeu Covas, Antonio Jorge Martins, Claudia Renata dos Santos Barros, David Schlesinger, Debora Botequio Moretti, Elaine Cristina Marqueze, Elaine Vieira Santos, Evandra Strazza Rodrigues, Heidge Fukumasu, Jayme Augusto de Souza-Neto, José Salvatore Leister Patané, Luiz Alcantara, Luiz Lehmann Coutinho, Maria Carolina Elias, Maurício Lacerda Nogueira, Rafael dos Santos Bezerra, Raul Machado Neto, Rejane Maria Tommasini Grotto, Ricardo Haddad, Sandra Coccuzzo Sampaio Vessoni, Simone Kashima, Svetoslav Nanev Slavov, Vincent Louis Viala |
| EPI_ISL_3374340, EPI_ISL_3374341, EPI_ISL_3374342, EPI_ISL_3374343, EPI_ISL_3374344, EPI_ISL_3374345, EPI_ISL_3374346, EPI_ISL_3374347, EPI_ISL_3374348, EPI_ISL_3374349, EPI_ISL_3374350, EPI_ISL_3374351, EPI_ISL_3374352, EPI_ISL_3374353, EPI_ISL_3374354, EPI_ISL_3374355, EPI_ISL_3374356, EPI_ISL_3374357, EPI_ISL_3374358 | | | |
| see above | BIOFAST | Instituto Butantan | Dimas Tadeu Covas, Antonio Jorge Martins, Claudia Renata dos Santos Barros, David Schlesinger, Debora Botequio Moretti, Elaine Cristina Marqueze, Elaine Vieira Santos, Evandra Strazza Rodrigues, Heidge Fukumasu, Jayme Augusto de Souza-Neto, José Salvatore Leister Patané, Luiz Alcantara, Luiz Lehmann Coutinho, Maria Carolina Elias, Maurício Lacerda Nogueira, Rafael dos Santos Bezerra, Raul Machado Neto, Rejane Maria Tommasini Grotto, Ricardo Haddad, Sandra Coccuzzo Sampaio Vessoni, Simone Kashima, Svetoslav Nanev Slavov, Vincent Louis Viala |
| EPI_ISL_3374359, EPI_ISL_3374360, EPI_ISL_3374361 | FACULDADE DE MEDICINA ABC | Instituto Butantan | Dimas Tadeu Covas, Antonio Jorge Martins, Claudia Renata dos Santos Barros, David Schlesinger, Debora Botequio Moretti, Elaine Cristina Marqueze, Elaine Vieira Santos, Evandra Strazza Rodrigues, Heidge Fukumasu, Jayme Augusto de Souza-Neto, José Salvatore Leister Patané, Luiz Alcantara, Luiz Lehmann Coutinho, Maria Carolina Elias, Maurício Lacerda Nogueira, Rafael dos Santos Bezerra, Raul Machado Neto, Rejane Maria Tommasini Grotto, Ricardo Haddad, Sandra Coccuzzo Sampaio Vessoni, Simone Kashima, Svetoslav Nanev Slavov, Vincent Louis Viala |
| EPI_ISL_3374362, EPI_ISL_3374363, EPI_ISL_3374364, EPI_ISL_3374365, EPI_ISL_3374366, EPI_ISL_3374367, EPI_ISL_3374368, EPI_ISL_3374369, EPI_ISL_3374370, EPI_ISL_3374371, EPI_ISL_3374372, EPI_ISL_3374373, EPI_ISL_3374374, EPI_ISL_3374375, EPI_ISL_3374376, EPI_ISL_3374377, EPI_ISL_3374378, EPI_ISL_3374379, EPI_ISL_3374380, EPI_ISL_3374381, EPI_ISL_3374382, EPI_ISL_3374383, EPI_ISL_3374384, EPI_ISL_3374385, EPI_ISL_3374386, EPI_ISL_3374387, EPI_ISL_3374388, EPI_ISL_3374389, EPI_ISL_3374390, EPI_ISL_3374391, EPI_ISL_3374392, EPI_ISL_3374393, EPI_ISL_3374394, EPI_ISL_3374395, EPI_ISL_3374396, EPI_ISL_3374397, EPI_ISL_3374398, EPI_ISL_3374399, EPI_ISL_3374400, EPI_ISL_3374401, EPI_ISL_3374402, EPI_ISL_3374403, EPI_ISL_3374404, EPI_ISL_3374405, EPI_ISL_3374406, EPI_ISL_3374407, EPI_ISL_3374408, EPI_ISL_3374409, EPI_ISL_3374410, EPI_ISL_3374411, EPI_ISL_3374412, EPI_ISL_3374413, EPI_ISL_3374414, EPI_ISL_3374415, EPI_ISL_3374416, EPI_ISL_3374417, EPI_ISL_3374418, EPI_ISL_3374419, EPI_ISL_3374420 | | | |
| see above | INSIDE DIAGNÓSTICOS | Instituto Butantan | Dimas Tadeu Covas, Antonio Jorge Martins, Claudia Renata dos Santos Barros, David Schlesinger, Debora Botequio Moretti, Elaine Cristina Marqueze, Elaine Vieira Santos, Evandra Strazza Rodrigues, Heidge Fukumasu, Jayme Augusto de Souza-Neto, José Salvatore Leister Patané, Luiz Alcantara, Luiz Lehmann Coutinho, Maria Carolina Elias, Maurício Lacerda Nogueira, Rafael dos Santos Bezerra, Raul Machado Neto, Rejane Maria Tommasini Grotto, Ricardo Haddad, Sandra Coccuzzo Sampaio Vessoni, Simone Kashima, Svetoslav Nanev Slavov, Vincent Louis Viala |
| EPI_ISL_3374426 | UiTM Sungai Buloh Hospital | Institute for Medical Research, Infectious Disease Research Centre, National Institutes of Health, Ministry of Health Malaysia | Suppiah J, Kamel K, Mohd Zawawi Z, Azizan MA, Ramly N, Robert F, Anasir MI, Thayan R |
| EPI_ISL_3374439, EPI_ISL_3374440, EPI_ISL_3374441, EPI_ISL_3374442, EPI_ISL_3374443, EPI_ISL_3374452, EPI_ISL_3374453, EPI_ISL_3374454, EPI_ISL_3374455, EPI_ISL_3374456, EPI_ISL_3374457, EPI_ISL_3374458, EPI_ISL_3374460, EPI_ISL_3374461, EPI_ISL_3374462, EPI_ISL_3374463, EPI_ISL_3374464, EPI_ISL_3374465, EPI_ISL_3374466, EPI_ISL_3374467, EPI_ISL_3374468, EPI_ISL_3374469 | | | |
| see above | Department of Virology I, National Institute of Infectious Diseases | Department of Veterinary Science, National Institute of Infectious Diseases | Yudai Kuroda, Tsukasa Yamamoto, Keita Ishijima, Tsuguto Fujimoto, Souichi Yamada, Shuetsu Fukushi, Ken Maeda |
| EPI_ISL_3374470 | Queen Elizabeth Hospital, Kota Kinabalu | Institute for Medical Research, Infectious Disease Research Centre, National Institutes of Health, Ministry of Health Malaysia | Suppiah J, Kamel K, Mohd Zawawi Z, Azizan MA, Ramly N, Robert F, Anasir MI, Thayan R |
| EPI_ISL_3374484 | Sultanah Bahiyah Hospital, Alor Setar | Institute for Medical Research, Infectious Disease Research Centre, National Institutes of Health, Ministry of Health Malaysia | Suppiah J, Kamel K, Mohd Zawawi Z, Azizan MA, Ramly N, Robert F, Anasir MI, Thayan R |
| EPI_ISL_3374485, EPI_ISL_3374486, EPI_ISL_3374487, EPI_ISL_3374488, EPI_ISL_3374489, EPI_ISL_3374490, EPI_ISL_3374491, EPI_ISL_3374492, EPI_ISL_3374493, EPI_ISL_3374494, EPI_ISL_3374495, EPI_ISL_3374496, EPI_ISL_3374497, EPI_ISL_3374498, EPI_ISL_3374499, EPI_ISL_3374500, EPI_ISL_3374501, EPI_ISL_3374502, EPI_ISL_3374503, EPI_ISL_3374504, EPI_ISL_3374505, EPI_ISL_3374506, EPI_ISL_3374507, EPI_ISL_3374508, EPI_ISL_3374509, EPI_ISL_3374510, EPI_ISL_3374511, EPI_ISL_3374512, EPI_ISL_3374513, EPI_ISL_3374514, EPI_ISL_3374515, EPI_ISL_3374516, EPI_ISL_3374517, EPI_ISL_3374518, EPI_ISL_3374519, EPI_ISL_3374520, EPI_ISL_3374521, EPI_ISL_3374522 | | | |
| see above | Tokyo Metropolitan Institute of Public Health | Tokyo Metropolitan Institute of Public Health | Masaki Hayashi, Takako Yamazaki, Akane Negishi, Hirofumi Miyake, Kenshirou Kuroki, Maya Isogai, Takushi Fujiwara, Mami Nagashima, Jun Suzuki, Kenji Sadamasu |
| EPI_ISL_3374523, EPI_ISL_3374524 | Lablink (M) Sdn. Bhd. | Institute for Medical Research, Infectious Disease Research Centre, National Institutes of Health, Ministry of Health Malaysia | Suppiah J, Kamel K, Mohd Zawawi Z, Azizan MA, Ramly N, Robert F, Anasir MI, Thayan R |
| EPI_ISL_3374529, EPI_ISL_3374530 | Diagnostic and Research Center of Infectious Diseases, Medical Faculty, Andalas University | Genomik Solidaritas Indonesia Laboratorium / Diagnostic and Research Center of Infectious Diseases, Medical Faculty, Andalas University | Himawan Masyhuri, Dhahlia Agustina Cahyono, Andani Eka Putra, Dede Rahman Agustian, Linosefa, Vania Gavrila Wikasa, Carissa Sintca Wijaya, Gracia Felias Enos Korompis, Annisa Muthiah Sukirman, Anuraj Shankar, Louisa Markus, Meutia Ayuputeri Kumaheri, Ariel Pradipta |
| EPI_ISL_3374532, EPI_ISL_3374533, EPI_ISL_3374534, EPI_ISL_3374535, EPI_ISL_3374536, EPI_ISL_3374537, EPI_ISL_3374538, EPI_ISL_3374539, EPI_ISL_3374540, EPI_ISL_3374541, EPI_ISL_3374542, EPI_ISL_3374543, EPI_ISL_3374544, EPI_ISL_3374545, EPI_ISL_3374546, EPI_ISL_3374547, EPI_ISL_3374548, EPI_ISL_3374549, EPI_ISL_3374550, EPI_ISL_3374551, EPI_ISL_3374552, EPI_ISL_3374553, EPI_ISL_3374555 | | | |
| see above | Zuckerberg San Francisco General Hospital | DeRisi Lab, University of California, San Francisco | Jeffery Whitman, Sabrina Mann, Anthea Mitchell, Matthias Hapte-Selassie, Jamin Liu, Sara Sunshine, Matthew Laurie, Genay Pilarowski, Amy Kistler, Manu Vanaerschot, Patrick Ayscue, Lucy Li, Eric Chow, IDseq Team, James Peng, Carina Marquez, Luis Rubio, Gabriel Chamie, Diane Jones, Jon Jacobo, Susana Rojas, Susy Rojas, Valerie Tulier-Laiwa, Douglas Black, Jackie Martinez, Maya Petersen, Diane Havlir, and Joseph DeRisi |
| EPI_ISL_3374556, EPI_ISL_3374557, EPI_ISL_3374558, EPI_ISL_3374559, EPI_ISL_3374560, EPI_ISL_3374561, EPI_ISL_3374562, EPI_ISL_3374563, EPI_ISL_3374564, EPI_ISL_3374565, EPI_ISL_3374566, EPI_ISL_3374567, EPI_ISL_3374568, EPI_ISL_3374569, EPI_ISL_3374570, EPI_ISL_3374571, EPI_ISL_3374572, EPI_ISL_3374573, EPI_ISL_3374574, EPI_ISL_3374575, EPI_ISL_3374576, EPI_ISL_3374577, EPI_ISL_3374578, EPI_ISL_3374579, EPI_ISL_3374580, EPI_ISL_3374581, EPI_ISL_3374582, EPI_ISL_3374583, EPI_ISL_3374584, EPI_ISL_3374585, EPI_ISL_3374586, EPI_ISL_3374587, EPI_ISL_3374588, EPI_ISL_3374589, EPI_ISL_3374590, EPI_ISL_3374591 | | | |
| see above | Dept. of Microbiology and Infection Control, Akershus University Hospital HF | Dept. of Microbiology and Infection Control, Akershus University Hospital HF | Hege Vangstein Aamot, Alexander Hesselberg Løvestad |
| EPI_ISL_3374594, EPI_ISL_3374601, EPI_ISL_3374602, EPI_ISL_3374603, EPI_ISL_3374604, EPI_ISL_3374605, EPI_ISL_3374606, EPI_ISL_3374610, EPI_ISL_3374611, EPI_ISL_3374612, EPI_ISL_3374613, EPI_ISL_3374618, EPI_ISL_3374619, EPI_ISL_3374625, EPI_ISL_3374626, EPI_ISL_3374627, EPI_ISL_3374632, EPI_ISL_3374633, EPI_ISL_3374634, EPI_ISL_3374636, EPI_ISL_3374637, EPI_ISL_3374638, EPI_ISL_3374639, EPI_ISL_3374640, EPI_ISL_3374641, EPI_ISL_3374642, EPI_ISL_3374643, EPI_ISL_3374644, EPI_ISL_3374645, EPI_ISL_3374646, EPI_ISL_3374647, EPI_ISL_3374648, EPI_ISL_3374649, EPI_ISL_3374650, EPI_ISL_3374651, EPI_ISL_3374653, EPI_ISL_3374656, EPI_ISL_3374662, EPI_ISL_3374663, EPI_ISL_3374664, EPI_ISL_3374670, EPI_ISL_3374672 | | | |
| see above | Virology Unit, Institut Pasteur du Cambodge | Virology Unit, Institut Pasteur du Cambodge | Leakhena Pum, Sophoannadedh Rath, Jurre Y Siegers, Cecile Troupin, Ly Sovann, Kraing Sidonn, Yi Sengdoeurn, Chin Savuth, Chau Darapheak, Veasna Duong, Erik A Karlsson |
| EPI_ISL_3374676, EPI_ISL_3374677 | Diagnostic and Research Center of Infectious Diseases, Medical Faculty, Andalas University | Genomik Solidaritas Indonesia Laboratorium / Diagnostic and Research Center of Infectious Diseases, Medical Faculty, Andalas University | Himawan Masyhuri, Carissa Sintca Wijaya,Andani Eka Putra, Dede Rahman Agustian, Linosefa, Vania Gavrila Wikasa, Dhahlia Agustina Cahyono, Gracia Felias Enos Korompis, Annisa Muthiah Sukirman, Anuraj Shankar, Louisa Markus, Meutia Ayuputeri Kumaheri, Ariel Pradipta |
| EPI_ISL_3374679 | Diagnostic and Research Center of Infectious Diseases, Medical Faculty, Andalas University | Genomik Solidaritas Indonesia Laboratorium / Diagnostic and Research Center of Infectious Diseases, Medical Faculty, Andalas University | Carissa Sintca Wijaya, Vania Gavrila Wikasa,Andani Eka Putra, Dede Rahman Agustian, Linosefa, Dhahlia Agustina Cahyono, Himawan Masyhuri, Gracia Felias Enos Korompis, Annisa Muthiah Sukirman, Anuraj Shankar, Louisa Markus, Meutia Ayuputeri Kumaheri, Ariel Pradipta |
| EPI_ISL_3374681 | Diagnostic and Research Center of Infectious Diseases, Medical Faculty, Andalas University | Genomik Solidaritas Indonesia Laboratorium / Diagnostic and Research Center of Infectious Diseases, Medical Faculty, Andalas University | Dhahlia Agustina Cahyono, Himawan Masyhuri,Andani Eka Putra, Dede Rahman Agustian, Linosefa, Carissa Sintca Wijaya, Vania Gavrila Wikasa, Annisa Muthiah Sukirman, Gracia Felias Enos Korompis, Anuraj Shankar, Louisa Markus, Meutia Ayuputeri Kumaheri, Ariel Pradipta |
| EPI_ISL_3374687 | Diagnostic and Research Center of Infectious Diseases, Medical Faculty, Andalas University | Genomik Solidaritas Indonesia Laboratorium / Diagnostic and Research Center of Infectious Diseases, Medical Faculty, Andalas University | Dhahlia Agustina Cahyono, Himawan Masyhuri, Andani Eka Putra, Dede Rahman Agustian, Linosefa, Carissa Sintca Wijaya, Vania Gavrila Wikasa, Annisa Muthiah Sukirman, Gracia Felias Enos Korompis, Anuraj Shankar, Louisa Markus, Meutia Ayuputeri Kumaheri, Ariel Pradipta |
| EPI_ISL_3374697 | Diagnostic and Research Center of Infectious Diseases, Medical Faculty, Andalas University | Genomik Solidaritas Indonesia Laboratorium / Diagnostic and Research Center of Infectious Diseases, Medical Faculty, Andalas University | Vania Gavrila Wikasa, Dhahlia Agustina Cahyono, Andani Eka Putra, Dede Rahman Agustian, Linosefa, Himawan Masyhuri, Carissa Sintca Wijaya, Annisa Muthiah Sukirman, Gracia Felias Enos Korompis, Anuraj Shankar, Louisa Markus, Meutia Ayuputeri Kumaheri, Ariel Pradipta |
| EPI_ISL_3374698, EPI_ISL_3374699 | Diagnostic and Research Center of Infectious Diseases, Medical Faculty, Andalas University | Genomik Solidaritas Indonesia Laboratorium / Diagnostic and Research Center of Infectious Diseases, Medical Faculty, Andalas University | Dhahlia Agustina Cahyono, Himawan Masyhuri, Andani Eka Putra, Dede Rahman Agustian, Linosefa, Carissa Sintca Wijaya, Vania Gavrila Wikasa, Annisa Muthiah Sukirman, Gracia Felias Enos Korompis, Anuraj Shankar, Louisa Markus, Meutia Ayuputeri Kumaheri, Ariel Pradipta |
| EPI_ISL_3374702 | RSUD KEBAYORAN BARU | Genomik Solidaritas Indonesia Laboratorium | Vania Gavrila Wikasa, Dhahlia Agustina Cahyono, Himawan Masyhuri, Carissa Sintca Wijaya, Annisa Muthiah Sukirman, Gracia Felias Enos Korompis, Anuraj Shankar, Louisa Markus, Meutia Ayuputeri Kumaheri, Ariel Pradipta |
| EPI_ISL_3374703 | RSUD KEBAYORAN BARU | Genomik Solidaritas Indonesia Laboratorium | Himawan Masyhuri, Carissa Sintca Wijaya, Vania Gavrila Wikasa, Dhahlia Agustina Cahyono, Gracia Felias Enos Korompis, Annisa Muthiah Sukirman, Anuraj Shankar, Louisa Markus, Meutia Ayuputeri Kumaheri, Ariel Pradipta |
| EPI_ISL_3374704 | RSU ADHYAKSA | Genomik Solidaritas Indonesia Laboratorium | Himawan Masyhuri, Carissa Sintca Wijaya, Vania Gavrila Wikasa, Dhahlia Agustina Cahyono, Gracia Felias Enos Korompis, Annisa Muthiah Sukirman, Anuraj Shankar, Louisa Markus, Meutia Ayuputeri Kumaheri, Ariel Pradipta |
| EPI_ISL_3374705 | LABKESDA DKI JAKARTA | Genomik Solidaritas Indonesia Laboratorium | Himawan Masyhuri, Carissa Sintca Wijaya, Vania Gavrila Wikasa, Dhahlia Agustina Cahyono, Gracia Felias Enos Korompis, Annisa Muthiah Sukirman, Anuraj Shankar, Louisa Markus, Meutia Ayuputeri Kumaheri, Ariel Pradipta |
| EPI_ISL_3374706, EPI_ISL_3374707, EPI_ISL_3374708 | Putrajaya Health Office | Institute for Medical Research, Infectious Disease Research Centre, National Institutes of Health, Ministry of Health Malaysia | Suppiah J, Kamel K, Mohd Zawawi Z, Azizan MA, Ramly N, Robert F, Anasir MI, Thayan R |
| EPI_ISL_3374709 | Cheras District Health Office | Institute for Medical Research, Infectious Disease Research Centre, National Institutes of Health, Ministry of Health Malaysia | Suppiah J, Kamel K, Mohd Zawawi Z, Azizan MA, Ramly N, Robert F, Anasir MI, Thayan R |
| EPI_ISL_3374710, EPI_ISL_3374711, EPI_ISL_3374712, EPI_ISL_3374713, EPI_ISL_3374714, EPI_ISL_3374715, EPI_ISL_3374716, EPI_ISL_3374717, EPI_ISL_3374718, EPI_ISL_3374719, EPI_ISL_3374720, EPI_ISL_3374721, EPI_ISL_3374723, EPI_ISL_3374724, EPI_ISL_3374725, EPI_ISL_3374726, EPI_ISL_3374727, EPI_ISL_3374728, EPI_ISL_3374729, EPI_ISL_3374730, EPI_ISL_3374731, EPI_ISL_3374732, EPI_ISL_3374733, EPI_ISL_3374734, EPI_ISL_3374735, EPI_ISL_3374736, EPI_ISL_3374737, EPI_ISL_3374738, EPI_ISL_3374739, EPI_ISL_3374740, EPI_ISL_3374741, EPI_ISL_3374742, EPI_ISL_3374743, EPI_ISL_3374744, EPI_ISL_3374745, EPI_ISL_3374746, EPI_ISL_3374747, EPI_ISL_3374748, EPI_ISL_3374749, EPI_ISL_3374751, EPI_ISL_3374752, EPI_ISL_3374753, EPI_ISL_3374754, EPI_ISL_3374755 | | | |
| see above | Hospital Universitari Vall d'Hebron - Vall d'Hebron Institut de Recerca | Hospital Universitari Vall d'Hebron - Vall d'Hebron Institut de Recerca | Cristina Andrés, Maria Piñana, Alejandra González-Sánchez, Damir Garcia-Cehic, Ariadna Rando, Juliana Esperalba, Maria Gema Codina, Carla Castillo, Karen García, Rodrigo Vásquez, Maria Carmen Martin, Tomàs Pumarola, Josep Quer, Andrés Antón |
| EPI_ISL_3374756, EPI_ISL_3374757, EPI_ISL_3374758, EPI_ISL_3374759, EPI_ISL_3374760, EPI_ISL_3374761, EPI_ISL_3374762, EPI_ISL_3374763, EPI_ISL_3374764, EPI_ISL_3374765, EPI_ISL_3374766, EPI_ISL_3374767, EPI_ISL_3374768, EPI_ISL_3374769, EPI_ISL_3374770, EPI_ISL_3374772, EPI_ISL_3374777, EPI_ISL_3374778, EPI_ISL_3374779, EPI_ISL_3374780, EPI_ISL_3374781, EPI_ISL_3374782, EPI_ISL_3374783, EPI_ISL_3374784, EPI_ISL_3374785, EPI_ISL_3374786, EPI_ISL_3374787 | | | |
| see above | Hospital Universitari Arnau de Vilanova | Hospital Universitari Vall d'Hebron - Vall d'Hebron Institut de Recerca | Cristina Andrés, Maria Piñana, Alejandra González-Sánchez, Damir Garcia-Cehic, Ariadna Rando, Juliana Esperalba, Maria Gema Codina, Carla Castillo, Karen García, Rodrigo Vásquez, Maria Carmen Martin, Tomàs Pumarola, Josep Quer, Andrés Antón |
| EPI_ISL_3374788, EPI_ISL_3374790, EPI_ISL_3374791, EPI_ISL_3374793 | Laboratory of Microbiology and Virology, Ospedale Amedeo di Savoia, ASL "Città di Torino" | Laboratory of Microbiology and Virology, Ospedale Amedeo di Savoia, ASL "Città di Torino" | Tiziano Allice, Francesco Cerutti, Maria Grazia Milia, Gabriella Gregori, Elisa Burdino, Valeria Ghisetti |
| EPI_ISL_3374801, | Laboratory of Microbiology and Virology, Ospedale Amedeo di | Laboratory of Microbiology and Virology, Ospedale Amedeo di | Francesco Cerutti, Tiziano Allice, Maria Grazia Milia, Gabriella Gregori, Elisa Burdino, Valeria Ghisetti |

| | | | |
|---|---|---|---|
| EPI_ISL_3374804, EPI_ISL_3374805, EPI_ISL_3374806, EPI_ISL_3374807, EPI_ISL_3374808, EPI_ISL_3374809, EPI_ISL_3374811, EPI_ISL_3374812 | Savoia, ASL "Città di Torino" | Savoia, ASL "Città di Torino" | |
| EPI_ISL_3374814, EPI_ISL_3374815, EPI_ISL_3374816, EPI_ISL_3374817, EPI_ISL_3374818, EPI_ISL_3374819, EPI_ISL_3374820, EPI_ISL_3374821, EPI_ISL_3374822, EPI_ISL_3374823, EPI_ISL_3374824, EPI_ISL_3374825, EPI_ISL_3374826, EPI_ISL_3374827, EPI_ISL_3374828, EPI_ISL_3374829, EPI_ISL_3374830, EPI_ISL_3374831, EPI_ISL_3374832, EPI_ISL_3374833, EPI_ISL_3374834, EPI_ISL_3374835, EPI_ISL_3374836, EPI_ISL_3374837, EPI_ISL_3374838, EPI_ISL_3374839, EPI_ISL_3374840, EPI_ISL_3374841, EPI_ISL_3374842, EPI_ISL_3374843, EPI_ISL_3374844, EPI_ISL_3374845, EPI_ISL_3374846, EPI_ISL_3374847, EPI_ISL_3374848, EPI_ISL_3374849, EPI_ISL_3374850, EPI_ISL_3374851, EPI_ISL_3374852, EPI_ISL_3374853, EPI_ISL_3374854, EPI_ISL_3374855, EPI_ISL_3374856, EPI_ISL_3374857, EPI_ISL_3374858, EPI_ISL_3374859, EPI_ISL_3374860, EPI_ISL_3374861, EPI_ISL_3374862, EPI_ISL_3374863, EPI_ISL_3374864, EPI_ISL_3374865, EPI_ISL_3374866, EPI_ISL_3374867, EPI_ISL_3374868, EPI_ISL_3374869, EPI_ISL_3374870, EPI_ISL_3374871, EPI_ISL_3374872, EPI_ISL_3374873, EPI_ISL_3374874, EPI_ISL_3374875, EPI_ISL_3374876, EPI_ISL_3374877, EPI_ISL_3374878, EPI_ISL_3374879, EPI_ISL_3374880, EPI_ISL_3374881, EPI_ISL_3374882, EPI_ISL_3374883, EPI_ISL_3374884, EPI_ISL_3374885, EPI_ISL_3374886, EPI_ISL_3374887, EPI_ISL_3374888, EPI_ISL_3374889, EPI_ISL_3374890, EPI_ISL_3374891, EPI_ISL_3374892, EPI_ISL_3374893, EPI_ISL_3374894, EPI_ISL_3374895, EPI_ISL_3374896, EPI_ISL_3374897, EPI_ISL_3374898, EPI_ISL_3374899, EPI_ISL_3374900, EPI_ISL_3374901, EPI_ISL_3374902, EPI_ISL_3374903, EPI_ISL_3374904, EPI_ISL_3374905, EPI_ISL_3374906, EPI_ISL_3374907, EPI_ISL_3374908, EPI_ISL_3374909, EPI_ISL_3374910, EPI_ISL_3374911, EPI_ISL_3374912, EPI_ISL_3374913, EPI_ISL_3374914, EPI_ISL_3374915, EPI_ISL_3374916, EPI_ISL_3374917, EPI_ISL_3374918, EPI_ISL_3374919, EPI_ISL_3374920, EPI_ISL_3374921, EPI_ISL_3374922, EPI_ISL_3374923, EPI_ISL_3374924, EPI_ISL_3374925, EPI_ISL_3374926, EPI_ISL_3374927, EPI_ISL_3374928, EPI_ISL_3374929, EPI_ISL_3374930, EPI_ISL_3374931, EPI_ISL_3374932, EPI_ISL_3374933, EPI_ISL_3374934, EPI_ISL_3374935, EPI_ISL_3374936, EPI_ISL_3374937, EPI_ISL_3374938, EPI_ISL_3374939, EPI_ISL_3374940, EPI_ISL_3374941, EPI_ISL_3374942, EPI_ISL_3374943, EPI_ISL_3374944, EPI_ISL_3374945, EPI_ISL_3374946, EPI_ISL_3374947, EPI_ISL_3374948, EPI_ISL_3374949, EPI_ISL_3374950, EPI_ISL_3374951, EPI_ISL_3374952, EPI_ISL_3374953, EPI_ISL_3374954, EPI_ISL_3374955, EPI_ISL_3374956, EPI_ISL_3374957, EPI_ISL_3374958, EPI_ISL_3374959, EPI_ISL_3374960, EPI_ISL_3374961, EPI_ISL_3374962, EPI_ISL_3374963, EPI_ISL_3374964, EPI_ISL_3374965, EPI_ISL_3374966, EPI_ISL_3374967, EPI_ISL_3374968, EPI_ISL_3374969, EPI_ISL_3374970, EPI_ISL_3374971, EPI_ISL_3374972, EPI_ISL_3374973, EPI_ISL_3374974, EPI_ISL_3374975, EPI_ISL_3374976, EPI_ISL_3374977, EPI_ISL_3374978, EPI_ISL_3374979, EPI_ISL_3374980, EPI_ISL_3374981, EPI_ISL_3374982, EPI_ISL_3374983, EPI_ISL_3374984, EPI_ISL_3374985, EPI_ISL_3374986, EPI_ISL_3374987, EPI_ISL_3374988, EPI_ISL_3374989, EPI_ISL_3374990, EPI_ISL_3374991, EPI_ISL_3374992, EPI_ISL_3374993, EPI_ISL_3374994, EPI_ISL_3374995, EPI_ISL_3374996, EPI_ISL_3374997, EPI_ISL_3374998, EPI_ISL_3374999, EPI_ISL_3375000, EPI_ISL_3375001, EPI_ISL_3375002, EPI_ISL_3375003, EPI_ISL_3375004, EPI_ISL_3375005, EPI_ISL_3375006, EPI_ISL_3375007, EPI_ISL_3375008, EPI_ISL_3375009, EPI_ISL_3375010, EPI_ISL_3375011, EPI_ISL_3375012, EPI_ISL_3375013, EPI_ISL_3375014, EPI_ISL_3375015, EPI_ISL_3375016, EPI_ISL_3375017, EPI_ISL_3375018, EPI_ISL_3375019, EPI_ISL_3375020, EPI_ISL_3375021, EPI_ISL_3375022, EPI_ISL_3375023, EPI_ISL_3375024, EPI_ISL_3375025, EPI_ISL_3375026, EPI_ISL_3375027, EPI_ISL_3375028, EPI_ISL_3375029, EPI_ISL_3375030, EPI_ISL_3375031, EPI_ISL_3375032, EPI_ISL_3375033, EPI_ISL_3375034, EPI_ISL_3375035, EPI_ISL_3375036, EPI_ISL_3375037, EPI_ISL_3375038, EPI_ISL_3375039, EPI_ISL_3375040, EPI_ISL_3375041, EPI_ISL_3375042, EPI_ISL_3375043, EPI_ISL_3375044, EPI_ISL_3375046, EPI_ISL_3375047, EPI_ISL_3375048, EPI_ISL_3375049, EPI_ISL_3375050, EPI_ISL_3375051, EPI_ISL_3375052, EPI_ISL_3375053, EPI_ISL_3375054, EPI_ISL_3375055, EPI_ISL_3375056, EPI_ISL_3375057, EPI_ISL_3375058, EPI_ISL_3375060, EPI_ISL_3375061, EPI_ISL_3375062, EPI_ISL_3375063, EPI_ISL_3375064, EPI_ISL_3375065, EPI_ISL_3375066, EPI_ISL_3375067, EPI_ISL_3375068, EPI_ISL_3375069, EPI_ISL_3375070, EPI_ISL_3375071, EPI_ISL_3375072, EPI_ISL_3375073, EPI_ISL_3375074, EPI_ISL_3375075, EPI_ISL_3375076, EPI_ISL_3375077, EPI_ISL_3375078, EPI_ISL_3375079, EPI_ISL_3375080, EPI_ISL_3375081, EPI_ISL_3375082, EPI_ISL_3375083 | | | |
| see above | National Virus Reference Laboratory | National Virus Reference Laboratory | Zoe Yandle, Charlene Bennett, Gabriel Gonzalez, Michael Carr, Jonathan Dean, Cillian F De Gascun |
| EPI_ISL_3375095, EPI_ISL_3375096, EPI_ISL_3375097, EPI_ISL_3375098, EPI_ISL_3375101, EPI_ISL_3375102, EPI_ISL_3375103, EPI_ISL_3375104, EPI_ISL_3375105, EPI_ISL_3375106, EPI_ISL_3375107, EPI_ISL_3375108 | | | |
| see above | Department of Microbiology, Yamanashi Institute for Public Health and Environment | Genome Analysis Center, Yamanashi Central Hospital | Yosuke Hirotsu |
| EPI_ISL_3375129, EPI_ISL_3375130, EPI_ISL_3375131 | Oddělení klinické mikrobiologi | Institute of Medical Microbiology and Virology, University Hospital Carl Gustav Carus, TU Dresden | Marlena Stadtmüller, Alexa Laubner, Johanna Beil, Fabian Rost, Sylvia Klemroth, Grit Mehnert, Montserrat Palau de Miquel, Susanne Reinhardt, Leo Büttner, Anett Zabzinski, Sylke Winkler, Alexander Dalpke, Eva Patrasová, Lenka Šimůnková, Lenka Dvořáková |
| EPI_ISL_3375132, EPI_ISL_3375133, EPI_ISL_3375134, EPI_ISL_3375135 | Zdravotní ústav Ústí nad Labem | Institute of Medical Microbiology and Virology, University Hospital Carl Gustav Carus, TU Dresden | Marlena Stadtmüller, Alexa Laubner, Johanna Beil, Fabian Rost, Sylvia Klemroth, Grit Mehnert, Montserrat Palau de Miquel, Susanne Reinhardt, Leo Büttner, Anett Zabzinski, Sylke Winkler, Alexander Dalpke, Romana Mikešová, Jitka Pohořská, Ivana Stiborová, Eva Patrasová, Lenka Šimůnková |
| EPI_ISL_3375136 | Pracownia Biologii Molekularnej Laboratorium Centralne SPZOZ w Krotoszynie | Wojewodzka Stacja Sanitarno-Epidemiologiczna w Lodzi, Oddzial Laboratoryjny Mikrobiologii i Parazytologii | Elian Terpo, Szymon Konkolewski, Marta Gawronska, Martyna Matera, Agnieszka Karnowska, Danuta Bartczak, Katarzyna Kosowska, Lucyna Kochanowska, Aleksandra Krawczyk |
| EPI_ISL_3375137, EPI_ISL_3375138, EPI_ISL_3375139, EPI_ISL_3375140, EPI_ISL_3375141 | Bialostockie Centrum Analiz Medycznych | Wojewodzka Stacja Sanitarno-Epidemiologiczna w Lodzi, Oddzial Laboratoryjny Mikrobiologii i Parazytologii | Elian Terpo, Szymon Konkolewski, Marta Gawronska, Martyna Matera, Agnieszka Karnowska, Danuta Bartczak, Katarzyna Kosowska, Lucyna Kochanowska, Aleksandra Krawczyk |
| EPI_ISL_3375142, EPI_ISL_3375143, EPI_ISL_3375144 | Centrum Medyczne MEDYK Sp. z o.o. Sp. k. Zaklad Diagnostyki Medycznej | Wojewodzka Stacja Sanitarno-Epidemiologiczna w Lodzi, Oddzial Laboratoryjny Mikrobiologii i Parazytologii | Elian Terpo, Szymon Konkolewski, Marta Gawronska, Martyna Matera, Agnieszka Karnowska, Danuta Bartczak, Katarzyna Kosowska, Lucyna Kochanowska, Aleksandra Krawczyk |
| EPI_ISL_3375145, EPI_ISL_3375146 | Szpital Uniwersytecki w Krakowie | Wojewodzka Stacja Sanitarno-Epidemiologiczna w Lodzi, Oddzial Laboratoryjny Mikrobiologii i Parazytologii | Elian Terpo, Szymon Konkolewski, Marta Gawronska, Martyna Matera, Agnieszka Karnowska, Danuta Bartczak, Katarzyna Kosowska, Lucyna Kochanowska, Aleksandra Krawczyk |
| EPI_ISL_3375147, EPI_ISL_3375148, EPI_ISL_3375149 | Vitomed Sp. z o.o. | Wojewodzka Stacja Sanitarno-Epidemiologiczna w Lodzi, Oddzial Laboratoryjny Mikrobiologii i Parazytologii | Elian Terpo, Szymon Konkolewski, Marta Gawronska, Martyna Matera, Agnieszka Karnowska, Danuta Bartczak, Katarzyna Kosowska, Lucyna Kochanowska, Aleksandra Krawczyk |
| EPI_ISL_3375150 | NZOZ Meditest Diagnostyka | Wojewodzka Stacja Sanitarno-Epidemiologiczna w Lodzi, Oddzial Laboratoryjny Mikrobiologii i Parazytologii | Elian Terpo, Szymon Konkolewski, Marta Gawronska, Martyna Matera, Agnieszka Karnowska, Danuta Bartczak, Katarzyna Kosowska, Lucyna Kochanowska, Aleksandra Krawczyk |
| EPI_ISL_3375151, EPI_ISL_3375152, EPI_ISL_3375153 | WSSE Gorzow Wlk. | Wojewodzka Stacja Sanitarno-Epidemiologiczna w Lodzi, Oddzial Laboratoryjny Mikrobiologii i Parazytologii | Elian Terpo, Szymon Konkolewski, Marta Gawronska, Martyna Matera, Agnieszka Karnowska, Danuta Bartczak, Katarzyna Kosowska, Lucyna Kochanowska, Aleksandra Krawczyk |
| EPI_ISL_3375154 | LAB-MED. | Wojewodzka Stacja Sanitarno-Epidemiologiczna w Lodzi, Oddzial Laboratoryjny Mikrobiologii i Parazytologii | Elian Terpo, Szymon Konkolewski, Marta Gawronska, Martyna Matera, Agnieszka Karnowska, Danuta Bartczak, Katarzyna Kosowska, Lucyna Kochanowska, Aleksandra Krawczyk |
| EPI_ISL_3375155, EPI_ISL_3375156, EPI_ISL_3375157 | Zaklad Diagnostyki Laboratoryjnej SPWSZ w Szczecinie | Wojewodzka Stacja Sanitarno-Epidemiologiczna w Lodzi, Oddzial Laboratoryjny Mikrobiologii i Parazytologii | Elian Terpo, Szymon Konkolewski, Marta Gawronska, Martyna Matera, Agnieszka Karnowska, Danuta Bartczak, Katarzyna Kosowska, Lucyna Kochanowska, Aleksandra Krawczyk |
| EPI_ISL_3375158, EPI_ISL_3375159, EPI_ISL_3375160 | Laboratorium Analiz Lekarskich ALAB. ul. Energetykow 2 Szczecin | Wojewodzka Stacja Sanitarno-Epidemiologiczna w Lodzi, Oddzial Laboratoryjny Mikrobiologii i Parazytologii | Elian Terpo, Szymon Konkolewski, Marta Gawronska, Martyna Matera, Agnieszka Karnowska, Danuta Bartczak, Katarzyna Kosowska, Lucyna Kochanowska, Aleksandra Krawczyk |
| EPI_ISL_3375161 | Centrum Medyczne MEDYK Sp. z o.o. Sp. k. Zaklad Diagnostyki Medycznej | Wojewodzka Stacja Sanitarno-Epidemiologiczna w Lodzi, Oddzial Laboratoryjny Mikrobiologii i Parazytologii | Elian Terpo, Szymon Konkolewski, Marta Gawronska, Martyna Matera, Agnieszka Karnowska, Danuta Bartczak, Katarzyna Kosowska, Lucyna Kochanowska, Aleksandra Krawczyk |
| EPI_ISL_3375162 | Laboratoria Medyczne Bruss grupa Alab sp. z o.o. Medyczne Laboratorium Diagnostyczne Pracownia Genetyki | Wojewodzka Stacja Sanitarno-Epidemiologiczna w Lodzi, Oddzial Laboratoryjny Mikrobiologii i Parazytologii | Elian Terpo, Szymon Konkolewski, Marta Gawronska, Martyna Matera, Agnieszka Karnowska, Danuta Bartczak, Katarzyna Kosowska, Lucyna Kochanowska, Aleksandra Krawczyk |
| EPI_ISL_3375163 | Zaklad Diagnostyki Laboratoryjnej SPWSZ w Szczecinie | Wojewodzka Stacja Sanitarno-Epidemiologiczna w Lodzi, Oddzial Laboratoryjny Mikrobiologii i Parazytologii | Elian Terpo, Szymon Konkolewski, Marta Gawronska, Martyna Matera, Agnieszka Karnowska, Danuta Bartczak, Katarzyna Kosowska, Lucyna Kochanowska, Aleksandra Krawczyk |
| EPI_ISL_3375167, EPI_ISL_3375168, EPI_ISL_3375169, EPI_ISL_3375170 | Department of Infectious Diseases and Immunology, National Hospital Organization Nagoya Medical Center | Clinical Research Center, National Hospital Organization Nagoya Medical Center | Yoshihiro Nakata, Hirotaka Ode, Mai Kubota, Masakazu Matsuda, Kazuhiro Matsuoka, Miho Nakasuji, Mikiko Mori, Mayumi Imahashi, Yoshiyuki Yokomaku, Yasumasa Iwatani |
| EPI_ISL_3375172, EPI_ISL_3375173 | Haukeland University Hospital, Dept. of Microbiology | Norwegian Institute of Public Health, Department of Virology | Kathrine Stene-Johansen, Kamilla Heddeland Instefjord, Hilde Elshaug, Garcia Llorente Ignacio, Jon Bråte, Engebretsen Serina Beate,Pedersen Benedikte Nevjen, Line Victoria Moen, Debech Nadia, Atiya R Ali,Marie Paulsen Madsen, Rasmus Riis Kopperud, Hilde Vollan, Karoline Bragstad, Olav Hungnes |
| EPI_ISL_3375175, EPI_ISL_3375176, EPI_ISL_3375178 | Foerde Hospital, Department of Microbiology | Norwegian Institute of Public Health, Department of Virology | Kathrine Stene-Johansen, Kamilla Heddeland Instefjord, Hilde Elshaug, Garcia Llorente Ignacio, Jon Bråte, Engebretsen Serina Beate,Pedersen Benedikte Nevjen, Line Victoria Moen, Debech Nadia, Atiya R Ali,Marie Paulsen Madsen, Rasmus Riis Kopperud, Hilde Vollan, Karoline Bragstad, Olav Hungnes |
| EPI_ISL_3375179, EPI_ISL_3375181, EPI_ISL_3375182 | Furst Medical Laboratory | Norwegian Institute of Public Health, Department of Virology | Kathrine Stene-Johansen, Kamilla Heddeland Instefjord, Hilde Elshaug, Garcia Llorente Ignacio, Jon Bråte, Engebretsen Serina Beate,Pedersen Benedikte Nevjen, Line Victoria Moen, Debech Nadia, Atiya R Ali,Marie Paulsen Madsen, Rasmus Riis Kopperud, Hilde Vollan, Karoline Bragstad, Olav Hungnes |
| EPI_ISL_3375183 | Dept. of Medical Microbiology, Stavanger University Hospital, Helse Stavanger HF | Norwegian Institute of Public Health, Department of Virology | Kathrine Stene-Johansen, Kamilla Heddeland Instefjord, Hilde Elshaug, Garcia Llorente Ignacio, Jon Bråte, Engebretsen Serina Beate,Pedersen Benedikte Nevjen, Line Victoria Moen, Debech Nadia, Atiya R Ali,Marie Paulsen Madsen, Rasmus Riis Kopperud, Hilde Vollan, Karoline Bragstad, Olav Hungnes |
| EPI_ISL_3375185, EPI_ISL_3375187 | Furst Medical Laboratory | Norwegian Institute of Public Health, Department of Virology | Kathrine Stene-Johansen, Kamilla Heddeland Instefjord, Hilde Elshaug, Garcia Llorente Ignacio, Jon Bråte, Engebretsen Serina Beate,Pedersen Benedikte Nevjen, Line Victoria Moen, Debech Nadia, Atiya R Ali,Marie Paulsen Madsen, Rasmus Riis Kopperud, Hilde Vollan, Karoline Bragstad, Olav Hungnes |
| EPI_ISL_3375188 | Foerde Hospital, Department of Microbiology | Norwegian Institute of Public Health, Department of Virology | Kathrine Stene-Johansen, Kamilla Heddeland Instefjord, Hilde Elshaug, Garcia Llorente Ignacio, Jon Bråte, Engebretsen Serina Beate,Pedersen Benedikte Nevjen, Line Victoria Moen, Debech Nadia, Atiya R Ali,Marie Paulsen Madsen, Rasmus Riis Kopperud, Hilde Vollan, Karoline Bragstad, Olav Hungnes |
| EPI_ISL_3375192, EPI_ISL_3375193, EPI_ISL_3375194 | Nordland Hospital - Bodo, Laboratory Department, Molecular Biology Unit | Norwegian Institute of Public Health, Department of Virology | Kathrine Stene-Johansen, Kamilla Heddeland Instefjord, Hilde Elshaug, Garcia Llorente Ignacio, Jon Bråte, Engebretsen Serina Beate,Pedersen Benedikte Nevjen, Line Victoria Moen, Debech Nadia, Atiya R Ali,Marie Paulsen Madsen, Rasmus Riis Kopperud, Hilde Vollan, Karoline Bragstad, Olav Hungnes |
| EPI_ISL_3375195 | Haukeland University Hospital, Dept. of Microbiology | Norwegian Institute of Public Health, Department of Virology | Kathrine Stene-Johansen, Kamilla Heddeland Instefjord, Hilde Elshaug, Garcia Llorente Ignacio, Jon Bråte, Engebretsen Serina Beate,Pedersen Benedikte Nevjen, Line Victoria Moen, Debech Nadia, Atiya R Ali,Marie Paulsen Madsen, Rasmus Riis Kopperud, Hilde Vollan, Karoline Bragstad, Olav Hungnes |
| EPI_ISL_3375196, EPI_ISL_3375197, EPI_ISL_3375198, EPI_ISL_3375199, EPI_ISL_3375200 | Furst Medical Laboratory | Norwegian Institute of Public Health, Department of Virology | Kathrine Stene-Johansen, Kamilla Heddeland Instefjord, Hilde Elshaug, Garcia Llorente Ignacio, Jon Bråte, Engebretsen Serina Beate,Pedersen Benedikte Nevjen, Line Victoria Moen, Debech Nadia, Atiya R Ali,Marie Paulsen Madsen, Rasmus Riis Kopperud, Hilde Vollan, Karoline Bragstad, Olav Hungnes |
| EPI_ISL_3375201 | Dept. of Medical Microbiology, Stavanger University Hospital, Helse Stavanger HF | Norwegian Institute of Public Health, Department of Virology | Kathrine Stene-Johansen, Kamilla Heddeland Instefjord, Hilde Elshaug, Garcia Llorente Ignacio, Jon Bråte, Engebretsen Serina Beate,Pedersen Benedikte Nevjen, Line Victoria Moen, Debech Nadia, Atiya R Ali,Marie Paulsen Madsen, Rasmus Riis Kopperud, Hilde Vollan, Karoline Bragstad, Olav Hungnes |

| | | | |
|---|---|---|---|
| EPI_ISL_3375202 | Furst Medical Laboratory | Norwegian Institute of Public Health, Department of Virology | Kathrine Stene-Johansen, Kamilla Heddeland Instefjord, Hilde Elshaug, Garcia Llorente Ignacio, Jon Bråte, Engebretsen Serina Beate,Pedersen Benedikte Nevjen, Line Victoria Moen, Debech Nadia, Atiya R Ali,Marie Paulsen Madsen, Rasmus Riis Kopperud, Hilde Vollan, Karoline Bragstad, Olav Hungnes |
| EPI_ISL_3375203 | Foerde Hospital, Department of Microbiology | Norwegian Institute of Public Health, Department of Virology | Kathrine Stene-Johansen, Kamilla Heddeland Instefjord, Hilde Elshaug, Garcia Llorente Ignacio, Jon Bråte, Engebretsen Serina Beate,Pedersen Benedikte Nevjen, Line Victoria Moen, Debech Nadia, Atiya R Ali,Marie Paulsen Madsen, Rasmus Riis Kopperud, Hilde Vollan, Karoline Bragstad, Olav Hungnes |
| EPI_ISL_3375204 | Dept. of Medical Microbiology, Stavanger University Hospital, Helse Stavanger HF | Norwegian Institute of Public Health, Department of Virology | Kathrine Stene-Johansen, Kamilla Heddeland Instefjord, Hilde Elshaug, Garcia Llorente Ignacio, Jon Bråte, Engebretsen Serina Beate,Pedersen Benedikte Nevjen, Line Victoria Moen, Debech Nadia, Atiya R Ali,Marie Paulsen Madsen, Rasmus Riis Kopperud, Hilde Vollan, Karoline Bragstad, Olav Hungnes |
| EPI_ISL_3375206 | Nordland Hospital - Bodo, Laboratory Department, Molecular Biology Unit | Norwegian Institute of Public Health, Department of Virology | Kathrine Stene-Johansen, Kamilla Heddeland Instefjord, Hilde Elshaug, Garcia Llorente Ignacio, Jon Bråte, Engebretsen Serina Beate,Pedersen Benedikte Nevjen, Line Victoria Moen, Debech Nadia, Atiya R Ali,Marie Paulsen Madsen, Rasmus Riis Kopperud, Hilde Vollan, Karoline Bragstad, Olav Hungnes |
| EPI_ISL_3375207 | University Hospital of Northern Norway, Department for Microbiology and Infectious Disease Control | Norwegian Institute of Public Health, Department of Virology | Kathrine Stene-Johansen, Kamilla Heddeland Instefjord, Hilde Elshaug, Garcia Llorente Ignacio, Jon Bråte, Engebretsen Serina Beate,Pedersen Benedikte Nevjen, Line Victoria Moen, Debech Nadia, Atiya R Ali,Marie Paulsen Madsen, Rasmus Riis Kopperud, Hilde Vollan, Karoline Bragstad, Olav Hungnes |
| EPI_ISL_3375208 | Haukeland University Hospital, Dept. of Microbiology | Norwegian Institute of Public Health, Department of Virology | Kathrine Stene-Johansen, Kamilla Heddeland Instefjord, Hilde Elshaug, Garcia Llorente Ignacio, Jon Bråte, Engebretsen Serina Beate,Pedersen Benedikte Nevjen, Line Victoria Moen, Debech Nadia, Atiya R Ali,Marie Paulsen Madsen, Rasmus Riis Kopperud, Hilde Vollan, Karoline Bragstad, Olav Hungnes |
| EPI_ISL_3375209 | Medical Microbiology Unit, Department for Laboratory Medicine, Drammen Hospital, Vestre Viken Health Trust, | Norwegian Institute of Public Health, Department of Virology | Kathrine Stene-Johansen, Kamilla Heddeland Instefjord, Hilde Elshaug, Garcia Llorente Ignacio, Jon Bråte, Engebretsen Serina Beate,Pedersen Benedikte Nevjen, Line Victoria Moen, Debech Nadia, Atiya R Ali,Marie Paulsen Madsen, Rasmus Riis Kopperud, Hilde Vollan, Karoline Bragstad, Olav Hungnes |
| EPI_ISL_3375210 | Ostfold Hospital Trust - Kalnes, Centre for Laboratory Medicine, Section for gene technology and infection serology | Norwegian Institute of Public Health, Department of Virology | Kathrine Stene-Johansen, Kamilla Heddeland Instefjord, Hilde Elshaug, Garcia Llorente Ignacio, Jon Bråte, Engebretsen Serina Beate,Pedersen Benedikte Nevjen, Line Victoria Moen, Debech Nadia, Atiya R Ali,Marie Paulsen Madsen, Rasmus Riis Kopperud, Hilde Vollan, Karoline Bragstad, Olav Hungnes |
| EPI_ISL_3375211, EPI_ISL_3375212 | Haukeland University Hospital, Dept. of Microbiology | Norwegian Institute of Public Health, Department of Virology | Kathrine Stene-Johansen, Kamilla Heddeland Instefjord, Hilde Elshaug, Garcia Llorente Ignacio, Jon Bråte, Engebretsen Serina Beate,Pedersen Benedikte Nevjen, Line Victoria Moen, Debech Nadia, Atiya R Ali,Marie Paulsen Madsen, Rasmus Riis Kopperud, Hilde Vollan, Karoline Bragstad, Olav Hungnes |
| EPI_ISL_3375213 | Furst Medical Laboratory | Norwegian Institute of Public Health, Department of Virology | Kathrine Stene-Johansen, Kamilla Heddeland Instefjord, Hilde Elshaug, Garcia Llorente Ignacio, Jon Bråte, Engebretsen Serina Beate,Pedersen Benedikte Nevjen, Line Victoria Moen, Debech Nadia, Atiya R Ali,Marie Paulsen Madsen, Rasmus Riis Kopperud, Hilde Vollan, Karoline Bragstad, Olav Hungnes |
| EPI_ISL_3375214 | Nordland Hospital - Bodo, Laboratory Department, Molecular Biology Unit | Norwegian Institute of Public Health, Department of Virology | Kathrine Stene-Johansen, Kamilla Heddeland Instefjord, Hilde Elshaug, Garcia Llorente Ignacio, Jon Bråte, Engebretsen Serina Beate,Pedersen Benedikte Nevjen, Line Victoria Moen, Debech Nadia, Atiya R Ali,Marie Paulsen Madsen, Rasmus Riis Kopperud, Hilde Vollan, Karoline Bragstad, Olav Hungnes |
| EPI_ISL_3375215 | Dept. of Medical Microbiology, Stavanger University Hospital, Helse Stavanger HF | Norwegian Institute of Public Health, Department of Virology | Kathrine Stene-Johansen, Kamilla Heddeland Instefjord, Hilde Elshaug, Garcia Llorente Ignacio, Jon Bråte, Engebretsen Serina Beate,Pedersen Benedikte Nevjen, Line Victoria Moen, Debech Nadia, Atiya R Ali,Marie Paulsen Madsen, Rasmus Riis Kopperud, Hilde Vollan, Karoline Bragstad, Olav Hungnes |
| EPI_ISL_3375216 | Foerde Hospital, Department of Microbiology | Norwegian Institute of Public Health, Department of Virology | Kathrine Stene-Johansen, Kamilla Heddeland Instefjord, Hilde Elshaug, Garcia Llorente Ignacio, Jon Bråte, Engebretsen Serina Beate,Pedersen Benedikte Nevjen, Line Victoria Moen, Debech Nadia, Atiya R Ali,Marie Paulsen Madsen, Rasmus Riis Kopperud, Hilde Vollan, Karoline Bragstad, Olav Hungnes |
| EPI_ISL_3375217 | Dept. of Medical Microbiology, Stavanger University Hospital, Helse Stavanger HF | Norwegian Institute of Public Health, Department of Virology | Kathrine Stene-Johansen, Kamilla Heddeland Instefjord, Hilde Elshaug, Garcia Llorente Ignacio, Jon Bråte, Engebretsen Serina Beate,Pedersen Benedikte Nevjen, Line Victoria Moen, Debech Nadia, Atiya R Ali,Marie Paulsen Madsen, Rasmus Riis Kopperud, Hilde Vollan, Karoline Bragstad, Olav Hungnes |
| EPI_ISL_3375218 | Foerde Hospital, Department of Microbiology | Norwegian Institute of Public Health, Department of Virology | Kathrine Stene-Johansen, Kamilla Heddeland Instefjord, Hilde Elshaug, Garcia Llorente Ignacio, Jon Bråte, Engebretsen Serina Beate,Pedersen Benedikte Nevjen, Line Victoria Moen, Debech Nadia, Atiya R Ali,Marie Paulsen Madsen, Rasmus Riis Kopperud, Hilde Vollan, Karoline Bragstad, Olav Hungnes |
| EPI_ISL_3375220 | Nordland Hospital - Bodo, Laboratory Department, Molecular Biology Unit | Norwegian Institute of Public Health, Department of Virology | Kathrine Stene-Johansen, Kamilla Heddeland Instefjord, Hilde Elshaug, Garcia Llorente Ignacio, Jon Bråte, Engebretsen Serina Beate,Pedersen Benedikte Nevjen, Line Victoria Moen, Debech Nadia, Atiya R Ali,Marie Paulsen Madsen, Rasmus Riis Kopperud, Hilde Vollan, Karoline Bragstad, Olav Hungnes |
| EPI_ISL_3375221 | Medical Microbiology Unit, Department for Laboratory Medicine, Drammen Hospital, Vestre Viken Health Trust, | Norwegian Institute of Public Health, Department of Virology | Kathrine Stene-Johansen, Kamilla Heddeland Instefjord, Hilde Elshaug, Garcia Llorente Ignacio, Jon Bråte, Engebretsen Serina Beate,Pedersen Benedikte Nevjen, Line Victoria Moen, Debech Nadia, Atiya R Ali,Marie Paulsen Madsen, Rasmus Riis Kopperud, Hilde Vollan, Karoline Bragstad, Olav Hungnes |
| EPI_ISL_3375222 | Nordland Hospital - Bodo, Laboratory Department, Molecular Biology Unit | Norwegian Institute of Public Health, Department of Virology | Kathrine Stene-Johansen, Kamilla Heddeland Instefjord, Hilde Elshaug, Garcia Llorente Ignacio, Jon Bråte, Engebretsen Serina Beate,Pedersen Benedikte Nevjen, Line Victoria Moen, Debech Nadia, Atiya R Ali,Marie Paulsen Madsen, Rasmus Riis Kopperud, Hilde Vollan, Karoline Bragstad, Olav Hungnes |
| EPI_ISL_3375223 | Dept. of Medical Microbiology, Stavanger University Hospital, Helse Stavanger HF | Norwegian Institute of Public Health, Department of Virology | Kathrine Stene-Johansen, Kamilla Heddeland Instefjord, Hilde Elshaug, Garcia Llorente Ignacio, Jon Bråte, Engebretsen Serina Beate,Pedersen Benedikte Nevjen, Line Victoria Moen, Debech Nadia, Atiya R Ali,Marie Paulsen Madsen, Rasmus Riis Kopperud, Hilde Vollan, Karoline Bragstad, Olav Hungnes |
| EPI_ISL_3375224 | Foerde Hospital, Department of Microbiology | Norwegian Institute of Public Health, Department of Virology | Kathrine Stene-Johansen, Kamilla Heddeland Instefjord, Hilde Elshaug, Garcia Llorente Ignacio, Jon Bråte, Engebretsen Serina Beate,Pedersen Benedikte Nevjen, Line Victoria Moen, Debech Nadia, Atiya R Ali,Marie Paulsen Madsen, Rasmus Riis Kopperud, Hilde Vollan, Karoline Bragstad, Olav Hungnes |
| EPI_ISL_3375225, EPI_ISL_3375226 | Nordland Hospital - Bodo, Laboratory Department, Molecular Biology Unit | Norwegian Institute of Public Health, Department of Virology | Kathrine Stene-Johansen, Kamilla Heddeland Instefjord, Hilde Elshaug, Garcia Llorente Ignacio, Jon Bråte, Engebretsen Serina Beate,Pedersen Benedikte Nevjen, Line Victoria Moen, Debech Nadia, Atiya R Ali,Marie Paulsen Madsen, Rasmus Riis Kopperud, Hilde Vollan, Karoline Bragstad, Olav Hungnes |
| EPI_ISL_3375227, EPI_ISL_3375228 | Foerde Hospital, Department of Microbiology | Norwegian Institute of Public Health, Department of Virology | Kathrine Stene-Johansen, Kamilla Heddeland Instefjord, Hilde Elshaug, Garcia Llorente Ignacio, Jon Bråte, Engebretsen Serina Beate,Pedersen Benedikte Nevjen, Line Victoria Moen, Debech Nadia, Atiya R Ali,Marie Paulsen Madsen, Rasmus Riis Kopperud, Hilde Vollan, Karoline Bragstad, Olav Hungnes |
| EPI_ISL_3375229 | Nordland Hospital - Bodo, Laboratory Department, Molecular Biology Unit | Norwegian Institute of Public Health, Department of Virology | Kathrine Stene-Johansen, Kamilla Heddeland Instefjord, Hilde Elshaug, Garcia Llorente Ignacio, Jon Bråte, Engebretsen Serina Beate,Pedersen Benedikte Nevjen, Line Victoria Moen, Debech Nadia, Atiya R Ali,Marie Paulsen Madsen, Rasmus Riis Kopperud, Hilde Vollan, Karoline Bragstad, Olav Hungnes |
| EPI_ISL_3375230 | Dept. of Medical Microbiology, Stavanger University Hospital, Helse Stavanger HF | Norwegian Institute of Public Health, Department of Virology | Kathrine Stene-Johansen, Kamilla Heddeland Instefjord, Hilde Elshaug, Garcia Llorente Ignacio, Jon Bråte, Engebretsen Serina Beate,Pedersen Benedikte Nevjen, Line Victoria Moen, Debech Nadia, Atiya R Ali,Marie Paulsen Madsen, Rasmus Riis Kopperud, Hilde Vollan, Karoline Bragstad, Olav Hungnes |
| EPI_ISL_3375231, EPI_ISL_3375232, EPI_ISL_3375233 | Nordland Hospital - Bodo, Laboratory Department, Molecular Biology Unit | Norwegian Institute of Public Health, Department of Virology | Kathrine Stene-Johansen, Kamilla Heddeland Instefjord, Hilde Elshaug, Garcia Llorente Ignacio, Jon Bråte, Engebretsen Serina Beate,Pedersen Benedikte Nevjen, Line Victoria Moen, Debech Nadia, Atiya R Ali,Marie Paulsen Madsen, Rasmus Riis Kopperud, Hilde Vollan, Karoline Bragstad, Olav Hungnes |
| EPI_ISL_3375234, EPI_ISL_3375235, EPI_ISL_3375236, EPI_ISL_3375237, EPI_ISL_3375238 | University Hospital of Northern Norway, Department for Microbiology and Infectious Disease Control | Norwegian Institute of Public Health, Department of Virology | Kathrine Stene-Johansen, Kamilla Heddeland Instefjord, Hilde Elshaug, Garcia Llorente Ignacio, Jon Bråte, Engebretsen Serina Beate,Pedersen Benedikte Nevjen, Line Victoria Moen, Debech Nadia, Atiya R Ali,Marie Paulsen Madsen, Rasmus Riis Kopperud, Hilde Vollan, Karoline Bragstad, Olav Hungnes |
| EPI_ISL_3375240, EPI_ISL_3375242, EPI_ISL_3375243 | Innlandet Hospital Trust, Division Lillehammer, Department for Medical Microbiology | Norwegian Institute of Public Health, Department of Virology | Kathrine Stene-Johansen, Kamilla Heddeland Instefjord, Hilde Elshaug, Garcia Llorente Ignacio, Jon Bråte, Engebretsen Serina Beate,Pedersen Benedikte Nevjen, Line Victoria Moen, Debech Nadia, Atiya R Ali,Marie Paulsen Madsen, Rasmus Riis Kopperud, Hilde Vollan, Karoline Bragstad, Olav Hungnes |
| EPI_ISL_3375244 | Vestfold Hospital, Toensberg Department of Microbiology | Norwegian Institute of Public Health, Department of Virology | Kathrine Stene-Johansen, Kamilla Heddeland Instefjord, Hilde Elshaug, Garcia Llorente Ignacio, Jon Bråte, Engebretsen Serina Beate,Pedersen Benedikte Nevjen, Line Victoria Moen, Debech Nadia, Atiya R Ali,Marie Paulsen Madsen, Rasmus Riis Kopperud, Hilde Vollan, Karoline Bragstad, Olav Hungnes |
| EPI_ISL_3375245, EPI_ISL_3375246, EPI_ISL_3375247 | Innlandet Hospital Trust, Division Lillehammer, Department for Medical Microbiology | Norwegian Institute of Public Health, Department of Virology | Kathrine Stene-Johansen, Kamilla Heddeland Instefjord, Hilde Elshaug, Garcia Llorente Ignacio, Jon Bråte, Engebretsen Serina Beate,Pedersen Benedikte Nevjen, Line Victoria Moen, Debech Nadia, Atiya R Ali,Marie Paulsen Madsen, Rasmus Riis Kopperud, Hilde Vollan, Karoline Bragstad, Olav Hungnes |
| EPI_ISL_3375248 | Department of Medical Microbiology, St. Olavs hospital | Norwegian Institute of Public Health, Department of Virology | Kathrine Stene-Johansen, Kamilla Heddeland Instefjord, Hilde Elshaug, Garcia Llorente Ignacio, Jon Bråte, Engebretsen Serina Beate,Pedersen Benedikte Nevjen, Line Victoria Moen, Debech Nadia, Atiya R Ali,Marie Paulsen Madsen, Rasmus Riis Kopperud, Hilde Vollan, Karoline Bragstad, Olav Hungnes |
| EPI_ISL_3375249 | Department of Medical Microbiology - section Molde, Molde Hospital | Norwegian Institute of Public Health, Department of Virology | Kathrine Stene-Johansen, Kamilla Heddeland Instefjord, Hilde Elshaug, Garcia Llorente Ignacio, Jon Bråte, Engebretsen Serina Beate,Pedersen Benedikte Nevjen, Line Victoria Moen, Debech Nadia, Atiya R Ali,Marie Paulsen Madsen, Rasmus Riis Kopperud, Hilde Vollan, Karoline Bragstad, Olav Hungnes |
| EPI_ISL_3375250 | Innlandet Hospital Trust, Division Lillehammer, Department for Medical Microbiology | Norwegian Institute of Public Health, Department of Virology | Kathrine Stene-Johansen, Kamilla Heddeland Instefjord, Hilde Elshaug, Garcia Llorente Ignacio, Jon Bråte, Engebretsen Serina Beate,Pedersen Benedikte Nevjen, Line Victoria Moen, Debech Nadia, Atiya R Ali,Marie Paulsen Madsen, Rasmus Riis Kopperud, Hilde Vollan, Karoline Bragstad, Olav Hungnes |
| EPI_ISL_3375251 | Department of Medical Microbiology, St. Olavs hospital | Norwegian Institute of Public Health, Department of Virology | Kathrine Stene-Johansen, Kamilla Heddeland Instefjord, Hilde Elshaug, Garcia Llorente Ignacio, Jon Bråte, Engebretsen Serina Beate,Pedersen Benedikte Nevjen, Line Victoria Moen, Debech Nadia, Atiya R Ali,Marie Paulsen Madsen, Rasmus Riis Kopperud, Hilde Vollan, Karoline Bragstad, Olav Hungnes |
| EPI_ISL_3375252 | University Hospital of Northern Norway, Department for Microbiology and Infectious Disease Control | Norwegian Institute of Public Health, Department of Virology | Kathrine Stene-Johansen, Kamilla Heddeland Instefjord, Hilde Elshaug, Garcia Llorente Ignacio, Jon Bråte, Engebretsen Serina Beate,Pedersen Benedikte Nevjen, Line Victoria Moen, Debech Nadia, Atiya R Ali,Marie Paulsen Madsen, Rasmus Riis Kopperud, Hilde Vollan, Karoline Bragstad, Olav Hungnes |
| EPI_ISL_3375253, EPI_ISL_3375254 | Department of Medical Microbiology, St. Olavs hospital | Norwegian Institute of Public Health, Department of Virology | Kathrine Stene-Johansen, Kamilla Heddeland Instefjord, Hilde Elshaug, Garcia Llorente Ignacio, Jon Bråte, Engebretsen Serina Beate,Pedersen Benedikte Nevjen, Line Victoria Moen, Debech Nadia, Atiya R Ali,Marie Paulsen Madsen, Rasmus Riis Kopperud, Hilde Vollan, Karoline Bragstad, Olav Hungnes |
| EPI_ISL_3375255 | Nordland Hospital - Bodo, Laboratory Department, Molecular Biology Unit | Norwegian Institute of Public Health, Department of Virology | Kathrine Stene-Johansen, Kamilla Heddeland Instefjord, Hilde Elshaug, Garcia Llorente Ignacio, Jon Bråte, Engebretsen Serina Beate,Pedersen Benedikte Nevjen, Line Victoria Moen, Debech Nadia, Atiya R Ali,Marie Paulsen Madsen, Rasmus Riis Kopperud, Hilde Vollan, Karoline Bragstad, Olav Hungnes |
| EPI_ISL_3375256 | Department of Medical Microbiology, St. Olavs hospital | Norwegian Institute of Public Health, Department of Virology | Kathrine Stene-Johansen, Kamilla Heddeland Instefjord, Hilde Elshaug, Garcia Llorente Ignacio, Jon Bråte, Engebretsen Serina Beate,Pedersen Benedikte Nevjen, Line Victoria Moen, Debech Nadia, Atiya R Ali,Marie Paulsen Madsen, Rasmus Riis Kopperud, Hilde Vollan, Karoline Bragstad, Olav Hungnes |
| EPI_ISL_3375259 | Collection of WHO National Influenza Center, N.F. Gamaleya National Research Center for Epidemiology and Microbiology of the Ministry of Health of the Russian Federation | Center for Precision Genome Editing and Genetic Technologies for Biomedicine, Pirogov Medical University Moscow Russian Federation | Yegor Botsmanov, Andrey Krivoy, Vera Belova, Svetlana Trushakova, Evgeniya Mukasheva, Kirill Krasnoslobodtsev, Alexandra Rosatkevich, Nataliya Breslav, Lyudmila Kolobukhina, Dmitriy Korostin, Anna Ignatieva, Elena Burtseva |
| EPI_ISL_3375261 | Collection of WHO National Influenza Center, N.F. Gamaleya National Research Center for Epidemiology and Microbiology of the Ministry of Health of the Russian Federation | Center for Precision Genome Editing and Genetic Technologies for Biomedicine, Pirogov Medical University Moscow Russian Federation | Yegor Botsmanov, Andrey Krivoy, Vera Belova, Svetlana Trushakova, Evgeniya Mukasheva, Kirill Krasnoslobodtsev, Alexandra Rosatkevich, Nataliya Breslav, Yuliya Levochkina, Lyudmila Kolobukhina, Dmitriy Korostin, Anna Ignatieva, Elena Burtseva |
| EPI_ISL_3375262, EPI_ISL_3375265 | Collection of WHO National Influenza Center, N.F. Gamaleya National Research Center for Epidemiology and Microbiology of the Ministry of Health of the Russian Federation | Center for Precision Genome Editing and Genetic Technologies for Biomedicine, Pirogov Medical University Moscow Russian Federation | Yegor Botsmanov, Andrey Krivoy, Vera Belova, Svetlana Trushakova, Evgeniya Mukasheva, Kirill Krasnoslobodtsev, Alexandra Rosatkevich, Nataliya Breslav, Lyudmila Kolobukhina, Dmitriy Korostin, Anna Ignatieva, Elena Burtseva |
| EPI_ISL_3375269 | Collection of WHO National Influenza Center, N.F. Gamaleya National Research Center for Epidemiology and Microbiology of the Ministry of Health of the Russian Federation | Center for Precision Genome Editing and Genetic Technologies for Biomedicine, Pirogov Medical University Moscow Russian Federation | Yegor Botsmanov, Andrey Krivoy, Vera Belova, Svetlana Trushakova, Evgeniya Mukasheva, Kirill Krasnoslobodtsev, Alexandra Rosatkevich, Nataliya Breslav, Yuliya Levochkina, Lyudmila Kolobukhina, Dmitriy Korostin, Anna Ignatieva, Elena Burtseva |

| | | | |
|---|---|---|---|
| EPI_ISL_3375297, EPI_ISL_3375298, EPI_ISL_3375299, EPI_ISL_3375301, EPI_ISL_3375302, EPI_ISL_3375304, EPI_ISL_3375305, EPI_ISL_3375307, EPI_ISL_3375308, EPI_ISL_3375310, EPI_ISL_3375311 | | | |
| see above | Dinkes Provinsi Sulawesi Tengah | National Institute of Health Research and Development | Vivi Setiawaty, Hana Apsari Pawestri, Hartanti Dian Ikawati, Subangkit, Kartika Dewi Puspa, Arie Ardiansyah Nugraha, Fajar Nur Sulistiyohadi, Triyani Soekarso, Krisna Pangesti, Nelly Puspandari |
| EPI_ISL_3375313, EPI_ISL_3375314, EPI_ISL_3375315, EPI_ISL_3375317, EPI_ISL_3375318, EPI_ISL_3375319, EPI_ISL_3375320, EPI_ISL_3375321, EPI_ISL_3375322 | Dinkes Provinsi Sulawesi Tengah | National Institute of Health Research and Development | Subangkit, Hana Apsari Pawestri, Hartanti Dian Ikawati, Kartika Dewi Puspa, Arie Ardiansyah Nugraha, Fajar Nur Sulistiyohadi, Triyani Soekarso, Krisna Pangesti, Nelly Puspandari, Vivi Setiawaty |
| EPI_ISL_3375327, EPI_ISL_3375329, EPI_ISL_3375330, EPI_ISL_3375332, EPI_ISL_3375333, EPI_ISL_3375334, EPI_ISL_3375335, EPI_ISL_3375336, EPI_ISL_3375338, EPI_ISL_3375339, EPI_ISL_3375341, EPI_ISL_3375342, EPI_ISL_3375345, EPI_ISL_3375346, EPI_ISL_3375348, EPI_ISL_3375349, EPI_ISL_3375350, EPI_ISL_3375351, EPI_ISL_3375352 | | | |
| see above | RSUP dr. M Djamil Padang | National Institute of Health Research and Development | Hana Apsari Pawestri, Hartanti Dian Ikawati, Subangkit, Kartika Dewi Puspa, Arie Ardiansyah Nugraha, Fajar Nur Sulistiyohadi, Triyani Soekarso, Krisna Pangesti, Nelly Puspandari, Deswita Sari, Vivi Setiawaty |
| EPI_ISL_3375354, EPI_ISL_3375355, EPI_ISL_3375357, EPI_ISL_3375358, EPI_ISL_3375359, EPI_ISL_3375360, EPI_ISL_3375361, EPI_ISL_3375362, EPI_ISL_3375363, EPI_ISL_3375364, EPI_ISL_3375365, EPI_ISL_3375366, EPI_ISL_3375367, EPI_ISL_3375368, EPI_ISL_3375370, EPI_ISL_3375372, EPI_ISL_3375373 | | | |
| see above | Rumah Sakit Universitas Tanjungpura | National Institute of Health Research and Development | Hartanti Dian Ikawati, Hana Apsari Pawestri, Subangkit, Kartika Dewi Puspa, Arie Ardiansyah Nugraha, Fajar Nur Sulistiyohadi, Triyani Soekarso, Krisna Pangesti, Nelly Puspandari, Andriani, Virhan Novianry, Delima Fajar Liana, Sofi Siti Shofiyah, Puji Astuti, Vivi Setiawaty |
| EPI_ISL_3375375, EPI_ISL_3375376, EPI_ISL_3375377, EPI_ISL_3375378, EPI_ISL_3375379, EPI_ISL_3375380, EPI_ISL_3375381, EPI_ISL_3375382, EPI_ISL_3375383, EPI_ISL_3375384, EPI_ISL_3375387 | | | |
| see above | Rumah Sakit Universitas Tanjungpura | National Institute of Health Research and Development | Kartika Dewi Puspa, Hana Apsari Pawestri, Subangkit, Hartanti Dian Ikawati, Arie Ardiansyah Nugraha, Fajar Nur Sulistiyohadi, Triyani Soekarso, Krisna Pangesti, Nelly Puspandari, Andriani, Virhan Novianry, Delima Fajar Liana, Sofi Siti Shofiyah, Puji Astuti Vivi Setiawaty |
| EPI_ISL_3375390, EPI_ISL_3375391, EPI_ISL_3375392, EPI_ISL_3375393, EPI_ISL_3375394, EPI_ISL_3375395, EPI_ISL_3375397, EPI_ISL_3375398, EPI_ISL_3375399 | CENTRE DE BIOLOGIE DU LANGUEDOC | CHU Purpan - Laboratoire de Virologie - Institut Fédératif de Biologie | Latour J., Jeanne N., Ranger N., Carcenac R., Harter A., Boyer P., Ferrer V., Tremeaux R., Izopet J. |
| EPI_ISL_3375400, EPI_ISL_3375401, EPI_ISL_3375402, EPI_ISL_3375403 | LABO BIO DOC | CHU Purpan - Laboratoire de Virologie - Institut Fédératif de Biologie | Latour J., Jeanne N., Ranger N., Carcenac R., Harter A., Boyer P., Ferrer V., Tremeaux R., Izopet J. |
| EPI_ISL_3375404 | CENTRE HOSPITALIER DE BIGORRE | CHU Purpan - Laboratoire de Virologie - Institut Fédératif de Biologie | Latour J., Jeanne N., Ranger N., Carcenac R., Harter A., Boyer P., Ferrer V., Tremeaux R., Izopet J. |
| EPI_ISL_3375405, EPI_ISL_3375406, EPI_ISL_3375407, EPI_ISL_3375408, EPI_ISL_3375409, EPI_ISL_3375410, EPI_ISL_3375411, EPI_ISL_3375413 | CHU Purpan - Laboratoire de Virologie - Institut Fédératif de Biologie | CHU Purpan - Laboratoire de Virologie - Institut Fédératif de Biologie | Latour J., Jeanne N., Ranger N., Carcenac R., Harter A., Boyer P., Ferrer V., Tremeaux R., Izopet J. |
| EPI_ISL_3375415, EPI_ISL_3375416, EPI_ISL_3375417, EPI_ISL_3375418, EPI_ISL_3375419, EPI_ISL_3375420, EPI_ISL_3375421, EPI_ISL_3375422 | LABO BIO DOC | CHU Purpan - Laboratoire de Virologie - Institut Fédératif de Biologie | Latour J., Jeanne N., Ranger N., Carcenac R., Harter A., Boyer P., Ferrer V., Tremeaux R., Izopet J. |
| EPI_ISL_3375426 | Hospital | National Reference Center for Viruses of Respiratory Infections, Institut Pasteur, Paris | Marion Barbet, Sylvie Behillil, Méline Bizard, Angela Brisebarre, Camille Capel, Vincent Enouf, Louise Lefrançois, Frédéric Lemoine, Christophe Malabat, Corinne Maufrais, Hub de bioinformatique et Biostatistique, Etienne Simon-Lorière, Maud Vanpeene, Sylvie Van der Werf ,Laura Djamdjian |
| EPI_ISL_3375427 | Hospital | National Reference Center for Viruses of Respiratory Infections, Institut Pasteur, Paris | Marion Barbet, Sylvie Behillil, Méline Bizard, Angela Brisebarre, Camille Capel, Vincent Enouf, Louise Lefrançois, Frédéric Lemoine, Christophe Malabat, Corinne Maufrais, Hub de bioinformatique et Biostatistique, Etienne Simon-Lorière, Maud Vanpeene, Sylvie Van der Werf ,Isabelle Gros |
| EPI_ISL_3375428, EPI_ISL_3375429 | Hospital | National Reference Center for Viruses of Respiratory Infections, Institut Pasteur, Paris | Marion Barbet, Sylvie Behillil, Méline Bizard, Angela Brisebarre, Camille Capel, Vincent Enouf, Louise Lefrançois, Frédéric Lemoine, Christophe Malabat, Corinne Maufrais, Hub de bioinformatique et Biostatistique, Etienne Simon-Lorière, Maud Vanpeene, Sylvie Van der Werf ,Eric Farfour |
| EPI_ISL_3375430 | Hospital | National Reference Center for Viruses of Respiratory Infections, Institut Pasteur, Paris | Marion Barbet, Sylvie Behillil, Méline Bizard, Angela Brisebarre, Camille Capel, Vincent Enouf, Louise Lefrançois, Frédéric Lemoine, Christophe Malabat, Corinne Maufrais, Hub de bioinformatique et Biostatistique, Etienne Simon-Lorière, Maud Vanpeene, Sylvie Van der Werf ,Benjamin Maneglie |
| EPI_ISL_3375431, EPI_ISL_3375432 | Labo Analyses Med | National Reference Center for Viruses of Respiratory Infections, Institut Pasteur, Paris | Marion Barbet, Sylvie Behillil, Méline Bizard, Angela Brisebarre, Camille Capel, Vincent Enouf, Louise Lefrançois, Frédéric Lemoine, Christophe Malabat, Corinne Maufrais, Hub de bioinformatique et Biostatistique, Etienne Simon-Lorière, Maud Vanpeene, Sylvie Van der Werf ,Nouredine Sadeg |
| EPI_ISL_3375433 | Hospital | National Reference Center for Viruses of Respiratory Infections, Institut Pasteur, Paris | Marion Barbet, Sylvie Behillil, Méline Bizard, Angela Brisebarre, Camille Capel, Vincent Enouf, Louise Lefrançois, Frédéric Lemoine, Christophe Malabat, Corinne Maufrais, Hub de bioinformatique et Biostatistique, Etienne Simon-Lorière, Maud Vanpeene, Sylvie Van der Werf ,Guy Galeazzi |
| EPI_ISL_3375434, EPI_ISL_3375435, EPI_ISL_3375436 | Hospital | National Reference Center for Viruses of Respiratory Infections, Institut Pasteur, Paris | Marion Barbet, Sylvie Behillil, Méline Bizard, Angela Brisebarre, Camille Capel, Vincent Enouf, Louise Lefrançois, Frédéric Lemoine, Christophe Malabat, Corinne Maufrais, Hub de bioinformatique et Biostatistique, Etienne Simon-Lorière, Maud Vanpeene, Sylvie Van der Werf ,Catherine FranÇOis |
| EPI_ISL_3375437, EPI_ISL_3375438, EPI_ISL_3375439 | Hospital | National Reference Center for Viruses of Respiratory Infections, Institut Pasteur, Paris | Marion Barbet, Sylvie Behillil, Méline Bizard, Angela Brisebarre, Camille Capel, Vincent Enouf, Louise Lefrançois, Frédéric Lemoine, Christophe Malabat, Corinne Maufrais, Hub de bioinformatique et Biostatistique, Etienne Simon-Lorière, Maud Vanpeene, Sylvie Van der Werf ,Guy Galeazzi |
| EPI_ISL_3375440, EPI_ISL_3375441, EPI_ISL_3375442 | Labo Analyses Med | National Reference Center for Viruses of Respiratory Infections, Institut Pasteur, Paris | Marion Barbet, Sylvie Behillil, Méline Bizard, Angela Brisebarre, Camille Capel, Vincent Enouf, Louise Lefrançois, Frédéric Lemoine, Christophe Malabat, Corinne Maufrais, Hub de bioinformatique et Biostatistique, Etienne Simon-Lorière, Maud Vanpeene, Sylvie Van der Werf ,Philippe Miara |
| EPI_ISL_3375443, EPI_ISL_3375444 | Hospital | National Reference Center for Viruses of Respiratory Infections, Institut Pasteur, Paris | Marion Barbet, Sylvie Behillil, Méline Bizard, Angela Brisebarre, Camille Capel, Vincent Enouf, Louise Lefrançois, Frédéric Lemoine, Christophe Malabat, Corinne Maufrais, Hub de bioinformatique et Biostatistique, Etienne Simon-Lorière, Maud Vanpeene, Sylvie Van der Werf ,Martine Escure |
| EPI_ISL_3375445, EPI_ISL_3375446, EPI_ISL_3375447 | Hospital | National Reference Center for Viruses of Respiratory Infections, Institut Pasteur, Paris | Marion Barbet, Sylvie Behillil, Méline Bizard, Angela Brisebarre, Camille Capel, Vincent Enouf, Louise Lefrançois, Frédéric Lemoine, Christophe Malabat, Corinne Maufrais, Hub de bioinformatique et Biostatistique, Etienne Simon-Lorière, Maud Vanpeene, Sylvie Van der Werf ,Pierre Patoz |
| EPI_ISL_3375448, EPI_ISL_3375449, EPI_ISL_3375450, EPI_ISL_3375451, EPI_ISL_3375452, EPI_ISL_3375453, EPI_ISL_3375454, EPI_ISL_3375455, EPI_ISL_3375456, EPI_ISL_3375457, EPI_ISL_3375458, EPI_ISL_3375459, EPI_ISL_3375460, EPI_ISL_3375461, EPI_ISL_3375462, EPI_ISL_3375463, EPI_ISL_3375464, EPI_ISL_3375465, EPI_ISL_3375466, EPI_ISL_3375467, EPI_ISL_3375468, EPI_ISL_3375469, EPI_ISL_3375470, EPI_ISL_3375471, EPI_ISL_3375472, EPI_ISL_3375473 | | | |
| see above | Labo Analyses Med | National Reference Center for Viruses of Respiratory Infections, Institut Pasteur, Paris | Marion Barbet, Sylvie Behillil, Méline Bizard, Angela Brisebarre, Camille Capel, Vincent Enouf, Louise Lefrançois, Frédéric Lemoine, Christophe Malabat, Corinne Maufrais, Hub de bioinformatique et Biostatistique, Etienne Simon-Lorière, Maud Vanpeene, Sylvie Van der Werf ,GréGoire Potiron |
| EPI_ISL_3375474, EPI_ISL_3375475 | Labo Analyses Med | National Reference Center for Viruses of Respiratory Infections, Institut Pasteur, Paris | Marion Barbet, Sylvie Behillil, Méline Bizard, Angela Brisebarre, Camille Capel, Vincent Enouf, Louise Lefrançois, Frédéric Lemoine, Christophe Malabat, Corinne Maufrais, Hub de bioinformatique et Biostatistique, Etienne Simon-Lorière, Maud Vanpeene, Sylvie Van der Werf ,Domitille Leman |
| EPI_ISL_3375476 | Hospital | National Reference Center for Viruses of Respiratory Infections, Institut Pasteur, Paris | Marion Barbet, Sylvie Behillil, Méline Bizard, Angela Brisebarre, Camille Capel, Vincent Enouf, Louise Lefrançois, Frédéric Lemoine, Christophe Malabat, Corinne Maufrais, Hub de bioinformatique et Biostatistique, Etienne Simon-Lorière, Maud Vanpeene, Sylvie Van der Werf ,Haciba Moudjahed |
| EPI_ISL_3375477, EPI_ISL_3375478, EPI_ISL_3375479, EPI_ISL_3375480, EPI_ISL_3375481, EPI_ISL_3375482, EPI_ISL_3375483, EPI_ISL_3375484, EPI_ISL_3375485, EPI_ISL_3375486, EPI_ISL_3375487, EPI_ISL_3375489, EPI_ISL_3375490, EPI_ISL_3375491, EPI_ISL_3375492, EPI_ISL_3375493, EPI_ISL_3375494, EPI_ISL_3375495, EPI_ISL_3375496, EPI_ISL_3375497, EPI_ISL_3375498, EPI_ISL_3375499, EPI_ISL_3375500, EPI_ISL_3375502, EPI_ISL_3375503, EPI_ISL_3375504, EPI_ISL_3375505, EPI_ISL_3375506, EPI_ISL_3375507, EPI_ISL_3375508, EPI_ISL_3375509, EPI_ISL_3375510, EPI_ISL_3375511, EPI_ISL_3375512, EPI_ISL_3375513, EPI_ISL_3375514, EPI_ISL_3375515, EPI_ISL_3375516, EPI_ISL_3375517, EPI_ISL_3375518, EPI_ISL_3375519, EPI_ISL_3375520, EPI_ISL_3375521, EPI_ISL_3375522, EPI_ISL_3375523, EPI_ISL_3375524, EPI_ISL_3375525, EPI_ISL_3375526, EPI_ISL_3375527, EPI_ISL_3375528 | | | |
| see above | Labo Analyses Med | National Reference Center for Viruses of Respiratory Infections, Institut Pasteur, Paris | Marion Barbet, Sylvie Behillil, Méline Bizard, Angela Brisebarre, Camille Capel, Vincent Enouf, Louise Lefrançois, Frédéric Lemoine, Christophe Malabat, Corinne Maufrais, Hub de bioinformatique et Biostatistique, Etienne Simon-Lorière, Maud Vanpeene, Sylvie Van der Werf ,Melanie Caron |
| EPI_ISL_3375529, EPI_ISL_3375530, EPI_ISL_3375531, EPI_ISL_3375532, EPI_ISL_3375533, EPI_ISL_3375534, EPI_ISL_3375535, EPI_ISL_3375536 | Labo Analyses Med | National Reference Center for Viruses of Respiratory Infections, Institut Pasteur, Paris | Marion Barbet, Sylvie Behillil, Méline Bizard, Angela Brisebarre, Camille Capel, Vincent Enouf, Louise Lefrançois, Frédéric Lemoine, Christophe Malabat, Corinne Maufrais, Hub de bioinformatique et Biostatistique, Etienne Simon-Lorière, Maud Vanpeene, Sylvie Van der Werf ,ValéRie Weber |

| | | | |
|---|---|---|---|
| EPI_ISL_3375537, EPI_ISL_3375538, EPI_ISL_3375539 | Labo Analyses Med | National Reference Center for Viruses of Respiratory Infections, Institut Pasteur, Paris | Marion Barbet, Sylvie Behillil, Méline Bizard, Angela Brisebarre, Camille Capel, Vincent Enouf, Louise Lefrançois, Frédéric Lemoine, Christophe Malabat, Corinne Maufrais, Hub de bioinformatique et Biostatistique, Etienne Simon-Lorière, Maud Vanpeene, Sylvie Van der Werf ,SéVerine Haouisee |
| EPI_ISL_3375540, EPI_ISL_3375541, EPI_ISL_3375543 | Labo Analyses Med | National Reference Center for Viruses of Respiratory Infections, Institut Pasteur, Paris | Marion Barbet, Sylvie Behillil, Méline Bizard, Angela Brisebarre, Camille Capel, Vincent Enouf, Louise Lefrançois, Frédéric Lemoine, Christophe Malabat, Corinne Maufrais, Hub de bioinformatique et Biostatistique, Etienne Simon-Lorière, Maud Vanpeene, Sylvie Van der Werf ,Xavier Guettier |
| EPI_ISL_3375544, EPI_ISL_3375545, EPI_ISL_3375546, EPI_ISL_3375547, EPI_ISL_3375548, EPI_ISL_3375549 | Labo Analyses Med | National Reference Center for Viruses of Respiratory Infections, Institut Pasteur, Paris | Marion Barbet, Sylvie Behillil, Méline Bizard, Angela Brisebarre, Camille Capel, Vincent Enouf, Louise Lefrançois, Frédéric Lemoine, Christophe Malabat, Corinne Maufrais, Hub de bioinformatique et Biostatistique, Etienne Simon-Lorière, Maud Vanpeene, Sylvie Van der Werf ,Beate Heym |
| EPI_ISL_3375550, EPI_ISL_3375551, EPI_ISL_3375552, EPI_ISL_3375553, EPI_ISL_3375554, EPI_ISL_3375555 | Labo Analyses Med | National Reference Center for Viruses of Respiratory Infections, Institut Pasteur, Paris | Marion Barbet, Sylvie Behillil, Méline Bizard, Angela Brisebarre, Camille Capel, Vincent Enouf, Louise Lefrançois, Frédéric Lemoine, Christophe Malabat, Corinne Maufrais, Hub de bioinformatique et Biostatistique, Etienne Simon-Lorière, Maud Vanpeene, Sylvie Van der Werf ,Eve Haguenoer |
| EPI_ISL_3375556, EPI_ISL_3375557, EPI_ISL_3375558, EPI_ISL_3375559 | Hospital | National Reference Center for Viruses of Respiratory Infections, Institut Pasteur, Paris | Marion Barbet, Sylvie Behillil, Méline Bizard, Angela Brisebarre, Camille Capel, Vincent Enouf, Louise Lefrançois, Frédéric Lemoine, Christophe Malabat, Corinne Maufrais, Hub de bioinformatique et Biostatistique, Etienne Simon-Lorière, Maud Vanpeene, Sylvie Van der Werf ,Alix Irimia |
| EPI_ISL_3375560, EPI_ISL_3375561, EPI_ISL_3375562, EPI_ISL_3375563, EPI_ISL_3375565, EPI_ISL_3375566, EPI_ISL_3375567, EPI_ISL_3375568, EPI_ISL_3375569, EPI_ISL_3375570, EPI_ISL_3375571, EPI_ISL_3375572, EPI_ISL_3375573, EPI_ISL_3375574, EPI_ISL_3375575, EPI_ISL_3375576 | | | |
| see above | Labo Analyses Med | National Reference Center for Viruses of Respiratory Infections, Institut Pasteur, Paris | Marion Barbet, Sylvie Behillil, Méline Bizard, Angela Brisebarre, Camille Capel, Vincent Enouf, Louise Lefrançois, Frédéric Lemoine, Christophe Malabat, Corinne Maufrais, Hub de bioinformatique et Biostatistique, Etienne Simon-Lorière, Maud Vanpeene, Sylvie Van der Werf ,Denis Le Minous |
| EPI_ISL_3375577, EPI_ISL_3375578 | Hospital | National Reference Center for Viruses of Respiratory Infections, Institut Pasteur, Paris | Marion Barbet, Sylvie Behillil, Méline Bizard, Angela Brisebarre, Camille Capel, Vincent Enouf, Louise Lefrançois, Frédéric Lemoine, Christophe Malabat, Corinne Maufrais, Hub de bioinformatique et Biostatistique, Etienne Simon-Lorière, Maud Vanpeene, Sylvie Van der Werf ,Julien Berthaud |
| EPI_ISL_3375579, EPI_ISL_3375580, EPI_ISL_3375581 | Hospital | National Reference Center for Viruses of Respiratory Infections, Institut Pasteur, Paris | Marion Barbet, Sylvie Behillil, Méline Bizard, Angela Brisebarre, Camille Capel, Vincent Enouf, Louise Lefrançois, Frédéric Lemoine, Christophe Malabat, Corinne Maufrais, Hub de bioinformatique et Biostatistique, Etienne Simon-Lorière, Maud Vanpeene, Sylvie Van der Werf ,Esther Gyde |
| EPI_ISL_3375582 | Hospital | National Reference Center for Viruses of Respiratory Infections, Institut Pasteur, Paris | Marion Barbet, Sylvie Behillil, Méline Bizard, Angela Brisebarre, Camille Capel, Vincent Enouf, Louise Lefrançois, Frédéric Lemoine, Christophe Malabat, Corinne Maufrais, Hub de bioinformatique et Biostatistique, Etienne Simon-Lorière, Maud Vanpeene, Sylvie Van der Werf ,Anne-CéCile Hochart |
| EPI_ISL_3375583, EPI_ISL_3375584, EPI_ISL_3375585, EPI_ISL_3375586, EPI_ISL_3375587 | Hospital | National Reference Center for Viruses of Respiratory Infections, Institut Pasteur, Paris | Marion Barbet, Sylvie Behillil, Méline Bizard, Angela Brisebarre, Camille Capel, Vincent Enouf, Louise Lefrançois, Frédéric Lemoine, Christophe Malabat, Corinne Maufrais, Hub de bioinformatique et Biostatistique, Etienne Simon-Lorière, Maud Vanpeene, Sylvie Van der Werf ,L. Courdavault |
| EPI_ISL_3375588, EPI_ISL_3375589, EPI_ISL_3375590, EPI_ISL_3375591 | Labo Analyses Med | National Reference Center for Viruses of Respiratory Infections, Institut Pasteur, Paris | Marion Barbet, Sylvie Behillil, Méline Bizard, Angela Brisebarre, Camille Capel, Vincent Enouf, Louise Lefrançois, Frédéric Lemoine, Christophe Malabat, Corinne Maufrais, Hub de bioinformatique et Biostatistique, Etienne Simon-Lorière, Maud Vanpeene, Sylvie Van der Werf |
| EPI_ISL_3375592, EPI_ISL_3375593, EPI_ISL_3375594, EPI_ISL_3375595, EPI_ISL_3375596, EPI_ISL_3375597, EPI_ISL_3375598, EPI_ISL_3375599 | Labo Analyses Med | National Reference Center for Viruses of Respiratory Infections, Institut Pasteur, Paris | Marion Barbet, Sylvie Behillil, Méline Bizard, Angela Brisebarre, Camille Capel, Vincent Enouf, Louise Lefrançois, Frédéric Lemoine, Christophe Malabat, Corinne Maufrais, Hub de bioinformatique et Biostatistique, Etienne Simon-Lorière, Maud Vanpeene, Sylvie Van der Werf ,Isabelle Aubin |
| EPI_ISL_3375600 | Labo Analyses Med | National Reference Center for Viruses of Respiratory Infections, Institut Pasteur, Paris | Marion Barbet, Sylvie Behillil, Méline Bizard, Angela Brisebarre, Camille Capel, Vincent Enouf, Louise Lefrançois, Frédéric Lemoine, Christophe Malabat, Corinne Maufrais, Hub de bioinformatique et Biostatistique, Etienne Simon-Lorière, Maud Vanpeene, Sylvie Van der Werf ,Sandrine Simon |
| EPI_ISL_3375602, EPI_ISL_3375603, EPI_ISL_3375604, EPI_ISL_3375605, EPI_ISL_3375606, EPI_ISL_3375607, EPI_ISL_3375608, EPI_ISL_3375609, EPI_ISL_3375610, EPI_ISL_3375611, EPI_ISL_3375612, EPI_ISL_3375613, EPI_ISL_3375614, EPI_ISL_3375615, EPI_ISL_3375616, EPI_ISL_3375617, EPI_ISL_3375618, EPI_ISL_3375619, EPI_ISL_3375620, EPI_ISL_3375621, EPI_ISL_3375622, EPI_ISL_3375623 | | | |
| see above | Kansas Health and Environmental Lab | Kansas Health and Environmental Lab | Mike Grose, Jonathan Barnell, Amanda Bradley, and Phil Adam |
| EPI_ISL_3375624, EPI_ISL_3375625, EPI_ISL_3375626, EPI_ISL_3375627, EPI_ISL_3375628, EPI_ISL_3375629, EPI_ISL_3375630, EPI_ISL_3375631, EPI_ISL_3375632, EPI_ISL_3375633, EPI_ISL_3375634 | | | |
| see above | NIC, Viral Respiratory Unit | Virology Departement | Fayez Ahmed Khardine, Amel Benyahia, Mohamed Amine Beloufa, Aissam Hachid, Aicha Bensalem, Fawzi Derrar |
| EPI_ISL_3375635 | Vitomed Sp. z o.o. | Wojewodzka Stacja Sanitarno-Epidemiologiczna w Lodzi, Oddzial Laboratoryjny Mikrobiologii i Parazytologii | Elian Terpo, Szymon Konkolewski, Marta Gawronska, Martyna Matera, Agnieszka Karnowska, Danuta Bartczak, Katarzyna Kosowska, Lucyna Kochanowska, Aleksandra Krawczyk |
| EPI_ISL_3375636, EPI_ISL_3375637, EPI_ISL_3375639, EPI_ISL_3375641 | DC Public Health Lab/ Dept. of Forensic Sciences | DC Public Health Lab/ Dept. of Forensic Sciences | Janis Doss, Scott Nguyen, Elizabeth Zelaya, Sarah Scott, Connie Maza, Monica Mann, Brittany Hamilton, David Payne, Jocelyn Hauser |
| EPI_ISL_3375662, EPI_ISL_3375663, EPI_ISL_3375664, EPI_ISL_3375665, EPI_ISL_3375666, EPI_ISL_3375667, EPI_ISL_3375668, EPI_ISL_3375669, EPI_ISL_3375670, EPI_ISL_3375671, EPI_ISL_3375672, EPI_ISL_3375673, EPI_ISL_3375674, EPI_ISL_3375675, EPI_ISL_3375676, EPI_ISL_3375677, EPI_ISL_3375678, EPI_ISL_3375679, EPI_ISL_3375680, EPI_ISL_3375681, EPI_ISL_3375682, EPI_ISL_3375683, EPI_ISL_3375684, EPI_ISL_3375685, EPI_ISL_3375686, EPI_ISL_3375687, EPI_ISL_3375688, EPI_ISL_3375689, EPI_ISL_3375690, EPI_ISL_3375691 | | | |
| see above | Kansas Health and Environmental Lab | Kansas Health and Environmental Lab | Mike Grose, Jonathan Barnell, Amanda Bradley, and Phil Adam |
| EPI_ISL_3375693 | Department of Public Health Bucharest | National Institute of Infectious Diseases-Prof. Dr. Matei Bals Molecular Diagnostics Laboratory | Corina Casangiu, Leontina Banica, Marius Surleac, Ovidiu Vlaicu, Andreea Tudor, Simona Paraschiv, Dan Otelea |
| EPI_ISL_3375694, EPI_ISL_3375695, EPI_ISL_3375696 | National Institute of Infectious Diseases-Prof. Dr. Matei Bals Molecular Diagnostics Laboratory | National Institute of Infectious Diseases-Prof. Dr. Matei Bals Molecular Diagnostics Laboratory | Corina Casangiu, Leontina Banica, Marius Surleac, Ovidiu Vlaicu, Andreea Tudor, Simona Paraschiv, Dan Otelea |
| EPI_ISL_3375702, EPI_ISL_3375703, EPI_ISL_3375704, EPI_ISL_3375705, EPI_ISL_3375706, EPI_ISL_3375707, EPI_ISL_3375708, EPI_ISL_3375709, EPI_ISL_3375710, EPI_ISL_3375711, EPI_ISL_3375712, EPI_ISL_3375713, EPI_ISL_3375714, EPI_ISL_3375715, EPI_ISL_3375716, EPI_ISL_3375717, EPI_ISL_3375718, EPI_ISL_3375719, EPI_ISL_3375720, EPI_ISL_3375721, EPI_ISL_3375722, EPI_ISL_3375723, EPI_ISL_3375724, EPI_ISL_3375725, EPI_ISL_3375726, EPI_ISL_3375728, EPI_ISL_3375729, EPI_ISL_3375731, EPI_ISL_3375732, EPI_ISL_3375733, EPI_ISL_3375734, EPI_ISL_3375735, EPI_ISL_3375736, EPI_ISL_3375737, EPI_ISL_3375738, EPI_ISL_3375739, EPI_ISL_3375740, EPI_ISL_3375741, EPI_ISL_3375742, EPI_ISL_3375743, EPI_ISL_3375744, EPI_ISL_3375745, EPI_ISL_3375746, EPI_ISL_3375747, EPI_ISL_3375748, EPI_ISL_3375749, EPI_ISL_3375750, EPI_ISL_3375751, EPI_ISL_3375752, EPI_ISL_3375753, EPI_ISL_3375754, EPI_ISL_3375755, EPI_ISL_3375756, EPI_ISL_3375757, EPI_ISL_3375758, EPI_ISL_3375759, EPI_ISL_3375760, EPI_ISL_3375761, EPI_ISL_3375762, EPI_ISL_3375763, EPI_ISL_3375764, EPI_ISL_3375765, EPI_ISL_3375766, EPI_ISL_3375767, EPI_ISL_3375768, EPI_ISL_3375769, EPI_ISL_3375770, EPI_ISL_3375771, EPI_ISL_3375772, EPI_ISL_3375773, EPI_ISL_3375774, EPI_ISL_3375775, EPI_ISL_3375776, EPI_ISL_3375777, EPI_ISL_3375778, EPI_ISL_3375779, EPI_ISL_3375780, EPI_ISL_3375781, EPI_ISL_3375783, EPI_ISL_3375784, EPI_ISL_3375785, EPI_ISL_3375786, EPI_ISL_3375787, EPI_ISL_3375788, EPI_ISL_3375789, EPI_ISL_3375790, EPI_ISL_3375791, EPI_ISL_3375792, EPI_ISL_3375793, EPI_ISL_3375794, EPI_ISL_3375795, EPI_ISL_3375796, EPI_ISL_3375797, EPI_ISL_3375799, EPI_ISL_3375800 | | | |
| see above | Laboratorio CQRC | CQRC_QUALITY CONTROL CHEMICAL BIOLOGICAL RISK_AOOR Villa Sofia Cervello Palermo | Di Gaudio F., Contino F., Brunacci G. |
| EPI_ISL_3375801, EPI_ISL_3375803, EPI_ISL_3375804, EPI_ISL_3375805, EPI_ISL_3375807, EPI_ISL_3375808, EPI_ISL_3375810, EPI_ISL_3375811, EPI_ISL_3375812, EPI_ISL_3375813, EPI_ISL_3375814, EPI_ISL_3375815, EPI_ISL_3375816, EPI_ISL_3375817, EPI_ISL_3375818, EPI_ISL_3375820, EPI_ISL_3375821, EPI_ISL_3375822, EPI_ISL_3375824, EPI_ISL_3375825, EPI_ISL_3375826, EPI_ISL_3375827, EPI_ISL_3375828, EPI_ISL_3375829, EPI_ISL_3375830, EPI_ISL_3375831, EPI_ISL_3375832, EPI_ISL_3375833, EPI_ISL_3375834, EPI_ISL_3375835, EPI_ISL_3375836, EPI_ISL_3375837, EPI_ISL_3375838, EPI_ISL_3375840, EPI_ISL_3375841, EPI_ISL_3375842, EPI_ISL_3375843, EPI_ISL_3375844, EPI_ISL_3375845, EPI_ISL_3375847, EPI_ISL_3375848, EPI_ISL_3375850, EPI_ISL_3375851, EPI_ISL_3375852 | | | |
| see above | US Air Force School of Aerospace Medicine | US Air Force School of Aerospace Medicine | Anthony Fries, Jennifer Meyer, William Gruner, Amanda Javorina, Carol Garrett, Sarah Purves, Clarise Starr, Elizabeth Macias |
| EPI_ISL_3375853, EPI_ISL_3375854 | US Air Force School of Aerospace Medicine | US Air Force School of Aerospace Medicine | Anthony Fries, Jennifer Meyer, William Gruner, Amanda Javorina, Carol Garrett, Sarah Purves, Clarise Starr, Elizabeth Macias, Fritz Castillo, Jennifer Black, Cole Anderson |
| EPI_ISL_3376055, EPI_ISL_3376057, EPI_ISL_3376059, EPI_ISL_3376064, EPI_ISL_3376066, EPI_ISL_3376072, EPI_ISL_3376073, EPI_ISL_3376077, EPI_ISL_3376082, EPI_ISL_3376083, EPI_ISL_3376085, EPI_ISL_3376087, EPI_ISL_3376089, EPI_ISL_3376094, EPI_ISL_3376101, EPI_ISL_3376106, EPI_ISL_3376109, EPI_ISL_3376110, EPI_ISL_3376114, EPI_ISL_3376115, EPI_ISL_3376118, EPI_ISL_3376120, EPI_ISL_3376124, EPI_ISL_3376128, EPI_ISL_3376131, EPI_ISL_3376135, EPI_ISL_3376137, EPI_ISL_3376139, EPI_ISL_3376143, EPI_ISL_3376144, EPI_ISL_3376414, EPI_ISL_3376415, EPI_ISL_3376416, EPI_ISL_3376417, EPI_ISL_3376418, EPI_ISL_3376419, EPI_ISL_3376420, EPI_ISL_3376421, EPI_ISL_3376423, EPI_ISL_3376424, EPI_ISL_3376426, EPI_ISL_3376427, EPI_ISL_3376428, EPI_ISL_3376430, EPI_ISL_3376431, EPI_ISL_3376432, EPI_ISL_3376433, EPI_ISL_3376434, EPI_ISL_3376435, EPI_ISL_3376436, EPI_ISL_3376437, EPI_ISL_3376438, EPI_ISL_3376439, EPI_ISL_3376441, EPI_ISL_3376442, EPI_ISL_3376443, EPI_ISL_3376444, EPI_ISL_3376445, EPI_ISL_3376446, EPI_ISL_3376447, EPI_ISL_3376448, EPI_ISL_3376449, EPI_ISL_3376450, EPI_ISL_3376451, EPI_ISL_3376452, EPI_ISL_3376453, EPI_ISL_3376454, EPI_ISL_3376457, EPI_ISL_3376458, EPI_ISL_3376459, EPI_ISL_3376460, EPI_ISL_3376461, EPI_ISL_3376462, EPI_ISL_3376463, EPI_ISL_3376464, EPI_ISL_3376465, EPI_ISL_3376466, EPI_ISL_3376467, EPI_ISL_3376468, EPI_ISL_3376469, EPI_ISL_3376470, EPI_ISL_3376471, EPI_ISL_3376472, EPI_ISL_3376473, EPI_ISL_3376474, EPI_ISL_3376476, EPI_ISL_3376477, EPI_ISL_3376478, EPI_ISL_3376479, EPI_ISL_3376480, EPI_ISL_3376481, EPI_ISL_3376482, EPI_ISL_3376483, EPI_ISL_3376484, EPI_ISL_3376485, EPI_ISL_3376486, EPI_ISL_3376487, EPI_ISL_3376488, EPI_ISL_3376489, EPI_ISL_3376490, EPI_ISL_3376491, EPI_ISL_3376492, EPI_ISL_3376493, EPI_ISL_3376494, EPI_ISL_3376495, EPI_ISL_3376496, EPI_ISL_3376497, EPI_ISL_3376499, EPI_ISL_3376501, EPI_ISL_3376502, EPI_ISL_3376503, EPI_ISL_3376504, EPI_ISL_3376505, EPI_ISL_3376506, EPI_ISL_3376507, EPI_ISL_3376508, EPI_ISL_3376509, EPI_ISL_3376510, EPI_ISL_3376511, EPI_ISL_3376512, EPI_ISL_3376517, EPI_ISL_3376518, EPI_ISL_3376520, EPI_ISL_3376522, EPI_ISL_3376523, EPI_ISL_3376524, EPI_ISL_3376525, EPI_ISL_3376527, EPI_ISL_3376528, EPI_ISL_3376529, EPI_ISL_3376530, EPI_ISL_3376531, EPI_ISL_3376533, EPI_ISL_3376534, EPI_ISL_3376535, EPI_ISL_3376537, EPI_ISL_3376538, EPI_ISL_3376539, EPI_ISL_3376540, EPI_ISL_3376542, EPI_ISL_3376543, EPI_ISL_3376544, EPI_ISL_3376545, EPI_ISL_3376546, EPI_ISL_3376548, EPI_ISL_3376549, EPI_ISL_3376550, EPI_ISL_3376552, EPI_ISL_3376553, EPI_ISL_3376555, EPI_ISL_3376556, EPI_ISL_3376557, EPI_ISL_3376558, EPI_ISL_3376559, EPI_ISL_3376560, EPI_ISL_3376562, EPI_ISL_3376563, EPI_ISL_3376564, EPI_ISL_3376565, EPI_ISL_3376568, EPI_ISL_3376570, EPI_ISL_3376572, EPI_ISL_3376573, EPI_ISL_3376574, EPI_ISL_3376578 | | | |
| see above | Laboratorio CQRC | CQRC_QUALITY CONTROL CHEMICAL BIOLOGICAL RISK_AOOR Villa Sofia Cervello Palermo | Di Gaudio F., Contino F., Brunacci G. |
| EPI_ISL_3376581, EPI_ISL_3376583, EPI_ISL_3376584, EPI_ISL_3376585, EPI_ISL_3376587, EPI_ISL_3376588, EPI_ISL_3376589, EPI_ISL_3376590, EPI_ISL_3376591, EPI_ISL_3376592, EPI_ISL_3376593, EPI_ISL_3376594, EPI_ISL_3376595, EPI_ISL_3376597, EPI_ISL_3376598 | | | |
| see above | A.S.L. CUNEO 2 | Fondazione del Piemonte per l'Oncologia IRCCS | Antonino Sottile, Silvia Brossa, Paola Marino, Giorgio Giardina |
| EPI_ISL_3376599, EPI_ISL_3376600, EPI_ISL_3376601, EPI_ISL_3376602, EPI_ISL_3376603, EPI_ISL_3376604, EPI_ISL_3376605, EPI_ISL_3376606, EPI_ISL_3376607, EPI_ISL_3376610, EPI_ISL_3376612, EPI_ISL_3376613, EPI_ISL_3376615, EPI_ISL_3376616, EPI_ISL_3376617, EPI_ISL_3376618, EPI_ISL_3376619, EPI_ISL_3376620, EPI_ISL_3376621, EPI_ISL_3376622, EPI_ISL_3376623, EPI_ISL_3376624, EPI_ISL_3376625, EPI_ISL_3376626, EPI_ISL_3376627, EPI_ISL_3376628, EPI_ISL_3376629, EPI_ISL_3376630, EPI_ISL_3376631, EPI_ISL_3376632, EPI_ISL_3376633, EPI_ISL_3376634, EPI_ISL_3376635, EPI_ISL_3376637, EPI_ISL_3376638, EPI_ISL_3376639, EPI_ISL_3376640, EPI_ISL_3376641, EPI_ISL_3376642, EPI_ISL_3376644, EPI_ISL_3376645 | | | |

| | | | |
|---|---|---|---|
| see above | A.S.L. NOVARA | Fondazione del Piemonte per l'Oncologia IRCCS | Antonino Sottile, Silvia Brossa, Paola Marino, Giorgio Giardina |
| EPI_ISL_3376646 | A.S.L. CUNEO 1 | Fondazione del Piemonte per l'Oncologia IRCCS | Antonino Sottile, Silvia Brossa, Paola Marino, Giorgio Giardina |
| EPI_ISL_3376647 | A.S.L. NOVARA | Fondazione del Piemonte per l'Oncologia IRCCS | Antonino Sottile, Silvia Brossa, Paola Marino, Giorgio Giardina |
| EPI_ISL_3376648, EPI_ISL_3376649, EPI_ISL_3376650, EPI_ISL_3376651, EPI_ISL_3376652, EPI_ISL_3376654, EPI_ISL_3376655, EPI_ISL_3376656, EPI_ISL_3376657, EPI_ISL_3376658, EPI_ISL_3376659, EPI_ISL_3376660, EPI_ISL_3376661, EPI_ISL_3376662, EPI_ISL_3376663, EPI_ISL_3376664, EPI_ISL_3376665 | | | |
| see above | A.S.L. CUNEO 1 | Fondazione del Piemonte per l'Oncologia IRCCS | Antonino Sottile, Silvia Brossa, Paola Marino, Giorgio Giardina |
| EPI_ISL_3376666, EPI_ISL_3376667, EPI_ISL_3376668, EPI_ISL_3376669, EPI_ISL_3376671, EPI_ISL_3376673, EPI_ISL_3376674, EPI_ISL_3376675 | A.S.L. VERCELLI | Fondazione del Piemonte per l'Oncologia IRCCS | Antonino Sottile, Silvia Brossa, Paola Marino, Giorgio Giardina |
| EPI_ISL_3376677 | A.S.L. TO4 | Fondazione del Piemonte per l'Oncologia IRCCS | Antonino Sottile, Silvia Brossa, Paola Marino, Giorgio Giardina |
| EPI_ISL_3376732, EPI_ISL_3376733, EPI_ISL_3376734, EPI_ISL_3376735, EPI_ISL_3376736, EPI_ISL_3376737, EPI_ISL_3376738, EPI_ISL_3376739, EPI_ISL_3376740, EPI_ISL_3376743, EPI_ISL_3376744, EPI_ISL_3376745, EPI_ISL_3376746, EPI_ISL_3376747, EPI_ISL_3376748, EPI_ISL_3376749, EPI_ISL_3376750, EPI_ISL_3376751, EPI_ISL_3376752, EPI_ISL_3376754, EPI_ISL_3376755, EPI_ISL_3376756, EPI_ISL_3376757, EPI_ISL_3376758, EPI_ISL_3376759, EPI_ISL_3376760, EPI_ISL_3376761, EPI_ISL_3376762, EPI_ISL_3376763, EPI_ISL_3376765, EPI_ISL_3376766, EPI_ISL_3376767, EPI_ISL_3376768, EPI_ISL_3376769, EPI_ISL_3376770, EPI_ISL_3376771, EPI_ISL_3376772, EPI_ISL_3376773, EPI_ISL_3376774, EPI_ISL_3376775, EPI_ISL_3376776, EPI_ISL_3376777, EPI_ISL_3376778, EPI_ISL_3376779, EPI_ISL_3376781, EPI_ISL_3376782, EPI_ISL_3376787, EPI_ISL_3376788, EPI_ISL_3376789, EPI_ISL_3376790, EPI_ISL_3376795, EPI_ISL_3376799, EPI_ISL_3376805, EPI_ISL_3376806, EPI_ISL_3376807, EPI_ISL_3376810, EPI_ISL_3376811, EPI_ISL_3376812, EPI_ISL_3376813, EPI_ISL_3376814, EPI_ISL_3376815, EPI_ISL_3376816, EPI_ISL_3376817, EPI_ISL_3376819, EPI_ISL_3376821, EPI_ISL_3376822, EPI_ISL_3376824, EPI_ISL_3376825, EPI_ISL_3376826, EPI_ISL_3376827, EPI_ISL_3376828, EPI_ISL_3376829, EPI_ISL_3376832, EPI_ISL_3376835, EPI_ISL_3376836, EPI_ISL_3376837, EPI_ISL_3376838, EPI_ISL_3376839, EPI_ISL_3376840, EPI_ISL_3376853, EPI_ISL_3376854, EPI_ISL_3376855, EPI_ISL_3376856, EPI_ISL_3376857, EPI_ISL_3376858, EPI_ISL_3376859, EPI_ISL_3376860, EPI_ISL_3376862, EPI_ISL_3376864, EPI_ISL_3376865, EPI_ISL_3376866, EPI_ISL_3376867, EPI_ISL_3376868, EPI_ISL_3376869, EPI_ISL_3376871, EPI_ISL_3376872, EPI_ISL_3376873, EPI_ISL_3376874, EPI_ISL_3376875, EPI_ISL_3376876, EPI_ISL_3376877, EPI_ISL_3376878, EPI_ISL_3376879, EPI_ISL_3376880, EPI_ISL_3376881, EPI_ISL_3376882, EPI_ISL_3376883 | | | |
| see above | Laboratorio CQRC | CQRC_QUALITY CONTROL CHEMICAL BIOLOGICAL RISK_AOOR Villa Sofia Cervello Palermo | Di Gaudio F., Contino F., Brunacci G. |
| EPI_ISL_3376887, EPI_ISL_3376888, EPI_ISL_3376890, EPI_ISL_3376891 | A.S.L. ASTI | Fondazione del Piemonte per l'Oncologia IRCCS | Antonino Sottile, Silvia Brossa, Paola Marino, Giorgio Giardina |
| EPI_ISL_3376892, EPI_ISL_3376893, EPI_ISL_3376894 | A.S.L. CITTA' DI TORINO - OSPEDALE MAURIZIANO | Fondazione del Piemonte per l'Oncologia IRCCS | Antonino Sottile, Silvia Brossa, Paola Marino, Giorgio Giardina |
| EPI_ISL_3376895, EPI_ISL_3376896, EPI_ISL_3376897, EPI_ISL_3376898, EPI_ISL_3376899, EPI_ISL_3376900, EPI_ISL_3376901, EPI_ISL_3376902, EPI_ISL_3376903, EPI_ISL_3376904, EPI_ISL_3376905, EPI_ISL_3376906, EPI_ISL_3376907, EPI_ISL_3376908, EPI_ISL_3376909, EPI_ISL_3376910 | | | |
| see above | A.S.L. CUNEO 1 | Fondazione del Piemonte per l'Oncologia IRCCS | Antonino Sottile, Silvia Brossa, Paola Marino, Giorgio Giardina |
| EPI_ISL_3376911 | A.S.L. ALESSANDRIA | Fondazione del Piemonte per l'Oncologia IRCCS | Antonino Sottile, Silvia Brossa, Paola Marino, Giorgio Giardina |
| EPI_ISL_3376912, EPI_ISL_3376913, EPI_ISL_3376914 | A.S.L. NOVARA | Fondazione del Piemonte per l'Oncologia IRCCS | Antonino Sottile, Silvia Brossa, Paola Marino, Giorgio Giardina |
| EPI_ISL_3376915 | A.S.L. CUNEO 1 | Fondazione del Piemonte per l'Oncologia IRCCS | Antonino Sottile, Silvia Brossa, Paola Marino, Giorgio Giardina |
| EPI_ISL_3376916, EPI_ISL_3376917, EPI_ISL_3376918, EPI_ISL_3376919, EPI_ISL_3376920, EPI_ISL_3376921, EPI_ISL_3376922, EPI_ISL_3376923, EPI_ISL_3376924, EPI_ISL_3376925, EPI_ISL_3376926, EPI_ISL_3376927, EPI_ISL_3376928, EPI_ISL_3376929, EPI_ISL_3376930, EPI_ISL_3376931, EPI_ISL_3376932, EPI_ISL_3376933, EPI_ISL_3376934, EPI_ISL_3376935, EPI_ISL_3376936, EPI_ISL_3376937, EPI_ISL_3376938, EPI_ISL_3376939, EPI_ISL_3376940, EPI_ISL_3376941, EPI_ISL_3376942, EPI_ISL_3376943, EPI_ISL_3376944, EPI_ISL_3376945, EPI_ISL_3376946, EPI_ISL_3376947, EPI_ISL_3376948 | | | |
| see above | A.S.L. NOVARA | Fondazione del Piemonte per l'Oncologia IRCCS | Antonino Sottile, Silvia Brossa, Paola Marino, Giorgio Giardina |
| EPI_ISL_3376949 | A.S.L. CUNEO 1 | Fondazione del Piemonte per l'Oncologia IRCCS | Antonino Sottile, Silvia Brossa, Paola Marino, Giorgio Giardina |
| EPI_ISL_3376951, EPI_ISL_3376952, EPI_ISL_3376953 | A.S.L. NOVARA | Fondazione del Piemonte per l'Oncologia IRCCS | Antonino Sottile, Silvia Brossa, Paola Marino, Giorgio Giardina |
| EPI_ISL_3376954, EPI_ISL_3376955, EPI_ISL_3376956, EPI_ISL_3376957, EPI_ISL_3376958, EPI_ISL_3376959, EPI_ISL_3376960, EPI_ISL_3376961, EPI_ISL_3376962, EPI_ISL_3376963, EPI_ISL_3376964, EPI_ISL_3376965, EPI_ISL_3376966, EPI_ISL_3376967, EPI_ISL_3376968, EPI_ISL_3376969, EPI_ISL_3376970, EPI_ISL_3376971, EPI_ISL_3376972, EPI_ISL_3376973, EPI_ISL_3376974, EPI_ISL_3376975, EPI_ISL_3376976, EPI_ISL_3376977, EPI_ISL_3376978, EPI_ISL_3376979, EPI_ISL_3376980 | | | |
| see above | A.S.L. CUNEO 1 | Fondazione del Piemonte per l'Oncologia IRCCS | Antonino Sottile, Silvia Brossa, Paola Marino, Giorgio Giardina |
| EPI_ISL_3376981, EPI_ISL_3376982, EPI_ISL_3376984 | A.S.L. NOVARA | Fondazione del Piemonte per l'Oncologia IRCCS | Antonino Sottile, Silvia Brossa, Paola Marino, Giorgio Giardina |
| EPI_ISL_3376997, EPI_ISL_3376998 | Lighthouse Lab in Alderley Park | Wellcome Sanger Institute for the COVID-19 Genomics UK (COG-UK) Consortium | Jacquelyn Wynn, Mairead Hyland, The Lighthouse Lab in Alderley Park and Alex Alderton, Roberto Amato, Jeffrey Barrett, Sonia Goncalves, Ewan Harrison, David K. Jackson, Ian Johnston, Dominic Kwiatkowski, Cordelia Langford, John Sillitoe on behalf of the Wellcome Sanger Institute COVID-19 Surveillance Team |
| EPI_ISL_3376999 | Berkshire and Surrey Pathology Services Lighthouse Laboratory | Wellcome Sanger Institute for the COVID-19 Genomics UK (COG-UK) Consortium | Berkshire and Surrey Pathology Services Lighthouse Laboratory and Alex Alderton, Roberto Amato, Jeffrey Barrett, Sonia Goncalves, Ewan Harrison, David K. Jackson, Ian Johnston, Dominic Kwiatkowski, Cordelia Langford, John Sillitoe on behalf of the Wellcome Sanger Institute COVID-19 Surveillance Team |
| EPI_ISL_3377000 | Lighthouse Lab in Alderley Park | Wellcome Sanger Institute for the COVID-19 Genomics UK (COG-UK) Consortium | Jacquelyn Wynn, Mairead Hyland, The Lighthouse Lab in Alderley Park and Alex Alderton, Roberto Amato, Jeffrey Barrett, Sonia Goncalves, Ewan Harrison, David K. Jackson, Ian Johnston, Dominic Kwiatkowski, Cordelia Langford, John Sillitoe on behalf of the Wellcome Sanger Institute COVID-19 Surveillance Team |
| EPI_ISL_3377001 | Berkshire and Surrey Pathology Services Lighthouse Laboratory | Wellcome Sanger Institute for the COVID-19 Genomics UK (COG-UK) Consortium | Berkshire and Surrey Pathology Services Lighthouse Laboratory and Alex Alderton, Roberto Amato, Jeffrey Barrett, Sonia Goncalves, Ewan Harrison, David K. Jackson, Ian Johnston, Dominic Kwiatkowski, Cordelia Langford, John Sillitoe on behalf of the Wellcome Sanger Institute COVID-19 Surveillance Team |
| EPI_ISL_3377002 | Lighthouse Lab in Milton Keynes | Wellcome Sanger Institute for the COVID-19 Genomics UK (COG-UK) Consortium | The Lighthouse Lab in Milton Keynes and Alex Alderton, Roberto Amato, Jeffrey Barrett, Sonia Goncalves, Ewan Harrison, David K. Jackson, Ian Johnston, Dominic Kwiatkowski, Cordelia Langford, John Sillitoe on behalf of the Wellcome Sanger Institute COVID-19 Surveillance Team |
| EPI_ISL_3377003 | Lighthouse Lab in Glasgow | Wellcome Sanger Institute for the COVID-19 Genomics UK (COG-UK) Consortium | Harper VanSteenhouse, Yumi Kasai, David Gray, Carol Clugston, Anna Dominiczak and Alex Alderton, Roberto Amato, Jeffrey Barrett, Sonia Goncalves, Ewan Harrison, David K. Jackson, Ian Johnston, Dominic Kwiatkowski, Cordelia Langford, John Sillitoe on behalf of the Wellcome Sanger Institute COVID-19 Surveillance Team |
| EPI_ISL_3377004 | Berkshire and Surrey Pathology Services Lighthouse Laboratory | Wellcome Sanger Institute for the COVID-19 Genomics UK (COG-UK) Consortium | Berkshire and Surrey Pathology Services Lighthouse Laboratory and Alex Alderton, Roberto Amato, Jeffrey Barrett, Sonia Goncalves, Ewan Harrison, David K. Jackson, Ian Johnston, Dominic Kwiatkowski, Cordelia Langford, John Sillitoe on behalf of the Wellcome Sanger Institute COVID-19 Surveillance Team |
| EPI_ISL_3377005, EPI_ISL_3377006 | Lighthouse Lab in Glasgow | Wellcome Sanger Institute for the COVID-19 Genomics UK (COG-UK) Consortium | Harper VanSteenhouse, Yumi Kasai, David Gray, Carol Clugston, Anna Dominiczak and Alex Alderton, Roberto Amato, Jeffrey Barrett, Sonia Goncalves, Ewan Harrison, David K. Jackson, Ian Johnston, Dominic Kwiatkowski, Cordelia Langford, John Sillitoe on behalf of the Wellcome Sanger Institute COVID-19 Surveillance Team |
| EPI_ISL_3377007 | Lighthouse Lab in Alderley Park | Wellcome Sanger Institute for the COVID-19 Genomics UK (COG-UK) Consortium | Jacquelyn Wynn, Mairead Hyland, The Lighthouse Lab in Alderley Park and Alex Alderton, Roberto Amato, Jeffrey Barrett, Sonia Goncalves, Ewan Harrison, David K. Jackson, Ian Johnston, Dominic Kwiatkowski, Cordelia Langford, John Sillitoe on behalf of the Wellcome Sanger Institute COVID-19 Surveillance Team |
| EPI_ISL_3377008 | Lighthouse Lab in Milton Keynes | Wellcome Sanger Institute for the COVID-19 Genomics UK (COG-UK) Consortium | The Lighthouse Lab in Milton Keynes and Alex Alderton, Roberto Amato, Jeffrey Barrett, Sonia Goncalves, Ewan Harrison, David K. Jackson, Ian Johnston, Dominic Kwiatkowski, Cordelia Langford, John Sillitoe on behalf of the Wellcome Sanger Institute COVID-19 Surveillance Team |
| EPI_ISL_3377009, EPI_ISL_3377010 | Lighthouse Lab in Alderley Park | Wellcome Sanger Institute for the COVID-19 Genomics UK (COG-UK) Consortium | Jacquelyn Wynn, Mairead Hyland, The Lighthouse Lab in Alderley Park and Alex Alderton, Roberto Amato, Jeffrey Barrett, Sonia Goncalves, Ewan Harrison, David K. Jackson, Ian Johnston, Dominic Kwiatkowski, Cordelia Langford, John Sillitoe on behalf of the Wellcome Sanger Institute COVID-19 Surveillance Team |
| EPI_ISL_3377011 | Berkshire and Surrey Pathology Services Lighthouse Laboratory | Wellcome Sanger Institute for the COVID-19 Genomics UK (COG-UK) Consortium | Berkshire and Surrey Pathology Services Lighthouse Laboratory and Alex Alderton, Roberto Amato, Jeffrey Barrett, Sonia Goncalves, Ewan Harrison, David K. Jackson, Ian Johnston, Dominic Kwiatkowski, Cordelia Langford, John Sillitoe on behalf of the Wellcome Sanger Institute COVID-19 Surveillance Team |
| EPI_ISL_3377012 | Lighthouse Lab in Glasgow | Wellcome Sanger Institute for the COVID-19 Genomics UK (COG-UK) Consortium | Harper VanSteenhouse, Yumi Kasai, David Gray, Carol Clugston, Anna Dominiczak and Alex Alderton, Roberto Amato, Jeffrey Barrett, Sonia Goncalves, Ewan Harrison, David K. Jackson, Ian Johnston, Dominic Kwiatkowski, Cordelia Langford, John Sillitoe on behalf of the Wellcome Sanger Institute COVID-19 Surveillance Team |
| EPI_ISL_3377013 | Berkshire and Surrey Pathology Services Lighthouse Laboratory | Wellcome Sanger Institute for the COVID-19 Genomics UK (COG-UK) Consortium | Berkshire and Surrey Pathology Services Lighthouse Laboratory and Alex Alderton, Roberto Amato, Jeffrey Barrett, Sonia Goncalves, Ewan Harrison, David K. Jackson, Ian Johnston, Dominic Kwiatkowski, Cordelia Langford, John Sillitoe on behalf of the Wellcome Sanger Institute COVID-19 Surveillance Team |
| EPI_ISL_3377014, EPI_ISL_3377015, EPI_ISL_3377016 | Lighthouse Lab in Alderley Park | Wellcome Sanger Institute for the COVID-19 Genomics UK (COG-UK) Consortium | Jacquelyn Wynn, Mairead Hyland, The Lighthouse Lab in Alderley Park and Alex Alderton, Roberto Amato, Jeffrey Barrett, Sonia Goncalves, Ewan Harrison, David K. Jackson, Ian Johnston, Dominic Kwiatkowski, Cordelia Langford, John Sillitoe on behalf of the Wellcome Sanger Institute COVID-19 Surveillance Team |
| EPI_ISL_3377017 | Lighthouse Lab in Milton Keynes | Wellcome Sanger Institute for the COVID-19 Genomics UK (COG-UK) Consortium | The Lighthouse Lab in Milton Keynes and Alex Alderton, Roberto Amato, Jeffrey Barrett, Sonia Goncalves, Ewan Harrison, David K. Jackson, Ian Johnston, Dominic Kwiatkowski, Cordelia Langford, John Sillitoe on behalf of the Wellcome Sanger Institute COVID-19 Surveillance Team |
| EPI_ISL_3377018 | Berkshire and Surrey Pathology Services Lighthouse Laboratory | Wellcome Sanger Institute for the COVID-19 Genomics UK (COG-UK) Consortium | Berkshire and Surrey Pathology Services Lighthouse Laboratory and Alex Alderton, Roberto Amato, Jeffrey Barrett, Sonia Goncalves, Ewan Harrison, David K. Jackson, Ian Johnston, Dominic Kwiatkowski, Cordelia Langford, John Sillitoe on behalf of the Wellcome Sanger Institute COVID-19 Surveillance Team |
| EPI_ISL_3377019 | Lighthouse Lab in Alderley Park | Wellcome Sanger Institute for the COVID-19 Genomics UK (COG-UK) Consortium | Jacquelyn Wynn, Mairead Hyland, The Lighthouse Lab in Alderley Park and Alex Alderton, Roberto Amato, Jeffrey Barrett, Sonia Goncalves, Ewan Harrison, David K. Jackson, Ian Johnston, Dominic Kwiatkowski, Cordelia Langford, John Sillitoe on behalf of the Wellcome Sanger Institute COVID-19 Surveillance Team |
| EPI_ISL_3377020 | Lighthouse Lab in Milton Keynes | Wellcome Sanger Institute for the COVID-19 Genomics UK (COG-UK) Consortium | The Lighthouse Lab in Milton Keynes and Alex Alderton, Roberto Amato, Jeffrey Barrett, Sonia Goncalves, Ewan Harrison, David K. Jackson, Ian Johnston, Dominic Kwiatkowski, Cordelia Langford, John Sillitoe on behalf of the Wellcome Sanger Institute COVID-19 Surveillance Team |
| EPI_ISL_3377021, EPI_ISL_3377022 | Lighthouse Lab in Alderley Park | Wellcome Sanger Institute for the COVID-19 Genomics UK (COG-UK) Consortium | Jacquelyn Wynn, Mairead Hyland, The Lighthouse Lab in Alderley Park and Alex Alderton, Roberto Amato, Jeffrey Barrett, Sonia Goncalves, Ewan Harrison, David K. Jackson, Ian Johnston, Dominic Kwiatkowski, Cordelia Langford, John Sillitoe on behalf of the Wellcome Sanger Institute COVID-19 Surveillance Team |
| EPI_ISL_3377023 | Berkshire and Surrey Pathology Services Lighthouse Laboratory | Wellcome Sanger Institute for the COVID-19 Genomics UK (COG-UK) Consortium | Berkshire and Surrey Pathology Services Lighthouse Laboratory and Alex Alderton, Roberto Amato, Jeffrey Barrett, Sonia Goncalves, Ewan Harrison, David K. Jackson, Ian Johnston, Dominic Kwiatkowski, Cordelia Langford, John Sillitoe on behalf of the Wellcome Sanger Institute COVID-19 Surveillance Team |

| | | | |
|---|---|---|---|
| EPI_ISL_3377024 | Lighthouse Lab in Alderley Park | Wellcome Sanger Institute for the COVID-19 Genomics UK (COG-UK) Consortium | Jacquelyn Wynn, Mairead Hyland, The Lighthouse Lab in Alderley Park and Alex Alderton, Roberto Amato, Jeffrey Barrett, Sonia Goncalves, Ewan Harrison, David K. Jackson, Ian Johnston, Dominic Kwiatkowski, Cordelia Langford, John Sillitoe on behalf of the Wellcome Sanger Institute COVID-19 Surveillance Team |
| EPI_ISL_3377025 | Lighthouse Lab in Milton Keynes | Wellcome Sanger Institute for the COVID-19 Genomics UK (COG-UK) Consortium | The Lighthouse Lab in Milton Keynes and Alex Alderton, Roberto Amato, Jeffrey Barrett, Sonia Goncalves, Ewan Harrison, David K. Jackson, Ian Johnston, Dominic Kwiatkowski, Cordelia Langford, John Sillitoe on behalf of the Wellcome Sanger Institute COVID-19 Surveillance Team |
| EPI_ISL_3377026 | Lighthouse Lab in Alderley Park | Wellcome Sanger Institute for the COVID-19 Genomics UK (COG-UK) Consortium | Jacquelyn Wynn, Mairead Hyland, The Lighthouse Lab in Alderley Park and Alex Alderton, Roberto Amato, Jeffrey Barrett, Sonia Goncalves, Ewan Harrison, David K. Jackson, Ian Johnston, Dominic Kwiatkowski, Cordelia Langford, John Sillitoe on behalf of the Wellcome Sanger Institute COVID-19 Surveillance Team |
| EPI_ISL_3377027 | Lighthouse Lab in Milton Keynes | Wellcome Sanger Institute for the COVID-19 Genomics UK (COG-UK) Consortium | The Lighthouse Lab in Milton Keynes and Alex Alderton, Roberto Amato, Jeffrey Barrett, Sonia Goncalves, Ewan Harrison, David K. Jackson, Ian Johnston, Dominic Kwiatkowski, Cordelia Langford, John Sillitoe on behalf of the Wellcome Sanger Institute COVID-19 Surveillance Team |
| EPI_ISL_3377028 | Lighthouse Lab in Alderley Park | Wellcome Sanger Institute for the COVID-19 Genomics UK (COG-UK) Consortium | Jacquelyn Wynn, Mairead Hyland, The Lighthouse Lab in Alderley Park and Alex Alderton, Roberto Amato, Jeffrey Barrett, Sonia Goncalves, Ewan Harrison, David K. Jackson, Ian Johnston, Dominic Kwiatkowski, Cordelia Langford, John Sillitoe on behalf of the Wellcome Sanger Institute COVID-19 Surveillance Team |
| EPI_ISL_3377029 | Lighthouse Lab in Milton Keynes | Wellcome Sanger Institute for the COVID-19 Genomics UK (COG-UK) Consortium | The Lighthouse Lab in Milton Keynes and Alex Alderton, Roberto Amato, Jeffrey Barrett, Sonia Goncalves, Ewan Harrison, David K. Jackson, Ian Johnston, Dominic Kwiatkowski, Cordelia Langford, John Sillitoe on behalf of the Wellcome Sanger Institute COVID-19 Surveillance Team |
| EPI_ISL_3377030 | Lighthouse Lab in Glasgow | Wellcome Sanger Institute for the COVID-19 Genomics UK (COG-UK) Consortium | Harper VanSteenhouse, Yumi Kasai, David Gray, Carol Clugston, Anna Dominiczak and Alex Alderton, Roberto Amato, Jeffrey Barrett, Sonia Goncalves, Ewan Harrison, David K. Jackson, Ian Johnston, Dominic Kwiatkowski, Cordelia Langford, John Sillitoe on behalf of the Wellcome Sanger Institute COVID-19 Surveillance Team |
| EPI_ISL_3377031 | Berkshire and Surrey Pathology Services Lighthouse Laboratory | Wellcome Sanger Institute for the COVID-19 Genomics UK (COG-UK) Consortium | Berkshire and Surrey Pathology Services Lighthouse Laboratory and Alex Alderton, Roberto Amato, Jeffrey Barrett, Sonia Goncalves, Ewan Harrison, David K. Jackson, Ian Johnston, Dominic Kwiatkowski, Cordelia Langford, John Sillitoe on behalf of the Wellcome Sanger Institute COVID-19 Surveillance Team |
| EPI_ISL_3377032 | Lighthouse Lab in Glasgow | Wellcome Sanger Institute for the COVID-19 Genomics UK (COG-UK) Consortium | Harper VanSteenhouse, Yumi Kasai, David Gray, Carol Clugston, Anna Dominiczak and Alex Alderton, Roberto Amato, Jeffrey Barrett, Sonia Goncalves, Ewan Harrison, David K. Jackson, Ian Johnston, Dominic Kwiatkowski, Cordelia Langford, John Sillitoe on behalf of the Wellcome Sanger Institute COVID-19 Surveillance Team |
| EPI_ISL_3377033 | Lighthouse Lab in Milton Keynes | Wellcome Sanger Institute for the COVID-19 Genomics UK (COG-UK) Consortium | The Lighthouse Lab in Milton Keynes and Alex Alderton, Roberto Amato, Jeffrey Barrett, Sonia Goncalves, Ewan Harrison, David K. Jackson, Ian Johnston, Dominic Kwiatkowski, Cordelia Langford, John Sillitoe on behalf of the Wellcome Sanger Institute COVID-19 Surveillance Team |
| EPI_ISL_3377034 | Lighthouse Lab in Glasgow | Wellcome Sanger Institute for the COVID-19 Genomics UK (COG-UK) Consortium | Harper VanSteenhouse, Yumi Kasai, David Gray, Carol Clugston, Anna Dominiczak and Alex Alderton, Roberto Amato, Jeffrey Barrett, Sonia Goncalves, Ewan Harrison, David K. Jackson, Ian Johnston, Dominic Kwiatkowski, Cordelia Langford, John Sillitoe on behalf of the Wellcome Sanger Institute COVID-19 Surveillance Team |
| EPI_ISL_3377035 | Lighthouse Lab in Alderley Park | Wellcome Sanger Institute for the COVID-19 Genomics UK (COG-UK) Consortium | Jacquelyn Wynn, Mairead Hyland, The Lighthouse Lab in Alderley Park and Alex Alderton, Roberto Amato, Jeffrey Barrett, Sonia Goncalves, Ewan Harrison, David K. Jackson, Ian Johnston, Dominic Kwiatkowski, Cordelia Langford, John Sillitoe on behalf of the Wellcome Sanger Institute COVID-19 Surveillance Team |
| EPI_ISL_3377036, EPI_ISL_3377037 | Lighthouse Lab in Milton Keynes | Wellcome Sanger Institute for the COVID-19 Genomics UK (COG-UK) Consortium | The Lighthouse Lab in Milton Keynes and Alex Alderton, Roberto Amato, Jeffrey Barrett, Sonia Goncalves, Ewan Harrison, David K. Jackson, Ian Johnston, Dominic Kwiatkowski, Cordelia Langford, John Sillitoe on behalf of the Wellcome Sanger Institute COVID-19 Surveillance Team |
| EPI_ISL_3377038 | Lighthouse Lab in Alderley Park | Wellcome Sanger Institute for the COVID-19 Genomics UK (COG-UK) Consortium | Jacquelyn Wynn, Mairead Hyland, The Lighthouse Lab in Alderley Park and Alex Alderton, Roberto Amato, Jeffrey Barrett, Sonia Goncalves, Ewan Harrison, David K. Jackson, Ian Johnston, Dominic Kwiatkowski, Cordelia Langford, John Sillitoe on behalf of the Wellcome Sanger Institute COVID-19 Surveillance Team |
| EPI_ISL_3377039 | Berkshire and Surrey Pathology Services Lighthouse Laboratory | Wellcome Sanger Institute for the COVID-19 Genomics UK (COG-UK) Consortium | Berkshire and Surrey Pathology Services Lighthouse Laboratory and Alex Alderton, Roberto Amato, Jeffrey Barrett, Sonia Goncalves, Ewan Harrison, David K. Jackson, Ian Johnston, Dominic Kwiatkowski, Cordelia Langford, John Sillitoe on behalf of the Wellcome Sanger Institute COVID-19 Surveillance Team |
| EPI_ISL_3377040 | Lighthouse Lab in Glasgow | Wellcome Sanger Institute for the COVID-19 Genomics UK (COG-UK) Consortium | Harper VanSteenhouse, Yumi Kasai, David Gray, Carol Clugston, Anna Dominiczak and Alex Alderton, Roberto Amato, Jeffrey Barrett, Sonia Goncalves, Ewan Harrison, David K. Jackson, Ian Johnston, Dominic Kwiatkowski, Cordelia Langford, John Sillitoe on behalf of the Wellcome Sanger Institute COVID-19 Surveillance Team |
| EPI_ISL_3377041, EPI_ISL_3377042 | Berkshire and Surrey Pathology Services Lighthouse Laboratory | Wellcome Sanger Institute for the COVID-19 Genomics UK (COG-UK) Consortium | Berkshire and Surrey Pathology Services Lighthouse Laboratory and Alex Alderton, Roberto Amato, Jeffrey Barrett, Sonia Goncalves, Ewan Harrison, David K. Jackson, Ian Johnston, Dominic Kwiatkowski, Cordelia Langford, John Sillitoe on behalf of the Wellcome Sanger Institute COVID-19 Surveillance Team |
| EPI_ISL_3377043 | Lighthouse Lab in Milton Keynes | Wellcome Sanger Institute for the COVID-19 Genomics UK (COG-UK) Consortium | The Lighthouse Lab in Milton Keynes and Alex Alderton, Roberto Amato, Jeffrey Barrett, Sonia Goncalves, Ewan Harrison, David K. Jackson, Ian Johnston, Dominic Kwiatkowski, Cordelia Langford, John Sillitoe on behalf of the Wellcome Sanger Institute COVID-19 Surveillance Team |
| EPI_ISL_3377044, EPI_ISL_3377045 | Berkshire and Surrey Pathology Services Lighthouse Laboratory | Wellcome Sanger Institute for the COVID-19 Genomics UK (COG-UK) Consortium | Berkshire and Surrey Pathology Services Lighthouse Laboratory and Alex Alderton, Roberto Amato, Jeffrey Barrett, Sonia Goncalves, Ewan Harrison, David K. Jackson, Ian Johnston, Dominic Kwiatkowski, Cordelia Langford, John Sillitoe on behalf of the Wellcome Sanger Institute COVID-19 Surveillance Team |
| EPI_ISL_3377046 | Lighthouse Lab in Milton Keynes | Wellcome Sanger Institute for the COVID-19 Genomics UK (COG-UK) Consortium | The Lighthouse Lab in Milton Keynes and Alex Alderton, Roberto Amato, Jeffrey Barrett, Sonia Goncalves, Ewan Harrison, David K. Jackson, Ian Johnston, Dominic Kwiatkowski, Cordelia Langford, John Sillitoe on behalf of the Wellcome Sanger Institute COVID-19 Surveillance Team |
| EPI_ISL_3377047 | Lighthouse Lab in Glasgow | Wellcome Sanger Institute for the COVID-19 Genomics UK (COG-UK) Consortium | Harper VanSteenhouse, Yumi Kasai, David Gray, Carol Clugston, Anna Dominiczak and Alex Alderton, Roberto Amato, Jeffrey Barrett, Sonia Goncalves, Ewan Harrison, David K. Jackson, Ian Johnston, Dominic Kwiatkowski, Cordelia Langford, John Sillitoe on behalf of the Wellcome Sanger Institute COVID-19 Surveillance Team |
| EPI_ISL_3377048 | Berkshire and Surrey Pathology Services Lighthouse Laboratory | Wellcome Sanger Institute for the COVID-19 Genomics UK (COG-UK) Consortium | Berkshire and Surrey Pathology Services Lighthouse Laboratory and Alex Alderton, Roberto Amato, Jeffrey Barrett, Sonia Goncalves, Ewan Harrison, David K. Jackson, Ian Johnston, Dominic Kwiatkowski, Cordelia Langford, John Sillitoe on behalf of the Wellcome Sanger Institute COVID-19 Surveillance Team |
| EPI_ISL_3377049 | Lighthouse Lab in Glasgow | Wellcome Sanger Institute for the COVID-19 Genomics UK (COG-UK) Consortium | Harper VanSteenhouse, Yumi Kasai, David Gray, Carol Clugston, Anna Dominiczak and Alex Alderton, Roberto Amato, Jeffrey Barrett, Sonia Goncalves, Ewan Harrison, David K. Jackson, Ian Johnston, Dominic Kwiatkowski, Cordelia Langford, John Sillitoe on behalf of the Wellcome Sanger Institute COVID-19 Surveillance Team |
| EPI_ISL_3377050 | Lighthouse Lab in Milton Keynes | Wellcome Sanger Institute for the COVID-19 Genomics UK (COG-UK) Consortium | The Lighthouse Lab in Milton Keynes and Alex Alderton, Roberto Amato, Jeffrey Barrett, Sonia Goncalves, Ewan Harrison, David K. Jackson, Ian Johnston, Dominic Kwiatkowski, Cordelia Langford, John Sillitoe on behalf of the Wellcome Sanger Institute COVID-19 Surveillance Team |
| EPI_ISL_3377051, EPI_ISL_3377052 | Lighthouse Lab in Alderley Park | Wellcome Sanger Institute for the COVID-19 Genomics UK (COG-UK) Consortium | Jacquelyn Wynn, Mairead Hyland, The Lighthouse Lab in Alderley Park and Alex Alderton, Roberto Amato, Jeffrey Barrett, Sonia Goncalves, Ewan Harrison, David K. Jackson, Ian Johnston, Dominic Kwiatkowski, Cordelia Langford, John Sillitoe on behalf of the Wellcome Sanger Institute COVID-19 Surveillance Team |
| EPI_ISL_3377053 | Lighthouse Lab in Glasgow | Wellcome Sanger Institute for the COVID-19 Genomics UK (COG-UK) Consortium | Harper VanSteenhouse, Yumi Kasai, David Gray, Carol Clugston, Anna Dominiczak and Alex Alderton, Roberto Amato, Jeffrey Barrett, Sonia Goncalves, Ewan Harrison, David K. Jackson, Ian Johnston, Dominic Kwiatkowski, Cordelia Langford, John Sillitoe on behalf of the Wellcome Sanger Institute COVID-19 Surveillance Team |
| EPI_ISL_3377054 | Lighthouse Lab in Alderley Park | Wellcome Sanger Institute for the COVID-19 Genomics UK (COG-UK) Consortium | Jacquelyn Wynn, Mairead Hyland, The Lighthouse Lab in Alderley Park and Alex Alderton, Roberto Amato, Jeffrey Barrett, Sonia Goncalves, Ewan Harrison, David K. Jackson, Ian Johnston, Dominic Kwiatkowski, Cordelia Langford, John Sillitoe on behalf of the Wellcome Sanger Institute COVID-19 Surveillance Team |
| EPI_ISL_3377055 | Lighthouse Lab in Milton Keynes | Wellcome Sanger Institute for the COVID-19 Genomics UK (COG-UK) Consortium | The Lighthouse Lab in Milton Keynes and Alex Alderton, Roberto Amato, Jeffrey Barrett, Sonia Goncalves, Ewan Harrison, David K. Jackson, Ian Johnston, Dominic Kwiatkowski, Cordelia Langford, John Sillitoe on behalf of the Wellcome Sanger Institute COVID-19 Surveillance Team |
| EPI_ISL_3377056 | Lighthouse Lab in Alderley Park | Wellcome Sanger Institute for the COVID-19 Genomics UK (COG-UK) Consortium | Jacquelyn Wynn, Mairead Hyland, The Lighthouse Lab in Alderley Park and Alex Alderton, Roberto Amato, Jeffrey Barrett, Sonia Goncalves, Ewan Harrison, David K. Jackson, Ian Johnston, Dominic Kwiatkowski, Cordelia Langford, John Sillitoe on behalf of the Wellcome Sanger Institute COVID-19 Surveillance Team |
| EPI_ISL_3377057 | Lighthouse Lab in Milton Keynes | Wellcome Sanger Institute for the COVID-19 Genomics UK (COG-UK) Consortium | The Lighthouse Lab in Milton Keynes and Alex Alderton, Roberto Amato, Jeffrey Barrett, Sonia Goncalves, Ewan Harrison, David K. Jackson, Ian Johnston, Dominic Kwiatkowski, Cordelia Langford, John Sillitoe on behalf of the Wellcome Sanger Institute COVID-19 Surveillance Team |
| EPI_ISL_3377058, EPI_ISL_3377059, EPI_ISL_3377060, EPI_ISL_3377061 | Lighthouse Lab in Alderley Park | Wellcome Sanger Institute for the COVID-19 Genomics UK (COG-UK) Consortium | Jacquelyn Wynn, Mairead Hyland, The Lighthouse Lab in Alderley Park and Alex Alderton, Roberto Amato, Jeffrey Barrett, Sonia Goncalves, Ewan Harrison, David K. Jackson, Ian Johnston, Dominic Kwiatkowski, Cordelia Langford, John Sillitoe on behalf of the Wellcome Sanger Institute COVID-19 Surveillance Team |
| EPI_ISL_3377062 | Lighthouse Lab in Milton Keynes | Wellcome Sanger Institute for the COVID-19 Genomics UK (COG-UK) Consortium | The Lighthouse Lab in Milton Keynes and Alex Alderton, Roberto Amato, Jeffrey Barrett, Sonia Goncalves, Ewan Harrison, David K. Jackson, Ian Johnston, Dominic Kwiatkowski, Cordelia Langford, John Sillitoe on behalf of the Wellcome Sanger Institute COVID-19 Surveillance Team |
| EPI_ISL_3377063, EPI_ISL_3377064, EPI_ISL_3377065 | Lighthouse Lab in Alderley Park | Wellcome Sanger Institute for the COVID-19 Genomics UK (COG-UK) Consortium | Jacquelyn Wynn, Mairead Hyland, The Lighthouse Lab in Alderley Park and Alex Alderton, Roberto Amato, Jeffrey Barrett, Sonia Goncalves, Ewan Harrison, David K. Jackson, Ian Johnston, Dominic Kwiatkowski, Cordelia Langford, John Sillitoe on behalf of the Wellcome Sanger Institute COVID-19 Surveillance Team |
| EPI_ISL_3377066, EPI_ISL_3377067 | Berkshire and Surrey Pathology Services Lighthouse Laboratory | Wellcome Sanger Institute for the COVID-19 Genomics UK (COG-UK) Consortium | Berkshire and Surrey Pathology Services Lighthouse Laboratory and Alex Alderton, Roberto Amato, Jeffrey Barrett, Sonia Goncalves, Ewan Harrison, David K. Jackson, Ian Johnston, Dominic Kwiatkowski, Cordelia Langford, John Sillitoe on behalf of the Wellcome Sanger Institute COVID-19 Surveillance Team |
| EPI_ISL_3377068 | Lighthouse Lab in Milton Keynes | Wellcome Sanger Institute for the COVID-19 Genomics UK (COG-UK) Consortium | The Lighthouse Lab in Milton Keynes and Alex Alderton, Roberto Amato, Jeffrey Barrett, Sonia Goncalves, Ewan Harrison, David K. Jackson, Ian Johnston, Dominic Kwiatkowski, Cordelia Langford, John Sillitoe on behalf of the Wellcome Sanger Institute COVID-19 Surveillance Team |
| EPI_ISL_3377069 | Lighthouse Lab in Glasgow | Wellcome Sanger Institute for the COVID-19 Genomics UK (COG-UK) Consortium | Harper VanSteenhouse, Yumi Kasai, David Gray, Carol Clugston, Anna Dominiczak and Alex Alderton, Roberto Amato, Jeffrey Barrett, Sonia Goncalves, Ewan Harrison, David K. Jackson, Ian Johnston, Dominic Kwiatkowski, Cordelia Langford, John Sillitoe on behalf of the Wellcome Sanger Institute COVID-19 Surveillance Team |
| EPI_ISL_3377070 | Lighthouse Lab in Alderley Park | Wellcome Sanger Institute for the COVID-19 Genomics UK (COG-UK) Consortium | Jacquelyn Wynn, Mairead Hyland, The Lighthouse Lab in Alderley Park and Alex Alderton, Roberto Amato, Jeffrey Barrett, Sonia Goncalves, Ewan Harrison, David K. Jackson, Ian Johnston, Dominic Kwiatkowski, Cordelia Langford, John Sillitoe on behalf of the Wellcome Sanger Institute COVID-19 Surveillance Team |
| EPI_ISL_3377071 | Berkshire and Surrey Pathology Services Lighthouse Laboratory | Wellcome Sanger Institute for the COVID-19 Genomics UK (COG-UK) Consortium | Berkshire and Surrey Pathology Services Lighthouse Laboratory and Alex Alderton, Roberto Amato, Jeffrey Barrett, Sonia Goncalves, Ewan Harrison, David K. Jackson, Ian Johnston, Dominic Kwiatkowski, Cordelia Langford, John Sillitoe on behalf of the Wellcome Sanger Institute COVID-19 Surveillance Team |
| EPI_ISL_3377072 | Lighthouse Lab in Milton Keynes | Wellcome Sanger Institute for the COVID-19 Genomics UK (COG-UK) Consortium | The Lighthouse Lab in Milton Keynes and Alex Alderton, Roberto Amato, Jeffrey Barrett, Sonia Goncalves, Ewan Harrison, David K. Jackson, Ian Johnston, Dominic Kwiatkowski, Cordelia Langford, John Sillitoe on behalf of the Wellcome Sanger Institute COVID-19 Surveillance Team |
| EPI_ISL_3377073, EPI_ISL_3377074 | Lighthouse Lab in Alderley Park | Wellcome Sanger Institute for the COVID-19 Genomics UK (COG-UK) Consortium | Jacquelyn Wynn, Mairead Hyland, The Lighthouse Lab in Alderley Park and Alex Alderton, Roberto Amato, Jeffrey Barrett, Sonia Goncalves, Ewan Harrison, David K. Jackson, Ian Johnston, Dominic Kwiatkowski, Cordelia Langford, John Sillitoe on behalf of the Wellcome Sanger Institute COVID-19 Surveillance Team |
| EPI_ISL_3377075, EPI_ISL_3377076, EPI_ISL_3377077 | Berkshire and Surrey Pathology Services Lighthouse Laboratory | Wellcome Sanger Institute for the COVID-19 Genomics UK (COG-UK) Consortium | Berkshire and Surrey Pathology Services Lighthouse Laboratory and Alex Alderton, Roberto Amato, Jeffrey Barrett, Sonia Goncalves, Ewan Harrison, David K. Jackson, Ian Johnston, Dominic Kwiatkowski, Cordelia Langford, John Sillitoe on behalf of the Wellcome Sanger Institute COVID-19 Surveillance Team |
| EPI_ISL_3377078 | Lighthouse Lab in Milton Keynes | Wellcome Sanger Institute for the COVID-19 Genomics UK (COG-UK) Consortium | The Lighthouse Lab in Milton Keynes and Alex Alderton, Roberto Amato, Jeffrey Barrett, Sonia Goncalves, Ewan Harrison, David K. Jackson, Ian Johnston, Dominic Kwiatkowski, Cordelia Langford, John Sillitoe on behalf of the Wellcome Sanger Institute COVID-19 Surveillance Team |
| EPI_ISL_3377079 | Berkshire and Surrey Pathology Services Lighthouse Laboratory | Wellcome Sanger Institute for the COVID-19 Genomics UK (COG-UK) Consortium | Berkshire and Surrey Pathology Services Lighthouse Laboratory and Alex Alderton, Roberto Amato, Jeffrey Barrett, Sonia Goncalves, Ewan Harrison, David K. Jackson, Ian Johnston, Dominic Kwiatkowski, Cordelia Langford, John Sillitoe on behalf of the Wellcome Sanger Institute COVID-19 Surveillance Team |
| EPI_ISL_3377080 | Lighthouse Lab in Glasgow | Wellcome Sanger Institute for the COVID-19 Genomics UK (COG-UK) | Harper VanSteenhouse, Yumi Kasai, David Gray, Carol Clugston, Anna Dominiczak and Alex Alderton, Roberto Amato, Jeffrey Barrett, Sonia Goncalves, Ewan Harrison, David K. Jackson, Ian Johnston, Dominic Kwiatkowski, Cordelia Langford, |

| | | | |
|---|---|---|---|
| | | Consortium | John Sillitoe on behalf of the Wellcome Sanger Institute COVID-19 Surveillance Team |
| EPI_ISL_3377081, EPI_ISL_3377082 | Lighthouse Lab in Alderley Park | Wellcome Sanger Institute for the COVID-19 Genomics UK (COG-UK) Consortium | Jacquelyn Wynn, Mairead Hyland, The Lighthouse Lab in Alderley Park and Alex Alderton, Roberto Amato, Jeffrey Barrett, Sonia Goncalves, Ewan Harrison, David K. Jackson, Ian Johnston, Dominic Kwiatkowski, Cordelia Langford, John Sillitoe on behalf of the Wellcome Sanger Institute COVID-19 Surveillance Team |
| EPI_ISL_3377083, EPI_ISL_3377084, EPI_ISL_3377085 | Lighthouse Lab in Milton Keynes | Wellcome Sanger Institute for the COVID-19 Genomics UK (COG-UK) Consortium | The Lighthouse Lab in Milton Keynes and Alex Alderton, Roberto Amato, Jeffrey Barrett, Sonia Goncalves, Ewan Harrison, David K. Jackson, Ian Johnston, Dominic Kwiatkowski, Cordelia Langford, John Sillitoe on behalf of the Wellcome Sanger Institute COVID-19 Surveillance Team |
| EPI_ISL_3377086 | Lighthouse Lab in Alderley Park | Wellcome Sanger Institute for the COVID-19 Genomics UK (COG-UK) Consortium | Jacquelyn Wynn, Mairead Hyland, The Lighthouse Lab in Alderley Park and Alex Alderton, Roberto Amato, Jeffrey Barrett, Sonia Goncalves, Ewan Harrison, David K. Jackson, Ian Johnston, Dominic Kwiatkowski, Cordelia Langford, John Sillitoe on behalf of the Wellcome Sanger Institute COVID-19 Surveillance Team |
| EPI_ISL_3377087 | Berkshire and Surrey Pathology Services Lighthouse Laboratory | Wellcome Sanger Institute for the COVID-19 Genomics UK (COG-UK) Consortium | Berkshire and Surrey Pathology Services Lighthouse Laboratory and Alex Alderton, Roberto Amato, Jeffrey Barrett, Sonia Goncalves, Ewan Harrison, David K. Jackson, Ian Johnston, Dominic Kwiatkowski, Cordelia Langford, John Sillitoe on behalf of the Wellcome Sanger Institute COVID-19 Surveillance Team |
| EPI_ISL_3377088 | Lighthouse Lab in Alderley Park | Wellcome Sanger Institute for the COVID-19 Genomics UK (COG-UK) Consortium | Jacquelyn Wynn, Mairead Hyland, The Lighthouse Lab in Alderley Park and Alex Alderton, Roberto Amato, Jeffrey Barrett, Sonia Goncalves, Ewan Harrison, David K. Jackson, Ian Johnston, Dominic Kwiatkowski, Cordelia Langford, John Sillitoe on behalf of the Wellcome Sanger Institute COVID-19 Surveillance Team |
| EPI_ISL_3377089 | Lighthouse Lab in Glasgow | Wellcome Sanger Institute for the COVID-19 Genomics UK (COG-UK) Consortium | Harper VanSteenhouse, Yumi Kasai, David Gray, Carol Clugston, Anna Dominiczak and Alex Alderton, Roberto Amato, Jeffrey Barrett, Sonia Goncalves, Ewan Harrison, David K. Jackson, Ian Johnston, Dominic Kwiatkowski, Cordelia Langford, John Sillitoe on behalf of the Wellcome Sanger Institute COVID-19 Surveillance Team |
| EPI_ISL_3377090 | Lighthouse Lab in Milton Keynes | Wellcome Sanger Institute for the COVID-19 Genomics UK (COG-UK) Consortium | The Lighthouse Lab in Milton Keynes and Alex Alderton, Roberto Amato, Jeffrey Barrett, Sonia Goncalves, Ewan Harrison, David K. Jackson, Ian Johnston, Dominic Kwiatkowski, Cordelia Langford, John Sillitoe on behalf of the Wellcome Sanger Institute COVID-19 Surveillance Team |
| EPI_ISL_3377091 | Berkshire and Surrey Pathology Services Lighthouse Laboratory | Wellcome Sanger Institute for the COVID-19 Genomics UK (COG-UK) Consortium | Berkshire and Surrey Pathology Services Lighthouse Laboratory and Alex Alderton, Roberto Amato, Jeffrey Barrett, Sonia Goncalves, Ewan Harrison, David K. Jackson, Ian Johnston, Dominic Kwiatkowski, Cordelia Langford, John Sillitoe on behalf of the Wellcome Sanger Institute COVID-19 Surveillance Team |
| EPI_ISL_3377092 | Lighthouse Lab in Milton Keynes | Wellcome Sanger Institute for the COVID-19 Genomics UK (COG-UK) Consortium | The Lighthouse Lab in Milton Keynes and Alex Alderton, Roberto Amato, Jeffrey Barrett, Sonia Goncalves, Ewan Harrison, David K. Jackson, Ian Johnston, Dominic Kwiatkowski, Cordelia Langford, John Sillitoe on behalf of the Wellcome Sanger Institute COVID-19 Surveillance Team |
| EPI_ISL_3377093 | Berkshire and Surrey Pathology Services Lighthouse Laboratory | Wellcome Sanger Institute for the COVID-19 Genomics UK (COG-UK) Consortium | Berkshire and Surrey Pathology Services Lighthouse Laboratory and Alex Alderton, Roberto Amato, Jeffrey Barrett, Sonia Goncalves, Ewan Harrison, David K. Jackson, Ian Johnston, Dominic Kwiatkowski, Cordelia Langford, John Sillitoe on behalf of the Wellcome Sanger Institute COVID-19 Surveillance Team |
| EPI_ISL_3377094 | Lighthouse Lab in Alderley Park | Wellcome Sanger Institute for the COVID-19 Genomics UK (COG-UK) Consortium | Jacquelyn Wynn, Mairead Hyland, The Lighthouse Lab in Alderley Park and Alex Alderton, Roberto Amato, Jeffrey Barrett, Sonia Goncalves, Ewan Harrison, David K. Jackson, Ian Johnston, Dominic Kwiatkowski, Cordelia Langford, John Sillitoe on behalf of the Wellcome Sanger Institute COVID-19 Surveillance Team |
| EPI_ISL_3377095 | Lighthouse Lab in Milton Keynes | Wellcome Sanger Institute for the COVID-19 Genomics UK (COG-UK) Consortium | The Lighthouse Lab in Milton Keynes and Alex Alderton, Roberto Amato, Jeffrey Barrett, Sonia Goncalves, Ewan Harrison, David K. Jackson, Ian Johnston, Dominic Kwiatkowski, Cordelia Langford, John Sillitoe on behalf of the Wellcome Sanger Institute COVID-19 Surveillance Team |
| EPI_ISL_3377096 | Lighthouse Lab in Glasgow | Wellcome Sanger Institute for the COVID-19 Genomics UK (COG-UK) Consortium | Harper VanSteenhouse, Yumi Kasai, David Gray, Carol Clugston, Anna Dominiczak and Alex Alderton, Roberto Amato, Jeffrey Barrett, Sonia Goncalves, Ewan Harrison, David K. Jackson, Ian Johnston, Dominic Kwiatkowski, Cordelia Langford, John Sillitoe on behalf of the Wellcome Sanger Institute COVID-19 Surveillance Team |
| EPI_ISL_3377097 | Lighthouse Lab in Milton Keynes | Wellcome Sanger Institute for the COVID-19 Genomics UK (COG-UK) Consortium | The Lighthouse Lab in Milton Keynes and Alex Alderton, Roberto Amato, Jeffrey Barrett, Sonia Goncalves, Ewan Harrison, David K. Jackson, Ian Johnston, Dominic Kwiatkowski, Cordelia Langford, John Sillitoe on behalf of the Wellcome Sanger Institute COVID-19 Surveillance Team |
| EPI_ISL_3377098 | Lighthouse Lab in Alderley Park | Wellcome Sanger Institute for the COVID-19 Genomics UK (COG-UK) Consortium | Jacquelyn Wynn, Mairead Hyland, The Lighthouse Lab in Alderley Park and Alex Alderton, Roberto Amato, Jeffrey Barrett, Sonia Goncalves, Ewan Harrison, David K. Jackson, Ian Johnston, Dominic Kwiatkowski, Cordelia Langford, John Sillitoe on behalf of the Wellcome Sanger Institute COVID-19 Surveillance Team |
| EPI_ISL_3377099 | Lighthouse Lab in Milton Keynes | Wellcome Sanger Institute for the COVID-19 Genomics UK (COG-UK) Consortium | The Lighthouse Lab in Milton Keynes and Alex Alderton, Roberto Amato, Jeffrey Barrett, Sonia Goncalves, Ewan Harrison, David K. Jackson, Ian Johnston, Dominic Kwiatkowski, Cordelia Langford, John Sillitoe on behalf of the Wellcome Sanger Institute COVID-19 Surveillance Team |
| EPI_ISL_3377100, EPI_ISL_3377101 | Lighthouse Lab in Glasgow | Wellcome Sanger Institute for the COVID-19 Genomics UK (COG-UK) Consortium | Harper VanSteenhouse, Yumi Kasai, David Gray, Carol Clugston, Anna Dominiczak and Alex Alderton, Roberto Amato, Jeffrey Barrett, Sonia Goncalves, Ewan Harrison, David K. Jackson, Ian Johnston, Dominic Kwiatkowski, Cordelia Langford, John Sillitoe on behalf of the Wellcome Sanger Institute COVID-19 Surveillance Team |
| EPI_ISL_3377102 | Lighthouse Lab in Alderley Park | Wellcome Sanger Institute for the COVID-19 Genomics UK (COG-UK) Consortium | Jacquelyn Wynn, Mairead Hyland, The Lighthouse Lab in Alderley Park and Alex Alderton, Roberto Amato, Jeffrey Barrett, Sonia Goncalves, Ewan Harrison, David K. Jackson, Ian Johnston, Dominic Kwiatkowski, Cordelia Langford, John Sillitoe on behalf of the Wellcome Sanger Institute COVID-19 Surveillance Team |
| EPI_ISL_3377103, EPI_ISL_3377104 | Lighthouse Lab in Milton Keynes | Wellcome Sanger Institute for the COVID-19 Genomics UK (COG-UK) Consortium | The Lighthouse Lab in Milton Keynes and Alex Alderton, Roberto Amato, Jeffrey Barrett, Sonia Goncalves, Ewan Harrison, David K. Jackson, Ian Johnston, Dominic Kwiatkowski, Cordelia Langford, John Sillitoe on behalf of the Wellcome Sanger Institute COVID-19 Surveillance Team |
| EPI_ISL_3377105 | Berkshire and Surrey Pathology Services Lighthouse Laboratory | Wellcome Sanger Institute for the COVID-19 Genomics UK (COG-UK) Consortium | Berkshire and Surrey Pathology Services Lighthouse Laboratory and Alex Alderton, Roberto Amato, Jeffrey Barrett, Sonia Goncalves, Ewan Harrison, David K. Jackson, Ian Johnston, Dominic Kwiatkowski, Cordelia Langford, John Sillitoe on behalf of the Wellcome Sanger Institute COVID-19 Surveillance Team |
| EPI_ISL_3377106 | Lighthouse Lab in Milton Keynes | Wellcome Sanger Institute for the COVID-19 Genomics UK (COG-UK) Consortium | The Lighthouse Lab in Milton Keynes and Alex Alderton, Roberto Amato, Jeffrey Barrett, Sonia Goncalves, Ewan Harrison, David K. Jackson, Ian Johnston, Dominic Kwiatkowski, Cordelia Langford, John Sillitoe on behalf of the Wellcome Sanger Institute COVID-19 Surveillance Team |
| EPI_ISL_3377107 | Lighthouse Lab in Alderley Park | Wellcome Sanger Institute for the COVID-19 Genomics UK (COG-UK) Consortium | Jacquelyn Wynn, Mairead Hyland, The Lighthouse Lab in Alderley Park and Alex Alderton, Roberto Amato, Jeffrey Barrett, Sonia Goncalves, Ewan Harrison, David K. Jackson, Ian Johnston, Dominic Kwiatkowski, Cordelia Langford, John Sillitoe on behalf of the Wellcome Sanger Institute COVID-19 Surveillance Team |
| EPI_ISL_3377108 | Lighthouse Lab in Glasgow | Wellcome Sanger Institute for the COVID-19 Genomics UK (COG-UK) Consortium | Harper VanSteenhouse, Yumi Kasai, David Gray, Carol Clugston, Anna Dominiczak and Alex Alderton, Roberto Amato, Jeffrey Barrett, Sonia Goncalves, Ewan Harrison, David K. Jackson, Ian Johnston, Dominic Kwiatkowski, Cordelia Langford, John Sillitoe on behalf of the Wellcome Sanger Institute COVID-19 Surveillance Team |
| EPI_ISL_3377109, EPI_ISL_3377110, EPI_ISL_3377111 | Berkshire and Surrey Pathology Services Lighthouse Laboratory | Wellcome Sanger Institute for the COVID-19 Genomics UK (COG-UK) Consortium | Berkshire and Surrey Pathology Services Lighthouse Laboratory and Alex Alderton, Roberto Amato, Jeffrey Barrett, Sonia Goncalves, Ewan Harrison, David K. Jackson, Ian Johnston, Dominic Kwiatkowski, Cordelia Langford, John Sillitoe on behalf of the Wellcome Sanger Institute COVID-19 Surveillance Team |
| EPI_ISL_3377112 | Lighthouse Lab in Milton Keynes | Wellcome Sanger Institute for the COVID-19 Genomics UK (COG-UK) Consortium | The Lighthouse Lab in Milton Keynes and Alex Alderton, Roberto Amato, Jeffrey Barrett, Sonia Goncalves, Ewan Harrison, David K. Jackson, Ian Johnston, Dominic Kwiatkowski, Cordelia Langford, John Sillitoe on behalf of the Wellcome Sanger Institute COVID-19 Surveillance Team |
| EPI_ISL_3377113, EPI_ISL_3377114 | Lighthouse Lab in Glasgow | Wellcome Sanger Institute for the COVID-19 Genomics UK (COG-UK) Consortium | Harper VanSteenhouse, Yumi Kasai, David Gray, Carol Clugston, Anna Dominiczak and Alex Alderton, Roberto Amato, Jeffrey Barrett, Sonia Goncalves, Ewan Harrison, David K. Jackson, Ian Johnston, Dominic Kwiatkowski, Cordelia Langford, John Sillitoe on behalf of the Wellcome Sanger Institute COVID-19 Surveillance Team |
| EPI_ISL_3377115, EPI_ISL_3377116 | Lighthouse Lab in Alderley Park | Wellcome Sanger Institute for the COVID-19 Genomics UK (COG-UK) Consortium | Jacquelyn Wynn, Mairead Hyland, The Lighthouse Lab in Alderley Park and Alex Alderton, Roberto Amato, Jeffrey Barrett, Sonia Goncalves, Ewan Harrison, David K. Jackson, Ian Johnston, Dominic Kwiatkowski, Cordelia Langford, John Sillitoe on behalf of the Wellcome Sanger Institute COVID-19 Surveillance Team |
| EPI_ISL_3377117 | Lighthouse Lab in Milton Keynes | Wellcome Sanger Institute for the COVID-19 Genomics UK (COG-UK) Consortium | The Lighthouse Lab in Milton Keynes and Alex Alderton, Roberto Amato, Jeffrey Barrett, Sonia Goncalves, Ewan Harrison, David K. Jackson, Ian Johnston, Dominic Kwiatkowski, Cordelia Langford, John Sillitoe on behalf of the Wellcome Sanger Institute COVID-19 Surveillance Team |
| EPI_ISL_3377118, EPI_ISL_3377119, EPI_ISL_3377120, EPI_ISL_3377121 | Lighthouse Lab in Alderley Park | Wellcome Sanger Institute for the COVID-19 Genomics UK (COG-UK) Consortium | Jacquelyn Wynn, Mairead Hyland, The Lighthouse Lab in Alderley Park and Alex Alderton, Roberto Amato, Jeffrey Barrett, Sonia Goncalves, Ewan Harrison, David K. Jackson, Ian Johnston, Dominic Kwiatkowski, Cordelia Langford, John Sillitoe on behalf of the Wellcome Sanger Institute COVID-19 Surveillance Team |
| EPI_ISL_3377122 | Lighthouse Lab in Milton Keynes | Wellcome Sanger Institute for the COVID-19 Genomics UK (COG-UK) Consortium | The Lighthouse Lab in Milton Keynes and Alex Alderton, Roberto Amato, Jeffrey Barrett, Sonia Goncalves, Ewan Harrison, David K. Jackson, Ian Johnston, Dominic Kwiatkowski, Cordelia Langford, John Sillitoe on behalf of the Wellcome Sanger Institute COVID-19 Surveillance Team |
| EPI_ISL_3377123 | Lighthouse Lab in Alderley Park | Wellcome Sanger Institute for the COVID-19 Genomics UK (COG-UK) Consortium | Jacquelyn Wynn, Mairead Hyland, The Lighthouse Lab in Alderley Park and Alex Alderton, Roberto Amato, Jeffrey Barrett, Sonia Goncalves, Ewan Harrison, David K. Jackson, Ian Johnston, Dominic Kwiatkowski, Cordelia Langford, John Sillitoe on behalf of the Wellcome Sanger Institute COVID-19 Surveillance Team |
| EPI_ISL_3377124 | Berkshire and Surrey Pathology Services Lighthouse Laboratory | Wellcome Sanger Institute for the COVID-19 Genomics UK (COG-UK) Consortium | Berkshire and Surrey Pathology Services Lighthouse Laboratory and Alex Alderton, Roberto Amato, Jeffrey Barrett, Sonia Goncalves, Ewan Harrison, David K. Jackson, Ian Johnston, Dominic Kwiatkowski, Cordelia Langford, John Sillitoe on behalf of the Wellcome Sanger Institute COVID-19 Surveillance Team |
| EPI_ISL_3377125 | Lighthouse Lab in Alderley Park | Wellcome Sanger Institute for the COVID-19 Genomics UK (COG-UK) Consortium | Jacquelyn Wynn, Mairead Hyland, The Lighthouse Lab in Alderley Park and Alex Alderton, Roberto Amato, Jeffrey Barrett, Sonia Goncalves, Ewan Harrison, David K. Jackson, Ian Johnston, Dominic Kwiatkowski, Cordelia Langford, John Sillitoe on behalf of the Wellcome Sanger Institute COVID-19 Surveillance Team |
| EPI_ISL_3377126 | Lighthouse Lab in Milton Keynes | Wellcome Sanger Institute for the COVID-19 Genomics UK (COG-UK) Consortium | The Lighthouse Lab in Milton Keynes and Alex Alderton, Roberto Amato, Jeffrey Barrett, Sonia Goncalves, Ewan Harrison, David K. Jackson, Ian Johnston, Dominic Kwiatkowski, Cordelia Langford, John Sillitoe on behalf of the Wellcome Sanger Institute COVID-19 Surveillance Team |
| EPI_ISL_3377127 | Lighthouse Lab in Glasgow | Wellcome Sanger Institute for the COVID-19 Genomics UK (COG-UK) Consortium | Harper VanSteenhouse, Yumi Kasai, David Gray, Carol Clugston, Anna Dominiczak and Alex Alderton, Roberto Amato, Jeffrey Barrett, Sonia Goncalves, Ewan Harrison, David K. Jackson, Ian Johnston, Dominic Kwiatkowski, Cordelia Langford, John Sillitoe on behalf of the Wellcome Sanger Institute COVID-19 Surveillance Team |
| EPI_ISL_3377128 | Lighthouse Lab in Alderley Park | Wellcome Sanger Institute for the COVID-19 Genomics UK (COG-UK) Consortium | Jacquelyn Wynn, Mairead Hyland, The Lighthouse Lab in Alderley Park and Alex Alderton, Roberto Amato, Jeffrey Barrett, Sonia Goncalves, Ewan Harrison, David K. Jackson, Ian Johnston, Dominic Kwiatkowski, Cordelia Langford, John Sillitoe on behalf of the Wellcome Sanger Institute COVID-19 Surveillance Team |
| EPI_ISL_3377129 | Lighthouse Lab in Milton Keynes | Wellcome Sanger Institute for the COVID-19 Genomics UK (COG-UK) Consortium | The Lighthouse Lab in Milton Keynes and Alex Alderton, Roberto Amato, Jeffrey Barrett, Sonia Goncalves, Ewan Harrison, David K. Jackson, Ian Johnston, Dominic Kwiatkowski, Cordelia Langford, John Sillitoe on behalf of the Wellcome Sanger Institute COVID-19 Surveillance Team |
| EPI_ISL_3377130 | Berkshire and Surrey Pathology Services Lighthouse Laboratory | Wellcome Sanger Institute for the COVID-19 Genomics UK (COG-UK) Consortium | Berkshire and Surrey Pathology Services Lighthouse Laboratory and Alex Alderton, Roberto Amato, Jeffrey Barrett, Sonia Goncalves, Ewan Harrison, David K. Jackson, Ian Johnston, Dominic Kwiatkowski, Cordelia Langford, John Sillitoe on behalf of the Wellcome Sanger Institute COVID-19 Surveillance Team |
| EPI_ISL_3377131 | Lighthouse Lab in Alderley Park | Wellcome Sanger Institute for the COVID-19 Genomics UK (COG-UK) Consortium | Jacquelyn Wynn, Mairead Hyland, The Lighthouse Lab in Alderley Park and Alex Alderton, Roberto Amato, Jeffrey Barrett, Sonia Goncalves, Ewan Harrison, David K. Jackson, Ian Johnston, Dominic Kwiatkowski, Cordelia Langford, John Sillitoe on behalf of the Wellcome Sanger Institute COVID-19 Surveillance Team |
| EPI_ISL_3377132, EPI_ISL_3377133, EPI_ISL_3377134, EPI_ISL_3377135, EPI_ISL_3377136 | Lighthouse Lab in Milton Keynes | Wellcome Sanger Institute for the COVID-19 Genomics UK (COG-UK) Consortium | The Lighthouse Lab in Milton Keynes and Alex Alderton, Roberto Amato, Jeffrey Barrett, Sonia Goncalves, Ewan Harrison, David K. Jackson, Ian Johnston, Dominic Kwiatkowski, Cordelia Langford, John Sillitoe on behalf of the Wellcome Sanger Institute COVID-19 Surveillance Team |
| EPI_ISL_3377137, EPI_ISL_3377138, EPI_ISL_3377139 | Lighthouse Lab in Alderley Park | Wellcome Sanger Institute for the COVID-19 Genomics UK (COG-UK) Consortium | Jacquelyn Wynn, Mairead Hyland, The Lighthouse Lab in Alderley Park and Alex Alderton, Roberto Amato, Jeffrey Barrett, Sonia Goncalves, Ewan Harrison, David K. Jackson, Ian Johnston, Dominic Kwiatkowski, Cordelia Langford, John Sillitoe on behalf of the Wellcome Sanger Institute COVID-19 Surveillance Team |
| EPI_ISL_3377140, | Lighthouse Lab in Glasgow | Wellcome Sanger Institute for the COVID-19 Genomics UK (COG-UK) | Harper VanSteenhouse, Yumi Kasai, David Gray, Carol Clugston, Anna Dominiczak and Alex Alderton, Roberto Amato, Jeffrey Barrett, Sonia Goncalves, Ewan Harrison, David K. Jackson, Ian Johnston, Dominic Kwiatkowski, Cordelia Langford, |

| | | | |
|---|---|---|---|
| EPI_ISL_3377141 | | Consortium | John Sillitoe on behalf of the Wellcome Sanger Institute COVID-19 Surveillance Team |
| EPI_ISL_3377142 | Berkshire and Surrey Pathology Services Lighthouse Laboratory | Wellcome Sanger Institute for the COVID-19 Genomics UK (COG-UK) Consortium | Berkshire and Surrey Pathology Services Lighthouse Laboratory and Alex Alderton, Roberto Amato, Jeffrey Barrett, Sonia Goncalves, Ewan Harrison, David K. Jackson, Ian Johnston, Dominic Kwiatkowski, Cordelia Langford, John Sillitoe on behalf of the Wellcome Sanger Institute COVID-19 Surveillance Team |
| EPI_ISL_3377143, EPI_ISL_3377144, EPI_ISL_3377145, EPI_ISL_3377146 | Lighthouse Lab in Milton Keynes | Wellcome Sanger Institute for the COVID-19 Genomics UK (COG-UK) Consortium | The Lighthouse Lab in Milton Keynes and Alex Alderton, Roberto Amato, Jeffrey Barrett, Sonia Goncalves, Ewan Harrison, David K. Jackson, Ian Johnston, Dominic Kwiatkowski, Cordelia Langford, John Sillitoe on behalf of the Wellcome Sanger Institute COVID-19 Surveillance Team |
| EPI_ISL_3377147 | Lighthouse Lab in Alderley Park | Wellcome Sanger Institute for the COVID-19 Genomics UK (COG-UK) Consortium | Jacquelyn Wynn, Mairead Hyland, The Lighthouse Lab in Alderley Park and Alex Alderton, Roberto Amato, Jeffrey Barrett, Sonia Goncalves, Ewan Harrison, David K. Jackson, Ian Johnston, Dominic Kwiatkowski, Cordelia Langford, John Sillitoe on behalf of the Wellcome Sanger Institute COVID-19 Surveillance Team |
| EPI_ISL_3377148 | Berkshire and Surrey Pathology Services Lighthouse Laboratory | Wellcome Sanger Institute for the COVID-19 Genomics UK (COG-UK) Consortium | Berkshire and Surrey Pathology Services Lighthouse Laboratory and Alex Alderton, Roberto Amato, Jeffrey Barrett, Sonia Goncalves, Ewan Harrison, David K. Jackson, Ian Johnston, Dominic Kwiatkowski, Cordelia Langford, John Sillitoe on behalf of the Wellcome Sanger Institute COVID-19 Surveillance Team |
| EPI_ISL_3377149 | Lighthouse Lab in Milton Keynes | Wellcome Sanger Institute for the COVID-19 Genomics UK (COG-UK) Consortium | The Lighthouse Lab in Milton Keynes and Alex Alderton, Roberto Amato, Jeffrey Barrett, Sonia Goncalves, Ewan Harrison, David K. Jackson, Ian Johnston, Dominic Kwiatkowski, Cordelia Langford, John Sillitoe on behalf of the Wellcome Sanger Institute COVID-19 Surveillance Team |
| EPI_ISL_3377150, EPI_ISL_3377151 | Lighthouse Lab in Alderley Park | Wellcome Sanger Institute for the COVID-19 Genomics UK (COG-UK) Consortium | Jacquelyn Wynn, Mairead Hyland, The Lighthouse Lab in Alderley Park and Alex Alderton, Roberto Amato, Jeffrey Barrett, Sonia Goncalves, Ewan Harrison, David K. Jackson, Ian Johnston, Dominic Kwiatkowski, Cordelia Langford, John Sillitoe on behalf of the Wellcome Sanger Institute COVID-19 Surveillance Team |
| EPI_ISL_3377152, EPI_ISL_3377153 | Lighthouse Lab in Milton Keynes | Wellcome Sanger Institute for the COVID-19 Genomics UK (COG-UK) Consortium | The Lighthouse Lab in Milton Keynes and Alex Alderton, Roberto Amato, Jeffrey Barrett, Sonia Goncalves, Ewan Harrison, David K. Jackson, Ian Johnston, Dominic Kwiatkowski, Cordelia Langford, John Sillitoe on behalf of the Wellcome Sanger Institute COVID-19 Surveillance Team |
| EPI_ISL_3377154 | Berkshire and Surrey Pathology Services Lighthouse Laboratory | Wellcome Sanger Institute for the COVID-19 Genomics UK (COG-UK) Consortium | Berkshire and Surrey Pathology Services Lighthouse Laboratory and Alex Alderton, Roberto Amato, Jeffrey Barrett, Sonia Goncalves, Ewan Harrison, David K. Jackson, Ian Johnston, Dominic Kwiatkowski, Cordelia Langford, John Sillitoe on behalf of the Wellcome Sanger Institute COVID-19 Surveillance Team |
| EPI_ISL_3377155, EPI_ISL_3377156 | Lighthouse Lab in Milton Keynes | Wellcome Sanger Institute for the COVID-19 Genomics UK (COG-UK) Consortium | The Lighthouse Lab in Milton Keynes and Alex Alderton, Roberto Amato, Jeffrey Barrett, Sonia Goncalves, Ewan Harrison, David K. Jackson, Ian Johnston, Dominic Kwiatkowski, Cordelia Langford, John Sillitoe on behalf of the Wellcome Sanger Institute COVID-19 Surveillance Team |
| EPI_ISL_3377157 | Lighthouse Lab in Alderley Park | Wellcome Sanger Institute for the COVID-19 Genomics UK (COG-UK) Consortium | Jacquelyn Wynn, Mairead Hyland, The Lighthouse Lab in Alderley Park and Alex Alderton, Roberto Amato, Jeffrey Barrett, Sonia Goncalves, Ewan Harrison, David K. Jackson, Ian Johnston, Dominic Kwiatkowski, Cordelia Langford, John Sillitoe on behalf of the Wellcome Sanger Institute COVID-19 Surveillance Team |
| EPI_ISL_3377158 | Berkshire and Surrey Pathology Services Lighthouse Laboratory | Wellcome Sanger Institute for the COVID-19 Genomics UK (COG-UK) Consortium | Berkshire and Surrey Pathology Services Lighthouse Laboratory and Alex Alderton, Roberto Amato, Jeffrey Barrett, Sonia Goncalves, Ewan Harrison, David K. Jackson, Ian Johnston, Dominic Kwiatkowski, Cordelia Langford, John Sillitoe on behalf of the Wellcome Sanger Institute COVID-19 Surveillance Team |
| EPI_ISL_3377159 | Lighthouse Lab in Glasgow | Wellcome Sanger Institute for the COVID-19 Genomics UK (COG-UK) Consortium | Harper VanSteenhouse, Yumi Kasai, David Gray, Carol Clugston, Anna Dominiczak and Alex Alderton, Roberto Amato, Jeffrey Barrett, Sonia Goncalves, Ewan Harrison, David K. Jackson, Ian Johnston, Dominic Kwiatkowski, Cordelia Langford, John Sillitoe on behalf of the Wellcome Sanger Institute COVID-19 Surveillance Team |
| EPI_ISL_3377160 | Lighthouse Lab in Alderley Park | Wellcome Sanger Institute for the COVID-19 Genomics UK (COG-UK) Consortium | Jacquelyn Wynn, Mairead Hyland, The Lighthouse Lab in Alderley Park and Alex Alderton, Roberto Amato, Jeffrey Barrett, Sonia Goncalves, Ewan Harrison, David K. Jackson, Ian Johnston, Dominic Kwiatkowski, Cordelia Langford, John Sillitoe on behalf of the Wellcome Sanger Institute COVID-19 Surveillance Team |
| EPI_ISL_3377161 | Lighthouse Lab in Milton Keynes | Wellcome Sanger Institute for the COVID-19 Genomics UK (COG-UK) Consortium | The Lighthouse Lab in Milton Keynes and Alex Alderton, Roberto Amato, Jeffrey Barrett, Sonia Goncalves, Ewan Harrison, David K. Jackson, Ian Johnston, Dominic Kwiatkowski, Cordelia Langford, John Sillitoe on behalf of the Wellcome Sanger Institute COVID-19 Surveillance Team |
| EPI_ISL_3377162, EPI_ISL_3377163 | Lighthouse Lab in Glasgow | Wellcome Sanger Institute for the COVID-19 Genomics UK (COG-UK) Consortium | Harper VanSteenhouse, Yumi Kasai, David Gray, Carol Clugston, Anna Dominiczak and Alex Alderton, Roberto Amato, Jeffrey Barrett, Sonia Goncalves, Ewan Harrison, David K. Jackson, Ian Johnston, Dominic Kwiatkowski, Cordelia Langford, John Sillitoe on behalf of the Wellcome Sanger Institute COVID-19 Surveillance Team |
| EPI_ISL_3377164, EPI_ISL_3377165 | Lighthouse Lab in Milton Keynes | Wellcome Sanger Institute for the COVID-19 Genomics UK (COG-UK) Consortium | The Lighthouse Lab in Milton Keynes and Alex Alderton, Roberto Amato, Jeffrey Barrett, Sonia Goncalves, Ewan Harrison, David K. Jackson, Ian Johnston, Dominic Kwiatkowski, Cordelia Langford, John Sillitoe on behalf of the Wellcome Sanger Institute COVID-19 Surveillance Team |
| EPI_ISL_3377166 | Lighthouse Lab in Glasgow | Wellcome Sanger Institute for the COVID-19 Genomics UK (COG-UK) Consortium | Harper VanSteenhouse, Yumi Kasai, David Gray, Carol Clugston, Anna Dominiczak and Alex Alderton, Roberto Amato, Jeffrey Barrett, Sonia Goncalves, Ewan Harrison, David K. Jackson, Ian Johnston, Dominic Kwiatkowski, Cordelia Langford, John Sillitoe on behalf of the Wellcome Sanger Institute COVID-19 Surveillance Team |
| EPI_ISL_3377167 | Lighthouse Lab in Alderley Park | Wellcome Sanger Institute for the COVID-19 Genomics UK (COG-UK) Consortium | Jacquelyn Wynn, Mairead Hyland, The Lighthouse Lab in Alderley Park and Alex Alderton, Roberto Amato, Jeffrey Barrett, Sonia Goncalves, Ewan Harrison, David K. Jackson, Ian Johnston, Dominic Kwiatkowski, Cordelia Langford, John Sillitoe on behalf of the Wellcome Sanger Institute COVID-19 Surveillance Team |
| EPI_ISL_3377168 | Lighthouse Lab in Glasgow | Wellcome Sanger Institute for the COVID-19 Genomics UK (COG-UK) Consortium | Harper VanSteenhouse, Yumi Kasai, David Gray, Carol Clugston, Anna Dominiczak and Alex Alderton, Roberto Amato, Jeffrey Barrett, Sonia Goncalves, Ewan Harrison, David K. Jackson, Ian Johnston, Dominic Kwiatkowski, Cordelia Langford, John Sillitoe on behalf of the Wellcome Sanger Institute COVID-19 Surveillance Team |
| EPI_ISL_3377169, EPI_ISL_3377170 | Lighthouse Lab in Alderley Park | Wellcome Sanger Institute for the COVID-19 Genomics UK (COG-UK) Consortium | Jacquelyn Wynn, Mairead Hyland, The Lighthouse Lab in Alderley Park and Alex Alderton, Roberto Amato, Jeffrey Barrett, Sonia Goncalves, Ewan Harrison, David K. Jackson, Ian Johnston, Dominic Kwiatkowski, Cordelia Langford, John Sillitoe on behalf of the Wellcome Sanger Institute COVID-19 Surveillance Team |
| EPI_ISL_3377171 | Lighthouse Lab in Glasgow | Wellcome Sanger Institute for the COVID-19 Genomics UK (COG-UK) Consortium | Harper VanSteenhouse, Yumi Kasai, David Gray, Carol Clugston, Anna Dominiczak and Alex Alderton, Roberto Amato, Jeffrey Barrett, Sonia Goncalves, Ewan Harrison, David K. Jackson, Ian Johnston, Dominic Kwiatkowski, Cordelia Langford, John Sillitoe on behalf of the Wellcome Sanger Institute COVID-19 Surveillance Team |
| EPI_ISL_3377172 | Lighthouse Lab in Milton Keynes | Wellcome Sanger Institute for the COVID-19 Genomics UK (COG-UK) Consortium | The Lighthouse Lab in Milton Keynes and Alex Alderton, Roberto Amato, Jeffrey Barrett, Sonia Goncalves, Ewan Harrison, David K. Jackson, Ian Johnston, Dominic Kwiatkowski, Cordelia Langford, John Sillitoe on behalf of the Wellcome Sanger Institute COVID-19 Surveillance Team |
| EPI_ISL_3377173 | Lighthouse Lab in Alderley Park | Wellcome Sanger Institute for the COVID-19 Genomics UK (COG-UK) Consortium | Jacquelyn Wynn, Mairead Hyland, The Lighthouse Lab in Alderley Park and Alex Alderton, Roberto Amato, Jeffrey Barrett, Sonia Goncalves, Ewan Harrison, David K. Jackson, Ian Johnston, Dominic Kwiatkowski, Cordelia Langford, John Sillitoe on behalf of the Wellcome Sanger Institute COVID-19 Surveillance Team |
| EPI_ISL_3377174 | Lighthouse Lab in Glasgow | Wellcome Sanger Institute for the COVID-19 Genomics UK (COG-UK) Consortium | Harper VanSteenhouse, Yumi Kasai, David Gray, Carol Clugston, Anna Dominiczak and Alex Alderton, Roberto Amato, Jeffrey Barrett, Sonia Goncalves, Ewan Harrison, David K. Jackson, Ian Johnston, Dominic Kwiatkowski, Cordelia Langford, John Sillitoe on behalf of the Wellcome Sanger Institute COVID-19 Surveillance Team |
| EPI_ISL_3377175, EPI_ISL_3377176 | Lighthouse Lab in Alderley Park | Wellcome Sanger Institute for the COVID-19 Genomics UK (COG-UK) Consortium | Jacquelyn Wynn, Mairead Hyland, The Lighthouse Lab in Alderley Park and Alex Alderton, Roberto Amato, Jeffrey Barrett, Sonia Goncalves, Ewan Harrison, David K. Jackson, Ian Johnston, Dominic Kwiatkowski, Cordelia Langford, John Sillitoe on behalf of the Wellcome Sanger Institute COVID-19 Surveillance Team |
| EPI_ISL_3377177, EPI_ISL_3377178, EPI_ISL_3377179, EPI_ISL_3377180 | Lighthouse Lab in Milton Keynes | Wellcome Sanger Institute for the COVID-19 Genomics UK (COG-UK) Consortium | The Lighthouse Lab in Milton Keynes and Alex Alderton, Roberto Amato, Jeffrey Barrett, Sonia Goncalves, Ewan Harrison, David K. Jackson, Ian Johnston, Dominic Kwiatkowski, Cordelia Langford, John Sillitoe on behalf of the Wellcome Sanger Institute COVID-19 Surveillance Team |
| EPI_ISL_3377181 | Berkshire and Surrey Pathology Services Lighthouse Laboratory | Wellcome Sanger Institute for the COVID-19 Genomics UK (COG-UK) Consortium | Berkshire and Surrey Pathology Services Lighthouse Laboratory and Alex Alderton, Roberto Amato, Jeffrey Barrett, Sonia Goncalves, Ewan Harrison, David K. Jackson, Ian Johnston, Dominic Kwiatkowski, Cordelia Langford, John Sillitoe on behalf of the Wellcome Sanger Institute COVID-19 Surveillance Team |
| EPI_ISL_3377182 | Lighthouse Lab in Milton Keynes | Wellcome Sanger Institute for the COVID-19 Genomics UK (COG-UK) Consortium | The Lighthouse Lab in Milton Keynes and Alex Alderton, Roberto Amato, Jeffrey Barrett, Sonia Goncalves, Ewan Harrison, David K. Jackson, Ian Johnston, Dominic Kwiatkowski, Cordelia Langford, John Sillitoe on behalf of the Wellcome Sanger Institute COVID-19 Surveillance Team |
| EPI_ISL_3377183 | Berkshire and Surrey Pathology Services Lighthouse Laboratory | Wellcome Sanger Institute for the COVID-19 Genomics UK (COG-UK) Consortium | Berkshire and Surrey Pathology Services Lighthouse Laboratory and Alex Alderton, Roberto Amato, Jeffrey Barrett, Sonia Goncalves, Ewan Harrison, David K. Jackson, Ian Johnston, Dominic Kwiatkowski, Cordelia Langford, John Sillitoe on behalf of the Wellcome Sanger Institute COVID-19 Surveillance Team |
| EPI_ISL_3377184 | Lighthouse Lab in Milton Keynes | Wellcome Sanger Institute for the COVID-19 Genomics UK (COG-UK) Consortium | The Lighthouse Lab in Milton Keynes and Alex Alderton, Roberto Amato, Jeffrey Barrett, Sonia Goncalves, Ewan Harrison, David K. Jackson, Ian Johnston, Dominic Kwiatkowski, Cordelia Langford, John Sillitoe on behalf of the Wellcome Sanger Institute COVID-19 Surveillance Team |
| EPI_ISL_3377185 | Berkshire and Surrey Pathology Services Lighthouse Laboratory | Wellcome Sanger Institute for the COVID-19 Genomics UK (COG-UK) Consortium | Berkshire and Surrey Pathology Services Lighthouse Laboratory and Alex Alderton, Roberto Amato, Jeffrey Barrett, Sonia Goncalves, Ewan Harrison, David K. Jackson, Ian Johnston, Dominic Kwiatkowski, Cordelia Langford, John Sillitoe on behalf of the Wellcome Sanger Institute COVID-19 Surveillance Team |
| EPI_ISL_3377186 | Lighthouse Lab in Alderley Park | Wellcome Sanger Institute for the COVID-19 Genomics UK (COG-UK) Consortium | Jacquelyn Wynn, Mairead Hyland, The Lighthouse Lab in Alderley Park and Alex Alderton, Roberto Amato, Jeffrey Barrett, Sonia Goncalves, Ewan Harrison, David K. Jackson, Ian Johnston, Dominic Kwiatkowski, Cordelia Langford, John Sillitoe on behalf of the Wellcome Sanger Institute COVID-19 Surveillance Team |
| EPI_ISL_3377187 | Lighthouse Lab in Milton Keynes | Wellcome Sanger Institute for the COVID-19 Genomics UK (COG-UK) Consortium | The Lighthouse Lab in Milton Keynes and Alex Alderton, Roberto Amato, Jeffrey Barrett, Sonia Goncalves, Ewan Harrison, David K. Jackson, Ian Johnston, Dominic Kwiatkowski, Cordelia Langford, John Sillitoe on behalf of the Wellcome Sanger Institute COVID-19 Surveillance Team |
| EPI_ISL_3377188 | Lighthouse Lab in Glasgow | Wellcome Sanger Institute for the COVID-19 Genomics UK (COG-UK) Consortium | Harper VanSteenhouse, Yumi Kasai, David Gray, Carol Clugston, Anna Dominiczak and Alex Alderton, Roberto Amato, Jeffrey Barrett, Sonia Goncalves, Ewan Harrison, David K. Jackson, Ian Johnston, Dominic Kwiatkowski, Cordelia Langford, John Sillitoe on behalf of the Wellcome Sanger Institute COVID-19 Surveillance Team |
| EPI_ISL_3377189, EPI_ISL_3377190, EPI_ISL_3377191 | Berkshire and Surrey Pathology Services Lighthouse Laboratory | Wellcome Sanger Institute for the COVID-19 Genomics UK (COG-UK) Consortium | Berkshire and Surrey Pathology Services Lighthouse Laboratory and Alex Alderton, Roberto Amato, Jeffrey Barrett, Sonia Goncalves, Ewan Harrison, David K. Jackson, Ian Johnston, Dominic Kwiatkowski, Cordelia Langford, John Sillitoe on behalf of the Wellcome Sanger Institute COVID-19 Surveillance Team |
| EPI_ISL_3377192 | Lighthouse Lab in Alderley Park | Wellcome Sanger Institute for the COVID-19 Genomics UK (COG-UK) Consortium | Jacquelyn Wynn, Mairead Hyland, The Lighthouse Lab in Alderley Park and Alex Alderton, Roberto Amato, Jeffrey Barrett, Sonia Goncalves, Ewan Harrison, David K. Jackson, Ian Johnston, Dominic Kwiatkowski, Cordelia Langford, John Sillitoe on behalf of the Wellcome Sanger Institute COVID-19 Surveillance Team |
| EPI_ISL_3377193 | Berkshire and Surrey Pathology Services Lighthouse Laboratory | Wellcome Sanger Institute for the COVID-19 Genomics UK (COG-UK) Consortium | Berkshire and Surrey Pathology Services Lighthouse Laboratory and Alex Alderton, Roberto Amato, Jeffrey Barrett, Sonia Goncalves, Ewan Harrison, David K. Jackson, Ian Johnston, Dominic Kwiatkowski, Cordelia Langford, John Sillitoe on behalf of the Wellcome Sanger Institute COVID-19 Surveillance Team |
| EPI_ISL_3377194 | Lighthouse Lab in Milton Keynes | Wellcome Sanger Institute for the COVID-19 Genomics UK (COG-UK) Consortium | The Lighthouse Lab in Milton Keynes and Alex Alderton, Roberto Amato, Jeffrey Barrett, Sonia Goncalves, Ewan Harrison, David K. Jackson, Ian Johnston, Dominic Kwiatkowski, Cordelia Langford, John Sillitoe on behalf of the Wellcome Sanger Institute COVID-19 Surveillance Team |
| EPI_ISL_3377195 | Lighthouse Lab in Glasgow | Wellcome Sanger Institute for the COVID-19 Genomics UK (COG-UK) Consortium | Harper VanSteenhouse, Yumi Kasai, David Gray, Carol Clugston, Anna Dominiczak and Alex Alderton, Roberto Amato, Jeffrey Barrett, Sonia Goncalves, Ewan Harrison, David K. Jackson, Ian Johnston, Dominic Kwiatkowski, Cordelia Langford, John Sillitoe on behalf of the Wellcome Sanger Institute COVID-19 Surveillance Team |
| EPI_ISL_3377196, EPI_ISL_3377197 | Lighthouse Lab in Alderley Park | Wellcome Sanger Institute for the COVID-19 Genomics UK (COG-UK) Consortium | Jacquelyn Wynn, Mairead Hyland, The Lighthouse Lab in Alderley Park and Alex Alderton, Roberto Amato, Jeffrey Barrett, Sonia Goncalves, Ewan Harrison, David K. Jackson, Ian Johnston, Dominic Kwiatkowski, Cordelia Langford, John Sillitoe on behalf of the Wellcome Sanger Institute COVID-19 Surveillance Team |
| EPI_ISL_3377198 | Lighthouse Lab in Milton Keynes | Wellcome Sanger Institute for the COVID-19 Genomics UK (COG-UK) Consortium | The Lighthouse Lab in Milton Keynes and Alex Alderton, Roberto Amato, Jeffrey Barrett, Sonia Goncalves, Ewan Harrison, David K. Jackson, Ian Johnston, Dominic Kwiatkowski, Cordelia Langford, John Sillitoe on behalf of the Wellcome Sanger Institute COVID-19 Surveillance Team |
| EPI_ISL_3377199 | Lighthouse Lab in Glasgow | Wellcome Sanger Institute for the COVID-19 Genomics UK (COG-UK) Consortium | Harper VanSteenhouse, Yumi Kasai, David Gray, Carol Clugston, Anna Dominiczak and Alex Alderton, Roberto Amato, Jeffrey Barrett, Sonia Goncalves, Ewan Harrison, David K. Jackson, Ian Johnston, Dominic Kwiatkowski, Cordelia Langford, John Sillitoe on behalf of the Wellcome Sanger Institute COVID-19 Surveillance Team |
| EPI_ISL_3377200 | Berkshire and Surrey Pathology Services Lighthouse Laboratory | Wellcome Sanger Institute for the COVID-19 Genomics UK (COG-UK) | Berkshire and Surrey Pathology Services Lighthouse Laboratory and Alex Alderton, Roberto Amato, Jeffrey Barrett, Sonia Goncalves, Ewan Harrison, David K. Jackson, Ian Johnston, Dominic Kwiatkowski, Cordelia Langford, John Sillitoe on behalf |

| | | | |
|---|---|---|---|
| | | Consortium | of the Wellcome Sanger Institute COVID-19 Surveillance Team |
| EPI_ISL_3377201 | Lighthouse Lab in Milton Keynes | Wellcome Sanger Institute for the COVID-19 Genomics UK (COG-UK) Consortium | The Lighthouse Lab in Milton Keynes and Alex Alderton, Roberto Amato, Jeffrey Barrett, Sonia Goncalves, Ewan Harrison, David K. Jackson, Ian Johnston, Dominic Kwiatkowski, Cordelia Langford, John Sillitoe on behalf of the Wellcome Sanger Institute COVID-19 Surveillance Team |
| EPI_ISL_3377202 | Lighthouse Lab in Alderley Park | Wellcome Sanger Institute for the COVID-19 Genomics UK (COG-UK) Consortium | Jacquelyn Wynn, Mairead Hyland, The Lighthouse Lab in Alderley Park and Alex Alderton, Roberto Amato, Jeffrey Barrett, Sonia Goncalves, Ewan Harrison, David K. Jackson, Ian Johnston, Dominic Kwiatkowski, Cordelia Langford, John Sillitoe on behalf of the Wellcome Sanger Institute COVID-19 Surveillance Team |
| EPI_ISL_3377203, EPI_ISL_3377204 | Lighthouse Lab in Milton Keynes | Wellcome Sanger Institute for the COVID-19 Genomics UK (COG-UK) Consortium | The Lighthouse Lab in Milton Keynes and Alex Alderton, Roberto Amato, Jeffrey Barrett, Sonia Goncalves, Ewan Harrison, David K. Jackson, Ian Johnston, Dominic Kwiatkowski, Cordelia Langford, John Sillitoe on behalf of the Wellcome Sanger Institute COVID-19 Surveillance Team |
| EPI_ISL_3377205 | Lighthouse Lab in Alderley Park | Wellcome Sanger Institute for the COVID-19 Genomics UK (COG-UK) Consortium | Jacquelyn Wynn, Mairead Hyland, The Lighthouse Lab in Alderley Park and Alex Alderton, Roberto Amato, Jeffrey Barrett, Sonia Goncalves, Ewan Harrison, David K. Jackson, Ian Johnston, Dominic Kwiatkowski, Cordelia Langford, John Sillitoe on behalf of the Wellcome Sanger Institute COVID-19 Surveillance Team |
| EPI_ISL_3377206 | Lighthouse Lab in Glasgow | Wellcome Sanger Institute for the COVID-19 Genomics UK (COG-UK) Consortium | Harper VanSteenhouse, Yumi Kasai, David Gray, Carol Clugston, Anna Dominiczak and Alex Alderton, Roberto Amato, Jeffrey Barrett, Sonia Goncalves, Ewan Harrison, David K. Jackson, Ian Johnston, Dominic Kwiatkowski, Cordelia Langford, John Sillitoe on behalf of the Wellcome Sanger Institute COVID-19 Surveillance Team |
| EPI_ISL_3377207 | Berkshire and Surrey Pathology Services Lighthouse Laboratory | Wellcome Sanger Institute for the COVID-19 Genomics UK (COG-UK) Consortium | Berkshire and Surrey Pathology Services Lighthouse Laboratory and Alex Alderton, Roberto Amato, Jeffrey Barrett, Sonia Goncalves, Ewan Harrison, David K. Jackson, Ian Johnston, Dominic Kwiatkowski, Cordelia Langford, John Sillitoe on behalf of the Wellcome Sanger Institute COVID-19 Surveillance Team |
| EPI_ISL_3377208, EPI_ISL_3377209 | Lighthouse Lab in Alderley Park | Wellcome Sanger Institute for the COVID-19 Genomics UK (COG-UK) Consortium | Jacquelyn Wynn, Mairead Hyland, The Lighthouse Lab in Alderley Park and Alex Alderton, Roberto Amato, Jeffrey Barrett, Sonia Goncalves, Ewan Harrison, David K. Jackson, Ian Johnston, Dominic Kwiatkowski, Cordelia Langford, John Sillitoe on behalf of the Wellcome Sanger Institute COVID-19 Surveillance Team |
| EPI_ISL_3377210, EPI_ISL_3377211, EPI_ISL_3377212, EPI_ISL_3377213 | Berkshire and Surrey Pathology Services Lighthouse Laboratory | Wellcome Sanger Institute for the COVID-19 Genomics UK (COG-UK) Consortium | Berkshire and Surrey Pathology Services Lighthouse Laboratory and Alex Alderton, Roberto Amato, Jeffrey Barrett, Sonia Goncalves, Ewan Harrison, David K. Jackson, Ian Johnston, Dominic Kwiatkowski, Cordelia Langford, John Sillitoe on behalf of the Wellcome Sanger Institute COVID-19 Surveillance Team |
| EPI_ISL_3377214 | Lighthouse Lab in Glasgow | Wellcome Sanger Institute for the COVID-19 Genomics UK (COG-UK) Consortium | Harper VanSteenhouse, Yumi Kasai, David Gray, Carol Clugston, Anna Dominiczak and Alex Alderton, Roberto Amato, Jeffrey Barrett, Sonia Goncalves, Ewan Harrison, David K. Jackson, Ian Johnston, Dominic Kwiatkowski, Cordelia Langford, John Sillitoe on behalf of the Wellcome Sanger Institute COVID-19 Surveillance Team |
| EPI_ISL_3377215 | Lighthouse Lab in Milton Keynes | Wellcome Sanger Institute for the COVID-19 Genomics UK (COG-UK) Consortium | The Lighthouse Lab in Milton Keynes and Alex Alderton, Roberto Amato, Jeffrey Barrett, Sonia Goncalves, Ewan Harrison, David K. Jackson, Ian Johnston, Dominic Kwiatkowski, Cordelia Langford, John Sillitoe on behalf of the Wellcome Sanger Institute COVID-19 Surveillance Team |
| EPI_ISL_3377216 | Lighthouse Lab in Glasgow | Wellcome Sanger Institute for the COVID-19 Genomics UK (COG-UK) Consortium | Harper VanSteenhouse, Yumi Kasai, David Gray, Carol Clugston, Anna Dominiczak and Alex Alderton, Roberto Amato, Jeffrey Barrett, Sonia Goncalves, Ewan Harrison, David K. Jackson, Ian Johnston, Dominic Kwiatkowski, Cordelia Langford, John Sillitoe on behalf of the Wellcome Sanger Institute COVID-19 Surveillance Team |
| EPI_ISL_3377217 | Lighthouse Lab in Alderley Park | Wellcome Sanger Institute for the COVID-19 Genomics UK (COG-UK) Consortium | Jacquelyn Wynn, Mairead Hyland, The Lighthouse Lab in Alderley Park and Alex Alderton, Roberto Amato, Jeffrey Barrett, Sonia Goncalves, Ewan Harrison, David K. Jackson, Ian Johnston, Dominic Kwiatkowski, Cordelia Langford, John Sillitoe on behalf of the Wellcome Sanger Institute COVID-19 Surveillance Team |
| EPI_ISL_3377218, EPI_ISL_3377219 | Berkshire and Surrey Pathology Services Lighthouse Laboratory | Wellcome Sanger Institute for the COVID-19 Genomics UK (COG-UK) Consortium | Berkshire and Surrey Pathology Services Lighthouse Laboratory and Alex Alderton, Roberto Amato, Jeffrey Barrett, Sonia Goncalves, Ewan Harrison, David K. Jackson, Ian Johnston, Dominic Kwiatkowski, Cordelia Langford, John Sillitoe on behalf of the Wellcome Sanger Institute COVID-19 Surveillance Team |
| EPI_ISL_3377220 | Lighthouse Lab in Milton Keynes | Wellcome Sanger Institute for the COVID-19 Genomics UK (COG-UK) Consortium | The Lighthouse Lab in Milton Keynes and Alex Alderton, Roberto Amato, Jeffrey Barrett, Sonia Goncalves, Ewan Harrison, David K. Jackson, Ian Johnston, Dominic Kwiatkowski, Cordelia Langford, John Sillitoe on behalf of the Wellcome Sanger Institute COVID-19 Surveillance Team |
| EPI_ISL_3377221 | Lighthouse Lab in Alderley Park | Wellcome Sanger Institute for the COVID-19 Genomics UK (COG-UK) Consortium | Jacquelyn Wynn, Mairead Hyland, The Lighthouse Lab in Alderley Park and Alex Alderton, Roberto Amato, Jeffrey Barrett, Sonia Goncalves, Ewan Harrison, David K. Jackson, Ian Johnston, Dominic Kwiatkowski, Cordelia Langford, John Sillitoe on behalf of the Wellcome Sanger Institute COVID-19 Surveillance Team |
| EPI_ISL_3377222 | Lighthouse Lab in Glasgow | Wellcome Sanger Institute for the COVID-19 Genomics UK (COG-UK) Consortium | Harper VanSteenhouse, Yumi Kasai, David Gray, Carol Clugston, Anna Dominiczak and Alex Alderton, Roberto Amato, Jeffrey Barrett, Sonia Goncalves, Ewan Harrison, David K. Jackson, Ian Johnston, Dominic Kwiatkowski, Cordelia Langford, John Sillitoe on behalf of the Wellcome Sanger Institute COVID-19 Surveillance Team |
| EPI_ISL_3377223 | Lighthouse Lab in Alderley Park | Wellcome Sanger Institute for the COVID-19 Genomics UK (COG-UK) Consortium | Jacquelyn Wynn, Mairead Hyland, The Lighthouse Lab in Alderley Park and Alex Alderton, Roberto Amato, Jeffrey Barrett, Sonia Goncalves, Ewan Harrison, David K. Jackson, Ian Johnston, Dominic Kwiatkowski, Cordelia Langford, John Sillitoe on behalf of the Wellcome Sanger Institute COVID-19 Surveillance Team |
| EPI_ISL_3377224, EPI_ISL_3377225 | Berkshire and Surrey Pathology Services Lighthouse Laboratory | Wellcome Sanger Institute for the COVID-19 Genomics UK (COG-UK) Consortium | Berkshire and Surrey Pathology Services Lighthouse Laboratory and Alex Alderton, Roberto Amato, Jeffrey Barrett, Sonia Goncalves, Ewan Harrison, David K. Jackson, Ian Johnston, Dominic Kwiatkowski, Cordelia Langford, John Sillitoe on behalf of the Wellcome Sanger Institute COVID-19 Surveillance Team |
| EPI_ISL_3377226 | Lighthouse Lab in Milton Keynes | Wellcome Sanger Institute for the COVID-19 Genomics UK (COG-UK) Consortium | The Lighthouse Lab in Milton Keynes and Alex Alderton, Roberto Amato, Jeffrey Barrett, Sonia Goncalves, Ewan Harrison, David K. Jackson, Ian Johnston, Dominic Kwiatkowski, Cordelia Langford, John Sillitoe on behalf of the Wellcome Sanger Institute COVID-19 Surveillance Team |
| EPI_ISL_3377227 | Berkshire and Surrey Pathology Services Lighthouse Laboratory | Wellcome Sanger Institute for the COVID-19 Genomics UK (COG-UK) Consortium | Berkshire and Surrey Pathology Services Lighthouse Laboratory and Alex Alderton, Roberto Amato, Jeffrey Barrett, Sonia Goncalves, Ewan Harrison, David K. Jackson, Ian Johnston, Dominic Kwiatkowski, Cordelia Langford, John Sillitoe on behalf of the Wellcome Sanger Institute COVID-19 Surveillance Team |
| EPI_ISL_3377228 | Lighthouse Lab in Alderley Park | Wellcome Sanger Institute for the COVID-19 Genomics UK (COG-UK) Consortium | Jacquelyn Wynn, Mairead Hyland, The Lighthouse Lab in Alderley Park and Alex Alderton, Roberto Amato, Jeffrey Barrett, Sonia Goncalves, Ewan Harrison, David K. Jackson, Ian Johnston, Dominic Kwiatkowski, Cordelia Langford, John Sillitoe on behalf of the Wellcome Sanger Institute COVID-19 Surveillance Team |
| EPI_ISL_3377229 | Lighthouse Lab in Glasgow | Wellcome Sanger Institute for the COVID-19 Genomics UK (COG-UK) Consortium | Harper VanSteenhouse, Yumi Kasai, David Gray, Carol Clugston, Anna Dominiczak and Alex Alderton, Roberto Amato, Jeffrey Barrett, Sonia Goncalves, Ewan Harrison, David K. Jackson, Ian Johnston, Dominic Kwiatkowski, Cordelia Langford, John Sillitoe on behalf of the Wellcome Sanger Institute COVID-19 Surveillance Team |
| EPI_ISL_3377230 | Lighthouse Lab in Milton Keynes | Wellcome Sanger Institute for the COVID-19 Genomics UK (COG-UK) Consortium | The Lighthouse Lab in Milton Keynes and Alex Alderton, Roberto Amato, Jeffrey Barrett, Sonia Goncalves, Ewan Harrison, David K. Jackson, Ian Johnston, Dominic Kwiatkowski, Cordelia Langford, John Sillitoe on behalf of the Wellcome Sanger Institute COVID-19 Surveillance Team |
| EPI_ISL_3377231, EPI_ISL_3377232 | Berkshire and Surrey Pathology Services Lighthouse Laboratory | Wellcome Sanger Institute for the COVID-19 Genomics UK (COG-UK) Consortium | Berkshire and Surrey Pathology Services Lighthouse Laboratory and Alex Alderton, Roberto Amato, Jeffrey Barrett, Sonia Goncalves, Ewan Harrison, David K. Jackson, Ian Johnston, Dominic Kwiatkowski, Cordelia Langford, John Sillitoe on behalf of the Wellcome Sanger Institute COVID-19 Surveillance Team |
| EPI_ISL_3377233 | Lighthouse Lab in Alderley Park | Wellcome Sanger Institute for the COVID-19 Genomics UK (COG-UK) Consortium | Jacquelyn Wynn, Mairead Hyland, The Lighthouse Lab in Alderley Park and Alex Alderton, Roberto Amato, Jeffrey Barrett, Sonia Goncalves, Ewan Harrison, David K. Jackson, Ian Johnston, Dominic Kwiatkowski, Cordelia Langford, John Sillitoe on behalf of the Wellcome Sanger Institute COVID-19 Surveillance Team |
| EPI_ISL_3377234 | Lighthouse Lab in Glasgow | Wellcome Sanger Institute for the COVID-19 Genomics UK (COG-UK) Consortium | Harper VanSteenhouse, Yumi Kasai, David Gray, Carol Clugston, Anna Dominiczak and Alex Alderton, Roberto Amato, Jeffrey Barrett, Sonia Goncalves, Ewan Harrison, David K. Jackson, Ian Johnston, Dominic Kwiatkowski, Cordelia Langford, John Sillitoe on behalf of the Wellcome Sanger Institute COVID-19 Surveillance Team |
| EPI_ISL_3377235 | Berkshire and Surrey Pathology Services Lighthouse Laboratory | Wellcome Sanger Institute for the COVID-19 Genomics UK (COG-UK) Consortium | Berkshire and Surrey Pathology Services Lighthouse Laboratory and Alex Alderton, Roberto Amato, Jeffrey Barrett, Sonia Goncalves, Ewan Harrison, David K. Jackson, Ian Johnston, Dominic Kwiatkowski, Cordelia Langford, John Sillitoe on behalf of the Wellcome Sanger Institute COVID-19 Surveillance Team |
| EPI_ISL_3377236 | Lighthouse Lab in Milton Keynes | Wellcome Sanger Institute for the COVID-19 Genomics UK (COG-UK) Consortium | The Lighthouse Lab in Milton Keynes and Alex Alderton, Roberto Amato, Jeffrey Barrett, Sonia Goncalves, Ewan Harrison, David K. Jackson, Ian Johnston, Dominic Kwiatkowski, Cordelia Langford, John Sillitoe on behalf of the Wellcome Sanger Institute COVID-19 Surveillance Team |
| EPI_ISL_3377237 | Berkshire and Surrey Pathology Services Lighthouse Laboratory | Wellcome Sanger Institute for the COVID-19 Genomics UK (COG-UK) Consortium | Berkshire and Surrey Pathology Services Lighthouse Laboratory and Alex Alderton, Roberto Amato, Jeffrey Barrett, Sonia Goncalves, Ewan Harrison, David K. Jackson, Ian Johnston, Dominic Kwiatkowski, Cordelia Langford, John Sillitoe on behalf of the Wellcome Sanger Institute COVID-19 Surveillance Team |
| EPI_ISL_3377238 | Lighthouse Lab in Alderley Park | Wellcome Sanger Institute for the COVID-19 Genomics UK (COG-UK) Consortium | Jacquelyn Wynn, Mairead Hyland, The Lighthouse Lab in Alderley Park and Alex Alderton, Roberto Amato, Jeffrey Barrett, Sonia Goncalves, Ewan Harrison, David K. Jackson, Ian Johnston, Dominic Kwiatkowski, Cordelia Langford, John Sillitoe on behalf of the Wellcome Sanger Institute COVID-19 Surveillance Team |
| EPI_ISL_3377239 | Lighthouse Lab in Milton Keynes | Wellcome Sanger Institute for the COVID-19 Genomics UK (COG-UK) Consortium | The Lighthouse Lab in Milton Keynes and Alex Alderton, Roberto Amato, Jeffrey Barrett, Sonia Goncalves, Ewan Harrison, David K. Jackson, Ian Johnston, Dominic Kwiatkowski, Cordelia Langford, John Sillitoe on behalf of the Wellcome Sanger Institute COVID-19 Surveillance Team |
| EPI_ISL_3377240 | Lighthouse Lab in Alderley Park | Wellcome Sanger Institute for the COVID-19 Genomics UK (COG-UK) Consortium | Jacquelyn Wynn, Mairead Hyland, The Lighthouse Lab in Alderley Park and Alex Alderton, Roberto Amato, Jeffrey Barrett, Sonia Goncalves, Ewan Harrison, David K. Jackson, Ian Johnston, Dominic Kwiatkowski, Cordelia Langford, John Sillitoe on behalf of the Wellcome Sanger Institute COVID-19 Surveillance Team |
| EPI_ISL_3377241 | Lighthouse Lab in Milton Keynes | Wellcome Sanger Institute for the COVID-19 Genomics UK (COG-UK) Consortium | The Lighthouse Lab in Milton Keynes and Alex Alderton, Roberto Amato, Jeffrey Barrett, Sonia Goncalves, Ewan Harrison, David K. Jackson, Ian Johnston, Dominic Kwiatkowski, Cordelia Langford, John Sillitoe on behalf of the Wellcome Sanger Institute COVID-19 Surveillance Team |
| EPI_ISL_3377242 | Lighthouse Lab in Glasgow | Wellcome Sanger Institute for the COVID-19 Genomics UK (COG-UK) Consortium | Harper VanSteenhouse, Yumi Kasai, David Gray, Carol Clugston, Anna Dominiczak and Alex Alderton, Roberto Amato, Jeffrey Barrett, Sonia Goncalves, Ewan Harrison, David K. Jackson, Ian Johnston, Dominic Kwiatkowski, Cordelia Langford, John Sillitoe on behalf of the Wellcome Sanger Institute COVID-19 Surveillance Team |
| EPI_ISL_3377243 | Berkshire and Surrey Pathology Services Lighthouse Laboratory | Wellcome Sanger Institute for the COVID-19 Genomics UK (COG-UK) Consortium | Berkshire and Surrey Pathology Services Lighthouse Laboratory and Alex Alderton, Roberto Amato, Jeffrey Barrett, Sonia Goncalves, Ewan Harrison, David K. Jackson, Ian Johnston, Dominic Kwiatkowski, Cordelia Langford, John Sillitoe on behalf of the Wellcome Sanger Institute COVID-19 Surveillance Team |
| EPI_ISL_3377244 | Lighthouse Lab in Alderley Park | Wellcome Sanger Institute for the COVID-19 Genomics UK (COG-UK) Consortium | Jacquelyn Wynn, Mairead Hyland, The Lighthouse Lab in Alderley Park and Alex Alderton, Roberto Amato, Jeffrey Barrett, Sonia Goncalves, Ewan Harrison, David K. Jackson, Ian Johnston, Dominic Kwiatkowski, Cordelia Langford, John Sillitoe on behalf of the Wellcome Sanger Institute COVID-19 Surveillance Team |
| EPI_ISL_3377245 | Berkshire and Surrey Pathology Services Lighthouse Laboratory | Wellcome Sanger Institute for the COVID-19 Genomics UK (COG-UK) Consortium | Berkshire and Surrey Pathology Services Lighthouse Laboratory and Alex Alderton, Roberto Amato, Jeffrey Barrett, Sonia Goncalves, Ewan Harrison, David K. Jackson, Ian Johnston, Dominic Kwiatkowski, Cordelia Langford, John Sillitoe on behalf of the Wellcome Sanger Institute COVID-19 Surveillance Team |
| EPI_ISL_3377246 | Lighthouse Lab in Alderley Park | Wellcome Sanger Institute for the COVID-19 Genomics UK (COG-UK) Consortium | Jacquelyn Wynn, Mairead Hyland, The Lighthouse Lab in Alderley Park and Alex Alderton, Roberto Amato, Jeffrey Barrett, Sonia Goncalves, Ewan Harrison, David K. Jackson, Ian Johnston, Dominic Kwiatkowski, Cordelia Langford, John Sillitoe on behalf of the Wellcome Sanger Institute COVID-19 Surveillance Team |
| EPI_ISL_3377247 | Berkshire and Surrey Pathology Services Lighthouse Laboratory | Wellcome Sanger Institute for the COVID-19 Genomics UK (COG-UK) Consortium | Berkshire and Surrey Pathology Services Lighthouse Laboratory and Alex Alderton, Roberto Amato, Jeffrey Barrett, Sonia Goncalves, Ewan Harrison, David K. Jackson, Ian Johnston, Dominic Kwiatkowski, Cordelia Langford, John Sillitoe on behalf of the Wellcome Sanger Institute COVID-19 Surveillance Team |
| EPI_ISL_3377248 | Lighthouse Lab in Milton Keynes | Wellcome Sanger Institute for the COVID-19 Genomics UK (COG-UK) Consortium | The Lighthouse Lab in Milton Keynes and Alex Alderton, Roberto Amato, Jeffrey Barrett, Sonia Goncalves, Ewan Harrison, David K. Jackson, Ian Johnston, Dominic Kwiatkowski, Cordelia Langford, John Sillitoe on behalf of the Wellcome Sanger Institute COVID-19 Surveillance Team |
| EPI_ISL_3377249, EPI_ISL_3377250 | Berkshire and Surrey Pathology Services Lighthouse Laboratory | Wellcome Sanger Institute for the COVID-19 Genomics UK (COG-UK) Consortium | Berkshire and Surrey Pathology Services Lighthouse Laboratory and Alex Alderton, Roberto Amato, Jeffrey Barrett, Sonia Goncalves, Ewan Harrison, David K. Jackson, Ian Johnston, Dominic Kwiatkowski, Cordelia Langford, John Sillitoe on behalf of the Wellcome Sanger Institute COVID-19 Surveillance Team |
| EPI_ISL_3377251 | Lighthouse Lab in Milton Keynes | Wellcome Sanger Institute for the COVID-19 Genomics UK (COG-UK) Consortium | The Lighthouse Lab in Milton Keynes and Alex Alderton, Roberto Amato, Jeffrey Barrett, Sonia Goncalves, Ewan Harrison, David K. Jackson, Ian Johnston, Dominic Kwiatkowski, Cordelia Langford, John Sillitoe on behalf of the Wellcome Sanger Institute COVID-19 Surveillance Team |
| EPI_ISL_3377252 | Berkshire and Surrey Pathology Services Lighthouse Laboratory | Wellcome Sanger Institute for the COVID-19 Genomics UK (COG-UK) Consortium | Berkshire and Surrey Pathology Services Lighthouse Laboratory and Alex Alderton, Roberto Amato, Jeffrey Barrett, Sonia Goncalves, Ewan Harrison, David K. Jackson, Ian Johnston, Dominic Kwiatkowski, Cordelia Langford, John Sillitoe on behalf of the Wellcome Sanger Institute COVID-19 Surveillance Team |
| EPI_ISL_3377253, EPI_ISL_3377254 | Lighthouse Lab in Milton Keynes | Wellcome Sanger Institute for the COVID-19 Genomics UK (COG-UK) Consortium | The Lighthouse Lab in Milton Keynes and Alex Alderton, Roberto Amato, Jeffrey Barrett, Sonia Goncalves, Ewan Harrison, David K. Jackson, Ian Johnston, Dominic Kwiatkowski, Cordelia Langford, John Sillitoe on behalf of the Wellcome Sanger Institute COVID-19 Surveillance Team |

| | | | |
|---|---|---|---|
| EPI_ISL_3377255 | Lighthouse Lab in Glasgow | Wellcome Sanger Institute for the COVID-19 Genomics UK (COG-UK) Consortium | Harper VanSteenhouse, Yumi Kasai, David Gray, Carol Clugston, Anna Dominiczak and Alex Alderton, Roberto Amato, Jeffrey Barrett, Sonia Goncalves, Ewan Harrison, David K. Jackson, Ian Johnston, Dominic Kwiatkowski, Cordelia Langford, John Sillitoe on behalf of the Wellcome Sanger Institute COVID-19 Surveillance Team |
| EPI_ISL_3377256 | Lighthouse Lab in Milton Keynes | Wellcome Sanger Institute for the COVID-19 Genomics UK (COG-UK) Consortium | The Lighthouse Lab in Milton Keynes and Alex Alderton, Roberto Amato, Jeffrey Barrett, Sonia Goncalves, Ewan Harrison, David K. Jackson, Ian Johnston, Dominic Kwiatkowski, Cordelia Langford, John Sillitoe on behalf of the Wellcome Sanger Institute COVID-19 Surveillance Team |
| EPI_ISL_3377257, EPI_ISL_3377258 | Lighthouse Lab in Glasgow | Wellcome Sanger Institute for the COVID-19 Genomics UK (COG-UK) Consortium | Harper VanSteenhouse, Yumi Kasai, David Gray, Carol Clugston, Anna Dominiczak and Alex Alderton, Roberto Amato, Jeffrey Barrett, Sonia Goncalves, Ewan Harrison, David K. Jackson, Ian Johnston, Dominic Kwiatkowski, Cordelia Langford, John Sillitoe on behalf of the Wellcome Sanger Institute COVID-19 Surveillance Team |
| EPI_ISL_3377259, EPI_ISL_3377260 | Berkshire and Surrey Pathology Services Lighthouse Laboratory | Wellcome Sanger Institute for the COVID-19 Genomics UK (COG-UK) Consortium | Berkshire and Surrey Pathology Services Lighthouse Laboratory and Alex Alderton, Roberto Amato, Jeffrey Barrett, Sonia Goncalves, Ewan Harrison, David K. Jackson, Ian Johnston, Dominic Kwiatkowski, Cordelia Langford, John Sillitoe on behalf of the Wellcome Sanger Institute COVID-19 Surveillance Team |
| EPI_ISL_3377261 | Lighthouse Lab in Alderley Park | Wellcome Sanger Institute for the COVID-19 Genomics UK (COG-UK) Consortium | Jacquelyn Wynn, Mairead Hyland, The Lighthouse Lab in Alderley Park and Alex Alderton, Roberto Amato, Jeffrey Barrett, Sonia Goncalves, Ewan Harrison, David K. Jackson, Ian Johnston, Dominic Kwiatkowski, Cordelia Langford, John Sillitoe on behalf of the Wellcome Sanger Institute COVID-19 Surveillance Team |
| EPI_ISL_3377262 | Lighthouse Lab in Glasgow | Wellcome Sanger Institute for the COVID-19 Genomics UK (COG-UK) Consortium | Harper VanSteenhouse, Yumi Kasai, David Gray, Carol Clugston, Anna Dominiczak and Alex Alderton, Roberto Amato, Jeffrey Barrett, Sonia Goncalves, Ewan Harrison, David K. Jackson, Ian Johnston, Dominic Kwiatkowski, Cordelia Langford, John Sillitoe on behalf of the Wellcome Sanger Institute COVID-19 Surveillance Team |
| EPI_ISL_3377263 | Lighthouse Lab in Milton Keynes | Wellcome Sanger Institute for the COVID-19 Genomics UK (COG-UK) Consortium | The Lighthouse Lab in Milton Keynes and Alex Alderton, Roberto Amato, Jeffrey Barrett, Sonia Goncalves, Ewan Harrison, David K. Jackson, Ian Johnston, Dominic Kwiatkowski, Cordelia Langford, John Sillitoe on behalf of the Wellcome Sanger Institute COVID-19 Surveillance Team |
| EPI_ISL_3377264 | Berkshire and Surrey Pathology Services Lighthouse Laboratory | Wellcome Sanger Institute for the COVID-19 Genomics UK (COG-UK) Consortium | Berkshire and Surrey Pathology Services Lighthouse Laboratory and Alex Alderton, Roberto Amato, Jeffrey Barrett, Sonia Goncalves, Ewan Harrison, David K. Jackson, Ian Johnston, Dominic Kwiatkowski, Cordelia Langford, John Sillitoe on behalf of the Wellcome Sanger Institute COVID-19 Surveillance Team |
| EPI_ISL_3377265 | Lighthouse Lab in Milton Keynes | Wellcome Sanger Institute for the COVID-19 Genomics UK (COG-UK) Consortium | The Lighthouse Lab in Milton Keynes and Alex Alderton, Roberto Amato, Jeffrey Barrett, Sonia Goncalves, Ewan Harrison, David K. Jackson, Ian Johnston, Dominic Kwiatkowski, Cordelia Langford, John Sillitoe on behalf of the Wellcome Sanger Institute COVID-19 Surveillance Team |
| EPI_ISL_3377266 | Berkshire and Surrey Pathology Services Lighthouse Laboratory | Wellcome Sanger Institute for the COVID-19 Genomics UK (COG-UK) Consortium | Berkshire and Surrey Pathology Services Lighthouse Laboratory and Alex Alderton, Roberto Amato, Jeffrey Barrett, Sonia Goncalves, Ewan Harrison, David K. Jackson, Ian Johnston, Dominic Kwiatkowski, Cordelia Langford, John Sillitoe on behalf of the Wellcome Sanger Institute COVID-19 Surveillance Team |
| EPI_ISL_3377267 | Lighthouse Lab in Alderley Park | Wellcome Sanger Institute for the COVID-19 Genomics UK (COG-UK) Consortium | Jacquelyn Wynn, Mairead Hyland, The Lighthouse Lab in Alderley Park and Alex Alderton, Roberto Amato, Jeffrey Barrett, Sonia Goncalves, Ewan Harrison, David K. Jackson, Ian Johnston, Dominic Kwiatkowski, Cordelia Langford, John Sillitoe on behalf of the Wellcome Sanger Institute COVID-19 Surveillance Team |
| EPI_ISL_3377268 | Lighthouse Lab in Glasgow | Wellcome Sanger Institute for the COVID-19 Genomics UK (COG-UK) Consortium | Harper VanSteenhouse, Yumi Kasai, David Gray, Carol Clugston, Anna Dominiczak and Alex Alderton, Roberto Amato, Jeffrey Barrett, Sonia Goncalves, Ewan Harrison, David K. Jackson, Ian Johnston, Dominic Kwiatkowski, Cordelia Langford, John Sillitoe on behalf of the Wellcome Sanger Institute COVID-19 Surveillance Team |
| EPI_ISL_3377269 | Lighthouse Lab in Alderley Park | Wellcome Sanger Institute for the COVID-19 Genomics UK (COG-UK) Consortium | Jacquelyn Wynn, Mairead Hyland, The Lighthouse Lab in Alderley Park and Alex Alderton, Roberto Amato, Jeffrey Barrett, Sonia Goncalves, Ewan Harrison, David K. Jackson, Ian Johnston, Dominic Kwiatkowski, Cordelia Langford, John Sillitoe on behalf of the Wellcome Sanger Institute COVID-19 Surveillance Team |
| EPI_ISL_3377270 | Lighthouse Lab in Milton Keynes | Wellcome Sanger Institute for the COVID-19 Genomics UK (COG-UK) Consortium | The Lighthouse Lab in Milton Keynes and Alex Alderton, Roberto Amato, Jeffrey Barrett, Sonia Goncalves, Ewan Harrison, David K. Jackson, Ian Johnston, Dominic Kwiatkowski, Cordelia Langford, John Sillitoe on behalf of the Wellcome Sanger Institute COVID-19 Surveillance Team |
| EPI_ISL_3377271 | Lighthouse Lab in Glasgow | Wellcome Sanger Institute for the COVID-19 Genomics UK (COG-UK) Consortium | Harper VanSteenhouse, Yumi Kasai, David Gray, Carol Clugston, Anna Dominiczak and Alex Alderton, Roberto Amato, Jeffrey Barrett, Sonia Goncalves, Ewan Harrison, David K. Jackson, Ian Johnston, Dominic Kwiatkowski, Cordelia Langford, John Sillitoe on behalf of the Wellcome Sanger Institute COVID-19 Surveillance Team |
| EPI_ISL_3377272 | Lighthouse Lab in Alderley Park | Wellcome Sanger Institute for the COVID-19 Genomics UK (COG-UK) Consortium | Jacquelyn Wynn, Mairead Hyland, The Lighthouse Lab in Alderley Park and Alex Alderton, Roberto Amato, Jeffrey Barrett, Sonia Goncalves, Ewan Harrison, David K. Jackson, Ian Johnston, Dominic Kwiatkowski, Cordelia Langford, John Sillitoe on behalf of the Wellcome Sanger Institute COVID-19 Surveillance Team |
| EPI_ISL_3377273 | Berkshire and Surrey Pathology Services Lighthouse Laboratory | Wellcome Sanger Institute for the COVID-19 Genomics UK (COG-UK) Consortium | Berkshire and Surrey Pathology Services Lighthouse Laboratory and Alex Alderton, Roberto Amato, Jeffrey Barrett, Sonia Goncalves, Ewan Harrison, David K. Jackson, Ian Johnston, Dominic Kwiatkowski, Cordelia Langford, John Sillitoe on behalf of the Wellcome Sanger Institute COVID-19 Surveillance Team |
| EPI_ISL_3377274, EPI_ISL_3377275 | Lighthouse Lab in Milton Keynes | Wellcome Sanger Institute for the COVID-19 Genomics UK (COG-UK) Consortium | The Lighthouse Lab in Milton Keynes and Alex Alderton, Roberto Amato, Jeffrey Barrett, Sonia Goncalves, Ewan Harrison, David K. Jackson, Ian Johnston, Dominic Kwiatkowski, Cordelia Langford, John Sillitoe on behalf of the Wellcome Sanger Institute COVID-19 Surveillance Team |
| EPI_ISL_3377276 | Lighthouse Lab in Alderley Park | Wellcome Sanger Institute for the COVID-19 Genomics UK (COG-UK) Consortium | Jacquelyn Wynn, Mairead Hyland, The Lighthouse Lab in Alderley Park and Alex Alderton, Roberto Amato, Jeffrey Barrett, Sonia Goncalves, Ewan Harrison, David K. Jackson, Ian Johnston, Dominic Kwiatkowski, Cordelia Langford, John Sillitoe on behalf of the Wellcome Sanger Institute COVID-19 Surveillance Team |
| EPI_ISL_3377277 | Berkshire and Surrey Pathology Services Lighthouse Laboratory | Wellcome Sanger Institute for the COVID-19 Genomics UK (COG-UK) Consortium | Berkshire and Surrey Pathology Services Lighthouse Laboratory and Alex Alderton, Roberto Amato, Jeffrey Barrett, Sonia Goncalves, Ewan Harrison, David K. Jackson, Ian Johnston, Dominic Kwiatkowski, Cordelia Langford, John Sillitoe on behalf of the Wellcome Sanger Institute COVID-19 Surveillance Team |
| EPI_ISL_3377278 | Lighthouse Lab in Alderley Park | Wellcome Sanger Institute for the COVID-19 Genomics UK (COG-UK) Consortium | Jacquelyn Wynn, Mairead Hyland, The Lighthouse Lab in Alderley Park and Alex Alderton, Roberto Amato, Jeffrey Barrett, Sonia Goncalves, Ewan Harrison, David K. Jackson, Ian Johnston, Dominic Kwiatkowski, Cordelia Langford, John Sillitoe on behalf of the Wellcome Sanger Institute COVID-19 Surveillance Team |
| EPI_ISL_3377279, EPI_ISL_3377280, EPI_ISL_3377281, EPI_ISL_3377282 | Lighthouse Lab in Milton Keynes | Wellcome Sanger Institute for the COVID-19 Genomics UK (COG-UK) Consortium | The Lighthouse Lab in Milton Keynes and Alex Alderton, Roberto Amato, Jeffrey Barrett, Sonia Goncalves, Ewan Harrison, David K. Jackson, Ian Johnston, Dominic Kwiatkowski, Cordelia Langford, John Sillitoe on behalf of the Wellcome Sanger Institute COVID-19 Surveillance Team |
| EPI_ISL_3377283 | Lighthouse Lab in Glasgow | Wellcome Sanger Institute for the COVID-19 Genomics UK (COG-UK) Consortium | Harper VanSteenhouse, Yumi Kasai, David Gray, Carol Clugston, Anna Dominiczak and Alex Alderton, Roberto Amato, Jeffrey Barrett, Sonia Goncalves, Ewan Harrison, David K. Jackson, Ian Johnston, Dominic Kwiatkowski, Cordelia Langford, John Sillitoe on behalf of the Wellcome Sanger Institute COVID-19 Surveillance Team |
| EPI_ISL_3377284, EPI_ISL_3377285 | Berkshire and Surrey Pathology Services Lighthouse Laboratory | Wellcome Sanger Institute for the COVID-19 Genomics UK (COG-UK) Consortium | Berkshire and Surrey Pathology Services Lighthouse Laboratory and Alex Alderton, Roberto Amato, Jeffrey Barrett, Sonia Goncalves, Ewan Harrison, David K. Jackson, Ian Johnston, Dominic Kwiatkowski, Cordelia Langford, John Sillitoe on behalf of the Wellcome Sanger Institute COVID-19 Surveillance Team |
| EPI_ISL_3377286 | Lighthouse Lab in Glasgow | Wellcome Sanger Institute for the COVID-19 Genomics UK (COG-UK) Consortium | Harper VanSteenhouse, Yumi Kasai, David Gray, Carol Clugston, Anna Dominiczak and Alex Alderton, Roberto Amato, Jeffrey Barrett, Sonia Goncalves, Ewan Harrison, David K. Jackson, Ian Johnston, Dominic Kwiatkowski, Cordelia Langford, John Sillitoe on behalf of the Wellcome Sanger Institute COVID-19 Surveillance Team |
| EPI_ISL_3377287 | Berkshire and Surrey Pathology Services Lighthouse Laboratory | Wellcome Sanger Institute for the COVID-19 Genomics UK (COG-UK) Consortium | Berkshire and Surrey Pathology Services Lighthouse Laboratory and Alex Alderton, Roberto Amato, Jeffrey Barrett, Sonia Goncalves, Ewan Harrison, David K. Jackson, Ian Johnston, Dominic Kwiatkowski, Cordelia Langford, John Sillitoe on behalf of the Wellcome Sanger Institute COVID-19 Surveillance Team |
| EPI_ISL_3377288 | Lighthouse Lab in Milton Keynes | Wellcome Sanger Institute for the COVID-19 Genomics UK (COG-UK) Consortium | The Lighthouse Lab in Milton Keynes and Alex Alderton, Roberto Amato, Jeffrey Barrett, Sonia Goncalves, Ewan Harrison, David K. Jackson, Ian Johnston, Dominic Kwiatkowski, Cordelia Langford, John Sillitoe on behalf of the Wellcome Sanger Institute COVID-19 Surveillance Team |
| EPI_ISL_3377289 | Berkshire and Surrey Pathology Services Lighthouse Laboratory | Wellcome Sanger Institute for the COVID-19 Genomics UK (COG-UK) Consortium | Berkshire and Surrey Pathology Services Lighthouse Laboratory and Alex Alderton, Roberto Amato, Jeffrey Barrett, Sonia Goncalves, Ewan Harrison, David K. Jackson, Ian Johnston, Dominic Kwiatkowski, Cordelia Langford, John Sillitoe on behalf of the Wellcome Sanger Institute COVID-19 Surveillance Team |
| EPI_ISL_3377290 | Lighthouse Laboratory Plymouth | Wellcome Sanger Institute for the COVID-19 Genomics UK (COG-UK) Consortium | Lighthouse Laboratory Plymouth and Alex Alderton, Roberto Amato, Jeffrey Barrett, Sonia Goncalves, Ewan Harrison, David K. Jackson, Ian Johnston, Dominic Kwiatkowski, Cordelia Langford, John Sillitoe on behalf of the Wellcome Sanger Institute COVID-19 Surveillance Team |
| EPI_ISL_3377291, EPI_ISL_3377292, EPI_ISL_3377293 | Lighthouse Lab in Milton Keynes | Wellcome Sanger Institute for the COVID-19 Genomics UK (COG-UK) Consortium | The Lighthouse Lab in Milton Keynes and Alex Alderton, Roberto Amato, Jeffrey Barrett, Sonia Goncalves, Ewan Harrison, David K. Jackson, Ian Johnston, Dominic Kwiatkowski, Cordelia Langford, John Sillitoe on behalf of the Wellcome Sanger Institute COVID-19 Surveillance Team |
| EPI_ISL_3377294 | Lighthouse Laboratory Plymouth | Wellcome Sanger Institute for the COVID-19 Genomics UK (COG-UK) Consortium | Lighthouse Laboratory Plymouth and Alex Alderton, Roberto Amato, Jeffrey Barrett, Sonia Goncalves, Ewan Harrison, David K. Jackson, Ian Johnston, Dominic Kwiatkowski, Cordelia Langford, John Sillitoe on behalf of the Wellcome Sanger Institute COVID-19 Surveillance Team |
| EPI_ISL_3377295 | Lighthouse Lab in Milton Keynes | Wellcome Sanger Institute for the COVID-19 Genomics UK (COG-UK) Consortium | The Lighthouse Lab in Milton Keynes and Alex Alderton, Roberto Amato, Jeffrey Barrett, Sonia Goncalves, Ewan Harrison, David K. Jackson, Ian Johnston, Dominic Kwiatkowski, Cordelia Langford, John Sillitoe on behalf of the Wellcome Sanger Institute COVID-19 Surveillance Team |
| EPI_ISL_3377296 | Berkshire and Surrey Pathology Services Lighthouse Laboratory | Wellcome Sanger Institute for the COVID-19 Genomics UK (COG-UK) Consortium | Berkshire and Surrey Pathology Services Lighthouse Laboratory and Alex Alderton, Roberto Amato, Jeffrey Barrett, Sonia Goncalves, Ewan Harrison, David K. Jackson, Ian Johnston, Dominic Kwiatkowski, Cordelia Langford, John Sillitoe on behalf of the Wellcome Sanger Institute COVID-19 Surveillance Team |
| EPI_ISL_3377297, EPI_ISL_3377298 | Lighthouse Lab in Milton Keynes | Wellcome Sanger Institute for the COVID-19 Genomics UK (COG-UK) Consortium | The Lighthouse Lab in Milton Keynes and Alex Alderton, Roberto Amato, Jeffrey Barrett, Sonia Goncalves, Ewan Harrison, David K. Jackson, Ian Johnston, Dominic Kwiatkowski, Cordelia Langford, John Sillitoe on behalf of the Wellcome Sanger Institute COVID-19 Surveillance Team |
| EPI_ISL_3377299 | Lighthouse Laboratory Plymouth | Wellcome Sanger Institute for the COVID-19 Genomics UK (COG-UK) Consortium | Lighthouse Laboratory Plymouth and Alex Alderton, Roberto Amato, Jeffrey Barrett, Sonia Goncalves, Ewan Harrison, David K. Jackson, Ian Johnston, Dominic Kwiatkowski, Cordelia Langford, John Sillitoe on behalf of the Wellcome Sanger Institute COVID-19 Surveillance Team |
| EPI_ISL_3377300 | Lighthouse Lab in Milton Keynes | Wellcome Sanger Institute for the COVID-19 Genomics UK (COG-UK) Consortium | The Lighthouse Lab in Milton Keynes and Alex Alderton, Roberto Amato, Jeffrey Barrett, Sonia Goncalves, Ewan Harrison, David K. Jackson, Ian Johnston, Dominic Kwiatkowski, Cordelia Langford, John Sillitoe on behalf of the Wellcome Sanger Institute COVID-19 Surveillance Team |
| EPI_ISL_3377301 | Berkshire and Surrey Pathology Services Lighthouse Laboratory | Wellcome Sanger Institute for the COVID-19 Genomics UK (COG-UK) Consortium | Berkshire and Surrey Pathology Services Lighthouse Laboratory and Alex Alderton, Roberto Amato, Jeffrey Barrett, Sonia Goncalves, Ewan Harrison, David K. Jackson, Ian Johnston, Dominic Kwiatkowski, Cordelia Langford, John Sillitoe on behalf of the Wellcome Sanger Institute COVID-19 Surveillance Team |
| EPI_ISL_3377302, EPI_ISL_3377303 | Lighthouse Lab in Milton Keynes | Wellcome Sanger Institute for the COVID-19 Genomics UK (COG-UK) Consortium | The Lighthouse Lab in Milton Keynes and Alex Alderton, Roberto Amato, Jeffrey Barrett, Sonia Goncalves, Ewan Harrison, David K. Jackson, Ian Johnston, Dominic Kwiatkowski, Cordelia Langford, John Sillitoe on behalf of the Wellcome Sanger Institute COVID-19 Surveillance Team |
| EPI_ISL_3377304, EPI_ISL_3377305 | Lighthouse Laboratory Plymouth | Wellcome Sanger Institute for the COVID-19 Genomics UK (COG-UK) Consortium | Lighthouse Laboratory Plymouth and Alex Alderton, Roberto Amato, Jeffrey Barrett, Sonia Goncalves, Ewan Harrison, David K. Jackson, Ian Johnston, Dominic Kwiatkowski, Cordelia Langford, John Sillitoe on behalf of the Wellcome Sanger Institute COVID-19 Surveillance Team |
| EPI_ISL_3377306 | Lighthouse Lab in Milton Keynes | Wellcome Sanger Institute for the COVID-19 Genomics UK (COG-UK) Consortium | The Lighthouse Lab in Milton Keynes and Alex Alderton, Roberto Amato, Jeffrey Barrett, Sonia Goncalves, Ewan Harrison, David K. Jackson, Ian Johnston, Dominic Kwiatkowski, Cordelia Langford, John Sillitoe on behalf of the Wellcome Sanger Institute COVID-19 Surveillance Team |
| EPI_ISL_3377307, EPI_ISL_3377308 | Lighthouse Laboratory Plymouth | Wellcome Sanger Institute for the COVID-19 Genomics UK (COG-UK) Consortium | Lighthouse Laboratory Plymouth and Alex Alderton, Roberto Amato, Jeffrey Barrett, Sonia Goncalves, Ewan Harrison, David K. Jackson, Ian Johnston, Dominic Kwiatkowski, Cordelia Langford, John Sillitoe on behalf of the Wellcome Sanger Institute COVID-19 Surveillance Team |
| EPI_ISL_3377309, EPI_ISL_3377310 | Lighthouse Lab in Milton Keynes | Wellcome Sanger Institute for the COVID-19 Genomics UK (COG-UK) Consortium | The Lighthouse Lab in Milton Keynes and Alex Alderton, Roberto Amato, Jeffrey Barrett, Sonia Goncalves, Ewan Harrison, David K. Jackson, Ian Johnston, Dominic Kwiatkowski, Cordelia Langford, John Sillitoe on behalf of the Wellcome Sanger Institute COVID-19 Surveillance Team |
| EPI_ISL_3377311 | Lighthouse Laboratory Plymouth | Wellcome Sanger Institute for the COVID-19 Genomics UK (COG-UK) Consortium | Lighthouse Laboratory Plymouth and Alex Alderton, Roberto Amato, Jeffrey Barrett, Sonia Goncalves, Ewan Harrison, David K. Jackson, Ian Johnston, Dominic Kwiatkowski, Cordelia Langford, John Sillitoe on behalf of the Wellcome Sanger Institute COVID-19 Surveillance Team |
| EPI_ISL_3377312, | Lighthouse Lab in Milton Keynes | Wellcome Sanger Institute for the COVID-19 Genomics UK (COG-UK) | The Lighthouse Lab in Milton Keynes and Alex Alderton, Roberto Amato, Jeffrey Barrett, Sonia Goncalves, Ewan Harrison, David K. Jackson, Ian Johnston, Dominic Kwiatkowski, Cordelia Langford, John Sillitoe on behalf of the Wellcome Sanger |

| | | | |
|---|---|---|---|
| EPI_ISL_3377313 | | Consortium | Institute COVID-19 Surveillance Team |
| EPI_ISL_3377314 | Lighthouse Laboratory Plymouth | Wellcome Sanger Institute for the COVID-19 Genomics UK (COG-UK) Consortium | Lighthouse Laboratory Plymouth and Alex Alderton, Roberto Amato, Jeffrey Barrett, Sonia Goncalves, Ewan Harrison, David K. Jackson, Ian Johnston, Dominic Kwiatkowski, Cordelia Langford, John Sillitoe on behalf of the Wellcome Sanger Institute COVID-19 Surveillance Team |
| EPI_ISL_3377315 | Lighthouse Lab in Milton Keynes | Wellcome Sanger Institute for the COVID-19 Genomics UK (COG-UK) Consortium | The Lighthouse Lab in Milton Keynes and Alex Alderton, Roberto Amato, Jeffrey Barrett, Sonia Goncalves, Ewan Harrison, David K. Jackson, Ian Johnston, Dominic Kwiatkowski, Cordelia Langford, John Sillitoe on behalf of the Wellcome Sanger Institute COVID-19 Surveillance Team |
| EPI_ISL_3377316, EPI_ISL_3377317, EPI_ISL_3377318 | Lighthouse Laboratory Plymouth | Wellcome Sanger Institute for the COVID-19 Genomics UK (COG-UK) Consortium | Lighthouse Laboratory Plymouth and Alex Alderton, Roberto Amato, Jeffrey Barrett, Sonia Goncalves, Ewan Harrison, David K. Jackson, Ian Johnston, Dominic Kwiatkowski, Cordelia Langford, John Sillitoe on behalf of the Wellcome Sanger Institute COVID-19 Surveillance Team |
| EPI_ISL_3377319, EPI_ISL_3377320 | Lighthouse Lab in Milton Keynes | Wellcome Sanger Institute for the COVID-19 Genomics UK (COG-UK) Consortium | The Lighthouse Lab in Milton Keynes and Alex Alderton, Roberto Amato, Jeffrey Barrett, Sonia Goncalves, Ewan Harrison, David K. Jackson, Ian Johnston, Dominic Kwiatkowski, Cordelia Langford, John Sillitoe on behalf of the Wellcome Sanger Institute COVID-19 Surveillance Team |
| EPI_ISL_3377321 | Lighthouse Laboratory Plymouth | Wellcome Sanger Institute for the COVID-19 Genomics UK (COG-UK) Consortium | Lighthouse Laboratory Plymouth and Alex Alderton, Roberto Amato, Jeffrey Barrett, Sonia Goncalves, Ewan Harrison, David K. Jackson, Ian Johnston, Dominic Kwiatkowski, Cordelia Langford, John Sillitoe on behalf of the Wellcome Sanger Institute COVID-19 Surveillance Team |
| EPI_ISL_3377322 | Lighthouse Lab in Milton Keynes | Wellcome Sanger Institute for the COVID-19 Genomics UK (COG-UK) Consortium | The Lighthouse Lab in Milton Keynes and Alex Alderton, Roberto Amato, Jeffrey Barrett, Sonia Goncalves, Ewan Harrison, David K. Jackson, Ian Johnston, Dominic Kwiatkowski, Cordelia Langford, John Sillitoe on behalf of the Wellcome Sanger Institute COVID-19 Surveillance Team |
| EPI_ISL_3377323 | Berkshire and Surrey Pathology Services Lighthouse Laboratory | Wellcome Sanger Institute for the COVID-19 Genomics UK (COG-UK) Consortium | Berkshire and Surrey Pathology Services Lighthouse Laboratory and Alex Alderton, Roberto Amato, Jeffrey Barrett, Sonia Goncalves, Ewan Harrison, David K. Jackson, Ian Johnston, Dominic Kwiatkowski, Cordelia Langford, John Sillitoe on behalf of the Wellcome Sanger Institute COVID-19 Surveillance Team |
| EPI_ISL_3377324, EPI_ISL_3377325 | Lighthouse Lab in Milton Keynes | Wellcome Sanger Institute for the COVID-19 Genomics UK (COG-UK) Consortium | The Lighthouse Lab in Milton Keynes and Alex Alderton, Roberto Amato, Jeffrey Barrett, Sonia Goncalves, Ewan Harrison, David K. Jackson, Ian Johnston, Dominic Kwiatkowski, Cordelia Langford, John Sillitoe on behalf of the Wellcome Sanger Institute COVID-19 Surveillance Team |
| EPI_ISL_3377326 | Lighthouse Laboratory Plymouth | Wellcome Sanger Institute for the COVID-19 Genomics UK (COG-UK) Consortium | Lighthouse Laboratory Plymouth and Alex Alderton, Roberto Amato, Jeffrey Barrett, Sonia Goncalves, Ewan Harrison, David K. Jackson, Ian Johnston, Dominic Kwiatkowski, Cordelia Langford, John Sillitoe on behalf of the Wellcome Sanger Institute COVID-19 Surveillance Team |
| EPI_ISL_3377327, EPI_ISL_3377328 | Lighthouse Lab in Milton Keynes | Wellcome Sanger Institute for the COVID-19 Genomics UK (COG-UK) Consortium | The Lighthouse Lab in Milton Keynes and Alex Alderton, Roberto Amato, Jeffrey Barrett, Sonia Goncalves, Ewan Harrison, David K. Jackson, Ian Johnston, Dominic Kwiatkowski, Cordelia Langford, John Sillitoe on behalf of the Wellcome Sanger Institute COVID-19 Surveillance Team |
| EPI_ISL_3377329, EPI_ISL_3377330 | Berkshire and Surrey Pathology Services Lighthouse Laboratory | Wellcome Sanger Institute for the COVID-19 Genomics UK (COG-UK) Consortium | Berkshire and Surrey Pathology Services Lighthouse Laboratory and Alex Alderton, Roberto Amato, Jeffrey Barrett, Sonia Goncalves, Ewan Harrison, David K. Jackson, Ian Johnston, Dominic Kwiatkowski, Cordelia Langford, John Sillitoe on behalf of the Wellcome Sanger Institute COVID-19 Surveillance Team |
| EPI_ISL_3377331 | Lighthouse Laboratory Plymouth | Wellcome Sanger Institute for the COVID-19 Genomics UK (COG-UK) Consortium | Lighthouse Laboratory Plymouth and Alex Alderton, Roberto Amato, Jeffrey Barrett, Sonia Goncalves, Ewan Harrison, David K. Jackson, Ian Johnston, Dominic Kwiatkowski, Cordelia Langford, John Sillitoe on behalf of the Wellcome Sanger Institute COVID-19 Surveillance Team |
| EPI_ISL_3377332, EPI_ISL_3377333 | Lighthouse Lab in Milton Keynes | Wellcome Sanger Institute for the COVID-19 Genomics UK (COG-UK) Consortium | The Lighthouse Lab in Milton Keynes and Alex Alderton, Roberto Amato, Jeffrey Barrett, Sonia Goncalves, Ewan Harrison, David K. Jackson, Ian Johnston, Dominic Kwiatkowski, Cordelia Langford, John Sillitoe on behalf of the Wellcome Sanger Institute COVID-19 Surveillance Team |
| EPI_ISL_3377334 | Lighthouse Laboratory Plymouth | Wellcome Sanger Institute for the COVID-19 Genomics UK (COG-UK) Consortium | Lighthouse Laboratory Plymouth and Alex Alderton, Roberto Amato, Jeffrey Barrett, Sonia Goncalves, Ewan Harrison, David K. Jackson, Ian Johnston, Dominic Kwiatkowski, Cordelia Langford, John Sillitoe on behalf of the Wellcome Sanger Institute COVID-19 Surveillance Team |
| EPI_ISL_3377335 | Lighthouse Lab in Milton Keynes | Wellcome Sanger Institute for the COVID-19 Genomics UK (COG-UK) Consortium | The Lighthouse Lab in Milton Keynes and Alex Alderton, Roberto Amato, Jeffrey Barrett, Sonia Goncalves, Ewan Harrison, David K. Jackson, Ian Johnston, Dominic Kwiatkowski, Cordelia Langford, John Sillitoe on behalf of the Wellcome Sanger Institute COVID-19 Surveillance Team |
| EPI_ISL_3377336 | Lighthouse Laboratory Plymouth | Wellcome Sanger Institute for the COVID-19 Genomics UK (COG-UK) Consortium | Lighthouse Laboratory Plymouth and Alex Alderton, Roberto Amato, Jeffrey Barrett, Sonia Goncalves, Ewan Harrison, David K. Jackson, Ian Johnston, Dominic Kwiatkowski, Cordelia Langford, John Sillitoe on behalf of the Wellcome Sanger Institute COVID-19 Surveillance Team |
| EPI_ISL_3377337, EPI_ISL_3377338, EPI_ISL_3377339 | Lighthouse Lab in Milton Keynes | Wellcome Sanger Institute for the COVID-19 Genomics UK (COG-UK) Consortium | The Lighthouse Lab in Milton Keynes and Alex Alderton, Roberto Amato, Jeffrey Barrett, Sonia Goncalves, Ewan Harrison, David K. Jackson, Ian Johnston, Dominic Kwiatkowski, Cordelia Langford, John Sillitoe on behalf of the Wellcome Sanger Institute COVID-19 Surveillance Team |
| EPI_ISL_3377340 | Berkshire and Surrey Pathology Services Lighthouse Laboratory | Wellcome Sanger Institute for the COVID-19 Genomics UK (COG-UK) Consortium | Berkshire and Surrey Pathology Services Lighthouse Laboratory and Alex Alderton, Roberto Amato, Jeffrey Barrett, Sonia Goncalves, Ewan Harrison, David K. Jackson, Ian Johnston, Dominic Kwiatkowski, Cordelia Langford, John Sillitoe on behalf of the Wellcome Sanger Institute COVID-19 Surveillance Team |
| EPI_ISL_3377341 | Lighthouse Lab in Milton Keynes | Wellcome Sanger Institute for the COVID-19 Genomics UK (COG-UK) Consortium | The Lighthouse Lab in Milton Keynes and Alex Alderton, Roberto Amato, Jeffrey Barrett, Sonia Goncalves, Ewan Harrison, David K. Jackson, Ian Johnston, Dominic Kwiatkowski, Cordelia Langford, John Sillitoe on behalf of the Wellcome Sanger Institute COVID-19 Surveillance Team |
| EPI_ISL_3377342 | Lighthouse Laboratory Plymouth | Wellcome Sanger Institute for the COVID-19 Genomics UK (COG-UK) Consortium | Lighthouse Laboratory Plymouth and Alex Alderton, Roberto Amato, Jeffrey Barrett, Sonia Goncalves, Ewan Harrison, David K. Jackson, Ian Johnston, Dominic Kwiatkowski, Cordelia Langford, John Sillitoe on behalf of the Wellcome Sanger Institute COVID-19 Surveillance Team |
| EPI_ISL_3377343, EPI_ISL_3377344 | Lighthouse Lab in Milton Keynes | Wellcome Sanger Institute for the COVID-19 Genomics UK (COG-UK) Consortium | The Lighthouse Lab in Milton Keynes and Alex Alderton, Roberto Amato, Jeffrey Barrett, Sonia Goncalves, Ewan Harrison, David K. Jackson, Ian Johnston, Dominic Kwiatkowski, Cordelia Langford, John Sillitoe on behalf of the Wellcome Sanger Institute COVID-19 Surveillance Team |
| EPI_ISL_3377345 | Lighthouse Laboratory Plymouth | Wellcome Sanger Institute for the COVID-19 Genomics UK (COG-UK) Consortium | Lighthouse Laboratory Plymouth and Alex Alderton, Roberto Amato, Jeffrey Barrett, Sonia Goncalves, Ewan Harrison, David K. Jackson, Ian Johnston, Dominic Kwiatkowski, Cordelia Langford, John Sillitoe on behalf of the Wellcome Sanger Institute COVID-19 Surveillance Team |
| EPI_ISL_3377346 | Berkshire and Surrey Pathology Services Lighthouse Laboratory | Wellcome Sanger Institute for the COVID-19 Genomics UK (COG-UK) Consortium | Berkshire and Surrey Pathology Services Lighthouse Laboratory and Alex Alderton, Roberto Amato, Jeffrey Barrett, Sonia Goncalves, Ewan Harrison, David K. Jackson, Ian Johnston, Dominic Kwiatkowski, Cordelia Langford, John Sillitoe on behalf of the Wellcome Sanger Institute COVID-19 Surveillance Team |
| EPI_ISL_3377347, EPI_ISL_3377348 | Lighthouse Lab in Milton Keynes | Wellcome Sanger Institute for the COVID-19 Genomics UK (COG-UK) Consortium | The Lighthouse Lab in Milton Keynes and Alex Alderton, Roberto Amato, Jeffrey Barrett, Sonia Goncalves, Ewan Harrison, David K. Jackson, Ian Johnston, Dominic Kwiatkowski, Cordelia Langford, John Sillitoe on behalf of the Wellcome Sanger Institute COVID-19 Surveillance Team |
| EPI_ISL_3377349 | Lighthouse Laboratory Plymouth | Wellcome Sanger Institute for the COVID-19 Genomics UK (COG-UK) Consortium | Lighthouse Laboratory Plymouth and Alex Alderton, Roberto Amato, Jeffrey Barrett, Sonia Goncalves, Ewan Harrison, David K. Jackson, Ian Johnston, Dominic Kwiatkowski, Cordelia Langford, John Sillitoe on behalf of the Wellcome Sanger Institute COVID-19 Surveillance Team |
| EPI_ISL_3377350, EPI_ISL_3377351, EPI_ISL_3377352, EPI_ISL_3377353 | Lighthouse Lab in Milton Keynes | Wellcome Sanger Institute for the COVID-19 Genomics UK (COG-UK) Consortium | The Lighthouse Lab in Milton Keynes and Alex Alderton, Roberto Amato, Jeffrey Barrett, Sonia Goncalves, Ewan Harrison, David K. Jackson, Ian Johnston, Dominic Kwiatkowski, Cordelia Langford, John Sillitoe on behalf of the Wellcome Sanger Institute COVID-19 Surveillance Team |
| EPI_ISL_3377354, EPI_ISL_3377355 | Berkshire and Surrey Pathology Services Lighthouse Laboratory | Wellcome Sanger Institute for the COVID-19 Genomics UK (COG-UK) Consortium | Berkshire and Surrey Pathology Services Lighthouse Laboratory and Alex Alderton, Roberto Amato, Jeffrey Barrett, Sonia Goncalves, Ewan Harrison, David K. Jackson, Ian Johnston, Dominic Kwiatkowski, Cordelia Langford, John Sillitoe on behalf of the Wellcome Sanger Institute COVID-19 Surveillance Team |
| EPI_ISL_3377356 | Lighthouse Lab in Milton Keynes | Wellcome Sanger Institute for the COVID-19 Genomics UK (COG-UK) Consortium | The Lighthouse Lab in Milton Keynes and Alex Alderton, Roberto Amato, Jeffrey Barrett, Sonia Goncalves, Ewan Harrison, David K. Jackson, Ian Johnston, Dominic Kwiatkowski, Cordelia Langford, John Sillitoe on behalf of the Wellcome Sanger Institute COVID-19 Surveillance Team |
| EPI_ISL_3377357 | Lighthouse Laboratory Plymouth | Wellcome Sanger Institute for the COVID-19 Genomics UK (COG-UK) Consortium | Lighthouse Laboratory Plymouth and Alex Alderton, Roberto Amato, Jeffrey Barrett, Sonia Goncalves, Ewan Harrison, David K. Jackson, Ian Johnston, Dominic Kwiatkowski, Cordelia Langford, John Sillitoe on behalf of the Wellcome Sanger Institute COVID-19 Surveillance Team |
| EPI_ISL_3377358, EPI_ISL_3377359 | Lighthouse Lab in Milton Keynes | Wellcome Sanger Institute for the COVID-19 Genomics UK (COG-UK) Consortium | The Lighthouse Lab in Milton Keynes and Alex Alderton, Roberto Amato, Jeffrey Barrett, Sonia Goncalves, Ewan Harrison, David K. Jackson, Ian Johnston, Dominic Kwiatkowski, Cordelia Langford, John Sillitoe on behalf of the Wellcome Sanger Institute COVID-19 Surveillance Team |
| EPI_ISL_3377360, EPI_ISL_3377361, EPI_ISL_3377362, EPI_ISL_3377363 | Berkshire and Surrey Pathology Services Lighthouse Laboratory | Wellcome Sanger Institute for the COVID-19 Genomics UK (COG-UK) Consortium | Berkshire and Surrey Pathology Services Lighthouse Laboratory and Alex Alderton, Roberto Amato, Jeffrey Barrett, Sonia Goncalves, Ewan Harrison, David K. Jackson, Ian Johnston, Dominic Kwiatkowski, Cordelia Langford, John Sillitoe on behalf of the Wellcome Sanger Institute COVID-19 Surveillance Team |
| EPI_ISL_3377364 | Lighthouse Laboratory Plymouth | Wellcome Sanger Institute for the COVID-19 Genomics UK (COG-UK) Consortium | Lighthouse Laboratory Plymouth and Alex Alderton, Roberto Amato, Jeffrey Barrett, Sonia Goncalves, Ewan Harrison, David K. Jackson, Ian Johnston, Dominic Kwiatkowski, Cordelia Langford, John Sillitoe on behalf of the Wellcome Sanger Institute COVID-19 Surveillance Team |
| EPI_ISL_3377365 | Lighthouse Lab in Milton Keynes | Wellcome Sanger Institute for the COVID-19 Genomics UK (COG-UK) Consortium | The Lighthouse Lab in Milton Keynes and Alex Alderton, Roberto Amato, Jeffrey Barrett, Sonia Goncalves, Ewan Harrison, David K. Jackson, Ian Johnston, Dominic Kwiatkowski, Cordelia Langford, John Sillitoe on behalf of the Wellcome Sanger Institute COVID-19 Surveillance Team |
| EPI_ISL_3377366, EPI_ISL_3377367, EPI_ISL_3377368, EPI_ISL_3377369, EPI_ISL_3377370 | Berkshire and Surrey Pathology Services Lighthouse Laboratory | Wellcome Sanger Institute for the COVID-19 Genomics UK (COG-UK) Consortium | Berkshire and Surrey Pathology Services Lighthouse Laboratory and Alex Alderton, Roberto Amato, Jeffrey Barrett, Sonia Goncalves, Ewan Harrison, David K. Jackson, Ian Johnston, Dominic Kwiatkowski, Cordelia Langford, John Sillitoe on behalf of the Wellcome Sanger Institute COVID-19 Surveillance Team |
| EPI_ISL_3377371, EPI_ISL_3377372 | Lighthouse Laboratory Plymouth | Wellcome Sanger Institute for the COVID-19 Genomics UK (COG-UK) Consortium | Lighthouse Laboratory Plymouth and Alex Alderton, Roberto Amato, Jeffrey Barrett, Sonia Goncalves, Ewan Harrison, David K. Jackson, Ian Johnston, Dominic Kwiatkowski, Cordelia Langford, John Sillitoe on behalf of the Wellcome Sanger Institute COVID-19 Surveillance Team |
| EPI_ISL_3377373 | Lighthouse Lab in Milton Keynes | Wellcome Sanger Institute for the COVID-19 Genomics UK (COG-UK) Consortium | The Lighthouse Lab in Milton Keynes and Alex Alderton, Roberto Amato, Jeffrey Barrett, Sonia Goncalves, Ewan Harrison, David K. Jackson, Ian Johnston, Dominic Kwiatkowski, Cordelia Langford, John Sillitoe on behalf of the Wellcome Sanger Institute COVID-19 Surveillance Team |
| EPI_ISL_3377374, EPI_ISL_3377375, EPI_ISL_3377376 | Berkshire and Surrey Pathology Services Lighthouse Laboratory | Wellcome Sanger Institute for the COVID-19 Genomics UK (COG-UK) Consortium | Berkshire and Surrey Pathology Services Lighthouse Laboratory and Alex Alderton, Roberto Amato, Jeffrey Barrett, Sonia Goncalves, Ewan Harrison, David K. Jackson, Ian Johnston, Dominic Kwiatkowski, Cordelia Langford, John Sillitoe on behalf of the Wellcome Sanger Institute COVID-19 Surveillance Team |
| EPI_ISL_3377377 | Lighthouse Laboratory Plymouth | Wellcome Sanger Institute for the COVID-19 Genomics UK (COG-UK) Consortium | Lighthouse Laboratory Plymouth and Alex Alderton, Roberto Amato, Jeffrey Barrett, Sonia Goncalves, Ewan Harrison, David K. Jackson, Ian Johnston, Dominic Kwiatkowski, Cordelia Langford, John Sillitoe on behalf of the Wellcome Sanger Institute COVID-19 Surveillance Team |
| EPI_ISL_3377378 | Berkshire and Surrey Pathology Services Lighthouse Laboratory | Wellcome Sanger Institute for the COVID-19 Genomics UK (COG-UK) Consortium | Berkshire and Surrey Pathology Services Lighthouse Laboratory and Alex Alderton, Roberto Amato, Jeffrey Barrett, Sonia Goncalves, Ewan Harrison, David K. Jackson, Ian Johnston, Dominic Kwiatkowski, Cordelia Langford, John Sillitoe on behalf of the Wellcome Sanger Institute COVID-19 Surveillance Team |
| EPI_ISL_3377379 | Lighthouse Laboratory Plymouth | Wellcome Sanger Institute for the COVID-19 Genomics UK (COG-UK) Consortium | Lighthouse Laboratory Plymouth and Alex Alderton, Roberto Amato, Jeffrey Barrett, Sonia Goncalves, Ewan Harrison, David K. Jackson, Ian Johnston, Dominic Kwiatkowski, Cordelia Langford, John Sillitoe on behalf of the Wellcome Sanger Institute COVID-19 Surveillance Team |
| EPI_ISL_3377380 | Berkshire and Surrey Pathology Services Lighthouse Laboratory | Wellcome Sanger Institute for the COVID-19 Genomics UK (COG-UK) | Berkshire and Surrey Pathology Services Lighthouse Laboratory and Alex Alderton, Roberto Amato, Jeffrey Barrett, Sonia Goncalves, Ewan Harrison, David K. Jackson, Ian Johnston, Dominic Kwiatkowski, Cordelia Langford, John Sillitoe on behalf |

| | | | |
|---|---|---|---|
| | | Consortium | of the Wellcome Sanger Institute COVID-19 Surveillance Team |
| EPI_ISL_3377381, EPI_ISL_3377382, EPI_ISL_3377383 | Lighthouse Laboratory Plymouth | Wellcome Sanger Institute for the COVID-19 Genomics UK (COG-UK) Consortium | Lighthouse Laboratory Plymouth and Alex Alderton, Roberto Amato, Jeffrey Barrett, Sonia Goncalves, Ewan Harrison, David K. Jackson, Ian Johnston, Dominic Kwiatkowski, Cordelia Langford, John Sillitoe on behalf of the Wellcome Sanger Institute COVID-19 Surveillance Team |
| EPI_ISL_3377384 | Lighthouse Lab in Milton Keynes | Wellcome Sanger Institute for the COVID-19 Genomics UK (COG-UK) Consortium | The Lighthouse Lab in Milton Keynes and Alex Alderton, Roberto Amato, Jeffrey Barrett, Sonia Goncalves, Ewan Harrison, David K. Jackson, Ian Johnston, Dominic Kwiatkowski, Cordelia Langford, John Sillitoe on behalf of the Wellcome Sanger Institute COVID-19 Surveillance Team |
| EPI_ISL_3377385 | Lighthouse Laboratory Plymouth | Wellcome Sanger Institute for the COVID-19 Genomics UK (COG-UK) Consortium | Lighthouse Laboratory Plymouth and Alex Alderton, Roberto Amato, Jeffrey Barrett, Sonia Goncalves, Ewan Harrison, David K. Jackson, Ian Johnston, Dominic Kwiatkowski, Cordelia Langford, John Sillitoe on behalf of the Wellcome Sanger Institute COVID-19 Surveillance Team |
| EPI_ISL_3377386 | Lighthouse Lab in Milton Keynes | Wellcome Sanger Institute for the COVID-19 Genomics UK (COG-UK) Consortium | The Lighthouse Lab in Milton Keynes and Alex Alderton, Roberto Amato, Jeffrey Barrett, Sonia Goncalves, Ewan Harrison, David K. Jackson, Ian Johnston, Dominic Kwiatkowski, Cordelia Langford, John Sillitoe on behalf of the Wellcome Sanger Institute COVID-19 Surveillance Team |
| EPI_ISL_3377387 | Berkshire and Surrey Pathology Services Lighthouse Laboratory | Wellcome Sanger Institute for the COVID-19 Genomics UK (COG-UK) Consortium | Berkshire and Surrey Pathology Services Lighthouse Laboratory and Alex Alderton, Roberto Amato, Jeffrey Barrett, Sonia Goncalves, Ewan Harrison, David K. Jackson, Ian Johnston, Dominic Kwiatkowski, Cordelia Langford, John Sillitoe on behalf of the Wellcome Sanger Institute COVID-19 Surveillance Team |
| EPI_ISL_3377388 | Lighthouse Laboratory Plymouth | Wellcome Sanger Institute for the COVID-19 Genomics UK (COG-UK) Consortium | Lighthouse Laboratory Plymouth and Alex Alderton, Roberto Amato, Jeffrey Barrett, Sonia Goncalves, Ewan Harrison, David K. Jackson, Ian Johnston, Dominic Kwiatkowski, Cordelia Langford, John Sillitoe on behalf of the Wellcome Sanger Institute COVID-19 Surveillance Team |
| EPI_ISL_3377389 | Berkshire and Surrey Pathology Services Lighthouse Laboratory | Wellcome Sanger Institute for the COVID-19 Genomics UK (COG-UK) Consortium | Berkshire and Surrey Pathology Services Lighthouse Laboratory and Alex Alderton, Roberto Amato, Jeffrey Barrett, Sonia Goncalves, Ewan Harrison, David K. Jackson, Ian Johnston, Dominic Kwiatkowski, Cordelia Langford, John Sillitoe on behalf of the Wellcome Sanger Institute COVID-19 Surveillance Team |
| EPI_ISL_3377390 | Lighthouse Lab in Milton Keynes | Wellcome Sanger Institute for the COVID-19 Genomics UK (COG-UK) Consortium | The Lighthouse Lab in Milton Keynes and Alex Alderton, Roberto Amato, Jeffrey Barrett, Sonia Goncalves, Ewan Harrison, David K. Jackson, Ian Johnston, Dominic Kwiatkowski, Cordelia Langford, John Sillitoe on behalf of the Wellcome Sanger Institute COVID-19 Surveillance Team |
| EPI_ISL_3377391 | Berkshire and Surrey Pathology Services Lighthouse Laboratory | Wellcome Sanger Institute for the COVID-19 Genomics UK (COG-UK) Consortium | Berkshire and Surrey Pathology Services Lighthouse Laboratory and Alex Alderton, Roberto Amato, Jeffrey Barrett, Sonia Goncalves, Ewan Harrison, David K. Jackson, Ian Johnston, Dominic Kwiatkowski, Cordelia Langford, John Sillitoe on behalf of the Wellcome Sanger Institute COVID-19 Surveillance Team |
| EPI_ISL_3377392 | Lighthouse Lab in Milton Keynes | Wellcome Sanger Institute for the COVID-19 Genomics UK (COG-UK) Consortium | The Lighthouse Lab in Milton Keynes and Alex Alderton, Roberto Amato, Jeffrey Barrett, Sonia Goncalves, Ewan Harrison, David K. Jackson, Ian Johnston, Dominic Kwiatkowski, Cordelia Langford, John Sillitoe on behalf of the Wellcome Sanger Institute COVID-19 Surveillance Team |
| EPI_ISL_3377393 | Lighthouse Laboratory Plymouth | Wellcome Sanger Institute for the COVID-19 Genomics UK (COG-UK) Consortium | Lighthouse Laboratory Plymouth and Alex Alderton, Roberto Amato, Jeffrey Barrett, Sonia Goncalves, Ewan Harrison, David K. Jackson, Ian Johnston, Dominic Kwiatkowski, Cordelia Langford, John Sillitoe on behalf of the Wellcome Sanger Institute COVID-19 Surveillance Team |
| EPI_ISL_3377394 | Lighthouse Lab in Milton Keynes | Wellcome Sanger Institute for the COVID-19 Genomics UK (COG-UK) Consortium | The Lighthouse Lab in Milton Keynes and Alex Alderton, Roberto Amato, Jeffrey Barrett, Sonia Goncalves, Ewan Harrison, David K. Jackson, Ian Johnston, Dominic Kwiatkowski, Cordelia Langford, John Sillitoe on behalf of the Wellcome Sanger Institute COVID-19 Surveillance Team |
| EPI_ISL_3377395 | Berkshire and Surrey Pathology Services Lighthouse Laboratory | Wellcome Sanger Institute for the COVID-19 Genomics UK (COG-UK) Consortium | Berkshire and Surrey Pathology Services Lighthouse Laboratory and Alex Alderton, Roberto Amato, Jeffrey Barrett, Sonia Goncalves, Ewan Harrison, David K. Jackson, Ian Johnston, Dominic Kwiatkowski, Cordelia Langford, John Sillitoe on behalf of the Wellcome Sanger Institute COVID-19 Surveillance Team |
| EPI_ISL_3377396 | Lighthouse Lab in Milton Keynes | Wellcome Sanger Institute for the COVID-19 Genomics UK (COG-UK) Consortium | The Lighthouse Lab in Milton Keynes and Alex Alderton, Roberto Amato, Jeffrey Barrett, Sonia Goncalves, Ewan Harrison, David K. Jackson, Ian Johnston, Dominic Kwiatkowski, Cordelia Langford, John Sillitoe on behalf of the Wellcome Sanger Institute COVID-19 Surveillance Team |
| EPI_ISL_3377397, EPI_ISL_3377398 | Berkshire and Surrey Pathology Services Lighthouse Laboratory | Wellcome Sanger Institute for the COVID-19 Genomics UK (COG-UK) Consortium | Berkshire and Surrey Pathology Services Lighthouse Laboratory and Alex Alderton, Roberto Amato, Jeffrey Barrett, Sonia Goncalves, Ewan Harrison, David K. Jackson, Ian Johnston, Dominic Kwiatkowski, Cordelia Langford, John Sillitoe on behalf of the Wellcome Sanger Institute COVID-19 Surveillance Team |
| EPI_ISL_3377399 | Lighthouse Lab in Milton Keynes | Wellcome Sanger Institute for the COVID-19 Genomics UK (COG-UK) Consortium | The Lighthouse Lab in Milton Keynes and Alex Alderton, Roberto Amato, Jeffrey Barrett, Sonia Goncalves, Ewan Harrison, David K. Jackson, Ian Johnston, Dominic Kwiatkowski, Cordelia Langford, John Sillitoe on behalf of the Wellcome Sanger Institute COVID-19 Surveillance Team |
| EPI_ISL_3377400, EPI_ISL_3377401 | Berkshire and Surrey Pathology Services Lighthouse Laboratory | Wellcome Sanger Institute for the COVID-19 Genomics UK (COG-UK) Consortium | Berkshire and Surrey Pathology Services Lighthouse Laboratory and Alex Alderton, Roberto Amato, Jeffrey Barrett, Sonia Goncalves, Ewan Harrison, David K. Jackson, Ian Johnston, Dominic Kwiatkowski, Cordelia Langford, John Sillitoe on behalf of the Wellcome Sanger Institute COVID-19 Surveillance Team |
| EPI_ISL_3377402, EPI_ISL_3377403, EPI_ISL_3377404, EPI_ISL_3377405, EPI_ISL_3377406 | Lighthouse Lab in Milton Keynes | Wellcome Sanger Institute for the COVID-19 Genomics UK (COG-UK) Consortium | The Lighthouse Lab in Milton Keynes and Alex Alderton, Roberto Amato, Jeffrey Barrett, Sonia Goncalves, Ewan Harrison, David K. Jackson, Ian Johnston, Dominic Kwiatkowski, Cordelia Langford, John Sillitoe on behalf of the Wellcome Sanger Institute COVID-19 Surveillance Team |
| EPI_ISL_3377407 | Lighthouse Laboratory Plymouth | Wellcome Sanger Institute for the COVID-19 Genomics UK (COG-UK) Consortium | Lighthouse Laboratory Plymouth and Alex Alderton, Roberto Amato, Jeffrey Barrett, Sonia Goncalves, Ewan Harrison, David K. Jackson, Ian Johnston, Dominic Kwiatkowski, Cordelia Langford, John Sillitoe on behalf of the Wellcome Sanger Institute COVID-19 Surveillance Team |
| EPI_ISL_3377408 | Berkshire and Surrey Pathology Services Lighthouse Laboratory | Wellcome Sanger Institute for the COVID-19 Genomics UK (COG-UK) Consortium | Berkshire and Surrey Pathology Services Lighthouse Laboratory and Alex Alderton, Roberto Amato, Jeffrey Barrett, Sonia Goncalves, Ewan Harrison, David K. Jackson, Ian Johnston, Dominic Kwiatkowski, Cordelia Langford, John Sillitoe on behalf of the Wellcome Sanger Institute COVID-19 Surveillance Team |
| EPI_ISL_3377409 | Lighthouse Laboratory Plymouth | Wellcome Sanger Institute for the COVID-19 Genomics UK (COG-UK) Consortium | Lighthouse Laboratory Plymouth and Alex Alderton, Roberto Amato, Jeffrey Barrett, Sonia Goncalves, Ewan Harrison, David K. Jackson, Ian Johnston, Dominic Kwiatkowski, Cordelia Langford, John Sillitoe on behalf of the Wellcome Sanger Institute COVID-19 Surveillance Team |
| EPI_ISL_3377410 | Lighthouse Lab in Milton Keynes | Wellcome Sanger Institute for the COVID-19 Genomics UK (COG-UK) Consortium | The Lighthouse Lab in Milton Keynes and Alex Alderton, Roberto Amato, Jeffrey Barrett, Sonia Goncalves, Ewan Harrison, David K. Jackson, Ian Johnston, Dominic Kwiatkowski, Cordelia Langford, John Sillitoe on behalf of the Wellcome Sanger Institute COVID-19 Surveillance Team |
| EPI_ISL_3377411, EPI_ISL_3377412 | Lighthouse Laboratory Plymouth | Wellcome Sanger Institute for the COVID-19 Genomics UK (COG-UK) Consortium | Lighthouse Laboratory Plymouth and Alex Alderton, Roberto Amato, Jeffrey Barrett, Sonia Goncalves, Ewan Harrison, David K. Jackson, Ian Johnston, Dominic Kwiatkowski, Cordelia Langford, John Sillitoe on behalf of the Wellcome Sanger Institute COVID-19 Surveillance Team |
| EPI_ISL_3377413 | Berkshire and Surrey Pathology Services Lighthouse Laboratory | Wellcome Sanger Institute for the COVID-19 Genomics UK (COG-UK) Consortium | Berkshire and Surrey Pathology Services Lighthouse Laboratory and Alex Alderton, Roberto Amato, Jeffrey Barrett, Sonia Goncalves, Ewan Harrison, David K. Jackson, Ian Johnston, Dominic Kwiatkowski, Cordelia Langford, John Sillitoe on behalf of the Wellcome Sanger Institute COVID-19 Surveillance Team |
| EPI_ISL_3377414 | Lighthouse Lab in Milton Keynes | Wellcome Sanger Institute for the COVID-19 Genomics UK (COG-UK) Consortium | The Lighthouse Lab in Milton Keynes and Alex Alderton, Roberto Amato, Jeffrey Barrett, Sonia Goncalves, Ewan Harrison, David K. Jackson, Ian Johnston, Dominic Kwiatkowski, Cordelia Langford, John Sillitoe on behalf of the Wellcome Sanger Institute COVID-19 Surveillance Team |
| EPI_ISL_3377415 | Lighthouse Laboratory Plymouth | Wellcome Sanger Institute for the COVID-19 Genomics UK (COG-UK) Consortium | Lighthouse Laboratory Plymouth and Alex Alderton, Roberto Amato, Jeffrey Barrett, Sonia Goncalves, Ewan Harrison, David K. Jackson, Ian Johnston, Dominic Kwiatkowski, Cordelia Langford, John Sillitoe on behalf of the Wellcome Sanger Institute COVID-19 Surveillance Team |
| EPI_ISL_3377416, EPI_ISL_3377417, EPI_ISL_3377418, EPI_ISL_3377419, EPI_ISL_3377420 | Lighthouse Lab in Milton Keynes | Wellcome Sanger Institute for the COVID-19 Genomics UK (COG-UK) Consortium | The Lighthouse Lab in Milton Keynes and Alex Alderton, Roberto Amato, Jeffrey Barrett, Sonia Goncalves, Ewan Harrison, David K. Jackson, Ian Johnston, Dominic Kwiatkowski, Cordelia Langford, John Sillitoe on behalf of the Wellcome Sanger Institute COVID-19 Surveillance Team |
| EPI_ISL_3377421 | Lighthouse Laboratory Plymouth | Wellcome Sanger Institute for the COVID-19 Genomics UK (COG-UK) Consortium | Lighthouse Laboratory Plymouth and Alex Alderton, Roberto Amato, Jeffrey Barrett, Sonia Goncalves, Ewan Harrison, David K. Jackson, Ian Johnston, Dominic Kwiatkowski, Cordelia Langford, John Sillitoe on behalf of the Wellcome Sanger Institute COVID-19 Surveillance Team |
| EPI_ISL_3377422, EPI_ISL_3377423, EPI_ISL_3377424, EPI_ISL_3377425 | Lighthouse Lab in Milton Keynes | Wellcome Sanger Institute for the COVID-19 Genomics UK (COG-UK) Consortium | The Lighthouse Lab in Milton Keynes and Alex Alderton, Roberto Amato, Jeffrey Barrett, Sonia Goncalves, Ewan Harrison, David K. Jackson, Ian Johnston, Dominic Kwiatkowski, Cordelia Langford, John Sillitoe on behalf of the Wellcome Sanger Institute COVID-19 Surveillance Team |
| EPI_ISL_3377426 | Berkshire and Surrey Pathology Services Lighthouse Laboratory | Wellcome Sanger Institute for the COVID-19 Genomics UK (COG-UK) Consortium | Berkshire and Surrey Pathology Services Lighthouse Laboratory and Alex Alderton, Roberto Amato, Jeffrey Barrett, Sonia Goncalves, Ewan Harrison, David K. Jackson, Ian Johnston, Dominic Kwiatkowski, Cordelia Langford, John Sillitoe on behalf of the Wellcome Sanger Institute COVID-19 Surveillance Team |
| EPI_ISL_3377427, EPI_ISL_3377428, EPI_ISL_3377429, EPI_ISL_3377430 | Lighthouse Lab in Milton Keynes | Wellcome Sanger Institute for the COVID-19 Genomics UK (COG-UK) Consortium | The Lighthouse Lab in Milton Keynes and Alex Alderton, Roberto Amato, Jeffrey Barrett, Sonia Goncalves, Ewan Harrison, David K. Jackson, Ian Johnston, Dominic Kwiatkowski, Cordelia Langford, John Sillitoe on behalf of the Wellcome Sanger Institute COVID-19 Surveillance Team |
| EPI_ISL_3377431, EPI_ISL_3377432 | Berkshire and Surrey Pathology Services Lighthouse Laboratory | Wellcome Sanger Institute for the COVID-19 Genomics UK (COG-UK) Consortium | Berkshire and Surrey Pathology Services Lighthouse Laboratory and Alex Alderton, Roberto Amato, Jeffrey Barrett, Sonia Goncalves, Ewan Harrison, David K. Jackson, Ian Johnston, Dominic Kwiatkowski, Cordelia Langford, John Sillitoe on behalf of the Wellcome Sanger Institute COVID-19 Surveillance Team |
| EPI_ISL_3377433 | Lighthouse Laboratory Plymouth | Wellcome Sanger Institute for the COVID-19 Genomics UK (COG-UK) Consortium | Lighthouse Laboratory Plymouth and Alex Alderton, Roberto Amato, Jeffrey Barrett, Sonia Goncalves, Ewan Harrison, David K. Jackson, Ian Johnston, Dominic Kwiatkowski, Cordelia Langford, John Sillitoe on behalf of the Wellcome Sanger Institute COVID-19 Surveillance Team |
| EPI_ISL_3377434, EPI_ISL_3377435, EPI_ISL_3377436 | Lighthouse Lab in Milton Keynes | Wellcome Sanger Institute for the COVID-19 Genomics UK (COG-UK) Consortium | The Lighthouse Lab in Milton Keynes and Alex Alderton, Roberto Amato, Jeffrey Barrett, Sonia Goncalves, Ewan Harrison, David K. Jackson, Ian Johnston, Dominic Kwiatkowski, Cordelia Langford, John Sillitoe on behalf of the Wellcome Sanger Institute COVID-19 Surveillance Team |
| EPI_ISL_3377437 | Berkshire and Surrey Pathology Services Lighthouse Laboratory | Wellcome Sanger Institute for the COVID-19 Genomics UK (COG-UK) Consortium | Berkshire and Surrey Pathology Services Lighthouse Laboratory and Alex Alderton, Roberto Amato, Jeffrey Barrett, Sonia Goncalves, Ewan Harrison, David K. Jackson, Ian Johnston, Dominic Kwiatkowski, Cordelia Langford, John Sillitoe on behalf of the Wellcome Sanger Institute COVID-19 Surveillance Team |
| EPI_ISL_3377438, EPI_ISL_3377439 | Lighthouse Lab in Milton Keynes | Wellcome Sanger Institute for the COVID-19 Genomics UK (COG-UK) Consortium | The Lighthouse Lab in Milton Keynes and Alex Alderton, Roberto Amato, Jeffrey Barrett, Sonia Goncalves, Ewan Harrison, David K. Jackson, Ian Johnston, Dominic Kwiatkowski, Cordelia Langford, John Sillitoe on behalf of the Wellcome Sanger Institute COVID-19 Surveillance Team |
| EPI_ISL_3377440, EPI_ISL_3377441 | Berkshire and Surrey Pathology Services Lighthouse Laboratory | Wellcome Sanger Institute for the COVID-19 Genomics UK (COG-UK) Consortium | Berkshire and Surrey Pathology Services Lighthouse Laboratory and Alex Alderton, Roberto Amato, Jeffrey Barrett, Sonia Goncalves, Ewan Harrison, David K. Jackson, Ian Johnston, Dominic Kwiatkowski, Cordelia Langford, John Sillitoe on behalf of the Wellcome Sanger Institute COVID-19 Surveillance Team |
| EPI_ISL_3377442, EPI_ISL_3377443 | Lighthouse Lab in Milton Keynes | Wellcome Sanger Institute for the COVID-19 Genomics UK (COG-UK) Consortium | The Lighthouse Lab in Milton Keynes and Alex Alderton, Roberto Amato, Jeffrey Barrett, Sonia Goncalves, Ewan Harrison, David K. Jackson, Ian Johnston, Dominic Kwiatkowski, Cordelia Langford, John Sillitoe on behalf of the Wellcome Sanger Institute COVID-19 Surveillance Team |
| EPI_ISL_3377444 | Berkshire and Surrey Pathology Services Lighthouse Laboratory | Wellcome Sanger Institute for the COVID-19 Genomics UK (COG-UK) Consortium | Berkshire and Surrey Pathology Services Lighthouse Laboratory and Alex Alderton, Roberto Amato, Jeffrey Barrett, Sonia Goncalves, Ewan Harrison, David K. Jackson, Ian Johnston, Dominic Kwiatkowski, Cordelia Langford, John Sillitoe on behalf of the Wellcome Sanger Institute COVID-19 Surveillance Team |
| EPI_ISL_3377445 | Lighthouse Laboratory Plymouth | Wellcome Sanger Institute for the COVID-19 Genomics UK (COG-UK) | Lighthouse Laboratory Plymouth and Alex Alderton, Roberto Amato, Jeffrey Barrett, Sonia Goncalves, Ewan Harrison, David K. Jackson, Ian Johnston, Dominic Kwiatkowski, Cordelia Langford, John Sillitoe on behalf of the Wellcome Sanger |

| | | | |
|---|---|---|---|
| | | Consortium | Institute COVID-19 Surveillance Team |
| EPI_ISL_3377446, EPI_ISL_3377447, EPI_ISL_3377448, EPI_ISL_3377449, EPI_ISL_3377450 | Lighthouse Lab in Milton Keynes | Wellcome Sanger Institute for the COVID-19 Genomics UK (COG-UK) Consortium | The Lighthouse Lab in Milton Keynes and Alex Alderton, Roberto Amato, Jeffrey Barrett, Sonia Goncalves, Ewan Harrison, David K. Jackson, Ian Johnston, Dominic Kwiatkowski, Cordelia Langford, John Sillitoe on behalf of the Wellcome Sanger Institute COVID-19 Surveillance Team |
| EPI_ISL_3377451 | Lighthouse Laboratory Plymouth | Wellcome Sanger Institute for the COVID-19 Genomics UK (COG-UK) Consortium | Lighthouse Laboratory Plymouth and Alex Alderton, Roberto Amato, Jeffrey Barrett, Sonia Goncalves, Ewan Harrison, David K. Jackson, Ian Johnston, Dominic Kwiatkowski, Cordelia Langford, John Sillitoe on behalf of the Wellcome Sanger Institute COVID-19 Surveillance Team |
| EPI_ISL_3377452 | Lighthouse Lab in Milton Keynes | Wellcome Sanger Institute for the COVID-19 Genomics UK (COG-UK) Consortium | The Lighthouse Lab in Milton Keynes and Alex Alderton, Roberto Amato, Jeffrey Barrett, Sonia Goncalves, Ewan Harrison, David K. Jackson, Ian Johnston, Dominic Kwiatkowski, Cordelia Langford, John Sillitoe on behalf of the Wellcome Sanger Institute COVID-19 Surveillance Team |
| EPI_ISL_3377453 | Lighthouse Laboratory Plymouth | Wellcome Sanger Institute for the COVID-19 Genomics UK (COG-UK) Consortium | Lighthouse Laboratory Plymouth and Alex Alderton, Roberto Amato, Jeffrey Barrett, Sonia Goncalves, Ewan Harrison, David K. Jackson, Ian Johnston, Dominic Kwiatkowski, Cordelia Langford, John Sillitoe on behalf of the Wellcome Sanger Institute COVID-19 Surveillance Team |
| EPI_ISL_3377454, EPI_ISL_3377455, EPI_ISL_3377456 | Lighthouse Lab in Milton Keynes | Wellcome Sanger Institute for the COVID-19 Genomics UK (COG-UK) Consortium | The Lighthouse Lab in Milton Keynes and Alex Alderton, Roberto Amato, Jeffrey Barrett, Sonia Goncalves, Ewan Harrison, David K. Jackson, Ian Johnston, Dominic Kwiatkowski, Cordelia Langford, John Sillitoe on behalf of the Wellcome Sanger Institute COVID-19 Surveillance Team |
| EPI_ISL_3377457, EPI_ISL_3377458, EPI_ISL_3377459 | Lighthouse Laboratory Plymouth | Wellcome Sanger Institute for the COVID-19 Genomics UK (COG-UK) Consortium | Lighthouse Laboratory Plymouth and Alex Alderton, Roberto Amato, Jeffrey Barrett, Sonia Goncalves, Ewan Harrison, David K. Jackson, Ian Johnston, Dominic Kwiatkowski, Cordelia Langford, John Sillitoe on behalf of the Wellcome Sanger Institute COVID-19 Surveillance Team |
| EPI_ISL_3377460, EPI_ISL_3377461 | Lighthouse Lab in Milton Keynes | Wellcome Sanger Institute for the COVID-19 Genomics UK (COG-UK) Consortium | The Lighthouse Lab in Milton Keynes and Alex Alderton, Roberto Amato, Jeffrey Barrett, Sonia Goncalves, Ewan Harrison, David K. Jackson, Ian Johnston, Dominic Kwiatkowski, Cordelia Langford, John Sillitoe on behalf of the Wellcome Sanger Institute COVID-19 Surveillance Team |
| EPI_ISL_3377462 | Berkshire and Surrey Pathology Services Lighthouse Laboratory | Wellcome Sanger Institute for the COVID-19 Genomics UK (COG-UK) Consortium | Berkshire and Surrey Pathology Services Lighthouse Laboratory and Alex Alderton, Roberto Amato, Jeffrey Barrett, Sonia Goncalves, Ewan Harrison, David K. Jackson, Ian Johnston, Dominic Kwiatkowski, Cordelia Langford, John Sillitoe on behalf of the Wellcome Sanger Institute COVID-19 Surveillance Team |
| EPI_ISL_3377463 | Lighthouse Laboratory Plymouth | Wellcome Sanger Institute for the COVID-19 Genomics UK (COG-UK) Consortium | Lighthouse Laboratory Plymouth and Alex Alderton, Roberto Amato, Jeffrey Barrett, Sonia Goncalves, Ewan Harrison, David K. Jackson, Ian Johnston, Dominic Kwiatkowski, Cordelia Langford, John Sillitoe on behalf of the Wellcome Sanger Institute COVID-19 Surveillance Team |
| EPI_ISL_3377464 | Lighthouse Lab in Milton Keynes | Wellcome Sanger Institute for the COVID-19 Genomics UK (COG-UK) Consortium | The Lighthouse Lab in Milton Keynes and Alex Alderton, Roberto Amato, Jeffrey Barrett, Sonia Goncalves, Ewan Harrison, David K. Jackson, Ian Johnston, Dominic Kwiatkowski, Cordelia Langford, John Sillitoe on behalf of the Wellcome Sanger Institute COVID-19 Surveillance Team |
| EPI_ISL_3377465 | Berkshire and Surrey Pathology Services Lighthouse Laboratory | Wellcome Sanger Institute for the COVID-19 Genomics UK (COG-UK) Consortium | Berkshire and Surrey Pathology Services Lighthouse Laboratory and Alex Alderton, Roberto Amato, Jeffrey Barrett, Sonia Goncalves, Ewan Harrison, David K. Jackson, Ian Johnston, Dominic Kwiatkowski, Cordelia Langford, John Sillitoe on behalf of the Wellcome Sanger Institute COVID-19 Surveillance Team |
| EPI_ISL_3377466 | Lighthouse Lab in Milton Keynes | Wellcome Sanger Institute for the COVID-19 Genomics UK (COG-UK) Consortium | The Lighthouse Lab in Milton Keynes and Alex Alderton, Roberto Amato, Jeffrey Barrett, Sonia Goncalves, Ewan Harrison, David K. Jackson, Ian Johnston, Dominic Kwiatkowski, Cordelia Langford, John Sillitoe on behalf of the Wellcome Sanger Institute COVID-19 Surveillance Team |
| EPI_ISL_3377467 | Lighthouse Laboratory Plymouth | Wellcome Sanger Institute for the COVID-19 Genomics UK (COG-UK) Consortium | Lighthouse Laboratory Plymouth and Alex Alderton, Roberto Amato, Jeffrey Barrett, Sonia Goncalves, Ewan Harrison, David K. Jackson, Ian Johnston, Dominic Kwiatkowski, Cordelia Langford, John Sillitoe on behalf of the Wellcome Sanger Institute COVID-19 Surveillance Team |
| EPI_ISL_3377468, EPI_ISL_3377469, EPI_ISL_3377470, EPI_ISL_3377471 | Lighthouse Lab in Milton Keynes | Wellcome Sanger Institute for the COVID-19 Genomics UK (COG-UK) Consortium | The Lighthouse Lab in Milton Keynes and Alex Alderton, Roberto Amato, Jeffrey Barrett, Sonia Goncalves, Ewan Harrison, David K. Jackson, Ian Johnston, Dominic Kwiatkowski, Cordelia Langford, John Sillitoe on behalf of the Wellcome Sanger Institute COVID-19 Surveillance Team |
| EPI_ISL_3377472 | Lighthouse Laboratory Plymouth | Wellcome Sanger Institute for the COVID-19 Genomics UK (COG-UK) Consortium | Lighthouse Laboratory Plymouth and Alex Alderton, Roberto Amato, Jeffrey Barrett, Sonia Goncalves, Ewan Harrison, David K. Jackson, Ian Johnston, Dominic Kwiatkowski, Cordelia Langford, John Sillitoe on behalf of the Wellcome Sanger Institute COVID-19 Surveillance Team |
| EPI_ISL_3377473 | Lighthouse Lab in Milton Keynes | Wellcome Sanger Institute for the COVID-19 Genomics UK (COG-UK) Consortium | The Lighthouse Lab in Milton Keynes and Alex Alderton, Roberto Amato, Jeffrey Barrett, Sonia Goncalves, Ewan Harrison, David K. Jackson, Ian Johnston, Dominic Kwiatkowski, Cordelia Langford, John Sillitoe on behalf of the Wellcome Sanger Institute COVID-19 Surveillance Team |
| EPI_ISL_3377474, EPI_ISL_3377475, EPI_ISL_3377476 | Berkshire and Surrey Pathology Services Lighthouse Laboratory | Wellcome Sanger Institute for the COVID-19 Genomics UK (COG-UK) Consortium | Berkshire and Surrey Pathology Services Lighthouse Laboratory and Alex Alderton, Roberto Amato, Jeffrey Barrett, Sonia Goncalves, Ewan Harrison, David K. Jackson, Ian Johnston, Dominic Kwiatkowski, Cordelia Langford, John Sillitoe on behalf of the Wellcome Sanger Institute COVID-19 Surveillance Team |
| EPI_ISL_3377477, EPI_ISL_3377478 | Lighthouse Lab in Milton Keynes | Wellcome Sanger Institute for the COVID-19 Genomics UK (COG-UK) Consortium | The Lighthouse Lab in Milton Keynes and Alex Alderton, Roberto Amato, Jeffrey Barrett, Sonia Goncalves, Ewan Harrison, David K. Jackson, Ian Johnston, Dominic Kwiatkowski, Cordelia Langford, John Sillitoe on behalf of the Wellcome Sanger Institute COVID-19 Surveillance Team |
| EPI_ISL_3377479 | Lighthouse Laboratory Plymouth | Wellcome Sanger Institute for the COVID-19 Genomics UK (COG-UK) Consortium | Lighthouse Laboratory Plymouth and Alex Alderton, Roberto Amato, Jeffrey Barrett, Sonia Goncalves, Ewan Harrison, David K. Jackson, Ian Johnston, Dominic Kwiatkowski, Cordelia Langford, John Sillitoe on behalf of the Wellcome Sanger Institute COVID-19 Surveillance Team |
| EPI_ISL_3377480 | Lighthouse Lab in Milton Keynes | Wellcome Sanger Institute for the COVID-19 Genomics UK (COG-UK) Consortium | The Lighthouse Lab in Milton Keynes and Alex Alderton, Roberto Amato, Jeffrey Barrett, Sonia Goncalves, Ewan Harrison, David K. Jackson, Ian Johnston, Dominic Kwiatkowski, Cordelia Langford, John Sillitoe on behalf of the Wellcome Sanger Institute COVID-19 Surveillance Team |
| EPI_ISL_3377481 | Berkshire and Surrey Pathology Services Lighthouse Laboratory | Wellcome Sanger Institute for the COVID-19 Genomics UK (COG-UK) Consortium | Berkshire and Surrey Pathology Services Lighthouse Laboratory and Alex Alderton, Roberto Amato, Jeffrey Barrett, Sonia Goncalves, Ewan Harrison, David K. Jackson, Ian Johnston, Dominic Kwiatkowski, Cordelia Langford, John Sillitoe on behalf of the Wellcome Sanger Institute COVID-19 Surveillance Team |
| EPI_ISL_3377482, EPI_ISL_3377483, EPI_ISL_3377484, EPI_ISL_3377485, EPI_ISL_3377486 | Lighthouse Lab in Milton Keynes | Wellcome Sanger Institute for the COVID-19 Genomics UK (COG-UK) Consortium | The Lighthouse Lab in Milton Keynes and Alex Alderton, Roberto Amato, Jeffrey Barrett, Sonia Goncalves, Ewan Harrison, David K. Jackson, Ian Johnston, Dominic Kwiatkowski, Cordelia Langford, John Sillitoe on behalf of the Wellcome Sanger Institute COVID-19 Surveillance Team |
| EPI_ISL_3377487, EPI_ISL_3377488 | Lighthouse Laboratory Plymouth | Wellcome Sanger Institute for the COVID-19 Genomics UK (COG-UK) Consortium | Lighthouse Laboratory Plymouth and Alex Alderton, Roberto Amato, Jeffrey Barrett, Sonia Goncalves, Ewan Harrison, David K. Jackson, Ian Johnston, Dominic Kwiatkowski, Cordelia Langford, John Sillitoe on behalf of the Wellcome Sanger Institute COVID-19 Surveillance Team |
| EPI_ISL_3377489 | Berkshire and Surrey Pathology Services Lighthouse Laboratory | Wellcome Sanger Institute for the COVID-19 Genomics UK (COG-UK) Consortium | Berkshire and Surrey Pathology Services Lighthouse Laboratory and Alex Alderton, Roberto Amato, Jeffrey Barrett, Sonia Goncalves, Ewan Harrison, David K. Jackson, Ian Johnston, Dominic Kwiatkowski, Cordelia Langford, John Sillitoe on behalf of the Wellcome Sanger Institute COVID-19 Surveillance Team |
| EPI_ISL_3377490, EPI_ISL_3377491 | Lighthouse Laboratory Plymouth | Wellcome Sanger Institute for the COVID-19 Genomics UK (COG-UK) Consortium | Lighthouse Laboratory Plymouth and Alex Alderton, Roberto Amato, Jeffrey Barrett, Sonia Goncalves, Ewan Harrison, David K. Jackson, Ian Johnston, Dominic Kwiatkowski, Cordelia Langford, John Sillitoe on behalf of the Wellcome Sanger Institute COVID-19 Surveillance Team |
| EPI_ISL_3377492, EPI_ISL_3377493, EPI_ISL_3377494, EPI_ISL_3377495 | Lighthouse Lab in Milton Keynes | Wellcome Sanger Institute for the COVID-19 Genomics UK (COG-UK) Consortium | The Lighthouse Lab in Milton Keynes and Alex Alderton, Roberto Amato, Jeffrey Barrett, Sonia Goncalves, Ewan Harrison, David K. Jackson, Ian Johnston, Dominic Kwiatkowski, Cordelia Langford, John Sillitoe on behalf of the Wellcome Sanger Institute COVID-19 Surveillance Team |
| EPI_ISL_3377496 | Berkshire and Surrey Pathology Services Lighthouse Laboratory | Wellcome Sanger Institute for the COVID-19 Genomics UK (COG-UK) Consortium | Berkshire and Surrey Pathology Services Lighthouse Laboratory and Alex Alderton, Roberto Amato, Jeffrey Barrett, Sonia Goncalves, Ewan Harrison, David K. Jackson, Ian Johnston, Dominic Kwiatkowski, Cordelia Langford, John Sillitoe on behalf of the Wellcome Sanger Institute COVID-19 Surveillance Team |
| EPI_ISL_3377497, EPI_ISL_3377498, EPI_ISL_3377499, EPI_ISL_3377500 | Lighthouse Lab in Milton Keynes | Wellcome Sanger Institute for the COVID-19 Genomics UK (COG-UK) Consortium | The Lighthouse Lab in Milton Keynes and Alex Alderton, Roberto Amato, Jeffrey Barrett, Sonia Goncalves, Ewan Harrison, David K. Jackson, Ian Johnston, Dominic Kwiatkowski, Cordelia Langford, John Sillitoe on behalf of the Wellcome Sanger Institute COVID-19 Surveillance Team |
| EPI_ISL_3377501, EPI_ISL_3377502 | Lighthouse Laboratory Plymouth | Wellcome Sanger Institute for the COVID-19 Genomics UK (COG-UK) Consortium | Lighthouse Laboratory Plymouth and Alex Alderton, Roberto Amato, Jeffrey Barrett, Sonia Goncalves, Ewan Harrison, David K. Jackson, Ian Johnston, Dominic Kwiatkowski, Cordelia Langford, John Sillitoe on behalf of the Wellcome Sanger Institute COVID-19 Surveillance Team |
| EPI_ISL_3377503 | Berkshire and Surrey Pathology Services Lighthouse Laboratory | Wellcome Sanger Institute for the COVID-19 Genomics UK (COG-UK) Consortium | Berkshire and Surrey Pathology Services Lighthouse Laboratory and Alex Alderton, Roberto Amato, Jeffrey Barrett, Sonia Goncalves, Ewan Harrison, David K. Jackson, Ian Johnston, Dominic Kwiatkowski, Cordelia Langford, John Sillitoe on behalf of the Wellcome Sanger Institute COVID-19 Surveillance Team |
| EPI_ISL_3377504 | Lighthouse Lab in Milton Keynes | Wellcome Sanger Institute for the COVID-19 Genomics UK (COG-UK) Consortium | The Lighthouse Lab in Milton Keynes and Alex Alderton, Roberto Amato, Jeffrey Barrett, Sonia Goncalves, Ewan Harrison, David K. Jackson, Ian Johnston, Dominic Kwiatkowski, Cordelia Langford, John Sillitoe on behalf of the Wellcome Sanger Institute COVID-19 Surveillance Team |
| EPI_ISL_3377505 | Lighthouse Laboratory Plymouth | Wellcome Sanger Institute for the COVID-19 Genomics UK (COG-UK) Consortium | Lighthouse Laboratory Plymouth and Alex Alderton, Roberto Amato, Jeffrey Barrett, Sonia Goncalves, Ewan Harrison, David K. Jackson, Ian Johnston, Dominic Kwiatkowski, Cordelia Langford, John Sillitoe on behalf of the Wellcome Sanger Institute COVID-19 Surveillance Team |
| EPI_ISL_3377506 | Lighthouse Lab in Milton Keynes | Wellcome Sanger Institute for the COVID-19 Genomics UK (COG-UK) Consortium | The Lighthouse Lab in Milton Keynes and Alex Alderton, Roberto Amato, Jeffrey Barrett, Sonia Goncalves, Ewan Harrison, David K. Jackson, Ian Johnston, Dominic Kwiatkowski, Cordelia Langford, John Sillitoe on behalf of the Wellcome Sanger Institute COVID-19 Surveillance Team |
| EPI_ISL_3377507 | Lighthouse Laboratory Plymouth | Wellcome Sanger Institute for the COVID-19 Genomics UK (COG-UK) Consortium | Lighthouse Laboratory Plymouth and Alex Alderton, Roberto Amato, Jeffrey Barrett, Sonia Goncalves, Ewan Harrison, David K. Jackson, Ian Johnston, Dominic Kwiatkowski, Cordelia Langford, John Sillitoe on behalf of the Wellcome Sanger Institute COVID-19 Surveillance Team |
| EPI_ISL_3377508 | Berkshire and Surrey Pathology Services Lighthouse Laboratory | Wellcome Sanger Institute for the COVID-19 Genomics UK (COG-UK) Consortium | Berkshire and Surrey Pathology Services Lighthouse Laboratory and Alex Alderton, Roberto Amato, Jeffrey Barrett, Sonia Goncalves, Ewan Harrison, David K. Jackson, Ian Johnston, Dominic Kwiatkowski, Cordelia Langford, John Sillitoe on behalf of the Wellcome Sanger Institute COVID-19 Surveillance Team |
| EPI_ISL_3377509, EPI_ISL_3377510 | Lighthouse Lab in Milton Keynes | Wellcome Sanger Institute for the COVID-19 Genomics UK (COG-UK) Consortium | The Lighthouse Lab in Milton Keynes and Alex Alderton, Roberto Amato, Jeffrey Barrett, Sonia Goncalves, Ewan Harrison, David K. Jackson, Ian Johnston, Dominic Kwiatkowski, Cordelia Langford, John Sillitoe on behalf of the Wellcome Sanger Institute COVID-19 Surveillance Team |
| EPI_ISL_3377511, EPI_ISL_3377512 | Lighthouse Laboratory Plymouth | Wellcome Sanger Institute for the COVID-19 Genomics UK (COG-UK) Consortium | Lighthouse Laboratory Plymouth and Alex Alderton, Roberto Amato, Jeffrey Barrett, Sonia Goncalves, Ewan Harrison, David K. Jackson, Ian Johnston, Dominic Kwiatkowski, Cordelia Langford, John Sillitoe on behalf of the Wellcome Sanger Institute COVID-19 Surveillance Team |
| EPI_ISL_3377513 | Lighthouse Lab in Milton Keynes | Wellcome Sanger Institute for the COVID-19 Genomics UK (COG-UK) Consortium | The Lighthouse Lab in Milton Keynes and Alex Alderton, Roberto Amato, Jeffrey Barrett, Sonia Goncalves, Ewan Harrison, David K. Jackson, Ian Johnston, Dominic Kwiatkowski, Cordelia Langford, John Sillitoe on behalf of the Wellcome Sanger Institute COVID-19 Surveillance Team |
| EPI_ISL_3377514, | Lighthouse Laboratory Plymouth | Wellcome Sanger Institute for the COVID-19 Genomics UK (COG-UK) | Lighthouse Laboratory Plymouth and Alex Alderton, Roberto Amato, Jeffrey Barrett, Sonia Goncalves, Ewan Harrison, David K. Jackson, Ian Johnston, Dominic Kwiatkowski, Cordelia Langford, John Sillitoe on behalf of the Wellcome Sanger |

| | | | |
|---|---|---|---|
| EPI_ISL_3377515 | | Consortium | Institute COVID-19 Surveillance Team |
| EPI_ISL_3377516 | Lighthouse Lab in Milton Keynes | Wellcome Sanger Institute for the COVID-19 Genomics UK (COG-UK) Consortium | The Lighthouse Lab in Milton Keynes and Alex Alderton, Roberto Amato, Jeffrey Barrett, Sonia Goncalves, Ewan Harrison, David K. Jackson, Ian Johnston, Dominic Kwiatkowski, Cordelia Langford, John Sillitoe on behalf of the Wellcome Sanger Institute COVID-19 Surveillance Team |
| EPI_ISL_3377517, EPI_ISL_3377518 | Berkshire and Surrey Pathology Services Lighthouse Laboratory | Wellcome Sanger Institute for the COVID-19 Genomics UK (COG-UK) Consortium | Berkshire and Surrey Pathology Services Lighthouse Laboratory and Alex Alderton, Roberto Amato, Jeffrey Barrett, Sonia Goncalves, Ewan Harrison, David K. Jackson, Ian Johnston, Dominic Kwiatkowski, Cordelia Langford, John Sillitoe on behalf of the Wellcome Sanger Institute COVID-19 Surveillance Team |
| EPI_ISL_3377519 | Lighthouse Laboratory Plymouth | Wellcome Sanger Institute for the COVID-19 Genomics UK (COG-UK) Consortium | Lighthouse Laboratory Plymouth and Alex Alderton, Roberto Amato, Jeffrey Barrett, Sonia Goncalves, Ewan Harrison, David K. Jackson, Ian Johnston, Dominic Kwiatkowski, Cordelia Langford, John Sillitoe on behalf of the Wellcome Sanger Institute COVID-19 Surveillance Team |
| EPI_ISL_3377520 | Berkshire and Surrey Pathology Services Lighthouse Laboratory | Wellcome Sanger Institute for the COVID-19 Genomics UK (COG-UK) Consortium | Berkshire and Surrey Pathology Services Lighthouse Laboratory and Alex Alderton, Roberto Amato, Jeffrey Barrett, Sonia Goncalves, Ewan Harrison, David K. Jackson, Ian Johnston, Dominic Kwiatkowski, Cordelia Langford, John Sillitoe on behalf of the Wellcome Sanger Institute COVID-19 Surveillance Team |
| EPI_ISL_3377521, EPI_ISL_3377522 | Lighthouse Lab in Milton Keynes | Wellcome Sanger Institute for the COVID-19 Genomics UK (COG-UK) Consortium | The Lighthouse Lab in Milton Keynes and Alex Alderton, Roberto Amato, Jeffrey Barrett, Sonia Goncalves, Ewan Harrison, David K. Jackson, Ian Johnston, Dominic Kwiatkowski, Cordelia Langford, John Sillitoe on behalf of the Wellcome Sanger Institute COVID-19 Surveillance Team |
| EPI_ISL_3377523 | Lighthouse Laboratory Plymouth | Wellcome Sanger Institute for the COVID-19 Genomics UK (COG-UK) Consortium | Lighthouse Laboratory Plymouth and Alex Alderton, Roberto Amato, Jeffrey Barrett, Sonia Goncalves, Ewan Harrison, David K. Jackson, Ian Johnston, Dominic Kwiatkowski, Cordelia Langford, John Sillitoe on behalf of the Wellcome Sanger Institute COVID-19 Surveillance Team |
| EPI_ISL_3377524 | Lighthouse Lab in Milton Keynes | Wellcome Sanger Institute for the COVID-19 Genomics UK (COG-UK) Consortium | The Lighthouse Lab in Milton Keynes and Alex Alderton, Roberto Amato, Jeffrey Barrett, Sonia Goncalves, Ewan Harrison, David K. Jackson, Ian Johnston, Dominic Kwiatkowski, Cordelia Langford, John Sillitoe on behalf of the Wellcome Sanger Institute COVID-19 Surveillance Team |
| EPI_ISL_3377525 | Lighthouse Laboratory Plymouth | Wellcome Sanger Institute for the COVID-19 Genomics UK (COG-UK) Consortium | Lighthouse Laboratory Plymouth and Alex Alderton, Roberto Amato, Jeffrey Barrett, Sonia Goncalves, Ewan Harrison, David K. Jackson, Ian Johnston, Dominic Kwiatkowski, Cordelia Langford, John Sillitoe on behalf of the Wellcome Sanger Institute COVID-19 Surveillance Team |
| EPI_ISL_3377526 | Berkshire and Surrey Pathology Services Lighthouse Laboratory | Wellcome Sanger Institute for the COVID-19 Genomics UK (COG-UK) Consortium | Berkshire and Surrey Pathology Services Lighthouse Laboratory and Alex Alderton, Roberto Amato, Jeffrey Barrett, Sonia Goncalves, Ewan Harrison, David K. Jackson, Ian Johnston, Dominic Kwiatkowski, Cordelia Langford, John Sillitoe on behalf of the Wellcome Sanger Institute COVID-19 Surveillance Team |
| EPI_ISL_3377527 | Lighthouse Laboratory Plymouth | Wellcome Sanger Institute for the COVID-19 Genomics UK (COG-UK) Consortium | Lighthouse Laboratory Plymouth and Alex Alderton, Roberto Amato, Jeffrey Barrett, Sonia Goncalves, Ewan Harrison, David K. Jackson, Ian Johnston, Dominic Kwiatkowski, Cordelia Langford, John Sillitoe on behalf of the Wellcome Sanger Institute COVID-19 Surveillance Team |
| EPI_ISL_3377528, EPI_ISL_3377529, EPI_ISL_3377530, EPI_ISL_3377531, EPI_ISL_3377532 | Lighthouse Lab in Milton Keynes | Wellcome Sanger Institute for the COVID-19 Genomics UK (COG-UK) Consortium | The Lighthouse Lab in Milton Keynes and Alex Alderton, Roberto Amato, Jeffrey Barrett, Sonia Goncalves, Ewan Harrison, David K. Jackson, Ian Johnston, Dominic Kwiatkowski, Cordelia Langford, John Sillitoe on behalf of the Wellcome Sanger Institute COVID-19 Surveillance Team |
| EPI_ISL_3377533, EPI_ISL_3377534 | Berkshire and Surrey Pathology Services Lighthouse Laboratory | Wellcome Sanger Institute for the COVID-19 Genomics UK (COG-UK) Consortium | Berkshire and Surrey Pathology Services Lighthouse Laboratory and Alex Alderton, Roberto Amato, Jeffrey Barrett, Sonia Goncalves, Ewan Harrison, David K. Jackson, Ian Johnston, Dominic Kwiatkowski, Cordelia Langford, John Sillitoe on behalf of the Wellcome Sanger Institute COVID-19 Surveillance Team |
| EPI_ISL_3377535, EPI_ISL_3377536 | Lighthouse Lab in Milton Keynes | Wellcome Sanger Institute for the COVID-19 Genomics UK (COG-UK) Consortium | The Lighthouse Lab in Milton Keynes and Alex Alderton, Roberto Amato, Jeffrey Barrett, Sonia Goncalves, Ewan Harrison, David K. Jackson, Ian Johnston, Dominic Kwiatkowski, Cordelia Langford, John Sillitoe on behalf of the Wellcome Sanger Institute COVID-19 Surveillance Team |
| EPI_ISL_3377537 | Lighthouse Laboratory Plymouth | Wellcome Sanger Institute for the COVID-19 Genomics UK (COG-UK) Consortium | Lighthouse Laboratory Plymouth and Alex Alderton, Roberto Amato, Jeffrey Barrett, Sonia Goncalves, Ewan Harrison, David K. Jackson, Ian Johnston, Dominic Kwiatkowski, Cordelia Langford, John Sillitoe on behalf of the Wellcome Sanger Institute COVID-19 Surveillance Team |
| EPI_ISL_3377538, EPI_ISL_3377539, EPI_ISL_3377540, EPI_ISL_3377541, EPI_ISL_3377542, EPI_ISL_3377543 | Lighthouse Lab in Milton Keynes | Wellcome Sanger Institute for the COVID-19 Genomics UK (COG-UK) Consortium | The Lighthouse Lab in Milton Keynes and Alex Alderton, Roberto Amato, Jeffrey Barrett, Sonia Goncalves, Ewan Harrison, David K. Jackson, Ian Johnston, Dominic Kwiatkowski, Cordelia Langford, John Sillitoe on behalf of the Wellcome Sanger Institute COVID-19 Surveillance Team |
| EPI_ISL_3377544 | Lighthouse Laboratory Plymouth | Wellcome Sanger Institute for the COVID-19 Genomics UK (COG-UK) Consortium | Lighthouse Laboratory Plymouth and Alex Alderton, Roberto Amato, Jeffrey Barrett, Sonia Goncalves, Ewan Harrison, David K. Jackson, Ian Johnston, Dominic Kwiatkowski, Cordelia Langford, John Sillitoe on behalf of the Wellcome Sanger Institute COVID-19 Surveillance Team |
| EPI_ISL_3377545, EPI_ISL_3377546, EPI_ISL_3377547, EPI_ISL_3377548, EPI_ISL_3377549 | Lighthouse Lab in Milton Keynes | Wellcome Sanger Institute for the COVID-19 Genomics UK (COG-UK) Consortium | The Lighthouse Lab in Milton Keynes and Alex Alderton, Roberto Amato, Jeffrey Barrett, Sonia Goncalves, Ewan Harrison, David K. Jackson, Ian Johnston, Dominic Kwiatkowski, Cordelia Langford, John Sillitoe on behalf of the Wellcome Sanger Institute COVID-19 Surveillance Team |
| EPI_ISL_3377550, EPI_ISL_3377551 | Lighthouse Laboratory Plymouth | Wellcome Sanger Institute for the COVID-19 Genomics UK (COG-UK) Consortium | Lighthouse Laboratory Plymouth and Alex Alderton, Roberto Amato, Jeffrey Barrett, Sonia Goncalves, Ewan Harrison, David K. Jackson, Ian Johnston, Dominic Kwiatkowski, Cordelia Langford, John Sillitoe on behalf of the Wellcome Sanger Institute COVID-19 Surveillance Team |
| EPI_ISL_3377552, EPI_ISL_3377553 | Lighthouse Lab in Milton Keynes | Wellcome Sanger Institute for the COVID-19 Genomics UK (COG-UK) Consortium | The Lighthouse Lab in Milton Keynes and Alex Alderton, Roberto Amato, Jeffrey Barrett, Sonia Goncalves, Ewan Harrison, David K. Jackson, Ian Johnston, Dominic Kwiatkowski, Cordelia Langford, John Sillitoe on behalf of the Wellcome Sanger Institute COVID-19 Surveillance Team |
| EPI_ISL_3377554 | Lighthouse Laboratory Plymouth | Wellcome Sanger Institute for the COVID-19 Genomics UK (COG-UK) Consortium | Lighthouse Laboratory Plymouth and Alex Alderton, Roberto Amato, Jeffrey Barrett, Sonia Goncalves, Ewan Harrison, David K. Jackson, Ian Johnston, Dominic Kwiatkowski, Cordelia Langford, John Sillitoe on behalf of the Wellcome Sanger Institute COVID-19 Surveillance Team |
| EPI_ISL_3377555 | Berkshire and Surrey Pathology Services Lighthouse Laboratory | Wellcome Sanger Institute for the COVID-19 Genomics UK (COG-UK) Consortium | Berkshire and Surrey Pathology Services Lighthouse Laboratory and Alex Alderton, Roberto Amato, Jeffrey Barrett, Sonia Goncalves, Ewan Harrison, David K. Jackson, Ian Johnston, Dominic Kwiatkowski, Cordelia Langford, John Sillitoe on behalf of the Wellcome Sanger Institute COVID-19 Surveillance Team |
| EPI_ISL_3377556, EPI_ISL_3377557 | Lighthouse Lab in Milton Keynes | Wellcome Sanger Institute for the COVID-19 Genomics UK (COG-UK) Consortium | The Lighthouse Lab in Milton Keynes and Alex Alderton, Roberto Amato, Jeffrey Barrett, Sonia Goncalves, Ewan Harrison, David K. Jackson, Ian Johnston, Dominic Kwiatkowski, Cordelia Langford, John Sillitoe on behalf of the Wellcome Sanger Institute COVID-19 Surveillance Team |
| EPI_ISL_3377558 | Berkshire and Surrey Pathology Services Lighthouse Laboratory | Wellcome Sanger Institute for the COVID-19 Genomics UK (COG-UK) Consortium | Berkshire and Surrey Pathology Services Lighthouse Laboratory and Alex Alderton, Roberto Amato, Jeffrey Barrett, Sonia Goncalves, Ewan Harrison, David K. Jackson, Ian Johnston, Dominic Kwiatkowski, Cordelia Langford, John Sillitoe on behalf of the Wellcome Sanger Institute COVID-19 Surveillance Team |
| EPI_ISL_3377560 | Lighthouse Laboratory Plymouth | Wellcome Sanger Institute for the COVID-19 Genomics UK (COG-UK) Consortium | Lighthouse Laboratory Plymouth and Alex Alderton, Roberto Amato, Jeffrey Barrett, Sonia Goncalves, Ewan Harrison, David K. Jackson, Ian Johnston, Dominic Kwiatkowski, Cordelia Langford, John Sillitoe on behalf of the Wellcome Sanger Institute COVID-19 Surveillance Team |
| EPI_ISL_3377561 | Lighthouse Lab in Milton Keynes | Wellcome Sanger Institute for the COVID-19 Genomics UK (COG-UK) Consortium | The Lighthouse Lab in Milton Keynes and Alex Alderton, Roberto Amato, Jeffrey Barrett, Sonia Goncalves, Ewan Harrison, David K. Jackson, Ian Johnston, Dominic Kwiatkowski, Cordelia Langford, John Sillitoe on behalf of the Wellcome Sanger Institute COVID-19 Surveillance Team |
| EPI_ISL_3377562 | Berkshire and Surrey Pathology Services Lighthouse Laboratory | Wellcome Sanger Institute for the COVID-19 Genomics UK (COG-UK) Consortium | Berkshire and Surrey Pathology Services Lighthouse Laboratory and Alex Alderton, Roberto Amato, Jeffrey Barrett, Sonia Goncalves, Ewan Harrison, David K. Jackson, Ian Johnston, Dominic Kwiatkowski, Cordelia Langford, John Sillitoe on behalf of the Wellcome Sanger Institute COVID-19 Surveillance Team |
| EPI_ISL_3377563, EPI_ISL_3377564, EPI_ISL_3377565 | Lighthouse Lab in Milton Keynes | Wellcome Sanger Institute for the COVID-19 Genomics UK (COG-UK) Consortium | The Lighthouse Lab in Milton Keynes and Alex Alderton, Roberto Amato, Jeffrey Barrett, Sonia Goncalves, Ewan Harrison, David K. Jackson, Ian Johnston, Dominic Kwiatkowski, Cordelia Langford, John Sillitoe on behalf of the Wellcome Sanger Institute COVID-19 Surveillance Team |
| EPI_ISL_3377566 | Lighthouse Laboratory Plymouth | Wellcome Sanger Institute for the COVID-19 Genomics UK (COG-UK) Consortium | Lighthouse Laboratory Plymouth and Alex Alderton, Roberto Amato, Jeffrey Barrett, Sonia Goncalves, Ewan Harrison, David K. Jackson, Ian Johnston, Dominic Kwiatkowski, Cordelia Langford, John Sillitoe on behalf of the Wellcome Sanger Institute COVID-19 Surveillance Team |
| EPI_ISL_3377567, EPI_ISL_3377568 | Berkshire and Surrey Pathology Services Lighthouse Laboratory | Wellcome Sanger Institute for the COVID-19 Genomics UK (COG-UK) Consortium | Berkshire and Surrey Pathology Services Lighthouse Laboratory and Alex Alderton, Roberto Amato, Jeffrey Barrett, Sonia Goncalves, Ewan Harrison, David K. Jackson, Ian Johnston, Dominic Kwiatkowski, Cordelia Langford, John Sillitoe on behalf of the Wellcome Sanger Institute COVID-19 Surveillance Team |
| EPI_ISL_3377569 | Lighthouse Lab in Milton Keynes | Wellcome Sanger Institute for the COVID-19 Genomics UK (COG-UK) Consortium | The Lighthouse Lab in Milton Keynes and Alex Alderton, Roberto Amato, Jeffrey Barrett, Sonia Goncalves, Ewan Harrison, David K. Jackson, Ian Johnston, Dominic Kwiatkowski, Cordelia Langford, John Sillitoe on behalf of the Wellcome Sanger Institute COVID-19 Surveillance Team |
| EPI_ISL_3377570 | Lighthouse Laboratory Plymouth | Wellcome Sanger Institute for the COVID-19 Genomics UK (COG-UK) Consortium | Lighthouse Laboratory Plymouth and Alex Alderton, Roberto Amato, Jeffrey Barrett, Sonia Goncalves, Ewan Harrison, David K. Jackson, Ian Johnston, Dominic Kwiatkowski, Cordelia Langford, John Sillitoe on behalf of the Wellcome Sanger Institute COVID-19 Surveillance Team |
| EPI_ISL_3377571 | Berkshire and Surrey Pathology Services Lighthouse Laboratory | Wellcome Sanger Institute for the COVID-19 Genomics UK (COG-UK) Consortium | Berkshire and Surrey Pathology Services Lighthouse Laboratory and Alex Alderton, Roberto Amato, Jeffrey Barrett, Sonia Goncalves, Ewan Harrison, David K. Jackson, Ian Johnston, Dominic Kwiatkowski, Cordelia Langford, John Sillitoe on behalf of the Wellcome Sanger Institute COVID-19 Surveillance Team |
| EPI_ISL_3377572, EPI_ISL_3377573, EPI_ISL_3377574, EPI_ISL_3377575 | Lighthouse Lab in Milton Keynes | Wellcome Sanger Institute for the COVID-19 Genomics UK (COG-UK) Consortium | The Lighthouse Lab in Milton Keynes and Alex Alderton, Roberto Amato, Jeffrey Barrett, Sonia Goncalves, Ewan Harrison, David K. Jackson, Ian Johnston, Dominic Kwiatkowski, Cordelia Langford, John Sillitoe on behalf of the Wellcome Sanger Institute COVID-19 Surveillance Team |
| EPI_ISL_3377576, EPI_ISL_3377577, EPI_ISL_3377578 | Lighthouse Laboratory Plymouth | Wellcome Sanger Institute for the COVID-19 Genomics UK (COG-UK) Consortium | Lighthouse Laboratory Plymouth and Alex Alderton, Roberto Amato, Jeffrey Barrett, Sonia Goncalves, Ewan Harrison, David K. Jackson, Ian Johnston, Dominic Kwiatkowski, Cordelia Langford, John Sillitoe on behalf of the Wellcome Sanger Institute COVID-19 Surveillance Team |
| EPI_ISL_3377579 | Lighthouse Lab in Milton Keynes | Wellcome Sanger Institute for the COVID-19 Genomics UK (COG-UK) Consortium | The Lighthouse Lab in Milton Keynes and Alex Alderton, Roberto Amato, Jeffrey Barrett, Sonia Goncalves, Ewan Harrison, David K. Jackson, Ian Johnston, Dominic Kwiatkowski, Cordelia Langford, John Sillitoe on behalf of the Wellcome Sanger Institute COVID-19 Surveillance Team |
| EPI_ISL_3377580 | Lighthouse Laboratory Plymouth | Wellcome Sanger Institute for the COVID-19 Genomics UK (COG-UK) Consortium | Lighthouse Laboratory Plymouth and Alex Alderton, Roberto Amato, Jeffrey Barrett, Sonia Goncalves, Ewan Harrison, David K. Jackson, Ian Johnston, Dominic Kwiatkowski, Cordelia Langford, John Sillitoe on behalf of the Wellcome Sanger Institute COVID-19 Surveillance Team |
| EPI_ISL_3377581 | Berkshire and Surrey Pathology Services Lighthouse Laboratory | Wellcome Sanger Institute for the COVID-19 Genomics UK (COG-UK) Consortium | Berkshire and Surrey Pathology Services Lighthouse Laboratory and Alex Alderton, Roberto Amato, Jeffrey Barrett, Sonia Goncalves, Ewan Harrison, David K. Jackson, Ian Johnston, Dominic Kwiatkowski, Cordelia Langford, John Sillitoe on behalf of the Wellcome Sanger Institute COVID-19 Surveillance Team |
| EPI_ISL_3377582 | Lighthouse Lab in Milton Keynes | Wellcome Sanger Institute for the COVID-19 Genomics UK (COG-UK) Consortium | The Lighthouse Lab in Milton Keynes and Alex Alderton, Roberto Amato, Jeffrey Barrett, Sonia Goncalves, Ewan Harrison, David K. Jackson, Ian Johnston, Dominic Kwiatkowski, Cordelia Langford, John Sillitoe on behalf of the Wellcome Sanger Institute COVID-19 Surveillance Team |
| EPI_ISL_3377583 | Lighthouse Laboratory Plymouth | Wellcome Sanger Institute for the COVID-19 Genomics UK (COG-UK) Consortium | Lighthouse Laboratory Plymouth and Alex Alderton, Roberto Amato, Jeffrey Barrett, Sonia Goncalves, Ewan Harrison, David K. Jackson, Ian Johnston, Dominic Kwiatkowski, Cordelia Langford, John Sillitoe on behalf of the Wellcome Sanger Institute COVID-19 Surveillance Team |

| | | | |
|---|---|---|---|
| EPI_ISL_3377584, EPI_ISL_3377585 | Lighthouse Lab in Milton Keynes | Wellcome Sanger Institute for the COVID-19 Genomics UK (COG-UK) Consortium | The Lighthouse Lab in Milton Keynes and Alex Alderton, Roberto Amato, Jeffrey Barrett, Sonia Goncalves, Ewan Harrison, David K. Jackson, Ian Johnston, Dominic Kwiatkowski, Cordelia Langford, John Sillitoe on behalf of the Wellcome Sanger Institute COVID-19 Surveillance Team |
| EPI_ISL_3377586 | Lighthouse Laboratory Plymouth | Wellcome Sanger Institute for the COVID-19 Genomics UK (COG-UK) Consortium | Lighthouse Laboratory Plymouth and Alex Alderton, Roberto Amato, Jeffrey Barrett, Sonia Goncalves, Ewan Harrison, David K. Jackson, Ian Johnston, Dominic Kwiatkowski, Cordelia Langford, John Sillitoe on behalf of the Wellcome Sanger Institute COVID-19 Surveillance Team |
| EPI_ISL_3377587 | Berkshire and Surrey Pathology Services Lighthouse Laboratory | Wellcome Sanger Institute for the COVID-19 Genomics UK (COG-UK) Consortium | Berkshire and Surrey Pathology Services Lighthouse Laboratory and Alex Alderton, Roberto Amato, Jeffrey Barrett, Sonia Goncalves, Ewan Harrison, David K. Jackson, Ian Johnston, Dominic Kwiatkowski, Cordelia Langford, John Sillitoe on behalf of the Wellcome Sanger Institute COVID-19 Surveillance Team |
| EPI_ISL_3377588 | Lighthouse Laboratory Plymouth | Wellcome Sanger Institute for the COVID-19 Genomics UK (COG-UK) Consortium | Lighthouse Laboratory Plymouth and Alex Alderton, Roberto Amato, Jeffrey Barrett, Sonia Goncalves, Ewan Harrison, David K. Jackson, Ian Johnston, Dominic Kwiatkowski, Cordelia Langford, John Sillitoe on behalf of the Wellcome Sanger Institute COVID-19 Surveillance Team |
| EPI_ISL_3377589 | Lighthouse Lab in Milton Keynes | Wellcome Sanger Institute for the COVID-19 Genomics UK (COG-UK) Consortium | The Lighthouse Lab in Milton Keynes and Alex Alderton, Roberto Amato, Jeffrey Barrett, Sonia Goncalves, Ewan Harrison, David K. Jackson, Ian Johnston, Dominic Kwiatkowski, Cordelia Langford, John Sillitoe on behalf of the Wellcome Sanger Institute COVID-19 Surveillance Team |
| EPI_ISL_3377590 | Berkshire and Surrey Pathology Services Lighthouse Laboratory | Wellcome Sanger Institute for the COVID-19 Genomics UK (COG-UK) Consortium | Berkshire and Surrey Pathology Services Lighthouse Laboratory and Alex Alderton, Roberto Amato, Jeffrey Barrett, Sonia Goncalves, Ewan Harrison, David K. Jackson, Ian Johnston, Dominic Kwiatkowski, Cordelia Langford, John Sillitoe on behalf of the Wellcome Sanger Institute COVID-19 Surveillance Team |
| EPI_ISL_3377591 | Lighthouse Lab in Milton Keynes | Wellcome Sanger Institute for the COVID-19 Genomics UK (COG-UK) Consortium | The Lighthouse Lab in Milton Keynes and Alex Alderton, Roberto Amato, Jeffrey Barrett, Sonia Goncalves, Ewan Harrison, David K. Jackson, Ian Johnston, Dominic Kwiatkowski, Cordelia Langford, John Sillitoe on behalf of the Wellcome Sanger Institute COVID-19 Surveillance Team |
| EPI_ISL_3377592 | Lighthouse Laboratory Plymouth | Wellcome Sanger Institute for the COVID-19 Genomics UK (COG-UK) Consortium | Lighthouse Laboratory Plymouth and Alex Alderton, Roberto Amato, Jeffrey Barrett, Sonia Goncalves, Ewan Harrison, David K. Jackson, Ian Johnston, Dominic Kwiatkowski, Cordelia Langford, John Sillitoe on behalf of the Wellcome Sanger Institute COVID-19 Surveillance Team |
| EPI_ISL_3377593 | Berkshire and Surrey Pathology Services Lighthouse Laboratory | Wellcome Sanger Institute for the COVID-19 Genomics UK (COG-UK) Consortium | Berkshire and Surrey Pathology Services Lighthouse Laboratory and Alex Alderton, Roberto Amato, Jeffrey Barrett, Sonia Goncalves, Ewan Harrison, David K. Jackson, Ian Johnston, Dominic Kwiatkowski, Cordelia Langford, John Sillitoe on behalf of the Wellcome Sanger Institute COVID-19 Surveillance Team |
| EPI_ISL_3377594 | Lighthouse Lab in Milton Keynes | Wellcome Sanger Institute for the COVID-19 Genomics UK (COG-UK) Consortium | The Lighthouse Lab in Milton Keynes and Alex Alderton, Roberto Amato, Jeffrey Barrett, Sonia Goncalves, Ewan Harrison, David K. Jackson, Ian Johnston, Dominic Kwiatkowski, Cordelia Langford, John Sillitoe on behalf of the Wellcome Sanger Institute COVID-19 Surveillance Team |
| EPI_ISL_3377595, EPI_ISL_3377596 | Lighthouse Laboratory Plymouth | Wellcome Sanger Institute for the COVID-19 Genomics UK (COG-UK) Consortium | Lighthouse Laboratory Plymouth and Alex Alderton, Roberto Amato, Jeffrey Barrett, Sonia Goncalves, Ewan Harrison, David K. Jackson, Ian Johnston, Dominic Kwiatkowski, Cordelia Langford, John Sillitoe on behalf of the Wellcome Sanger Institute COVID-19 Surveillance Team |
| EPI_ISL_3377597 | Lighthouse Lab in Milton Keynes | Wellcome Sanger Institute for the COVID-19 Genomics UK (COG-UK) Consortium | The Lighthouse Lab in Milton Keynes and Alex Alderton, Roberto Amato, Jeffrey Barrett, Sonia Goncalves, Ewan Harrison, David K. Jackson, Ian Johnston, Dominic Kwiatkowski, Cordelia Langford, John Sillitoe on behalf of the Wellcome Sanger Institute COVID-19 Surveillance Team |
| EPI_ISL_3377598 | Berkshire and Surrey Pathology Services Lighthouse Laboratory | Wellcome Sanger Institute for the COVID-19 Genomics UK (COG-UK) Consortium | Berkshire and Surrey Pathology Services Lighthouse Laboratory and Alex Alderton, Roberto Amato, Jeffrey Barrett, Sonia Goncalves, Ewan Harrison, David K. Jackson, Ian Johnston, Dominic Kwiatkowski, Cordelia Langford, John Sillitoe on behalf of the Wellcome Sanger Institute COVID-19 Surveillance Team |
| EPI_ISL_3377599 | Lighthouse Lab in Milton Keynes | Wellcome Sanger Institute for the COVID-19 Genomics UK (COG-UK) Consortium | The Lighthouse Lab in Milton Keynes and Alex Alderton, Roberto Amato, Jeffrey Barrett, Sonia Goncalves, Ewan Harrison, David K. Jackson, Ian Johnston, Dominic Kwiatkowski, Cordelia Langford, John Sillitoe on behalf of the Wellcome Sanger Institute COVID-19 Surveillance Team |
| EPI_ISL_3377600, EPI_ISL_3377601 | Berkshire and Surrey Pathology Services Lighthouse Laboratory | Wellcome Sanger Institute for the COVID-19 Genomics UK (COG-UK) Consortium | Berkshire and Surrey Pathology Services Lighthouse Laboratory and Alex Alderton, Roberto Amato, Jeffrey Barrett, Sonia Goncalves, Ewan Harrison, David K. Jackson, Ian Johnston, Dominic Kwiatkowski, Cordelia Langford, John Sillitoe on behalf of the Wellcome Sanger Institute COVID-19 Surveillance Team |
| EPI_ISL_3377602 | Lighthouse Laboratory Plymouth | Wellcome Sanger Institute for the COVID-19 Genomics UK (COG-UK) Consortium | Lighthouse Laboratory Plymouth and Alex Alderton, Roberto Amato, Jeffrey Barrett, Sonia Goncalves, Ewan Harrison, David K. Jackson, Ian Johnston, Dominic Kwiatkowski, Cordelia Langford, John Sillitoe on behalf of the Wellcome Sanger Institute COVID-19 Surveillance Team |
| EPI_ISL_3377603, EPI_ISL_3377604 | Lighthouse Lab in Milton Keynes | Wellcome Sanger Institute for the COVID-19 Genomics UK (COG-UK) Consortium | The Lighthouse Lab in Milton Keynes and Alex Alderton, Roberto Amato, Jeffrey Barrett, Sonia Goncalves, Ewan Harrison, David K. Jackson, Ian Johnston, Dominic Kwiatkowski, Cordelia Langford, John Sillitoe on behalf of the Wellcome Sanger Institute COVID-19 Surveillance Team |
| EPI_ISL_3377605 | Lighthouse Laboratory Plymouth | Wellcome Sanger Institute for the COVID-19 Genomics UK (COG-UK) Consortium | Lighthouse Laboratory Plymouth and Alex Alderton, Roberto Amato, Jeffrey Barrett, Sonia Goncalves, Ewan Harrison, David K. Jackson, Ian Johnston, Dominic Kwiatkowski, Cordelia Langford, John Sillitoe on behalf of the Wellcome Sanger Institute COVID-19 Surveillance Team |
| EPI_ISL_3377606 | Lighthouse Lab in Milton Keynes | Wellcome Sanger Institute for the COVID-19 Genomics UK (COG-UK) Consortium | The Lighthouse Lab in Milton Keynes and Alex Alderton, Roberto Amato, Jeffrey Barrett, Sonia Goncalves, Ewan Harrison, David K. Jackson, Ian Johnston, Dominic Kwiatkowski, Cordelia Langford, John Sillitoe on behalf of the Wellcome Sanger Institute COVID-19 Surveillance Team |
| EPI_ISL_3377607 | Berkshire and Surrey Pathology Services Lighthouse Laboratory | Wellcome Sanger Institute for the COVID-19 Genomics UK (COG-UK) Consortium | Berkshire and Surrey Pathology Services Lighthouse Laboratory and Alex Alderton, Roberto Amato, Jeffrey Barrett, Sonia Goncalves, Ewan Harrison, David K. Jackson, Ian Johnston, Dominic Kwiatkowski, Cordelia Langford, John Sillitoe on behalf of the Wellcome Sanger Institute COVID-19 Surveillance Team |
| EPI_ISL_3377608, EPI_ISL_3377609 | Lighthouse Lab in Milton Keynes | Wellcome Sanger Institute for the COVID-19 Genomics UK (COG-UK) Consortium | The Lighthouse Lab in Milton Keynes and Alex Alderton, Roberto Amato, Jeffrey Barrett, Sonia Goncalves, Ewan Harrison, David K. Jackson, Ian Johnston, Dominic Kwiatkowski, Cordelia Langford, John Sillitoe on behalf of the Wellcome Sanger Institute COVID-19 Surveillance Team |
| EPI_ISL_3377610 | Berkshire and Surrey Pathology Services Lighthouse Laboratory | Wellcome Sanger Institute for the COVID-19 Genomics UK (COG-UK) Consortium | Berkshire and Surrey Pathology Services Lighthouse Laboratory and Alex Alderton, Roberto Amato, Jeffrey Barrett, Sonia Goncalves, Ewan Harrison, David K. Jackson, Ian Johnston, Dominic Kwiatkowski, Cordelia Langford, John Sillitoe on behalf of the Wellcome Sanger Institute COVID-19 Surveillance Team |
| EPI_ISL_3377611, EPI_ISL_3377612, EPI_ISL_3377613, EPI_ISL_3377614 | Lighthouse Lab in Milton Keynes | Wellcome Sanger Institute for the COVID-19 Genomics UK (COG-UK) Consortium | The Lighthouse Lab in Milton Keynes and Alex Alderton, Roberto Amato, Jeffrey Barrett, Sonia Goncalves, Ewan Harrison, David K. Jackson, Ian Johnston, Dominic Kwiatkowski, Cordelia Langford, John Sillitoe on behalf of the Wellcome Sanger Institute COVID-19 Surveillance Team |
| EPI_ISL_3377615 | Lighthouse Laboratory Plymouth | Wellcome Sanger Institute for the COVID-19 Genomics UK (COG-UK) Consortium | Lighthouse Laboratory Plymouth and Alex Alderton, Roberto Amato, Jeffrey Barrett, Sonia Goncalves, Ewan Harrison, David K. Jackson, Ian Johnston, Dominic Kwiatkowski, Cordelia Langford, John Sillitoe on behalf of the Wellcome Sanger Institute COVID-19 Surveillance Team |
| EPI_ISL_3377616 | Berkshire and Surrey Pathology Services Lighthouse Laboratory | Wellcome Sanger Institute for the COVID-19 Genomics UK (COG-UK) Consortium | Berkshire and Surrey Pathology Services Lighthouse Laboratory and Alex Alderton, Roberto Amato, Jeffrey Barrett, Sonia Goncalves, Ewan Harrison, David K. Jackson, Ian Johnston, Dominic Kwiatkowski, Cordelia Langford, John Sillitoe on behalf of the Wellcome Sanger Institute COVID-19 Surveillance Team |
| EPI_ISL_3377617 | Lighthouse Laboratory Plymouth | Wellcome Sanger Institute for the COVID-19 Genomics UK (COG-UK) Consortium | Lighthouse Laboratory Plymouth and Alex Alderton, Roberto Amato, Jeffrey Barrett, Sonia Goncalves, Ewan Harrison, David K. Jackson, Ian Johnston, Dominic Kwiatkowski, Cordelia Langford, John Sillitoe on behalf of the Wellcome Sanger Institute COVID-19 Surveillance Team |
| EPI_ISL_3377618, EPI_ISL_3377619, EPI_ISL_3377620, EPI_ISL_3377621, EPI_ISL_3377622 | Lighthouse Lab in Milton Keynes | Wellcome Sanger Institute for the COVID-19 Genomics UK (COG-UK) Consortium | The Lighthouse Lab in Milton Keynes and Alex Alderton, Roberto Amato, Jeffrey Barrett, Sonia Goncalves, Ewan Harrison, David K. Jackson, Ian Johnston, Dominic Kwiatkowski, Cordelia Langford, John Sillitoe on behalf of the Wellcome Sanger Institute COVID-19 Surveillance Team |
| EPI_ISL_3377623 | Lighthouse Laboratory Plymouth | Wellcome Sanger Institute for the COVID-19 Genomics UK (COG-UK) Consortium | Lighthouse Laboratory Plymouth and Alex Alderton, Roberto Amato, Jeffrey Barrett, Sonia Goncalves, Ewan Harrison, David K. Jackson, Ian Johnston, Dominic Kwiatkowski, Cordelia Langford, John Sillitoe on behalf of the Wellcome Sanger Institute COVID-19 Surveillance Team |
| EPI_ISL_3377624 | Lighthouse Lab in Milton Keynes | Wellcome Sanger Institute for the COVID-19 Genomics UK (COG-UK) Consortium | The Lighthouse Lab in Milton Keynes and Alex Alderton, Roberto Amato, Jeffrey Barrett, Sonia Goncalves, Ewan Harrison, David K. Jackson, Ian Johnston, Dominic Kwiatkowski, Cordelia Langford, John Sillitoe on behalf of the Wellcome Sanger Institute COVID-19 Surveillance Team |
| EPI_ISL_3377625 | Lighthouse Laboratory Plymouth | Wellcome Sanger Institute for the COVID-19 Genomics UK (COG-UK) Consortium | Lighthouse Laboratory Plymouth and Alex Alderton, Roberto Amato, Jeffrey Barrett, Sonia Goncalves, Ewan Harrison, David K. Jackson, Ian Johnston, Dominic Kwiatkowski, Cordelia Langford, John Sillitoe on behalf of the Wellcome Sanger Institute COVID-19 Surveillance Team |
| EPI_ISL_3377626 | Lighthouse Lab in Glasgow | Wellcome Sanger Institute for the COVID-19 Genomics UK (COG-UK) Consortium | Harper VanSteenhouse, Yumi Kasai, David Gray, Carol Clugston, Anna Dominiczak and Alex Alderton, Roberto Amato, Jeffrey Barrett, Sonia Goncalves, Ewan Harrison, David K. Jackson, Ian Johnston, Dominic Kwiatkowski, Cordelia Langford, John Sillitoe on behalf of the Wellcome Sanger Institute COVID-19 Surveillance Team |
| EPI_ISL_3377627 | Lighthouse Lab in Alderley Park | Wellcome Sanger Institute for the COVID-19 Genomics UK (COG-UK) Consortium | Jacquelyn Wynn, Mairead Hyland, The Lighthouse Lab in Alderley Park and Alex Alderton, Roberto Amato, Jeffrey Barrett, Sonia Goncalves, Ewan Harrison, David K. Jackson, Ian Johnston, Dominic Kwiatkowski, Cordelia Langford, John Sillitoe on behalf of the Wellcome Sanger Institute COVID-19 Surveillance Team |
| EPI_ISL_3377628 | Lighthouse Laboratory Plymouth | Wellcome Sanger Institute for the COVID-19 Genomics UK (COG-UK) Consortium | Lighthouse Laboratory Plymouth and Alex Alderton, Roberto Amato, Jeffrey Barrett, Sonia Goncalves, Ewan Harrison, David K. Jackson, Ian Johnston, Dominic Kwiatkowski, Cordelia Langford, John Sillitoe on behalf of the Wellcome Sanger Institute COVID-19 Surveillance Team |
| EPI_ISL_3377630 | Lighthouse Lab in Alderley Park | Wellcome Sanger Institute for the COVID-19 Genomics UK (COG-UK) Consortium | Jacquelyn Wynn, Mairead Hyland, The Lighthouse Lab in Alderley Park and Alex Alderton, Roberto Amato, Jeffrey Barrett, Sonia Goncalves, Ewan Harrison, David K. Jackson, Ian Johnston, Dominic Kwiatkowski, Cordelia Langford, John Sillitoe on behalf of the Wellcome Sanger Institute COVID-19 Surveillance Team |
| EPI_ISL_3377631 | Lighthouse Lab in Glasgow | Wellcome Sanger Institute for the COVID-19 Genomics UK (COG-UK) Consortium | Harper VanSteenhouse, Yumi Kasai, David Gray, Carol Clugston, Anna Dominiczak and Alex Alderton, Roberto Amato, Jeffrey Barrett, Sonia Goncalves, Ewan Harrison, David K. Jackson, Ian Johnston, Dominic Kwiatkowski, Cordelia Langford, John Sillitoe on behalf of the Wellcome Sanger Institute COVID-19 Surveillance Team |
| EPI_ISL_3377632, EPI_ISL_3377633, EPI_ISL_3377634 | Lighthouse Lab in Alderley Park | Wellcome Sanger Institute for the COVID-19 Genomics UK (COG-UK) Consortium | Jacquelyn Wynn, Mairead Hyland, The Lighthouse Lab in Alderley Park and Alex Alderton, Roberto Amato, Jeffrey Barrett, Sonia Goncalves, Ewan Harrison, David K. Jackson, Ian Johnston, Dominic Kwiatkowski, Cordelia Langford, John Sillitoe on behalf of the Wellcome Sanger Institute COVID-19 Surveillance Team |
| EPI_ISL_3377635 | Lighthouse Laboratory Plymouth | Wellcome Sanger Institute for the COVID-19 Genomics UK (COG-UK) Consortium | Lighthouse Laboratory Plymouth and Alex Alderton, Roberto Amato, Jeffrey Barrett, Sonia Goncalves, Ewan Harrison, David K. Jackson, Ian Johnston, Dominic Kwiatkowski, Cordelia Langford, John Sillitoe on behalf of the Wellcome Sanger Institute COVID-19 Surveillance Team |
| EPI_ISL_3377636, EPI_ISL_3377637, EPI_ISL_3377638, EPI_ISL_3377639, EPI_ISL_3377640 | Lighthouse Lab in Alderley Park | Wellcome Sanger Institute for the COVID-19 Genomics UK (COG-UK) Consortium | Jacquelyn Wynn, Mairead Hyland, The Lighthouse Lab in Alderley Park and Alex Alderton, Roberto Amato, Jeffrey Barrett, Sonia Goncalves, Ewan Harrison, David K. Jackson, Ian Johnston, Dominic Kwiatkowski, Cordelia Langford, John Sillitoe on behalf of the Wellcome Sanger Institute COVID-19 Surveillance Team |
| EPI_ISL_3377641 | Lighthouse Lab in Glasgow | Wellcome Sanger Institute for the COVID-19 Genomics UK (COG-UK) Consortium | Harper VanSteenhouse, Yumi Kasai, David Gray, Carol Clugston, Anna Dominiczak and Alex Alderton, Roberto Amato, Jeffrey Barrett, Sonia Goncalves, Ewan Harrison, David K. Jackson, Ian Johnston, Dominic Kwiatkowski, Cordelia Langford, John Sillitoe on behalf of the Wellcome Sanger Institute COVID-19 Surveillance Team |
| EPI_ISL_3377642 | Lighthouse Lab in Alderley Park | Wellcome Sanger Institute for the COVID-19 Genomics UK (COG-UK) Consortium | Jacquelyn Wynn, Mairead Hyland, The Lighthouse Lab in Alderley Park and Alex Alderton, Roberto Amato, Jeffrey Barrett, Sonia Goncalves, Ewan Harrison, David K. Jackson, Ian Johnston, Dominic Kwiatkowski, Cordelia Langford, John Sillitoe on behalf of the Wellcome Sanger Institute COVID-19 Surveillance Team |
| EPI_ISL_3377643 | Lighthouse Laboratory Plymouth | Wellcome Sanger Institute for the COVID-19 Genomics UK (COG-UK) Consortium | Lighthouse Laboratory Plymouth and Alex Alderton, Roberto Amato, Jeffrey Barrett, Sonia Goncalves, Ewan Harrison, David K. Jackson, Ian Johnston, Dominic Kwiatkowski, Cordelia Langford, John Sillitoe on behalf of the Wellcome Sanger Institute COVID-19 Surveillance Team |
| EPI_ISL_3377644 | Lighthouse Lab in Alderley Park | Wellcome Sanger Institute for the COVID-19 Genomics UK (COG-UK) | Jacquelyn Wynn, Mairead Hyland, The Lighthouse Lab in Alderley Park and Alex Alderton, Roberto Amato, Jeffrey Barrett, Sonia Goncalves, Ewan Harrison, David K. Jackson, Ian Johnston, Dominic Kwiatkowski, Cordelia Langford, John Sillitoe on |

| | | | |
|---|---|---|---|
| | | Consortium | behalf of the Wellcome Sanger Institute COVID-19 Surveillance Team |
| EPI_ISL_3377645 | Lighthouse Laboratory Plymouth | Wellcome Sanger Institute for the COVID-19 Genomics UK (COG-UK) Consortium | Lighthouse Laboratory Plymouth and Alex Alderton, Roberto Amato, Jeffrey Barrett, Sonia Goncalves, Ewan Harrison, David K. Jackson, Ian Johnston, Dominic Kwiatkowski, Cordelia Langford, John Sillitoe on behalf of the Wellcome Sanger Institute COVID-19 Surveillance Team |
| EPI_ISL_3377646, EPI_ISL_3377647 | Lighthouse Lab in Alderley Park | Wellcome Sanger Institute for the COVID-19 Genomics UK (COG-UK) Consortium | Jacquelyn Wynn, Mairead Hyland, The Lighthouse Lab in Alderley Park and Alex Alderton, Roberto Amato, Jeffrey Barrett, Sonia Goncalves, Ewan Harrison, David K. Jackson, Ian Johnston, Dominic Kwiatkowski, Cordelia Langford, John Sillitoe on behalf of the Wellcome Sanger Institute COVID-19 Surveillance Team |
| EPI_ISL_3377648 | Lighthouse Laboratory Plymouth | Wellcome Sanger Institute for the COVID-19 Genomics UK (COG-UK) Consortium | Lighthouse Laboratory Plymouth and Alex Alderton, Roberto Amato, Jeffrey Barrett, Sonia Goncalves, Ewan Harrison, David K. Jackson, Ian Johnston, Dominic Kwiatkowski, Cordelia Langford, John Sillitoe on behalf of the Wellcome Sanger Institute COVID-19 Surveillance Team |
| EPI_ISL_3377649 | Lighthouse Lab in Glasgow | Wellcome Sanger Institute for the COVID-19 Genomics UK (COG-UK) Consortium | Harper VanSteenhouse, Yumi Kasai, David Gray, Carol Clugston, Anna Dominiczak and Alex Alderton, Roberto Amato, Jeffrey Barrett, Sonia Goncalves, Ewan Harrison, David K. Jackson, Ian Johnston, Dominic Kwiatkowski, Cordelia Langford, John Sillitoe on behalf of the Wellcome Sanger Institute COVID-19 Surveillance Team |
| EPI_ISL_3377650, EPI_ISL_3377651 | Lighthouse Laboratory Plymouth | Wellcome Sanger Institute for the COVID-19 Genomics UK (COG-UK) Consortium | Lighthouse Laboratory Plymouth and Alex Alderton, Roberto Amato, Jeffrey Barrett, Sonia Goncalves, Ewan Harrison, David K. Jackson, Ian Johnston, Dominic Kwiatkowski, Cordelia Langford, John Sillitoe on behalf of the Wellcome Sanger Institute COVID-19 Surveillance Team |
| EPI_ISL_3377652 | Lighthouse Lab in Glasgow | Wellcome Sanger Institute for the COVID-19 Genomics UK (COG-UK) Consortium | Harper VanSteenhouse, Yumi Kasai, David Gray, Carol Clugston, Anna Dominiczak and Alex Alderton, Roberto Amato, Jeffrey Barrett, Sonia Goncalves, Ewan Harrison, David K. Jackson, Ian Johnston, Dominic Kwiatkowski, Cordelia Langford, John Sillitoe on behalf of the Wellcome Sanger Institute COVID-19 Surveillance Team |
| EPI_ISL_3377653, EPI_ISL_3377654, EPI_ISL_3377655, EPI_ISL_3377656 | Lighthouse Laboratory Plymouth | Wellcome Sanger Institute for the COVID-19 Genomics UK (COG-UK) Consortium | Lighthouse Laboratory Plymouth and Alex Alderton, Roberto Amato, Jeffrey Barrett, Sonia Goncalves, Ewan Harrison, David K. Jackson, Ian Johnston, Dominic Kwiatkowski, Cordelia Langford, John Sillitoe on behalf of the Wellcome Sanger Institute COVID-19 Surveillance Team |
| EPI_ISL_3377657 | Lighthouse Lab in Glasgow | Wellcome Sanger Institute for the COVID-19 Genomics UK (COG-UK) Consortium | Harper VanSteenhouse, Yumi Kasai, David Gray, Carol Clugston, Anna Dominiczak and Alex Alderton, Roberto Amato, Jeffrey Barrett, Sonia Goncalves, Ewan Harrison, David K. Jackson, Ian Johnston, Dominic Kwiatkowski, Cordelia Langford, John Sillitoe on behalf of the Wellcome Sanger Institute COVID-19 Surveillance Team |
| EPI_ISL_3377658 | Lighthouse Laboratory Plymouth | Wellcome Sanger Institute for the COVID-19 Genomics UK (COG-UK) Consortium | Lighthouse Laboratory Plymouth and Alex Alderton, Roberto Amato, Jeffrey Barrett, Sonia Goncalves, Ewan Harrison, David K. Jackson, Ian Johnston, Dominic Kwiatkowski, Cordelia Langford, John Sillitoe on behalf of the Wellcome Sanger Institute COVID-19 Surveillance Team |
| EPI_ISL_3377659 | Lighthouse Lab in Alderley Park | Wellcome Sanger Institute for the COVID-19 Genomics UK (COG-UK) Consortium | Jacquelyn Wynn, Mairead Hyland, The Lighthouse Lab in Alderley Park and Alex Alderton, Roberto Amato, Jeffrey Barrett, Sonia Goncalves, Ewan Harrison, David K. Jackson, Ian Johnston, Dominic Kwiatkowski, Cordelia Langford, John Sillitoe on behalf of the Wellcome Sanger Institute COVID-19 Surveillance Team |
| EPI_ISL_3377660 | Lighthouse Laboratory Plymouth | Wellcome Sanger Institute for the COVID-19 Genomics UK (COG-UK) Consortium | Lighthouse Laboratory Plymouth and Alex Alderton, Roberto Amato, Jeffrey Barrett, Sonia Goncalves, Ewan Harrison, David K. Jackson, Ian Johnston, Dominic Kwiatkowski, Cordelia Langford, John Sillitoe on behalf of the Wellcome Sanger Institute COVID-19 Surveillance Team |
| EPI_ISL_3377661 | Lighthouse Lab in Alderley Park | Wellcome Sanger Institute for the COVID-19 Genomics UK (COG-UK) Consortium | Jacquelyn Wynn, Mairead Hyland, The Lighthouse Lab in Alderley Park and Alex Alderton, Roberto Amato, Jeffrey Barrett, Sonia Goncalves, Ewan Harrison, David K. Jackson, Ian Johnston, Dominic Kwiatkowski, Cordelia Langford, John Sillitoe on behalf of the Wellcome Sanger Institute COVID-19 Surveillance Team |
| EPI_ISL_3377662 | Lighthouse Lab in Glasgow | Wellcome Sanger Institute for the COVID-19 Genomics UK (COG-UK) Consortium | Harper VanSteenhouse, Yumi Kasai, David Gray, Carol Clugston, Anna Dominiczak and Alex Alderton, Roberto Amato, Jeffrey Barrett, Sonia Goncalves, Ewan Harrison, David K. Jackson, Ian Johnston, Dominic Kwiatkowski, Cordelia Langford, John Sillitoe on behalf of the Wellcome Sanger Institute COVID-19 Surveillance Team |
| EPI_ISL_3377663, EPI_ISL_3377664 | Lighthouse Laboratory Plymouth | Wellcome Sanger Institute for the COVID-19 Genomics UK (COG-UK) Consortium | Lighthouse Laboratory Plymouth and Alex Alderton, Roberto Amato, Jeffrey Barrett, Sonia Goncalves, Ewan Harrison, David K. Jackson, Ian Johnston, Dominic Kwiatkowski, Cordelia Langford, John Sillitoe on behalf of the Wellcome Sanger Institute COVID-19 Surveillance Team |
| EPI_ISL_3377665, EPI_ISL_3377666 | Lighthouse Lab in Glasgow | Wellcome Sanger Institute for the COVID-19 Genomics UK (COG-UK) Consortium | Harper VanSteenhouse, Yumi Kasai, David Gray, Carol Clugston, Anna Dominiczak and Alex Alderton, Roberto Amato, Jeffrey Barrett, Sonia Goncalves, Ewan Harrison, David K. Jackson, Ian Johnston, Dominic Kwiatkowski, Cordelia Langford, John Sillitoe on behalf of the Wellcome Sanger Institute COVID-19 Surveillance Team |
| EPI_ISL_3377667 | Lighthouse Lab in Alderley Park | Wellcome Sanger Institute for the COVID-19 Genomics UK (COG-UK) Consortium | Jacquelyn Wynn, Mairead Hyland, The Lighthouse Lab in Alderley Park and Alex Alderton, Roberto Amato, Jeffrey Barrett, Sonia Goncalves, Ewan Harrison, David K. Jackson, Ian Johnston, Dominic Kwiatkowski, Cordelia Langford, John Sillitoe on behalf of the Wellcome Sanger Institute COVID-19 Surveillance Team |
| EPI_ISL_3377668 | Lighthouse Lab in Glasgow | Wellcome Sanger Institute for the COVID-19 Genomics UK (COG-UK) Consortium | Harper VanSteenhouse, Yumi Kasai, David Gray, Carol Clugston, Anna Dominiczak and Alex Alderton, Roberto Amato, Jeffrey Barrett, Sonia Goncalves, Ewan Harrison, David K. Jackson, Ian Johnston, Dominic Kwiatkowski, Cordelia Langford, John Sillitoe on behalf of the Wellcome Sanger Institute COVID-19 Surveillance Team |
| EPI_ISL_3377669, EPI_ISL_3377670 | Lighthouse Lab in Alderley Park | Wellcome Sanger Institute for the COVID-19 Genomics UK (COG-UK) Consortium | Jacquelyn Wynn, Mairead Hyland, The Lighthouse Lab in Alderley Park and Alex Alderton, Roberto Amato, Jeffrey Barrett, Sonia Goncalves, Ewan Harrison, David K. Jackson, Ian Johnston, Dominic Kwiatkowski, Cordelia Langford, John Sillitoe on behalf of the Wellcome Sanger Institute COVID-19 Surveillance Team |
| EPI_ISL_3377671 | Lighthouse Lab in Glasgow | Wellcome Sanger Institute for the COVID-19 Genomics UK (COG-UK) Consortium | Harper VanSteenhouse, Yumi Kasai, David Gray, Carol Clugston, Anna Dominiczak and Alex Alderton, Roberto Amato, Jeffrey Barrett, Sonia Goncalves, Ewan Harrison, David K. Jackson, Ian Johnston, Dominic Kwiatkowski, Cordelia Langford, John Sillitoe on behalf of the Wellcome Sanger Institute COVID-19 Surveillance Team |
| EPI_ISL_3377672 | Lighthouse Laboratory Plymouth | Wellcome Sanger Institute for the COVID-19 Genomics UK (COG-UK) Consortium | Lighthouse Laboratory Plymouth and Alex Alderton, Roberto Amato, Jeffrey Barrett, Sonia Goncalves, Ewan Harrison, David K. Jackson, Ian Johnston, Dominic Kwiatkowski, Cordelia Langford, John Sillitoe on behalf of the Wellcome Sanger Institute COVID-19 Surveillance Team |
| EPI_ISL_3377673, EPI_ISL_3377674 | Lighthouse Lab in Glasgow | Wellcome Sanger Institute for the COVID-19 Genomics UK (COG-UK) Consortium | Harper VanSteenhouse, Yumi Kasai, David Gray, Carol Clugston, Anna Dominiczak and Alex Alderton, Roberto Amato, Jeffrey Barrett, Sonia Goncalves, Ewan Harrison, David K. Jackson, Ian Johnston, Dominic Kwiatkowski, Cordelia Langford, John Sillitoe on behalf of the Wellcome Sanger Institute COVID-19 Surveillance Team |
| EPI_ISL_3377675 | Lighthouse Laboratory Plymouth | Wellcome Sanger Institute for the COVID-19 Genomics UK (COG-UK) Consortium | Lighthouse Laboratory Plymouth and Alex Alderton, Roberto Amato, Jeffrey Barrett, Sonia Goncalves, Ewan Harrison, David K. Jackson, Ian Johnston, Dominic Kwiatkowski, Cordelia Langford, John Sillitoe on behalf of the Wellcome Sanger Institute COVID-19 Surveillance Team |
| EPI_ISL_3377676 | Lighthouse Lab in Alderley Park | Wellcome Sanger Institute for the COVID-19 Genomics UK (COG-UK) Consortium | Jacquelyn Wynn, Mairead Hyland, The Lighthouse Lab in Alderley Park and Alex Alderton, Roberto Amato, Jeffrey Barrett, Sonia Goncalves, Ewan Harrison, David K. Jackson, Ian Johnston, Dominic Kwiatkowski, Cordelia Langford, John Sillitoe on behalf of the Wellcome Sanger Institute COVID-19 Surveillance Team |
| EPI_ISL_3377677, EPI_ISL_3377678 | Lighthouse Lab in Glasgow | Wellcome Sanger Institute for the COVID-19 Genomics UK (COG-UK) Consortium | Harper VanSteenhouse, Yumi Kasai, David Gray, Carol Clugston, Anna Dominiczak and Alex Alderton, Roberto Amato, Jeffrey Barrett, Sonia Goncalves, Ewan Harrison, David K. Jackson, Ian Johnston, Dominic Kwiatkowski, Cordelia Langford, John Sillitoe on behalf of the Wellcome Sanger Institute COVID-19 Surveillance Team |
| EPI_ISL_3377679, EPI_ISL_3377680 | Lighthouse Lab in Alderley Park | Wellcome Sanger Institute for the COVID-19 Genomics UK (COG-UK) Consortium | Jacquelyn Wynn, Mairead Hyland, The Lighthouse Lab in Alderley Park and Alex Alderton, Roberto Amato, Jeffrey Barrett, Sonia Goncalves, Ewan Harrison, David K. Jackson, Ian Johnston, Dominic Kwiatkowski, Cordelia Langford, John Sillitoe on behalf of the Wellcome Sanger Institute COVID-19 Surveillance Team |
| EPI_ISL_3377681, EPI_ISL_3377682 | Lighthouse Laboratory Plymouth | Wellcome Sanger Institute for the COVID-19 Genomics UK (COG-UK) Consortium | Lighthouse Laboratory Plymouth and Alex Alderton, Roberto Amato, Jeffrey Barrett, Sonia Goncalves, Ewan Harrison, David K. Jackson, Ian Johnston, Dominic Kwiatkowski, Cordelia Langford, John Sillitoe on behalf of the Wellcome Sanger Institute COVID-19 Surveillance Team |
| EPI_ISL_3377683 | Lighthouse Lab in Alderley Park | Wellcome Sanger Institute for the COVID-19 Genomics UK (COG-UK) Consortium | Jacquelyn Wynn, Mairead Hyland, The Lighthouse Lab in Alderley Park and Alex Alderton, Roberto Amato, Jeffrey Barrett, Sonia Goncalves, Ewan Harrison, David K. Jackson, Ian Johnston, Dominic Kwiatkowski, Cordelia Langford, John Sillitoe on behalf of the Wellcome Sanger Institute COVID-19 Surveillance Team |
| EPI_ISL_3377684 | Lighthouse Lab in Glasgow | Wellcome Sanger Institute for the COVID-19 Genomics UK (COG-UK) Consortium | Harper VanSteenhouse, Yumi Kasai, David Gray, Carol Clugston, Anna Dominiczak and Alex Alderton, Roberto Amato, Jeffrey Barrett, Sonia Goncalves, Ewan Harrison, David K. Jackson, Ian Johnston, Dominic Kwiatkowski, Cordelia Langford, John Sillitoe on behalf of the Wellcome Sanger Institute COVID-19 Surveillance Team |
| EPI_ISL_3377685, EPI_ISL_3377686 | Lighthouse Lab in Alderley Park | Wellcome Sanger Institute for the COVID-19 Genomics UK (COG-UK) Consortium | Jacquelyn Wynn, Mairead Hyland, The Lighthouse Lab in Alderley Park and Alex Alderton, Roberto Amato, Jeffrey Barrett, Sonia Goncalves, Ewan Harrison, David K. Jackson, Ian Johnston, Dominic Kwiatkowski, Cordelia Langford, John Sillitoe on behalf of the Wellcome Sanger Institute COVID-19 Surveillance Team |
| EPI_ISL_3377687, EPI_ISL_3377688 | Lighthouse Laboratory Plymouth | Wellcome Sanger Institute for the COVID-19 Genomics UK (COG-UK) Consortium | Lighthouse Laboratory Plymouth and Alex Alderton, Roberto Amato, Jeffrey Barrett, Sonia Goncalves, Ewan Harrison, David K. Jackson, Ian Johnston, Dominic Kwiatkowski, Cordelia Langford, John Sillitoe on behalf of the Wellcome Sanger Institute COVID-19 Surveillance Team |
| EPI_ISL_3377689, EPI_ISL_3377690 | Lighthouse Lab in Alderley Park | Wellcome Sanger Institute for the COVID-19 Genomics UK (COG-UK) Consortium | Jacquelyn Wynn, Mairead Hyland, The Lighthouse Lab in Alderley Park and Alex Alderton, Roberto Amato, Jeffrey Barrett, Sonia Goncalves, Ewan Harrison, David K. Jackson, Ian Johnston, Dominic Kwiatkowski, Cordelia Langford, John Sillitoe on behalf of the Wellcome Sanger Institute COVID-19 Surveillance Team |
| EPI_ISL_3377691, EPI_ISL_3377692 | Lighthouse Lab in Glasgow | Wellcome Sanger Institute for the COVID-19 Genomics UK (COG-UK) Consortium | Harper VanSteenhouse, Yumi Kasai, David Gray, Carol Clugston, Anna Dominiczak and Alex Alderton, Roberto Amato, Jeffrey Barrett, Sonia Goncalves, Ewan Harrison, David K. Jackson, Ian Johnston, Dominic Kwiatkowski, Cordelia Langford, John Sillitoe on behalf of the Wellcome Sanger Institute COVID-19 Surveillance Team |
| EPI_ISL_3377693, EPI_ISL_3377694 | Lighthouse Lab in Alderley Park | Wellcome Sanger Institute for the COVID-19 Genomics UK (COG-UK) Consortium | Jacquelyn Wynn, Mairead Hyland, The Lighthouse Lab in Alderley Park and Alex Alderton, Roberto Amato, Jeffrey Barrett, Sonia Goncalves, Ewan Harrison, David K. Jackson, Ian Johnston, Dominic Kwiatkowski, Cordelia Langford, John Sillitoe on behalf of the Wellcome Sanger Institute COVID-19 Surveillance Team |
| EPI_ISL_3377695 | Lighthouse Lab in Glasgow | Wellcome Sanger Institute for the COVID-19 Genomics UK (COG-UK) Consortium | Harper VanSteenhouse, Yumi Kasai, David Gray, Carol Clugston, Anna Dominiczak and Alex Alderton, Roberto Amato, Jeffrey Barrett, Sonia Goncalves, Ewan Harrison, David K. Jackson, Ian Johnston, Dominic Kwiatkowski, Cordelia Langford, John Sillitoe on behalf of the Wellcome Sanger Institute COVID-19 Surveillance Team |
| EPI_ISL_3377696 | Lighthouse Lab in Alderley Park | Wellcome Sanger Institute for the COVID-19 Genomics UK (COG-UK) Consortium | Jacquelyn Wynn, Mairead Hyland, The Lighthouse Lab in Alderley Park and Alex Alderton, Roberto Amato, Jeffrey Barrett, Sonia Goncalves, Ewan Harrison, David K. Jackson, Ian Johnston, Dominic Kwiatkowski, Cordelia Langford, John Sillitoe on behalf of the Wellcome Sanger Institute COVID-19 Surveillance Team |
| EPI_ISL_3377697 | Lighthouse Lab in Glasgow | Wellcome Sanger Institute for the COVID-19 Genomics UK (COG-UK) Consortium | Harper VanSteenhouse, Yumi Kasai, David Gray, Carol Clugston, Anna Dominiczak and Alex Alderton, Roberto Amato, Jeffrey Barrett, Sonia Goncalves, Ewan Harrison, David K. Jackson, Ian Johnston, Dominic Kwiatkowski, Cordelia Langford, John Sillitoe on behalf of the Wellcome Sanger Institute COVID-19 Surveillance Team |
| EPI_ISL_3377698 | Lighthouse Laboratory Plymouth | Wellcome Sanger Institute for the COVID-19 Genomics UK (COG-UK) Consortium | Lighthouse Laboratory Plymouth and Alex Alderton, Roberto Amato, Jeffrey Barrett, Sonia Goncalves, Ewan Harrison, David K. Jackson, Ian Johnston, Dominic Kwiatkowski, Cordelia Langford, John Sillitoe on behalf of the Wellcome Sanger Institute COVID-19 Surveillance Team |
| EPI_ISL_3377699, EPI_ISL_3377700 | Lighthouse Lab in Alderley Park | Wellcome Sanger Institute for the COVID-19 Genomics UK (COG-UK) Consortium | Jacquelyn Wynn, Mairead Hyland, The Lighthouse Lab in Alderley Park and Alex Alderton, Roberto Amato, Jeffrey Barrett, Sonia Goncalves, Ewan Harrison, David K. Jackson, Ian Johnston, Dominic Kwiatkowski, Cordelia Langford, John Sillitoe on behalf of the Wellcome Sanger Institute COVID-19 Surveillance Team |
| EPI_ISL_3377701 | Lighthouse Laboratory Plymouth | Wellcome Sanger Institute for the COVID-19 Genomics UK (COG-UK) Consortium | Lighthouse Laboratory Plymouth and Alex Alderton, Roberto Amato, Jeffrey Barrett, Sonia Goncalves, Ewan Harrison, David K. Jackson, Ian Johnston, Dominic Kwiatkowski, Cordelia Langford, John Sillitoe on behalf of the Wellcome Sanger Institute COVID-19 Surveillance Team |
| EPI_ISL_3377702, EPI_ISL_3377703, EPI_ISL_3377704 | Lighthouse Lab in Glasgow | Wellcome Sanger Institute for the COVID-19 Genomics UK (COG-UK) Consortium | Harper VanSteenhouse, Yumi Kasai, David Gray, Carol Clugston, Anna Dominiczak and Alex Alderton, Roberto Amato, Jeffrey Barrett, Sonia Goncalves, Ewan Harrison, David K. Jackson, Ian Johnston, Dominic Kwiatkowski, Cordelia Langford, John Sillitoe on behalf of the Wellcome Sanger Institute COVID-19 Surveillance Team |
| EPI_ISL_3377705 | Lighthouse Lab in Alderley Park | Wellcome Sanger Institute for the COVID-19 Genomics UK (COG-UK) Consortium | Jacquelyn Wynn, Mairead Hyland, The Lighthouse Lab in Alderley Park and Alex Alderton, Roberto Amato, Jeffrey Barrett, Sonia Goncalves, Ewan Harrison, David K. Jackson, Ian Johnston, Dominic Kwiatkowski, Cordelia Langford, John Sillitoe on behalf of the Wellcome Sanger Institute COVID-19 Surveillance Team |
| EPI_ISL_3377706, EPI_ISL_3377707 | Lighthouse Lab in Glasgow | Wellcome Sanger Institute for the COVID-19 Genomics UK (COG-UK) Consortium | Harper VanSteenhouse, Yumi Kasai, David Gray, Carol Clugston, Anna Dominiczak and Alex Alderton, Roberto Amato, Jeffrey Barrett, Sonia Goncalves, Ewan Harrison, David K. Jackson, Ian Johnston, Dominic Kwiatkowski, Cordelia Langford, John Sillitoe on behalf of the Wellcome Sanger Institute COVID-19 Surveillance Team |
| EPI_ISL_3377708, EPI_ISL_3377709, EPI_ISL_3377710 | Lighthouse Lab in Alderley Park | Wellcome Sanger Institute for the COVID-19 Genomics UK (COG-UK) Consortium | Jacquelyn Wynn, Mairead Hyland, The Lighthouse Lab in Alderley Park and Alex Alderton, Roberto Amato, Jeffrey Barrett, Sonia Goncalves, Ewan Harrison, David K. Jackson, Ian Johnston, Dominic Kwiatkowski, Cordelia Langford, John Sillitoe on behalf of the Wellcome Sanger Institute COVID-19 Surveillance Team |

| | | | |
|---|---|---|---|
| EPI_ISL_3377711 | Lighthouse Laboratory Plymouth | Wellcome Sanger Institute for the COVID-19 Genomics UK (COG-UK) Consortium | Lighthouse Laboratory Plymouth and Alex Alderton, Roberto Amato, Jeffrey Barrett, Sonia Goncalves, Ewan Harrison, David K. Jackson, Ian Johnston, Dominic Kwiatkowski, Cordelia Langford, John Sillitoe on behalf of the Wellcome Sanger Institute COVID-19 Surveillance Team |
| EPI_ISL_3377712 | Lighthouse Lab in Glasgow | Wellcome Sanger Institute for the COVID-19 Genomics UK (COG-UK) Consortium | Harper VanSteenhouse, Yumi Kasai, David Gray, Carol Clugston, Anna Dominiczak and Alex Alderton, Roberto Amato, Jeffrey Barrett, Sonia Goncalves, Ewan Harrison, David K. Jackson, Ian Johnston, Dominic Kwiatkowski, Cordelia Langford, John Sillitoe on behalf of the Wellcome Sanger Institute COVID-19 Surveillance Team |
| EPI_ISL_3377713 | Lighthouse Lab in Alderley Park | Wellcome Sanger Institute for the COVID-19 Genomics UK (COG-UK) Consortium | Jacquelyn Wynn, Mairead Hyland, The Lighthouse Lab in Alderley Park and Alex Alderton, Roberto Amato, Jeffrey Barrett, Sonia Goncalves, Ewan Harrison, David K. Jackson, Ian Johnston, Dominic Kwiatkowski, Cordelia Langford, John Sillitoe on behalf of the Wellcome Sanger Institute COVID-19 Surveillance Team |
| EPI_ISL_3377714 | Lighthouse Lab in Glasgow | Wellcome Sanger Institute for the COVID-19 Genomics UK (COG-UK) Consortium | Harper VanSteenhouse, Yumi Kasai, David Gray, Carol Clugston, Anna Dominiczak and Alex Alderton, Roberto Amato, Jeffrey Barrett, Sonia Goncalves, Ewan Harrison, David K. Jackson, Ian Johnston, Dominic Kwiatkowski, Cordelia Langford, John Sillitoe on behalf of the Wellcome Sanger Institute COVID-19 Surveillance Team |
| EPI_ISL_3377715 | Lighthouse Lab in Alderley Park | Wellcome Sanger Institute for the COVID-19 Genomics UK (COG-UK) Consortium | Jacquelyn Wynn, Mairead Hyland, The Lighthouse Lab in Alderley Park and Alex Alderton, Roberto Amato, Jeffrey Barrett, Sonia Goncalves, Ewan Harrison, David K. Jackson, Ian Johnston, Dominic Kwiatkowski, Cordelia Langford, John Sillitoe on behalf of the Wellcome Sanger Institute COVID-19 Surveillance Team |
| EPI_ISL_3377716 | Lighthouse Lab in Glasgow | Wellcome Sanger Institute for the COVID-19 Genomics UK (COG-UK) Consortium | Harper VanSteenhouse, Yumi Kasai, David Gray, Carol Clugston, Anna Dominiczak and Alex Alderton, Roberto Amato, Jeffrey Barrett, Sonia Goncalves, Ewan Harrison, David K. Jackson, Ian Johnston, Dominic Kwiatkowski, Cordelia Langford, John Sillitoe on behalf of the Wellcome Sanger Institute COVID-19 Surveillance Team |
| EPI_ISL_3377717, EPI_ISL_3377718 | Lighthouse Laboratory Plymouth | Wellcome Sanger Institute for the COVID-19 Genomics UK (COG-UK) Consortium | Lighthouse Laboratory Plymouth and Alex Alderton, Roberto Amato, Jeffrey Barrett, Sonia Goncalves, Ewan Harrison, David K. Jackson, Ian Johnston, Dominic Kwiatkowski, Cordelia Langford, John Sillitoe on behalf of the Wellcome Sanger Institute COVID-19 Surveillance Team |
| EPI_ISL_3377719 | Lighthouse Lab in Alderley Park | Wellcome Sanger Institute for the COVID-19 Genomics UK (COG-UK) Consortium | Jacquelyn Wynn, Mairead Hyland, The Lighthouse Lab in Alderley Park and Alex Alderton, Roberto Amato, Jeffrey Barrett, Sonia Goncalves, Ewan Harrison, David K. Jackson, Ian Johnston, Dominic Kwiatkowski, Cordelia Langford, John Sillitoe on behalf of the Wellcome Sanger Institute COVID-19 Surveillance Team |
| EPI_ISL_3377720 | Lighthouse Lab in Glasgow | Wellcome Sanger Institute for the COVID-19 Genomics UK (COG-UK) Consortium | Harper VanSteenhouse, Yumi Kasai, David Gray, Carol Clugston, Anna Dominiczak and Alex Alderton, Roberto Amato, Jeffrey Barrett, Sonia Goncalves, Ewan Harrison, David K. Jackson, Ian Johnston, Dominic Kwiatkowski, Cordelia Langford, John Sillitoe on behalf of the Wellcome Sanger Institute COVID-19 Surveillance Team |
| EPI_ISL_3377721 | Lighthouse Lab in Alderley Park | Wellcome Sanger Institute for the COVID-19 Genomics UK (COG-UK) Consortium | Jacquelyn Wynn, Mairead Hyland, The Lighthouse Lab in Alderley Park and Alex Alderton, Roberto Amato, Jeffrey Barrett, Sonia Goncalves, Ewan Harrison, David K. Jackson, Ian Johnston, Dominic Kwiatkowski, Cordelia Langford, John Sillitoe on behalf of the Wellcome Sanger Institute COVID-19 Surveillance Team |
| EPI_ISL_3377722 | Lighthouse Laboratory Plymouth | Wellcome Sanger Institute for the COVID-19 Genomics UK (COG-UK) Consortium | Lighthouse Laboratory Plymouth and Alex Alderton, Roberto Amato, Jeffrey Barrett, Sonia Goncalves, Ewan Harrison, David K. Jackson, Ian Johnston, Dominic Kwiatkowski, Cordelia Langford, John Sillitoe on behalf of the Wellcome Sanger Institute COVID-19 Surveillance Team |
| EPI_ISL_3377723, EPI_ISL_3377724 | Lighthouse Lab in Glasgow | Wellcome Sanger Institute for the COVID-19 Genomics UK (COG-UK) Consortium | Harper VanSteenhouse, Yumi Kasai, David Gray, Carol Clugston, Anna Dominiczak and Alex Alderton, Roberto Amato, Jeffrey Barrett, Sonia Goncalves, Ewan Harrison, David K. Jackson, Ian Johnston, Dominic Kwiatkowski, Cordelia Langford, John Sillitoe on behalf of the Wellcome Sanger Institute COVID-19 Surveillance Team |
| EPI_ISL_3377725 | Lighthouse Lab in Alderley Park | Wellcome Sanger Institute for the COVID-19 Genomics UK (COG-UK) Consortium | Jacquelyn Wynn, Mairead Hyland, The Lighthouse Lab in Alderley Park and Alex Alderton, Roberto Amato, Jeffrey Barrett, Sonia Goncalves, Ewan Harrison, David K. Jackson, Ian Johnston, Dominic Kwiatkowski, Cordelia Langford, John Sillitoe on behalf of the Wellcome Sanger Institute COVID-19 Surveillance Team |
| EPI_ISL_3377726, EPI_ISL_3377727, EPI_ISL_3377728 | Lighthouse Laboratory Plymouth | Wellcome Sanger Institute for the COVID-19 Genomics UK (COG-UK) Consortium | Lighthouse Laboratory Plymouth and Alex Alderton, Roberto Amato, Jeffrey Barrett, Sonia Goncalves, Ewan Harrison, David K. Jackson, Ian Johnston, Dominic Kwiatkowski, Cordelia Langford, John Sillitoe on behalf of the Wellcome Sanger Institute COVID-19 Surveillance Team |
| EPI_ISL_3377729 | Lighthouse Lab in Glasgow | Wellcome Sanger Institute for the COVID-19 Genomics UK (COG-UK) Consortium | Harper VanSteenhouse, Yumi Kasai, David Gray, Carol Clugston, Anna Dominiczak and Alex Alderton, Roberto Amato, Jeffrey Barrett, Sonia Goncalves, Ewan Harrison, David K. Jackson, Ian Johnston, Dominic Kwiatkowski, Cordelia Langford, John Sillitoe on behalf of the Wellcome Sanger Institute COVID-19 Surveillance Team |
| EPI_ISL_3377730 | Lighthouse Lab in Alderley Park | Wellcome Sanger Institute for the COVID-19 Genomics UK (COG-UK) Consortium | Jacquelyn Wynn, Mairead Hyland, The Lighthouse Lab in Alderley Park and Alex Alderton, Roberto Amato, Jeffrey Barrett, Sonia Goncalves, Ewan Harrison, David K. Jackson, Ian Johnston, Dominic Kwiatkowski, Cordelia Langford, John Sillitoe on behalf of the Wellcome Sanger Institute COVID-19 Surveillance Team |
| EPI_ISL_3377731, EPI_ISL_3377732 | Lighthouse Laboratory Plymouth | Wellcome Sanger Institute for the COVID-19 Genomics UK (COG-UK) Consortium | Lighthouse Laboratory Plymouth and Alex Alderton, Roberto Amato, Jeffrey Barrett, Sonia Goncalves, Ewan Harrison, David K. Jackson, Ian Johnston, Dominic Kwiatkowski, Cordelia Langford, John Sillitoe on behalf of the Wellcome Sanger Institute COVID-19 Surveillance Team |
| EPI_ISL_3377733, EPI_ISL_3377734, EPI_ISL_3377735 | Lighthouse Lab in Alderley Park | Wellcome Sanger Institute for the COVID-19 Genomics UK (COG-UK) Consortium | Jacquelyn Wynn, Mairead Hyland, The Lighthouse Lab in Alderley Park and Alex Alderton, Roberto Amato, Jeffrey Barrett, Sonia Goncalves, Ewan Harrison, David K. Jackson, Ian Johnston, Dominic Kwiatkowski, Cordelia Langford, John Sillitoe on behalf of the Wellcome Sanger Institute COVID-19 Surveillance Team |
| EPI_ISL_3377736 | Lighthouse Laboratory Plymouth | Wellcome Sanger Institute for the COVID-19 Genomics UK (COG-UK) Consortium | Lighthouse Laboratory Plymouth and Alex Alderton, Roberto Amato, Jeffrey Barrett, Sonia Goncalves, Ewan Harrison, David K. Jackson, Ian Johnston, Dominic Kwiatkowski, Cordelia Langford, John Sillitoe on behalf of the Wellcome Sanger Institute COVID-19 Surveillance Team |
| EPI_ISL_3377737 | Lighthouse Lab in Alderley Park | Wellcome Sanger Institute for the COVID-19 Genomics UK (COG-UK) Consortium | Jacquelyn Wynn, Mairead Hyland, The Lighthouse Lab in Alderley Park and Alex Alderton, Roberto Amato, Jeffrey Barrett, Sonia Goncalves, Ewan Harrison, David K. Jackson, Ian Johnston, Dominic Kwiatkowski, Cordelia Langford, John Sillitoe on behalf of the Wellcome Sanger Institute COVID-19 Surveillance Team |
| EPI_ISL_3377738 | Lighthouse Lab in Glasgow | Wellcome Sanger Institute for the COVID-19 Genomics UK (COG-UK) Consortium | Harper VanSteenhouse, Yumi Kasai, David Gray, Carol Clugston, Anna Dominiczak and Alex Alderton, Roberto Amato, Jeffrey Barrett, Sonia Goncalves, Ewan Harrison, David K. Jackson, Ian Johnston, Dominic Kwiatkowski, Cordelia Langford, John Sillitoe on behalf of the Wellcome Sanger Institute COVID-19 Surveillance Team |
| EPI_ISL_3377739 | Lighthouse Lab in Alderley Park | Wellcome Sanger Institute for the COVID-19 Genomics UK (COG-UK) Consortium | Jacquelyn Wynn, Mairead Hyland, The Lighthouse Lab in Alderley Park and Alex Alderton, Roberto Amato, Jeffrey Barrett, Sonia Goncalves, Ewan Harrison, David K. Jackson, Ian Johnston, Dominic Kwiatkowski, Cordelia Langford, John Sillitoe on behalf of the Wellcome Sanger Institute COVID-19 Surveillance Team |
| EPI_ISL_3377740 | Lighthouse Laboratory Plymouth | Wellcome Sanger Institute for the COVID-19 Genomics UK (COG-UK) Consortium | Lighthouse Laboratory Plymouth and Alex Alderton, Roberto Amato, Jeffrey Barrett, Sonia Goncalves, Ewan Harrison, David K. Jackson, Ian Johnston, Dominic Kwiatkowski, Cordelia Langford, John Sillitoe on behalf of the Wellcome Sanger Institute COVID-19 Surveillance Team |
| EPI_ISL_3377741, EPI_ISL_3377742 | Lighthouse Lab in Alderley Park | Wellcome Sanger Institute for the COVID-19 Genomics UK (COG-UK) Consortium | Jacquelyn Wynn, Mairead Hyland, The Lighthouse Lab in Alderley Park and Alex Alderton, Roberto Amato, Jeffrey Barrett, Sonia Goncalves, Ewan Harrison, David K. Jackson, Ian Johnston, Dominic Kwiatkowski, Cordelia Langford, John Sillitoe on behalf of the Wellcome Sanger Institute COVID-19 Surveillance Team |
| EPI_ISL_3377743 | Lighthouse Laboratory Plymouth | Wellcome Sanger Institute for the COVID-19 Genomics UK (COG-UK) Consortium | Lighthouse Laboratory Plymouth and Alex Alderton, Roberto Amato, Jeffrey Barrett, Sonia Goncalves, Ewan Harrison, David K. Jackson, Ian Johnston, Dominic Kwiatkowski, Cordelia Langford, John Sillitoe on behalf of the Wellcome Sanger Institute COVID-19 Surveillance Team |
| EPI_ISL_3377744 | Lighthouse Lab in Glasgow | Wellcome Sanger Institute for the COVID-19 Genomics UK (COG-UK) Consortium | Harper VanSteenhouse, Yumi Kasai, David Gray, Carol Clugston, Anna Dominiczak and Alex Alderton, Roberto Amato, Jeffrey Barrett, Sonia Goncalves, Ewan Harrison, David K. Jackson, Ian Johnston, Dominic Kwiatkowski, Cordelia Langford, John Sillitoe on behalf of the Wellcome Sanger Institute COVID-19 Surveillance Team |
| EPI_ISL_3377745, EPI_ISL_3377746 | Lighthouse Laboratory Plymouth | Wellcome Sanger Institute for the COVID-19 Genomics UK (COG-UK) Consortium | Lighthouse Laboratory Plymouth and Alex Alderton, Roberto Amato, Jeffrey Barrett, Sonia Goncalves, Ewan Harrison, David K. Jackson, Ian Johnston, Dominic Kwiatkowski, Cordelia Langford, John Sillitoe on behalf of the Wellcome Sanger Institute COVID-19 Surveillance Team |
| EPI_ISL_3377747 | Lighthouse Lab in Alderley Park | Wellcome Sanger Institute for the COVID-19 Genomics UK (COG-UK) Consortium | Jacquelyn Wynn, Mairead Hyland, The Lighthouse Lab in Alderley Park and Alex Alderton, Roberto Amato, Jeffrey Barrett, Sonia Goncalves, Ewan Harrison, David K. Jackson, Ian Johnston, Dominic Kwiatkowski, Cordelia Langford, John Sillitoe on behalf of the Wellcome Sanger Institute COVID-19 Surveillance Team |
| EPI_ISL_3377748 | Lighthouse Lab in Glasgow | Wellcome Sanger Institute for the COVID-19 Genomics UK (COG-UK) Consortium | Harper VanSteenhouse, Yumi Kasai, David Gray, Carol Clugston, Anna Dominiczak and Alex Alderton, Roberto Amato, Jeffrey Barrett, Sonia Goncalves, Ewan Harrison, David K. Jackson, Ian Johnston, Dominic Kwiatkowski, Cordelia Langford, John Sillitoe on behalf of the Wellcome Sanger Institute COVID-19 Surveillance Team |
| EPI_ISL_3377749 | Lighthouse Laboratory Plymouth | Wellcome Sanger Institute for the COVID-19 Genomics UK (COG-UK) Consortium | Lighthouse Laboratory Plymouth and Alex Alderton, Roberto Amato, Jeffrey Barrett, Sonia Goncalves, Ewan Harrison, David K. Jackson, Ian Johnston, Dominic Kwiatkowski, Cordelia Langford, John Sillitoe on behalf of the Wellcome Sanger Institute COVID-19 Surveillance Team |
| EPI_ISL_3377750, EPI_ISL_3377751 | Lighthouse Lab in Glasgow | Wellcome Sanger Institute for the COVID-19 Genomics UK (COG-UK) Consortium | Harper VanSteenhouse, Yumi Kasai, David Gray, Carol Clugston, Anna Dominiczak and Alex Alderton, Roberto Amato, Jeffrey Barrett, Sonia Goncalves, Ewan Harrison, David K. Jackson, Ian Johnston, Dominic Kwiatkowski, Cordelia Langford, John Sillitoe on behalf of the Wellcome Sanger Institute COVID-19 Surveillance Team |
| EPI_ISL_3377752 | Lighthouse Lab in Alderley Park | Wellcome Sanger Institute for the COVID-19 Genomics UK (COG-UK) Consortium | Jacquelyn Wynn, Mairead Hyland, The Lighthouse Lab in Alderley Park and Alex Alderton, Roberto Amato, Jeffrey Barrett, Sonia Goncalves, Ewan Harrison, David K. Jackson, Ian Johnston, Dominic Kwiatkowski, Cordelia Langford, John Sillitoe on behalf of the Wellcome Sanger Institute COVID-19 Surveillance Team |
| EPI_ISL_3377753, EPI_ISL_3377754 | Lighthouse Laboratory Plymouth | Wellcome Sanger Institute for the COVID-19 Genomics UK (COG-UK) Consortium | Lighthouse Laboratory Plymouth and Alex Alderton, Roberto Amato, Jeffrey Barrett, Sonia Goncalves, Ewan Harrison, David K. Jackson, Ian Johnston, Dominic Kwiatkowski, Cordelia Langford, John Sillitoe on behalf of the Wellcome Sanger Institute COVID-19 Surveillance Team |
| EPI_ISL_3377755, EPI_ISL_3377756 | Lighthouse Lab in Alderley Park | Wellcome Sanger Institute for the COVID-19 Genomics UK (COG-UK) Consortium | Jacquelyn Wynn, Mairead Hyland, The Lighthouse Lab in Alderley Park and Alex Alderton, Roberto Amato, Jeffrey Barrett, Sonia Goncalves, Ewan Harrison, David K. Jackson, Ian Johnston, Dominic Kwiatkowski, Cordelia Langford, John Sillitoe on behalf of the Wellcome Sanger Institute COVID-19 Surveillance Team |
| EPI_ISL_3377757 | Lighthouse Laboratory Plymouth | Wellcome Sanger Institute for the COVID-19 Genomics UK (COG-UK) Consortium | Lighthouse Laboratory Plymouth and Alex Alderton, Roberto Amato, Jeffrey Barrett, Sonia Goncalves, Ewan Harrison, David K. Jackson, Ian Johnston, Dominic Kwiatkowski, Cordelia Langford, John Sillitoe on behalf of the Wellcome Sanger Institute COVID-19 Surveillance Team |
| EPI_ISL_3377758, EPI_ISL_3377759, EPI_ISL_3377760 | Lighthouse Lab in Glasgow | Wellcome Sanger Institute for the COVID-19 Genomics UK (COG-UK) Consortium | Harper VanSteenhouse, Yumi Kasai, David Gray, Carol Clugston, Anna Dominiczak and Alex Alderton, Roberto Amato, Jeffrey Barrett, Sonia Goncalves, Ewan Harrison, David K. Jackson, Ian Johnston, Dominic Kwiatkowski, Cordelia Langford, John Sillitoe on behalf of the Wellcome Sanger Institute COVID-19 Surveillance Team |
| EPI_ISL_3377761 | Lighthouse Laboratory Plymouth | Wellcome Sanger Institute for the COVID-19 Genomics UK (COG-UK) Consortium | Lighthouse Laboratory Plymouth and Alex Alderton, Roberto Amato, Jeffrey Barrett, Sonia Goncalves, Ewan Harrison, David K. Jackson, Ian Johnston, Dominic Kwiatkowski, Cordelia Langford, John Sillitoe on behalf of the Wellcome Sanger Institute COVID-19 Surveillance Team |
| EPI_ISL_3377762, EPI_ISL_3377763 | Lighthouse Lab in Glasgow | Wellcome Sanger Institute for the COVID-19 Genomics UK (COG-UK) Consortium | Harper VanSteenhouse, Yumi Kasai, David Gray, Carol Clugston, Anna Dominiczak and Alex Alderton, Roberto Amato, Jeffrey Barrett, Sonia Goncalves, Ewan Harrison, David K. Jackson, Ian Johnston, Dominic Kwiatkowski, Cordelia Langford, John Sillitoe on behalf of the Wellcome Sanger Institute COVID-19 Surveillance Team |
| EPI_ISL_3377764 | Lighthouse Laboratory Plymouth | Wellcome Sanger Institute for the COVID-19 Genomics UK (COG-UK) Consortium | Lighthouse Laboratory Plymouth and Alex Alderton, Roberto Amato, Jeffrey Barrett, Sonia Goncalves, Ewan Harrison, David K. Jackson, Ian Johnston, Dominic Kwiatkowski, Cordelia Langford, John Sillitoe on behalf of the Wellcome Sanger Institute COVID-19 Surveillance Team |
| EPI_ISL_3377765 | Lighthouse Lab in Glasgow | Wellcome Sanger Institute for the COVID-19 Genomics UK (COG-UK) Consortium | Harper VanSteenhouse, Yumi Kasai, David Gray, Carol Clugston, Anna Dominiczak and Alex Alderton, Roberto Amato, Jeffrey Barrett, Sonia Goncalves, Ewan Harrison, David K. Jackson, Ian Johnston, Dominic Kwiatkowski, Cordelia Langford, John Sillitoe on behalf of the Wellcome Sanger Institute COVID-19 Surveillance Team |
| EPI_ISL_3377766 | Lighthouse Laboratory Plymouth | Wellcome Sanger Institute for the COVID-19 Genomics UK (COG-UK) Consortium | Lighthouse Laboratory Plymouth and Alex Alderton, Roberto Amato, Jeffrey Barrett, Sonia Goncalves, Ewan Harrison, David K. Jackson, Ian Johnston, Dominic Kwiatkowski, Cordelia Langford, John Sillitoe on behalf of the Wellcome Sanger Institute COVID-19 Surveillance Team |
| EPI_ISL_3377767, EPI_ISL_3377768 | Lighthouse Lab in Glasgow | Wellcome Sanger Institute for the COVID-19 Genomics UK (COG-UK) Consortium | Harper VanSteenhouse, Yumi Kasai, David Gray, Carol Clugston, Anna Dominiczak and Alex Alderton, Roberto Amato, Jeffrey Barrett, Sonia Goncalves, Ewan Harrison, David K. Jackson, Ian Johnston, Dominic Kwiatkowski, Cordelia Langford, John Sillitoe on behalf of the Wellcome Sanger Institute COVID-19 Surveillance Team |
| EPI_ISL_3377769 | Lighthouse Lab in Alderley Park | Wellcome Sanger Institute for the COVID-19 Genomics UK (COG-UK) Consortium | Jacquelyn Wynn, Mairead Hyland, The Lighthouse Lab in Alderley Park and Alex Alderton, Roberto Amato, Jeffrey Barrett, Sonia Goncalves, Ewan Harrison, David K. Jackson, Ian Johnston, Dominic Kwiatkowski, Cordelia Langford, John Sillitoe on behalf of the Wellcome Sanger Institute COVID-19 Surveillance Team |
| EPI_ISL_3377770 | Lighthouse Lab in Glasgow | Wellcome Sanger Institute for the COVID-19 Genomics UK (COG-UK) Consortium | Harper VanSteenhouse, Yumi Kasai, David Gray, Carol Clugston, Anna Dominiczak and Alex Alderton, Roberto Amato, Jeffrey Barrett, Sonia Goncalves, Ewan Harrison, David K. Jackson, Ian Johnston, Dominic Kwiatkowski, Cordelia Langford, John Sillitoe on behalf of the Wellcome Sanger Institute COVID-19 Surveillance Team |

| | | | |
|---|---|---|---|
| EPI_ISL_3377771 | Lighthouse Lab in Alderley Park | Wellcome Sanger Institute for the COVID-19 Genomics UK (COG-UK) Consortium | Jacquelyn Wynn, Mairead Hyland, The Lighthouse Lab in Alderley Park and Alex Alderton, Roberto Amato, Jeffrey Barrett, Sonia Goncalves, Ewan Harrison, David K. Jackson, Ian Johnston, Dominic Kwiatkowski, Cordelia Langford, John Sillitoe on behalf of the Wellcome Sanger Institute COVID-19 Surveillance Team |
| EPI_ISL_3377772 | Lighthouse Laboratory Plymouth | Wellcome Sanger Institute for the COVID-19 Genomics UK (COG-UK) Consortium | Lighthouse Laboratory Plymouth and Alex Alderton, Roberto Amato, Jeffrey Barrett, Sonia Goncalves, Ewan Harrison, David K. Jackson, Ian Johnston, Dominic Kwiatkowski, Cordelia Langford, John Sillitoe on behalf of the Wellcome Sanger Institute COVID-19 Surveillance Team |
| EPI_ISL_3377773 | Lighthouse Lab in Alderley Park | Wellcome Sanger Institute for the COVID-19 Genomics UK (COG-UK) Consortium | Jacquelyn Wynn, Mairead Hyland, The Lighthouse Lab in Alderley Park and Alex Alderton, Roberto Amato, Jeffrey Barrett, Sonia Goncalves, Ewan Harrison, David K. Jackson, Ian Johnston, Dominic Kwiatkowski, Cordelia Langford, John Sillitoe on behalf of the Wellcome Sanger Institute COVID-19 Surveillance Team |
| EPI_ISL_3377774 | Lighthouse Laboratory Plymouth | Wellcome Sanger Institute for the COVID-19 Genomics UK (COG-UK) Consortium | Lighthouse Laboratory Plymouth and Alex Alderton, Roberto Amato, Jeffrey Barrett, Sonia Goncalves, Ewan Harrison, David K. Jackson, Ian Johnston, Dominic Kwiatkowski, Cordelia Langford, John Sillitoe on behalf of the Wellcome Sanger Institute COVID-19 Surveillance Team |
| EPI_ISL_3377775 | Lighthouse Lab in Glasgow | Wellcome Sanger Institute for the COVID-19 Genomics UK (COG-UK) Consortium | Harper VanSteenhouse, Yumi Kasai, David Gray, Carol Clugston, Anna Dominiczak and Alex Alderton, Roberto Amato, Jeffrey Barrett, Sonia Goncalves, Ewan Harrison, David K. Jackson, Ian Johnston, Dominic Kwiatkowski, Cordelia Langford, John Sillitoe on behalf of the Wellcome Sanger Institute COVID-19 Surveillance Team |
| EPI_ISL_3377776 | Lighthouse Lab in Alderley Park | Wellcome Sanger Institute for the COVID-19 Genomics UK (COG-UK) Consortium | Jacquelyn Wynn, Mairead Hyland, The Lighthouse Lab in Alderley Park and Alex Alderton, Roberto Amato, Jeffrey Barrett, Sonia Goncalves, Ewan Harrison, David K. Jackson, Ian Johnston, Dominic Kwiatkowski, Cordelia Langford, John Sillitoe on behalf of the Wellcome Sanger Institute COVID-19 Surveillance Team |
| EPI_ISL_3377777 | Lighthouse Lab in Glasgow | Wellcome Sanger Institute for the COVID-19 Genomics UK (COG-UK) Consortium | Harper VanSteenhouse, Yumi Kasai, David Gray, Carol Clugston, Anna Dominiczak and Alex Alderton, Roberto Amato, Jeffrey Barrett, Sonia Goncalves, Ewan Harrison, David K. Jackson, Ian Johnston, Dominic Kwiatkowski, Cordelia Langford, John Sillitoe on behalf of the Wellcome Sanger Institute COVID-19 Surveillance Team |
| EPI_ISL_3377778 | Lighthouse Laboratory Plymouth | Wellcome Sanger Institute for the COVID-19 Genomics UK (COG-UK) Consortium | Lighthouse Laboratory Plymouth and Alex Alderton, Roberto Amato, Jeffrey Barrett, Sonia Goncalves, Ewan Harrison, David K. Jackson, Ian Johnston, Dominic Kwiatkowski, Cordelia Langford, John Sillitoe on behalf of the Wellcome Sanger Institute COVID-19 Surveillance Team |
| EPI_ISL_3377779, EPI_ISL_3377780 | Lighthouse Lab in Glasgow | Wellcome Sanger Institute for the COVID-19 Genomics UK (COG-UK) Consortium | Harper VanSteenhouse, Yumi Kasai, David Gray, Carol Clugston, Anna Dominiczak and Alex Alderton, Roberto Amato, Jeffrey Barrett, Sonia Goncalves, Ewan Harrison, David K. Jackson, Ian Johnston, Dominic Kwiatkowski, Cordelia Langford, John Sillitoe on behalf of the Wellcome Sanger Institute COVID-19 Surveillance Team |
| EPI_ISL_3377781, EPI_ISL_3377782 | Lighthouse Laboratory Plymouth | Wellcome Sanger Institute for the COVID-19 Genomics UK (COG-UK) Consortium | Lighthouse Laboratory Plymouth and Alex Alderton, Roberto Amato, Jeffrey Barrett, Sonia Goncalves, Ewan Harrison, David K. Jackson, Ian Johnston, Dominic Kwiatkowski, Cordelia Langford, John Sillitoe on behalf of the Wellcome Sanger Institute COVID-19 Surveillance Team |
| EPI_ISL_3377783 | Lighthouse Lab in Glasgow | Wellcome Sanger Institute for the COVID-19 Genomics UK (COG-UK) Consortium | Harper VanSteenhouse, Yumi Kasai, David Gray, Carol Clugston, Anna Dominiczak and Alex Alderton, Roberto Amato, Jeffrey Barrett, Sonia Goncalves, Ewan Harrison, David K. Jackson, Ian Johnston, Dominic Kwiatkowski, Cordelia Langford, John Sillitoe on behalf of the Wellcome Sanger Institute COVID-19 Surveillance Team |
| EPI_ISL_3377784 | Lighthouse Laboratory Plymouth | Wellcome Sanger Institute for the COVID-19 Genomics UK (COG-UK) Consortium | Lighthouse Laboratory Plymouth and Alex Alderton, Roberto Amato, Jeffrey Barrett, Sonia Goncalves, Ewan Harrison, David K. Jackson, Ian Johnston, Dominic Kwiatkowski, Cordelia Langford, John Sillitoe on behalf of the Wellcome Sanger Institute COVID-19 Surveillance Team |
| EPI_ISL_3377785, EPI_ISL_3377786, EPI_ISL_3377787 | Lighthouse Lab in Glasgow | Wellcome Sanger Institute for the COVID-19 Genomics UK (COG-UK) Consortium | Harper VanSteenhouse, Yumi Kasai, David Gray, Carol Clugston, Anna Dominiczak and Alex Alderton, Roberto Amato, Jeffrey Barrett, Sonia Goncalves, Ewan Harrison, David K. Jackson, Ian Johnston, Dominic Kwiatkowski, Cordelia Langford, John Sillitoe on behalf of the Wellcome Sanger Institute COVID-19 Surveillance Team |
| EPI_ISL_3377788, EPI_ISL_3377789 | Lighthouse Laboratory Plymouth | Wellcome Sanger Institute for the COVID-19 Genomics UK (COG-UK) Consortium | Lighthouse Laboratory Plymouth and Alex Alderton, Roberto Amato, Jeffrey Barrett, Sonia Goncalves, Ewan Harrison, David K. Jackson, Ian Johnston, Dominic Kwiatkowski, Cordelia Langford, John Sillitoe on behalf of the Wellcome Sanger Institute COVID-19 Surveillance Team |
| EPI_ISL_3377790, EPI_ISL_3377791, EPI_ISL_3377792, EPI_ISL_3377793 | Lighthouse Lab in Alderley Park | Wellcome Sanger Institute for the COVID-19 Genomics UK (COG-UK) Consortium | Jacquelyn Wynn, Mairead Hyland, The Lighthouse Lab in Alderley Park and Alex Alderton, Roberto Amato, Jeffrey Barrett, Sonia Goncalves, Ewan Harrison, David K. Jackson, Ian Johnston, Dominic Kwiatkowski, Cordelia Langford, John Sillitoe on behalf of the Wellcome Sanger Institute COVID-19 Surveillance Team |
| EPI_ISL_3377794 | Lighthouse Laboratory Plymouth | Wellcome Sanger Institute for the COVID-19 Genomics UK (COG-UK) Consortium | Lighthouse Laboratory Plymouth and Alex Alderton, Roberto Amato, Jeffrey Barrett, Sonia Goncalves, Ewan Harrison, David K. Jackson, Ian Johnston, Dominic Kwiatkowski, Cordelia Langford, John Sillitoe on behalf of the Wellcome Sanger Institute COVID-19 Surveillance Team |
| EPI_ISL_3377795 | Lighthouse Lab in Glasgow | Wellcome Sanger Institute for the COVID-19 Genomics UK (COG-UK) Consortium | Harper VanSteenhouse, Yumi Kasai, David Gray, Carol Clugston, Anna Dominiczak and Alex Alderton, Roberto Amato, Jeffrey Barrett, Sonia Goncalves, Ewan Harrison, David K. Jackson, Ian Johnston, Dominic Kwiatkowski, Cordelia Langford, John Sillitoe on behalf of the Wellcome Sanger Institute COVID-19 Surveillance Team |
| EPI_ISL_3377796 | Lighthouse Laboratory Plymouth | Wellcome Sanger Institute for the COVID-19 Genomics UK (COG-UK) Consortium | Lighthouse Laboratory Plymouth and Alex Alderton, Roberto Amato, Jeffrey Barrett, Sonia Goncalves, Ewan Harrison, David K. Jackson, Ian Johnston, Dominic Kwiatkowski, Cordelia Langford, John Sillitoe on behalf of the Wellcome Sanger Institute COVID-19 Surveillance Team |
| EPI_ISL_3377797, EPI_ISL_3377798 | Lighthouse Lab in Glasgow | Wellcome Sanger Institute for the COVID-19 Genomics UK (COG-UK) Consortium | Harper VanSteenhouse, Yumi Kasai, David Gray, Carol Clugston, Anna Dominiczak and Alex Alderton, Roberto Amato, Jeffrey Barrett, Sonia Goncalves, Ewan Harrison, David K. Jackson, Ian Johnston, Dominic Kwiatkowski, Cordelia Langford, John Sillitoe on behalf of the Wellcome Sanger Institute COVID-19 Surveillance Team |
| EPI_ISL_3377799 | Lighthouse Laboratory Plymouth | Wellcome Sanger Institute for the COVID-19 Genomics UK (COG-UK) Consortium | Lighthouse Laboratory Plymouth and Alex Alderton, Roberto Amato, Jeffrey Barrett, Sonia Goncalves, Ewan Harrison, David K. Jackson, Ian Johnston, Dominic Kwiatkowski, Cordelia Langford, John Sillitoe on behalf of the Wellcome Sanger Institute COVID-19 Surveillance Team |
| EPI_ISL_3377800, EPI_ISL_3377801 | Lighthouse Lab in Glasgow | Wellcome Sanger Institute for the COVID-19 Genomics UK (COG-UK) Consortium | Harper VanSteenhouse, Yumi Kasai, David Gray, Carol Clugston, Anna Dominiczak and Alex Alderton, Roberto Amato, Jeffrey Barrett, Sonia Goncalves, Ewan Harrison, David K. Jackson, Ian Johnston, Dominic Kwiatkowski, Cordelia Langford, John Sillitoe on behalf of the Wellcome Sanger Institute COVID-19 Surveillance Team |
| EPI_ISL_3377802, EPI_ISL_3377803 | Lighthouse Lab in Alderley Park | Wellcome Sanger Institute for the COVID-19 Genomics UK (COG-UK) Consortium | Jacquelyn Wynn, Mairead Hyland, The Lighthouse Lab in Alderley Park and Alex Alderton, Roberto Amato, Jeffrey Barrett, Sonia Goncalves, Ewan Harrison, David K. Jackson, Ian Johnston, Dominic Kwiatkowski, Cordelia Langford, John Sillitoe on behalf of the Wellcome Sanger Institute COVID-19 Surveillance Team |
| EPI_ISL_3377804, EPI_ISL_3377805 | Lighthouse Lab in Glasgow | Wellcome Sanger Institute for the COVID-19 Genomics UK (COG-UK) Consortium | Harper VanSteenhouse, Yumi Kasai, David Gray, Carol Clugston, Anna Dominiczak and Alex Alderton, Roberto Amato, Jeffrey Barrett, Sonia Goncalves, Ewan Harrison, David K. Jackson, Ian Johnston, Dominic Kwiatkowski, Cordelia Langford, John Sillitoe on behalf of the Wellcome Sanger Institute COVID-19 Surveillance Team |
| EPI_ISL_3377806, EPI_ISL_3377807 | Lighthouse Lab in Alderley Park | Wellcome Sanger Institute for the COVID-19 Genomics UK (COG-UK) Consortium | Jacquelyn Wynn, Mairead Hyland, The Lighthouse Lab in Alderley Park and Alex Alderton, Roberto Amato, Jeffrey Barrett, Sonia Goncalves, Ewan Harrison, David K. Jackson, Ian Johnston, Dominic Kwiatkowski, Cordelia Langford, John Sillitoe on behalf of the Wellcome Sanger Institute COVID-19 Surveillance Team |
| EPI_ISL_3377808, EPI_ISL_3377809 | Lighthouse Lab in Glasgow | Wellcome Sanger Institute for the COVID-19 Genomics UK (COG-UK) Consortium | Harper VanSteenhouse, Yumi Kasai, David Gray, Carol Clugston, Anna Dominiczak and Alex Alderton, Roberto Amato, Jeffrey Barrett, Sonia Goncalves, Ewan Harrison, David K. Jackson, Ian Johnston, Dominic Kwiatkowski, Cordelia Langford, John Sillitoe on behalf of the Wellcome Sanger Institute COVID-19 Surveillance Team |
| EPI_ISL_3377810 | Lighthouse Laboratory Plymouth | Wellcome Sanger Institute for the COVID-19 Genomics UK (COG-UK) Consortium | Lighthouse Laboratory Plymouth and Alex Alderton, Roberto Amato, Jeffrey Barrett, Sonia Goncalves, Ewan Harrison, David K. Jackson, Ian Johnston, Dominic Kwiatkowski, Cordelia Langford, John Sillitoe on behalf of the Wellcome Sanger Institute COVID-19 Surveillance Team |
| EPI_ISL_3377811 | Lighthouse Lab in Alderley Park | Wellcome Sanger Institute for the COVID-19 Genomics UK (COG-UK) Consortium | Jacquelyn Wynn, Mairead Hyland, The Lighthouse Lab in Alderley Park and Alex Alderton, Roberto Amato, Jeffrey Barrett, Sonia Goncalves, Ewan Harrison, David K. Jackson, Ian Johnston, Dominic Kwiatkowski, Cordelia Langford, John Sillitoe on behalf of the Wellcome Sanger Institute COVID-19 Surveillance Team |
| EPI_ISL_3377812 | Lighthouse Lab in Glasgow | Wellcome Sanger Institute for the COVID-19 Genomics UK (COG-UK) Consortium | Harper VanSteenhouse, Yumi Kasai, David Gray, Carol Clugston, Anna Dominiczak and Alex Alderton, Roberto Amato, Jeffrey Barrett, Sonia Goncalves, Ewan Harrison, David K. Jackson, Ian Johnston, Dominic Kwiatkowski, Cordelia Langford, John Sillitoe on behalf of the Wellcome Sanger Institute COVID-19 Surveillance Team |
| EPI_ISL_3377813, EPI_ISL_3377814 | Lighthouse Laboratory Plymouth | Wellcome Sanger Institute for the COVID-19 Genomics UK (COG-UK) Consortium | Lighthouse Laboratory Plymouth and Alex Alderton, Roberto Amato, Jeffrey Barrett, Sonia Goncalves, Ewan Harrison, David K. Jackson, Ian Johnston, Dominic Kwiatkowski, Cordelia Langford, John Sillitoe on behalf of the Wellcome Sanger Institute COVID-19 Surveillance Team |
| EPI_ISL_3377815, EPI_ISL_3377816, EPI_ISL_3377817, EPI_ISL_3377818 | Lighthouse Lab in Glasgow | Wellcome Sanger Institute for the COVID-19 Genomics UK (COG-UK) Consortium | Harper VanSteenhouse, Yumi Kasai, David Gray, Carol Clugston, Anna Dominiczak and Alex Alderton, Roberto Amato, Jeffrey Barrett, Sonia Goncalves, Ewan Harrison, David K. Jackson, Ian Johnston, Dominic Kwiatkowski, Cordelia Langford, John Sillitoe on behalf of the Wellcome Sanger Institute COVID-19 Surveillance Team |
| EPI_ISL_3377819 | Lighthouse Lab in Alderley Park | Wellcome Sanger Institute for the COVID-19 Genomics UK (COG-UK) Consortium | Jacquelyn Wynn, Mairead Hyland, The Lighthouse Lab in Alderley Park and Alex Alderton, Roberto Amato, Jeffrey Barrett, Sonia Goncalves, Ewan Harrison, David K. Jackson, Ian Johnston, Dominic Kwiatkowski, Cordelia Langford, John Sillitoe on behalf of the Wellcome Sanger Institute COVID-19 Surveillance Team |
| EPI_ISL_3377820, EPI_ISL_3377821 | Lighthouse Lab in Glasgow | Wellcome Sanger Institute for the COVID-19 Genomics UK (COG-UK) Consortium | Harper VanSteenhouse, Yumi Kasai, David Gray, Carol Clugston, Anna Dominiczak and Alex Alderton, Roberto Amato, Jeffrey Barrett, Sonia Goncalves, Ewan Harrison, David K. Jackson, Ian Johnston, Dominic Kwiatkowski, Cordelia Langford, John Sillitoe on behalf of the Wellcome Sanger Institute COVID-19 Surveillance Team |
| EPI_ISL_3377822, EPI_ISL_3377823 | Lighthouse Lab in Alderley Park | Wellcome Sanger Institute for the COVID-19 Genomics UK (COG-UK) Consortium | Jacquelyn Wynn, Mairead Hyland, The Lighthouse Lab in Alderley Park and Alex Alderton, Roberto Amato, Jeffrey Barrett, Sonia Goncalves, Ewan Harrison, David K. Jackson, Ian Johnston, Dominic Kwiatkowski, Cordelia Langford, John Sillitoe on behalf of the Wellcome Sanger Institute COVID-19 Surveillance Team |
| EPI_ISL_3377824 | Lighthouse Laboratory Plymouth | Wellcome Sanger Institute for the COVID-19 Genomics UK (COG-UK) Consortium | Lighthouse Laboratory Plymouth and Alex Alderton, Roberto Amato, Jeffrey Barrett, Sonia Goncalves, Ewan Harrison, David K. Jackson, Ian Johnston, Dominic Kwiatkowski, Cordelia Langford, John Sillitoe on behalf of the Wellcome Sanger Institute COVID-19 Surveillance Team |
| EPI_ISL_3377825 | Lighthouse Lab in Glasgow | Wellcome Sanger Institute for the COVID-19 Genomics UK (COG-UK) Consortium | Harper VanSteenhouse, Yumi Kasai, David Gray, Carol Clugston, Anna Dominiczak and Alex Alderton, Roberto Amato, Jeffrey Barrett, Sonia Goncalves, Ewan Harrison, David K. Jackson, Ian Johnston, Dominic Kwiatkowski, Cordelia Langford, John Sillitoe on behalf of the Wellcome Sanger Institute COVID-19 Surveillance Team |
| EPI_ISL_3377826 | Lighthouse Lab in Alderley Park | Wellcome Sanger Institute for the COVID-19 Genomics UK (COG-UK) Consortium | Jacquelyn Wynn, Mairead Hyland, The Lighthouse Lab in Alderley Park and Alex Alderton, Roberto Amato, Jeffrey Barrett, Sonia Goncalves, Ewan Harrison, David K. Jackson, Ian Johnston, Dominic Kwiatkowski, Cordelia Langford, John Sillitoe on behalf of the Wellcome Sanger Institute COVID-19 Surveillance Team |
| EPI_ISL_3377827, EPI_ISL_3377828 | Lighthouse Lab in Glasgow | Wellcome Sanger Institute for the COVID-19 Genomics UK (COG-UK) Consortium | Harper VanSteenhouse, Yumi Kasai, David Gray, Carol Clugston, Anna Dominiczak and Alex Alderton, Roberto Amato, Jeffrey Barrett, Sonia Goncalves, Ewan Harrison, David K. Jackson, Ian Johnston, Dominic Kwiatkowski, Cordelia Langford, John Sillitoe on behalf of the Wellcome Sanger Institute COVID-19 Surveillance Team |
| EPI_ISL_3377829 | Lighthouse Laboratory Plymouth | Wellcome Sanger Institute for the COVID-19 Genomics UK (COG-UK) Consortium | Lighthouse Laboratory Plymouth and Alex Alderton, Roberto Amato, Jeffrey Barrett, Sonia Goncalves, Ewan Harrison, David K. Jackson, Ian Johnston, Dominic Kwiatkowski, Cordelia Langford, John Sillitoe on behalf of the Wellcome Sanger Institute COVID-19 Surveillance Team |
| EPI_ISL_3377830, EPI_ISL_3377831 | Lighthouse Lab in Glasgow | Wellcome Sanger Institute for the COVID-19 Genomics UK (COG-UK) Consortium | Harper VanSteenhouse, Yumi Kasai, David Gray, Carol Clugston, Anna Dominiczak and Alex Alderton, Roberto Amato, Jeffrey Barrett, Sonia Goncalves, Ewan Harrison, David K. Jackson, Ian Johnston, Dominic Kwiatkowski, Cordelia Langford, John Sillitoe on behalf of the Wellcome Sanger Institute COVID-19 Surveillance Team |
| EPI_ISL_3377832 | Lighthouse Laboratory Plymouth | Wellcome Sanger Institute for the COVID-19 Genomics UK (COG-UK) Consortium | Lighthouse Laboratory Plymouth and Alex Alderton, Roberto Amato, Jeffrey Barrett, Sonia Goncalves, Ewan Harrison, David K. Jackson, Ian Johnston, Dominic Kwiatkowski, Cordelia Langford, John Sillitoe on behalf of the Wellcome Sanger Institute COVID-19 Surveillance Team |
| EPI_ISL_3377833 | Lighthouse Lab in Alderley Park | Wellcome Sanger Institute for the COVID-19 Genomics UK (COG-UK) Consortium | Jacquelyn Wynn, Mairead Hyland, The Lighthouse Lab in Alderley Park and Alex Alderton, Roberto Amato, Jeffrey Barrett, Sonia Goncalves, Ewan Harrison, David K. Jackson, Ian Johnston, Dominic Kwiatkowski, Cordelia Langford, John Sillitoe on behalf of the Wellcome Sanger Institute COVID-19 Surveillance Team |
| EPI_ISL_3377834 | Lighthouse Lab in Glasgow | Wellcome Sanger Institute for the COVID-19 Genomics UK (COG-UK) Consortium | Harper VanSteenhouse, Yumi Kasai, David Gray, Carol Clugston, Anna Dominiczak and Alex Alderton, Roberto Amato, Jeffrey Barrett, Sonia Goncalves, Ewan Harrison, David K. Jackson, Ian Johnston, Dominic Kwiatkowski, Cordelia Langford, John Sillitoe on behalf of the Wellcome Sanger Institute COVID-19 Surveillance Team |
| EPI_ISL_3377835 | Lighthouse Laboratory Plymouth | Wellcome Sanger Institute for the COVID-19 Genomics UK (COG-UK) | Lighthouse Laboratory Plymouth and Alex Alderton, Roberto Amato, Jeffrey Barrett, Sonia Goncalves, Ewan Harrison, David K. Jackson, Ian Johnston, Dominic Kwiatkowski, Cordelia Langford, John Sillitoe on behalf of the Wellcome Sanger |

| | | | |
|---|---|---|---|
| | | Consortium | Institute COVID-19 Surveillance Team |
| EPI_ISL_3377836 | Lighthouse Lab in Alderley Park | Wellcome Sanger Institute for the COVID-19 Genomics UK (COG-UK) Consortium | Jacquelyn Wynn, Mairead Hyland, The Lighthouse Lab in Alderley Park and Alex Alderton, Roberto Amato, Jeffrey Barrett, Sonia Goncalves, Ewan Harrison, David K. Jackson, Ian Johnston, Dominic Kwiatkowski, Cordelia Langford, John Sillitoe on behalf of the Wellcome Sanger Institute COVID-19 Surveillance Team |
| EPI_ISL_3377837 | Lighthouse Lab in Glasgow | Wellcome Sanger Institute for the COVID-19 Genomics UK (COG-UK) Consortium | Harper VanSteenhouse, Yumi Kasai, David Gray, Carol Clugston, Anna Dominiczak and Alex Alderton, Roberto Amato, Jeffrey Barrett, Sonia Goncalves, Ewan Harrison, David K. Jackson, Ian Johnston, Dominic Kwiatkowski, Cordelia Langford, John Sillitoe on behalf of the Wellcome Sanger Institute COVID-19 Surveillance Team |
| EPI_ISL_3377838 | Lighthouse Laboratory Plymouth | Wellcome Sanger Institute for the COVID-19 Genomics UK (COG-UK) Consortium | Lighthouse Laboratory Plymouth and Alex Alderton, Roberto Amato, Jeffrey Barrett, Sonia Goncalves, Ewan Harrison, David K. Jackson, Ian Johnston, Dominic Kwiatkowski, Cordelia Langford, John Sillitoe on behalf of the Wellcome Sanger Institute COVID-19 Surveillance Team |
| EPI_ISL_3377839, EPI_ISL_3377840 | Lighthouse Lab in Glasgow | Wellcome Sanger Institute for the COVID-19 Genomics UK (COG-UK) Consortium | Harper VanSteenhouse, Yumi Kasai, David Gray, Carol Clugston, Anna Dominiczak and Alex Alderton, Roberto Amato, Jeffrey Barrett, Sonia Goncalves, Ewan Harrison, David K. Jackson, Ian Johnston, Dominic Kwiatkowski, Cordelia Langford, John Sillitoe on behalf of the Wellcome Sanger Institute COVID-19 Surveillance Team |
| EPI_ISL_3377841 | Lighthouse Lab in Alderley Park | Wellcome Sanger Institute for the COVID-19 Genomics UK (COG-UK) Consortium | Jacquelyn Wynn, Mairead Hyland, The Lighthouse Lab in Alderley Park and Alex Alderton, Roberto Amato, Jeffrey Barrett, Sonia Goncalves, Ewan Harrison, David K. Jackson, Ian Johnston, Dominic Kwiatkowski, Cordelia Langford, John Sillitoe on behalf of the Wellcome Sanger Institute COVID-19 Surveillance Team |
| EPI_ISL_3377842 | Lighthouse Lab in Glasgow | Wellcome Sanger Institute for the COVID-19 Genomics UK (COG-UK) Consortium | Harper VanSteenhouse, Yumi Kasai, David Gray, Carol Clugston, Anna Dominiczak and Alex Alderton, Roberto Amato, Jeffrey Barrett, Sonia Goncalves, Ewan Harrison, David K. Jackson, Ian Johnston, Dominic Kwiatkowski, Cordelia Langford, John Sillitoe on behalf of the Wellcome Sanger Institute COVID-19 Surveillance Team |
| EPI_ISL_3377843 | Lighthouse Lab in Alderley Park | Wellcome Sanger Institute for the COVID-19 Genomics UK (COG-UK) Consortium | Jacquelyn Wynn, Mairead Hyland, The Lighthouse Lab in Alderley Park and Alex Alderton, Roberto Amato, Jeffrey Barrett, Sonia Goncalves, Ewan Harrison, David K. Jackson, Ian Johnston, Dominic Kwiatkowski, Cordelia Langford, John Sillitoe on behalf of the Wellcome Sanger Institute COVID-19 Surveillance Team |
| EPI_ISL_3377844 | Lighthouse Laboratory Plymouth | Wellcome Sanger Institute for the COVID-19 Genomics UK (COG-UK) Consortium | Lighthouse Laboratory Plymouth and Alex Alderton, Roberto Amato, Jeffrey Barrett, Sonia Goncalves, Ewan Harrison, David K. Jackson, Ian Johnston, Dominic Kwiatkowski, Cordelia Langford, John Sillitoe on behalf of the Wellcome Sanger Institute COVID-19 Surveillance Team |
| EPI_ISL_3377845 | Lighthouse Lab in Glasgow | Wellcome Sanger Institute for the COVID-19 Genomics UK (COG-UK) Consortium | Harper VanSteenhouse, Yumi Kasai, David Gray, Carol Clugston, Anna Dominiczak and Alex Alderton, Roberto Amato, Jeffrey Barrett, Sonia Goncalves, Ewan Harrison, David K. Jackson, Ian Johnston, Dominic Kwiatkowski, Cordelia Langford, John Sillitoe on behalf of the Wellcome Sanger Institute COVID-19 Surveillance Team |
| EPI_ISL_3377846 | Lighthouse Laboratory Plymouth | Wellcome Sanger Institute for the COVID-19 Genomics UK (COG-UK) Consortium | Lighthouse Laboratory Plymouth and Alex Alderton, Roberto Amato, Jeffrey Barrett, Sonia Goncalves, Ewan Harrison, David K. Jackson, Ian Johnston, Dominic Kwiatkowski, Cordelia Langford, John Sillitoe on behalf of the Wellcome Sanger Institute COVID-19 Surveillance Team |
| EPI_ISL_3377847 | Lighthouse Lab in Alderley Park | Wellcome Sanger Institute for the COVID-19 Genomics UK (COG-UK) Consortium | Jacquelyn Wynn, Mairead Hyland, The Lighthouse Lab in Alderley Park and Alex Alderton, Roberto Amato, Jeffrey Barrett, Sonia Goncalves, Ewan Harrison, David K. Jackson, Ian Johnston, Dominic Kwiatkowski, Cordelia Langford, John Sillitoe on behalf of the Wellcome Sanger Institute COVID-19 Surveillance Team |
| EPI_ISL_3377848, EPI_ISL_3377849 | Lighthouse Lab in Glasgow | Wellcome Sanger Institute for the COVID-19 Genomics UK (COG-UK) Consortium | Harper VanSteenhouse, Yumi Kasai, David Gray, Carol Clugston, Anna Dominiczak and Alex Alderton, Roberto Amato, Jeffrey Barrett, Sonia Goncalves, Ewan Harrison, David K. Jackson, Ian Johnston, Dominic Kwiatkowski, Cordelia Langford, John Sillitoe on behalf of the Wellcome Sanger Institute COVID-19 Surveillance Team |
| EPI_ISL_3377850 | Lighthouse Laboratory Plymouth | Wellcome Sanger Institute for the COVID-19 Genomics UK (COG-UK) Consortium | Lighthouse Laboratory Plymouth and Alex Alderton, Roberto Amato, Jeffrey Barrett, Sonia Goncalves, Ewan Harrison, David K. Jackson, Ian Johnston, Dominic Kwiatkowski, Cordelia Langford, John Sillitoe on behalf of the Wellcome Sanger Institute COVID-19 Surveillance Team |
| EPI_ISL_3377851 | Lighthouse Lab in Alderley Park | Wellcome Sanger Institute for the COVID-19 Genomics UK (COG-UK) Consortium | Jacquelyn Wynn, Mairead Hyland, The Lighthouse Lab in Alderley Park and Alex Alderton, Roberto Amato, Jeffrey Barrett, Sonia Goncalves, Ewan Harrison, David K. Jackson, Ian Johnston, Dominic Kwiatkowski, Cordelia Langford, John Sillitoe on behalf of the Wellcome Sanger Institute COVID-19 Surveillance Team |
| EPI_ISL_3377852 | Lighthouse Laboratory Plymouth | Wellcome Sanger Institute for the COVID-19 Genomics UK (COG-UK) Consortium | Lighthouse Laboratory Plymouth and Alex Alderton, Roberto Amato, Jeffrey Barrett, Sonia Goncalves, Ewan Harrison, David K. Jackson, Ian Johnston, Dominic Kwiatkowski, Cordelia Langford, John Sillitoe on behalf of the Wellcome Sanger Institute COVID-19 Surveillance Team |
| EPI_ISL_3377853, EPI_ISL_3377854 | Lighthouse Lab in Glasgow | Wellcome Sanger Institute for the COVID-19 Genomics UK (COG-UK) Consortium | Harper VanSteenhouse, Yumi Kasai, David Gray, Carol Clugston, Anna Dominiczak and Alex Alderton, Roberto Amato, Jeffrey Barrett, Sonia Goncalves, Ewan Harrison, David K. Jackson, Ian Johnston, Dominic Kwiatkowski, Cordelia Langford, John Sillitoe on behalf of the Wellcome Sanger Institute COVID-19 Surveillance Team |
| EPI_ISL_3377855 | Lighthouse Lab in Alderley Park | Wellcome Sanger Institute for the COVID-19 Genomics UK (COG-UK) Consortium | Jacquelyn Wynn, Mairead Hyland, The Lighthouse Lab in Alderley Park and Alex Alderton, Roberto Amato, Jeffrey Barrett, Sonia Goncalves, Ewan Harrison, David K. Jackson, Ian Johnston, Dominic Kwiatkowski, Cordelia Langford, John Sillitoe on behalf of the Wellcome Sanger Institute COVID-19 Surveillance Team |
| EPI_ISL_3377856 | Lighthouse Lab in Glasgow | Wellcome Sanger Institute for the COVID-19 Genomics UK (COG-UK) Consortium | Harper VanSteenhouse, Yumi Kasai, David Gray, Carol Clugston, Anna Dominiczak and Alex Alderton, Roberto Amato, Jeffrey Barrett, Sonia Goncalves, Ewan Harrison, David K. Jackson, Ian Johnston, Dominic Kwiatkowski, Cordelia Langford, John Sillitoe on behalf of the Wellcome Sanger Institute COVID-19 Surveillance Team |
| EPI_ISL_3377857 | Lighthouse Lab in Alderley Park | Wellcome Sanger Institute for the COVID-19 Genomics UK (COG-UK) Consortium | Jacquelyn Wynn, Mairead Hyland, The Lighthouse Lab in Alderley Park and Alex Alderton, Roberto Amato, Jeffrey Barrett, Sonia Goncalves, Ewan Harrison, David K. Jackson, Ian Johnston, Dominic Kwiatkowski, Cordelia Langford, John Sillitoe on behalf of the Wellcome Sanger Institute COVID-19 Surveillance Team |
| EPI_ISL_3377858, EPI_ISL_3377859, EPI_ISL_3377860 | Lighthouse Laboratory Plymouth | Wellcome Sanger Institute for the COVID-19 Genomics UK (COG-UK) Consortium | Lighthouse Laboratory Plymouth and Alex Alderton, Roberto Amato, Jeffrey Barrett, Sonia Goncalves, Ewan Harrison, David K. Jackson, Ian Johnston, Dominic Kwiatkowski, Cordelia Langford, John Sillitoe on behalf of the Wellcome Sanger Institute COVID-19 Surveillance Team |
| EPI_ISL_3377861, EPI_ISL_3377862, EPI_ISL_3377863, EPI_ISL_3377864, EPI_ISL_3377865, EPI_ISL_3377866 | Lighthouse Lab in Alderley Park | Wellcome Sanger Institute for the COVID-19 Genomics UK (COG-UK) Consortium | Jacquelyn Wynn, Mairead Hyland, The Lighthouse Lab in Alderley Park and Alex Alderton, Roberto Amato, Jeffrey Barrett, Sonia Goncalves, Ewan Harrison, David K. Jackson, Ian Johnston, Dominic Kwiatkowski, Cordelia Langford, John Sillitoe on behalf of the Wellcome Sanger Institute COVID-19 Surveillance Team |
| EPI_ISL_3377867 | Lighthouse Laboratory Plymouth | Wellcome Sanger Institute for the COVID-19 Genomics UK (COG-UK) Consortium | Lighthouse Laboratory Plymouth and Alex Alderton, Roberto Amato, Jeffrey Barrett, Sonia Goncalves, Ewan Harrison, David K. Jackson, Ian Johnston, Dominic Kwiatkowski, Cordelia Langford, John Sillitoe on behalf of the Wellcome Sanger Institute COVID-19 Surveillance Team |
| EPI_ISL_3377868 | Lighthouse Lab in Alderley Park | Wellcome Sanger Institute for the COVID-19 Genomics UK (COG-UK) Consortium | Jacquelyn Wynn, Mairead Hyland, The Lighthouse Lab in Alderley Park and Alex Alderton, Roberto Amato, Jeffrey Barrett, Sonia Goncalves, Ewan Harrison, David K. Jackson, Ian Johnston, Dominic Kwiatkowski, Cordelia Langford, John Sillitoe on behalf of the Wellcome Sanger Institute COVID-19 Surveillance Team |
| EPI_ISL_3377869 | Lighthouse Laboratory Plymouth | Wellcome Sanger Institute for the COVID-19 Genomics UK (COG-UK) Consortium | Lighthouse Laboratory Plymouth and Alex Alderton, Roberto Amato, Jeffrey Barrett, Sonia Goncalves, Ewan Harrison, David K. Jackson, Ian Johnston, Dominic Kwiatkowski, Cordelia Langford, John Sillitoe on behalf of the Wellcome Sanger Institute COVID-19 Surveillance Team |
| EPI_ISL_3377870, EPI_ISL_3377871 | Lighthouse Lab in Alderley Park | Wellcome Sanger Institute for the COVID-19 Genomics UK (COG-UK) Consortium | Jacquelyn Wynn, Mairead Hyland, The Lighthouse Lab in Alderley Park and Alex Alderton, Roberto Amato, Jeffrey Barrett, Sonia Goncalves, Ewan Harrison, David K. Jackson, Ian Johnston, Dominic Kwiatkowski, Cordelia Langford, John Sillitoe on behalf of the Wellcome Sanger Institute COVID-19 Surveillance Team |
| EPI_ISL_3377872, EPI_ISL_3377873 | Lighthouse Laboratory Plymouth | Wellcome Sanger Institute for the COVID-19 Genomics UK (COG-UK) Consortium | Lighthouse Laboratory Plymouth and Alex Alderton, Roberto Amato, Jeffrey Barrett, Sonia Goncalves, Ewan Harrison, David K. Jackson, Ian Johnston, Dominic Kwiatkowski, Cordelia Langford, John Sillitoe on behalf of the Wellcome Sanger Institute COVID-19 Surveillance Team |
| EPI_ISL_3377874, EPI_ISL_3377875 | Lighthouse Lab in Alderley Park | Wellcome Sanger Institute for the COVID-19 Genomics UK (COG-UK) Consortium | Jacquelyn Wynn, Mairead Hyland, The Lighthouse Lab in Alderley Park and Alex Alderton, Roberto Amato, Jeffrey Barrett, Sonia Goncalves, Ewan Harrison, David K. Jackson, Ian Johnston, Dominic Kwiatkowski, Cordelia Langford, John Sillitoe on behalf of the Wellcome Sanger Institute COVID-19 Surveillance Team |
| EPI_ISL_3377876 | Lighthouse Laboratory Plymouth | Wellcome Sanger Institute for the COVID-19 Genomics UK (COG-UK) Consortium | Lighthouse Laboratory Plymouth and Alex Alderton, Roberto Amato, Jeffrey Barrett, Sonia Goncalves, Ewan Harrison, David K. Jackson, Ian Johnston, Dominic Kwiatkowski, Cordelia Langford, John Sillitoe on behalf of the Wellcome Sanger Institute COVID-19 Surveillance Team |
| EPI_ISL_3377877 | Lighthouse Lab in Alderley Park | Wellcome Sanger Institute for the COVID-19 Genomics UK (COG-UK) Consortium | Jacquelyn Wynn, Mairead Hyland, The Lighthouse Lab in Alderley Park and Alex Alderton, Roberto Amato, Jeffrey Barrett, Sonia Goncalves, Ewan Harrison, David K. Jackson, Ian Johnston, Dominic Kwiatkowski, Cordelia Langford, John Sillitoe on behalf of the Wellcome Sanger Institute COVID-19 Surveillance Team |
| EPI_ISL_3377878 | Lighthouse Laboratory Plymouth | Wellcome Sanger Institute for the COVID-19 Genomics UK (COG-UK) Consortium | Lighthouse Laboratory Plymouth and Alex Alderton, Roberto Amato, Jeffrey Barrett, Sonia Goncalves, Ewan Harrison, David K. Jackson, Ian Johnston, Dominic Kwiatkowski, Cordelia Langford, John Sillitoe on behalf of the Wellcome Sanger Institute COVID-19 Surveillance Team |
| EPI_ISL_3377879, EPI_ISL_3377880, EPI_ISL_3377881, EPI_ISL_3377882, EPI_ISL_3377883, EPI_ISL_3377884 | Lighthouse Lab in Alderley Park | Wellcome Sanger Institute for the COVID-19 Genomics UK (COG-UK) Consortium | Jacquelyn Wynn, Mairead Hyland, The Lighthouse Lab in Alderley Park and Alex Alderton, Roberto Amato, Jeffrey Barrett, Sonia Goncalves, Ewan Harrison, David K. Jackson, Ian Johnston, Dominic Kwiatkowski, Cordelia Langford, John Sillitoe on behalf of the Wellcome Sanger Institute COVID-19 Surveillance Team |
| EPI_ISL_3377885 | Lighthouse Laboratory Plymouth | Wellcome Sanger Institute for the COVID-19 Genomics UK (COG-UK) Consortium | Lighthouse Laboratory Plymouth and Alex Alderton, Roberto Amato, Jeffrey Barrett, Sonia Goncalves, Ewan Harrison, David K. Jackson, Ian Johnston, Dominic Kwiatkowski, Cordelia Langford, John Sillitoe on behalf of the Wellcome Sanger Institute COVID-19 Surveillance Team |
| EPI_ISL_3377886, EPI_ISL_3377887, EPI_ISL_3377888, EPI_ISL_3377889, EPI_ISL_3377890 | Lighthouse Lab in Alderley Park | Wellcome Sanger Institute for the COVID-19 Genomics UK (COG-UK) Consortium | Jacquelyn Wynn, Mairead Hyland, The Lighthouse Lab in Alderley Park and Alex Alderton, Roberto Amato, Jeffrey Barrett, Sonia Goncalves, Ewan Harrison, David K. Jackson, Ian Johnston, Dominic Kwiatkowski, Cordelia Langford, John Sillitoe on behalf of the Wellcome Sanger Institute COVID-19 Surveillance Team |
| EPI_ISL_3377891 | Lighthouse Laboratory Plymouth | Wellcome Sanger Institute for the COVID-19 Genomics UK (COG-UK) Consortium | Lighthouse Laboratory Plymouth and Alex Alderton, Roberto Amato, Jeffrey Barrett, Sonia Goncalves, Ewan Harrison, David K. Jackson, Ian Johnston, Dominic Kwiatkowski, Cordelia Langford, John Sillitoe on behalf of the Wellcome Sanger Institute COVID-19 Surveillance Team |
| EPI_ISL_3377892, EPI_ISL_3377893 | Lighthouse Lab in Alderley Park | Wellcome Sanger Institute for the COVID-19 Genomics UK (COG-UK) Consortium | Jacquelyn Wynn, Mairead Hyland, The Lighthouse Lab in Alderley Park and Alex Alderton, Roberto Amato, Jeffrey Barrett, Sonia Goncalves, Ewan Harrison, David K. Jackson, Ian Johnston, Dominic Kwiatkowski, Cordelia Langford, John Sillitoe on behalf of the Wellcome Sanger Institute COVID-19 Surveillance Team |
| EPI_ISL_3377894, EPI_ISL_3377895 | Lighthouse Laboratory Plymouth | Wellcome Sanger Institute for the COVID-19 Genomics UK (COG-UK) Consortium | Lighthouse Laboratory Plymouth and Alex Alderton, Roberto Amato, Jeffrey Barrett, Sonia Goncalves, Ewan Harrison, David K. Jackson, Ian Johnston, Dominic Kwiatkowski, Cordelia Langford, John Sillitoe on behalf of the Wellcome Sanger Institute COVID-19 Surveillance Team |
| EPI_ISL_3377896, EPI_ISL_3377897, EPI_ISL_3377898, EPI_ISL_3377899, EPI_ISL_3377900, EPI_ISL_3377901, EPI_ISL_3377902 | Lighthouse Lab in Alderley Park | Wellcome Sanger Institute for the COVID-19 Genomics UK (COG-UK) Consortium | Jacquelyn Wynn, Mairead Hyland, The Lighthouse Lab in Alderley Park and Alex Alderton, Roberto Amato, Jeffrey Barrett, Sonia Goncalves, Ewan Harrison, David K. Jackson, Ian Johnston, Dominic Kwiatkowski, Cordelia Langford, John Sillitoe on behalf of the Wellcome Sanger Institute COVID-19 Surveillance Team |
| EPI_ISL_3377903, | Lighthouse Laboratory Plymouth | Wellcome Sanger Institute for the COVID-19 Genomics UK (COG-UK) | Lighthouse Laboratory Plymouth and Alex Alderton, Roberto Amato, Jeffrey Barrett, Sonia Goncalves, Ewan Harrison, David K. Jackson, Ian Johnston, Dominic Kwiatkowski, Cordelia Langford, John Sillitoe on behalf of the Wellcome Sanger |

| | | | |
|---|---|---|---|
| EPI_ISL_3377904 | | Consortium | Institute COVID-19 Surveillance Team |
| EPI_ISL_3377905, EPI_ISL_3377906, EPI_ISL_3377907 | Lighthouse Lab in Alderley Park | Wellcome Sanger Institute for the COVID-19 Genomics UK (COG-UK) Consortium | Jacquelyn Wynn, Mairead Hyland, The Lighthouse Lab in Alderley Park and Alex Alderton, Roberto Amato, Jeffrey Barrett, Sonia Goncalves, Ewan Harrison, David K. Jackson, Ian Johnston, Dominic Kwiatkowski, Cordelia Langford, John Sillitoe on behalf of the Wellcome Sanger Institute COVID-19 Surveillance Team |
| EPI_ISL_3377908, EPI_ISL_3377909 | Lighthouse Laboratory Plymouth | Wellcome Sanger Institute for the COVID-19 Genomics UK (COG-UK) Consortium | Lighthouse Laboratory Plymouth and Alex Alderton, Roberto Amato, Jeffrey Barrett, Sonia Goncalves, Ewan Harrison, David K. Jackson, Ian Johnston, Dominic Kwiatkowski, Cordelia Langford, John Sillitoe on behalf of the Wellcome Sanger Institute COVID-19 Surveillance Team |
| EPI_ISL_3377910 | Lighthouse Lab in Alderley Park | Wellcome Sanger Institute for the COVID-19 Genomics UK (COG-UK) Consortium | Jacquelyn Wynn, Mairead Hyland, The Lighthouse Lab in Alderley Park and Alex Alderton, Roberto Amato, Jeffrey Barrett, Sonia Goncalves, Ewan Harrison, David K. Jackson, Ian Johnston, Dominic Kwiatkowski, Cordelia Langford, John Sillitoe on behalf of the Wellcome Sanger Institute COVID-19 Surveillance Team |
| EPI_ISL_3377911, EPI_ISL_3377912 | Lighthouse Laboratory Plymouth | Wellcome Sanger Institute for the COVID-19 Genomics UK (COG-UK) Consortium | Lighthouse Laboratory Plymouth and Alex Alderton, Roberto Amato, Jeffrey Barrett, Sonia Goncalves, Ewan Harrison, David K. Jackson, Ian Johnston, Dominic Kwiatkowski, Cordelia Langford, John Sillitoe on behalf of the Wellcome Sanger Institute COVID-19 Surveillance Team |
| EPI_ISL_3377913 | Lighthouse Lab in Alderley Park | Wellcome Sanger Institute for the COVID-19 Genomics UK (COG-UK) Consortium | Jacquelyn Wynn, Mairead Hyland, The Lighthouse Lab in Alderley Park and Alex Alderton, Roberto Amato, Jeffrey Barrett, Sonia Goncalves, Ewan Harrison, David K. Jackson, Ian Johnston, Dominic Kwiatkowski, Cordelia Langford, John Sillitoe on behalf of the Wellcome Sanger Institute COVID-19 Surveillance Team |
| EPI_ISL_3377914 | Lighthouse Laboratory Plymouth | Wellcome Sanger Institute for the COVID-19 Genomics UK (COG-UK) Consortium | Lighthouse Laboratory Plymouth and Alex Alderton, Roberto Amato, Jeffrey Barrett, Sonia Goncalves, Ewan Harrison, David K. Jackson, Ian Johnston, Dominic Kwiatkowski, Cordelia Langford, John Sillitoe on behalf of the Wellcome Sanger Institute COVID-19 Surveillance Team |
| EPI_ISL_3377915 | Lighthouse Lab in Alderley Park | Wellcome Sanger Institute for the COVID-19 Genomics UK (COG-UK) Consortium | Jacquelyn Wynn, Mairead Hyland, The Lighthouse Lab in Alderley Park and Alex Alderton, Roberto Amato, Jeffrey Barrett, Sonia Goncalves, Ewan Harrison, David K. Jackson, Ian Johnston, Dominic Kwiatkowski, Cordelia Langford, John Sillitoe on behalf of the Wellcome Sanger Institute COVID-19 Surveillance Team |
| EPI_ISL_3377916 | Lighthouse Laboratory Plymouth | Wellcome Sanger Institute for the COVID-19 Genomics UK (COG-UK) Consortium | Lighthouse Laboratory Plymouth and Alex Alderton, Roberto Amato, Jeffrey Barrett, Sonia Goncalves, Ewan Harrison, David K. Jackson, Ian Johnston, Dominic Kwiatkowski, Cordelia Langford, John Sillitoe on behalf of the Wellcome Sanger Institute COVID-19 Surveillance Team |
| EPI_ISL_3377917 | Lighthouse Lab in Alderley Park | Wellcome Sanger Institute for the COVID-19 Genomics UK (COG-UK) Consortium | Jacquelyn Wynn, Mairead Hyland, The Lighthouse Lab in Alderley Park and Alex Alderton, Roberto Amato, Jeffrey Barrett, Sonia Goncalves, Ewan Harrison, David K. Jackson, Ian Johnston, Dominic Kwiatkowski, Cordelia Langford, John Sillitoe on behalf of the Wellcome Sanger Institute COVID-19 Surveillance Team |
| EPI_ISL_3377918, EPI_ISL_3377919, EPI_ISL_3377920 | Lighthouse Laboratory Plymouth | Wellcome Sanger Institute for the COVID-19 Genomics UK (COG-UK) Consortium | Lighthouse Laboratory Plymouth and Alex Alderton, Roberto Amato, Jeffrey Barrett, Sonia Goncalves, Ewan Harrison, David K. Jackson, Ian Johnston, Dominic Kwiatkowski, Cordelia Langford, John Sillitoe on behalf of the Wellcome Sanger Institute COVID-19 Surveillance Team |
| EPI_ISL_3377921 | Lighthouse Lab in Alderley Park | Wellcome Sanger Institute for the COVID-19 Genomics UK (COG-UK) Consortium | Jacquelyn Wynn, Mairead Hyland, The Lighthouse Lab in Alderley Park and Alex Alderton, Roberto Amato, Jeffrey Barrett, Sonia Goncalves, Ewan Harrison, David K. Jackson, Ian Johnston, Dominic Kwiatkowski, Cordelia Langford, John Sillitoe on behalf of the Wellcome Sanger Institute COVID-19 Surveillance Team |
| EPI_ISL_3377922 | Lighthouse Laboratory Plymouth | Wellcome Sanger Institute for the COVID-19 Genomics UK (COG-UK) Consortium | Lighthouse Laboratory Plymouth and Alex Alderton, Roberto Amato, Jeffrey Barrett, Sonia Goncalves, Ewan Harrison, David K. Jackson, Ian Johnston, Dominic Kwiatkowski, Cordelia Langford, John Sillitoe on behalf of the Wellcome Sanger Institute COVID-19 Surveillance Team |
| EPI_ISL_3377923, EPI_ISL_3377924, EPI_ISL_3377925 | Lighthouse Lab in Alderley Park | Wellcome Sanger Institute for the COVID-19 Genomics UK (COG-UK) Consortium | Jacquelyn Wynn, Mairead Hyland, The Lighthouse Lab in Alderley Park and Alex Alderton, Roberto Amato, Jeffrey Barrett, Sonia Goncalves, Ewan Harrison, David K. Jackson, Ian Johnston, Dominic Kwiatkowski, Cordelia Langford, John Sillitoe on behalf of the Wellcome Sanger Institute COVID-19 Surveillance Team |
| EPI_ISL_3377926 | Lighthouse Laboratory Plymouth | Wellcome Sanger Institute for the COVID-19 Genomics UK (COG-UK) Consortium | Lighthouse Laboratory Plymouth and Alex Alderton, Roberto Amato, Jeffrey Barrett, Sonia Goncalves, Ewan Harrison, David K. Jackson, Ian Johnston, Dominic Kwiatkowski, Cordelia Langford, John Sillitoe on behalf of the Wellcome Sanger Institute COVID-19 Surveillance Team |
| EPI_ISL_3377927, EPI_ISL_3377928, EPI_ISL_3377929 | Lighthouse Lab in Alderley Park | Wellcome Sanger Institute for the COVID-19 Genomics UK (COG-UK) Consortium | Jacquelyn Wynn, Mairead Hyland, The Lighthouse Lab in Alderley Park and Alex Alderton, Roberto Amato, Jeffrey Barrett, Sonia Goncalves, Ewan Harrison, David K. Jackson, Ian Johnston, Dominic Kwiatkowski, Cordelia Langford, John Sillitoe on behalf of the Wellcome Sanger Institute COVID-19 Surveillance Team |
| EPI_ISL_3377930 | Lighthouse Laboratory Plymouth | Wellcome Sanger Institute for the COVID-19 Genomics UK (COG-UK) Consortium | Lighthouse Laboratory Plymouth and Alex Alderton, Roberto Amato, Jeffrey Barrett, Sonia Goncalves, Ewan Harrison, David K. Jackson, Ian Johnston, Dominic Kwiatkowski, Cordelia Langford, John Sillitoe on behalf of the Wellcome Sanger Institute COVID-19 Surveillance Team |
| EPI_ISL_3377931, EPI_ISL_3377932, EPI_ISL_3377933, EPI_ISL_3377934 | Lighthouse Lab in Alderley Park | Wellcome Sanger Institute for the COVID-19 Genomics UK (COG-UK) Consortium | Jacquelyn Wynn, Mairead Hyland, The Lighthouse Lab in Alderley Park and Alex Alderton, Roberto Amato, Jeffrey Barrett, Sonia Goncalves, Ewan Harrison, David K. Jackson, Ian Johnston, Dominic Kwiatkowski, Cordelia Langford, John Sillitoe on behalf of the Wellcome Sanger Institute COVID-19 Surveillance Team |
| EPI_ISL_3377935 | Lighthouse Laboratory Plymouth | Wellcome Sanger Institute for the COVID-19 Genomics UK (COG-UK) Consortium | Lighthouse Laboratory Plymouth and Alex Alderton, Roberto Amato, Jeffrey Barrett, Sonia Goncalves, Ewan Harrison, David K. Jackson, Ian Johnston, Dominic Kwiatkowski, Cordelia Langford, John Sillitoe on behalf of the Wellcome Sanger Institute COVID-19 Surveillance Team |
| EPI_ISL_3377936, EPI_ISL_3377937, EPI_ISL_3377938, EPI_ISL_3377939, EPI_ISL_3377940, EPI_ISL_3377941 | Lighthouse Lab in Alderley Park | Wellcome Sanger Institute for the COVID-19 Genomics UK (COG-UK) Consortium | Jacquelyn Wynn, Mairead Hyland, The Lighthouse Lab in Alderley Park and Alex Alderton, Roberto Amato, Jeffrey Barrett, Sonia Goncalves, Ewan Harrison, David K. Jackson, Ian Johnston, Dominic Kwiatkowski, Cordelia Langford, John Sillitoe on behalf of the Wellcome Sanger Institute COVID-19 Surveillance Team |
| EPI_ISL_3377942, EPI_ISL_3377943 | Lighthouse Laboratory Plymouth | Wellcome Sanger Institute for the COVID-19 Genomics UK (COG-UK) Consortium | Lighthouse Laboratory Plymouth and Alex Alderton, Roberto Amato, Jeffrey Barrett, Sonia Goncalves, Ewan Harrison, David K. Jackson, Ian Johnston, Dominic Kwiatkowski, Cordelia Langford, John Sillitoe on behalf of the Wellcome Sanger Institute COVID-19 Surveillance Team |
| EPI_ISL_3377944 | Lighthouse Lab in Alderley Park | Wellcome Sanger Institute for the COVID-19 Genomics UK (COG-UK) Consortium | Jacquelyn Wynn, Mairead Hyland, The Lighthouse Lab in Alderley Park and Alex Alderton, Roberto Amato, Jeffrey Barrett, Sonia Goncalves, Ewan Harrison, David K. Jackson, Ian Johnston, Dominic Kwiatkowski, Cordelia Langford, John Sillitoe on behalf of the Wellcome Sanger Institute COVID-19 Surveillance Team |
| EPI_ISL_3377945 | Lighthouse Laboratory Plymouth | Wellcome Sanger Institute for the COVID-19 Genomics UK (COG-UK) Consortium | Lighthouse Laboratory Plymouth and Alex Alderton, Roberto Amato, Jeffrey Barrett, Sonia Goncalves, Ewan Harrison, David K. Jackson, Ian Johnston, Dominic Kwiatkowski, Cordelia Langford, John Sillitoe on behalf of the Wellcome Sanger Institute COVID-19 Surveillance Team |
| EPI_ISL_3377946, EPI_ISL_3377947, EPI_ISL_3377948, EPI_ISL_3377949 | Lighthouse Lab in Alderley Park | Wellcome Sanger Institute for the COVID-19 Genomics UK (COG-UK) Consortium | Jacquelyn Wynn, Mairead Hyland, The Lighthouse Lab in Alderley Park and Alex Alderton, Roberto Amato, Jeffrey Barrett, Sonia Goncalves, Ewan Harrison, David K. Jackson, Ian Johnston, Dominic Kwiatkowski, Cordelia Langford, John Sillitoe on behalf of the Wellcome Sanger Institute COVID-19 Surveillance Team |
| EPI_ISL_3377950, EPI_ISL_3377951, EPI_ISL_3377952, EPI_ISL_3377953 | Lighthouse Laboratory Plymouth | Wellcome Sanger Institute for the COVID-19 Genomics UK (COG-UK) Consortium | Lighthouse Laboratory Plymouth and Alex Alderton, Roberto Amato, Jeffrey Barrett, Sonia Goncalves, Ewan Harrison, David K. Jackson, Ian Johnston, Dominic Kwiatkowski, Cordelia Langford, John Sillitoe on behalf of the Wellcome Sanger Institute COVID-19 Surveillance Team |
| EPI_ISL_3377954, EPI_ISL_3377955, EPI_ISL_3377956, EPI_ISL_3377957 | Lighthouse Lab in Alderley Park | Wellcome Sanger Institute for the COVID-19 Genomics UK (COG-UK) Consortium | Jacquelyn Wynn, Mairead Hyland, The Lighthouse Lab in Alderley Park and Alex Alderton, Roberto Amato, Jeffrey Barrett, Sonia Goncalves, Ewan Harrison, David K. Jackson, Ian Johnston, Dominic Kwiatkowski, Cordelia Langford, John Sillitoe on behalf of the Wellcome Sanger Institute COVID-19 Surveillance Team |
| EPI_ISL_3377958 | Lighthouse Laboratory Plymouth | Wellcome Sanger Institute for the COVID-19 Genomics UK (COG-UK) Consortium | Lighthouse Laboratory Plymouth and Alex Alderton, Roberto Amato, Jeffrey Barrett, Sonia Goncalves, Ewan Harrison, David K. Jackson, Ian Johnston, Dominic Kwiatkowski, Cordelia Langford, John Sillitoe on behalf of the Wellcome Sanger Institute COVID-19 Surveillance Team |
| EPI_ISL_3377959, EPI_ISL_3377960, EPI_ISL_3377961, EPI_ISL_3377962 | Lighthouse Lab in Alderley Park | Wellcome Sanger Institute for the COVID-19 Genomics UK (COG-UK) Consortium | Jacquelyn Wynn, Mairead Hyland, The Lighthouse Lab in Alderley Park and Alex Alderton, Roberto Amato, Jeffrey Barrett, Sonia Goncalves, Ewan Harrison, David K. Jackson, Ian Johnston, Dominic Kwiatkowski, Cordelia Langford, John Sillitoe on behalf of the Wellcome Sanger Institute COVID-19 Surveillance Team |
| EPI_ISL_3377963, EPI_ISL_3377964, EPI_ISL_3377965 | Lighthouse Laboratory Plymouth | Wellcome Sanger Institute for the COVID-19 Genomics UK (COG-UK) Consortium | Lighthouse Laboratory Plymouth and Alex Alderton, Roberto Amato, Jeffrey Barrett, Sonia Goncalves, Ewan Harrison, David K. Jackson, Ian Johnston, Dominic Kwiatkowski, Cordelia Langford, John Sillitoe on behalf of the Wellcome Sanger Institute COVID-19 Surveillance Team |
| EPI_ISL_3377966, EPI_ISL_3377967 | Lighthouse Lab in Alderley Park | Wellcome Sanger Institute for the COVID-19 Genomics UK (COG-UK) Consortium | Jacquelyn Wynn, Mairead Hyland, The Lighthouse Lab in Alderley Park and Alex Alderton, Roberto Amato, Jeffrey Barrett, Sonia Goncalves, Ewan Harrison, David K. Jackson, Ian Johnston, Dominic Kwiatkowski, Cordelia Langford, John Sillitoe on behalf of the Wellcome Sanger Institute COVID-19 Surveillance Team |
| EPI_ISL_3377968 | Lighthouse Laboratory Plymouth | Wellcome Sanger Institute for the COVID-19 Genomics UK (COG-UK) Consortium | Lighthouse Laboratory Plymouth and Alex Alderton, Roberto Amato, Jeffrey Barrett, Sonia Goncalves, Ewan Harrison, David K. Jackson, Ian Johnston, Dominic Kwiatkowski, Cordelia Langford, John Sillitoe on behalf of the Wellcome Sanger Institute COVID-19 Surveillance Team |
| EPI_ISL_3377969 | Lighthouse Lab in Alderley Park | Wellcome Sanger Institute for the COVID-19 Genomics UK (COG-UK) Consortium | Jacquelyn Wynn, Mairead Hyland, The Lighthouse Lab in Alderley Park and Alex Alderton, Roberto Amato, Jeffrey Barrett, Sonia Goncalves, Ewan Harrison, David K. Jackson, Ian Johnston, Dominic Kwiatkowski, Cordelia Langford, John Sillitoe on behalf of the Wellcome Sanger Institute COVID-19 Surveillance Team |
| EPI_ISL_3377970 | Lighthouse Laboratory Plymouth | Wellcome Sanger Institute for the COVID-19 Genomics UK (COG-UK) Consortium | Lighthouse Laboratory Plymouth and Alex Alderton, Roberto Amato, Jeffrey Barrett, Sonia Goncalves, Ewan Harrison, David K. Jackson, Ian Johnston, Dominic Kwiatkowski, Cordelia Langford, John Sillitoe on behalf of the Wellcome Sanger Institute COVID-19 Surveillance Team |
| EPI_ISL_3377971 | Lighthouse Lab in Alderley Park | Wellcome Sanger Institute for the COVID-19 Genomics UK (COG-UK) Consortium | Jacquelyn Wynn, Mairead Hyland, The Lighthouse Lab in Alderley Park and Alex Alderton, Roberto Amato, Jeffrey Barrett, Sonia Goncalves, Ewan Harrison, David K. Jackson, Ian Johnston, Dominic Kwiatkowski, Cordelia Langford, John Sillitoe on behalf of the Wellcome Sanger Institute COVID-19 Surveillance Team |
| EPI_ISL_3377972 | Lighthouse Laboratory Plymouth | Wellcome Sanger Institute for the COVID-19 Genomics UK (COG-UK) Consortium | Lighthouse Laboratory Plymouth and Alex Alderton, Roberto Amato, Jeffrey Barrett, Sonia Goncalves, Ewan Harrison, David K. Jackson, Ian Johnston, Dominic Kwiatkowski, Cordelia Langford, John Sillitoe on behalf of the Wellcome Sanger Institute COVID-19 Surveillance Team |
| EPI_ISL_3377973, EPI_ISL_3377974, EPI_ISL_3377975 | Lighthouse Lab in Alderley Park | Wellcome Sanger Institute for the COVID-19 Genomics UK (COG-UK) Consortium | Jacquelyn Wynn, Mairead Hyland, The Lighthouse Lab in Alderley Park and Alex Alderton, Roberto Amato, Jeffrey Barrett, Sonia Goncalves, Ewan Harrison, David K. Jackson, Ian Johnston, Dominic Kwiatkowski, Cordelia Langford, John Sillitoe on behalf of the Wellcome Sanger Institute COVID-19 Surveillance Team |
| EPI_ISL_3377976 | Lighthouse Laboratory Plymouth | Wellcome Sanger Institute for the COVID-19 Genomics UK (COG-UK) Consortium | Lighthouse Laboratory Plymouth and Alex Alderton, Roberto Amato, Jeffrey Barrett, Sonia Goncalves, Ewan Harrison, David K. Jackson, Ian Johnston, Dominic Kwiatkowski, Cordelia Langford, John Sillitoe on behalf of the Wellcome Sanger Institute COVID-19 Surveillance Team |

| | | | |
|---|---|---|---|
| EPI_ISL_3377977 | Lighthouse Lab in Alderley Park | Wellcome Sanger Institute for the COVID-19 Genomics UK (COG-UK) Consortium | Jacquelyn Wynn, Mairead Hyland, The Lighthouse Lab in Alderley Park and Alex Alderton, Roberto Amato, Jeffrey Barrett, Sonia Goncalves, Ewan Harrison, David K. Jackson, Ian Johnston, Dominic Kwiatkowski, Cordelia Langford, John Sillitoe on behalf of the Wellcome Sanger Institute COVID-19 Surveillance Team |
| EPI_ISL_3377978 | Lighthouse Laboratory Plymouth | Wellcome Sanger Institute for the COVID-19 Genomics UK (COG-UK) Consortium | Lighthouse Laboratory Plymouth and Alex Alderton, Roberto Amato, Jeffrey Barrett, Sonia Goncalves, Ewan Harrison, David K. Jackson, Ian Johnston, Dominic Kwiatkowski, Cordelia Langford, John Sillitoe on behalf of the Wellcome Sanger Institute COVID-19 Surveillance Team |
| EPI_ISL_3377979, EPI_ISL_3377980, EPI_ISL_3377981, EPI_ISL_3377982 | Lighthouse Lab in Alderley Park | Wellcome Sanger Institute for the COVID-19 Genomics UK (COG-UK) Consortium | Jacquelyn Wynn, Mairead Hyland, The Lighthouse Lab in Alderley Park and Alex Alderton, Roberto Amato, Jeffrey Barrett, Sonia Goncalves, Ewan Harrison, David K. Jackson, Ian Johnston, Dominic Kwiatkowski, Cordelia Langford, John Sillitoe on behalf of the Wellcome Sanger Institute COVID-19 Surveillance Team |
| EPI_ISL_3377983 | Lighthouse Laboratory Plymouth | Wellcome Sanger Institute for the COVID-19 Genomics UK (COG-UK) Consortium | Lighthouse Laboratory Plymouth and Alex Alderton, Roberto Amato, Jeffrey Barrett, Sonia Goncalves, Ewan Harrison, David K. Jackson, Ian Johnston, Dominic Kwiatkowski, Cordelia Langford, John Sillitoe on behalf of the Wellcome Sanger Institute COVID-19 Surveillance Team |
| EPI_ISL_3377984, EPI_ISL_3377985, EPI_ISL_3377986, EPI_ISL_3377987, EPI_ISL_3377988, EPI_ISL_3377989 | Lighthouse Lab in Alderley Park | Wellcome Sanger Institute for the COVID-19 Genomics UK (COG-UK) Consortium | Jacquelyn Wynn, Mairead Hyland, The Lighthouse Lab in Alderley Park and Alex Alderton, Roberto Amato, Jeffrey Barrett, Sonia Goncalves, Ewan Harrison, David K. Jackson, Ian Johnston, Dominic Kwiatkowski, Cordelia Langford, John Sillitoe on behalf of the Wellcome Sanger Institute COVID-19 Surveillance Team |
| EPI_ISL_3377990 | Lighthouse Laboratory Plymouth | Wellcome Sanger Institute for the COVID-19 Genomics UK (COG-UK) Consortium | Lighthouse Laboratory Plymouth and Alex Alderton, Roberto Amato, Jeffrey Barrett, Sonia Goncalves, Ewan Harrison, David K. Jackson, Ian Johnston, Dominic Kwiatkowski, Cordelia Langford, John Sillitoe on behalf of the Wellcome Sanger Institute COVID-19 Surveillance Team |
| EPI_ISL_3377991, EPI_ISL_3377992, EPI_ISL_3377993, EPI_ISL_3377994 | Lighthouse Lab in Alderley Park | Wellcome Sanger Institute for the COVID-19 Genomics UK (COG-UK) Consortium | Jacquelyn Wynn, Mairead Hyland, The Lighthouse Lab in Alderley Park and Alex Alderton, Roberto Amato, Jeffrey Barrett, Sonia Goncalves, Ewan Harrison, David K. Jackson, Ian Johnston, Dominic Kwiatkowski, Cordelia Langford, John Sillitoe on behalf of the Wellcome Sanger Institute COVID-19 Surveillance Team |
| EPI_ISL_3377995 | LAM CERBALLIANCE | CNR Virus des Infections Respiratoires - France SUD | Antonin Bal, Gregory Destras, Gwendolyne Burfin, Hadrien Regue, Quentin Semanas, Martine Valette, Bruno Lina, Laurence Josset |
| EPI_ISL_3377996 | Lighthouse Lab in Alderley Park | Wellcome Sanger Institute for the COVID-19 Genomics UK (COG-UK) Consortium | Jacquelyn Wynn, Mairead Hyland, The Lighthouse Lab in Alderley Park and Alex Alderton, Roberto Amato, Jeffrey Barrett, Sonia Goncalves, Ewan Harrison, David K. Jackson, Ian Johnston, Dominic Kwiatkowski, Cordelia Langford, John Sillitoe on behalf of the Wellcome Sanger Institute COVID-19 Surveillance Team |
| EPI_ISL_3377997 | Lighthouse Laboratory Plymouth | Wellcome Sanger Institute for the COVID-19 Genomics UK (COG-UK) Consortium | Lighthouse Laboratory Plymouth and Alex Alderton, Roberto Amato, Jeffrey Barrett, Sonia Goncalves, Ewan Harrison, David K. Jackson, Ian Johnston, Dominic Kwiatkowski, Cordelia Langford, John Sillitoe on behalf of the Wellcome Sanger Institute COVID-19 Surveillance Team |
| EPI_ISL_3377998, EPI_ISL_3377999 | Lighthouse Lab in Alderley Park | Wellcome Sanger Institute for the COVID-19 Genomics UK (COG-UK) Consortium | Jacquelyn Wynn, Mairead Hyland, The Lighthouse Lab in Alderley Park and Alex Alderton, Roberto Amato, Jeffrey Barrett, Sonia Goncalves, Ewan Harrison, David K. Jackson, Ian Johnston, Dominic Kwiatkowski, Cordelia Langford, John Sillitoe on behalf of the Wellcome Sanger Institute COVID-19 Surveillance Team |
| EPI_ISL_3378000, EPI_ISL_3378001, EPI_ISL_3378002, EPI_ISL_3378003, EPI_ISL_3378004, EPI_ISL_3378005 | LAM CERBALLIANCE | CNR Virus des Infections Respiratoires - France SUD | Antonin Bal, Gregory Destras, Gwendolyne Burfin, Hadrien Regue, Quentin Semanas, Martine Valette, Bruno Lina, Laurence Josset |
| EPI_ISL_3378006 | LABORATOIRE UNILIANS DECINES | CNR Virus des Infections Respiratoires - France SUD | Antonin Bal, Gregory Destras, Gwendolyne Burfin, Hadrien Regue, Quentin Semanas, Martine Valette, Bruno Lina, Laurence Josset |
| EPI_ISL_3378007, EPI_ISL_3378008, EPI_ISL_3378009, EPI_ISL_3378010, EPI_ISL_3378011, EPI_ISL_3378012, EPI_ISL_3378013, EPI_ISL_3378014, EPI_ISL_3378015, EPI_ISL_3378016, EPI_ISL_3378017, EPI_ISL_3378018, EPI_ISL_3378019, EPI_ISL_3378020, EPI_ISL_3378021, EPI_ISL_3378022, EPI_ISL_3378023, EPI_ISL_3378024 | | | |
| see above | LAM CERBALLIANCE | CNR Virus des Infections Respiratoires - France SUD | Antonin Bal, Gregory Destras, Gwendolyne Burfin, Hadrien Regue, Quentin Semanas, Martine Valette, Bruno Lina, Laurence Josset |
| EPI_ISL_3378025 | LABORATOIRE UNILIANS DECINES | CNR Virus des Infections Respiratoires - France SUD | Antonin Bal, Gregory Destras, Gwendolyne Burfin, Hadrien Regue, Quentin Semanas, Martine Valette, Bruno Lina, Laurence Josset |
| EPI_ISL_3378026, EPI_ISL_3378027, EPI_ISL_3378029, EPI_ISL_3378030, EPI_ISL_3378031, EPI_ISL_3378032, EPI_ISL_3378033, EPI_ISL_3378034, EPI_ISL_3378035, EPI_ISL_3378036, EPI_ISL_3378037, EPI_ISL_3378038, EPI_ISL_3378039 | | | |
| see above | LAM CERBALLIANCE | CNR Virus des Infections Respiratoires - France SUD | Antonin Bal, Gregory Destras, Gwendolyne Burfin, Hadrien Regue, Quentin Semanas, Martine Valette, Bruno Lina, Laurence Josset |
| EPI_ISL_3378040 | Lighthouse Lab in Alderley Park | Wellcome Sanger Institute for the COVID-19 Genomics UK (COG-UK) Consortium | Jacquelyn Wynn, Mairead Hyland, The Lighthouse Lab in Alderley Park and Alex Alderton, Roberto Amato, Jeffrey Barrett, Sonia Goncalves, Ewan Harrison, David K. Jackson, Ian Johnston, Dominic Kwiatkowski, Cordelia Langford, John Sillitoe on behalf of the Wellcome Sanger Institute COVID-19 Surveillance Team |
| EPI_ISL_3378041 | LABORATOIRE ACCOLAB SUD OUEST | CNR Virus des Infections Respiratoires - France SUD | Antonin Bal, Gregory Destras, Gwendolyne Burfin, Hadrien Regue, Quentin Semanas, Martine Valette, Bruno Lina, Laurence Josset |
| EPI_ISL_3378042 | Lighthouse Lab in Alderley Park | Wellcome Sanger Institute for the COVID-19 Genomics UK (COG-UK) Consortium | Jacquelyn Wynn, Mairead Hyland, The Lighthouse Lab in Alderley Park and Alex Alderton, Roberto Amato, Jeffrey Barrett, Sonia Goncalves, Ewan Harrison, David K. Jackson, Ian Johnston, Dominic Kwiatkowski, Cordelia Langford, John Sillitoe on behalf of the Wellcome Sanger Institute COVID-19 Surveillance Team |
| EPI_ISL_3378043 | Lighthouse Laboratory Plymouth | Wellcome Sanger Institute for the COVID-19 Genomics UK (COG-UK) Consortium | Lighthouse Laboratory Plymouth and Alex Alderton, Roberto Amato, Jeffrey Barrett, Sonia Goncalves, Ewan Harrison, David K. Jackson, Ian Johnston, Dominic Kwiatkowski, Cordelia Langford, John Sillitoe on behalf of the Wellcome Sanger Institute COVID-19 Surveillance Team |
| EPI_ISL_3378044 | LABORATOIRE ACCOLAB SUD OUEST | CNR Virus des Infections Respiratoires - France SUD | Antonin Bal, Gregory Destras, Gwendolyne Burfin, Hadrien Regue, Quentin Semanas, Martine Valette, Bruno Lina, Laurence Josset |
| EPI_ISL_3378045 | Lighthouse Lab in Alderley Park | Wellcome Sanger Institute for the COVID-19 Genomics UK (COG-UK) Consortium | Jacquelyn Wynn, Mairead Hyland, The Lighthouse Lab in Alderley Park and Alex Alderton, Roberto Amato, Jeffrey Barrett, Sonia Goncalves, Ewan Harrison, David K. Jackson, Ian Johnston, Dominic Kwiatkowski, Cordelia Langford, John Sillitoe on behalf of the Wellcome Sanger Institute COVID-19 Surveillance Team |
| EPI_ISL_3378046, EPI_ISL_3378047 | Lighthouse Laboratory Plymouth | Wellcome Sanger Institute for the COVID-19 Genomics UK (COG-UK) Consortium | Lighthouse Laboratory Plymouth and Alex Alderton, Roberto Amato, Jeffrey Barrett, Sonia Goncalves, Ewan Harrison, David K. Jackson, Ian Johnston, Dominic Kwiatkowski, Cordelia Langford, John Sillitoe on behalf of the Wellcome Sanger Institute COVID-19 Surveillance Team |
| EPI_ISL_3378048 | CNR Virus des Infections Respiratoires - France SUD | CNR Virus des Infections Respiratoires - France SUD | Antonin Bal, Gregory Destras, Gwendolyne Burfin, Hadrien Regue, Quentin Semanas, Martine Valette, Bruno Lina, Laurence Josset |
| EPI_ISL_3378049 | Lighthouse Laboratory Plymouth | Wellcome Sanger Institute for the COVID-19 Genomics UK (COG-UK) Consortium | Lighthouse Laboratory Plymouth and Alex Alderton, Roberto Amato, Jeffrey Barrett, Sonia Goncalves, Ewan Harrison, David K. Jackson, Ian Johnston, Dominic Kwiatkowski, Cordelia Langford, John Sillitoe on behalf of the Wellcome Sanger Institute COVID-19 Surveillance Team |
| EPI_ISL_3378050 | Lighthouse Lab in Alderley Park | Wellcome Sanger Institute for the COVID-19 Genomics UK (COG-UK) Consortium | Jacquelyn Wynn, Mairead Hyland, The Lighthouse Lab in Alderley Park and Alex Alderton, Roberto Amato, Jeffrey Barrett, Sonia Goncalves, Ewan Harrison, David K. Jackson, Ian Johnston, Dominic Kwiatkowski, Cordelia Langford, John Sillitoe on behalf of the Wellcome Sanger Institute COVID-19 Surveillance Team |
| EPI_ISL_3378051 | LABORATOIRE ACCOLAB SUD OUEST | CNR Virus des Infections Respiratoires - France SUD | Antonin Bal, Gregory Destras, Gwendolyne Burfin, Hadrien Regue, Quentin Semanas, Martine Valette, Bruno Lina, Laurence Josset |
| EPI_ISL_3378052 | Lighthouse Lab in Alderley Park | Wellcome Sanger Institute for the COVID-19 Genomics UK (COG-UK) Consortium | Jacquelyn Wynn, Mairead Hyland, The Lighthouse Lab in Alderley Park and Alex Alderton, Roberto Amato, Jeffrey Barrett, Sonia Goncalves, Ewan Harrison, David K. Jackson, Ian Johnston, Dominic Kwiatkowski, Cordelia Langford, John Sillitoe on behalf of the Wellcome Sanger Institute COVID-19 Surveillance Team |
| EPI_ISL_3378053 | Lighthouse Laboratory Plymouth | Wellcome Sanger Institute for the COVID-19 Genomics UK (COG-UK) Consortium | Lighthouse Laboratory Plymouth and Alex Alderton, Roberto Amato, Jeffrey Barrett, Sonia Goncalves, Ewan Harrison, David K. Jackson, Ian Johnston, Dominic Kwiatkowski, Cordelia Langford, John Sillitoe on behalf of the Wellcome Sanger Institute COVID-19 Surveillance Team |
| EPI_ISL_3378054 | Lighthouse Lab in Alderley Park | Wellcome Sanger Institute for the COVID-19 Genomics UK (COG-UK) Consortium | Jacquelyn Wynn, Mairead Hyland, The Lighthouse Lab in Alderley Park and Alex Alderton, Roberto Amato, Jeffrey Barrett, Sonia Goncalves, Ewan Harrison, David K. Jackson, Ian Johnston, Dominic Kwiatkowski, Cordelia Langford, John Sillitoe on behalf of the Wellcome Sanger Institute COVID-19 Surveillance Team |
| EPI_ISL_3378055 | LABORATOIRE ACCOLAB SUD OUEST | CNR Virus des Infections Respiratoires - France SUD | Antonin Bal, Gregory Destras, Gwendolyne Burfin, Hadrien Regue, Quentin Semanas, Martine Valette, Bruno Lina, Laurence Josset |
| EPI_ISL_3378056 | Lighthouse Lab in Alderley Park | Wellcome Sanger Institute for the COVID-19 Genomics UK (COG-UK) Consortium | Jacquelyn Wynn, Mairead Hyland, The Lighthouse Lab in Alderley Park and Alex Alderton, Roberto Amato, Jeffrey Barrett, Sonia Goncalves, Ewan Harrison, David K. Jackson, Ian Johnston, Dominic Kwiatkowski, Cordelia Langford, John Sillitoe on behalf of the Wellcome Sanger Institute COVID-19 Surveillance Team |
| EPI_ISL_3378057 | Lighthouse Laboratory Plymouth | Wellcome Sanger Institute for the COVID-19 Genomics UK (COG-UK) Consortium | Lighthouse Laboratory Plymouth and Alex Alderton, Roberto Amato, Jeffrey Barrett, Sonia Goncalves, Ewan Harrison, David K. Jackson, Ian Johnston, Dominic Kwiatkowski, Cordelia Langford, John Sillitoe on behalf of the Wellcome Sanger Institute COVID-19 Surveillance Team |
| EPI_ISL_3378058 | Lighthouse Lab in Alderley Park | Wellcome Sanger Institute for the COVID-19 Genomics UK (COG-UK) Consortium | Jacquelyn Wynn, Mairead Hyland, The Lighthouse Lab in Alderley Park and Alex Alderton, Roberto Amato, Jeffrey Barrett, Sonia Goncalves, Ewan Harrison, David K. Jackson, Ian Johnston, Dominic Kwiatkowski, Cordelia Langford, John Sillitoe on behalf of the Wellcome Sanger Institute COVID-19 Surveillance Team |
| EPI_ISL_3378059 | LABORATOIRE ACCOLAB SUD OUEST | CNR Virus des Infections Respiratoires - France SUD | Antonin Bal, Gregory Destras, Gwendolyne Burfin, Hadrien Regue, Quentin Semanas, Martine Valette, Bruno Lina, Laurence Josset |
| EPI_ISL_3378060 | Lighthouse Lab in Alderley Park | Wellcome Sanger Institute for the COVID-19 Genomics UK (COG-UK) Consortium | Jacquelyn Wynn, Mairead Hyland, The Lighthouse Lab in Alderley Park and Alex Alderton, Roberto Amato, Jeffrey Barrett, Sonia Goncalves, Ewan Harrison, David K. Jackson, Ian Johnston, Dominic Kwiatkowski, Cordelia Langford, John Sillitoe on behalf of the Wellcome Sanger Institute COVID-19 Surveillance Team |
| EPI_ISL_3378061 | Lighthouse Laboratory Plymouth | Wellcome Sanger Institute for the COVID-19 Genomics UK (COG-UK) Consortium | Lighthouse Laboratory Plymouth and Alex Alderton, Roberto Amato, Jeffrey Barrett, Sonia Goncalves, Ewan Harrison, David K. Jackson, Ian Johnston, Dominic Kwiatkowski, Cordelia Langford, John Sillitoe on behalf of the Wellcome Sanger Institute COVID-19 Surveillance Team |
| EPI_ISL_3378062 | Lighthouse Lab in Alderley Park | Wellcome Sanger Institute for the COVID-19 Genomics UK (COG-UK) Consortium | Jacquelyn Wynn, Mairead Hyland, The Lighthouse Lab in Alderley Park and Alex Alderton, Roberto Amato, Jeffrey Barrett, Sonia Goncalves, Ewan Harrison, David K. Jackson, Ian Johnston, Dominic Kwiatkowski, Cordelia Langford, John Sillitoe on behalf of the Wellcome Sanger Institute COVID-19 Surveillance Team |
| EPI_ISL_3378063 | LABORATOIRE ACCOLAB SUD OUEST | CNR Virus des Infections Respiratoires - France SUD | Antonin Bal, Gregory Destras, Gwendolyne Burfin, Hadrien Regue, Quentin Semanas, Martine Valette, Bruno Lina, Laurence Josset |
| EPI_ISL_3378064, EPI_ISL_3378065 | Lighthouse Lab in Alderley Park | Wellcome Sanger Institute for the COVID-19 Genomics UK (COG-UK) Consortium | Jacquelyn Wynn, Mairead Hyland, The Lighthouse Lab in Alderley Park and Alex Alderton, Roberto Amato, Jeffrey Barrett, Sonia Goncalves, Ewan Harrison, David K. Jackson, Ian Johnston, Dominic Kwiatkowski, Cordelia Langford, John Sillitoe on behalf of the Wellcome Sanger Institute COVID-19 Surveillance Team |
| EPI_ISL_3378066 | LABORATOIRE ACCOLAB SUD OUEST | CNR Virus des Infections Respiratoires - France SUD | Antonin Bal, Gregory Destras, Gwendolyne Burfin, Hadrien Regue, Quentin Semanas, Martine Valette, Bruno Lina, Laurence Josset |
| EPI_ISL_3378067, EPI_ISL_3378068 | Lighthouse Laboratory Plymouth | Wellcome Sanger Institute for the COVID-19 Genomics UK (COG-UK) Consortium | Lighthouse Laboratory Plymouth and Alex Alderton, Roberto Amato, Jeffrey Barrett, Sonia Goncalves, Ewan Harrison, David K. Jackson, Ian Johnston, Dominic Kwiatkowski, Cordelia Langford, John Sillitoe on behalf of the Wellcome Sanger Institute COVID-19 Surveillance Team |
| EPI_ISL_3378069 | Lighthouse Lab in Alderley Park | Wellcome Sanger Institute for the COVID-19 Genomics UK (COG-UK) Consortium | Jacquelyn Wynn, Mairead Hyland, The Lighthouse Lab in Alderley Park and Alex Alderton, Roberto Amato, Jeffrey Barrett, Sonia Goncalves, Ewan Harrison, David K. Jackson, Ian Johnston, Dominic Kwiatkowski, Cordelia Langford, John Sillitoe on behalf of the Wellcome Sanger Institute COVID-19 Surveillance Team |
| EPI_ISL_3378070 | LABORATOIRE ACCOLAB SUD OUEST | CNR Virus des Infections Respiratoires - France SUD | Antonin Bal, Gregory Destras, Gwendolyne Burfin, Hadrien Regue, Quentin Semanas, Martine Valette, Bruno Lina, Laurence Josset |
| EPI_ISL_3378071 | Lighthouse Laboratory Plymouth | Wellcome Sanger Institute for the COVID-19 Genomics UK (COG-UK) Consortium | Lighthouse Laboratory Plymouth and Alex Alderton, Roberto Amato, Jeffrey Barrett, Sonia Goncalves, Ewan Harrison, David K. Jackson, Ian Johnston, Dominic Kwiatkowski, Cordelia Langford, John Sillitoe on behalf of the Wellcome Sanger Institute COVID-19 Surveillance Team |

| | | | |
|---|---|---|---|
| EPI_ISL_3378072 | Lighthouse Lab in Alderley Park | Wellcome Sanger Institute for the COVID-19 Genomics UK (COG-UK) Consortium | Jacquelyn Wynn, Mairead Hyland, The Lighthouse Lab in Alderley Park and Alex Alderton, Roberto Amato, Jeffrey Barrett, Sonia Goncalves, Ewan Harrison, David K. Jackson, Ian Johnston, Dominic Kwiatkowski, Cordelia Langford, John Sillitoe on behalf of the Wellcome Sanger Institute COVID-19 Surveillance Team |
| EPI_ISL_3378073 | LABORATOIRE ACCOLAB SUD OUEST | CNR Virus des Infections Respiratoires - France SUD | Antonin Bal, Gregory Destras, Gwendolyne Burfin, Hadrien Regue, Quentin Semanas, Martine Valette, Bruno Lina, Laurence Josset |
| EPI_ISL_3378074 | Lighthouse Laboratory Plymouth | Wellcome Sanger Institute for the COVID-19 Genomics UK (COG-UK) Consortium | Lighthouse Laboratory Plymouth and Alex Alderton, Roberto Amato, Jeffrey Barrett, Sonia Goncalves, Ewan Harrison, David K. Jackson, Ian Johnston, Dominic Kwiatkowski, Cordelia Langford, John Sillitoe on behalf of the Wellcome Sanger Institute COVID-19 Surveillance Team |
| EPI_ISL_3378075 | Lighthouse Lab in Alderley Park | Wellcome Sanger Institute for the COVID-19 Genomics UK (COG-UK) Consortium | Jacquelyn Wynn, Mairead Hyland, The Lighthouse Lab in Alderley Park and Alex Alderton, Roberto Amato, Jeffrey Barrett, Sonia Goncalves, Ewan Harrison, David K. Jackson, Ian Johnston, Dominic Kwiatkowski, Cordelia Langford, John Sillitoe on behalf of the Wellcome Sanger Institute COVID-19 Surveillance Team |
| EPI_ISL_3378076 | Lighthouse Laboratory Plymouth | Wellcome Sanger Institute for the COVID-19 Genomics UK (COG-UK) Consortium | Lighthouse Laboratory Plymouth and Alex Alderton, Roberto Amato, Jeffrey Barrett, Sonia Goncalves, Ewan Harrison, David K. Jackson, Ian Johnston, Dominic Kwiatkowski, Cordelia Langford, John Sillitoe on behalf of the Wellcome Sanger Institute COVID-19 Surveillance Team |
| EPI_ISL_3378077 | Lighthouse Lab in Alderley Park | Wellcome Sanger Institute for the COVID-19 Genomics UK (COG-UK) Consortium | Jacquelyn Wynn, Mairead Hyland, The Lighthouse Lab in Alderley Park and Alex Alderton, Roberto Amato, Jeffrey Barrett, Sonia Goncalves, Ewan Harrison, David K. Jackson, Ian Johnston, Dominic Kwiatkowski, Cordelia Langford, John Sillitoe on behalf of the Wellcome Sanger Institute COVID-19 Surveillance Team |
| EPI_ISL_3378078 | LABORATOIRE ACCOLAB SUD OUEST | CNR Virus des Infections Respiratoires - France SUD | Antonin Bal, Gregory Destras, Gwendolyne Burfin, Hadrien Regue, Quentin Semanas, Martine Valette, Bruno Lina, Laurence Josset |
| EPI_ISL_3378079 | Lighthouse Laboratory Plymouth | Wellcome Sanger Institute for the COVID-19 Genomics UK (COG-UK) Consortium | Lighthouse Laboratory Plymouth and Alex Alderton, Roberto Amato, Jeffrey Barrett, Sonia Goncalves, Ewan Harrison, David K. Jackson, Ian Johnston, Dominic Kwiatkowski, Cordelia Langford, John Sillitoe on behalf of the Wellcome Sanger Institute COVID-19 Surveillance Team |
| EPI_ISL_3378080, EPI_ISL_3378081 | Lighthouse Lab in Alderley Park | Wellcome Sanger Institute for the COVID-19 Genomics UK (COG-UK) Consortium | Jacquelyn Wynn, Mairead Hyland, The Lighthouse Lab in Alderley Park and Alex Alderton, Roberto Amato, Jeffrey Barrett, Sonia Goncalves, Ewan Harrison, David K. Jackson, Ian Johnston, Dominic Kwiatkowski, Cordelia Langford, John Sillitoe on behalf of the Wellcome Sanger Institute COVID-19 Surveillance Team |
| EPI_ISL_3378082 | Lighthouse Laboratory Plymouth | Wellcome Sanger Institute for the COVID-19 Genomics UK (COG-UK) Consortium | Lighthouse Laboratory Plymouth and Alex Alderton, Roberto Amato, Jeffrey Barrett, Sonia Goncalves, Ewan Harrison, David K. Jackson, Ian Johnston, Dominic Kwiatkowski, Cordelia Langford, John Sillitoe on behalf of the Wellcome Sanger Institute COVID-19 Surveillance Team |
| EPI_ISL_3378083 | LABORATOIRE ACCOLAB SUD OUEST | CNR Virus des Infections Respiratoires - France SUD | Antonin Bal, Gregory Destras, Gwendolyne Burfin, Hadrien Regue, Quentin Semanas, Martine Valette, Bruno Lina, Laurence Josset |
| EPI_ISL_3378084, EPI_ISL_3378085 | Lighthouse Lab in Alderley Park | Wellcome Sanger Institute for the COVID-19 Genomics UK (COG-UK) Consortium | Jacquelyn Wynn, Mairead Hyland, The Lighthouse Lab in Alderley Park and Alex Alderton, Roberto Amato, Jeffrey Barrett, Sonia Goncalves, Ewan Harrison, David K. Jackson, Ian Johnston, Dominic Kwiatkowski, Cordelia Langford, John Sillitoe on behalf of the Wellcome Sanger Institute COVID-19 Surveillance Team |
| EPI_ISL_3378086 | Lighthouse Laboratory Plymouth | Wellcome Sanger Institute for the COVID-19 Genomics UK (COG-UK) Consortium | Lighthouse Laboratory Plymouth and Alex Alderton, Roberto Amato, Jeffrey Barrett, Sonia Goncalves, Ewan Harrison, David K. Jackson, Ian Johnston, Dominic Kwiatkowski, Cordelia Langford, John Sillitoe on behalf of the Wellcome Sanger Institute COVID-19 Surveillance Team |
| EPI_ISL_3378087 | CNR Virus des Infections Respiratoires - France SUD | CNR Virus des Infections Respiratoires - France SUD | Antonin Bal, Gregory Destras, Gwendolyne Burfin, Hadrien Regue, Quentin Semanas, Martine Valette, Bruno Lina, Laurence Josset |
| EPI_ISL_3378088, EPI_ISL_3378089, EPI_ISL_3378090 | Lighthouse Lab in Alderley Park | Wellcome Sanger Institute for the COVID-19 Genomics UK (COG-UK) Consortium | Jacquelyn Wynn, Mairead Hyland, The Lighthouse Lab in Alderley Park and Alex Alderton, Roberto Amato, Jeffrey Barrett, Sonia Goncalves, Ewan Harrison, David K. Jackson, Ian Johnston, Dominic Kwiatkowski, Cordelia Langford, John Sillitoe on behalf of the Wellcome Sanger Institute COVID-19 Surveillance Team |
| EPI_ISL_3378091 | CNR Virus des Infections Respiratoires - France SUD | CNR Virus des Infections Respiratoires - France SUD | Antonin Bal, Gregory Destras, Gwendolyne Burfin, Hadrien Regue, Quentin Semanas, Martine Valette, Bruno Lina, Laurence Josset |
| EPI_ISL_3378092, EPI_ISL_3378093 | Lighthouse Lab in Alderley Park | Wellcome Sanger Institute for the COVID-19 Genomics UK (COG-UK) Consortium | Jacquelyn Wynn, Mairead Hyland, The Lighthouse Lab in Alderley Park and Alex Alderton, Roberto Amato, Jeffrey Barrett, Sonia Goncalves, Ewan Harrison, David K. Jackson, Ian Johnston, Dominic Kwiatkowski, Cordelia Langford, John Sillitoe on behalf of the Wellcome Sanger Institute COVID-19 Surveillance Team |
| EPI_ISL_3378094 | LABORATOIRE DE L'OSPEDALE - 2A 2B | CNR Virus des Infections Respiratoires - France SUD | Antonin Bal, Gregory Destras, Gwendolyne Burfin, Hadrien Regue, Quentin Semanas, Martine Valette, Bruno Lina, Laurence Josset |
| EPI_ISL_3378095 | Lighthouse Laboratory Plymouth | Wellcome Sanger Institute for the COVID-19 Genomics UK (COG-UK) Consortium | Lighthouse Laboratory Plymouth and Alex Alderton, Roberto Amato, Jeffrey Barrett, Sonia Goncalves, Ewan Harrison, David K. Jackson, Ian Johnston, Dominic Kwiatkowski, Cordelia Langford, John Sillitoe on behalf of the Wellcome Sanger Institute COVID-19 Surveillance Team |
| EPI_ISL_3378096, EPI_ISL_3378097 | Lighthouse Lab in Alderley Park | Wellcome Sanger Institute for the COVID-19 Genomics UK (COG-UK) Consortium | Jacquelyn Wynn, Mairead Hyland, The Lighthouse Lab in Alderley Park and Alex Alderton, Roberto Amato, Jeffrey Barrett, Sonia Goncalves, Ewan Harrison, David K. Jackson, Ian Johnston, Dominic Kwiatkowski, Cordelia Langford, John Sillitoe on behalf of the Wellcome Sanger Institute COVID-19 Surveillance Team |
| EPI_ISL_3378098 | Lighthouse Laboratory Plymouth | Wellcome Sanger Institute for the COVID-19 Genomics UK (COG-UK) Consortium | Lighthouse Laboratory Plymouth and Alex Alderton, Roberto Amato, Jeffrey Barrett, Sonia Goncalves, Ewan Harrison, David K. Jackson, Ian Johnston, Dominic Kwiatkowski, Cordelia Langford, John Sillitoe on behalf of the Wellcome Sanger Institute COVID-19 Surveillance Team |
| EPI_ISL_3378099 | LABORATOIRE DE L'OSPEDALE - 2A 2B | CNR Virus des Infections Respiratoires - France SUD | Antonin Bal, Gregory Destras, Gwendolyne Burfin, Hadrien Regue, Quentin Semanas, Martine Valette, Bruno Lina, Laurence Josset |
| EPI_ISL_3378100, EPI_ISL_3378101, EPI_ISL_3378102 | Lighthouse Lab in Alderley Park | Wellcome Sanger Institute for the COVID-19 Genomics UK (COG-UK) Consortium | Jacquelyn Wynn, Mairead Hyland, The Lighthouse Lab in Alderley Park and Alex Alderton, Roberto Amato, Jeffrey Barrett, Sonia Goncalves, Ewan Harrison, David K. Jackson, Ian Johnston, Dominic Kwiatkowski, Cordelia Langford, John Sillitoe on behalf of the Wellcome Sanger Institute COVID-19 Surveillance Team |
| EPI_ISL_3378103 | LAM DE BIO HD | CNR Virus des Infections Respiratoires - France SUD | Antonin Bal, Gregory Destras, Gwendolyne Burfin, Hadrien Regue, Quentin Semanas, Martine Valette, Bruno Lina, Laurence Josset |
| EPI_ISL_3378104 | Lighthouse Laboratory Plymouth | Wellcome Sanger Institute for the COVID-19 Genomics UK (COG-UK) Consortium | Lighthouse Laboratory Plymouth and Alex Alderton, Roberto Amato, Jeffrey Barrett, Sonia Goncalves, Ewan Harrison, David K. Jackson, Ian Johnston, Dominic Kwiatkowski, Cordelia Langford, John Sillitoe on behalf of the Wellcome Sanger Institute COVID-19 Surveillance Team |
| EPI_ISL_3378105, EPI_ISL_3378106 | Lighthouse Lab in Alderley Park | Wellcome Sanger Institute for the COVID-19 Genomics UK (COG-UK) Consortium | Jacquelyn Wynn, Mairead Hyland, The Lighthouse Lab in Alderley Park and Alex Alderton, Roberto Amato, Jeffrey Barrett, Sonia Goncalves, Ewan Harrison, David K. Jackson, Ian Johnston, Dominic Kwiatkowski, Cordelia Langford, John Sillitoe on behalf of the Wellcome Sanger Institute COVID-19 Surveillance Team |
| EPI_ISL_3378107 | LAM DE BIO HD | CNR Virus des Infections Respiratoires - France SUD | Antonin Bal, Gregory Destras, Gwendolyne Burfin, Hadrien Regue, Quentin Semanas, Martine Valette, Bruno Lina, Laurence Josset |
| EPI_ISL_3378108 | Lighthouse Lab in Alderley Park | Wellcome Sanger Institute for the COVID-19 Genomics UK (COG-UK) Consortium | Jacquelyn Wynn, Mairead Hyland, The Lighthouse Lab in Alderley Park and Alex Alderton, Roberto Amato, Jeffrey Barrett, Sonia Goncalves, Ewan Harrison, David K. Jackson, Ian Johnston, Dominic Kwiatkowski, Cordelia Langford, John Sillitoe on behalf of the Wellcome Sanger Institute COVID-19 Surveillance Team |
| EPI_ISL_3378109, EPI_ISL_3378110 | Lighthouse Laboratory Plymouth | Wellcome Sanger Institute for the COVID-19 Genomics UK (COG-UK) Consortium | Lighthouse Laboratory Plymouth and Alex Alderton, Roberto Amato, Jeffrey Barrett, Sonia Goncalves, Ewan Harrison, David K. Jackson, Ian Johnston, Dominic Kwiatkowski, Cordelia Langford, John Sillitoe on behalf of the Wellcome Sanger Institute COVID-19 Surveillance Team |
| EPI_ISL_3378111 | CNR Virus des Infections Respiratoires - France SUD | CNR Virus des Infections Respiratoires - France SUD | Antonin Bal, Gregory Destras, Gwendolyne Burfin, Hadrien Regue, Quentin Semanas, Martine Valette, Bruno Lina, Laurence Josset |
| EPI_ISL_3378112, EPI_ISL_3378113, EPI_ISL_3378114 | Lighthouse Lab in Alderley Park | Wellcome Sanger Institute for the COVID-19 Genomics UK (COG-UK) Consortium | Jacquelyn Wynn, Mairead Hyland, The Lighthouse Lab in Alderley Park and Alex Alderton, Roberto Amato, Jeffrey Barrett, Sonia Goncalves, Ewan Harrison, David K. Jackson, Ian Johnston, Dominic Kwiatkowski, Cordelia Langford, John Sillitoe on behalf of the Wellcome Sanger Institute COVID-19 Surveillance Team |
| EPI_ISL_3378115 | LAM DE BIO HD | CNR Virus des Infections Respiratoires - France SUD | Antonin Bal, Gregory Destras, Gwendolyne Burfin, Hadrien Regue, Quentin Semanas, Martine Valette, Bruno Lina, Laurence Josset |
| EPI_ISL_3378116, EPI_ISL_3378117, EPI_ISL_3378118 | Lighthouse Lab in Alderley Park | Wellcome Sanger Institute for the COVID-19 Genomics UK (COG-UK) Consortium | Jacquelyn Wynn, Mairead Hyland, The Lighthouse Lab in Alderley Park and Alex Alderton, Roberto Amato, Jeffrey Barrett, Sonia Goncalves, Ewan Harrison, David K. Jackson, Ian Johnston, Dominic Kwiatkowski, Cordelia Langford, John Sillitoe on behalf of the Wellcome Sanger Institute COVID-19 Surveillance Team |
| EPI_ISL_3378119 | LAM DE BIO HD | CNR Virus des Infections Respiratoires - France SUD | Antonin Bal, Gregory Destras, Gwendolyne Burfin, Hadrien Regue, Quentin Semanas, Martine Valette, Bruno Lina, Laurence Josset |
| EPI_ISL_3378120, EPI_ISL_3378121, EPI_ISL_3378122 | Lighthouse Lab in Alderley Park | Wellcome Sanger Institute for the COVID-19 Genomics UK (COG-UK) Consortium | Jacquelyn Wynn, Mairead Hyland, The Lighthouse Lab in Alderley Park and Alex Alderton, Roberto Amato, Jeffrey Barrett, Sonia Goncalves, Ewan Harrison, David K. Jackson, Ian Johnston, Dominic Kwiatkowski, Cordelia Langford, John Sillitoe on behalf of the Wellcome Sanger Institute COVID-19 Surveillance Team |
| EPI_ISL_3378123 | LAM DE BIO HD | CNR Virus des Infections Respiratoires - France SUD | Antonin Bal, Gregory Destras, Gwendolyne Burfin, Hadrien Regue, Quentin Semanas, Martine Valette, Bruno Lina, Laurence Josset |
| EPI_ISL_3378124, EPI_ISL_3378125, EPI_ISL_3378127, EPI_ISL_3378128, EPI_ISL_3378129, EPI_ISL_3378130 | Lighthouse Lab in Alderley Park | Wellcome Sanger Institute for the COVID-19 Genomics UK (COG-UK) Consortium | Jacquelyn Wynn, Mairead Hyland, The Lighthouse Lab in Alderley Park and Alex Alderton, Roberto Amato, Jeffrey Barrett, Sonia Goncalves, Ewan Harrison, David K. Jackson, Ian Johnston, Dominic Kwiatkowski, Cordelia Langford, John Sillitoe on behalf of the Wellcome Sanger Institute COVID-19 Surveillance Team |
| EPI_ISL_3378131 | DYOMEDEA-LABORATOIRE DE LA SAUVEGARDE | CNR Virus des Infections Respiratoires - France SUD | Antonin Bal, Gregory Destras, Gwendolyne Burfin, Hadrien Regue, Quentin Semanas, Martine Valette, Bruno Lina, Laurence Josset |
| EPI_ISL_3378132 | Lighthouse Lab in Alderley Park | Wellcome Sanger Institute for the COVID-19 Genomics UK (COG-UK) Consortium | Jacquelyn Wynn, Mairead Hyland, The Lighthouse Lab in Alderley Park and Alex Alderton, Roberto Amato, Jeffrey Barrett, Sonia Goncalves, Ewan Harrison, David K. Jackson, Ian Johnston, Dominic Kwiatkowski, Cordelia Langford, John Sillitoe on behalf of the Wellcome Sanger Institute COVID-19 Surveillance Team |
| EPI_ISL_3378133 | Lighthouse Laboratory Plymouth | Wellcome Sanger Institute for the COVID-19 Genomics UK (COG-UK) Consortium | Lighthouse Laboratory Plymouth and Alex Alderton, Roberto Amato, Jeffrey Barrett, Sonia Goncalves, Ewan Harrison, David K. Jackson, Ian Johnston, Dominic Kwiatkowski, Cordelia Langford, John Sillitoe on behalf of the Wellcome Sanger Institute COVID-19 Surveillance Team |
| EPI_ISL_3378134 | Lighthouse Lab in Alderley Park | Wellcome Sanger Institute for the COVID-19 Genomics UK (COG-UK) Consortium | Jacquelyn Wynn, Mairead Hyland, The Lighthouse Lab in Alderley Park and Alex Alderton, Roberto Amato, Jeffrey Barrett, Sonia Goncalves, Ewan Harrison, David K. Jackson, Ian Johnston, Dominic Kwiatkowski, Cordelia Langford, John Sillitoe on behalf of the Wellcome Sanger Institute COVID-19 Surveillance Team |
| EPI_ISL_3378135 | CH GIVORS | CNR Virus des Infections Respiratoires - France SUD | Antonin Bal, Gregory Destras, Gwendolyne Burfin, Hadrien Regue, Quentin Semanas, Martine Valette, Bruno Lina, Laurence Josset |
| EPI_ISL_3378136 | Lighthouse Laboratory Plymouth | Wellcome Sanger Institute for the COVID-19 Genomics UK (COG-UK) Consortium | Lighthouse Laboratory Plymouth and Alex Alderton, Roberto Amato, Jeffrey Barrett, Sonia Goncalves, Ewan Harrison, David K. Jackson, Ian Johnston, Dominic Kwiatkowski, Cordelia Langford, John Sillitoe on behalf of the Wellcome Sanger Institute COVID-19 Surveillance Team |
| EPI_ISL_3378137 | Lighthouse Lab in Alderley Park | Wellcome Sanger Institute for the COVID-19 Genomics UK (COG-UK) Consortium | Jacquelyn Wynn, Mairead Hyland, The Lighthouse Lab in Alderley Park and Alex Alderton, Roberto Amato, Jeffrey Barrett, Sonia Goncalves, Ewan Harrison, David K. Jackson, Ian Johnston, Dominic Kwiatkowski, Cordelia Langford, John Sillitoe on behalf of the Wellcome Sanger Institute COVID-19 Surveillance Team |
| EPI_ISL_3378138 | Lighthouse Laboratory Plymouth | Wellcome Sanger Institute for the COVID-19 Genomics UK (COG-UK) Consortium | Lighthouse Laboratory Plymouth and Alex Alderton, Roberto Amato, Jeffrey Barrett, Sonia Goncalves, Ewan Harrison, David K. Jackson, Ian Johnston, Dominic Kwiatkowski, Cordelia Langford, John Sillitoe on behalf of the Wellcome Sanger Institute COVID-19 Surveillance Team |
| EPI_ISL_3378139 | CNR Virus des Infections Respiratoires - France SUD | CNR Virus des Infections Respiratoires - France SUD | Antonin Bal, Gregory Destras, Gwendolyne Burfin, Hadrien Regue, Quentin Semanas, Martine Valette, Bruno Lina, Laurence Josset |
| EPI_ISL_3378140, EPI_ISL_3378141, EPI_ISL_3378142 | Lighthouse Lab in Alderley Park | Wellcome Sanger Institute for the COVID-19 Genomics UK (COG-UK) Consortium | Jacquelyn Wynn, Mairead Hyland, The Lighthouse Lab in Alderley Park and Alex Alderton, Roberto Amato, Jeffrey Barrett, Sonia Goncalves, Ewan Harrison, David K. Jackson, Ian Johnston, Dominic Kwiatkowski, Cordelia Langford, John Sillitoe on behalf of the Wellcome Sanger Institute COVID-19 Surveillance Team |

| | | | |
|---|---|---|---|
| EPI_ISL_3378143 | CNR Virus des Infections Respiratoires - France SUD | CNR Virus des Infections Respiratoires - France SUD | Antonin Bal, Gregory Destras, Gwendolyne Burfin, Hadrien Regue, Quentin Semanas, Martine Valette, Bruno Lina, Laurence Josset |
| EPI_ISL_3378144, EPI_ISL_3378145, EPI_ISL_3378146 | Lighthouse Lab in Alderley Park | Wellcome Sanger Institute for the COVID-19 Genomics UK (COG-UK) Consortium | Jacquelyn Wynn, Mairead Hyland, The Lighthouse Lab in Alderley Park and Alex Alderton, Roberto Amato, Jeffrey Barrett, Sonia Goncalves, Ewan Harrison, David K. Jackson, Ian Johnston, Dominic Kwiatkowski, Cordelia Langford, John Sillitoe on behalf of the Wellcome Sanger Institute COVID-19 Surveillance Team |
| EPI_ISL_3378147 | CNR Virus des Infections Respiratoires - France SUD | CNR Virus des Infections Respiratoires - France SUD | Antonin Bal, Gregory Destras, Gwendolyne Burfin, Hadrien Regue, Quentin Semanas, Martine Valette, Bruno Lina, Laurence Josset |
| EPI_ISL_3378148, EPI_ISL_3378149 | Lighthouse Laboratory Plymouth | Wellcome Sanger Institute for the COVID-19 Genomics UK (COG-UK) Consortium | Lighthouse Laboratory Plymouth and Alex Alderton, Roberto Amato, Jeffrey Barrett, Sonia Goncalves, Ewan Harrison, David K. Jackson, Ian Johnston, Dominic Kwiatkowski, Cordelia Langford, John Sillitoe on behalf of the Wellcome Sanger Institute COVID-19 Surveillance Team |
| EPI_ISL_3378150 | Lighthouse Lab in Alderley Park | Wellcome Sanger Institute for the COVID-19 Genomics UK (COG-UK) Consortium | Jacquelyn Wynn, Mairead Hyland, The Lighthouse Lab in Alderley Park and Alex Alderton, Roberto Amato, Jeffrey Barrett, Sonia Goncalves, Ewan Harrison, David K. Jackson, Ian Johnston, Dominic Kwiatkowski, Cordelia Langford, John Sillitoe on behalf of the Wellcome Sanger Institute COVID-19 Surveillance Team |
| EPI_ISL_3378151 | LABORATOIRE UNILIANS DECINES | CNR Virus des Infections Respiratoires - France SUD | Antonin Bal, Gregory Destras, Gwendolyne Burfin, Hadrien Regue, Quentin Semanas, Martine Valette, Bruno Lina, Laurence Josset |
| EPI_ISL_3378152 | Lighthouse Laboratory Plymouth | Wellcome Sanger Institute for the COVID-19 Genomics UK (COG-UK) Consortium | Lighthouse Laboratory Plymouth and Alex Alderton, Roberto Amato, Jeffrey Barrett, Sonia Goncalves, Ewan Harrison, David K. Jackson, Ian Johnston, Dominic Kwiatkowski, Cordelia Langford, John Sillitoe on behalf of the Wellcome Sanger Institute COVID-19 Surveillance Team |
| EPI_ISL_3378153, EPI_ISL_3378154 | Lighthouse Lab in Alderley Park | Wellcome Sanger Institute for the COVID-19 Genomics UK (COG-UK) Consortium | Jacquelyn Wynn, Mairead Hyland, The Lighthouse Lab in Alderley Park and Alex Alderton, Roberto Amato, Jeffrey Barrett, Sonia Goncalves, Ewan Harrison, David K. Jackson, Ian Johnston, Dominic Kwiatkowski, Cordelia Langford, John Sillitoe on behalf of the Wellcome Sanger Institute COVID-19 Surveillance Team |
| EPI_ISL_3378155 | LABORATOIRE UNILIANS DECINES | CNR Virus des Infections Respiratoires - France SUD | Antonin Bal, Gregory Destras, Gwendolyne Burfin, Hadrien Regue, Quentin Semanas, Martine Valette, Bruno Lina, Laurence Josset |
| EPI_ISL_3378156, EPI_ISL_3378157, EPI_ISL_3378158 | Lighthouse Laboratory Plymouth | Wellcome Sanger Institute for the COVID-19 Genomics UK (COG-UK) Consortium | Lighthouse Laboratory Plymouth and Alex Alderton, Roberto Amato, Jeffrey Barrett, Sonia Goncalves, Ewan Harrison, David K. Jackson, Ian Johnston, Dominic Kwiatkowski, Cordelia Langford, John Sillitoe on behalf of the Wellcome Sanger Institute COVID-19 Surveillance Team |
| EPI_ISL_3378159 | LABORATOIRE UNILIANS DECINES | CNR Virus des Infections Respiratoires - France SUD | Antonin Bal, Gregory Destras, Gwendolyne Burfin, Hadrien Regue, Quentin Semanas, Martine Valette, Bruno Lina, Laurence Josset |
| EPI_ISL_3378160, EPI_ISL_3378161 | Lighthouse Lab in Alderley Park | Wellcome Sanger Institute for the COVID-19 Genomics UK (COG-UK) Consortium | Jacquelyn Wynn, Mairead Hyland, The Lighthouse Lab in Alderley Park and Alex Alderton, Roberto Amato, Jeffrey Barrett, Sonia Goncalves, Ewan Harrison, David K. Jackson, Ian Johnston, Dominic Kwiatkowski, Cordelia Langford, John Sillitoe on behalf of the Wellcome Sanger Institute COVID-19 Surveillance Team |
| EPI_ISL_3378162 | Lighthouse Laboratory Plymouth | Wellcome Sanger Institute for the COVID-19 Genomics UK (COG-UK) Consortium | Lighthouse Laboratory Plymouth and Alex Alderton, Roberto Amato, Jeffrey Barrett, Sonia Goncalves, Ewan Harrison, David K. Jackson, Ian Johnston, Dominic Kwiatkowski, Cordelia Langford, John Sillitoe on behalf of the Wellcome Sanger Institute COVID-19 Surveillance Team |
| EPI_ISL_3378163 | LABORATOIRE UNILIANS DECINES | CNR Virus des Infections Respiratoires - France SUD | Antonin Bal, Gregory Destras, Gwendolyne Burfin, Hadrien Regue, Quentin Semanas, Martine Valette, Bruno Lina, Laurence Josset |
| EPI_ISL_3378164, EPI_ISL_3378165 | Lighthouse Lab in Alderley Park | Wellcome Sanger Institute for the COVID-19 Genomics UK (COG-UK) Consortium | Jacquelyn Wynn, Mairead Hyland, The Lighthouse Lab in Alderley Park and Alex Alderton, Roberto Amato, Jeffrey Barrett, Sonia Goncalves, Ewan Harrison, David K. Jackson, Ian Johnston, Dominic Kwiatkowski, Cordelia Langford, John Sillitoe on behalf of the Wellcome Sanger Institute COVID-19 Surveillance Team |
| EPI_ISL_3378166 | LABORATOIRE UNILIANS DECINES | CNR Virus des Infections Respiratoires - France SUD | Antonin Bal, Gregory Destras, Gwendolyne Burfin, Hadrien Regue, Quentin Semanas, Martine Valette, Bruno Lina, Laurence Josset |
| EPI_ISL_3378167 | Lighthouse Lab in Alderley Park | Wellcome Sanger Institute for the COVID-19 Genomics UK (COG-UK) Consortium | Jacquelyn Wynn, Mairead Hyland, The Lighthouse Lab in Alderley Park and Alex Alderton, Roberto Amato, Jeffrey Barrett, Sonia Goncalves, Ewan Harrison, David K. Jackson, Ian Johnston, Dominic Kwiatkowski, Cordelia Langford, John Sillitoe on behalf of the Wellcome Sanger Institute COVID-19 Surveillance Team |
| EPI_ISL_3378168 | Lighthouse Laboratory Plymouth | Wellcome Sanger Institute for the COVID-19 Genomics UK (COG-UK) Consortium | Lighthouse Laboratory Plymouth and Alex Alderton, Roberto Amato, Jeffrey Barrett, Sonia Goncalves, Ewan Harrison, David K. Jackson, Ian Johnston, Dominic Kwiatkowski, Cordelia Langford, John Sillitoe on behalf of the Wellcome Sanger Institute COVID-19 Surveillance Team |
| EPI_ISL_3378169 | Lighthouse Lab in Alderley Park | Wellcome Sanger Institute for the COVID-19 Genomics UK (COG-UK) Consortium | Jacquelyn Wynn, Mairead Hyland, The Lighthouse Lab in Alderley Park and Alex Alderton, Roberto Amato, Jeffrey Barrett, Sonia Goncalves, Ewan Harrison, David K. Jackson, Ian Johnston, Dominic Kwiatkowski, Cordelia Langford, John Sillitoe on behalf of the Wellcome Sanger Institute COVID-19 Surveillance Team |
| EPI_ISL_3378170 | LABORATOIRE UNILIANS DECINES | CNR Virus des Infections Respiratoires - France SUD | Antonin Bal, Gregory Destras, Gwendolyne Burfin, Hadrien Regue, Quentin Semanas, Martine Valette, Bruno Lina, Laurence Josset |
| EPI_ISL_3378171, EPI_ISL_3378172, EPI_ISL_3378173 | Lighthouse Lab in Alderley Park | Wellcome Sanger Institute for the COVID-19 Genomics UK (COG-UK) Consortium | Jacquelyn Wynn, Mairead Hyland, The Lighthouse Lab in Alderley Park and Alex Alderton, Roberto Amato, Jeffrey Barrett, Sonia Goncalves, Ewan Harrison, David K. Jackson, Ian Johnston, Dominic Kwiatkowski, Cordelia Langford, John Sillitoe on behalf of the Wellcome Sanger Institute COVID-19 Surveillance Team |
| EPI_ISL_3378174 | Lighthouse Laboratory Plymouth | Wellcome Sanger Institute for the COVID-19 Genomics UK (COG-UK) Consortium | Lighthouse Laboratory Plymouth and Alex Alderton, Roberto Amato, Jeffrey Barrett, Sonia Goncalves, Ewan Harrison, David K. Jackson, Ian Johnston, Dominic Kwiatkowski, Cordelia Langford, John Sillitoe on behalf of the Wellcome Sanger Institute COVID-19 Surveillance Team |
| EPI_ISL_3378175 | LABORATOIRE UNILIANS DECINES | CNR Virus des Infections Respiratoires - France SUD | Antonin Bal, Gregory Destras, Gwendolyne Burfin, Hadrien Regue, Quentin Semanas, Martine Valette, Bruno Lina, Laurence Josset |
| EPI_ISL_3378176 | Lighthouse Lab in Alderley Park | Wellcome Sanger Institute for the COVID-19 Genomics UK (COG-UK) Consortium | Jacquelyn Wynn, Mairead Hyland, The Lighthouse Lab in Alderley Park and Alex Alderton, Roberto Amato, Jeffrey Barrett, Sonia Goncalves, Ewan Harrison, David K. Jackson, Ian Johnston, Dominic Kwiatkowski, Cordelia Langford, John Sillitoe on behalf of the Wellcome Sanger Institute COVID-19 Surveillance Team |
| EPI_ISL_3378177, EPI_ISL_3378178 | Berkshire and Surrey Pathology Services Lighthouse Laboratory | Wellcome Sanger Institute for the COVID-19 Genomics UK (COG-UK) Consortium | Berkshire and Surrey Pathology Services Lighthouse Laboratory and Alex Alderton, Roberto Amato, Jeffrey Barrett, Sonia Goncalves, Ewan Harrison, David K. Jackson, Ian Johnston, Dominic Kwiatkowski, Cordelia Langford, John Sillitoe on behalf of the Wellcome Sanger Institute COVID-19 Surveillance Team |
| EPI_ISL_3378179 | LABORATOIRE UNILIANS DECINES | CNR Virus des Infections Respiratoires - France SUD | Antonin Bal, Gregory Destras, Gwendolyne Burfin, Hadrien Regue, Quentin Semanas, Martine Valette, Bruno Lina, Laurence Josset |
| EPI_ISL_3378180 | Lighthouse Lab in Alderley Park | Wellcome Sanger Institute for the COVID-19 Genomics UK (COG-UK) Consortium | Jacquelyn Wynn, Mairead Hyland, The Lighthouse Lab in Alderley Park and Alex Alderton, Roberto Amato, Jeffrey Barrett, Sonia Goncalves, Ewan Harrison, David K. Jackson, Ian Johnston, Dominic Kwiatkowski, Cordelia Langford, John Sillitoe on behalf of the Wellcome Sanger Institute COVID-19 Surveillance Team |
| EPI_ISL_3378181 | Berkshire and Surrey Pathology Services Lighthouse Laboratory | Wellcome Sanger Institute for the COVID-19 Genomics UK (COG-UK) Consortium | Berkshire and Surrey Pathology Services Lighthouse Laboratory and Alex Alderton, Roberto Amato, Jeffrey Barrett, Sonia Goncalves, Ewan Harrison, David K. Jackson, Ian Johnston, Dominic Kwiatkowski, Cordelia Langford, John Sillitoe on behalf of the Wellcome Sanger Institute COVID-19 Surveillance Team |
| EPI_ISL_3378182 | Lighthouse Lab in Milton Keynes | Wellcome Sanger Institute for the COVID-19 Genomics UK (COG-UK) Consortium | The Lighthouse Lab in Milton Keynes and Alex Alderton, Roberto Amato, Jeffrey Barrett, Sonia Goncalves, Ewan Harrison, David K. Jackson, Ian Johnston, Dominic Kwiatkowski, Cordelia Langford, John Sillitoe on behalf of the Wellcome Sanger Institute COVID-19 Surveillance Team |
| EPI_ISL_3378183 | LABORATOIRE UNILIANS DECINES | CNR Virus des Infections Respiratoires - France SUD | Antonin Bal, Gregory Destras, Gwendolyne Burfin, Hadrien Regue, Quentin Semanas, Martine Valette, Bruno Lina, Laurence Josset |
| EPI_ISL_3378184, EPI_ISL_3378185 | Lighthouse Lab in Milton Keynes | Wellcome Sanger Institute for the COVID-19 Genomics UK (COG-UK) Consortium | The Lighthouse Lab in Milton Keynes and Alex Alderton, Roberto Amato, Jeffrey Barrett, Sonia Goncalves, Ewan Harrison, David K. Jackson, Ian Johnston, Dominic Kwiatkowski, Cordelia Langford, John Sillitoe on behalf of the Wellcome Sanger Institute COVID-19 Surveillance Team |
| EPI_ISL_3378186 | Lighthouse Lab in Alderley Park | Wellcome Sanger Institute for the COVID-19 Genomics UK (COG-UK) Consortium | Jacquelyn Wynn, Mairead Hyland, The Lighthouse Lab in Alderley Park and Alex Alderton, Roberto Amato, Jeffrey Barrett, Sonia Goncalves, Ewan Harrison, David K. Jackson, Ian Johnston, Dominic Kwiatkowski, Cordelia Langford, John Sillitoe on behalf of the Wellcome Sanger Institute COVID-19 Surveillance Team |
| EPI_ISL_3378187 | LABORATOIRE UNILIANS DECINES | CNR Virus des Infections Respiratoires - France SUD | Antonin Bal, Gregory Destras, Gwendolyne Burfin, Hadrien Regue, Quentin Semanas, Martine Valette, Bruno Lina, Laurence Josset |
| EPI_ISL_3378188, EPI_ISL_3378189, EPI_ISL_3378190 | Lighthouse Lab in Milton Keynes | Wellcome Sanger Institute for the COVID-19 Genomics UK (COG-UK) Consortium | The Lighthouse Lab in Milton Keynes and Alex Alderton, Roberto Amato, Jeffrey Barrett, Sonia Goncalves, Ewan Harrison, David K. Jackson, Ian Johnston, Dominic Kwiatkowski, Cordelia Langford, John Sillitoe on behalf of the Wellcome Sanger Institute COVID-19 Surveillance Team |
| EPI_ISL_3378191 | LABORATOIRE UNILIANS DECINES | CNR Virus des Infections Respiratoires - France SUD | Antonin Bal, Gregory Destras, Gwendolyne Burfin, Hadrien Regue, Quentin Semanas, Martine Valette, Bruno Lina, Laurence Josset |
| EPI_ISL_3378192 | Berkshire and Surrey Pathology Services Lighthouse Laboratory | Wellcome Sanger Institute for the COVID-19 Genomics UK (COG-UK) Consortium | Berkshire and Surrey Pathology Services Lighthouse Laboratory and Alex Alderton, Roberto Amato, Jeffrey Barrett, Sonia Goncalves, Ewan Harrison, David K. Jackson, Ian Johnston, Dominic Kwiatkowski, Cordelia Langford, John Sillitoe on behalf of the Wellcome Sanger Institute COVID-19 Surveillance Team |
| EPI_ISL_3378193 | Lighthouse Lab in Alderley Park | Wellcome Sanger Institute for the COVID-19 Genomics UK (COG-UK) Consortium | Jacquelyn Wynn, Mairead Hyland, The Lighthouse Lab in Alderley Park and Alex Alderton, Roberto Amato, Jeffrey Barrett, Sonia Goncalves, Ewan Harrison, David K. Jackson, Ian Johnston, Dominic Kwiatkowski, Cordelia Langford, John Sillitoe on behalf of the Wellcome Sanger Institute COVID-19 Surveillance Team |
| EPI_ISL_3378194 | Berkshire and Surrey Pathology Services Lighthouse Laboratory | Wellcome Sanger Institute for the COVID-19 Genomics UK (COG-UK) Consortium | Berkshire and Surrey Pathology Services Lighthouse Laboratory and Alex Alderton, Roberto Amato, Jeffrey Barrett, Sonia Goncalves, Ewan Harrison, David K. Jackson, Ian Johnston, Dominic Kwiatkowski, Cordelia Langford, John Sillitoe on behalf of the Wellcome Sanger Institute COVID-19 Surveillance Team |
| EPI_ISL_3378195 | LABORATOIRE UNILIANS DECINES | CNR Virus des Infections Respiratoires - France SUD | Antonin Bal, Gregory Destras, Gwendolyne Burfin, Hadrien Regue, Quentin Semanas, Martine Valette, Bruno Lina, Laurence Josset |
| EPI_ISL_3378196, EPI_ISL_3378197, EPI_ISL_3378198 | Berkshire and Surrey Pathology Services Lighthouse Laboratory | Wellcome Sanger Institute for the COVID-19 Genomics UK (COG-UK) Consortium | Berkshire and Surrey Pathology Services Lighthouse Laboratory and Alex Alderton, Roberto Amato, Jeffrey Barrett, Sonia Goncalves, Ewan Harrison, David K. Jackson, Ian Johnston, Dominic Kwiatkowski, Cordelia Langford, John Sillitoe on behalf of the Wellcome Sanger Institute COVID-19 Surveillance Team |
| EPI_ISL_3378199 | LABORATOIRE UNILIANS DECINES | CNR Virus des Infections Respiratoires - France SUD | Antonin Bal, Gregory Destras, Gwendolyne Burfin, Hadrien Regue, Quentin Semanas, Martine Valette, Bruno Lina, Laurence Josset |
| EPI_ISL_3378200, EPI_ISL_3378201 | Lighthouse Lab in Milton Keynes | Wellcome Sanger Institute for the COVID-19 Genomics UK (COG-UK) Consortium | The Lighthouse Lab in Milton Keynes and Alex Alderton, Roberto Amato, Jeffrey Barrett, Sonia Goncalves, Ewan Harrison, David K. Jackson, Ian Johnston, Dominic Kwiatkowski, Cordelia Langford, John Sillitoe on behalf of the Wellcome Sanger Institute COVID-19 Surveillance Team |
| EPI_ISL_3378202 | LABORATOIRE UNILIANS DECINES | CNR Virus des Infections Respiratoires - France SUD | Antonin Bal, Gregory Destras, Gwendolyne Burfin, Hadrien Regue, Quentin Semanas, Martine Valette, Bruno Lina, Laurence Josset |
| EPI_ISL_3378203 | Lighthouse Lab in Milton Keynes | Wellcome Sanger Institute for the COVID-19 Genomics UK (COG-UK) Consortium | The Lighthouse Lab in Milton Keynes and Alex Alderton, Roberto Amato, Jeffrey Barrett, Sonia Goncalves, Ewan Harrison, David K. Jackson, Ian Johnston, Dominic Kwiatkowski, Cordelia Langford, John Sillitoe on behalf of the Wellcome Sanger Institute COVID-19 Surveillance Team |
| EPI_ISL_3378204, EPI_ISL_3378205 | Berkshire and Surrey Pathology Services Lighthouse Laboratory | Wellcome Sanger Institute for the COVID-19 Genomics UK (COG-UK) Consortium | Berkshire and Surrey Pathology Services Lighthouse Laboratory and Alex Alderton, Roberto Amato, Jeffrey Barrett, Sonia Goncalves, Ewan Harrison, David K. Jackson, Ian Johnston, Dominic Kwiatkowski, Cordelia Langford, John Sillitoe on behalf of the Wellcome Sanger Institute COVID-19 Surveillance Team |
| EPI_ISL_3378206, EPI_ISL_3378207, EPI_ISL_3378208, EPI_ISL_3378209 | Lighthouse Lab in Milton Keynes | Wellcome Sanger Institute for the COVID-19 Genomics UK (COG-UK) Consortium | The Lighthouse Lab in Milton Keynes and Alex Alderton, Roberto Amato, Jeffrey Barrett, Sonia Goncalves, Ewan Harrison, David K. Jackson, Ian Johnston, Dominic Kwiatkowski, Cordelia Langford, John Sillitoe on behalf of the Wellcome Sanger Institute COVID-19 Surveillance Team |
| EPI_ISL_3378210 | Lighthouse Lab in Alderley Park | Wellcome Sanger Institute for the COVID-19 Genomics UK (COG-UK) Consortium | Jacquelyn Wynn, Mairead Hyland, The Lighthouse Lab in Alderley Park and Alex Alderton, Roberto Amato, Jeffrey Barrett, Sonia Goncalves, Ewan Harrison, David K. Jackson, Ian Johnston, Dominic Kwiatkowski, Cordelia Langford, John Sillitoe on behalf of the Wellcome Sanger Institute COVID-19 Surveillance Team |
| EPI_ISL_3378211 | LABORATOIRE UNILIANS DECINES | CNR Virus des Infections Respiratoires - France SUD | Antonin Bal, Gregory Destras, Gwendolyne Burfin, Hadrien Regue, Quentin Semanas, Martine Valette, Bruno Lina, Laurence Josset |
| EPI_ISL_3378212 | Lighthouse Lab in Milton Keynes | Wellcome Sanger Institute for the COVID-19 Genomics UK (COG-UK) Consortium | The Lighthouse Lab in Milton Keynes and Alex Alderton, Roberto Amato, Jeffrey Barrett, Sonia Goncalves, Ewan Harrison, David K. Jackson, Ian Johnston, Dominic Kwiatkowski, Cordelia Langford, John Sillitoe on behalf of the Wellcome Sanger Institute COVID-19 Surveillance Team |
| EPI_ISL_3378213, | Berkshire and Surrey Pathology Services Lighthouse Laboratory | Wellcome Sanger Institute for the COVID-19 Genomics UK (COG-UK) | Berkshire and Surrey Pathology Services Lighthouse Laboratory and Alex Alderton, Roberto Amato, Jeffrey Barrett, Sonia Goncalves, Ewan Harrison, David K. Jackson, Ian Johnston, Dominic Kwiatkowski, Cordelia Langford, John Sillitoe on behalf |

| | | | |
|---|---|---|---|
| EPI_ISL_3378214 | | Consortium | of the Wellcome Sanger Institute COVID-19 Surveillance Team |
| EPI_ISL_3378215, EPI_ISL_3378216, EPI_ISL_3378217, EPI_ISL_3378218 | Lighthouse Lab in Milton Keynes | Wellcome Sanger Institute for the COVID-19 Genomics UK (COG-UK) Consortium | The Lighthouse Lab in Milton Keynes and Alex Alderton, Roberto Amato, Jeffrey Barrett, Sonia Goncalves, Ewan Harrison, David K. Jackson, Ian Johnston, Dominic Kwiatkowski, Cordelia Langford, John Sillitoe on behalf of the Wellcome Sanger Institute COVID-19 Surveillance Team |
| EPI_ISL_3378219 | LABORATOIRE UNILIANS DECINES | CNR Virus des Infections Respiratoires - France SUD | Antonin Bal, Gregory Destras, Gwendolyne Burfin, Hadrien Regue, Quentin Semanas, Martine Valette, Bruno Lina, Laurence Josset |
| EPI_ISL_3378220, EPI_ISL_3378221 | Lighthouse Lab in Alderley Park | Wellcome Sanger Institute for the COVID-19 Genomics UK (COG-UK) Consortium | Jacquelyn Wynn, Mairead Hyland, The Lighthouse Lab in Alderley Park and Alex Alderton, Roberto Amato, Jeffrey Barrett, Sonia Goncalves, Ewan Harrison, David K. Jackson, Ian Johnston, Dominic Kwiatkowski, Cordelia Langford, John Sillitoe on behalf of the Wellcome Sanger Institute COVID-19 Surveillance Team |
| EPI_ISL_3378222 | Lighthouse Lab in Milton Keynes | Wellcome Sanger Institute for the COVID-19 Genomics UK (COG-UK) Consortium | The Lighthouse Lab in Milton Keynes and Alex Alderton, Roberto Amato, Jeffrey Barrett, Sonia Goncalves, Ewan Harrison, David K. Jackson, Ian Johnston, Dominic Kwiatkowski, Cordelia Langford, John Sillitoe on behalf of the Wellcome Sanger Institute COVID-19 Surveillance Team |
| EPI_ISL_3378223 | LABORATOIRE UNILIANS DECINES | CNR Virus des Infections Respiratoires - France SUD | Antonin Bal, Gregory Destras, Gwendolyne Burfin, Hadrien Regue, Quentin Semanas, Martine Valette, Bruno Lina, Laurence Josset |
| EPI_ISL_3378224 | Berkshire and Surrey Pathology Services Lighthouse Laboratory | Wellcome Sanger Institute for the COVID-19 Genomics UK (COG-UK) Consortium | Berkshire and Surrey Pathology Services Lighthouse Laboratory and Alex Alderton, Roberto Amato, Jeffrey Barrett, Sonia Goncalves, Ewan Harrison, David K. Jackson, Ian Johnston, Dominic Kwiatkowski, Cordelia Langford, John Sillitoe on behalf of the Wellcome Sanger Institute COVID-19 Surveillance Team |
| EPI_ISL_3378225, EPI_ISL_3378226 | Lighthouse Lab in Alderley Park | Wellcome Sanger Institute for the COVID-19 Genomics UK (COG-UK) Consortium | Jacquelyn Wynn, Mairead Hyland, The Lighthouse Lab in Alderley Park and Alex Alderton, Roberto Amato, Jeffrey Barrett, Sonia Goncalves, Ewan Harrison, David K. Jackson, Ian Johnston, Dominic Kwiatkowski, Cordelia Langford, John Sillitoe on behalf of the Wellcome Sanger Institute COVID-19 Surveillance Team |
| EPI_ISL_3378227 | LABORATOIRE UNILIANS DECINES | CNR Virus des Infections Respiratoires - France SUD | Antonin Bal, Gregory Destras, Gwendolyne Burfin, Hadrien Regue, Quentin Semanas, Martine Valette, Bruno Lina, Laurence Josset |
| EPI_ISL_3378228 | Lighthouse Lab in Alderley Park | Wellcome Sanger Institute for the COVID-19 Genomics UK (COG-UK) Consortium | Jacquelyn Wynn, Mairead Hyland, The Lighthouse Lab in Alderley Park and Alex Alderton, Roberto Amato, Jeffrey Barrett, Sonia Goncalves, Ewan Harrison, David K. Jackson, Ian Johnston, Dominic Kwiatkowski, Cordelia Langford, John Sillitoe on behalf of the Wellcome Sanger Institute COVID-19 Surveillance Team |
| EPI_ISL_3378229 | Lighthouse Lab in Milton Keynes | Wellcome Sanger Institute for the COVID-19 Genomics UK (COG-UK) Consortium | The Lighthouse Lab in Milton Keynes and Alex Alderton, Roberto Amato, Jeffrey Barrett, Sonia Goncalves, Ewan Harrison, David K. Jackson, Ian Johnston, Dominic Kwiatkowski, Cordelia Langford, John Sillitoe on behalf of the Wellcome Sanger Institute COVID-19 Surveillance Team |
| EPI_ISL_3378230 | Lighthouse Lab in Alderley Park | Wellcome Sanger Institute for the COVID-19 Genomics UK (COG-UK) Consortium | Jacquelyn Wynn, Mairead Hyland, The Lighthouse Lab in Alderley Park and Alex Alderton, Roberto Amato, Jeffrey Barrett, Sonia Goncalves, Ewan Harrison, David K. Jackson, Ian Johnston, Dominic Kwiatkowski, Cordelia Langford, John Sillitoe on behalf of the Wellcome Sanger Institute COVID-19 Surveillance Team |
| EPI_ISL_3378231 | CNR Virus des Infections Respiratoires - France SUD | CNR Virus des Infections Respiratoires - France SUD | Antonin Bal, Gregory Destras, Gwendolyne Burfin, Hadrien Regue, Quentin Semanas, Martine Valette, Bruno Lina, Laurence Josset |
| EPI_ISL_3378232 | Berkshire and Surrey Pathology Services Lighthouse Laboratory | Wellcome Sanger Institute for the COVID-19 Genomics UK (COG-UK) Consortium | Berkshire and Surrey Pathology Services Lighthouse Laboratory and Alex Alderton, Roberto Amato, Jeffrey Barrett, Sonia Goncalves, Ewan Harrison, David K. Jackson, Ian Johnston, Dominic Kwiatkowski, Cordelia Langford, John Sillitoe on behalf of the Wellcome Sanger Institute COVID-19 Surveillance Team |
| EPI_ISL_3378233 | Lighthouse Lab in Alderley Park | Wellcome Sanger Institute for the COVID-19 Genomics UK (COG-UK) Consortium | Jacquelyn Wynn, Mairead Hyland, The Lighthouse Lab in Alderley Park and Alex Alderton, Roberto Amato, Jeffrey Barrett, Sonia Goncalves, Ewan Harrison, David K. Jackson, Ian Johnston, Dominic Kwiatkowski, Cordelia Langford, John Sillitoe on behalf of the Wellcome Sanger Institute COVID-19 Surveillance Team |
| EPI_ISL_3378234 | Berkshire and Surrey Pathology Services Lighthouse Laboratory | Wellcome Sanger Institute for the COVID-19 Genomics UK (COG-UK) Consortium | Berkshire and Surrey Pathology Services Lighthouse Laboratory and Alex Alderton, Roberto Amato, Jeffrey Barrett, Sonia Goncalves, Ewan Harrison, David K. Jackson, Ian Johnston, Dominic Kwiatkowski, Cordelia Langford, John Sillitoe on behalf of the Wellcome Sanger Institute COVID-19 Surveillance Team |
| EPI_ISL_3378235 | LABORATOIRE UNILIANS DECINES | CNR Virus des Infections Respiratoires - France SUD | Antonin Bal, Gregory Destras, Gwendolyne Burfin, Hadrien Regue, Quentin Semanas, Martine Valette, Bruno Lina, Laurence Josset |
| EPI_ISL_3378236 | Berkshire and Surrey Pathology Services Lighthouse Laboratory | Wellcome Sanger Institute for the COVID-19 Genomics UK (COG-UK) Consortium | Berkshire and Surrey Pathology Services Lighthouse Laboratory and Alex Alderton, Roberto Amato, Jeffrey Barrett, Sonia Goncalves, Ewan Harrison, David K. Jackson, Ian Johnston, Dominic Kwiatkowski, Cordelia Langford, John Sillitoe on behalf of the Wellcome Sanger Institute COVID-19 Surveillance Team |
| EPI_ISL_3378237, EPI_ISL_3378238 | Lighthouse Lab in Alderley Park | Wellcome Sanger Institute for the COVID-19 Genomics UK (COG-UK) Consortium | Jacquelyn Wynn, Mairead Hyland, The Lighthouse Lab in Alderley Park and Alex Alderton, Roberto Amato, Jeffrey Barrett, Sonia Goncalves, Ewan Harrison, David K. Jackson, Ian Johnston, Dominic Kwiatkowski, Cordelia Langford, John Sillitoe on behalf of the Wellcome Sanger Institute COVID-19 Surveillance Team |
| EPI_ISL_3378239 | LABORATOIRE UNILIANS DECINES | CNR Virus des Infections Respiratoires - France SUD | Antonin Bal, Gregory Destras, Gwendolyne Burfin, Hadrien Regue, Quentin Semanas, Martine Valette, Bruno Lina, Laurence Josset |
| EPI_ISL_3378240 | Lighthouse Lab in Alderley Park | Wellcome Sanger Institute for the COVID-19 Genomics UK (COG-UK) Consortium | Jacquelyn Wynn, Mairead Hyland, The Lighthouse Lab in Alderley Park and Alex Alderton, Roberto Amato, Jeffrey Barrett, Sonia Goncalves, Ewan Harrison, David K. Jackson, Ian Johnston, Dominic Kwiatkowski, Cordelia Langford, John Sillitoe on behalf of the Wellcome Sanger Institute COVID-19 Surveillance Team |
| EPI_ISL_3378241, EPI_ISL_3378242 | Lighthouse Lab in Milton Keynes | Wellcome Sanger Institute for the COVID-19 Genomics UK (COG-UK) Consortium | The Lighthouse Lab in Milton Keynes and Alex Alderton, Roberto Amato, Jeffrey Barrett, Sonia Goncalves, Ewan Harrison, David K. Jackson, Ian Johnston, Dominic Kwiatkowski, Cordelia Langford, John Sillitoe on behalf of the Wellcome Sanger Institute COVID-19 Surveillance Team |
| EPI_ISL_3378243 | LABORATOIRE UNILIANS DECINES | CNR Virus des Infections Respiratoires - France SUD | Antonin Bal, Gregory Destras, Gwendolyne Burfin, Hadrien Regue, Quentin Semanas, Martine Valette, Bruno Lina, Laurence Josset |
| EPI_ISL_3378244, EPI_ISL_3378245 | Lighthouse Lab in Milton Keynes | Wellcome Sanger Institute for the COVID-19 Genomics UK (COG-UK) Consortium | The Lighthouse Lab in Milton Keynes and Alex Alderton, Roberto Amato, Jeffrey Barrett, Sonia Goncalves, Ewan Harrison, David K. Jackson, Ian Johnston, Dominic Kwiatkowski, Cordelia Langford, John Sillitoe on behalf of the Wellcome Sanger Institute COVID-19 Surveillance Team |
| EPI_ISL_3378246 | Berkshire and Surrey Pathology Services Lighthouse Laboratory | Wellcome Sanger Institute for the COVID-19 Genomics UK (COG-UK) Consortium | Berkshire and Surrey Pathology Services Lighthouse Laboratory and Alex Alderton, Roberto Amato, Jeffrey Barrett, Sonia Goncalves, Ewan Harrison, David K. Jackson, Ian Johnston, Dominic Kwiatkowski, Cordelia Langford, John Sillitoe on behalf of the Wellcome Sanger Institute COVID-19 Surveillance Team |
| EPI_ISL_3378247 | LABORATOIRE UNILIANS DECINES | CNR Virus des Infections Respiratoires - France SUD | Antonin Bal, Gregory Destras, Gwendolyne Burfin, Hadrien Regue, Quentin Semanas, Martine Valette, Bruno Lina, Laurence Josset |
| EPI_ISL_3378248, EPI_ISL_3378249, EPI_ISL_3378250 | Lighthouse Lab in Milton Keynes | Wellcome Sanger Institute for the COVID-19 Genomics UK (COG-UK) Consortium | The Lighthouse Lab in Milton Keynes and Alex Alderton, Roberto Amato, Jeffrey Barrett, Sonia Goncalves, Ewan Harrison, David K. Jackson, Ian Johnston, Dominic Kwiatkowski, Cordelia Langford, John Sillitoe on behalf of the Wellcome Sanger Institute COVID-19 Surveillance Team |
| EPI_ISL_3378251 | LABORATOIRE UNILIANS DECINES | CNR Virus des Infections Respiratoires - France SUD | Antonin Bal, Gregory Destras, Gwendolyne Burfin, Hadrien Regue, Quentin Semanas, Martine Valette, Bruno Lina, Laurence Josset |
| EPI_ISL_3378252 | Lighthouse Lab in Alderley Park | Wellcome Sanger Institute for the COVID-19 Genomics UK (COG-UK) Consortium | Jacquelyn Wynn, Mairead Hyland, The Lighthouse Lab in Alderley Park and Alex Alderton, Roberto Amato, Jeffrey Barrett, Sonia Goncalves, Ewan Harrison, David K. Jackson, Ian Johnston, Dominic Kwiatkowski, Cordelia Langford, John Sillitoe on behalf of the Wellcome Sanger Institute COVID-19 Surveillance Team |
| EPI_ISL_3378253 | Lighthouse Lab in Milton Keynes | Wellcome Sanger Institute for the COVID-19 Genomics UK (COG-UK) Consortium | The Lighthouse Lab in Milton Keynes and Alex Alderton, Roberto Amato, Jeffrey Barrett, Sonia Goncalves, Ewan Harrison, David K. Jackson, Ian Johnston, Dominic Kwiatkowski, Cordelia Langford, John Sillitoe on behalf of the Wellcome Sanger Institute COVID-19 Surveillance Team |
| EPI_ISL_3378254 | Berkshire and Surrey Pathology Services Lighthouse Laboratory | Wellcome Sanger Institute for the COVID-19 Genomics UK (COG-UK) Consortium | Berkshire and Surrey Pathology Services Lighthouse Laboratory and Alex Alderton, Roberto Amato, Jeffrey Barrett, Sonia Goncalves, Ewan Harrison, David K. Jackson, Ian Johnston, Dominic Kwiatkowski, Cordelia Langford, John Sillitoe on behalf of the Wellcome Sanger Institute COVID-19 Surveillance Team |
| EPI_ISL_3378255 | CH GIVORS | CNR Virus des Infections Respiratoires - France SUD | Antonin Bal, Gregory Destras, Gwendolyne Burfin, Hadrien Regue, Quentin Semanas, Martine Valette, Bruno Lina, Laurence Josset |
| EPI_ISL_3378256 | Lighthouse Lab in Milton Keynes | Wellcome Sanger Institute for the COVID-19 Genomics UK (COG-UK) Consortium | The Lighthouse Lab in Milton Keynes and Alex Alderton, Roberto Amato, Jeffrey Barrett, Sonia Goncalves, Ewan Harrison, David K. Jackson, Ian Johnston, Dominic Kwiatkowski, Cordelia Langford, John Sillitoe on behalf of the Wellcome Sanger Institute COVID-19 Surveillance Team |
| EPI_ISL_3378257 | Berkshire and Surrey Pathology Services Lighthouse Laboratory | Wellcome Sanger Institute for the COVID-19 Genomics UK (COG-UK) Consortium | Berkshire and Surrey Pathology Services Lighthouse Laboratory and Alex Alderton, Roberto Amato, Jeffrey Barrett, Sonia Goncalves, Ewan Harrison, David K. Jackson, Ian Johnston, Dominic Kwiatkowski, Cordelia Langford, John Sillitoe on behalf of the Wellcome Sanger Institute COVID-19 Surveillance Team |
| EPI_ISL_3378258 | Lighthouse Lab in Milton Keynes | Wellcome Sanger Institute for the COVID-19 Genomics UK (COG-UK) Consortium | The Lighthouse Lab in Milton Keynes and Alex Alderton, Roberto Amato, Jeffrey Barrett, Sonia Goncalves, Ewan Harrison, David K. Jackson, Ian Johnston, Dominic Kwiatkowski, Cordelia Langford, John Sillitoe on behalf of the Wellcome Sanger Institute COVID-19 Surveillance Team |
| EPI_ISL_3378259 | LABORATOIRE UNILIANS DECINES | CNR Virus des Infections Respiratoires - France SUD | Antonin Bal, Gregory Destras, Gwendolyne Burfin, Hadrien Regue, Quentin Semanas, Martine Valette, Bruno Lina, Laurence Josset |
| EPI_ISL_3378260, EPI_ISL_3378261 | Lighthouse Lab in Alderley Park | Wellcome Sanger Institute for the COVID-19 Genomics UK (COG-UK) Consortium | Jacquelyn Wynn, Mairead Hyland, The Lighthouse Lab in Alderley Park and Alex Alderton, Roberto Amato, Jeffrey Barrett, Sonia Goncalves, Ewan Harrison, David K. Jackson, Ian Johnston, Dominic Kwiatkowski, Cordelia Langford, John Sillitoe on behalf of the Wellcome Sanger Institute COVID-19 Surveillance Team |
| EPI_ISL_3378262, EPI_ISL_3378263 | Lighthouse Lab in Milton Keynes | Wellcome Sanger Institute for the COVID-19 Genomics UK (COG-UK) Consortium | The Lighthouse Lab in Milton Keynes and Alex Alderton, Roberto Amato, Jeffrey Barrett, Sonia Goncalves, Ewan Harrison, David K. Jackson, Ian Johnston, Dominic Kwiatkowski, Cordelia Langford, John Sillitoe on behalf of the Wellcome Sanger Institute COVID-19 Surveillance Team |
| EPI_ISL_3378264 | LABORATOIRE UNILIANS DECINES | CNR Virus des Infections Respiratoires - France SUD | Antonin Bal, Gregory Destras, Gwendolyne Burfin, Hadrien Regue, Quentin Semanas, Martine Valette, Bruno Lina, Laurence Josset |
| EPI_ISL_3378265, EPI_ISL_3378266 | Lighthouse Lab in Alderley Park | Wellcome Sanger Institute for the COVID-19 Genomics UK (COG-UK) Consortium | Jacquelyn Wynn, Mairead Hyland, The Lighthouse Lab in Alderley Park and Alex Alderton, Roberto Amato, Jeffrey Barrett, Sonia Goncalves, Ewan Harrison, David K. Jackson, Ian Johnston, Dominic Kwiatkowski, Cordelia Langford, John Sillitoe on behalf of the Wellcome Sanger Institute COVID-19 Surveillance Team |
| EPI_ISL_3378267 | Berkshire and Surrey Pathology Services Lighthouse Laboratory | Wellcome Sanger Institute for the COVID-19 Genomics UK (COG-UK) Consortium | Berkshire and Surrey Pathology Services Lighthouse Laboratory and Alex Alderton, Roberto Amato, Jeffrey Barrett, Sonia Goncalves, Ewan Harrison, David K. Jackson, Ian Johnston, Dominic Kwiatkowski, Cordelia Langford, John Sillitoe on behalf of the Wellcome Sanger Institute COVID-19 Surveillance Team |
| EPI_ISL_3378268 | CNR Virus des Infections Respiratoires - France SUD | CNR Virus des Infections Respiratoires - France SUD | Antonin Bal, Gregory Destras, Gwendolyne Burfin, Hadrien Regue, Quentin Semanas, Martine Valette, Bruno Lina, Laurence Josset |
| EPI_ISL_3378269 | Lighthouse Lab in Alderley Park | Wellcome Sanger Institute for the COVID-19 Genomics UK (COG-UK) Consortium | Jacquelyn Wynn, Mairead Hyland, The Lighthouse Lab in Alderley Park and Alex Alderton, Roberto Amato, Jeffrey Barrett, Sonia Goncalves, Ewan Harrison, David K. Jackson, Ian Johnston, Dominic Kwiatkowski, Cordelia Langford, John Sillitoe on behalf of the Wellcome Sanger Institute COVID-19 Surveillance Team |
| EPI_ISL_3378270 | Lighthouse Lab in Milton Keynes | Wellcome Sanger Institute for the COVID-19 Genomics UK (COG-UK) Consortium | The Lighthouse Lab in Milton Keynes and Alex Alderton, Roberto Amato, Jeffrey Barrett, Sonia Goncalves, Ewan Harrison, David K. Jackson, Ian Johnston, Dominic Kwiatkowski, Cordelia Langford, John Sillitoe on behalf of the Wellcome Sanger Institute COVID-19 Surveillance Team |
| EPI_ISL_3378271 | Lighthouse Lab in Alderley Park | Wellcome Sanger Institute for the COVID-19 Genomics UK (COG-UK) Consortium | Jacquelyn Wynn, Mairead Hyland, The Lighthouse Lab in Alderley Park and Alex Alderton, Roberto Amato, Jeffrey Barrett, Sonia Goncalves, Ewan Harrison, David K. Jackson, Ian Johnston, Dominic Kwiatkowski, Cordelia Langford, John Sillitoe on behalf of the Wellcome Sanger Institute COVID-19 Surveillance Team |
| EPI_ISL_3378272 | Lighthouse Lab in Milton Keynes | Wellcome Sanger Institute for the COVID-19 Genomics UK (COG-UK) Consortium | The Lighthouse Lab in Milton Keynes and Alex Alderton, Roberto Amato, Jeffrey Barrett, Sonia Goncalves, Ewan Harrison, David K. Jackson, Ian Johnston, Dominic Kwiatkowski, Cordelia Langford, John Sillitoe on behalf of the Wellcome Sanger Institute COVID-19 Surveillance Team |
| EPI_ISL_3378273 | LABORATOIRE ACCOLAB SUD OUEST | CNR Virus des Infections Respiratoires - France SUD | Antonin Bal, Gregory Destras, Gwendolyne Burfin, Hadrien Regue, Quentin Semanas, Martine Valette, Bruno Lina, Laurence Josset |
| EPI_ISL_3378274 | Lighthouse Lab in Alderley Park | Wellcome Sanger Institute for the COVID-19 Genomics UK (COG-UK) Consortium | Jacquelyn Wynn, Mairead Hyland, The Lighthouse Lab in Alderley Park and Alex Alderton, Roberto Amato, Jeffrey Barrett, Sonia Goncalves, Ewan Harrison, David K. Jackson, Ian Johnston, Dominic Kwiatkowski, Cordelia Langford, John Sillitoe on behalf of the Wellcome Sanger Institute COVID-19 Surveillance Team |
| EPI_ISL_3378275, EPI_ISL_3378276 | Lighthouse Lab in Milton Keynes | Wellcome Sanger Institute for the COVID-19 Genomics UK (COG-UK) Consortium | The Lighthouse Lab in Milton Keynes and Alex Alderton, Roberto Amato, Jeffrey Barrett, Sonia Goncalves, Ewan Harrison, David K. Jackson, Ian Johnston, Dominic Kwiatkowski, Cordelia Langford, John Sillitoe on behalf of the Wellcome Sanger Institute COVID-19 Surveillance Team |
| EPI_ISL_3378277 | LABORATOIRE ACCOLAB SUD OUEST | CNR Virus des Infections Respiratoires - France SUD | Antonin Bal, Gregory Destras, Gwendolyne Burfin, Hadrien Regue, Quentin Semanas, Martine Valette, Bruno Lina, Laurence Josset |

| | | | |
|---|---|---|---|
| EPI_ISL_3378278, EPI_ISL_3378279 | Berkshire and Surrey Pathology Services Lighthouse Laboratory | Wellcome Sanger Institute for the COVID-19 Genomics UK (COG-UK) Consortium | Berkshire and Surrey Pathology Services Lighthouse Laboratory and Alex Alderton, Roberto Amato, Jeffrey Barrett, Sonia Goncalves, Ewan Harrison, David K. Jackson, Ian Johnston, Dominic Kwiatkowski, Cordelia Langford, John Sillitoe on behalf of the Wellcome Sanger Institute COVID-19 Surveillance Team |
| EPI_ISL_3378280 | Lighthouse Lab in Milton Keynes | Wellcome Sanger Institute for the COVID-19 Genomics UK (COG-UK) Consortium | The Lighthouse Lab in Milton Keynes and Alex Alderton, Roberto Amato, Jeffrey Barrett, Sonia Goncalves, Ewan Harrison, David K. Jackson, Ian Johnston, Dominic Kwiatkowski, Cordelia Langford, John Sillitoe on behalf of the Wellcome Sanger Institute COVID-19 Surveillance Team |
| EPI_ISL_3378281 | LABORATOIRE ACCOLAB SUD OUEST | CNR Virus des Infections Respiratoires - France SUD | Antonin Bal, Gregory Destras, Gwendolyne Burfin, Hadrien Regue, Quentin Semanas, Martine Valette, Bruno Lina, Laurence Josset |
| EPI_ISL_3378282 | Lighthouse Lab in Milton Keynes | Wellcome Sanger Institute for the COVID-19 Genomics UK (COG-UK) Consortium | The Lighthouse Lab in Milton Keynes and Alex Alderton, Roberto Amato, Jeffrey Barrett, Sonia Goncalves, Ewan Harrison, David K. Jackson, Ian Johnston, Dominic Kwiatkowski, Cordelia Langford, John Sillitoe on behalf of the Wellcome Sanger Institute COVID-19 Surveillance Team |
| EPI_ISL_3378283 | Berkshire and Surrey Pathology Services Lighthouse Laboratory | Wellcome Sanger Institute for the COVID-19 Genomics UK (COG-UK) Consortium | Berkshire and Surrey Pathology Services Lighthouse Laboratory and Alex Alderton, Roberto Amato, Jeffrey Barrett, Sonia Goncalves, Ewan Harrison, David K. Jackson, Ian Johnston, Dominic Kwiatkowski, Cordelia Langford, John Sillitoe on behalf of the Wellcome Sanger Institute COVID-19 Surveillance Team |
| EPI_ISL_3378284 | Lighthouse Lab in Milton Keynes | Wellcome Sanger Institute for the COVID-19 Genomics UK (COG-UK) Consortium | The Lighthouse Lab in Milton Keynes and Alex Alderton, Roberto Amato, Jeffrey Barrett, Sonia Goncalves, Ewan Harrison, David K. Jackson, Ian Johnston, Dominic Kwiatkowski, Cordelia Langford, John Sillitoe on behalf of the Wellcome Sanger Institute COVID-19 Surveillance Team |
| EPI_ISL_3378285 | LABORATOIRE ACCOLAB SUD OUEST | CNR Virus des Infections Respiratoires - France SUD | Antonin Bal, Gregory Destras, Gwendolyne Burfin, Hadrien Regue, Quentin Semanas, Martine Valette, Bruno Lina, Laurence Josset |
| EPI_ISL_3378286, EPI_ISL_3378287 | Lighthouse Lab in Milton Keynes | Wellcome Sanger Institute for the COVID-19 Genomics UK (COG-UK) Consortium | The Lighthouse Lab in Milton Keynes and Alex Alderton, Roberto Amato, Jeffrey Barrett, Sonia Goncalves, Ewan Harrison, David K. Jackson, Ian Johnston, Dominic Kwiatkowski, Cordelia Langford, John Sillitoe on behalf of the Wellcome Sanger Institute COVID-19 Surveillance Team |
| EPI_ISL_3378288 | Lighthouse Lab in Alderley Park | Wellcome Sanger Institute for the COVID-19 Genomics UK (COG-UK) Consortium | Jacquelyn Wynn, Mairead Hyland, The Lighthouse Lab in Alderley Park and Alex Alderton, Roberto Amato, Jeffrey Barrett, Sonia Goncalves, Ewan Harrison, David K. Jackson, Ian Johnston, Dominic Kwiatkowski, Cordelia Langford, John Sillitoe on behalf of the Wellcome Sanger Institute COVID-19 Surveillance Team |
| EPI_ISL_3378289 | LABORATOIRE ACCOLAB SUD OUEST | CNR Virus des Infections Respiratoires - France SUD | Antonin Bal, Gregory Destras, Gwendolyne Burfin, Hadrien Regue, Quentin Semanas, Martine Valette, Bruno Lina, Laurence Josset |
| EPI_ISL_3378290 | Berkshire and Surrey Pathology Services Lighthouse Laboratory | Wellcome Sanger Institute for the COVID-19 Genomics UK (COG-UK) Consortium | Berkshire and Surrey Pathology Services Lighthouse Laboratory and Alex Alderton, Roberto Amato, Jeffrey Barrett, Sonia Goncalves, Ewan Harrison, David K. Jackson, Ian Johnston, Dominic Kwiatkowski, Cordelia Langford, John Sillitoe on behalf of the Wellcome Sanger Institute COVID-19 Surveillance Team |
| EPI_ISL_3378291, EPI_ISL_3378292 | Lighthouse Lab in Milton Keynes | Wellcome Sanger Institute for the COVID-19 Genomics UK (COG-UK) Consortium | The Lighthouse Lab in Milton Keynes and Alex Alderton, Roberto Amato, Jeffrey Barrett, Sonia Goncalves, Ewan Harrison, David K. Jackson, Ian Johnston, Dominic Kwiatkowski, Cordelia Langford, John Sillitoe on behalf of the Wellcome Sanger Institute COVID-19 Surveillance Team |
| EPI_ISL_3378293 | LABORATOIRE ACCOLAB SUD OUEST | CNR Virus des Infections Respiratoires - France SUD | Antonin Bal, Gregory Destras, Gwendolyne Burfin, Hadrien Regue, Quentin Semanas, Martine Valette, Bruno Lina, Laurence Josset |
| EPI_ISL_3378294, EPI_ISL_3378295 | Lighthouse Lab in Milton Keynes | Wellcome Sanger Institute for the COVID-19 Genomics UK (COG-UK) Consortium | The Lighthouse Lab in Milton Keynes and Alex Alderton, Roberto Amato, Jeffrey Barrett, Sonia Goncalves, Ewan Harrison, David K. Jackson, Ian Johnston, Dominic Kwiatkowski, Cordelia Langford, John Sillitoe on behalf of the Wellcome Sanger Institute COVID-19 Surveillance Team |
| EPI_ISL_3378296 | Berkshire and Surrey Pathology Services Lighthouse Laboratory | Wellcome Sanger Institute for the COVID-19 Genomics UK (COG-UK) Consortium | Berkshire and Surrey Pathology Services Lighthouse Laboratory and Alex Alderton, Roberto Amato, Jeffrey Barrett, Sonia Goncalves, Ewan Harrison, David K. Jackson, Ian Johnston, Dominic Kwiatkowski, Cordelia Langford, John Sillitoe on behalf of the Wellcome Sanger Institute COVID-19 Surveillance Team |
| EPI_ISL_3378298 | Lighthouse Lab in Milton Keynes | Wellcome Sanger Institute for the COVID-19 Genomics UK (COG-UK) Consortium | The Lighthouse Lab in Milton Keynes and Alex Alderton, Roberto Amato, Jeffrey Barrett, Sonia Goncalves, Ewan Harrison, David K. Jackson, Ian Johnston, Dominic Kwiatkowski, Cordelia Langford, John Sillitoe on behalf of the Wellcome Sanger Institute COVID-19 Surveillance Team |
| EPI_ISL_3378299 | Berkshire and Surrey Pathology Services Lighthouse Laboratory | Wellcome Sanger Institute for the COVID-19 Genomics UK (COG-UK) Consortium | Berkshire and Surrey Pathology Services Lighthouse Laboratory and Alex Alderton, Roberto Amato, Jeffrey Barrett, Sonia Goncalves, Ewan Harrison, David K. Jackson, Ian Johnston, Dominic Kwiatkowski, Cordelia Langford, John Sillitoe on behalf of the Wellcome Sanger Institute COVID-19 Surveillance Team |
| EPI_ISL_3378300 | Lighthouse Lab in Alderley Park | Wellcome Sanger Institute for the COVID-19 Genomics UK (COG-UK) Consortium | Jacquelyn Wynn, Mairead Hyland, The Lighthouse Lab in Alderley Park and Alex Alderton, Roberto Amato, Jeffrey Barrett, Sonia Goncalves, Ewan Harrison, David K. Jackson, Ian Johnston, Dominic Kwiatkowski, Cordelia Langford, John Sillitoe on behalf of the Wellcome Sanger Institute COVID-19 Surveillance Team |
| EPI_ISL_3378302 | Berkshire and Surrey Pathology Services Lighthouse Laboratory | Wellcome Sanger Institute for the COVID-19 Genomics UK (COG-UK) Consortium | Berkshire and Surrey Pathology Services Lighthouse Laboratory and Alex Alderton, Roberto Amato, Jeffrey Barrett, Sonia Goncalves, Ewan Harrison, David K. Jackson, Ian Johnston, Dominic Kwiatkowski, Cordelia Langford, John Sillitoe on behalf of the Wellcome Sanger Institute COVID-19 Surveillance Team |
| EPI_ISL_3378303, EPI_ISL_3378304 | Lighthouse Lab in Milton Keynes | Wellcome Sanger Institute for the COVID-19 Genomics UK (COG-UK) Consortium | The Lighthouse Lab in Milton Keynes and Alex Alderton, Roberto Amato, Jeffrey Barrett, Sonia Goncalves, Ewan Harrison, David K. Jackson, Ian Johnston, Dominic Kwiatkowski, Cordelia Langford, John Sillitoe on behalf of the Wellcome Sanger Institute COVID-19 Surveillance Team |
| EPI_ISL_3378305 | LABORATOIRE ACCOLAB SUD OUEST | CNR Virus des Infections Respiratoires - France SUD | Antonin Bal, Gregory Destras, Gwendolyne Burfin, Hadrien Regue, Quentin Semanas, Martine Valette, Bruno Lina, Laurence Josset |
| EPI_ISL_3378306, EPI_ISL_3378307 | Berkshire and Surrey Pathology Services Lighthouse Laboratory | Wellcome Sanger Institute for the COVID-19 Genomics UK (COG-UK) Consortium | Berkshire and Surrey Pathology Services Lighthouse Laboratory and Alex Alderton, Roberto Amato, Jeffrey Barrett, Sonia Goncalves, Ewan Harrison, David K. Jackson, Ian Johnston, Dominic Kwiatkowski, Cordelia Langford, John Sillitoe on behalf of the Wellcome Sanger Institute COVID-19 Surveillance Team |
| EPI_ISL_3378308 | Lighthouse Lab in Milton Keynes | Wellcome Sanger Institute for the COVID-19 Genomics UK (COG-UK) Consortium | The Lighthouse Lab in Milton Keynes and Alex Alderton, Roberto Amato, Jeffrey Barrett, Sonia Goncalves, Ewan Harrison, David K. Jackson, Ian Johnston, Dominic Kwiatkowski, Cordelia Langford, John Sillitoe on behalf of the Wellcome Sanger Institute COVID-19 Surveillance Team |
| EPI_ISL_3378309 | LABORATOIRE ACCOLAB SUD OUEST | CNR Virus des Infections Respiratoires - France SUD | Antonin Bal, Gregory Destras, Gwendolyne Burfin, Hadrien Regue, Quentin Semanas, Martine Valette, Bruno Lina, Laurence Josset |
| EPI_ISL_3378310 | Lighthouse Lab in Milton Keynes | Wellcome Sanger Institute for the COVID-19 Genomics UK (COG-UK) Consortium | The Lighthouse Lab in Milton Keynes and Alex Alderton, Roberto Amato, Jeffrey Barrett, Sonia Goncalves, Ewan Harrison, David K. Jackson, Ian Johnston, Dominic Kwiatkowski, Cordelia Langford, John Sillitoe on behalf of the Wellcome Sanger Institute COVID-19 Surveillance Team |
| EPI_ISL_3378311 | Lighthouse Lab in Alderley Park | Wellcome Sanger Institute for the COVID-19 Genomics UK (COG-UK) Consortium | Jacquelyn Wynn, Mairead Hyland, The Lighthouse Lab in Alderley Park and Alex Alderton, Roberto Amato, Jeffrey Barrett, Sonia Goncalves, Ewan Harrison, David K. Jackson, Ian Johnston, Dominic Kwiatkowski, Cordelia Langford, John Sillitoe on behalf of the Wellcome Sanger Institute COVID-19 Surveillance Team |
| EPI_ISL_3378312 | ORIAPOLE | CNR Virus des Infections Respiratoires - France SUD | Antonin Bal, Gregory Destras, Gwendolyne Burfin, Hadrien Regue, Quentin Semanas, Martine Valette, Bruno Lina, Laurence Josset |
| EPI_ISL_3378313, EPI_ISL_3378314, EPI_ISL_3378315 | Lighthouse Lab in Milton Keynes | Wellcome Sanger Institute for the COVID-19 Genomics UK (COG-UK) Consortium | The Lighthouse Lab in Milton Keynes and Alex Alderton, Roberto Amato, Jeffrey Barrett, Sonia Goncalves, Ewan Harrison, David K. Jackson, Ian Johnston, Dominic Kwiatkowski, Cordelia Langford, John Sillitoe on behalf of the Wellcome Sanger Institute COVID-19 Surveillance Team |
| EPI_ISL_3378316 | ORIAPOLE | CNR Virus des Infections Respiratoires - France SUD | Antonin Bal, Gregory Destras, Gwendolyne Burfin, Hadrien Regue, Quentin Semanas, Martine Valette, Bruno Lina, Laurence Josset |
| EPI_ISL_3378317 | Berkshire and Surrey Pathology Services Lighthouse Laboratory | Wellcome Sanger Institute for the COVID-19 Genomics UK (COG-UK) Consortium | Berkshire and Surrey Pathology Services Lighthouse Laboratory and Alex Alderton, Roberto Amato, Jeffrey Barrett, Sonia Goncalves, Ewan Harrison, David K. Jackson, Ian Johnston, Dominic Kwiatkowski, Cordelia Langford, John Sillitoe on behalf of the Wellcome Sanger Institute COVID-19 Surveillance Team |
| EPI_ISL_3378318, EPI_ISL_3378319 | Lighthouse Lab in Milton Keynes | Wellcome Sanger Institute for the COVID-19 Genomics UK (COG-UK) Consortium | The Lighthouse Lab in Milton Keynes and Alex Alderton, Roberto Amato, Jeffrey Barrett, Sonia Goncalves, Ewan Harrison, David K. Jackson, Ian Johnston, Dominic Kwiatkowski, Cordelia Langford, John Sillitoe on behalf of the Wellcome Sanger Institute COVID-19 Surveillance Team |
| EPI_ISL_3378320 | LABORATOIRE ACCOLAB SUD OUEST | CNR Virus des Infections Respiratoires - France SUD | Antonin Bal, Gregory Destras, Gwendolyne Burfin, Hadrien Regue, Quentin Semanas, Martine Valette, Bruno Lina, Laurence Josset |
| EPI_ISL_3378321, EPI_ISL_3378322 | Lighthouse Lab in Milton Keynes | Wellcome Sanger Institute for the COVID-19 Genomics UK (COG-UK) Consortium | The Lighthouse Lab in Milton Keynes and Alex Alderton, Roberto Amato, Jeffrey Barrett, Sonia Goncalves, Ewan Harrison, David K. Jackson, Ian Johnston, Dominic Kwiatkowski, Cordelia Langford, John Sillitoe on behalf of the Wellcome Sanger Institute COVID-19 Surveillance Team |
| EPI_ISL_3378323 | Berkshire and Surrey Pathology Services Lighthouse Laboratory | Wellcome Sanger Institute for the COVID-19 Genomics UK (COG-UK) Consortium | Berkshire and Surrey Pathology Services Lighthouse Laboratory and Alex Alderton, Roberto Amato, Jeffrey Barrett, Sonia Goncalves, Ewan Harrison, David K. Jackson, Ian Johnston, Dominic Kwiatkowski, Cordelia Langford, John Sillitoe on behalf of the Wellcome Sanger Institute COVID-19 Surveillance Team |
| EPI_ISL_3378324 | Lighthouse Lab in Alderley Park | Wellcome Sanger Institute for the COVID-19 Genomics UK (COG-UK) Consortium | Jacquelyn Wynn, Mairead Hyland, The Lighthouse Lab in Alderley Park and Alex Alderton, Roberto Amato, Jeffrey Barrett, Sonia Goncalves, Ewan Harrison, David K. Jackson, Ian Johnston, Dominic Kwiatkowski, Cordelia Langford, John Sillitoe on behalf of the Wellcome Sanger Institute COVID-19 Surveillance Team |
| EPI_ISL_3378325 | ORIAPOLE | CNR Virus des Infections Respiratoires - France SUD | Antonin Bal, Gregory Destras, Gwendolyne Burfin, Hadrien Regue, Quentin Semanas, Martine Valette, Bruno Lina, Laurence Josset |
| EPI_ISL_3378326 | Lighthouse Lab in Milton Keynes | Wellcome Sanger Institute for the COVID-19 Genomics UK (COG-UK) Consortium | The Lighthouse Lab in Milton Keynes and Alex Alderton, Roberto Amato, Jeffrey Barrett, Sonia Goncalves, Ewan Harrison, David K. Jackson, Ian Johnston, Dominic Kwiatkowski, Cordelia Langford, John Sillitoe on behalf of the Wellcome Sanger Institute COVID-19 Surveillance Team |
| EPI_ISL_3378327 | Lighthouse Lab in Alderley Park | Wellcome Sanger Institute for the COVID-19 Genomics UK (COG-UK) Consortium | Jacquelyn Wynn, Mairead Hyland, The Lighthouse Lab in Alderley Park and Alex Alderton, Roberto Amato, Jeffrey Barrett, Sonia Goncalves, Ewan Harrison, David K. Jackson, Ian Johnston, Dominic Kwiatkowski, Cordelia Langford, John Sillitoe on behalf of the Wellcome Sanger Institute COVID-19 Surveillance Team |
| EPI_ISL_3378328 | Lighthouse Lab in Milton Keynes | Wellcome Sanger Institute for the COVID-19 Genomics UK (COG-UK) Consortium | The Lighthouse Lab in Milton Keynes and Alex Alderton, Roberto Amato, Jeffrey Barrett, Sonia Goncalves, Ewan Harrison, David K. Jackson, Ian Johnston, Dominic Kwiatkowski, Cordelia Langford, John Sillitoe on behalf of the Wellcome Sanger Institute COVID-19 Surveillance Team |
| EPI_ISL_3378329 | ORIAPOLE | CNR Virus des Infections Respiratoires - France SUD | Antonin Bal, Gregory Destras, Gwendolyne Burfin, Hadrien Regue, Quentin Semanas, Martine Valette, Bruno Lina, Laurence Josset |
| EPI_ISL_3378330 | Lighthouse Lab in Alderley Park | Wellcome Sanger Institute for the COVID-19 Genomics UK (COG-UK) Consortium | Jacquelyn Wynn, Mairead Hyland, The Lighthouse Lab in Alderley Park and Alex Alderton, Roberto Amato, Jeffrey Barrett, Sonia Goncalves, Ewan Harrison, David K. Jackson, Ian Johnston, Dominic Kwiatkowski, Cordelia Langford, John Sillitoe on behalf of the Wellcome Sanger Institute COVID-19 Surveillance Team |
| EPI_ISL_3378331 | Berkshire and Surrey Pathology Services Lighthouse Laboratory | Wellcome Sanger Institute for the COVID-19 Genomics UK (COG-UK) Consortium | Berkshire and Surrey Pathology Services Lighthouse Laboratory and Alex Alderton, Roberto Amato, Jeffrey Barrett, Sonia Goncalves, Ewan Harrison, David K. Jackson, Ian Johnston, Dominic Kwiatkowski, Cordelia Langford, John Sillitoe on behalf of the Wellcome Sanger Institute COVID-19 Surveillance Team |
| EPI_ISL_3378332 | Lighthouse Lab in Milton Keynes | Wellcome Sanger Institute for the COVID-19 Genomics UK (COG-UK) Consortium | The Lighthouse Lab in Milton Keynes and Alex Alderton, Roberto Amato, Jeffrey Barrett, Sonia Goncalves, Ewan Harrison, David K. Jackson, Ian Johnston, Dominic Kwiatkowski, Cordelia Langford, John Sillitoe on behalf of the Wellcome Sanger Institute COVID-19 Surveillance Team |
| EPI_ISL_3378333 | ORIAPOLE | CNR Virus des Infections Respiratoires - France SUD | Antonin Bal, Gregory Destras, Gwendolyne Burfin, Hadrien Regue, Quentin Semanas, Martine Valette, Bruno Lina, Laurence Josset |
| EPI_ISL_3378334 | Lighthouse Lab in Milton Keynes | Wellcome Sanger Institute for the COVID-19 Genomics UK (COG-UK) Consortium | The Lighthouse Lab in Milton Keynes and Alex Alderton, Roberto Amato, Jeffrey Barrett, Sonia Goncalves, Ewan Harrison, David K. Jackson, Ian Johnston, Dominic Kwiatkowski, Cordelia Langford, John Sillitoe on behalf of the Wellcome Sanger Institute COVID-19 Surveillance Team |
| EPI_ISL_3378335 | Berkshire and Surrey Pathology Services Lighthouse Laboratory | Wellcome Sanger Institute for the COVID-19 Genomics UK (COG-UK) Consortium | Berkshire and Surrey Pathology Services Lighthouse Laboratory and Alex Alderton, Roberto Amato, Jeffrey Barrett, Sonia Goncalves, Ewan Harrison, David K. Jackson, Ian Johnston, Dominic Kwiatkowski, Cordelia Langford, John Sillitoe on behalf of the Wellcome Sanger Institute COVID-19 Surveillance Team |
| EPI_ISL_3378336 | Lighthouse Lab in Milton Keynes | Wellcome Sanger Institute for the COVID-19 Genomics UK (COG-UK) Consortium | The Lighthouse Lab in Milton Keynes and Alex Alderton, Roberto Amato, Jeffrey Barrett, Sonia Goncalves, Ewan Harrison, David K. Jackson, Ian Johnston, Dominic Kwiatkowski, Cordelia Langford, John Sillitoe on behalf of the Wellcome Sanger Institute COVID-19 Surveillance Team |
| EPI_ISL_3378337 | LABORATOIRE ACCOLAB SUD OUEST | CNR Virus des Infections Respiratoires - France SUD | Antonin Bal, Gregory Destras, Gwendolyne Burfin, Hadrien Regue, Quentin Semanas, Martine Valette, Bruno Lina, Laurence Josset |
| EPI_ISL_3378338 | Lighthouse Lab in Milton Keynes | Wellcome Sanger Institute for the COVID-19 Genomics UK (COG-UK) Consortium | The Lighthouse Lab in Milton Keynes and Alex Alderton, Roberto Amato, Jeffrey Barrett, Sonia Goncalves, Ewan Harrison, David K. Jackson, Ian Johnston, Dominic Kwiatkowski, Cordelia Langford, John Sillitoe on behalf of the Wellcome Sanger Institute COVID-19 Surveillance Team |
| EPI_ISL_3378339 | Lighthouse Lab in Alderley Park | Wellcome Sanger Institute for the COVID-19 Genomics UK (COG-UK) Consortium | Jacquelyn Wynn, Mairead Hyland, The Lighthouse Lab in Alderley Park and Alex Alderton, Roberto Amato, Jeffrey Barrett, Sonia Goncalves, Ewan Harrison, David K. Jackson, Ian Johnston, Dominic Kwiatkowski, Cordelia Langford, John Sillitoe on behalf of the Wellcome Sanger Institute COVID-19 Surveillance Team |

| | | | |
|---|---|---|---|
| EPI_ISL_3378340, EPI_ISL_3378341 | Lighthouse Lab in Milton Keynes | Wellcome Sanger Institute for the COVID-19 Genomics UK (COG-UK) Consortium | The Lighthouse Lab in Milton Keynes and Alex Alderton, Roberto Amato, Jeffrey Barrett, Sonia Goncalves, Ewan Harrison, David K. Jackson, Ian Johnston, Dominic Kwiatkowski, Cordelia Langford, John Sillitoe on behalf of the Wellcome Sanger Institute COVID-19 Surveillance Team |
| EPI_ISL_3378342 | ORIAPOLE | CNR Virus des Infections Respiratoires - France SUD | Antonin Bal, Gregory Destras, Gwendolyne Burfin, Hadrien Regue, Quentin Semanas, Martine Valette, Bruno Lina, Laurence Josset |
| EPI_ISL_3378343 | Lighthouse Lab in Alderley Park | Wellcome Sanger Institute for the COVID-19 Genomics UK (COG-UK) Consortium | Jacquelyn Wynn, Mairead Hyland, The Lighthouse Lab in Alderley Park and Alex Alderton, Roberto Amato, Jeffrey Barrett, Sonia Goncalves, Ewan Harrison, David K. Jackson, Ian Johnston, Dominic Kwiatkowski, Cordelia Langford, John Sillitoe on behalf of the Wellcome Sanger Institute COVID-19 Surveillance Team |
| EPI_ISL_3378344 | Lighthouse Lab in Milton Keynes | Wellcome Sanger Institute for the COVID-19 Genomics UK (COG-UK) Consortium | The Lighthouse Lab in Milton Keynes and Alex Alderton, Roberto Amato, Jeffrey Barrett, Sonia Goncalves, Ewan Harrison, David K. Jackson, Ian Johnston, Dominic Kwiatkowski, Cordelia Langford, John Sillitoe on behalf of the Wellcome Sanger Institute COVID-19 Surveillance Team |
| EPI_ISL_3378345 | Berkshire and Surrey Pathology Services Lighthouse Laboratory | Wellcome Sanger Institute for the COVID-19 Genomics UK (COG-UK) Consortium | Berkshire and Surrey Pathology Services Lighthouse Laboratory and Alex Alderton, Roberto Amato, Jeffrey Barrett, Sonia Goncalves, Ewan Harrison, David K. Jackson, Ian Johnston, Dominic Kwiatkowski, Cordelia Langford, John Sillitoe on behalf of the Wellcome Sanger Institute COVID-19 Surveillance Team |
| EPI_ISL_3378346 | CNR Virus des Infections Respiratoires - France SUD | CNR Virus des Infections Respiratoires - France SUD | Antonin Bal, Gregory Destras, Gwendolyne Burfin, Hadrien Regue, Quentin Semanas, Martine Valette, Bruno Lina, Laurence Josset |
| EPI_ISL_3378347, EPI_ISL_3378348 | Lighthouse Lab in Milton Keynes | Wellcome Sanger Institute for the COVID-19 Genomics UK (COG-UK) Consortium | The Lighthouse Lab in Milton Keynes and Alex Alderton, Roberto Amato, Jeffrey Barrett, Sonia Goncalves, Ewan Harrison, David K. Jackson, Ian Johnston, Dominic Kwiatkowski, Cordelia Langford, John Sillitoe on behalf of the Wellcome Sanger Institute COVID-19 Surveillance Team |
| EPI_ISL_3378349 | Berkshire and Surrey Pathology Services Lighthouse Laboratory | Wellcome Sanger Institute for the COVID-19 Genomics UK (COG-UK) Consortium | Berkshire and Surrey Pathology Services Lighthouse Laboratory and Alex Alderton, Roberto Amato, Jeffrey Barrett, Sonia Goncalves, Ewan Harrison, David K. Jackson, Ian Johnston, Dominic Kwiatkowski, Cordelia Langford, John Sillitoe on behalf of the Wellcome Sanger Institute COVID-19 Surveillance Team |
| EPI_ISL_3378350 | CENTRE HOSPITALIER DE VALENCE | CNR Virus des Infections Respiratoires - France SUD | Antonin Bal, Gregory Destras, Gwendolyne Burfin, Hadrien Regue, Quentin Semanas, Martine Valette, Bruno Lina, Laurence Josset |
| EPI_ISL_3378351, EPI_ISL_3378352 | Berkshire and Surrey Pathology Services Lighthouse Laboratory | Wellcome Sanger Institute for the COVID-19 Genomics UK (COG-UK) Consortium | Berkshire and Surrey Pathology Services Lighthouse Laboratory and Alex Alderton, Roberto Amato, Jeffrey Barrett, Sonia Goncalves, Ewan Harrison, David K. Jackson, Ian Johnston, Dominic Kwiatkowski, Cordelia Langford, John Sillitoe on behalf of the Wellcome Sanger Institute COVID-19 Surveillance Team |
| EPI_ISL_3378353 | Lighthouse Lab in Milton Keynes | Wellcome Sanger Institute for the COVID-19 Genomics UK (COG-UK) Consortium | The Lighthouse Lab in Milton Keynes and Alex Alderton, Roberto Amato, Jeffrey Barrett, Sonia Goncalves, Ewan Harrison, David K. Jackson, Ian Johnston, Dominic Kwiatkowski, Cordelia Langford, John Sillitoe on behalf of the Wellcome Sanger Institute COVID-19 Surveillance Team |
| EPI_ISL_3378354 | Lighthouse Lab in Alderley Park | Wellcome Sanger Institute for the COVID-19 Genomics UK (COG-UK) Consortium | Jacquelyn Wynn, Mairead Hyland, The Lighthouse Lab in Alderley Park and Alex Alderton, Roberto Amato, Jeffrey Barrett, Sonia Goncalves, Ewan Harrison, David K. Jackson, Ian Johnston, Dominic Kwiatkowski, Cordelia Langford, John Sillitoe on behalf of the Wellcome Sanger Institute COVID-19 Surveillance Team |
| EPI_ISL_3378355 | LABORATOIRE BIOESTEREL | CNR Virus des Infections Respiratoires - France SUD | Antonin Bal, Gregory Destras, Gwendolyne Burfin, Hadrien Regue, Quentin Semanas, Martine Valette, Bruno Lina, Laurence Josset |
| EPI_ISL_3378356 | Lighthouse Lab in Milton Keynes | Wellcome Sanger Institute for the COVID-19 Genomics UK (COG-UK) Consortium | The Lighthouse Lab in Milton Keynes and Alex Alderton, Roberto Amato, Jeffrey Barrett, Sonia Goncalves, Ewan Harrison, David K. Jackson, Ian Johnston, Dominic Kwiatkowski, Cordelia Langford, John Sillitoe on behalf of the Wellcome Sanger Institute COVID-19 Surveillance Team |
| EPI_ISL_3378357 | Berkshire and Surrey Pathology Services Lighthouse Laboratory | Wellcome Sanger Institute for the COVID-19 Genomics UK (COG-UK) Consortium | Berkshire and Surrey Pathology Services Lighthouse Laboratory and Alex Alderton, Roberto Amato, Jeffrey Barrett, Sonia Goncalves, Ewan Harrison, David K. Jackson, Ian Johnston, Dominic Kwiatkowski, Cordelia Langford, John Sillitoe on behalf of the Wellcome Sanger Institute COVID-19 Surveillance Team |
| EPI_ISL_3378358 | Lighthouse Lab in Alderley Park | Wellcome Sanger Institute for the COVID-19 Genomics UK (COG-UK) Consortium | Jacquelyn Wynn, Mairead Hyland, The Lighthouse Lab in Alderley Park and Alex Alderton, Roberto Amato, Jeffrey Barrett, Sonia Goncalves, Ewan Harrison, David K. Jackson, Ian Johnston, Dominic Kwiatkowski, Cordelia Langford, John Sillitoe on behalf of the Wellcome Sanger Institute COVID-19 Surveillance Team |
| EPI_ISL_3378359 | LABORATOIRE BIOESTEREL | CNR Virus des Infections Respiratoires - France SUD | Antonin Bal, Gregory Destras, Gwendolyne Burfin, Hadrien Regue, Quentin Semanas, Martine Valette, Bruno Lina, Laurence Josset |
| EPI_ISL_3378360 | Lighthouse Lab in Milton Keynes | Wellcome Sanger Institute for the COVID-19 Genomics UK (COG-UK) Consortium | The Lighthouse Lab in Milton Keynes and Alex Alderton, Roberto Amato, Jeffrey Barrett, Sonia Goncalves, Ewan Harrison, David K. Jackson, Ian Johnston, Dominic Kwiatkowski, Cordelia Langford, John Sillitoe on behalf of the Wellcome Sanger Institute COVID-19 Surveillance Team |
| EPI_ISL_3378361 | Berkshire and Surrey Pathology Services Lighthouse Laboratory | Wellcome Sanger Institute for the COVID-19 Genomics UK (COG-UK) Consortium | Berkshire and Surrey Pathology Services Lighthouse Laboratory and Alex Alderton, Roberto Amato, Jeffrey Barrett, Sonia Goncalves, Ewan Harrison, David K. Jackson, Ian Johnston, Dominic Kwiatkowski, Cordelia Langford, John Sillitoe on behalf of the Wellcome Sanger Institute COVID-19 Surveillance Team |
| EPI_ISL_3378362 | Lighthouse Lab in Milton Keynes | Wellcome Sanger Institute for the COVID-19 Genomics UK (COG-UK) Consortium | The Lighthouse Lab in Milton Keynes and Alex Alderton, Roberto Amato, Jeffrey Barrett, Sonia Goncalves, Ewan Harrison, David K. Jackson, Ian Johnston, Dominic Kwiatkowski, Cordelia Langford, John Sillitoe on behalf of the Wellcome Sanger Institute COVID-19 Surveillance Team |
| EPI_ISL_3378363 | LABORATOIRE BIOESTEREL | CNR Virus des Infections Respiratoires - France SUD | Antonin Bal, Gregory Destras, Gwendolyne Burfin, Hadrien Regue, Quentin Semanas, Martine Valette, Bruno Lina, Laurence Josset |
| EPI_ISL_3378364 | Lighthouse Lab in Milton Keynes | Wellcome Sanger Institute for the COVID-19 Genomics UK (COG-UK) Consortium | The Lighthouse Lab in Milton Keynes and Alex Alderton, Roberto Amato, Jeffrey Barrett, Sonia Goncalves, Ewan Harrison, David K. Jackson, Ian Johnston, Dominic Kwiatkowski, Cordelia Langford, John Sillitoe on behalf of the Wellcome Sanger Institute COVID-19 Surveillance Team |
| EPI_ISL_3378365 | Berkshire and Surrey Pathology Services Lighthouse Laboratory | Wellcome Sanger Institute for the COVID-19 Genomics UK (COG-UK) Consortium | Berkshire and Surrey Pathology Services Lighthouse Laboratory and Alex Alderton, Roberto Amato, Jeffrey Barrett, Sonia Goncalves, Ewan Harrison, David K. Jackson, Ian Johnston, Dominic Kwiatkowski, Cordelia Langford, John Sillitoe on behalf of the Wellcome Sanger Institute COVID-19 Surveillance Team |
| EPI_ISL_3378366 | Lighthouse Lab in Milton Keynes | Wellcome Sanger Institute for the COVID-19 Genomics UK (COG-UK) Consortium | The Lighthouse Lab in Milton Keynes and Alex Alderton, Roberto Amato, Jeffrey Barrett, Sonia Goncalves, Ewan Harrison, David K. Jackson, Ian Johnston, Dominic Kwiatkowski, Cordelia Langford, John Sillitoe on behalf of the Wellcome Sanger Institute COVID-19 Surveillance Team |
| EPI_ISL_3378367 | LABORATOIRE BIOESTEREL | CNR Virus des Infections Respiratoires - France SUD | Antonin Bal, Gregory Destras, Gwendolyne Burfin, Hadrien Regue, Quentin Semanas, Martine Valette, Bruno Lina, Laurence Josset |
| EPI_ISL_3378368, EPI_ISL_3378369 | Berkshire and Surrey Pathology Services Lighthouse Laboratory | Wellcome Sanger Institute for the COVID-19 Genomics UK (COG-UK) Consortium | Berkshire and Surrey Pathology Services Lighthouse Laboratory and Alex Alderton, Roberto Amato, Jeffrey Barrett, Sonia Goncalves, Ewan Harrison, David K. Jackson, Ian Johnston, Dominic Kwiatkowski, Cordelia Langford, John Sillitoe on behalf of the Wellcome Sanger Institute COVID-19 Surveillance Team |
| EPI_ISL_3378370 | Lighthouse Lab in Alderley Park | Wellcome Sanger Institute for the COVID-19 Genomics UK (COG-UK) Consortium | Jacquelyn Wynn, Mairead Hyland, The Lighthouse Lab in Alderley Park and Alex Alderton, Roberto Amato, Jeffrey Barrett, Sonia Goncalves, Ewan Harrison, David K. Jackson, Ian Johnston, Dominic Kwiatkowski, Cordelia Langford, John Sillitoe on behalf of the Wellcome Sanger Institute COVID-19 Surveillance Team |
| EPI_ISL_3378371 | CENTRE HOSPITALIER ST JOSEPH ST LUC | CNR Virus des Infections Respiratoires - France SUD | Antonin Bal, Gregory Destras, Gwendolyne Burfin, Hadrien Regue, Quentin Semanas, Martine Valette, Bruno Lina, Laurence Josset |
| EPI_ISL_3378372, EPI_ISL_3378373 | Lighthouse Lab in Alderley Park | Wellcome Sanger Institute for the COVID-19 Genomics UK (COG-UK) Consortium | Jacquelyn Wynn, Mairead Hyland, The Lighthouse Lab in Alderley Park and Alex Alderton, Roberto Amato, Jeffrey Barrett, Sonia Goncalves, Ewan Harrison, David K. Jackson, Ian Johnston, Dominic Kwiatkowski, Cordelia Langford, John Sillitoe on behalf of the Wellcome Sanger Institute COVID-19 Surveillance Team |
| EPI_ISL_3378374 | Lighthouse Lab in Milton Keynes | Wellcome Sanger Institute for the COVID-19 Genomics UK (COG-UK) Consortium | The Lighthouse Lab in Milton Keynes and Alex Alderton, Roberto Amato, Jeffrey Barrett, Sonia Goncalves, Ewan Harrison, David K. Jackson, Ian Johnston, Dominic Kwiatkowski, Cordelia Langford, John Sillitoe on behalf of the Wellcome Sanger Institute COVID-19 Surveillance Team |
| EPI_ISL_3378375 | LABORATOIRE BIOESTEREL | CNR Virus des Infections Respiratoires - France SUD | Antonin Bal, Gregory Destras, Gwendolyne Burfin, Hadrien Regue, Quentin Semanas, Martine Valette, Bruno Lina, Laurence Josset |
| EPI_ISL_3378376 | Lighthouse Lab in Alderley Park | Wellcome Sanger Institute for the COVID-19 Genomics UK (COG-UK) Consortium | Jacquelyn Wynn, Mairead Hyland, The Lighthouse Lab in Alderley Park and Alex Alderton, Roberto Amato, Jeffrey Barrett, Sonia Goncalves, Ewan Harrison, David K. Jackson, Ian Johnston, Dominic Kwiatkowski, Cordelia Langford, John Sillitoe on behalf of the Wellcome Sanger Institute COVID-19 Surveillance Team |
| EPI_ISL_3378377 | Lighthouse Lab in Milton Keynes | Wellcome Sanger Institute for the COVID-19 Genomics UK (COG-UK) Consortium | The Lighthouse Lab in Milton Keynes and Alex Alderton, Roberto Amato, Jeffrey Barrett, Sonia Goncalves, Ewan Harrison, David K. Jackson, Ian Johnston, Dominic Kwiatkowski, Cordelia Langford, John Sillitoe on behalf of the Wellcome Sanger Institute COVID-19 Surveillance Team |
| EPI_ISL_3378378 | Berkshire and Surrey Pathology Services Lighthouse Laboratory | Wellcome Sanger Institute for the COVID-19 Genomics UK (COG-UK) Consortium | Berkshire and Surrey Pathology Services Lighthouse Laboratory and Alex Alderton, Roberto Amato, Jeffrey Barrett, Sonia Goncalves, Ewan Harrison, David K. Jackson, Ian Johnston, Dominic Kwiatkowski, Cordelia Langford, John Sillitoe on behalf of the Wellcome Sanger Institute COVID-19 Surveillance Team |
| EPI_ISL_3378379 | Lighthouse Lab in Milton Keynes | Wellcome Sanger Institute for the COVID-19 Genomics UK (COG-UK) Consortium | The Lighthouse Lab in Milton Keynes and Alex Alderton, Roberto Amato, Jeffrey Barrett, Sonia Goncalves, Ewan Harrison, David K. Jackson, Ian Johnston, Dominic Kwiatkowski, Cordelia Langford, John Sillitoe on behalf of the Wellcome Sanger Institute COVID-19 Surveillance Team |
| EPI_ISL_3378380 | LABORATOIRE BIOESTEREL | CNR Virus des Infections Respiratoires - France SUD | Antonin Bal, Gregory Destras, Gwendolyne Burfin, Hadrien Regue, Quentin Semanas, Martine Valette, Bruno Lina, Laurence Josset |
| EPI_ISL_3378381, EPI_ISL_3378382 | Lighthouse Lab in Milton Keynes | Wellcome Sanger Institute for the COVID-19 Genomics UK (COG-UK) Consortium | The Lighthouse Lab in Milton Keynes and Alex Alderton, Roberto Amato, Jeffrey Barrett, Sonia Goncalves, Ewan Harrison, David K. Jackson, Ian Johnston, Dominic Kwiatkowski, Cordelia Langford, John Sillitoe on behalf of the Wellcome Sanger Institute COVID-19 Surveillance Team |
| EPI_ISL_3378383 | Berkshire and Surrey Pathology Services Lighthouse Laboratory | Wellcome Sanger Institute for the COVID-19 Genomics UK (COG-UK) Consortium | Berkshire and Surrey Pathology Services Lighthouse Laboratory and Alex Alderton, Roberto Amato, Jeffrey Barrett, Sonia Goncalves, Ewan Harrison, David K. Jackson, Ian Johnston, Dominic Kwiatkowski, Cordelia Langford, John Sillitoe on behalf of the Wellcome Sanger Institute COVID-19 Surveillance Team |
| EPI_ISL_3378384 | LABORATOIRE BIOESTEREL | CNR Virus des Infections Respiratoires - France SUD | Antonin Bal, Gregory Destras, Gwendolyne Burfin, Hadrien Regue, Quentin Semanas, Martine Valette, Bruno Lina, Laurence Josset |
| EPI_ISL_3378385, EPI_ISL_3378386 | Lighthouse Lab in Milton Keynes | Wellcome Sanger Institute for the COVID-19 Genomics UK (COG-UK) Consortium | The Lighthouse Lab in Milton Keynes and Alex Alderton, Roberto Amato, Jeffrey Barrett, Sonia Goncalves, Ewan Harrison, David K. Jackson, Ian Johnston, Dominic Kwiatkowski, Cordelia Langford, John Sillitoe on behalf of the Wellcome Sanger Institute COVID-19 Surveillance Team |
| EPI_ISL_3378387 | Lighthouse Lab in Alderley Park | Wellcome Sanger Institute for the COVID-19 Genomics UK (COG-UK) Consortium | Jacquelyn Wynn, Mairead Hyland, The Lighthouse Lab in Alderley Park and Alex Alderton, Roberto Amato, Jeffrey Barrett, Sonia Goncalves, Ewan Harrison, David K. Jackson, Ian Johnston, Dominic Kwiatkowski, Cordelia Langford, John Sillitoe on behalf of the Wellcome Sanger Institute COVID-19 Surveillance Team |
| EPI_ISL_3378388 | Lighthouse Lab in Milton Keynes | Wellcome Sanger Institute for the COVID-19 Genomics UK (COG-UK) Consortium | The Lighthouse Lab in Milton Keynes and Alex Alderton, Roberto Amato, Jeffrey Barrett, Sonia Goncalves, Ewan Harrison, David K. Jackson, Ian Johnston, Dominic Kwiatkowski, Cordelia Langford, John Sillitoe on behalf of the Wellcome Sanger Institute COVID-19 Surveillance Team |
| EPI_ISL_3378389 | LABORATOIRE BIOESTEREL | CNR Virus des Infections Respiratoires - France SUD | Antonin Bal, Gregory Destras, Gwendolyne Burfin, Hadrien Regue, Quentin Semanas, Martine Valette, Bruno Lina, Laurence Josset |
| EPI_ISL_3378390, EPI_ISL_3378391, EPI_ISL_3378392 | Lighthouse Lab in Milton Keynes | Wellcome Sanger Institute for the COVID-19 Genomics UK (COG-UK) Consortium | The Lighthouse Lab in Milton Keynes and Alex Alderton, Roberto Amato, Jeffrey Barrett, Sonia Goncalves, Ewan Harrison, David K. Jackson, Ian Johnston, Dominic Kwiatkowski, Cordelia Langford, John Sillitoe on behalf of the Wellcome Sanger Institute COVID-19 Surveillance Team |
| EPI_ISL_3378393 | LABORATOIRE BIOESTEREL | CNR Virus des Infections Respiratoires - France SUD | Antonin Bal, Gregory Destras, Gwendolyne Burfin, Hadrien Regue, Quentin Semanas, Martine Valette, Bruno Lina, Laurence Josset |
| EPI_ISL_3378394 | Lighthouse Lab in Alderley Park | Wellcome Sanger Institute for the COVID-19 Genomics UK (COG-UK) Consortium | Jacquelyn Wynn, Mairead Hyland, The Lighthouse Lab in Alderley Park and Alex Alderton, Roberto Amato, Jeffrey Barrett, Sonia Goncalves, Ewan Harrison, David K. Jackson, Ian Johnston, Dominic Kwiatkowski, Cordelia Langford, John Sillitoe on behalf of the Wellcome Sanger Institute COVID-19 Surveillance Team |
| EPI_ISL_3378395 | Berkshire and Surrey Pathology Services Lighthouse Laboratory | Wellcome Sanger Institute for the COVID-19 Genomics UK (COG-UK) Consortium | Berkshire and Surrey Pathology Services Lighthouse Laboratory and Alex Alderton, Roberto Amato, Jeffrey Barrett, Sonia Goncalves, Ewan Harrison, David K. Jackson, Ian Johnston, Dominic Kwiatkowski, Cordelia Langford, John Sillitoe on behalf of the Wellcome Sanger Institute COVID-19 Surveillance Team |
| EPI_ISL_3378396 | Lighthouse Lab in Alderley Park | Wellcome Sanger Institute for the COVID-19 Genomics UK (COG-UK) Consortium | Jacquelyn Wynn, Mairead Hyland, The Lighthouse Lab in Alderley Park and Alex Alderton, Roberto Amato, Jeffrey Barrett, Sonia Goncalves, Ewan Harrison, David K. Jackson, Ian Johnston, Dominic Kwiatkowski, Cordelia Langford, John Sillitoe on behalf of the Wellcome Sanger Institute COVID-19 Surveillance Team |
| EPI_ISL_3378397 | LABORATOIRE ACCOLAB SUD OUEST | CNR Virus des Infections Respiratoires - France SUD | Antonin Bal, Gregory Destras, Gwendolyne Burfin, Hadrien Regue, Quentin Semanas, Martine Valette, Bruno Lina, Laurence Josset |
| EPI_ISL_3378398, EPI_ISL_3378399, EPI_ISL_3378400 | Lighthouse Lab in Milton Keynes | Wellcome Sanger Institute for the COVID-19 Genomics UK (COG-UK) Consortium | The Lighthouse Lab in Milton Keynes and Alex Alderton, Roberto Amato, Jeffrey Barrett, Sonia Goncalves, Ewan Harrison, David K. Jackson, Ian Johnston, Dominic Kwiatkowski, Cordelia Langford, John Sillitoe on behalf of the Wellcome Sanger Institute COVID-19 Surveillance Team |

| | | | |
|---|---|---|---|
| EPI_ISL_3378401 | LABORATOIRE BIOESTEREL | CNR Virus des Infections Respiratoires - France SUD | Antonin Bal, Gregory Destras, Gwendolyne Burfin, Hadrien Regue, Quentin Semanas, Martine Valette, Bruno Lina, Laurence Josset |
| EPI_ISL_3378402, EPI_ISL_3378403 | Lighthouse Lab in Alderley Park | Wellcome Sanger Institute for the COVID-19 Genomics UK (COG-UK) Consortium | Jacquelyn Wynn, Mairead Hyland, The Lighthouse Lab in Alderley Park and Alex Alderton, Roberto Amato, Jeffrey Barrett, Sonia Goncalves, Ewan Harrison, David K. Jackson, Ian Johnston, Dominic Kwiatkowski, Cordelia Langford, John Sillitoe on behalf of the Wellcome Sanger Institute COVID-19 Surveillance Team |
| EPI_ISL_3378404 | Berkshire and Surrey Pathology Services Lighthouse Laboratory | Wellcome Sanger Institute for the COVID-19 Genomics UK (COG-UK) Consortium | Berkshire and Surrey Pathology Services Lighthouse Laboratory and Alex Alderton, Roberto Amato, Jeffrey Barrett, Sonia Goncalves, Ewan Harrison, David K. Jackson, Ian Johnston, Dominic Kwiatkowski, Cordelia Langford, John Sillitoe on behalf of the Wellcome Sanger Institute COVID-19 Surveillance Team |
| EPI_ISL_3378405 | LABORATOIRE BIOESTEREL | CNR Virus des Infections Respiratoires - France SUD | Antonin Bal, Gregory Destras, Gwendolyne Burfin, Hadrien Regue, Quentin Semanas, Martine Valette, Bruno Lina, Laurence Josset |
| EPI_ISL_3378406 | Berkshire and Surrey Pathology Services Lighthouse Laboratory | Wellcome Sanger Institute for the COVID-19 Genomics UK (COG-UK) Consortium | Berkshire and Surrey Pathology Services Lighthouse Laboratory and Alex Alderton, Roberto Amato, Jeffrey Barrett, Sonia Goncalves, Ewan Harrison, David K. Jackson, Ian Johnston, Dominic Kwiatkowski, Cordelia Langford, John Sillitoe on behalf of the Wellcome Sanger Institute COVID-19 Surveillance Team |
| EPI_ISL_3378407 | Lighthouse Lab in Milton Keynes | Wellcome Sanger Institute for the COVID-19 Genomics UK (COG-UK) Consortium | The Lighthouse Lab in Milton Keynes and Alex Alderton, Roberto Amato, Jeffrey Barrett, Sonia Goncalves, Ewan Harrison, David K. Jackson, Ian Johnston, Dominic Kwiatkowski, Cordelia Langford, John Sillitoe on behalf of the Wellcome Sanger Institute COVID-19 Surveillance Team |
| EPI_ISL_3378408 | Lighthouse Lab in Alderley Park | Wellcome Sanger Institute for the COVID-19 Genomics UK (COG-UK) Consortium | Jacquelyn Wynn, Mairead Hyland, The Lighthouse Lab in Alderley Park and Alex Alderton, Roberto Amato, Jeffrey Barrett, Sonia Goncalves, Ewan Harrison, David K. Jackson, Ian Johnston, Dominic Kwiatkowski, Cordelia Langford, John Sillitoe on behalf of the Wellcome Sanger Institute COVID-19 Surveillance Team |
| EPI_ISL_3378409 | CNR Virus des Infections Respiratoires - France SUD | CNR Virus des Infections Respiratoires - France SUD | Antonin Bal, Gregory Destras, Gwendolyne Burfin, Hadrien Regue, Quentin Semanas, Martine Valette, Bruno Lina, Laurence Josset |
| EPI_ISL_3378410 | Lighthouse Lab in Milton Keynes | Wellcome Sanger Institute for the COVID-19 Genomics UK (COG-UK) Consortium | The Lighthouse Lab in Milton Keynes and Alex Alderton, Roberto Amato, Jeffrey Barrett, Sonia Goncalves, Ewan Harrison, David K. Jackson, Ian Johnston, Dominic Kwiatkowski, Cordelia Langford, John Sillitoe on behalf of the Wellcome Sanger Institute COVID-19 Surveillance Team |
| EPI_ISL_3378411 | Berkshire and Surrey Pathology Services Lighthouse Laboratory | Wellcome Sanger Institute for the COVID-19 Genomics UK (COG-UK) Consortium | Berkshire and Surrey Pathology Services Lighthouse Laboratory and Alex Alderton, Roberto Amato, Jeffrey Barrett, Sonia Goncalves, Ewan Harrison, David K. Jackson, Ian Johnston, Dominic Kwiatkowski, Cordelia Langford, John Sillitoe on behalf of the Wellcome Sanger Institute COVID-19 Surveillance Team |
| EPI_ISL_3378412 | Lighthouse Lab in Alderley Park | Wellcome Sanger Institute for the COVID-19 Genomics UK (COG-UK) Consortium | Jacquelyn Wynn, Mairead Hyland, The Lighthouse Lab in Alderley Park and Alex Alderton, Roberto Amato, Jeffrey Barrett, Sonia Goncalves, Ewan Harrison, David K. Jackson, Ian Johnston, Dominic Kwiatkowski, Cordelia Langford, John Sillitoe on behalf of the Wellcome Sanger Institute COVID-19 Surveillance Team |
| EPI_ISL_3378413 | LABORATOIRE BIOESTEREL | CNR Virus des Infections Respiratoires - France SUD | Antonin Bal, Gregory Destras, Gwendolyne Burfin, Hadrien Regue, Quentin Semanas, Martine Valette, Bruno Lina, Laurence Josset |
| EPI_ISL_3378414 | Lighthouse Lab in Milton Keynes | Wellcome Sanger Institute for the COVID-19 Genomics UK (COG-UK) Consortium | The Lighthouse Lab in Milton Keynes and Alex Alderton, Roberto Amato, Jeffrey Barrett, Sonia Goncalves, Ewan Harrison, David K. Jackson, Ian Johnston, Dominic Kwiatkowski, Cordelia Langford, John Sillitoe on behalf of the Wellcome Sanger Institute COVID-19 Surveillance Team |
| EPI_ISL_3378415 | Lighthouse Lab in Alderley Park | Wellcome Sanger Institute for the COVID-19 Genomics UK (COG-UK) Consortium | Jacquelyn Wynn, Mairead Hyland, The Lighthouse Lab in Alderley Park and Alex Alderton, Roberto Amato, Jeffrey Barrett, Sonia Goncalves, Ewan Harrison, David K. Jackson, Ian Johnston, Dominic Kwiatkowski, Cordelia Langford, John Sillitoe on behalf of the Wellcome Sanger Institute COVID-19 Surveillance Team |
| EPI_ISL_3378416 | Lighthouse Lab in Milton Keynes | Wellcome Sanger Institute for the COVID-19 Genomics UK (COG-UK) Consortium | The Lighthouse Lab in Milton Keynes and Alex Alderton, Roberto Amato, Jeffrey Barrett, Sonia Goncalves, Ewan Harrison, David K. Jackson, Ian Johnston, Dominic Kwiatkowski, Cordelia Langford, John Sillitoe on behalf of the Wellcome Sanger Institute COVID-19 Surveillance Team |
| EPI_ISL_3378417 | LABORATOIRE BIOESTEREL | CNR Virus des Infections Respiratoires - France SUD | Antonin Bal, Gregory Destras, Gwendolyne Burfin, Hadrien Regue, Quentin Semanas, Martine Valette, Bruno Lina, Laurence Josset |
| EPI_ISL_3378418, EPI_ISL_3378419, EPI_ISL_3378420 | Lighthouse Lab in Milton Keynes | Wellcome Sanger Institute for the COVID-19 Genomics UK (COG-UK) Consortium | The Lighthouse Lab in Milton Keynes and Alex Alderton, Roberto Amato, Jeffrey Barrett, Sonia Goncalves, Ewan Harrison, David K. Jackson, Ian Johnston, Dominic Kwiatkowski, Cordelia Langford, John Sillitoe on behalf of the Wellcome Sanger Institute COVID-19 Surveillance Team |
| EPI_ISL_3378421 | LABORATOIRE BIOESTEREL | CNR Virus des Infections Respiratoires - France SUD | Antonin Bal, Gregory Destras, Gwendolyne Burfin, Hadrien Regue, Quentin Semanas, Martine Valette, Bruno Lina, Laurence Josset |
| EPI_ISL_3378422 | Lighthouse Lab in Milton Keynes | Wellcome Sanger Institute for the COVID-19 Genomics UK (COG-UK) Consortium | The Lighthouse Lab in Milton Keynes and Alex Alderton, Roberto Amato, Jeffrey Barrett, Sonia Goncalves, Ewan Harrison, David K. Jackson, Ian Johnston, Dominic Kwiatkowski, Cordelia Langford, John Sillitoe on behalf of the Wellcome Sanger Institute COVID-19 Surveillance Team |
| EPI_ISL_3378423 | Berkshire and Surrey Pathology Services Lighthouse Laboratory | Wellcome Sanger Institute for the COVID-19 Genomics UK (COG-UK) Consortium | Berkshire and Surrey Pathology Services Lighthouse Laboratory and Alex Alderton, Roberto Amato, Jeffrey Barrett, Sonia Goncalves, Ewan Harrison, David K. Jackson, Ian Johnston, Dominic Kwiatkowski, Cordelia Langford, John Sillitoe on behalf of the Wellcome Sanger Institute COVID-19 Surveillance Team |
| EPI_ISL_3378424 | Lighthouse Lab in Alderley Park | Wellcome Sanger Institute for the COVID-19 Genomics UK (COG-UK) Consortium | Jacquelyn Wynn, Mairead Hyland, The Lighthouse Lab in Alderley Park and Alex Alderton, Roberto Amato, Jeffrey Barrett, Sonia Goncalves, Ewan Harrison, David K. Jackson, Ian Johnston, Dominic Kwiatkowski, Cordelia Langford, John Sillitoe on behalf of the Wellcome Sanger Institute COVID-19 Surveillance Team |
| EPI_ISL_3378425 | LABORATOIRE BIOESTEREL | CNR Virus des Infections Respiratoires - France SUD | Antonin Bal, Gregory Destras, Gwendolyne Burfin, Hadrien Regue, Quentin Semanas, Martine Valette, Bruno Lina, Laurence Josset |
| EPI_ISL_3378426 | Lighthouse Lab in Milton Keynes | Wellcome Sanger Institute for the COVID-19 Genomics UK (COG-UK) Consortium | The Lighthouse Lab in Milton Keynes and Alex Alderton, Roberto Amato, Jeffrey Barrett, Sonia Goncalves, Ewan Harrison, David K. Jackson, Ian Johnston, Dominic Kwiatkowski, Cordelia Langford, John Sillitoe on behalf of the Wellcome Sanger Institute COVID-19 Surveillance Team |
| EPI_ISL_3378427 | Lighthouse Lab in Alderley Park | Wellcome Sanger Institute for the COVID-19 Genomics UK (COG-UK) Consortium | Jacquelyn Wynn, Mairead Hyland, The Lighthouse Lab in Alderley Park and Alex Alderton, Roberto Amato, Jeffrey Barrett, Sonia Goncalves, Ewan Harrison, David K. Jackson, Ian Johnston, Dominic Kwiatkowski, Cordelia Langford, John Sillitoe on behalf of the Wellcome Sanger Institute COVID-19 Surveillance Team |
| EPI_ISL_3378428 | Lighthouse Lab in Milton Keynes | Wellcome Sanger Institute for the COVID-19 Genomics UK (COG-UK) Consortium | The Lighthouse Lab in Milton Keynes and Alex Alderton, Roberto Amato, Jeffrey Barrett, Sonia Goncalves, Ewan Harrison, David K. Jackson, Ian Johnston, Dominic Kwiatkowski, Cordelia Langford, John Sillitoe on behalf of the Wellcome Sanger Institute COVID-19 Surveillance Team |
| EPI_ISL_3378429 | Lighthouse Lab in Alderley Park | Wellcome Sanger Institute for the COVID-19 Genomics UK (COG-UK) Consortium | Jacquelyn Wynn, Mairead Hyland, The Lighthouse Lab in Alderley Park and Alex Alderton, Roberto Amato, Jeffrey Barrett, Sonia Goncalves, Ewan Harrison, David K. Jackson, Ian Johnston, Dominic Kwiatkowski, Cordelia Langford, John Sillitoe on behalf of the Wellcome Sanger Institute COVID-19 Surveillance Team |
| EPI_ISL_3378430 | Lighthouse Lab in Milton Keynes | Wellcome Sanger Institute for the COVID-19 Genomics UK (COG-UK) Consortium | The Lighthouse Lab in Milton Keynes and Alex Alderton, Roberto Amato, Jeffrey Barrett, Sonia Goncalves, Ewan Harrison, David K. Jackson, Ian Johnston, Dominic Kwiatkowski, Cordelia Langford, John Sillitoe on behalf of the Wellcome Sanger Institute COVID-19 Surveillance Team |
| EPI_ISL_3378431 | LABORATOIRE BIOESTEREL | CNR Virus des Infections Respiratoires - France SUD | Antonin Bal, Gregory Destras, Gwendolyne Burfin, Hadrien Regue, Quentin Semanas, Martine Valette, Bruno Lina, Laurence Josset |
| EPI_ISL_3378432, EPI_ISL_3378433 | Lighthouse Lab in Milton Keynes | Wellcome Sanger Institute for the COVID-19 Genomics UK (COG-UK) Consortium | The Lighthouse Lab in Milton Keynes and Alex Alderton, Roberto Amato, Jeffrey Barrett, Sonia Goncalves, Ewan Harrison, David K. Jackson, Ian Johnston, Dominic Kwiatkowski, Cordelia Langford, John Sillitoe on behalf of the Wellcome Sanger Institute COVID-19 Surveillance Team |
| EPI_ISL_3378434 | LABORATOIRE BIOESTEREL | CNR Virus des Infections Respiratoires - France SUD | Antonin Bal, Gregory Destras, Gwendolyne Burfin, Hadrien Regue, Quentin Semanas, Martine Valette, Bruno Lina, Laurence Josset |
| EPI_ISL_3378435 | Berkshire and Surrey Pathology Services Lighthouse Laboratory | Wellcome Sanger Institute for the COVID-19 Genomics UK (COG-UK) Consortium | Berkshire and Surrey Pathology Services Lighthouse Laboratory and Alex Alderton, Roberto Amato, Jeffrey Barrett, Sonia Goncalves, Ewan Harrison, David K. Jackson, Ian Johnston, Dominic Kwiatkowski, Cordelia Langford, John Sillitoe on behalf of the Wellcome Sanger Institute COVID-19 Surveillance Team |
| EPI_ISL_3378436 | Lighthouse Lab in Alderley Park | Wellcome Sanger Institute for the COVID-19 Genomics UK (COG-UK) Consortium | Jacquelyn Wynn, Mairead Hyland, The Lighthouse Lab in Alderley Park and Alex Alderton, Roberto Amato, Jeffrey Barrett, Sonia Goncalves, Ewan Harrison, David K. Jackson, Ian Johnston, Dominic Kwiatkowski, Cordelia Langford, John Sillitoe on behalf of the Wellcome Sanger Institute COVID-19 Surveillance Team |
| EPI_ISL_3378437 | Lighthouse Lab in Milton Keynes | Wellcome Sanger Institute for the COVID-19 Genomics UK (COG-UK) Consortium | The Lighthouse Lab in Milton Keynes and Alex Alderton, Roberto Amato, Jeffrey Barrett, Sonia Goncalves, Ewan Harrison, David K. Jackson, Ian Johnston, Dominic Kwiatkowski, Cordelia Langford, John Sillitoe on behalf of the Wellcome Sanger Institute COVID-19 Surveillance Team |
| EPI_ISL_3378438 | LABORATOIRE BIOESTEREL | CNR Virus des Infections Respiratoires - France SUD | Antonin Bal, Gregory Destras, Gwendolyne Burfin, Hadrien Regue, Quentin Semanas, Martine Valette, Bruno Lina, Laurence Josset |
| EPI_ISL_3378439 | Lighthouse Lab in Alderley Park | Wellcome Sanger Institute for the COVID-19 Genomics UK (COG-UK) Consortium | Jacquelyn Wynn, Mairead Hyland, The Lighthouse Lab in Alderley Park and Alex Alderton, Roberto Amato, Jeffrey Barrett, Sonia Goncalves, Ewan Harrison, David K. Jackson, Ian Johnston, Dominic Kwiatkowski, Cordelia Langford, John Sillitoe on behalf of the Wellcome Sanger Institute COVID-19 Surveillance Team |
| EPI_ISL_3378440 | Lighthouse Lab in Milton Keynes | Wellcome Sanger Institute for the COVID-19 Genomics UK (COG-UK) Consortium | The Lighthouse Lab in Milton Keynes and Alex Alderton, Roberto Amato, Jeffrey Barrett, Sonia Goncalves, Ewan Harrison, David K. Jackson, Ian Johnston, Dominic Kwiatkowski, Cordelia Langford, John Sillitoe on behalf of the Wellcome Sanger Institute COVID-19 Surveillance Team |
| EPI_ISL_3378441 | Lighthouse Lab in Alderley Park | Wellcome Sanger Institute for the COVID-19 Genomics UK (COG-UK) Consortium | Jacquelyn Wynn, Mairead Hyland, The Lighthouse Lab in Alderley Park and Alex Alderton, Roberto Amato, Jeffrey Barrett, Sonia Goncalves, Ewan Harrison, David K. Jackson, Ian Johnston, Dominic Kwiatkowski, Cordelia Langford, John Sillitoe on behalf of the Wellcome Sanger Institute COVID-19 Surveillance Team |
| EPI_ISL_3378442 | LABORATOIRE BIOESTEREL | CNR Virus des Infections Respiratoires - France SUD | Antonin Bal, Gregory Destras, Gwendolyne Burfin, Hadrien Regue, Quentin Semanas, Martine Valette, Bruno Lina, Laurence Josset |
| EPI_ISL_3378443, EPI_ISL_3378444 | Lighthouse Lab in Milton Keynes | Wellcome Sanger Institute for the COVID-19 Genomics UK (COG-UK) Consortium | The Lighthouse Lab in Milton Keynes and Alex Alderton, Roberto Amato, Jeffrey Barrett, Sonia Goncalves, Ewan Harrison, David K. Jackson, Ian Johnston, Dominic Kwiatkowski, Cordelia Langford, John Sillitoe on behalf of the Wellcome Sanger Institute COVID-19 Surveillance Team |
| EPI_ISL_3378445 | Lighthouse Lab in Alderley Park | Wellcome Sanger Institute for the COVID-19 Genomics UK (COG-UK) Consortium | Jacquelyn Wynn, Mairead Hyland, The Lighthouse Lab in Alderley Park and Alex Alderton, Roberto Amato, Jeffrey Barrett, Sonia Goncalves, Ewan Harrison, David K. Jackson, Ian Johnston, Dominic Kwiatkowski, Cordelia Langford, John Sillitoe on behalf of the Wellcome Sanger Institute COVID-19 Surveillance Team |
| EPI_ISL_3378446 | LABORATOIRE BIOESTEREL | CNR Virus des Infections Respiratoires - France SUD | Antonin Bal, Gregory Destras, Gwendolyne Burfin, Hadrien Regue, Quentin Semanas, Martine Valette, Bruno Lina, Laurence Josset |
| EPI_ISL_3378447, EPI_ISL_3378448, EPI_ISL_3378449 | Lighthouse Lab in Milton Keynes | Wellcome Sanger Institute for the COVID-19 Genomics UK (COG-UK) Consortium | The Lighthouse Lab in Milton Keynes and Alex Alderton, Roberto Amato, Jeffrey Barrett, Sonia Goncalves, Ewan Harrison, David K. Jackson, Ian Johnston, Dominic Kwiatkowski, Cordelia Langford, John Sillitoe on behalf of the Wellcome Sanger Institute COVID-19 Surveillance Team |
| EPI_ISL_3378450 | LABORATOIRE BIOESTEREL | CNR Virus des Infections Respiratoires - France SUD | Antonin Bal, Gregory Destras, Gwendolyne Burfin, Hadrien Regue, Quentin Semanas, Martine Valette, Bruno Lina, Laurence Josset |
| EPI_ISL_3378451, EPI_ISL_3378452 | Lighthouse Lab in Milton Keynes | Wellcome Sanger Institute for the COVID-19 Genomics UK (COG-UK) Consortium | The Lighthouse Lab in Milton Keynes and Alex Alderton, Roberto Amato, Jeffrey Barrett, Sonia Goncalves, Ewan Harrison, David K. Jackson, Ian Johnston, Dominic Kwiatkowski, Cordelia Langford, John Sillitoe on behalf of the Wellcome Sanger Institute COVID-19 Surveillance Team |
| EPI_ISL_3378453 | Berkshire and Surrey Pathology Services Lighthouse Laboratory | Wellcome Sanger Institute for the COVID-19 Genomics UK (COG-UK) Consortium | Berkshire and Surrey Pathology Services Lighthouse Laboratory and Alex Alderton, Roberto Amato, Jeffrey Barrett, Sonia Goncalves, Ewan Harrison, David K. Jackson, Ian Johnston, Dominic Kwiatkowski, Cordelia Langford, John Sillitoe on behalf of the Wellcome Sanger Institute COVID-19 Surveillance Team |
| EPI_ISL_3378454 | LABORATOIRE BIOESTEREL | CNR Virus des Infections Respiratoires - France SUD | Antonin Bal, Gregory Destras, Gwendolyne Burfin, Hadrien Regue, Quentin Semanas, Martine Valette, Bruno Lina, Laurence Josset |
| EPI_ISL_3378455 | Lighthouse Lab in Milton Keynes | Wellcome Sanger Institute for the COVID-19 Genomics UK (COG-UK) Consortium | The Lighthouse Lab in Milton Keynes and Alex Alderton, Roberto Amato, Jeffrey Barrett, Sonia Goncalves, Ewan Harrison, David K. Jackson, Ian Johnston, Dominic Kwiatkowski, Cordelia Langford, John Sillitoe on behalf of the Wellcome Sanger Institute COVID-19 Surveillance Team |
| EPI_ISL_3378456 | Berkshire and Surrey Pathology Services Lighthouse Laboratory | Wellcome Sanger Institute for the COVID-19 Genomics UK (COG-UK) Consortium | Berkshire and Surrey Pathology Services Lighthouse Laboratory and Alex Alderton, Roberto Amato, Jeffrey Barrett, Sonia Goncalves, Ewan Harrison, David K. Jackson, Ian Johnston, Dominic Kwiatkowski, Cordelia Langford, John Sillitoe on behalf of the Wellcome Sanger Institute COVID-19 Surveillance Team |
| EPI_ISL_3378457 | Lighthouse Lab in Alderley Park | Wellcome Sanger Institute for the COVID-19 Genomics UK (COG-UK) Consortium | Jacquelyn Wynn, Mairead Hyland, The Lighthouse Lab in Alderley Park and Alex Alderton, Roberto Amato, Jeffrey Barrett, Sonia Goncalves, Ewan Harrison, David K. Jackson, Ian Johnston, Dominic Kwiatkowski, Cordelia Langford, John Sillitoe on behalf of the Wellcome Sanger Institute COVID-19 Surveillance Team |
| EPI_ISL_3378458 | LABORATOIRE BIOESTEREL | CNR Virus des Infections Respiratoires - France SUD | Antonin Bal, Gregory Destras, Gwendolyne Burfin, Hadrien Regue, Quentin Semanas, Martine Valette, Bruno Lina, Laurence Josset |
| EPI_ISL_3378459 | Berkshire and Surrey Pathology Services Lighthouse Laboratory | Wellcome Sanger Institute for the COVID-19 Genomics UK (COG-UK) | Berkshire and Surrey Pathology Services Lighthouse Laboratory and Alex Alderton, Roberto Amato, Jeffrey Barrett, Sonia Goncalves, Ewan Harrison, David K. Jackson, Ian Johnston, Dominic Kwiatkowski, Cordelia Langford, John Sillitoe on behalf |

| | | | |
|---|---|---|---|
| | | Consortium | of the Wellcome Sanger Institute COVID-19 Surveillance Team |
| EPI_ISL_3378460 | Lighthouse Lab in Milton Keynes | Wellcome Sanger Institute for the COVID-19 Genomics UK (COG-UK) Consortium | The Lighthouse Lab in Milton Keynes and Alex Alderton, Roberto Amato, Jeffrey Barrett, Sonia Goncalves, Ewan Harrison, David K. Jackson, Ian Johnston, Dominic Kwiatkowski, Cordelia Langford, John Sillitoe on behalf of the Wellcome Sanger Institute COVID-19 Surveillance Team |
| EPI_ISL_3378461 | LABORATOIRE BIOESTEREL | CNR Virus des Infections Respiratoires - France SUD | Antonin Bal, Gregory Destras, Gwendolyne Burfin, Hadrien Regue, Quentin Semanas, Martine Valette, Bruno Lina, Laurence Josset |
| EPI_ISL_3378462 | Berkshire and Surrey Pathology Services Lighthouse Laboratory | Wellcome Sanger Institute for the COVID-19 Genomics UK (COG-UK) Consortium | Berkshire and Surrey Pathology Services Lighthouse Laboratory and Alex Alderton, Roberto Amato, Jeffrey Barrett, Sonia Goncalves, Ewan Harrison, David K. Jackson, Ian Johnston, Dominic Kwiatkowski, Cordelia Langford, John Sillitoe on behalf of the Wellcome Sanger Institute COVID-19 Surveillance Team |
| EPI_ISL_3378463 | Lighthouse Lab in Alderley Park | Wellcome Sanger Institute for the COVID-19 Genomics UK (COG-UK) Consortium | Jacquelyn Wynn, Mairead Hyland, The Lighthouse Lab in Alderley Park and Alex Alderton, Roberto Amato, Jeffrey Barrett, Sonia Goncalves, Ewan Harrison, David K. Jackson, Ian Johnston, Dominic Kwiatkowski, Cordelia Langford, John Sillitoe on behalf of the Wellcome Sanger Institute COVID-19 Surveillance Team |
| EPI_ISL_3378464 | Lighthouse Lab in Milton Keynes | Wellcome Sanger Institute for the COVID-19 Genomics UK (COG-UK) Consortium | The Lighthouse Lab in Milton Keynes and Alex Alderton, Roberto Amato, Jeffrey Barrett, Sonia Goncalves, Ewan Harrison, David K. Jackson, Ian Johnston, Dominic Kwiatkowski, Cordelia Langford, John Sillitoe on behalf of the Wellcome Sanger Institute COVID-19 Surveillance Team |
| EPI_ISL_3378465 | Berkshire and Surrey Pathology Services Lighthouse Laboratory | Wellcome Sanger Institute for the COVID-19 Genomics UK (COG-UK) Consortium | Berkshire and Surrey Pathology Services Lighthouse Laboratory and Alex Alderton, Roberto Amato, Jeffrey Barrett, Sonia Goncalves, Ewan Harrison, David K. Jackson, Ian Johnston, Dominic Kwiatkowski, Cordelia Langford, John Sillitoe on behalf of the Wellcome Sanger Institute COVID-19 Surveillance Team |
| EPI_ISL_3378466 | LABORATOIRE BIOESTEREL | CNR Virus des Infections Respiratoires - France SUD | Antonin Bal, Gregory Destras, Gwendolyne Burfin, Hadrien Regue, Quentin Semanas, Martine Valette, Bruno Lina, Laurence Josset |
| EPI_ISL_3378467 | Berkshire and Surrey Pathology Services Lighthouse Laboratory | Wellcome Sanger Institute for the COVID-19 Genomics UK (COG-UK) Consortium | Berkshire and Surrey Pathology Services Lighthouse Laboratory and Alex Alderton, Roberto Amato, Jeffrey Barrett, Sonia Goncalves, Ewan Harrison, David K. Jackson, Ian Johnston, Dominic Kwiatkowski, Cordelia Langford, John Sillitoe on behalf of the Wellcome Sanger Institute COVID-19 Surveillance Team |
| EPI_ISL_3378468 | Lighthouse Lab in Milton Keynes | Wellcome Sanger Institute for the COVID-19 Genomics UK (COG-UK) Consortium | The Lighthouse Lab in Milton Keynes and Alex Alderton, Roberto Amato, Jeffrey Barrett, Sonia Goncalves, Ewan Harrison, David K. Jackson, Ian Johnston, Dominic Kwiatkowski, Cordelia Langford, John Sillitoe on behalf of the Wellcome Sanger Institute COVID-19 Surveillance Team |
| EPI_ISL_3378469 | Berkshire and Surrey Pathology Services Lighthouse Laboratory | Wellcome Sanger Institute for the COVID-19 Genomics UK (COG-UK) Consortium | Berkshire and Surrey Pathology Services Lighthouse Laboratory and Alex Alderton, Roberto Amato, Jeffrey Barrett, Sonia Goncalves, Ewan Harrison, David K. Jackson, Ian Johnston, Dominic Kwiatkowski, Cordelia Langford, John Sillitoe on behalf of the Wellcome Sanger Institute COVID-19 Surveillance Team |
| EPI_ISL_3378470 | LABORATOIRE BIOESTEREL | CNR Virus des Infections Respiratoires - France SUD | Antonin Bal, Gregory Destras, Gwendolyne Burfin, Hadrien Regue, Quentin Semanas, Martine Valette, Bruno Lina, Laurence Josset |
| EPI_ISL_3378471 | Lighthouse Lab in Alderley Park | Wellcome Sanger Institute for the COVID-19 Genomics UK (COG-UK) Consortium | Jacquelyn Wynn, Mairead Hyland, The Lighthouse Lab in Alderley Park and Alex Alderton, Roberto Amato, Jeffrey Barrett, Sonia Goncalves, Ewan Harrison, David K. Jackson, Ian Johnston, Dominic Kwiatkowski, Cordelia Langford, John Sillitoe on behalf of the Wellcome Sanger Institute COVID-19 Surveillance Team |
| EPI_ISL_3378472 | Lighthouse Lab in Milton Keynes | Wellcome Sanger Institute for the COVID-19 Genomics UK (COG-UK) Consortium | The Lighthouse Lab in Milton Keynes and Alex Alderton, Roberto Amato, Jeffrey Barrett, Sonia Goncalves, Ewan Harrison, David K. Jackson, Ian Johnston, Dominic Kwiatkowski, Cordelia Langford, John Sillitoe on behalf of the Wellcome Sanger Institute COVID-19 Surveillance Team |
| EPI_ISL_3378473 | LABORATOIRE BIOESTEREL | CNR Virus des Infections Respiratoires - France SUD | Antonin Bal, Gregory Destras, Gwendolyne Burfin, Hadrien Regue, Quentin Semanas, Martine Valette, Bruno Lina, Laurence Josset |
| EPI_ISL_3378474, EPI_ISL_3378475 | Lighthouse Lab in Alderley Park | Wellcome Sanger Institute for the COVID-19 Genomics UK (COG-UK) Consortium | Jacquelyn Wynn, Mairead Hyland, The Lighthouse Lab in Alderley Park and Alex Alderton, Roberto Amato, Jeffrey Barrett, Sonia Goncalves, Ewan Harrison, David K. Jackson, Ian Johnston, Dominic Kwiatkowski, Cordelia Langford, John Sillitoe on behalf of the Wellcome Sanger Institute COVID-19 Surveillance Team |
| EPI_ISL_3378476, EPI_ISL_3378477 | Berkshire and Surrey Pathology Services Lighthouse Laboratory | Wellcome Sanger Institute for the COVID-19 Genomics UK (COG-UK) Consortium | Berkshire and Surrey Pathology Services Lighthouse Laboratory and Alex Alderton, Roberto Amato, Jeffrey Barrett, Sonia Goncalves, Ewan Harrison, David K. Jackson, Ian Johnston, Dominic Kwiatkowski, Cordelia Langford, John Sillitoe on behalf of the Wellcome Sanger Institute COVID-19 Surveillance Team |
| EPI_ISL_3378478 | LABORATOIRE BIOESTEREL | CNR Virus des Infections Respiratoires - France SUD | Antonin Bal, Gregory Destras, Gwendolyne Burfin, Hadrien Regue, Quentin Semanas, Martine Valette, Bruno Lina, Laurence Josset |
| EPI_ISL_3378479 | Lighthouse Lab in Alderley Park | Wellcome Sanger Institute for the COVID-19 Genomics UK (COG-UK) Consortium | Jacquelyn Wynn, Mairead Hyland, The Lighthouse Lab in Alderley Park and Alex Alderton, Roberto Amato, Jeffrey Barrett, Sonia Goncalves, Ewan Harrison, David K. Jackson, Ian Johnston, Dominic Kwiatkowski, Cordelia Langford, John Sillitoe on behalf of the Wellcome Sanger Institute COVID-19 Surveillance Team |
| EPI_ISL_3378480 | Lighthouse Lab in Milton Keynes | Wellcome Sanger Institute for the COVID-19 Genomics UK (COG-UK) Consortium | The Lighthouse Lab in Milton Keynes and Alex Alderton, Roberto Amato, Jeffrey Barrett, Sonia Goncalves, Ewan Harrison, David K. Jackson, Ian Johnston, Dominic Kwiatkowski, Cordelia Langford, John Sillitoe on behalf of the Wellcome Sanger Institute COVID-19 Surveillance Team |
| EPI_ISL_3378481 | LABORATOIRE BIOESTEREL | CNR Virus des Infections Respiratoires - France SUD | Antonin Bal, Gregory Destras, Gwendolyne Burfin, Hadrien Regue, Quentin Semanas, Martine Valette, Bruno Lina, Laurence Josset |
| EPI_ISL_3378482 | Lighthouse Lab in Milton Keynes | Wellcome Sanger Institute for the COVID-19 Genomics UK (COG-UK) Consortium | The Lighthouse Lab in Milton Keynes and Alex Alderton, Roberto Amato, Jeffrey Barrett, Sonia Goncalves, Ewan Harrison, David K. Jackson, Ian Johnston, Dominic Kwiatkowski, Cordelia Langford, John Sillitoe on behalf of the Wellcome Sanger Institute COVID-19 Surveillance Team |
| EPI_ISL_3378483, EPI_ISL_3378484 | Lighthouse Lab in Alderley Park | Wellcome Sanger Institute for the COVID-19 Genomics UK (COG-UK) Consortium | Jacquelyn Wynn, Mairead Hyland, The Lighthouse Lab in Alderley Park and Alex Alderton, Roberto Amato, Jeffrey Barrett, Sonia Goncalves, Ewan Harrison, David K. Jackson, Ian Johnston, Dominic Kwiatkowski, Cordelia Langford, John Sillitoe on behalf of the Wellcome Sanger Institute COVID-19 Surveillance Team |
| EPI_ISL_3378485 | LABORATOIRE BIOESTEREL | CNR Virus des Infections Respiratoires - France SUD | Antonin Bal, Gregory Destras, Gwendolyne Burfin, Hadrien Regue, Quentin Semanas, Martine Valette, Bruno Lina, Laurence Josset |
| EPI_ISL_3378486, EPI_ISL_3378487, EPI_ISL_3378488 | Lighthouse Lab in Milton Keynes | Wellcome Sanger Institute for the COVID-19 Genomics UK (COG-UK) Consortium | The Lighthouse Lab in Milton Keynes and Alex Alderton, Roberto Amato, Jeffrey Barrett, Sonia Goncalves, Ewan Harrison, David K. Jackson, Ian Johnston, Dominic Kwiatkowski, Cordelia Langford, John Sillitoe on behalf of the Wellcome Sanger Institute COVID-19 Surveillance Team |
| EPI_ISL_3378489 | LABORATOIRE BIOESTEREL | CNR Virus des Infections Respiratoires - France SUD | Antonin Bal, Gregory Destras, Gwendolyne Burfin, Hadrien Regue, Quentin Semanas, Martine Valette, Bruno Lina, Laurence Josset |
| EPI_ISL_3378490 | Berkshire and Surrey Pathology Services Lighthouse Laboratory | Wellcome Sanger Institute for the COVID-19 Genomics UK (COG-UK) Consortium | Berkshire and Surrey Pathology Services Lighthouse Laboratory and Alex Alderton, Roberto Amato, Jeffrey Barrett, Sonia Goncalves, Ewan Harrison, David K. Jackson, Ian Johnston, Dominic Kwiatkowski, Cordelia Langford, John Sillitoe on behalf of the Wellcome Sanger Institute COVID-19 Surveillance Team |
| EPI_ISL_3378491 | Lighthouse Lab in Alderley Park | Wellcome Sanger Institute for the COVID-19 Genomics UK (COG-UK) Consortium | Jacquelyn Wynn, Mairead Hyland, The Lighthouse Lab in Alderley Park and Alex Alderton, Roberto Amato, Jeffrey Barrett, Sonia Goncalves, Ewan Harrison, David K. Jackson, Ian Johnston, Dominic Kwiatkowski, Cordelia Langford, John Sillitoe on behalf of the Wellcome Sanger Institute COVID-19 Surveillance Team |
| EPI_ISL_3378492 | LABORATOIRE BIOESTEREL | CNR Virus des Infections Respiratoires - France SUD | Antonin Bal, Gregory Destras, Gwendolyne Burfin, Hadrien Regue, Quentin Semanas, Martine Valette, Bruno Lina, Laurence Josset |
| EPI_ISL_3378493, EPI_ISL_3378494 | Lighthouse Lab in Milton Keynes | Wellcome Sanger Institute for the COVID-19 Genomics UK (COG-UK) Consortium | The Lighthouse Lab in Milton Keynes and Alex Alderton, Roberto Amato, Jeffrey Barrett, Sonia Goncalves, Ewan Harrison, David K. Jackson, Ian Johnston, Dominic Kwiatkowski, Cordelia Langford, John Sillitoe on behalf of the Wellcome Sanger Institute COVID-19 Surveillance Team |
| EPI_ISL_3378495 | Lighthouse Lab in Alderley Park | Wellcome Sanger Institute for the COVID-19 Genomics UK (COG-UK) Consortium | Jacquelyn Wynn, Mairead Hyland, The Lighthouse Lab in Alderley Park and Alex Alderton, Roberto Amato, Jeffrey Barrett, Sonia Goncalves, Ewan Harrison, David K. Jackson, Ian Johnston, Dominic Kwiatkowski, Cordelia Langford, John Sillitoe on behalf of the Wellcome Sanger Institute COVID-19 Surveillance Team |
| EPI_ISL_3378496 | LABORATOIRE BIOESTEREL | CNR Virus des Infections Respiratoires - France SUD | Antonin Bal, Gregory Destras, Gwendolyne Burfin, Hadrien Regue, Quentin Semanas, Martine Valette, Bruno Lina, Laurence Josset |
| EPI_ISL_3378497, EPI_ISL_3378498, EPI_ISL_3378499 | Lighthouse Lab in Milton Keynes | Wellcome Sanger Institute for the COVID-19 Genomics UK (COG-UK) Consortium | The Lighthouse Lab in Milton Keynes and Alex Alderton, Roberto Amato, Jeffrey Barrett, Sonia Goncalves, Ewan Harrison, David K. Jackson, Ian Johnston, Dominic Kwiatkowski, Cordelia Langford, John Sillitoe on behalf of the Wellcome Sanger Institute COVID-19 Surveillance Team |
| EPI_ISL_3378500 | Lighthouse Lab in Alderley Park | Wellcome Sanger Institute for the COVID-19 Genomics UK (COG-UK) Consortium | Jacquelyn Wynn, Mairead Hyland, The Lighthouse Lab in Alderley Park and Alex Alderton, Roberto Amato, Jeffrey Barrett, Sonia Goncalves, Ewan Harrison, David K. Jackson, Ian Johnston, Dominic Kwiatkowski, Cordelia Langford, John Sillitoe on behalf of the Wellcome Sanger Institute COVID-19 Surveillance Team |
| EPI_ISL_3378501 | LABORATOIRE BIOESTEREL | CNR Virus des Infections Respiratoires - France SUD | Antonin Bal, Gregory Destras, Gwendolyne Burfin, Hadrien Regue, Quentin Semanas, Martine Valette, Bruno Lina, Laurence Josset |
| EPI_ISL_3378502 | Lighthouse Lab in Milton Keynes | Wellcome Sanger Institute for the COVID-19 Genomics UK (COG-UK) Consortium | The Lighthouse Lab in Milton Keynes and Alex Alderton, Roberto Amato, Jeffrey Barrett, Sonia Goncalves, Ewan Harrison, David K. Jackson, Ian Johnston, Dominic Kwiatkowski, Cordelia Langford, John Sillitoe on behalf of the Wellcome Sanger Institute COVID-19 Surveillance Team |
| EPI_ISL_3378503 | Lighthouse Lab in Alderley Park | Wellcome Sanger Institute for the COVID-19 Genomics UK (COG-UK) Consortium | Jacquelyn Wynn, Mairead Hyland, The Lighthouse Lab in Alderley Park and Alex Alderton, Roberto Amato, Jeffrey Barrett, Sonia Goncalves, Ewan Harrison, David K. Jackson, Ian Johnston, Dominic Kwiatkowski, Cordelia Langford, John Sillitoe on behalf of the Wellcome Sanger Institute COVID-19 Surveillance Team |
| EPI_ISL_3378504 | Lighthouse Lab in Milton Keynes | Wellcome Sanger Institute for the COVID-19 Genomics UK (COG-UK) Consortium | The Lighthouse Lab in Milton Keynes and Alex Alderton, Roberto Amato, Jeffrey Barrett, Sonia Goncalves, Ewan Harrison, David K. Jackson, Ian Johnston, Dominic Kwiatkowski, Cordelia Langford, John Sillitoe on behalf of the Wellcome Sanger Institute COVID-19 Surveillance Team |
| EPI_ISL_3378505 | LABORATOIRE BIOESTEREL | CNR Virus des Infections Respiratoires - France SUD | Antonin Bal, Gregory Destras, Gwendolyne Burfin, Hadrien Regue, Quentin Semanas, Martine Valette, Bruno Lina, Laurence Josset |
| EPI_ISL_3378506 | Lighthouse Lab in Milton Keynes | Wellcome Sanger Institute for the COVID-19 Genomics UK (COG-UK) Consortium | The Lighthouse Lab in Milton Keynes and Alex Alderton, Roberto Amato, Jeffrey Barrett, Sonia Goncalves, Ewan Harrison, David K. Jackson, Ian Johnston, Dominic Kwiatkowski, Cordelia Langford, John Sillitoe on behalf of the Wellcome Sanger Institute COVID-19 Surveillance Team |
| EPI_ISL_3378507 | Berkshire and Surrey Pathology Services Lighthouse Laboratory | Wellcome Sanger Institute for the COVID-19 Genomics UK (COG-UK) Consortium | Berkshire and Surrey Pathology Services Lighthouse Laboratory and Alex Alderton, Roberto Amato, Jeffrey Barrett, Sonia Goncalves, Ewan Harrison, David K. Jackson, Ian Johnston, Dominic Kwiatkowski, Cordelia Langford, John Sillitoe on behalf of the Wellcome Sanger Institute COVID-19 Surveillance Team |
| EPI_ISL_3378508 | LABORATOIRE ACCOLAB SUD OUEST | CNR Virus des Infections Respiratoires - France SUD | Antonin Bal, Gregory Destras, Gwendolyne Burfin, Hadrien Regue, Quentin Semanas, Martine Valette, Bruno Lina, Laurence Josset |
| EPI_ISL_3378509 | Berkshire and Surrey Pathology Services Lighthouse Laboratory | Wellcome Sanger Institute for the COVID-19 Genomics UK (COG-UK) Consortium | Berkshire and Surrey Pathology Services Lighthouse Laboratory and Alex Alderton, Roberto Amato, Jeffrey Barrett, Sonia Goncalves, Ewan Harrison, David K. Jackson, Ian Johnston, Dominic Kwiatkowski, Cordelia Langford, John Sillitoe on behalf of the Wellcome Sanger Institute COVID-19 Surveillance Team |
| EPI_ISL_3378510 | Lighthouse Lab in Milton Keynes | Wellcome Sanger Institute for the COVID-19 Genomics UK (COG-UK) Consortium | The Lighthouse Lab in Milton Keynes and Alex Alderton, Roberto Amato, Jeffrey Barrett, Sonia Goncalves, Ewan Harrison, David K. Jackson, Ian Johnston, Dominic Kwiatkowski, Cordelia Langford, John Sillitoe on behalf of the Wellcome Sanger Institute COVID-19 Surveillance Team |
| EPI_ISL_3378511 | Berkshire and Surrey Pathology Services Lighthouse Laboratory | Wellcome Sanger Institute for the COVID-19 Genomics UK (COG-UK) Consortium | Berkshire and Surrey Pathology Services Lighthouse Laboratory and Alex Alderton, Roberto Amato, Jeffrey Barrett, Sonia Goncalves, Ewan Harrison, David K. Jackson, Ian Johnston, Dominic Kwiatkowski, Cordelia Langford, John Sillitoe on behalf of the Wellcome Sanger Institute COVID-19 Surveillance Team |
| EPI_ISL_3378512 | LABORATOIRE BIOESTEREL | CNR Virus des Infections Respiratoires - France SUD | Antonin Bal, Gregory Destras, Gwendolyne Burfin, Hadrien Regue, Quentin Semanas, Martine Valette, Bruno Lina, Laurence Josset |
| EPI_ISL_3378513, EPI_ISL_3378514, EPI_ISL_3378515 | Lighthouse Lab in Milton Keynes | Wellcome Sanger Institute for the COVID-19 Genomics UK (COG-UK) Consortium | The Lighthouse Lab in Milton Keynes and Alex Alderton, Roberto Amato, Jeffrey Barrett, Sonia Goncalves, Ewan Harrison, David K. Jackson, Ian Johnston, Dominic Kwiatkowski, Cordelia Langford, John Sillitoe on behalf of the Wellcome Sanger Institute COVID-19 Surveillance Team |
| EPI_ISL_3378516 | LABORATOIRE ACCOLAB SUD OUEST | CNR Virus des Infections Respiratoires - France SUD | Antonin Bal, Gregory Destras, Gwendolyne Burfin, Hadrien Regue, Quentin Semanas, Martine Valette, Bruno Lina, Laurence Josset |
| EPI_ISL_3378517 | Berkshire and Surrey Pathology Services Lighthouse Laboratory | Wellcome Sanger Institute for the COVID-19 Genomics UK (COG-UK) Consortium | Berkshire and Surrey Pathology Services Lighthouse Laboratory and Alex Alderton, Roberto Amato, Jeffrey Barrett, Sonia Goncalves, Ewan Harrison, David K. Jackson, Ian Johnston, Dominic Kwiatkowski, Cordelia Langford, John Sillitoe on behalf of the Wellcome Sanger Institute COVID-19 Surveillance Team |
| EPI_ISL_3378518, EPI_ISL_3378519 | Lighthouse Lab in Milton Keynes | Wellcome Sanger Institute for the COVID-19 Genomics UK (COG-UK) Consortium | The Lighthouse Lab in Milton Keynes and Alex Alderton, Roberto Amato, Jeffrey Barrett, Sonia Goncalves, Ewan Harrison, David K. Jackson, Ian Johnston, Dominic Kwiatkowski, Cordelia Langford, John Sillitoe on behalf of the Wellcome Sanger Institute COVID-19 Surveillance Team |
| EPI_ISL_3378520 | LABORATOIRE BIOESTEREL | CNR Virus des Infections Respiratoires - France SUD | Antonin Bal, Gregory Destras, Gwendolyne Burfin, Hadrien Regue, Quentin Semanas, Martine Valette, Bruno Lina, Laurence Josset |

| | | | |
|---|---|---|---|
| EPI_ISL_3378521, EPI_ISL_3378522 | Lighthouse Lab in Milton Keynes | Wellcome Sanger Institute for the COVID-19 Genomics UK (COG-UK) Consortium | The Lighthouse Lab in Milton Keynes and Alex Alderton, Roberto Amato, Jeffrey Barrett, Sonia Goncalves, Ewan Harrison, David K. Jackson, Ian Johnston, Dominic Kwiatkowski, Cordelia Langford, John Sillitoe on behalf of the Wellcome Sanger Institute COVID-19 Surveillance Team |
| EPI_ISL_3378523 | LABORATOIRE BIOESTEREL | CNR Virus des Infections Respiratoires - France SUD | Antonin Bal, Gregory Destras, Gwendolyne Burfin, Hadrien Regue, Quentin Semanas, Martine Valette, Bruno Lina, Laurence Josset |
| EPI_ISL_3378524, EPI_ISL_3378525 | Lighthouse Lab in Milton Keynes | Wellcome Sanger Institute for the COVID-19 Genomics UK (COG-UK) Consortium | The Lighthouse Lab in Milton Keynes and Alex Alderton, Roberto Amato, Jeffrey Barrett, Sonia Goncalves, Ewan Harrison, David K. Jackson, Ian Johnston, Dominic Kwiatkowski, Cordelia Langford, John Sillitoe on behalf of the Wellcome Sanger Institute COVID-19 Surveillance Team |
| EPI_ISL_3378526 | LABORATOIRE ACCOLAB SUD OUEST | CNR Virus des Infections Respiratoires - France SUD | Antonin Bal, Gregory Destras, Gwendolyne Burfin, Hadrien Regue, Quentin Semanas, Martine Valette, Bruno Lina, Laurence Josset |
| EPI_ISL_3378527, EPI_ISL_3378528, EPI_ISL_3378529 | Lighthouse Lab in Milton Keynes | Wellcome Sanger Institute for the COVID-19 Genomics UK (COG-UK) Consortium | The Lighthouse Lab in Milton Keynes and Alex Alderton, Roberto Amato, Jeffrey Barrett, Sonia Goncalves, Ewan Harrison, David K. Jackson, Ian Johnston, Dominic Kwiatkowski, Cordelia Langford, John Sillitoe on behalf of the Wellcome Sanger Institute COVID-19 Surveillance Team |
| EPI_ISL_3378530 | LABORATOIRE BIOESTEREL | CNR Virus des Infections Respiratoires - France SUD | Antonin Bal, Gregory Destras, Gwendolyne Burfin, Hadrien Regue, Quentin Semanas, Martine Valette, Bruno Lina, Laurence Josset |
| EPI_ISL_3378531, EPI_ISL_3378532 | Lighthouse Lab in Milton Keynes | Wellcome Sanger Institute for the COVID-19 Genomics UK (COG-UK) Consortium | The Lighthouse Lab in Milton Keynes and Alex Alderton, Roberto Amato, Jeffrey Barrett, Sonia Goncalves, Ewan Harrison, David K. Jackson, Ian Johnston, Dominic Kwiatkowski, Cordelia Langford, John Sillitoe on behalf of the Wellcome Sanger Institute COVID-19 Surveillance Team |
| EPI_ISL_3378533 | Lighthouse Lab in Alderley Park | Wellcome Sanger Institute for the COVID-19 Genomics UK (COG-UK) Consortium | Jacquelyn Wynn, Mairead Hyland, The Lighthouse Lab in Alderley Park and Alex Alderton, Roberto Amato, Jeffrey Barrett, Sonia Goncalves, Ewan Harrison, David K. Jackson, Ian Johnston, Dominic Kwiatkowski, Cordelia Langford, John Sillitoe on behalf of the Wellcome Sanger Institute COVID-19 Surveillance Team |
| EPI_ISL_3378534 | Berkshire and Surrey Pathology Services Lighthouse Laboratory | Wellcome Sanger Institute for the COVID-19 Genomics UK (COG-UK) Consortium | Berkshire and Surrey Pathology Services Lighthouse Laboratory and Alex Alderton, Roberto Amato, Jeffrey Barrett, Sonia Goncalves, Ewan Harrison, David K. Jackson, Ian Johnston, Dominic Kwiatkowski, Cordelia Langford, John Sillitoe on behalf of the Wellcome Sanger Institute COVID-19 Surveillance Team |
| EPI_ISL_3378535 | LABORATOIRE ACCOLAB SUD OUEST | CNR Virus des Infections Respiratoires - France SUD | Antonin Bal, Gregory Destras, Gwendolyne Burfin, Hadrien Regue, Quentin Semanas, Martine Valette, Bruno Lina, Laurence Josset |
| EPI_ISL_3378536, EPI_ISL_3378537 | Lighthouse Lab in Milton Keynes | Wellcome Sanger Institute for the COVID-19 Genomics UK (COG-UK) Consortium | The Lighthouse Lab in Milton Keynes and Alex Alderton, Roberto Amato, Jeffrey Barrett, Sonia Goncalves, Ewan Harrison, David K. Jackson, Ian Johnston, Dominic Kwiatkowski, Cordelia Langford, John Sillitoe on behalf of the Wellcome Sanger Institute COVID-19 Surveillance Team |
| EPI_ISL_3378538 | Lighthouse Lab in Alderley Park | Wellcome Sanger Institute for the COVID-19 Genomics UK (COG-UK) Consortium | Jacquelyn Wynn, Mairead Hyland, The Lighthouse Lab in Alderley Park and Alex Alderton, Roberto Amato, Jeffrey Barrett, Sonia Goncalves, Ewan Harrison, David K. Jackson, Ian Johnston, Dominic Kwiatkowski, Cordelia Langford, John Sillitoe on behalf of the Wellcome Sanger Institute COVID-19 Surveillance Team |
| EPI_ISL_3378539 | LABORATOIRE ACCOLAB SUD OUEST | CNR Virus des Infections Respiratoires - France SUD | Antonin Bal, Gregory Destras, Gwendolyne Burfin, Hadrien Regue, Quentin Semanas, Martine Valette, Bruno Lina, Laurence Josset |
| EPI_ISL_3378540 | Lighthouse Lab in Alderley Park | Wellcome Sanger Institute for the COVID-19 Genomics UK (COG-UK) Consortium | Jacquelyn Wynn, Mairead Hyland, The Lighthouse Lab in Alderley Park and Alex Alderton, Roberto Amato, Jeffrey Barrett, Sonia Goncalves, Ewan Harrison, David K. Jackson, Ian Johnston, Dominic Kwiatkowski, Cordelia Langford, John Sillitoe on behalf of the Wellcome Sanger Institute COVID-19 Surveillance Team |
| EPI_ISL_3378541, EPI_ISL_3378542 | Lighthouse Lab in Milton Keynes | Wellcome Sanger Institute for the COVID-19 Genomics UK (COG-UK) Consortium | The Lighthouse Lab in Milton Keynes and Alex Alderton, Roberto Amato, Jeffrey Barrett, Sonia Goncalves, Ewan Harrison, David K. Jackson, Ian Johnston, Dominic Kwiatkowski, Cordelia Langford, John Sillitoe on behalf of the Wellcome Sanger Institute COVID-19 Surveillance Team |
| EPI_ISL_3378543 | LABORATOIRE BIOESTEREL | CNR Virus des Infections Respiratoires - France SUD | Antonin Bal, Gregory Destras, Gwendolyne Burfin, Hadrien Regue, Quentin Semanas, Martine Valette, Bruno Lina, Laurence Josset |
| EPI_ISL_3378544 | Lighthouse Lab in Alderley Park | Wellcome Sanger Institute for the COVID-19 Genomics UK (COG-UK) Consortium | Jacquelyn Wynn, Mairead Hyland, The Lighthouse Lab in Alderley Park and Alex Alderton, Roberto Amato, Jeffrey Barrett, Sonia Goncalves, Ewan Harrison, David K. Jackson, Ian Johnston, Dominic Kwiatkowski, Cordelia Langford, John Sillitoe on behalf of the Wellcome Sanger Institute COVID-19 Surveillance Team |
| EPI_ISL_3378545 | Lighthouse Lab in Milton Keynes | Wellcome Sanger Institute for the COVID-19 Genomics UK (COG-UK) Consortium | The Lighthouse Lab in Milton Keynes and Alex Alderton, Roberto Amato, Jeffrey Barrett, Sonia Goncalves, Ewan Harrison, David K. Jackson, Ian Johnston, Dominic Kwiatkowski, Cordelia Langford, John Sillitoe on behalf of the Wellcome Sanger Institute COVID-19 Surveillance Team |
| EPI_ISL_3378546 | LABORATOIRE ACCOLAB SUD OUEST | CNR Virus des Infections Respiratoires - France SUD | Antonin Bal, Gregory Destras, Gwendolyne Burfin, Hadrien Regue, Quentin Semanas, Martine Valette, Bruno Lina, Laurence Josset |
| EPI_ISL_3378547, EPI_ISL_3378548, EPI_ISL_3378549 | Lighthouse Lab in Alderley Park | Wellcome Sanger Institute for the COVID-19 Genomics UK (COG-UK) Consortium | Jacquelyn Wynn, Mairead Hyland, The Lighthouse Lab in Alderley Park and Alex Alderton, Roberto Amato, Jeffrey Barrett, Sonia Goncalves, Ewan Harrison, David K. Jackson, Ian Johnston, Dominic Kwiatkowski, Cordelia Langford, John Sillitoe on behalf of the Wellcome Sanger Institute COVID-19 Surveillance Team |
| EPI_ISL_3378550 | LABORATOIRE ACCOLAB SUD OUEST | CNR Virus des Infections Respiratoires - France SUD | Antonin Bal, Gregory Destras, Gwendolyne Burfin, Hadrien Regue, Quentin Semanas, Martine Valette, Bruno Lina, Laurence Josset |
| EPI_ISL_3378551 | Lighthouse Lab in Alderley Park | Wellcome Sanger Institute for the COVID-19 Genomics UK (COG-UK) Consortium | Jacquelyn Wynn, Mairead Hyland, The Lighthouse Lab in Alderley Park and Alex Alderton, Roberto Amato, Jeffrey Barrett, Sonia Goncalves, Ewan Harrison, David K. Jackson, Ian Johnston, Dominic Kwiatkowski, Cordelia Langford, John Sillitoe on behalf of the Wellcome Sanger Institute COVID-19 Surveillance Team |
| EPI_ISL_3378552, EPI_ISL_3378553 | Lighthouse Lab in Glasgow | Wellcome Sanger Institute for the COVID-19 Genomics UK (COG-UK) Consortium | Harper VanSteenhouse, Yumi Kasai, David Gray, Carol Clugston, Anna Dominiczak and Alex Alderton, Roberto Amato, Jeffrey Barrett, Sonia Goncalves, Ewan Harrison, David K. Jackson, Ian Johnston, Dominic Kwiatkowski, Cordelia Langford, John Sillitoe on behalf of the Wellcome Sanger Institute COVID-19 Surveillance Team |
| EPI_ISL_3378554 | CNR Virus des Infections Respiratoires - France SUD | CNR Virus des Infections Respiratoires - France SUD | Antonin Bal, Gregory Destras, Gwendolyne Burfin, Hadrien Regue, Quentin Semanas, Martine Valette, Bruno Lina, Laurence Josset |
| EPI_ISL_3378555 | Lighthouse Lab in Glasgow | Wellcome Sanger Institute for the COVID-19 Genomics UK (COG-UK) Consortium | Harper VanSteenhouse, Yumi Kasai, David Gray, Carol Clugston, Anna Dominiczak and Alex Alderton, Roberto Amato, Jeffrey Barrett, Sonia Goncalves, Ewan Harrison, David K. Jackson, Ian Johnston, Dominic Kwiatkowski, Cordelia Langford, John Sillitoe on behalf of the Wellcome Sanger Institute COVID-19 Surveillance Team |
| EPI_ISL_3378556, EPI_ISL_3378557 | Lighthouse Lab in Alderley Park | Wellcome Sanger Institute for the COVID-19 Genomics UK (COG-UK) Consortium | Jacquelyn Wynn, Mairead Hyland, The Lighthouse Lab in Alderley Park and Alex Alderton, Roberto Amato, Jeffrey Barrett, Sonia Goncalves, Ewan Harrison, David K. Jackson, Ian Johnston, Dominic Kwiatkowski, Cordelia Langford, John Sillitoe on behalf of the Wellcome Sanger Institute COVID-19 Surveillance Team |
| EPI_ISL_3378558 | LABORATOIRE ACCOLAB SUD OUEST | CNR Virus des Infections Respiratoires - France SUD | Antonin Bal, Gregory Destras, Gwendolyne Burfin, Hadrien Regue, Quentin Semanas, Martine Valette, Bruno Lina, Laurence Josset |
| EPI_ISL_3378559, EPI_ISL_3378560 | Lighthouse Lab in Alderley Park | Wellcome Sanger Institute for the COVID-19 Genomics UK (COG-UK) Consortium | Jacquelyn Wynn, Mairead Hyland, The Lighthouse Lab in Alderley Park and Alex Alderton, Roberto Amato, Jeffrey Barrett, Sonia Goncalves, Ewan Harrison, David K. Jackson, Ian Johnston, Dominic Kwiatkowski, Cordelia Langford, John Sillitoe on behalf of the Wellcome Sanger Institute COVID-19 Surveillance Team |
| EPI_ISL_3378561 | Lighthouse Lab in Glasgow | Wellcome Sanger Institute for the COVID-19 Genomics UK (COG-UK) Consortium | Harper VanSteenhouse, Yumi Kasai, David Gray, Carol Clugston, Anna Dominiczak and Alex Alderton, Roberto Amato, Jeffrey Barrett, Sonia Goncalves, Ewan Harrison, David K. Jackson, Ian Johnston, Dominic Kwiatkowski, Cordelia Langford, John Sillitoe on behalf of the Wellcome Sanger Institute COVID-19 Surveillance Team |
| EPI_ISL_3378562 | LABORATOIRE ACCOLAB SUD OUEST | CNR Virus des Infections Respiratoires - France SUD | Antonin Bal, Gregory Destras, Gwendolyne Burfin, Hadrien Regue, Quentin Semanas, Martine Valette, Bruno Lina, Laurence Josset |
| EPI_ISL_3378563, EPI_ISL_3378564 | Lighthouse Lab in Alderley Park | Wellcome Sanger Institute for the COVID-19 Genomics UK (COG-UK) Consortium | Jacquelyn Wynn, Mairead Hyland, The Lighthouse Lab in Alderley Park and Alex Alderton, Roberto Amato, Jeffrey Barrett, Sonia Goncalves, Ewan Harrison, David K. Jackson, Ian Johnston, Dominic Kwiatkowski, Cordelia Langford, John Sillitoe on behalf of the Wellcome Sanger Institute COVID-19 Surveillance Team |
| EPI_ISL_3378565 | LABORATOIRE BIOESTEREL | CNR Virus des Infections Respiratoires - France SUD | Antonin Bal, Gregory Destras, Gwendolyne Burfin, Hadrien Regue, Quentin Semanas, Martine Valette, Bruno Lina, Laurence Josset |
| EPI_ISL_3378566 | Lighthouse Lab in Glasgow | Wellcome Sanger Institute for the COVID-19 Genomics UK (COG-UK) Consortium | Harper VanSteenhouse, Yumi Kasai, David Gray, Carol Clugston, Anna Dominiczak and Alex Alderton, Roberto Amato, Jeffrey Barrett, Sonia Goncalves, Ewan Harrison, David K. Jackson, Ian Johnston, Dominic Kwiatkowski, Cordelia Langford, John Sillitoe on behalf of the Wellcome Sanger Institute COVID-19 Surveillance Team |
| EPI_ISL_3378567, EPI_ISL_3378568 | Lighthouse Lab in Alderley Park | Wellcome Sanger Institute for the COVID-19 Genomics UK (COG-UK) Consortium | Jacquelyn Wynn, Mairead Hyland, The Lighthouse Lab in Alderley Park and Alex Alderton, Roberto Amato, Jeffrey Barrett, Sonia Goncalves, Ewan Harrison, David K. Jackson, Ian Johnston, Dominic Kwiatkowski, Cordelia Langford, John Sillitoe on behalf of the Wellcome Sanger Institute COVID-19 Surveillance Team |
| EPI_ISL_3378569, EPI_ISL_3378570 | Lighthouse Lab in Glasgow | Wellcome Sanger Institute for the COVID-19 Genomics UK (COG-UK) Consortium | Harper VanSteenhouse, Yumi Kasai, David Gray, Carol Clugston, Anna Dominiczak and Alex Alderton, Roberto Amato, Jeffrey Barrett, Sonia Goncalves, Ewan Harrison, David K. Jackson, Ian Johnston, Dominic Kwiatkowski, Cordelia Langford, John Sillitoe on behalf of the Wellcome Sanger Institute COVID-19 Surveillance Team |
| EPI_ISL_3378571 | LABORATOIRE ACCOLAB SUD OUEST | CNR Virus des Infections Respiratoires - France SUD | Antonin Bal, Gregory Destras, Gwendolyne Burfin, Hadrien Regue, Quentin Semanas, Martine Valette, Bruno Lina, Laurence Josset |
| EPI_ISL_3378572 | Lighthouse Lab in Glasgow | Wellcome Sanger Institute for the COVID-19 Genomics UK (COG-UK) Consortium | Harper VanSteenhouse, Yumi Kasai, David Gray, Carol Clugston, Anna Dominiczak and Alex Alderton, Roberto Amato, Jeffrey Barrett, Sonia Goncalves, Ewan Harrison, David K. Jackson, Ian Johnston, Dominic Kwiatkowski, Cordelia Langford, John Sillitoe on behalf of the Wellcome Sanger Institute COVID-19 Surveillance Team |
| EPI_ISL_3378573 | Lighthouse Lab in Alderley Park | Wellcome Sanger Institute for the COVID-19 Genomics UK (COG-UK) Consortium | Jacquelyn Wynn, Mairead Hyland, The Lighthouse Lab in Alderley Park and Alex Alderton, Roberto Amato, Jeffrey Barrett, Sonia Goncalves, Ewan Harrison, David K. Jackson, Ian Johnston, Dominic Kwiatkowski, Cordelia Langford, John Sillitoe on behalf of the Wellcome Sanger Institute COVID-19 Surveillance Team |
| EPI_ISL_3378574 | Lighthouse Lab in Glasgow | Wellcome Sanger Institute for the COVID-19 Genomics UK (COG-UK) Consortium | Harper VanSteenhouse, Yumi Kasai, David Gray, Carol Clugston, Anna Dominiczak and Alex Alderton, Roberto Amato, Jeffrey Barrett, Sonia Goncalves, Ewan Harrison, David K. Jackson, Ian Johnston, Dominic Kwiatkowski, Cordelia Langford, John Sillitoe on behalf of the Wellcome Sanger Institute COVID-19 Surveillance Team |
| EPI_ISL_3378575 | LABORATOIRE BIOESTEREL | CNR Virus des Infections Respiratoires - France SUD | Antonin Bal, Gregory Destras, Gwendolyne Burfin, Hadrien Regue, Quentin Semanas, Martine Valette, Bruno Lina, Laurence Josset |
| EPI_ISL_3378576, EPI_ISL_3378577, EPI_ISL_3378578 | Lighthouse Lab in Alderley Park | Wellcome Sanger Institute for the COVID-19 Genomics UK (COG-UK) Consortium | Jacquelyn Wynn, Mairead Hyland, The Lighthouse Lab in Alderley Park and Alex Alderton, Roberto Amato, Jeffrey Barrett, Sonia Goncalves, Ewan Harrison, David K. Jackson, Ian Johnston, Dominic Kwiatkowski, Cordelia Langford, John Sillitoe on behalf of the Wellcome Sanger Institute COVID-19 Surveillance Team |
| EPI_ISL_3378579 | LABORATOIRE ACCOLAB SUD OUEST | CNR Virus des Infections Respiratoires - France SUD | Antonin Bal, Gregory Destras, Gwendolyne Burfin, Hadrien Regue, Quentin Semanas, Martine Valette, Bruno Lina, Laurence Josset |
| EPI_ISL_3378580, EPI_ISL_3378581, EPI_ISL_3378582 | Lighthouse Lab in Alderley Park | Wellcome Sanger Institute for the COVID-19 Genomics UK (COG-UK) Consortium | Jacquelyn Wynn, Mairead Hyland, The Lighthouse Lab in Alderley Park and Alex Alderton, Roberto Amato, Jeffrey Barrett, Sonia Goncalves, Ewan Harrison, David K. Jackson, Ian Johnston, Dominic Kwiatkowski, Cordelia Langford, John Sillitoe on behalf of the Wellcome Sanger Institute COVID-19 Surveillance Team |
| EPI_ISL_3378583 | LABORATOIRE BIOESTEREL | CNR Virus des Infections Respiratoires - France SUD | Antonin Bal, Gregory Destras, Gwendolyne Burfin, Hadrien Regue, Quentin Semanas, Martine Valette, Bruno Lina, Laurence Josset |
| EPI_ISL_3378584 | Lighthouse Lab in Glasgow | Wellcome Sanger Institute for the COVID-19 Genomics UK (COG-UK) Consortium | Harper VanSteenhouse, Yumi Kasai, David Gray, Carol Clugston, Anna Dominiczak and Alex Alderton, Roberto Amato, Jeffrey Barrett, Sonia Goncalves, Ewan Harrison, David K. Jackson, Ian Johnston, Dominic Kwiatkowski, Cordelia Langford, John Sillitoe on behalf of the Wellcome Sanger Institute COVID-19 Surveillance Team |
| EPI_ISL_3378585 | Lighthouse Lab in Alderley Park | Wellcome Sanger Institute for the COVID-19 Genomics UK (COG-UK) Consortium | Jacquelyn Wynn, Mairead Hyland, The Lighthouse Lab in Alderley Park and Alex Alderton, Roberto Amato, Jeffrey Barrett, Sonia Goncalves, Ewan Harrison, David K. Jackson, Ian Johnston, Dominic Kwiatkowski, Cordelia Langford, John Sillitoe on behalf of the Wellcome Sanger Institute COVID-19 Surveillance Team |
| EPI_ISL_3378586 | LABORATOIRE BIOESTEREL | CNR Virus des Infections Respiratoires - France SUD | Antonin Bal, Gregory Destras, Gwendolyne Burfin, Hadrien Regue, Quentin Semanas, Martine Valette, Bruno Lina, Laurence Josset |
| EPI_ISL_3378587, EPI_ISL_3378588 | Lighthouse Lab in Alderley Park | Wellcome Sanger Institute for the COVID-19 Genomics UK (COG-UK) Consortium | Jacquelyn Wynn, Mairead Hyland, The Lighthouse Lab in Alderley Park and Alex Alderton, Roberto Amato, Jeffrey Barrett, Sonia Goncalves, Ewan Harrison, David K. Jackson, Ian Johnston, Dominic Kwiatkowski, Cordelia Langford, John Sillitoe on behalf of the Wellcome Sanger Institute COVID-19 Surveillance Team |
| EPI_ISL_3378589 | Lighthouse Lab in Glasgow | Wellcome Sanger Institute for the COVID-19 Genomics UK (COG-UK) Consortium | Harper VanSteenhouse, Yumi Kasai, David Gray, Carol Clugston, Anna Dominiczak and Alex Alderton, Roberto Amato, Jeffrey Barrett, Sonia Goncalves, Ewan Harrison, David K. Jackson, Ian Johnston, Dominic Kwiatkowski, Cordelia Langford, John Sillitoe on behalf of the Wellcome Sanger Institute COVID-19 Surveillance Team |
| EPI_ISL_3378590 | Lighthouse Lab in Alderley Park | Wellcome Sanger Institute for the COVID-19 Genomics UK (COG-UK) Consortium | Jacquelyn Wynn, Mairead Hyland, The Lighthouse Lab in Alderley Park and Alex Alderton, Roberto Amato, Jeffrey Barrett, Sonia Goncalves, Ewan Harrison, David K. Jackson, Ian Johnston, Dominic Kwiatkowski, Cordelia Langford, John Sillitoe on behalf of the Wellcome Sanger Institute COVID-19 Surveillance Team |
| EPI_ISL_3378591 | LABORATOIRE BIOESTEREL | CNR Virus des Infections Respiratoires - France SUD | Antonin Bal, Gregory Destras, Gwendolyne Burfin, Hadrien Regue, Quentin Semanas, Martine Valette, Bruno Lina, Laurence Josset |

| | | | |
|---|---|---|---|
| EPI_ISL_3378592 | Lighthouse Lab in Alderley Park | Wellcome Sanger Institute for the COVID-19 Genomics UK (COG-UK) Consortium | Jacquelyn Wynn, Mairead Hyland, The Lighthouse Lab in Alderley Park and Alex Alderton, Roberto Amato, Jeffrey Barrett, Sonia Goncalves, Ewan Harrison, David K. Jackson, Ian Johnston, Dominic Kwiatkowski, Cordelia Langford, John Sillitoe on behalf of the Wellcome Sanger Institute COVID-19 Surveillance Team |
| EPI_ISL_3378593 | Lighthouse Lab in Glasgow | Wellcome Sanger Institute for the COVID-19 Genomics UK (COG-UK) Consortium | Harper VanSteenhouse, Yumi Kasai, David Gray, Carol Clugston, Anna Dominiczak and Alex Alderton, Roberto Amato, Jeffrey Barrett, Sonia Goncalves, Ewan Harrison, David K. Jackson, Ian Johnston, Dominic Kwiatkowski, Cordelia Langford, John Sillitoe on behalf of the Wellcome Sanger Institute COVID-19 Surveillance Team |
| EPI_ISL_3378594 | LABORATOIRE ACCOLAB SUD OUEST | CNR Virus des Infections Respiratoires - France SUD | Antonin Bal, Gregory Destras, Gwendolyne Burfin, Hadrien Regue, Quentin Semanas, Martine Valette, Bruno Lina, Laurence Josset |
| EPI_ISL_3378595, EPI_ISL_3378596, EPI_ISL_3378597 | Health Services Laboratories | Wellcome Sanger Institute for the COVID-19 Genomics UK (COG-UK) Consortium | Health Services Laboratories and Alex Alderton, Roberto Amato, Jeffrey Barrett, Sonia Goncalves, Ewan Harrison, David K. Jackson, Ian Johnston, Dominic Kwiatkowski, Cordelia Langford, John Sillitoe on behalf of the Wellcome Sanger Institute COVID-19 Surveillance Team |
| EPI_ISL_3378598 | Lighthouse Lab in Glasgow | Wellcome Sanger Institute for the COVID-19 Genomics UK (COG-UK) Consortium | Harper VanSteenhouse, Yumi Kasai, David Gray, Carol Clugston, Anna Dominiczak and Alex Alderton, Roberto Amato, Jeffrey Barrett, Sonia Goncalves, Ewan Harrison, David K. Jackson, Ian Johnston, Dominic Kwiatkowski, Cordelia Langford, John Sillitoe on behalf of the Wellcome Sanger Institute COVID-19 Surveillance Team |
| EPI_ISL_3378599 | LABORATOIRE BIOESTEREL | CNR Virus des Infections Respiratoires - France SUD | Antonin Bal, Gregory Destras, Gwendolyne Burfin, Hadrien Regue, Quentin Semanas, Martine Valette, Bruno Lina, Laurence Josset |
| EPI_ISL_3378600 | Health Services Laboratories | Wellcome Sanger Institute for the COVID-19 Genomics UK (COG-UK) Consortium | Health Services Laboratories and Alex Alderton, Roberto Amato, Jeffrey Barrett, Sonia Goncalves, Ewan Harrison, David K. Jackson, Ian Johnston, Dominic Kwiatkowski, Cordelia Langford, John Sillitoe on behalf of the Wellcome Sanger Institute COVID-19 Surveillance Team |
| EPI_ISL_3378601 | Lighthouse Lab in Glasgow | Wellcome Sanger Institute for the COVID-19 Genomics UK (COG-UK) Consortium | Harper VanSteenhouse, Yumi Kasai, David Gray, Carol Clugston, Anna Dominiczak and Alex Alderton, Roberto Amato, Jeffrey Barrett, Sonia Goncalves, Ewan Harrison, David K. Jackson, Ian Johnston, Dominic Kwiatkowski, Cordelia Langford, John Sillitoe on behalf of the Wellcome Sanger Institute COVID-19 Surveillance Team |
| EPI_ISL_3378603 | Health Services Laboratories | Wellcome Sanger Institute for the COVID-19 Genomics UK (COG-UK) Consortium | Health Services Laboratories and Alex Alderton, Roberto Amato, Jeffrey Barrett, Sonia Goncalves, Ewan Harrison, David K. Jackson, Ian Johnston, Dominic Kwiatkowski, Cordelia Langford, John Sillitoe on behalf of the Wellcome Sanger Institute COVID-19 Surveillance Team |
| EPI_ISL_3378604 | Lighthouse Lab in Glasgow | Wellcome Sanger Institute for the COVID-19 Genomics UK (COG-UK) Consortium | Harper VanSteenhouse, Yumi Kasai, David Gray, Carol Clugston, Anna Dominiczak and Alex Alderton, Roberto Amato, Jeffrey Barrett, Sonia Goncalves, Ewan Harrison, David K. Jackson, Ian Johnston, Dominic Kwiatkowski, Cordelia Langford, John Sillitoe on behalf of the Wellcome Sanger Institute COVID-19 Surveillance Team |
| EPI_ISL_3378605 | Health Services Laboratories | Wellcome Sanger Institute for the COVID-19 Genomics UK (COG-UK) Consortium | Health Services Laboratories and Alex Alderton, Roberto Amato, Jeffrey Barrett, Sonia Goncalves, Ewan Harrison, David K. Jackson, Ian Johnston, Dominic Kwiatkowski, Cordelia Langford, John Sillitoe on behalf of the Wellcome Sanger Institute COVID-19 Surveillance Team |
| EPI_ISL_3378606 | LABORATOIRE BIOESTEREL | CNR Virus des Infections Respiratoires - France SUD | Antonin Bal, Gregory Destras, Gwendolyne Burfin, Hadrien Regue, Quentin Semanas, Martine Valette, Bruno Lina, Laurence Josset |
| EPI_ISL_3378607 | Health Services Laboratories | Wellcome Sanger Institute for the COVID-19 Genomics UK (COG-UK) Consortium | Health Services Laboratories and Alex Alderton, Roberto Amato, Jeffrey Barrett, Sonia Goncalves, Ewan Harrison, David K. Jackson, Ian Johnston, Dominic Kwiatkowski, Cordelia Langford, John Sillitoe on behalf of the Wellcome Sanger Institute COVID-19 Surveillance Team |
| EPI_ISL_3378608, EPI_ISL_3378609 | Lighthouse Lab in Glasgow | Wellcome Sanger Institute for the COVID-19 Genomics UK (COG-UK) Consortium | Harper VanSteenhouse, Yumi Kasai, David Gray, Carol Clugston, Anna Dominiczak and Alex Alderton, Roberto Amato, Jeffrey Barrett, Sonia Goncalves, Ewan Harrison, David K. Jackson, Ian Johnston, Dominic Kwiatkowski, Cordelia Langford, John Sillitoe on behalf of the Wellcome Sanger Institute COVID-19 Surveillance Team |
| EPI_ISL_3378610 | LABORATOIRE BIOESTEREL | CNR Virus des Infections Respiratoires - France SUD | Antonin Bal, Gregory Destras, Gwendolyne Burfin, Hadrien Regue, Quentin Semanas, Martine Valette, Bruno Lina, Laurence Josset |
| EPI_ISL_3378611 | Lighthouse Lab in Glasgow | Wellcome Sanger Institute for the COVID-19 Genomics UK (COG-UK) Consortium | Harper VanSteenhouse, Yumi Kasai, David Gray, Carol Clugston, Anna Dominiczak and Alex Alderton, Roberto Amato, Jeffrey Barrett, Sonia Goncalves, Ewan Harrison, David K. Jackson, Ian Johnston, Dominic Kwiatkowski, Cordelia Langford, John Sillitoe on behalf of the Wellcome Sanger Institute COVID-19 Surveillance Team |
| EPI_ISL_3378612 | Lighthouse Lab in Alderley Park | Wellcome Sanger Institute for the COVID-19 Genomics UK (COG-UK) Consortium | Jacquelyn Wynn, Mairead Hyland, The Lighthouse Lab in Alderley Park and Alex Alderton, Roberto Amato, Jeffrey Barrett, Sonia Goncalves, Ewan Harrison, David K. Jackson, Ian Johnston, Dominic Kwiatkowski, Cordelia Langford, John Sillitoe on behalf of the Wellcome Sanger Institute COVID-19 Surveillance Team |
| EPI_ISL_3378613 | Health Services Laboratories | Wellcome Sanger Institute for the COVID-19 Genomics UK (COG-UK) Consortium | Health Services Laboratories and Alex Alderton, Roberto Amato, Jeffrey Barrett, Sonia Goncalves, Ewan Harrison, David K. Jackson, Ian Johnston, Dominic Kwiatkowski, Cordelia Langford, John Sillitoe on behalf of the Wellcome Sanger Institute COVID-19 Surveillance Team |
| EPI_ISL_3378614 | LABORATOIRE BIOESTEREL | CNR Virus des Infections Respiratoires - France SUD | Antonin Bal, Gregory Destras, Gwendolyne Burfin, Hadrien Regue, Quentin Semanas, Martine Valette, Bruno Lina, Laurence Josset |
| EPI_ISL_3378615 | Lighthouse Lab in Alderley Park | Wellcome Sanger Institute for the COVID-19 Genomics UK (COG-UK) Consortium | Jacquelyn Wynn, Mairead Hyland, The Lighthouse Lab in Alderley Park and Alex Alderton, Roberto Amato, Jeffrey Barrett, Sonia Goncalves, Ewan Harrison, David K. Jackson, Ian Johnston, Dominic Kwiatkowski, Cordelia Langford, John Sillitoe on behalf of the Wellcome Sanger Institute COVID-19 Surveillance Team |
| EPI_ISL_3378616 | Lighthouse Lab in Glasgow | Wellcome Sanger Institute for the COVID-19 Genomics UK (COG-UK) Consortium | Harper VanSteenhouse, Yumi Kasai, David Gray, Carol Clugston, Anna Dominiczak and Alex Alderton, Roberto Amato, Jeffrey Barrett, Sonia Goncalves, Ewan Harrison, David K. Jackson, Ian Johnston, Dominic Kwiatkowski, Cordelia Langford, John Sillitoe on behalf of the Wellcome Sanger Institute COVID-19 Surveillance Team |
| EPI_ISL_3378617 | LABORATOIRE BIOESTEREL | CNR Virus des Infections Respiratoires - France SUD | Antonin Bal, Gregory Destras, Gwendolyne Burfin, Hadrien Regue, Quentin Semanas, Martine Valette, Bruno Lina, Laurence Josset |
| EPI_ISL_3378618, EPI_ISL_3378619, EPI_ISL_3378620 | Health Services Laboratories | Wellcome Sanger Institute for the COVID-19 Genomics UK (COG-UK) Consortium | Health Services Laboratories and Alex Alderton, Roberto Amato, Jeffrey Barrett, Sonia Goncalves, Ewan Harrison, David K. Jackson, Ian Johnston, Dominic Kwiatkowski, Cordelia Langford, John Sillitoe on behalf of the Wellcome Sanger Institute COVID-19 Surveillance Team |
| EPI_ISL_3378621 | LABORATOIRE ACCOLAB SUD OUEST | CNR Virus des Infections Respiratoires - France SUD | Antonin Bal, Gregory Destras, Gwendolyne Burfin, Hadrien Regue, Quentin Semanas, Martine Valette, Bruno Lina, Laurence Josset |
| EPI_ISL_3378622, EPI_ISL_3378623 | Lighthouse Lab in Glasgow | Wellcome Sanger Institute for the COVID-19 Genomics UK (COG-UK) Consortium | Harper VanSteenhouse, Yumi Kasai, David Gray, Carol Clugston, Anna Dominiczak and Alex Alderton, Roberto Amato, Jeffrey Barrett, Sonia Goncalves, Ewan Harrison, David K. Jackson, Ian Johnston, Dominic Kwiatkowski, Cordelia Langford, John Sillitoe on behalf of the Wellcome Sanger Institute COVID-19 Surveillance Team |
| EPI_ISL_3378624 | Health Services Laboratories | Wellcome Sanger Institute for the COVID-19 Genomics UK (COG-UK) Consortium | Health Services Laboratories and Alex Alderton, Roberto Amato, Jeffrey Barrett, Sonia Goncalves, Ewan Harrison, David K. Jackson, Ian Johnston, Dominic Kwiatkowski, Cordelia Langford, John Sillitoe on behalf of the Wellcome Sanger Institute COVID-19 Surveillance Team |
| EPI_ISL_3378625 | LABORATOIRE BIOESTEREL | CNR Virus des Infections Respiratoires - France SUD | Antonin Bal, Gregory Destras, Gwendolyne Burfin, Hadrien Regue, Quentin Semanas, Martine Valette, Bruno Lina, Laurence Josset |
| EPI_ISL_3378626, EPI_ISL_3378627 | Lighthouse Lab in Alderley Park | Wellcome Sanger Institute for the COVID-19 Genomics UK (COG-UK) Consortium | Jacquelyn Wynn, Mairead Hyland, The Lighthouse Lab in Alderley Park and Alex Alderton, Roberto Amato, Jeffrey Barrett, Sonia Goncalves, Ewan Harrison, David K. Jackson, Ian Johnston, Dominic Kwiatkowski, Cordelia Langford, John Sillitoe on behalf of the Wellcome Sanger Institute COVID-19 Surveillance Team |
| EPI_ISL_3378628 | Health Services Laboratories | Wellcome Sanger Institute for the COVID-19 Genomics UK (COG-UK) Consortium | Health Services Laboratories and Alex Alderton, Roberto Amato, Jeffrey Barrett, Sonia Goncalves, Ewan Harrison, David K. Jackson, Ian Johnston, Dominic Kwiatkowski, Cordelia Langford, John Sillitoe on behalf of the Wellcome Sanger Institute COVID-19 Surveillance Team |
| EPI_ISL_3378629 | LABORATOIRE ACCOLAB SUD OUEST | CNR Virus des Infections Respiratoires - France SUD | Antonin Bal, Gregory Destras, Gwendolyne Burfin, Hadrien Regue, Quentin Semanas, Martine Valette, Bruno Lina, Laurence Josset |
| EPI_ISL_3378630 | Lighthouse Lab in Alderley Park | Wellcome Sanger Institute for the COVID-19 Genomics UK (COG-UK) Consortium | Jacquelyn Wynn, Mairead Hyland, The Lighthouse Lab in Alderley Park and Alex Alderton, Roberto Amato, Jeffrey Barrett, Sonia Goncalves, Ewan Harrison, David K. Jackson, Ian Johnston, Dominic Kwiatkowski, Cordelia Langford, John Sillitoe on behalf of the Wellcome Sanger Institute COVID-19 Surveillance Team |
| EPI_ISL_3378631 | Lighthouse Lab in Glasgow | Wellcome Sanger Institute for the COVID-19 Genomics UK (COG-UK) Consortium | Harper VanSteenhouse, Yumi Kasai, David Gray, Carol Clugston, Anna Dominiczak and Alex Alderton, Roberto Amato, Jeffrey Barrett, Sonia Goncalves, Ewan Harrison, David K. Jackson, Ian Johnston, Dominic Kwiatkowski, Cordelia Langford, John Sillitoe on behalf of the Wellcome Sanger Institute COVID-19 Surveillance Team |
| EPI_ISL_3378632 | LABORATOIRE BIOESTEREL | CNR Virus des Infections Respiratoires - France SUD | Antonin Bal, Gregory Destras, Gwendolyne Burfin, Hadrien Regue, Quentin Semanas, Martine Valette, Bruno Lina, Laurence Josset |
| EPI_ISL_3378633 | Health Services Laboratories | Wellcome Sanger Institute for the COVID-19 Genomics UK (COG-UK) Consortium | Health Services Laboratories and Alex Alderton, Roberto Amato, Jeffrey Barrett, Sonia Goncalves, Ewan Harrison, David K. Jackson, Ian Johnston, Dominic Kwiatkowski, Cordelia Langford, John Sillitoe on behalf of the Wellcome Sanger Institute COVID-19 Surveillance Team |
| EPI_ISL_3378634 | Lighthouse Lab in Glasgow | Wellcome Sanger Institute for the COVID-19 Genomics UK (COG-UK) Consortium | Harper VanSteenhouse, Yumi Kasai, David Gray, Carol Clugston, Anna Dominiczak and Alex Alderton, Roberto Amato, Jeffrey Barrett, Sonia Goncalves, Ewan Harrison, David K. Jackson, Ian Johnston, Dominic Kwiatkowski, Cordelia Langford, John Sillitoe on behalf of the Wellcome Sanger Institute COVID-19 Surveillance Team |
| EPI_ISL_3378635 | Lighthouse Lab in Alderley Park | Wellcome Sanger Institute for the COVID-19 Genomics UK (COG-UK) Consortium | Jacquelyn Wynn, Mairead Hyland, The Lighthouse Lab in Alderley Park and Alex Alderton, Roberto Amato, Jeffrey Barrett, Sonia Goncalves, Ewan Harrison, David K. Jackson, Ian Johnston, Dominic Kwiatkowski, Cordelia Langford, John Sillitoe on behalf of the Wellcome Sanger Institute COVID-19 Surveillance Team |
| EPI_ISL_3378636 | LABORATOIRE BIOESTEREL | CNR Virus des Infections Respiratoires - France SUD | Antonin Bal, Gregory Destras, Gwendolyne Burfin, Hadrien Regue, Quentin Semanas, Martine Valette, Bruno Lina, Laurence Josset |
| EPI_ISL_3378637, EPI_ISL_3378638 | Lighthouse Lab in Glasgow | Wellcome Sanger Institute for the COVID-19 Genomics UK (COG-UK) Consortium | Harper VanSteenhouse, Yumi Kasai, David Gray, Carol Clugston, Anna Dominiczak and Alex Alderton, Roberto Amato, Jeffrey Barrett, Sonia Goncalves, Ewan Harrison, David K. Jackson, Ian Johnston, Dominic Kwiatkowski, Cordelia Langford, John Sillitoe on behalf of the Wellcome Sanger Institute COVID-19 Surveillance Team |
| EPI_ISL_3378639 | LABORATOIRE ACCOLAB SUD OUEST | CNR Virus des Infections Respiratoires - France SUD | Antonin Bal, Gregory Destras, Gwendolyne Burfin, Hadrien Regue, Quentin Semanas, Martine Valette, Bruno Lina, Laurence Josset |
| EPI_ISL_3378640, EPI_ISL_3378641 | Health Services Laboratories | Wellcome Sanger Institute for the COVID-19 Genomics UK (COG-UK) Consortium | Health Services Laboratories and Alex Alderton, Roberto Amato, Jeffrey Barrett, Sonia Goncalves, Ewan Harrison, David K. Jackson, Ian Johnston, Dominic Kwiatkowski, Cordelia Langford, John Sillitoe on behalf of the Wellcome Sanger Institute COVID-19 Surveillance Team |
| EPI_ISL_3378642 | Lighthouse Lab in Glasgow | Wellcome Sanger Institute for the COVID-19 Genomics UK (COG-UK) Consortium | Harper VanSteenhouse, Yumi Kasai, David Gray, Carol Clugston, Anna Dominiczak and Alex Alderton, Roberto Amato, Jeffrey Barrett, Sonia Goncalves, Ewan Harrison, David K. Jackson, Ian Johnston, Dominic Kwiatkowski, Cordelia Langford, John Sillitoe on behalf of the Wellcome Sanger Institute COVID-19 Surveillance Team |
| EPI_ISL_3378643 | LABORATOIRE BIOESTEREL | CNR Virus des Infections Respiratoires - France SUD | Antonin Bal, Gregory Destras, Gwendolyne Burfin, Hadrien Regue, Quentin Semanas, Martine Valette, Bruno Lina, Laurence Josset |
| EPI_ISL_3378644 | Lighthouse Lab in Milton Keynes | Wellcome Sanger Institute for the COVID-19 Genomics UK (COG-UK) Consortium | The Lighthouse Lab in Milton Keynes and Alex Alderton, Roberto Amato, Jeffrey Barrett, Sonia Goncalves, Ewan Harrison, David K. Jackson, Ian Johnston, Dominic Kwiatkowski, Cordelia Langford, John Sillitoe on behalf of the Wellcome Sanger Institute COVID-19 Surveillance Team |
| EPI_ISL_3378645, EPI_ISL_3378646 | Lighthouse Lab in Glasgow | Wellcome Sanger Institute for the COVID-19 Genomics UK (COG-UK) Consortium | Harper VanSteenhouse, Yumi Kasai, David Gray, Carol Clugston, Anna Dominiczak and Alex Alderton, Roberto Amato, Jeffrey Barrett, Sonia Goncalves, Ewan Harrison, David K. Jackson, Ian Johnston, Dominic Kwiatkowski, Cordelia Langford, John Sillitoe on behalf of the Wellcome Sanger Institute COVID-19 Surveillance Team |
| EPI_ISL_3378647 | LABORATOIRE ACCOLAB SUD OUEST | CNR Virus des Infections Respiratoires - France SUD | Antonin Bal, Gregory Destras, Gwendolyne Burfin, Hadrien Regue, Quentin Semanas, Martine Valette, Bruno Lina, Laurence Josset |
| EPI_ISL_3378648 | Lighthouse Lab in Glasgow | Wellcome Sanger Institute for the COVID-19 Genomics UK (COG-UK) Consortium | Harper VanSteenhouse, Yumi Kasai, David Gray, Carol Clugston, Anna Dominiczak and Alex Alderton, Roberto Amato, Jeffrey Barrett, Sonia Goncalves, Ewan Harrison, David K. Jackson, Ian Johnston, Dominic Kwiatkowski, Cordelia Langford, John Sillitoe on behalf of the Wellcome Sanger Institute COVID-19 Surveillance Team |
| EPI_ISL_3378649 | Health Services Laboratories | Wellcome Sanger Institute for the COVID-19 Genomics UK (COG-UK) Consortium | Health Services Laboratories and Alex Alderton, Roberto Amato, Jeffrey Barrett, Sonia Goncalves, Ewan Harrison, David K. Jackson, Ian Johnston, Dominic Kwiatkowski, Cordelia Langford, John Sillitoe on behalf of the Wellcome Sanger Institute COVID-19 Surveillance Team |
| EPI_ISL_3378650 | Lighthouse Lab in Glasgow | Wellcome Sanger Institute for the COVID-19 Genomics UK (COG-UK) Consortium | Harper VanSteenhouse, Yumi Kasai, David Gray, Carol Clugston, Anna Dominiczak and Alex Alderton, Roberto Amato, Jeffrey Barrett, Sonia Goncalves, Ewan Harrison, David K. Jackson, Ian Johnston, Dominic Kwiatkowski, Cordelia Langford, John Sillitoe on behalf of the Wellcome Sanger Institute COVID-19 Surveillance Team |
| EPI_ISL_3378651 | LABORATOIRE BIOESTEREL | CNR Virus des Infections Respiratoires - France SUD | Antonin Bal, Gregory Destras, Gwendolyne Burfin, Hadrien Regue, Quentin Semanas, Martine Valette, Bruno Lina, Laurence Josset |
| EPI_ISL_3378652 | Health Services Laboratories | Wellcome Sanger Institute for the COVID-19 Genomics UK (COG-UK) Consortium | Health Services Laboratories and Alex Alderton, Roberto Amato, Jeffrey Barrett, Sonia Goncalves, Ewan Harrison, David K. Jackson, Ian Johnston, Dominic Kwiatkowski, Cordelia Langford, John Sillitoe on behalf of the Wellcome Sanger Institute COVID-19 Surveillance Team |
| EPI_ISL_3378653 | Lighthouse Lab in Glasgow | Wellcome Sanger Institute for the COVID-19 Genomics UK (COG-UK) | Harper VanSteenhouse, Yumi Kasai, David Gray, Carol Clugston, Anna Dominiczak and Alex Alderton, Roberto Amato, Jeffrey Barrett, Sonia Goncalves, Ewan Harrison, David K. Jackson, Ian Johnston, Dominic Kwiatkowski, Cordelia Langford, |

| | | | |
|---|---|---|---|
| | | Consortium | John Sillitoe on behalf of the Wellcome Sanger Institute COVID-19 Surveillance Team |
| EPI_ISL_3378654 | Health Services Laboratories | Wellcome Sanger Institute for the COVID-19 Genomics UK (COG-UK) Consortium | Health Services Laboratories and Alex Alderton, Roberto Amato, Jeffrey Barrett, Sonia Goncalves, Ewan Harrison, David K. Jackson, Ian Johnston, Dominic Kwiatkowski, Cordelia Langford, John Sillitoe on behalf of the Wellcome Sanger Institute COVID-19 Surveillance Team |
| EPI_ISL_3378655 | LABORATOIRE ACCOLAB SUD OUEST | CNR Virus des Infections Respiratoires - France SUD | Antonin Bal, Gregory Destras, Gwendolyne Burfin, Hadrien Regue, Quentin Semanas, Martine Valette, Bruno Lina, Laurence Josset |
| EPI_ISL_3378656 | Health Services Laboratories | Wellcome Sanger Institute for the COVID-19 Genomics UK (COG-UK) Consortium | Health Services Laboratories and Alex Alderton, Roberto Amato, Jeffrey Barrett, Sonia Goncalves, Ewan Harrison, David K. Jackson, Ian Johnston, Dominic Kwiatkowski, Cordelia Langford, John Sillitoe on behalf of the Wellcome Sanger Institute COVID-19 Surveillance Team |
| EPI_ISL_3378657, EPI_ISL_3378658 | Lighthouse Lab in Glasgow | Wellcome Sanger Institute for the COVID-19 Genomics UK (COG-UK) Consortium | Harper VanSteenhouse, Yumi Kasai, David Gray, Carol Clugston, Anna Dominiczak and Alex Alderton, Roberto Amato, Jeffrey Barrett, Sonia Goncalves, Ewan Harrison, David K. Jackson, Ian Johnston, Dominic Kwiatkowski, Cordelia Langford, John Sillitoe on behalf of the Wellcome Sanger Institute COVID-19 Surveillance Team |
| EPI_ISL_3378659 | LABORATOIRE BIOESTEREL | CNR Virus des Infections Respiratoires - France SUD | Antonin Bal, Gregory Destras, Gwendolyne Burfin, Hadrien Regue, Quentin Semanas, Martine Valette, Bruno Lina, Laurence Josset |
| EPI_ISL_3378660 | Health Services Laboratories | Wellcome Sanger Institute for the COVID-19 Genomics UK (COG-UK) Consortium | Health Services Laboratories and Alex Alderton, Roberto Amato, Jeffrey Barrett, Sonia Goncalves, Ewan Harrison, David K. Jackson, Ian Johnston, Dominic Kwiatkowski, Cordelia Langford, John Sillitoe on behalf of the Wellcome Sanger Institute COVID-19 Surveillance Team |
| EPI_ISL_3378661, EPI_ISL_3378662 | Lighthouse Lab in Glasgow | Wellcome Sanger Institute for the COVID-19 Genomics UK (COG-UK) Consortium | Harper VanSteenhouse, Yumi Kasai, David Gray, Carol Clugston, Anna Dominiczak and Alex Alderton, Roberto Amato, Jeffrey Barrett, Sonia Goncalves, Ewan Harrison, David K. Jackson, Ian Johnston, Dominic Kwiatkowski, Cordelia Langford, John Sillitoe on behalf of the Wellcome Sanger Institute COVID-19 Surveillance Team |
| EPI_ISL_3378663 | LABORATOIRE ACCOLAB SUD OUEST | CNR Virus des Infections Respiratoires - France SUD | Antonin Bal, Gregory Destras, Gwendolyne Burfin, Hadrien Regue, Quentin Semanas, Martine Valette, Bruno Lina, Laurence Josset |
| EPI_ISL_3378664, EPI_ISL_3378665 | Lighthouse Lab in Alderley Park | Wellcome Sanger Institute for the COVID-19 Genomics UK (COG-UK) Consortium | Jacquelyn Wynn, Mairead Hyland, The Lighthouse Lab in Alderley Park and Alex Alderton, Roberto Amato, Jeffrey Barrett, Sonia Goncalves, Ewan Harrison, David K. Jackson, Ian Johnston, Dominic Kwiatkowski, Cordelia Langford, John Sillitoe on behalf of the Wellcome Sanger Institute COVID-19 Surveillance Team |
| EPI_ISL_3378666 | Lighthouse Lab in Glasgow | Wellcome Sanger Institute for the COVID-19 Genomics UK (COG-UK) Consortium | Harper VanSteenhouse, Yumi Kasai, David Gray, Carol Clugston, Anna Dominiczak and Alex Alderton, Roberto Amato, Jeffrey Barrett, Sonia Goncalves, Ewan Harrison, David K. Jackson, Ian Johnston, Dominic Kwiatkowski, Cordelia Langford, John Sillitoe on behalf of the Wellcome Sanger Institute COVID-19 Surveillance Team |
| EPI_ISL_3378667 | LABORATOIRE BIOESTEREL | CNR Virus des Infections Respiratoires - France SUD | Antonin Bal, Gregory Destras, Gwendolyne Burfin, Hadrien Regue, Quentin Semanas, Martine Valette, Bruno Lina, Laurence Josset |
| EPI_ISL_3378668 | Lighthouse Lab in Alderley Park | Wellcome Sanger Institute for the COVID-19 Genomics UK (COG-UK) Consortium | Jacquelyn Wynn, Mairead Hyland, The Lighthouse Lab in Alderley Park and Alex Alderton, Roberto Amato, Jeffrey Barrett, Sonia Goncalves, Ewan Harrison, David K. Jackson, Ian Johnston, Dominic Kwiatkowski, Cordelia Langford, John Sillitoe on behalf of the Wellcome Sanger Institute COVID-19 Surveillance Team |
| EPI_ISL_3378669 | Lighthouse Lab in Glasgow | Wellcome Sanger Institute for the COVID-19 Genomics UK (COG-UK) Consortium | Harper VanSteenhouse, Yumi Kasai, David Gray, Carol Clugston, Anna Dominiczak and Alex Alderton, Roberto Amato, Jeffrey Barrett, Sonia Goncalves, Ewan Harrison, David K. Jackson, Ian Johnston, Dominic Kwiatkowski, Cordelia Langford, John Sillitoe on behalf of the Wellcome Sanger Institute COVID-19 Surveillance Team |
| EPI_ISL_3378670 | LABORATOIRE BIOESTEREL | CNR Virus des Infections Respiratoires - France SUD | Antonin Bal, Gregory Destras, Gwendolyne Burfin, Hadrien Regue, Quentin Semanas, Martine Valette, Bruno Lina, Laurence Josset |
| EPI_ISL_3378671 | Lighthouse Lab in Glasgow | Wellcome Sanger Institute for the COVID-19 Genomics UK (COG-UK) Consortium | Harper VanSteenhouse, Yumi Kasai, David Gray, Carol Clugston, Anna Dominiczak and Alex Alderton, Roberto Amato, Jeffrey Barrett, Sonia Goncalves, Ewan Harrison, David K. Jackson, Ian Johnston, Dominic Kwiatkowski, Cordelia Langford, John Sillitoe on behalf of the Wellcome Sanger Institute COVID-19 Surveillance Team |
| EPI_ISL_3378672 | Lighthouse Lab in Milton Keynes | Wellcome Sanger Institute for the COVID-19 Genomics UK (COG-UK) Consortium | The Lighthouse Lab in Milton Keynes and Alex Alderton, Roberto Amato, Jeffrey Barrett, Sonia Goncalves, Ewan Harrison, David K. Jackson, Ian Johnston, Dominic Kwiatkowski, Cordelia Langford, John Sillitoe on behalf of the Wellcome Sanger Institute COVID-19 Surveillance Team |
| EPI_ISL_3378673 | Lighthouse Lab in Alderley Park | Wellcome Sanger Institute for the COVID-19 Genomics UK (COG-UK) Consortium | Jacquelyn Wynn, Mairead Hyland, The Lighthouse Lab in Alderley Park and Alex Alderton, Roberto Amato, Jeffrey Barrett, Sonia Goncalves, Ewan Harrison, David K. Jackson, Ian Johnston, Dominic Kwiatkowski, Cordelia Langford, John Sillitoe on behalf of the Wellcome Sanger Institute COVID-19 Surveillance Team |
| EPI_ISL_3378674 | CNR Virus des Infections Respiratoires - France SUD | CNR Virus des Infections Respiratoires - France SUD | Antonin Bal, Gregory Destras, Gwendolyne Burfin, Hadrien Regue, Quentin Semanas, Martine Valette, Bruno Lina, Laurence Josset |
| EPI_ISL_3378675 | Lighthouse Lab in Alderley Park | Wellcome Sanger Institute for the COVID-19 Genomics UK (COG-UK) Consortium | Jacquelyn Wynn, Mairead Hyland, The Lighthouse Lab in Alderley Park and Alex Alderton, Roberto Amato, Jeffrey Barrett, Sonia Goncalves, Ewan Harrison, David K. Jackson, Ian Johnston, Dominic Kwiatkowski, Cordelia Langford, John Sillitoe on behalf of the Wellcome Sanger Institute COVID-19 Surveillance Team |
| EPI_ISL_3378676 | Lighthouse Lab in Glasgow | Wellcome Sanger Institute for the COVID-19 Genomics UK (COG-UK) Consortium | Harper VanSteenhouse, Yumi Kasai, David Gray, Carol Clugston, Anna Dominiczak and Alex Alderton, Roberto Amato, Jeffrey Barrett, Sonia Goncalves, Ewan Harrison, David K. Jackson, Ian Johnston, Dominic Kwiatkowski, Cordelia Langford, John Sillitoe on behalf of the Wellcome Sanger Institute COVID-19 Surveillance Team |
| EPI_ISL_3378677 | Lighthouse Lab in Alderley Park | Wellcome Sanger Institute for the COVID-19 Genomics UK (COG-UK) Consortium | Jacquelyn Wynn, Mairead Hyland, The Lighthouse Lab in Alderley Park and Alex Alderton, Roberto Amato, Jeffrey Barrett, Sonia Goncalves, Ewan Harrison, David K. Jackson, Ian Johnston, Dominic Kwiatkowski, Cordelia Langford, John Sillitoe on behalf of the Wellcome Sanger Institute COVID-19 Surveillance Team |
| EPI_ISL_3378678 | LABORATOIRE BIOESTEREL | CNR Virus des Infections Respiratoires - France SUD | Antonin Bal, Gregory Destras, Gwendolyne Burfin, Hadrien Regue, Quentin Semanas, Martine Valette, Bruno Lina, Laurence Josset |
| EPI_ISL_3378679 | Lighthouse Lab in Glasgow | Wellcome Sanger Institute for the COVID-19 Genomics UK (COG-UK) Consortium | Harper VanSteenhouse, Yumi Kasai, David Gray, Carol Clugston, Anna Dominiczak and Alex Alderton, Roberto Amato, Jeffrey Barrett, Sonia Goncalves, Ewan Harrison, David K. Jackson, Ian Johnston, Dominic Kwiatkowski, Cordelia Langford, John Sillitoe on behalf of the Wellcome Sanger Institute COVID-19 Surveillance Team |
| EPI_ISL_3378680 | Lighthouse Lab in Alderley Park | Wellcome Sanger Institute for the COVID-19 Genomics UK (COG-UK) Consortium | Jacquelyn Wynn, Mairead Hyland, The Lighthouse Lab in Alderley Park and Alex Alderton, Roberto Amato, Jeffrey Barrett, Sonia Goncalves, Ewan Harrison, David K. Jackson, Ian Johnston, Dominic Kwiatkowski, Cordelia Langford, John Sillitoe on behalf of the Wellcome Sanger Institute COVID-19 Surveillance Team |
| EPI_ISL_3378681 | Lighthouse Lab in Glasgow | Wellcome Sanger Institute for the COVID-19 Genomics UK (COG-UK) Consortium | Harper VanSteenhouse, Yumi Kasai, David Gray, Carol Clugston, Anna Dominiczak and Alex Alderton, Roberto Amato, Jeffrey Barrett, Sonia Goncalves, Ewan Harrison, David K. Jackson, Ian Johnston, Dominic Kwiatkowski, Cordelia Langford, John Sillitoe on behalf of the Wellcome Sanger Institute COVID-19 Surveillance Team |
| EPI_ISL_3378682 | LABORATOIRE BIOESTEREL | CNR Virus des Infections Respiratoires - France SUD | Antonin Bal, Gregory Destras, Gwendolyne Burfin, Hadrien Regue, Quentin Semanas, Martine Valette, Bruno Lina, Laurence Josset |
| EPI_ISL_3378683, EPI_ISL_3378684 | Lighthouse Lab in Alderley Park | Wellcome Sanger Institute for the COVID-19 Genomics UK (COG-UK) Consortium | Jacquelyn Wynn, Mairead Hyland, The Lighthouse Lab in Alderley Park and Alex Alderton, Roberto Amato, Jeffrey Barrett, Sonia Goncalves, Ewan Harrison, David K. Jackson, Ian Johnston, Dominic Kwiatkowski, Cordelia Langford, John Sillitoe on behalf of the Wellcome Sanger Institute COVID-19 Surveillance Team |
| EPI_ISL_3378685 | Lighthouse Lab in Glasgow | Wellcome Sanger Institute for the COVID-19 Genomics UK (COG-UK) Consortium | Harper VanSteenhouse, Yumi Kasai, David Gray, Carol Clugston, Anna Dominiczak and Alex Alderton, Roberto Amato, Jeffrey Barrett, Sonia Goncalves, Ewan Harrison, David K. Jackson, Ian Johnston, Dominic Kwiatkowski, Cordelia Langford, John Sillitoe on behalf of the Wellcome Sanger Institute COVID-19 Surveillance Team |
| EPI_ISL_3378686 | LABORATOIRE BIOESTEREL | CNR Virus des Infections Respiratoires - France SUD | Antonin Bal, Gregory Destras, Gwendolyne Burfin, Hadrien Regue, Quentin Semanas, Martine Valette, Bruno Lina, Laurence Josset |
| EPI_ISL_3378687 | Lighthouse Lab in Alderley Park | Wellcome Sanger Institute for the COVID-19 Genomics UK (COG-UK) Consortium | Jacquelyn Wynn, Mairead Hyland, The Lighthouse Lab in Alderley Park and Alex Alderton, Roberto Amato, Jeffrey Barrett, Sonia Goncalves, Ewan Harrison, David K. Jackson, Ian Johnston, Dominic Kwiatkowski, Cordelia Langford, John Sillitoe on behalf of the Wellcome Sanger Institute COVID-19 Surveillance Team |
| EPI_ISL_3378688 | Lighthouse Lab in Glasgow | Wellcome Sanger Institute for the COVID-19 Genomics UK (COG-UK) Consortium | Harper VanSteenhouse, Yumi Kasai, David Gray, Carol Clugston, Anna Dominiczak and Alex Alderton, Roberto Amato, Jeffrey Barrett, Sonia Goncalves, Ewan Harrison, David K. Jackson, Ian Johnston, Dominic Kwiatkowski, Cordelia Langford, John Sillitoe on behalf of the Wellcome Sanger Institute COVID-19 Surveillance Team |
| EPI_ISL_3378689 | Lighthouse Lab in Alderley Park | Wellcome Sanger Institute for the COVID-19 Genomics UK (COG-UK) Consortium | Jacquelyn Wynn, Mairead Hyland, The Lighthouse Lab in Alderley Park and Alex Alderton, Roberto Amato, Jeffrey Barrett, Sonia Goncalves, Ewan Harrison, David K. Jackson, Ian Johnston, Dominic Kwiatkowski, Cordelia Langford, John Sillitoe on behalf of the Wellcome Sanger Institute COVID-19 Surveillance Team |
| EPI_ISL_3378690 | LABORATOIRE BIOESTEREL | CNR Virus des Infections Respiratoires - France SUD | Antonin Bal, Gregory Destras, Gwendolyne Burfin, Hadrien Regue, Quentin Semanas, Martine Valette, Bruno Lina, Laurence Josset |
| EPI_ISL_3378691, EPI_ISL_3378692 | Lighthouse Lab in Glasgow | Wellcome Sanger Institute for the COVID-19 Genomics UK (COG-UK) Consortium | Harper VanSteenhouse, Yumi Kasai, David Gray, Carol Clugston, Anna Dominiczak and Alex Alderton, Roberto Amato, Jeffrey Barrett, Sonia Goncalves, Ewan Harrison, David K. Jackson, Ian Johnston, Dominic Kwiatkowski, Cordelia Langford, John Sillitoe on behalf of the Wellcome Sanger Institute COVID-19 Surveillance Team |
| EPI_ISL_3378693 | Lighthouse Lab in Alderley Park | Wellcome Sanger Institute for the COVID-19 Genomics UK (COG-UK) Consortium | Jacquelyn Wynn, Mairead Hyland, The Lighthouse Lab in Alderley Park and Alex Alderton, Roberto Amato, Jeffrey Barrett, Sonia Goncalves, Ewan Harrison, David K. Jackson, Ian Johnston, Dominic Kwiatkowski, Cordelia Langford, John Sillitoe on behalf of the Wellcome Sanger Institute COVID-19 Surveillance Team |
| EPI_ISL_3378694 | LABORATOIRE BIOESTEREL | CNR Virus des Infections Respiratoires - France SUD | Antonin Bal, Gregory Destras, Gwendolyne Burfin, Hadrien Regue, Quentin Semanas, Martine Valette, Bruno Lina, Laurence Josset |
| EPI_ISL_3378695, EPI_ISL_3378696 | Lighthouse Lab in Glasgow | Wellcome Sanger Institute for the COVID-19 Genomics UK (COG-UK) Consortium | Harper VanSteenhouse, Yumi Kasai, David Gray, Carol Clugston, Anna Dominiczak and Alex Alderton, Roberto Amato, Jeffrey Barrett, Sonia Goncalves, Ewan Harrison, David K. Jackson, Ian Johnston, Dominic Kwiatkowski, Cordelia Langford, John Sillitoe on behalf of the Wellcome Sanger Institute COVID-19 Surveillance Team |
| EPI_ISL_3378697 | CNR Virus des Infections Respiratoires - France SUD | CNR Virus des Infections Respiratoires - France SUD | Antonin Bal, Gregory Destras, Gwendolyne Burfin, Hadrien Regue, Quentin Semanas, Martine Valette, Bruno Lina, Laurence Josset |
| EPI_ISL_3378698 | Lighthouse Lab in Alderley Park | Wellcome Sanger Institute for the COVID-19 Genomics UK (COG-UK) Consortium | Jacquelyn Wynn, Mairead Hyland, The Lighthouse Lab in Alderley Park and Alex Alderton, Roberto Amato, Jeffrey Barrett, Sonia Goncalves, Ewan Harrison, David K. Jackson, Ian Johnston, Dominic Kwiatkowski, Cordelia Langford, John Sillitoe on behalf of the Wellcome Sanger Institute COVID-19 Surveillance Team |
| EPI_ISL_3378699, EPI_ISL_3378700 | Lighthouse Lab in Glasgow | Wellcome Sanger Institute for the COVID-19 Genomics UK (COG-UK) Consortium | Harper VanSteenhouse, Yumi Kasai, David Gray, Carol Clugston, Anna Dominiczak and Alex Alderton, Roberto Amato, Jeffrey Barrett, Sonia Goncalves, Ewan Harrison, David K. Jackson, Ian Johnston, Dominic Kwiatkowski, Cordelia Langford, John Sillitoe on behalf of the Wellcome Sanger Institute COVID-19 Surveillance Team |
| EPI_ISL_3378701 | CNR Virus des Infections Respiratoires - France SUD | CNR Virus des Infections Respiratoires - France SUD | Antonin Bal, Gregory Destras, Gwendolyne Burfin, Hadrien Regue, Quentin Semanas, Martine Valette, Bruno Lina, Laurence Josset |
| EPI_ISL_3378702 | Lighthouse Lab in Glasgow | Wellcome Sanger Institute for the COVID-19 Genomics UK (COG-UK) Consortium | Harper VanSteenhouse, Yumi Kasai, David Gray, Carol Clugston, Anna Dominiczak and Alex Alderton, Roberto Amato, Jeffrey Barrett, Sonia Goncalves, Ewan Harrison, David K. Jackson, Ian Johnston, Dominic Kwiatkowski, Cordelia Langford, John Sillitoe on behalf of the Wellcome Sanger Institute COVID-19 Surveillance Team |
| EPI_ISL_3378703 | Lighthouse Lab in Alderley Park | Wellcome Sanger Institute for the COVID-19 Genomics UK (COG-UK) Consortium | Jacquelyn Wynn, Mairead Hyland, The Lighthouse Lab in Alderley Park and Alex Alderton, Roberto Amato, Jeffrey Barrett, Sonia Goncalves, Ewan Harrison, David K. Jackson, Ian Johnston, Dominic Kwiatkowski, Cordelia Langford, John Sillitoe on behalf of the Wellcome Sanger Institute COVID-19 Surveillance Team |
| EPI_ISL_3378704 | CNR Virus des Infections Respiratoires - France SUD | CNR Virus des Infections Respiratoires - France SUD | Antonin Bal, Gregory Destras, Gwendolyne Burfin, Hadrien Regue, Quentin Semanas, Martine Valette, Bruno Lina, Laurence Josset |
| EPI_ISL_3378705 | Lighthouse Lab in Glasgow | Wellcome Sanger Institute for the COVID-19 Genomics UK (COG-UK) Consortium | Harper VanSteenhouse, Yumi Kasai, David Gray, Carol Clugston, Anna Dominiczak and Alex Alderton, Roberto Amato, Jeffrey Barrett, Sonia Goncalves, Ewan Harrison, David K. Jackson, Ian Johnston, Dominic Kwiatkowski, Cordelia Langford, John Sillitoe on behalf of the Wellcome Sanger Institute COVID-19 Surveillance Team |
| EPI_ISL_3378706, EPI_ISL_3378707, EPI_ISL_3378708, EPI_ISL_3378709 | CNR Virus des Infections Respiratoires - France SUD | CNR Virus des Infections Respiratoires - France SUD | Antonin Bal, Gregory Destras, Gwendolyne Burfin, Hadrien Regue, Quentin Semanas, Martine Valette, Bruno Lina, Laurence Josset |
| EPI_ISL_3378710, EPI_ISL_3378711, EPI_ISL_3378712, EPI_ISL_3378713, EPI_ISL_3378714, EPI_ISL_3378715, EPI_ISL_3378716, EPI_ISL_3378717, EPI_ISL_3378718, EPI_ISL_3378719, EPI_ISL_3378720, EPI_ISL_3378721, EPI_ISL_3378722, EPI_ISL_3378723, EPI_ISL_3378724, EPI_ISL_3378725, EPI_ISL_3378726, EPI_ISL_3378727, EPI_ISL_3378728, EPI_ISL_3378729, EPI_ISL_3378730, EPI_ISL_3378731, EPI_ISL_3378732, EPI_ISL_3378733, EPI_ISL_3378734, EPI_ISL_3378735, EPI_ISL_3378736, EPI_ISL_3378737, EPI_ISL_3378738, EPI_ISL_3378739 | | | |
| see above | SYNLAB GASCOGNE | CNR Virus des Infections Respiratoires - France SUD | Antonin Bal, Gregory Destras, Gwendolyne Burfin, Hadrien Regue, Quentin Semanas, Martine Valette, Bruno Lina, Laurence Josset |
| EPI_ISL_3378740, EPI_ISL_3378741 | LABORATOIRE UNILIANS DECINES | CNR Virus des Infections Respiratoires - France SUD | Antonin Bal, Gregory Destras, Gwendolyne Burfin, Hadrien Regue, Quentin Semanas, Martine Valette, Bruno Lina, Laurence Josset |
| EPI_ISL_3378742, EPI_ISL_3378743 | Lighthouse Lab in Glasgow | Wellcome Sanger Institute for the COVID-19 Genomics UK (COG-UK) Consortium | Harper VanSteenhouse, Yumi Kasai, David Gray, Carol Clugston, Anna Dominiczak and Alex Alderton, Roberto Amato, Jeffrey Barrett, Sonia Goncalves, Ewan Harrison, David K. Jackson, Ian Johnston, Dominic Kwiatkowski, Cordelia Langford, John Sillitoe on behalf of the Wellcome Sanger Institute COVID-19 Surveillance Team |

| | | | |
|---|---|---|---|
| EPI_ISL_3378744 | LABORATOIRE UNILIANS DECINES | CNR Virus des Infections Respiratoires - France SUD | Antonin Bal, Gregory Destras, Gwendolyne Burfin, Hadrien Regue, Quentin Semanas, Martine Valette, Bruno Lina, Laurence Josset |
| EPI_ISL_3378745, EPI_ISL_3378746 | Lighthouse Lab in Alderley Park | Wellcome Sanger Institute for the COVID-19 Genomics UK (COG-UK) Consortium | Jacquelyn Wynn, Mairead Hyland, The Lighthouse Lab in Alderley Park and Alex Alderton, Roberto Amato, Jeffrey Barrett, Sonia Goncalves, Ewan Harrison, David K. Jackson, Ian Johnston, Dominic Kwiatkowski, Cordelia Langford, John Sillitoe on behalf of the Wellcome Sanger Institute COVID-19 Surveillance Team |
| EPI_ISL_3378747 | LABORATOIRE UNILIANS DECINES | CNR Virus des Infections Respiratoires - France SUD | Antonin Bal, Gregory Destras, Gwendolyne Burfin, Hadrien Regue, Quentin Semanas, Martine Valette, Bruno Lina, Laurence Josset |
| EPI_ISL_3378748, EPI_ISL_3378749 | Lighthouse Lab in Alderley Park | Wellcome Sanger Institute for the COVID-19 Genomics UK (COG-UK) Consortium | Jacquelyn Wynn, Mairead Hyland, The Lighthouse Lab in Alderley Park and Alex Alderton, Roberto Amato, Jeffrey Barrett, Sonia Goncalves, Ewan Harrison, David K. Jackson, Ian Johnston, Dominic Kwiatkowski, Cordelia Langford, John Sillitoe on behalf of the Wellcome Sanger Institute COVID-19 Surveillance Team |
| EPI_ISL_3378750 | LABORATOIRE UNILIANS DECINES | CNR Virus des Infections Respiratoires - France SUD | Antonin Bal, Gregory Destras, Gwendolyne Burfin, Hadrien Regue, Quentin Semanas, Martine Valette, Bruno Lina, Laurence Josset |
| EPI_ISL_3378751 | Lighthouse Lab in Alderley Park | Wellcome Sanger Institute for the COVID-19 Genomics UK (COG-UK) Consortium | Jacquelyn Wynn, Mairead Hyland, The Lighthouse Lab in Alderley Park and Alex Alderton, Roberto Amato, Jeffrey Barrett, Sonia Goncalves, Ewan Harrison, David K. Jackson, Ian Johnston, Dominic Kwiatkowski, Cordelia Langford, John Sillitoe on behalf of the Wellcome Sanger Institute COVID-19 Surveillance Team |
| EPI_ISL_3378752, EPI_ISL_3378753 | Lighthouse Lab in Glasgow | Wellcome Sanger Institute for the COVID-19 Genomics UK (COG-UK) Consortium | Harper VanSteenhouse, Yumi Kasai, David Gray, Carol Clugston, Anna Dominiczak and Alex Alderton, Roberto Amato, Jeffrey Barrett, Sonia Goncalves, Ewan Harrison, David K. Jackson, Ian Johnston, Dominic Kwiatkowski, Cordelia Langford, John Sillitoe on behalf of the Wellcome Sanger Institute COVID-19 Surveillance Team |
| EPI_ISL_3378754 | DYOMEDEA-LABORATOIRE DE LA SAUVEGARDE | CNR Virus des Infections Respiratoires - France SUD | Antonin Bal, Gregory Destras, Gwendolyne Burfin, Hadrien Regue, Quentin Semanas, Martine Valette, Bruno Lina, Laurence Josset |
| EPI_ISL_3378755, EPI_ISL_3378756 | Lighthouse Lab in Glasgow | Wellcome Sanger Institute for the COVID-19 Genomics UK (COG-UK) Consortium | Harper VanSteenhouse, Yumi Kasai, David Gray, Carol Clugston, Anna Dominiczak and Alex Alderton, Roberto Amato, Jeffrey Barrett, Sonia Goncalves, Ewan Harrison, David K. Jackson, Ian Johnston, Dominic Kwiatkowski, Cordelia Langford, John Sillitoe on behalf of the Wellcome Sanger Institute COVID-19 Surveillance Team |
| EPI_ISL_3378757 | DYOMEDEA-LABORATOIRE DE LA SAUVEGARDE | CNR Virus des Infections Respiratoires - France SUD | Antonin Bal, Gregory Destras, Gwendolyne Burfin, Hadrien Regue, Quentin Semanas, Martine Valette, Bruno Lina, Laurence Josset |
| EPI_ISL_3378758, EPI_ISL_3378759 | Lighthouse Lab in Glasgow | Wellcome Sanger Institute for the COVID-19 Genomics UK (COG-UK) Consortium | Harper VanSteenhouse, Yumi Kasai, David Gray, Carol Clugston, Anna Dominiczak and Alex Alderton, Roberto Amato, Jeffrey Barrett, Sonia Goncalves, Ewan Harrison, David K. Jackson, Ian Johnston, Dominic Kwiatkowski, Cordelia Langford, John Sillitoe on behalf of the Wellcome Sanger Institute COVID-19 Surveillance Team |
| EPI_ISL_3378760 | Lighthouse Lab in Alderley Park | Wellcome Sanger Institute for the COVID-19 Genomics UK (COG-UK) Consortium | Jacquelyn Wynn, Mairead Hyland, The Lighthouse Lab in Alderley Park and Alex Alderton, Roberto Amato, Jeffrey Barrett, Sonia Goncalves, Ewan Harrison, David K. Jackson, Ian Johnston, Dominic Kwiatkowski, Cordelia Langford, John Sillitoe on behalf of the Wellcome Sanger Institute COVID-19 Surveillance Team |
| EPI_ISL_3378761 | DYOMEDEA-LABORATOIRE DE LA SAUVEGARDE | CNR Virus des Infections Respiratoires - France SUD | Antonin Bal, Gregory Destras, Gwendolyne Burfin, Hadrien Regue, Quentin Semanas, Martine Valette, Bruno Lina, Laurence Josset |
| EPI_ISL_3378762, EPI_ISL_3378763, EPI_ISL_3378764 | Lighthouse Lab in Glasgow | Wellcome Sanger Institute for the COVID-19 Genomics UK (COG-UK) Consortium | Harper VanSteenhouse, Yumi Kasai, David Gray, Carol Clugston, Anna Dominiczak and Alex Alderton, Roberto Amato, Jeffrey Barrett, Sonia Goncalves, Ewan Harrison, David K. Jackson, Ian Johnston, Dominic Kwiatkowski, Cordelia Langford, John Sillitoe on behalf of the Wellcome Sanger Institute COVID-19 Surveillance Team |
| EPI_ISL_3378765 | DYOMEDEA-LABORATOIRE DE LA SAUVEGARDE | CNR Virus des Infections Respiratoires - France SUD | Antonin Bal, Gregory Destras, Gwendolyne Burfin, Hadrien Regue, Quentin Semanas, Martine Valette, Bruno Lina, Laurence Josset |
| EPI_ISL_3378766 | Lighthouse Lab in Milton Keynes | Wellcome Sanger Institute for the COVID-19 Genomics UK (COG-UK) Consortium | The Lighthouse Lab in Milton Keynes and Alex Alderton, Roberto Amato, Jeffrey Barrett, Sonia Goncalves, Ewan Harrison, David K. Jackson, Ian Johnston, Dominic Kwiatkowski, Cordelia Langford, John Sillitoe on behalf of the Wellcome Sanger Institute COVID-19 Surveillance Team |
| EPI_ISL_3378767, EPI_ISL_3378768 | Lighthouse Lab in Alderley Park | Wellcome Sanger Institute for the COVID-19 Genomics UK (COG-UK) Consortium | Jacquelyn Wynn, Mairead Hyland, The Lighthouse Lab in Alderley Park and Alex Alderton, Roberto Amato, Jeffrey Barrett, Sonia Goncalves, Ewan Harrison, David K. Jackson, Ian Johnston, Dominic Kwiatkowski, Cordelia Langford, John Sillitoe on behalf of the Wellcome Sanger Institute COVID-19 Surveillance Team |
| EPI_ISL_3378769 | DYOMEDEA-LABORATOIRE DE LA SAUVEGARDE | CNR Virus des Infections Respiratoires - France SUD | Antonin Bal, Gregory Destras, Gwendolyne Burfin, Hadrien Regue, Quentin Semanas, Martine Valette, Bruno Lina, Laurence Josset |
| EPI_ISL_3378770 | Lighthouse Lab in Alderley Park | Wellcome Sanger Institute for the COVID-19 Genomics UK (COG-UK) Consortium | Jacquelyn Wynn, Mairead Hyland, The Lighthouse Lab in Alderley Park and Alex Alderton, Roberto Amato, Jeffrey Barrett, Sonia Goncalves, Ewan Harrison, David K. Jackson, Ian Johnston, Dominic Kwiatkowski, Cordelia Langford, John Sillitoe on behalf of the Wellcome Sanger Institute COVID-19 Surveillance Team |
| EPI_ISL_3378771 | Lighthouse Lab in Glasgow | Wellcome Sanger Institute for the COVID-19 Genomics UK (COG-UK) Consortium | Harper VanSteenhouse, Yumi Kasai, David Gray, Carol Clugston, Anna Dominiczak and Alex Alderton, Roberto Amato, Jeffrey Barrett, Sonia Goncalves, Ewan Harrison, David K. Jackson, Ian Johnston, Dominic Kwiatkowski, Cordelia Langford, John Sillitoe on behalf of the Wellcome Sanger Institute COVID-19 Surveillance Team |
| EPI_ISL_3378772 | Lighthouse Lab in Alderley Park | Wellcome Sanger Institute for the COVID-19 Genomics UK (COG-UK) Consortium | Jacquelyn Wynn, Mairead Hyland, The Lighthouse Lab in Alderley Park and Alex Alderton, Roberto Amato, Jeffrey Barrett, Sonia Goncalves, Ewan Harrison, David K. Jackson, Ian Johnston, Dominic Kwiatkowski, Cordelia Langford, John Sillitoe on behalf of the Wellcome Sanger Institute COVID-19 Surveillance Team |
| EPI_ISL_3378773 | DYOMEDEA-LABORATOIRE DE LA SAUVEGARDE | CNR Virus des Infections Respiratoires - France SUD | Antonin Bal, Gregory Destras, Gwendolyne Burfin, Hadrien Regue, Quentin Semanas, Martine Valette, Bruno Lina, Laurence Josset |
| EPI_ISL_3378774 | Lighthouse Lab in Glasgow | Wellcome Sanger Institute for the COVID-19 Genomics UK (COG-UK) Consortium | Harper VanSteenhouse, Yumi Kasai, David Gray, Carol Clugston, Anna Dominiczak and Alex Alderton, Roberto Amato, Jeffrey Barrett, Sonia Goncalves, Ewan Harrison, David K. Jackson, Ian Johnston, Dominic Kwiatkowski, Cordelia Langford, John Sillitoe on behalf of the Wellcome Sanger Institute COVID-19 Surveillance Team |
| EPI_ISL_3378775 | Lighthouse Lab in Alderley Park | Wellcome Sanger Institute for the COVID-19 Genomics UK (COG-UK) Consortium | Jacquelyn Wynn, Mairead Hyland, The Lighthouse Lab in Alderley Park and Alex Alderton, Roberto Amato, Jeffrey Barrett, Sonia Goncalves, Ewan Harrison, David K. Jackson, Ian Johnston, Dominic Kwiatkowski, Cordelia Langford, John Sillitoe on behalf of the Wellcome Sanger Institute COVID-19 Surveillance Team |
| EPI_ISL_3378777 | Lighthouse Lab in Glasgow | Wellcome Sanger Institute for the COVID-19 Genomics UK (COG-UK) Consortium | Harper VanSteenhouse, Yumi Kasai, David Gray, Carol Clugston, Anna Dominiczak and Alex Alderton, Roberto Amato, Jeffrey Barrett, Sonia Goncalves, Ewan Harrison, David K. Jackson, Ian Johnston, Dominic Kwiatkowski, Cordelia Langford, John Sillitoe on behalf of the Wellcome Sanger Institute COVID-19 Surveillance Team |
| EPI_ISL_3378778 | DYOMEDEA-LABORATOIRE DE LA SAUVEGARDE | CNR Virus des Infections Respiratoires - France SUD | Antonin Bal, Gregory Destras, Gwendolyne Burfin, Hadrien Regue, Quentin Semanas, Martine Valette, Bruno Lina, Laurence Josset |
| EPI_ISL_3378779 | Lighthouse Lab in Alderley Park | Wellcome Sanger Institute for the COVID-19 Genomics UK (COG-UK) Consortium | Jacquelyn Wynn, Mairead Hyland, The Lighthouse Lab in Alderley Park and Alex Alderton, Roberto Amato, Jeffrey Barrett, Sonia Goncalves, Ewan Harrison, David K. Jackson, Ian Johnston, Dominic Kwiatkowski, Cordelia Langford, John Sillitoe on behalf of the Wellcome Sanger Institute COVID-19 Surveillance Team |
| EPI_ISL_3378780 | Lighthouse Lab in Glasgow | Wellcome Sanger Institute for the COVID-19 Genomics UK (COG-UK) Consortium | Harper VanSteenhouse, Yumi Kasai, David Gray, Carol Clugston, Anna Dominiczak and Alex Alderton, Roberto Amato, Jeffrey Barrett, Sonia Goncalves, Ewan Harrison, David K. Jackson, Ian Johnston, Dominic Kwiatkowski, Cordelia Langford, John Sillitoe on behalf of the Wellcome Sanger Institute COVID-19 Surveillance Team |
| EPI_ISL_3378781 | DYOMEDEA-LABORATOIRE DE LA SAUVEGARDE | CNR Virus des Infections Respiratoires - France SUD | Antonin Bal, Gregory Destras, Gwendolyne Burfin, Hadrien Regue, Quentin Semanas, Martine Valette, Bruno Lina, Laurence Josset |
| EPI_ISL_3378782, EPI_ISL_3378783 | Lighthouse Lab in Alderley Park | Wellcome Sanger Institute for the COVID-19 Genomics UK (COG-UK) Consortium | Jacquelyn Wynn, Mairead Hyland, The Lighthouse Lab in Alderley Park and Alex Alderton, Roberto Amato, Jeffrey Barrett, Sonia Goncalves, Ewan Harrison, David K. Jackson, Ian Johnston, Dominic Kwiatkowski, Cordelia Langford, John Sillitoe on behalf of the Wellcome Sanger Institute COVID-19 Surveillance Team |
| EPI_ISL_3378784 | Lighthouse Lab in Glasgow | Wellcome Sanger Institute for the COVID-19 Genomics UK (COG-UK) Consortium | Harper VanSteenhouse, Yumi Kasai, David Gray, Carol Clugston, Anna Dominiczak and Alex Alderton, Roberto Amato, Jeffrey Barrett, Sonia Goncalves, Ewan Harrison, David K. Jackson, Ian Johnston, Dominic Kwiatkowski, Cordelia Langford, John Sillitoe on behalf of the Wellcome Sanger Institute COVID-19 Surveillance Team |
| EPI_ISL_3378785 | Lighthouse Lab in Alderley Park | Wellcome Sanger Institute for the COVID-19 Genomics UK (COG-UK) Consortium | Jacquelyn Wynn, Mairead Hyland, The Lighthouse Lab in Alderley Park and Alex Alderton, Roberto Amato, Jeffrey Barrett, Sonia Goncalves, Ewan Harrison, David K. Jackson, Ian Johnston, Dominic Kwiatkowski, Cordelia Langford, John Sillitoe on behalf of the Wellcome Sanger Institute COVID-19 Surveillance Team |
| EPI_ISL_3378786 | DYOMEDEA-LABORATOIRE DE LA SAUVEGARDE | CNR Virus des Infections Respiratoires - France SUD | Antonin Bal, Gregory Destras, Gwendolyne Burfin, Hadrien Regue, Quentin Semanas, Martine Valette, Bruno Lina, Laurence Josset |
| EPI_ISL_3378787 | Lighthouse Lab in Glasgow | Wellcome Sanger Institute for the COVID-19 Genomics UK (COG-UK) Consortium | Harper VanSteenhouse, Yumi Kasai, David Gray, Carol Clugston, Anna Dominiczak and Alex Alderton, Roberto Amato, Jeffrey Barrett, Sonia Goncalves, Ewan Harrison, David K. Jackson, Ian Johnston, Dominic Kwiatkowski, Cordelia Langford, John Sillitoe on behalf of the Wellcome Sanger Institute COVID-19 Surveillance Team |
| EPI_ISL_3378788, EPI_ISL_3378789 | Lighthouse Lab in Alderley Park | Wellcome Sanger Institute for the COVID-19 Genomics UK (COG-UK) Consortium | Jacquelyn Wynn, Mairead Hyland, The Lighthouse Lab in Alderley Park and Alex Alderton, Roberto Amato, Jeffrey Barrett, Sonia Goncalves, Ewan Harrison, David K. Jackson, Ian Johnston, Dominic Kwiatkowski, Cordelia Langford, John Sillitoe on behalf of the Wellcome Sanger Institute COVID-19 Surveillance Team |
| EPI_ISL_3378790 | DYOMEDEA-LABORATOIRE DE LA SAUVEGARDE | CNR Virus des Infections Respiratoires - France SUD | Antonin Bal, Gregory Destras, Gwendolyne Burfin, Hadrien Regue, Quentin Semanas, Martine Valette, Bruno Lina, Laurence Josset |
| EPI_ISL_3378791, EPI_ISL_3378792 | Lighthouse Lab in Alderley Park | Wellcome Sanger Institute for the COVID-19 Genomics UK (COG-UK) Consortium | Jacquelyn Wynn, Mairead Hyland, The Lighthouse Lab in Alderley Park and Alex Alderton, Roberto Amato, Jeffrey Barrett, Sonia Goncalves, Ewan Harrison, David K. Jackson, Ian Johnston, Dominic Kwiatkowski, Cordelia Langford, John Sillitoe on behalf of the Wellcome Sanger Institute COVID-19 Surveillance Team |
| EPI_ISL_3378793 | DYOMEDEA-LABORATOIRE DE LA SAUVEGARDE | CNR Virus des Infections Respiratoires - France SUD | Antonin Bal, Gregory Destras, Gwendolyne Burfin, Hadrien Regue, Quentin Semanas, Martine Valette, Bruno Lina, Laurence Josset |
| EPI_ISL_3378794, EPI_ISL_3378795, EPI_ISL_3378796 | Lighthouse Lab in Glasgow | Wellcome Sanger Institute for the COVID-19 Genomics UK (COG-UK) Consortium | Harper VanSteenhouse, Yumi Kasai, David Gray, Carol Clugston, Anna Dominiczak and Alex Alderton, Roberto Amato, Jeffrey Barrett, Sonia Goncalves, Ewan Harrison, David K. Jackson, Ian Johnston, Dominic Kwiatkowski, Cordelia Langford, John Sillitoe on behalf of the Wellcome Sanger Institute COVID-19 Surveillance Team |
| EPI_ISL_3378797 | DYOMEDEA-LABORATOIRE DE LA SAUVEGARDE | CNR Virus des Infections Respiratoires - France SUD | Antonin Bal, Gregory Destras, Gwendolyne Burfin, Hadrien Regue, Quentin Semanas, Martine Valette, Bruno Lina, Laurence Josset |
| EPI_ISL_3378798, EPI_ISL_3378799, EPI_ISL_3378800 | Lighthouse Lab in Alderley Park | Wellcome Sanger Institute for the COVID-19 Genomics UK (COG-UK) Consortium | Jacquelyn Wynn, Mairead Hyland, The Lighthouse Lab in Alderley Park and Alex Alderton, Roberto Amato, Jeffrey Barrett, Sonia Goncalves, Ewan Harrison, David K. Jackson, Ian Johnston, Dominic Kwiatkowski, Cordelia Langford, John Sillitoe on behalf of the Wellcome Sanger Institute COVID-19 Surveillance Team |
| EPI_ISL_3378801 | LAM BIORYLIS | CNR Virus des Infections Respiratoires - France SUD | Antonin Bal, Gregory Destras, Gwendolyne Burfin, Hadrien Regue, Quentin Semanas, Martine Valette, Bruno Lina, Laurence Josset |
| EPI_ISL_3378802 | Lighthouse Lab in Alderley Park | Wellcome Sanger Institute for the COVID-19 Genomics UK (COG-UK) Consortium | Jacquelyn Wynn, Mairead Hyland, The Lighthouse Lab in Alderley Park and Alex Alderton, Roberto Amato, Jeffrey Barrett, Sonia Goncalves, Ewan Harrison, David K. Jackson, Ian Johnston, Dominic Kwiatkowski, Cordelia Langford, John Sillitoe on behalf of the Wellcome Sanger Institute COVID-19 Surveillance Team |
| EPI_ISL_3378803 | Lighthouse Lab in Glasgow | Wellcome Sanger Institute for the COVID-19 Genomics UK (COG-UK) Consortium | Harper VanSteenhouse, Yumi Kasai, David Gray, Carol Clugston, Anna Dominiczak and Alex Alderton, Roberto Amato, Jeffrey Barrett, Sonia Goncalves, Ewan Harrison, David K. Jackson, Ian Johnston, Dominic Kwiatkowski, Cordelia Langford, John Sillitoe on behalf of the Wellcome Sanger Institute COVID-19 Surveillance Team |
| EPI_ISL_3378804 | Lighthouse Lab in Alderley Park | Wellcome Sanger Institute for the COVID-19 Genomics UK (COG-UK) Consortium | Jacquelyn Wynn, Mairead Hyland, The Lighthouse Lab in Alderley Park and Alex Alderton, Roberto Amato, Jeffrey Barrett, Sonia Goncalves, Ewan Harrison, David K. Jackson, Ian Johnston, Dominic Kwiatkowski, Cordelia Langford, John Sillitoe on behalf of the Wellcome Sanger Institute COVID-19 Surveillance Team |
| EPI_ISL_3378805 | LAM BIORYLIS | CNR Virus des Infections Respiratoires - France SUD | Antonin Bal, Gregory Destras, Gwendolyne Burfin, Hadrien Regue, Quentin Semanas, Martine Valette, Bruno Lina, Laurence Josset |
| EPI_ISL_3378806, EPI_ISL_3378807 | Lighthouse Lab in Glasgow | Wellcome Sanger Institute for the COVID-19 Genomics UK (COG-UK) Consortium | Harper VanSteenhouse, Yumi Kasai, David Gray, Carol Clugston, Anna Dominiczak and Alex Alderton, Roberto Amato, Jeffrey Barrett, Sonia Goncalves, Ewan Harrison, David K. Jackson, Ian Johnston, Dominic Kwiatkowski, Cordelia Langford, John Sillitoe on behalf of the Wellcome Sanger Institute COVID-19 Surveillance Team |
| EPI_ISL_3378808 | Lighthouse Lab in Alderley Park | Wellcome Sanger Institute for the COVID-19 Genomics UK (COG-UK) Consortium | Jacquelyn Wynn, Mairead Hyland, The Lighthouse Lab in Alderley Park and Alex Alderton, Roberto Amato, Jeffrey Barrett, Sonia Goncalves, Ewan Harrison, David K. Jackson, Ian Johnston, Dominic Kwiatkowski, Cordelia Langford, John Sillitoe on behalf of the Wellcome Sanger Institute COVID-19 Surveillance Team |
| EPI_ISL_3378809 | MIRIALIS CLUSES BECHET | CNR Virus des Infections Respiratoires - France SUD | Antonin Bal, Gregory Destras, Gwendolyne Burfin, Hadrien Regue, Quentin Semanas, Martine Valette, Bruno Lina, Laurence Josset |
| EPI_ISL_3378810, EPI_ISL_3378811, EPI_ISL_3378812 | Lighthouse Lab in Alderley Park | Wellcome Sanger Institute for the COVID-19 Genomics UK (COG-UK) Consortium | Jacquelyn Wynn, Mairead Hyland, The Lighthouse Lab in Alderley Park and Alex Alderton, Roberto Amato, Jeffrey Barrett, Sonia Goncalves, Ewan Harrison, David K. Jackson, Ian Johnston, Dominic Kwiatkowski, Cordelia Langford, John Sillitoe on behalf of the Wellcome Sanger Institute COVID-19 Surveillance Team |
| EPI_ISL_3378813 | MIRIALIS CLUSES BECHET | CNR Virus des Infections Respiratoires - France SUD | Antonin Bal, Gregory Destras, Gwendolyne Burfin, Hadrien Regue, Quentin Semanas, Martine Valette, Bruno Lina, Laurence Josset |

| | | | |
|---|---|---|---|
| EPI_ISL_3378814, EPI_ISL_3378815 | Lighthouse Lab in Alderley Park | Wellcome Sanger Institute for the COVID-19 Genomics UK (COG-UK) Consortium | Jacquelyn Wynn, Mairead Hyland, The Lighthouse Lab in Alderley Park and Alex Alderton, Roberto Amato, Jeffrey Barrett, Sonia Goncalves, Ewan Harrison, David K. Jackson, Ian Johnston, Dominic Kwiatkowski, Cordelia Langford, John Sillitoe on behalf of the Wellcome Sanger Institute COVID-19 Surveillance Team |
| EPI_ISL_3378816 | MIRIALIS CLUSES BECHET | CNR Virus des Infections Respiratoires - France SUD | Antonin Bal, Gregory Destras, Gwendolyne Burfin, Hadrien Regue, Quentin Semanas, Martine Valette, Bruno Lina, Laurence Josset |
| EPI_ISL_3378817 | Lighthouse Lab in Glasgow | Wellcome Sanger Institute for the COVID-19 Genomics UK (COG-UK) Consortium | Harper VanSteenhouse, Yumi Kasai, David Gray, Carol Clugston, Anna Dominiczak and Alex Alderton, Roberto Amato, Jeffrey Barrett, Sonia Goncalves, Ewan Harrison, David K. Jackson, Ian Johnston, Dominic Kwiatkowski, Cordelia Langford, John Sillitoe on behalf of the Wellcome Sanger Institute COVID-19 Surveillance Team |
| EPI_ISL_3378818 | Lighthouse Lab in Alderley Park | Wellcome Sanger Institute for the COVID-19 Genomics UK (COG-UK) Consortium | Jacquelyn Wynn, Mairead Hyland, The Lighthouse Lab in Alderley Park and Alex Alderton, Roberto Amato, Jeffrey Barrett, Sonia Goncalves, Ewan Harrison, David K. Jackson, Ian Johnston, Dominic Kwiatkowski, Cordelia Langford, John Sillitoe on behalf of the Wellcome Sanger Institute COVID-19 Surveillance Team |
| EPI_ISL_3378819 | Lighthouse Lab in Glasgow | Wellcome Sanger Institute for the COVID-19 Genomics UK (COG-UK) Consortium | Harper VanSteenhouse, Yumi Kasai, David Gray, Carol Clugston, Anna Dominiczak and Alex Alderton, Roberto Amato, Jeffrey Barrett, Sonia Goncalves, Ewan Harrison, David K. Jackson, Ian Johnston, Dominic Kwiatkowski, Cordelia Langford, John Sillitoe on behalf of the Wellcome Sanger Institute COVID-19 Surveillance Team |
| EPI_ISL_3378820 | MIRIALIS CLUSES BECHET | CNR Virus des Infections Respiratoires - France SUD | Antonin Bal, Gregory Destras, Gwendolyne Burfin, Hadrien Regue, Quentin Semanas, Martine Valette, Bruno Lina, Laurence Josset |
| EPI_ISL_3378821 | Lighthouse Lab in Alderley Park | Wellcome Sanger Institute for the COVID-19 Genomics UK (COG-UK) Consortium | Jacquelyn Wynn, Mairead Hyland, The Lighthouse Lab in Alderley Park and Alex Alderton, Roberto Amato, Jeffrey Barrett, Sonia Goncalves, Ewan Harrison, David K. Jackson, Ian Johnston, Dominic Kwiatkowski, Cordelia Langford, John Sillitoe on behalf of the Wellcome Sanger Institute COVID-19 Surveillance Team |
| EPI_ISL_3378822, EPI_ISL_3378823 | Lighthouse Lab in Glasgow | Wellcome Sanger Institute for the COVID-19 Genomics UK (COG-UK) Consortium | Harper VanSteenhouse, Yumi Kasai, David Gray, Carol Clugston, Anna Dominiczak and Alex Alderton, Roberto Amato, Jeffrey Barrett, Sonia Goncalves, Ewan Harrison, David K. Jackson, Ian Johnston, Dominic Kwiatkowski, Cordelia Langford, John Sillitoe on behalf of the Wellcome Sanger Institute COVID-19 Surveillance Team |
| EPI_ISL_3378825 | Lighthouse Lab in Alderley Park | Wellcome Sanger Institute for the COVID-19 Genomics UK (COG-UK) Consortium | Jacquelyn Wynn, Mairead Hyland, The Lighthouse Lab in Alderley Park and Alex Alderton, Roberto Amato, Jeffrey Barrett, Sonia Goncalves, Ewan Harrison, David K. Jackson, Ian Johnston, Dominic Kwiatkowski, Cordelia Langford, John Sillitoe on behalf of the Wellcome Sanger Institute COVID-19 Surveillance Team |
| EPI_ISL_3378826, EPI_ISL_3378827 | Lighthouse Lab in Glasgow | Wellcome Sanger Institute for the COVID-19 Genomics UK (COG-UK) Consortium | Harper VanSteenhouse, Yumi Kasai, David Gray, Carol Clugston, Anna Dominiczak and Alex Alderton, Roberto Amato, Jeffrey Barrett, Sonia Goncalves, Ewan Harrison, David K. Jackson, Ian Johnston, Dominic Kwiatkowski, Cordelia Langford, John Sillitoe on behalf of the Wellcome Sanger Institute COVID-19 Surveillance Team |
| EPI_ISL_3378828 | MIRIALIS CLUSES BECHET | CNR Virus des Infections Respiratoires - France SUD | Antonin Bal, Gregory Destras, Gwendolyne Burfin, Hadrien Regue, Quentin Semanas, Martine Valette, Bruno Lina, Laurence Josset |
| EPI_ISL_3378829, EPI_ISL_3378830, EPI_ISL_3378831 | Lighthouse Lab in Glasgow | Wellcome Sanger Institute for the COVID-19 Genomics UK (COG-UK) Consortium | Harper VanSteenhouse, Yumi Kasai, David Gray, Carol Clugston, Anna Dominiczak and Alex Alderton, Roberto Amato, Jeffrey Barrett, Sonia Goncalves, Ewan Harrison, David K. Jackson, Ian Johnston, Dominic Kwiatkowski, Cordelia Langford, John Sillitoe on behalf of the Wellcome Sanger Institute COVID-19 Surveillance Team |
| EPI_ISL_3378832 | MIRIALIS CLUSES BECHET | CNR Virus des Infections Respiratoires - France SUD | Antonin Bal, Gregory Destras, Gwendolyne Burfin, Hadrien Regue, Quentin Semanas, Martine Valette, Bruno Lina, Laurence Josset |
| EPI_ISL_3378833, EPI_ISL_3378834 | Lighthouse Lab in Alderley Park | Wellcome Sanger Institute for the COVID-19 Genomics UK (COG-UK) Consortium | Jacquelyn Wynn, Mairead Hyland, The Lighthouse Lab in Alderley Park and Alex Alderton, Roberto Amato, Jeffrey Barrett, Sonia Goncalves, Ewan Harrison, David K. Jackson, Ian Johnston, Dominic Kwiatkowski, Cordelia Langford, John Sillitoe on behalf of the Wellcome Sanger Institute COVID-19 Surveillance Team |
| EPI_ISL_3378835, EPI_ISL_3378837, EPI_ISL_3378838 | Lighthouse Lab in Glasgow | Wellcome Sanger Institute for the COVID-19 Genomics UK (COG-UK) Consortium | Harper VanSteenhouse, Yumi Kasai, David Gray, Carol Clugston, Anna Dominiczak and Alex Alderton, Roberto Amato, Jeffrey Barrett, Sonia Goncalves, Ewan Harrison, David K. Jackson, Ian Johnston, Dominic Kwiatkowski, Cordelia Langford, John Sillitoe on behalf of the Wellcome Sanger Institute COVID-19 Surveillance Team |
| EPI_ISL_3378839 | MIRIALIS CLUSES BECHET | CNR Virus des Infections Respiratoires - France SUD | Antonin Bal, Gregory Destras, Gwendolyne Burfin, Hadrien Regue, Quentin Semanas, Martine Valette, Bruno Lina, Laurence Josset |
| EPI_ISL_3378840 | Lighthouse Lab in Alderley Park | Wellcome Sanger Institute for the COVID-19 Genomics UK (COG-UK) Consortium | Jacquelyn Wynn, Mairead Hyland, The Lighthouse Lab in Alderley Park and Alex Alderton, Roberto Amato, Jeffrey Barrett, Sonia Goncalves, Ewan Harrison, David K. Jackson, Ian Johnston, Dominic Kwiatkowski, Cordelia Langford, John Sillitoe on behalf of the Wellcome Sanger Institute COVID-19 Surveillance Team |
| EPI_ISL_3378841, EPI_ISL_3378842 | Lighthouse Lab in Glasgow | Wellcome Sanger Institute for the COVID-19 Genomics UK (COG-UK) Consortium | Harper VanSteenhouse, Yumi Kasai, David Gray, Carol Clugston, Anna Dominiczak and Alex Alderton, Roberto Amato, Jeffrey Barrett, Sonia Goncalves, Ewan Harrison, David K. Jackson, Ian Johnston, Dominic Kwiatkowski, Cordelia Langford, John Sillitoe on behalf of the Wellcome Sanger Institute COVID-19 Surveillance Team |
| EPI_ISL_3378843 | MIRIALIS CLUSES BECHET | CNR Virus des Infections Respiratoires - France SUD | Antonin Bal, Gregory Destras, Gwendolyne Burfin, Hadrien Regue, Quentin Semanas, Martine Valette, Bruno Lina, Laurence Josset |
| EPI_ISL_3378844 | Lighthouse Lab in Glasgow | Wellcome Sanger Institute for the COVID-19 Genomics UK (COG-UK) Consortium | Harper VanSteenhouse, Yumi Kasai, David Gray, Carol Clugston, Anna Dominiczak and Alex Alderton, Roberto Amato, Jeffrey Barrett, Sonia Goncalves, Ewan Harrison, David K. Jackson, Ian Johnston, Dominic Kwiatkowski, Cordelia Langford, John Sillitoe on behalf of the Wellcome Sanger Institute COVID-19 Surveillance Team |
| EPI_ISL_3378845, EPI_ISL_3378846 | Lighthouse Lab in Alderley Park | Wellcome Sanger Institute for the COVID-19 Genomics UK (COG-UK) Consortium | Jacquelyn Wynn, Mairead Hyland, The Lighthouse Lab in Alderley Park and Alex Alderton, Roberto Amato, Jeffrey Barrett, Sonia Goncalves, Ewan Harrison, David K. Jackson, Ian Johnston, Dominic Kwiatkowski, Cordelia Langford, John Sillitoe on behalf of the Wellcome Sanger Institute COVID-19 Surveillance Team |
| EPI_ISL_3378847 | MIRIALIS CLUSES BECHET | CNR Virus des Infections Respiratoires - France SUD | Antonin Bal, Gregory Destras, Gwendolyne Burfin, Hadrien Regue, Quentin Semanas, Martine Valette, Bruno Lina, Laurence Josset |
| EPI_ISL_3378848 | Lighthouse Lab in Glasgow | Wellcome Sanger Institute for the COVID-19 Genomics UK (COG-UK) Consortium | Harper VanSteenhouse, Yumi Kasai, David Gray, Carol Clugston, Anna Dominiczak and Alex Alderton, Roberto Amato, Jeffrey Barrett, Sonia Goncalves, Ewan Harrison, David K. Jackson, Ian Johnston, Dominic Kwiatkowski, Cordelia Langford, John Sillitoe on behalf of the Wellcome Sanger Institute COVID-19 Surveillance Team |
| EPI_ISL_3378849 | Lighthouse Lab in Alderley Park | Wellcome Sanger Institute for the COVID-19 Genomics UK (COG-UK) Consortium | Jacquelyn Wynn, Mairead Hyland, The Lighthouse Lab in Alderley Park and Alex Alderton, Roberto Amato, Jeffrey Barrett, Sonia Goncalves, Ewan Harrison, David K. Jackson, Ian Johnston, Dominic Kwiatkowski, Cordelia Langford, John Sillitoe on behalf of the Wellcome Sanger Institute COVID-19 Surveillance Team |
| EPI_ISL_3378850 | CNR Virus des Infections Respiratoires - France SUD | CNR Virus des Infections Respiratoires - France SUD | Antonin Bal, Gregory Destras, Gwendolyne Burfin, Hadrien Regue, Quentin Semanas, Martine Valette, Bruno Lina, Laurence Josset |
| EPI_ISL_3378851 | Lighthouse Lab in Glasgow | Wellcome Sanger Institute for the COVID-19 Genomics UK (COG-UK) Consortium | Harper VanSteenhouse, Yumi Kasai, David Gray, Carol Clugston, Anna Dominiczak and Alex Alderton, Roberto Amato, Jeffrey Barrett, Sonia Goncalves, Ewan Harrison, David K. Jackson, Ian Johnston, Dominic Kwiatkowski, Cordelia Langford, John Sillitoe on behalf of the Wellcome Sanger Institute COVID-19 Surveillance Team |
| EPI_ISL_3378852 | Lighthouse Lab in Alderley Park | Wellcome Sanger Institute for the COVID-19 Genomics UK (COG-UK) Consortium | Jacquelyn Wynn, Mairead Hyland, The Lighthouse Lab in Alderley Park and Alex Alderton, Roberto Amato, Jeffrey Barrett, Sonia Goncalves, Ewan Harrison, David K. Jackson, Ian Johnston, Dominic Kwiatkowski, Cordelia Langford, John Sillitoe on behalf of the Wellcome Sanger Institute COVID-19 Surveillance Team |
| EPI_ISL_3378853 | Lighthouse Lab in Glasgow | Wellcome Sanger Institute for the COVID-19 Genomics UK (COG-UK) Consortium | Harper VanSteenhouse, Yumi Kasai, David Gray, Carol Clugston, Anna Dominiczak and Alex Alderton, Roberto Amato, Jeffrey Barrett, Sonia Goncalves, Ewan Harrison, David K. Jackson, Ian Johnston, Dominic Kwiatkowski, Cordelia Langford, John Sillitoe on behalf of the Wellcome Sanger Institute COVID-19 Surveillance Team |
| EPI_ISL_3378854 | MIRIALIS CLUSES BECHET | CNR Virus des Infections Respiratoires - France SUD | Antonin Bal, Gregory Destras, Gwendolyne Burfin, Hadrien Regue, Quentin Semanas, Martine Valette, Bruno Lina, Laurence Josset |
| EPI_ISL_3378855 | Lighthouse Lab in Alderley Park | Wellcome Sanger Institute for the COVID-19 Genomics UK (COG-UK) Consortium | Jacquelyn Wynn, Mairead Hyland, The Lighthouse Lab in Alderley Park and Alex Alderton, Roberto Amato, Jeffrey Barrett, Sonia Goncalves, Ewan Harrison, David K. Jackson, Ian Johnston, Dominic Kwiatkowski, Cordelia Langford, John Sillitoe on behalf of the Wellcome Sanger Institute COVID-19 Surveillance Team |
| EPI_ISL_3378856 | Lighthouse Lab in Glasgow | Wellcome Sanger Institute for the COVID-19 Genomics UK (COG-UK) Consortium | Harper VanSteenhouse, Yumi Kasai, David Gray, Carol Clugston, Anna Dominiczak and Alex Alderton, Roberto Amato, Jeffrey Barrett, Sonia Goncalves, Ewan Harrison, David K. Jackson, Ian Johnston, Dominic Kwiatkowski, Cordelia Langford, John Sillitoe on behalf of the Wellcome Sanger Institute COVID-19 Surveillance Team |
| EPI_ISL_3378857 | Lighthouse Lab in Alderley Park | Wellcome Sanger Institute for the COVID-19 Genomics UK (COG-UK) Consortium | Jacquelyn Wynn, Mairead Hyland, The Lighthouse Lab in Alderley Park and Alex Alderton, Roberto Amato, Jeffrey Barrett, Sonia Goncalves, Ewan Harrison, David K. Jackson, Ian Johnston, Dominic Kwiatkowski, Cordelia Langford, John Sillitoe on behalf of the Wellcome Sanger Institute COVID-19 Surveillance Team |
| EPI_ISL_3378858 | MIRIALIS CLUSES BECHET | CNR Virus des Infections Respiratoires - France SUD | Antonin Bal, Gregory Destras, Gwendolyne Burfin, Hadrien Regue, Quentin Semanas, Martine Valette, Bruno Lina, Laurence Josset |
| EPI_ISL_3378859 | Lighthouse Lab in Glasgow | Wellcome Sanger Institute for the COVID-19 Genomics UK (COG-UK) Consortium | Harper VanSteenhouse, Yumi Kasai, David Gray, Carol Clugston, Anna Dominiczak and Alex Alderton, Roberto Amato, Jeffrey Barrett, Sonia Goncalves, Ewan Harrison, David K. Jackson, Ian Johnston, Dominic Kwiatkowski, Cordelia Langford, John Sillitoe on behalf of the Wellcome Sanger Institute COVID-19 Surveillance Team |
| EPI_ISL_3378860, EPI_ISL_3378861 | Lighthouse Lab in Alderley Park | Wellcome Sanger Institute for the COVID-19 Genomics UK (COG-UK) Consortium | Jacquelyn Wynn, Mairead Hyland, The Lighthouse Lab in Alderley Park and Alex Alderton, Roberto Amato, Jeffrey Barrett, Sonia Goncalves, Ewan Harrison, David K. Jackson, Ian Johnston, Dominic Kwiatkowski, Cordelia Langford, John Sillitoe on behalf of the Wellcome Sanger Institute COVID-19 Surveillance Team |
| EPI_ISL_3378862 | MIRIALIS CLUSES BECHET | CNR Virus des Infections Respiratoires - France SUD | Antonin Bal, Gregory Destras, Gwendolyne Burfin, Hadrien Regue, Quentin Semanas, Martine Valette, Bruno Lina, Laurence Josset |
| EPI_ISL_3378863 | Lighthouse Lab in Alderley Park | Wellcome Sanger Institute for the COVID-19 Genomics UK (COG-UK) Consortium | Jacquelyn Wynn, Mairead Hyland, The Lighthouse Lab in Alderley Park and Alex Alderton, Roberto Amato, Jeffrey Barrett, Sonia Goncalves, Ewan Harrison, David K. Jackson, Ian Johnston, Dominic Kwiatkowski, Cordelia Langford, John Sillitoe on behalf of the Wellcome Sanger Institute COVID-19 Surveillance Team |
| EPI_ISL_3378864 | Lighthouse Lab in Glasgow | Wellcome Sanger Institute for the COVID-19 Genomics UK (COG-UK) Consortium | Harper VanSteenhouse, Yumi Kasai, David Gray, Carol Clugston, Anna Dominiczak and Alex Alderton, Roberto Amato, Jeffrey Barrett, Sonia Goncalves, Ewan Harrison, David K. Jackson, Ian Johnston, Dominic Kwiatkowski, Cordelia Langford, John Sillitoe on behalf of the Wellcome Sanger Institute COVID-19 Surveillance Team |
| EPI_ISL_3378865 | Lighthouse Lab in Alderley Park | Wellcome Sanger Institute for the COVID-19 Genomics UK (COG-UK) Consortium | Jacquelyn Wynn, Mairead Hyland, The Lighthouse Lab in Alderley Park and Alex Alderton, Roberto Amato, Jeffrey Barrett, Sonia Goncalves, Ewan Harrison, David K. Jackson, Ian Johnston, Dominic Kwiatkowski, Cordelia Langford, John Sillitoe on behalf of the Wellcome Sanger Institute COVID-19 Surveillance Team |
| EPI_ISL_3378866 | MIRIALIS CLUSES BECHET | CNR Virus des Infections Respiratoires - France SUD | Antonin Bal, Gregory Destras, Gwendolyne Burfin, Hadrien Regue, Quentin Semanas, Martine Valette, Bruno Lina, Laurence Josset |
| EPI_ISL_3378867, EPI_ISL_3378868, EPI_ISL_3378869 | Lighthouse Lab in Glasgow | Wellcome Sanger Institute for the COVID-19 Genomics UK (COG-UK) Consortium | Harper VanSteenhouse, Yumi Kasai, David Gray, Carol Clugston, Anna Dominiczak and Alex Alderton, Roberto Amato, Jeffrey Barrett, Sonia Goncalves, Ewan Harrison, David K. Jackson, Ian Johnston, Dominic Kwiatkowski, Cordelia Langford, John Sillitoe on behalf of the Wellcome Sanger Institute COVID-19 Surveillance Team |
| EPI_ISL_3378870 | ORIAPOLE | CNR Virus des Infections Respiratoires - France SUD | Antonin Bal, Gregory Destras, Gwendolyne Burfin, Hadrien Regue, Quentin Semanas, Martine Valette, Bruno Lina, Laurence Josset |
| EPI_ISL_3378871, EPI_ISL_3378872 | Lighthouse Lab in Glasgow | Wellcome Sanger Institute for the COVID-19 Genomics UK (COG-UK) Consortium | Harper VanSteenhouse, Yumi Kasai, David Gray, Carol Clugston, Anna Dominiczak and Alex Alderton, Roberto Amato, Jeffrey Barrett, Sonia Goncalves, Ewan Harrison, David K. Jackson, Ian Johnston, Dominic Kwiatkowski, Cordelia Langford, John Sillitoe on behalf of the Wellcome Sanger Institute COVID-19 Surveillance Team |
| EPI_ISL_3378873 | ORIAPOLE | CNR Virus des Infections Respiratoires - France SUD | Antonin Bal, Gregory Destras, Gwendolyne Burfin, Hadrien Regue, Quentin Semanas, Martine Valette, Bruno Lina, Laurence Josset |
| EPI_ISL_3378874 | Lighthouse Lab in Glasgow | Wellcome Sanger Institute for the COVID-19 Genomics UK (COG-UK) Consortium | Harper VanSteenhouse, Yumi Kasai, David Gray, Carol Clugston, Anna Dominiczak and Alex Alderton, Roberto Amato, Jeffrey Barrett, Sonia Goncalves, Ewan Harrison, David K. Jackson, Ian Johnston, Dominic Kwiatkowski, Cordelia Langford, John Sillitoe on behalf of the Wellcome Sanger Institute COVID-19 Surveillance Team |
| EPI_ISL_3378875 | Lighthouse Lab in Milton Keynes | Wellcome Sanger Institute for the COVID-19 Genomics UK (COG-UK) Consortium | The Lighthouse Lab in Milton Keynes and Alex Alderton, Roberto Amato, Jeffrey Barrett, Sonia Goncalves, Ewan Harrison, David K. Jackson, Ian Johnston, Dominic Kwiatkowski, Cordelia Langford, John Sillitoe on behalf of the Wellcome Sanger Institute COVID-19 Surveillance Team |
| EPI_ISL_3378876 | Lighthouse Lab in Glasgow | Wellcome Sanger Institute for the COVID-19 Genomics UK (COG-UK) Consortium | Harper VanSteenhouse, Yumi Kasai, David Gray, Carol Clugston, Anna Dominiczak and Alex Alderton, Roberto Amato, Jeffrey Barrett, Sonia Goncalves, Ewan Harrison, David K. Jackson, Ian Johnston, Dominic Kwiatkowski, Cordelia Langford, John Sillitoe on behalf of the Wellcome Sanger Institute COVID-19 Surveillance Team |
| EPI_ISL_3378877 | ORIAPOLE | CNR Virus des Infections Respiratoires - France SUD | Antonin Bal, Gregory Destras, Gwendolyne Burfin, Hadrien Regue, Quentin Semanas, Martine Valette, Bruno Lina, Laurence Josset |
| EPI_ISL_3378878 | Lighthouse Lab in Alderley Park | Wellcome Sanger Institute for the COVID-19 Genomics UK (COG-UK) Consortium | Jacquelyn Wynn, Mairead Hyland, The Lighthouse Lab in Alderley Park and Alex Alderton, Roberto Amato, Jeffrey Barrett, Sonia Goncalves, Ewan Harrison, David K. Jackson, Ian Johnston, Dominic Kwiatkowski, Cordelia Langford, John Sillitoe on behalf of the Wellcome Sanger Institute COVID-19 Surveillance Team |
| EPI_ISL_3378879, EPI_ISL_3378880 | Lighthouse Lab in Glasgow | Wellcome Sanger Institute for the COVID-19 Genomics UK (COG-UK) Consortium | Harper VanSteenhouse, Yumi Kasai, David Gray, Carol Clugston, Anna Dominiczak and Alex Alderton, Roberto Amato, Jeffrey Barrett, Sonia Goncalves, Ewan Harrison, David K. Jackson, Ian Johnston, Dominic Kwiatkowski, Cordelia Langford, John Sillitoe on behalf of the Wellcome Sanger Institute COVID-19 Surveillance Team |

| | | | |
|---|---|---|---|
| EPI_ISL_3378881 | ORIAPOLE | CNR Virus des Infections Respiratoires - France SUD | Antonin Bal, Gregory Destras, Gwendolyne Burfin, Hadrien Regue, Quentin Semanas, Martine Valette, Bruno Lina, Laurence Josset |
| EPI_ISL_3378882 | Lighthouse Lab in Alderley Park | Wellcome Sanger Institute for the COVID-19 Genomics UK (COG-UK) Consortium | Jacquelyn Wynn, Mairead Hyland, The Lighthouse Lab in Alderley Park and Alex Alderton, Roberto Amato, Jeffrey Barrett, Sonia Goncalves, Ewan Harrison, David K. Jackson, Ian Johnston, Dominic Kwiatkowski, Cordelia Langford, John Sillitoe on behalf of the Wellcome Sanger Institute COVID-19 Surveillance Team |
| EPI_ISL_3378883, EPI_ISL_3378884, EPI_ISL_3378885 | Lighthouse Lab in Glasgow | Wellcome Sanger Institute for the COVID-19 Genomics UK (COG-UK) Consortium | Harper VanSteenhouse, Yumi Kasai, David Gray, Carol Clugston, Anna Dominiczak and Alex Alderton, Roberto Amato, Jeffrey Barrett, Sonia Goncalves, Ewan Harrison, David K. Jackson, Ian Johnston, Dominic Kwiatkowski, Cordelia Langford, John Sillitoe on behalf of the Wellcome Sanger Institute COVID-19 Surveillance Team |
| EPI_ISL_3378886 | Lighthouse Lab in Alderley Park | Wellcome Sanger Institute for the COVID-19 Genomics UK (COG-UK) Consortium | Jacquelyn Wynn, Mairead Hyland, The Lighthouse Lab in Alderley Park and Alex Alderton, Roberto Amato, Jeffrey Barrett, Sonia Goncalves, Ewan Harrison, David K. Jackson, Ian Johnston, Dominic Kwiatkowski, Cordelia Langford, John Sillitoe on behalf of the Wellcome Sanger Institute COVID-19 Surveillance Team |
| EPI_ISL_3378887 | Lighthouse Lab in Glasgow | Wellcome Sanger Institute for the COVID-19 Genomics UK (COG-UK) Consortium | Harper VanSteenhouse, Yumi Kasai, David Gray, Carol Clugston, Anna Dominiczak and Alex Alderton, Roberto Amato, Jeffrey Barrett, Sonia Goncalves, Ewan Harrison, David K. Jackson, Ian Johnston, Dominic Kwiatkowski, Cordelia Langford, John Sillitoe on behalf of the Wellcome Sanger Institute COVID-19 Surveillance Team |
| EPI_ISL_3378888 | Lighthouse Lab in Alderley Park | Wellcome Sanger Institute for the COVID-19 Genomics UK (COG-UK) Consortium | Jacquelyn Wynn, Mairead Hyland, The Lighthouse Lab in Alderley Park and Alex Alderton, Roberto Amato, Jeffrey Barrett, Sonia Goncalves, Ewan Harrison, David K. Jackson, Ian Johnston, Dominic Kwiatkowski, Cordelia Langford, John Sillitoe on behalf of the Wellcome Sanger Institute COVID-19 Surveillance Team |
| EPI_ISL_3378889 | Lighthouse Lab in Glasgow | Wellcome Sanger Institute for the COVID-19 Genomics UK (COG-UK) Consortium | Harper VanSteenhouse, Yumi Kasai, David Gray, Carol Clugston, Anna Dominiczak and Alex Alderton, Roberto Amato, Jeffrey Barrett, Sonia Goncalves, Ewan Harrison, David K. Jackson, Ian Johnston, Dominic Kwiatkowski, Cordelia Langford, John Sillitoe on behalf of the Wellcome Sanger Institute COVID-19 Surveillance Team |
| EPI_ISL_3378890 | ORIAPOLE | CNR Virus des Infections Respiratoires - France SUD | Antonin Bal, Gregory Destras, Gwendolyne Burfin, Hadrien Regue, Quentin Semanas, Martine Valette, Bruno Lina, Laurence Josset |
| EPI_ISL_3378891 | Lighthouse Lab in Glasgow | Wellcome Sanger Institute for the COVID-19 Genomics UK (COG-UK) Consortium | Harper VanSteenhouse, Yumi Kasai, David Gray, Carol Clugston, Anna Dominiczak and Alex Alderton, Roberto Amato, Jeffrey Barrett, Sonia Goncalves, Ewan Harrison, David K. Jackson, Ian Johnston, Dominic Kwiatkowski, Cordelia Langford, John Sillitoe on behalf of the Wellcome Sanger Institute COVID-19 Surveillance Team |
| EPI_ISL_3378892, EPI_ISL_3378893, EPI_ISL_3378894, EPI_ISL_3378895, EPI_ISL_3378896 | Lighthouse Lab in Alderley Park | Wellcome Sanger Institute for the COVID-19 Genomics UK (COG-UK) Consortium | Jacquelyn Wynn, Mairead Hyland, The Lighthouse Lab in Alderley Park and Alex Alderton, Roberto Amato, Jeffrey Barrett, Sonia Goncalves, Ewan Harrison, David K. Jackson, Ian Johnston, Dominic Kwiatkowski, Cordelia Langford, John Sillitoe on behalf of the Wellcome Sanger Institute COVID-19 Surveillance Team |
| EPI_ISL_3378897 | Lighthouse Lab in Glasgow | Wellcome Sanger Institute for the COVID-19 Genomics UK (COG-UK) Consortium | Harper VanSteenhouse, Yumi Kasai, David Gray, Carol Clugston, Anna Dominiczak and Alex Alderton, Roberto Amato, Jeffrey Barrett, Sonia Goncalves, Ewan Harrison, David K. Jackson, Ian Johnston, Dominic Kwiatkowski, Cordelia Langford, John Sillitoe on behalf of the Wellcome Sanger Institute COVID-19 Surveillance Team |
| EPI_ISL_3378898 | CNR Virus des Infections Respiratoires - France SUD | CNR Virus des Infections Respiratoires - France SUD | Antonin Bal, Gregory Destras, Gwendolyne Burfin, Hadrien Regue, Quentin Semanas, Martine Valette, Bruno Lina, Laurence Josset |
| EPI_ISL_3378899 | Lighthouse Lab in Glasgow | Wellcome Sanger Institute for the COVID-19 Genomics UK (COG-UK) Consortium | Harper VanSteenhouse, Yumi Kasai, David Gray, Carol Clugston, Anna Dominiczak and Alex Alderton, Roberto Amato, Jeffrey Barrett, Sonia Goncalves, Ewan Harrison, David K. Jackson, Ian Johnston, Dominic Kwiatkowski, Cordelia Langford, John Sillitoe on behalf of the Wellcome Sanger Institute COVID-19 Surveillance Team |
| EPI_ISL_3378900 | Lighthouse Lab in Alderley Park | Wellcome Sanger Institute for the COVID-19 Genomics UK (COG-UK) Consortium | Jacquelyn Wynn, Mairead Hyland, The Lighthouse Lab in Alderley Park and Alex Alderton, Roberto Amato, Jeffrey Barrett, Sonia Goncalves, Ewan Harrison, David K. Jackson, Ian Johnston, Dominic Kwiatkowski, Cordelia Langford, John Sillitoe on behalf of the Wellcome Sanger Institute COVID-19 Surveillance Team |
| EPI_ISL_3378901 | Lighthouse Lab in Glasgow | Wellcome Sanger Institute for the COVID-19 Genomics UK (COG-UK) Consortium | Harper VanSteenhouse, Yumi Kasai, David Gray, Carol Clugston, Anna Dominiczak and Alex Alderton, Roberto Amato, Jeffrey Barrett, Sonia Goncalves, Ewan Harrison, David K. Jackson, Ian Johnston, Dominic Kwiatkowski, Cordelia Langford, John Sillitoe on behalf of the Wellcome Sanger Institute COVID-19 Surveillance Team |
| EPI_ISL_3378902 | Lighthouse Lab in Alderley Park | Wellcome Sanger Institute for the COVID-19 Genomics UK (COG-UK) Consortium | Jacquelyn Wynn, Mairead Hyland, The Lighthouse Lab in Alderley Park and Alex Alderton, Roberto Amato, Jeffrey Barrett, Sonia Goncalves, Ewan Harrison, David K. Jackson, Ian Johnston, Dominic Kwiatkowski, Cordelia Langford, John Sillitoe on behalf of the Wellcome Sanger Institute COVID-19 Surveillance Team |
| EPI_ISL_3378903 | Lighthouse Lab in Glasgow | Wellcome Sanger Institute for the COVID-19 Genomics UK (COG-UK) Consortium | Harper VanSteenhouse, Yumi Kasai, David Gray, Carol Clugston, Anna Dominiczak and Alex Alderton, Roberto Amato, Jeffrey Barrett, Sonia Goncalves, Ewan Harrison, David K. Jackson, Ian Johnston, Dominic Kwiatkowski, Cordelia Langford, John Sillitoe on behalf of the Wellcome Sanger Institute COVID-19 Surveillance Team |
| EPI_ISL_3378904 | Lighthouse Lab in Alderley Park | Wellcome Sanger Institute for the COVID-19 Genomics UK (COG-UK) Consortium | Jacquelyn Wynn, Mairead Hyland, The Lighthouse Lab in Alderley Park and Alex Alderton, Roberto Amato, Jeffrey Barrett, Sonia Goncalves, Ewan Harrison, David K. Jackson, Ian Johnston, Dominic Kwiatkowski, Cordelia Langford, John Sillitoe on behalf of the Wellcome Sanger Institute COVID-19 Surveillance Team |
| EPI_ISL_3378905 | ORIAPOLE | CNR Virus des Infections Respiratoires - France SUD | Antonin Bal, Gregory Destras, Gwendolyne Burfin, Hadrien Regue, Quentin Semanas, Martine Valette, Bruno Lina, Laurence Josset |
| EPI_ISL_3378906, EPI_ISL_3378907, EPI_ISL_3378908, EPI_ISL_3378909, EPI_ISL_3378910 | Lighthouse Lab in Alderley Park | Wellcome Sanger Institute for the COVID-19 Genomics UK (COG-UK) Consortium | Jacquelyn Wynn, Mairead Hyland, The Lighthouse Lab in Alderley Park and Alex Alderton, Roberto Amato, Jeffrey Barrett, Sonia Goncalves, Ewan Harrison, David K. Jackson, Ian Johnston, Dominic Kwiatkowski, Cordelia Langford, John Sillitoe on behalf of the Wellcome Sanger Institute COVID-19 Surveillance Team |
| EPI_ISL_3378911 | ORIAPOLE | CNR Virus des Infections Respiratoires - France SUD | Antonin Bal, Gregory Destras, Gwendolyne Burfin, Hadrien Regue, Quentin Semanas, Martine Valette, Bruno Lina, Laurence Josset |
| EPI_ISL_3378912 | Lighthouse Lab in Glasgow | Wellcome Sanger Institute for the COVID-19 Genomics UK (COG-UK) Consortium | Harper VanSteenhouse, Yumi Kasai, David Gray, Carol Clugston, Anna Dominiczak and Alex Alderton, Roberto Amato, Jeffrey Barrett, Sonia Goncalves, Ewan Harrison, David K. Jackson, Ian Johnston, Dominic Kwiatkowski, Cordelia Langford, John Sillitoe on behalf of the Wellcome Sanger Institute COVID-19 Surveillance Team |
| EPI_ISL_3378913 | Lighthouse Lab in Alderley Park | Wellcome Sanger Institute for the COVID-19 Genomics UK (COG-UK) Consortium | Jacquelyn Wynn, Mairead Hyland, The Lighthouse Lab in Alderley Park and Alex Alderton, Roberto Amato, Jeffrey Barrett, Sonia Goncalves, Ewan Harrison, David K. Jackson, Ian Johnston, Dominic Kwiatkowski, Cordelia Langford, John Sillitoe on behalf of the Wellcome Sanger Institute COVID-19 Surveillance Team |
| EPI_ISL_3378914 | Lighthouse Lab in Glasgow | Wellcome Sanger Institute for the COVID-19 Genomics UK (COG-UK) Consortium | Harper VanSteenhouse, Yumi Kasai, David Gray, Carol Clugston, Anna Dominiczak and Alex Alderton, Roberto Amato, Jeffrey Barrett, Sonia Goncalves, Ewan Harrison, David K. Jackson, Ian Johnston, Dominic Kwiatkowski, Cordelia Langford, John Sillitoe on behalf of the Wellcome Sanger Institute COVID-19 Surveillance Team |
| EPI_ISL_3378915 | ORIAPOLE | CNR Virus des Infections Respiratoires - France SUD | Antonin Bal, Gregory Destras, Gwendolyne Burfin, Hadrien Regue, Quentin Semanas, Martine Valette, Bruno Lina, Laurence Josset |
| EPI_ISL_3378916, EPI_ISL_3378917 | Lighthouse Lab in Alderley Park | Wellcome Sanger Institute for the COVID-19 Genomics UK (COG-UK) Consortium | Jacquelyn Wynn, Mairead Hyland, The Lighthouse Lab in Alderley Park and Alex Alderton, Roberto Amato, Jeffrey Barrett, Sonia Goncalves, Ewan Harrison, David K. Jackson, Ian Johnston, Dominic Kwiatkowski, Cordelia Langford, John Sillitoe on behalf of the Wellcome Sanger Institute COVID-19 Surveillance Team |
| EPI_ISL_3378918 | Lighthouse Lab in Milton Keynes | Wellcome Sanger Institute for the COVID-19 Genomics UK (COG-UK) Consortium | The Lighthouse Lab in Milton Keynes and Alex Alderton, Roberto Amato, Jeffrey Barrett, Sonia Goncalves, Ewan Harrison, David K. Jackson, Ian Johnston, Dominic Kwiatkowski, Cordelia Langford, John Sillitoe on behalf of the Wellcome Sanger Institute COVID-19 Surveillance Team |
| EPI_ISL_3378919 | ORIAPOLE | CNR Virus des Infections Respiratoires - France SUD | Antonin Bal, Gregory Destras, Gwendolyne Burfin, Hadrien Regue, Quentin Semanas, Martine Valette, Bruno Lina, Laurence Josset |
| EPI_ISL_3378920, EPI_ISL_3378921 | Lighthouse Lab in Glasgow | Wellcome Sanger Institute for the COVID-19 Genomics UK (COG-UK) Consortium | Harper VanSteenhouse, Yumi Kasai, David Gray, Carol Clugston, Anna Dominiczak and Alex Alderton, Roberto Amato, Jeffrey Barrett, Sonia Goncalves, Ewan Harrison, David K. Jackson, Ian Johnston, Dominic Kwiatkowski, Cordelia Langford, John Sillitoe on behalf of the Wellcome Sanger Institute COVID-19 Surveillance Team |
| EPI_ISL_3378922 | ORIAPOLE | CNR Virus des Infections Respiratoires - France SUD | Antonin Bal, Gregory Destras, Gwendolyne Burfin, Hadrien Regue, Quentin Semanas, Martine Valette, Bruno Lina, Laurence Josset |
| EPI_ISL_3378923 | Berkshire and Surrey Pathology Services Lighthouse Laboratory | Wellcome Sanger Institute for the COVID-19 Genomics UK (COG-UK) Consortium | Berkshire and Surrey Pathology Services Lighthouse Laboratory and Alex Alderton, Roberto Amato, Jeffrey Barrett, Sonia Goncalves, Ewan Harrison, David K. Jackson, Ian Johnston, Dominic Kwiatkowski, Cordelia Langford, John Sillitoe on behalf of the Wellcome Sanger Institute COVID-19 Surveillance Team |
| EPI_ISL_3378924 | Lighthouse Lab in Alderley Park | Wellcome Sanger Institute for the COVID-19 Genomics UK (COG-UK) Consortium | Jacquelyn Wynn, Mairead Hyland, The Lighthouse Lab in Alderley Park and Alex Alderton, Roberto Amato, Jeffrey Barrett, Sonia Goncalves, Ewan Harrison, David K. Jackson, Ian Johnston, Dominic Kwiatkowski, Cordelia Langford, John Sillitoe on behalf of the Wellcome Sanger Institute COVID-19 Surveillance Team |
| EPI_ISL_3378925 | Lighthouse Lab in Glasgow | Wellcome Sanger Institute for the COVID-19 Genomics UK (COG-UK) Consortium | Harper VanSteenhouse, Yumi Kasai, David Gray, Carol Clugston, Anna Dominiczak and Alex Alderton, Roberto Amato, Jeffrey Barrett, Sonia Goncalves, Ewan Harrison, David K. Jackson, Ian Johnston, Dominic Kwiatkowski, Cordelia Langford, John Sillitoe on behalf of the Wellcome Sanger Institute COVID-19 Surveillance Team |
| EPI_ISL_3378926 | ORIAPOLE | CNR Virus des Infections Respiratoires - France SUD | Antonin Bal, Gregory Destras, Gwendolyne Burfin, Hadrien Regue, Quentin Semanas, Martine Valette, Bruno Lina, Laurence Josset |
| EPI_ISL_3378927 | Lighthouse Lab in Alderley Park | Wellcome Sanger Institute for the COVID-19 Genomics UK (COG-UK) Consortium | Jacquelyn Wynn, Mairead Hyland, The Lighthouse Lab in Alderley Park and Alex Alderton, Roberto Amato, Jeffrey Barrett, Sonia Goncalves, Ewan Harrison, David K. Jackson, Ian Johnston, Dominic Kwiatkowski, Cordelia Langford, John Sillitoe on behalf of the Wellcome Sanger Institute COVID-19 Surveillance Team |
| EPI_ISL_3378928 | Lighthouse Lab in Milton Keynes | Wellcome Sanger Institute for the COVID-19 Genomics UK (COG-UK) Consortium | The Lighthouse Lab in Milton Keynes and Alex Alderton, Roberto Amato, Jeffrey Barrett, Sonia Goncalves, Ewan Harrison, David K. Jackson, Ian Johnston, Dominic Kwiatkowski, Cordelia Langford, John Sillitoe on behalf of the Wellcome Sanger Institute COVID-19 Surveillance Team |
| EPI_ISL_3378929 | Lighthouse Lab in Glasgow | Wellcome Sanger Institute for the COVID-19 Genomics UK (COG-UK) Consortium | Harper VanSteenhouse, Yumi Kasai, David Gray, Carol Clugston, Anna Dominiczak and Alex Alderton, Roberto Amato, Jeffrey Barrett, Sonia Goncalves, Ewan Harrison, David K. Jackson, Ian Johnston, Dominic Kwiatkowski, Cordelia Langford, John Sillitoe on behalf of the Wellcome Sanger Institute COVID-19 Surveillance Team |
| EPI_ISL_3378930 | ORIAPOLE | CNR Virus des Infections Respiratoires - France SUD | Antonin Bal, Gregory Destras, Gwendolyne Burfin, Hadrien Regue, Quentin Semanas, Martine Valette, Bruno Lina, Laurence Josset |
| EPI_ISL_3378931 | Lighthouse Lab in Alderley Park | Wellcome Sanger Institute for the COVID-19 Genomics UK (COG-UK) Consortium | Jacquelyn Wynn, Mairead Hyland, The Lighthouse Lab in Alderley Park and Alex Alderton, Roberto Amato, Jeffrey Barrett, Sonia Goncalves, Ewan Harrison, David K. Jackson, Ian Johnston, Dominic Kwiatkowski, Cordelia Langford, John Sillitoe on behalf of the Wellcome Sanger Institute COVID-19 Surveillance Team |
| EPI_ISL_3378932 | Berkshire and Surrey Pathology Services Lighthouse Laboratory | Wellcome Sanger Institute for the COVID-19 Genomics UK (COG-UK) Consortium | Berkshire and Surrey Pathology Services Lighthouse Laboratory and Alex Alderton, Roberto Amato, Jeffrey Barrett, Sonia Goncalves, Ewan Harrison, David K. Jackson, Ian Johnston, Dominic Kwiatkowski, Cordelia Langford, John Sillitoe on behalf of the Wellcome Sanger Institute COVID-19 Surveillance Team |
| EPI_ISL_3378933 | Lighthouse Lab in Milton Keynes | Wellcome Sanger Institute for the COVID-19 Genomics UK (COG-UK) Consortium | The Lighthouse Lab in Milton Keynes and Alex Alderton, Roberto Amato, Jeffrey Barrett, Sonia Goncalves, Ewan Harrison, David K. Jackson, Ian Johnston, Dominic Kwiatkowski, Cordelia Langford, John Sillitoe on behalf of the Wellcome Sanger Institute COVID-19 Surveillance Team |
| EPI_ISL_3378934 | ORIAPOLE | CNR Virus des Infections Respiratoires - France SUD | Antonin Bal, Gregory Destras, Gwendolyne Burfin, Hadrien Regue, Quentin Semanas, Martine Valette, Bruno Lina, Laurence Josset |
| EPI_ISL_3378935 | Lighthouse Lab in Milton Keynes | Wellcome Sanger Institute for the COVID-19 Genomics UK (COG-UK) Consortium | The Lighthouse Lab in Milton Keynes and Alex Alderton, Roberto Amato, Jeffrey Barrett, Sonia Goncalves, Ewan Harrison, David K. Jackson, Ian Johnston, Dominic Kwiatkowski, Cordelia Langford, John Sillitoe on behalf of the Wellcome Sanger Institute COVID-19 Surveillance Team |
| EPI_ISL_3378936 | Lighthouse Lab in Alderley Park | Wellcome Sanger Institute for the COVID-19 Genomics UK (COG-UK) Consortium | Jacquelyn Wynn, Mairead Hyland, The Lighthouse Lab in Alderley Park and Alex Alderton, Roberto Amato, Jeffrey Barrett, Sonia Goncalves, Ewan Harrison, David K. Jackson, Ian Johnston, Dominic Kwiatkowski, Cordelia Langford, John Sillitoe on behalf of the Wellcome Sanger Institute COVID-19 Surveillance Team |
| EPI_ISL_3378937 | ORIAPOLE | CNR Virus des Infections Respiratoires - France SUD | Antonin Bal, Gregory Destras, Gwendolyne Burfin, Hadrien Regue, Quentin Semanas, Martine Valette, Bruno Lina, Laurence Josset |
| EPI_ISL_3378938 | Lighthouse Lab in Alderley Park | Wellcome Sanger Institute for the COVID-19 Genomics UK (COG-UK) Consortium | Jacquelyn Wynn, Mairead Hyland, The Lighthouse Lab in Alderley Park and Alex Alderton, Roberto Amato, Jeffrey Barrett, Sonia Goncalves, Ewan Harrison, David K. Jackson, Ian Johnston, Dominic Kwiatkowski, Cordelia Langford, John Sillitoe on behalf of the Wellcome Sanger Institute COVID-19 Surveillance Team |
| EPI_ISL_3378939 | Lighthouse Lab in Milton Keynes | Wellcome Sanger Institute for the COVID-19 Genomics UK (COG-UK) Consortium | The Lighthouse Lab in Milton Keynes and Alex Alderton, Roberto Amato, Jeffrey Barrett, Sonia Goncalves, Ewan Harrison, David K. Jackson, Ian Johnston, Dominic Kwiatkowski, Cordelia Langford, John Sillitoe on behalf of the Wellcome Sanger Institute COVID-19 Surveillance Team |

| | | | |
|---|---|---|---|
| EPI_ISL_3378940 | Berkshire and Surrey Pathology Services Lighthouse Laboratory | Wellcome Sanger Institute for the COVID-19 Genomics UK (COG-UK) Consortium | Berkshire and Surrey Pathology Services Lighthouse Laboratory and Alex Alderton, Roberto Amato, Jeffrey Barrett, Sonia Goncalves, Ewan Harrison, David K. Jackson, Ian Johnston, Dominic Kwiatkowski, Cordelia Langford, John Sillitoe on behalf of the Wellcome Sanger Institute COVID-19 Surveillance Team |
| EPI_ISL_3378941 | ORIAPOLE | CNR Virus des Infections Respiratoires - France SUD | Antonin Bal, Gregory Destras, Gwendolyne Burfin, Hadrien Regue, Quentin Semanas, Martine Valette, Bruno Lina, Laurence Josset |
| EPI_ISL_3378942 | Lighthouse Lab in Glasgow | Wellcome Sanger Institute for the COVID-19 Genomics UK (COG-UK) Consortium | Harper VanSteenhouse, Yumi Kasai, David Gray, Carol Clugston, Anna Dominiczak and Alex Alderton, Roberto Amato, Jeffrey Barrett, Sonia Goncalves, Ewan Harrison, David K. Jackson, Ian Johnston, Dominic Kwiatkowski, Cordelia Langford, John Sillitoe on behalf of the Wellcome Sanger Institute COVID-19 Surveillance Team |
| EPI_ISL_3378943 | Lighthouse Lab in Alderley Park | Wellcome Sanger Institute for the COVID-19 Genomics UK (COG-UK) Consortium | Jacquelyn Wynn, Mairead Hyland, The Lighthouse Lab in Alderley Park and Alex Alderton, Roberto Amato, Jeffrey Barrett, Sonia Goncalves, Ewan Harrison, David K. Jackson, Ian Johnston, Dominic Kwiatkowski, Cordelia Langford, John Sillitoe on behalf of the Wellcome Sanger Institute COVID-19 Surveillance Team |
| EPI_ISL_3378944 | Lighthouse Lab in Milton Keynes | Wellcome Sanger Institute for the COVID-19 Genomics UK (COG-UK) Consortium | The Lighthouse Lab in Milton Keynes and Alex Alderton, Roberto Amato, Jeffrey Barrett, Sonia Goncalves, Ewan Harrison, David K. Jackson, Ian Johnston, Dominic Kwiatkowski, Cordelia Langford, John Sillitoe on behalf of the Wellcome Sanger Institute COVID-19 Surveillance Team |
| EPI_ISL_3378945 | ORIAPOLE | CNR Virus des Infections Respiratoires - France SUD | Antonin Bal, Gregory Destras, Gwendolyne Burfin, Hadrien Regue, Quentin Semanas, Martine Valette, Bruno Lina, Laurence Josset |
| EPI_ISL_3378946, EPI_ISL_3378947 | Lighthouse Lab in Milton Keynes | Wellcome Sanger Institute for the COVID-19 Genomics UK (COG-UK) Consortium | The Lighthouse Lab in Milton Keynes and Alex Alderton, Roberto Amato, Jeffrey Barrett, Sonia Goncalves, Ewan Harrison, David K. Jackson, Ian Johnston, Dominic Kwiatkowski, Cordelia Langford, John Sillitoe on behalf of the Wellcome Sanger Institute COVID-19 Surveillance Team |
| EPI_ISL_3378948 | ORIAPOLE | CNR Virus des Infections Respiratoires - France SUD | Antonin Bal, Gregory Destras, Gwendolyne Burfin, Hadrien Regue, Quentin Semanas, Martine Valette, Bruno Lina, Laurence Josset |
| EPI_ISL_3378949 | Lighthouse Lab in Glasgow | Wellcome Sanger Institute for the COVID-19 Genomics UK (COG-UK) Consortium | Harper VanSteenhouse, Yumi Kasai, David Gray, Carol Clugston, Anna Dominiczak and Alex Alderton, Roberto Amato, Jeffrey Barrett, Sonia Goncalves, Ewan Harrison, David K. Jackson, Ian Johnston, Dominic Kwiatkowski, Cordelia Langford, John Sillitoe on behalf of the Wellcome Sanger Institute COVID-19 Surveillance Team |
| EPI_ISL_3378950, EPI_ISL_3378951 | Lighthouse Lab in Alderley Park | Wellcome Sanger Institute for the COVID-19 Genomics UK (COG-UK) Consortium | Jacquelyn Wynn, Mairead Hyland, The Lighthouse Lab in Alderley Park and Alex Alderton, Roberto Amato, Jeffrey Barrett, Sonia Goncalves, Ewan Harrison, David K. Jackson, Ian Johnston, Dominic Kwiatkowski, Cordelia Langford, John Sillitoe on behalf of the Wellcome Sanger Institute COVID-19 Surveillance Team |
| EPI_ISL_3378952 | ORIAPOLE | CNR Virus des Infections Respiratoires - France SUD | Antonin Bal, Gregory Destras, Gwendolyne Burfin, Hadrien Regue, Quentin Semanas, Martine Valette, Bruno Lina, Laurence Josset |
| EPI_ISL_3378953 | Lighthouse Lab in Alderley Park | Wellcome Sanger Institute for the COVID-19 Genomics UK (COG-UK) Consortium | Jacquelyn Wynn, Mairead Hyland, The Lighthouse Lab in Alderley Park and Alex Alderton, Roberto Amato, Jeffrey Barrett, Sonia Goncalves, Ewan Harrison, David K. Jackson, Ian Johnston, Dominic Kwiatkowski, Cordelia Langford, John Sillitoe on behalf of the Wellcome Sanger Institute COVID-19 Surveillance Team |
| EPI_ISL_3378954 | Berkshire and Surrey Pathology Services Lighthouse Laboratory | Wellcome Sanger Institute for the COVID-19 Genomics UK (COG-UK) Consortium | Berkshire and Surrey Pathology Services Lighthouse Laboratory and Alex Alderton, Roberto Amato, Jeffrey Barrett, Sonia Goncalves, Ewan Harrison, David K. Jackson, Ian Johnston, Dominic Kwiatkowski, Cordelia Langford, John Sillitoe on behalf of the Wellcome Sanger Institute COVID-19 Surveillance Team |
| EPI_ISL_3378955 | ORIAPOLE | CNR Virus des Infections Respiratoires - France SUD | Antonin Bal, Gregory Destras, Gwendolyne Burfin, Hadrien Regue, Quentin Semanas, Martine Valette, Bruno Lina, Laurence Josset |
| EPI_ISL_3378956, EPI_ISL_3378957 | Lighthouse Lab in Milton Keynes | Wellcome Sanger Institute for the COVID-19 Genomics UK (COG-UK) Consortium | The Lighthouse Lab in Milton Keynes and Alex Alderton, Roberto Amato, Jeffrey Barrett, Sonia Goncalves, Ewan Harrison, David K. Jackson, Ian Johnston, Dominic Kwiatkowski, Cordelia Langford, John Sillitoe on behalf of the Wellcome Sanger Institute COVID-19 Surveillance Team |
| EPI_ISL_3378958 | Berkshire and Surrey Pathology Services Lighthouse Laboratory | Wellcome Sanger Institute for the COVID-19 Genomics UK (COG-UK) Consortium | Berkshire and Surrey Pathology Services Lighthouse Laboratory and Alex Alderton, Roberto Amato, Jeffrey Barrett, Sonia Goncalves, Ewan Harrison, David K. Jackson, Ian Johnston, Dominic Kwiatkowski, Cordelia Langford, John Sillitoe on behalf of the Wellcome Sanger Institute COVID-19 Surveillance Team |
| EPI_ISL_3378959 | Lighthouse Lab in Glasgow | Wellcome Sanger Institute for the COVID-19 Genomics UK (COG-UK) Consortium | Harper VanSteenhouse, Yumi Kasai, David Gray, Carol Clugston, Anna Dominiczak and Alex Alderton, Roberto Amato, Jeffrey Barrett, Sonia Goncalves, Ewan Harrison, David K. Jackson, Ian Johnston, Dominic Kwiatkowski, Cordelia Langford, John Sillitoe on behalf of the Wellcome Sanger Institute COVID-19 Surveillance Team |
| EPI_ISL_3378960 | ORIAPOLE | CNR Virus des Infections Respiratoires - France SUD | Antonin Bal, Gregory Destras, Gwendolyne Burfin, Hadrien Regue, Quentin Semanas, Martine Valette, Bruno Lina, Laurence Josset |
| EPI_ISL_3378961 | Lighthouse Lab in Alderley Park | Wellcome Sanger Institute for the COVID-19 Genomics UK (COG-UK) Consortium | Jacquelyn Wynn, Mairead Hyland, The Lighthouse Lab in Alderley Park and Alex Alderton, Roberto Amato, Jeffrey Barrett, Sonia Goncalves, Ewan Harrison, David K. Jackson, Ian Johnston, Dominic Kwiatkowski, Cordelia Langford, John Sillitoe on behalf of the Wellcome Sanger Institute COVID-19 Surveillance Team |
| EPI_ISL_3378962 | Lighthouse Lab in Glasgow | Wellcome Sanger Institute for the COVID-19 Genomics UK (COG-UK) Consortium | Harper VanSteenhouse, Yumi Kasai, David Gray, Carol Clugston, Anna Dominiczak and Alex Alderton, Roberto Amato, Jeffrey Barrett, Sonia Goncalves, Ewan Harrison, David K. Jackson, Ian Johnston, Dominic Kwiatkowski, Cordelia Langford, John Sillitoe on behalf of the Wellcome Sanger Institute COVID-19 Surveillance Team |
| EPI_ISL_3378963 | Lighthouse Lab in Milton Keynes | Wellcome Sanger Institute for the COVID-19 Genomics UK (COG-UK) Consortium | The Lighthouse Lab in Milton Keynes and Alex Alderton, Roberto Amato, Jeffrey Barrett, Sonia Goncalves, Ewan Harrison, David K. Jackson, Ian Johnston, Dominic Kwiatkowski, Cordelia Langford, John Sillitoe on behalf of the Wellcome Sanger Institute COVID-19 Surveillance Team |
| EPI_ISL_3378964 | ORIAPOLE | CNR Virus des Infections Respiratoires - France SUD | Antonin Bal, Gregory Destras, Gwendolyne Burfin, Hadrien Regue, Quentin Semanas, Martine Valette, Bruno Lina, Laurence Josset |
| EPI_ISL_3378965, EPI_ISL_3378966 | Lighthouse Lab in Milton Keynes | Wellcome Sanger Institute for the COVID-19 Genomics UK (COG-UK) Consortium | The Lighthouse Lab in Milton Keynes and Alex Alderton, Roberto Amato, Jeffrey Barrett, Sonia Goncalves, Ewan Harrison, David K. Jackson, Ian Johnston, Dominic Kwiatkowski, Cordelia Langford, John Sillitoe on behalf of the Wellcome Sanger Institute COVID-19 Surveillance Team |
| EPI_ISL_3378967 | Lighthouse Lab in Glasgow | Wellcome Sanger Institute for the COVID-19 Genomics UK (COG-UK) Consortium | Harper VanSteenhouse, Yumi Kasai, David Gray, Carol Clugston, Anna Dominiczak and Alex Alderton, Roberto Amato, Jeffrey Barrett, Sonia Goncalves, Ewan Harrison, David K. Jackson, Ian Johnston, Dominic Kwiatkowski, Cordelia Langford, John Sillitoe on behalf of the Wellcome Sanger Institute COVID-19 Surveillance Team |
| EPI_ISL_3378968 | ORIAPOLE | CNR Virus des Infections Respiratoires - France SUD | Antonin Bal, Gregory Destras, Gwendolyne Burfin, Hadrien Regue, Quentin Semanas, Martine Valette, Bruno Lina, Laurence Josset |
| EPI_ISL_3378969 | Lighthouse Lab in Milton Keynes | Wellcome Sanger Institute for the COVID-19 Genomics UK (COG-UK) Consortium | The Lighthouse Lab in Milton Keynes and Alex Alderton, Roberto Amato, Jeffrey Barrett, Sonia Goncalves, Ewan Harrison, David K. Jackson, Ian Johnston, Dominic Kwiatkowski, Cordelia Langford, John Sillitoe on behalf of the Wellcome Sanger Institute COVID-19 Surveillance Team |
| EPI_ISL_3378970 | Lighthouse Lab in Alderley Park | Wellcome Sanger Institute for the COVID-19 Genomics UK (COG-UK) Consortium | Jacquelyn Wynn, Mairead Hyland, The Lighthouse Lab in Alderley Park and Alex Alderton, Roberto Amato, Jeffrey Barrett, Sonia Goncalves, Ewan Harrison, David K. Jackson, Ian Johnston, Dominic Kwiatkowski, Cordelia Langford, John Sillitoe on behalf of the Wellcome Sanger Institute COVID-19 Surveillance Team |
| EPI_ISL_3378971 | Lighthouse Lab in Milton Keynes | Wellcome Sanger Institute for the COVID-19 Genomics UK (COG-UK) Consortium | The Lighthouse Lab in Milton Keynes and Alex Alderton, Roberto Amato, Jeffrey Barrett, Sonia Goncalves, Ewan Harrison, David K. Jackson, Ian Johnston, Dominic Kwiatkowski, Cordelia Langford, John Sillitoe on behalf of the Wellcome Sanger Institute COVID-19 Surveillance Team |
| EPI_ISL_3378972 | LABORATOIRE ACCOLAB SUD OUEST | CNR Virus des Infections Respiratoires - France SUD | Antonin Bal, Gregory Destras, Gwendolyne Burfin, Hadrien Regue, Quentin Semanas, Martine Valette, Bruno Lina, Laurence Josset |
| EPI_ISL_3378973 | Lighthouse Lab in Glasgow | Wellcome Sanger Institute for the COVID-19 Genomics UK (COG-UK) Consortium | Harper VanSteenhouse, Yumi Kasai, David Gray, Carol Clugston, Anna Dominiczak and Alex Alderton, Roberto Amato, Jeffrey Barrett, Sonia Goncalves, Ewan Harrison, David K. Jackson, Ian Johnston, Dominic Kwiatkowski, Cordelia Langford, John Sillitoe on behalf of the Wellcome Sanger Institute COVID-19 Surveillance Team |
| EPI_ISL_3378974 | Lighthouse Lab in Milton Keynes | Wellcome Sanger Institute for the COVID-19 Genomics UK (COG-UK) Consortium | The Lighthouse Lab in Milton Keynes and Alex Alderton, Roberto Amato, Jeffrey Barrett, Sonia Goncalves, Ewan Harrison, David K. Jackson, Ian Johnston, Dominic Kwiatkowski, Cordelia Langford, John Sillitoe on behalf of the Wellcome Sanger Institute COVID-19 Surveillance Team |
| EPI_ISL_3378975 | ORIAPOLE | CNR Virus des Infections Respiratoires - France SUD | Antonin Bal, Gregory Destras, Gwendolyne Burfin, Hadrien Regue, Quentin Semanas, Martine Valette, Bruno Lina, Laurence Josset |
| EPI_ISL_3378976 | Lighthouse Lab in Glasgow | Wellcome Sanger Institute for the COVID-19 Genomics UK (COG-UK) Consortium | Harper VanSteenhouse, Yumi Kasai, David Gray, Carol Clugston, Anna Dominiczak and Alex Alderton, Roberto Amato, Jeffrey Barrett, Sonia Goncalves, Ewan Harrison, David K. Jackson, Ian Johnston, Dominic Kwiatkowski, Cordelia Langford, John Sillitoe on behalf of the Wellcome Sanger Institute COVID-19 Surveillance Team |
| EPI_ISL_3378977 | Berkshire and Surrey Pathology Services Lighthouse Laboratory | Wellcome Sanger Institute for the COVID-19 Genomics UK (COG-UK) Consortium | Berkshire and Surrey Pathology Services Lighthouse Laboratory and Alex Alderton, Roberto Amato, Jeffrey Barrett, Sonia Goncalves, Ewan Harrison, David K. Jackson, Ian Johnston, Dominic Kwiatkowski, Cordelia Langford, John Sillitoe on behalf of the Wellcome Sanger Institute COVID-19 Surveillance Team |
| EPI_ISL_3378978 | Lighthouse Lab in Milton Keynes | Wellcome Sanger Institute for the COVID-19 Genomics UK (COG-UK) Consortium | The Lighthouse Lab in Milton Keynes and Alex Alderton, Roberto Amato, Jeffrey Barrett, Sonia Goncalves, Ewan Harrison, David K. Jackson, Ian Johnston, Dominic Kwiatkowski, Cordelia Langford, John Sillitoe on behalf of the Wellcome Sanger Institute COVID-19 Surveillance Team |
| EPI_ISL_3378979 | Lighthouse Lab in Alderley Park | Wellcome Sanger Institute for the COVID-19 Genomics UK (COG-UK) Consortium | Jacquelyn Wynn, Mairead Hyland, The Lighthouse Lab in Alderley Park and Alex Alderton, Roberto Amato, Jeffrey Barrett, Sonia Goncalves, Ewan Harrison, David K. Jackson, Ian Johnston, Dominic Kwiatkowski, Cordelia Langford, John Sillitoe on behalf of the Wellcome Sanger Institute COVID-19 Surveillance Team |
| EPI_ISL_3378980 | ORIAPOLE | CNR Virus des Infections Respiratoires - France SUD | Antonin Bal, Gregory Destras, Gwendolyne Burfin, Hadrien Regue, Quentin Semanas, Martine Valette, Bruno Lina, Laurence Josset |
| EPI_ISL_3378981 | Lighthouse Lab in Milton Keynes | Wellcome Sanger Institute for the COVID-19 Genomics UK (COG-UK) Consortium | The Lighthouse Lab in Milton Keynes and Alex Alderton, Roberto Amato, Jeffrey Barrett, Sonia Goncalves, Ewan Harrison, David K. Jackson, Ian Johnston, Dominic Kwiatkowski, Cordelia Langford, John Sillitoe on behalf of the Wellcome Sanger Institute COVID-19 Surveillance Team |
| EPI_ISL_3378982 | Lighthouse Lab in Glasgow | Wellcome Sanger Institute for the COVID-19 Genomics UK (COG-UK) Consortium | Harper VanSteenhouse, Yumi Kasai, David Gray, Carol Clugston, Anna Dominiczak and Alex Alderton, Roberto Amato, Jeffrey Barrett, Sonia Goncalves, Ewan Harrison, David K. Jackson, Ian Johnston, Dominic Kwiatkowski, Cordelia Langford, John Sillitoe on behalf of the Wellcome Sanger Institute COVID-19 Surveillance Team |
| EPI_ISL_3378983 | Lighthouse Lab in Milton Keynes | Wellcome Sanger Institute for the COVID-19 Genomics UK (COG-UK) Consortium | The Lighthouse Lab in Milton Keynes and Alex Alderton, Roberto Amato, Jeffrey Barrett, Sonia Goncalves, Ewan Harrison, David K. Jackson, Ian Johnston, Dominic Kwiatkowski, Cordelia Langford, John Sillitoe on behalf of the Wellcome Sanger Institute COVID-19 Surveillance Team |
| EPI_ISL_3378984 | ORIAPOLE | CNR Virus des Infections Respiratoires - France SUD | Antonin Bal, Gregory Destras, Gwendolyne Burfin, Hadrien Regue, Quentin Semanas, Martine Valette, Bruno Lina, Laurence Josset |
| EPI_ISL_3378985 | Lighthouse Lab in Alderley Park | Wellcome Sanger Institute for the COVID-19 Genomics UK (COG-UK) Consortium | Jacquelyn Wynn, Mairead Hyland, The Lighthouse Lab in Alderley Park and Alex Alderton, Roberto Amato, Jeffrey Barrett, Sonia Goncalves, Ewan Harrison, David K. Jackson, Ian Johnston, Dominic Kwiatkowski, Cordelia Langford, John Sillitoe on behalf of the Wellcome Sanger Institute COVID-19 Surveillance Team |
| EPI_ISL_3378986 | Lighthouse Lab in Milton Keynes | Wellcome Sanger Institute for the COVID-19 Genomics UK (COG-UK) Consortium | The Lighthouse Lab in Milton Keynes and Alex Alderton, Roberto Amato, Jeffrey Barrett, Sonia Goncalves, Ewan Harrison, David K. Jackson, Ian Johnston, Dominic Kwiatkowski, Cordelia Langford, John Sillitoe on behalf of the Wellcome Sanger Institute COVID-19 Surveillance Team |
| EPI_ISL_3378987 | Lighthouse Lab in Glasgow | Wellcome Sanger Institute for the COVID-19 Genomics UK (COG-UK) Consortium | Harper VanSteenhouse, Yumi Kasai, David Gray, Carol Clugston, Anna Dominiczak and Alex Alderton, Roberto Amato, Jeffrey Barrett, Sonia Goncalves, Ewan Harrison, David K. Jackson, Ian Johnston, Dominic Kwiatkowski, Cordelia Langford, John Sillitoe on behalf of the Wellcome Sanger Institute COVID-19 Surveillance Team |
| EPI_ISL_3378988 | Berkshire and Surrey Pathology Services Lighthouse Laboratory | Wellcome Sanger Institute for the COVID-19 Genomics UK (COG-UK) Consortium | Berkshire and Surrey Pathology Services Lighthouse Laboratory and Alex Alderton, Roberto Amato, Jeffrey Barrett, Sonia Goncalves, Ewan Harrison, David K. Jackson, Ian Johnston, Dominic Kwiatkowski, Cordelia Langford, John Sillitoe on behalf of the Wellcome Sanger Institute COVID-19 Surveillance Team |
| EPI_ISL_3378989 | Lighthouse Lab in Milton Keynes | Wellcome Sanger Institute for the COVID-19 Genomics UK (COG-UK) Consortium | The Lighthouse Lab in Milton Keynes and Alex Alderton, Roberto Amato, Jeffrey Barrett, Sonia Goncalves, Ewan Harrison, David K. Jackson, Ian Johnston, Dominic Kwiatkowski, Cordelia Langford, John Sillitoe on behalf of the Wellcome Sanger Institute COVID-19 Surveillance Team |
| EPI_ISL_3378990 | Lighthouse Lab in Glasgow | Wellcome Sanger Institute for the COVID-19 Genomics UK (COG-UK) Consortium | Harper VanSteenhouse, Yumi Kasai, David Gray, Carol Clugston, Anna Dominiczak and Alex Alderton, Roberto Amato, Jeffrey Barrett, Sonia Goncalves, Ewan Harrison, David K. Jackson, Ian Johnston, Dominic Kwiatkowski, Cordelia Langford, John Sillitoe on behalf of the Wellcome Sanger Institute COVID-19 Surveillance Team |
| EPI_ISL_3378991 | Lighthouse Lab in Milton Keynes | Wellcome Sanger Institute for the COVID-19 Genomics UK (COG-UK) Consortium | The Lighthouse Lab in Milton Keynes and Alex Alderton, Roberto Amato, Jeffrey Barrett, Sonia Goncalves, Ewan Harrison, David K. Jackson, Ian Johnston, Dominic Kwiatkowski, Cordelia Langford, John Sillitoe on behalf of the Wellcome Sanger Institute COVID-19 Surveillance Team |
| EPI_ISL_3378992 | Lighthouse Lab in Alderley Park | Wellcome Sanger Institute for the COVID-19 Genomics UK (COG-UK) Consortium | Jacquelyn Wynn, Mairead Hyland, The Lighthouse Lab in Alderley Park and Alex Alderton, Roberto Amato, Jeffrey Barrett, Sonia Goncalves, Ewan Harrison, David K. Jackson, Ian Johnston, Dominic Kwiatkowski, Cordelia Langford, John Sillitoe on behalf of the Wellcome Sanger Institute COVID-19 Surveillance Team |
| EPI_ISL_3378993 | LABORATOIRE ACCOLAB SUD OUEST | CNR Virus des Infections Respiratoires - France SUD | Antonin Bal, Gregory Destras, Gwendolyne Burfin, Hadrien Regue, Quentin Semanas, Martine Valette, Bruno Lina, Laurence Josset |

| | | | |
|---|---|---|---|
| EPI_ISL_3378994 | Lighthouse Lab in Alderley Park | Wellcome Sanger Institute for the COVID-19 Genomics UK (COG-UK) Consortium | Jacquelyn Wynn, Mairead Hyland, The Lighthouse Lab in Alderley Park and Alex Alderton, Roberto Amato, Jeffrey Barrett, Sonia Goncalves, Ewan Harrison, David K. Jackson, Ian Johnston, Dominic Kwiatkowski, Cordelia Langford, John Sillitoe on behalf of the Wellcome Sanger Institute COVID-19 Surveillance Team |
| EPI_ISL_3378995 | Lighthouse Lab in Milton Keynes | Wellcome Sanger Institute for the COVID-19 Genomics UK (COG-UK) Consortium | The Lighthouse Lab in Milton Keynes and Alex Alderton, Roberto Amato, Jeffrey Barrett, Sonia Goncalves, Ewan Harrison, David K. Jackson, Ian Johnston, Dominic Kwiatkowski, Cordelia Langford, John Sillitoe on behalf of the Wellcome Sanger Institute COVID-19 Surveillance Team |
| EPI_ISL_3378996 | Lighthouse Lab in Alderley Park | Wellcome Sanger Institute for the COVID-19 Genomics UK (COG-UK) Consortium | Jacquelyn Wynn, Mairead Hyland, The Lighthouse Lab in Alderley Park and Alex Alderton, Roberto Amato, Jeffrey Barrett, Sonia Goncalves, Ewan Harrison, David K. Jackson, Ian Johnston, Dominic Kwiatkowski, Cordelia Langford, John Sillitoe on behalf of the Wellcome Sanger Institute COVID-19 Surveillance Team |
| EPI_ISL_3378997 | Berkshire and Surrey Pathology Services Lighthouse Laboratory | Wellcome Sanger Institute for the COVID-19 Genomics UK (COG-UK) Consortium | Berkshire and Surrey Pathology Services Lighthouse Laboratory and Alex Alderton, Roberto Amato, Jeffrey Barrett, Sonia Goncalves, Ewan Harrison, David K. Jackson, Ian Johnston, Dominic Kwiatkowski, Cordelia Langford, John Sillitoe on behalf of the Wellcome Sanger Institute COVID-19 Surveillance Team |
| EPI_ISL_3378998 | Lighthouse Lab in Glasgow | Wellcome Sanger Institute for the COVID-19 Genomics UK (COG-UK) Consortium | Harper VanSteenhouse, Yumi Kasai, David Gray, Carol Clugston, Anna Dominiczak and Alex Alderton, Roberto Amato, Jeffrey Barrett, Sonia Goncalves, Ewan Harrison, David K. Jackson, Ian Johnston, Dominic Kwiatkowski, Cordelia Langford, John Sillitoe on behalf of the Wellcome Sanger Institute COVID-19 Surveillance Team |
| EPI_ISL_3378999 | Lighthouse Lab in Milton Keynes | Wellcome Sanger Institute for the COVID-19 Genomics UK (COG-UK) Consortium | The Lighthouse Lab in Milton Keynes and Alex Alderton, Roberto Amato, Jeffrey Barrett, Sonia Goncalves, Ewan Harrison, David K. Jackson, Ian Johnston, Dominic Kwiatkowski, Cordelia Langford, John Sillitoe on behalf of the Wellcome Sanger Institute COVID-19 Surveillance Team |
| EPI_ISL_3379000 | Lighthouse Lab in Alderley Park | Wellcome Sanger Institute for the COVID-19 Genomics UK (COG-UK) Consortium | Jacquelyn Wynn, Mairead Hyland, The Lighthouse Lab in Alderley Park and Alex Alderton, Roberto Amato, Jeffrey Barrett, Sonia Goncalves, Ewan Harrison, David K. Jackson, Ian Johnston, Dominic Kwiatkowski, Cordelia Langford, John Sillitoe on behalf of the Wellcome Sanger Institute COVID-19 Surveillance Team |
| EPI_ISL_3379001 | LABORATOIRE ACCOLAB SUD OUEST | CNR Virus des Infections Respiratoires - France SUD | Antonin Bal, Gregory Destras, Gwendolyne Burfin, Hadrien Regue, Quentin Semanas, Martine Valette, Bruno Lina, Laurence Josset |
| EPI_ISL_3379002 | Lighthouse Lab in Glasgow | Wellcome Sanger Institute for the COVID-19 Genomics UK (COG-UK) Consortium | Harper VanSteenhouse, Yumi Kasai, David Gray, Carol Clugston, Anna Dominiczak and Alex Alderton, Roberto Amato, Jeffrey Barrett, Sonia Goncalves, Ewan Harrison, David K. Jackson, Ian Johnston, Dominic Kwiatkowski, Cordelia Langford, John Sillitoe on behalf of the Wellcome Sanger Institute COVID-19 Surveillance Team |
| EPI_ISL_3379003, EPI_ISL_3379004 | Lighthouse Lab in Alderley Park | Wellcome Sanger Institute for the COVID-19 Genomics UK (COG-UK) Consortium | Jacquelyn Wynn, Mairead Hyland, The Lighthouse Lab in Alderley Park and Alex Alderton, Roberto Amato, Jeffrey Barrett, Sonia Goncalves, Ewan Harrison, David K. Jackson, Ian Johnston, Dominic Kwiatkowski, Cordelia Langford, John Sillitoe on behalf of the Wellcome Sanger Institute COVID-19 Surveillance Team |
| EPI_ISL_3379005 | ORIAPOLE | CNR Virus des Infections Respiratoires - France SUD | Antonin Bal, Gregory Destras, Gwendolyne Burfin, Hadrien Regue, Quentin Semanas, Martine Valette, Bruno Lina, Laurence Josset |
| EPI_ISL_3379006 | Berkshire and Surrey Pathology Services Lighthouse Laboratory | Wellcome Sanger Institute for the COVID-19 Genomics UK (COG-UK) Consortium | Berkshire and Surrey Pathology Services Lighthouse Laboratory and Alex Alderton, Roberto Amato, Jeffrey Barrett, Sonia Goncalves, Ewan Harrison, David K. Jackson, Ian Johnston, Dominic Kwiatkowski, Cordelia Langford, John Sillitoe on behalf of the Wellcome Sanger Institute COVID-19 Surveillance Team |
| EPI_ISL_3379007 | Lighthouse Lab in Milton Keynes | Wellcome Sanger Institute for the COVID-19 Genomics UK (COG-UK) Consortium | The Lighthouse Lab in Milton Keynes and Alex Alderton, Roberto Amato, Jeffrey Barrett, Sonia Goncalves, Ewan Harrison, David K. Jackson, Ian Johnston, Dominic Kwiatkowski, Cordelia Langford, John Sillitoe on behalf of the Wellcome Sanger Institute COVID-19 Surveillance Team |
| EPI_ISL_3379008 | Lighthouse Lab in Alderley Park | Wellcome Sanger Institute for the COVID-19 Genomics UK (COG-UK) Consortium | Jacquelyn Wynn, Mairead Hyland, The Lighthouse Lab in Alderley Park and Alex Alderton, Roberto Amato, Jeffrey Barrett, Sonia Goncalves, Ewan Harrison, David K. Jackson, Ian Johnston, Dominic Kwiatkowski, Cordelia Langford, John Sillitoe on behalf of the Wellcome Sanger Institute COVID-19 Surveillance Team |
| EPI_ISL_3379009 | Lighthouse Lab in Glasgow | Wellcome Sanger Institute for the COVID-19 Genomics UK (COG-UK) Consortium | Harper VanSteenhouse, Yumi Kasai, David Gray, Carol Clugston, Anna Dominiczak and Alex Alderton, Roberto Amato, Jeffrey Barrett, Sonia Goncalves, Ewan Harrison, David K. Jackson, Ian Johnston, Dominic Kwiatkowski, Cordelia Langford, John Sillitoe on behalf of the Wellcome Sanger Institute COVID-19 Surveillance Team |
| EPI_ISL_3379010 | LABORATOIRE ACCOLAB SUD OUEST | CNR Virus des Infections Respiratoires - France SUD | Antonin Bal, Gregory Destras, Gwendolyne Burfin, Hadrien Regue, Quentin Semanas, Martine Valette, Bruno Lina, Laurence Josset |
| EPI_ISL_3379011, EPI_ISL_3379012 | Lighthouse Lab in Milton Keynes | Wellcome Sanger Institute for the COVID-19 Genomics UK (COG-UK) Consortium | The Lighthouse Lab in Milton Keynes and Alex Alderton, Roberto Amato, Jeffrey Barrett, Sonia Goncalves, Ewan Harrison, David K. Jackson, Ian Johnston, Dominic Kwiatkowski, Cordelia Langford, John Sillitoe on behalf of the Wellcome Sanger Institute COVID-19 Surveillance Team |
| EPI_ISL_3379013 | Lighthouse Lab in Alderley Park | Wellcome Sanger Institute for the COVID-19 Genomics UK (COG-UK) Consortium | Jacquelyn Wynn, Mairead Hyland, The Lighthouse Lab in Alderley Park and Alex Alderton, Roberto Amato, Jeffrey Barrett, Sonia Goncalves, Ewan Harrison, David K. Jackson, Ian Johnston, Dominic Kwiatkowski, Cordelia Langford, John Sillitoe on behalf of the Wellcome Sanger Institute COVID-19 Surveillance Team |
| EPI_ISL_3379014, EPI_ISL_3379015 | Lighthouse Lab in Glasgow | Wellcome Sanger Institute for the COVID-19 Genomics UK (COG-UK) Consortium | Harper VanSteenhouse, Yumi Kasai, David Gray, Carol Clugston, Anna Dominiczak and Alex Alderton, Roberto Amato, Jeffrey Barrett, Sonia Goncalves, Ewan Harrison, David K. Jackson, Ian Johnston, Dominic Kwiatkowski, Cordelia Langford, John Sillitoe on behalf of the Wellcome Sanger Institute COVID-19 Surveillance Team |
| EPI_ISL_3379016 | Lighthouse Lab in Alderley Park | Wellcome Sanger Institute for the COVID-19 Genomics UK (COG-UK) Consortium | Jacquelyn Wynn, Mairead Hyland, The Lighthouse Lab in Alderley Park and Alex Alderton, Roberto Amato, Jeffrey Barrett, Sonia Goncalves, Ewan Harrison, David K. Jackson, Ian Johnston, Dominic Kwiatkowski, Cordelia Langford, John Sillitoe on behalf of the Wellcome Sanger Institute COVID-19 Surveillance Team |
| EPI_ISL_3379017 | Lighthouse Lab in Glasgow | Wellcome Sanger Institute for the COVID-19 Genomics UK (COG-UK) Consortium | Harper VanSteenhouse, Yumi Kasai, David Gray, Carol Clugston, Anna Dominiczak and Alex Alderton, Roberto Amato, Jeffrey Barrett, Sonia Goncalves, Ewan Harrison, David K. Jackson, Ian Johnston, Dominic Kwiatkowski, Cordelia Langford, John Sillitoe on behalf of the Wellcome Sanger Institute COVID-19 Surveillance Team |
| EPI_ISL_3379018 | ORIAPOLE | CNR Virus des Infections Respiratoires - France SUD | Antonin Bal, Gregory Destras, Gwendolyne Burfin, Hadrien Regue, Quentin Semanas, Martine Valette, Bruno Lina, Laurence Josset |
| EPI_ISL_3379019 | Lighthouse Lab in Alderley Park | Wellcome Sanger Institute for the COVID-19 Genomics UK (COG-UK) Consortium | Jacquelyn Wynn, Mairead Hyland, The Lighthouse Lab in Alderley Park and Alex Alderton, Roberto Amato, Jeffrey Barrett, Sonia Goncalves, Ewan Harrison, David K. Jackson, Ian Johnston, Dominic Kwiatkowski, Cordelia Langford, John Sillitoe on behalf of the Wellcome Sanger Institute COVID-19 Surveillance Team |
| EPI_ISL_3379020, EPI_ISL_3379021 | Lighthouse Lab in Milton Keynes | Wellcome Sanger Institute for the COVID-19 Genomics UK (COG-UK) Consortium | The Lighthouse Lab in Milton Keynes and Alex Alderton, Roberto Amato, Jeffrey Barrett, Sonia Goncalves, Ewan Harrison, David K. Jackson, Ian Johnston, Dominic Kwiatkowski, Cordelia Langford, John Sillitoe on behalf of the Wellcome Sanger Institute COVID-19 Surveillance Team |
| EPI_ISL_3379022 | Lighthouse Lab in Alderley Park | Wellcome Sanger Institute for the COVID-19 Genomics UK (COG-UK) Consortium | Jacquelyn Wynn, Mairead Hyland, The Lighthouse Lab in Alderley Park and Alex Alderton, Roberto Amato, Jeffrey Barrett, Sonia Goncalves, Ewan Harrison, David K. Jackson, Ian Johnston, Dominic Kwiatkowski, Cordelia Langford, John Sillitoe on behalf of the Wellcome Sanger Institute COVID-19 Surveillance Team |
| EPI_ISL_3379023 | Lighthouse Lab in Milton Keynes | Wellcome Sanger Institute for the COVID-19 Genomics UK (COG-UK) Consortium | The Lighthouse Lab in Milton Keynes and Alex Alderton, Roberto Amato, Jeffrey Barrett, Sonia Goncalves, Ewan Harrison, David K. Jackson, Ian Johnston, Dominic Kwiatkowski, Cordelia Langford, John Sillitoe on behalf of the Wellcome Sanger Institute COVID-19 Surveillance Team |
| EPI_ISL_3379024 | Lighthouse Lab in Alderley Park | Wellcome Sanger Institute for the COVID-19 Genomics UK (COG-UK) Consortium | Jacquelyn Wynn, Mairead Hyland, The Lighthouse Lab in Alderley Park and Alex Alderton, Roberto Amato, Jeffrey Barrett, Sonia Goncalves, Ewan Harrison, David K. Jackson, Ian Johnston, Dominic Kwiatkowski, Cordelia Langford, John Sillitoe on behalf of the Wellcome Sanger Institute COVID-19 Surveillance Team |
| EPI_ISL_3379025 | Lighthouse Lab in Glasgow | Wellcome Sanger Institute for the COVID-19 Genomics UK (COG-UK) Consortium | Harper VanSteenhouse, Yumi Kasai, David Gray, Carol Clugston, Anna Dominiczak and Alex Alderton, Roberto Amato, Jeffrey Barrett, Sonia Goncalves, Ewan Harrison, David K. Jackson, Ian Johnston, Dominic Kwiatkowski, Cordelia Langford, John Sillitoe on behalf of the Wellcome Sanger Institute COVID-19 Surveillance Team |
| EPI_ISL_3379026 | ORIAPOLE | CNR Virus des Infections Respiratoires - France SUD | Antonin Bal, Gregory Destras, Gwendolyne Burfin, Hadrien Regue, Quentin Semanas, Martine Valette, Bruno Lina, Laurence Josset |
| EPI_ISL_3379027 | Lighthouse Lab in Milton Keynes | Wellcome Sanger Institute for the COVID-19 Genomics UK (COG-UK) Consortium | The Lighthouse Lab in Milton Keynes and Alex Alderton, Roberto Amato, Jeffrey Barrett, Sonia Goncalves, Ewan Harrison, David K. Jackson, Ian Johnston, Dominic Kwiatkowski, Cordelia Langford, John Sillitoe on behalf of the Wellcome Sanger Institute COVID-19 Surveillance Team |
| EPI_ISL_3379028 | Lighthouse Lab in Alderley Park | Wellcome Sanger Institute for the COVID-19 Genomics UK (COG-UK) Consortium | Jacquelyn Wynn, Mairead Hyland, The Lighthouse Lab in Alderley Park and Alex Alderton, Roberto Amato, Jeffrey Barrett, Sonia Goncalves, Ewan Harrison, David K. Jackson, Ian Johnston, Dominic Kwiatkowski, Cordelia Langford, John Sillitoe on behalf of the Wellcome Sanger Institute COVID-19 Surveillance Team |
| EPI_ISL_3379029 | Lighthouse Lab in Milton Keynes | Wellcome Sanger Institute for the COVID-19 Genomics UK (COG-UK) Consortium | The Lighthouse Lab in Milton Keynes and Alex Alderton, Roberto Amato, Jeffrey Barrett, Sonia Goncalves, Ewan Harrison, David K. Jackson, Ian Johnston, Dominic Kwiatkowski, Cordelia Langford, John Sillitoe on behalf of the Wellcome Sanger Institute COVID-19 Surveillance Team |
| EPI_ISL_3379030 | LABORATOIRE ACCOLAB SUD OUEST | CNR Virus des Infections Respiratoires - France SUD | Antonin Bal, Gregory Destras, Gwendolyne Burfin, Hadrien Regue, Quentin Semanas, Martine Valette, Bruno Lina, Laurence Josset |
| EPI_ISL_3379031 | Lighthouse Lab in Milton Keynes | Wellcome Sanger Institute for the COVID-19 Genomics UK (COG-UK) Consortium | The Lighthouse Lab in Milton Keynes and Alex Alderton, Roberto Amato, Jeffrey Barrett, Sonia Goncalves, Ewan Harrison, David K. Jackson, Ian Johnston, Dominic Kwiatkowski, Cordelia Langford, John Sillitoe on behalf of the Wellcome Sanger Institute COVID-19 Surveillance Team |
| EPI_ISL_3379032 | Lighthouse Lab in Glasgow | Wellcome Sanger Institute for the COVID-19 Genomics UK (COG-UK) Consortium | Harper VanSteenhouse, Yumi Kasai, David Gray, Carol Clugston, Anna Dominiczak and Alex Alderton, Roberto Amato, Jeffrey Barrett, Sonia Goncalves, Ewan Harrison, David K. Jackson, Ian Johnston, Dominic Kwiatkowski, Cordelia Langford, John Sillitoe on behalf of the Wellcome Sanger Institute COVID-19 Surveillance Team |
| EPI_ISL_3379033 | Lighthouse Lab in Milton Keynes | Wellcome Sanger Institute for the COVID-19 Genomics UK (COG-UK) Consortium | The Lighthouse Lab in Milton Keynes and Alex Alderton, Roberto Amato, Jeffrey Barrett, Sonia Goncalves, Ewan Harrison, David K. Jackson, Ian Johnston, Dominic Kwiatkowski, Cordelia Langford, John Sillitoe on behalf of the Wellcome Sanger Institute COVID-19 Surveillance Team |
| EPI_ISL_3379034 | LABORATOIRE ACCOLAB SUD OUEST | CNR Virus des Infections Respiratoires - France SUD | Antonin Bal, Gregory Destras, Gwendolyne Burfin, Hadrien Regue, Quentin Semanas, Martine Valette, Bruno Lina, Laurence Josset |
| EPI_ISL_3379035, EPI_ISL_3379036 | Lighthouse Lab in Milton Keynes | Wellcome Sanger Institute for the COVID-19 Genomics UK (COG-UK) Consortium | The Lighthouse Lab in Milton Keynes and Alex Alderton, Roberto Amato, Jeffrey Barrett, Sonia Goncalves, Ewan Harrison, David K. Jackson, Ian Johnston, Dominic Kwiatkowski, Cordelia Langford, John Sillitoe on behalf of the Wellcome Sanger Institute COVID-19 Surveillance Team |
| EPI_ISL_3379037 | Berkshire and Surrey Pathology Services Lighthouse Laboratory | Wellcome Sanger Institute for the COVID-19 Genomics UK (COG-UK) Consortium | Berkshire and Surrey Pathology Services Lighthouse Laboratory and Alex Alderton, Roberto Amato, Jeffrey Barrett, Sonia Goncalves, Ewan Harrison, David K. Jackson, Ian Johnston, Dominic Kwiatkowski, Cordelia Langford, John Sillitoe on behalf of the Wellcome Sanger Institute COVID-19 Surveillance Team |
| EPI_ISL_3379038 | ORIAPOLE | CNR Virus des Infections Respiratoires - France SUD | Antonin Bal, Gregory Destras, Gwendolyne Burfin, Hadrien Regue, Quentin Semanas, Martine Valette, Bruno Lina, Laurence Josset |
| EPI_ISL_3379039, EPI_ISL_3379040 | Lighthouse Lab in Milton Keynes | Wellcome Sanger Institute for the COVID-19 Genomics UK (COG-UK) Consortium | The Lighthouse Lab in Milton Keynes and Alex Alderton, Roberto Amato, Jeffrey Barrett, Sonia Goncalves, Ewan Harrison, David K. Jackson, Ian Johnston, Dominic Kwiatkowski, Cordelia Langford, John Sillitoe on behalf of the Wellcome Sanger Institute COVID-19 Surveillance Team |
| EPI_ISL_3379041 | Lighthouse Lab in Alderley Park | Wellcome Sanger Institute for the COVID-19 Genomics UK (COG-UK) Consortium | Jacquelyn Wynn, Mairead Hyland, The Lighthouse Lab in Alderley Park and Alex Alderton, Roberto Amato, Jeffrey Barrett, Sonia Goncalves, Ewan Harrison, David K. Jackson, Ian Johnston, Dominic Kwiatkowski, Cordelia Langford, John Sillitoe on behalf of the Wellcome Sanger Institute COVID-19 Surveillance Team |
| EPI_ISL_3379042 | ORIAPOLE | CNR Virus des Infections Respiratoires - France SUD | Antonin Bal, Gregory Destras, Gwendolyne Burfin, Hadrien Regue, Quentin Semanas, Martine Valette, Bruno Lina, Laurence Josset |
| EPI_ISL_3379043 | Lighthouse Lab in Milton Keynes | Wellcome Sanger Institute for the COVID-19 Genomics UK (COG-UK) Consortium | The Lighthouse Lab in Milton Keynes and Alex Alderton, Roberto Amato, Jeffrey Barrett, Sonia Goncalves, Ewan Harrison, David K. Jackson, Ian Johnston, Dominic Kwiatkowski, Cordelia Langford, John Sillitoe on behalf of the Wellcome Sanger Institute COVID-19 Surveillance Team |
| EPI_ISL_3379044, EPI_ISL_3379045 | Lighthouse Lab in Alderley Park | Wellcome Sanger Institute for the COVID-19 Genomics UK (COG-UK) Consortium | Jacquelyn Wynn, Mairead Hyland, The Lighthouse Lab in Alderley Park and Alex Alderton, Roberto Amato, Jeffrey Barrett, Sonia Goncalves, Ewan Harrison, David K. Jackson, Ian Johnston, Dominic Kwiatkowski, Cordelia Langford, John Sillitoe on behalf of the Wellcome Sanger Institute COVID-19 Surveillance Team |
| EPI_ISL_3379046 | Lighthouse Lab in Glasgow | Wellcome Sanger Institute for the COVID-19 Genomics UK (COG-UK) Consortium | Harper VanSteenhouse, Yumi Kasai, David Gray, Carol Clugston, Anna Dominiczak and Alex Alderton, Roberto Amato, Jeffrey Barrett, Sonia Goncalves, Ewan Harrison, David K. Jackson, Ian Johnston, Dominic Kwiatkowski, Cordelia Langford, John Sillitoe on behalf of the Wellcome Sanger Institute COVID-19 Surveillance Team |
| EPI_ISL_3379048 | Lighthouse Lab in Alderley Park | Wellcome Sanger Institute for the COVID-19 Genomics UK (COG-UK) Consortium | Jacquelyn Wynn, Mairead Hyland, The Lighthouse Lab in Alderley Park and Alex Alderton, Roberto Amato, Jeffrey Barrett, Sonia Goncalves, Ewan Harrison, David K. Jackson, Ian Johnston, Dominic Kwiatkowski, Cordelia Langford, John Sillitoe on behalf of the Wellcome Sanger Institute COVID-19 Surveillance Team |
| EPI_ISL_3379049 | Berkshire and Surrey Pathology Services Lighthouse Laboratory | Wellcome Sanger Institute for the COVID-19 Genomics UK (COG-UK) Consortium | Berkshire and Surrey Pathology Services Lighthouse Laboratory and Alex Alderton, Roberto Amato, Jeffrey Barrett, Sonia Goncalves, Ewan Harrison, David K. Jackson, Ian Johnston, Dominic Kwiatkowski, Cordelia Langford, John Sillitoe on behalf of the Wellcome Sanger Institute COVID-19 Surveillance Team |
| EPI_ISL_3379050 | Lighthouse Lab in Milton Keynes | Wellcome Sanger Institute for the COVID-19 Genomics UK (COG-UK) Consortium | The Lighthouse Lab in Milton Keynes and Alex Alderton, Roberto Amato, Jeffrey Barrett, Sonia Goncalves, Ewan Harrison, David K. Jackson, Ian Johnston, Dominic Kwiatkowski, Cordelia Langford, John Sillitoe on behalf of the Wellcome Sanger Institute COVID-19 Surveillance Team |

| | | | |
|---|---|---|---|
| EPI_ISL_3379051 | ORIAPOLE | CNR Virus des Infections Respiratoires - France SUD | Antonin Bal, Gregory Destras, Gwendolyne Burfin, Hadrien Regue, Quentin Semanas, Martine Valette, Bruno Lina, Laurence Josset |
| EPI_ISL_3379052, EPI_ISL_3379053 | Lighthouse Lab in Glasgow | Wellcome Sanger Institute for the COVID-19 Genomics UK (COG-UK) Consortium | Harper VanSteenhouse, Yumi Kasai, David Gray, Carol Clugston, Anna Dominiczak and Alex Alderton, Roberto Amato, Jeffrey Barrett, Sonia Goncalves, Ewan Harrison, David K. Jackson, Ian Johnston, Dominic Kwiatkowski, Cordelia Langford, John Sillitoe on behalf of the Wellcome Sanger Institute COVID-19 Surveillance Team |
| EPI_ISL_3379054 | Lighthouse Lab in Alderley Park | Wellcome Sanger Institute for the COVID-19 Genomics UK (COG-UK) Consortium | Jacquelyn Wynn, Mairead Hyland, The Lighthouse Lab in Alderley Park and Alex Alderton, Roberto Amato, Jeffrey Barrett, Sonia Goncalves, Ewan Harrison, David K. Jackson, Ian Johnston, Dominic Kwiatkowski, Cordelia Langford, John Sillitoe on behalf of the Wellcome Sanger Institute COVID-19 Surveillance Team |
| EPI_ISL_3379055 | Berkshire and Surrey Pathology Services Lighthouse Laboratory | Wellcome Sanger Institute for the COVID-19 Genomics UK (COG-UK) Consortium | Berkshire and Surrey Pathology Services Lighthouse Laboratory and Alex Alderton, Roberto Amato, Jeffrey Barrett, Sonia Goncalves, Ewan Harrison, David K. Jackson, Ian Johnston, Dominic Kwiatkowski, Cordelia Langford, John Sillitoe on behalf of the Wellcome Sanger Institute COVID-19 Surveillance Team |
| EPI_ISL_3379056 | Lighthouse Lab in Glasgow | Wellcome Sanger Institute for the COVID-19 Genomics UK (COG-UK) Consortium | Harper VanSteenhouse, Yumi Kasai, David Gray, Carol Clugston, Anna Dominiczak and Alex Alderton, Roberto Amato, Jeffrey Barrett, Sonia Goncalves, Ewan Harrison, David K. Jackson, Ian Johnston, Dominic Kwiatkowski, Cordelia Langford, John Sillitoe on behalf of the Wellcome Sanger Institute COVID-19 Surveillance Team |
| EPI_ISL_3379057 | Berkshire and Surrey Pathology Services Lighthouse Laboratory | Wellcome Sanger Institute for the COVID-19 Genomics UK (COG-UK) Consortium | Berkshire and Surrey Pathology Services Lighthouse Laboratory and Alex Alderton, Roberto Amato, Jeffrey Barrett, Sonia Goncalves, Ewan Harrison, David K. Jackson, Ian Johnston, Dominic Kwiatkowski, Cordelia Langford, John Sillitoe on behalf of the Wellcome Sanger Institute COVID-19 Surveillance Team |
| EPI_ISL_3379058 | Lighthouse Lab in Glasgow | Wellcome Sanger Institute for the COVID-19 Genomics UK (COG-UK) Consortium | Harper VanSteenhouse, Yumi Kasai, David Gray, Carol Clugston, Anna Dominiczak and Alex Alderton, Roberto Amato, Jeffrey Barrett, Sonia Goncalves, Ewan Harrison, David K. Jackson, Ian Johnston, Dominic Kwiatkowski, Cordelia Langford, John Sillitoe on behalf of the Wellcome Sanger Institute COVID-19 Surveillance Team |
| EPI_ISL_3379059 | Berkshire and Surrey Pathology Services Lighthouse Laboratory | Wellcome Sanger Institute for the COVID-19 Genomics UK (COG-UK) Consortium | Berkshire and Surrey Pathology Services Lighthouse Laboratory and Alex Alderton, Roberto Amato, Jeffrey Barrett, Sonia Goncalves, Ewan Harrison, David K. Jackson, Ian Johnston, Dominic Kwiatkowski, Cordelia Langford, John Sillitoe on behalf of the Wellcome Sanger Institute COVID-19 Surveillance Team |
| EPI_ISL_3379060 | Lighthouse Lab in Glasgow | Wellcome Sanger Institute for the COVID-19 Genomics UK (COG-UK) Consortium | Harper VanSteenhouse, Yumi Kasai, David Gray, Carol Clugston, Anna Dominiczak and Alex Alderton, Roberto Amato, Jeffrey Barrett, Sonia Goncalves, Ewan Harrison, David K. Jackson, Ian Johnston, Dominic Kwiatkowski, Cordelia Langford, John Sillitoe on behalf of the Wellcome Sanger Institute COVID-19 Surveillance Team |
| EPI_ISL_3379061 | Lighthouse Lab in Alderley Park | Wellcome Sanger Institute for the COVID-19 Genomics UK (COG-UK) Consortium | Jacquelyn Wynn, Mairead Hyland, The Lighthouse Lab in Alderley Park and Alex Alderton, Roberto Amato, Jeffrey Barrett, Sonia Goncalves, Ewan Harrison, David K. Jackson, Ian Johnston, Dominic Kwiatkowski, Cordelia Langford, John Sillitoe on behalf of the Wellcome Sanger Institute COVID-19 Surveillance Team |
| EPI_ISL_3379062 | ORIAPOLE | CNR Virus des Infections Respiratoires - France SUD | Antonin Bal, Gregory Destras, Gwendolyne Burfin, Hadrien Regue, Quentin Semanas, Martine Valette, Bruno Lina, Laurence Josset |
| EPI_ISL_3379063 | Lighthouse Lab in Milton Keynes | Wellcome Sanger Institute for the COVID-19 Genomics UK (COG-UK) Consortium | The Lighthouse Lab in Milton Keynes and Alex Alderton, Roberto Amato, Jeffrey Barrett, Sonia Goncalves, Ewan Harrison, David K. Jackson, Ian Johnston, Dominic Kwiatkowski, Cordelia Langford, John Sillitoe on behalf of the Wellcome Sanger Institute COVID-19 Surveillance Team |
| EPI_ISL_3379064 | Lighthouse Lab in Glasgow | Wellcome Sanger Institute for the COVID-19 Genomics UK (COG-UK) Consortium | Harper VanSteenhouse, Yumi Kasai, David Gray, Carol Clugston, Anna Dominiczak and Alex Alderton, Roberto Amato, Jeffrey Barrett, Sonia Goncalves, Ewan Harrison, David K. Jackson, Ian Johnston, Dominic Kwiatkowski, Cordelia Langford, John Sillitoe on behalf of the Wellcome Sanger Institute COVID-19 Surveillance Team |
| EPI_ISL_3379065 | Lighthouse Lab in Milton Keynes | Wellcome Sanger Institute for the COVID-19 Genomics UK (COG-UK) Consortium | The Lighthouse Lab in Milton Keynes and Alex Alderton, Roberto Amato, Jeffrey Barrett, Sonia Goncalves, Ewan Harrison, David K. Jackson, Ian Johnston, Dominic Kwiatkowski, Cordelia Langford, John Sillitoe on behalf of the Wellcome Sanger Institute COVID-19 Surveillance Team |
| EPI_ISL_3379066 | LABORATOIRE ACCOLAB SUD OUEST | CNR Virus des Infections Respiratoires - France SUD | Antonin Bal, Gregory Destras, Gwendolyne Burfin, Hadrien Regue, Quentin Semanas, Martine Valette, Bruno Lina, Laurence Josset |
| EPI_ISL_3379067 | Lighthouse Lab in Glasgow | Wellcome Sanger Institute for the COVID-19 Genomics UK (COG-UK) Consortium | Harper VanSteenhouse, Yumi Kasai, David Gray, Carol Clugston, Anna Dominiczak and Alex Alderton, Roberto Amato, Jeffrey Barrett, Sonia Goncalves, Ewan Harrison, David K. Jackson, Ian Johnston, Dominic Kwiatkowski, Cordelia Langford, John Sillitoe on behalf of the Wellcome Sanger Institute COVID-19 Surveillance Team |
| EPI_ISL_3379068 | Lighthouse Lab in Milton Keynes | Wellcome Sanger Institute for the COVID-19 Genomics UK (COG-UK) Consortium | The Lighthouse Lab in Milton Keynes and Alex Alderton, Roberto Amato, Jeffrey Barrett, Sonia Goncalves, Ewan Harrison, David K. Jackson, Ian Johnston, Dominic Kwiatkowski, Cordelia Langford, John Sillitoe on behalf of the Wellcome Sanger Institute COVID-19 Surveillance Team |
| EPI_ISL_3379069 | Lighthouse Lab in Alderley Park | Wellcome Sanger Institute for the COVID-19 Genomics UK (COG-UK) Consortium | Jacquelyn Wynn, Mairead Hyland, The Lighthouse Lab in Alderley Park and Alex Alderton, Roberto Amato, Jeffrey Barrett, Sonia Goncalves, Ewan Harrison, David K. Jackson, Ian Johnston, Dominic Kwiatkowski, Cordelia Langford, John Sillitoe on behalf of the Wellcome Sanger Institute COVID-19 Surveillance Team |
| EPI_ISL_3379070 | ORIAPOLE | CNR Virus des Infections Respiratoires - France SUD | Antonin Bal, Gregory Destras, Gwendolyne Burfin, Hadrien Regue, Quentin Semanas, Martine Valette, Bruno Lina, Laurence Josset |
| EPI_ISL_3379071 | Lighthouse Lab in Glasgow | Wellcome Sanger Institute for the COVID-19 Genomics UK (COG-UK) Consortium | Harper VanSteenhouse, Yumi Kasai, David Gray, Carol Clugston, Anna Dominiczak and Alex Alderton, Roberto Amato, Jeffrey Barrett, Sonia Goncalves, Ewan Harrison, David K. Jackson, Ian Johnston, Dominic Kwiatkowski, Cordelia Langford, John Sillitoe on behalf of the Wellcome Sanger Institute COVID-19 Surveillance Team |
| EPI_ISL_3379072, EPI_ISL_3379073 | Lighthouse Lab in Alderley Park | Wellcome Sanger Institute for the COVID-19 Genomics UK (COG-UK) Consortium | Jacquelyn Wynn, Mairead Hyland, The Lighthouse Lab in Alderley Park and Alex Alderton, Roberto Amato, Jeffrey Barrett, Sonia Goncalves, Ewan Harrison, David K. Jackson, Ian Johnston, Dominic Kwiatkowski, Cordelia Langford, John Sillitoe on behalf of the Wellcome Sanger Institute COVID-19 Surveillance Team |
| EPI_ISL_3379074 | ORIAPOLE | CNR Virus des Infections Respiratoires - France SUD | Antonin Bal, Gregory Destras, Gwendolyne Burfin, Hadrien Regue, Quentin Semanas, Martine Valette, Bruno Lina, Laurence Josset |
| EPI_ISL_3379075 | Lighthouse Lab in Glasgow | Wellcome Sanger Institute for the COVID-19 Genomics UK (COG-UK) Consortium | Harper VanSteenhouse, Yumi Kasai, David Gray, Carol Clugston, Anna Dominiczak and Alex Alderton, Roberto Amato, Jeffrey Barrett, Sonia Goncalves, Ewan Harrison, David K. Jackson, Ian Johnston, Dominic Kwiatkowski, Cordelia Langford, John Sillitoe on behalf of the Wellcome Sanger Institute COVID-19 Surveillance Team |
| EPI_ISL_3379076 | Lighthouse Lab in Milton Keynes | Wellcome Sanger Institute for the COVID-19 Genomics UK (COG-UK) Consortium | The Lighthouse Lab in Milton Keynes and Alex Alderton, Roberto Amato, Jeffrey Barrett, Sonia Goncalves, Ewan Harrison, David K. Jackson, Ian Johnston, Dominic Kwiatkowski, Cordelia Langford, John Sillitoe on behalf of the Wellcome Sanger Institute COVID-19 Surveillance Team |
| EPI_ISL_3379077 | Lighthouse Lab in Alderley Park | Wellcome Sanger Institute for the COVID-19 Genomics UK (COG-UK) Consortium | Jacquelyn Wynn, Mairead Hyland, The Lighthouse Lab in Alderley Park and Alex Alderton, Roberto Amato, Jeffrey Barrett, Sonia Goncalves, Ewan Harrison, David K. Jackson, Ian Johnston, Dominic Kwiatkowski, Cordelia Langford, John Sillitoe on behalf of the Wellcome Sanger Institute COVID-19 Surveillance Team |
| EPI_ISL_3379078 | LABORATOIRE ACCOLAB SUD OUEST | CNR Virus des Infections Respiratoires - France SUD | Antonin Bal, Gregory Destras, Gwendolyne Burfin, Hadrien Regue, Quentin Semanas, Martine Valette, Bruno Lina, Laurence Josset |
| EPI_ISL_3379079, EPI_ISL_3379080, EPI_ISL_3379081 | Lighthouse Lab in Milton Keynes | Wellcome Sanger Institute for the COVID-19 Genomics UK (COG-UK) Consortium | The Lighthouse Lab in Milton Keynes and Alex Alderton, Roberto Amato, Jeffrey Barrett, Sonia Goncalves, Ewan Harrison, David K. Jackson, Ian Johnston, Dominic Kwiatkowski, Cordelia Langford, John Sillitoe on behalf of the Wellcome Sanger Institute COVID-19 Surveillance Team |
| EPI_ISL_3379082 | ORIAPOLE | CNR Virus des Infections Respiratoires - France SUD | Antonin Bal, Gregory Destras, Gwendolyne Burfin, Hadrien Regue, Quentin Semanas, Martine Valette, Bruno Lina, Laurence Josset |
| EPI_ISL_3379083 | Lighthouse Lab in Milton Keynes | Wellcome Sanger Institute for the COVID-19 Genomics UK (COG-UK) Consortium | The Lighthouse Lab in Milton Keynes and Alex Alderton, Roberto Amato, Jeffrey Barrett, Sonia Goncalves, Ewan Harrison, David K. Jackson, Ian Johnston, Dominic Kwiatkowski, Cordelia Langford, John Sillitoe on behalf of the Wellcome Sanger Institute COVID-19 Surveillance Team |
| EPI_ISL_3379084 | Lighthouse Lab in Alderley Park | Wellcome Sanger Institute for the COVID-19 Genomics UK (COG-UK) Consortium | Jacquelyn Wynn, Mairead Hyland, The Lighthouse Lab in Alderley Park and Alex Alderton, Roberto Amato, Jeffrey Barrett, Sonia Goncalves, Ewan Harrison, David K. Jackson, Ian Johnston, Dominic Kwiatkowski, Cordelia Langford, John Sillitoe on behalf of the Wellcome Sanger Institute COVID-19 Surveillance Team |
| EPI_ISL_3379085 | LABORATOIRE ACCOLAB SUD OUEST | CNR Virus des Infections Respiratoires - France SUD | Antonin Bal, Gregory Destras, Gwendolyne Burfin, Hadrien Regue, Quentin Semanas, Martine Valette, Bruno Lina, Laurence Josset |
| EPI_ISL_3379086, EPI_ISL_3379087, EPI_ISL_3379088 | Lighthouse Lab in Milton Keynes | Wellcome Sanger Institute for the COVID-19 Genomics UK (COG-UK) Consortium | The Lighthouse Lab in Milton Keynes and Alex Alderton, Roberto Amato, Jeffrey Barrett, Sonia Goncalves, Ewan Harrison, David K. Jackson, Ian Johnston, Dominic Kwiatkowski, Cordelia Langford, John Sillitoe on behalf of the Wellcome Sanger Institute COVID-19 Surveillance Team |
| EPI_ISL_3379089 | LABORATOIRE ACCOLAB SUD OUEST | CNR Virus des Infections Respiratoires - France SUD | Antonin Bal, Gregory Destras, Gwendolyne Burfin, Hadrien Regue, Quentin Semanas, Martine Valette, Bruno Lina, Laurence Josset |
| EPI_ISL_3379090, EPI_ISL_3379091 | Lighthouse Lab in Alderley Park | Wellcome Sanger Institute for the COVID-19 Genomics UK (COG-UK) Consortium | Jacquelyn Wynn, Mairead Hyland, The Lighthouse Lab in Alderley Park and Alex Alderton, Roberto Amato, Jeffrey Barrett, Sonia Goncalves, Ewan Harrison, David K. Jackson, Ian Johnston, Dominic Kwiatkowski, Cordelia Langford, John Sillitoe on behalf of the Wellcome Sanger Institute COVID-19 Surveillance Team |
| EPI_ISL_3379092 | Lighthouse Lab in Milton Keynes | Wellcome Sanger Institute for the COVID-19 Genomics UK (COG-UK) Consortium | The Lighthouse Lab in Milton Keynes and Alex Alderton, Roberto Amato, Jeffrey Barrett, Sonia Goncalves, Ewan Harrison, David K. Jackson, Ian Johnston, Dominic Kwiatkowski, Cordelia Langford, John Sillitoe on behalf of the Wellcome Sanger Institute COVID-19 Surveillance Team |
| EPI_ISL_3379093 | Lighthouse Lab in Glasgow | Wellcome Sanger Institute for the COVID-19 Genomics UK (COG-UK) Consortium | Harper VanSteenhouse, Yumi Kasai, David Gray, Carol Clugston, Anna Dominiczak and Alex Alderton, Roberto Amato, Jeffrey Barrett, Sonia Goncalves, Ewan Harrison, David K. Jackson, Ian Johnston, Dominic Kwiatkowski, Cordelia Langford, John Sillitoe on behalf of the Wellcome Sanger Institute COVID-19 Surveillance Team |
| EPI_ISL_3379094 | Lighthouse Lab in Alderley Park | Wellcome Sanger Institute for the COVID-19 Genomics UK (COG-UK) Consortium | Jacquelyn Wynn, Mairead Hyland, The Lighthouse Lab in Alderley Park and Alex Alderton, Roberto Amato, Jeffrey Barrett, Sonia Goncalves, Ewan Harrison, David K. Jackson, Ian Johnston, Dominic Kwiatkowski, Cordelia Langford, John Sillitoe on behalf of the Wellcome Sanger Institute COVID-19 Surveillance Team |
| EPI_ISL_3379095 | Berkshire and Surrey Pathology Services Lighthouse Laboratory | Wellcome Sanger Institute for the COVID-19 Genomics UK (COG-UK) Consortium | Berkshire and Surrey Pathology Services Lighthouse Laboratory and Alex Alderton, Roberto Amato, Jeffrey Barrett, Sonia Goncalves, Ewan Harrison, David K. Jackson, Ian Johnston, Dominic Kwiatkowski, Cordelia Langford, John Sillitoe on behalf of the Wellcome Sanger Institute COVID-19 Surveillance Team |
| EPI_ISL_3379096 | Lighthouse Lab in Milton Keynes | Wellcome Sanger Institute for the COVID-19 Genomics UK (COG-UK) Consortium | The Lighthouse Lab in Milton Keynes and Alex Alderton, Roberto Amato, Jeffrey Barrett, Sonia Goncalves, Ewan Harrison, David K. Jackson, Ian Johnston, Dominic Kwiatkowski, Cordelia Langford, John Sillitoe on behalf of the Wellcome Sanger Institute COVID-19 Surveillance Team |
| EPI_ISL_3379097 | Berkshire and Surrey Pathology Services Lighthouse Laboratory | Wellcome Sanger Institute for the COVID-19 Genomics UK (COG-UK) Consortium | Berkshire and Surrey Pathology Services Lighthouse Laboratory and Alex Alderton, Roberto Amato, Jeffrey Barrett, Sonia Goncalves, Ewan Harrison, David K. Jackson, Ian Johnston, Dominic Kwiatkowski, Cordelia Langford, John Sillitoe on behalf of the Wellcome Sanger Institute COVID-19 Surveillance Team |
| EPI_ISL_3379098 | LABORATOIRE ACCOLAB SUD OUEST | CNR Virus des Infections Respiratoires - France SUD | Antonin Bal, Gregory Destras, Gwendolyne Burfin, Hadrien Regue, Quentin Semanas, Martine Valette, Bruno Lina, Laurence Josset |
| EPI_ISL_3379099 | Berkshire and Surrey Pathology Services Lighthouse Laboratory | Wellcome Sanger Institute for the COVID-19 Genomics UK (COG-UK) Consortium | Berkshire and Surrey Pathology Services Lighthouse Laboratory and Alex Alderton, Roberto Amato, Jeffrey Barrett, Sonia Goncalves, Ewan Harrison, David K. Jackson, Ian Johnston, Dominic Kwiatkowski, Cordelia Langford, John Sillitoe on behalf of the Wellcome Sanger Institute COVID-19 Surveillance Team |
| EPI_ISL_3379100 | Lighthouse Lab in Alderley Park | Wellcome Sanger Institute for the COVID-19 Genomics UK (COG-UK) Consortium | Jacquelyn Wynn, Mairead Hyland, The Lighthouse Lab in Alderley Park and Alex Alderton, Roberto Amato, Jeffrey Barrett, Sonia Goncalves, Ewan Harrison, David K. Jackson, Ian Johnston, Dominic Kwiatkowski, Cordelia Langford, John Sillitoe on behalf of the Wellcome Sanger Institute COVID-19 Surveillance Team |
| EPI_ISL_3379101 | LABORATOIRE ACCOLAB SUD OUEST | CNR Virus des Infections Respiratoires - France SUD | Antonin Bal, Gregory Destras, Gwendolyne Burfin, Hadrien Regue, Quentin Semanas, Martine Valette, Bruno Lina, Laurence Josset |
| EPI_ISL_3379102 | Lighthouse Lab in Alderley Park | Wellcome Sanger Institute for the COVID-19 Genomics UK (COG-UK) Consortium | Jacquelyn Wynn, Mairead Hyland, The Lighthouse Lab in Alderley Park and Alex Alderton, Roberto Amato, Jeffrey Barrett, Sonia Goncalves, Ewan Harrison, David K. Jackson, Ian Johnston, Dominic Kwiatkowski, Cordelia Langford, John Sillitoe on behalf of the Wellcome Sanger Institute COVID-19 Surveillance Team |
| EPI_ISL_3379103 | Lighthouse Lab in Glasgow | Wellcome Sanger Institute for the COVID-19 Genomics UK (COG-UK) Consortium | Harper VanSteenhouse, Yumi Kasai, David Gray, Carol Clugston, Anna Dominiczak and Alex Alderton, Roberto Amato, Jeffrey Barrett, Sonia Goncalves, Ewan Harrison, David K. Jackson, Ian Johnston, Dominic Kwiatkowski, Cordelia Langford, John Sillitoe on behalf of the Wellcome Sanger Institute COVID-19 Surveillance Team |
| EPI_ISL_3379104 | Lighthouse Lab in Milton Keynes | Wellcome Sanger Institute for the COVID-19 Genomics UK (COG-UK) Consortium | The Lighthouse Lab in Milton Keynes and Alex Alderton, Roberto Amato, Jeffrey Barrett, Sonia Goncalves, Ewan Harrison, David K. Jackson, Ian Johnston, Dominic Kwiatkowski, Cordelia Langford, John Sillitoe on behalf of the Wellcome Sanger Institute COVID-19 Surveillance Team |
| EPI_ISL_3379105 | LABORATOIRE ACCOLAB SUD OUEST | CNR Virus des Infections Respiratoires - France SUD | Antonin Bal, Gregory Destras, Gwendolyne Burfin, Hadrien Regue, Quentin Semanas, Martine Valette, Bruno Lina, Laurence Josset |
| EPI_ISL_3379106 | Lighthouse Lab in Milton Keynes | Wellcome Sanger Institute for the COVID-19 Genomics UK (COG-UK) Consortium | The Lighthouse Lab in Milton Keynes and Alex Alderton, Roberto Amato, Jeffrey Barrett, Sonia Goncalves, Ewan Harrison, David K. Jackson, Ian Johnston, Dominic Kwiatkowski, Cordelia Langford, John Sillitoe on behalf of the Wellcome Sanger Institute COVID-19 Surveillance Team |
| EPI_ISL_3379107 | Lighthouse Lab in Alderley Park | Wellcome Sanger Institute for the COVID-19 Genomics UK (COG-UK) | Jacquelyn Wynn, Mairead Hyland, The Lighthouse Lab in Alderley Park and Alex Alderton, Roberto Amato, Jeffrey Barrett, Sonia Goncalves, Ewan Harrison, David K. Jackson, Ian Johnston, Dominic Kwiatkowski, Cordelia Langford, John Sillitoe on |

| | | | |
|---|---|---|---|
| | | Consortium | behalf of the Wellcome Sanger Institute COVID-19 Surveillance Team |
| EPI_ISL_3379108 | Lighthouse Lab in Glasgow | Wellcome Sanger Institute for the COVID-19 Genomics UK (COG-UK) Consortium | Harper VanSteenhouse, Yumi Kasai, David Gray, Carol Clugston, Anna Dominiczak and Alex Alderton, Roberto Amato, Jeffrey Barrett, Sonia Goncalves, Ewan Harrison, David K. Jackson, Ian Johnston, Dominic Kwiatkowski, Cordelia Langford, John Sillitoe on behalf of the Wellcome Sanger Institute COVID-19 Surveillance Team |
| EPI_ISL_3379110 | Lighthouse Lab in Milton Keynes | Wellcome Sanger Institute for the COVID-19 Genomics UK (COG-UK) Consortium | The Lighthouse Lab in Milton Keynes and Alex Alderton, Roberto Amato, Jeffrey Barrett, Sonia Goncalves, Ewan Harrison, David K. Jackson, Ian Johnston, Dominic Kwiatkowski, Cordelia Langford, John Sillitoe on behalf of the Wellcome Sanger Institute COVID-19 Surveillance Team |
| EPI_ISL_3379111, EPI_ISL_3379112 | Lighthouse Lab in Alderley Park | Wellcome Sanger Institute for the COVID-19 Genomics UK (COG-UK) Consortium | Jacquelyn Wynn, Mairead Hyland, The Lighthouse Lab in Alderley Park and Alex Alderton, Roberto Amato, Jeffrey Barrett, Sonia Goncalves, Ewan Harrison, David K. Jackson, Ian Johnston, Dominic Kwiatkowski, Cordelia Langford, John Sillitoe on behalf of the Wellcome Sanger Institute COVID-19 Surveillance Team |
| EPI_ISL_3379113 | LABORATOIRE ACCOLAB SUD OUEST | CNR Virus des Infections Respiratoires - France SUD | Antonin Bal, Gregory Destras, Gwendolyne Burfin, Hadrien Regue, Quentin Semanas, Martine Valette, Bruno Lina, Laurence Josset |
| EPI_ISL_3379114 | Lighthouse Lab in Glasgow | Wellcome Sanger Institute for the COVID-19 Genomics UK (COG-UK) Consortium | Harper VanSteenhouse, Yumi Kasai, David Gray, Carol Clugston, Anna Dominiczak and Alex Alderton, Roberto Amato, Jeffrey Barrett, Sonia Goncalves, Ewan Harrison, David K. Jackson, Ian Johnston, Dominic Kwiatkowski, Cordelia Langford, John Sillitoe on behalf of the Wellcome Sanger Institute COVID-19 Surveillance Team |
| EPI_ISL_3379115 | Lighthouse Lab in Milton Keynes | Wellcome Sanger Institute for the COVID-19 Genomics UK (COG-UK) Consortium | The Lighthouse Lab in Milton Keynes and Alex Alderton, Roberto Amato, Jeffrey Barrett, Sonia Goncalves, Ewan Harrison, David K. Jackson, Ian Johnston, Dominic Kwiatkowski, Cordelia Langford, John Sillitoe on behalf of the Wellcome Sanger Institute COVID-19 Surveillance Team |
| EPI_ISL_3379116 | Lighthouse Lab in Alderley Park | Wellcome Sanger Institute for the COVID-19 Genomics UK (COG-UK) Consortium | Jacquelyn Wynn, Mairead Hyland, The Lighthouse Lab in Alderley Park and Alex Alderton, Roberto Amato, Jeffrey Barrett, Sonia Goncalves, Ewan Harrison, David K. Jackson, Ian Johnston, Dominic Kwiatkowski, Cordelia Langford, John Sillitoe on behalf of the Wellcome Sanger Institute COVID-19 Surveillance Team |
| EPI_ISL_3379117 | LABORATOIRE ACCOLAB SUD OUEST | CNR Virus des Infections Respiratoires - France SUD | Antonin Bal, Gregory Destras, Gwendolyne Burfin, Hadrien Regue, Quentin Semanas, Martine Valette, Bruno Lina, Laurence Josset |
| EPI_ISL_3379118 | Lighthouse Lab in Milton Keynes | Wellcome Sanger Institute for the COVID-19 Genomics UK (COG-UK) Consortium | The Lighthouse Lab in Milton Keynes and Alex Alderton, Roberto Amato, Jeffrey Barrett, Sonia Goncalves, Ewan Harrison, David K. Jackson, Ian Johnston, Dominic Kwiatkowski, Cordelia Langford, John Sillitoe on behalf of the Wellcome Sanger Institute COVID-19 Surveillance Team |
| EPI_ISL_3379119, EPI_ISL_3379120 | Lighthouse Lab in Glasgow | Wellcome Sanger Institute for the COVID-19 Genomics UK (COG-UK) Consortium | Harper VanSteenhouse, Yumi Kasai, David Gray, Carol Clugston, Anna Dominiczak and Alex Alderton, Roberto Amato, Jeffrey Barrett, Sonia Goncalves, Ewan Harrison, David K. Jackson, Ian Johnston, Dominic Kwiatkowski, Cordelia Langford, John Sillitoe on behalf of the Wellcome Sanger Institute COVID-19 Surveillance Team |
| EPI_ISL_3379121 | LABORATOIRE ACCOLAB SUD OUEST | CNR Virus des Infections Respiratoires - France SUD | Antonin Bal, Gregory Destras, Gwendolyne Burfin, Hadrien Regue, Quentin Semanas, Martine Valette, Bruno Lina, Laurence Josset |
| EPI_ISL_3379122 | Berkshire and Surrey Pathology Services Lighthouse Laboratory | Wellcome Sanger Institute for the COVID-19 Genomics UK (COG-UK) Consortium | Berkshire and Surrey Pathology Services Lighthouse Laboratory and Alex Alderton, Roberto Amato, Jeffrey Barrett, Sonia Goncalves, Ewan Harrison, David K. Jackson, Ian Johnston, Dominic Kwiatkowski, Cordelia Langford, John Sillitoe on behalf of the Wellcome Sanger Institute COVID-19 Surveillance Team |
| EPI_ISL_3379123 | Lighthouse Lab in Alderley Park | Wellcome Sanger Institute for the COVID-19 Genomics UK (COG-UK) Consortium | Jacquelyn Wynn, Mairead Hyland, The Lighthouse Lab in Alderley Park and Alex Alderton, Roberto Amato, Jeffrey Barrett, Sonia Goncalves, Ewan Harrison, David K. Jackson, Ian Johnston, Dominic Kwiatkowski, Cordelia Langford, John Sillitoe on behalf of the Wellcome Sanger Institute COVID-19 Surveillance Team |
| EPI_ISL_3379124 | ORIAPOLE | CNR Virus des Infections Respiratoires - France SUD | Antonin Bal, Gregory Destras, Gwendolyne Burfin, Hadrien Regue, Quentin Semanas, Martine Valette, Bruno Lina, Laurence Josset |
| EPI_ISL_3379125 | Berkshire and Surrey Pathology Services Lighthouse Laboratory | Wellcome Sanger Institute for the COVID-19 Genomics UK (COG-UK) Consortium | Berkshire and Surrey Pathology Services Lighthouse Laboratory and Alex Alderton, Roberto Amato, Jeffrey Barrett, Sonia Goncalves, Ewan Harrison, David K. Jackson, Ian Johnston, Dominic Kwiatkowski, Cordelia Langford, John Sillitoe on behalf of the Wellcome Sanger Institute COVID-19 Surveillance Team |
| EPI_ISL_3379126 | Lighthouse Lab in Alderley Park | Wellcome Sanger Institute for the COVID-19 Genomics UK (COG-UK) Consortium | Jacquelyn Wynn, Mairead Hyland, The Lighthouse Lab in Alderley Park and Alex Alderton, Roberto Amato, Jeffrey Barrett, Sonia Goncalves, Ewan Harrison, David K. Jackson, Ian Johnston, Dominic Kwiatkowski, Cordelia Langford, John Sillitoe on behalf of the Wellcome Sanger Institute COVID-19 Surveillance Team |
| EPI_ISL_3379127 | Lighthouse Lab in Glasgow | Wellcome Sanger Institute for the COVID-19 Genomics UK (COG-UK) Consortium | Harper VanSteenhouse, Yumi Kasai, David Gray, Carol Clugston, Anna Dominiczak and Alex Alderton, Roberto Amato, Jeffrey Barrett, Sonia Goncalves, Ewan Harrison, David K. Jackson, Ian Johnston, Dominic Kwiatkowski, Cordelia Langford, John Sillitoe on behalf of the Wellcome Sanger Institute COVID-19 Surveillance Team |
| EPI_ISL_3379128 | LABORATOIRE ACCOLAB SUD OUEST | CNR Virus des Infections Respiratoires - France SUD | Antonin Bal, Gregory Destras, Gwendolyne Burfin, Hadrien Regue, Quentin Semanas, Martine Valette, Bruno Lina, Laurence Josset |
| EPI_ISL_3379129, EPI_ISL_3379130 | Lighthouse Lab in Milton Keynes | Wellcome Sanger Institute for the COVID-19 Genomics UK (COG-UK) Consortium | The Lighthouse Lab in Milton Keynes and Alex Alderton, Roberto Amato, Jeffrey Barrett, Sonia Goncalves, Ewan Harrison, David K. Jackson, Ian Johnston, Dominic Kwiatkowski, Cordelia Langford, John Sillitoe on behalf of the Wellcome Sanger Institute COVID-19 Surveillance Team |
| EPI_ISL_3379131 | LABORATOIRE ACCOLAB SUD OUEST | CNR Virus des Infections Respiratoires - France SUD | Antonin Bal, Gregory Destras, Gwendolyne Burfin, Hadrien Regue, Quentin Semanas, Martine Valette, Bruno Lina, Laurence Josset |
| EPI_ISL_3379132 | Lighthouse Lab in Alderley Park | Wellcome Sanger Institute for the COVID-19 Genomics UK (COG-UK) Consortium | Jacquelyn Wynn, Mairead Hyland, The Lighthouse Lab in Alderley Park and Alex Alderton, Roberto Amato, Jeffrey Barrett, Sonia Goncalves, Ewan Harrison, David K. Jackson, Ian Johnston, Dominic Kwiatkowski, Cordelia Langford, John Sillitoe on behalf of the Wellcome Sanger Institute COVID-19 Surveillance Team |
| EPI_ISL_3379133 | Berkshire and Surrey Pathology Services Lighthouse Laboratory | Wellcome Sanger Institute for the COVID-19 Genomics UK (COG-UK) Consortium | Berkshire and Surrey Pathology Services Lighthouse Laboratory and Alex Alderton, Roberto Amato, Jeffrey Barrett, Sonia Goncalves, Ewan Harrison, David K. Jackson, Ian Johnston, Dominic Kwiatkowski, Cordelia Langford, John Sillitoe on behalf of the Wellcome Sanger Institute COVID-19 Surveillance Team |
| EPI_ISL_3379134 | Lighthouse Lab in Milton Keynes | Wellcome Sanger Institute for the COVID-19 Genomics UK (COG-UK) Consortium | The Lighthouse Lab in Milton Keynes and Alex Alderton, Roberto Amato, Jeffrey Barrett, Sonia Goncalves, Ewan Harrison, David K. Jackson, Ian Johnston, Dominic Kwiatkowski, Cordelia Langford, John Sillitoe on behalf of the Wellcome Sanger Institute COVID-19 Surveillance Team |
| EPI_ISL_3379135 | LABORATOIRE ACCOLAB SUD OUEST | CNR Virus des Infections Respiratoires - France SUD | Antonin Bal, Gregory Destras, Gwendolyne Burfin, Hadrien Regue, Quentin Semanas, Martine Valette, Bruno Lina, Laurence Josset |
| EPI_ISL_3379136 | Lighthouse Lab in Glasgow | Wellcome Sanger Institute for the COVID-19 Genomics UK (COG-UK) Consortium | Harper VanSteenhouse, Yumi Kasai, David Gray, Carol Clugston, Anna Dominiczak and Alex Alderton, Roberto Amato, Jeffrey Barrett, Sonia Goncalves, Ewan Harrison, David K. Jackson, Ian Johnston, Dominic Kwiatkowski, Cordelia Langford, John Sillitoe on behalf of the Wellcome Sanger Institute COVID-19 Surveillance Team |
| EPI_ISL_3379137 | Lighthouse Lab in Alderley Park | Wellcome Sanger Institute for the COVID-19 Genomics UK (COG-UK) Consortium | Jacquelyn Wynn, Mairead Hyland, The Lighthouse Lab in Alderley Park and Alex Alderton, Roberto Amato, Jeffrey Barrett, Sonia Goncalves, Ewan Harrison, David K. Jackson, Ian Johnston, Dominic Kwiatkowski, Cordelia Langford, John Sillitoe on behalf of the Wellcome Sanger Institute COVID-19 Surveillance Team |
| EPI_ISL_3379138 | Lighthouse Lab in Milton Keynes | Wellcome Sanger Institute for the COVID-19 Genomics UK (COG-UK) Consortium | The Lighthouse Lab in Milton Keynes and Alex Alderton, Roberto Amato, Jeffrey Barrett, Sonia Goncalves, Ewan Harrison, David K. Jackson, Ian Johnston, Dominic Kwiatkowski, Cordelia Langford, John Sillitoe on behalf of the Wellcome Sanger Institute COVID-19 Surveillance Team |
| EPI_ISL_3379139 | LABORATOIRE ACCOLAB SUD OUEST | CNR Virus des Infections Respiratoires - France SUD | Antonin Bal, Gregory Destras, Gwendolyne Burfin, Hadrien Regue, Quentin Semanas, Martine Valette, Bruno Lina, Laurence Josset |
| EPI_ISL_3379140 | Lighthouse Lab in Alderley Park | Wellcome Sanger Institute for the COVID-19 Genomics UK (COG-UK) Consortium | Jacquelyn Wynn, Mairead Hyland, The Lighthouse Lab in Alderley Park and Alex Alderton, Roberto Amato, Jeffrey Barrett, Sonia Goncalves, Ewan Harrison, David K. Jackson, Ian Johnston, Dominic Kwiatkowski, Cordelia Langford, John Sillitoe on behalf of the Wellcome Sanger Institute COVID-19 Surveillance Team |
| EPI_ISL_3379141 | Lighthouse Lab in Milton Keynes | Wellcome Sanger Institute for the COVID-19 Genomics UK (COG-UK) Consortium | The Lighthouse Lab in Milton Keynes and Alex Alderton, Roberto Amato, Jeffrey Barrett, Sonia Goncalves, Ewan Harrison, David K. Jackson, Ian Johnston, Dominic Kwiatkowski, Cordelia Langford, John Sillitoe on behalf of the Wellcome Sanger Institute COVID-19 Surveillance Team |
| EPI_ISL_3379142 | Lighthouse Lab in Glasgow | Wellcome Sanger Institute for the COVID-19 Genomics UK (COG-UK) Consortium | Harper VanSteenhouse, Yumi Kasai, David Gray, Carol Clugston, Anna Dominiczak and Alex Alderton, Roberto Amato, Jeffrey Barrett, Sonia Goncalves, Ewan Harrison, David K. Jackson, Ian Johnston, Dominic Kwiatkowski, Cordelia Langford, John Sillitoe on behalf of the Wellcome Sanger Institute COVID-19 Surveillance Team |
| EPI_ISL_3379143 | LABORATOIRE ACCOLAB SUD OUEST | CNR Virus des Infections Respiratoires - France SUD | Antonin Bal, Gregory Destras, Gwendolyne Burfin, Hadrien Regue, Quentin Semanas, Martine Valette, Bruno Lina, Laurence Josset |
| EPI_ISL_3379144 | Lighthouse Lab in Glasgow | Wellcome Sanger Institute for the COVID-19 Genomics UK (COG-UK) Consortium | Harper VanSteenhouse, Yumi Kasai, David Gray, Carol Clugston, Anna Dominiczak and Alex Alderton, Roberto Amato, Jeffrey Barrett, Sonia Goncalves, Ewan Harrison, David K. Jackson, Ian Johnston, Dominic Kwiatkowski, Cordelia Langford, John Sillitoe on behalf of the Wellcome Sanger Institute COVID-19 Surveillance Team |
| EPI_ISL_3379145, EPI_ISL_3379146 | Berkshire and Surrey Pathology Services Lighthouse Laboratory | Wellcome Sanger Institute for the COVID-19 Genomics UK (COG-UK) Consortium | Berkshire and Surrey Pathology Services Lighthouse Laboratory and Alex Alderton, Roberto Amato, Jeffrey Barrett, Sonia Goncalves, Ewan Harrison, David K. Jackson, Ian Johnston, Dominic Kwiatkowski, Cordelia Langford, John Sillitoe on behalf of the Wellcome Sanger Institute COVID-19 Surveillance Team |
| EPI_ISL_3379147, EPI_ISL_3379148, EPI_ISL_3379149 | Lighthouse Lab in Alderley Park | Wellcome Sanger Institute for the COVID-19 Genomics UK (COG-UK) Consortium | Jacquelyn Wynn, Mairead Hyland, The Lighthouse Lab in Alderley Park and Alex Alderton, Roberto Amato, Jeffrey Barrett, Sonia Goncalves, Ewan Harrison, David K. Jackson, Ian Johnston, Dominic Kwiatkowski, Cordelia Langford, John Sillitoe on behalf of the Wellcome Sanger Institute COVID-19 Surveillance Team |
| EPI_ISL_3379150 | Lighthouse Lab in Glasgow | Wellcome Sanger Institute for the COVID-19 Genomics UK (COG-UK) Consortium | Harper VanSteenhouse, Yumi Kasai, David Gray, Carol Clugston, Anna Dominiczak and Alex Alderton, Roberto Amato, Jeffrey Barrett, Sonia Goncalves, Ewan Harrison, David K. Jackson, Ian Johnston, Dominic Kwiatkowski, Cordelia Langford, John Sillitoe on behalf of the Wellcome Sanger Institute COVID-19 Surveillance Team |
| EPI_ISL_3379151 | LABORATOIRE ACCOLAB SUD OUEST | CNR Virus des Infections Respiratoires - France SUD | Antonin Bal, Gregory Destras, Gwendolyne Burfin, Hadrien Regue, Quentin Semanas, Martine Valette, Bruno Lina, Laurence Josset |
| EPI_ISL_3379152, EPI_ISL_3379153 | Lighthouse Lab in Alderley Park | Wellcome Sanger Institute for the COVID-19 Genomics UK (COG-UK) Consortium | Jacquelyn Wynn, Mairead Hyland, The Lighthouse Lab in Alderley Park and Alex Alderton, Roberto Amato, Jeffrey Barrett, Sonia Goncalves, Ewan Harrison, David K. Jackson, Ian Johnston, Dominic Kwiatkowski, Cordelia Langford, John Sillitoe on behalf of the Wellcome Sanger Institute COVID-19 Surveillance Team |
| EPI_ISL_3379154 | Lighthouse Lab in Glasgow | Wellcome Sanger Institute for the COVID-19 Genomics UK (COG-UK) Consortium | Harper VanSteenhouse, Yumi Kasai, David Gray, Carol Clugston, Anna Dominiczak and Alex Alderton, Roberto Amato, Jeffrey Barrett, Sonia Goncalves, Ewan Harrison, David K. Jackson, Ian Johnston, Dominic Kwiatkowski, Cordelia Langford, John Sillitoe on behalf of the Wellcome Sanger Institute COVID-19 Surveillance Team |
| EPI_ISL_3379155 | Lighthouse Lab in Milton Keynes | Wellcome Sanger Institute for the COVID-19 Genomics UK (COG-UK) Consortium | The Lighthouse Lab in Milton Keynes and Alex Alderton, Roberto Amato, Jeffrey Barrett, Sonia Goncalves, Ewan Harrison, David K. Jackson, Ian Johnston, Dominic Kwiatkowski, Cordelia Langford, John Sillitoe on behalf of the Wellcome Sanger Institute COVID-19 Surveillance Team |
| EPI_ISL_3379156 | Lighthouse Lab in Alderley Park | Wellcome Sanger Institute for the COVID-19 Genomics UK (COG-UK) Consortium | Jacquelyn Wynn, Mairead Hyland, The Lighthouse Lab in Alderley Park and Alex Alderton, Roberto Amato, Jeffrey Barrett, Sonia Goncalves, Ewan Harrison, David K. Jackson, Ian Johnston, Dominic Kwiatkowski, Cordelia Langford, John Sillitoe on behalf of the Wellcome Sanger Institute COVID-19 Surveillance Team |
| EPI_ISL_3379157, EPI_ISL_3379158 | Lighthouse Lab in Glasgow | Wellcome Sanger Institute for the COVID-19 Genomics UK (COG-UK) Consortium | Harper VanSteenhouse, Yumi Kasai, David Gray, Carol Clugston, Anna Dominiczak and Alex Alderton, Roberto Amato, Jeffrey Barrett, Sonia Goncalves, Ewan Harrison, David K. Jackson, Ian Johnston, Dominic Kwiatkowski, Cordelia Langford, John Sillitoe on behalf of the Wellcome Sanger Institute COVID-19 Surveillance Team |
| EPI_ISL_3379159 | LABORATOIRE ACCOLAB SUD OUEST | CNR Virus des Infections Respiratoires - France SUD | Antonin Bal, Gregory Destras, Gwendolyne Burfin, Hadrien Regue, Quentin Semanas, Martine Valette, Bruno Lina, Laurence Josset |
| EPI_ISL_3379160 | Lighthouse Lab in Alderley Park | Wellcome Sanger Institute for the COVID-19 Genomics UK (COG-UK) Consortium | Jacquelyn Wynn, Mairead Hyland, The Lighthouse Lab in Alderley Park and Alex Alderton, Roberto Amato, Jeffrey Barrett, Sonia Goncalves, Ewan Harrison, David K. Jackson, Ian Johnston, Dominic Kwiatkowski, Cordelia Langford, John Sillitoe on behalf of the Wellcome Sanger Institute COVID-19 Surveillance Team |
| EPI_ISL_3379161 | Lighthouse Lab in Milton Keynes | Wellcome Sanger Institute for the COVID-19 Genomics UK (COG-UK) Consortium | The Lighthouse Lab in Milton Keynes and Alex Alderton, Roberto Amato, Jeffrey Barrett, Sonia Goncalves, Ewan Harrison, David K. Jackson, Ian Johnston, Dominic Kwiatkowski, Cordelia Langford, John Sillitoe on behalf of the Wellcome Sanger Institute COVID-19 Surveillance Team |
| EPI_ISL_3379162, EPI_ISL_3379163 | Lighthouse Lab in Alderley Park | Wellcome Sanger Institute for the COVID-19 Genomics UK (COG-UK) Consortium | Jacquelyn Wynn, Mairead Hyland, The Lighthouse Lab in Alderley Park and Alex Alderton, Roberto Amato, Jeffrey Barrett, Sonia Goncalves, Ewan Harrison, David K. Jackson, Ian Johnston, Dominic Kwiatkowski, Cordelia Langford, John Sillitoe on behalf of the Wellcome Sanger Institute COVID-19 Surveillance Team |
| EPI_ISL_3379164 | Lighthouse Lab in Glasgow | Wellcome Sanger Institute for the COVID-19 Genomics UK (COG-UK) Consortium | Harper VanSteenhouse, Yumi Kasai, David Gray, Carol Clugston, Anna Dominiczak and Alex Alderton, Roberto Amato, Jeffrey Barrett, Sonia Goncalves, Ewan Harrison, David K. Jackson, Ian Johnston, Dominic Kwiatkowski, Cordelia Langford, John Sillitoe on behalf of the Wellcome Sanger Institute COVID-19 Surveillance Team |
| EPI_ISL_3379165 | Lighthouse Lab in Milton Keynes | Wellcome Sanger Institute for the COVID-19 Genomics UK (COG-UK) Consortium | The Lighthouse Lab in Milton Keynes and Alex Alderton, Roberto Amato, Jeffrey Barrett, Sonia Goncalves, Ewan Harrison, David K. Jackson, Ian Johnston, Dominic Kwiatkowski, Cordelia Langford, John Sillitoe on behalf of the Wellcome Sanger Institute COVID-19 Surveillance Team |
| EPI_ISL_3379166 | LABORATOIRE ACCOLAB SUD OUEST | CNR Virus des Infections Respiratoires - France SUD | Antonin Bal, Gregory Destras, Gwendolyne Burfin, Hadrien Regue, Quentin Semanas, Martine Valette, Bruno Lina, Laurence Josset |

| | | | |
|---|---|---|---|
| EPI_ISL_3379167 | Berkshire and Surrey Pathology Services Lighthouse Laboratory | Wellcome Sanger Institute for the COVID-19 Genomics UK (COG-UK) Consortium | Berkshire and Surrey Pathology Services Lighthouse Laboratory and Alex Alderton, Roberto Amato, Jeffrey Barrett, Sonia Goncalves, Ewan Harrison, David K. Jackson, Ian Johnston, Dominic Kwiatkowski, Cordelia Langford, John Sillitoe on behalf of the Wellcome Sanger Institute COVID-19 Surveillance Team |
| EPI_ISL_3379168 | Lighthouse Lab in Glasgow | Wellcome Sanger Institute for the COVID-19 Genomics UK (COG-UK) Consortium | Harper VanSteenhouse, Yumi Kasai, David Gray, Carol Clugston, Anna Dominiczak and Alex Alderton, Roberto Amato, Jeffrey Barrett, Sonia Goncalves, Ewan Harrison, David K. Jackson, Ian Johnston, Dominic Kwiatkowski, Cordelia Langford, John Sillitoe on behalf of the Wellcome Sanger Institute COVID-19 Surveillance Team |
| EPI_ISL_3379169 | Lighthouse Lab in Alderley Park | Wellcome Sanger Institute for the COVID-19 Genomics UK (COG-UK) Consortium | Jacquelyn Wynn, Mairead Hyland, The Lighthouse Lab in Alderley Park and Alex Alderton, Roberto Amato, Jeffrey Barrett, Sonia Goncalves, Ewan Harrison, David K. Jackson, Ian Johnston, Dominic Kwiatkowski, Cordelia Langford, John Sillitoe on behalf of the Wellcome Sanger Institute COVID-19 Surveillance Team |
| EPI_ISL_3379170 | LABORATOIRE ACCOLAB SUD OUEST | CNR Virus des Infections Respiratoires - France SUD | Antonin Bal, Gregory Destras, Gwendolyne Burfin, Hadrien Regue, Quentin Semanas, Martine Valette, Bruno Lina, Laurence Josset |
| EPI_ISL_3379171 | Berkshire and Surrey Pathology Services Lighthouse Laboratory | Wellcome Sanger Institute for the COVID-19 Genomics UK (COG-UK) Consortium | Berkshire and Surrey Pathology Services Lighthouse Laboratory and Alex Alderton, Roberto Amato, Jeffrey Barrett, Sonia Goncalves, Ewan Harrison, David K. Jackson, Ian Johnston, Dominic Kwiatkowski, Cordelia Langford, John Sillitoe on behalf of the Wellcome Sanger Institute COVID-19 Surveillance Team |
| EPI_ISL_3379172 | Lighthouse Lab in Alderley Park | Wellcome Sanger Institute for the COVID-19 Genomics UK (COG-UK) Consortium | Jacquelyn Wynn, Mairead Hyland, The Lighthouse Lab in Alderley Park and Alex Alderton, Roberto Amato, Jeffrey Barrett, Sonia Goncalves, Ewan Harrison, David K. Jackson, Ian Johnston, Dominic Kwiatkowski, Cordelia Langford, John Sillitoe on behalf of the Wellcome Sanger Institute COVID-19 Surveillance Team |
| EPI_ISL_3379173 | Berkshire and Surrey Pathology Services Lighthouse Laboratory | Wellcome Sanger Institute for the COVID-19 Genomics UK (COG-UK) Consortium | Berkshire and Surrey Pathology Services Lighthouse Laboratory and Alex Alderton, Roberto Amato, Jeffrey Barrett, Sonia Goncalves, Ewan Harrison, David K. Jackson, Ian Johnston, Dominic Kwiatkowski, Cordelia Langford, John Sillitoe on behalf of the Wellcome Sanger Institute COVID-19 Surveillance Team |
| EPI_ISL_3379174 | LABORATOIRE ACCOLAB SUD OUEST | CNR Virus des Infections Respiratoires - France SUD | Antonin Bal, Gregory Destras, Gwendolyne Burfin, Hadrien Regue, Quentin Semanas, Martine Valette, Bruno Lina, Laurence Josset |
| EPI_ISL_3379175 | Berkshire and Surrey Pathology Services Lighthouse Laboratory | Wellcome Sanger Institute for the COVID-19 Genomics UK (COG-UK) Consortium | Berkshire and Surrey Pathology Services Lighthouse Laboratory and Alex Alderton, Roberto Amato, Jeffrey Barrett, Sonia Goncalves, Ewan Harrison, David K. Jackson, Ian Johnston, Dominic Kwiatkowski, Cordelia Langford, John Sillitoe on behalf of the Wellcome Sanger Institute COVID-19 Surveillance Team |
| EPI_ISL_3379176, EPI_ISL_3379177 | Lighthouse Lab in Glasgow | Wellcome Sanger Institute for the COVID-19 Genomics UK (COG-UK) Consortium | Harper VanSteenhouse, Yumi Kasai, David Gray, Carol Clugston, Anna Dominiczak and Alex Alderton, Roberto Amato, Jeffrey Barrett, Sonia Goncalves, Ewan Harrison, David K. Jackson, Ian Johnston, Dominic Kwiatkowski, Cordelia Langford, John Sillitoe on behalf of the Wellcome Sanger Institute COVID-19 Surveillance Team |
| EPI_ISL_3379178 | LABORATOIRE ACCOLAB SUD OUEST | CNR Virus des Infections Respiratoires - France SUD | Antonin Bal, Gregory Destras, Gwendolyne Burfin, Hadrien Regue, Quentin Semanas, Martine Valette, Bruno Lina, Laurence Josset |
| EPI_ISL_3379179, EPI_ISL_3379180 | Berkshire and Surrey Pathology Services Lighthouse Laboratory | Wellcome Sanger Institute for the COVID-19 Genomics UK (COG-UK) Consortium | Berkshire and Surrey Pathology Services Lighthouse Laboratory and Alex Alderton, Roberto Amato, Jeffrey Barrett, Sonia Goncalves, Ewan Harrison, David K. Jackson, Ian Johnston, Dominic Kwiatkowski, Cordelia Langford, John Sillitoe on behalf of the Wellcome Sanger Institute COVID-19 Surveillance Team |
| EPI_ISL_3379181 | Lighthouse Lab in Milton Keynes | Wellcome Sanger Institute for the COVID-19 Genomics UK (COG-UK) Consortium | The Lighthouse Lab in Milton Keynes and Alex Alderton, Roberto Amato, Jeffrey Barrett, Sonia Goncalves, Ewan Harrison, David K. Jackson, Ian Johnston, Dominic Kwiatkowski, Cordelia Langford, John Sillitoe on behalf of the Wellcome Sanger Institute COVID-19 Surveillance Team |
| EPI_ISL_3379182 | LABORATOIRE ACCOLAB SUD OUEST | CNR Virus des Infections Respiratoires - France SUD | Antonin Bal, Gregory Destras, Gwendolyne Burfin, Hadrien Regue, Quentin Semanas, Martine Valette, Bruno Lina, Laurence Josset |
| EPI_ISL_3379183 | Lighthouse Lab in Alderley Park | Wellcome Sanger Institute for the COVID-19 Genomics UK (COG-UK) Consortium | Jacquelyn Wynn, Mairead Hyland, The Lighthouse Lab in Alderley Park and Alex Alderton, Roberto Amato, Jeffrey Barrett, Sonia Goncalves, Ewan Harrison, David K. Jackson, Ian Johnston, Dominic Kwiatkowski, Cordelia Langford, John Sillitoe on behalf of the Wellcome Sanger Institute COVID-19 Surveillance Team |
| EPI_ISL_3379184 | Berkshire and Surrey Pathology Services Lighthouse Laboratory | Wellcome Sanger Institute for the COVID-19 Genomics UK (COG-UK) Consortium | Berkshire and Surrey Pathology Services Lighthouse Laboratory and Alex Alderton, Roberto Amato, Jeffrey Barrett, Sonia Goncalves, Ewan Harrison, David K. Jackson, Ian Johnston, Dominic Kwiatkowski, Cordelia Langford, John Sillitoe on behalf of the Wellcome Sanger Institute COVID-19 Surveillance Team |
| EPI_ISL_3379185 | Lighthouse Lab in Milton Keynes | Wellcome Sanger Institute for the COVID-19 Genomics UK (COG-UK) Consortium | The Lighthouse Lab in Milton Keynes and Alex Alderton, Roberto Amato, Jeffrey Barrett, Sonia Goncalves, Ewan Harrison, David K. Jackson, Ian Johnston, Dominic Kwiatkowski, Cordelia Langford, John Sillitoe on behalf of the Wellcome Sanger Institute COVID-19 Surveillance Team |
| EPI_ISL_3379186 | LABORATOIRE ACCOLAB SUD OUEST | CNR Virus des Infections Respiratoires - France SUD | Antonin Bal, Gregory Destras, Gwendolyne Burfin, Hadrien Regue, Quentin Semanas, Martine Valette, Bruno Lina, Laurence Josset |
| EPI_ISL_3379187, EPI_ISL_3379188 | Berkshire and Surrey Pathology Services Lighthouse Laboratory | Wellcome Sanger Institute for the COVID-19 Genomics UK (COG-UK) Consortium | Berkshire and Surrey Pathology Services Lighthouse Laboratory and Alex Alderton, Roberto Amato, Jeffrey Barrett, Sonia Goncalves, Ewan Harrison, David K. Jackson, Ian Johnston, Dominic Kwiatkowski, Cordelia Langford, John Sillitoe on behalf of the Wellcome Sanger Institute COVID-19 Surveillance Team |
| EPI_ISL_3379189 | LABORATOIRE ACCOLAB SUD OUEST | CNR Virus des Infections Respiratoires - France SUD | Antonin Bal, Gregory Destras, Gwendolyne Burfin, Hadrien Regue, Quentin Semanas, Martine Valette, Bruno Lina, Laurence Josset |
| EPI_ISL_3379190 | Berkshire and Surrey Pathology Services Lighthouse Laboratory | Wellcome Sanger Institute for the COVID-19 Genomics UK (COG-UK) Consortium | Berkshire and Surrey Pathology Services Lighthouse Laboratory and Alex Alderton, Roberto Amato, Jeffrey Barrett, Sonia Goncalves, Ewan Harrison, David K. Jackson, Ian Johnston, Dominic Kwiatkowski, Cordelia Langford, John Sillitoe on behalf of the Wellcome Sanger Institute COVID-19 Surveillance Team |
| EPI_ISL_3379191, EPI_ISL_3379192 | Lighthouse Lab in Alderley Park | Wellcome Sanger Institute for the COVID-19 Genomics UK (COG-UK) Consortium | Jacquelyn Wynn, Mairead Hyland, The Lighthouse Lab in Alderley Park and Alex Alderton, Roberto Amato, Jeffrey Barrett, Sonia Goncalves, Ewan Harrison, David K. Jackson, Ian Johnston, Dominic Kwiatkowski, Cordelia Langford, John Sillitoe on behalf of the Wellcome Sanger Institute COVID-19 Surveillance Team |
| EPI_ISL_3379193 | LABORATOIRE ACCOLAB SUD OUEST | CNR Virus des Infections Respiratoires - France SUD | Antonin Bal, Gregory Destras, Gwendolyne Burfin, Hadrien Regue, Quentin Semanas, Martine Valette, Bruno Lina, Laurence Josset |
| EPI_ISL_3379194, EPI_ISL_3379195 | Lighthouse Lab in Milton Keynes | Wellcome Sanger Institute for the COVID-19 Genomics UK (COG-UK) Consortium | The Lighthouse Lab in Milton Keynes and Alex Alderton, Roberto Amato, Jeffrey Barrett, Sonia Goncalves, Ewan Harrison, David K. Jackson, Ian Johnston, Dominic Kwiatkowski, Cordelia Langford, John Sillitoe on behalf of the Wellcome Sanger Institute COVID-19 Surveillance Team |
| EPI_ISL_3379196 | Lighthouse Lab in Glasgow | Wellcome Sanger Institute for the COVID-19 Genomics UK (COG-UK) Consortium | Harper VanSteenhouse, Yumi Kasai, David Gray, Carol Clugston, Anna Dominiczak and Alex Alderton, Roberto Amato, Jeffrey Barrett, Sonia Goncalves, Ewan Harrison, David K. Jackson, Ian Johnston, Dominic Kwiatkowski, Cordelia Langford, John Sillitoe on behalf of the Wellcome Sanger Institute COVID-19 Surveillance Team |
| EPI_ISL_3379197, EPI_ISL_3379198 | Lighthouse Lab in Alderley Park | Wellcome Sanger Institute for the COVID-19 Genomics UK (COG-UK) Consortium | Jacquelyn Wynn, Mairead Hyland, The Lighthouse Lab in Alderley Park and Alex Alderton, Roberto Amato, Jeffrey Barrett, Sonia Goncalves, Ewan Harrison, David K. Jackson, Ian Johnston, Dominic Kwiatkowski, Cordelia Langford, John Sillitoe on behalf of the Wellcome Sanger Institute COVID-19 Surveillance Team |
| EPI_ISL_3379199 | Lighthouse Lab in Glasgow | Wellcome Sanger Institute for the COVID-19 Genomics UK (COG-UK) Consortium | Harper VanSteenhouse, Yumi Kasai, David Gray, Carol Clugston, Anna Dominiczak and Alex Alderton, Roberto Amato, Jeffrey Barrett, Sonia Goncalves, Ewan Harrison, David K. Jackson, Ian Johnston, Dominic Kwiatkowski, Cordelia Langford, John Sillitoe on behalf of the Wellcome Sanger Institute COVID-19 Surveillance Team |
| EPI_ISL_3379200 | Lighthouse Lab in Milton Keynes | Wellcome Sanger Institute for the COVID-19 Genomics UK (COG-UK) Consortium | The Lighthouse Lab in Milton Keynes and Alex Alderton, Roberto Amato, Jeffrey Barrett, Sonia Goncalves, Ewan Harrison, David K. Jackson, Ian Johnston, Dominic Kwiatkowski, Cordelia Langford, John Sillitoe on behalf of the Wellcome Sanger Institute COVID-19 Surveillance Team |
| EPI_ISL_3379201 | LABORATOIRE ACCOLAB SUD OUEST | CNR Virus des Infections Respiratoires - France SUD | Antonin Bal, Gregory Destras, Gwendolyne Burfin, Hadrien Regue, Quentin Semanas, Martine Valette, Bruno Lina, Laurence Josset |
| EPI_ISL_3379202 | Lighthouse Lab in Alderley Park | Wellcome Sanger Institute for the COVID-19 Genomics UK (COG-UK) Consortium | Jacquelyn Wynn, Mairead Hyland, The Lighthouse Lab in Alderley Park and Alex Alderton, Roberto Amato, Jeffrey Barrett, Sonia Goncalves, Ewan Harrison, David K. Jackson, Ian Johnston, Dominic Kwiatkowski, Cordelia Langford, John Sillitoe on behalf of the Wellcome Sanger Institute COVID-19 Surveillance Team |
| EPI_ISL_3379203 | Lighthouse Lab in Glasgow | Wellcome Sanger Institute for the COVID-19 Genomics UK (COG-UK) Consortium | Harper VanSteenhouse, Yumi Kasai, David Gray, Carol Clugston, Anna Dominiczak and Alex Alderton, Roberto Amato, Jeffrey Barrett, Sonia Goncalves, Ewan Harrison, David K. Jackson, Ian Johnston, Dominic Kwiatkowski, Cordelia Langford, John Sillitoe on behalf of the Wellcome Sanger Institute COVID-19 Surveillance Team |
| EPI_ISL_3379204 | Lighthouse Lab in Milton Keynes | Wellcome Sanger Institute for the COVID-19 Genomics UK (COG-UK) Consortium | The Lighthouse Lab in Milton Keynes and Alex Alderton, Roberto Amato, Jeffrey Barrett, Sonia Goncalves, Ewan Harrison, David K. Jackson, Ian Johnston, Dominic Kwiatkowski, Cordelia Langford, John Sillitoe on behalf of the Wellcome Sanger Institute COVID-19 Surveillance Team |
| EPI_ISL_3379205, EPI_ISL_3379206, EPI_ISL_3379207 | Lighthouse Lab in Glasgow | Wellcome Sanger Institute for the COVID-19 Genomics UK (COG-UK) Consortium | Harper VanSteenhouse, Yumi Kasai, David Gray, Carol Clugston, Anna Dominiczak and Alex Alderton, Roberto Amato, Jeffrey Barrett, Sonia Goncalves, Ewan Harrison, David K. Jackson, Ian Johnston, Dominic Kwiatkowski, Cordelia Langford, John Sillitoe on behalf of the Wellcome Sanger Institute COVID-19 Surveillance Team |
| EPI_ISL_3379208 | Lighthouse Lab in Alderley Park | Wellcome Sanger Institute for the COVID-19 Genomics UK (COG-UK) Consortium | Jacquelyn Wynn, Mairead Hyland, The Lighthouse Lab in Alderley Park and Alex Alderton, Roberto Amato, Jeffrey Barrett, Sonia Goncalves, Ewan Harrison, David K. Jackson, Ian Johnston, Dominic Kwiatkowski, Cordelia Langford, John Sillitoe on behalf of the Wellcome Sanger Institute COVID-19 Surveillance Team |
| EPI_ISL_3379209 | Lighthouse Lab in Milton Keynes | Wellcome Sanger Institute for the COVID-19 Genomics UK (COG-UK) Consortium | The Lighthouse Lab in Milton Keynes and Alex Alderton, Roberto Amato, Jeffrey Barrett, Sonia Goncalves, Ewan Harrison, David K. Jackson, Ian Johnston, Dominic Kwiatkowski, Cordelia Langford, John Sillitoe on behalf of the Wellcome Sanger Institute COVID-19 Surveillance Team |
| EPI_ISL_3379210 | LABORATOIRE ACCOLAB SUD OUEST | CNR Virus des Infections Respiratoires - France SUD | Antonin Bal, Gregory Destras, Gwendolyne Burfin, Hadrien Regue, Quentin Semanas, Martine Valette, Bruno Lina, Laurence Josset |
| EPI_ISL_3379211 | Berkshire and Surrey Pathology Services Lighthouse Laboratory | Wellcome Sanger Institute for the COVID-19 Genomics UK (COG-UK) Consortium | Berkshire and Surrey Pathology Services Lighthouse Laboratory and Alex Alderton, Roberto Amato, Jeffrey Barrett, Sonia Goncalves, Ewan Harrison, David K. Jackson, Ian Johnston, Dominic Kwiatkowski, Cordelia Langford, John Sillitoe on behalf of the Wellcome Sanger Institute COVID-19 Surveillance Team |
| EPI_ISL_3379212 | Lighthouse Lab in Glasgow | Wellcome Sanger Institute for the COVID-19 Genomics UK (COG-UK) Consortium | Harper VanSteenhouse, Yumi Kasai, David Gray, Carol Clugston, Anna Dominiczak and Alex Alderton, Roberto Amato, Jeffrey Barrett, Sonia Goncalves, Ewan Harrison, David K. Jackson, Ian Johnston, Dominic Kwiatkowski, Cordelia Langford, John Sillitoe on behalf of the Wellcome Sanger Institute COVID-19 Surveillance Team |
| EPI_ISL_3379213 | CENTRE HOSPITALIER DE VALENCE | CNR Virus des Infections Respiratoires - France SUD | Antonin Bal, Gregory Destras, Gwendolyne Burfin, Hadrien Regue, Quentin Semanas, Martine Valette, Bruno Lina, Laurence Josset |
| EPI_ISL_3379214 | Lighthouse Lab in Alderley Park | Wellcome Sanger Institute for the COVID-19 Genomics UK (COG-UK) Consortium | Jacquelyn Wynn, Mairead Hyland, The Lighthouse Lab in Alderley Park and Alex Alderton, Roberto Amato, Jeffrey Barrett, Sonia Goncalves, Ewan Harrison, David K. Jackson, Ian Johnston, Dominic Kwiatkowski, Cordelia Langford, John Sillitoe on behalf of the Wellcome Sanger Institute COVID-19 Surveillance Team |
| EPI_ISL_3379215, EPI_ISL_3379216 | Lighthouse Lab in Glasgow | Wellcome Sanger Institute for the COVID-19 Genomics UK (COG-UK) Consortium | Harper VanSteenhouse, Yumi Kasai, David Gray, Carol Clugston, Anna Dominiczak and Alex Alderton, Roberto Amato, Jeffrey Barrett, Sonia Goncalves, Ewan Harrison, David K. Jackson, Ian Johnston, Dominic Kwiatkowski, Cordelia Langford, John Sillitoe on behalf of the Wellcome Sanger Institute COVID-19 Surveillance Team |
| EPI_ISL_3379217 | CENTRE HOSPITALIER DE VALENCE | CNR Virus des Infections Respiratoires - France SUD | Antonin Bal, Gregory Destras, Gwendolyne Burfin, Hadrien Regue, Quentin Semanas, Martine Valette, Bruno Lina, Laurence Josset |
| EPI_ISL_3379218 | Lighthouse Lab in Glasgow | Wellcome Sanger Institute for the COVID-19 Genomics UK (COG-UK) Consortium | Harper VanSteenhouse, Yumi Kasai, David Gray, Carol Clugston, Anna Dominiczak and Alex Alderton, Roberto Amato, Jeffrey Barrett, Sonia Goncalves, Ewan Harrison, David K. Jackson, Ian Johnston, Dominic Kwiatkowski, Cordelia Langford, John Sillitoe on behalf of the Wellcome Sanger Institute COVID-19 Surveillance Team |
| EPI_ISL_3379219 | Lighthouse Lab in Milton Keynes | Wellcome Sanger Institute for the COVID-19 Genomics UK (COG-UK) Consortium | The Lighthouse Lab in Milton Keynes and Alex Alderton, Roberto Amato, Jeffrey Barrett, Sonia Goncalves, Ewan Harrison, David K. Jackson, Ian Johnston, Dominic Kwiatkowski, Cordelia Langford, John Sillitoe on behalf of the Wellcome Sanger Institute COVID-19 Surveillance Team |
| EPI_ISL_3379220 | Lighthouse Lab in Glasgow | Wellcome Sanger Institute for the COVID-19 Genomics UK (COG-UK) Consortium | Harper VanSteenhouse, Yumi Kasai, David Gray, Carol Clugston, Anna Dominiczak and Alex Alderton, Roberto Amato, Jeffrey Barrett, Sonia Goncalves, Ewan Harrison, David K. Jackson, Ian Johnston, Dominic Kwiatkowski, Cordelia Langford, John Sillitoe on behalf of the Wellcome Sanger Institute COVID-19 Surveillance Team |
| EPI_ISL_3379221 | CENTRE HOSPITALIER DE VALENCE | CNR Virus des Infections Respiratoires - France SUD | Antonin Bal, Gregory Destras, Gwendolyne Burfin, Hadrien Regue, Quentin Semanas, Martine Valette, Bruno Lina, Laurence Josset |
| EPI_ISL_3379222, EPI_ISL_3379223 | Lighthouse Lab in Alderley Park | Wellcome Sanger Institute for the COVID-19 Genomics UK (COG-UK) Consortium | Jacquelyn Wynn, Mairead Hyland, The Lighthouse Lab in Alderley Park and Alex Alderton, Roberto Amato, Jeffrey Barrett, Sonia Goncalves, Ewan Harrison, David K. Jackson, Ian Johnston, Dominic Kwiatkowski, Cordelia Langford, John Sillitoe on behalf of the Wellcome Sanger Institute COVID-19 Surveillance Team |
| EPI_ISL_3379224 | CENTRE HOSPITALIER DE VALENCE | CNR Virus des Infections Respiratoires - France SUD | Antonin Bal, Gregory Destras, Gwendolyne Burfin, Hadrien Regue, Quentin Semanas, Martine Valette, Bruno Lina, Laurence Josset |
| EPI_ISL_3379225 | Lighthouse Lab in Milton Keynes | Wellcome Sanger Institute for the COVID-19 Genomics UK (COG-UK) Consortium | The Lighthouse Lab in Milton Keynes and Alex Alderton, Roberto Amato, Jeffrey Barrett, Sonia Goncalves, Ewan Harrison, David K. Jackson, Ian Johnston, Dominic Kwiatkowski, Cordelia Langford, John Sillitoe on behalf of the Wellcome Sanger Institute COVID-19 Surveillance Team |
| EPI_ISL_3379226 | Berkshire and Surrey Pathology Services Lighthouse Laboratory | Wellcome Sanger Institute for the COVID-19 Genomics UK (COG-UK) Consortium | Berkshire and Surrey Pathology Services Lighthouse Laboratory and Alex Alderton, Roberto Amato, Jeffrey Barrett, Sonia Goncalves, Ewan Harrison, David K. Jackson, Ian Johnston, Dominic Kwiatkowski, Cordelia Langford, John Sillitoe on behalf of the Wellcome Sanger Institute COVID-19 Surveillance Team |

| | | | |
|---|---|---|---|
| EPI_ISL_3379227 | Lighthouse Lab in Alderley Park | Wellcome Sanger Institute for the COVID-19 Genomics UK (COG-UK) Consortium | Jacquelyn Wynn, Mairead Hyland, The Lighthouse Lab in Alderley Park and Alex Alderton, Roberto Amato, Jeffrey Barrett, Sonia Goncalves, Ewan Harrison, David K. Jackson, Ian Johnston, Dominic Kwiatkowski, Cordelia Langford, John Sillitoe on behalf of the Wellcome Sanger Institute COVID-19 Surveillance Team |
| EPI_ISL_3379228 | CNR Virus des Infections Respiratoires - France SUD | CNR Virus des Infections Respiratoires - France SUD | Antonin Bal, Gregory Destras, Gwendolyne Burfin, Hadrien Regue, Quentin Semanas, Martine Valette, Bruno Lina, Laurence Josset |
| EPI_ISL_3379229 | Lighthouse Lab in Milton Keynes | Wellcome Sanger Institute for the COVID-19 Genomics UK (COG-UK) Consortium | The Lighthouse Lab in Milton Keynes and Alex Alderton, Roberto Amato, Jeffrey Barrett, Sonia Goncalves, Ewan Harrison, David K. Jackson, Ian Johnston, Dominic Kwiatkowski, Cordelia Langford, John Sillitoe on behalf of the Wellcome Sanger Institute COVID-19 Surveillance Team |
| EPI_ISL_3379230 | Lighthouse Lab in Glasgow | Wellcome Sanger Institute for the COVID-19 Genomics UK (COG-UK) Consortium | Harper VanSteenhouse, Yumi Kasai, David Gray, Carol Clugston, Anna Dominiczak and Alex Alderton, Roberto Amato, Jeffrey Barrett, Sonia Goncalves, Ewan Harrison, David K. Jackson, Ian Johnston, Dominic Kwiatkowski, Cordelia Langford, John Sillitoe on behalf of the Wellcome Sanger Institute COVID-19 Surveillance Team |
| EPI_ISL_3379231 | Lighthouse Lab in Milton Keynes | Wellcome Sanger Institute for the COVID-19 Genomics UK (COG-UK) Consortium | The Lighthouse Lab in Milton Keynes and Alex Alderton, Roberto Amato, Jeffrey Barrett, Sonia Goncalves, Ewan Harrison, David K. Jackson, Ian Johnston, Dominic Kwiatkowski, Cordelia Langford, John Sillitoe on behalf of the Wellcome Sanger Institute COVID-19 Surveillance Team |
| EPI_ISL_3379232 | LABORATOIRE BIOESTEREL | CNR Virus des Infections Respiratoires - France SUD | Antonin Bal, Gregory Destras, Gwendolyne Burfin, Hadrien Regue, Quentin Semanas, Martine Valette, Bruno Lina, Laurence Josset |
| EPI_ISL_3379233 | Lighthouse Lab in Milton Keynes | Wellcome Sanger Institute for the COVID-19 Genomics UK (COG-UK) Consortium | The Lighthouse Lab in Milton Keynes and Alex Alderton, Roberto Amato, Jeffrey Barrett, Sonia Goncalves, Ewan Harrison, David K. Jackson, Ian Johnston, Dominic Kwiatkowski, Cordelia Langford, John Sillitoe on behalf of the Wellcome Sanger Institute COVID-19 Surveillance Team |
| EPI_ISL_3379234 | Lighthouse Lab in Glasgow | Wellcome Sanger Institute for the COVID-19 Genomics UK (COG-UK) Consortium | Harper VanSteenhouse, Yumi Kasai, David Gray, Carol Clugston, Anna Dominiczak and Alex Alderton, Roberto Amato, Jeffrey Barrett, Sonia Goncalves, Ewan Harrison, David K. Jackson, Ian Johnston, Dominic Kwiatkowski, Cordelia Langford, John Sillitoe on behalf of the Wellcome Sanger Institute COVID-19 Surveillance Team |
| EPI_ISL_3379235 | Lighthouse Lab in Milton Keynes | Wellcome Sanger Institute for the COVID-19 Genomics UK (COG-UK) Consortium | The Lighthouse Lab in Milton Keynes and Alex Alderton, Roberto Amato, Jeffrey Barrett, Sonia Goncalves, Ewan Harrison, David K. Jackson, Ian Johnston, Dominic Kwiatkowski, Cordelia Langford, John Sillitoe on behalf of the Wellcome Sanger Institute COVID-19 Surveillance Team |
| EPI_ISL_3379236 | LABORATOIRE BIOESTEREL | CNR Virus des Infections Respiratoires - France SUD | Antonin Bal, Gregory Destras, Gwendolyne Burfin, Hadrien Regue, Quentin Semanas, Martine Valette, Bruno Lina, Laurence Josset |
| EPI_ISL_3379237 | Lighthouse Lab in Glasgow | Wellcome Sanger Institute for the COVID-19 Genomics UK (COG-UK) Consortium | Harper VanSteenhouse, Yumi Kasai, David Gray, Carol Clugston, Anna Dominiczak and Alex Alderton, Roberto Amato, Jeffrey Barrett, Sonia Goncalves, Ewan Harrison, David K. Jackson, Ian Johnston, Dominic Kwiatkowski, Cordelia Langford, John Sillitoe on behalf of the Wellcome Sanger Institute COVID-19 Surveillance Team |
| EPI_ISL_3379238 | Berkshire and Surrey Pathology Services Lighthouse Laboratory | Wellcome Sanger Institute for the COVID-19 Genomics UK (COG-UK) Consortium | Berkshire and Surrey Pathology Services Lighthouse Laboratory and Alex Alderton, Roberto Amato, Jeffrey Barrett, Sonia Goncalves, Ewan Harrison, David K. Jackson, Ian Johnston, Dominic Kwiatkowski, Cordelia Langford, John Sillitoe on behalf of the Wellcome Sanger Institute COVID-19 Surveillance Team |
| EPI_ISL_3379239 | LABORATOIRE BIOESTEREL | CNR Virus des Infections Respiratoires - France SUD | Antonin Bal, Gregory Destras, Gwendolyne Burfin, Hadrien Regue, Quentin Semanas, Martine Valette, Bruno Lina, Laurence Josset |
| EPI_ISL_3379240 | Lighthouse Lab in Alderley Park | Wellcome Sanger Institute for the COVID-19 Genomics UK (COG-UK) Consortium | Jacquelyn Wynn, Mairead Hyland, The Lighthouse Lab in Alderley Park and Alex Alderton, Roberto Amato, Jeffrey Barrett, Sonia Goncalves, Ewan Harrison, David K. Jackson, Ian Johnston, Dominic Kwiatkowski, Cordelia Langford, John Sillitoe on behalf of the Wellcome Sanger Institute COVID-19 Surveillance Team |
| EPI_ISL_3379241 | Lighthouse Lab in Glasgow | Wellcome Sanger Institute for the COVID-19 Genomics UK (COG-UK) Consortium | Harper VanSteenhouse, Yumi Kasai, David Gray, Carol Clugston, Anna Dominiczak and Alex Alderton, Roberto Amato, Jeffrey Barrett, Sonia Goncalves, Ewan Harrison, David K. Jackson, Ian Johnston, Dominic Kwiatkowski, Cordelia Langford, John Sillitoe on behalf of the Wellcome Sanger Institute COVID-19 Surveillance Team |
| EPI_ISL_3379242 | Lighthouse Lab in Alderley Park | Wellcome Sanger Institute for the COVID-19 Genomics UK (COG-UK) Consortium | Jacquelyn Wynn, Mairead Hyland, The Lighthouse Lab in Alderley Park and Alex Alderton, Roberto Amato, Jeffrey Barrett, Sonia Goncalves, Ewan Harrison, David K. Jackson, Ian Johnston, Dominic Kwiatkowski, Cordelia Langford, John Sillitoe on behalf of the Wellcome Sanger Institute COVID-19 Surveillance Team |
| EPI_ISL_3379243 | CNR Virus des Infections Respiratoires - France SUD | CNR Virus des Infections Respiratoires - France SUD | Antonin Bal, Gregory Destras, Gwendolyne Burfin, Hadrien Regue, Quentin Semanas, Martine Valette, Bruno Lina, Laurence Josset |
| EPI_ISL_3379244 | Lighthouse Lab in Alderley Park | Wellcome Sanger Institute for the COVID-19 Genomics UK (COG-UK) Consortium | Jacquelyn Wynn, Mairead Hyland, The Lighthouse Lab in Alderley Park and Alex Alderton, Roberto Amato, Jeffrey Barrett, Sonia Goncalves, Ewan Harrison, David K. Jackson, Ian Johnston, Dominic Kwiatkowski, Cordelia Langford, John Sillitoe on behalf of the Wellcome Sanger Institute COVID-19 Surveillance Team |
| EPI_ISL_3379245, EPI_ISL_3379246 | Lighthouse Lab in Glasgow | Wellcome Sanger Institute for the COVID-19 Genomics UK (COG-UK) Consortium | Harper VanSteenhouse, Yumi Kasai, David Gray, Carol Clugston, Anna Dominiczak and Alex Alderton, Roberto Amato, Jeffrey Barrett, Sonia Goncalves, Ewan Harrison, David K. Jackson, Ian Johnston, Dominic Kwiatkowski, Cordelia Langford, John Sillitoe on behalf of the Wellcome Sanger Institute COVID-19 Surveillance Team |
| EPI_ISL_3379247 | LABORATOIRE BIOESTEREL | CNR Virus des Infections Respiratoires - France SUD | Antonin Bal, Gregory Destras, Gwendolyne Burfin, Hadrien Regue, Quentin Semanas, Martine Valette, Bruno Lina, Laurence Josset |
| EPI_ISL_3379248, EPI_ISL_3379249 | Lighthouse Lab in Glasgow | Wellcome Sanger Institute for the COVID-19 Genomics UK (COG-UK) Consortium | Harper VanSteenhouse, Yumi Kasai, David Gray, Carol Clugston, Anna Dominiczak and Alex Alderton, Roberto Amato, Jeffrey Barrett, Sonia Goncalves, Ewan Harrison, David K. Jackson, Ian Johnston, Dominic Kwiatkowski, Cordelia Langford, John Sillitoe on behalf of the Wellcome Sanger Institute COVID-19 Surveillance Team |
| EPI_ISL_3379250 | Lighthouse Lab in Alderley Park | Wellcome Sanger Institute for the COVID-19 Genomics UK (COG-UK) Consortium | Jacquelyn Wynn, Mairead Hyland, The Lighthouse Lab in Alderley Park and Alex Alderton, Roberto Amato, Jeffrey Barrett, Sonia Goncalves, Ewan Harrison, David K. Jackson, Ian Johnston, Dominic Kwiatkowski, Cordelia Langford, John Sillitoe on behalf of the Wellcome Sanger Institute COVID-19 Surveillance Team |
| EPI_ISL_3379251 | Lighthouse Lab in Milton Keynes | Wellcome Sanger Institute for the COVID-19 Genomics UK (COG-UK) Consortium | The Lighthouse Lab in Milton Keynes and Alex Alderton, Roberto Amato, Jeffrey Barrett, Sonia Goncalves, Ewan Harrison, David K. Jackson, Ian Johnston, Dominic Kwiatkowski, Cordelia Langford, John Sillitoe on behalf of the Wellcome Sanger Institute COVID-19 Surveillance Team |
| EPI_ISL_3379252 | Lighthouse Lab in Glasgow | Wellcome Sanger Institute for the COVID-19 Genomics UK (COG-UK) Consortium | Harper VanSteenhouse, Yumi Kasai, David Gray, Carol Clugston, Anna Dominiczak and Alex Alderton, Roberto Amato, Jeffrey Barrett, Sonia Goncalves, Ewan Harrison, David K. Jackson, Ian Johnston, Dominic Kwiatkowski, Cordelia Langford, John Sillitoe on behalf of the Wellcome Sanger Institute COVID-19 Surveillance Team |
| EPI_ISL_3379253 | Lighthouse Lab in Alderley Park | Wellcome Sanger Institute for the COVID-19 Genomics UK (COG-UK) Consortium | Jacquelyn Wynn, Mairead Hyland, The Lighthouse Lab in Alderley Park and Alex Alderton, Roberto Amato, Jeffrey Barrett, Sonia Goncalves, Ewan Harrison, David K. Jackson, Ian Johnston, Dominic Kwiatkowski, Cordelia Langford, John Sillitoe on behalf of the Wellcome Sanger Institute COVID-19 Surveillance Team |
| EPI_ISL_3379254 | Berkshire and Surrey Pathology Services Lighthouse Laboratory | Wellcome Sanger Institute for the COVID-19 Genomics UK (COG-UK) Consortium | Berkshire and Surrey Pathology Services Lighthouse Laboratory and Alex Alderton, Roberto Amato, Jeffrey Barrett, Sonia Goncalves, Ewan Harrison, David K. Jackson, Ian Johnston, Dominic Kwiatkowski, Cordelia Langford, John Sillitoe on behalf of the Wellcome Sanger Institute COVID-19 Surveillance Team |
| EPI_ISL_3379255 | LABORATOIRE BIOESTEREL | CNR Virus des Infections Respiratoires - France SUD | Antonin Bal, Gregory Destras, Gwendolyne Burfin, Hadrien Regue, Quentin Semanas, Martine Valette, Bruno Lina, Laurence Josset |
| EPI_ISL_3379256 | Lighthouse Lab in Glasgow | Wellcome Sanger Institute for the COVID-19 Genomics UK (COG-UK) Consortium | Harper VanSteenhouse, Yumi Kasai, David Gray, Carol Clugston, Anna Dominiczak and Alex Alderton, Roberto Amato, Jeffrey Barrett, Sonia Goncalves, Ewan Harrison, David K. Jackson, Ian Johnston, Dominic Kwiatkowski, Cordelia Langford, John Sillitoe on behalf of the Wellcome Sanger Institute COVID-19 Surveillance Team |
| EPI_ISL_3379257 | Lighthouse Lab in Alderley Park | Wellcome Sanger Institute for the COVID-19 Genomics UK (COG-UK) Consortium | Jacquelyn Wynn, Mairead Hyland, The Lighthouse Lab in Alderley Park and Alex Alderton, Roberto Amato, Jeffrey Barrett, Sonia Goncalves, Ewan Harrison, David K. Jackson, Ian Johnston, Dominic Kwiatkowski, Cordelia Langford, John Sillitoe on behalf of the Wellcome Sanger Institute COVID-19 Surveillance Team |
| EPI_ISL_3379258, EPI_ISL_3379259 | Lighthouse Lab in Milton Keynes | Wellcome Sanger Institute for the COVID-19 Genomics UK (COG-UK) Consortium | The Lighthouse Lab in Milton Keynes and Alex Alderton, Roberto Amato, Jeffrey Barrett, Sonia Goncalves, Ewan Harrison, David K. Jackson, Ian Johnston, Dominic Kwiatkowski, Cordelia Langford, John Sillitoe on behalf of the Wellcome Sanger Institute COVID-19 Surveillance Team |
| EPI_ISL_3379260 | Lighthouse Lab in Glasgow | Wellcome Sanger Institute for the COVID-19 Genomics UK (COG-UK) Consortium | Harper VanSteenhouse, Yumi Kasai, David Gray, Carol Clugston, Anna Dominiczak and Alex Alderton, Roberto Amato, Jeffrey Barrett, Sonia Goncalves, Ewan Harrison, David K. Jackson, Ian Johnston, Dominic Kwiatkowski, Cordelia Langford, John Sillitoe on behalf of the Wellcome Sanger Institute COVID-19 Surveillance Team |
| EPI_ISL_3379261 | Berkshire and Surrey Pathology Services Lighthouse Laboratory | Wellcome Sanger Institute for the COVID-19 Genomics UK (COG-UK) Consortium | Berkshire and Surrey Pathology Services Lighthouse Laboratory and Alex Alderton, Roberto Amato, Jeffrey Barrett, Sonia Goncalves, Ewan Harrison, David K. Jackson, Ian Johnston, Dominic Kwiatkowski, Cordelia Langford, John Sillitoe on behalf of the Wellcome Sanger Institute COVID-19 Surveillance Team |
| EPI_ISL_3379262 | Lighthouse Lab in Glasgow | Wellcome Sanger Institute for the COVID-19 Genomics UK (COG-UK) Consortium | Harper VanSteenhouse, Yumi Kasai, David Gray, Carol Clugston, Anna Dominiczak and Alex Alderton, Roberto Amato, Jeffrey Barrett, Sonia Goncalves, Ewan Harrison, David K. Jackson, Ian Johnston, Dominic Kwiatkowski, Cordelia Langford, John Sillitoe on behalf of the Wellcome Sanger Institute COVID-19 Surveillance Team |
| EPI_ISL_3379263 | HOPITAL ALBI | CNR Virus des Infections Respiratoires - France SUD | Antonin Bal, Gregory Destras, Gwendolyne Burfin, Hadrien Regue, Quentin Semanas, Martine Valette, Bruno Lina, Laurence Josset |
| EPI_ISL_3379264, EPI_ISL_3379265 | Lighthouse Lab in Glasgow | Wellcome Sanger Institute for the COVID-19 Genomics UK (COG-UK) Consortium | Harper VanSteenhouse, Yumi Kasai, David Gray, Carol Clugston, Anna Dominiczak and Alex Alderton, Roberto Amato, Jeffrey Barrett, Sonia Goncalves, Ewan Harrison, David K. Jackson, Ian Johnston, Dominic Kwiatkowski, Cordelia Langford, John Sillitoe on behalf of the Wellcome Sanger Institute COVID-19 Surveillance Team |
| EPI_ISL_3379266 | Lighthouse Lab in Alderley Park | Wellcome Sanger Institute for the COVID-19 Genomics UK (COG-UK) Consortium | Jacquelyn Wynn, Mairead Hyland, The Lighthouse Lab in Alderley Park and Alex Alderton, Roberto Amato, Jeffrey Barrett, Sonia Goncalves, Ewan Harrison, David K. Jackson, Ian Johnston, Dominic Kwiatkowski, Cordelia Langford, John Sillitoe on behalf of the Wellcome Sanger Institute COVID-19 Surveillance Team |
| EPI_ISL_3379267 | Lighthouse Lab in Glasgow | Wellcome Sanger Institute for the COVID-19 Genomics UK (COG-UK) Consortium | Harper VanSteenhouse, Yumi Kasai, David Gray, Carol Clugston, Anna Dominiczak and Alex Alderton, Roberto Amato, Jeffrey Barrett, Sonia Goncalves, Ewan Harrison, David K. Jackson, Ian Johnston, Dominic Kwiatkowski, Cordelia Langford, John Sillitoe on behalf of the Wellcome Sanger Institute COVID-19 Surveillance Team |
| EPI_ISL_3379268 | Lighthouse Lab in Milton Keynes | Wellcome Sanger Institute for the COVID-19 Genomics UK (COG-UK) Consortium | The Lighthouse Lab in Milton Keynes and Alex Alderton, Roberto Amato, Jeffrey Barrett, Sonia Goncalves, Ewan Harrison, David K. Jackson, Ian Johnston, Dominic Kwiatkowski, Cordelia Langford, John Sillitoe on behalf of the Wellcome Sanger Institute COVID-19 Surveillance Team |
| EPI_ISL_3379269, EPI_ISL_3379270 | Lighthouse Lab in Alderley Park | Wellcome Sanger Institute for the COVID-19 Genomics UK (COG-UK) Consortium | Jacquelyn Wynn, Mairead Hyland, The Lighthouse Lab in Alderley Park and Alex Alderton, Roberto Amato, Jeffrey Barrett, Sonia Goncalves, Ewan Harrison, David K. Jackson, Ian Johnston, Dominic Kwiatkowski, Cordelia Langford, John Sillitoe on behalf of the Wellcome Sanger Institute COVID-19 Surveillance Team |
| EPI_ISL_3379271 | Lighthouse Lab in Milton Keynes | Wellcome Sanger Institute for the COVID-19 Genomics UK (COG-UK) Consortium | The Lighthouse Lab in Milton Keynes and Alex Alderton, Roberto Amato, Jeffrey Barrett, Sonia Goncalves, Ewan Harrison, David K. Jackson, Ian Johnston, Dominic Kwiatkowski, Cordelia Langford, John Sillitoe on behalf of the Wellcome Sanger Institute COVID-19 Surveillance Team |
| EPI_ISL_3379272 | HOPITAL ALBI | CNR Virus des Infections Respiratoires - France SUD | Antonin Bal, Gregory Destras, Gwendolyne Burfin, Hadrien Regue, Quentin Semanas, Martine Valette, Bruno Lina, Laurence Josset |
| EPI_ISL_3379273 | Berkshire and Surrey Pathology Services Lighthouse Laboratory | Wellcome Sanger Institute for the COVID-19 Genomics UK (COG-UK) Consortium | Berkshire and Surrey Pathology Services Lighthouse Laboratory and Alex Alderton, Roberto Amato, Jeffrey Barrett, Sonia Goncalves, Ewan Harrison, David K. Jackson, Ian Johnston, Dominic Kwiatkowski, Cordelia Langford, John Sillitoe on behalf of the Wellcome Sanger Institute COVID-19 Surveillance Team |
| EPI_ISL_3379274 | Lighthouse Lab in Alderley Park | Wellcome Sanger Institute for the COVID-19 Genomics UK (COG-UK) Consortium | Jacquelyn Wynn, Mairead Hyland, The Lighthouse Lab in Alderley Park and Alex Alderton, Roberto Amato, Jeffrey Barrett, Sonia Goncalves, Ewan Harrison, David K. Jackson, Ian Johnston, Dominic Kwiatkowski, Cordelia Langford, John Sillitoe on behalf of the Wellcome Sanger Institute COVID-19 Surveillance Team |
| EPI_ISL_3379275 | Lighthouse Lab in Glasgow | Wellcome Sanger Institute for the COVID-19 Genomics UK (COG-UK) Consortium | Harper VanSteenhouse, Yumi Kasai, David Gray, Carol Clugston, Anna Dominiczak and Alex Alderton, Roberto Amato, Jeffrey Barrett, Sonia Goncalves, Ewan Harrison, David K. Jackson, Ian Johnston, Dominic Kwiatkowski, Cordelia Langford, John Sillitoe on behalf of the Wellcome Sanger Institute COVID-19 Surveillance Team |
| EPI_ISL_3379276, EPI_ISL_3379277 | Lighthouse Lab in Alderley Park | Wellcome Sanger Institute for the COVID-19 Genomics UK (COG-UK) Consortium | Jacquelyn Wynn, Mairead Hyland, The Lighthouse Lab in Alderley Park and Alex Alderton, Roberto Amato, Jeffrey Barrett, Sonia Goncalves, Ewan Harrison, David K. Jackson, Ian Johnston, Dominic Kwiatkowski, Cordelia Langford, John Sillitoe on behalf of the Wellcome Sanger Institute COVID-19 Surveillance Team |
| EPI_ISL_3379278, EPI_ISL_3379279 | Lighthouse Lab in Glasgow | Wellcome Sanger Institute for the COVID-19 Genomics UK (COG-UK) Consortium | Harper VanSteenhouse, Yumi Kasai, David Gray, Carol Clugston, Anna Dominiczak and Alex Alderton, Roberto Amato, Jeffrey Barrett, Sonia Goncalves, Ewan Harrison, David K. Jackson, Ian Johnston, Dominic Kwiatkowski, Cordelia Langford, John Sillitoe on behalf of the Wellcome Sanger Institute COVID-19 Surveillance Team |
| EPI_ISL_3379280 | HOPITAL ALBI | CNR Virus des Infections Respiratoires - France SUD | Antonin Bal, Gregory Destras, Gwendolyne Burfin, Hadrien Regue, Quentin Semanas, Martine Valette, Bruno Lina, Laurence Josset |
| EPI_ISL_3379281 | Lighthouse Lab in Glasgow | Wellcome Sanger Institute for the COVID-19 Genomics UK (COG-UK) Consortium | Harper VanSteenhouse, Yumi Kasai, David Gray, Carol Clugston, Anna Dominiczak and Alex Alderton, Roberto Amato, Jeffrey Barrett, Sonia Goncalves, Ewan Harrison, David K. Jackson, Ian Johnston, Dominic Kwiatkowski, Cordelia Langford, John Sillitoe on behalf of the Wellcome Sanger Institute COVID-19 Surveillance Team |
| EPI_ISL_3379282, EPI_ISL_3379283 | Lighthouse Lab in Milton Keynes | Wellcome Sanger Institute for the COVID-19 Genomics UK (COG-UK) Consortium | The Lighthouse Lab in Milton Keynes and Alex Alderton, Roberto Amato, Jeffrey Barrett, Sonia Goncalves, Ewan Harrison, David K. Jackson, Ian Johnston, Dominic Kwiatkowski, Cordelia Langford, John Sillitoe on behalf of the Wellcome Sanger Institute COVID-19 Surveillance Team |

| | | | |
|---|---|---|---|
| EPI_ISL_3379284 | Berkshire and Surrey Pathology Services Lighthouse Laboratory | Wellcome Sanger Institute for the COVID-19 Genomics UK (COG-UK) Consortium | Berkshire and Surrey Pathology Services Lighthouse Laboratory and Alex Alderton, Roberto Amato, Jeffrey Barrett, Sonia Goncalves, Ewan Harrison, David K. Jackson, Ian Johnston, Dominic Kwiatkowski, Cordelia Langford, John Sillitoe on behalf of the Wellcome Sanger Institute COVID-19 Surveillance Team |
| EPI_ISL_3379285 | Lighthouse Lab in Milton Keynes | Wellcome Sanger Institute for the COVID-19 Genomics UK (COG-UK) Consortium | The Lighthouse Lab in Milton Keynes and Alex Alderton, Roberto Amato, Jeffrey Barrett, Sonia Goncalves, Ewan Harrison, David K. Jackson, Ian Johnston, Dominic Kwiatkowski, Cordelia Langford, John Sillitoe on behalf of the Wellcome Sanger Institute COVID-19 Surveillance Team |
| EPI_ISL_3379286 | Lighthouse Lab in Alderley Park | Wellcome Sanger Institute for the COVID-19 Genomics UK (COG-UK) Consortium | Jacquelyn Wynn, Mairead Hyland, The Lighthouse Lab in Alderley Park and Alex Alderton, Roberto Amato, Jeffrey Barrett, Sonia Goncalves, Ewan Harrison, David K. Jackson, Ian Johnston, Dominic Kwiatkowski, Cordelia Langford, John Sillitoe on behalf of the Wellcome Sanger Institute COVID-19 Surveillance Team |
| EPI_ISL_3379287 | Lighthouse Lab in Milton Keynes | Wellcome Sanger Institute for the COVID-19 Genomics UK (COG-UK) Consortium | The Lighthouse Lab in Milton Keynes and Alex Alderton, Roberto Amato, Jeffrey Barrett, Sonia Goncalves, Ewan Harrison, David K. Jackson, Ian Johnston, Dominic Kwiatkowski, Cordelia Langford, John Sillitoe on behalf of the Wellcome Sanger Institute COVID-19 Surveillance Team |
| EPI_ISL_3379288 | HOPITAL ALBI | CNR Virus des Infections Respiratoires - France SUD | Antonin Bal, Gregory Destras, Gwendolyne Burfin, Hadrien Regue, Quentin Semanas, Martine Valette, Bruno Lina, Laurence Josset |
| EPI_ISL_3379289, EPI_ISL_3379290 | Lighthouse Lab in Milton Keynes | Wellcome Sanger Institute for the COVID-19 Genomics UK (COG-UK) Consortium | The Lighthouse Lab in Milton Keynes and Alex Alderton, Roberto Amato, Jeffrey Barrett, Sonia Goncalves, Ewan Harrison, David K. Jackson, Ian Johnston, Dominic Kwiatkowski, Cordelia Langford, John Sillitoe on behalf of the Wellcome Sanger Institute COVID-19 Surveillance Team |
| EPI_ISL_3379291 | Berkshire and Surrey Pathology Services Lighthouse Laboratory | Wellcome Sanger Institute for the COVID-19 Genomics UK (COG-UK) Consortium | Berkshire and Surrey Pathology Services Lighthouse Laboratory and Alex Alderton, Roberto Amato, Jeffrey Barrett, Sonia Goncalves, Ewan Harrison, David K. Jackson, Ian Johnston, Dominic Kwiatkowski, Cordelia Langford, John Sillitoe on behalf of the Wellcome Sanger Institute COVID-19 Surveillance Team |
| EPI_ISL_3379292 | HOPITAL ALBI | CNR Virus des Infections Respiratoires - France SUD | Antonin Bal, Gregory Destras, Gwendolyne Burfin, Hadrien Regue, Quentin Semanas, Martine Valette, Bruno Lina, Laurence Josset |
| EPI_ISL_3379293, EPI_ISL_3379294 | Lighthouse Lab in Milton Keynes | Wellcome Sanger Institute for the COVID-19 Genomics UK (COG-UK) Consortium | The Lighthouse Lab in Milton Keynes and Alex Alderton, Roberto Amato, Jeffrey Barrett, Sonia Goncalves, Ewan Harrison, David K. Jackson, Ian Johnston, Dominic Kwiatkowski, Cordelia Langford, John Sillitoe on behalf of the Wellcome Sanger Institute COVID-19 Surveillance Team |
| EPI_ISL_3379295 | CNR Virus des Infections Respiratoires - France SUD | CNR Virus des Infections Respiratoires - France SUD | Antonin Bal, Gregory Destras, Gwendolyne Burfin, Hadrien Regue, Quentin Semanas, Martine Valette, Bruno Lina, Laurence Josset |
| EPI_ISL_3379296 | Lighthouse Lab in Milton Keynes | Wellcome Sanger Institute for the COVID-19 Genomics UK (COG-UK) Consortium | The Lighthouse Lab in Milton Keynes and Alex Alderton, Roberto Amato, Jeffrey Barrett, Sonia Goncalves, Ewan Harrison, David K. Jackson, Ian Johnston, Dominic Kwiatkowski, Cordelia Langford, John Sillitoe on behalf of the Wellcome Sanger Institute COVID-19 Surveillance Team |
| EPI_ISL_3379297 | Lighthouse Lab in Alderley Park | Wellcome Sanger Institute for the COVID-19 Genomics UK (COG-UK) Consortium | Jacquelyn Wynn, Mairead Hyland, The Lighthouse Lab in Alderley Park and Alex Alderton, Roberto Amato, Jeffrey Barrett, Sonia Goncalves, Ewan Harrison, David K. Jackson, Ian Johnston, Dominic Kwiatkowski, Cordelia Langford, John Sillitoe on behalf of the Wellcome Sanger Institute COVID-19 Surveillance Team |
| EPI_ISL_3379298 | Berkshire and Surrey Pathology Services Lighthouse Laboratory | Wellcome Sanger Institute for the COVID-19 Genomics UK (COG-UK) Consortium | Berkshire and Surrey Pathology Services Lighthouse Laboratory and Alex Alderton, Roberto Amato, Jeffrey Barrett, Sonia Goncalves, Ewan Harrison, David K. Jackson, Ian Johnston, Dominic Kwiatkowski, Cordelia Langford, John Sillitoe on behalf of the Wellcome Sanger Institute COVID-19 Surveillance Team |
| EPI_ISL_3379299 | Lighthouse Lab in Alderley Park | Wellcome Sanger Institute for the COVID-19 Genomics UK (COG-UK) Consortium | Jacquelyn Wynn, Mairead Hyland, The Lighthouse Lab in Alderley Park and Alex Alderton, Roberto Amato, Jeffrey Barrett, Sonia Goncalves, Ewan Harrison, David K. Jackson, Ian Johnston, Dominic Kwiatkowski, Cordelia Langford, John Sillitoe on behalf of the Wellcome Sanger Institute COVID-19 Surveillance Team |
| EPI_ISL_3379300 | CNR Virus des Infections Respiratoires - France SUD | CNR Virus des Infections Respiratoires - France SUD | Antonin Bal, Gregory Destras, Gwendolyne Burfin, Hadrien Regue, Quentin Semanas, Martine Valette, Bruno Lina, Laurence Josset |
| EPI_ISL_3379301 | Berkshire and Surrey Pathology Services Lighthouse Laboratory | Wellcome Sanger Institute for the COVID-19 Genomics UK (COG-UK) Consortium | Berkshire and Surrey Pathology Services Lighthouse Laboratory and Alex Alderton, Roberto Amato, Jeffrey Barrett, Sonia Goncalves, Ewan Harrison, David K. Jackson, Ian Johnston, Dominic Kwiatkowski, Cordelia Langford, John Sillitoe on behalf of the Wellcome Sanger Institute COVID-19 Surveillance Team |
| EPI_ISL_3379302 | Lighthouse Lab in Milton Keynes | Wellcome Sanger Institute for the COVID-19 Genomics UK (COG-UK) Consortium | The Lighthouse Lab in Milton Keynes and Alex Alderton, Roberto Amato, Jeffrey Barrett, Sonia Goncalves, Ewan Harrison, David K. Jackson, Ian Johnston, Dominic Kwiatkowski, Cordelia Langford, John Sillitoe on behalf of the Wellcome Sanger Institute COVID-19 Surveillance Team |
| EPI_ISL_3379303 | CNR Virus des Infections Respiratoires - France SUD | CNR Virus des Infections Respiratoires - France SUD | Antonin Bal, Gregory Destras, Gwendolyne Burfin, Hadrien Regue, Quentin Semanas, Martine Valette, Bruno Lina, Laurence Josset |
| EPI_ISL_3379304, EPI_ISL_3379305, EPI_ISL_3379306 | Lighthouse Lab in Milton Keynes | Wellcome Sanger Institute for the COVID-19 Genomics UK (COG-UK) Consortium | The Lighthouse Lab in Milton Keynes and Alex Alderton, Roberto Amato, Jeffrey Barrett, Sonia Goncalves, Ewan Harrison, David K. Jackson, Ian Johnston, Dominic Kwiatkowski, Cordelia Langford, John Sillitoe on behalf of the Wellcome Sanger Institute COVID-19 Surveillance Team |
| EPI_ISL_3379307 | CNR Virus des Infections Respiratoires - France SUD | CNR Virus des Infections Respiratoires - France SUD | Antonin Bal, Gregory Destras, Gwendolyne Burfin, Hadrien Regue, Quentin Semanas, Martine Valette, Bruno Lina, Laurence Josset |
| EPI_ISL_3379308 | Lighthouse Lab in Milton Keynes | Wellcome Sanger Institute for the COVID-19 Genomics UK (COG-UK) Consortium | The Lighthouse Lab in Milton Keynes and Alex Alderton, Roberto Amato, Jeffrey Barrett, Sonia Goncalves, Ewan Harrison, David K. Jackson, Ian Johnston, Dominic Kwiatkowski, Cordelia Langford, John Sillitoe on behalf of the Wellcome Sanger Institute COVID-19 Surveillance Team |
| EPI_ISL_3379309 | Berkshire and Surrey Pathology Services Lighthouse Laboratory | Wellcome Sanger Institute for the COVID-19 Genomics UK (COG-UK) Consortium | Berkshire and Surrey Pathology Services Lighthouse Laboratory and Alex Alderton, Roberto Amato, Jeffrey Barrett, Sonia Goncalves, Ewan Harrison, David K. Jackson, Ian Johnston, Dominic Kwiatkowski, Cordelia Langford, John Sillitoe on behalf of the Wellcome Sanger Institute COVID-19 Surveillance Team |
| EPI_ISL_3379310 | Lighthouse Lab in Alderley Park | Wellcome Sanger Institute for the COVID-19 Genomics UK (COG-UK) Consortium | Jacquelyn Wynn, Mairead Hyland, The Lighthouse Lab in Alderley Park and Alex Alderton, Roberto Amato, Jeffrey Barrett, Sonia Goncalves, Ewan Harrison, David K. Jackson, Ian Johnston, Dominic Kwiatkowski, Cordelia Langford, John Sillitoe on behalf of the Wellcome Sanger Institute COVID-19 Surveillance Team |
| EPI_ISL_3379311 | LX BIO | CNR Virus des Infections Respiratoires - France SUD | Antonin Bal, Gregory Destras, Gwendolyne Burfin, Hadrien Regue, Quentin Semanas, Martine Valette, Bruno Lina, Laurence Josset |
| EPI_ISL_3379312, EPI_ISL_3379313 | Lighthouse Lab in Milton Keynes | Wellcome Sanger Institute for the COVID-19 Genomics UK (COG-UK) Consortium | The Lighthouse Lab in Milton Keynes and Alex Alderton, Roberto Amato, Jeffrey Barrett, Sonia Goncalves, Ewan Harrison, David K. Jackson, Ian Johnston, Dominic Kwiatkowski, Cordelia Langford, John Sillitoe on behalf of the Wellcome Sanger Institute COVID-19 Surveillance Team |
| EPI_ISL_3379314 | Lighthouse Lab in Alderley Park | Wellcome Sanger Institute for the COVID-19 Genomics UK (COG-UK) Consortium | Jacquelyn Wynn, Mairead Hyland, The Lighthouse Lab in Alderley Park and Alex Alderton, Roberto Amato, Jeffrey Barrett, Sonia Goncalves, Ewan Harrison, David K. Jackson, Ian Johnston, Dominic Kwiatkowski, Cordelia Langford, John Sillitoe on behalf of the Wellcome Sanger Institute COVID-19 Surveillance Team |
| EPI_ISL_3379315 | LX BIO | CNR Virus des Infections Respiratoires - France SUD | Antonin Bal, Gregory Destras, Gwendolyne Burfin, Hadrien Regue, Quentin Semanas, Martine Valette, Bruno Lina, Laurence Josset |
| EPI_ISL_3379316 | Lighthouse Lab in Alderley Park | Wellcome Sanger Institute for the COVID-19 Genomics UK (COG-UK) Consortium | Jacquelyn Wynn, Mairead Hyland, The Lighthouse Lab in Alderley Park and Alex Alderton, Roberto Amato, Jeffrey Barrett, Sonia Goncalves, Ewan Harrison, David K. Jackson, Ian Johnston, Dominic Kwiatkowski, Cordelia Langford, John Sillitoe on behalf of the Wellcome Sanger Institute COVID-19 Surveillance Team |
| EPI_ISL_3379317 | Lighthouse Lab in Milton Keynes | Wellcome Sanger Institute for the COVID-19 Genomics UK (COG-UK) Consortium | The Lighthouse Lab in Milton Keynes and Alex Alderton, Roberto Amato, Jeffrey Barrett, Sonia Goncalves, Ewan Harrison, David K. Jackson, Ian Johnston, Dominic Kwiatkowski, Cordelia Langford, John Sillitoe on behalf of the Wellcome Sanger Institute COVID-19 Surveillance Team |
| EPI_ISL_3379318 | LX BIO | CNR Virus des Infections Respiratoires - France SUD | Antonin Bal, Gregory Destras, Gwendolyne Burfin, Hadrien Regue, Quentin Semanas, Martine Valette, Bruno Lina, Laurence Josset |
| EPI_ISL_3379319 | Lighthouse Lab in Alderley Park | Wellcome Sanger Institute for the COVID-19 Genomics UK (COG-UK) Consortium | Jacquelyn Wynn, Mairead Hyland, The Lighthouse Lab in Alderley Park and Alex Alderton, Roberto Amato, Jeffrey Barrett, Sonia Goncalves, Ewan Harrison, David K. Jackson, Ian Johnston, Dominic Kwiatkowski, Cordelia Langford, John Sillitoe on behalf of the Wellcome Sanger Institute COVID-19 Surveillance Team |
| EPI_ISL_3379320 | Lighthouse Lab in Milton Keynes | Wellcome Sanger Institute for the COVID-19 Genomics UK (COG-UK) Consortium | The Lighthouse Lab in Milton Keynes and Alex Alderton, Roberto Amato, Jeffrey Barrett, Sonia Goncalves, Ewan Harrison, David K. Jackson, Ian Johnston, Dominic Kwiatkowski, Cordelia Langford, John Sillitoe on behalf of the Wellcome Sanger Institute COVID-19 Surveillance Team |
| EPI_ISL_3379321 | Lighthouse Lab in Alderley Park | Wellcome Sanger Institute for the COVID-19 Genomics UK (COG-UK) Consortium | Jacquelyn Wynn, Mairead Hyland, The Lighthouse Lab in Alderley Park and Alex Alderton, Roberto Amato, Jeffrey Barrett, Sonia Goncalves, Ewan Harrison, David K. Jackson, Ian Johnston, Dominic Kwiatkowski, Cordelia Langford, John Sillitoe on behalf of the Wellcome Sanger Institute COVID-19 Surveillance Team |
| EPI_ISL_3379322 | LX BIO | CNR Virus des Infections Respiratoires - France SUD | Antonin Bal, Gregory Destras, Gwendolyne Burfin, Hadrien Regue, Quentin Semanas, Martine Valette, Bruno Lina, Laurence Josset |
| EPI_ISL_3379323 | Berkshire and Surrey Pathology Services Lighthouse Laboratory | Wellcome Sanger Institute for the COVID-19 Genomics UK (COG-UK) Consortium | Berkshire and Surrey Pathology Services Lighthouse Laboratory and Alex Alderton, Roberto Amato, Jeffrey Barrett, Sonia Goncalves, Ewan Harrison, David K. Jackson, Ian Johnston, Dominic Kwiatkowski, Cordelia Langford, John Sillitoe on behalf of the Wellcome Sanger Institute COVID-19 Surveillance Team |
| EPI_ISL_3379324, EPI_ISL_3379325, EPI_ISL_3379326 | Lighthouse Lab in Milton Keynes | Wellcome Sanger Institute for the COVID-19 Genomics UK (COG-UK) Consortium | The Lighthouse Lab in Milton Keynes and Alex Alderton, Roberto Amato, Jeffrey Barrett, Sonia Goncalves, Ewan Harrison, David K. Jackson, Ian Johnston, Dominic Kwiatkowski, Cordelia Langford, John Sillitoe on behalf of the Wellcome Sanger Institute COVID-19 Surveillance Team |
| EPI_ISL_3379327, EPI_ISL_3379328, EPI_ISL_3379329 | Lighthouse Lab in Alderley Park | Wellcome Sanger Institute for the COVID-19 Genomics UK (COG-UK) Consortium | Jacquelyn Wynn, Mairead Hyland, The Lighthouse Lab in Alderley Park and Alex Alderton, Roberto Amato, Jeffrey Barrett, Sonia Goncalves, Ewan Harrison, David K. Jackson, Ian Johnston, Dominic Kwiatkowski, Cordelia Langford, John Sillitoe on behalf of the Wellcome Sanger Institute COVID-19 Surveillance Team |
| EPI_ISL_3379330, EPI_ISL_3379331 | Lighthouse Lab in Milton Keynes | Wellcome Sanger Institute for the COVID-19 Genomics UK (COG-UK) Consortium | The Lighthouse Lab in Milton Keynes and Alex Alderton, Roberto Amato, Jeffrey Barrett, Sonia Goncalves, Ewan Harrison, David K. Jackson, Ian Johnston, Dominic Kwiatkowski, Cordelia Langford, John Sillitoe on behalf of the Wellcome Sanger Institute COVID-19 Surveillance Team |
| EPI_ISL_3379332 | Lighthouse Lab in Alderley Park | Wellcome Sanger Institute for the COVID-19 Genomics UK (COG-UK) Consortium | Jacquelyn Wynn, Mairead Hyland, The Lighthouse Lab in Alderley Park and Alex Alderton, Roberto Amato, Jeffrey Barrett, Sonia Goncalves, Ewan Harrison, David K. Jackson, Ian Johnston, Dominic Kwiatkowski, Cordelia Langford, John Sillitoe on behalf of the Wellcome Sanger Institute COVID-19 Surveillance Team |
| EPI_ISL_3379333 | LX BIO | CNR Virus des Infections Respiratoires - France SUD | Antonin Bal, Gregory Destras, Gwendolyne Burfin, Hadrien Regue, Quentin Semanas, Martine Valette, Bruno Lina, Laurence Josset |
| EPI_ISL_3379334 | Lighthouse Lab in Milton Keynes | Wellcome Sanger Institute for the COVID-19 Genomics UK (COG-UK) Consortium | The Lighthouse Lab in Milton Keynes and Alex Alderton, Roberto Amato, Jeffrey Barrett, Sonia Goncalves, Ewan Harrison, David K. Jackson, Ian Johnston, Dominic Kwiatkowski, Cordelia Langford, John Sillitoe on behalf of the Wellcome Sanger Institute COVID-19 Surveillance Team |
| EPI_ISL_3379335 | Lighthouse Lab in Alderley Park | Wellcome Sanger Institute for the COVID-19 Genomics UK (COG-UK) Consortium | Jacquelyn Wynn, Mairead Hyland, The Lighthouse Lab in Alderley Park and Alex Alderton, Roberto Amato, Jeffrey Barrett, Sonia Goncalves, Ewan Harrison, David K. Jackson, Ian Johnston, Dominic Kwiatkowski, Cordelia Langford, John Sillitoe on behalf of the Wellcome Sanger Institute COVID-19 Surveillance Team |
| EPI_ISL_3379336, EPI_ISL_3379337 | Lighthouse Lab in Milton Keynes | Wellcome Sanger Institute for the COVID-19 Genomics UK (COG-UK) Consortium | The Lighthouse Lab in Milton Keynes and Alex Alderton, Roberto Amato, Jeffrey Barrett, Sonia Goncalves, Ewan Harrison, David K. Jackson, Ian Johnston, Dominic Kwiatkowski, Cordelia Langford, John Sillitoe on behalf of the Wellcome Sanger Institute COVID-19 Surveillance Team |
| EPI_ISL_3379338, EPI_ISL_3379339 | Berkshire and Surrey Pathology Services Lighthouse Laboratory | Wellcome Sanger Institute for the COVID-19 Genomics UK (COG-UK) Consortium | Berkshire and Surrey Pathology Services Lighthouse Laboratory and Alex Alderton, Roberto Amato, Jeffrey Barrett, Sonia Goncalves, Ewan Harrison, David K. Jackson, Ian Johnston, Dominic Kwiatkowski, Cordelia Langford, John Sillitoe on behalf of the Wellcome Sanger Institute COVID-19 Surveillance Team |
| EPI_ISL_3379340 | CNR Virus des Infections Respiratoires - France SUD | CNR Virus des Infections Respiratoires - France SUD | Antonin Bal, Gregory Destras, Gwendolyne Burfin, Hadrien Regue, Quentin Semanas, Martine Valette, Bruno Lina, Laurence Josset |
| EPI_ISL_3379341, EPI_ISL_3379342 | Berkshire and Surrey Pathology Services Lighthouse Laboratory | Wellcome Sanger Institute for the COVID-19 Genomics UK (COG-UK) Consortium | Berkshire and Surrey Pathology Services Lighthouse Laboratory and Alex Alderton, Roberto Amato, Jeffrey Barrett, Sonia Goncalves, Ewan Harrison, David K. Jackson, Ian Johnston, Dominic Kwiatkowski, Cordelia Langford, John Sillitoe on behalf of the Wellcome Sanger Institute COVID-19 Surveillance Team |
| EPI_ISL_3379343, EPI_ISL_3379344 | Lighthouse Lab in Milton Keynes | Wellcome Sanger Institute for the COVID-19 Genomics UK (COG-UK) Consortium | The Lighthouse Lab in Milton Keynes and Alex Alderton, Roberto Amato, Jeffrey Barrett, Sonia Goncalves, Ewan Harrison, David K. Jackson, Ian Johnston, Dominic Kwiatkowski, Cordelia Langford, John Sillitoe on behalf of the Wellcome Sanger Institute COVID-19 Surveillance Team |
| EPI_ISL_3379345, EPI_ISL_3379346, EPI_ISL_3379347 | Berkshire and Surrey Pathology Services Lighthouse Laboratory | Wellcome Sanger Institute for the COVID-19 Genomics UK (COG-UK) Consortium | Berkshire and Surrey Pathology Services Lighthouse Laboratory and Alex Alderton, Roberto Amato, Jeffrey Barrett, Sonia Goncalves, Ewan Harrison, David K. Jackson, Ian Johnston, Dominic Kwiatkowski, Cordelia Langford, John Sillitoe on behalf of the Wellcome Sanger Institute COVID-19 Surveillance Team |

| | | | |
|---|---|---|---|
| EPI_ISL_3379348 | LX BIO | CNR Virus des Infections Respiratoires - France SUD | Antonin Bal, Gregory Destras, Gwendolyne Burfin, Hadrien Regue, Quentin Semanas, Martine Valette, Bruno Lina, Laurence Josset |
| EPI_ISL_3379349, EPI_ISL_3379350 | Lighthouse Lab in Milton Keynes | Wellcome Sanger Institute for the COVID-19 Genomics UK (COG-UK) Consortium | The Lighthouse Lab in Milton Keynes and Alex Alderton, Roberto Amato, Jeffrey Barrett, Sonia Goncalves, Ewan Harrison, David K. Jackson, Ian Johnston, Dominic Kwiatkowski, Cordelia Langford, John Sillitoe on behalf of the Wellcome Sanger Institute COVID-19 Surveillance Team |
| EPI_ISL_3379351 | CNR Virus des Infections Respiratoires - France SUD | CNR Virus des Infections Respiratoires - France SUD | Antonin Bal, Gregory Destras, Gwendolyne Burfin, Hadrien Regue, Quentin Semanas, Martine Valette, Bruno Lina, Laurence Josset |
| EPI_ISL_3379352 | Berkshire and Surrey Pathology Services Lighthouse Laboratory | Wellcome Sanger Institute for the COVID-19 Genomics UK (COG-UK) Consortium | Berkshire and Surrey Pathology Services Lighthouse Laboratory and Alex Alderton, Roberto Amato, Jeffrey Barrett, Sonia Goncalves, Ewan Harrison, David K. Jackson, Ian Johnston, Dominic Kwiatkowski, Cordelia Langford, John Sillitoe on behalf of the Wellcome Sanger Institute COVID-19 Surveillance Team |
| EPI_ISL_3379353, EPI_ISL_3379354, EPI_ISL_3379355 | Lighthouse Lab in Milton Keynes | Wellcome Sanger Institute for the COVID-19 Genomics UK (COG-UK) Consortium | The Lighthouse Lab in Milton Keynes and Alex Alderton, Roberto Amato, Jeffrey Barrett, Sonia Goncalves, Ewan Harrison, David K. Jackson, Ian Johnston, Dominic Kwiatkowski, Cordelia Langford, John Sillitoe on behalf of the Wellcome Sanger Institute COVID-19 Surveillance Team |
| EPI_ISL_3379356 | LX BIO | CNR Virus des Infections Respiratoires - France SUD | Antonin Bal, Gregory Destras, Gwendolyne Burfin, Hadrien Regue, Quentin Semanas, Martine Valette, Bruno Lina, Laurence Josset |
| EPI_ISL_3379357, EPI_ISL_3379358 | Berkshire and Surrey Pathology Services Lighthouse Laboratory | Wellcome Sanger Institute for the COVID-19 Genomics UK (COG-UK) Consortium | Berkshire and Surrey Pathology Services Lighthouse Laboratory and Alex Alderton, Roberto Amato, Jeffrey Barrett, Sonia Goncalves, Ewan Harrison, David K. Jackson, Ian Johnston, Dominic Kwiatkowski, Cordelia Langford, John Sillitoe on behalf of the Wellcome Sanger Institute COVID-19 Surveillance Team |
| EPI_ISL_3379359 | LX BIO | CNR Virus des Infections Respiratoires - France SUD | Antonin Bal, Gregory Destras, Gwendolyne Burfin, Hadrien Regue, Quentin Semanas, Martine Valette, Bruno Lina, Laurence Josset |
| EPI_ISL_3379360, EPI_ISL_3379361 | Lighthouse Lab in Alderley Park | Wellcome Sanger Institute for the COVID-19 Genomics UK (COG-UK) Consortium | Jacquelyn Wynn, Mairead Hyland, The Lighthouse Lab in Alderley Park and Alex Alderton, Roberto Amato, Jeffrey Barrett, Sonia Goncalves, Ewan Harrison, David K. Jackson, Ian Johnston, Dominic Kwiatkowski, Cordelia Langford, John Sillitoe on behalf of the Wellcome Sanger Institute COVID-19 Surveillance Team |
| EPI_ISL_3379362 | Lighthouse Lab in Milton Keynes | Wellcome Sanger Institute for the COVID-19 Genomics UK (COG-UK) Consortium | The Lighthouse Lab in Milton Keynes and Alex Alderton, Roberto Amato, Jeffrey Barrett, Sonia Goncalves, Ewan Harrison, David K. Jackson, Ian Johnston, Dominic Kwiatkowski, Cordelia Langford, John Sillitoe on behalf of the Wellcome Sanger Institute COVID-19 Surveillance Team |
| EPI_ISL_3379363 | LX BIO | CNR Virus des Infections Respiratoires - France SUD | Antonin Bal, Gregory Destras, Gwendolyne Burfin, Hadrien Regue, Quentin Semanas, Martine Valette, Bruno Lina, Laurence Josset |
| EPI_ISL_3379364 | Lighthouse Lab in Alderley Park | Wellcome Sanger Institute for the COVID-19 Genomics UK (COG-UK) Consortium | Jacquelyn Wynn, Mairead Hyland, The Lighthouse Lab in Alderley Park and Alex Alderton, Roberto Amato, Jeffrey Barrett, Sonia Goncalves, Ewan Harrison, David K. Jackson, Ian Johnston, Dominic Kwiatkowski, Cordelia Langford, John Sillitoe on behalf of the Wellcome Sanger Institute COVID-19 Surveillance Team |
| EPI_ISL_3379365 | Lighthouse Lab in Milton Keynes | Wellcome Sanger Institute for the COVID-19 Genomics UK (COG-UK) Consortium | The Lighthouse Lab in Milton Keynes and Alex Alderton, Roberto Amato, Jeffrey Barrett, Sonia Goncalves, Ewan Harrison, David K. Jackson, Ian Johnston, Dominic Kwiatkowski, Cordelia Langford, John Sillitoe on behalf of the Wellcome Sanger Institute COVID-19 Surveillance Team |
| EPI_ISL_3379366 | Lighthouse Lab in Alderley Park | Wellcome Sanger Institute for the COVID-19 Genomics UK (COG-UK) Consortium | Jacquelyn Wynn, Mairead Hyland, The Lighthouse Lab in Alderley Park and Alex Alderton, Roberto Amato, Jeffrey Barrett, Sonia Goncalves, Ewan Harrison, David K. Jackson, Ian Johnston, Dominic Kwiatkowski, Cordelia Langford, John Sillitoe on behalf of the Wellcome Sanger Institute COVID-19 Surveillance Team |
| EPI_ISL_3379367 | LX BIO | CNR Virus des Infections Respiratoires - France SUD | Antonin Bal, Gregory Destras, Gwendolyne Burfin, Hadrien Regue, Quentin Semanas, Martine Valette, Bruno Lina, Laurence Josset |
| EPI_ISL_3379368, EPI_ISL_3379369, EPI_ISL_3379370 | Berkshire and Surrey Pathology Services Lighthouse Laboratory | Wellcome Sanger Institute for the COVID-19 Genomics UK (COG-UK) Consortium | Berkshire and Surrey Pathology Services Lighthouse Laboratory and Alex Alderton, Roberto Amato, Jeffrey Barrett, Sonia Goncalves, Ewan Harrison, David K. Jackson, Ian Johnston, Dominic Kwiatkowski, Cordelia Langford, John Sillitoe on behalf of the Wellcome Sanger Institute COVID-19 Surveillance Team |
| EPI_ISL_3379371 | LX BIO | CNR Virus des Infections Respiratoires - France SUD | Antonin Bal, Gregory Destras, Gwendolyne Burfin, Hadrien Regue, Quentin Semanas, Martine Valette, Bruno Lina, Laurence Josset |
| EPI_ISL_3379372 | Berkshire and Surrey Pathology Services Lighthouse Laboratory | Wellcome Sanger Institute for the COVID-19 Genomics UK (COG-UK) Consortium | Berkshire and Surrey Pathology Services Lighthouse Laboratory and Alex Alderton, Roberto Amato, Jeffrey Barrett, Sonia Goncalves, Ewan Harrison, David K. Jackson, Ian Johnston, Dominic Kwiatkowski, Cordelia Langford, John Sillitoe on behalf of the Wellcome Sanger Institute COVID-19 Surveillance Team |
| EPI_ISL_3379373 | Lighthouse Lab in Alderley Park | Wellcome Sanger Institute for the COVID-19 Genomics UK (COG-UK) Consortium | Jacquelyn Wynn, Mairead Hyland, The Lighthouse Lab in Alderley Park and Alex Alderton, Roberto Amato, Jeffrey Barrett, Sonia Goncalves, Ewan Harrison, David K. Jackson, Ian Johnston, Dominic Kwiatkowski, Cordelia Langford, John Sillitoe on behalf of the Wellcome Sanger Institute COVID-19 Surveillance Team |
| EPI_ISL_3379374, EPI_ISL_3379375 | Lighthouse Lab in Milton Keynes | Wellcome Sanger Institute for the COVID-19 Genomics UK (COG-UK) Consortium | The Lighthouse Lab in Milton Keynes and Alex Alderton, Roberto Amato, Jeffrey Barrett, Sonia Goncalves, Ewan Harrison, David K. Jackson, Ian Johnston, Dominic Kwiatkowski, Cordelia Langford, John Sillitoe on behalf of the Wellcome Sanger Institute COVID-19 Surveillance Team |
| EPI_ISL_3379376 | LX BIO | CNR Virus des Infections Respiratoires - France SUD | Antonin Bal, Gregory Destras, Gwendolyne Burfin, Hadrien Regue, Quentin Semanas, Martine Valette, Bruno Lina, Laurence Josset |
| EPI_ISL_3379377 | Lighthouse Lab in Milton Keynes | Wellcome Sanger Institute for the COVID-19 Genomics UK (COG-UK) Consortium | The Lighthouse Lab in Milton Keynes and Alex Alderton, Roberto Amato, Jeffrey Barrett, Sonia Goncalves, Ewan Harrison, David K. Jackson, Ian Johnston, Dominic Kwiatkowski, Cordelia Langford, John Sillitoe on behalf of the Wellcome Sanger Institute COVID-19 Surveillance Team |
| EPI_ISL_3379378, EPI_ISL_3379379 | Berkshire and Surrey Pathology Services Lighthouse Laboratory | Wellcome Sanger Institute for the COVID-19 Genomics UK (COG-UK) Consortium | Berkshire and Surrey Pathology Services Lighthouse Laboratory and Alex Alderton, Roberto Amato, Jeffrey Barrett, Sonia Goncalves, Ewan Harrison, David K. Jackson, Ian Johnston, Dominic Kwiatkowski, Cordelia Langford, John Sillitoe on behalf of the Wellcome Sanger Institute COVID-19 Surveillance Team |
| EPI_ISL_3379380 | LX BIO | CNR Virus des Infections Respiratoires - France SUD | Antonin Bal, Gregory Destras, Gwendolyne Burfin, Hadrien Regue, Quentin Semanas, Martine Valette, Bruno Lina, Laurence Josset |
| EPI_ISL_3379381 | Berkshire and Surrey Pathology Services Lighthouse Laboratory | Wellcome Sanger Institute for the COVID-19 Genomics UK (COG-UK) Consortium | Berkshire and Surrey Pathology Services Lighthouse Laboratory and Alex Alderton, Roberto Amato, Jeffrey Barrett, Sonia Goncalves, Ewan Harrison, David K. Jackson, Ian Johnston, Dominic Kwiatkowski, Cordelia Langford, John Sillitoe on behalf of the Wellcome Sanger Institute COVID-19 Surveillance Team |
| EPI_ISL_3379382 | Lighthouse Lab in Alderley Park | Wellcome Sanger Institute for the COVID-19 Genomics UK (COG-UK) Consortium | Jacquelyn Wynn, Mairead Hyland, The Lighthouse Lab in Alderley Park and Alex Alderton, Roberto Amato, Jeffrey Barrett, Sonia Goncalves, Ewan Harrison, David K. Jackson, Ian Johnston, Dominic Kwiatkowski, Cordelia Langford, John Sillitoe on behalf of the Wellcome Sanger Institute COVID-19 Surveillance Team |
| EPI_ISL_3379383 | LX BIO | CNR Virus des Infections Respiratoires - France SUD | Antonin Bal, Gregory Destras, Gwendolyne Burfin, Hadrien Regue, Quentin Semanas, Martine Valette, Bruno Lina, Laurence Josset |
| EPI_ISL_3379384, EPI_ISL_3379385, EPI_ISL_3379386 | C.H.R.U. MONTPIED | CNR Virus des Infections Respiratoires - France SUD | Antonin Bal, Gregory Destras, Gwendolyne Burfin, Hadrien Regue, Quentin Semanas, Martine Valette, Bruno Lina, Laurence Josset |
| EPI_ISL_3379387, EPI_ISL_3379388, EPI_ISL_3379389, EPI_ISL_3379390, EPI_ISL_3379391, EPI_ISL_3379392, EPI_ISL_3379393 | CNR Virus des Infections Respiratoires - France SUD | CNR Virus des Infections Respiratoires - France SUD | Antonin Bal, Gregory Destras, Gwendolyne Burfin, Hadrien Regue, Quentin Semanas, Martine Valette, Bruno Lina, Laurence Josset |
| EPI_ISL_3379394 | ORIAPOLE | CNR Virus des Infections Respiratoires - France SUD | Antonin Bal, Gregory Destras, Gwendolyne Burfin, Hadrien Regue, Quentin Semanas, Martine Valette, Bruno Lina, Laurence Josset |
| EPI_ISL_3379395 | LABORATOIRE ACCOLAB SUD OUEST | CNR Virus des Infections Respiratoires - France SUD | Antonin Bal, Gregory Destras, Gwendolyne Burfin, Hadrien Regue, Quentin Semanas, Martine Valette, Bruno Lina, Laurence Josset |
| EPI_ISL_3379396, EPI_ISL_3379397, EPI_ISL_3379398, EPI_ISL_3379399, EPI_ISL_3379400, EPI_ISL_3379401, EPI_ISL_3379402, EPI_ISL_3379403, EPI_ISL_3379404, EPI_ISL_3379405, EPI_ISL_3379406, EPI_ISL_3379407 | | | |
| see above | ORIAPOLE | CNR Virus des Infections Respiratoires - France SUD | Antonin Bal, Gregory Destras, Gwendolyne Burfin, Hadrien Regue, Quentin Semanas, Martine Valette, Bruno Lina, Laurence Josset |
| EPI_ISL_3379408 | LABORATOIRE ACCOLAB SUD OUEST | CNR Virus des Infections Respiratoires - France SUD | Antonin Bal, Gregory Destras, Gwendolyne Burfin, Hadrien Regue, Quentin Semanas, Martine Valette, Bruno Lina, Laurence Josset |
| EPI_ISL_3379409, EPI_ISL_3379410 | ORIAPOLE | CNR Virus des Infections Respiratoires - France SUD | Antonin Bal, Gregory Destras, Gwendolyne Burfin, Hadrien Regue, Quentin Semanas, Martine Valette, Bruno Lina, Laurence Josset |
| EPI_ISL_3379411 | LABORATOIRE ACCOLAB SUD OUEST | CNR Virus des Infections Respiratoires - France SUD | Antonin Bal, Gregory Destras, Gwendolyne Burfin, Hadrien Regue, Quentin Semanas, Martine Valette, Bruno Lina, Laurence Josset |
| EPI_ISL_3379412 | Berkshire and Surrey Pathology Services Lighthouse Laboratory | Wellcome Sanger Institute for the COVID-19 Genomics UK (COG-UK) Consortium | Berkshire and Surrey Pathology Services Lighthouse Laboratory and Alex Alderton, Roberto Amato, Jeffrey Barrett, Sonia Goncalves, Ewan Harrison, David K. Jackson, Ian Johnston, Dominic Kwiatkowski, Cordelia Langford, John Sillitoe on behalf of the Wellcome Sanger Institute COVID-19 Surveillance Team |
| EPI_ISL_3379413 | Lighthouse Lab in Milton Keynes | Wellcome Sanger Institute for the COVID-19 Genomics UK (COG-UK) Consortium | The Lighthouse Lab in Milton Keynes and Alex Alderton, Roberto Amato, Jeffrey Barrett, Sonia Goncalves, Ewan Harrison, David K. Jackson, Ian Johnston, Dominic Kwiatkowski, Cordelia Langford, John Sillitoe on behalf of the Wellcome Sanger Institute COVID-19 Surveillance Team |
| EPI_ISL_3379414 | ORIAPOLE | CNR Virus des Infections Respiratoires - France SUD | Antonin Bal, Gregory Destras, Gwendolyne Burfin, Hadrien Regue, Quentin Semanas, Martine Valette, Bruno Lina, Laurence Josset |
| EPI_ISL_3379415 | Lighthouse Lab in Milton Keynes | Wellcome Sanger Institute for the COVID-19 Genomics UK (COG-UK) Consortium | The Lighthouse Lab in Milton Keynes and Alex Alderton, Roberto Amato, Jeffrey Barrett, Sonia Goncalves, Ewan Harrison, David K. Jackson, Ian Johnston, Dominic Kwiatkowski, Cordelia Langford, John Sillitoe on behalf of the Wellcome Sanger Institute COVID-19 Surveillance Team |
| EPI_ISL_3379416 | Berkshire and Surrey Pathology Services Lighthouse Laboratory | Wellcome Sanger Institute for the COVID-19 Genomics UK (COG-UK) Consortium | Berkshire and Surrey Pathology Services Lighthouse Laboratory and Alex Alderton, Roberto Amato, Jeffrey Barrett, Sonia Goncalves, Ewan Harrison, David K. Jackson, Ian Johnston, Dominic Kwiatkowski, Cordelia Langford, John Sillitoe on behalf of the Wellcome Sanger Institute COVID-19 Surveillance Team |
| EPI_ISL_3379417 | LABORATOIRE ACCOLAB SUD OUEST | CNR Virus des Infections Respiratoires - France SUD | Antonin Bal, Gregory Destras, Gwendolyne Burfin, Hadrien Regue, Quentin Semanas, Martine Valette, Bruno Lina, Laurence Josset |
| EPI_ISL_3379418 | Lighthouse Lab in Milton Keynes | Wellcome Sanger Institute for the COVID-19 Genomics UK (COG-UK) Consortium | The Lighthouse Lab in Milton Keynes and Alex Alderton, Roberto Amato, Jeffrey Barrett, Sonia Goncalves, Ewan Harrison, David K. Jackson, Ian Johnston, Dominic Kwiatkowski, Cordelia Langford, John Sillitoe on behalf of the Wellcome Sanger Institute COVID-19 Surveillance Team |
| EPI_ISL_3379419 | Lighthouse Lab in Alderley Park | Wellcome Sanger Institute for the COVID-19 Genomics UK (COG-UK) Consortium | Jacquelyn Wynn, Mairead Hyland, The Lighthouse Lab in Alderley Park and Alex Alderton, Roberto Amato, Jeffrey Barrett, Sonia Goncalves, Ewan Harrison, David K. Jackson, Ian Johnston, Dominic Kwiatkowski, Cordelia Langford, John Sillitoe on behalf of the Wellcome Sanger Institute COVID-19 Surveillance Team |
| EPI_ISL_3379420 | Lighthouse Lab in Milton Keynes | Wellcome Sanger Institute for the COVID-19 Genomics UK (COG-UK) Consortium | The Lighthouse Lab in Milton Keynes and Alex Alderton, Roberto Amato, Jeffrey Barrett, Sonia Goncalves, Ewan Harrison, David K. Jackson, Ian Johnston, Dominic Kwiatkowski, Cordelia Langford, John Sillitoe on behalf of the Wellcome Sanger Institute COVID-19 Surveillance Team |
| EPI_ISL_3379421 | ORIAPOLE | CNR Virus des Infections Respiratoires - France SUD | Antonin Bal, Gregory Destras, Gwendolyne Burfin, Hadrien Regue, Quentin Semanas, Martine Valette, Bruno Lina, Laurence Josset |
| EPI_ISL_3379422 | Lighthouse Lab in Milton Keynes | Wellcome Sanger Institute for the COVID-19 Genomics UK (COG-UK) Consortium | The Lighthouse Lab in Milton Keynes and Alex Alderton, Roberto Amato, Jeffrey Barrett, Sonia Goncalves, Ewan Harrison, David K. Jackson, Ian Johnston, Dominic Kwiatkowski, Cordelia Langford, John Sillitoe on behalf of the Wellcome Sanger Institute COVID-19 Surveillance Team |
| EPI_ISL_3379423 | Lighthouse Lab in Alderley Park | Wellcome Sanger Institute for the COVID-19 Genomics UK (COG-UK) Consortium | Jacquelyn Wynn, Mairead Hyland, The Lighthouse Lab in Alderley Park and Alex Alderton, Roberto Amato, Jeffrey Barrett, Sonia Goncalves, Ewan Harrison, David K. Jackson, Ian Johnston, Dominic Kwiatkowski, Cordelia Langford, John Sillitoe on behalf of the Wellcome Sanger Institute COVID-19 Surveillance Team |
| EPI_ISL_3379424 | Lighthouse Lab in Milton Keynes | Wellcome Sanger Institute for the COVID-19 Genomics UK (COG-UK) Consortium | The Lighthouse Lab in Milton Keynes and Alex Alderton, Roberto Amato, Jeffrey Barrett, Sonia Goncalves, Ewan Harrison, David K. Jackson, Ian Johnston, Dominic Kwiatkowski, Cordelia Langford, John Sillitoe on behalf of the Wellcome Sanger Institute COVID-19 Surveillance Team |
| EPI_ISL_3379425 | ORIAPOLE | CNR Virus des Infections Respiratoires - France SUD | Antonin Bal, Gregory Destras, Gwendolyne Burfin, Hadrien Regue, Quentin Semanas, Martine Valette, Bruno Lina, Laurence Josset |

| | | | |
|---|---|---|---|
| EPI_ISL_3379426 | Lighthouse Lab in Milton Keynes | Wellcome Sanger Institute for the COVID-19 Genomics UK (COG-UK) Consortium | The Lighthouse Lab in Milton Keynes and Alex Alderton, Roberto Amato, Jeffrey Barrett, Sonia Goncalves, Ewan Harrison, David K. Jackson, Ian Johnston, Dominic Kwiatkowski, Cordelia Langford, John Sillitoe on behalf of the Wellcome Sanger Institute COVID-19 Surveillance Team |
| EPI_ISL_3379427 | Lighthouse Lab in Alderley Park | Wellcome Sanger Institute for the COVID-19 Genomics UK (COG-UK) Consortium | Jacquelyn Wynn, Mairead Hyland, The Lighthouse Lab in Alderley Park and Alex Alderton, Roberto Amato, Jeffrey Barrett, Sonia Goncalves, Ewan Harrison, David K. Jackson, Ian Johnston, Dominic Kwiatkowski, Cordelia Langford, John Sillitoe on behalf of the Wellcome Sanger Institute COVID-19 Surveillance Team |
| EPI_ISL_3379428 | Berkshire and Surrey Pathology Services Lighthouse Laboratory | Wellcome Sanger Institute for the COVID-19 Genomics UK (COG-UK) Consortium | Berkshire and Surrey Pathology Services Lighthouse Laboratory and Alex Alderton, Roberto Amato, Jeffrey Barrett, Sonia Goncalves, Ewan Harrison, David K. Jackson, Ian Johnston, Dominic Kwiatkowski, Cordelia Langford, John Sillitoe on behalf of the Wellcome Sanger Institute COVID-19 Surveillance Team |
| EPI_ISL_3379429 | Lighthouse Lab in Alderley Park | Wellcome Sanger Institute for the COVID-19 Genomics UK (COG-UK) Consortium | Jacquelyn Wynn, Mairead Hyland, The Lighthouse Lab in Alderley Park and Alex Alderton, Roberto Amato, Jeffrey Barrett, Sonia Goncalves, Ewan Harrison, David K. Jackson, Ian Johnston, Dominic Kwiatkowski, Cordelia Langford, John Sillitoe on behalf of the Wellcome Sanger Institute COVID-19 Surveillance Team |
| EPI_ISL_3379430, EPI_ISL_3379431, EPI_ISL_3379432 | Lighthouse Lab in Milton Keynes | Wellcome Sanger Institute for the COVID-19 Genomics UK (COG-UK) Consortium | The Lighthouse Lab in Milton Keynes and Alex Alderton, Roberto Amato, Jeffrey Barrett, Sonia Goncalves, Ewan Harrison, David K. Jackson, Ian Johnston, Dominic Kwiatkowski, Cordelia Langford, John Sillitoe on behalf of the Wellcome Sanger Institute COVID-19 Surveillance Team |
| EPI_ISL_3379433 | LABORATOIRE ACCOLAB SUD OUEST | CNR Virus des Infections Respiratoires - France SUD | Antonin Bal, Gregory Destras, Gwendolyne Burfin, Hadrien Regue, Quentin Semanas, Martine Valette, Bruno Lina, Laurence Josset |
| EPI_ISL_3379434 | Berkshire and Surrey Pathology Services Lighthouse Laboratory | Wellcome Sanger Institute for the COVID-19 Genomics UK (COG-UK) Consortium | Berkshire and Surrey Pathology Services Lighthouse Laboratory and Alex Alderton, Roberto Amato, Jeffrey Barrett, Sonia Goncalves, Ewan Harrison, David K. Jackson, Ian Johnston, Dominic Kwiatkowski, Cordelia Langford, John Sillitoe on behalf of the Wellcome Sanger Institute COVID-19 Surveillance Team |
| EPI_ISL_3379435, EPI_ISL_3379436 | Lighthouse Lab in Milton Keynes | Wellcome Sanger Institute for the COVID-19 Genomics UK (COG-UK) Consortium | The Lighthouse Lab in Milton Keynes and Alex Alderton, Roberto Amato, Jeffrey Barrett, Sonia Goncalves, Ewan Harrison, David K. Jackson, Ian Johnston, Dominic Kwiatkowski, Cordelia Langford, John Sillitoe on behalf of the Wellcome Sanger Institute COVID-19 Surveillance Team |
| EPI_ISL_3379437 | ORIAPOLE | CNR Virus des Infections Respiratoires - France SUD | Antonin Bal, Gregory Destras, Gwendolyne Burfin, Hadrien Regue, Quentin Semanas, Martine Valette, Bruno Lina, Laurence Josset |
| EPI_ISL_3379438 | Berkshire and Surrey Pathology Services Lighthouse Laboratory | Wellcome Sanger Institute for the COVID-19 Genomics UK (COG-UK) Consortium | Berkshire and Surrey Pathology Services Lighthouse Laboratory and Alex Alderton, Roberto Amato, Jeffrey Barrett, Sonia Goncalves, Ewan Harrison, David K. Jackson, Ian Johnston, Dominic Kwiatkowski, Cordelia Langford, John Sillitoe on behalf of the Wellcome Sanger Institute COVID-19 Surveillance Team |
| EPI_ISL_3379439 | Lighthouse Lab in Milton Keynes | Wellcome Sanger Institute for the COVID-19 Genomics UK (COG-UK) Consortium | The Lighthouse Lab in Milton Keynes and Alex Alderton, Roberto Amato, Jeffrey Barrett, Sonia Goncalves, Ewan Harrison, David K. Jackson, Ian Johnston, Dominic Kwiatkowski, Cordelia Langford, John Sillitoe on behalf of the Wellcome Sanger Institute COVID-19 Surveillance Team |
| EPI_ISL_3379440 | Lighthouse Lab in Alderley Park | Wellcome Sanger Institute for the COVID-19 Genomics UK (COG-UK) Consortium | Jacquelyn Wynn, Mairead Hyland, The Lighthouse Lab in Alderley Park and Alex Alderton, Roberto Amato, Jeffrey Barrett, Sonia Goncalves, Ewan Harrison, David K. Jackson, Ian Johnston, Dominic Kwiatkowski, Cordelia Langford, John Sillitoe on behalf of the Wellcome Sanger Institute COVID-19 Surveillance Team |
| EPI_ISL_3379441 | ORIAPOLE | CNR Virus des Infections Respiratoires - France SUD | Antonin Bal, Gregory Destras, Gwendolyne Burfin, Hadrien Regue, Quentin Semanas, Martine Valette, Bruno Lina, Laurence Josset |
| EPI_ISL_3379442 | Berkshire and Surrey Pathology Services Lighthouse Laboratory | Wellcome Sanger Institute for the COVID-19 Genomics UK (COG-UK) Consortium | Berkshire and Surrey Pathology Services Lighthouse Laboratory and Alex Alderton, Roberto Amato, Jeffrey Barrett, Sonia Goncalves, Ewan Harrison, David K. Jackson, Ian Johnston, Dominic Kwiatkowski, Cordelia Langford, John Sillitoe on behalf of the Wellcome Sanger Institute COVID-19 Surveillance Team |
| EPI_ISL_3379443, EPI_ISL_3379444, EPI_ISL_3379445, EPI_ISL_3379446, EPI_ISL_3379447, EPI_ISL_3379448 | Lighthouse Lab in Milton Keynes | Wellcome Sanger Institute for the COVID-19 Genomics UK (COG-UK) Consortium | The Lighthouse Lab in Milton Keynes and Alex Alderton, Roberto Amato, Jeffrey Barrett, Sonia Goncalves, Ewan Harrison, David K. Jackson, Ian Johnston, Dominic Kwiatkowski, Cordelia Langford, John Sillitoe on behalf of the Wellcome Sanger Institute COVID-19 Surveillance Team |
| EPI_ISL_3379449 | ORIAPOLE | CNR Virus des Infections Respiratoires - France SUD | Antonin Bal, Gregory Destras, Gwendolyne Burfin, Hadrien Regue, Quentin Semanas, Martine Valette, Bruno Lina, Laurence Josset |
| EPI_ISL_3379450, EPI_ISL_3379451 | Lighthouse Lab in Milton Keynes | Wellcome Sanger Institute for the COVID-19 Genomics UK (COG-UK) Consortium | The Lighthouse Lab in Milton Keynes and Alex Alderton, Roberto Amato, Jeffrey Barrett, Sonia Goncalves, Ewan Harrison, David K. Jackson, Ian Johnston, Dominic Kwiatkowski, Cordelia Langford, John Sillitoe on behalf of the Wellcome Sanger Institute COVID-19 Surveillance Team |
| EPI_ISL_3379452, EPI_ISL_3379453 | Berkshire and Surrey Pathology Services Lighthouse Laboratory | Wellcome Sanger Institute for the COVID-19 Genomics UK (COG-UK) Consortium | Berkshire and Surrey Pathology Services Lighthouse Laboratory and Alex Alderton, Roberto Amato, Jeffrey Barrett, Sonia Goncalves, Ewan Harrison, David K. Jackson, Ian Johnston, Dominic Kwiatkowski, Cordelia Langford, John Sillitoe on behalf of the Wellcome Sanger Institute COVID-19 Surveillance Team |
| EPI_ISL_3379454, EPI_ISL_3379455 | Lighthouse Lab in Milton Keynes | Wellcome Sanger Institute for the COVID-19 Genomics UK (COG-UK) Consortium | The Lighthouse Lab in Milton Keynes and Alex Alderton, Roberto Amato, Jeffrey Barrett, Sonia Goncalves, Ewan Harrison, David K. Jackson, Ian Johnston, Dominic Kwiatkowski, Cordelia Langford, John Sillitoe on behalf of the Wellcome Sanger Institute COVID-19 Surveillance Team |
| EPI_ISL_3379456 | Lighthouse Lab in Alderley Park | Wellcome Sanger Institute for the COVID-19 Genomics UK (COG-UK) Consortium | Jacquelyn Wynn, Mairead Hyland, The Lighthouse Lab in Alderley Park and Alex Alderton, Roberto Amato, Jeffrey Barrett, Sonia Goncalves, Ewan Harrison, David K. Jackson, Ian Johnston, Dominic Kwiatkowski, Cordelia Langford, John Sillitoe on behalf of the Wellcome Sanger Institute COVID-19 Surveillance Team |
| EPI_ISL_3379457 | LABORATOIRE ACCOLAB SUD OUEST | CNR Virus des Infections Respiratoires - France SUD | Antonin Bal, Gregory Destras, Gwendolyne Burfin, Hadrien Regue, Quentin Semanas, Martine Valette, Bruno Lina, Laurence Josset |
| EPI_ISL_3379458 | Lighthouse Lab in Alderley Park | Wellcome Sanger Institute for the COVID-19 Genomics UK (COG-UK) Consortium | Jacquelyn Wynn, Mairead Hyland, The Lighthouse Lab in Alderley Park and Alex Alderton, Roberto Amato, Jeffrey Barrett, Sonia Goncalves, Ewan Harrison, David K. Jackson, Ian Johnston, Dominic Kwiatkowski, Cordelia Langford, John Sillitoe on behalf of the Wellcome Sanger Institute COVID-19 Surveillance Team |
| EPI_ISL_3379459 | Berkshire and Surrey Pathology Services Lighthouse Laboratory | Wellcome Sanger Institute for the COVID-19 Genomics UK (COG-UK) Consortium | Berkshire and Surrey Pathology Services Lighthouse Laboratory and Alex Alderton, Roberto Amato, Jeffrey Barrett, Sonia Goncalves, Ewan Harrison, David K. Jackson, Ian Johnston, Dominic Kwiatkowski, Cordelia Langford, John Sillitoe on behalf of the Wellcome Sanger Institute COVID-19 Surveillance Team |
| EPI_ISL_3379460, EPI_ISL_3379461 | Lighthouse Lab in Milton Keynes | Wellcome Sanger Institute for the COVID-19 Genomics UK (COG-UK) Consortium | The Lighthouse Lab in Milton Keynes and Alex Alderton, Roberto Amato, Jeffrey Barrett, Sonia Goncalves, Ewan Harrison, David K. Jackson, Ian Johnston, Dominic Kwiatkowski, Cordelia Langford, John Sillitoe on behalf of the Wellcome Sanger Institute COVID-19 Surveillance Team |
| EPI_ISL_3379462 | Berkshire and Surrey Pathology Services Lighthouse Laboratory | Wellcome Sanger Institute for the COVID-19 Genomics UK (COG-UK) Consortium | Berkshire and Surrey Pathology Services Lighthouse Laboratory and Alex Alderton, Roberto Amato, Jeffrey Barrett, Sonia Goncalves, Ewan Harrison, David K. Jackson, Ian Johnston, Dominic Kwiatkowski, Cordelia Langford, John Sillitoe on behalf of the Wellcome Sanger Institute COVID-19 Surveillance Team |
| EPI_ISL_3379463, EPI_ISL_3379464, EPI_ISL_3379465, EPI_ISL_3379467 | Lighthouse Lab in Milton Keynes | Wellcome Sanger Institute for the COVID-19 Genomics UK (COG-UK) Consortium | The Lighthouse Lab in Milton Keynes and Alex Alderton, Roberto Amato, Jeffrey Barrett, Sonia Goncalves, Ewan Harrison, David K. Jackson, Ian Johnston, Dominic Kwiatkowski, Cordelia Langford, John Sillitoe on behalf of the Wellcome Sanger Institute COVID-19 Surveillance Team |
| EPI_ISL_3379468, EPI_ISL_3379469 | Berkshire and Surrey Pathology Services Lighthouse Laboratory | Wellcome Sanger Institute for the COVID-19 Genomics UK (COG-UK) Consortium | Berkshire and Surrey Pathology Services Lighthouse Laboratory and Alex Alderton, Roberto Amato, Jeffrey Barrett, Sonia Goncalves, Ewan Harrison, David K. Jackson, Ian Johnston, Dominic Kwiatkowski, Cordelia Langford, John Sillitoe on behalf of the Wellcome Sanger Institute COVID-19 Surveillance Team |
| EPI_ISL_3379470 | Lighthouse Lab in Milton Keynes | Wellcome Sanger Institute for the COVID-19 Genomics UK (COG-UK) Consortium | The Lighthouse Lab in Milton Keynes and Alex Alderton, Roberto Amato, Jeffrey Barrett, Sonia Goncalves, Ewan Harrison, David K. Jackson, Ian Johnston, Dominic Kwiatkowski, Cordelia Langford, John Sillitoe on behalf of the Wellcome Sanger Institute COVID-19 Surveillance Team |
| EPI_ISL_3379471 | Berkshire and Surrey Pathology Services Lighthouse Laboratory | Wellcome Sanger Institute for the COVID-19 Genomics UK (COG-UK) Consortium | Berkshire and Surrey Pathology Services Lighthouse Laboratory and Alex Alderton, Roberto Amato, Jeffrey Barrett, Sonia Goncalves, Ewan Harrison, David K. Jackson, Ian Johnston, Dominic Kwiatkowski, Cordelia Langford, John Sillitoe on behalf of the Wellcome Sanger Institute COVID-19 Surveillance Team |
| EPI_ISL_3379472 | Lighthouse Lab in Milton Keynes | Wellcome Sanger Institute for the COVID-19 Genomics UK (COG-UK) Consortium | The Lighthouse Lab in Milton Keynes and Alex Alderton, Roberto Amato, Jeffrey Barrett, Sonia Goncalves, Ewan Harrison, David K. Jackson, Ian Johnston, Dominic Kwiatkowski, Cordelia Langford, John Sillitoe on behalf of the Wellcome Sanger Institute COVID-19 Surveillance Team |
| EPI_ISL_3379473 | LABORATOIRE ACCOLAB SUD OUEST | CNR Virus des Infections Respiratoires - France SUD | Antonin Bal, Gregory Destras, Gwendolyne Burfin, Hadrien Regue, Quentin Semanas, Martine Valette, Bruno Lina, Laurence Josset |
| EPI_ISL_3379474 | Lighthouse Lab in Milton Keynes | Wellcome Sanger Institute for the COVID-19 Genomics UK (COG-UK) Consortium | The Lighthouse Lab in Milton Keynes and Alex Alderton, Roberto Amato, Jeffrey Barrett, Sonia Goncalves, Ewan Harrison, David K. Jackson, Ian Johnston, Dominic Kwiatkowski, Cordelia Langford, John Sillitoe on behalf of the Wellcome Sanger Institute COVID-19 Surveillance Team |
| EPI_ISL_3379475 | Lighthouse Lab in Alderley Park | Wellcome Sanger Institute for the COVID-19 Genomics UK (COG-UK) Consortium | Jacquelyn Wynn, Mairead Hyland, The Lighthouse Lab in Alderley Park and Alex Alderton, Roberto Amato, Jeffrey Barrett, Sonia Goncalves, Ewan Harrison, David K. Jackson, Ian Johnston, Dominic Kwiatkowski, Cordelia Langford, John Sillitoe on behalf of the Wellcome Sanger Institute COVID-19 Surveillance Team |
| EPI_ISL_3379476 | ORIAPOLE | CNR Virus des Infections Respiratoires - France SUD | Antonin Bal, Gregory Destras, Gwendolyne Burfin, Hadrien Regue, Quentin Semanas, Martine Valette, Bruno Lina, Laurence Josset |
| EPI_ISL_3379477, EPI_ISL_3379478, EPI_ISL_3379479 | Lighthouse Lab in Milton Keynes | Wellcome Sanger Institute for the COVID-19 Genomics UK (COG-UK) Consortium | The Lighthouse Lab in Milton Keynes and Alex Alderton, Roberto Amato, Jeffrey Barrett, Sonia Goncalves, Ewan Harrison, David K. Jackson, Ian Johnston, Dominic Kwiatkowski, Cordelia Langford, John Sillitoe on behalf of the Wellcome Sanger Institute COVID-19 Surveillance Team |
| EPI_ISL_3379480 | LABORATOIRE ACCOLAB SUD OUEST | CNR Virus des Infections Respiratoires - France SUD | Antonin Bal, Gregory Destras, Gwendolyne Burfin, Hadrien Regue, Quentin Semanas, Martine Valette, Bruno Lina, Laurence Josset |
| EPI_ISL_3379481 | Berkshire and Surrey Pathology Services Lighthouse Laboratory | Wellcome Sanger Institute for the COVID-19 Genomics UK (COG-UK) Consortium | Berkshire and Surrey Pathology Services Lighthouse Laboratory and Alex Alderton, Roberto Amato, Jeffrey Barrett, Sonia Goncalves, Ewan Harrison, David K. Jackson, Ian Johnston, Dominic Kwiatkowski, Cordelia Langford, John Sillitoe on behalf of the Wellcome Sanger Institute COVID-19 Surveillance Team |
| EPI_ISL_3379482 | Lighthouse Lab in Milton Keynes | Wellcome Sanger Institute for the COVID-19 Genomics UK (COG-UK) Consortium | The Lighthouse Lab in Milton Keynes and Alex Alderton, Roberto Amato, Jeffrey Barrett, Sonia Goncalves, Ewan Harrison, David K. Jackson, Ian Johnston, Dominic Kwiatkowski, Cordelia Langford, John Sillitoe on behalf of the Wellcome Sanger Institute COVID-19 Surveillance Team |
| EPI_ISL_3379483 | Lighthouse Lab in Alderley Park | Wellcome Sanger Institute for the COVID-19 Genomics UK (COG-UK) Consortium | Jacquelyn Wynn, Mairead Hyland, The Lighthouse Lab in Alderley Park and Alex Alderton, Roberto Amato, Jeffrey Barrett, Sonia Goncalves, Ewan Harrison, David K. Jackson, Ian Johnston, Dominic Kwiatkowski, Cordelia Langford, John Sillitoe on behalf of the Wellcome Sanger Institute COVID-19 Surveillance Team |
| EPI_ISL_3379484 | ORIAPOLE | CNR Virus des Infections Respiratoires - France SUD | Antonin Bal, Gregory Destras, Gwendolyne Burfin, Hadrien Regue, Quentin Semanas, Martine Valette, Bruno Lina, Laurence Josset |
| EPI_ISL_3379485 | Lighthouse Lab in Milton Keynes | Wellcome Sanger Institute for the COVID-19 Genomics UK (COG-UK) Consortium | The Lighthouse Lab in Milton Keynes and Alex Alderton, Roberto Amato, Jeffrey Barrett, Sonia Goncalves, Ewan Harrison, David K. Jackson, Ian Johnston, Dominic Kwiatkowski, Cordelia Langford, John Sillitoe on behalf of the Wellcome Sanger Institute COVID-19 Surveillance Team |
| EPI_ISL_3379486 | Lighthouse Lab in Alderley Park | Wellcome Sanger Institute for the COVID-19 Genomics UK (COG-UK) Consortium | Jacquelyn Wynn, Mairead Hyland, The Lighthouse Lab in Alderley Park and Alex Alderton, Roberto Amato, Jeffrey Barrett, Sonia Goncalves, Ewan Harrison, David K. Jackson, Ian Johnston, Dominic Kwiatkowski, Cordelia Langford, John Sillitoe on behalf of the Wellcome Sanger Institute COVID-19 Surveillance Team |
| EPI_ISL_3379487 | LABORATOIRE ACCOLAB SUD OUEST | CNR Virus des Infections Respiratoires - France SUD | Antonin Bal, Gregory Destras, Gwendolyne Burfin, Hadrien Regue, Quentin Semanas, Martine Valette, Bruno Lina, Laurence Josset |
| EPI_ISL_3379488, EPI_ISL_3379489 | Lighthouse Lab in Milton Keynes | Wellcome Sanger Institute for the COVID-19 Genomics UK (COG-UK) Consortium | The Lighthouse Lab in Milton Keynes and Alex Alderton, Roberto Amato, Jeffrey Barrett, Sonia Goncalves, Ewan Harrison, David K. Jackson, Ian Johnston, Dominic Kwiatkowski, Cordelia Langford, John Sillitoe on behalf of the Wellcome Sanger Institute COVID-19 Surveillance Team |
| EPI_ISL_3379490 | Berkshire and Surrey Pathology Services Lighthouse Laboratory | Wellcome Sanger Institute for the COVID-19 Genomics UK (COG-UK) Consortium | Berkshire and Surrey Pathology Services Lighthouse Laboratory and Alex Alderton, Roberto Amato, Jeffrey Barrett, Sonia Goncalves, Ewan Harrison, David K. Jackson, Ian Johnston, Dominic Kwiatkowski, Cordelia Langford, John Sillitoe on behalf of the Wellcome Sanger Institute COVID-19 Surveillance Team |
| EPI_ISL_3379491 | Lighthouse Lab in Milton Keynes | Wellcome Sanger Institute for the COVID-19 Genomics UK (COG-UK) Consortium | The Lighthouse Lab in Milton Keynes and Alex Alderton, Roberto Amato, Jeffrey Barrett, Sonia Goncalves, Ewan Harrison, David K. Jackson, Ian Johnston, Dominic Kwiatkowski, Cordelia Langford, John Sillitoe on behalf of the Wellcome Sanger Institute COVID-19 Surveillance Team |

| | | | |
|---|---|---|---|
| EPI_ISL_3379492 | ORIAPOLE | CNR Virus des Infections Respiratoires - France SUD | Antonin Bal, Gregory Destras, Gwendolyne Burfin, Hadrien Regue, Quentin Semanas, Martine Valette, Bruno Lina, Laurence Josset |
| EPI_ISL_3379493, EPI_ISL_3379494 | Lighthouse Lab in Alderley Park | Wellcome Sanger Institute for the COVID-19 Genomics UK (COG-UK) Consortium | Jacquelyn Wynn, Mairead Hyland, The Lighthouse Lab in Alderley Park and Alex Alderton, Roberto Amato, Jeffrey Barrett, Sonia Goncalves, Ewan Harrison, David K. Jackson, Ian Johnston, Dominic Kwiatkowski, Cordelia Langford, John Sillitoe on behalf of the Wellcome Sanger Institute COVID-19 Surveillance Team |
| EPI_ISL_3379495 | Lighthouse Lab in Milton Keynes | Wellcome Sanger Institute for the COVID-19 Genomics UK (COG-UK) Consortium | The Lighthouse Lab in Milton Keynes and Alex Alderton, Roberto Amato, Jeffrey Barrett, Sonia Goncalves, Ewan Harrison, David K. Jackson, Ian Johnston, Dominic Kwiatkowski, Cordelia Langford, John Sillitoe on behalf of the Wellcome Sanger Institute COVID-19 Surveillance Team |
| EPI_ISL_3379496 | ORIAPOLE | CNR Virus des Infections Respiratoires - France SUD | Antonin Bal, Gregory Destras, Gwendolyne Burfin, Hadrien Regue, Quentin Semanas, Martine Valette, Bruno Lina, Laurence Josset |
| EPI_ISL_3379497, EPI_ISL_3379498 | Lighthouse Lab in Alderley Park | Wellcome Sanger Institute for the COVID-19 Genomics UK (COG-UK) Consortium | Jacquelyn Wynn, Mairead Hyland, The Lighthouse Lab in Alderley Park and Alex Alderton, Roberto Amato, Jeffrey Barrett, Sonia Goncalves, Ewan Harrison, David K. Jackson, Ian Johnston, Dominic Kwiatkowski, Cordelia Langford, John Sillitoe on behalf of the Wellcome Sanger Institute COVID-19 Surveillance Team |
| EPI_ISL_3379499 | Berkshire and Surrey Pathology Services Lighthouse Laboratory | Wellcome Sanger Institute for the COVID-19 Genomics UK (COG-UK) Consortium | Berkshire and Surrey Pathology Services Lighthouse Laboratory and Alex Alderton, Roberto Amato, Jeffrey Barrett, Sonia Goncalves, Ewan Harrison, David K. Jackson, Ian Johnston, Dominic Kwiatkowski, Cordelia Langford, John Sillitoe on behalf of the Wellcome Sanger Institute COVID-19 Surveillance Team |
| EPI_ISL_3379500 | LABORATOIRE ACCOLAB SUD OUEST | CNR Virus des Infections Respiratoires - France SUD | Antonin Bal, Gregory Destras, Gwendolyne Burfin, Hadrien Regue, Quentin Semanas, Martine Valette, Bruno Lina, Laurence Josset |
| EPI_ISL_3379501 | Lighthouse Lab in Milton Keynes | Wellcome Sanger Institute for the COVID-19 Genomics UK (COG-UK) Consortium | The Lighthouse Lab in Milton Keynes and Alex Alderton, Roberto Amato, Jeffrey Barrett, Sonia Goncalves, Ewan Harrison, David K. Jackson, Ian Johnston, Dominic Kwiatkowski, Cordelia Langford, John Sillitoe on behalf of the Wellcome Sanger Institute COVID-19 Surveillance Team |
| EPI_ISL_3379502, EPI_ISL_3379503, EPI_ISL_3379504 | Berkshire and Surrey Pathology Services Lighthouse Laboratory | Wellcome Sanger Institute for the COVID-19 Genomics UK (COG-UK) Consortium | Berkshire and Surrey Pathology Services Lighthouse Laboratory and Alex Alderton, Roberto Amato, Jeffrey Barrett, Sonia Goncalves, Ewan Harrison, David K. Jackson, Ian Johnston, Dominic Kwiatkowski, Cordelia Langford, John Sillitoe on behalf of the Wellcome Sanger Institute COVID-19 Surveillance Team |
| EPI_ISL_3379505, EPI_ISL_3379506 | Lighthouse Lab in Milton Keynes | Wellcome Sanger Institute for the COVID-19 Genomics UK (COG-UK) Consortium | The Lighthouse Lab in Milton Keynes and Alex Alderton, Roberto Amato, Jeffrey Barrett, Sonia Goncalves, Ewan Harrison, David K. Jackson, Ian Johnston, Dominic Kwiatkowski, Cordelia Langford, John Sillitoe on behalf of the Wellcome Sanger Institute COVID-19 Surveillance Team |
| EPI_ISL_3379507 | Berkshire and Surrey Pathology Services Lighthouse Laboratory | Wellcome Sanger Institute for the COVID-19 Genomics UK (COG-UK) Consortium | Berkshire and Surrey Pathology Services Lighthouse Laboratory and Alex Alderton, Roberto Amato, Jeffrey Barrett, Sonia Goncalves, Ewan Harrison, David K. Jackson, Ian Johnston, Dominic Kwiatkowski, Cordelia Langford, John Sillitoe on behalf of the Wellcome Sanger Institute COVID-19 Surveillance Team |
| EPI_ISL_3379508 | Lighthouse Lab in Milton Keynes | Wellcome Sanger Institute for the COVID-19 Genomics UK (COG-UK) Consortium | The Lighthouse Lab in Milton Keynes and Alex Alderton, Roberto Amato, Jeffrey Barrett, Sonia Goncalves, Ewan Harrison, David K. Jackson, Ian Johnston, Dominic Kwiatkowski, Cordelia Langford, John Sillitoe on behalf of the Wellcome Sanger Institute COVID-19 Surveillance Team |
| EPI_ISL_3379509 | Lighthouse Lab in Alderley Park | Wellcome Sanger Institute for the COVID-19 Genomics UK (COG-UK) Consortium | Jacquelyn Wynn, Mairead Hyland, The Lighthouse Lab in Alderley Park and Alex Alderton, Roberto Amato, Jeffrey Barrett, Sonia Goncalves, Ewan Harrison, David K. Jackson, Ian Johnston, Dominic Kwiatkowski, Cordelia Langford, John Sillitoe on behalf of the Wellcome Sanger Institute COVID-19 Surveillance Team |
| EPI_ISL_3379510 | ORIAPOLE | CNR Virus des Infections Respiratoires - France SUD | Antonin Bal, Gregory Destras, Gwendolyne Burfin, Hadrien Regue, Quentin Semanas, Martine Valette, Bruno Lina, Laurence Josset |
| EPI_ISL_3379511, EPI_ISL_3379512 | Lighthouse Lab in Milton Keynes | Wellcome Sanger Institute for the COVID-19 Genomics UK (COG-UK) Consortium | The Lighthouse Lab in Milton Keynes and Alex Alderton, Roberto Amato, Jeffrey Barrett, Sonia Goncalves, Ewan Harrison, David K. Jackson, Ian Johnston, Dominic Kwiatkowski, Cordelia Langford, John Sillitoe on behalf of the Wellcome Sanger Institute COVID-19 Surveillance Team |
| EPI_ISL_3379513 | Berkshire and Surrey Pathology Services Lighthouse Laboratory | Wellcome Sanger Institute for the COVID-19 Genomics UK (COG-UK) Consortium | Berkshire and Surrey Pathology Services Lighthouse Laboratory and Alex Alderton, Roberto Amato, Jeffrey Barrett, Sonia Goncalves, Ewan Harrison, David K. Jackson, Ian Johnston, Dominic Kwiatkowski, Cordelia Langford, John Sillitoe on behalf of the Wellcome Sanger Institute COVID-19 Surveillance Team |
| EPI_ISL_3379514, EPI_ISL_3379515, EPI_ISL_3379516 | Lighthouse Lab in Milton Keynes | Wellcome Sanger Institute for the COVID-19 Genomics UK (COG-UK) Consortium | The Lighthouse Lab in Milton Keynes and Alex Alderton, Roberto Amato, Jeffrey Barrett, Sonia Goncalves, Ewan Harrison, David K. Jackson, Ian Johnston, Dominic Kwiatkowski, Cordelia Langford, John Sillitoe on behalf of the Wellcome Sanger Institute COVID-19 Surveillance Team |
| EPI_ISL_3379517, EPI_ISL_3379518 | Lighthouse Lab in Alderley Park | Wellcome Sanger Institute for the COVID-19 Genomics UK (COG-UK) Consortium | Jacquelyn Wynn, Mairead Hyland, The Lighthouse Lab in Alderley Park and Alex Alderton, Roberto Amato, Jeffrey Barrett, Sonia Goncalves, Ewan Harrison, David K. Jackson, Ian Johnston, Dominic Kwiatkowski, Cordelia Langford, John Sillitoe on behalf of the Wellcome Sanger Institute COVID-19 Surveillance Team |
| EPI_ISL_3379519 | ORIAPOLE | CNR Virus des Infections Respiratoires - France SUD | Antonin Bal, Gregory Destras, Gwendolyne Burfin, Hadrien Regue, Quentin Semanas, Martine Valette, Bruno Lina, Laurence Josset |
| EPI_ISL_3379520 | Berkshire and Surrey Pathology Services Lighthouse Laboratory | Wellcome Sanger Institute for the COVID-19 Genomics UK (COG-UK) Consortium | Berkshire and Surrey Pathology Services Lighthouse Laboratory and Alex Alderton, Roberto Amato, Jeffrey Barrett, Sonia Goncalves, Ewan Harrison, David K. Jackson, Ian Johnston, Dominic Kwiatkowski, Cordelia Langford, John Sillitoe on behalf of the Wellcome Sanger Institute COVID-19 Surveillance Team |
| EPI_ISL_3379521, EPI_ISL_3379522 | Lighthouse Lab in Milton Keynes | Wellcome Sanger Institute for the COVID-19 Genomics UK (COG-UK) Consortium | The Lighthouse Lab in Milton Keynes and Alex Alderton, Roberto Amato, Jeffrey Barrett, Sonia Goncalves, Ewan Harrison, David K. Jackson, Ian Johnston, Dominic Kwiatkowski, Cordelia Langford, John Sillitoe on behalf of the Wellcome Sanger Institute COVID-19 Surveillance Team |
| EPI_ISL_3379523 | LABORATOIRE ACCOLAB SUD OUEST | CNR Virus des Infections Respiratoires - France SUD | Antonin Bal, Gregory Destras, Gwendolyne Burfin, Hadrien Regue, Quentin Semanas, Martine Valette, Bruno Lina, Laurence Josset |
| EPI_ISL_3379524 | Lighthouse Lab in Alderley Park | Wellcome Sanger Institute for the COVID-19 Genomics UK (COG-UK) Consortium | Jacquelyn Wynn, Mairead Hyland, The Lighthouse Lab in Alderley Park and Alex Alderton, Roberto Amato, Jeffrey Barrett, Sonia Goncalves, Ewan Harrison, David K. Jackson, Ian Johnston, Dominic Kwiatkowski, Cordelia Langford, John Sillitoe on behalf of the Wellcome Sanger Institute COVID-19 Surveillance Team |
| EPI_ISL_3379525 | Berkshire and Surrey Pathology Services Lighthouse Laboratory | Wellcome Sanger Institute for the COVID-19 Genomics UK (COG-UK) Consortium | Berkshire and Surrey Pathology Services Lighthouse Laboratory and Alex Alderton, Roberto Amato, Jeffrey Barrett, Sonia Goncalves, Ewan Harrison, David K. Jackson, Ian Johnston, Dominic Kwiatkowski, Cordelia Langford, John Sillitoe on behalf of the Wellcome Sanger Institute COVID-19 Surveillance Team |
| EPI_ISL_3379526 | Lighthouse Lab in Alderley Park | Wellcome Sanger Institute for the COVID-19 Genomics UK (COG-UK) Consortium | Jacquelyn Wynn, Mairead Hyland, The Lighthouse Lab in Alderley Park and Alex Alderton, Roberto Amato, Jeffrey Barrett, Sonia Goncalves, Ewan Harrison, David K. Jackson, Ian Johnston, Dominic Kwiatkowski, Cordelia Langford, John Sillitoe on behalf of the Wellcome Sanger Institute COVID-19 Surveillance Team |
| EPI_ISL_3379527 | ORIAPOLE | CNR Virus des Infections Respiratoires - France SUD | Antonin Bal, Gregory Destras, Gwendolyne Burfin, Hadrien Regue, Quentin Semanas, Martine Valette, Bruno Lina, Laurence Josset |
| EPI_ISL_3379528 | Lighthouse Lab in Alderley Park | Wellcome Sanger Institute for the COVID-19 Genomics UK (COG-UK) Consortium | Jacquelyn Wynn, Mairead Hyland, The Lighthouse Lab in Alderley Park and Alex Alderton, Roberto Amato, Jeffrey Barrett, Sonia Goncalves, Ewan Harrison, David K. Jackson, Ian Johnston, Dominic Kwiatkowski, Cordelia Langford, John Sillitoe on behalf of the Wellcome Sanger Institute COVID-19 Surveillance Team |
| EPI_ISL_3379529 | Berkshire and Surrey Pathology Services Lighthouse Laboratory | Wellcome Sanger Institute for the COVID-19 Genomics UK (COG-UK) Consortium | Berkshire and Surrey Pathology Services Lighthouse Laboratory and Alex Alderton, Roberto Amato, Jeffrey Barrett, Sonia Goncalves, Ewan Harrison, David K. Jackson, Ian Johnston, Dominic Kwiatkowski, Cordelia Langford, John Sillitoe on behalf of the Wellcome Sanger Institute COVID-19 Surveillance Team |
| EPI_ISL_3379530 | Lighthouse Lab in Milton Keynes | Wellcome Sanger Institute for the COVID-19 Genomics UK (COG-UK) Consortium | The Lighthouse Lab in Milton Keynes and Alex Alderton, Roberto Amato, Jeffrey Barrett, Sonia Goncalves, Ewan Harrison, David K. Jackson, Ian Johnston, Dominic Kwiatkowski, Cordelia Langford, John Sillitoe on behalf of the Wellcome Sanger Institute COVID-19 Surveillance Team |
| EPI_ISL_3379531 | ORIAPOLE | CNR Virus des Infections Respiratoires - France SUD | Antonin Bal, Gregory Destras, Gwendolyne Burfin, Hadrien Regue, Quentin Semanas, Martine Valette, Bruno Lina, Laurence Josset |
| EPI_ISL_3379532, EPI_ISL_3379533 | Berkshire and Surrey Pathology Services Lighthouse Laboratory | Wellcome Sanger Institute for the COVID-19 Genomics UK (COG-UK) Consortium | Berkshire and Surrey Pathology Services Lighthouse Laboratory and Alex Alderton, Roberto Amato, Jeffrey Barrett, Sonia Goncalves, Ewan Harrison, David K. Jackson, Ian Johnston, Dominic Kwiatkowski, Cordelia Langford, John Sillitoe on behalf of the Wellcome Sanger Institute COVID-19 Surveillance Team |
| EPI_ISL_3379534 | LABORATOIRE ACCOLAB SUD OUEST | CNR Virus des Infections Respiratoires - France SUD | Antonin Bal, Gregory Destras, Gwendolyne Burfin, Hadrien Regue, Quentin Semanas, Martine Valette, Bruno Lina, Laurence Josset |
| EPI_ISL_3379535 | Lighthouse Lab in Alderley Park | Wellcome Sanger Institute for the COVID-19 Genomics UK (COG-UK) Consortium | Jacquelyn Wynn, Mairead Hyland, The Lighthouse Lab in Alderley Park and Alex Alderton, Roberto Amato, Jeffrey Barrett, Sonia Goncalves, Ewan Harrison, David K. Jackson, Ian Johnston, Dominic Kwiatkowski, Cordelia Langford, John Sillitoe on behalf of the Wellcome Sanger Institute COVID-19 Surveillance Team |
| EPI_ISL_3379536 | Lighthouse Lab in Milton Keynes | Wellcome Sanger Institute for the COVID-19 Genomics UK (COG-UK) Consortium | The Lighthouse Lab in Milton Keynes and Alex Alderton, Roberto Amato, Jeffrey Barrett, Sonia Goncalves, Ewan Harrison, David K. Jackson, Ian Johnston, Dominic Kwiatkowski, Cordelia Langford, John Sillitoe on behalf of the Wellcome Sanger Institute COVID-19 Surveillance Team |
| EPI_ISL_3379537 | Berkshire and Surrey Pathology Services Lighthouse Laboratory | Wellcome Sanger Institute for the COVID-19 Genomics UK (COG-UK) Consortium | Berkshire and Surrey Pathology Services Lighthouse Laboratory and Alex Alderton, Roberto Amato, Jeffrey Barrett, Sonia Goncalves, Ewan Harrison, David K. Jackson, Ian Johnston, Dominic Kwiatkowski, Cordelia Langford, John Sillitoe on behalf of the Wellcome Sanger Institute COVID-19 Surveillance Team |
| EPI_ISL_3379538 | ORIAPOLE | CNR Virus des Infections Respiratoires - France SUD | Antonin Bal, Gregory Destras, Gwendolyne Burfin, Hadrien Regue, Quentin Semanas, Martine Valette, Bruno Lina, Laurence Josset |
| EPI_ISL_3379539 | Lighthouse Lab in Alderley Park | Wellcome Sanger Institute for the COVID-19 Genomics UK (COG-UK) Consortium | Jacquelyn Wynn, Mairead Hyland, The Lighthouse Lab in Alderley Park and Alex Alderton, Roberto Amato, Jeffrey Barrett, Sonia Goncalves, Ewan Harrison, David K. Jackson, Ian Johnston, Dominic Kwiatkowski, Cordelia Langford, John Sillitoe on behalf of the Wellcome Sanger Institute COVID-19 Surveillance Team |
| EPI_ISL_3379540 | Lighthouse Lab in Milton Keynes | Wellcome Sanger Institute for the COVID-19 Genomics UK (COG-UK) Consortium | The Lighthouse Lab in Milton Keynes and Alex Alderton, Roberto Amato, Jeffrey Barrett, Sonia Goncalves, Ewan Harrison, David K. Jackson, Ian Johnston, Dominic Kwiatkowski, Cordelia Langford, John Sillitoe on behalf of the Wellcome Sanger Institute COVID-19 Surveillance Team |
| EPI_ISL_3379541 | Lighthouse Lab in Alderley Park | Wellcome Sanger Institute for the COVID-19 Genomics UK (COG-UK) Consortium | Jacquelyn Wynn, Mairead Hyland, The Lighthouse Lab in Alderley Park and Alex Alderton, Roberto Amato, Jeffrey Barrett, Sonia Goncalves, Ewan Harrison, David K. Jackson, Ian Johnston, Dominic Kwiatkowski, Cordelia Langford, John Sillitoe on behalf of the Wellcome Sanger Institute COVID-19 Surveillance Team |
| EPI_ISL_3379542, EPI_ISL_3379543, EPI_ISL_3379544, EPI_ISL_3379545, EPI_ISL_3379547, EPI_ISL_3379548, EPI_ISL_3379549 | Lighthouse Lab in Milton Keynes | Wellcome Sanger Institute for the COVID-19 Genomics UK (COG-UK) Consortium | The Lighthouse Lab in Milton Keynes and Alex Alderton, Roberto Amato, Jeffrey Barrett, Sonia Goncalves, Ewan Harrison, David K. Jackson, Ian Johnston, Dominic Kwiatkowski, Cordelia Langford, John Sillitoe on behalf of the Wellcome Sanger Institute COVID-19 Surveillance Team |
| EPI_ISL_3379550 | LABORATOIRE ACCOLAB SUD OUEST | CNR Virus des Infections Respiratoires - France SUD | Antonin Bal, Gregory Destras, Gwendolyne Burfin, Hadrien Regue, Quentin Semanas, Martine Valette, Bruno Lina, Laurence Josset |
| EPI_ISL_3379551 | Lighthouse Lab in Alderley Park | Wellcome Sanger Institute for the COVID-19 Genomics UK (COG-UK) Consortium | Jacquelyn Wynn, Mairead Hyland, The Lighthouse Lab in Alderley Park and Alex Alderton, Roberto Amato, Jeffrey Barrett, Sonia Goncalves, Ewan Harrison, David K. Jackson, Ian Johnston, Dominic Kwiatkowski, Cordelia Langford, John Sillitoe on behalf of the Wellcome Sanger Institute COVID-19 Surveillance Team |
| EPI_ISL_3379552 | Lighthouse Lab in Milton Keynes | Wellcome Sanger Institute for the COVID-19 Genomics UK (COG-UK) Consortium | The Lighthouse Lab in Milton Keynes and Alex Alderton, Roberto Amato, Jeffrey Barrett, Sonia Goncalves, Ewan Harrison, David K. Jackson, Ian Johnston, Dominic Kwiatkowski, Cordelia Langford, John Sillitoe on behalf of the Wellcome Sanger Institute COVID-19 Surveillance Team |
| EPI_ISL_3379553 | Berkshire and Surrey Pathology Services Lighthouse Laboratory | Wellcome Sanger Institute for the COVID-19 Genomics UK (COG-UK) Consortium | Berkshire and Surrey Pathology Services Lighthouse Laboratory and Alex Alderton, Roberto Amato, Jeffrey Barrett, Sonia Goncalves, Ewan Harrison, David K. Jackson, Ian Johnston, Dominic Kwiatkowski, Cordelia Langford, John Sillitoe on behalf of the Wellcome Sanger Institute COVID-19 Surveillance Team |
| EPI_ISL_3379554 | LABORATOIRE ACCOLAB SUD OUEST | CNR Virus des Infections Respiratoires - France SUD | Antonin Bal, Gregory Destras, Gwendolyne Burfin, Hadrien Regue, Quentin Semanas, Martine Valette, Bruno Lina, Laurence Josset |
| EPI_ISL_3379555 | Berkshire and Surrey Pathology Services Lighthouse Laboratory | Wellcome Sanger Institute for the COVID-19 Genomics UK (COG-UK) Consortium | Berkshire and Surrey Pathology Services Lighthouse Laboratory and Alex Alderton, Roberto Amato, Jeffrey Barrett, Sonia Goncalves, Ewan Harrison, David K. Jackson, Ian Johnston, Dominic Kwiatkowski, Cordelia Langford, John Sillitoe on behalf of the Wellcome Sanger Institute COVID-19 Surveillance Team |
| EPI_ISL_3379556 | Lighthouse Lab in Alderley Park | Wellcome Sanger Institute for the COVID-19 Genomics UK (COG-UK) Consortium | Jacquelyn Wynn, Mairead Hyland, The Lighthouse Lab in Alderley Park and Alex Alderton, Roberto Amato, Jeffrey Barrett, Sonia Goncalves, Ewan Harrison, David K. Jackson, Ian Johnston, Dominic Kwiatkowski, Cordelia Langford, John Sillitoe on behalf of the Wellcome Sanger Institute COVID-19 Surveillance Team |

| | | | |
|---|---|---|---|
| EPI_ISL_3379557 | Lighthouse Lab in Milton Keynes | Wellcome Sanger Institute for the COVID-19 Genomics UK (COG-UK) Consortium | The Lighthouse Lab in Milton Keynes and Alex Alderton, Roberto Amato, Jeffrey Barrett, Sonia Goncalves, Ewan Harrison, David K. Jackson, Ian Johnston, Dominic Kwiatkowski, Cordelia Langford, John Sillitoe on behalf of the Wellcome Sanger Institute COVID-19 Surveillance Team |
| EPI_ISL_3379558 | Lighthouse Lab in Alderley Park | Wellcome Sanger Institute for the COVID-19 Genomics UK (COG-UK) Consortium | Jacquelyn Wynn, Mairead Hyland, The Lighthouse Lab in Alderley Park and Alex Alderton, Roberto Amato, Jeffrey Barrett, Sonia Goncalves, Ewan Harrison, David K. Jackson, Ian Johnston, Dominic Kwiatkowski, Cordelia Langford, John Sillitoe on behalf of the Wellcome Sanger Institute COVID-19 Surveillance Team |
| EPI_ISL_3379559 | LABORATOIRE ACCOLAB SUD OUEST | CNR Virus des Infections Respiratoires - France SUD | Antonin Bal, Gregory Destras, Gwendolyne Burfin, Hadrien Regue, Quentin Semanas, Martine Valette, Bruno Lina, Laurence Josset |
| EPI_ISL_3379560 | Lighthouse Lab in Alderley Park | Wellcome Sanger Institute for the COVID-19 Genomics UK (COG-UK) Consortium | Jacquelyn Wynn, Mairead Hyland, The Lighthouse Lab in Alderley Park and Alex Alderton, Roberto Amato, Jeffrey Barrett, Sonia Goncalves, Ewan Harrison, David K. Jackson, Ian Johnston, Dominic Kwiatkowski, Cordelia Langford, John Sillitoe on behalf of the Wellcome Sanger Institute COVID-19 Surveillance Team |
| EPI_ISL_3379561 | Lighthouse Lab in Milton Keynes | Wellcome Sanger Institute for the COVID-19 Genomics UK (COG-UK) Consortium | The Lighthouse Lab in Milton Keynes and Alex Alderton, Roberto Amato, Jeffrey Barrett, Sonia Goncalves, Ewan Harrison, David K. Jackson, Ian Johnston, Dominic Kwiatkowski, Cordelia Langford, John Sillitoe on behalf of the Wellcome Sanger Institute COVID-19 Surveillance Team |
| EPI_ISL_3379562 | Berkshire and Surrey Pathology Services Lighthouse Laboratory | Wellcome Sanger Institute for the COVID-19 Genomics UK (COG-UK) Consortium | Berkshire and Surrey Pathology Services Lighthouse Laboratory and Alex Alderton, Roberto Amato, Jeffrey Barrett, Sonia Goncalves, Ewan Harrison, David K. Jackson, Ian Johnston, Dominic Kwiatkowski, Cordelia Langford, John Sillitoe on behalf of the Wellcome Sanger Institute COVID-19 Surveillance Team |
| EPI_ISL_3379563 | LABORATOIRE ACCOLAB SUD OUEST | CNR Virus des Infections Respiratoires - France SUD | Antonin Bal, Gregory Destras, Gwendolyne Burfin, Hadrien Regue, Quentin Semanas, Martine Valette, Bruno Lina, Laurence Josset |
| EPI_ISL_3379564 | Lighthouse Lab in Milton Keynes | Wellcome Sanger Institute for the COVID-19 Genomics UK (COG-UK) Consortium | The Lighthouse Lab in Milton Keynes and Alex Alderton, Roberto Amato, Jeffrey Barrett, Sonia Goncalves, Ewan Harrison, David K. Jackson, Ian Johnston, Dominic Kwiatkowski, Cordelia Langford, John Sillitoe on behalf of the Wellcome Sanger Institute COVID-19 Surveillance Team |
| EPI_ISL_3379565 | Berkshire and Surrey Pathology Services Lighthouse Laboratory | Wellcome Sanger Institute for the COVID-19 Genomics UK (COG-UK) Consortium | Berkshire and Surrey Pathology Services Lighthouse Laboratory and Alex Alderton, Roberto Amato, Jeffrey Barrett, Sonia Goncalves, Ewan Harrison, David K. Jackson, Ian Johnston, Dominic Kwiatkowski, Cordelia Langford, John Sillitoe on behalf of the Wellcome Sanger Institute COVID-19 Surveillance Team |
| EPI_ISL_3379566, EPI_ISL_3379567 | Lighthouse Lab in Milton Keynes | Wellcome Sanger Institute for the COVID-19 Genomics UK (COG-UK) Consortium | The Lighthouse Lab in Milton Keynes and Alex Alderton, Roberto Amato, Jeffrey Barrett, Sonia Goncalves, Ewan Harrison, David K. Jackson, Ian Johnston, Dominic Kwiatkowski, Cordelia Langford, John Sillitoe on behalf of the Wellcome Sanger Institute COVID-19 Surveillance Team |
| EPI_ISL_3379568, EPI_ISL_3379569, EPI_ISL_3379570, EPI_ISL_3379571 | Lighthouse Lab in Alderley Park | Wellcome Sanger Institute for the COVID-19 Genomics UK (COG-UK) Consortium | Jacquelyn Wynn, Mairead Hyland, The Lighthouse Lab in Alderley Park and Alex Alderton, Roberto Amato, Jeffrey Barrett, Sonia Goncalves, Ewan Harrison, David K. Jackson, Ian Johnston, Dominic Kwiatkowski, Cordelia Langford, John Sillitoe on behalf of the Wellcome Sanger Institute COVID-19 Surveillance Team |
| EPI_ISL_3379572 | LABORATOIRE ACCOLAB SUD OUEST | CNR Virus des Infections Respiratoires - France SUD | Antonin Bal, Gregory Destras, Gwendolyne Burfin, Hadrien Regue, Quentin Semanas, Martine Valette, Bruno Lina, Laurence Josset |
| EPI_ISL_3379573 | Lighthouse Lab in Alderley Park | Wellcome Sanger Institute for the COVID-19 Genomics UK (COG-UK) Consortium | Jacquelyn Wynn, Mairead Hyland, The Lighthouse Lab in Alderley Park and Alex Alderton, Roberto Amato, Jeffrey Barrett, Sonia Goncalves, Ewan Harrison, David K. Jackson, Ian Johnston, Dominic Kwiatkowski, Cordelia Langford, John Sillitoe on behalf of the Wellcome Sanger Institute COVID-19 Surveillance Team |
| EPI_ISL_3379574 | Lighthouse Lab in Milton Keynes | Wellcome Sanger Institute for the COVID-19 Genomics UK (COG-UK) Consortium | The Lighthouse Lab in Milton Keynes and Alex Alderton, Roberto Amato, Jeffrey Barrett, Sonia Goncalves, Ewan Harrison, David K. Jackson, Ian Johnston, Dominic Kwiatkowski, Cordelia Langford, John Sillitoe on behalf of the Wellcome Sanger Institute COVID-19 Surveillance Team |
| EPI_ISL_3379575 | Berkshire and Surrey Pathology Services Lighthouse Laboratory | Wellcome Sanger Institute for the COVID-19 Genomics UK (COG-UK) Consortium | Berkshire and Surrey Pathology Services Lighthouse Laboratory and Alex Alderton, Roberto Amato, Jeffrey Barrett, Sonia Goncalves, Ewan Harrison, David K. Jackson, Ian Johnston, Dominic Kwiatkowski, Cordelia Langford, John Sillitoe on behalf of the Wellcome Sanger Institute COVID-19 Surveillance Team |
| EPI_ISL_3379576, EPI_ISL_3379577 | Lighthouse Lab in Milton Keynes | Wellcome Sanger Institute for the COVID-19 Genomics UK (COG-UK) Consortium | The Lighthouse Lab in Milton Keynes and Alex Alderton, Roberto Amato, Jeffrey Barrett, Sonia Goncalves, Ewan Harrison, David K. Jackson, Ian Johnston, Dominic Kwiatkowski, Cordelia Langford, John Sillitoe on behalf of the Wellcome Sanger Institute COVID-19 Surveillance Team |
| EPI_ISL_3379578 | LABORATOIRE ACCOLAB SUD OUEST | CNR Virus des Infections Respiratoires - France SUD | Antonin Bal, Gregory Destras, Gwendolyne Burfin, Hadrien Regue, Quentin Semanas, Martine Valette, Bruno Lina, Laurence Josset |
| EPI_ISL_3379579 | Lighthouse Lab in Alderley Park | Wellcome Sanger Institute for the COVID-19 Genomics UK (COG-UK) Consortium | Jacquelyn Wynn, Mairead Hyland, The Lighthouse Lab in Alderley Park and Alex Alderton, Roberto Amato, Jeffrey Barrett, Sonia Goncalves, Ewan Harrison, David K. Jackson, Ian Johnston, Dominic Kwiatkowski, Cordelia Langford, John Sillitoe on behalf of the Wellcome Sanger Institute COVID-19 Surveillance Team |
| EPI_ISL_3379580, EPI_ISL_3379581 | Lighthouse Lab in Milton Keynes | Wellcome Sanger Institute for the COVID-19 Genomics UK (COG-UK) Consortium | The Lighthouse Lab in Milton Keynes and Alex Alderton, Roberto Amato, Jeffrey Barrett, Sonia Goncalves, Ewan Harrison, David K. Jackson, Ian Johnston, Dominic Kwiatkowski, Cordelia Langford, John Sillitoe on behalf of the Wellcome Sanger Institute COVID-19 Surveillance Team |
| EPI_ISL_3379582 | LABORATOIRE BIOFUSION | CNR Virus des Infections Respiratoires - France SUD | Antonin Bal, Gregory Destras, Gwendolyne Burfin, Hadrien Regue, Quentin Semanas, Martine Valette, Bruno Lina, Laurence Josset |
| EPI_ISL_3379583 | Lighthouse Lab in Milton Keynes | Wellcome Sanger Institute for the COVID-19 Genomics UK (COG-UK) Consortium | The Lighthouse Lab in Milton Keynes and Alex Alderton, Roberto Amato, Jeffrey Barrett, Sonia Goncalves, Ewan Harrison, David K. Jackson, Ian Johnston, Dominic Kwiatkowski, Cordelia Langford, John Sillitoe on behalf of the Wellcome Sanger Institute COVID-19 Surveillance Team |
| EPI_ISL_3379584 | Lighthouse Lab in Alderley Park | Wellcome Sanger Institute for the COVID-19 Genomics UK (COG-UK) Consortium | Jacquelyn Wynn, Mairead Hyland, The Lighthouse Lab in Alderley Park and Alex Alderton, Roberto Amato, Jeffrey Barrett, Sonia Goncalves, Ewan Harrison, David K. Jackson, Ian Johnston, Dominic Kwiatkowski, Cordelia Langford, John Sillitoe on behalf of the Wellcome Sanger Institute COVID-19 Surveillance Team |
| EPI_ISL_3379585 | Lighthouse Lab in Milton Keynes | Wellcome Sanger Institute for the COVID-19 Genomics UK (COG-UK) Consortium | The Lighthouse Lab in Milton Keynes and Alex Alderton, Roberto Amato, Jeffrey Barrett, Sonia Goncalves, Ewan Harrison, David K. Jackson, Ian Johnston, Dominic Kwiatkowski, Cordelia Langford, John Sillitoe on behalf of the Wellcome Sanger Institute COVID-19 Surveillance Team |
| EPI_ISL_3379586 | LABORATOIRE BIOFUSION | CNR Virus des Infections Respiratoires - France SUD | Antonin Bal, Gregory Destras, Gwendolyne Burfin, Hadrien Regue, Quentin Semanas, Martine Valette, Bruno Lina, Laurence Josset |
| EPI_ISL_3379587 | Berkshire and Surrey Pathology Services Lighthouse Laboratory | Wellcome Sanger Institute for the COVID-19 Genomics UK (COG-UK) Consortium | Berkshire and Surrey Pathology Services Lighthouse Laboratory and Alex Alderton, Roberto Amato, Jeffrey Barrett, Sonia Goncalves, Ewan Harrison, David K. Jackson, Ian Johnston, Dominic Kwiatkowski, Cordelia Langford, John Sillitoe on behalf of the Wellcome Sanger Institute COVID-19 Surveillance Team |
| EPI_ISL_3379588, EPI_ISL_3379589 | Lighthouse Lab in Alderley Park | Wellcome Sanger Institute for the COVID-19 Genomics UK (COG-UK) Consortium | Jacquelyn Wynn, Mairead Hyland, The Lighthouse Lab in Alderley Park and Alex Alderton, Roberto Amato, Jeffrey Barrett, Sonia Goncalves, Ewan Harrison, David K. Jackson, Ian Johnston, Dominic Kwiatkowski, Cordelia Langford, John Sillitoe on behalf of the Wellcome Sanger Institute COVID-19 Surveillance Team |
| EPI_ISL_3379590 | LABORATOIRE ACCOLAB SUD OUEST | CNR Virus des Infections Respiratoires - France SUD | Antonin Bal, Gregory Destras, Gwendolyne Burfin, Hadrien Regue, Quentin Semanas, Martine Valette, Bruno Lina, Laurence Josset |
| EPI_ISL_3379591 | Lighthouse Lab in Alderley Park | Wellcome Sanger Institute for the COVID-19 Genomics UK (COG-UK) Consortium | Jacquelyn Wynn, Mairead Hyland, The Lighthouse Lab in Alderley Park and Alex Alderton, Roberto Amato, Jeffrey Barrett, Sonia Goncalves, Ewan Harrison, David K. Jackson, Ian Johnston, Dominic Kwiatkowski, Cordelia Langford, John Sillitoe on behalf of the Wellcome Sanger Institute COVID-19 Surveillance Team |
| EPI_ISL_3379592 | Lighthouse Lab in Milton Keynes | Wellcome Sanger Institute for the COVID-19 Genomics UK (COG-UK) Consortium | The Lighthouse Lab in Milton Keynes and Alex Alderton, Roberto Amato, Jeffrey Barrett, Sonia Goncalves, Ewan Harrison, David K. Jackson, Ian Johnston, Dominic Kwiatkowski, Cordelia Langford, John Sillitoe on behalf of the Wellcome Sanger Institute COVID-19 Surveillance Team |
| EPI_ISL_3379593 | Lighthouse Lab in Alderley Park | Wellcome Sanger Institute for the COVID-19 Genomics UK (COG-UK) Consortium | Jacquelyn Wynn, Mairead Hyland, The Lighthouse Lab in Alderley Park and Alex Alderton, Roberto Amato, Jeffrey Barrett, Sonia Goncalves, Ewan Harrison, David K. Jackson, Ian Johnston, Dominic Kwiatkowski, Cordelia Langford, John Sillitoe on behalf of the Wellcome Sanger Institute COVID-19 Surveillance Team |
| EPI_ISL_3379594, EPI_ISL_3379595 | Lighthouse Lab in Milton Keynes | Wellcome Sanger Institute for the COVID-19 Genomics UK (COG-UK) Consortium | The Lighthouse Lab in Milton Keynes and Alex Alderton, Roberto Amato, Jeffrey Barrett, Sonia Goncalves, Ewan Harrison, David K. Jackson, Ian Johnston, Dominic Kwiatkowski, Cordelia Langford, John Sillitoe on behalf of the Wellcome Sanger Institute COVID-19 Surveillance Team |
| EPI_ISL_3379596 | Berkshire and Surrey Pathology Services Lighthouse Laboratory | Wellcome Sanger Institute for the COVID-19 Genomics UK (COG-UK) Consortium | Berkshire and Surrey Pathology Services Lighthouse Laboratory and Alex Alderton, Roberto Amato, Jeffrey Barrett, Sonia Goncalves, Ewan Harrison, David K. Jackson, Ian Johnston, Dominic Kwiatkowski, Cordelia Langford, John Sillitoe on behalf of the Wellcome Sanger Institute COVID-19 Surveillance Team |
| EPI_ISL_3379597 | Lighthouse Lab in Milton Keynes | Wellcome Sanger Institute for the COVID-19 Genomics UK (COG-UK) Consortium | The Lighthouse Lab in Milton Keynes and Alex Alderton, Roberto Amato, Jeffrey Barrett, Sonia Goncalves, Ewan Harrison, David K. Jackson, Ian Johnston, Dominic Kwiatkowski, Cordelia Langford, John Sillitoe on behalf of the Wellcome Sanger Institute COVID-19 Surveillance Team |
| EPI_ISL_3379598 | Lighthouse Lab in Alderley Park | Wellcome Sanger Institute for the COVID-19 Genomics UK (COG-UK) Consortium | Jacquelyn Wynn, Mairead Hyland, The Lighthouse Lab in Alderley Park and Alex Alderton, Roberto Amato, Jeffrey Barrett, Sonia Goncalves, Ewan Harrison, David K. Jackson, Ian Johnston, Dominic Kwiatkowski, Cordelia Langford, John Sillitoe on behalf of the Wellcome Sanger Institute COVID-19 Surveillance Team |
| EPI_ISL_3379599 | LABORATOIRE BIOFUSION | CNR Virus des Infections Respiratoires - France SUD | Antonin Bal, Gregory Destras, Gwendolyne Burfin, Hadrien Regue, Quentin Semanas, Martine Valette, Bruno Lina, Laurence Josset |
| EPI_ISL_3379600 | Lighthouse Lab in Milton Keynes | Wellcome Sanger Institute for the COVID-19 Genomics UK (COG-UK) Consortium | The Lighthouse Lab in Milton Keynes and Alex Alderton, Roberto Amato, Jeffrey Barrett, Sonia Goncalves, Ewan Harrison, David K. Jackson, Ian Johnston, Dominic Kwiatkowski, Cordelia Langford, John Sillitoe on behalf of the Wellcome Sanger Institute COVID-19 Surveillance Team |
| EPI_ISL_3379601 | Berkshire and Surrey Pathology Services Lighthouse Laboratory | Wellcome Sanger Institute for the COVID-19 Genomics UK (COG-UK) Consortium | Berkshire and Surrey Pathology Services Lighthouse Laboratory and Alex Alderton, Roberto Amato, Jeffrey Barrett, Sonia Goncalves, Ewan Harrison, David K. Jackson, Ian Johnston, Dominic Kwiatkowski, Cordelia Langford, John Sillitoe on behalf of the Wellcome Sanger Institute COVID-19 Surveillance Team |
| EPI_ISL_3379602, EPI_ISL_3379603, EPI_ISL_3379604, EPI_ISL_3379605 | Lighthouse Lab in Milton Keynes | Wellcome Sanger Institute for the COVID-19 Genomics UK (COG-UK) Consortium | The Lighthouse Lab in Milton Keynes and Alex Alderton, Roberto Amato, Jeffrey Barrett, Sonia Goncalves, Ewan Harrison, David K. Jackson, Ian Johnston, Dominic Kwiatkowski, Cordelia Langford, John Sillitoe on behalf of the Wellcome Sanger Institute COVID-19 Surveillance Team |
| EPI_ISL_3379606 | Berkshire and Surrey Pathology Services Lighthouse Laboratory | Wellcome Sanger Institute for the COVID-19 Genomics UK (COG-UK) Consortium | Berkshire and Surrey Pathology Services Lighthouse Laboratory and Alex Alderton, Roberto Amato, Jeffrey Barrett, Sonia Goncalves, Ewan Harrison, David K. Jackson, Ian Johnston, Dominic Kwiatkowski, Cordelia Langford, John Sillitoe on behalf of the Wellcome Sanger Institute COVID-19 Surveillance Team |
| EPI_ISL_3379607 | Lighthouse Lab in Milton Keynes | Wellcome Sanger Institute for the COVID-19 Genomics UK (COG-UK) Consortium | The Lighthouse Lab in Milton Keynes and Alex Alderton, Roberto Amato, Jeffrey Barrett, Sonia Goncalves, Ewan Harrison, David K. Jackson, Ian Johnston, Dominic Kwiatkowski, Cordelia Langford, John Sillitoe on behalf of the Wellcome Sanger Institute COVID-19 Surveillance Team |
| EPI_ISL_3379608 | LABORATOIRE BIOFUSION | CNR Virus des Infections Respiratoires - France SUD | Antonin Bal, Gregory Destras, Gwendolyne Burfin, Hadrien Regue, Quentin Semanas, Martine Valette, Bruno Lina, Laurence Josset |
| EPI_ISL_3379609, EPI_ISL_3379610, EPI_ISL_3379611 | Lighthouse Lab in Milton Keynes | Wellcome Sanger Institute for the COVID-19 Genomics UK (COG-UK) Consortium | The Lighthouse Lab in Milton Keynes and Alex Alderton, Roberto Amato, Jeffrey Barrett, Sonia Goncalves, Ewan Harrison, David K. Jackson, Ian Johnston, Dominic Kwiatkowski, Cordelia Langford, John Sillitoe on behalf of the Wellcome Sanger Institute COVID-19 Surveillance Team |
| EPI_ISL_3379612, EPI_ISL_3379613 | Berkshire and Surrey Pathology Services Lighthouse Laboratory | Wellcome Sanger Institute for the COVID-19 Genomics UK (COG-UK) Consortium | Berkshire and Surrey Pathology Services Lighthouse Laboratory and Alex Alderton, Roberto Amato, Jeffrey Barrett, Sonia Goncalves, Ewan Harrison, David K. Jackson, Ian Johnston, Dominic Kwiatkowski, Cordelia Langford, John Sillitoe on behalf of the Wellcome Sanger Institute COVID-19 Surveillance Team |
| EPI_ISL_3379614, EPI_ISL_3379615 | Lighthouse Lab in Milton Keynes | Wellcome Sanger Institute for the COVID-19 Genomics UK (COG-UK) Consortium | The Lighthouse Lab in Milton Keynes and Alex Alderton, Roberto Amato, Jeffrey Barrett, Sonia Goncalves, Ewan Harrison, David K. Jackson, Ian Johnston, Dominic Kwiatkowski, Cordelia Langford, John Sillitoe on behalf of the Wellcome Sanger Institute COVID-19 Surveillance Team |
| EPI_ISL_3379616 | LABORATOIRE BIOFUSION | CNR Virus des Infections Respiratoires - France SUD | Antonin Bal, Gregory Destras, Gwendolyne Burfin, Hadrien Regue, Quentin Semanas, Martine Valette, Bruno Lina, Laurence Josset |
| EPI_ISL_3379617 | Berkshire and Surrey Pathology Services Lighthouse Laboratory | Wellcome Sanger Institute for the COVID-19 Genomics UK (COG-UK) Consortium | Berkshire and Surrey Pathology Services Lighthouse Laboratory and Alex Alderton, Roberto Amato, Jeffrey Barrett, Sonia Goncalves, Ewan Harrison, David K. Jackson, Ian Johnston, Dominic Kwiatkowski, Cordelia Langford, John Sillitoe on behalf of the Wellcome Sanger Institute COVID-19 Surveillance Team |
| EPI_ISL_3379618 | Lighthouse Lab in Alderley Park | Wellcome Sanger Institute for the COVID-19 Genomics UK (COG-UK) Consortium | Jacquelyn Wynn, Mairead Hyland, The Lighthouse Lab in Alderley Park and Alex Alderton, Roberto Amato, Jeffrey Barrett, Sonia Goncalves, Ewan Harrison, David K. Jackson, Ian Johnston, Dominic Kwiatkowski, Cordelia Langford, John Sillitoe on behalf of the Wellcome Sanger Institute COVID-19 Surveillance Team |

| | | | |
|---|---|---|---|
| EPI_ISL_3379619 | Lighthouse Lab in Milton Keynes | Wellcome Sanger Institute for the COVID-19 Genomics UK (COG-UK) Consortium | The Lighthouse Lab in Milton Keynes and Alex Alderton, Roberto Amato, Jeffrey Barrett, Sonia Goncalves, Ewan Harrison, David K. Jackson, Ian Johnston, Dominic Kwiatkowski, Cordelia Langford, John Sillitoe on behalf of the Wellcome Sanger Institute COVID-19 Surveillance Team |
| EPI_ISL_3379620, EPI_ISL_3379621 | Lighthouse Lab in Alderley Park | Wellcome Sanger Institute for the COVID-19 Genomics UK (COG-UK) Consortium | Jacquelyn Wynn, Mairead Hyland, The Lighthouse Lab in Alderley Park and Alex Alderton, Roberto Amato, Jeffrey Barrett, Sonia Goncalves, Ewan Harrison, David K. Jackson, Ian Johnston, Dominic Kwiatkowski, Cordelia Langford, John Sillitoe on behalf of the Wellcome Sanger Institute COVID-19 Surveillance Team |
| EPI_ISL_3379622 | Lighthouse Lab in Milton Keynes | Wellcome Sanger Institute for the COVID-19 Genomics UK (COG-UK) Consortium | The Lighthouse Lab in Milton Keynes and Alex Alderton, Roberto Amato, Jeffrey Barrett, Sonia Goncalves, Ewan Harrison, David K. Jackson, Ian Johnston, Dominic Kwiatkowski, Cordelia Langford, John Sillitoe on behalf of the Wellcome Sanger Institute COVID-19 Surveillance Team |
| EPI_ISL_3379623 | Lighthouse Lab in Alderley Park | Wellcome Sanger Institute for the COVID-19 Genomics UK (COG-UK) Consortium | Jacquelyn Wynn, Mairead Hyland, The Lighthouse Lab in Alderley Park and Alex Alderton, Roberto Amato, Jeffrey Barrett, Sonia Goncalves, Ewan Harrison, David K. Jackson, Ian Johnston, Dominic Kwiatkowski, Cordelia Langford, John Sillitoe on behalf of the Wellcome Sanger Institute COVID-19 Surveillance Team |
| EPI_ISL_3379624 | LABORATOIRE ACCOLAB SUD OUEST | CNR Virus des Infections Respiratoires - France SUD | Antonin Bal, Gregory Destras, Gwendolyne Burfin, Hadrien Regue, Quentin Semanas, Martine Valette, Bruno Lina, Laurence Josset |
| EPI_ISL_3379625, EPI_ISL_3379626 | Lighthouse Lab in Milton Keynes | Wellcome Sanger Institute for the COVID-19 Genomics UK (COG-UK) Consortium | The Lighthouse Lab in Milton Keynes and Alex Alderton, Roberto Amato, Jeffrey Barrett, Sonia Goncalves, Ewan Harrison, David K. Jackson, Ian Johnston, Dominic Kwiatkowski, Cordelia Langford, John Sillitoe on behalf of the Wellcome Sanger Institute COVID-19 Surveillance Team |
| EPI_ISL_3379627 | Berkshire and Surrey Pathology Services Lighthouse Laboratory | Wellcome Sanger Institute for the COVID-19 Genomics UK (COG-UK) Consortium | Berkshire and Surrey Pathology Services Lighthouse Laboratory and Alex Alderton, Roberto Amato, Jeffrey Barrett, Sonia Goncalves, Ewan Harrison, David K. Jackson, Ian Johnston, Dominic Kwiatkowski, Cordelia Langford, John Sillitoe on behalf of the Wellcome Sanger Institute COVID-19 Surveillance Team |
| EPI_ISL_3379628 | Lighthouse Lab in Alderley Park | Wellcome Sanger Institute for the COVID-19 Genomics UK (COG-UK) Consortium | Jacquelyn Wynn, Mairead Hyland, The Lighthouse Lab in Alderley Park and Alex Alderton, Roberto Amato, Jeffrey Barrett, Sonia Goncalves, Ewan Harrison, David K. Jackson, Ian Johnston, Dominic Kwiatkowski, Cordelia Langford, John Sillitoe on behalf of the Wellcome Sanger Institute COVID-19 Surveillance Team |
| EPI_ISL_3379629 | Berkshire and Surrey Pathology Services Lighthouse Laboratory | Wellcome Sanger Institute for the COVID-19 Genomics UK (COG-UK) Consortium | Berkshire and Surrey Pathology Services Lighthouse Laboratory and Alex Alderton, Roberto Amato, Jeffrey Barrett, Sonia Goncalves, Ewan Harrison, David K. Jackson, Ian Johnston, Dominic Kwiatkowski, Cordelia Langford, John Sillitoe on behalf of the Wellcome Sanger Institute COVID-19 Surveillance Team |
| EPI_ISL_3379630 | Lighthouse Lab in Alderley Park | Wellcome Sanger Institute for the COVID-19 Genomics UK (COG-UK) Consortium | Jacquelyn Wynn, Mairead Hyland, The Lighthouse Lab in Alderley Park and Alex Alderton, Roberto Amato, Jeffrey Barrett, Sonia Goncalves, Ewan Harrison, David K. Jackson, Ian Johnston, Dominic Kwiatkowski, Cordelia Langford, John Sillitoe on behalf of the Wellcome Sanger Institute COVID-19 Surveillance Team |
| EPI_ISL_3379631, EPI_ISL_3379632 | Berkshire and Surrey Pathology Services Lighthouse Laboratory | Wellcome Sanger Institute for the COVID-19 Genomics UK (COG-UK) Consortium | Berkshire and Surrey Pathology Services Lighthouse Laboratory and Alex Alderton, Roberto Amato, Jeffrey Barrett, Sonia Goncalves, Ewan Harrison, David K. Jackson, Ian Johnston, Dominic Kwiatkowski, Cordelia Langford, John Sillitoe on behalf of the Wellcome Sanger Institute COVID-19 Surveillance Team |
| EPI_ISL_3379633 | LABORATOIRE BIOFUSION | CNR Virus des Infections Respiratoires - France SUD | Antonin Bal, Gregory Destras, Gwendolyne Burfin, Hadrien Regue, Quentin Semanas, Martine Valette, Bruno Lina, Laurence Josset |
| EPI_ISL_3379634 | Lighthouse Lab in Milton Keynes | Wellcome Sanger Institute for the COVID-19 Genomics UK (COG-UK) Consortium | The Lighthouse Lab in Milton Keynes and Alex Alderton, Roberto Amato, Jeffrey Barrett, Sonia Goncalves, Ewan Harrison, David K. Jackson, Ian Johnston, Dominic Kwiatkowski, Cordelia Langford, John Sillitoe on behalf of the Wellcome Sanger Institute COVID-19 Surveillance Team |
| EPI_ISL_3379635 | Lighthouse Lab in Alderley Park | Wellcome Sanger Institute for the COVID-19 Genomics UK (COG-UK) Consortium | Jacquelyn Wynn, Mairead Hyland, The Lighthouse Lab in Alderley Park and Alex Alderton, Roberto Amato, Jeffrey Barrett, Sonia Goncalves, Ewan Harrison, David K. Jackson, Ian Johnston, Dominic Kwiatkowski, Cordelia Langford, John Sillitoe on behalf of the Wellcome Sanger Institute COVID-19 Surveillance Team |
| EPI_ISL_3379636 | LABORATOIRE BIOFUSION | CNR Virus des Infections Respiratoires - France SUD | Antonin Bal, Gregory Destras, Gwendolyne Burfin, Hadrien Regue, Quentin Semanas, Martine Valette, Bruno Lina, Laurence Josset |
| EPI_ISL_3379637 | Lighthouse Lab in Alderley Park | Wellcome Sanger Institute for the COVID-19 Genomics UK (COG-UK) Consortium | Jacquelyn Wynn, Mairead Hyland, The Lighthouse Lab in Alderley Park and Alex Alderton, Roberto Amato, Jeffrey Barrett, Sonia Goncalves, Ewan Harrison, David K. Jackson, Ian Johnston, Dominic Kwiatkowski, Cordelia Langford, John Sillitoe on behalf of the Wellcome Sanger Institute COVID-19 Surveillance Team |
| EPI_ISL_3379638, EPI_ISL_3379639 | Berkshire and Surrey Pathology Services Lighthouse Laboratory | Wellcome Sanger Institute for the COVID-19 Genomics UK (COG-UK) Consortium | Berkshire and Surrey Pathology Services Lighthouse Laboratory and Alex Alderton, Roberto Amato, Jeffrey Barrett, Sonia Goncalves, Ewan Harrison, David K. Jackson, Ian Johnston, Dominic Kwiatkowski, Cordelia Langford, John Sillitoe on behalf of the Wellcome Sanger Institute COVID-19 Surveillance Team |
| EPI_ISL_3379640 | LABORATOIRE BIOFUSION | CNR Virus des Infections Respiratoires - France SUD | Antonin Bal, Gregory Destras, Gwendolyne Burfin, Hadrien Regue, Quentin Semanas, Martine Valette, Bruno Lina, Laurence Josset |
| EPI_ISL_3379641 | Lighthouse Lab in Alderley Park | Wellcome Sanger Institute for the COVID-19 Genomics UK (COG-UK) Consortium | Jacquelyn Wynn, Mairead Hyland, The Lighthouse Lab in Alderley Park and Alex Alderton, Roberto Amato, Jeffrey Barrett, Sonia Goncalves, Ewan Harrison, David K. Jackson, Ian Johnston, Dominic Kwiatkowski, Cordelia Langford, John Sillitoe on behalf of the Wellcome Sanger Institute COVID-19 Surveillance Team |
| EPI_ISL_3379642 | Lighthouse Lab in Milton Keynes | Wellcome Sanger Institute for the COVID-19 Genomics UK (COG-UK) Consortium | The Lighthouse Lab in Milton Keynes and Alex Alderton, Roberto Amato, Jeffrey Barrett, Sonia Goncalves, Ewan Harrison, David K. Jackson, Ian Johnston, Dominic Kwiatkowski, Cordelia Langford, John Sillitoe on behalf of the Wellcome Sanger Institute COVID-19 Surveillance Team |
| EPI_ISL_3379643 | Berkshire and Surrey Pathology Services Lighthouse Laboratory | Wellcome Sanger Institute for the COVID-19 Genomics UK (COG-UK) Consortium | Berkshire and Surrey Pathology Services Lighthouse Laboratory and Alex Alderton, Roberto Amato, Jeffrey Barrett, Sonia Goncalves, Ewan Harrison, David K. Jackson, Ian Johnston, Dominic Kwiatkowski, Cordelia Langford, John Sillitoe on behalf of the Wellcome Sanger Institute COVID-19 Surveillance Team |
| EPI_ISL_3379644 | Lighthouse Lab in Alderley Park | Wellcome Sanger Institute for the COVID-19 Genomics UK (COG-UK) Consortium | Jacquelyn Wynn, Mairead Hyland, The Lighthouse Lab in Alderley Park and Alex Alderton, Roberto Amato, Jeffrey Barrett, Sonia Goncalves, Ewan Harrison, David K. Jackson, Ian Johnston, Dominic Kwiatkowski, Cordelia Langford, John Sillitoe on behalf of the Wellcome Sanger Institute COVID-19 Surveillance Team |
| EPI_ISL_3379645 | Lighthouse Lab in Milton Keynes | Wellcome Sanger Institute for the COVID-19 Genomics UK (COG-UK) Consortium | The Lighthouse Lab in Milton Keynes and Alex Alderton, Roberto Amato, Jeffrey Barrett, Sonia Goncalves, Ewan Harrison, David K. Jackson, Ian Johnston, Dominic Kwiatkowski, Cordelia Langford, John Sillitoe on behalf of the Wellcome Sanger Institute COVID-19 Surveillance Team |
| EPI_ISL_3379646 | LABORATOIRE BIOFUSION | CNR Virus des Infections Respiratoires - France SUD | Antonin Bal, Gregory Destras, Gwendolyne Burfin, Hadrien Regue, Quentin Semanas, Martine Valette, Bruno Lina, Laurence Josset |
| EPI_ISL_3379647, EPI_ISL_3379648 | Lighthouse Lab in Milton Keynes | Wellcome Sanger Institute for the COVID-19 Genomics UK (COG-UK) Consortium | The Lighthouse Lab in Milton Keynes and Alex Alderton, Roberto Amato, Jeffrey Barrett, Sonia Goncalves, Ewan Harrison, David K. Jackson, Ian Johnston, Dominic Kwiatkowski, Cordelia Langford, John Sillitoe on behalf of the Wellcome Sanger Institute COVID-19 Surveillance Team |
| EPI_ISL_3379649 | Berkshire and Surrey Pathology Services Lighthouse Laboratory | Wellcome Sanger Institute for the COVID-19 Genomics UK (COG-UK) Consortium | Berkshire and Surrey Pathology Services Lighthouse Laboratory and Alex Alderton, Roberto Amato, Jeffrey Barrett, Sonia Goncalves, Ewan Harrison, David K. Jackson, Ian Johnston, Dominic Kwiatkowski, Cordelia Langford, John Sillitoe on behalf of the Wellcome Sanger Institute COVID-19 Surveillance Team |
| EPI_ISL_3379650 | LABORATOIRE ACCOLAB SUD OUEST | CNR Virus des Infections Respiratoires - France SUD | Antonin Bal, Gregory Destras, Gwendolyne Burfin, Hadrien Regue, Quentin Semanas, Martine Valette, Bruno Lina, Laurence Josset |
| EPI_ISL_3379651 | Lighthouse Lab in Milton Keynes | Wellcome Sanger Institute for the COVID-19 Genomics UK (COG-UK) Consortium | The Lighthouse Lab in Milton Keynes and Alex Alderton, Roberto Amato, Jeffrey Barrett, Sonia Goncalves, Ewan Harrison, David K. Jackson, Ian Johnston, Dominic Kwiatkowski, Cordelia Langford, John Sillitoe on behalf of the Wellcome Sanger Institute COVID-19 Surveillance Team |
| EPI_ISL_3379652 | Lighthouse Lab in Alderley Park | Wellcome Sanger Institute for the COVID-19 Genomics UK (COG-UK) Consortium | Jacquelyn Wynn, Mairead Hyland, The Lighthouse Lab in Alderley Park and Alex Alderton, Roberto Amato, Jeffrey Barrett, Sonia Goncalves, Ewan Harrison, David K. Jackson, Ian Johnston, Dominic Kwiatkowski, Cordelia Langford, John Sillitoe on behalf of the Wellcome Sanger Institute COVID-19 Surveillance Team |
| EPI_ISL_3379653 | Lighthouse Lab in Milton Keynes | Wellcome Sanger Institute for the COVID-19 Genomics UK (COG-UK) Consortium | The Lighthouse Lab in Milton Keynes and Alex Alderton, Roberto Amato, Jeffrey Barrett, Sonia Goncalves, Ewan Harrison, David K. Jackson, Ian Johnston, Dominic Kwiatkowski, Cordelia Langford, John Sillitoe on behalf of the Wellcome Sanger Institute COVID-19 Surveillance Team |
| EPI_ISL_3379654 | LABORATOIRE BIOFUSION | CNR Virus des Infections Respiratoires - France SUD | Antonin Bal, Gregory Destras, Gwendolyne Burfin, Hadrien Regue, Quentin Semanas, Martine Valette, Bruno Lina, Laurence Josset |
| EPI_ISL_3379655, EPI_ISL_3379656 | Lighthouse Lab in Alderley Park | Wellcome Sanger Institute for the COVID-19 Genomics UK (COG-UK) Consortium | Jacquelyn Wynn, Mairead Hyland, The Lighthouse Lab in Alderley Park and Alex Alderton, Roberto Amato, Jeffrey Barrett, Sonia Goncalves, Ewan Harrison, David K. Jackson, Ian Johnston, Dominic Kwiatkowski, Cordelia Langford, John Sillitoe on behalf of the Wellcome Sanger Institute COVID-19 Surveillance Team |
| EPI_ISL_3379657 | Lighthouse Lab in Milton Keynes | Wellcome Sanger Institute for the COVID-19 Genomics UK (COG-UK) Consortium | The Lighthouse Lab in Milton Keynes and Alex Alderton, Roberto Amato, Jeffrey Barrett, Sonia Goncalves, Ewan Harrison, David K. Jackson, Ian Johnston, Dominic Kwiatkowski, Cordelia Langford, John Sillitoe on behalf of the Wellcome Sanger Institute COVID-19 Surveillance Team |
| EPI_ISL_3379658 | LABORATOIRE BIOFUSION | CNR Virus des Infections Respiratoires - France SUD | Antonin Bal, Gregory Destras, Gwendolyne Burfin, Hadrien Regue, Quentin Semanas, Martine Valette, Bruno Lina, Laurence Josset |
| EPI_ISL_3379659 | Lighthouse Lab in Milton Keynes | Wellcome Sanger Institute for the COVID-19 Genomics UK (COG-UK) Consortium | The Lighthouse Lab in Milton Keynes and Alex Alderton, Roberto Amato, Jeffrey Barrett, Sonia Goncalves, Ewan Harrison, David K. Jackson, Ian Johnston, Dominic Kwiatkowski, Cordelia Langford, John Sillitoe on behalf of the Wellcome Sanger Institute COVID-19 Surveillance Team |
| EPI_ISL_3379660 | Lighthouse Lab in Alderley Park | Wellcome Sanger Institute for the COVID-19 Genomics UK (COG-UK) Consortium | Jacquelyn Wynn, Mairead Hyland, The Lighthouse Lab in Alderley Park and Alex Alderton, Roberto Amato, Jeffrey Barrett, Sonia Goncalves, Ewan Harrison, David K. Jackson, Ian Johnston, Dominic Kwiatkowski, Cordelia Langford, John Sillitoe on behalf of the Wellcome Sanger Institute COVID-19 Surveillance Team |
| EPI_ISL_3379661 | Lighthouse Lab in Milton Keynes | Wellcome Sanger Institute for the COVID-19 Genomics UK (COG-UK) Consortium | The Lighthouse Lab in Milton Keynes and Alex Alderton, Roberto Amato, Jeffrey Barrett, Sonia Goncalves, Ewan Harrison, David K. Jackson, Ian Johnston, Dominic Kwiatkowski, Cordelia Langford, John Sillitoe on behalf of the Wellcome Sanger Institute COVID-19 Surveillance Team |
| EPI_ISL_3379662 | LABORATOIRE BIOFUSION | CNR Virus des Infections Respiratoires - France SUD | Antonin Bal, Gregory Destras, Gwendolyne Burfin, Hadrien Regue, Quentin Semanas, Martine Valette, Bruno Lina, Laurence Josset |
| EPI_ISL_3379663, EPI_ISL_3379664 | Lighthouse Lab in Milton Keynes | Wellcome Sanger Institute for the COVID-19 Genomics UK (COG-UK) Consortium | The Lighthouse Lab in Milton Keynes and Alex Alderton, Roberto Amato, Jeffrey Barrett, Sonia Goncalves, Ewan Harrison, David K. Jackson, Ian Johnston, Dominic Kwiatkowski, Cordelia Langford, John Sillitoe on behalf of the Wellcome Sanger Institute COVID-19 Surveillance Team |
| EPI_ISL_3379665 | Lighthouse Lab in Alderley Park | Wellcome Sanger Institute for the COVID-19 Genomics UK (COG-UK) Consortium | Jacquelyn Wynn, Mairead Hyland, The Lighthouse Lab in Alderley Park and Alex Alderton, Roberto Amato, Jeffrey Barrett, Sonia Goncalves, Ewan Harrison, David K. Jackson, Ian Johnston, Dominic Kwiatkowski, Cordelia Langford, John Sillitoe on behalf of the Wellcome Sanger Institute COVID-19 Surveillance Team |
| EPI_ISL_3379666 | LABORATOIRE ACCOLAB SUD OUEST | CNR Virus des Infections Respiratoires - France SUD | Antonin Bal, Gregory Destras, Gwendolyne Burfin, Hadrien Regue, Quentin Semanas, Martine Valette, Bruno Lina, Laurence Josset |
| EPI_ISL_3379667 | Lighthouse Lab in Milton Keynes | Wellcome Sanger Institute for the COVID-19 Genomics UK (COG-UK) Consortium | The Lighthouse Lab in Milton Keynes and Alex Alderton, Roberto Amato, Jeffrey Barrett, Sonia Goncalves, Ewan Harrison, David K. Jackson, Ian Johnston, Dominic Kwiatkowski, Cordelia Langford, John Sillitoe on behalf of the Wellcome Sanger Institute COVID-19 Surveillance Team |
| EPI_ISL_3379668 | Berkshire and Surrey Pathology Services Lighthouse Laboratory | Wellcome Sanger Institute for the COVID-19 Genomics UK (COG-UK) Consortium | Berkshire and Surrey Pathology Services Lighthouse Laboratory and Alex Alderton, Roberto Amato, Jeffrey Barrett, Sonia Goncalves, Ewan Harrison, David K. Jackson, Ian Johnston, Dominic Kwiatkowski, Cordelia Langford, John Sillitoe on behalf of the Wellcome Sanger Institute COVID-19 Surveillance Team |
| EPI_ISL_3379669 | Lighthouse Lab in Milton Keynes | Wellcome Sanger Institute for the COVID-19 Genomics UK (COG-UK) Consortium | The Lighthouse Lab in Milton Keynes and Alex Alderton, Roberto Amato, Jeffrey Barrett, Sonia Goncalves, Ewan Harrison, David K. Jackson, Ian Johnston, Dominic Kwiatkowski, Cordelia Langford, John Sillitoe on behalf of the Wellcome Sanger Institute COVID-19 Surveillance Team |
| EPI_ISL_3379671 | Lighthouse Lab in Alderley Park | Wellcome Sanger Institute for the COVID-19 Genomics UK (COG-UK) Consortium | Jacquelyn Wynn, Mairead Hyland, The Lighthouse Lab in Alderley Park and Alex Alderton, Roberto Amato, Jeffrey Barrett, Sonia Goncalves, Ewan Harrison, David K. Jackson, Ian Johnston, Dominic Kwiatkowski, Cordelia Langford, John Sillitoe on behalf of the Wellcome Sanger Institute COVID-19 Surveillance Team |
| EPI_ISL_3379672 | Berkshire and Surrey Pathology Services Lighthouse Laboratory | Wellcome Sanger Institute for the COVID-19 Genomics UK (COG-UK) Consortium | Berkshire and Surrey Pathology Services Lighthouse Laboratory and Alex Alderton, Roberto Amato, Jeffrey Barrett, Sonia Goncalves, Ewan Harrison, David K. Jackson, Ian Johnston, Dominic Kwiatkowski, Cordelia Langford, John Sillitoe on behalf of the Wellcome Sanger Institute COVID-19 Surveillance Team |
| EPI_ISL_3379673 | Lighthouse Lab in Alderley Park | Wellcome Sanger Institute for the COVID-19 Genomics UK (COG-UK) Consortium | Jacquelyn Wynn, Mairead Hyland, The Lighthouse Lab in Alderley Park and Alex Alderton, Roberto Amato, Jeffrey Barrett, Sonia Goncalves, Ewan Harrison, David K. Jackson, Ian Johnston, Dominic Kwiatkowski, Cordelia Langford, John Sillitoe on behalf of the Wellcome Sanger Institute COVID-19 Surveillance Team |
| EPI_ISL_3379674 | LABORATOIRE ACCOLAB SUD OUEST | CNR Virus des Infections Respiratoires - France SUD | Antonin Bal, Gregory Destras, Gwendolyne Burfin, Hadrien Regue, Quentin Semanas, Martine Valette, Bruno Lina, Laurence Josset |
| EPI_ISL_3379675 | Lighthouse Lab in Alderley Park | Wellcome Sanger Institute for the COVID-19 Genomics UK (COG-UK) Consortium | Jacquelyn Wynn, Mairead Hyland, The Lighthouse Lab in Alderley Park and Alex Alderton, Roberto Amato, Jeffrey Barrett, Sonia Goncalves, Ewan Harrison, David K. Jackson, Ian Johnston, Dominic Kwiatkowski, Cordelia Langford, John Sillitoe on behalf of the Wellcome Sanger Institute COVID-19 Surveillance Team |

| | | | |
|---|---|---|---|
| EPI_ISL_3379676, EPI_ISL_3379677 | Lighthouse Lab in Milton Keynes | Wellcome Sanger Institute for the COVID-19 Genomics UK (COG-UK) Consortium | The Lighthouse Lab in Milton Keynes and Alex Alderton, Roberto Amato, Jeffrey Barrett, Sonia Goncalves, Ewan Harrison, David K. Jackson, Ian Johnston, Dominic Kwiatkowski, Cordelia Langford, John Sillitoe on behalf of the Wellcome Sanger Institute COVID-19 Surveillance Team |
| EPI_ISL_3379678 | LABORATOIRE BIOFUSION | CNR Virus des Infections Respiratoires - France SUD | Antonin Bal, Gregory Destras, Gwendolyne Burfin, Hadrien Regue, Quentin Semanas, Martine Valette, Bruno Lina, Laurence Josset |
| EPI_ISL_3379679 | Lighthouse Lab in Milton Keynes | Wellcome Sanger Institute for the COVID-19 Genomics UK (COG-UK) Consortium | The Lighthouse Lab in Milton Keynes and Alex Alderton, Roberto Amato, Jeffrey Barrett, Sonia Goncalves, Ewan Harrison, David K. Jackson, Ian Johnston, Dominic Kwiatkowski, Cordelia Langford, John Sillitoe on behalf of the Wellcome Sanger Institute COVID-19 Surveillance Team |
| EPI_ISL_3379680 | Berkshire and Surrey Pathology Services Lighthouse Laboratory | Wellcome Sanger Institute for the COVID-19 Genomics UK (COG-UK) Consortium | Berkshire and Surrey Pathology Services Lighthouse Laboratory and Alex Alderton, Roberto Amato, Jeffrey Barrett, Sonia Goncalves, Ewan Harrison, David K. Jackson, Ian Johnston, Dominic Kwiatkowski, Cordelia Langford, John Sillitoe on behalf of the Wellcome Sanger Institute COVID-19 Surveillance Team |
| EPI_ISL_3379681 | LABORATOIRE BIOFUSION | CNR Virus des Infections Respiratoires - France SUD | Antonin Bal, Gregory Destras, Gwendolyne Burfin, Hadrien Regue, Quentin Semanas, Martine Valette, Bruno Lina, Laurence Josset |
| EPI_ISL_3379682 | Lighthouse Lab in Milton Keynes | Wellcome Sanger Institute for the COVID-19 Genomics UK (COG-UK) Consortium | The Lighthouse Lab in Milton Keynes and Alex Alderton, Roberto Amato, Jeffrey Barrett, Sonia Goncalves, Ewan Harrison, David K. Jackson, Ian Johnston, Dominic Kwiatkowski, Cordelia Langford, John Sillitoe on behalf of the Wellcome Sanger Institute COVID-19 Surveillance Team |
| EPI_ISL_3379683, EPI_ISL_3379684 | Lighthouse Lab in Alderley Park | Wellcome Sanger Institute for the COVID-19 Genomics UK (COG-UK) Consortium | Jacquelyn Wynn, Mairead Hyland, The Lighthouse Lab in Alderley Park and Alex Alderton, Roberto Amato, Jeffrey Barrett, Sonia Goncalves, Ewan Harrison, David K. Jackson, Ian Johnston, Dominic Kwiatkowski, Cordelia Langford, John Sillitoe on behalf of the Wellcome Sanger Institute COVID-19 Surveillance Team |
| EPI_ISL_3379685 | LABORATOIRE ACCOLAB SUD OUEST | CNR Virus des Infections Respiratoires - France SUD | Antonin Bal, Gregory Destras, Gwendolyne Burfin, Hadrien Regue, Quentin Semanas, Martine Valette, Bruno Lina, Laurence Josset |
| EPI_ISL_3379686 | Berkshire and Surrey Pathology Services Lighthouse Laboratory | Wellcome Sanger Institute for the COVID-19 Genomics UK (COG-UK) Consortium | Berkshire and Surrey Pathology Services Lighthouse Laboratory and Alex Alderton, Roberto Amato, Jeffrey Barrett, Sonia Goncalves, Ewan Harrison, David K. Jackson, Ian Johnston, Dominic Kwiatkowski, Cordelia Langford, John Sillitoe on behalf of the Wellcome Sanger Institute COVID-19 Surveillance Team |
| EPI_ISL_3379687 | Lighthouse Lab in Alderley Park | Wellcome Sanger Institute for the COVID-19 Genomics UK (COG-UK) Consortium | Jacquelyn Wynn, Mairead Hyland, The Lighthouse Lab in Alderley Park and Alex Alderton, Roberto Amato, Jeffrey Barrett, Sonia Goncalves, Ewan Harrison, David K. Jackson, Ian Johnston, Dominic Kwiatkowski, Cordelia Langford, John Sillitoe on behalf of the Wellcome Sanger Institute COVID-19 Surveillance Team |
| EPI_ISL_3379688 | LABORATOIRE BIOFUSION | CNR Virus des Infections Respiratoires - France SUD | Antonin Bal, Gregory Destras, Gwendolyne Burfin, Hadrien Regue, Quentin Semanas, Martine Valette, Bruno Lina, Laurence Josset |
| EPI_ISL_3379689, EPI_ISL_3379690 | Lighthouse Lab in Alderley Park | Wellcome Sanger Institute for the COVID-19 Genomics UK (COG-UK) Consortium | Jacquelyn Wynn, Mairead Hyland, The Lighthouse Lab in Alderley Park and Alex Alderton, Roberto Amato, Jeffrey Barrett, Sonia Goncalves, Ewan Harrison, David K. Jackson, Ian Johnston, Dominic Kwiatkowski, Cordelia Langford, John Sillitoe on behalf of the Wellcome Sanger Institute COVID-19 Surveillance Team |
| EPI_ISL_3379691 | Lighthouse Lab in Milton Keynes | Wellcome Sanger Institute for the COVID-19 Genomics UK (COG-UK) Consortium | The Lighthouse Lab in Milton Keynes and Alex Alderton, Roberto Amato, Jeffrey Barrett, Sonia Goncalves, Ewan Harrison, David K. Jackson, Ian Johnston, Dominic Kwiatkowski, Cordelia Langford, John Sillitoe on behalf of the Wellcome Sanger Institute COVID-19 Surveillance Team |
| EPI_ISL_3379692 | LABORATOIRE ACCOLAB SUD OUEST | CNR Virus des Infections Respiratoires - France SUD | Antonin Bal, Gregory Destras, Gwendolyne Burfin, Hadrien Regue, Quentin Semanas, Martine Valette, Bruno Lina, Laurence Josset |
| EPI_ISL_3379693 | Lighthouse Lab in Alderley Park | Wellcome Sanger Institute for the COVID-19 Genomics UK (COG-UK) Consortium | Jacquelyn Wynn, Mairead Hyland, The Lighthouse Lab in Alderley Park and Alex Alderton, Roberto Amato, Jeffrey Barrett, Sonia Goncalves, Ewan Harrison, David K. Jackson, Ian Johnston, Dominic Kwiatkowski, Cordelia Langford, John Sillitoe on behalf of the Wellcome Sanger Institute COVID-19 Surveillance Team |
| EPI_ISL_3379694 | Berkshire and Surrey Pathology Services Lighthouse Laboratory | Wellcome Sanger Institute for the COVID-19 Genomics UK (COG-UK) Consortium | Berkshire and Surrey Pathology Services Lighthouse Laboratory and Alex Alderton, Roberto Amato, Jeffrey Barrett, Sonia Goncalves, Ewan Harrison, David K. Jackson, Ian Johnston, Dominic Kwiatkowski, Cordelia Langford, John Sillitoe on behalf of the Wellcome Sanger Institute COVID-19 Surveillance Team |
| EPI_ISL_3379695 | Lighthouse Lab in Milton Keynes | Wellcome Sanger Institute for the COVID-19 Genomics UK (COG-UK) Consortium | The Lighthouse Lab in Milton Keynes and Alex Alderton, Roberto Amato, Jeffrey Barrett, Sonia Goncalves, Ewan Harrison, David K. Jackson, Ian Johnston, Dominic Kwiatkowski, Cordelia Langford, John Sillitoe on behalf of the Wellcome Sanger Institute COVID-19 Surveillance Team |
| EPI_ISL_3379696 | LABORATOIRE BIOFUSION | CNR Virus des Infections Respiratoires - France SUD | Antonin Bal, Gregory Destras, Gwendolyne Burfin, Hadrien Regue, Quentin Semanas, Martine Valette, Bruno Lina, Laurence Josset |
| EPI_ISL_3379697 | Berkshire and Surrey Pathology Services Lighthouse Laboratory | Wellcome Sanger Institute for the COVID-19 Genomics UK (COG-UK) Consortium | Berkshire and Surrey Pathology Services Lighthouse Laboratory and Alex Alderton, Roberto Amato, Jeffrey Barrett, Sonia Goncalves, Ewan Harrison, David K. Jackson, Ian Johnston, Dominic Kwiatkowski, Cordelia Langford, John Sillitoe on behalf of the Wellcome Sanger Institute COVID-19 Surveillance Team |
| EPI_ISL_3379698 | Lighthouse Lab in Milton Keynes | Wellcome Sanger Institute for the COVID-19 Genomics UK (COG-UK) Consortium | The Lighthouse Lab in Milton Keynes and Alex Alderton, Roberto Amato, Jeffrey Barrett, Sonia Goncalves, Ewan Harrison, David K. Jackson, Ian Johnston, Dominic Kwiatkowski, Cordelia Langford, John Sillitoe on behalf of the Wellcome Sanger Institute COVID-19 Surveillance Team |
| EPI_ISL_3379699 | Lighthouse Lab in Glasgow | Wellcome Sanger Institute for the COVID-19 Genomics UK (COG-UK) Consortium | Harper VanSteenhouse, Yumi Kasai, David Gray, Carol Clugston, Anna Dominiczak and Alex Alderton, Roberto Amato, Jeffrey Barrett, Sonia Goncalves, Ewan Harrison, David K. Jackson, Ian Johnston, Dominic Kwiatkowski, Cordelia Langford, John Sillitoe on behalf of the Wellcome Sanger Institute COVID-19 Surveillance Team |
| EPI_ISL_3379700 | LABORATOIRE BIOFUSION | CNR Virus des Infections Respiratoires - France SUD | Antonin Bal, Gregory Destras, Gwendolyne Burfin, Hadrien Regue, Quentin Semanas, Martine Valette, Bruno Lina, Laurence Josset |
| EPI_ISL_3379701, EPI_ISL_3379702 | Lighthouse Lab in Glasgow | Wellcome Sanger Institute for the COVID-19 Genomics UK (COG-UK) Consortium | Harper VanSteenhouse, Yumi Kasai, David Gray, Carol Clugston, Anna Dominiczak and Alex Alderton, Roberto Amato, Jeffrey Barrett, Sonia Goncalves, Ewan Harrison, David K. Jackson, Ian Johnston, Dominic Kwiatkowski, Cordelia Langford, John Sillitoe on behalf of the Wellcome Sanger Institute COVID-19 Surveillance Team |
| EPI_ISL_3379703 | Lighthouse Lab in Alderley Park | Wellcome Sanger Institute for the COVID-19 Genomics UK (COG-UK) Consortium | Jacquelyn Wynn, Mairead Hyland, The Lighthouse Lab in Alderley Park and Alex Alderton, Roberto Amato, Jeffrey Barrett, Sonia Goncalves, Ewan Harrison, David K. Jackson, Ian Johnston, Dominic Kwiatkowski, Cordelia Langford, John Sillitoe on behalf of the Wellcome Sanger Institute COVID-19 Surveillance Team |
| EPI_ISL_3379704 | LABORATOIRE BIOFUSION | CNR Virus des Infections Respiratoires - France SUD | Antonin Bal, Gregory Destras, Gwendolyne Burfin, Hadrien Regue, Quentin Semanas, Martine Valette, Bruno Lina, Laurence Josset |
| EPI_ISL_3379705, EPI_ISL_3379706, EPI_ISL_3379707, EPI_ISL_3379708 | Lighthouse Lab in Glasgow | Wellcome Sanger Institute for the COVID-19 Genomics UK (COG-UK) Consortium | Harper VanSteenhouse, Yumi Kasai, David Gray, Carol Clugston, Anna Dominiczak and Alex Alderton, Roberto Amato, Jeffrey Barrett, Sonia Goncalves, Ewan Harrison, David K. Jackson, Ian Johnston, Dominic Kwiatkowski, Cordelia Langford, John Sillitoe on behalf of the Wellcome Sanger Institute COVID-19 Surveillance Team |
| EPI_ISL_3379709 | LABORATOIRE ACCOLAB SUD OUEST | CNR Virus des Infections Respiratoires - France SUD | Antonin Bal, Gregory Destras, Gwendolyne Burfin, Hadrien Regue, Quentin Semanas, Martine Valette, Bruno Lina, Laurence Josset |
| EPI_ISL_3379710 | Lighthouse Lab in Glasgow | Wellcome Sanger Institute for the COVID-19 Genomics UK (COG-UK) Consortium | Harper VanSteenhouse, Yumi Kasai, David Gray, Carol Clugston, Anna Dominiczak and Alex Alderton, Roberto Amato, Jeffrey Barrett, Sonia Goncalves, Ewan Harrison, David K. Jackson, Ian Johnston, Dominic Kwiatkowski, Cordelia Langford, John Sillitoe on behalf of the Wellcome Sanger Institute COVID-19 Surveillance Team |
| EPI_ISL_3379711, EPI_ISL_3379712 | Lighthouse Lab in Milton Keynes | Wellcome Sanger Institute for the COVID-19 Genomics UK (COG-UK) Consortium | The Lighthouse Lab in Milton Keynes and Alex Alderton, Roberto Amato, Jeffrey Barrett, Sonia Goncalves, Ewan Harrison, David K. Jackson, Ian Johnston, Dominic Kwiatkowski, Cordelia Langford, John Sillitoe on behalf of the Wellcome Sanger Institute COVID-19 Surveillance Team |
| EPI_ISL_3379713 | LABORATOIRE BIOFUSION | CNR Virus des Infections Respiratoires - France SUD | Antonin Bal, Gregory Destras, Gwendolyne Burfin, Hadrien Regue, Quentin Semanas, Martine Valette, Bruno Lina, Laurence Josset |
| EPI_ISL_3379714, EPI_ISL_3379715 | Lighthouse Lab in Milton Keynes | Wellcome Sanger Institute for the COVID-19 Genomics UK (COG-UK) Consortium | The Lighthouse Lab in Milton Keynes and Alex Alderton, Roberto Amato, Jeffrey Barrett, Sonia Goncalves, Ewan Harrison, David K. Jackson, Ian Johnston, Dominic Kwiatkowski, Cordelia Langford, John Sillitoe on behalf of the Wellcome Sanger Institute COVID-19 Surveillance Team |
| EPI_ISL_3379716 | Lighthouse Lab in Alderley Park | Wellcome Sanger Institute for the COVID-19 Genomics UK (COG-UK) Consortium | Jacquelyn Wynn, Mairead Hyland, The Lighthouse Lab in Alderley Park and Alex Alderton, Roberto Amato, Jeffrey Barrett, Sonia Goncalves, Ewan Harrison, David K. Jackson, Ian Johnston, Dominic Kwiatkowski, Cordelia Langford, John Sillitoe on behalf of the Wellcome Sanger Institute COVID-19 Surveillance Team |
| EPI_ISL_3379717 | LABORATOIRE BIOFUSION | CNR Virus des Infections Respiratoires - France SUD | Antonin Bal, Gregory Destras, Gwendolyne Burfin, Hadrien Regue, Quentin Semanas, Martine Valette, Bruno Lina, Laurence Josset |
| EPI_ISL_3379718, EPI_ISL_3379719 | Lighthouse Lab in Alderley Park | Wellcome Sanger Institute for the COVID-19 Genomics UK (COG-UK) Consortium | Jacquelyn Wynn, Mairead Hyland, The Lighthouse Lab in Alderley Park and Alex Alderton, Roberto Amato, Jeffrey Barrett, Sonia Goncalves, Ewan Harrison, David K. Jackson, Ian Johnston, Dominic Kwiatkowski, Cordelia Langford, John Sillitoe on behalf of the Wellcome Sanger Institute COVID-19 Surveillance Team |
| EPI_ISL_3379720 | LABORATOIRE BIOFUSION | CNR Virus des Infections Respiratoires - France SUD | Antonin Bal, Gregory Destras, Gwendolyne Burfin, Hadrien Regue, Quentin Semanas, Martine Valette, Bruno Lina, Laurence Josset |
| EPI_ISL_3379721, EPI_ISL_3379722 | Lighthouse Lab in Glasgow | Wellcome Sanger Institute for the COVID-19 Genomics UK (COG-UK) Consortium | Harper VanSteenhouse, Yumi Kasai, David Gray, Carol Clugston, Anna Dominiczak and Alex Alderton, Roberto Amato, Jeffrey Barrett, Sonia Goncalves, Ewan Harrison, David K. Jackson, Ian Johnston, Dominic Kwiatkowski, Cordelia Langford, John Sillitoe on behalf of the Wellcome Sanger Institute COVID-19 Surveillance Team |
| EPI_ISL_3379723 | Lighthouse Lab in Alderley Park | Wellcome Sanger Institute for the COVID-19 Genomics UK (COG-UK) Consortium | Jacquelyn Wynn, Mairead Hyland, The Lighthouse Lab in Alderley Park and Alex Alderton, Roberto Amato, Jeffrey Barrett, Sonia Goncalves, Ewan Harrison, David K. Jackson, Ian Johnston, Dominic Kwiatkowski, Cordelia Langford, John Sillitoe on behalf of the Wellcome Sanger Institute COVID-19 Surveillance Team |
| EPI_ISL_3379724 | LABORATOIRE BIOFUSION | CNR Virus des Infections Respiratoires - France SUD | Antonin Bal, Gregory Destras, Gwendolyne Burfin, Hadrien Regue, Quentin Semanas, Martine Valette, Bruno Lina, Laurence Josset |
| EPI_ISL_3379725 | Lighthouse Lab in Alderley Park | Wellcome Sanger Institute for the COVID-19 Genomics UK (COG-UK) Consortium | Jacquelyn Wynn, Mairead Hyland, The Lighthouse Lab in Alderley Park and Alex Alderton, Roberto Amato, Jeffrey Barrett, Sonia Goncalves, Ewan Harrison, David K. Jackson, Ian Johnston, Dominic Kwiatkowski, Cordelia Langford, John Sillitoe on behalf of the Wellcome Sanger Institute COVID-19 Surveillance Team |
| EPI_ISL_3379726, EPI_ISL_3379727 | Lighthouse Lab in Milton Keynes | Wellcome Sanger Institute for the COVID-19 Genomics UK (COG-UK) Consortium | The Lighthouse Lab in Milton Keynes and Alex Alderton, Roberto Amato, Jeffrey Barrett, Sonia Goncalves, Ewan Harrison, David K. Jackson, Ian Johnston, Dominic Kwiatkowski, Cordelia Langford, John Sillitoe on behalf of the Wellcome Sanger Institute COVID-19 Surveillance Team |
| EPI_ISL_3379728 | LABORATOIRE BIOFUSION | CNR Virus des Infections Respiratoires - France SUD | Antonin Bal, Gregory Destras, Gwendolyne Burfin, Hadrien Regue, Quentin Semanas, Martine Valette, Bruno Lina, Laurence Josset |
| EPI_ISL_3379729 | Lighthouse Lab in Alderley Park | Wellcome Sanger Institute for the COVID-19 Genomics UK (COG-UK) Consortium | Jacquelyn Wynn, Mairead Hyland, The Lighthouse Lab in Alderley Park and Alex Alderton, Roberto Amato, Jeffrey Barrett, Sonia Goncalves, Ewan Harrison, David K. Jackson, Ian Johnston, Dominic Kwiatkowski, Cordelia Langford, John Sillitoe on behalf of the Wellcome Sanger Institute COVID-19 Surveillance Team |
| EPI_ISL_3379730, EPI_ISL_3379731 | Lighthouse Lab in Glasgow | Wellcome Sanger Institute for the COVID-19 Genomics UK (COG-UK) Consortium | Harper VanSteenhouse, Yumi Kasai, David Gray, Carol Clugston, Anna Dominiczak and Alex Alderton, Roberto Amato, Jeffrey Barrett, Sonia Goncalves, Ewan Harrison, David K. Jackson, Ian Johnston, Dominic Kwiatkowski, Cordelia Langford, John Sillitoe on behalf of the Wellcome Sanger Institute COVID-19 Surveillance Team |
| EPI_ISL_3379732 | LABORATOIRE ACCOLAB SUD OUEST | CNR Virus des Infections Respiratoires - France SUD | Antonin Bal, Gregory Destras, Gwendolyne Burfin, Hadrien Regue, Quentin Semanas, Martine Valette, Bruno Lina, Laurence Josset |
| EPI_ISL_3379733 | Lighthouse Lab in Glasgow | Wellcome Sanger Institute for the COVID-19 Genomics UK (COG-UK) Consortium | Harper VanSteenhouse, Yumi Kasai, David Gray, Carol Clugston, Anna Dominiczak and Alex Alderton, Roberto Amato, Jeffrey Barrett, Sonia Goncalves, Ewan Harrison, David K. Jackson, Ian Johnston, Dominic Kwiatkowski, Cordelia Langford, John Sillitoe on behalf of the Wellcome Sanger Institute COVID-19 Surveillance Team |
| EPI_ISL_3379734 | Lighthouse Lab in Milton Keynes | Wellcome Sanger Institute for the COVID-19 Genomics UK (COG-UK) Consortium | The Lighthouse Lab in Milton Keynes and Alex Alderton, Roberto Amato, Jeffrey Barrett, Sonia Goncalves, Ewan Harrison, David K. Jackson, Ian Johnston, Dominic Kwiatkowski, Cordelia Langford, John Sillitoe on behalf of the Wellcome Sanger Institute COVID-19 Surveillance Team |
| EPI_ISL_3379735 | Lighthouse Lab in Glasgow | Wellcome Sanger Institute for the COVID-19 Genomics UK (COG-UK) Consortium | Harper VanSteenhouse, Yumi Kasai, David Gray, Carol Clugston, Anna Dominiczak and Alex Alderton, Roberto Amato, Jeffrey Barrett, Sonia Goncalves, Ewan Harrison, David K. Jackson, Ian Johnston, Dominic Kwiatkowski, Cordelia Langford, John Sillitoe on behalf of the Wellcome Sanger Institute COVID-19 Surveillance Team |
| EPI_ISL_3379736 | LABORATOIRE BIOFUSION | CNR Virus des Infections Respiratoires - France SUD | Antonin Bal, Gregory Destras, Gwendolyne Burfin, Hadrien Regue, Quentin Semanas, Martine Valette, Bruno Lina, Laurence Josset |
| EPI_ISL_3379737 | Lighthouse Lab in Alderley Park | Wellcome Sanger Institute for the COVID-19 Genomics UK (COG-UK) Consortium | Jacquelyn Wynn, Mairead Hyland, The Lighthouse Lab in Alderley Park and Alex Alderton, Roberto Amato, Jeffrey Barrett, Sonia Goncalves, Ewan Harrison, David K. Jackson, Ian Johnston, Dominic Kwiatkowski, Cordelia Langford, John Sillitoe on behalf of the Wellcome Sanger Institute COVID-19 Surveillance Team |
| EPI_ISL_3379738 | Lighthouse Lab in Milton Keynes | Wellcome Sanger Institute for the COVID-19 Genomics UK (COG-UK) Consortium | The Lighthouse Lab in Milton Keynes and Alex Alderton, Roberto Amato, Jeffrey Barrett, Sonia Goncalves, Ewan Harrison, David K. Jackson, Ian Johnston, Dominic Kwiatkowski, Cordelia Langford, John Sillitoe on behalf of the Wellcome Sanger Institute COVID-19 Surveillance Team |
| EPI_ISL_3379739 | Lighthouse Lab in Alderley Park | Wellcome Sanger Institute for the COVID-19 Genomics UK (COG-UK) | Jacquelyn Wynn, Mairead Hyland, The Lighthouse Lab in Alderley Park and Alex Alderton, Roberto Amato, Jeffrey Barrett, Sonia Goncalves, Ewan Harrison, David K. Jackson, Ian Johnston, Dominic Kwiatkowski, Cordelia Langford, John Sillitoe on |

| | | | |
|---|---|---|---|
| | | Consortium | behalf of the Wellcome Sanger Institute COVID-19 Surveillance Team |
| EPI_ISL_3379740 | LABORATOIRE ACCOLAB SUD OUEST | CNR Virus des Infections Respiratoires - France SUD | Antonin Bal, Gregory Destras, Gwendolyne Burfin, Hadrien Regue, Quentin Semanas, Martine Valette, Bruno Lina, Laurence Josset |
| EPI_ISL_3379741 | Lighthouse Lab in Alderley Park | Wellcome Sanger Institute for the COVID-19 Genomics UK (COG-UK) Consortium | Jacquelyn Wynn, Mairead Hyland, The Lighthouse Lab in Alderley Park and Alex Alderton, Roberto Amato, Jeffrey Barrett, Sonia Goncalves, Ewan Harrison, David K. Jackson, Ian Johnston, Dominic Kwiatkowski, Cordelia Langford, John Sillitoe on behalf of the Wellcome Sanger Institute COVID-19 Surveillance Team |
| EPI_ISL_3379742 | Lighthouse Lab in Glasgow | Wellcome Sanger Institute for the COVID-19 Genomics UK (COG-UK) Consortium | Harper VanSteenhouse, Yumi Kasai, David Gray, Carol Clugston, Anna Dominiczak and Alex Alderton, Roberto Amato, Jeffrey Barrett, Sonia Goncalves, Ewan Harrison, David K. Jackson, Ian Johnston, Dominic Kwiatkowski, Cordelia Langford, John Sillitoe on behalf of the Wellcome Sanger Institute COVID-19 Surveillance Team |
| EPI_ISL_3379743 | LABORATOIRE ACCOLAB SUD OUEST | CNR Virus des Infections Respiratoires - France SUD | Antonin Bal, Gregory Destras, Gwendolyne Burfin, Hadrien Regue, Quentin Semanas, Martine Valette, Bruno Lina, Laurence Josset |
| EPI_ISL_3379744, EPI_ISL_3379745, EPI_ISL_3379746 | Lighthouse Lab in Glasgow | Wellcome Sanger Institute for the COVID-19 Genomics UK (COG-UK) Consortium | Harper VanSteenhouse, Yumi Kasai, David Gray, Carol Clugston, Anna Dominiczak and Alex Alderton, Roberto Amato, Jeffrey Barrett, Sonia Goncalves, Ewan Harrison, David K. Jackson, Ian Johnston, Dominic Kwiatkowski, Cordelia Langford, John Sillitoe on behalf of the Wellcome Sanger Institute COVID-19 Surveillance Team |
| EPI_ISL_3379747 | LABORATOIRE BIOESTEREL | CNR Virus des Infections Respiratoires - France SUD | Antonin Bal, Gregory Destras, Gwendolyne Burfin, Hadrien Regue, Quentin Semanas, Martine Valette, Bruno Lina, Laurence Josset |
| EPI_ISL_3379748, EPI_ISL_3379749, EPI_ISL_3379750 | Lighthouse Lab in Glasgow | Wellcome Sanger Institute for the COVID-19 Genomics UK (COG-UK) Consortium | Harper VanSteenhouse, Yumi Kasai, David Gray, Carol Clugston, Anna Dominiczak and Alex Alderton, Roberto Amato, Jeffrey Barrett, Sonia Goncalves, Ewan Harrison, David K. Jackson, Ian Johnston, Dominic Kwiatkowski, Cordelia Langford, John Sillitoe on behalf of the Wellcome Sanger Institute COVID-19 Surveillance Team |
| EPI_ISL_3379751 | CENTRE HOSPITALIER DE VALENCE | CNR Virus des Infections Respiratoires - France SUD | Antonin Bal, Gregory Destras, Gwendolyne Burfin, Hadrien Regue, Quentin Semanas, Martine Valette, Bruno Lina, Laurence Josset |
| EPI_ISL_3379752 | Lighthouse Lab in Milton Keynes | Wellcome Sanger Institute for the COVID-19 Genomics UK (COG-UK) Consortium | The Lighthouse Lab in Milton Keynes and Alex Alderton, Roberto Amato, Jeffrey Barrett, Sonia Goncalves, Ewan Harrison, David K. Jackson, Ian Johnston, Dominic Kwiatkowski, Cordelia Langford, John Sillitoe on behalf of the Wellcome Sanger Institute COVID-19 Surveillance Team |
| EPI_ISL_3379753 | Lighthouse Lab in Glasgow | Wellcome Sanger Institute for the COVID-19 Genomics UK (COG-UK) Consortium | Harper VanSteenhouse, Yumi Kasai, David Gray, Carol Clugston, Anna Dominiczak and Alex Alderton, Roberto Amato, Jeffrey Barrett, Sonia Goncalves, Ewan Harrison, David K. Jackson, Ian Johnston, Dominic Kwiatkowski, Cordelia Langford, John Sillitoe on behalf of the Wellcome Sanger Institute COVID-19 Surveillance Team |
| EPI_ISL_3379754 | BIOLITTORAL BIOGROUP PLATEAU TECH | CNR Virus des Infections Respiratoires - France SUD | Antonin Bal, Gregory Destras, Gwendolyne Burfin, Hadrien Regue, Quentin Semanas, Martine Valette, Bruno Lina, Laurence Josset |
| EPI_ISL_3379755, EPI_ISL_3379756 | Lighthouse Lab in Glasgow | Wellcome Sanger Institute for the COVID-19 Genomics UK (COG-UK) Consortium | Harper VanSteenhouse, Yumi Kasai, David Gray, Carol Clugston, Anna Dominiczak and Alex Alderton, Roberto Amato, Jeffrey Barrett, Sonia Goncalves, Ewan Harrison, David K. Jackson, Ian Johnston, Dominic Kwiatkowski, Cordelia Langford, John Sillitoe on behalf of the Wellcome Sanger Institute COVID-19 Surveillance Team |
| EPI_ISL_3379757 | Lighthouse Lab in Alderley Park | Wellcome Sanger Institute for the COVID-19 Genomics UK (COG-UK) Consortium | Jacquelyn Wynn, Mairead Hyland, The Lighthouse Lab in Alderley Park and Alex Alderton, Roberto Amato, Jeffrey Barrett, Sonia Goncalves, Ewan Harrison, David K. Jackson, Ian Johnston, Dominic Kwiatkowski, Cordelia Langford, John Sillitoe on behalf of the Wellcome Sanger Institute COVID-19 Surveillance Team |
| EPI_ISL_3379758 | BIOLITTORAL BIOGROUP PLATEAU TECH | CNR Virus des Infections Respiratoires - France SUD | Antonin Bal, Gregory Destras, Gwendolyne Burfin, Hadrien Regue, Quentin Semanas, Martine Valette, Bruno Lina, Laurence Josset |
| EPI_ISL_3379759 | Lighthouse Lab in Alderley Park | Wellcome Sanger Institute for the COVID-19 Genomics UK (COG-UK) Consortium | Jacquelyn Wynn, Mairead Hyland, The Lighthouse Lab in Alderley Park and Alex Alderton, Roberto Amato, Jeffrey Barrett, Sonia Goncalves, Ewan Harrison, David K. Jackson, Ian Johnston, Dominic Kwiatkowski, Cordelia Langford, John Sillitoe on behalf of the Wellcome Sanger Institute COVID-19 Surveillance Team |
| EPI_ISL_3379760 | Lighthouse Lab in Glasgow | Wellcome Sanger Institute for the COVID-19 Genomics UK (COG-UK) Consortium | Harper VanSteenhouse, Yumi Kasai, David Gray, Carol Clugston, Anna Dominiczak and Alex Alderton, Roberto Amato, Jeffrey Barrett, Sonia Goncalves, Ewan Harrison, David K. Jackson, Ian Johnston, Dominic Kwiatkowski, Cordelia Langford, John Sillitoe on behalf of the Wellcome Sanger Institute COVID-19 Surveillance Team |
| EPI_ISL_3379761 | Lighthouse Lab in Milton Keynes | Wellcome Sanger Institute for the COVID-19 Genomics UK (COG-UK) Consortium | The Lighthouse Lab in Milton Keynes and Alex Alderton, Roberto Amato, Jeffrey Barrett, Sonia Goncalves, Ewan Harrison, David K. Jackson, Ian Johnston, Dominic Kwiatkowski, Cordelia Langford, John Sillitoe on behalf of the Wellcome Sanger Institute COVID-19 Surveillance Team |
| EPI_ISL_3379762 | BIOLITTORAL BIOGROUP PLATEAU TECH | CNR Virus des Infections Respiratoires - France SUD | Antonin Bal, Gregory Destras, Gwendolyne Burfin, Hadrien Regue, Quentin Semanas, Martine Valette, Bruno Lina, Laurence Josset |
| EPI_ISL_3379763 | Lighthouse Lab in Glasgow | Wellcome Sanger Institute for the COVID-19 Genomics UK (COG-UK) Consortium | Harper VanSteenhouse, Yumi Kasai, David Gray, Carol Clugston, Anna Dominiczak and Alex Alderton, Roberto Amato, Jeffrey Barrett, Sonia Goncalves, Ewan Harrison, David K. Jackson, Ian Johnston, Dominic Kwiatkowski, Cordelia Langford, John Sillitoe on behalf of the Wellcome Sanger Institute COVID-19 Surveillance Team |
| EPI_ISL_3379764 | Lighthouse Lab in Alderley Park | Wellcome Sanger Institute for the COVID-19 Genomics UK (COG-UK) Consortium | Jacquelyn Wynn, Mairead Hyland, The Lighthouse Lab in Alderley Park and Alex Alderton, Roberto Amato, Jeffrey Barrett, Sonia Goncalves, Ewan Harrison, David K. Jackson, Ian Johnston, Dominic Kwiatkowski, Cordelia Langford, John Sillitoe on behalf of the Wellcome Sanger Institute COVID-19 Surveillance Team |
| EPI_ISL_3379765 | Lighthouse Lab in Milton Keynes | Wellcome Sanger Institute for the COVID-19 Genomics UK (COG-UK) Consortium | The Lighthouse Lab in Milton Keynes and Alex Alderton, Roberto Amato, Jeffrey Barrett, Sonia Goncalves, Ewan Harrison, David K. Jackson, Ian Johnston, Dominic Kwiatkowski, Cordelia Langford, John Sillitoe on behalf of the Wellcome Sanger Institute COVID-19 Surveillance Team |
| EPI_ISL_3379766 | BIOLITTORAL BIOGROUP PLATEAU TECH | CNR Virus des Infections Respiratoires - France SUD | Antonin Bal, Gregory Destras, Gwendolyne Burfin, Hadrien Regue, Quentin Semanas, Martine Valette, Bruno Lina, Laurence Josset |
| EPI_ISL_3379767 | Lighthouse Lab in Glasgow | Wellcome Sanger Institute for the COVID-19 Genomics UK (COG-UK) Consortium | Harper VanSteenhouse, Yumi Kasai, David Gray, Carol Clugston, Anna Dominiczak and Alex Alderton, Roberto Amato, Jeffrey Barrett, Sonia Goncalves, Ewan Harrison, David K. Jackson, Ian Johnston, Dominic Kwiatkowski, Cordelia Langford, John Sillitoe on behalf of the Wellcome Sanger Institute COVID-19 Surveillance Team |
| EPI_ISL_3379768 | Lighthouse Lab in Milton Keynes | Wellcome Sanger Institute for the COVID-19 Genomics UK (COG-UK) Consortium | The Lighthouse Lab in Milton Keynes and Alex Alderton, Roberto Amato, Jeffrey Barrett, Sonia Goncalves, Ewan Harrison, David K. Jackson, Ian Johnston, Dominic Kwiatkowski, Cordelia Langford, John Sillitoe on behalf of the Wellcome Sanger Institute COVID-19 Surveillance Team |
| EPI_ISL_3379769 | BIOLITTORAL BIOGROUP PLATEAU TECH | CNR Virus des Infections Respiratoires - France SUD | Antonin Bal, Gregory Destras, Gwendolyne Burfin, Hadrien Regue, Quentin Semanas, Martine Valette, Bruno Lina, Laurence Josset |
| EPI_ISL_3379770 | Lighthouse Lab in Alderley Park | Wellcome Sanger Institute for the COVID-19 Genomics UK (COG-UK) Consortium | Jacquelyn Wynn, Mairead Hyland, The Lighthouse Lab in Alderley Park and Alex Alderton, Roberto Amato, Jeffrey Barrett, Sonia Goncalves, Ewan Harrison, David K. Jackson, Ian Johnston, Dominic Kwiatkowski, Cordelia Langford, John Sillitoe on behalf of the Wellcome Sanger Institute COVID-19 Surveillance Team |
| EPI_ISL_3379771, EPI_ISL_3379772 | Lighthouse Lab in Glasgow | Wellcome Sanger Institute for the COVID-19 Genomics UK (COG-UK) Consortium | Harper VanSteenhouse, Yumi Kasai, David Gray, Carol Clugston, Anna Dominiczak and Alex Alderton, Roberto Amato, Jeffrey Barrett, Sonia Goncalves, Ewan Harrison, David K. Jackson, Ian Johnston, Dominic Kwiatkowski, Cordelia Langford, John Sillitoe on behalf of the Wellcome Sanger Institute COVID-19 Surveillance Team |
| EPI_ISL_3379773 | LABORATOIRE BIOESTEREL | CNR Virus des Infections Respiratoires - France SUD | Antonin Bal, Gregory Destras, Gwendolyne Burfin, Hadrien Regue, Quentin Semanas, Martine Valette, Bruno Lina, Laurence Josset |
| EPI_ISL_3379774 | Lighthouse Lab in Glasgow | Wellcome Sanger Institute for the COVID-19 Genomics UK (COG-UK) Consortium | Harper VanSteenhouse, Yumi Kasai, David Gray, Carol Clugston, Anna Dominiczak and Alex Alderton, Roberto Amato, Jeffrey Barrett, Sonia Goncalves, Ewan Harrison, David K. Jackson, Ian Johnston, Dominic Kwiatkowski, Cordelia Langford, John Sillitoe on behalf of the Wellcome Sanger Institute COVID-19 Surveillance Team |
| EPI_ISL_3379775 | Lighthouse Lab in Milton Keynes | Wellcome Sanger Institute for the COVID-19 Genomics UK (COG-UK) Consortium | The Lighthouse Lab in Milton Keynes and Alex Alderton, Roberto Amato, Jeffrey Barrett, Sonia Goncalves, Ewan Harrison, David K. Jackson, Ian Johnston, Dominic Kwiatkowski, Cordelia Langford, John Sillitoe on behalf of the Wellcome Sanger Institute COVID-19 Surveillance Team |
| EPI_ISL_3379776 | Lighthouse Lab in Alderley Park | Wellcome Sanger Institute for the COVID-19 Genomics UK (COG-UK) Consortium | Jacquelyn Wynn, Mairead Hyland, The Lighthouse Lab in Alderley Park and Alex Alderton, Roberto Amato, Jeffrey Barrett, Sonia Goncalves, Ewan Harrison, David K. Jackson, Ian Johnston, Dominic Kwiatkowski, Cordelia Langford, John Sillitoe on behalf of the Wellcome Sanger Institute COVID-19 Surveillance Team |
| EPI_ISL_3379777 | BIOLITTORAL BIOGROUP PLATEAU TECH | CNR Virus des Infections Respiratoires - France SUD | Antonin Bal, Gregory Destras, Gwendolyne Burfin, Hadrien Regue, Quentin Semanas, Martine Valette, Bruno Lina, Laurence Josset |
| EPI_ISL_3379778, EPI_ISL_3379779, EPI_ISL_3379780 | Lighthouse Lab in Glasgow | Wellcome Sanger Institute for the COVID-19 Genomics UK (COG-UK) Consortium | Harper VanSteenhouse, Yumi Kasai, David Gray, Carol Clugston, Anna Dominiczak and Alex Alderton, Roberto Amato, Jeffrey Barrett, Sonia Goncalves, Ewan Harrison, David K. Jackson, Ian Johnston, Dominic Kwiatkowski, Cordelia Langford, John Sillitoe on behalf of the Wellcome Sanger Institute COVID-19 Surveillance Team |
| EPI_ISL_3379781 | BIOLITTORAL BIOGROUP PLATEAU TECH | CNR Virus des Infections Respiratoires - France SUD | Antonin Bal, Gregory Destras, Gwendolyne Burfin, Hadrien Regue, Quentin Semanas, Martine Valette, Bruno Lina, Laurence Josset |
| EPI_ISL_3379782 | Lighthouse Lab in Milton Keynes | Wellcome Sanger Institute for the COVID-19 Genomics UK (COG-UK) Consortium | The Lighthouse Lab in Milton Keynes and Alex Alderton, Roberto Amato, Jeffrey Barrett, Sonia Goncalves, Ewan Harrison, David K. Jackson, Ian Johnston, Dominic Kwiatkowski, Cordelia Langford, John Sillitoe on behalf of the Wellcome Sanger Institute COVID-19 Surveillance Team |
| EPI_ISL_3379783, EPI_ISL_3379784 | Lighthouse Lab in Alderley Park | Wellcome Sanger Institute for the COVID-19 Genomics UK (COG-UK) Consortium | Jacquelyn Wynn, Mairead Hyland, The Lighthouse Lab in Alderley Park and Alex Alderton, Roberto Amato, Jeffrey Barrett, Sonia Goncalves, Ewan Harrison, David K. Jackson, Ian Johnston, Dominic Kwiatkowski, Cordelia Langford, John Sillitoe on behalf of the Wellcome Sanger Institute COVID-19 Surveillance Team |
| EPI_ISL_3379785 | BIOLITTORAL BIOGROUP PLATEAU TECH | CNR Virus des Infections Respiratoires - France SUD | Antonin Bal, Gregory Destras, Gwendolyne Burfin, Hadrien Regue, Quentin Semanas, Martine Valette, Bruno Lina, Laurence Josset |
| EPI_ISL_3379786, EPI_ISL_3379787, EPI_ISL_3379788 | Lighthouse Lab in Milton Keynes | Wellcome Sanger Institute for the COVID-19 Genomics UK (COG-UK) Consortium | The Lighthouse Lab in Milton Keynes and Alex Alderton, Roberto Amato, Jeffrey Barrett, Sonia Goncalves, Ewan Harrison, David K. Jackson, Ian Johnston, Dominic Kwiatkowski, Cordelia Langford, John Sillitoe on behalf of the Wellcome Sanger Institute COVID-19 Surveillance Team |
| EPI_ISL_3379789 | BIOLITTORAL BIOGROUP PLATEAU TECH | CNR Virus des Infections Respiratoires - France SUD | Antonin Bal, Gregory Destras, Gwendolyne Burfin, Hadrien Regue, Quentin Semanas, Martine Valette, Bruno Lina, Laurence Josset |
| EPI_ISL_3379790, EPI_ISL_3379791 | Lighthouse Lab in Milton Keynes | Wellcome Sanger Institute for the COVID-19 Genomics UK (COG-UK) Consortium | The Lighthouse Lab in Milton Keynes and Alex Alderton, Roberto Amato, Jeffrey Barrett, Sonia Goncalves, Ewan Harrison, David K. Jackson, Ian Johnston, Dominic Kwiatkowski, Cordelia Langford, John Sillitoe on behalf of the Wellcome Sanger Institute COVID-19 Surveillance Team |
| EPI_ISL_3379792 | BIOLITTORAL BIOGROUP PLATEAU TECH | CNR Virus des Infections Respiratoires - France SUD | Antonin Bal, Gregory Destras, Gwendolyne Burfin, Hadrien Regue, Quentin Semanas, Martine Valette, Bruno Lina, Laurence Josset |
| EPI_ISL_3379793 | Lighthouse Lab in Alderley Park | Wellcome Sanger Institute for the COVID-19 Genomics UK (COG-UK) Consortium | Jacquelyn Wynn, Mairead Hyland, The Lighthouse Lab in Alderley Park and Alex Alderton, Roberto Amato, Jeffrey Barrett, Sonia Goncalves, Ewan Harrison, David K. Jackson, Ian Johnston, Dominic Kwiatkowski, Cordelia Langford, John Sillitoe on behalf of the Wellcome Sanger Institute COVID-19 Surveillance Team |
| EPI_ISL_3379794 | Lighthouse Lab in Milton Keynes | Wellcome Sanger Institute for the COVID-19 Genomics UK (COG-UK) Consortium | The Lighthouse Lab in Milton Keynes and Alex Alderton, Roberto Amato, Jeffrey Barrett, Sonia Goncalves, Ewan Harrison, David K. Jackson, Ian Johnston, Dominic Kwiatkowski, Cordelia Langford, John Sillitoe on behalf of the Wellcome Sanger Institute COVID-19 Surveillance Team |
| EPI_ISL_3379795 | Lighthouse Lab in Glasgow | Wellcome Sanger Institute for the COVID-19 Genomics UK (COG-UK) Consortium | Harper VanSteenhouse, Yumi Kasai, David Gray, Carol Clugston, Anna Dominiczak and Alex Alderton, Roberto Amato, Jeffrey Barrett, Sonia Goncalves, Ewan Harrison, David K. Jackson, Ian Johnston, Dominic Kwiatkowski, Cordelia Langford, John Sillitoe on behalf of the Wellcome Sanger Institute COVID-19 Surveillance Team |
| EPI_ISL_3379796 | BIOLITTORAL BIOGROUP PLATEAU TECH | CNR Virus des Infections Respiratoires - France SUD | Antonin Bal, Gregory Destras, Gwendolyne Burfin, Hadrien Regue, Quentin Semanas, Martine Valette, Bruno Lina, Laurence Josset |
| EPI_ISL_3379797, EPI_ISL_3379798 | Lighthouse Lab in Glasgow | Wellcome Sanger Institute for the COVID-19 Genomics UK (COG-UK) Consortium | Harper VanSteenhouse, Yumi Kasai, David Gray, Carol Clugston, Anna Dominiczak and Alex Alderton, Roberto Amato, Jeffrey Barrett, Sonia Goncalves, Ewan Harrison, David K. Jackson, Ian Johnston, Dominic Kwiatkowski, Cordelia Langford, John Sillitoe on behalf of the Wellcome Sanger Institute COVID-19 Surveillance Team |
| EPI_ISL_3379799 | BIOLITTORAL BIOGROUP PLATEAU TECH | CNR Virus des Infections Respiratoires - France SUD | Antonin Bal, Gregory Destras, Gwendolyne Burfin, Hadrien Regue, Quentin Semanas, Martine Valette, Bruno Lina, Laurence Josset |
| EPI_ISL_3379800, EPI_ISL_3379801 | Lighthouse Lab in Glasgow | Wellcome Sanger Institute for the COVID-19 Genomics UK (COG-UK) Consortium | Harper VanSteenhouse, Yumi Kasai, David Gray, Carol Clugston, Anna Dominiczak and Alex Alderton, Roberto Amato, Jeffrey Barrett, Sonia Goncalves, Ewan Harrison, David K. Jackson, Ian Johnston, Dominic Kwiatkowski, Cordelia Langford, John Sillitoe on behalf of the Wellcome Sanger Institute COVID-19 Surveillance Team |
| EPI_ISL_3379802, EPI_ISL_3379803 | Lighthouse Lab in Alderley Park | Wellcome Sanger Institute for the COVID-19 Genomics UK (COG-UK) Consortium | Jacquelyn Wynn, Mairead Hyland, The Lighthouse Lab in Alderley Park and Alex Alderton, Roberto Amato, Jeffrey Barrett, Sonia Goncalves, Ewan Harrison, David K. Jackson, Ian Johnston, Dominic Kwiatkowski, Cordelia Langford, John Sillitoe on behalf of the Wellcome Sanger Institute COVID-19 Surveillance Team |
| EPI_ISL_3379804 | BIOLITTORAL BIOGROUP PLATEAU TECH | CNR Virus des Infections Respiratoires - France SUD | Antonin Bal, Gregory Destras, Gwendolyne Burfin, Hadrien Regue, Quentin Semanas, Martine Valette, Bruno Lina, Laurence Josset |

| | | | |
|---|---|---|---|
| EPI_ISL_3379805 | Lighthouse Lab in Alderley Park | Wellcome Sanger Institute for the COVID-19 Genomics UK (COG-UK) Consortium | Jacquelyn Wynn, Mairead Hyland, The Lighthouse Lab in Alderley Park and Alex Alderton, Roberto Amato, Jeffrey Barrett, Sonia Goncalves, Ewan Harrison, David K. Jackson, Ian Johnston, Dominic Kwiatkowski, Cordelia Langford, John Sillitoe on behalf of the Wellcome Sanger Institute COVID-19 Surveillance Team |
| EPI_ISL_3379806 | Lighthouse Lab in Glasgow | Wellcome Sanger Institute for the COVID-19 Genomics UK (COG-UK) Consortium | Harper VanSteenhouse, Yumi Kasai, David Gray, Carol Clugston, Anna Dominiczak and Alex Alderton, Roberto Amato, Jeffrey Barrett, Sonia Goncalves, Ewan Harrison, David K. Jackson, Ian Johnston, Dominic Kwiatkowski, Cordelia Langford, John Sillitoe on behalf of the Wellcome Sanger Institute COVID-19 Surveillance Team |
| EPI_ISL_3379807 | Lighthouse Lab in Alderley Park | Wellcome Sanger Institute for the COVID-19 Genomics UK (COG-UK) Consortium | Jacquelyn Wynn, Mairead Hyland, The Lighthouse Lab in Alderley Park and Alex Alderton, Roberto Amato, Jeffrey Barrett, Sonia Goncalves, Ewan Harrison, David K. Jackson, Ian Johnston, Dominic Kwiatkowski, Cordelia Langford, John Sillitoe on behalf of the Wellcome Sanger Institute COVID-19 Surveillance Team |
| EPI_ISL_3379808, EPI_ISL_3379809 | Lighthouse Lab in Glasgow | Wellcome Sanger Institute for the COVID-19 Genomics UK (COG-UK) Consortium | Harper VanSteenhouse, Yumi Kasai, David Gray, Carol Clugston, Anna Dominiczak and Alex Alderton, Roberto Amato, Jeffrey Barrett, Sonia Goncalves, Ewan Harrison, David K. Jackson, Ian Johnston, Dominic Kwiatkowski, Cordelia Langford, John Sillitoe on behalf of the Wellcome Sanger Institute COVID-19 Surveillance Team |
| EPI_ISL_3379810 | BIOLITTORAL BIOGROUP PLATEAU TECH | CNR Virus des Infections Respiratoires - France SUD | Antonin Bal, Gregory Destras, Gwendolyne Burfin, Hadrien Regue, Quentin Semanas, Martine Valette, Bruno Lina, Laurence Josset |
| EPI_ISL_3379811 | Lighthouse Lab in Milton Keynes | Wellcome Sanger Institute for the COVID-19 Genomics UK (COG-UK) Consortium | The Lighthouse Lab in Milton Keynes and Alex Alderton, Roberto Amato, Jeffrey Barrett, Sonia Goncalves, Ewan Harrison, David K. Jackson, Ian Johnston, Dominic Kwiatkowski, Cordelia Langford, John Sillitoe on behalf of the Wellcome Sanger Institute COVID-19 Surveillance Team |
| EPI_ISL_3379812 | Lighthouse Lab in Glasgow | Wellcome Sanger Institute for the COVID-19 Genomics UK (COG-UK) Consortium | Harper VanSteenhouse, Yumi Kasai, David Gray, Carol Clugston, Anna Dominiczak and Alex Alderton, Roberto Amato, Jeffrey Barrett, Sonia Goncalves, Ewan Harrison, David K. Jackson, Ian Johnston, Dominic Kwiatkowski, Cordelia Langford, John Sillitoe on behalf of the Wellcome Sanger Institute COVID-19 Surveillance Team |
| EPI_ISL_3379813 | BIOLITTORAL BIOGROUP PLATEAU TECH | CNR Virus des Infections Respiratoires - France SUD | Antonin Bal, Gregory Destras, Gwendolyne Burfin, Hadrien Regue, Quentin Semanas, Martine Valette, Bruno Lina, Laurence Josset |
| EPI_ISL_3379814, EPI_ISL_3379815 | Lighthouse Lab in Glasgow | Wellcome Sanger Institute for the COVID-19 Genomics UK (COG-UK) Consortium | Harper VanSteenhouse, Yumi Kasai, David Gray, Carol Clugston, Anna Dominiczak and Alex Alderton, Roberto Amato, Jeffrey Barrett, Sonia Goncalves, Ewan Harrison, David K. Jackson, Ian Johnston, Dominic Kwiatkowski, Cordelia Langford, John Sillitoe on behalf of the Wellcome Sanger Institute COVID-19 Surveillance Team |
| EPI_ISL_3379816 | BIOLITTORAL BIOGROUP PLATEAU TECH | CNR Virus des Infections Respiratoires - France SUD | Antonin Bal, Gregory Destras, Gwendolyne Burfin, Hadrien Regue, Quentin Semanas, Martine Valette, Bruno Lina, Laurence Josset |
| EPI_ISL_3379817, EPI_ISL_3379818 | Lighthouse Lab in Glasgow | Wellcome Sanger Institute for the COVID-19 Genomics UK (COG-UK) Consortium | Harper VanSteenhouse, Yumi Kasai, David Gray, Carol Clugston, Anna Dominiczak and Alex Alderton, Roberto Amato, Jeffrey Barrett, Sonia Goncalves, Ewan Harrison, David K. Jackson, Ian Johnston, Dominic Kwiatkowski, Cordelia Langford, John Sillitoe on behalf of the Wellcome Sanger Institute COVID-19 Surveillance Team |
| EPI_ISL_3379819 | Lighthouse Lab in Alderley Park | Wellcome Sanger Institute for the COVID-19 Genomics UK (COG-UK) Consortium | Jacquelyn Wynn, Mairead Hyland, The Lighthouse Lab in Alderley Park and Alex Alderton, Roberto Amato, Jeffrey Barrett, Sonia Goncalves, Ewan Harrison, David K. Jackson, Ian Johnston, Dominic Kwiatkowski, Cordelia Langford, John Sillitoe on behalf of the Wellcome Sanger Institute COVID-19 Surveillance Team |
| EPI_ISL_3379820 | Lighthouse Lab in Glasgow | Wellcome Sanger Institute for the COVID-19 Genomics UK (COG-UK) Consortium | Harper VanSteenhouse, Yumi Kasai, David Gray, Carol Clugston, Anna Dominiczak and Alex Alderton, Roberto Amato, Jeffrey Barrett, Sonia Goncalves, Ewan Harrison, David K. Jackson, Ian Johnston, Dominic Kwiatkowski, Cordelia Langford, John Sillitoe on behalf of the Wellcome Sanger Institute COVID-19 Surveillance Team |
| EPI_ISL_3379821 | Lighthouse Lab in Milton Keynes | Wellcome Sanger Institute for the COVID-19 Genomics UK (COG-UK) Consortium | The Lighthouse Lab in Milton Keynes and Alex Alderton, Roberto Amato, Jeffrey Barrett, Sonia Goncalves, Ewan Harrison, David K. Jackson, Ian Johnston, Dominic Kwiatkowski, Cordelia Langford, John Sillitoe on behalf of the Wellcome Sanger Institute COVID-19 Surveillance Team |
| EPI_ISL_3379822, EPI_ISL_3379823 | Lighthouse Lab in Glasgow | Wellcome Sanger Institute for the COVID-19 Genomics UK (COG-UK) Consortium | Harper VanSteenhouse, Yumi Kasai, David Gray, Carol Clugston, Anna Dominiczak and Alex Alderton, Roberto Amato, Jeffrey Barrett, Sonia Goncalves, Ewan Harrison, David K. Jackson, Ian Johnston, Dominic Kwiatkowski, Cordelia Langford, John Sillitoe on behalf of the Wellcome Sanger Institute COVID-19 Surveillance Team |
| EPI_ISL_3379824 | BIOLITTORAL BIOGROUP PLATEAU TECH | CNR Virus des Infections Respiratoires - France SUD | Antonin Bal, Gregory Destras, Gwendolyne Burfin, Hadrien Regue, Quentin Semanas, Martine Valette, Bruno Lina, Laurence Josset |
| EPI_ISL_3379825 | Lighthouse Lab in Milton Keynes | Wellcome Sanger Institute for the COVID-19 Genomics UK (COG-UK) Consortium | The Lighthouse Lab in Milton Keynes and Alex Alderton, Roberto Amato, Jeffrey Barrett, Sonia Goncalves, Ewan Harrison, David K. Jackson, Ian Johnston, Dominic Kwiatkowski, Cordelia Langford, John Sillitoe on behalf of the Wellcome Sanger Institute COVID-19 Surveillance Team |
| EPI_ISL_3379826, EPI_ISL_3379827 | Lighthouse Lab in Glasgow | Wellcome Sanger Institute for the COVID-19 Genomics UK (COG-UK) Consortium | Harper VanSteenhouse, Yumi Kasai, David Gray, Carol Clugston, Anna Dominiczak and Alex Alderton, Roberto Amato, Jeffrey Barrett, Sonia Goncalves, Ewan Harrison, David K. Jackson, Ian Johnston, Dominic Kwiatkowski, Cordelia Langford, John Sillitoe on behalf of the Wellcome Sanger Institute COVID-19 Surveillance Team |
| EPI_ISL_3379828 | BIOLITTORAL BIOGROUP PLATEAU TECH | CNR Virus des Infections Respiratoires - France SUD | Antonin Bal, Gregory Destras, Gwendolyne Burfin, Hadrien Regue, Quentin Semanas, Martine Valette, Bruno Lina, Laurence Josset |
| EPI_ISL_3379829 | Lighthouse Lab in Milton Keynes | Wellcome Sanger Institute for the COVID-19 Genomics UK (COG-UK) Consortium | The Lighthouse Lab in Milton Keynes and Alex Alderton, Roberto Amato, Jeffrey Barrett, Sonia Goncalves, Ewan Harrison, David K. Jackson, Ian Johnston, Dominic Kwiatkowski, Cordelia Langford, John Sillitoe on behalf of the Wellcome Sanger Institute COVID-19 Surveillance Team |
| EPI_ISL_3379830, EPI_ISL_3379831 | Lighthouse Lab in Glasgow | Wellcome Sanger Institute for the COVID-19 Genomics UK (COG-UK) Consortium | Harper VanSteenhouse, Yumi Kasai, David Gray, Carol Clugston, Anna Dominiczak and Alex Alderton, Roberto Amato, Jeffrey Barrett, Sonia Goncalves, Ewan Harrison, David K. Jackson, Ian Johnston, Dominic Kwiatkowski, Cordelia Langford, John Sillitoe on behalf of the Wellcome Sanger Institute COVID-19 Surveillance Team |
| EPI_ISL_3379832 | BIOLITTORAL BIOGROUP PLATEAU TECH | CNR Virus des Infections Respiratoires - France SUD | Antonin Bal, Gregory Destras, Gwendolyne Burfin, Hadrien Regue, Quentin Semanas, Martine Valette, Bruno Lina, Laurence Josset |
| EPI_ISL_3379833 | Lighthouse Lab in Alderley Park | Wellcome Sanger Institute for the COVID-19 Genomics UK (COG-UK) Consortium | Jacquelyn Wynn, Mairead Hyland, The Lighthouse Lab in Alderley Park and Alex Alderton, Roberto Amato, Jeffrey Barrett, Sonia Goncalves, Ewan Harrison, David K. Jackson, Ian Johnston, Dominic Kwiatkowski, Cordelia Langford, John Sillitoe on behalf of the Wellcome Sanger Institute COVID-19 Surveillance Team |
| EPI_ISL_3379834 | Lighthouse Lab in Milton Keynes | Wellcome Sanger Institute for the COVID-19 Genomics UK (COG-UK) Consortium | The Lighthouse Lab in Milton Keynes and Alex Alderton, Roberto Amato, Jeffrey Barrett, Sonia Goncalves, Ewan Harrison, David K. Jackson, Ian Johnston, Dominic Kwiatkowski, Cordelia Langford, John Sillitoe on behalf of the Wellcome Sanger Institute COVID-19 Surveillance Team |
| EPI_ISL_3379835 | BIOLITTORAL BIOGROUP PLATEAU TECH | CNR Virus des Infections Respiratoires - France SUD | Antonin Bal, Gregory Destras, Gwendolyne Burfin, Hadrien Regue, Quentin Semanas, Martine Valette, Bruno Lina, Laurence Josset |
| EPI_ISL_3379836 | Lighthouse Lab in Milton Keynes | Wellcome Sanger Institute for the COVID-19 Genomics UK (COG-UK) Consortium | The Lighthouse Lab in Milton Keynes and Alex Alderton, Roberto Amato, Jeffrey Barrett, Sonia Goncalves, Ewan Harrison, David K. Jackson, Ian Johnston, Dominic Kwiatkowski, Cordelia Langford, John Sillitoe on behalf of the Wellcome Sanger Institute COVID-19 Surveillance Team |
| EPI_ISL_3379837, EPI_ISL_3379838 | Lighthouse Lab in Glasgow | Wellcome Sanger Institute for the COVID-19 Genomics UK (COG-UK) Consortium | Harper VanSteenhouse, Yumi Kasai, David Gray, Carol Clugston, Anna Dominiczak and Alex Alderton, Roberto Amato, Jeffrey Barrett, Sonia Goncalves, Ewan Harrison, David K. Jackson, Ian Johnston, Dominic Kwiatkowski, Cordelia Langford, John Sillitoe on behalf of the Wellcome Sanger Institute COVID-19 Surveillance Team |
| EPI_ISL_3379839 | BIOLITTORAL BIOGROUP PLATEAU TECH | CNR Virus des Infections Respiratoires - France SUD | Antonin Bal, Gregory Destras, Gwendolyne Burfin, Hadrien Regue, Quentin Semanas, Martine Valette, Bruno Lina, Laurence Josset |
| EPI_ISL_3379840 | Lighthouse Lab in Glasgow | Wellcome Sanger Institute for the COVID-19 Genomics UK (COG-UK) Consortium | Harper VanSteenhouse, Yumi Kasai, David Gray, Carol Clugston, Anna Dominiczak and Alex Alderton, Roberto Amato, Jeffrey Barrett, Sonia Goncalves, Ewan Harrison, David K. Jackson, Ian Johnston, Dominic Kwiatkowski, Cordelia Langford, John Sillitoe on behalf of the Wellcome Sanger Institute COVID-19 Surveillance Team |
| EPI_ISL_3379841 | Lighthouse Lab in Alderley Park | Wellcome Sanger Institute for the COVID-19 Genomics UK (COG-UK) Consortium | Jacquelyn Wynn, Mairead Hyland, The Lighthouse Lab in Alderley Park and Alex Alderton, Roberto Amato, Jeffrey Barrett, Sonia Goncalves, Ewan Harrison, David K. Jackson, Ian Johnston, Dominic Kwiatkowski, Cordelia Langford, John Sillitoe on behalf of the Wellcome Sanger Institute COVID-19 Surveillance Team |
| EPI_ISL_3379842 | Lighthouse Lab in Milton Keynes | Wellcome Sanger Institute for the COVID-19 Genomics UK (COG-UK) Consortium | The Lighthouse Lab in Milton Keynes and Alex Alderton, Roberto Amato, Jeffrey Barrett, Sonia Goncalves, Ewan Harrison, David K. Jackson, Ian Johnston, Dominic Kwiatkowski, Cordelia Langford, John Sillitoe on behalf of the Wellcome Sanger Institute COVID-19 Surveillance Team |
| EPI_ISL_3379843 | ORIAPOLE | CNR Virus des Infections Respiratoires - France SUD | Antonin Bal, Gregory Destras, Gwendolyne Burfin, Hadrien Regue, Quentin Semanas, Martine Valette, Bruno Lina, Laurence Josset |
| EPI_ISL_3379844, EPI_ISL_3379845 | Lighthouse Lab in Milton Keynes | Wellcome Sanger Institute for the COVID-19 Genomics UK (COG-UK) Consortium | The Lighthouse Lab in Milton Keynes and Alex Alderton, Roberto Amato, Jeffrey Barrett, Sonia Goncalves, Ewan Harrison, David K. Jackson, Ian Johnston, Dominic Kwiatkowski, Cordelia Langford, John Sillitoe on behalf of the Wellcome Sanger Institute COVID-19 Surveillance Team |
| EPI_ISL_3379846 | Lighthouse Lab in Alderley Park | Wellcome Sanger Institute for the COVID-19 Genomics UK (COG-UK) Consortium | Jacquelyn Wynn, Mairead Hyland, The Lighthouse Lab in Alderley Park and Alex Alderton, Roberto Amato, Jeffrey Barrett, Sonia Goncalves, Ewan Harrison, David K. Jackson, Ian Johnston, Dominic Kwiatkowski, Cordelia Langford, John Sillitoe on behalf of the Wellcome Sanger Institute COVID-19 Surveillance Team |
| EPI_ISL_3379847 | ORIAPOLE | CNR Virus des Infections Respiratoires - France SUD | Antonin Bal, Gregory Destras, Gwendolyne Burfin, Hadrien Regue, Quentin Semanas, Martine Valette, Bruno Lina, Laurence Josset |
| EPI_ISL_3379848, EPI_ISL_3379849, EPI_ISL_3379850 | Lighthouse Lab in Glasgow | Wellcome Sanger Institute for the COVID-19 Genomics UK (COG-UK) Consortium | Harper VanSteenhouse, Yumi Kasai, David Gray, Carol Clugston, Anna Dominiczak and Alex Alderton, Roberto Amato, Jeffrey Barrett, Sonia Goncalves, Ewan Harrison, David K. Jackson, Ian Johnston, Dominic Kwiatkowski, Cordelia Langford, John Sillitoe on behalf of the Wellcome Sanger Institute COVID-19 Surveillance Team |
| EPI_ISL_3379851 | ORIAPOLE | CNR Virus des Infections Respiratoires - France SUD | Antonin Bal, Gregory Destras, Gwendolyne Burfin, Hadrien Regue, Quentin Semanas, Martine Valette, Bruno Lina, Laurence Josset |
| EPI_ISL_3379852 | Lighthouse Lab in Alderley Park | Wellcome Sanger Institute for the COVID-19 Genomics UK (COG-UK) Consortium | Jacquelyn Wynn, Mairead Hyland, The Lighthouse Lab in Alderley Park and Alex Alderton, Roberto Amato, Jeffrey Barrett, Sonia Goncalves, Ewan Harrison, David K. Jackson, Ian Johnston, Dominic Kwiatkowski, Cordelia Langford, John Sillitoe on behalf of the Wellcome Sanger Institute COVID-19 Surveillance Team |
| EPI_ISL_3379853, EPI_ISL_3379854, EPI_ISL_3379855 | Lighthouse Lab in Glasgow | Wellcome Sanger Institute for the COVID-19 Genomics UK (COG-UK) Consortium | Harper VanSteenhouse, Yumi Kasai, David Gray, Carol Clugston, Anna Dominiczak and Alex Alderton, Roberto Amato, Jeffrey Barrett, Sonia Goncalves, Ewan Harrison, David K. Jackson, Ian Johnston, Dominic Kwiatkowski, Cordelia Langford, John Sillitoe on behalf of the Wellcome Sanger Institute COVID-19 Surveillance Team |
| EPI_ISL_3379856 | CENTRE HOSPITALIER PIERRE OUDOT | CNR Virus des Infections Respiratoires - France SUD | Antonin Bal, Gregory Destras, Gwendolyne Burfin, Hadrien Regue, Quentin Semanas, Martine Valette, Bruno Lina, Laurence Josset |
| EPI_ISL_3379857, EPI_ISL_3379858 | Lighthouse Lab in Glasgow | Wellcome Sanger Institute for the COVID-19 Genomics UK (COG-UK) Consortium | Harper VanSteenhouse, Yumi Kasai, David Gray, Carol Clugston, Anna Dominiczak and Alex Alderton, Roberto Amato, Jeffrey Barrett, Sonia Goncalves, Ewan Harrison, David K. Jackson, Ian Johnston, Dominic Kwiatkowski, Cordelia Langford, John Sillitoe on behalf of the Wellcome Sanger Institute COVID-19 Surveillance Team |
| EPI_ISL_3379859 | BIOLITTORAL BIOGROUP PLATEAU TECH | CNR Virus des Infections Respiratoires - France SUD | Antonin Bal, Gregory Destras, Gwendolyne Burfin, Hadrien Regue, Quentin Semanas, Martine Valette, Bruno Lina, Laurence Josset |
| EPI_ISL_3379860 | Lighthouse Lab in Alderley Park | Wellcome Sanger Institute for the COVID-19 Genomics UK (COG-UK) Consortium | Jacquelyn Wynn, Mairead Hyland, The Lighthouse Lab in Alderley Park and Alex Alderton, Roberto Amato, Jeffrey Barrett, Sonia Goncalves, Ewan Harrison, David K. Jackson, Ian Johnston, Dominic Kwiatkowski, Cordelia Langford, John Sillitoe on behalf of the Wellcome Sanger Institute COVID-19 Surveillance Team |
| EPI_ISL_3379861 | Lighthouse Lab in Milton Keynes | Wellcome Sanger Institute for the COVID-19 Genomics UK (COG-UK) Consortium | The Lighthouse Lab in Milton Keynes and Alex Alderton, Roberto Amato, Jeffrey Barrett, Sonia Goncalves, Ewan Harrison, David K. Jackson, Ian Johnston, Dominic Kwiatkowski, Cordelia Langford, John Sillitoe on behalf of the Wellcome Sanger Institute COVID-19 Surveillance Team |
| EPI_ISL_3379862 | Lighthouse Lab in Alderley Park | Wellcome Sanger Institute for the COVID-19 Genomics UK (COG-UK) Consortium | Jacquelyn Wynn, Mairead Hyland, The Lighthouse Lab in Alderley Park and Alex Alderton, Roberto Amato, Jeffrey Barrett, Sonia Goncalves, Ewan Harrison, David K. Jackson, Ian Johnston, Dominic Kwiatkowski, Cordelia Langford, John Sillitoe on behalf of the Wellcome Sanger Institute COVID-19 Surveillance Team |
| EPI_ISL_3379863 | BIOLITTORAL BIOGROUP PLATEAU TECH | CNR Virus des Infections Respiratoires - France SUD | Antonin Bal, Gregory Destras, Gwendolyne Burfin, Hadrien Regue, Quentin Semanas, Martine Valette, Bruno Lina, Laurence Josset |
| EPI_ISL_3379864, EPI_ISL_3379865 | Lighthouse Lab in Glasgow | Wellcome Sanger Institute for the COVID-19 Genomics UK (COG-UK) Consortium | Harper VanSteenhouse, Yumi Kasai, David Gray, Carol Clugston, Anna Dominiczak and Alex Alderton, Roberto Amato, Jeffrey Barrett, Sonia Goncalves, Ewan Harrison, David K. Jackson, Ian Johnston, Dominic Kwiatkowski, Cordelia Langford, John Sillitoe on behalf of the Wellcome Sanger Institute COVID-19 Surveillance Team |
| EPI_ISL_3379866 | Lighthouse Lab in Milton Keynes | Wellcome Sanger Institute for the COVID-19 Genomics UK (COG-UK) Consortium | The Lighthouse Lab in Milton Keynes and Alex Alderton, Roberto Amato, Jeffrey Barrett, Sonia Goncalves, Ewan Harrison, David K. Jackson, Ian Johnston, Dominic Kwiatkowski, Cordelia Langford, John Sillitoe on behalf of the Wellcome Sanger Institute COVID-19 Surveillance Team |
| EPI_ISL_3379867 | BIOLITTORAL BIOGROUP PLATEAU TECH | CNR Virus des Infections Respiratoires - France SUD | Antonin Bal, Gregory Destras, Gwendolyne Burfin, Hadrien Regue, Quentin Semanas, Martine Valette, Bruno Lina, Laurence Josset |
| EPI_ISL_3379868 | Lighthouse Lab in Alderley Park | Wellcome Sanger Institute for the COVID-19 Genomics UK (COG-UK) Consortium | Jacquelyn Wynn, Mairead Hyland, The Lighthouse Lab in Alderley Park and Alex Alderton, Roberto Amato, Jeffrey Barrett, Sonia Goncalves, Ewan Harrison, David K. Jackson, Ian Johnston, Dominic Kwiatkowski, Cordelia Langford, John Sillitoe on behalf of the Wellcome Sanger Institute COVID-19 Surveillance Team |

| | | | |
|---|---|---|---|
| EPI_ISL_3379869, EPI_ISL_3379870, EPI_ISL_3379871 | Lighthouse Lab in Glasgow | Wellcome Sanger Institute for the COVID-19 Genomics UK (COG-UK) Consortium | Harper VanSteenhouse, Yumi Kasai, David Gray, Carol Clugston, Anna Dominiczak and Alex Alderton, Roberto Amato, Jeffrey Barrett, Sonia Goncalves, Ewan Harrison, David K. Jackson, Ian Johnston, Dominic Kwiatkowski, Cordelia Langford, John Sillitoe on behalf of the Wellcome Sanger Institute COVID-19 Surveillance Team |
| EPI_ISL_3379872 | LAM BIORYLIS | CNR Virus des Infections Respiratoires - France SUD | Antonin Bal, Gregory Destras, Gwendolyne Burfin, Hadrien Regue, Quentin Semanas, Martine Valette, Bruno Lina, Laurence Josset |
| EPI_ISL_3379873 | Lighthouse Lab in Glasgow | Wellcome Sanger Institute for the COVID-19 Genomics UK (COG-UK) Consortium | Harper VanSteenhouse, Yumi Kasai, David Gray, Carol Clugston, Anna Dominiczak and Alex Alderton, Roberto Amato, Jeffrey Barrett, Sonia Goncalves, Ewan Harrison, David K. Jackson, Ian Johnston, Dominic Kwiatkowski, Cordelia Langford, John Sillitoe on behalf of the Wellcome Sanger Institute COVID-19 Surveillance Team |
| EPI_ISL_3379874 | Lighthouse Lab in Alderley Park | Wellcome Sanger Institute for the COVID-19 Genomics UK (COG-UK) Consortium | Jacquelyn Wynn, Mairead Hyland, The Lighthouse Lab in Alderley Park and Alex Alderton, Roberto Amato, Jeffrey Barrett, Sonia Goncalves, Ewan Harrison, David K. Jackson, Ian Johnston, Dominic Kwiatkowski, Cordelia Langford, John Sillitoe on behalf of the Wellcome Sanger Institute COVID-19 Surveillance Team |
| EPI_ISL_3379875 | Lighthouse Lab in Milton Keynes | Wellcome Sanger Institute for the COVID-19 Genomics UK (COG-UK) Consortium | The Lighthouse Lab in Milton Keynes and Alex Alderton, Roberto Amato, Jeffrey Barrett, Sonia Goncalves, Ewan Harrison, David K. Jackson, Ian Johnston, Dominic Kwiatkowski, Cordelia Langford, John Sillitoe on behalf of the Wellcome Sanger Institute COVID-19 Surveillance Team |
| EPI_ISL_3379876 | BIOLITTORAL BIOGROUP PLATEAU TECH | CNR Virus des Infections Respiratoires - France SUD | Antonin Bal, Gregory Destras, Gwendolyne Burfin, Hadrien Regue, Quentin Semanas, Martine Valette, Bruno Lina, Laurence Josset |
| EPI_ISL_3379877, EPI_ISL_3379878 | Lighthouse Lab in Glasgow | Wellcome Sanger Institute for the COVID-19 Genomics UK (COG-UK) Consortium | Harper VanSteenhouse, Yumi Kasai, David Gray, Carol Clugston, Anna Dominiczak and Alex Alderton, Roberto Amato, Jeffrey Barrett, Sonia Goncalves, Ewan Harrison, David K. Jackson, Ian Johnston, Dominic Kwiatkowski, Cordelia Langford, John Sillitoe on behalf of the Wellcome Sanger Institute COVID-19 Surveillance Team |
| EPI_ISL_3379879 | BIOLITTORAL BIOGROUP PLATEAU TECH | CNR Virus des Infections Respiratoires - France SUD | Antonin Bal, Gregory Destras, Gwendolyne Burfin, Hadrien Regue, Quentin Semanas, Martine Valette, Bruno Lina, Laurence Josset |
| EPI_ISL_3379880, EPI_ISL_3379881, EPI_ISL_3379882 | Lighthouse Lab in Glasgow | Wellcome Sanger Institute for the COVID-19 Genomics UK (COG-UK) Consortium | Harper VanSteenhouse, Yumi Kasai, David Gray, Carol Clugston, Anna Dominiczak and Alex Alderton, Roberto Amato, Jeffrey Barrett, Sonia Goncalves, Ewan Harrison, David K. Jackson, Ian Johnston, Dominic Kwiatkowski, Cordelia Langford, John Sillitoe on behalf of the Wellcome Sanger Institute COVID-19 Surveillance Team |
| EPI_ISL_3379883 | LAM BIORYLIS | CNR Virus des Infections Respiratoires - France SUD | Antonin Bal, Gregory Destras, Gwendolyne Burfin, Hadrien Regue, Quentin Semanas, Martine Valette, Bruno Lina, Laurence Josset |
| EPI_ISL_3379884 | Lighthouse Lab in Alderley Park | Wellcome Sanger Institute for the COVID-19 Genomics UK (COG-UK) Consortium | Jacquelyn Wynn, Mairead Hyland, The Lighthouse Lab in Alderley Park and Alex Alderton, Roberto Amato, Jeffrey Barrett, Sonia Goncalves, Ewan Harrison, David K. Jackson, Ian Johnston, Dominic Kwiatkowski, Cordelia Langford, John Sillitoe on behalf of the Wellcome Sanger Institute COVID-19 Surveillance Team |
| EPI_ISL_3379885 | Lighthouse Lab in Glasgow | Wellcome Sanger Institute for the COVID-19 Genomics UK (COG-UK) Consortium | Harper VanSteenhouse, Yumi Kasai, David Gray, Carol Clugston, Anna Dominiczak and Alex Alderton, Roberto Amato, Jeffrey Barrett, Sonia Goncalves, Ewan Harrison, David K. Jackson, Ian Johnston, Dominic Kwiatkowski, Cordelia Langford, John Sillitoe on behalf of the Wellcome Sanger Institute COVID-19 Surveillance Team |
| EPI_ISL_3379886 | Lighthouse Lab in Alderley Park | Wellcome Sanger Institute for the COVID-19 Genomics UK (COG-UK) Consortium | Jacquelyn Wynn, Mairead Hyland, The Lighthouse Lab in Alderley Park and Alex Alderton, Roberto Amato, Jeffrey Barrett, Sonia Goncalves, Ewan Harrison, David K. Jackson, Ian Johnston, Dominic Kwiatkowski, Cordelia Langford, John Sillitoe on behalf of the Wellcome Sanger Institute COVID-19 Surveillance Team |
| EPI_ISL_3379887 | BIOLITTORAL BIOGROUP PLATEAU TECH | CNR Virus des Infections Respiratoires - France SUD | Antonin Bal, Gregory Destras, Gwendolyne Burfin, Hadrien Regue, Quentin Semanas, Martine Valette, Bruno Lina, Laurence Josset |
| EPI_ISL_3379888, EPI_ISL_3379889 | Lighthouse Lab in Alderley Park | Wellcome Sanger Institute for the COVID-19 Genomics UK (COG-UK) Consortium | Jacquelyn Wynn, Mairead Hyland, The Lighthouse Lab in Alderley Park and Alex Alderton, Roberto Amato, Jeffrey Barrett, Sonia Goncalves, Ewan Harrison, David K. Jackson, Ian Johnston, Dominic Kwiatkowski, Cordelia Langford, John Sillitoe on behalf of the Wellcome Sanger Institute COVID-19 Surveillance Team |
| EPI_ISL_3379890 | Lighthouse Lab in Milton Keynes | Wellcome Sanger Institute for the COVID-19 Genomics UK (COG-UK) Consortium | The Lighthouse Lab in Milton Keynes and Alex Alderton, Roberto Amato, Jeffrey Barrett, Sonia Goncalves, Ewan Harrison, David K. Jackson, Ian Johnston, Dominic Kwiatkowski, Cordelia Langford, John Sillitoe on behalf of the Wellcome Sanger Institute COVID-19 Surveillance Team |
| EPI_ISL_3379891 | BIOLITTORAL BIOGROUP PLATEAU TECH | CNR Virus des Infections Respiratoires - France SUD | Antonin Bal, Gregory Destras, Gwendolyne Burfin, Hadrien Regue, Quentin Semanas, Martine Valette, Bruno Lina, Laurence Josset |
| EPI_ISL_3379892, EPI_ISL_3379893 | Lighthouse Lab in Glasgow | Wellcome Sanger Institute for the COVID-19 Genomics UK (COG-UK) Consortium | Harper VanSteenhouse, Yumi Kasai, David Gray, Carol Clugston, Anna Dominiczak and Alex Alderton, Roberto Amato, Jeffrey Barrett, Sonia Goncalves, Ewan Harrison, David K. Jackson, Ian Johnston, Dominic Kwiatkowski, Cordelia Langford, John Sillitoe on behalf of the Wellcome Sanger Institute COVID-19 Surveillance Team |
| EPI_ISL_3379894 | Lighthouse Lab in Alderley Park | Wellcome Sanger Institute for the COVID-19 Genomics UK (COG-UK) Consortium | Jacquelyn Wynn, Mairead Hyland, The Lighthouse Lab in Alderley Park and Alex Alderton, Roberto Amato, Jeffrey Barrett, Sonia Goncalves, Ewan Harrison, David K. Jackson, Ian Johnston, Dominic Kwiatkowski, Cordelia Langford, John Sillitoe on behalf of the Wellcome Sanger Institute COVID-19 Surveillance Team |
| EPI_ISL_3379895 | BIOLITTORAL BIOGROUP PLATEAU TECH | CNR Virus des Infections Respiratoires - France SUD | Antonin Bal, Gregory Destras, Gwendolyne Burfin, Hadrien Regue, Quentin Semanas, Martine Valette, Bruno Lina, Laurence Josset |
| EPI_ISL_3379896, EPI_ISL_3379897, EPI_ISL_3379898 | Lighthouse Lab in Glasgow | Wellcome Sanger Institute for the COVID-19 Genomics UK (COG-UK) Consortium | Harper VanSteenhouse, Yumi Kasai, David Gray, Carol Clugston, Anna Dominiczak and Alex Alderton, Roberto Amato, Jeffrey Barrett, Sonia Goncalves, Ewan Harrison, David K. Jackson, Ian Johnston, Dominic Kwiatkowski, Cordelia Langford, John Sillitoe on behalf of the Wellcome Sanger Institute COVID-19 Surveillance Team |
| EPI_ISL_3379899 | Lighthouse Lab in Milton Keynes | Wellcome Sanger Institute for the COVID-19 Genomics UK (COG-UK) Consortium | The Lighthouse Lab in Milton Keynes and Alex Alderton, Roberto Amato, Jeffrey Barrett, Sonia Goncalves, Ewan Harrison, David K. Jackson, Ian Johnston, Dominic Kwiatkowski, Cordelia Langford, John Sillitoe on behalf of the Wellcome Sanger Institute COVID-19 Surveillance Team |
| EPI_ISL_3379900 | LAM BIORYLIS | CNR Virus des Infections Respiratoires - France SUD | Antonin Bal, Gregory Destras, Gwendolyne Burfin, Hadrien Regue, Quentin Semanas, Martine Valette, Bruno Lina, Laurence Josset |
| EPI_ISL_3379901 | Lighthouse Lab in Glasgow | Wellcome Sanger Institute for the COVID-19 Genomics UK (COG-UK) Consortium | Harper VanSteenhouse, Yumi Kasai, David Gray, Carol Clugston, Anna Dominiczak and Alex Alderton, Roberto Amato, Jeffrey Barrett, Sonia Goncalves, Ewan Harrison, David K. Jackson, Ian Johnston, Dominic Kwiatkowski, Cordelia Langford, John Sillitoe on behalf of the Wellcome Sanger Institute COVID-19 Surveillance Team |
| EPI_ISL_3379902 | Lighthouse Lab in Alderley Park | Wellcome Sanger Institute for the COVID-19 Genomics UK (COG-UK) Consortium | Jacquelyn Wynn, Mairead Hyland, The Lighthouse Lab in Alderley Park and Alex Alderton, Roberto Amato, Jeffrey Barrett, Sonia Goncalves, Ewan Harrison, David K. Jackson, Ian Johnston, Dominic Kwiatkowski, Cordelia Langford, John Sillitoe on behalf of the Wellcome Sanger Institute COVID-19 Surveillance Team |
| EPI_ISL_3379903 | BIOLITTORAL BIOGROUP PLATEAU TECH | CNR Virus des Infections Respiratoires - France SUD | Antonin Bal, Gregory Destras, Gwendolyne Burfin, Hadrien Regue, Quentin Semanas, Martine Valette, Bruno Lina, Laurence Josset |
| EPI_ISL_3379904, EPI_ISL_3379905, EPI_ISL_3379906 | Lighthouse Lab in Glasgow | Wellcome Sanger Institute for the COVID-19 Genomics UK (COG-UK) Consortium | Harper VanSteenhouse, Yumi Kasai, David Gray, Carol Clugston, Anna Dominiczak and Alex Alderton, Roberto Amato, Jeffrey Barrett, Sonia Goncalves, Ewan Harrison, David K. Jackson, Ian Johnston, Dominic Kwiatkowski, Cordelia Langford, John Sillitoe on behalf of the Wellcome Sanger Institute COVID-19 Surveillance Team |
| EPI_ISL_3379907 | LAM BIORYLIS | CNR Virus des Infections Respiratoires - France SUD | Antonin Bal, Gregory Destras, Gwendolyne Burfin, Hadrien Regue, Quentin Semanas, Martine Valette, Bruno Lina, Laurence Josset |
| EPI_ISL_3379908, EPI_ISL_3379909, EPI_ISL_3379910 | Lighthouse Lab in Glasgow | Wellcome Sanger Institute for the COVID-19 Genomics UK (COG-UK) Consortium | Harper VanSteenhouse, Yumi Kasai, David Gray, Carol Clugston, Anna Dominiczak and Alex Alderton, Roberto Amato, Jeffrey Barrett, Sonia Goncalves, Ewan Harrison, David K. Jackson, Ian Johnston, Dominic Kwiatkowski, Cordelia Langford, John Sillitoe on behalf of the Wellcome Sanger Institute COVID-19 Surveillance Team |
| EPI_ISL_3379911 | BIOLITTORAL BIOGROUP PLATEAU TECH | CNR Virus des Infections Respiratoires - France SUD | Antonin Bal, Gregory Destras, Gwendolyne Burfin, Hadrien Regue, Quentin Semanas, Martine Valette, Bruno Lina, Laurence Josset |
| EPI_ISL_3379912 | Lighthouse Lab in Glasgow | Wellcome Sanger Institute for the COVID-19 Genomics UK (COG-UK) Consortium | Harper VanSteenhouse, Yumi Kasai, David Gray, Carol Clugston, Anna Dominiczak and Alex Alderton, Roberto Amato, Jeffrey Barrett, Sonia Goncalves, Ewan Harrison, David K. Jackson, Ian Johnston, Dominic Kwiatkowski, Cordelia Langford, John Sillitoe on behalf of the Wellcome Sanger Institute COVID-19 Surveillance Team |
| EPI_ISL_3379913 | Lighthouse Lab in Alderley Park | Wellcome Sanger Institute for the COVID-19 Genomics UK (COG-UK) Consortium | Jacquelyn Wynn, Mairead Hyland, The Lighthouse Lab in Alderley Park and Alex Alderton, Roberto Amato, Jeffrey Barrett, Sonia Goncalves, Ewan Harrison, David K. Jackson, Ian Johnston, Dominic Kwiatkowski, Cordelia Langford, John Sillitoe on behalf of the Wellcome Sanger Institute COVID-19 Surveillance Team |
| EPI_ISL_3379914 | Lighthouse Lab in Glasgow | Wellcome Sanger Institute for the COVID-19 Genomics UK (COG-UK) Consortium | Harper VanSteenhouse, Yumi Kasai, David Gray, Carol Clugston, Anna Dominiczak and Alex Alderton, Roberto Amato, Jeffrey Barrett, Sonia Goncalves, Ewan Harrison, David K. Jackson, Ian Johnston, Dominic Kwiatkowski, Cordelia Langford, John Sillitoe on behalf of the Wellcome Sanger Institute COVID-19 Surveillance Team |
| EPI_ISL_3379915 | BIOLITTORAL BIOGROUP PLATEAU TECH | CNR Virus des Infections Respiratoires - France SUD | Antonin Bal, Gregory Destras, Gwendolyne Burfin, Hadrien Regue, Quentin Semanas, Martine Valette, Bruno Lina, Laurence Josset |
| EPI_ISL_3379916, EPI_ISL_3379917 | Lighthouse Lab in Glasgow | Wellcome Sanger Institute for the COVID-19 Genomics UK (COG-UK) Consortium | Harper VanSteenhouse, Yumi Kasai, David Gray, Carol Clugston, Anna Dominiczak and Alex Alderton, Roberto Amato, Jeffrey Barrett, Sonia Goncalves, Ewan Harrison, David K. Jackson, Ian Johnston, Dominic Kwiatkowski, Cordelia Langford, John Sillitoe on behalf of the Wellcome Sanger Institute COVID-19 Surveillance Team |
| EPI_ISL_3379918 | BIOLITTORAL BIOGROUP PLATEAU TECH | CNR Virus des Infections Respiratoires - France SUD | Antonin Bal, Gregory Destras, Gwendolyne Burfin, Hadrien Regue, Quentin Semanas, Martine Valette, Bruno Lina, Laurence Josset |
| EPI_ISL_3379919 | Lighthouse Lab in Alderley Park | Wellcome Sanger Institute for the COVID-19 Genomics UK (COG-UK) Consortium | Jacquelyn Wynn, Mairead Hyland, The Lighthouse Lab in Alderley Park and Alex Alderton, Roberto Amato, Jeffrey Barrett, Sonia Goncalves, Ewan Harrison, David K. Jackson, Ian Johnston, Dominic Kwiatkowski, Cordelia Langford, John Sillitoe on behalf of the Wellcome Sanger Institute COVID-19 Surveillance Team |
| EPI_ISL_3379920, EPI_ISL_3379921 | Lighthouse Lab in Glasgow | Wellcome Sanger Institute for the COVID-19 Genomics UK (COG-UK) Consortium | Harper VanSteenhouse, Yumi Kasai, David Gray, Carol Clugston, Anna Dominiczak and Alex Alderton, Roberto Amato, Jeffrey Barrett, Sonia Goncalves, Ewan Harrison, David K. Jackson, Ian Johnston, Dominic Kwiatkowski, Cordelia Langford, John Sillitoe on behalf of the Wellcome Sanger Institute COVID-19 Surveillance Team |
| EPI_ISL_3379922 | BIOLITTORAL BIOGROUP PLATEAU TECH | CNR Virus des Infections Respiratoires - France SUD | Antonin Bal, Gregory Destras, Gwendolyne Burfin, Hadrien Regue, Quentin Semanas, Martine Valette, Bruno Lina, Laurence Josset |
| EPI_ISL_3379923 | Lighthouse Lab in Glasgow | Wellcome Sanger Institute for the COVID-19 Genomics UK (COG-UK) Consortium | Harper VanSteenhouse, Yumi Kasai, David Gray, Carol Clugston, Anna Dominiczak and Alex Alderton, Roberto Amato, Jeffrey Barrett, Sonia Goncalves, Ewan Harrison, David K. Jackson, Ian Johnston, Dominic Kwiatkowski, Cordelia Langford, John Sillitoe on behalf of the Wellcome Sanger Institute COVID-19 Surveillance Team |
| EPI_ISL_3379924 | Lighthouse Lab in Milton Keynes | Wellcome Sanger Institute for the COVID-19 Genomics UK (COG-UK) Consortium | The Lighthouse Lab in Milton Keynes and Alex Alderton, Roberto Amato, Jeffrey Barrett, Sonia Goncalves, Ewan Harrison, David K. Jackson, Ian Johnston, Dominic Kwiatkowski, Cordelia Langford, John Sillitoe on behalf of the Wellcome Sanger Institute COVID-19 Surveillance Team |
| EPI_ISL_3379925 | LAM BIORYLIS | CNR Virus des Infections Respiratoires - France SUD | Antonin Bal, Gregory Destras, Gwendolyne Burfin, Hadrien Regue, Quentin Semanas, Martine Valette, Bruno Lina, Laurence Josset |
| EPI_ISL_3379926 | Lighthouse Lab in Milton Keynes | Wellcome Sanger Institute for the COVID-19 Genomics UK (COG-UK) Consortium | The Lighthouse Lab in Milton Keynes and Alex Alderton, Roberto Amato, Jeffrey Barrett, Sonia Goncalves, Ewan Harrison, David K. Jackson, Ian Johnston, Dominic Kwiatkowski, Cordelia Langford, John Sillitoe on behalf of the Wellcome Sanger Institute COVID-19 Surveillance Team |
| EPI_ISL_3379927 | Lighthouse Lab in Alderley Park | Wellcome Sanger Institute for the COVID-19 Genomics UK (COG-UK) Consortium | Jacquelyn Wynn, Mairead Hyland, The Lighthouse Lab in Alderley Park and Alex Alderton, Roberto Amato, Jeffrey Barrett, Sonia Goncalves, Ewan Harrison, David K. Jackson, Ian Johnston, Dominic Kwiatkowski, Cordelia Langford, John Sillitoe on behalf of the Wellcome Sanger Institute COVID-19 Surveillance Team |
| EPI_ISL_3379928 | Lighthouse Lab in Glasgow | Wellcome Sanger Institute for the COVID-19 Genomics UK (COG-UK) Consortium | Harper VanSteenhouse, Yumi Kasai, David Gray, Carol Clugston, Anna Dominiczak and Alex Alderton, Roberto Amato, Jeffrey Barrett, Sonia Goncalves, Ewan Harrison, David K. Jackson, Ian Johnston, Dominic Kwiatkowski, Cordelia Langford, John Sillitoe on behalf of the Wellcome Sanger Institute COVID-19 Surveillance Team |
| EPI_ISL_3379929 | BIOLITTORAL BIOGROUP PLATEAU TECH | CNR Virus des Infections Respiratoires - France SUD | Antonin Bal, Gregory Destras, Gwendolyne Burfin, Hadrien Regue, Quentin Semanas, Martine Valette, Bruno Lina, Laurence Josset |
| EPI_ISL_3379930 | Lighthouse Lab in Glasgow | Wellcome Sanger Institute for the COVID-19 Genomics UK (COG-UK) Consortium | Harper VanSteenhouse, Yumi Kasai, David Gray, Carol Clugston, Anna Dominiczak and Alex Alderton, Roberto Amato, Jeffrey Barrett, Sonia Goncalves, Ewan Harrison, David K. Jackson, Ian Johnston, Dominic Kwiatkowski, Cordelia Langford, John Sillitoe on behalf of the Wellcome Sanger Institute COVID-19 Surveillance Team |
| EPI_ISL_3379931, EPI_ISL_3379932 | Lighthouse Lab in Alderley Park | Wellcome Sanger Institute for the COVID-19 Genomics UK (COG-UK) Consortium | Jacquelyn Wynn, Mairead Hyland, The Lighthouse Lab in Alderley Park and Alex Alderton, Roberto Amato, Jeffrey Barrett, Sonia Goncalves, Ewan Harrison, David K. Jackson, Ian Johnston, Dominic Kwiatkowski, Cordelia Langford, John Sillitoe on behalf of the Wellcome Sanger Institute COVID-19 Surveillance Team |
| EPI_ISL_3379933 | BIOLITTORAL BIOGROUP PLATEAU TECH | CNR Virus des Infections Respiratoires - France SUD | Antonin Bal, Gregory Destras, Gwendolyne Burfin, Hadrien Regue, Quentin Semanas, Martine Valette, Bruno Lina, Laurence Josset |
| EPI_ISL_3379934, EPI_ISL_3379935, | Lighthouse Lab in Alderley Park | Wellcome Sanger Institute for the COVID-19 Genomics UK (COG-UK) Consortium | Jacquelyn Wynn, Mairead Hyland, The Lighthouse Lab in Alderley Park and Alex Alderton, Roberto Amato, Jeffrey Barrett, Sonia Goncalves, Ewan Harrison, David K. Jackson, Ian Johnston, Dominic Kwiatkowski, Cordelia Langford, John Sillitoe on behalf of the Wellcome Sanger Institute COVID-19 Surveillance Team |

| | | | |
|---|---|---|---|
| EPI_ISL_3379936 | | | |
| EPI_ISL_3379937 | LAM BIORYLIS | CNR Virus des Infections Respiratoires - France SUD | Antonin Bal, Gregory Destras, Gwendolyne Burfin, Hadrien Regue, Quentin Semanas, Martine Valette, Bruno Lina, Laurence Josset |
| EPI_ISL_3379938 | Lighthouse Lab in Milton Keynes | Wellcome Sanger Institute for the COVID-19 Genomics UK (COG-UK) Consortium | The Lighthouse Lab in Milton Keynes and Alex Alderton, Roberto Amato, Jeffrey Barrett, Sonia Goncalves, Ewan Harrison, David K. Jackson, Ian Johnston, Dominic Kwiatkowski, Cordelia Langford, John Sillitoe on behalf of the Wellcome Sanger Institute COVID-19 Surveillance Team |
| EPI_ISL_3379939, EPI_ISL_3379940 | Lighthouse Lab in Glasgow | Wellcome Sanger Institute for the COVID-19 Genomics UK (COG-UK) Consortium | Harper VanSteenhouse, Yumi Kasai, David Gray, Carol Clugston, Anna Dominiczak and Alex Alderton, Roberto Amato, Jeffrey Barrett, Sonia Goncalves, Ewan Harrison, David K. Jackson, Ian Johnston, Dominic Kwiatkowski, Cordelia Langford, John Sillitoe on behalf of the Wellcome Sanger Institute COVID-19 Surveillance Team |
| EPI_ISL_3379941 | LAM BIORYLIS | CNR Virus des Infections Respiratoires - France SUD | Antonin Bal, Gregory Destras, Gwendolyne Burfin, Hadrien Regue, Quentin Semanas, Martine Valette, Bruno Lina, Laurence Josset |
| EPI_ISL_3379942 | Lighthouse Lab in Alderley Park | Wellcome Sanger Institute for the COVID-19 Genomics UK (COG-UK) Consortium | Jacquelyn Wynn, Mairead Hyland, The Lighthouse Lab in Alderley Park and Alex Alderton, Roberto Amato, Jeffrey Barrett, Sonia Goncalves, Ewan Harrison, David K. Jackson, Ian Johnston, Dominic Kwiatkowski, Cordelia Langford, John Sillitoe on behalf of the Wellcome Sanger Institute COVID-19 Surveillance Team |
| EPI_ISL_3379943 | Lighthouse Lab in Milton Keynes | Wellcome Sanger Institute for the COVID-19 Genomics UK (COG-UK) Consortium | The Lighthouse Lab in Milton Keynes and Alex Alderton, Roberto Amato, Jeffrey Barrett, Sonia Goncalves, Ewan Harrison, David K. Jackson, Ian Johnston, Dominic Kwiatkowski, Cordelia Langford, John Sillitoe on behalf of the Wellcome Sanger Institute COVID-19 Surveillance Team |
| EPI_ISL_3379944 | Lighthouse Lab in Alderley Park | Wellcome Sanger Institute for the COVID-19 Genomics UK (COG-UK) Consortium | Jacquelyn Wynn, Mairead Hyland, The Lighthouse Lab in Alderley Park and Alex Alderton, Roberto Amato, Jeffrey Barrett, Sonia Goncalves, Ewan Harrison, David K. Jackson, Ian Johnston, Dominic Kwiatkowski, Cordelia Langford, John Sillitoe on behalf of the Wellcome Sanger Institute COVID-19 Surveillance Team |
| EPI_ISL_3379945 | LAM BIORYLIS | CNR Virus des Infections Respiratoires - France SUD | Antonin Bal, Gregory Destras, Gwendolyne Burfin, Hadrien Regue, Quentin Semanas, Martine Valette, Bruno Lina, Laurence Josset |
| EPI_ISL_3379946 | Lighthouse Lab in Glasgow | Wellcome Sanger Institute for the COVID-19 Genomics UK (COG-UK) Consortium | Harper VanSteenhouse, Yumi Kasai, David Gray, Carol Clugston, Anna Dominiczak and Alex Alderton, Roberto Amato, Jeffrey Barrett, Sonia Goncalves, Ewan Harrison, David K. Jackson, Ian Johnston, Dominic Kwiatkowski, Cordelia Langford, John Sillitoe on behalf of the Wellcome Sanger Institute COVID-19 Surveillance Team |
| EPI_ISL_3379947 | Lighthouse Lab in Alderley Park | Wellcome Sanger Institute for the COVID-19 Genomics UK (COG-UK) Consortium | Jacquelyn Wynn, Mairead Hyland, The Lighthouse Lab in Alderley Park and Alex Alderton, Roberto Amato, Jeffrey Barrett, Sonia Goncalves, Ewan Harrison, David K. Jackson, Ian Johnston, Dominic Kwiatkowski, Cordelia Langford, John Sillitoe on behalf of the Wellcome Sanger Institute COVID-19 Surveillance Team |
| EPI_ISL_3379948 | Lighthouse Lab in Glasgow | Wellcome Sanger Institute for the COVID-19 Genomics UK (COG-UK) Consortium | Harper VanSteenhouse, Yumi Kasai, David Gray, Carol Clugston, Anna Dominiczak and Alex Alderton, Roberto Amato, Jeffrey Barrett, Sonia Goncalves, Ewan Harrison, David K. Jackson, Ian Johnston, Dominic Kwiatkowski, Cordelia Langford, John Sillitoe on behalf of the Wellcome Sanger Institute COVID-19 Surveillance Team |
| EPI_ISL_3379949 | LAM BIORYLIS | CNR Virus des Infections Respiratoires - France SUD | Antonin Bal, Gregory Destras, Gwendolyne Burfin, Hadrien Regue, Quentin Semanas, Martine Valette, Bruno Lina, Laurence Josset |
| EPI_ISL_3379950 | Lighthouse Lab in Milton Keynes | Wellcome Sanger Institute for the COVID-19 Genomics UK (COG-UK) Consortium | The Lighthouse Lab in Milton Keynes and Alex Alderton, Roberto Amato, Jeffrey Barrett, Sonia Goncalves, Ewan Harrison, David K. Jackson, Ian Johnston, Dominic Kwiatkowski, Cordelia Langford, John Sillitoe on behalf of the Wellcome Sanger Institute COVID-19 Surveillance Team |
| EPI_ISL_3379951 | Lighthouse Lab in Glasgow | Wellcome Sanger Institute for the COVID-19 Genomics UK (COG-UK) Consortium | Harper VanSteenhouse, Yumi Kasai, David Gray, Carol Clugston, Anna Dominiczak and Alex Alderton, Roberto Amato, Jeffrey Barrett, Sonia Goncalves, Ewan Harrison, David K. Jackson, Ian Johnston, Dominic Kwiatkowski, Cordelia Langford, John Sillitoe on behalf of the Wellcome Sanger Institute COVID-19 Surveillance Team |
| EPI_ISL_3379952 | LAM BIORYLIS | CNR Virus des Infections Respiratoires - France SUD | Antonin Bal, Gregory Destras, Gwendolyne Burfin, Hadrien Regue, Quentin Semanas, Martine Valette, Bruno Lina, Laurence Josset |
| EPI_ISL_3379953 | Lighthouse Lab in Glasgow | Wellcome Sanger Institute for the COVID-19 Genomics UK (COG-UK) Consortium | Harper VanSteenhouse, Yumi Kasai, David Gray, Carol Clugston, Anna Dominiczak and Alex Alderton, Roberto Amato, Jeffrey Barrett, Sonia Goncalves, Ewan Harrison, David K. Jackson, Ian Johnston, Dominic Kwiatkowski, Cordelia Langford, John Sillitoe on behalf of the Wellcome Sanger Institute COVID-19 Surveillance Team |
| EPI_ISL_3379954 | Lighthouse Lab in Alderley Park | Wellcome Sanger Institute for the COVID-19 Genomics UK (COG-UK) Consortium | Jacquelyn Wynn, Mairead Hyland, The Lighthouse Lab in Alderley Park and Alex Alderton, Roberto Amato, Jeffrey Barrett, Sonia Goncalves, Ewan Harrison, David K. Jackson, Ian Johnston, Dominic Kwiatkowski, Cordelia Langford, John Sillitoe on behalf of the Wellcome Sanger Institute COVID-19 Surveillance Team |
| EPI_ISL_3379955 | Lighthouse Lab in Glasgow | Wellcome Sanger Institute for the COVID-19 Genomics UK (COG-UK) Consortium | Harper VanSteenhouse, Yumi Kasai, David Gray, Carol Clugston, Anna Dominiczak and Alex Alderton, Roberto Amato, Jeffrey Barrett, Sonia Goncalves, Ewan Harrison, David K. Jackson, Ian Johnston, Dominic Kwiatkowski, Cordelia Langford, John Sillitoe on behalf of the Wellcome Sanger Institute COVID-19 Surveillance Team |
| EPI_ISL_3379956 | LAM DE BIO HD | CNR Virus des Infections Respiratoires - France SUD | Antonin Bal, Gregory Destras, Gwendolyne Burfin, Hadrien Regue, Quentin Semanas, Martine Valette, Bruno Lina, Laurence Josset |
| EPI_ISL_3379957 | Lighthouse Lab in Alderley Park | Wellcome Sanger Institute for the COVID-19 Genomics UK (COG-UK) Consortium | Jacquelyn Wynn, Mairead Hyland, The Lighthouse Lab in Alderley Park and Alex Alderton, Roberto Amato, Jeffrey Barrett, Sonia Goncalves, Ewan Harrison, David K. Jackson, Ian Johnston, Dominic Kwiatkowski, Cordelia Langford, John Sillitoe on behalf of the Wellcome Sanger Institute COVID-19 Surveillance Team |
| EPI_ISL_3379958 | Lighthouse Lab in Milton Keynes | Wellcome Sanger Institute for the COVID-19 Genomics UK (COG-UK) Consortium | The Lighthouse Lab in Milton Keynes and Alex Alderton, Roberto Amato, Jeffrey Barrett, Sonia Goncalves, Ewan Harrison, David K. Jackson, Ian Johnston, Dominic Kwiatkowski, Cordelia Langford, John Sillitoe on behalf of the Wellcome Sanger Institute COVID-19 Surveillance Team |
| EPI_ISL_3379959 | Lighthouse Lab in Alderley Park | Wellcome Sanger Institute for the COVID-19 Genomics UK (COG-UK) Consortium | Jacquelyn Wynn, Mairead Hyland, The Lighthouse Lab in Alderley Park and Alex Alderton, Roberto Amato, Jeffrey Barrett, Sonia Goncalves, Ewan Harrison, David K. Jackson, Ian Johnston, Dominic Kwiatkowski, Cordelia Langford, John Sillitoe on behalf of the Wellcome Sanger Institute COVID-19 Surveillance Team |
| EPI_ISL_3379960 | LAM DE BIO HD | CNR Virus des Infections Respiratoires - France SUD | Antonin Bal, Gregory Destras, Gwendolyne Burfin, Hadrien Regue, Quentin Semanas, Martine Valette, Bruno Lina, Laurence Josset |
| EPI_ISL_3379961, EPI_ISL_3379962, EPI_ISL_3379963 | Lighthouse Lab in Glasgow | Wellcome Sanger Institute for the COVID-19 Genomics UK (COG-UK) Consortium | Harper VanSteenhouse, Yumi Kasai, David Gray, Carol Clugston, Anna Dominiczak and Alex Alderton, Roberto Amato, Jeffrey Barrett, Sonia Goncalves, Ewan Harrison, David K. Jackson, Ian Johnston, Dominic Kwiatkowski, Cordelia Langford, John Sillitoe on behalf of the Wellcome Sanger Institute COVID-19 Surveillance Team |
| EPI_ISL_3379964 | BIOLITTORAL BIOGROUP PLATEAU TECH | CNR Virus des Infections Respiratoires - France SUD | Antonin Bal, Gregory Destras, Gwendolyne Burfin, Hadrien Regue, Quentin Semanas, Martine Valette, Bruno Lina, Laurence Josset |
| EPI_ISL_3379965 | Lighthouse Lab in Glasgow | Wellcome Sanger Institute for the COVID-19 Genomics UK (COG-UK) Consortium | Harper VanSteenhouse, Yumi Kasai, David Gray, Carol Clugston, Anna Dominiczak and Alex Alderton, Roberto Amato, Jeffrey Barrett, Sonia Goncalves, Ewan Harrison, David K. Jackson, Ian Johnston, Dominic Kwiatkowski, Cordelia Langford, John Sillitoe on behalf of the Wellcome Sanger Institute COVID-19 Surveillance Team |
| EPI_ISL_3379966 | Lighthouse Lab in Alderley Park | Wellcome Sanger Institute for the COVID-19 Genomics UK (COG-UK) Consortium | Jacquelyn Wynn, Mairead Hyland, The Lighthouse Lab in Alderley Park and Alex Alderton, Roberto Amato, Jeffrey Barrett, Sonia Goncalves, Ewan Harrison, David K. Jackson, Ian Johnston, Dominic Kwiatkowski, Cordelia Langford, John Sillitoe on behalf of the Wellcome Sanger Institute COVID-19 Surveillance Team |
| EPI_ISL_3379967 | Lighthouse Lab in Glasgow | Wellcome Sanger Institute for the COVID-19 Genomics UK (COG-UK) Consortium | Harper VanSteenhouse, Yumi Kasai, David Gray, Carol Clugston, Anna Dominiczak and Alex Alderton, Roberto Amato, Jeffrey Barrett, Sonia Goncalves, Ewan Harrison, David K. Jackson, Ian Johnston, Dominic Kwiatkowski, Cordelia Langford, John Sillitoe on behalf of the Wellcome Sanger Institute COVID-19 Surveillance Team |
| EPI_ISL_3379968 | BIOLITTORAL BIOGROUP PLATEAU TECH | CNR Virus des Infections Respiratoires - France SUD | Antonin Bal, Gregory Destras, Gwendolyne Burfin, Hadrien Regue, Quentin Semanas, Martine Valette, Bruno Lina, Laurence Josset |
| EPI_ISL_3379969 | Lighthouse Lab in Milton Keynes | Wellcome Sanger Institute for the COVID-19 Genomics UK (COG-UK) Consortium | The Lighthouse Lab in Milton Keynes and Alex Alderton, Roberto Amato, Jeffrey Barrett, Sonia Goncalves, Ewan Harrison, David K. Jackson, Ian Johnston, Dominic Kwiatkowski, Cordelia Langford, John Sillitoe on behalf of the Wellcome Sanger Institute COVID-19 Surveillance Team |
| EPI_ISL_3379970 | Lighthouse Lab in Glasgow | Wellcome Sanger Institute for the COVID-19 Genomics UK (COG-UK) Consortium | Harper VanSteenhouse, Yumi Kasai, David Gray, Carol Clugston, Anna Dominiczak and Alex Alderton, Roberto Amato, Jeffrey Barrett, Sonia Goncalves, Ewan Harrison, David K. Jackson, Ian Johnston, Dominic Kwiatkowski, Cordelia Langford, John Sillitoe on behalf of the Wellcome Sanger Institute COVID-19 Surveillance Team |
| EPI_ISL_3379971 | Lighthouse Lab in Milton Keynes | Wellcome Sanger Institute for the COVID-19 Genomics UK (COG-UK) Consortium | The Lighthouse Lab in Milton Keynes and Alex Alderton, Roberto Amato, Jeffrey Barrett, Sonia Goncalves, Ewan Harrison, David K. Jackson, Ian Johnston, Dominic Kwiatkowski, Cordelia Langford, John Sillitoe on behalf of the Wellcome Sanger Institute COVID-19 Surveillance Team |
| EPI_ISL_3379972 | BIOLITTORAL BIOGROUP PLATEAU TECH | CNR Virus des Infections Respiratoires - France SUD | Antonin Bal, Gregory Destras, Gwendolyne Burfin, Hadrien Regue, Quentin Semanas, Martine Valette, Bruno Lina, Laurence Josset |
| EPI_ISL_3379973 | Lighthouse Lab in Alderley Park | Wellcome Sanger Institute for the COVID-19 Genomics UK (COG-UK) Consortium | Jacquelyn Wynn, Mairead Hyland, The Lighthouse Lab in Alderley Park and Alex Alderton, Roberto Amato, Jeffrey Barrett, Sonia Goncalves, Ewan Harrison, David K. Jackson, Ian Johnston, Dominic Kwiatkowski, Cordelia Langford, John Sillitoe on behalf of the Wellcome Sanger Institute COVID-19 Surveillance Team |
| EPI_ISL_3379974 | Lighthouse Lab in Milton Keynes | Wellcome Sanger Institute for the COVID-19 Genomics UK (COG-UK) Consortium | The Lighthouse Lab in Milton Keynes and Alex Alderton, Roberto Amato, Jeffrey Barrett, Sonia Goncalves, Ewan Harrison, David K. Jackson, Ian Johnston, Dominic Kwiatkowski, Cordelia Langford, John Sillitoe on behalf of the Wellcome Sanger Institute COVID-19 Surveillance Team |
| EPI_ISL_3379975 | Lighthouse Lab in Alderley Park | Wellcome Sanger Institute for the COVID-19 Genomics UK (COG-UK) Consortium | Jacquelyn Wynn, Mairead Hyland, The Lighthouse Lab in Alderley Park and Alex Alderton, Roberto Amato, Jeffrey Barrett, Sonia Goncalves, Ewan Harrison, David K. Jackson, Ian Johnston, Dominic Kwiatkowski, Cordelia Langford, John Sillitoe on behalf of the Wellcome Sanger Institute COVID-19 Surveillance Team |
| EPI_ISL_3379976 | BIOLITTORAL BIOGROUP PLATEAU TECH | CNR Virus des Infections Respiratoires - France SUD | Antonin Bal, Gregory Destras, Gwendolyne Burfin, Hadrien Regue, Quentin Semanas, Martine Valette, Bruno Lina, Laurence Josset |
| EPI_ISL_3379977 | Lighthouse Lab in Alderley Park | Wellcome Sanger Institute for the COVID-19 Genomics UK (COG-UK) Consortium | Jacquelyn Wynn, Mairead Hyland, The Lighthouse Lab in Alderley Park and Alex Alderton, Roberto Amato, Jeffrey Barrett, Sonia Goncalves, Ewan Harrison, David K. Jackson, Ian Johnston, Dominic Kwiatkowski, Cordelia Langford, John Sillitoe on behalf of the Wellcome Sanger Institute COVID-19 Surveillance Team |
| EPI_ISL_3379978 | Lighthouse Lab in Glasgow | Wellcome Sanger Institute for the COVID-19 Genomics UK (COG-UK) Consortium | Harper VanSteenhouse, Yumi Kasai, David Gray, Carol Clugston, Anna Dominiczak and Alex Alderton, Roberto Amato, Jeffrey Barrett, Sonia Goncalves, Ewan Harrison, David K. Jackson, Ian Johnston, Dominic Kwiatkowski, Cordelia Langford, John Sillitoe on behalf of the Wellcome Sanger Institute COVID-19 Surveillance Team |
| EPI_ISL_3379979 | LAM DE BIO HD | CNR Virus des Infections Respiratoires - France SUD | Antonin Bal, Gregory Destras, Gwendolyne Burfin, Hadrien Regue, Quentin Semanas, Martine Valette, Bruno Lina, Laurence Josset |
| EPI_ISL_3379980 | Lighthouse Lab in Alderley Park | Wellcome Sanger Institute for the COVID-19 Genomics UK (COG-UK) Consortium | Jacquelyn Wynn, Mairead Hyland, The Lighthouse Lab in Alderley Park and Alex Alderton, Roberto Amato, Jeffrey Barrett, Sonia Goncalves, Ewan Harrison, David K. Jackson, Ian Johnston, Dominic Kwiatkowski, Cordelia Langford, John Sillitoe on behalf of the Wellcome Sanger Institute COVID-19 Surveillance Team |
| EPI_ISL_3379981 | Lighthouse Lab in Glasgow | Wellcome Sanger Institute for the COVID-19 Genomics UK (COG-UK) Consortium | Harper VanSteenhouse, Yumi Kasai, David Gray, Carol Clugston, Anna Dominiczak and Alex Alderton, Roberto Amato, Jeffrey Barrett, Sonia Goncalves, Ewan Harrison, David K. Jackson, Ian Johnston, Dominic Kwiatkowski, Cordelia Langford, John Sillitoe on behalf of the Wellcome Sanger Institute COVID-19 Surveillance Team |
| EPI_ISL_3379982 | Lighthouse Lab in Alderley Park | Wellcome Sanger Institute for the COVID-19 Genomics UK (COG-UK) Consortium | Jacquelyn Wynn, Mairead Hyland, The Lighthouse Lab in Alderley Park and Alex Alderton, Roberto Amato, Jeffrey Barrett, Sonia Goncalves, Ewan Harrison, David K. Jackson, Ian Johnston, Dominic Kwiatkowski, Cordelia Langford, John Sillitoe on behalf of the Wellcome Sanger Institute COVID-19 Surveillance Team |
| EPI_ISL_3379983 | BIOLITTORAL BIOGROUP PLATEAU TECH | CNR Virus des Infections Respiratoires - France SUD | Antonin Bal, Gregory Destras, Gwendolyne Burfin, Hadrien Regue, Quentin Semanas, Martine Valette, Bruno Lina, Laurence Josset |
| EPI_ISL_3379984 | Lighthouse Lab in Glasgow | Wellcome Sanger Institute for the COVID-19 Genomics UK (COG-UK) Consortium | Harper VanSteenhouse, Yumi Kasai, David Gray, Carol Clugston, Anna Dominiczak and Alex Alderton, Roberto Amato, Jeffrey Barrett, Sonia Goncalves, Ewan Harrison, David K. Jackson, Ian Johnston, Dominic Kwiatkowski, Cordelia Langford, John Sillitoe on behalf of the Wellcome Sanger Institute COVID-19 Surveillance Team |
| EPI_ISL_3379985 | Lighthouse Lab in Milton Keynes | Wellcome Sanger Institute for the COVID-19 Genomics UK (COG-UK) Consortium | The Lighthouse Lab in Milton Keynes and Alex Alderton, Roberto Amato, Jeffrey Barrett, Sonia Goncalves, Ewan Harrison, David K. Jackson, Ian Johnston, Dominic Kwiatkowski, Cordelia Langford, John Sillitoe on behalf of the Wellcome Sanger Institute COVID-19 Surveillance Team |
| EPI_ISL_3379986 | LAM DE BIO HD | CNR Virus des Infections Respiratoires - France SUD | Antonin Bal, Gregory Destras, Gwendolyne Burfin, Hadrien Regue, Quentin Semanas, Martine Valette, Bruno Lina, Laurence Josset |
| EPI_ISL_3379987 | Lighthouse Lab in Glasgow | Wellcome Sanger Institute for the COVID-19 Genomics UK (COG-UK) Consortium | Harper VanSteenhouse, Yumi Kasai, David Gray, Carol Clugston, Anna Dominiczak and Alex Alderton, Roberto Amato, Jeffrey Barrett, Sonia Goncalves, Ewan Harrison, David K. Jackson, Ian Johnston, Dominic Kwiatkowski, Cordelia Langford, John Sillitoe on behalf of the Wellcome Sanger Institute COVID-19 Surveillance Team |
| EPI_ISL_3379988, EPI_ISL_3379989 | Lighthouse Lab in Alderley Park | Wellcome Sanger Institute for the COVID-19 Genomics UK (COG-UK) Consortium | Jacquelyn Wynn, Mairead Hyland, The Lighthouse Lab in Alderley Park and Alex Alderton, Roberto Amato, Jeffrey Barrett, Sonia Goncalves, Ewan Harrison, David K. Jackson, Ian Johnston, Dominic Kwiatkowski, Cordelia Langford, John Sillitoe on behalf of the Wellcome Sanger Institute COVID-19 Surveillance Team |
| EPI_ISL_3379990 | BIOLITTORAL BIOGROUP PLATEAU TECH | CNR Virus des Infections Respiratoires - France SUD | Antonin Bal, Gregory Destras, Gwendolyne Burfin, Hadrien Regue, Quentin Semanas, Martine Valette, Bruno Lina, Laurence Josset |
| EPI_ISL_3379991 | Lighthouse Lab in Glasgow | Wellcome Sanger Institute for the COVID-19 Genomics UK (COG-UK) | Harper VanSteenhouse, Yumi Kasai, David Gray, Carol Clugston, Anna Dominiczak and Alex Alderton, Roberto Amato, Jeffrey Barrett, Sonia Goncalves, Ewan Harrison, David K. Jackson, Ian Johnston, Dominic Kwiatkowski, Cordelia Langford, |

| | | | |
|---|---|---|---|
| | | Consortium | John Sillitoe on behalf of the Wellcome Sanger Institute COVID-19 Surveillance Team |
| EPI_ISL_3379992 | Lighthouse Lab in Alderley Park | Wellcome Sanger Institute for the COVID-19 Genomics UK (COG-UK) Consortium | Jacquelyn Wynn, Mairead Hyland, The Lighthouse Lab in Alderley Park and Alex Alderton, Roberto Amato, Jeffrey Barrett, Sonia Goncalves, Ewan Harrison, David K. Jackson, Ian Johnston, Dominic Kwiatkowski, Cordelia Langford, John Sillitoe on behalf of the Wellcome Sanger Institute COVID-19 Surveillance Team |
| EPI_ISL_3379993 | Lighthouse Lab in Glasgow | Wellcome Sanger Institute for the COVID-19 Genomics UK (COG-UK) Consortium | Harper VanSteenhouse, Yumi Kasai, David Gray, Carol Clugston, Anna Dominiczak and Alex Alderton, Roberto Amato, Jeffrey Barrett, Sonia Goncalves, Ewan Harrison, David K. Jackson, Ian Johnston, Dominic Kwiatkowski, Cordelia Langford, John Sillitoe on behalf of the Wellcome Sanger Institute COVID-19 Surveillance Team |
| EPI_ISL_3379994 | LAM DE BIO HD | CNR Virus des Infections Respiratoires - France SUD | Antonin Bal, Gregory Destras, Gwendolyne Burfin, Hadrien Regue, Quentin Semanas, Martine Valette, Bruno Lina, Laurence Josset |
| EPI_ISL_3379995, EPI_ISL_3379996 | Lighthouse Lab in Glasgow | Wellcome Sanger Institute for the COVID-19 Genomics UK (COG-UK) Consortium | Harper VanSteenhouse, Yumi Kasai, David Gray, Carol Clugston, Anna Dominiczak and Alex Alderton, Roberto Amato, Jeffrey Barrett, Sonia Goncalves, Ewan Harrison, David K. Jackson, Ian Johnston, Dominic Kwiatkowski, Cordelia Langford, John Sillitoe on behalf of the Wellcome Sanger Institute COVID-19 Surveillance Team |
| EPI_ISL_3379997 | Lighthouse Lab in Milton Keynes | Wellcome Sanger Institute for the COVID-19 Genomics UK (COG-UK) Consortium | The Lighthouse Lab in Milton Keynes and Alex Alderton, Roberto Amato, Jeffrey Barrett, Sonia Goncalves, Ewan Harrison, David K. Jackson, Ian Johnston, Dominic Kwiatkowski, Cordelia Langford, John Sillitoe on behalf of the Wellcome Sanger Institute COVID-19 Surveillance Team |
| EPI_ISL_3379998 | BIOLITTORAL BIOGROUP PLATEAU TECH | CNR Virus des Infections Respiratoires - France SUD | Antonin Bal, Gregory Destras, Gwendolyne Burfin, Hadrien Regue, Quentin Semanas, Martine Valette, Bruno Lina, Laurence Josset |
| EPI_ISL_3379999 | Lighthouse Lab in Milton Keynes | Wellcome Sanger Institute for the COVID-19 Genomics UK (COG-UK) Consortium | The Lighthouse Lab in Milton Keynes and Alex Alderton, Roberto Amato, Jeffrey Barrett, Sonia Goncalves, Ewan Harrison, David K. Jackson, Ian Johnston, Dominic Kwiatkowski, Cordelia Langford, John Sillitoe on behalf of the Wellcome Sanger Institute COVID-19 Surveillance Team |
| EPI_ISL_3380000 | Lighthouse Lab in Glasgow | Wellcome Sanger Institute for the COVID-19 Genomics UK (COG-UK) Consortium | Harper VanSteenhouse, Yumi Kasai, David Gray, Carol Clugston, Anna Dominiczak and Alex Alderton, Roberto Amato, Jeffrey Barrett, Sonia Goncalves, Ewan Harrison, David K. Jackson, Ian Johnston, Dominic Kwiatkowski, Cordelia Langford, John Sillitoe on behalf of the Wellcome Sanger Institute COVID-19 Surveillance Team |
| EPI_ISL_3380001, EPI_ISL_3380003 | Lighthouse Lab in Milton Keynes | Wellcome Sanger Institute for the COVID-19 Genomics UK (COG-UK) Consortium | The Lighthouse Lab in Milton Keynes and Alex Alderton, Roberto Amato, Jeffrey Barrett, Sonia Goncalves, Ewan Harrison, David K. Jackson, Ian Johnston, Dominic Kwiatkowski, Cordelia Langford, John Sillitoe on behalf of the Wellcome Sanger Institute COVID-19 Surveillance Team |
| EPI_ISL_3380004, EPI_ISL_3380005 | Lighthouse Lab in Glasgow | Wellcome Sanger Institute for the COVID-19 Genomics UK (COG-UK) Consortium | Harper VanSteenhouse, Yumi Kasai, David Gray, Carol Clugston, Anna Dominiczak and Alex Alderton, Roberto Amato, Jeffrey Barrett, Sonia Goncalves, Ewan Harrison, David K. Jackson, Ian Johnston, Dominic Kwiatkowski, Cordelia Langford, John Sillitoe on behalf of the Wellcome Sanger Institute COVID-19 Surveillance Team |
| EPI_ISL_3380006 | BIOLITTORAL BIOGROUP PLATEAU TECH | CNR Virus des Infections Respiratoires - France SUD | Antonin Bal, Gregory Destras, Gwendolyne Burfin, Hadrien Regue, Quentin Semanas, Martine Valette, Bruno Lina, Laurence Josset |
| EPI_ISL_3380007 | Lighthouse Lab in Alderley Park | Wellcome Sanger Institute for the COVID-19 Genomics UK (COG-UK) Consortium | Jacquelyn Wynn, Mairead Hyland, The Lighthouse Lab in Alderley Park and Alex Alderton, Roberto Amato, Jeffrey Barrett, Sonia Goncalves, Ewan Harrison, David K. Jackson, Ian Johnston, Dominic Kwiatkowski, Cordelia Langford, John Sillitoe on behalf of the Wellcome Sanger Institute COVID-19 Surveillance Team |
| EPI_ISL_3380008 | Lighthouse Lab in Glasgow | Wellcome Sanger Institute for the COVID-19 Genomics UK (COG-UK) Consortium | Harper VanSteenhouse, Yumi Kasai, David Gray, Carol Clugston, Anna Dominiczak and Alex Alderton, Roberto Amato, Jeffrey Barrett, Sonia Goncalves, Ewan Harrison, David K. Jackson, Ian Johnston, Dominic Kwiatkowski, Cordelia Langford, John Sillitoe on behalf of the Wellcome Sanger Institute COVID-19 Surveillance Team |
| EPI_ISL_3380009 | LAM DE TERRE ROUGE | CNR Virus des Infections Respiratoires - France SUD | Antonin Bal, Gregory Destras, Gwendolyne Burfin, Hadrien Regue, Quentin Semanas, Martine Valette, Bruno Lina, Laurence Josset |
| EPI_ISL_3380010, EPI_ISL_3380011 | Lighthouse Lab in Milton Keynes | Wellcome Sanger Institute for the COVID-19 Genomics UK (COG-UK) Consortium | The Lighthouse Lab in Milton Keynes and Alex Alderton, Roberto Amato, Jeffrey Barrett, Sonia Goncalves, Ewan Harrison, David K. Jackson, Ian Johnston, Dominic Kwiatkowski, Cordelia Langford, John Sillitoe on behalf of the Wellcome Sanger Institute COVID-19 Surveillance Team |
| EPI_ISL_3380012 | Lighthouse Lab in Glasgow | Wellcome Sanger Institute for the COVID-19 Genomics UK (COG-UK) Consortium | Harper VanSteenhouse, Yumi Kasai, David Gray, Carol Clugston, Anna Dominiczak and Alex Alderton, Roberto Amato, Jeffrey Barrett, Sonia Goncalves, Ewan Harrison, David K. Jackson, Ian Johnston, Dominic Kwiatkowski, Cordelia Langford, John Sillitoe on behalf of the Wellcome Sanger Institute COVID-19 Surveillance Team |
| EPI_ISL_3380013 | Lighthouse Lab in Alderley Park | Wellcome Sanger Institute for the COVID-19 Genomics UK (COG-UK) Consortium | Jacquelyn Wynn, Mairead Hyland, The Lighthouse Lab in Alderley Park and Alex Alderton, Roberto Amato, Jeffrey Barrett, Sonia Goncalves, Ewan Harrison, David K. Jackson, Ian Johnston, Dominic Kwiatkowski, Cordelia Langford, John Sillitoe on behalf of the Wellcome Sanger Institute COVID-19 Surveillance Team |
| EPI_ISL_3380014 | LAM DE TERRE ROUGE | CNR Virus des Infections Respiratoires - France SUD | Antonin Bal, Gregory Destras, Gwendolyne Burfin, Hadrien Regue, Quentin Semanas, Martine Valette, Bruno Lina, Laurence Josset |
| EPI_ISL_3380015 | Lighthouse Lab in Glasgow | Wellcome Sanger Institute for the COVID-19 Genomics UK (COG-UK) Consortium | Harper VanSteenhouse, Yumi Kasai, David Gray, Carol Clugston, Anna Dominiczak and Alex Alderton, Roberto Amato, Jeffrey Barrett, Sonia Goncalves, Ewan Harrison, David K. Jackson, Ian Johnston, Dominic Kwiatkowski, Cordelia Langford, John Sillitoe on behalf of the Wellcome Sanger Institute COVID-19 Surveillance Team |
| EPI_ISL_3380016 | Lighthouse Lab in Alderley Park | Wellcome Sanger Institute for the COVID-19 Genomics UK (COG-UK) Consortium | Jacquelyn Wynn, Mairead Hyland, The Lighthouse Lab in Alderley Park and Alex Alderton, Roberto Amato, Jeffrey Barrett, Sonia Goncalves, Ewan Harrison, David K. Jackson, Ian Johnston, Dominic Kwiatkowski, Cordelia Langford, John Sillitoe on behalf of the Wellcome Sanger Institute COVID-19 Surveillance Team |
| EPI_ISL_3380017 | LAM DE TERRE ROUGE | CNR Virus des Infections Respiratoires - France SUD | Antonin Bal, Gregory Destras, Gwendolyne Burfin, Hadrien Regue, Quentin Semanas, Martine Valette, Bruno Lina, Laurence Josset |
| EPI_ISL_3380018, EPI_ISL_3380019, EPI_ISL_3380020 | Lighthouse Lab in Glasgow | Wellcome Sanger Institute for the COVID-19 Genomics UK (COG-UK) Consortium | Harper VanSteenhouse, Yumi Kasai, David Gray, Carol Clugston, Anna Dominiczak and Alex Alderton, Roberto Amato, Jeffrey Barrett, Sonia Goncalves, Ewan Harrison, David K. Jackson, Ian Johnston, Dominic Kwiatkowski, Cordelia Langford, John Sillitoe on behalf of the Wellcome Sanger Institute COVID-19 Surveillance Team |
| EPI_ISL_3380021 | LAM DE TERRE ROUGE | CNR Virus des Infections Respiratoires - France SUD | Antonin Bal, Gregory Destras, Gwendolyne Burfin, Hadrien Regue, Quentin Semanas, Martine Valette, Bruno Lina, Laurence Josset |
| EPI_ISL_3380022, EPI_ISL_3380023, EPI_ISL_3380024 | Lighthouse Lab in Glasgow | Wellcome Sanger Institute for the COVID-19 Genomics UK (COG-UK) Consortium | Harper VanSteenhouse, Yumi Kasai, David Gray, Carol Clugston, Anna Dominiczak and Alex Alderton, Roberto Amato, Jeffrey Barrett, Sonia Goncalves, Ewan Harrison, David K. Jackson, Ian Johnston, Dominic Kwiatkowski, Cordelia Langford, John Sillitoe on behalf of the Wellcome Sanger Institute COVID-19 Surveillance Team |
| EPI_ISL_3380025 | LAM DE TERRE ROUGE | CNR Virus des Infections Respiratoires - France SUD | Antonin Bal, Gregory Destras, Gwendolyne Burfin, Hadrien Regue, Quentin Semanas, Martine Valette, Bruno Lina, Laurence Josset |
| EPI_ISL_3380026 | Lighthouse Lab in Glasgow | Wellcome Sanger Institute for the COVID-19 Genomics UK (COG-UK) Consortium | Harper VanSteenhouse, Yumi Kasai, David Gray, Carol Clugston, Anna Dominiczak and Alex Alderton, Roberto Amato, Jeffrey Barrett, Sonia Goncalves, Ewan Harrison, David K. Jackson, Ian Johnston, Dominic Kwiatkowski, Cordelia Langford, John Sillitoe on behalf of the Wellcome Sanger Institute COVID-19 Surveillance Team |
| EPI_ISL_3380027 | Lighthouse Lab in Alderley Park | Wellcome Sanger Institute for the COVID-19 Genomics UK (COG-UK) Consortium | Jacquelyn Wynn, Mairead Hyland, The Lighthouse Lab in Alderley Park and Alex Alderton, Roberto Amato, Jeffrey Barrett, Sonia Goncalves, Ewan Harrison, David K. Jackson, Ian Johnston, Dominic Kwiatkowski, Cordelia Langford, John Sillitoe on behalf of the Wellcome Sanger Institute COVID-19 Surveillance Team |
| EPI_ISL_3380028 | Lighthouse Lab in Glasgow | Wellcome Sanger Institute for the COVID-19 Genomics UK (COG-UK) Consortium | Harper VanSteenhouse, Yumi Kasai, David Gray, Carol Clugston, Anna Dominiczak and Alex Alderton, Roberto Amato, Jeffrey Barrett, Sonia Goncalves, Ewan Harrison, David K. Jackson, Ian Johnston, Dominic Kwiatkowski, Cordelia Langford, John Sillitoe on behalf of the Wellcome Sanger Institute COVID-19 Surveillance Team |
| EPI_ISL_3380029 | LAM DE TERRE ROUGE | CNR Virus des Infections Respiratoires - France SUD | Antonin Bal, Gregory Destras, Gwendolyne Burfin, Hadrien Regue, Quentin Semanas, Martine Valette, Bruno Lina, Laurence Josset |
| EPI_ISL_3380030 | Lighthouse Lab in Glasgow | Wellcome Sanger Institute for the COVID-19 Genomics UK (COG-UK) Consortium | Harper VanSteenhouse, Yumi Kasai, David Gray, Carol Clugston, Anna Dominiczak and Alex Alderton, Roberto Amato, Jeffrey Barrett, Sonia Goncalves, Ewan Harrison, David K. Jackson, Ian Johnston, Dominic Kwiatkowski, Cordelia Langford, John Sillitoe on behalf of the Wellcome Sanger Institute COVID-19 Surveillance Team |
| EPI_ISL_3380031 | Lighthouse Lab in Milton Keynes | Wellcome Sanger Institute for the COVID-19 Genomics UK (COG-UK) Consortium | The Lighthouse Lab in Milton Keynes and Alex Alderton, Roberto Amato, Jeffrey Barrett, Sonia Goncalves, Ewan Harrison, David K. Jackson, Ian Johnston, Dominic Kwiatkowski, Cordelia Langford, John Sillitoe on behalf of the Wellcome Sanger Institute COVID-19 Surveillance Team |
| EPI_ISL_3380032 | LAM DE TERRE ROUGE | CNR Virus des Infections Respiratoires - France SUD | Antonin Bal, Gregory Destras, Gwendolyne Burfin, Hadrien Regue, Quentin Semanas, Martine Valette, Bruno Lina, Laurence Josset |
| EPI_ISL_3380033, EPI_ISL_3380034 | Lighthouse Lab in Glasgow | Wellcome Sanger Institute for the COVID-19 Genomics UK (COG-UK) Consortium | Harper VanSteenhouse, Yumi Kasai, David Gray, Carol Clugston, Anna Dominiczak and Alex Alderton, Roberto Amato, Jeffrey Barrett, Sonia Goncalves, Ewan Harrison, David K. Jackson, Ian Johnston, Dominic Kwiatkowski, Cordelia Langford, John Sillitoe on behalf of the Wellcome Sanger Institute COVID-19 Surveillance Team |
| EPI_ISL_3380035 | Lighthouse Lab in Alderley Park | Wellcome Sanger Institute for the COVID-19 Genomics UK (COG-UK) Consortium | Jacquelyn Wynn, Mairead Hyland, The Lighthouse Lab in Alderley Park and Alex Alderton, Roberto Amato, Jeffrey Barrett, Sonia Goncalves, Ewan Harrison, David K. Jackson, Ian Johnston, Dominic Kwiatkowski, Cordelia Langford, John Sillitoe on behalf of the Wellcome Sanger Institute COVID-19 Surveillance Team |
| EPI_ISL_3380036 | LAM DE TERRE ROUGE | CNR Virus des Infections Respiratoires - France SUD | Antonin Bal, Gregory Destras, Gwendolyne Burfin, Hadrien Regue, Quentin Semanas, Martine Valette, Bruno Lina, Laurence Josset |
| EPI_ISL_3380037 | Lighthouse Lab in Glasgow | Wellcome Sanger Institute for the COVID-19 Genomics UK (COG-UK) Consortium | Harper VanSteenhouse, Yumi Kasai, David Gray, Carol Clugston, Anna Dominiczak and Alex Alderton, Roberto Amato, Jeffrey Barrett, Sonia Goncalves, Ewan Harrison, David K. Jackson, Ian Johnston, Dominic Kwiatkowski, Cordelia Langford, John Sillitoe on behalf of the Wellcome Sanger Institute COVID-19 Surveillance Team |
| EPI_ISL_3380038 | Lighthouse Lab in Milton Keynes | Wellcome Sanger Institute for the COVID-19 Genomics UK (COG-UK) Consortium | The Lighthouse Lab in Milton Keynes and Alex Alderton, Roberto Amato, Jeffrey Barrett, Sonia Goncalves, Ewan Harrison, David K. Jackson, Ian Johnston, Dominic Kwiatkowski, Cordelia Langford, John Sillitoe on behalf of the Wellcome Sanger Institute COVID-19 Surveillance Team |
| EPI_ISL_3380039 | LABORATOIRE UNILIANS DECINES | CNR Virus des Infections Respiratoires - France SUD | Antonin Bal, Gregory Destras, Gwendolyne Burfin, Hadrien Regue, Quentin Semanas, Martine Valette, Bruno Lina, Laurence Josset |
| EPI_ISL_3380040 | Lighthouse Lab in Alderley Park | Wellcome Sanger Institute for the COVID-19 Genomics UK (COG-UK) Consortium | Jacquelyn Wynn, Mairead Hyland, The Lighthouse Lab in Alderley Park and Alex Alderton, Roberto Amato, Jeffrey Barrett, Sonia Goncalves, Ewan Harrison, David K. Jackson, Ian Johnston, Dominic Kwiatkowski, Cordelia Langford, John Sillitoe on behalf of the Wellcome Sanger Institute COVID-19 Surveillance Team |
| EPI_ISL_3380041 | Lighthouse Lab in Glasgow | Wellcome Sanger Institute for the COVID-19 Genomics UK (COG-UK) Consortium | Harper VanSteenhouse, Yumi Kasai, David Gray, Carol Clugston, Anna Dominiczak and Alex Alderton, Roberto Amato, Jeffrey Barrett, Sonia Goncalves, Ewan Harrison, David K. Jackson, Ian Johnston, Dominic Kwiatkowski, Cordelia Langford, John Sillitoe on behalf of the Wellcome Sanger Institute COVID-19 Surveillance Team |
| EPI_ISL_3380042 | Lighthouse Lab in Alderley Park | Wellcome Sanger Institute for the COVID-19 Genomics UK (COG-UK) Consortium | Jacquelyn Wynn, Mairead Hyland, The Lighthouse Lab in Alderley Park and Alex Alderton, Roberto Amato, Jeffrey Barrett, Sonia Goncalves, Ewan Harrison, David K. Jackson, Ian Johnston, Dominic Kwiatkowski, Cordelia Langford, John Sillitoe on behalf of the Wellcome Sanger Institute COVID-19 Surveillance Team |
| EPI_ISL_3380043 | LABORATOIRE UNILIANS DECINES | CNR Virus des Infections Respiratoires - France SUD | Antonin Bal, Gregory Destras, Gwendolyne Burfin, Hadrien Regue, Quentin Semanas, Martine Valette, Bruno Lina, Laurence Josset |
| EPI_ISL_3380044 | Lighthouse Lab in Milton Keynes | Wellcome Sanger Institute for the COVID-19 Genomics UK (COG-UK) Consortium | The Lighthouse Lab in Milton Keynes and Alex Alderton, Roberto Amato, Jeffrey Barrett, Sonia Goncalves, Ewan Harrison, David K. Jackson, Ian Johnston, Dominic Kwiatkowski, Cordelia Langford, John Sillitoe on behalf of the Wellcome Sanger Institute COVID-19 Surveillance Team |
| EPI_ISL_3380045, EPI_ISL_3380046 | Lighthouse Lab in Glasgow | Wellcome Sanger Institute for the COVID-19 Genomics UK (COG-UK) Consortium | Harper VanSteenhouse, Yumi Kasai, David Gray, Carol Clugston, Anna Dominiczak and Alex Alderton, Roberto Amato, Jeffrey Barrett, Sonia Goncalves, Ewan Harrison, David K. Jackson, Ian Johnston, Dominic Kwiatkowski, Cordelia Langford, John Sillitoe on behalf of the Wellcome Sanger Institute COVID-19 Surveillance Team |
| EPI_ISL_3380047 | LABORATOIRE UNILIANS DECINES | CNR Virus des Infections Respiratoires - France SUD | Antonin Bal, Gregory Destras, Gwendolyne Burfin, Hadrien Regue, Quentin Semanas, Martine Valette, Bruno Lina, Laurence Josset |
| EPI_ISL_3380048 | Lighthouse Lab in Glasgow | Wellcome Sanger Institute for the COVID-19 Genomics UK (COG-UK) Consortium | Harper VanSteenhouse, Yumi Kasai, David Gray, Carol Clugston, Anna Dominiczak and Alex Alderton, Roberto Amato, Jeffrey Barrett, Sonia Goncalves, Ewan Harrison, David K. Jackson, Ian Johnston, Dominic Kwiatkowski, Cordelia Langford, John Sillitoe on behalf of the Wellcome Sanger Institute COVID-19 Surveillance Team |
| EPI_ISL_3380049 | Lighthouse Lab in Milton Keynes | Wellcome Sanger Institute for the COVID-19 Genomics UK (COG-UK) Consortium | The Lighthouse Lab in Milton Keynes and Alex Alderton, Roberto Amato, Jeffrey Barrett, Sonia Goncalves, Ewan Harrison, David K. Jackson, Ian Johnston, Dominic Kwiatkowski, Cordelia Langford, John Sillitoe on behalf of the Wellcome Sanger Institute COVID-19 Surveillance Team |
| EPI_ISL_3380050 | LABORATOIRE UNILIANS DECINES | CNR Virus des Infections Respiratoires - France SUD | Antonin Bal, Gregory Destras, Gwendolyne Burfin, Hadrien Regue, Quentin Semanas, Martine Valette, Bruno Lina, Laurence Josset |
| EPI_ISL_3380051 | Lighthouse Lab in Milton Keynes | Wellcome Sanger Institute for the COVID-19 Genomics UK (COG-UK) Consortium | The Lighthouse Lab in Milton Keynes and Alex Alderton, Roberto Amato, Jeffrey Barrett, Sonia Goncalves, Ewan Harrison, David K. Jackson, Ian Johnston, Dominic Kwiatkowski, Cordelia Langford, John Sillitoe on behalf of the Wellcome Sanger Institute COVID-19 Surveillance Team |
| EPI_ISL_3380052, EPI_ISL_3380053 | Lighthouse Lab in Glasgow | Wellcome Sanger Institute for the COVID-19 Genomics UK (COG-UK) Consortium | Harper VanSteenhouse, Yumi Kasai, David Gray, Carol Clugston, Anna Dominiczak and Alex Alderton, Roberto Amato, Jeffrey Barrett, Sonia Goncalves, Ewan Harrison, David K. Jackson, Ian Johnston, Dominic Kwiatkowski, Cordelia Langford, John Sillitoe on behalf of the Wellcome Sanger Institute COVID-19 Surveillance Team |

| | | | |
|---|---|---|---|
| EPI_ISL_3380054 | LABORATOIRE UNILIANS DECINES | CNR Virus des Infections Respiratoires - France SUD | Antonin Bal, Gregory Destras, Gwendolyne Burfin, Hadrien Regue, Quentin Semanas, Martine Valette, Bruno Lina, Laurence Josset |
| EPI_ISL_3380055, EPI_ISL_3380056 | Lighthouse Lab in Glasgow | Wellcome Sanger Institute for the COVID-19 Genomics UK (COG-UK) Consortium | Harper VanSteenhouse, Yumi Kasai, David Gray, Carol Clugston, Anna Dominiczak and Alex Alderton, Roberto Amato, Jeffrey Barrett, Sonia Goncalves, Ewan Harrison, David K. Jackson, Ian Johnston, Dominic Kwiatkowski, Cordelia Langford, John Sillitoe on behalf of the Wellcome Sanger Institute COVID-19 Surveillance Team |
| EPI_ISL_3380057 | Lighthouse Lab in Alderley Park | Wellcome Sanger Institute for the COVID-19 Genomics UK (COG-UK) Consortium | Jacquelyn Wynn, Mairead Hyland, The Lighthouse Lab in Alderley Park and Alex Alderton, Roberto Amato, Jeffrey Barrett, Sonia Goncalves, Ewan Harrison, David K. Jackson, Ian Johnston, Dominic Kwiatkowski, Cordelia Langford, John Sillitoe on behalf of the Wellcome Sanger Institute COVID-19 Surveillance Team |
| EPI_ISL_3380058 | LABORATOIRE UNILIANS DECINES | CNR Virus des Infections Respiratoires - France SUD | Antonin Bal, Gregory Destras, Gwendolyne Burfin, Hadrien Regue, Quentin Semanas, Martine Valette, Bruno Lina, Laurence Josset |
| EPI_ISL_3380059 | Lighthouse Lab in Alderley Park | Wellcome Sanger Institute for the COVID-19 Genomics UK (COG-UK) Consortium | Jacquelyn Wynn, Mairead Hyland, The Lighthouse Lab in Alderley Park and Alex Alderton, Roberto Amato, Jeffrey Barrett, Sonia Goncalves, Ewan Harrison, David K. Jackson, Ian Johnston, Dominic Kwiatkowski, Cordelia Langford, John Sillitoe on behalf of the Wellcome Sanger Institute COVID-19 Surveillance Team |
| EPI_ISL_3380060 | Lighthouse Lab in Glasgow | Wellcome Sanger Institute for the COVID-19 Genomics UK (COG-UK) Consortium | Harper VanSteenhouse, Yumi Kasai, David Gray, Carol Clugston, Anna Dominiczak and Alex Alderton, Roberto Amato, Jeffrey Barrett, Sonia Goncalves, Ewan Harrison, David K. Jackson, Ian Johnston, Dominic Kwiatkowski, Cordelia Langford, John Sillitoe on behalf of the Wellcome Sanger Institute COVID-19 Surveillance Team |
| EPI_ISL_3380061, EPI_ISL_3380062, EPI_ISL_3380063, EPI_ISL_3380064, EPI_ISL_3380065, EPI_ISL_3380066, EPI_ISL_3380067, EPI_ISL_3380068, EPI_ISL_3380069, EPI_ISL_3380070, EPI_ISL_3380071, EPI_ISL_3380072, EPI_ISL_3380074, EPI_ISL_3380075, EPI_ISL_3380076, EPI_ISL_3380077, EPI_ISL_3380078, EPI_ISL_3380079, EPI_ISL_3380080, EPI_ISL_3380081, EPI_ISL_3380082, EPI_ISL_3380083, EPI_ISL_3380084, EPI_ISL_3380085, EPI_ISL_3380086, EPI_ISL_3380087, EPI_ISL_3380088, EPI_ISL_3380089, EPI_ISL_3380090, EPI_ISL_3380091, EPI_ISL_3380092, EPI_ISL_3380093, EPI_ISL_3380094, EPI_ISL_3380095, EPI_ISL_3380096, EPI_ISL_3380097, EPI_ISL_3380098, EPI_ISL_3380099, EPI_ISL_3380100 | | | |
| see above | LABORATOIRE UNILIANS DECINES | CNR Virus des Infections Respiratoires - France SUD | Antonin Bal, Gregory Destras, Gwendolyne Burfin, Hadrien Regue, Quentin Semanas, Martine Valette, Bruno Lina, Laurence Josset |
| EPI_ISL_3380101 | Lighthouse Lab in Milton Keynes | Wellcome Sanger Institute for the COVID-19 Genomics UK (COG-UK) Consortium | The Lighthouse Lab in Milton Keynes and Alex Alderton, Roberto Amato, Jeffrey Barrett, Sonia Goncalves, Ewan Harrison, David K. Jackson, Ian Johnston, Dominic Kwiatkowski, Cordelia Langford, John Sillitoe on behalf of the Wellcome Sanger Institute COVID-19 Surveillance Team |
| EPI_ISL_3380102 | LABORATOIRE UNILIANS DECINES | CNR Virus des Infections Respiratoires - France SUD | Antonin Bal, Gregory Destras, Gwendolyne Burfin, Hadrien Regue, Quentin Semanas, Martine Valette, Bruno Lina, Laurence Josset |
| EPI_ISL_3380103, EPI_ISL_3380104, EPI_ISL_3380105 | Lighthouse Lab in Glasgow | Wellcome Sanger Institute for the COVID-19 Genomics UK (COG-UK) Consortium | Harper VanSteenhouse, Yumi Kasai, David Gray, Carol Clugston, Anna Dominiczak and Alex Alderton, Roberto Amato, Jeffrey Barrett, Sonia Goncalves, Ewan Harrison, David K. Jackson, Ian Johnston, Dominic Kwiatkowski, Cordelia Langford, John Sillitoe on behalf of the Wellcome Sanger Institute COVID-19 Surveillance Team |
| EPI_ISL_3380106 | LABORATOIRE BIOMEDICA | CNR Virus des Infections Respiratoires - France SUD | Antonin Bal, Gregory Destras, Gwendolyne Burfin, Hadrien Regue, Quentin Semanas, Martine Valette, Bruno Lina, Laurence Josset |
| EPI_ISL_3380107, EPI_ISL_3380108 | Lighthouse Lab in Alderley Park | Wellcome Sanger Institute for the COVID-19 Genomics UK (COG-UK) Consortium | Jacquelyn Wynn, Mairead Hyland, The Lighthouse Lab in Alderley Park and Alex Alderton, Roberto Amato, Jeffrey Barrett, Sonia Goncalves, Ewan Harrison, David K. Jackson, Ian Johnston, Dominic Kwiatkowski, Cordelia Langford, John Sillitoe on behalf of the Wellcome Sanger Institute COVID-19 Surveillance Team |
| EPI_ISL_3380109 | LABORATOIRE BIOMEDICA | CNR Virus des Infections Respiratoires - France SUD | Antonin Bal, Gregory Destras, Gwendolyne Burfin, Hadrien Regue, Quentin Semanas, Martine Valette, Bruno Lina, Laurence Josset |
| EPI_ISL_3380110 | Lighthouse Lab in Milton Keynes | Wellcome Sanger Institute for the COVID-19 Genomics UK (COG-UK) Consortium | The Lighthouse Lab in Milton Keynes and Alex Alderton, Roberto Amato, Jeffrey Barrett, Sonia Goncalves, Ewan Harrison, David K. Jackson, Ian Johnston, Dominic Kwiatkowski, Cordelia Langford, John Sillitoe on behalf of the Wellcome Sanger Institute COVID-19 Surveillance Team |
| EPI_ISL_3380111 | Lighthouse Lab in Glasgow | Wellcome Sanger Institute for the COVID-19 Genomics UK (COG-UK) Consortium | Harper VanSteenhouse, Yumi Kasai, David Gray, Carol Clugston, Anna Dominiczak and Alex Alderton, Roberto Amato, Jeffrey Barrett, Sonia Goncalves, Ewan Harrison, David K. Jackson, Ian Johnston, Dominic Kwiatkowski, Cordelia Langford, John Sillitoe on behalf of the Wellcome Sanger Institute COVID-19 Surveillance Team |
| EPI_ISL_3380112 | Lighthouse Lab in Milton Keynes | Wellcome Sanger Institute for the COVID-19 Genomics UK (COG-UK) Consortium | The Lighthouse Lab in Milton Keynes and Alex Alderton, Roberto Amato, Jeffrey Barrett, Sonia Goncalves, Ewan Harrison, David K. Jackson, Ian Johnston, Dominic Kwiatkowski, Cordelia Langford, John Sillitoe on behalf of the Wellcome Sanger Institute COVID-19 Surveillance Team |
| EPI_ISL_3380113 | LABORATOIRE BIOMEDICA | CNR Virus des Infections Respiratoires - France SUD | Antonin Bal, Gregory Destras, Gwendolyne Burfin, Hadrien Regue, Quentin Semanas, Martine Valette, Bruno Lina, Laurence Josset |
| EPI_ISL_3380114 | Lighthouse Lab in Glasgow | Wellcome Sanger Institute for the COVID-19 Genomics UK (COG-UK) Consortium | Harper VanSteenhouse, Yumi Kasai, David Gray, Carol Clugston, Anna Dominiczak and Alex Alderton, Roberto Amato, Jeffrey Barrett, Sonia Goncalves, Ewan Harrison, David K. Jackson, Ian Johnston, Dominic Kwiatkowski, Cordelia Langford, John Sillitoe on behalf of the Wellcome Sanger Institute COVID-19 Surveillance Team |
| EPI_ISL_3380115, EPI_ISL_3380116 | Lighthouse Lab in Milton Keynes | Wellcome Sanger Institute for the COVID-19 Genomics UK (COG-UK) Consortium | The Lighthouse Lab in Milton Keynes and Alex Alderton, Roberto Amato, Jeffrey Barrett, Sonia Goncalves, Ewan Harrison, David K. Jackson, Ian Johnston, Dominic Kwiatkowski, Cordelia Langford, John Sillitoe on behalf of the Wellcome Sanger Institute COVID-19 Surveillance Team |
| EPI_ISL_3380117 | LABORATOIRE BIOMEDICA | CNR Virus des Infections Respiratoires - France SUD | Antonin Bal, Gregory Destras, Gwendolyne Burfin, Hadrien Regue, Quentin Semanas, Martine Valette, Bruno Lina, Laurence Josset |
| EPI_ISL_3380118 | Lighthouse Lab in Glasgow | Wellcome Sanger Institute for the COVID-19 Genomics UK (COG-UK) Consortium | Harper VanSteenhouse, Yumi Kasai, David Gray, Carol Clugston, Anna Dominiczak and Alex Alderton, Roberto Amato, Jeffrey Barrett, Sonia Goncalves, Ewan Harrison, David K. Jackson, Ian Johnston, Dominic Kwiatkowski, Cordelia Langford, John Sillitoe on behalf of the Wellcome Sanger Institute COVID-19 Surveillance Team |
| EPI_ISL_3380119 | Lighthouse Lab in Milton Keynes | Wellcome Sanger Institute for the COVID-19 Genomics UK (COG-UK) Consortium | The Lighthouse Lab in Milton Keynes and Alex Alderton, Roberto Amato, Jeffrey Barrett, Sonia Goncalves, Ewan Harrison, David K. Jackson, Ian Johnston, Dominic Kwiatkowski, Cordelia Langford, John Sillitoe on behalf of the Wellcome Sanger Institute COVID-19 Surveillance Team |
| EPI_ISL_3380120 | LABORATOIRE BIOMEDICA | CNR Virus des Infections Respiratoires - France SUD | Antonin Bal, Gregory Destras, Gwendolyne Burfin, Hadrien Regue, Quentin Semanas, Martine Valette, Bruno Lina, Laurence Josset |
| EPI_ISL_3380121 | Lighthouse Lab in Milton Keynes | Wellcome Sanger Institute for the COVID-19 Genomics UK (COG-UK) Consortium | The Lighthouse Lab in Milton Keynes and Alex Alderton, Roberto Amato, Jeffrey Barrett, Sonia Goncalves, Ewan Harrison, David K. Jackson, Ian Johnston, Dominic Kwiatkowski, Cordelia Langford, John Sillitoe on behalf of the Wellcome Sanger Institute COVID-19 Surveillance Team |
| EPI_ISL_3380122 | Lighthouse Lab in Glasgow | Wellcome Sanger Institute for the COVID-19 Genomics UK (COG-UK) Consortium | Harper VanSteenhouse, Yumi Kasai, David Gray, Carol Clugston, Anna Dominiczak and Alex Alderton, Roberto Amato, Jeffrey Barrett, Sonia Goncalves, Ewan Harrison, David K. Jackson, Ian Johnston, Dominic Kwiatkowski, Cordelia Langford, John Sillitoe on behalf of the Wellcome Sanger Institute COVID-19 Surveillance Team |
| EPI_ISL_3380123 | LABORATOIRE BIOMEDICA | CNR Virus des Infections Respiratoires - France SUD | Antonin Bal, Gregory Destras, Gwendolyne Burfin, Hadrien Regue, Quentin Semanas, Martine Valette, Bruno Lina, Laurence Josset |
| EPI_ISL_3380124, EPI_ISL_3380125, EPI_ISL_3380126 | Lighthouse Lab in Glasgow | Wellcome Sanger Institute for the COVID-19 Genomics UK (COG-UK) Consortium | Harper VanSteenhouse, Yumi Kasai, David Gray, Carol Clugston, Anna Dominiczak and Alex Alderton, Roberto Amato, Jeffrey Barrett, Sonia Goncalves, Ewan Harrison, David K. Jackson, Ian Johnston, Dominic Kwiatkowski, Cordelia Langford, John Sillitoe on behalf of the Wellcome Sanger Institute COVID-19 Surveillance Team |
| EPI_ISL_3380127 | LABORATOIRE BIOMEDICA | CNR Virus des Infections Respiratoires - France SUD | Antonin Bal, Gregory Destras, Gwendolyne Burfin, Hadrien Regue, Quentin Semanas, Martine Valette, Bruno Lina, Laurence Josset |
| EPI_ISL_3380128 | Lighthouse Lab in Glasgow | Wellcome Sanger Institute for the COVID-19 Genomics UK (COG-UK) Consortium | Harper VanSteenhouse, Yumi Kasai, David Gray, Carol Clugston, Anna Dominiczak and Alex Alderton, Roberto Amato, Jeffrey Barrett, Sonia Goncalves, Ewan Harrison, David K. Jackson, Ian Johnston, Dominic Kwiatkowski, Cordelia Langford, John Sillitoe on behalf of the Wellcome Sanger Institute COVID-19 Surveillance Team |
| EPI_ISL_3380129 | Lighthouse Lab in Alderley Park | Wellcome Sanger Institute for the COVID-19 Genomics UK (COG-UK) Consortium | Jacquelyn Wynn, Mairead Hyland, The Lighthouse Lab in Alderley Park and Alex Alderton, Roberto Amato, Jeffrey Barrett, Sonia Goncalves, Ewan Harrison, David K. Jackson, Ian Johnston, Dominic Kwiatkowski, Cordelia Langford, John Sillitoe on behalf of the Wellcome Sanger Institute COVID-19 Surveillance Team |
| EPI_ISL_3380130 | LABORATOIRE BIOMEDICA | CNR Virus des Infections Respiratoires - France SUD | Antonin Bal, Gregory Destras, Gwendolyne Burfin, Hadrien Regue, Quentin Semanas, Martine Valette, Bruno Lina, Laurence Josset |
| EPI_ISL_3380131 | Lighthouse Lab in Alderley Park | Wellcome Sanger Institute for the COVID-19 Genomics UK (COG-UK) Consortium | Jacquelyn Wynn, Mairead Hyland, The Lighthouse Lab in Alderley Park and Alex Alderton, Roberto Amato, Jeffrey Barrett, Sonia Goncalves, Ewan Harrison, David K. Jackson, Ian Johnston, Dominic Kwiatkowski, Cordelia Langford, John Sillitoe on behalf of the Wellcome Sanger Institute COVID-19 Surveillance Team |
| EPI_ISL_3380132, EPI_ISL_3380133 | Lighthouse Lab in Glasgow | Wellcome Sanger Institute for the COVID-19 Genomics UK (COG-UK) Consortium | Harper VanSteenhouse, Yumi Kasai, David Gray, Carol Clugston, Anna Dominiczak and Alex Alderton, Roberto Amato, Jeffrey Barrett, Sonia Goncalves, Ewan Harrison, David K. Jackson, Ian Johnston, Dominic Kwiatkowski, Cordelia Langford, John Sillitoe on behalf of the Wellcome Sanger Institute COVID-19 Surveillance Team |
| EPI_ISL_3380135, EPI_ISL_3380136 | Lighthouse Lab in Alderley Park | Wellcome Sanger Institute for the COVID-19 Genomics UK (COG-UK) Consortium | Jacquelyn Wynn, Mairead Hyland, The Lighthouse Lab in Alderley Park and Alex Alderton, Roberto Amato, Jeffrey Barrett, Sonia Goncalves, Ewan Harrison, David K. Jackson, Ian Johnston, Dominic Kwiatkowski, Cordelia Langford, John Sillitoe on behalf of the Wellcome Sanger Institute COVID-19 Surveillance Team |
| EPI_ISL_3380137 | Lighthouse Lab in Glasgow | Wellcome Sanger Institute for the COVID-19 Genomics UK (COG-UK) Consortium | Harper VanSteenhouse, Yumi Kasai, David Gray, Carol Clugston, Anna Dominiczak and Alex Alderton, Roberto Amato, Jeffrey Barrett, Sonia Goncalves, Ewan Harrison, David K. Jackson, Ian Johnston, Dominic Kwiatkowski, Cordelia Langford, John Sillitoe on behalf of the Wellcome Sanger Institute COVID-19 Surveillance Team |
| EPI_ISL_3380138 | LABORATOIRE BIOMEDICA | CNR Virus des Infections Respiratoires - France SUD | Antonin Bal, Gregory Destras, Gwendolyne Burfin, Hadrien Regue, Quentin Semanas, Martine Valette, Bruno Lina, Laurence Josset |
| EPI_ISL_3380139 | Lighthouse Lab in Alderley Park | Wellcome Sanger Institute for the COVID-19 Genomics UK (COG-UK) Consortium | Jacquelyn Wynn, Mairead Hyland, The Lighthouse Lab in Alderley Park and Alex Alderton, Roberto Amato, Jeffrey Barrett, Sonia Goncalves, Ewan Harrison, David K. Jackson, Ian Johnston, Dominic Kwiatkowski, Cordelia Langford, John Sillitoe on behalf of the Wellcome Sanger Institute COVID-19 Surveillance Team |
| EPI_ISL_3380140 | Lighthouse Lab in Glasgow | Wellcome Sanger Institute for the COVID-19 Genomics UK (COG-UK) Consortium | Harper VanSteenhouse, Yumi Kasai, David Gray, Carol Clugston, Anna Dominiczak and Alex Alderton, Roberto Amato, Jeffrey Barrett, Sonia Goncalves, Ewan Harrison, David K. Jackson, Ian Johnston, Dominic Kwiatkowski, Cordelia Langford, John Sillitoe on behalf of the Wellcome Sanger Institute COVID-19 Surveillance Team |
| EPI_ISL_3380141 | LABORATOIRE BIOMEDICA | CNR Virus des Infections Respiratoires - France SUD | Antonin Bal, Gregory Destras, Gwendolyne Burfin, Hadrien Regue, Quentin Semanas, Martine Valette, Bruno Lina, Laurence Josset |
| EPI_ISL_3380142, EPI_ISL_3380143, EPI_ISL_3380144 | Lighthouse Lab in Glasgow | Wellcome Sanger Institute for the COVID-19 Genomics UK (COG-UK) Consortium | Harper VanSteenhouse, Yumi Kasai, David Gray, Carol Clugston, Anna Dominiczak and Alex Alderton, Roberto Amato, Jeffrey Barrett, Sonia Goncalves, Ewan Harrison, David K. Jackson, Ian Johnston, Dominic Kwiatkowski, Cordelia Langford, John Sillitoe on behalf of the Wellcome Sanger Institute COVID-19 Surveillance Team |
| EPI_ISL_3380145 | LABORATOIRE BIOMEDICA | CNR Virus des Infections Respiratoires - France SUD | Antonin Bal, Gregory Destras, Gwendolyne Burfin, Hadrien Regue, Quentin Semanas, Martine Valette, Bruno Lina, Laurence Josset |
| EPI_ISL_3380146 | Lighthouse Lab in Milton Keynes | Wellcome Sanger Institute for the COVID-19 Genomics UK (COG-UK) Consortium | The Lighthouse Lab in Milton Keynes and Alex Alderton, Roberto Amato, Jeffrey Barrett, Sonia Goncalves, Ewan Harrison, David K. Jackson, Ian Johnston, Dominic Kwiatkowski, Cordelia Langford, John Sillitoe on behalf of the Wellcome Sanger Institute COVID-19 Surveillance Team |
| EPI_ISL_3380147, EPI_ISL_3380148, EPI_ISL_3380150 | Lighthouse Lab in Glasgow | Wellcome Sanger Institute for the COVID-19 Genomics UK (COG-UK) Consortium | Harper VanSteenhouse, Yumi Kasai, David Gray, Carol Clugston, Anna Dominiczak and Alex Alderton, Roberto Amato, Jeffrey Barrett, Sonia Goncalves, Ewan Harrison, David K. Jackson, Ian Johnston, Dominic Kwiatkowski, Cordelia Langford, John Sillitoe on behalf of the Wellcome Sanger Institute COVID-19 Surveillance Team |
| EPI_ISL_3380151 | Lighthouse Lab in Alderley Park | Wellcome Sanger Institute for the COVID-19 Genomics UK (COG-UK) Consortium | Jacquelyn Wynn, Mairead Hyland, The Lighthouse Lab in Alderley Park and Alex Alderton, Roberto Amato, Jeffrey Barrett, Sonia Goncalves, Ewan Harrison, David K. Jackson, Ian Johnston, Dominic Kwiatkowski, Cordelia Langford, John Sillitoe on behalf of the Wellcome Sanger Institute COVID-19 Surveillance Team |
| EPI_ISL_3380152 | LABORATOIRE BIOMEDICA | CNR Virus des Infections Respiratoires - France SUD | Antonin Bal, Gregory Destras, Gwendolyne Burfin, Hadrien Regue, Quentin Semanas, Martine Valette, Bruno Lina, Laurence Josset |
| EPI_ISL_3380153 | Lighthouse Lab in Glasgow | Wellcome Sanger Institute for the COVID-19 Genomics UK (COG-UK) Consortium | Harper VanSteenhouse, Yumi Kasai, David Gray, Carol Clugston, Anna Dominiczak and Alex Alderton, Roberto Amato, Jeffrey Barrett, Sonia Goncalves, Ewan Harrison, David K. Jackson, Ian Johnston, Dominic Kwiatkowski, Cordelia Langford, John Sillitoe on behalf of the Wellcome Sanger Institute COVID-19 Surveillance Team |
| EPI_ISL_3380154, EPI_ISL_3380155 | Lighthouse Lab in Alderley Park | Wellcome Sanger Institute for the COVID-19 Genomics UK (COG-UK) Consortium | Jacquelyn Wynn, Mairead Hyland, The Lighthouse Lab in Alderley Park and Alex Alderton, Roberto Amato, Jeffrey Barrett, Sonia Goncalves, Ewan Harrison, David K. Jackson, Ian Johnston, Dominic Kwiatkowski, Cordelia Langford, John Sillitoe on behalf of the Wellcome Sanger Institute COVID-19 Surveillance Team |
| EPI_ISL_3380156 | LABORATOIRE BIOMEDICA | CNR Virus des Infections Respiratoires - France SUD | Antonin Bal, Gregory Destras, Gwendolyne Burfin, Hadrien Regue, Quentin Semanas, Martine Valette, Bruno Lina, Laurence Josset |
| EPI_ISL_3380157 | Lighthouse Lab in Alderley Park | Wellcome Sanger Institute for the COVID-19 Genomics UK (COG-UK) Consortium | Jacquelyn Wynn, Mairead Hyland, The Lighthouse Lab in Alderley Park and Alex Alderton, Roberto Amato, Jeffrey Barrett, Sonia Goncalves, Ewan Harrison, David K. Jackson, Ian Johnston, Dominic Kwiatkowski, Cordelia Langford, John Sillitoe on behalf of the Wellcome Sanger Institute COVID-19 Surveillance Team |
| EPI_ISL_3380158 | Lighthouse Lab in Milton Keynes | Wellcome Sanger Institute for the COVID-19 Genomics UK (COG-UK) | The Lighthouse Lab in Milton Keynes and Alex Alderton, Roberto Amato, Jeffrey Barrett, Sonia Goncalves, Ewan Harrison, David K. Jackson, Ian Johnston, Dominic Kwiatkowski, Cordelia Langford, John Sillitoe on behalf of the Wellcome Sanger |

| | | | |
|---|---|---|---|
| | | Consortium | Institute COVID-19 Surveillance Team |
| EPI_ISL_3380159 | Lighthouse Lab in Alderley Park | Wellcome Sanger Institute for the COVID-19 Genomics UK (COG-UK) Consortium | Jacquelyn Wynn, Mairead Hyland, The Lighthouse Lab in Alderley Park and Alex Alderton, Roberto Amato, Jeffrey Barrett, Sonia Goncalves, Ewan Harrison, David K. Jackson, Ian Johnston, Dominic Kwiatkowski, Cordelia Langford, John Sillitoe on behalf of the Wellcome Sanger Institute COVID-19 Surveillance Team |
| EPI_ISL_3380160 | LABORATOIRE BIOMEDICA | CNR Virus des Infections Respiratoires - France SUD | Antonin Bal, Gregory Destras, Gwendolyne Burfin, Hadrien Regue, Quentin Semanas, Martine Valette, Bruno Lina, Laurence Josset |
| EPI_ISL_3380161, EPI_ISL_3380162, EPI_ISL_3380163 | Lighthouse Lab in Glasgow | Wellcome Sanger Institute for the COVID-19 Genomics UK (COG-UK) Consortium | Harper VanSteenhouse, Yumi Kasai, David Gray, Carol Clugston, Anna Dominiczak and Alex Alderton, Roberto Amato, Jeffrey Barrett, Sonia Goncalves, Ewan Harrison, David K. Jackson, Ian Johnston, Dominic Kwiatkowski, Cordelia Langford, John Sillitoe on behalf of the Wellcome Sanger Institute COVID-19 Surveillance Team |
| EPI_ISL_3380164 | LABORATOIRE BIOMEDICA | CNR Virus des Infections Respiratoires - France SUD | Antonin Bal, Gregory Destras, Gwendolyne Burfin, Hadrien Regue, Quentin Semanas, Martine Valette, Bruno Lina, Laurence Josset |
| EPI_ISL_3380165 | Lighthouse Lab in Alderley Park | Wellcome Sanger Institute for the COVID-19 Genomics UK (COG-UK) Consortium | Jacquelyn Wynn, Mairead Hyland, The Lighthouse Lab in Alderley Park and Alex Alderton, Roberto Amato, Jeffrey Barrett, Sonia Goncalves, Ewan Harrison, David K. Jackson, Ian Johnston, Dominic Kwiatkowski, Cordelia Langford, John Sillitoe on behalf of the Wellcome Sanger Institute COVID-19 Surveillance Team |
| EPI_ISL_3380166 | Lighthouse Lab in Milton Keynes | Wellcome Sanger Institute for the COVID-19 Genomics UK (COG-UK) Consortium | The Lighthouse Lab in Milton Keynes and Alex Alderton, Roberto Amato, Jeffrey Barrett, Sonia Goncalves, Ewan Harrison, David K. Jackson, Ian Johnston, Dominic Kwiatkowski, Cordelia Langford, John Sillitoe on behalf of the Wellcome Sanger Institute COVID-19 Surveillance Team |
| EPI_ISL_3380167 | LABORATOIRE BIOMEDICA | CNR Virus des Infections Respiratoires - France SUD | Antonin Bal, Gregory Destras, Gwendolyne Burfin, Hadrien Regue, Quentin Semanas, Martine Valette, Bruno Lina, Laurence Josset |
| EPI_ISL_3380168, EPI_ISL_3380169 | Lighthouse Lab in Milton Keynes | Wellcome Sanger Institute for the COVID-19 Genomics UK (COG-UK) Consortium | The Lighthouse Lab in Milton Keynes and Alex Alderton, Roberto Amato, Jeffrey Barrett, Sonia Goncalves, Ewan Harrison, David K. Jackson, Ian Johnston, Dominic Kwiatkowski, Cordelia Langford, John Sillitoe on behalf of the Wellcome Sanger Institute COVID-19 Surveillance Team |
| EPI_ISL_3380170 | Lighthouse Lab in Alderley Park | Wellcome Sanger Institute for the COVID-19 Genomics UK (COG-UK) Consortium | Jacquelyn Wynn, Mairead Hyland, The Lighthouse Lab in Alderley Park and Alex Alderton, Roberto Amato, Jeffrey Barrett, Sonia Goncalves, Ewan Harrison, David K. Jackson, Ian Johnston, Dominic Kwiatkowski, Cordelia Langford, John Sillitoe on behalf of the Wellcome Sanger Institute COVID-19 Surveillance Team |
| EPI_ISL_3380171 | LABORATOIRE BIOMEDICA | CNR Virus des Infections Respiratoires - France SUD | Antonin Bal, Gregory Destras, Gwendolyne Burfin, Hadrien Regue, Quentin Semanas, Martine Valette, Bruno Lina, Laurence Josset |
| EPI_ISL_3380172 | Lighthouse Lab in Alderley Park | Wellcome Sanger Institute for the COVID-19 Genomics UK (COG-UK) Consortium | Jacquelyn Wynn, Mairead Hyland, The Lighthouse Lab in Alderley Park and Alex Alderton, Roberto Amato, Jeffrey Barrett, Sonia Goncalves, Ewan Harrison, David K. Jackson, Ian Johnston, Dominic Kwiatkowski, Cordelia Langford, John Sillitoe on behalf of the Wellcome Sanger Institute COVID-19 Surveillance Team |
| EPI_ISL_3380173 | Lighthouse Lab in Milton Keynes | Wellcome Sanger Institute for the COVID-19 Genomics UK (COG-UK) Consortium | The Lighthouse Lab in Milton Keynes and Alex Alderton, Roberto Amato, Jeffrey Barrett, Sonia Goncalves, Ewan Harrison, David K. Jackson, Ian Johnston, Dominic Kwiatkowski, Cordelia Langford, John Sillitoe on behalf of the Wellcome Sanger Institute COVID-19 Surveillance Team |
| EPI_ISL_3380174 | Lighthouse Lab in Alderley Park | Wellcome Sanger Institute for the COVID-19 Genomics UK (COG-UK) Consortium | Jacquelyn Wynn, Mairead Hyland, The Lighthouse Lab in Alderley Park and Alex Alderton, Roberto Amato, Jeffrey Barrett, Sonia Goncalves, Ewan Harrison, David K. Jackson, Ian Johnston, Dominic Kwiatkowski, Cordelia Langford, John Sillitoe on behalf of the Wellcome Sanger Institute COVID-19 Surveillance Team |
| EPI_ISL_3380175 | LABORATOIRE BIOMEDICA | CNR Virus des Infections Respiratoires - France SUD | Antonin Bal, Gregory Destras, Gwendolyne Burfin, Hadrien Regue, Quentin Semanas, Martine Valette, Bruno Lina, Laurence Josset |
| EPI_ISL_3380176 | Lighthouse Lab in Milton Keynes | Wellcome Sanger Institute for the COVID-19 Genomics UK (COG-UK) Consortium | The Lighthouse Lab in Milton Keynes and Alex Alderton, Roberto Amato, Jeffrey Barrett, Sonia Goncalves, Ewan Harrison, David K. Jackson, Ian Johnston, Dominic Kwiatkowski, Cordelia Langford, John Sillitoe on behalf of the Wellcome Sanger Institute COVID-19 Surveillance Team |
| EPI_ISL_3380177 | Lighthouse Lab in Alderley Park | Wellcome Sanger Institute for the COVID-19 Genomics UK (COG-UK) Consortium | Jacquelyn Wynn, Mairead Hyland, The Lighthouse Lab in Alderley Park and Alex Alderton, Roberto Amato, Jeffrey Barrett, Sonia Goncalves, Ewan Harrison, David K. Jackson, Ian Johnston, Dominic Kwiatkowski, Cordelia Langford, John Sillitoe on behalf of the Wellcome Sanger Institute COVID-19 Surveillance Team |
| EPI_ISL_3380178 | LABORATOIRE BIOMEDICA | CNR Virus des Infections Respiratoires - France SUD | Antonin Bal, Gregory Destras, Gwendolyne Burfin, Hadrien Regue, Quentin Semanas, Martine Valette, Bruno Lina, Laurence Josset |
| EPI_ISL_3380179, EPI_ISL_3380180 | Lighthouse Lab in Alderley Park | Wellcome Sanger Institute for the COVID-19 Genomics UK (COG-UK) Consortium | Jacquelyn Wynn, Mairead Hyland, The Lighthouse Lab in Alderley Park and Alex Alderton, Roberto Amato, Jeffrey Barrett, Sonia Goncalves, Ewan Harrison, David K. Jackson, Ian Johnston, Dominic Kwiatkowski, Cordelia Langford, John Sillitoe on behalf of the Wellcome Sanger Institute COVID-19 Surveillance Team |
| EPI_ISL_3380181 | Lighthouse Lab in Milton Keynes | Wellcome Sanger Institute for the COVID-19 Genomics UK (COG-UK) Consortium | The Lighthouse Lab in Milton Keynes and Alex Alderton, Roberto Amato, Jeffrey Barrett, Sonia Goncalves, Ewan Harrison, David K. Jackson, Ian Johnston, Dominic Kwiatkowski, Cordelia Langford, John Sillitoe on behalf of the Wellcome Sanger Institute COVID-19 Surveillance Team |
| EPI_ISL_3380182 | LABORATOIRE BIOMEDICA | CNR Virus des Infections Respiratoires - France SUD | Antonin Bal, Gregory Destras, Gwendolyne Burfin, Hadrien Regue, Quentin Semanas, Martine Valette, Bruno Lina, Laurence Josset |
| EPI_ISL_3380183, EPI_ISL_3380184, EPI_ISL_3380185 | Lighthouse Lab in Milton Keynes | Wellcome Sanger Institute for the COVID-19 Genomics UK (COG-UK) Consortium | The Lighthouse Lab in Milton Keynes and Alex Alderton, Roberto Amato, Jeffrey Barrett, Sonia Goncalves, Ewan Harrison, David K. Jackson, Ian Johnston, Dominic Kwiatkowski, Cordelia Langford, John Sillitoe on behalf of the Wellcome Sanger Institute COVID-19 Surveillance Team |
| EPI_ISL_3380186 | LABORATOIRE BIOMEDICA | CNR Virus des Infections Respiratoires - France SUD | Antonin Bal, Gregory Destras, Gwendolyne Burfin, Hadrien Regue, Quentin Semanas, Martine Valette, Bruno Lina, Laurence Josset |
| EPI_ISL_3380187, EPI_ISL_3380188 | Lighthouse Lab in Alderley Park | Wellcome Sanger Institute for the COVID-19 Genomics UK (COG-UK) Consortium | Jacquelyn Wynn, Mairead Hyland, The Lighthouse Lab in Alderley Park and Alex Alderton, Roberto Amato, Jeffrey Barrett, Sonia Goncalves, Ewan Harrison, David K. Jackson, Ian Johnston, Dominic Kwiatkowski, Cordelia Langford, John Sillitoe on behalf of the Wellcome Sanger Institute COVID-19 Surveillance Team |
| EPI_ISL_3380189 | Lighthouse Lab in Milton Keynes | Wellcome Sanger Institute for the COVID-19 Genomics UK (COG-UK) Consortium | The Lighthouse Lab in Milton Keynes and Alex Alderton, Roberto Amato, Jeffrey Barrett, Sonia Goncalves, Ewan Harrison, David K. Jackson, Ian Johnston, Dominic Kwiatkowski, Cordelia Langford, John Sillitoe on behalf of the Wellcome Sanger Institute COVID-19 Surveillance Team |
| EPI_ISL_3380190 | LABORATOIRE BIOMEDICA | CNR Virus des Infections Respiratoires - France SUD | Antonin Bal, Gregory Destras, Gwendolyne Burfin, Hadrien Regue, Quentin Semanas, Martine Valette, Bruno Lina, Laurence Josset |
| EPI_ISL_3380191 | Lighthouse Lab in Alderley Park | Wellcome Sanger Institute for the COVID-19 Genomics UK (COG-UK) Consortium | Jacquelyn Wynn, Mairead Hyland, The Lighthouse Lab in Alderley Park and Alex Alderton, Roberto Amato, Jeffrey Barrett, Sonia Goncalves, Ewan Harrison, David K. Jackson, Ian Johnston, Dominic Kwiatkowski, Cordelia Langford, John Sillitoe on behalf of the Wellcome Sanger Institute COVID-19 Surveillance Team |
| EPI_ISL_3380192, EPI_ISL_3380193 | Lighthouse Lab in Milton Keynes | Wellcome Sanger Institute for the COVID-19 Genomics UK (COG-UK) Consortium | The Lighthouse Lab in Milton Keynes and Alex Alderton, Roberto Amato, Jeffrey Barrett, Sonia Goncalves, Ewan Harrison, David K. Jackson, Ian Johnston, Dominic Kwiatkowski, Cordelia Langford, John Sillitoe on behalf of the Wellcome Sanger Institute COVID-19 Surveillance Team |
| EPI_ISL_3380194 | LABORATOIRE BIOMEDICA | CNR Virus des Infections Respiratoires - France SUD | Antonin Bal, Gregory Destras, Gwendolyne Burfin, Hadrien Regue, Quentin Semanas, Martine Valette, Bruno Lina, Laurence Josset |
| EPI_ISL_3380195, EPI_ISL_3380196, EPI_ISL_3380197 | Lighthouse Lab in Milton Keynes | Wellcome Sanger Institute for the COVID-19 Genomics UK (COG-UK) Consortium | The Lighthouse Lab in Milton Keynes and Alex Alderton, Roberto Amato, Jeffrey Barrett, Sonia Goncalves, Ewan Harrison, David K. Jackson, Ian Johnston, Dominic Kwiatkowski, Cordelia Langford, John Sillitoe on behalf of the Wellcome Sanger Institute COVID-19 Surveillance Team |
| EPI_ISL_3380198 | LABORATOIRE BIOMEDICA | CNR Virus des Infections Respiratoires - France SUD | Antonin Bal, Gregory Destras, Gwendolyne Burfin, Hadrien Regue, Quentin Semanas, Martine Valette, Bruno Lina, Laurence Josset |
| EPI_ISL_3380199 | Lighthouse Lab in Milton Keynes | Wellcome Sanger Institute for the COVID-19 Genomics UK (COG-UK) Consortium | The Lighthouse Lab in Milton Keynes and Alex Alderton, Roberto Amato, Jeffrey Barrett, Sonia Goncalves, Ewan Harrison, David K. Jackson, Ian Johnston, Dominic Kwiatkowski, Cordelia Langford, John Sillitoe on behalf of the Wellcome Sanger Institute COVID-19 Surveillance Team |
| EPI_ISL_3380200 | Lighthouse Lab in Alderley Park | Wellcome Sanger Institute for the COVID-19 Genomics UK (COG-UK) Consortium | Jacquelyn Wynn, Mairead Hyland, The Lighthouse Lab in Alderley Park and Alex Alderton, Roberto Amato, Jeffrey Barrett, Sonia Goncalves, Ewan Harrison, David K. Jackson, Ian Johnston, Dominic Kwiatkowski, Cordelia Langford, John Sillitoe on behalf of the Wellcome Sanger Institute COVID-19 Surveillance Team |
| EPI_ISL_3380201 | Lighthouse Lab in Milton Keynes | Wellcome Sanger Institute for the COVID-19 Genomics UK (COG-UK) Consortium | The Lighthouse Lab in Milton Keynes and Alex Alderton, Roberto Amato, Jeffrey Barrett, Sonia Goncalves, Ewan Harrison, David K. Jackson, Ian Johnston, Dominic Kwiatkowski, Cordelia Langford, John Sillitoe on behalf of the Wellcome Sanger Institute COVID-19 Surveillance Team |
| EPI_ISL_3380202 | LABORATOIRE BIOMEDICA | CNR Virus des Infections Respiratoires - France SUD | Antonin Bal, Gregory Destras, Gwendolyne Burfin, Hadrien Regue, Quentin Semanas, Martine Valette, Bruno Lina, Laurence Josset |
| EPI_ISL_3380203, EPI_ISL_3380204, EPI_ISL_3380205 | Lighthouse Lab in Milton Keynes | Wellcome Sanger Institute for the COVID-19 Genomics UK (COG-UK) Consortium | The Lighthouse Lab in Milton Keynes and Alex Alderton, Roberto Amato, Jeffrey Barrett, Sonia Goncalves, Ewan Harrison, David K. Jackson, Ian Johnston, Dominic Kwiatkowski, Cordelia Langford, John Sillitoe on behalf of the Wellcome Sanger Institute COVID-19 Surveillance Team |
| EPI_ISL_3380206 | LABORATOIRE BIOMEDICA | CNR Virus des Infections Respiratoires - France SUD | Antonin Bal, Gregory Destras, Gwendolyne Burfin, Hadrien Regue, Quentin Semanas, Martine Valette, Bruno Lina, Laurence Josset |
| EPI_ISL_3380207 | Lighthouse Lab in Alderley Park | Wellcome Sanger Institute for the COVID-19 Genomics UK (COG-UK) Consortium | Jacquelyn Wynn, Mairead Hyland, The Lighthouse Lab in Alderley Park and Alex Alderton, Roberto Amato, Jeffrey Barrett, Sonia Goncalves, Ewan Harrison, David K. Jackson, Ian Johnston, Dominic Kwiatkowski, Cordelia Langford, John Sillitoe on behalf of the Wellcome Sanger Institute COVID-19 Surveillance Team |
| EPI_ISL_3380208, EPI_ISL_3380209 | Lighthouse Lab in Milton Keynes | Wellcome Sanger Institute for the COVID-19 Genomics UK (COG-UK) Consortium | The Lighthouse Lab in Milton Keynes and Alex Alderton, Roberto Amato, Jeffrey Barrett, Sonia Goncalves, Ewan Harrison, David K. Jackson, Ian Johnston, Dominic Kwiatkowski, Cordelia Langford, John Sillitoe on behalf of the Wellcome Sanger Institute COVID-19 Surveillance Team |
| EPI_ISL_3380210 | LABORATOIRE BIOMEDICA | CNR Virus des Infections Respiratoires - France SUD | Antonin Bal, Gregory Destras, Gwendolyne Burfin, Hadrien Regue, Quentin Semanas, Martine Valette, Bruno Lina, Laurence Josset |
| EPI_ISL_3380211, EPI_ISL_3380212, EPI_ISL_3380213 | Lighthouse Lab in Milton Keynes | Wellcome Sanger Institute for the COVID-19 Genomics UK (COG-UK) Consortium | The Lighthouse Lab in Milton Keynes and Alex Alderton, Roberto Amato, Jeffrey Barrett, Sonia Goncalves, Ewan Harrison, David K. Jackson, Ian Johnston, Dominic Kwiatkowski, Cordelia Langford, John Sillitoe on behalf of the Wellcome Sanger Institute COVID-19 Surveillance Team |
| EPI_ISL_3380214 | LABORATOIRE BIOMEDICA | CNR Virus des Infections Respiratoires - France SUD | Antonin Bal, Gregory Destras, Gwendolyne Burfin, Hadrien Regue, Quentin Semanas, Martine Valette, Bruno Lina, Laurence Josset |
| EPI_ISL_3380215, EPI_ISL_3380216 | Lighthouse Lab in Alderley Park | Wellcome Sanger Institute for the COVID-19 Genomics UK (COG-UK) Consortium | Jacquelyn Wynn, Mairead Hyland, The Lighthouse Lab in Alderley Park and Alex Alderton, Roberto Amato, Jeffrey Barrett, Sonia Goncalves, Ewan Harrison, David K. Jackson, Ian Johnston, Dominic Kwiatkowski, Cordelia Langford, John Sillitoe on behalf of the Wellcome Sanger Institute COVID-19 Surveillance Team |
| EPI_ISL_3380217 | BIOLITTORAL BIOGROUP PLATEAU TECH | CNR Virus des Infections Respiratoires - France SUD | Antonin Bal, Gregory Destras, Gwendolyne Burfin, Hadrien Regue, Quentin Semanas, Martine Valette, Bruno Lina, Laurence Josset |
| EPI_ISL_3380218, EPI_ISL_3380219, EPI_ISL_3380220 | Lighthouse Lab in Milton Keynes | Wellcome Sanger Institute for the COVID-19 Genomics UK (COG-UK) Consortium | The Lighthouse Lab in Milton Keynes and Alex Alderton, Roberto Amato, Jeffrey Barrett, Sonia Goncalves, Ewan Harrison, David K. Jackson, Ian Johnston, Dominic Kwiatkowski, Cordelia Langford, John Sillitoe on behalf of the Wellcome Sanger Institute COVID-19 Surveillance Team |
| EPI_ISL_3380221 | BIOLITTORAL BIOGROUP PLATEAU TECH | CNR Virus des Infections Respiratoires - France SUD | Antonin Bal, Gregory Destras, Gwendolyne Burfin, Hadrien Regue, Quentin Semanas, Martine Valette, Bruno Lina, Laurence Josset |
| EPI_ISL_3380222 | Lighthouse Lab in Milton Keynes | Wellcome Sanger Institute for the COVID-19 Genomics UK (COG-UK) Consortium | The Lighthouse Lab in Milton Keynes and Alex Alderton, Roberto Amato, Jeffrey Barrett, Sonia Goncalves, Ewan Harrison, David K. Jackson, Ian Johnston, Dominic Kwiatkowski, Cordelia Langford, John Sillitoe on behalf of the Wellcome Sanger Institute COVID-19 Surveillance Team |
| EPI_ISL_3380223, EPI_ISL_3380224 | Lighthouse Lab in Alderley Park | Wellcome Sanger Institute for the COVID-19 Genomics UK (COG-UK) Consortium | Jacquelyn Wynn, Mairead Hyland, The Lighthouse Lab in Alderley Park and Alex Alderton, Roberto Amato, Jeffrey Barrett, Sonia Goncalves, Ewan Harrison, David K. Jackson, Ian Johnston, Dominic Kwiatkowski, Cordelia Langford, John Sillitoe on behalf of the Wellcome Sanger Institute COVID-19 Surveillance Team |
| EPI_ISL_3380225 | BIOLITTORAL BIOGROUP PLATEAU TECH | CNR Virus des Infections Respiratoires - France SUD | Antonin Bal, Gregory Destras, Gwendolyne Burfin, Hadrien Regue, Quentin Semanas, Martine Valette, Bruno Lina, Laurence Josset |
| EPI_ISL_3380226 | Lighthouse Lab in Milton Keynes | Wellcome Sanger Institute for the COVID-19 Genomics UK (COG-UK) Consortium | The Lighthouse Lab in Milton Keynes and Alex Alderton, Roberto Amato, Jeffrey Barrett, Sonia Goncalves, Ewan Harrison, David K. Jackson, Ian Johnston, Dominic Kwiatkowski, Cordelia Langford, John Sillitoe on behalf of the Wellcome Sanger Institute COVID-19 Surveillance Team |
| EPI_ISL_3380227 | Lighthouse Lab in Alderley Park | Wellcome Sanger Institute for the COVID-19 Genomics UK (COG-UK) | Jacquelyn Wynn, Mairead Hyland, The Lighthouse Lab in Alderley Park and Alex Alderton, Roberto Amato, Jeffrey Barrett, Sonia Goncalves, Ewan Harrison, David K. Jackson, Ian Johnston, Dominic Kwiatkowski, Cordelia Langford, John Sillitoe on |

| | | | |
|---|---|---|---|
| | | Consortium | behalf of the Wellcome Sanger Institute COVID-19 Surveillance Team |
| EPI_ISL_3380228 | BIOLITTORAL BIOGROUP PLATEAU TECH | CNR Virus des Infections Respiratoires - France SUD | Antonin Bal, Gregory Destras, Gwendolyne Burfin, Hadrien Regue, Quentin Semanas, Martine Valette, Bruno Lina, Laurence Josset |
| EPI_ISL_3380229 | Lighthouse Lab in Alderley Park | Wellcome Sanger Institute for the COVID-19 Genomics UK (COG-UK) Consortium | Jacquelyn Wynn, Mairead Hyland, The Lighthouse Lab in Alderley Park and Alex Alderton, Roberto Amato, Jeffrey Barrett, Sonia Goncalves, Ewan Harrison, David K. Jackson, Ian Johnston, Dominic Kwiatkowski, Cordelia Langford, John Sillitoe on behalf of the Wellcome Sanger Institute COVID-19 Surveillance Team |
| EPI_ISL_3380230 | Lighthouse Lab in Milton Keynes | Wellcome Sanger Institute for the COVID-19 Genomics UK (COG-UK) Consortium | The Lighthouse Lab in Milton Keynes and Alex Alderton, Roberto Amato, Jeffrey Barrett, Sonia Goncalves, Ewan Harrison, David K. Jackson, Ian Johnston, Dominic Kwiatkowski, Cordelia Langford, John Sillitoe on behalf of the Wellcome Sanger Institute COVID-19 Surveillance Team |
| EPI_ISL_3380231 | Lighthouse Lab in Alderley Park | Wellcome Sanger Institute for the COVID-19 Genomics UK (COG-UK) Consortium | Jacquelyn Wynn, Mairead Hyland, The Lighthouse Lab in Alderley Park and Alex Alderton, Roberto Amato, Jeffrey Barrett, Sonia Goncalves, Ewan Harrison, David K. Jackson, Ian Johnston, Dominic Kwiatkowski, Cordelia Langford, John Sillitoe on behalf of the Wellcome Sanger Institute COVID-19 Surveillance Team |
| EPI_ISL_3380232 | BIOLITTORAL BIOGROUP PLATEAU TECH | CNR Virus des Infections Respiratoires - France SUD | Antonin Bal, Gregory Destras, Gwendolyne Burfin, Hadrien Regue, Quentin Semanas, Martine Valette, Bruno Lina, Laurence Josset |
| EPI_ISL_3380233 | Lighthouse Lab in Milton Keynes | Wellcome Sanger Institute for the COVID-19 Genomics UK (COG-UK) Consortium | The Lighthouse Lab in Milton Keynes and Alex Alderton, Roberto Amato, Jeffrey Barrett, Sonia Goncalves, Ewan Harrison, David K. Jackson, Ian Johnston, Dominic Kwiatkowski, Cordelia Langford, John Sillitoe on behalf of the Wellcome Sanger Institute COVID-19 Surveillance Team |
| EPI_ISL_3380234 | Lighthouse Lab in Alderley Park | Wellcome Sanger Institute for the COVID-19 Genomics UK (COG-UK) Consortium | Jacquelyn Wynn, Mairead Hyland, The Lighthouse Lab in Alderley Park and Alex Alderton, Roberto Amato, Jeffrey Barrett, Sonia Goncalves, Ewan Harrison, David K. Jackson, Ian Johnston, Dominic Kwiatkowski, Cordelia Langford, John Sillitoe on behalf of the Wellcome Sanger Institute COVID-19 Surveillance Team |
| EPI_ISL_3380235 | Lighthouse Lab in Milton Keynes | Wellcome Sanger Institute for the COVID-19 Genomics UK (COG-UK) Consortium | The Lighthouse Lab in Milton Keynes and Alex Alderton, Roberto Amato, Jeffrey Barrett, Sonia Goncalves, Ewan Harrison, David K. Jackson, Ian Johnston, Dominic Kwiatkowski, Cordelia Langford, John Sillitoe on behalf of the Wellcome Sanger Institute COVID-19 Surveillance Team |
| EPI_ISL_3380236 | BIOLITTORAL BIOGROUP PLATEAU TECH | CNR Virus des Infections Respiratoires - France SUD | Antonin Bal, Gregory Destras, Gwendolyne Burfin, Hadrien Regue, Quentin Semanas, Martine Valette, Bruno Lina, Laurence Josset |
| EPI_ISL_3380237 | Lighthouse Lab in Alderley Park | Wellcome Sanger Institute for the COVID-19 Genomics UK (COG-UK) Consortium | Jacquelyn Wynn, Mairead Hyland, The Lighthouse Lab in Alderley Park and Alex Alderton, Roberto Amato, Jeffrey Barrett, Sonia Goncalves, Ewan Harrison, David K. Jackson, Ian Johnston, Dominic Kwiatkowski, Cordelia Langford, John Sillitoe on behalf of the Wellcome Sanger Institute COVID-19 Surveillance Team |
| EPI_ISL_3380238 | Lighthouse Lab in Milton Keynes | Wellcome Sanger Institute for the COVID-19 Genomics UK (COG-UK) Consortium | The Lighthouse Lab in Milton Keynes and Alex Alderton, Roberto Amato, Jeffrey Barrett, Sonia Goncalves, Ewan Harrison, David K. Jackson, Ian Johnston, Dominic Kwiatkowski, Cordelia Langford, John Sillitoe on behalf of the Wellcome Sanger Institute COVID-19 Surveillance Team |
| EPI_ISL_3380239 | BIOLITTORAL BIOGROUP PLATEAU TECH | CNR Virus des Infections Respiratoires - France SUD | Antonin Bal, Gregory Destras, Gwendolyne Burfin, Hadrien Regue, Quentin Semanas, Martine Valette, Bruno Lina, Laurence Josset |
| EPI_ISL_3380240 | Lighthouse Lab in Milton Keynes | Wellcome Sanger Institute for the COVID-19 Genomics UK (COG-UK) Consortium | The Lighthouse Lab in Milton Keynes and Alex Alderton, Roberto Amato, Jeffrey Barrett, Sonia Goncalves, Ewan Harrison, David K. Jackson, Ian Johnston, Dominic Kwiatkowski, Cordelia Langford, John Sillitoe on behalf of the Wellcome Sanger Institute COVID-19 Surveillance Team |
| EPI_ISL_3380241, EPI_ISL_3380242 | Lighthouse Lab in Alderley Park | Wellcome Sanger Institute for the COVID-19 Genomics UK (COG-UK) Consortium | Jacquelyn Wynn, Mairead Hyland, The Lighthouse Lab in Alderley Park and Alex Alderton, Roberto Amato, Jeffrey Barrett, Sonia Goncalves, Ewan Harrison, David K. Jackson, Ian Johnston, Dominic Kwiatkowski, Cordelia Langford, John Sillitoe on behalf of the Wellcome Sanger Institute COVID-19 Surveillance Team |
| EPI_ISL_3380243 | BIOLITTORAL BIOGROUP PLATEAU TECH | CNR Virus des Infections Respiratoires - France SUD | Antonin Bal, Gregory Destras, Gwendolyne Burfin, Hadrien Regue, Quentin Semanas, Martine Valette, Bruno Lina, Laurence Josset |
| EPI_ISL_3380244, EPI_ISL_3380245, EPI_ISL_3380246 | Lighthouse Lab in Milton Keynes | Wellcome Sanger Institute for the COVID-19 Genomics UK (COG-UK) Consortium | The Lighthouse Lab in Milton Keynes and Alex Alderton, Roberto Amato, Jeffrey Barrett, Sonia Goncalves, Ewan Harrison, David K. Jackson, Ian Johnston, Dominic Kwiatkowski, Cordelia Langford, John Sillitoe on behalf of the Wellcome Sanger Institute COVID-19 Surveillance Team |
| EPI_ISL_3380247 | CERBAILLIANCE LA REUNION | CNR Virus des Infections Respiratoires - France SUD | Antonin Bal, Gregory Destras, Gwendolyne Burfin, Hadrien Regue, Quentin Semanas, Martine Valette, Bruno Lina, Laurence Josset |
| EPI_ISL_3380248 | Lighthouse Lab in Alderley Park | Wellcome Sanger Institute for the COVID-19 Genomics UK (COG-UK) Consortium | Jacquelyn Wynn, Mairead Hyland, The Lighthouse Lab in Alderley Park and Alex Alderton, Roberto Amato, Jeffrey Barrett, Sonia Goncalves, Ewan Harrison, David K. Jackson, Ian Johnston, Dominic Kwiatkowski, Cordelia Langford, John Sillitoe on behalf of the Wellcome Sanger Institute COVID-19 Surveillance Team |
| EPI_ISL_3380249, EPI_ISL_3380250 | Lighthouse Lab in Milton Keynes | Wellcome Sanger Institute for the COVID-19 Genomics UK (COG-UK) Consortium | The Lighthouse Lab in Milton Keynes and Alex Alderton, Roberto Amato, Jeffrey Barrett, Sonia Goncalves, Ewan Harrison, David K. Jackson, Ian Johnston, Dominic Kwiatkowski, Cordelia Langford, John Sillitoe on behalf of the Wellcome Sanger Institute COVID-19 Surveillance Team |
| EPI_ISL_3380251 | BIOLITTORAL BIOGROUP PLATEAU TECH | CNR Virus des Infections Respiratoires - France SUD | Antonin Bal, Gregory Destras, Gwendolyne Burfin, Hadrien Regue, Quentin Semanas, Martine Valette, Bruno Lina, Laurence Josset |
| EPI_ISL_3380252, EPI_ISL_3380253, EPI_ISL_3380254 | Lighthouse Lab in Milton Keynes | Wellcome Sanger Institute for the COVID-19 Genomics UK (COG-UK) Consortium | The Lighthouse Lab in Milton Keynes and Alex Alderton, Roberto Amato, Jeffrey Barrett, Sonia Goncalves, Ewan Harrison, David K. Jackson, Ian Johnston, Dominic Kwiatkowski, Cordelia Langford, John Sillitoe on behalf of the Wellcome Sanger Institute COVID-19 Surveillance Team |
| EPI_ISL_3380255 | BIOLITTORAL BIOGROUP PLATEAU TECH | CNR Virus des Infections Respiratoires - France SUD | Antonin Bal, Gregory Destras, Gwendolyne Burfin, Hadrien Regue, Quentin Semanas, Martine Valette, Bruno Lina, Laurence Josset |
| EPI_ISL_3380256, EPI_ISL_3380257 | Lighthouse Lab in Milton Keynes | Wellcome Sanger Institute for the COVID-19 Genomics UK (COG-UK) Consortium | The Lighthouse Lab in Milton Keynes and Alex Alderton, Roberto Amato, Jeffrey Barrett, Sonia Goncalves, Ewan Harrison, David K. Jackson, Ian Johnston, Dominic Kwiatkowski, Cordelia Langford, John Sillitoe on behalf of the Wellcome Sanger Institute COVID-19 Surveillance Team |
| EPI_ISL_3380258 | Lighthouse Lab in Alderley Park | Wellcome Sanger Institute for the COVID-19 Genomics UK (COG-UK) Consortium | Jacquelyn Wynn, Mairead Hyland, The Lighthouse Lab in Alderley Park and Alex Alderton, Roberto Amato, Jeffrey Barrett, Sonia Goncalves, Ewan Harrison, David K. Jackson, Ian Johnston, Dominic Kwiatkowski, Cordelia Langford, John Sillitoe on behalf of the Wellcome Sanger Institute COVID-19 Surveillance Team |
| EPI_ISL_3380260, EPI_ISL_3380261, EPI_ISL_3380262 | Lighthouse Lab in Milton Keynes | Wellcome Sanger Institute for the COVID-19 Genomics UK (COG-UK) Consortium | The Lighthouse Lab in Milton Keynes and Alex Alderton, Roberto Amato, Jeffrey Barrett, Sonia Goncalves, Ewan Harrison, David K. Jackson, Ian Johnston, Dominic Kwiatkowski, Cordelia Langford, John Sillitoe on behalf of the Wellcome Sanger Institute COVID-19 Surveillance Team |
| EPI_ISL_3380263 | BIOLITTORAL BIOGROUP PLATEAU TECH | CNR Virus des Infections Respiratoires - France SUD | Antonin Bal, Gregory Destras, Gwendolyne Burfin, Hadrien Regue, Quentin Semanas, Martine Valette, Bruno Lina, Laurence Josset |
| EPI_ISL_3380264, EPI_ISL_3380265 | Lighthouse Lab in Alderley Park | Wellcome Sanger Institute for the COVID-19 Genomics UK (COG-UK) Consortium | Jacquelyn Wynn, Mairead Hyland, The Lighthouse Lab in Alderley Park and Alex Alderton, Roberto Amato, Jeffrey Barrett, Sonia Goncalves, Ewan Harrison, David K. Jackson, Ian Johnston, Dominic Kwiatkowski, Cordelia Langford, John Sillitoe on behalf of the Wellcome Sanger Institute COVID-19 Surveillance Team |
| EPI_ISL_3380267 | Lighthouse Lab in Milton Keynes | Wellcome Sanger Institute for the COVID-19 Genomics UK (COG-UK) Consortium | The Lighthouse Lab in Milton Keynes and Alex Alderton, Roberto Amato, Jeffrey Barrett, Sonia Goncalves, Ewan Harrison, David K. Jackson, Ian Johnston, Dominic Kwiatkowski, Cordelia Langford, John Sillitoe on behalf of the Wellcome Sanger Institute COVID-19 Surveillance Team |
| EPI_ISL_3380268 | Lighthouse Lab in Alderley Park | Wellcome Sanger Institute for the COVID-19 Genomics UK (COG-UK) Consortium | Jacquelyn Wynn, Mairead Hyland, The Lighthouse Lab in Alderley Park and Alex Alderton, Roberto Amato, Jeffrey Barrett, Sonia Goncalves, Ewan Harrison, David K. Jackson, Ian Johnston, Dominic Kwiatkowski, Cordelia Langford, John Sillitoe on behalf of the Wellcome Sanger Institute COVID-19 Surveillance Team |
| EPI_ISL_3380269 | Lighthouse Lab in Milton Keynes | Wellcome Sanger Institute for the COVID-19 Genomics UK (COG-UK) Consortium | The Lighthouse Lab in Milton Keynes and Alex Alderton, Roberto Amato, Jeffrey Barrett, Sonia Goncalves, Ewan Harrison, David K. Jackson, Ian Johnston, Dominic Kwiatkowski, Cordelia Langford, John Sillitoe on behalf of the Wellcome Sanger Institute COVID-19 Surveillance Team |
| EPI_ISL_3380270 | BIOLITTORAL BIOGROUP PLATEAU TECH | CNR Virus des Infections Respiratoires - France SUD | Antonin Bal, Gregory Destras, Gwendolyne Burfin, Hadrien Regue, Quentin Semanas, Martine Valette, Bruno Lina, Laurence Josset |
| EPI_ISL_3380271 | Lighthouse Lab in Alderley Park | Wellcome Sanger Institute for the COVID-19 Genomics UK (COG-UK) Consortium | Jacquelyn Wynn, Mairead Hyland, The Lighthouse Lab in Alderley Park and Alex Alderton, Roberto Amato, Jeffrey Barrett, Sonia Goncalves, Ewan Harrison, David K. Jackson, Ian Johnston, Dominic Kwiatkowski, Cordelia Langford, John Sillitoe on behalf of the Wellcome Sanger Institute COVID-19 Surveillance Team |
| EPI_ISL_3380272, EPI_ISL_3380273 | Lighthouse Lab in Milton Keynes | Wellcome Sanger Institute for the COVID-19 Genomics UK (COG-UK) Consortium | The Lighthouse Lab in Milton Keynes and Alex Alderton, Roberto Amato, Jeffrey Barrett, Sonia Goncalves, Ewan Harrison, David K. Jackson, Ian Johnston, Dominic Kwiatkowski, Cordelia Langford, John Sillitoe on behalf of the Wellcome Sanger Institute COVID-19 Surveillance Team |
| EPI_ISL_3380274 | BIOLITTORAL BIOGROUP PLATEAU TECH | CNR Virus des Infections Respiratoires - France SUD | Antonin Bal, Gregory Destras, Gwendolyne Burfin, Hadrien Regue, Quentin Semanas, Martine Valette, Bruno Lina, Laurence Josset |
| EPI_ISL_3380275 | Lighthouse Lab in Alderley Park | Wellcome Sanger Institute for the COVID-19 Genomics UK (COG-UK) Consortium | Jacquelyn Wynn, Mairead Hyland, The Lighthouse Lab in Alderley Park and Alex Alderton, Roberto Amato, Jeffrey Barrett, Sonia Goncalves, Ewan Harrison, David K. Jackson, Ian Johnston, Dominic Kwiatkowski, Cordelia Langford, John Sillitoe on behalf of the Wellcome Sanger Institute COVID-19 Surveillance Team |
| EPI_ISL_3380276, EPI_ISL_3380277 | Lighthouse Lab in Milton Keynes | Wellcome Sanger Institute for the COVID-19 Genomics UK (COG-UK) Consortium | The Lighthouse Lab in Milton Keynes and Alex Alderton, Roberto Amato, Jeffrey Barrett, Sonia Goncalves, Ewan Harrison, David K. Jackson, Ian Johnston, Dominic Kwiatkowski, Cordelia Langford, John Sillitoe on behalf of the Wellcome Sanger Institute COVID-19 Surveillance Team |
| EPI_ISL_3380278 | CERBAILLIANCE LA REUNION | CNR Virus des Infections Respiratoires - France SUD | Antonin Bal, Gregory Destras, Gwendolyne Burfin, Hadrien Regue, Quentin Semanas, Martine Valette, Bruno Lina, Laurence Josset |
| EPI_ISL_3380279, EPI_ISL_3380280, EPI_ISL_3380281 | Lighthouse Lab in Milton Keynes | Wellcome Sanger Institute for the COVID-19 Genomics UK (COG-UK) Consortium | The Lighthouse Lab in Milton Keynes and Alex Alderton, Roberto Amato, Jeffrey Barrett, Sonia Goncalves, Ewan Harrison, David K. Jackson, Ian Johnston, Dominic Kwiatkowski, Cordelia Langford, John Sillitoe on behalf of the Wellcome Sanger Institute COVID-19 Surveillance Team |
| EPI_ISL_3380282 | BIOLITTORAL BIOGROUP PLATEAU TECH | CNR Virus des Infections Respiratoires - France SUD | Antonin Bal, Gregory Destras, Gwendolyne Burfin, Hadrien Regue, Quentin Semanas, Martine Valette, Bruno Lina, Laurence Josset |
| EPI_ISL_3380283 | Lighthouse Lab in Milton Keynes | Wellcome Sanger Institute for the COVID-19 Genomics UK (COG-UK) Consortium | The Lighthouse Lab in Milton Keynes and Alex Alderton, Roberto Amato, Jeffrey Barrett, Sonia Goncalves, Ewan Harrison, David K. Jackson, Ian Johnston, Dominic Kwiatkowski, Cordelia Langford, John Sillitoe on behalf of the Wellcome Sanger Institute COVID-19 Surveillance Team |
| EPI_ISL_3380284 | Lighthouse Lab in Alderley Park | Wellcome Sanger Institute for the COVID-19 Genomics UK (COG-UK) Consortium | Jacquelyn Wynn, Mairead Hyland, The Lighthouse Lab in Alderley Park and Alex Alderton, Roberto Amato, Jeffrey Barrett, Sonia Goncalves, Ewan Harrison, David K. Jackson, Ian Johnston, Dominic Kwiatkowski, Cordelia Langford, John Sillitoe on behalf of the Wellcome Sanger Institute COVID-19 Surveillance Team |
| EPI_ISL_3380285 | Lighthouse Lab in Milton Keynes | Wellcome Sanger Institute for the COVID-19 Genomics UK (COG-UK) Consortium | The Lighthouse Lab in Milton Keynes and Alex Alderton, Roberto Amato, Jeffrey Barrett, Sonia Goncalves, Ewan Harrison, David K. Jackson, Ian Johnston, Dominic Kwiatkowski, Cordelia Langford, John Sillitoe on behalf of the Wellcome Sanger Institute COVID-19 Surveillance Team |
| EPI_ISL_3380286 | CERBAILLIANCE LA REUNION | CNR Virus des Infections Respiratoires - France SUD | Antonin Bal, Gregory Destras, Gwendolyne Burfin, Hadrien Regue, Quentin Semanas, Martine Valette, Bruno Lina, Laurence Josset |
| EPI_ISL_3380287, EPI_ISL_3380288 | Lighthouse Lab in Alderley Park | Wellcome Sanger Institute for the COVID-19 Genomics UK (COG-UK) Consortium | Jacquelyn Wynn, Mairead Hyland, The Lighthouse Lab in Alderley Park and Alex Alderton, Roberto Amato, Jeffrey Barrett, Sonia Goncalves, Ewan Harrison, David K. Jackson, Ian Johnston, Dominic Kwiatkowski, Cordelia Langford, John Sillitoe on behalf of the Wellcome Sanger Institute COVID-19 Surveillance Team |
| EPI_ISL_3380289 | Lighthouse Lab in Milton Keynes | Wellcome Sanger Institute for the COVID-19 Genomics UK (COG-UK) Consortium | The Lighthouse Lab in Milton Keynes and Alex Alderton, Roberto Amato, Jeffrey Barrett, Sonia Goncalves, Ewan Harrison, David K. Jackson, Ian Johnston, Dominic Kwiatkowski, Cordelia Langford, John Sillitoe on behalf of the Wellcome Sanger Institute COVID-19 Surveillance Team |
| EPI_ISL_3380290 | BIOLITTORAL BIOGROUP PLATEAU TECH | CNR Virus des Infections Respiratoires - France SUD | Antonin Bal, Gregory Destras, Gwendolyne Burfin, Hadrien Regue, Quentin Semanas, Martine Valette, Bruno Lina, Laurence Josset |
| EPI_ISL_3380291, EPI_ISL_3380292 | Lighthouse Lab in Alderley Park | Wellcome Sanger Institute for the COVID-19 Genomics UK (COG-UK) Consortium | Jacquelyn Wynn, Mairead Hyland, The Lighthouse Lab in Alderley Park and Alex Alderton, Roberto Amato, Jeffrey Barrett, Sonia Goncalves, Ewan Harrison, David K. Jackson, Ian Johnston, Dominic Kwiatkowski, Cordelia Langford, John Sillitoe on behalf of the Wellcome Sanger Institute COVID-19 Surveillance Team |
| EPI_ISL_3380293 | Lighthouse Lab in Milton Keynes | Wellcome Sanger Institute for the COVID-19 Genomics UK (COG-UK) Consortium | The Lighthouse Lab in Milton Keynes and Alex Alderton, Roberto Amato, Jeffrey Barrett, Sonia Goncalves, Ewan Harrison, David K. Jackson, Ian Johnston, Dominic Kwiatkowski, Cordelia Langford, John Sillitoe on behalf of the Wellcome Sanger Institute COVID-19 Surveillance Team |
| EPI_ISL_3380294 | BIOLITTORAL BIOGROUP PLATEAU TECH | CNR Virus des Infections Respiratoires - France SUD | Antonin Bal, Gregory Destras, Gwendolyne Burfin, Hadrien Regue, Quentin Semanas, Martine Valette, Bruno Lina, Laurence Josset |

| | | | |
|---|---|---|---|
| EPI_ISL_3380295, EPI_ISL_3380296, EPI_ISL_3380297 | Lighthouse Lab in Milton Keynes | Wellcome Sanger Institute for the COVID-19 Genomics UK (COG-UK) Consortium | The Lighthouse Lab in Milton Keynes and Alex Alderton, Roberto Amato, Jeffrey Barrett, Sonia Goncalves, Ewan Harrison, David K. Jackson, Ian Johnston, Dominic Kwiatkowski, Cordelia Langford, John Sillitoe on behalf of the Wellcome Sanger Institute COVID-19 Surveillance Team |
| EPI_ISL_3380298 | BIOLITTORAL BIOGROUP PLATEAU TECH | CNR Virus des Infections Respiratoires - France SUD | Antonin Bal, Gregory Destras, Gwendolyne Burfin, Hadrien Regue, Quentin Semanas, Martine Valette, Bruno Lina, Laurence Josset |
| EPI_ISL_3380299, EPI_ISL_3380300 | Lighthouse Lab in Milton Keynes | Wellcome Sanger Institute for the COVID-19 Genomics UK (COG-UK) Consortium | The Lighthouse Lab in Milton Keynes and Alex Alderton, Roberto Amato, Jeffrey Barrett, Sonia Goncalves, Ewan Harrison, David K. Jackson, Ian Johnston, Dominic Kwiatkowski, Cordelia Langford, John Sillitoe on behalf of the Wellcome Sanger Institute COVID-19 Surveillance Team |
| EPI_ISL_3380302 | BIOLITTORAL BIOGROUP PLATEAU TECH | CNR Virus des Infections Respiratoires - France SUD | Antonin Bal, Gregory Destras, Gwendolyne Burfin, Hadrien Regue, Quentin Semanas, Martine Valette, Bruno Lina, Laurence Josset |
| EPI_ISL_3380303, EPI_ISL_3380304, EPI_ISL_3380305, EPI_ISL_3380307 | Lighthouse Lab in Milton Keynes | Wellcome Sanger Institute for the COVID-19 Genomics UK (COG-UK) Consortium | The Lighthouse Lab in Milton Keynes and Alex Alderton, Roberto Amato, Jeffrey Barrett, Sonia Goncalves, Ewan Harrison, David K. Jackson, Ian Johnston, Dominic Kwiatkowski, Cordelia Langford, John Sillitoe on behalf of the Wellcome Sanger Institute COVID-19 Surveillance Team |
| EPI_ISL_3380308, EPI_ISL_3380309 | Lighthouse Lab in Alderley Park | Wellcome Sanger Institute for the COVID-19 Genomics UK (COG-UK) Consortium | Jacquelyn Wynn, Mairead Hyland, The Lighthouse Lab in Alderley Park and Alex Alderton, Roberto Amato, Jeffrey Barrett, Sonia Goncalves, Ewan Harrison, David K. Jackson, Ian Johnston, Dominic Kwiatkowski, Cordelia Langford, John Sillitoe on behalf of the Wellcome Sanger Institute COVID-19 Surveillance Team |
| EPI_ISL_3380310 | BIOLITTORAL BIOGROUP PLATEAU TECH | CNR Virus des Infections Respiratoires - France SUD | Antonin Bal, Gregory Destras, Gwendolyne Burfin, Hadrien Regue, Quentin Semanas, Martine Valette, Bruno Lina, Laurence Josset |
| EPI_ISL_3380311, EPI_ISL_3380312 | Lighthouse Lab in Milton Keynes | Wellcome Sanger Institute for the COVID-19 Genomics UK (COG-UK) Consortium | The Lighthouse Lab in Milton Keynes and Alex Alderton, Roberto Amato, Jeffrey Barrett, Sonia Goncalves, Ewan Harrison, David K. Jackson, Ian Johnston, Dominic Kwiatkowski, Cordelia Langford, John Sillitoe on behalf of the Wellcome Sanger Institute COVID-19 Surveillance Team |
| EPI_ISL_3380313 | BIOLITTORAL BIOGROUP PLATEAU TECH | CNR Virus des Infections Respiratoires - France SUD | Antonin Bal, Gregory Destras, Gwendolyne Burfin, Hadrien Regue, Quentin Semanas, Martine Valette, Bruno Lina, Laurence Josset |
| EPI_ISL_3380314, EPI_ISL_3380315 | Lighthouse Lab in Alderley Park | Wellcome Sanger Institute for the COVID-19 Genomics UK (COG-UK) Consortium | Jacquelyn Wynn, Mairead Hyland, The Lighthouse Lab in Alderley Park and Alex Alderton, Roberto Amato, Jeffrey Barrett, Sonia Goncalves, Ewan Harrison, David K. Jackson, Ian Johnston, Dominic Kwiatkowski, Cordelia Langford, John Sillitoe on behalf of the Wellcome Sanger Institute COVID-19 Surveillance Team |
| EPI_ISL_3380316, EPI_ISL_3380318, EPI_ISL_3380319, EPI_ISL_3380320 | Lighthouse Lab in Milton Keynes | Wellcome Sanger Institute for the COVID-19 Genomics UK (COG-UK) Consortium | The Lighthouse Lab in Milton Keynes and Alex Alderton, Roberto Amato, Jeffrey Barrett, Sonia Goncalves, Ewan Harrison, David K. Jackson, Ian Johnston, Dominic Kwiatkowski, Cordelia Langford, John Sillitoe on behalf of the Wellcome Sanger Institute COVID-19 Surveillance Team |
| EPI_ISL_3380321 | BIOLITTORAL BIOGROUP PLATEAU TECH | CNR Virus des Infections Respiratoires - France SUD | Antonin Bal, Gregory Destras, Gwendolyne Burfin, Hadrien Regue, Quentin Semanas, Martine Valette, Bruno Lina, Laurence Josset |
| EPI_ISL_3380322, EPI_ISL_3380323, EPI_ISL_3380324 | Lighthouse Lab in Milton Keynes | Wellcome Sanger Institute for the COVID-19 Genomics UK (COG-UK) Consortium | The Lighthouse Lab in Milton Keynes and Alex Alderton, Roberto Amato, Jeffrey Barrett, Sonia Goncalves, Ewan Harrison, David K. Jackson, Ian Johnston, Dominic Kwiatkowski, Cordelia Langford, John Sillitoe on behalf of the Wellcome Sanger Institute COVID-19 Surveillance Team |
| EPI_ISL_3380325 | BIOLITTORAL BIOGROUP PLATEAU TECH | CNR Virus des Infections Respiratoires - France SUD | Antonin Bal, Gregory Destras, Gwendolyne Burfin, Hadrien Regue, Quentin Semanas, Martine Valette, Bruno Lina, Laurence Josset |
| EPI_ISL_3380326, EPI_ISL_3380327, EPI_ISL_3380328 | Lighthouse Lab in Milton Keynes | Wellcome Sanger Institute for the COVID-19 Genomics UK (COG-UK) Consortium | The Lighthouse Lab in Milton Keynes and Alex Alderton, Roberto Amato, Jeffrey Barrett, Sonia Goncalves, Ewan Harrison, David K. Jackson, Ian Johnston, Dominic Kwiatkowski, Cordelia Langford, John Sillitoe on behalf of the Wellcome Sanger Institute COVID-19 Surveillance Team |
| EPI_ISL_3380329 | Lighthouse Lab in Alderley Park | Wellcome Sanger Institute for the COVID-19 Genomics UK (COG-UK) Consortium | Jacquelyn Wynn, Mairead Hyland, The Lighthouse Lab in Alderley Park and Alex Alderton, Roberto Amato, Jeffrey Barrett, Sonia Goncalves, Ewan Harrison, David K. Jackson, Ian Johnston, Dominic Kwiatkowski, Cordelia Langford, John Sillitoe on behalf of the Wellcome Sanger Institute COVID-19 Surveillance Team |
| EPI_ISL_3380330 | BIOLITTORAL BIOGROUP PLATEAU TECH | CNR Virus des Infections Respiratoires - France SUD | Antonin Bal, Gregory Destras, Gwendolyne Burfin, Hadrien Regue, Quentin Semanas, Martine Valette, Bruno Lina, Laurence Josset |
| EPI_ISL_3380331, EPI_ISL_3380332, EPI_ISL_3380333 | Lighthouse Lab in Milton Keynes | Wellcome Sanger Institute for the COVID-19 Genomics UK (COG-UK) Consortium | The Lighthouse Lab in Milton Keynes and Alex Alderton, Roberto Amato, Jeffrey Barrett, Sonia Goncalves, Ewan Harrison, David K. Jackson, Ian Johnston, Dominic Kwiatkowski, Cordelia Langford, John Sillitoe on behalf of the Wellcome Sanger Institute COVID-19 Surveillance Team |
| EPI_ISL_3380334 | BIOLITTORAL BIOGROUP PLATEAU TECH | CNR Virus des Infections Respiratoires - France SUD | Antonin Bal, Gregory Destras, Gwendolyne Burfin, Hadrien Regue, Quentin Semanas, Martine Valette, Bruno Lina, Laurence Josset |
| EPI_ISL_3380335, EPI_ISL_3380336, EPI_ISL_3380337 | Lighthouse Lab in Milton Keynes | Wellcome Sanger Institute for the COVID-19 Genomics UK (COG-UK) Consortium | The Lighthouse Lab in Milton Keynes and Alex Alderton, Roberto Amato, Jeffrey Barrett, Sonia Goncalves, Ewan Harrison, David K. Jackson, Ian Johnston, Dominic Kwiatkowski, Cordelia Langford, John Sillitoe on behalf of the Wellcome Sanger Institute COVID-19 Surveillance Team |
| EPI_ISL_3380339 | Lighthouse Lab in Alderley Park | Wellcome Sanger Institute for the COVID-19 Genomics UK (COG-UK) Consortium | Jacquelyn Wynn, Mairead Hyland, The Lighthouse Lab in Alderley Park and Alex Alderton, Roberto Amato, Jeffrey Barrett, Sonia Goncalves, Ewan Harrison, David K. Jackson, Ian Johnston, Dominic Kwiatkowski, Cordelia Langford, John Sillitoe on behalf of the Wellcome Sanger Institute COVID-19 Surveillance Team |
| EPI_ISL_3380340 | Lighthouse Lab in Milton Keynes | Wellcome Sanger Institute for the COVID-19 Genomics UK (COG-UK) Consortium | The Lighthouse Lab in Milton Keynes and Alex Alderton, Roberto Amato, Jeffrey Barrett, Sonia Goncalves, Ewan Harrison, David K. Jackson, Ian Johnston, Dominic Kwiatkowski, Cordelia Langford, John Sillitoe on behalf of the Wellcome Sanger Institute COVID-19 Surveillance Team |
| EPI_ISL_3380341 | Lighthouse Lab in Alderley Park | Wellcome Sanger Institute for the COVID-19 Genomics UK (COG-UK) Consortium | Jacquelyn Wynn, Mairead Hyland, The Lighthouse Lab in Alderley Park and Alex Alderton, Roberto Amato, Jeffrey Barrett, Sonia Goncalves, Ewan Harrison, David K. Jackson, Ian Johnston, Dominic Kwiatkowski, Cordelia Langford, John Sillitoe on behalf of the Wellcome Sanger Institute COVID-19 Surveillance Team |
| EPI_ISL_3380342 | Lighthouse Lab in Milton Keynes | Wellcome Sanger Institute for the COVID-19 Genomics UK (COG-UK) Consortium | The Lighthouse Lab in Milton Keynes and Alex Alderton, Roberto Amato, Jeffrey Barrett, Sonia Goncalves, Ewan Harrison, David K. Jackson, Ian Johnston, Dominic Kwiatkowski, Cordelia Langford, John Sillitoe on behalf of the Wellcome Sanger Institute COVID-19 Surveillance Team |
| EPI_ISL_3380343 | BIOLITTORAL BIOGROUP PLATEAU TECH | CNR Virus des Infections Respiratoires - France SUD | Antonin Bal, Gregory Destras, Gwendolyne Burfin, Hadrien Regue, Quentin Semanas, Martine Valette, Bruno Lina, Laurence Josset |
| EPI_ISL_3380344, EPI_ISL_3380345, EPI_ISL_3380346 | Lighthouse Lab in Alderley Park | Wellcome Sanger Institute for the COVID-19 Genomics UK (COG-UK) Consortium | Jacquelyn Wynn, Mairead Hyland, The Lighthouse Lab in Alderley Park and Alex Alderton, Roberto Amato, Jeffrey Barrett, Sonia Goncalves, Ewan Harrison, David K. Jackson, Ian Johnston, Dominic Kwiatkowski, Cordelia Langford, John Sillitoe on behalf of the Wellcome Sanger Institute COVID-19 Surveillance Team |
| EPI_ISL_3380347 | BIOLITTORAL BIOGROUP PLATEAU TECH | CNR Virus des Infections Respiratoires - France SUD | Antonin Bal, Gregory Destras, Gwendolyne Burfin, Hadrien Regue, Quentin Semanas, Martine Valette, Bruno Lina, Laurence Josset |
| EPI_ISL_3380348, EPI_ISL_3380349, EPI_ISL_3380350 | Lighthouse Lab in Milton Keynes | Wellcome Sanger Institute for the COVID-19 Genomics UK (COG-UK) Consortium | The Lighthouse Lab in Milton Keynes and Alex Alderton, Roberto Amato, Jeffrey Barrett, Sonia Goncalves, Ewan Harrison, David K. Jackson, Ian Johnston, Dominic Kwiatkowski, Cordelia Langford, John Sillitoe on behalf of the Wellcome Sanger Institute COVID-19 Surveillance Team |
| EPI_ISL_3380352, EPI_ISL_3380353 | Lighthouse Lab in Alderley Park | Wellcome Sanger Institute for the COVID-19 Genomics UK (COG-UK) Consortium | Jacquelyn Wynn, Mairead Hyland, The Lighthouse Lab in Alderley Park and Alex Alderton, Roberto Amato, Jeffrey Barrett, Sonia Goncalves, Ewan Harrison, David K. Jackson, Ian Johnston, Dominic Kwiatkowski, Cordelia Langford, John Sillitoe on behalf of the Wellcome Sanger Institute COVID-19 Surveillance Team |
| EPI_ISL_3380354, EPI_ISL_3380355 | Lighthouse Lab in Milton Keynes | Wellcome Sanger Institute for the COVID-19 Genomics UK (COG-UK) Consortium | The Lighthouse Lab in Milton Keynes and Alex Alderton, Roberto Amato, Jeffrey Barrett, Sonia Goncalves, Ewan Harrison, David K. Jackson, Ian Johnston, Dominic Kwiatkowski, Cordelia Langford, John Sillitoe on behalf of the Wellcome Sanger Institute COVID-19 Surveillance Team |
| EPI_ISL_3380356 | BIOLITTORAL BIOGROUP PLATEAU TECH | CNR Virus des Infections Respiratoires - France SUD | Antonin Bal, Gregory Destras, Gwendolyne Burfin, Hadrien Regue, Quentin Semanas, Martine Valette, Bruno Lina, Laurence Josset |
| EPI_ISL_3380357, EPI_ISL_3380358, EPI_ISL_3380359 | Lighthouse Lab in Milton Keynes | Wellcome Sanger Institute for the COVID-19 Genomics UK (COG-UK) Consortium | The Lighthouse Lab in Milton Keynes and Alex Alderton, Roberto Amato, Jeffrey Barrett, Sonia Goncalves, Ewan Harrison, David K. Jackson, Ian Johnston, Dominic Kwiatkowski, Cordelia Langford, John Sillitoe on behalf of the Wellcome Sanger Institute COVID-19 Surveillance Team |
| EPI_ISL_3380360 | BIOLITTORAL BIOGROUP PLATEAU TECH | CNR Virus des Infections Respiratoires - France SUD | Antonin Bal, Gregory Destras, Gwendolyne Burfin, Hadrien Regue, Quentin Semanas, Martine Valette, Bruno Lina, Laurence Josset |
| EPI_ISL_3380361 | Lighthouse Lab in Alderley Park | Wellcome Sanger Institute for the COVID-19 Genomics UK (COG-UK) Consortium | Jacquelyn Wynn, Mairead Hyland, The Lighthouse Lab in Alderley Park and Alex Alderton, Roberto Amato, Jeffrey Barrett, Sonia Goncalves, Ewan Harrison, David K. Jackson, Ian Johnston, Dominic Kwiatkowski, Cordelia Langford, John Sillitoe on behalf of the Wellcome Sanger Institute COVID-19 Surveillance Team |
| EPI_ISL_3380362, EPI_ISL_3380363, EPI_ISL_3380365, EPI_ISL_3380366 | Lighthouse Lab in Milton Keynes | Wellcome Sanger Institute for the COVID-19 Genomics UK (COG-UK) Consortium | The Lighthouse Lab in Milton Keynes and Alex Alderton, Roberto Amato, Jeffrey Barrett, Sonia Goncalves, Ewan Harrison, David K. Jackson, Ian Johnston, Dominic Kwiatkowski, Cordelia Langford, John Sillitoe on behalf of the Wellcome Sanger Institute COVID-19 Surveillance Team |
| EPI_ISL_3380368 | Lighthouse Lab in Alderley Park | Wellcome Sanger Institute for the COVID-19 Genomics UK (COG-UK) Consortium | Jacquelyn Wynn, Mairead Hyland, The Lighthouse Lab in Alderley Park and Alex Alderton, Roberto Amato, Jeffrey Barrett, Sonia Goncalves, Ewan Harrison, David K. Jackson, Ian Johnston, Dominic Kwiatkowski, Cordelia Langford, John Sillitoe on behalf of the Wellcome Sanger Institute COVID-19 Surveillance Team |
| EPI_ISL_3380369, EPI_ISL_3380370 | Lighthouse Lab in Milton Keynes | Wellcome Sanger Institute for the COVID-19 Genomics UK (COG-UK) Consortium | The Lighthouse Lab in Milton Keynes and Alex Alderton, Roberto Amato, Jeffrey Barrett, Sonia Goncalves, Ewan Harrison, David K. Jackson, Ian Johnston, Dominic Kwiatkowski, Cordelia Langford, John Sillitoe on behalf of the Wellcome Sanger Institute COVID-19 Surveillance Team |
| EPI_ISL_3380371 | BIOLITTORAL BIOGROUP PLATEAU TECH | CNR Virus des Infections Respiratoires - France SUD | Antonin Bal, Gregory Destras, Gwendolyne Burfin, Hadrien Regue, Quentin Semanas, Martine Valette, Bruno Lina, Laurence Josset |
| EPI_ISL_3380372, EPI_ISL_3380373, EPI_ISL_3380375, EPI_ISL_3380376, EPI_ISL_3380377, EPI_ISL_3380378, EPI_ISL_3380380, EPI_ISL_3380381 | Lighthouse Lab in Milton Keynes | Wellcome Sanger Institute for the COVID-19 Genomics UK (COG-UK) Consortium | The Lighthouse Lab in Milton Keynes and Alex Alderton, Roberto Amato, Jeffrey Barrett, Sonia Goncalves, Ewan Harrison, David K. Jackson, Ian Johnston, Dominic Kwiatkowski, Cordelia Langford, John Sillitoe on behalf of the Wellcome Sanger Institute COVID-19 Surveillance Team |
| EPI_ISL_3380382 | BIOLITTORAL BIOGROUP PLATEAU TECH | CNR Virus des Infections Respiratoires - France SUD | Antonin Bal, Gregory Destras, Gwendolyne Burfin, Hadrien Regue, Quentin Semanas, Martine Valette, Bruno Lina, Laurence Josset |
| EPI_ISL_3380383, EPI_ISL_3380384, EPI_ISL_3380385, EPI_ISL_3380387, EPI_ISL_3380388, EPI_ISL_3380389 | Lighthouse Lab in Milton Keynes | Wellcome Sanger Institute for the COVID-19 Genomics UK (COG-UK) Consortium | The Lighthouse Lab in Milton Keynes and Alex Alderton, Roberto Amato, Jeffrey Barrett, Sonia Goncalves, Ewan Harrison, David K. Jackson, Ian Johnston, Dominic Kwiatkowski, Cordelia Langford, John Sillitoe on behalf of the Wellcome Sanger Institute COVID-19 Surveillance Team |

| | | | |
|---|---|---|---|
| EPI_ISL_3380390 | BIOLITTORAL BIOGROUP PLATEAU TECH | CNR Virus des Infections Respiratoires - France SUD | Antonin Bal, Gregory Destras, Gwendolyne Burfin, Hadrien Regue, Quentin Semanas, Martine Valette, Bruno Lina, Laurence Josset |
| EPI_ISL_3380391, EPI_ISL_3380392, EPI_ISL_3380393, EPI_ISL_3380395, EPI_ISL_3380396, EPI_ISL_3380397 | Lighthouse Lab in Milton Keynes | Wellcome Sanger Institute for the COVID-19 Genomics UK (COG-UK) Consortium | The Lighthouse Lab in Milton Keynes and Alex Alderton, Roberto Amato, Jeffrey Barrett, Sonia Goncalves, Ewan Harrison, David K. Jackson, Ian Johnston, Dominic Kwiatkowski, Cordelia Langford, John Sillitoe on behalf of the Wellcome Sanger Institute COVID-19 Surveillance Team |
| EPI_ISL_3380398 | BIOLITTORAL BIOGROUP PLATEAU TECH | CNR Virus des Infections Respiratoires - France SUD | Antonin Bal, Gregory Destras, Gwendolyne Burfin, Hadrien Regue, Quentin Semanas, Martine Valette, Bruno Lina, Laurence Josset |
| EPI_ISL_3380399, EPI_ISL_3380400, EPI_ISL_3380401, EPI_ISL_3380403, EPI_ISL_3380404, EPI_ISL_3380405, EPI_ISL_3380407, EPI_ISL_3380408, EPI_ISL_3380409, EPI_ISL_3380411, EPI_ISL_3380412, EPI_ISL_3380414, EPI_ISL_3380415, EPI_ISL_3380416 | | | |
| see above | Lighthouse Lab in Milton Keynes | Wellcome Sanger Institute for the COVID-19 Genomics UK (COG-UK) Consortium | The Lighthouse Lab in Milton Keynes and Alex Alderton, Roberto Amato, Jeffrey Barrett, Sonia Goncalves, Ewan Harrison, David K. Jackson, Ian Johnston, Dominic Kwiatkowski, Cordelia Langford, John Sillitoe on behalf of the Wellcome Sanger Institute COVID-19 Surveillance Team |
| EPI_ISL_3380417 | CERBAILLIANCE LA REUNION | CNR Virus des Infections Respiratoires - France SUD | Antonin Bal, Gregory Destras, Gwendolyne Burfin, Hadrien Regue, Quentin Semanas, Martine Valette, Bruno Lina, Laurence Josset |
| EPI_ISL_3380418, EPI_ISL_3380419 | Lighthouse Lab in Milton Keynes | Wellcome Sanger Institute for the COVID-19 Genomics UK (COG-UK) Consortium | The Lighthouse Lab in Milton Keynes and Alex Alderton, Roberto Amato, Jeffrey Barrett, Sonia Goncalves, Ewan Harrison, David K. Jackson, Ian Johnston, Dominic Kwiatkowski, Cordelia Langford, John Sillitoe on behalf of the Wellcome Sanger Institute COVID-19 Surveillance Team |
| EPI_ISL_3380420 | Lighthouse Lab in Alderley Park | Wellcome Sanger Institute for the COVID-19 Genomics UK (COG-UK) Consortium | Jacquelyn Wynn, Mairead Hyland, The Lighthouse Lab in Alderley Park and Alex Alderton, Roberto Amato, Jeffrey Barrett, Sonia Goncalves, Ewan Harrison, David K. Jackson, Ian Johnston, Dominic Kwiatkowski, Cordelia Langford, John Sillitoe on behalf of the Wellcome Sanger Institute COVID-19 Surveillance Team |
| EPI_ISL_3380421 | CERBAILLIANCE LA REUNION | CNR Virus des Infections Respiratoires - France SUD | Antonin Bal, Gregory Destras, Gwendolyne Burfin, Hadrien Regue, Quentin Semanas, Martine Valette, Bruno Lina, Laurence Josset |
| EPI_ISL_3380422, EPI_ISL_3380423 | Lighthouse Lab in Alderley Park | Wellcome Sanger Institute for the COVID-19 Genomics UK (COG-UK) Consortium | Jacquelyn Wynn, Mairead Hyland, The Lighthouse Lab in Alderley Park and Alex Alderton, Roberto Amato, Jeffrey Barrett, Sonia Goncalves, Ewan Harrison, David K. Jackson, Ian Johnston, Dominic Kwiatkowski, Cordelia Langford, John Sillitoe on behalf of the Wellcome Sanger Institute COVID-19 Surveillance Team |
| EPI_ISL_3380424, EPI_ISL_3380426 | Lighthouse Lab in Milton Keynes | Wellcome Sanger Institute for the COVID-19 Genomics UK (COG-UK) Consortium | The Lighthouse Lab in Milton Keynes and Alex Alderton, Roberto Amato, Jeffrey Barrett, Sonia Goncalves, Ewan Harrison, David K. Jackson, Ian Johnston, Dominic Kwiatkowski, Cordelia Langford, John Sillitoe on behalf of the Wellcome Sanger Institute COVID-19 Surveillance Team |
| EPI_ISL_3380427 | Lighthouse Lab in Alderley Park | Wellcome Sanger Institute for the COVID-19 Genomics UK (COG-UK) Consortium | Jacquelyn Wynn, Mairead Hyland, The Lighthouse Lab in Alderley Park and Alex Alderton, Roberto Amato, Jeffrey Barrett, Sonia Goncalves, Ewan Harrison, David K. Jackson, Ian Johnston, Dominic Kwiatkowski, Cordelia Langford, John Sillitoe on behalf of the Wellcome Sanger Institute COVID-19 Surveillance Team |
| EPI_ISL_3380428, EPI_ISL_3380430, EPI_ISL_3380431, EPI_ISL_3380432 | Lighthouse Lab in Milton Keynes | Wellcome Sanger Institute for the COVID-19 Genomics UK (COG-UK) Consortium | The Lighthouse Lab in Milton Keynes and Alex Alderton, Roberto Amato, Jeffrey Barrett, Sonia Goncalves, Ewan Harrison, David K. Jackson, Ian Johnston, Dominic Kwiatkowski, Cordelia Langford, John Sillitoe on behalf of the Wellcome Sanger Institute COVID-19 Surveillance Team |
| EPI_ISL_3380433 | CERBAILLIANCE LA REUNION | CNR Virus des Infections Respiratoires - France SUD | Antonin Bal, Gregory Destras, Gwendolyne Burfin, Hadrien Regue, Quentin Semanas, Martine Valette, Bruno Lina, Laurence Josset |
| EPI_ISL_3380434 | Lighthouse Lab in Milton Keynes | Wellcome Sanger Institute for the COVID-19 Genomics UK (COG-UK) Consortium | The Lighthouse Lab in Milton Keynes and Alex Alderton, Roberto Amato, Jeffrey Barrett, Sonia Goncalves, Ewan Harrison, David K. Jackson, Ian Johnston, Dominic Kwiatkowski, Cordelia Langford, John Sillitoe on behalf of the Wellcome Sanger Institute COVID-19 Surveillance Team |
| EPI_ISL_3380435 | Lighthouse Lab in Alderley Park | Wellcome Sanger Institute for the COVID-19 Genomics UK (COG-UK) Consortium | Jacquelyn Wynn, Mairead Hyland, The Lighthouse Lab in Alderley Park and Alex Alderton, Roberto Amato, Jeffrey Barrett, Sonia Goncalves, Ewan Harrison, David K. Jackson, Ian Johnston, Dominic Kwiatkowski, Cordelia Langford, John Sillitoe on behalf of the Wellcome Sanger Institute COVID-19 Surveillance Team |
| EPI_ISL_3380436, EPI_ISL_3380437 | Lighthouse Lab in Milton Keynes | Wellcome Sanger Institute for the COVID-19 Genomics UK (COG-UK) Consortium | The Lighthouse Lab in Milton Keynes and Alex Alderton, Roberto Amato, Jeffrey Barrett, Sonia Goncalves, Ewan Harrison, David K. Jackson, Ian Johnston, Dominic Kwiatkowski, Cordelia Langford, John Sillitoe on behalf of the Wellcome Sanger Institute COVID-19 Surveillance Team |
| EPI_ISL_3380438 | CERBAILLIANCE LA REUNION | CNR Virus des Infections Respiratoires - France SUD | Antonin Bal, Gregory Destras, Gwendolyne Burfin, Hadrien Regue, Quentin Semanas, Martine Valette, Bruno Lina, Laurence Josset |
| EPI_ISL_3380439, EPI_ISL_3380440 | Lighthouse Lab in Milton Keynes | Wellcome Sanger Institute for the COVID-19 Genomics UK (COG-UK) Consortium | The Lighthouse Lab in Milton Keynes and Alex Alderton, Roberto Amato, Jeffrey Barrett, Sonia Goncalves, Ewan Harrison, David K. Jackson, Ian Johnston, Dominic Kwiatkowski, Cordelia Langford, John Sillitoe on behalf of the Wellcome Sanger Institute COVID-19 Surveillance Team |
| EPI_ISL_3380441 | Lighthouse Lab in Alderley Park | Wellcome Sanger Institute for the COVID-19 Genomics UK (COG-UK) Consortium | Jacquelyn Wynn, Mairead Hyland, The Lighthouse Lab in Alderley Park and Alex Alderton, Roberto Amato, Jeffrey Barrett, Sonia Goncalves, Ewan Harrison, David K. Jackson, Ian Johnston, Dominic Kwiatkowski, Cordelia Langford, John Sillitoe on behalf of the Wellcome Sanger Institute COVID-19 Surveillance Team |
| EPI_ISL_3380442 | LABORATOIRE BIOESTEREL | CNR Virus des Infections Respiratoires - France SUD | Antonin Bal, Gregory Destras, Gwendolyne Burfin, Hadrien Regue, Quentin Semanas, Martine Valette, Bruno Lina, Laurence Josset |
| EPI_ISL_3380443, EPI_ISL_3380444 | Lighthouse Lab in Milton Keynes | Wellcome Sanger Institute for the COVID-19 Genomics UK (COG-UK) Consortium | The Lighthouse Lab in Milton Keynes and Alex Alderton, Roberto Amato, Jeffrey Barrett, Sonia Goncalves, Ewan Harrison, David K. Jackson, Ian Johnston, Dominic Kwiatkowski, Cordelia Langford, John Sillitoe on behalf of the Wellcome Sanger Institute COVID-19 Surveillance Team |
| EPI_ISL_3380445 | Lighthouse Lab in Alderley Park | Wellcome Sanger Institute for the COVID-19 Genomics UK (COG-UK) Consortium | Jacquelyn Wynn, Mairead Hyland, The Lighthouse Lab in Alderley Park and Alex Alderton, Roberto Amato, Jeffrey Barrett, Sonia Goncalves, Ewan Harrison, David K. Jackson, Ian Johnston, Dominic Kwiatkowski, Cordelia Langford, John Sillitoe on behalf of the Wellcome Sanger Institute COVID-19 Surveillance Team |
| EPI_ISL_3380446 | LABORATOIRE BIOESTEREL | CNR Virus des Infections Respiratoires - France SUD | Antonin Bal, Gregory Destras, Gwendolyne Burfin, Hadrien Regue, Quentin Semanas, Martine Valette, Bruno Lina, Laurence Josset |
| EPI_ISL_3380447, EPI_ISL_3380448, EPI_ISL_3380449 | Lighthouse Lab in Milton Keynes | Wellcome Sanger Institute for the COVID-19 Genomics UK (COG-UK) Consortium | The Lighthouse Lab in Milton Keynes and Alex Alderton, Roberto Amato, Jeffrey Barrett, Sonia Goncalves, Ewan Harrison, David K. Jackson, Ian Johnston, Dominic Kwiatkowski, Cordelia Langford, John Sillitoe on behalf of the Wellcome Sanger Institute COVID-19 Surveillance Team |
| EPI_ISL_3380450 | LABORATOIRE BIOESTEREL | CNR Virus des Infections Respiratoires - France SUD | Antonin Bal, Gregory Destras, Gwendolyne Burfin, Hadrien Regue, Quentin Semanas, Martine Valette, Bruno Lina, Laurence Josset |
| EPI_ISL_3380451, EPI_ISL_3380452 | Lighthouse Lab in Alderley Park | Wellcome Sanger Institute for the COVID-19 Genomics UK (COG-UK) Consortium | Jacquelyn Wynn, Mairead Hyland, The Lighthouse Lab in Alderley Park and Alex Alderton, Roberto Amato, Jeffrey Barrett, Sonia Goncalves, Ewan Harrison, David K. Jackson, Ian Johnston, Dominic Kwiatkowski, Cordelia Langford, John Sillitoe on behalf of the Wellcome Sanger Institute COVID-19 Surveillance Team |
| EPI_ISL_3380453 | Lighthouse Lab in Milton Keynes | Wellcome Sanger Institute for the COVID-19 Genomics UK (COG-UK) Consortium | The Lighthouse Lab in Milton Keynes and Alex Alderton, Roberto Amato, Jeffrey Barrett, Sonia Goncalves, Ewan Harrison, David K. Jackson, Ian Johnston, Dominic Kwiatkowski, Cordelia Langford, John Sillitoe on behalf of the Wellcome Sanger Institute COVID-19 Surveillance Team |
| EPI_ISL_3380454 | LABORATOIRE BIOESTEREL | CNR Virus des Infections Respiratoires - France SUD | Antonin Bal, Gregory Destras, Gwendolyne Burfin, Hadrien Regue, Quentin Semanas, Martine Valette, Bruno Lina, Laurence Josset |
| EPI_ISL_3380455, EPI_ISL_3380456, EPI_ISL_3380457 | Lighthouse Lab in Milton Keynes | Wellcome Sanger Institute for the COVID-19 Genomics UK (COG-UK) Consortium | The Lighthouse Lab in Milton Keynes and Alex Alderton, Roberto Amato, Jeffrey Barrett, Sonia Goncalves, Ewan Harrison, David K. Jackson, Ian Johnston, Dominic Kwiatkowski, Cordelia Langford, John Sillitoe on behalf of the Wellcome Sanger Institute COVID-19 Surveillance Team |
| EPI_ISL_3380458 | LABORATOIRE BIOESTEREL | CNR Virus des Infections Respiratoires - France SUD | Antonin Bal, Gregory Destras, Gwendolyne Burfin, Hadrien Regue, Quentin Semanas, Martine Valette, Bruno Lina, Laurence Josset |
| EPI_ISL_3380459 | Lighthouse Lab in Alderley Park | Wellcome Sanger Institute for the COVID-19 Genomics UK (COG-UK) Consortium | Jacquelyn Wynn, Mairead Hyland, The Lighthouse Lab in Alderley Park and Alex Alderton, Roberto Amato, Jeffrey Barrett, Sonia Goncalves, Ewan Harrison, David K. Jackson, Ian Johnston, Dominic Kwiatkowski, Cordelia Langford, John Sillitoe on behalf of the Wellcome Sanger Institute COVID-19 Surveillance Team |
| EPI_ISL_3380460 | Lighthouse Lab in Milton Keynes | Wellcome Sanger Institute for the COVID-19 Genomics UK (COG-UK) Consortium | The Lighthouse Lab in Milton Keynes and Alex Alderton, Roberto Amato, Jeffrey Barrett, Sonia Goncalves, Ewan Harrison, David K. Jackson, Ian Johnston, Dominic Kwiatkowski, Cordelia Langford, John Sillitoe on behalf of the Wellcome Sanger Institute COVID-19 Surveillance Team |
| EPI_ISL_3380461 | Lighthouse Lab in Alderley Park | Wellcome Sanger Institute for the COVID-19 Genomics UK (COG-UK) Consortium | Jacquelyn Wynn, Mairead Hyland, The Lighthouse Lab in Alderley Park and Alex Alderton, Roberto Amato, Jeffrey Barrett, Sonia Goncalves, Ewan Harrison, David K. Jackson, Ian Johnston, Dominic Kwiatkowski, Cordelia Langford, John Sillitoe on behalf of the Wellcome Sanger Institute COVID-19 Surveillance Team |
| EPI_ISL_3380462 | LABORATOIRE BIOESTEREL | CNR Virus des Infections Respiratoires - France SUD | Antonin Bal, Gregory Destras, Gwendolyne Burfin, Hadrien Regue, Quentin Semanas, Martine Valette, Bruno Lina, Laurence Josset |
| EPI_ISL_3380463, EPI_ISL_3380464 | Lighthouse Lab in Milton Keynes | Wellcome Sanger Institute for the COVID-19 Genomics UK (COG-UK) Consortium | The Lighthouse Lab in Milton Keynes and Alex Alderton, Roberto Amato, Jeffrey Barrett, Sonia Goncalves, Ewan Harrison, David K. Jackson, Ian Johnston, Dominic Kwiatkowski, Cordelia Langford, John Sillitoe on behalf of the Wellcome Sanger Institute COVID-19 Surveillance Team |
| EPI_ISL_3380465 | Lighthouse Lab in Alderley Park | Wellcome Sanger Institute for the COVID-19 Genomics UK (COG-UK) Consortium | Jacquelyn Wynn, Mairead Hyland, The Lighthouse Lab in Alderley Park and Alex Alderton, Roberto Amato, Jeffrey Barrett, Sonia Goncalves, Ewan Harrison, David K. Jackson, Ian Johnston, Dominic Kwiatkowski, Cordelia Langford, John Sillitoe on behalf of the Wellcome Sanger Institute COVID-19 Surveillance Team |
| EPI_ISL_3380466 | LABORATOIRE BIOESTEREL | CNR Virus des Infections Respiratoires - France SUD | Antonin Bal, Gregory Destras, Gwendolyne Burfin, Hadrien Regue, Quentin Semanas, Martine Valette, Bruno Lina, Laurence Josset |
| EPI_ISL_3380467, EPI_ISL_3380468, EPI_ISL_3380469 | Lighthouse Lab in Milton Keynes | Wellcome Sanger Institute for the COVID-19 Genomics UK (COG-UK) Consortium | The Lighthouse Lab in Milton Keynes and Alex Alderton, Roberto Amato, Jeffrey Barrett, Sonia Goncalves, Ewan Harrison, David K. Jackson, Ian Johnston, Dominic Kwiatkowski, Cordelia Langford, John Sillitoe on behalf of the Wellcome Sanger Institute COVID-19 Surveillance Team |
| EPI_ISL_3380470 | LABORATOIRE BIOESTEREL | CNR Virus des Infections Respiratoires - France SUD | Antonin Bal, Gregory Destras, Gwendolyne Burfin, Hadrien Regue, Quentin Semanas, Martine Valette, Bruno Lina, Laurence Josset |
| EPI_ISL_3380471, EPI_ISL_3380472 | Lighthouse Lab in Milton Keynes | Wellcome Sanger Institute for the COVID-19 Genomics UK (COG-UK) Consortium | The Lighthouse Lab in Milton Keynes and Alex Alderton, Roberto Amato, Jeffrey Barrett, Sonia Goncalves, Ewan Harrison, David K. Jackson, Ian Johnston, Dominic Kwiatkowski, Cordelia Langford, John Sillitoe on behalf of the Wellcome Sanger Institute COVID-19 Surveillance Team |
| EPI_ISL_3380473 | Lighthouse Lab in Alderley Park | Wellcome Sanger Institute for the COVID-19 Genomics UK (COG-UK) Consortium | Jacquelyn Wynn, Mairead Hyland, The Lighthouse Lab in Alderley Park and Alex Alderton, Roberto Amato, Jeffrey Barrett, Sonia Goncalves, Ewan Harrison, David K. Jackson, Ian Johnston, Dominic Kwiatkowski, Cordelia Langford, John Sillitoe on behalf of the Wellcome Sanger Institute COVID-19 Surveillance Team |
| EPI_ISL_3380474 | CERBAILLIANCE LA REUNION | CNR Virus des Infections Respiratoires - France SUD | Antonin Bal, Gregory Destras, Gwendolyne Burfin, Hadrien Regue, Quentin Semanas, Martine Valette, Bruno Lina, Laurence Josset |
| EPI_ISL_3380475, EPI_ISL_3380476, EPI_ISL_3380477 | Lighthouse Lab in Milton Keynes | Wellcome Sanger Institute for the COVID-19 Genomics UK (COG-UK) Consortium | The Lighthouse Lab in Milton Keynes and Alex Alderton, Roberto Amato, Jeffrey Barrett, Sonia Goncalves, Ewan Harrison, David K. Jackson, Ian Johnston, Dominic Kwiatkowski, Cordelia Langford, John Sillitoe on behalf of the Wellcome Sanger Institute COVID-19 Surveillance Team |
| EPI_ISL_3380478 | LABORATOIRE BIOESTEREL | CNR Virus des Infections Respiratoires - France SUD | Antonin Bal, Gregory Destras, Gwendolyne Burfin, Hadrien Regue, Quentin Semanas, Martine Valette, Bruno Lina, Laurence Josset |
| EPI_ISL_3380479, EPI_ISL_3380480 | Lighthouse Lab in Milton Keynes | Wellcome Sanger Institute for the COVID-19 Genomics UK (COG-UK) Consortium | The Lighthouse Lab in Milton Keynes and Alex Alderton, Roberto Amato, Jeffrey Barrett, Sonia Goncalves, Ewan Harrison, David K. Jackson, Ian Johnston, Dominic Kwiatkowski, Cordelia Langford, John Sillitoe on behalf of the Wellcome Sanger Institute COVID-19 Surveillance Team |
| EPI_ISL_3380481 | LABORATOIRE BIOESTEREL | CNR Virus des Infections Respiratoires - France SUD | Antonin Bal, Gregory Destras, Gwendolyne Burfin, Hadrien Regue, Quentin Semanas, Martine Valette, Bruno Lina, Laurence Josset |
| EPI_ISL_3380482 | Lighthouse Lab in Alderley Park | Wellcome Sanger Institute for the COVID-19 Genomics UK (COG-UK) Consortium | Jacquelyn Wynn, Mairead Hyland, The Lighthouse Lab in Alderley Park and Alex Alderton, Roberto Amato, Jeffrey Barrett, Sonia Goncalves, Ewan Harrison, David K. Jackson, Ian Johnston, Dominic Kwiatkowski, Cordelia Langford, John Sillitoe on behalf of the Wellcome Sanger Institute COVID-19 Surveillance Team |

| | | | |
|---|---|---|---|
| EPI_ISL_3380483, EPI_ISL_3380484 | Lighthouse Lab in Milton Keynes | Wellcome Sanger Institute for the COVID-19 Genomics UK (COG-UK) Consortium | The Lighthouse Lab in Milton Keynes and Alex Alderton, Roberto Amato, Jeffrey Barrett, Sonia Goncalves, Ewan Harrison, David K. Jackson, Ian Johnston, Dominic Kwiatkowski, Cordelia Langford, John Sillitoe on behalf of the Wellcome Sanger Institute COVID-19 Surveillance Team |
| EPI_ISL_3380485 | LABORATOIRE BIOESTEREL | CNR Virus des Infections Respiratoires - France SUD | Antonin Bal, Gregory Destras, Gwendolyne Burfin, Hadrien Regue, Quentin Semanas, Martine Valette, Bruno Lina, Laurence Josset |
| EPI_ISL_3380486 | Lighthouse Lab in Milton Keynes | Wellcome Sanger Institute for the COVID-19 Genomics UK (COG-UK) Consortium | The Lighthouse Lab in Milton Keynes and Alex Alderton, Roberto Amato, Jeffrey Barrett, Sonia Goncalves, Ewan Harrison, David K. Jackson, Ian Johnston, Dominic Kwiatkowski, Cordelia Langford, John Sillitoe on behalf of the Wellcome Sanger Institute COVID-19 Surveillance Team |
| EPI_ISL_3380487 | Lighthouse Lab in Alderley Park | Wellcome Sanger Institute for the COVID-19 Genomics UK (COG-UK) Consortium | Jacquelyn Wynn, Mairead Hyland, The Lighthouse Lab in Alderley Park and Alex Alderton, Roberto Amato, Jeffrey Barrett, Sonia Goncalves, Ewan Harrison, David K. Jackson, Ian Johnston, Dominic Kwiatkowski, Cordelia Langford, John Sillitoe on behalf of the Wellcome Sanger Institute COVID-19 Surveillance Team |
| EPI_ISL_3380488 | Lighthouse Lab in Milton Keynes | Wellcome Sanger Institute for the COVID-19 Genomics UK (COG-UK) Consortium | The Lighthouse Lab in Milton Keynes and Alex Alderton, Roberto Amato, Jeffrey Barrett, Sonia Goncalves, Ewan Harrison, David K. Jackson, Ian Johnston, Dominic Kwiatkowski, Cordelia Langford, John Sillitoe on behalf of the Wellcome Sanger Institute COVID-19 Surveillance Team |
| EPI_ISL_3380490, EPI_ISL_3380491 | Lighthouse Lab in Alderley Park | Wellcome Sanger Institute for the COVID-19 Genomics UK (COG-UK) Consortium | Jacquelyn Wynn, Mairead Hyland, The Lighthouse Lab in Alderley Park and Alex Alderton, Roberto Amato, Jeffrey Barrett, Sonia Goncalves, Ewan Harrison, David K. Jackson, Ian Johnston, Dominic Kwiatkowski, Cordelia Langford, John Sillitoe on behalf of the Wellcome Sanger Institute COVID-19 Surveillance Team |
| EPI_ISL_3380492, EPI_ISL_3380493 | Lighthouse Lab in Milton Keynes | Wellcome Sanger Institute for the COVID-19 Genomics UK (COG-UK) Consortium | The Lighthouse Lab in Milton Keynes and Alex Alderton, Roberto Amato, Jeffrey Barrett, Sonia Goncalves, Ewan Harrison, David K. Jackson, Ian Johnston, Dominic Kwiatkowski, Cordelia Langford, John Sillitoe on behalf of the Wellcome Sanger Institute COVID-19 Surveillance Team |
| EPI_ISL_3380494 | LABORATOIRE BIOESTEREL | CNR Virus des Infections Respiratoires - France SUD | Antonin Bal, Gregory Destras, Gwendolyne Burfin, Hadrien Regue, Quentin Semanas, Martine Valette, Bruno Lina, Laurence Josset |
| EPI_ISL_3380495 | Lighthouse Lab in Milton Keynes | Wellcome Sanger Institute for the COVID-19 Genomics UK (COG-UK) Consortium | The Lighthouse Lab in Milton Keynes and Alex Alderton, Roberto Amato, Jeffrey Barrett, Sonia Goncalves, Ewan Harrison, David K. Jackson, Ian Johnston, Dominic Kwiatkowski, Cordelia Langford, John Sillitoe on behalf of the Wellcome Sanger Institute COVID-19 Surveillance Team |
| EPI_ISL_3380496 | Lighthouse Lab in Alderley Park | Wellcome Sanger Institute for the COVID-19 Genomics UK (COG-UK) Consortium | Jacquelyn Wynn, Mairead Hyland, The Lighthouse Lab in Alderley Park and Alex Alderton, Roberto Amato, Jeffrey Barrett, Sonia Goncalves, Ewan Harrison, David K. Jackson, Ian Johnston, Dominic Kwiatkowski, Cordelia Langford, John Sillitoe on behalf of the Wellcome Sanger Institute COVID-19 Surveillance Team |
| EPI_ISL_3380497 | Lighthouse Lab in Milton Keynes | Wellcome Sanger Institute for the COVID-19 Genomics UK (COG-UK) Consortium | The Lighthouse Lab in Milton Keynes and Alex Alderton, Roberto Amato, Jeffrey Barrett, Sonia Goncalves, Ewan Harrison, David K. Jackson, Ian Johnston, Dominic Kwiatkowski, Cordelia Langford, John Sillitoe on behalf of the Wellcome Sanger Institute COVID-19 Surveillance Team |
| EPI_ISL_3380498 | LABORATOIRE BIOESTEREL | CNR Virus des Infections Respiratoires - France SUD | Antonin Bal, Gregory Destras, Gwendolyne Burfin, Hadrien Regue, Quentin Semanas, Martine Valette, Bruno Lina, Laurence Josset |
| EPI_ISL_3380499, EPI_ISL_3380500, EPI_ISL_3380501, EPI_ISL_3380503, EPI_ISL_3380504 | Lighthouse Lab in Milton Keynes | Wellcome Sanger Institute for the COVID-19 Genomics UK (COG-UK) Consortium | The Lighthouse Lab in Milton Keynes and Alex Alderton, Roberto Amato, Jeffrey Barrett, Sonia Goncalves, Ewan Harrison, David K. Jackson, Ian Johnston, Dominic Kwiatkowski, Cordelia Langford, John Sillitoe on behalf of the Wellcome Sanger Institute COVID-19 Surveillance Team |
| EPI_ISL_3380505 | CERBAILLIANCE LA REUNION | CNR Virus des Infections Respiratoires - France SUD | Antonin Bal, Gregory Destras, Gwendolyne Burfin, Hadrien Regue, Quentin Semanas, Martine Valette, Bruno Lina, Laurence Josset |
| EPI_ISL_3380506 | Lighthouse Lab in Milton Keynes | Wellcome Sanger Institute for the COVID-19 Genomics UK (COG-UK) Consortium | The Lighthouse Lab in Milton Keynes and Alex Alderton, Roberto Amato, Jeffrey Barrett, Sonia Goncalves, Ewan Harrison, David K. Jackson, Ian Johnston, Dominic Kwiatkowski, Cordelia Langford, John Sillitoe on behalf of the Wellcome Sanger Institute COVID-19 Surveillance Team |
| EPI_ISL_3380507 | Lighthouse Lab in Alderley Park | Wellcome Sanger Institute for the COVID-19 Genomics UK (COG-UK) Consortium | Jacquelyn Wynn, Mairead Hyland, The Lighthouse Lab in Alderley Park and Alex Alderton, Roberto Amato, Jeffrey Barrett, Sonia Goncalves, Ewan Harrison, David K. Jackson, Ian Johnston, Dominic Kwiatkowski, Cordelia Langford, John Sillitoe on behalf of the Wellcome Sanger Institute COVID-19 Surveillance Team |
| EPI_ISL_3380508, EPI_ISL_3380509 | Lighthouse Lab in Milton Keynes | Wellcome Sanger Institute for the COVID-19 Genomics UK (COG-UK) Consortium | The Lighthouse Lab in Milton Keynes and Alex Alderton, Roberto Amato, Jeffrey Barrett, Sonia Goncalves, Ewan Harrison, David K. Jackson, Ian Johnston, Dominic Kwiatkowski, Cordelia Langford, John Sillitoe on behalf of the Wellcome Sanger Institute COVID-19 Surveillance Team |
| EPI_ISL_3380511 | Lighthouse Lab in Alderley Park | Wellcome Sanger Institute for the COVID-19 Genomics UK (COG-UK) Consortium | Jacquelyn Wynn, Mairead Hyland, The Lighthouse Lab in Alderley Park and Alex Alderton, Roberto Amato, Jeffrey Barrett, Sonia Goncalves, Ewan Harrison, David K. Jackson, Ian Johnston, Dominic Kwiatkowski, Cordelia Langford, John Sillitoe on behalf of the Wellcome Sanger Institute COVID-19 Surveillance Team |
| EPI_ISL_3380512 | Lighthouse Lab in Milton Keynes | Wellcome Sanger Institute for the COVID-19 Genomics UK (COG-UK) Consortium | The Lighthouse Lab in Milton Keynes and Alex Alderton, Roberto Amato, Jeffrey Barrett, Sonia Goncalves, Ewan Harrison, David K. Jackson, Ian Johnston, Dominic Kwiatkowski, Cordelia Langford, John Sillitoe on behalf of the Wellcome Sanger Institute COVID-19 Surveillance Team |
| EPI_ISL_3380513 | Lighthouse Lab in Alderley Park | Wellcome Sanger Institute for the COVID-19 Genomics UK (COG-UK) Consortium | Jacquelyn Wynn, Mairead Hyland, The Lighthouse Lab in Alderley Park and Alex Alderton, Roberto Amato, Jeffrey Barrett, Sonia Goncalves, Ewan Harrison, David K. Jackson, Ian Johnston, Dominic Kwiatkowski, Cordelia Langford, John Sillitoe on behalf of the Wellcome Sanger Institute COVID-19 Surveillance Team |
| EPI_ISL_3380514 | LABORATOIRE BIOESTEREL | CNR Virus des Infections Respiratoires - France SUD | Antonin Bal, Gregory Destras, Gwendolyne Burfin, Hadrien Regue, Quentin Semanas, Martine Valette, Bruno Lina, Laurence Josset |
| EPI_ISL_3380515, EPI_ISL_3380516 | Lighthouse Lab in Milton Keynes | Wellcome Sanger Institute for the COVID-19 Genomics UK (COG-UK) Consortium | The Lighthouse Lab in Milton Keynes and Alex Alderton, Roberto Amato, Jeffrey Barrett, Sonia Goncalves, Ewan Harrison, David K. Jackson, Ian Johnston, Dominic Kwiatkowski, Cordelia Langford, John Sillitoe on behalf of the Wellcome Sanger Institute COVID-19 Surveillance Team |
| EPI_ISL_3380517 | CERBAILLIANCE LA REUNION | CNR Virus des Infections Respiratoires - France SUD | Antonin Bal, Gregory Destras, Gwendolyne Burfin, Hadrien Regue, Quentin Semanas, Martine Valette, Bruno Lina, Laurence Josset |
| EPI_ISL_3380518, EPI_ISL_3380519, EPI_ISL_3380520 | Lighthouse Lab in Milton Keynes | Wellcome Sanger Institute for the COVID-19 Genomics UK (COG-UK) Consortium | The Lighthouse Lab in Milton Keynes and Alex Alderton, Roberto Amato, Jeffrey Barrett, Sonia Goncalves, Ewan Harrison, David K. Jackson, Ian Johnston, Dominic Kwiatkowski, Cordelia Langford, John Sillitoe on behalf of the Wellcome Sanger Institute COVID-19 Surveillance Team |
| EPI_ISL_3380521 | LABORATOIRE BIOESTEREL | CNR Virus des Infections Respiratoires - France SUD | Antonin Bal, Gregory Destras, Gwendolyne Burfin, Hadrien Regue, Quentin Semanas, Martine Valette, Bruno Lina, Laurence Josset |
| EPI_ISL_3380522, EPI_ISL_3380523, EPI_ISL_3380524 | Lighthouse Lab in Milton Keynes | Wellcome Sanger Institute for the COVID-19 Genomics UK (COG-UK) Consortium | The Lighthouse Lab in Milton Keynes and Alex Alderton, Roberto Amato, Jeffrey Barrett, Sonia Goncalves, Ewan Harrison, David K. Jackson, Ian Johnston, Dominic Kwiatkowski, Cordelia Langford, John Sillitoe on behalf of the Wellcome Sanger Institute COVID-19 Surveillance Team |
| EPI_ISL_3380525 | LABORATOIRE BIOESTEREL | CNR Virus des Infections Respiratoires - France SUD | Antonin Bal, Gregory Destras, Gwendolyne Burfin, Hadrien Regue, Quentin Semanas, Martine Valette, Bruno Lina, Laurence Josset |
| EPI_ISL_3380526 | Lighthouse Lab in Alderley Park | Wellcome Sanger Institute for the COVID-19 Genomics UK (COG-UK) Consortium | Jacquelyn Wynn, Mairead Hyland, The Lighthouse Lab in Alderley Park and Alex Alderton, Roberto Amato, Jeffrey Barrett, Sonia Goncalves, Ewan Harrison, David K. Jackson, Ian Johnston, Dominic Kwiatkowski, Cordelia Langford, John Sillitoe on behalf of the Wellcome Sanger Institute COVID-19 Surveillance Team |
| EPI_ISL_3380527, EPI_ISL_3380528 | Lighthouse Lab in Milton Keynes | Wellcome Sanger Institute for the COVID-19 Genomics UK (COG-UK) Consortium | The Lighthouse Lab in Milton Keynes and Alex Alderton, Roberto Amato, Jeffrey Barrett, Sonia Goncalves, Ewan Harrison, David K. Jackson, Ian Johnston, Dominic Kwiatkowski, Cordelia Langford, John Sillitoe on behalf of the Wellcome Sanger Institute COVID-19 Surveillance Team |
| EPI_ISL_3380529 | LABORATOIRE BIOESTEREL | CNR Virus des Infections Respiratoires - France SUD | Antonin Bal, Gregory Destras, Gwendolyne Burfin, Hadrien Regue, Quentin Semanas, Martine Valette, Bruno Lina, Laurence Josset |
| EPI_ISL_3380530, EPI_ISL_3380531 | Lighthouse Lab in Milton Keynes | Wellcome Sanger Institute for the COVID-19 Genomics UK (COG-UK) Consortium | The Lighthouse Lab in Milton Keynes and Alex Alderton, Roberto Amato, Jeffrey Barrett, Sonia Goncalves, Ewan Harrison, David K. Jackson, Ian Johnston, Dominic Kwiatkowski, Cordelia Langford, John Sillitoe on behalf of the Wellcome Sanger Institute COVID-19 Surveillance Team |
| EPI_ISL_3380532 | LABORATOIRE BIOESTEREL | CNR Virus des Infections Respiratoires - France SUD | Antonin Bal, Gregory Destras, Gwendolyne Burfin, Hadrien Regue, Quentin Semanas, Martine Valette, Bruno Lina, Laurence Josset |
| EPI_ISL_3380533, EPI_ISL_3380534, EPI_ISL_3380535 | Lighthouse Lab in Milton Keynes | Wellcome Sanger Institute for the COVID-19 Genomics UK (COG-UK) Consortium | The Lighthouse Lab in Milton Keynes and Alex Alderton, Roberto Amato, Jeffrey Barrett, Sonia Goncalves, Ewan Harrison, David K. Jackson, Ian Johnston, Dominic Kwiatkowski, Cordelia Langford, John Sillitoe on behalf of the Wellcome Sanger Institute COVID-19 Surveillance Team |
| EPI_ISL_3380536 | LABORATOIRE BIOESTEREL | CNR Virus des Infections Respiratoires - France SUD | Antonin Bal, Gregory Destras, Gwendolyne Burfin, Hadrien Regue, Quentin Semanas, Martine Valette, Bruno Lina, Laurence Josset |
| EPI_ISL_3380537 | Lighthouse Lab in Alderley Park | Wellcome Sanger Institute for the COVID-19 Genomics UK (COG-UK) Consortium | Jacquelyn Wynn, Mairead Hyland, The Lighthouse Lab in Alderley Park and Alex Alderton, Roberto Amato, Jeffrey Barrett, Sonia Goncalves, Ewan Harrison, David K. Jackson, Ian Johnston, Dominic Kwiatkowski, Cordelia Langford, John Sillitoe on behalf of the Wellcome Sanger Institute COVID-19 Surveillance Team |
| EPI_ISL_3380538, EPI_ISL_3380539 | Lighthouse Lab in Milton Keynes | Wellcome Sanger Institute for the COVID-19 Genomics UK (COG-UK) Consortium | The Lighthouse Lab in Milton Keynes and Alex Alderton, Roberto Amato, Jeffrey Barrett, Sonia Goncalves, Ewan Harrison, David K. Jackson, Ian Johnston, Dominic Kwiatkowski, Cordelia Langford, John Sillitoe on behalf of the Wellcome Sanger Institute COVID-19 Surveillance Team |
| EPI_ISL_3380540 | LABORATOIRE BIOESTEREL | CNR Virus des Infections Respiratoires - France SUD | Antonin Bal, Gregory Destras, Gwendolyne Burfin, Hadrien Regue, Quentin Semanas, Martine Valette, Bruno Lina, Laurence Josset |
| EPI_ISL_3380541, EPI_ISL_3380542 | Lighthouse Lab in Milton Keynes | Wellcome Sanger Institute for the COVID-19 Genomics UK (COG-UK) Consortium | The Lighthouse Lab in Milton Keynes and Alex Alderton, Roberto Amato, Jeffrey Barrett, Sonia Goncalves, Ewan Harrison, David K. Jackson, Ian Johnston, Dominic Kwiatkowski, Cordelia Langford, John Sillitoe on behalf of the Wellcome Sanger Institute COVID-19 Surveillance Team |
| EPI_ISL_3380543 | LABORATOIRE BIOESTEREL | CNR Virus des Infections Respiratoires - France SUD | Antonin Bal, Gregory Destras, Gwendolyne Burfin, Hadrien Regue, Quentin Semanas, Martine Valette, Bruno Lina, Laurence Josset |
| EPI_ISL_3380544 | Lighthouse Lab in Milton Keynes | Wellcome Sanger Institute for the COVID-19 Genomics UK (COG-UK) Consortium | The Lighthouse Lab in Milton Keynes and Alex Alderton, Roberto Amato, Jeffrey Barrett, Sonia Goncalves, Ewan Harrison, David K. Jackson, Ian Johnston, Dominic Kwiatkowski, Cordelia Langford, John Sillitoe on behalf of the Wellcome Sanger Institute COVID-19 Surveillance Team |
| EPI_ISL_3380545 | Lighthouse Lab in Alderley Park | Wellcome Sanger Institute for the COVID-19 Genomics UK (COG-UK) Consortium | Jacquelyn Wynn, Mairead Hyland, The Lighthouse Lab in Alderley Park and Alex Alderton, Roberto Amato, Jeffrey Barrett, Sonia Goncalves, Ewan Harrison, David K. Jackson, Ian Johnston, Dominic Kwiatkowski, Cordelia Langford, John Sillitoe on behalf of the Wellcome Sanger Institute COVID-19 Surveillance Team |
| EPI_ISL_3380546 | Lighthouse Lab in Milton Keynes | Wellcome Sanger Institute for the COVID-19 Genomics UK (COG-UK) Consortium | The Lighthouse Lab in Milton Keynes and Alex Alderton, Roberto Amato, Jeffrey Barrett, Sonia Goncalves, Ewan Harrison, David K. Jackson, Ian Johnston, Dominic Kwiatkowski, Cordelia Langford, John Sillitoe on behalf of the Wellcome Sanger Institute COVID-19 Surveillance Team |
| EPI_ISL_3380547 | CERBAILLIANCE LA REUNION | CNR Virus des Infections Respiratoires - France SUD | Antonin Bal, Gregory Destras, Gwendolyne Burfin, Hadrien Regue, Quentin Semanas, Martine Valette, Bruno Lina, Laurence Josset |
| EPI_ISL_3380548, EPI_ISL_3380549, EPI_ISL_3380550 | Lighthouse Lab in Milton Keynes | Wellcome Sanger Institute for the COVID-19 Genomics UK (COG-UK) Consortium | The Lighthouse Lab in Milton Keynes and Alex Alderton, Roberto Amato, Jeffrey Barrett, Sonia Goncalves, Ewan Harrison, David K. Jackson, Ian Johnston, Dominic Kwiatkowski, Cordelia Langford, John Sillitoe on behalf of the Wellcome Sanger Institute COVID-19 Surveillance Team |
| EPI_ISL_3380551 | LABORATOIRE BIOESTEREL | CNR Virus des Infections Respiratoires - France SUD | Antonin Bal, Gregory Destras, Gwendolyne Burfin, Hadrien Regue, Quentin Semanas, Martine Valette, Bruno Lina, Laurence Josset |
| EPI_ISL_3380552, EPI_ISL_3380553 | Lighthouse Lab in Milton Keynes | Wellcome Sanger Institute for the COVID-19 Genomics UK (COG-UK) Consortium | The Lighthouse Lab in Milton Keynes and Alex Alderton, Roberto Amato, Jeffrey Barrett, Sonia Goncalves, Ewan Harrison, David K. Jackson, Ian Johnston, Dominic Kwiatkowski, Cordelia Langford, John Sillitoe on behalf of the Wellcome Sanger Institute COVID-19 Surveillance Team |
| EPI_ISL_3380554 | LABORATOIRE BIOESTEREL | CNR Virus des Infections Respiratoires - France SUD | Antonin Bal, Gregory Destras, Gwendolyne Burfin, Hadrien Regue, Quentin Semanas, Martine Valette, Bruno Lina, Laurence Josset |
| EPI_ISL_3380555, EPI_ISL_3380556 | Lighthouse Lab in Milton Keynes | Wellcome Sanger Institute for the COVID-19 Genomics UK (COG-UK) Consortium | The Lighthouse Lab in Milton Keynes and Alex Alderton, Roberto Amato, Jeffrey Barrett, Sonia Goncalves, Ewan Harrison, David K. Jackson, Ian Johnston, Dominic Kwiatkowski, Cordelia Langford, John Sillitoe on behalf of the Wellcome Sanger Institute COVID-19 Surveillance Team |
| EPI_ISL_3380557 | Lighthouse Lab in Alderley Park | Wellcome Sanger Institute for the COVID-19 Genomics UK (COG-UK) Consortium | Jacquelyn Wynn, Mairead Hyland, The Lighthouse Lab in Alderley Park and Alex Alderton, Roberto Amato, Jeffrey Barrett, Sonia Goncalves, Ewan Harrison, David K. Jackson, Ian Johnston, Dominic Kwiatkowski, Cordelia Langford, John Sillitoe on behalf of the Wellcome Sanger Institute COVID-19 Surveillance Team |

| | | | |
|---|---|---|---|
| EPI_ISL_3380558 | LABORATOIRE BIOESTEREL | CNR Virus des Infections Respiratoires - France SUD | Antonin Bal, Gregory Destras, Gwendolyne Burfin, Hadrien Regue, Quentin Semanas, Martine Valette, Bruno Lina, Laurence Josset |
| EPI_ISL_3380559 | Lighthouse Lab in Alderley Park | Wellcome Sanger Institute for the COVID-19 Genomics UK (COG-UK) Consortium | Jacquelyn Wynn, Mairead Hyland, The Lighthouse Lab in Alderley Park and Alex Alderton, Roberto Amato, Jeffrey Barrett, Sonia Goncalves, Ewan Harrison, David K. Jackson, Ian Johnston, Dominic Kwiatkowski, Cordelia Langford, John Sillitoe on behalf of the Wellcome Sanger Institute COVID-19 Surveillance Team |
| EPI_ISL_3380560, EPI_ISL_3380561 | Lighthouse Lab in Milton Keynes | Wellcome Sanger Institute for the COVID-19 Genomics UK (COG-UK) Consortium | The Lighthouse Lab in Milton Keynes and Alex Alderton, Roberto Amato, Jeffrey Barrett, Sonia Goncalves, Ewan Harrison, David K. Jackson, Ian Johnston, Dominic Kwiatkowski, Cordelia Langford, John Sillitoe on behalf of the Wellcome Sanger Institute COVID-19 Surveillance Team |
| EPI_ISL_3380563, EPI_ISL_3380564, EPI_ISL_3380565 | Lighthouse Lab in Alderley Park | Wellcome Sanger Institute for the COVID-19 Genomics UK (COG-UK) Consortium | Jacquelyn Wynn, Mairead Hyland, The Lighthouse Lab in Alderley Park and Alex Alderton, Roberto Amato, Jeffrey Barrett, Sonia Goncalves, Ewan Harrison, David K. Jackson, Ian Johnston, Dominic Kwiatkowski, Cordelia Langford, John Sillitoe on behalf of the Wellcome Sanger Institute COVID-19 Surveillance Team |
| EPI_ISL_3380566 | LABORATOIRE BIOESTEREL | CNR Virus des Infections Respiratoires - France SUD | Antonin Bal, Gregory Destras, Gwendolyne Burfin, Hadrien Regue, Quentin Semanas, Martine Valette, Bruno Lina, Laurence Josset |
| EPI_ISL_3380567, EPI_ISL_3380568, EPI_ISL_3380569 | Lighthouse Lab in Milton Keynes | Wellcome Sanger Institute for the COVID-19 Genomics UK (COG-UK) Consortium | The Lighthouse Lab in Milton Keynes and Alex Alderton, Roberto Amato, Jeffrey Barrett, Sonia Goncalves, Ewan Harrison, David K. Jackson, Ian Johnston, Dominic Kwiatkowski, Cordelia Langford, John Sillitoe on behalf of the Wellcome Sanger Institute COVID-19 Surveillance Team |
| EPI_ISL_3380570 | LABORATOIRE BIOESTEREL | CNR Virus des Infections Respiratoires - France SUD | Antonin Bal, Gregory Destras, Gwendolyne Burfin, Hadrien Regue, Quentin Semanas, Martine Valette, Bruno Lina, Laurence Josset |
| EPI_ISL_3380571 | Lighthouse Lab in Milton Keynes | Wellcome Sanger Institute for the COVID-19 Genomics UK (COG-UK) Consortium | The Lighthouse Lab in Milton Keynes and Alex Alderton, Roberto Amato, Jeffrey Barrett, Sonia Goncalves, Ewan Harrison, David K. Jackson, Ian Johnston, Dominic Kwiatkowski, Cordelia Langford, John Sillitoe on behalf of the Wellcome Sanger Institute COVID-19 Surveillance Team |
| EPI_ISL_3380572 | Lighthouse Lab in Alderley Park | Wellcome Sanger Institute for the COVID-19 Genomics UK (COG-UK) Consortium | Jacquelyn Wynn, Mairead Hyland, The Lighthouse Lab in Alderley Park and Alex Alderton, Roberto Amato, Jeffrey Barrett, Sonia Goncalves, Ewan Harrison, David K. Jackson, Ian Johnston, Dominic Kwiatkowski, Cordelia Langford, John Sillitoe on behalf of the Wellcome Sanger Institute COVID-19 Surveillance Team |
| EPI_ISL_3380573 | Lighthouse Lab in Milton Keynes | Wellcome Sanger Institute for the COVID-19 Genomics UK (COG-UK) Consortium | The Lighthouse Lab in Milton Keynes and Alex Alderton, Roberto Amato, Jeffrey Barrett, Sonia Goncalves, Ewan Harrison, David K. Jackson, Ian Johnston, Dominic Kwiatkowski, Cordelia Langford, John Sillitoe on behalf of the Wellcome Sanger Institute COVID-19 Surveillance Team |
| EPI_ISL_3380574 | LABORATOIRE BIOESTEREL | CNR Virus des Infections Respiratoires - France SUD | Antonin Bal, Gregory Destras, Gwendolyne Burfin, Hadrien Regue, Quentin Semanas, Martine Valette, Bruno Lina, Laurence Josset |
| EPI_ISL_3380575, EPI_ISL_3380576 | Lighthouse Lab in Milton Keynes | Wellcome Sanger Institute for the COVID-19 Genomics UK (COG-UK) Consortium | The Lighthouse Lab in Milton Keynes and Alex Alderton, Roberto Amato, Jeffrey Barrett, Sonia Goncalves, Ewan Harrison, David K. Jackson, Ian Johnston, Dominic Kwiatkowski, Cordelia Langford, John Sillitoe on behalf of the Wellcome Sanger Institute COVID-19 Surveillance Team |
| EPI_ISL_3380577 | LABORATOIRE BIOESTEREL | CNR Virus des Infections Respiratoires - France SUD | Antonin Bal, Gregory Destras, Gwendolyne Burfin, Hadrien Regue, Quentin Semanas, Martine Valette, Bruno Lina, Laurence Josset |
| EPI_ISL_3380578 | Lighthouse Lab in Alderley Park | Wellcome Sanger Institute for the COVID-19 Genomics UK (COG-UK) Consortium | Jacquelyn Wynn, Mairead Hyland, The Lighthouse Lab in Alderley Park and Alex Alderton, Roberto Amato, Jeffrey Barrett, Sonia Goncalves, Ewan Harrison, David K. Jackson, Ian Johnston, Dominic Kwiatkowski, Cordelia Langford, John Sillitoe on behalf of the Wellcome Sanger Institute COVID-19 Surveillance Team |
| EPI_ISL_3380579 | Lighthouse Lab in Milton Keynes | Wellcome Sanger Institute for the COVID-19 Genomics UK (COG-UK) Consortium | The Lighthouse Lab in Milton Keynes and Alex Alderton, Roberto Amato, Jeffrey Barrett, Sonia Goncalves, Ewan Harrison, David K. Jackson, Ian Johnston, Dominic Kwiatkowski, Cordelia Langford, John Sillitoe on behalf of the Wellcome Sanger Institute COVID-19 Surveillance Team |
| EPI_ISL_3380580 | Lighthouse Lab in Alderley Park | Wellcome Sanger Institute for the COVID-19 Genomics UK (COG-UK) Consortium | Jacquelyn Wynn, Mairead Hyland, The Lighthouse Lab in Alderley Park and Alex Alderton, Roberto Amato, Jeffrey Barrett, Sonia Goncalves, Ewan Harrison, David K. Jackson, Ian Johnston, Dominic Kwiatkowski, Cordelia Langford, John Sillitoe on behalf of the Wellcome Sanger Institute COVID-19 Surveillance Team |
| EPI_ISL_3380581 | LABORATOIRE BIOESTEREL | CNR Virus des Infections Respiratoires - France SUD | Antonin Bal, Gregory Destras, Gwendolyne Burfin, Hadrien Regue, Quentin Semanas, Martine Valette, Bruno Lina, Laurence Josset |
| EPI_ISL_3380582, EPI_ISL_3380583, EPI_ISL_3380585, EPI_ISL_3380586 | Lighthouse Lab in Milton Keynes | Wellcome Sanger Institute for the COVID-19 Genomics UK (COG-UK) Consortium | The Lighthouse Lab in Milton Keynes and Alex Alderton, Roberto Amato, Jeffrey Barrett, Sonia Goncalves, Ewan Harrison, David K. Jackson, Ian Johnston, Dominic Kwiatkowski, Cordelia Langford, John Sillitoe on behalf of the Wellcome Sanger Institute COVID-19 Surveillance Team |
| EPI_ISL_3380587 | Lighthouse Lab in Alderley Park | Wellcome Sanger Institute for the COVID-19 Genomics UK (COG-UK) Consortium | Jacquelyn Wynn, Mairead Hyland, The Lighthouse Lab in Alderley Park and Alex Alderton, Roberto Amato, Jeffrey Barrett, Sonia Goncalves, Ewan Harrison, David K. Jackson, Ian Johnston, Dominic Kwiatkowski, Cordelia Langford, John Sillitoe on behalf of the Wellcome Sanger Institute COVID-19 Surveillance Team |
| EPI_ISL_3380588 | LABORATOIRE BIOESTEREL | CNR Virus des Infections Respiratoires - France SUD | Antonin Bal, Gregory Destras, Gwendolyne Burfin, Hadrien Regue, Quentin Semanas, Martine Valette, Bruno Lina, Laurence Josset |
| EPI_ISL_3380589, EPI_ISL_3380590, EPI_ISL_3380591 | Lighthouse Lab in Milton Keynes | Wellcome Sanger Institute for the COVID-19 Genomics UK (COG-UK) Consortium | The Lighthouse Lab in Milton Keynes and Alex Alderton, Roberto Amato, Jeffrey Barrett, Sonia Goncalves, Ewan Harrison, David K. Jackson, Ian Johnston, Dominic Kwiatkowski, Cordelia Langford, John Sillitoe on behalf of the Wellcome Sanger Institute COVID-19 Surveillance Team |
| EPI_ISL_3380592 | LABORATOIRE BIOESTEREL | CNR Virus des Infections Respiratoires - France SUD | Antonin Bal, Gregory Destras, Gwendolyne Burfin, Hadrien Regue, Quentin Semanas, Martine Valette, Bruno Lina, Laurence Josset |
| EPI_ISL_3380593, EPI_ISL_3380594, EPI_ISL_3380595 | Lighthouse Lab in Milton Keynes | Wellcome Sanger Institute for the COVID-19 Genomics UK (COG-UK) Consortium | The Lighthouse Lab in Milton Keynes and Alex Alderton, Roberto Amato, Jeffrey Barrett, Sonia Goncalves, Ewan Harrison, David K. Jackson, Ian Johnston, Dominic Kwiatkowski, Cordelia Langford, John Sillitoe on behalf of the Wellcome Sanger Institute COVID-19 Surveillance Team |
| EPI_ISL_3380596 | LABORATOIRE BIOESTEREL | CNR Virus des Infections Respiratoires - France SUD | Antonin Bal, Gregory Destras, Gwendolyne Burfin, Hadrien Regue, Quentin Semanas, Martine Valette, Bruno Lina, Laurence Josset |
| EPI_ISL_3380597, EPI_ISL_3380598 | Lighthouse Lab in Alderley Park | Wellcome Sanger Institute for the COVID-19 Genomics UK (COG-UK) Consortium | Jacquelyn Wynn, Mairead Hyland, The Lighthouse Lab in Alderley Park and Alex Alderton, Roberto Amato, Jeffrey Barrett, Sonia Goncalves, Ewan Harrison, David K. Jackson, Ian Johnston, Dominic Kwiatkowski, Cordelia Langford, John Sillitoe on behalf of the Wellcome Sanger Institute COVID-19 Surveillance Team |
| EPI_ISL_3380599 | CERBAILLIANCE LA REUNION | CNR Virus des Infections Respiratoires - France SUD | Antonin Bal, Gregory Destras, Gwendolyne Burfin, Hadrien Regue, Quentin Semanas, Martine Valette, Bruno Lina, Laurence Josset |
| EPI_ISL_3380600 | Lighthouse Lab in Alderley Park | Wellcome Sanger Institute for the COVID-19 Genomics UK (COG-UK) Consortium | Jacquelyn Wynn, Mairead Hyland, The Lighthouse Lab in Alderley Park and Alex Alderton, Roberto Amato, Jeffrey Barrett, Sonia Goncalves, Ewan Harrison, David K. Jackson, Ian Johnston, Dominic Kwiatkowski, Cordelia Langford, John Sillitoe on behalf of the Wellcome Sanger Institute COVID-19 Surveillance Team |
| EPI_ISL_3380601 | Lighthouse Lab in Milton Keynes | Wellcome Sanger Institute for the COVID-19 Genomics UK (COG-UK) Consortium | The Lighthouse Lab in Milton Keynes and Alex Alderton, Roberto Amato, Jeffrey Barrett, Sonia Goncalves, Ewan Harrison, David K. Jackson, Ian Johnston, Dominic Kwiatkowski, Cordelia Langford, John Sillitoe on behalf of the Wellcome Sanger Institute COVID-19 Surveillance Team |
| EPI_ISL_3380602 | Lighthouse Lab in Alderley Park | Wellcome Sanger Institute for the COVID-19 Genomics UK (COG-UK) Consortium | Jacquelyn Wynn, Mairead Hyland, The Lighthouse Lab in Alderley Park and Alex Alderton, Roberto Amato, Jeffrey Barrett, Sonia Goncalves, Ewan Harrison, David K. Jackson, Ian Johnston, Dominic Kwiatkowski, Cordelia Langford, John Sillitoe on behalf of the Wellcome Sanger Institute COVID-19 Surveillance Team |
| EPI_ISL_3380603 | LABORATOIRE BIOESTEREL | CNR Virus des Infections Respiratoires - France SUD | Antonin Bal, Gregory Destras, Gwendolyne Burfin, Hadrien Regue, Quentin Semanas, Martine Valette, Bruno Lina, Laurence Josset |
| EPI_ISL_3380604 | Lighthouse Lab in Milton Keynes | Wellcome Sanger Institute for the COVID-19 Genomics UK (COG-UK) Consortium | The Lighthouse Lab in Milton Keynes and Alex Alderton, Roberto Amato, Jeffrey Barrett, Sonia Goncalves, Ewan Harrison, David K. Jackson, Ian Johnston, Dominic Kwiatkowski, Cordelia Langford, John Sillitoe on behalf of the Wellcome Sanger Institute COVID-19 Surveillance Team |
| EPI_ISL_3380605 | Lighthouse Lab in Alderley Park | Wellcome Sanger Institute for the COVID-19 Genomics UK (COG-UK) Consortium | Jacquelyn Wynn, Mairead Hyland, The Lighthouse Lab in Alderley Park and Alex Alderton, Roberto Amato, Jeffrey Barrett, Sonia Goncalves, Ewan Harrison, David K. Jackson, Ian Johnston, Dominic Kwiatkowski, Cordelia Langford, John Sillitoe on behalf of the Wellcome Sanger Institute COVID-19 Surveillance Team |
| EPI_ISL_3380606 | Lighthouse Lab in Milton Keynes | Wellcome Sanger Institute for the COVID-19 Genomics UK (COG-UK) Consortium | The Lighthouse Lab in Milton Keynes and Alex Alderton, Roberto Amato, Jeffrey Barrett, Sonia Goncalves, Ewan Harrison, David K. Jackson, Ian Johnston, Dominic Kwiatkowski, Cordelia Langford, John Sillitoe on behalf of the Wellcome Sanger Institute COVID-19 Surveillance Team |
| EPI_ISL_3380607 | LABORATOIRE BIOESTEREL | CNR Virus des Infections Respiratoires - France SUD | Antonin Bal, Gregory Destras, Gwendolyne Burfin, Hadrien Regue, Quentin Semanas, Martine Valette, Bruno Lina, Laurence Josset |
| EPI_ISL_3380608, EPI_ISL_3380609, EPI_ISL_3380610 | Lighthouse Lab in Milton Keynes | Wellcome Sanger Institute for the COVID-19 Genomics UK (COG-UK) Consortium | The Lighthouse Lab in Milton Keynes and Alex Alderton, Roberto Amato, Jeffrey Barrett, Sonia Goncalves, Ewan Harrison, David K. Jackson, Ian Johnston, Dominic Kwiatkowski, Cordelia Langford, John Sillitoe on behalf of the Wellcome Sanger Institute COVID-19 Surveillance Team |
| EPI_ISL_3380611 | LABORATOIRE BIOESTEREL | CNR Virus des Infections Respiratoires - France SUD | Antonin Bal, Gregory Destras, Gwendolyne Burfin, Hadrien Regue, Quentin Semanas, Martine Valette, Bruno Lina, Laurence Josset |
| EPI_ISL_3380612, EPI_ISL_3380613 | Berkshire and Surrey Pathology Services Lighthouse Laboratory | Wellcome Sanger Institute for the COVID-19 Genomics UK (COG-UK) Consortium | Berkshire and Surrey Pathology Services Lighthouse Laboratory and Alex Alderton, Roberto Amato, Jeffrey Barrett, Sonia Goncalves, Ewan Harrison, David K. Jackson, Ian Johnston, Dominic Kwiatkowski, Cordelia Langford, John Sillitoe on behalf of the Wellcome Sanger Institute COVID-19 Surveillance Team |
| EPI_ISL_3380614 | LABORATOIRE BIOESTEREL | CNR Virus des Infections Respiratoires - France SUD | Antonin Bal, Gregory Destras, Gwendolyne Burfin, Hadrien Regue, Quentin Semanas, Martine Valette, Bruno Lina, Laurence Josset |
| EPI_ISL_3380615 | Berkshire and Surrey Pathology Services Lighthouse Laboratory | Wellcome Sanger Institute for the COVID-19 Genomics UK (COG-UK) Consortium | Berkshire and Surrey Pathology Services Lighthouse Laboratory and Alex Alderton, Roberto Amato, Jeffrey Barrett, Sonia Goncalves, Ewan Harrison, David K. Jackson, Ian Johnston, Dominic Kwiatkowski, Cordelia Langford, John Sillitoe on behalf of the Wellcome Sanger Institute COVID-19 Surveillance Team |
| EPI_ISL_3380616 | Lighthouse Lab in Milton Keynes | Wellcome Sanger Institute for the COVID-19 Genomics UK (COG-UK) Consortium | The Lighthouse Lab in Milton Keynes and Alex Alderton, Roberto Amato, Jeffrey Barrett, Sonia Goncalves, Ewan Harrison, David K. Jackson, Ian Johnston, Dominic Kwiatkowski, Cordelia Langford, John Sillitoe on behalf of the Wellcome Sanger Institute COVID-19 Surveillance Team |
| EPI_ISL_3380617 | Berkshire and Surrey Pathology Services Lighthouse Laboratory | Wellcome Sanger Institute for the COVID-19 Genomics UK (COG-UK) Consortium | Berkshire and Surrey Pathology Services Lighthouse Laboratory and Alex Alderton, Roberto Amato, Jeffrey Barrett, Sonia Goncalves, Ewan Harrison, David K. Jackson, Ian Johnston, Dominic Kwiatkowski, Cordelia Langford, John Sillitoe on behalf of the Wellcome Sanger Institute COVID-19 Surveillance Team |
| EPI_ISL_3380619, EPI_ISL_3380620 | Health Services Laboratories | Wellcome Sanger Institute for the COVID-19 Genomics UK (COG-UK) Consortium | Health Services Laboratories and Alex Alderton, Roberto Amato, Jeffrey Barrett, Sonia Goncalves, Ewan Harrison, David K. Jackson, Ian Johnston, Dominic Kwiatkowski, Cordelia Langford, John Sillitoe on behalf of the Wellcome Sanger Institute COVID-19 Surveillance Team |
| EPI_ISL_3380621 | Berkshire and Surrey Pathology Services Lighthouse Laboratory | Wellcome Sanger Institute for the COVID-19 Genomics UK (COG-UK) Consortium | Berkshire and Surrey Pathology Services Lighthouse Laboratory and Alex Alderton, Roberto Amato, Jeffrey Barrett, Sonia Goncalves, Ewan Harrison, David K. Jackson, Ian Johnston, Dominic Kwiatkowski, Cordelia Langford, John Sillitoe on behalf of the Wellcome Sanger Institute COVID-19 Surveillance Team |
| EPI_ISL_3380622 | LABORATOIRE BIOESTEREL | CNR Virus des Infections Respiratoires - France SUD | Antonin Bal, Gregory Destras, Gwendolyne Burfin, Hadrien Regue, Quentin Semanas, Martine Valette, Bruno Lina, Laurence Josset |
| EPI_ISL_3380623 | Berkshire and Surrey Pathology Services Lighthouse Laboratory | Wellcome Sanger Institute for the COVID-19 Genomics UK (COG-UK) Consortium | Berkshire and Surrey Pathology Services Lighthouse Laboratory and Alex Alderton, Roberto Amato, Jeffrey Barrett, Sonia Goncalves, Ewan Harrison, David K. Jackson, Ian Johnston, Dominic Kwiatkowski, Cordelia Langford, John Sillitoe on behalf of the Wellcome Sanger Institute COVID-19 Surveillance Team |
| EPI_ISL_3380624, EPI_ISL_3380625 | Health Services Laboratories | Wellcome Sanger Institute for the COVID-19 Genomics UK (COG-UK) Consortium | Health Services Laboratories and Alex Alderton, Roberto Amato, Jeffrey Barrett, Sonia Goncalves, Ewan Harrison, David K. Jackson, Ian Johnston, Dominic Kwiatkowski, Cordelia Langford, John Sillitoe on behalf of the Wellcome Sanger Institute COVID-19 Surveillance Team |
| EPI_ISL_3380626 | LABORATOIRE BIOESTEREL | CNR Virus des Infections Respiratoires - France SUD | Antonin Bal, Gregory Destras, Gwendolyne Burfin, Hadrien Regue, Quentin Semanas, Martine Valette, Bruno Lina, Laurence Josset |
| EPI_ISL_3380627 | Berkshire and Surrey Pathology Services Lighthouse Laboratory | Wellcome Sanger Institute for the COVID-19 Genomics UK (COG-UK) Consortium | Berkshire and Surrey Pathology Services Lighthouse Laboratory and Alex Alderton, Roberto Amato, Jeffrey Barrett, Sonia Goncalves, Ewan Harrison, David K. Jackson, Ian Johnston, Dominic Kwiatkowski, Cordelia Langford, John Sillitoe on behalf of the Wellcome Sanger Institute COVID-19 Surveillance Team |
| EPI_ISL_3380628 | Lighthouse Lab in Milton Keynes | Wellcome Sanger Institute for the COVID-19 Genomics UK (COG-UK) Consortium | The Lighthouse Lab in Milton Keynes and Alex Alderton, Roberto Amato, Jeffrey Barrett, Sonia Goncalves, Ewan Harrison, David K. Jackson, Ian Johnston, Dominic Kwiatkowski, Cordelia Langford, John Sillitoe on behalf of the Wellcome Sanger Institute COVID-19 Surveillance Team |

| | | | |
|---|---|---|---|
| EPI_ISL_3380629 | LABORATOIRE BIOESTEREL | CNR Virus des Infections Respiratoires - France SUD | Antonin Bal, Gregory Destras, Gwendolyne Burfin, Hadrien Regue, Quentin Semanas, Martine Valette, Bruno Lina, Laurence Josset |
| EPI_ISL_3380630, EPI_ISL_3380631 | Berkshire and Surrey Pathology Services Lighthouse Laboratory | Wellcome Sanger Institute for the COVID-19 Genomics UK (COG-UK) Consortium | Berkshire and Surrey Pathology Services Lighthouse Laboratory and Alex Alderton, Roberto Amato, Jeffrey Barrett, Sonia Goncalves, Ewan Harrison, David K. Jackson, Ian Johnston, Dominic Kwiatkowski, Cordelia Langford, John Sillitoe on behalf of the Wellcome Sanger Institute COVID-19 Surveillance Team |
| EPI_ISL_3380632, EPI_ISL_3380633 | Lighthouse Lab in Milton Keynes | Wellcome Sanger Institute for the COVID-19 Genomics UK (COG-UK) Consortium | The Lighthouse Lab in Milton Keynes and Alex Alderton, Roberto Amato, Jeffrey Barrett, Sonia Goncalves, Ewan Harrison, David K. Jackson, Ian Johnston, Dominic Kwiatkowski, Cordelia Langford, John Sillitoe on behalf of the Wellcome Sanger Institute COVID-19 Surveillance Team |
| EPI_ISL_3380634 | LABORATOIRE BIOESTEREL | CNR Virus des Infections Respiratoires - France SUD | Antonin Bal, Gregory Destras, Gwendolyne Burfin, Hadrien Regue, Quentin Semanas, Martine Valette, Bruno Lina, Laurence Josset |
| EPI_ISL_3380635, EPI_ISL_3380636 | Berkshire and Surrey Pathology Services Lighthouse Laboratory | Wellcome Sanger Institute for the COVID-19 Genomics UK (COG-UK) Consortium | Berkshire and Surrey Pathology Services Lighthouse Laboratory and Alex Alderton, Roberto Amato, Jeffrey Barrett, Sonia Goncalves, Ewan Harrison, David K. Jackson, Ian Johnston, Dominic Kwiatkowski, Cordelia Langford, John Sillitoe on behalf of the Wellcome Sanger Institute COVID-19 Surveillance Team |
| EPI_ISL_3380637 | Lighthouse Laboratory Plymouth | Wellcome Sanger Institute for the COVID-19 Genomics UK (COG-UK) Consortium | Lighthouse Laboratory Plymouth and Alex Alderton, Roberto Amato, Jeffrey Barrett, Sonia Goncalves, Ewan Harrison, David K. Jackson, Ian Johnston, Dominic Kwiatkowski, Cordelia Langford, John Sillitoe on behalf of the Wellcome Sanger Institute COVID-19 Surveillance Team |
| EPI_ISL_3380638 | CERBAILLIANCE LA REUNION | CNR Virus des Infections Respiratoires - France SUD | Antonin Bal, Gregory Destras, Gwendolyne Burfin, Hadrien Regue, Quentin Semanas, Martine Valette, Bruno Lina, Laurence Josset |
| EPI_ISL_3380639 | Lighthouse Laboratory Plymouth | Wellcome Sanger Institute for the COVID-19 Genomics UK (COG-UK) Consortium | Lighthouse Laboratory Plymouth and Alex Alderton, Roberto Amato, Jeffrey Barrett, Sonia Goncalves, Ewan Harrison, David K. Jackson, Ian Johnston, Dominic Kwiatkowski, Cordelia Langford, John Sillitoe on behalf of the Wellcome Sanger Institute COVID-19 Surveillance Team |
| EPI_ISL_3380640 | Berkshire and Surrey Pathology Services Lighthouse Laboratory | Wellcome Sanger Institute for the COVID-19 Genomics UK (COG-UK) Consortium | Berkshire and Surrey Pathology Services Lighthouse Laboratory and Alex Alderton, Roberto Amato, Jeffrey Barrett, Sonia Goncalves, Ewan Harrison, David K. Jackson, Ian Johnston, Dominic Kwiatkowski, Cordelia Langford, John Sillitoe on behalf of the Wellcome Sanger Institute COVID-19 Surveillance Team |
| EPI_ISL_3380641 | Health Services Laboratories | Wellcome Sanger Institute for the COVID-19 Genomics UK (COG-UK) Consortium | Health Services Laboratories and Alex Alderton, Roberto Amato, Jeffrey Barrett, Sonia Goncalves, Ewan Harrison, David K. Jackson, Ian Johnston, Dominic Kwiatkowski, Cordelia Langford, John Sillitoe on behalf of the Wellcome Sanger Institute COVID-19 Surveillance Team |
| EPI_ISL_3380642 | LABORATOIRE BIOESTEREL | CNR Virus des Infections Respiratoires - France SUD | Antonin Bal, Gregory Destras, Gwendolyne Burfin, Hadrien Regue, Quentin Semanas, Martine Valette, Bruno Lina, Laurence Josset |
| EPI_ISL_3380643 | Health Services Laboratories | Wellcome Sanger Institute for the COVID-19 Genomics UK (COG-UK) Consortium | Health Services Laboratories and Alex Alderton, Roberto Amato, Jeffrey Barrett, Sonia Goncalves, Ewan Harrison, David K. Jackson, Ian Johnston, Dominic Kwiatkowski, Cordelia Langford, John Sillitoe on behalf of the Wellcome Sanger Institute COVID-19 Surveillance Team |
| EPI_ISL_3380644, EPI_ISL_3380645 | Lighthouse Lab in Milton Keynes | Wellcome Sanger Institute for the COVID-19 Genomics UK (COG-UK) Consortium | The Lighthouse Lab in Milton Keynes and Alex Alderton, Roberto Amato, Jeffrey Barrett, Sonia Goncalves, Ewan Harrison, David K. Jackson, Ian Johnston, Dominic Kwiatkowski, Cordelia Langford, John Sillitoe on behalf of the Wellcome Sanger Institute COVID-19 Surveillance Team |
| EPI_ISL_3380646 | LABORATOIRE BIOESTEREL | CNR Virus des Infections Respiratoires - France SUD | Antonin Bal, Gregory Destras, Gwendolyne Burfin, Hadrien Regue, Quentin Semanas, Martine Valette, Bruno Lina, Laurence Josset |
| EPI_ISL_3380647 | Lighthouse Lab in Milton Keynes | Wellcome Sanger Institute for the COVID-19 Genomics UK (COG-UK) Consortium | The Lighthouse Lab in Milton Keynes and Alex Alderton, Roberto Amato, Jeffrey Barrett, Sonia Goncalves, Ewan Harrison, David K. Jackson, Ian Johnston, Dominic Kwiatkowski, Cordelia Langford, John Sillitoe on behalf of the Wellcome Sanger Institute COVID-19 Surveillance Team |
| EPI_ISL_3380648 | Health Services Laboratories | Wellcome Sanger Institute for the COVID-19 Genomics UK (COG-UK) Consortium | Health Services Laboratories and Alex Alderton, Roberto Amato, Jeffrey Barrett, Sonia Goncalves, Ewan Harrison, David K. Jackson, Ian Johnston, Dominic Kwiatkowski, Cordelia Langford, John Sillitoe on behalf of the Wellcome Sanger Institute COVID-19 Surveillance Team |
| EPI_ISL_3380649 | Berkshire and Surrey Pathology Services Lighthouse Laboratory | Wellcome Sanger Institute for the COVID-19 Genomics UK (COG-UK) Consortium | Berkshire and Surrey Pathology Services Lighthouse Laboratory and Alex Alderton, Roberto Amato, Jeffrey Barrett, Sonia Goncalves, Ewan Harrison, David K. Jackson, Ian Johnston, Dominic Kwiatkowski, Cordelia Langford, John Sillitoe on behalf of the Wellcome Sanger Institute COVID-19 Surveillance Team |
| EPI_ISL_3380650 | LABORATOIRE BIOESTEREL | CNR Virus des Infections Respiratoires - France SUD | Antonin Bal, Gregory Destras, Gwendolyne Burfin, Hadrien Regue, Quentin Semanas, Martine Valette, Bruno Lina, Laurence Josset |
| EPI_ISL_3380651 | Berkshire and Surrey Pathology Services Lighthouse Laboratory | Wellcome Sanger Institute for the COVID-19 Genomics UK (COG-UK) Consortium | Berkshire and Surrey Pathology Services Lighthouse Laboratory and Alex Alderton, Roberto Amato, Jeffrey Barrett, Sonia Goncalves, Ewan Harrison, David K. Jackson, Ian Johnston, Dominic Kwiatkowski, Cordelia Langford, John Sillitoe on behalf of the Wellcome Sanger Institute COVID-19 Surveillance Team |
| EPI_ISL_3380652 | Lighthouse Lab in Milton Keynes | Wellcome Sanger Institute for the COVID-19 Genomics UK (COG-UK) Consortium | The Lighthouse Lab in Milton Keynes and Alex Alderton, Roberto Amato, Jeffrey Barrett, Sonia Goncalves, Ewan Harrison, David K. Jackson, Ian Johnston, Dominic Kwiatkowski, Cordelia Langford, John Sillitoe on behalf of the Wellcome Sanger Institute COVID-19 Surveillance Team |
| EPI_ISL_3380653 | Berkshire and Surrey Pathology Services Lighthouse Laboratory | Wellcome Sanger Institute for the COVID-19 Genomics UK (COG-UK) Consortium | Berkshire and Surrey Pathology Services Lighthouse Laboratory and Alex Alderton, Roberto Amato, Jeffrey Barrett, Sonia Goncalves, Ewan Harrison, David K. Jackson, Ian Johnston, Dominic Kwiatkowski, Cordelia Langford, John Sillitoe on behalf of the Wellcome Sanger Institute COVID-19 Surveillance Team |
| EPI_ISL_3380654 | LABORATOIRE BIOESTEREL | CNR Virus des Infections Respiratoires - France SUD | Antonin Bal, Gregory Destras, Gwendolyne Burfin, Hadrien Regue, Quentin Semanas, Martine Valette, Bruno Lina, Laurence Josset |
| EPI_ISL_3380655 | Lighthouse Lab in Milton Keynes | Wellcome Sanger Institute for the COVID-19 Genomics UK (COG-UK) Consortium | The Lighthouse Lab in Milton Keynes and Alex Alderton, Roberto Amato, Jeffrey Barrett, Sonia Goncalves, Ewan Harrison, David K. Jackson, Ian Johnston, Dominic Kwiatkowski, Cordelia Langford, John Sillitoe on behalf of the Wellcome Sanger Institute COVID-19 Surveillance Team |
| EPI_ISL_3380656, EPI_ISL_3380657 | Berkshire and Surrey Pathology Services Lighthouse Laboratory | Wellcome Sanger Institute for the COVID-19 Genomics UK (COG-UK) Consortium | Berkshire and Surrey Pathology Services Lighthouse Laboratory and Alex Alderton, Roberto Amato, Jeffrey Barrett, Sonia Goncalves, Ewan Harrison, David K. Jackson, Ian Johnston, Dominic Kwiatkowski, Cordelia Langford, John Sillitoe on behalf of the Wellcome Sanger Institute COVID-19 Surveillance Team |
| EPI_ISL_3380658 | LABORATOIRE BIOESTEREL | CNR Virus des Infections Respiratoires - France SUD | Antonin Bal, Gregory Destras, Gwendolyne Burfin, Hadrien Regue, Quentin Semanas, Martine Valette, Bruno Lina, Laurence Josset |
| EPI_ISL_3380659 | Lighthouse Lab in Milton Keynes | Wellcome Sanger Institute for the COVID-19 Genomics UK (COG-UK) Consortium | The Lighthouse Lab in Milton Keynes and Alex Alderton, Roberto Amato, Jeffrey Barrett, Sonia Goncalves, Ewan Harrison, David K. Jackson, Ian Johnston, Dominic Kwiatkowski, Cordelia Langford, John Sillitoe on behalf of the Wellcome Sanger Institute COVID-19 Surveillance Team |
| EPI_ISL_3380660, EPI_ISL_3380661 | Health Services Laboratories | Wellcome Sanger Institute for the COVID-19 Genomics UK (COG-UK) Consortium | Health Services Laboratories and Alex Alderton, Roberto Amato, Jeffrey Barrett, Sonia Goncalves, Ewan Harrison, David K. Jackson, Ian Johnston, Dominic Kwiatkowski, Cordelia Langford, John Sillitoe on behalf of the Wellcome Sanger Institute COVID-19 Surveillance Team |
| EPI_ISL_3380662 | LABORATOIRE BIOESTEREL | CNR Virus des Infections Respiratoires - France SUD | Antonin Bal, Gregory Destras, Gwendolyne Burfin, Hadrien Regue, Quentin Semanas, Martine Valette, Bruno Lina, Laurence Josset |
| EPI_ISL_3380663, EPI_ISL_3380664 | Berkshire and Surrey Pathology Services Lighthouse Laboratory | Wellcome Sanger Institute for the COVID-19 Genomics UK (COG-UK) Consortium | Berkshire and Surrey Pathology Services Lighthouse Laboratory and Alex Alderton, Roberto Amato, Jeffrey Barrett, Sonia Goncalves, Ewan Harrison, David K. Jackson, Ian Johnston, Dominic Kwiatkowski, Cordelia Langford, John Sillitoe on behalf of the Wellcome Sanger Institute COVID-19 Surveillance Team |
| EPI_ISL_3380665 | CENTRE HOSPITALIER DE LA COTE BASQUE | CNR Virus des Infections Respiratoires - France SUD | Antonin Bal, Gregory Destras, Gwendolyne Burfin, Hadrien Regue, Quentin Semanas, Martine Valette, Bruno Lina, Laurence Josset |
| EPI_ISL_3380666, EPI_ISL_3380667, EPI_ISL_3380668 | Lighthouse Lab in Milton Keynes | Wellcome Sanger Institute for the COVID-19 Genomics UK (COG-UK) Consortium | The Lighthouse Lab in Milton Keynes and Alex Alderton, Roberto Amato, Jeffrey Barrett, Sonia Goncalves, Ewan Harrison, David K. Jackson, Ian Johnston, Dominic Kwiatkowski, Cordelia Langford, John Sillitoe on behalf of the Wellcome Sanger Institute COVID-19 Surveillance Team |
| EPI_ISL_3380669 | CNR Virus des Infections Respiratoires - France SUD | CNR Virus des Infections Respiratoires - France SUD | Antonin Bal, Gregory Destras, Gwendolyne Burfin, Hadrien Regue, Quentin Semanas, Martine Valette, Bruno Lina, Laurence Josset |
| EPI_ISL_3380670 | Lighthouse Lab in Milton Keynes | Wellcome Sanger Institute for the COVID-19 Genomics UK (COG-UK) Consortium | The Lighthouse Lab in Milton Keynes and Alex Alderton, Roberto Amato, Jeffrey Barrett, Sonia Goncalves, Ewan Harrison, David K. Jackson, Ian Johnston, Dominic Kwiatkowski, Cordelia Langford, John Sillitoe on behalf of the Wellcome Sanger Institute COVID-19 Surveillance Team |
| EPI_ISL_3380671 | Health Services Laboratories | Wellcome Sanger Institute for the COVID-19 Genomics UK (COG-UK) Consortium | Health Services Laboratories and Alex Alderton, Roberto Amato, Jeffrey Barrett, Sonia Goncalves, Ewan Harrison, David K. Jackson, Ian Johnston, Dominic Kwiatkowski, Cordelia Langford, John Sillitoe on behalf of the Wellcome Sanger Institute COVID-19 Surveillance Team |
| EPI_ISL_3380672 | Lighthouse Lab in Milton Keynes | Wellcome Sanger Institute for the COVID-19 Genomics UK (COG-UK) Consortium | The Lighthouse Lab in Milton Keynes and Alex Alderton, Roberto Amato, Jeffrey Barrett, Sonia Goncalves, Ewan Harrison, David K. Jackson, Ian Johnston, Dominic Kwiatkowski, Cordelia Langford, John Sillitoe on behalf of the Wellcome Sanger Institute COVID-19 Surveillance Team |
| EPI_ISL_3380673 | CNR Virus des Infections Respiratoires - France SUD | CNR Virus des Infections Respiratoires - France SUD | Antonin Bal, Gregory Destras, Gwendolyne Burfin, Hadrien Regue, Quentin Semanas, Martine Valette, Bruno Lina, Laurence Josset |
| EPI_ISL_3380674 | Berkshire and Surrey Pathology Services Lighthouse Laboratory | Wellcome Sanger Institute for the COVID-19 Genomics UK (COG-UK) Consortium | Berkshire and Surrey Pathology Services Lighthouse Laboratory and Alex Alderton, Roberto Amato, Jeffrey Barrett, Sonia Goncalves, Ewan Harrison, David K. Jackson, Ian Johnston, Dominic Kwiatkowski, Cordelia Langford, John Sillitoe on behalf of the Wellcome Sanger Institute COVID-19 Surveillance Team |
| EPI_ISL_3380675 | Lighthouse Lab in Milton Keynes | Wellcome Sanger Institute for the COVID-19 Genomics UK (COG-UK) Consortium | The Lighthouse Lab in Milton Keynes and Alex Alderton, Roberto Amato, Jeffrey Barrett, Sonia Goncalves, Ewan Harrison, David K. Jackson, Ian Johnston, Dominic Kwiatkowski, Cordelia Langford, John Sillitoe on behalf of the Wellcome Sanger Institute COVID-19 Surveillance Team |
| EPI_ISL_3380676 | Berkshire and Surrey Pathology Services Lighthouse Laboratory | Wellcome Sanger Institute for the COVID-19 Genomics UK (COG-UK) Consortium | Berkshire and Surrey Pathology Services Lighthouse Laboratory and Alex Alderton, Roberto Amato, Jeffrey Barrett, Sonia Goncalves, Ewan Harrison, David K. Jackson, Ian Johnston, Dominic Kwiatkowski, Cordelia Langford, John Sillitoe on behalf of the Wellcome Sanger Institute COVID-19 Surveillance Team |
| EPI_ISL_3380677 | CNR Virus des Infections Respiratoires - France SUD | CNR Virus des Infections Respiratoires - France SUD | Antonin Bal, Gregory Destras, Gwendolyne Burfin, Hadrien Regue, Quentin Semanas, Martine Valette, Bruno Lina, Laurence Josset |
| EPI_ISL_3380678 | Berkshire and Surrey Pathology Services Lighthouse Laboratory | Wellcome Sanger Institute for the COVID-19 Genomics UK (COG-UK) Consortium | Berkshire and Surrey Pathology Services Lighthouse Laboratory and Alex Alderton, Roberto Amato, Jeffrey Barrett, Sonia Goncalves, Ewan Harrison, David K. Jackson, Ian Johnston, Dominic Kwiatkowski, Cordelia Langford, John Sillitoe on behalf of the Wellcome Sanger Institute COVID-19 Surveillance Team |
| EPI_ISL_3380679 | Lighthouse Lab in Milton Keynes | Wellcome Sanger Institute for the COVID-19 Genomics UK (COG-UK) Consortium | The Lighthouse Lab in Milton Keynes and Alex Alderton, Roberto Amato, Jeffrey Barrett, Sonia Goncalves, Ewan Harrison, David K. Jackson, Ian Johnston, Dominic Kwiatkowski, Cordelia Langford, John Sillitoe on behalf of the Wellcome Sanger Institute COVID-19 Surveillance Team |
| EPI_ISL_3380680 | Berkshire and Surrey Pathology Services Lighthouse Laboratory | Wellcome Sanger Institute for the COVID-19 Genomics UK (COG-UK) Consortium | Berkshire and Surrey Pathology Services Lighthouse Laboratory and Alex Alderton, Roberto Amato, Jeffrey Barrett, Sonia Goncalves, Ewan Harrison, David K. Jackson, Ian Johnston, Dominic Kwiatkowski, Cordelia Langford, John Sillitoe on behalf of the Wellcome Sanger Institute COVID-19 Surveillance Team |
| EPI_ISL_3380681, EPI_ISL_3380682, EPI_ISL_3380683 | CNR Virus des Infections Respiratoires - France SUD | CNR Virus des Infections Respiratoires - France SUD | Antonin Bal, Gregory Destras, Gwendolyne Burfin, Hadrien Regue, Quentin Semanas, Martine Valette, Bruno Lina, Laurence Josset |
| EPI_ISL_3380684 | Health Services Laboratories | Wellcome Sanger Institute for the COVID-19 Genomics UK (COG-UK) Consortium | Health Services Laboratories and Alex Alderton, Roberto Amato, Jeffrey Barrett, Sonia Goncalves, Ewan Harrison, David K. Jackson, Ian Johnston, Dominic Kwiatkowski, Cordelia Langford, John Sillitoe on behalf of the Wellcome Sanger Institute COVID-19 Surveillance Team |
| EPI_ISL_3380685 | Lighthouse Lab in Milton Keynes | Wellcome Sanger Institute for the COVID-19 Genomics UK (COG-UK) Consortium | The Lighthouse Lab in Milton Keynes and Alex Alderton, Roberto Amato, Jeffrey Barrett, Sonia Goncalves, Ewan Harrison, David K. Jackson, Ian Johnston, Dominic Kwiatkowski, Cordelia Langford, John Sillitoe on behalf of the Wellcome Sanger Institute COVID-19 Surveillance Team |
| EPI_ISL_3380686 | CNR Virus des Infections Respiratoires - France SUD | CNR Virus des Infections Respiratoires - France SUD | Antonin Bal, Gregory Destras, Gwendolyne Burfin, Hadrien Regue, Quentin Semanas, Martine Valette, Bruno Lina, Laurence Josset |
| EPI_ISL_3380687 | Lighthouse Laboratory Plymouth | Wellcome Sanger Institute for the COVID-19 Genomics UK (COG-UK) Consortium | Lighthouse Laboratory Plymouth and Alex Alderton, Roberto Amato, Jeffrey Barrett, Sonia Goncalves, Ewan Harrison, David K. Jackson, Ian Johnston, Dominic Kwiatkowski, Cordelia Langford, John Sillitoe on behalf of the Wellcome Sanger Institute COVID-19 Surveillance Team |
| EPI_ISL_3380688, EPI_ISL_3380689 | Lighthouse Lab in Milton Keynes | Wellcome Sanger Institute for the COVID-19 Genomics UK (COG-UK) Consortium | The Lighthouse Lab in Milton Keynes and Alex Alderton, Roberto Amato, Jeffrey Barrett, Sonia Goncalves, Ewan Harrison, David K. Jackson, Ian Johnston, Dominic Kwiatkowski, Cordelia Langford, John Sillitoe on behalf of the Wellcome Sanger Institute COVID-19 Surveillance Team |
| EPI_ISL_3380690 | CNR Virus des Infections Respiratoires - France SUD | CNR Virus des Infections Respiratoires - France SUD | Antonin Bal, Gregory Destras, Gwendolyne Burfin, Hadrien Regue, Quentin Semanas, Martine Valette, Bruno Lina, Laurence Josset |
| EPI_ISL_3380691 | Berkshire and Surrey Pathology Services Lighthouse Laboratory | Wellcome Sanger Institute for the COVID-19 Genomics UK (COG-UK) | Berkshire and Surrey Pathology Services Lighthouse Laboratory and Alex Alderton, Roberto Amato, Jeffrey Barrett, Sonia Goncalves, Ewan Harrison, David K. Jackson, Ian Johnston, Dominic Kwiatkowski, Cordelia Langford, John Sillitoe on behalf |

| | | | |
|---|---|---|---|
| | | Consortium | of the Wellcome Sanger Institute COVID-19 Surveillance Team |
| EPI_ISL_3380692 | Lighthouse Lab in Milton Keynes | Wellcome Sanger Institute for the COVID-19 Genomics UK (COG-UK) Consortium | The Lighthouse Lab in Milton Keynes and Alex Alderton, Roberto Amato, Jeffrey Barrett, Sonia Goncalves, Ewan Harrison, David K. Jackson, Ian Johnston, Dominic Kwiatkowski, Cordelia Langford, John Sillitoe on behalf of the Wellcome Sanger Institute COVID-19 Surveillance Team |
| EPI_ISL_3380693 | CNR Virus des Infections Respiratoires - France SUD | CNR Virus des Infections Respiratoires - France SUD | Antonin Bal, Gregory Destras, Gwendolyne Burfin, Hadrien Regue, Quentin Semanas, Martine Valette, Bruno Lina, Laurence Josset |
| EPI_ISL_3380694 | Health Services Laboratories | Wellcome Sanger Institute for the COVID-19 Genomics UK (COG-UK) Consortium | Health Services Laboratories and Alex Alderton, Roberto Amato, Jeffrey Barrett, Sonia Goncalves, Ewan Harrison, David K. Jackson, Ian Johnston, Dominic Kwiatkowski, Cordelia Langford, John Sillitoe on behalf of the Wellcome Sanger Institute COVID-19 Surveillance Team |
| EPI_ISL_3380695 | Lighthouse Laboratory Plymouth | Wellcome Sanger Institute for the COVID-19 Genomics UK (COG-UK) Consortium | Lighthouse Laboratory Plymouth and Alex Alderton, Roberto Amato, Jeffrey Barrett, Sonia Goncalves, Ewan Harrison, David K. Jackson, Ian Johnston, Dominic Kwiatkowski, Cordelia Langford, John Sillitoe on behalf of the Wellcome Sanger Institute COVID-19 Surveillance Team |
| EPI_ISL_3380696 | Berkshire and Surrey Pathology Services Lighthouse Laboratory | Wellcome Sanger Institute for the COVID-19 Genomics UK (COG-UK) Consortium | Berkshire and Surrey Pathology Services Lighthouse Laboratory and Alex Alderton, Roberto Amato, Jeffrey Barrett, Sonia Goncalves, Ewan Harrison, David K. Jackson, Ian Johnston, Dominic Kwiatkowski, Cordelia Langford, John Sillitoe on behalf of the Wellcome Sanger Institute COVID-19 Surveillance Team |
| EPI_ISL_3380697 | CNR Virus des Infections Respiratoires - France SUD | CNR Virus des Infections Respiratoires - France SUD | Antonin Bal, Gregory Destras, Gwendolyne Burfin, Hadrien Regue, Quentin Semanas, Martine Valette, Bruno Lina, Laurence Josset |
| EPI_ISL_3380698 | Health Services Laboratories | Wellcome Sanger Institute for the COVID-19 Genomics UK (COG-UK) Consortium | Health Services Laboratories and Alex Alderton, Roberto Amato, Jeffrey Barrett, Sonia Goncalves, Ewan Harrison, David K. Jackson, Ian Johnston, Dominic Kwiatkowski, Cordelia Langford, John Sillitoe on behalf of the Wellcome Sanger Institute COVID-19 Surveillance Team |
| EPI_ISL_3380699 | Lighthouse Laboratory Plymouth | Wellcome Sanger Institute for the COVID-19 Genomics UK (COG-UK) Consortium | Lighthouse Laboratory Plymouth and Alex Alderton, Roberto Amato, Jeffrey Barrett, Sonia Goncalves, Ewan Harrison, David K. Jackson, Ian Johnston, Dominic Kwiatkowski, Cordelia Langford, John Sillitoe on behalf of the Wellcome Sanger Institute COVID-19 Surveillance Team |
| EPI_ISL_3380700 | Lighthouse Lab in Milton Keynes | Wellcome Sanger Institute for the COVID-19 Genomics UK (COG-UK) Consortium | The Lighthouse Lab in Milton Keynes and Alex Alderton, Roberto Amato, Jeffrey Barrett, Sonia Goncalves, Ewan Harrison, David K. Jackson, Ian Johnston, Dominic Kwiatkowski, Cordelia Langford, John Sillitoe on behalf of the Wellcome Sanger Institute COVID-19 Surveillance Team |
| EPI_ISL_3380701 | Health Services Laboratories | Wellcome Sanger Institute for the COVID-19 Genomics UK (COG-UK) Consortium | Health Services Laboratories and Alex Alderton, Roberto Amato, Jeffrey Barrett, Sonia Goncalves, Ewan Harrison, David K. Jackson, Ian Johnston, Dominic Kwiatkowski, Cordelia Langford, John Sillitoe on behalf of the Wellcome Sanger Institute COVID-19 Surveillance Team |
| EPI_ISL_3380702 | Berkshire and Surrey Pathology Services Lighthouse Laboratory | Wellcome Sanger Institute for the COVID-19 Genomics UK (COG-UK) Consortium | Berkshire and Surrey Pathology Services Lighthouse Laboratory and Alex Alderton, Roberto Amato, Jeffrey Barrett, Sonia Goncalves, Ewan Harrison, David K. Jackson, Ian Johnston, Dominic Kwiatkowski, Cordelia Langford, John Sillitoe on behalf of the Wellcome Sanger Institute COVID-19 Surveillance Team |
| EPI_ISL_3380703 | CNR Virus des Infections Respiratoires - France SUD | CNR Virus des Infections Respiratoires - France SUD | Antonin Bal, Gregory Destras, Gwendolyne Burfin, Hadrien Regue, Quentin Semanas, Martine Valette, Bruno Lina, Laurence Josset |
| EPI_ISL_3380704, EPI_ISL_3380705 | Health Services Laboratories | Wellcome Sanger Institute for the COVID-19 Genomics UK (COG-UK) Consortium | Health Services Laboratories and Alex Alderton, Roberto Amato, Jeffrey Barrett, Sonia Goncalves, Ewan Harrison, David K. Jackson, Ian Johnston, Dominic Kwiatkowski, Cordelia Langford, John Sillitoe on behalf of the Wellcome Sanger Institute COVID-19 Surveillance Team |
| EPI_ISL_3380706 | CNR Virus des Infections Respiratoires - France SUD | CNR Virus des Infections Respiratoires - France SUD | Antonin Bal, Gregory Destras, Gwendolyne Burfin, Hadrien Regue, Quentin Semanas, Martine Valette, Bruno Lina, Laurence Josset |
| EPI_ISL_3380707 | Lighthouse Lab in Milton Keynes | Wellcome Sanger Institute for the COVID-19 Genomics UK (COG-UK) Consortium | The Lighthouse Lab in Milton Keynes and Alex Alderton, Roberto Amato, Jeffrey Barrett, Sonia Goncalves, Ewan Harrison, David K. Jackson, Ian Johnston, Dominic Kwiatkowski, Cordelia Langford, John Sillitoe on behalf of the Wellcome Sanger Institute COVID-19 Surveillance Team |
| EPI_ISL_3380708 | Lighthouse Laboratory Plymouth | Wellcome Sanger Institute for the COVID-19 Genomics UK (COG-UK) Consortium | Lighthouse Laboratory Plymouth and Alex Alderton, Roberto Amato, Jeffrey Barrett, Sonia Goncalves, Ewan Harrison, David K. Jackson, Ian Johnston, Dominic Kwiatkowski, Cordelia Langford, John Sillitoe on behalf of the Wellcome Sanger Institute COVID-19 Surveillance Team |
| EPI_ISL_3380709 | CNR Virus des Infections Respiratoires - France SUD | CNR Virus des Infections Respiratoires - France SUD | Antonin Bal, Gregory Destras, Gwendolyne Burfin, Hadrien Regue, Quentin Semanas, Martine Valette, Bruno Lina, Laurence Josset |
| EPI_ISL_3380710 | Lighthouse Lab in Milton Keynes | Wellcome Sanger Institute for the COVID-19 Genomics UK (COG-UK) Consortium | The Lighthouse Lab in Milton Keynes and Alex Alderton, Roberto Amato, Jeffrey Barrett, Sonia Goncalves, Ewan Harrison, David K. Jackson, Ian Johnston, Dominic Kwiatkowski, Cordelia Langford, John Sillitoe on behalf of the Wellcome Sanger Institute COVID-19 Surveillance Team |
| EPI_ISL_3380711 | Berkshire and Surrey Pathology Services Lighthouse Laboratory | Wellcome Sanger Institute for the COVID-19 Genomics UK (COG-UK) Consortium | Berkshire and Surrey Pathology Services Lighthouse Laboratory and Alex Alderton, Roberto Amato, Jeffrey Barrett, Sonia Goncalves, Ewan Harrison, David K. Jackson, Ian Johnston, Dominic Kwiatkowski, Cordelia Langford, John Sillitoe on behalf of the Wellcome Sanger Institute COVID-19 Surveillance Team |
| EPI_ISL_3380712 | Lighthouse Lab in Milton Keynes | Wellcome Sanger Institute for the COVID-19 Genomics UK (COG-UK) Consortium | The Lighthouse Lab in Milton Keynes and Alex Alderton, Roberto Amato, Jeffrey Barrett, Sonia Goncalves, Ewan Harrison, David K. Jackson, Ian Johnston, Dominic Kwiatkowski, Cordelia Langford, John Sillitoe on behalf of the Wellcome Sanger Institute COVID-19 Surveillance Team |
| EPI_ISL_3380713 | CNR Virus des Infections Respiratoires - France SUD | CNR Virus des Infections Respiratoires - France SUD | Antonin Bal, Gregory Destras, Gwendolyne Burfin, Hadrien Regue, Quentin Semanas, Martine Valette, Bruno Lina, Laurence Josset |
| EPI_ISL_3380714, EPI_ISL_3380715 | Lighthouse Lab in Milton Keynes | Wellcome Sanger Institute for the COVID-19 Genomics UK (COG-UK) Consortium | The Lighthouse Lab in Milton Keynes and Alex Alderton, Roberto Amato, Jeffrey Barrett, Sonia Goncalves, Ewan Harrison, David K. Jackson, Ian Johnston, Dominic Kwiatkowski, Cordelia Langford, John Sillitoe on behalf of the Wellcome Sanger Institute COVID-19 Surveillance Team |
| EPI_ISL_3380716 | Berkshire and Surrey Pathology Services Lighthouse Laboratory | Wellcome Sanger Institute for the COVID-19 Genomics UK (COG-UK) Consortium | Berkshire and Surrey Pathology Services Lighthouse Laboratory and Alex Alderton, Roberto Amato, Jeffrey Barrett, Sonia Goncalves, Ewan Harrison, David K. Jackson, Ian Johnston, Dominic Kwiatkowski, Cordelia Langford, John Sillitoe on behalf of the Wellcome Sanger Institute COVID-19 Surveillance Team |
| EPI_ISL_3380717 | CNR Virus des Infections Respiratoires - France SUD | CNR Virus des Infections Respiratoires - France SUD | Antonin Bal, Gregory Destras, Gwendolyne Burfin, Hadrien Regue, Quentin Semanas, Martine Valette, Bruno Lina, Laurence Josset |
| EPI_ISL_3380718 | Lighthouse Lab in Milton Keynes | Wellcome Sanger Institute for the COVID-19 Genomics UK (COG-UK) Consortium | The Lighthouse Lab in Milton Keynes and Alex Alderton, Roberto Amato, Jeffrey Barrett, Sonia Goncalves, Ewan Harrison, David K. Jackson, Ian Johnston, Dominic Kwiatkowski, Cordelia Langford, John Sillitoe on behalf of the Wellcome Sanger Institute COVID-19 Surveillance Team |
| EPI_ISL_3380719 | Berkshire and Surrey Pathology Services Lighthouse Laboratory | Wellcome Sanger Institute for the COVID-19 Genomics UK (COG-UK) Consortium | Berkshire and Surrey Pathology Services Lighthouse Laboratory and Alex Alderton, Roberto Amato, Jeffrey Barrett, Sonia Goncalves, Ewan Harrison, David K. Jackson, Ian Johnston, Dominic Kwiatkowski, Cordelia Langford, John Sillitoe on behalf of the Wellcome Sanger Institute COVID-19 Surveillance Team |
| EPI_ISL_3380720 | Lighthouse Lab in Milton Keynes | Wellcome Sanger Institute for the COVID-19 Genomics UK (COG-UK) Consortium | The Lighthouse Lab in Milton Keynes and Alex Alderton, Roberto Amato, Jeffrey Barrett, Sonia Goncalves, Ewan Harrison, David K. Jackson, Ian Johnston, Dominic Kwiatkowski, Cordelia Langford, John Sillitoe on behalf of the Wellcome Sanger Institute COVID-19 Surveillance Team |
| EPI_ISL_3380721 | CNR Virus des Infections Respiratoires - France SUD | CNR Virus des Infections Respiratoires - France SUD | Antonin Bal, Gregory Destras, Gwendolyne Burfin, Hadrien Regue, Quentin Semanas, Martine Valette, Bruno Lina, Laurence Josset |
| EPI_ISL_3380722 | Lighthouse Lab in Milton Keynes | Wellcome Sanger Institute for the COVID-19 Genomics UK (COG-UK) Consortium | The Lighthouse Lab in Milton Keynes and Alex Alderton, Roberto Amato, Jeffrey Barrett, Sonia Goncalves, Ewan Harrison, David K. Jackson, Ian Johnston, Dominic Kwiatkowski, Cordelia Langford, John Sillitoe on behalf of the Wellcome Sanger Institute COVID-19 Surveillance Team |
| EPI_ISL_3380723 | Berkshire and Surrey Pathology Services Lighthouse Laboratory | Wellcome Sanger Institute for the COVID-19 Genomics UK (COG-UK) Consortium | Berkshire and Surrey Pathology Services Lighthouse Laboratory and Alex Alderton, Roberto Amato, Jeffrey Barrett, Sonia Goncalves, Ewan Harrison, David K. Jackson, Ian Johnston, Dominic Kwiatkowski, Cordelia Langford, John Sillitoe on behalf of the Wellcome Sanger Institute COVID-19 Surveillance Team |
| EPI_ISL_3380724 | Lighthouse Laboratory Plymouth | Wellcome Sanger Institute for the COVID-19 Genomics UK (COG-UK) Consortium | Lighthouse Laboratory Plymouth and Alex Alderton, Roberto Amato, Jeffrey Barrett, Sonia Goncalves, Ewan Harrison, David K. Jackson, Ian Johnston, Dominic Kwiatkowski, Cordelia Langford, John Sillitoe on behalf of the Wellcome Sanger Institute COVID-19 Surveillance Team |
| EPI_ISL_3380725 | CNR Virus des Infections Respiratoires - France SUD | CNR Virus des Infections Respiratoires - France SUD | Antonin Bal, Gregory Destras, Gwendolyne Burfin, Hadrien Regue, Quentin Semanas, Martine Valette, Bruno Lina, Laurence Josset |
| EPI_ISL_3380726 | Lighthouse Laboratory Plymouth | Wellcome Sanger Institute for the COVID-19 Genomics UK (COG-UK) Consortium | Lighthouse Laboratory Plymouth and Alex Alderton, Roberto Amato, Jeffrey Barrett, Sonia Goncalves, Ewan Harrison, David K. Jackson, Ian Johnston, Dominic Kwiatkowski, Cordelia Langford, John Sillitoe on behalf of the Wellcome Sanger Institute COVID-19 Surveillance Team |
| EPI_ISL_3380727 | Lighthouse Lab in Milton Keynes | Wellcome Sanger Institute for the COVID-19 Genomics UK (COG-UK) Consortium | The Lighthouse Lab in Milton Keynes and Alex Alderton, Roberto Amato, Jeffrey Barrett, Sonia Goncalves, Ewan Harrison, David K. Jackson, Ian Johnston, Dominic Kwiatkowski, Cordelia Langford, John Sillitoe on behalf of the Wellcome Sanger Institute COVID-19 Surveillance Team |
| EPI_ISL_3380728 | Berkshire and Surrey Pathology Services Lighthouse Laboratory | Wellcome Sanger Institute for the COVID-19 Genomics UK (COG-UK) Consortium | Berkshire and Surrey Pathology Services Lighthouse Laboratory and Alex Alderton, Roberto Amato, Jeffrey Barrett, Sonia Goncalves, Ewan Harrison, David K. Jackson, Ian Johnston, Dominic Kwiatkowski, Cordelia Langford, John Sillitoe on behalf of the Wellcome Sanger Institute COVID-19 Surveillance Team |
| EPI_ISL_3380729 | CNR Virus des Infections Respiratoires - France SUD | CNR Virus des Infections Respiratoires - France SUD | Antonin Bal, Gregory Destras, Gwendolyne Burfin, Hadrien Regue, Quentin Semanas, Martine Valette, Bruno Lina, Laurence Josset |
| EPI_ISL_3380730 | Health Services Laboratories | Wellcome Sanger Institute for the COVID-19 Genomics UK (COG-UK) Consortium | Health Services Laboratories and Alex Alderton, Roberto Amato, Jeffrey Barrett, Sonia Goncalves, Ewan Harrison, David K. Jackson, Ian Johnston, Dominic Kwiatkowski, Cordelia Langford, John Sillitoe on behalf of the Wellcome Sanger Institute COVID-19 Surveillance Team |
| EPI_ISL_3380731 | Lighthouse Lab in Milton Keynes | Wellcome Sanger Institute for the COVID-19 Genomics UK (COG-UK) Consortium | The Lighthouse Lab in Milton Keynes and Alex Alderton, Roberto Amato, Jeffrey Barrett, Sonia Goncalves, Ewan Harrison, David K. Jackson, Ian Johnston, Dominic Kwiatkowski, Cordelia Langford, John Sillitoe on behalf of the Wellcome Sanger Institute COVID-19 Surveillance Team |
| EPI_ISL_3380732 | Health Services Laboratories | Wellcome Sanger Institute for the COVID-19 Genomics UK (COG-UK) Consortium | Health Services Laboratories and Alex Alderton, Roberto Amato, Jeffrey Barrett, Sonia Goncalves, Ewan Harrison, David K. Jackson, Ian Johnston, Dominic Kwiatkowski, Cordelia Langford, John Sillitoe on behalf of the Wellcome Sanger Institute COVID-19 Surveillance Team |
| EPI_ISL_3380733 | Lighthouse Laboratory Plymouth | Wellcome Sanger Institute for the COVID-19 Genomics UK (COG-UK) Consortium | Lighthouse Laboratory Plymouth and Alex Alderton, Roberto Amato, Jeffrey Barrett, Sonia Goncalves, Ewan Harrison, David K. Jackson, Ian Johnston, Dominic Kwiatkowski, Cordelia Langford, John Sillitoe on behalf of the Wellcome Sanger Institute COVID-19 Surveillance Team |
| EPI_ISL_3380734 | CNR Virus des Infections Respiratoires - France SUD | CNR Virus des Infections Respiratoires - France SUD | Antonin Bal, Gregory Destras, Gwendolyne Burfin, Hadrien Regue, Quentin Semanas, Martine Valette, Bruno Lina, Laurence Josset |
| EPI_ISL_3380735 | Lighthouse Laboratory Plymouth | Wellcome Sanger Institute for the COVID-19 Genomics UK (COG-UK) Consortium | Lighthouse Laboratory Plymouth and Alex Alderton, Roberto Amato, Jeffrey Barrett, Sonia Goncalves, Ewan Harrison, David K. Jackson, Ian Johnston, Dominic Kwiatkowski, Cordelia Langford, John Sillitoe on behalf of the Wellcome Sanger Institute COVID-19 Surveillance Team |
| EPI_ISL_3380736 | Health Services Laboratories | Wellcome Sanger Institute for the COVID-19 Genomics UK (COG-UK) Consortium | Health Services Laboratories and Alex Alderton, Roberto Amato, Jeffrey Barrett, Sonia Goncalves, Ewan Harrison, David K. Jackson, Ian Johnston, Dominic Kwiatkowski, Cordelia Langford, John Sillitoe on behalf of the Wellcome Sanger Institute COVID-19 Surveillance Team |
| EPI_ISL_3380737 | Lighthouse Laboratory Plymouth | Wellcome Sanger Institute for the COVID-19 Genomics UK (COG-UK) Consortium | Lighthouse Laboratory Plymouth and Alex Alderton, Roberto Amato, Jeffrey Barrett, Sonia Goncalves, Ewan Harrison, David K. Jackson, Ian Johnston, Dominic Kwiatkowski, Cordelia Langford, John Sillitoe on behalf of the Wellcome Sanger Institute COVID-19 Surveillance Team |
| EPI_ISL_3380738 | CNR Virus des Infections Respiratoires - France SUD | CNR Virus des Infections Respiratoires - France SUD | Antonin Bal, Gregory Destras, Gwendolyne Burfin, Hadrien Regue, Quentin Semanas, Martine Valette, Bruno Lina, Laurence Josset |
| EPI_ISL_3380739 | Lighthouse Laboratory Plymouth | Wellcome Sanger Institute for the COVID-19 Genomics UK (COG-UK) Consortium | Lighthouse Laboratory Plymouth and Alex Alderton, Roberto Amato, Jeffrey Barrett, Sonia Goncalves, Ewan Harrison, David K. Jackson, Ian Johnston, Dominic Kwiatkowski, Cordelia Langford, John Sillitoe on behalf of the Wellcome Sanger Institute COVID-19 Surveillance Team |
| EPI_ISL_3380740 | Berkshire and Surrey Pathology Services Lighthouse Laboratory | Wellcome Sanger Institute for the COVID-19 Genomics UK (COG-UK) Consortium | Berkshire and Surrey Pathology Services Lighthouse Laboratory and Alex Alderton, Roberto Amato, Jeffrey Barrett, Sonia Goncalves, Ewan Harrison, David K. Jackson, Ian Johnston, Dominic Kwiatkowski, Cordelia Langford, John Sillitoe on behalf of the Wellcome Sanger Institute COVID-19 Surveillance Team |
| EPI_ISL_3380741 | Lighthouse Lab in Milton Keynes | Wellcome Sanger Institute for the COVID-19 Genomics UK (COG-UK) Consortium | The Lighthouse Lab in Milton Keynes and Alex Alderton, Roberto Amato, Jeffrey Barrett, Sonia Goncalves, Ewan Harrison, David K. Jackson, Ian Johnston, Dominic Kwiatkowski, Cordelia Langford, John Sillitoe on behalf of the Wellcome Sanger Institute COVID-19 Surveillance Team |
| EPI_ISL_3380742 | CNR Virus des Infections Respiratoires - France SUD | CNR Virus des Infections Respiratoires - France SUD | Antonin Bal, Gregory Destras, Gwendolyne Burfin, Hadrien Regue, Quentin Semanas, Martine Valette, Bruno Lina, Laurence Josset |
| EPI_ISL_3380743 | Berkshire and Surrey Pathology Services Lighthouse Laboratory | Wellcome Sanger Institute for the COVID-19 Genomics UK (COG-UK) Consortium | Berkshire and Surrey Pathology Services Lighthouse Laboratory and Alex Alderton, Roberto Amato, Jeffrey Barrett, Sonia Goncalves, Ewan Harrison, David K. Jackson, Ian Johnston, Dominic Kwiatkowski, Cordelia Langford, John Sillitoe on behalf of the Wellcome Sanger Institute COVID-19 Surveillance Team |

| | | | |
|---|---|---|---|
| EPI_ISL_3380744 | Lighthouse Laboratory Plymouth | Wellcome Sanger Institute for the COVID-19 Genomics UK (COG-UK) Consortium | Lighthouse Laboratory Plymouth and Alex Alderton, Roberto Amato, Jeffrey Barrett, Sonia Goncalves, Ewan Harrison, David K. Jackson, Ian Johnston, Dominic Kwiatkowski, Cordelia Langford, John Sillitoe on behalf of the Wellcome Sanger Institute COVID-19 Surveillance Team |
| EPI_ISL_3380745 | Berkshire and Surrey Pathology Services Lighthouse Laboratory | Wellcome Sanger Institute for the COVID-19 Genomics UK (COG-UK) Consortium | Berkshire and Surrey Pathology Services Lighthouse Laboratory and Alex Alderton, Roberto Amato, Jeffrey Barrett, Sonia Goncalves, Ewan Harrison, David K. Jackson, Ian Johnston, Dominic Kwiatkowski, Cordelia Langford, John Sillitoe on behalf of the Wellcome Sanger Institute COVID-19 Surveillance Team |
| EPI_ISL_3380746 | CNR Virus des Infections Respiratoires - France SUD | CNR Virus des Infections Respiratoires - France SUD | Antonin Bal, Gregory Destras, Gwendolyne Burfin, Hadrien Regue, Quentin Semanas, Martine Valette, Bruno Lina, Laurence Josset |
| EPI_ISL_3380747 | Lighthouse Laboratory Plymouth | Wellcome Sanger Institute for the COVID-19 Genomics UK (COG-UK) Consortium | Lighthouse Laboratory Plymouth and Alex Alderton, Roberto Amato, Jeffrey Barrett, Sonia Goncalves, Ewan Harrison, David K. Jackson, Ian Johnston, Dominic Kwiatkowski, Cordelia Langford, John Sillitoe on behalf of the Wellcome Sanger Institute COVID-19 Surveillance Team |
| EPI_ISL_3380748 | Health Services Laboratories | Wellcome Sanger Institute for the COVID-19 Genomics UK (COG-UK) Consortium | Health Services Laboratories and Alex Alderton, Roberto Amato, Jeffrey Barrett, Sonia Goncalves, Ewan Harrison, David K. Jackson, Ian Johnston, Dominic Kwiatkowski, Cordelia Langford, John Sillitoe on behalf of the Wellcome Sanger Institute COVID-19 Surveillance Team |
| EPI_ISL_3380749 | CNR Virus des Infections Respiratoires - France SUD | CNR Virus des Infections Respiratoires - France SUD | Antonin Bal, Gregory Destras, Gwendolyne Burfin, Hadrien Regue, Quentin Semanas, Martine Valette, Bruno Lina, Laurence Josset |
| EPI_ISL_3380750 | Berkshire and Surrey Pathology Services Lighthouse Laboratory | Wellcome Sanger Institute for the COVID-19 Genomics UK (COG-UK) Consortium | Berkshire and Surrey Pathology Services Lighthouse Laboratory and Alex Alderton, Roberto Amato, Jeffrey Barrett, Sonia Goncalves, Ewan Harrison, David K. Jackson, Ian Johnston, Dominic Kwiatkowski, Cordelia Langford, John Sillitoe on behalf of the Wellcome Sanger Institute COVID-19 Surveillance Team |
| EPI_ISL_3380751 | Health Services Laboratories | Wellcome Sanger Institute for the COVID-19 Genomics UK (COG-UK) Consortium | Health Services Laboratories and Alex Alderton, Roberto Amato, Jeffrey Barrett, Sonia Goncalves, Ewan Harrison, David K. Jackson, Ian Johnston, Dominic Kwiatkowski, Cordelia Langford, John Sillitoe on behalf of the Wellcome Sanger Institute COVID-19 Surveillance Team |
| EPI_ISL_3380752 | Lighthouse Laboratory Plymouth | Wellcome Sanger Institute for the COVID-19 Genomics UK (COG-UK) Consortium | Lighthouse Laboratory Plymouth and Alex Alderton, Roberto Amato, Jeffrey Barrett, Sonia Goncalves, Ewan Harrison, David K. Jackson, Ian Johnston, Dominic Kwiatkowski, Cordelia Langford, John Sillitoe on behalf of the Wellcome Sanger Institute COVID-19 Surveillance Team |
| EPI_ISL_3380753 | Lighthouse Lab in Milton Keynes | Wellcome Sanger Institute for the COVID-19 Genomics UK (COG-UK) Consortium | The Lighthouse Lab in Milton Keynes and Alex Alderton, Roberto Amato, Jeffrey Barrett, Sonia Goncalves, Ewan Harrison, David K. Jackson, Ian Johnston, Dominic Kwiatkowski, Cordelia Langford, John Sillitoe on behalf of the Wellcome Sanger Institute COVID-19 Surveillance Team |
| EPI_ISL_3380754 | CNR Virus des Infections Respiratoires - France SUD | CNR Virus des Infections Respiratoires - France SUD | Antonin Bal, Gregory Destras, Gwendolyne Burfin, Hadrien Regue, Quentin Semanas, Martine Valette, Bruno Lina, Laurence Josset |
| EPI_ISL_3380755 | Berkshire and Surrey Pathology Services Lighthouse Laboratory | Wellcome Sanger Institute for the COVID-19 Genomics UK (COG-UK) Consortium | Berkshire and Surrey Pathology Services Lighthouse Laboratory and Alex Alderton, Roberto Amato, Jeffrey Barrett, Sonia Goncalves, Ewan Harrison, David K. Jackson, Ian Johnston, Dominic Kwiatkowski, Cordelia Langford, John Sillitoe on behalf of the Wellcome Sanger Institute COVID-19 Surveillance Team |
| EPI_ISL_3380756 | Lighthouse Laboratory Plymouth | Wellcome Sanger Institute for the COVID-19 Genomics UK (COG-UK) Consortium | Lighthouse Laboratory Plymouth and Alex Alderton, Roberto Amato, Jeffrey Barrett, Sonia Goncalves, Ewan Harrison, David K. Jackson, Ian Johnston, Dominic Kwiatkowski, Cordelia Langford, John Sillitoe on behalf of the Wellcome Sanger Institute COVID-19 Surveillance Team |
| EPI_ISL_3380757 | CNR Virus des Infections Respiratoires - France SUD | CNR Virus des Infections Respiratoires - France SUD | Antonin Bal, Gregory Destras, Gwendolyne Burfin, Hadrien Regue, Quentin Semanas, Martine Valette, Bruno Lina, Laurence Josset |
| EPI_ISL_3380758, EPI_ISL_3380759 | Lighthouse Lab in Milton Keynes | Wellcome Sanger Institute for the COVID-19 Genomics UK (COG-UK) Consortium | The Lighthouse Lab in Milton Keynes and Alex Alderton, Roberto Amato, Jeffrey Barrett, Sonia Goncalves, Ewan Harrison, David K. Jackson, Ian Johnston, Dominic Kwiatkowski, Cordelia Langford, John Sillitoe on behalf of the Wellcome Sanger Institute COVID-19 Surveillance Team |
| EPI_ISL_3380760 | CNR Virus des Infections Respiratoires - France SUD | CNR Virus des Infections Respiratoires - France SUD | Antonin Bal, Gregory Destras, Gwendolyne Burfin, Hadrien Regue, Quentin Semanas, Martine Valette, Bruno Lina, Laurence Josset |
| EPI_ISL_3380761 | Lighthouse Laboratory Plymouth | Wellcome Sanger Institute for the COVID-19 Genomics UK (COG-UK) Consortium | Lighthouse Laboratory Plymouth and Alex Alderton, Roberto Amato, Jeffrey Barrett, Sonia Goncalves, Ewan Harrison, David K. Jackson, Ian Johnston, Dominic Kwiatkowski, Cordelia Langford, John Sillitoe on behalf of the Wellcome Sanger Institute COVID-19 Surveillance Team |
| EPI_ISL_3380762 | Berkshire and Surrey Pathology Services Lighthouse Laboratory | Wellcome Sanger Institute for the COVID-19 Genomics UK (COG-UK) Consortium | Berkshire and Surrey Pathology Services Lighthouse Laboratory and Alex Alderton, Roberto Amato, Jeffrey Barrett, Sonia Goncalves, Ewan Harrison, David K. Jackson, Ian Johnston, Dominic Kwiatkowski, Cordelia Langford, John Sillitoe on behalf of the Wellcome Sanger Institute COVID-19 Surveillance Team |
| EPI_ISL_3380763 | Health Services Laboratories | Wellcome Sanger Institute for the COVID-19 Genomics UK (COG-UK) Consortium | Health Services Laboratories and Alex Alderton, Roberto Amato, Jeffrey Barrett, Sonia Goncalves, Ewan Harrison, David K. Jackson, Ian Johnston, Dominic Kwiatkowski, Cordelia Langford, John Sillitoe on behalf of the Wellcome Sanger Institute COVID-19 Surveillance Team |
| EPI_ISL_3380764 | Lighthouse Lab in Milton Keynes | Wellcome Sanger Institute for the COVID-19 Genomics UK (COG-UK) Consortium | The Lighthouse Lab in Milton Keynes and Alex Alderton, Roberto Amato, Jeffrey Barrett, Sonia Goncalves, Ewan Harrison, David K. Jackson, Ian Johnston, Dominic Kwiatkowski, Cordelia Langford, John Sillitoe on behalf of the Wellcome Sanger Institute COVID-19 Surveillance Team |
| EPI_ISL_3380765 | Health Services Laboratories | Wellcome Sanger Institute for the COVID-19 Genomics UK (COG-UK) Consortium | Health Services Laboratories and Alex Alderton, Roberto Amato, Jeffrey Barrett, Sonia Goncalves, Ewan Harrison, David K. Jackson, Ian Johnston, Dominic Kwiatkowski, Cordelia Langford, John Sillitoe on behalf of the Wellcome Sanger Institute COVID-19 Surveillance Team |
| EPI_ISL_3380766 | Lighthouse Lab in Milton Keynes | Wellcome Sanger Institute for the COVID-19 Genomics UK (COG-UK) Consortium | The Lighthouse Lab in Milton Keynes and Alex Alderton, Roberto Amato, Jeffrey Barrett, Sonia Goncalves, Ewan Harrison, David K. Jackson, Ian Johnston, Dominic Kwiatkowski, Cordelia Langford, John Sillitoe on behalf of the Wellcome Sanger Institute COVID-19 Surveillance Team |
| EPI_ISL_3380767 | CNR Virus des Infections Respiratoires - France SUD | CNR Virus des Infections Respiratoires - France SUD | Antonin Bal, Gregory Destras, Gwendolyne Burfin, Hadrien Regue, Quentin Semanas, Martine Valette, Bruno Lina, Laurence Josset |
| EPI_ISL_3380768 | Health Services Laboratories | Wellcome Sanger Institute for the COVID-19 Genomics UK (COG-UK) Consortium | Health Services Laboratories and Alex Alderton, Roberto Amato, Jeffrey Barrett, Sonia Goncalves, Ewan Harrison, David K. Jackson, Ian Johnston, Dominic Kwiatkowski, Cordelia Langford, John Sillitoe on behalf of the Wellcome Sanger Institute COVID-19 Surveillance Team |
| EPI_ISL_3380769 | Lighthouse Lab in Milton Keynes | Wellcome Sanger Institute for the COVID-19 Genomics UK (COG-UK) Consortium | The Lighthouse Lab in Milton Keynes and Alex Alderton, Roberto Amato, Jeffrey Barrett, Sonia Goncalves, Ewan Harrison, David K. Jackson, Ian Johnston, Dominic Kwiatkowski, Cordelia Langford, John Sillitoe on behalf of the Wellcome Sanger Institute COVID-19 Surveillance Team |
| EPI_ISL_3380770 | CNR Virus des Infections Respiratoires - France SUD | CNR Virus des Infections Respiratoires - France SUD | Antonin Bal, Gregory Destras, Gwendolyne Burfin, Hadrien Regue, Quentin Semanas, Martine Valette, Bruno Lina, Laurence Josset |
| EPI_ISL_3380771 | Berkshire and Surrey Pathology Services Lighthouse Laboratory | Wellcome Sanger Institute for the COVID-19 Genomics UK (COG-UK) Consortium | Berkshire and Surrey Pathology Services Lighthouse Laboratory and Alex Alderton, Roberto Amato, Jeffrey Barrett, Sonia Goncalves, Ewan Harrison, David K. Jackson, Ian Johnston, Dominic Kwiatkowski, Cordelia Langford, John Sillitoe on behalf of the Wellcome Sanger Institute COVID-19 Surveillance Team |
| EPI_ISL_3380772 | Lighthouse Lab in Milton Keynes | Wellcome Sanger Institute for the COVID-19 Genomics UK (COG-UK) Consortium | The Lighthouse Lab in Milton Keynes and Alex Alderton, Roberto Amato, Jeffrey Barrett, Sonia Goncalves, Ewan Harrison, David K. Jackson, Ian Johnston, Dominic Kwiatkowski, Cordelia Langford, John Sillitoe on behalf of the Wellcome Sanger Institute COVID-19 Surveillance Team |
| EPI_ISL_3380773 | Berkshire and Surrey Pathology Services Lighthouse Laboratory | Wellcome Sanger Institute for the COVID-19 Genomics UK (COG-UK) Consortium | Berkshire and Surrey Pathology Services Lighthouse Laboratory and Alex Alderton, Roberto Amato, Jeffrey Barrett, Sonia Goncalves, Ewan Harrison, David K. Jackson, Ian Johnston, Dominic Kwiatkowski, Cordelia Langford, John Sillitoe on behalf of the Wellcome Sanger Institute COVID-19 Surveillance Team |
| EPI_ISL_3380774 | CNR Virus des Infections Respiratoires - France SUD | CNR Virus des Infections Respiratoires - France SUD | Antonin Bal, Gregory Destras, Gwendolyne Burfin, Hadrien Regue, Quentin Semanas, Martine Valette, Bruno Lina, Laurence Josset |
| EPI_ISL_3380775 | Health Services Laboratories | Wellcome Sanger Institute for the COVID-19 Genomics UK (COG-UK) Consortium | Health Services Laboratories and Alex Alderton, Roberto Amato, Jeffrey Barrett, Sonia Goncalves, Ewan Harrison, David K. Jackson, Ian Johnston, Dominic Kwiatkowski, Cordelia Langford, John Sillitoe on behalf of the Wellcome Sanger Institute COVID-19 Surveillance Team |
| EPI_ISL_3380776, EPI_ISL_3380777 | Lighthouse Lab in Milton Keynes | Wellcome Sanger Institute for the COVID-19 Genomics UK (COG-UK) Consortium | The Lighthouse Lab in Milton Keynes and Alex Alderton, Roberto Amato, Jeffrey Barrett, Sonia Goncalves, Ewan Harrison, David K. Jackson, Ian Johnston, Dominic Kwiatkowski, Cordelia Langford, John Sillitoe on behalf of the Wellcome Sanger Institute COVID-19 Surveillance Team |
| EPI_ISL_3380778, EPI_ISL_3380779, EPI_ISL_3380780, EPI_ISL_3380781, EPI_ISL_3380782, EPI_ISL_3380783, EPI_ISL_3380784, EPI_ISL_3380785, EPI_ISL_3380786, EPI_ISL_3380787, EPI_ISL_3380788, EPI_ISL_3380789, EPI_ISL_3380790, EPI_ISL_3380791, EPI_ISL_3380792, EPI_ISL_3380793, EPI_ISL_3380794, EPI_ISL_3380795, EPI_ISL_3380796, EPI_ISL_3380797, EPI_ISL_3380798, EPI_ISL_3380799, EPI_ISL_3380800, EPI_ISL_3380801, EPI_ISL_3380802, EPI_ISL_3380803, EPI_ISL_3380804, EPI_ISL_3380805, EPI_ISL_3380806, EPI_ISL_3380807, EPI_ISL_3380808, EPI_ISL_3380809, EPI_ISL_3380810, EPI_ISL_3380811, EPI_ISL_3380812, EPI_ISL_3380813, EPI_ISL_3380814, EPI_ISL_3380815, EPI_ISL_3380816, EPI_ISL_3380817, EPI_ISL_3380818 | | | |
| see above | CNR Virus des Infections Respiratoires - France SUD | CNR Virus des Infections Respiratoires - France SUD | Antonin Bal, Gregory Destras, Gwendolyne Burfin, Hadrien Regue, Quentin Semanas, Martine Valette, Bruno Lina, Laurence Josset |
| EPI_ISL_3380819 | Lighthouse Laboratory Plymouth | Wellcome Sanger Institute for the COVID-19 Genomics UK (COG-UK) Consortium | Lighthouse Laboratory Plymouth and Alex Alderton, Roberto Amato, Jeffrey Barrett, Sonia Goncalves, Ewan Harrison, David K. Jackson, Ian Johnston, Dominic Kwiatkowski, Cordelia Langford, John Sillitoe on behalf of the Wellcome Sanger Institute COVID-19 Surveillance Team |
| EPI_ISL_3380820 | CNR Virus des Infections Respiratoires - France SUD | CNR Virus des Infections Respiratoires - France SUD | Antonin Bal, Gregory Destras, Gwendolyne Burfin, Hadrien Regue, Quentin Semanas, Martine Valette, Bruno Lina, Laurence Josset |
| EPI_ISL_3380821, EPI_ISL_3380822 | Berkshire and Surrey Pathology Services Lighthouse Laboratory | Wellcome Sanger Institute for the COVID-19 Genomics UK (COG-UK) Consortium | Berkshire and Surrey Pathology Services Lighthouse Laboratory and Alex Alderton, Roberto Amato, Jeffrey Barrett, Sonia Goncalves, Ewan Harrison, David K. Jackson, Ian Johnston, Dominic Kwiatkowski, Cordelia Langford, John Sillitoe on behalf of the Wellcome Sanger Institute COVID-19 Surveillance Team |
| EPI_ISL_3380823 | Health Services Laboratories | Wellcome Sanger Institute for the COVID-19 Genomics UK (COG-UK) Consortium | Health Services Laboratories and Alex Alderton, Roberto Amato, Jeffrey Barrett, Sonia Goncalves, Ewan Harrison, David K. Jackson, Ian Johnston, Dominic Kwiatkowski, Cordelia Langford, John Sillitoe on behalf of the Wellcome Sanger Institute COVID-19 Surveillance Team |
| EPI_ISL_3380824 | CNR Virus des Infections Respiratoires - France SUD | CNR Virus des Infections Respiratoires - France SUD | Antonin Bal, Gregory Destras, Gwendolyne Burfin, Hadrien Regue, Quentin Semanas, Martine Valette, Bruno Lina, Laurence Josset |
| EPI_ISL_3380825 | Lighthouse Laboratory Plymouth | Wellcome Sanger Institute for the COVID-19 Genomics UK (COG-UK) Consortium | Lighthouse Laboratory Plymouth and Alex Alderton, Roberto Amato, Jeffrey Barrett, Sonia Goncalves, Ewan Harrison, David K. Jackson, Ian Johnston, Dominic Kwiatkowski, Cordelia Langford, John Sillitoe on behalf of the Wellcome Sanger Institute COVID-19 Surveillance Team |
| EPI_ISL_3380826 | Berkshire and Surrey Pathology Services Lighthouse Laboratory | Wellcome Sanger Institute for the COVID-19 Genomics UK (COG-UK) Consortium | Berkshire and Surrey Pathology Services Lighthouse Laboratory and Alex Alderton, Roberto Amato, Jeffrey Barrett, Sonia Goncalves, Ewan Harrison, David K. Jackson, Ian Johnston, Dominic Kwiatkowski, Cordelia Langford, John Sillitoe on behalf of the Wellcome Sanger Institute COVID-19 Surveillance Team |
| EPI_ISL_3380827 | Lighthouse Laboratory Plymouth | Wellcome Sanger Institute for the COVID-19 Genomics UK (COG-UK) Consortium | Lighthouse Laboratory Plymouth and Alex Alderton, Roberto Amato, Jeffrey Barrett, Sonia Goncalves, Ewan Harrison, David K. Jackson, Ian Johnston, Dominic Kwiatkowski, Cordelia Langford, John Sillitoe on behalf of the Wellcome Sanger Institute COVID-19 Surveillance Team |
| EPI_ISL_3380828 | CNR Virus des Infections Respiratoires - France SUD | CNR Virus des Infections Respiratoires - France SUD | Antonin Bal, Gregory Destras, Gwendolyne Burfin, Hadrien Regue, Quentin Semanas, Martine Valette, Bruno Lina, Laurence Josset |
| EPI_ISL_3380829 | Lighthouse Lab in Milton Keynes | Wellcome Sanger Institute for the COVID-19 Genomics UK (COG-UK) Consortium | The Lighthouse Lab in Milton Keynes and Alex Alderton, Roberto Amato, Jeffrey Barrett, Sonia Goncalves, Ewan Harrison, David K. Jackson, Ian Johnston, Dominic Kwiatkowski, Cordelia Langford, John Sillitoe on behalf of the Wellcome Sanger Institute COVID-19 Surveillance Team |
| EPI_ISL_3380830, EPI_ISL_3380831 | Berkshire and Surrey Pathology Services Lighthouse Laboratory | Wellcome Sanger Institute for the COVID-19 Genomics UK (COG-UK) Consortium | Berkshire and Surrey Pathology Services Lighthouse Laboratory and Alex Alderton, Roberto Amato, Jeffrey Barrett, Sonia Goncalves, Ewan Harrison, David K. Jackson, Ian Johnston, Dominic Kwiatkowski, Cordelia Langford, John Sillitoe on behalf of the Wellcome Sanger Institute COVID-19 Surveillance Team |
| EPI_ISL_3380832, EPI_ISL_3380833 | Health Services Laboratories | Wellcome Sanger Institute for the COVID-19 Genomics UK (COG-UK) Consortium | Health Services Laboratories and Alex Alderton, Roberto Amato, Jeffrey Barrett, Sonia Goncalves, Ewan Harrison, David K. Jackson, Ian Johnston, Dominic Kwiatkowski, Cordelia Langford, John Sillitoe on behalf of the Wellcome Sanger Institute COVID-19 Surveillance Team |
| EPI_ISL_3380834 | CNR Virus des Infections Respiratoires - France SUD | CNR Virus des Infections Respiratoires - France SUD | Antonin Bal, Gregory Destras, Gwendolyne Burfin, Hadrien Regue, Quentin Semanas, Martine Valette, Bruno Lina, Laurence Josset |
| EPI_ISL_3380835 | Berkshire and Surrey Pathology Services Lighthouse Laboratory | Wellcome Sanger Institute for the COVID-19 Genomics UK (COG-UK) Consortium | Berkshire and Surrey Pathology Services Lighthouse Laboratory and Alex Alderton, Roberto Amato, Jeffrey Barrett, Sonia Goncalves, Ewan Harrison, David K. Jackson, Ian Johnston, Dominic Kwiatkowski, Cordelia Langford, John Sillitoe on behalf of the Wellcome Sanger Institute COVID-19 Surveillance Team |
| EPI_ISL_3380836 | Health Services Laboratories | Wellcome Sanger Institute for the COVID-19 Genomics UK (COG-UK) Consortium | Health Services Laboratories and Alex Alderton, Roberto Amato, Jeffrey Barrett, Sonia Goncalves, Ewan Harrison, David K. Jackson, Ian Johnston, Dominic Kwiatkowski, Cordelia Langford, John Sillitoe on behalf of the Wellcome Sanger Institute COVID-19 Surveillance Team |
| EPI_ISL_3380837 | Lighthouse Lab in Milton Keynes | Wellcome Sanger Institute for the COVID-19 Genomics UK (COG-UK) Consortium | The Lighthouse Lab in Milton Keynes and Alex Alderton, Roberto Amato, Jeffrey Barrett, Sonia Goncalves, Ewan Harrison, David K. Jackson, Ian Johnston, Dominic Kwiatkowski, Cordelia Langford, John Sillitoe on behalf of the Wellcome Sanger Institute COVID-19 Surveillance Team |

| | | | |
|---|---|---|---|
| EPI_ISL_3380838 | Berkshire and Surrey Pathology Services Lighthouse Laboratory | Wellcome Sanger Institute for the COVID-19 Genomics UK (COG-UK) Consortium | Berkshire and Surrey Pathology Services Lighthouse Laboratory and Alex Alderton, Roberto Amato, Jeffrey Barrett, Sonia Goncalves, Ewan Harrison, David K. Jackson, Ian Johnston, Dominic Kwiatkowski, Cordelia Langford, John Sillitoe on behalf of the Wellcome Sanger Institute COVID-19 Surveillance Team |
| EPI_ISL_3380839 | Lighthouse Laboratory Plymouth | Wellcome Sanger Institute for the COVID-19 Genomics UK (COG-UK) Consortium | Lighthouse Laboratory Plymouth and Alex Alderton, Roberto Amato, Jeffrey Barrett, Sonia Goncalves, Ewan Harrison, David K. Jackson, Ian Johnston, Dominic Kwiatkowski, Cordelia Langford, John Sillitoe on behalf of the Wellcome Sanger Institute COVID-19 Surveillance Team |
| EPI_ISL_3380840 | CNR Virus des Infections Respiratoires - France SUD | CNR Virus des Infections Respiratoires - France SUD | Antonin Bal, Gregory Destras, Gwendolyne Burfin, Hadrien Regue, Quentin Semanas, Martine Valette, Bruno Lina, Laurence Josset |
| EPI_ISL_3380841 | Lighthouse Laboratory Plymouth | Wellcome Sanger Institute for the COVID-19 Genomics UK (COG-UK) Consortium | Lighthouse Laboratory Plymouth and Alex Alderton, Roberto Amato, Jeffrey Barrett, Sonia Goncalves, Ewan Harrison, David K. Jackson, Ian Johnston, Dominic Kwiatkowski, Cordelia Langford, John Sillitoe on behalf of the Wellcome Sanger Institute COVID-19 Surveillance Team |
| EPI_ISL_3380842 | Berkshire and Surrey Pathology Services Lighthouse Laboratory | Wellcome Sanger Institute for the COVID-19 Genomics UK (COG-UK) Consortium | Berkshire and Surrey Pathology Services Lighthouse Laboratory and Alex Alderton, Roberto Amato, Jeffrey Barrett, Sonia Goncalves, Ewan Harrison, David K. Jackson, Ian Johnston, Dominic Kwiatkowski, Cordelia Langford, John Sillitoe on behalf of the Wellcome Sanger Institute COVID-19 Surveillance Team |
| EPI_ISL_3380843, EPI_ISL_3380844 | Lighthouse Laboratory Plymouth | Wellcome Sanger Institute for the COVID-19 Genomics UK (COG-UK) Consortium | Lighthouse Laboratory Plymouth and Alex Alderton, Roberto Amato, Jeffrey Barrett, Sonia Goncalves, Ewan Harrison, David K. Jackson, Ian Johnston, Dominic Kwiatkowski, Cordelia Langford, John Sillitoe on behalf of the Wellcome Sanger Institute COVID-19 Surveillance Team |
| EPI_ISL_3380845 | CNR Virus des Infections Respiratoires - France SUD | CNR Virus des Infections Respiratoires - France SUD | Antonin Bal, Gregory Destras, Gwendolyne Burfin, Hadrien Regue, Quentin Semanas, Martine Valette, Bruno Lina, Laurence Josset |
| EPI_ISL_3380846 | Berkshire and Surrey Pathology Services Lighthouse Laboratory | Wellcome Sanger Institute for the COVID-19 Genomics UK (COG-UK) Consortium | Berkshire and Surrey Pathology Services Lighthouse Laboratory and Alex Alderton, Roberto Amato, Jeffrey Barrett, Sonia Goncalves, Ewan Harrison, David K. Jackson, Ian Johnston, Dominic Kwiatkowski, Cordelia Langford, John Sillitoe on behalf of the Wellcome Sanger Institute COVID-19 Surveillance Team |
| EPI_ISL_3380847 | Lighthouse Lab in Milton Keynes | Wellcome Sanger Institute for the COVID-19 Genomics UK (COG-UK) Consortium | The Lighthouse Lab in Milton Keynes and Alex Alderton, Roberto Amato, Jeffrey Barrett, Sonia Goncalves, Ewan Harrison, David K. Jackson, Ian Johnston, Dominic Kwiatkowski, Cordelia Langford, John Sillitoe on behalf of the Wellcome Sanger Institute COVID-19 Surveillance Team |
| EPI_ISL_3380848 | Health Services Laboratories | Wellcome Sanger Institute for the COVID-19 Genomics UK (COG-UK) Consortium | Health Services Laboratories and Alex Alderton, Roberto Amato, Jeffrey Barrett, Sonia Goncalves, Ewan Harrison, David K. Jackson, Ian Johnston, Dominic Kwiatkowski, Cordelia Langford, John Sillitoe on behalf of the Wellcome Sanger Institute COVID-19 Surveillance Team |
| EPI_ISL_3380849 | Berkshire and Surrey Pathology Services Lighthouse Laboratory | Wellcome Sanger Institute for the COVID-19 Genomics UK (COG-UK) Consortium | Berkshire and Surrey Pathology Services Lighthouse Laboratory and Alex Alderton, Roberto Amato, Jeffrey Barrett, Sonia Goncalves, Ewan Harrison, David K. Jackson, Ian Johnston, Dominic Kwiatkowski, Cordelia Langford, John Sillitoe on behalf of the Wellcome Sanger Institute COVID-19 Surveillance Team |
| EPI_ISL_3380850 | Lighthouse Lab in Milton Keynes | Wellcome Sanger Institute for the COVID-19 Genomics UK (COG-UK) Consortium | The Lighthouse Lab in Milton Keynes and Alex Alderton, Roberto Amato, Jeffrey Barrett, Sonia Goncalves, Ewan Harrison, David K. Jackson, Ian Johnston, Dominic Kwiatkowski, Cordelia Langford, John Sillitoe on behalf of the Wellcome Sanger Institute COVID-19 Surveillance Team |
| EPI_ISL_3380851 | CNR Virus des Infections Respiratoires - France SUD | CNR Virus des Infections Respiratoires - France SUD | Antonin Bal, Gregory Destras, Gwendolyne Burfin, Hadrien Regue, Quentin Semanas, Martine Valette, Bruno Lina, Laurence Josset |
| EPI_ISL_3380852 | Health Services Laboratories | Wellcome Sanger Institute for the COVID-19 Genomics UK (COG-UK) Consortium | Health Services Laboratories and Alex Alderton, Roberto Amato, Jeffrey Barrett, Sonia Goncalves, Ewan Harrison, David K. Jackson, Ian Johnston, Dominic Kwiatkowski, Cordelia Langford, John Sillitoe on behalf of the Wellcome Sanger Institute COVID-19 Surveillance Team |
| EPI_ISL_3380853 | Lighthouse Laboratory Plymouth | Wellcome Sanger Institute for the COVID-19 Genomics UK (COG-UK) Consortium | Lighthouse Laboratory Plymouth and Alex Alderton, Roberto Amato, Jeffrey Barrett, Sonia Goncalves, Ewan Harrison, David K. Jackson, Ian Johnston, Dominic Kwiatkowski, Cordelia Langford, John Sillitoe on behalf of the Wellcome Sanger Institute COVID-19 Surveillance Team |
| EPI_ISL_3380854 | Berkshire and Surrey Pathology Services Lighthouse Laboratory | Wellcome Sanger Institute for the COVID-19 Genomics UK (COG-UK) Consortium | Berkshire and Surrey Pathology Services Lighthouse Laboratory and Alex Alderton, Roberto Amato, Jeffrey Barrett, Sonia Goncalves, Ewan Harrison, David K. Jackson, Ian Johnston, Dominic Kwiatkowski, Cordelia Langford, John Sillitoe on behalf of the Wellcome Sanger Institute COVID-19 Surveillance Team |
| EPI_ISL_3380855 | Lighthouse Laboratory Plymouth | Wellcome Sanger Institute for the COVID-19 Genomics UK (COG-UK) Consortium | Lighthouse Laboratory Plymouth and Alex Alderton, Roberto Amato, Jeffrey Barrett, Sonia Goncalves, Ewan Harrison, David K. Jackson, Ian Johnston, Dominic Kwiatkowski, Cordelia Langford, John Sillitoe on behalf of the Wellcome Sanger Institute COVID-19 Surveillance Team |
| EPI_ISL_3380856 | CNR Virus des Infections Respiratoires - France SUD | CNR Virus des Infections Respiratoires - France SUD | Antonin Bal, Gregory Destras, Gwendolyne Burfin, Hadrien Regue, Quentin Semanas, Martine Valette, Bruno Lina, Laurence Josset |
| EPI_ISL_3380857 | Health Services Laboratories | Wellcome Sanger Institute for the COVID-19 Genomics UK (COG-UK) Consortium | Health Services Laboratories and Alex Alderton, Roberto Amato, Jeffrey Barrett, Sonia Goncalves, Ewan Harrison, David K. Jackson, Ian Johnston, Dominic Kwiatkowski, Cordelia Langford, John Sillitoe on behalf of the Wellcome Sanger Institute COVID-19 Surveillance Team |
| EPI_ISL_3380858, EPI_ISL_3380859 | Lighthouse Lab in Milton Keynes | Wellcome Sanger Institute for the COVID-19 Genomics UK (COG-UK) Consortium | The Lighthouse Lab in Milton Keynes and Alex Alderton, Roberto Amato, Jeffrey Barrett, Sonia Goncalves, Ewan Harrison, David K. Jackson, Ian Johnston, Dominic Kwiatkowski, Cordelia Langford, John Sillitoe on behalf of the Wellcome Sanger Institute COVID-19 Surveillance Team |
| EPI_ISL_3380860 | Lighthouse Laboratory Plymouth | Wellcome Sanger Institute for the COVID-19 Genomics UK (COG-UK) Consortium | Lighthouse Laboratory Plymouth and Alex Alderton, Roberto Amato, Jeffrey Barrett, Sonia Goncalves, Ewan Harrison, David K. Jackson, Ian Johnston, Dominic Kwiatkowski, Cordelia Langford, John Sillitoe on behalf of the Wellcome Sanger Institute COVID-19 Surveillance Team |
| EPI_ISL_3380861 | CNR Virus des Infections Respiratoires - France SUD | CNR Virus des Infections Respiratoires - France SUD | Antonin Bal, Gregory Destras, Gwendolyne Burfin, Hadrien Regue, Quentin Semanas, Martine Valette, Bruno Lina, Laurence Josset |
| EPI_ISL_3380862 | Berkshire and Surrey Pathology Services Lighthouse Laboratory | Wellcome Sanger Institute for the COVID-19 Genomics UK (COG-UK) Consortium | Berkshire and Surrey Pathology Services Lighthouse Laboratory and Alex Alderton, Roberto Amato, Jeffrey Barrett, Sonia Goncalves, Ewan Harrison, David K. Jackson, Ian Johnston, Dominic Kwiatkowski, Cordelia Langford, John Sillitoe on behalf of the Wellcome Sanger Institute COVID-19 Surveillance Team |
| EPI_ISL_3380863 | Lighthouse Laboratory Plymouth | Wellcome Sanger Institute for the COVID-19 Genomics UK (COG-UK) Consortium | Lighthouse Laboratory Plymouth and Alex Alderton, Roberto Amato, Jeffrey Barrett, Sonia Goncalves, Ewan Harrison, David K. Jackson, Ian Johnston, Dominic Kwiatkowski, Cordelia Langford, John Sillitoe on behalf of the Wellcome Sanger Institute COVID-19 Surveillance Team |
| EPI_ISL_3380864 | Berkshire and Surrey Pathology Services Lighthouse Laboratory | Wellcome Sanger Institute for the COVID-19 Genomics UK (COG-UK) Consortium | Berkshire and Surrey Pathology Services Lighthouse Laboratory and Alex Alderton, Roberto Amato, Jeffrey Barrett, Sonia Goncalves, Ewan Harrison, David K. Jackson, Ian Johnston, Dominic Kwiatkowski, Cordelia Langford, John Sillitoe on behalf of the Wellcome Sanger Institute COVID-19 Surveillance Team |
| EPI_ISL_3380865 | Lighthouse Laboratory Plymouth | Wellcome Sanger Institute for the COVID-19 Genomics UK (COG-UK) Consortium | Lighthouse Laboratory Plymouth and Alex Alderton, Roberto Amato, Jeffrey Barrett, Sonia Goncalves, Ewan Harrison, David K. Jackson, Ian Johnston, Dominic Kwiatkowski, Cordelia Langford, John Sillitoe on behalf of the Wellcome Sanger Institute COVID-19 Surveillance Team |
| EPI_ISL_3380866 | Berkshire and Surrey Pathology Services Lighthouse Laboratory | Wellcome Sanger Institute for the COVID-19 Genomics UK (COG-UK) Consortium | Berkshire and Surrey Pathology Services Lighthouse Laboratory and Alex Alderton, Roberto Amato, Jeffrey Barrett, Sonia Goncalves, Ewan Harrison, David K. Jackson, Ian Johnston, Dominic Kwiatkowski, Cordelia Langford, John Sillitoe on behalf of the Wellcome Sanger Institute COVID-19 Surveillance Team |
| EPI_ISL_3380867 | CNR Virus des Infections Respiratoires - France SUD | CNR Virus des Infections Respiratoires - France SUD | Antonin Bal, Gregory Destras, Gwendolyne Burfin, Hadrien Regue, Quentin Semanas, Martine Valette, Bruno Lina, Laurence Josset |
| EPI_ISL_3380868 | Lighthouse Lab in Milton Keynes | Wellcome Sanger Institute for the COVID-19 Genomics UK (COG-UK) Consortium | The Lighthouse Lab in Milton Keynes and Alex Alderton, Roberto Amato, Jeffrey Barrett, Sonia Goncalves, Ewan Harrison, David K. Jackson, Ian Johnston, Dominic Kwiatkowski, Cordelia Langford, John Sillitoe on behalf of the Wellcome Sanger Institute COVID-19 Surveillance Team |
| EPI_ISL_3380869 | Lighthouse Laboratory Plymouth | Wellcome Sanger Institute for the COVID-19 Genomics UK (COG-UK) Consortium | Lighthouse Laboratory Plymouth and Alex Alderton, Roberto Amato, Jeffrey Barrett, Sonia Goncalves, Ewan Harrison, David K. Jackson, Ian Johnston, Dominic Kwiatkowski, Cordelia Langford, John Sillitoe on behalf of the Wellcome Sanger Institute COVID-19 Surveillance Team |
| EPI_ISL_3380870 | Berkshire and Surrey Pathology Services Lighthouse Laboratory | Wellcome Sanger Institute for the COVID-19 Genomics UK (COG-UK) Consortium | Berkshire and Surrey Pathology Services Lighthouse Laboratory and Alex Alderton, Roberto Amato, Jeffrey Barrett, Sonia Goncalves, Ewan Harrison, David K. Jackson, Ian Johnston, Dominic Kwiatkowski, Cordelia Langford, John Sillitoe on behalf of the Wellcome Sanger Institute COVID-19 Surveillance Team |
| EPI_ISL_3380871 | CNR Virus des Infections Respiratoires - France SUD | CNR Virus des Infections Respiratoires - France SUD | Antonin Bal, Gregory Destras, Gwendolyne Burfin, Hadrien Regue, Quentin Semanas, Martine Valette, Bruno Lina, Laurence Josset |
| EPI_ISL_3380872 | Lighthouse Lab in Milton Keynes | Wellcome Sanger Institute for the COVID-19 Genomics UK (COG-UK) Consortium | The Lighthouse Lab in Milton Keynes and Alex Alderton, Roberto Amato, Jeffrey Barrett, Sonia Goncalves, Ewan Harrison, David K. Jackson, Ian Johnston, Dominic Kwiatkowski, Cordelia Langford, John Sillitoe on behalf of the Wellcome Sanger Institute COVID-19 Surveillance Team |
| EPI_ISL_3380873 | Lighthouse Laboratory Plymouth | Wellcome Sanger Institute for the COVID-19 Genomics UK (COG-UK) Consortium | Lighthouse Laboratory Plymouth and Alex Alderton, Roberto Amato, Jeffrey Barrett, Sonia Goncalves, Ewan Harrison, David K. Jackson, Ian Johnston, Dominic Kwiatkowski, Cordelia Langford, John Sillitoe on behalf of the Wellcome Sanger Institute COVID-19 Surveillance Team |
| EPI_ISL_3380874 | Health Services Laboratories | Wellcome Sanger Institute for the COVID-19 Genomics UK (COG-UK) Consortium | Health Services Laboratories and Alex Alderton, Roberto Amato, Jeffrey Barrett, Sonia Goncalves, Ewan Harrison, David K. Jackson, Ian Johnston, Dominic Kwiatkowski, Cordelia Langford, John Sillitoe on behalf of the Wellcome Sanger Institute COVID-19 Surveillance Team |
| EPI_ISL_3380876 | Lighthouse Lab in Milton Keynes | Wellcome Sanger Institute for the COVID-19 Genomics UK (COG-UK) Consortium | The Lighthouse Lab in Milton Keynes and Alex Alderton, Roberto Amato, Jeffrey Barrett, Sonia Goncalves, Ewan Harrison, David K. Jackson, Ian Johnston, Dominic Kwiatkowski, Cordelia Langford, John Sillitoe on behalf of the Wellcome Sanger Institute COVID-19 Surveillance Team |
| EPI_ISL_3380877 | Health Services Laboratories | Wellcome Sanger Institute for the COVID-19 Genomics UK (COG-UK) Consortium | Health Services Laboratories and Alex Alderton, Roberto Amato, Jeffrey Barrett, Sonia Goncalves, Ewan Harrison, David K. Jackson, Ian Johnston, Dominic Kwiatkowski, Cordelia Langford, John Sillitoe on behalf of the Wellcome Sanger Institute COVID-19 Surveillance Team |
| EPI_ISL_3380878 | Lighthouse Lab in Milton Keynes | Wellcome Sanger Institute for the COVID-19 Genomics UK (COG-UK) Consortium | The Lighthouse Lab in Milton Keynes and Alex Alderton, Roberto Amato, Jeffrey Barrett, Sonia Goncalves, Ewan Harrison, David K. Jackson, Ian Johnston, Dominic Kwiatkowski, Cordelia Langford, John Sillitoe on behalf of the Wellcome Sanger Institute COVID-19 Surveillance Team |
| EPI_ISL_3380879 | CNR Virus des Infections Respiratoires - France SUD | CNR Virus des Infections Respiratoires - France SUD | Antonin Bal, Gregory Destras, Gwendolyne Burfin, Hadrien Regue, Quentin Semanas, Martine Valette, Bruno Lina, Laurence Josset |
| EPI_ISL_3380880, EPI_ISL_3380881 | Health Services Laboratories | Wellcome Sanger Institute for the COVID-19 Genomics UK (COG-UK) Consortium | Health Services Laboratories and Alex Alderton, Roberto Amato, Jeffrey Barrett, Sonia Goncalves, Ewan Harrison, David K. Jackson, Ian Johnston, Dominic Kwiatkowski, Cordelia Langford, John Sillitoe on behalf of the Wellcome Sanger Institute COVID-19 Surveillance Team |
| EPI_ISL_3380882 | CNR Virus des Infections Respiratoires - France SUD | CNR Virus des Infections Respiratoires - France SUD | Antonin Bal, Gregory Destras, Gwendolyne Burfin, Hadrien Regue, Quentin Semanas, Martine Valette, Bruno Lina, Laurence Josset |
| EPI_ISL_3380883 | Lighthouse Lab in Milton Keynes | Wellcome Sanger Institute for the COVID-19 Genomics UK (COG-UK) Consortium | The Lighthouse Lab in Milton Keynes and Alex Alderton, Roberto Amato, Jeffrey Barrett, Sonia Goncalves, Ewan Harrison, David K. Jackson, Ian Johnston, Dominic Kwiatkowski, Cordelia Langford, John Sillitoe on behalf of the Wellcome Sanger Institute COVID-19 Surveillance Team |
| EPI_ISL_3380884 | Lighthouse Laboratory Plymouth | Wellcome Sanger Institute for the COVID-19 Genomics UK (COG-UK) Consortium | Lighthouse Laboratory Plymouth and Alex Alderton, Roberto Amato, Jeffrey Barrett, Sonia Goncalves, Ewan Harrison, David K. Jackson, Ian Johnston, Dominic Kwiatkowski, Cordelia Langford, John Sillitoe on behalf of the Wellcome Sanger Institute COVID-19 Surveillance Team |
| EPI_ISL_3380885 | Health Services Laboratories | Wellcome Sanger Institute for the COVID-19 Genomics UK (COG-UK) Consortium | Health Services Laboratories and Alex Alderton, Roberto Amato, Jeffrey Barrett, Sonia Goncalves, Ewan Harrison, David K. Jackson, Ian Johnston, Dominic Kwiatkowski, Cordelia Langford, John Sillitoe on behalf of the Wellcome Sanger Institute COVID-19 Surveillance Team |
| EPI_ISL_3380886 | CNR Virus des Infections Respiratoires - France SUD | CNR Virus des Infections Respiratoires - France SUD | Antonin Bal, Gregory Destras, Gwendolyne Burfin, Hadrien Regue, Quentin Semanas, Martine Valette, Bruno Lina, Laurence Josset |
| EPI_ISL_3380887 | Health Services Laboratories | Wellcome Sanger Institute for the COVID-19 Genomics UK (COG-UK) Consortium | Health Services Laboratories and Alex Alderton, Roberto Amato, Jeffrey Barrett, Sonia Goncalves, Ewan Harrison, David K. Jackson, Ian Johnston, Dominic Kwiatkowski, Cordelia Langford, John Sillitoe on behalf of the Wellcome Sanger Institute COVID-19 Surveillance Team |
| EPI_ISL_3380888 | Lighthouse Lab in Milton Keynes | Wellcome Sanger Institute for the COVID-19 Genomics UK (COG-UK) Consortium | The Lighthouse Lab in Milton Keynes and Alex Alderton, Roberto Amato, Jeffrey Barrett, Sonia Goncalves, Ewan Harrison, David K. Jackson, Ian Johnston, Dominic Kwiatkowski, Cordelia Langford, John Sillitoe on behalf of the Wellcome Sanger Institute COVID-19 Surveillance Team |
| EPI_ISL_3380889 | CNR Virus des Infections Respiratoires - France SUD | CNR Virus des Infections Respiratoires - France SUD | Antonin Bal, Gregory Destras, Gwendolyne Burfin, Hadrien Regue, Quentin Semanas, Martine Valette, Bruno Lina, Laurence Josset |
| EPI_ISL_3380890 | Berkshire and Surrey Pathology Services Lighthouse Laboratory | Wellcome Sanger Institute for the COVID-19 Genomics UK (COG-UK) Consortium | Berkshire and Surrey Pathology Services Lighthouse Laboratory and Alex Alderton, Roberto Amato, Jeffrey Barrett, Sonia Goncalves, Ewan Harrison, David K. Jackson, Ian Johnston, Dominic Kwiatkowski, Cordelia Langford, John Sillitoe on behalf of the Wellcome Sanger Institute COVID-19 Surveillance Team |
| EPI_ISL_3380891 | Health Services Laboratories | Wellcome Sanger Institute for the COVID-19 Genomics UK (COG-UK) | Health Services Laboratories and Alex Alderton, Roberto Amato, Jeffrey Barrett, Sonia Goncalves, Ewan Harrison, David K. Jackson, Ian Johnston, Dominic Kwiatkowski, Cordelia Langford, John Sillitoe on behalf of the Wellcome Sanger Institute |

| | | | |
|---|---|---|---|
| | | Consortium | COVID-19 Surveillance Team |
| EPI_ISL_3380892 | Lighthouse Laboratory Plymouth | Wellcome Sanger Institute for the COVID-19 Genomics UK (COG-UK) Consortium | Lighthouse Laboratory Plymouth and Alex Alderton, Roberto Amato, Jeffrey Barrett, Sonia Goncalves, Ewan Harrison, David K. Jackson, Ian Johnston, Dominic Kwiatkowski, Cordelia Langford, John Sillitoe on behalf of the Wellcome Sanger Institute COVID-19 Surveillance Team |
| EPI_ISL_3380893 | CNR Virus des Infections Respiratoires - France SUD | CNR Virus des Infections Respiratoires - France SUD | Antonin Bal, Gregory Destras, Gwendolyne Burfin, Hadrien Regue, Quentin Semanas, Martine Valette, Bruno Lina, Laurence Josset |
| EPI_ISL_3380894 | Lighthouse Lab in Milton Keynes | Wellcome Sanger Institute for the COVID-19 Genomics UK (COG-UK) Consortium | The Lighthouse Lab in Milton Keynes and Alex Alderton, Roberto Amato, Jeffrey Barrett, Sonia Goncalves, Ewan Harrison, David K. Jackson, Ian Johnston, Dominic Kwiatkowski, Cordelia Langford, John Sillitoe on behalf of the Wellcome Sanger Institute COVID-19 Surveillance Team |
| EPI_ISL_3380895 | Health Services Laboratories | Wellcome Sanger Institute for the COVID-19 Genomics UK (COG-UK) Consortium | Health Services Laboratories and Alex Alderton, Roberto Amato, Jeffrey Barrett, Sonia Goncalves, Ewan Harrison, David K. Jackson, Ian Johnston, Dominic Kwiatkowski, Cordelia Langford, John Sillitoe on behalf of the Wellcome Sanger Institute COVID-19 Surveillance Team |
| EPI_ISL_3380896 | Berkshire and Surrey Pathology Services Lighthouse Laboratory | Wellcome Sanger Institute for the COVID-19 Genomics UK (COG-UK) Consortium | Berkshire and Surrey Pathology Services Lighthouse Laboratory and Alex Alderton, Roberto Amato, Jeffrey Barrett, Sonia Goncalves, Ewan Harrison, David K. Jackson, Ian Johnston, Dominic Kwiatkowski, Cordelia Langford, John Sillitoe on behalf of the Wellcome Sanger Institute COVID-19 Surveillance Team |
| EPI_ISL_3380897 | CNR Virus des Infections Respiratoires - France SUD | CNR Virus des Infections Respiratoires - France SUD | Antonin Bal, Gregory Destras, Gwendolyne Burfin, Hadrien Regue, Quentin Semanas, Martine Valette, Bruno Lina, Laurence Josset |
| EPI_ISL_3380898 | Lighthouse Lab in Milton Keynes | Wellcome Sanger Institute for the COVID-19 Genomics UK (COG-UK) Consortium | The Lighthouse Lab in Milton Keynes and Alex Alderton, Roberto Amato, Jeffrey Barrett, Sonia Goncalves, Ewan Harrison, David K. Jackson, Ian Johnston, Dominic Kwiatkowski, Cordelia Langford, John Sillitoe on behalf of the Wellcome Sanger Institute COVID-19 Surveillance Team |
| EPI_ISL_3380899, EPI_ISL_3380900 | Lighthouse Laboratory Plymouth | Wellcome Sanger Institute for the COVID-19 Genomics UK (COG-UK) Consortium | Lighthouse Laboratory Plymouth and Alex Alderton, Roberto Amato, Jeffrey Barrett, Sonia Goncalves, Ewan Harrison, David K. Jackson, Ian Johnston, Dominic Kwiatkowski, Cordelia Langford, John Sillitoe on behalf of the Wellcome Sanger Institute COVID-19 Surveillance Team |
| EPI_ISL_3380901 | CNR Virus des Infections Respiratoires - France SUD | CNR Virus des Infections Respiratoires - France SUD | Antonin Bal, Gregory Destras, Gwendolyne Burfin, Hadrien Regue, Quentin Semanas, Martine Valette, Bruno Lina, Laurence Josset |
| EPI_ISL_3380902 | Lighthouse Laboratory Plymouth | Wellcome Sanger Institute for the COVID-19 Genomics UK (COG-UK) Consortium | Lighthouse Laboratory Plymouth and Alex Alderton, Roberto Amato, Jeffrey Barrett, Sonia Goncalves, Ewan Harrison, David K. Jackson, Ian Johnston, Dominic Kwiatkowski, Cordelia Langford, John Sillitoe on behalf of the Wellcome Sanger Institute COVID-19 Surveillance Team |
| EPI_ISL_3380903 | Lighthouse Lab in Milton Keynes | Wellcome Sanger Institute for the COVID-19 Genomics UK (COG-UK) Consortium | The Lighthouse Lab in Milton Keynes and Alex Alderton, Roberto Amato, Jeffrey Barrett, Sonia Goncalves, Ewan Harrison, David K. Jackson, Ian Johnston, Dominic Kwiatkowski, Cordelia Langford, John Sillitoe on behalf of the Wellcome Sanger Institute COVID-19 Surveillance Team |
| EPI_ISL_3380904 | CNR Virus des Infections Respiratoires - France SUD | CNR Virus des Infections Respiratoires - France SUD | Antonin Bal, Gregory Destras, Gwendolyne Burfin, Hadrien Regue, Quentin Semanas, Martine Valette, Bruno Lina, Laurence Josset |
| EPI_ISL_3380905 | Berkshire and Surrey Pathology Services Lighthouse Laboratory | Wellcome Sanger Institute for the COVID-19 Genomics UK (COG-UK) Consortium | Berkshire and Surrey Pathology Services Lighthouse Laboratory and Alex Alderton, Roberto Amato, Jeffrey Barrett, Sonia Goncalves, Ewan Harrison, David K. Jackson, Ian Johnston, Dominic Kwiatkowski, Cordelia Langford, John Sillitoe on behalf of the Wellcome Sanger Institute COVID-19 Surveillance Team |
| EPI_ISL_3380906 | Lighthouse Laboratory Plymouth | Wellcome Sanger Institute for the COVID-19 Genomics UK (COG-UK) Consortium | Lighthouse Laboratory Plymouth and Alex Alderton, Roberto Amato, Jeffrey Barrett, Sonia Goncalves, Ewan Harrison, David K. Jackson, Ian Johnston, Dominic Kwiatkowski, Cordelia Langford, John Sillitoe on behalf of the Wellcome Sanger Institute COVID-19 Surveillance Team |
| EPI_ISL_3380907 | Lighthouse Lab in Milton Keynes | Wellcome Sanger Institute for the COVID-19 Genomics UK (COG-UK) Consortium | The Lighthouse Lab in Milton Keynes and Alex Alderton, Roberto Amato, Jeffrey Barrett, Sonia Goncalves, Ewan Harrison, David K. Jackson, Ian Johnston, Dominic Kwiatkowski, Cordelia Langford, John Sillitoe on behalf of the Wellcome Sanger Institute COVID-19 Surveillance Team |
| EPI_ISL_3380908 | CNR Virus des Infections Respiratoires - France SUD | CNR Virus des Infections Respiratoires - France SUD | Antonin Bal, Gregory Destras, Gwendolyne Burfin, Hadrien Regue, Quentin Semanas, Martine Valette, Bruno Lina, Laurence Josset |
| EPI_ISL_3380909, EPI_ISL_3380910, EPI_ISL_3380911 | Lighthouse Lab in Milton Keynes | Wellcome Sanger Institute for the COVID-19 Genomics UK (COG-UK) Consortium | The Lighthouse Lab in Milton Keynes and Alex Alderton, Roberto Amato, Jeffrey Barrett, Sonia Goncalves, Ewan Harrison, David K. Jackson, Ian Johnston, Dominic Kwiatkowski, Cordelia Langford, John Sillitoe on behalf of the Wellcome Sanger Institute COVID-19 Surveillance Team |
| EPI_ISL_3380912 | CNR Virus des Infections Respiratoires - France SUD | CNR Virus des Infections Respiratoires - France SUD | Antonin Bal, Gregory Destras, Gwendolyne Burfin, Hadrien Regue, Quentin Semanas, Martine Valette, Bruno Lina, Laurence Josset |
| EPI_ISL_3380913 | Berkshire and Surrey Pathology Services Lighthouse Laboratory | Wellcome Sanger Institute for the COVID-19 Genomics UK (COG-UK) Consortium | Berkshire and Surrey Pathology Services Lighthouse Laboratory and Alex Alderton, Roberto Amato, Jeffrey Barrett, Sonia Goncalves, Ewan Harrison, David K. Jackson, Ian Johnston, Dominic Kwiatkowski, Cordelia Langford, John Sillitoe on behalf of the Wellcome Sanger Institute COVID-19 Surveillance Team |
| EPI_ISL_3380914 | Lighthouse Laboratory Plymouth | Wellcome Sanger Institute for the COVID-19 Genomics UK (COG-UK) Consortium | Lighthouse Laboratory Plymouth and Alex Alderton, Roberto Amato, Jeffrey Barrett, Sonia Goncalves, Ewan Harrison, David K. Jackson, Ian Johnston, Dominic Kwiatkowski, Cordelia Langford, John Sillitoe on behalf of the Wellcome Sanger Institute COVID-19 Surveillance Team |
| EPI_ISL_3380915 | CNR Virus des Infections Respiratoires - France SUD | CNR Virus des Infections Respiratoires - France SUD | Antonin Bal, Gregory Destras, Gwendolyne Burfin, Hadrien Regue, Quentin Semanas, Martine Valette, Bruno Lina, Laurence Josset |
| EPI_ISL_3380916, EPI_ISL_3380917, EPI_ISL_3380918 | Lighthouse Lab in Milton Keynes | Wellcome Sanger Institute for the COVID-19 Genomics UK (COG-UK) Consortium | The Lighthouse Lab in Milton Keynes and Alex Alderton, Roberto Amato, Jeffrey Barrett, Sonia Goncalves, Ewan Harrison, David K. Jackson, Ian Johnston, Dominic Kwiatkowski, Cordelia Langford, John Sillitoe on behalf of the Wellcome Sanger Institute COVID-19 Surveillance Team |
| EPI_ISL_3380919 | CNR Virus des Infections Respiratoires - France SUD | CNR Virus des Infections Respiratoires - France SUD | Antonin Bal, Gregory Destras, Gwendolyne Burfin, Hadrien Regue, Quentin Semanas, Martine Valette, Bruno Lina, Laurence Josset |
| EPI_ISL_3380920 | Lighthouse Lab in Milton Keynes | Wellcome Sanger Institute for the COVID-19 Genomics UK (COG-UK) Consortium | The Lighthouse Lab in Milton Keynes and Alex Alderton, Roberto Amato, Jeffrey Barrett, Sonia Goncalves, Ewan Harrison, David K. Jackson, Ian Johnston, Dominic Kwiatkowski, Cordelia Langford, John Sillitoe on behalf of the Wellcome Sanger Institute COVID-19 Surveillance Team |
| EPI_ISL_3380921 | Health Services Laboratories | Wellcome Sanger Institute for the COVID-19 Genomics UK (COG-UK) Consortium | Health Services Laboratories and Alex Alderton, Roberto Amato, Jeffrey Barrett, Sonia Goncalves, Ewan Harrison, David K. Jackson, Ian Johnston, Dominic Kwiatkowski, Cordelia Langford, John Sillitoe on behalf of the Wellcome Sanger Institute COVID-19 Surveillance Team |
| EPI_ISL_3380922 | Berkshire and Surrey Pathology Services Lighthouse Laboratory | Wellcome Sanger Institute for the COVID-19 Genomics UK (COG-UK) Consortium | Berkshire and Surrey Pathology Services Lighthouse Laboratory and Alex Alderton, Roberto Amato, Jeffrey Barrett, Sonia Goncalves, Ewan Harrison, David K. Jackson, Ian Johnston, Dominic Kwiatkowski, Cordelia Langford, John Sillitoe on behalf of the Wellcome Sanger Institute COVID-19 Surveillance Team |
| EPI_ISL_3380923 | CNR Virus des Infections Respiratoires - France SUD | CNR Virus des Infections Respiratoires - France SUD | Antonin Bal, Gregory Destras, Gwendolyne Burfin, Hadrien Regue, Quentin Semanas, Martine Valette, Bruno Lina, Laurence Josset |
| EPI_ISL_3380924 | Health Services Laboratories | Wellcome Sanger Institute for the COVID-19 Genomics UK (COG-UK) Consortium | Health Services Laboratories and Alex Alderton, Roberto Amato, Jeffrey Barrett, Sonia Goncalves, Ewan Harrison, David K. Jackson, Ian Johnston, Dominic Kwiatkowski, Cordelia Langford, John Sillitoe on behalf of the Wellcome Sanger Institute COVID-19 Surveillance Team |
| EPI_ISL_3380925 | Lighthouse Lab in Milton Keynes | Wellcome Sanger Institute for the COVID-19 Genomics UK (COG-UK) Consortium | The Lighthouse Lab in Milton Keynes and Alex Alderton, Roberto Amato, Jeffrey Barrett, Sonia Goncalves, Ewan Harrison, David K. Jackson, Ian Johnston, Dominic Kwiatkowski, Cordelia Langford, John Sillitoe on behalf of the Wellcome Sanger Institute COVID-19 Surveillance Team |
| EPI_ISL_3380926 | CNR Virus des Infections Respiratoires - France SUD | CNR Virus des Infections Respiratoires - France SUD | Antonin Bal, Gregory Destras, Gwendolyne Burfin, Hadrien Regue, Quentin Semanas, Martine Valette, Bruno Lina, Laurence Josset |
| EPI_ISL_3380927 | Health Services Laboratories | Wellcome Sanger Institute for the COVID-19 Genomics UK (COG-UK) Consortium | Health Services Laboratories and Alex Alderton, Roberto Amato, Jeffrey Barrett, Sonia Goncalves, Ewan Harrison, David K. Jackson, Ian Johnston, Dominic Kwiatkowski, Cordelia Langford, John Sillitoe on behalf of the Wellcome Sanger Institute COVID-19 Surveillance Team |
| EPI_ISL_3380928 | Lighthouse Laboratory Plymouth | Wellcome Sanger Institute for the COVID-19 Genomics UK (COG-UK) Consortium | Lighthouse Laboratory Plymouth and Alex Alderton, Roberto Amato, Jeffrey Barrett, Sonia Goncalves, Ewan Harrison, David K. Jackson, Ian Johnston, Dominic Kwiatkowski, Cordelia Langford, John Sillitoe on behalf of the Wellcome Sanger Institute COVID-19 Surveillance Team |
| EPI_ISL_3380929 | Berkshire and Surrey Pathology Services Lighthouse Laboratory | Wellcome Sanger Institute for the COVID-19 Genomics UK (COG-UK) Consortium | Berkshire and Surrey Pathology Services Lighthouse Laboratory and Alex Alderton, Roberto Amato, Jeffrey Barrett, Sonia Goncalves, Ewan Harrison, David K. Jackson, Ian Johnston, Dominic Kwiatkowski, Cordelia Langford, John Sillitoe on behalf of the Wellcome Sanger Institute COVID-19 Surveillance Team |
| EPI_ISL_3380930 | CNR Virus des Infections Respiratoires - France SUD | CNR Virus des Infections Respiratoires - France SUD | Antonin Bal, Gregory Destras, Gwendolyne Burfin, Hadrien Regue, Quentin Semanas, Martine Valette, Bruno Lina, Laurence Josset |
| EPI_ISL_3380931 | Berkshire and Surrey Pathology Services Lighthouse Laboratory | Wellcome Sanger Institute for the COVID-19 Genomics UK (COG-UK) Consortium | Berkshire and Surrey Pathology Services Lighthouse Laboratory and Alex Alderton, Roberto Amato, Jeffrey Barrett, Sonia Goncalves, Ewan Harrison, David K. Jackson, Ian Johnston, Dominic Kwiatkowski, Cordelia Langford, John Sillitoe on behalf of the Wellcome Sanger Institute COVID-19 Surveillance Team |
| EPI_ISL_3380932, EPI_ISL_3380933 | Lighthouse Laboratory Plymouth | Wellcome Sanger Institute for the COVID-19 Genomics UK (COG-UK) Consortium | Lighthouse Laboratory Plymouth and Alex Alderton, Roberto Amato, Jeffrey Barrett, Sonia Goncalves, Ewan Harrison, David K. Jackson, Ian Johnston, Dominic Kwiatkowski, Cordelia Langford, John Sillitoe on behalf of the Wellcome Sanger Institute COVID-19 Surveillance Team |
| EPI_ISL_3380934 | CNR Virus des Infections Respiratoires - France SUD | CNR Virus des Infections Respiratoires - France SUD | Antonin Bal, Gregory Destras, Gwendolyne Burfin, Hadrien Regue, Quentin Semanas, Martine Valette, Bruno Lina, Laurence Josset |
| EPI_ISL_3380935, EPI_ISL_3380936 | Health Services Laboratories | Wellcome Sanger Institute for the COVID-19 Genomics UK (COG-UK) Consortium | Health Services Laboratories and Alex Alderton, Roberto Amato, Jeffrey Barrett, Sonia Goncalves, Ewan Harrison, David K. Jackson, Ian Johnston, Dominic Kwiatkowski, Cordelia Langford, John Sillitoe on behalf of the Wellcome Sanger Institute COVID-19 Surveillance Team |
| EPI_ISL_3380937 | Lighthouse Lab in Milton Keynes | Wellcome Sanger Institute for the COVID-19 Genomics UK (COG-UK) Consortium | The Lighthouse Lab in Milton Keynes and Alex Alderton, Roberto Amato, Jeffrey Barrett, Sonia Goncalves, Ewan Harrison, David K. Jackson, Ian Johnston, Dominic Kwiatkowski, Cordelia Langford, John Sillitoe on behalf of the Wellcome Sanger Institute COVID-19 Surveillance Team |
| EPI_ISL_3380938 | CNR Virus des Infections Respiratoires - France SUD | CNR Virus des Infections Respiratoires - France SUD | Antonin Bal, Gregory Destras, Gwendolyne Burfin, Hadrien Regue, Quentin Semanas, Martine Valette, Bruno Lina, Laurence Josset |
| EPI_ISL_3380939, EPI_ISL_3380940 | Berkshire and Surrey Pathology Services Lighthouse Laboratory | Wellcome Sanger Institute for the COVID-19 Genomics UK (COG-UK) Consortium | Berkshire and Surrey Pathology Services Lighthouse Laboratory and Alex Alderton, Roberto Amato, Jeffrey Barrett, Sonia Goncalves, Ewan Harrison, David K. Jackson, Ian Johnston, Dominic Kwiatkowski, Cordelia Langford, John Sillitoe on behalf of the Wellcome Sanger Institute COVID-19 Surveillance Team |
| EPI_ISL_3380941 | Lighthouse Lab in Milton Keynes | Wellcome Sanger Institute for the COVID-19 Genomics UK (COG-UK) Consortium | The Lighthouse Lab in Milton Keynes and Alex Alderton, Roberto Amato, Jeffrey Barrett, Sonia Goncalves, Ewan Harrison, David K. Jackson, Ian Johnston, Dominic Kwiatkowski, Cordelia Langford, John Sillitoe on behalf of the Wellcome Sanger Institute COVID-19 Surveillance Team |
| EPI_ISL_3380942 | CNR Virus des Infections Respiratoires - France SUD | CNR Virus des Infections Respiratoires - France SUD | Antonin Bal, Gregory Destras, Gwendolyne Burfin, Hadrien Regue, Quentin Semanas, Martine Valette, Bruno Lina, Laurence Josset |
| EPI_ISL_3380943 | Lighthouse Lab in Milton Keynes | Wellcome Sanger Institute for the COVID-19 Genomics UK (COG-UK) Consortium | The Lighthouse Lab in Milton Keynes and Alex Alderton, Roberto Amato, Jeffrey Barrett, Sonia Goncalves, Ewan Harrison, David K. Jackson, Ian Johnston, Dominic Kwiatkowski, Cordelia Langford, John Sillitoe on behalf of the Wellcome Sanger Institute COVID-19 Surveillance Team |
| EPI_ISL_3380944 | Berkshire and Surrey Pathology Services Lighthouse Laboratory | Wellcome Sanger Institute for the COVID-19 Genomics UK (COG-UK) Consortium | Berkshire and Surrey Pathology Services Lighthouse Laboratory and Alex Alderton, Roberto Amato, Jeffrey Barrett, Sonia Goncalves, Ewan Harrison, David K. Jackson, Ian Johnston, Dominic Kwiatkowski, Cordelia Langford, John Sillitoe on behalf of the Wellcome Sanger Institute COVID-19 Surveillance Team |
| EPI_ISL_3380945 | CNR Virus des Infections Respiratoires - France SUD | CNR Virus des Infections Respiratoires - France SUD | Antonin Bal, Gregory Destras, Gwendolyne Burfin, Hadrien Regue, Quentin Semanas, Martine Valette, Bruno Lina, Laurence Josset |
| EPI_ISL_3380946, EPI_ISL_3380947 | Berkshire and Surrey Pathology Services Lighthouse Laboratory | Wellcome Sanger Institute for the COVID-19 Genomics UK (COG-UK) Consortium | Berkshire and Surrey Pathology Services Lighthouse Laboratory and Alex Alderton, Roberto Amato, Jeffrey Barrett, Sonia Goncalves, Ewan Harrison, David K. Jackson, Ian Johnston, Dominic Kwiatkowski, Cordelia Langford, John Sillitoe on behalf of the Wellcome Sanger Institute COVID-19 Surveillance Team |
| EPI_ISL_3380948 | Health Services Laboratories | Wellcome Sanger Institute for the COVID-19 Genomics UK (COG-UK) Consortium | Health Services Laboratories and Alex Alderton, Roberto Amato, Jeffrey Barrett, Sonia Goncalves, Ewan Harrison, David K. Jackson, Ian Johnston, Dominic Kwiatkowski, Cordelia Langford, John Sillitoe on behalf of the Wellcome Sanger Institute COVID-19 Surveillance Team |
| EPI_ISL_3380949 | CNR Virus des Infections Respiratoires - France SUD | CNR Virus des Infections Respiratoires - France SUD | Antonin Bal, Gregory Destras, Gwendolyne Burfin, Hadrien Regue, Quentin Semanas, Martine Valette, Bruno Lina, Laurence Josset |
| EPI_ISL_3380950 | Lighthouse Lab in Milton Keynes | Wellcome Sanger Institute for the COVID-19 Genomics UK (COG-UK) Consortium | The Lighthouse Lab in Milton Keynes and Alex Alderton, Roberto Amato, Jeffrey Barrett, Sonia Goncalves, Ewan Harrison, David K. Jackson, Ian Johnston, Dominic Kwiatkowski, Cordelia Langford, John Sillitoe on behalf of the Wellcome Sanger Institute COVID-19 Surveillance Team |

| | | | |
|---|---|---|---|
| EPI_ISL_3380951 | Health Services Laboratories | Wellcome Sanger Institute for the COVID-19 Genomics UK (COG-UK) Consortium | Health Services Laboratories and Alex Alderton, Roberto Amato, Jeffrey Barrett, Sonia Goncalves, Ewan Harrison, David K. Jackson, Ian Johnston, Dominic Kwiatkowski, Cordelia Langford, John Sillitoe on behalf of the Wellcome Sanger Institute COVID-19 Surveillance Team |
| EPI_ISL_3380952 | Berkshire and Surrey Pathology Services Lighthouse Laboratory | Wellcome Sanger Institute for the COVID-19 Genomics UK (COG-UK) Consortium | Berkshire and Surrey Pathology Services Lighthouse Laboratory and Alex Alderton, Roberto Amato, Jeffrey Barrett, Sonia Goncalves, Ewan Harrison, David K. Jackson, Ian Johnston, Dominic Kwiatkowski, Cordelia Langford, John Sillitoe on behalf of the Wellcome Sanger Institute COVID-19 Surveillance Team |
| EPI_ISL_3380953 | CNR Virus des Infections Respiratoires - France SUD | CNR Virus des Infections Respiratoires - France SUD | Antonin Bal, Gregory Destras, Gwendolyne Burfin, Hadrien Regue, Quentin Semanas, Martine Valette, Bruno Lina, Laurence Josset |
| EPI_ISL_3380954 | Berkshire and Surrey Pathology Services Lighthouse Laboratory | Wellcome Sanger Institute for the COVID-19 Genomics UK (COG-UK) Consortium | Berkshire and Surrey Pathology Services Lighthouse Laboratory and Alex Alderton, Roberto Amato, Jeffrey Barrett, Sonia Goncalves, Ewan Harrison, David K. Jackson, Ian Johnston, Dominic Kwiatkowski, Cordelia Langford, John Sillitoe on behalf of the Wellcome Sanger Institute COVID-19 Surveillance Team |
| EPI_ISL_3380955 | Lighthouse Laboratory Plymouth | Wellcome Sanger Institute for the COVID-19 Genomics UK (COG-UK) Consortium | Lighthouse Laboratory Plymouth and Alex Alderton, Roberto Amato, Jeffrey Barrett, Sonia Goncalves, Ewan Harrison, David K. Jackson, Ian Johnston, Dominic Kwiatkowski, Cordelia Langford, John Sillitoe on behalf of the Wellcome Sanger Institute COVID-19 Surveillance Team |
| EPI_ISL_3380956 | Lighthouse Lab in Milton Keynes | Wellcome Sanger Institute for the COVID-19 Genomics UK (COG-UK) Consortium | The Lighthouse Lab in Milton Keynes and Alex Alderton, Roberto Amato, Jeffrey Barrett, Sonia Goncalves, Ewan Harrison, David K. Jackson, Ian Johnston, Dominic Kwiatkowski, Cordelia Langford, John Sillitoe on behalf of the Wellcome Sanger Institute COVID-19 Surveillance Team |
| EPI_ISL_3380957 | CNR Virus des Infections Respiratoires - France SUD | CNR Virus des Infections Respiratoires - France SUD | Antonin Bal, Gregory Destras, Gwendolyne Burfin, Hadrien Regue, Quentin Semanas, Martine Valette, Bruno Lina, Laurence Josset |
| EPI_ISL_3380958 | Lighthouse Laboratory Plymouth | Wellcome Sanger Institute for the COVID-19 Genomics UK (COG-UK) Consortium | Lighthouse Laboratory Plymouth and Alex Alderton, Roberto Amato, Jeffrey Barrett, Sonia Goncalves, Ewan Harrison, David K. Jackson, Ian Johnston, Dominic Kwiatkowski, Cordelia Langford, John Sillitoe on behalf of the Wellcome Sanger Institute COVID-19 Surveillance Team |
| EPI_ISL_3380959 | Health Services Laboratories | Wellcome Sanger Institute for the COVID-19 Genomics UK (COG-UK) Consortium | Health Services Laboratories and Alex Alderton, Roberto Amato, Jeffrey Barrett, Sonia Goncalves, Ewan Harrison, David K. Jackson, Ian Johnston, Dominic Kwiatkowski, Cordelia Langford, John Sillitoe on behalf of the Wellcome Sanger Institute COVID-19 Surveillance Team |
| EPI_ISL_3380960, EPI_ISL_3380961 | Lighthouse Laboratory Plymouth | Wellcome Sanger Institute for the COVID-19 Genomics UK (COG-UK) Consortium | Lighthouse Laboratory Plymouth and Alex Alderton, Roberto Amato, Jeffrey Barrett, Sonia Goncalves, Ewan Harrison, David K. Jackson, Ian Johnston, Dominic Kwiatkowski, Cordelia Langford, John Sillitoe on behalf of the Wellcome Sanger Institute COVID-19 Surveillance Team |
| EPI_ISL_3380962 | Berkshire and Surrey Pathology Services Lighthouse Laboratory | Wellcome Sanger Institute for the COVID-19 Genomics UK (COG-UK) Consortium | Berkshire and Surrey Pathology Services Lighthouse Laboratory and Alex Alderton, Roberto Amato, Jeffrey Barrett, Sonia Goncalves, Ewan Harrison, David K. Jackson, Ian Johnston, Dominic Kwiatkowski, Cordelia Langford, John Sillitoe on behalf of the Wellcome Sanger Institute COVID-19 Surveillance Team |
| EPI_ISL_3380963 | Health Services Laboratories | Wellcome Sanger Institute for the COVID-19 Genomics UK (COG-UK) Consortium | Health Services Laboratories and Alex Alderton, Roberto Amato, Jeffrey Barrett, Sonia Goncalves, Ewan Harrison, David K. Jackson, Ian Johnston, Dominic Kwiatkowski, Cordelia Langford, John Sillitoe on behalf of the Wellcome Sanger Institute COVID-19 Surveillance Team |
| EPI_ISL_3380964 | Berkshire and Surrey Pathology Services Lighthouse Laboratory | Wellcome Sanger Institute for the COVID-19 Genomics UK (COG-UK) Consortium | Berkshire and Surrey Pathology Services Lighthouse Laboratory and Alex Alderton, Roberto Amato, Jeffrey Barrett, Sonia Goncalves, Ewan Harrison, David K. Jackson, Ian Johnston, Dominic Kwiatkowski, Cordelia Langford, John Sillitoe on behalf of the Wellcome Sanger Institute COVID-19 Surveillance Team |
| EPI_ISL_3380965 | Lighthouse Laboratory Plymouth | Wellcome Sanger Institute for the COVID-19 Genomics UK (COG-UK) Consortium | Lighthouse Laboratory Plymouth and Alex Alderton, Roberto Amato, Jeffrey Barrett, Sonia Goncalves, Ewan Harrison, David K. Jackson, Ian Johnston, Dominic Kwiatkowski, Cordelia Langford, John Sillitoe on behalf of the Wellcome Sanger Institute COVID-19 Surveillance Team |
| EPI_ISL_3380966, EPI_ISL_3380967, EPI_ISL_3380968 | Lighthouse Lab in Milton Keynes | Wellcome Sanger Institute for the COVID-19 Genomics UK (COG-UK) Consortium | The Lighthouse Lab in Milton Keynes and Alex Alderton, Roberto Amato, Jeffrey Barrett, Sonia Goncalves, Ewan Harrison, David K. Jackson, Ian Johnston, Dominic Kwiatkowski, Cordelia Langford, John Sillitoe on behalf of the Wellcome Sanger Institute COVID-19 Surveillance Team |
| EPI_ISL_3380969 | Berkshire and Surrey Pathology Services Lighthouse Laboratory | Wellcome Sanger Institute for the COVID-19 Genomics UK (COG-UK) Consortium | Berkshire and Surrey Pathology Services Lighthouse Laboratory and Alex Alderton, Roberto Amato, Jeffrey Barrett, Sonia Goncalves, Ewan Harrison, David K. Jackson, Ian Johnston, Dominic Kwiatkowski, Cordelia Langford, John Sillitoe on behalf of the Wellcome Sanger Institute COVID-19 Surveillance Team |
| EPI_ISL_3380970, EPI_ISL_3380971 | Lighthouse Lab in Milton Keynes | Wellcome Sanger Institute for the COVID-19 Genomics UK (COG-UK) Consortium | The Lighthouse Lab in Milton Keynes and Alex Alderton, Roberto Amato, Jeffrey Barrett, Sonia Goncalves, Ewan Harrison, David K. Jackson, Ian Johnston, Dominic Kwiatkowski, Cordelia Langford, John Sillitoe on behalf of the Wellcome Sanger Institute COVID-19 Surveillance Team |
| EPI_ISL_3380972 | Health Services Laboratories | Wellcome Sanger Institute for the COVID-19 Genomics UK (COG-UK) Consortium | Health Services Laboratories and Alex Alderton, Roberto Amato, Jeffrey Barrett, Sonia Goncalves, Ewan Harrison, David K. Jackson, Ian Johnston, Dominic Kwiatkowski, Cordelia Langford, John Sillitoe on behalf of the Wellcome Sanger Institute COVID-19 Surveillance Team |
| EPI_ISL_3380973 | Lighthouse Laboratory Plymouth | Wellcome Sanger Institute for the COVID-19 Genomics UK (COG-UK) Consortium | Lighthouse Laboratory Plymouth and Alex Alderton, Roberto Amato, Jeffrey Barrett, Sonia Goncalves, Ewan Harrison, David K. Jackson, Ian Johnston, Dominic Kwiatkowski, Cordelia Langford, John Sillitoe on behalf of the Wellcome Sanger Institute COVID-19 Surveillance Team |
| EPI_ISL_3380974 | Lighthouse Lab in Milton Keynes | Wellcome Sanger Institute for the COVID-19 Genomics UK (COG-UK) Consortium | The Lighthouse Lab in Milton Keynes and Alex Alderton, Roberto Amato, Jeffrey Barrett, Sonia Goncalves, Ewan Harrison, David K. Jackson, Ian Johnston, Dominic Kwiatkowski, Cordelia Langford, John Sillitoe on behalf of the Wellcome Sanger Institute COVID-19 Surveillance Team |
| EPI_ISL_3380975 | Lighthouse Laboratory Plymouth | Wellcome Sanger Institute for the COVID-19 Genomics UK (COG-UK) Consortium | Lighthouse Laboratory Plymouth and Alex Alderton, Roberto Amato, Jeffrey Barrett, Sonia Goncalves, Ewan Harrison, David K. Jackson, Ian Johnston, Dominic Kwiatkowski, Cordelia Langford, John Sillitoe on behalf of the Wellcome Sanger Institute COVID-19 Surveillance Team |
| EPI_ISL_3380976 | Health Services Laboratories | Wellcome Sanger Institute for the COVID-19 Genomics UK (COG-UK) Consortium | Health Services Laboratories and Alex Alderton, Roberto Amato, Jeffrey Barrett, Sonia Goncalves, Ewan Harrison, David K. Jackson, Ian Johnston, Dominic Kwiatkowski, Cordelia Langford, John Sillitoe on behalf of the Wellcome Sanger Institute COVID-19 Surveillance Team |
| EPI_ISL_3380977 | Lighthouse Laboratory Plymouth | Wellcome Sanger Institute for the COVID-19 Genomics UK (COG-UK) Consortium | Lighthouse Laboratory Plymouth and Alex Alderton, Roberto Amato, Jeffrey Barrett, Sonia Goncalves, Ewan Harrison, David K. Jackson, Ian Johnston, Dominic Kwiatkowski, Cordelia Langford, John Sillitoe on behalf of the Wellcome Sanger Institute COVID-19 Surveillance Team |
| EPI_ISL_3380978, EPI_ISL_3380979 | Lighthouse Lab in Milton Keynes | Wellcome Sanger Institute for the COVID-19 Genomics UK (COG-UK) Consortium | The Lighthouse Lab in Milton Keynes and Alex Alderton, Roberto Amato, Jeffrey Barrett, Sonia Goncalves, Ewan Harrison, David K. Jackson, Ian Johnston, Dominic Kwiatkowski, Cordelia Langford, John Sillitoe on behalf of the Wellcome Sanger Institute COVID-19 Surveillance Team |
| EPI_ISL_3380980 | Berkshire and Surrey Pathology Services Lighthouse Laboratory | Wellcome Sanger Institute for the COVID-19 Genomics UK (COG-UK) Consortium | Berkshire and Surrey Pathology Services Lighthouse Laboratory and Alex Alderton, Roberto Amato, Jeffrey Barrett, Sonia Goncalves, Ewan Harrison, David K. Jackson, Ian Johnston, Dominic Kwiatkowski, Cordelia Langford, John Sillitoe on behalf of the Wellcome Sanger Institute COVID-19 Surveillance Team |
| EPI_ISL_3380981 | Health Services Laboratories | Wellcome Sanger Institute for the COVID-19 Genomics UK (COG-UK) Consortium | Health Services Laboratories and Alex Alderton, Roberto Amato, Jeffrey Barrett, Sonia Goncalves, Ewan Harrison, David K. Jackson, Ian Johnston, Dominic Kwiatkowski, Cordelia Langford, John Sillitoe on behalf of the Wellcome Sanger Institute COVID-19 Surveillance Team |
| EPI_ISL_3380982, EPI_ISL_3380983 | Lighthouse Laboratory Plymouth | Wellcome Sanger Institute for the COVID-19 Genomics UK (COG-UK) Consortium | Lighthouse Laboratory Plymouth and Alex Alderton, Roberto Amato, Jeffrey Barrett, Sonia Goncalves, Ewan Harrison, David K. Jackson, Ian Johnston, Dominic Kwiatkowski, Cordelia Langford, John Sillitoe on behalf of the Wellcome Sanger Institute COVID-19 Surveillance Team |
| EPI_ISL_3380984 | Berkshire and Surrey Pathology Services Lighthouse Laboratory | Wellcome Sanger Institute for the COVID-19 Genomics UK (COG-UK) Consortium | Berkshire and Surrey Pathology Services Lighthouse Laboratory and Alex Alderton, Roberto Amato, Jeffrey Barrett, Sonia Goncalves, Ewan Harrison, David K. Jackson, Ian Johnston, Dominic Kwiatkowski, Cordelia Langford, John Sillitoe on behalf of the Wellcome Sanger Institute COVID-19 Surveillance Team |
| EPI_ISL_3380985 | Health Services Laboratories | Wellcome Sanger Institute for the COVID-19 Genomics UK (COG-UK) Consortium | Health Services Laboratories and Alex Alderton, Roberto Amato, Jeffrey Barrett, Sonia Goncalves, Ewan Harrison, David K. Jackson, Ian Johnston, Dominic Kwiatkowski, Cordelia Langford, John Sillitoe on behalf of the Wellcome Sanger Institute COVID-19 Surveillance Team |
| EPI_ISL_3380986 | Lighthouse Lab in Milton Keynes | Wellcome Sanger Institute for the COVID-19 Genomics UK (COG-UK) Consortium | The Lighthouse Lab in Milton Keynes and Alex Alderton, Roberto Amato, Jeffrey Barrett, Sonia Goncalves, Ewan Harrison, David K. Jackson, Ian Johnston, Dominic Kwiatkowski, Cordelia Langford, John Sillitoe on behalf of the Wellcome Sanger Institute COVID-19 Surveillance Team |
| EPI_ISL_3380987 | Lighthouse Laboratory Plymouth | Wellcome Sanger Institute for the COVID-19 Genomics UK (COG-UK) Consortium | Lighthouse Laboratory Plymouth and Alex Alderton, Roberto Amato, Jeffrey Barrett, Sonia Goncalves, Ewan Harrison, David K. Jackson, Ian Johnston, Dominic Kwiatkowski, Cordelia Langford, John Sillitoe on behalf of the Wellcome Sanger Institute COVID-19 Surveillance Team |
| EPI_ISL_3380988 | Lighthouse Lab in Milton Keynes | Wellcome Sanger Institute for the COVID-19 Genomics UK (COG-UK) Consortium | The Lighthouse Lab in Milton Keynes and Alex Alderton, Roberto Amato, Jeffrey Barrett, Sonia Goncalves, Ewan Harrison, David K. Jackson, Ian Johnston, Dominic Kwiatkowski, Cordelia Langford, John Sillitoe on behalf of the Wellcome Sanger Institute COVID-19 Surveillance Team |
| EPI_ISL_3380989 | Lighthouse Laboratory Plymouth | Wellcome Sanger Institute for the COVID-19 Genomics UK (COG-UK) Consortium | Lighthouse Laboratory Plymouth and Alex Alderton, Roberto Amato, Jeffrey Barrett, Sonia Goncalves, Ewan Harrison, David K. Jackson, Ian Johnston, Dominic Kwiatkowski, Cordelia Langford, John Sillitoe on behalf of the Wellcome Sanger Institute COVID-19 Surveillance Team |
| EPI_ISL_3380990, EPI_ISL_3380991 | Health Services Laboratories | Wellcome Sanger Institute for the COVID-19 Genomics UK (COG-UK) Consortium | Health Services Laboratories and Alex Alderton, Roberto Amato, Jeffrey Barrett, Sonia Goncalves, Ewan Harrison, David K. Jackson, Ian Johnston, Dominic Kwiatkowski, Cordelia Langford, John Sillitoe on behalf of the Wellcome Sanger Institute COVID-19 Surveillance Team |
| EPI_ISL_3380992 | Lighthouse Lab in Milton Keynes | Wellcome Sanger Institute for the COVID-19 Genomics UK (COG-UK) Consortium | The Lighthouse Lab in Milton Keynes and Alex Alderton, Roberto Amato, Jeffrey Barrett, Sonia Goncalves, Ewan Harrison, David K. Jackson, Ian Johnston, Dominic Kwiatkowski, Cordelia Langford, John Sillitoe on behalf of the Wellcome Sanger Institute COVID-19 Surveillance Team |
| EPI_ISL_3380993 | Berkshire and Surrey Pathology Services Lighthouse Laboratory | Wellcome Sanger Institute for the COVID-19 Genomics UK (COG-UK) Consortium | Berkshire and Surrey Pathology Services Lighthouse Laboratory and Alex Alderton, Roberto Amato, Jeffrey Barrett, Sonia Goncalves, Ewan Harrison, David K. Jackson, Ian Johnston, Dominic Kwiatkowski, Cordelia Langford, John Sillitoe on behalf of the Wellcome Sanger Institute COVID-19 Surveillance Team |
| EPI_ISL_3380994 | Lighthouse Lab in Milton Keynes | Wellcome Sanger Institute for the COVID-19 Genomics UK (COG-UK) Consortium | The Lighthouse Lab in Milton Keynes and Alex Alderton, Roberto Amato, Jeffrey Barrett, Sonia Goncalves, Ewan Harrison, David K. Jackson, Ian Johnston, Dominic Kwiatkowski, Cordelia Langford, John Sillitoe on behalf of the Wellcome Sanger Institute COVID-19 Surveillance Team |
| EPI_ISL_3380995, EPI_ISL_3380996 | Berkshire and Surrey Pathology Services Lighthouse Laboratory | Wellcome Sanger Institute for the COVID-19 Genomics UK (COG-UK) Consortium | Berkshire and Surrey Pathology Services Lighthouse Laboratory and Alex Alderton, Roberto Amato, Jeffrey Barrett, Sonia Goncalves, Ewan Harrison, David K. Jackson, Ian Johnston, Dominic Kwiatkowski, Cordelia Langford, John Sillitoe on behalf of the Wellcome Sanger Institute COVID-19 Surveillance Team |
| EPI_ISL_3380997 | Lighthouse Laboratory Plymouth | Wellcome Sanger Institute for the COVID-19 Genomics UK (COG-UK) Consortium | Lighthouse Laboratory Plymouth and Alex Alderton, Roberto Amato, Jeffrey Barrett, Sonia Goncalves, Ewan Harrison, David K. Jackson, Ian Johnston, Dominic Kwiatkowski, Cordelia Langford, John Sillitoe on behalf of the Wellcome Sanger Institute COVID-19 Surveillance Team |
| EPI_ISL_3380998 | Berkshire and Surrey Pathology Services Lighthouse Laboratory | Wellcome Sanger Institute for the COVID-19 Genomics UK (COG-UK) Consortium | Berkshire and Surrey Pathology Services Lighthouse Laboratory and Alex Alderton, Roberto Amato, Jeffrey Barrett, Sonia Goncalves, Ewan Harrison, David K. Jackson, Ian Johnston, Dominic Kwiatkowski, Cordelia Langford, John Sillitoe on behalf of the Wellcome Sanger Institute COVID-19 Surveillance Team |
| EPI_ISL_3380999 | Lighthouse Laboratory Plymouth | Wellcome Sanger Institute for the COVID-19 Genomics UK (COG-UK) Consortium | Lighthouse Laboratory Plymouth and Alex Alderton, Roberto Amato, Jeffrey Barrett, Sonia Goncalves, Ewan Harrison, David K. Jackson, Ian Johnston, Dominic Kwiatkowski, Cordelia Langford, John Sillitoe on behalf of the Wellcome Sanger Institute COVID-19 Surveillance Team |
| EPI_ISL_3381000 | Health Services Laboratories | Wellcome Sanger Institute for the COVID-19 Genomics UK (COG-UK) Consortium | Health Services Laboratories and Alex Alderton, Roberto Amato, Jeffrey Barrett, Sonia Goncalves, Ewan Harrison, David K. Jackson, Ian Johnston, Dominic Kwiatkowski, Cordelia Langford, John Sillitoe on behalf of the Wellcome Sanger Institute COVID-19 Surveillance Team |
| EPI_ISL_3381001 | Berkshire and Surrey Pathology Services Lighthouse Laboratory | Wellcome Sanger Institute for the COVID-19 Genomics UK (COG-UK) Consortium | Berkshire and Surrey Pathology Services Lighthouse Laboratory and Alex Alderton, Roberto Amato, Jeffrey Barrett, Sonia Goncalves, Ewan Harrison, David K. Jackson, Ian Johnston, Dominic Kwiatkowski, Cordelia Langford, John Sillitoe on behalf of the Wellcome Sanger Institute COVID-19 Surveillance Team |
| EPI_ISL_3381002 | Health Services Laboratories | Wellcome Sanger Institute for the COVID-19 Genomics UK (COG-UK) Consortium | Health Services Laboratories and Alex Alderton, Roberto Amato, Jeffrey Barrett, Sonia Goncalves, Ewan Harrison, David K. Jackson, Ian Johnston, Dominic Kwiatkowski, Cordelia Langford, John Sillitoe on behalf of the Wellcome Sanger Institute COVID-19 Surveillance Team |
| EPI_ISL_3381003 | Berkshire and Surrey Pathology Services Lighthouse Laboratory | Wellcome Sanger Institute for the COVID-19 Genomics UK (COG-UK) Consortium | Berkshire and Surrey Pathology Services Lighthouse Laboratory and Alex Alderton, Roberto Amato, Jeffrey Barrett, Sonia Goncalves, Ewan Harrison, David K. Jackson, Ian Johnston, Dominic Kwiatkowski, Cordelia Langford, John Sillitoe on behalf of the Wellcome Sanger Institute COVID-19 Surveillance Team |
| EPI_ISL_3381004, | Lighthouse Laboratory Plymouth | Wellcome Sanger Institute for the COVID-19 Genomics UK (COG-UK) | Lighthouse Laboratory Plymouth and Alex Alderton, Roberto Amato, Jeffrey Barrett, Sonia Goncalves, Ewan Harrison, David K. Jackson, Ian Johnston, Dominic Kwiatkowski, Cordelia Langford, John Sillitoe on behalf of the Wellcome Sanger |

| | | | |
|---|---|---|---|
| EPI_ISL_3381005 | | Consortium | Institute COVID-19 Surveillance Team |
| EPI_ISL_3381006 | Berkshire and Surrey Pathology Services Lighthouse Laboratory | Wellcome Sanger Institute for the COVID-19 Genomics UK (COG-UK) Consortium | Berkshire and Surrey Pathology Services Lighthouse Laboratory and Alex Alderton, Roberto Amato, Jeffrey Barrett, Sonia Goncalves, Ewan Harrison, David K. Jackson, Ian Johnston, Dominic Kwiatkowski, Cordelia Langford, John Sillitoe on behalf of the Wellcome Sanger Institute COVID-19 Surveillance Team |
| EPI_ISL_3381007 | Lighthouse Lab in Milton Keynes | Wellcome Sanger Institute for the COVID-19 Genomics UK (COG-UK) Consortium | The Lighthouse Lab in Milton Keynes and Alex Alderton, Roberto Amato, Jeffrey Barrett, Sonia Goncalves, Ewan Harrison, David K. Jackson, Ian Johnston, Dominic Kwiatkowski, Cordelia Langford, John Sillitoe on behalf of the Wellcome Sanger Institute COVID-19 Surveillance Team |
| EPI_ISL_3381008 | Health Services Laboratories | Wellcome Sanger Institute for the COVID-19 Genomics UK (COG-UK) Consortium | Health Services Laboratories and Alex Alderton, Roberto Amato, Jeffrey Barrett, Sonia Goncalves, Ewan Harrison, David K. Jackson, Ian Johnston, Dominic Kwiatkowski, Cordelia Langford, John Sillitoe on behalf of the Wellcome Sanger Institute COVID-19 Surveillance Team |
| EPI_ISL_3381009 | Lighthouse Lab in Milton Keynes | Wellcome Sanger Institute for the COVID-19 Genomics UK (COG-UK) Consortium | The Lighthouse Lab in Milton Keynes and Alex Alderton, Roberto Amato, Jeffrey Barrett, Sonia Goncalves, Ewan Harrison, David K. Jackson, Ian Johnston, Dominic Kwiatkowski, Cordelia Langford, John Sillitoe on behalf of the Wellcome Sanger Institute COVID-19 Surveillance Team |
| EPI_ISL_3381010 | Health Services Laboratories | Wellcome Sanger Institute for the COVID-19 Genomics UK (COG-UK) Consortium | Health Services Laboratories and Alex Alderton, Roberto Amato, Jeffrey Barrett, Sonia Goncalves, Ewan Harrison, David K. Jackson, Ian Johnston, Dominic Kwiatkowski, Cordelia Langford, John Sillitoe on behalf of the Wellcome Sanger Institute COVID-19 Surveillance Team |
| EPI_ISL_3381011 | Lighthouse Lab in Milton Keynes | Wellcome Sanger Institute for the COVID-19 Genomics UK (COG-UK) Consortium | The Lighthouse Lab in Milton Keynes and Alex Alderton, Roberto Amato, Jeffrey Barrett, Sonia Goncalves, Ewan Harrison, David K. Jackson, Ian Johnston, Dominic Kwiatkowski, Cordelia Langford, John Sillitoe on behalf of the Wellcome Sanger Institute COVID-19 Surveillance Team |
| EPI_ISL_3381012 | Health Services Laboratories | Wellcome Sanger Institute for the COVID-19 Genomics UK (COG-UK) Consortium | Health Services Laboratories and Alex Alderton, Roberto Amato, Jeffrey Barrett, Sonia Goncalves, Ewan Harrison, David K. Jackson, Ian Johnston, Dominic Kwiatkowski, Cordelia Langford, John Sillitoe on behalf of the Wellcome Sanger Institute COVID-19 Surveillance Team |
| EPI_ISL_3381013, EPI_ISL_3381014, EPI_ISL_3381015 | Berkshire and Surrey Pathology Services Lighthouse Laboratory | Wellcome Sanger Institute for the COVID-19 Genomics UK (COG-UK) Consortium | Berkshire and Surrey Pathology Services Lighthouse Laboratory and Alex Alderton, Roberto Amato, Jeffrey Barrett, Sonia Goncalves, Ewan Harrison, David K. Jackson, Ian Johnston, Dominic Kwiatkowski, Cordelia Langford, John Sillitoe on behalf of the Wellcome Sanger Institute COVID-19 Surveillance Team |
| EPI_ISL_3381016 | Lighthouse Lab in Milton Keynes | Wellcome Sanger Institute for the COVID-19 Genomics UK (COG-UK) Consortium | The Lighthouse Lab in Milton Keynes and Alex Alderton, Roberto Amato, Jeffrey Barrett, Sonia Goncalves, Ewan Harrison, David K. Jackson, Ian Johnston, Dominic Kwiatkowski, Cordelia Langford, John Sillitoe on behalf of the Wellcome Sanger Institute COVID-19 Surveillance Team |
| EPI_ISL_3381017, EPI_ISL_3381018, EPI_ISL_3381019 | Lighthouse Lab in Glasgow | Wellcome Sanger Institute for the COVID-19 Genomics UK (COG-UK) Consortium | Harper VanSteenhouse, Yumi Kasai, David Gray, Carol Clugston, Anna Dominiczak and Alex Alderton, Roberto Amato, Jeffrey Barrett, Sonia Goncalves, Ewan Harrison, David K. Jackson, Ian Johnston, Dominic Kwiatkowski, Cordelia Langford, John Sillitoe on behalf of the Wellcome Sanger Institute COVID-19 Surveillance Team |
| EPI_ISL_3381020 | Lighthouse Lab in Alderley Park | Wellcome Sanger Institute for the COVID-19 Genomics UK (COG-UK) Consortium | Jacquelyn Wynn, Mairead Hyland, The Lighthouse Lab in Alderley Park and Alex Alderton, Roberto Amato, Jeffrey Barrett, Sonia Goncalves, Ewan Harrison, David K. Jackson, Ian Johnston, Dominic Kwiatkowski, Cordelia Langford, John Sillitoe on behalf of the Wellcome Sanger Institute COVID-19 Surveillance Team |
| EPI_ISL_3381021 | Lighthouse Lab in Glasgow | Wellcome Sanger Institute for the COVID-19 Genomics UK (COG-UK) Consortium | Harper VanSteenhouse, Yumi Kasai, David Gray, Carol Clugston, Anna Dominiczak and Alex Alderton, Roberto Amato, Jeffrey Barrett, Sonia Goncalves, Ewan Harrison, David K. Jackson, Ian Johnston, Dominic Kwiatkowski, Cordelia Langford, John Sillitoe on behalf of the Wellcome Sanger Institute COVID-19 Surveillance Team |
| EPI_ISL_3381022 | Lighthouse Lab in Alderley Park | Wellcome Sanger Institute for the COVID-19 Genomics UK (COG-UK) Consortium | Jacquelyn Wynn, Mairead Hyland, The Lighthouse Lab in Alderley Park and Alex Alderton, Roberto Amato, Jeffrey Barrett, Sonia Goncalves, Ewan Harrison, David K. Jackson, Ian Johnston, Dominic Kwiatkowski, Cordelia Langford, John Sillitoe on behalf of the Wellcome Sanger Institute COVID-19 Surveillance Team |
| EPI_ISL_3381023 | Lighthouse Lab in Glasgow | Wellcome Sanger Institute for the COVID-19 Genomics UK (COG-UK) Consortium | Harper VanSteenhouse, Yumi Kasai, David Gray, Carol Clugston, Anna Dominiczak and Alex Alderton, Roberto Amato, Jeffrey Barrett, Sonia Goncalves, Ewan Harrison, David K. Jackson, Ian Johnston, Dominic Kwiatkowski, Cordelia Langford, John Sillitoe on behalf of the Wellcome Sanger Institute COVID-19 Surveillance Team |
| EPI_ISL_3381024, EPI_ISL_3381025 | Lighthouse Lab in Milton Keynes | Wellcome Sanger Institute for the COVID-19 Genomics UK (COG-UK) Consortium | The Lighthouse Lab in Milton Keynes and Alex Alderton, Roberto Amato, Jeffrey Barrett, Sonia Goncalves, Ewan Harrison, David K. Jackson, Ian Johnston, Dominic Kwiatkowski, Cordelia Langford, John Sillitoe on behalf of the Wellcome Sanger Institute COVID-19 Surveillance Team |
| EPI_ISL_3381026 | Lighthouse Lab in Glasgow | Wellcome Sanger Institute for the COVID-19 Genomics UK (COG-UK) Consortium | Harper VanSteenhouse, Yumi Kasai, David Gray, Carol Clugston, Anna Dominiczak and Alex Alderton, Roberto Amato, Jeffrey Barrett, Sonia Goncalves, Ewan Harrison, David K. Jackson, Ian Johnston, Dominic Kwiatkowski, Cordelia Langford, John Sillitoe on behalf of the Wellcome Sanger Institute COVID-19 Surveillance Team |
| EPI_ISL_3381027 | Lighthouse Lab in Milton Keynes | Wellcome Sanger Institute for the COVID-19 Genomics UK (COG-UK) Consortium | The Lighthouse Lab in Milton Keynes and Alex Alderton, Roberto Amato, Jeffrey Barrett, Sonia Goncalves, Ewan Harrison, David K. Jackson, Ian Johnston, Dominic Kwiatkowski, Cordelia Langford, John Sillitoe on behalf of the Wellcome Sanger Institute COVID-19 Surveillance Team |
| EPI_ISL_3381028, EPI_ISL_3381029, EPI_ISL_3381030, EPI_ISL_3381031, EPI_ISL_3381032, EPI_ISL_3381033, EPI_ISL_3381034, EPI_ISL_3381035, EPI_ISL_3381036, EPI_ISL_3381037, EPI_ISL_3381038, EPI_ISL_3381039, EPI_ISL_3381040, EPI_ISL_3381041, EPI_ISL_3381042, EPI_ISL_3381043, EPI_ISL_3381044 | | | |
| see above | Lighthouse Lab in Glasgow | Wellcome Sanger Institute for the COVID-19 Genomics UK (COG-UK) Consortium | Harper VanSteenhouse, Yumi Kasai, David Gray, Carol Clugston, Anna Dominiczak and Alex Alderton, Roberto Amato, Jeffrey Barrett, Sonia Goncalves, Ewan Harrison, David K. Jackson, Ian Johnston, Dominic Kwiatkowski, Cordelia Langford, John Sillitoe on behalf of the Wellcome Sanger Institute COVID-19 Surveillance Team |
| EPI_ISL_3381045 | Lighthouse Lab in Milton Keynes | Wellcome Sanger Institute for the COVID-19 Genomics UK (COG-UK) Consortium | The Lighthouse Lab in Milton Keynes and Alex Alderton, Roberto Amato, Jeffrey Barrett, Sonia Goncalves, Ewan Harrison, David K. Jackson, Ian Johnston, Dominic Kwiatkowski, Cordelia Langford, John Sillitoe on behalf of the Wellcome Sanger Institute COVID-19 Surveillance Team |
| EPI_ISL_3381046 | Berkshire and Surrey Pathology Services Lighthouse Laboratory | Wellcome Sanger Institute for the COVID-19 Genomics UK (COG-UK) Consortium | Berkshire and Surrey Pathology Services Lighthouse Laboratory and Alex Alderton, Roberto Amato, Jeffrey Barrett, Sonia Goncalves, Ewan Harrison, David K. Jackson, Ian Johnston, Dominic Kwiatkowski, Cordelia Langford, John Sillitoe on behalf of the Wellcome Sanger Institute COVID-19 Surveillance Team |
| EPI_ISL_3381047 | Lighthouse Lab in Glasgow | Wellcome Sanger Institute for the COVID-19 Genomics UK (COG-UK) Consortium | Harper VanSteenhouse, Yumi Kasai, David Gray, Carol Clugston, Anna Dominiczak and Alex Alderton, Roberto Amato, Jeffrey Barrett, Sonia Goncalves, Ewan Harrison, David K. Jackson, Ian Johnston, Dominic Kwiatkowski, Cordelia Langford, John Sillitoe on behalf of the Wellcome Sanger Institute COVID-19 Surveillance Team |
| EPI_ISL_3381048, EPI_ISL_3381049 | Lighthouse Lab in Alderley Park | Wellcome Sanger Institute for the COVID-19 Genomics UK (COG-UK) Consortium | Jacquelyn Wynn, Mairead Hyland, The Lighthouse Lab in Alderley Park and Alex Alderton, Roberto Amato, Jeffrey Barrett, Sonia Goncalves, Ewan Harrison, David K. Jackson, Ian Johnston, Dominic Kwiatkowski, Cordelia Langford, John Sillitoe on behalf of the Wellcome Sanger Institute COVID-19 Surveillance Team |
| EPI_ISL_3381050 | Lighthouse Lab in Glasgow | Wellcome Sanger Institute for the COVID-19 Genomics UK (COG-UK) Consortium | Harper VanSteenhouse, Yumi Kasai, David Gray, Carol Clugston, Anna Dominiczak and Alex Alderton, Roberto Amato, Jeffrey Barrett, Sonia Goncalves, Ewan Harrison, David K. Jackson, Ian Johnston, Dominic Kwiatkowski, Cordelia Langford, John Sillitoe on behalf of the Wellcome Sanger Institute COVID-19 Surveillance Team |
| EPI_ISL_3381051, EPI_ISL_3381052, EPI_ISL_3381053 | Lighthouse Lab in Alderley Park | Wellcome Sanger Institute for the COVID-19 Genomics UK (COG-UK) Consortium | Jacquelyn Wynn, Mairead Hyland, The Lighthouse Lab in Alderley Park and Alex Alderton, Roberto Amato, Jeffrey Barrett, Sonia Goncalves, Ewan Harrison, David K. Jackson, Ian Johnston, Dominic Kwiatkowski, Cordelia Langford, John Sillitoe on behalf of the Wellcome Sanger Institute COVID-19 Surveillance Team |
| EPI_ISL_3381054, EPI_ISL_3381055, EPI_ISL_3381056 | Lighthouse Lab in Glasgow | Wellcome Sanger Institute for the COVID-19 Genomics UK (COG-UK) Consortium | Harper VanSteenhouse, Yumi Kasai, David Gray, Carol Clugston, Anna Dominiczak and Alex Alderton, Roberto Amato, Jeffrey Barrett, Sonia Goncalves, Ewan Harrison, David K. Jackson, Ian Johnston, Dominic Kwiatkowski, Cordelia Langford, John Sillitoe on behalf of the Wellcome Sanger Institute COVID-19 Surveillance Team |
| EPI_ISL_3381057 | Lighthouse Lab in Alderley Park | Wellcome Sanger Institute for the COVID-19 Genomics UK (COG-UK) Consortium | Jacquelyn Wynn, Mairead Hyland, The Lighthouse Lab in Alderley Park and Alex Alderton, Roberto Amato, Jeffrey Barrett, Sonia Goncalves, Ewan Harrison, David K. Jackson, Ian Johnston, Dominic Kwiatkowski, Cordelia Langford, John Sillitoe on behalf of the Wellcome Sanger Institute COVID-19 Surveillance Team |
| EPI_ISL_3381058 | Lighthouse Lab in Glasgow | Wellcome Sanger Institute for the COVID-19 Genomics UK (COG-UK) Consortium | Harper VanSteenhouse, Yumi Kasai, David Gray, Carol Clugston, Anna Dominiczak and Alex Alderton, Roberto Amato, Jeffrey Barrett, Sonia Goncalves, Ewan Harrison, David K. Jackson, Ian Johnston, Dominic Kwiatkowski, Cordelia Langford, John Sillitoe on behalf of the Wellcome Sanger Institute COVID-19 Surveillance Team |
| EPI_ISL_3381059 | Lighthouse Lab in Alderley Park | Wellcome Sanger Institute for the COVID-19 Genomics UK (COG-UK) Consortium | Jacquelyn Wynn, Mairead Hyland, The Lighthouse Lab in Alderley Park and Alex Alderton, Roberto Amato, Jeffrey Barrett, Sonia Goncalves, Ewan Harrison, David K. Jackson, Ian Johnston, Dominic Kwiatkowski, Cordelia Langford, John Sillitoe on behalf of the Wellcome Sanger Institute COVID-19 Surveillance Team |
| EPI_ISL_3381060 | Lighthouse Lab in Glasgow | Wellcome Sanger Institute for the COVID-19 Genomics UK (COG-UK) Consortium | Harper VanSteenhouse, Yumi Kasai, David Gray, Carol Clugston, Anna Dominiczak and Alex Alderton, Roberto Amato, Jeffrey Barrett, Sonia Goncalves, Ewan Harrison, David K. Jackson, Ian Johnston, Dominic Kwiatkowski, Cordelia Langford, John Sillitoe on behalf of the Wellcome Sanger Institute COVID-19 Surveillance Team |
| EPI_ISL_3381061 | Lighthouse Lab in Alderley Park | Wellcome Sanger Institute for the COVID-19 Genomics UK (COG-UK) Consortium | Jacquelyn Wynn, Mairead Hyland, The Lighthouse Lab in Alderley Park and Alex Alderton, Roberto Amato, Jeffrey Barrett, Sonia Goncalves, Ewan Harrison, David K. Jackson, Ian Johnston, Dominic Kwiatkowski, Cordelia Langford, John Sillitoe on behalf of the Wellcome Sanger Institute COVID-19 Surveillance Team |
| EPI_ISL_3381062, EPI_ISL_3381063 | Lighthouse Lab in Glasgow | Wellcome Sanger Institute for the COVID-19 Genomics UK (COG-UK) Consortium | Harper VanSteenhouse, Yumi Kasai, David Gray, Carol Clugston, Anna Dominiczak and Alex Alderton, Roberto Amato, Jeffrey Barrett, Sonia Goncalves, Ewan Harrison, David K. Jackson, Ian Johnston, Dominic Kwiatkowski, Cordelia Langford, John Sillitoe on behalf of the Wellcome Sanger Institute COVID-19 Surveillance Team |
| EPI_ISL_3381064 | Lighthouse Laboratory Plymouth | Wellcome Sanger Institute for the COVID-19 Genomics UK (COG-UK) Consortium | Lighthouse Laboratory Plymouth and Alex Alderton, Roberto Amato, Jeffrey Barrett, Sonia Goncalves, Ewan Harrison, David K. Jackson, Ian Johnston, Dominic Kwiatkowski, Cordelia Langford, John Sillitoe on behalf of the Wellcome Sanger Institute COVID-19 Surveillance Team |
| EPI_ISL_3381065, EPI_ISL_3381066 | Lighthouse Lab in Alderley Park | Wellcome Sanger Institute for the COVID-19 Genomics UK (COG-UK) Consortium | Jacquelyn Wynn, Mairead Hyland, The Lighthouse Lab in Alderley Park and Alex Alderton, Roberto Amato, Jeffrey Barrett, Sonia Goncalves, Ewan Harrison, David K. Jackson, Ian Johnston, Dominic Kwiatkowski, Cordelia Langford, John Sillitoe on behalf of the Wellcome Sanger Institute COVID-19 Surveillance Team |
| EPI_ISL_3381067, EPI_ISL_3381068 | Lighthouse Lab in Glasgow | Wellcome Sanger Institute for the COVID-19 Genomics UK (COG-UK) Consortium | Harper VanSteenhouse, Yumi Kasai, David Gray, Carol Clugston, Anna Dominiczak and Alex Alderton, Roberto Amato, Jeffrey Barrett, Sonia Goncalves, Ewan Harrison, David K. Jackson, Ian Johnston, Dominic Kwiatkowski, Cordelia Langford, John Sillitoe on behalf of the Wellcome Sanger Institute COVID-19 Surveillance Team |
| EPI_ISL_3381069 | Lighthouse Lab in Alderley Park | Wellcome Sanger Institute for the COVID-19 Genomics UK (COG-UK) Consortium | Jacquelyn Wynn, Mairead Hyland, The Lighthouse Lab in Alderley Park and Alex Alderton, Roberto Amato, Jeffrey Barrett, Sonia Goncalves, Ewan Harrison, David K. Jackson, Ian Johnston, Dominic Kwiatkowski, Cordelia Langford, John Sillitoe on behalf of the Wellcome Sanger Institute COVID-19 Surveillance Team |
| EPI_ISL_3381070, EPI_ISL_3381071, EPI_ISL_3381072, EPI_ISL_3381073, EPI_ISL_3381074 | Lighthouse Lab in Glasgow | Wellcome Sanger Institute for the COVID-19 Genomics UK (COG-UK) Consortium | Harper VanSteenhouse, Yumi Kasai, David Gray, Carol Clugston, Anna Dominiczak and Alex Alderton, Roberto Amato, Jeffrey Barrett, Sonia Goncalves, Ewan Harrison, David K. Jackson, Ian Johnston, Dominic Kwiatkowski, Cordelia Langford, John Sillitoe on behalf of the Wellcome Sanger Institute COVID-19 Surveillance Team |
| EPI_ISL_3381075 | Lighthouse Lab in Alderley Park | Wellcome Sanger Institute for the COVID-19 Genomics UK (COG-UK) Consortium | Jacquelyn Wynn, Mairead Hyland, The Lighthouse Lab in Alderley Park and Alex Alderton, Roberto Amato, Jeffrey Barrett, Sonia Goncalves, Ewan Harrison, David K. Jackson, Ian Johnston, Dominic Kwiatkowski, Cordelia Langford, John Sillitoe on behalf of the Wellcome Sanger Institute COVID-19 Surveillance Team |
| EPI_ISL_3381076 | Lighthouse Lab in Glasgow | Wellcome Sanger Institute for the COVID-19 Genomics UK (COG-UK) Consortium | Harper VanSteenhouse, Yumi Kasai, David Gray, Carol Clugston, Anna Dominiczak and Alex Alderton, Roberto Amato, Jeffrey Barrett, Sonia Goncalves, Ewan Harrison, David K. Jackson, Ian Johnston, Dominic Kwiatkowski, Cordelia Langford, John Sillitoe on behalf of the Wellcome Sanger Institute COVID-19 Surveillance Team |
| EPI_ISL_3381077, EPI_ISL_3381078 | Lighthouse Lab in Alderley Park | Wellcome Sanger Institute for the COVID-19 Genomics UK (COG-UK) Consortium | Jacquelyn Wynn, Mairead Hyland, The Lighthouse Lab in Alderley Park and Alex Alderton, Roberto Amato, Jeffrey Barrett, Sonia Goncalves, Ewan Harrison, David K. Jackson, Ian Johnston, Dominic Kwiatkowski, Cordelia Langford, John Sillitoe on behalf of the Wellcome Sanger Institute COVID-19 Surveillance Team |
| EPI_ISL_3381079 | Lighthouse Lab in Glasgow | Wellcome Sanger Institute for the COVID-19 Genomics UK (COG-UK) Consortium | Harper VanSteenhouse, Yumi Kasai, David Gray, Carol Clugston, Anna Dominiczak and Alex Alderton, Roberto Amato, Jeffrey Barrett, Sonia Goncalves, Ewan Harrison, David K. Jackson, Ian Johnston, Dominic Kwiatkowski, Cordelia Langford, John Sillitoe on behalf of the Wellcome Sanger Institute COVID-19 Surveillance Team |
| EPI_ISL_3381080, EPI_ISL_3381081 | Lighthouse Lab in Alderley Park | Wellcome Sanger Institute for the COVID-19 Genomics UK (COG-UK) Consortium | Jacquelyn Wynn, Mairead Hyland, The Lighthouse Lab in Alderley Park and Alex Alderton, Roberto Amato, Jeffrey Barrett, Sonia Goncalves, Ewan Harrison, David K. Jackson, Ian Johnston, Dominic Kwiatkowski, Cordelia Langford, John Sillitoe on behalf of the Wellcome Sanger Institute COVID-19 Surveillance Team |

| | | | |
|---|---|---|---|
| EPI_ISL_3381082 | Lighthouse Lab in Glasgow | Wellcome Sanger Institute for the COVID-19 Genomics UK (COG-UK) Consortium | Harper VanSteenhouse, Yumi Kasai, David Gray, Carol Clugston, Anna Dominiczak and Alex Alderton, Roberto Amato, Jeffrey Barrett, Sonia Goncalves, Ewan Harrison, David K. Jackson, Ian Johnston, Dominic Kwiatkowski, Cordelia Langford, John Sillitoe on behalf of the Wellcome Sanger Institute COVID-19 Surveillance Team |
| EPI_ISL_3381083 | Lighthouse Lab in Alderley Park | Wellcome Sanger Institute for the COVID-19 Genomics UK (COG-UK) Consortium | Jacquelyn Wynn, Mairead Hyland, The Lighthouse Lab in Alderley Park and Alex Alderton, Roberto Amato, Jeffrey Barrett, Sonia Goncalves, Ewan Harrison, David K. Jackson, Ian Johnston, Dominic Kwiatkowski, Cordelia Langford, John Sillitoe on behalf of the Wellcome Sanger Institute COVID-19 Surveillance Team |
| EPI_ISL_3381084, EPI_ISL_3381085, EPI_ISL_3381086 | Lighthouse Lab in Glasgow | Wellcome Sanger Institute for the COVID-19 Genomics UK (COG-UK) Consortium | Harper VanSteenhouse, Yumi Kasai, David Gray, Carol Clugston, Anna Dominiczak and Alex Alderton, Roberto Amato, Jeffrey Barrett, Sonia Goncalves, Ewan Harrison, David K. Jackson, Ian Johnston, Dominic Kwiatkowski, Cordelia Langford, John Sillitoe on behalf of the Wellcome Sanger Institute COVID-19 Surveillance Team |
| EPI_ISL_3381087 | Lighthouse Lab in Alderley Park | Wellcome Sanger Institute for the COVID-19 Genomics UK (COG-UK) Consortium | Jacquelyn Wynn, Mairead Hyland, The Lighthouse Lab in Alderley Park and Alex Alderton, Roberto Amato, Jeffrey Barrett, Sonia Goncalves, Ewan Harrison, David K. Jackson, Ian Johnston, Dominic Kwiatkowski, Cordelia Langford, John Sillitoe on behalf of the Wellcome Sanger Institute COVID-19 Surveillance Team |
| EPI_ISL_3381088, EPI_ISL_3381089 | Lighthouse Lab in Glasgow | Wellcome Sanger Institute for the COVID-19 Genomics UK (COG-UK) Consortium | Harper VanSteenhouse, Yumi Kasai, David Gray, Carol Clugston, Anna Dominiczak and Alex Alderton, Roberto Amato, Jeffrey Barrett, Sonia Goncalves, Ewan Harrison, David K. Jackson, Ian Johnston, Dominic Kwiatkowski, Cordelia Langford, John Sillitoe on behalf of the Wellcome Sanger Institute COVID-19 Surveillance Team |
| EPI_ISL_3381090, EPI_ISL_3381091, EPI_ISL_3381092, EPI_ISL_3381093 | Lighthouse Lab in Alderley Park | Wellcome Sanger Institute for the COVID-19 Genomics UK (COG-UK) Consortium | Jacquelyn Wynn, Mairead Hyland, The Lighthouse Lab in Alderley Park and Alex Alderton, Roberto Amato, Jeffrey Barrett, Sonia Goncalves, Ewan Harrison, David K. Jackson, Ian Johnston, Dominic Kwiatkowski, Cordelia Langford, John Sillitoe on behalf of the Wellcome Sanger Institute COVID-19 Surveillance Team |
| EPI_ISL_3381094, EPI_ISL_3381095 | Lighthouse Lab in Glasgow | Wellcome Sanger Institute for the COVID-19 Genomics UK (COG-UK) Consortium | Harper VanSteenhouse, Yumi Kasai, David Gray, Carol Clugston, Anna Dominiczak and Alex Alderton, Roberto Amato, Jeffrey Barrett, Sonia Goncalves, Ewan Harrison, David K. Jackson, Ian Johnston, Dominic Kwiatkowski, Cordelia Langford, John Sillitoe on behalf of the Wellcome Sanger Institute COVID-19 Surveillance Team |
| EPI_ISL_3381096, EPI_ISL_3381097, EPI_ISL_3381098 | Lighthouse Lab in Alderley Park | Wellcome Sanger Institute for the COVID-19 Genomics UK (COG-UK) Consortium | Jacquelyn Wynn, Mairead Hyland, The Lighthouse Lab in Alderley Park and Alex Alderton, Roberto Amato, Jeffrey Barrett, Sonia Goncalves, Ewan Harrison, David K. Jackson, Ian Johnston, Dominic Kwiatkowski, Cordelia Langford, John Sillitoe on behalf of the Wellcome Sanger Institute COVID-19 Surveillance Team |
| EPI_ISL_3381099, EPI_ISL_3381100, EPI_ISL_3381101 | Lighthouse Lab in Glasgow | Wellcome Sanger Institute for the COVID-19 Genomics UK (COG-UK) Consortium | Harper VanSteenhouse, Yumi Kasai, David Gray, Carol Clugston, Anna Dominiczak and Alex Alderton, Roberto Amato, Jeffrey Barrett, Sonia Goncalves, Ewan Harrison, David K. Jackson, Ian Johnston, Dominic Kwiatkowski, Cordelia Langford, John Sillitoe on behalf of the Wellcome Sanger Institute COVID-19 Surveillance Team |
| EPI_ISL_3381102 | Lighthouse Lab in Alderley Park | Wellcome Sanger Institute for the COVID-19 Genomics UK (COG-UK) Consortium | Jacquelyn Wynn, Mairead Hyland, The Lighthouse Lab in Alderley Park and Alex Alderton, Roberto Amato, Jeffrey Barrett, Sonia Goncalves, Ewan Harrison, David K. Jackson, Ian Johnston, Dominic Kwiatkowski, Cordelia Langford, John Sillitoe on behalf of the Wellcome Sanger Institute COVID-19 Surveillance Team |
| EPI_ISL_3381103, EPI_ISL_3381104 | Lighthouse Lab in Glasgow | Wellcome Sanger Institute for the COVID-19 Genomics UK (COG-UK) Consortium | Harper VanSteenhouse, Yumi Kasai, David Gray, Carol Clugston, Anna Dominiczak and Alex Alderton, Roberto Amato, Jeffrey Barrett, Sonia Goncalves, Ewan Harrison, David K. Jackson, Ian Johnston, Dominic Kwiatkowski, Cordelia Langford, John Sillitoe on behalf of the Wellcome Sanger Institute COVID-19 Surveillance Team |
| EPI_ISL_3381105, EPI_ISL_3381106 | Lighthouse Lab in Alderley Park | Wellcome Sanger Institute for the COVID-19 Genomics UK (COG-UK) Consortium | Jacquelyn Wynn, Mairead Hyland, The Lighthouse Lab in Alderley Park and Alex Alderton, Roberto Amato, Jeffrey Barrett, Sonia Goncalves, Ewan Harrison, David K. Jackson, Ian Johnston, Dominic Kwiatkowski, Cordelia Langford, John Sillitoe on behalf of the Wellcome Sanger Institute COVID-19 Surveillance Team |
| EPI_ISL_3381107 | Lighthouse Lab in Glasgow | Wellcome Sanger Institute for the COVID-19 Genomics UK (COG-UK) Consortium | Harper VanSteenhouse, Yumi Kasai, David Gray, Carol Clugston, Anna Dominiczak and Alex Alderton, Roberto Amato, Jeffrey Barrett, Sonia Goncalves, Ewan Harrison, David K. Jackson, Ian Johnston, Dominic Kwiatkowski, Cordelia Langford, John Sillitoe on behalf of the Wellcome Sanger Institute COVID-19 Surveillance Team |
| EPI_ISL_3381108, EPI_ISL_3381109, EPI_ISL_3381110, EPI_ISL_3381111 | Lighthouse Lab in Alderley Park | Wellcome Sanger Institute for the COVID-19 Genomics UK (COG-UK) Consortium | Jacquelyn Wynn, Mairead Hyland, The Lighthouse Lab in Alderley Park and Alex Alderton, Roberto Amato, Jeffrey Barrett, Sonia Goncalves, Ewan Harrison, David K. Jackson, Ian Johnston, Dominic Kwiatkowski, Cordelia Langford, John Sillitoe on behalf of the Wellcome Sanger Institute COVID-19 Surveillance Team |
| EPI_ISL_3381112 | Lighthouse Lab in Glasgow | Wellcome Sanger Institute for the COVID-19 Genomics UK (COG-UK) Consortium | Harper VanSteenhouse, Yumi Kasai, David Gray, Carol Clugston, Anna Dominiczak and Alex Alderton, Roberto Amato, Jeffrey Barrett, Sonia Goncalves, Ewan Harrison, David K. Jackson, Ian Johnston, Dominic Kwiatkowski, Cordelia Langford, John Sillitoe on behalf of the Wellcome Sanger Institute COVID-19 Surveillance Team |
| EPI_ISL_3381113 | Lighthouse Lab in Alderley Park | Wellcome Sanger Institute for the COVID-19 Genomics UK (COG-UK) Consortium | Jacquelyn Wynn, Mairead Hyland, The Lighthouse Lab in Alderley Park and Alex Alderton, Roberto Amato, Jeffrey Barrett, Sonia Goncalves, Ewan Harrison, David K. Jackson, Ian Johnston, Dominic Kwiatkowski, Cordelia Langford, John Sillitoe on behalf of the Wellcome Sanger Institute COVID-19 Surveillance Team |
| EPI_ISL_3381114, EPI_ISL_3381115, EPI_ISL_3381116, EPI_ISL_3381117, EPI_ISL_3381118 | Lighthouse Lab in Glasgow | Wellcome Sanger Institute for the COVID-19 Genomics UK (COG-UK) Consortium | Harper VanSteenhouse, Yumi Kasai, David Gray, Carol Clugston, Anna Dominiczak and Alex Alderton, Roberto Amato, Jeffrey Barrett, Sonia Goncalves, Ewan Harrison, David K. Jackson, Ian Johnston, Dominic Kwiatkowski, Cordelia Langford, John Sillitoe on behalf of the Wellcome Sanger Institute COVID-19 Surveillance Team |
| EPI_ISL_3381119 | Lighthouse Laboratory Plymouth | Wellcome Sanger Institute for the COVID-19 Genomics UK (COG-UK) Consortium | Lighthouse Laboratory Plymouth and Alex Alderton, Roberto Amato, Jeffrey Barrett, Sonia Goncalves, Ewan Harrison, David K. Jackson, Ian Johnston, Dominic Kwiatkowski, Cordelia Langford, John Sillitoe on behalf of the Wellcome Sanger Institute COVID-19 Surveillance Team |
| EPI_ISL_3381120 | Lighthouse Lab in Glasgow | Wellcome Sanger Institute for the COVID-19 Genomics UK (COG-UK) Consortium | Harper VanSteenhouse, Yumi Kasai, David Gray, Carol Clugston, Anna Dominiczak and Alex Alderton, Roberto Amato, Jeffrey Barrett, Sonia Goncalves, Ewan Harrison, David K. Jackson, Ian Johnston, Dominic Kwiatkowski, Cordelia Langford, John Sillitoe on behalf of the Wellcome Sanger Institute COVID-19 Surveillance Team |
| EPI_ISL_3381121 | Lighthouse Laboratory Plymouth | Wellcome Sanger Institute for the COVID-19 Genomics UK (COG-UK) Consortium | Lighthouse Laboratory Plymouth and Alex Alderton, Roberto Amato, Jeffrey Barrett, Sonia Goncalves, Ewan Harrison, David K. Jackson, Ian Johnston, Dominic Kwiatkowski, Cordelia Langford, John Sillitoe on behalf of the Wellcome Sanger Institute COVID-19 Surveillance Team |
| EPI_ISL_3381122, EPI_ISL_3381123 | Lighthouse Lab in Glasgow | Wellcome Sanger Institute for the COVID-19 Genomics UK (COG-UK) Consortium | Harper VanSteenhouse, Yumi Kasai, David Gray, Carol Clugston, Anna Dominiczak and Alex Alderton, Roberto Amato, Jeffrey Barrett, Sonia Goncalves, Ewan Harrison, David K. Jackson, Ian Johnston, Dominic Kwiatkowski, Cordelia Langford, John Sillitoe on behalf of the Wellcome Sanger Institute COVID-19 Surveillance Team |
| EPI_ISL_3381124, EPI_ISL_3381125 | Lighthouse Lab in Alderley Park | Wellcome Sanger Institute for the COVID-19 Genomics UK (COG-UK) Consortium | Jacquelyn Wynn, Mairead Hyland, The Lighthouse Lab in Alderley Park and Alex Alderton, Roberto Amato, Jeffrey Barrett, Sonia Goncalves, Ewan Harrison, David K. Jackson, Ian Johnston, Dominic Kwiatkowski, Cordelia Langford, John Sillitoe on behalf of the Wellcome Sanger Institute COVID-19 Surveillance Team |
| EPI_ISL_3381126 | Lighthouse Lab in Glasgow | Wellcome Sanger Institute for the COVID-19 Genomics UK (COG-UK) Consortium | Harper VanSteenhouse, Yumi Kasai, David Gray, Carol Clugston, Anna Dominiczak and Alex Alderton, Roberto Amato, Jeffrey Barrett, Sonia Goncalves, Ewan Harrison, David K. Jackson, Ian Johnston, Dominic Kwiatkowski, Cordelia Langford, John Sillitoe on behalf of the Wellcome Sanger Institute COVID-19 Surveillance Team |
| EPI_ISL_3381127 | Lighthouse Lab in Alderley Park | Wellcome Sanger Institute for the COVID-19 Genomics UK (COG-UK) Consortium | Jacquelyn Wynn, Mairead Hyland, The Lighthouse Lab in Alderley Park and Alex Alderton, Roberto Amato, Jeffrey Barrett, Sonia Goncalves, Ewan Harrison, David K. Jackson, Ian Johnston, Dominic Kwiatkowski, Cordelia Langford, John Sillitoe on behalf of the Wellcome Sanger Institute COVID-19 Surveillance Team |
| EPI_ISL_3381128 | Lighthouse Lab in Glasgow | Wellcome Sanger Institute for the COVID-19 Genomics UK (COG-UK) Consortium | Harper VanSteenhouse, Yumi Kasai, David Gray, Carol Clugston, Anna Dominiczak and Alex Alderton, Roberto Amato, Jeffrey Barrett, Sonia Goncalves, Ewan Harrison, David K. Jackson, Ian Johnston, Dominic Kwiatkowski, Cordelia Langford, John Sillitoe on behalf of the Wellcome Sanger Institute COVID-19 Surveillance Team |
| EPI_ISL_3381129, EPI_ISL_3381130 | Lighthouse Lab in Alderley Park | Wellcome Sanger Institute for the COVID-19 Genomics UK (COG-UK) Consortium | Jacquelyn Wynn, Mairead Hyland, The Lighthouse Lab in Alderley Park and Alex Alderton, Roberto Amato, Jeffrey Barrett, Sonia Goncalves, Ewan Harrison, David K. Jackson, Ian Johnston, Dominic Kwiatkowski, Cordelia Langford, John Sillitoe on behalf of the Wellcome Sanger Institute COVID-19 Surveillance Team |
| EPI_ISL_3381131 | Lighthouse Lab in Glasgow | Wellcome Sanger Institute for the COVID-19 Genomics UK (COG-UK) Consortium | Harper VanSteenhouse, Yumi Kasai, David Gray, Carol Clugston, Anna Dominiczak and Alex Alderton, Roberto Amato, Jeffrey Barrett, Sonia Goncalves, Ewan Harrison, David K. Jackson, Ian Johnston, Dominic Kwiatkowski, Cordelia Langford, John Sillitoe on behalf of the Wellcome Sanger Institute COVID-19 Surveillance Team |
| EPI_ISL_3381132 | Lighthouse Lab in Alderley Park | Wellcome Sanger Institute for the COVID-19 Genomics UK (COG-UK) Consortium | Jacquelyn Wynn, Mairead Hyland, The Lighthouse Lab in Alderley Park and Alex Alderton, Roberto Amato, Jeffrey Barrett, Sonia Goncalves, Ewan Harrison, David K. Jackson, Ian Johnston, Dominic Kwiatkowski, Cordelia Langford, John Sillitoe on behalf of the Wellcome Sanger Institute COVID-19 Surveillance Team |
| EPI_ISL_3381133 | Lighthouse Lab in Glasgow | Wellcome Sanger Institute for the COVID-19 Genomics UK (COG-UK) Consortium | Harper VanSteenhouse, Yumi Kasai, David Gray, Carol Clugston, Anna Dominiczak and Alex Alderton, Roberto Amato, Jeffrey Barrett, Sonia Goncalves, Ewan Harrison, David K. Jackson, Ian Johnston, Dominic Kwiatkowski, Cordelia Langford, John Sillitoe on behalf of the Wellcome Sanger Institute COVID-19 Surveillance Team |
| EPI_ISL_3381134 | Lighthouse Laboratory Plymouth | Wellcome Sanger Institute for the COVID-19 Genomics UK (COG-UK) Consortium | Lighthouse Laboratory Plymouth and Alex Alderton, Roberto Amato, Jeffrey Barrett, Sonia Goncalves, Ewan Harrison, David K. Jackson, Ian Johnston, Dominic Kwiatkowski, Cordelia Langford, John Sillitoe on behalf of the Wellcome Sanger Institute COVID-19 Surveillance Team |
| EPI_ISL_3381135 | Lighthouse Lab in Alderley Park | Wellcome Sanger Institute for the COVID-19 Genomics UK (COG-UK) Consortium | Jacquelyn Wynn, Mairead Hyland, The Lighthouse Lab in Alderley Park and Alex Alderton, Roberto Amato, Jeffrey Barrett, Sonia Goncalves, Ewan Harrison, David K. Jackson, Ian Johnston, Dominic Kwiatkowski, Cordelia Langford, John Sillitoe on behalf of the Wellcome Sanger Institute COVID-19 Surveillance Team |
| EPI_ISL_3381136, EPI_ISL_3381137, EPI_ISL_3381138 | Lighthouse Lab in Glasgow | Wellcome Sanger Institute for the COVID-19 Genomics UK (COG-UK) Consortium | Harper VanSteenhouse, Yumi Kasai, David Gray, Carol Clugston, Anna Dominiczak and Alex Alderton, Roberto Amato, Jeffrey Barrett, Sonia Goncalves, Ewan Harrison, David K. Jackson, Ian Johnston, Dominic Kwiatkowski, Cordelia Langford, John Sillitoe on behalf of the Wellcome Sanger Institute COVID-19 Surveillance Team |
| EPI_ISL_3381139 | Lighthouse Laboratory Plymouth | Wellcome Sanger Institute for the COVID-19 Genomics UK (COG-UK) Consortium | Lighthouse Laboratory Plymouth and Alex Alderton, Roberto Amato, Jeffrey Barrett, Sonia Goncalves, Ewan Harrison, David K. Jackson, Ian Johnston, Dominic Kwiatkowski, Cordelia Langford, John Sillitoe on behalf of the Wellcome Sanger Institute COVID-19 Surveillance Team |
| EPI_ISL_3381140 | Lighthouse Lab in Glasgow | Wellcome Sanger Institute for the COVID-19 Genomics UK (COG-UK) Consortium | Harper VanSteenhouse, Yumi Kasai, David Gray, Carol Clugston, Anna Dominiczak and Alex Alderton, Roberto Amato, Jeffrey Barrett, Sonia Goncalves, Ewan Harrison, David K. Jackson, Ian Johnston, Dominic Kwiatkowski, Cordelia Langford, John Sillitoe on behalf of the Wellcome Sanger Institute COVID-19 Surveillance Team |
| EPI_ISL_3381141, EPI_ISL_3381142, EPI_ISL_3381143 | Lighthouse Laboratory Plymouth | Wellcome Sanger Institute for the COVID-19 Genomics UK (COG-UK) Consortium | Lighthouse Laboratory Plymouth and Alex Alderton, Roberto Amato, Jeffrey Barrett, Sonia Goncalves, Ewan Harrison, David K. Jackson, Ian Johnston, Dominic Kwiatkowski, Cordelia Langford, John Sillitoe on behalf of the Wellcome Sanger Institute COVID-19 Surveillance Team |
| EPI_ISL_3381144, EPI_ISL_3381145 | Lighthouse Lab in Glasgow | Wellcome Sanger Institute for the COVID-19 Genomics UK (COG-UK) Consortium | Harper VanSteenhouse, Yumi Kasai, David Gray, Carol Clugston, Anna Dominiczak and Alex Alderton, Roberto Amato, Jeffrey Barrett, Sonia Goncalves, Ewan Harrison, David K. Jackson, Ian Johnston, Dominic Kwiatkowski, Cordelia Langford, John Sillitoe on behalf of the Wellcome Sanger Institute COVID-19 Surveillance Team |
| EPI_ISL_3381146 | Lighthouse Laboratory Plymouth | Wellcome Sanger Institute for the COVID-19 Genomics UK (COG-UK) Consortium | Lighthouse Laboratory Plymouth and Alex Alderton, Roberto Amato, Jeffrey Barrett, Sonia Goncalves, Ewan Harrison, David K. Jackson, Ian Johnston, Dominic Kwiatkowski, Cordelia Langford, John Sillitoe on behalf of the Wellcome Sanger Institute COVID-19 Surveillance Team |
| EPI_ISL_3381147 | Lighthouse Lab in Glasgow | Wellcome Sanger Institute for the COVID-19 Genomics UK (COG-UK) Consortium | Harper VanSteenhouse, Yumi Kasai, David Gray, Carol Clugston, Anna Dominiczak and Alex Alderton, Roberto Amato, Jeffrey Barrett, Sonia Goncalves, Ewan Harrison, David K. Jackson, Ian Johnston, Dominic Kwiatkowski, Cordelia Langford, John Sillitoe on behalf of the Wellcome Sanger Institute COVID-19 Surveillance Team |
| EPI_ISL_3381148 | Lighthouse Laboratory Plymouth | Wellcome Sanger Institute for the COVID-19 Genomics UK (COG-UK) Consortium | Lighthouse Laboratory Plymouth and Alex Alderton, Roberto Amato, Jeffrey Barrett, Sonia Goncalves, Ewan Harrison, David K. Jackson, Ian Johnston, Dominic Kwiatkowski, Cordelia Langford, John Sillitoe on behalf of the Wellcome Sanger Institute COVID-19 Surveillance Team |
| EPI_ISL_3381149 | Lighthouse Lab in Alderley Park | Wellcome Sanger Institute for the COVID-19 Genomics UK (COG-UK) Consortium | Jacquelyn Wynn, Mairead Hyland, The Lighthouse Lab in Alderley Park and Alex Alderton, Roberto Amato, Jeffrey Barrett, Sonia Goncalves, Ewan Harrison, David K. Jackson, Ian Johnston, Dominic Kwiatkowski, Cordelia Langford, John Sillitoe on behalf of the Wellcome Sanger Institute COVID-19 Surveillance Team |
| EPI_ISL_3381150 | Lighthouse Lab in Glasgow | Wellcome Sanger Institute for the COVID-19 Genomics UK (COG-UK) | Harper VanSteenhouse, Yumi Kasai, David Gray, Carol Clugston, Anna Dominiczak and Alex Alderton, Roberto Amato, Jeffrey Barrett, Sonia Goncalves, Ewan Harrison, David K. Jackson, Ian Johnston, Dominic Kwiatkowski, Cordelia Langford, |

| | | | |
|---|---|---|---|
| | | Consortium | John Sillitoe on behalf of the Wellcome Sanger Institute COVID-19 Surveillance Team |
| EPI_ISL_3381151, EPI_ISL_3381152, EPI_ISL_3381153, EPI_ISL_3381154 | Lighthouse Laboratory Plymouth | Wellcome Sanger Institute for the COVID-19 Genomics UK (COG-UK) Consortium | Lighthouse Laboratory Plymouth and Alex Alderton, Roberto Amato, Jeffrey Barrett, Sonia Goncalves, Ewan Harrison, David K. Jackson, Ian Johnston, Dominic Kwiatkowski, Cordelia Langford, John Sillitoe on behalf of the Wellcome Sanger Institute COVID-19 Surveillance Team |
| EPI_ISL_3381155, EPI_ISL_3381156, EPI_ISL_3381157 | Lighthouse Lab in Glasgow | Wellcome Sanger Institute for the COVID-19 Genomics UK (COG-UK) Consortium | Harper VanSteenhouse, Yumi Kasai, David Gray, Carol Clugston, Anna Dominiczak and Alex Alderton, Roberto Amato, Jeffrey Barrett, Sonia Goncalves, Ewan Harrison, David K. Jackson, Ian Johnston, Dominic Kwiatkowski, Cordelia Langford, John Sillitoe on behalf of the Wellcome Sanger Institute COVID-19 Surveillance Team |
| EPI_ISL_3381158 | Lighthouse Laboratory Plymouth | Wellcome Sanger Institute for the COVID-19 Genomics UK (COG-UK) Consortium | Lighthouse Laboratory Plymouth and Alex Alderton, Roberto Amato, Jeffrey Barrett, Sonia Goncalves, Ewan Harrison, David K. Jackson, Ian Johnston, Dominic Kwiatkowski, Cordelia Langford, John Sillitoe on behalf of the Wellcome Sanger Institute COVID-19 Surveillance Team |
| EPI_ISL_3381159 | Lighthouse Lab in Alderley Park | Wellcome Sanger Institute for the COVID-19 Genomics UK (COG-UK) Consortium | Jacquelyn Wynn, Mairead Hyland, The Lighthouse Lab in Alderley Park and Alex Alderton, Roberto Amato, Jeffrey Barrett, Sonia Goncalves, Ewan Harrison, David K. Jackson, Ian Johnston, Dominic Kwiatkowski, Cordelia Langford, John Sillitoe on behalf of the Wellcome Sanger Institute COVID-19 Surveillance Team |
| EPI_ISL_3381160, EPI_ISL_3381161, EPI_ISL_3381162 | Lighthouse Lab in Glasgow | Wellcome Sanger Institute for the COVID-19 Genomics UK (COG-UK) Consortium | Harper VanSteenhouse, Yumi Kasai, David Gray, Carol Clugston, Anna Dominiczak and Alex Alderton, Roberto Amato, Jeffrey Barrett, Sonia Goncalves, Ewan Harrison, David K. Jackson, Ian Johnston, Dominic Kwiatkowski, Cordelia Langford, John Sillitoe on behalf of the Wellcome Sanger Institute COVID-19 Surveillance Team |
| EPI_ISL_3381163, EPI_ISL_3381164, EPI_ISL_3381165 | Lighthouse Laboratory Plymouth | Wellcome Sanger Institute for the COVID-19 Genomics UK (COG-UK) Consortium | Lighthouse Laboratory Plymouth and Alex Alderton, Roberto Amato, Jeffrey Barrett, Sonia Goncalves, Ewan Harrison, David K. Jackson, Ian Johnston, Dominic Kwiatkowski, Cordelia Langford, John Sillitoe on behalf of the Wellcome Sanger Institute COVID-19 Surveillance Team |
| EPI_ISL_3381166, EPI_ISL_3381167 | Lighthouse Lab in Alderley Park | Wellcome Sanger Institute for the COVID-19 Genomics UK (COG-UK) Consortium | Jacquelyn Wynn, Mairead Hyland, The Lighthouse Lab in Alderley Park and Alex Alderton, Roberto Amato, Jeffrey Barrett, Sonia Goncalves, Ewan Harrison, David K. Jackson, Ian Johnston, Dominic Kwiatkowski, Cordelia Langford, John Sillitoe on behalf of the Wellcome Sanger Institute COVID-19 Surveillance Team |
| EPI_ISL_3381168 | Lighthouse Laboratory Plymouth | Wellcome Sanger Institute for the COVID-19 Genomics UK (COG-UK) Consortium | Lighthouse Laboratory Plymouth and Alex Alderton, Roberto Amato, Jeffrey Barrett, Sonia Goncalves, Ewan Harrison, David K. Jackson, Ian Johnston, Dominic Kwiatkowski, Cordelia Langford, John Sillitoe on behalf of the Wellcome Sanger Institute COVID-19 Surveillance Team |
| EPI_ISL_3381169, EPI_ISL_3381170, EPI_ISL_3381171, EPI_ISL_3381172 | Lighthouse Lab in Glasgow | Wellcome Sanger Institute for the COVID-19 Genomics UK (COG-UK) Consortium | Harper VanSteenhouse, Yumi Kasai, David Gray, Carol Clugston, Anna Dominiczak and Alex Alderton, Roberto Amato, Jeffrey Barrett, Sonia Goncalves, Ewan Harrison, David K. Jackson, Ian Johnston, Dominic Kwiatkowski, Cordelia Langford, John Sillitoe on behalf of the Wellcome Sanger Institute COVID-19 Surveillance Team |
| EPI_ISL_3381173, EPI_ISL_3381174 | Lighthouse Laboratory Plymouth | Wellcome Sanger Institute for the COVID-19 Genomics UK (COG-UK) Consortium | Lighthouse Laboratory Plymouth and Alex Alderton, Roberto Amato, Jeffrey Barrett, Sonia Goncalves, Ewan Harrison, David K. Jackson, Ian Johnston, Dominic Kwiatkowski, Cordelia Langford, John Sillitoe on behalf of the Wellcome Sanger Institute COVID-19 Surveillance Team |
| EPI_ISL_3381175 | Lighthouse Lab in Alderley Park | Wellcome Sanger Institute for the COVID-19 Genomics UK (COG-UK) Consortium | Jacquelyn Wynn, Mairead Hyland, The Lighthouse Lab in Alderley Park and Alex Alderton, Roberto Amato, Jeffrey Barrett, Sonia Goncalves, Ewan Harrison, David K. Jackson, Ian Johnston, Dominic Kwiatkowski, Cordelia Langford, John Sillitoe on behalf of the Wellcome Sanger Institute COVID-19 Surveillance Team |
| EPI_ISL_3381176 | Lighthouse Laboratory Plymouth | Wellcome Sanger Institute for the COVID-19 Genomics UK (COG-UK) Consortium | Lighthouse Laboratory Plymouth and Alex Alderton, Roberto Amato, Jeffrey Barrett, Sonia Goncalves, Ewan Harrison, David K. Jackson, Ian Johnston, Dominic Kwiatkowski, Cordelia Langford, John Sillitoe on behalf of the Wellcome Sanger Institute COVID-19 Surveillance Team |
| EPI_ISL_3381177 | Lighthouse Lab in Glasgow | Wellcome Sanger Institute for the COVID-19 Genomics UK (COG-UK) Consortium | Harper VanSteenhouse, Yumi Kasai, David Gray, Carol Clugston, Anna Dominiczak and Alex Alderton, Roberto Amato, Jeffrey Barrett, Sonia Goncalves, Ewan Harrison, David K. Jackson, Ian Johnston, Dominic Kwiatkowski, Cordelia Langford, John Sillitoe on behalf of the Wellcome Sanger Institute COVID-19 Surveillance Team |
| EPI_ISL_3381178 | Lighthouse Lab in Alderley Park | Wellcome Sanger Institute for the COVID-19 Genomics UK (COG-UK) Consortium | Jacquelyn Wynn, Mairead Hyland, The Lighthouse Lab in Alderley Park and Alex Alderton, Roberto Amato, Jeffrey Barrett, Sonia Goncalves, Ewan Harrison, David K. Jackson, Ian Johnston, Dominic Kwiatkowski, Cordelia Langford, John Sillitoe on behalf of the Wellcome Sanger Institute COVID-19 Surveillance Team |
| EPI_ISL_3381179, EPI_ISL_3381180, EPI_ISL_3381181, EPI_ISL_3381182, EPI_ISL_3381183 | Lighthouse Lab in Glasgow | Wellcome Sanger Institute for the COVID-19 Genomics UK (COG-UK) Consortium | Harper VanSteenhouse, Yumi Kasai, David Gray, Carol Clugston, Anna Dominiczak and Alex Alderton, Roberto Amato, Jeffrey Barrett, Sonia Goncalves, Ewan Harrison, David K. Jackson, Ian Johnston, Dominic Kwiatkowski, Cordelia Langford, John Sillitoe on behalf of the Wellcome Sanger Institute COVID-19 Surveillance Team |
| EPI_ISL_3381184 | Lighthouse Laboratory Plymouth | Wellcome Sanger Institute for the COVID-19 Genomics UK (COG-UK) Consortium | Lighthouse Laboratory Plymouth and Alex Alderton, Roberto Amato, Jeffrey Barrett, Sonia Goncalves, Ewan Harrison, David K. Jackson, Ian Johnston, Dominic Kwiatkowski, Cordelia Langford, John Sillitoe on behalf of the Wellcome Sanger Institute COVID-19 Surveillance Team |
| EPI_ISL_3381185, EPI_ISL_3381186, EPI_ISL_3381187 | Lighthouse Lab in Glasgow | Wellcome Sanger Institute for the COVID-19 Genomics UK (COG-UK) Consortium | Harper VanSteenhouse, Yumi Kasai, David Gray, Carol Clugston, Anna Dominiczak and Alex Alderton, Roberto Amato, Jeffrey Barrett, Sonia Goncalves, Ewan Harrison, David K. Jackson, Ian Johnston, Dominic Kwiatkowski, Cordelia Langford, John Sillitoe on behalf of the Wellcome Sanger Institute COVID-19 Surveillance Team |
| EPI_ISL_3381188, EPI_ISL_3381189, EPI_ISL_3381190, EPI_ISL_3381191 | Lighthouse Lab in Alderley Park | Wellcome Sanger Institute for the COVID-19 Genomics UK (COG-UK) Consortium | Jacquelyn Wynn, Mairead Hyland, The Lighthouse Lab in Alderley Park and Alex Alderton, Roberto Amato, Jeffrey Barrett, Sonia Goncalves, Ewan Harrison, David K. Jackson, Ian Johnston, Dominic Kwiatkowski, Cordelia Langford, John Sillitoe on behalf of the Wellcome Sanger Institute COVID-19 Surveillance Team |
| EPI_ISL_3381192, EPI_ISL_3381193 | Lighthouse Lab in Glasgow | Wellcome Sanger Institute for the COVID-19 Genomics UK (COG-UK) Consortium | Harper VanSteenhouse, Yumi Kasai, David Gray, Carol Clugston, Anna Dominiczak and Alex Alderton, Roberto Amato, Jeffrey Barrett, Sonia Goncalves, Ewan Harrison, David K. Jackson, Ian Johnston, Dominic Kwiatkowski, Cordelia Langford, John Sillitoe on behalf of the Wellcome Sanger Institute COVID-19 Surveillance Team |
| EPI_ISL_3381194 | Lighthouse Laboratory Plymouth | Wellcome Sanger Institute for the COVID-19 Genomics UK (COG-UK) Consortium | Lighthouse Laboratory Plymouth and Alex Alderton, Roberto Amato, Jeffrey Barrett, Sonia Goncalves, Ewan Harrison, David K. Jackson, Ian Johnston, Dominic Kwiatkowski, Cordelia Langford, John Sillitoe on behalf of the Wellcome Sanger Institute COVID-19 Surveillance Team |
| EPI_ISL_3381195, EPI_ISL_3381196 | Lighthouse Lab in Glasgow | Wellcome Sanger Institute for the COVID-19 Genomics UK (COG-UK) Consortium | Harper VanSteenhouse, Yumi Kasai, David Gray, Carol Clugston, Anna Dominiczak and Alex Alderton, Roberto Amato, Jeffrey Barrett, Sonia Goncalves, Ewan Harrison, David K. Jackson, Ian Johnston, Dominic Kwiatkowski, Cordelia Langford, John Sillitoe on behalf of the Wellcome Sanger Institute COVID-19 Surveillance Team |
| EPI_ISL_3381197 | Lighthouse Lab in Alderley Park | Wellcome Sanger Institute for the COVID-19 Genomics UK (COG-UK) Consortium | Jacquelyn Wynn, Mairead Hyland, The Lighthouse Lab in Alderley Park and Alex Alderton, Roberto Amato, Jeffrey Barrett, Sonia Goncalves, Ewan Harrison, David K. Jackson, Ian Johnston, Dominic Kwiatkowski, Cordelia Langford, John Sillitoe on behalf of the Wellcome Sanger Institute COVID-19 Surveillance Team |
| EPI_ISL_3381198 | Lighthouse Lab in Glasgow | Wellcome Sanger Institute for the COVID-19 Genomics UK (COG-UK) Consortium | Harper VanSteenhouse, Yumi Kasai, David Gray, Carol Clugston, Anna Dominiczak and Alex Alderton, Roberto Amato, Jeffrey Barrett, Sonia Goncalves, Ewan Harrison, David K. Jackson, Ian Johnston, Dominic Kwiatkowski, Cordelia Langford, John Sillitoe on behalf of the Wellcome Sanger Institute COVID-19 Surveillance Team |
| EPI_ISL_3381199 | Lighthouse Lab in Alderley Park | Wellcome Sanger Institute for the COVID-19 Genomics UK (COG-UK) Consortium | Jacquelyn Wynn, Mairead Hyland, The Lighthouse Lab in Alderley Park and Alex Alderton, Roberto Amato, Jeffrey Barrett, Sonia Goncalves, Ewan Harrison, David K. Jackson, Ian Johnston, Dominic Kwiatkowski, Cordelia Langford, John Sillitoe on behalf of the Wellcome Sanger Institute COVID-19 Surveillance Team |
| EPI_ISL_3381200 | Lighthouse Lab in Glasgow | Wellcome Sanger Institute for the COVID-19 Genomics UK (COG-UK) Consortium | Harper VanSteenhouse, Yumi Kasai, David Gray, Carol Clugston, Anna Dominiczak and Alex Alderton, Roberto Amato, Jeffrey Barrett, Sonia Goncalves, Ewan Harrison, David K. Jackson, Ian Johnston, Dominic Kwiatkowski, Cordelia Langford, John Sillitoe on behalf of the Wellcome Sanger Institute COVID-19 Surveillance Team |
| EPI_ISL_3381201 | Lighthouse Laboratory Plymouth | Wellcome Sanger Institute for the COVID-19 Genomics UK (COG-UK) Consortium | Lighthouse Laboratory Plymouth and Alex Alderton, Roberto Amato, Jeffrey Barrett, Sonia Goncalves, Ewan Harrison, David K. Jackson, Ian Johnston, Dominic Kwiatkowski, Cordelia Langford, John Sillitoe on behalf of the Wellcome Sanger Institute COVID-19 Surveillance Team |
| EPI_ISL_3381202, EPI_ISL_3381203, EPI_ISL_3381204 | Lighthouse Lab in Glasgow | Wellcome Sanger Institute for the COVID-19 Genomics UK (COG-UK) Consortium | Harper VanSteenhouse, Yumi Kasai, David Gray, Carol Clugston, Anna Dominiczak and Alex Alderton, Roberto Amato, Jeffrey Barrett, Sonia Goncalves, Ewan Harrison, David K. Jackson, Ian Johnston, Dominic Kwiatkowski, Cordelia Langford, John Sillitoe on behalf of the Wellcome Sanger Institute COVID-19 Surveillance Team |
| EPI_ISL_3381205 | Lighthouse Laboratory Plymouth | Wellcome Sanger Institute for the COVID-19 Genomics UK (COG-UK) Consortium | Lighthouse Laboratory Plymouth and Alex Alderton, Roberto Amato, Jeffrey Barrett, Sonia Goncalves, Ewan Harrison, David K. Jackson, Ian Johnston, Dominic Kwiatkowski, Cordelia Langford, John Sillitoe on behalf of the Wellcome Sanger Institute COVID-19 Surveillance Team |
| EPI_ISL_3381206, EPI_ISL_3381207 | Lighthouse Lab in Alderley Park | Wellcome Sanger Institute for the COVID-19 Genomics UK (COG-UK) Consortium | Jacquelyn Wynn, Mairead Hyland, The Lighthouse Lab in Alderley Park and Alex Alderton, Roberto Amato, Jeffrey Barrett, Sonia Goncalves, Ewan Harrison, David K. Jackson, Ian Johnston, Dominic Kwiatkowski, Cordelia Langford, John Sillitoe on behalf of the Wellcome Sanger Institute COVID-19 Surveillance Team |
| EPI_ISL_3381208, EPI_ISL_3381209 | Lighthouse Lab in Glasgow | Wellcome Sanger Institute for the COVID-19 Genomics UK (COG-UK) Consortium | Harper VanSteenhouse, Yumi Kasai, David Gray, Carol Clugston, Anna Dominiczak and Alex Alderton, Roberto Amato, Jeffrey Barrett, Sonia Goncalves, Ewan Harrison, David K. Jackson, Ian Johnston, Dominic Kwiatkowski, Cordelia Langford, John Sillitoe on behalf of the Wellcome Sanger Institute COVID-19 Surveillance Team |
| EPI_ISL_3381210 | Lighthouse Laboratory Plymouth | Wellcome Sanger Institute for the COVID-19 Genomics UK (COG-UK) Consortium | Lighthouse Laboratory Plymouth and Alex Alderton, Roberto Amato, Jeffrey Barrett, Sonia Goncalves, Ewan Harrison, David K. Jackson, Ian Johnston, Dominic Kwiatkowski, Cordelia Langford, John Sillitoe on behalf of the Wellcome Sanger Institute COVID-19 Surveillance Team |
| EPI_ISL_3381211 | Lighthouse Lab in Alderley Park | Wellcome Sanger Institute for the COVID-19 Genomics UK (COG-UK) Consortium | Jacquelyn Wynn, Mairead Hyland, The Lighthouse Lab in Alderley Park and Alex Alderton, Roberto Amato, Jeffrey Barrett, Sonia Goncalves, Ewan Harrison, David K. Jackson, Ian Johnston, Dominic Kwiatkowski, Cordelia Langford, John Sillitoe on behalf of the Wellcome Sanger Institute COVID-19 Surveillance Team |
| EPI_ISL_3381212, EPI_ISL_3381213, EPI_ISL_3381214 | Lighthouse Lab in Glasgow | Wellcome Sanger Institute for the COVID-19 Genomics UK (COG-UK) Consortium | Harper VanSteenhouse, Yumi Kasai, David Gray, Carol Clugston, Anna Dominiczak and Alex Alderton, Roberto Amato, Jeffrey Barrett, Sonia Goncalves, Ewan Harrison, David K. Jackson, Ian Johnston, Dominic Kwiatkowski, Cordelia Langford, John Sillitoe on behalf of the Wellcome Sanger Institute COVID-19 Surveillance Team |
| EPI_ISL_3381215 | Lighthouse Laboratory Plymouth | Wellcome Sanger Institute for the COVID-19 Genomics UK (COG-UK) Consortium | Lighthouse Laboratory Plymouth and Alex Alderton, Roberto Amato, Jeffrey Barrett, Sonia Goncalves, Ewan Harrison, David K. Jackson, Ian Johnston, Dominic Kwiatkowski, Cordelia Langford, John Sillitoe on behalf of the Wellcome Sanger Institute COVID-19 Surveillance Team |
| EPI_ISL_3381216, EPI_ISL_3381217, EPI_ISL_3381218 | Lighthouse Lab in Alderley Park | Wellcome Sanger Institute for the COVID-19 Genomics UK (COG-UK) Consortium | Jacquelyn Wynn, Mairead Hyland, The Lighthouse Lab in Alderley Park and Alex Alderton, Roberto Amato, Jeffrey Barrett, Sonia Goncalves, Ewan Harrison, David K. Jackson, Ian Johnston, Dominic Kwiatkowski, Cordelia Langford, John Sillitoe on behalf of the Wellcome Sanger Institute COVID-19 Surveillance Team |
| EPI_ISL_3381219, EPI_ISL_3381220 | Lighthouse Lab in Glasgow | Wellcome Sanger Institute for the COVID-19 Genomics UK (COG-UK) Consortium | Harper VanSteenhouse, Yumi Kasai, David Gray, Carol Clugston, Anna Dominiczak and Alex Alderton, Roberto Amato, Jeffrey Barrett, Sonia Goncalves, Ewan Harrison, David K. Jackson, Ian Johnston, Dominic Kwiatkowski, Cordelia Langford, John Sillitoe on behalf of the Wellcome Sanger Institute COVID-19 Surveillance Team |
| EPI_ISL_3381221 | Lighthouse Laboratory Plymouth | Wellcome Sanger Institute for the COVID-19 Genomics UK (COG-UK) Consortium | Lighthouse Laboratory Plymouth and Alex Alderton, Roberto Amato, Jeffrey Barrett, Sonia Goncalves, Ewan Harrison, David K. Jackson, Ian Johnston, Dominic Kwiatkowski, Cordelia Langford, John Sillitoe on behalf of the Wellcome Sanger Institute COVID-19 Surveillance Team |
| EPI_ISL_3381222, | Lighthouse Lab in Alderley Park | Wellcome Sanger Institute for the COVID-19 Genomics UK (COG-UK) | Jacquelyn Wynn, Mairead Hyland, The Lighthouse Lab in Alderley Park and Alex Alderton, Roberto Amato, Jeffrey Barrett, Sonia Goncalves, Ewan Harrison, David K. Jackson, Ian Johnston, Dominic Kwiatkowski, Cordelia Langford, John Sillitoe on |

| | | | |
|---|---|---|---|
| EPI_ISL_3381223 | | Consortium | behalf of the Wellcome Sanger Institute COVID-19 Surveillance Team |
| EPI_ISL_3381224, EPI_ISL_3381225, EPI_ISL_3381226 | Lighthouse Laboratory Plymouth | Wellcome Sanger Institute for the COVID-19 Genomics UK (COG-UK) Consortium | Lighthouse Laboratory Plymouth and Alex Alderton, Roberto Amato, Jeffrey Barrett, Sonia Goncalves, Ewan Harrison, David K. Jackson, Ian Johnston, Dominic Kwiatkowski, Cordelia Langford, John Sillitoe on behalf of the Wellcome Sanger Institute COVID-19 Surveillance Team |
| EPI_ISL_3381227 | Lighthouse Lab in Alderley Park | Wellcome Sanger Institute for the COVID-19 Genomics UK (COG-UK) Consortium | Jacquelyn Wynn, Mairead Hyland, The Lighthouse Lab in Alderley Park and Alex Alderton, Roberto Amato, Jeffrey Barrett, Sonia Goncalves, Ewan Harrison, David K. Jackson, Ian Johnston, Dominic Kwiatkowski, Cordelia Langford, John Sillitoe on behalf of the Wellcome Sanger Institute COVID-19 Surveillance Team |
| EPI_ISL_3381228, EPI_ISL_3381229, EPI_ISL_3381230 | Lighthouse Lab in Glasgow | Wellcome Sanger Institute for the COVID-19 Genomics UK (COG-UK) Consortium | Harper VanSteenhouse, Yumi Kasai, David Gray, Carol Clugston, Anna Dominiczak and Alex Alderton, Roberto Amato, Jeffrey Barrett, Sonia Goncalves, Ewan Harrison, David K. Jackson, Ian Johnston, Dominic Kwiatkowski, Cordelia Langford, John Sillitoe on behalf of the Wellcome Sanger Institute COVID-19 Surveillance Team |
| EPI_ISL_3381231, EPI_ISL_3381232 | Lighthouse Laboratory Plymouth | Wellcome Sanger Institute for the COVID-19 Genomics UK (COG-UK) Consortium | Lighthouse Laboratory Plymouth and Alex Alderton, Roberto Amato, Jeffrey Barrett, Sonia Goncalves, Ewan Harrison, David K. Jackson, Ian Johnston, Dominic Kwiatkowski, Cordelia Langford, John Sillitoe on behalf of the Wellcome Sanger Institute COVID-19 Surveillance Team |
| EPI_ISL_3381233 | Lighthouse Lab in Glasgow | Wellcome Sanger Institute for the COVID-19 Genomics UK (COG-UK) Consortium | Harper VanSteenhouse, Yumi Kasai, David Gray, Carol Clugston, Anna Dominiczak and Alex Alderton, Roberto Amato, Jeffrey Barrett, Sonia Goncalves, Ewan Harrison, David K. Jackson, Ian Johnston, Dominic Kwiatkowski, Cordelia Langford, John Sillitoe on behalf of the Wellcome Sanger Institute COVID-19 Surveillance Team |
| EPI_ISL_3381234 | Lighthouse Laboratory Plymouth | Wellcome Sanger Institute for the COVID-19 Genomics UK (COG-UK) Consortium | Lighthouse Laboratory Plymouth and Alex Alderton, Roberto Amato, Jeffrey Barrett, Sonia Goncalves, Ewan Harrison, David K. Jackson, Ian Johnston, Dominic Kwiatkowski, Cordelia Langford, John Sillitoe on behalf of the Wellcome Sanger Institute COVID-19 Surveillance Team |
| EPI_ISL_3381235, EPI_ISL_3381236, EPI_ISL_3381237, EPI_ISL_3381238 | Lighthouse Lab in Glasgow | Wellcome Sanger Institute for the COVID-19 Genomics UK (COG-UK) Consortium | Harper VanSteenhouse, Yumi Kasai, David Gray, Carol Clugston, Anna Dominiczak and Alex Alderton, Roberto Amato, Jeffrey Barrett, Sonia Goncalves, Ewan Harrison, David K. Jackson, Ian Johnston, Dominic Kwiatkowski, Cordelia Langford, John Sillitoe on behalf of the Wellcome Sanger Institute COVID-19 Surveillance Team |
| EPI_ISL_3381239 | Lighthouse Lab in Alderley Park | Wellcome Sanger Institute for the COVID-19 Genomics UK (COG-UK) Consortium | Jacquelyn Wynn, Mairead Hyland, The Lighthouse Lab in Alderley Park and Alex Alderton, Roberto Amato, Jeffrey Barrett, Sonia Goncalves, Ewan Harrison, David K. Jackson, Ian Johnston, Dominic Kwiatkowski, Cordelia Langford, John Sillitoe on behalf of the Wellcome Sanger Institute COVID-19 Surveillance Team |
| EPI_ISL_3381240, EPI_ISL_3381241, EPI_ISL_3381242 | Lighthouse Lab in Glasgow | Wellcome Sanger Institute for the COVID-19 Genomics UK (COG-UK) Consortium | Harper VanSteenhouse, Yumi Kasai, David Gray, Carol Clugston, Anna Dominiczak and Alex Alderton, Roberto Amato, Jeffrey Barrett, Sonia Goncalves, Ewan Harrison, David K. Jackson, Ian Johnston, Dominic Kwiatkowski, Cordelia Langford, John Sillitoe on behalf of the Wellcome Sanger Institute COVID-19 Surveillance Team |
| EPI_ISL_3381243, EPI_ISL_3381244 | Lighthouse Lab in Alderley Park | Wellcome Sanger Institute for the COVID-19 Genomics UK (COG-UK) Consortium | Jacquelyn Wynn, Mairead Hyland, The Lighthouse Lab in Alderley Park and Alex Alderton, Roberto Amato, Jeffrey Barrett, Sonia Goncalves, Ewan Harrison, David K. Jackson, Ian Johnston, Dominic Kwiatkowski, Cordelia Langford, John Sillitoe on behalf of the Wellcome Sanger Institute COVID-19 Surveillance Team |
| EPI_ISL_3381245, EPI_ISL_3381246, EPI_ISL_3381247 | Lighthouse Lab in Glasgow | Wellcome Sanger Institute for the COVID-19 Genomics UK (COG-UK) Consortium | Harper VanSteenhouse, Yumi Kasai, David Gray, Carol Clugston, Anna Dominiczak and Alex Alderton, Roberto Amato, Jeffrey Barrett, Sonia Goncalves, Ewan Harrison, David K. Jackson, Ian Johnston, Dominic Kwiatkowski, Cordelia Langford, John Sillitoe on behalf of the Wellcome Sanger Institute COVID-19 Surveillance Team |
| EPI_ISL_3381248, EPI_ISL_3381249, EPI_ISL_3381250, EPI_ISL_3381251 | Lighthouse Lab in Alderley Park | Wellcome Sanger Institute for the COVID-19 Genomics UK (COG-UK) Consortium | Jacquelyn Wynn, Mairead Hyland, The Lighthouse Lab in Alderley Park and Alex Alderton, Roberto Amato, Jeffrey Barrett, Sonia Goncalves, Ewan Harrison, David K. Jackson, Ian Johnston, Dominic Kwiatkowski, Cordelia Langford, John Sillitoe on behalf of the Wellcome Sanger Institute COVID-19 Surveillance Team |
| EPI_ISL_3381252, EPI_ISL_3381253, EPI_ISL_3381254, EPI_ISL_3381255 | Lighthouse Lab in Glasgow | Wellcome Sanger Institute for the COVID-19 Genomics UK (COG-UK) Consortium | Harper VanSteenhouse, Yumi Kasai, David Gray, Carol Clugston, Anna Dominiczak and Alex Alderton, Roberto Amato, Jeffrey Barrett, Sonia Goncalves, Ewan Harrison, David K. Jackson, Ian Johnston, Dominic Kwiatkowski, Cordelia Langford, John Sillitoe on behalf of the Wellcome Sanger Institute COVID-19 Surveillance Team |
| EPI_ISL_3381256 | Lighthouse Laboratory Plymouth | Wellcome Sanger Institute for the COVID-19 Genomics UK (COG-UK) Consortium | Lighthouse Laboratory Plymouth and Alex Alderton, Roberto Amato, Jeffrey Barrett, Sonia Goncalves, Ewan Harrison, David K. Jackson, Ian Johnston, Dominic Kwiatkowski, Cordelia Langford, John Sillitoe on behalf of the Wellcome Sanger Institute COVID-19 Surveillance Team |
| EPI_ISL_3381257 | Lighthouse Lab in Alderley Park | Wellcome Sanger Institute for the COVID-19 Genomics UK (COG-UK) Consortium | Jacquelyn Wynn, Mairead Hyland, The Lighthouse Lab in Alderley Park and Alex Alderton, Roberto Amato, Jeffrey Barrett, Sonia Goncalves, Ewan Harrison, David K. Jackson, Ian Johnston, Dominic Kwiatkowski, Cordelia Langford, John Sillitoe on behalf of the Wellcome Sanger Institute COVID-19 Surveillance Team |
| EPI_ISL_3381258 | Lighthouse Lab in Glasgow | Wellcome Sanger Institute for the COVID-19 Genomics UK (COG-UK) Consortium | Harper VanSteenhouse, Yumi Kasai, David Gray, Carol Clugston, Anna Dominiczak and Alex Alderton, Roberto Amato, Jeffrey Barrett, Sonia Goncalves, Ewan Harrison, David K. Jackson, Ian Johnston, Dominic Kwiatkowski, Cordelia Langford, John Sillitoe on behalf of the Wellcome Sanger Institute COVID-19 Surveillance Team |
| EPI_ISL_3381259 | Lighthouse Lab in Alderley Park | Wellcome Sanger Institute for the COVID-19 Genomics UK (COG-UK) Consortium | Jacquelyn Wynn, Mairead Hyland, The Lighthouse Lab in Alderley Park and Alex Alderton, Roberto Amato, Jeffrey Barrett, Sonia Goncalves, Ewan Harrison, David K. Jackson, Ian Johnston, Dominic Kwiatkowski, Cordelia Langford, John Sillitoe on behalf of the Wellcome Sanger Institute COVID-19 Surveillance Team |
| EPI_ISL_3381260, EPI_ISL_3381261, EPI_ISL_3381262 | Lighthouse Lab in Glasgow | Wellcome Sanger Institute for the COVID-19 Genomics UK (COG-UK) Consortium | Harper VanSteenhouse, Yumi Kasai, David Gray, Carol Clugston, Anna Dominiczak and Alex Alderton, Roberto Amato, Jeffrey Barrett, Sonia Goncalves, Ewan Harrison, David K. Jackson, Ian Johnston, Dominic Kwiatkowski, Cordelia Langford, John Sillitoe on behalf of the Wellcome Sanger Institute COVID-19 Surveillance Team |
| EPI_ISL_3381263 | Lighthouse Lab in Alderley Park | Wellcome Sanger Institute for the COVID-19 Genomics UK (COG-UK) Consortium | Jacquelyn Wynn, Mairead Hyland, The Lighthouse Lab in Alderley Park and Alex Alderton, Roberto Amato, Jeffrey Barrett, Sonia Goncalves, Ewan Harrison, David K. Jackson, Ian Johnston, Dominic Kwiatkowski, Cordelia Langford, John Sillitoe on behalf of the Wellcome Sanger Institute COVID-19 Surveillance Team |
| EPI_ISL_3381264, EPI_ISL_3381265, EPI_ISL_3381266 | Lighthouse Lab in Glasgow | Wellcome Sanger Institute for the COVID-19 Genomics UK (COG-UK) Consortium | Harper VanSteenhouse, Yumi Kasai, David Gray, Carol Clugston, Anna Dominiczak and Alex Alderton, Roberto Amato, Jeffrey Barrett, Sonia Goncalves, Ewan Harrison, David K. Jackson, Ian Johnston, Dominic Kwiatkowski, Cordelia Langford, John Sillitoe on behalf of the Wellcome Sanger Institute COVID-19 Surveillance Team |
| EPI_ISL_3381267, EPI_ISL_3381268, EPI_ISL_3381269, EPI_ISL_3381270, EPI_ISL_3381271 | Lighthouse Laboratory Plymouth | Wellcome Sanger Institute for the COVID-19 Genomics UK (COG-UK) Consortium | Lighthouse Laboratory Plymouth and Alex Alderton, Roberto Amato, Jeffrey Barrett, Sonia Goncalves, Ewan Harrison, David K. Jackson, Ian Johnston, Dominic Kwiatkowski, Cordelia Langford, John Sillitoe on behalf of the Wellcome Sanger Institute COVID-19 Surveillance Team |
| EPI_ISL_3381272 | Lighthouse Lab in Alderley Park | Wellcome Sanger Institute for the COVID-19 Genomics UK (COG-UK) Consortium | Jacquelyn Wynn, Mairead Hyland, The Lighthouse Lab in Alderley Park and Alex Alderton, Roberto Amato, Jeffrey Barrett, Sonia Goncalves, Ewan Harrison, David K. Jackson, Ian Johnston, Dominic Kwiatkowski, Cordelia Langford, John Sillitoe on behalf of the Wellcome Sanger Institute COVID-19 Surveillance Team |
| EPI_ISL_3381273 | Lighthouse Laboratory Plymouth | Wellcome Sanger Institute for the COVID-19 Genomics UK (COG-UK) Consortium | Lighthouse Laboratory Plymouth and Alex Alderton, Roberto Amato, Jeffrey Barrett, Sonia Goncalves, Ewan Harrison, David K. Jackson, Ian Johnston, Dominic Kwiatkowski, Cordelia Langford, John Sillitoe on behalf of the Wellcome Sanger Institute COVID-19 Surveillance Team |
| EPI_ISL_3381274, EPI_ISL_3381275 | Lighthouse Lab in Glasgow | Wellcome Sanger Institute for the COVID-19 Genomics UK (COG-UK) Consortium | Harper VanSteenhouse, Yumi Kasai, David Gray, Carol Clugston, Anna Dominiczak and Alex Alderton, Roberto Amato, Jeffrey Barrett, Sonia Goncalves, Ewan Harrison, David K. Jackson, Ian Johnston, Dominic Kwiatkowski, Cordelia Langford, John Sillitoe on behalf of the Wellcome Sanger Institute COVID-19 Surveillance Team |
| EPI_ISL_3381276 | Lighthouse Laboratory Plymouth | Wellcome Sanger Institute for the COVID-19 Genomics UK (COG-UK) Consortium | Lighthouse Laboratory Plymouth and Alex Alderton, Roberto Amato, Jeffrey Barrett, Sonia Goncalves, Ewan Harrison, David K. Jackson, Ian Johnston, Dominic Kwiatkowski, Cordelia Langford, John Sillitoe on behalf of the Wellcome Sanger Institute COVID-19 Surveillance Team |
| EPI_ISL_3381277 | Lighthouse Lab in Alderley Park | Wellcome Sanger Institute for the COVID-19 Genomics UK (COG-UK) Consortium | Jacquelyn Wynn, Mairead Hyland, The Lighthouse Lab in Alderley Park and Alex Alderton, Roberto Amato, Jeffrey Barrett, Sonia Goncalves, Ewan Harrison, David K. Jackson, Ian Johnston, Dominic Kwiatkowski, Cordelia Langford, John Sillitoe on behalf of the Wellcome Sanger Institute COVID-19 Surveillance Team |
| EPI_ISL_3381278, EPI_ISL_3381279, EPI_ISL_3381280, EPI_ISL_3381281, EPI_ISL_3381282, EPI_ISL_3381283 | Lighthouse Lab in Glasgow | Wellcome Sanger Institute for the COVID-19 Genomics UK (COG-UK) Consortium | Harper VanSteenhouse, Yumi Kasai, David Gray, Carol Clugston, Anna Dominiczak and Alex Alderton, Roberto Amato, Jeffrey Barrett, Sonia Goncalves, Ewan Harrison, David K. Jackson, Ian Johnston, Dominic Kwiatkowski, Cordelia Langford, John Sillitoe on behalf of the Wellcome Sanger Institute COVID-19 Surveillance Team |
| EPI_ISL_3381284, EPI_ISL_3381285, EPI_ISL_3381286, EPI_ISL_3381287 | Lighthouse Laboratory Plymouth | Wellcome Sanger Institute for the COVID-19 Genomics UK (COG-UK) Consortium | Lighthouse Laboratory Plymouth and Alex Alderton, Roberto Amato, Jeffrey Barrett, Sonia Goncalves, Ewan Harrison, David K. Jackson, Ian Johnston, Dominic Kwiatkowski, Cordelia Langford, John Sillitoe on behalf of the Wellcome Sanger Institute COVID-19 Surveillance Team |
| EPI_ISL_3381288 | Lighthouse Lab in Glasgow | Wellcome Sanger Institute for the COVID-19 Genomics UK (COG-UK) Consortium | Harper VanSteenhouse, Yumi Kasai, David Gray, Carol Clugston, Anna Dominiczak and Alex Alderton, Roberto Amato, Jeffrey Barrett, Sonia Goncalves, Ewan Harrison, David K. Jackson, Ian Johnston, Dominic Kwiatkowski, Cordelia Langford, John Sillitoe on behalf of the Wellcome Sanger Institute COVID-19 Surveillance Team |
| EPI_ISL_3381289 | Lighthouse Lab in Alderley Park | Wellcome Sanger Institute for the COVID-19 Genomics UK (COG-UK) Consortium | Jacquelyn Wynn, Mairead Hyland, The Lighthouse Lab in Alderley Park and Alex Alderton, Roberto Amato, Jeffrey Barrett, Sonia Goncalves, Ewan Harrison, David K. Jackson, Ian Johnston, Dominic Kwiatkowski, Cordelia Langford, John Sillitoe on behalf of the Wellcome Sanger Institute COVID-19 Surveillance Team |
| EPI_ISL_3381290 | Lighthouse Lab in Glasgow | Wellcome Sanger Institute for the COVID-19 Genomics UK (COG-UK) Consortium | Harper VanSteenhouse, Yumi Kasai, David Gray, Carol Clugston, Anna Dominiczak and Alex Alderton, Roberto Amato, Jeffrey Barrett, Sonia Goncalves, Ewan Harrison, David K. Jackson, Ian Johnston, Dominic Kwiatkowski, Cordelia Langford, John Sillitoe on behalf of the Wellcome Sanger Institute COVID-19 Surveillance Team |
| EPI_ISL_3381291 | Lighthouse Lab in Alderley Park | Wellcome Sanger Institute for the COVID-19 Genomics UK (COG-UK) Consortium | Jacquelyn Wynn, Mairead Hyland, The Lighthouse Lab in Alderley Park and Alex Alderton, Roberto Amato, Jeffrey Barrett, Sonia Goncalves, Ewan Harrison, David K. Jackson, Ian Johnston, Dominic Kwiatkowski, Cordelia Langford, John Sillitoe on behalf of the Wellcome Sanger Institute COVID-19 Surveillance Team |
| EPI_ISL_3381292 | Lighthouse Lab in Glasgow | Wellcome Sanger Institute for the COVID-19 Genomics UK (COG-UK) Consortium | Harper VanSteenhouse, Yumi Kasai, David Gray, Carol Clugston, Anna Dominiczak and Alex Alderton, Roberto Amato, Jeffrey Barrett, Sonia Goncalves, Ewan Harrison, David K. Jackson, Ian Johnston, Dominic Kwiatkowski, Cordelia Langford, John Sillitoe on behalf of the Wellcome Sanger Institute COVID-19 Surveillance Team |
| EPI_ISL_3381293, EPI_ISL_3381294 | Lighthouse Laboratory Plymouth | Wellcome Sanger Institute for the COVID-19 Genomics UK (COG-UK) Consortium | Lighthouse Laboratory Plymouth and Alex Alderton, Roberto Amato, Jeffrey Barrett, Sonia Goncalves, Ewan Harrison, David K. Jackson, Ian Johnston, Dominic Kwiatkowski, Cordelia Langford, John Sillitoe on behalf of the Wellcome Sanger Institute COVID-19 Surveillance Team |
| EPI_ISL_3381295 | Lighthouse Lab in Glasgow | Wellcome Sanger Institute for the COVID-19 Genomics UK (COG-UK) Consortium | Harper VanSteenhouse, Yumi Kasai, David Gray, Carol Clugston, Anna Dominiczak and Alex Alderton, Roberto Amato, Jeffrey Barrett, Sonia Goncalves, Ewan Harrison, David K. Jackson, Ian Johnston, Dominic Kwiatkowski, Cordelia Langford, John Sillitoe on behalf of the Wellcome Sanger Institute COVID-19 Surveillance Team |
| EPI_ISL_3381296 | Lighthouse Lab in Alderley Park | Wellcome Sanger Institute for the COVID-19 Genomics UK (COG-UK) | Jacquelyn Wynn, Mairead Hyland, The Lighthouse Lab in Alderley Park and Alex Alderton, Roberto Amato, Jeffrey Barrett, Sonia Goncalves, Ewan Harrison, David K. Jackson, Ian Johnston, Dominic Kwiatkowski, Cordelia Langford, John Sillitoe on |

| | | | |
|---|---|---|---|
| | | Consortium | behalf of the Wellcome Sanger Institute COVID-19 Surveillance Team |
| EPI_ISL_3381297 | Lighthouse Lab in Glasgow | Wellcome Sanger Institute for the COVID-19 Genomics UK (COG-UK) Consortium | Harper VanSteenhouse, Yumi Kasai, David Gray, Carol Clugston, Anna Dominiczak and Alex Alderton, Roberto Amato, Jeffrey Barrett, Sonia Goncalves, Ewan Harrison, David K. Jackson, Ian Johnston, Dominic Kwiatkowski, Cordelia Langford, John Sillitoe on behalf of the Wellcome Sanger Institute COVID-19 Surveillance Team |
| EPI_ISL_3381298 | Lighthouse Lab in Alderley Park | Wellcome Sanger Institute for the COVID-19 Genomics UK (COG-UK) Consortium | Jacquelyn Wynn, Mairead Hyland, The Lighthouse Lab in Alderley Park and Alex Alderton, Roberto Amato, Jeffrey Barrett, Sonia Goncalves, Ewan Harrison, David K. Jackson, Ian Johnston, Dominic Kwiatkowski, Cordelia Langford, John Sillitoe on behalf of the Wellcome Sanger Institute COVID-19 Surveillance Team |
| EPI_ISL_3381299, EPI_ISL_3381300 | Lighthouse Lab in Glasgow | Wellcome Sanger Institute for the COVID-19 Genomics UK (COG-UK) Consortium | Harper VanSteenhouse, Yumi Kasai, David Gray, Carol Clugston, Anna Dominiczak and Alex Alderton, Roberto Amato, Jeffrey Barrett, Sonia Goncalves, Ewan Harrison, David K. Jackson, Ian Johnston, Dominic Kwiatkowski, Cordelia Langford, John Sillitoe on behalf of the Wellcome Sanger Institute COVID-19 Surveillance Team |
| EPI_ISL_3381301, EPI_ISL_3381302 | Lighthouse Laboratory Plymouth | Wellcome Sanger Institute for the COVID-19 Genomics UK (COG-UK) Consortium | Lighthouse Laboratory Plymouth and Alex Alderton, Roberto Amato, Jeffrey Barrett, Sonia Goncalves, Ewan Harrison, David K. Jackson, Ian Johnston, Dominic Kwiatkowski, Cordelia Langford, John Sillitoe on behalf of the Wellcome Sanger Institute COVID-19 Surveillance Team |
| EPI_ISL_3381303, EPI_ISL_3381304 | Lighthouse Lab in Alderley Park | Wellcome Sanger Institute for the COVID-19 Genomics UK (COG-UK) Consortium | Jacquelyn Wynn, Mairead Hyland, The Lighthouse Lab in Alderley Park and Alex Alderton, Roberto Amato, Jeffrey Barrett, Sonia Goncalves, Ewan Harrison, David K. Jackson, Ian Johnston, Dominic Kwiatkowski, Cordelia Langford, John Sillitoe on behalf of the Wellcome Sanger Institute COVID-19 Surveillance Team |
| EPI_ISL_3381305, EPI_ISL_3381306, EPI_ISL_3381307, EPI_ISL_3381308, EPI_ISL_3381309 | Lighthouse Lab in Glasgow | Wellcome Sanger Institute for the COVID-19 Genomics UK (COG-UK) Consortium | Harper VanSteenhouse, Yumi Kasai, David Gray, Carol Clugston, Anna Dominiczak and Alex Alderton, Roberto Amato, Jeffrey Barrett, Sonia Goncalves, Ewan Harrison, David K. Jackson, Ian Johnston, Dominic Kwiatkowski, Cordelia Langford, John Sillitoe on behalf of the Wellcome Sanger Institute COVID-19 Surveillance Team |
| EPI_ISL_3381310 | Lighthouse Laboratory Plymouth | Wellcome Sanger Institute for the COVID-19 Genomics UK (COG-UK) Consortium | Lighthouse Laboratory Plymouth and Alex Alderton, Roberto Amato, Jeffrey Barrett, Sonia Goncalves, Ewan Harrison, David K. Jackson, Ian Johnston, Dominic Kwiatkowski, Cordelia Langford, John Sillitoe on behalf of the Wellcome Sanger Institute COVID-19 Surveillance Team |
| EPI_ISL_3381311, EPI_ISL_3381312 | Lighthouse Lab in Glasgow | Wellcome Sanger Institute for the COVID-19 Genomics UK (COG-UK) Consortium | Harper VanSteenhouse, Yumi Kasai, David Gray, Carol Clugston, Anna Dominiczak and Alex Alderton, Roberto Amato, Jeffrey Barrett, Sonia Goncalves, Ewan Harrison, David K. Jackson, Ian Johnston, Dominic Kwiatkowski, Cordelia Langford, John Sillitoe on behalf of the Wellcome Sanger Institute COVID-19 Surveillance Team |
| EPI_ISL_3381313 | Lighthouse Lab in Alderley Park | Wellcome Sanger Institute for the COVID-19 Genomics UK (COG-UK) Consortium | Jacquelyn Wynn, Mairead Hyland, The Lighthouse Lab in Alderley Park and Alex Alderton, Roberto Amato, Jeffrey Barrett, Sonia Goncalves, Ewan Harrison, David K. Jackson, Ian Johnston, Dominic Kwiatkowski, Cordelia Langford, John Sillitoe on behalf of the Wellcome Sanger Institute COVID-19 Surveillance Team |
| EPI_ISL_3381314 | Lighthouse Lab in Glasgow | Wellcome Sanger Institute for the COVID-19 Genomics UK (COG-UK) Consortium | Harper VanSteenhouse, Yumi Kasai, David Gray, Carol Clugston, Anna Dominiczak and Alex Alderton, Roberto Amato, Jeffrey Barrett, Sonia Goncalves, Ewan Harrison, David K. Jackson, Ian Johnston, Dominic Kwiatkowski, Cordelia Langford, John Sillitoe on behalf of the Wellcome Sanger Institute COVID-19 Surveillance Team |
| EPI_ISL_3381315, EPI_ISL_3381316 | Lighthouse Lab in Alderley Park | Wellcome Sanger Institute for the COVID-19 Genomics UK (COG-UK) Consortium | Jacquelyn Wynn, Mairead Hyland, The Lighthouse Lab in Alderley Park and Alex Alderton, Roberto Amato, Jeffrey Barrett, Sonia Goncalves, Ewan Harrison, David K. Jackson, Ian Johnston, Dominic Kwiatkowski, Cordelia Langford, John Sillitoe on behalf of the Wellcome Sanger Institute COVID-19 Surveillance Team |
| EPI_ISL_3381317, EPI_ISL_3381318, EPI_ISL_3381319, EPI_ISL_3381320 | Lighthouse Lab in Glasgow | Wellcome Sanger Institute for the COVID-19 Genomics UK (COG-UK) Consortium | Harper VanSteenhouse, Yumi Kasai, David Gray, Carol Clugston, Anna Dominiczak and Alex Alderton, Roberto Amato, Jeffrey Barrett, Sonia Goncalves, Ewan Harrison, David K. Jackson, Ian Johnston, Dominic Kwiatkowski, Cordelia Langford, John Sillitoe on behalf of the Wellcome Sanger Institute COVID-19 Surveillance Team |
| EPI_ISL_3381321 | Lighthouse Laboratory Plymouth | Wellcome Sanger Institute for the COVID-19 Genomics UK (COG-UK) Consortium | Lighthouse Laboratory Plymouth and Alex Alderton, Roberto Amato, Jeffrey Barrett, Sonia Goncalves, Ewan Harrison, David K. Jackson, Ian Johnston, Dominic Kwiatkowski, Cordelia Langford, John Sillitoe on behalf of the Wellcome Sanger Institute COVID-19 Surveillance Team |
| EPI_ISL_3381322 | Lighthouse Lab in Glasgow | Wellcome Sanger Institute for the COVID-19 Genomics UK (COG-UK) Consortium | Harper VanSteenhouse, Yumi Kasai, David Gray, Carol Clugston, Anna Dominiczak and Alex Alderton, Roberto Amato, Jeffrey Barrett, Sonia Goncalves, Ewan Harrison, David K. Jackson, Ian Johnston, Dominic Kwiatkowski, Cordelia Langford, John Sillitoe on behalf of the Wellcome Sanger Institute COVID-19 Surveillance Team |
| EPI_ISL_3381323 | Lighthouse Lab in Alderley Park | Wellcome Sanger Institute for the COVID-19 Genomics UK (COG-UK) Consortium | Jacquelyn Wynn, Mairead Hyland, The Lighthouse Lab in Alderley Park and Alex Alderton, Roberto Amato, Jeffrey Barrett, Sonia Goncalves, Ewan Harrison, David K. Jackson, Ian Johnston, Dominic Kwiatkowski, Cordelia Langford, John Sillitoe on behalf of the Wellcome Sanger Institute COVID-19 Surveillance Team |
| EPI_ISL_3381324, EPI_ISL_3381325, EPI_ISL_3381326, EPI_ISL_3381327, EPI_ISL_3381328, EPI_ISL_3381329 | Lighthouse Lab in Glasgow | Wellcome Sanger Institute for the COVID-19 Genomics UK (COG-UK) Consortium | Harper VanSteenhouse, Yumi Kasai, David Gray, Carol Clugston, Anna Dominiczak and Alex Alderton, Roberto Amato, Jeffrey Barrett, Sonia Goncalves, Ewan Harrison, David K. Jackson, Ian Johnston, Dominic Kwiatkowski, Cordelia Langford, John Sillitoe on behalf of the Wellcome Sanger Institute COVID-19 Surveillance Team |
| EPI_ISL_3381330 | Lighthouse Laboratory Plymouth | Wellcome Sanger Institute for the COVID-19 Genomics UK (COG-UK) Consortium | Lighthouse Laboratory Plymouth and Alex Alderton, Roberto Amato, Jeffrey Barrett, Sonia Goncalves, Ewan Harrison, David K. Jackson, Ian Johnston, Dominic Kwiatkowski, Cordelia Langford, John Sillitoe on behalf of the Wellcome Sanger Institute COVID-19 Surveillance Team |
| EPI_ISL_3381331, EPI_ISL_3381332 | Lighthouse Lab in Alderley Park | Wellcome Sanger Institute for the COVID-19 Genomics UK (COG-UK) Consortium | Jacquelyn Wynn, Mairead Hyland, The Lighthouse Lab in Alderley Park and Alex Alderton, Roberto Amato, Jeffrey Barrett, Sonia Goncalves, Ewan Harrison, David K. Jackson, Ian Johnston, Dominic Kwiatkowski, Cordelia Langford, John Sillitoe on behalf of the Wellcome Sanger Institute COVID-19 Surveillance Team |
| EPI_ISL_3381333 | Lighthouse Lab in Glasgow | Wellcome Sanger Institute for the COVID-19 Genomics UK (COG-UK) Consortium | Harper VanSteenhouse, Yumi Kasai, David Gray, Carol Clugston, Anna Dominiczak and Alex Alderton, Roberto Amato, Jeffrey Barrett, Sonia Goncalves, Ewan Harrison, David K. Jackson, Ian Johnston, Dominic Kwiatkowski, Cordelia Langford, John Sillitoe on behalf of the Wellcome Sanger Institute COVID-19 Surveillance Team |
| EPI_ISL_3381334 | Lighthouse Lab in Alderley Park | Wellcome Sanger Institute for the COVID-19 Genomics UK (COG-UK) Consortium | Jacquelyn Wynn, Mairead Hyland, The Lighthouse Lab in Alderley Park and Alex Alderton, Roberto Amato, Jeffrey Barrett, Sonia Goncalves, Ewan Harrison, David K. Jackson, Ian Johnston, Dominic Kwiatkowski, Cordelia Langford, John Sillitoe on behalf of the Wellcome Sanger Institute COVID-19 Surveillance Team |
| EPI_ISL_3381335 | Lighthouse Lab in Glasgow | Wellcome Sanger Institute for the COVID-19 Genomics UK (COG-UK) Consortium | Harper VanSteenhouse, Yumi Kasai, David Gray, Carol Clugston, Anna Dominiczak and Alex Alderton, Roberto Amato, Jeffrey Barrett, Sonia Goncalves, Ewan Harrison, David K. Jackson, Ian Johnston, Dominic Kwiatkowski, Cordelia Langford, John Sillitoe on behalf of the Wellcome Sanger Institute COVID-19 Surveillance Team |
| EPI_ISL_3381336 | Lighthouse Lab in Alderley Park | Wellcome Sanger Institute for the COVID-19 Genomics UK (COG-UK) Consortium | Jacquelyn Wynn, Mairead Hyland, The Lighthouse Lab in Alderley Park and Alex Alderton, Roberto Amato, Jeffrey Barrett, Sonia Goncalves, Ewan Harrison, David K. Jackson, Ian Johnston, Dominic Kwiatkowski, Cordelia Langford, John Sillitoe on behalf of the Wellcome Sanger Institute COVID-19 Surveillance Team |
| EPI_ISL_3381337 | Lighthouse Lab in Glasgow | Wellcome Sanger Institute for the COVID-19 Genomics UK (COG-UK) Consortium | Harper VanSteenhouse, Yumi Kasai, David Gray, Carol Clugston, Anna Dominiczak and Alex Alderton, Roberto Amato, Jeffrey Barrett, Sonia Goncalves, Ewan Harrison, David K. Jackson, Ian Johnston, Dominic Kwiatkowski, Cordelia Langford, John Sillitoe on behalf of the Wellcome Sanger Institute COVID-19 Surveillance Team |
| EPI_ISL_3381338 | Lighthouse Laboratory Plymouth | Wellcome Sanger Institute for the COVID-19 Genomics UK (COG-UK) Consortium | Lighthouse Laboratory Plymouth and Alex Alderton, Roberto Amato, Jeffrey Barrett, Sonia Goncalves, Ewan Harrison, David K. Jackson, Ian Johnston, Dominic Kwiatkowski, Cordelia Langford, John Sillitoe on behalf of the Wellcome Sanger Institute COVID-19 Surveillance Team |
| EPI_ISL_3381339, EPI_ISL_3381340 | Lighthouse Lab in Glasgow | Wellcome Sanger Institute for the COVID-19 Genomics UK (COG-UK) Consortium | Harper VanSteenhouse, Yumi Kasai, David Gray, Carol Clugston, Anna Dominiczak and Alex Alderton, Roberto Amato, Jeffrey Barrett, Sonia Goncalves, Ewan Harrison, David K. Jackson, Ian Johnston, Dominic Kwiatkowski, Cordelia Langford, John Sillitoe on behalf of the Wellcome Sanger Institute COVID-19 Surveillance Team |
| EPI_ISL_3381341, EPI_ISL_3381342 | Lighthouse Lab in Alderley Park | Wellcome Sanger Institute for the COVID-19 Genomics UK (COG-UK) Consortium | Jacquelyn Wynn, Mairead Hyland, The Lighthouse Lab in Alderley Park and Alex Alderton, Roberto Amato, Jeffrey Barrett, Sonia Goncalves, Ewan Harrison, David K. Jackson, Ian Johnston, Dominic Kwiatkowski, Cordelia Langford, John Sillitoe on behalf of the Wellcome Sanger Institute COVID-19 Surveillance Team |
| EPI_ISL_3381343, EPI_ISL_3381344 | Lighthouse Lab in Glasgow | Wellcome Sanger Institute for the COVID-19 Genomics UK (COG-UK) Consortium | Harper VanSteenhouse, Yumi Kasai, David Gray, Carol Clugston, Anna Dominiczak and Alex Alderton, Roberto Amato, Jeffrey Barrett, Sonia Goncalves, Ewan Harrison, David K. Jackson, Ian Johnston, Dominic Kwiatkowski, Cordelia Langford, John Sillitoe on behalf of the Wellcome Sanger Institute COVID-19 Surveillance Team |
| EPI_ISL_3381345 | Lighthouse Laboratory Plymouth | Wellcome Sanger Institute for the COVID-19 Genomics UK (COG-UK) Consortium | Lighthouse Laboratory Plymouth and Alex Alderton, Roberto Amato, Jeffrey Barrett, Sonia Goncalves, Ewan Harrison, David K. Jackson, Ian Johnston, Dominic Kwiatkowski, Cordelia Langford, John Sillitoe on behalf of the Wellcome Sanger Institute COVID-19 Surveillance Team |
| EPI_ISL_3381346 | Lighthouse Lab in Alderley Park | Wellcome Sanger Institute for the COVID-19 Genomics UK (COG-UK) Consortium | Jacquelyn Wynn, Mairead Hyland, The Lighthouse Lab in Alderley Park and Alex Alderton, Roberto Amato, Jeffrey Barrett, Sonia Goncalves, Ewan Harrison, David K. Jackson, Ian Johnston, Dominic Kwiatkowski, Cordelia Langford, John Sillitoe on behalf of the Wellcome Sanger Institute COVID-19 Surveillance Team |
| EPI_ISL_3381347 | Lighthouse Laboratory Plymouth | Wellcome Sanger Institute for the COVID-19 Genomics UK (COG-UK) Consortium | Lighthouse Laboratory Plymouth and Alex Alderton, Roberto Amato, Jeffrey Barrett, Sonia Goncalves, Ewan Harrison, David K. Jackson, Ian Johnston, Dominic Kwiatkowski, Cordelia Langford, John Sillitoe on behalf of the Wellcome Sanger Institute COVID-19 Surveillance Team |
| EPI_ISL_3381348 | Lighthouse Lab in Glasgow | Wellcome Sanger Institute for the COVID-19 Genomics UK (COG-UK) Consortium | Harper VanSteenhouse, Yumi Kasai, David Gray, Carol Clugston, Anna Dominiczak and Alex Alderton, Roberto Amato, Jeffrey Barrett, Sonia Goncalves, Ewan Harrison, David K. Jackson, Ian Johnston, Dominic Kwiatkowski, Cordelia Langford, John Sillitoe on behalf of the Wellcome Sanger Institute COVID-19 Surveillance Team |
| EPI_ISL_3381349 | Lighthouse Laboratory Plymouth | Wellcome Sanger Institute for the COVID-19 Genomics UK (COG-UK) Consortium | Lighthouse Laboratory Plymouth and Alex Alderton, Roberto Amato, Jeffrey Barrett, Sonia Goncalves, Ewan Harrison, David K. Jackson, Ian Johnston, Dominic Kwiatkowski, Cordelia Langford, John Sillitoe on behalf of the Wellcome Sanger Institute COVID-19 Surveillance Team |
| EPI_ISL_3381350, EPI_ISL_3381351 | Lighthouse Lab in Alderley Park | Wellcome Sanger Institute for the COVID-19 Genomics UK (COG-UK) Consortium | Jacquelyn Wynn, Mairead Hyland, The Lighthouse Lab in Alderley Park and Alex Alderton, Roberto Amato, Jeffrey Barrett, Sonia Goncalves, Ewan Harrison, David K. Jackson, Ian Johnston, Dominic Kwiatkowski, Cordelia Langford, John Sillitoe on behalf of the Wellcome Sanger Institute COVID-19 Surveillance Team |
| EPI_ISL_3381352, EPI_ISL_3381353 | Lighthouse Laboratory Plymouth | Wellcome Sanger Institute for the COVID-19 Genomics UK (COG-UK) Consortium | Lighthouse Laboratory Plymouth and Alex Alderton, Roberto Amato, Jeffrey Barrett, Sonia Goncalves, Ewan Harrison, David K. Jackson, Ian Johnston, Dominic Kwiatkowski, Cordelia Langford, John Sillitoe on behalf of the Wellcome Sanger Institute COVID-19 Surveillance Team |
| EPI_ISL_3381354, EPI_ISL_3381355 | Lighthouse Lab in Glasgow | Wellcome Sanger Institute for the COVID-19 Genomics UK (COG-UK) Consortium | Harper VanSteenhouse, Yumi Kasai, David Gray, Carol Clugston, Anna Dominiczak and Alex Alderton, Roberto Amato, Jeffrey Barrett, Sonia Goncalves, Ewan Harrison, David K. Jackson, Ian Johnston, Dominic Kwiatkowski, Cordelia Langford, John Sillitoe on behalf of the Wellcome Sanger Institute COVID-19 Surveillance Team |
| EPI_ISL_3381356 | Lighthouse Lab in Alderley Park | Wellcome Sanger Institute for the COVID-19 Genomics UK (COG-UK) Consortium | Jacquelyn Wynn, Mairead Hyland, The Lighthouse Lab in Alderley Park and Alex Alderton, Roberto Amato, Jeffrey Barrett, Sonia Goncalves, Ewan Harrison, David K. Jackson, Ian Johnston, Dominic Kwiatkowski, Cordelia Langford, John Sillitoe on behalf of the Wellcome Sanger Institute COVID-19 Surveillance Team |
| EPI_ISL_3381357, EPI_ISL_3381358 | Lighthouse Lab in Glasgow | Wellcome Sanger Institute for the COVID-19 Genomics UK (COG-UK) Consortium | Harper VanSteenhouse, Yumi Kasai, David Gray, Carol Clugston, Anna Dominiczak and Alex Alderton, Roberto Amato, Jeffrey Barrett, Sonia Goncalves, Ewan Harrison, David K. Jackson, Ian Johnston, Dominic Kwiatkowski, Cordelia Langford, John Sillitoe on behalf of the Wellcome Sanger Institute COVID-19 Surveillance Team |
| EPI_ISL_3381359 | Lighthouse Lab in Alderley Park | Wellcome Sanger Institute for the COVID-19 Genomics UK (COG-UK) Consortium | Jacquelyn Wynn, Mairead Hyland, The Lighthouse Lab in Alderley Park and Alex Alderton, Roberto Amato, Jeffrey Barrett, Sonia Goncalves, Ewan Harrison, David K. Jackson, Ian Johnston, Dominic Kwiatkowski, Cordelia Langford, John Sillitoe on behalf of the Wellcome Sanger Institute COVID-19 Surveillance Team |
| EPI_ISL_3381360, EPI_ISL_3381361, EPI_ISL_3381362, EPI_ISL_3381363 | Lighthouse Lab in Glasgow | Wellcome Sanger Institute for the COVID-19 Genomics UK (COG-UK) Consortium | Harper VanSteenhouse, Yumi Kasai, David Gray, Carol Clugston, Anna Dominiczak and Alex Alderton, Roberto Amato, Jeffrey Barrett, Sonia Goncalves, Ewan Harrison, David K. Jackson, Ian Johnston, Dominic Kwiatkowski, Cordelia Langford, John Sillitoe on behalf of the Wellcome Sanger Institute COVID-19 Surveillance Team |
| EPI_ISL_3381364 | Lighthouse Lab in Alderley Park | Wellcome Sanger Institute for the COVID-19 Genomics UK (COG-UK) Consortium | Jacquelyn Wynn, Mairead Hyland, The Lighthouse Lab in Alderley Park and Alex Alderton, Roberto Amato, Jeffrey Barrett, Sonia Goncalves, Ewan Harrison, David K. Jackson, Ian Johnston, Dominic Kwiatkowski, Cordelia Langford, John Sillitoe on behalf of the Wellcome Sanger Institute COVID-19 Surveillance Team |

| | | | |
|---|---|---|---|
| EPI_ISL_3381365 | Lighthouse Laboratory Plymouth | Wellcome Sanger Institute for the COVID-19 Genomics UK (COG-UK) Consortium | Lighthouse Laboratory Plymouth and Alex Alderton, Roberto Amato, Jeffrey Barrett, Sonia Goncalves, Ewan Harrison, David K. Jackson, Ian Johnston, Dominic Kwiatkowski, Cordelia Langford, John Sillitoe on behalf of the Wellcome Sanger Institute COVID-19 Surveillance Team |
| EPI_ISL_3381366 | Lighthouse Lab in Alderley Park | Wellcome Sanger Institute for the COVID-19 Genomics UK (COG-UK) Consortium | Jacquelyn Wynn, Mairead Hyland, The Lighthouse Lab in Alderley Park and Alex Alderton, Roberto Amato, Jeffrey Barrett, Sonia Goncalves, Ewan Harrison, David K. Jackson, Ian Johnston, Dominic Kwiatkowski, Cordelia Langford, John Sillitoe on behalf of the Wellcome Sanger Institute COVID-19 Surveillance Team |
| EPI_ISL_3381367, EPI_ISL_3381368, EPI_ISL_3381369, EPI_ISL_3381370 | Lighthouse Lab in Glasgow | Wellcome Sanger Institute for the COVID-19 Genomics UK (COG-UK) Consortium | Harper VanSteenhouse, Yumi Kasai, David Gray, Carol Clugston, Anna Dominiczak and Alex Alderton, Roberto Amato, Jeffrey Barrett, Sonia Goncalves, Ewan Harrison, David K. Jackson, Ian Johnston, Dominic Kwiatkowski, Cordelia Langford, John Sillitoe on behalf of the Wellcome Sanger Institute COVID-19 Surveillance Team |
| EPI_ISL_3381371 | Lighthouse Laboratory Plymouth | Wellcome Sanger Institute for the COVID-19 Genomics UK (COG-UK) Consortium | Lighthouse Laboratory Plymouth and Alex Alderton, Roberto Amato, Jeffrey Barrett, Sonia Goncalves, Ewan Harrison, David K. Jackson, Ian Johnston, Dominic Kwiatkowski, Cordelia Langford, John Sillitoe on behalf of the Wellcome Sanger Institute COVID-19 Surveillance Team |
| EPI_ISL_3381372, EPI_ISL_3381373, EPI_ISL_3381374 | Lighthouse Lab in Glasgow | Wellcome Sanger Institute for the COVID-19 Genomics UK (COG-UK) Consortium | Harper VanSteenhouse, Yumi Kasai, David Gray, Carol Clugston, Anna Dominiczak and Alex Alderton, Roberto Amato, Jeffrey Barrett, Sonia Goncalves, Ewan Harrison, David K. Jackson, Ian Johnston, Dominic Kwiatkowski, Cordelia Langford, John Sillitoe on behalf of the Wellcome Sanger Institute COVID-19 Surveillance Team |
| EPI_ISL_3381375, EPI_ISL_3381376 | Lighthouse Laboratory Plymouth | Wellcome Sanger Institute for the COVID-19 Genomics UK (COG-UK) Consortium | Lighthouse Laboratory Plymouth and Alex Alderton, Roberto Amato, Jeffrey Barrett, Sonia Goncalves, Ewan Harrison, David K. Jackson, Ian Johnston, Dominic Kwiatkowski, Cordelia Langford, John Sillitoe on behalf of the Wellcome Sanger Institute COVID-19 Surveillance Team |
| EPI_ISL_3381377 | Lighthouse Lab in Glasgow | Wellcome Sanger Institute for the COVID-19 Genomics UK (COG-UK) Consortium | Harper VanSteenhouse, Yumi Kasai, David Gray, Carol Clugston, Anna Dominiczak and Alex Alderton, Roberto Amato, Jeffrey Barrett, Sonia Goncalves, Ewan Harrison, David K. Jackson, Ian Johnston, Dominic Kwiatkowski, Cordelia Langford, John Sillitoe on behalf of the Wellcome Sanger Institute COVID-19 Surveillance Team |
| EPI_ISL_3381378 | Lighthouse Laboratory Plymouth | Wellcome Sanger Institute for the COVID-19 Genomics UK (COG-UK) Consortium | Lighthouse Laboratory Plymouth and Alex Alderton, Roberto Amato, Jeffrey Barrett, Sonia Goncalves, Ewan Harrison, David K. Jackson, Ian Johnston, Dominic Kwiatkowski, Cordelia Langford, John Sillitoe on behalf of the Wellcome Sanger Institute COVID-19 Surveillance Team |
| EPI_ISL_3381379 | Lighthouse Lab in Alderley Park | Wellcome Sanger Institute for the COVID-19 Genomics UK (COG-UK) Consortium | Jacquelyn Wynn, Mairead Hyland, The Lighthouse Lab in Alderley Park and Alex Alderton, Roberto Amato, Jeffrey Barrett, Sonia Goncalves, Ewan Harrison, David K. Jackson, Ian Johnston, Dominic Kwiatkowski, Cordelia Langford, John Sillitoe on behalf of the Wellcome Sanger Institute COVID-19 Surveillance Team |
| EPI_ISL_3381380, EPI_ISL_3381381 | Lighthouse Laboratory Plymouth | Wellcome Sanger Institute for the COVID-19 Genomics UK (COG-UK) Consortium | Lighthouse Laboratory Plymouth and Alex Alderton, Roberto Amato, Jeffrey Barrett, Sonia Goncalves, Ewan Harrison, David K. Jackson, Ian Johnston, Dominic Kwiatkowski, Cordelia Langford, John Sillitoe on behalf of the Wellcome Sanger Institute COVID-19 Surveillance Team |
| EPI_ISL_3381382 | Lighthouse Lab in Glasgow | Wellcome Sanger Institute for the COVID-19 Genomics UK (COG-UK) Consortium | Harper VanSteenhouse, Yumi Kasai, David Gray, Carol Clugston, Anna Dominiczak and Alex Alderton, Roberto Amato, Jeffrey Barrett, Sonia Goncalves, Ewan Harrison, David K. Jackson, Ian Johnston, Dominic Kwiatkowski, Cordelia Langford, John Sillitoe on behalf of the Wellcome Sanger Institute COVID-19 Surveillance Team |
| EPI_ISL_3381383, EPI_ISL_3381384 | Lighthouse Laboratory Plymouth | Wellcome Sanger Institute for the COVID-19 Genomics UK (COG-UK) Consortium | Lighthouse Laboratory Plymouth and Alex Alderton, Roberto Amato, Jeffrey Barrett, Sonia Goncalves, Ewan Harrison, David K. Jackson, Ian Johnston, Dominic Kwiatkowski, Cordelia Langford, John Sillitoe on behalf of the Wellcome Sanger Institute COVID-19 Surveillance Team |
| EPI_ISL_3381385, EPI_ISL_3381386 | Lighthouse Lab in Glasgow | Wellcome Sanger Institute for the COVID-19 Genomics UK (COG-UK) Consortium | Harper VanSteenhouse, Yumi Kasai, David Gray, Carol Clugston, Anna Dominiczak and Alex Alderton, Roberto Amato, Jeffrey Barrett, Sonia Goncalves, Ewan Harrison, David K. Jackson, Ian Johnston, Dominic Kwiatkowski, Cordelia Langford, John Sillitoe on behalf of the Wellcome Sanger Institute COVID-19 Surveillance Team |
| EPI_ISL_3381387, EPI_ISL_3381388 | Lighthouse Lab in Alderley Park | Wellcome Sanger Institute for the COVID-19 Genomics UK (COG-UK) Consortium | Jacquelyn Wynn, Mairead Hyland, The Lighthouse Lab in Alderley Park and Alex Alderton, Roberto Amato, Jeffrey Barrett, Sonia Goncalves, Ewan Harrison, David K. Jackson, Ian Johnston, Dominic Kwiatkowski, Cordelia Langford, John Sillitoe on behalf of the Wellcome Sanger Institute COVID-19 Surveillance Team |
| EPI_ISL_3381389 | Lighthouse Laboratory Plymouth | Wellcome Sanger Institute for the COVID-19 Genomics UK (COG-UK) Consortium | Lighthouse Laboratory Plymouth and Alex Alderton, Roberto Amato, Jeffrey Barrett, Sonia Goncalves, Ewan Harrison, David K. Jackson, Ian Johnston, Dominic Kwiatkowski, Cordelia Langford, John Sillitoe on behalf of the Wellcome Sanger Institute COVID-19 Surveillance Team |
| EPI_ISL_3381390 | Lighthouse Lab in Glasgow | Wellcome Sanger Institute for the COVID-19 Genomics UK (COG-UK) Consortium | Harper VanSteenhouse, Yumi Kasai, David Gray, Carol Clugston, Anna Dominiczak and Alex Alderton, Roberto Amato, Jeffrey Barrett, Sonia Goncalves, Ewan Harrison, David K. Jackson, Ian Johnston, Dominic Kwiatkowski, Cordelia Langford, John Sillitoe on behalf of the Wellcome Sanger Institute COVID-19 Surveillance Team |
| EPI_ISL_3381391, EPI_ISL_3381392 | Lighthouse Laboratory Plymouth | Wellcome Sanger Institute for the COVID-19 Genomics UK (COG-UK) Consortium | Lighthouse Laboratory Plymouth and Alex Alderton, Roberto Amato, Jeffrey Barrett, Sonia Goncalves, Ewan Harrison, David K. Jackson, Ian Johnston, Dominic Kwiatkowski, Cordelia Langford, John Sillitoe on behalf of the Wellcome Sanger Institute COVID-19 Surveillance Team |
| EPI_ISL_3381393 | Lighthouse Lab in Alderley Park | Wellcome Sanger Institute for the COVID-19 Genomics UK (COG-UK) Consortium | Jacquelyn Wynn, Mairead Hyland, The Lighthouse Lab in Alderley Park and Alex Alderton, Roberto Amato, Jeffrey Barrett, Sonia Goncalves, Ewan Harrison, David K. Jackson, Ian Johnston, Dominic Kwiatkowski, Cordelia Langford, John Sillitoe on behalf of the Wellcome Sanger Institute COVID-19 Surveillance Team |
| EPI_ISL_3381394, EPI_ISL_3381395 | Lighthouse Lab in Glasgow | Wellcome Sanger Institute for the COVID-19 Genomics UK (COG-UK) Consortium | Harper VanSteenhouse, Yumi Kasai, David Gray, Carol Clugston, Anna Dominiczak and Alex Alderton, Roberto Amato, Jeffrey Barrett, Sonia Goncalves, Ewan Harrison, David K. Jackson, Ian Johnston, Dominic Kwiatkowski, Cordelia Langford, John Sillitoe on behalf of the Wellcome Sanger Institute COVID-19 Surveillance Team |
| EPI_ISL_3381396 | Lighthouse Lab in Alderley Park | Wellcome Sanger Institute for the COVID-19 Genomics UK (COG-UK) Consortium | Jacquelyn Wynn, Mairead Hyland, The Lighthouse Lab in Alderley Park and Alex Alderton, Roberto Amato, Jeffrey Barrett, Sonia Goncalves, Ewan Harrison, David K. Jackson, Ian Johnston, Dominic Kwiatkowski, Cordelia Langford, John Sillitoe on behalf of the Wellcome Sanger Institute COVID-19 Surveillance Team |
| EPI_ISL_3381397, EPI_ISL_3381398, EPI_ISL_3381399, EPI_ISL_3381400 | Lighthouse Lab in Glasgow | Wellcome Sanger Institute for the COVID-19 Genomics UK (COG-UK) Consortium | Harper VanSteenhouse, Yumi Kasai, David Gray, Carol Clugston, Anna Dominiczak and Alex Alderton, Roberto Amato, Jeffrey Barrett, Sonia Goncalves, Ewan Harrison, David K. Jackson, Ian Johnston, Dominic Kwiatkowski, Cordelia Langford, John Sillitoe on behalf of the Wellcome Sanger Institute COVID-19 Surveillance Team |
| EPI_ISL_3381401, EPI_ISL_3381402 | Lighthouse Laboratory Plymouth | Wellcome Sanger Institute for the COVID-19 Genomics UK (COG-UK) Consortium | Lighthouse Laboratory Plymouth and Alex Alderton, Roberto Amato, Jeffrey Barrett, Sonia Goncalves, Ewan Harrison, David K. Jackson, Ian Johnston, Dominic Kwiatkowski, Cordelia Langford, John Sillitoe on behalf of the Wellcome Sanger Institute COVID-19 Surveillance Team |
| EPI_ISL_3381403, EPI_ISL_3381404, EPI_ISL_3381405 | Lighthouse Lab in Glasgow | Wellcome Sanger Institute for the COVID-19 Genomics UK (COG-UK) Consortium | Harper VanSteenhouse, Yumi Kasai, David Gray, Carol Clugston, Anna Dominiczak and Alex Alderton, Roberto Amato, Jeffrey Barrett, Sonia Goncalves, Ewan Harrison, David K. Jackson, Ian Johnston, Dominic Kwiatkowski, Cordelia Langford, John Sillitoe on behalf of the Wellcome Sanger Institute COVID-19 Surveillance Team |
| EPI_ISL_3381406 | Lighthouse Laboratory Plymouth | Wellcome Sanger Institute for the COVID-19 Genomics UK (COG-UK) Consortium | Lighthouse Laboratory Plymouth and Alex Alderton, Roberto Amato, Jeffrey Barrett, Sonia Goncalves, Ewan Harrison, David K. Jackson, Ian Johnston, Dominic Kwiatkowski, Cordelia Langford, John Sillitoe on behalf of the Wellcome Sanger Institute COVID-19 Surveillance Team |
| EPI_ISL_3381407 | Lighthouse Lab in Glasgow | Wellcome Sanger Institute for the COVID-19 Genomics UK (COG-UK) Consortium | Harper VanSteenhouse, Yumi Kasai, David Gray, Carol Clugston, Anna Dominiczak and Alex Alderton, Roberto Amato, Jeffrey Barrett, Sonia Goncalves, Ewan Harrison, David K. Jackson, Ian Johnston, Dominic Kwiatkowski, Cordelia Langford, John Sillitoe on behalf of the Wellcome Sanger Institute COVID-19 Surveillance Team |
| EPI_ISL_3381408 | Lighthouse Laboratory Plymouth | Wellcome Sanger Institute for the COVID-19 Genomics UK (COG-UK) Consortium | Lighthouse Laboratory Plymouth and Alex Alderton, Roberto Amato, Jeffrey Barrett, Sonia Goncalves, Ewan Harrison, David K. Jackson, Ian Johnston, Dominic Kwiatkowski, Cordelia Langford, John Sillitoe on behalf of the Wellcome Sanger Institute COVID-19 Surveillance Team |
| EPI_ISL_3381409 | Lighthouse Lab in Alderley Park | Wellcome Sanger Institute for the COVID-19 Genomics UK (COG-UK) Consortium | Jacquelyn Wynn, Mairead Hyland, The Lighthouse Lab in Alderley Park and Alex Alderton, Roberto Amato, Jeffrey Barrett, Sonia Goncalves, Ewan Harrison, David K. Jackson, Ian Johnston, Dominic Kwiatkowski, Cordelia Langford, John Sillitoe on behalf of the Wellcome Sanger Institute COVID-19 Surveillance Team |
| EPI_ISL_3381410 | Lighthouse Lab in Glasgow | Wellcome Sanger Institute for the COVID-19 Genomics UK (COG-UK) Consortium | Harper VanSteenhouse, Yumi Kasai, David Gray, Carol Clugston, Anna Dominiczak and Alex Alderton, Roberto Amato, Jeffrey Barrett, Sonia Goncalves, Ewan Harrison, David K. Jackson, Ian Johnston, Dominic Kwiatkowski, Cordelia Langford, John Sillitoe on behalf of the Wellcome Sanger Institute COVID-19 Surveillance Team |
| EPI_ISL_3381411 | Lighthouse Laboratory Plymouth | Wellcome Sanger Institute for the COVID-19 Genomics UK (COG-UK) Consortium | Lighthouse Laboratory Plymouth and Alex Alderton, Roberto Amato, Jeffrey Barrett, Sonia Goncalves, Ewan Harrison, David K. Jackson, Ian Johnston, Dominic Kwiatkowski, Cordelia Langford, John Sillitoe on behalf of the Wellcome Sanger Institute COVID-19 Surveillance Team |
| EPI_ISL_3381412 | Lighthouse Lab in Glasgow | Wellcome Sanger Institute for the COVID-19 Genomics UK (COG-UK) Consortium | Harper VanSteenhouse, Yumi Kasai, David Gray, Carol Clugston, Anna Dominiczak and Alex Alderton, Roberto Amato, Jeffrey Barrett, Sonia Goncalves, Ewan Harrison, David K. Jackson, Ian Johnston, Dominic Kwiatkowski, Cordelia Langford, John Sillitoe on behalf of the Wellcome Sanger Institute COVID-19 Surveillance Team |
| EPI_ISL_3381413 | Lighthouse Laboratory Plymouth | Wellcome Sanger Institute for the COVID-19 Genomics UK (COG-UK) Consortium | Lighthouse Laboratory Plymouth and Alex Alderton, Roberto Amato, Jeffrey Barrett, Sonia Goncalves, Ewan Harrison, David K. Jackson, Ian Johnston, Dominic Kwiatkowski, Cordelia Langford, John Sillitoe on behalf of the Wellcome Sanger Institute COVID-19 Surveillance Team |
| EPI_ISL_3381414 | Lighthouse Lab in Glasgow | Wellcome Sanger Institute for the COVID-19 Genomics UK (COG-UK) Consortium | Harper VanSteenhouse, Yumi Kasai, David Gray, Carol Clugston, Anna Dominiczak and Alex Alderton, Roberto Amato, Jeffrey Barrett, Sonia Goncalves, Ewan Harrison, David K. Jackson, Ian Johnston, Dominic Kwiatkowski, Cordelia Langford, John Sillitoe on behalf of the Wellcome Sanger Institute COVID-19 Surveillance Team |
| EPI_ISL_3381415, EPI_ISL_3381416 | Lighthouse Laboratory Plymouth | Wellcome Sanger Institute for the COVID-19 Genomics UK (COG-UK) Consortium | Lighthouse Laboratory Plymouth and Alex Alderton, Roberto Amato, Jeffrey Barrett, Sonia Goncalves, Ewan Harrison, David K. Jackson, Ian Johnston, Dominic Kwiatkowski, Cordelia Langford, John Sillitoe on behalf of the Wellcome Sanger Institute COVID-19 Surveillance Team |
| EPI_ISL_3381417, EPI_ISL_3381418 | Lighthouse Lab in Alderley Park | Wellcome Sanger Institute for the COVID-19 Genomics UK (COG-UK) Consortium | Jacquelyn Wynn, Mairead Hyland, The Lighthouse Lab in Alderley Park and Alex Alderton, Roberto Amato, Jeffrey Barrett, Sonia Goncalves, Ewan Harrison, David K. Jackson, Ian Johnston, Dominic Kwiatkowski, Cordelia Langford, John Sillitoe on behalf of the Wellcome Sanger Institute COVID-19 Surveillance Team |
| EPI_ISL_3381419, EPI_ISL_3381420 | Lighthouse Laboratory Plymouth | Wellcome Sanger Institute for the COVID-19 Genomics UK (COG-UK) Consortium | Lighthouse Laboratory Plymouth and Alex Alderton, Roberto Amato, Jeffrey Barrett, Sonia Goncalves, Ewan Harrison, David K. Jackson, Ian Johnston, Dominic Kwiatkowski, Cordelia Langford, John Sillitoe on behalf of the Wellcome Sanger Institute COVID-19 Surveillance Team |
| EPI_ISL_3381421 | Lighthouse Lab in Glasgow | Wellcome Sanger Institute for the COVID-19 Genomics UK (COG-UK) Consortium | Harper VanSteenhouse, Yumi Kasai, David Gray, Carol Clugston, Anna Dominiczak and Alex Alderton, Roberto Amato, Jeffrey Barrett, Sonia Goncalves, Ewan Harrison, David K. Jackson, Ian Johnston, Dominic Kwiatkowski, Cordelia Langford, John Sillitoe on behalf of the Wellcome Sanger Institute COVID-19 Surveillance Team |
| EPI_ISL_3381422, EPI_ISL_3381423 | Lighthouse Laboratory Plymouth | Wellcome Sanger Institute for the COVID-19 Genomics UK (COG-UK) Consortium | Lighthouse Laboratory Plymouth and Alex Alderton, Roberto Amato, Jeffrey Barrett, Sonia Goncalves, Ewan Harrison, David K. Jackson, Ian Johnston, Dominic Kwiatkowski, Cordelia Langford, John Sillitoe on behalf of the Wellcome Sanger Institute COVID-19 Surveillance Team |
| EPI_ISL_3381424 | Lighthouse Lab in Glasgow | Wellcome Sanger Institute for the COVID-19 Genomics UK (COG-UK) Consortium | Harper VanSteenhouse, Yumi Kasai, David Gray, Carol Clugston, Anna Dominiczak and Alex Alderton, Roberto Amato, Jeffrey Barrett, Sonia Goncalves, Ewan Harrison, David K. Jackson, Ian Johnston, Dominic Kwiatkowski, Cordelia Langford, John Sillitoe on behalf of the Wellcome Sanger Institute COVID-19 Surveillance Team |
| EPI_ISL_3381425 | Lighthouse Laboratory Plymouth | Wellcome Sanger Institute for the COVID-19 Genomics UK (COG-UK) Consortium | Lighthouse Laboratory Plymouth and Alex Alderton, Roberto Amato, Jeffrey Barrett, Sonia Goncalves, Ewan Harrison, David K. Jackson, Ian Johnston, Dominic Kwiatkowski, Cordelia Langford, John Sillitoe on behalf of the Wellcome Sanger Institute COVID-19 Surveillance Team |
| EPI_ISL_3381426 | Lighthouse Lab in Alderley Park | Wellcome Sanger Institute for the COVID-19 Genomics UK (COG-UK) Consortium | Jacquelyn Wynn, Mairead Hyland, The Lighthouse Lab in Alderley Park and Alex Alderton, Roberto Amato, Jeffrey Barrett, Sonia Goncalves, Ewan Harrison, David K. Jackson, Ian Johnston, Dominic Kwiatkowski, Cordelia Langford, John Sillitoe on behalf of the Wellcome Sanger Institute COVID-19 Surveillance Team |
| EPI_ISL_3381427 | Lighthouse Lab in Glasgow | Wellcome Sanger Institute for the COVID-19 Genomics UK (COG-UK) Consortium | Harper VanSteenhouse, Yumi Kasai, David Gray, Carol Clugston, Anna Dominiczak and Alex Alderton, Roberto Amato, Jeffrey Barrett, Sonia Goncalves, Ewan Harrison, David K. Jackson, Ian Johnston, Dominic Kwiatkowski, Cordelia Langford, John Sillitoe on behalf of the Wellcome Sanger Institute COVID-19 Surveillance Team |
| EPI_ISL_3381428 | Lighthouse Laboratory Plymouth | Wellcome Sanger Institute for the COVID-19 Genomics UK (COG-UK) Consortium | Lighthouse Laboratory Plymouth and Alex Alderton, Roberto Amato, Jeffrey Barrett, Sonia Goncalves, Ewan Harrison, David K. Jackson, Ian Johnston, Dominic Kwiatkowski, Cordelia Langford, John Sillitoe on behalf of the Wellcome Sanger Institute COVID-19 Surveillance Team |

EPI_ISL_3381429, EPI_ISL_3381430, EPI_ISL_3381432, EPI_ISL_3381433, EPI_ISL_3381434, EPI_ISL_3381435, EPI_ISL_3381436, EPI_ISL_3381437, EPI_ISL_3381438, EPI_ISL_3381439, EPI_ISL_3381440, EPI_ISL_3381441, EPI_ISL_3381442, EPI_ISL_3381443, EPI_ISL_3381444, EPI_ISL_3381445, EPI_ISL_3381446, EPI_ISL_3381447, EPI_ISL_3381448, EPI_ISL_3381449, EPI_ISL_3381450, EPI_ISL_3381451,

| | | | |
|---|---|---|---|
| EPI_ISL_3381452, EPI_ISL_3381453, EPI_ISL_3381454, EPI_ISL_3381455, EPI_ISL_3381457, EPI_ISL_3381458, EPI_ISL_3381459, EPI_ISL_3381460, EPI_ISL_3381461, EPI_ISL_3381462, EPI_ISL_3381463, EPI_ISL_3381464, EPI_ISL_3381465, EPI_ISL_3381466, EPI_ISL_3381467, EPI_ISL_3381468, EPI_ISL_3381469, EPI_ISL_3381470, EPI_ISL_3381471, EPI_ISL_3381472, EPI_ISL_3381473, EPI_ISL_3381474, EPI_ISL_3381475, EPI_ISL_3381476, EPI_ISL_3381477, EPI_ISL_3381478, EPI_ISL_3381479, EPI_ISL_3381480, EPI_ISL_3381481, EPI_ISL_3381482, EPI_ISL_3381483, EPI_ISL_3381484, EPI_ISL_3381485, EPI_ISL_3381486, EPI_ISL_3381487, EPI_ISL_3381488, EPI_ISL_3381489, EPI_ISL_3381490, EPI_ISL_3381491, EPI_ISL_3381492, EPI_ISL_3381493, EPI_ISL_3381494, EPI_ISL_3381495, EPI_ISL_3381496, EPI_ISL_3381497, EPI_ISL_3381498, EPI_ISL_3381499, EPI_ISL_3381500, EPI_ISL_3381501, EPI_ISL_3381502, EPI_ISL_3381503, EPI_ISL_3381504, EPI_ISL_3381505, EPI_ISL_3381506, EPI_ISL_3381507, EPI_ISL_3381508, EPI_ISL_3381509, EPI_ISL_3381510, EPI_ISL_3381511, EPI_ISL_3381512, EPI_ISL_3381513, EPI_ISL_3381514, EPI_ISL_3381515, EPI_ISL_3381516, EPI_ISL_3381517, EPI_ISL_3381518, EPI_ISL_3381519, EPI_ISL_3381520, EPI_ISL_3381521, EPI_ISL_3381522, EPI_ISL_3381523, EPI_ISL_3381524, EPI_ISL_3381525, EPI_ISL_3381526, EPI_ISL_3381527, EPI_ISL_3381528, EPI_ISL_3381529, EPI_ISL_3381530, EPI_ISL_3381531, EPI_ISL_3381532, EPI_ISL_3381533, EPI_ISL_3381534, EPI_ISL_3381535, EPI_ISL_3381536, EPI_ISL_3381537, EPI_ISL_3381538, EPI_ISL_3381539, EPI_ISL_3381540, EPI_ISL_3381541, EPI_ISL_3381542, EPI_ISL_3381543, EPI_ISL_3381544, EPI_ISL_3381545, EPI_ISL_3381546, EPI_ISL_3381547, EPI_ISL_3381548, EPI_ISL_3381549, EPI_ISL_3381550, EPI_ISL_3381551, EPI_ISL_3381552, EPI_ISL_3381553, EPI_ISL_3381554, EPI_ISL_3381555, EPI_ISL_3381556, EPI_ISL_3381557, EPI_ISL_3381558, EPI_ISL_3381559, EPI_ISL_3381560, EPI_ISL_3381561, EPI_ISL_3381562, EPI_ISL_3381563, EPI_ISL_3381564, EPI_ISL_3381565, EPI_ISL_3381566, EPI_ISL_3381567, EPI_ISL_3381568, EPI_ISL_3381569, EPI_ISL_3381570, EPI_ISL_3381571, EPI_ISL_3381572, EPI_ISL_3381573, EPI_ISL_3381574, EPI_ISL_3381575, EPI_ISL_3381576, EPI_ISL_3381577, EPI_ISL_3381578, EPI_ISL_3381579, EPI_ISL_3381580, EPI_ISL_3381581, EPI_ISL_3381582, EPI_ISL_3381583, EPI_ISL_3381584, EPI_ISL_3381585, EPI_ISL_3381586, EPI_ISL_3381587, EPI_ISL_3381588, EPI_ISL_3381589, EPI_ISL_3381590, EPI_ISL_3381591, EPI_ISL_3381592, EPI_ISL_3381593, EPI_ISL_3381594 | | | |
| see above | Randox Laboratories | Wellcome Sanger Institute for the COVID-19 Genomics UK (COG-UK) Consortium | Randox Laboratories and Alex Alderton, Roberto Amato, Jeffrey Barrett, Sonia Goncalves, Ewan Harrison, David K. Jackson, Ian Johnston, Dominic Kwiatkowski, Cordelia Langford, John Sillitoe on behalf of the Wellcome Sanger Institute COVID-19 Surveillance Team |
| EPI_ISL_3381595 | Lighthouse Laboratory Plymouth | Wellcome Sanger Institute for the COVID-19 Genomics UK (COG-UK) Consortium | Lighthouse Laboratory Plymouth and Alex Alderton, Roberto Amato, Jeffrey Barrett, Sonia Goncalves, Ewan Harrison, David K. Jackson, Ian Johnston, Dominic Kwiatkowski, Cordelia Langford, John Sillitoe on behalf of the Wellcome Sanger Institute COVID-19 Surveillance Team |
| EPI_ISL_3381596, EPI_ISL_3381597, EPI_ISL_3381598, EPI_ISL_3381599, EPI_ISL_3381600, EPI_ISL_3381601, EPI_ISL_3381602, EPI_ISL_3381603, EPI_ISL_3381604, EPI_ISL_3381605, EPI_ISL_3381606, EPI_ISL_3381607, EPI_ISL_3381608, EPI_ISL_3381609, EPI_ISL_3381610, EPI_ISL_3381611, EPI_ISL_3381612, EPI_ISL_3381613, EPI_ISL_3381614, EPI_ISL_3381615, EPI_ISL_3381616, EPI_ISL_3381617, EPI_ISL_3381618, EPI_ISL_3381619, EPI_ISL_3381620, EPI_ISL_3381621, EPI_ISL_3381622, EPI_ISL_3381623, EPI_ISL_3381624, EPI_ISL_3381625, EPI_ISL_3381626, EPI_ISL_3381627, EPI_ISL_3381628, EPI_ISL_3381629, EPI_ISL_3381630, EPI_ISL_3381631, EPI_ISL_3381632, EPI_ISL_3381633, EPI_ISL_3381634, EPI_ISL_3381635, EPI_ISL_3381636, EPI_ISL_3381637, EPI_ISL_3381638, EPI_ISL_3381639, EPI_ISL_3381640, EPI_ISL_3381641, EPI_ISL_3381642, EPI_ISL_3381643, EPI_ISL_3381644, EPI_ISL_3381645, EPI_ISL_3381646, EPI_ISL_3381647, EPI_ISL_3381648, EPI_ISL_3381649, EPI_ISL_3381650, EPI_ISL_3381651, EPI_ISL_3381652, EPI_ISL_3381653, EPI_ISL_3381654, EPI_ISL_3381655, EPI_ISL_3381656, EPI_ISL_3381657, EPI_ISL_3381658, EPI_ISL_3381659, EPI_ISL_3381660, EPI_ISL_3381661, EPI_ISL_3381662, EPI_ISL_3381663, EPI_ISL_3381664, EPI_ISL_3381665, EPI_ISL_3381666, EPI_ISL_3381667, EPI_ISL_3381668, EPI_ISL_3381669, EPI_ISL_3381670, EPI_ISL_3381671, EPI_ISL_3381672, EPI_ISL_3381673, EPI_ISL_3381674, EPI_ISL_3381675, EPI_ISL_3381676, EPI_ISL_3381677, EPI_ISL_3381678, EPI_ISL_3381679, EPI_ISL_3381680, EPI_ISL_3381681, EPI_ISL_3381682, EPI_ISL_3381683, EPI_ISL_3381684, EPI_ISL_3381685, EPI_ISL_3381686, EPI_ISL_3381687, EPI_ISL_3381688, EPI_ISL_3381689, EPI_ISL_3381690, EPI_ISL_3381691, EPI_ISL_3381692, EPI_ISL_3381693, EPI_ISL_3381694, EPI_ISL_3381695, EPI_ISL_3381696, EPI_ISL_3381697, EPI_ISL_3381698, EPI_ISL_3381699, EPI_ISL_3381700, EPI_ISL_3381701, EPI_ISL_3381702, EPI_ISL_3381703, EPI_ISL_3381704, EPI_ISL_3381705, EPI_ISL_3381706, EPI_ISL_3381707, EPI_ISL_3381708, EPI_ISL_3381709, EPI_ISL_3381710, EPI_ISL_3381711, EPI_ISL_3381712, EPI_ISL_3381713, EPI_ISL_3381714, EPI_ISL_3381715, EPI_ISL_3381716, EPI_ISL_3381717, EPI_ISL_3381718, EPI_ISL_3381719, EPI_ISL_3381720, EPI_ISL_3381721, EPI_ISL_3381722, EPI_ISL_3381723, EPI_ISL_3381724, EPI_ISL_3381725, EPI_ISL_3381726, EPI_ISL_3381727, EPI_ISL_3381728, EPI_ISL_3381729, EPI_ISL_3381730, EPI_ISL_3381731, EPI_ISL_3381732, EPI_ISL_3381733, EPI_ISL_3381734, EPI_ISL_3381735, EPI_ISL_3381736, EPI_ISL_3381737, EPI_ISL_3381738, EPI_ISL_3381739, EPI_ISL_3381740, EPI_ISL_3381741, EPI_ISL_3381742, EPI_ISL_3381743, EPI_ISL_3381744, EPI_ISL_3381745, EPI_ISL_3381746, EPI_ISL_3381747, EPI_ISL_3381748, EPI_ISL_3381749, EPI_ISL_3381750, EPI_ISL_3381751, EPI_ISL_3381752, EPI_ISL_3381753, EPI_ISL_3381754, EPI_ISL_3381755, EPI_ISL_3381756, EPI_ISL_3381757, EPI_ISL_3381758, EPI_ISL_3381759, EPI_ISL_3381760, EPI_ISL_3381761, EPI_ISL_3381762, EPI_ISL_3381763, EPI_ISL_3381764, EPI_ISL_3381765, EPI_ISL_3381766, EPI_ISL_3381767, EPI_ISL_3381768, EPI_ISL_3381769, EPI_ISL_3381770, EPI_ISL_3381771, EPI_ISL_3381772, EPI_ISL_3381773, EPI_ISL_3381774, EPI_ISL_3381775, EPI_ISL_3381776, EPI_ISL_3381777, EPI_ISL_3381778, EPI_ISL_3381779, EPI_ISL_3381780, EPI_ISL_3381781, EPI_ISL_3381782, EPI_ISL_3381783, EPI_ISL_3381784, EPI_ISL_3381785, EPI_ISL_3381786, EPI_ISL_3381787, EPI_ISL_3381788, EPI_ISL_3381789, EPI_ISL_3381790, EPI_ISL_3381791, EPI_ISL_3381792, EPI_ISL_3381793, EPI_ISL_3381794, EPI_ISL_3381795, EPI_ISL_3381796, EPI_ISL_3381797, EPI_ISL_3381798, EPI_ISL_3381799, EPI_ISL_3381800, EPI_ISL_3381801, EPI_ISL_3381802, EPI_ISL_3381803, EPI_ISL_3381804, EPI_ISL_3381805, EPI_ISL_3381806, EPI_ISL_3381807, EPI_ISL_3381808, EPI_ISL_3381809, EPI_ISL_3381810, EPI_ISL_3381811, EPI_ISL_3381812, EPI_ISL_3381813, EPI_ISL_3381814, EPI_ISL_3381815, EPI_ISL_3381816, EPI_ISL_3381817, EPI_ISL_3381818, EPI_ISL_3381819, EPI_ISL_3381820, EPI_ISL_3381821, EPI_ISL_3381822, EPI_ISL_3381823, EPI_ISL_3381824, EPI_ISL_3381825, EPI_ISL_3381826, EPI_ISL_3381827, EPI_ISL_3381828, EPI_ISL_3381829, EPI_ISL_3381830, EPI_ISL_3381831, EPI_ISL_3381832, EPI_ISL_3381833, EPI_ISL_3381834, EPI_ISL_3381835, EPI_ISL_3381836, EPI_ISL_3381837, EPI_ISL_3381838, EPI_ISL_3381839, EPI_ISL_3381840, EPI_ISL_3381841, EPI_ISL_3381842 | | | |
| see above | Randox Laboratories | Wellcome Sanger Institute for the COVID-19 Genomics UK (COG-UK) Consortium | Randox Laboratories and Alex Alderton, Roberto Amato, Jeffrey Barrett, Sonia Goncalves, Ewan Harrison, David K. Jackson, Ian Johnston, Dominic Kwiatkowski, Cordelia Langford, John Sillitoe on behalf of the Wellcome Sanger Institute COVID-19 Surveillance Team |
| EPI_ISL_3381843 | Lighthouse Laboratory Plymouth | Wellcome Sanger Institute for the COVID-19 Genomics UK (COG-UK) Consortium | Lighthouse Laboratory Plymouth and Alex Alderton, Roberto Amato, Jeffrey Barrett, Sonia Goncalves, Ewan Harrison, David K. Jackson, Ian Johnston, Dominic Kwiatkowski, Cordelia Langford, John Sillitoe on behalf of the Wellcome Sanger Institute COVID-19 Surveillance Team |
| EPI_ISL_3381844, EPI_ISL_3381845, EPI_ISL_3381846, EPI_ISL_3381847, EPI_ISL_3381848, EPI_ISL_3381849, EPI_ISL_3381850, EPI_ISL_3381851, EPI_ISL_3381852, EPI_ISL_3381853, EPI_ISL_3381854, EPI_ISL_3381855, EPI_ISL_3381856, EPI_ISL_3381857, EPI_ISL_3381858, EPI_ISL_3381859, EPI_ISL_3381860, EPI_ISL_3381861, EPI_ISL_3381862, EPI_ISL_3381863, EPI_ISL_3381864, EPI_ISL_3381865, EPI_ISL_3381866, EPI_ISL_3381867 | | | |
| see above | Randox Laboratories | Wellcome Sanger Institute for the COVID-19 Genomics UK (COG-UK) Consortium | Randox Laboratories and Alex Alderton, Roberto Amato, Jeffrey Barrett, Sonia Goncalves, Ewan Harrison, David K. Jackson, Ian Johnston, Dominic Kwiatkowski, Cordelia Langford, John Sillitoe on behalf of the Wellcome Sanger Institute COVID-19 Surveillance Team |
| EPI_ISL_3381868 | Lighthouse Lab in Glasgow | Wellcome Sanger Institute for the COVID-19 Genomics UK (COG-UK) Consortium | Harper VanSteenhouse, Yumi Kasai, David Gray, Carol Clugston, Anna Dominiczak and Alex Alderton, Roberto Amato, Jeffrey Barrett, Sonia Goncalves, Ewan Harrison, David K. Jackson, Ian Johnston, Dominic Kwiatkowski, Cordelia Langford, John Sillitoe on behalf of the Wellcome Sanger Institute COVID-19 Surveillance Team |
| EPI_ISL_3381869 | Lighthouse Lab in Alderley Park | Wellcome Sanger Institute for the COVID-19 Genomics UK (COG-UK) Consortium | Jacquelyn Wynn, Mairead Hyland, The Lighthouse Lab in Alderley Park and Alex Alderton, Roberto Amato, Jeffrey Barrett, Sonia Goncalves, Ewan Harrison, David K. Jackson, Ian Johnston, Dominic Kwiatkowski, Cordelia Langford, John Sillitoe on behalf of the Wellcome Sanger Institute COVID-19 Surveillance Team |
| EPI_ISL_3381870 | Lighthouse Lab in Glasgow | Wellcome Sanger Institute for the COVID-19 Genomics UK (COG-UK) Consortium | Harper VanSteenhouse, Yumi Kasai, David Gray, Carol Clugston, Anna Dominiczak and Alex Alderton, Roberto Amato, Jeffrey Barrett, Sonia Goncalves, Ewan Harrison, David K. Jackson, Ian Johnston, Dominic Kwiatkowski, Cordelia Langford, John Sillitoe on behalf of the Wellcome Sanger Institute COVID-19 Surveillance Team |
| EPI_ISL_3381871 | Lighthouse Lab in Alderley Park | Wellcome Sanger Institute for the COVID-19 Genomics UK (COG-UK) Consortium | Jacquelyn Wynn, Mairead Hyland, The Lighthouse Lab in Alderley Park and Alex Alderton, Roberto Amato, Jeffrey Barrett, Sonia Goncalves, Ewan Harrison, David K. Jackson, Ian Johnston, Dominic Kwiatkowski, Cordelia Langford, John Sillitoe on behalf of the Wellcome Sanger Institute COVID-19 Surveillance Team |
| EPI_ISL_3381872, EPI_ISL_3381873 | Lighthouse Lab in Glasgow | Wellcome Sanger Institute for the COVID-19 Genomics UK (COG-UK) Consortium | Harper VanSteenhouse, Yumi Kasai, David Gray, Carol Clugston, Anna Dominiczak and Alex Alderton, Roberto Amato, Jeffrey Barrett, Sonia Goncalves, Ewan Harrison, David K. Jackson, Ian Johnston, Dominic Kwiatkowski, Cordelia Langford, John Sillitoe on behalf of the Wellcome Sanger Institute COVID-19 Surveillance Team |
| EPI_ISL_3381874, EPI_ISL_3381875 | Lighthouse Lab in Alderley Park | Wellcome Sanger Institute for the COVID-19 Genomics UK (COG-UK) Consortium | Jacquelyn Wynn, Mairead Hyland, The Lighthouse Lab in Alderley Park and Alex Alderton, Roberto Amato, Jeffrey Barrett, Sonia Goncalves, Ewan Harrison, David K. Jackson, Ian Johnston, Dominic Kwiatkowski, Cordelia Langford, John Sillitoe on behalf of the Wellcome Sanger Institute COVID-19 Surveillance Team |
| EPI_ISL_3381876 | Lighthouse Lab in Glasgow | Wellcome Sanger Institute for the COVID-19 Genomics UK (COG-UK) Consortium | Harper VanSteenhouse, Yumi Kasai, David Gray, Carol Clugston, Anna Dominiczak and Alex Alderton, Roberto Amato, Jeffrey Barrett, Sonia Goncalves, Ewan Harrison, David K. Jackson, Ian Johnston, Dominic Kwiatkowski, Cordelia Langford, John Sillitoe on behalf of the Wellcome Sanger Institute COVID-19 Surveillance Team |
| EPI_ISL_3381877, EPI_ISL_3381878 | Lighthouse Lab in Alderley Park | Wellcome Sanger Institute for the COVID-19 Genomics UK (COG-UK) Consortium | Jacquelyn Wynn, Mairead Hyland, The Lighthouse Lab in Alderley Park and Alex Alderton, Roberto Amato, Jeffrey Barrett, Sonia Goncalves, Ewan Harrison, David K. Jackson, Ian Johnston, Dominic Kwiatkowski, Cordelia Langford, John Sillitoe on behalf of the Wellcome Sanger Institute COVID-19 Surveillance Team |
| EPI_ISL_3381879 | Berkshire and Surrey Pathology Services Lighthouse Laboratory | Wellcome Sanger Institute for the COVID-19 Genomics UK (COG-UK) Consortium | Berkshire and Surrey Pathology Services Lighthouse Laboratory and Alex Alderton, Roberto Amato, Jeffrey Barrett, Sonia Goncalves, Ewan Harrison, David K. Jackson, Ian Johnston, Dominic Kwiatkowski, Cordelia Langford, John Sillitoe on behalf of the Wellcome Sanger Institute COVID-19 Surveillance Team |
| EPI_ISL_3381880, EPI_ISL_3381881, EPI_ISL_3381882 | Lighthouse Lab in Glasgow | Wellcome Sanger Institute for the COVID-19 Genomics UK (COG-UK) Consortium | Harper VanSteenhouse, Yumi Kasai, David Gray, Carol Clugston, Anna Dominiczak and Alex Alderton, Roberto Amato, Jeffrey Barrett, Sonia Goncalves, Ewan Harrison, David K. Jackson, Ian Johnston, Dominic Kwiatkowski, Cordelia Langford, John Sillitoe on behalf of the Wellcome Sanger Institute COVID-19 Surveillance Team |
| EPI_ISL_3381883 | Lighthouse Lab in Alderley Park | Wellcome Sanger Institute for the COVID-19 Genomics UK (COG-UK) Consortium | Jacquelyn Wynn, Mairead Hyland, The Lighthouse Lab in Alderley Park and Alex Alderton, Roberto Amato, Jeffrey Barrett, Sonia Goncalves, Ewan Harrison, David K. Jackson, Ian Johnston, Dominic Kwiatkowski, Cordelia Langford, John Sillitoe on behalf of the Wellcome Sanger Institute COVID-19 Surveillance Team |
| EPI_ISL_3381884 | Lighthouse Lab in Glasgow | Wellcome Sanger Institute for the COVID-19 Genomics UK (COG-UK) Consortium | Harper VanSteenhouse, Yumi Kasai, David Gray, Carol Clugston, Anna Dominiczak and Alex Alderton, Roberto Amato, Jeffrey Barrett, Sonia Goncalves, Ewan Harrison, David K. Jackson, Ian Johnston, Dominic Kwiatkowski, Cordelia Langford, John Sillitoe on behalf of the Wellcome Sanger Institute COVID-19 Surveillance Team |
| EPI_ISL_3381885 | Lighthouse Lab in Alderley Park | Wellcome Sanger Institute for the COVID-19 Genomics UK (COG-UK) Consortium | Jacquelyn Wynn, Mairead Hyland, The Lighthouse Lab in Alderley Park and Alex Alderton, Roberto Amato, Jeffrey Barrett, Sonia Goncalves, Ewan Harrison, David K. Jackson, Ian Johnston, Dominic Kwiatkowski, Cordelia Langford, John Sillitoe on behalf of the Wellcome Sanger Institute COVID-19 Surveillance Team |
| EPI_ISL_3381886 | Lighthouse Lab in Glasgow | Wellcome Sanger Institute for the COVID-19 Genomics UK (COG-UK) Consortium | Harper VanSteenhouse, Yumi Kasai, David Gray, Carol Clugston, Anna Dominiczak and Alex Alderton, Roberto Amato, Jeffrey Barrett, Sonia Goncalves, Ewan Harrison, David K. Jackson, Ian Johnston, Dominic Kwiatkowski, Cordelia Langford, John Sillitoe on behalf of the Wellcome Sanger Institute COVID-19 Surveillance Team |
| EPI_ISL_3381887 | Lighthouse Lab in Alderley Park | Wellcome Sanger Institute for the COVID-19 Genomics UK (COG-UK) Consortium | Jacquelyn Wynn, Mairead Hyland, The Lighthouse Lab in Alderley Park and Alex Alderton, Roberto Amato, Jeffrey Barrett, Sonia Goncalves, Ewan Harrison, David K. Jackson, Ian Johnston, Dominic Kwiatkowski, Cordelia Langford, John Sillitoe on behalf of the Wellcome Sanger Institute COVID-19 Surveillance Team |
| EPI_ISL_3381888 | Berkshire and Surrey Pathology Services Lighthouse Laboratory | Wellcome Sanger Institute for the COVID-19 Genomics UK (COG-UK) Consortium | Berkshire and Surrey Pathology Services Lighthouse Laboratory and Alex Alderton, Roberto Amato, Jeffrey Barrett, Sonia Goncalves, Ewan Harrison, David K. Jackson, Ian Johnston, Dominic Kwiatkowski, Cordelia Langford, John Sillitoe on behalf of the Wellcome Sanger Institute COVID-19 Surveillance Team |
| EPI_ISL_3381889 | Lighthouse Lab in Alderley Park | Wellcome Sanger Institute for the COVID-19 Genomics UK (COG-UK) Consortium | Jacquelyn Wynn, Mairead Hyland, The Lighthouse Lab in Alderley Park and Alex Alderton, Roberto Amato, Jeffrey Barrett, Sonia Goncalves, Ewan Harrison, David K. Jackson, Ian Johnston, Dominic Kwiatkowski, Cordelia Langford, John Sillitoe on behalf of the Wellcome Sanger Institute COVID-19 Surveillance Team |
| EPI_ISL_3381890 | Berkshire and Surrey Pathology Services Lighthouse Laboratory | Wellcome Sanger Institute for the COVID-19 Genomics UK (COG-UK) Consortium | Berkshire and Surrey Pathology Services Lighthouse Laboratory and Alex Alderton, Roberto Amato, Jeffrey Barrett, Sonia Goncalves, Ewan Harrison, David K. Jackson, Ian Johnston, Dominic Kwiatkowski, Cordelia Langford, John Sillitoe on behalf of the Wellcome Sanger Institute COVID-19 Surveillance Team |
| EPI_ISL_3381891 | Lighthouse Lab in Milton Keynes | Wellcome Sanger Institute for the COVID-19 Genomics UK (COG-UK) Consortium | The Lighthouse Lab in Milton Keynes and Alex Alderton, Roberto Amato, Jeffrey Barrett, Sonia Goncalves, Ewan Harrison, David K. Jackson, Ian Johnston, Dominic Kwiatkowski, Cordelia Langford, John Sillitoe on behalf of the Wellcome Sanger Institute COVID-19 Surveillance Team |
| EPI_ISL_3381892 | Berkshire and Surrey Pathology Services Lighthouse Laboratory | Wellcome Sanger Institute for the COVID-19 Genomics UK (COG-UK) Consortium | Berkshire and Surrey Pathology Services Lighthouse Laboratory and Alex Alderton, Roberto Amato, Jeffrey Barrett, Sonia Goncalves, Ewan Harrison, David K. Jackson, Ian Johnston, Dominic Kwiatkowski, Cordelia Langford, John Sillitoe on behalf of the Wellcome Sanger Institute COVID-19 Surveillance Team |
| EPI_ISL_3381893, EPI_ISL_3381894 | Lighthouse Lab in Alderley Park | Wellcome Sanger Institute for the COVID-19 Genomics UK (COG-UK) Consortium | Jacquelyn Wynn, Mairead Hyland, The Lighthouse Lab in Alderley Park and Alex Alderton, Roberto Amato, Jeffrey Barrett, Sonia Goncalves, Ewan Harrison, David K. Jackson, Ian Johnston, Dominic Kwiatkowski, Cordelia Langford, John Sillitoe on behalf of the Wellcome Sanger Institute COVID-19 Surveillance Team |
| EPI_ISL_3381895 | Berkshire and Surrey Pathology Services Lighthouse Laboratory | Wellcome Sanger Institute for the COVID-19 Genomics UK (COG-UK) Consortium | Berkshire and Surrey Pathology Services Lighthouse Laboratory and Alex Alderton, Roberto Amato, Jeffrey Barrett, Sonia Goncalves, Ewan Harrison, David K. Jackson, Ian Johnston, Dominic Kwiatkowski, Cordelia Langford, John Sillitoe on behalf of the Wellcome Sanger Institute COVID-19 Surveillance Team |
| EPI_ISL_3381896, EPI_ISL_3381897, EPI_ISL_3381898, EPI_ISL_3381899, EPI_ISL_3381900, EPI_ISL_3381901, EPI_ISL_3381902, EPI_ISL_3381903, EPI_ISL_3381904, EPI_ISL_3381905, EPI_ISL_3381906, EPI_ISL_3381907, EPI_ISL_3381908, EPI_ISL_3381909, EPI_ISL_3381910 | | | |
| see above | Lighthouse Lab in Alderley Park | Wellcome Sanger Institute for the COVID-19 Genomics UK (COG-UK) Consortium | Jacquelyn Wynn, Mairead Hyland, The Lighthouse Lab in Alderley Park and Alex Alderton, Roberto Amato, Jeffrey Barrett, Sonia Goncalves, Ewan Harrison, David K. Jackson, Ian Johnston, Dominic Kwiatkowski, Cordelia Langford, John Sillitoe on behalf of the Wellcome Sanger Institute COVID-19 Surveillance Team |
| EPI_ISL_3381911 | Health Services Laboratories | Wellcome Sanger Institute for the COVID-19 Genomics UK (COG-UK) Consortium | Health Services Laboratories and Alex Alderton, Roberto Amato, Jeffrey Barrett, Sonia Goncalves, Ewan Harrison, David K. Jackson, Ian Johnston, Dominic Kwiatkowski, Cordelia Langford, John Sillitoe on behalf of the Wellcome Sanger Institute COVID-19 Surveillance Team |
| EPI_ISL_3381912 | Lighthouse Lab in Glasgow | Wellcome Sanger Institute for the COVID-19 Genomics UK (COG-UK) Consortium | Harper VanSteenhouse, Yumi Kasai, David Gray, Carol Clugston, Anna Dominiczak and Alex Alderton, Roberto Amato, Jeffrey Barrett, Sonia Goncalves, Ewan Harrison, David K. Jackson, Ian Johnston, Dominic Kwiatkowski, Cordelia Langford, John Sillitoe on behalf of the Wellcome Sanger Institute COVID-19 Surveillance Team |
| EPI_ISL_3381913, EPI_ISL_3381914 | Lighthouse Lab in Milton Keynes | Wellcome Sanger Institute for the COVID-19 Genomics UK (COG-UK) Consortium | The Lighthouse Lab in Milton Keynes and Alex Alderton, Roberto Amato, Jeffrey Barrett, Sonia Goncalves, Ewan Harrison, David K. Jackson, Ian Johnston, Dominic Kwiatkowski, Cordelia Langford, John Sillitoe on behalf of the Wellcome Sanger Institute COVID-19 Surveillance Team |
| EPI_ISL_3381915, EPI_ISL_3381916, EPI_ISL_3381917 | Health Services Laboratories | Wellcome Sanger Institute for the COVID-19 Genomics UK (COG-UK) Consortium | Health Services Laboratories and Alex Alderton, Roberto Amato, Jeffrey Barrett, Sonia Goncalves, Ewan Harrison, David K. Jackson, Ian Johnston, Dominic Kwiatkowski, Cordelia Langford, John Sillitoe on behalf of the Wellcome Sanger Institute COVID-19 Surveillance Team |
| EPI_ISL_3381918, EPI_ISL_3381919, EPI_ISL_3381920, EPI_ISL_3381921, EPI_ISL_3381922 | Lighthouse Lab in Glasgow | Wellcome Sanger Institute for the COVID-19 Genomics UK (COG-UK) Consortium | Harper VanSteenhouse, Yumi Kasai, David Gray, Carol Clugston, Anna Dominiczak and Alex Alderton, Roberto Amato, Jeffrey Barrett, Sonia Goncalves, Ewan Harrison, David K. Jackson, Ian Johnston, Dominic Kwiatkowski, Cordelia Langford, John Sillitoe on behalf of the Wellcome Sanger Institute COVID-19 Surveillance Team |

| | | | |
|---|---|---|---|
| EPI_ISL_3381923 | Health Services Laboratories | Wellcome Sanger Institute for the COVID-19 Genomics UK (COG-UK) Consortium | Health Services Laboratories and Alex Alderton, Roberto Amato, Jeffrey Barrett, Sonia Goncalves, Ewan Harrison, David K. Jackson, Ian Johnston, Dominic Kwiatkowski, Cordelia Langford, John Sillitoe on behalf of the Wellcome Sanger Institute COVID-19 Surveillance Team |
| EPI_ISL_3381924, EPI_ISL_3381925 | Lighthouse Lab in Glasgow | Wellcome Sanger Institute for the COVID-19 Genomics UK (COG-UK) Consortium | Harper VanSteenhouse, Yumi Kasai, David Gray, Carol Clugston, Anna Dominiczak and Alex Alderton, Roberto Amato, Jeffrey Barrett, Sonia Goncalves, Ewan Harrison, David K. Jackson, Ian Johnston, Dominic Kwiatkowski, Cordelia Langford, John Sillitoe on behalf of the Wellcome Sanger Institute COVID-19 Surveillance Team |
| EPI_ISL_3381926 | Lighthouse Lab in Alderley Park | Wellcome Sanger Institute for the COVID-19 Genomics UK (COG-UK) Consortium | Jacquelyn Wynn, Mairead Hyland, The Lighthouse Lab in Alderley Park and Alex Alderton, Roberto Amato, Jeffrey Barrett, Sonia Goncalves, Ewan Harrison, David K. Jackson, Ian Johnston, Dominic Kwiatkowski, Cordelia Langford, John Sillitoe on behalf of the Wellcome Sanger Institute COVID-19 Surveillance Team |
| EPI_ISL_3381927, EPI_ISL_3381928 | Health Services Laboratories | Wellcome Sanger Institute for the COVID-19 Genomics UK (COG-UK) Consortium | Health Services Laboratories and Alex Alderton, Roberto Amato, Jeffrey Barrett, Sonia Goncalves, Ewan Harrison, David K. Jackson, Ian Johnston, Dominic Kwiatkowski, Cordelia Langford, John Sillitoe on behalf of the Wellcome Sanger Institute COVID-19 Surveillance Team |
| EPI_ISL_3381929 | Lighthouse Lab in Glasgow | Wellcome Sanger Institute for the COVID-19 Genomics UK (COG-UK) Consortium | Harper VanSteenhouse, Yumi Kasai, David Gray, Carol Clugston, Anna Dominiczak and Alex Alderton, Roberto Amato, Jeffrey Barrett, Sonia Goncalves, Ewan Harrison, David K. Jackson, Ian Johnston, Dominic Kwiatkowski, Cordelia Langford, John Sillitoe on behalf of the Wellcome Sanger Institute COVID-19 Surveillance Team |
| EPI_ISL_3381930 | Health Services Laboratories | Wellcome Sanger Institute for the COVID-19 Genomics UK (COG-UK) Consortium | Health Services Laboratories and Alex Alderton, Roberto Amato, Jeffrey Barrett, Sonia Goncalves, Ewan Harrison, David K. Jackson, Ian Johnston, Dominic Kwiatkowski, Cordelia Langford, John Sillitoe on behalf of the Wellcome Sanger Institute COVID-19 Surveillance Team |
| EPI_ISL_3381931, EPI_ISL_3381932 | Lighthouse Lab in Glasgow | Wellcome Sanger Institute for the COVID-19 Genomics UK (COG-UK) Consortium | Harper VanSteenhouse, Yumi Kasai, David Gray, Carol Clugston, Anna Dominiczak and Alex Alderton, Roberto Amato, Jeffrey Barrett, Sonia Goncalves, Ewan Harrison, David K. Jackson, Ian Johnston, Dominic Kwiatkowski, Cordelia Langford, John Sillitoe on behalf of the Wellcome Sanger Institute COVID-19 Surveillance Team |
| EPI_ISL_3381933, EPI_ISL_3381934, EPI_ISL_3381935, EPI_ISL_3381936, EPI_ISL_3381937 | Health Services Laboratories | Wellcome Sanger Institute for the COVID-19 Genomics UK (COG-UK) Consortium | Health Services Laboratories and Alex Alderton, Roberto Amato, Jeffrey Barrett, Sonia Goncalves, Ewan Harrison, David K. Jackson, Ian Johnston, Dominic Kwiatkowski, Cordelia Langford, John Sillitoe on behalf of the Wellcome Sanger Institute COVID-19 Surveillance Team |
| EPI_ISL_3381938, EPI_ISL_3381939, EPI_ISL_3381940, EPI_ISL_3381941, EPI_ISL_3381942, EPI_ISL_3381943 | Lighthouse Lab in Glasgow | Wellcome Sanger Institute for the COVID-19 Genomics UK (COG-UK) Consortium | Harper VanSteenhouse, Yumi Kasai, David Gray, Carol Clugston, Anna Dominiczak and Alex Alderton, Roberto Amato, Jeffrey Barrett, Sonia Goncalves, Ewan Harrison, David K. Jackson, Ian Johnston, Dominic Kwiatkowski, Cordelia Langford, John Sillitoe on behalf of the Wellcome Sanger Institute COVID-19 Surveillance Team |
| EPI_ISL_3381944 | Lighthouse Lab in Milton Keynes | Wellcome Sanger Institute for the COVID-19 Genomics UK (COG-UK) Consortium | The Lighthouse Lab in Milton Keynes and Alex Alderton, Roberto Amato, Jeffrey Barrett, Sonia Goncalves, Ewan Harrison, David K. Jackson, Ian Johnston, Dominic Kwiatkowski, Cordelia Langford, John Sillitoe on behalf of the Wellcome Sanger Institute COVID-19 Surveillance Team |
| EPI_ISL_3381945, EPI_ISL_3381946, EPI_ISL_3381947, EPI_ISL_3381948, EPI_ISL_3381949 | Lighthouse Lab in Glasgow | Wellcome Sanger Institute for the COVID-19 Genomics UK (COG-UK) Consortium | Harper VanSteenhouse, Yumi Kasai, David Gray, Carol Clugston, Anna Dominiczak and Alex Alderton, Roberto Amato, Jeffrey Barrett, Sonia Goncalves, Ewan Harrison, David K. Jackson, Ian Johnston, Dominic Kwiatkowski, Cordelia Langford, John Sillitoe on behalf of the Wellcome Sanger Institute COVID-19 Surveillance Team |
| EPI_ISL_3381950, EPI_ISL_3381951, EPI_ISL_3381952 | Health Services Laboratories | Wellcome Sanger Institute for the COVID-19 Genomics UK (COG-UK) Consortium | Health Services Laboratories and Alex Alderton, Roberto Amato, Jeffrey Barrett, Sonia Goncalves, Ewan Harrison, David K. Jackson, Ian Johnston, Dominic Kwiatkowski, Cordelia Langford, John Sillitoe on behalf of the Wellcome Sanger Institute COVID-19 Surveillance Team |
| EPI_ISL_3381953 | Lighthouse Lab in Glasgow | Wellcome Sanger Institute for the COVID-19 Genomics UK (COG-UK) Consortium | Harper VanSteenhouse, Yumi Kasai, David Gray, Carol Clugston, Anna Dominiczak and Alex Alderton, Roberto Amato, Jeffrey Barrett, Sonia Goncalves, Ewan Harrison, David K. Jackson, Ian Johnston, Dominic Kwiatkowski, Cordelia Langford, John Sillitoe on behalf of the Wellcome Sanger Institute COVID-19 Surveillance Team |
| EPI_ISL_3381954, EPI_ISL_3381955 | Health Services Laboratories | Wellcome Sanger Institute for the COVID-19 Genomics UK (COG-UK) Consortium | Health Services Laboratories and Alex Alderton, Roberto Amato, Jeffrey Barrett, Sonia Goncalves, Ewan Harrison, David K. Jackson, Ian Johnston, Dominic Kwiatkowski, Cordelia Langford, John Sillitoe on behalf of the Wellcome Sanger Institute COVID-19 Surveillance Team |
| EPI_ISL_3381956 | Lighthouse Lab in Glasgow | Wellcome Sanger Institute for the COVID-19 Genomics UK (COG-UK) Consortium | Harper VanSteenhouse, Yumi Kasai, David Gray, Carol Clugston, Anna Dominiczak and Alex Alderton, Roberto Amato, Jeffrey Barrett, Sonia Goncalves, Ewan Harrison, David K. Jackson, Ian Johnston, Dominic Kwiatkowski, Cordelia Langford, John Sillitoe on behalf of the Wellcome Sanger Institute COVID-19 Surveillance Team |
| EPI_ISL_3381957 | Health Services Laboratories | Wellcome Sanger Institute for the COVID-19 Genomics UK (COG-UK) Consortium | Health Services Laboratories and Alex Alderton, Roberto Amato, Jeffrey Barrett, Sonia Goncalves, Ewan Harrison, David K. Jackson, Ian Johnston, Dominic Kwiatkowski, Cordelia Langford, John Sillitoe on behalf of the Wellcome Sanger Institute COVID-19 Surveillance Team |
| EPI_ISL_3381958 | Lighthouse Lab in Milton Keynes | Wellcome Sanger Institute for the COVID-19 Genomics UK (COG-UK) Consortium | The Lighthouse Lab in Milton Keynes and Alex Alderton, Roberto Amato, Jeffrey Barrett, Sonia Goncalves, Ewan Harrison, David K. Jackson, Ian Johnston, Dominic Kwiatkowski, Cordelia Langford, John Sillitoe on behalf of the Wellcome Sanger Institute COVID-19 Surveillance Team |
| EPI_ISL_3381959, EPI_ISL_3381960 | Lighthouse Lab in Alderley Park | Wellcome Sanger Institute for the COVID-19 Genomics UK (COG-UK) Consortium | Jacquelyn Wynn, Mairead Hyland, The Lighthouse Lab in Alderley Park and Alex Alderton, Roberto Amato, Jeffrey Barrett, Sonia Goncalves, Ewan Harrison, David K. Jackson, Ian Johnston, Dominic Kwiatkowski, Cordelia Langford, John Sillitoe on behalf of the Wellcome Sanger Institute COVID-19 Surveillance Team |
| EPI_ISL_3381961 | Lighthouse Laboratory Plymouth | Wellcome Sanger Institute for the COVID-19 Genomics UK (COG-UK) Consortium | Lighthouse Laboratory Plymouth and Alex Alderton, Roberto Amato, Jeffrey Barrett, Sonia Goncalves, Ewan Harrison, David K. Jackson, Ian Johnston, Dominic Kwiatkowski, Cordelia Langford, John Sillitoe on behalf of the Wellcome Sanger Institute COVID-19 Surveillance Team |
| EPI_ISL_3381962, EPI_ISL_3381963 | Lighthouse Lab in Milton Keynes | Wellcome Sanger Institute for the COVID-19 Genomics UK (COG-UK) Consortium | The Lighthouse Lab in Milton Keynes and Alex Alderton, Roberto Amato, Jeffrey Barrett, Sonia Goncalves, Ewan Harrison, David K. Jackson, Ian Johnston, Dominic Kwiatkowski, Cordelia Langford, John Sillitoe on behalf of the Wellcome Sanger Institute COVID-19 Surveillance Team |
| EPI_ISL_3381964 | Lighthouse Lab in Alderley Park | Wellcome Sanger Institute for the COVID-19 Genomics UK (COG-UK) Consortium | Jacquelyn Wynn, Mairead Hyland, The Lighthouse Lab in Alderley Park and Alex Alderton, Roberto Amato, Jeffrey Barrett, Sonia Goncalves, Ewan Harrison, David K. Jackson, Ian Johnston, Dominic Kwiatkowski, Cordelia Langford, John Sillitoe on behalf of the Wellcome Sanger Institute COVID-19 Surveillance Team |
| EPI_ISL_3381965, EPI_ISL_3381966, EPI_ISL_3381967 | Lighthouse Lab in Milton Keynes | Wellcome Sanger Institute for the COVID-19 Genomics UK (COG-UK) Consortium | The Lighthouse Lab in Milton Keynes and Alex Alderton, Roberto Amato, Jeffrey Barrett, Sonia Goncalves, Ewan Harrison, David K. Jackson, Ian Johnston, Dominic Kwiatkowski, Cordelia Langford, John Sillitoe on behalf of the Wellcome Sanger Institute COVID-19 Surveillance Team |
| EPI_ISL_3381968, EPI_ISL_3381969, EPI_ISL_3381970 | Lighthouse Lab in Alderley Park | Wellcome Sanger Institute for the COVID-19 Genomics UK (COG-UK) Consortium | Jacquelyn Wynn, Mairead Hyland, The Lighthouse Lab in Alderley Park and Alex Alderton, Roberto Amato, Jeffrey Barrett, Sonia Goncalves, Ewan Harrison, David K. Jackson, Ian Johnston, Dominic Kwiatkowski, Cordelia Langford, John Sillitoe on behalf of the Wellcome Sanger Institute COVID-19 Surveillance Team |
| EPI_ISL_3381971, EPI_ISL_3381972, EPI_ISL_3381973, EPI_ISL_3381974, EPI_ISL_3381975, EPI_ISL_3381976, EPI_ISL_3381977, EPI_ISL_3381978, EPI_ISL_3381979, EPI_ISL_3381980, EPI_ISL_3381981, EPI_ISL_3381982, EPI_ISL_3381983, EPI_ISL_3381984, EPI_ISL_3381985, EPI_ISL_3381986, EPI_ISL_3381987, EPI_ISL_3381988, EPI_ISL_3381989, EPI_ISL_3381990, EPI_ISL_3381991, EPI_ISL_3381992, EPI_ISL_3381993, EPI_ISL_3381994, EPI_ISL_3381995, EPI_ISL_3381996, EPI_ISL_3381997, EPI_ISL_3381998, EPI_ISL_3381999, EPI_ISL_3382000, EPI_ISL_3382001, EPI_ISL_3382002, EPI_ISL_3382003, EPI_ISL_3382004, EPI_ISL_3382005, EPI_ISL_3382006, EPI_ISL_3382007, EPI_ISL_3382008, EPI_ISL_3382009, EPI_ISL_3382010, EPI_ISL_3382011, EPI_ISL_3382012, EPI_ISL_3382013, EPI_ISL_3382014, EPI_ISL_3382015, EPI_ISL_3382016, EPI_ISL_3382017, EPI_ISL_3382018, EPI_ISL_3382019, EPI_ISL_3382020, EPI_ISL_3382021, EPI_ISL_3382022, EPI_ISL_3382023, EPI_ISL_3382024, EPI_ISL_3382025, EPI_ISL_3382026, EPI_ISL_3382027, EPI_ISL_3382028, EPI_ISL_3382029, EPI_ISL_3382030, EPI_ISL_3382031, EPI_ISL_3382032, EPI_ISL_3382033, EPI_ISL_3382034, EPI_ISL_3382035, EPI_ISL_3382036, EPI_ISL_3382037, EPI_ISL_3382038, EPI_ISL_3382039, EPI_ISL_3382040, EPI_ISL_3382041, EPI_ISL_3382042, EPI_ISL_3382043, EPI_ISL_3382044, EPI_ISL_3382045, EPI_ISL_3382046, EPI_ISL_3382047, EPI_ISL_3382048, EPI_ISL_3382049, EPI_ISL_3382050, EPI_ISL_3382051, EPI_ISL_3382052, EPI_ISL_3382053, EPI_ISL_3382054, EPI_ISL_3382055, EPI_ISL_3382056, EPI_ISL_3382057, EPI_ISL_3382058, EPI_ISL_3382059, EPI_ISL_3382060, EPI_ISL_3382061, EPI_ISL_3382062, EPI_ISL_3382063, EPI_ISL_3382064, EPI_ISL_3382065, EPI_ISL_3382066 | | | |
| see above | Lighthouse Lab in Glasgow | Wellcome Sanger Institute for the COVID-19 Genomics UK (COG-UK) Consortium | Harper VanSteenhouse, Yumi Kasai, David Gray, Carol Clugston, Anna Dominiczak and Alex Alderton, Roberto Amato, Jeffrey Barrett, Sonia Goncalves, Ewan Harrison, David K. Jackson, Ian Johnston, Dominic Kwiatkowski, Cordelia Langford, John Sillitoe on behalf of the Wellcome Sanger Institute COVID-19 Surveillance Team |
| EPI_ISL_3382067 | Lighthouse Lab in Milton Keynes | Wellcome Sanger Institute for the COVID-19 Genomics UK (COG-UK) Consortium | The Lighthouse Lab in Milton Keynes and Alex Alderton, Roberto Amato, Jeffrey Barrett, Sonia Goncalves, Ewan Harrison, David K. Jackson, Ian Johnston, Dominic Kwiatkowski, Cordelia Langford, John Sillitoe on behalf of the Wellcome Sanger Institute COVID-19 Surveillance Team |
| EPI_ISL_3382068, EPI_ISL_3382069, EPI_ISL_3382070, EPI_ISL_3382071, EPI_ISL_3382072, EPI_ISL_3382073, EPI_ISL_3382074, EPI_ISL_3382075, EPI_ISL_3382076, EPI_ISL_3382077, EPI_ISL_3382078, EPI_ISL_3382079, EPI_ISL_3382080, EPI_ISL_3382081, EPI_ISL_3382082, EPI_ISL_3382083, EPI_ISL_3382084, EPI_ISL_3382085, EPI_ISL_3382086, EPI_ISL_3382087, EPI_ISL_3382088, EPI_ISL_3382089, EPI_ISL_3382090, EPI_ISL_3382091, EPI_ISL_3382092, EPI_ISL_3382093, EPI_ISL_3382094, EPI_ISL_3382095, EPI_ISL_3382096, EPI_ISL_3382097, EPI_ISL_3382098, EPI_ISL_3382099, EPI_ISL_3382100, EPI_ISL_3382101, EPI_ISL_3382102 | | | |
| see above | Lighthouse Lab in Glasgow | Wellcome Sanger Institute for the COVID-19 Genomics UK (COG-UK) Consortium | Harper VanSteenhouse, Yumi Kasai, David Gray, Carol Clugston, Anna Dominiczak and Alex Alderton, Roberto Amato, Jeffrey Barrett, Sonia Goncalves, Ewan Harrison, David K. Jackson, Ian Johnston, Dominic Kwiatkowski, Cordelia Langford, John Sillitoe on behalf of the Wellcome Sanger Institute COVID-19 Surveillance Team |
| EPI_ISL_3382103 | Lighthouse Lab in Alderley Park | Wellcome Sanger Institute for the COVID-19 Genomics UK (COG-UK) Consortium | Jacquelyn Wynn, Mairead Hyland, The Lighthouse Lab in Alderley Park and Alex Alderton, Roberto Amato, Jeffrey Barrett, Sonia Goncalves, Ewan Harrison, David K. Jackson, Ian Johnston, Dominic Kwiatkowski, Cordelia Langford, John Sillitoe on behalf of the Wellcome Sanger Institute COVID-19 Surveillance Team |
| EPI_ISL_3382104, EPI_ISL_3382105, EPI_ISL_3382106, EPI_ISL_3382107, EPI_ISL_3382108, EPI_ISL_3382109, EPI_ISL_3382110, EPI_ISL_3382111, EPI_ISL_3382112, EPI_ISL_3382113, EPI_ISL_3382114, EPI_ISL_3382115, EPI_ISL_3382116, EPI_ISL_3382117, EPI_ISL_3382118, EPI_ISL_3382119, EPI_ISL_3382120, EPI_ISL_3382121, EPI_ISL_3382122, EPI_ISL_3382123, EPI_ISL_3382124, EPI_ISL_3382125 | | | |
| see above | Lighthouse Lab in Glasgow | Wellcome Sanger Institute for the COVID-19 Genomics UK (COG-UK) Consortium | Harper VanSteenhouse, Yumi Kasai, David Gray, Carol Clugston, Anna Dominiczak and Alex Alderton, Roberto Amato, Jeffrey Barrett, Sonia Goncalves, Ewan Harrison, David K. Jackson, Ian Johnston, Dominic Kwiatkowski, Cordelia Langford, John Sillitoe on behalf of the Wellcome Sanger Institute COVID-19 Surveillance Team |
| EPI_ISL_3382126 | Lighthouse Lab in Alderley Park | Wellcome Sanger Institute for the COVID-19 Genomics UK (COG-UK) Consortium | Jacquelyn Wynn, Mairead Hyland, The Lighthouse Lab in Alderley Park and Alex Alderton, Roberto Amato, Jeffrey Barrett, Sonia Goncalves, Ewan Harrison, David K. Jackson, Ian Johnston, Dominic Kwiatkowski, Cordelia Langford, John Sillitoe on behalf of the Wellcome Sanger Institute COVID-19 Surveillance Team |
| EPI_ISL_3382127, EPI_ISL_3382128, EPI_ISL_3382129, EPI_ISL_3382130, EPI_ISL_3382131, EPI_ISL_3382132, EPI_ISL_3382133, EPI_ISL_3382134, EPI_ISL_3382135, EPI_ISL_3382136, EPI_ISL_3382137, EPI_ISL_3382138, EPI_ISL_3382139 | | | |
| see above | Lighthouse Lab in Glasgow | Wellcome Sanger Institute for the COVID-19 Genomics UK (COG-UK) Consortium | Harper VanSteenhouse, Yumi Kasai, David Gray, Carol Clugston, Anna Dominiczak and Alex Alderton, Roberto Amato, Jeffrey Barrett, Sonia Goncalves, Ewan Harrison, David K. Jackson, Ian Johnston, Dominic Kwiatkowski, Cordelia Langford, John Sillitoe on behalf of the Wellcome Sanger Institute COVID-19 Surveillance Team |
| EPI_ISL_3382140 | Lighthouse Lab in Alderley Park | Wellcome Sanger Institute for the COVID-19 Genomics UK (COG-UK) Consortium | Jacquelyn Wynn, Mairead Hyland, The Lighthouse Lab in Alderley Park and Alex Alderton, Roberto Amato, Jeffrey Barrett, Sonia Goncalves, Ewan Harrison, David K. Jackson, Ian Johnston, Dominic Kwiatkowski, Cordelia Langford, John Sillitoe on behalf of the Wellcome Sanger Institute COVID-19 Surveillance Team |
| EPI_ISL_3382141, EPI_ISL_3382142, EPI_ISL_3382143, EPI_ISL_3382144, EPI_ISL_3382145, EPI_ISL_3382146, EPI_ISL_3382147, EPI_ISL_3382148, EPI_ISL_3382149, EPI_ISL_3382150, EPI_ISL_3382151, EPI_ISL_3382152, EPI_ISL_3382153, EPI_ISL_3382154, EPI_ISL_3382155, EPI_ISL_3382156, EPI_ISL_3382157, EPI_ISL_3382158, EPI_ISL_3382159, EPI_ISL_3382160, EPI_ISL_3382161, EPI_ISL_3382162, EPI_ISL_3382163, EPI_ISL_3382164, EPI_ISL_3382165, EPI_ISL_3382166, EPI_ISL_3382167, EPI_ISL_3382168, EPI_ISL_3382169, EPI_ISL_3382170, EPI_ISL_3382171 | | | |
| see above | Randox Laboratories | Wellcome Sanger Institute for the COVID-19 Genomics UK (COG-UK) Consortium | Randox Laboratories and Alex Alderton, Roberto Amato, Jeffrey Barrett, Sonia Goncalves, Ewan Harrison, David K. Jackson, Ian Johnston, Dominic Kwiatkowski, Cordelia Langford, John Sillitoe on behalf of the Wellcome Sanger Institute COVID-19 Surveillance Team |
| EPI_ISL_3382172 | Lighthouse Lab in Alderley Park | Wellcome Sanger Institute for the COVID-19 Genomics UK (COG-UK) Consortium | Jacquelyn Wynn, Mairead Hyland, The Lighthouse Lab in Alderley Park and Alex Alderton, Roberto Amato, Jeffrey Barrett, Sonia Goncalves, Ewan Harrison, David K. Jackson, Ian Johnston, Dominic Kwiatkowski, Cordelia Langford, John Sillitoe on behalf of the Wellcome Sanger Institute COVID-19 Surveillance Team |
| EPI_ISL_3382173, EPI_ISL_3382174, EPI_ISL_3382175, EPI_ISL_3382176, EPI_ISL_3382177, EPI_ISL_3382178, EPI_ISL_3382179, EPI_ISL_3382180, EPI_ISL_3382181, EPI_ISL_3382182, EPI_ISL_3382183, EPI_ISL_3382184, EPI_ISL_3382185, EPI_ISL_3382186, EPI_ISL_3382187, EPI_ISL_3382188, EPI_ISL_3382189, EPI_ISL_3382190, EPI_ISL_3382191, EPI_ISL_3382192, EPI_ISL_3382193, EPI_ISL_3382194, EPI_ISL_3382195, EPI_ISL_3382196, EPI_ISL_3382197, EPI_ISL_3382198, EPI_ISL_3382199, EPI_ISL_3382200, EPI_ISL_3382201, EPI_ISL_3382202, EPI_ISL_3382203, EPI_ISL_3382204, EPI_ISL_3382205, EPI_ISL_3382206, EPI_ISL_3382207, EPI_ISL_3382208, EPI_ISL_3382209, EPI_ISL_3382210, EPI_ISL_3382211, EPI_ISL_3382212, EPI_ISL_3382213, EPI_ISL_3382214, EPI_ISL_3382215, EPI_ISL_3382216, EPI_ISL_3382217, EPI_ISL_3382218, EPI_ISL_3382219, EPI_ISL_3382220, EPI_ISL_3382221, EPI_ISL_3382222, EPI_ISL_3382223, EPI_ISL_3382224, EPI_ISL_3382225, EPI_ISL_3382226, EPI_ISL_3382227, EPI_ISL_3382228, EPI_ISL_3382229, EPI_ISL_3382230, EPI_ISL_3382231, EPI_ISL_3382232, EPI_ISL_3382233, EPI_ISL_3382234, EPI_ISL_3382235, EPI_ISL_3382236, EPI_ISL_3382237, EPI_ISL_3382238, | | | |

| | | | |
|---|---|---|---|
| EPI_ISL_3382239, EPI_ISL_3382240, EPI_ISL_3382241, EPI_ISL_3382242, EPI_ISL_3382243, EPI_ISL_3382244, EPI_ISL_3382245, EPI_ISL_3382246, EPI_ISL_3382247, EPI_ISL_3382248, EPI_ISL_3382249, EPI_ISL_3382250, EPI_ISL_3382251, EPI_ISL_3382252, EPI_ISL_3382253, EPI_ISL_3382254, EPI_ISL_3382255, EPI_ISL_3382256, EPI_ISL_3382257, EPI_ISL_3382258 | | | |
| see above | Randox Laboratories | Wellcome Sanger Institute for the COVID-19 Genomics UK (COG-UK) Consortium | Randox Laboratories and Alex Alderton, Roberto Amato, Jeffrey Barrett, Sonia Goncalves, Ewan Harrison, David K. Jackson, Ian Johnston, Dominic Kwiatkowski, Cordelia Langford, John Sillitoe on behalf of the Wellcome Sanger Institute COVID-19 Surveillance Team |
| EPI_ISL_3382259, EPI_ISL_3382260 | Berkshire and Surrey Pathology Services Lighthouse Laboratory | Wellcome Sanger Institute for the COVID-19 Genomics UK (COG-UK) Consortium | Berkshire and Surrey Pathology Services Lighthouse Laboratory and Alex Alderton, Roberto Amato, Jeffrey Barrett, Sonia Goncalves, Ewan Harrison, David K. Jackson, Ian Johnston, Dominic Kwiatkowski, Cordelia Langford, John Sillitoe on behalf of the Wellcome Sanger Institute COVID-19 Surveillance Team |
| EPI_ISL_3382261, EPI_ISL_3382262, EPI_ISL_3382263, EPI_ISL_3382264, EPI_ISL_3382265 | Lighthouse Lab in Glasgow | Wellcome Sanger Institute for the COVID-19 Genomics UK (COG-UK) Consortium | Harper VanSteenhouse, Yumi Kasai, David Gray, Carol Clugston, Anna Dominiczak and Alex Alderton, Roberto Amato, Jeffrey Barrett, Sonia Goncalves, Ewan Harrison, David K. Jackson, Ian Johnston, Dominic Kwiatkowski, Cordelia Langford, John Sillitoe on behalf of the Wellcome Sanger Institute COVID-19 Surveillance Team |
| EPI_ISL_3382266 | Berkshire and Surrey Pathology Services Lighthouse Laboratory | Wellcome Sanger Institute for the COVID-19 Genomics UK (COG-UK) Consortium | Berkshire and Surrey Pathology Services Lighthouse Laboratory and Alex Alderton, Roberto Amato, Jeffrey Barrett, Sonia Goncalves, Ewan Harrison, David K. Jackson, Ian Johnston, Dominic Kwiatkowski, Cordelia Langford, John Sillitoe on behalf of the Wellcome Sanger Institute COVID-19 Surveillance Team |
| EPI_ISL_3382267 | Lighthouse Lab in Glasgow | Wellcome Sanger Institute for the COVID-19 Genomics UK (COG-UK) Consortium | Harper VanSteenhouse, Yumi Kasai, David Gray, Carol Clugston, Anna Dominiczak and Alex Alderton, Roberto Amato, Jeffrey Barrett, Sonia Goncalves, Ewan Harrison, David K. Jackson, Ian Johnston, Dominic Kwiatkowski, Cordelia Langford, John Sillitoe on behalf of the Wellcome Sanger Institute COVID-19 Surveillance Team |
| EPI_ISL_3382268 | Berkshire and Surrey Pathology Services Lighthouse Laboratory | Wellcome Sanger Institute for the COVID-19 Genomics UK (COG-UK) Consortium | Berkshire and Surrey Pathology Services Lighthouse Laboratory and Alex Alderton, Roberto Amato, Jeffrey Barrett, Sonia Goncalves, Ewan Harrison, David K. Jackson, Ian Johnston, Dominic Kwiatkowski, Cordelia Langford, John Sillitoe on behalf of the Wellcome Sanger Institute COVID-19 Surveillance Team |
| EPI_ISL_3382269, EPI_ISL_3382270, EPI_ISL_3382271, EPI_ISL_3382272, EPI_ISL_3382273 | Lighthouse Lab in Glasgow | Wellcome Sanger Institute for the COVID-19 Genomics UK (COG-UK) Consortium | Harper VanSteenhouse, Yumi Kasai, David Gray, Carol Clugston, Anna Dominiczak and Alex Alderton, Roberto Amato, Jeffrey Barrett, Sonia Goncalves, Ewan Harrison, David K. Jackson, Ian Johnston, Dominic Kwiatkowski, Cordelia Langford, John Sillitoe on behalf of the Wellcome Sanger Institute COVID-19 Surveillance Team |
| EPI_ISL_3382274 | Berkshire and Surrey Pathology Services Lighthouse Laboratory | Wellcome Sanger Institute for the COVID-19 Genomics UK (COG-UK) Consortium | Berkshire and Surrey Pathology Services Lighthouse Laboratory and Alex Alderton, Roberto Amato, Jeffrey Barrett, Sonia Goncalves, Ewan Harrison, David K. Jackson, Ian Johnston, Dominic Kwiatkowski, Cordelia Langford, John Sillitoe on behalf of the Wellcome Sanger Institute COVID-19 Surveillance Team |
| EPI_ISL_3382275, EPI_ISL_3382276, EPI_ISL_3382277, EPI_ISL_3382278, EPI_ISL_3382279, EPI_ISL_3382280, EPI_ISL_3382281, EPI_ISL_3382282, EPI_ISL_3382283, EPI_ISL_3382284, EPI_ISL_3382285, EPI_ISL_3382286, EPI_ISL_3382287, EPI_ISL_3382288, EPI_ISL_3382289, EPI_ISL_3382290, EPI_ISL_3382291, EPI_ISL_3382292, EPI_ISL_3382293, EPI_ISL_3382294, EPI_ISL_3382295, EPI_ISL_3382296, EPI_ISL_3382297, EPI_ISL_3382298, EPI_ISL_3382299, EPI_ISL_3382300, EPI_ISL_3382301, EPI_ISL_3382302, EPI_ISL_3382303, EPI_ISL_3382304, EPI_ISL_3382305, EPI_ISL_3382306, EPI_ISL_3382307, EPI_ISL_3382308, EPI_ISL_3382309, EPI_ISL_3382310, EPI_ISL_3382311, EPI_ISL_3382312, EPI_ISL_3382313, EPI_ISL_3382314, EPI_ISL_3382315, EPI_ISL_3382316, EPI_ISL_3382317, EPI_ISL_3382318, EPI_ISL_3382319, EPI_ISL_3382320, EPI_ISL_3382321, EPI_ISL_3382322, EPI_ISL_3382323, EPI_ISL_3382324, EPI_ISL_3382325, EPI_ISL_3382326, EPI_ISL_3382327, EPI_ISL_3382328, EPI_ISL_3382329, EPI_ISL_3382330, EPI_ISL_3382331, EPI_ISL_3382332, EPI_ISL_3382333, EPI_ISL_3382334 | | | |
| see above | Lighthouse Lab in Glasgow | Wellcome Sanger Institute for the COVID-19 Genomics UK (COG-UK) Consortium | Harper VanSteenhouse, Yumi Kasai, David Gray, Carol Clugston, Anna Dominiczak and Alex Alderton, Roberto Amato, Jeffrey Barrett, Sonia Goncalves, Ewan Harrison, David K. Jackson, Ian Johnston, Dominic Kwiatkowski, Cordelia Langford, John Sillitoe on behalf of the Wellcome Sanger Institute COVID-19 Surveillance Team |
| EPI_ISL_3382335, EPI_ISL_3382336, EPI_ISL_3382337, EPI_ISL_3382338, EPI_ISL_3382339, EPI_ISL_3382340, EPI_ISL_3382341, EPI_ISL_3382342, EPI_ISL_3382343, EPI_ISL_3382344, EPI_ISL_3382345, EPI_ISL_3382346, EPI_ISL_3382347, EPI_ISL_3382348, EPI_ISL_3382349, EPI_ISL_3382350, EPI_ISL_3382351, EPI_ISL_3382352, EPI_ISL_3382353, EPI_ISL_3382354, EPI_ISL_3382355, EPI_ISL_3382356, EPI_ISL_3382357, EPI_ISL_3382358, EPI_ISL_3382359, EPI_ISL_3382360, EPI_ISL_3382361, EPI_ISL_3382362, EPI_ISL_3382363, EPI_ISL_3382364, EPI_ISL_3382365, EPI_ISL_3382366, EPI_ISL_3382367, EPI_ISL_3382368, EPI_ISL_3382369, EPI_ISL_3382370, EPI_ISL_3382371, EPI_ISL_3382372, EPI_ISL_3382373, EPI_ISL_3382374, EPI_ISL_3382375, EPI_ISL_3382376, EPI_ISL_3382377, EPI_ISL_3382378, EPI_ISL_3382379, EPI_ISL_3382380, EPI_ISL_3382381, EPI_ISL_3382382, EPI_ISL_3382383, EPI_ISL_3382384, EPI_ISL_3382385, EPI_ISL_3382386, EPI_ISL_3382387, EPI_ISL_3382388, EPI_ISL_3382389, EPI_ISL_3382390, EPI_ISL_3382391, EPI_ISL_3382392, EPI_ISL_3382393, EPI_ISL_3382394, EPI_ISL_3382395, EPI_ISL_3382396, EPI_ISL_3382397, EPI_ISL_3382398, EPI_ISL_3382399, EPI_ISL_3382400, EPI_ISL_3382401, EPI_ISL_3382402, EPI_ISL_3382403, EPI_ISL_3382404, EPI_ISL_3382405, EPI_ISL_3382406, EPI_ISL_3382407, EPI_ISL_3382408, EPI_ISL_3382409, EPI_ISL_3382410, EPI_ISL_3382411, EPI_ISL_3382412, EPI_ISL_3382413, EPI_ISL_3382414, EPI_ISL_3382415, EPI_ISL_3382416, EPI_ISL_3382417, EPI_ISL_3382418, EPI_ISL_3382419, EPI_ISL_3382420, EPI_ISL_3382421, EPI_ISL_3382422, EPI_ISL_3382423, EPI_ISL_3382424, EPI_ISL_3382425, EPI_ISL_3382426, EPI_ISL_3382427 | | | |
| see above | Randox Laboratories | Wellcome Sanger Institute for the COVID-19 Genomics UK (COG-UK) Consortium | Randox Laboratories and Alex Alderton, Roberto Amato, Jeffrey Barrett, Sonia Goncalves, Ewan Harrison, David K. Jackson, Ian Johnston, Dominic Kwiatkowski, Cordelia Langford, John Sillitoe on behalf of the Wellcome Sanger Institute COVID-19 Surveillance Team |
| EPI_ISL_3382434, EPI_ISL_3382435 | LABORATOIRE UNILIANS DECINES | CNR Virus des Infections Respiratoires - France SUD | Antonin Bal, Gregory Destras, Gwendolyne Burfin, Hadrien Regue, Quentin Semanas, Martine Valette, Bruno Lina, Laurence Josset |
| EPI_ISL_3382436, EPI_ISL_3382437, EPI_ISL_3382438 | CNR Virus des Infections Respiratoires - France SUD | CNR Virus des Infections Respiratoires - France SUD | Antonin Bal, Gregory Destras, Gwendolyne Burfin, Hadrien Regue, Quentin Semanas, Martine Valette, Bruno Lina, Laurence Josset |
| EPI_ISL_3382439 | LABORATOIRE UNILIANS DECINES | CNR Virus des Infections Respiratoires - France SUD | Antonin Bal, Gregory Destras, Gwendolyne Burfin, Hadrien Regue, Quentin Semanas, Martine Valette, Bruno Lina, Laurence Josset |
| EPI_ISL_3382440 | LABORATOIRE d'ANALYSES de BIOLOGIE MEDICALES PUILBOREAU | CNR Virus des Infections Respiratoires - France SUD | Antonin Bal, Gregory Destras, Gwendolyne Burfin, Hadrien Regue, Quentin Semanas, Martine Valette, Bruno Lina, Laurence Josset |
| EPI_ISL_3382441 | CNR Virus des Infections Respiratoires - France SUD | CNR Virus des Infections Respiratoires - France SUD | Antonin Bal, Gregory Destras, Gwendolyne Burfin, Hadrien Regue, Quentin Semanas, Martine Valette, Bruno Lina, Laurence Josset |
| EPI_ISL_3382442 | LABORATOIRE d'ANALYSES de BIOLOGIE MEDICALES PUILBOREAU | CNR Virus des Infections Respiratoires - France SUD | Antonin Bal, Gregory Destras, Gwendolyne Burfin, Hadrien Regue, Quentin Semanas, Martine Valette, Bruno Lina, Laurence Josset |
| EPI_ISL_3382443, EPI_ISL_3382444 | LABORATOIRE MAYMAT | CNR Virus des Infections Respiratoires - France SUD | Antonin Bal, Gregory Destras, Gwendolyne Burfin, Hadrien Regue, Quentin Semanas, Martine Valette, Bruno Lina, Laurence Josset |
| EPI_ISL_3382445, EPI_ISL_3382446 | CNR Virus des Infections Respiratoires - France SUD | CNR Virus des Infections Respiratoires - France SUD | Antonin Bal, Gregory Destras, Gwendolyne Burfin, Hadrien Regue, Quentin Semanas, Martine Valette, Bruno Lina, Laurence Josset |
| EPI_ISL_3382447 | DYOMEDEA-LABORATOIRE DE LA SAUVEGARDE | CNR Virus des Infections Respiratoires - France SUD | Antonin Bal, Gregory Destras, Gwendolyne Burfin, Hadrien Regue, Quentin Semanas, Martine Valette, Bruno Lina, Laurence Josset |
| EPI_ISL_3382448 | LABORATOIRE d'ANALYSES de BIOLOGIE MEDICALES PUILBOREAU | CNR Virus des Infections Respiratoires - France SUD | Antonin Bal, Gregory Destras, Gwendolyne Burfin, Hadrien Regue, Quentin Semanas, Martine Valette, Bruno Lina, Laurence Josset |
| EPI_ISL_3382449 | CNR Virus des Infections Respiratoires - France SUD | CNR Virus des Infections Respiratoires - France SUD | Antonin Bal, Gregory Destras, Gwendolyne Burfin, Hadrien Regue, Quentin Semanas, Martine Valette, Bruno Lina, Laurence Josset |
| EPI_ISL_3382450, EPI_ISL_3382451 | LABORATOIRE CERBALLIANCE PLT VILLON | CNR Virus des Infections Respiratoires - France SUD | Antonin Bal, Gregory Destras, Gwendolyne Burfin, Hadrien Regue, Quentin Semanas, Martine Valette, Bruno Lina, Laurence Josset |
| EPI_ISL_3382452, EPI_ISL_3382453, EPI_ISL_3382454, EPI_ISL_3382455, EPI_ISL_3382456 | LABORATOIRE d'ANALYSES de BIOLOGIE MEDICALES PUILBOREAU | CNR Virus des Infections Respiratoires - France SUD | Antonin Bal, Gregory Destras, Gwendolyne Burfin, Hadrien Regue, Quentin Semanas, Martine Valette, Bruno Lina, Laurence Josset |
| EPI_ISL_3382459 | REUNILAB | CNR Virus des Infections Respiratoires - France SUD | Antonin Bal, Gregory Destras, Gwendolyne Burfin, Hadrien Regue, Quentin Semanas, Martine Valette, Bruno Lina, Laurence Josset |
| EPI_ISL_3382461, EPI_ISL_3382462, EPI_ISL_3382463, EPI_ISL_3382464, EPI_ISL_3382465, EPI_ISL_3382466, EPI_ISL_3382467, EPI_ISL_3382468, EPI_ISL_3382469, EPI_ISL_3382470, EPI_ISL_3382471, EPI_ISL_3382472, EPI_ISL_3382473, EPI_ISL_3382474 | | | |
| see above | LABORATOIRE d'ANALYSES de BIOLOGIE MEDICALES PUILBOREAU | CNR Virus des Infections Respiratoires - France SUD | Antonin Bal, Gregory Destras, Gwendolyne Burfin, Hadrien Regue, Quentin Semanas, Martine Valette, Bruno Lina, Laurence Josset |
| EPI_ISL_3382476 | LABORATOIRE BOUVIER | CNR Virus des Infections Respiratoires - France SUD | Antonin Bal, Gregory Destras, Gwendolyne Burfin, Hadrien Regue, Quentin Semanas, Martine Valette, Bruno Lina, Laurence Josset |
| EPI_ISL_3382477, EPI_ISL_3382478, EPI_ISL_3382479, EPI_ISL_3382480, EPI_ISL_3382481, EPI_ISL_3382482, EPI_ISL_3382483, EPI_ISL_3382484, EPI_ISL_3382485, EPI_ISL_3382486, EPI_ISL_3382487, EPI_ISL_3382488, EPI_ISL_3382489, EPI_ISL_3382490, EPI_ISL_3382491, EPI_ISL_3382492 | | | |
| see above | LABORATOIRE d'ANALYSES de BIOLOGIE MEDICALES PUILBOREAU | CNR Virus des Infections Respiratoires - France SUD | Antonin Bal, Gregory Destras, Gwendolyne Burfin, Hadrien Regue, Quentin Semanas, Martine Valette, Bruno Lina, Laurence Josset |
| EPI_ISL_3382493, EPI_ISL_3382494 | DYOMEDEA-LABORATOIRE DE LA SAUVEGARDE | CNR Virus des Infections Respiratoires - France SUD | Antonin Bal, Gregory Destras, Gwendolyne Burfin, Hadrien Regue, Quentin Semanas, Martine Valette, Bruno Lina, Laurence Josset |
| EPI_ISL_3382495 | LABORATOIRE d'ANALYSES de BIOLOGIE MEDICALES PUILBOREAU | CNR Virus des Infections Respiratoires - France SUD | Antonin Bal, Gregory Destras, Gwendolyne Burfin, Hadrien Regue, Quentin Semanas, Martine Valette, Bruno Lina, Laurence Josset |
| EPI_ISL_3382496 | DYOMEDEA-LABORATOIRE DE LA SAUVEGARDE | CNR Virus des Infections Respiratoires - France SUD | Antonin Bal, Gregory Destras, Gwendolyne Burfin, Hadrien Regue, Quentin Semanas, Martine Valette, Bruno Lina, Laurence Josset |
| EPI_ISL_3382497, EPI_ISL_3382498, EPI_ISL_3382499, EPI_ISL_3382500, EPI_ISL_3382501, EPI_ISL_3382502 | LABORATOIRE d'ANALYSES de BIOLOGIE MEDICALES PUILBOREAU | CNR Virus des Infections Respiratoires - France SUD | Antonin Bal, Gregory Destras, Gwendolyne Burfin, Hadrien Regue, Quentin Semanas, Martine Valette, Bruno Lina, Laurence Josset |
| EPI_ISL_3382503, EPI_ISL_3382504, EPI_ISL_3382505 | DYOMEDEA-LABORATOIRE DE LA SAUVEGARDE | CNR Virus des Infections Respiratoires - France SUD | Antonin Bal, Gregory Destras, Gwendolyne Burfin, Hadrien Regue, Quentin Semanas, Martine Valette, Bruno Lina, Laurence Josset |
| EPI_ISL_3382506, EPI_ISL_3382507, EPI_ISL_3382508, EPI_ISL_3382509, EPI_ISL_3382510, EPI_ISL_3382511 | LABORATOIRE d'ANALYSES de BIOLOGIE MEDICALES PUILBOREAU | CNR Virus des Infections Respiratoires - France SUD | Antonin Bal, Gregory Destras, Gwendolyne Burfin, Hadrien Regue, Quentin Semanas, Martine Valette, Bruno Lina, Laurence Josset |
| EPI_ISL_3382512, EPI_ISL_3382513 | DYOMEDEA-LABORATOIRE DE LA SAUVEGARDE | CNR Virus des Infections Respiratoires - France SUD | Antonin Bal, Gregory Destras, Gwendolyne Burfin, Hadrien Regue, Quentin Semanas, Martine Valette, Bruno Lina, Laurence Josset |
| EPI_ISL_3382514, EPI_ISL_3382515 | LABORATOIRE d'ANALYSES de BIOLOGIE MEDICALES PUILBOREAU | CNR Virus des Infections Respiratoires - France SUD | Antonin Bal, Gregory Destras, Gwendolyne Burfin, Hadrien Regue, Quentin Semanas, Martine Valette, Bruno Lina, Laurence Josset |

| | | | |
|---|---|---|---|
| EPI_ISL_3382516 | DYOMEDEA-LABORATOIRE DE LA SAUVEGARDE | CNR Virus des Infections Respiratoires - France SUD | Antonin Bal, Gregory Destras, Gwendolyne Burfin, Hadrien Regue, Quentin Semanas, Martine Valette, Bruno Lina, Laurence Josset |
| EPI_ISL_3382517, EPI_ISL_3382518 | LABORATOIRE d'ANALYSES de BIOLOGIE MEDICALES PUILBOREAU | CNR Virus des Infections Respiratoires - France SUD | Antonin Bal, Gregory Destras, Gwendolyne Burfin, Hadrien Regue, Quentin Semanas, Martine Valette, Bruno Lina, Laurence Josset |
| EPI_ISL_3382519 | DYOMEDEA-LABORATOIRE DE LA SAUVEGARDE | CNR Virus des Infections Respiratoires - France SUD | Antonin Bal, Gregory Destras, Gwendolyne Burfin, Hadrien Regue, Quentin Semanas, Martine Valette, Bruno Lina, Laurence Josset |
| EPI_ISL_3382520 | LABORATOIRE d'ANALYSES de BIOLOGIE MEDICALES PUILBOREAU | CNR Virus des Infections Respiratoires - France SUD | Antonin Bal, Gregory Destras, Gwendolyne Burfin, Hadrien Regue, Quentin Semanas, Martine Valette, Bruno Lina, Laurence Josset |
| EPI_ISL_3382521, EPI_ISL_3382522, EPI_ISL_3382523 | DYOMEDEA-LABORATOIRE DE LA SAUVEGARDE | CNR Virus des Infections Respiratoires - France SUD | Antonin Bal, Gregory Destras, Gwendolyne Burfin, Hadrien Regue, Quentin Semanas, Martine Valette, Bruno Lina, Laurence Josset |
| EPI_ISL_3382524 | LABORATOIRE d'ANALYSES de BIOLOGIE MEDICALES PUILBOREAU | CNR Virus des Infections Respiratoires - France SUD | Antonin Bal, Gregory Destras, Gwendolyne Burfin, Hadrien Regue, Quentin Semanas, Martine Valette, Bruno Lina, Laurence Josset |
| EPI_ISL_3382525 | DYOMEDEA-LABORATOIRE DE LA SAUVEGARDE | CNR Virus des Infections Respiratoires - France SUD | Antonin Bal, Gregory Destras, Gwendolyne Burfin, Hadrien Regue, Quentin Semanas, Martine Valette, Bruno Lina, Laurence Josset |
| EPI_ISL_3382526 | LABORATOIRE d'ANALYSES de BIOLOGIE MEDICALES PUILBOREAU | CNR Virus des Infections Respiratoires - France SUD | Antonin Bal, Gregory Destras, Gwendolyne Burfin, Hadrien Regue, Quentin Semanas, Martine Valette, Bruno Lina, Laurence Josset |
| EPI_ISL_3382527, EPI_ISL_3382528, EPI_ISL_3382530, EPI_ISL_3382531 | DYOMEDEA-LABORATOIRE DE LA SAUVEGARDE | CNR Virus des Infections Respiratoires - France SUD | Antonin Bal, Gregory Destras, Gwendolyne Burfin, Hadrien Regue, Quentin Semanas, Martine Valette, Bruno Lina, Laurence Josset |
| EPI_ISL_3382532 | LABORATOIRE d'ANALYSES de BIOLOGIE MEDICALES PUILBOREAU | CNR Virus des Infections Respiratoires - France SUD | Antonin Bal, Gregory Destras, Gwendolyne Burfin, Hadrien Regue, Quentin Semanas, Martine Valette, Bruno Lina, Laurence Josset |
| EPI_ISL_3382533, EPI_ISL_3382534 | DYOMEDEA-LABORATOIRE DE LA SAUVEGARDE | CNR Virus des Infections Respiratoires - France SUD | Antonin Bal, Gregory Destras, Gwendolyne Burfin, Hadrien Regue, Quentin Semanas, Martine Valette, Bruno Lina, Laurence Josset |
| EPI_ISL_3382535 | LABORATOIRE d'ANALYSES de BIOLOGIE MEDICALES PUILBOREAU | CNR Virus des Infections Respiratoires - France SUD | Antonin Bal, Gregory Destras, Gwendolyne Burfin, Hadrien Regue, Quentin Semanas, Martine Valette, Bruno Lina, Laurence Josset |
| EPI_ISL_3382536, EPI_ISL_3382537 | DYOMEDEA-LABORATOIRE DE LA SAUVEGARDE | CNR Virus des Infections Respiratoires - France SUD | Antonin Bal, Gregory Destras, Gwendolyne Burfin, Hadrien Regue, Quentin Semanas, Martine Valette, Bruno Lina, Laurence Josset |
| EPI_ISL_3382538, EPI_ISL_3382539 | LABORATOIRE d'ANALYSES de BIOLOGIE MEDICALES PUILBOREAU | CNR Virus des Infections Respiratoires - France SUD | Antonin Bal, Gregory Destras, Gwendolyne Burfin, Hadrien Regue, Quentin Semanas, Martine Valette, Bruno Lina, Laurence Josset |
| EPI_ISL_3382540 | CENTRE HOSPITALIER DE VALENCE | CNR Virus des Infections Respiratoires - France SUD | Antonin Bal, Gregory Destras, Gwendolyne Burfin, Hadrien Regue, Quentin Semanas, Martine Valette, Bruno Lina, Laurence Josset |
| EPI_ISL_3382541 | LABORATOIRE d'ANALYSES de BIOLOGIE MEDICALES PUILBOREAU | CNR Virus des Infections Respiratoires - France SUD | Antonin Bal, Gregory Destras, Gwendolyne Burfin, Hadrien Regue, Quentin Semanas, Martine Valette, Bruno Lina, Laurence Josset |
| EPI_ISL_3382542 | CENTRE HOSPITALIER DE VALENCE | CNR Virus des Infections Respiratoires - France SUD | Antonin Bal, Gregory Destras, Gwendolyne Burfin, Hadrien Regue, Quentin Semanas, Martine Valette, Bruno Lina, Laurence Josset |
| EPI_ISL_3382543 | LABORATOIRE d'ANALYSES de BIOLOGIE MEDICALES PUILBOREAU | CNR Virus des Infections Respiratoires - France SUD | Antonin Bal, Gregory Destras, Gwendolyne Burfin, Hadrien Regue, Quentin Semanas, Martine Valette, Bruno Lina, Laurence Josset |
| EPI_ISL_3382544, EPI_ISL_3382545 | CENTRE HOSPITALIER DE VALENCE | CNR Virus des Infections Respiratoires - France SUD | Antonin Bal, Gregory Destras, Gwendolyne Burfin, Hadrien Regue, Quentin Semanas, Martine Valette, Bruno Lina, Laurence Josset |
| EPI_ISL_3382546 | DYOMEDEA-LABORATOIRE DE LA SAUVEGARDE | CNR Virus des Infections Respiratoires - France SUD | Antonin Bal, Gregory Destras, Gwendolyne Burfin, Hadrien Regue, Quentin Semanas, Martine Valette, Bruno Lina, Laurence Josset |
| EPI_ISL_3382547, EPI_ISL_3382548 | CENTRE HOSPITALIER DE VALENCE | CNR Virus des Infections Respiratoires - France SUD | Antonin Bal, Gregory Destras, Gwendolyne Burfin, Hadrien Regue, Quentin Semanas, Martine Valette, Bruno Lina, Laurence Josset |
| EPI_ISL_3382549 | LABORATOIRE d'ANALYSES de BIOLOGIE MEDICALES PUILBOREAU | CNR Virus des Infections Respiratoires - France SUD | Antonin Bal, Gregory Destras, Gwendolyne Burfin, Hadrien Regue, Quentin Semanas, Martine Valette, Bruno Lina, Laurence Josset |
| EPI_ISL_3382550, EPI_ISL_3382551 | CENTRE HOSPITALIER ST JOSEPH ST LUC | CNR Virus des Infections Respiratoires - France SUD | Antonin Bal, Gregory Destras, Gwendolyne Burfin, Hadrien Regue, Quentin Semanas, Martine Valette, Bruno Lina, Laurence Josset |
| EPI_ISL_3382552, EPI_ISL_3382553, EPI_ISL_3382554, EPI_ISL_3382555, EPI_ISL_3382556, EPI_ISL_3382557, EPI_ISL_3382558, EPI_ISL_3382559, EPI_ISL_3382560, EPI_ISL_3382561, EPI_ISL_3382562 | | | |
| see above | DYOMEDEA-LABORATOIRE DE LA SAUVEGARDE | CNR Virus des Infections Respiratoires - France SUD | Antonin Bal, Gregory Destras, Gwendolyne Burfin, Hadrien Regue, Quentin Semanas, Martine Valette, Bruno Lina, Laurence Josset |
| EPI_ISL_3382563, EPI_ISL_3382564 | LABORATOIRE d'ANALYSES de BIOLOGIE MEDICALES PUILBOREAU | CNR Virus des Infections Respiratoires - France SUD | Antonin Bal, Gregory Destras, Gwendolyne Burfin, Hadrien Regue, Quentin Semanas, Martine Valette, Bruno Lina, Laurence Josset |
| EPI_ISL_3382565 | DYOMEDEA-LABORATOIRE DE LA SAUVEGARDE | CNR Virus des Infections Respiratoires - France SUD | Antonin Bal, Gregory Destras, Gwendolyne Burfin, Hadrien Regue, Quentin Semanas, Martine Valette, Bruno Lina, Laurence Josset |
| EPI_ISL_3382566 | LABORATOIRE d'ANALYSES de BIOLOGIE MEDICALES PUILBOREAU | CNR Virus des Infections Respiratoires - France SUD | Antonin Bal, Gregory Destras, Gwendolyne Burfin, Hadrien Regue, Quentin Semanas, Martine Valette, Bruno Lina, Laurence Josset |
| EPI_ISL_3382567 | DYOMEDEA-LABORATOIRE DE LA SAUVEGARDE | CNR Virus des Infections Respiratoires - France SUD | Antonin Bal, Gregory Destras, Gwendolyne Burfin, Hadrien Regue, Quentin Semanas, Martine Valette, Bruno Lina, Laurence Josset |
| EPI_ISL_3382568, EPI_ISL_3382569, EPI_ISL_3382570 | LABORATOIRE d'ANALYSES de BIOLOGIE MEDICALES PUILBOREAU | CNR Virus des Infections Respiratoires - France SUD | Antonin Bal, Gregory Destras, Gwendolyne Burfin, Hadrien Regue, Quentin Semanas, Martine Valette, Bruno Lina, Laurence Josset |
| EPI_ISL_3382571 | DYOMEDEA-LABORATOIRE DE LA SAUVEGARDE | CNR Virus des Infections Respiratoires - France SUD | Antonin Bal, Gregory Destras, Gwendolyne Burfin, Hadrien Regue, Quentin Semanas, Martine Valette, Bruno Lina, Laurence Josset |
| EPI_ISL_3382572 | LABORATOIRE d'ANALYSES de BIOLOGIE MEDICALES PUILBOREAU | CNR Virus des Infections Respiratoires - France SUD | Antonin Bal, Gregory Destras, Gwendolyne Burfin, Hadrien Regue, Quentin Semanas, Martine Valette, Bruno Lina, Laurence Josset |
| EPI_ISL_3382573, EPI_ISL_3382574 | DYOMEDEA-LABORATOIRE DE LA SAUVEGARDE | CNR Virus des Infections Respiratoires - France SUD | Antonin Bal, Gregory Destras, Gwendolyne Burfin, Hadrien Regue, Quentin Semanas, Martine Valette, Bruno Lina, Laurence Josset |
| EPI_ISL_3382575, EPI_ISL_3382576, EPI_ISL_3382577, EPI_ISL_3382578 | LABORATOIRE d'ANALYSES de BIOLOGIE MEDICALES PUILBOREAU | CNR Virus des Infections Respiratoires - France SUD | Antonin Bal, Gregory Destras, Gwendolyne Burfin, Hadrien Regue, Quentin Semanas, Martine Valette, Bruno Lina, Laurence Josset |
| EPI_ISL_3382579, EPI_ISL_3382580, EPI_ISL_3382581 | DYOMEDEA-LABORATOIRE DE LA SAUVEGARDE | CNR Virus des Infections Respiratoires - France SUD | Antonin Bal, Gregory Destras, Gwendolyne Burfin, Hadrien Regue, Quentin Semanas, Martine Valette, Bruno Lina, Laurence Josset |
| EPI_ISL_3382582, EPI_ISL_3382583 | LBM DYNABIO CARREAU | CNR Virus des Infections Respiratoires - France SUD | Antonin Bal, Gregory Destras, Gwendolyne Burfin, Hadrien Regue, Quentin Semanas, Martine Valette, Bruno Lina, Laurence Josset |
| EPI_ISL_3382584, EPI_ISL_3382585, EPI_ISL_3382586, EPI_ISL_3382587, EPI_ISL_3382588, EPI_ISL_3382589 | DYOMEDEA-LABORATOIRE DE LA SAUVEGARDE | CNR Virus des Infections Respiratoires - France SUD | Antonin Bal, Gregory Destras, Gwendolyne Burfin, Hadrien Regue, Quentin Semanas, Martine Valette, Bruno Lina, Laurence Josset |
| EPI_ISL_3382590 | LAM DU MARAIS | CNR Virus des Infections Respiratoires - France SUD | Antonin Bal, Gregory Destras, Gwendolyne Burfin, Hadrien Regue, Quentin Semanas, Martine Valette, Bruno Lina, Laurence Josset |
| EPI_ISL_3382591 | DYOMEDEA-LABORATOIRE DE LA SAUVEGARDE | CNR Virus des Infections Respiratoires - France SUD | Antonin Bal, Gregory Destras, Gwendolyne Burfin, Hadrien Regue, Quentin Semanas, Martine Valette, Bruno Lina, Laurence Josset |
| EPI_ISL_3382592 | LAM DU MARAIS | CNR Virus des Infections Respiratoires - France SUD | Antonin Bal, Gregory Destras, Gwendolyne Burfin, Hadrien Regue, Quentin Semanas, Martine Valette, Bruno Lina, Laurence Josset |
| EPI_ISL_3382593 | DYOMEDEA-LABORATOIRE DE LA SAUVEGARDE | CNR Virus des Infections Respiratoires - France SUD | Antonin Bal, Gregory Destras, Gwendolyne Burfin, Hadrien Regue, Quentin Semanas, Martine Valette, Bruno Lina, Laurence Josset |
| EPI_ISL_3382594, EPI_ISL_3382595, EPI_ISL_3382596 | LBM DYNABIO CARREAU | CNR Virus des Infections Respiratoires - France SUD | Antonin Bal, Gregory Destras, Gwendolyne Burfin, Hadrien Regue, Quentin Semanas, Martine Valette, Bruno Lina, Laurence Josset |
| EPI_ISL_3382597 | MIRIALIS CLUSES BECHET | CNR Virus des Infections Respiratoires - France SUD | Antonin Bal, Gregory Destras, Gwendolyne Burfin, Hadrien Regue, Quentin Semanas, Martine Valette, Bruno Lina, Laurence Josset |
| EPI_ISL_3382598 | LAM DU MARAIS | CNR Virus des Infections Respiratoires - France SUD | Antonin Bal, Gregory Destras, Gwendolyne Burfin, Hadrien Regue, Quentin Semanas, Martine Valette, Bruno Lina, Laurence Josset |
| EPI_ISL_3382599 | DYOMEDEA-LABORATOIRE DE LA SAUVEGARDE | CNR Virus des Infections Respiratoires - France SUD | Antonin Bal, Gregory Destras, Gwendolyne Burfin, Hadrien Regue, Quentin Semanas, Martine Valette, Bruno Lina, Laurence Josset |
| EPI_ISL_3382600, EPI_ISL_3382601 | LAM DU MARAIS | CNR Virus des Infections Respiratoires - France SUD | Antonin Bal, Gregory Destras, Gwendolyne Burfin, Hadrien Regue, Quentin Semanas, Martine Valette, Bruno Lina, Laurence Josset |
| EPI_ISL_3382602, EPI_ISL_3382603 | MIRIALIS CLUSES BECHET | CNR Virus des Infections Respiratoires - France SUD | Antonin Bal, Gregory Destras, Gwendolyne Burfin, Hadrien Regue, Quentin Semanas, Martine Valette, Bruno Lina, Laurence Josset |
| EPI_ISL_3382608 | REUNILAB | CNR Virus des Infections Respiratoires - France SUD | Antonin Bal, Gregory Destras, Gwendolyne Burfin, Hadrien Regue, Quentin Semanas, Martine Valette, Bruno Lina, Laurence Josset |

| | | | |
|---|---|---|---|
| EPI_ISL_3382614 | CNR Virus des Infections Respiratoires - France SUD | CNR Virus des Infections Respiratoires - France SUD | Antonin Bal, Gregory Destras, Gwendolyne Burfin, Hadrien Regue, Quentin Semanas, Martine Valette, Bruno Lina, Laurence Josset |
| EPI_ISL_3382619 | REUNILAB | CNR Virus des Infections Respiratoires - France SUD | Antonin Bal, Gregory Destras, Gwendolyne Burfin, Hadrien Regue, Quentin Semanas, Martine Valette, Bruno Lina, Laurence Josset |
| EPI_ISL_3382624, EPI_ISL_3382625, EPI_ISL_3382626, EPI_ISL_3382627, EPI_ISL_3382628 | LABORATOIRE BIOGROUP | CNR Virus des Infections Respiratoires - France SUD | Antonin Bal, Gregory Destras, Gwendolyne Burfin, Hadrien Regue, Quentin Semanas, Martine Valette, Bruno Lina, Laurence Josset |
| EPI_ISL_3382629 | CENTRE HOSPITALIER DE VALENCE | CNR Virus des Infections Respiratoires - France SUD | Antonin Bal, Gregory Destras, Gwendolyne Burfin, Hadrien Regue, Quentin Semanas, Martine Valette, Bruno Lina, Laurence Josset |
| EPI_ISL_3382630 | CNR Virus des Infections Respiratoires - France SUD | CNR Virus des Infections Respiratoires - France SUD | Antonin Bal, Gregory Destras, Gwendolyne Burfin, Hadrien Regue, Quentin Semanas, Martine Valette, Bruno Lina, Laurence Josset |
| EPI_ISL_3382631, EPI_ISL_3382632 | MIRIALIS CLUSES BECHET | CNR Virus des Infections Respiratoires - France SUD | Antonin Bal, Gregory Destras, Gwendolyne Burfin, Hadrien Regue, Quentin Semanas, Martine Valette, Bruno Lina, Laurence Josset |
| EPI_ISL_3382633, EPI_ISL_3382634 | AX BIO OCEAN | CNR Virus des Infections Respiratoires - France SUD | Antonin Bal, Gregory Destras, Gwendolyne Burfin, Hadrien Regue, Quentin Semanas, Martine Valette, Bruno Lina, Laurence Josset |
| EPI_ISL_3382635, EPI_ISL_3382636 | MIRIALIS CLUSES BECHET | CNR Virus des Infections Respiratoires - France SUD | Antonin Bal, Gregory Destras, Gwendolyne Burfin, Hadrien Regue, Quentin Semanas, Martine Valette, Bruno Lina, Laurence Josset |
| EPI_ISL_3382637, EPI_ISL_3382638 | AX BIO OCEAN | CNR Virus des Infections Respiratoires - France SUD | Antonin Bal, Gregory Destras, Gwendolyne Burfin, Hadrien Regue, Quentin Semanas, Martine Valette, Bruno Lina, Laurence Josset |
| EPI_ISL_3382639 | MIRIALIS CLUSES BECHET | CNR Virus des Infections Respiratoires - France SUD | Antonin Bal, Gregory Destras, Gwendolyne Burfin, Hadrien Regue, Quentin Semanas, Martine Valette, Bruno Lina, Laurence Josset |
| EPI_ISL_3382640 | LABORATOIRE LABAZUR | CNR Virus des Infections Respiratoires - France SUD | Antonin Bal, Gregory Destras, Gwendolyne Burfin, Hadrien Regue, Quentin Semanas, Martine Valette, Bruno Lina, Laurence Josset |
| EPI_ISL_3382641 | MIRIALIS CLUSES BECHET | CNR Virus des Infections Respiratoires - France SUD | Antonin Bal, Gregory Destras, Gwendolyne Burfin, Hadrien Regue, Quentin Semanas, Martine Valette, Bruno Lina, Laurence Josset |
| EPI_ISL_3382642 | LABORATOIRE LABAZUR | CNR Virus des Infections Respiratoires - France SUD | Antonin Bal, Gregory Destras, Gwendolyne Burfin, Hadrien Regue, Quentin Semanas, Martine Valette, Bruno Lina, Laurence Josset |
| EPI_ISL_3382643 | CNR Virus des Infections Respiratoires - France SUD | CNR Virus des Infections Respiratoires - France SUD | Antonin Bal, Gregory Destras, Gwendolyne Burfin, Hadrien Regue, Quentin Semanas, Martine Valette, Bruno Lina, Laurence Josset |
| EPI_ISL_3382644, EPI_ISL_3382645 | AX BIO OCEAN | CNR Virus des Infections Respiratoires - France SUD | Antonin Bal, Gregory Destras, Gwendolyne Burfin, Hadrien Regue, Quentin Semanas, Martine Valette, Bruno Lina, Laurence Josset |
| EPI_ISL_3382646 | MIRIALIS CLUSES BECHET | CNR Virus des Infections Respiratoires - France SUD | Antonin Bal, Gregory Destras, Gwendolyne Burfin, Hadrien Regue, Quentin Semanas, Martine Valette, Bruno Lina, Laurence Josset |
| EPI_ISL_3382647, EPI_ISL_3382648, EPI_ISL_3382649 | AX BIO OCEAN | CNR Virus des Infections Respiratoires - France SUD | Antonin Bal, Gregory Destras, Gwendolyne Burfin, Hadrien Regue, Quentin Semanas, Martine Valette, Bruno Lina, Laurence Josset |
| EPI_ISL_3382651 | CNR Virus des Infections Respiratoires - France SUD | CNR Virus des Infections Respiratoires - France SUD | Antonin Bal, Gregory Destras, Gwendolyne Burfin, Hadrien Regue, Quentin Semanas, Martine Valette, Bruno Lina, Laurence Josset |
| EPI_ISL_3382652 | AX BIO OCEAN | CNR Virus des Infections Respiratoires - France SUD | Antonin Bal, Gregory Destras, Gwendolyne Burfin, Hadrien Regue, Quentin Semanas, Martine Valette, Bruno Lina, Laurence Josset |
| EPI_ISL_3382653 | MIRIALIS CLUSES BECHET | CNR Virus des Infections Respiratoires - France SUD | Antonin Bal, Gregory Destras, Gwendolyne Burfin, Hadrien Regue, Quentin Semanas, Martine Valette, Bruno Lina, Laurence Josset |
| EPI_ISL_3382654 | AX BIO OCEAN | CNR Virus des Infections Respiratoires - France SUD | Antonin Bal, Gregory Destras, Gwendolyne Burfin, Hadrien Regue, Quentin Semanas, Martine Valette, Bruno Lina, Laurence Josset |
| EPI_ISL_3382655 | MIRIALIS CLUSES BECHET | CNR Virus des Infections Respiratoires - France SUD | Antonin Bal, Gregory Destras, Gwendolyne Burfin, Hadrien Regue, Quentin Semanas, Martine Valette, Bruno Lina, Laurence Josset |
| EPI_ISL_3382656, EPI_ISL_3382657, EPI_ISL_3382658 | AX BIO OCEAN | CNR Virus des Infections Respiratoires - France SUD | Antonin Bal, Gregory Destras, Gwendolyne Burfin, Hadrien Regue, Quentin Semanas, Martine Valette, Bruno Lina, Laurence Josset |
| EPI_ISL_3382659 | MIRIALIS CLUSES BECHET | CNR Virus des Infections Respiratoires - France SUD | Antonin Bal, Gregory Destras, Gwendolyne Burfin, Hadrien Regue, Quentin Semanas, Martine Valette, Bruno Lina, Laurence Josset |
| EPI_ISL_3382660, EPI_ISL_3382661 | AX BIO OCEAN | CNR Virus des Infections Respiratoires - France SUD | Antonin Bal, Gregory Destras, Gwendolyne Burfin, Hadrien Regue, Quentin Semanas, Martine Valette, Bruno Lina, Laurence Josset |
| EPI_ISL_3382662 | MIRIALIS CLUSES BECHET | CNR Virus des Infections Respiratoires - France SUD | Antonin Bal, Gregory Destras, Gwendolyne Burfin, Hadrien Regue, Quentin Semanas, Martine Valette, Bruno Lina, Laurence Josset |
| EPI_ISL_3382663, EPI_ISL_3382664, EPI_ISL_3382665 | AX BIO OCEAN | CNR Virus des Infections Respiratoires - France SUD | Antonin Bal, Gregory Destras, Gwendolyne Burfin, Hadrien Regue, Quentin Semanas, Martine Valette, Bruno Lina, Laurence Josset |
| EPI_ISL_3382666 | MIRIALIS CLUSES BECHET | CNR Virus des Infections Respiratoires - France SUD | Antonin Bal, Gregory Destras, Gwendolyne Burfin, Hadrien Regue, Quentin Semanas, Martine Valette, Bruno Lina, Laurence Josset |
| EPI_ISL_3382667, EPI_ISL_3382668, EPI_ISL_3382669, EPI_ISL_3382670, EPI_ISL_3382671 | AX BIO OCEAN | CNR Virus des Infections Respiratoires - France SUD | Antonin Bal, Gregory Destras, Gwendolyne Burfin, Hadrien Regue, Quentin Semanas, Martine Valette, Bruno Lina, Laurence Josset |
| EPI_ISL_3382672, EPI_ISL_3382673 | ORIAPOLE | CNR Virus des Infections Respiratoires - France SUD | Antonin Bal, Gregory Destras, Gwendolyne Burfin, Hadrien Regue, Quentin Semanas, Martine Valette, Bruno Lina, Laurence Josset |
| EPI_ISL_3382674 | MIRIALIS CLUSES BECHET | CNR Virus des Infections Respiratoires - France SUD | Antonin Bal, Gregory Destras, Gwendolyne Burfin, Hadrien Regue, Quentin Semanas, Martine Valette, Bruno Lina, Laurence Josset |
| EPI_ISL_3382675 | AX BIO OCEAN | CNR Virus des Infections Respiratoires - France SUD | Antonin Bal, Gregory Destras, Gwendolyne Burfin, Hadrien Regue, Quentin Semanas, Martine Valette, Bruno Lina, Laurence Josset |
| EPI_ISL_3382676 | ORIAPOLE | CNR Virus des Infections Respiratoires - France SUD | Antonin Bal, Gregory Destras, Gwendolyne Burfin, Hadrien Regue, Quentin Semanas, Martine Valette, Bruno Lina, Laurence Josset |
| EPI_ISL_3382677 | AX BIO OCEAN | CNR Virus des Infections Respiratoires - France SUD | Antonin Bal, Gregory Destras, Gwendolyne Burfin, Hadrien Regue, Quentin Semanas, Martine Valette, Bruno Lina, Laurence Josset |
| EPI_ISL_3382678 | ORIAPOLE | CNR Virus des Infections Respiratoires - France SUD | Antonin Bal, Gregory Destras, Gwendolyne Burfin, Hadrien Regue, Quentin Semanas, Martine Valette, Bruno Lina, Laurence Josset |
| EPI_ISL_3382679, EPI_ISL_3382680 | AX BIO OCEAN | CNR Virus des Infections Respiratoires - France SUD | Antonin Bal, Gregory Destras, Gwendolyne Burfin, Hadrien Regue, Quentin Semanas, Martine Valette, Bruno Lina, Laurence Josset |
| EPI_ISL_3382681 | LABORATOIRE d'ANALYSES de BIOLOGIE MEDICALES PUILBOREAU | CNR Virus des Infections Respiratoires - France SUD | Antonin Bal, Gregory Destras, Gwendolyne Burfin, Hadrien Regue, Quentin Semanas, Martine Valette, Bruno Lina, Laurence Josset |
| EPI_ISL_3382682 | AX BIO OCEAN | CNR Virus des Infections Respiratoires - France SUD | Antonin Bal, Gregory Destras, Gwendolyne Burfin, Hadrien Regue, Quentin Semanas, Martine Valette, Bruno Lina, Laurence Josset |
| EPI_ISL_3382683 | CH METROPOLE SAVOIE | CNR Virus des Infections Respiratoires - France SUD | Antonin Bal, Gregory Destras, Gwendolyne Burfin, Hadrien Regue, Quentin Semanas, Martine Valette, Bruno Lina, Laurence Josset |
| EPI_ISL_3382684, EPI_ISL_3382685 | AX BIO OCEAN | CNR Virus des Infections Respiratoires - France SUD | Antonin Bal, Gregory Destras, Gwendolyne Burfin, Hadrien Regue, Quentin Semanas, Martine Valette, Bruno Lina, Laurence Josset |
| EPI_ISL_3382686 | CENTRE HOSPITALIER LUCIEN HUSSEL | CNR Virus des Infections Respiratoires - France SUD | Antonin Bal, Gregory Destras, Gwendolyne Burfin, Hadrien Regue, Quentin Semanas, Martine Valette, Bruno Lina, Laurence Josset |
| EPI_ISL_3382687, EPI_ISL_3382689, EPI_ISL_3382690, EPI_ISL_3382691, EPI_ISL_3382692, EPI_ISL_3382693, EPI_ISL_3382694, EPI_ISL_3382695, EPI_ISL_3382696, EPI_ISL_3382697 | AX BIO OCEAN | CNR Virus des Infections Respiratoires - France SUD | Antonin Bal, Gregory Destras, Gwendolyne Burfin, Hadrien Regue, Quentin Semanas, Martine Valette, Bruno Lina, Laurence Josset |
| EPI_ISL_3382698, EPI_ISL_3382699, EPI_ISL_3382700, EPI_ISL_3382701, EPI_ISL_3382702, EPI_ISL_3382703, EPI_ISL_3382704 | MIRIALIS CLUSES BECHET | CNR Virus des Infections Respiratoires - France SUD | Antonin Bal, Gregory Destras, Gwendolyne Burfin, Hadrien Regue, Quentin Semanas, Martine Valette, Bruno Lina, Laurence Josset |
| EPI_ISL_3382705, EPI_ISL_3382706, EPI_ISL_3382707, EPI_ISL_3382708 | CNR Virus des Infections Respiratoires - France SUD | CNR Virus des Infections Respiratoires - France SUD | Antonin Bal, Gregory Destras, Gwendolyne Burfin, Hadrien Regue, Quentin Semanas, Martine Valette, Bruno Lina, Laurence Josset |
| EPI_ISL_3382709, EPI_ISL_3382710, EPI_ISL_3382711, EPI_ISL_3382712, EPI_ISL_3382713, EPI_ISL_3382715, EPI_ISL_3382716, EPI_ISL_3382717, EPI_ISL_3382718, EPI_ISL_3382719, EPI_ISL_3382720, EPI_ISL_3382721, EPI_ISL_3382722, EPI_ISL_3382723, EPI_ISL_3382724, EPI_ISL_3382725, EPI_ISL_3382726, EPI_ISL_3382727, EPI_ISL_3382728, EPI_ISL_3382729, EPI_ISL_3382730, EPI_ISL_3382731 | | | |
| see above | AX BIO OCEAN | CNR Virus des Infections Respiratoires - France SUD | Antonin Bal, Gregory Destras, Gwendolyne Burfin, Hadrien Regue, Quentin Semanas, Martine Valette, Bruno Lina, Laurence Josset |
| EPI_ISL_3382735, EPI_ISL_3382736 | CHR LA REUNION FELIX GUYON | CNR Virus des Infections Respiratoires - France SUD | Antonin Bal, Gregory Destras, Gwendolyne Burfin, Hadrien Regue, Quentin Semanas, Martine Valette, Bruno Lina, Laurence Josset |

| | | | |
|---|---|---|---|
| EPI_ISL_3382740 | LABORATOIRE CBM DE MURET | CNR Virus des Infections Respiratoires - France SUD | Antonin Bal, Gregory Destras, Gwendolyne Burfin, Hadrien Regue, Quentin Semanas, Martine Valette, Bruno Lina, Laurence Josset |
| EPI_ISL_3382741 | LABORATOIRE UNILIANS DECINES | CNR Virus des Infections Respiratoires - France SUD | Antonin Bal, Gregory Destras, Gwendolyne Burfin, Hadrien Regue, Quentin Semanas, Martine Valette, Bruno Lina, Laurence Josset |
| EPI_ISL_3382742 | CHR LA REUNION FELIX GUYON | CNR Virus des Infections Respiratoires - France SUD | Antonin Bal, Gregory Destras, Gwendolyne Burfin, Hadrien Regue, Quentin Semanas, Martine Valette, Bruno Lina, Laurence Josset |
| EPI_ISL_3382744 | LABORATOIRE CBM DE MURET | CNR Virus des Infections Respiratoires - France SUD | Antonin Bal, Gregory Destras, Gwendolyne Burfin, Hadrien Regue, Quentin Semanas, Martine Valette, Bruno Lina, Laurence Josset |
| EPI_ISL_3382745 | LABORATOIRE UNILIANS DECINES | CNR Virus des Infections Respiratoires - France SUD | Antonin Bal, Gregory Destras, Gwendolyne Burfin, Hadrien Regue, Quentin Semanas, Martine Valette, Bruno Lina, Laurence Josset |
| EPI_ISL_3382748, EPI_ISL_3382749, EPI_ISL_3382750 | LABORATOIRE CBM DE MURET | CNR Virus des Infections Respiratoires - France SUD | Antonin Bal, Gregory Destras, Gwendolyne Burfin, Hadrien Regue, Quentin Semanas, Martine Valette, Bruno Lina, Laurence Josset |
| EPI_ISL_3382751 | LAM BAYARD-BIOPYRENEES | CNR Virus des Infections Respiratoires - France SUD | Antonin Bal, Gregory Destras, Gwendolyne Burfin, Hadrien Regue, Quentin Semanas, Martine Valette, Bruno Lina, Laurence Josset |
| EPI_ISL_3382752 | LABORATOIRE CBM DE MURET | CNR Virus des Infections Respiratoires - France SUD | Antonin Bal, Gregory Destras, Gwendolyne Burfin, Hadrien Regue, Quentin Semanas, Martine Valette, Bruno Lina, Laurence Josset |
| EPI_ISL_3382753 | LABORATOIRE UNILIANS DECINES | CNR Virus des Infections Respiratoires - France SUD | Antonin Bal, Gregory Destras, Gwendolyne Burfin, Hadrien Regue, Quentin Semanas, Martine Valette, Bruno Lina, Laurence Josset |
| EPI_ISL_3382754 | LABORATOIRE CBM DE MURET | CNR Virus des Infections Respiratoires - France SUD | Antonin Bal, Gregory Destras, Gwendolyne Burfin, Hadrien Regue, Quentin Semanas, Martine Valette, Bruno Lina, Laurence Josset |
| EPI_ISL_3382755 | LAM BAYARD-BIOPYRENEES | CNR Virus des Infections Respiratoires - France SUD | Antonin Bal, Gregory Destras, Gwendolyne Burfin, Hadrien Regue, Quentin Semanas, Martine Valette, Bruno Lina, Laurence Josset |
| EPI_ISL_3382756 | LABORATOIRE CBM DE MURET | CNR Virus des Infections Respiratoires - France SUD | Antonin Bal, Gregory Destras, Gwendolyne Burfin, Hadrien Regue, Quentin Semanas, Martine Valette, Bruno Lina, Laurence Josset |
| EPI_ISL_3382757 | LAM BAYARD-BIOPYRENEES | CNR Virus des Infections Respiratoires - France SUD | Antonin Bal, Gregory Destras, Gwendolyne Burfin, Hadrien Regue, Quentin Semanas, Martine Valette, Bruno Lina, Laurence Josset |
| EPI_ISL_3382758 | LABORATOIRE UNILIANS DECINES | CNR Virus des Infections Respiratoires - France SUD | Antonin Bal, Gregory Destras, Gwendolyne Burfin, Hadrien Regue, Quentin Semanas, Martine Valette, Bruno Lina, Laurence Josset |
| EPI_ISL_3382759 | LABORATOIRE CBM DE MURET | CNR Virus des Infections Respiratoires - France SUD | Antonin Bal, Gregory Destras, Gwendolyne Burfin, Hadrien Regue, Quentin Semanas, Martine Valette, Bruno Lina, Laurence Josset |
| EPI_ISL_3382760 | LAM BAYARD-BIOPYRENEES | CNR Virus des Infections Respiratoires - France SUD | Antonin Bal, Gregory Destras, Gwendolyne Burfin, Hadrien Regue, Quentin Semanas, Martine Valette, Bruno Lina, Laurence Josset |
| EPI_ISL_3382761 | LABORATOIRE CBM DE MURET | CNR Virus des Infections Respiratoires - France SUD | Antonin Bal, Gregory Destras, Gwendolyne Burfin, Hadrien Regue, Quentin Semanas, Martine Valette, Bruno Lina, Laurence Josset |
| EPI_ISL_3382762 | LABORATOIRE d'ANALYSES de BIOLOGIE MEDICALES PUILBOREAU | CNR Virus des Infections Respiratoires - France SUD | Antonin Bal, Gregory Destras, Gwendolyne Burfin, Hadrien Regue, Quentin Semanas, Martine Valette, Bruno Lina, Laurence Josset |
| EPI_ISL_3382764 | LABORATOIRE CBM DE MURET | CNR Virus des Infections Respiratoires - France SUD | Antonin Bal, Gregory Destras, Gwendolyne Burfin, Hadrien Regue, Quentin Semanas, Martine Valette, Bruno Lina, Laurence Josset |
| EPI_ISL_3382765 | LABORATOIRE UNILIANS DECINES | CNR Virus des Infections Respiratoires - France SUD | Antonin Bal, Gregory Destras, Gwendolyne Burfin, Hadrien Regue, Quentin Semanas, Martine Valette, Bruno Lina, Laurence Josset |
| EPI_ISL_3382766, EPI_ISL_3382767, EPI_ISL_3382768 | LABORATOIRE d'ANALYSES de BIOLOGIE MEDICALES PUILBOREAU | CNR Virus des Infections Respiratoires - France SUD | Antonin Bal, Gregory Destras, Gwendolyne Burfin, Hadrien Regue, Quentin Semanas, Martine Valette, Bruno Lina, Laurence Josset |
| EPI_ISL_3382769 | LABORATOIRE CBM DE MURET | CNR Virus des Infections Respiratoires - France SUD | Antonin Bal, Gregory Destras, Gwendolyne Burfin, Hadrien Regue, Quentin Semanas, Martine Valette, Bruno Lina, Laurence Josset |
| EPI_ISL_3382770 | LABORATOIRE UNILIANS DECINES | CNR Virus des Infections Respiratoires - France SUD | Antonin Bal, Gregory Destras, Gwendolyne Burfin, Hadrien Regue, Quentin Semanas, Martine Valette, Bruno Lina, Laurence Josset |
| EPI_ISL_3382772 | LABORATOIRE d'ANALYSES de BIOLOGIE MEDICALES PUILBOREAU | CNR Virus des Infections Respiratoires - France SUD | Antonin Bal, Gregory Destras, Gwendolyne Burfin, Hadrien Regue, Quentin Semanas, Martine Valette, Bruno Lina, Laurence Josset |
| EPI_ISL_3382773 | LABORATOIRE CBM DE MURET | CNR Virus des Infections Respiratoires - France SUD | Antonin Bal, Gregory Destras, Gwendolyne Burfin, Hadrien Regue, Quentin Semanas, Martine Valette, Bruno Lina, Laurence Josset |
| EPI_ISL_3382774 | LABORATOIRE UNILIANS DECINES | CNR Virus des Infections Respiratoires - France SUD | Antonin Bal, Gregory Destras, Gwendolyne Burfin, Hadrien Regue, Quentin Semanas, Martine Valette, Bruno Lina, Laurence Josset |
| EPI_ISL_3382775, EPI_ISL_3382776, EPI_ISL_3382777 | LABORATOIRE d'ANALYSES de BIOLOGIE MEDICALES PUILBOREAU | CNR Virus des Infections Respiratoires - France SUD | Antonin Bal, Gregory Destras, Gwendolyne Burfin, Hadrien Regue, Quentin Semanas, Martine Valette, Bruno Lina, Laurence Josset |
| EPI_ISL_3382778 | LABORATOIRE CBM DE MURET | CNR Virus des Infections Respiratoires - France SUD | Antonin Bal, Gregory Destras, Gwendolyne Burfin, Hadrien Regue, Quentin Semanas, Martine Valette, Bruno Lina, Laurence Josset |
| EPI_ISL_3382779 | LABORATOIRE UNILIANS DECINES | CNR Virus des Infections Respiratoires - France SUD | Antonin Bal, Gregory Destras, Gwendolyne Burfin, Hadrien Regue, Quentin Semanas, Martine Valette, Bruno Lina, Laurence Josset |
| EPI_ISL_3382780 | LABORATOIRE d'ANALYSES de BIOLOGIE MEDICALES PUILBOREAU | CNR Virus des Infections Respiratoires - France SUD | Antonin Bal, Gregory Destras, Gwendolyne Burfin, Hadrien Regue, Quentin Semanas, Martine Valette, Bruno Lina, Laurence Josset |
| EPI_ISL_3382781 | LABORATOIRE CBM DE MURET | CNR Virus des Infections Respiratoires - France SUD | Antonin Bal, Gregory Destras, Gwendolyne Burfin, Hadrien Regue, Quentin Semanas, Martine Valette, Bruno Lina, Laurence Josset |
| EPI_ISL_3382782, EPI_ISL_3382783 | LABORATOIRE d'ANALYSES de BIOLOGIE MEDICALES PUILBOREAU | CNR Virus des Infections Respiratoires - France SUD | Antonin Bal, Gregory Destras, Gwendolyne Burfin, Hadrien Regue, Quentin Semanas, Martine Valette, Bruno Lina, Laurence Josset |
| EPI_ISL_3382784 | LABORATOIRE CBM DE MURET | CNR Virus des Infections Respiratoires - France SUD | Antonin Bal, Gregory Destras, Gwendolyne Burfin, Hadrien Regue, Quentin Semanas, Martine Valette, Bruno Lina, Laurence Josset |
| EPI_ISL_3382786, EPI_ISL_3382787 | LABORATOIRE d'ANALYSES de BIOLOGIE MEDICALES PUILBOREAU | CNR Virus des Infections Respiratoires - France SUD | Antonin Bal, Gregory Destras, Gwendolyne Burfin, Hadrien Regue, Quentin Semanas, Martine Valette, Bruno Lina, Laurence Josset |
| EPI_ISL_3382788 | LABORATOIRE CBM DE MURET | CNR Virus des Infections Respiratoires - France SUD | Antonin Bal, Gregory Destras, Gwendolyne Burfin, Hadrien Regue, Quentin Semanas, Martine Valette, Bruno Lina, Laurence Josset |
| EPI_ISL_3382789 | LAM BIO 86 SITE DE CHAUMONT | CNR Virus des Infections Respiratoires - France SUD | Antonin Bal, Gregory Destras, Gwendolyne Burfin, Hadrien Regue, Quentin Semanas, Martine Valette, Bruno Lina, Laurence Josset |
| EPI_ISL_3382790 | LABORATOIRE d'ANALYSES de BIOLOGIE MEDICALES PUILBOREAU | CNR Virus des Infections Respiratoires - France SUD | Antonin Bal, Gregory Destras, Gwendolyne Burfin, Hadrien Regue, Quentin Semanas, Martine Valette, Bruno Lina, Laurence Josset |
| EPI_ISL_3382791 | LAM BAYARD-BIOPYRENEES | CNR Virus des Infections Respiratoires - France SUD | Antonin Bal, Gregory Destras, Gwendolyne Burfin, Hadrien Regue, Quentin Semanas, Martine Valette, Bruno Lina, Laurence Josset |
| EPI_ISL_3382792 | LABORATOIRE UNILIANS DECINES | CNR Virus des Infections Respiratoires - France SUD | Antonin Bal, Gregory Destras, Gwendolyne Burfin, Hadrien Regue, Quentin Semanas, Martine Valette, Bruno Lina, Laurence Josset |
| EPI_ISL_3382793 | LABORATOIRE d'ANALYSES de BIOLOGIE MEDICALES PUILBOREAU | CNR Virus des Infections Respiratoires - France SUD | Antonin Bal, Gregory Destras, Gwendolyne Burfin, Hadrien Regue, Quentin Semanas, Martine Valette, Bruno Lina, Laurence Josset |
| EPI_ISL_3382794 | LAM BIO 86 SITE DE CHAUMONT | CNR Virus des Infections Respiratoires - France SUD | Antonin Bal, Gregory Destras, Gwendolyne Burfin, Hadrien Regue, Quentin Semanas, Martine Valette, Bruno Lina, Laurence Josset |
| EPI_ISL_3382795 | LABORATOIRE d'ANALYSES de BIOLOGIE MEDICALES PUILBOREAU | CNR Virus des Infections Respiratoires - France SUD | Antonin Bal, Gregory Destras, Gwendolyne Burfin, Hadrien Regue, Quentin Semanas, Martine Valette, Bruno Lina, Laurence Josset |
| EPI_ISL_3382796 | LABORATOIRE CBM DE MURET | CNR Virus des Infections Respiratoires - France SUD | Antonin Bal, Gregory Destras, Gwendolyne Burfin, Hadrien Regue, Quentin Semanas, Martine Valette, Bruno Lina, Laurence Josset |
| EPI_ISL_3382797, EPI_ISL_3382798 | LABORATOIRE d'ANALYSES de BIOLOGIE MEDICALES PUILBOREAU | CNR Virus des Infections Respiratoires - France SUD | Antonin Bal, Gregory Destras, Gwendolyne Burfin, Hadrien Regue, Quentin Semanas, Martine Valette, Bruno Lina, Laurence Josset |
| EPI_ISL_3382799 | LAM BAYARD-BIOPYRENEES | CNR Virus des Infections Respiratoires - France SUD | Antonin Bal, Gregory Destras, Gwendolyne Burfin, Hadrien Regue, Quentin Semanas, Martine Valette, Bruno Lina, Laurence Josset |
| EPI_ISL_3382800 | LABORATOIRE UNILIANS DECINES | CNR Virus des Infections Respiratoires - France SUD | Antonin Bal, Gregory Destras, Gwendolyne Burfin, Hadrien Regue, Quentin Semanas, Martine Valette, Bruno Lina, Laurence Josset |
| EPI_ISL_3382801 | LAM BIO 86 SITE DE CHAUMONT | CNR Virus des Infections Respiratoires - France SUD | Antonin Bal, Gregory Destras, Gwendolyne Burfin, Hadrien Regue, Quentin Semanas, Martine Valette, Bruno Lina, Laurence Josset |
| EPI_ISL_3382802 | LABORATOIRE CBM DE MURET | CNR Virus des Infections Respiratoires - France SUD | Antonin Bal, Gregory Destras, Gwendolyne Burfin, Hadrien Regue, Quentin Semanas, Martine Valette, Bruno Lina, Laurence Josset |
| EPI_ISL_3382803 | LAM BIO 86 SITE DE CHAUMONT | CNR Virus des Infections Respiratoires - France SUD | Antonin Bal, Gregory Destras, Gwendolyne Burfin, Hadrien Regue, Quentin Semanas, Martine Valette, Bruno Lina, Laurence Josset |
| EPI_ISL_3382804 | LAM BAYARD-BIOPYRENEES | CNR Virus des Infections Respiratoires - France SUD | Antonin Bal, Gregory Destras, Gwendolyne Burfin, Hadrien Regue, Quentin Semanas, Martine Valette, Bruno Lina, Laurence Josset |
| EPI_ISL_3382805 | LAM BIO 86 SITE DE CHAUMONT | CNR Virus des Infections Respiratoires - France SUD | Antonin Bal, Gregory Destras, Gwendolyne Burfin, Hadrien Regue, Quentin Semanas, Martine Valette, Bruno Lina, Laurence Josset |
| EPI_ISL_3382806 | LAM BAYARD-BIOPYRENEES | CNR Virus des Infections Respiratoires - France SUD | Antonin Bal, Gregory Destras, Gwendolyne Burfin, Hadrien Regue, Quentin Semanas, Martine Valette, Bruno Lina, Laurence Josset |
| EPI_ISL_3382807 | LABORATOIRE UNILIANS DECINES | CNR Virus des Infections Respiratoires - France SUD | Antonin Bal, Gregory Destras, Gwendolyne Burfin, Hadrien Regue, Quentin Semanas, Martine Valette, Bruno Lina, Laurence Josset |
| EPI_ISL_3382808 | LAM BAYARD-BIOPYRENEES | CNR Virus des Infections Respiratoires - France SUD | Antonin Bal, Gregory Destras, Gwendolyne Burfin, Hadrien Regue, Quentin Semanas, Martine Valette, Bruno Lina, Laurence Josset |
| EPI_ISL_3382809, EPI_ISL_3382810 | LABORATOIRE UNILIANS DECINES | CNR Virus des Infections Respiratoires - France SUD | Antonin Bal, Gregory Destras, Gwendolyne Burfin, Hadrien Regue, Quentin Semanas, Martine Valette, Bruno Lina, Laurence Josset |
| EPI_ISL_3382811 | LAM BAYARD-BIOPYRENEES | CNR Virus des Infections Respiratoires - France SUD | Antonin Bal, Gregory Destras, Gwendolyne Burfin, Hadrien Regue, Quentin Semanas, Martine Valette, Bruno Lina, Laurence Josset |
| EPI_ISL_3382812 | LAM BIO 86 SITE DE CHAUMONT | CNR Virus des Infections Respiratoires - France SUD | Antonin Bal, Gregory Destras, Gwendolyne Burfin, Hadrien Regue, Quentin Semanas, Martine Valette, Bruno Lina, Laurence Josset |
| EPI_ISL_3382813 | LABORATOIRE UNILIANS DECINES | CNR Virus des Infections Respiratoires - France SUD | Antonin Bal, Gregory Destras, Gwendolyne Burfin, Hadrien Regue, Quentin Semanas, Martine Valette, Bruno Lina, Laurence Josset |
| EPI_ISL_3382814 | LAM BAYARD-BIOPYRENEES | CNR Virus des Infections Respiratoires - France SUD | Antonin Bal, Gregory Destras, Gwendolyne Burfin, Hadrien Regue, Quentin Semanas, Martine Valette, Bruno Lina, Laurence Josset |
| EPI_ISL_3382815 | LABORATOIRE d'ANALYSES de BIOLOGIE MEDICALES PUILBOREAU | CNR Virus des Infections Respiratoires - France SUD | Antonin Bal, Gregory Destras, Gwendolyne Burfin, Hadrien Regue, Quentin Semanas, Martine Valette, Bruno Lina, Laurence Josset |
| EPI_ISL_3382816 | LAM BIO 86 SITE DE CHAUMONT | CNR Virus des Infections Respiratoires - France SUD | Antonin Bal, Gregory Destras, Gwendolyne Burfin, Hadrien Regue, Quentin Semanas, Martine Valette, Bruno Lina, Laurence Josset |
| EPI_ISL_3382817 | LAM BAYARD-BIOPYRENEES | CNR Virus des Infections Respiratoires - France SUD | Antonin Bal, Gregory Destras, Gwendolyne Burfin, Hadrien Regue, Quentin Semanas, Martine Valette, Bruno Lina, Laurence Josset |
| EPI_ISL_3382818 | LABORATOIRE UNILIANS DECINES | CNR Virus des Infections Respiratoires - France SUD | Antonin Bal, Gregory Destras, Gwendolyne Burfin, Hadrien Regue, Quentin Semanas, Martine Valette, Bruno Lina, Laurence Josset |
| EPI_ISL_3382819 | LAM BIO 86 SITE DE CHAUMONT | CNR Virus des Infections Respiratoires - France SUD | Antonin Bal, Gregory Destras, Gwendolyne Burfin, Hadrien Regue, Quentin Semanas, Martine Valette, Bruno Lina, Laurence Josset |

| | | | |
|---|---|---|---|
| EPI_ISL_3382820 | LABORATOIRE d'ANALYSES de BIOLOGIE MEDICALES PUILBOREAU | CNR Virus des Infections Respiratoires - France SUD | Antonin Bal, Gregory Destras, Gwendolyne Burfin, Hadrien Regue, Quentin Semanas, Martine Valette, Bruno Lina, Laurence Josset |
| EPI_ISL_3382821 | LAM BAYARD-BIOPYRENEES | CNR Virus des Infections Respiratoires - France SUD | Antonin Bal, Gregory Destras, Gwendolyne Burfin, Hadrien Regue, Quentin Semanas, Martine Valette, Bruno Lina, Laurence Josset |
| EPI_ISL_3382822 | LAM BIO 86 SITE DE CHAUMONT | CNR Virus des Infections Respiratoires - France SUD | Antonin Bal, Gregory Destras, Gwendolyne Burfin, Hadrien Regue, Quentin Semanas, Martine Valette, Bruno Lina, Laurence Josset |
| EPI_ISL_3382823, EPI_ISL_3382824 | LABORATOIRE d'ANALYSES de BIOLOGIE MEDICALES PUILBOREAU | CNR Virus des Infections Respiratoires - France SUD | Antonin Bal, Gregory Destras, Gwendolyne Burfin, Hadrien Regue, Quentin Semanas, Martine Valette, Bruno Lina, Laurence Josset |
| EPI_ISL_3382825 | LAM BIO 86 SITE DE CHAUMONT | CNR Virus des Infections Respiratoires - France SUD | Antonin Bal, Gregory Destras, Gwendolyne Burfin, Hadrien Regue, Quentin Semanas, Martine Valette, Bruno Lina, Laurence Josset |
| EPI_ISL_3382826 | LABORATOIRE d'ANALYSES de BIOLOGIE MEDICALES PUILBOREAU | CNR Virus des Infections Respiratoires - France SUD | Antonin Bal, Gregory Destras, Gwendolyne Burfin, Hadrien Regue, Quentin Semanas, Martine Valette, Bruno Lina, Laurence Josset |
| EPI_ISL_3382827 | LABORATOIRE UNILIANS DECINES | CNR Virus des Infections Respiratoires - France SUD | Antonin Bal, Gregory Destras, Gwendolyne Burfin, Hadrien Regue, Quentin Semanas, Martine Valette, Bruno Lina, Laurence Josset |
| EPI_ISL_3382828 | LAM BAYARD-BIOPYRENEES | CNR Virus des Infections Respiratoires - France SUD | Antonin Bal, Gregory Destras, Gwendolyne Burfin, Hadrien Regue, Quentin Semanas, Martine Valette, Bruno Lina, Laurence Josset |
| EPI_ISL_3382829 | LABORATOIRE d'ANALYSES de BIOLOGIE MEDICALES PUILBOREAU | CNR Virus des Infections Respiratoires - France SUD | Antonin Bal, Gregory Destras, Gwendolyne Burfin, Hadrien Regue, Quentin Semanas, Martine Valette, Bruno Lina, Laurence Josset |
| EPI_ISL_3382830 | LABORATOIRE UNILIANS DECINES | CNR Virus des Infections Respiratoires - France SUD | Antonin Bal, Gregory Destras, Gwendolyne Burfin, Hadrien Regue, Quentin Semanas, Martine Valette, Bruno Lina, Laurence Josset |
| EPI_ISL_3382831, EPI_ISL_3382832, EPI_ISL_3382833, EPI_ISL_3382834, EPI_ISL_3382835, EPI_ISL_3382836, EPI_ISL_3382837 | LABORATOIRE d'ANALYSES de BIOLOGIE MEDICALES PUILBOREAU | CNR Virus des Infections Respiratoires - France SUD | Antonin Bal, Gregory Destras, Gwendolyne Burfin, Hadrien Regue, Quentin Semanas, Martine Valette, Bruno Lina, Laurence Josset |
| EPI_ISL_3382838 | LABORATOIRE UNILIANS DECINES | CNR Virus des Infections Respiratoires - France SUD | Antonin Bal, Gregory Destras, Gwendolyne Burfin, Hadrien Regue, Quentin Semanas, Martine Valette, Bruno Lina, Laurence Josset |
| EPI_ISL_3382839, EPI_ISL_3382840 | LABORATOIRE d'ANALYSES de BIOLOGIE MEDICALES PUILBOREAU | CNR Virus des Infections Respiratoires - France SUD | Antonin Bal, Gregory Destras, Gwendolyne Burfin, Hadrien Regue, Quentin Semanas, Martine Valette, Bruno Lina, Laurence Josset |
| EPI_ISL_3382841 | LABORATOIRE UNILIANS DECINES | CNR Virus des Infections Respiratoires - France SUD | Antonin Bal, Gregory Destras, Gwendolyne Burfin, Hadrien Regue, Quentin Semanas, Martine Valette, Bruno Lina, Laurence Josset |
| EPI_ISL_3382842 | LABORATOIRE d'ANALYSES de BIOLOGIE MEDICALES PUILBOREAU | CNR Virus des Infections Respiratoires - France SUD | Antonin Bal, Gregory Destras, Gwendolyne Burfin, Hadrien Regue, Quentin Semanas, Martine Valette, Bruno Lina, Laurence Josset |
| EPI_ISL_3382843, EPI_ISL_3382844, EPI_ISL_3382845, EPI_ISL_3382846, EPI_ISL_3382847, EPI_ISL_3382848, EPI_ISL_3382849, EPI_ISL_3382851, EPI_ISL_3382853, EPI_ISL_3382854, EPI_ISL_3382855 | | | |
| see above | SELAS BIO 86 SITE DU PARC | CNR Virus des Infections Respiratoires - France SUD | Antonin Bal, Gregory Destras, Gwendolyne Burfin, Hadrien Regue, Quentin Semanas, Martine Valette, Bruno Lina, Laurence Josset |
| EPI_ISL_3382856 | LAM EIMER LENYS | CNR Virus des Infections Respiratoires - France SUD | Antonin Bal, Gregory Destras, Gwendolyne Burfin, Hadrien Regue, Quentin Semanas, Martine Valette, Bruno Lina, Laurence Josset |
| EPI_ISL_3382857 | SELAS BIO 86 SITE DU PARC | CNR Virus des Infections Respiratoires - France SUD | Antonin Bal, Gregory Destras, Gwendolyne Burfin, Hadrien Regue, Quentin Semanas, Martine Valette, Bruno Lina, Laurence Josset |
| EPI_ISL_3382858 | LAM EIMER LENYS | CNR Virus des Infections Respiratoires - France SUD | Antonin Bal, Gregory Destras, Gwendolyne Burfin, Hadrien Regue, Quentin Semanas, Martine Valette, Bruno Lina, Laurence Josset |
| EPI_ISL_3382859 | SELAS BIO 86 SITE DU PARC | CNR Virus des Infections Respiratoires - France SUD | Antonin Bal, Gregory Destras, Gwendolyne Burfin, Hadrien Regue, Quentin Semanas, Martine Valette, Bruno Lina, Laurence Josset |
| EPI_ISL_3382860 | LAM EIMER LENYS | CNR Virus des Infections Respiratoires - France SUD | Antonin Bal, Gregory Destras, Gwendolyne Burfin, Hadrien Regue, Quentin Semanas, Martine Valette, Bruno Lina, Laurence Josset |
| EPI_ISL_3382861 | SELAS BIO 86 SITE DU PARC | CNR Virus des Infections Respiratoires - France SUD | Antonin Bal, Gregory Destras, Gwendolyne Burfin, Hadrien Regue, Quentin Semanas, Martine Valette, Bruno Lina, Laurence Josset |
| EPI_ISL_3382862, EPI_ISL_3382863, EPI_ISL_3382864, EPI_ISL_3382865, EPI_ISL_3382866, EPI_ISL_3382867, EPI_ISL_3382868 | LAM EIMER LENYS | CNR Virus des Infections Respiratoires - France SUD | Antonin Bal, Gregory Destras, Gwendolyne Burfin, Hadrien Regue, Quentin Semanas, Martine Valette, Bruno Lina, Laurence Josset |
| EPI_ISL_3382869 | CENTRE HOSPITALIER DE BASTIA | CNR Virus des Infections Respiratoires - France SUD | Antonin Bal, Gregory Destras, Gwendolyne Burfin, Hadrien Regue, Quentin Semanas, Martine Valette, Bruno Lina, Laurence Josset |
| EPI_ISL_3382870 | HOPITAL NOTRE-DAME LA MISERICORDE | CNR Virus des Infections Respiratoires - France SUD | Antonin Bal, Gregory Destras, Gwendolyne Burfin, Hadrien Regue, Quentin Semanas, Martine Valette, Bruno Lina, Laurence Josset |
| EPI_ISL_3382871, EPI_ISL_3382872, EPI_ISL_3382873 | LAM EIMER LENYS | CNR Virus des Infections Respiratoires - France SUD | Antonin Bal, Gregory Destras, Gwendolyne Burfin, Hadrien Regue, Quentin Semanas, Martine Valette, Bruno Lina, Laurence Josset |
| EPI_ISL_3382874 | HOPITAL NOTRE-DAME LA MISERICORDE | CNR Virus des Infections Respiratoires - France SUD | Antonin Bal, Gregory Destras, Gwendolyne Burfin, Hadrien Regue, Quentin Semanas, Martine Valette, Bruno Lina, Laurence Josset |
| EPI_ISL_3382875, EPI_ISL_3382876, EPI_ISL_3382877, EPI_ISL_3382878, EPI_ISL_3382879, EPI_ISL_3382880, EPI_ISL_3382881, EPI_ISL_3382882, EPI_ISL_3382883, EPI_ISL_3382884, EPI_ISL_3382885, EPI_ISL_3382886, EPI_ISL_3382887, EPI_ISL_3382888, EPI_ISL_3382889, EPI_ISL_3382890, EPI_ISL_3382891, EPI_ISL_3382892, EPI_ISL_3382893, EPI_ISL_3382894, EPI_ISL_3382895, EPI_ISL_3382896 | | | |
| see above | LAM EIMER LENYS | CNR Virus des Infections Respiratoires - France SUD | Antonin Bal, Gregory Destras, Gwendolyne Burfin, Hadrien Regue, Quentin Semanas, Martine Valette, Bruno Lina, Laurence Josset |
| EPI_ISL_3382897, EPI_ISL_3382898, EPI_ISL_3382899 | LABORATOIRE d'ANALYSES de BIOLOGIE MEDICALES PUILBOREAU | CNR Virus des Infections Respiratoires - France SUD | Antonin Bal, Gregory Destras, Gwendolyne Burfin, Hadrien Regue, Quentin Semanas, Martine Valette, Bruno Lina, Laurence Josset |
| EPI_ISL_3382900, EPI_ISL_3382901 | LAM EIMER LENYS | CNR Virus des Infections Respiratoires - France SUD | Antonin Bal, Gregory Destras, Gwendolyne Burfin, Hadrien Regue, Quentin Semanas, Martine Valette, Bruno Lina, Laurence Josset |
| EPI_ISL_3382902 | LABORATOIRE d'ANALYSES de BIOLOGIE MEDICALES PUILBOREAU | CNR Virus des Infections Respiratoires - France SUD | Antonin Bal, Gregory Destras, Gwendolyne Burfin, Hadrien Regue, Quentin Semanas, Martine Valette, Bruno Lina, Laurence Josset |
| EPI_ISL_3382903 | LAM EIMER LENYS | CNR Virus des Infections Respiratoires - France SUD | Antonin Bal, Gregory Destras, Gwendolyne Burfin, Hadrien Regue, Quentin Semanas, Martine Valette, Bruno Lina, Laurence Josset |
| EPI_ISL_3382904 | LABORATOIRE d'ANALYSES de BIOLOGIE MEDICALES PUILBOREAU | CNR Virus des Infections Respiratoires - France SUD | Antonin Bal, Gregory Destras, Gwendolyne Burfin, Hadrien Regue, Quentin Semanas, Martine Valette, Bruno Lina, Laurence Josset |
| EPI_ISL_3382905 | LAM EIMER LENYS | CNR Virus des Infections Respiratoires - France SUD | Antonin Bal, Gregory Destras, Gwendolyne Burfin, Hadrien Regue, Quentin Semanas, Martine Valette, Bruno Lina, Laurence Josset |
| EPI_ISL_3382906, EPI_ISL_3382907 | LABORATOIRE d'ANALYSES de BIOLOGIE MEDICALES PUILBOREAU | CNR Virus des Infections Respiratoires - France SUD | Antonin Bal, Gregory Destras, Gwendolyne Burfin, Hadrien Regue, Quentin Semanas, Martine Valette, Bruno Lina, Laurence Josset |
| EPI_ISL_3382908 | LAM EIMER LENYS | CNR Virus des Infections Respiratoires - France SUD | Antonin Bal, Gregory Destras, Gwendolyne Burfin, Hadrien Regue, Quentin Semanas, Martine Valette, Bruno Lina, Laurence Josset |
| EPI_ISL_3382909 | LABORATOIRE d'ANALYSES de BIOLOGIE MEDICALES PUILBOREAU | CNR Virus des Infections Respiratoires - France SUD | Antonin Bal, Gregory Destras, Gwendolyne Burfin, Hadrien Regue, Quentin Semanas, Martine Valette, Bruno Lina, Laurence Josset |
| EPI_ISL_3382910, EPI_ISL_3382911, EPI_ISL_3382912, EPI_ISL_3382913, EPI_ISL_3382914, EPI_ISL_3382915, EPI_ISL_3382916 | LAM EIMER LENYS | CNR Virus des Infections Respiratoires - France SUD | Antonin Bal, Gregory Destras, Gwendolyne Burfin, Hadrien Regue, Quentin Semanas, Martine Valette, Bruno Lina, Laurence Josset |
| EPI_ISL_3382917, EPI_ISL_3382918, EPI_ISL_3382919, EPI_ISL_3382920, EPI_ISL_3382921, EPI_ISL_3382922, EPI_ISL_3382924 | CNR Virus des Infections Respiratoires - France SUD | CNR Virus des Infections Respiratoires - France SUD | Antonin Bal, Gregory Destras, Gwendolyne Burfin, Hadrien Regue, Quentin Semanas, Martine Valette, Bruno Lina, Laurence Josset |
| EPI_ISL_3382925, EPI_ISL_3382927, EPI_ISL_3382928 | CHR LA REUNION FELIX GUYON | CNR Virus des Infections Respiratoires - France SUD | Antonin Bal, Gregory Destras, Gwendolyne Burfin, Hadrien Regue, Quentin Semanas, Martine Valette, Bruno Lina, Laurence Josset |
| EPI_ISL_3382930 | CENTRE HOSPITALIER POITIERS | CNR Virus des Infections Respiratoires - France SUD | Antonin Bal, Gregory Destras, Gwendolyne Burfin, Hadrien Regue, Quentin Semanas, Martine Valette, Bruno Lina, Laurence Josset |
| EPI_ISL_3382931 | LAM EIMER LENYS | CNR Virus des Infections Respiratoires - France SUD | Antonin Bal, Gregory Destras, Gwendolyne Burfin, Hadrien Regue, Quentin Semanas, Martine Valette, Bruno Lina, Laurence Josset |
| EPI_ISL_3382932 | LAM DE TERRE ROUGE | CNR Virus des Infections Respiratoires - France SUD | Antonin Bal, Gregory Destras, Gwendolyne Burfin, Hadrien Regue, Quentin Semanas, Martine Valette, Bruno Lina, Laurence Josset |
| EPI_ISL_3382933, EPI_ISL_3382934 | LAM EIMER LENYS | CNR Virus des Infections Respiratoires - France SUD | Antonin Bal, Gregory Destras, Gwendolyne Burfin, Hadrien Regue, Quentin Semanas, Martine Valette, Bruno Lina, Laurence Josset |
| EPI_ISL_3382935, | LAM DE TERRE ROUGE | CNR Virus des Infections Respiratoires - France SUD | Antonin Bal, Gregory Destras, Gwendolyne Burfin, Hadrien Regue, Quentin Semanas, Martine Valette, Bruno Lina, Laurence Josset |

| | | | |
|---|---|---|---|
| EPI_ISL_3382936, EPI_ISL_3382937 | | | |
| EPI_ISL_3382938 | LAM EIMER LENYS | CNR Virus des Infections Respiratoires - France SUD | Antonin Bal, Gregory Destras, Gwendolyne Burfin, Hadrien Regue, Quentin Semanas, Martine Valette, Bruno Lina, Laurence Josset |
| EPI_ISL_3382939 | LAM DE TERRE ROUGE | CNR Virus des Infections Respiratoires - France SUD | Antonin Bal, Gregory Destras, Gwendolyne Burfin, Hadrien Regue, Quentin Semanas, Martine Valette, Bruno Lina, Laurence Josset |
| EPI_ISL_3382940 | LAM EIMER LENYS | CNR Virus des Infections Respiratoires - France SUD | Antonin Bal, Gregory Destras, Gwendolyne Burfin, Hadrien Regue, Quentin Semanas, Martine Valette, Bruno Lina, Laurence Josset |
| EPI_ISL_3382941 | LAM DE TERRE ROUGE | CNR Virus des Infections Respiratoires - France SUD | Antonin Bal, Gregory Destras, Gwendolyne Burfin, Hadrien Regue, Quentin Semanas, Martine Valette, Bruno Lina, Laurence Josset |
| EPI_ISL_3382942, EPI_ISL_3382943, EPI_ISL_3382944, EPI_ISL_3382945, EPI_ISL_3382946, EPI_ISL_3382947, EPI_ISL_3382948, EPI_ISL_3382949, EPI_ISL_3382950, EPI_ISL_3382951 | LAM EIMER LENYS | CNR Virus des Infections Respiratoires - France SUD | Antonin Bal, Gregory Destras, Gwendolyne Burfin, Hadrien Regue, Quentin Semanas, Martine Valette, Bruno Lina, Laurence Josset |
| EPI_ISL_3382952 | DYOMEDEA-LABORATOIRE DE LA SAUVEGARDE | CNR Virus des Infections Respiratoires - France SUD | Antonin Bal, Gregory Destras, Gwendolyne Burfin, Hadrien Regue, Quentin Semanas, Martine Valette, Bruno Lina, Laurence Josset |
| EPI_ISL_3382953 | CNR Virus des Infections Respiratoires - France SUD | CNR Virus des Infections Respiratoires - France SUD | Antonin Bal, Gregory Destras, Gwendolyne Burfin, Hadrien Regue, Quentin Semanas, Martine Valette, Bruno Lina, Laurence Josset |
| EPI_ISL_3382954, EPI_ISL_3382955 | DYOMEDEA-LABORATOIRE DE LA SAUVEGARDE | CNR Virus des Infections Respiratoires - France SUD | Antonin Bal, Gregory Destras, Gwendolyne Burfin, Hadrien Regue, Quentin Semanas, Martine Valette, Bruno Lina, Laurence Josset |
| EPI_ISL_3382956 | CNR Virus des Infections Respiratoires - France SUD | CNR Virus des Infections Respiratoires - France SUD | Antonin Bal, Gregory Destras, Gwendolyne Burfin, Hadrien Regue, Quentin Semanas, Martine Valette, Bruno Lina, Laurence Josset |
| EPI_ISL_3382957, EPI_ISL_3382958, EPI_ISL_3382959, EPI_ISL_3382960, EPI_ISL_3382961, EPI_ISL_3382962, EPI_ISL_3382963 | DYOMEDEA-LABORATOIRE DE LA SAUVEGARDE | CNR Virus des Infections Respiratoires - France SUD | Antonin Bal, Gregory Destras, Gwendolyne Burfin, Hadrien Regue, Quentin Semanas, Martine Valette, Bruno Lina, Laurence Josset |
| EPI_ISL_3382964 | LAM EIMER LENYS | CNR Virus des Infections Respiratoires - France SUD | Antonin Bal, Gregory Destras, Gwendolyne Burfin, Hadrien Regue, Quentin Semanas, Martine Valette, Bruno Lina, Laurence Josset |
| EPI_ISL_3382965 | DYOMEDEA-LABORATOIRE DE LA SAUVEGARDE | CNR Virus des Infections Respiratoires - France SUD | Antonin Bal, Gregory Destras, Gwendolyne Burfin, Hadrien Regue, Quentin Semanas, Martine Valette, Bruno Lina, Laurence Josset |
| EPI_ISL_3382966 | LAM EIMER LENYS | CNR Virus des Infections Respiratoires - France SUD | Antonin Bal, Gregory Destras, Gwendolyne Burfin, Hadrien Regue, Quentin Semanas, Martine Valette, Bruno Lina, Laurence Josset |
| EPI_ISL_3382967 | DYOMEDEA-LABORATOIRE DE LA SAUVEGARDE | CNR Virus des Infections Respiratoires - France SUD | Antonin Bal, Gregory Destras, Gwendolyne Burfin, Hadrien Regue, Quentin Semanas, Martine Valette, Bruno Lina, Laurence Josset |
| EPI_ISL_3382968 | LAM EIMER LENYS | CNR Virus des Infections Respiratoires - France SUD | Antonin Bal, Gregory Destras, Gwendolyne Burfin, Hadrien Regue, Quentin Semanas, Martine Valette, Bruno Lina, Laurence Josset |
| EPI_ISL_3382969 | DYOMEDEA-LABORATOIRE DE LA SAUVEGARDE | CNR Virus des Infections Respiratoires - France SUD | Antonin Bal, Gregory Destras, Gwendolyne Burfin, Hadrien Regue, Quentin Semanas, Martine Valette, Bruno Lina, Laurence Josset |
| EPI_ISL_3382970 | LAM EIMER LENYS | CNR Virus des Infections Respiratoires - France SUD | Antonin Bal, Gregory Destras, Gwendolyne Burfin, Hadrien Regue, Quentin Semanas, Martine Valette, Bruno Lina, Laurence Josset |
| EPI_ISL_3382971, EPI_ISL_3382972 | DYOMEDEA-LABORATOIRE DE LA SAUVEGARDE | CNR Virus des Infections Respiratoires - France SUD | Antonin Bal, Gregory Destras, Gwendolyne Burfin, Hadrien Regue, Quentin Semanas, Martine Valette, Bruno Lina, Laurence Josset |
| EPI_ISL_3382973 | LAM EIMER LENYS | CNR Virus des Infections Respiratoires - France SUD | Antonin Bal, Gregory Destras, Gwendolyne Burfin, Hadrien Regue, Quentin Semanas, Martine Valette, Bruno Lina, Laurence Josset |
| EPI_ISL_3382974 | DYOMEDEA-LABORATOIRE DE LA SAUVEGARDE | CNR Virus des Infections Respiratoires - France SUD | Antonin Bal, Gregory Destras, Gwendolyne Burfin, Hadrien Regue, Quentin Semanas, Martine Valette, Bruno Lina, Laurence Josset |
| EPI_ISL_3382975 | LAM EIMER LENYS | CNR Virus des Infections Respiratoires - France SUD | Antonin Bal, Gregory Destras, Gwendolyne Burfin, Hadrien Regue, Quentin Semanas, Martine Valette, Bruno Lina, Laurence Josset |
| EPI_ISL_3382976, EPI_ISL_3382977 | DYOMEDEA-LABORATOIRE DE LA SAUVEGARDE | CNR Virus des Infections Respiratoires - France SUD | Antonin Bal, Gregory Destras, Gwendolyne Burfin, Hadrien Regue, Quentin Semanas, Martine Valette, Bruno Lina, Laurence Josset |
| EPI_ISL_3382978 | LAM EIMER LENYS | CNR Virus des Infections Respiratoires - France SUD | Antonin Bal, Gregory Destras, Gwendolyne Burfin, Hadrien Regue, Quentin Semanas, Martine Valette, Bruno Lina, Laurence Josset |
| EPI_ISL_3382979, EPI_ISL_3382980, EPI_ISL_3382981 | DYOMEDEA-LABORATOIRE DE LA SAUVEGARDE | CNR Virus des Infections Respiratoires - France SUD | Antonin Bal, Gregory Destras, Gwendolyne Burfin, Hadrien Regue, Quentin Semanas, Martine Valette, Bruno Lina, Laurence Josset |
| EPI_ISL_3382982 | CH GIVORS | CNR Virus des Infections Respiratoires - France SUD | Antonin Bal, Gregory Destras, Gwendolyne Burfin, Hadrien Regue, Quentin Semanas, Martine Valette, Bruno Lina, Laurence Josset |
| EPI_ISL_3382983, EPI_ISL_3382984, EPI_ISL_3382986, EPI_ISL_3382987 | LAM EIMER LENYS | CNR Virus des Infections Respiratoires - France SUD | Antonin Bal, Gregory Destras, Gwendolyne Burfin, Hadrien Regue, Quentin Semanas, Martine Valette, Bruno Lina, Laurence Josset |
| EPI_ISL_3382988, EPI_ISL_3382990 | CNR Virus des Infections Respiratoires - France SUD | CNR Virus des Infections Respiratoires - France SUD | Antonin Bal, Gregory Destras, Gwendolyne Burfin, Hadrien Regue, Quentin Semanas, Martine Valette, Bruno Lina, Laurence Josset |
| EPI_ISL_3382993, EPI_ISL_3382994, EPI_ISL_3382995, EPI_ISL_3382996, EPI_ISL_3382997, EPI_ISL_3382998, EPI_ISL_3383000, EPI_ISL_3383001, EPI_ISL_3383002, EPI_ISL_3383003, EPI_ISL_3383004, EPI_ISL_3383005, EPI_ISL_3383006, EPI_ISL_3383007, EPI_ISL_3383008, EPI_ISL_3383009, EPI_ISL_3383010, EPI_ISL_3383011, EPI_ISL_3383012 | | | |
| see above | DYOMEDEA-LABORATOIRE DE LA SAUVEGARDE | CNR Virus des Infections Respiratoires - France SUD | Antonin Bal, Gregory Destras, Gwendolyne Burfin, Hadrien Regue, Quentin Semanas, Martine Valette, Bruno Lina, Laurence Josset |
| EPI_ISL_3383013, EPI_ISL_3383014 | CNR Virus des Infections Respiratoires - France SUD | CNR Virus des Infections Respiratoires - France SUD | Antonin Bal, Gregory Destras, Gwendolyne Burfin, Hadrien Regue, Quentin Semanas, Martine Valette, Bruno Lina, Laurence Josset |
| EPI_ISL_3383015 | CH GIVORS | CNR Virus des Infections Respiratoires - France SUD | Antonin Bal, Gregory Destras, Gwendolyne Burfin, Hadrien Regue, Quentin Semanas, Martine Valette, Bruno Lina, Laurence Josset |
| EPI_ISL_3383016, EPI_ISL_3383017 | CNR Virus des Infections Respiratoires - France SUD | CNR Virus des Infections Respiratoires - France SUD | Antonin Bal, Gregory Destras, Gwendolyne Burfin, Hadrien Regue, Quentin Semanas, Martine Valette, Bruno Lina, Laurence Josset |
| EPI_ISL_3383018, EPI_ISL_3383019, EPI_ISL_3383020, EPI_ISL_3383021, EPI_ISL_3383022, EPI_ISL_3383023, EPI_ISL_3383024, EPI_ISL_3383025, EPI_ISL_3383026, EPI_ISL_3383027, EPI_ISL_3383028 | | | |
| see above | LABORATOIRE UNILIANS DECINES | CNR Virus des Infections Respiratoires - France SUD | Antonin Bal, Gregory Destras, Gwendolyne Burfin, Hadrien Regue, Quentin Semanas, Martine Valette, Bruno Lina, Laurence Josset |
| EPI_ISL_3383029, EPI_ISL_3383030 | CNR Virus des Infections Respiratoires - France SUD | CNR Virus des Infections Respiratoires - France SUD | Antonin Bal, Gregory Destras, Gwendolyne Burfin, Hadrien Regue, Quentin Semanas, Martine Valette, Bruno Lina, Laurence Josset |
| EPI_ISL_3383031, EPI_ISL_3383032, EPI_ISL_3383033, EPI_ISL_3383034, EPI_ISL_3383035 | CENTRE HOSPITALIER DE VALENCE | CNR Virus des Infections Respiratoires - France SUD | Antonin Bal, Gregory Destras, Gwendolyne Burfin, Hadrien Regue, Quentin Semanas, Martine Valette, Bruno Lina, Laurence Josset |
| EPI_ISL_3383036 | LABORATOIRE MAYMAT | CNR Virus des Infections Respiratoires - France SUD | Antonin Bal, Gregory Destras, Gwendolyne Burfin, Hadrien Regue, Quentin Semanas, Martine Valette, Bruno Lina, Laurence Josset |
| EPI_ISL_3383037 | CENTRE HOSPITALIER DE VALENCE | CNR Virus des Infections Respiratoires - France SUD | Antonin Bal, Gregory Destras, Gwendolyne Burfin, Hadrien Regue, Quentin Semanas, Martine Valette, Bruno Lina, Laurence Josset |
| EPI_ISL_3383038 | LABORATOIRE MAYMAT | CNR Virus des Infections Respiratoires - France SUD | Antonin Bal, Gregory Destras, Gwendolyne Burfin, Hadrien Regue, Quentin Semanas, Martine Valette, Bruno Lina, Laurence Josset |
| EPI_ISL_3383039, EPI_ISL_3383041, EPI_ISL_3383042 | CENTRE HOSPITALIER DE VALENCE | CNR Virus des Infections Respiratoires - France SUD | Antonin Bal, Gregory Destras, Gwendolyne Burfin, Hadrien Regue, Quentin Semanas, Martine Valette, Bruno Lina, Laurence Josset |
| EPI_ISL_3383043 | CH GIVORS | CNR Virus des Infections Respiratoires - France SUD | Antonin Bal, Gregory Destras, Gwendolyne Burfin, Hadrien Regue, Quentin Semanas, Martine Valette, Bruno Lina, Laurence Josset |
| EPI_ISL_3383044, EPI_ISL_3383045 | CENTRE HOSPITALIER DE VALENCE | CNR Virus des Infections Respiratoires - France SUD | Antonin Bal, Gregory Destras, Gwendolyne Burfin, Hadrien Regue, Quentin Semanas, Martine Valette, Bruno Lina, Laurence Josset |
| EPI_ISL_3383046 | CENTRE HOSPITALIER EMILE ROUX | CNR Virus des Infections Respiratoires - France SUD | Antonin Bal, Gregory Destras, Gwendolyne Burfin, Hadrien Regue, Quentin Semanas, Martine Valette, Bruno Lina, Laurence Josset |
| EPI_ISL_3383047 | LABO BIOSEVRES BRESSUIRE | CNR Virus des Infections Respiratoires - France SUD | Antonin Bal, Gregory Destras, Gwendolyne Burfin, Hadrien Regue, Quentin Semanas, Martine Valette, Bruno Lina, Laurence Josset |
| EPI_ISL_3383048 | CNR Virus des Infections Respiratoires - France SUD | CNR Virus des Infections Respiratoires - France SUD | Antonin Bal, Gregory Destras, Gwendolyne Burfin, Hadrien Regue, Quentin Semanas, Martine Valette, Bruno Lina, Laurence Josset |
| EPI_ISL_3383049 | CENTRE HOSPITALIER DE TULLE | CNR Virus des Infections Respiratoires - France SUD | Antonin Bal, Gregory Destras, Gwendolyne Burfin, Hadrien Regue, Quentin Semanas, Martine Valette, Bruno Lina, Laurence Josset |
| EPI_ISL_3383050 | DYOMEDEA-LABORATOIRE DE LA SAUVEGARDE | CNR Virus des Infections Respiratoires - France SUD | Antonin Bal, Gregory Destras, Gwendolyne Burfin, Hadrien Regue, Quentin Semanas, Martine Valette, Bruno Lina, Laurence Josset |
| EPI_ISL_3383051 | CENTRE HOSPITALIER PIERRE OUDOT | CNR Virus des Infections Respiratoires - France SUD | Antonin Bal, Gregory Destras, Gwendolyne Burfin, Hadrien Regue, Quentin Semanas, Martine Valette, Bruno Lina, Laurence Josset |
| EPI_ISL_3383053, EPI_ISL_3383054, EPI_ISL_3383055 | DYOMEDEA-LABORATOIRE DE LA SAUVEGARDE | CNR Virus des Infections Respiratoires - France SUD | Antonin Bal, Gregory Destras, Gwendolyne Burfin, Hadrien Regue, Quentin Semanas, Martine Valette, Bruno Lina, Laurence Josset |
| EPI_ISL_3383056 | CERBALLIANCE MIDI-PYRENEES | CNR Virus des Infections Respiratoires - France SUD | Antonin Bal, Gregory Destras, Gwendolyne Burfin, Hadrien Regue, Quentin Semanas, Martine Valette, Bruno Lina, Laurence Josset |

| | | | |
|---|---|---|---|
| EPI_ISL_3383057 | DYOMEDEA-LABORATOIRE DE LA SAUVEGARDE | CNR Virus des Infections Respiratoires - France SUD | Antonin Bal, Gregory Destras, Gwendolyne Burfin, Hadrien Regue, Quentin Semanas, Martine Valette, Bruno Lina, Laurence Josset |
| EPI_ISL_3383058 | ASTRALAB TASSIGNY | CNR Virus des Infections Respiratoires - France SUD | Antonin Bal, Gregory Destras, Gwendolyne Burfin, Hadrien Regue, Quentin Semanas, Martine Valette, Bruno Lina, Laurence Josset |
| EPI_ISL_3383059 | DYOMEDEA-LABORATOIRE DE LA SAUVEGARDE | CNR Virus des Infections Respiratoires - France SUD | Antonin Bal, Gregory Destras, Gwendolyne Burfin, Hadrien Regue, Quentin Semanas, Martine Valette, Bruno Lina, Laurence Josset |
| EPI_ISL_3383060 | ASTRALAB TASSIGNY | CNR Virus des Infections Respiratoires - France SUD | Antonin Bal, Gregory Destras, Gwendolyne Burfin, Hadrien Regue, Quentin Semanas, Martine Valette, Bruno Lina, Laurence Josset |
| EPI_ISL_3383061 | CERBALLIANCE MIDI-PYRENEES | CNR Virus des Infections Respiratoires - France SUD | Antonin Bal, Gregory Destras, Gwendolyne Burfin, Hadrien Regue, Quentin Semanas, Martine Valette, Bruno Lina, Laurence Josset |
| EPI_ISL_3383062, EPI_ISL_3383063 | DYOMEDEA-LABORATOIRE DE LA SAUVEGARDE | CNR Virus des Infections Respiratoires - France SUD | Antonin Bal, Gregory Destras, Gwendolyne Burfin, Hadrien Regue, Quentin Semanas, Martine Valette, Bruno Lina, Laurence Josset |
| EPI_ISL_3383064 | ASTRALAB TASSIGNY | CNR Virus des Infections Respiratoires - France SUD | Antonin Bal, Gregory Destras, Gwendolyne Burfin, Hadrien Regue, Quentin Semanas, Martine Valette, Bruno Lina, Laurence Josset |
| EPI_ISL_3383065 | CERBALLIANCE MIDI-PYRENEES | CNR Virus des Infections Respiratoires - France SUD | Antonin Bal, Gregory Destras, Gwendolyne Burfin, Hadrien Regue, Quentin Semanas, Martine Valette, Bruno Lina, Laurence Josset |
| EPI_ISL_3383066 | DYOMEDEA-LABORATOIRE DE LA SAUVEGARDE | CNR Virus des Infections Respiratoires - France SUD | Antonin Bal, Gregory Destras, Gwendolyne Burfin, Hadrien Regue, Quentin Semanas, Martine Valette, Bruno Lina, Laurence Josset |
| EPI_ISL_3383067 | ASTRALAB TASSIGNY | CNR Virus des Infections Respiratoires - France SUD | Antonin Bal, Gregory Destras, Gwendolyne Burfin, Hadrien Regue, Quentin Semanas, Martine Valette, Bruno Lina, Laurence Josset |
| EPI_ISL_3383068 | CERBALLIANCE MIDI-PYRENEES | CNR Virus des Infections Respiratoires - France SUD | Antonin Bal, Gregory Destras, Gwendolyne Burfin, Hadrien Regue, Quentin Semanas, Martine Valette, Bruno Lina, Laurence Josset |
| EPI_ISL_3383069 | ASTRALAB TASSIGNY | CNR Virus des Infections Respiratoires - France SUD | Antonin Bal, Gregory Destras, Gwendolyne Burfin, Hadrien Regue, Quentin Semanas, Martine Valette, Bruno Lina, Laurence Josset |
| EPI_ISL_3383070 | CERBALLIANCE MIDI-PYRENEES | CNR Virus des Infections Respiratoires - France SUD | Antonin Bal, Gregory Destras, Gwendolyne Burfin, Hadrien Regue, Quentin Semanas, Martine Valette, Bruno Lina, Laurence Josset |
| EPI_ISL_3383071 | DYOMEDEA-LABORATOIRE DE LA SAUVEGARDE | CNR Virus des Infections Respiratoires - France SUD | Antonin Bal, Gregory Destras, Gwendolyne Burfin, Hadrien Regue, Quentin Semanas, Martine Valette, Bruno Lina, Laurence Josset |
| EPI_ISL_3383072 | ASTRALAB TASSIGNY | CNR Virus des Infections Respiratoires - France SUD | Antonin Bal, Gregory Destras, Gwendolyne Burfin, Hadrien Regue, Quentin Semanas, Martine Valette, Bruno Lina, Laurence Josset |
| EPI_ISL_3383073 | DYOMEDEA-LABORATOIRE DE LA SAUVEGARDE | CNR Virus des Infections Respiratoires - France SUD | Antonin Bal, Gregory Destras, Gwendolyne Burfin, Hadrien Regue, Quentin Semanas, Martine Valette, Bruno Lina, Laurence Josset |
| EPI_ISL_3383074, EPI_ISL_3383075 | ASTRALAB TASSIGNY | CNR Virus des Infections Respiratoires - France SUD | Antonin Bal, Gregory Destras, Gwendolyne Burfin, Hadrien Regue, Quentin Semanas, Martine Valette, Bruno Lina, Laurence Josset |
| EPI_ISL_3383076 | DYOMEDEA-LABORATOIRE DE LA SAUVEGARDE | CNR Virus des Infections Respiratoires - France SUD | Antonin Bal, Gregory Destras, Gwendolyne Burfin, Hadrien Regue, Quentin Semanas, Martine Valette, Bruno Lina, Laurence Josset |
| EPI_ISL_3383077 | ASTRALAB TASSIGNY | CNR Virus des Infections Respiratoires - France SUD | Antonin Bal, Gregory Destras, Gwendolyne Burfin, Hadrien Regue, Quentin Semanas, Martine Valette, Bruno Lina, Laurence Josset |
| EPI_ISL_3383078, EPI_ISL_3383079, EPI_ISL_3383080 | DYOMEDEA-LABORATOIRE DE LA SAUVEGARDE | CNR Virus des Infections Respiratoires - France SUD | Antonin Bal, Gregory Destras, Gwendolyne Burfin, Hadrien Regue, Quentin Semanas, Martine Valette, Bruno Lina, Laurence Josset |
| EPI_ISL_3383081, EPI_ISL_3383082 | ASTRALAB TASSIGNY | CNR Virus des Infections Respiratoires - France SUD | Antonin Bal, Gregory Destras, Gwendolyne Burfin, Hadrien Regue, Quentin Semanas, Martine Valette, Bruno Lina, Laurence Josset |
| EPI_ISL_3383083 | DYOMEDEA-LABORATOIRE DE LA SAUVEGARDE | CNR Virus des Infections Respiratoires - France SUD | Antonin Bal, Gregory Destras, Gwendolyne Burfin, Hadrien Regue, Quentin Semanas, Martine Valette, Bruno Lina, Laurence Josset |
| EPI_ISL_3383084, EPI_ISL_3383085 | ASTRALAB TASSIGNY | CNR Virus des Infections Respiratoires - France SUD | Antonin Bal, Gregory Destras, Gwendolyne Burfin, Hadrien Regue, Quentin Semanas, Martine Valette, Bruno Lina, Laurence Josset |
| EPI_ISL_3383086 | CERBALLIANCE MIDI-PYRENEES | CNR Virus des Infections Respiratoires - France SUD | Antonin Bal, Gregory Destras, Gwendolyne Burfin, Hadrien Regue, Quentin Semanas, Martine Valette, Bruno Lina, Laurence Josset |
| EPI_ISL_3383087 | DYOMEDEA-LABORATOIRE DE LA SAUVEGARDE | CNR Virus des Infections Respiratoires - France SUD | Antonin Bal, Gregory Destras, Gwendolyne Burfin, Hadrien Regue, Quentin Semanas, Martine Valette, Bruno Lina, Laurence Josset |
| EPI_ISL_3383088 | ASTRALAB TASSIGNY | CNR Virus des Infections Respiratoires - France SUD | Antonin Bal, Gregory Destras, Gwendolyne Burfin, Hadrien Regue, Quentin Semanas, Martine Valette, Bruno Lina, Laurence Josset |
| EPI_ISL_3383089 | DYOMEDEA-LABORATOIRE DE LA SAUVEGARDE | CNR Virus des Infections Respiratoires - France SUD | Antonin Bal, Gregory Destras, Gwendolyne Burfin, Hadrien Regue, Quentin Semanas, Martine Valette, Bruno Lina, Laurence Josset |
| EPI_ISL_3383090 | CERBALLIANCE MIDI-PYRENEES | CNR Virus des Infections Respiratoires - France SUD | Antonin Bal, Gregory Destras, Gwendolyne Burfin, Hadrien Regue, Quentin Semanas, Martine Valette, Bruno Lina, Laurence Josset |
| EPI_ISL_3383091 | ASTRALAB TASSIGNY | CNR Virus des Infections Respiratoires - France SUD | Antonin Bal, Gregory Destras, Gwendolyne Burfin, Hadrien Regue, Quentin Semanas, Martine Valette, Bruno Lina, Laurence Josset |
| EPI_ISL_3383092 | CNR Virus des Infections Respiratoires - France SUD | CNR Virus des Infections Respiratoires - France SUD | Antonin Bal, Gregory Destras, Gwendolyne Burfin, Hadrien Regue, Quentin Semanas, Martine Valette, Bruno Lina, Laurence Josset |
| EPI_ISL_3383093, EPI_ISL_3383094 | CERBALLIANCE MIDI-PYRENEES | CNR Virus des Infections Respiratoires - France SUD | Antonin Bal, Gregory Destras, Gwendolyne Burfin, Hadrien Regue, Quentin Semanas, Martine Valette, Bruno Lina, Laurence Josset |
| EPI_ISL_3383095 | DYOMEDEA-LABORATOIRE DE LA SAUVEGARDE | CNR Virus des Infections Respiratoires - France SUD | Antonin Bal, Gregory Destras, Gwendolyne Burfin, Hadrien Regue, Quentin Semanas, Martine Valette, Bruno Lina, Laurence Josset |
| EPI_ISL_3383096 | ASTRALAB TASSIGNY | CNR Virus des Infections Respiratoires - France SUD | Antonin Bal, Gregory Destras, Gwendolyne Burfin, Hadrien Regue, Quentin Semanas, Martine Valette, Bruno Lina, Laurence Josset |
| EPI_ISL_3383097 | CERBALLIANCE MIDI-PYRENEES | CNR Virus des Infections Respiratoires - France SUD | Antonin Bal, Gregory Destras, Gwendolyne Burfin, Hadrien Regue, Quentin Semanas, Martine Valette, Bruno Lina, Laurence Josset |
| EPI_ISL_3383098 | DYOMEDEA-LABORATOIRE DE LA SAUVEGARDE | CNR Virus des Infections Respiratoires - France SUD | Antonin Bal, Gregory Destras, Gwendolyne Burfin, Hadrien Regue, Quentin Semanas, Martine Valette, Bruno Lina, Laurence Josset |
| EPI_ISL_3383100, EPI_ISL_3383101, EPI_ISL_3383102 | CERBALLIANCE MIDI-PYRENEES | CNR Virus des Infections Respiratoires - France SUD | Antonin Bal, Gregory Destras, Gwendolyne Burfin, Hadrien Regue, Quentin Semanas, Martine Valette, Bruno Lina, Laurence Josset |
| EPI_ISL_3383103, EPI_ISL_3383104, EPI_ISL_3383106 | DYOMEDEA-LABORATOIRE DE LA SAUVEGARDE | CNR Virus des Infections Respiratoires - France SUD | Antonin Bal, Gregory Destras, Gwendolyne Burfin, Hadrien Regue, Quentin Semanas, Martine Valette, Bruno Lina, Laurence Josset |
| EPI_ISL_3383107 | CNR Virus des Infections Respiratoires - France SUD | CNR Virus des Infections Respiratoires - France SUD | Antonin Bal, Gregory Destras, Gwendolyne Burfin, Hadrien Regue, Quentin Semanas, Martine Valette, Bruno Lina, Laurence Josset |
| EPI_ISL_3383108, EPI_ISL_3383109 | DYOMEDEA-LABORATOIRE DE LA SAUVEGARDE | CNR Virus des Infections Respiratoires - France SUD | Antonin Bal, Gregory Destras, Gwendolyne Burfin, Hadrien Regue, Quentin Semanas, Martine Valette, Bruno Lina, Laurence Josset |
| EPI_ISL_3383110 | CERBALLIANCE MIDI-PYRENEES | CNR Virus des Infections Respiratoires - France SUD | Antonin Bal, Gregory Destras, Gwendolyne Burfin, Hadrien Regue, Quentin Semanas, Martine Valette, Bruno Lina, Laurence Josset |
| EPI_ISL_3383111 | DYOMEDEA-LABORATOIRE DE LA SAUVEGARDE | CNR Virus des Infections Respiratoires - France SUD | Antonin Bal, Gregory Destras, Gwendolyne Burfin, Hadrien Regue, Quentin Semanas, Martine Valette, Bruno Lina, Laurence Josset |
| EPI_ISL_3383112, EPI_ISL_3383113 | CERBALLIANCE MIDI-PYRENEES | CNR Virus des Infections Respiratoires - France SUD | Antonin Bal, Gregory Destras, Gwendolyne Burfin, Hadrien Regue, Quentin Semanas, Martine Valette, Bruno Lina, Laurence Josset |
| EPI_ISL_3383114 | DYOMEDEA-LABORATOIRE DE LA SAUVEGARDE | CNR Virus des Infections Respiratoires - France SUD | Antonin Bal, Gregory Destras, Gwendolyne Burfin, Hadrien Regue, Quentin Semanas, Martine Valette, Bruno Lina, Laurence Josset |
| EPI_ISL_3383115, EPI_ISL_3383116, EPI_ISL_3383117 | CERBALLIANCE MIDI-PYRENEES | CNR Virus des Infections Respiratoires - France SUD | Antonin Bal, Gregory Destras, Gwendolyne Burfin, Hadrien Regue, Quentin Semanas, Martine Valette, Bruno Lina, Laurence Josset |
| EPI_ISL_3383118 | DYOMEDEA-LABORATOIRE DE LA SAUVEGARDE | CNR Virus des Infections Respiratoires - France SUD | Antonin Bal, Gregory Destras, Gwendolyne Burfin, Hadrien Regue, Quentin Semanas, Martine Valette, Bruno Lina, Laurence Josset |
| EPI_ISL_3383119, EPI_ISL_3383120 | CERBALLIANCE MIDI-PYRENEES | CNR Virus des Infections Respiratoires - France SUD | Antonin Bal, Gregory Destras, Gwendolyne Burfin, Hadrien Regue, Quentin Semanas, Martine Valette, Bruno Lina, Laurence Josset |
| EPI_ISL_3383121 | DYOMEDEA-LABORATOIRE DE LA SAUVEGARDE | CNR Virus des Infections Respiratoires - France SUD | Antonin Bal, Gregory Destras, Gwendolyne Burfin, Hadrien Regue, Quentin Semanas, Martine Valette, Bruno Lina, Laurence Josset |
| EPI_ISL_3383122 | CERBALLIANCE MIDI-PYRENEES | CNR Virus des Infections Respiratoires - France SUD | Antonin Bal, Gregory Destras, Gwendolyne Burfin, Hadrien Regue, Quentin Semanas, Martine Valette, Bruno Lina, Laurence Josset |
| EPI_ISL_3383123 | DYOMEDEA-LABORATOIRE DE LA SAUVEGARDE | CNR Virus des Infections Respiratoires - France SUD | Antonin Bal, Gregory Destras, Gwendolyne Burfin, Hadrien Regue, Quentin Semanas, Martine Valette, Bruno Lina, Laurence Josset |
| EPI_ISL_3383124 | CERBALLIANCE MIDI-PYRENEES | CNR Virus des Infections Respiratoires - France SUD | Antonin Bal, Gregory Destras, Gwendolyne Burfin, Hadrien Regue, Quentin Semanas, Martine Valette, Bruno Lina, Laurence Josset |
| EPI_ISL_3383125 | DYOMEDEA-LABORATOIRE DE LA SAUVEGARDE | CNR Virus des Infections Respiratoires - France SUD | Antonin Bal, Gregory Destras, Gwendolyne Burfin, Hadrien Regue, Quentin Semanas, Martine Valette, Bruno Lina, Laurence Josset |
| EPI_ISL_3383126 | CERBALLIANCE MIDI-PYRENEES | CNR Virus des Infections Respiratoires - France SUD | Antonin Bal, Gregory Destras, Gwendolyne Burfin, Hadrien Regue, Quentin Semanas, Martine Valette, Bruno Lina, Laurence Josset |
| EPI_ISL_3383127 | DYOMEDEA-LABORATOIRE DE LA SAUVEGARDE | CNR Virus des Infections Respiratoires - France SUD | Antonin Bal, Gregory Destras, Gwendolyne Burfin, Hadrien Regue, Quentin Semanas, Martine Valette, Bruno Lina, Laurence Josset |
| EPI_ISL_3383128, EPI_ISL_3383129 | CERBALLIANCE MIDI-PYRENEES | CNR Virus des Infections Respiratoires - France SUD | Antonin Bal, Gregory Destras, Gwendolyne Burfin, Hadrien Regue, Quentin Semanas, Martine Valette, Bruno Lina, Laurence Josset |
| EPI_ISL_3383130 | DYOMEDEA-LABORATOIRE DE LA SAUVEGARDE | CNR Virus des Infections Respiratoires - France SUD | Antonin Bal, Gregory Destras, Gwendolyne Burfin, Hadrien Regue, Quentin Semanas, Martine Valette, Bruno Lina, Laurence Josset |
| EPI_ISL_3383131, EPI_ISL_3383132, EPI_ISL_3383133, EPI_ISL_3383134 | CERBALLIANCE MIDI-PYRENEES | CNR Virus des Infections Respiratoires - France SUD | Antonin Bal, Gregory Destras, Gwendolyne Burfin, Hadrien Regue, Quentin Semanas, Martine Valette, Bruno Lina, Laurence Josset |
| EPI_ISL_3383135 | DYOMEDEA-LABORATOIRE DE LA SAUVEGARDE | CNR Virus des Infections Respiratoires - France SUD | Antonin Bal, Gregory Destras, Gwendolyne Burfin, Hadrien Regue, Quentin Semanas, Martine Valette, Bruno Lina, Laurence Josset |
| EPI_ISL_3383136, EPI_ISL_3383137 | CERBALLIANCE MIDI-PYRENEES | CNR Virus des Infections Respiratoires - France SUD | Antonin Bal, Gregory Destras, Gwendolyne Burfin, Hadrien Regue, Quentin Semanas, Martine Valette, Bruno Lina, Laurence Josset |
| EPI_ISL_3383138 | DYOMEDEA-LABORATOIRE DE LA SAUVEGARDE | CNR Virus des Infections Respiratoires - France SUD | Antonin Bal, Gregory Destras, Gwendolyne Burfin, Hadrien Regue, Quentin Semanas, Martine Valette, Bruno Lina, Laurence Josset |
| EPI_ISL_3383139, EPI_ISL_3383140 | CERBALLIANCE MIDI-PYRENEES | CNR Virus des Infections Respiratoires - France SUD | Antonin Bal, Gregory Destras, Gwendolyne Burfin, Hadrien Regue, Quentin Semanas, Martine Valette, Bruno Lina, Laurence Josset |
| EPI_ISL_3383141 | DYOMEDEA-LABORATOIRE DE LA SAUVEGARDE | CNR Virus des Infections Respiratoires - France SUD | Antonin Bal, Gregory Destras, Gwendolyne Burfin, Hadrien Regue, Quentin Semanas, Martine Valette, Bruno Lina, Laurence Josset |

| | | | |
|---|---|---|---|
| EPI_ISL_3383142 | CERBALLIANCE MIDI-PYRENEES | CNR Virus des Infections Respiratoires - France SUD | Antonin Bal, Gregory Destras, Gwendolyne Burfin, Hadrien Regue, Quentin Semanas, Martine Valette, Bruno Lina, Laurence Josset |
| EPI_ISL_3383143 | DYOMEDEA-LABORATOIRE DE LA SAUVEGARDE | CNR Virus des Infections Respiratoires - France SUD | Antonin Bal, Gregory Destras, Gwendolyne Burfin, Hadrien Regue, Quentin Semanas, Martine Valette, Bruno Lina, Laurence Josset |
| EPI_ISL_3383144, EPI_ISL_3383145 | CERBALLIANCE MIDI-PYRENEES | CNR Virus des Infections Respiratoires - France SUD | Antonin Bal, Gregory Destras, Gwendolyne Burfin, Hadrien Regue, Quentin Semanas, Martine Valette, Bruno Lina, Laurence Josset |
| EPI_ISL_3383146 | DYOMEDEA-LABORATOIRE DE LA SAUVEGARDE | CNR Virus des Infections Respiratoires - France SUD | Antonin Bal, Gregory Destras, Gwendolyne Burfin, Hadrien Regue, Quentin Semanas, Martine Valette, Bruno Lina, Laurence Josset |
| EPI_ISL_3383147, EPI_ISL_3383148, EPI_ISL_3383149 | CERBALLIANCE MIDI-PYRENEES | CNR Virus des Infections Respiratoires - France SUD | Antonin Bal, Gregory Destras, Gwendolyne Burfin, Hadrien Regue, Quentin Semanas, Martine Valette, Bruno Lina, Laurence Josset |
| EPI_ISL_3383150, EPI_ISL_3383151, EPI_ISL_3383152, EPI_ISL_3383154, EPI_ISL_3383155, EPI_ISL_3383157, EPI_ISL_3383158, EPI_ISL_3383159 | LBM SEALAB DARRASSE ET ASSOCIES | CNR Virus des Infections Respiratoires - France SUD | Antonin Bal, Gregory Destras, Gwendolyne Burfin, Hadrien Regue, Quentin Semanas, Martine Valette, Bruno Lina, Laurence Josset |
| EPI_ISL_3383160 | DYOMEDEA-LABORATOIRE DE LA SAUVEGARDE | CNR Virus des Infections Respiratoires - France SUD | Antonin Bal, Gregory Destras, Gwendolyne Burfin, Hadrien Regue, Quentin Semanas, Martine Valette, Bruno Lina, Laurence Josset |
| EPI_ISL_3383161 | CNR Virus des Infections Respiratoires - France SUD | CNR Virus des Infections Respiratoires - France SUD | Antonin Bal, Gregory Destras, Gwendolyne Burfin, Hadrien Regue, Quentin Semanas, Martine Valette, Bruno Lina, Laurence Josset |
| EPI_ISL_3383167 | CENTRE HOSPITALIER PIERRE OUDOT | CNR Virus des Infections Respiratoires - France SUD | Antonin Bal, Gregory Destras, Gwendolyne Burfin, Hadrien Regue, Quentin Semanas, Martine Valette, Bruno Lina, Laurence Josset |
| EPI_ISL_3383168, EPI_ISL_3383169, EPI_ISL_3383170 | DYOMEDEA-LABORATOIRE DE LA SAUVEGARDE | CNR Virus des Infections Respiratoires - France SUD | Antonin Bal, Gregory Destras, Gwendolyne Burfin, Hadrien Regue, Quentin Semanas, Martine Valette, Bruno Lina, Laurence Josset |
| EPI_ISL_3383171, EPI_ISL_3383172 | LABORATOIRE LBA JAYAN AGEN | CNR Virus des Infections Respiratoires - France SUD | Antonin Bal, Gregory Destras, Gwendolyne Burfin, Hadrien Regue, Quentin Semanas, Martine Valette, Bruno Lina, Laurence Josset |
| EPI_ISL_3383173, EPI_ISL_3383174, EPI_ISL_3383175 | CNR Virus des Infections Respiratoires - France SUD | CNR Virus des Infections Respiratoires - France SUD | Antonin Bal, Gregory Destras, Gwendolyne Burfin, Hadrien Regue, Quentin Semanas, Martine Valette, Bruno Lina, Laurence Josset |
| EPI_ISL_3383176, EPI_ISL_3383177, EPI_ISL_3383178, EPI_ISL_3383179, EPI_ISL_3383180, EPI_ISL_3383181, EPI_ISL_3383182, EPI_ISL_3383183, EPI_ISL_3383184, EPI_ISL_3383185, EPI_ISL_3383186, EPI_ISL_3383187, EPI_ISL_3383188, EPI_ISL_3383189 | | | |
| see above | LABORATOIRE UNILIANS DECINES | CNR Virus des Infections Respiratoires - France SUD | Antonin Bal, Gregory Destras, Gwendolyne Burfin, Hadrien Regue, Quentin Semanas, Martine Valette, Bruno Lina, Laurence Josset |
| EPI_ISL_3383199, EPI_ISL_3383200, EPI_ISL_3383201 | CENTRE HOSPITALIER DE VALENCE | CNR Virus des Infections Respiratoires - France SUD | Antonin Bal, Gregory Destras, Gwendolyne Burfin, Hadrien Regue, Quentin Semanas, Martine Valette, Bruno Lina, Laurence Josset |
| EPI_ISL_3383202 | CNR Virus des Infections Respiratoires - France SUD | CNR Virus des Infections Respiratoires - France SUD | Antonin Bal, Gregory Destras, Gwendolyne Burfin, Hadrien Regue, Quentin Semanas, Martine Valette, Bruno Lina, Laurence Josset |
| EPI_ISL_3383203, EPI_ISL_3383204, EPI_ISL_3383205, EPI_ISL_3383206, EPI_ISL_3383207, EPI_ISL_3383208, EPI_ISL_3383209, EPI_ISL_3383210, EPI_ISL_3383211, EPI_ISL_3383212, EPI_ISL_3383213, EPI_ISL_3383214, EPI_ISL_3383215, EPI_ISL_3383216, EPI_ISL_3383217, EPI_ISL_3383218, EPI_ISL_3383219, EPI_ISL_3383220 | | | |
| see above | LBM SEALAB DARRASSE ET ASSOCIES | CNR Virus des Infections Respiratoires - France SUD | Antonin Bal, Gregory Destras, Gwendolyne Burfin, Hadrien Regue, Quentin Semanas, Martine Valette, Bruno Lina, Laurence Josset |
| EPI_ISL_3383221 | CNR Virus des Infections Respiratoires - France SUD | CNR Virus des Infections Respiratoires - France SUD | Antonin Bal, Gregory Destras, Gwendolyne Burfin, Hadrien Regue, Quentin Semanas, Martine Valette, Bruno Lina, Laurence Josset |
| EPI_ISL_3383222, EPI_ISL_3383223, EPI_ISL_3383224, EPI_ISL_3383226, EPI_ISL_3383227, EPI_ISL_3383228 | LBM SEALAB DARRASSE ET ASSOCIES | CNR Virus des Infections Respiratoires - France SUD | Antonin Bal, Gregory Destras, Gwendolyne Burfin, Hadrien Regue, Quentin Semanas, Martine Valette, Bruno Lina, Laurence Josset |
| EPI_ISL_3383229, EPI_ISL_3383230 | LABORATOIRE ANALYSEO | CNR Virus des Infections Respiratoires - France SUD | Antonin Bal, Gregory Destras, Gwendolyne Burfin, Hadrien Regue, Quentin Semanas, Martine Valette, Bruno Lina, Laurence Josset |
| EPI_ISL_3383231, EPI_ISL_3383232 | LAM CERBALLIANCE | CNR Virus des Infections Respiratoires - France SUD | Antonin Bal, Gregory Destras, Gwendolyne Burfin, Hadrien Regue, Quentin Semanas, Martine Valette, Bruno Lina, Laurence Josset |
| EPI_ISL_3383233 | LABORATOIRE ANALYSEO | CNR Virus des Infections Respiratoires - France SUD | Antonin Bal, Gregory Destras, Gwendolyne Burfin, Hadrien Regue, Quentin Semanas, Martine Valette, Bruno Lina, Laurence Josset |
| EPI_ISL_3383235, EPI_ISL_3383236 | LAM CERBALLIANCE | CNR Virus des Infections Respiratoires - France SUD | Antonin Bal, Gregory Destras, Gwendolyne Burfin, Hadrien Regue, Quentin Semanas, Martine Valette, Bruno Lina, Laurence Josset |
| EPI_ISL_3383237, EPI_ISL_3383238 | LABORATOIRE ANALYSEO | CNR Virus des Infections Respiratoires - France SUD | Antonin Bal, Gregory Destras, Gwendolyne Burfin, Hadrien Regue, Quentin Semanas, Martine Valette, Bruno Lina, Laurence Josset |
| EPI_ISL_3383239 | CNR Virus des Infections Respiratoires - France SUD | CNR Virus des Infections Respiratoires - France SUD | Antonin Bal, Gregory Destras, Gwendolyne Burfin, Hadrien Regue, Quentin Semanas, Martine Valette, Bruno Lina, Laurence Josset |
| EPI_ISL_3383240, EPI_ISL_3383241, EPI_ISL_3383242 | LABORATOIRE ANALYSEO | CNR Virus des Infections Respiratoires - France SUD | Antonin Bal, Gregory Destras, Gwendolyne Burfin, Hadrien Regue, Quentin Semanas, Martine Valette, Bruno Lina, Laurence Josset |
| EPI_ISL_3383243 | LABORATOIRE LABAZUR | CNR Virus des Infections Respiratoires - France SUD | Antonin Bal, Gregory Destras, Gwendolyne Burfin, Hadrien Regue, Quentin Semanas, Martine Valette, Bruno Lina, Laurence Josset |
| EPI_ISL_3383245 | LAM BIORYLIS | CNR Virus des Infections Respiratoires - France SUD | Antonin Bal, Gregory Destras, Gwendolyne Burfin, Hadrien Regue, Quentin Semanas, Martine Valette, Bruno Lina, Laurence Josset |
| EPI_ISL_3383246 | LABORATOIRE LABAZUR | CNR Virus des Infections Respiratoires - France SUD | Antonin Bal, Gregory Destras, Gwendolyne Burfin, Hadrien Regue, Quentin Semanas, Martine Valette, Bruno Lina, Laurence Josset |
| EPI_ISL_3383247 | LABO HOSPILAB 47 SITE D AGEN | CNR Virus des Infections Respiratoires - France SUD | Antonin Bal, Gregory Destras, Gwendolyne Burfin, Hadrien Regue, Quentin Semanas, Martine Valette, Bruno Lina, Laurence Josset |
| EPI_ISL_3383248 | LABORATOIRE LABAZUR | CNR Virus des Infections Respiratoires - France SUD | Antonin Bal, Gregory Destras, Gwendolyne Burfin, Hadrien Regue, Quentin Semanas, Martine Valette, Bruno Lina, Laurence Josset |
| EPI_ISL_3383249 | LAM BIORYLIS | CNR Virus des Infections Respiratoires - France SUD | Antonin Bal, Gregory Destras, Gwendolyne Burfin, Hadrien Regue, Quentin Semanas, Martine Valette, Bruno Lina, Laurence Josset |
| EPI_ISL_3383250 | LABO HOSPILAB 47 SITE D AGEN | CNR Virus des Infections Respiratoires - France SUD | Antonin Bal, Gregory Destras, Gwendolyne Burfin, Hadrien Regue, Quentin Semanas, Martine Valette, Bruno Lina, Laurence Josset |
| EPI_ISL_3383251 | LABORATOIRE BOUVIER | CNR Virus des Infections Respiratoires - France SUD | Antonin Bal, Gregory Destras, Gwendolyne Burfin, Hadrien Regue, Quentin Semanas, Martine Valette, Bruno Lina, Laurence Josset |
| EPI_ISL_3383252, EPI_ISL_3383253 | LAM BIORYLIS | CNR Virus des Infections Respiratoires - France SUD | Antonin Bal, Gregory Destras, Gwendolyne Burfin, Hadrien Regue, Quentin Semanas, Martine Valette, Bruno Lina, Laurence Josset |
| EPI_ISL_3383254 | LAM CERBALLIANCE | CNR Virus des Infections Respiratoires - France SUD | Antonin Bal, Gregory Destras, Gwendolyne Burfin, Hadrien Regue, Quentin Semanas, Martine Valette, Bruno Lina, Laurence Josset |
| EPI_ISL_3383255 | LAM BIORYLIS | CNR Virus des Infections Respiratoires - France SUD | Antonin Bal, Gregory Destras, Gwendolyne Burfin, Hadrien Regue, Quentin Semanas, Martine Valette, Bruno Lina, Laurence Josset |
| EPI_ISL_3383256, EPI_ISL_3383257, EPI_ISL_3383258, EPI_ISL_3383259, EPI_ISL_3383260, EPI_ISL_3383261, EPI_ISL_3383262 | LAM CERBALLIANCE | CNR Virus des Infections Respiratoires - France SUD | Antonin Bal, Gregory Destras, Gwendolyne Burfin, Hadrien Regue, Quentin Semanas, Martine Valette, Bruno Lina, Laurence Josset |
| EPI_ISL_3383263 | LAM BIORYLIS | CNR Virus des Infections Respiratoires - France SUD | Antonin Bal, Gregory Destras, Gwendolyne Burfin, Hadrien Regue, Quentin Semanas, Martine Valette, Bruno Lina, Laurence Josset |
| EPI_ISL_3383264, EPI_ISL_3383265 | LAM CERBALLIANCE | CNR Virus des Infections Respiratoires - France SUD | Antonin Bal, Gregory Destras, Gwendolyne Burfin, Hadrien Regue, Quentin Semanas, Martine Valette, Bruno Lina, Laurence Josset |
| EPI_ISL_3383266 | LAM BIORYLIS | CNR Virus des Infections Respiratoires - France SUD | Antonin Bal, Gregory Destras, Gwendolyne Burfin, Hadrien Regue, Quentin Semanas, Martine Valette, Bruno Lina, Laurence Josset |
| EPI_ISL_3383267 | LAM CERBALLIANCE | CNR Virus des Infections Respiratoires - France SUD | Antonin Bal, Gregory Destras, Gwendolyne Burfin, Hadrien Regue, Quentin Semanas, Martine Valette, Bruno Lina, Laurence Josset |
| EPI_ISL_3383268 | LAM BIORYLIS | CNR Virus des Infections Respiratoires - France SUD | Antonin Bal, Gregory Destras, Gwendolyne Burfin, Hadrien Regue, Quentin Semanas, Martine Valette, Bruno Lina, Laurence Josset |
| EPI_ISL_3383269 | LAM CERBALLIANCE | CNR Virus des Infections Respiratoires - France SUD | Antonin Bal, Gregory Destras, Gwendolyne Burfin, Hadrien Regue, Quentin Semanas, Martine Valette, Bruno Lina, Laurence Josset |
| EPI_ISL_3383270, EPI_ISL_3383271 | LAM BIORYLIS | CNR Virus des Infections Respiratoires - France SUD | Antonin Bal, Gregory Destras, Gwendolyne Burfin, Hadrien Regue, Quentin Semanas, Martine Valette, Bruno Lina, Laurence Josset |
| EPI_ISL_3383272 | LAM CERBALLIANCE | CNR Virus des Infections Respiratoires - France SUD | Antonin Bal, Gregory Destras, Gwendolyne Burfin, Hadrien Regue, Quentin Semanas, Martine Valette, Bruno Lina, Laurence Josset |
| EPI_ISL_3383273 | CNR Virus des Infections Respiratoires - France SUD | CNR Virus des Infections Respiratoires - France SUD | Antonin Bal, Gregory Destras, Gwendolyne Burfin, Hadrien Regue, Quentin Semanas, Martine Valette, Bruno Lina, Laurence Josset |
| EPI_ISL_3383274 | LAM CERBALLIANCE | CNR Virus des Infections Respiratoires - France SUD | Antonin Bal, Gregory Destras, Gwendolyne Burfin, Hadrien Regue, Quentin Semanas, Martine Valette, Bruno Lina, Laurence Josset |
| EPI_ISL_3383275 | LAM BIORYLIS | CNR Virus des Infections Respiratoires - France SUD | Antonin Bal, Gregory Destras, Gwendolyne Burfin, Hadrien Regue, Quentin Semanas, Martine Valette, Bruno Lina, Laurence Josset |
| EPI_ISL_3383276 | LAM CERBALLIANCE | CNR Virus des Infections Respiratoires - France SUD | Antonin Bal, Gregory Destras, Gwendolyne Burfin, Hadrien Regue, Quentin Semanas, Martine Valette, Bruno Lina, Laurence Josset |

| | | | |
|---|---|---|---|
| EPI_ISL_3383277 | CNR Virus des Infections Respiratoires - France SUD | CNR Virus des Infections Respiratoires - France SUD | Antonin Bal, Gregory Destras, Gwendolyne Burfin, Hadrien Regue, Quentin Semanas, Martine Valette, Bruno Lina, Laurence Josset |
| EPI_ISL_3383278 | LAM BIORYLIS | CNR Virus des Infections Respiratoires - France SUD | Antonin Bal, Gregory Destras, Gwendolyne Burfin, Hadrien Regue, Quentin Semanas, Martine Valette, Bruno Lina, Laurence Josset |
| EPI_ISL_3383279, EPI_ISL_3383280, EPI_ISL_3383281 | LAM CERBALLIANCE | CNR Virus des Infections Respiratoires - France SUD | Antonin Bal, Gregory Destras, Gwendolyne Burfin, Hadrien Regue, Quentin Semanas, Martine Valette, Bruno Lina, Laurence Josset |
| EPI_ISL_3383282 | LABORATOIRE UNILIANS DECINES | CNR Virus des Infections Respiratoires - France SUD | Antonin Bal, Gregory Destras, Gwendolyne Burfin, Hadrien Regue, Quentin Semanas, Martine Valette, Bruno Lina, Laurence Josset |
| EPI_ISL_3383283, EPI_ISL_3383284, EPI_ISL_3383285, EPI_ISL_3383286 | LAM CERBALLIANCE | CNR Virus des Infections Respiratoires - France SUD | Antonin Bal, Gregory Destras, Gwendolyne Burfin, Hadrien Regue, Quentin Semanas, Martine Valette, Bruno Lina, Laurence Josset |
| EPI_ISL_3383288 | LABORATOIRE UNILIANS DECINES | CNR Virus des Infections Respiratoires - France SUD | Antonin Bal, Gregory Destras, Gwendolyne Burfin, Hadrien Regue, Quentin Semanas, Martine Valette, Bruno Lina, Laurence Josset |
| EPI_ISL_3383289, EPI_ISL_3383290 | LAM CERBALLIANCE | CNR Virus des Infections Respiratoires - France SUD | Antonin Bal, Gregory Destras, Gwendolyne Burfin, Hadrien Regue, Quentin Semanas, Martine Valette, Bruno Lina, Laurence Josset |
| EPI_ISL_3383291, EPI_ISL_3383292 | LABORATOIRE UNILIANS DECINES | CNR Virus des Infections Respiratoires - France SUD | Antonin Bal, Gregory Destras, Gwendolyne Burfin, Hadrien Regue, Quentin Semanas, Martine Valette, Bruno Lina, Laurence Josset |
| EPI_ISL_3383293 | LAM CERBALLIANCE | CNR Virus des Infections Respiratoires - France SUD | Antonin Bal, Gregory Destras, Gwendolyne Burfin, Hadrien Regue, Quentin Semanas, Martine Valette, Bruno Lina, Laurence Josset |
| EPI_ISL_3383296, EPI_ISL_3383298, EPI_ISL_3383299, EPI_ISL_3383300, EPI_ISL_3383301, EPI_ISL_3383302, EPI_ISL_3383303, EPI_ISL_3383304, EPI_ISL_3383305, EPI_ISL_3383306, EPI_ISL_3383307, EPI_ISL_3383308, EPI_ISL_3383309, EPI_ISL_3383310, EPI_ISL_3383311, EPI_ISL_3383312, EPI_ISL_3383313, EPI_ISL_3383314, EPI_ISL_3383315, EPI_ISL_3383316, EPI_ISL_3383317, EPI_ISL_3383318, EPI_ISL_3383319, EPI_ISL_3383320, EPI_ISL_3383321, EPI_ISL_3383322, EPI_ISL_3383323, EPI_ISL_3383324, EPI_ISL_3383325, EPI_ISL_3383326 | | | |
| see above | National Virus Reference Laboratory | National Virus Reference Laboratory | Guerrino Macori, Gabriel Gonzalez, Michael Carr, Zoe Yandle, Charlene Bennett, Jonathan Dean, Seamus Fanning, Cillian F De Gascun |
| EPI_ISL_3383355, EPI_ISL_3383356, EPI_ISL_3383357, EPI_ISL_3383358, EPI_ISL_3383359, EPI_ISL_3383360, EPI_ISL_3383361, EPI_ISL_3383362, EPI_ISL_3383363, EPI_ISL_3383364 | Wyoming Public Health Laboratory | Wyoming Public Health Laboratory | Jim Mildenberger, Taylor Fearing, Channing Weber, Ashley Norberg, Chayse Rowley, Marley Goetz, Brian Dominguez, Elliot Thomasson, Sam Britz, Cari Sloma, Robert Petit, and Rob Christensen |
| EPI_ISL_3383365, EPI_ISL_3383366, EPI_ISL_3383367, EPI_ISL_3383368, EPI_ISL_3383369, EPI_ISL_3383370, EPI_ISL_3383371, EPI_ISL_3383373, EPI_ISL_3383374, EPI_ISL_3383375, EPI_ISL_3383376, EPI_ISL_3383377, EPI_ISL_3383378, EPI_ISL_3383379, EPI_ISL_3383380, EPI_ISL_3383381, EPI_ISL_3383382, EPI_ISL_3383383, EPI_ISL_3383384, EPI_ISL_3383385, EPI_ISL_3383386, EPI_ISL_3383387, EPI_ISL_3383388, EPI_ISL_3383389, EPI_ISL_3383390, EPI_ISL_3383391, EPI_ISL_3383392, EPI_ISL_3383393, EPI_ISL_3383394, EPI_ISL_3383395, EPI_ISL_3383396, EPI_ISL_3383397, EPI_ISL_3383398, EPI_ISL_3383399, EPI_ISL_3383400, EPI_ISL_3383401, EPI_ISL_3383402 | | | |
| see above | Fondazione IRCCS Ca' Granda Ospedale Maggiore Policlinico | Fondazione IRCCS Ca' Granda Ospedale Maggiore Policlinico | Ferruccio Ceriotti, Sara Uceda Renteria |
| EPI_ISL_3383405 | CEDIBIO UNILABS | CHU Purpan - Laboratoire de Virologie - Institut Fédératif de Biologie | Latour J., Jeanne N., Ranger N., Carcenac R., Harter A., Boyer P., Ferrer V., Tremeaux R., Izopet J. |
| EPI_ISL_3383410 | Diagnostic and Research Center of Infectious Diseases, Medical Faculty, Andalas University | Genomik Solidaritas Indonesia Laboratorium / Diagnostic and Research Center of Infectious Diseases, Medical Faculty, Andalas University | Vania Gavrila Wikasa, Carissa Sintca Wijaya, Andani Eka Putra, Dede Rahman Agustian, Linosefa, Himawan Masyhuri, Dhahlia Agustina Cahyono, Gracia Felias Enos Korompis, Annisa Muthiah Sukirman, Anuraj Shankar, Louisa Markus, Meutia Ayuputeri Kumaheri, Ariel Pradipta |
| EPI_ISL_3383420, EPI_ISL_3383421 | Diagnostic and Research Center of Infectious Diseases, Medical Faculty, Andalas University | Genomik Solidaritas Indonesia Laboratorium / Diagnostic and Research Center of Infectious Diseases, Medical Faculty, Andalas University | Carissa Sintca Wijaya, Vania Gavrila Wikasa, Andani Eka Putra, Dede Rahman Agustian, Linosefa, Himawan Masyhuri, Dhahlia Agustina Cahyono, Gracia Felias Enos Korompis, Annisa Muthiah Sukirman, Anuraj Shankar, Louisa Markus, Meutia Ayuputeri Kumaheri, Ariel Pradipta |
| EPI_ISL_3383424, EPI_ISL_3383426, EPI_ISL_3383427, EPI_ISL_3383428, EPI_ISL_3383430, EPI_ISL_3383431 | Diagnostic and Research Center of Infectious Diseases, Medical Faculty, Andalas University | Genomik Solidaritas Indonesia Laboratorium / Diagnostic and Research Center of Infectious Diseases, Medical Faculty, Andalas University | Dhahlia Agustina Cahyono, Himawan Masyhuri, Andani Eka Putra, Dede Rahman Agustian, Linosefa, Vania Gavrila Wikasa, Carissa Sintca Wijaya, Gracia Felias Enos Korompis, Annisa Muthiah Sukirman, Anuraj Shankar, Louisa Markus, Meutia Ayuputeri Kumaheri, Ariel Pradipta |
| EPI_ISL_3383434, EPI_ISL_3383435, EPI_ISL_3383436, EPI_ISL_3383439 | Diagnostic and Research Center of Infectious Diseases, Medical Faculty, Andalas University | Genomik Solidaritas Indonesia Laboratorium / Diagnostic and Research Center of Infectious Diseases, Medical Faculty, Andalas University | Himawan Masyhuri, Dhahlia Agustina Cahyono, Andani Eka Putra, Dede Rahman Agustian, Linosefa, Vania Gavrila Wikasa, Carissa Sintca Wijaya, Gracia Felias Enos Korompis, Annisa Muthiah Sukirman, Anuraj Shankar, Louisa Markus, Meutia Ayuputeri Kumaheri, Ariel Pradipta |
| EPI_ISL_3383441, EPI_ISL_3383442, EPI_ISL_3383443, EPI_ISL_3383444, EPI_ISL_3383445, EPI_ISL_3383446, EPI_ISL_3383447, EPI_ISL_3383448, EPI_ISL_3383449, EPI_ISL_3383450, EPI_ISL_3383451, EPI_ISL_3383452, EPI_ISL_3383453, EPI_ISL_3383454, EPI_ISL_3383455, EPI_ISL_3383456, EPI_ISL_3383457, EPI_ISL_3383458, EPI_ISL_3383459, EPI_ISL_3383460, EPI_ISL_3383461, EPI_ISL_3383462, EPI_ISL_3383463, EPI_ISL_3383464, EPI_ISL_3383465, EPI_ISL_3383466, EPI_ISL_3383467, EPI_ISL_3383468, EPI_ISL_3383469, EPI_ISL_3383470, EPI_ISL_3383471, EPI_ISL_3383472, EPI_ISL_3383473, EPI_ISL_3383474, EPI_ISL_3383475, EPI_ISL_3383476, EPI_ISL_3383477, EPI_ISL_3383478, EPI_ISL_3383479, EPI_ISL_3383480, EPI_ISL_3383481, EPI_ISL_3383482, EPI_ISL_3383483, EPI_ISL_3383484, EPI_ISL_3383485, EPI_ISL_3383486, EPI_ISL_3383487, EPI_ISL_3383488, EPI_ISL_3383489, EPI_ISL_3383490, EPI_ISL_3383491, EPI_ISL_3383492 | | | |
| see above | Department of Bacteria, Parasites and Fungi, Statens Serum Institut, Copenhagen, Denmark | Statens Serum Institut Bioinformatics and Microbial Genomics | Danish Covid-19 Genome Consortium |
| EPI_ISL_3383493 | Molekylær Medicinsk Afdeling, Aarhus University Hospital, Aarhus, Denmark | Statens Serum Institut Bioinformatics and Microbial Genomics | Danish Covid-19 Genome Consortium |
| EPI_ISL_3383494, EPI_ISL_3383495, EPI_ISL_3383497, EPI_ISL_3383498, EPI_ISL_3383499, EPI_ISL_3383500, EPI_ISL_3383501, EPI_ISL_3383502, EPI_ISL_3383503, EPI_ISL_3383504, EPI_ISL_3383505, EPI_ISL_3383506, EPI_ISL_3383507, EPI_ISL_3383508, EPI_ISL_3383509, EPI_ISL_3383510, EPI_ISL_3383511, EPI_ISL_3383512, EPI_ISL_3383513, EPI_ISL_3383514, EPI_ISL_3383515, EPI_ISL_3383516, EPI_ISL_3383517, EPI_ISL_3383518, EPI_ISL_3383519, EPI_ISL_3383520, EPI_ISL_3383521, EPI_ISL_3383522, EPI_ISL_3383523, EPI_ISL_3383524, EPI_ISL_3383525, EPI_ISL_3383526 | | | |
| see above | Department of Bacteria, Parasites and Fungi, Statens Serum Institut, Copenhagen, Denmark | Statens Serum Institut Bioinformatics and Microbial Genomics | Danish Covid-19 Genome Consortium |
| EPI_ISL_3383527 | Molekylær Medicinsk Afdeling, Aarhus University Hospital, Aarhus, Denmark | Statens Serum Institut Bioinformatics and Microbial Genomics | Danish Covid-19 Genome Consortium |
| EPI_ISL_3383528, EPI_ISL_3383529, EPI_ISL_3383530, EPI_ISL_3383531, EPI_ISL_3383532, EPI_ISL_3383533, EPI_ISL_3383534, EPI_ISL_3383535, EPI_ISL_3383536, EPI_ISL_3383537, EPI_ISL_3383538, EPI_ISL_3383539, EPI_ISL_3383540, EPI_ISL_3383541, EPI_ISL_3383542, EPI_ISL_3383543, EPI_ISL_3383544, EPI_ISL_3383545, EPI_ISL_3383546, EPI_ISL_3383547, EPI_ISL_3383548, EPI_ISL_3383549, EPI_ISL_3383550, EPI_ISL_3383551, EPI_ISL_3383552, EPI_ISL_3383553, EPI_ISL_3383554, EPI_ISL_3383555, EPI_ISL_3383556, EPI_ISL_3383557, EPI_ISL_3383558, EPI_ISL_3383559, EPI_ISL_3383560, EPI_ISL_3383561, EPI_ISL_3383562, EPI_ISL_3383563, EPI_ISL_3383564 | | | |
| see above | Department of Bacteria, Parasites and Fungi, Statens Serum Institut, Copenhagen, Denmark | Statens Serum Institut Bioinformatics and Microbial Genomics | Danish Covid-19 Genome Consortium |
| EPI_ISL_3383565 | Molekylær Medicinsk Afdeling, Aarhus University Hospital, Aarhus, Denmark | Statens Serum Institut Bioinformatics and Microbial Genomics | Danish Covid-19 Genome Consortium |
| EPI_ISL_3383566, EPI_ISL_3383567, EPI_ISL_3383568, EPI_ISL_3383569, EPI_ISL_3383570, EPI_ISL_3383571, EPI_ISL_3383572, EPI_ISL_3383573, EPI_ISL_3383574, EPI_ISL_3383575, EPI_ISL_3383576, EPI_ISL_3383577, EPI_ISL_3383578, EPI_ISL_3383579, EPI_ISL_3383580, EPI_ISL_3383581, EPI_ISL_3383582, EPI_ISL_3383583, EPI_ISL_3383584, EPI_ISL_3383585, EPI_ISL_3383586, EPI_ISL_3383587, EPI_ISL_3383588, EPI_ISL_3383589, EPI_ISL_3383590, EPI_ISL_3383591, EPI_ISL_3383592, EPI_ISL_3383593, EPI_ISL_3383594, EPI_ISL_3383595, EPI_ISL_3383596, EPI_ISL_3383597, EPI_ISL_3383599, EPI_ISL_3383600, EPI_ISL_3383601, EPI_ISL_3383602, EPI_ISL_3383603, EPI_ISL_3383604, EPI_ISL_3383605, EPI_ISL_3383606, EPI_ISL_3383607, EPI_ISL_3383608, EPI_ISL_3383609, EPI_ISL_3383610, EPI_ISL_3383611, EPI_ISL_3383612, EPI_ISL_3383613, EPI_ISL_3383614, EPI_ISL_3383615, EPI_ISL_3383616, EPI_ISL_3383617, EPI_ISL_3383618, EPI_ISL_3383619, EPI_ISL_3383620, EPI_ISL_3383621, EPI_ISL_3383622, EPI_ISL_3383623, EPI_ISL_3383624, EPI_ISL_3383625, EPI_ISL_3383626, EPI_ISL_3383627, EPI_ISL_3383628, EPI_ISL_3383629, EPI_ISL_3383630, EPI_ISL_3383631, EPI_ISL_3383632, EPI_ISL_3383633, EPI_ISL_3383634, EPI_ISL_3383635, EPI_ISL_3383636, EPI_ISL_3383637, EPI_ISL_3383638, EPI_ISL_3383639, EPI_ISL_3383640, EPI_ISL_3383642, EPI_ISL_3383643, EPI_ISL_3383645, EPI_ISL_3383647, EPI_ISL_3383648, EPI_ISL_3383649, EPI_ISL_3383650, EPI_ISL_3383651, EPI_ISL_3383652, EPI_ISL_3383653, EPI_ISL_3383654, EPI_ISL_3383655, EPI_ISL_3383656, EPI_ISL_3383657, EPI_ISL_3383658, EPI_ISL_3383659, EPI_ISL_3383660, EPI_ISL_3383661, EPI_ISL_3383662, EPI_ISL_3383663, EPI_ISL_3383664, EPI_ISL_3383665, EPI_ISL_3383666, EPI_ISL_3383667, EPI_ISL_3383668, EPI_ISL_3383669, EPI_ISL_3383670, EPI_ISL_3383671, EPI_ISL_3383672, EPI_ISL_3383673, EPI_ISL_3383674, EPI_ISL_3383675, EPI_ISL_3383676, EPI_ISL_3383677, EPI_ISL_3383678, EPI_ISL_3383679, EPI_ISL_3383680, EPI_ISL_3383681, EPI_ISL_3383682, EPI_ISL_3383683, EPI_ISL_3383684, EPI_ISL_3383685, EPI_ISL_3383686, EPI_ISL_3383687, EPI_ISL_3383688, EPI_ISL_3383689, EPI_ISL_3383690, EPI_ISL_3383691, EPI_ISL_3383692, EPI_ISL_3383693, EPI_ISL_3383694, EPI_ISL_3383695, EPI_ISL_3383696, EPI_ISL_3383697, EPI_ISL_3383698 | | | |
| see above | Department of Bacteria, Parasites and Fungi, Statens Serum Institut, Copenhagen, Denmark | Statens Serum Institut Bioinformatics and Microbial Genomics | Danish Covid-19 Genome Consortium |
| EPI_ISL_3383699 | Department of Clinical Microbiology and Center for Genomic Medicine, Rigshospitalet, Copenhagen, Denmark | Statens Serum Institut Bioinformatics and Microbial Genomics | Danish Covid-19 Genome Consortium |
| EPI_ISL_3383700, EPI_ISL_3383701, EPI_ISL_3383702, EPI_ISL_3383703, EPI_ISL_3383704, EPI_ISL_3383705, EPI_ISL_3383706, EPI_ISL_3383707, EPI_ISL_3383708, EPI_ISL_3383709, EPI_ISL_3383710, EPI_ISL_3383711, EPI_ISL_3383712, EPI_ISL_3383713, EPI_ISL_3383714, EPI_ISL_3383715, EPI_ISL_3383716, EPI_ISL_3383717, EPI_ISL_3383718, EPI_ISL_3383719, EPI_ISL_3383720, EPI_ISL_3383721, EPI_ISL_3383722, EPI_ISL_3383723, EPI_ISL_3383724, EPI_ISL_3383725, EPI_ISL_3383726, EPI_ISL_3383727, EPI_ISL_3383728, EPI_ISL_3383729, EPI_ISL_3383730, EPI_ISL_3383731, EPI_ISL_3383732, EPI_ISL_3383733, EPI_ISL_3383734, EPI_ISL_3383735, EPI_ISL_3383736, EPI_ISL_3383737, EPI_ISL_3383738, EPI_ISL_3383739, EPI_ISL_3383740, EPI_ISL_3383741, EPI_ISL_3383742, EPI_ISL_3383743, EPI_ISL_3383744, EPI_ISL_3383745, EPI_ISL_3383746, EPI_ISL_3383747, EPI_ISL_3383748, EPI_ISL_3383749, EPI_ISL_3383750, EPI_ISL_3383751, EPI_ISL_3383752, EPI_ISL_3383753, EPI_ISL_3383754, EPI_ISL_3383755, EPI_ISL_3383756, EPI_ISL_3383757, EPI_ISL_3383758, EPI_ISL_3383759, EPI_ISL_3383760, EPI_ISL_3383761, EPI_ISL_3383762, EPI_ISL_3383763, EPI_ISL_3383764, EPI_ISL_3383765, EPI_ISL_3383766, EPI_ISL_3383767, EPI_ISL_3383768, EPI_ISL_3383769, EPI_ISL_3383770, EPI_ISL_3383771, EPI_ISL_3383772, EPI_ISL_3383773, EPI_ISL_3383774, EPI_ISL_3383775, EPI_ISL_3383776, EPI_ISL_3383777, EPI_ISL_3383778, EPI_ISL_3383779, EPI_ISL_3383780, EPI_ISL_3383781, EPI_ISL_3383782 | | | |
| see above | Department of Bacteria, Parasites and Fungi, Statens Serum Institut, Copenhagen, Denmark | Statens Serum Institut Bioinformatics and Microbial Genomics | Danish Covid-19 Genome Consortium |
| EPI_ISL_3383783 | Molekylær Medicinsk Afdeling, Aarhus University Hospital, Aarhus, Denmark | Statens Serum Institut Bioinformatics and Microbial Genomics | Danish Covid-19 Genome Consortium |
| EPI_ISL_3383784, EPI_ISL_3383785, EPI_ISL_3383786, EPI_ISL_3383787, EPI_ISL_3383788, EPI_ISL_3383789, EPI_ISL_3383790, EPI_ISL_3383791, | Department of Bacteria, Parasites and Fungi, Statens Serum Institut, Copenhagen, Denmark | Statens Serum Institut Bioinformatics and Microbial Genomics | Danish Covid-19 Genome Consortium |

| | | | |
|---|---|---|---|
| EPI_ISL_3383792, EPI_ISL_3383793 | | | |
| EPI_ISL_3383794 | Molekylær Medicinsk Afdeling, Aarhus University Hospital, Aarhus, Denmark | Statens Serum Institut Bioinformatics and Microbial Genomics | Danish Covid-19 Genome Consortium |
| EPI_ISL_3383795, EPI_ISL_3383796, EPI_ISL_3383797, EPI_ISL_3383798, EPI_ISL_3383799, EPI_ISL_3383800, EPI_ISL_3383801, EPI_ISL_3383802, EPI_ISL_3383803, EPI_ISL_3383804, EPI_ISL_3383805, EPI_ISL_3383806, EPI_ISL_3383807, EPI_ISL_3383808, EPI_ISL_3383809, EPI_ISL_3383810, EPI_ISL_3383811, EPI_ISL_3383812, EPI_ISL_3383813, EPI_ISL_3383814, EPI_ISL_3383815, EPI_ISL_3383816, EPI_ISL_3383817, EPI_ISL_3383818, EPI_ISL_3383819, EPI_ISL_3383820, EPI_ISL_3383821, EPI_ISL_3383822, EPI_ISL_3383823, EPI_ISL_3383824, EPI_ISL_3383825, EPI_ISL_3383826, EPI_ISL_3383827, EPI_ISL_3383828, EPI_ISL_3383829, EPI_ISL_3383830, EPI_ISL_3383831, EPI_ISL_3383832, EPI_ISL_3383833, EPI_ISL_3383834, EPI_ISL_3383835, EPI_ISL_3383836, EPI_ISL_3383837, EPI_ISL_3383838, EPI_ISL_3383839, EPI_ISL_3383840, EPI_ISL_3383841, EPI_ISL_3383842, EPI_ISL_3383843, EPI_ISL_3383844, EPI_ISL_3383845, EPI_ISL_3383846, EPI_ISL_3383847, EPI_ISL_3383848, EPI_ISL_3383849, EPI_ISL_3383851, EPI_ISL_3383852, EPI_ISL_3383853, EPI_ISL_3383854, EPI_ISL_3383855, EPI_ISL_3383856, EPI_ISL_3383857, EPI_ISL_3383858, EPI_ISL_3383859, EPI_ISL_3383860, EPI_ISL_3383861, EPI_ISL_3383862, EPI_ISL_3383863, EPI_ISL_3383864, EPI_ISL_3383865, EPI_ISL_3383866, EPI_ISL_3383867, EPI_ISL_3383868, EPI_ISL_3383869, EPI_ISL_3383870, EPI_ISL_3383871, EPI_ISL_3383872, EPI_ISL_3383873, EPI_ISL_3383874, EPI_ISL_3383875 | | | |
| see above | Department of Bacteria, Parasites and Fungi, Statens Serum Institut, Copenhagen, Denmark | Statens Serum Institut Bioinformatics and Microbial Genomics | Danish Covid-19 Genome Consortium |
| EPI_ISL_3383876 | Molekylær Medicinsk Afdeling, Aarhus University Hospital, Aarhus, Denmark | Statens Serum Institut Bioinformatics and Microbial Genomics | Danish Covid-19 Genome Consortium |
| EPI_ISL_3383877, EPI_ISL_3383878, EPI_ISL_3383879, EPI_ISL_3383880, EPI_ISL_3383881, EPI_ISL_3383882, EPI_ISL_3383883, EPI_ISL_3383884, EPI_ISL_3383885, EPI_ISL_3383886, EPI_ISL_3383887, EPI_ISL_3383888, EPI_ISL_3383889, EPI_ISL_3383890, EPI_ISL_3383892, EPI_ISL_3383893, EPI_ISL_3383894, EPI_ISL_3383895, EPI_ISL_3383896, EPI_ISL_3383897, EPI_ISL_3383898, EPI_ISL_3383899, EPI_ISL_3383900, EPI_ISL_3383901, EPI_ISL_3383902, EPI_ISL_3383903, EPI_ISL_3383904, EPI_ISL_3383905 | | | |
| see above | Department of Bacteria, Parasites and Fungi, Statens Serum Institut, Copenhagen, Denmark | Statens Serum Institut Bioinformatics and Microbial Genomics | Danish Covid-19 Genome Consortium |
| EPI_ISL_3383906 | Molekylær Medicinsk Afdeling, Aarhus University Hospital, Aarhus, Denmark | Statens Serum Institut Bioinformatics and Microbial Genomics | Danish Covid-19 Genome Consortium |
| EPI_ISL_3383907, EPI_ISL_3383908 | Department of Bacteria, Parasites and Fungi, Statens Serum Institut, Copenhagen, Denmark | Statens Serum Institut Bioinformatics and Microbial Genomics | Danish Covid-19 Genome Consortium |
| EPI_ISL_3383909 | Molekylær Medicinsk Afdeling, Aarhus University Hospital, Aarhus, Denmark | Statens Serum Institut Bioinformatics and Microbial Genomics | Danish Covid-19 Genome Consortium |
| EPI_ISL_3383910, EPI_ISL_3383911, EPI_ISL_3383912, EPI_ISL_3383913, EPI_ISL_3383914, EPI_ISL_3383915, EPI_ISL_3383916, EPI_ISL_3383917, EPI_ISL_3383918, EPI_ISL_3383919, EPI_ISL_3383920, EPI_ISL_3383921, EPI_ISL_3383922, EPI_ISL_3383923, EPI_ISL_3383924, EPI_ISL_3383925, EPI_ISL_3383926, EPI_ISL_3383927, EPI_ISL_3383928, EPI_ISL_3383929, EPI_ISL_3383930, EPI_ISL_3383931, EPI_ISL_3383932, EPI_ISL_3383933, EPI_ISL_3383934, EPI_ISL_3383935, EPI_ISL_3383936, EPI_ISL_3383937, EPI_ISL_3383938, EPI_ISL_3383939, EPI_ISL_3383940, EPI_ISL_3383941, EPI_ISL_3383942, EPI_ISL_3383943, EPI_ISL_3383944, EPI_ISL_3383945, EPI_ISL_3383946, EPI_ISL_3383947, EPI_ISL_3383948, EPI_ISL_3383949, EPI_ISL_3383950, EPI_ISL_3383951, EPI_ISL_3383952, EPI_ISL_3383953, EPI_ISL_3383954, EPI_ISL_3383955, EPI_ISL_3383956, EPI_ISL_3383957, EPI_ISL_3383958, EPI_ISL_3383959, EPI_ISL_3383960, EPI_ISL_3383961, EPI_ISL_3383962, EPI_ISL_3383963, EPI_ISL_3383964, EPI_ISL_3383966, EPI_ISL_3383967, EPI_ISL_3383968, EPI_ISL_3383969, EPI_ISL_3383970, EPI_ISL_3383971, EPI_ISL_3383972, EPI_ISL_3383973 | | | |
| see above | Department of Bacteria, Parasites and Fungi, Statens Serum Institut, Copenhagen, Denmark | Statens Serum Institut Bioinformatics and Microbial Genomics | Danish Covid-19 Genome Consortium |
| EPI_ISL_3383974 | Molekylær Medicinsk Afdeling, Aarhus University Hospital, Aarhus, Denmark | Statens Serum Institut Bioinformatics and Microbial Genomics | Danish Covid-19 Genome Consortium |
| EPI_ISL_3383975, EPI_ISL_3383976, EPI_ISL_3383978, EPI_ISL_3383979, EPI_ISL_3383981, EPI_ISL_3383983, EPI_ISL_3383984, EPI_ISL_3383986, EPI_ISL_3383987, EPI_ISL_3383988, EPI_ISL_3383989, EPI_ISL_3383991, EPI_ISL_3383992, EPI_ISL_3383993, EPI_ISL_3383995, EPI_ISL_3383996, EPI_ISL_3383997, EPI_ISL_3383999, EPI_ISL_3384001, EPI_ISL_3384003, EPI_ISL_3384005, EPI_ISL_3384006, EPI_ISL_3384008, EPI_ISL_3384010, EPI_ISL_3384011, EPI_ISL_3384013, EPI_ISL_3384015, EPI_ISL_3384017, EPI_ISL_3384019, EPI_ISL_3384021 | | | |
| see above | Department of Bacteria, Parasites and Fungi, Statens Serum Institut, Copenhagen, Denmark | Statens Serum Institut Bioinformatics and Microbial Genomics | Danish Covid-19 Genome Consortium |
| EPI_ISL_3384023 | Molekylær Medicinsk Afdeling, Aarhus University Hospital, Aarhus, Denmark | Statens Serum Institut Bioinformatics and Microbial Genomics | Danish Covid-19 Genome Consortium |
| EPI_ISL_3384024, EPI_ISL_3384026, EPI_ISL_3384027, EPI_ISL_3384028, EPI_ISL_3384029, EPI_ISL_3384031, EPI_ISL_3384033, EPI_ISL_3384034, EPI_ISL_3384035, EPI_ISL_3384037, EPI_ISL_3384038, EPI_ISL_3384040, EPI_ISL_3384042, EPI_ISL_3384044, EPI_ISL_3384045, EPI_ISL_3384047, EPI_ISL_3384049, EPI_ISL_3384050, EPI_ISL_3384051, EPI_ISL_3384053 | | | |
| see above | Department of Bacteria, Parasites and Fungi, Statens Serum Institut, Copenhagen, Denmark | Statens Serum Institut Bioinformatics and Microbial Genomics | Danish Covid-19 Genome Consortium |
| EPI_ISL_3384054 | Department of Clinical Microbiology and Center for Genomic Medicine, Rigshospitalet, Copenhagen, Denmark | Statens Serum Institut Bioinformatics and Microbial Genomics | Danish Covid-19 Genome Consortium |
| EPI_ISL_3384055, EPI_ISL_3384056, EPI_ISL_3384058, EPI_ISL_3384060 | Department of Bacteria, Parasites and Fungi, Statens Serum Institut, Copenhagen, Denmark | Statens Serum Institut Bioinformatics and Microbial Genomics | Danish Covid-19 Genome Consortium |
| EPI_ISL_3384061 | Molekylær Medicinsk Afdeling, Aarhus University Hospital, Aarhus, Denmark | Statens Serum Institut Bioinformatics and Microbial Genomics | Danish Covid-19 Genome Consortium |
| EPI_ISL_3384065, EPI_ISL_3384067, EPI_ISL_3384069, EPI_ISL_3384071, EPI_ISL_3384072, EPI_ISL_3384074, EPI_ISL_3384076, EPI_ISL_3384078, EPI_ISL_3384080, EPI_ISL_3384081, EPI_ISL_3384083, EPI_ISL_3384085, EPI_ISL_3384087, EPI_ISL_3384089 | | | |
| see above | Department of Bacteria, Parasites and Fungi, Statens Serum Institut, Copenhagen, Denmark | Statens Serum Institut Bioinformatics and Microbial Genomics | Danish Covid-19 Genome Consortium |
| EPI_ISL_3384090 | Molekylær Medicinsk Afdeling, Aarhus University Hospital, Aarhus, Denmark | Statens Serum Institut Bioinformatics and Microbial Genomics | Danish Covid-19 Genome Consortium |
| EPI_ISL_3384092, EPI_ISL_3384094, EPI_ISL_3384096, EPI_ISL_3384098, EPI_ISL_3384099, EPI_ISL_3384101, EPI_ISL_3384103, EPI_ISL_3384105, EPI_ISL_3384106, EPI_ISL_3384108, EPI_ISL_3384110, EPI_ISL_3384112, EPI_ISL_3384114, EPI_ISL_3384115, EPI_ISL_3384117, EPI_ISL_3384119, EPI_ISL_3384121, EPI_ISL_3384123, EPI_ISL_3384124, EPI_ISL_3384126, EPI_ISL_3384128, EPI_ISL_3384130, EPI_ISL_3384132, EPI_ISL_3384134, EPI_ISL_3384136, EPI_ISL_3384137, EPI_ISL_3384140, EPI_ISL_3384141, EPI_ISL_3384143, EPI_ISL_3384145, EPI_ISL_3384147, EPI_ISL_3384148, EPI_ISL_3384150, EPI_ISL_3384153, EPI_ISL_3384154, EPI_ISL_3384155, EPI_ISL_3384156, EPI_ISL_3384157, EPI_ISL_3384158, EPI_ISL_3384159, EPI_ISL_3384160, EPI_ISL_3384161, EPI_ISL_3384162, EPI_ISL_3384163, EPI_ISL_3384164, EPI_ISL_3384165, EPI_ISL_3384166, EPI_ISL_3384167, EPI_ISL_3384169, EPI_ISL_3384170, EPI_ISL_3384171, EPI_ISL_3384172, EPI_ISL_3384173, EPI_ISL_3384174, EPI_ISL_3384175, EPI_ISL_3384176, EPI_ISL_3384178, EPI_ISL_3384179, EPI_ISL_3384180, EPI_ISL_3384181, EPI_ISL_3384182, EPI_ISL_3384183, EPI_ISL_3384184, EPI_ISL_3384185, EPI_ISL_3384186, EPI_ISL_3384187, EPI_ISL_3384188, EPI_ISL_3384189, EPI_ISL_3384190, EPI_ISL_3384191, EPI_ISL_3384192, EPI_ISL_3384193, EPI_ISL_3384194, EPI_ISL_3384195, EPI_ISL_3384196, EPI_ISL_3384197, EPI_ISL_3384198, EPI_ISL_3384199, EPI_ISL_3384200, EPI_ISL_3384201, EPI_ISL_3384202, EPI_ISL_3384203, EPI_ISL_3384204, EPI_ISL_3384205, EPI_ISL_3384206, EPI_ISL_3384207, EPI_ISL_3384208, EPI_ISL_3384209, EPI_ISL_3384210, EPI_ISL_3384211, EPI_ISL_3384212, EPI_ISL_3384213, EPI_ISL_3384214, EPI_ISL_3384215, EPI_ISL_3384216, EPI_ISL_3384217, EPI_ISL_3384218, EPI_ISL_3384219, EPI_ISL_3384220, EPI_ISL_3384221, EPI_ISL_3384222, EPI_ISL_3384223, EPI_ISL_3384224, EPI_ISL_3384225, EPI_ISL_3384226, EPI_ISL_3384227, EPI_ISL_3384228, EPI_ISL_3384229, EPI_ISL_3384230, EPI_ISL_3384231, EPI_ISL_3384232, EPI_ISL_3384233, EPI_ISL_3384234, EPI_ISL_3384235, EPI_ISL_3384236, EPI_ISL_3384237, EPI_ISL_3384238, EPI_ISL_3384239, EPI_ISL_3384240, EPI_ISL_3384241, EPI_ISL_3384242, EPI_ISL_3384243, EPI_ISL_3384244, EPI_ISL_3384245, EPI_ISL_3384246, EPI_ISL_3384247, EPI_ISL_3384248, EPI_ISL_3384249, EPI_ISL_3384250, EPI_ISL_3384251, EPI_ISL_3384252 | | | |
| see above | Department of Bacteria, Parasites and Fungi, Statens Serum Institut, Copenhagen, Denmark | Statens Serum Institut Bioinformatics and Microbial Genomics | Danish Covid-19 Genome Consortium |
| EPI_ISL_3384253 | Molekylær Medicinsk Afdeling, Aarhus University Hospital, Aarhus, Denmark | Statens Serum Institut Bioinformatics and Microbial Genomics | Danish Covid-19 Genome Consortium |
| EPI_ISL_3384254, EPI_ISL_3384255, EPI_ISL_3384256, EPI_ISL_3384257, EPI_ISL_3384258, EPI_ISL_3384259, EPI_ISL_3384260, EPI_ISL_3384261, EPI_ISL_3384262, EPI_ISL_3384263, EPI_ISL_3384264, EPI_ISL_3384265, EPI_ISL_3384266, EPI_ISL_3384267, EPI_ISL_3384268, EPI_ISL_3384269, EPI_ISL_3384270, EPI_ISL_3384271, EPI_ISL_3384272, EPI_ISL_3384273, EPI_ISL_3384274, EPI_ISL_3384275, EPI_ISL_3384276, EPI_ISL_3384277, EPI_ISL_3384278, EPI_ISL_3384279, EPI_ISL_3384280, EPI_ISL_3384281, EPI_ISL_3384282, EPI_ISL_3384283, EPI_ISL_3384284, EPI_ISL_3384285, EPI_ISL_3384286, EPI_ISL_3384287, EPI_ISL_3384288, EPI_ISL_3384289, EPI_ISL_3384290, EPI_ISL_3384291, EPI_ISL_3384292, EPI_ISL_3384293, EPI_ISL_3384294, EPI_ISL_3384295, EPI_ISL_3384296 | | | |
| see above | Department of Bacteria, Parasites and Fungi, Statens Serum Institut, Copenhagen, Denmark | Statens Serum Institut Bioinformatics and Microbial Genomics | Danish Covid-19 Genome Consortium |
| EPI_ISL_3384297 | Department of Clinical Microbiology and Center for Genomic Medicine, Rigshospitalet, Copenhagen, Denmark | Statens Serum Institut Bioinformatics and Microbial Genomics | Danish Covid-19 Genome Consortium |
| EPI_ISL_3384298, EPI_ISL_3384299, EPI_ISL_3384300, EPI_ISL_3384301, EPI_ISL_3384302, EPI_ISL_3384303, EPI_ISL_3384304, EPI_ISL_3384305, EPI_ISL_3384306, EPI_ISL_3384307, EPI_ISL_3384308, EPI_ISL_3384309, EPI_ISL_3384310, EPI_ISL_3384311, EPI_ISL_3384312, EPI_ISL_3384313, EPI_ISL_3384314, EPI_ISL_3384315, EPI_ISL_3384316, EPI_ISL_3384317, EPI_ISL_3384318, EPI_ISL_3384319, EPI_ISL_3384320, EPI_ISL_3384321, EPI_ISL_3384322, EPI_ISL_3384323, EPI_ISL_3384324, EPI_ISL_3384325, EPI_ISL_3384326, EPI_ISL_3384328 | | | |
| see above | Department of Bacteria, Parasites and Fungi, Statens Serum Institut, Copenhagen, Denmark | Statens Serum Institut Bioinformatics and Microbial Genomics | Danish Covid-19 Genome Consortium |
| EPI_ISL_3384329 | Department of Clinical Microbiology and Center for Genomic Medicine, Rigshospitalet, Copenhagen, Denmark | Statens Serum Institut Bioinformatics and Microbial Genomics | Danish Covid-19 Genome Consortium |
| EPI_ISL_3384330, EPI_ISL_3384331, EPI_ISL_3384332, EPI_ISL_3384333, EPI_ISL_3384334, EPI_ISL_3384335, EPI_ISL_3384336, EPI_ISL_3384337, EPI_ISL_3384338, EPI_ISL_3384339, EPI_ISL_3384340, EPI_ISL_3384341, EPI_ISL_3384342, EPI_ISL_3384343, EPI_ISL_3384344, EPI_ISL_3384345, EPI_ISL_3384346, EPI_ISL_3384347, EPI_ISL_3384348, EPI_ISL_3384349, EPI_ISL_3384350, EPI_ISL_3384351, EPI_ISL_3384352, EPI_ISL_3384353, EPI_ISL_3384354, EPI_ISL_3384355, EPI_ISL_3384356, EPI_ISL_3384357, EPI_ISL_3384358, EPI_ISL_3384359, EPI_ISL_3384360, EPI_ISL_3384361, EPI_ISL_3384362, EPI_ISL_3384363, EPI_ISL_3384364, EPI_ISL_3384365, EPI_ISL_3384366, EPI_ISL_3384367, EPI_ISL_3384368, EPI_ISL_3384369, EPI_ISL_3384370, EPI_ISL_3384371 | | | |
| see above | Department of Bacteria, Parasites and Fungi, Statens Serum Institut, Copenhagen, Denmark | Statens Serum Institut Bioinformatics and Microbial Genomics | Danish Covid-19 Genome Consortium |
| EPI_ISL_3384372 | Molekylær Medicinsk Afdeling, Aarhus University Hospital, Aarhus, Denmark | Statens Serum Institut Bioinformatics and Microbial Genomics | Danish Covid-19 Genome Consortium |
| EPI_ISL_3384373, EPI_ISL_3384374, EPI_ISL_3384375, EPI_ISL_3384376, EPI_ISL_3384377, EPI_ISL_3384378, EPI_ISL_3384380, EPI_ISL_3384381, EPI_ISL_3384382, EPI_ISL_3384383, EPI_ISL_3384384 | | | |
| see above | Department of Bacteria, Parasites and Fungi, Statens Serum Institut, Copenhagen, Denmark | Statens Serum Institut Bioinformatics and Microbial Genomics | Danish Covid-19 Genome Consortium |
| EPI_ISL_3384385 | Molekylær Medicinsk Afdeling, Aarhus University Hospital, Aarhus, Denmark | Statens Serum Institut Bioinformatics and Microbial Genomics | Danish Covid-19 Genome Consortium |
| EPI_ISL_3384386, EPI_ISL_3384387, EPI_ISL_3384388, EPI_ISL_3384389, EPI_ISL_3384390, EPI_ISL_3384391, EPI_ISL_3384392, EPI_ISL_3384393, EPI_ISL_3384394, EPI_ISL_3384395, EPI_ISL_3384396, EPI_ISL_3384397, EPI_ISL_3384398, EPI_ISL_3384399, EPI_ISL_3384400, EPI_ISL_3384401, EPI_ISL_3384402, EPI_ISL_3384403, EPI_ISL_3384404, EPI_ISL_3384405, EPI_ISL_3384406, EPI_ISL_3384407, EPI_ISL_3384408, EPI_ISL_3384409, EPI_ISL_3384410, EPI_ISL_3384411, EPI_ISL_3384412, EPI_ISL_3384413, EPI_ISL_3384414, EPI_ISL_3384415, EPI_ISL_3384416, EPI_ISL_3384417, EPI_ISL_3384418, EPI_ISL_3384419, EPI_ISL_3384420, EPI_ISL_3384421, EPI_ISL_3384422, EPI_ISL_3384423, EPI_ISL_3384424, EPI_ISL_3384425, EPI_ISL_3384426, EPI_ISL_3384427, EPI_ISL_3384428, EPI_ISL_3384429, EPI_ISL_3384430, EPI_ISL_3384431, EPI_ISL_3384432, EPI_ISL_3384433, EPI_ISL_3384434, EPI_ISL_3384435, EPI_ISL_3384436, EPI_ISL_3384437, EPI_ISL_3384438, EPI_ISL_3384439, EPI_ISL_3384440, EPI_ISL_3384441, EPI_ISL_3384442, EPI_ISL_3384443, EPI_ISL_3384444, EPI_ISL_3384445, EPI_ISL_3384446 | | | |
| see above | Department of Bacteria, Parasites and Fungi, Statens Serum Institut, Copenhagen, Denmark | Statens Serum Institut Bioinformatics and Microbial Genomics | Danish Covid-19 Genome Consortium |
| EPI_ISL_3384447 | Molekylær Medicinsk Afdeling, Aarhus University Hospital, Aarhus, Denmark | Statens Serum Institut Bioinformatics and Microbial Genomics | Danish Covid-19 Genome Consortium |
| EPI_ISL_3384448, EPI_ISL_3384449, EPI_ISL_3384450, EPI_ISL_3384451, EPI_ISL_3384452, EPI_ISL_3384453, EPI_ISL_3384454, EPI_ISL_3384455, EPI_ISL_3384456, EPI_ISL_3384457, EPI_ISL_3384459, EPI_ISL_3384460, EPI_ISL_3384461, EPI_ISL_3384462, EPI_ISL_3384464, EPI_ISL_3384465, EPI_ISL_3384466, EPI_ISL_3384467, EPI_ISL_3384468, EPI_ISL_3384469, EPI_ISL_3384470, EPI_ISL_3384471, | | | |

| | | | |
|---|---|---|---|
| EPI_ISL_3384472, EPI_ISL_3384473, EPI_ISL_3384474, EPI_ISL_3384475, EPI_ISL_3384476, EPI_ISL_3384477, EPI_ISL_3384478, EPI_ISL_3384479, EPI_ISL_3384480, EPI_ISL_3384481, EPI_ISL_3384483, EPI_ISL_3384484, EPI_ISL_3384485, EPI_ISL_3384486, EPI_ISL_3384487, EPI_ISL_3384488, EPI_ISL_3384489, EPI_ISL_3384490, EPI_ISL_3384491, EPI_ISL_3384492, EPI_ISL_3384493, EPI_ISL_3384494, EPI_ISL_3384495, EPI_ISL_3384496, EPI_ISL_3384497, EPI_ISL_3384498, EPI_ISL_3384499, EPI_ISL_3384500, EPI_ISL_3384501, EPI_ISL_3384502, EPI_ISL_3384503, EPI_ISL_3384504, EPI_ISL_3384505, EPI_ISL_3384506, EPI_ISL_3384507, EPI_ISL_3384508, EPI_ISL_3384509, EPI_ISL_3384510, EPI_ISL_3384511, EPI_ISL_3384512, EPI_ISL_3384513, EPI_ISL_3384514, EPI_ISL_3384515, EPI_ISL_3384516, EPI_ISL_3384517, EPI_ISL_3384518, EPI_ISL_3384519, EPI_ISL_3384520, EPI_ISL_3384521, EPI_ISL_3384522, EPI_ISL_3384523, EPI_ISL_3384524, EPI_ISL_3384525, EPI_ISL_3384526, EPI_ISL_3384527, EPI_ISL_3384528, EPI_ISL_3384529, EPI_ISL_3384530, EPI_ISL_3384532, EPI_ISL_3384533, EPI_ISL_3384534, EPI_ISL_3384535, EPI_ISL_3384536, EPI_ISL_3384537, EPI_ISL_3384538, EPI_ISL_3384539, EPI_ISL_3384541, EPI_ISL_3384542, EPI_ISL_3384543, EPI_ISL_3384544, EPI_ISL_3384545, EPI_ISL_3384546, EPI_ISL_3384547, EPI_ISL_3384548, EPI_ISL_3384549, EPI_ISL_3384550, EPI_ISL_3384551, EPI_ISL_3384552, EPI_ISL_3384553, EPI_ISL_3384554, EPI_ISL_3384555, EPI_ISL_3384556, EPI_ISL_3384557, EPI_ISL_3384558, EPI_ISL_3384559, EPI_ISL_3384560, EPI_ISL_3384561, EPI_ISL_3384562, EPI_ISL_3384563, EPI_ISL_3384564, EPI_ISL_3384565, EPI_ISL_3384566, EPI_ISL_3384567, EPI_ISL_3384568, EPI_ISL_3384569, EPI_ISL_3384570, EPI_ISL_3384571, EPI_ISL_3384572, EPI_ISL_3384573, EPI_ISL_3384574, EPI_ISL_3384575, EPI_ISL_3384576, EPI_ISL_3384577, EPI_ISL_3384578 | | | |
| see above | Department of Bacteria, Parasites and Fungi, Statens Serum Institut, Copenhagen, Denmark | Statens Serum Institut Bioinformatics and Microbial Genomics | Danish Covid-19 Genome Consortium |
| EPI_ISL_3384580 | Molekylær Medicinsk Afdeling, Aarhus University Hospital, Aarhus, Denmark | Statens Serum Institut Bioinformatics and Microbial Genomics | Danish Covid-19 Genome Consortium |
| EPI_ISL_3384581, EPI_ISL_3384582, EPI_ISL_3384583, EPI_ISL_3384584, EPI_ISL_3384585, EPI_ISL_3384586, EPI_ISL_3384587, EPI_ISL_3384588, EPI_ISL_3384589, EPI_ISL_3384590, EPI_ISL_3384591, EPI_ISL_3384592, EPI_ISL_3384593, EPI_ISL_3384594, EPI_ISL_3384595, EPI_ISL_3384596, EPI_ISL_3384597 | | | |
| see above | Department of Bacteria, Parasites and Fungi, Statens Serum Institut, Copenhagen, Denmark | Statens Serum Institut Bioinformatics and Microbial Genomics | Danish Covid-19 Genome Consortium |
| EPI_ISL_3384598 | Molekylær Medicinsk Afdeling, Aarhus University Hospital, Aarhus, Denmark | Statens Serum Institut Bioinformatics and Microbial Genomics | Danish Covid-19 Genome Consortium |
| EPI_ISL_3384599, EPI_ISL_3384600, EPI_ISL_3384601, EPI_ISL_3384602, EPI_ISL_3384603, EPI_ISL_3384604, EPI_ISL_3384605, EPI_ISL_3384606, EPI_ISL_3384607, EPI_ISL_3384608, EPI_ISL_3384609, EPI_ISL_3384610, EPI_ISL_3384611, EPI_ISL_3384612, EPI_ISL_3384613, EPI_ISL_3384614, EPI_ISL_3384615, EPI_ISL_3384616, EPI_ISL_3384617, EPI_ISL_3384618 | | | |
| see above | Department of Bacteria, Parasites and Fungi, Statens Serum Institut, Copenhagen, Denmark | Statens Serum Institut Bioinformatics and Microbial Genomics | Danish Covid-19 Genome Consortium |
| EPI_ISL_3384619 | Molekylær Medicinsk Afdeling, Aarhus University Hospital, Aarhus, Denmark | Statens Serum Institut Bioinformatics and Microbial Genomics | Danish Covid-19 Genome Consortium |
| EPI_ISL_3384620, EPI_ISL_3384621, EPI_ISL_3384622, EPI_ISL_3384623, EPI_ISL_3384624, EPI_ISL_3384625, EPI_ISL_3384626, EPI_ISL_3384627, EPI_ISL_3384628, EPI_ISL_3384629, EPI_ISL_3384630, EPI_ISL_3384631, EPI_ISL_3384632, EPI_ISL_3384633, EPI_ISL_3384634, EPI_ISL_3384636, EPI_ISL_3384637, EPI_ISL_3384638, EPI_ISL_3384639, EPI_ISL_3384640, EPI_ISL_3384641, EPI_ISL_3384642, EPI_ISL_3384643, EPI_ISL_3384644, EPI_ISL_3384645, EPI_ISL_3384646, EPI_ISL_3384647, EPI_ISL_3384648, EPI_ISL_3384649, EPI_ISL_3384650, EPI_ISL_3384651, EPI_ISL_3384652, EPI_ISL_3384653, EPI_ISL_3384654, EPI_ISL_3384655, EPI_ISL_3384656, EPI_ISL_3384657, EPI_ISL_3384658, EPI_ISL_3384659 | | | |
| see above | Department of Bacteria, Parasites and Fungi, Statens Serum Institut, Copenhagen, Denmark | Statens Serum Institut Bioinformatics and Microbial Genomics | Danish Covid-19 Genome Consortium |
| EPI_ISL_3384660 | Department of Clinical Microbiology and Center for Genomic Medicine, Rigshospitalet, Copenhagen, Denmark | Statens Serum Institut Bioinformatics and Microbial Genomics | Danish Covid-19 Genome Consortium |
| EPI_ISL_3384661, EPI_ISL_3384662, EPI_ISL_3384663, EPI_ISL_3384664, EPI_ISL_3384665, EPI_ISL_3384666, EPI_ISL_3384667, EPI_ISL_3384668, EPI_ISL_3384669, EPI_ISL_3384670, EPI_ISL_3384671, EPI_ISL_3384673, EPI_ISL_3384675, EPI_ISL_3384676, EPI_ISL_3384677, EPI_ISL_3384678, EPI_ISL_3384680, EPI_ISL_3384681, EPI_ISL_3384682, EPI_ISL_3384683, EPI_ISL_3384684, EPI_ISL_3384685, EPI_ISL_3384686 | | | |
| see above | Department of Bacteria, Parasites and Fungi, Statens Serum Institut, Copenhagen, Denmark | Statens Serum Institut Bioinformatics and Microbial Genomics | Danish Covid-19 Genome Consortium |
| EPI_ISL_3384687 | Department of Clinical Microbiology and Center for Genomic Medicine, Rigshospitalet, Copenhagen, Denmark | Statens Serum Institut Bioinformatics and Microbial Genomics | Danish Covid-19 Genome Consortium |
| EPI_ISL_3384688, EPI_ISL_3384689, EPI_ISL_3384690, EPI_ISL_3384691, EPI_ISL_3384692, EPI_ISL_3384693, EPI_ISL_3384694, EPI_ISL_3384695, EPI_ISL_3384696, EPI_ISL_3384697, EPI_ISL_3384698, EPI_ISL_3384699, EPI_ISL_3384700, EPI_ISL_3384701, EPI_ISL_3384702, EPI_ISL_3384703, EPI_ISL_3384704, EPI_ISL_3384705, EPI_ISL_3384706, EPI_ISL_3384707, EPI_ISL_3384709, EPI_ISL_3384710, EPI_ISL_3384711, EPI_ISL_3384712, EPI_ISL_3384713, EPI_ISL_3384714, EPI_ISL_3384715, EPI_ISL_3384716, EPI_ISL_3384717, EPI_ISL_3384718, EPI_ISL_3384719, EPI_ISL_3384720, EPI_ISL_3384721, EPI_ISL_3384722, EPI_ISL_3384723, EPI_ISL_3384724, EPI_ISL_3384725, EPI_ISL_3384726, EPI_ISL_3384727, EPI_ISL_3384728, EPI_ISL_3384729, EPI_ISL_3384730, EPI_ISL_3384731, EPI_ISL_3384732, EPI_ISL_3384733, EPI_ISL_3384734, EPI_ISL_3384735, EPI_ISL_3384736, EPI_ISL_3384737, EPI_ISL_3384738, EPI_ISL_3384739, EPI_ISL_3384740, EPI_ISL_3384741, EPI_ISL_3384742, EPI_ISL_3384743 | | | |
| see above | Department of Bacteria, Parasites and Fungi, Statens Serum Institut, Copenhagen, Denmark | Statens Serum Institut Bioinformatics and Microbial Genomics | Danish Covid-19 Genome Consortium |
| EPI_ISL_3384744 | Molekylær Medicinsk Afdeling, Aarhus University Hospital, Aarhus, Denmark | Statens Serum Institut Bioinformatics and Microbial Genomics | Danish Covid-19 Genome Consortium |
| EPI_ISL_3384745, EPI_ISL_3384746, EPI_ISL_3384747, EPI_ISL_3384748, EPI_ISL_3384749, EPI_ISL_3384750, EPI_ISL_3384751, EPI_ISL_3384752, EPI_ISL_3384753, EPI_ISL_3384754, EPI_ISL_3384755, EPI_ISL_3384756, EPI_ISL_3384757, EPI_ISL_3384758, EPI_ISL_3384759, EPI_ISL_3384761, EPI_ISL_3384762, EPI_ISL_3384763, EPI_ISL_3384764, EPI_ISL_3384765, EPI_ISL_3384766, EPI_ISL_3384767, EPI_ISL_3384768, EPI_ISL_3384769 | | | |
| see above | Department of Bacteria, Parasites and Fungi, Statens Serum Institut, Copenhagen, Denmark | Statens Serum Institut Bioinformatics and Microbial Genomics | Danish Covid-19 Genome Consortium |
| EPI_ISL_3384770 | RSUD PROVINSI NTB | Genomik Solidaritas Indonesia Laboratorium | Vania Gavrila Wikasa, Dhahlia Agustina Cahyono, Himawan Masyhuri, Carissa Sintca Wijaya, Annisa Muthiah Sukirman, Gracia Felias Enos Korompis, Anuraj Shankar, Louisa Markus, Meutia Ayuputeri Kumaheri, Ariel Pradipta |
| EPI_ISL_3384771, EPI_ISL_3384772 | RSUD PROVINSI NTB | Genomik Solidaritas Indonesia Laboratorium | Dhahlia Agustina Cahyono, Himawan Masyhuri, Carissa Sintca Wijaya, Vania Gavrila Wikasa, Annisa Muthiah Sukirman, Gracia Felias Enos Korompis, Anuraj Shankar, Louisa Markus, Meutia Ayuputeri Kumaheri, Ariel Pradipta |
| EPI_ISL_3384773 | RSUD PROVINSI NTB | Genomik Solidaritas Indonesia Laboratorium | Himawan Masyhuri, Carissa Sintca Wijaya, Vania Gavrila Wikasa, Dhahlia Agustina Cahyono, Gracia Felias Enos Korompis, Annisa Muthiah Sukirman, Anuraj Shankar, Louisa Markus, Meutia Ayuputeri Kumaheri, Ariel Pradipta |
| EPI_ISL_3384774 | RSUD PROVINSI NTB | Genomik Solidaritas Indonesia Laboratorium | Carissa Sintca Wijaya, Vania Gavrila Wikasa, Dhahlia Agustina Cahyono, Himawan Masyhuri, Gracia Felias Enos Korompis, Annisa Muthiah Sukirman, Anuraj Shankar, Louisa Markus, Meutia Ayuputeri Kumaheri, Ariel Pradipta |
| EPI_ISL_3384775 | RSUD PROVINSI NTB | Genomik Solidaritas Indonesia Laboratorium | Vania Gavrila Wikasa, Dhahlia Agustina Cahyono, Himawan Masyhuri, Carissa Sintca Wijaya, Gracia Felias Enos Korompis, Annisa Muthiah Sukirman, Anuraj Shankar, Louisa Markus, Meutia Ayuputeri Kumaheri, Ariel Pradipta |
| EPI_ISL_3384776 | RSUD PROVINSI NTB | Genomik Solidaritas Indonesia Laboratorium | Dhahlia Agustina Cahyono, Himawan Masyhuri, Carissa Sintca Wijaya, Vania Gavrila Wikasa, Annisa Muthiah Sukirman, Gracia Felias Enos Korompis, Anuraj Shankar, Louisa Markus, Meutia Ayuputeri Kumaheri, Ariel Pradipta |
| EPI_ISL_3384777 | RSUD PROVINSI NTB | Genomik Solidaritas Indonesia Laboratorium | Himawan Masyhuri, Carissa Sintca Wijaya, Vania Gavrila Wikasa, Dhahlia Agustina Cahyono, Gracia Felias Enos Korompis, Annisa Muthiah Sukirman, Anuraj Shankar, Louisa Markus, Meutia Ayuputeri Kumaheri, Ariel Pradipta |
| EPI_ISL_3384778 | RSUD PROVINSI NTB | Genomik Solidaritas Indonesia Laboratorium | Carissa Sintca Wijaya, Vania Gavrila Wikasa, Dhahlia Agustina Cahyono, Himawan Masyhuri, Gracia Felias Enos Korompis, Annisa Muthiah Sukirman, Anuraj Shankar, Louisa Markus, Meutia Ayuputeri Kumaheri, Ariel Pradipta |
| EPI_ISL_3384779 | RSUD PROVINSI NTB | Genomik Solidaritas Indonesia Laboratorium | Himawan Masyhuri, Carissa Sintca Wijaya, Vania Gavrila Wikasa, Dhahlia Agustina Cahyono, Gracia Felias Enos Korompis, Annisa Muthiah Sukirman, Anuraj Shankar, Louisa Markus, Meutia Ayuputeri Kumaheri, Ariel Pradipta |
| EPI_ISL_3384780 | RSUD PROVINSI NTB | Genomik Solidaritas Indonesia Laboratorium | Vania Gavrila Wikasa, Dhahlia Agustina Cahyono, Himawan Masyhuri, Carissa Sintca Wijaya, Annisa Muthiah Sukirman, Gracia Felias Enos Korompis, Anuraj Shankar, Louisa Markus, Meutia Ayuputeri Kumaheri, Ariel Pradipta |
| EPI_ISL_3384781 | RSUD PROVINSI NTB | Genomik Solidaritas Indonesia Laboratorium | Carissa Sintca Wijaya, Vania Gavrila Wikasa, Dhahlia Agustina Cahyono, Himawan Masyhuri, Gracia Felias Enos Korompis, Annisa Muthiah Sukirman, Anuraj Shankar, Louisa Markus, Meutia Ayuputeri Kumaheri, Ariel Pradipta |
| EPI_ISL_3384782 | RSUD PROVINSI NTB | Genomik Solidaritas Indonesia Laboratorium | Himawan Masyhuri, Carissa Sintca Wijaya, Vania Gavrila Wikasa, Dhahlia Agustina Cahyono, Gracia Felias Enos Korompis, Annisa Muthiah Sukirman, Anuraj Shankar, Louisa Markus, Meutia Ayuputeri Kumaheri, Ariel Pradipta |
| EPI_ISL_3384783 | RSUD PROVINSI NTB | Genomik Solidaritas Indonesia Laboratorium | Dhahlia Agustina Cahyono, Himawan Masyhuri, Carissa Sintca Wijaya, Vania Gavrila Wikasa, Annisa Muthiah Sukirman, Gracia Felias Enos Korompis, Anuraj Shankar, Louisa Markus, Meutia Ayuputeri Kumaheri, Ariel Pradipta |
| EPI_ISL_3384805, EPI_ISL_3384806, EPI_ISL_3384807, EPI_ISL_3384808, EPI_ISL_3384809 | US Air Force School of Aerospace Medicine | US Air Force School of Aerospace Medicine | Anthony Fries, Jennifer Meyer, William Gruner, Amanda Javorina, Carol Garrett, Sarah Purves, Clarise Starr, Elizabeth Macias |
| EPI_ISL_3384810 | US Air Force School of Aerospace Medicine | US Air Force School of Aerospace Medicine | Anthony Fries, Jennifer Meyer, William Gruner, Amanda Javorina, Carol Garrett, Sarah Purves, Clarise Starr, Elizabeth Macias, Fritz Castillo, Jennifer Black, Cole Anderson |
| EPI_ISL_3384811, EPI_ISL_3384812, EPI_ISL_3384813, EPI_ISL_3384814, EPI_ISL_3384818, EPI_ISL_3384819, EPI_ISL_3384820, EPI_ISL_3384821, EPI_ISL_3384822, EPI_ISL_3384823, EPI_ISL_3384824, EPI_ISL_3384826, EPI_ISL_3384827, EPI_ISL_3384830, EPI_ISL_3384831, EPI_ISL_3384832, EPI_ISL_3384833, EPI_ISL_3384834, EPI_ISL_3384835, EPI_ISL_3384836, EPI_ISL_3384837, EPI_ISL_3384838, EPI_ISL_3384839, EPI_ISL_3384840, EPI_ISL_3384841, EPI_ISL_3384842, EPI_ISL_3384843, EPI_ISL_3384844, EPI_ISL_3384845, EPI_ISL_3384846, EPI_ISL_3384847, EPI_ISL_3384849, EPI_ISL_3384850, EPI_ISL_3384851, EPI_ISL_3384852, EPI_ISL_3384853, EPI_ISL_3384854, EPI_ISL_3384855, EPI_ISL_3384856, EPI_ISL_3384857, EPI_ISL_3384858, EPI_ISL_3384859, EPI_ISL_3384860, EPI_ISL_3384861, EPI_ISL_3384862, EPI_ISL_3384863, EPI_ISL_3384864, EPI_ISL_3384865, EPI_ISL_3384866, EPI_ISL_3384867, EPI_ISL_3384868, EPI_ISL_3384869, EPI_ISL_3384870, EPI_ISL_3384871, EPI_ISL_3384872, EPI_ISL_3384873, EPI_ISL_3384874, EPI_ISL_3384875, EPI_ISL_3384876 | | | |
| see above | US Air Force School of Aerospace Medicine | US Air Force School of Aerospace Medicine | Anthony Fries, Jennifer Meyer, William Gruner, Amanda Javorina, Carol Garrett, Sarah Purves, Clarise Starr, Elizabeth Macias |
| EPI_ISL_3384877, EPI_ISL_3384878, EPI_ISL_3384879, EPI_ISL_3384880, EPI_ISL_3384881, EPI_ISL_3384882, EPI_ISL_3384883, EPI_ISL_3384884 | US Air Force School of Aerospace Medicine | US Air Force School of Aerospace Medicine | Anthony Fries, Jennifer Meyer, William Gruner, Amanda Javorina, Carol Garrett, Sarah Purves, Clarise Starr, Elizabeth Macias, Fritz Castillo, Jennifer Black, Cole Anderson |
| EPI_ISL_3384886, EPI_ISL_3384890, EPI_ISL_3384891, EPI_ISL_3384892 | Laboratori d'anàlisis clíniques, Hospital Nostra Senyora de Meritxell | LBM de CHU de Toulouse, Hôpitaux de Toulouse | M,Rendon et Al |
| EPI_ISL_3384897, EPI_ISL_3384898, EPI_ISL_3384899, EPI_ISL_3384900, EPI_ISL_3384901 | KAISER SAN JOSE | Santa Clara County Public Health Laboratory | Santa Clara County Public Health Department |
| EPI_ISL_3384902, EPI_ISL_3384903, EPI_ISL_3384904, EPI_ISL_3384905, EPI_ISL_3384906, EPI_ISL_3384907, EPI_ISL_3384908, EPI_ISL_3384909, EPI_ISL_3384910, EPI_ISL_3384911 | KAISER SANTA CLARA | Santa Clara County Public Health Laboratory | Santa Clara County Public Health Department |

EPI_ISL_3384912, EPI_ISL_3384913, EPI_ISL_3384914, EPI_ISL_3384915, EPI_ISL_3384916, EPI_ISL_3384917, EPI_ISL_3384918, EPI_ISL_3384919, EPI_ISL_3384920, EPI_ISL_3384921, EPI_ISL_3384922, EPI_ISL_3384923, EPI_ISL_3384924, EPI_ISL_3384925, EPI_ISL_3384926, EPI_ISL_3384927, EPI_ISL_3384928, EPI_ISL_3384929, EPI_ISL_3384930, EPI_ISL_3384931, EPI_ISL_3384932

| | | | |
|---|---|---|---|
| see above | PAMF PALO ALTO | Santa Clara County Public Health Laboratory | Santa Clara County Public Health Department |
| EPI_ISL_3384933, EPI_ISL_3384934, EPI_ISL_3384935 | Santa Clara Valley Medical Center | Santa Clara County Public Health Laboratory | |
| EPI_ISL_3384936 | KAISER SAN JOSE | Santa Clara County Public Health Laboratory | Santa Clara County Public Health Department |
| EPI_ISL_3384938, EPI_ISL_3384939 | KAISER SANTA CLARA | Santa Clara County Public Health Laboratory | Santa Clara County Public Health Department |
| EPI_ISL_3384940 | Santa Clara Valley Medical Center | Santa Clara County Public Health Laboratory | Santa Clara County Public Health Department |
| EPI_ISL_3384941, EPI_ISL_3384942, EPI_ISL_3384943 | PAMF PALO ALTO | Santa Clara County Public Health Laboratory | Santa Clara County Public Health Department |
| EPI_ISL_3384944 | KAISER SAN JOSE | Santa Clara County Public Health Laboratory | Santa Clara County Public Health Department |
| EPI_ISL_3384945 | PAMF PALO ALTO | Santa Clara County Public Health Laboratory | Santa Clara County Public Health Department |
| EPI_ISL_3384947 | A.S.L. NOVARA | Fondazione del Piemonte per l'Oncologia IRCCS | Antonino Sottile, Silvia Brossa, Paola Marino, Giorgio Giardina |
| EPI_ISL_3384948 | Santa Clara Valley Medical Center | Santa Clara County Public Health Laboratory | Santa Clara County Public Health Department |
| EPI_ISL_3384949 | A.S.L. NOVARA | Fondazione del Piemonte per l'Oncologia IRCCS | Antonino Sottile, Silvia Brossa, Paola Marino, Giorgio Giardina |
| EPI_ISL_3384950 | PAMF PALO ALTO | Santa Clara County Public Health Laboratory | Santa Clara County Public Health Department |
| EPI_ISL_3384951 | A.S.L. NOVARA | Fondazione del Piemonte per l'Oncologia IRCCS | Antonino Sottile, Silvia Brossa, Paola Marino, Giorgio Giardina |
| EPI_ISL_3384952 | PAMF PALO ALTO | Santa Clara County Public Health Laboratory | Santa Clara County Public Health Department |
| EPI_ISL_3384953 | A.S.L. NOVARA | Fondazione del Piemonte per l'Oncologia IRCCS | Antonino Sottile, Silvia Brossa, Paola Marino, Giorgio Giardina |
| EPI_ISL_3384954 | PAMF PALO ALTO | Santa Clara County Public Health Laboratory | Santa Clara County Public Health Department |
| EPI_ISL_3384956 | Santa Clara Valley Medical Center | Santa Clara County Public Health Laboratory | Santa Clara County Public Health Department |
| EPI_ISL_3384957 | A.S.L. NOVARA | Fondazione del Piemonte per l'Oncologia IRCCS | Antonino Sottile, Silvia Brossa, Paola Marino, Giorgio Giardina |
| EPI_ISL_3384958 | Santa Clara Valley Medical Center | Santa Clara County Public Health Laboratory | Santa Clara County Public Health Department |
| EPI_ISL_3384959 | A.S.L. NOVARA | Fondazione del Piemonte per l'Oncologia IRCCS | Antonino Sottile, Silvia Brossa, Paola Marino, Giorgio Giardina |
| EPI_ISL_3384960 | PAMF PALO ALTO | Santa Clara County Public Health Laboratory | Santa Clara County Public Health Department |
| EPI_ISL_3384961 | A.S.L. NOVARA | Fondazione del Piemonte per l'Oncologia IRCCS | Antonino Sottile, Silvia Brossa, Paola Marino, Giorgio Giardina |
| EPI_ISL_3384962 | Santa Clara Valley Medical Center | Santa Clara County Public Health Laboratory | Santa Clara County Public Health Department |
| EPI_ISL_3384963 | A.S.L. NOVARA | Fondazione del Piemonte per l'Oncologia IRCCS | Antonino Sottile, Silvia Brossa, Paola Marino, Giorgio Giardina |
| EPI_ISL_3384964 | Santa Clara Valley Medical Center | Santa Clara County Public Health Laboratory | Santa Clara County Public Health Department |
| EPI_ISL_3384966 | PAMF PALO ALTO | Santa Clara County Public Health Laboratory | Santa Clara County Public Health Department |
| EPI_ISL_3384967 | A.S.L. NOVARA | Fondazione del Piemonte per l'Oncologia IRCCS | Antonino Sottile, Silvia Brossa, Paola Marino, Giorgio Giardina |
| EPI_ISL_3384968, EPI_ISL_3384970 | PAMF PALO ALTO | Santa Clara County Public Health Laboratory | Santa Clara County Public Health Department |
| EPI_ISL_3384971 | A.S.L. CUNEO 1 | Fondazione del Piemonte per l'Oncologia IRCCS | Antonino Sottile, Silvia Brossa, Paola Marino, Giorgio Giardina |
| EPI_ISL_3384972 | PAMF PALO ALTO | Santa Clara County Public Health Laboratory | Santa Clara County Public Health Department |
| EPI_ISL_3384973 | A.S.L. BIELLA | Fondazione del Piemonte per l'Oncologia IRCCS | Antonino Sottile, Silvia Brossa, Paola Marino, Giorgio Giardina |
| EPI_ISL_3384974 | KAISER SANTA CLARA | Santa Clara County Public Health Laboratory | Santa Clara County Public Health Department |
| EPI_ISL_3384975 | A.S.L. CUNEO 1 | Fondazione del Piemonte per l'Oncologia IRCCS | Antonino Sottile, Silvia Brossa, Paola Marino, Giorgio Giardina |
| EPI_ISL_3384976 | PAMF PALO ALTO | Santa Clara County Public Health Laboratory | Santa Clara County Public Health Department |
| EPI_ISL_3384977 | A.S.L. CUNEO 1 | Fondazione del Piemonte per l'Oncologia IRCCS | Antonino Sottile, Silvia Brossa, Paola Marino, Giorgio Giardina |
| EPI_ISL_3384978 | Santa Clara Valley Medical Center | Santa Clara County Public Health Laboratory | Santa Clara County Public Health Department |
| EPI_ISL_3384979 | A.S.L. CUNEO 1 | Fondazione del Piemonte per l'Oncologia IRCCS | Antonino Sottile, Silvia Brossa, Paola Marino, Giorgio Giardina |
| EPI_ISL_3384980 | KAISER SAN JOSE | Santa Clara County Public Health Laboratory | Santa Clara County Public Health Department |
| EPI_ISL_3384981 | A.S.L. CUNEO 1 | Fondazione del Piemonte per l'Oncologia IRCCS | Antonino Sottile, Silvia Brossa, Paola Marino, Giorgio Giardina |
| EPI_ISL_3384982 | KAISER SANTA CLARA | Santa Clara County Public Health Laboratory | Santa Clara County Public Health Department |
| EPI_ISL_3384983, EPI_ISL_3384985 | A.S.L. CUNEO 1 | Fondazione del Piemonte per l'Oncologia IRCCS | Antonino Sottile, Silvia Brossa, Paola Marino, Giorgio Giardina |
| EPI_ISL_3384986 | PAMF PALO ALTO | Santa Clara County Public Health Laboratory | Santa Clara County Public Health Department |
| EPI_ISL_3384987 | A.S.L. CUNEO 1 | Fondazione del Piemonte per l'Oncologia IRCCS | Antonino Sottile, Silvia Brossa, Paola Marino, Giorgio Giardina |
| EPI_ISL_3384988 | KAISER SANTA CLARA | Santa Clara County Public Health Laboratory | Santa Clara County Public Health Department |
| EPI_ISL_3384989 | A.S.L. CUNEO 1 | Fondazione del Piemonte per l'Oncologia IRCCS | Antonino Sottile, Silvia Brossa, Paola Marino, Giorgio Giardina |
| EPI_ISL_3384990 | PAMF PALO ALTO | Santa Clara County Public Health Laboratory | Santa Clara County Public Health Department |
| EPI_ISL_3384991 | A.S.L. CUNEO 1 | Fondazione del Piemonte per l'Oncologia IRCCS | Antonino Sottile, Silvia Brossa, Paola Marino, Giorgio Giardina |
| EPI_ISL_3384992 | PAMF PALO ALTO | Santa Clara County Public Health Laboratory | Santa Clara County Public Health Department |
| EPI_ISL_3384993 | A.S.L. CUNEO 1 | Fondazione del Piemonte per l'Oncologia IRCCS | Antonino Sottile, Silvia Brossa, Paola Marino, Giorgio Giardina |
| EPI_ISL_3384994 | KAISER SANTA CLARA | Santa Clara County Public Health Laboratory | Santa Clara County Public Health Department |
| EPI_ISL_3384995, EPI_ISL_3384996 | A.S.L. CUNEO 1 | Fondazione del Piemonte per l'Oncologia IRCCS | Antonino Sottile, Silvia Brossa, Paola Marino, Giorgio Giardina |
| EPI_ISL_3384997 | PAMF PALO ALTO | Santa Clara County Public Health Laboratory | Santa Clara County Public Health Department |
| EPI_ISL_3384998 | A.S.L. CUNEO 1 | Fondazione del Piemonte per l'Oncologia IRCCS | Antonino Sottile, Silvia Brossa, Paola Marino, Giorgio Giardina |
| EPI_ISL_3384999 | PAMF PALO ALTO | Santa Clara County Public Health Laboratory | Santa Clara County Public Health Department |
| EPI_ISL_3385000 | A.S.L. CUNEO 1 | Fondazione del Piemonte per l'Oncologia IRCCS | Antonino Sottile, Silvia Brossa, Paola Marino, Giorgio Giardina |
| EPI_ISL_3385001 | Santa Clara Valley Medical Center | Santa Clara County Public Health Laboratory | Santa Clara County Public Health Department |
| EPI_ISL_3385002 | A.S.L. CUNEO 1 | Fondazione del Piemonte per l'Oncologia IRCCS | Antonino Sottile, Silvia Brossa, Paola Marino, Giorgio Giardina |
| EPI_ISL_3385003 | KAISER SAN JOSE | Santa Clara County Public Health Laboratory | Santa Clara County Public Health Department |
| EPI_ISL_3385004 | A.S.L. CUNEO 1 | Fondazione del Piemonte per l'Oncologia IRCCS | Antonino Sottile, Silvia Brossa, Paola Marino, Giorgio Giardina |
| EPI_ISL_3385005 | PAMF PALO ALTO | Santa Clara County Public Health Laboratory | Santa Clara County Public Health Department |
| EPI_ISL_3385006 | A.S.L. CUNEO 1 | Fondazione del Piemonte per l'Oncologia IRCCS | Antonino Sottile, Silvia Brossa, Paola Marino, Giorgio Giardina |
| EPI_ISL_3385007 | KAISER SAN JOSE | Santa Clara County Public Health Laboratory | Santa Clara County Public Health Department |
| EPI_ISL_3385008 | A.S.L. CUNEO 1 | Fondazione del Piemonte per l'Oncologia IRCCS | Antonino Sottile, Silvia Brossa, Paola Marino, Giorgio Giardina |
| EPI_ISL_3385009 | KAISER SAN JOSE | Santa Clara County Public Health Laboratory | Santa Clara County Public Health Department |
| EPI_ISL_3385010 | A.S.L. CUNEO 1 | Fondazione del Piemonte per l'Oncologia IRCCS | Antonino Sottile, Silvia Brossa, Paola Marino, Giorgio Giardina |
| EPI_ISL_3385011 | PAMF PALO ALTO | Santa Clara County Public Health Laboratory | Santa Clara County Public Health Department |
| EPI_ISL_3385012 | A.S.L. CUNEO 1 | Fondazione del Piemonte per l'Oncologia IRCCS | Antonino Sottile, Silvia Brossa, Paola Marino, Giorgio Giardina |
| EPI_ISL_3385013 | PAMF PALO ALTO | Santa Clara County Public Health Laboratory | Santa Clara County Public Health Department |
| EPI_ISL_3385014 | A.S.L. CUNEO 1 | Fondazione del Piemonte per l'Oncologia IRCCS | Antonino Sottile, Silvia Brossa, Paola Marino, Giorgio Giardina |
| EPI_ISL_3385015 | Santa Clara Valley Medical Center | Santa Clara County Public Health Laboratory | Santa Clara County Public Health Department |

| | | | |
|---|---|---|---|
| EPI_ISL_3385016 | A.S.L. CUNEO 1 | Fondazione del Piemonte per l'Oncologia IRCCS | Antonino Sottile, Silvia Brossa, Paola Marino, Giorgio Giardina |
| EPI_ISL_3385017 | PAMF PALO ALTO | Santa Clara County Public Health Laboratory | Santa Clara County Public Health Department |
| EPI_ISL_3385018, EPI_ISL_3385019 | A.S.L. CUNEO 1 | Fondazione del Piemonte per l'Oncologia IRCCS | Antonino Sottile, Silvia Brossa, Paola Marino, Giorgio Giardina |
| EPI_ISL_3385020 | KAISER SAN JOSE | Santa Clara County Public Health Laboratory | Santa Clara County Public Health Department |
| EPI_ISL_3385021 | A.S.L. CUNEO 1 | Fondazione del Piemonte per l'Oncologia IRCCS | Antonino Sottile, Silvia Brossa, Paola Marino, Giorgio Giardina |
| EPI_ISL_3385022 | PAMF PALO ALTO | Santa Clara County Public Health Laboratory | Santa Clara County Public Health Department |
| EPI_ISL_3385023 | A.S.L. CUNEO 1 | Fondazione del Piemonte per l'Oncologia IRCCS | Antonino Sottile, Silvia Brossa, Paola Marino, Giorgio Giardina |
| EPI_ISL_3385024 | PAMF PALO ALTO | Santa Clara County Public Health Laboratory | Santa Clara County Public Health Department |
| EPI_ISL_3385025 | A.S.L. TO5 | Fondazione del Piemonte per l'Oncologia IRCCS | Antonino Sottile, Silvia Brossa, Paola Marino, Giorgio Giardina |
| EPI_ISL_3385026 | KAISER SANTA CLARA | Santa Clara County Public Health Laboratory | Santa Clara County Public Health Department |
| EPI_ISL_3385029 | A.S.L. TO5 | Fondazione del Piemonte per l'Oncologia IRCCS | Antonino Sottile, Silvia Brossa, Paola Marino, Giorgio Giardina |
| EPI_ISL_3385030 | PAMF PALO ALTO | Santa Clara County Public Health Laboratory | Santa Clara County Public Health Department |
| EPI_ISL_3385089, EPI_ISL_3385090, EPI_ISL_3385091, EPI_ISL_3385092, EPI_ISL_3385093, EPI_ISL_3385094, EPI_ISL_3385095, EPI_ISL_3385096, EPI_ISL_3385097, EPI_ISL_3385098, EPI_ISL_3385099, EPI_ISL_3385100, EPI_ISL_3385101, EPI_ISL_3385102, EPI_ISL_3385103, EPI_ISL_3385104, EPI_ISL_3385105, EPI_ISL_3385106, EPI_ISL_3385107, EPI_ISL_3385109, EPI_ISL_3385110, EPI_ISL_3385111, EPI_ISL_3385112, EPI_ISL_3385113, EPI_ISL_3385114, EPI_ISL_3385115, EPI_ISL_3385116, EPI_ISL_3385117, EPI_ISL_3385118, EPI_ISL_3385119, EPI_ISL_3385120, EPI_ISL_3385121, EPI_ISL_3385122, EPI_ISL_3385123, EPI_ISL_3385124, EPI_ISL_3385125, EPI_ISL_3385126, EPI_ISL_3385127 | | | |
| see above | AZDelta | AZ Delta Medical Laboratories in Roeselare, Belgium | Geert Martens, Dieter De Smet, Frederik Van Hoecke, on behalf of AZ Delta COVID-19 Genomics core (member of Genomic surveillance of SARS-CoV-2 in Belgium network) |
| EPI_ISL_3385128, EPI_ISL_3385129, EPI_ISL_3385130, EPI_ISL_3385131, EPI_ISL_3385132, EPI_ISL_3385133, EPI_ISL_3385134, EPI_ISL_3385135, EPI_ISL_3385136, EPI_ISL_3385137, EPI_ISL_3385138, EPI_ISL_3385139 | | | |
| see above | LATE - Laboratório de Técnicas Especiais - Hospital Israelita Albert Einstein | LATE - Laboratório de Técnicas Especiais - Hospital Israelita Albert Einstein | Deyvid Amgarten, Fernanda de Mello Malta, Raquel Riyuzo, Ana Paula Moreira Salles, Pedro Henrique Sebe Rodrigues, Alexandre Hideaki Takara, Anelisie da Silva Santos, Erick Gustavo Dorlass, Marcio Anunciacao Menezes, João Renato Rebello Pinho |
| EPI_ISL_3385155, EPI_ISL_3385156, EPI_ISL_3385157, EPI_ISL_3385158, EPI_ISL_3385159, EPI_ISL_3385160, EPI_ISL_3385161, EPI_ISL_3385162, EPI_ISL_3385163, EPI_ISL_3385164, EPI_ISL_3385165, EPI_ISL_3385166, EPI_ISL_3385167, EPI_ISL_3385168, EPI_ISL_3385169, EPI_ISL_3385170, EPI_ISL_3385171, EPI_ISL_3385172, EPI_ISL_3385173, EPI_ISL_3385174, EPI_ISL_3385175, EPI_ISL_3385176, EPI_ISL_3385177, EPI_ISL_3385178, EPI_ISL_3385179, EPI_ISL_3385180, EPI_ISL_3385181, EPI_ISL_3385182, EPI_ISL_3385183, EPI_ISL_3385184, EPI_ISL_3385185, EPI_ISL_3385186, EPI_ISL_3385187, EPI_ISL_3385188, EPI_ISL_3385189, EPI_ISL_3385190, EPI_ISL_3385191 | | | |
| see above | Dipartimento di Medicina di Laboratorio, Azienda sanitaria universitaria Friuli Centrale (ASU FC) | Dipartimento di Medicina di Laboratorio, Azienda sanitaria universitaria Friuli Centrale (ASU FC) | Corrado Pipan, Catia Mio, Chiara Dal Secco, Stefania Marzinotto, Francesco Curcio |
| EPI_ISL_3385192, EPI_ISL_3385193, EPI_ISL_3385194, EPI_ISL_3385195, EPI_ISL_3385196, EPI_ISL_3385197, EPI_ISL_3385198, EPI_ISL_3385199, EPI_ISL_3385200, EPI_ISL_3385201, EPI_ISL_3385202, EPI_ISL_3385203, EPI_ISL_3385204, EPI_ISL_3385205, EPI_ISL_3385206, EPI_ISL_3385207, EPI_ISL_3385208, EPI_ISL_3385209, EPI_ISL_3385210, EPI_ISL_3385211, EPI_ISL_3385212, EPI_ISL_3385213, EPI_ISL_3385214, EPI_ISL_3385215, EPI_ISL_3385216, EPI_ISL_3385217, EPI_ISL_3385218, EPI_ISL_3385219 | | | |
| see above | Nebraska Public Health Laboratory | NPHL COVID-19 Response Team | NPHL COVID-19 Response Team |
| EPI_ISL_3385220, EPI_ISL_3385221, EPI_ISL_3385222, EPI_ISL_3385223, EPI_ISL_3385224, EPI_ISL_3385225, EPI_ISL_3385227, EPI_ISL_3385229, EPI_ISL_3385230, EPI_ISL_3385231, EPI_ISL_3385232, EPI_ISL_3385233, EPI_ISL_3385234, EPI_ISL_3385235, EPI_ISL_3385236, EPI_ISL_3385237 | | | |
| see above | TXDSHS | TXDSHS | Rashmi Tuladhar, Bonnie Oh, Jenny Zhang, Maliha Rahman, Mayela Pedrueza, Anita Pokharel, Karen Bobier, Lorraine Rodriguez, Myong Koag, Chun Wang, Rachel Lee, Grace Kubin |
| EPI_ISL_3385238, EPI_ISL_3385239, EPI_ISL_3385240, EPI_ISL_3385241, EPI_ISL_3385242, EPI_ISL_3385243, EPI_ISL_3385244, EPI_ISL_3385245, EPI_ISL_3385246, EPI_ISL_3385247, EPI_ISL_3385248, EPI_ISL_3385249, EPI_ISL_3385250, EPI_ISL_3385251, EPI_ISL_3385252, EPI_ISL_3385253, EPI_ISL_3385254, EPI_ISL_3385255, EPI_ISL_3385256, EPI_ISL_3385257, EPI_ISL_3385258, EPI_ISL_3385259, EPI_ISL_3385260, EPI_ISL_3385261, EPI_ISL_3385262, EPI_ISL_3385263, EPI_ISL_3385264, EPI_ISL_3385265, EPI_ISL_3385266, EPI_ISL_3385267, EPI_ISL_3385268, EPI_ISL_3385269, EPI_ISL_3385270, EPI_ISL_3385271, EPI_ISL_3385272, EPI_ISL_3385273, EPI_ISL_3385274, EPI_ISL_3385275, EPI_ISL_3385276, EPI_ISL_3385277, EPI_ISL_3385278, EPI_ISL_3385279, EPI_ISL_3385280, EPI_ISL_3385281, EPI_ISL_3385282, EPI_ISL_3385283, EPI_ISL_3385284, EPI_ISL_3385285, EPI_ISL_3385286, EPI_ISL_3385287, EPI_ISL_3385288, EPI_ISL_3385289, EPI_ISL_3385290, EPI_ISL_3385291, EPI_ISL_3385292, EPI_ISL_3385293, EPI_ISL_3385294, EPI_ISL_3385295, EPI_ISL_3385296, EPI_ISL_3385297, EPI_ISL_3385298, EPI_ISL_3385299 | | | |
| see above | NJDOH, Public Health and Environmental Laboratories | NJ_PHEL | Lindsey Bodnar, Shiv K. Verma, Ryan Pachucki, Chelsea San Filippo, Jacquelyn Deverell, Dana Woell, Allison Roder, Byeong Jeong |
| EPI_ISL_3385301 | AULSS 8 Berica | Istituto Zooprofilattico Sperimentale delle Venezie | Adelaide Milani, Alessia Schivo, Annalisa Salviato, Elisa Palumbo, Erika Giorgia Quaranta, Luca Tassoni, Ambra Pastori, Edoardo Giussani, Alice Fusaro, Isabella Monne, Calogero Terregino, Antonia Ricci |
| EPI_ISL_3385303, EPI_ISL_3385304, EPI_ISL_3385305, EPI_ISL_3385306, EPI_ISL_3385307, EPI_ISL_3385308, EPI_ISL_3385309, EPI_ISL_3385310, EPI_ISL_3385311 | AOUIVR | Istituto Zooprofilattico Sperimentale delle Venezie | Adelaide Milani, Alessia Schivo, Annalisa Salviato, Elisa Palumbo, Erika Giorgia Quaranta, Luca Tassoni, Ambra Pastori, Edoardo Giussani, Alice Fusaro, Isabella Monne, Calogero Terregino, Antonia Ricci |
| EPI_ISL_3385312 | AULSS 9 Scaligera - Legnago | Istituto Zooprofilattico Sperimentale delle Venezie | Adelaide Milani, Alessia Schivo, Annalisa Salviato, Elisa Palumbo, Erika Giorgia Quaranta, Luca Tassoni, Ambra Pastori, Edoardo Giussani, Alice Fusaro, Isabella Monne, Calogero Terregino, Antonia Ricci |
| EPI_ISL_3385313 | AOUIVR | Istituto Zooprofilattico Sperimentale delle Venezie | Adelaide Milani, Alessia Schivo, Annalisa Salviato, Elisa Palumbo, Erika Giorgia Quaranta, Luca Tassoni, Ambra Pastori, Edoardo Giussani, Alice Fusaro, Isabella Monne, Calogero Terregino, Antonia Ricci |
| EPI_ISL_3385314, EPI_ISL_3385315 | AULSS 9 Scaligera - Legnago | Istituto Zooprofilattico Sperimentale delle Venezie | Adelaide Milani, Alessia Schivo, Annalisa Salviato, Elisa Palumbo, Erika Giorgia Quaranta, Luca Tassoni, Ambra Pastori, Edoardo Giussani, Alice Fusaro, Isabella Monne, Calogero Terregino, Antonia Ricci |
| EPI_ISL_3385316 | AULSS 5 Polesana | Istituto Zooprofilattico Sperimentale delle Venezie | Adelaide Milani, Alessia Schivo, Annalisa Salviato, Elisa Palumbo, Erika Giorgia Quaranta, Luca Tassoni, Ambra Pastori, Edoardo Giussani, Alice Fusaro, Isabella Monne, Calogero Terregino, Antonia Ricci |
| EPI_ISL_3385317, EPI_ISL_3385318, EPI_ISL_3385319 | AULSS 9 Scaligera - Legnago | Istituto Zooprofilattico Sperimentale delle Venezie | Adelaide Milani, Alessia Schivo, Annalisa Salviato, Elisa Palumbo, Erika Giorgia Quaranta, Luca Tassoni, Ambra Pastori, Edoardo Giussani, Alice Fusaro, Isabella Monne, Calogero Terregino, Antonia Ricci |
| EPI_ISL_3385320, EPI_ISL_3385321, EPI_ISL_3385322, EPI_ISL_3385323 | AULSS 5 Polesana | Istituto Zooprofilattico Sperimentale delle Venezie | Adelaide Milani, Alessia Schivo, Annalisa Salviato, Elisa Palumbo, Erika Giorgia Quaranta, Luca Tassoni, Ambra Pastori, Edoardo Giussani, Alice Fusaro, Isabella Monne, Calogero Terregino, Antonia Ricci |
| EPI_ISL_3385324, EPI_ISL_3385325, EPI_ISL_3385326, EPI_ISL_3385327, EPI_ISL_3385328, EPI_ISL_3385329, EPI_ISL_3385330, EPI_ISL_3385331, EPI_ISL_3385332, EPI_ISL_3385333, EPI_ISL_3385334, EPI_ISL_3385335, EPI_ISL_3385336, EPI_ISL_3385337, EPI_ISL_3385338, EPI_ISL_3385339, EPI_ISL_3385340, EPI_ISL_3385341, EPI_ISL_3385342, EPI_ISL_3385343, EPI_ISL_3385344, EPI_ISL_3385345, EPI_ISL_3385346, EPI_ISL_3385347, EPI_ISL_3385348, EPI_ISL_3385349, EPI_ISL_3385350, EPI_ISL_3385351, EPI_ISL_3385352, EPI_ISL_3385354, EPI_ISL_3385355, EPI_ISL_3385356, EPI_ISL_3385357, EPI_ISL_3385358, EPI_ISL_3385359, EPI_ISL_3385360, EPI_ISL_3385361, EPI_ISL_3385362, EPI_ISL_3385363, EPI_ISL_3385364, EPI_ISL_3385365, EPI_ISL_3385366, EPI_ISL_3385367, EPI_ISL_3385368, EPI_ISL_3385369, EPI_ISL_3385371, EPI_ISL_3385372, EPI_ISL_3385373, EPI_ISL_3385374, EPI_ISL_3385375, EPI_ISL_3385376, EPI_ISL_3385377, EPI_ISL_3385378, EPI_ISL_3385379, EPI_ISL_3385380, EPI_ISL_3385382, EPI_ISL_3385383, EPI_ISL_3385384, EPI_ISL_3385385, EPI_ISL_3385386, EPI_ISL_3385387, EPI_ISL_3385388, EPI_ISL_3385389 | | | |
| see above | TXDSHS | TXDSHS | Rashmi Tuladhar, Bonnie Oh, Jenny Zhang, Maliha Rahman, Mayela Pedrueza, Anita Pokharel, Karen Bobier, Lorraine Rodriguez, Myong Koag, Chun Wang, Rachel Lee, Grace Kubin |
| EPI_ISL_3385390, EPI_ISL_3385391, EPI_ISL_3385392, EPI_ISL_3385393, EPI_ISL_3385394, EPI_ISL_3385395, EPI_ISL_3385396, EPI_ISL_3385397 | Texas Children's Hospital | Texas Children's Microbiome Center | Ruth Ann Luna, Jennifer K. Spinler, James Dunn, James Versalovic, Ila Singh |
| EPI_ISL_3385398, EPI_ISL_3385399 | RSUD PROVINSI NTB | Genomik Solidaritas Indonesia Laboratorium | Himawan Masyhuri, Carissa Sintca Wijaya, Vania Gavrila Wikasa, Dhahlia Agustina Cahyono, Gracia Felias Enos Korompis, Annisa Muthiah Sukirman, Anuraj Shankar, Louisa Markus, Meutia Ayuputeri Kumaheri, Ariel Pradipta |
| EPI_ISL_3385400, EPI_ISL_3385401 | RSUD PROVINSI NTB | Genomik Solidaritas Indonesia Laboratorium | Carissa Sintca Wijaya, Vania Gavrila Wikasa, Dhahlia Agustina Cahyono, Himawan Masyhuri, Gracia Felias Enos Korompis, Annisa Muthiah Sukirman, Anuraj Shankar, Louisa Markus, Meutia Ayuputeri Kumaheri, Ariel Pradipta |
| EPI_ISL_3385402 | Laboratorium Khusus Speedlab Jakarta | Genomik Solidaritas Indonesia Laboratorium | Carissa Sintca Wijaya, Vania Gavrila Wikasa, Dhahlia Agustina Cahyono, Himawan Masyhuri, Gracia Felias Enos Korompis, Annisa Muthiah Sukirman, Anuraj Shankar, Louisa Markus, Meutia Ayuputeri Kumaheri, Ariel Pradipta |
| EPI_ISL_3385403 | Laboratorium Khusus Speedlab Jakarta | Genomik Solidaritas Indonesia Laboratorium | Dhahlia Agustina Cahyono, Himawan Masyhuri, Carissa Sintca Wijaya, Vania Gavrila Wikasa, Annisa Muthiah Sukirman, Gracia Felias Enos Korompis, Anuraj Shankar, Louisa Markus, Meutia Ayuputeri Kumaheri, Ariel Pradipta |
| EPI_ISL_3385404, EPI_ISL_3385405, EPI_ISL_3385406 | Laboratorium Khusus Speedlab Jakarta | Genomik Solidaritas Indonesia Laboratorium | Vania Gavrila Wikasa, Dhahlia Agustina Cahyono, Himawan Masyhuri, Carissa Sintca Wijaya, Annisa Muthiah Sukirman, Gracia Felias Enos Korompis, Anuraj Shankar, Louisa Markus, Meutia Ayuputeri Kumaheri, Ariel Pradipta |
| EPI_ISL_3385407, EPI_ISL_3385408 | Laboratorium Khusus Speedlab Jakarta | Genomik Solidaritas Indonesia Laboratorium | Dhahlia Agustina Cahyono, Himawan Masyhuri, Carissa Sintca Wijaya, Vania Gavrila Wikasa, Annisa Muthiah Sukirman, Gracia Felias Enos Korompis, Anuraj Shankar, Louisa Markus, Meutia Ayuputeri Kumaheri, Ariel Pradipta |
| EPI_ISL_3385409 | Laboratorium Khusus Speedlab Jakarta | Genomik Solidaritas Indonesia Laboratorium | Himawan Masyhuri, Carissa Sintca Wijaya, Vania Gavrila Wikasa, Dhahlia Agustina Cahyono, Gracia Felias Enos Korompis, Annisa Muthiah Sukirman, Anuraj Shankar, Louisa Markus, Meutia Ayuputeri Kumaheri, Ariel Pradipta |
| EPI_ISL_3385410 | Laboratorium Khusus Speedlab Jakarta | Genomik Solidaritas Indonesia Laboratorium | Vania Gavrila Wikasa, Dhahlia Agustina Cahyono, Himawan Masyhuri, Carissa Sintca Wijaya, Annisa Muthiah Sukirman, Gracia Felias Enos Korompis, Anuraj Shankar, Louisa Markus, Meutia Ayuputeri Kumaheri, Ariel Pradipta |
| EPI_ISL_3385411, EPI_ISL_3385412 | Laboratorium Khusus Speedlab Jakarta | Genomik Solidaritas Indonesia Laboratorium | Himawan Masyhuri, Carissa Sintca Wijaya, Vania Gavrila Wikasa, Dhahlia Agustina Cahyono, Gracia Felias Enos Korompis, Annisa Muthiah Sukirman, Anuraj Shankar, Louisa Markus, Meutia Ayuputeri Kumaheri, Ariel Pradipta |
| EPI_ISL_3385413, EPI_ISL_3385414, EPI_ISL_3385415, EPI_ISL_3385416 | Laboratorium Khusus Speedlab Jakarta | Genomik Solidaritas Indonesia Laboratorium | Carissa Sintca Wijaya, Vania Gavrila Wikasa, Dhahlia Agustina Cahyono, Himawan Masyhuri, Gracia Felias Enos Korompis, Annisa Muthiah Sukirman, Anuraj Shankar, Louisa Markus, Meutia Ayuputeri Kumaheri, Ariel Pradipta |
| EPI_ISL_3385417 | LABORATORIUM RSUD PROF DR. W. Z. JOHANNES KUPANG | Genomik Solidaritas Indonesia Laboratorium | Vania Gavrila Wikasa, Dhahlia Agustina Cahyono, Hermi Indita Malewa, Chandra Apriadi Panduwal, Don Bosko Joni Dumbaris, Himawan Masyhuri, Carissa Sintca Wijaya, Annisa Muthiah Sukirman, Gracia Felias Enos Korompis, Anuraj Shankar, Louisa Markus, Meutia Ayuputeri Kumaheri, Ariel Pradipta |
| EPI_ISL_3385418, | LABORATORIUM RSUD PROF DR. W. Z. JOHANNES KUPANG | Genomik Solidaritas Indonesia Laboratorium | Dhahlia Agustina Cahyono, Himawan Masyhuri, Hermi Indita Malewa, Chandra Apriadi Panduwal, Don Bosko Joni Dumbaris, Carissa Sintca Wijaya, Vania Gavrila Wikasa, Annisa Muthiah Sukirman, Gracia Felias Enos Korompis, Anuraj Shankar, |

| | | | |
|---|---|---|---|
| EPI_ISL_3385419, EPI_ISL_3385420 | | | Louisa Markus, Meutia Ayuputeri Kumaheri, Ariel Pradipta |
| EPI_ISL_3385421 | LABORATORIUM RSUD PROF DR. W. Z. JOHANNES KUPANG | Genomik Solidaritas Indonesia Laboratorium | Himawan Masyhuri, Carissa Sintca Wijaya, Hermi Indita Malewa, Chandra Apriadi Panduwal, Don Bosko Joni Dumbaris, Vania Gavrila Wikasa, Dhahlia Agustina Cahyono, Gracia Felias Enos Korompis, Annisa Muthiah Sukirman, Anuraj Shankar, Louisa Markus, Meutia Ayuputeri Kumaheri, Ariel Pradipta |
| EPI_ISL_3385422 | LABORATORIUM RSUD PROF DR. W. Z. JOHANNES KUPANG | Genomik Solidaritas Indonesia Laboratorium | Vania Gavrila Wikasa, Dhahlia Agustina Cahyono, Hermi Indita Malewa, Chandra Apriadi Panduwal, Don Bosko Joni Dumbaris, Himawan Masyhuri, Carissa Sintca Wijaya, Annisa Muthiah Sukirman, Gracia Felias Enos Korompis, Anuraj Shankar, Louisa Markus, Meutia Ayuputeri Kumaheri, Ariel Pradipta |
| EPI_ISL_3385423, EPI_ISL_3385424, EPI_ISL_3385425, EPI_ISL_3385426, EPI_ISL_3385427, EPI_ISL_3385428, EPI_ISL_3385429, EPI_ISL_3385430, EPI_ISL_3385431, EPI_ISL_3385432, EPI_ISL_3385433, EPI_ISL_3385434 | | | |
| see above | LABORATORIUM RSUD PROF DR. W. Z. JOHANNES KUPANG | Genomik Solidaritas Indonesia Laboratorium | Himawan Masyhuri, Carissa Sintca Wijaya, Hermi Indita Malewa, Chandra Apriadi Panduwal, Don Bosko Joni Dumbaris, Vania Gavrila Wikasa, Dhahlia Agustina Cahyono, Gracia Felias Enos Korompis, Annisa Muthiah Sukirman, Anuraj Shankar, Louisa Markus, Meutia Ayuputeri Kumaheri, Ariel Pradipta |
| EPI_ISL_3385435, EPI_ISL_3385436, EPI_ISL_3385437, EPI_ISL_3385438, EPI_ISL_3385439, EPI_ISL_3385440, EPI_ISL_3385441 | LABORATORIUM RSUD PROF DR. W. Z. JOHANNES KUPANG | Genomik Solidaritas Indonesia Laboratorium | Vania Gavrila Wikasa, Dhahlia Agustina Cahyono, Hermi Indita Malewa, Chandra Apriadi Panduwal, Don Bosko Joni Dumbaris, Himawan Masyhuri, Carissa Sintca Wijaya, Annisa Muthiah Sukirman, Gracia Felias Enos Korompis, Anuraj Shankar, Louisa Markus, Meutia Ayuputeri Kumaheri, Ariel Pradipta |
| EPI_ISL_3385442, EPI_ISL_3385443 | Genomik Solidaritas Indonesia Laboratorium | Genomik Solidaritas Indonesia Laboratorium | Carissa Sintca Wijaya, Vania Gavrila Wikasa, Dhahlia Agustina Cahyono, Himawan Masyhuri, Gracia Felias Enos Korompis, Annisa Muthiah Sukirman, Anuraj Shankar, Louisa Markus, Meutia Ayuputeri Kumaheri, Ariel Pradipta |
| EPI_ISL_3385444, EPI_ISL_3385445, EPI_ISL_3385446, EPI_ISL_3385447, EPI_ISL_3385448 | LABORATORIUM RSUD PROF DR. W. Z. JOHANNES KUPANG | Genomik Solidaritas Indonesia Laboratorium | Vania Gavrila Wikasa, Dhahlia Agustina Cahyono, Hermi Indita Malewa, Chandra Apriadi Panduwal, Don Bosko Joni Dumbaris, Himawan Masyhuri, Carissa Sintca Wijaya, Annisa Muthiah Sukirman, Gracia Felias Enos Korompis, Anuraj Shankar, Louisa Markus, Meutia Ayuputeri Kumaheri, Ariel Pradipta |
| EPI_ISL_3385449, EPI_ISL_3385450, EPI_ISL_3385451, EPI_ISL_3385452, EPI_ISL_3385453, EPI_ISL_3385454, EPI_ISL_3385455, EPI_ISL_3385456, EPI_ISL_3385457, EPI_ISL_3385458, EPI_ISL_3385459, EPI_ISL_3385460, EPI_ISL_3385461 | | | |
| see above | LABORATORIUM RSUD PROF DR. W. Z. JOHANNES KUPANG | Genomik Solidaritas Indonesia Laboratorium | Dhahlia Agustina Cahyono, Himawan Masyhuri, Hermi Indita Malewa, Chandra Apriadi Panduwal, Don Bosko Joni Dumbaris, Carissa Sintca Wijaya, Vania Gavrila Wikasa, Annisa Muthiah Sukirman, Gracia Felias Enos Korompis, Anuraj Shankar, Louisa Markus, Meutia Ayuputeri Kumaheri, Ariel Pradipta |
| EPI_ISL_3385462, EPI_ISL_3385463, EPI_ISL_3385464, EPI_ISL_3385465, EPI_ISL_3385466 | LABORATORIUM RSUD PROF DR. W. Z. JOHANNES KUPANG | Genomik Solidaritas Indonesia Laboratorium | Carissa Sintca Wijaya, Vania Gavrila Wikasa, Hermi Indita Malewa, Chandra Apriadi Panduwal, Don Bosko Joni Dumbaris, Dhahlia Agustina Cahyono, Himawan Masyhuri, Gracia Felias Enos Korompis, Annisa Muthiah Sukirman, Anuraj Shankar, Louisa Markus, Meutia Ayuputeri Kumaheri, Ariel Pradipta |
| EPI_ISL_3385467, EPI_ISL_3385468 | Genomik Solidaritas Indonesia Laboratorium | Genomik Solidaritas Indonesia Laboratorium | Himawan Masyhuri, Carissa Sintca Wijaya, Vania Gavrila Wikasa, Dhahlia Agustina Cahyono, Gracia Felias Enos Korompis, Annisa Muthiah Sukirman, Anuraj Shankar, Louisa Markus, Meutia Ayuputeri Kumaheri, Ariel Pradipta |
| EPI_ISL_3385469, EPI_ISL_3385470, EPI_ISL_3385471 | Genomik Solidaritas Indonesia Laboratorium | Genomik Solidaritas Indonesia Laboratorium | Carissa Sintca Wijaya, Vania Gavrila Wikasa, Dhahlia Agustina Cahyono, Himawan Masyhuri, Gracia Felias Enos Korompis, Annisa Muthiah Sukirman, Anuraj Shankar, Louisa Markus, Meutia Ayuputeri Kumaheri, Ariel Pradipta |
| EPI_ISL_3385472 | Genomik Solidaritas Indonesia Laboratorium | Genomik Solidaritas Indonesia Laboratorium | Dhahlia Agustina Cahyono, Himawan Masyhuri, Carissa Sintca Wijaya, Vania Gavrila Wikasa, Annisa Muthiah Sukirman, Gracia Felias Enos Korompis, Anuraj Shankar, Louisa Markus, Meutia Ayuputeri Kumaheri, Ariel Pradipta |
| EPI_ISL_3385473, EPI_ISL_3385474, EPI_ISL_3385475 | Genomik Solidaritas Indonesia Laboratorium | Genomik Solidaritas Indonesia Laboratorium | Vania Gavrila Wikasa, Dhahlia Agustina Cahyono, Himawan Masyhuri, Carissa Sintca Wijaya, Annisa Muthiah Sukirman, Gracia Felias Enos Korompis, Anuraj Shankar, Louisa Markus, Meutia Ayuputeri Kumaheri, Ariel Pradipta |
| EPI_ISL_3385476, EPI_ISL_3385477, EPI_ISL_3385478 | Genomik Solidaritas Indonesia Laboratorium | Genomik Solidaritas Indonesia Laboratorium | Dhahlia Agustina Cahyono, Himawan Masyhuri, Carissa Sintca Wijaya, Vania Gavrila Wikasa, Annisa Muthiah Sukirman, Gracia Felias Enos Korompis, Anuraj Shankar, Louisa Markus, Meutia Ayuputeri Kumaheri, Ariel Pradipta |
| EPI_ISL_3385479 | Genomik Solidaritas Indonesia Laboratorium | Genomik Solidaritas Indonesia Laboratorium | Himawan Masyhuri, Carissa Sintca Wijaya, Vania Gavrila Wikasa, Dhahlia Agustina Cahyono, Gracia Felias Enos Korompis, Annisa Muthiah Sukirman, Anuraj Shankar, Louisa Markus, Meutia Ayuputeri Kumaheri, Ariel Pradipta |
| EPI_ISL_3385480, EPI_ISL_3385481 | Genomik Solidaritas Indonesia Laboratorium | Genomik Solidaritas Indonesia Laboratorium | Carissa Sintca Wijaya, Vania Gavrila Wikasa, Dhahlia Agustina Cahyono, Himawan Masyhuri, Gracia Felias Enos Korompis, Annisa Muthiah Sukirman, Anuraj Shankar, Louisa Markus, Meutia Ayuputeri Kumaheri, Ariel Pradipta |
| EPI_ISL_3385482 | Genomik Solidaritas Indonesia Laboratorium | Genomik Solidaritas Indonesia Laboratorium | Dhahlia Agustina Cahyono, Himawan Masyhuri, Carissa Sintca Wijaya, Vania Gavrila Wikasa, Annisa Muthiah Sukirman, Gracia Felias Enos Korompis, Anuraj Shankar, Louisa Markus, Meutia Ayuputeri Kumaheri, Ariel Pradipta |
| EPI_ISL_3385483, EPI_ISL_3385484, EPI_ISL_3385485 | Genomik Solidaritas Indonesia Laboratorium | Genomik Solidaritas Indonesia Laboratorium | Vania Gavrila Wikasa, Dhahlia Agustina Cahyono, Himawan Masyhuri, Carissa Sintca Wijaya, Annisa Muthiah Sukirman, Gracia Felias Enos Korompis, Anuraj Shankar, Louisa Markus, Meutia Ayuputeri Kumaheri, Ariel Pradipta |
| EPI_ISL_3385486, EPI_ISL_3385487 | Genomik Solidaritas Indonesia Laboratorium | Genomik Solidaritas Indonesia Laboratorium | Dhahlia Agustina Cahyono, Himawan Masyhuri, Carissa Sintca Wijaya, Vania Gavrila Wikasa, Annisa Muthiah Sukirman, Gracia Felias Enos Korompis, Anuraj Shankar, Louisa Markus, Meutia Ayuputeri Kumaheri, Ariel Pradipta |
| EPI_ISL_3385488, EPI_ISL_3385489, EPI_ISL_3385490, EPI_ISL_3385491, EPI_ISL_3385492, EPI_ISL_3385493, EPI_ISL_3385494, EPI_ISL_3385495, EPI_ISL_3385496, EPI_ISL_3385497, EPI_ISL_3385498, EPI_ISL_3385499, EPI_ISL_3385500, EPI_ISL_3385501, EPI_ISL_3385502, EPI_ISL_3385503, EPI_ISL_3385504, EPI_ISL_3385505, EPI_ISL_3385507, EPI_ISL_3385508, EPI_ISL_3385509, EPI_ISL_3385510, EPI_ISL_3385511, EPI_ISL_3385512, EPI_ISL_3385513, EPI_ISL_3385514, EPI_ISL_3385515, EPI_ISL_3385516, EPI_ISL_3385518, EPI_ISL_3385519, EPI_ISL_3385520, EPI_ISL_3385521, EPI_ISL_3385522, EPI_ISL_3385523, EPI_ISL_3385524, EPI_ISL_3385525, EPI_ISL_3385526, EPI_ISL_3385527, EPI_ISL_3385528, EPI_ISL_3385529, EPI_ISL_3385530, EPI_ISL_3385531, EPI_ISL_3385532, EPI_ISL_3385533, EPI_ISL_3385534, EPI_ISL_3385535, EPI_ISL_3385536, EPI_ISL_3385537, EPI_ISL_3385538, EPI_ISL_3385539, EPI_ISL_3385540, EPI_ISL_3385541, EPI_ISL_3385542, EPI_ISL_3385543, EPI_ISL_3385544, EPI_ISL_3385545, EPI_ISL_3385546, EPI_ISL_3385547, EPI_ISL_3385548, EPI_ISL_3385549, EPI_ISL_3385550, EPI_ISL_3385551, EPI_ISL_3385552, EPI_ISL_3385553, EPI_ISL_3385554, EPI_ISL_3385555, EPI_ISL_3385556, EPI_ISL_3385557, EPI_ISL_3385558, EPI_ISL_3385559, EPI_ISL_3385560, EPI_ISL_3385561, EPI_ISL_3385562, EPI_ISL_3385563, EPI_ISL_3385564, EPI_ISL_3385565, EPI_ISL_3385567, EPI_ISL_3385568, EPI_ISL_3385569, EPI_ISL_3385570, EPI_ISL_3385571, EPI_ISL_3385572, EPI_ISL_3385573, EPI_ISL_3385574, EPI_ISL_3385575, EPI_ISL_3385576, EPI_ISL_3385577, EPI_ISL_3385578, EPI_ISL_3385579, EPI_ISL_3385580, EPI_ISL_3385581, EPI_ISL_3385582, EPI_ISL_3385583, EPI_ISL_3385584, EPI_ISL_3385585, EPI_ISL_3385586, EPI_ISL_3385587, EPI_ISL_3385588, EPI_ISL_3385589, EPI_ISL_3385590, EPI_ISL_3385591, EPI_ISL_3385592, EPI_ISL_3385593, EPI_ISL_3385594, EPI_ISL_3385595, EPI_ISL_3385596, EPI_ISL_3385597, EPI_ISL_3385598, EPI_ISL_3385599, EPI_ISL_3385600, EPI_ISL_3385601, EPI_ISL_3385602, EPI_ISL_3385603, EPI_ISL_3385604, EPI_ISL_3385605, EPI_ISL_3385606, EPI_ISL_3385607, EPI_ISL_3385608, EPI_ISL_3385609, EPI_ISL_3385610, EPI_ISL_3385611, EPI_ISL_3385612, EPI_ISL_3385613, EPI_ISL_3385614, EPI_ISL_3385615, EPI_ISL_3385616, EPI_ISL_3385617, EPI_ISL_3385618, EPI_ISL_3385619, EPI_ISL_3385620, EPI_ISL_3385621, EPI_ISL_3385622, EPI_ISL_3385623, EPI_ISL_3385624, EPI_ISL_3385625, EPI_ISL_3385626, EPI_ISL_3385627, EPI_ISL_3385628, EPI_ISL_3385629, EPI_ISL_3385630, EPI_ISL_3385631, EPI_ISL_3385632, EPI_ISL_3385633, EPI_ISL_3385634, EPI_ISL_3385635, EPI_ISL_3385636, EPI_ISL_3385637, EPI_ISL_3385638, EPI_ISL_3385639, EPI_ISL_3385640, EPI_ISL_3385641, EPI_ISL_3385642, EPI_ISL_3385643, EPI_ISL_3385644, EPI_ISL_3385645, EPI_ISL_3385646, EPI_ISL_3385647, EPI_ISL_3385648, EPI_ISL_3385649, EPI_ISL_3385650, EPI_ISL_3385651, EPI_ISL_3385652, EPI_ISL_3385653, EPI_ISL_3385654, EPI_ISL_3385655, EPI_ISL_3385656, EPI_ISL_3385657, EPI_ISL_3385658, EPI_ISL_3385659, EPI_ISL_3385660, EPI_ISL_3385661, EPI_ISL_3385662, EPI_ISL_3385663, EPI_ISL_3385664, EPI_ISL_3385665, EPI_ISL_3385666, EPI_ISL_3385667, EPI_ISL_3385669, EPI_ISL_3385670, EPI_ISL_3385671, EPI_ISL_3385672, EPI_ISL_3385673, EPI_ISL_3385674, EPI_ISL_3385675, EPI_ISL_3385676, EPI_ISL_3385677, EPI_ISL_3385678, EPI_ISL_3385679, EPI_ISL_3385680, EPI_ISL_3385681, EPI_ISL_3385682, EPI_ISL_3385683, EPI_ISL_3385684, EPI_ISL_3385685, EPI_ISL_3385686, EPI_ISL_3385687, EPI_ISL_3385688, EPI_ISL_3385689, EPI_ISL_3385690, EPI_ISL_3385691, EPI_ISL_3385692, EPI_ISL_3385693, EPI_ISL_3385694, EPI_ISL_3385695, EPI_ISL_3385696, EPI_ISL_3385697, EPI_ISL_3385698, EPI_ISL_3385699, EPI_ISL_3385700, EPI_ISL_3385701, EPI_ISL_3385702, EPI_ISL_3385703, EPI_ISL_3385704, EPI_ISL_3385705, EPI_ISL_3385706, EPI_ISL_3385707, EPI_ISL_3385708, EPI_ISL_3385709, EPI_ISL_3385710, EPI_ISL_3385711, EPI_ISL_3385712, EPI_ISL_3385713, EPI_ISL_3385714, EPI_ISL_3385715, EPI_ISL_3385716, EPI_ISL_3385717, EPI_ISL_3385718, EPI_ISL_3385719, EPI_ISL_3385720, EPI_ISL_3385721, EPI_ISL_3385722, EPI_ISL_3385723, EPI_ISL_3385724, EPI_ISL_3385725, EPI_ISL_3385726, EPI_ISL_3385727, EPI_ISL_3385729, EPI_ISL_3385730, EPI_ISL_3385731, EPI_ISL_3385732, EPI_ISL_3385733, EPI_ISL_3385734, EPI_ISL_3385735, EPI_ISL_3385736, EPI_ISL_3385737, EPI_ISL_3385738, EPI_ISL_3385739, EPI_ISL_3385740, EPI_ISL_3385741, EPI_ISL_3385742, EPI_ISL_3385743, EPI_ISL_3385745, EPI_ISL_3385746, EPI_ISL_3385747, EPI_ISL_3385748, EPI_ISL_3385749, EPI_ISL_3385750, EPI_ISL_3385751, EPI_ISL_3385752, EPI_ISL_3385753, EPI_ISL_3385754, EPI_ISL_3385755, EPI_ISL_3385756, EPI_ISL_3385757, EPI_ISL_3385758, EPI_ISL_3385759, EPI_ISL_3385760, EPI_ISL_3385761, EPI_ISL_3385762, EPI_ISL_3385763, EPI_ISL_3385764, EPI_ISL_3385765, EPI_ISL_3385766, EPI_ISL_3385767 | | | |
| see above | Arizona State University | Arizona State University | Ajeet Bains, LaRinda A. Holland, Matthew F. Smith, Peter T. Skidmore, Rabia Maqsood, Nicholas J. Mellor, Nathaniel Johnson, Joshua LaBaer, Vel Murugan, Efrem S. Lim |
| EPI_ISL_3385856, EPI_ISL_3385857, EPI_ISL_3385858, EPI_ISL_3385859, EPI_ISL_3385860, EPI_ISL_3385861, EPI_ISL_3385862, EPI_ISL_3385863, EPI_ISL_3385864, EPI_ISL_3385865, EPI_ISL_3385866, EPI_ISL_3385867, EPI_ISL_3385868, EPI_ISL_3385869, EPI_ISL_3385870, EPI_ISL_3385871, EPI_ISL_3385872, EPI_ISL_3385873, EPI_ISL_3385874, EPI_ISL_3385875, EPI_ISL_3385876, EPI_ISL_3385877, EPI_ISL_3385878, EPI_ISL_3385879, EPI_ISL_3385880, EPI_ISL_3385881, EPI_ISL_3385882, EPI_ISL_3385883, EPI_ISL_3385884, EPI_ISL_3385885, EPI_ISL_3385886, EPI_ISL_3385887, EPI_ISL_3385888, EPI_ISL_3385889, EPI_ISL_3385890, EPI_ISL_3385891, EPI_ISL_3385892, EPI_ISL_3385893, EPI_ISL_3385894, EPI_ISL_3385895, EPI_ISL_3385896, EPI_ISL_3385897, EPI_ISL_3385898, EPI_ISL_3385899, EPI_ISL_3385900, EPI_ISL_3385901, EPI_ISL_3385902, EPI_ISL_3385903, EPI_ISL_3385905, EPI_ISL_3385906, EPI_ISL_3385907, EPI_ISL_3385908, EPI_ISL_3385909, EPI_ISL_3385910, EPI_ISL_3385911, EPI_ISL_3385912, EPI_ISL_3385913, EPI_ISL_3385914, EPI_ISL_3385915, EPI_ISL_3385916, EPI_ISL_3385917, EPI_ISL_3385918, EPI_ISL_3385919, EPI_ISL_3385920, EPI_ISL_3385921, EPI_ISL_3385922, EPI_ISL_3385923, EPI_ISL_3385924, EPI_ISL_3385925, EPI_ISL_3385926, EPI_ISL_3385927, EPI_ISL_3385928, EPI_ISL_3385929, EPI_ISL_3385930, EPI_ISL_3385931, EPI_ISL_3385932, EPI_ISL_3385933, EPI_ISL_3385934, EPI_ISL_3385935, EPI_ISL_3385936, EPI_ISL_3385937, EPI_ISL_3385938, EPI_ISL_3385939, EPI_ISL_3385940, EPI_ISL_3385941 | | | |
| see above | Idaho Bureau of Laboratories | Idaho Bureau of Laboratories | "R. Beukelman, Matthew Charles Burns, Aimee Ceniseros, Christian Loera, Robert L. Voermans, Christopher Ball" |
| EPI_ISL_3385942 | AULSS 8 Berica | Istituto Zooprofilattico Sperimentale delle Venezie | Adelaide Milani, Alessia Schivo, Annalisa Salviato, Elisa Palumbo, Erika Giorgia Quaranta, Luca Tassoni, Ambra Pastori, Edoardo Giussani, Alice Fusaro, Isabella Monne, Calogero Terregino, Antonia Ricci |
| EPI_ISL_3385943 | AULSS 7 Pedemontana - Santorso | Istituto Zooprofilattico Sperimentale delle Venezie | Adelaide Milani, Alessia Schivo, Annalisa Salviato, Elisa Palumbo, Erika Giorgia Quaranta, Luca Tassoni, Ambra Pastori, Edoardo Giussani, Alice Fusaro, Isabella Monne, Calogero Terregino, Antonia Ricci |
| EPI_ISL_3385944, EPI_ISL_3385945, EPI_ISL_3385947, EPI_ISL_3385948, EPI_ISL_3385953, EPI_ISL_3385955, EPI_ISL_3385956 | AULSS 8 Berica | Istituto Zooprofilattico Sperimentale delle Venezie | Adelaide Milani, Alessia Schivo, Annalisa Salviato, Elisa Palumbo, Erika Giorgia Quaranta, Luca Tassoni, Ambra Pastori, Edoardo Giussani, Alice Fusaro, Isabella Monne, Calogero Terregino, Antonia Ricci |
| EPI_ISL_3385957, EPI_ISL_3385958, EPI_ISL_3385959, EPI_ISL_3385960, EPI_ISL_3385961, EPI_ISL_3385962, EPI_ISL_3385963 | AOUIVR | Istituto Zooprofilattico Sperimentale delle Venezie | Adelaide Milani, Alessia Schivo, Annalisa Salviato, Elisa Palumbo, Erika Giorgia Quaranta, Luca Tassoni, Ambra Pastori, Edoardo Giussani, Alice Fusaro, Isabella Monne, Calogero Terregino, Antonia Ricci |
| EPI_ISL_3385967, EPI_ISL_3385968, EPI_ISL_3385970 | AULSS 9 Scaligera - Legnago | Istituto Zooprofilattico Sperimentale delle Venezie | Adelaide Milani, Alessia Schivo, Annalisa Salviato, Elisa Palumbo, Erika Giorgia Quaranta, Luca Tassoni, Ambra Pastori, Edoardo Giussani, Alice Fusaro, Isabella Monne, Calogero Terregino, Antonia Ricci |
| EPI_ISL_3385971 | AULSS 1 Dolomiti | Istituto Zooprofilattico Sperimentale delle Venezie | Adelaide Milani, Alessia Schivo, Annalisa Salviato, Elisa Palumbo, Erika Giorgia Quaranta, Luca Tassoni, Ambra Pastori, Edoardo Giussani, Alice Fusaro, Isabella Monne, Calogero Terregino, Antonia Ricci |
| EPI_ISL_3385972 | AULSS 9 Scaligera - Legnago | Istituto Zooprofilattico Sperimentale delle Venezie | Adelaide Milani, Alessia Schivo, Annalisa Salviato, Elisa Palumbo, Erika Giorgia Quaranta, Luca Tassoni, Ambra Pastori, Edoardo Giussani, Alice Fusaro, Isabella Monne, Calogero Terregino, Antonia Ricci |
| EPI_ISL_3385973, EPI_ISL_3385974, EPI_ISL_3385975, EPI_ISL_3385976, | AULSS 1 Dolomiti | Istituto Zooprofilattico Sperimentale delle Venezie | Adelaide Milani, Alessia Schivo, Annalisa Salviato, Elisa Palumbo, Erika Giorgia Quaranta, Luca Tassoni, Ambra Pastori, Edoardo Giussani, Alice Fusaro, Isabella Monne, Calogero Terregino, Antonia Ricci |

| | | | |
|---|---|---|---|
| EPI_ISL_3385977, EPI_ISL_3385980, EPI_ISL_3385981 | | | |
| EPI_ISL_3385982, EPI_ISL_3385983, EPI_ISL_3385984, EPI_ISL_3385985, EPI_ISL_3385986, EPI_ISL_3385987, EPI_ISL_3385988, EPI_ISL_3385989, EPI_ISL_3385990, EPI_ISL_3385991, EPI_ISL_3385992, EPI_ISL_3385993, EPI_ISL_3385994 | | | |
| see above | AULSS 9 Scaligera - Villafranca | Istituto Zooprofilattico Sperimentale delle Venezie | Adelaide Milani, Alessia Schivo, Annalisa Salviato, Elisa Palumbo, Erika Giorgia Quaranta, Luca Tassoni, Ambra Pastori, Edoardo Giussani, Alice Fusaro, Isabella Monne, Calogero Terregino, Antonia Ricci |
| EPI_ISL_3385995 | AULSS 7 Pedemontana - Santorso | Istituto Zooprofilattico Sperimentale delle Venezie | Adelaide Milani, Alessia Schivo, Annalisa Salviato, Elisa Palumbo, Erika Giorgia Quaranta, Luca Tassoni, Ambra Pastori, Edoardo Giussani, Alice Fusaro, Isabella Monne, Calogero Terregino, Antonia Ricci |
| EPI_ISL_3385996 | AULSS 8 Berica | Istituto Zooprofilattico Sperimentale delle Venezie | Adelaide Milani, Alessia Schivo, Annalisa Salviato, Elisa Palumbo, Erika Giorgia Quaranta, Luca Tassoni, Ambra Pastori, Edoardo Giussani, Alice Fusaro, Isabella Monne, Calogero Terregino, Antonia Ricci |
| EPI_ISL_3385997, EPI_ISL_3385998 | AULSS 7 Pedemontana - Santorso | Istituto Zooprofilattico Sperimentale delle Venezie | Adelaide Milani, Alessia Schivo, Annalisa Salviato, Elisa Palumbo, Erika Giorgia Quaranta, Luca Tassoni, Ambra Pastori, Edoardo Giussani, Alice Fusaro, Isabella Monne, Calogero Terregino, Antonia Ricci |
| EPI_ISL_3386000, EPI_ISL_3386001, EPI_ISL_3386002, EPI_ISL_3386004, EPI_ISL_3386006 | AULSS 8 Berica | Istituto Zooprofilattico Sperimentale delle Venezie | Adelaide Milani, Alessia Schivo, Annalisa Salviato, Elisa Palumbo, Erika Giorgia Quaranta, Luca Tassoni, Ambra Pastori, Edoardo Giussani, Alice Fusaro, Isabella Monne, Calogero Terregino, Antonia Ricci |
| EPI_ISL_3386008 | Dipartimento di Medicina di Laboratorio, Azienda sanitaria universitaria Friuli Centrale (ASU FC) | Dipartimento di Medicina di Laboratorio, Azienda sanitaria universitaria Friuli Centrale (ASU FC) | Corrado Pipan, Catia Mio, Chiara Dal Secco, Stefania Marzinotto, Francesco Curcio |
| EPI_ISL_3386009 | AULSS 7 Pedemontana - Santorso | Istituto Zooprofilattico Sperimentale delle Venezie | Adelaide Milani, Alessia Schivo, Annalisa Salviato, Elisa Palumbo, Erika Giorgia Quaranta, Luca Tassoni, Ambra Pastori, Edoardo Giussani, Alice Fusaro, Isabella Monne, Calogero Terregino, Antonia Ricci |
| EPI_ISL_3386014, EPI_ISL_3386016, EPI_ISL_3386019, EPI_ISL_3386021, EPI_ISL_3386022, EPI_ISL_3386029, EPI_ISL_3386037, EPI_ISL_3386038, EPI_ISL_3386039, EPI_ISL_3386040, EPI_ISL_3386045, EPI_ISL_3386048, EPI_ISL_3386049, EPI_ISL_3386050, EPI_ISL_3386053, EPI_ISL_3386054, EPI_ISL_3386055, EPI_ISL_3386057, EPI_ISL_3386058, EPI_ISL_3386059, EPI_ISL_3386072, EPI_ISL_3386073, EPI_ISL_3386074, EPI_ISL_3386075, EPI_ISL_3386077, EPI_ISL_3386083, EPI_ISL_3386085, EPI_ISL_3386086, EPI_ISL_3386087, EPI_ISL_3386088, EPI_ISL_3386090, EPI_ISL_3386091, EPI_ISL_3386092 | | | |
| see above | Colorado Department of Public Health and Environment | Colorado Department of Public Health and Environment | Laura Bankers, Molly C. Hetherington-Rauth, Diana Ir, Alexandria Rossheim, Michael A. Martin, Mandy Waters, Shannon R. Matzinger, Sarah Elizabeth Totten, Emily A. Travanty |
| EPI_ISL_3386093 | Laboratorium Khusus Speedlab Jakarta | Genomik Solidaritas Indonesia Laboratorium | Dhahlia Agustina Cahyono, Himawan Masyhuri, Carissa Sintca Wijaya, Vania Gavrila Wikasa, Annisa Muthiah Sukirman, Gracia Felias Enos Korompis, Anuraj Shankar, Louisa Markus, Meutia Ayuputeri Kumaheri, Ariel Pradipta |
| EPI_ISL_3386094, EPI_ISL_3386095 | LABORATORIUM RSUD PROF DR. W. Z. JOHANNES KUPANG | Genomik Solidaritas Indonesia Laboratorium | Vania Gavrila Wikasa, Dhahlia Agustina Cahyono, Hermi Indita Malewa, Chandra Apriadi Panduwal, Don Bosko Joni Dumbaris, Himawan Masyhuri, Carissa Sintca Wijaya, Annisa Muthiah Sukirman, Gracia Felias Enos Korompis, Anuraj Shankar, Louisa Markus, Meutia Ayuputeri Kumaheri, Ariel Pradipta |
| EPI_ISL_3386143 | HLAGYN - Laboratorio de Imunologia de Transplantes de Goias | HLAGYN - Laboratorio de Imunologia de Transplantes de Goias | Fernando Antonio Vinhal dos Santos, Erika Lopes Rocha Batista, Alessandro Leonardo Alvares Magalhaes, Frederico Rodrigues Vinhal, Sabrina Sara Moreira Duarte, Lucas Carlos Gomes Pereira, Kamila Oliveira Reis De Freitas. |
| EPI_ISL_3386158, EPI_ISL_3386159, EPI_ISL_3386160, EPI_ISL_3386161, EPI_ISL_3386162, EPI_ISL_3386163, EPI_ISL_3386164, EPI_ISL_3386165, EPI_ISL_3386166, EPI_ISL_3386167, EPI_ISL_3386168, EPI_ISL_3386169, EPI_ISL_3386170, EPI_ISL_3386171, EPI_ISL_3386177, EPI_ISL_3386179, EPI_ISL_3386180, EPI_ISL_3386181, EPI_ISL_3386183, EPI_ISL_3386185, EPI_ISL_3386186, EPI_ISL_3386188, EPI_ISL_3386189, EPI_ISL_3386190, EPI_ISL_3386193, EPI_ISL_3386195, EPI_ISL_3386200, EPI_ISL_3386201, EPI_ISL_3386203, EPI_ISL_3386204, EPI_ISL_3386205, EPI_ISL_3386206, EPI_ISL_3386208, EPI_ISL_3386209, EPI_ISL_3386212, EPI_ISL_3386215, EPI_ISL_3386216, EPI_ISL_3386217, EPI_ISL_3386218, EPI_ISL_3386219, EPI_ISL_3386220, EPI_ISL_3386222, EPI_ISL_3386223, EPI_ISL_3386224, EPI_ISL_3386225, EPI_ISL_3386226, EPI_ISL_3386227, EPI_ISL_3386228, EPI_ISL_3386229, EPI_ISL_3386231, EPI_ISL_3386232, EPI_ISL_3386233, EPI_ISL_3386234, EPI_ISL_3386235, EPI_ISL_3386236, EPI_ISL_3386237, EPI_ISL_3386238, EPI_ISL_3386239, EPI_ISL_3386240, EPI_ISL_3386241, EPI_ISL_3386242, EPI_ISL_3386243, EPI_ISL_3386245, EPI_ISL_3386246, EPI_ISL_3386247, EPI_ISL_3386248, EPI_ISL_3386249, EPI_ISL_3386250, EPI_ISL_3386251, EPI_ISL_3386252, EPI_ISL_3386253, EPI_ISL_3386254, EPI_ISL_3386255, EPI_ISL_3386256, EPI_ISL_3386257, EPI_ISL_3386258, EPI_ISL_3386259, EPI_ISL_3386260, EPI_ISL_3386261, EPI_ISL_3386262, EPI_ISL_3386263, EPI_ISL_3386264, EPI_ISL_3386265, EPI_ISL_3386266, EPI_ISL_3386267, EPI_ISL_3386269, EPI_ISL_3386271, EPI_ISL_3386272, EPI_ISL_3386273, EPI_ISL_3386274, EPI_ISL_3386275, EPI_ISL_3386277, EPI_ISL_3386278, EPI_ISL_3386279, EPI_ISL_3386280, EPI_ISL_3386282, EPI_ISL_3386283, EPI_ISL_3386284, EPI_ISL_3386285, EPI_ISL_3386286, EPI_ISL_3386287, EPI_ISL_3386288, EPI_ISL_3386289, EPI_ISL_3386291, EPI_ISL_3386292, EPI_ISL_3386293, EPI_ISL_3386294, EPI_ISL_3386295, EPI_ISL_3386296, EPI_ISL_3386297, EPI_ISL_3386298, EPI_ISL_3386300, EPI_ISL_3386301, EPI_ISL_3386302, EPI_ISL_3386303, EPI_ISL_3386304, EPI_ISL_3386305, EPI_ISL_3386306, EPI_ISL_3386307, EPI_ISL_3386308, EPI_ISL_3386309, EPI_ISL_3386312, EPI_ISL_3386313, EPI_ISL_3386314, EPI_ISL_3386315, EPI_ISL_3386316, EPI_ISL_3386317, EPI_ISL_3386319, EPI_ISL_3386320, EPI_ISL_3386321, EPI_ISL_3386322, EPI_ISL_3386323, EPI_ISL_3386324, EPI_ISL_3386325, EPI_ISL_3386326, EPI_ISL_3386327, EPI_ISL_3386328, EPI_ISL_3386329, EPI_ISL_3386330, EPI_ISL_3386331, EPI_ISL_3386332, EPI_ISL_3386333, EPI_ISL_3386334, EPI_ISL_3386335, EPI_ISL_3386336, EPI_ISL_3386337, EPI_ISL_3386338, EPI_ISL_3386340, EPI_ISL_3386341, EPI_ISL_3386342, EPI_ISL_3386344, EPI_ISL_3386346, EPI_ISL_3386348, EPI_ISL_3386349, EPI_ISL_3386350, EPI_ISL_3386352, EPI_ISL_3386354, EPI_ISL_3386355, EPI_ISL_3386356, EPI_ISL_3386357, EPI_ISL_3386358, EPI_ISL_3386360, EPI_ISL_3386362, EPI_ISL_3386363, EPI_ISL_3386364, EPI_ISL_3386365, EPI_ISL_3386366, EPI_ISL_3386367, EPI_ISL_3386368, EPI_ISL_3386369, EPI_ISL_3386370, EPI_ISL_3386372, EPI_ISL_3386373, EPI_ISL_3386375, EPI_ISL_3386376, EPI_ISL_3386377, EPI_ISL_3386378, EPI_ISL_3386379, EPI_ISL_3386380, EPI_ISL_3386381, EPI_ISL_3386382, EPI_ISL_3386383, EPI_ISL_3386384, EPI_ISL_3386385, EPI_ISL_3386386, EPI_ISL_3386387, EPI_ISL_3386388, EPI_ISL_3386389, EPI_ISL_3386390, EPI_ISL_3386391, EPI_ISL_3386392, EPI_ISL_3386393, EPI_ISL_3386395 | | | |
| see above | Utah Public Health Laboratory | Utah Public Health Laboratory | Erin L. Young, Kelly F. Oakeson, Pooja Gupta, Olinto Linares-Perdomo |
| EPI_ISL_3386396, EPI_ISL_3386398, EPI_ISL_3386399 | Arizona State University | Arizona State University | Ajeet Bains, LaRinda A. Holland, Matthew F. Smith, Peter T. Skidmore, Rabia Maqsood, Nicholas J. Mellor, Nathaniel Johnson, Joshua LaBaer, Vel Murugan, Efrem S. Lim |
| EPI_ISL_3386401 | CAP El Remei | Banc de Sang i Teixits | Francisco Vidal, Irene Corrales, Nina Borràs, Carlos Hobeich, Natália Comes, Lorena Ramírez, Noemí Gonzalez, Silvia Sauleda, Maria Glòria Soria |
| EPI_ISL_3386403 | CAP Vic Nord | Banc de Sang i Teixits | Francisco Vidal, Irene Corrales, Nina Borràs, Carlos Hobeich, Natália Comes, Lorena Ramírez, Noemí Gonzalez, Silvia Sauleda, Maria Glòria Soria |
| EPI_ISL_3386404 | CAP Manlleu | Banc de Sang i Teixits | Francisco Vidal, Irene Corrales, Nina Borràs, Carlos Hobeich, Natália Comes, Lorena Ramírez, Noemí Gonzalez, Silvia Sauleda, Maria Glòria Soria |
| EPI_ISL_3386406, EPI_ISL_3386407 | CAP El Remei | Banc de Sang i Teixits | Francisco Vidal, Irene Corrales, Nina Borràs, Carlos Hobeich, Natália Comes, Lorena Ramírez, Noemí Gonzalez, Silvia Sauleda, Maria Glòria Soria |
| EPI_ISL_3386408 | CAP Sant Hipolit de Voltrega | Banc de Sang i Teixits | Francisco Vidal, Irene Corrales, Nina Borràs, Carlos Hobeich, Natália Comes, Lorena Ramírez, Noemí Gonzalez, Silvia Sauleda, Maria Glòria Soria |
| EPI_ISL_3386409 | CAP Sarria | Banc de Sang i Teixits | Francisco Vidal, Irene Corrales, Nina Borràs, Carlos Hobeich, Natália Comes, Lorena Ramírez, Noemí Gonzalez, Silvia Sauleda, Maria Glòria Soria |
| EPI_ISL_3386410 | CAP Manlleu | Banc de Sang i Teixits | Francisco Vidal, Irene Corrales, Nina Borràs, Carlos Hobeich, Natália Comes, Lorena Ramírez, Noemí Gonzalez, Silvia Sauleda, Maria Glòria Soria |
| EPI_ISL_3386411 | Fundacio Althaia-Manresa | Banc de Sang i Teixits | Francisco Vidal, Irene Corrales, Nina Borràs, Carlos Hobeich, Natália Comes, Lorena Ramírez, Noemí Gonzalez, Silvia Sauleda, Maria Glòria Soria |
| EPI_ISL_3386412 | CAP El Remei | Banc de Sang i Teixits | Francisco Vidal, Irene Corrales, Nina Borràs, Carlos Hobeich, Natália Comes, Lorena Ramírez, Noemí Gonzalez, Silvia Sauleda, Maria Glòria Soria |
| EPI_ISL_3386413 | Fundacio Althaia-Manresa | Banc de Sang i Teixits | Francisco Vidal, Irene Corrales, Nina Borràs, Carlos Hobeich, Natália Comes, Lorena Ramírez, Noemí Gonzalez, Silvia Sauleda, Maria Glòria Soria |
| EPI_ISL_3386414 | CAP Sarria | Banc de Sang i Teixits | Francisco Vidal, Irene Corrales, Nina Borràs, Carlos Hobeich, Natália Comes, Lorena Ramírez, Noemí Gonzalez, Silvia Sauleda, Maria Glòria Soria |
| EPI_ISL_3386415 | CAP Vic Nord | Banc de Sang i Teixits | Francisco Vidal, Irene Corrales, Nina Borràs, Carlos Hobeich, Natália Comes, Lorena Ramírez, Noemí Gonzalez, Silvia Sauleda, Maria Glòria Soria |
| EPI_ISL_3386416 | CAP El Remei | Banc de Sang i Teixits | Francisco Vidal, Irene Corrales, Nina Borràs, Carlos Hobeich, Natália Comes, Lorena Ramírez, Noemí Gonzalez, Silvia Sauleda, Maria Glòria Soria |
| EPI_ISL_3386417, EPI_ISL_3386418 | Fundacio Althaia-Manresa | Banc de Sang i Teixits | Francisco Vidal, Irene Corrales, Nina Borràs, Carlos Hobeich, Natália Comes, Lorena Ramírez, Noemí Gonzalez, Silvia Sauleda, Maria Glòria Soria |
| EPI_ISL_3386419 | CAP El Remei | Banc de Sang i Teixits | Francisco Vidal, Irene Corrales, Nina Borràs, Carlos Hobeich, Natália Comes, Lorena Ramírez, Noemí Gonzalez, Silvia Sauleda, Maria Glòria Soria |
| EPI_ISL_3386420 | Fundacio Althaia-Manresa | Banc de Sang i Teixits | Francisco Vidal, Irene Corrales, Nina Borràs, Carlos Hobeich, Natália Comes, Lorena Ramírez, Noemí Gonzalez, Silvia Sauleda, Maria Glòria Soria |
| EPI_ISL_3386421, EPI_ISL_3386422 | CAP El Remei | Banc de Sang i Teixits | Francisco Vidal, Irene Corrales, Nina Borràs, Carlos Hobeich, Natália Comes, Lorena Ramírez, Noemí Gonzalez, Silvia Sauleda, Maria Glòria Soria |
| EPI_ISL_3386423, EPI_ISL_3386424, EPI_ISL_3386425 | Fundacio Althaia-Manresa | Banc de Sang i Teixits | Francisco Vidal, Irene Corrales, Nina Borràs, Carlos Hobeich, Natália Comes, Lorena Ramírez, Noemí Gonzalez, Silvia Sauleda, Maria Glòria Soria |
| EPI_ISL_3386426 | CAP Manlleu | Banc de Sang i Teixits | Francisco Vidal, Irene Corrales, Nina Borràs, Carlos Hobeich, Natália Comes, Lorena Ramírez, Noemí Gonzalez, Silvia Sauleda, Maria Glòria Soria |
| EPI_ISL_3386427 | CAP Vic Nord | Banc de Sang i Teixits | Francisco Vidal, Irene Corrales, Nina Borràs, Carlos Hobeich, Natália Comes, Lorena Ramírez, Noemí Gonzalez, Silvia Sauleda, Maria Glòria Soria |
| EPI_ISL_3386428 | CAP Manlleu | Banc de Sang i Teixits | Francisco Vidal, Irene Corrales, Nina Borràs, Carlos Hobeich, Natália Comes, Lorena Ramírez, Noemí Gonzalez, Silvia Sauleda, Maria Glòria Soria |
| EPI_ISL_3386429 | CAP Olost de Llucanes | Banc de Sang i Teixits | Francisco Vidal, Irene Corrales, Nina Borràs, Carlos Hobeich, Natália Comes, Lorena Ramírez, Noemí Gonzalez, Silvia Sauleda, Maria Glòria Soria |
| EPI_ISL_3386430 | CAP Sant Quirze de Besora | Banc de Sang i Teixits | Francisco Vidal, Irene Corrales, Nina Borràs, Carlos Hobeich, Natália Comes, Lorena Ramírez, Noemí Gonzalez, Silvia Sauleda, Maria Glòria Soria |
| EPI_ISL_3386431 | CAP Tona | Banc de Sang i Teixits | Francisco Vidal, Irene Corrales, Nina Borràs, Carlos Hobeich, Natália Comes, Lorena Ramírez, Noemí Gonzalez, Silvia Sauleda, Maria Glòria Soria |
| EPI_ISL_3386432, EPI_ISL_3386433 | Fundacio Althaia-Manresa | Banc de Sang i Teixits | Francisco Vidal, Irene Corrales, Nina Borràs, Carlos Hobeich, Natália Comes, Lorena Ramírez, Noemí Gonzalez, Silvia Sauleda, Maria Glòria Soria |
| EPI_ISL_3386434 | CAP Manlleu | Banc de Sang i Teixits | Francisco Vidal, Irene Corrales, Nina Borràs, Carlos Hobeich, Natália Comes, Lorena Ramírez, Noemí Gonzalez, Silvia Sauleda, Maria Glòria Soria |
| EPI_ISL_3386435 | CAP Vic Nord | Banc de Sang i Teixits | Francisco Vidal, Irene Corrales, Nina Borràs, Carlos Hobeich, Natália Comes, Lorena Ramírez, Noemí Gonzalez, Silvia Sauleda, Maria Glòria Soria |
| EPI_ISL_3386436 | Fundacio Althaia-Manresa | Banc de Sang i Teixits | Francisco Vidal, Irene Corrales, Nina Borràs, Carlos Hobeich, Natália Comes, Lorena Ramírez, Noemí Gonzalez, Silvia Sauleda, Maria Glòria Soria |
| EPI_ISL_3386437 | CAP Tona | Banc de Sang i Teixits | Francisco Vidal, Irene Corrales, Nina Borràs, Carlos Hobeich, Natália Comes, Lorena Ramírez, Noemí Gonzalez, Silvia Sauleda, Maria Glòria Soria |
| EPI_ISL_3386439 | Fundacio Althaia-Manresa | Banc de Sang i Teixits | Francisco Vidal, Irene Corrales, Nina Borràs, Carlos Hobeich, Natália Comes, Lorena Ramírez, Noemí Gonzalez, Silvia Sauleda, Maria Glòria Soria |
| EPI_ISL_3386440 | CAP Sarria | Banc de Sang i Teixits | Francisco Vidal, Irene Corrales, Nina Borràs, Carlos Hobeich, Natália Comes, Lorena Ramírez, Noemí Gonzalez, Silvia Sauleda, Maria Glòria Soria |
| EPI_ISL_3386441 | CAP El Remei | Banc de Sang i Teixits | Francisco Vidal, Irene Corrales, Nina Borràs, Carlos Hobeich, Natália Comes, Lorena Ramírez, Noemí Gonzalez, Silvia Sauleda, Maria Glòria Soria |
| EPI_ISL_3386442 | CAP Tona | Banc de Sang i Teixits | Francisco Vidal, Irene Corrales, Nina Borràs, Carlos Hobeich, Natália Comes, Lorena Ramírez, Noemí Gonzalez, Silvia Sauleda, Maria Glòria Soria |
| EPI_ISL_3386443 | CAP El Remei | Banc de Sang i Teixits | Francisco Vidal, Irene Corrales, Nina Borràs, Carlos Hobeich, Natália Comes, Lorena Ramírez, Noemí Gonzalez, Silvia Sauleda, Maria Glòria Soria |
| EPI_ISL_3386444 | Fundacio Althaia-Manresa | Banc de Sang i Teixits | Francisco Vidal, Irene Corrales, Nina Borràs, Carlos Hobeich, Natália Comes, Lorena Ramírez, Noemí Gonzalez, Silvia Sauleda, Maria Glòria Soria |
| EPI_ISL_3386445 | CAP Santa Eugenia de Berga | Banc de Sang i Teixits | Francisco Vidal, Irene Corrales, Nina Borràs, Carlos Hobeich, Natália Comes, Lorena Ramírez, Noemí Gonzalez, Silvia Sauleda, Maria Glòria Soria |
| EPI_ISL_3386450, EPI_ISL_3386454, EPI_ISL_3386455, EPI_ISL_3386456, EPI_ISL_3386457 | Arizona State University | Arizona State University | Ajeet Bains, LaRinda A. Holland, Matthew F. Smith, Peter T. Skidmore, Rabia Maqsood, Nicholas J. Mellor, Nathaniel Johnson, Joshua LaBaer, Vel Murugan, Efrem S. Lim |
| EPI_ISL_3386458 | Serrana | Hemocentro de Ribeirao Preto | Sao Paulo State Network for Pandemic Alert of Emerging SARS-CoV-2 |
| EPI_ISL_3386460, EPI_ISL_3386461, EPI_ISL_3386462, EPI_ISL_3386464, EPI_ISL_3386465, EPI_ISL_3386466, EPI_ISL_3386467, EPI_ISL_3386468, EPI_ISL_3386469, EPI_ISL_3386470, EPI_ISL_3386471, EPI_ISL_3386472, EPI_ISL_3386473, EPI_ISL_3386474, EPI_ISL_3386475, EPI_ISL_3386476, EPI_ISL_3386477, EPI_ISL_3386478, EPI_ISL_3386479, EPI_ISL_3386480, EPI_ISL_3386481, EPI_ISL_3386482, | | | |

| | | | |
|---|---|---|---|
| EPI_ISL_3386483, EPI_ISL_3386484, EPI_ISL_3386485, EPI_ISL_3386486, EPI_ISL_3386487, EPI_ISL_3386488, EPI_ISL_3386489, EPI_ISL_3386490, EPI_ISL_3386491, EPI_ISL_3386492, EPI_ISL_3386493, EPI_ISL_3386494, EPI_ISL_3386495, EPI_ISL_3386496, EPI_ISL_3386497, EPI_ISL_3386498, EPI_ISL_3386500, EPI_ISL_3386501, EPI_ISL_3386502, EPI_ISL_3386503, EPI_ISL_3386505, EPI_ISL_3386506, EPI_ISL_3386507, EPI_ISL_3386508, EPI_ISL_3386509, EPI_ISL_3386510, EPI_ISL_3386512, EPI_ISL_3386513, EPI_ISL_3386514, EPI_ISL_3386515, EPI_ISL_3386516, EPI_ISL_3386517, EPI_ISL_3386518, EPI_ISL_3386519, EPI_ISL_3386520, EPI_ISL_3386521, EPI_ISL_3386523, EPI_ISL_3386524, EPI_ISL_3386525, EPI_ISL_3386526, EPI_ISL_3386527, EPI_ISL_3386528, EPI_ISL_3386530, EPI_ISL_3386531, EPI_ISL_3386532, EPI_ISL_3386533, EPI_ISL_3386534, EPI_ISL_3386535, EPI_ISL_3386536, EPI_ISL_3386537, EPI_ISL_3386538, EPI_ISL_3386539, EPI_ISL_3386540, EPI_ISL_3386541, EPI_ISL_3386542, EPI_ISL_3386543, EPI_ISL_3386544, EPI_ISL_3386545, EPI_ISL_3386546, EPI_ISL_3386547, EPI_ISL_3386548, EPI_ISL_3386549, EPI_ISL_3386550, EPI_ISL_3386552, EPI_ISL_3386553, EPI_ISL_3386554, EPI_ISL_3386556 | | | |
| see above | Ministry of Health Turkey | Ministry of Health Turkey | Fatma Bayrakdar, Yasemin Coşgun, Süleyman Yalcin, Gülay Korukluoğlu |
| EPI_ISL_3386557, EPI_ISL_3386558, EPI_ISL_3386559, EPI_ISL_3386560, EPI_ISL_3386561, EPI_ISL_3386562, EPI_ISL_3386563, EPI_ISL_3386564, EPI_ISL_3386565, EPI_ISL_3386566 | Klinika za infektivne bolesti "Dr. Fran Mihaljević" | Hrvatski zavod za javno zdravstvo | Irena Tabain, Ivana Ferenčak |
| EPI_ISL_3386567, EPI_ISL_3386570, EPI_ISL_3386571 | Klinički bolnički centar Split | Hrvatski zavod za javno zdravstvo | Irena Tabain, Ivana Ferenčak |
| EPI_ISL_3386572, EPI_ISL_3386573 | Nastavni zavod za javno zdravstvo Splitsko- Dalmatinske Županije | Hrvatski zavod za javno zdravstvo | Irena Tabain, Ivana Ferenčak |
| EPI_ISL_3386574 | Zavod za javno zdravstvo Dubrovačko-Neretvanske Županije | Hrvatski zavod za javno zdravstvo | Irena Tabain, Ivana Ferenčak |
| EPI_ISL_3386575, EPI_ISL_3386576, EPI_ISL_3386577, EPI_ISL_3386578 | Zavod za javno zdravstvo Zagrebačke Županije | Hrvatski zavod za javno zdravstvo | Irena Tabain, Ivana Ferenčak |
| EPI_ISL_3386579, EPI_ISL_3386580, EPI_ISL_3386581, EPI_ISL_3386582, EPI_ISL_3386583, EPI_ISL_3386584, EPI_ISL_3386585 | Zavod za javno zdravstvo Dubrovačko-Neretvanske Županije | Hrvatski zavod za javno zdravstvo | Irena Tabain, Ivana Ferenčak |
| EPI_ISL_3386586 | Klinički bolnički centar Zagreb | Hrvatski zavod za javno zdravstvo | Irena Tabain, Ivana Ferenčak |
| EPI_ISL_3386590, EPI_ISL_3386591, EPI_ISL_3386592, EPI_ISL_3386593, EPI_ISL_3386594, EPI_ISL_3386595 | Klinika za infektivne bolesti "Dr. Fran Mihaljević" | Hrvatski zavod za javno zdravstvo | Irena Tabain, Ivana Ferenčak |
| EPI_ISL_3386596, EPI_ISL_3386597 | Zavod za javno zdravstvo Varaždinske županije | Hrvatski zavod za javno zdravstvo | Irena Tabain, Ivana Ferenčak |
| EPI_ISL_3386600, EPI_ISL_3386601 | Zavod za javno zdravstvo Zagrebačke Županije | Hrvatski zavod za javno zdravstvo | Irena Tabain, Ivana Ferenčak |
| EPI_ISL_3386603, EPI_ISL_3386604, EPI_ISL_3386606, EPI_ISL_3386607, EPI_ISL_3386608, EPI_ISL_3386609, EPI_ISL_3386610, EPI_ISL_3386611, EPI_ISL_3386612, EPI_ISL_3386614, EPI_ISL_3386615, EPI_ISL_3386616, EPI_ISL_3386617, EPI_ISL_3386619, EPI_ISL_3386620, EPI_ISL_3386621 | | | |
| see above | Klinika za infektivne bolesti "Dr. Fran Mihaljević" | Hrvatski zavod za javno zdravstvo | Irena Tabain, Ivana Ferenčak |
| EPI_ISL_3386622 | Opća bolnica "Dr. Josip Benčević" Slavonski Brod | Hrvatski zavod za javno zdravstvo | Irena Tabain, Ivana Ferenčak |
| EPI_ISL_3386623, EPI_ISL_3386624 | Zavod za javno zdravstvo Međimurske Županije | Hrvatski zavod za javno zdravstvo | Irena Tabain, Ivana Ferenčak |
| EPI_ISL_3386636 | Zavod za javno zdravstvo Šibensko- Kninske Županije | Hrvatski zavod za javno zdravstvo | Irena Tabain, Ivana Ferenčak |
| EPI_ISL_3386639 | Zavod za javno zdravstvo Međimurske Županije | Hrvatski zavod za javno zdravstvo | Irena Tabain, Ivana Ferenčak |
| EPI_ISL_3386648, EPI_ISL_3386650 | National Institute of Infectious Diseases-Prof. Dr. Matei Bals Molecular Diagnostics Laboratory | National Institute of Infectious Diseases-Prof. Dr. Matei Bals Molecular Diagnostics Laboratory | Corina Casangiu, Leontina Banica, Marius Surleac, Ovidiu Vlaicu, Petre Milu, Robert Hohan, Simona Paraschiv, Dan Otelea |
| EPI_ISL_3386651, EPI_ISL_3386652, EPI_ISL_3386653, EPI_ISL_3386654, EPI_ISL_3386655 | Department of Public Health Ilfov | National Institute of Infectious Diseases-Prof. Dr. Matei Bals Molecular Diagnostics Laboratory | Corina Casangiu, Leontina Banica, Marius Surleac, Ovidiu Vlaicu, Petre Milu, Robert Hohan, Simona Paraschiv, Dan Otelea |
| EPI_ISL_3386656 | National Institute of Infectious Diseases-Prof. Dr. Matei Bals Molecular Diagnostics Laboratory | National Institute of Infectious Diseases-Prof. Dr. Matei Bals Molecular Diagnostics Laboratory | Corina Casangiu, Leontina Banica, Marius Surleac, Ovidiu Vlaicu, Petre Milu, Robert Hohan, Simona Paraschiv, Dan Otelea |
| EPI_ISL_3386657, EPI_ISL_3386658, EPI_ISL_3386659, EPI_ISL_3386660, EPI_ISL_3386661, EPI_ISL_3386662 | Department of Public Health Buzau | National Institute of Infectious Diseases-Prof. Dr. Matei Bals Molecular Diagnostics Laboratory | Corina Casangiu, Leontina Banica, Marius Surleac, Ovidiu Vlaicu, Petre Milu, Robert Hohan, Simona Paraschiv, Dan Otelea |
| EPI_ISL_3386663, EPI_ISL_3386664 | Department of Public Health Vrancea | National Institute of Infectious Diseases-Prof. Dr. Matei Bals Molecular Diagnostics Laboratory | Corina Casangiu, Leontina Banica, Marius Surleac, Ovidiu Vlaicu, Petre Milu, Robert Hohan, Simona Paraschiv, Dan Otelea |
| EPI_ISL_3386665, EPI_ISL_3386666, EPI_ISL_3386667, EPI_ISL_3386668, EPI_ISL_3386669 | Department of Public Health Neamt | National Institute of Infectious Diseases-Prof. Dr. Matei Bals Molecular Diagnostics Laboratory | Corina Casangiu, Leontina Banica, Marius Surleac, Ovidiu Vlaicu, Petre Milu, Robert Hohan, Simona Paraschiv, Dan Otelea |
| EPI_ISL_3386670, EPI_ISL_3386671 | National Institute of Infectious Diseases-Prof. Dr. Matei Bals Molecular Diagnostics Laboratory | National Institute of Infectious Diseases-Prof. Dr. Matei Bals Molecular Diagnostics Laboratory | Corina Casangiu, Leontina Banica, Marius Surleac, Ovidiu Vlaicu, Petre Milu, Robert Hohan, Simona Paraschiv, Dan Otelea |
| EPI_ISL_3386672, EPI_ISL_3386673, EPI_ISL_3386674, EPI_ISL_3386675 | Department of Public Health Constanta | National Institute of Infectious Diseases-Prof. Dr. Matei Bals Molecular Diagnostics Laboratory | Corina Casangiu, Leontina Banica, Marius Surleac, Ovidiu Vlaicu, Petre Milu, Robert Hohan, Simona Paraschiv, Dan Otelea |
| EPI_ISL_3386676 | Department of Public Health Covasna | National Institute of Infectious Diseases-Prof. Dr. Matei Bals Molecular Diagnostics Laboratory | Corina Casangiu, Leontina Banica, Marius Surleac, Ovidiu Vlaicu, Petre Milu, Robert Hohan, Simona Paraschiv, Dan Otelea |
| EPI_ISL_3386677, EPI_ISL_3386678, EPI_ISL_3386679, EPI_ISL_3386680, EPI_ISL_3386681 | Department of Public Health Iasi | National Institute of Infectious Diseases-Prof. Dr. Matei Bals Molecular Diagnostics Laboratory | Corina Casangiu, Leontina Banica, Marius Surleac, Ovidiu Vlaicu, Petre Milu, Robert Hohan, Simona Paraschiv, Dan Otelea |
| EPI_ISL_3386682 | Institute of Infectious Diseases-Cuvioasa Paraschiva, Iasi | National Institute of Infectious Diseases-Prof. Dr. Matei Bals Molecular Diagnostics Laboratory | Corina Casangiu, Leontina Banica, Marius Surleac, Ovidiu Vlaicu, Petre Milu, Robert Hohan, Simona Paraschiv, Dan Otelea |
| EPI_ISL_3386683 | Department of Public Health Botosani | National Institute of Infectious Diseases-Prof. Dr. Matei Bals Molecular Diagnostics Laboratory | Corina Casangiu, Leontina Banica, Marius Surleac, Ovidiu Vlaicu, Petre Milu, Robert Hohan, Simona Paraschiv, Dan Otelea |
| EPI_ISL_3386684, EPI_ISL_3386685 | Department of Public Health Bacau | National Institute of Infectious Diseases-Prof. Dr. Matei Bals Molecular Diagnostics Laboratory | Corina Casangiu, Leontina Banica, Marius Surleac, Ovidiu Vlaicu, Petre Milu, Robert Hohan, Simona Paraschiv, Dan Otelea |
| EPI_ISL_3386686, EPI_ISL_3386687, EPI_ISL_3386688, EPI_ISL_3386689, EPI_ISL_3386690 | Department of Public Health Suceava | National Institute of Infectious Diseases-Prof. Dr. Matei Bals Molecular Diagnostics Laboratory | Corina Casangiu, Leontina Banica, Marius Surleac, Ovidiu Vlaicu, Petre Milu, Robert Hohan, Simona Paraschiv, Dan Otelea |

| | | | |
|---|---|---|---|
| EPI_ISL_3386691 | Department of Public Health Neamt | National Institute of Infectious Diseases-Prof. Dr. Matei Bals Molecular Diagnostics Laboratory | Corina Casangiu, Leontina Banica, Marius Surleac, Ovidiu Vlaicu, Petre Milu, Robert Hohan, Simona Paraschiv, Dan Otelea |
| EPI_ISL_3386692, EPI_ISL_3386693 | Department of Public Health Brasov | National Institute of Infectious Diseases-Prof. Dr. Matei Bals Molecular Diagnostics Laboratory | Corina Casangiu, Leontina Banica, Marius Surleac, Ovidiu Vlaicu, Petre Milu, Robert Hohan, Simona Paraschiv, Dan Otelea |
| EPI_ISL_3386694 | Department of Public Health Alba | National Institute of Infectious Diseases-Prof. Dr. Matei Bals Molecular Diagnostics Laboratory | Corina Casangiu, Leontina Banica, Marius Surleac, Ovidiu Vlaicu, Petre Milu, Robert Hohan, Simona Paraschiv, Dan Otelea |
| EPI_ISL_3386820, EPI_ISL_3386821, EPI_ISL_3386822, EPI_ISL_3386823, EPI_ISL_3386824, EPI_ISL_3386825, EPI_ISL_3386826, EPI_ISL_3386827, EPI_ISL_3386828, EPI_ISL_3386829, EPI_ISL_3386830, EPI_ISL_3386831, EPI_ISL_3386832, EPI_ISL_3386833, EPI_ISL_3386834, EPI_ISL_3386835, EPI_ISL_3386836, EPI_ISL_3386837, EPI_ISL_3386838, EPI_ISL_3386839, EPI_ISL_3386840, EPI_ISL_3386841, EPI_ISL_3386842, EPI_ISL_3386843, EPI_ISL_3386844, EPI_ISL_3386845, EPI_ISL_3386846, EPI_ISL_3386847, EPI_ISL_3386848, EPI_ISL_3386849, EPI_ISL_3386850, EPI_ISL_3386851, EPI_ISL_3386852, EPI_ISL_3386853, EPI_ISL_3386854, EPI_ISL_3386855, EPI_ISL_3386856, EPI_ISL_3386857, EPI_ISL_3386858, EPI_ISL_3386859, EPI_ISL_3386860, EPI_ISL_3386861, EPI_ISL_3386862, EPI_ISL_3386863, EPI_ISL_3386864, EPI_ISL_3386865, EPI_ISL_3386866, EPI_ISL_3386867, EPI_ISL_3386868, EPI_ISL_3386869, EPI_ISL_3386870, EPI_ISL_3386871, EPI_ISL_3386872, EPI_ISL_3386873, EPI_ISL_3386874, EPI_ISL_3386875, EPI_ISL_3386876, EPI_ISL_3386877, EPI_ISL_3386878, EPI_ISL_3386879, EPI_ISL_3386880, EPI_ISL_3386882, EPI_ISL_3386883, EPI_ISL_3386884, EPI_ISL_3386885, EPI_ISL_3386886 | | | |
| see above | Cliniques universitaires Saint-Luc | UCLouvain/IREC/MBLG | Jean Ruelle, Bertrand Bearzatto, Benoit Kabamba Mukadi |
| EPI_ISL_3386887, EPI_ISL_3386888, EPI_ISL_3386889, EPI_ISL_3386890, EPI_ISL_3386891, EPI_ISL_3386892, EPI_ISL_3386893, EPI_ISL_3386894, EPI_ISL_3386895, EPI_ISL_3386896, EPI_ISL_3386897, EPI_ISL_3386898, EPI_ISL_3386899, EPI_ISL_3386900, EPI_ISL_3386901, EPI_ISL_3386902, EPI_ISL_3386903, EPI_ISL_3386904, EPI_ISL_3386905, EPI_ISL_3386906 | | | |
| see above | Clinique Saint-Pierre Ottignies | UCLouvain/IREC/MBLG | Jean Ruelle, Bertrand Bearzatto, Benoit Kabamba Mukadi |
| EPI_ISL_3386907, EPI_ISL_3386908 | Centre Hospitalier Interrégional Edith Cavell | UCLouvain/IREC/MBLG | Jean Ruelle, Bertrand Bearzatto, Benoit Kabamba Mukadi |
| EPI_ISL_3386909 | Cliniques universitaires Saint-Luc | UCLouvain/IREC/MBLG | Jean Ruelle, Bertrand Bearzatto, Benoit Kabamba Mukadi |
| EPI_ISL_3386910 | Centre Hospitalier Interrégional Edith Cavell | UCLouvain/IREC/MBLG | Jean Ruelle, Bertrand Bearzatto, Benoit Kabamba Mukadi |
| EPI_ISL_3386911 | Cliniques universitaires Saint-Luc | UCLouvain/IREC/MBLG | Jean Ruelle, Bertrand Bearzatto, Benoit Kabamba Mukadi |
| EPI_ISL_3386912, EPI_ISL_3386913, EPI_ISL_3386914, EPI_ISL_3386915, EPI_ISL_3386916, EPI_ISL_3386917, EPI_ISL_3386918, EPI_ISL_3386919, EPI_ISL_3386920, EPI_ISL_3386921, EPI_ISL_3386922, EPI_ISL_3386923, EPI_ISL_3386924, EPI_ISL_3386925, EPI_ISL_3386926, EPI_ISL_3386927, EPI_ISL_3386928, EPI_ISL_3386929, EPI_ISL_3386930, EPI_ISL_3386931, EPI_ISL_3386932, EPI_ISL_3386933, EPI_ISL_3386934, EPI_ISL_3386936, EPI_ISL_3386937, EPI_ISL_3386938, EPI_ISL_3386939, EPI_ISL_3386940, EPI_ISL_3386941, EPI_ISL_3386942, EPI_ISL_3386943, EPI_ISL_3386944, EPI_ISL_3386945, EPI_ISL_3386946, EPI_ISL_3386947, EPI_ISL_3386948, EPI_ISL_3386949, EPI_ISL_3386950, EPI_ISL_3386951, EPI_ISL_3386952, EPI_ISL_3386953, EPI_ISL_3386954, EPI_ISL_3386955, EPI_ISL_3386956, EPI_ISL_3386957, EPI_ISL_3386958, EPI_ISL_3386959, EPI_ISL_3386960, EPI_ISL_3386961, EPI_ISL_3386962, EPI_ISL_3386963, EPI_ISL_3386964, EPI_ISL_3386965, EPI_ISL_3386966, EPI_ISL_3386967, EPI_ISL_3386968 | | | |
| see above | Hospital General Universitario Gregorio Marañón | Hospital General Universitario Gregorio Marañón | Sergio Buenestado Serrano, Pedro Sola Campoy, Laura Pérez-Lago, Cristina Rodriguez-Grande, Marta Herranz Martin, Victor Manuel de la Cueva, Julia Suárez, Pilar Catalán, Patricia Muñoz, Darío García de Viedma |
| EPI_ISL_3386969, EPI_ISL_3386970, EPI_ISL_3386971, EPI_ISL_3386972, EPI_ISL_3386973, EPI_ISL_3386974, EPI_ISL_3386975, EPI_ISL_3386976, EPI_ISL_3386977, EPI_ISL_3386978, EPI_ISL_3386979, EPI_ISL_3386980, EPI_ISL_3386981, EPI_ISL_3386982, EPI_ISL_3386983, EPI_ISL_3386984, EPI_ISL_3386985, EPI_ISL_3386986, EPI_ISL_3386987, EPI_ISL_3386988, EPI_ISL_3386989, EPI_ISL_3386990, EPI_ISL_3386991, EPI_ISL_3386992, EPI_ISL_3386993, EPI_ISL_3386994, EPI_ISL_3386995, EPI_ISL_3386996, EPI_ISL_3386997 | | | |
| see above | AZDelta | AZ Delta Medical Laboratories in Roeselare, Belgium | Geert Martens, Dieter De Smet, Frederik Van Hoecke, on behalf of AZ Delta COVID-19 Genomics core (member of Genomic surveillance of SARS-CoV-2 in Belgium network) |
| EPI_ISL_3386999, EPI_ISL_3387000, EPI_ISL_3387001, EPI_ISL_3387002, EPI_ISL_3387003, EPI_ISL_3387004, EPI_ISL_3387005, EPI_ISL_3387006, EPI_ISL_3387007, EPI_ISL_3387008, EPI_ISL_3387009, EPI_ISL_3387010, EPI_ISL_3387011, EPI_ISL_3387012, EPI_ISL_3387013, EPI_ISL_3387014, EPI_ISL_3387015, EPI_ISL_3387016, EPI_ISL_3387017, EPI_ISL_3387018, EPI_ISL_3387019, EPI_ISL_3387020, EPI_ISL_3387021, EPI_ISL_3387022, EPI_ISL_3387023, EPI_ISL_3387024, EPI_ISL_3387025, EPI_ISL_3387026, EPI_ISL_3387027, EPI_ISL_3387028, EPI_ISL_3387029, EPI_ISL_3387030, EPI_ISL_3387031, EPI_ISL_3387032, EPI_ISL_3387033, EPI_ISL_3387035, EPI_ISL_3387036, EPI_ISL_3387040, EPI_ISL_3387041, EPI_ISL_3387042, EPI_ISL_3387043, EPI_ISL_3387044, EPI_ISL_3387045, EPI_ISL_3387046, EPI_ISL_3387047, EPI_ISL_3387048, EPI_ISL_3387049, EPI_ISL_3387050, EPI_ISL_3387051, EPI_ISL_3387052, EPI_ISL_3387053, EPI_ISL_3387054, EPI_ISL_3387055, EPI_ISL_3387056, EPI_ISL_3387057, EPI_ISL_3387058, EPI_ISL_3387059 | | | |
| see above | Alaska State Virology Laboratory | Alaska State Virology Laboratory | Stephanie DeRonde, Elva House, Jacob Zidek, Lisa Smith, Ph.D., Jack Chen, Ph.D. |
| EPI_ISL_3387060, EPI_ISL_3387061, EPI_ISL_3387062, EPI_ISL_3387063, EPI_ISL_3387064, EPI_ISL_3387065, EPI_ISL_3387066, EPI_ISL_3387067, EPI_ISL_3387068, EPI_ISL_3387069, EPI_ISL_3387070, EPI_ISL_3387071, EPI_ISL_3387072, EPI_ISL_3387073, EPI_ISL_3387074, EPI_ISL_3387075 | | | |
| see above | Hospital Universitari de Bellvitge | Microbiology Department | Sara Marti, Aida Gonzalez-Diaz, Yolanda Hernandez, Laura Calatayud, Jordi Niubó, Carmen Ardanuy, Jordi Camara, M Angeles Domínguez |
| EPI_ISL_3387076, EPI_ISL_3387078, EPI_ISL_3387079, EPI_ISL_3387080, EPI_ISL_3387081 | CLILAB | Microbiology Department | Sara Marti, Aida Gonzalez-Diaz, Yolanda Hernandez, Laura Calatayud, Jordi Niubó, Carmen Ardanuy, Jordi Camara, M Angeles Domínguez |
| EPI_ISL_3387082 | Hospital Universitari de Bellvitge | Microbiology Department | Sara Marti, Aida Gonzalez-Diaz, Yolanda Hernandez, Laura Calatayud, Jordi Niubó, Carmen Ardanuy, Jordi Camara, M Angeles Domínguez |
| EPI_ISL_3387083, EPI_ISL_3387084, EPI_ISL_3387085, EPI_ISL_3387086, EPI_ISL_3387087, EPI_ISL_3387088, EPI_ISL_3387089 | CLILAB | Microbiology Department | Sara Marti, Aida Gonzalez-Diaz, Yolanda Hernandez, Laura Calatayud, Jordi Niubó, Carmen Ardanuy, Jordi Camara, M Angeles Domínguez |
| EPI_ISL_3387091, EPI_ISL_3387092, EPI_ISL_3387093, EPI_ISL_3387094, EPI_ISL_3387095, EPI_ISL_3387096, EPI_ISL_3387097, EPI_ISL_3387098, EPI_ISL_3387099, EPI_ISL_3387100, EPI_ISL_3387101, EPI_ISL_3387102, EPI_ISL_3387103, EPI_ISL_3387104, EPI_ISL_3387105, EPI_ISL_3387106, EPI_ISL_3387107, EPI_ISL_3387108, EPI_ISL_3387109, EPI_ISL_3387110, EPI_ISL_3387111, EPI_ISL_3387112, EPI_ISL_3387113, EPI_ISL_3387114, EPI_ISL_3387115, EPI_ISL_3387116, EPI_ISL_3387117, EPI_ISL_3387118, EPI_ISL_3387119, EPI_ISL_3387120, EPI_ISL_3387121, EPI_ISL_3387122, EPI_ISL_3387123, EPI_ISL_3387124, EPI_ISL_3387125, EPI_ISL_3387126, EPI_ISL_3387127, EPI_ISL_3387128, EPI_ISL_3387129, EPI_ISL_3387130, EPI_ISL_3387131, EPI_ISL_3387132, EPI_ISL_3387133, EPI_ISL_3387134, EPI_ISL_3387135, EPI_ISL_3387136, EPI_ISL_3387137, EPI_ISL_3387138, EPI_ISL_3387139, EPI_ISL_3387140, EPI_ISL_3387141, EPI_ISL_3387142, EPI_ISL_3387143, EPI_ISL_3387144, EPI_ISL_3387145, EPI_ISL_3387146, EPI_ISL_3387147 | | | |
| see above | Hospital Universitari de Bellvitge | Microbiology Department | Sara Marti, Aida Gonzalez-Diaz, Yolanda Hernandez, Laura Calatayud, Jordi Niubó, Carmen Ardanuy, Jordi Camara, M Angeles Domínguez |
| EPI_ISL_3387148, EPI_ISL_3387149, EPI_ISL_3387151 | Instituto Nacional de Rehabilitación | Centro de Investigación en Enfermedades Infecciosas (CIENI), Instituto Nacional de Enfermedades Respiratorias (INER) | Margarita Matias-Florentino, Claudia Adriana Colín-Castro, Luis Esaú López-Jácome, María Guadalupe Martínez-Zavaleta, Guillermo Cerón-González, Noé Becerra-Lobato, Arturo Mondragón-Eguiluz, Edgar Samuel Vanegas-Rodríguez, Rafael Franco-Cendejas, Santiago Avila-Rios, Celia Boukadida |
| EPI_ISL_3387152, EPI_ISL_3387153, EPI_ISL_3387154, EPI_ISL_3387155, EPI_ISL_3387156, EPI_ISL_3387157, EPI_ISL_3387158, EPI_ISL_3387159, EPI_ISL_3387160, EPI_ISL_3387161, EPI_ISL_3387162, EPI_ISL_3387163, EPI_ISL_3387164, EPI_ISL_3387165, EPI_ISL_3387166, EPI_ISL_3387167, EPI_ISL_3387168, EPI_ISL_3387169, EPI_ISL_3387170, EPI_ISL_3387171, EPI_ISL_3387172, EPI_ISL_3387173, EPI_ISL_3387174, EPI_ISL_3387175, EPI_ISL_3387176, EPI_ISL_3387177, EPI_ISL_3387178, EPI_ISL_3387179, EPI_ISL_3387180, EPI_ISL_3387181, EPI_ISL_3387182, EPI_ISL_3387183, EPI_ISL_3387184, EPI_ISL_3387185, EPI_ISL_3387186, EPI_ISL_3387187, EPI_ISL_3387188, EPI_ISL_3387189, EPI_ISL_3387190, EPI_ISL_3387191, EPI_ISL_3387192, EPI_ISL_3387193, EPI_ISL_3387194, EPI_ISL_3387195, EPI_ISL_3387196, EPI_ISL_3387197, EPI_ISL_3387198, EPI_ISL_3387199, EPI_ISL_3387200, EPI_ISL_3387201, EPI_ISL_3387202, EPI_ISL_3387203, EPI_ISL_3387204, EPI_ISL_3387205, EPI_ISL_3387206, EPI_ISL_3387207, EPI_ISL_3387208, EPI_ISL_3387209, EPI_ISL_3387210, EPI_ISL_3387211, EPI_ISL_3387212, EPI_ISL_3387213, EPI_ISL_3387214, EPI_ISL_3387215, EPI_ISL_3387216, EPI_ISL_3387217, EPI_ISL_3387218, EPI_ISL_3387219, EPI_ISL_3387220, EPI_ISL_3387221, EPI_ISL_3387222, EPI_ISL_3387223, EPI_ISL_3387224, EPI_ISL_3387225, EPI_ISL_3387226, EPI_ISL_3387227, EPI_ISL_3387228, EPI_ISL_3387229, EPI_ISL_3387230 | | | |
| see above | National Public Health Laboratory, National Centre for Infectious Diseases | National Public Health Laboratory, National Centre for Infectious Diseases | Tze Minn Mak, Katherine Ching, Grace Ngan, Zhenyang Zhou, Royce Ang, Lin Cui, Raymond Tzer Pin Lin |
| EPI_ISL_3387231, EPI_ISL_3387232 | Jessa | Jessa | Severine Berden et al. on behalf of the Jessa_cmdLab |
| EPI_ISL_3387233, EPI_ISL_3387234, EPI_ISL_3387235, EPI_ISL_3387236, EPI_ISL_3387237, EPI_ISL_3387238, EPI_ISL_3387239, EPI_ISL_3387240, EPI_ISL_3387241, EPI_ISL_3387242, EPI_ISL_3387243, EPI_ISL_3387244, EPI_ISL_3387245, EPI_ISL_3387246, EPI_ISL_3387247, EPI_ISL_3387248, EPI_ISL_3387249, EPI_ISL_3387250, EPI_ISL_3387251 | | | |
| see above | ZOL | Jessa | Severine Berden et al. on behalf of the Jessa_cmdLab |
| EPI_ISL_3387252, EPI_ISL_3387253 | Jessa | Jessa | Severine Berden et al. on behalf of the Jessa_cmdLab |
| EPI_ISL_3387254, EPI_ISL_3387255 | LKO | Jessa | Severine Berden et al. on behalf of the Jessa_cmdLab |
| EPI_ISL_3387256, EPI_ISL_3387257, EPI_ISL_3387258, EPI_ISL_3387259, EPI_ISL_3387260, EPI_ISL_3387261, EPI_ISL_3387262, EPI_ISL_3387263, EPI_ISL_3387264, EPI_ISL_3387265, EPI_ISL_3387266, EPI_ISL_3387267, EPI_ISL_3387268, EPI_ISL_3387269, EPI_ISL_3387270, EPI_ISL_3387271, EPI_ISL_3387272, EPI_ISL_3387273 | | | |
| see above | Jessa | Jessa | Severine Berden et al. on behalf of the Jessa_cmdLab |
| EPI_ISL_3387274, EPI_ISL_3387275, EPI_ISL_3387277, EPI_ISL_3387278, EPI_ISL_3387279, EPI_ISL_3387280, EPI_ISL_3387281, EPI_ISL_3387282, EPI_ISL_3387283, EPI_ISL_3387284, EPI_ISL_3387285, EPI_ISL_3387286, EPI_ISL_3387287, EPI_ISL_3387288, EPI_ISL_3387289, EPI_ISL_3387290, EPI_ISL_3387291, EPI_ISL_3387292, EPI_ISL_3387293, EPI_ISL_3387294, EPI_ISL_3387295, EPI_ISL_3387296, EPI_ISL_3387297, EPI_ISL_3387298, EPI_ISL_3387299, EPI_ISL_3387300, EPI_ISL_3387301, EPI_ISL_3387302, EPI_ISL_3387304 | | | |
| see above | State Hygienic Laboratory at the University of Iowa | State Hygienic Laboratory at the University of Iowa | Valerie Reeb, Erik Twait, Anna Yakos, Mariah Knutson, Kris Eveland, Kristen Zanon, Jeff, Benfer, Wes Hottel, Alankar Kampoowale |
| EPI_ISL_3387350 | MS PHL | MS PHL | Beata Karowlewicz |
| EPI_ISL_3387378, EPI_ISL_3387379, EPI_ISL_3387380, EPI_ISL_3387382, EPI_ISL_3387386 | Virology Unit, Institut Pasteur du Cambodge | Virology Unit, Institut Pasteur du Cambodge | Leakhena Pum, Sophoannadedh Rath, Jurre Y Siegers, Cecile Troupin, Ly Sovann, Kraing Sidonn, Yi Sengdoeurn, Chin Savuth, Chau Darapheak, Veasna Duong, Erik A Karlsson |
| EPI_ISL_3387417, EPI_ISL_3387418, EPI_ISL_3387419, EPI_ISL_3387421, EPI_ISL_3387422, EPI_ISL_3387423, EPI_ISL_3387424, EPI_ISL_3387430, EPI_ISL_3387431, EPI_ISL_3387433, EPI_ISL_3387434, EPI_ISL_3387435 | | | |
| see above | Azienda Sanitaria dell'Alto Adige - Laboratorio Aziendale di Microbiologia e Virologia | Azienda Sanitaria dell'Alto Adige | Irene Bianconi |
| EPI_ISL_3387459 | Sonic Reference Laboratory | Sonic Reference Laboratory | Robert Moller, Bijaya Dhakal, Krishna Maddula, John Woody, Cora Lahey, Ellen Pizarek |
| EPI_ISL_3387460 | MS PHL | MS PHL | Beata Karowlewicz |
| EPI_ISL_3387473, EPI_ISL_3387474, EPI_ISL_3387499, EPI_ISL_3387502, EPI_ISL_3387503, EPI_ISL_3387504, EPI_ISL_3387505, EPI_ISL_3387506, EPI_ISL_3387507, EPI_ISL_3387508, EPI_ISL_3387509, EPI_ISL_3387510, EPI_ISL_3387511, EPI_ISL_3387512, EPI_ISL_3387513 | | | |
| see above | NJDOH, Public Health and Environmental Laboratories | NJ_PHEL | Lindsey Bodnar, Shiv K. Verma, Ryan Pachucki, Chelsea San Filippo, Jacquelyn Deverell, Dana Woell, Allison Roder, Byeong Jeong |
| EPI_ISL_3387525 | Arizona State University | Arizona State University | Ajeet Bains, LaRinda A. Holland, Matthew F. Smith, Peter T. Skidmore, Rabia Maqsood, Nicholas J. Mellor, Nathaniel Johnson, Joshua LaBaer, Vel Murugan, Efrem S. Lim |
| EPI_ISL_3388708, EPI_ISL_3388729, EPI_ISL_3388730, EPI_ISL_3388733, EPI_ISL_3388734, EPI_ISL_3388735, EPI_ISL_3388736, | Rush University Medical Center | RIPHL at Rush University Medical Center | Stefan Green, Kevin Kunstman, Max Kolton, Marieta Hyde, Laura Furtado, Felix Araujo Perez, Edith Perez, Mary Hayden, Joyce Houlihan, Diane Springer |

| | | | |
|---|---|---|---|
| EPI_ISL_3388737, EPI_ISL_3388738, EPI_ISL_3388739 | | | |
| EPI_ISL_3388740 | Alverno Labs | RIPHL at Rush University Medical Center | Stefan Green, Kevin Kunstman, Max Kolton, Marieta Hyde, Laura Furtado, Felix Araujo Perez, Edith Perez, Melissa Mace |
| EPI_ISL_3388741, EPI_ISL_3388742, EPI_ISL_3388743, EPI_ISL_3388744, EPI_ISL_3388745, EPI_ISL_3388746, EPI_ISL_3388747, EPI_ISL_3388748 | Rush University Medical Center | RIPHL at Rush University Medical Center | Stefan Green, Kevin Kunstman, Max Kolton, Marieta Hyde, Laura Furtado, Felix Araujo Perez, Edith Perez, Mary Hayden, Joyce Houlihan, Diane Springer |
| EPI_ISL_3388749, EPI_ISL_3388750, EPI_ISL_3388751, EPI_ISL_3388752 | University of Chicago Medicine | RIPHL at Rush University Medical Center | Stefan Green, Kevin Kunstman, Max Kolton, Marieta Hyde, Laura Furtado, Felix Araujo Perez, Edith Perez, Lauren Megger |
| EPI_ISL_3388753, EPI_ISL_3388754, EPI_ISL_3388755 | John H Stroger Jr Hospital of Cook County | RIPHL at Rush University Medical Center | Stefan Green, Kevin Kunstman, Max Kolton, Marieta Hyde, Laura Furtado, Felix Araujo Perez, Edith Perez, Ah Young Sung, Sharon Welbel |
| EPI_ISL_3388756 | University of Illinois at Chicago | RIPHL at Rush University Medical Center | Stefan Green, Kevin Kunstman, Max Kolton, Marieta Hyde, Laura Furtado, Felix Araujo Perez, Edith Perez, Iuliana Bentea, Omar Perez |
| EPI_ISL_3388757, EPI_ISL_3388758, EPI_ISL_3388759, EPI_ISL_3388760, EPI_ISL_3388761, EPI_ISL_3388762 | Alverno Labs | RIPHL at Rush University Medical Center | Stefan Green, Kevin Kunstman, Max Kolton, Marieta Hyde, Laura Furtado, Felix Araujo Perez, Edith Perez, Melissa Mace |
| EPI_ISL_3388763, EPI_ISL_3388764, EPI_ISL_3388765, EPI_ISL_3388766 | Rush University Medical Center | RIPHL at Rush University Medical Center | Stefan Green, Kevin Kunstman, Max Kolton, Marieta Hyde, Laura Furtado, Felix Araujo Perez, Edith Perez, Mary Hayden, Joyce Houlihan, Diane Springer |
| EPI_ISL_3388767 | University of Chicago Medicine | RIPHL at Rush University Medical Center | Stefan Green, Kevin Kunstman, Max Kolton, Marieta Hyde, Laura Furtado, Felix Araujo Perez, Edith Perez, Lauren Megger |
| EPI_ISL_3388768, EPI_ISL_3388769, EPI_ISL_3388770, EPI_ISL_3388771 | Northwestern Memorial Hospital | RIPHL at Rush University Medical Center | Stefan Green, Kevin Kunstman, Max Kolton, Marieta Hyde, Laura Furtado, Felix Araujo Perez, Edith Perez, Amber Kimble, Chao Qi |
| EPI_ISL_3388772, EPI_ISL_3388773, EPI_ISL_3388774, EPI_ISL_3388775, EPI_ISL_3388776, EPI_ISL_3388777, EPI_ISL_3388778, EPI_ISL_3388779, EPI_ISL_3388780, EPI_ISL_3388781 | University of Illinois at Chicago | RIPHL at Rush University Medical Center | Stefan Green, Kevin Kunstman, Max Kolton, Marieta Hyde, Laura Furtado, Felix Araujo Perez, Edith Perez, Iuliana Bentea, Omar Perez |
| EPI_ISL_3388782, EPI_ISL_3388783, EPI_ISL_3388784, EPI_ISL_3388785, EPI_ISL_3388786, EPI_ISL_3388787, EPI_ISL_3388788, EPI_ISL_3388789, EPI_ISL_3388790, EPI_ISL_3388791, EPI_ISL_3388792, EPI_ISL_3388793, EPI_ISL_3388804, EPI_ISL_3388809 | | | |
| see above | Alverno Labs | RIPHL at Rush University Medical Center | Stefan Green, Kevin Kunstman, Max Kolton, Marieta Hyde, Laura Furtado, Felix Araujo Perez, Edith Perez, Melissa Mace |
| EPI_ISL_3388810, EPI_ISL_3388811, EPI_ISL_3388812 | University of Illinois at Chicago | RIPHL at Rush University Medical Center | Stefan Green, Kevin Kunstman, Max Kolton, Marieta Hyde, Laura Furtado, Felix Araujo Perez, Edith Perez, Iuliana Bentea, Omar Perez |
| EPI_ISL_3388813 | Alverno Labs | RIPHL at Rush University Medical Center | Stefan Green, Kevin Kunstman, Max Kolton, Marieta Hyde, Laura Furtado, Felix Araujo Perez, Edith Perez, Melissa Mace |
| EPI_ISL_3388814, EPI_ISL_3388815 | Rush University Medical Center | RIPHL at Rush University Medical Center | Stefan Green, Kevin Kunstman, Max Kolton, Marieta Hyde, Laura Furtado, Felix Araujo Perez, Edith Perez, Mary Hayden, Joyce Houlihan, Diane Springer |
| EPI_ISL_3388816 | University of Illinois at Chicago | RIPHL at Rush University Medical Center | Stefan Green, Kevin Kunstman, Max Kolton, Marieta Hyde, Laura Furtado, Felix Araujo Perez, Edith Perez, Iuliana Bentea, Omar Perez |
| EPI_ISL_3388817, EPI_ISL_3388818 | Alverno Labs | RIPHL at Rush University Medical Center | Stefan Green, Kevin Kunstman, Max Kolton, Marieta Hyde, Laura Furtado, Felix Araujo Perez, Edith Perez, Melissa Mace |
| EPI_ISL_3388819 | University of Chicago Medicine | RIPHL at Rush University Medical Center | Stefan Green, Kevin Kunstman, Max Kolton, Marieta Hyde, Laura Furtado, Felix Araujo Perez, Edith Perez, Lauren Megger |
| EPI_ISL_3388820 | University of Illinois at Chicago | RIPHL at Rush University Medical Center | Stefan Green, Kevin Kunstman, Max Kolton, Marieta Hyde, Laura Furtado, Felix Araujo Perez, Edith Perez, Iuliana Bentea, Omar Perez |
| EPI_ISL_3388821, EPI_ISL_3388822, EPI_ISL_3388823, EPI_ISL_3388824 | Alverno Labs | RIPHL at Rush University Medical Center | Stefan Green, Kevin Kunstman, Max Kolton, Marieta Hyde, Laura Furtado, Felix Araujo Perez, Edith Perez, Melissa Mace |
| EPI_ISL_3388835, EPI_ISL_3388836, EPI_ISL_3388837 | Hospital General Universitario de Ciudad Real | Hospital General Universitario de Ciudad Real | José Martínez-Alarcón, Soledad Illescas Fernández-Bermejo, Marta Torres-Narbona, Cristina Colmenarejo, Lidia García-Agudo |
| EPI_ISL_3388838 | Hospital General de Valdepeñas | Hospital General Universitario de Ciudad Real | José Martínez-Alarcón, Soledad Illescas Fernández-Bermejo, Marta Torres-Narbona, Cristina Colmenarejo, Lidia García-Agudo |
| EPI_ISL_3388839 | Hospital General Universitario de Ciudad Real | Hospital General Universitario de Ciudad Real | José Martínez-Alarcón, Soledad Illescas Fernández-Bermejo, Marta Torres-Narbona, Cristina Colmenarejo, Lidia García-Agudo |
| EPI_ISL_3388840 | Hospital Virgen De Altagracia | Hospital General Universitario de Ciudad Real | José Martínez-Alarcón, Soledad Illescas Fernández-Bermejo, Marta Torres-Narbona, Cristina Colmenarejo, Lidia García-Agudo |
| EPI_ISL_3388841 | Hospital General de Valdepeñas | Hospital General Universitario de Ciudad Real | José Martínez-Alarcón, Soledad Illescas Fernández-Bermejo, Marta Torres-Narbona, Cristina Colmenarejo, Lidia García-Agudo |
| EPI_ISL_3388848, EPI_ISL_3388849, EPI_ISL_3388850, EPI_ISL_3388851, EPI_ISL_3388852, EPI_ISL_3388853, EPI_ISL_3388854, EPI_ISL_3388855, EPI_ISL_3388856, EPI_ISL_3388857, EPI_ISL_3388859 | | | |
| see above | Boise State University Campus School | Boise State University Genetics and Infectious Disease Laboratory | Stephanie Hudon, Joe Christianson, Adriana Rodriguez, Milan Sengthep |
| EPI_ISL_3388863, EPI_ISL_3388864, EPI_ISL_3388869, EPI_ISL_3388870, EPI_ISL_3388871, EPI_ISL_3388872, EPI_ISL_3388873, EPI_ISL_3388874, EPI_ISL_3388875, EPI_ISL_3388876, EPI_ISL_3388877, EPI_ISL_3388878, EPI_ISL_3388879, EPI_ISL_3388880, EPI_ISL_3388881, EPI_ISL_3388882, EPI_ISL_3388883, EPI_ISL_3388884 | | | |
| see above | San Diego County Public Health Laboratory | San Diego County Public Health Laboratory | Ashleigh Murphy Schafer, Brett Austin |
| EPI_ISL_3388891, EPI_ISL_3388892, EPI_ISL_3388893, EPI_ISL_3388894, EPI_ISL_3388895, EPI_ISL_3388896, EPI_ISL_3388897, EPI_ISL_3388898, EPI_ISL_3388899, EPI_ISL_3388900, EPI_ISL_3388901, EPI_ISL_3388902, EPI_ISL_3388903 | | | |
| see above | Dr. Risch Ostschweiz AG | Microbiology, Dr. Risch | Nadia Wohlwend |
| EPI_ISL_3388904, EPI_ISL_3388905, EPI_ISL_3388907, EPI_ISL_3388908, EPI_ISL_3388909, EPI_ISL_3388910, EPI_ISL_3388911, EPI_ISL_3388912, EPI_ISL_3388913, EPI_ISL_3388914, EPI_ISL_3388915, EPI_ISL_3388916, EPI_ISL_3388917, EPI_ISL_3388918, EPI_ISL_3388920, EPI_ISL_3388921, EPI_ISL_3388922, EPI_ISL_3388923, EPI_ISL_3388924, EPI_ISL_3388925, EPI_ISL_3388926, EPI_ISL_3388927, EPI_ISL_3388929, EPI_ISL_3388930, EPI_ISL_3388931, EPI_ISL_3388932, EPI_ISL_3388933, EPI_ISL_3388934, EPI_ISL_3388935, EPI_ISL_3388936, EPI_ISL_3388937, EPI_ISL_3388938, EPI_ISL_3388939, EPI_ISL_3388940, EPI_ISL_3388941, EPI_ISL_3388942, EPI_ISL_3388943, EPI_ISL_3388944, EPI_ISL_3388945, EPI_ISL_3388946, EPI_ISL_3388947, EPI_ISL_3388948, EPI_ISL_3388949, EPI_ISL_3388950, EPI_ISL_3388951, EPI_ISL_3388952, EPI_ISL_3388953, EPI_ISL_3388954 | | | |
| see above | Dutch COVID-19 response team | National Institute for Public Health and the Environment (RIVM) | Adam Meijer, Harry Vennema, Dirk Eggink, Jeroen Cremer, Sharon van den Brink, Bas van der Veer, AnneMarie van den Brandt, Lisa Wijsman, Kim Freriks, Ryanne Jaarsma, Euníce Then, Lynn Aarts, Sanne Bos, Stijn van Rossum, Florian Zwagemaker, Dennis Schmitz, Annelies Kroneman, Karim Hajji, Chantal Reusken, on behalf of the national COVID-19 response team |
| EPI_ISL_3388983, EPI_ISL_3388984, EPI_ISL_3388986 | Ender Diagnostics AG | Microbiology, Dr. Risch | Nadia Wohlwend |
| EPI_ISL_3388988, EPI_ISL_3388990, EPI_ISL_3388992, EPI_ISL_3388993 | Dr. Risch Ostschweiz AG | Microbiology, Dr. Risch | Nadia Wohlwend |
| EPI_ISL_3388995 | Ender Diagnostics AG | Microbiology, Dr. Risch | Nadia Wohlwend |
| EPI_ISL_3388997 | ZLC Zentrallabor Chur | Microbiology, Dr. Risch | Nadia Wohlwend |
| EPI_ISL_3388999, EPI_ISL_3389000, EPI_ISL_3389002 | Dr. Risch Ostschweiz AG | Microbiology, Dr. Risch | Nadia Wohlwend |
| EPI_ISL_3389004 | ZLC Zentrallabor Chur | Microbiology, Dr. Risch | Nadia Wohlwend |

| | | | |
|---|---|---|---|
| EPI_ISL_3389005 | Dr. Risch Ostschweiz AG | Microbiology, Dr. Risch | Nadia Wohlwend |
| EPI_ISL_3389007, EPI_ISL_3389009, EPI_ISL_3389011, EPI_ISL_3389013 | Ender Diagnostics AG | Microbiology, Dr. Risch | Nadia Wohlwend |
| EPI_ISL_3389015, EPI_ISL_3389022, EPI_ISL_3389029, EPI_ISL_3389030, EPI_ISL_3389031, EPI_ISL_3389032, EPI_ISL_3389033, EPI_ISL_3389034, EPI_ISL_3389035, EPI_ISL_3389036, EPI_ISL_3389037, EPI_ISL_3389039, EPI_ISL_3389040, EPI_ISL_3389041, EPI_ISL_3389042, EPI_ISL_3389043, EPI_ISL_3389044, EPI_ISL_3389045, EPI_ISL_3389046, EPI_ISL_3389047, EPI_ISL_3389048, EPI_ISL_3389049, EPI_ISL_3389050, EPI_ISL_3389051, EPI_ISL_3389052, EPI_ISL_3389053, EPI_ISL_3389054, EPI_ISL_3389055, EPI_ISL_3389056, EPI_ISL_3389058, EPI_ISL_3389060, EPI_ISL_3389061, EPI_ISL_3389062, EPI_ISL_3389063, EPI_ISL_3389064, EPI_ISL_3389065, EPI_ISL_3389066, EPI_ISL_3389068, EPI_ISL_3389069, EPI_ISL_3389070, EPI_ISL_3389071, EPI_ISL_3389072, EPI_ISL_3389083, EPI_ISL_3389084, EPI_ISL_3389085, EPI_ISL_3389086, EPI_ISL_3389087, EPI_ISL_3389088, EPI_ISL_3389089, EPI_ISL_3389090, EPI_ISL_3389091, EPI_ISL_3389092, EPI_ISL_3389093, EPI_ISL_3389094, EPI_ISL_3389095, EPI_ISL_3389096, EPI_ISL_3389097, EPI_ISL_3389098, EPI_ISL_3389099, EPI_ISL_3389100, EPI_ISL_3389101, EPI_ISL_3389102, EPI_ISL_3389103, EPI_ISL_3389104, EPI_ISL_3389105, EPI_ISL_3389106, EPI_ISL_3389107, EPI_ISL_3389108, EPI_ISL_3389109, EPI_ISL_3389110, EPI_ISL_3389112, EPI_ISL_3389113, EPI_ISL_3389114, EPI_ISL_3389115, EPI_ISL_3389116, EPI_ISL_3389117, EPI_ISL_3389118, EPI_ISL_3389119, EPI_ISL_3389120, EPI_ISL_3389121, EPI_ISL_3389122, EPI_ISL_3389123, EPI_ISL_3389124, EPI_ISL_3389125, EPI_ISL_3389126, EPI_ISL_3389127, EPI_ISL_3389128, EPI_ISL_3389129, EPI_ISL_3389130, EPI_ISL_3389131, EPI_ISL_3389132, EPI_ISL_3389133, EPI_ISL_3389134, EPI_ISL_3389135, EPI_ISL_3389136, EPI_ISL_3389137, EPI_ISL_3389138, EPI_ISL_3389139, EPI_ISL_3389140, EPI_ISL_3389141, EPI_ISL_3389142, EPI_ISL_3389143, EPI_ISL_3389144, EPI_ISL_3389145, EPI_ISL_3389146, EPI_ISL_3389147, EPI_ISL_3389148, EPI_ISL_3389149, EPI_ISL_3389150, EPI_ISL_3389151, EPI_ISL_3389152, EPI_ISL_3389153, EPI_ISL_3389154, EPI_ISL_3389155, EPI_ISL_3389156, EPI_ISL_3389157, EPI_ISL_3389158, EPI_ISL_3389159, EPI_ISL_3389160, EPI_ISL_3389161, EPI_ISL_3389162, EPI_ISL_3389163, EPI_ISL_3389164, EPI_ISL_3389165, EPI_ISL_3389166, EPI_ISL_3389167, EPI_ISL_3389168, EPI_ISL_3389169, EPI_ISL_3389170, EPI_ISL_3389171, EPI_ISL_3389172, EPI_ISL_3389173, EPI_ISL_3389174, EPI_ISL_3389175, EPI_ISL_3389176, EPI_ISL_3389177, EPI_ISL_3389178, EPI_ISL_3389179, EPI_ISL_3389180, EPI_ISL_3389181, EPI_ISL_3389182, EPI_ISL_3389183, EPI_ISL_3389184, EPI_ISL_3389185, EPI_ISL_3389186, EPI_ISL_3389187, EPI_ISL_3389188, EPI_ISL_3389189, EPI_ISL_3389190, EPI_ISL_3389191, EPI_ISL_3389192, EPI_ISL_3389193, EPI_ISL_3389194, EPI_ISL_3389197, EPI_ISL_3389199, EPI_ISL_3389201, EPI_ISL_3389204, EPI_ISL_3389206 | | | |
| see above | Dutch COVID-19 response team | National Institute for Public Health and the Environment (RIVM) | Adam Meijer, Harry Vennema, Dirk Eggink, Jeroen Cremer, Sharon van den Brink, Bas van der Veer, AnneMarie van den Brandt, Lisa Wijsman, Kim Freriks, Ryanne Jaarsma, Euníce Then, Lynn Aarts, Sanne Bos, Stijn van Rossum, Florian Zwagemaker, Dennis Schmitz, Annelies Kroneman, Karim Hajji, Chantal Reusken, on behalf of the national COVID-19 response team |
| EPI_ISL_3389207 | Ender Diagnostics AG | Microbiology, Dr. Risch | Nadia Wohlwend |
| EPI_ISL_3389208, EPI_ISL_3389211, EPI_ISL_3389213, EPI_ISL_3389215, EPI_ISL_3389217, EPI_ISL_3389220, EPI_ISL_3389222, EPI_ISL_3389224, EPI_ISL_3389225, EPI_ISL_3389226, EPI_ISL_3389227, EPI_ISL_3389228, EPI_ISL_3389229, EPI_ISL_3389230, EPI_ISL_3389231, EPI_ISL_3389232, EPI_ISL_3389233, EPI_ISL_3389234, EPI_ISL_3389235, EPI_ISL_3389236, EPI_ISL_3389237, EPI_ISL_3389238, EPI_ISL_3389239, EPI_ISL_3389240, EPI_ISL_3389241, EPI_ISL_3389242, EPI_ISL_3389243, EPI_ISL_3389244, EPI_ISL_3389245, EPI_ISL_3389246, EPI_ISL_3389247, EPI_ISL_3389248, EPI_ISL_3389249, EPI_ISL_3389250, EPI_ISL_3389251, EPI_ISL_3389252, EPI_ISL_3389253, EPI_ISL_3389254, EPI_ISL_3389255, EPI_ISL_3389256, EPI_ISL_3389257, EPI_ISL_3389258, EPI_ISL_3389259, EPI_ISL_3389260, EPI_ISL_3389261, EPI_ISL_3389262, EPI_ISL_3389263, EPI_ISL_3389264, EPI_ISL_3389265, EPI_ISL_3389266, EPI_ISL_3389267, EPI_ISL_3389268, EPI_ISL_3389269, EPI_ISL_3389270, EPI_ISL_3389271, EPI_ISL_3389272, EPI_ISL_3389273, EPI_ISL_3389274, EPI_ISL_3389275, EPI_ISL_3389276, EPI_ISL_3389277, EPI_ISL_3389278, EPI_ISL_3389279, EPI_ISL_3389280, EPI_ISL_3389281, EPI_ISL_3389282, EPI_ISL_3389283, EPI_ISL_3389284, EPI_ISL_3389285, EPI_ISL_3389286, EPI_ISL_3389287, EPI_ISL_3389288, EPI_ISL_3389289, EPI_ISL_3389290, EPI_ISL_3389291, EPI_ISL_3389292, EPI_ISL_3389293, EPI_ISL_3389294, EPI_ISL_3389295, EPI_ISL_3389296, EPI_ISL_3389297, EPI_ISL_3389298, EPI_ISL_3389299, EPI_ISL_3389300, EPI_ISL_3389301, EPI_ISL_3389302, EPI_ISL_3389303, EPI_ISL_3389304, EPI_ISL_3389305, EPI_ISL_3389306, EPI_ISL_3389307, EPI_ISL_3389308, EPI_ISL_3389309, EPI_ISL_3389310, EPI_ISL_3389311, EPI_ISL_3389312, EPI_ISL_3389313, EPI_ISL_3389314, EPI_ISL_3389315, EPI_ISL_3389316, EPI_ISL_3389317, EPI_ISL_3389318, EPI_ISL_3389319, EPI_ISL_3389320, EPI_ISL_3389321, EPI_ISL_3389322, EPI_ISL_3389323, EPI_ISL_3389324, EPI_ISL_3389325, EPI_ISL_3389326, EPI_ISL_3389327, EPI_ISL_3389328, EPI_ISL_3389329, EPI_ISL_3389330, EPI_ISL_3389331, EPI_ISL_3389332, EPI_ISL_3389333, EPI_ISL_3389334, EPI_ISL_3389335, EPI_ISL_3389336, EPI_ISL_3389337, EPI_ISL_3389338, EPI_ISL_3389339, EPI_ISL_3389340, EPI_ISL_3389341, EPI_ISL_3389342, EPI_ISL_3389343, EPI_ISL_3389344, EPI_ISL_3389345, EPI_ISL_3389346, EPI_ISL_3389347, EPI_ISL_3389348, EPI_ISL_3389349, EPI_ISL_3389350, EPI_ISL_3389351, EPI_ISL_3389352, EPI_ISL_3389353, EPI_ISL_3389354, EPI_ISL_3389355, EPI_ISL_3389356, EPI_ISL_3389357, EPI_ISL_3389358, EPI_ISL_3389359, EPI_ISL_3389360, EPI_ISL_3389361, EPI_ISL_3389362, EPI_ISL_3389363, EPI_ISL_3389364, EPI_ISL_3389365, EPI_ISL_3389366, EPI_ISL_3389367, EPI_ISL_3389368, EPI_ISL_3389369, EPI_ISL_3389370, EPI_ISL_3389371, EPI_ISL_3389372, EPI_ISL_3389373, EPI_ISL_3389374, EPI_ISL_3389375, EPI_ISL_3389376, EPI_ISL_3389377, EPI_ISL_3389378, EPI_ISL_3389379, EPI_ISL_3389380, EPI_ISL_3389381, EPI_ISL_3389382, EPI_ISL_3389383, EPI_ISL_3389384, EPI_ISL_3389385, EPI_ISL_3389386, EPI_ISL_3389387, EPI_ISL_3389388, EPI_ISL_3389389, EPI_ISL_3389390, EPI_ISL_3389391, EPI_ISL_3389392, EPI_ISL_3389393, EPI_ISL_3389394, EPI_ISL_3389395, EPI_ISL_3389396, EPI_ISL_3389397, EPI_ISL_3389398, EPI_ISL_3389399, EPI_ISL_3389400, EPI_ISL_3389401, EPI_ISL_3389402, EPI_ISL_3389403, EPI_ISL_3389404, EPI_ISL_3389405, EPI_ISL_3389406, EPI_ISL_3389407, EPI_ISL_3389408, EPI_ISL_3389409, EPI_ISL_3389410, EPI_ISL_3389411, EPI_ISL_3389412, EPI_ISL_3389413, EPI_ISL_3389414, EPI_ISL_3389415, EPI_ISL_3389416, EPI_ISL_3389417, EPI_ISL_3389418, EPI_ISL_3389419, EPI_ISL_3389420, EPI_ISL_3389421, EPI_ISL_3389422, EPI_ISL_3389423, EPI_ISL_3389424, EPI_ISL_3389425, EPI_ISL_3389426, EPI_ISL_3389427, EPI_ISL_3389428, EPI_ISL_3389429, EPI_ISL_3389430, EPI_ISL_3389431, EPI_ISL_3389432, EPI_ISL_3389433, EPI_ISL_3389434, EPI_ISL_3389435, EPI_ISL_3389436, EPI_ISL_3389437, EPI_ISL_3389438, EPI_ISL_3389439, EPI_ISL_3389440, EPI_ISL_3389441, EPI_ISL_3389442, EPI_ISL_3389443, EPI_ISL_3389444, EPI_ISL_3389445, EPI_ISL_3389446, EPI_ISL_3389447, EPI_ISL_3389448, EPI_ISL_3389449, EPI_ISL_3389450, EPI_ISL_3389451, EPI_ISL_3389452, EPI_ISL_3389453, EPI_ISL_3389454, EPI_ISL_3389455, EPI_ISL_3389456, EPI_ISL_3389457, EPI_ISL_3389458, EPI_ISL_3389459, EPI_ISL_3389460, EPI_ISL_3389461, EPI_ISL_3389462, EPI_ISL_3389463, EPI_ISL_3389464, EPI_ISL_3389465, EPI_ISL_3389466, EPI_ISL_3389467, EPI_ISL_3389468, EPI_ISL_3389469, EPI_ISL_3389470, EPI_ISL_3389471, EPI_ISL_3389472, EPI_ISL_3389473, EPI_ISL_3389474, EPI_ISL_3389475, EPI_ISL_3389476, EPI_ISL_3389477, EPI_ISL_3389478, EPI_ISL_3389479, EPI_ISL_3389480, EPI_ISL_3389481, EPI_ISL_3389482, EPI_ISL_3389483, EPI_ISL_3389484, EPI_ISL_3389485, EPI_ISL_3389486, EPI_ISL_3389487, EPI_ISL_3389488, EPI_ISL_3389489, EPI_ISL_3389490, EPI_ISL_3389491, EPI_ISL_3389492, EPI_ISL_3389493, EPI_ISL_3389494, EPI_ISL_3389495, EPI_ISL_3389496, EPI_ISL_3389497, EPI_ISL_3389498, EPI_ISL_3389499, EPI_ISL_3389500, EPI_ISL_3389501, EPI_ISL_3389502, EPI_ISL_3389503, EPI_ISL_3389504, EPI_ISL_3389505, EPI_ISL_3389506, EPI_ISL_3389507, EPI_ISL_3389508, EPI_ISL_3389509, EPI_ISL_3389510, EPI_ISL_3389511, EPI_ISL_3389512, EPI_ISL_3389513, EPI_ISL_3389514, EPI_ISL_3389515, EPI_ISL_3389516, EPI_ISL_3389517, EPI_ISL_3389518, EPI_ISL_3389519, EPI_ISL_3389520, EPI_ISL_3389521, EPI_ISL_3389522, EPI_ISL_3389523, EPI_ISL_3389524, EPI_ISL_3389525, EPI_ISL_3389526, EPI_ISL_3389527, EPI_ISL_3389528, EPI_ISL_3389529, EPI_ISL_3389530, EPI_ISL_3389531, EPI_ISL_3389532, EPI_ISL_3389533, EPI_ISL_3389534, EPI_ISL_3389535, EPI_ISL_3389536, EPI_ISL_3389537, EPI_ISL_3389538, EPI_ISL_3389539, EPI_ISL_3389540, EPI_ISL_3389541, EPI_ISL_3389542, EPI_ISL_3389543, EPI_ISL_3389544, EPI_ISL_3389545, EPI_ISL_3389546, EPI_ISL_3389547, EPI_ISL_3389548, EPI_ISL_3389549, EPI_ISL_3389550, EPI_ISL_3389551, EPI_ISL_3389552, EPI_ISL_3389553, EPI_ISL_3389554, EPI_ISL_3389555, EPI_ISL_3389556, EPI_ISL_3389557, EPI_ISL_3389558, EPI_ISL_3389559, EPI_ISL_3389560, EPI_ISL_3389561, EPI_ISL_3389562, EPI_ISL_3389563, EPI_ISL_3389564, EPI_ISL_3389565, EPI_ISL_3389566, EPI_ISL_3389567, EPI_ISL_3389568, EPI_ISL_3389569, EPI_ISL_3389570, EPI_ISL_3389571, EPI_ISL_3389572, EPI_ISL_3389573, EPI_ISL_3389574, EPI_ISL_3389575, EPI_ISL_3389576, EPI_ISL_3389577, EPI_ISL_3389578, EPI_ISL_3389579, EPI_ISL_3389580, EPI_ISL_3389581, EPI_ISL_3389582, EPI_ISL_3389583, EPI_ISL_3389584, EPI_ISL_3389585, EPI_ISL_3389586, EPI_ISL_3389587, EPI_ISL_3389588, EPI_ISL_3389589, EPI_ISL_3389590, EPI_ISL_3389591, EPI_ISL_3389592, EPI_ISL_3389593, EPI_ISL_3389594, EPI_ISL_3389595, EPI_ISL_3389596, EPI_ISL_3389597, EPI_ISL_3389598, EPI_ISL_3389599, EPI_ISL_3389600, EPI_ISL_3389601, EPI_ISL_3389602, EPI_ISL_3389603, EPI_ISL_3389604, EPI_ISL_3389605, EPI_ISL_3389606, EPI_ISL_3389607, EPI_ISL_3389608, EPI_ISL_3389609, EPI_ISL_3389610, EPI_ISL_3389611, EPI_ISL_3389612, EPI_ISL_3389613, EPI_ISL_3389614, EPI_ISL_3389615, EPI_ISL_3389616, EPI_ISL_3389617, EPI_ISL_3389618, EPI_ISL_3389619, EPI_ISL_3389620, EPI_ISL_3389621, EPI_ISL_3389622, EPI_ISL_3389623, EPI_ISL_3389624, EPI_ISL_3389625, EPI_ISL_3389626, EPI_ISL_3389627, EPI_ISL_3389628, EPI_ISL_3389629, EPI_ISL_3389630, EPI_ISL_3389631, EPI_ISL_3389632, EPI_ISL_3389633, EPI_ISL_3389634, EPI_ISL_3389635, EPI_ISL_3389636, EPI_ISL_3389637, EPI_ISL_3389638, EPI_ISL_3389639, EPI_ISL_3389640, EPI_ISL_3389641, EPI_ISL_3389642, EPI_ISL_3389643, EPI_ISL_3389644, EPI_ISL_3389645, EPI_ISL_3389646, EPI_ISL_3389647, EPI_ISL_3389648, EPI_ISL_3389649, EPI_ISL_3389650, EPI_ISL_3389651, EPI_ISL_3389652, EPI_ISL_3389653, EPI_ISL_3389654, EPI_ISL_3389655, EPI_ISL_3389656, EPI_ISL_3389657, EPI_ISL_3389658, EPI_ISL_3389659, EPI_ISL_3389660, EPI_ISL_3389661, EPI_ISL_3389662, EPI_ISL_3389663, EPI_ISL_3389664, EPI_ISL_3389665, EPI_ISL_3389666, EPI_ISL_3389667, EPI_ISL_3389668, EPI_ISL_3389669, EPI_ISL_3389670, EPI_ISL_3389671, EPI_ISL_3389672, EPI_ISL_3389673, EPI_ISL_3389674, EPI_ISL_3389675, EPI_ISL_3389676, EPI_ISL_3389677, EPI_ISL_3389678, EPI_ISL_3389679, EPI_ISL_3389680, EPI_ISL_3389681, EPI_ISL_3389682, EPI_ISL_3389683, EPI_ISL_3389684, EPI_ISL_3389685, EPI_ISL_3389686, EPI_ISL_3389687, EPI_ISL_3389688, EPI_ISL_3389689, EPI_ISL_3389690, EPI_ISL_3389691, EPI_ISL_3389692, EPI_ISL_3389693, EPI_ISL_3389694, EPI_ISL_3389695, EPI_ISL_3389696, EPI_ISL_3389697, EPI_ISL_3389698, EPI_ISL_3389699, EPI_ISL_3389700, EPI_ISL_3389701, EPI_ISL_3389702, EPI_ISL_3389703, EPI_ISL_3389704, EPI_ISL_3389705, EPI_ISL_3389706, EPI_ISL_3389707, EPI_ISL_3389708, EPI_ISL_3389709, EPI_ISL_3389710, EPI_ISL_3389711, EPI_ISL_3389712, EPI_ISL_3389713, EPI_ISL_3389714, EPI_ISL_3389715, EPI_ISL_3389716, EPI_ISL_3389717, EPI_ISL_3389718, EPI_ISL_3389719, EPI_ISL_3389720, EPI_ISL_3389721, EPI_ISL_3389722, EPI_ISL_3389723, EPI_ISL_3389724, EPI_ISL_3389725, EPI_ISL_3389726, EPI_ISL_3389727, EPI_ISL_3389728, EPI_ISL_3389729, EPI_ISL_3389730, EPI_ISL_3389731, EPI_ISL_3389732, EPI_ISL_3389733, EPI_ISL_3389734, EPI_ISL_3389735, EPI_ISL_3389736, EPI_ISL_3389737, EPI_ISL_3389738, EPI_ISL_3389739, EPI_ISL_3389740, EPI_ISL_3389741, EPI_ISL_3389742, EPI_ISL_3389743, EPI_ISL_3389744, EPI_ISL_3389745, EPI_ISL_3389746, EPI_ISL_3389747, EPI_ISL_3389748, EPI_ISL_3389749, EPI_ISL_3389750, EPI_ISL_3389751, EPI_ISL_3389752, EPI_ISL_3389753, EPI_ISL_3389754, EPI_ISL_3389755, EPI_ISL_3389756, EPI_ISL_3389757, EPI_ISL_3389758, EPI_ISL_3389759, EPI_ISL_3389760, EPI_ISL_3389761, EPI_ISL_3389762, EPI_ISL_3389763, EPI_ISL_3389764, EPI_ISL_3389765, EPI_ISL_3389766, EPI_ISL_3389767, EPI_ISL_3389768, EPI_ISL_3389769, EPI_ISL_3389770, EPI_ISL_3389771, EPI_ISL_3389772, EPI_ISL_3389773, EPI_ISL_3389774, EPI_ISL_3389775, EPI_ISL_3389776, EPI_ISL_3389777, EPI_ISL_3389778, EPI_ISL_3389779, EPI_ISL_3389780, EPI_ISL_3389781, EPI_ISL_3389782, EPI_ISL_3389783, EPI_ISL_3389784, EPI_ISL_3389785, EPI_ISL_3389786, EPI_ISL_3389787, EPI_ISL_3389788, EPI_ISL_3389789, EPI_ISL_3389790, EPI_ISL_3389791, EPI_ISL_3389792, EPI_ISL_3389793, EPI_ISL_3389794, EPI_ISL_3389795, EPI_ISL_3389796, EPI_ISL_3389797, EPI_ISL_3389798, EPI_ISL_3389799, EPI_ISL_3389800, EPI_ISL_3389801, EPI_ISL_3389802, EPI_ISL_3389803, EPI_ISL_3389804, EPI_ISL_3389805, EPI_ISL_3389806, EPI_ISL_3389807, EPI_ISL_3389808, EPI_ISL_3389809, EPI_ISL_3389810, EPI_ISL_3389811, EPI_ISL_3389812, EPI_ISL_3389813, EPI_ISL_3389814, EPI_ISL_3389815, EPI_ISL_3389816, EPI_ISL_3389817, EPI_ISL_3389818, EPI_ISL_3389819, EPI_ISL_3389820, EPI_ISL_3389821, EPI_ISL_3389822, EPI_ISL_3389823, EPI_ISL_3389824, EPI_ISL_3389825, EPI_ISL_3389826, EPI_ISL_3389827, EPI_ISL_3389828, EPI_ISL_3389829, EPI_ISL_3389830, EPI_ISL_3389831, EPI_ISL_3389832, EPI_ISL_3389833, EPI_ISL_3389834, EPI_ISL_3389835, EPI_ISL_3389836, EPI_ISL_3389837, EPI_ISL_3389838, EPI_ISL_3389839, EPI_ISL_3389840, EPI_ISL_3389841, EPI_ISL_3389842, EPI_ISL_3389843, EPI_ISL_3389844, EPI_ISL_3389845, EPI_ISL_3389846, EPI_ISL_3389847, EPI_ISL_3389848, EPI_ISL_3389849, EPI_ISL_3389850, EPI_ISL_3389851, EPI_ISL_3389852, EPI_ISL_3389853, EPI_ISL_3389854, EPI_ISL_3389855, EPI_ISL_3389856, EPI_ISL_3389857, EPI_ISL_3389858, EPI_ISL_3389859, EPI_ISL_3389860, EPI_ISL_3389861, EPI_ISL_3389862, EPI_ISL_3389863, EPI_ISL_3389864, EPI_ISL_3389865, EPI_ISL_3389866, EPI_ISL_3389867, EPI_ISL_3389868, EPI_ISL_3389869, EPI_ISL_3389870, EPI_ISL_3389871, EPI_ISL_3389872, EPI_ISL_3389873, EPI_ISL_3389874, EPI_ISL_3389875, EPI_ISL_3389876, EPI_ISL_3389877, EPI_ISL_3389878, EPI_ISL_3389879, EPI_ISL_3389880, EPI_ISL_3389881, EPI_ISL_3389882, EPI_ISL_3389883, EPI_ISL_3389884, EPI_ISL_3389885, EPI_ISL_3389886, EPI_ISL_3389887, EPI_ISL_3389888, EPI_ISL_3389889, EPI_ISL_3389890, EPI_ISL_3389891, EPI_ISL_3389892, EPI_ISL_3389893, EPI_ISL_3389894, EPI_ISL_3389895, EPI_ISL_3389896, EPI_ISL_3389897, EPI_ISL_3389898, EPI_ISL_3389899, EPI_ISL_3389900, EPI_ISL_3389901, EPI_ISL_3389902, EPI_ISL_3389903, EPI_ISL_3389904, EPI_ISL_3389905, EPI_ISL_3389906, EPI_ISL_3389907, EPI_ISL_3389908, EPI_ISL_3389909, EPI_ISL_3389910, EPI_ISL_3389911, EPI_ISL_3389912, EPI_ISL_3389913, EPI_ISL_3389914, EPI_ISL_3389915, EPI_ISL_3389916, EPI_ISL_3389917, EPI_ISL_3389918, EPI_ISL_3389919, EPI_ISL_3389920, EPI_ISL_3389921, EPI_ISL_3389922, EPI_ISL_3389923, EPI_ISL_3389924, EPI_ISL_3389925, EPI_ISL_3389926, EPI_ISL_3389927, EPI_ISL_3389928, EPI_ISL_3389929, EPI_ISL_3389930, EPI_ISL_3389931, EPI_ISL_3389932, EPI_ISL_3389933, EPI_ISL_3389934, EPI_ISL_3389935, EPI_ISL_3389936, EPI_ISL_3389937, EPI_ISL_3389938, EPI_ISL_3389939, EPI_ISL_3389940, EPI_ISL_3389941, EPI_ISL_3389942, EPI_ISL_3389943, EPI_ISL_3389944, EPI_ISL_3389945, EPI_ISL_3389946, EPI_ISL_3389947, EPI_ISL_3389948, EPI_ISL_3389949, EPI_ISL_3389950, EPI_ISL_3389951, EPI_ISL_3389952, EPI_ISL_3389953, EPI_ISL_3389954, EPI_ISL_3389955, EPI_ISL_3389956, EPI_ISL_3389957, EPI_ISL_3389958, EPI_ISL_3389959, EPI_ISL_3389960, EPI_ISL_3389961, EPI_ISL_3389962, EPI_ISL_3389963, EPI_ISL_3389964, EPI_ISL_3389965, EPI_ISL_3389966, EPI_ISL_3389967, EPI_ISL_3389968, EPI_ISL_3389969, EPI_ISL_3389970, EPI_ISL_3389971, EPI_ISL_3389972, EPI_ISL_3389973, EPI_ISL_3389974, EPI_ISL_3389975, EPI_ISL_3389976, EPI_ISL_3389977, EPI_ISL_3389978, EPI_ISL_3389979, EPI_ISL_3389980, EPI_ISL_3389981, EPI_ISL_3389982, EPI_ISL_3389983, EPI_ISL_3389984, EPI_ISL_3389985, EPI_ISL_3389986, EPI_ISL_3389987, EPI_ISL_3389988, EPI_ISL_3389989, EPI_ISL_3389990, EPI_ISL_3389991, EPI_ISL_3389992, EPI_ISL_3389993, EPI_ISL_3389994, EPI_ISL_3389995, EPI_ISL_3389996, EPI_ISL_3389997, EPI_ISL_3389998, EPI_ISL_3389999, EPI_ISL_3390000, EPI_ISL_3390001, EPI_ISL_3390002, EPI_ISL_3390003, EPI_ISL_3390004, EPI_ISL_3390005, EPI_ISL_3390006, EPI_ISL_3390007, EPI_ISL_3390008, EPI_ISL_3390009, EPI_ISL_3390010, EPI_ISL_3390011, EPI_ISL_3390012, EPI_ISL_3390013, EPI_ISL_3390014, EPI_ISL_3390015, EPI_ISL_3390016, EPI_ISL_3390017, EPI_ISL_3390018, EPI_ISL_3390019, EPI_ISL_3390020, EPI_ISL_3390021, EPI_ISL_3390022, EPI_ISL_3390023, EPI_ISL_3390024, EPI_ISL_3390025, EPI_ISL_3390026, EPI_ISL_3390027, EPI_ISL_3390028, EPI_ISL_3390029, EPI_ISL_3390030, EPI_ISL_3390031, EPI_ISL_3390032, EPI_ISL_3390033, EPI_ISL_3390034, EPI_ISL_3390035, EPI_ISL_3390036, EPI_ISL_3390037, EPI_ISL_3390038, EPI_ISL_3390039, EPI_ISL_3390040, EPI_ISL_3390041, EPI_ISL_3390042, EPI_ISL_3390043, EPI_ISL_3390044, EPI_ISL_3390045, EPI_ISL_3390046, EPI_ISL_3390047, EPI_ISL_3390048, EPI_ISL_3390049, EPI_ISL_3390050, EPI_ISL_3390051, EPI_ISL_3390052, EPI_ISL_3390053, EPI_ISL_3390054, EPI_ISL_3390055, EPI_ISL_3390056, EPI_ISL_3390057, EPI_ISL_3390058, EPI_ISL_3390059, EPI_ISL_3390060, EPI_ISL_3390061, EPI_ISL_3390062, EPI_ISL_3390063, EPI_ISL_3390064, EPI_ISL_3390065, EPI_ISL_3390066, EPI_ISL_3390067, EPI_ISL_3390068, EPI_ISL_3390069, EPI_ISL_3390070, EPI_ISL_3390071, EPI_ISL_3390072, EPI_ISL_3390073, EPI_ISL_3390074, EPI_ISL_3390075, EPI_ISL_3390076, EPI_ISL_3390077, EPI_ISL_3390078, EPI_ISL_3390079, EPI_ISL_3390080, EPI_ISL_3390081, EPI_ISL_3390082, EPI_ISL_3390083, EPI_ISL_3390084, EPI_ISL_3390085, EPI_ISL_3390086, EPI_ISL_3390087, EPI_ISL_3390088, EPI_ISL_3390089, EPI_ISL_3390090, EPI_ISL_3390091, EPI_ISL_3390092, EPI_ISL_3390093, EPI_ISL_3390094, EPI_ISL_3390095, EPI_ISL_3390096, EPI_ISL_3390097, EPI_ISL_3390098, EPI_ISL_3390099, EPI_ISL_3390100, EPI_ISL_3390101, EPI_ISL_3390102, EPI_ISL_3390103, EPI_ISL_3390104, EPI_ISL_3390105, EPI_ISL_3390106, EPI_ISL_3390107, EPI_ISL_3390108, EPI_ISL_3390109, EPI_ISL_3390110, EPI_ISL_3390111, EPI_ISL_3390112, EPI_ISL_3390113, EPI_ISL_3390114, EPI_ISL_3390115, EPI_ISL_3390116, EPI_ISL_3390117, EPI_ISL_3390118, EPI_ISL_3390119, EPI_ISL_3390120, EPI_ISL_3390121, EPI_ISL_3390122, EPI_ISL_3390123, EPI_ISL_3390124, EPI_ISL_3390125, EPI_ISL_3390126, EPI_ISL_3390127, EPI_ISL_3390128, EPI_ISL_3390129, EPI_ISL_3390130, EPI_ISL_3390131, EPI_ISL_3390132, EPI_ISL_3390133, EPI_ISL_3390134, EPI_ISL_3390135, EPI_ISL_3390136, EPI_ISL_3390137, EPI_ISL_3390138, EPI_ISL_3390139, EPI_ISL_3390140, EPI_ISL_3390141, EPI_ISL_3390142, EPI_ISL_3390143, EPI_ISL_3390144, EPI_ISL_3390145, EPI_ISL_3390146, EPI_ISL_3390147, EPI_ISL_3390148, EPI_ISL_3390149, EPI_ISL_3390150, EPI_ISL_3390151, EPI_ISL_3390152, EPI_ISL_3390153, EPI_ISL_3390154, EPI_ISL_3390155, EPI_ISL_3390156, EPI_ISL_3390157, EPI_ISL_3390158, EPI_ISL_3390159, EPI_ISL_3390160, EPI_ISL_3390161, EPI_ISL_3390162, EPI_ISL_3390163, EPI_ISL_3390164, EPI_ISL_3390165, EPI_ISL_3390166, EPI_ISL_3390167, EPI_ISL_3390168, EPI_ISL_3390169, EPI_ISL_3390170, EPI_ISL_3390171, EPI_ISL_3390172, EPI_ISL_3390173, EPI_ISL_3390174, EPI_ISL_3390175, EPI_ISL_3390176, EPI_ISL_3390177, EPI_ISL_3390178, EPI_ISL_3390179, EPI_ISL_3390180, EPI_ISL_3390181, EPI_ISL_3390182, EPI_ISL_3390183, EPI_ISL_3390184, EPI_ISL_3390185, EPI_ISL_3390186, EPI_ISL_3390187, EPI_ISL_3390188, EPI_ISL_3390189, EPI_ISL_3390190, EPI_ISL_3390191, EPI_ISL_3390192, EPI_ISL_3390193, EPI_ISL_3390194, EPI_ISL_3390195, EPI_ISL_3390196, EPI_ISL_3390197, EPI_ISL_3390198, EPI_ISL_3390199, EPI_ISL_3390200, EPI_ISL_3390201, EPI_ISL_3390202, EPI_ISL_3390203, EPI_ISL_3390204, EPI_ISL_3390205, EPI_ISL_3390206, EPI_ISL_3390207, EPI_ISL_3390208, EPI_ISL_3390209, EPI_ISL_3390210, EPI_ISL_3390211, EPI_ISL_3390212, EPI_ISL_3390213, EPI_ISL_3390214, EPI_ISL_3390215, EPI_ISL_3390216, EPI_ISL_3390217, EPI_ISL_3390218, EPI_ISL_3390219, EPI_ISL_3390220, EPI_ISL_3390221, EPI_ISL_3390222, EPI_ISL_3390223, EPI_ISL_3390224, EPI_ISL_3390225, EPI_ISL_3390226, EPI_ISL_3390227, EPI_ISL_3390228, EPI_ISL_3390229, EPI_ISL_3390230, EPI_ISL_3390231, EPI_ISL_3390232, EPI_ISL_3390233, EPI_ISL_3390234, EPI_ISL_3390235, EPI_ISL_3390236, EPI_ISL_3390237, EPI_ISL_3390238, EPI_ISL_3390239, EPI_ISL_3390240, EPI_ISL_3390241, EPI_ISL_3390242, EPI_ISL_3390243, EPI_ISL_3390244, EPI_ISL_3390245, EPI_ISL_3390246, EPI_ISL_3390247, EPI_ISL_3390248, EPI_ISL_3390249, EPI_ISL_3390250, EPI_ISL_3390251, EPI_ISL_3390252, EPI_ISL_3390253, EPI_ISL_3390254, EPI_ISL_3390255, EPI_ISL_3390256, EPI_ISL_3390257, EPI_ISL_3390258, EPI_ISL_3390259, EPI_ISL_3390260, EPI_ISL_3390261, EPI_ISL_3390262, EPI_ISL_3390263, EPI_ISL_3390264, EPI_ISL_3390265, EPI_ISL_3390266, EPI_ISL_3390267, EPI_ISL_3390268, EPI_ISL_3390269, EPI_ISL_3390270, EPI_ISL_3390271, EPI_ISL_3390272, EPI_ISL_3390273, EPI_ISL_3390274, EPI_ISL_3390275, EPI_ISL_3390276, EPI_ISL_3390277, EPI_ISL_3390278, EPI_ISL_3390279, EPI_ISL_3390280, EPI_ISL_3390281, EPI_ISL_3390282, EPI_ISL_3390283, EPI_ISL_3390284, EPI_ISL_3390285, EPI_ISL_3390286, EPI_ISL_3390287, EPI_ISL_3390288, EPI_ISL_3390289, EPI_ISL_3390290, EPI_ISL_3390291, EPI_ISL_3390292, EPI_ISL_3390293, EPI_ISL_3390294, EPI_ISL_3390295, EPI_ISL_3390296, EPI_ISL_3390297, EPI_ISL_3390298, EPI_ISL_3390299, EPI_ISL_3390300, EPI_ISL_3390301, EPI_ISL_3390302, EPI_ISL_3390303, EPI_ISL_3390304, EPI_ISL_3390305, EPI_ISL_3390306, EPI_ISL_3390307, EPI_ISL_3390308, EPI_ISL_3390309, EPI_ISL_3390310, EPI_ISL_3390311, EPI_ISL_3390312, EPI_ISL_3390313, EPI_ISL_3390314, EPI_ISL_3390315, EPI_ISL_3390316, EPI_ISL_3390317, EPI_ISL_3390318, EPI_ISL_3390319, EPI_ISL_3390320, EPI_ISL_3390321, EPI_ISL_3390322, EPI_ISL_3390323, EPI_ISL_3390324, EPI_ISL_3390325, EPI_ISL_3390326, EPI_ISL_3390327, EPI_ISL_3390328, EPI_ISL_3390329, EPI_ISL_3390330, EPI_ISL_3390331, EPI_ISL_3390332, EPI_ISL_3390333, EPI_ISL_3390334, EPI_ISL_3390335, EPI_ISL_3390336, EPI_ISL_3390337, EPI_ISL_3390338, EPI_ISL_3390339, EPI_ISL_3390340, EPI_ISL_3390341, EPI_ISL_3390342, EPI_ISL_3390343, EPI_ISL_3390344, EPI_ISL_3390345, EPI_ISL_3390346, EPI_ISL_3390347, EPI_ISL_3390348, EPI_ISL_3390349, EPI_ISL_3390350, EPI_ISL_3390351, EPI_ISL_3390352, EPI_ISL_3390353, EPI_ISL_3390354, EPI_ISL_3390355, EPI_ISL_3390356, EPI_ISL_3390357, EPI_ISL_3390358, EPI_ISL_3390359, EPI_ISL_3390360, EPI_ISL_3390361, EPI_ISL_3390362, EPI_ISL_3390363, EPI_ISL_3390364, EPI_ISL_3390365, EPI_ISL_3390366, EPI_ISL_3390367, EPI_ISL_3390368, EPI_ISL_3390369, EPI_ISL_3390370, EPI_ISL_3390371, EPI_ISL_3390372, EPI_ISL_3390373, EPI_ISL_3390374, EPI_ISL_3390375, EPI_ISL_3390376, EPI_ISL_3390377, EPI_ISL_3390378, EPI_ISL_3390379, EPI_ISL_3390380, EPI_ISL_3390381, EPI_ISL_3390382, EPI_ISL_3390383, EPI_ISL_3390384, EPI_ISL_3390385, EPI_ISL_3390386, EPI_ISL_3390387, EPI_ISL_3390388, EPI_ISL_3390389, EPI_ISL_3390390, EPI_ISL_3390391, EPI_ISL_3390392, EPI_ISL_3390393, EPI_ISL_3390394, EPI_ISL_3390395, EPI_ISL_3390396, EPI_ISL_3390397, EPI_ISL_3390398, EPI_ISL_3390399, EPI_ISL_3390400, EPI_ISL_3390401, EPI_ISL_3390402, EPI_ISL_3390403, EPI_ISL_3390404, EPI_ISL_3390405, EPI_ISL_3390406, EPI_ISL_3390407, EPI_ISL_3390408, EPI_ISL_3390409, EPI_ISL_3390410, EPI_ISL_3390411, EPI_ISL_3390412, EPI_ISL_3390413, EPI_ISL_3390414, EPI_ISL_3390415, EPI_ISL_3390416, EPI_ISL_3390417, EPI_ISL_3390418, EPI_ISL_3390419, EPI_ISL_3390420, EPI_ISL_3390421, EPI_ISL_3390422, EPI_ISL_3390423, EPI_ISL_3390424, EPI_ISL_3390425, EPI_ISL_3390426, EPI_ISL_3390427, EPI_ISL_3390428, EPI_ISL_3390429, EPI_ISL_3390430, EPI_ISL_3390431, EPI_ISL_3390432, EPI_ISL_3390433, EPI_ISL_3390434, EPI_ISL_3390435, EPI_ISL_3390436, EPI_ISL_3390437, EPI_ISL_3390438, EPI_ISL_3390439, EPI_ISL_3390440, EPI_ISL_3390441, EPI_ISL_3390442, EPI_ISL_3390443, EPI_ISL_3390444, EPI_ISL_3390445, EPI_ISL_3390446, EPI_ISL_3390447, EPI_ISL_3390448, EPI_ISL_3390449, EPI_ISL_3390450, EPI_ISL_3390451, EPI_ISL_3390452, EPI_ISL_3390453, EPI_ISL_3390454, EPI_ISL_3390455, EPI_ISL_3390456, EPI_ISL_3390457, EPI_ISL_3390458, EPI_ISL_3390459, EPI_ISL_3390460, EPI_ISL_3390461, EPI_ISL_3390462, EPI_ISL_3390463, EPI_ISL_3390464, EPI_ISL_3390465, EPI_ISL_3390466, EPI_ISL_3390467, EPI_ISL_3390468, EPI_ISL_3390469, EPI_ISL_3390470, EPI_ISL_3390471, EPI_ISL_3390472, EPI_ISL_3390473, EPI_ISL_3390474, EPI_ISL_3390475, EPI_ISL_3390476, EPI_ISL_3390477, EPI_ISL_3390478, EPI_ISL_3390479, EPI_ISL_3390480, EPI_ISL_3390481, EPI_ISL_3390482, EPI_ISL_3390483, EPI_ISL_3390484, EPI_ISL_3390485, EPI_ISL_3390486, EPI_ISL_3390487, EPI_ISL_3390488, EPI_ISL_3390489, EPI_ISL_3390490, EPI_ISL_3390491, EPI_ISL_3390492, EPI_ISL_3390493, EPI_ISL_3390494, EPI_ISL_3390495, EPI_ISL_3390496, EPI_ISL_3390497, EPI_ISL_3390498, EPI_ISL_3390499, EPI_ISL_3390500, EPI_ISL_3390501, EPI_ISL_3390502, EPI_ISL_3390503, EPI_ISL_3390504, EPI_ISL_3390505, EPI_ISL_3390506, EPI_ISL_3390507, EPI_ISL_3390508, EPI_ISL_3390509, EPI_ISL_3390510, EPI_ISL_3390511, EPI_ISL_3390512, EPI_ISL_3390513, EPI_ISL_3390514, EPI_ISL_3390515, EPI_ISL_3390516, EPI_ISL_3390517, EPI_ISL_3390518, EPI_ISL_3390519, EPI_ISL_3390520, EPI_ISL_3390521, EPI_ISL_3390522, EPI_ISL_3390523, EPI_ISL_3390524, EPI_ISL_3390525, EPI_ISL_3390526, EPI_ISL_3390527, EPI_ISL_3390528, EPI_ISL_3390529, EPI_ISL_3390530, EPI_ISL_3390531, EPI_ISL_3390532, EPI_ISL_3390533, EPI_ISL_3390534, EPI_ISL_3390535, EPI_ISL_3390536, EPI_ISL_3390537, EPI_ISL_3390538, EPI_ISL_3390539, EPI_ISL_3390540, EPI_ISL_3390541, EPI_ISL_3390542, EPI_ISL_3390543, EPI_ISL_3390544, EPI_ISL_3390545, EPI_ISL_3390546, EPI_ISL_3390547, EPI_ISL_3390548, EPI_ISL_3390549, EPI_ISL_3390550, EPI_ISL_3390551, EPI_ISL_3390552, EPI_ISL_3390553, EPI_ISL_3390554, EPI_ISL_3390555, EPI_ISL_3390556, EPI_ISL_3390557, EPI_ISL_3390558, EPI_ISL_3390559, EPI_ISL_3390560, EPI_ISL_3390561, EPI_ISL_3390562, EPI_ISL_3390563, EPI_ISL_3390564, EPI_ISL_3390565, EPI_ISL_3390566, EPI_ISL_3390567, EPI_ISL_3390568, EPI_ISL_3390569, EPI_ISL_3390570, EPI_ISL_3390571, EPI_ISL_3390572, EPI_ISL_3390573, EPI_ISL_3390574, EPI_ISL_3390575, EPI_ISL_3390576, EPI_ISL_3390577, EPI_ISL_3390578, EPI_ISL_3390579, EPI_ISL_3390580, EPI_ISL_3390581, EPI_ISL_3390582, EPI_ISL_3390583, EPI_ISL_3390584, EPI_ISL_3390585, EPI_ISL_3390586, EPI_ISL_3390587, EPI_ISL_3390588, EPI_ISL_3390589, EPI_ISL_3390590, EPI_ISL_3390591, EPI_ISL_3390592, EPI_ISL_3390593, EPI_ISL_3390594, EPI_ISL_3390595, EPI_ISL_3390596, EPI_ISL_3390597, EPI_ISL_3390598, EPI_ISL_3390599, EPI_ISL_3390600, EPI_ISL_3390601, EPI_ISL_3390602, EPI_ISL_3390603, EPI_ISL_3390604, EPI_ISL_3390605, EPI_ISL_3390606, EPI_ISL_3390607, EPI_ISL_3390608, EPI_ISL_3390609, EPI_ISL_3390610, EPI_ISL_3390611, EPI_ISL_3390612, EPI_ISL_3390613, EPI_ISL_3390614, EPI_ISL_3390615, EPI_ISL_3390616, EPI_ISL_3390617, EPI_ISL_3390618, EPI_ISL_3390619, EPI_ISL_3390620, EPI_ISL_3390621, EPI_ISL_3390622, EPI_ISL_3390623, EPI_ISL_3390624, EPI_ISL_3390625, EPI_ISL_3390626, EPI_ISL_3390627, EPI_ISL_3390628, EPI_ISL_3390629, EPI_ISL_3390630, EPI_ISL_3390631, EPI_ISL_3390632, EPI_ISL_3390633, EPI_ISL_3390634, EPI_ISL_3390635, EPI_ISL_3390636, EPI_ISL_3390637, EPI_ISL_3390638, EPI_ISL_3390639, EPI_ISL_3390640, EPI_ISL_3390641, EPI_ISL_3390642, EPI_ISL_3390643, EPI_ISL_3390644, EPI_ISL_3390645, EPI_ISL_3390646, EPI_ISL_3390647, EPI_ISL_3390648, EPI_ISL_3390649, EPI_ISL_3390650, EPI_ISL_3390651, EPI_ISL_3390652, EPI_ISL_3390653, EPI_ISL_3390654, EPI_ISL_3390655, EPI_ISL_3390656, EPI_ISL_3390657, EPI_ISL_3390658, EPI_ISL_3390659, EPI_ISL_3390660, EPI_ISL_3390661, EPI_ISL_3390662, EPI_ISL_3390663, EPI_ISL_3390664, EPI_ISL_3390665, EPI_ISL_3390666, EPI_ISL_3390667, EPI_ISL_3390668, EPI_ISL_3390669, EPI_ISL_3390670, EPI_ISL_3390671, EPI_ISL_3390672, EPI_ISL_3390673, EPI_ISL_3390674, EPI_ISL_3390675, EPI_ISL_3390676, EPI_ISL_3390677, EPI_ISL_3390678, EPI_ISL_3390679, EPI_ISL_3390680, EPI_ISL_3390681, EPI_ISL_3390682, EPI_ISL_3390683, EPI_ISL_3390684, EPI_ISL_3390685, EPI_ISL_3390686, EPI_ISL_3390687, EPI_ISL_3390688, EPI_ISL_3390689, EPI_ISL_3390690, EPI_ISL_3390691, EPI_ISL_3390692, EPI_ISL_3390693, EPI_ISL_3390694, EPI_ISL_3390695, EPI_ISL_3390696, EPI_ISL_3390697, EPI_ISL_3390698, EPI_ISL_3390699, EPI_ISL_3390700, EPI_ISL_3390701, EPI_ISL_3390702, EPI_ISL_3390703, EPI_ISL_3390704, EPI_ISL_3390705, EPI_ISL_3390706, EPI_ISL_3390707, EPI_ISL_3390708, EPI_ISL_3390709, EPI_ISL_3390710, EPI_ISL_3390711, EPI_ISL_3390712, EPI_ISL_3390713, EPI_ISL_3390714, EPI_ISL_3390715, EPI_ISL_3390716, EPI_ISL_3390717, EPI_ISL_3390718, EPI_ISL_3390719 | | | |
| see above | Dutch COVID-19 response team | National Institute for Public Health and the Environment (RIVM) | Adam Meijer, Harry Vennema, Dirk Eggink, Jeroen Cremer, Sharon van den Brink, Bas van der Veer, AnneMarie van den Brandt, Lisa Wijsman, Kim Freriks, Ryanne Jaarsma, Euníce Then, Lynn Aarts, Sanne Bos, Stijn van Rossum, Florian Zwagemaker, Dennis Schmitz, Annelies Kroneman, Karim Hajji, Chantal Reusken, on behalf of the national COVID-19 response team |
| EPI_ISL_3390724, EPI_ISL_3390725, EPI_ISL_3390726, EPI_ISL_3390727, EPI_ISL_3390728, EPI_ISL_3390729, EPI_ISL_3390730, EPI_ISL_3390731, EPI_ISL_3390732, EPI_ISL_3390733 | Stanford Health Care | Stanford University School of Medicine, Clinical Virology Laboratory | Bernadette Troung, Malaya K. Sahoo, Becky Jiang, Nathan Hammond, James Zehnder, and Benjamin A. Pinsky |
| EPI_ISL_3390734, EPI_ISL_3390735, EPI_ISL_3390736, EPI_ISL_3390737, EPI_ISL_3390738, EPI_ISL_3390739, EPI_ISL_3390740, EPI_ISL_3390741, EPI_ISL_3390742, | Stanford Health Care | Stanford University School of Medicine, Clinical Virology Laboratory | Becky Jiang, Malaya K. Sahoo, Bernadette Troung, Nathan Hammond, James Zehnder, and Benjamin A. Pinsky |

| | | | |
|---|---|---|---|
| EPI_ISL_3390743 | | | |
| EPI_ISL_3390744, EPI_ISL_3390745, EPI_ISL_3390746, EPI_ISL_3390747, EPI_ISL_3390748, EPI_ISL_3390749, EPI_ISL_3390750, EPI_ISL_3390751, EPI_ISL_3390752 | Stanford Health Care | Stanford University School of Medicine, Clinical Virology Laboratory | Malaya K. Sahoo, Becky Jiang, Bernadette Troung, Nathan Hammond, James Zehnder, and Benjamin A. Pinsky |
| EPI_ISL_3390753 | Stanford Health Care | Stanford University School of Medicine, Clinical Virology Laboratory | Bernadette Troung, Malaya K. Sahoo, Becky Jiang, Nathan Hammond, James Zehnder, and Benjamin A. Pinsky |
| EPI_ISL_3390754, EPI_ISL_3390755 | Stanford Health Care | Stanford University School of Medicine, Clinical Virology Laboratory | Becky Jiang, Malaya K. Sahoo, Bernadette Troung, Nathan Hammond, James Zehnder, and Benjamin A. Pinsky |
| EPI_ISL_3390756 | Stanford Health Care | Stanford University School of Medicine, Clinical Virology Laboratory | Bernadette Troung, Malaya K. Sahoo, Becky Jiang, Nathan Hammond, James Zehnder, and Benjamin A. Pinsky |
| EPI_ISL_3390758 | Stanford Health Care | Stanford University School of Medicine, Clinical Virology Laboratory | Malaya K. Sahoo, Becky Jiang, Bernadette Troung, Nathan Hammond, James Zehnder, and Benjamin A. Pinsky |
| EPI_ISL_3390759, EPI_ISL_3390760, EPI_ISL_3390761, EPI_ISL_3390762, EPI_ISL_3390763, EPI_ISL_3390764, EPI_ISL_3390765, EPI_ISL_3390766, EPI_ISL_3390767, EPI_ISL_3390768, EPI_ISL_3390769, EPI_ISL_3390770, EPI_ISL_3390771, EPI_ISL_3390772, EPI_ISL_3390773, EPI_ISL_3390774, EPI_ISL_3390775, EPI_ISL_3390776, EPI_ISL_3390777, EPI_ISL_3390778, EPI_ISL_3390779, EPI_ISL_3390780, EPI_ISL_3390781, EPI_ISL_3390782, EPI_ISL_3390783, EPI_ISL_3390784, EPI_ISL_3390785, EPI_ISL_3390786 | | | |
| see above | State Hygienic Laboratory at the University of Iowa | State Hygienic Laboratory at the University of Iowa | Valerie Reeb, Erik Twait, Anna Yakos, Mariah Knutson, Kris Eveland, Kristen Zanon, Jeff Benfer, Wes Hottel, Alankar Kampoowale |
| EPI_ISL_3390908 | Sonic Reference Laboratory | Sonic Reference Laboratory | Robert Moller, Bijaya Dhakal, Krishna Maddula, John Woody, Cora Lahey, Ellen Pizarek |
| EPI_ISL_3390909, EPI_ISL_3390910, EPI_ISL_3390911, EPI_ISL_3390912, EPI_ISL_3390913, EPI_ISL_3390914, EPI_ISL_3390916, EPI_ISL_3390917, EPI_ISL_3390918, EPI_ISL_3390919, EPI_ISL_3390920, EPI_ISL_3390921, EPI_ISL_3390922, EPI_ISL_3390923, EPI_ISL_3390924, EPI_ISL_3390925, EPI_ISL_3390926, EPI_ISL_3390927, EPI_ISL_3390928, EPI_ISL_3390929, EPI_ISL_3390930, EPI_ISL_3390931, EPI_ISL_3390932, EPI_ISL_3390933, EPI_ISL_3390934, EPI_ISL_3390935, EPI_ISL_3390936, EPI_ISL_3390937, EPI_ISL_3390938, EPI_ISL_3390939, EPI_ISL_3390940, EPI_ISL_3390941, EPI_ISL_3390942, EPI_ISL_3390943, EPI_ISL_3390944, EPI_ISL_3390945 | | | |
| see above | Institute for Infectious Diseases | Institute for Infectious Diseases | Stefan Neuenschwander, Christian Baumann, Miguel A Terrazos Miani, Cora SÃ¤gesser, Pascal Bittel, Peter Keller, Franziska Suter-Riniker, Stephen L Leib, Alban Ramette |
| EPI_ISL_3390946, EPI_ISL_3390947, EPI_ISL_3390948 | Biopole Antilles | Department of Virology, Henri Mondor University Hospital, Assistance Publique Hôpitaux de Paris, Université Paris-Est Créteil, INSERM U955 | Christophe Rodriguez, Slim Fourati, Vanessa Demontant, Guillaume Gricourt, Melissa N'Debi, Alexandre Soulier, Elisabeth Trawinski, Jean-Michel Pawlotsky |
| EPI_ISL_3390949, EPI_ISL_3390950 | LISIEUX Cerballiance | Department of Virology, Henri Mondor University Hospital, Assistance Publique Hôpitaux de Paris, Université Paris-Est Créteil, INSERM U955 | Christophe Rodriguez, Slim Fourati, Vanessa Demontant, Guillaume Gricourt, Melissa N'Debi, Alexandre Soulier, Elisabeth Trawinski, Jean-Michel Pawlotsky |
| EPI_ISL_3390951 | CERBALLIANCE SAINT MARTIN | Department of Virology, Henri Mondor University Hospital, Assistance Publique Hôpitaux de Paris, Université Paris-Est Créteil, INSERM U955 | Christophe Rodriguez, Slim Fourati, Vanessa Demontant, Guillaume Gricourt, Melissa N'Debi, Alexandre Soulier, Elisabeth Trawinski, Jean-Michel Pawlotsky |
| EPI_ISL_3390953 | CH Mémorial F.E.U Laboratoire | Department of Virology, Henri Mondor University Hospital, Assistance Publique Hôpitaux de Paris, Université Paris-Est Créteil, INSERM U955 | Christophe Rodriguez, Slim Fourati, Vanessa Demontant, Guillaume Gricourt, Melissa N'Debi, Alexandre Soulier, Elisabeth Trawinski, Jean-Michel Pawlotsky |
| EPI_ISL_3390954, EPI_ISL_3390955, EPI_ISL_3390956 | LISIEUX Cerballiance | Department of Virology, Henri Mondor University Hospital, Assistance Publique Hôpitaux de Paris, Université Paris-Est Créteil, INSERM U955 | Christophe Rodriguez, Slim Fourati, Vanessa Demontant, Guillaume Gricourt, Melissa N'Debi, Alexandre Soulier, Elisabeth Trawinski, Jean-Michel Pawlotsky |
| EPI_ISL_3390957 | CERBALLIANCE SAINT MARTIN | Department of Virology, Henri Mondor University Hospital, Assistance Publique Hôpitaux de Paris, Université Paris-Est Créteil, INSERM U955 | Christophe Rodriguez, Slim Fourati, Vanessa Demontant, Guillaume Gricourt, Melissa N'Debi, Alexandre Soulier, Elisabeth Trawinski, Jean-Michel Pawlotsky |
| EPI_ISL_3390958 | LISIEUX Cerballiance | Department of Virology, Henri Mondor University Hospital, Assistance Publique Hôpitaux de Paris, Université Paris-Est Créteil, INSERM U955 | Christophe Rodriguez, Slim Fourati, Vanessa Demontant, Guillaume Gricourt, Melissa N'Debi, Alexandre Soulier, Elisabeth Trawinski, Jean-Michel Pawlotsky |
| EPI_ISL_3390959 | CH. ANDRE GREGOIRE | Department of Virology, Henri Mondor University Hospital, Assistance Publique Hôpitaux de Paris, Université Paris-Est Créteil, INSERM U955 | Christophe Rodriguez, Slim Fourati, Vanessa Demontant, Guillaume Gricourt, Melissa N'Debi, Alexandre Soulier, Elisabeth Trawinski, Jean-Michel Pawlotsky |
| EPI_ISL_3390960, EPI_ISL_3390961 | Biogroup Bio Lam-LCD Saint-Denis | Department of Virology, Henri Mondor University Hospital, Assistance Publique Hôpitaux de Paris, Université Paris-Est Créteil, INSERM U955 | Christophe Rodriguez, Slim Fourati, Vanessa Demontant, Guillaume Gricourt, Melissa N'Debi, Alexandre Soulier, Elisabeth Trawinski, Jean-Michel Pawlotsky |
| EPI_ISL_3390962 | GH de l'Est Francilien | Department of Virology, Henri Mondor University Hospital, Assistance Publique Hôpitaux de Paris, Université Paris-Est Créteil, INSERM U955 | Christophe Rodriguez, Slim Fourati, Vanessa Demontant, Guillaume Gricourt, Melissa N'Debi, Alexandre Soulier, Elisabeth Trawinski, Jean-Michel Pawlotsky |
| EPI_ISL_3390965 | CH. ANDRE GREGOIRE | Department of Virology, Henri Mondor University Hospital, Assistance Publique Hôpitaux de Paris, Université Paris-Est Créteil, INSERM U955 | Christophe Rodriguez, Slim Fourati, Vanessa Demontant, Guillaume Gricourt, Melissa N'Debi, Alexandre Soulier, Elisabeth Trawinski, Jean-Michel Pawlotsky |
| EPI_ISL_3390967, EPI_ISL_3390969 | Biogroup Bio Lam-LCD Saint-Denis | Department of Virology, Henri Mondor University Hospital, Assistance Publique Hôpitaux de Paris, Université Paris-Est Créteil, INSERM U955 | Christophe Rodriguez, Slim Fourati, Vanessa Demontant, Guillaume Gricourt, Melissa N'Debi, Alexandre Soulier, Elisabeth Trawinski, Jean-Michel Pawlotsky |
| EPI_ISL_3390970 | CH. ANDRE GREGOIRE | Department of Virology, Henri Mondor University Hospital, Assistance Publique Hôpitaux de Paris, Université Paris-Est Créteil, INSERM U955 | Christophe Rodriguez, Slim Fourati, Vanessa Demontant, Guillaume Gricourt, Melissa N'Debi, Alexandre Soulier, Elisabeth Trawinski, Jean-Michel Pawlotsky |
| EPI_ISL_3390971, EPI_ISL_3390972, EPI_ISL_3390973, EPI_ISL_3390974, EPI_ISL_3390975 | Biogroup Bio Lam-LCD Saint-Denis | Department of Virology, Henri Mondor University Hospital, Assistance Publique Hôpitaux de Paris, Université Paris-Est Créteil, INSERM U955 | Christophe Rodriguez, Slim Fourati, Vanessa Demontant, Guillaume Gricourt, Melissa N'Debi, Alexandre Soulier, Elisabeth Trawinski, Jean-Michel Pawlotsky |
| EPI_ISL_3390976 | CH. ANDRE GREGOIRE | Department of Virology, Henri Mondor University Hospital, Assistance Publique Hôpitaux de Paris, Université Paris-Est Créteil, INSERM U955 | Christophe Rodriguez, Slim Fourati, Vanessa Demontant, Guillaume Gricourt, Melissa N'Debi, Alexandre Soulier, Elisabeth Trawinski, Jean-Michel Pawlotsky |
| EPI_ISL_3390977, EPI_ISL_3390978, EPI_ISL_3390979 | Biogroup Bio Lam-LCD Saint-Denis | Department of Virology, Henri Mondor University Hospital, Assistance Publique Hôpitaux de Paris, Université Paris-Est Créteil, INSERM U955 | Christophe Rodriguez, Slim Fourati, Vanessa Demontant, Guillaume Gricourt, Melissa N'Debi, Alexandre Soulier, Elisabeth Trawinski, Jean-Michel Pawlotsky |
| EPI_ISL_3390980 | GH de l'Est Francilien | Department of Virology, Henri Mondor University Hospital, Assistance Publique Hôpitaux de Paris, Université Paris-Est Créteil, INSERM U955 | Christophe Rodriguez, Slim Fourati, Vanessa Demontant, Guillaume Gricourt, Melissa N'Debi, Alexandre Soulier, Elisabeth Trawinski, Jean-Michel Pawlotsky |
| EPI_ISL_3390981, EPI_ISL_3390982, EPI_ISL_3390983, EPI_ISL_3390984, EPI_ISL_3390985, EPI_ISL_3390987 | Biogroup Bio Lam-LCD Saint-Denis | Department of Virology, Henri Mondor University Hospital, Assistance Publique Hôpitaux de Paris, Université Paris-Est Créteil, INSERM U955 | Christophe Rodriguez, Slim Fourati, Vanessa Demontant, Guillaume Gricourt, Melissa N'Debi, Alexandre Soulier, Elisabeth Trawinski, Jean-Michel Pawlotsky |
| EPI_ISL_3390988 | CH. ANDRE GREGOIRE | Department of Virology, Henri Mondor University Hospital, Assistance Publique Hôpitaux de Paris, Université Paris-Est Créteil, INSERM U955 | Christophe Rodriguez, Slim Fourati, Vanessa Demontant, Guillaume Gricourt, Melissa N'Debi, Alexandre Soulier, Elisabeth Trawinski, Jean-Michel Pawlotsky |
| EPI_ISL_3390989 | Biogroup Bio Lam-LCD Saint-Denis | Department of Virology, Henri Mondor University Hospital, Assistance Publique Hôpitaux de Paris, Université Paris-Est Créteil, INSERM U955 | Christophe Rodriguez, Slim Fourati, Vanessa Demontant, Guillaume Gricourt, Melissa N'Debi, Alexandre Soulier, Elisabeth Trawinski, Jean-Michel Pawlotsky |
| EPI_ISL_3390990 | CH. BEGIN | Department of Virology, Henri Mondor University Hospital, Assistance Publique Hôpitaux de Paris, Université Paris-Est Créteil, INSERM U955 | Christophe Rodriguez, Slim Fourati, Vanessa Demontant, Guillaume Gricourt, Melissa N'Debi, Alexandre Soulier, Elisabeth Trawinski, Jean-Michel Pawlotsky |
| EPI_ISL_3390991, EPI_ISL_3390992, EPI_ISL_3390993, EPI_ISL_3390994, EPI_ISL_3390995, EPI_ISL_3390996, EPI_ISL_3390998, EPI_ISL_3390999, EPI_ISL_3391000, EPI_ISL_3391001, EPI_ISL_3391002, EPI_ISL_3391003, EPI_ISL_3391004, EPI_ISL_3391006, EPI_ISL_3391007, EPI_ISL_3391008, EPI_ISL_3391010, EPI_ISL_3391011 | | | |
| see above | Biogroup Bio Lam-LCD Saint-Denis | Department of Virology, Henri Mondor University Hospital, Assistance Publique Hôpitaux de Paris, Université Paris-Est Créteil, INSERM U955 | Christophe Rodriguez, Slim Fourati, Vanessa Demontant, Guillaume Gricourt, Melissa N'Debi, Alexandre Soulier, Elisabeth Trawinski, Jean-Michel Pawlotsky |
| EPI_ISL_3391012, | CH. ANDRE GREGOIRE | Department of Virology, Henri Mondor University Hospital, | Christophe Rodriguez, Slim Fourati, Vanessa Demontant, Guillaume Gricourt, Melissa N'Debi, Alexandre Soulier, Elisabeth Trawinski, Jean-Michel Pawlotsky |

| | | | |
|---|---|---|---|
| EPI_ISL_3391013 | | Assistance Publique Hôpitaux de Paris, Université Paris-Est Créteil, INSERM U955 | |
| EPI_ISL_3391014, EPI_ISL_3391015, EPI_ISL_3391016, EPI_ISL_3391017, EPI_ISL_3391018, EPI_ISL_3391019, EPI_ISL_3391020, EPI_ISL_3391021, EPI_ISL_3391022, EPI_ISL_3391023, EPI_ISL_3391024, EPI_ISL_3391025, EPI_ISL_3391026, EPI_ISL_3391027, EPI_ISL_3391028, EPI_ISL_3391029, EPI_ISL_3391030, EPI_ISL_3391031, EPI_ISL_3391034, EPI_ISL_3391035, EPI_ISL_3391036, EPI_ISL_3391037, EPI_ISL_3391040 | | | |
| see above | Biogroup Bio Lam-LCD Saint-Denis | Department of Virology, Henri Mondor University Hospital, Assistance Publique Hôpitaux de Paris, Université Paris-Est Créteil, INSERM U955 | Christophe Rodriguez, Slim Fourati, Vanessa Demontant, Guillaume Gricourt, Melissa N'Debi, Alexandre Soulier, Elisabeth Trawinski, Jean-Michel Pawlotsky |
| EPI_ISL_3391041 | GH de l'Est Francilien | Department of Virology, Henri Mondor University Hospital, Assistance Publique Hôpitaux de Paris, Université Paris-Est Créteil, INSERM U955 | Christophe Rodriguez, Slim Fourati, Vanessa Demontant, Guillaume Gricourt, Melissa N'Debi, Alexandre Soulier, Elisabeth Trawinski, Jean-Michel Pawlotsky |
| EPI_ISL_3391042 | Biogroup Bio Lam-LCD Saint-Denis | Department of Virology, Henri Mondor University Hospital, Assistance Publique Hôpitaux de Paris, Université Paris-Est Créteil, INSERM U955 | Christophe Rodriguez, Slim Fourati, Vanessa Demontant, Guillaume Gricourt, Melissa N'Debi, Alexandre Soulier, Elisabeth Trawinski, Jean-Michel Pawlotsky |
| EPI_ISL_3391043 | GH de l'Est Francilien | Department of Virology, Henri Mondor University Hospital, Assistance Publique Hôpitaux de Paris, Université Paris-Est Créteil, INSERM U955 | Christophe Rodriguez, Slim Fourati, Vanessa Demontant, Guillaume Gricourt, Melissa N'Debi, Alexandre Soulier, Elisabeth Trawinski, Jean-Michel Pawlotsky |
| EPI_ISL_3391044, EPI_ISL_3391048, EPI_ISL_3391049, EPI_ISL_3391050, EPI_ISL_3391051, EPI_ISL_3391054, EPI_ISL_3391056 | Biogroup Bio Lam-LCD Saint-Denis | Department of Virology, Henri Mondor University Hospital, Assistance Publique Hôpitaux de Paris, Université Paris-Est Créteil, INSERM U955 | Christophe Rodriguez, Slim Fourati, Vanessa Demontant, Guillaume Gricourt, Melissa N'Debi, Alexandre Soulier, Elisabeth Trawinski, Jean-Michel Pawlotsky |
| EPI_ISL_3391057 | GH de l'Est Francilien | Department of Virology, Henri Mondor University Hospital, Assistance Publique Hôpitaux de Paris, Université Paris-Est Créteil, INSERM U955 | Christophe Rodriguez, Slim Fourati, Vanessa Demontant, Guillaume Gricourt, Melissa N'Debi, Alexandre Soulier, Elisabeth Trawinski, Jean-Michel Pawlotsky |
| EPI_ISL_3391058, EPI_ISL_3391059, EPI_ISL_3391061, EPI_ISL_3391063 | Biogroup Bio Lam-LCD Saint-Denis | Department of Virology, Henri Mondor University Hospital, Assistance Publique Hôpitaux de Paris, Université Paris-Est Créteil, INSERM U955 | Christophe Rodriguez, Slim Fourati, Vanessa Demontant, Guillaume Gricourt, Melissa N'Debi, Alexandre Soulier, Elisabeth Trawinski, Jean-Michel Pawlotsky |
| EPI_ISL_3391064 | CH. ANDRE GREGOIRE | Department of Virology, Henri Mondor University Hospital, Assistance Publique Hôpitaux de Paris, Université Paris-Est Créteil, INSERM U955 | Christophe Rodriguez, Slim Fourati, Vanessa Demontant, Guillaume Gricourt, Melissa N'Debi, Alexandre Soulier, Elisabeth Trawinski, Jean-Michel Pawlotsky |
| EPI_ISL_3391065, EPI_ISL_3391066, EPI_ISL_3391067 | Biogroup Bio Lam-LCD Saint-Denis | Department of Virology, Henri Mondor University Hospital, Assistance Publique Hôpitaux de Paris, Université Paris-Est Créteil, INSERM U955 | Christophe Rodriguez, Slim Fourati, Vanessa Demontant, Guillaume Gricourt, Melissa N'Debi, Alexandre Soulier, Elisabeth Trawinski, Jean-Michel Pawlotsky |
| EPI_ISL_3391068 | GH de l'Est Francilien | Department of Virology, Henri Mondor University Hospital, Assistance Publique Hôpitaux de Paris, Université Paris-Est Créteil, INSERM U955 | Christophe Rodriguez, Slim Fourati, Vanessa Demontant, Guillaume Gricourt, Melissa N'Debi, Alexandre Soulier, Elisabeth Trawinski, Jean-Michel Pawlotsky |
| EPI_ISL_3391069 | Biogroup Bio Lam-LCD Saint-Denis | Department of Virology, Henri Mondor University Hospital, Assistance Publique Hôpitaux de Paris, Université Paris-Est Créteil, INSERM U955 | Christophe Rodriguez, Slim Fourati, Vanessa Demontant, Guillaume Gricourt, Melissa N'Debi, Alexandre Soulier, Elisabeth Trawinski, Jean-Michel Pawlotsky |
| EPI_ISL_3391070 | BIO ARD'AISNE | Department of Virology, Henri Mondor University Hospital, Assistance Publique Hôpitaux de Paris, Université Paris-Est Créteil, INSERM U955 | Christophe Rodriguez, Slim Fourati, Vanessa Demontant, Guillaume Gricourt, Melissa N'Debi, Alexandre Soulier, Elisabeth Trawinski, Jean-Michel Pawlotsky |
| EPI_ISL_3391072, EPI_ISL_3391075 | BIO67-BIOSPHERE | Department of Virology, Henri Mondor University Hospital, Assistance Publique Hôpitaux de Paris, Université Paris-Est Créteil, INSERM U955 | Christophe Rodriguez, Slim Fourati, Vanessa Demontant, Guillaume Gricourt, Melissa N'Debi, Alexandre Soulier, Elisabeth Trawinski, Jean-Michel Pawlotsky |
| EPI_ISL_3391076 | Plateau technique Biomer Metz Verriers | Department of Virology, Henri Mondor University Hospital, Assistance Publique Hôpitaux de Paris, Université Paris-Est Créteil, INSERM U955 | Christophe Rodriguez, Slim Fourati, Vanessa Demontant, Guillaume Gricourt, Melissa N'Debi, Alexandre Soulier, Elisabeth Trawinski, Jean-Michel Pawlotsky |
| EPI_ISL_3391078, EPI_ISL_3391079 | BIO67-BIOSPHERE | Department of Virology, Henri Mondor University Hospital, Assistance Publique Hôpitaux de Paris, Université Paris-Est Créteil, INSERM U955 | Christophe Rodriguez, Slim Fourati, Vanessa Demontant, Guillaume Gricourt, Melissa N'Debi, Alexandre Soulier, Elisabeth Trawinski, Jean-Michel Pawlotsky |
| EPI_ISL_3391080 | Laboratoire LENYS | Department of Virology, Henri Mondor University Hospital, Assistance Publique Hôpitaux de Paris, Université Paris-Est Créteil, INSERM U955 | Christophe Rodriguez, Slim Fourati, Vanessa Demontant, Guillaume Gricourt, Melissa N'Debi, Alexandre Soulier, Elisabeth Trawinski, Jean-Michel Pawlotsky |
| EPI_ISL_3391081, EPI_ISL_3391083 | BIO67-BIOSPHERE | Department of Virology, Henri Mondor University Hospital, Assistance Publique Hôpitaux de Paris, Université Paris-Est Créteil, INSERM U955 | Christophe Rodriguez, Slim Fourati, Vanessa Demontant, Guillaume Gricourt, Melissa N'Debi, Alexandre Soulier, Elisabeth Trawinski, Jean-Michel Pawlotsky |
| EPI_ISL_3391084 | Laboratoire LENYS | Department of Virology, Henri Mondor University Hospital, Assistance Publique Hôpitaux de Paris, Université Paris-Est Créteil, INSERM U955 | Christophe Rodriguez, Slim Fourati, Vanessa Demontant, Guillaume Gricourt, Melissa N'Debi, Alexandre Soulier, Elisabeth Trawinski, Jean-Michel Pawlotsky |
| EPI_ISL_3391085, EPI_ISL_3391086 | Plateau technique Biomer Metz Verriers | Department of Virology, Henri Mondor University Hospital, Assistance Publique Hôpitaux de Paris, Université Paris-Est Créteil, INSERM U955 | Christophe Rodriguez, Slim Fourati, Vanessa Demontant, Guillaume Gricourt, Melissa N'Debi, Alexandre Soulier, Elisabeth Trawinski, Jean-Michel Pawlotsky |
| EPI_ISL_3391087 | BIO67-BIOSPHERE | Department of Virology, Henri Mondor University Hospital, Assistance Publique Hôpitaux de Paris, Université Paris-Est Créteil, INSERM U955 | Christophe Rodriguez, Slim Fourati, Vanessa Demontant, Guillaume Gricourt, Melissa N'Debi, Alexandre Soulier, Elisabeth Trawinski, Jean-Michel Pawlotsky |
| EPI_ISL_3391088 | Laboratoire LENYS | Department of Virology, Henri Mondor University Hospital, Assistance Publique Hôpitaux de Paris, Université Paris-Est Créteil, INSERM U955 | Christophe Rodriguez, Slim Fourati, Vanessa Demontant, Guillaume Gricourt, Melissa N'Debi, Alexandre Soulier, Elisabeth Trawinski, Jean-Michel Pawlotsky |
| EPI_ISL_3391089, EPI_ISL_3391090 | BIO67-BIOSPHERE | Department of Virology, Henri Mondor University Hospital, Assistance Publique Hôpitaux de Paris, Université Paris-Est Créteil, INSERM U955 | Christophe Rodriguez, Slim Fourati, Vanessa Demontant, Guillaume Gricourt, Melissa N'Debi, Alexandre Soulier, Elisabeth Trawinski, Jean-Michel Pawlotsky |
| EPI_ISL_3391091 | Plateau technique Biomer Metz Verriers | Department of Virology, Henri Mondor University Hospital, Assistance Publique Hôpitaux de Paris, Université Paris-Est Créteil, INSERM U955 | Christophe Rodriguez, Slim Fourati, Vanessa Demontant, Guillaume Gricourt, Melissa N'Debi, Alexandre Soulier, Elisabeth Trawinski, Jean-Michel Pawlotsky |
| EPI_ISL_3391092 | Laboratoire LENYS | Department of Virology, Henri Mondor University Hospital, Assistance Publique Hôpitaux de Paris, Université Paris-Est Créteil, INSERM U955 | Christophe Rodriguez, Slim Fourati, Vanessa Demontant, Guillaume Gricourt, Melissa N'Debi, Alexandre Soulier, Elisabeth Trawinski, Jean-Michel Pawlotsky |
| EPI_ISL_3391093 | BIO67-BIOSPHERE | Department of Virology, Henri Mondor University Hospital, Assistance Publique Hôpitaux de Paris, Université Paris-Est Créteil, INSERM U955 | Christophe Rodriguez, Slim Fourati, Vanessa Demontant, Guillaume Gricourt, Melissa N'Debi, Alexandre Soulier, Elisabeth Trawinski, Jean-Michel Pawlotsky |
| EPI_ISL_3391094 | Plateau technique Biomer Metz Verriers | Department of Virology, Henri Mondor University Hospital, Assistance Publique Hôpitaux de Paris, Université Paris-Est Créteil, INSERM U955 | Christophe Rodriguez, Slim Fourati, Vanessa Demontant, Guillaume Gricourt, Melissa N'Debi, Alexandre Soulier, Elisabeth Trawinski, Jean-Michel Pawlotsky |
| EPI_ISL_3391096, EPI_ISL_3391097 | BIO67-BIOSPHERE | Department of Virology, Henri Mondor University Hospital, Assistance Publique Hôpitaux de Paris, Université Paris-Est Créteil, INSERM U955 | Christophe Rodriguez, Slim Fourati, Vanessa Demontant, Guillaume Gricourt, Melissa N'Debi, Alexandre Soulier, Elisabeth Trawinski, Jean-Michel Pawlotsky |
| EPI_ISL_3391098 | Laboratoire LENYS | Department of Virology, Henri Mondor University Hospital, Assistance Publique Hôpitaux de Paris, Université Paris-Est Créteil, INSERM U955 | Christophe Rodriguez, Slim Fourati, Vanessa Demontant, Guillaume Gricourt, Melissa N'Debi, Alexandre Soulier, Elisabeth Trawinski, Jean-Michel Pawlotsky |
| EPI_ISL_3391099 | Plateau technique Biomer Metz Verriers | Department of Virology, Henri Mondor University Hospital, Assistance Publique Hôpitaux de Paris, Université Paris-Est Créteil, INSERM U955 | Christophe Rodriguez, Slim Fourati, Vanessa Demontant, Guillaume Gricourt, Melissa N'Debi, Alexandre Soulier, Elisabeth Trawinski, Jean-Michel Pawlotsky |

| | | | |
|---|---|---|---|
| EPI_ISL_3391100, EPI_ISL_3391101, EPI_ISL_3391102 | BIO67-BIOSPHERE | Department of Virology, Henri Mondor University Hospital, Assistance Publique Hôpitaux de Paris, Université Paris-Est Créteil, INSERM U955 | Christophe Rodriguez, Slim Fourati, Vanessa Demontant, Guillaume Gricourt, Melissa N'Debi, Alexandre Soulier, Elisabeth Trawinski, Jean-Michel Pawlotsky |
| EPI_ISL_3391103, EPI_ISL_3391106 | Plateau technique Biomer Metz Verriers | Department of Virology, Henri Mondor University Hospital, Assistance Publique Hôpitaux de Paris, Université Paris-Est Créteil, INSERM U955 | Christophe Rodriguez, Slim Fourati, Vanessa Demontant, Guillaume Gricourt, Melissa N'Debi, Alexandre Soulier, Elisabeth Trawinski, Jean-Michel Pawlotsky |
| EPI_ISL_3391107 | BIO67-BIOSPHERE | Department of Virology, Henri Mondor University Hospital, Assistance Publique Hôpitaux de Paris, Université Paris-Est Créteil, INSERM U955 | Christophe Rodriguez, Slim Fourati, Vanessa Demontant, Guillaume Gricourt, Melissa N'Debi, Alexandre Soulier, Elisabeth Trawinski, Jean-Michel Pawlotsky |
| EPI_ISL_3391108 | Laboratoire LENYS | Department of Virology, Henri Mondor University Hospital, Assistance Publique Hôpitaux de Paris, Université Paris-Est Créteil, INSERM U955 | Christophe Rodriguez, Slim Fourati, Vanessa Demontant, Guillaume Gricourt, Melissa N'Debi, Alexandre Soulier, Elisabeth Trawinski, Jean-Michel Pawlotsky |
| EPI_ISL_3391109 | BIO67-BIOSPHERE | Department of Virology, Henri Mondor University Hospital, Assistance Publique Hôpitaux de Paris, Université Paris-Est Créteil, INSERM U955 | Christophe Rodriguez, Slim Fourati, Vanessa Demontant, Guillaume Gricourt, Melissa N'Debi, Alexandre Soulier, Elisabeth Trawinski, Jean-Michel Pawlotsky |
| EPI_ISL_3391111, EPI_ISL_3391112 | Plateau technique Biomer Metz Verriers | Department of Virology, Henri Mondor University Hospital, Assistance Publique Hôpitaux de Paris, Université Paris-Est Créteil, INSERM U955 | Christophe Rodriguez, Slim Fourati, Vanessa Demontant, Guillaume Gricourt, Melissa N'Debi, Alexandre Soulier, Elisabeth Trawinski, Jean-Michel Pawlotsky |
| EPI_ISL_3391114, EPI_ISL_3391115, EPI_ISL_3391116, EPI_ISL_3391118 | BIO67-BIOSPHERE | Department of Virology, Henri Mondor University Hospital, Assistance Publique Hôpitaux de Paris, Université Paris-Est Créteil, INSERM U955 | Christophe Rodriguez, Slim Fourati, Vanessa Demontant, Guillaume Gricourt, Melissa N'Debi, Alexandre Soulier, Elisabeth Trawinski, Jean-Michel Pawlotsky |
| EPI_ISL_3391119 | Laboratoire multi-site du Diaconat | Department of Virology, Henri Mondor University Hospital, Assistance Publique Hôpitaux de Paris, Université Paris-Est Créteil, INSERM U955 | Christophe Rodriguez, Slim Fourati, Vanessa Demontant, Guillaume Gricourt, Melissa N'Debi, Alexandre Soulier, Elisabeth Trawinski, Jean-Michel Pawlotsky |
| EPI_ISL_3391121 | BIO67-BIOSPHERE | Department of Virology, Henri Mondor University Hospital, Assistance Publique Hôpitaux de Paris, Université Paris-Est Créteil, INSERM U955 | Christophe Rodriguez, Slim Fourati, Vanessa Demontant, Guillaume Gricourt, Melissa N'Debi, Alexandre Soulier, Elisabeth Trawinski, Jean-Michel Pawlotsky |
| EPI_ISL_3391124 | Plateau technique Biomer Metz Verriers | Department of Virology, Henri Mondor University Hospital, Assistance Publique Hôpitaux de Paris, Université Paris-Est Créteil, INSERM U955 | Christophe Rodriguez, Slim Fourati, Vanessa Demontant, Guillaume Gricourt, Melissa N'Debi, Alexandre Soulier, Elisabeth Trawinski, Jean-Michel Pawlotsky |
| EPI_ISL_3391125 | Laboratoire LENYS | Department of Virology, Henri Mondor University Hospital, Assistance Publique Hôpitaux de Paris, Université Paris-Est Créteil, INSERM U955 | Christophe Rodriguez, Slim Fourati, Vanessa Demontant, Guillaume Gricourt, Melissa N'Debi, Alexandre Soulier, Elisabeth Trawinski, Jean-Michel Pawlotsky |
| EPI_ISL_3391126, EPI_ISL_3391127 | BIO67-BIOSPHERE | Department of Virology, Henri Mondor University Hospital, Assistance Publique Hôpitaux de Paris, Université Paris-Est Créteil, INSERM U955 | Christophe Rodriguez, Slim Fourati, Vanessa Demontant, Guillaume Gricourt, Melissa N'Debi, Alexandre Soulier, Elisabeth Trawinski, Jean-Michel Pawlotsky |
| EPI_ISL_3391128 | Laboratoire LENYS | Department of Virology, Henri Mondor University Hospital, Assistance Publique Hôpitaux de Paris, Université Paris-Est Créteil, INSERM U955 | Christophe Rodriguez, Slim Fourati, Vanessa Demontant, Guillaume Gricourt, Melissa N'Debi, Alexandre Soulier, Elisabeth Trawinski, Jean-Michel Pawlotsky |
| EPI_ISL_3391129 | Plateau technique Biomer Metz Verriers | Department of Virology, Henri Mondor University Hospital, Assistance Publique Hôpitaux de Paris, Université Paris-Est Créteil, INSERM U955 | Christophe Rodriguez, Slim Fourati, Vanessa Demontant, Guillaume Gricourt, Melissa N'Debi, Alexandre Soulier, Elisabeth Trawinski, Jean-Michel Pawlotsky |
| EPI_ISL_3391131, EPI_ISL_3391132 | BIO67-BIOSPHERE | Department of Virology, Henri Mondor University Hospital, Assistance Publique Hôpitaux de Paris, Université Paris-Est Créteil, INSERM U955 | Christophe Rodriguez, Slim Fourati, Vanessa Demontant, Guillaume Gricourt, Melissa N'Debi, Alexandre Soulier, Elisabeth Trawinski, Jean-Michel Pawlotsky |
| EPI_ISL_3391133 | Plateau technique Biomer Metz Verriers | Department of Virology, Henri Mondor University Hospital, Assistance Publique Hôpitaux de Paris, Université Paris-Est Créteil, INSERM U955 | Christophe Rodriguez, Slim Fourati, Vanessa Demontant, Guillaume Gricourt, Melissa N'Debi, Alexandre Soulier, Elisabeth Trawinski, Jean-Michel Pawlotsky |
| EPI_ISL_3391135 | Laboratoire LENYS | Department of Virology, Henri Mondor University Hospital, Assistance Publique Hôpitaux de Paris, Université Paris-Est Créteil, INSERM U955 | Christophe Rodriguez, Slim Fourati, Vanessa Demontant, Guillaume Gricourt, Melissa N'Debi, Alexandre Soulier, Elisabeth Trawinski, Jean-Michel Pawlotsky |
| EPI_ISL_3391137 | BIO67-BIOSPHERE | Department of Virology, Henri Mondor University Hospital, Assistance Publique Hôpitaux de Paris, Université Paris-Est Créteil, INSERM U955 | Christophe Rodriguez, Slim Fourati, Vanessa Demontant, Guillaume Gricourt, Melissa N'Debi, Alexandre Soulier, Elisabeth Trawinski, Jean-Michel Pawlotsky |
| EPI_ISL_3391138 | Plateau technique Biomer Metz Verriers | Department of Virology, Henri Mondor University Hospital, Assistance Publique Hôpitaux de Paris, Université Paris-Est Créteil, INSERM U955 | Christophe Rodriguez, Slim Fourati, Vanessa Demontant, Guillaume Gricourt, Melissa N'Debi, Alexandre Soulier, Elisabeth Trawinski, Jean-Michel Pawlotsky |
| EPI_ISL_3391139 | BIO67-BIOSPHERE | Department of Virology, Henri Mondor University Hospital, Assistance Publique Hôpitaux de Paris, Université Paris-Est Créteil, INSERM U955 | Christophe Rodriguez, Slim Fourati, Vanessa Demontant, Guillaume Gricourt, Melissa N'Debi, Alexandre Soulier, Elisabeth Trawinski, Jean-Michel Pawlotsky |
| EPI_ISL_3391141 | Laboratoire LENYS | Department of Virology, Henri Mondor University Hospital, Assistance Publique Hôpitaux de Paris, Université Paris-Est Créteil, INSERM U955 | Christophe Rodriguez, Slim Fourati, Vanessa Demontant, Guillaume Gricourt, Melissa N'Debi, Alexandre Soulier, Elisabeth Trawinski, Jean-Michel Pawlotsky |
| EPI_ISL_3391142, EPI_ISL_3391143 | BIO67-BIOSPHERE | Department of Virology, Henri Mondor University Hospital, Assistance Publique Hôpitaux de Paris, Université Paris-Est Créteil, INSERM U955 | Christophe Rodriguez, Slim Fourati, Vanessa Demontant, Guillaume Gricourt, Melissa N'Debi, Alexandre Soulier, Elisabeth Trawinski, Jean-Michel Pawlotsky |
| EPI_ISL_3391144 | Laboratoire LENYS | Department of Virology, Henri Mondor University Hospital, Assistance Publique Hôpitaux de Paris, Université Paris-Est Créteil, INSERM U955 | Christophe Rodriguez, Slim Fourati, Vanessa Demontant, Guillaume Gricourt, Melissa N'Debi, Alexandre Soulier, Elisabeth Trawinski, Jean-Michel Pawlotsky |
| EPI_ISL_3391145, EPI_ISL_3391146 | BIO67-BIOSPHERE | Department of Virology, Henri Mondor University Hospital, Assistance Publique Hôpitaux de Paris, Université Paris-Est Créteil, INSERM U955 | Christophe Rodriguez, Slim Fourati, Vanessa Demontant, Guillaume Gricourt, Melissa N'Debi, Alexandre Soulier, Elisabeth Trawinski, Jean-Michel Pawlotsky |
| EPI_ISL_3391147, EPI_ISL_3391148 | Plateau technique Biomer Metz Verriers | Department of Virology, Henri Mondor University Hospital, Assistance Publique Hôpitaux de Paris, Université Paris-Est Créteil, INSERM U955 | Christophe Rodriguez, Slim Fourati, Vanessa Demontant, Guillaume Gricourt, Melissa N'Debi, Alexandre Soulier, Elisabeth Trawinski, Jean-Michel Pawlotsky |
| EPI_ISL_3391149, EPI_ISL_3391150, EPI_ISL_3391151 | BIO67-BIOSPHERE | Department of Virology, Henri Mondor University Hospital, Assistance Publique Hôpitaux de Paris, Université Paris-Est Créteil, INSERM U955 | Christophe Rodriguez, Slim Fourati, Vanessa Demontant, Guillaume Gricourt, Melissa N'Debi, Alexandre Soulier, Elisabeth Trawinski, Jean-Michel Pawlotsky |
| EPI_ISL_3391152, EPI_ISL_3391153, EPI_ISL_3391154 | Plateau technique Biomer Metz Verriers | Department of Virology, Henri Mondor University Hospital, Assistance Publique Hôpitaux de Paris, Université Paris-Est Créteil, INSERM U955 | Christophe Rodriguez, Slim Fourati, Vanessa Demontant, Guillaume Gricourt, Melissa N'Debi, Alexandre Soulier, Elisabeth Trawinski, Jean-Michel Pawlotsky |
| EPI_ISL_3391155 | BIO67-BIOSPHERE | Department of Virology, Henri Mondor University Hospital, Assistance Publique Hôpitaux de Paris, Université Paris-Est Créteil, INSERM U955 | Christophe Rodriguez, Slim Fourati, Vanessa Demontant, Guillaume Gricourt, Melissa N'Debi, Alexandre Soulier, Elisabeth Trawinski, Jean-Michel Pawlotsky |
| EPI_ISL_3391157, EPI_ISL_3391158 | Plateau technique Biomer Metz Verriers | Department of Virology, Henri Mondor University Hospital, Assistance Publique Hôpitaux de Paris, Université Paris-Est Créteil, INSERM U955 | Christophe Rodriguez, Slim Fourati, Vanessa Demontant, Guillaume Gricourt, Melissa N'Debi, Alexandre Soulier, Elisabeth Trawinski, Jean-Michel Pawlotsky |
| EPI_ISL_3391159 | BIO67-BIOSPHERE | Department of Virology, Henri Mondor University Hospital, Assistance Publique Hôpitaux de Paris, Université Paris-Est Créteil, INSERM U955 | Christophe Rodriguez, Slim Fourati, Vanessa Demontant, Guillaume Gricourt, Melissa N'Debi, Alexandre Soulier, Elisabeth Trawinski, Jean-Michel Pawlotsky |
| EPI_ISL_3391160 | Plateau technique Biomer Metz Verriers | Department of Virology, Henri Mondor University Hospital, Assistance Publique Hôpitaux de Paris, Université Paris-Est Créteil, INSERM U955 | Christophe Rodriguez, Slim Fourati, Vanessa Demontant, Guillaume Gricourt, Melissa N'Debi, Alexandre Soulier, Elisabeth Trawinski, Jean-Michel Pawlotsky |
| EPI_ISL_3391161 | BIO67-BIOSPHERE | Department of Virology, Henri Mondor University Hospital, Assistance Publique Hôpitaux de Paris, Université Paris-Est Créteil, INSERM U955 | Christophe Rodriguez, Slim Fourati, Vanessa Demontant, Guillaume Gricourt, Melissa N'Debi, Alexandre Soulier, Elisabeth Trawinski, Jean-Michel Pawlotsky |

| | | | |
|---|---|---|---|
| EPI_ISL_3391162 | Laboratoire LENYS | Department of Virology, Henri Mondor University Hospital, Assistance Publique Hôpitaux de Paris, Université Paris-Est Créteil, INSERM U955 | Christophe Rodriguez, Slim Fourati, Vanessa Demontant, Guillaume Gricourt, Melissa N'Debi, Alexandre Soulier, Elisabeth Trawinski, Jean-Michel Pawlotsky |
| EPI_ISL_3391163, EPI_ISL_3391164, EPI_ISL_3391165 | Plateau technique Biomer Metz Verriers | Department of Virology, Henri Mondor University Hospital, Assistance Publique Hôpitaux de Paris, Université Paris-Est Créteil, INSERM U955 | Christophe Rodriguez, Slim Fourati, Vanessa Demontant, Guillaume Gricourt, Melissa N'Debi, Alexandre Soulier, Elisabeth Trawinski, Jean-Michel Pawlotsky |
| EPI_ISL_3391166, EPI_ISL_3391167 | Laboratoire LENYS | Department of Virology, Henri Mondor University Hospital, Assistance Publique Hôpitaux de Paris, Université Paris-Est Créteil, INSERM U955 | Christophe Rodriguez, Slim Fourati, Vanessa Demontant, Guillaume Gricourt, Melissa N'Debi, Alexandre Soulier, Elisabeth Trawinski, Jean-Michel Pawlotsky |
| EPI_ISL_3391168 | BIO67-BIOSPHERE | Department of Virology, Henri Mondor University Hospital, Assistance Publique Hôpitaux de Paris, Université Paris-Est Créteil, INSERM U955 | Christophe Rodriguez, Slim Fourati, Vanessa Demontant, Guillaume Gricourt, Melissa N'Debi, Alexandre Soulier, Elisabeth Trawinski, Jean-Michel Pawlotsky |
| EPI_ISL_3391169 | Plateau technique Biomer Metz Verriers | Department of Virology, Henri Mondor University Hospital, Assistance Publique Hôpitaux de Paris, Université Paris-Est Créteil, INSERM U955 | Christophe Rodriguez, Slim Fourati, Vanessa Demontant, Guillaume Gricourt, Melissa N'Debi, Alexandre Soulier, Elisabeth Trawinski, Jean-Michel Pawlotsky |
| EPI_ISL_3391170 | BIO67-BIOSPHERE | Department of Virology, Henri Mondor University Hospital, Assistance Publique Hôpitaux de Paris, Université Paris-Est Créteil, INSERM U955 | Christophe Rodriguez, Slim Fourati, Vanessa Demontant, Guillaume Gricourt, Melissa N'Debi, Alexandre Soulier, Elisabeth Trawinski, Jean-Michel Pawlotsky |
| EPI_ISL_3391171 | Laboratoire LENYS | Department of Virology, Henri Mondor University Hospital, Assistance Publique Hôpitaux de Paris, Université Paris-Est Créteil, INSERM U955 | Christophe Rodriguez, Slim Fourati, Vanessa Demontant, Guillaume Gricourt, Melissa N'Debi, Alexandre Soulier, Elisabeth Trawinski, Jean-Michel Pawlotsky |
| EPI_ISL_3391172, EPI_ISL_3391173, EPI_ISL_3391174 | Plateau technique Biomer Metz Verriers | Department of Virology, Henri Mondor University Hospital, Assistance Publique Hôpitaux de Paris, Université Paris-Est Créteil, INSERM U955 | Christophe Rodriguez, Slim Fourati, Vanessa Demontant, Guillaume Gricourt, Melissa N'Debi, Alexandre Soulier, Elisabeth Trawinski, Jean-Michel Pawlotsky |
| EPI_ISL_3391175, EPI_ISL_3391178 | BIO67-BIOSPHERE | Department of Virology, Henri Mondor University Hospital, Assistance Publique Hôpitaux de Paris, Université Paris-Est Créteil, INSERM U955 | Christophe Rodriguez, Slim Fourati, Vanessa Demontant, Guillaume Gricourt, Melissa N'Debi, Alexandre Soulier, Elisabeth Trawinski, Jean-Michel Pawlotsky |
| EPI_ISL_3391179 | Plateau technique Biomer Metz Verriers | Department of Virology, Henri Mondor University Hospital, Assistance Publique Hôpitaux de Paris, Université Paris-Est Créteil, INSERM U955 | Christophe Rodriguez, Slim Fourati, Vanessa Demontant, Guillaume Gricourt, Melissa N'Debi, Alexandre Soulier, Elisabeth Trawinski, Jean-Michel Pawlotsky |
| EPI_ISL_3391180, EPI_ISL_3391181 | BIO67-BIOSPHERE | Department of Virology, Henri Mondor University Hospital, Assistance Publique Hôpitaux de Paris, Université Paris-Est Créteil, INSERM U955 | Christophe Rodriguez, Slim Fourati, Vanessa Demontant, Guillaume Gricourt, Melissa N'Debi, Alexandre Soulier, Elisabeth Trawinski, Jean-Michel Pawlotsky |
| EPI_ISL_3391182 | Plateau technique Biomer Metz Verriers | Department of Virology, Henri Mondor University Hospital, Assistance Publique Hôpitaux de Paris, Université Paris-Est Créteil, INSERM U955 | Christophe Rodriguez, Slim Fourati, Vanessa Demontant, Guillaume Gricourt, Melissa N'Debi, Alexandre Soulier, Elisabeth Trawinski, Jean-Michel Pawlotsky |
| EPI_ISL_3391183 | BIO67-BIOSPHERE | Department of Virology, Henri Mondor University Hospital, Assistance Publique Hôpitaux de Paris, Université Paris-Est Créteil, INSERM U955 | Christophe Rodriguez, Slim Fourati, Vanessa Demontant, Guillaume Gricourt, Melissa N'Debi, Alexandre Soulier, Elisabeth Trawinski, Jean-Michel Pawlotsky |
| EPI_ISL_3391184 | Laboratoire LENYS | Department of Virology, Henri Mondor University Hospital, Assistance Publique Hôpitaux de Paris, Université Paris-Est Créteil, INSERM U955 | Christophe Rodriguez, Slim Fourati, Vanessa Demontant, Guillaume Gricourt, Melissa N'Debi, Alexandre Soulier, Elisabeth Trawinski, Jean-Michel Pawlotsky |
| EPI_ISL_3391185 | Plateau technique Biomer Metz Verriers | Department of Virology, Henri Mondor University Hospital, Assistance Publique Hôpitaux de Paris, Université Paris-Est Créteil, INSERM U955 | Christophe Rodriguez, Slim Fourati, Vanessa Demontant, Guillaume Gricourt, Melissa N'Debi, Alexandre Soulier, Elisabeth Trawinski, Jean-Michel Pawlotsky |
| EPI_ISL_3391186 | BIO67-BIOSPHERE | Department of Virology, Henri Mondor University Hospital, Assistance Publique Hôpitaux de Paris, Université Paris-Est Créteil, INSERM U955 | Christophe Rodriguez, Slim Fourati, Vanessa Demontant, Guillaume Gricourt, Melissa N'Debi, Alexandre Soulier, Elisabeth Trawinski, Jean-Michel Pawlotsky |
| EPI_ISL_3391187 | Laboratoire LENYS | Department of Virology, Henri Mondor University Hospital, Assistance Publique Hôpitaux de Paris, Université Paris-Est Créteil, INSERM U955 | Christophe Rodriguez, Slim Fourati, Vanessa Demontant, Guillaume Gricourt, Melissa N'Debi, Alexandre Soulier, Elisabeth Trawinski, Jean-Michel Pawlotsky |
| EPI_ISL_3391188, EPI_ISL_3391189 | BIO67-BIOSPHERE | Department of Virology, Henri Mondor University Hospital, Assistance Publique Hôpitaux de Paris, Université Paris-Est Créteil, INSERM U955 | Christophe Rodriguez, Slim Fourati, Vanessa Demontant, Guillaume Gricourt, Melissa N'Debi, Alexandre Soulier, Elisabeth Trawinski, Jean-Michel Pawlotsky |
| EPI_ISL_3391190 | Laboratoire LENYS | Department of Virology, Henri Mondor University Hospital, Assistance Publique Hôpitaux de Paris, Université Paris-Est Créteil, INSERM U955 | Christophe Rodriguez, Slim Fourati, Vanessa Demontant, Guillaume Gricourt, Melissa N'Debi, Alexandre Soulier, Elisabeth Trawinski, Jean-Michel Pawlotsky |
| EPI_ISL_3391191, EPI_ISL_3391193 | BIO67-BIOSPHERE | Department of Virology, Henri Mondor University Hospital, Assistance Publique Hôpitaux de Paris, Université Paris-Est Créteil, INSERM U955 | Christophe Rodriguez, Slim Fourati, Vanessa Demontant, Guillaume Gricourt, Melissa N'Debi, Alexandre Soulier, Elisabeth Trawinski, Jean-Michel Pawlotsky |
| EPI_ISL_3391194, EPI_ISL_3391195 | Plateau technique Biomer Metz Verriers | Department of Virology, Henri Mondor University Hospital, Assistance Publique Hôpitaux de Paris, Université Paris-Est Créteil, INSERM U955 | Christophe Rodriguez, Slim Fourati, Vanessa Demontant, Guillaume Gricourt, Melissa N'Debi, Alexandre Soulier, Elisabeth Trawinski, Jean-Michel Pawlotsky |
| EPI_ISL_3391197 | SELAS BC-LAB | Department of Virology, Henri Mondor University Hospital, Assistance Publique Hôpitaux de Paris, Université Paris-Est Créteil, INSERM U955 | Christophe Rodriguez, Slim Fourati, Vanessa Demontant, Guillaume Gricourt, Melissa N'Debi, Alexandre Soulier, Elisabeth Trawinski, Jean-Michel Pawlotsky |
| EPI_ISL_3391199, EPI_ISL_3391200, EPI_ISL_3391204, EPI_ISL_3391206, EPI_ISL_3391207, EPI_ISL_3391208, EPI_ISL_3391209 | SELAS MEDILYS | Department of Virology, Henri Mondor University Hospital, Assistance Publique Hôpitaux de Paris, Université Paris-Est Créteil, INSERM U955 | Christophe Rodriguez, Slim Fourati, Vanessa Demontant, Guillaume Gricourt, Melissa N'Debi, Alexandre Soulier, Elisabeth Trawinski, Jean-Michel Pawlotsky |
| EPI_ISL_3391212 | SELAS BC-LAB | Department of Virology, Henri Mondor University Hospital, Assistance Publique Hôpitaux de Paris, Université Paris-Est Créteil, INSERM U955 | Christophe Rodriguez, Slim Fourati, Vanessa Demontant, Guillaume Gricourt, Melissa N'Debi, Alexandre Soulier, Elisabeth Trawinski, Jean-Michel Pawlotsky |
| EPI_ISL_3391213, EPI_ISL_3391214, EPI_ISL_3391215, EPI_ISL_3391216, EPI_ISL_3391218 | SELAS MEDILYS | Department of Virology, Henri Mondor University Hospital, Assistance Publique Hôpitaux de Paris, Université Paris-Est Créteil, INSERM U955 | Christophe Rodriguez, Slim Fourati, Vanessa Demontant, Guillaume Gricourt, Melissa N'Debi, Alexandre Soulier, Elisabeth Trawinski, Jean-Michel Pawlotsky |
| EPI_ISL_3391219 | SELAS BC-LAB | Department of Virology, Henri Mondor University Hospital, Assistance Publique Hôpitaux de Paris, Université Paris-Est Créteil, INSERM U955 | Christophe Rodriguez, Slim Fourati, Vanessa Demontant, Guillaume Gricourt, Melissa N'Debi, Alexandre Soulier, Elisabeth Trawinski, Jean-Michel Pawlotsky |
| EPI_ISL_3391220 | SELAS MEDILYS | Department of Virology, Henri Mondor University Hospital, Assistance Publique Hôpitaux de Paris, Université Paris-Est Créteil, INSERM U955 | Christophe Rodriguez, Slim Fourati, Vanessa Demontant, Guillaume Gricourt, Melissa N'Debi, Alexandre Soulier, Elisabeth Trawinski, Jean-Michel Pawlotsky |
| EPI_ISL_3391221 | SELAS BC-LAB | Department of Virology, Henri Mondor University Hospital, Assistance Publique Hôpitaux de Paris, Université Paris-Est Créteil, INSERM U955 | Christophe Rodriguez, Slim Fourati, Vanessa Demontant, Guillaume Gricourt, Melissa N'Debi, Alexandre Soulier, Elisabeth Trawinski, Jean-Michel Pawlotsky |
| EPI_ISL_3391222, EPI_ISL_3391223 | SELAS MEDILYS | Department of Virology, Henri Mondor University Hospital, Assistance Publique Hôpitaux de Paris, Université Paris-Est Créteil, INSERM U955 | Christophe Rodriguez, Slim Fourati, Vanessa Demontant, Guillaume Gricourt, Melissa N'Debi, Alexandre Soulier, Elisabeth Trawinski, Jean-Michel Pawlotsky |
| EPI_ISL_3391224 | SELAS BC-LAB | Department of Virology, Henri Mondor University Hospital, Assistance Publique Hôpitaux de Paris, Université Paris-Est Créteil, INSERM U955 | Christophe Rodriguez, Slim Fourati, Vanessa Demontant, Guillaume Gricourt, Melissa N'Debi, Alexandre Soulier, Elisabeth Trawinski, Jean-Michel Pawlotsky |

| | | | |
|---|---|---|---|
| EPI_ISL_3391225, EPI_ISL_3391226, EPI_ISL_3391227, EPI_ISL_3391228, EPI_ISL_3391229 | SELAS MEDILYS | Department of Virology, Henri Mondor University Hospital, Assistance Publique Hôpitaux de Paris, Université Paris-Est Créteil, INSERM U955 | Christophe Rodriguez, Slim Fourati, Vanessa Demontant, Guillaume Gricourt, Melissa N'Debi, Alexandre Soulier, Elisabeth Trawinski, Jean-Michel Pawlotsky |
| EPI_ISL_3391230, EPI_ISL_3391232 | SELAS BC-LAB | Department of Virology, Henri Mondor University Hospital, Assistance Publique Hôpitaux de Paris, Université Paris-Est Créteil, INSERM U955 | Christophe Rodriguez, Slim Fourati, Vanessa Demontant, Guillaume Gricourt, Melissa N'Debi, Alexandre Soulier, Elisabeth Trawinski, Jean-Michel Pawlotsky |
| EPI_ISL_3391233, EPI_ISL_3391235, EPI_ISL_3391236 | SELAS MEDILYS | Department of Virology, Henri Mondor University Hospital, Assistance Publique Hôpitaux de Paris, Université Paris-Est Créteil, INSERM U955 | Christophe Rodriguez, Slim Fourati, Vanessa Demontant, Guillaume Gricourt, Melissa N'Debi, Alexandre Soulier, Elisabeth Trawinski, Jean-Michel Pawlotsky |
| EPI_ISL_3391237 | SELAS BC-LAB | Department of Virology, Henri Mondor University Hospital, Assistance Publique Hôpitaux de Paris, Université Paris-Est Créteil, INSERM U955 | Christophe Rodriguez, Slim Fourati, Vanessa Demontant, Guillaume Gricourt, Melissa N'Debi, Alexandre Soulier, Elisabeth Trawinski, Jean-Michel Pawlotsky |
| EPI_ISL_3391238, EPI_ISL_3391239, EPI_ISL_3391240, EPI_ISL_3391241 | SELAS MEDILYS | Department of Virology, Henri Mondor University Hospital, Assistance Publique Hôpitaux de Paris, Université Paris-Est Créteil, INSERM U955 | Christophe Rodriguez, Slim Fourati, Vanessa Demontant, Guillaume Gricourt, Melissa N'Debi, Alexandre Soulier, Elisabeth Trawinski, Jean-Michel Pawlotsky |
| EPI_ISL_3391245 | SELAS BC-LAB | Department of Virology, Henri Mondor University Hospital, Assistance Publique Hôpitaux de Paris, Université Paris-Est Créteil, INSERM U955 | Christophe Rodriguez, Slim Fourati, Vanessa Demontant, Guillaume Gricourt, Melissa N'Debi, Alexandre Soulier, Elisabeth Trawinski, Jean-Michel Pawlotsky |
| EPI_ISL_3391246, EPI_ISL_3391247, EPI_ISL_3391248, EPI_ISL_3391249, EPI_ISL_3391252, EPI_ISL_3391253 | SELAS MEDILYS | Department of Virology, Henri Mondor University Hospital, Assistance Publique Hôpitaux de Paris, Université Paris-Est Créteil, INSERM U955 | Christophe Rodriguez, Slim Fourati, Vanessa Demontant, Guillaume Gricourt, Melissa N'Debi, Alexandre Soulier, Elisabeth Trawinski, Jean-Michel Pawlotsky |
| EPI_ISL_3391254, EPI_ISL_3391255 | SELAS BC-LAB | Department of Virology, Henri Mondor University Hospital, Assistance Publique Hôpitaux de Paris, Université Paris-Est Créteil, INSERM U955 | Christophe Rodriguez, Slim Fourati, Vanessa Demontant, Guillaume Gricourt, Melissa N'Debi, Alexandre Soulier, Elisabeth Trawinski, Jean-Michel Pawlotsky |
| EPI_ISL_3391256, EPI_ISL_3391257, EPI_ISL_3391258, EPI_ISL_3391259, EPI_ISL_3391260, EPI_ISL_3391261 | SELAS MEDILYS | Department of Virology, Henri Mondor University Hospital, Assistance Publique Hôpitaux de Paris, Université Paris-Est Créteil, INSERM U955 | Christophe Rodriguez, Slim Fourati, Vanessa Demontant, Guillaume Gricourt, Melissa N'Debi, Alexandre Soulier, Elisabeth Trawinski, Jean-Michel Pawlotsky |
| EPI_ISL_3391263 | SELAS BC-LAB | Department of Virology, Henri Mondor University Hospital, Assistance Publique Hôpitaux de Paris, Université Paris-Est Créteil, INSERM U955 | Christophe Rodriguez, Slim Fourati, Vanessa Demontant, Guillaume Gricourt, Melissa N'Debi, Alexandre Soulier, Elisabeth Trawinski, Jean-Michel Pawlotsky |
| EPI_ISL_3391264, EPI_ISL_3391266 | SELAS MEDILYS | Department of Virology, Henri Mondor University Hospital, Assistance Publique Hôpitaux de Paris, Université Paris-Est Créteil, INSERM U955 | Christophe Rodriguez, Slim Fourati, Vanessa Demontant, Guillaume Gricourt, Melissa N'Debi, Alexandre Soulier, Elisabeth Trawinski, Jean-Michel Pawlotsky |
| EPI_ISL_3391269, EPI_ISL_3391270 | SELAS BC-LAB | Department of Virology, Henri Mondor University Hospital, Assistance Publique Hôpitaux de Paris, Université Paris-Est Créteil, INSERM U955 | Christophe Rodriguez, Slim Fourati, Vanessa Demontant, Guillaume Gricourt, Melissa N'Debi, Alexandre Soulier, Elisabeth Trawinski, Jean-Michel Pawlotsky |
| EPI_ISL_3391271, EPI_ISL_3391273 | SELAS MEDILYS | Department of Virology, Henri Mondor University Hospital, Assistance Publique Hôpitaux de Paris, Université Paris-Est Créteil, INSERM U955 | Christophe Rodriguez, Slim Fourati, Vanessa Demontant, Guillaume Gricourt, Melissa N'Debi, Alexandre Soulier, Elisabeth Trawinski, Jean-Michel Pawlotsky |
| EPI_ISL_3391274 | SELAS BC-LAB | Department of Virology, Henri Mondor University Hospital, Assistance Publique Hôpitaux de Paris, Université Paris-Est Créteil, INSERM U955 | Christophe Rodriguez, Slim Fourati, Vanessa Demontant, Guillaume Gricourt, Melissa N'Debi, Alexandre Soulier, Elisabeth Trawinski, Jean-Michel Pawlotsky |
| EPI_ISL_3391275, EPI_ISL_3391276 | SELAS MEDILYS | Department of Virology, Henri Mondor University Hospital, Assistance Publique Hôpitaux de Paris, Université Paris-Est Créteil, INSERM U955 | Christophe Rodriguez, Slim Fourati, Vanessa Demontant, Guillaume Gricourt, Melissa N'Debi, Alexandre Soulier, Elisabeth Trawinski, Jean-Michel Pawlotsky |
| EPI_ISL_3391277, EPI_ISL_3391278, EPI_ISL_3391279, EPI_ISL_3391280, EPI_ISL_3391281, EPI_ISL_3391282, EPI_ISL_3391283, EPI_ISL_3391284, EPI_ISL_3391285, EPI_ISL_3391286, EPI_ISL_3391287, EPI_ISL_3391288, EPI_ISL_3391289, EPI_ISL_3391290, EPI_ISL_3391291, EPI_ISL_3391292, EPI_ISL_3391293, EPI_ISL_3391294, EPI_ISL_3391295, EPI_ISL_3391296, EPI_ISL_3391297, EPI_ISL_3391298, EPI_ISL_3391299, EPI_ISL_3391300, EPI_ISL_3391301, EPI_ISL_3391302, EPI_ISL_3391303, EPI_ISL_3391304 | | | |
| see above | State Hygienic Laboratory at the University of Iowa | State Hygienic Laboratory at the University of Iowa | Valerie Reeb, Erik Twait, Anna Yakos, Mariah Knutson, Kris Eveland, Kristen Zanon, Jeff Benfer, Wes Hottel, Alankar Kampoowale |
| EPI_ISL_3391305 | BIOMNIS EUROFINS IVRY | Department of Virology, Henri Mondor University Hospital, Assistance Publique Hôpitaux de Paris, Université Paris-Est Créteil, INSERM U955 | Christophe Rodriguez, Slim Fourati, Vanessa Demontant, Guillaume Gricourt, Melissa N'Debi, Alexandre Soulier, Elisabeth Trawinski, Jean-Michel Pawlotsky |
| EPI_ISL_3391306 | Biogroup Bio Lam-LCD Saint-Denis | Department of Virology, Henri Mondor University Hospital, Assistance Publique Hôpitaux de Paris, Université Paris-Est Créteil, INSERM U955 | Christophe Rodriguez, Slim Fourati, Vanessa Demontant, Guillaume Gricourt, Melissa N'Debi, Alexandre Soulier, Elisabeth Trawinski, Jean-Michel Pawlotsky |
| EPI_ISL_3391307 | BIOMNIS EUROFINS IVRY | Department of Virology, Henri Mondor University Hospital, Assistance Publique Hôpitaux de Paris, Université Paris-Est Créteil, INSERM U955 | Christophe Rodriguez, Slim Fourati, Vanessa Demontant, Guillaume Gricourt, Melissa N'Debi, Alexandre Soulier, Elisabeth Trawinski, Jean-Michel Pawlotsky |
| EPI_ISL_3391308 | Biogroup Bio Lam-LCD Saint-Denis | Department of Virology, Henri Mondor University Hospital, Assistance Publique Hôpitaux de Paris, Université Paris-Est Créteil, INSERM U955 | Christophe Rodriguez, Slim Fourati, Vanessa Demontant, Guillaume Gricourt, Melissa N'Debi, Alexandre Soulier, Elisabeth Trawinski, Jean-Michel Pawlotsky |
| EPI_ISL_3391309 | BIO67-BIOSPHERE | Department of Virology, Henri Mondor University Hospital, Assistance Publique Hôpitaux de Paris, Université Paris-Est Créteil, INSERM U955 | Christophe Rodriguez, Slim Fourati, Vanessa Demontant, Guillaume Gricourt, Melissa N'Debi, Alexandre Soulier, Elisabeth Trawinski, Jean-Michel Pawlotsky |
| EPI_ISL_3391310 | Biogroup Bio Lam-LCD Saint-Denis | Department of Virology, Henri Mondor University Hospital, Assistance Publique Hôpitaux de Paris, Université Paris-Est Créteil, INSERM U955 | Christophe Rodriguez, Slim Fourati, Vanessa Demontant, Guillaume Gricourt, Melissa N'Debi, Alexandre Soulier, Elisabeth Trawinski, Jean-Michel Pawlotsky |
| EPI_ISL_3391311, EPI_ISL_3391312 | BIOMNIS EUROFINS IVRY | Department of Virology, Henri Mondor University Hospital, Assistance Publique Hôpitaux de Paris, Université Paris-Est Créteil, INSERM U955 | Christophe Rodriguez, Slim Fourati, Vanessa Demontant, Guillaume Gricourt, Melissa N'Debi, Alexandre Soulier, Elisabeth Trawinski, Jean-Michel Pawlotsky |
| EPI_ISL_3391313 | Biogroup Bio Lam-LCD Saint-Denis | Department of Virology, Henri Mondor University Hospital, Assistance Publique Hôpitaux de Paris, Université Paris-Est Créteil, INSERM U955 | Christophe Rodriguez, Slim Fourati, Vanessa Demontant, Guillaume Gricourt, Melissa N'Debi, Alexandre Soulier, Elisabeth Trawinski, Jean-Michel Pawlotsky |
| EPI_ISL_3391314, EPI_ISL_3391315 | BIO67-BIOSPHERE | Department of Virology, Henri Mondor University Hospital, Assistance Publique Hôpitaux de Paris, Université Paris-Est Créteil, INSERM U955 | Christophe Rodriguez, Slim Fourati, Vanessa Demontant, Guillaume Gricourt, Melissa N'Debi, Alexandre Soulier, Elisabeth Trawinski, Jean-Michel Pawlotsky |
| EPI_ISL_3391316 | BIOMNIS EUROFINS IVRY | Department of Virology, Henri Mondor University Hospital, Assistance Publique Hôpitaux de Paris, Université Paris-Est Créteil, INSERM U955 | Christophe Rodriguez, Slim Fourati, Vanessa Demontant, Guillaume Gricourt, Melissa N'Debi, Alexandre Soulier, Elisabeth Trawinski, Jean-Michel Pawlotsky |
| EPI_ISL_3391317 | Biogroup Bio Lam-LCD Saint-Denis | Department of Virology, Henri Mondor University Hospital, Assistance Publique Hôpitaux de Paris, Université Paris-Est Créteil, INSERM U955 | Christophe Rodriguez, Slim Fourati, Vanessa Demontant, Guillaume Gricourt, Melissa N'Debi, Alexandre Soulier, Elisabeth Trawinski, Jean-Michel Pawlotsky |
| EPI_ISL_3391318 | BIOMNIS EUROFINS IVRY | Department of Virology, Henri Mondor University Hospital, Assistance Publique Hôpitaux de Paris, Université Paris-Est Créteil, INSERM U955 | Christophe Rodriguez, Slim Fourati, Vanessa Demontant, Guillaume Gricourt, Melissa N'Debi, Alexandre Soulier, Elisabeth Trawinski, Jean-Michel Pawlotsky |
| EPI_ISL_3391321 | Biogroup Bio Lam-LCD Saint-Denis | Department of Virology, Henri Mondor University Hospital, Assistance Publique Hôpitaux de Paris, Université Paris-Est Créteil, INSERM U955 | Christophe Rodriguez, Slim Fourati, Vanessa Demontant, Guillaume Gricourt, Melissa N'Debi, Alexandre Soulier, Elisabeth Trawinski, Jean-Michel Pawlotsky |
| EPI_ISL_3391322 | BIO67-BIOSPHERE | Department of Virology, Henri Mondor University Hospital, | Christophe Rodriguez, Slim Fourati, Vanessa Demontant, Guillaume Gricourt, Melissa N'Debi, Alexandre Soulier, Elisabeth Trawinski, Jean-Michel Pawlotsky |

| | | | |
|---|---|---|---|
| | | Assistance Publique Hôpitaux de Paris, Université Paris-Est Créteil, INSERM U955 | |
| EPI_ISL_3391323, EPI_ISL_3391324, EPI_ISL_3391325, EPI_ISL_3391326, EPI_ISL_3391327, EPI_ISL_3391328 | Biogroup Bio Lam-LCD Saint-Denis | Department of Virology, Henri Mondor University Hospital, Assistance Publique Hôpitaux de Paris, Université Paris-Est Créteil, INSERM U955 | Christophe Rodriguez, Slim Fourati, Vanessa Demontant, Guillaume Gricourt, Melissa N'Debi, Alexandre Soulier, Elisabeth Trawinski, Jean-Michel Pawlotsky |
| EPI_ISL_3391329 | BIOMNIS EUROFINS IVRY | Department of Virology, Henri Mondor University Hospital, Assistance Publique Hôpitaux de Paris, Université Paris-Est Créteil, INSERM U955 | Christophe Rodriguez, Slim Fourati, Vanessa Demontant, Guillaume Gricourt, Melissa N'Debi, Alexandre Soulier, Elisabeth Trawinski, Jean-Michel Pawlotsky |
| EPI_ISL_3391330, EPI_ISL_3391331 | Biogroup Bio Lam-LCD Saint-Denis | Department of Virology, Henri Mondor University Hospital, Assistance Publique Hôpitaux de Paris, Université Paris-Est Créteil, INSERM U955 | Christophe Rodriguez, Slim Fourati, Vanessa Demontant, Guillaume Gricourt, Melissa N'Debi, Alexandre Soulier, Elisabeth Trawinski, Jean-Michel Pawlotsky |
| EPI_ISL_3391332 | LBM CARRON SELAS | Department of Virology, Henri Mondor University Hospital, Assistance Publique Hôpitaux de Paris, Université Paris-Est Créteil, INSERM U955 | Christophe Rodriguez, Slim Fourati, Vanessa Demontant, Guillaume Gricourt, Melissa N'Debi, Alexandre Soulier, Elisabeth Trawinski, Jean-Michel Pawlotsky |
| EPI_ISL_3391333 | BIOMNIS EUROFINS IVRY | Department of Virology, Henri Mondor University Hospital, Assistance Publique Hôpitaux de Paris, Université Paris-Est Créteil, INSERM U955 | Christophe Rodriguez, Slim Fourati, Vanessa Demontant, Guillaume Gricourt, Melissa N'Debi, Alexandre Soulier, Elisabeth Trawinski, Jean-Michel Pawlotsky |
| EPI_ISL_3391334, EPI_ISL_3391335, EPI_ISL_3391336 | Biogroup Bio Lam-LCD Saint-Denis | Department of Virology, Henri Mondor University Hospital, Assistance Publique Hôpitaux de Paris, Université Paris-Est Créteil, INSERM U955 | Christophe Rodriguez, Slim Fourati, Vanessa Demontant, Guillaume Gricourt, Melissa N'Debi, Alexandre Soulier, Elisabeth Trawinski, Jean-Michel Pawlotsky |
| EPI_ISL_3391337, EPI_ISL_3391338 | Laboratoire Ana-L | Department of Virology, Henri Mondor University Hospital, Assistance Publique Hôpitaux de Paris, Université Paris-Est Créteil, INSERM U955 | Christophe Rodriguez, Slim Fourati, Vanessa Demontant, Guillaume Gricourt, Melissa N'Debi, Alexandre Soulier, Elisabeth Trawinski, Jean-Michel Pawlotsky |
| EPI_ISL_3391339 | BIOMNIS EUROFINS IVRY | Department of Virology, Henri Mondor University Hospital, Assistance Publique Hôpitaux de Paris, Université Paris-Est Créteil, INSERM U955 | Christophe Rodriguez, Slim Fourati, Vanessa Demontant, Guillaume Gricourt, Melissa N'Debi, Alexandre Soulier, Elisabeth Trawinski, Jean-Michel Pawlotsky |
| EPI_ISL_3391340, EPI_ISL_3391341 | Biogroup Bio Lam-LCD Saint-Denis | Department of Virology, Henri Mondor University Hospital, Assistance Publique Hôpitaux de Paris, Université Paris-Est Créteil, INSERM U955 | Christophe Rodriguez, Slim Fourati, Vanessa Demontant, Guillaume Gricourt, Melissa N'Debi, Alexandre Soulier, Elisabeth Trawinski, Jean-Michel Pawlotsky |
| EPI_ISL_3391342, EPI_ISL_3391344 | Laboratoire Ana-L | Department of Virology, Henri Mondor University Hospital, Assistance Publique Hôpitaux de Paris, Université Paris-Est Créteil, INSERM U955 | Christophe Rodriguez, Slim Fourati, Vanessa Demontant, Guillaume Gricourt, Melissa N'Debi, Alexandre Soulier, Elisabeth Trawinski, Jean-Michel Pawlotsky |
| EPI_ISL_3391345 | BIO67-BIOSPHERE | Department of Virology, Henri Mondor University Hospital, Assistance Publique Hôpitaux de Paris, Université Paris-Est Créteil, INSERM U955 | Christophe Rodriguez, Slim Fourati, Vanessa Demontant, Guillaume Gricourt, Melissa N'Debi, Alexandre Soulier, Elisabeth Trawinski, Jean-Michel Pawlotsky |
| EPI_ISL_3391346 | DYNABIO UNILABS | Department of Virology, Henri Mondor University Hospital, Assistance Publique Hôpitaux de Paris, Université Paris-Est Créteil, INSERM U955 | Christophe Rodriguez, Slim Fourati, Vanessa Demontant, Guillaume Gricourt, Melissa N'Debi, Alexandre Soulier, Elisabeth Trawinski, Jean-Michel Pawlotsky |
| EPI_ISL_3391347 | SELAS BC-LAB | Department of Virology, Henri Mondor University Hospital, Assistance Publique Hôpitaux de Paris, Université Paris-Est Créteil, INSERM U955 | Christophe Rodriguez, Slim Fourati, Vanessa Demontant, Guillaume Gricourt, Melissa N'Debi, Alexandre Soulier, Elisabeth Trawinski, Jean-Michel Pawlotsky |
| EPI_ISL_3391348 | Biogroup Bio Lam-LCD Saint-Denis | Department of Virology, Henri Mondor University Hospital, Assistance Publique Hôpitaux de Paris, Université Paris-Est Créteil, INSERM U955 | Christophe Rodriguez, Slim Fourati, Vanessa Demontant, Guillaume Gricourt, Melissa N'Debi, Alexandre Soulier, Elisabeth Trawinski, Jean-Michel Pawlotsky |
| EPI_ISL_3391349 | Laboratoire Ana-L | Department of Virology, Henri Mondor University Hospital, Assistance Publique Hôpitaux de Paris, Université Paris-Est Créteil, INSERM U955 | Christophe Rodriguez, Slim Fourati, Vanessa Demontant, Guillaume Gricourt, Melissa N'Debi, Alexandre Soulier, Elisabeth Trawinski, Jean-Michel Pawlotsky |
| EPI_ISL_3391350, EPI_ISL_3391351 | Biogroup Bio Lam-LCD Saint-Denis | Department of Virology, Henri Mondor University Hospital, Assistance Publique Hôpitaux de Paris, Université Paris-Est Créteil, INSERM U955 | Christophe Rodriguez, Slim Fourati, Vanessa Demontant, Guillaume Gricourt, Melissa N'Debi, Alexandre Soulier, Elisabeth Trawinski, Jean-Michel Pawlotsky |
| EPI_ISL_3391352 | BIOMNIS EUROFINS IVRY | Department of Virology, Henri Mondor University Hospital, Assistance Publique Hôpitaux de Paris, Université Paris-Est Créteil, INSERM U955 | Christophe Rodriguez, Slim Fourati, Vanessa Demontant, Guillaume Gricourt, Melissa N'Debi, Alexandre Soulier, Elisabeth Trawinski, Jean-Michel Pawlotsky |
| EPI_ISL_3391353 | Biogroup Bio Lam-LCD Saint-Denis | Department of Virology, Henri Mondor University Hospital, Assistance Publique Hôpitaux de Paris, Université Paris-Est Créteil, INSERM U955 | Christophe Rodriguez, Slim Fourati, Vanessa Demontant, Guillaume Gricourt, Melissa N'Debi, Alexandre Soulier, Elisabeth Trawinski, Jean-Michel Pawlotsky |
| EPI_ISL_3391354 | BIO67-BIOSPHERE | Department of Virology, Henri Mondor University Hospital, Assistance Publique Hôpitaux de Paris, Université Paris-Est Créteil, INSERM U955 | Christophe Rodriguez, Slim Fourati, Vanessa Demontant, Guillaume Gricourt, Melissa N'Debi, Alexandre Soulier, Elisabeth Trawinski, Jean-Michel Pawlotsky |
| EPI_ISL_3391355 | Biogroup Bio Lam-LCD Saint-Denis | Department of Virology, Henri Mondor University Hospital, Assistance Publique Hôpitaux de Paris, Université Paris-Est Créteil, INSERM U955 | Christophe Rodriguez, Slim Fourati, Vanessa Demontant, Guillaume Gricourt, Melissa N'Debi, Alexandre Soulier, Elisabeth Trawinski, Jean-Michel Pawlotsky |
| EPI_ISL_3391356 | BIO67-BIOSPHERE | Department of Virology, Henri Mondor University Hospital, Assistance Publique Hôpitaux de Paris, Université Paris-Est Créteil, INSERM U955 | Christophe Rodriguez, Slim Fourati, Vanessa Demontant, Guillaume Gricourt, Melissa N'Debi, Alexandre Soulier, Elisabeth Trawinski, Jean-Michel Pawlotsky |
| EPI_ISL_3391357 | Biogroup Bio Lam-LCD Saint-Denis | Department of Virology, Henri Mondor University Hospital, Assistance Publique Hôpitaux de Paris, Université Paris-Est Créteil, INSERM U955 | Christophe Rodriguez, Slim Fourati, Vanessa Demontant, Guillaume Gricourt, Melissa N'Debi, Alexandre Soulier, Elisabeth Trawinski, Jean-Michel Pawlotsky |
| EPI_ISL_3391359 | Laboratoire Ana-L | Department of Virology, Henri Mondor University Hospital, Assistance Publique Hôpitaux de Paris, Université Paris-Est Créteil, INSERM U955 | Christophe Rodriguez, Slim Fourati, Vanessa Demontant, Guillaume Gricourt, Melissa N'Debi, Alexandre Soulier, Elisabeth Trawinski, Jean-Michel Pawlotsky |
| EPI_ISL_3391360 | Biogroup Bio Lam-LCD Saint-Denis | Department of Virology, Henri Mondor University Hospital, Assistance Publique Hôpitaux de Paris, Université Paris-Est Créteil, INSERM U955 | Christophe Rodriguez, Slim Fourati, Vanessa Demontant, Guillaume Gricourt, Melissa N'Debi, Alexandre Soulier, Elisabeth Trawinski, Jean-Michel Pawlotsky |
| EPI_ISL_3391361 | BIOMNIS EUROFINS IVRY | Department of Virology, Henri Mondor University Hospital, Assistance Publique Hôpitaux de Paris, Université Paris-Est Créteil, INSERM U955 | Christophe Rodriguez, Slim Fourati, Vanessa Demontant, Guillaume Gricourt, Melissa N'Debi, Alexandre Soulier, Elisabeth Trawinski, Jean-Michel Pawlotsky |
| EPI_ISL_3391362 | Biogroup Bio Lam-LCD Saint-Denis | Department of Virology, Henri Mondor University Hospital, Assistance Publique Hôpitaux de Paris, Université Paris-Est Créteil, INSERM U955 | Christophe Rodriguez, Slim Fourati, Vanessa Demontant, Guillaume Gricourt, Melissa N'Debi, Alexandre Soulier, Elisabeth Trawinski, Jean-Michel Pawlotsky |
| EPI_ISL_3391363 | DYNABIO UNILABS | Department of Virology, Henri Mondor University Hospital, Assistance Publique Hôpitaux de Paris, Université Paris-Est Créteil, INSERM U955 | Christophe Rodriguez, Slim Fourati, Vanessa Demontant, Guillaume Gricourt, Melissa N'Debi, Alexandre Soulier, Elisabeth Trawinski, Jean-Michel Pawlotsky |
| EPI_ISL_3391364, EPI_ISL_3391366 | Biogroup Bio Lam-LCD Saint-Denis | Department of Virology, Henri Mondor University Hospital, Assistance Publique Hôpitaux de Paris, Université Paris-Est Créteil, INSERM U955 | Christophe Rodriguez, Slim Fourati, Vanessa Demontant, Guillaume Gricourt, Melissa N'Debi, Alexandre Soulier, Elisabeth Trawinski, Jean-Michel Pawlotsky |
| EPI_ISL_3391367 | SELAS BC-LAB | Department of Virology, Henri Mondor University Hospital, Assistance Publique Hôpitaux de Paris, Université Paris-Est Créteil, INSERM U955 | Christophe Rodriguez, Slim Fourati, Vanessa Demontant, Guillaume Gricourt, Melissa N'Debi, Alexandre Soulier, Elisabeth Trawinski, Jean-Michel Pawlotsky |
| EPI_ISL_3391368 | Biogroup Bio Lam-LCD Saint-Denis | Department of Virology, Henri Mondor University Hospital, Assistance Publique Hôpitaux de Paris, Université Paris-Est Créteil, INSERM U955 | Christophe Rodriguez, Slim Fourati, Vanessa Demontant, Guillaume Gricourt, Melissa N'Debi, Alexandre Soulier, Elisabeth Trawinski, Jean-Michel Pawlotsky |
| EPI_ISL_3391370 | BIO67-BIOSPHERE | Department of Virology, Henri Mondor University Hospital, | Christophe Rodriguez, Slim Fourati, Vanessa Demontant, Guillaume Gricourt, Melissa N'Debi, Alexandre Soulier, Elisabeth Trawinski, Jean-Michel Pawlotsky |

| | | | |
|---|---|---|---|
| | | Assistance Publique Hôpitaux de Paris, Université Paris-Est Créteil, INSERM U955 | |
| EPI_ISL_3391371 | Biogroup Bio Lam-LCD Saint-Denis | Department of Virology, Henri Mondor University Hospital, Assistance Publique Hôpitaux de Paris, Université Paris-Est Créteil, INSERM U955 | Christophe Rodriguez, Slim Fourati, Vanessa Demontant, Guillaume Gricourt, Melissa N'Debi, Alexandre Soulier, Elisabeth Trawinski, Jean-Michel Pawlotsky |
| EPI_ISL_3391372 | Laboratoire Ana-L | Department of Virology, Henri Mondor University Hospital, Assistance Publique Hôpitaux de Paris, Université Paris-Est Créteil, INSERM U955 | Christophe Rodriguez, Slim Fourati, Vanessa Demontant, Guillaume Gricourt, Melissa N'Debi, Alexandre Soulier, Elisabeth Trawinski, Jean-Michel Pawlotsky |
| EPI_ISL_3391373 | BIOMNIS EUROFINS IVRY | Department of Virology, Henri Mondor University Hospital, Assistance Publique Hôpitaux de Paris, Université Paris-Est Créteil, INSERM U955 | Christophe Rodriguez, Slim Fourati, Vanessa Demontant, Guillaume Gricourt, Melissa N'Debi, Alexandre Soulier, Elisabeth Trawinski, Jean-Michel Pawlotsky |
| EPI_ISL_3391374 | Laboratoire Ana-L | Department of Virology, Henri Mondor University Hospital, Assistance Publique Hôpitaux de Paris, Université Paris-Est Créteil, INSERM U955 | Christophe Rodriguez, Slim Fourati, Vanessa Demontant, Guillaume Gricourt, Melissa N'Debi, Alexandre Soulier, Elisabeth Trawinski, Jean-Michel Pawlotsky |
| EPI_ISL_3391375 | BIOMNIS EUROFINS IVRY | Department of Virology, Henri Mondor University Hospital, Assistance Publique Hôpitaux de Paris, Université Paris-Est Créteil, INSERM U955 | Christophe Rodriguez, Slim Fourati, Vanessa Demontant, Guillaume Gricourt, Melissa N'Debi, Alexandre Soulier, Elisabeth Trawinski, Jean-Michel Pawlotsky |
| EPI_ISL_3391377 | Biogroup Bio Lam-LCD Saint-Denis | Department of Virology, Henri Mondor University Hospital, Assistance Publique Hôpitaux de Paris, Université Paris-Est Créteil, INSERM U955 | Christophe Rodriguez, Slim Fourati, Vanessa Demontant, Guillaume Gricourt, Melissa N'Debi, Alexandre Soulier, Elisabeth Trawinski, Jean-Michel Pawlotsky |
| EPI_ISL_3391379 | BIOMNIS EUROFINS IVRY | Department of Virology, Henri Mondor University Hospital, Assistance Publique Hôpitaux de Paris, Université Paris-Est Créteil, INSERM U955 | Christophe Rodriguez, Slim Fourati, Vanessa Demontant, Guillaume Gricourt, Melissa N'Debi, Alexandre Soulier, Elisabeth Trawinski, Jean-Michel Pawlotsky |
| EPI_ISL_3391381 | DYNABIO UNILABS | Department of Virology, Henri Mondor University Hospital, Assistance Publique Hôpitaux de Paris, Université Paris-Est Créteil, INSERM U955 | Christophe Rodriguez, Slim Fourati, Vanessa Demontant, Guillaume Gricourt, Melissa N'Debi, Alexandre Soulier, Elisabeth Trawinski, Jean-Michel Pawlotsky |
| EPI_ISL_3391382 | Laboratoire Ana-L | Department of Virology, Henri Mondor University Hospital, Assistance Publique Hôpitaux de Paris, Université Paris-Est Créteil, INSERM U955 | Christophe Rodriguez, Slim Fourati, Vanessa Demontant, Guillaume Gricourt, Melissa N'Debi, Alexandre Soulier, Elisabeth Trawinski, Jean-Michel Pawlotsky |
| EPI_ISL_3391383 | BIO67-BIOSPHERE | Department of Virology, Henri Mondor University Hospital, Assistance Publique Hôpitaux de Paris, Université Paris-Est Créteil, INSERM U955 | Christophe Rodriguez, Slim Fourati, Vanessa Demontant, Guillaume Gricourt, Melissa N'Debi, Alexandre Soulier, Elisabeth Trawinski, Jean-Michel Pawlotsky |
| EPI_ISL_3391384, EPI_ISL_3391385, EPI_ISL_3391386 | Biogroup Bio Lam-LCD Saint-Denis | Department of Virology, Henri Mondor University Hospital, Assistance Publique Hôpitaux de Paris, Université Paris-Est Créteil, INSERM U955 | Christophe Rodriguez, Slim Fourati, Vanessa Demontant, Guillaume Gricourt, Melissa N'Debi, Alexandre Soulier, Elisabeth Trawinski, Jean-Michel Pawlotsky |
| EPI_ISL_3391387 | Laboratoire Ana-L | Department of Virology, Henri Mondor University Hospital, Assistance Publique Hôpitaux de Paris, Université Paris-Est Créteil, INSERM U955 | Christophe Rodriguez, Slim Fourati, Vanessa Demontant, Guillaume Gricourt, Melissa N'Debi, Alexandre Soulier, Elisabeth Trawinski, Jean-Michel Pawlotsky |
| EPI_ISL_3391388, EPI_ISL_3391389 | Biogroup Bio Lam-LCD Saint-Denis | Department of Virology, Henri Mondor University Hospital, Assistance Publique Hôpitaux de Paris, Université Paris-Est Créteil, INSERM U955 | Christophe Rodriguez, Slim Fourati, Vanessa Demontant, Guillaume Gricourt, Melissa N'Debi, Alexandre Soulier, Elisabeth Trawinski, Jean-Michel Pawlotsky |
| EPI_ISL_3391390 | GROUPE HOSPITALIER SUD ILE DE FRANCE | Department of Virology, Henri Mondor University Hospital, Assistance Publique Hôpitaux de Paris, Université Paris-Est Créteil, INSERM U955 | Christophe Rodriguez, Slim Fourati, Vanessa Demontant, Guillaume Gricourt, Melissa N'Debi, Alexandre Soulier, Elisabeth Trawinski, Jean-Michel Pawlotsky |
| EPI_ISL_3391391 | DYNABIO UNILABS | Department of Virology, Henri Mondor University Hospital, Assistance Publique Hôpitaux de Paris, Université Paris-Est Créteil, INSERM U955 | Christophe Rodriguez, Slim Fourati, Vanessa Demontant, Guillaume Gricourt, Melissa N'Debi, Alexandre Soulier, Elisabeth Trawinski, Jean-Michel Pawlotsky |
| EPI_ISL_3391392, EPI_ISL_3391393, EPI_ISL_3391394 | Biogroup Bio Lam-LCD Saint-Denis | Department of Virology, Henri Mondor University Hospital, Assistance Publique Hôpitaux de Paris, Université Paris-Est Créteil, INSERM U955 | Christophe Rodriguez, Slim Fourati, Vanessa Demontant, Guillaume Gricourt, Melissa N'Debi, Alexandre Soulier, Elisabeth Trawinski, Jean-Michel Pawlotsky |
| EPI_ISL_3391395 | BIO67-BIOSPHERE | Department of Virology, Henri Mondor University Hospital, Assistance Publique Hôpitaux de Paris, Université Paris-Est Créteil, INSERM U955 | Christophe Rodriguez, Slim Fourati, Vanessa Demontant, Guillaume Gricourt, Melissa N'Debi, Alexandre Soulier, Elisabeth Trawinski, Jean-Michel Pawlotsky |
| EPI_ISL_3391396, EPI_ISL_3391397, EPI_ISL_3391398 | Biogroup Bio Lam-LCD Saint-Denis | Department of Virology, Henri Mondor University Hospital, Assistance Publique Hôpitaux de Paris, Université Paris-Est Créteil, INSERM U955 | Christophe Rodriguez, Slim Fourati, Vanessa Demontant, Guillaume Gricourt, Melissa N'Debi, Alexandre Soulier, Elisabeth Trawinski, Jean-Michel Pawlotsky |
| EPI_ISL_3391399 | DYNABIO UNILABS | Department of Virology, Henri Mondor University Hospital, Assistance Publique Hôpitaux de Paris, Université Paris-Est Créteil, INSERM U955 | Christophe Rodriguez, Slim Fourati, Vanessa Demontant, Guillaume Gricourt, Melissa N'Debi, Alexandre Soulier, Elisabeth Trawinski, Jean-Michel Pawlotsky |
| EPI_ISL_3391400 | BIOMNIS EUROFINS IVRY | Department of Virology, Henri Mondor University Hospital, Assistance Publique Hôpitaux de Paris, Université Paris-Est Créteil, INSERM U955 | Christophe Rodriguez, Slim Fourati, Vanessa Demontant, Guillaume Gricourt, Melissa N'Debi, Alexandre Soulier, Elisabeth Trawinski, Jean-Michel Pawlotsky |
| EPI_ISL_3391401 | CHI VILLENEUVE ST GEORGES | Department of Virology, Henri Mondor University Hospital, Assistance Publique Hôpitaux de Paris, Université Paris-Est Créteil, INSERM U955 | Christophe Rodriguez, Slim Fourati, Vanessa Demontant, Guillaume Gricourt, Melissa N'Debi, Alexandre Soulier, Elisabeth Trawinski, Jean-Michel Pawlotsky |
| EPI_ISL_3391402 | CH.LE RAINCY_MONTFERMEIL | Department of Virology, Henri Mondor University Hospital, Assistance Publique Hôpitaux de Paris, Université Paris-Est Créteil, INSERM U955 | Christophe Rodriguez, Slim Fourati, Vanessa Demontant, Guillaume Gricourt, Melissa N'Debi, Alexandre Soulier, Elisabeth Trawinski, Jean-Michel Pawlotsky |
| EPI_ISL_3391403, EPI_ISL_3391404, EPI_ISL_3391405 | Biogroup Bio Lam-LCD Saint-Denis | Department of Virology, Henri Mondor University Hospital, Assistance Publique Hôpitaux de Paris, Université Paris-Est Créteil, INSERM U955 | Christophe Rodriguez, Slim Fourati, Vanessa Demontant, Guillaume Gricourt, Melissa N'Debi, Alexandre Soulier, Elisabeth Trawinski, Jean-Michel Pawlotsky |
| EPI_ISL_3391406 | DYNABIO UNILABS | Department of Virology, Henri Mondor University Hospital, Assistance Publique Hôpitaux de Paris, Université Paris-Est Créteil, INSERM U955 | Christophe Rodriguez, Slim Fourati, Vanessa Demontant, Guillaume Gricourt, Melissa N'Debi, Alexandre Soulier, Elisabeth Trawinski, Jean-Michel Pawlotsky |
| EPI_ISL_3391407, EPI_ISL_3391408 | BIO67-BIOSPHERE | Department of Virology, Henri Mondor University Hospital, Assistance Publique Hôpitaux de Paris, Université Paris-Est Créteil, INSERM U955 | Christophe Rodriguez, Slim Fourati, Vanessa Demontant, Guillaume Gricourt, Melissa N'Debi, Alexandre Soulier, Elisabeth Trawinski, Jean-Michel Pawlotsky |
| EPI_ISL_3391409, EPI_ISL_3391411 | Biogroup Bio Lam-LCD Saint-Denis | Department of Virology, Henri Mondor University Hospital, Assistance Publique Hôpitaux de Paris, Université Paris-Est Créteil, INSERM U955 | Christophe Rodriguez, Slim Fourati, Vanessa Demontant, Guillaume Gricourt, Melissa N'Debi, Alexandre Soulier, Elisabeth Trawinski, Jean-Michel Pawlotsky |
| EPI_ISL_3391414 | Laboratoire Ana-L | Department of Virology, Henri Mondor University Hospital, Assistance Publique Hôpitaux de Paris, Université Paris-Est Créteil, INSERM U955 | Christophe Rodriguez, Slim Fourati, Vanessa Demontant, Guillaume Gricourt, Melissa N'Debi, Alexandre Soulier, Elisabeth Trawinski, Jean-Michel Pawlotsky |
| EPI_ISL_3391415, EPI_ISL_3391416 | BIOMNIS EUROFINS IVRY | Department of Virology, Henri Mondor University Hospital, Assistance Publique Hôpitaux de Paris, Université Paris-Est Créteil, INSERM U955 | Christophe Rodriguez, Slim Fourati, Vanessa Demontant, Guillaume Gricourt, Melissa N'Debi, Alexandre Soulier, Elisabeth Trawinski, Jean-Michel Pawlotsky |
| EPI_ISL_3391417 | DYNALAB | Department of Virology, Henri Mondor University Hospital, Assistance Publique Hôpitaux de Paris, Université Paris-Est Créteil, INSERM U955 | Christophe Rodriguez, Slim Fourati, Vanessa Demontant, Guillaume Gricourt, Melissa N'Debi, Alexandre Soulier, Elisabeth Trawinski, Jean-Michel Pawlotsky |
| EPI_ISL_3391418, EPI_ISL_3391419 | Laboratoire Ana-L | Department of Virology, Henri Mondor University Hospital, Assistance Publique Hôpitaux de Paris, Université Paris-Est Créteil, INSERM U955 | Christophe Rodriguez, Slim Fourati, Vanessa Demontant, Guillaume Gricourt, Melissa N'Debi, Alexandre Soulier, Elisabeth Trawinski, Jean-Michel Pawlotsky |
| EPI_ISL_3391421, EPI_ISL_3391422 | Biogroup Bio Lam-LCD Saint-Denis | Department of Virology, Henri Mondor University Hospital, Assistance Publique Hôpitaux de Paris, Université Paris-Est Créteil, INSERM U955 | Christophe Rodriguez, Slim Fourati, Vanessa Demontant, Guillaume Gricourt, Melissa N'Debi, Alexandre Soulier, Elisabeth Trawinski, Jean-Michel Pawlotsky |
| EPI_ISL_3391423 | BIOMNIS EUROFINS IVRY | Department of Virology, Henri Mondor University Hospital, | Christophe Rodriguez, Slim Fourati, Vanessa Demontant, Guillaume Gricourt, Melissa N'Debi, Alexandre Soulier, Elisabeth Trawinski, Jean-Michel Pawlotsky |

| | | | |
|---|---|---|---|
| | | Assistance Publique Hôpitaux de Paris, Université Paris-Est Créteil, INSERM U955 | |
| EPI_ISL_3391424 | Laboratoire Ana-L | Department of Virology, Henri Mondor University Hospital, Assistance Publique Hôpitaux de Paris, Université Paris-Est Créteil, INSERM U955 | Christophe Rodriguez, Slim Fourati, Vanessa Demontant, Guillaume Gricourt, Melissa N'Debi, Alexandre Soulier, Elisabeth Trawinski, Jean-Michel Pawlotsky |
| EPI_ISL_3391425, EPI_ISL_3391427 | Biogroup Bio Lam-LCD Saint-Denis | Department of Virology, Henri Mondor University Hospital, Assistance Publique Hôpitaux de Paris, Université Paris-Est Créteil, INSERM U955 | Christophe Rodriguez, Slim Fourati, Vanessa Demontant, Guillaume Gricourt, Melissa N'Debi, Alexandre Soulier, Elisabeth Trawinski, Jean-Michel Pawlotsky |
| EPI_ISL_3391428 | BIO67-BIOSPHERE | Department of Virology, Henri Mondor University Hospital, Assistance Publique Hôpitaux de Paris, Université Paris-Est Créteil, INSERM U955 | Christophe Rodriguez, Slim Fourati, Vanessa Demontant, Guillaume Gricourt, Melissa N'Debi, Alexandre Soulier, Elisabeth Trawinski, Jean-Michel Pawlotsky |
| EPI_ISL_3391429, EPI_ISL_3391430, EPI_ISL_3391431 | Biogroup Bio Lam-LCD Saint-Denis | Department of Virology, Henri Mondor University Hospital, Assistance Publique Hôpitaux de Paris, Université Paris-Est Créteil, INSERM U955 | Christophe Rodriguez, Slim Fourati, Vanessa Demontant, Guillaume Gricourt, Melissa N'Debi, Alexandre Soulier, Elisabeth Trawinski, Jean-Michel Pawlotsky |
| EPI_ISL_3391432 | Laboratoire Ana-L | Department of Virology, Henri Mondor University Hospital, Assistance Publique Hôpitaux de Paris, Université Paris-Est Créteil, INSERM U955 | Christophe Rodriguez, Slim Fourati, Vanessa Demontant, Guillaume Gricourt, Melissa N'Debi, Alexandre Soulier, Elisabeth Trawinski, Jean-Michel Pawlotsky |
| EPI_ISL_3391433 | BIOMNIS EUROFINS IVRY | Department of Virology, Henri Mondor University Hospital, Assistance Publique Hôpitaux de Paris, Université Paris-Est Créteil, INSERM U955 | Christophe Rodriguez, Slim Fourati, Vanessa Demontant, Guillaume Gricourt, Melissa N'Debi, Alexandre Soulier, Elisabeth Trawinski, Jean-Michel Pawlotsky |
| EPI_ISL_3391434, EPI_ISL_3391435 | Biogroup Bio Lam-LCD Saint-Denis | Department of Virology, Henri Mondor University Hospital, Assistance Publique Hôpitaux de Paris, Université Paris-Est Créteil, INSERM U955 | Christophe Rodriguez, Slim Fourati, Vanessa Demontant, Guillaume Gricourt, Melissa N'Debi, Alexandre Soulier, Elisabeth Trawinski, Jean-Michel Pawlotsky |
| EPI_ISL_3391436 | CH.LE RAINCY_MONTFERMEIL | Department of Virology, Henri Mondor University Hospital, Assistance Publique Hôpitaux de Paris, Université Paris-Est Créteil, INSERM U955 | Christophe Rodriguez, Slim Fourati, Vanessa Demontant, Guillaume Gricourt, Melissa N'Debi, Alexandre Soulier, Elisabeth Trawinski, Jean-Michel Pawlotsky |
| EPI_ISL_3391437 | Biogroup Bio Lam-LCD Saint-Denis | Department of Virology, Henri Mondor University Hospital, Assistance Publique Hôpitaux de Paris, Université Paris-Est Créteil, INSERM U955 | Christophe Rodriguez, Slim Fourati, Vanessa Demontant, Guillaume Gricourt, Melissa N'Debi, Alexandre Soulier, Elisabeth Trawinski, Jean-Michel Pawlotsky |
| EPI_ISL_3391438 | DYNABIO UNILABS | Department of Virology, Henri Mondor University Hospital, Assistance Publique Hôpitaux de Paris, Université Paris-Est Créteil, INSERM U955 | Christophe Rodriguez, Slim Fourati, Vanessa Demontant, Guillaume Gricourt, Melissa N'Debi, Alexandre Soulier, Elisabeth Trawinski, Jean-Michel Pawlotsky |
| EPI_ISL_3391439, EPI_ISL_3391440, EPI_ISL_3391441, EPI_ISL_3391442, EPI_ISL_3391443, EPI_ISL_3391446 | Biogroup Bio Lam-LCD Saint-Denis | Department of Virology, Henri Mondor University Hospital, Assistance Publique Hôpitaux de Paris, Université Paris-Est Créteil, INSERM U955 | Christophe Rodriguez, Slim Fourati, Vanessa Demontant, Guillaume Gricourt, Melissa N'Debi, Alexandre Soulier, Elisabeth Trawinski, Jean-Michel Pawlotsky |
| EPI_ISL_3391447 | CH.DIEPPE laboratoire | Department of Virology, Henri Mondor University Hospital, Assistance Publique Hôpitaux de Paris, Université Paris-Est Créteil, INSERM U955 | Christophe Rodriguez, Slim Fourati, Vanessa Demontant, Guillaume Gricourt, Melissa N'Debi, Alexandre Soulier, Elisabeth Trawinski, Jean-Michel Pawlotsky |
| EPI_ISL_3391448 | BIOMNIS EUROFINS IVRY | Department of Virology, Henri Mondor University Hospital, Assistance Publique Hôpitaux de Paris, Université Paris-Est Créteil, INSERM U955 | Christophe Rodriguez, Slim Fourati, Vanessa Demontant, Guillaume Gricourt, Melissa N'Debi, Alexandre Soulier, Elisabeth Trawinski, Jean-Michel Pawlotsky |
| EPI_ISL_3391449 | DYNALAB | Department of Virology, Henri Mondor University Hospital, Assistance Publique Hôpitaux de Paris, Université Paris-Est Créteil, INSERM U955 | Christophe Rodriguez, Slim Fourati, Vanessa Demontant, Guillaume Gricourt, Melissa N'Debi, Alexandre Soulier, Elisabeth Trawinski, Jean-Michel Pawlotsky |
| EPI_ISL_3391450, EPI_ISL_3391451 | Laboratoire Ana-L | Department of Virology, Henri Mondor University Hospital, Assistance Publique Hôpitaux de Paris, Université Paris-Est Créteil, INSERM U955 | Christophe Rodriguez, Slim Fourati, Vanessa Demontant, Guillaume Gricourt, Melissa N'Debi, Alexandre Soulier, Elisabeth Trawinski, Jean-Michel Pawlotsky |
| EPI_ISL_3391452 | BIO67-BIOSPHERE | Department of Virology, Henri Mondor University Hospital, Assistance Publique Hôpitaux de Paris, Université Paris-Est Créteil, INSERM U955 | Christophe Rodriguez, Slim Fourati, Vanessa Demontant, Guillaume Gricourt, Melissa N'Debi, Alexandre Soulier, Elisabeth Trawinski, Jean-Michel Pawlotsky |
| EPI_ISL_3391453 | Biogroup Bio Lam-LCD Saint-Denis | Department of Virology, Henri Mondor University Hospital, Assistance Publique Hôpitaux de Paris, Université Paris-Est Créteil, INSERM U955 | Christophe Rodriguez, Slim Fourati, Vanessa Demontant, Guillaume Gricourt, Melissa N'Debi, Alexandre Soulier, Elisabeth Trawinski, Jean-Michel Pawlotsky |
| EPI_ISL_3391454 | GROUPE HOSPITALIER SUD ILE DE FRANCE | Department of Virology, Henri Mondor University Hospital, Assistance Publique Hôpitaux de Paris, Université Paris-Est Créteil, INSERM U955 | Christophe Rodriguez, Slim Fourati, Vanessa Demontant, Guillaume Gricourt, Melissa N'Debi, Alexandre Soulier, Elisabeth Trawinski, Jean-Michel Pawlotsky |
| EPI_ISL_3391455, EPI_ISL_3391456, EPI_ISL_3391457, EPI_ISL_3391458 | Biogroup Bio Lam-LCD Saint-Denis | Department of Virology, Henri Mondor University Hospital, Assistance Publique Hôpitaux de Paris, Université Paris-Est Créteil, INSERM U955 | Christophe Rodriguez, Slim Fourati, Vanessa Demontant, Guillaume Gricourt, Melissa N'Debi, Alexandre Soulier, Elisabeth Trawinski, Jean-Michel Pawlotsky |
| EPI_ISL_3391459 | BIO67-BIOSPHERE | Department of Virology, Henri Mondor University Hospital, Assistance Publique Hôpitaux de Paris, Université Paris-Est Créteil, INSERM U955 | Christophe Rodriguez, Slim Fourati, Vanessa Demontant, Guillaume Gricourt, Melissa N'Debi, Alexandre Soulier, Elisabeth Trawinski, Jean-Michel Pawlotsky |
| EPI_ISL_3391460 | BIOMNIS EUROFINS IVRY | Department of Virology, Henri Mondor University Hospital, Assistance Publique Hôpitaux de Paris, Université Paris-Est Créteil, INSERM U955 | Christophe Rodriguez, Slim Fourati, Vanessa Demontant, Guillaume Gricourt, Melissa N'Debi, Alexandre Soulier, Elisabeth Trawinski, Jean-Michel Pawlotsky |
| EPI_ISL_3391461, EPI_ISL_3391462 | Biogroup Bio Lam-LCD Saint-Denis | Department of Virology, Henri Mondor University Hospital, Assistance Publique Hôpitaux de Paris, Université Paris-Est Créteil, INSERM U955 | Christophe Rodriguez, Slim Fourati, Vanessa Demontant, Guillaume Gricourt, Melissa N'Debi, Alexandre Soulier, Elisabeth Trawinski, Jean-Michel Pawlotsky |
| EPI_ISL_3391463 | Laboratoire Ana-L | Department of Virology, Henri Mondor University Hospital, Assistance Publique Hôpitaux de Paris, Université Paris-Est Créteil, INSERM U955 | Christophe Rodriguez, Slim Fourati, Vanessa Demontant, Guillaume Gricourt, Melissa N'Debi, Alexandre Soulier, Elisabeth Trawinski, Jean-Michel Pawlotsky |
| EPI_ISL_3391464 | Biogroup Bio Lam-LCD Saint-Denis | Department of Virology, Henri Mondor University Hospital, Assistance Publique Hôpitaux de Paris, Université Paris-Est Créteil, INSERM U955 | Christophe Rodriguez, Slim Fourati, Vanessa Demontant, Guillaume Gricourt, Melissa N'Debi, Alexandre Soulier, Elisabeth Trawinski, Jean-Michel Pawlotsky |
| EPI_ISL_3391465 | BIOMNIS EUROFINS IVRY | Department of Virology, Henri Mondor University Hospital, Assistance Publique Hôpitaux de Paris, Université Paris-Est Créteil, INSERM U955 | Christophe Rodriguez, Slim Fourati, Vanessa Demontant, Guillaume Gricourt, Melissa N'Debi, Alexandre Soulier, Elisabeth Trawinski, Jean-Michel Pawlotsky |
| EPI_ISL_3391466 | BIO67-BIOSPHERE | Department of Virology, Henri Mondor University Hospital, Assistance Publique Hôpitaux de Paris, Université Paris-Est Créteil, INSERM U955 | Christophe Rodriguez, Slim Fourati, Vanessa Demontant, Guillaume Gricourt, Melissa N'Debi, Alexandre Soulier, Elisabeth Trawinski, Jean-Michel Pawlotsky |
| EPI_ISL_3391468 | Biogroup Bio Lam-LCD Saint-Denis | Department of Virology, Henri Mondor University Hospital, Assistance Publique Hôpitaux de Paris, Université Paris-Est Créteil, INSERM U955 | Christophe Rodriguez, Slim Fourati, Vanessa Demontant, Guillaume Gricourt, Melissa N'Debi, Alexandre Soulier, Elisabeth Trawinski, Jean-Michel Pawlotsky |
| EPI_ISL_3391469 | BIO67-BIOSPHERE | Department of Virology, Henri Mondor University Hospital, Assistance Publique Hôpitaux de Paris, Université Paris-Est Créteil, INSERM U955 | Christophe Rodriguez, Slim Fourati, Vanessa Demontant, Guillaume Gricourt, Melissa N'Debi, Alexandre Soulier, Elisabeth Trawinski, Jean-Michel Pawlotsky |
| EPI_ISL_3391470, EPI_ISL_3391471, EPI_ISL_3391472, EPI_ISL_3391473 | Biogroup Bio Lam-LCD Saint-Denis | Department of Virology, Henri Mondor University Hospital, Assistance Publique Hôpitaux de Paris, Université Paris-Est Créteil, INSERM U955 | Christophe Rodriguez, Slim Fourati, Vanessa Demontant, Guillaume Gricourt, Melissa N'Debi, Alexandre Soulier, Elisabeth Trawinski, Jean-Michel Pawlotsky |
| EPI_ISL_3391474 | SELAS BC-LAB | Department of Virology, Henri Mondor University Hospital, Assistance Publique Hôpitaux de Paris, Université Paris-Est Créteil, | Christophe Rodriguez, Slim Fourati, Vanessa Demontant, Guillaume Gricourt, Melissa N'Debi, Alexandre Soulier, Elisabeth Trawinski, Jean-Michel Pawlotsky |

| | | | |
|---|---|---|---|
| | | INSERM U955 | |
| EPI_ISL_3391475 | Laboratoire Ana-L | Department of Virology, Henri Mondor University Hospital, Assistance Publique Hôpitaux de Paris, Université Paris-Est Créteil, INSERM U955 | Christophe Rodriguez, Slim Fourati, Vanessa Demontant, Guillaume Gricourt, Melissa N'Debi, Alexandre Soulier, Elisabeth Trawinski, Jean-Michel Pawlotsky |
| EPI_ISL_3391476 | BIO67-BIOSPHERE | Department of Virology, Henri Mondor University Hospital, Assistance Publique Hôpitaux de Paris, Université Paris-Est Créteil, INSERM U955 | Christophe Rodriguez, Slim Fourati, Vanessa Demontant, Guillaume Gricourt, Melissa N'Debi, Alexandre Soulier, Elisabeth Trawinski, Jean-Michel Pawlotsky |
| EPI_ISL_3391477 | Biogroup Bio Lam-LCD Saint-Denis | Department of Virology, Henri Mondor University Hospital, Assistance Publique Hôpitaux de Paris, Université Paris-Est Créteil, INSERM U955 | Christophe Rodriguez, Slim Fourati, Vanessa Demontant, Guillaume Gricourt, Melissa N'Debi, Alexandre Soulier, Elisabeth Trawinski, Jean-Michel Pawlotsky |
| EPI_ISL_3391478 | BIOMNIS EUROFINS IVRY | Department of Virology, Henri Mondor University Hospital, Assistance Publique Hôpitaux de Paris, Université Paris-Est Créteil, INSERM U955 | Christophe Rodriguez, Slim Fourati, Vanessa Demontant, Guillaume Gricourt, Melissa N'Debi, Alexandre Soulier, Elisabeth Trawinski, Jean-Michel Pawlotsky |
| EPI_ISL_3391479 | Biogroup Bio Lam-LCD Saint-Denis | Department of Virology, Henri Mondor University Hospital, Assistance Publique Hôpitaux de Paris, Université Paris-Est Créteil, INSERM U955 | Christophe Rodriguez, Slim Fourati, Vanessa Demontant, Guillaume Gricourt, Melissa N'Debi, Alexandre Soulier, Elisabeth Trawinski, Jean-Michel Pawlotsky |
| EPI_ISL_3391480 | BIO67-BIOSPHERE | Department of Virology, Henri Mondor University Hospital, Assistance Publique Hôpitaux de Paris, Université Paris-Est Créteil, INSERM U955 | Christophe Rodriguez, Slim Fourati, Vanessa Demontant, Guillaume Gricourt, Melissa N'Debi, Alexandre Soulier, Elisabeth Trawinski, Jean-Michel Pawlotsky |
| EPI_ISL_3391481 | BIOMNIS EUROFINS IVRY | Department of Virology, Henri Mondor University Hospital, Assistance Publique Hôpitaux de Paris, Université Paris-Est Créteil, INSERM U955 | Christophe Rodriguez, Slim Fourati, Vanessa Demontant, Guillaume Gricourt, Melissa N'Debi, Alexandre Soulier, Elisabeth Trawinski, Jean-Michel Pawlotsky |
| EPI_ISL_3391482 | Laboratoire Ana-L | Department of Virology, Henri Mondor University Hospital, Assistance Publique Hôpitaux de Paris, Université Paris-Est Créteil, INSERM U955 | Christophe Rodriguez, Slim Fourati, Vanessa Demontant, Guillaume Gricourt, Melissa N'Debi, Alexandre Soulier, Elisabeth Trawinski, Jean-Michel Pawlotsky |
| EPI_ISL_3391483, EPI_ISL_3391484 | Biogroup Bio Lam-LCD Saint-Denis | Department of Virology, Henri Mondor University Hospital, Assistance Publique Hôpitaux de Paris, Université Paris-Est Créteil, INSERM U955 | Christophe Rodriguez, Slim Fourati, Vanessa Demontant, Guillaume Gricourt, Melissa N'Debi, Alexandre Soulier, Elisabeth Trawinski, Jean-Michel Pawlotsky |
| EPI_ISL_3391486 | DYNALAB | Department of Virology, Henri Mondor University Hospital, Assistance Publique Hôpitaux de Paris, Université Paris-Est Créteil, INSERM U955 | Christophe Rodriguez, Slim Fourati, Vanessa Demontant, Guillaume Gricourt, Melissa N'Debi, Alexandre Soulier, Elisabeth Trawinski, Jean-Michel Pawlotsky |
| EPI_ISL_3391487 | BIOMNIS EUROFINS IVRY | Department of Virology, Henri Mondor University Hospital, Assistance Publique Hôpitaux de Paris, Université Paris-Est Créteil, INSERM U955 | Christophe Rodriguez, Slim Fourati, Vanessa Demontant, Guillaume Gricourt, Melissa N'Debi, Alexandre Soulier, Elisabeth Trawinski, Jean-Michel Pawlotsky |
| EPI_ISL_3391488 | Biogroup Bio Lam-LCD Saint-Denis | Department of Virology, Henri Mondor University Hospital, Assistance Publique Hôpitaux de Paris, Université Paris-Est Créteil, INSERM U955 | Christophe Rodriguez, Slim Fourati, Vanessa Demontant, Guillaume Gricourt, Melissa N'Debi, Alexandre Soulier, Elisabeth Trawinski, Jean-Michel Pawlotsky |
| EPI_ISL_3391489 | BIO67-BIOSPHERE | Department of Virology, Henri Mondor University Hospital, Assistance Publique Hôpitaux de Paris, Université Paris-Est Créteil, INSERM U955 | Christophe Rodriguez, Slim Fourati, Vanessa Demontant, Guillaume Gricourt, Melissa N'Debi, Alexandre Soulier, Elisabeth Trawinski, Jean-Michel Pawlotsky |
| EPI_ISL_3391491 | Biogroup Bio Lam-LCD Saint-Denis | Department of Virology, Henri Mondor University Hospital, Assistance Publique Hôpitaux de Paris, Université Paris-Est Créteil, INSERM U955 | Christophe Rodriguez, Slim Fourati, Vanessa Demontant, Guillaume Gricourt, Melissa N'Debi, Alexandre Soulier, Elisabeth Trawinski, Jean-Michel Pawlotsky |
| EPI_ISL_3391492 | BIO67-BIOSPHERE | Department of Virology, Henri Mondor University Hospital, Assistance Publique Hôpitaux de Paris, Université Paris-Est Créteil, INSERM U955 | Christophe Rodriguez, Slim Fourati, Vanessa Demontant, Guillaume Gricourt, Melissa N'Debi, Alexandre Soulier, Elisabeth Trawinski, Jean-Michel Pawlotsky |
| EPI_ISL_3391493, EPI_ISL_3391494 | Biogroup Bio Lam-LCD Saint-Denis | Department of Virology, Henri Mondor University Hospital, Assistance Publique Hôpitaux de Paris, Université Paris-Est Créteil, INSERM U955 | Christophe Rodriguez, Slim Fourati, Vanessa Demontant, Guillaume Gricourt, Melissa N'Debi, Alexandre Soulier, Elisabeth Trawinski, Jean-Michel Pawlotsky |
| EPI_ISL_3391495 | BIOMNIS EUROFINS IVRY | Department of Virology, Henri Mondor University Hospital, Assistance Publique Hôpitaux de Paris, Université Paris-Est Créteil, INSERM U955 | Christophe Rodriguez, Slim Fourati, Vanessa Demontant, Guillaume Gricourt, Melissa N'Debi, Alexandre Soulier, Elisabeth Trawinski, Jean-Michel Pawlotsky |
| EPI_ISL_3391496, EPI_ISL_3391497 | BIO67-BIOSPHERE | Department of Virology, Henri Mondor University Hospital, Assistance Publique Hôpitaux de Paris, Université Paris-Est Créteil, INSERM U955 | Christophe Rodriguez, Slim Fourati, Vanessa Demontant, Guillaume Gricourt, Melissa N'Debi, Alexandre Soulier, Elisabeth Trawinski, Jean-Michel Pawlotsky |
| EPI_ISL_3391498 | Biogroup Bio Lam-LCD Saint-Denis | Department of Virology, Henri Mondor University Hospital, Assistance Publique Hôpitaux de Paris, Université Paris-Est Créteil, INSERM U955 | Christophe Rodriguez, Slim Fourati, Vanessa Demontant, Guillaume Gricourt, Melissa N'Debi, Alexandre Soulier, Elisabeth Trawinski, Jean-Michel Pawlotsky |
| EPI_ISL_3391499 | Laboratoire Ana-L | Department of Virology, Henri Mondor University Hospital, Assistance Publique Hôpitaux de Paris, Université Paris-Est Créteil, INSERM U955 | Christophe Rodriguez, Slim Fourati, Vanessa Demontant, Guillaume Gricourt, Melissa N'Debi, Alexandre Soulier, Elisabeth Trawinski, Jean-Michel Pawlotsky |
| EPI_ISL_3391500 | Biogroup Bio Lam-LCD Saint-Denis | Department of Virology, Henri Mondor University Hospital, Assistance Publique Hôpitaux de Paris, Université Paris-Est Créteil, INSERM U955 | Christophe Rodriguez, Slim Fourati, Vanessa Demontant, Guillaume Gricourt, Melissa N'Debi, Alexandre Soulier, Elisabeth Trawinski, Jean-Michel Pawlotsky |
| EPI_ISL_3391501 | Laboratoire Ana-L | Department of Virology, Henri Mondor University Hospital, Assistance Publique Hôpitaux de Paris, Université Paris-Est Créteil, INSERM U955 | Christophe Rodriguez, Slim Fourati, Vanessa Demontant, Guillaume Gricourt, Melissa N'Debi, Alexandre Soulier, Elisabeth Trawinski, Jean-Michel Pawlotsky |
| EPI_ISL_3391502 | BIOMNIS EUROFINS IVRY | Department of Virology, Henri Mondor University Hospital, Assistance Publique Hôpitaux de Paris, Université Paris-Est Créteil, INSERM U955 | Christophe Rodriguez, Slim Fourati, Vanessa Demontant, Guillaume Gricourt, Melissa N'Debi, Alexandre Soulier, Elisabeth Trawinski, Jean-Michel Pawlotsky |
| EPI_ISL_3391503 | Biogroup Bio Lam-LCD Saint-Denis | Department of Virology, Henri Mondor University Hospital, Assistance Publique Hôpitaux de Paris, Université Paris-Est Créteil, INSERM U955 | Christophe Rodriguez, Slim Fourati, Vanessa Demontant, Guillaume Gricourt, Melissa N'Debi, Alexandre Soulier, Elisabeth Trawinski, Jean-Michel Pawlotsky |
| EPI_ISL_3391504 | Laboratoire Ana-L | Department of Virology, Henri Mondor University Hospital, Assistance Publique Hôpitaux de Paris, Université Paris-Est Créteil, INSERM U955 | Christophe Rodriguez, Slim Fourati, Vanessa Demontant, Guillaume Gricourt, Melissa N'Debi, Alexandre Soulier, Elisabeth Trawinski, Jean-Michel Pawlotsky |
| EPI_ISL_3391506 | BIO67-BIOSPHERE | Department of Virology, Henri Mondor University Hospital, Assistance Publique Hôpitaux de Paris, Université Paris-Est Créteil, INSERM U955 | Christophe Rodriguez, Slim Fourati, Vanessa Demontant, Guillaume Gricourt, Melissa N'Debi, Alexandre Soulier, Elisabeth Trawinski, Jean-Michel Pawlotsky |
| EPI_ISL_3391508 | Biogroup Bio Lam-LCD Saint-Denis | Department of Virology, Henri Mondor University Hospital, Assistance Publique Hôpitaux de Paris, Université Paris-Est Créteil, INSERM U955 | Christophe Rodriguez, Slim Fourati, Vanessa Demontant, Guillaume Gricourt, Melissa N'Debi, Alexandre Soulier, Elisabeth Trawinski, Jean-Michel Pawlotsky |
| EPI_ISL_3391509 | Laboratoire Ana-L | Department of Virology, Henri Mondor University Hospital, Assistance Publique Hôpitaux de Paris, Université Paris-Est Créteil, INSERM U955 | Christophe Rodriguez, Slim Fourati, Vanessa Demontant, Guillaume Gricourt, Melissa N'Debi, Alexandre Soulier, Elisabeth Trawinski, Jean-Michel Pawlotsky |
| EPI_ISL_3391510 | Biogroup Bio Lam-LCD Saint-Denis | Department of Virology, Henri Mondor University Hospital, Assistance Publique Hôpitaux de Paris, Université Paris-Est Créteil, INSERM U955 | Christophe Rodriguez, Slim Fourati, Vanessa Demontant, Guillaume Gricourt, Melissa N'Debi, Alexandre Soulier, Elisabeth Trawinski, Jean-Michel Pawlotsky |
| EPI_ISL_3391511 | BIO67-BIOSPHERE | Department of Virology, Henri Mondor University Hospital, Assistance Publique Hôpitaux de Paris, Université Paris-Est Créteil, INSERM U955 | Christophe Rodriguez, Slim Fourati, Vanessa Demontant, Guillaume Gricourt, Melissa N'Debi, Alexandre Soulier, Elisabeth Trawinski, Jean-Michel Pawlotsky |
| EPI_ISL_3391512 | BIOMNIS EUROFINS IVRY | Department of Virology, Henri Mondor University Hospital, Assistance Publique Hôpitaux de Paris, Université Paris-Est Créteil, INSERM U955 | Christophe Rodriguez, Slim Fourati, Vanessa Demontant, Guillaume Gricourt, Melissa N'Debi, Alexandre Soulier, Elisabeth Trawinski, Jean-Michel Pawlotsky |
| EPI_ISL_3391513 | Laboratoire Ana-L | Department of Virology, Henri Mondor University Hospital, | Christophe Rodriguez, Slim Fourati, Vanessa Demontant, Guillaume Gricourt, Melissa N'Debi, Alexandre Soulier, Elisabeth Trawinski, Jean-Michel Pawlotsky |

| | | | |
|---|---|---|---|
| | | Assistance Publique Hôpitaux de Paris, Université Paris-Est Créteil, INSERM U955 | |
| EPI_ISL_3391514 | BIOMNIS EUROFINS IVRY | Department of Virology, Henri Mondor University Hospital, Assistance Publique Hôpitaux de Paris, Université Paris-Est Créteil, INSERM U955 | Christophe Rodriguez, Slim Fourati, Vanessa Demontant, Guillaume Gricourt, Melissa N'Debi, Alexandre Soulier, Elisabeth Trawinski, Jean-Michel Pawlotsky |
| EPI_ISL_3391515, EPI_ISL_3391516 | Biogroup Bio Lam-LCD Saint-Denis | Department of Virology, Henri Mondor University Hospital, Assistance Publique Hôpitaux de Paris, Université Paris-Est Créteil, INSERM U955 | Christophe Rodriguez, Slim Fourati, Vanessa Demontant, Guillaume Gricourt, Melissa N'Debi, Alexandre Soulier, Elisabeth Trawinski, Jean-Michel Pawlotsky |
| EPI_ISL_3391517, EPI_ISL_3391518 | BIOMNIS EUROFINS IVRY | Department of Virology, Henri Mondor University Hospital, Assistance Publique Hôpitaux de Paris, Université Paris-Est Créteil, INSERM U955 | Christophe Rodriguez, Slim Fourati, Vanessa Demontant, Guillaume Gricourt, Melissa N'Debi, Alexandre Soulier, Elisabeth Trawinski, Jean-Michel Pawlotsky |
| EPI_ISL_3391519 | CH.LE RAINCY_MONTFERMEIL | Department of Virology, Henri Mondor University Hospital, Assistance Publique Hôpitaux de Paris, Université Paris-Est Créteil, INSERM U955 | Christophe Rodriguez, Slim Fourati, Vanessa Demontant, Guillaume Gricourt, Melissa N'Debi, Alexandre Soulier, Elisabeth Trawinski, Jean-Michel Pawlotsky |
| EPI_ISL_3391520 | SELAS BC-LAB | Department of Virology, Henri Mondor University Hospital, Assistance Publique Hôpitaux de Paris, Université Paris-Est Créteil, INSERM U955 | Christophe Rodriguez, Slim Fourati, Vanessa Demontant, Guillaume Gricourt, Melissa N'Debi, Alexandre Soulier, Elisabeth Trawinski, Jean-Michel Pawlotsky |
| EPI_ISL_3391522 | BIOMNIS EUROFINS IVRY | Department of Virology, Henri Mondor University Hospital, Assistance Publique Hôpitaux de Paris, Université Paris-Est Créteil, INSERM U955 | Christophe Rodriguez, Slim Fourati, Vanessa Demontant, Guillaume Gricourt, Melissa N'Debi, Alexandre Soulier, Elisabeth Trawinski, Jean-Michel Pawlotsky |
| EPI_ISL_3391523 | Biogroup Bio Lam-LCD Saint-Denis | Department of Virology, Henri Mondor University Hospital, Assistance Publique Hôpitaux de Paris, Université Paris-Est Créteil, INSERM U955 | Christophe Rodriguez, Slim Fourati, Vanessa Demontant, Guillaume Gricourt, Melissa N'Debi, Alexandre Soulier, Elisabeth Trawinski, Jean-Michel Pawlotsky |
| EPI_ISL_3391524 | SELAS BC-LAB | Department of Virology, Henri Mondor University Hospital, Assistance Publique Hôpitaux de Paris, Université Paris-Est Créteil, INSERM U955 | Christophe Rodriguez, Slim Fourati, Vanessa Demontant, Guillaume Gricourt, Melissa N'Debi, Alexandre Soulier, Elisabeth Trawinski, Jean-Michel Pawlotsky |
| EPI_ISL_3391525 | Biogroup Bio Lam-LCD Saint-Denis | Department of Virology, Henri Mondor University Hospital, Assistance Publique Hôpitaux de Paris, Université Paris-Est Créteil, INSERM U955 | Christophe Rodriguez, Slim Fourati, Vanessa Demontant, Guillaume Gricourt, Melissa N'Debi, Alexandre Soulier, Elisabeth Trawinski, Jean-Michel Pawlotsky |
| EPI_ISL_3391526 | Laboratoire Ana-L | Department of Virology, Henri Mondor University Hospital, Assistance Publique Hôpitaux de Paris, Université Paris-Est Créteil, INSERM U955 | Christophe Rodriguez, Slim Fourati, Vanessa Demontant, Guillaume Gricourt, Melissa N'Debi, Alexandre Soulier, Elisabeth Trawinski, Jean-Michel Pawlotsky |
| EPI_ISL_3391527 | BIO67-BIOSPHERE | Department of Virology, Henri Mondor University Hospital, Assistance Publique Hôpitaux de Paris, Université Paris-Est Créteil, INSERM U955 | Christophe Rodriguez, Slim Fourati, Vanessa Demontant, Guillaume Gricourt, Melissa N'Debi, Alexandre Soulier, Elisabeth Trawinski, Jean-Michel Pawlotsky |
| EPI_ISL_3391528 | Biogroup Bio Lam-LCD Saint-Denis | Department of Virology, Henri Mondor University Hospital, Assistance Publique Hôpitaux de Paris, Université Paris-Est Créteil, INSERM U955 | Christophe Rodriguez, Slim Fourati, Vanessa Demontant, Guillaume Gricourt, Melissa N'Debi, Alexandre Soulier, Elisabeth Trawinski, Jean-Michel Pawlotsky |
| EPI_ISL_3391529 | BIO67-BIOSPHERE | Department of Virology, Henri Mondor University Hospital, Assistance Publique Hôpitaux de Paris, Université Paris-Est Créteil, INSERM U955 | Christophe Rodriguez, Slim Fourati, Vanessa Demontant, Guillaume Gricourt, Melissa N'Debi, Alexandre Soulier, Elisabeth Trawinski, Jean-Michel Pawlotsky |
| EPI_ISL_3391530, EPI_ISL_3391531, EPI_ISL_3391532, EPI_ISL_3391533 | Biogroup Bio Lam-LCD Saint-Denis | Department of Virology, Henri Mondor University Hospital, Assistance Publique Hôpitaux de Paris, Université Paris-Est Créteil, INSERM U955 | Christophe Rodriguez, Slim Fourati, Vanessa Demontant, Guillaume Gricourt, Melissa N'Debi, Alexandre Soulier, Elisabeth Trawinski, Jean-Michel Pawlotsky |
| EPI_ISL_3391534 | BIO67-BIOSPHERE | Department of Virology, Henri Mondor University Hospital, Assistance Publique Hôpitaux de Paris, Université Paris-Est Créteil, INSERM U955 | Christophe Rodriguez, Slim Fourati, Vanessa Demontant, Guillaume Gricourt, Melissa N'Debi, Alexandre Soulier, Elisabeth Trawinski, Jean-Michel Pawlotsky |
| EPI_ISL_3391537 | Biogroup Bio Lam-LCD Saint-Denis | Department of Virology, Henri Mondor University Hospital, Assistance Publique Hôpitaux de Paris, Université Paris-Est Créteil, INSERM U955 | Christophe Rodriguez, Slim Fourati, Vanessa Demontant, Guillaume Gricourt, Melissa N'Debi, Alexandre Soulier, Elisabeth Trawinski, Jean-Michel Pawlotsky |
| EPI_ISL_3391538 | BIOMNIS EUROFINS IVRY | Department of Virology, Henri Mondor University Hospital, Assistance Publique Hôpitaux de Paris, Université Paris-Est Créteil, INSERM U955 | Christophe Rodriguez, Slim Fourati, Vanessa Demontant, Guillaume Gricourt, Melissa N'Debi, Alexandre Soulier, Elisabeth Trawinski, Jean-Michel Pawlotsky |
| EPI_ISL_3391539 | Laboratoire Ana-L | Department of Virology, Henri Mondor University Hospital, Assistance Publique Hôpitaux de Paris, Université Paris-Est Créteil, INSERM U955 | Christophe Rodriguez, Slim Fourati, Vanessa Demontant, Guillaume Gricourt, Melissa N'Debi, Alexandre Soulier, Elisabeth Trawinski, Jean-Michel Pawlotsky |
| EPI_ISL_3391542 | CH.LE RAINCY_MONTFERMEIL | Department of Virology, Henri Mondor University Hospital, Assistance Publique Hôpitaux de Paris, Université Paris-Est Créteil, INSERM U955 | Christophe Rodriguez, Slim Fourati, Vanessa Demontant, Guillaume Gricourt, Melissa N'Debi, Alexandre Soulier, Elisabeth Trawinski, Jean-Michel Pawlotsky |
| EPI_ISL_3391543 | BIOMNIS EUROFINS IVRY | Department of Virology, Henri Mondor University Hospital, Assistance Publique Hôpitaux de Paris, Université Paris-Est Créteil, INSERM U955 | Christophe Rodriguez, Slim Fourati, Vanessa Demontant, Guillaume Gricourt, Melissa N'Debi, Alexandre Soulier, Elisabeth Trawinski, Jean-Michel Pawlotsky |
| EPI_ISL_3391544 | BIO VSM LAB | Department of Virology, Henri Mondor University Hospital, Assistance Publique Hôpitaux de Paris, Université Paris-Est Créteil, INSERM U955 | Christophe Rodriguez, Slim Fourati, Vanessa Demontant, Guillaume Gricourt, Melissa N'Debi, Alexandre Soulier, Elisabeth Trawinski, Jean-Michel Pawlotsky |
| EPI_ISL_3391545 | BIOMNIS EUROFINS IVRY | Department of Virology, Henri Mondor University Hospital, Assistance Publique Hôpitaux de Paris, Université Paris-Est Créteil, INSERM U955 | Christophe Rodriguez, Slim Fourati, Vanessa Demontant, Guillaume Gricourt, Melissa N'Debi, Alexandre Soulier, Elisabeth Trawinski, Jean-Michel Pawlotsky |
| EPI_ISL_3391546, EPI_ISL_3391547 | Laboratoire Ana-L | Department of Virology, Henri Mondor University Hospital, Assistance Publique Hôpitaux de Paris, Université Paris-Est Créteil, INSERM U955 | Christophe Rodriguez, Slim Fourati, Vanessa Demontant, Guillaume Gricourt, Melissa N'Debi, Alexandre Soulier, Elisabeth Trawinski, Jean-Michel Pawlotsky |
| EPI_ISL_3391548 | CHI VILLENEUVE ST GEORGES | Department of Virology, Henri Mondor University Hospital, Assistance Publique Hôpitaux de Paris, Université Paris-Est Créteil, INSERM U955 | Christophe Rodriguez, Slim Fourati, Vanessa Demontant, Guillaume Gricourt, Melissa N'Debi, Alexandre Soulier, Elisabeth Trawinski, Jean-Michel Pawlotsky |
| EPI_ISL_3391549, EPI_ISL_3391550 | Biogroup Bio Lam-LCD Saint-Denis | Department of Virology, Henri Mondor University Hospital, Assistance Publique Hôpitaux de Paris, Université Paris-Est Créteil, INSERM U955 | Christophe Rodriguez, Slim Fourati, Vanessa Demontant, Guillaume Gricourt, Melissa N'Debi, Alexandre Soulier, Elisabeth Trawinski, Jean-Michel Pawlotsky |
| EPI_ISL_3391551 | Laboratoire Bioconcorde | Department of Virology, Henri Mondor University Hospital, Assistance Publique Hôpitaux de Paris, Université Paris-Est Créteil, INSERM U955 | Christophe Rodriguez, Slim Fourati, Vanessa Demontant, Guillaume Gricourt, Melissa N'Debi, Alexandre Soulier, Elisabeth Trawinski, Jean-Michel Pawlotsky |
| EPI_ISL_3391552 | CH.LE RAINCY_MONTFERMEIL | Department of Virology, Henri Mondor University Hospital, Assistance Publique Hôpitaux de Paris, Université Paris-Est Créteil, INSERM U955 | Christophe Rodriguez, Slim Fourati, Vanessa Demontant, Guillaume Gricourt, Melissa N'Debi, Alexandre Soulier, Elisabeth Trawinski, Jean-Michel Pawlotsky |
| EPI_ISL_3391555, EPI_ISL_3391556, EPI_ISL_3391557 | Biogroup Bio Lam-LCD Saint-Denis | Department of Virology, Henri Mondor University Hospital, Assistance Publique Hôpitaux de Paris, Université Paris-Est Créteil, INSERM U955 | Christophe Rodriguez, Slim Fourati, Vanessa Demontant, Guillaume Gricourt, Melissa N'Debi, Alexandre Soulier, Elisabeth Trawinski, Jean-Michel Pawlotsky |
| EPI_ISL_3391558 | BIOMNIS EUROFINS IVRY | Department of Virology, Henri Mondor University Hospital, Assistance Publique Hôpitaux de Paris, Université Paris-Est Créteil, INSERM U955 | Christophe Rodriguez, Slim Fourati, Vanessa Demontant, Guillaume Gricourt, Melissa N'Debi, Alexandre Soulier, Elisabeth Trawinski, Jean-Michel Pawlotsky |
| EPI_ISL_3391559 | Biogroup Bio Lam-LCD Saint-Denis | Department of Virology, Henri Mondor University Hospital, Assistance Publique Hôpitaux de Paris, Université Paris-Est Créteil, INSERM U955 | Christophe Rodriguez, Slim Fourati, Vanessa Demontant, Guillaume Gricourt, Melissa N'Debi, Alexandre Soulier, Elisabeth Trawinski, Jean-Michel Pawlotsky |
| EPI_ISL_3391560 | BIOMNIS EUROFINS IVRY | Department of Virology, Henri Mondor University Hospital, Assistance Publique Hôpitaux de Paris, Université Paris-Est Créteil, INSERM U955 | Christophe Rodriguez, Slim Fourati, Vanessa Demontant, Guillaume Gricourt, Melissa N'Debi, Alexandre Soulier, Elisabeth Trawinski, Jean-Michel Pawlotsky |

| | | | |
|---|---|---|---|
| EPI_ISL_3391561 | Biogroup Bio Lam-LCD Saint-Denis | Department of Virology, Henri Mondor University Hospital, Assistance Publique Hôpitaux de Paris, Université Paris-Est Créteil, INSERM U955 | Christophe Rodriguez, Slim Fourati, Vanessa Demontant, Guillaume Gricourt, Melissa N'Debi, Alexandre Soulier, Elisabeth Trawinski, Jean-Michel Pawlotsky |
| EPI_ISL_3391562 | BIOMNIS EUROFINS IVRY | Department of Virology, Henri Mondor University Hospital, Assistance Publique Hôpitaux de Paris, Université Paris-Est Créteil, INSERM U955 | Christophe Rodriguez, Slim Fourati, Vanessa Demontant, Guillaume Gricourt, Melissa N'Debi, Alexandre Soulier, Elisabeth Trawinski, Jean-Michel Pawlotsky |
| EPI_ISL_3391563 | Laboratoire Ana-L | Department of Virology, Henri Mondor University Hospital, Assistance Publique Hôpitaux de Paris, Université Paris-Est Créteil, INSERM U955 | Christophe Rodriguez, Slim Fourati, Vanessa Demontant, Guillaume Gricourt, Melissa N'Debi, Alexandre Soulier, Elisabeth Trawinski, Jean-Michel Pawlotsky |
| EPI_ISL_3391564 | CH.LE RAINCY_MONTFERMEIL | Department of Virology, Henri Mondor University Hospital, Assistance Publique Hôpitaux de Paris, Université Paris-Est Créteil, INSERM U955 | Christophe Rodriguez, Slim Fourati, Vanessa Demontant, Guillaume Gricourt, Melissa N'Debi, Alexandre Soulier, Elisabeth Trawinski, Jean-Michel Pawlotsky |
| EPI_ISL_3391565 | Laboratoire Ana-L | Department of Virology, Henri Mondor University Hospital, Assistance Publique Hôpitaux de Paris, Université Paris-Est Créteil, INSERM U955 | Christophe Rodriguez, Slim Fourati, Vanessa Demontant, Guillaume Gricourt, Melissa N'Debi, Alexandre Soulier, Elisabeth Trawinski, Jean-Michel Pawlotsky |
| EPI_ISL_3391566 | Biogroup Bio Lam-LCD Saint-Denis | Department of Virology, Henri Mondor University Hospital, Assistance Publique Hôpitaux de Paris, Université Paris-Est Créteil, INSERM U955 | Christophe Rodriguez, Slim Fourati, Vanessa Demontant, Guillaume Gricourt, Melissa N'Debi, Alexandre Soulier, Elisabeth Trawinski, Jean-Michel Pawlotsky |
| EPI_ISL_3391567, EPI_ISL_3391568 | Laboratoire Ana-L | Department of Virology, Henri Mondor University Hospital, Assistance Publique Hôpitaux de Paris, Université Paris-Est Créteil, INSERM U955 | Christophe Rodriguez, Slim Fourati, Vanessa Demontant, Guillaume Gricourt, Melissa N'Debi, Alexandre Soulier, Elisabeth Trawinski, Jean-Michel Pawlotsky |
| EPI_ISL_3391569, EPI_ISL_3391570, EPI_ISL_3391571 | Biogroup Bio Lam-LCD Saint-Denis | Department of Virology, Henri Mondor University Hospital, Assistance Publique Hôpitaux de Paris, Université Paris-Est Créteil, INSERM U955 | Christophe Rodriguez, Slim Fourati, Vanessa Demontant, Guillaume Gricourt, Melissa N'Debi, Alexandre Soulier, Elisabeth Trawinski, Jean-Michel Pawlotsky |
| EPI_ISL_3391572, EPI_ISL_3391573 | Laboratoire Ana-L | Department of Virology, Henri Mondor University Hospital, Assistance Publique Hôpitaux de Paris, Université Paris-Est Créteil, INSERM U955 | Christophe Rodriguez, Slim Fourati, Vanessa Demontant, Guillaume Gricourt, Melissa N'Debi, Alexandre Soulier, Elisabeth Trawinski, Jean-Michel Pawlotsky |
| EPI_ISL_3391574, EPI_ISL_3391575, EPI_ISL_3391576 | Biogroup Bio Lam-LCD Saint-Denis | Department of Virology, Henri Mondor University Hospital, Assistance Publique Hôpitaux de Paris, Université Paris-Est Créteil, INSERM U955 | Christophe Rodriguez, Slim Fourati, Vanessa Demontant, Guillaume Gricourt, Melissa N'Debi, Alexandre Soulier, Elisabeth Trawinski, Jean-Michel Pawlotsky |
| EPI_ISL_3391577 | BIOMNIS EUROFINS IVRY | Department of Virology, Henri Mondor University Hospital, Assistance Publique Hôpitaux de Paris, Université Paris-Est Créteil, INSERM U955 | Christophe Rodriguez, Slim Fourati, Vanessa Demontant, Guillaume Gricourt, Melissa N'Debi, Alexandre Soulier, Elisabeth Trawinski, Jean-Michel Pawlotsky |
| EPI_ISL_3391578, EPI_ISL_3391579 | Biogroup Bio Lam-LCD Saint-Denis | Department of Virology, Henri Mondor University Hospital, Assistance Publique Hôpitaux de Paris, Université Paris-Est Créteil, INSERM U955 | Christophe Rodriguez, Slim Fourati, Vanessa Demontant, Guillaume Gricourt, Melissa N'Debi, Alexandre Soulier, Elisabeth Trawinski, Jean-Michel Pawlotsky |
| EPI_ISL_3391580 | DYNALAB | Department of Virology, Henri Mondor University Hospital, Assistance Publique Hôpitaux de Paris, Université Paris-Est Créteil, INSERM U955 | Christophe Rodriguez, Slim Fourati, Vanessa Demontant, Guillaume Gricourt, Melissa N'Debi, Alexandre Soulier, Elisabeth Trawinski, Jean-Michel Pawlotsky |
| EPI_ISL_3391581 | BIO VSM LAB | Department of Virology, Henri Mondor University Hospital, Assistance Publique Hôpitaux de Paris, Université Paris-Est Créteil, INSERM U955 | Christophe Rodriguez, Slim Fourati, Vanessa Demontant, Guillaume Gricourt, Melissa N'Debi, Alexandre Soulier, Elisabeth Trawinski, Jean-Michel Pawlotsky |
| EPI_ISL_3391582 | Biogroup Bio Lam-LCD Saint-Denis | Department of Virology, Henri Mondor University Hospital, Assistance Publique Hôpitaux de Paris, Université Paris-Est Créteil, INSERM U955 | Christophe Rodriguez, Slim Fourati, Vanessa Demontant, Guillaume Gricourt, Melissa N'Debi, Alexandre Soulier, Elisabeth Trawinski, Jean-Michel Pawlotsky |
| EPI_ISL_3391583 | BIOMNIS EUROFINS IVRY | Department of Virology, Henri Mondor University Hospital, Assistance Publique Hôpitaux de Paris, Université Paris-Est Créteil, INSERM U955 | Christophe Rodriguez, Slim Fourati, Vanessa Demontant, Guillaume Gricourt, Melissa N'Debi, Alexandre Soulier, Elisabeth Trawinski, Jean-Michel Pawlotsky |
| EPI_ISL_3391584 | Biogroup Bio Lam-LCD Saint-Denis | Department of Virology, Henri Mondor University Hospital, Assistance Publique Hôpitaux de Paris, Université Paris-Est Créteil, INSERM U955 | Christophe Rodriguez, Slim Fourati, Vanessa Demontant, Guillaume Gricourt, Melissa N'Debi, Alexandre Soulier, Elisabeth Trawinski, Jean-Michel Pawlotsky |
| EPI_ISL_3391585, EPI_ISL_3391586 | Laboratoire Ana-L | Department of Virology, Henri Mondor University Hospital, Assistance Publique Hôpitaux de Paris, Université Paris-Est Créteil, INSERM U955 | Christophe Rodriguez, Slim Fourati, Vanessa Demontant, Guillaume Gricourt, Melissa N'Debi, Alexandre Soulier, Elisabeth Trawinski, Jean-Michel Pawlotsky |
| EPI_ISL_3391587, EPI_ISL_3391588, EPI_ISL_3391589, EPI_ISL_3391590 | Biogroup Bio Lam-LCD Saint-Denis | Department of Virology, Henri Mondor University Hospital, Assistance Publique Hôpitaux de Paris, Université Paris-Est Créteil, INSERM U955 | Christophe Rodriguez, Slim Fourati, Vanessa Demontant, Guillaume Gricourt, Melissa N'Debi, Alexandre Soulier, Elisabeth Trawinski, Jean-Michel Pawlotsky |
| EPI_ISL_3391591 | SELAS BC-LAB | Department of Virology, Henri Mondor University Hospital, Assistance Publique Hôpitaux de Paris, Université Paris-Est Créteil, INSERM U955 | Christophe Rodriguez, Slim Fourati, Vanessa Demontant, Guillaume Gricourt, Melissa N'Debi, Alexandre Soulier, Elisabeth Trawinski, Jean-Michel Pawlotsky |
| EPI_ISL_3391592 | CH.LE RAINCY_MONTFERMEIL | Department of Virology, Henri Mondor University Hospital, Assistance Publique Hôpitaux de Paris, Université Paris-Est Créteil, INSERM U955 | Christophe Rodriguez, Slim Fourati, Vanessa Demontant, Guillaume Gricourt, Melissa N'Debi, Alexandre Soulier, Elisabeth Trawinski, Jean-Michel Pawlotsky |
| EPI_ISL_3391594 | Biogroup Bio Lam-LCD Saint-Denis | Department of Virology, Henri Mondor University Hospital, Assistance Publique Hôpitaux de Paris, Université Paris-Est Créteil, INSERM U955 | Christophe Rodriguez, Slim Fourati, Vanessa Demontant, Guillaume Gricourt, Melissa N'Debi, Alexandre Soulier, Elisabeth Trawinski, Jean-Michel Pawlotsky |
| EPI_ISL_3391595 | BIO67-BIOSPHERE | Department of Virology, Henri Mondor University Hospital, Assistance Publique Hôpitaux de Paris, Université Paris-Est Créteil, INSERM U955 | Christophe Rodriguez, Slim Fourati, Vanessa Demontant, Guillaume Gricourt, Melissa N'Debi, Alexandre Soulier, Elisabeth Trawinski, Jean-Michel Pawlotsky |
| EPI_ISL_3391596 | Biogroup Bio Lam-LCD Saint-Denis | Department of Virology, Henri Mondor University Hospital, Assistance Publique Hôpitaux de Paris, Université Paris-Est Créteil, INSERM U955 | Christophe Rodriguez, Slim Fourati, Vanessa Demontant, Guillaume Gricourt, Melissa N'Debi, Alexandre Soulier, Elisabeth Trawinski, Jean-Michel Pawlotsky |
| EPI_ISL_3391597, EPI_ISL_3391598 | BIOMNIS EUROFINS IVRY | Department of Virology, Henri Mondor University Hospital, Assistance Publique Hôpitaux de Paris, Université Paris-Est Créteil, INSERM U955 | Christophe Rodriguez, Slim Fourati, Vanessa Demontant, Guillaume Gricourt, Melissa N'Debi, Alexandre Soulier, Elisabeth Trawinski, Jean-Michel Pawlotsky |
| EPI_ISL_3391599, EPI_ISL_3391600, EPI_ISL_3391601 | Biogroup Bio Lam-LCD Saint-Denis | Department of Virology, Henri Mondor University Hospital, Assistance Publique Hôpitaux de Paris, Université Paris-Est Créteil, INSERM U955 | Christophe Rodriguez, Slim Fourati, Vanessa Demontant, Guillaume Gricourt, Melissa N'Debi, Alexandre Soulier, Elisabeth Trawinski, Jean-Michel Pawlotsky |
| EPI_ISL_3391602 | BIOMNIS EUROFINS IVRY | Department of Virology, Henri Mondor University Hospital, Assistance Publique Hôpitaux de Paris, Université Paris-Est Créteil, INSERM U955 | Christophe Rodriguez, Slim Fourati, Vanessa Demontant, Guillaume Gricourt, Melissa N'Debi, Alexandre Soulier, Elisabeth Trawinski, Jean-Michel Pawlotsky |
| EPI_ISL_3391603 | Biogroup Bio Lam-LCD Saint-Denis | Department of Virology, Henri Mondor University Hospital, Assistance Publique Hôpitaux de Paris, Université Paris-Est Créteil, INSERM U955 | Christophe Rodriguez, Slim Fourati, Vanessa Demontant, Guillaume Gricourt, Melissa N'Debi, Alexandre Soulier, Elisabeth Trawinski, Jean-Michel Pawlotsky |
| EPI_ISL_3391604 | Laboratoire Ana-L | Department of Virology, Henri Mondor University Hospital, Assistance Publique Hôpitaux de Paris, Université Paris-Est Créteil, INSERM U955 | Christophe Rodriguez, Slim Fourati, Vanessa Demontant, Guillaume Gricourt, Melissa N'Debi, Alexandre Soulier, Elisabeth Trawinski, Jean-Michel Pawlotsky |
| EPI_ISL_3391605, EPI_ISL_3391606, EPI_ISL_3391607, EPI_ISL_3391608 | Biogroup Bio Lam-LCD Saint-Denis | Department of Virology, Henri Mondor University Hospital, Assistance Publique Hôpitaux de Paris, Université Paris-Est Créteil, INSERM U955 | Christophe Rodriguez, Slim Fourati, Vanessa Demontant, Guillaume Gricourt, Melissa N'Debi, Alexandre Soulier, Elisabeth Trawinski, Jean-Michel Pawlotsky |
| EPI_ISL_3391609 | BIOMNIS EUROFINS IVRY | Department of Virology, Henri Mondor University Hospital, Assistance Publique Hôpitaux de Paris, Université Paris-Est Créteil, INSERM U955 | Christophe Rodriguez, Slim Fourati, Vanessa Demontant, Guillaume Gricourt, Melissa N'Debi, Alexandre Soulier, Elisabeth Trawinski, Jean-Michel Pawlotsky |
| EPI_ISL_3391610 | Biogroup Bio Lam-LCD Saint-Denis | Department of Virology, Henri Mondor University Hospital, Assistance Publique Hôpitaux de Paris, Université Paris-Est Créteil, | Christophe Rodriguez, Slim Fourati, Vanessa Demontant, Guillaume Gricourt, Melissa N'Debi, Alexandre Soulier, Elisabeth Trawinski, Jean-Michel Pawlotsky |

| | | | |
|---|---|---|---|
| | | INSERM U955 | |
| EPI_ISL_3391611 | Laboratoire Ana-L | Department of Virology, Henri Mondor University Hospital, Assistance Publique Hôpitaux de Paris, Université Paris-Est Créteil, INSERM U955 | Christophe Rodriguez, Slim Fourati, Vanessa Demontant, Guillaume Gricourt, Melissa N'Debi, Alexandre Soulier, Elisabeth Trawinski, Jean-Michel Pawlotsky |
| EPI_ISL_3391612, EPI_ISL_3391613, EPI_ISL_3391614, EPI_ISL_3391615, EPI_ISL_3391616, EPI_ISL_3391617 | Biogroup Bio Lam-LCD Saint-Denis | Department of Virology, Henri Mondor University Hospital, Assistance Publique Hôpitaux de Paris, Université Paris-Est Créteil, INSERM U955 | Christophe Rodriguez, Slim Fourati, Vanessa Demontant, Guillaume Gricourt, Melissa N'Debi, Alexandre Soulier, Elisabeth Trawinski, Jean-Michel Pawlotsky |
| EPI_ISL_3391618 | BIO67-BIOSPHERE | Department of Virology, Henri Mondor University Hospital, Assistance Publique Hôpitaux de Paris, Université Paris-Est Créteil, INSERM U955 | Christophe Rodriguez, Slim Fourati, Vanessa Demontant, Guillaume Gricourt, Melissa N'Debi, Alexandre Soulier, Elisabeth Trawinski, Jean-Michel Pawlotsky |
| EPI_ISL_3391619, EPI_ISL_3391620, EPI_ISL_3391621, EPI_ISL_3391622, EPI_ISL_3391623, EPI_ISL_3391624, EPI_ISL_3391625 | Biogroup Bio Lam-LCD Saint-Denis | Department of Virology, Henri Mondor University Hospital, Assistance Publique Hôpitaux de Paris, Université Paris-Est Créteil, INSERM U955 | Christophe Rodriguez, Slim Fourati, Vanessa Demontant, Guillaume Gricourt, Melissa N'Debi, Alexandre Soulier, Elisabeth Trawinski, Jean-Michel Pawlotsky |
| EPI_ISL_3391626 | Laboratoire Ana-L | Department of Virology, Henri Mondor University Hospital, Assistance Publique Hôpitaux de Paris, Université Paris-Est Créteil, INSERM U955 | Christophe Rodriguez, Slim Fourati, Vanessa Demontant, Guillaume Gricourt, Melissa N'Debi, Alexandre Soulier, Elisabeth Trawinski, Jean-Michel Pawlotsky |
| EPI_ISL_3391628 | SELAS BC-LAB | Department of Virology, Henri Mondor University Hospital, Assistance Publique Hôpitaux de Paris, Université Paris-Est Créteil, INSERM U955 | Christophe Rodriguez, Slim Fourati, Vanessa Demontant, Guillaume Gricourt, Melissa N'Debi, Alexandre Soulier, Elisabeth Trawinski, Jean-Michel Pawlotsky |
| EPI_ISL_3391629, EPI_ISL_3391630 | Biogroup Bio Lam-LCD Saint-Denis | Department of Virology, Henri Mondor University Hospital, Assistance Publique Hôpitaux de Paris, Université Paris-Est Créteil, INSERM U955 | Christophe Rodriguez, Slim Fourati, Vanessa Demontant, Guillaume Gricourt, Melissa N'Debi, Alexandre Soulier, Elisabeth Trawinski, Jean-Michel Pawlotsky |
| EPI_ISL_3391631 | BIOMNIS EUROFINS IVRY | Department of Virology, Henri Mondor University Hospital, Assistance Publique Hôpitaux de Paris, Université Paris-Est Créteil, INSERM U955 | Christophe Rodriguez, Slim Fourati, Vanessa Demontant, Guillaume Gricourt, Melissa N'Debi, Alexandre Soulier, Elisabeth Trawinski, Jean-Michel Pawlotsky |
| EPI_ISL_3391632 | Biogroup Bio Lam-LCD Saint-Denis | Department of Virology, Henri Mondor University Hospital, Assistance Publique Hôpitaux de Paris, Université Paris-Est Créteil, INSERM U955 | Christophe Rodriguez, Slim Fourati, Vanessa Demontant, Guillaume Gricourt, Melissa N'Debi, Alexandre Soulier, Elisabeth Trawinski, Jean-Michel Pawlotsky |
| EPI_ISL_3391633 | BIO67-BIOSPHERE | Department of Virology, Henri Mondor University Hospital, Assistance Publique Hôpitaux de Paris, Université Paris-Est Créteil, INSERM U955 | Christophe Rodriguez, Slim Fourati, Vanessa Demontant, Guillaume Gricourt, Melissa N'Debi, Alexandre Soulier, Elisabeth Trawinski, Jean-Michel Pawlotsky |
| EPI_ISL_3391634 | Biogroup Bio Lam-LCD Saint-Denis | Department of Virology, Henri Mondor University Hospital, Assistance Publique Hôpitaux de Paris, Université Paris-Est Créteil, INSERM U955 | Christophe Rodriguez, Slim Fourati, Vanessa Demontant, Guillaume Gricourt, Melissa N'Debi, Alexandre Soulier, Elisabeth Trawinski, Jean-Michel Pawlotsky |
| EPI_ISL_3391635 | BIO67-BIOSPHERE | Department of Virology, Henri Mondor University Hospital, Assistance Publique Hôpitaux de Paris, Université Paris-Est Créteil, INSERM U955 | Christophe Rodriguez, Slim Fourati, Vanessa Demontant, Guillaume Gricourt, Melissa N'Debi, Alexandre Soulier, Elisabeth Trawinski, Jean-Michel Pawlotsky |
| EPI_ISL_3391636, EPI_ISL_3391637 | Biogroup Bio Lam-LCD Saint-Denis | Department of Virology, Henri Mondor University Hospital, Assistance Publique Hôpitaux de Paris, Université Paris-Est Créteil, INSERM U955 | Christophe Rodriguez, Slim Fourati, Vanessa Demontant, Guillaume Gricourt, Melissa N'Debi, Alexandre Soulier, Elisabeth Trawinski, Jean-Michel Pawlotsky |
| EPI_ISL_3391639 | BIO VSM LAB | Department of Virology, Henri Mondor University Hospital, Assistance Publique Hôpitaux de Paris, Université Paris-Est Créteil, INSERM U955 | Christophe Rodriguez, Slim Fourati, Vanessa Demontant, Guillaume Gricourt, Melissa N'Debi, Alexandre Soulier, Elisabeth Trawinski, Jean-Michel Pawlotsky |
| EPI_ISL_3391640 | Biogroup Bio Lam-LCD Saint-Denis | Department of Virology, Henri Mondor University Hospital, Assistance Publique Hôpitaux de Paris, Université Paris-Est Créteil, INSERM U955 | Christophe Rodriguez, Slim Fourati, Vanessa Demontant, Guillaume Gricourt, Melissa N'Debi, Alexandre Soulier, Elisabeth Trawinski, Jean-Michel Pawlotsky |
| EPI_ISL_3391641 | BIOMNIS EUROFINS IVRY | Department of Virology, Henri Mondor University Hospital, Assistance Publique Hôpitaux de Paris, Université Paris-Est Créteil, INSERM U955 | Christophe Rodriguez, Slim Fourati, Vanessa Demontant, Guillaume Gricourt, Melissa N'Debi, Alexandre Soulier, Elisabeth Trawinski, Jean-Michel Pawlotsky |
| EPI_ISL_3391642, EPI_ISL_3391643 | Laboratoire Ana-L | Department of Virology, Henri Mondor University Hospital, Assistance Publique Hôpitaux de Paris, Université Paris-Est Créteil, INSERM U955 | Christophe Rodriguez, Slim Fourati, Vanessa Demontant, Guillaume Gricourt, Melissa N'Debi, Alexandre Soulier, Elisabeth Trawinski, Jean-Michel Pawlotsky |
| EPI_ISL_3391644 | GROUPE HOSPITALIER SUD ILE DE FRANCE | Department of Virology, Henri Mondor University Hospital, Assistance Publique Hôpitaux de Paris, Université Paris-Est Créteil, INSERM U955 | Christophe Rodriguez, Slim Fourati, Vanessa Demontant, Guillaume Gricourt, Melissa N'Debi, Alexandre Soulier, Elisabeth Trawinski, Jean-Michel Pawlotsky |
| EPI_ISL_3391645, EPI_ISL_3391646, EPI_ISL_3391647, EPI_ISL_3391648, EPI_ISL_3391649, EPI_ISL_3391654 | Biogroup Bio Lam-LCD Saint-Denis | Department of Virology, Henri Mondor University Hospital, Assistance Publique Hôpitaux de Paris, Université Paris-Est Créteil, INSERM U955 | Christophe Rodriguez, Slim Fourati, Vanessa Demontant, Guillaume Gricourt, Melissa N'Debi, Alexandre Soulier, Elisabeth Trawinski, Jean-Michel Pawlotsky |
| EPI_ISL_3391655 | SELAS CERBALLIANCE Normandie ouest | Department of Virology, Henri Mondor University Hospital, Assistance Publique Hôpitaux de Paris, Université Paris-Est Créteil, INSERM U955 | Christophe Rodriguez, Slim Fourati, Vanessa Demontant, Guillaume Gricourt, Melissa N'Debi, Alexandre Soulier, Elisabeth Trawinski, Jean-Michel Pawlotsky |
| EPI_ISL_3391656, EPI_ISL_3391657 | Biogroup Bio Lam-LCD Saint-Denis | Department of Virology, Henri Mondor University Hospital, Assistance Publique Hôpitaux de Paris, Université Paris-Est Créteil, INSERM U955 | Christophe Rodriguez, Slim Fourati, Vanessa Demontant, Guillaume Gricourt, Melissa N'Debi, Alexandre Soulier, Elisabeth Trawinski, Jean-Michel Pawlotsky |
| EPI_ISL_3391658 | LAM BARRAND | Department of Virology, Henri Mondor University Hospital, Assistance Publique Hôpitaux de Paris, Université Paris-Est Créteil, INSERM U955 | Christophe Rodriguez, Slim Fourati, Vanessa Demontant, Guillaume Gricourt, Melissa N'Debi, Alexandre Soulier, Elisabeth Trawinski, Jean-Michel Pawlotsky |
| EPI_ISL_3391659 | Biogroup Bio Lam-LCD Saint-Denis | Department of Virology, Henri Mondor University Hospital, Assistance Publique Hôpitaux de Paris, Université Paris-Est Créteil, INSERM U955 | Christophe Rodriguez, Slim Fourati, Vanessa Demontant, Guillaume Gricourt, Melissa N'Debi, Alexandre Soulier, Elisabeth Trawinski, Jean-Michel Pawlotsky |
| EPI_ISL_3391660 | SELAS MEDILYS | Department of Virology, Henri Mondor University Hospital, Assistance Publique Hôpitaux de Paris, Université Paris-Est Créteil, INSERM U955 | Christophe Rodriguez, Slim Fourati, Vanessa Demontant, Guillaume Gricourt, Melissa N'Debi, Alexandre Soulier, Elisabeth Trawinski, Jean-Michel Pawlotsky |
| EPI_ISL_3391661 | CHI VILLENEUVE ST GEORGES | Department of Virology, Henri Mondor University Hospital, Assistance Publique Hôpitaux de Paris, Université Paris-Est Créteil, INSERM U955 | Christophe Rodriguez, Slim Fourati, Vanessa Demontant, Guillaume Gricourt, Melissa N'Debi, Alexandre Soulier, Elisabeth Trawinski, Jean-Michel Pawlotsky |
| EPI_ISL_3391662 | BPR Analyses spécialisées | Department of Virology, Henri Mondor University Hospital, Assistance Publique Hôpitaux de Paris, Université Paris-Est Créteil, INSERM U955 | Christophe Rodriguez, Slim Fourati, Vanessa Demontant, Guillaume Gricourt, Melissa N'Debi, Alexandre Soulier, Elisabeth Trawinski, Jean-Michel Pawlotsky |
| EPI_ISL_3391663, EPI_ISL_3391664, EPI_ISL_3391665 | SELAS MEDILYS | Department of Virology, Henri Mondor University Hospital, Assistance Publique Hôpitaux de Paris, Université Paris-Est Créteil, INSERM U955 | Christophe Rodriguez, Slim Fourati, Vanessa Demontant, Guillaume Gricourt, Melissa N'Debi, Alexandre Soulier, Elisabeth Trawinski, Jean-Michel Pawlotsky |
| EPI_ISL_3391666 | Biogroup Bio Lam-LCD Saint-Denis | Department of Virology, Henri Mondor University Hospital, Assistance Publique Hôpitaux de Paris, Université Paris-Est Créteil, INSERM U955 | Christophe Rodriguez, Slim Fourati, Vanessa Demontant, Guillaume Gricourt, Melissa N'Debi, Alexandre Soulier, Elisabeth Trawinski, Jean-Michel Pawlotsky |
| EPI_ISL_3391667 | BIO ARD'AISNE | Department of Virology, Henri Mondor University Hospital, Assistance Publique Hôpitaux de Paris, Université Paris-Est Créteil, | Christophe Rodriguez, Slim Fourati, Vanessa Demontant, Guillaume Gricourt, Melissa N'Debi, Alexandre Soulier, Elisabeth Trawinski, Jean-Michel Pawlotsky |

| | | | |
|---|---|---|---|
| | | INSERM U955 | |
| EPI_ISL_3391668, EPI_ISL_3391669 | SELAS MEDILYS | Department of Virology, Henri Mondor University Hospital, Assistance Publique Hôpitaux de Paris, Université Paris-Est Créteil, INSERM U955 | Christophe Rodriguez, Slim Fourati, Vanessa Demontant, Guillaume Gricourt, Melissa N'Debi, Alexandre Soulier, Elisabeth Trawinski, Jean-Michel Pawlotsky |
| EPI_ISL_3391670, EPI_ISL_3391671, EPI_ISL_3391672, EPI_ISL_3391673, EPI_ISL_3391674, EPI_ISL_3391675 | Biogroup Bio Lam-LCD Saint-Denis | Department of Virology, Henri Mondor University Hospital, Assistance Publique Hôpitaux de Paris, Université Paris-Est Créteil, INSERM U955 | Christophe Rodriguez, Slim Fourati, Vanessa Demontant, Guillaume Gricourt, Melissa N'Debi, Alexandre Soulier, Elisabeth Trawinski, Jean-Michel Pawlotsky |
| EPI_ISL_3391676 | BPR Analyses spécialisées | Department of Virology, Henri Mondor University Hospital, Assistance Publique Hôpitaux de Paris, Université Paris-Est Créteil, INSERM U955 | Christophe Rodriguez, Slim Fourati, Vanessa Demontant, Guillaume Gricourt, Melissa N'Debi, Alexandre Soulier, Elisabeth Trawinski, Jean-Michel Pawlotsky |
| EPI_ISL_3391677 | Biogroup Bio Lam-LCD Saint-Denis | Department of Virology, Henri Mondor University Hospital, Assistance Publique Hôpitaux de Paris, Université Paris-Est Créteil, INSERM U955 | Christophe Rodriguez, Slim Fourati, Vanessa Demontant, Guillaume Gricourt, Melissa N'Debi, Alexandre Soulier, Elisabeth Trawinski, Jean-Michel Pawlotsky |
| EPI_ISL_3391678 | GH de l'Est Francilien | Department of Virology, Henri Mondor University Hospital, Assistance Publique Hôpitaux de Paris, Université Paris-Est Créteil, INSERM U955 | Christophe Rodriguez, Slim Fourati, Vanessa Demontant, Guillaume Gricourt, Melissa N'Debi, Alexandre Soulier, Elisabeth Trawinski, Jean-Michel Pawlotsky |
| EPI_ISL_3391679 | SELAS MEDILYS | Department of Virology, Henri Mondor University Hospital, Assistance Publique Hôpitaux de Paris, Université Paris-Est Créteil, INSERM U955 | Christophe Rodriguez, Slim Fourati, Vanessa Demontant, Guillaume Gricourt, Melissa N'Debi, Alexandre Soulier, Elisabeth Trawinski, Jean-Michel Pawlotsky |
| EPI_ISL_3391680 | CH. CLEMENCEAU | Department of Virology, Henri Mondor University Hospital, Assistance Publique Hôpitaux de Paris, Université Paris-Est Créteil, INSERM U955 | Christophe Rodriguez, Slim Fourati, Vanessa Demontant, Guillaume Gricourt, Melissa N'Debi, Alexandre Soulier, Elisabeth Trawinski, Jean-Michel Pawlotsky |
| EPI_ISL_3391681 | BPR Analyses spécialisées | Department of Virology, Henri Mondor University Hospital, Assistance Publique Hôpitaux de Paris, Université Paris-Est Créteil, INSERM U955 | Christophe Rodriguez, Slim Fourati, Vanessa Demontant, Guillaume Gricourt, Melissa N'Debi, Alexandre Soulier, Elisabeth Trawinski, Jean-Michel Pawlotsky |
| EPI_ISL_3391682 | SELAS MEDILYS | Department of Virology, Henri Mondor University Hospital, Assistance Publique Hôpitaux de Paris, Université Paris-Est Créteil, INSERM U955 | Christophe Rodriguez, Slim Fourati, Vanessa Demontant, Guillaume Gricourt, Melissa N'Debi, Alexandre Soulier, Elisabeth Trawinski, Jean-Michel Pawlotsky |
| EPI_ISL_3391683 | SELAS CERBALLIANCE Normandie ouest | Department of Virology, Henri Mondor University Hospital, Assistance Publique Hôpitaux de Paris, Université Paris-Est Créteil, INSERM U955 | Christophe Rodriguez, Slim Fourati, Vanessa Demontant, Guillaume Gricourt, Melissa N'Debi, Alexandre Soulier, Elisabeth Trawinski, Jean-Michel Pawlotsky |
| EPI_ISL_3391684 | Biogroup Bio Lam-LCD Saint-Denis | Department of Virology, Henri Mondor University Hospital, Assistance Publique Hôpitaux de Paris, Université Paris-Est Créteil, INSERM U955 | Christophe Rodriguez, Slim Fourati, Vanessa Demontant, Guillaume Gricourt, Melissa N'Debi, Alexandre Soulier, Elisabeth Trawinski, Jean-Michel Pawlotsky |
| EPI_ISL_3391686 | CH. CLEMENCEAU | Department of Virology, Henri Mondor University Hospital, Assistance Publique Hôpitaux de Paris, Université Paris-Est Créteil, INSERM U955 | Christophe Rodriguez, Slim Fourati, Vanessa Demontant, Guillaume Gricourt, Melissa N'Debi, Alexandre Soulier, Elisabeth Trawinski, Jean-Michel Pawlotsky |
| EPI_ISL_3391687 | CH.LE RAINCY_MONTFERMEIL | Department of Virology, Henri Mondor University Hospital, Assistance Publique Hôpitaux de Paris, Université Paris-Est Créteil, INSERM U955 | Christophe Rodriguez, Slim Fourati, Vanessa Demontant, Guillaume Gricourt, Melissa N'Debi, Alexandre Soulier, Elisabeth Trawinski, Jean-Michel Pawlotsky |
| EPI_ISL_3391688 | SELAS MEDILYS | Department of Virology, Henri Mondor University Hospital, Assistance Publique Hôpitaux de Paris, Université Paris-Est Créteil, INSERM U955 | Christophe Rodriguez, Slim Fourati, Vanessa Demontant, Guillaume Gricourt, Melissa N'Debi, Alexandre Soulier, Elisabeth Trawinski, Jean-Michel Pawlotsky |
| EPI_ISL_3391689 | Biogroup Bio Lam-LCD Saint-Denis | Department of Virology, Henri Mondor University Hospital, Assistance Publique Hôpitaux de Paris, Université Paris-Est Créteil, INSERM U955 | Christophe Rodriguez, Slim Fourati, Vanessa Demontant, Guillaume Gricourt, Melissa N'Debi, Alexandre Soulier, Elisabeth Trawinski, Jean-Michel Pawlotsky |
| EPI_ISL_3391690 | BIO ARD'AISNE | Department of Virology, Henri Mondor University Hospital, Assistance Publique Hôpitaux de Paris, Université Paris-Est Créteil, INSERM U955 | Christophe Rodriguez, Slim Fourati, Vanessa Demontant, Guillaume Gricourt, Melissa N'Debi, Alexandre Soulier, Elisabeth Trawinski, Jean-Michel Pawlotsky |
| EPI_ISL_3391691 | Biogroup Bio Lam-LCD Saint-Denis | Department of Virology, Henri Mondor University Hospital, Assistance Publique Hôpitaux de Paris, Université Paris-Est Créteil, INSERM U955 | Christophe Rodriguez, Slim Fourati, Vanessa Demontant, Guillaume Gricourt, Melissa N'Debi, Alexandre Soulier, Elisabeth Trawinski, Jean-Michel Pawlotsky |
| EPI_ISL_3391692 | BIO ARD'AISNE | Department of Virology, Henri Mondor University Hospital, Assistance Publique Hôpitaux de Paris, Université Paris-Est Créteil, INSERM U955 | Christophe Rodriguez, Slim Fourati, Vanessa Demontant, Guillaume Gricourt, Melissa N'Debi, Alexandre Soulier, Elisabeth Trawinski, Jean-Michel Pawlotsky |
| EPI_ISL_3391693 | SELAS MEDILYS | Department of Virology, Henri Mondor University Hospital, Assistance Publique Hôpitaux de Paris, Université Paris-Est Créteil, INSERM U955 | Christophe Rodriguez, Slim Fourati, Vanessa Demontant, Guillaume Gricourt, Melissa N'Debi, Alexandre Soulier, Elisabeth Trawinski, Jean-Michel Pawlotsky |
| EPI_ISL_3391694, EPI_ISL_3391695 | Biogroup Bio Lam-LCD Saint-Denis | Department of Virology, Henri Mondor University Hospital, Assistance Publique Hôpitaux de Paris, Université Paris-Est Créteil, INSERM U955 | Christophe Rodriguez, Slim Fourati, Vanessa Demontant, Guillaume Gricourt, Melissa N'Debi, Alexandre Soulier, Elisabeth Trawinski, Jean-Michel Pawlotsky |
| EPI_ISL_3391696 | CHI VILLENEUVE ST GEORGES | Department of Virology, Henri Mondor University Hospital, Assistance Publique Hôpitaux de Paris, Université Paris-Est Créteil, INSERM U955 | Christophe Rodriguez, Slim Fourati, Vanessa Demontant, Guillaume Gricourt, Melissa N'Debi, Alexandre Soulier, Elisabeth Trawinski, Jean-Michel Pawlotsky |
| EPI_ISL_3391697, EPI_ISL_3391698 | Biogroup Bio Lam-LCD Saint-Denis | Department of Virology, Henri Mondor University Hospital, Assistance Publique Hôpitaux de Paris, Université Paris-Est Créteil, INSERM U955 | Christophe Rodriguez, Slim Fourati, Vanessa Demontant, Guillaume Gricourt, Melissa N'Debi, Alexandre Soulier, Elisabeth Trawinski, Jean-Michel Pawlotsky |
| EPI_ISL_3391699 | Laboratoire multi-site du Diaconat | Department of Virology, Henri Mondor University Hospital, Assistance Publique Hôpitaux de Paris, Université Paris-Est Créteil, INSERM U955 | Christophe Rodriguez, Slim Fourati, Vanessa Demontant, Guillaume Gricourt, Melissa N'Debi, Alexandre Soulier, Elisabeth Trawinski, Jean-Michel Pawlotsky |
| EPI_ISL_3391701 | Biogroup Bio Lam-LCD Saint-Denis | Department of Virology, Henri Mondor University Hospital, Assistance Publique Hôpitaux de Paris, Université Paris-Est Créteil, INSERM U955 | Christophe Rodriguez, Slim Fourati, Vanessa Demontant, Guillaume Gricourt, Melissa N'Debi, Alexandre Soulier, Elisabeth Trawinski, Jean-Michel Pawlotsky |
| EPI_ISL_3391702 | BIOLAB UNILABS | Department of Virology, Henri Mondor University Hospital, Assistance Publique Hôpitaux de Paris, Université Paris-Est Créteil, INSERM U955 | Christophe Rodriguez, Slim Fourati, Vanessa Demontant, Guillaume Gricourt, Melissa N'Debi, Alexandre Soulier, Elisabeth Trawinski, Jean-Michel Pawlotsky |
| EPI_ISL_3391703 | SELAS MEDILYS | Department of Virology, Henri Mondor University Hospital, Assistance Publique Hôpitaux de Paris, Université Paris-Est Créteil, INSERM U955 | Christophe Rodriguez, Slim Fourati, Vanessa Demontant, Guillaume Gricourt, Melissa N'Debi, Alexandre Soulier, Elisabeth Trawinski, Jean-Michel Pawlotsky |
| EPI_ISL_3391705 | Biogroup Bio Lam-LCD Saint-Denis | Department of Virology, Henri Mondor University Hospital, Assistance Publique Hôpitaux de Paris, Université Paris-Est Créteil, INSERM U955 | Christophe Rodriguez, Slim Fourati, Vanessa Demontant, Guillaume Gricourt, Melissa N'Debi, Alexandre Soulier, Elisabeth Trawinski, Jean-Michel Pawlotsky |
| EPI_ISL_3391706 | SELAS MEDILYS | Department of Virology, Henri Mondor University Hospital, Assistance Publique Hôpitaux de Paris, Université Paris-Est Créteil, INSERM U955 | Christophe Rodriguez, Slim Fourati, Vanessa Demontant, Guillaume Gricourt, Melissa N'Debi, Alexandre Soulier, Elisabeth Trawinski, Jean-Michel Pawlotsky |
| EPI_ISL_3391707, EPI_ISL_3391708 | BIO ARD'AISNE | Department of Virology, Henri Mondor University Hospital, Assistance Publique Hôpitaux de Paris, Université Paris-Est Créteil, INSERM U955 | Christophe Rodriguez, Slim Fourati, Vanessa Demontant, Guillaume Gricourt, Melissa N'Debi, Alexandre Soulier, Elisabeth Trawinski, Jean-Michel Pawlotsky |
| EPI_ISL_3391709 | SELAS MEDILYS | Department of Virology, Henri Mondor University Hospital, Assistance Publique Hôpitaux de Paris, Université Paris-Est Créteil, INSERM U955 | Christophe Rodriguez, Slim Fourati, Vanessa Demontant, Guillaume Gricourt, Melissa N'Debi, Alexandre Soulier, Elisabeth Trawinski, Jean-Michel Pawlotsky |
| EPI_ISL_3391710 | Biogroup Bio Lam-LCD Saint-Denis | Department of Virology, Henri Mondor University Hospital, Assistance Publique Hôpitaux de Paris, Université Paris-Est Créteil, | Christophe Rodriguez, Slim Fourati, Vanessa Demontant, Guillaume Gricourt, Melissa N'Debi, Alexandre Soulier, Elisabeth Trawinski, Jean-Michel Pawlotsky |

| | | | |
|---|---|---|---|
| | | INSERM U955 | |
| EPI_ISL_3391711 | SELAS MEDILYS | Department of Virology, Henri Mondor University Hospital, Assistance Publique Hôpitaux de Paris, Université Paris-Est Créteil, INSERM U955 | Christophe Rodriguez, Slim Fourati, Vanessa Demontant, Guillaume Gricourt, Melissa N'Debi, Alexandre Soulier, Elisabeth Trawinski, Jean-Michel Pawlotsky |
| EPI_ISL_3391713 | LAM BARRAND | Department of Virology, Henri Mondor University Hospital, Assistance Publique Hôpitaux de Paris, Université Paris-Est Créteil, INSERM U955 | Christophe Rodriguez, Slim Fourati, Vanessa Demontant, Guillaume Gricourt, Melissa N'Debi, Alexandre Soulier, Elisabeth Trawinski, Jean-Michel Pawlotsky |
| EPI_ISL_3391714 | Biogroup Bio Lam-LCD Saint-Denis | Department of Virology, Henri Mondor University Hospital, Assistance Publique Hôpitaux de Paris, Université Paris-Est Créteil, INSERM U955 | Christophe Rodriguez, Slim Fourati, Vanessa Demontant, Guillaume Gricourt, Melissa N'Debi, Alexandre Soulier, Elisabeth Trawinski, Jean-Michel Pawlotsky |
| EPI_ISL_3391715 | BIO ARD'AISNE | Department of Virology, Henri Mondor University Hospital, Assistance Publique Hôpitaux de Paris, Université Paris-Est Créteil, INSERM U955 | Christophe Rodriguez, Slim Fourati, Vanessa Demontant, Guillaume Gricourt, Melissa N'Debi, Alexandre Soulier, Elisabeth Trawinski, Jean-Michel Pawlotsky |
| EPI_ISL_3391716 | SELAS MEDILYS | Department of Virology, Henri Mondor University Hospital, Assistance Publique Hôpitaux de Paris, Université Paris-Est Créteil, INSERM U955 | Christophe Rodriguez, Slim Fourati, Vanessa Demontant, Guillaume Gricourt, Melissa N'Debi, Alexandre Soulier, Elisabeth Trawinski, Jean-Michel Pawlotsky |
| EPI_ISL_3391717 | BPR Analyses spécialisées | Department of Virology, Henri Mondor University Hospital, Assistance Publique Hôpitaux de Paris, Université Paris-Est Créteil, INSERM U955 | Christophe Rodriguez, Slim Fourati, Vanessa Demontant, Guillaume Gricourt, Melissa N'Debi, Alexandre Soulier, Elisabeth Trawinski, Jean-Michel Pawlotsky |
| EPI_ISL_3391718 | SELAS MEDILYS | Department of Virology, Henri Mondor University Hospital, Assistance Publique Hôpitaux de Paris, Université Paris-Est Créteil, INSERM U955 | Christophe Rodriguez, Slim Fourati, Vanessa Demontant, Guillaume Gricourt, Melissa N'Debi, Alexandre Soulier, Elisabeth Trawinski, Jean-Michel Pawlotsky |
| EPI_ISL_3391719 | SELAS BC-LAB | Department of Virology, Henri Mondor University Hospital, Assistance Publique Hôpitaux de Paris, Université Paris-Est Créteil, INSERM U955 | Christophe Rodriguez, Slim Fourati, Vanessa Demontant, Guillaume Gricourt, Melissa N'Debi, Alexandre Soulier, Elisabeth Trawinski, Jean-Michel Pawlotsky |
| EPI_ISL_3391720 | SELAS MEDILYS | Department of Virology, Henri Mondor University Hospital, Assistance Publique Hôpitaux de Paris, Université Paris-Est Créteil, INSERM U955 | Christophe Rodriguez, Slim Fourati, Vanessa Demontant, Guillaume Gricourt, Melissa N'Debi, Alexandre Soulier, Elisabeth Trawinski, Jean-Michel Pawlotsky |
| EPI_ISL_3391721 | Biogroup Bio Lam-LCD Saint-Denis | Department of Virology, Henri Mondor University Hospital, Assistance Publique Hôpitaux de Paris, Université Paris-Est Créteil, INSERM U955 | Christophe Rodriguez, Slim Fourati, Vanessa Demontant, Guillaume Gricourt, Melissa N'Debi, Alexandre Soulier, Elisabeth Trawinski, Jean-Michel Pawlotsky |
| EPI_ISL_3391722 | CH AUXERRE | Department of Virology, Henri Mondor University Hospital, Assistance Publique Hôpitaux de Paris, Université Paris-Est Créteil, INSERM U955 | Christophe Rodriguez, Slim Fourati, Vanessa Demontant, Guillaume Gricourt, Melissa N'Debi, Alexandre Soulier, Elisabeth Trawinski, Jean-Michel Pawlotsky |
| EPI_ISL_3391723, EPI_ISL_3391724 | Biogroup Bio Lam-LCD Saint-Denis | Department of Virology, Henri Mondor University Hospital, Assistance Publique Hôpitaux de Paris, Université Paris-Est Créteil, INSERM U955 | Christophe Rodriguez, Slim Fourati, Vanessa Demontant, Guillaume Gricourt, Melissa N'Debi, Alexandre Soulier, Elisabeth Trawinski, Jean-Michel Pawlotsky |
| EPI_ISL_3391725 | LAM BARRAND | Department of Virology, Henri Mondor University Hospital, Assistance Publique Hôpitaux de Paris, Université Paris-Est Créteil, INSERM U955 | Christophe Rodriguez, Slim Fourati, Vanessa Demontant, Guillaume Gricourt, Melissa N'Debi, Alexandre Soulier, Elisabeth Trawinski, Jean-Michel Pawlotsky |
| EPI_ISL_3391726, EPI_ISL_3391727 | SELAS MEDILYS | Department of Virology, Henri Mondor University Hospital, Assistance Publique Hôpitaux de Paris, Université Paris-Est Créteil, INSERM U955 | Christophe Rodriguez, Slim Fourati, Vanessa Demontant, Guillaume Gricourt, Melissa N'Debi, Alexandre Soulier, Elisabeth Trawinski, Jean-Michel Pawlotsky |
| EPI_ISL_3391728 | Biogroup Bio Lam-LCD Saint-Denis | Department of Virology, Henri Mondor University Hospital, Assistance Publique Hôpitaux de Paris, Université Paris-Est Créteil, INSERM U955 | Christophe Rodriguez, Slim Fourati, Vanessa Demontant, Guillaume Gricourt, Melissa N'Debi, Alexandre Soulier, Elisabeth Trawinski, Jean-Michel Pawlotsky |
| EPI_ISL_3391729 | SELAS MEDILYS | Department of Virology, Henri Mondor University Hospital, Assistance Publique Hôpitaux de Paris, Université Paris-Est Créteil, INSERM U955 | Christophe Rodriguez, Slim Fourati, Vanessa Demontant, Guillaume Gricourt, Melissa N'Debi, Alexandre Soulier, Elisabeth Trawinski, Jean-Michel Pawlotsky |
| EPI_ISL_3391730 | CH Mémorial F.E.U Laboratoire | Department of Virology, Henri Mondor University Hospital, Assistance Publique Hôpitaux de Paris, Université Paris-Est Créteil, INSERM U955 | Christophe Rodriguez, Slim Fourati, Vanessa Demontant, Guillaume Gricourt, Melissa N'Debi, Alexandre Soulier, Elisabeth Trawinski, Jean-Michel Pawlotsky |
| EPI_ISL_3391731 | CH. CLEMENCEAU | Department of Virology, Henri Mondor University Hospital, Assistance Publique Hôpitaux de Paris, Université Paris-Est Créteil, INSERM U955 | Christophe Rodriguez, Slim Fourati, Vanessa Demontant, Guillaume Gricourt, Melissa N'Debi, Alexandre Soulier, Elisabeth Trawinski, Jean-Michel Pawlotsky |
| EPI_ISL_3391732 | SELAS MEDILYS | Department of Virology, Henri Mondor University Hospital, Assistance Publique Hôpitaux de Paris, Université Paris-Est Créteil, INSERM U955 | Christophe Rodriguez, Slim Fourati, Vanessa Demontant, Guillaume Gricourt, Melissa N'Debi, Alexandre Soulier, Elisabeth Trawinski, Jean-Michel Pawlotsky |
| EPI_ISL_3391733, EPI_ISL_3391734 | BIO ARD'AISNE | Department of Virology, Henri Mondor University Hospital, Assistance Publique Hôpitaux de Paris, Université Paris-Est Créteil, INSERM U955 | Christophe Rodriguez, Slim Fourati, Vanessa Demontant, Guillaume Gricourt, Melissa N'Debi, Alexandre Soulier, Elisabeth Trawinski, Jean-Michel Pawlotsky |
| EPI_ISL_3391735 | SELAS CERBALLIANCE Normandie ouest | Department of Virology, Henri Mondor University Hospital, Assistance Publique Hôpitaux de Paris, Université Paris-Est Créteil, INSERM U955 | Christophe Rodriguez, Slim Fourati, Vanessa Demontant, Guillaume Gricourt, Melissa N'Debi, Alexandre Soulier, Elisabeth Trawinski, Jean-Michel Pawlotsky |
| EPI_ISL_3391736 | LABM DEFRANCE | Department of Virology, Henri Mondor University Hospital, Assistance Publique Hôpitaux de Paris, Université Paris-Est Créteil, INSERM U955 | Christophe Rodriguez, Slim Fourati, Vanessa Demontant, Guillaume Gricourt, Melissa N'Debi, Alexandre Soulier, Elisabeth Trawinski, Jean-Michel Pawlotsky |
| EPI_ISL_3391737 | Biogroup Bio Lam-LCD Saint-Denis | Department of Virology, Henri Mondor University Hospital, Assistance Publique Hôpitaux de Paris, Université Paris-Est Créteil, INSERM U955 | Christophe Rodriguez, Slim Fourati, Vanessa Demontant, Guillaume Gricourt, Melissa N'Debi, Alexandre Soulier, Elisabeth Trawinski, Jean-Michel Pawlotsky |
| EPI_ISL_3391738 | BIO ARD'AISNE | Department of Virology, Henri Mondor University Hospital, Assistance Publique Hôpitaux de Paris, Université Paris-Est Créteil, INSERM U955 | Christophe Rodriguez, Slim Fourati, Vanessa Demontant, Guillaume Gricourt, Melissa N'Debi, Alexandre Soulier, Elisabeth Trawinski, Jean-Michel Pawlotsky |
| EPI_ISL_3391739, EPI_ISL_3391740 | SELAS MEDILYS | Department of Virology, Henri Mondor University Hospital, Assistance Publique Hôpitaux de Paris, Université Paris-Est Créteil, INSERM U955 | Christophe Rodriguez, Slim Fourati, Vanessa Demontant, Guillaume Gricourt, Melissa N'Debi, Alexandre Soulier, Elisabeth Trawinski, Jean-Michel Pawlotsky |
| EPI_ISL_3391741 | Biogroup Bio Lam-LCD Saint-Denis | Department of Virology, Henri Mondor University Hospital, Assistance Publique Hôpitaux de Paris, Université Paris-Est Créteil, INSERM U955 | Christophe Rodriguez, Slim Fourati, Vanessa Demontant, Guillaume Gricourt, Melissa N'Debi, Alexandre Soulier, Elisabeth Trawinski, Jean-Michel Pawlotsky |
| EPI_ISL_3391742 | CH. CLEMENCEAU | Department of Virology, Henri Mondor University Hospital, Assistance Publique Hôpitaux de Paris, Université Paris-Est Créteil, INSERM U955 | Christophe Rodriguez, Slim Fourati, Vanessa Demontant, Guillaume Gricourt, Melissa N'Debi, Alexandre Soulier, Elisabeth Trawinski, Jean-Michel Pawlotsky |
| EPI_ISL_3391743 | BIOLBS Rouen | Department of Virology, Henri Mondor University Hospital, Assistance Publique Hôpitaux de Paris, Université Paris-Est Créteil, INSERM U955 | Christophe Rodriguez, Slim Fourati, Vanessa Demontant, Guillaume Gricourt, Melissa N'Debi, Alexandre Soulier, Elisabeth Trawinski, Jean-Michel Pawlotsky |
| EPI_ISL_3391744 | BIO ARD'AISNE | Department of Virology, Henri Mondor University Hospital, Assistance Publique Hôpitaux de Paris, Université Paris-Est Créteil, INSERM U955 | Christophe Rodriguez, Slim Fourati, Vanessa Demontant, Guillaume Gricourt, Melissa N'Debi, Alexandre Soulier, Elisabeth Trawinski, Jean-Michel Pawlotsky |
| EPI_ISL_3391745 | BPR Analyses spécialisées | Department of Virology, Henri Mondor University Hospital, Assistance Publique Hôpitaux de Paris, Université Paris-Est Créteil, INSERM U955 | Christophe Rodriguez, Slim Fourati, Vanessa Demontant, Guillaume Gricourt, Melissa N'Debi, Alexandre Soulier, Elisabeth Trawinski, Jean-Michel Pawlotsky |
| EPI_ISL_3391746 | Biogroup Bio Lam-LCD Saint-Denis | Department of Virology, Henri Mondor University Hospital, Assistance Publique Hôpitaux de Paris, Université Paris-Est Créteil, INSERM U955 | Christophe Rodriguez, Slim Fourati, Vanessa Demontant, Guillaume Gricourt, Melissa N'Debi, Alexandre Soulier, Elisabeth Trawinski, Jean-Michel Pawlotsky |
| EPI_ISL_3391748 | SELAS BC-LAB | Department of Virology, Henri Mondor University Hospital, Assistance Publique Hôpitaux de Paris, Université Paris-Est Créteil, | Christophe Rodriguez, Slim Fourati, Vanessa Demontant, Guillaume Gricourt, Melissa N'Debi, Alexandre Soulier, Elisabeth Trawinski, Jean-Michel Pawlotsky |

| | | | |
|---|---|---|---|
| | | INSERM U955 | |
| EPI_ISL_3391749 | BPR Analyses spécialisées | Department of Virology, Henri Mondor University Hospital, Assistance Publique Hôpitaux de Paris, Université Paris-Est Créteil, INSERM U955 | Christophe Rodriguez, Slim Fourati, Vanessa Demontant, Guillaume Gricourt, Melissa N'Debi, Alexandre Soulier, Elisabeth Trawinski, Jean-Michel Pawlotsky |
| EPI_ISL_3391750 | SELAS D LAB | Department of Virology, Henri Mondor University Hospital, Assistance Publique Hôpitaux de Paris, Université Paris-Est Créteil, INSERM U955 | Christophe Rodriguez, Slim Fourati, Vanessa Demontant, Guillaume Gricourt, Melissa N'Debi, Alexandre Soulier, Elisabeth Trawinski, Jean-Michel Pawlotsky |
| EPI_ISL_3391751, EPI_ISL_3391752, EPI_ISL_3391753 | Biogroup Bio Lam-LCD Saint-Denis | Department of Virology, Henri Mondor University Hospital, Assistance Publique Hôpitaux de Paris, Université Paris-Est Créteil, INSERM U955 | Christophe Rodriguez, Slim Fourati, Vanessa Demontant, Guillaume Gricourt, Melissa N'Debi, Alexandre Soulier, Elisabeth Trawinski, Jean-Michel Pawlotsky |
| EPI_ISL_3391754 | BIO ARD'AISNE | Department of Virology, Henri Mondor University Hospital, Assistance Publique Hôpitaux de Paris, Université Paris-Est Créteil, INSERM U955 | Christophe Rodriguez, Slim Fourati, Vanessa Demontant, Guillaume Gricourt, Melissa N'Debi, Alexandre Soulier, Elisabeth Trawinski, Jean-Michel Pawlotsky |
| EPI_ISL_3391755 | Biogroup Bio Lam-LCD Saint-Denis | Department of Virology, Henri Mondor University Hospital, Assistance Publique Hôpitaux de Paris, Université Paris-Est Créteil, INSERM U955 | Christophe Rodriguez, Slim Fourati, Vanessa Demontant, Guillaume Gricourt, Melissa N'Debi, Alexandre Soulier, Elisabeth Trawinski, Jean-Michel Pawlotsky |
| EPI_ISL_3391757 | SELAS MEDILYS | Department of Virology, Henri Mondor University Hospital, Assistance Publique Hôpitaux de Paris, Université Paris-Est Créteil, INSERM U955 | Christophe Rodriguez, Slim Fourati, Vanessa Demontant, Guillaume Gricourt, Melissa N'Debi, Alexandre Soulier, Elisabeth Trawinski, Jean-Michel Pawlotsky |
| EPI_ISL_3391758, EPI_ISL_3391759 | Biogroup Bio Lam-LCD Saint-Denis | Department of Virology, Henri Mondor University Hospital, Assistance Publique Hôpitaux de Paris, Université Paris-Est Créteil, INSERM U955 | Christophe Rodriguez, Slim Fourati, Vanessa Demontant, Guillaume Gricourt, Melissa N'Debi, Alexandre Soulier, Elisabeth Trawinski, Jean-Michel Pawlotsky |
| EPI_ISL_3391760 | GH de l'Est Francilien | Department of Virology, Henri Mondor University Hospital, Assistance Publique Hôpitaux de Paris, Université Paris-Est Créteil, INSERM U955 | Christophe Rodriguez, Slim Fourati, Vanessa Demontant, Guillaume Gricourt, Melissa N'Debi, Alexandre Soulier, Elisabeth Trawinski, Jean-Michel Pawlotsky |
| EPI_ISL_3391761 | Biogroup Bio Lam-LCD Saint-Denis | Department of Virology, Henri Mondor University Hospital, Assistance Publique Hôpitaux de Paris, Université Paris-Est Créteil, INSERM U955 | Christophe Rodriguez, Slim Fourati, Vanessa Demontant, Guillaume Gricourt, Melissa N'Debi, Alexandre Soulier, Elisabeth Trawinski, Jean-Michel Pawlotsky |
| EPI_ISL_3391762 | SELAS MEDILYS | Department of Virology, Henri Mondor University Hospital, Assistance Publique Hôpitaux de Paris, Université Paris-Est Créteil, INSERM U955 | Christophe Rodriguez, Slim Fourati, Vanessa Demontant, Guillaume Gricourt, Melissa N'Debi, Alexandre Soulier, Elisabeth Trawinski, Jean-Michel Pawlotsky |
| EPI_ISL_3391763 | BPR Analyses spécialisées | Department of Virology, Henri Mondor University Hospital, Assistance Publique Hôpitaux de Paris, Université Paris-Est Créteil, INSERM U955 | Christophe Rodriguez, Slim Fourati, Vanessa Demontant, Guillaume Gricourt, Melissa N'Debi, Alexandre Soulier, Elisabeth Trawinski, Jean-Michel Pawlotsky |
| EPI_ISL_3391764, EPI_ISL_3391765 | Biogroup Bio Lam-LCD Saint-Denis | Department of Virology, Henri Mondor University Hospital, Assistance Publique Hôpitaux de Paris, Université Paris-Est Créteil, INSERM U955 | Christophe Rodriguez, Slim Fourati, Vanessa Demontant, Guillaume Gricourt, Melissa N'Debi, Alexandre Soulier, Elisabeth Trawinski, Jean-Michel Pawlotsky |
| EPI_ISL_3391766, EPI_ISL_3391767 | SELAS MEDILYS | Department of Virology, Henri Mondor University Hospital, Assistance Publique Hôpitaux de Paris, Université Paris-Est Créteil, INSERM U955 | Christophe Rodriguez, Slim Fourati, Vanessa Demontant, Guillaume Gricourt, Melissa N'Debi, Alexandre Soulier, Elisabeth Trawinski, Jean-Michel Pawlotsky |
| EPI_ISL_3391768 | Laboratoire EspaceBio | Department of Virology, Henri Mondor University Hospital, Assistance Publique Hôpitaux de Paris, Université Paris-Est Créteil, INSERM U955 | Christophe Rodriguez, Slim Fourati, Vanessa Demontant, Guillaume Gricourt, Melissa N'Debi, Alexandre Soulier, Elisabeth Trawinski, Jean-Michel Pawlotsky |
| EPI_ISL_3391769 | BIO ARD'AISNE | Department of Virology, Henri Mondor University Hospital, Assistance Publique Hôpitaux de Paris, Université Paris-Est Créteil, INSERM U955 | Christophe Rodriguez, Slim Fourati, Vanessa Demontant, Guillaume Gricourt, Melissa N'Debi, Alexandre Soulier, Elisabeth Trawinski, Jean-Michel Pawlotsky |
| EPI_ISL_3391770 | Biogroup Bio Lam-LCD Saint-Denis | Department of Virology, Henri Mondor University Hospital, Assistance Publique Hôpitaux de Paris, Université Paris-Est Créteil, INSERM U955 | Christophe Rodriguez, Slim Fourati, Vanessa Demontant, Guillaume Gricourt, Melissa N'Debi, Alexandre Soulier, Elisabeth Trawinski, Jean-Michel Pawlotsky |
| EPI_ISL_3391772 | SELAS MEDILYS | Department of Virology, Henri Mondor University Hospital, Assistance Publique Hôpitaux de Paris, Université Paris-Est Créteil, INSERM U955 | Christophe Rodriguez, Slim Fourati, Vanessa Demontant, Guillaume Gricourt, Melissa N'Debi, Alexandre Soulier, Elisabeth Trawinski, Jean-Michel Pawlotsky |
| EPI_ISL_3391773, EPI_ISL_3391774, EPI_ISL_3391775, EPI_ISL_3391776, EPI_ISL_3391777 | Biogroup Bio Lam-LCD Saint-Denis | Department of Virology, Henri Mondor University Hospital, Assistance Publique Hôpitaux de Paris, Université Paris-Est Créteil, INSERM U955 | Christophe Rodriguez, Slim Fourati, Vanessa Demontant, Guillaume Gricourt, Melissa N'Debi, Alexandre Soulier, Elisabeth Trawinski, Jean-Michel Pawlotsky |
| EPI_ISL_3391778 | SELAS MEDILYS | Department of Virology, Henri Mondor University Hospital, Assistance Publique Hôpitaux de Paris, Université Paris-Est Créteil, INSERM U955 | Christophe Rodriguez, Slim Fourati, Vanessa Demontant, Guillaume Gricourt, Melissa N'Debi, Alexandre Soulier, Elisabeth Trawinski, Jean-Michel Pawlotsky |
| EPI_ISL_3391779, EPI_ISL_3391780 | BPR Analyses spécialisées | Department of Virology, Henri Mondor University Hospital, Assistance Publique Hôpitaux de Paris, Université Paris-Est Créteil, INSERM U955 | Christophe Rodriguez, Slim Fourati, Vanessa Demontant, Guillaume Gricourt, Melissa N'Debi, Alexandre Soulier, Elisabeth Trawinski, Jean-Michel Pawlotsky |
| EPI_ISL_3391781, EPI_ISL_3391782 | BIO ARD'AISNE | Department of Virology, Henri Mondor University Hospital, Assistance Publique Hôpitaux de Paris, Université Paris-Est Créteil, INSERM U955 | Christophe Rodriguez, Slim Fourati, Vanessa Demontant, Guillaume Gricourt, Melissa N'Debi, Alexandre Soulier, Elisabeth Trawinski, Jean-Michel Pawlotsky |
| EPI_ISL_3391783, EPI_ISL_3391784, EPI_ISL_3391785 | Biogroup Bio Lam-LCD Saint-Denis | Department of Virology, Henri Mondor University Hospital, Assistance Publique Hôpitaux de Paris, Université Paris-Est Créteil, INSERM U955 | Christophe Rodriguez, Slim Fourati, Vanessa Demontant, Guillaume Gricourt, Melissa N'Debi, Alexandre Soulier, Elisabeth Trawinski, Jean-Michel Pawlotsky |
| EPI_ISL_3391786 | BIO ARD'AISNE | Department of Virology, Henri Mondor University Hospital, Assistance Publique Hôpitaux de Paris, Université Paris-Est Créteil, INSERM U955 | Christophe Rodriguez, Slim Fourati, Vanessa Demontant, Guillaume Gricourt, Melissa N'Debi, Alexandre Soulier, Elisabeth Trawinski, Jean-Michel Pawlotsky |
| EPI_ISL_3391787 | Biogroup Bio Lam-LCD Saint-Denis | Department of Virology, Henri Mondor University Hospital, Assistance Publique Hôpitaux de Paris, Université Paris-Est Créteil, INSERM U955 | Christophe Rodriguez, Slim Fourati, Vanessa Demontant, Guillaume Gricourt, Melissa N'Debi, Alexandre Soulier, Elisabeth Trawinski, Jean-Michel Pawlotsky |
| EPI_ISL_3391788 | BPR Analyses spécialisées | Department of Virology, Henri Mondor University Hospital, Assistance Publique Hôpitaux de Paris, Université Paris-Est Créteil, INSERM U955 | Christophe Rodriguez, Slim Fourati, Vanessa Demontant, Guillaume Gricourt, Melissa N'Debi, Alexandre Soulier, Elisabeth Trawinski, Jean-Michel Pawlotsky |
| EPI_ISL_3391789 | Biogroup Bio Lam-LCD Saint-Denis | Department of Virology, Henri Mondor University Hospital, Assistance Publique Hôpitaux de Paris, Université Paris-Est Créteil, INSERM U955 | Christophe Rodriguez, Slim Fourati, Vanessa Demontant, Guillaume Gricourt, Melissa N'Debi, Alexandre Soulier, Elisabeth Trawinski, Jean-Michel Pawlotsky |
| EPI_ISL_3391790 | CH. CLEMENCEAU | Department of Virology, Henri Mondor University Hospital, Assistance Publique Hôpitaux de Paris, Université Paris-Est Créteil, INSERM U955 | Christophe Rodriguez, Slim Fourati, Vanessa Demontant, Guillaume Gricourt, Melissa N'Debi, Alexandre Soulier, Elisabeth Trawinski, Jean-Michel Pawlotsky |
| EPI_ISL_3391791, EPI_ISL_3391792 | Biogroup Bio Lam-LCD Saint-Denis | Department of Virology, Henri Mondor University Hospital, Assistance Publique Hôpitaux de Paris, Université Paris-Est Créteil, INSERM U955 | Christophe Rodriguez, Slim Fourati, Vanessa Demontant, Guillaume Gricourt, Melissa N'Debi, Alexandre Soulier, Elisabeth Trawinski, Jean-Michel Pawlotsky |
| EPI_ISL_3391793, EPI_ISL_3391794, EPI_ISL_3391795 | SELAS MEDILYS | Department of Virology, Henri Mondor University Hospital, Assistance Publique Hôpitaux de Paris, Université Paris-Est Créteil, INSERM U955 | Christophe Rodriguez, Slim Fourati, Vanessa Demontant, Guillaume Gricourt, Melissa N'Debi, Alexandre Soulier, Elisabeth Trawinski, Jean-Michel Pawlotsky |
| EPI_ISL_3391796 | GH de l'Est Francilien | Department of Virology, Henri Mondor University Hospital, Assistance Publique Hôpitaux de Paris, Université Paris-Est Créteil, INSERM U955 | Christophe Rodriguez, Slim Fourati, Vanessa Demontant, Guillaume Gricourt, Melissa N'Debi, Alexandre Soulier, Elisabeth Trawinski, Jean-Michel Pawlotsky |
| EPI_ISL_3391797 | BIO ARD'AISNE | Department of Virology, Henri Mondor University Hospital, Assistance Publique Hôpitaux de Paris, Université Paris-Est Créteil, INSERM U955 | Christophe Rodriguez, Slim Fourati, Vanessa Demontant, Guillaume Gricourt, Melissa N'Debi, Alexandre Soulier, Elisabeth Trawinski, Jean-Michel Pawlotsky |

| | | | |
|---|---|---|---|
| EPI_ISL_3391798, EPI_ISL_3391799 | Biogroup Bio Lam-LCD Saint-Denis | Department of Virology, Henri Mondor University Hospital, Assistance Publique Hôpitaux de Paris, Université Paris-Est Créteil, INSERM U955 | Christophe Rodriguez, Slim Fourati, Vanessa Demontant, Guillaume Gricourt, Melissa N'Debi, Alexandre Soulier, Elisabeth Trawinski, Jean-Michel Pawlotsky |
| EPI_ISL_3391800, EPI_ISL_3391801 | SELAS MEDILYS | Department of Virology, Henri Mondor University Hospital, Assistance Publique Hôpitaux de Paris, Université Paris-Est Créteil, INSERM U955 | Christophe Rodriguez, Slim Fourati, Vanessa Demontant, Guillaume Gricourt, Melissa N'Debi, Alexandre Soulier, Elisabeth Trawinski, Jean-Michel Pawlotsky |
| EPI_ISL_3391802 | CHU Rouen Normandie | Department of Virology, Henri Mondor University Hospital, Assistance Publique Hôpitaux de Paris, Université Paris-Est Créteil, INSERM U955 | Christophe Rodriguez, Slim Fourati, Vanessa Demontant, Guillaume Gricourt, Melissa N'Debi, Alexandre Soulier, Elisabeth Trawinski, Jean-Michel Pawlotsky |
| EPI_ISL_3391803 | Biogroup Bio Lam-LCD Saint-Denis | Department of Virology, Henri Mondor University Hospital, Assistance Publique Hôpitaux de Paris, Université Paris-Est Créteil, INSERM U955 | Christophe Rodriguez, Slim Fourati, Vanessa Demontant, Guillaume Gricourt, Melissa N'Debi, Alexandre Soulier, Elisabeth Trawinski, Jean-Michel Pawlotsky |
| EPI_ISL_3391804 | SELAS MEDILYS | Department of Virology, Henri Mondor University Hospital, Assistance Publique Hôpitaux de Paris, Université Paris-Est Créteil, INSERM U955 | Christophe Rodriguez, Slim Fourati, Vanessa Demontant, Guillaume Gricourt, Melissa N'Debi, Alexandre Soulier, Elisabeth Trawinski, Jean-Michel Pawlotsky |
| EPI_ISL_3391805 | BPR Analyses spécialisées | Department of Virology, Henri Mondor University Hospital, Assistance Publique Hôpitaux de Paris, Université Paris-Est Créteil, INSERM U955 | Christophe Rodriguez, Slim Fourati, Vanessa Demontant, Guillaume Gricourt, Melissa N'Debi, Alexandre Soulier, Elisabeth Trawinski, Jean-Michel Pawlotsky |
| EPI_ISL_3391806 | SELAS MEDILYS | Department of Virology, Henri Mondor University Hospital, Assistance Publique Hôpitaux de Paris, Université Paris-Est Créteil, INSERM U955 | Christophe Rodriguez, Slim Fourati, Vanessa Demontant, Guillaume Gricourt, Melissa N'Debi, Alexandre Soulier, Elisabeth Trawinski, Jean-Michel Pawlotsky |
| EPI_ISL_3391807 | Biogroup Bio Lam-LCD Saint-Denis | Department of Virology, Henri Mondor University Hospital, Assistance Publique Hôpitaux de Paris, Université Paris-Est Créteil, INSERM U955 | Christophe Rodriguez, Slim Fourati, Vanessa Demontant, Guillaume Gricourt, Melissa N'Debi, Alexandre Soulier, Elisabeth Trawinski, Jean-Michel Pawlotsky |
| EPI_ISL_3391808 | BIO ARD'AISNE | Department of Virology, Henri Mondor University Hospital, Assistance Publique Hôpitaux de Paris, Université Paris-Est Créteil, INSERM U955 | Christophe Rodriguez, Slim Fourati, Vanessa Demontant, Guillaume Gricourt, Melissa N'Debi, Alexandre Soulier, Elisabeth Trawinski, Jean-Michel Pawlotsky |
| EPI_ISL_3391810 | SELAS MEDILYS | Department of Virology, Henri Mondor University Hospital, Assistance Publique Hôpitaux de Paris, Université Paris-Est Créteil, INSERM U955 | Christophe Rodriguez, Slim Fourati, Vanessa Demontant, Guillaume Gricourt, Melissa N'Debi, Alexandre Soulier, Elisabeth Trawinski, Jean-Michel Pawlotsky |
| EPI_ISL_3391811 | SELAS CERBALLIANCE Normandie ouest | Department of Virology, Henri Mondor University Hospital, Assistance Publique Hôpitaux de Paris, Université Paris-Est Créteil, INSERM U955 | Christophe Rodriguez, Slim Fourati, Vanessa Demontant, Guillaume Gricourt, Melissa N'Debi, Alexandre Soulier, Elisabeth Trawinski, Jean-Michel Pawlotsky |
| EPI_ISL_3391812 | SELAS MEDILYS | Department of Virology, Henri Mondor University Hospital, Assistance Publique Hôpitaux de Paris, Université Paris-Est Créteil, INSERM U955 | Christophe Rodriguez, Slim Fourati, Vanessa Demontant, Guillaume Gricourt, Melissa N'Debi, Alexandre Soulier, Elisabeth Trawinski, Jean-Michel Pawlotsky |
| EPI_ISL_3391813 | Biogroup Bio Lam-LCD Saint-Denis | Department of Virology, Henri Mondor University Hospital, Assistance Publique Hôpitaux de Paris, Université Paris-Est Créteil, INSERM U955 | Christophe Rodriguez, Slim Fourati, Vanessa Demontant, Guillaume Gricourt, Melissa N'Debi, Alexandre Soulier, Elisabeth Trawinski, Jean-Michel Pawlotsky |
| EPI_ISL_3391814 | SELAS MEDILYS | Department of Virology, Henri Mondor University Hospital, Assistance Publique Hôpitaux de Paris, Université Paris-Est Créteil, INSERM U955 | Christophe Rodriguez, Slim Fourati, Vanessa Demontant, Guillaume Gricourt, Melissa N'Debi, Alexandre Soulier, Elisabeth Trawinski, Jean-Michel Pawlotsky |
| EPI_ISL_3391815 | SELAS CERBALLIANCE Normandie ouest | Department of Virology, Henri Mondor University Hospital, Assistance Publique Hôpitaux de Paris, Université Paris-Est Créteil, INSERM U955 | Christophe Rodriguez, Slim Fourati, Vanessa Demontant, Guillaume Gricourt, Melissa N'Debi, Alexandre Soulier, Elisabeth Trawinski, Jean-Michel Pawlotsky |
| EPI_ISL_3391816 | SELAS MEDILYS | Department of Virology, Henri Mondor University Hospital, Assistance Publique Hôpitaux de Paris, Université Paris-Est Créteil, INSERM U955 | Christophe Rodriguez, Slim Fourati, Vanessa Demontant, Guillaume Gricourt, Melissa N'Debi, Alexandre Soulier, Elisabeth Trawinski, Jean-Michel Pawlotsky |
| EPI_ISL_3391817 | Biogroup Bio Lam-LCD Saint-Denis | Department of Virology, Henri Mondor University Hospital, Assistance Publique Hôpitaux de Paris, Université Paris-Est Créteil, INSERM U955 | Christophe Rodriguez, Slim Fourati, Vanessa Demontant, Guillaume Gricourt, Melissa N'Debi, Alexandre Soulier, Elisabeth Trawinski, Jean-Michel Pawlotsky |
| EPI_ISL_3391818, EPI_ISL_3391820 | BIO ARD'AISNE | Department of Virology, Henri Mondor University Hospital, Assistance Publique Hôpitaux de Paris, Université Paris-Est Créteil, INSERM U955 | Christophe Rodriguez, Slim Fourati, Vanessa Demontant, Guillaume Gricourt, Melissa N'Debi, Alexandre Soulier, Elisabeth Trawinski, Jean-Michel Pawlotsky |
| EPI_ISL_3391822 | CHU Rouen Normandie | Department of Virology, Henri Mondor University Hospital, Assistance Publique Hôpitaux de Paris, Université Paris-Est Créteil, INSERM U955 | Christophe Rodriguez, Slim Fourati, Vanessa Demontant, Guillaume Gricourt, Melissa N'Debi, Alexandre Soulier, Elisabeth Trawinski, Jean-Michel Pawlotsky |
| EPI_ISL_3391823 | Biogroup Bio Lam-LCD Saint-Denis | Department of Virology, Henri Mondor University Hospital, Assistance Publique Hôpitaux de Paris, Université Paris-Est Créteil, INSERM U955 | Christophe Rodriguez, Slim Fourati, Vanessa Demontant, Guillaume Gricourt, Melissa N'Debi, Alexandre Soulier, Elisabeth Trawinski, Jean-Michel Pawlotsky |
| EPI_ISL_3391824 | SELAS CERBALLIANCE Normandie ouest | Department of Virology, Henri Mondor University Hospital, Assistance Publique Hôpitaux de Paris, Université Paris-Est Créteil, INSERM U955 | Christophe Rodriguez, Slim Fourati, Vanessa Demontant, Guillaume Gricourt, Melissa N'Debi, Alexandre Soulier, Elisabeth Trawinski, Jean-Michel Pawlotsky |
| EPI_ISL_3391825 | Biogroup Bio Lam-LCD Saint-Denis | Department of Virology, Henri Mondor University Hospital, Assistance Publique Hôpitaux de Paris, Université Paris-Est Créteil, INSERM U955 | Christophe Rodriguez, Slim Fourati, Vanessa Demontant, Guillaume Gricourt, Melissa N'Debi, Alexandre Soulier, Elisabeth Trawinski, Jean-Michel Pawlotsky |
| EPI_ISL_3391826 | SELAS MEDILYS | Department of Virology, Henri Mondor University Hospital, Assistance Publique Hôpitaux de Paris, Université Paris-Est Créteil, INSERM U955 | Christophe Rodriguez, Slim Fourati, Vanessa Demontant, Guillaume Gricourt, Melissa N'Debi, Alexandre Soulier, Elisabeth Trawinski, Jean-Michel Pawlotsky |
| EPI_ISL_3391827, EPI_ISL_3391828 | Biogroup Bio Lam-LCD Saint-Denis | Department of Virology, Henri Mondor University Hospital, Assistance Publique Hôpitaux de Paris, Université Paris-Est Créteil, INSERM U955 | Christophe Rodriguez, Slim Fourati, Vanessa Demontant, Guillaume Gricourt, Melissa N'Debi, Alexandre Soulier, Elisabeth Trawinski, Jean-Michel Pawlotsky |
| EPI_ISL_3391829 | BPR Analyses spécialisées | Department of Virology, Henri Mondor University Hospital, Assistance Publique Hôpitaux de Paris, Université Paris-Est Créteil, INSERM U955 | Christophe Rodriguez, Slim Fourati, Vanessa Demontant, Guillaume Gricourt, Melissa N'Debi, Alexandre Soulier, Elisabeth Trawinski, Jean-Michel Pawlotsky |
| EPI_ISL_3391831, EPI_ISL_3391832 | Biogroup Bio Lam-LCD Saint-Denis | Department of Virology, Henri Mondor University Hospital, Assistance Publique Hôpitaux de Paris, Université Paris-Est Créteil, INSERM U955 | Christophe Rodriguez, Slim Fourati, Vanessa Demontant, Guillaume Gricourt, Melissa N'Debi, Alexandre Soulier, Elisabeth Trawinski, Jean-Michel Pawlotsky |
| EPI_ISL_3391833 | SELAS MEDILYS | Department of Virology, Henri Mondor University Hospital, Assistance Publique Hôpitaux de Paris, Université Paris-Est Créteil, INSERM U955 | Christophe Rodriguez, Slim Fourati, Vanessa Demontant, Guillaume Gricourt, Melissa N'Debi, Alexandre Soulier, Elisabeth Trawinski, Jean-Michel Pawlotsky |
| EPI_ISL_3391834 | CH. ROBERT BALLANGER | Department of Virology, Henri Mondor University Hospital, Assistance Publique Hôpitaux de Paris, Université Paris-Est Créteil, INSERM U955 | Christophe Rodriguez, Slim Fourati, Vanessa Demontant, Guillaume Gricourt, Melissa N'Debi, Alexandre Soulier, Elisabeth Trawinski, Jean-Michel Pawlotsky |
| EPI_ISL_3391835 | SELAS MEDILYS | Department of Virology, Henri Mondor University Hospital, Assistance Publique Hôpitaux de Paris, Université Paris-Est Créteil, INSERM U955 | Christophe Rodriguez, Slim Fourati, Vanessa Demontant, Guillaume Gricourt, Melissa N'Debi, Alexandre Soulier, Elisabeth Trawinski, Jean-Michel Pawlotsky |
| EPI_ISL_3391836 | BIO ARD'AISNE | Department of Virology, Henri Mondor University Hospital, Assistance Publique Hôpitaux de Paris, Université Paris-Est Créteil, INSERM U955 | Christophe Rodriguez, Slim Fourati, Vanessa Demontant, Guillaume Gricourt, Melissa N'Debi, Alexandre Soulier, Elisabeth Trawinski, Jean-Michel Pawlotsky |
| EPI_ISL_3391837 | BPR Analyses spécialisées | Department of Virology, Henri Mondor University Hospital, Assistance Publique Hôpitaux de Paris, Université Paris-Est Créteil, INSERM U955 | Christophe Rodriguez, Slim Fourati, Vanessa Demontant, Guillaume Gricourt, Melissa N'Debi, Alexandre Soulier, Elisabeth Trawinski, Jean-Michel Pawlotsky |
| EPI_ISL_3391838 | Biogroup Bio Lam-LCD Saint-Denis | Department of Virology, Henri Mondor University Hospital, Assistance Publique Hôpitaux de Paris, Université Paris-Est Créteil, INSERM U955 | Christophe Rodriguez, Slim Fourati, Vanessa Demontant, Guillaume Gricourt, Melissa N'Debi, Alexandre Soulier, Elisabeth Trawinski, Jean-Michel Pawlotsky |

| | | | |
|---|---|---|---|
| EPI_ISL_3391839 | SELAS MEDILYS | Department of Virology, Henri Mondor University Hospital, Assistance Publique Hôpitaux de Paris, Université Paris-Est Créteil, INSERM U955 | Christophe Rodriguez, Slim Fourati, Vanessa Demontant, Guillaume Gricourt, Melissa N'Debi, Alexandre Soulier, Elisabeth Trawinski, Jean-Michel Pawlotsky |
| EPI_ISL_3391840 | LAM BARRAND | Department of Virology, Henri Mondor University Hospital, Assistance Publique Hôpitaux de Paris, Université Paris-Est Créteil, INSERM U955 | Christophe Rodriguez, Slim Fourati, Vanessa Demontant, Guillaume Gricourt, Melissa N'Debi, Alexandre Soulier, Elisabeth Trawinski, Jean-Michel Pawlotsky |
| EPI_ISL_3391841, EPI_ISL_3391842 | SELAS MEDILYS | Department of Virology, Henri Mondor University Hospital, Assistance Publique Hôpitaux de Paris, Université Paris-Est Créteil, INSERM U955 | Christophe Rodriguez, Slim Fourati, Vanessa Demontant, Guillaume Gricourt, Melissa N'Debi, Alexandre Soulier, Elisabeth Trawinski, Jean-Michel Pawlotsky |
| EPI_ISL_3391843, EPI_ISL_3391844 | Biogroup Bio Lam-LCD Saint-Denis | Department of Virology, Henri Mondor University Hospital, Assistance Publique Hôpitaux de Paris, Université Paris-Est Créteil, INSERM U955 | Christophe Rodriguez, Slim Fourati, Vanessa Demontant, Guillaume Gricourt, Melissa N'Debi, Alexandre Soulier, Elisabeth Trawinski, Jean-Michel Pawlotsky |
| EPI_ISL_3391845 | SELAS MEDILYS | Department of Virology, Henri Mondor University Hospital, Assistance Publique Hôpitaux de Paris, Université Paris-Est Créteil, INSERM U955 | Christophe Rodriguez, Slim Fourati, Vanessa Demontant, Guillaume Gricourt, Melissa N'Debi, Alexandre Soulier, Elisabeth Trawinski, Jean-Michel Pawlotsky |
| EPI_ISL_3391846 | CHU Rouen Normandie | Department of Virology, Henri Mondor University Hospital, Assistance Publique Hôpitaux de Paris, Université Paris-Est Créteil, INSERM U955 | Christophe Rodriguez, Slim Fourati, Vanessa Demontant, Guillaume Gricourt, Melissa N'Debi, Alexandre Soulier, Elisabeth Trawinski, Jean-Michel Pawlotsky |
| EPI_ISL_3391847, EPI_ISL_3391848, EPI_ISL_3391849 | SELAS MEDILYS | Department of Virology, Henri Mondor University Hospital, Assistance Publique Hôpitaux de Paris, Université Paris-Est Créteil, INSERM U955 | Christophe Rodriguez, Slim Fourati, Vanessa Demontant, Guillaume Gricourt, Melissa N'Debi, Alexandre Soulier, Elisabeth Trawinski, Jean-Michel Pawlotsky |
| EPI_ISL_3391850, EPI_ISL_3391851, EPI_ISL_3391852 | Biogroup Bio Lam-LCD Saint-Denis | Department of Virology, Henri Mondor University Hospital, Assistance Publique Hôpitaux de Paris, Université Paris-Est Créteil, INSERM U955 | Christophe Rodriguez, Slim Fourati, Vanessa Demontant, Guillaume Gricourt, Melissa N'Debi, Alexandre Soulier, Elisabeth Trawinski, Jean-Michel Pawlotsky |
| EPI_ISL_3391854 | BIO ARD'AISNE | Department of Virology, Henri Mondor University Hospital, Assistance Publique Hôpitaux de Paris, Université Paris-Est Créteil, INSERM U955 | Christophe Rodriguez, Slim Fourati, Vanessa Demontant, Guillaume Gricourt, Melissa N'Debi, Alexandre Soulier, Elisabeth Trawinski, Jean-Michel Pawlotsky |
| EPI_ISL_3391855 | SELAS MEDILYS | Department of Virology, Henri Mondor University Hospital, Assistance Publique Hôpitaux de Paris, Université Paris-Est Créteil, INSERM U955 | Christophe Rodriguez, Slim Fourati, Vanessa Demontant, Guillaume Gricourt, Melissa N'Debi, Alexandre Soulier, Elisabeth Trawinski, Jean-Michel Pawlotsky |
| EPI_ISL_3391856 | Biogroup Bio Lam-LCD Saint-Denis | Department of Virology, Henri Mondor University Hospital, Assistance Publique Hôpitaux de Paris, Université Paris-Est Créteil, INSERM U955 | Christophe Rodriguez, Slim Fourati, Vanessa Demontant, Guillaume Gricourt, Melissa N'Debi, Alexandre Soulier, Elisabeth Trawinski, Jean-Michel Pawlotsky |
| EPI_ISL_3391857 | CH. ROBERT BALLANGER | Department of Virology, Henri Mondor University Hospital, Assistance Publique Hôpitaux de Paris, Université Paris-Est Créteil, INSERM U955 | Christophe Rodriguez, Slim Fourati, Vanessa Demontant, Guillaume Gricourt, Melissa N'Debi, Alexandre Soulier, Elisabeth Trawinski, Jean-Michel Pawlotsky |
| EPI_ISL_3391858 | SELAS MEDILYS | Department of Virology, Henri Mondor University Hospital, Assistance Publique Hôpitaux de Paris, Université Paris-Est Créteil, INSERM U955 | Christophe Rodriguez, Slim Fourati, Vanessa Demontant, Guillaume Gricourt, Melissa N'Debi, Alexandre Soulier, Elisabeth Trawinski, Jean-Michel Pawlotsky |
| EPI_ISL_3391859 | BIO ARD'AISNE | Department of Virology, Henri Mondor University Hospital, Assistance Publique Hôpitaux de Paris, Université Paris-Est Créteil, INSERM U955 | Christophe Rodriguez, Slim Fourati, Vanessa Demontant, Guillaume Gricourt, Melissa N'Debi, Alexandre Soulier, Elisabeth Trawinski, Jean-Michel Pawlotsky |
| EPI_ISL_3391860 | Biogroup Bio Lam-LCD Saint-Denis | Department of Virology, Henri Mondor University Hospital, Assistance Publique Hôpitaux de Paris, Université Paris-Est Créteil, INSERM U955 | Christophe Rodriguez, Slim Fourati, Vanessa Demontant, Guillaume Gricourt, Melissa N'Debi, Alexandre Soulier, Elisabeth Trawinski, Jean-Michel Pawlotsky |
| EPI_ISL_3391861 | SELAS MEDILYS | Department of Virology, Henri Mondor University Hospital, Assistance Publique Hôpitaux de Paris, Université Paris-Est Créteil, INSERM U955 | Christophe Rodriguez, Slim Fourati, Vanessa Demontant, Guillaume Gricourt, Melissa N'Debi, Alexandre Soulier, Elisabeth Trawinski, Jean-Michel Pawlotsky |
| EPI_ISL_3391862 | BIO ARD'AISNE | Department of Virology, Henri Mondor University Hospital, Assistance Publique Hôpitaux de Paris, Université Paris-Est Créteil, INSERM U955 | Christophe Rodriguez, Slim Fourati, Vanessa Demontant, Guillaume Gricourt, Melissa N'Debi, Alexandre Soulier, Elisabeth Trawinski, Jean-Michel Pawlotsky |
| EPI_ISL_3391863 | Laboratoire EspaceBio | Department of Virology, Henri Mondor University Hospital, Assistance Publique Hôpitaux de Paris, Université Paris-Est Créteil, INSERM U955 | Christophe Rodriguez, Slim Fourati, Vanessa Demontant, Guillaume Gricourt, Melissa N'Debi, Alexandre Soulier, Elisabeth Trawinski, Jean-Michel Pawlotsky |
| EPI_ISL_3391864 | SELAS BC-LAB | Department of Virology, Henri Mondor University Hospital, Assistance Publique Hôpitaux de Paris, Université Paris-Est Créteil, INSERM U955 | Christophe Rodriguez, Slim Fourati, Vanessa Demontant, Guillaume Gricourt, Melissa N'Debi, Alexandre Soulier, Elisabeth Trawinski, Jean-Michel Pawlotsky |
| EPI_ISL_3391865, EPI_ISL_3391866 | SELAS CERBALLIANCE Normandie ouest | Department of Virology, Henri Mondor University Hospital, Assistance Publique Hôpitaux de Paris, Université Paris-Est Créteil, INSERM U955 | Christophe Rodriguez, Slim Fourati, Vanessa Demontant, Guillaume Gricourt, Melissa N'Debi, Alexandre Soulier, Elisabeth Trawinski, Jean-Michel Pawlotsky |
| EPI_ISL_3391867 | SELAS MEDILYS | Department of Virology, Henri Mondor University Hospital, Assistance Publique Hôpitaux de Paris, Université Paris-Est Créteil, INSERM U955 | Christophe Rodriguez, Slim Fourati, Vanessa Demontant, Guillaume Gricourt, Melissa N'Debi, Alexandre Soulier, Elisabeth Trawinski, Jean-Michel Pawlotsky |
| EPI_ISL_3391868 | SELAS BC-LAB | Department of Virology, Henri Mondor University Hospital, Assistance Publique Hôpitaux de Paris, Université Paris-Est Créteil, INSERM U955 | Christophe Rodriguez, Slim Fourati, Vanessa Demontant, Guillaume Gricourt, Melissa N'Debi, Alexandre Soulier, Elisabeth Trawinski, Jean-Michel Pawlotsky |
| EPI_ISL_3391869 | Biogroup Bio Lam-LCD Saint-Denis | Department of Virology, Henri Mondor University Hospital, Assistance Publique Hôpitaux de Paris, Université Paris-Est Créteil, INSERM U955 | Christophe Rodriguez, Slim Fourati, Vanessa Demontant, Guillaume Gricourt, Melissa N'Debi, Alexandre Soulier, Elisabeth Trawinski, Jean-Michel Pawlotsky |
| EPI_ISL_3391870 | SELAS MEDILYS | Department of Virology, Henri Mondor University Hospital, Assistance Publique Hôpitaux de Paris, Université Paris-Est Créteil, INSERM U955 | Christophe Rodriguez, Slim Fourati, Vanessa Demontant, Guillaume Gricourt, Melissa N'Debi, Alexandre Soulier, Elisabeth Trawinski, Jean-Michel Pawlotsky |
| EPI_ISL_3391871 | Biogroup Bio Lam-LCD Saint-Denis | Department of Virology, Henri Mondor University Hospital, Assistance Publique Hôpitaux de Paris, Université Paris-Est Créteil, INSERM U955 | Christophe Rodriguez, Slim Fourati, Vanessa Demontant, Guillaume Gricourt, Melissa N'Debi, Alexandre Soulier, Elisabeth Trawinski, Jean-Michel Pawlotsky |
| EPI_ISL_3391872 | SELAS MEDILYS | Department of Virology, Henri Mondor University Hospital, Assistance Publique Hôpitaux de Paris, Université Paris-Est Créteil, INSERM U955 | Christophe Rodriguez, Slim Fourati, Vanessa Demontant, Guillaume Gricourt, Melissa N'Debi, Alexandre Soulier, Elisabeth Trawinski, Jean-Michel Pawlotsky |
| EPI_ISL_3391873 | GH de l'Est Francilien | Department of Virology, Henri Mondor University Hospital, Assistance Publique Hôpitaux de Paris, Université Paris-Est Créteil, INSERM U955 | Christophe Rodriguez, Slim Fourati, Vanessa Demontant, Guillaume Gricourt, Melissa N'Debi, Alexandre Soulier, Elisabeth Trawinski, Jean-Michel Pawlotsky |
| EPI_ISL_3391874 | CH.LE RAINCY_MONTFERMEIL | Department of Virology, Henri Mondor University Hospital, Assistance Publique Hôpitaux de Paris, Université Paris-Est Créteil, INSERM U955 | Christophe Rodriguez, Slim Fourati, Vanessa Demontant, Guillaume Gricourt, Melissa N'Debi, Alexandre Soulier, Elisabeth Trawinski, Jean-Michel Pawlotsky |
| EPI_ISL_3391875 | SELAS MEDILYS | Department of Virology, Henri Mondor University Hospital, Assistance Publique Hôpitaux de Paris, Université Paris-Est Créteil, INSERM U955 | Christophe Rodriguez, Slim Fourati, Vanessa Demontant, Guillaume Gricourt, Melissa N'Debi, Alexandre Soulier, Elisabeth Trawinski, Jean-Michel Pawlotsky |
| EPI_ISL_3391876 | BIO ARD'AISNE | Department of Virology, Henri Mondor University Hospital, Assistance Publique Hôpitaux de Paris, Université Paris-Est Créteil, INSERM U955 | Christophe Rodriguez, Slim Fourati, Vanessa Demontant, Guillaume Gricourt, Melissa N'Debi, Alexandre Soulier, Elisabeth Trawinski, Jean-Michel Pawlotsky |
| EPI_ISL_3391877 | SELAS MEDILYS | Department of Virology, Henri Mondor University Hospital, Assistance Publique Hôpitaux de Paris, Université Paris-Est Créteil, INSERM U955 | Christophe Rodriguez, Slim Fourati, Vanessa Demontant, Guillaume Gricourt, Melissa N'Debi, Alexandre Soulier, Elisabeth Trawinski, Jean-Michel Pawlotsky |
| EPI_ISL_3391878 | CH Mémorial F.E.U Laboratoire | Department of Virology, Henri Mondor University Hospital, Assistance Publique Hôpitaux de Paris, Université Paris-Est Créteil, INSERM U955 | Christophe Rodriguez, Slim Fourati, Vanessa Demontant, Guillaume Gricourt, Melissa N'Debi, Alexandre Soulier, Elisabeth Trawinski, Jean-Michel Pawlotsky |

| | | | |
|---|---|---|---|
| EPI_ISL_3391879 | Biogroup Bio Lam-LCD Saint-Denis | Department of Virology, Henri Mondor University Hospital, Assistance Publique Hôpitaux de Paris, Université Paris-Est Créteil, INSERM U955 | Christophe Rodriguez, Slim Fourati, Vanessa Demontant, Guillaume Gricourt, Melissa N'Debi, Alexandre Soulier, Elisabeth Trawinski, Jean-Michel Pawlotsky |
| EPI_ISL_3391880 | LAM BARRAND | Department of Virology, Henri Mondor University Hospital, Assistance Publique Hôpitaux de Paris, Université Paris-Est Créteil, INSERM U955 | Christophe Rodriguez, Slim Fourati, Vanessa Demontant, Guillaume Gricourt, Melissa N'Debi, Alexandre Soulier, Elisabeth Trawinski, Jean-Michel Pawlotsky |
| EPI_ISL_3391881 | SELAS MEDILYS | Department of Virology, Henri Mondor University Hospital, Assistance Publique Hôpitaux de Paris, Université Paris-Est Créteil, INSERM U955 | Christophe Rodriguez, Slim Fourati, Vanessa Demontant, Guillaume Gricourt, Melissa N'Debi, Alexandre Soulier, Elisabeth Trawinski, Jean-Michel Pawlotsky |
| EPI_ISL_3391882 | GH de l'Est Francilien | Department of Virology, Henri Mondor University Hospital, Assistance Publique Hôpitaux de Paris, Université Paris-Est Créteil, INSERM U955 | Christophe Rodriguez, Slim Fourati, Vanessa Demontant, Guillaume Gricourt, Melissa N'Debi, Alexandre Soulier, Elisabeth Trawinski, Jean-Michel Pawlotsky |
| EPI_ISL_3391883 | Biogroup Bio Lam-LCD Saint-Denis | Department of Virology, Henri Mondor University Hospital, Assistance Publique Hôpitaux de Paris, Université Paris-Est Créteil, INSERM U955 | Christophe Rodriguez, Slim Fourati, Vanessa Demontant, Guillaume Gricourt, Melissa N'Debi, Alexandre Soulier, Elisabeth Trawinski, Jean-Michel Pawlotsky |
| EPI_ISL_3391884 | CH.LE RAINCY_MONTFERMEIL | Department of Virology, Henri Mondor University Hospital, Assistance Publique Hôpitaux de Paris, Université Paris-Est Créteil, INSERM U955 | Christophe Rodriguez, Slim Fourati, Vanessa Demontant, Guillaume Gricourt, Melissa N'Debi, Alexandre Soulier, Elisabeth Trawinski, Jean-Michel Pawlotsky |
| EPI_ISL_3391885 | BIO ARD'AISNE | Department of Virology, Henri Mondor University Hospital, Assistance Publique Hôpitaux de Paris, Université Paris-Est Créteil, INSERM U955 | Christophe Rodriguez, Slim Fourati, Vanessa Demontant, Guillaume Gricourt, Melissa N'Debi, Alexandre Soulier, Elisabeth Trawinski, Jean-Michel Pawlotsky |
| EPI_ISL_3391886 | GH de l'Est Francilien | Department of Virology, Henri Mondor University Hospital, Assistance Publique Hôpitaux de Paris, Université Paris-Est Créteil, INSERM U955 | Christophe Rodriguez, Slim Fourati, Vanessa Demontant, Guillaume Gricourt, Melissa N'Debi, Alexandre Soulier, Elisabeth Trawinski, Jean-Michel Pawlotsky |
| EPI_ISL_3391887 | CH. CLEMENCEAU | Department of Virology, Henri Mondor University Hospital, Assistance Publique Hôpitaux de Paris, Université Paris-Est Créteil, INSERM U955 | Christophe Rodriguez, Slim Fourati, Vanessa Demontant, Guillaume Gricourt, Melissa N'Debi, Alexandre Soulier, Elisabeth Trawinski, Jean-Michel Pawlotsky |
| EPI_ISL_3391888, EPI_ISL_3391890 | SELAS MEDILYS | Department of Virology, Henri Mondor University Hospital, Assistance Publique Hôpitaux de Paris, Université Paris-Est Créteil, INSERM U955 | Christophe Rodriguez, Slim Fourati, Vanessa Demontant, Guillaume Gricourt, Melissa N'Debi, Alexandre Soulier, Elisabeth Trawinski, Jean-Michel Pawlotsky |
| EPI_ISL_3391893 | Laboratoire multi-site du Diaconat | Department of Virology, Henri Mondor University Hospital, Assistance Publique Hôpitaux de Paris, Université Paris-Est Créteil, INSERM U955 | Christophe Rodriguez, Slim Fourati, Vanessa Demontant, Guillaume Gricourt, Melissa N'Debi, Alexandre Soulier, Elisabeth Trawinski, Jean-Michel Pawlotsky |
| EPI_ISL_3391894 | Biogroup Bio Lam-LCD Saint-Denis | Department of Virology, Henri Mondor University Hospital, Assistance Publique Hôpitaux de Paris, Université Paris-Est Créteil, INSERM U955 | Christophe Rodriguez, Slim Fourati, Vanessa Demontant, Guillaume Gricourt, Melissa N'Debi, Alexandre Soulier, Elisabeth Trawinski, Jean-Michel Pawlotsky |
| EPI_ISL_3391895 | SELAS MEDILYS | Department of Virology, Henri Mondor University Hospital, Assistance Publique Hôpitaux de Paris, Université Paris-Est Créteil, INSERM U955 | Christophe Rodriguez, Slim Fourati, Vanessa Demontant, Guillaume Gricourt, Melissa N'Debi, Alexandre Soulier, Elisabeth Trawinski, Jean-Michel Pawlotsky |
| EPI_ISL_3391896 | GH de l'Est Francilien | Department of Virology, Henri Mondor University Hospital, Assistance Publique Hôpitaux de Paris, Université Paris-Est Créteil, INSERM U955 | Christophe Rodriguez, Slim Fourati, Vanessa Demontant, Guillaume Gricourt, Melissa N'Debi, Alexandre Soulier, Elisabeth Trawinski, Jean-Michel Pawlotsky |
| EPI_ISL_3391897 | Biogroup Bio Lam-LCD Saint-Denis | Department of Virology, Henri Mondor University Hospital, Assistance Publique Hôpitaux de Paris, Université Paris-Est Créteil, INSERM U955 | Christophe Rodriguez, Slim Fourati, Vanessa Demontant, Guillaume Gricourt, Melissa N'Debi, Alexandre Soulier, Elisabeth Trawinski, Jean-Michel Pawlotsky |
| EPI_ISL_3391899 | CH.LE RAINCY_MONTFERMEIL | Department of Virology, Henri Mondor University Hospital, Assistance Publique Hôpitaux de Paris, Université Paris-Est Créteil, INSERM U955 | Christophe Rodriguez, Slim Fourati, Vanessa Demontant, Guillaume Gricourt, Melissa N'Debi, Alexandre Soulier, Elisabeth Trawinski, Jean-Michel Pawlotsky |
| EPI_ISL_3391900 | SELAS MEDILYS | Department of Virology, Henri Mondor University Hospital, Assistance Publique Hôpitaux de Paris, Université Paris-Est Créteil, INSERM U955 | Christophe Rodriguez, Slim Fourati, Vanessa Demontant, Guillaume Gricourt, Melissa N'Debi, Alexandre Soulier, Elisabeth Trawinski, Jean-Michel Pawlotsky |
| EPI_ISL_3391901 | Laboratoire multi-site du Diaconat | Department of Virology, Henri Mondor University Hospital, Assistance Publique Hôpitaux de Paris, Université Paris-Est Créteil, INSERM U955 | Christophe Rodriguez, Slim Fourati, Vanessa Demontant, Guillaume Gricourt, Melissa N'Debi, Alexandre Soulier, Elisabeth Trawinski, Jean-Michel Pawlotsky |
| EPI_ISL_3391902 | SELAS MEDILYS | Department of Virology, Henri Mondor University Hospital, Assistance Publique Hôpitaux de Paris, Université Paris-Est Créteil, INSERM U955 | Christophe Rodriguez, Slim Fourati, Vanessa Demontant, Guillaume Gricourt, Melissa N'Debi, Alexandre Soulier, Elisabeth Trawinski, Jean-Michel Pawlotsky |
| EPI_ISL_3391903 | BIO ARD'AISNE | Department of Virology, Henri Mondor University Hospital, Assistance Publique Hôpitaux de Paris, Université Paris-Est Créteil, INSERM U955 | Christophe Rodriguez, Slim Fourati, Vanessa Demontant, Guillaume Gricourt, Melissa N'Debi, Alexandre Soulier, Elisabeth Trawinski, Jean-Michel Pawlotsky |
| EPI_ISL_3391904, EPI_ISL_3391905 | BPR Analyses spécialisées | Department of Virology, Henri Mondor University Hospital, Assistance Publique Hôpitaux de Paris, Université Paris-Est Créteil, INSERM U955 | Christophe Rodriguez, Slim Fourati, Vanessa Demontant, Guillaume Gricourt, Melissa N'Debi, Alexandre Soulier, Elisabeth Trawinski, Jean-Michel Pawlotsky |
| EPI_ISL_3391906 | Biogroup Bio Lam-LCD Saint-Denis | Department of Virology, Henri Mondor University Hospital, Assistance Publique Hôpitaux de Paris, Université Paris-Est Créteil, INSERM U955 | Christophe Rodriguez, Slim Fourati, Vanessa Demontant, Guillaume Gricourt, Melissa N'Debi, Alexandre Soulier, Elisabeth Trawinski, Jean-Michel Pawlotsky |
| EPI_ISL_3391907 | SELAS BC-LAB | Department of Virology, Henri Mondor University Hospital, Assistance Publique Hôpitaux de Paris, Université Paris-Est Créteil, INSERM U955 | Christophe Rodriguez, Slim Fourati, Vanessa Demontant, Guillaume Gricourt, Melissa N'Debi, Alexandre Soulier, Elisabeth Trawinski, Jean-Michel Pawlotsky |
| EPI_ISL_3391908 | SELAS CERBALLIANCE Normandie ouest | Department of Virology, Henri Mondor University Hospital, Assistance Publique Hôpitaux de Paris, Université Paris-Est Créteil, INSERM U955 | Christophe Rodriguez, Slim Fourati, Vanessa Demontant, Guillaume Gricourt, Melissa N'Debi, Alexandre Soulier, Elisabeth Trawinski, Jean-Michel Pawlotsky |
| EPI_ISL_3391910, EPI_ISL_3391911 | LABM DEFRANCE | Department of Virology, Henri Mondor University Hospital, Assistance Publique Hôpitaux de Paris, Université Paris-Est Créteil, INSERM U955 | Christophe Rodriguez, Slim Fourati, Vanessa Demontant, Guillaume Gricourt, Melissa N'Debi, Alexandre Soulier, Elisabeth Trawinski, Jean-Michel Pawlotsky |
| EPI_ISL_3391912 | SELAS MEDILYS | Department of Virology, Henri Mondor University Hospital, Assistance Publique Hôpitaux de Paris, Université Paris-Est Créteil, INSERM U955 | Christophe Rodriguez, Slim Fourati, Vanessa Demontant, Guillaume Gricourt, Melissa N'Debi, Alexandre Soulier, Elisabeth Trawinski, Jean-Michel Pawlotsky |
| EPI_ISL_3391913 | GH de l'Est Francilien | Department of Virology, Henri Mondor University Hospital, Assistance Publique Hôpitaux de Paris, Université Paris-Est Créteil, INSERM U955 | Christophe Rodriguez, Slim Fourati, Vanessa Demontant, Guillaume Gricourt, Melissa N'Debi, Alexandre Soulier, Elisabeth Trawinski, Jean-Michel Pawlotsky |
| EPI_ISL_3391914 | Laboratoire multi-site du Diaconat | Department of Virology, Henri Mondor University Hospital, Assistance Publique Hôpitaux de Paris, Université Paris-Est Créteil, INSERM U955 | Christophe Rodriguez, Slim Fourati, Vanessa Demontant, Guillaume Gricourt, Melissa N'Debi, Alexandre Soulier, Elisabeth Trawinski, Jean-Michel Pawlotsky |
| EPI_ISL_3391916, EPI_ISL_3391917 | Biogroup Bio Lam-LCD Saint-Denis | Department of Virology, Henri Mondor University Hospital, Assistance Publique Hôpitaux de Paris, Université Paris-Est Créteil, INSERM U955 | Christophe Rodriguez, Slim Fourati, Vanessa Demontant, Guillaume Gricourt, Melissa N'Debi, Alexandre Soulier, Elisabeth Trawinski, Jean-Michel Pawlotsky |
| EPI_ISL_3391918 | BPR Analyses spécialisées | Department of Virology, Henri Mondor University Hospital, Assistance Publique Hôpitaux de Paris, Université Paris-Est Créteil, INSERM U955 | Christophe Rodriguez, Slim Fourati, Vanessa Demontant, Guillaume Gricourt, Melissa N'Debi, Alexandre Soulier, Elisabeth Trawinski, Jean-Michel Pawlotsky |
| EPI_ISL_3391919 | SELAS BC-LAB | Department of Virology, Henri Mondor University Hospital, Assistance Publique Hôpitaux de Paris, Université Paris-Est Créteil, INSERM U955 | Christophe Rodriguez, Slim Fourati, Vanessa Demontant, Guillaume Gricourt, Melissa N'Debi, Alexandre Soulier, Elisabeth Trawinski, Jean-Michel Pawlotsky |
| EPI_ISL_3391920 | SELAS MEDILYS | Department of Virology, Henri Mondor University Hospital, Assistance Publique Hôpitaux de Paris, Université Paris-Est Créteil, INSERM U955 | Christophe Rodriguez, Slim Fourati, Vanessa Demontant, Guillaume Gricourt, Melissa N'Debi, Alexandre Soulier, Elisabeth Trawinski, Jean-Michel Pawlotsky |

| | | | |
|---|---|---|---|
| EPI_ISL_3391921 | BIOLAB UNILABS | Department of Virology, Henri Mondor University Hospital, Assistance Publique Hôpitaux de Paris, Université Paris-Est Créteil, INSERM U955 | Christophe Rodriguez, Slim Fourati, Vanessa Demontant, Guillaume Gricourt, Melissa N'Debi, Alexandre Soulier, Elisabeth Trawinski, Jean-Michel Pawlotsky |
| EPI_ISL_3391922 | Biogroup Bio Lam-LCD Saint-Denis | Department of Virology, Henri Mondor University Hospital, Assistance Publique Hôpitaux de Paris, Université Paris-Est Créteil, INSERM U955 | Christophe Rodriguez, Slim Fourati, Vanessa Demontant, Guillaume Gricourt, Melissa N'Debi, Alexandre Soulier, Elisabeth Trawinski, Jean-Michel Pawlotsky |
| EPI_ISL_3391923 | GH de l'Est Francilien | Department of Virology, Henri Mondor University Hospital, Assistance Publique Hôpitaux de Paris, Université Paris-Est Créteil, INSERM U955 | Christophe Rodriguez, Slim Fourati, Vanessa Demontant, Guillaume Gricourt, Melissa N'Debi, Alexandre Soulier, Elisabeth Trawinski, Jean-Michel Pawlotsky |
| EPI_ISL_3391924, EPI_ISL_3391925, EPI_ISL_3391926, EPI_ISL_3391927 | Biogroup Bio Lam-LCD Saint-Denis | Department of Virology, Henri Mondor University Hospital, Assistance Publique Hôpitaux de Paris, Université Paris-Est Créteil, INSERM U955 | Christophe Rodriguez, Slim Fourati, Vanessa Demontant, Guillaume Gricourt, Melissa N'Debi, Alexandre Soulier, Elisabeth Trawinski, Jean-Michel Pawlotsky |
| EPI_ISL_3391928 | GH de l'Est Francilien | Department of Virology, Henri Mondor University Hospital, Assistance Publique Hôpitaux de Paris, Université Paris-Est Créteil, INSERM U955 | Christophe Rodriguez, Slim Fourati, Vanessa Demontant, Guillaume Gricourt, Melissa N'Debi, Alexandre Soulier, Elisabeth Trawinski, Jean-Michel Pawlotsky |
| EPI_ISL_3391929 | Biogroup Bio Lam-LCD Saint-Denis | Department of Virology, Henri Mondor University Hospital, Assistance Publique Hôpitaux de Paris, Université Paris-Est Créteil, INSERM U955 | Christophe Rodriguez, Slim Fourati, Vanessa Demontant, Guillaume Gricourt, Melissa N'Debi, Alexandre Soulier, Elisabeth Trawinski, Jean-Michel Pawlotsky |
| EPI_ISL_3391930 | BIOLBS Rouen | Department of Virology, Henri Mondor University Hospital, Assistance Publique Hôpitaux de Paris, Université Paris-Est Créteil, INSERM U955 | Christophe Rodriguez, Slim Fourati, Vanessa Demontant, Guillaume Gricourt, Melissa N'Debi, Alexandre Soulier, Elisabeth Trawinski, Jean-Michel Pawlotsky |
| EPI_ISL_3391931 | Biogroup Bio Lam-LCD Saint-Denis | Department of Virology, Henri Mondor University Hospital, Assistance Publique Hôpitaux de Paris, Université Paris-Est Créteil, INSERM U955 | Christophe Rodriguez, Slim Fourati, Vanessa Demontant, Guillaume Gricourt, Melissa N'Debi, Alexandre Soulier, Elisabeth Trawinski, Jean-Michel Pawlotsky |
| EPI_ISL_3391932 | CHU Rouen Normandie | Department of Virology, Henri Mondor University Hospital, Assistance Publique Hôpitaux de Paris, Université Paris-Est Créteil, INSERM U955 | Christophe Rodriguez, Slim Fourati, Vanessa Demontant, Guillaume Gricourt, Melissa N'Debi, Alexandre Soulier, Elisabeth Trawinski, Jean-Michel Pawlotsky |
| EPI_ISL_3391933 | SELAS CERBALLIANCE Normandie ouest | Department of Virology, Henri Mondor University Hospital, Assistance Publique Hôpitaux de Paris, Université Paris-Est Créteil, INSERM U955 | Christophe Rodriguez, Slim Fourati, Vanessa Demontant, Guillaume Gricourt, Melissa N'Debi, Alexandre Soulier, Elisabeth Trawinski, Jean-Michel Pawlotsky |
| EPI_ISL_3391934 | Biogroup Bio Lam-LCD Saint-Denis | Department of Virology, Henri Mondor University Hospital, Assistance Publique Hôpitaux de Paris, Université Paris-Est Créteil, INSERM U955 | Christophe Rodriguez, Slim Fourati, Vanessa Demontant, Guillaume Gricourt, Melissa N'Debi, Alexandre Soulier, Elisabeth Trawinski, Jean-Michel Pawlotsky |
| EPI_ISL_3391935 | SELAS BC-LAB | Department of Virology, Henri Mondor University Hospital, Assistance Publique Hôpitaux de Paris, Université Paris-Est Créteil, INSERM U955 | Christophe Rodriguez, Slim Fourati, Vanessa Demontant, Guillaume Gricourt, Melissa N'Debi, Alexandre Soulier, Elisabeth Trawinski, Jean-Michel Pawlotsky |
| EPI_ISL_3391936 | SELAS MEDILYS | Department of Virology, Henri Mondor University Hospital, Assistance Publique Hôpitaux de Paris, Université Paris-Est Créteil, INSERM U955 | Christophe Rodriguez, Slim Fourati, Vanessa Demontant, Guillaume Gricourt, Melissa N'Debi, Alexandre Soulier, Elisabeth Trawinski, Jean-Michel Pawlotsky |
| EPI_ISL_3391937 | Biogroup Bio Lam-LCD Saint-Denis | Department of Virology, Henri Mondor University Hospital, Assistance Publique Hôpitaux de Paris, Université Paris-Est Créteil, INSERM U955 | Christophe Rodriguez, Slim Fourati, Vanessa Demontant, Guillaume Gricourt, Melissa N'Debi, Alexandre Soulier, Elisabeth Trawinski, Jean-Michel Pawlotsky |
| EPI_ISL_3391939 | BIO ARD'AISNE | Department of Virology, Henri Mondor University Hospital, Assistance Publique Hôpitaux de Paris, Université Paris-Est Créteil, INSERM U955 | Christophe Rodriguez, Slim Fourati, Vanessa Demontant, Guillaume Gricourt, Melissa N'Debi, Alexandre Soulier, Elisabeth Trawinski, Jean-Michel Pawlotsky |
| EPI_ISL_3391940 | GH de l'Est Francilien | Department of Virology, Henri Mondor University Hospital, Assistance Publique Hôpitaux de Paris, Université Paris-Est Créteil, INSERM U955 | Christophe Rodriguez, Slim Fourati, Vanessa Demontant, Guillaume Gricourt, Melissa N'Debi, Alexandre Soulier, Elisabeth Trawinski, Jean-Michel Pawlotsky |
| EPI_ISL_3391941 | Biogroup Bio Lam-LCD Saint-Denis | Department of Virology, Henri Mondor University Hospital, Assistance Publique Hôpitaux de Paris, Université Paris-Est Créteil, INSERM U955 | Christophe Rodriguez, Slim Fourati, Vanessa Demontant, Guillaume Gricourt, Melissa N'Debi, Alexandre Soulier, Elisabeth Trawinski, Jean-Michel Pawlotsky |
| EPI_ISL_3391942 | BPR Analyses spécialisées | Department of Virology, Henri Mondor University Hospital, Assistance Publique Hôpitaux de Paris, Université Paris-Est Créteil, INSERM U955 | Christophe Rodriguez, Slim Fourati, Vanessa Demontant, Guillaume Gricourt, Melissa N'Debi, Alexandre Soulier, Elisabeth Trawinski, Jean-Michel Pawlotsky |
| EPI_ISL_3391943 | Biogroup Bio Lam-LCD Saint-Denis | Department of Virology, Henri Mondor University Hospital, Assistance Publique Hôpitaux de Paris, Université Paris-Est Créteil, INSERM U955 | Christophe Rodriguez, Slim Fourati, Vanessa Demontant, Guillaume Gricourt, Melissa N'Debi, Alexandre Soulier, Elisabeth Trawinski, Jean-Michel Pawlotsky |
| EPI_ISL_3391945 | SELAS MEDILYS | Department of Virology, Henri Mondor University Hospital, Assistance Publique Hôpitaux de Paris, Université Paris-Est Créteil, INSERM U955 | Christophe Rodriguez, Slim Fourati, Vanessa Demontant, Guillaume Gricourt, Melissa N'Debi, Alexandre Soulier, Elisabeth Trawinski, Jean-Michel Pawlotsky |
| EPI_ISL_3391946 | Biogroup Bio Lam-LCD Saint-Denis | Department of Virology, Henri Mondor University Hospital, Assistance Publique Hôpitaux de Paris, Université Paris-Est Créteil, INSERM U955 | Christophe Rodriguez, Slim Fourati, Vanessa Demontant, Guillaume Gricourt, Melissa N'Debi, Alexandre Soulier, Elisabeth Trawinski, Jean-Michel Pawlotsky |
| EPI_ISL_3391948 | SELAS MEDILYS | Department of Virology, Henri Mondor University Hospital, Assistance Publique Hôpitaux de Paris, Université Paris-Est Créteil, INSERM U955 | Christophe Rodriguez, Slim Fourati, Vanessa Demontant, Guillaume Gricourt, Melissa N'Debi, Alexandre Soulier, Elisabeth Trawinski, Jean-Michel Pawlotsky |
| EPI_ISL_3391949 | SELAS BC-LAB | Department of Virology, Henri Mondor University Hospital, Assistance Publique Hôpitaux de Paris, Université Paris-Est Créteil, INSERM U955 | Christophe Rodriguez, Slim Fourati, Vanessa Demontant, Guillaume Gricourt, Melissa N'Debi, Alexandre Soulier, Elisabeth Trawinski, Jean-Michel Pawlotsky |
| EPI_ISL_3391950, EPI_ISL_3391951 | SELAS MEDILYS | Department of Virology, Henri Mondor University Hospital, Assistance Publique Hôpitaux de Paris, Université Paris-Est Créteil, INSERM U955 | Christophe Rodriguez, Slim Fourati, Vanessa Demontant, Guillaume Gricourt, Melissa N'Debi, Alexandre Soulier, Elisabeth Trawinski, Jean-Michel Pawlotsky |
| EPI_ISL_3391952, EPI_ISL_3391953, EPI_ISL_3391954 | Biogroup Bio Lam-LCD Saint-Denis | Department of Virology, Henri Mondor University Hospital, Assistance Publique Hôpitaux de Paris, Université Paris-Est Créteil, INSERM U955 | Christophe Rodriguez, Slim Fourati, Vanessa Demontant, Guillaume Gricourt, Melissa N'Debi, Alexandre Soulier, Elisabeth Trawinski, Jean-Michel Pawlotsky |
| EPI_ISL_3391955 | LABM DEFRANCE | Department of Virology, Henri Mondor University Hospital, Assistance Publique Hôpitaux de Paris, Université Paris-Est Créteil, INSERM U955 | Christophe Rodriguez, Slim Fourati, Vanessa Demontant, Guillaume Gricourt, Melissa N'Debi, Alexandre Soulier, Elisabeth Trawinski, Jean-Michel Pawlotsky |
| EPI_ISL_3391956 | SELAS MEDILYS | Department of Virology, Henri Mondor University Hospital, Assistance Publique Hôpitaux de Paris, Université Paris-Est Créteil, INSERM U955 | Christophe Rodriguez, Slim Fourati, Vanessa Demontant, Guillaume Gricourt, Melissa N'Debi, Alexandre Soulier, Elisabeth Trawinski, Jean-Michel Pawlotsky |
| EPI_ISL_3391957 | Biogroup Bio Lam-LCD Saint-Denis | Department of Virology, Henri Mondor University Hospital, Assistance Publique Hôpitaux de Paris, Université Paris-Est Créteil, INSERM U955 | Christophe Rodriguez, Slim Fourati, Vanessa Demontant, Guillaume Gricourt, Melissa N'Debi, Alexandre Soulier, Elisabeth Trawinski, Jean-Michel Pawlotsky |
| EPI_ISL_3391958 | SELAS MEDILYS | Department of Virology, Henri Mondor University Hospital, Assistance Publique Hôpitaux de Paris, Université Paris-Est Créteil, INSERM U955 | Christophe Rodriguez, Slim Fourati, Vanessa Demontant, Guillaume Gricourt, Melissa N'Debi, Alexandre Soulier, Elisabeth Trawinski, Jean-Michel Pawlotsky |
| EPI_ISL_3391959 | Biogroup Bio Lam-LCD Saint-Denis | Department of Virology, Henri Mondor University Hospital, Assistance Publique Hôpitaux de Paris, Université Paris-Est Créteil, INSERM U955 | Christophe Rodriguez, Slim Fourati, Vanessa Demontant, Guillaume Gricourt, Melissa N'Debi, Alexandre Soulier, Elisabeth Trawinski, Jean-Michel Pawlotsky |
| EPI_ISL_3391960 | Laboratoire multi-site du Diaconat | Department of Virology, Henri Mondor University Hospital, Assistance Publique Hôpitaux de Paris, Université Paris-Est Créteil, INSERM U955 | Christophe Rodriguez, Slim Fourati, Vanessa Demontant, Guillaume Gricourt, Melissa N'Debi, Alexandre Soulier, Elisabeth Trawinski, Jean-Michel Pawlotsky |
| EPI_ISL_3391961 | SELAS MEDILYS | Department of Virology, Henri Mondor University Hospital, Assistance Publique Hôpitaux de Paris, Université Paris-Est Créteil, | Christophe Rodriguez, Slim Fourati, Vanessa Demontant, Guillaume Gricourt, Melissa N'Debi, Alexandre Soulier, Elisabeth Trawinski, Jean-Michel Pawlotsky |

| | | | |
|---|---|---|---|
| | | INSERM U955 | |
| EPI_ISL_3391962, EPI_ISL_3391963 | Biogroup Bio Lam-LCD Saint-Denis | Department of Virology, Henri Mondor University Hospital, Assistance Publique Hôpitaux de Paris, Université Paris-Est Créteil, INSERM U955 | Christophe Rodriguez, Slim Fourati, Vanessa Demontant, Guillaume Gricourt, Melissa N'Debi, Alexandre Soulier, Elisabeth Trawinski, Jean-Michel Pawlotsky |
| EPI_ISL_3391964 | LABM DEFRANCE | Department of Virology, Henri Mondor University Hospital, Assistance Publique Hôpitaux de Paris, Université Paris-Est Créteil, INSERM U955 | Christophe Rodriguez, Slim Fourati, Vanessa Demontant, Guillaume Gricourt, Melissa N'Debi, Alexandre Soulier, Elisabeth Trawinski, Jean-Michel Pawlotsky |
| EPI_ISL_3391965 | Biogroup Bio Lam-LCD Saint-Denis | Department of Virology, Henri Mondor University Hospital, Assistance Publique Hôpitaux de Paris, Université Paris-Est Créteil, INSERM U955 | Christophe Rodriguez, Slim Fourati, Vanessa Demontant, Guillaume Gricourt, Melissa N'Debi, Alexandre Soulier, Elisabeth Trawinski, Jean-Michel Pawlotsky |
| EPI_ISL_3391966 | SELAS MEDILYS | Department of Virology, Henri Mondor University Hospital, Assistance Publique Hôpitaux de Paris, Université Paris-Est Créteil, INSERM U955 | Christophe Rodriguez, Slim Fourati, Vanessa Demontant, Guillaume Gricourt, Melissa N'Debi, Alexandre Soulier, Elisabeth Trawinski, Jean-Michel Pawlotsky |
| EPI_ISL_3391967 | CH. CLEMENCEAU | Department of Virology, Henri Mondor University Hospital, Assistance Publique Hôpitaux de Paris, Université Paris-Est Créteil, INSERM U955 | Christophe Rodriguez, Slim Fourati, Vanessa Demontant, Guillaume Gricourt, Melissa N'Debi, Alexandre Soulier, Elisabeth Trawinski, Jean-Michel Pawlotsky |
| EPI_ISL_3391968, EPI_ISL_3391969 | SELAS BC-LAB | Department of Virology, Henri Mondor University Hospital, Assistance Publique Hôpitaux de Paris, Université Paris-Est Créteil, INSERM U955 | Christophe Rodriguez, Slim Fourati, Vanessa Demontant, Guillaume Gricourt, Melissa N'Debi, Alexandre Soulier, Elisabeth Trawinski, Jean-Michel Pawlotsky |
| EPI_ISL_3391970 | SELAS MEDILYS | Department of Virology, Henri Mondor University Hospital, Assistance Publique Hôpitaux de Paris, Université Paris-Est Créteil, INSERM U955 | Christophe Rodriguez, Slim Fourati, Vanessa Demontant, Guillaume Gricourt, Melissa N'Debi, Alexandre Soulier, Elisabeth Trawinski, Jean-Michel Pawlotsky |
| EPI_ISL_3391971 | LAM BARRAND | Department of Virology, Henri Mondor University Hospital, Assistance Publique Hôpitaux de Paris, Université Paris-Est Créteil, INSERM U955 | Christophe Rodriguez, Slim Fourati, Vanessa Demontant, Guillaume Gricourt, Melissa N'Debi, Alexandre Soulier, Elisabeth Trawinski, Jean-Michel Pawlotsky |
| EPI_ISL_3391972 | BIO ARD'AISNE | Department of Virology, Henri Mondor University Hospital, Assistance Publique Hôpitaux de Paris, Université Paris-Est Créteil, INSERM U955 | Christophe Rodriguez, Slim Fourati, Vanessa Demontant, Guillaume Gricourt, Melissa N'Debi, Alexandre Soulier, Elisabeth Trawinski, Jean-Michel Pawlotsky |
| EPI_ISL_3391974 | Biogroup Bio Lam-LCD Saint-Denis | Department of Virology, Henri Mondor University Hospital, Assistance Publique Hôpitaux de Paris, Université Paris-Est Créteil, INSERM U955 | Christophe Rodriguez, Slim Fourati, Vanessa Demontant, Guillaume Gricourt, Melissa N'Debi, Alexandre Soulier, Elisabeth Trawinski, Jean-Michel Pawlotsky |
| EPI_ISL_3391975 | CH.LE RAINCY_MONTFERMEIL | Department of Virology, Henri Mondor University Hospital, Assistance Publique Hôpitaux de Paris, Université Paris-Est Créteil, INSERM U955 | Christophe Rodriguez, Slim Fourati, Vanessa Demontant, Guillaume Gricourt, Melissa N'Debi, Alexandre Soulier, Elisabeth Trawinski, Jean-Michel Pawlotsky |
| EPI_ISL_3391976 | BIO ARD'AISNE | Department of Virology, Henri Mondor University Hospital, Assistance Publique Hôpitaux de Paris, Université Paris-Est Créteil, INSERM U955 | Christophe Rodriguez, Slim Fourati, Vanessa Demontant, Guillaume Gricourt, Melissa N'Debi, Alexandre Soulier, Elisabeth Trawinski, Jean-Michel Pawlotsky |
| EPI_ISL_3391988, EPI_ISL_3391989, EPI_ISL_3391990, EPI_ISL_3391991, EPI_ISL_3391992 | SA Pathology | SA Pathology | Caitlin Selway, Lex Leong, Chuan Kok Lim, Mark Turra, Ivan Bastian |
| EPI_ISL_3391993 | SELAS MEDILYS | Department of Virology, Henri Mondor University Hospital, Assistance Publique Hôpitaux de Paris, Université Paris-Est Créteil, INSERM U955 | Christophe Rodriguez, Slim Fourati, Vanessa Demontant, Guillaume Gricourt, Melissa N'Debi, Alexandre Soulier, Elisabeth Trawinski, Jean-Michel Pawlotsky |
| EPI_ISL_3391994 | Laboratoire ATOUTBIO JDC | Department of Virology, Henri Mondor University Hospital, Assistance Publique Hôpitaux de Paris, Université Paris-Est Créteil, INSERM U955 | Christophe Rodriguez, Slim Fourati, Vanessa Demontant, Guillaume Gricourt, Melissa N'Debi, Alexandre Soulier, Elisabeth Trawinski, Jean-Michel Pawlotsky |
| EPI_ISL_3391995 | SELAS MEDILYS | Department of Virology, Henri Mondor University Hospital, Assistance Publique Hôpitaux de Paris, Université Paris-Est Créteil, INSERM U955 | Christophe Rodriguez, Slim Fourati, Vanessa Demontant, Guillaume Gricourt, Melissa N'Debi, Alexandre Soulier, Elisabeth Trawinski, Jean-Michel Pawlotsky |
| EPI_ISL_3391996 | GH A.CHENEVIER-H.MONDOR | Department of Virology, Henri Mondor University Hospital, Assistance Publique Hôpitaux de Paris, Université Paris-Est Créteil, INSERM U955 | Christophe Rodriguez, Slim Fourati, Vanessa Demontant, Guillaume Gricourt, Melissa N'Debi, Alexandre Soulier, Elisabeth Trawinski, Jean-Michel Pawlotsky |
| EPI_ISL_3391997 | BROUSSAIS | Department of Virology, Henri Mondor University Hospital, Assistance Publique Hôpitaux de Paris, Université Paris-Est Créteil, INSERM U955 | Christophe Rodriguez, Slim Fourati, Vanessa Demontant, Guillaume Gricourt, Melissa N'Debi, Alexandre Soulier, Elisabeth Trawinski, Jean-Michel Pawlotsky |
| EPI_ISL_3392000 | BIO67-BIOSPHERE | Department of Virology, Henri Mondor University Hospital, Assistance Publique Hôpitaux de Paris, Université Paris-Est Créteil, INSERM U955 | Christophe Rodriguez, Slim Fourati, Vanessa Demontant, Guillaume Gricourt, Melissa N'Debi, Alexandre Soulier, Elisabeth Trawinski, Jean-Michel Pawlotsky |
| EPI_ISL_3392001 | SELAS MEDILYS | Department of Virology, Henri Mondor University Hospital, Assistance Publique Hôpitaux de Paris, Université Paris-Est Créteil, INSERM U955 | Christophe Rodriguez, Slim Fourati, Vanessa Demontant, Guillaume Gricourt, Melissa N'Debi, Alexandre Soulier, Elisabeth Trawinski, Jean-Michel Pawlotsky |
| EPI_ISL_3392003 | BIO67-BIOSPHERE | Department of Virology, Henri Mondor University Hospital, Assistance Publique Hôpitaux de Paris, Université Paris-Est Créteil, INSERM U955 | Christophe Rodriguez, Slim Fourati, Vanessa Demontant, Guillaume Gricourt, Melissa N'Debi, Alexandre Soulier, Elisabeth Trawinski, Jean-Michel Pawlotsky |
| EPI_ISL_3392004, EPI_ISL_3392006 | SELAS MEDILYS | Department of Virology, Henri Mondor University Hospital, Assistance Publique Hôpitaux de Paris, Université Paris-Est Créteil, INSERM U955 | Christophe Rodriguez, Slim Fourati, Vanessa Demontant, Guillaume Gricourt, Melissa N'Debi, Alexandre Soulier, Elisabeth Trawinski, Jean-Michel Pawlotsky |
| EPI_ISL_3392008 | LBM du Val d'Orne | Department of Virology, Henri Mondor University Hospital, Assistance Publique Hôpitaux de Paris, Université Paris-Est Créteil, INSERM U955 | Christophe Rodriguez, Slim Fourati, Vanessa Demontant, Guillaume Gricourt, Melissa N'Debi, Alexandre Soulier, Elisabeth Trawinski, Jean-Michel Pawlotsky |
| EPI_ISL_3392009 | SELAS MEDILYS | Department of Virology, Henri Mondor University Hospital, Assistance Publique Hôpitaux de Paris, Université Paris-Est Créteil, INSERM U955 | Christophe Rodriguez, Slim Fourati, Vanessa Demontant, Guillaume Gricourt, Melissa N'Debi, Alexandre Soulier, Elisabeth Trawinski, Jean-Michel Pawlotsky |
| EPI_ISL_3392011 | GH A.CHENEVIER-H.MONDOR | Department of Virology, Henri Mondor University Hospital, Assistance Publique Hôpitaux de Paris, Université Paris-Est Créteil, INSERM U955 | Christophe Rodriguez, Slim Fourati, Vanessa Demontant, Guillaume Gricourt, Melissa N'Debi, Alexandre Soulier, Elisabeth Trawinski, Jean-Michel Pawlotsky |
| EPI_ISL_3392012 | BROUSSAIS | Department of Virology, Henri Mondor University Hospital, Assistance Publique Hôpitaux de Paris, Université Paris-Est Créteil, INSERM U955 | Christophe Rodriguez, Slim Fourati, Vanessa Demontant, Guillaume Gricourt, Melissa N'Debi, Alexandre Soulier, Elisabeth Trawinski, Jean-Michel Pawlotsky |
| EPI_ISL_3392013 | SELAS MEDILYS | Department of Virology, Henri Mondor University Hospital, Assistance Publique Hôpitaux de Paris, Université Paris-Est Créteil, INSERM U955 | Christophe Rodriguez, Slim Fourati, Vanessa Demontant, Guillaume Gricourt, Melissa N'Debi, Alexandre Soulier, Elisabeth Trawinski, Jean-Michel Pawlotsky |
| EPI_ISL_3392015 | Laboratoire CBM 25 TERRE ROUGE | Department of Virology, Henri Mondor University Hospital, Assistance Publique Hôpitaux de Paris, Université Paris-Est Créteil, INSERM U955 | Christophe Rodriguez, Slim Fourati, Vanessa Demontant, Guillaume Gricourt, Melissa N'Debi, Alexandre Soulier, Elisabeth Trawinski, Jean-Michel Pawlotsky |
| EPI_ISL_3392016, EPI_ISL_3392017 | SELAS MEDILYS | Department of Virology, Henri Mondor University Hospital, Assistance Publique Hôpitaux de Paris, Université Paris-Est Créteil, INSERM U955 | Christophe Rodriguez, Slim Fourati, Vanessa Demontant, Guillaume Gricourt, Melissa N'Debi, Alexandre Soulier, Elisabeth Trawinski, Jean-Michel Pawlotsky |
| EPI_ISL_3392020, EPI_ISL_3392021 | BROUSSAIS | Department of Virology, Henri Mondor University Hospital, Assistance Publique Hôpitaux de Paris, Université Paris-Est Créteil, INSERM U955 | Christophe Rodriguez, Slim Fourati, Vanessa Demontant, Guillaume Gricourt, Melissa N'Debi, Alexandre Soulier, Elisabeth Trawinski, Jean-Michel Pawlotsky |
| EPI_ISL_3392022 | SELAS MEDILYS | Department of Virology, Henri Mondor University Hospital, Assistance Publique Hôpitaux de Paris, Université Paris-Est Créteil, INSERM U955 | Christophe Rodriguez, Slim Fourati, Vanessa Demontant, Guillaume Gricourt, Melissa N'Debi, Alexandre Soulier, Elisabeth Trawinski, Jean-Michel Pawlotsky |

| | | | |
|---|---|---|---|
| EPI_ISL_3392023, EPI_ISL_3392024, EPI_ISL_3392025 | BIO67-BIOSPHERE | Department of Virology, Henri Mondor University Hospital, Assistance Publique Hôpitaux de Paris, Université Paris-Est Créteil, INSERM U955 | Christophe Rodriguez, Slim Fourati, Vanessa Demontant, Guillaume Gricourt, Melissa N'Debi, Alexandre Soulier, Elisabeth Trawinski, Jean-Michel Pawlotsky |
| EPI_ISL_3392026, EPI_ISL_3392027 | SELAS MEDILYS | Department of Virology, Henri Mondor University Hospital, Assistance Publique Hôpitaux de Paris, Université Paris-Est Créteil, INSERM U955 | Christophe Rodriguez, Slim Fourati, Vanessa Demontant, Guillaume Gricourt, Melissa N'Debi, Alexandre Soulier, Elisabeth Trawinski, Jean-Michel Pawlotsky |
| EPI_ISL_3392028 | BROUSSAIS | Department of Virology, Henri Mondor University Hospital, Assistance Publique Hôpitaux de Paris, Université Paris-Est Créteil, INSERM U955 | Christophe Rodriguez, Slim Fourati, Vanessa Demontant, Guillaume Gricourt, Melissa N'Debi, Alexandre Soulier, Elisabeth Trawinski, Jean-Michel Pawlotsky |
| EPI_ISL_3392029 | BIO67-BIOSPHERE | Department of Virology, Henri Mondor University Hospital, Assistance Publique Hôpitaux de Paris, Université Paris-Est Créteil, INSERM U955 | Christophe Rodriguez, Slim Fourati, Vanessa Demontant, Guillaume Gricourt, Melissa N'Debi, Alexandre Soulier, Elisabeth Trawinski, Jean-Michel Pawlotsky |
| EPI_ISL_3392031 | SELAS MEDILYS | Department of Virology, Henri Mondor University Hospital, Assistance Publique Hôpitaux de Paris, Université Paris-Est Créteil, INSERM U955 | Christophe Rodriguez, Slim Fourati, Vanessa Demontant, Guillaume Gricourt, Melissa N'Debi, Alexandre Soulier, Elisabeth Trawinski, Jean-Michel Pawlotsky |
| EPI_ISL_3392032 | BIO67-BIOSPHERE | Department of Virology, Henri Mondor University Hospital, Assistance Publique Hôpitaux de Paris, Université Paris-Est Créteil, INSERM U955 | Christophe Rodriguez, Slim Fourati, Vanessa Demontant, Guillaume Gricourt, Melissa N'Debi, Alexandre Soulier, Elisabeth Trawinski, Jean-Michel Pawlotsky |
| EPI_ISL_3392033 | SELAFA SANTE LABO | Department of Virology, Henri Mondor University Hospital, Assistance Publique Hôpitaux de Paris, Université Paris-Est Créteil, INSERM U955 | Christophe Rodriguez, Slim Fourati, Vanessa Demontant, Guillaume Gricourt, Melissa N'Debi, Alexandre Soulier, Elisabeth Trawinski, Jean-Michel Pawlotsky |
| EPI_ISL_3392035 | SELAS MEDILYS | Department of Virology, Henri Mondor University Hospital, Assistance Publique Hôpitaux de Paris, Université Paris-Est Créteil, INSERM U955 | Christophe Rodriguez, Slim Fourati, Vanessa Demontant, Guillaume Gricourt, Melissa N'Debi, Alexandre Soulier, Elisabeth Trawinski, Jean-Michel Pawlotsky |
| EPI_ISL_3392036 | Laboratoire CBM 25 TERRE ROUGE | Department of Virology, Henri Mondor University Hospital, Assistance Publique Hôpitaux de Paris, Université Paris-Est Créteil, INSERM U955 | Christophe Rodriguez, Slim Fourati, Vanessa Demontant, Guillaume Gricourt, Melissa N'Debi, Alexandre Soulier, Elisabeth Trawinski, Jean-Michel Pawlotsky |
| EPI_ISL_3392037, EPI_ISL_3392039, EPI_ISL_3392041 | BIO67-BIOSPHERE | Department of Virology, Henri Mondor University Hospital, Assistance Publique Hôpitaux de Paris, Université Paris-Est Créteil, INSERM U955 | Christophe Rodriguez, Slim Fourati, Vanessa Demontant, Guillaume Gricourt, Melissa N'Debi, Alexandre Soulier, Elisabeth Trawinski, Jean-Michel Pawlotsky |
| EPI_ISL_3392042, EPI_ISL_3392044 | SELAS MEDILYS | Department of Virology, Henri Mondor University Hospital, Assistance Publique Hôpitaux de Paris, Université Paris-Est Créteil, INSERM U955 | Christophe Rodriguez, Slim Fourati, Vanessa Demontant, Guillaume Gricourt, Melissa N'Debi, Alexandre Soulier, Elisabeth Trawinski, Jean-Michel Pawlotsky |
| EPI_ISL_3392045 | SELAFA SANTE LABO | Department of Virology, Henri Mondor University Hospital, Assistance Publique Hôpitaux de Paris, Université Paris-Est Créteil, INSERM U955 | Christophe Rodriguez, Slim Fourati, Vanessa Demontant, Guillaume Gricourt, Melissa N'Debi, Alexandre Soulier, Elisabeth Trawinski, Jean-Michel Pawlotsky |
| EPI_ISL_3392047 | BIO67-BIOSPHERE | Department of Virology, Henri Mondor University Hospital, Assistance Publique Hôpitaux de Paris, Université Paris-Est Créteil, INSERM U955 | Christophe Rodriguez, Slim Fourati, Vanessa Demontant, Guillaume Gricourt, Melissa N'Debi, Alexandre Soulier, Elisabeth Trawinski, Jean-Michel Pawlotsky |
| EPI_ISL_3392048 | SELAS MEDILYS | Department of Virology, Henri Mondor University Hospital, Assistance Publique Hôpitaux de Paris, Université Paris-Est Créteil, INSERM U955 | Christophe Rodriguez, Slim Fourati, Vanessa Demontant, Guillaume Gricourt, Melissa N'Debi, Alexandre Soulier, Elisabeth Trawinski, Jean-Michel Pawlotsky |
| EPI_ISL_3392052 | BIO67-BIOSPHERE | Department of Virology, Henri Mondor University Hospital, Assistance Publique Hôpitaux de Paris, Université Paris-Est Créteil, INSERM U955 | Christophe Rodriguez, Slim Fourati, Vanessa Demontant, Guillaume Gricourt, Melissa N'Debi, Alexandre Soulier, Elisabeth Trawinski, Jean-Michel Pawlotsky |
| EPI_ISL_3392053 | GH A.CHENEVIER-H.MONDOR | Department of Virology, Henri Mondor University Hospital, Assistance Publique Hôpitaux de Paris, Université Paris-Est Créteil, INSERM U955 | Christophe Rodriguez, Slim Fourati, Vanessa Demontant, Guillaume Gricourt, Melissa N'Debi, Alexandre Soulier, Elisabeth Trawinski, Jean-Michel Pawlotsky |
| EPI_ISL_3392055 | Hôpital Cochin | Department of Virology, Henri Mondor University Hospital, Assistance Publique Hôpitaux de Paris, Université Paris-Est Créteil, INSERM U955 | Christophe Rodriguez, Slim Fourati, Vanessa Demontant, Guillaume Gricourt, Melissa N'Debi, Alexandre Soulier, Elisabeth Trawinski, Jean-Michel Pawlotsky |
| EPI_ISL_3392056 | SELAS MEDILYS | Department of Virology, Henri Mondor University Hospital, Assistance Publique Hôpitaux de Paris, Université Paris-Est Créteil, INSERM U955 | Christophe Rodriguez, Slim Fourati, Vanessa Demontant, Guillaume Gricourt, Melissa N'Debi, Alexandre Soulier, Elisabeth Trawinski, Jean-Michel Pawlotsky |
| EPI_ISL_3392058 | Laboratoire ATOUTBIO JDC | Department of Virology, Henri Mondor University Hospital, Assistance Publique Hôpitaux de Paris, Université Paris-Est Créteil, INSERM U955 | Christophe Rodriguez, Slim Fourati, Vanessa Demontant, Guillaume Gricourt, Melissa N'Debi, Alexandre Soulier, Elisabeth Trawinski, Jean-Michel Pawlotsky |
| EPI_ISL_3392059 | Hôpital Cochin | Department of Virology, Henri Mondor University Hospital, Assistance Publique Hôpitaux de Paris, Université Paris-Est Créteil, INSERM U955 | Christophe Rodriguez, Slim Fourati, Vanessa Demontant, Guillaume Gricourt, Melissa N'Debi, Alexandre Soulier, Elisabeth Trawinski, Jean-Michel Pawlotsky |
| EPI_ISL_3392060 | SELAS MEDILYS | Department of Virology, Henri Mondor University Hospital, Assistance Publique Hôpitaux de Paris, Université Paris-Est Créteil, INSERM U955 | Christophe Rodriguez, Slim Fourati, Vanessa Demontant, Guillaume Gricourt, Melissa N'Debi, Alexandre Soulier, Elisabeth Trawinski, Jean-Michel Pawlotsky |
| EPI_ISL_3392061 | BROUSSAIS | Department of Virology, Henri Mondor University Hospital, Assistance Publique Hôpitaux de Paris, Université Paris-Est Créteil, INSERM U955 | Christophe Rodriguez, Slim Fourati, Vanessa Demontant, Guillaume Gricourt, Melissa N'Debi, Alexandre Soulier, Elisabeth Trawinski, Jean-Michel Pawlotsky |
| EPI_ISL_3392062 | BIO67-BIOSPHERE | Department of Virology, Henri Mondor University Hospital, Assistance Publique Hôpitaux de Paris, Université Paris-Est Créteil, INSERM U955 | Christophe Rodriguez, Slim Fourati, Vanessa Demontant, Guillaume Gricourt, Melissa N'Debi, Alexandre Soulier, Elisabeth Trawinski, Jean-Michel Pawlotsky |
| EPI_ISL_3392063 | Laboratoire CBM 25 TERRE ROUGE | Department of Virology, Henri Mondor University Hospital, Assistance Publique Hôpitaux de Paris, Université Paris-Est Créteil, INSERM U955 | Christophe Rodriguez, Slim Fourati, Vanessa Demontant, Guillaume Gricourt, Melissa N'Debi, Alexandre Soulier, Elisabeth Trawinski, Jean-Michel Pawlotsky |
| EPI_ISL_3392064 | BIOMNIS EUROFINS IVRY | Department of Virology, Henri Mondor University Hospital, Assistance Publique Hôpitaux de Paris, Université Paris-Est Créteil, INSERM U955 | Christophe Rodriguez, Slim Fourati, Vanessa Demontant, Guillaume Gricourt, Melissa N'Debi, Alexandre Soulier, Elisabeth Trawinski, Jean-Michel Pawlotsky |
| EPI_ISL_3392065 | SELAS MEDILYS | Department of Virology, Henri Mondor University Hospital, Assistance Publique Hôpitaux de Paris, Université Paris-Est Créteil, INSERM U955 | Christophe Rodriguez, Slim Fourati, Vanessa Demontant, Guillaume Gricourt, Melissa N'Debi, Alexandre Soulier, Elisabeth Trawinski, Jean-Michel Pawlotsky |
| EPI_ISL_3392066, EPI_ISL_3392067 | BIO67-BIOSPHERE | Department of Virology, Henri Mondor University Hospital, Assistance Publique Hôpitaux de Paris, Université Paris-Est Créteil, INSERM U955 | Christophe Rodriguez, Slim Fourati, Vanessa Demontant, Guillaume Gricourt, Melissa N'Debi, Alexandre Soulier, Elisabeth Trawinski, Jean-Michel Pawlotsky |
| EPI_ISL_3392068 | GH A.CHENEVIER-H.MONDOR | Department of Virology, Henri Mondor University Hospital, Assistance Publique Hôpitaux de Paris, Université Paris-Est Créteil, INSERM U955 | Christophe Rodriguez, Slim Fourati, Vanessa Demontant, Guillaume Gricourt, Melissa N'Debi, Alexandre Soulier, Elisabeth Trawinski, Jean-Michel Pawlotsky |
| EPI_ISL_3392069 | SELAS MEDILYS | Department of Virology, Henri Mondor University Hospital, Assistance Publique Hôpitaux de Paris, Université Paris-Est Créteil, INSERM U955 | Christophe Rodriguez, Slim Fourati, Vanessa Demontant, Guillaume Gricourt, Melissa N'Debi, Alexandre Soulier, Elisabeth Trawinski, Jean-Michel Pawlotsky |
| EPI_ISL_3392070 | BIO67-BIOSPHERE | Department of Virology, Henri Mondor University Hospital, Assistance Publique Hôpitaux de Paris, Université Paris-Est Créteil, INSERM U955 | Christophe Rodriguez, Slim Fourati, Vanessa Demontant, Guillaume Gricourt, Melissa N'Debi, Alexandre Soulier, Elisabeth Trawinski, Jean-Michel Pawlotsky |
| EPI_ISL_3392072 | SELAS MEDILYS | Department of Virology, Henri Mondor University Hospital, Assistance Publique Hôpitaux de Paris, Université Paris-Est Créteil, INSERM U955 | Christophe Rodriguez, Slim Fourati, Vanessa Demontant, Guillaume Gricourt, Melissa N'Debi, Alexandre Soulier, Elisabeth Trawinski, Jean-Michel Pawlotsky |
| EPI_ISL_3392074 | BROUSSAIS | Department of Virology, Henri Mondor University Hospital, Assistance Publique Hôpitaux de Paris, Université Paris-Est Créteil, INSERM U955 | Christophe Rodriguez, Slim Fourati, Vanessa Demontant, Guillaume Gricourt, Melissa N'Debi, Alexandre Soulier, Elisabeth Trawinski, Jean-Michel Pawlotsky |

| | | | |
|---|---|---|---|
| EPI_ISL_3392075 | SELAS MEDILYS | Department of Virology, Henri Mondor University Hospital, Assistance Publique Hôpitaux de Paris, Université Paris-Est Créteil, INSERM U955 | Christophe Rodriguez, Slim Fourati, Vanessa Demontant, Guillaume Gricourt, Melissa N'Debi, Alexandre Soulier, Elisabeth Trawinski, Jean-Michel Pawlotsky |
| EPI_ISL_3392076 | BROUSSAIS | Department of Virology, Henri Mondor University Hospital, Assistance Publique Hôpitaux de Paris, Université Paris-Est Créteil, INSERM U955 | Christophe Rodriguez, Slim Fourati, Vanessa Demontant, Guillaume Gricourt, Melissa N'Debi, Alexandre Soulier, Elisabeth Trawinski, Jean-Michel Pawlotsky |
| EPI_ISL_3392079, EPI_ISL_3392080 | SELAS MEDILYS | Department of Virology, Henri Mondor University Hospital, Assistance Publique Hôpitaux de Paris, Université Paris-Est Créteil, INSERM U955 | Christophe Rodriguez, Slim Fourati, Vanessa Demontant, Guillaume Gricourt, Melissa N'Debi, Alexandre Soulier, Elisabeth Trawinski, Jean-Michel Pawlotsky |
| EPI_ISL_3392081 | BROUSSAIS | Department of Virology, Henri Mondor University Hospital, Assistance Publique Hôpitaux de Paris, Université Paris-Est Créteil, INSERM U955 | Christophe Rodriguez, Slim Fourati, Vanessa Demontant, Guillaume Gricourt, Melissa N'Debi, Alexandre Soulier, Elisabeth Trawinski, Jean-Michel Pawlotsky |
| EPI_ISL_3392082 | BIOXA | Department of Virology, Henri Mondor University Hospital, Assistance Publique Hôpitaux de Paris, Université Paris-Est Créteil, INSERM U955 | Christophe Rodriguez, Slim Fourati, Vanessa Demontant, Guillaume Gricourt, Melissa N'Debi, Alexandre Soulier, Elisabeth Trawinski, Jean-Michel Pawlotsky |
| EPI_ISL_3392083 | SELAS MEDILYS | Department of Virology, Henri Mondor University Hospital, Assistance Publique Hôpitaux de Paris, Université Paris-Est Créteil, INSERM U955 | Christophe Rodriguez, Slim Fourati, Vanessa Demontant, Guillaume Gricourt, Melissa N'Debi, Alexandre Soulier, Elisabeth Trawinski, Jean-Michel Pawlotsky |
| EPI_ISL_3392085 | Laboratoire LENYS | Department of Virology, Henri Mondor University Hospital, Assistance Publique Hôpitaux de Paris, Université Paris-Est Créteil, INSERM U955 | Christophe Rodriguez, Slim Fourati, Vanessa Demontant, Guillaume Gricourt, Melissa N'Debi, Alexandre Soulier, Elisabeth Trawinski, Jean-Michel Pawlotsky |
| EPI_ISL_3392088 | BIO67-BIOSPHERE | Department of Virology, Henri Mondor University Hospital, Assistance Publique Hôpitaux de Paris, Université Paris-Est Créteil, INSERM U955 | Christophe Rodriguez, Slim Fourati, Vanessa Demontant, Guillaume Gricourt, Melissa N'Debi, Alexandre Soulier, Elisabeth Trawinski, Jean-Michel Pawlotsky |
| EPI_ISL_3392089 | BROUSSAIS | Department of Virology, Henri Mondor University Hospital, Assistance Publique Hôpitaux de Paris, Université Paris-Est Créteil, INSERM U955 | Christophe Rodriguez, Slim Fourati, Vanessa Demontant, Guillaume Gricourt, Melissa N'Debi, Alexandre Soulier, Elisabeth Trawinski, Jean-Michel Pawlotsky |
| EPI_ISL_3392091 | BIO67-BIOSPHERE | Department of Virology, Henri Mondor University Hospital, Assistance Publique Hôpitaux de Paris, Université Paris-Est Créteil, INSERM U955 | Christophe Rodriguez, Slim Fourati, Vanessa Demontant, Guillaume Gricourt, Melissa N'Debi, Alexandre Soulier, Elisabeth Trawinski, Jean-Michel Pawlotsky |
| EPI_ISL_3392092 | SELAS MEDILYS | Department of Virology, Henri Mondor University Hospital, Assistance Publique Hôpitaux de Paris, Université Paris-Est Créteil, INSERM U955 | Christophe Rodriguez, Slim Fourati, Vanessa Demontant, Guillaume Gricourt, Melissa N'Debi, Alexandre Soulier, Elisabeth Trawinski, Jean-Michel Pawlotsky |
| EPI_ISL_3392093 | Laboratoire ATOUTBIO JDC | Department of Virology, Henri Mondor University Hospital, Assistance Publique Hôpitaux de Paris, Université Paris-Est Créteil, INSERM U955 | Christophe Rodriguez, Slim Fourati, Vanessa Demontant, Guillaume Gricourt, Melissa N'Debi, Alexandre Soulier, Elisabeth Trawinski, Jean-Michel Pawlotsky |
| EPI_ISL_3392094 | SELAS MEDILYS | Department of Virology, Henri Mondor University Hospital, Assistance Publique Hôpitaux de Paris, Université Paris-Est Créteil, INSERM U955 | Christophe Rodriguez, Slim Fourati, Vanessa Demontant, Guillaume Gricourt, Melissa N'Debi, Alexandre Soulier, Elisabeth Trawinski, Jean-Michel Pawlotsky |
| EPI_ISL_3392097, EPI_ISL_3392099 | BIO67-BIOSPHERE | Department of Virology, Henri Mondor University Hospital, Assistance Publique Hôpitaux de Paris, Université Paris-Est Créteil, INSERM U955 | Christophe Rodriguez, Slim Fourati, Vanessa Demontant, Guillaume Gricourt, Melissa N'Debi, Alexandre Soulier, Elisabeth Trawinski, Jean-Michel Pawlotsky |
| EPI_ISL_3392100 | SELAS MEDILYS | Department of Virology, Henri Mondor University Hospital, Assistance Publique Hôpitaux de Paris, Université Paris-Est Créteil, INSERM U955 | Christophe Rodriguez, Slim Fourati, Vanessa Demontant, Guillaume Gricourt, Melissa N'Debi, Alexandre Soulier, Elisabeth Trawinski, Jean-Michel Pawlotsky |
| EPI_ISL_3392101 | Laboratoire ATOUTBIO JDC | Department of Virology, Henri Mondor University Hospital, Assistance Publique Hôpitaux de Paris, Université Paris-Est Créteil, INSERM U955 | Christophe Rodriguez, Slim Fourati, Vanessa Demontant, Guillaume Gricourt, Melissa N'Debi, Alexandre Soulier, Elisabeth Trawinski, Jean-Michel Pawlotsky |
| EPI_ISL_3392104 | BIO67-BIOSPHERE | Department of Virology, Henri Mondor University Hospital, Assistance Publique Hôpitaux de Paris, Université Paris-Est Créteil, INSERM U955 | Christophe Rodriguez, Slim Fourati, Vanessa Demontant, Guillaume Gricourt, Melissa N'Debi, Alexandre Soulier, Elisabeth Trawinski, Jean-Michel Pawlotsky |
| EPI_ISL_3392108 | SELAS MEDILYS | Department of Virology, Henri Mondor University Hospital, Assistance Publique Hôpitaux de Paris, Université Paris-Est Créteil, INSERM U955 | Christophe Rodriguez, Slim Fourati, Vanessa Demontant, Guillaume Gricourt, Melissa N'Debi, Alexandre Soulier, Elisabeth Trawinski, Jean-Michel Pawlotsky |
| EPI_ISL_3392109 | BIO67-BIOSPHERE | Department of Virology, Henri Mondor University Hospital, Assistance Publique Hôpitaux de Paris, Université Paris-Est Créteil, INSERM U955 | Christophe Rodriguez, Slim Fourati, Vanessa Demontant, Guillaume Gricourt, Melissa N'Debi, Alexandre Soulier, Elisabeth Trawinski, Jean-Michel Pawlotsky |
| EPI_ISL_3392110 | SELAS MEDILYS | Department of Virology, Henri Mondor University Hospital, Assistance Publique Hôpitaux de Paris, Université Paris-Est Créteil, INSERM U955 | Christophe Rodriguez, Slim Fourati, Vanessa Demontant, Guillaume Gricourt, Melissa N'Debi, Alexandre Soulier, Elisabeth Trawinski, Jean-Michel Pawlotsky |
| EPI_ISL_3392111 | GH A.CHENEVIER-H.MONDOR | Department of Virology, Henri Mondor University Hospital, Assistance Publique Hôpitaux de Paris, Université Paris-Est Créteil, INSERM U955 | Christophe Rodriguez, Slim Fourati, Vanessa Demontant, Guillaume Gricourt, Melissa N'Debi, Alexandre Soulier, Elisabeth Trawinski, Jean-Michel Pawlotsky |
| EPI_ISL_3392113 | SELAS MEDILYS | Department of Virology, Henri Mondor University Hospital, Assistance Publique Hôpitaux de Paris, Université Paris-Est Créteil, INSERM U955 | Christophe Rodriguez, Slim Fourati, Vanessa Demontant, Guillaume Gricourt, Melissa N'Debi, Alexandre Soulier, Elisabeth Trawinski, Jean-Michel Pawlotsky |
| EPI_ISL_3392114 | GH A.CHENEVIER-H.MONDOR | Department of Virology, Henri Mondor University Hospital, Assistance Publique Hôpitaux de Paris, Université Paris-Est Créteil, INSERM U955 | Christophe Rodriguez, Slim Fourati, Vanessa Demontant, Guillaume Gricourt, Melissa N'Debi, Alexandre Soulier, Elisabeth Trawinski, Jean-Michel Pawlotsky |
| EPI_ISL_3392115 | Laboratoire LENYS | Department of Virology, Henri Mondor University Hospital, Assistance Publique Hôpitaux de Paris, Université Paris-Est Créteil, INSERM U955 | Christophe Rodriguez, Slim Fourati, Vanessa Demontant, Guillaume Gricourt, Melissa N'Debi, Alexandre Soulier, Elisabeth Trawinski, Jean-Michel Pawlotsky |
| EPI_ISL_3392116, EPI_ISL_3392117 | SELAS MEDILYS | Department of Virology, Henri Mondor University Hospital, Assistance Publique Hôpitaux de Paris, Université Paris-Est Créteil, INSERM U955 | Christophe Rodriguez, Slim Fourati, Vanessa Demontant, Guillaume Gricourt, Melissa N'Debi, Alexandre Soulier, Elisabeth Trawinski, Jean-Michel Pawlotsky |
| EPI_ISL_3392118, EPI_ISL_3392119, EPI_ISL_3392120, EPI_ISL_3392121 | BIO67-BIOSPHERE | Department of Virology, Henri Mondor University Hospital, Assistance Publique Hôpitaux de Paris, Université Paris-Est Créteil, INSERM U955 | Christophe Rodriguez, Slim Fourati, Vanessa Demontant, Guillaume Gricourt, Melissa N'Debi, Alexandre Soulier, Elisabeth Trawinski, Jean-Michel Pawlotsky |
| EPI_ISL_3392123, EPI_ISL_3392124 | SELAS MEDILYS | Department of Virology, Henri Mondor University Hospital, Assistance Publique Hôpitaux de Paris, Université Paris-Est Créteil, INSERM U955 | Christophe Rodriguez, Slim Fourati, Vanessa Demontant, Guillaume Gricourt, Melissa N'Debi, Alexandre Soulier, Elisabeth Trawinski, Jean-Michel Pawlotsky |
| EPI_ISL_3392127 | Hôpital Cochin | Department of Virology, Henri Mondor University Hospital, Assistance Publique Hôpitaux de Paris, Université Paris-Est Créteil, INSERM U955 | Christophe Rodriguez, Slim Fourati, Vanessa Demontant, Guillaume Gricourt, Melissa N'Debi, Alexandre Soulier, Elisabeth Trawinski, Jean-Michel Pawlotsky |
| EPI_ISL_3392129, EPI_ISL_3392131 | SELAS MEDILYS | Department of Virology, Henri Mondor University Hospital, Assistance Publique Hôpitaux de Paris, Université Paris-Est Créteil, INSERM U955 | Christophe Rodriguez, Slim Fourati, Vanessa Demontant, Guillaume Gricourt, Melissa N'Debi, Alexandre Soulier, Elisabeth Trawinski, Jean-Michel Pawlotsky |
| EPI_ISL_3392132 | BIOXA | Department of Virology, Henri Mondor University Hospital, Assistance Publique Hôpitaux de Paris, Université Paris-Est Créteil, INSERM U955 | Christophe Rodriguez, Slim Fourati, Vanessa Demontant, Guillaume Gricourt, Melissa N'Debi, Alexandre Soulier, Elisabeth Trawinski, Jean-Michel Pawlotsky |
| EPI_ISL_3392133 | SELAS MEDILYS | Department of Virology, Henri Mondor University Hospital, Assistance Publique Hôpitaux de Paris, Université Paris-Est Créteil, INSERM U955 | Christophe Rodriguez, Slim Fourati, Vanessa Demontant, Guillaume Gricourt, Melissa N'Debi, Alexandre Soulier, Elisabeth Trawinski, Jean-Michel Pawlotsky |
| EPI_ISL_3392134 | Laboratoire ATOUTBIO JDC | Department of Virology, Henri Mondor University Hospital, Assistance Publique Hôpitaux de Paris, Université Paris-Est Créteil, | Christophe Rodriguez, Slim Fourati, Vanessa Demontant, Guillaume Gricourt, Melissa N'Debi, Alexandre Soulier, Elisabeth Trawinski, Jean-Michel Pawlotsky |

| | | | |
|---|---|---|---|
| | | INSERM U955 | |
| EPI_ISL_3392135 | BIO67-BIOSPHERE | Department of Virology, Henri Mondor University Hospital, Assistance Publique Hôpitaux de Paris, Université Paris-Est Créteil, INSERM U955 | Christophe Rodriguez, Slim Fourati, Vanessa Demontant, Guillaume Gricourt, Melissa N'Debi, Alexandre Soulier, Elisabeth Trawinski, Jean-Michel Pawlotsky |
| EPI_ISL_3392136 | BROUSSAIS | Department of Virology, Henri Mondor University Hospital, Assistance Publique Hôpitaux de Paris, Université Paris-Est Créteil, INSERM U955 | Christophe Rodriguez, Slim Fourati, Vanessa Demontant, Guillaume Gricourt, Melissa N'Debi, Alexandre Soulier, Elisabeth Trawinski, Jean-Michel Pawlotsky |
| EPI_ISL_3392139, EPI_ISL_3392140, EPI_ISL_3392141 | SELAS MEDILYS | Department of Virology, Henri Mondor University Hospital, Assistance Publique Hôpitaux de Paris, Université Paris-Est Créteil, INSERM U955 | Christophe Rodriguez, Slim Fourati, Vanessa Demontant, Guillaume Gricourt, Melissa N'Debi, Alexandre Soulier, Elisabeth Trawinski, Jean-Michel Pawlotsky |
| EPI_ISL_3392142 | BIO67-BIOSPHERE | Department of Virology, Henri Mondor University Hospital, Assistance Publique Hôpitaux de Paris, Université Paris-Est Créteil, INSERM U955 | Christophe Rodriguez, Slim Fourati, Vanessa Demontant, Guillaume Gricourt, Melissa N'Debi, Alexandre Soulier, Elisabeth Trawinski, Jean-Michel Pawlotsky |
| EPI_ISL_3392143 | GH A.CHENEVIER-H.MONDOR | Department of Virology, Henri Mondor University Hospital, Assistance Publique Hôpitaux de Paris, Université Paris-Est Créteil, INSERM U955 | Christophe Rodriguez, Slim Fourati, Vanessa Demontant, Guillaume Gricourt, Melissa N'Debi, Alexandre Soulier, Elisabeth Trawinski, Jean-Michel Pawlotsky |
| EPI_ISL_3392145 | BROUSSAIS | Department of Virology, Henri Mondor University Hospital, Assistance Publique Hôpitaux de Paris, Université Paris-Est Créteil, INSERM U955 | Christophe Rodriguez, Slim Fourati, Vanessa Demontant, Guillaume Gricourt, Melissa N'Debi, Alexandre Soulier, Elisabeth Trawinski, Jean-Michel Pawlotsky |
| EPI_ISL_3392147, EPI_ISL_3392149, EPI_ISL_3392150, EPI_ISL_3392151 | SELAS MEDILYS | Department of Virology, Henri Mondor University Hospital, Assistance Publique Hôpitaux de Paris, Université Paris-Est Créteil, INSERM U955 | Christophe Rodriguez, Slim Fourati, Vanessa Demontant, Guillaume Gricourt, Melissa N'Debi, Alexandre Soulier, Elisabeth Trawinski, Jean-Michel Pawlotsky |
| EPI_ISL_3392152 | BIO67-BIOSPHERE | Department of Virology, Henri Mondor University Hospital, Assistance Publique Hôpitaux de Paris, Université Paris-Est Créteil, INSERM U955 | Christophe Rodriguez, Slim Fourati, Vanessa Demontant, Guillaume Gricourt, Melissa N'Debi, Alexandre Soulier, Elisabeth Trawinski, Jean-Michel Pawlotsky |
| EPI_ISL_3392153, EPI_ISL_3392154, EPI_ISL_3392157 | SELAS MEDILYS | Department of Virology, Henri Mondor University Hospital, Assistance Publique Hôpitaux de Paris, Université Paris-Est Créteil, INSERM U955 | Christophe Rodriguez, Slim Fourati, Vanessa Demontant, Guillaume Gricourt, Melissa N'Debi, Alexandre Soulier, Elisabeth Trawinski, Jean-Michel Pawlotsky |
| EPI_ISL_3392158 | Hôpital Cochin | Department of Virology, Henri Mondor University Hospital, Assistance Publique Hôpitaux de Paris, Université Paris-Est Créteil, INSERM U955 | Christophe Rodriguez, Slim Fourati, Vanessa Demontant, Guillaume Gricourt, Melissa N'Debi, Alexandre Soulier, Elisabeth Trawinski, Jean-Michel Pawlotsky |
| EPI_ISL_3392159 | BIOCEANE | Department of Virology, Henri Mondor University Hospital, Assistance Publique Hôpitaux de Paris, Université Paris-Est Créteil, INSERM U955 | Christophe Rodriguez, Slim Fourati, Vanessa Demontant, Guillaume Gricourt, Melissa N'Debi, Alexandre Soulier, Elisabeth Trawinski, Jean-Michel Pawlotsky |
| EPI_ISL_3392160 | BIO67-BIOSPHERE | Department of Virology, Henri Mondor University Hospital, Assistance Publique Hôpitaux de Paris, Université Paris-Est Créteil, INSERM U955 | Christophe Rodriguez, Slim Fourati, Vanessa Demontant, Guillaume Gricourt, Melissa N'Debi, Alexandre Soulier, Elisabeth Trawinski, Jean-Michel Pawlotsky |
| EPI_ISL_3392161 | GH A.CHENEVIER-H.MONDOR | Department of Virology, Henri Mondor University Hospital, Assistance Publique Hôpitaux de Paris, Université Paris-Est Créteil, INSERM U955 | Christophe Rodriguez, Slim Fourati, Vanessa Demontant, Guillaume Gricourt, Melissa N'Debi, Alexandre Soulier, Elisabeth Trawinski, Jean-Michel Pawlotsky |
| EPI_ISL_3392162 | BROUSSAIS | Department of Virology, Henri Mondor University Hospital, Assistance Publique Hôpitaux de Paris, Université Paris-Est Créteil, INSERM U955 | Christophe Rodriguez, Slim Fourati, Vanessa Demontant, Guillaume Gricourt, Melissa N'Debi, Alexandre Soulier, Elisabeth Trawinski, Jean-Michel Pawlotsky |
| EPI_ISL_3392164 | GH A.CHENEVIER-H.MONDOR | Department of Virology, Henri Mondor University Hospital, Assistance Publique Hôpitaux de Paris, Université Paris-Est Créteil, INSERM U955 | Christophe Rodriguez, Slim Fourati, Vanessa Demontant, Guillaume Gricourt, Melissa N'Debi, Alexandre Soulier, Elisabeth Trawinski, Jean-Michel Pawlotsky |
| EPI_ISL_3392165 | BIO67-BIOSPHERE | Department of Virology, Henri Mondor University Hospital, Assistance Publique Hôpitaux de Paris, Université Paris-Est Créteil, INSERM U955 | Christophe Rodriguez, Slim Fourati, Vanessa Demontant, Guillaume Gricourt, Melissa N'Debi, Alexandre Soulier, Elisabeth Trawinski, Jean-Michel Pawlotsky |
| EPI_ISL_3392166 | SELAS MEDILYS | Department of Virology, Henri Mondor University Hospital, Assistance Publique Hôpitaux de Paris, Université Paris-Est Créteil, INSERM U955 | Christophe Rodriguez, Slim Fourati, Vanessa Demontant, Guillaume Gricourt, Melissa N'Debi, Alexandre Soulier, Elisabeth Trawinski, Jean-Michel Pawlotsky |
| EPI_ISL_3392167 | Laboratoire LENYS | Department of Virology, Henri Mondor University Hospital, Assistance Publique Hôpitaux de Paris, Université Paris-Est Créteil, INSERM U955 | Christophe Rodriguez, Slim Fourati, Vanessa Demontant, Guillaume Gricourt, Melissa N'Debi, Alexandre Soulier, Elisabeth Trawinski, Jean-Michel Pawlotsky |
| EPI_ISL_3392168, EPI_ISL_3392170, EPI_ISL_3392171 | SELAS MEDILYS | Department of Virology, Henri Mondor University Hospital, Assistance Publique Hôpitaux de Paris, Université Paris-Est Créteil, INSERM U955 | Christophe Rodriguez, Slim Fourati, Vanessa Demontant, Guillaume Gricourt, Melissa N'Debi, Alexandre Soulier, Elisabeth Trawinski, Jean-Michel Pawlotsky |
| EPI_ISL_3392172 | Laboratoire ATOUTBIO JDC | Department of Virology, Henri Mondor University Hospital, Assistance Publique Hôpitaux de Paris, Université Paris-Est Créteil, INSERM U955 | Christophe Rodriguez, Slim Fourati, Vanessa Demontant, Guillaume Gricourt, Melissa N'Debi, Alexandre Soulier, Elisabeth Trawinski, Jean-Michel Pawlotsky |
| EPI_ISL_3392174 | SELAS MEDILYS | Department of Virology, Henri Mondor University Hospital, Assistance Publique Hôpitaux de Paris, Université Paris-Est Créteil, INSERM U955 | Christophe Rodriguez, Slim Fourati, Vanessa Demontant, Guillaume Gricourt, Melissa N'Debi, Alexandre Soulier, Elisabeth Trawinski, Jean-Michel Pawlotsky |
| EPI_ISL_3392175 | Laboratoire CBM 25 TERRE ROUGE | Department of Virology, Henri Mondor University Hospital, Assistance Publique Hôpitaux de Paris, Université Paris-Est Créteil, INSERM U955 | Christophe Rodriguez, Slim Fourati, Vanessa Demontant, Guillaume Gricourt, Melissa N'Debi, Alexandre Soulier, Elisabeth Trawinski, Jean-Michel Pawlotsky |
| EPI_ISL_3392176, EPI_ISL_3392178 | BIOXA | Department of Virology, Henri Mondor University Hospital, Assistance Publique Hôpitaux de Paris, Université Paris-Est Créteil, INSERM U955 | Christophe Rodriguez, Slim Fourati, Vanessa Demontant, Guillaume Gricourt, Melissa N'Debi, Alexandre Soulier, Elisabeth Trawinski, Jean-Michel Pawlotsky |
| EPI_ISL_3392180 | Laboratoire ATOUTBIO JDC | Department of Virology, Henri Mondor University Hospital, Assistance Publique Hôpitaux de Paris, Université Paris-Est Créteil, INSERM U955 | Christophe Rodriguez, Slim Fourati, Vanessa Demontant, Guillaume Gricourt, Melissa N'Debi, Alexandre Soulier, Elisabeth Trawinski, Jean-Michel Pawlotsky |
| EPI_ISL_3392182, EPI_ISL_3392183, EPI_ISL_3392184 | SELAS MEDILYS | Department of Virology, Henri Mondor University Hospital, Assistance Publique Hôpitaux de Paris, Université Paris-Est Créteil, INSERM U955 | Christophe Rodriguez, Slim Fourati, Vanessa Demontant, Guillaume Gricourt, Melissa N'Debi, Alexandre Soulier, Elisabeth Trawinski, Jean-Michel Pawlotsky |
| EPI_ISL_3392185 | BIO67-BIOSPHERE | Department of Virology, Henri Mondor University Hospital, Assistance Publique Hôpitaux de Paris, Université Paris-Est Créteil, INSERM U955 | Christophe Rodriguez, Slim Fourati, Vanessa Demontant, Guillaume Gricourt, Melissa N'Debi, Alexandre Soulier, Elisabeth Trawinski, Jean-Michel Pawlotsky |
| EPI_ISL_3392186 | SELAS MEDILYS | Department of Virology, Henri Mondor University Hospital, Assistance Publique Hôpitaux de Paris, Université Paris-Est Créteil, INSERM U955 | Christophe Rodriguez, Slim Fourati, Vanessa Demontant, Guillaume Gricourt, Melissa N'Debi, Alexandre Soulier, Elisabeth Trawinski, Jean-Michel Pawlotsky |
| EPI_ISL_3392187 | BIO67-BIOSPHERE | Department of Virology, Henri Mondor University Hospital, Assistance Publique Hôpitaux de Paris, Université Paris-Est Créteil, INSERM U955 | Christophe Rodriguez, Slim Fourati, Vanessa Demontant, Guillaume Gricourt, Melissa N'Debi, Alexandre Soulier, Elisabeth Trawinski, Jean-Michel Pawlotsky |
| EPI_ISL_3392188 | BROUSSAIS | Department of Virology, Henri Mondor University Hospital, Assistance Publique Hôpitaux de Paris, Université Paris-Est Créteil, INSERM U955 | Christophe Rodriguez, Slim Fourati, Vanessa Demontant, Guillaume Gricourt, Melissa N'Debi, Alexandre Soulier, Elisabeth Trawinski, Jean-Michel Pawlotsky |
| EPI_ISL_3392189 | Laboratoire ATOUTBIO JDC | Department of Virology, Henri Mondor University Hospital, Assistance Publique Hôpitaux de Paris, Université Paris-Est Créteil, INSERM U955 | Christophe Rodriguez, Slim Fourati, Vanessa Demontant, Guillaume Gricourt, Melissa N'Debi, Alexandre Soulier, Elisabeth Trawinski, Jean-Michel Pawlotsky |
| EPI_ISL_3392190 | BROUSSAIS | Department of Virology, Henri Mondor University Hospital, Assistance Publique Hôpitaux de Paris, Université Paris-Est Créteil, INSERM U955 | Christophe Rodriguez, Slim Fourati, Vanessa Demontant, Guillaume Gricourt, Melissa N'Debi, Alexandre Soulier, Elisabeth Trawinski, Jean-Michel Pawlotsky |
| EPI_ISL_3392191 | GH A.CHENEVIER-H.MONDOR | Department of Virology, Henri Mondor University Hospital, | Christophe Rodriguez, Slim Fourati, Vanessa Demontant, Guillaume Gricourt, Melissa N'Debi, Alexandre Soulier, Elisabeth Trawinski, Jean-Michel Pawlotsky |

| | | | |
|---|---|---|---|
| | | Assistance Publique Hôpitaux de Paris, Université Paris-Est Créteil, INSERM U955 | |
| EPI_ISL_3392192, EPI_ISL_3392193 | SELAS MEDILYS | Department of Virology, Henri Mondor University Hospital, Assistance Publique Hôpitaux de Paris, Université Paris-Est Créteil, INSERM U955 | Christophe Rodriguez, Slim Fourati, Vanessa Demontant, Guillaume Gricourt, Melissa N'Debi, Alexandre Soulier, Elisabeth Trawinski, Jean-Michel Pawlotsky |
| EPI_ISL_3392194 | BIOMNIS EUROFINS IVRY | Department of Virology, Henri Mondor University Hospital, Assistance Publique Hôpitaux de Paris, Université Paris-Est Créteil, INSERM U955 | Christophe Rodriguez, Slim Fourati, Vanessa Demontant, Guillaume Gricourt, Melissa N'Debi, Alexandre Soulier, Elisabeth Trawinski, Jean-Michel Pawlotsky |
| EPI_ISL_3392195 | SELAS MEDILYS | Department of Virology, Henri Mondor University Hospital, Assistance Publique Hôpitaux de Paris, Université Paris-Est Créteil, INSERM U955 | Christophe Rodriguez, Slim Fourati, Vanessa Demontant, Guillaume Gricourt, Melissa N'Debi, Alexandre Soulier, Elisabeth Trawinski, Jean-Michel Pawlotsky |
| EPI_ISL_3392196 | GH A.CHENEVIER-H.MONDOR | Department of Virology, Henri Mondor University Hospital, Assistance Publique Hôpitaux de Paris, Université Paris-Est Créteil, INSERM U955 | Christophe Rodriguez, Slim Fourati, Vanessa Demontant, Guillaume Gricourt, Melissa N'Debi, Alexandre Soulier, Elisabeth Trawinski, Jean-Michel Pawlotsky |
| EPI_ISL_3392197 | SELAS MEDILYS | Department of Virology, Henri Mondor University Hospital, Assistance Publique Hôpitaux de Paris, Université Paris-Est Créteil, INSERM U955 | Christophe Rodriguez, Slim Fourati, Vanessa Demontant, Guillaume Gricourt, Melissa N'Debi, Alexandre Soulier, Elisabeth Trawinski, Jean-Michel Pawlotsky |
| EPI_ISL_3392199 | BROUSSAIS | Department of Virology, Henri Mondor University Hospital, Assistance Publique Hôpitaux de Paris, Université Paris-Est Créteil, INSERM U955 | Christophe Rodriguez, Slim Fourati, Vanessa Demontant, Guillaume Gricourt, Melissa N'Debi, Alexandre Soulier, Elisabeth Trawinski, Jean-Michel Pawlotsky |
| EPI_ISL_3392200 | SELAS MEDILYS | Department of Virology, Henri Mondor University Hospital, Assistance Publique Hôpitaux de Paris, Université Paris-Est Créteil, INSERM U955 | Christophe Rodriguez, Slim Fourati, Vanessa Demontant, Guillaume Gricourt, Melissa N'Debi, Alexandre Soulier, Elisabeth Trawinski, Jean-Michel Pawlotsky |
| EPI_ISL_3392201 | BIO67-BIOSPHERE | Department of Virology, Henri Mondor University Hospital, Assistance Publique Hôpitaux de Paris, Université Paris-Est Créteil, INSERM U955 | Christophe Rodriguez, Slim Fourati, Vanessa Demontant, Guillaume Gricourt, Melissa N'Debi, Alexandre Soulier, Elisabeth Trawinski, Jean-Michel Pawlotsky |
| EPI_ISL_3392203 | Laboratoire ATOUTBIO JDC | Department of Virology, Henri Mondor University Hospital, Assistance Publique Hôpitaux de Paris, Université Paris-Est Créteil, INSERM U955 | Christophe Rodriguez, Slim Fourati, Vanessa Demontant, Guillaume Gricourt, Melissa N'Debi, Alexandre Soulier, Elisabeth Trawinski, Jean-Michel Pawlotsky |
| EPI_ISL_3392204 | SELAS MEDILYS | Department of Virology, Henri Mondor University Hospital, Assistance Publique Hôpitaux de Paris, Université Paris-Est Créteil, INSERM U955 | Christophe Rodriguez, Slim Fourati, Vanessa Demontant, Guillaume Gricourt, Melissa N'Debi, Alexandre Soulier, Elisabeth Trawinski, Jean-Michel Pawlotsky |
| EPI_ISL_3392207 | GH A.CHENEVIER-H.MONDOR | Department of Virology, Henri Mondor University Hospital, Assistance Publique Hôpitaux de Paris, Université Paris-Est Créteil, INSERM U955 | Christophe Rodriguez, Slim Fourati, Vanessa Demontant, Guillaume Gricourt, Melissa N'Debi, Alexandre Soulier, Elisabeth Trawinski, Jean-Michel Pawlotsky |
| EPI_ISL_3392210 | SELAS MEDILYS | Department of Virology, Henri Mondor University Hospital, Assistance Publique Hôpitaux de Paris, Université Paris-Est Créteil, INSERM U955 | Christophe Rodriguez, Slim Fourati, Vanessa Demontant, Guillaume Gricourt, Melissa N'Debi, Alexandre Soulier, Elisabeth Trawinski, Jean-Michel Pawlotsky |
| EPI_ISL_3392211, EPI_ISL_3392212 | BIO67-BIOSPHERE | Department of Virology, Henri Mondor University Hospital, Assistance Publique Hôpitaux de Paris, Université Paris-Est Créteil, INSERM U955 | Christophe Rodriguez, Slim Fourati, Vanessa Demontant, Guillaume Gricourt, Melissa N'Debi, Alexandre Soulier, Elisabeth Trawinski, Jean-Michel Pawlotsky |
| EPI_ISL_3392213 | SELAS MEDILYS | Department of Virology, Henri Mondor University Hospital, Assistance Publique Hôpitaux de Paris, Université Paris-Est Créteil, INSERM U955 | Christophe Rodriguez, Slim Fourati, Vanessa Demontant, Guillaume Gricourt, Melissa N'Debi, Alexandre Soulier, Elisabeth Trawinski, Jean-Michel Pawlotsky |
| EPI_ISL_3392216 | Hôpital Cochin | Department of Virology, Henri Mondor University Hospital, Assistance Publique Hôpitaux de Paris, Université Paris-Est Créteil, INSERM U955 | Christophe Rodriguez, Slim Fourati, Vanessa Demontant, Guillaume Gricourt, Melissa N'Debi, Alexandre Soulier, Elisabeth Trawinski, Jean-Michel Pawlotsky |
| EPI_ISL_3392217 | BROUSSAIS | Department of Virology, Henri Mondor University Hospital, Assistance Publique Hôpitaux de Paris, Université Paris-Est Créteil, INSERM U955 | Christophe Rodriguez, Slim Fourati, Vanessa Demontant, Guillaume Gricourt, Melissa N'Debi, Alexandre Soulier, Elisabeth Trawinski, Jean-Michel Pawlotsky |
| EPI_ISL_3392218 | SELAS MEDILYS | Department of Virology, Henri Mondor University Hospital, Assistance Publique Hôpitaux de Paris, Université Paris-Est Créteil, INSERM U955 | Christophe Rodriguez, Slim Fourati, Vanessa Demontant, Guillaume Gricourt, Melissa N'Debi, Alexandre Soulier, Elisabeth Trawinski, Jean-Michel Pawlotsky |
| EPI_ISL_3392219 | Laboratoire ATOUTBIO JDC | Department of Virology, Henri Mondor University Hospital, Assistance Publique Hôpitaux de Paris, Université Paris-Est Créteil, INSERM U955 | Christophe Rodriguez, Slim Fourati, Vanessa Demontant, Guillaume Gricourt, Melissa N'Debi, Alexandre Soulier, Elisabeth Trawinski, Jean-Michel Pawlotsky |
| EPI_ISL_3392220 | BIO67-BIOSPHERE | Department of Virology, Henri Mondor University Hospital, Assistance Publique Hôpitaux de Paris, Université Paris-Est Créteil, INSERM U955 | Christophe Rodriguez, Slim Fourati, Vanessa Demontant, Guillaume Gricourt, Melissa N'Debi, Alexandre Soulier, Elisabeth Trawinski, Jean-Michel Pawlotsky |
| EPI_ISL_3392221 | SELAS MEDILYS | Department of Virology, Henri Mondor University Hospital, Assistance Publique Hôpitaux de Paris, Université Paris-Est Créteil, INSERM U955 | Christophe Rodriguez, Slim Fourati, Vanessa Demontant, Guillaume Gricourt, Melissa N'Debi, Alexandre Soulier, Elisabeth Trawinski, Jean-Michel Pawlotsky |
| EPI_ISL_3392222 | BIO67-BIOSPHERE | Department of Virology, Henri Mondor University Hospital, Assistance Publique Hôpitaux de Paris, Université Paris-Est Créteil, INSERM U955 | Christophe Rodriguez, Slim Fourati, Vanessa Demontant, Guillaume Gricourt, Melissa N'Debi, Alexandre Soulier, Elisabeth Trawinski, Jean-Michel Pawlotsky |
| EPI_ISL_3392223 | SELAS MEDILYS | Department of Virology, Henri Mondor University Hospital, Assistance Publique Hôpitaux de Paris, Université Paris-Est Créteil, INSERM U955 | Christophe Rodriguez, Slim Fourati, Vanessa Demontant, Guillaume Gricourt, Melissa N'Debi, Alexandre Soulier, Elisabeth Trawinski, Jean-Michel Pawlotsky |
| EPI_ISL_3392226, EPI_ISL_3392227, EPI_ISL_3392228 | BIO67-BIOSPHERE | Department of Virology, Henri Mondor University Hospital, Assistance Publique Hôpitaux de Paris, Université Paris-Est Créteil, INSERM U955 | Christophe Rodriguez, Slim Fourati, Vanessa Demontant, Guillaume Gricourt, Melissa N'Debi, Alexandre Soulier, Elisabeth Trawinski, Jean-Michel Pawlotsky |
| EPI_ISL_3392229 | SELAS MEDILYS | Department of Virology, Henri Mondor University Hospital, Assistance Publique Hôpitaux de Paris, Université Paris-Est Créteil, INSERM U955 | Christophe Rodriguez, Slim Fourati, Vanessa Demontant, Guillaume Gricourt, Melissa N'Debi, Alexandre Soulier, Elisabeth Trawinski, Jean-Michel Pawlotsky |
| EPI_ISL_3392230 | Laboratoire CBM 25 TERRE ROUGE | Department of Virology, Henri Mondor University Hospital, Assistance Publique Hôpitaux de Paris, Université Paris-Est Créteil, INSERM U955 | Christophe Rodriguez, Slim Fourati, Vanessa Demontant, Guillaume Gricourt, Melissa N'Debi, Alexandre Soulier, Elisabeth Trawinski, Jean-Michel Pawlotsky |
| EPI_ISL_3392231 | SELAS MEDILYS | Department of Virology, Henri Mondor University Hospital, Assistance Publique Hôpitaux de Paris, Université Paris-Est Créteil, INSERM U955 | Christophe Rodriguez, Slim Fourati, Vanessa Demontant, Guillaume Gricourt, Melissa N'Debi, Alexandre Soulier, Elisabeth Trawinski, Jean-Michel Pawlotsky |
| EPI_ISL_3392232 | Laboratoire ATOUTBIO JDC | Department of Virology, Henri Mondor University Hospital, Assistance Publique Hôpitaux de Paris, Université Paris-Est Créteil, INSERM U955 | Christophe Rodriguez, Slim Fourati, Vanessa Demontant, Guillaume Gricourt, Melissa N'Debi, Alexandre Soulier, Elisabeth Trawinski, Jean-Michel Pawlotsky |
| EPI_ISL_3392233 | SELAS MEDILYS | Department of Virology, Henri Mondor University Hospital, Assistance Publique Hôpitaux de Paris, Université Paris-Est Créteil, INSERM U955 | Christophe Rodriguez, Slim Fourati, Vanessa Demontant, Guillaume Gricourt, Melissa N'Debi, Alexandre Soulier, Elisabeth Trawinski, Jean-Michel Pawlotsky |
| EPI_ISL_3392234 | BIOMNIS EUROFINS IVRY | Department of Virology, Henri Mondor University Hospital, Assistance Publique Hôpitaux de Paris, Université Paris-Est Créteil, INSERM U955 | Christophe Rodriguez, Slim Fourati, Vanessa Demontant, Guillaume Gricourt, Melissa N'Debi, Alexandre Soulier, Elisabeth Trawinski, Jean-Michel Pawlotsky |
| EPI_ISL_3392235 | SELAS MEDILYS | Department of Virology, Henri Mondor University Hospital, Assistance Publique Hôpitaux de Paris, Université Paris-Est Créteil, INSERM U955 | Christophe Rodriguez, Slim Fourati, Vanessa Demontant, Guillaume Gricourt, Melissa N'Debi, Alexandre Soulier, Elisabeth Trawinski, Jean-Michel Pawlotsky |
| EPI_ISL_3392236 | BIO67-BIOSPHERE | Department of Virology, Henri Mondor University Hospital, Assistance Publique Hôpitaux de Paris, Université Paris-Est Créteil, INSERM U955 | Christophe Rodriguez, Slim Fourati, Vanessa Demontant, Guillaume Gricourt, Melissa N'Debi, Alexandre Soulier, Elisabeth Trawinski, Jean-Michel Pawlotsky |
| EPI_ISL_3392237 | BROUSSAIS | Department of Virology, Henri Mondor University Hospital, | Christophe Rodriguez, Slim Fourati, Vanessa Demontant, Guillaume Gricourt, Melissa N'Debi, Alexandre Soulier, Elisabeth Trawinski, Jean-Michel Pawlotsky |

| | | | |
|---|---|---|---|
| | | Assistance Publique Hôpitaux de Paris, Université Paris-Est Créteil, INSERM U955 | |
| EPI_ISL_3392238 | SELAS MEDILYS | Department of Virology, Henri Mondor University Hospital, Assistance Publique Hôpitaux de Paris, Université Paris-Est Créteil, INSERM U955 | Christophe Rodriguez, Slim Fourati, Vanessa Demontant, Guillaume Gricourt, Melissa N'Debi, Alexandre Soulier, Elisabeth Trawinski, Jean-Michel Pawlotsky |
| EPI_ISL_3392239, EPI_ISL_3392240 | BIO67-BIOSPHERE | Department of Virology, Henri Mondor University Hospital, Assistance Publique Hôpitaux de Paris, Université Paris-Est Créteil, INSERM U955 | Christophe Rodriguez, Slim Fourati, Vanessa Demontant, Guillaume Gricourt, Melissa N'Debi, Alexandre Soulier, Elisabeth Trawinski, Jean-Michel Pawlotsky |
| EPI_ISL_3392241, EPI_ISL_3392242 | SELAS MEDILYS | Department of Virology, Henri Mondor University Hospital, Assistance Publique Hôpitaux de Paris, Université Paris-Est Créteil, INSERM U955 | Christophe Rodriguez, Slim Fourati, Vanessa Demontant, Guillaume Gricourt, Melissa N'Debi, Alexandre Soulier, Elisabeth Trawinski, Jean-Michel Pawlotsky |
| EPI_ISL_3392244, EPI_ISL_3392249 | BIO67-BIOSPHERE | Department of Virology, Henri Mondor University Hospital, Assistance Publique Hôpitaux de Paris, Université Paris-Est Créteil, INSERM U955 | Christophe Rodriguez, Slim Fourati, Vanessa Demontant, Guillaume Gricourt, Melissa N'Debi, Alexandre Soulier, Elisabeth Trawinski, Jean-Michel Pawlotsky |
| EPI_ISL_3392250 | SELAS MEDILYS | Department of Virology, Henri Mondor University Hospital, Assistance Publique Hôpitaux de Paris, Université Paris-Est Créteil, INSERM U955 | Christophe Rodriguez, Slim Fourati, Vanessa Demontant, Guillaume Gricourt, Melissa N'Debi, Alexandre Soulier, Elisabeth Trawinski, Jean-Michel Pawlotsky |
| EPI_ISL_3392251 | BROUSSAIS | Department of Virology, Henri Mondor University Hospital, Assistance Publique Hôpitaux de Paris, Université Paris-Est Créteil, INSERM U955 | Christophe Rodriguez, Slim Fourati, Vanessa Demontant, Guillaume Gricourt, Melissa N'Debi, Alexandre Soulier, Elisabeth Trawinski, Jean-Michel Pawlotsky |
| EPI_ISL_3392253 | BIO67-BIOSPHERE | Department of Virology, Henri Mondor University Hospital, Assistance Publique Hôpitaux de Paris, Université Paris-Est Créteil, INSERM U955 | Christophe Rodriguez, Slim Fourati, Vanessa Demontant, Guillaume Gricourt, Melissa N'Debi, Alexandre Soulier, Elisabeth Trawinski, Jean-Michel Pawlotsky |
| EPI_ISL_3392255 | BROUSSAIS | Department of Virology, Henri Mondor University Hospital, Assistance Publique Hôpitaux de Paris, Université Paris-Est Créteil, INSERM U955 | Christophe Rodriguez, Slim Fourati, Vanessa Demontant, Guillaume Gricourt, Melissa N'Debi, Alexandre Soulier, Elisabeth Trawinski, Jean-Michel Pawlotsky |
| EPI_ISL_3392256, EPI_ISL_3392257 | SELAS MEDILYS | Department of Virology, Henri Mondor University Hospital, Assistance Publique Hôpitaux de Paris, Université Paris-Est Créteil, INSERM U955 | Christophe Rodriguez, Slim Fourati, Vanessa Demontant, Guillaume Gricourt, Melissa N'Debi, Alexandre Soulier, Elisabeth Trawinski, Jean-Michel Pawlotsky |
| EPI_ISL_3392258 | BROUSSAIS | Department of Virology, Henri Mondor University Hospital, Assistance Publique Hôpitaux de Paris, Université Paris-Est Créteil, INSERM U955 | Christophe Rodriguez, Slim Fourati, Vanessa Demontant, Guillaume Gricourt, Melissa N'Debi, Alexandre Soulier, Elisabeth Trawinski, Jean-Michel Pawlotsky |
| EPI_ISL_3392259, EPI_ISL_3392260 | BIO67-BIOSPHERE | Department of Virology, Henri Mondor University Hospital, Assistance Publique Hôpitaux de Paris, Université Paris-Est Créteil, INSERM U955 | Christophe Rodriguez, Slim Fourati, Vanessa Demontant, Guillaume Gricourt, Melissa N'Debi, Alexandre Soulier, Elisabeth Trawinski, Jean-Michel Pawlotsky |
| EPI_ISL_3392261 | SELAS MEDILYS | Department of Virology, Henri Mondor University Hospital, Assistance Publique Hôpitaux de Paris, Université Paris-Est Créteil, INSERM U955 | Christophe Rodriguez, Slim Fourati, Vanessa Demontant, Guillaume Gricourt, Melissa N'Debi, Alexandre Soulier, Elisabeth Trawinski, Jean-Michel Pawlotsky |
| EPI_ISL_3392262 | Hôpital Cochin | Department of Virology, Henri Mondor University Hospital, Assistance Publique Hôpitaux de Paris, Université Paris-Est Créteil, INSERM U955 | Christophe Rodriguez, Slim Fourati, Vanessa Demontant, Guillaume Gricourt, Melissa N'Debi, Alexandre Soulier, Elisabeth Trawinski, Jean-Michel Pawlotsky |
| EPI_ISL_3392263 | SELAS MEDILYS | Department of Virology, Henri Mondor University Hospital, Assistance Publique Hôpitaux de Paris, Université Paris-Est Créteil, INSERM U955 | Christophe Rodriguez, Slim Fourati, Vanessa Demontant, Guillaume Gricourt, Melissa N'Debi, Alexandre Soulier, Elisabeth Trawinski, Jean-Michel Pawlotsky |
| EPI_ISL_3392264 | Laboratoire LENYS | Department of Virology, Henri Mondor University Hospital, Assistance Publique Hôpitaux de Paris, Université Paris-Est Créteil, INSERM U955 | Christophe Rodriguez, Slim Fourati, Vanessa Demontant, Guillaume Gricourt, Melissa N'Debi, Alexandre Soulier, Elisabeth Trawinski, Jean-Michel Pawlotsky |
| EPI_ISL_3392268, EPI_ISL_3392269, EPI_ISL_3392270, EPI_ISL_3392271, EPI_ISL_3392272, EPI_ISL_3392273, EPI_ISL_3392275, EPI_ISL_3392276, EPI_ISL_3392278, EPI_ISL_3392279, EPI_ISL_3392280, EPI_ISL_3392281, EPI_ISL_3392282, EPI_ISL_3392283, EPI_ISL_3392284, EPI_ISL_3392285, EPI_ISL_3392286, EPI_ISL_3392287, EPI_ISL_3392288, EPI_ISL_3392289, EPI_ISL_3392290, EPI_ISL_3392291, EPI_ISL_3392292, EPI_ISL_3392293, EPI_ISL_3392294, EPI_ISL_3392295, EPI_ISL_3392296, EPI_ISL_3392297, EPI_ISL_3392298, EPI_ISL_3392299, EPI_ISL_3392300, EPI_ISL_3392301, EPI_ISL_3392302, EPI_ISL_3392303, EPI_ISL_3392304, EPI_ISL_3392305, EPI_ISL_3392306, EPI_ISL_3392307, EPI_ISL_3392308, EPI_ISL_3392309, EPI_ISL_3392310, EPI_ISL_3392311, EPI_ISL_3392312, EPI_ISL_3392313, EPI_ISL_3392314, EPI_ISL_3392315, EPI_ISL_3392316, EPI_ISL_3392317, EPI_ISL_3392318, EPI_ISL_3392319, EPI_ISL_3392320, EPI_ISL_3392321, EPI_ISL_3392322, EPI_ISL_3392323, EPI_ISL_3392324, EPI_ISL_3392325, EPI_ISL_3392326, EPI_ISL_3392327, EPI_ISL_3392328, EPI_ISL_3392329, EPI_ISL_3392330, EPI_ISL_3392331, EPI_ISL_3392332, EPI_ISL_3392333, EPI_ISL_3392334, EPI_ISL_3392335, EPI_ISL_3392336, EPI_ISL_3392337, EPI_ISL_3392338, EPI_ISL_3392339, EPI_ISL_3392340, EPI_ISL_3392341, EPI_ISL_3392342, EPI_ISL_3392343, EPI_ISL_3392344, EPI_ISL_3392345, EPI_ISL_3392346, EPI_ISL_3392347, EPI_ISL_3392348, EPI_ISL_3392349, EPI_ISL_3392350, EPI_ISL_3392351, EPI_ISL_3392352, EPI_ISL_3392353, EPI_ISL_3392354 | | | |
| see above | SARS-CoV-2 testing team, National Institute of Infectious Diseases | Pathogen Genomics Center, National Institute of Infectious Diseases | Tsuyoshi Sekizuka, Kentaro Itokawa, Rina Tanaka, Masanori Hashino, Nozomu Hanaoka, Masumichi Saito, Naomi Nojiri, Hazuka Y Furihata, Hiromizu Takahashi, Tsuguto Fujimoto, Makoto Kuroda |
| EPI_ISL_3392355, EPI_ISL_3392356, EPI_ISL_3392357, EPI_ISL_3392358, EPI_ISL_3392359, EPI_ISL_3392360, EPI_ISL_3392361, EPI_ISL_3392362, EPI_ISL_3392363, EPI_ISL_3392364, EPI_ISL_3392365, EPI_ISL_3392366, EPI_ISL_3392367, EPI_ISL_3392368, EPI_ISL_3392369, EPI_ISL_3392370, EPI_ISL_3392371, EPI_ISL_3392372, EPI_ISL_3392373, EPI_ISL_3392374, EPI_ISL_3392375, EPI_ISL_3392376, EPI_ISL_3392377, EPI_ISL_3392378, EPI_ISL_3392379, EPI_ISL_3392380, EPI_ISL_3392381, EPI_ISL_3392382, EPI_ISL_3392383, EPI_ISL_3392384, EPI_ISL_3392385, EPI_ISL_3392386, EPI_ISL_3392387, EPI_ISL_3392388, EPI_ISL_3392389, EPI_ISL_3392390, EPI_ISL_3392391, EPI_ISL_3392392, EPI_ISL_3392393, EPI_ISL_3392394, EPI_ISL_3392395, EPI_ISL_3392396, EPI_ISL_3392397, EPI_ISL_3392398, EPI_ISL_3392399, EPI_ISL_3392400, EPI_ISL_3392401, EPI_ISL_3392402, EPI_ISL_3392403, EPI_ISL_3392404, EPI_ISL_3392405, EPI_ISL_3392406, EPI_ISL_3392407, EPI_ISL_3392408, EPI_ISL_3392409, EPI_ISL_3392410, EPI_ISL_3392411, EPI_ISL_3392412, EPI_ISL_3392413, EPI_ISL_3392414, EPI_ISL_3392415, EPI_ISL_3392416, EPI_ISL_3392417, EPI_ISL_3392418, EPI_ISL_3392419, EPI_ISL_3392420, EPI_ISL_3392421, EPI_ISL_3392422, EPI_ISL_3392423, EPI_ISL_3392424, EPI_ISL_3392425, EPI_ISL_3392426, EPI_ISL_3392427, EPI_ISL_3392428, EPI_ISL_3392429, EPI_ISL_3392430, EPI_ISL_3392431, EPI_ISL_3392432, EPI_ISL_3392433, EPI_ISL_3392434, EPI_ISL_3392435, EPI_ISL_3392436, EPI_ISL_3392437, EPI_ISL_3392438, EPI_ISL_3392439, EPI_ISL_3392440, EPI_ISL_3392441, EPI_ISL_3392442, EPI_ISL_3392443, EPI_ISL_3392444, EPI_ISL_3392445, EPI_ISL_3392446, EPI_ISL_3392447, EPI_ISL_3392448, EPI_ISL_3392449, EPI_ISL_3392450, EPI_ISL_3392451, EPI_ISL_3392452, EPI_ISL_3392453, EPI_ISL_3392454, EPI_ISL_3392455, EPI_ISL_3392456, EPI_ISL_3392457, EPI_ISL_3392458, EPI_ISL_3392459, EPI_ISL_3392460, EPI_ISL_3392461, EPI_ISL_3392462, EPI_ISL_3392463, EPI_ISL_3392464, EPI_ISL_3392465, EPI_ISL_3392466, EPI_ISL_3392467, EPI_ISL_3392468, EPI_ISL_3392469, EPI_ISL_3392470, EPI_ISL_3392471, EPI_ISL_3392472, EPI_ISL_3392473, EPI_ISL_3392474, EPI_ISL_3392475, EPI_ISL_3392476, EPI_ISL_3392477, EPI_ISL_3392478, EPI_ISL_3392479, EPI_ISL_3392480, EPI_ISL_3392481, EPI_ISL_3392482, EPI_ISL_3392483, EPI_ISL_3392484, EPI_ISL_3392485, EPI_ISL_3392486, EPI_ISL_3392487, EPI_ISL_3392488, EPI_ISL_3392489, EPI_ISL_3392490, EPI_ISL_3392491, EPI_ISL_3392492, EPI_ISL_3392493, EPI_ISL_3392494, EPI_ISL_3392495, EPI_ISL_3392496, EPI_ISL_3392497, EPI_ISL_3392498, EPI_ISL_3392499, EPI_ISL_3392500, EPI_ISL_3392501, EPI_ISL_3392502, EPI_ISL_3392503, EPI_ISL_3392504, EPI_ISL_3392505, EPI_ISL_3392506, EPI_ISL_3392507, EPI_ISL_3392508, EPI_ISL_3392509, EPI_ISL_3392510, EPI_ISL_3392511, EPI_ISL_3392512, EPI_ISL_3392513, EPI_ISL_3392514, EPI_ISL_3392515, EPI_ISL_3392516, EPI_ISL_3392517, EPI_ISL_3392518, EPI_ISL_3392519, EPI_ISL_3392520, EPI_ISL_3392521, EPI_ISL_3392522, EPI_ISL_3392523, EPI_ISL_3392524, EPI_ISL_3392525, EPI_ISL_3392526, EPI_ISL_3392527, EPI_ISL_3392528, EPI_ISL_3392529, EPI_ISL_3392530, EPI_ISL_3392531, EPI_ISL_3392532, EPI_ISL_3392533, EPI_ISL_3392534, EPI_ISL_3392535, EPI_ISL_3392536, EPI_ISL_3392537, EPI_ISL_3392538, EPI_ISL_3392539, EPI_ISL_3392540, EPI_ISL_3392541, EPI_ISL_3392542, EPI_ISL_3392543, EPI_ISL_3392544, EPI_ISL_3392545, EPI_ISL_3392546, EPI_ISL_3392547, EPI_ISL_3392548, EPI_ISL_3392549, EPI_ISL_3392550, EPI_ISL_3392551, EPI_ISL_3392552, EPI_ISL_3392553, EPI_ISL_3392554, EPI_ISL_3392555, EPI_ISL_3392556, EPI_ISL_3392557, EPI_ISL_3392558, EPI_ISL_3392559, EPI_ISL_3392560, EPI_ISL_3392561, EPI_ISL_3392562, EPI_ISL_3392563, EPI_ISL_3392564, EPI_ISL_3392565, EPI_ISL_3392566, EPI_ISL_3392567, EPI_ISL_3392568, EPI_ISL_3392569, EPI_ISL_3392570, EPI_ISL_3392571, EPI_ISL_3392572, EPI_ISL_3392573, EPI_ISL_3392574, EPI_ISL_3392575, EPI_ISL_3392576, EPI_ISL_3392577, EPI_ISL_3392578, EPI_ISL_3392579, EPI_ISL_3392580, EPI_ISL_3392581, EPI_ISL_3392582, EPI_ISL_3392583, EPI_ISL_3392584, EPI_ISL_3392585, EPI_ISL_3392586, EPI_ISL_3392587, EPI_ISL_3392588, EPI_ISL_3392589, EPI_ISL_3392590, EPI_ISL_3392591, EPI_ISL_3392592, EPI_ISL_3392593, EPI_ISL_3392594, EPI_ISL_3392595, EPI_ISL_3392596 | | | |
| see above | University Hospitals of Geneva, Laboratory of Virology | HUG, Laboratory of Virology and the Health2030 Genome Center | Samuel Cordey, Aline Mamin, Francisco Perez, Ana Rita Goncalves, Laurent Kaiser, Lorenzo Cerutti, Cedric Howald, Henri Pegeot, Melyssa Elies, Deborah Penet, Keith Harshman, Ioannis Xenarios |
| EPI_ISL_3392597 | Hospital | National Reference Center for Viruses of Respiratory Infections, Institut Pasteur, Paris | Marion Barbet, Sylvie Behillil, Méline Bizard, Angela Brisebarre, Camille Capel, Vincent Enouf, Louise Lefrançois, Frédéric Lemoine, Christophe Malabat, Corinne Maufrais, Hub de bioinformatique et Biostatistique, Etienne Simon-Lorière, Maud Vanpeene, Sylvie Van der Werf ,Marianne Burgard |
| EPI_ISL_3392598, EPI_ISL_3392600 | Labo Analyses Med | National Reference Center for Viruses of Respiratory Infections, Institut Pasteur, Paris | Marion Barbet, Sylvie Behillil, Méline Bizard, Angela Brisebarre, Camille Capel, Vincent Enouf, Louise Lefrançois, Frédéric Lemoine, Christophe Malabat, Corinne Maufrais, Hub de bioinformatique et Biostatistique, Etienne Simon-Lorière, Maud Vanpeene, Sylvie Van der Werf ,Valentin Robbe |
| EPI_ISL_3392601 | Hospital | National Reference Center for Viruses of Respiratory Infections, Institut Pasteur, Paris | Marion Barbet, Sylvie Behillil, Méline Bizard, Angela Brisebarre, Camille Capel, Vincent Enouf, Louise Lefrançois, Frédéric Lemoine, Christophe Malabat, Corinne Maufrais, Hub de bioinformatique et Biostatistique, Etienne Simon-Lorière, Maud Vanpeene, Sylvie Van der Werf ,Marine Desroches |
| EPI_ISL_3392602 | Labo Analyses Med | National Reference Center for Viruses of Respiratory Infections, Institut Pasteur, Paris | Marion Barbet, Sylvie Behillil, Méline Bizard, Angela Brisebarre, Camille Capel, Vincent Enouf, Louise Lefrançois, Frédéric Lemoine, Christophe Malabat, Corinne Maufrais, Hub de bioinformatique et Biostatistique, Etienne Simon-Lorière, Maud Vanpeene, Sylvie Van der Werf ,Valentin Robbe |
| EPI_ISL_3392604 | Hospital | National Reference Center for Viruses of Respiratory Infections, Institut Pasteur, Paris | Marion Barbet, Sylvie Behillil, Méline Bizard, Angela Brisebarre, Camille Capel, Vincent Enouf, Louise Lefrançois, Frédéric Lemoine, Christophe Malabat, Corinne Maufrais, Hub de bioinformatique et Biostatistique, Etienne Simon-Lorière, Maud Vanpeene, Sylvie Van der Werf ,Pascale Martres |
| EPI_ISL_3392605 | Labo Analyses Med | National Reference Center for Viruses of Respiratory Infections, Institut Pasteur, Paris | Marion Barbet, Sylvie Behillil, Méline Bizard, Angela Brisebarre, Camille Capel, Vincent Enouf, Louise Lefrançois, Frédéric Lemoine, Christophe Malabat, Corinne Maufrais, Hub de bioinformatique et Biostatistique, Etienne Simon-Lorière, Maud Vanpeene, Sylvie Van der Werf ,Sabine Trombert |
| EPI_ISL_3392606 | Hospital | National Reference Center for Viruses of Respiratory Infections, Institut Pasteur, Paris | Marion Barbet, Sylvie Behillil, Méline Bizard, Angela Brisebarre, Camille Capel, Vincent Enouf, Louise Lefrançois, Frédéric Lemoine, Christophe Malabat, Corinne Maufrais, Hub de bioinformatique et Biostatistique, Etienne Simon-Lorière, Maud Vanpeene, Sylvie Van der Werf ,ValéRie Serazin |
| EPI_ISL_3392607 | Hospital | National Reference Center for Viruses of Respiratory Infections, Institut Pasteur, Paris | Marion Barbet, Sylvie Behillil, Méline Bizard, Angela Brisebarre, Camille Capel, Vincent Enouf, Louise Lefrançois, Frédéric Lemoine, Christophe Malabat, Corinne Maufrais, Hub de bioinformatique et Biostatistique, Etienne Simon-Lorière, Maud Vanpeene, Sylvie Van der Werf ,Laurence Fagour |
| EPI_ISL_3392608 | Labo Analyses Med | National Reference Center for Viruses of Respiratory Infections, Institut Pasteur, Paris | Marion Barbet, Sylvie Behillil, Méline Bizard, Angela Brisebarre, Camille Capel, Vincent Enouf, Louise Lefrançois, Frédéric Lemoine, Christophe Malabat, Corinne Maufrais, Hub de bioinformatique et Biostatistique, Etienne Simon-Lorière, Maud Vanpeene, Sylvie Van der Werf |
| EPI_ISL_3392610 | Hospital | National Reference Center for Viruses of Respiratory Infections, Institut Pasteur, Paris | Marion Barbet, Sylvie Behillil, Méline Bizard, Angela Brisebarre, Camille Capel, Vincent Enouf, Louise Lefrançois, Frédéric Lemoine, Christophe Malabat, Corinne Maufrais, Hub de bioinformatique et Biostatistique, Etienne Simon-Lorière, Maud Vanpeene, Sylvie Van der Werf ,Sylvaine Bastian |
| EPI_ISL_3392628, EPI_ISL_3392639 | Labo Analyses Med | National Reference Center for Viruses of Respiratory Infections, Institut Pasteur, Paris | Marion Barbet, Sylvie Behillil, Méline Bizard, Angela Brisebarre, Camille Capel, Vincent Enouf, Louise Lefrançois, Frédéric Lemoine, Christophe Malabat, Corinne Maufrais, Hub de bioinformatique et Biostatistique, Etienne Simon-Lorière, Maud Vanpeene, Sylvie Van der Werf |
| EPI_ISL_3392640 | Hospital | National Reference Center for Viruses of Respiratory Infections, Institut Pasteur, Paris | Marion Barbet, Sylvie Behillil, Méline Bizard, Angela Brisebarre, Camille Capel, Vincent Enouf, Louise Lefrançois, Frédéric Lemoine, Christophe Malabat, Corinne Maufrais, Hub de bioinformatique et Biostatistique, Etienne Simon-Lorière, Maud Vanpeene, Sylvie Van der Werf ,ValéRie Serazin |
| EPI_ISL_3392641 | Hospital | National Reference Center for Viruses of Respiratory Infections, Institut Pasteur, Paris | Marion Barbet, Sylvie Behillil, Méline Bizard, Angela Brisebarre, Camille Capel, Vincent Enouf, Louise Lefrançois, Frédéric Lemoine, Christophe Malabat, Corinne Maufrais, Hub de bioinformatique et Biostatistique, Etienne Simon-Lorière, Maud Vanpeene, Sylvie Van der Werf ,Laurence Fagour |
| EPI_ISL_3392643, EPI_ISL_3392644 | Labo Analyses Med | National Reference Center for Viruses of Respiratory Infections, Institut Pasteur, Paris | Marion Barbet, Sylvie Behillil, Méline Bizard, Angela Brisebarre, Camille Capel, Vincent Enouf, Louise Lefrançois, Frédéric Lemoine, Christophe Malabat, Corinne Maufrais, Hub de bioinformatique et Biostatistique, Etienne Simon-Lorière, Maud Vanpeene, Sylvie Van der Werf |
| EPI_ISL_3392645 | Labo Analyses Med | National Reference Center for Viruses of Respiratory Infections, | Marion Barbet, Sylvie Behillil, Méline Bizard, Angela Brisebarre, Camille Capel, Vincent Enouf, Louise Lefrançois, Frédéric Lemoine, Christophe Malabat, Corinne Maufrais, Hub de bioinformatique et Biostatistique, Etienne Simon-Lorière, Maud |

| | | | |
|---|---|---|---|
| | | Institut Pasteur, Paris | Vanpeene, Sylvie Van der Werf ,Isabelle Souron |
| EPI_ISL_3392646 | Hospital | National Reference Center for Viruses of Respiratory Infections, Institut Pasteur, Paris | Marion Barbet, Sylvie Behillil, Méline Bizard, Angela Brisebarre, Camille Capel, Vincent Enouf, Louise Lefrançois, Frédéric Lemoine, Christophe Malabat, Corinne Maufrais, Hub de bioinformatique et Biostatistique, Etienne Simon-Lorière, Maud Vanpeene, Sylvie Van der Werf ,Laura Djamdjian |
| EPI_ISL_3392648, EPI_ISL_3392649, EPI_ISL_3392650, EPI_ISL_3392651, EPI_ISL_3392652, EPI_ISL_3392653, EPI_ISL_3392654, EPI_ISL_3392655, EPI_ISL_3392656, EPI_ISL_3392657, EPI_ISL_3392658, EPI_ISL_3392659, EPI_ISL_3392660, EPI_ISL_3392661, EPI_ISL_3392662, EPI_ISL_3392663, EPI_ISL_3392664, EPI_ISL_3392665 | | | |
| see above | Labo Analyses Med | National Reference Center for Viruses of Respiratory Infections, Institut Pasteur, Paris | Marion Barbet, Sylvie Behillil, Méline Bizard, Angela Brisebarre, Camille Capel, Vincent Enouf, Louise Lefrançois, Frédéric Lemoine, Christophe Malabat, Corinne Maufrais, Hub de bioinformatique et Biostatistique, Etienne Simon-Lorière, Maud Vanpeene, Sylvie Van der Werf |
| EPI_ISL_3392666, EPI_ISL_3392667, EPI_ISL_3392668 | Labo Analyses Med | National Reference Center for Viruses of Respiratory Infections, Institut Pasteur, Paris | Marion Barbet, Sylvie Behillil, Méline Bizard, Angela Brisebarre, Camille Capel, Vincent Enouf, Louise Lefrançois, Frédéric Lemoine, Christophe Malabat, Corinne Maufrais, Hub de bioinformatique et Biostatistique, Etienne Simon-Lorière, Maud Vanpeene, Sylvie Van der Werf ,Valentin Robbe |
| EPI_ISL_3392669, EPI_ISL_3392670 | Hospital | National Reference Center for Viruses of Respiratory Infections, Institut Pasteur, Paris | Marion Barbet, Sylvie Behillil, Méline Bizard, Angela Brisebarre, Camille Capel, Vincent Enouf, Louise Lefrançois, Frédéric Lemoine, Christophe Malabat, Corinne Maufrais, Hub de bioinformatique et Biostatistique, Etienne Simon-Lorière, Maud Vanpeene, Sylvie Van der Werf ,ValéRie Serazin |
| EPI_ISL_3392671, EPI_ISL_3392672 | Hospital | National Reference Center for Viruses of Respiratory Infections, Institut Pasteur, Paris | Marion Barbet, Sylvie Behillil, Méline Bizard, Angela Brisebarre, Camille Capel, Vincent Enouf, Louise Lefrançois, Frédéric Lemoine, Christophe Malabat, Corinne Maufrais, Hub de bioinformatique et Biostatistique, Etienne Simon-Lorière, Maud Vanpeene, Sylvie Van der Werf ,Eric Farfour |
| EPI_ISL_3392673, EPI_ISL_3392674 | Labo Analyses Med | National Reference Center for Viruses of Respiratory Infections, Institut Pasteur, Paris | Marion Barbet, Sylvie Behillil, Méline Bizard, Angela Brisebarre, Camille Capel, Vincent Enouf, Louise Lefrançois, Frédéric Lemoine, Christophe Malabat, Corinne Maufrais, Hub de bioinformatique et Biostatistique, Etienne Simon-Lorière, Maud Vanpeene, Sylvie Van der Werf ,Caroline Malderet |
| EPI_ISL_3392675 | Hospital | National Reference Center for Viruses of Respiratory Infections, Institut Pasteur, Paris | Marion Barbet, Sylvie Behillil, Méline Bizard, Angela Brisebarre, Camille Capel, Vincent Enouf, Louise Lefrançois, Frédéric Lemoine, Christophe Malabat, Corinne Maufrais, Hub de bioinformatique et Biostatistique, Etienne Simon-Lorière, Maud Vanpeene, Sylvie Van der Werf ,Marianne Burgard |
| EPI_ISL_3392676 | Hospital | National Reference Center for Viruses of Respiratory Infections, Institut Pasteur, Paris | Marion Barbet, Sylvie Behillil, Méline Bizard, Angela Brisebarre, Camille Capel, Vincent Enouf, Louise Lefrançois, Frédéric Lemoine, Christophe Malabat, Corinne Maufrais, Hub de bioinformatique et Biostatistique, Etienne Simon-Lorière, Maud Vanpeene, Sylvie Van der Werf ,Olivia Raulin |
| EPI_ISL_3392677, EPI_ISL_3392678, EPI_ISL_3392679 | Hospital | National Reference Center for Viruses of Respiratory Infections, Institut Pasteur, Paris | Marion Barbet, Sylvie Behillil, Méline Bizard, Angela Brisebarre, Camille Capel, Vincent Enouf, Louise Lefrançois, Frédéric Lemoine, Christophe Malabat, Corinne Maufrais, Hub de bioinformatique et Biostatistique, Etienne Simon-Lorière, Maud Vanpeene, Sylvie Van der Werf ,Pascale Martres |
| EPI_ISL_3392680, EPI_ISL_3392681 | Hospital | National Reference Center for Viruses of Respiratory Infections, Institut Pasteur, Paris | Marion Barbet, Sylvie Behillil, Méline Bizard, Angela Brisebarre, Camille Capel, Vincent Enouf, Louise Lefrançois, Frédéric Lemoine, Christophe Malabat, Corinne Maufrais, Hub de bioinformatique et Biostatistique, Etienne Simon-Lorière, Maud Vanpeene, Sylvie Van der Werf ,Marianne Burgard |
| EPI_ISL_3392682, EPI_ISL_3392683 | Hospital | National Reference Center for Viruses of Respiratory Infections, Institut Pasteur, Paris | Marion Barbet, Sylvie Behillil, Méline Bizard, Angela Brisebarre, Camille Capel, Vincent Enouf, Louise Lefrançois, Frédéric Lemoine, Christophe Malabat, Corinne Maufrais, Hub de bioinformatique et Biostatistique, Etienne Simon-Lorière, Maud Vanpeene, Sylvie Van der Werf ,Sylvaine Bastian |
| EPI_ISL_3392684, EPI_ISL_3392685, EPI_ISL_3392686, EPI_ISL_3392687, EPI_ISL_3392688, EPI_ISL_3392689 | Labo Analyses Med | National Reference Center for Viruses of Respiratory Infections, Institut Pasteur, Paris | Marion Barbet, Sylvie Behillil, Méline Bizard, Angela Brisebarre, Camille Capel, Vincent Enouf, Louise Lefrançois, Frédéric Lemoine, Christophe Malabat, Corinne Maufrais, Hub de bioinformatique et Biostatistique, Etienne Simon-Lorière, Maud Vanpeene, Sylvie Van der Werf ,Sabine Trombert |
| EPI_ISL_3392690 | Hospital | National Reference Center for Viruses of Respiratory Infections, Institut Pasteur, Paris | Marion Barbet, Sylvie Behillil, Méline Bizard, Angela Brisebarre, Camille Capel, Vincent Enouf, Louise Lefrançois, Frédéric Lemoine, Christophe Malabat, Corinne Maufrais, Hub de bioinformatique et Biostatistique, Etienne Simon-Lorière, Maud Vanpeene, Sylvie Van der Werf ,Marianne Burgard |
| EPI_ISL_3392691, EPI_ISL_3392692, EPI_ISL_3392693, EPI_ISL_3392694, EPI_ISL_3392695, EPI_ISL_3392696, EPI_ISL_3392697, EPI_ISL_3392698, EPI_ISL_3392699, EPI_ISL_3392700, EPI_ISL_3392701, EPI_ISL_3392702, EPI_ISL_3392703, EPI_ISL_3392704, EPI_ISL_3392705 | | | |
| see above | Labo Analyses Med | National Reference Center for Viruses of Respiratory Infections, Institut Pasteur, Paris | Marion Barbet, Sylvie Behillil, Méline Bizard, Angela Brisebarre, Camille Capel, Vincent Enouf, Louise Lefrançois, Frédéric Lemoine, Christophe Malabat, Corinne Maufrais, Hub de bioinformatique et Biostatistique, Etienne Simon-Lorière, Maud Vanpeene, Sylvie Van der Werf ,Isabelle Souron |
| EPI_ISL_3392706, EPI_ISL_3392707 | Hospital | National Reference Center for Viruses of Respiratory Infections, Institut Pasteur, Paris | Marion Barbet, Sylvie Behillil, Méline Bizard, Angela Brisebarre, Camille Capel, Vincent Enouf, Louise Lefrançois, Frédéric Lemoine, Christophe Malabat, Corinne Maufrais, Hub de bioinformatique et Biostatistique, Etienne Simon-Lorière, Maud Vanpeene, Sylvie Van der Werf ,Marine Desroches |
| EPI_ISL_3392708 | Hospital | National Reference Center for Viruses of Respiratory Infections, Institut Pasteur, Paris | Marion Barbet, Sylvie Behillil, Méline Bizard, Angela Brisebarre, Camille Capel, Vincent Enouf, Louise Lefrançois, Frédéric Lemoine, Christophe Malabat, Corinne Maufrais, Hub de bioinformatique et Biostatistique, Etienne Simon-Lorière, Maud Vanpeene, Sylvie Van der Werf ,Laura Djamdjian |
| EPI_ISL_3392709, EPI_ISL_3392710, EPI_ISL_3392712, EPI_ISL_3392713, EPI_ISL_3392714, EPI_ISL_3392715, EPI_ISL_3392716, EPI_ISL_3392717, EPI_ISL_3392718, EPI_ISL_3392719, EPI_ISL_3392720, EPI_ISL_3392721, EPI_ISL_3392722, EPI_ISL_3392723, EPI_ISL_3392724, EPI_ISL_3392725, EPI_ISL_3392726, EPI_ISL_3392727, EPI_ISL_3392728, EPI_ISL_3392729, EPI_ISL_3392730, EPI_ISL_3392731, EPI_ISL_3392732, EPI_ISL_3392733, EPI_ISL_3392734, EPI_ISL_3392735, EPI_ISL_3392736, EPI_ISL_3392737, EPI_ISL_3392738, EPI_ISL_3392739, EPI_ISL_3392740, EPI_ISL_3392741, EPI_ISL_3392742, EPI_ISL_3392743, EPI_ISL_3392744, EPI_ISL_3392745, EPI_ISL_3392746, EPI_ISL_3392747, EPI_ISL_3392748, EPI_ISL_3392749, EPI_ISL_3392750, EPI_ISL_3392751, EPI_ISL_3392752, EPI_ISL_3392753, EPI_ISL_3392754, EPI_ISL_3392755, EPI_ISL_3392756, EPI_ISL_3392757, EPI_ISL_3392758, EPI_ISL_3392759, EPI_ISL_3392760, EPI_ISL_3392761, EPI_ISL_3392762, EPI_ISL_3392763, EPI_ISL_3392764, EPI_ISL_3392765, EPI_ISL_3392766, EPI_ISL_3392767, EPI_ISL_3392768, EPI_ISL_3392769, EPI_ISL_3392770, EPI_ISL_3392771, EPI_ISL_3392772, EPI_ISL_3392773, EPI_ISL_3392774, EPI_ISL_3392775, EPI_ISL_3392776, EPI_ISL_3392777, EPI_ISL_3392778, EPI_ISL_3392779, EPI_ISL_3392780, EPI_ISL_3392781, EPI_ISL_3392782, EPI_ISL_3392783, EPI_ISL_3392784, EPI_ISL_3392785, EPI_ISL_3392786, EPI_ISL_3392787, EPI_ISL_3392788 | | | |
| see above | Minnesota Department of Health, Public Health Laboratory | Minnesota Department of Health, Public Health Laboratory | Alexandra Lorentz, Jacob Garfin, Matt Plumb, and Xiong Wang |
| EPI_ISL_3392789, EPI_ISL_3392790, EPI_ISL_3392791, EPI_ISL_3392792, EPI_ISL_3392793, EPI_ISL_3392794, EPI_ISL_3392795, EPI_ISL_3392796, EPI_ISL_3392797, EPI_ISL_3392798, EPI_ISL_3392799, EPI_ISL_3392800, EPI_ISL_3392801, EPI_ISL_3392802, EPI_ISL_3392803, EPI_ISL_3392804, EPI_ISL_3392805, EPI_ISL_3392806, EPI_ISL_3392807, EPI_ISL_3392808, EPI_ISL_3392809, EPI_ISL_3392810, EPI_ISL_3392811, EPI_ISL_3392812, EPI_ISL_3392813, EPI_ISL_3392814, EPI_ISL_3392815, EPI_ISL_3392816, EPI_ISL_3392817, EPI_ISL_3392818, EPI_ISL_3392819, EPI_ISL_3392820, EPI_ISL_3392821, EPI_ISL_3392822, EPI_ISL_3392823, EPI_ISL_3392824, EPI_ISL_3392825, EPI_ISL_3392826, EPI_ISL_3392827, EPI_ISL_3392828, EPI_ISL_3392829, EPI_ISL_3392830, EPI_ISL_3392831, EPI_ISL_3392832, EPI_ISL_3392833, EPI_ISL_3392834, EPI_ISL_3392835, EPI_ISL_3392836, EPI_ISL_3392837, EPI_ISL_3392838, EPI_ISL_3392839, EPI_ISL_3392840, EPI_ISL_3392841, EPI_ISL_3392842, EPI_ISL_3392843, EPI_ISL_3392844, EPI_ISL_3392845, EPI_ISL_3392846, EPI_ISL_3392847, EPI_ISL_3392848, EPI_ISL_3392849, EPI_ISL_3392850, EPI_ISL_3392851, EPI_ISL_3392852, EPI_ISL_3392853, EPI_ISL_3392854, EPI_ISL_3392855, EPI_ISL_3392856, EPI_ISL_3392857, EPI_ISL_3392858, EPI_ISL_3392859, EPI_ISL_3392860, EPI_ISL_3392861, EPI_ISL_3392862, EPI_ISL_3392863, EPI_ISL_3392864, EPI_ISL_3392865, EPI_ISL_3392866, EPI_ISL_3392867, EPI_ISL_3392868, EPI_ISL_3392869, EPI_ISL_3392870, EPI_ISL_3392871, EPI_ISL_3392872, EPI_ISL_3392873, EPI_ISL_3392874, EPI_ISL_3392875, EPI_ISL_3392876, EPI_ISL_3392877, EPI_ISL_3392878, EPI_ISL_3392879, EPI_ISL_3392880, EPI_ISL_3392881, EPI_ISL_3392882, EPI_ISL_3392883, EPI_ISL_3392884, EPI_ISL_3392885, EPI_ISL_3392886, EPI_ISL_3392887, EPI_ISL_3392888, EPI_ISL_3392889, EPI_ISL_3392890, EPI_ISL_3392891, EPI_ISL_3392892, EPI_ISL_3392893, EPI_ISL_3392894, EPI_ISL_3392895, EPI_ISL_3392896, EPI_ISL_3392897, EPI_ISL_3392898, EPI_ISL_3392899, EPI_ISL_3392900, EPI_ISL_3392901, EPI_ISL_3392902, EPI_ISL_3392903, EPI_ISL_3392904, EPI_ISL_3392905, EPI_ISL_3392906, EPI_ISL_3392907, EPI_ISL_3392908, EPI_ISL_3392909, EPI_ISL_3392910, EPI_ISL_3392911, EPI_ISL_3392912, EPI_ISL_3392913, EPI_ISL_3392914, EPI_ISL_3392915, EPI_ISL_3392916, EPI_ISL_3392917, EPI_ISL_3392918, EPI_ISL_3392919, EPI_ISL_3392920, EPI_ISL_3392921, EPI_ISL_3392922, EPI_ISL_3392923, EPI_ISL_3392924, EPI_ISL_3392925, EPI_ISL_3392926, EPI_ISL_3392927, EPI_ISL_3392928, EPI_ISL_3392929, EPI_ISL_3392930, EPI_ISL_3392931, EPI_ISL_3392932, EPI_ISL_3392933, EPI_ISL_3392934, EPI_ISL_3392935, EPI_ISL_3392936, EPI_ISL_3392937, EPI_ISL_3392938, EPI_ISL_3392939, EPI_ISL_3392940, EPI_ISL_3392941, EPI_ISL_3392942, EPI_ISL_3392943, EPI_ISL_3392944, EPI_ISL_3392945, EPI_ISL_3392946, EPI_ISL_3392947, EPI_ISL_3392948, EPI_ISL_3392949, EPI_ISL_3392950, EPI_ISL_3392951, EPI_ISL_3392952, EPI_ISL_3392953, EPI_ISL_3392954, EPI_ISL_3392955, EPI_ISL_3392956, EPI_ISL_3392957, EPI_ISL_3392958, EPI_ISL_3392959, EPI_ISL_3392968, EPI_ISL_3392971 | | | |
| see above | Pathology and Laboratory Medicine Institute, Cleveland Clinic, Ohio, USA | Pathology and Laboratory Medicine Institute, Cleveland Clinic, Ohio, USA | Jessica Spildener, Concetta Peck, Jennifer Starbuck, Joy Nakitandwe, Kristen McDonnell, David Plunkett, Zheng Jin Tu, Jay Brock, Yu-Wei Cheng, Gary Procop, Daniel Rhoads, Daniel H. Farkas, David Bosler |
| EPI_ISL_3392972, EPI_ISL_3392973, EPI_ISL_3392974, EPI_ISL_3392975, EPI_ISL_3392976, EPI_ISL_3392977, EPI_ISL_3392978, EPI_ISL_3392979, EPI_ISL_3392980, EPI_ISL_3392981, EPI_ISL_3392982, EPI_ISL_3392983, EPI_ISL_3392984, EPI_ISL_3392985, EPI_ISL_3392986, EPI_ISL_3392987, EPI_ISL_3392988, EPI_ISL_3392989, EPI_ISL_3392990, EPI_ISL_3392991, EPI_ISL_3392992, EPI_ISL_3392993, EPI_ISL_3392994, EPI_ISL_3392995, EPI_ISL_3392996, EPI_ISL_3392997, EPI_ISL_3392998, EPI_ISL_3392999, EPI_ISL_3393000, EPI_ISL_3393001, EPI_ISL_3393002, EPI_ISL_3393004, EPI_ISL_3393005, EPI_ISL_3393006, EPI_ISL_3393007, EPI_ISL_3393008, EPI_ISL_3393009, EPI_ISL_3393010, EPI_ISL_3393011, EPI_ISL_3393012, EPI_ISL_3393013, EPI_ISL_3393014, EPI_ISL_3393015, EPI_ISL_3393016, EPI_ISL_3393017, EPI_ISL_3393018, EPI_ISL_3393019, EPI_ISL_3393021, EPI_ISL_3393022, EPI_ISL_3393023, EPI_ISL_3393024, EPI_ISL_3393025, EPI_ISL_3393026, EPI_ISL_3393027, EPI_ISL_3393028, EPI_ISL_3393029, EPI_ISL_3393030, EPI_ISL_3393031, EPI_ISL_3393032, EPI_ISL_3393033, EPI_ISL_3393034, EPI_ISL_3393035, EPI_ISL_3393036, EPI_ISL_3393037, EPI_ISL_3393038, EPI_ISL_3393039, EPI_ISL_3393040, EPI_ISL_3393042, EPI_ISL_3393043, EPI_ISL_3393044, EPI_ISL_3393045, EPI_ISL_3393046, EPI_ISL_3393047, EPI_ISL_3393048, EPI_ISL_3393049, EPI_ISL_3393050, EPI_ISL_3393051, EPI_ISL_3393052, EPI_ISL_3393053, EPI_ISL_3393054, EPI_ISL_3393055, EPI_ISL_3393056, EPI_ISL_3393057, EPI_ISL_3393059, EPI_ISL_3393060, EPI_ISL_3393061, EPI_ISL_3393062, EPI_ISL_3393063, EPI_ISL_3393064, EPI_ISL_3393065, EPI_ISL_3393066, EPI_ISL_3393067, EPI_ISL_3393068, EPI_ISL_3393069, EPI_ISL_3393070, EPI_ISL_3393071, EPI_ISL_3393072, EPI_ISL_3393073, EPI_ISL_3393074, EPI_ISL_3393075, EPI_ISL_3393076, EPI_ISL_3393077, EPI_ISL_3393078, EPI_ISL_3393080, EPI_ISL_3393081, EPI_ISL_3393082, EPI_ISL_3393083, EPI_ISL_3393084, EPI_ISL_3393085, EPI_ISL_3393086, EPI_ISL_3393087, EPI_ISL_3393088, EPI_ISL_3393089, EPI_ISL_3393090, EPI_ISL_3393091, EPI_ISL_3393092, EPI_ISL_3393093, EPI_ISL_3393094, EPI_ISL_3393095, EPI_ISL_3393097, EPI_ISL_3393098, EPI_ISL_3393099, EPI_ISL_3393100, EPI_ISL_3393101, EPI_ISL_3393102, EPI_ISL_3393104, EPI_ISL_3393105, EPI_ISL_3393106, EPI_ISL_3393107, EPI_ISL_3393108, EPI_ISL_3393109, EPI_ISL_3393111, EPI_ISL_3393112, EPI_ISL_3393113, EPI_ISL_3393114, EPI_ISL_3393115, EPI_ISL_3393116, EPI_ISL_3393118, EPI_ISL_3393119, EPI_ISL_3393120, EPI_ISL_3393122, EPI_ISL_3393123, EPI_ISL_3393124, EPI_ISL_3393125, EPI_ISL_3393126, EPI_ISL_3393127, EPI_ISL_3393128, EPI_ISL_3393129, EPI_ISL_3393131, EPI_ISL_3393132, EPI_ISL_3393133, EPI_ISL_3393134, EPI_ISL_3393135, EPI_ISL_3393136, EPI_ISL_3393137, EPI_ISL_3393138, EPI_ISL_3393139, EPI_ISL_3393141, EPI_ISL_3393142, EPI_ISL_3393143, EPI_ISL_3393144, EPI_ISL_3393145, EPI_ISL_3393146, EPI_ISL_3393148, EPI_ISL_3393149, EPI_ISL_3393150, EPI_ISL_3393151, EPI_ISL_3393152, EPI_ISL_3393153, EPI_ISL_3393155, EPI_ISL_3393156, EPI_ISL_3393157, EPI_ISL_3393159, EPI_ISL_3393160, EPI_ISL_3393161, EPI_ISL_3393162, EPI_ISL_3393163, EPI_ISL_3393164, EPI_ISL_3393165, EPI_ISL_3393166, EPI_ISL_3393168, EPI_ISL_3393169, EPI_ISL_3393170, EPI_ISL_3393171, EPI_ISL_3393172, EPI_ISL_3393173, EPI_ISL_3393174, EPI_ISL_3393175, EPI_ISL_3393176, EPI_ISL_3393178, EPI_ISL_3393179, EPI_ISL_3393180, EPI_ISL_3393181, EPI_ISL_3393182, EPI_ISL_3393183, EPI_ISL_3393185, EPI_ISL_3393186, EPI_ISL_3393187, EPI_ISL_3393188, EPI_ISL_3393189, EPI_ISL_3393190, EPI_ISL_3393192, EPI_ISL_3393193, EPI_ISL_3393194, EPI_ISL_3393196, EPI_ISL_3393197, EPI_ISL_3393198, EPI_ISL_3393199, EPI_ISL_3393200, EPI_ISL_3393201, EPI_ISL_3393202, EPI_ISL_3393203, EPI_ISL_3393205, EPI_ISL_3393206, EPI_ISL_3393207, EPI_ISL_3393208, EPI_ISL_3393209, EPI_ISL_3393210, EPI_ISL_3393211, EPI_ISL_3393212, EPI_ISL_3393214, EPI_ISL_3393215, EPI_ISL_3393216, EPI_ISL_3393217, EPI_ISL_3393218, EPI_ISL_3393219, EPI_ISL_3393221, EPI_ISL_3393222, EPI_ISL_3393223, EPI_ISL_3393224, EPI_ISL_3393225, EPI_ISL_3393226, EPI_ISL_3393228, EPI_ISL_3393229, EPI_ISL_3393230, EPI_ISL_3393232, EPI_ISL_3393233, EPI_ISL_3393234, EPI_ISL_3393235, EPI_ISL_3393236, EPI_ISL_3393237, EPI_ISL_3393238, EPI_ISL_3393239, EPI_ISL_3393241, EPI_ISL_3393242, EPI_ISL_3393243, EPI_ISL_3393244, EPI_ISL_3393245, EPI_ISL_3393246, EPI_ISL_3393247, EPI_ISL_3393248, EPI_ISL_3393249, EPI_ISL_3393251, EPI_ISL_3393252, EPI_ISL_3393253, EPI_ISL_3393254, EPI_ISL_3393255, EPI_ISL_3393256, EPI_ISL_3393258, EPI_ISL_3393259, EPI_ISL_3393260, EPI_ISL_3393261, EPI_ISL_3393262, EPI_ISL_3393263, EPI_ISL_3393265, EPI_ISL_3393266, EPI_ISL_3393267, EPI_ISL_3393269, EPI_ISL_3393270, EPI_ISL_3393271, EPI_ISL_3393272, EPI_ISL_3393273, EPI_ISL_3393274, EPI_ISL_3393275, EPI_ISL_3393276, EPI_ISL_3393278, EPI_ISL_3393279, EPI_ISL_3393280, EPI_ISL_3393281, EPI_ISL_3393282, EPI_ISL_3393283, EPI_ISL_3393284, EPI_ISL_3393285, EPI_ISL_3393286, EPI_ISL_3393287, EPI_ISL_3393288, EPI_ISL_3393289, EPI_ISL_3393290, EPI_ISL_3393291, EPI_ISL_3393292, EPI_ISL_3393293, EPI_ISL_3393294, EPI_ISL_3393295, EPI_ISL_3393296, EPI_ISL_3393297, EPI_ISL_3393298, EPI_ISL_3393299, EPI_ISL_3393300, EPI_ISL_3393301, EPI_ISL_3393302, EPI_ISL_3393303, EPI_ISL_3393304, EPI_ISL_3393305, EPI_ISL_3393306, EPI_ISL_3393307, EPI_ISL_3393308, EPI_ISL_3393309, EPI_ISL_3393310, EPI_ISL_3393311, EPI_ISL_3393312, EPI_ISL_3393313, EPI_ISL_3393314, EPI_ISL_3393315, EPI_ISL_3393316, EPI_ISL_3393317, EPI_ISL_3393318, EPI_ISL_3393319, EPI_ISL_3393320, EPI_ISL_3393321, EPI_ISL_3393322, EPI_ISL_3393323, EPI_ISL_3393324, EPI_ISL_3393325, EPI_ISL_3393326, EPI_ISL_3393327 | | | |
| see above | Ministry of Health Turkey | Ministry of Health Turkey | Fatma Bayrakdar, Yasemin Coşgun, Süleyman Yalcin, Gülay Korukluoğlu |
| EPI_ISL_3393328 | LISIEUX Cerballiance | Department of Virology, Henri Mondor University Hospital, Assistance Publique Hôpitaux de Paris, Université Paris-Est Créteil, INSERM U955 | Christophe Rodriguez, Slim Fourati, Vanessa Demontant, Guillaume Gricourt, Melissa N'Debi, Alexandre Soulier, Elisabeth Trawinski, Jean-Michel Pawlotsky |
| EPI_ISL_3393330 | Hôpital Avicenne | Department of Virology, Henri Mondor University Hospital, Assistance Publique Hôpitaux de Paris, Université Paris-Est Créteil, INSERM U955 | Christophe Rodriguez, Slim Fourati, Vanessa Demontant, Guillaume Gricourt, Melissa N'Debi, Alexandre Soulier, Elisabeth Trawinski, Jean-Michel Pawlotsky |
| EPI_ISL_3393331 | BIO ARD'AISNE | Department of Virology, Henri Mondor University Hospital, Assistance Publique Hôpitaux de Paris, Université Paris-Est Créteil, INSERM U955 | Christophe Rodriguez, Slim Fourati, Vanessa Demontant, Guillaume Gricourt, Melissa N'Debi, Alexandre Soulier, Elisabeth Trawinski, Jean-Michel Pawlotsky |
| EPI_ISL_3393332 | Biogroup Bio Lam-LCD Saint-Denis | Department of Virology, Henri Mondor University Hospital, Assistance Publique Hôpitaux de Paris, Université Paris-Est Créteil, INSERM U955 | Christophe Rodriguez, Slim Fourati, Vanessa Demontant, Guillaume Gricourt, Melissa N'Debi, Alexandre Soulier, Elisabeth Trawinski, Jean-Michel Pawlotsky |
| EPI_ISL_3393333 | BIO67-BIOSPHERE | Department of Virology, Henri Mondor University Hospital, Assistance Publique Hôpitaux de Paris, Université Paris-Est Créteil, INSERM U955 | Christophe Rodriguez, Slim Fourati, Vanessa Demontant, Guillaume Gricourt, Melissa N'Debi, Alexandre Soulier, Elisabeth Trawinski, Jean-Michel Pawlotsky |
| EPI_ISL_3393334 | BROUSSAIS | Department of Virology, Henri Mondor University Hospital, Assistance Publique Hôpitaux de Paris, Université Paris-Est Créteil, INSERM U955 | Christophe Rodriguez, Slim Fourati, Vanessa Demontant, Guillaume Gricourt, Melissa N'Debi, Alexandre Soulier, Elisabeth Trawinski, Jean-Michel Pawlotsky |
| EPI_ISL_3393336, EPI_ISL_3393337 | Biogroup Bio Lam-LCD Saint-Denis | Department of Virology, Henri Mondor University Hospital, Assistance Publique Hôpitaux de Paris, Université Paris-Est Créteil, INSERM U955 | Christophe Rodriguez, Slim Fourati, Vanessa Demontant, Guillaume Gricourt, Melissa N'Debi, Alexandre Soulier, Elisabeth Trawinski, Jean-Michel Pawlotsky |
| EPI_ISL_3393338 | Cerballiance Paris IDF est Saint-Denis | Department of Virology, Henri Mondor University Hospital, Assistance Publique Hôpitaux de Paris, Université Paris-Est Créteil, INSERM U955 | Christophe Rodriguez, Slim Fourati, Vanessa Demontant, Guillaume Gricourt, Melissa N'Debi, Alexandre Soulier, Elisabeth Trawinski, Jean-Michel Pawlotsky |

| | | | |
|---|---|---|---|
| EPI_ISL_3393339 | SELAS MEDILYS | Department of Virology, Henri Mondor University Hospital, Assistance Publique Hôpitaux de Paris, Université Paris-Est Créteil, INSERM U955 | Christophe Rodriguez, Slim Fourati, Vanessa Demontant, Guillaume Gricourt, Melissa N'Debi, Alexandre Soulier, Elisabeth Trawinski, Jean-Michel Pawlotsky |
| EPI_ISL_3393340 | laboratoire Belle Epine | Department of Virology, Henri Mondor University Hospital, Assistance Publique Hôpitaux de Paris, Université Paris-Est Créteil, INSERM U955 | Christophe Rodriguez, Slim Fourati, Vanessa Demontant, Guillaume Gricourt, Melissa N'Debi, Alexandre Soulier, Elisabeth Trawinski, Jean-Michel Pawlotsky |
| EPI_ISL_3393342 | Biogroup Bio Lam-LCD Saint-Denis | Department of Virology, Henri Mondor University Hospital, Assistance Publique Hôpitaux de Paris, Université Paris-Est Créteil, INSERM U955 | Christophe Rodriguez, Slim Fourati, Vanessa Demontant, Guillaume Gricourt, Melissa N'Debi, Alexandre Soulier, Elisabeth Trawinski, Jean-Michel Pawlotsky |
| EPI_ISL_3393343 | BIO67-BIOSPHERE | Department of Virology, Henri Mondor University Hospital, Assistance Publique Hôpitaux de Paris, Université Paris-Est Créteil, INSERM U955 | Christophe Rodriguez, Slim Fourati, Vanessa Demontant, Guillaume Gricourt, Melissa N'Debi, Alexandre Soulier, Elisabeth Trawinski, Jean-Michel Pawlotsky |
| EPI_ISL_3393344 | BROUSSAIS | Department of Virology, Henri Mondor University Hospital, Assistance Publique Hôpitaux de Paris, Université Paris-Est Créteil, INSERM U955 | Christophe Rodriguez, Slim Fourati, Vanessa Demontant, Guillaume Gricourt, Melissa N'Debi, Alexandre Soulier, Elisabeth Trawinski, Jean-Michel Pawlotsky |
| EPI_ISL_3393346, EPI_ISL_3393349 | SELAS MEDILYS | Department of Virology, Henri Mondor University Hospital, Assistance Publique Hôpitaux de Paris, Université Paris-Est Créteil, INSERM U955 | Christophe Rodriguez, Slim Fourati, Vanessa Demontant, Guillaume Gricourt, Melissa N'Debi, Alexandre Soulier, Elisabeth Trawinski, Jean-Michel Pawlotsky |
| EPI_ISL_3393350 | BROUSSAIS | Department of Virology, Henri Mondor University Hospital, Assistance Publique Hôpitaux de Paris, Université Paris-Est Créteil, INSERM U955 | Christophe Rodriguez, Slim Fourati, Vanessa Demontant, Guillaume Gricourt, Melissa N'Debi, Alexandre Soulier, Elisabeth Trawinski, Jean-Michel Pawlotsky |
| EPI_ISL_3393351 | BIO ARD'AISNE | Department of Virology, Henri Mondor University Hospital, Assistance Publique Hôpitaux de Paris, Université Paris-Est Créteil, INSERM U955 | Christophe Rodriguez, Slim Fourati, Vanessa Demontant, Guillaume Gricourt, Melissa N'Debi, Alexandre Soulier, Elisabeth Trawinski, Jean-Michel Pawlotsky |
| EPI_ISL_3393353 | CHU Rouen Normandie | Department of Virology, Henri Mondor University Hospital, Assistance Publique Hôpitaux de Paris, Université Paris-Est Créteil, INSERM U955 | Christophe Rodriguez, Slim Fourati, Vanessa Demontant, Guillaume Gricourt, Melissa N'Debi, Alexandre Soulier, Elisabeth Trawinski, Jean-Michel Pawlotsky |
| EPI_ISL_3393354 | Cerballiance Paris IDF est Saint-Denis | Department of Virology, Henri Mondor University Hospital, Assistance Publique Hôpitaux de Paris, Université Paris-Est Créteil, INSERM U955 | Christophe Rodriguez, Slim Fourati, Vanessa Demontant, Guillaume Gricourt, Melissa N'Debi, Alexandre Soulier, Elisabeth Trawinski, Jean-Michel Pawlotsky |
| EPI_ISL_3393356 | Hôp Civils de COLMAR | Department of Virology, Henri Mondor University Hospital, Assistance Publique Hôpitaux de Paris, Université Paris-Est Créteil, INSERM U955 | Christophe Rodriguez, Slim Fourati, Vanessa Demontant, Guillaume Gricourt, Melissa N'Debi, Alexandre Soulier, Elisabeth Trawinski, Jean-Michel Pawlotsky |
| EPI_ISL_3393357 | Biogroup Bio Lam-LCD Saint-Denis | Department of Virology, Henri Mondor University Hospital, Assistance Publique Hôpitaux de Paris, Université Paris-Est Créteil, INSERM U955 | Christophe Rodriguez, Slim Fourati, Vanessa Demontant, Guillaume Gricourt, Melissa N'Debi, Alexandre Soulier, Elisabeth Trawinski, Jean-Michel Pawlotsky |
| EPI_ISL_3393358 | BROUSSAIS | Department of Virology, Henri Mondor University Hospital, Assistance Publique Hôpitaux de Paris, Université Paris-Est Créteil, INSERM U955 | Christophe Rodriguez, Slim Fourati, Vanessa Demontant, Guillaume Gricourt, Melissa N'Debi, Alexandre Soulier, Elisabeth Trawinski, Jean-Michel Pawlotsky |
| EPI_ISL_3393359 | CHU Rouen Normandie | Department of Virology, Henri Mondor University Hospital, Assistance Publique Hôpitaux de Paris, Université Paris-Est Créteil, INSERM U955 | Christophe Rodriguez, Slim Fourati, Vanessa Demontant, Guillaume Gricourt, Melissa N'Debi, Alexandre Soulier, Elisabeth Trawinski, Jean-Michel Pawlotsky |
| EPI_ISL_3393362 | BROUSSAIS | Department of Virology, Henri Mondor University Hospital, Assistance Publique Hôpitaux de Paris, Université Paris-Est Créteil, INSERM U955 | Christophe Rodriguez, Slim Fourati, Vanessa Demontant, Guillaume Gricourt, Melissa N'Debi, Alexandre Soulier, Elisabeth Trawinski, Jean-Michel Pawlotsky |
| EPI_ISL_3393363 | SELAS MEDILYS | Department of Virology, Henri Mondor University Hospital, Assistance Publique Hôpitaux de Paris, Université Paris-Est Créteil, INSERM U955 | Christophe Rodriguez, Slim Fourati, Vanessa Demontant, Guillaume Gricourt, Melissa N'Debi, Alexandre Soulier, Elisabeth Trawinski, Jean-Michel Pawlotsky |
| EPI_ISL_3393364 | BROUSSAIS | Department of Virology, Henri Mondor University Hospital, Assistance Publique Hôpitaux de Paris, Université Paris-Est Créteil, INSERM U955 | Christophe Rodriguez, Slim Fourati, Vanessa Demontant, Guillaume Gricourt, Melissa N'Debi, Alexandre Soulier, Elisabeth Trawinski, Jean-Michel Pawlotsky |
| EPI_ISL_3393366 | BIO ARD'AISNE | Department of Virology, Henri Mondor University Hospital, Assistance Publique Hôpitaux de Paris, Université Paris-Est Créteil, INSERM U955 | Christophe Rodriguez, Slim Fourati, Vanessa Demontant, Guillaume Gricourt, Melissa N'Debi, Alexandre Soulier, Elisabeth Trawinski, Jean-Michel Pawlotsky |
| EPI_ISL_3393367 | Hôpital Pitié-Salpêtrière | Department of Virology, Henri Mondor University Hospital, Assistance Publique Hôpitaux de Paris, Université Paris-Est Créteil, INSERM U955 | Christophe Rodriguez, Slim Fourati, Vanessa Demontant, Guillaume Gricourt, Melissa N'Debi, Alexandre Soulier, Elisabeth Trawinski, Jean-Michel Pawlotsky |
| EPI_ISL_3393368 | BROUSSAIS | Department of Virology, Henri Mondor University Hospital, Assistance Publique Hôpitaux de Paris, Université Paris-Est Créteil, INSERM U955 | Christophe Rodriguez, Slim Fourati, Vanessa Demontant, Guillaume Gricourt, Melissa N'Debi, Alexandre Soulier, Elisabeth Trawinski, Jean-Michel Pawlotsky |
| EPI_ISL_3393370 | laboratoire Belle Epine | Department of Virology, Henri Mondor University Hospital, Assistance Publique Hôpitaux de Paris, Université Paris-Est Créteil, INSERM U955 | Christophe Rodriguez, Slim Fourati, Vanessa Demontant, Guillaume Gricourt, Melissa N'Debi, Alexandre Soulier, Elisabeth Trawinski, Jean-Michel Pawlotsky |
| EPI_ISL_3393371 | BROUSSAIS | Department of Virology, Henri Mondor University Hospital, Assistance Publique Hôpitaux de Paris, Université Paris-Est Créteil, INSERM U955 | Christophe Rodriguez, Slim Fourati, Vanessa Demontant, Guillaume Gricourt, Melissa N'Debi, Alexandre Soulier, Elisabeth Trawinski, Jean-Michel Pawlotsky |
| EPI_ISL_3393372 | SELAS MEDILYS | Department of Virology, Henri Mondor University Hospital, Assistance Publique Hôpitaux de Paris, Université Paris-Est Créteil, INSERM U955 | Christophe Rodriguez, Slim Fourati, Vanessa Demontant, Guillaume Gricourt, Melissa N'Debi, Alexandre Soulier, Elisabeth Trawinski, Jean-Michel Pawlotsky |
| EPI_ISL_3393373 | CHU Rouen Normandie | Department of Virology, Henri Mondor University Hospital, Assistance Publique Hôpitaux de Paris, Université Paris-Est Créteil, INSERM U955 | Christophe Rodriguez, Slim Fourati, Vanessa Demontant, Guillaume Gricourt, Melissa N'Debi, Alexandre Soulier, Elisabeth Trawinski, Jean-Michel Pawlotsky |
| EPI_ISL_3393375 | BIO67-BIOSPHERE | Department of Virology, Henri Mondor University Hospital, Assistance Publique Hôpitaux de Paris, Université Paris-Est Créteil, INSERM U955 | Christophe Rodriguez, Slim Fourati, Vanessa Demontant, Guillaume Gricourt, Melissa N'Debi, Alexandre Soulier, Elisabeth Trawinski, Jean-Michel Pawlotsky |
| EPI_ISL_3393376 | GH A.CHENEVIER-H.MONDOR | Department of Virology, Henri Mondor University Hospital, Assistance Publique Hôpitaux de Paris, Université Paris-Est Créteil, INSERM U955 | Christophe Rodriguez, Slim Fourati, Vanessa Demontant, Guillaume Gricourt, Melissa N'Debi, Alexandre Soulier, Elisabeth Trawinski, Jean-Michel Pawlotsky |
| EPI_ISL_3393377 | BROUSSAIS | Department of Virology, Henri Mondor University Hospital, Assistance Publique Hôpitaux de Paris, Université Paris-Est Créteil, INSERM U955 | Christophe Rodriguez, Slim Fourati, Vanessa Demontant, Guillaume Gricourt, Melissa N'Debi, Alexandre Soulier, Elisabeth Trawinski, Jean-Michel Pawlotsky |
| EPI_ISL_3393378 | SELAS BC-LAB | Department of Virology, Henri Mondor University Hospital, Assistance Publique Hôpitaux de Paris, Université Paris-Est Créteil, INSERM U955 | Christophe Rodriguez, Slim Fourati, Vanessa Demontant, Guillaume Gricourt, Melissa N'Debi, Alexandre Soulier, Elisabeth Trawinski, Jean-Michel Pawlotsky |
| EPI_ISL_3393379 | Cerballiance Paris IDF est Saint-Denis | Department of Virology, Henri Mondor University Hospital, Assistance Publique Hôpitaux de Paris, Université Paris-Est Créteil, INSERM U955 | Christophe Rodriguez, Slim Fourati, Vanessa Demontant, Guillaume Gricourt, Melissa N'Debi, Alexandre Soulier, Elisabeth Trawinski, Jean-Michel Pawlotsky |
| EPI_ISL_3393381 | GH A.CHENEVIER-H.MONDOR | Department of Virology, Henri Mondor University Hospital, Assistance Publique Hôpitaux de Paris, Université Paris-Est Créteil, INSERM U955 | Christophe Rodriguez, Slim Fourati, Vanessa Demontant, Guillaume Gricourt, Melissa N'Debi, Alexandre Soulier, Elisabeth Trawinski, Jean-Michel Pawlotsky |
| EPI_ISL_3393383, EPI_ISL_3393386 | BROUSSAIS | Department of Virology, Henri Mondor University Hospital, Assistance Publique Hôpitaux de Paris, Université Paris-Est Créteil, INSERM U955 | Christophe Rodriguez, Slim Fourati, Vanessa Demontant, Guillaume Gricourt, Melissa N'Debi, Alexandre Soulier, Elisabeth Trawinski, Jean-Michel Pawlotsky |
| EPI_ISL_3393388 | SELAS BC-LAB | Department of Virology, Henri Mondor University Hospital, Assistance Publique Hôpitaux de Paris, Université Paris-Est Créteil, | Christophe Rodriguez, Slim Fourati, Vanessa Demontant, Guillaume Gricourt, Melissa N'Debi, Alexandre Soulier, Elisabeth Trawinski, Jean-Michel Pawlotsky |

| | | | |
|---|---|---|---|
| | | INSERM U955 | |
| EPI_ISL_3393389 | laboratoire Belle Epine | Department of Virology, Henri Mondor University Hospital, Assistance Publique Hôpitaux de Paris, Université Paris-Est Créteil, INSERM U955 | Christophe Rodriguez, Slim Fourati, Vanessa Demontant, Guillaume Gricourt, Melissa N'Debi, Alexandre Soulier, Elisabeth Trawinski, Jean-Michel Pawlotsky |
| EPI_ISL_3393390 | LISIEUX Cerballiance | Department of Virology, Henri Mondor University Hospital, Assistance Publique Hôpitaux de Paris, Université Paris-Est Créteil, INSERM U955 | Christophe Rodriguez, Slim Fourati, Vanessa Demontant, Guillaume Gricourt, Melissa N'Debi, Alexandre Soulier, Elisabeth Trawinski, Jean-Michel Pawlotsky |
| EPI_ISL_3393391 | SELAS MEDILYS | Department of Virology, Henri Mondor University Hospital, Assistance Publique Hôpitaux de Paris, Université Paris-Est Créteil, INSERM U955 | Christophe Rodriguez, Slim Fourati, Vanessa Demontant, Guillaume Gricourt, Melissa N'Debi, Alexandre Soulier, Elisabeth Trawinski, Jean-Michel Pawlotsky |
| EPI_ISL_3393392 | BROUSSAIS | Department of Virology, Henri Mondor University Hospital, Assistance Publique Hôpitaux de Paris, Université Paris-Est Créteil, INSERM U955 | Christophe Rodriguez, Slim Fourati, Vanessa Demontant, Guillaume Gricourt, Melissa N'Debi, Alexandre Soulier, Elisabeth Trawinski, Jean-Michel Pawlotsky |
| EPI_ISL_3393394 | GH A.CHENEVIER-H.MONDOR | Department of Virology, Henri Mondor University Hospital, Assistance Publique Hôpitaux de Paris, Université Paris-Est Créteil, INSERM U955 | Christophe Rodriguez, Slim Fourati, Vanessa Demontant, Guillaume Gricourt, Melissa N'Debi, Alexandre Soulier, Elisabeth Trawinski, Jean-Michel Pawlotsky |
| EPI_ISL_3393395 | BIOCEANE | Department of Virology, Henri Mondor University Hospital, Assistance Publique Hôpitaux de Paris, Université Paris-Est Créteil, INSERM U955 | Christophe Rodriguez, Slim Fourati, Vanessa Demontant, Guillaume Gricourt, Melissa N'Debi, Alexandre Soulier, Elisabeth Trawinski, Jean-Michel Pawlotsky |
| EPI_ISL_3393396, EPI_ISL_3393397 | laboratoire Belle Epine | Department of Virology, Henri Mondor University Hospital, Assistance Publique Hôpitaux de Paris, Université Paris-Est Créteil, INSERM U955 | Christophe Rodriguez, Slim Fourati, Vanessa Demontant, Guillaume Gricourt, Melissa N'Debi, Alexandre Soulier, Elisabeth Trawinski, Jean-Michel Pawlotsky |
| EPI_ISL_3393398 | CH. CHU DE REIMS | Department of Virology, Henri Mondor University Hospital, Assistance Publique Hôpitaux de Paris, Université Paris-Est Créteil, INSERM U955 | Christophe Rodriguez, Slim Fourati, Vanessa Demontant, Guillaume Gricourt, Melissa N'Debi, Alexandre Soulier, Elisabeth Trawinski, Jean-Michel Pawlotsky |
| EPI_ISL_3393399 | SELAS MEDILYS | Department of Virology, Henri Mondor University Hospital, Assistance Publique Hôpitaux de Paris, Université Paris-Est Créteil, INSERM U955 | Christophe Rodriguez, Slim Fourati, Vanessa Demontant, Guillaume Gricourt, Melissa N'Debi, Alexandre Soulier, Elisabeth Trawinski, Jean-Michel Pawlotsky |
| EPI_ISL_3393400 | SELAS BC-LAB | Department of Virology, Henri Mondor University Hospital, Assistance Publique Hôpitaux de Paris, Université Paris-Est Créteil, INSERM U955 | Christophe Rodriguez, Slim Fourati, Vanessa Demontant, Guillaume Gricourt, Melissa N'Debi, Alexandre Soulier, Elisabeth Trawinski, Jean-Michel Pawlotsky |
| EPI_ISL_3393401 | BIOMNIS EUROFINS IVRY | Department of Virology, Henri Mondor University Hospital, Assistance Publique Hôpitaux de Paris, Université Paris-Est Créteil, INSERM U955 | Christophe Rodriguez, Slim Fourati, Vanessa Demontant, Guillaume Gricourt, Melissa N'Debi, Alexandre Soulier, Elisabeth Trawinski, Jean-Michel Pawlotsky |
| EPI_ISL_3393402 | BROUSSAIS | Department of Virology, Henri Mondor University Hospital, Assistance Publique Hôpitaux de Paris, Université Paris-Est Créteil, INSERM U955 | Christophe Rodriguez, Slim Fourati, Vanessa Demontant, Guillaume Gricourt, Melissa N'Debi, Alexandre Soulier, Elisabeth Trawinski, Jean-Michel Pawlotsky |
| EPI_ISL_3393404 | laboratoire Belle Epine | Department of Virology, Henri Mondor University Hospital, Assistance Publique Hôpitaux de Paris, Université Paris-Est Créteil, INSERM U955 | Christophe Rodriguez, Slim Fourati, Vanessa Demontant, Guillaume Gricourt, Melissa N'Debi, Alexandre Soulier, Elisabeth Trawinski, Jean-Michel Pawlotsky |
| EPI_ISL_3393405 | BROUSSAIS | Department of Virology, Henri Mondor University Hospital, Assistance Publique Hôpitaux de Paris, Université Paris-Est Créteil, INSERM U955 | Christophe Rodriguez, Slim Fourati, Vanessa Demontant, Guillaume Gricourt, Melissa N'Debi, Alexandre Soulier, Elisabeth Trawinski, Jean-Michel Pawlotsky |
| EPI_ISL_3393406 | SELARL BIO | Department of Virology, Henri Mondor University Hospital, Assistance Publique Hôpitaux de Paris, Université Paris-Est Créteil, INSERM U955 | Christophe Rodriguez, Slim Fourati, Vanessa Demontant, Guillaume Gricourt, Melissa N'Debi, Alexandre Soulier, Elisabeth Trawinski, Jean-Michel Pawlotsky |
| EPI_ISL_3393407 | BIO ARD'AISNE | Department of Virology, Henri Mondor University Hospital, Assistance Publique Hôpitaux de Paris, Université Paris-Est Créteil, INSERM U955 | Christophe Rodriguez, Slim Fourati, Vanessa Demontant, Guillaume Gricourt, Melissa N'Debi, Alexandre Soulier, Elisabeth Trawinski, Jean-Michel Pawlotsky |
| EPI_ISL_3393408 | BIOCEANE | Department of Virology, Henri Mondor University Hospital, Assistance Publique Hôpitaux de Paris, Université Paris-Est Créteil, INSERM U955 | Christophe Rodriguez, Slim Fourati, Vanessa Demontant, Guillaume Gricourt, Melissa N'Debi, Alexandre Soulier, Elisabeth Trawinski, Jean-Michel Pawlotsky |
| EPI_ISL_3393410 | BIOMNIS EUROFINS IVRY | Department of Virology, Henri Mondor University Hospital, Assistance Publique Hôpitaux de Paris, Université Paris-Est Créteil, INSERM U955 | Christophe Rodriguez, Slim Fourati, Vanessa Demontant, Guillaume Gricourt, Melissa N'Debi, Alexandre Soulier, Elisabeth Trawinski, Jean-Michel Pawlotsky |
| EPI_ISL_3393411 | CH. CLEMENCEAU | Department of Virology, Henri Mondor University Hospital, Assistance Publique Hôpitaux de Paris, Université Paris-Est Créteil, INSERM U955 | Christophe Rodriguez, Slim Fourati, Vanessa Demontant, Guillaume Gricourt, Melissa N'Debi, Alexandre Soulier, Elisabeth Trawinski, Jean-Michel Pawlotsky |
| EPI_ISL_3393412 | BROUSSAIS | Department of Virology, Henri Mondor University Hospital, Assistance Publique Hôpitaux de Paris, Université Paris-Est Créteil, INSERM U955 | Christophe Rodriguez, Slim Fourati, Vanessa Demontant, Guillaume Gricourt, Melissa N'Debi, Alexandre Soulier, Elisabeth Trawinski, Jean-Michel Pawlotsky |
| EPI_ISL_3393413 | BIOMNIS EUROFINS IVRY | Department of Virology, Henri Mondor University Hospital, Assistance Publique Hôpitaux de Paris, Université Paris-Est Créteil, INSERM U955 | Christophe Rodriguez, Slim Fourati, Vanessa Demontant, Guillaume Gricourt, Melissa N'Debi, Alexandre Soulier, Elisabeth Trawinski, Jean-Michel Pawlotsky |
| EPI_ISL_3393414 | BIO67-BIOSPHERE | Department of Virology, Henri Mondor University Hospital, Assistance Publique Hôpitaux de Paris, Université Paris-Est Créteil, INSERM U955 | Christophe Rodriguez, Slim Fourati, Vanessa Demontant, Guillaume Gricourt, Melissa N'Debi, Alexandre Soulier, Elisabeth Trawinski, Jean-Michel Pawlotsky |
| EPI_ISL_3393415 | SELAS MEDILYS | Department of Virology, Henri Mondor University Hospital, Assistance Publique Hôpitaux de Paris, Université Paris-Est Créteil, INSERM U955 | Christophe Rodriguez, Slim Fourati, Vanessa Demontant, Guillaume Gricourt, Melissa N'Debi, Alexandre Soulier, Elisabeth Trawinski, Jean-Michel Pawlotsky |
| EPI_ISL_3393416 | BROUSSAIS | Department of Virology, Henri Mondor University Hospital, Assistance Publique Hôpitaux de Paris, Université Paris-Est Créteil, INSERM U955 | Christophe Rodriguez, Slim Fourati, Vanessa Demontant, Guillaume Gricourt, Melissa N'Debi, Alexandre Soulier, Elisabeth Trawinski, Jean-Michel Pawlotsky |
| EPI_ISL_3393417 | SELARL BIO | Department of Virology, Henri Mondor University Hospital, Assistance Publique Hôpitaux de Paris, Université Paris-Est Créteil, INSERM U955 | Christophe Rodriguez, Slim Fourati, Vanessa Demontant, Guillaume Gricourt, Melissa N'Debi, Alexandre Soulier, Elisabeth Trawinski, Jean-Michel Pawlotsky |
| EPI_ISL_3393419 | laboratoire Belle Epine | Department of Virology, Henri Mondor University Hospital, Assistance Publique Hôpitaux de Paris, Université Paris-Est Créteil, INSERM U955 | Christophe Rodriguez, Slim Fourati, Vanessa Demontant, Guillaume Gricourt, Melissa N'Debi, Alexandre Soulier, Elisabeth Trawinski, Jean-Michel Pawlotsky |
| EPI_ISL_3393420 | SELAS BC-LAB | Department of Virology, Henri Mondor University Hospital, Assistance Publique Hôpitaux de Paris, Université Paris-Est Créteil, INSERM U955 | Christophe Rodriguez, Slim Fourati, Vanessa Demontant, Guillaume Gricourt, Melissa N'Debi, Alexandre Soulier, Elisabeth Trawinski, Jean-Michel Pawlotsky |
| EPI_ISL_3393422, EPI_ISL_3393423 | BROUSSAIS | Department of Virology, Henri Mondor University Hospital, Assistance Publique Hôpitaux de Paris, Université Paris-Est Créteil, INSERM U955 | Christophe Rodriguez, Slim Fourati, Vanessa Demontant, Guillaume Gricourt, Melissa N'Debi, Alexandre Soulier, Elisabeth Trawinski, Jean-Michel Pawlotsky |
| EPI_ISL_3393424 | CH. BEGIN | Department of Virology, Henri Mondor University Hospital, Assistance Publique Hôpitaux de Paris, Université Paris-Est Créteil, INSERM U955 | Christophe Rodriguez, Slim Fourati, Vanessa Demontant, Guillaume Gricourt, Melissa N'Debi, Alexandre Soulier, Elisabeth Trawinski, Jean-Michel Pawlotsky |
| EPI_ISL_3393425 | laboratoire Belle Epine | Department of Virology, Henri Mondor University Hospital, Assistance Publique Hôpitaux de Paris, Université Paris-Est Créteil, INSERM U955 | Christophe Rodriguez, Slim Fourati, Vanessa Demontant, Guillaume Gricourt, Melissa N'Debi, Alexandre Soulier, Elisabeth Trawinski, Jean-Michel Pawlotsky |
| EPI_ISL_3393427 | Biogroup Bio Lam-LCD Saint-Denis | Department of Virology, Henri Mondor University Hospital, Assistance Publique Hôpitaux de Paris, Université Paris-Est Créteil, INSERM U955 | Christophe Rodriguez, Slim Fourati, Vanessa Demontant, Guillaume Gricourt, Melissa N'Debi, Alexandre Soulier, Elisabeth Trawinski, Jean-Michel Pawlotsky |
| EPI_ISL_3393428, EPI_ISL_3393429 | laboratoire Belle Epine | Department of Virology, Henri Mondor University Hospital, Assistance Publique Hôpitaux de Paris, Université Paris-Est Créteil, | Christophe Rodriguez, Slim Fourati, Vanessa Demontant, Guillaume Gricourt, Melissa N'Debi, Alexandre Soulier, Elisabeth Trawinski, Jean-Michel Pawlotsky |

| | | | |
|---|---|---|---|
| | | INSERM U955 | |
| EPI_ISL_3393430 | CH. CLEMENCEAU | Department of Virology, Henri Mondor University Hospital, Assistance Publique Hôpitaux de Paris, Université Paris-Est Créteil, INSERM U955 | Christophe Rodriguez, Slim Fourati, Vanessa Demontant, Guillaume Gricourt, Melissa N'Debi, Alexandre Soulier, Elisabeth Trawinski, Jean-Michel Pawlotsky |
| EPI_ISL_3393431 | BROUSSAIS | Department of Virology, Henri Mondor University Hospital, Assistance Publique Hôpitaux de Paris, Université Paris-Est Créteil, INSERM U955 | Christophe Rodriguez, Slim Fourati, Vanessa Demontant, Guillaume Gricourt, Melissa N'Debi, Alexandre Soulier, Elisabeth Trawinski, Jean-Michel Pawlotsky |
| EPI_ISL_3393433 | BIO67-BIOSPHERE | Department of Virology, Henri Mondor University Hospital, Assistance Publique Hôpitaux de Paris, Université Paris-Est Créteil, INSERM U955 | Christophe Rodriguez, Slim Fourati, Vanessa Demontant, Guillaume Gricourt, Melissa N'Debi, Alexandre Soulier, Elisabeth Trawinski, Jean-Michel Pawlotsky |
| EPI_ISL_3393434 | Cerballiance Paris IDF est Saint-Denis | Department of Virology, Henri Mondor University Hospital, Assistance Publique Hôpitaux de Paris, Université Paris-Est Créteil, INSERM U955 | Christophe Rodriguez, Slim Fourati, Vanessa Demontant, Guillaume Gricourt, Melissa N'Debi, Alexandre Soulier, Elisabeth Trawinski, Jean-Michel Pawlotsky |
| EPI_ISL_3393435 | BROUSSAIS | Department of Virology, Henri Mondor University Hospital, Assistance Publique Hôpitaux de Paris, Université Paris-Est Créteil, INSERM U955 | Christophe Rodriguez, Slim Fourati, Vanessa Demontant, Guillaume Gricourt, Melissa N'Debi, Alexandre Soulier, Elisabeth Trawinski, Jean-Michel Pawlotsky |
| EPI_ISL_3393436 | LISIEUX Cerballiance | Department of Virology, Henri Mondor University Hospital, Assistance Publique Hôpitaux de Paris, Université Paris-Est Créteil, INSERM U955 | Christophe Rodriguez, Slim Fourati, Vanessa Demontant, Guillaume Gricourt, Melissa N'Debi, Alexandre Soulier, Elisabeth Trawinski, Jean-Michel Pawlotsky |
| EPI_ISL_3393437 | GH A.CHENEVIER-H.MONDOR | Department of Virology, Henri Mondor University Hospital, Assistance Publique Hôpitaux de Paris, Université Paris-Est Créteil, INSERM U955 | Christophe Rodriguez, Slim Fourati, Vanessa Demontant, Guillaume Gricourt, Melissa N'Debi, Alexandre Soulier, Elisabeth Trawinski, Jean-Michel Pawlotsky |
| EPI_ISL_3393438 | Biogroup Bio Lam-LCD Saint-Denis | Department of Virology, Henri Mondor University Hospital, Assistance Publique Hôpitaux de Paris, Université Paris-Est Créteil, INSERM U955 | Christophe Rodriguez, Slim Fourati, Vanessa Demontant, Guillaume Gricourt, Melissa N'Debi, Alexandre Soulier, Elisabeth Trawinski, Jean-Michel Pawlotsky |
| EPI_ISL_3393439 | Hôpital Avicenne | Department of Virology, Henri Mondor University Hospital, Assistance Publique Hôpitaux de Paris, Université Paris-Est Créteil, INSERM U955 | Christophe Rodriguez, Slim Fourati, Vanessa Demontant, Guillaume Gricourt, Melissa N'Debi, Alexandre Soulier, Elisabeth Trawinski, Jean-Michel Pawlotsky |
| EPI_ISL_3393440 | Biogroup Bio Lam-LCD Saint-Denis | Department of Virology, Henri Mondor University Hospital, Assistance Publique Hôpitaux de Paris, Université Paris-Est Créteil, INSERM U955 | Christophe Rodriguez, Slim Fourati, Vanessa Demontant, Guillaume Gricourt, Melissa N'Debi, Alexandre Soulier, Elisabeth Trawinski, Jean-Michel Pawlotsky |
| EPI_ISL_3393442 | CHU Rouen Normandie | Department of Virology, Henri Mondor University Hospital, Assistance Publique Hôpitaux de Paris, Université Paris-Est Créteil, INSERM U955 | Christophe Rodriguez, Slim Fourati, Vanessa Demontant, Guillaume Gricourt, Melissa N'Debi, Alexandre Soulier, Elisabeth Trawinski, Jean-Michel Pawlotsky |
| EPI_ISL_3393443 | Hôpital Bichat-Claude Bernard | Department of Virology, Henri Mondor University Hospital, Assistance Publique Hôpitaux de Paris, Université Paris-Est Créteil, INSERM U955 | Christophe Rodriguez, Slim Fourati, Vanessa Demontant, Guillaume Gricourt, Melissa N'Debi, Alexandre Soulier, Elisabeth Trawinski, Jean-Michel Pawlotsky |
| EPI_ISL_3393445 | BIO67-BIOSPHERE | Department of Virology, Henri Mondor University Hospital, Assistance Publique Hôpitaux de Paris, Université Paris-Est Créteil, INSERM U955 | Christophe Rodriguez, Slim Fourati, Vanessa Demontant, Guillaume Gricourt, Melissa N'Debi, Alexandre Soulier, Elisabeth Trawinski, Jean-Michel Pawlotsky |
| EPI_ISL_3393446 | CERBALLIANCE Valmy | Department of Virology, Henri Mondor University Hospital, Assistance Publique Hôpitaux de Paris, Université Paris-Est Créteil, INSERM U955 | Christophe Rodriguez, Slim Fourati, Vanessa Demontant, Guillaume Gricourt, Melissa N'Debi, Alexandre Soulier, Elisabeth Trawinski, Jean-Michel Pawlotsky |
| EPI_ISL_3393451 | laboratoire Belle Epine | Department of Virology, Henri Mondor University Hospital, Assistance Publique Hôpitaux de Paris, Université Paris-Est Créteil, INSERM U955 | Christophe Rodriguez, Slim Fourati, Vanessa Demontant, Guillaume Gricourt, Melissa N'Debi, Alexandre Soulier, Elisabeth Trawinski, Jean-Michel Pawlotsky |
| EPI_ISL_3393452 | BIO67-BIOSPHERE | Department of Virology, Henri Mondor University Hospital, Assistance Publique Hôpitaux de Paris, Université Paris-Est Créteil, INSERM U955 | Christophe Rodriguez, Slim Fourati, Vanessa Demontant, Guillaume Gricourt, Melissa N'Debi, Alexandre Soulier, Elisabeth Trawinski, Jean-Michel Pawlotsky |
| EPI_ISL_3393453 | BROUSSAIS | Department of Virology, Henri Mondor University Hospital, Assistance Publique Hôpitaux de Paris, Université Paris-Est Créteil, INSERM U955 | Christophe Rodriguez, Slim Fourati, Vanessa Demontant, Guillaume Gricourt, Melissa N'Debi, Alexandre Soulier, Elisabeth Trawinski, Jean-Michel Pawlotsky |
| EPI_ISL_3393454, EPI_ISL_3393455 | SELAS MEDILYS | Department of Virology, Henri Mondor University Hospital, Assistance Publique Hôpitaux de Paris, Université Paris-Est Créteil, INSERM U955 | Christophe Rodriguez, Slim Fourati, Vanessa Demontant, Guillaume Gricourt, Melissa N'Debi, Alexandre Soulier, Elisabeth Trawinski, Jean-Michel Pawlotsky |
| EPI_ISL_3393456 | BIOLBS Rouen | Department of Virology, Henri Mondor University Hospital, Assistance Publique Hôpitaux de Paris, Université Paris-Est Créteil, INSERM U955 | Christophe Rodriguez, Slim Fourati, Vanessa Demontant, Guillaume Gricourt, Melissa N'Debi, Alexandre Soulier, Elisabeth Trawinski, Jean-Michel Pawlotsky |
| EPI_ISL_3393458 | BIO67-BIOSPHERE | Department of Virology, Henri Mondor University Hospital, Assistance Publique Hôpitaux de Paris, Université Paris-Est Créteil, INSERM U955 | Christophe Rodriguez, Slim Fourati, Vanessa Demontant, Guillaume Gricourt, Melissa N'Debi, Alexandre Soulier, Elisabeth Trawinski, Jean-Michel Pawlotsky |
| EPI_ISL_3393460 | LISIEUX Cerballiance | Department of Virology, Henri Mondor University Hospital, Assistance Publique Hôpitaux de Paris, Université Paris-Est Créteil, INSERM U955 | Christophe Rodriguez, Slim Fourati, Vanessa Demontant, Guillaume Gricourt, Melissa N'Debi, Alexandre Soulier, Elisabeth Trawinski, Jean-Michel Pawlotsky |
| EPI_ISL_3393461 | laboratoire Belle Epine | Department of Virology, Henri Mondor University Hospital, Assistance Publique Hôpitaux de Paris, Université Paris-Est Créteil, INSERM U955 | Christophe Rodriguez, Slim Fourati, Vanessa Demontant, Guillaume Gricourt, Melissa N'Debi, Alexandre Soulier, Elisabeth Trawinski, Jean-Michel Pawlotsky |
| EPI_ISL_3393463 | BIO67-BIOSPHERE | Department of Virology, Henri Mondor University Hospital, Assistance Publique Hôpitaux de Paris, Université Paris-Est Créteil, INSERM U955 | Christophe Rodriguez, Slim Fourati, Vanessa Demontant, Guillaume Gricourt, Melissa N'Debi, Alexandre Soulier, Elisabeth Trawinski, Jean-Michel Pawlotsky |
| EPI_ISL_3393464 | SELAS MEDILYS | Department of Virology, Henri Mondor University Hospital, Assistance Publique Hôpitaux de Paris, Université Paris-Est Créteil, INSERM U955 | Christophe Rodriguez, Slim Fourati, Vanessa Demontant, Guillaume Gricourt, Melissa N'Debi, Alexandre Soulier, Elisabeth Trawinski, Jean-Michel Pawlotsky |
| EPI_ISL_3393465 | BIOCEANE | Department of Virology, Henri Mondor University Hospital, Assistance Publique Hôpitaux de Paris, Université Paris-Est Créteil, INSERM U955 | Christophe Rodriguez, Slim Fourati, Vanessa Demontant, Guillaume Gricourt, Melissa N'Debi, Alexandre Soulier, Elisabeth Trawinski, Jean-Michel Pawlotsky |
| EPI_ISL_3393466 | laboratoire Belle Epine | Department of Virology, Henri Mondor University Hospital, Assistance Publique Hôpitaux de Paris, Université Paris-Est Créteil, INSERM U955 | Christophe Rodriguez, Slim Fourati, Vanessa Demontant, Guillaume Gricourt, Melissa N'Debi, Alexandre Soulier, Elisabeth Trawinski, Jean-Michel Pawlotsky |
| EPI_ISL_3393467, EPI_ISL_3393468 | BROUSSAIS | Department of Virology, Henri Mondor University Hospital, Assistance Publique Hôpitaux de Paris, Université Paris-Est Créteil, INSERM U955 | Christophe Rodriguez, Slim Fourati, Vanessa Demontant, Guillaume Gricourt, Melissa N'Debi, Alexandre Soulier, Elisabeth Trawinski, Jean-Michel Pawlotsky |
| EPI_ISL_3393469 | BIO67-BIOSPHERE | Department of Virology, Henri Mondor University Hospital, Assistance Publique Hôpitaux de Paris, Université Paris-Est Créteil, INSERM U955 | Christophe Rodriguez, Slim Fourati, Vanessa Demontant, Guillaume Gricourt, Melissa N'Debi, Alexandre Soulier, Elisabeth Trawinski, Jean-Michel Pawlotsky |
| EPI_ISL_3393470 | CH. BEGIN | Department of Virology, Henri Mondor University Hospital, Assistance Publique Hôpitaux de Paris, Université Paris-Est Créteil, INSERM U955 | Christophe Rodriguez, Slim Fourati, Vanessa Demontant, Guillaume Gricourt, Melissa N'Debi, Alexandre Soulier, Elisabeth Trawinski, Jean-Michel Pawlotsky |
| EPI_ISL_3393472, EPI_ISL_3393473 | Cerballiance Paris IDF est Saint-Denis | Department of Virology, Henri Mondor University Hospital, Assistance Publique Hôpitaux de Paris, Université Paris-Est Créteil, INSERM U955 | Christophe Rodriguez, Slim Fourati, Vanessa Demontant, Guillaume Gricourt, Melissa N'Debi, Alexandre Soulier, Elisabeth Trawinski, Jean-Michel Pawlotsky |
| EPI_ISL_3393474 | SELAS MEDILYS | Department of Virology, Henri Mondor University Hospital, Assistance Publique Hôpitaux de Paris, Université Paris-Est Créteil, INSERM U955 | Christophe Rodriguez, Slim Fourati, Vanessa Demontant, Guillaume Gricourt, Melissa N'Debi, Alexandre Soulier, Elisabeth Trawinski, Jean-Michel Pawlotsky |
| EPI_ISL_3393475 | laboratoire Belle Epine | Department of Virology, Henri Mondor University Hospital, Assistance Publique Hôpitaux de Paris, Université Paris-Est Créteil, | Christophe Rodriguez, Slim Fourati, Vanessa Demontant, Guillaume Gricourt, Melissa N'Debi, Alexandre Soulier, Elisabeth Trawinski, Jean-Michel Pawlotsky |

| | | | |
|---|---|---|---|
| | | INSERM U955 | |
| EPI_ISL_3393476, EPI_ISL_3393477 | BROUSSAIS | Department of Virology, Henri Mondor University Hospital, Assistance Publique Hôpitaux de Paris, Université Paris-Est Créteil, INSERM U955 | Christophe Rodriguez, Slim Fourati, Vanessa Demontant, Guillaume Gricourt, Melissa N'Debi, Alexandre Soulier, Elisabeth Trawinski, Jean-Michel Pawlotsky |
| EPI_ISL_3393478 | CH. CLEMENCEAU | Department of Virology, Henri Mondor University Hospital, Assistance Publique Hôpitaux de Paris, Université Paris-Est Créteil, INSERM U955 | Christophe Rodriguez, Slim Fourati, Vanessa Demontant, Guillaume Gricourt, Melissa N'Debi, Alexandre Soulier, Elisabeth Trawinski, Jean-Michel Pawlotsky |
| EPI_ISL_3393479 | SELAS MEDILYS | Department of Virology, Henri Mondor University Hospital, Assistance Publique Hôpitaux de Paris, Université Paris-Est Créteil, INSERM U955 | Christophe Rodriguez, Slim Fourati, Vanessa Demontant, Guillaume Gricourt, Melissa N'Debi, Alexandre Soulier, Elisabeth Trawinski, Jean-Michel Pawlotsky |
| EPI_ISL_3393480 | BIO67-BIOSPHERE | Department of Virology, Henri Mondor University Hospital, Assistance Publique Hôpitaux de Paris, Université Paris-Est Créteil, INSERM U955 | Christophe Rodriguez, Slim Fourati, Vanessa Demontant, Guillaume Gricourt, Melissa N'Debi, Alexandre Soulier, Elisabeth Trawinski, Jean-Michel Pawlotsky |
| EPI_ISL_3393481 | BROUSSAIS | Department of Virology, Henri Mondor University Hospital, Assistance Publique Hôpitaux de Paris, Université Paris-Est Créteil, INSERM U955 | Christophe Rodriguez, Slim Fourati, Vanessa Demontant, Guillaume Gricourt, Melissa N'Debi, Alexandre Soulier, Elisabeth Trawinski, Jean-Michel Pawlotsky |
| EPI_ISL_3393482 | Biogroup Bio Lam-LCD Saint-Denis | Department of Virology, Henri Mondor University Hospital, Assistance Publique Hôpitaux de Paris, Université Paris-Est Créteil, INSERM U955 | Christophe Rodriguez, Slim Fourati, Vanessa Demontant, Guillaume Gricourt, Melissa N'Debi, Alexandre Soulier, Elisabeth Trawinski, Jean-Michel Pawlotsky |
| EPI_ISL_3393483 | BROUSSAIS | Department of Virology, Henri Mondor University Hospital, Assistance Publique Hôpitaux de Paris, Université Paris-Est Créteil, INSERM U955 | Christophe Rodriguez, Slim Fourati, Vanessa Demontant, Guillaume Gricourt, Melissa N'Debi, Alexandre Soulier, Elisabeth Trawinski, Jean-Michel Pawlotsky |
| EPI_ISL_3393486 | SELAS MEDILYS | Department of Virology, Henri Mondor University Hospital, Assistance Publique Hôpitaux de Paris, Université Paris-Est Créteil, INSERM U955 | Christophe Rodriguez, Slim Fourati, Vanessa Demontant, Guillaume Gricourt, Melissa N'Debi, Alexandre Soulier, Elisabeth Trawinski, Jean-Michel Pawlotsky |
| EPI_ISL_3393487 | Biogroup Bio Lam-LCD Saint-Denis | Department of Virology, Henri Mondor University Hospital, Assistance Publique Hôpitaux de Paris, Université Paris-Est Créteil, INSERM U955 | Christophe Rodriguez, Slim Fourati, Vanessa Demontant, Guillaume Gricourt, Melissa N'Debi, Alexandre Soulier, Elisabeth Trawinski, Jean-Michel Pawlotsky |
| EPI_ISL_3393488 | Cerballiance Paris IDF est Saint-Denis | Department of Virology, Henri Mondor University Hospital, Assistance Publique Hôpitaux de Paris, Université Paris-Est Créteil, INSERM U955 | Christophe Rodriguez, Slim Fourati, Vanessa Demontant, Guillaume Gricourt, Melissa N'Debi, Alexandre Soulier, Elisabeth Trawinski, Jean-Michel Pawlotsky |
| EPI_ISL_3393489 | Biogroup Bio Lam-LCD Saint-Denis | Department of Virology, Henri Mondor University Hospital, Assistance Publique Hôpitaux de Paris, Université Paris-Est Créteil, INSERM U955 | Christophe Rodriguez, Slim Fourati, Vanessa Demontant, Guillaume Gricourt, Melissa N'Debi, Alexandre Soulier, Elisabeth Trawinski, Jean-Michel Pawlotsky |
| EPI_ISL_3393490 | LISIEUX Cerballiance | Department of Virology, Henri Mondor University Hospital, Assistance Publique Hôpitaux de Paris, Université Paris-Est Créteil, INSERM U955 | Christophe Rodriguez, Slim Fourati, Vanessa Demontant, Guillaume Gricourt, Melissa N'Debi, Alexandre Soulier, Elisabeth Trawinski, Jean-Michel Pawlotsky |
| EPI_ISL_3393492 | BROUSSAIS | Department of Virology, Henri Mondor University Hospital, Assistance Publique Hôpitaux de Paris, Université Paris-Est Créteil, INSERM U955 | Christophe Rodriguez, Slim Fourati, Vanessa Demontant, Guillaume Gricourt, Melissa N'Debi, Alexandre Soulier, Elisabeth Trawinski, Jean-Michel Pawlotsky |
| EPI_ISL_3393493 | Hôpital Avicenne | Department of Virology, Henri Mondor University Hospital, Assistance Publique Hôpitaux de Paris, Université Paris-Est Créteil, INSERM U955 | Christophe Rodriguez, Slim Fourati, Vanessa Demontant, Guillaume Gricourt, Melissa N'Debi, Alexandre Soulier, Elisabeth Trawinski, Jean-Michel Pawlotsky |
| EPI_ISL_3393494 | laboratoire Belle Epine | Department of Virology, Henri Mondor University Hospital, Assistance Publique Hôpitaux de Paris, Université Paris-Est Créteil, INSERM U955 | Christophe Rodriguez, Slim Fourati, Vanessa Demontant, Guillaume Gricourt, Melissa N'Debi, Alexandre Soulier, Elisabeth Trawinski, Jean-Michel Pawlotsky |
| EPI_ISL_3393495 | BIO67-BIOSPHERE | Department of Virology, Henri Mondor University Hospital, Assistance Publique Hôpitaux de Paris, Université Paris-Est Créteil, INSERM U955 | Christophe Rodriguez, Slim Fourati, Vanessa Demontant, Guillaume Gricourt, Melissa N'Debi, Alexandre Soulier, Elisabeth Trawinski, Jean-Michel Pawlotsky |
| EPI_ISL_3393496 | laboratoire Belle Epine | Department of Virology, Henri Mondor University Hospital, Assistance Publique Hôpitaux de Paris, Université Paris-Est Créteil, INSERM U955 | Christophe Rodriguez, Slim Fourati, Vanessa Demontant, Guillaume Gricourt, Melissa N'Debi, Alexandre Soulier, Elisabeth Trawinski, Jean-Michel Pawlotsky |
| EPI_ISL_3393497 | BIO67-BIOSPHERE | Department of Virology, Henri Mondor University Hospital, Assistance Publique Hôpitaux de Paris, Université Paris-Est Créteil, INSERM U955 | Christophe Rodriguez, Slim Fourati, Vanessa Demontant, Guillaume Gricourt, Melissa N'Debi, Alexandre Soulier, Elisabeth Trawinski, Jean-Michel Pawlotsky |
| EPI_ISL_3393498 | BROUSSAIS | Department of Virology, Henri Mondor University Hospital, Assistance Publique Hôpitaux de Paris, Université Paris-Est Créteil, INSERM U955 | Christophe Rodriguez, Slim Fourati, Vanessa Demontant, Guillaume Gricourt, Melissa N'Debi, Alexandre Soulier, Elisabeth Trawinski, Jean-Michel Pawlotsky |
| EPI_ISL_3393499 | BIO67-BIOSPHERE | Department of Virology, Henri Mondor University Hospital, Assistance Publique Hôpitaux de Paris, Université Paris-Est Créteil, INSERM U955 | Christophe Rodriguez, Slim Fourati, Vanessa Demontant, Guillaume Gricourt, Melissa N'Debi, Alexandre Soulier, Elisabeth Trawinski, Jean-Michel Pawlotsky |
| EPI_ISL_3393500 | SELARL BIO | Department of Virology, Henri Mondor University Hospital, Assistance Publique Hôpitaux de Paris, Université Paris-Est Créteil, INSERM U955 | Christophe Rodriguez, Slim Fourati, Vanessa Demontant, Guillaume Gricourt, Melissa N'Debi, Alexandre Soulier, Elisabeth Trawinski, Jean-Michel Pawlotsky |
| EPI_ISL_3393501 | BIO ARD'AISNE | Department of Virology, Henri Mondor University Hospital, Assistance Publique Hôpitaux de Paris, Université Paris-Est Créteil, INSERM U955 | Christophe Rodriguez, Slim Fourati, Vanessa Demontant, Guillaume Gricourt, Melissa N'Debi, Alexandre Soulier, Elisabeth Trawinski, Jean-Michel Pawlotsky |
| EPI_ISL_3393502 | BIO67-BIOSPHERE | Department of Virology, Henri Mondor University Hospital, Assistance Publique Hôpitaux de Paris, Université Paris-Est Créteil, INSERM U955 | Christophe Rodriguez, Slim Fourati, Vanessa Demontant, Guillaume Gricourt, Melissa N'Debi, Alexandre Soulier, Elisabeth Trawinski, Jean-Michel Pawlotsky |
| EPI_ISL_3393503 | laboratoire Belle Epine | Department of Virology, Henri Mondor University Hospital, Assistance Publique Hôpitaux de Paris, Université Paris-Est Créteil, INSERM U955 | Christophe Rodriguez, Slim Fourati, Vanessa Demontant, Guillaume Gricourt, Melissa N'Debi, Alexandre Soulier, Elisabeth Trawinski, Jean-Michel Pawlotsky |
| EPI_ISL_3393504 | CH de Saint Dié des vosges | Department of Virology, Henri Mondor University Hospital, Assistance Publique Hôpitaux de Paris, Université Paris-Est Créteil, INSERM U955 | Christophe Rodriguez, Slim Fourati, Vanessa Demontant, Guillaume Gricourt, Melissa N'Debi, Alexandre Soulier, Elisabeth Trawinski, Jean-Michel Pawlotsky |
| EPI_ISL_3393505 | SELAS MEDILYS | Department of Virology, Henri Mondor University Hospital, Assistance Publique Hôpitaux de Paris, Université Paris-Est Créteil, INSERM U955 | Christophe Rodriguez, Slim Fourati, Vanessa Demontant, Guillaume Gricourt, Melissa N'Debi, Alexandre Soulier, Elisabeth Trawinski, Jean-Michel Pawlotsky |
| EPI_ISL_3393507 | SELAS BC-LAB | Department of Virology, Henri Mondor University Hospital, Assistance Publique Hôpitaux de Paris, Université Paris-Est Créteil, INSERM U955 | Christophe Rodriguez, Slim Fourati, Vanessa Demontant, Guillaume Gricourt, Melissa N'Debi, Alexandre Soulier, Elisabeth Trawinski, Jean-Michel Pawlotsky |
| EPI_ISL_3393508 | DYNABIO UNILABS | Department of Virology, Henri Mondor University Hospital, Assistance Publique Hôpitaux de Paris, Université Paris-Est Créteil, INSERM U955 | Christophe Rodriguez, Slim Fourati, Vanessa Demontant, Guillaume Gricourt, Melissa N'Debi, Alexandre Soulier, Elisabeth Trawinski, Jean-Michel Pawlotsky |
| EPI_ISL_3393510 | Biogroup Bio Lam-LCD Saint-Denis | Department of Virology, Henri Mondor University Hospital, Assistance Publique Hôpitaux de Paris, Université Paris-Est Créteil, INSERM U955 | Christophe Rodriguez, Slim Fourati, Vanessa Demontant, Guillaume Gricourt, Melissa N'Debi, Alexandre Soulier, Elisabeth Trawinski, Jean-Michel Pawlotsky |
| EPI_ISL_3393511 | SELARL BIO | Department of Virology, Henri Mondor University Hospital, Assistance Publique Hôpitaux de Paris, Université Paris-Est Créteil, INSERM U955 | Christophe Rodriguez, Slim Fourati, Vanessa Demontant, Guillaume Gricourt, Melissa N'Debi, Alexandre Soulier, Elisabeth Trawinski, Jean-Michel Pawlotsky |
| EPI_ISL_3393512 | SELAS MEDILYS | Department of Virology, Henri Mondor University Hospital, Assistance Publique Hôpitaux de Paris, Université Paris-Est Créteil, INSERM U955 | Christophe Rodriguez, Slim Fourati, Vanessa Demontant, Guillaume Gricourt, Melissa N'Debi, Alexandre Soulier, Elisabeth Trawinski, Jean-Michel Pawlotsky |
| EPI_ISL_3393513 | Biogroup Bio Lam-LCD Saint-Denis | Department of Virology, Henri Mondor University Hospital, Assistance Publique Hôpitaux de Paris, Université Paris-Est Créteil, | Christophe Rodriguez, Slim Fourati, Vanessa Demontant, Guillaume Gricourt, Melissa N'Debi, Alexandre Soulier, Elisabeth Trawinski, Jean-Michel Pawlotsky |

| | | | |
|---|---|---|---|
| | | INSERM U955 | |
| EPI_ISL_3393516, EPI_ISL_3393517 | SELAS MEDILYS | Department of Virology, Henri Mondor University Hospital, Assistance Publique Hôpitaux de Paris, Université Paris-Est Créteil, INSERM U955 | Christophe Rodriguez, Slim Fourati, Vanessa Demontant, Guillaume Gricourt, Melissa N'Debi, Alexandre Soulier, Elisabeth Trawinski, Jean-Michel Pawlotsky |
| EPI_ISL_3393518 | Hôpital Cochin | Department of Virology, Henri Mondor University Hospital, Assistance Publique Hôpitaux de Paris, Université Paris-Est Créteil, INSERM U955 | Christophe Rodriguez, Slim Fourati, Vanessa Demontant, Guillaume Gricourt, Melissa N'Debi, Alexandre Soulier, Elisabeth Trawinski, Jean-Michel Pawlotsky |
| EPI_ISL_3393519 | Biogroup Bio Lam-LCD Saint-Denis | Department of Virology, Henri Mondor University Hospital, Assistance Publique Hôpitaux de Paris, Université Paris-Est Créteil, INSERM U955 | Christophe Rodriguez, Slim Fourati, Vanessa Demontant, Guillaume Gricourt, Melissa N'Debi, Alexandre Soulier, Elisabeth Trawinski, Jean-Michel Pawlotsky |
| EPI_ISL_3393520 | SELAS MEDILYS | Department of Virology, Henri Mondor University Hospital, Assistance Publique Hôpitaux de Paris, Université Paris-Est Créteil, INSERM U955 | Christophe Rodriguez, Slim Fourati, Vanessa Demontant, Guillaume Gricourt, Melissa N'Debi, Alexandre Soulier, Elisabeth Trawinski, Jean-Michel Pawlotsky |
| EPI_ISL_3393521 | BIO67-BIOSPHERE | Department of Virology, Henri Mondor University Hospital, Assistance Publique Hôpitaux de Paris, Université Paris-Est Créteil, INSERM U955 | Christophe Rodriguez, Slim Fourati, Vanessa Demontant, Guillaume Gricourt, Melissa N'Debi, Alexandre Soulier, Elisabeth Trawinski, Jean-Michel Pawlotsky |
| EPI_ISL_3393522 | SELAS MEDILYS | Department of Virology, Henri Mondor University Hospital, Assistance Publique Hôpitaux de Paris, Université Paris-Est Créteil, INSERM U955 | Christophe Rodriguez, Slim Fourati, Vanessa Demontant, Guillaume Gricourt, Melissa N'Debi, Alexandre Soulier, Elisabeth Trawinski, Jean-Michel Pawlotsky |
| EPI_ISL_3393523 | CH de Saint Dié des vosges | Department of Virology, Henri Mondor University Hospital, Assistance Publique Hôpitaux de Paris, Université Paris-Est Créteil, INSERM U955 | Christophe Rodriguez, Slim Fourati, Vanessa Demontant, Guillaume Gricourt, Melissa N'Debi, Alexandre Soulier, Elisabeth Trawinski, Jean-Michel Pawlotsky |
| EPI_ISL_3393525 | BIO67-BIOSPHERE | Department of Virology, Henri Mondor University Hospital, Assistance Publique Hôpitaux de Paris, Université Paris-Est Créteil, INSERM U955 | Christophe Rodriguez, Slim Fourati, Vanessa Demontant, Guillaume Gricourt, Melissa N'Debi, Alexandre Soulier, Elisabeth Trawinski, Jean-Michel Pawlotsky |
| EPI_ISL_3393526, EPI_ISL_3393527 | GH A.CHENEVIER-H.MONDOR | Department of Virology, Henri Mondor University Hospital, Assistance Publique Hôpitaux de Paris, Université Paris-Est Créteil, INSERM U955 | Christophe Rodriguez, Slim Fourati, Vanessa Demontant, Guillaume Gricourt, Melissa N'Debi, Alexandre Soulier, Elisabeth Trawinski, Jean-Michel Pawlotsky |
| EPI_ISL_3393528 | Hôpital Avicenne | Department of Virology, Henri Mondor University Hospital, Assistance Publique Hôpitaux de Paris, Université Paris-Est Créteil, INSERM U955 | Christophe Rodriguez, Slim Fourati, Vanessa Demontant, Guillaume Gricourt, Melissa N'Debi, Alexandre Soulier, Elisabeth Trawinski, Jean-Michel Pawlotsky |
| EPI_ISL_3393529 | BIO67-BIOSPHERE | Department of Virology, Henri Mondor University Hospital, Assistance Publique Hôpitaux de Paris, Université Paris-Est Créteil, INSERM U955 | Christophe Rodriguez, Slim Fourati, Vanessa Demontant, Guillaume Gricourt, Melissa N'Debi, Alexandre Soulier, Elisabeth Trawinski, Jean-Michel Pawlotsky |
| EPI_ISL_3393530 | laboratoire Belle Epine | Department of Virology, Henri Mondor University Hospital, Assistance Publique Hôpitaux de Paris, Université Paris-Est Créteil, INSERM U955 | Christophe Rodriguez, Slim Fourati, Vanessa Demontant, Guillaume Gricourt, Melissa N'Debi, Alexandre Soulier, Elisabeth Trawinski, Jean-Michel Pawlotsky |
| EPI_ISL_3393531, EPI_ISL_3393532, EPI_ISL_3393533 | SELAS MEDILYS | Department of Virology, Henri Mondor University Hospital, Assistance Publique Hôpitaux de Paris, Université Paris-Est Créteil, INSERM U955 | Christophe Rodriguez, Slim Fourati, Vanessa Demontant, Guillaume Gricourt, Melissa N'Debi, Alexandre Soulier, Elisabeth Trawinski, Jean-Michel Pawlotsky |
| EPI_ISL_3393535, EPI_ISL_3393536 | BROUSSAIS | Department of Virology, Henri Mondor University Hospital, Assistance Publique Hôpitaux de Paris, Université Paris-Est Créteil, INSERM U955 | Christophe Rodriguez, Slim Fourati, Vanessa Demontant, Guillaume Gricourt, Melissa N'Debi, Alexandre Soulier, Elisabeth Trawinski, Jean-Michel Pawlotsky |
| EPI_ISL_3393537, EPI_ISL_3393538 | laboratoire Belle Epine | Department of Virology, Henri Mondor University Hospital, Assistance Publique Hôpitaux de Paris, Université Paris-Est Créteil, INSERM U955 | Christophe Rodriguez, Slim Fourati, Vanessa Demontant, Guillaume Gricourt, Melissa N'Debi, Alexandre Soulier, Elisabeth Trawinski, Jean-Michel Pawlotsky |
| EPI_ISL_3393539 | Hôpital Avicenne | Department of Virology, Henri Mondor University Hospital, Assistance Publique Hôpitaux de Paris, Université Paris-Est Créteil, INSERM U955 | Christophe Rodriguez, Slim Fourati, Vanessa Demontant, Guillaume Gricourt, Melissa N'Debi, Alexandre Soulier, Elisabeth Trawinski, Jean-Michel Pawlotsky |
| EPI_ISL_3393540 | Cerballiance Paris IDF est Saint-Denis | Department of Virology, Henri Mondor University Hospital, Assistance Publique Hôpitaux de Paris, Université Paris-Est Créteil, INSERM U955 | Christophe Rodriguez, Slim Fourati, Vanessa Demontant, Guillaume Gricourt, Melissa N'Debi, Alexandre Soulier, Elisabeth Trawinski, Jean-Michel Pawlotsky |
| EPI_ISL_3393541 | laboratoire Belle Epine | Department of Virology, Henri Mondor University Hospital, Assistance Publique Hôpitaux de Paris, Université Paris-Est Créteil, INSERM U955 | Christophe Rodriguez, Slim Fourati, Vanessa Demontant, Guillaume Gricourt, Melissa N'Debi, Alexandre Soulier, Elisabeth Trawinski, Jean-Michel Pawlotsky |
| EPI_ISL_3393542 | BIO67-BIOSPHERE | Department of Virology, Henri Mondor University Hospital, Assistance Publique Hôpitaux de Paris, Université Paris-Est Créteil, INSERM U955 | Christophe Rodriguez, Slim Fourati, Vanessa Demontant, Guillaume Gricourt, Melissa N'Debi, Alexandre Soulier, Elisabeth Trawinski, Jean-Michel Pawlotsky |
| EPI_ISL_3393543 | GH A.CHENEVIER-H.MONDOR | Department of Virology, Henri Mondor University Hospital, Assistance Publique Hôpitaux de Paris, Université Paris-Est Créteil, INSERM U955 | Christophe Rodriguez, Slim Fourati, Vanessa Demontant, Guillaume Gricourt, Melissa N'Debi, Alexandre Soulier, Elisabeth Trawinski, Jean-Michel Pawlotsky |
| EPI_ISL_3393544, EPI_ISL_3393545, EPI_ISL_3393546, EPI_ISL_3393547, EPI_ISL_3393548, EPI_ISL_3393549, EPI_ISL_3393550, EPI_ISL_3393551, EPI_ISL_3393552, EPI_ISL_3393553, EPI_ISL_3393554, EPI_ISL_3393555, EPI_ISL_3393556, EPI_ISL_3393557, EPI_ISL_3393558, EPI_ISL_3393559, EPI_ISL_3393560, EPI_ISL_3393561, EPI_ISL_3393562, EPI_ISL_3393563, EPI_ISL_3393564, EPI_ISL_3393565, EPI_ISL_3393566, EPI_ISL_3393567, EPI_ISL_3393568, EPI_ISL_3393569, EPI_ISL_3393570, EPI_ISL_3393571, EPI_ISL_3393572, EPI_ISL_3393573, EPI_ISL_3393574, EPI_ISL_3393575, EPI_ISL_3393576, EPI_ISL_3393577, EPI_ISL_3393578, EPI_ISL_3393579, EPI_ISL_3393580, EPI_ISL_3393581, EPI_ISL_3393582, EPI_ISL_3393583, EPI_ISL_3393584, EPI_ISL_3393585, EPI_ISL_3393586, EPI_ISL_3393587, EPI_ISL_3393588, EPI_ISL_3393589, EPI_ISL_3393590, EPI_ISL_3393591, EPI_ISL_3393592, EPI_ISL_3393593, EPI_ISL_3393594, EPI_ISL_3393595, EPI_ISL_3393596, EPI_ISL_3393597, EPI_ISL_3393598, EPI_ISL_3393599, EPI_ISL_3393600, EPI_ISL_3393601, EPI_ISL_3393602, EPI_ISL_3393603, EPI_ISL_3393604, EPI_ISL_3393605, EPI_ISL_3393606, EPI_ISL_3393607, EPI_ISL_3393608, EPI_ISL_3393609, EPI_ISL_3393610, EPI_ISL_3393611, EPI_ISL_3393612, EPI_ISL_3393613, EPI_ISL_3393614, EPI_ISL_3393615, EPI_ISL_3393616, EPI_ISL_3393617, EPI_ISL_3393618, EPI_ISL_3393619, EPI_ISL_3393620, EPI_ISL_3393621, EPI_ISL_3393622, EPI_ISL_3393623, EPI_ISL_3393624, EPI_ISL_3393625, EPI_ISL_3393626, EPI_ISL_3393627, EPI_ISL_3393628, EPI_ISL_3393629, EPI_ISL_3393630, EPI_ISL_3393631, EPI_ISL_3393632, EPI_ISL_3393633, EPI_ISL_3393634, EPI_ISL_3393635, EPI_ISL_3393636, EPI_ISL_3393637, EPI_ISL_3393638, EPI_ISL_3393639, EPI_ISL_3393640, EPI_ISL_3393641, EPI_ISL_3393642, EPI_ISL_3393643, EPI_ISL_3393644, EPI_ISL_3393645, EPI_ISL_3393646, EPI_ISL_3393647, EPI_ISL_3393648, EPI_ISL_3393649, EPI_ISL_3393650, EPI_ISL_3393651, EPI_ISL_3393652, EPI_ISL_3393653, EPI_ISL_3393654 | | | |
| see above | National Public Health Laboratory, National Centre for Infectious Diseases | National Public Health Laboratory, National Centre for Infectious Diseases | Tze Minn Mak, Katherine Ching, Grace Ngan, Zhenyang Zhou, Royce Ang, Lin Cui, Raymond Tzer Pin Lin |
| EPI_ISL_3393971, EPI_ISL_3393974, EPI_ISL_3393977, EPI_ISL_3393981, EPI_ISL_3393983, EPI_ISL_3393987, EPI_ISL_3393991, EPI_ISL_3393994, EPI_ISL_3393997, EPI_ISL_3394000, EPI_ISL_3394004, EPI_ISL_3394006, EPI_ISL_3394009, EPI_ISL_3394011, EPI_ISL_3394015, EPI_ISL_3394017, EPI_ISL_3394021, EPI_ISL_3394024, EPI_ISL_3394026, EPI_ISL_3394030, EPI_ISL_3394034, EPI_ISL_3394040, EPI_ISL_3394043, EPI_ISL_3394047, EPI_ISL_3394049, EPI_ISL_3394053, EPI_ISL_3394056, EPI_ISL_3394059, EPI_ISL_3394062, EPI_ISL_3394069 | | | |
| see above | IN State Department of Health Laboratory Services | IN State Department of Health Laboratory Services | Cassandra Campion, Jamie Yeadon, Brian Pope, Lixia Liu, Kyle Brownlee, Melissa Hindenlang, Mark Glazier |
| EPI_ISL_3394075, EPI_ISL_3394082, EPI_ISL_3394088, EPI_ISL_3394101, EPI_ISL_3394129 | Sonic Reference Laboratory | Sonic Reference Laboratory | "Robert Moller, Bijaya Dhakal, Krishna Maddula, John Woody, Cora Lahey, Ellen Pizarek" |
| EPI_ISL_3394162, EPI_ISL_3394163, EPI_ISL_3394164, EPI_ISL_3394165, EPI_ISL_3394166, EPI_ISL_3394167, EPI_ISL_3394168, EPI_ISL_3394169, EPI_ISL_3394170, EPI_ISL_3394171, EPI_ISL_3394172, EPI_ISL_3394173, EPI_ISL_3394174, EPI_ISL_3394175, EPI_ISL_3394176, EPI_ISL_3394177, EPI_ISL_3394178, EPI_ISL_3394179, EPI_ISL_3394180, EPI_ISL_3394181, EPI_ISL_3394182, EPI_ISL_3394183, EPI_ISL_3394184, EPI_ISL_3394185, EPI_ISL_3394186, EPI_ISL_3394187, EPI_ISL_3394188, EPI_ISL_3394189, EPI_ISL_3394192, EPI_ISL_3394193, EPI_ISL_3394194, EPI_ISL_3394196, EPI_ISL_3394197, EPI_ISL_3394198, EPI_ISL_3394199, EPI_ISL_3394200, EPI_ISL_3394201, EPI_ISL_3394202, EPI_ISL_3394203, EPI_ISL_3394204, EPI_ISL_3394205, EPI_ISL_3394206, EPI_ISL_3394208, EPI_ISL_3394209, EPI_ISL_3394210, EPI_ISL_3394211, EPI_ISL_3394212, EPI_ISL_3394213, EPI_ISL_3394214, EPI_ISL_3394215, EPI_ISL_3394216, EPI_ISL_3394217, EPI_ISL_3394218, EPI_ISL_3394219, EPI_ISL_3394220, EPI_ISL_3394221, EPI_ISL_3394222, EPI_ISL_3394223, EPI_ISL_3394224, EPI_ISL_3394225, EPI_ISL_3394226, EPI_ISL_3394227, EPI_ISL_3394228, EPI_ISL_3394230, EPI_ISL_3394231, EPI_ISL_3394232, EPI_ISL_3394234, EPI_ISL_3394235, EPI_ISL_3394236, EPI_ISL_3394237, EPI_ISL_3394239, EPI_ISL_3394240, EPI_ISL_3394241, EPI_ISL_3394242, EPI_ISL_3394243, EPI_ISL_3394244, EPI_ISL_3394245, EPI_ISL_3394246, EPI_ISL_3394247, EPI_ISL_3394248, EPI_ISL_3394249, EPI_ISL_3394250, EPI_ISL_3394251, EPI_ISL_3394252, EPI_ISL_3394253, EPI_ISL_3394254, EPI_ISL_3394255, EPI_ISL_3394256, EPI_ISL_3394257, EPI_ISL_3394258, EPI_ISL_3394259, EPI_ISL_3394260, EPI_ISL_3394261, EPI_ISL_3394262, EPI_ISL_3394263, EPI_ISL_3394264, EPI_ISL_3394265, EPI_ISL_3394266, EPI_ISL_3394267, EPI_ISL_3394268, EPI_ISL_3394269, EPI_ISL_3394270, EPI_ISL_3394271, EPI_ISL_3394272, EPI_ISL_3394273, EPI_ISL_3394275, EPI_ISL_3394276, EPI_ISL_3394277, EPI_ISL_3394278, EPI_ISL_3394279, EPI_ISL_3394280, EPI_ISL_3394281, EPI_ISL_3394282, EPI_ISL_3394283, EPI_ISL_3394284, EPI_ISL_3394285, EPI_ISL_3394286, EPI_ISL_3394287, EPI_ISL_3394288, EPI_ISL_3394289, EPI_ISL_3394290, EPI_ISL_3394291, EPI_ISL_3394292, EPI_ISL_3394293, EPI_ISL_3394294, EPI_ISL_3394295, EPI_ISL_3394296, EPI_ISL_3394297, EPI_ISL_3394298, EPI_ISL_3394299, EPI_ISL_3394301, EPI_ISL_3394302, EPI_ISL_3394303, EPI_ISL_3394304, EPI_ISL_3394305, EPI_ISL_3394306, EPI_ISL_3394307, EPI_ISL_3394311, EPI_ISL_3394312, EPI_ISL_3394313, EPI_ISL_3394314, EPI_ISL_3394315, EPI_ISL_3394316, EPI_ISL_3394317, EPI_ISL_3394318, EPI_ISL_3394319, EPI_ISL_3394320, EPI_ISL_3394324, EPI_ISL_3394325, EPI_ISL_3394326, EPI_ISL_3394327, EPI_ISL_3394328, EPI_ISL_3394329, EPI_ISL_3394330, EPI_ISL_3394331, EPI_ISL_3394332, EPI_ISL_3394333, EPI_ISL_3394334, EPI_ISL_3394335, EPI_ISL_3394336, EPI_ISL_3394337, EPI_ISL_3394338, EPI_ISL_3394339, EPI_ISL_3394340, EPI_ISL_3394341, EPI_ISL_3394342, EPI_ISL_3394343, EPI_ISL_3394344, EPI_ISL_3394345, EPI_ISL_3394346, EPI_ISL_3394347, EPI_ISL_3394348, EPI_ISL_3394349, EPI_ISL_3394350, EPI_ISL_3394351, EPI_ISL_3394352, EPI_ISL_3394353, EPI_ISL_3394354, EPI_ISL_3394355, EPI_ISL_3394356, EPI_ISL_3394357, EPI_ISL_3394358, EPI_ISL_3394359, EPI_ISL_3394360, EPI_ISL_3394361, EPI_ISL_3394362, EPI_ISL_3394363, EPI_ISL_3394364, EPI_ISL_3394365, EPI_ISL_3394366, EPI_ISL_3394367, EPI_ISL_3394368, EPI_ISL_3394369, EPI_ISL_3394370, EPI_ISL_3394371, EPI_ISL_3394372, EPI_ISL_3394373, EPI_ISL_3394374, EPI_ISL_3394375, EPI_ISL_3394376, EPI_ISL_3394377, EPI_ISL_3394378, EPI_ISL_3394379, EPI_ISL_3394380, EPI_ISL_3394381, EPI_ISL_3394382, EPI_ISL_3394383, EPI_ISL_3394384, EPI_ISL_3394385, EPI_ISL_3394386, EPI_ISL_3394387, EPI_ISL_3394388, EPI_ISL_3394389, EPI_ISL_3394390, EPI_ISL_3394391, EPI_ISL_3394392, EPI_ISL_3394393, EPI_ISL_3394394, EPI_ISL_3394395, EPI_ISL_3394396, EPI_ISL_3394397, EPI_ISL_3394398, EPI_ISL_3394399, EPI_ISL_3394400, EPI_ISL_3394401, EPI_ISL_3394402, EPI_ISL_3394403, EPI_ISL_3394404, EPI_ISL_3394405, EPI_ISL_3394406, EPI_ISL_3394407, EPI_ISL_3394408, EPI_ISL_3394409, EPI_ISL_3394410, EPI_ISL_3394411, EPI_ISL_3394412, EPI_ISL_3394413, EPI_ISL_3394414, EPI_ISL_3394415, EPI_ISL_3394416, EPI_ISL_3394417, EPI_ISL_3394418, EPI_ISL_3394419, EPI_ISL_3394420, EPI_ISL_3394421, EPI_ISL_3394422, EPI_ISL_3394423, EPI_ISL_3394424, EPI_ISL_3394425, EPI_ISL_3394426, EPI_ISL_3394427, EPI_ISL_3394428, EPI_ISL_3394429, EPI_ISL_3394430, EPI_ISL_3394431, EPI_ISL_3394432, EPI_ISL_3394433, EPI_ISL_3394434, EPI_ISL_3394435, EPI_ISL_3394436, EPI_ISL_3394437, EPI_ISL_3394438, EPI_ISL_3394439, EPI_ISL_3394440, EPI_ISL_3394441, EPI_ISL_3394442, EPI_ISL_3394443, EPI_ISL_3394444, EPI_ISL_3394445, EPI_ISL_3394446, EPI_ISL_3394447, EPI_ISL_3394448, EPI_ISL_3394449, EPI_ISL_3394450, EPI_ISL_3394451, EPI_ISL_3394452, EPI_ISL_3394453, EPI_ISL_3394454, EPI_ISL_3394455, EPI_ISL_3394456, EPI_ISL_3394457, EPI_ISL_3394458, EPI_ISL_3394459, EPI_ISL_3394460, EPI_ISL_3394461, EPI_ISL_3394462, EPI_ISL_3394463, EPI_ISL_3394464, EPI_ISL_3394465, EPI_ISL_3394466, EPI_ISL_3394467, EPI_ISL_3394468, EPI_ISL_3394469, EPI_ISL_3394470, EPI_ISL_3394471, EPI_ISL_3394472, EPI_ISL_3394473, EPI_ISL_3394474, EPI_ISL_3394475, EPI_ISL_3394476, EPI_ISL_3394477, EPI_ISL_3394478, EPI_ISL_3394479, EPI_ISL_3394480, EPI_ISL_3394481, EPI_ISL_3394482, EPI_ISL_3394483, EPI_ISL_3394484, EPI_ISL_3394485, EPI_ISL_3394486, EPI_ISL_3394487, EPI_ISL_3394488, EPI_ISL_3394489, EPI_ISL_3394490, EPI_ISL_3394491, EPI_ISL_3394492, EPI_ISL_3394493, EPI_ISL_3394494, EPI_ISL_3394495, EPI_ISL_3394496, EPI_ISL_3394497, EPI_ISL_3394498, EPI_ISL_3394499, EPI_ISL_3394500, EPI_ISL_3394501, EPI_ISL_3394502, EPI_ISL_3394503, EPI_ISL_3394504, EPI_ISL_3394505, EPI_ISL_3394506, EPI_ISL_3394507, EPI_ISL_3394508, EPI_ISL_3394509, EPI_ISL_3394510, EPI_ISL_3394511, EPI_ISL_3394512, EPI_ISL_3394513, EPI_ISL_3394514, EPI_ISL_3394515, EPI_ISL_3394516, EPI_ISL_3394517, EPI_ISL_3394518, EPI_ISL_3394519, EPI_ISL_3394520, EPI_ISL_3394521, EPI_ISL_3394522, EPI_ISL_3394523, EPI_ISL_3394524, EPI_ISL_3394525, EPI_ISL_3394526, EPI_ISL_3394527, EPI_ISL_3394528, EPI_ISL_3394529, EPI_ISL_3394530, EPI_ISL_3394531, EPI_ISL_3394532, EPI_ISL_3394533, EPI_ISL_3394534, EPI_ISL_3394535, EPI_ISL_3394536, EPI_ISL_3394537, EPI_ISL_3394538, EPI_ISL_3394539, EPI_ISL_3394540, EPI_ISL_3394541, EPI_ISL_3394542, EPI_ISL_3394543, EPI_ISL_3394544, EPI_ISL_3394545, EPI_ISL_3394546, EPI_ISL_3394547, EPI_ISL_3394548, EPI_ISL_3394549, EPI_ISL_3394550, EPI_ISL_3394551, EPI_ISL_3394552, EPI_ISL_3394553, EPI_ISL_3394554, EPI_ISL_3394555, EPI_ISL_3394556, EPI_ISL_3394557, EPI_ISL_3394558, EPI_ISL_3394559, EPI_ISL_3394560, EPI_ISL_3394561, EPI_ISL_3394562, EPI_ISL_3394563, EPI_ISL_3394564, EPI_ISL_3394565, EPI_ISL_3394566, EPI_ISL_3394567, EPI_ISL_3394568, EPI_ISL_3394569, EPI_ISL_3394570, EPI_ISL_3394571, EPI_ISL_3394572, EPI_ISL_3394573, EPI_ISL_3394574, EPI_ISL_3394575, EPI_ISL_3394576, EPI_ISL_3394577, EPI_ISL_3394578, EPI_ISL_3394579, EPI_ISL_3394580, EPI_ISL_3394581, EPI_ISL_3394582, EPI_ISL_3394583, EPI_ISL_3394584, EPI_ISL_3394585, EPI_ISL_3394586, EPI_ISL_3394587, EPI_ISL_3394588, EPI_ISL_3394589, EPI_ISL_3394590, EPI_ISL_3394591, EPI_ISL_3394592, EPI_ISL_3394593, EPI_ISL_3394594, EPI_ISL_3394595, EPI_ISL_3394596, EPI_ISL_3394597, EPI_ISL_3394598, EPI_ISL_3394599, EPI_ISL_3394600, EPI_ISL_3394601, EPI_ISL_3394602, EPI_ISL_3394603, EPI_ISL_3394604, EPI_ISL_3394605, EPI_ISL_3394606, EPI_ISL_3394607, EPI_ISL_3394608, EPI_ISL_3394609, EPI_ISL_3394610, EPI_ISL_3394611, EPI_ISL_3394612, EPI_ISL_3394613, EPI_ISL_3394614, EPI_ISL_3394615, EPI_ISL_3394616, EPI_ISL_3394617, EPI_ISL_3394618, EPI_ISL_3394619, EPI_ISL_3394620, EPI_ISL_3394621, EPI_ISL_3394622, EPI_ISL_3394623, EPI_ISL_3394624, EPI_ISL_3394625, EPI_ISL_3394626, EPI_ISL_3394627, EPI_ISL_3394628, EPI_ISL_3394629, EPI_ISL_3394630, EPI_ISL_3394631, EPI_ISL_3394632, EPI_ISL_3394633, EPI_ISL_3394634, EPI_ISL_3394635, EPI_ISL_3394636, EPI_ISL_3394637, EPI_ISL_3394638, EPI_ISL_3394639, EPI_ISL_3394640, EPI_ISL_3394641, EPI_ISL_3394642, EPI_ISL_3394643, EPI_ISL_3394644, EPI_ISL_3394645, EPI_ISL_3394646, EPI_ISL_3394647, EPI_ISL_3394648, EPI_ISL_3394649, EPI_ISL_3394650, EPI_ISL_3394651, EPI_ISL_3394652, EPI_ISL_3394653, EPI_ISL_3394654, EPI_ISL_3394655, EPI_ISL_3394656, EPI_ISL_3394657, EPI_ISL_3394658, EPI_ISL_3394659, EPI_ISL_3394660, | | | |

EPI_ISL_3394661, EPI_ISL_3394662, EPI_ISL_3394663, EPI_ISL_3394664, EPI_ISL_3394665, EPI_ISL_3394666, EPI_ISL_3394667, EPI_ISL_3394668, EPI_ISL_3394669, EPI_ISL_3394670, EPI_ISL_3394671, EPI_ISL_3394672, EPI_ISL_3394673, EPI_ISL_3394674, EPI_ISL_3394675, EPI_ISL_3394676, EPI_ISL_3394677, EPI_ISL_3394678, EPI_ISL_3394679, EPI_ISL_3394680, EPI_ISL_3394681, EPI_ISL_3394682, EPI_ISL_3394683, EPI_ISL_3394684, EPI_ISL_3394685, EPI_ISL_3394686, EPI_ISL_3394687, EPI_ISL_3394688, EPI_ISL_3394689, EPI_ISL_3394690, EPI_ISL_3394691, EPI_ISL_3394692, EPI_ISL_3394693, EPI_ISL_3394694, EPI_ISL_3394695, EPI_ISL_3394696, EPI_ISL_3394697, EPI_ISL_3394698, EPI_ISL_3394699, EPI_ISL_3394700, EPI_ISL_3394701, EPI_ISL_3394702, EPI_ISL_3394703, EPI_ISL_3394704, EPI_ISL_3394705, EPI_ISL_3394706, EPI_ISL_3394707, EPI_ISL_3394708, EPI_ISL_3394709, EPI_ISL_3394710, EPI_ISL_3394711, EPI_ISL_3394712, EPI_ISL_3394713, EPI_ISL_3394714, EPI_ISL_3394715, EPI_ISL_3394716, EPI_ISL_3394717, EPI_ISL_3394718, EPI_ISL_3394719, EPI_ISL_3394720, EPI_ISL_3394721, EPI_ISL_3394722, EPI_ISL_3394723, EPI_ISL_3394724, EPI_ISL_3394725, EPI_ISL_3394726, EPI_ISL_3394727, EPI_ISL_3394728, EPI_ISL_3394729, EPI_ISL_3394730, EPI_ISL_3394731, EPI_ISL_3394732, EPI_ISL_3394733, EPI_ISL_3394734, EPI_ISL_3394735, EPI_ISL_3394736, EPI_ISL_3394737, EPI_ISL_3394738, EPI_ISL_3394739, EPI_ISL_3394740, EPI_ISL_3394741, EPI_ISL_3394742, EPI_ISL_3394743, EPI_ISL_3394744, EPI_ISL_3394745, EPI_ISL_3394746, EPI_ISL_3394747, EPI_ISL_3394748, EPI_ISL_3394749, EPI_ISL_3394750, EPI_ISL_3394751, EPI_ISL_3394752, EPI_ISL_3394753, EPI_ISL_3394754, EPI_ISL_3394755, EPI_ISL_3394756, EPI_ISL_3394757, EPI_ISL_3394758, EPI_ISL_3394759, EPI_ISL_3394760, EPI_ISL_3394761, EPI_ISL_3394762, EPI_ISL_3394763, EPI_ISL_3394764, EPI_ISL_3394765, EPI_ISL_3394766, EPI_ISL_3394767, EPI_ISL_3394768, EPI_ISL_3394769, EPI_ISL_3394770, EPI_ISL_3394771, EPI_ISL_3394772, EPI_ISL_3394773, EPI_ISL_3394774, EPI_ISL_3394775, EPI_ISL_3394776, EPI_ISL_3394777, EPI_ISL_3394778, EPI_ISL_3394779, EPI_ISL_3394780, EPI_ISL_3394781, EPI_ISL_3394782, EPI_ISL_3394783, EPI_ISL_3394784, EPI_ISL_3394785, EPI_ISL_3394786, EPI_ISL_3394787, EPI_ISL_3394788, EPI_ISL_3394789, EPI_ISL_3394790, EPI_ISL_3394791, EPI_ISL_3394792, EPI_ISL_3394793, EPI_ISL_3394794, EPI_ISL_3394795, EPI_ISL_3394796, EPI_ISL_3394797, EPI_ISL_3394798, EPI_ISL_3394799, EPI_ISL_3394800, EPI_ISL_3394801, EPI_ISL_3394802, EPI_ISL_3394803, EPI_ISL_3394804, EPI_ISL_3394805, EPI_ISL_3394806, EPI_ISL_3394807, EPI_ISL_3394808, EPI_ISL_3394809, EPI_ISL_3394810, EPI_ISL_3394811, EPI_ISL_3394812, EPI_ISL_3394813, EPI_ISL_3394814, EPI_ISL_3394815, EPI_ISL_3394816, EPI_ISL_3394817, EPI_ISL_3394818, EPI_ISL_3394819, EPI_ISL_3394820, EPI_ISL_3394821, EPI_ISL_3394822, EPI_ISL_3394823, EPI_ISL_3394824, EPI_ISL_3394825, EPI_ISL_3394826, EPI_ISL_3394827, EPI_ISL_3394828, EPI_ISL_3394829, EPI_ISL_3394830, EPI_ISL_3394831, EPI_ISL_3394832, EPI_ISL_3394833, EPI_ISL_3394834, EPI_ISL_3394835, EPI_ISL_3394836, EPI_ISL_3394837, EPI_ISL_3394838, EPI_ISL_3394839, EPI_ISL_3394840, EPI_ISL_3394841, EPI_ISL_3394842, EPI_ISL_3394843, EPI_ISL_3394844, EPI_ISL_3394845, EPI_ISL_3394846, EPI_ISL_3394847, EPI_ISL_3394848, EPI_ISL_3394849, EPI_ISL_3394850, EPI_ISL_3394851, EPI_ISL_3394852, EPI_ISL_3394853, EPI_ISL_3394854, EPI_ISL_3394855, EPI_ISL_3394856, EPI_ISL_3394857, EPI_ISL_3394858, EPI_ISL_3394859, EPI_ISL_3394860

| | | | |
|---|---|---|---|
| see above | Pandemic Response Lab - NYC | Pandemic Response Lab, R&D | Henry Lee, Michael Hammerling, Melissa Hopkins, Cybill del Castillo, Shinyoung Clair Kang, William Ward, Pradeep Bugga, Sol Rey, Dylan Law, Alex Carpio, Isabel Fernandez Escapa, Haiping Hao, Jon Laurent |

EPI_ISL_3394861, EPI_ISL_3394862, EPI_ISL_3394864, EPI_ISL_3394865, EPI_ISL_3394866, EPI_ISL_3394867, EPI_ISL_3394868, EPI_ISL_3394869, EPI_ISL_3394870, EPI_ISL_3394871, EPI_ISL_3394872, EPI_ISL_3394873, EPI_ISL_3394874, EPI_ISL_3394875, EPI_ISL_3394876, EPI_ISL_3394878, EPI_ISL_3394888, EPI_ISL_3394890, EPI_ISL_3394893, EPI_ISL_3394894, EPI_ISL_3394895, EPI_ISL_3394897, EPI_ISL_3394898, EPI_ISL_3394899, EPI_ISL_3394900, EPI_ISL_3394901, EPI_ISL_3394902, EPI_ISL_3394903, EPI_ISL_3394904, EPI_ISL_3394905, EPI_ISL_3394906, EPI_ISL_3394907, EPI_ISL_3394908, EPI_ISL_3394909, EPI_ISL_3394910, EPI_ISL_3394911, EPI_ISL_3394912, EPI_ISL_3394913, EPI_ISL_3394914, EPI_ISL_3394915, EPI_ISL_3394916

| | | | |
|---|---|---|---|
| see above | Devki Devi Foundation, a unit of Max Healthcare | CSIR-IGIB | Rajesh Pandey, Priti Devi, Sheeba Saifi, Aparna S, Ranjeet Maurya, Partha Chattopadhyay, Akshay Kanakan, Priyanka Mehta, Bansidhar Das, Sandeep Budhiraja |

EPI_ISL_3394921, EPI_ISL_3394923, EPI_ISL_3394924, EPI_ISL_3394925, EPI_ISL_3394926, EPI_ISL_3394927, EPI_ISL_3394928, EPI_ISL_3394929, EPI_ISL_3394930, EPI_ISL_3394931, EPI_ISL_3394932, EPI_ISL_3394933, EPI_ISL_3394934, EPI_ISL_3394935, EPI_ISL_3394936, EPI_ISL_3394937, EPI_ISL_3394938, EPI_ISL_3394939, EPI_ISL_3394940, EPI_ISL_3394941, EPI_ISL_3394942, EPI_ISL_3394943, EPI_ISL_3394944, EPI_ISL_3394945, EPI_ISL_3394946, EPI_ISL_3394947, EPI_ISL_3394948, EPI_ISL_3394950, EPI_ISL_3394951, EPI_ISL_3394952, EPI_ISL_3394953, EPI_ISL_3394954, EPI_ISL_3394955

| | | | |
|---|---|---|---|
| see above | Dept. of Microbiology and Infection Control, Akershus University Hospital HF | Dept. of Microbiology and Infection Control, Akershus University Hospital HF | Hege Vangstein Aamot, Alexander Hesselberg Løvestad |

EPI_ISL_3394956, EPI_ISL_3394957, EPI_ISL_3394958, EPI_ISL_3394959, EPI_ISL_3394960, EPI_ISL_3394961, EPI_ISL_3394962, EPI_ISL_3394963, EPI_ISL_3394964, EPI_ISL_3394965, EPI_ISL_3394966, EPI_ISL_3394967, EPI_ISL_3394968, EPI_ISL_3394969, EPI_ISL_3394970, EPI_ISL_3394971, EPI_ISL_3394972, EPI_ISL_3394973, EPI_ISL_3394974, EPI_ISL_3394975, EPI_ISL_3394976, EPI_ISL_3394977

| | | | |
|---|---|---|---|
| see above | Dept. of Laboratory Medicine | Dept. of Laboratory Medicine | Harald Esterbauer, Anna Gschaider, Petra Jurkowitsch, Fabian Konig, Sabina Plumer, Robert Strassl, Claudia Weber, Oswald Wagner |
| EPI_ISL_3394979, EPI_ISL_3394980, EPI_ISL_3394981, EPI_ISL_3394982, EPI_ISL_3394985, EPI_ISL_3394986, EPI_ISL_3394987, EPI_ISL_3394988, EPI_ISL_3394989 | Balai Penelitian dan Pengembangan Aceh | National Institute of Health Research and Development | Vivi Setiawaty, Hana Apsari Pawestri, Subangkit, Kartika Dewi Puspa, Arie Ardiansyah Nugraha, Hartanti Dian Ikawati, Krisna Pangesti, Triyani Soekarso, Fahmi Ichwansyah, Nelly Puspandari |
| EPI_ISL_3394992, EPI_ISL_3394994, EPI_ISL_3394995, EPI_ISL_3394996, EPI_ISL_3395000, EPI_ISL_3395001, EPI_ISL_3395002, EPI_ISL_3395005, EPI_ISL_3395006 | Balai Penelitian dan Pengembangan Aceh | National Institute of Health Research and Development | Hana Apsari Pawestri, Subangkit, Kartika Dewi Puspa, Arie Ardiansyah Nugraha, Hartanti Dian Ikawati, Krisna Pangesti, Triyani Soekarso, Nelly Puspandari, Fahmi Ichwansyah, Vivi Setiawaty |

EPI_ISL_3395011, EPI_ISL_3395013, EPI_ISL_3395014, EPI_ISL_3395015, EPI_ISL_3395016, EPI_ISL_3395017, EPI_ISL_3395018, EPI_ISL_3395019, EPI_ISL_3395020, EPI_ISL_3395021, EPI_ISL_3395022, EPI_ISL_3395023, EPI_ISL_3395024, EPI_ISL_3395025, EPI_ISL_3395026, EPI_ISL_3395027, EPI_ISL_3395028, EPI_ISL_3395029, EPI_ISL_3395030, EPI_ISL_3395031, EPI_ISL_3395032, EPI_ISL_3395033, EPI_ISL_3395034, EPI_ISL_3395035, EPI_ISL_3395036, EPI_ISL_3395037, EPI_ISL_3395038, EPI_ISL_3395039, EPI_ISL_3395040, EPI_ISL_3395041, EPI_ISL_3395042, EPI_ISL_3395043, EPI_ISL_3395044, EPI_ISL_3395045, EPI_ISL_3395047, EPI_ISL_3395048, EPI_ISL_3395049, EPI_ISL_3395050, EPI_ISL_3395051, EPI_ISL_3395052, EPI_ISL_3395053, EPI_ISL_3395054, EPI_ISL_3395055, EPI_ISL_3395056, EPI_ISL_3395057, EPI_ISL_3395058, EPI_ISL_3395059, EPI_ISL_3395060, EPI_ISL_3395061, EPI_ISL_3395062, EPI_ISL_3395063, EPI_ISL_3395064, EPI_ISL_3395065, EPI_ISL_3395066, EPI_ISL_3395067, EPI_ISL_3395068, EPI_ISL_3395069, EPI_ISL_3395070, EPI_ISL_3395071, EPI_ISL_3395072, EPI_ISL_3395073, EPI_ISL_3395074, EPI_ISL_3395075, EPI_ISL_3395076, EPI_ISL_3395077, EPI_ISL_3395078, EPI_ISL_3395079, EPI_ISL_3395080, EPI_ISL_3395081, EPI_ISL_3395082, EPI_ISL_3395083, EPI_ISL_3395084, EPI_ISL_3395085, EPI_ISL_3395086, EPI_ISL_3395087, EPI_ISL_3395088, EPI_ISL_3395089, EPI_ISL_3395090, EPI_ISL_3395091, EPI_ISL_3395092, EPI_ISL_3395093, EPI_ISL_3395094, EPI_ISL_3395095, EPI_ISL_3395096, EPI_ISL_3395097, EPI_ISL_3395098, EPI_ISL_3395099, EPI_ISL_3395100, EPI_ISL_3395101, EPI_ISL_3395102, EPI_ISL_3395103, EPI_ISL_3395104, EPI_ISL_3395105, EPI_ISL_3395106, EPI_ISL_3395107, EPI_ISL_3395108, EPI_ISL_3395109, EPI_ISL_3395111, EPI_ISL_3395112, EPI_ISL_3395114, EPI_ISL_3395115, EPI_ISL_3395116, EPI_ISL_3395117, EPI_ISL_3395118, EPI_ISL_3395119, EPI_ISL_3395120, EPI_ISL_3395121, EPI_ISL_3395122, EPI_ISL_3395123, EPI_ISL_3395124, EPI_ISL_3395125, EPI_ISL_3395127, EPI_ISL_3395128, EPI_ISL_3395129, EPI_ISL_3395130, EPI_ISL_3395131, EPI_ISL_3395132, EPI_ISL_3395133, EPI_ISL_3395134, EPI_ISL_3395135, EPI_ISL_3395136, EPI_ISL_3395137, EPI_ISL_3395139, EPI_ISL_3395140, EPI_ISL_3395141, EPI_ISL_3395142, EPI_ISL_3395143, EPI_ISL_3395144, EPI_ISL_3395145, EPI_ISL_3395146, EPI_ISL_3395147, EPI_ISL_3395148, EPI_ISL_3395149, EPI_ISL_3395150, EPI_ISL_3395151, EPI_ISL_3395152, EPI_ISL_3395153, EPI_ISL_3395154, EPI_ISL_3395155, EPI_ISL_3395156, EPI_ISL_3395157, EPI_ISL_3395158, EPI_ISL_3395159, EPI_ISL_3395160, EPI_ISL_3395161, EPI_ISL_3395162, EPI_ISL_3395163, EPI_ISL_3395165, EPI_ISL_3395166, EPI_ISL_3395167, EPI_ISL_3395168, EPI_ISL_3395169, EPI_ISL_3395170, EPI_ISL_3395171, EPI_ISL_3395172, EPI_ISL_3395173, EPI_ISL_3395174, EPI_ISL_3395175, EPI_ISL_3395176, EPI_ISL_3395177, EPI_ISL_3395178, EPI_ISL_3395179, EPI_ISL_3395180, EPI_ISL_3395182, EPI_ISL_3395183, EPI_ISL_3395184, EPI_ISL_3395185, EPI_ISL_3395186, EPI_ISL_3395187, EPI_ISL_3395188, EPI_ISL_3395189, EPI_ISL_3395190, EPI_ISL_3395191, EPI_ISL_3395192, EPI_ISL_3395193, EPI_ISL_3395194, EPI_ISL_3395195, EPI_ISL_3395196, EPI_ISL_3395197, EPI_ISL_3395198, EPI_ISL_3395199, EPI_ISL_3395200, EPI_ISL_3395201, EPI_ISL_3395202, EPI_ISL_3395203, EPI_ISL_3395204, EPI_ISL_3395205, EPI_ISL_3395206, EPI_ISL_3395208, EPI_ISL_3395209, EPI_ISL_3395210, EPI_ISL_3395211, EPI_ISL_3395212, EPI_ISL_3395214, EPI_ISL_3395215, EPI_ISL_3395216, EPI_ISL_3395217, EPI_ISL_3395218, EPI_ISL_3395219, EPI_ISL_3395220, EPI_ISL_3395221, EPI_ISL_3395222, EPI_ISL_3395223, EPI_ISL_3395225, EPI_ISL_3395226, EPI_ISL_3395227, EPI_ISL_3395228, EPI_ISL_3395229, EPI_ISL_3395230, EPI_ISL_3395231, EPI_ISL_3395232, EPI_ISL_3395233, EPI_ISL_3395234, EPI_ISL_3395235, EPI_ISL_3395236, EPI_ISL_3395237, EPI_ISL_3395238, EPI_ISL_3395240, EPI_ISL_3395241, EPI_ISL_3395242, EPI_ISL_3395243, EPI_ISL_3395244, EPI_ISL_3395245, EPI_ISL_3395246, EPI_ISL_3395247, EPI_ISL_3395248, EPI_ISL_3395249, EPI_ISL_3395250, EPI_ISL_3395251, EPI_ISL_3395252, EPI_ISL_3395253, EPI_ISL_3395254, EPI_ISL_3395255, EPI_ISL_3395256, EPI_ISL_3395257, EPI_ISL_3395258, EPI_ISL_3395259, EPI_ISL_3395260, EPI_ISL_3395261, EPI_ISL_3395262, EPI_ISL_3395263, EPI_ISL_3395264, EPI_ISL_3395265, EPI_ISL_3395266, EPI_ISL_3395267, EPI_ISL_3395268, EPI_ISL_3395269, EPI_ISL_3395270, EPI_ISL_3395271, EPI_ISL_3395272, EPI_ISL_3395273, EPI_ISL_3395274, EPI_ISL_3395275, EPI_ISL_3395276, EPI_ISL_3395277, EPI_ISL_3395278, EPI_ISL_3395279, EPI_ISL_3395280, EPI_ISL_3395282, EPI_ISL_3395283, EPI_ISL_3395284, EPI_ISL_3395285, EPI_ISL_3395286, EPI_ISL_3395287, EPI_ISL_3395288, EPI_ISL_3395289, EPI_ISL_3395290, EPI_ISL_3395291, EPI_ISL_3395292, EPI_ISL_3395293, EPI_ISL_3395294, EPI_ISL_3395295, EPI_ISL_3395297, EPI_ISL_3395298, EPI_ISL_3395299, EPI_ISL_3395301, EPI_ISL_3395302, EPI_ISL_3395303, EPI_ISL_3395304, EPI_ISL_3395306, EPI_ISL_3395307, EPI_ISL_3395308, EPI_ISL_3395309, EPI_ISL_3395310, EPI_ISL_3395311, EPI_ISL_3395312, EPI_ISL_3395314, EPI_ISL_3395315, EPI_ISL_3395316, EPI_ISL_3395317, EPI_ISL_3395318, EPI_ISL_3395319, EPI_ISL_3395320, EPI_ISL_3395321, EPI_ISL_3395322, EPI_ISL_3395323, EPI_ISL_3395324, EPI_ISL_3395325, EPI_ISL_3395326, EPI_ISL_3395327, EPI_ISL_3395328, EPI_ISL_3395329, EPI_ISL_3395330, EPI_ISL_3395331, EPI_ISL_3395332, EPI_ISL_3395333, EPI_ISL_3395334, EPI_ISL_3395335, EPI_ISL_3395336, EPI_ISL_3395337, EPI_ISL_3395338, EPI_ISL_3395339, EPI_ISL_3395340, EPI_ISL_3395341, EPI_ISL_3395342, EPI_ISL_3395343, EPI_ISL_3395344, EPI_ISL_3395345, EPI_ISL_3395346, EPI_ISL_3395347, EPI_ISL_3395348, EPI_ISL_3395349, EPI_ISL_3395350, EPI_ISL_3395351, EPI_ISL_3395352, EPI_ISL_3395353, EPI_ISL_3395354, EPI_ISL_3395355, EPI_ISL_3395356, EPI_ISL_3395357, EPI_ISL_3395358, EPI_ISL_3395359, EPI_ISL_3395360, EPI_ISL_3395361, EPI_ISL_3395362, EPI_ISL_3395363, EPI_ISL_3395364, EPI_ISL_3395365, EPI_ISL_3395366, EPI_ISL_3395367, EPI_ISL_3395368, EPI_ISL_3395369, EPI_ISL_3395370, EPI_ISL_3395371, EPI_ISL_3395372, EPI_ISL_3395373, EPI_ISL_3395374, EPI_ISL_3395375, EPI_ISL_3395376, EPI_ISL_3395377, EPI_ISL_3395378, EPI_ISL_3395379, EPI_ISL_3395380, EPI_ISL_3395381, EPI_ISL_3395382, EPI_ISL_3395383, EPI_ISL_3395384, EPI_ISL_3395385, EPI_ISL_3395386, EPI_ISL_3395387, EPI_ISL_3395388, EPI_ISL_3395389, EPI_ISL_3395391, EPI_ISL_3395392, EPI_ISL_3395394, EPI_ISL_3395395, EPI_ISL_3395396, EPI_ISL_3395397, EPI_ISL_3395398, EPI_ISL_3395399, EPI_ISL_3395400, EPI_ISL_3395401, EPI_ISL_3395402, EPI_ISL_3395403, EPI_ISL_3395404, EPI_ISL_3395405, EPI_ISL_3395406, EPI_ISL_3395407, EPI_ISL_3395408, EPI_ISL_3395409, EPI_ISL_3395410, EPI_ISL_3395411, EPI_ISL_3395412, EPI_ISL_3395413, EPI_ISL_3395414, EPI_ISL_3395415, EPI_ISL_3395416, EPI_ISL_3395417, EPI_ISL_3395418, EPI_ISL_3395419, EPI_ISL_3395420, EPI_ISL_3395421, EPI_ISL_3395422, EPI_ISL_3395423, EPI_ISL_3395424, EPI_ISL_3395425, EPI_ISL_3395426, EPI_ISL_3395427, EPI_ISL_3395428, EPI_ISL_3395429, EPI_ISL_3395430, EPI_ISL_3395431, EPI_ISL_3395432, EPI_ISL_3395433, EPI_ISL_3395434, EPI_ISL_3395435, EPI_ISL_3395436, EPI_ISL_3395437, EPI_ISL_3395439, EPI_ISL_3395440, EPI_ISL_3395441, EPI_ISL_3395442, EPI_ISL_3395443, EPI_ISL_3395444, EPI_ISL_3395445, EPI_ISL_3395446, EPI_ISL_3395447, EPI_ISL_3395448, EPI_ISL_3395449, EPI_ISL_3395450, EPI_ISL_3395451, EPI_ISL_3395452, EPI_ISL_3395453, EPI_ISL_3395454, EPI_ISL_3395456, EPI_ISL_3395457, EPI_ISL_3395458, EPI_ISL_3395459, EPI_ISL_3395460, EPI_ISL_3395461, EPI_ISL_3395462, EPI_ISL_3395463, EPI_ISL_3395464, EPI_ISL_3395465, EPI_ISL_3395466, EPI_ISL_3395467, EPI_ISL_3395468, EPI_ISL_3395469, EPI_ISL_3395470, EPI_ISL_3395471, EPI_ISL_3395472, EPI_ISL_3395473, EPI_ISL_3395474, EPI_ISL_3395475, EPI_ISL_3395477, EPI_ISL_3395478, EPI_ISL_3395479, EPI_ISL_3395480, EPI_ISL_3395481, EPI_ISL_3395482, EPI_ISL_3395483, EPI_ISL_3395484, EPI_ISL_3395485, EPI_ISL_3395486, EPI_ISL_3395487, EPI_ISL_3395488, EPI_ISL_3395489, EPI_ISL_3395490, EPI_ISL_3395491, EPI_ISL_3395492, EPI_ISL_3395493, EPI_ISL_3395494, EPI_ISL_3395495, EPI_ISL_3395496, EPI_ISL_3395497, EPI_ISL_3395498, EPI_ISL_3395499, EPI_ISL_3395500, EPI_ISL_3395501, EPI_ISL_3395502, EPI_ISL_3395503, EPI_ISL_3395504, EPI_ISL_3395505, EPI_ISL_3395506, EPI_ISL_3395508, EPI_ISL_3395509, EPI_ISL_3395510, EPI_ISL_3395511, EPI_ISL_3395512, EPI_ISL_3395513, EPI_ISL_3395515, EPI_ISL_3395516, EPI_ISL_3395517, EPI_ISL_3395518, EPI_ISL_3395519, EPI_ISL_3395520, EPI_ISL_3395521, EPI_ISL_3395522, EPI_ISL_3395523, EPI_ISL_3395524, EPI_ISL_3395525, EPI_ISL_3395526, EPI_ISL_3395527, EPI_ISL_3395528, EPI_ISL_3395529, EPI_ISL_3395530, EPI_ISL_3395531, EPI_ISL_3395532, EPI_ISL_3395533, EPI_ISL_3395534, EPI_ISL_3395535, EPI_ISL_3395536, EPI_ISL_3395537, EPI_ISL_3395538, EPI_ISL_3395539, EPI_ISL_3395540, EPI_ISL_3395541, EPI_ISL_3395542, EPI_ISL_3395543, EPI_ISL_3395544, EPI_ISL_3395545, EPI_ISL_3395546, EPI_ISL_3395547, EPI_ISL_3395548, EPI_ISL_3395549, EPI_ISL_3395550, EPI_ISL_3395552, EPI_ISL_3395553, EPI_ISL_3395554, EPI_ISL_3395555, EPI_ISL_3395556, EPI_ISL_3395557, EPI_ISL_3395558, EPI_ISL_3395559, EPI_ISL_3395560, EPI_ISL_3395561, EPI_ISL_3395562, EPI_ISL_3395563, EPI_ISL_3395564, EPI_ISL_3395565, EPI_ISL_3395566, EPI_ISL_3395567, EPI_ISL_3395568, EPI_ISL_3395569, EPI_ISL_3395570, EPI_ISL_3395571, EPI_ISL_3395572, EPI_ISL_3395573, EPI_ISL_3395574, EPI_ISL_3395575, EPI_ISL_3395576, EPI_ISL_3395577, EPI_ISL_3395578, EPI_ISL_3395579, EPI_ISL_3395580, EPI_ISL_3395581, EPI_ISL_3395582, EPI_ISL_3395583, EPI_ISL_3395584, EPI_ISL_3395585, EPI_ISL_3395586, EPI_ISL_3395587, EPI_ISL_3395589, EPI_ISL_3395590, EPI_ISL_3395591, EPI_ISL_3395592, EPI_ISL_3395593, EPI_ISL_3395594, EPI_ISL_3395595, EPI_ISL_3395596, EPI_ISL_3395597, EPI_ISL_3395598, EPI_ISL_3395599, EPI_ISL_3395600, EPI_ISL_3395601, EPI_ISL_3395602, EPI_ISL_3395603, EPI_ISL_3395604, EPI_ISL_3395605, EPI_ISL_3395606, EPI_ISL_3395607, EPI_ISL_3395608, EPI_ISL_3395609, EPI_ISL_3395610, EPI_ISL_3395611, EPI_ISL_3395612, EPI_ISL_3395613, EPI_ISL_3395614, EPI_ISL_3395615, EPI_ISL_3395616, EPI_ISL_3395617, EPI_ISL_3395618, EPI_ISL_3395619, EPI_ISL_3395620, EPI_ISL_3395621, EPI_ISL_3395622, EPI_ISL_3395623, EPI_ISL_3395624, EPI_ISL_3395625, EPI_ISL_3395626, EPI_ISL_3395627, EPI_ISL_3395628, EPI_ISL_3395629, EPI_ISL_3395630, EPI_ISL_3395632, EPI_ISL_3395633, EPI_ISL_3395634, EPI_ISL_3395635, EPI_ISL_3395636, EPI_ISL_3395637, EPI_ISL_3395638, EPI_ISL_3395639, EPI_ISL_3395640, EPI_ISL_3395641, EPI_ISL_3395642, EPI_ISL_3395643, EPI_ISL_3395644, EPI_ISL_3395645, EPI_ISL_3395646, EPI_ISL_3395647, EPI_ISL_3395648, EPI_ISL_3395651, EPI_ISL_3395652, EPI_ISL_3395653, EPI_ISL_3395654, EPI_ISL_3395655, EPI_ISL_3395656, EPI_ISL_3395657, EPI_ISL_3395658, EPI_ISL_3395659, EPI_ISL_3395660, EPI_ISL_3395661, EPI_ISL_3395662, EPI_ISL_3395663, EPI_ISL_3395664, EPI_ISL_3395665, EPI_ISL_3395666, EPI_ISL_3395667, EPI_ISL_3395668, EPI_ISL_3395669, EPI_ISL_3395670, EPI_ISL_3395671, EPI_ISL_3395672, EPI_ISL_3395673, EPI_ISL_3395674, EPI_ISL_3395675, EPI_ISL_3395676, EPI_ISL_3395677, EPI_ISL_3395678, EPI_ISL_3395679, EPI_ISL_3395682, EPI_ISL_3395683, EPI_ISL_3395684, EPI_ISL_3395686, EPI_ISL_3395687, EPI_ISL_3395688, EPI_ISL_3395689, EPI_ISL_3395690, EPI_ISL_3395691, EPI_ISL_3395692, EPI_ISL_3395693, EPI_ISL_3395694, EPI_ISL_3395695, EPI_ISL_3395696, EPI_ISL_3395697, EPI_ISL_3395698, EPI_ISL_3395699, EPI_ISL_3395700, EPI_ISL_3395701, EPI_ISL_3395702, EPI_ISL_3395703, EPI_ISL_3395704, EPI_ISL_3395705, EPI_ISL_3395706, EPI_ISL_3395707, EPI_ISL_3395708, EPI_ISL_3395709, EPI_ISL_3395710, EPI_ISL_3395711, EPI_ISL_3395712, EPI_ISL_3395713, EPI_ISL_3395714, EPI_ISL_3395715, EPI_ISL_3395716, EPI_ISL_3395717, EPI_ISL_3395718, EPI_ISL_3395719, EPI_ISL_3395720, EPI_ISL_3395721, EPI_ISL_3395722, EPI_ISL_3395723, EPI_ISL_3395724, EPI_ISL_3395725, EPI_ISL_3395726, EPI_ISL_3395727, EPI_ISL_3395728, EPI_ISL_3395730, EPI_ISL_3395731, EPI_ISL_3395732, EPI_ISL_3395733, EPI_ISL_3395734, EPI_ISL_3395735, EPI_ISL_3395736, EPI_ISL_3395737, EPI_ISL_3395738, EPI_ISL_3395739, EPI_ISL_3395740, EPI_ISL_3395741, EPI_ISL_3395742, EPI_ISL_3395743, EPI_ISL_3395744, EPI_ISL_3395745, EPI_ISL_3395746, EPI_ISL_3395747, EPI_ISL_3395748, EPI_ISL_3395749, EPI_ISL_3395750, EPI_ISL_3395751, EPI_ISL_3395752, EPI_ISL_3395753, EPI_ISL_3395754, EPI_ISL_3395755, EPI_ISL_3395756, EPI_ISL_3395757, EPI_ISL_3395758, EPI_ISL_3395760, EPI_ISL_3395761, EPI_ISL_3395762, EPI_ISL_3395763, EPI_ISL_3395764, EPI_ISL_3395765, EPI_ISL_3395766, EPI_ISL_3395767, EPI_ISL_3395768, EPI_ISL_3395769, EPI_ISL_3395770, EPI_ISL_3395771, EPI_ISL_3395772, EPI_ISL_3395773, EPI_ISL_3395774, EPI_ISL_3395775, EPI_ISL_3395776, EPI_ISL_3395777, EPI_ISL_3395778, EPI_ISL_3395779, EPI_ISL_3395780, EPI_ISL_3395781, EPI_ISL_3395782, EPI_ISL_3395783, EPI_ISL_3395784, EPI_ISL_3395785, EPI_ISL_3395787, EPI_ISL_3395788, EPI_ISL_3395789, EPI_ISL_3395790, EPI_ISL_3395791, EPI_ISL_3395792, EPI_ISL_3395793, EPI_ISL_3395794, EPI_ISL_3395795, EPI_ISL_3395796, EPI_ISL_3395797, EPI_ISL_3395798, EPI_ISL_3395799, EPI_ISL_3395800, EPI_ISL_3395801, EPI_ISL_3395802, EPI_ISL_3395803, EPI_ISL_3395804, EPI_ISL_3395805, EPI_ISL_3395806, EPI_ISL_3395807, EPI_ISL_3395808, EPI_ISL_3395809, EPI_ISL_3395810, EPI_ISL_3395811, EPI_ISL_3395812, EPI_ISL_3395813, EPI_ISL_3395814, EPI_ISL_3395815, EPI_ISL_3395816, EPI_ISL_3395817, EPI_ISL_3395818, EPI_ISL_3395819, EPI_ISL_3395820, EPI_ISL_3395821, EPI_ISL_3395822, EPI_ISL_3395823, EPI_ISL_3395824, EPI_ISL_3395826, EPI_ISL_3395828

| | | | |
|---|---|---|---|
| see above | Quest Diagnostics Incorporated | Centers for Disease Control and Prevention Division of Viral Diseases, Pathogen Discovery | Dakota Howard, Dhwani Batra, Peter W. Cook, Kara Moser, Adrian Paskey, Jason Caravas, Benjamin Rambo-Martin, Shatavia Morrison, Christopher Gulvick, Scott Sammons, Yvette Unoarumhi, Darlene Wagner, Matthew Schmerer, S. H. Rosenthal, A. Gerasimova, R. M. Kagan, B. Anderson, M. Hua, Y. Liu, L.E. Bernstein, K.E. Livingston, A. Perez, I. A. Shlyakhter, R. V. Rolando, R. Owen, P. Tanpaiboon, F. Lacbawan, Clinton R. Paden, Duncan MacCannell |
| EPI_ISL_3395829, EPI_ISL_3395830 | OLVZ Aalst | OLVZ Aalst | Astrid Holderbeke |
| EPI_ISL_3395831 | Quest Diagnostics Incorporated | Centers for Disease Control and Prevention Division of Viral Diseases, Pathogen Discovery | Dakota Howard, Dhwani Batra, Peter W. Cook, Kara Moser, Adrian Paskey, Jason Caravas, Benjamin Rambo-Martin, Shatavia Morrison, Christopher Gulvick, Scott Sammons, Yvette Unoarumhi, Darlene Wagner, Matthew Schmerer, S. H. Rosenthal, A. Gerasimova, R. M. Kagan, B. Anderson, M. Hua, Y. Liu, L.E. Bernstein, K.E. Livingston, A. Perez, I. A. Shlyakhter, R. V. Rolando, R. Owen, P. Tanpaiboon, F. Lacbawan, Clinton R. Paden, Duncan MacCannell |
| EPI_ISL_3395832 | OLVZ Aalst | OLVZ Aalst | Astrid Holderbeke |
| EPI_ISL_3395833 | Quest Diagnostics Incorporated | Centers for Disease Control and Prevention Division of Viral Diseases, Pathogen Discovery | Dakota Howard, Dhwani Batra, Peter W. Cook, Kara Moser, Adrian Paskey, Jason Caravas, Benjamin Rambo-Martin, Shatavia Morrison, Christopher Gulvick, Scott Sammons, Yvette Unoarumhi, Darlene Wagner, Matthew Schmerer, S. H. Rosenthal, A. Gerasimova, R. M. Kagan, B. Anderson, M. Hua, Y. Liu, L.E. Bernstein, K.E. Livingston, A. Perez, I. A. Shlyakhter, R. V. Rolando, R. Owen, P. Tanpaiboon, F. Lacbawan, Clinton R. Paden, Duncan MacCannell |
| EPI_ISL_3395834, EPI_ISL_3395835 | OLVZ Aalst | OLVZ Aalst | Astrid Holderbeke |
| EPI_ISL_3395836 | Quest Diagnostics Incorporated | Centers for Disease Control and Prevention Division of Viral Diseases, Pathogen Discovery | Dakota Howard, Dhwani Batra, Peter W. Cook, Kara Moser, Adrian Paskey, Jason Caravas, Benjamin Rambo-Martin, Shatavia Morrison, Christopher Gulvick, Scott Sammons, Yvette Unoarumhi, Darlene Wagner, Matthew Schmerer, S. H. Rosenthal, A. Gerasimova, R. M. Kagan, B. Anderson, M. Hua, Y. Liu, L.E. Bernstein, K.E. Livingston, A. Perez, I. A. Shlyakhter, R. V. Rolando, R. Owen, P. Tanpaiboon, F. Lacbawan, Clinton R. Paden, Duncan MacCannell |
| EPI_ISL_3395837 | OLVZ Aalst | OLVZ Aalst | Astrid Holderbeke |
| EPI_ISL_3395838 | Quest Diagnostics Incorporated | Centers for Disease Control and Prevention Division of Viral Diseases, Pathogen Discovery | Dakota Howard, Dhwani Batra, Peter W. Cook, Kara Moser, Adrian Paskey, Jason Caravas, Benjamin Rambo-Martin, Shatavia Morrison, Christopher Gulvick, Scott Sammons, Yvette Unoarumhi, Darlene Wagner, Matthew Schmerer, S. H. Rosenthal, A. Gerasimova, R. M. Kagan, B. Anderson, M. Hua, Y. Liu, L.E. Bernstein, K.E. Livingston, A. Perez, I. A. Shlyakhter, R. V. Rolando, R. Owen, P. Tanpaiboon, F. Lacbawan, Clinton R. Paden, Duncan MacCannell |
| EPI_ISL_3395839, EPI_ISL_3395840 | OLVZ Aalst | OLVZ Aalst | Astrid Holderbeke |
| EPI_ISL_3395841 | Quest Diagnostics Incorporated | Centers for Disease Control and Prevention Division of Viral Diseases, Pathogen Discovery | Dakota Howard, Dhwani Batra, Peter W. Cook, Kara Moser, Adrian Paskey, Jason Caravas, Benjamin Rambo-Martin, Shatavia Morrison, Christopher Gulvick, Scott Sammons, Yvette Unoarumhi, Darlene Wagner, Matthew Schmerer, S. H. Rosenthal, A. Gerasimova, R. M. Kagan, B. Anderson, M. Hua, Y. Liu, L.E. Bernstein, K.E. Livingston, A. Perez, I. A. Shlyakhter, R. V. Rolando, R. Owen, P. Tanpaiboon, F. Lacbawan, Clinton R. Paden, Duncan MacCannell |
| EPI_ISL_3395842 | OLVZ Aalst | OLVZ Aalst | Astrid Holderbeke |
| EPI_ISL_3395843 | Quest Diagnostics Incorporated | Centers for Disease Control and Prevention Division of Viral Diseases, Pathogen Discovery | Dakota Howard, Dhwani Batra, Peter W. Cook, Kara Moser, Adrian Paskey, Jason Caravas, Benjamin Rambo-Martin, Shatavia Morrison, Christopher Gulvick, Scott Sammons, Yvette Unoarumhi, Darlene Wagner, Matthew Schmerer, S. H. Rosenthal, A. Gerasimova, R. M. Kagan, B. Anderson, M. Hua, Y. Liu, L.E. Bernstein, K.E. Livingston, A. Perez, I. A. Shlyakhter, R. V. Rolando, R. Owen, P. Tanpaiboon, F. Lacbawan, Clinton R. Paden, Duncan MacCannell |
| EPI_ISL_3395844, EPI_ISL_3395845 | OLVZ Aalst | OLVZ Aalst | Astrid Holderbeke |

| | | | |
|---|---|---|---|
| EPI_ISL_3395846 | Quest Diagnostics Incorporated | Centers for Disease Control and Prevention Division of Viral Diseases, Pathogen Discovery | Dakota Howard, Dhwani Batra, Peter W. Cook, Kara Moser, Adrian Paskey, Jason Caravas, Benjamin Rambo-Martin, Shatavia Morrison, Christopher Gulvick, Scott Sammons, Yvette Unoarumhi, Darlene Wagner, Matthew Schmerer, S. H. Rosenthal, A. Gerasimova, R. M. Kagan, B. Anderson, M. Hua, Y. Liu, L.E. Bernstein, K.E. Livingston, A. Perez, I. A. Shlyakhter, R. V. Rolando, R. Owen, P. Tanpaiboon, F. Lacbawan, Clinton R. Paden, Duncan MacCannell |
| EPI_ISL_3395847 | OLVZ Aalst | OLVZ Aalst | Astrid Holderbeke |
| EPI_ISL_3395848 | Quest Diagnostics Incorporated | Centers for Disease Control and Prevention Division of Viral Diseases, Pathogen Discovery | Dakota Howard, Dhwani Batra, Peter W. Cook, Kara Moser, Adrian Paskey, Jason Caravas, Benjamin Rambo-Martin, Shatavia Morrison, Christopher Gulvick, Scott Sammons, Yvette Unoarumhi, Darlene Wagner, Matthew Schmerer, S. H. Rosenthal, A. Gerasimova, R. M. Kagan, B. Anderson, M. Hua, Y. Liu, L.E. Bernstein, K.E. Livingston, A. Perez, I. A. Shlyakhter, R. V. Rolando, R. Owen, P. Tanpaiboon, F. Lacbawan, Clinton R. Paden, Duncan MacCannell |
| EPI_ISL_3395849, EPI_ISL_3395850 | OLVZ Aalst | OLVZ Aalst | Astrid Holderbeke |
| EPI_ISL_3395851 | Quest Diagnostics Incorporated | Centers for Disease Control and Prevention Division of Viral Diseases, Pathogen Discovery | Dakota Howard, Dhwani Batra, Peter W. Cook, Kara Moser, Adrian Paskey, Jason Caravas, Benjamin Rambo-Martin, Shatavia Morrison, Christopher Gulvick, Scott Sammons, Yvette Unoarumhi, Darlene Wagner, Matthew Schmerer, S. H. Rosenthal, A. Gerasimova, R. M. Kagan, B. Anderson, M. Hua, Y. Liu, L.E. Bernstein, K.E. Livingston, A. Perez, I. A. Shlyakhter, R. V. Rolando, R. Owen, P. Tanpaiboon, F. Lacbawan, Clinton R. Paden, Duncan MacCannell |
| EPI_ISL_3395852 | OLVZ Aalst | OLVZ Aalst | Astrid Holderbeke |
| EPI_ISL_3395853 | Quest Diagnostics Incorporated | Centers for Disease Control and Prevention Division of Viral Diseases, Pathogen Discovery | Dakota Howard, Dhwani Batra, Peter W. Cook, Kara Moser, Adrian Paskey, Jason Caravas, Benjamin Rambo-Martin, Shatavia Morrison, Christopher Gulvick, Scott Sammons, Yvette Unoarumhi, Darlene Wagner, Matthew Schmerer, S. H. Rosenthal, A. Gerasimova, R. M. Kagan, B. Anderson, M. Hua, Y. Liu, L.E. Bernstein, K.E. Livingston, A. Perez, I. A. Shlyakhter, R. V. Rolando, R. Owen, P. Tanpaiboon, F. Lacbawan, Clinton R. Paden, Duncan MacCannell |
| EPI_ISL_3395854, EPI_ISL_3395855 | OLVZ Aalst | OLVZ Aalst | Astrid Holderbeke |
| EPI_ISL_3395856 | Quest Diagnostics Incorporated | Centers for Disease Control and Prevention Division of Viral Diseases, Pathogen Discovery | Dakota Howard, Dhwani Batra, Peter W. Cook, Kara Moser, Adrian Paskey, Jason Caravas, Benjamin Rambo-Martin, Shatavia Morrison, Christopher Gulvick, Scott Sammons, Yvette Unoarumhi, Darlene Wagner, Matthew Schmerer, S. H. Rosenthal, A. Gerasimova, R. M. Kagan, B. Anderson, M. Hua, Y. Liu, L.E. Bernstein, K.E. Livingston, A. Perez, I. A. Shlyakhter, R. V. Rolando, R. Owen, P. Tanpaiboon, F. Lacbawan, Clinton R. Paden, Duncan MacCannell |
| EPI_ISL_3395857 | OLVZ Aalst | OLVZ Aalst | Astrid Holderbeke |
| EPI_ISL_3395858 | Quest Diagnostics Incorporated | Centers for Disease Control and Prevention Division of Viral Diseases, Pathogen Discovery | Dakota Howard, Dhwani Batra, Peter W. Cook, Kara Moser, Adrian Paskey, Jason Caravas, Benjamin Rambo-Martin, Shatavia Morrison, Christopher Gulvick, Scott Sammons, Yvette Unoarumhi, Darlene Wagner, Matthew Schmerer, S. H. Rosenthal, A. Gerasimova, R. M. Kagan, B. Anderson, M. Hua, Y. Liu, L.E. Bernstein, K.E. Livingston, A. Perez, I. A. Shlyakhter, R. V. Rolando, R. Owen, P. Tanpaiboon, F. Lacbawan, Clinton R. Paden, Duncan MacCannell |
| EPI_ISL_3395859 | OLVZ Aalst | OLVZ Aalst | Astrid Holderbeke |
| EPI_ISL_3395860 | Quest Diagnostics Incorporated | Centers for Disease Control and Prevention Division of Viral Diseases, Pathogen Discovery | Dakota Howard, Dhwani Batra, Peter W. Cook, Kara Moser, Adrian Paskey, Jason Caravas, Benjamin Rambo-Martin, Shatavia Morrison, Christopher Gulvick, Scott Sammons, Yvette Unoarumhi, Darlene Wagner, Matthew Schmerer, S. H. Rosenthal, A. Gerasimova, R. M. Kagan, B. Anderson, M. Hua, Y. Liu, L.E. Bernstein, K.E. Livingston, A. Perez, I. A. Shlyakhter, R. V. Rolando, R. Owen, P. Tanpaiboon, F. Lacbawan, Clinton R. Paden, Duncan MacCannell |
| EPI_ISL_3395861, EPI_ISL_3395862 | OLVZ Aalst | OLVZ Aalst | Astrid Holderbeke |
| EPI_ISL_3395863 | Quest Diagnostics Incorporated | Centers for Disease Control and Prevention Division of Viral Diseases, Pathogen Discovery | Dakota Howard, Dhwani Batra, Peter W. Cook, Kara Moser, Adrian Paskey, Jason Caravas, Benjamin Rambo-Martin, Shatavia Morrison, Christopher Gulvick, Scott Sammons, Yvette Unoarumhi, Darlene Wagner, Matthew Schmerer, S. H. Rosenthal, A. Gerasimova, R. M. Kagan, B. Anderson, M. Hua, Y. Liu, L.E. Bernstein, K.E. Livingston, A. Perez, I. A. Shlyakhter, R. V. Rolando, R. Owen, P. Tanpaiboon, F. Lacbawan, Clinton R. Paden, Duncan MacCannell |
| EPI_ISL_3395864 | OLVZ Aalst | OLVZ Aalst | Astrid Holderbeke |
| EPI_ISL_3395865 | Quest Diagnostics Incorporated | Centers for Disease Control and Prevention Division of Viral Diseases, Pathogen Discovery | Dakota Howard, Dhwani Batra, Peter W. Cook, Kara Moser, Adrian Paskey, Jason Caravas, Benjamin Rambo-Martin, Shatavia Morrison, Christopher Gulvick, Scott Sammons, Yvette Unoarumhi, Darlene Wagner, Matthew Schmerer, S. H. Rosenthal, A. Gerasimova, R. M. Kagan, B. Anderson, M. Hua, Y. Liu, L.E. Bernstein, K.E. Livingston, A. Perez, I. A. Shlyakhter, R. V. Rolando, R. Owen, P. Tanpaiboon, F. Lacbawan, Clinton R. Paden, Duncan MacCannell |
| EPI_ISL_3395866, EPI_ISL_3395867 | OLVZ Aalst | OLVZ Aalst | Astrid Holderbeke |
| EPI_ISL_3395868, EPI_ISL_3395869, EPI_ISL_3395870, EPI_ISL_3395871, EPI_ISL_3395872, EPI_ISL_3395873, EPI_ISL_3395874, EPI_ISL_3395875, EPI_ISL_3395876, EPI_ISL_3395877, EPI_ISL_3395878, EPI_ISL_3395879, EPI_ISL_3395880, EPI_ISL_3395881, EPI_ISL_3395882, EPI_ISL_3395883, EPI_ISL_3395884, EPI_ISL_3395885, EPI_ISL_3395886, EPI_ISL_3395887, EPI_ISL_3395888, EPI_ISL_3395889, EPI_ISL_3395890, EPI_ISL_3395891, EPI_ISL_3395892, EPI_ISL_3395893, EPI_ISL_3395894, EPI_ISL_3395895, EPI_ISL_3395896, EPI_ISL_3395897, EPI_ISL_3395898, EPI_ISL_3395899, EPI_ISL_3395900, EPI_ISL_3395901, EPI_ISL_3395902, EPI_ISL_3395903, EPI_ISL_3395904, EPI_ISL_3395905, EPI_ISL_3395906, EPI_ISL_3395907, EPI_ISL_3395908, EPI_ISL_3395909, EPI_ISL_3395910, EPI_ISL_3395911, EPI_ISL_3395912, EPI_ISL_3395913, EPI_ISL_3395914, EPI_ISL_3395915, EPI_ISL_3395916, EPI_ISL_3395917, EPI_ISL_3395918, EPI_ISL_3395919, EPI_ISL_3395920, EPI_ISL_3395921, EPI_ISL_3395922, EPI_ISL_3395923, EPI_ISL_3395924, EPI_ISL_3395925, EPI_ISL_3395926, EPI_ISL_3395927, EPI_ISL_3395928, EPI_ISL_3395929, EPI_ISL_3395930, EPI_ISL_3395931, EPI_ISL_3395933, EPI_ISL_3395934, EPI_ISL_3395935, EPI_ISL_3395936, EPI_ISL_3395937, EPI_ISL_3395938, EPI_ISL_3395939, EPI_ISL_3395940, EPI_ISL_3395941, EPI_ISL_3395942, EPI_ISL_3395943, EPI_ISL_3395944, EPI_ISL_3395945, EPI_ISL_3395946, EPI_ISL_3395947, EPI_ISL_3395948, EPI_ISL_3395949, EPI_ISL_3395950, EPI_ISL_3395951, EPI_ISL_3395952, EPI_ISL_3395953, EPI_ISL_3395954, EPI_ISL_3395955, EPI_ISL_3395956, EPI_ISL_3395958, EPI_ISL_3395959, EPI_ISL_3395960, EPI_ISL_3395961, EPI_ISL_3395962, EPI_ISL_3395963, EPI_ISL_3395964, EPI_ISL_3395965, EPI_ISL_3395966, EPI_ISL_3395967, EPI_ISL_3395968, EPI_ISL_3395969, EPI_ISL_3395970, EPI_ISL_3395971, EPI_ISL_3395972, EPI_ISL_3395973, EPI_ISL_3395975, EPI_ISL_3395976, EPI_ISL_3395977, EPI_ISL_3395978, EPI_ISL_3395979, EPI_ISL_3395980, EPI_ISL_3395981, EPI_ISL_3395982, EPI_ISL_3395983, EPI_ISL_3395984, EPI_ISL_3395985, EPI_ISL_3395986, EPI_ISL_3395987, EPI_ISL_3395988, EPI_ISL_3395989, EPI_ISL_3395990, EPI_ISL_3395991, EPI_ISL_3395992, EPI_ISL_3395993, EPI_ISL_3395994, EPI_ISL_3395995, EPI_ISL_3395996, EPI_ISL_3395997, EPI_ISL_3395998, EPI_ISL_3395999, EPI_ISL_3396000, EPI_ISL_3396001, EPI_ISL_3396002, EPI_ISL_3396003, EPI_ISL_3396004, EPI_ISL_3396005, EPI_ISL_3396006, EPI_ISL_3396007, EPI_ISL_3396008, EPI_ISL_3396009, EPI_ISL_3396010, EPI_ISL_3396011, EPI_ISL_3396012, EPI_ISL_3396013, EPI_ISL_3396014, EPI_ISL_3396015, EPI_ISL_3396016, EPI_ISL_3396017, EPI_ISL_3396018, EPI_ISL_3396019, EPI_ISL_3396020, EPI_ISL_3396021, EPI_ISL_3396022, EPI_ISL_3396023, EPI_ISL_3396024, EPI_ISL_3396025, EPI_ISL_3396026, EPI_ISL_3396027, EPI_ISL_3396028, EPI_ISL_3396029, EPI_ISL_3396030, EPI_ISL_3396031, EPI_ISL_3396032, EPI_ISL_3396033, EPI_ISL_3396034, EPI_ISL_3396035, EPI_ISL_3396037, EPI_ISL_3396042, EPI_ISL_3396045, EPI_ISL_3396047, EPI_ISL_3396049, EPI_ISL_3396052, EPI_ISL_3396054, EPI_ISL_3396057, EPI_ISL_3396059, EPI_ISL_3396061, EPI_ISL_3396063, EPI_ISL_3396065, EPI_ISL_3396068, EPI_ISL_3396069, EPI_ISL_3396070, EPI_ISL_3396071, EPI_ISL_3396072, EPI_ISL_3396073, EPI_ISL_3396074, EPI_ISL_3396075, EPI_ISL_3396076, EPI_ISL_3396077, EPI_ISL_3396078, EPI_ISL_3396079, EPI_ISL_3396080, EPI_ISL_3396081, EPI_ISL_3396082, EPI_ISL_3396083, EPI_ISL_3396084, EPI_ISL_3396085, EPI_ISL_3396086, EPI_ISL_3396087, EPI_ISL_3396088, EPI_ISL_3396090, EPI_ISL_3396091, EPI_ISL_3396092, EPI_ISL_3396093, EPI_ISL_3396094, EPI_ISL_3396095, EPI_ISL_3396096, EPI_ISL_3396097, EPI_ISL_3396098, EPI_ISL_3396099, EPI_ISL_3396100, EPI_ISL_3396101, EPI_ISL_3396102, EPI_ISL_3396103, EPI_ISL_3396105, EPI_ISL_3396106, EPI_ISL_3396107, EPI_ISL_3396108, EPI_ISL_3396109, EPI_ISL_3396110, EPI_ISL_3396111, EPI_ISL_3396112, EPI_ISL_3396113, EPI_ISL_3396114, EPI_ISL_3396115, EPI_ISL_3396116, EPI_ISL_3396117, EPI_ISL_3396118, EPI_ISL_3396119, EPI_ISL_3396120, EPI_ISL_3396121, EPI_ISL_3396122, EPI_ISL_3396123, EPI_ISL_3396124, EPI_ISL_3396125, EPI_ISL_3396126, EPI_ISL_3396127, EPI_ISL_3396128, EPI_ISL_3396129, EPI_ISL_3396130, EPI_ISL_3396131, EPI_ISL_3396132, EPI_ISL_3396133, EPI_ISL_3396134, EPI_ISL_3396135, EPI_ISL_3396136, EPI_ISL_3396137, EPI_ISL_3396138, EPI_ISL_3396139, EPI_ISL_3396140, EPI_ISL_3396141, EPI_ISL_3396142, EPI_ISL_3396143, EPI_ISL_3396145, EPI_ISL_3396146, EPI_ISL_3396147, EPI_ISL_3396148, EPI_ISL_3396149, EPI_ISL_3396150, EPI_ISL_3396151, EPI_ISL_3396152, EPI_ISL_3396153, EPI_ISL_3396154, EPI_ISL_3396155, EPI_ISL_3396156, EPI_ISL_3396157, EPI_ISL_3396158, EPI_ISL_3396159, EPI_ISL_3396160, EPI_ISL_3396161, EPI_ISL_3396162, EPI_ISL_3396163, EPI_ISL_3396164, EPI_ISL_3396165, EPI_ISL_3396166, EPI_ISL_3396167, EPI_ISL_3396168, EPI_ISL_3396169, EPI_ISL_3396170, EPI_ISL_3396171, EPI_ISL_3396172, EPI_ISL_3396173, EPI_ISL_3396174, EPI_ISL_3396175, EPI_ISL_3396176, EPI_ISL_3396177, EPI_ISL_3396178, EPI_ISL_3396179, EPI_ISL_3396180, EPI_ISL_3396181, EPI_ISL_3396182 | | | |
| see above | Quest Diagnostics Incorporated | Centers for Disease Control and Prevention Division of Viral Diseases, Pathogen Discovery | Dakota Howard, Dhwani Batra, Peter W. Cook, Kara Moser, Adrian Paskey, Jason Caravas, Benjamin Rambo-Martin, Shatavia Morrison, Christopher Gulvick, Scott Sammons, Yvette Unoarumhi, Darlene Wagner, Matthew Schmerer, S. H. Rosenthal, A. Gerasimova, R. M. Kagan, B. Anderson, M. Hua, Y. Liu, L.E. Bernstein, K.E. Livingston, A. Perez, I. A. Shlyakhter, R. V. Rolando, R. Owen, P. Tanpaiboon, F. Lacbawan, Clinton R. Paden, Duncan MacCannell |
| EPI_ISL_3396183 | MD PHL | Maryland Department of Health Laboratories Administration | Eric N. Keller, Jillian Loomis, Kwang Low, Terence L. Moore, Ami Patel, and Robert Myers |
| EPI_ISL_3396184 | Quest Diagnostics Incorporated | Centers for Disease Control and Prevention Division of Viral Diseases, Pathogen Discovery | Dakota Howard, Dhwani Batra, Peter W. Cook, Kara Moser, Adrian Paskey, Jason Caravas, Benjamin Rambo-Martin, Shatavia Morrison, Christopher Gulvick, Scott Sammons, Yvette Unoarumhi, Darlene Wagner, Matthew Schmerer, S. H. Rosenthal, A. Gerasimova, R. M. Kagan, B. Anderson, M. Hua, Y. Liu, L.E. Bernstein, K.E. Livingston, A. Perez, I. A. Shlyakhter, R. V. Rolando, R. Owen, P. Tanpaiboon, F. Lacbawan, Clinton R. Paden, Duncan MacCannell |
| EPI_ISL_3396185, EPI_ISL_3396186 | MD PHL | Maryland Department of Health Laboratories Administration | Eric N. Keller, Jillian Loomis, Kwang Low, Terence L. Moore, Ami Patel, and Robert Myers |
| EPI_ISL_3396187 | Quest Diagnostics Incorporated | Centers for Disease Control and Prevention Division of Viral Diseases, Pathogen Discovery | Dakota Howard, Dhwani Batra, Peter W. Cook, Kara Moser, Adrian Paskey, Jason Caravas, Benjamin Rambo-Martin, Shatavia Morrison, Christopher Gulvick, Scott Sammons, Yvette Unoarumhi, Darlene Wagner, Matthew Schmerer, S. H. Rosenthal, A. Gerasimova, R. M. Kagan, B. Anderson, M. Hua, Y. Liu, L.E. Bernstein, K.E. Livingston, A. Perez, I. A. Shlyakhter, R. V. Rolando, R. Owen, P. Tanpaiboon, F. Lacbawan, Clinton R. Paden, Duncan MacCannell |
| EPI_ISL_3396188 | MD PHL | Maryland Department of Health Laboratories Administration | Eric N. Keller, Jillian Loomis, Kwang Low, Terence L. Moore, Ami Patel, and Robert Myers |
| EPI_ISL_3396189 | Quest Diagnostics Incorporated | Centers for Disease Control and Prevention Division of Viral Diseases, Pathogen Discovery | Dakota Howard, Dhwani Batra, Peter W. Cook, Kara Moser, Adrian Paskey, Jason Caravas, Benjamin Rambo-Martin, Shatavia Morrison, Christopher Gulvick, Scott Sammons, Yvette Unoarumhi, Darlene Wagner, Matthew Schmerer, S. H. Rosenthal, A. Gerasimova, R. M. Kagan, B. Anderson, M. Hua, Y. Liu, L.E. Bernstein, K.E. Livingston, A. Perez, I. A. Shlyakhter, R. V. Rolando, R. Owen, P. Tanpaiboon, F. Lacbawan, Clinton R. Paden, Duncan MacCannell |
| EPI_ISL_3396190, EPI_ISL_3396191 | MD PHL | Maryland Department of Health Laboratories Administration | Eric N. Keller, Jillian Loomis, Kwang Low, Terence L. Moore, Ami Patel, and Robert Myers |
| EPI_ISL_3396192 | Quest Diagnostics Incorporated | Centers for Disease Control and Prevention Division of Viral Diseases, Pathogen Discovery | Dakota Howard, Dhwani Batra, Peter W. Cook, Kara Moser, Adrian Paskey, Jason Caravas, Benjamin Rambo-Martin, Shatavia Morrison, Christopher Gulvick, Scott Sammons, Yvette Unoarumhi, Darlene Wagner, Matthew Schmerer, S. H. Rosenthal, A. Gerasimova, R. M. Kagan, B. Anderson, M. Hua, Y. Liu, L.E. Bernstein, K.E. Livingston, A. Perez, I. A. Shlyakhter, R. V. Rolando, R. Owen, P. Tanpaiboon, F. Lacbawan, Clinton R. Paden, Duncan MacCannell |
| EPI_ISL_3396193 | MD PHL | Maryland Department of Health Laboratories Administration | Eric N. Keller, Jillian Loomis, Kwang Low, Terence L. Moore, Ami Patel, and Robert Myers |
| EPI_ISL_3396194 | Quest Diagnostics Incorporated | Centers for Disease Control and Prevention Division of Viral Diseases, Pathogen Discovery | Dakota Howard, Dhwani Batra, Peter W. Cook, Kara Moser, Adrian Paskey, Jason Caravas, Benjamin Rambo-Martin, Shatavia Morrison, Christopher Gulvick, Scott Sammons, Yvette Unoarumhi, Darlene Wagner, Matthew Schmerer, S. H. Rosenthal, A. Gerasimova, R. M. Kagan, B. Anderson, M. Hua, Y. Liu, L.E. Bernstein, K.E. Livingston, A. Perez, I. A. Shlyakhter, R. V. Rolando, R. Owen, P. Tanpaiboon, F. Lacbawan, Clinton R. Paden, Duncan MacCannell |
| EPI_ISL_3396195, EPI_ISL_3396196 | MD PHL | Maryland Department of Health Laboratories Administration | Eric N. Keller, Jillian Loomis, Kwang Low, Terence L. Moore, Ami Patel, and Robert Myers |
| EPI_ISL_3396197 | Quest Diagnostics Incorporated | Centers for Disease Control and Prevention Division of Viral Diseases, Pathogen Discovery | Dakota Howard, Dhwani Batra, Peter W. Cook, Kara Moser, Adrian Paskey, Jason Caravas, Benjamin Rambo-Martin, Shatavia Morrison, Christopher Gulvick, Scott Sammons, Yvette Unoarumhi, Darlene Wagner, Matthew Schmerer, S. H. Rosenthal, A. Gerasimova, R. M. Kagan, B. Anderson, M. Hua, Y. Liu, L.E. Bernstein, K.E. Livingston, A. Perez, I. A. Shlyakhter, R. V. Rolando, R. Owen, P. Tanpaiboon, F. Lacbawan, Clinton R. Paden, Duncan MacCannell |
| EPI_ISL_3396198, EPI_ISL_3396199 | MD PHL | Maryland Department of Health Laboratories Administration | Eric N. Keller, Jillian Loomis, Kwang Low, Terence L. Moore, Ami Patel, and Robert Myers |
| EPI_ISL_3396200 | Quest Diagnostics Incorporated | Centers for Disease Control and Prevention Division of Viral Diseases, Pathogen Discovery | Dakota Howard, Dhwani Batra, Peter W. Cook, Kara Moser, Adrian Paskey, Jason Caravas, Benjamin Rambo-Martin, Shatavia Morrison, Christopher Gulvick, Scott Sammons, Yvette Unoarumhi, Darlene Wagner, Matthew Schmerer, S. H. Rosenthal, A. Gerasimova, R. M. Kagan, B. Anderson, M. Hua, Y. Liu, L.E. Bernstein, K.E. Livingston, A. Perez, I. A. Shlyakhter, R. V. Rolando, R. Owen, P. Tanpaiboon, F. Lacbawan, Clinton R. Paden, Duncan MacCannell |
| EPI_ISL_3396201, EPI_ISL_3396202 | MD PHL | Maryland Department of Health Laboratories Administration | Eric N. Keller, Jillian Loomis, Kwang Low, Terence L. Moore, Ami Patel, and Robert Myers |
| EPI_ISL_3396203 | Quest Diagnostics Incorporated | Centers for Disease Control and Prevention Division of Viral Diseases, Pathogen Discovery | Dakota Howard, Dhwani Batra, Peter W. Cook, Kara Moser, Adrian Paskey, Jason Caravas, Benjamin Rambo-Martin, Shatavia Morrison, Christopher Gulvick, Scott Sammons, Yvette Unoarumhi, Darlene Wagner, Matthew Schmerer, S. H. Rosenthal, A. Gerasimova, R. M. Kagan, B. Anderson, M. Hua, Y. Liu, L.E. Bernstein, K.E. Livingston, A. Perez, I. A. Shlyakhter, R. V. Rolando, R. Owen, P. Tanpaiboon, F. Lacbawan, Clinton R. Paden, Duncan MacCannell |
| EPI_ISL_3396204 | MD PHL | Maryland Department of Health Laboratories Administration | Eric N. Keller, Jillian Loomis, Kwang Low, Terence L. Moore, Ami Patel, and Robert Myers |
| EPI_ISL_3396205 | Quest Diagnostics Incorporated | Centers for Disease Control and Prevention Division of Viral Diseases, Pathogen Discovery | Dakota Howard, Dhwani Batra, Peter W. Cook, Kara Moser, Adrian Paskey, Jason Caravas, Benjamin Rambo-Martin, Shatavia Morrison, Christopher Gulvick, Scott Sammons, Yvette Unoarumhi, Darlene Wagner, Matthew Schmerer, S. H. Rosenthal, A. Gerasimova, R. M. Kagan, B. Anderson, M. Hua, Y. Liu, L.E. Bernstein, K.E. Livingston, A. Perez, I. A. Shlyakhter, R. V. Rolando, R. Owen, P. Tanpaiboon, F. Lacbawan, Clinton R. Paden, Duncan MacCannell |
| EPI_ISL_3396206 | MD PHL | Maryland Department of Health Laboratories Administration | Eric N. Keller, Jillian Loomis, Kwang Low, Terence L. Moore, Ami Patel, and Robert Myers |
| EPI_ISL_3396207 | Quest Diagnostics Incorporated | Centers for Disease Control and Prevention Division of Viral Diseases, Pathogen Discovery | Dakota Howard, Dhwani Batra, Peter W. Cook, Kara Moser, Adrian Paskey, Jason Caravas, Benjamin Rambo-Martin, Shatavia Morrison, Christopher Gulvick, Scott Sammons, Yvette Unoarumhi, Darlene Wagner, Matthew Schmerer, S. H. Rosenthal, A. Gerasimova, R. M. Kagan, B. Anderson, M. Hua, Y. Liu, L.E. Bernstein, K.E. Livingston, A. Perez, I. A. Shlyakhter, R. V. Rolando, R. Owen, P. Tanpaiboon, F. Lacbawan, Clinton R. Paden, Duncan MacCannell |
| EPI_ISL_3396208, EPI_ISL_3396209 | MD PHL | Maryland Department of Health Laboratories Administration | Eric N. Keller, Jillian Loomis, Kwang Low, Terence L. Moore, Ami Patel, and Robert Myers |
| EPI_ISL_3396210 | Quest Diagnostics Incorporated | Centers for Disease Control and Prevention Division of Viral Diseases, Pathogen Discovery | Dakota Howard, Dhwani Batra, Peter W. Cook, Kara Moser, Adrian Paskey, Jason Caravas, Benjamin Rambo-Martin, Shatavia Morrison, Christopher Gulvick, Scott Sammons, Yvette Unoarumhi, Darlene Wagner, Matthew Schmerer, S. H. Rosenthal, A. Gerasimova, R. M. Kagan, B. Anderson, M. Hua, Y. Liu, L.E. Bernstein, K.E. Livingston, A. Perez, I. A. Shlyakhter, R. V. Rolando, R. Owen, P. Tanpaiboon, F. Lacbawan, Clinton R. Paden, Duncan MacCannell |
| EPI_ISL_3396211, EPI_ISL_3396212 | MD PHL | Maryland Department of Health Laboratories Administration | Eric N. Keller, Jillian Loomis, Kwang Low, Terence L. Moore, Ami Patel, and Robert Myers |
| EPI_ISL_3396213 | Quest Diagnostics Incorporated | Centers for Disease Control and Prevention Division of Viral Diseases, Pathogen Discovery | Dakota Howard, Dhwani Batra, Peter W. Cook, Kara Moser, Adrian Paskey, Jason Caravas, Benjamin Rambo-Martin, Shatavia Morrison, Christopher Gulvick, Scott Sammons, Yvette Unoarumhi, Darlene Wagner, Matthew Schmerer, S. H. Rosenthal, A. Gerasimova, R. M. Kagan, B. Anderson, M. Hua, Y. Liu, L.E. Bernstein, K.E. Livingston, A. Perez, I. A. Shlyakhter, R. V. Rolando, R. Owen, P. Tanpaiboon, F. Lacbawan, Clinton R. Paden, Duncan MacCannell |

| | | | |
|---|---|---|---|
| EPI_ISL_3396214 | MD PHL | Maryland Department of Health Laboratories Administration | Eric N. Keller, Jillian Loomis, Kwang Low, Terence L. Moore, Ami Patel, and Robert Myers |
| EPI_ISL_3396215 | Quest Diagnostics Incorporated | Centers for Disease Control and Prevention Division of Viral Diseases, Pathogen Discovery | Dakota Howard, Dhwani Batra, Peter W. Cook, Kara Moser, Adrian Paskey, Jason Caravas, Benjamin Rambo-Martin, Shatavia Morrison, Christopher Gulvick, Scott Sammons, Yvette Unoarumhi, Darlene Wagner, Matthew Schmerer, S. H. Rosenthal, A. Gerasimova, R. M. Kagan, B. Anderson, M. Hua, Y. Liu, L.E. Bernstein, K.E. Livingston, A. Perez, I. A. Shlyakhter, R. V. Rolando, R. Owen, P. Tanpaiboon, F. Lacbawan, Clinton R. Paden, Duncan MacCannell |
| EPI_ISL_3396216, EPI_ISL_3396217 | MD PHL | Maryland Department of Health Laboratories Administration | Eric N. Keller, Jillian Loomis, Kwang Low, Terence L. Moore, Ami Patel, and Robert Myers |
| EPI_ISL_3396218, EPI_ISL_3396220 | Quest Diagnostics Incorporated | Centers for Disease Control and Prevention Division of Viral Diseases, Pathogen Discovery | Dakota Howard, Dhwani Batra, Peter W. Cook, Kara Moser, Adrian Paskey, Jason Caravas, Benjamin Rambo-Martin, Shatavia Morrison, Christopher Gulvick, Scott Sammons, Yvette Unoarumhi, Darlene Wagner, Matthew Schmerer, S. H. Rosenthal, A. Gerasimova, R. M. Kagan, B. Anderson, M. Hua, Y. Liu, L.E. Bernstein, K.E. Livingston, A. Perez, I. A. Shlyakhter, R. V. Rolando, R. Owen, P. Tanpaiboon, F. Lacbawan, Clinton R. Paden, Duncan MacCannell |
| EPI_ISL_3396221, EPI_ISL_3396222 | MD PHL | Maryland Department of Health Laboratories Administration | Eric N. Keller, Jillian Loomis, Kwang Low, Terence L. Moore, Ami Patel, and Robert Myers |
| EPI_ISL_3396223 | Quest Diagnostics Incorporated | Centers for Disease Control and Prevention Division of Viral Diseases, Pathogen Discovery | Dakota Howard, Dhwani Batra, Peter W. Cook, Kara Moser, Adrian Paskey, Jason Caravas, Benjamin Rambo-Martin, Shatavia Morrison, Christopher Gulvick, Scott Sammons, Yvette Unoarumhi, Darlene Wagner, Matthew Schmerer, S. H. Rosenthal, A. Gerasimova, R. M. Kagan, B. Anderson, M. Hua, Y. Liu, L.E. Bernstein, K.E. Livingston, A. Perez, I. A. Shlyakhter, R. V. Rolando, R. Owen, P. Tanpaiboon, F. Lacbawan, Clinton R. Paden, Duncan MacCannell |
| EPI_ISL_3396224 | MD PHL | Maryland Department of Health Laboratories Administration | Eric N. Keller, Jillian Loomis, Kwang Low, Terence L. Moore, Ami Patel, and Robert Myers |
| EPI_ISL_3396225 | Quest Diagnostics Incorporated | Centers for Disease Control and Prevention Division of Viral Diseases, Pathogen Discovery | Dakota Howard, Dhwani Batra, Peter W. Cook, Kara Moser, Adrian Paskey, Jason Caravas, Benjamin Rambo-Martin, Shatavia Morrison, Christopher Gulvick, Scott Sammons, Yvette Unoarumhi, Darlene Wagner, Matthew Schmerer, S. H. Rosenthal, A. Gerasimova, R. M. Kagan, B. Anderson, M. Hua, Y. Liu, L.E. Bernstein, K.E. Livingston, A. Perez, I. A. Shlyakhter, R. V. Rolando, R. Owen, P. Tanpaiboon, F. Lacbawan, Clinton R. Paden, Duncan MacCannell |
| EPI_ISL_3396226, EPI_ISL_3396227 | MD PHL | Maryland Department of Health Laboratories Administration | Eric N. Keller, Jillian Loomis, Kwang Low, Terence L. Moore, Ami Patel, and Robert Myers |
| EPI_ISL_3396228 | Quest Diagnostics Incorporated | Centers for Disease Control and Prevention Division of Viral Diseases, Pathogen Discovery | Dakota Howard, Dhwani Batra, Peter W. Cook, Kara Moser, Adrian Paskey, Jason Caravas, Benjamin Rambo-Martin, Shatavia Morrison, Christopher Gulvick, Scott Sammons, Yvette Unoarumhi, Darlene Wagner, Matthew Schmerer, S. H. Rosenthal, A. Gerasimova, R. M. Kagan, B. Anderson, M. Hua, Y. Liu, L.E. Bernstein, K.E. Livingston, A. Perez, I. A. Shlyakhter, R. V. Rolando, R. Owen, P. Tanpaiboon, F. Lacbawan, Clinton R. Paden, Duncan MacCannell |
| EPI_ISL_3396229 | MD PHL | Maryland Department of Health Laboratories Administration | Eric N. Keller, Jillian Loomis, Kwang Low, Terence L. Moore, Ami Patel, and Robert Myers |
| EPI_ISL_3396230 | Quest Diagnostics Incorporated | Centers for Disease Control and Prevention Division of Viral Diseases, Pathogen Discovery | Dakota Howard, Dhwani Batra, Peter W. Cook, Kara Moser, Adrian Paskey, Jason Caravas, Benjamin Rambo-Martin, Shatavia Morrison, Christopher Gulvick, Scott Sammons, Yvette Unoarumhi, Darlene Wagner, Matthew Schmerer, S. H. Rosenthal, A. Gerasimova, R. M. Kagan, B. Anderson, M. Hua, Y. Liu, L.E. Bernstein, K.E. Livingston, A. Perez, I. A. Shlyakhter, R. V. Rolando, R. Owen, P. Tanpaiboon, F. Lacbawan, Clinton R. Paden, Duncan MacCannell |
| EPI_ISL_3396231, EPI_ISL_3396232 | MD PHL | Maryland Department of Health Laboratories Administration | Eric N. Keller, Jillian Loomis, Kwang Low, Terence L. Moore, Ami Patel, and Robert Myers |
| EPI_ISL_3396233 | Quest Diagnostics Incorporated | Centers for Disease Control and Prevention Division of Viral Diseases, Pathogen Discovery | Dakota Howard, Dhwani Batra, Peter W. Cook, Kara Moser, Adrian Paskey, Jason Caravas, Benjamin Rambo-Martin, Shatavia Morrison, Christopher Gulvick, Scott Sammons, Yvette Unoarumhi, Darlene Wagner, Matthew Schmerer, S. H. Rosenthal, A. Gerasimova, R. M. Kagan, B. Anderson, M. Hua, Y. Liu, L.E. Bernstein, K.E. Livingston, A. Perez, I. A. Shlyakhter, R. V. Rolando, R. Owen, P. Tanpaiboon, F. Lacbawan, Clinton R. Paden, Duncan MacCannell |
| EPI_ISL_3396234 | MD PHL | Maryland Department of Health Laboratories Administration | Eric N. Keller, Jillian Loomis, Kwang Low, Terence L. Moore, Ami Patel, and Robert Myers |
| EPI_ISL_3396235 | Quest Diagnostics Incorporated | Centers for Disease Control and Prevention Division of Viral Diseases, Pathogen Discovery | Dakota Howard, Dhwani Batra, Peter W. Cook, Kara Moser, Adrian Paskey, Jason Caravas, Benjamin Rambo-Martin, Shatavia Morrison, Christopher Gulvick, Scott Sammons, Yvette Unoarumhi, Darlene Wagner, Matthew Schmerer, S. H. Rosenthal, A. Gerasimova, R. M. Kagan, B. Anderson, M. Hua, Y. Liu, L.E. Bernstein, K.E. Livingston, A. Perez, I. A. Shlyakhter, R. V. Rolando, R. Owen, P. Tanpaiboon, F. Lacbawan, Clinton R. Paden, Duncan MacCannell |
| EPI_ISL_3396236, EPI_ISL_3396237 | MD PHL | Maryland Department of Health Laboratories Administration | Eric N. Keller, Jillian Loomis, Kwang Low, Terence L. Moore, Ami Patel, and Robert Myers |
| EPI_ISL_3396238 | Quest Diagnostics Incorporated | Centers for Disease Control and Prevention Division of Viral Diseases, Pathogen Discovery | Dakota Howard, Dhwani Batra, Peter W. Cook, Kara Moser, Adrian Paskey, Jason Caravas, Benjamin Rambo-Martin, Shatavia Morrison, Christopher Gulvick, Scott Sammons, Yvette Unoarumhi, Darlene Wagner, Matthew Schmerer, S. H. Rosenthal, A. Gerasimova, R. M. Kagan, B. Anderson, M. Hua, Y. Liu, L.E. Bernstein, K.E. Livingston, A. Perez, I. A. Shlyakhter, R. V. Rolando, R. Owen, P. Tanpaiboon, F. Lacbawan, Clinton R. Paden, Duncan MacCannell |
| EPI_ISL_3396239 | MD PHL | Maryland Department of Health Laboratories Administration | Eric N. Keller, Jillian Loomis, Kwang Low, Terence L. Moore, Ami Patel, and Robert Myers |
| EPI_ISL_3396240 | Quest Diagnostics Incorporated | Centers for Disease Control and Prevention Division of Viral Diseases, Pathogen Discovery | Dakota Howard, Dhwani Batra, Peter W. Cook, Kara Moser, Adrian Paskey, Jason Caravas, Benjamin Rambo-Martin, Shatavia Morrison, Christopher Gulvick, Scott Sammons, Yvette Unoarumhi, Darlene Wagner, Matthew Schmerer, S. H. Rosenthal, A. Gerasimova, R. M. Kagan, B. Anderson, M. Hua, Y. Liu, L.E. Bernstein, K.E. Livingston, A. Perez, I. A. Shlyakhter, R. V. Rolando, R. Owen, P. Tanpaiboon, F. Lacbawan, Clinton R. Paden, Duncan MacCannell |
| EPI_ISL_3396241 | MD PHL | Maryland Department of Health Laboratories Administration | Eric N. Keller, Jillian Loomis, Kwang Low, Terence L. Moore, Ami Patel, and Robert Myers |
| EPI_ISL_3396242, EPI_ISL_3396243 | Quest Diagnostics Incorporated | Centers for Disease Control and Prevention Division of Viral Diseases, Pathogen Discovery | Dakota Howard, Dhwani Batra, Peter W. Cook, Kara Moser, Adrian Paskey, Jason Caravas, Benjamin Rambo-Martin, Shatavia Morrison, Christopher Gulvick, Scott Sammons, Yvette Unoarumhi, Darlene Wagner, Matthew Schmerer, S. H. Rosenthal, A. Gerasimova, R. M. Kagan, B. Anderson, M. Hua, Y. Liu, L.E. Bernstein, K.E. Livingston, A. Perez, I. A. Shlyakhter, R. V. Rolando, R. Owen, P. Tanpaiboon, F. Lacbawan, Clinton R. Paden, Duncan MacCannell |
| EPI_ISL_3396244 | MD PHL | Maryland Department of Health Laboratories Administration | Eric N. Keller, Jillian Loomis, Kwang Low, Terence L. Moore, Ami Patel, and Robert Myers |
| EPI_ISL_3396245, EPI_ISL_3396246 | Quest Diagnostics Incorporated | Centers for Disease Control and Prevention Division of Viral Diseases, Pathogen Discovery | Dakota Howard, Dhwani Batra, Peter W. Cook, Kara Moser, Adrian Paskey, Jason Caravas, Benjamin Rambo-Martin, Shatavia Morrison, Christopher Gulvick, Scott Sammons, Yvette Unoarumhi, Darlene Wagner, Matthew Schmerer, S. H. Rosenthal, A. Gerasimova, R. M. Kagan, B. Anderson, M. Hua, Y. Liu, L.E. Bernstein, K.E. Livingston, A. Perez, I. A. Shlyakhter, R. V. Rolando, R. Owen, P. Tanpaiboon, F. Lacbawan, Clinton R. Paden, Duncan MacCannell |
| EPI_ISL_3396247 | MD PHL | Maryland Department of Health Laboratories Administration | Eric N. Keller, Jillian Loomis, Kwang Low, Terence L. Moore, Ami Patel, and Robert Myers |
| EPI_ISL_3396248 | Quest Diagnostics Incorporated | Centers for Disease Control and Prevention Division of Viral Diseases, Pathogen Discovery | Dakota Howard, Dhwani Batra, Peter W. Cook, Kara Moser, Adrian Paskey, Jason Caravas, Benjamin Rambo-Martin, Shatavia Morrison, Christopher Gulvick, Scott Sammons, Yvette Unoarumhi, Darlene Wagner, Matthew Schmerer, S. H. Rosenthal, A. Gerasimova, R. M. Kagan, B. Anderson, M. Hua, Y. Liu, L.E. Bernstein, K.E. Livingston, A. Perez, I. A. Shlyakhter, R. V. Rolando, R. Owen, P. Tanpaiboon, F. Lacbawan, Clinton R. Paden, Duncan MacCannell |
| EPI_ISL_3396249 | MD PHL | Maryland Department of Health Laboratories Administration | Eric N. Keller, Jillian Loomis, Kwang Low, Terence L. Moore, Ami Patel, and Robert Myers |
| EPI_ISL_3396250 | Quest Diagnostics Incorporated | Centers for Disease Control and Prevention Division of Viral Diseases, Pathogen Discovery | Dakota Howard, Dhwani Batra, Peter W. Cook, Kara Moser, Adrian Paskey, Jason Caravas, Benjamin Rambo-Martin, Shatavia Morrison, Christopher Gulvick, Scott Sammons, Yvette Unoarumhi, Darlene Wagner, Matthew Schmerer, S. H. Rosenthal, A. Gerasimova, R. M. Kagan, B. Anderson, M. Hua, Y. Liu, L.E. Bernstein, K.E. Livingston, A. Perez, I. A. Shlyakhter, R. V. Rolando, R. Owen, P. Tanpaiboon, F. Lacbawan, Clinton R. Paden, Duncan MacCannell |
| EPI_ISL_3396252 | MD PHL | Maryland Department of Health Laboratories Administration | Eric N. Keller, Jillian Loomis, Kwang Low, Terence L. Moore, Ami Patel, and Robert Myers |
| EPI_ISL_3396253 | Quest Diagnostics Incorporated | Centers for Disease Control and Prevention Division of Viral Diseases, Pathogen Discovery | Dakota Howard, Dhwani Batra, Peter W. Cook, Kara Moser, Adrian Paskey, Jason Caravas, Benjamin Rambo-Martin, Shatavia Morrison, Christopher Gulvick, Scott Sammons, Yvette Unoarumhi, Darlene Wagner, Matthew Schmerer, S. H. Rosenthal, A. Gerasimova, R. M. Kagan, B. Anderson, M. Hua, Y. Liu, L.E. Bernstein, K.E. Livingston, A. Perez, I. A. Shlyakhter, R. V. Rolando, R. Owen, P. Tanpaiboon, F. Lacbawan, Clinton R. Paden, Duncan MacCannell |
| EPI_ISL_3396254 | MD PHL | Maryland Department of Health Laboratories Administration | Eric N. Keller, Jillian Loomis, Kwang Low, Terence L. Moore, Ami Patel, and Robert Myers |
| EPI_ISL_3396255, EPI_ISL_3396256 | Quest Diagnostics Incorporated | Centers for Disease Control and Prevention Division of Viral Diseases, Pathogen Discovery | Dakota Howard, Dhwani Batra, Peter W. Cook, Kara Moser, Adrian Paskey, Jason Caravas, Benjamin Rambo-Martin, Shatavia Morrison, Christopher Gulvick, Scott Sammons, Yvette Unoarumhi, Darlene Wagner, Matthew Schmerer, S. H. Rosenthal, A. Gerasimova, R. M. Kagan, B. Anderson, M. Hua, Y. Liu, L.E. Bernstein, K.E. Livingston, A. Perez, I. A. Shlyakhter, R. V. Rolando, R. Owen, P. Tanpaiboon, F. Lacbawan, Clinton R. Paden, Duncan MacCannell |
| EPI_ISL_3396257 | MD PHL | Maryland Department of Health Laboratories Administration | Eric N. Keller, Jillian Loomis, Kwang Low, Terence L. Moore, Ami Patel, and Robert Myers |
| EPI_ISL_3396258, EPI_ISL_3396259 | Quest Diagnostics Incorporated | Centers for Disease Control and Prevention Division of Viral Diseases, Pathogen Discovery | Dakota Howard, Dhwani Batra, Peter W. Cook, Kara Moser, Adrian Paskey, Jason Caravas, Benjamin Rambo-Martin, Shatavia Morrison, Christopher Gulvick, Scott Sammons, Yvette Unoarumhi, Darlene Wagner, Matthew Schmerer, S. H. Rosenthal, A. Gerasimova, R. M. Kagan, B. Anderson, M. Hua, Y. Liu, L.E. Bernstein, K.E. Livingston, A. Perez, I. A. Shlyakhter, R. V. Rolando, R. Owen, P. Tanpaiboon, F. Lacbawan, Clinton R. Paden, Duncan MacCannell |
| EPI_ISL_3396260 | MD PHL | Maryland Department of Health Laboratories Administration | Eric N. Keller, Jillian Loomis, Kwang Low, Terence L. Moore, Ami Patel, and Robert Myers |
| EPI_ISL_3396261 | Quest Diagnostics Incorporated | Centers for Disease Control and Prevention Division of Viral Diseases, Pathogen Discovery | Dakota Howard, Dhwani Batra, Peter W. Cook, Kara Moser, Adrian Paskey, Jason Caravas, Benjamin Rambo-Martin, Shatavia Morrison, Christopher Gulvick, Scott Sammons, Yvette Unoarumhi, Darlene Wagner, Matthew Schmerer, S. H. Rosenthal, A. Gerasimova, R. M. Kagan, B. Anderson, M. Hua, Y. Liu, L.E. Bernstein, K.E. Livingston, A. Perez, I. A. Shlyakhter, R. V. Rolando, R. Owen, P. Tanpaiboon, F. Lacbawan, Clinton R. Paden, Duncan MacCannell |
| EPI_ISL_3396262 | MD PHL | Maryland Department of Health Laboratories Administration | Eric N. Keller, Jillian Loomis, Kwang Low, Terence L. Moore, Ami Patel, and Robert Myers |
| EPI_ISL_3396263, EPI_ISL_3396264 | Quest Diagnostics Incorporated | Centers for Disease Control and Prevention Division of Viral Diseases, Pathogen Discovery | Dakota Howard, Dhwani Batra, Peter W. Cook, Kara Moser, Adrian Paskey, Jason Caravas, Benjamin Rambo-Martin, Shatavia Morrison, Christopher Gulvick, Scott Sammons, Yvette Unoarumhi, Darlene Wagner, Matthew Schmerer, S. H. Rosenthal, A. Gerasimova, R. M. Kagan, B. Anderson, M. Hua, Y. Liu, L.E. Bernstein, K.E. Livingston, A. Perez, I. A. Shlyakhter, R. V. Rolando, R. Owen, P. Tanpaiboon, F. Lacbawan, Clinton R. Paden, Duncan MacCannell |
| EPI_ISL_3396265 | MD PHL | Maryland Department of Health Laboratories Administration | Eric N. Keller, Jillian Loomis, Kwang Low, Terence L. Moore, Ami Patel, and Robert Myers |
| EPI_ISL_3396266, EPI_ISL_3396267 | Quest Diagnostics Incorporated | Centers for Disease Control and Prevention Division of Viral Diseases, Pathogen Discovery | Dakota Howard, Dhwani Batra, Peter W. Cook, Kara Moser, Adrian Paskey, Jason Caravas, Benjamin Rambo-Martin, Shatavia Morrison, Christopher Gulvick, Scott Sammons, Yvette Unoarumhi, Darlene Wagner, Matthew Schmerer, S. H. Rosenthal, A. Gerasimova, R. M. Kagan, B. Anderson, M. Hua, Y. Liu, L.E. Bernstein, K.E. Livingston, A. Perez, I. A. Shlyakhter, R. V. Rolando, R. Owen, P. Tanpaiboon, F. Lacbawan, Clinton R. Paden, Duncan MacCannell |
| EPI_ISL_3396268 | MD PHL | Maryland Department of Health Laboratories Administration | Eric N. Keller, Jillian Loomis, Kwang Low, Terence L. Moore, Ami Patel, and Robert Myers |
| EPI_ISL_3396269 | Quest Diagnostics Incorporated | Centers for Disease Control and Prevention Division of Viral Diseases, Pathogen Discovery | Dakota Howard, Dhwani Batra, Peter W. Cook, Kara Moser, Adrian Paskey, Jason Caravas, Benjamin Rambo-Martin, Shatavia Morrison, Christopher Gulvick, Scott Sammons, Yvette Unoarumhi, Darlene Wagner, Matthew Schmerer, S. H. Rosenthal, A. Gerasimova, R. M. Kagan, B. Anderson, M. Hua, Y. Liu, L.E. Bernstein, K.E. Livingston, A. Perez, I. A. Shlyakhter, R. V. Rolando, R. Owen, P. Tanpaiboon, F. Lacbawan, Clinton R. Paden, Duncan MacCannell |
| EPI_ISL_3396270 | MD PHL | Maryland Department of Health Laboratories Administration | Eric N. Keller, Jillian Loomis, Kwang Low, Terence L. Moore, Ami Patel, and Robert Myers |
| EPI_ISL_3396271, EPI_ISL_3396272 | Quest Diagnostics Incorporated | Centers for Disease Control and Prevention Division of Viral Diseases, Pathogen Discovery | Dakota Howard, Dhwani Batra, Peter W. Cook, Kara Moser, Adrian Paskey, Jason Caravas, Benjamin Rambo-Martin, Shatavia Morrison, Christopher Gulvick, Scott Sammons, Yvette Unoarumhi, Darlene Wagner, Matthew Schmerer, S. H. Rosenthal, A. Gerasimova, R. M. Kagan, B. Anderson, M. Hua, Y. Liu, L.E. Bernstein, K.E. Livingston, A. Perez, I. A. Shlyakhter, R. V. Rolando, R. Owen, P. Tanpaiboon, F. Lacbawan, Clinton R. Paden, Duncan MacCannell |
| EPI_ISL_3396273 | MD PHL | Maryland Department of Health Laboratories Administration | Eric N. Keller, Jillian Loomis, Kwang Low, Terence L. Moore, Ami Patel, and Robert Myers |
| EPI_ISL_3396274 | Quest Diagnostics Incorporated | Centers for Disease Control and Prevention Division of Viral Diseases, Pathogen Discovery | Dakota Howard, Dhwani Batra, Peter W. Cook, Kara Moser, Adrian Paskey, Jason Caravas, Benjamin Rambo-Martin, Shatavia Morrison, Christopher Gulvick, Scott Sammons, Yvette Unoarumhi, Darlene Wagner, Matthew Schmerer, S. H. Rosenthal, A. Gerasimova, R. M. Kagan, B. Anderson, M. Hua, Y. Liu, L.E. Bernstein, K.E. Livingston, A. Perez, I. A. Shlyakhter, R. V. Rolando, R. Owen, P. Tanpaiboon, F. Lacbawan, Clinton R. Paden, Duncan MacCannell |
| EPI_ISL_3396275 | MD PHL | Maryland Department of Health Laboratories Administration | Eric N. Keller, Jillian Loomis, Kwang Low, Terence L. Moore, Ami Patel, and Robert Myers |
| EPI_ISL_3396276, EPI_ISL_3396277 | Quest Diagnostics Incorporated | Centers for Disease Control and Prevention Division of Viral Diseases, Pathogen Discovery | Dakota Howard, Dhwani Batra, Peter W. Cook, Kara Moser, Adrian Paskey, Jason Caravas, Benjamin Rambo-Martin, Shatavia Morrison, Christopher Gulvick, Scott Sammons, Yvette Unoarumhi, Darlene Wagner, Matthew Schmerer, S. H. Rosenthal, A. Gerasimova, R. M. Kagan, B. Anderson, M. Hua, Y. Liu, L.E. Bernstein, K.E. Livingston, A. Perez, I. A. Shlyakhter, R. V. Rolando, R. Owen, P. Tanpaiboon, F. Lacbawan, Clinton R. Paden, Duncan MacCannell |
| EPI_ISL_3396278 | MD PHL | Maryland Department of Health Laboratories Administration | Eric N. Keller, Jillian Loomis, Kwang Low, Terence L. Moore, Ami Patel, and Robert Myers |
| EPI_ISL_3396279, EPI_ISL_3396280 | Quest Diagnostics Incorporated | Centers for Disease Control and Prevention Division of Viral Diseases, Pathogen Discovery | Dakota Howard, Dhwani Batra, Peter W. Cook, Kara Moser, Adrian Paskey, Jason Caravas, Benjamin Rambo-Martin, Shatavia Morrison, Christopher Gulvick, Scott Sammons, Yvette Unoarumhi, Darlene Wagner, Matthew Schmerer, S. H. Rosenthal, A. Gerasimova, R. M. Kagan, B. Anderson, M. Hua, Y. Liu, L.E. Bernstein, K.E. Livingston, A. Perez, I. A. Shlyakhter, R. V. Rolando, R. Owen, P. Tanpaiboon, F. Lacbawan, Clinton R. Paden, Duncan MacCannell |
| EPI_ISL_3396281 | MD PHL | Maryland Department of Health Laboratories Administration | Eric N. Keller, Jillian Loomis, Kwang Low, Terence L. Moore, Ami Patel, and Robert Myers |
| EPI_ISL_3396282 | Quest Diagnostics Incorporated | Centers for Disease Control and Prevention Division of Viral Diseases, Pathogen Discovery | Dakota Howard, Dhwani Batra, Peter W. Cook, Kara Moser, Adrian Paskey, Jason Caravas, Benjamin Rambo-Martin, Shatavia Morrison, Christopher Gulvick, Scott Sammons, Yvette Unoarumhi, Darlene Wagner, Matthew Schmerer, S. H. Rosenthal, A. Gerasimova, R. M. Kagan, B. Anderson, M. Hua, Y. Liu, L.E. Bernstein, K.E. Livingston, A. Perez, I. A. Shlyakhter, R. V. Rolando, R. Owen, P. Tanpaiboon, F. Lacbawan, Clinton R. Paden, Duncan MacCannell |
| EPI_ISL_3396283 | MD PHL | Maryland Department of Health Laboratories Administration | Eric N. Keller, Jillian Loomis, Kwang Low, Terence L. Moore, Ami Patel, and Robert Myers |
| EPI_ISL_3396284 | Quest Diagnostics Incorporated | Centers for Disease Control and Prevention Division of Viral Diseases, Pathogen Discovery | Dakota Howard, Dhwani Batra, Peter W. Cook, Kara Moser, Adrian Paskey, Jason Caravas, Benjamin Rambo-Martin, Shatavia Morrison, Christopher Gulvick, Scott Sammons, Yvette Unoarumhi, Darlene Wagner, Matthew Schmerer, S. H. Rosenthal, A. Gerasimova, R. M. Kagan, B. Anderson, M. Hua, Y. Liu, L.E. Bernstein, K.E. Livingston, A. Perez, I. A. Shlyakhter, R. V. Rolando, R. Owen, P. Tanpaiboon, F. Lacbawan, Clinton R. Paden, Duncan MacCannell |

| | | | |
|---|---|---|---|
| EPI_ISL_3396286 | MD PHL | Maryland Department of Health Laboratories Administration | Eric N. Keller, Jillian Loomis, Kwang Low, Terence L. Moore, Ami Patel, and Robert Myers |
| EPI_ISL_3396287 | Quest Diagnostics Incorporated | Centers for Disease Control and Prevention Division of Viral Diseases, Pathogen Discovery | Dakota Howard, Dhwani Batra, Peter W. Cook, Kara Moser, Adrian Paskey, Jason Caravas, Benjamin Rambo-Martin, Shatavia Morrison, Christopher Gulvick, Scott Sammons, Yvette Unoarumhi, Darlene Wagner, Matthew Schmerer, S. H. Rosenthal, A. Gerasimova, R. M. Kagan, B. Anderson, M. Hua, Y. Liu, L.E. Bernstein, K.E. Livingston, A. Perez, I. A. Shlyakhter, R. V. Rolando, R. Owen, P. Tanpaiboon, F. Lacbawan, Clinton R. Paden, Duncan MacCannell |
| EPI_ISL_3396288, EPI_ISL_3396289 | MD PHL | Maryland Department of Health Laboratories Administration | Eric N. Keller, Jillian Loomis, Kwang Low, Terence L. Moore, Ami Patel, and Robert Myers |
| EPI_ISL_3396290 | Quest Diagnostics Incorporated | Centers for Disease Control and Prevention Division of Viral Diseases, Pathogen Discovery | Dakota Howard, Dhwani Batra, Peter W. Cook, Kara Moser, Adrian Paskey, Jason Caravas, Benjamin Rambo-Martin, Shatavia Morrison, Christopher Gulvick, Scott Sammons, Yvette Unoarumhi, Darlene Wagner, Matthew Schmerer, S. H. Rosenthal, A. Gerasimova, R. M. Kagan, B. Anderson, M. Hua, Y. Liu, L.E. Bernstein, K.E. Livingston, A. Perez, I. A. Shlyakhter, R. V. Rolando, R. Owen, P. Tanpaiboon, F. Lacbawan, Clinton R. Paden, Duncan MacCannell |
| EPI_ISL_3396291 | MD PHL | Maryland Department of Health Laboratories Administration | Eric N. Keller, Jillian Loomis, Kwang Low, Terence L. Moore, Ami Patel, and Robert Myers |
| EPI_ISL_3396293 | Quest Diagnostics Incorporated | Centers for Disease Control and Prevention Division of Viral Diseases, Pathogen Discovery | Dakota Howard, Dhwani Batra, Peter W. Cook, Kara Moser, Adrian Paskey, Jason Caravas, Benjamin Rambo-Martin, Shatavia Morrison, Christopher Gulvick, Scott Sammons, Yvette Unoarumhi, Darlene Wagner, Matthew Schmerer, S. H. Rosenthal, A. Gerasimova, R. M. Kagan, B. Anderson, M. Hua, Y. Liu, L.E. Bernstein, K.E. Livingston, A. Perez, I. A. Shlyakhter, R. V. Rolando, R. Owen, P. Tanpaiboon, F. Lacbawan, Clinton R. Paden, Duncan MacCannell |
| EPI_ISL_3396294 | MD PHL | Maryland Department of Health Laboratories Administration | Eric N. Keller, Jillian Loomis, Kwang Low, Terence L. Moore, Ami Patel, and Robert Myers |
| EPI_ISL_3396295 | Quest Diagnostics Incorporated | Centers for Disease Control and Prevention Division of Viral Diseases, Pathogen Discovery | Dakota Howard, Dhwani Batra, Peter W. Cook, Kara Moser, Adrian Paskey, Jason Caravas, Benjamin Rambo-Martin, Shatavia Morrison, Christopher Gulvick, Scott Sammons, Yvette Unoarumhi, Darlene Wagner, Matthew Schmerer, S. H. Rosenthal, A. Gerasimova, R. M. Kagan, B. Anderson, M. Hua, Y. Liu, L.E. Bernstein, K.E. Livingston, A. Perez, I. A. Shlyakhter, R. V. Rolando, R. Owen, P. Tanpaiboon, F. Lacbawan, Clinton R. Paden, Duncan MacCannell |
| EPI_ISL_3396296, EPI_ISL_3396297 | MD PHL | Maryland Department of Health Laboratories Administration | Eric N. Keller, Jillian Loomis, Kwang Low, Terence L. Moore, Ami Patel, and Robert Myers |
| EPI_ISL_3396298 | Quest Diagnostics Incorporated | Centers for Disease Control and Prevention Division of Viral Diseases, Pathogen Discovery | Dakota Howard, Dhwani Batra, Peter W. Cook, Kara Moser, Adrian Paskey, Jason Caravas, Benjamin Rambo-Martin, Shatavia Morrison, Christopher Gulvick, Scott Sammons, Yvette Unoarumhi, Darlene Wagner, Matthew Schmerer, S. H. Rosenthal, A. Gerasimova, R. M. Kagan, B. Anderson, M. Hua, Y. Liu, L.E. Bernstein, K.E. Livingston, A. Perez, I. A. Shlyakhter, R. V. Rolando, R. Owen, P. Tanpaiboon, F. Lacbawan, Clinton R. Paden, Duncan MacCannell |
| EPI_ISL_3396299, EPI_ISL_3396300 | MD PHL | Maryland Department of Health Laboratories Administration | Eric N. Keller, Jillian Loomis, Kwang Low, Terence L. Moore, Ami Patel, and Robert Myers |
| EPI_ISL_3396301 | Quest Diagnostics Incorporated | Centers for Disease Control and Prevention Division of Viral Diseases, Pathogen Discovery | Dakota Howard, Dhwani Batra, Peter W. Cook, Kara Moser, Adrian Paskey, Jason Caravas, Benjamin Rambo-Martin, Shatavia Morrison, Christopher Gulvick, Scott Sammons, Yvette Unoarumhi, Darlene Wagner, Matthew Schmerer, S. H. Rosenthal, A. Gerasimova, R. M. Kagan, B. Anderson, M. Hua, Y. Liu, L.E. Bernstein, K.E. Livingston, A. Perez, I. A. Shlyakhter, R. V. Rolando, R. Owen, P. Tanpaiboon, F. Lacbawan, Clinton R. Paden, Duncan MacCannell |
| EPI_ISL_3396302 | MD PHL | Maryland Department of Health Laboratories Administration | Eric N. Keller, Jillian Loomis, Kwang Low, Terence L. Moore, Ami Patel, and Robert Myers |
| EPI_ISL_3396303 | Quest Diagnostics Incorporated | Centers for Disease Control and Prevention Division of Viral Diseases, Pathogen Discovery | Dakota Howard, Dhwani Batra, Peter W. Cook, Kara Moser, Adrian Paskey, Jason Caravas, Benjamin Rambo-Martin, Shatavia Morrison, Christopher Gulvick, Scott Sammons, Yvette Unoarumhi, Darlene Wagner, Matthew Schmerer, S. H. Rosenthal, A. Gerasimova, R. M. Kagan, B. Anderson, M. Hua, Y. Liu, L.E. Bernstein, K.E. Livingston, A. Perez, I. A. Shlyakhter, R. V. Rolando, R. Owen, P. Tanpaiboon, F. Lacbawan, Clinton R. Paden, Duncan MacCannell |
| EPI_ISL_3396304, EPI_ISL_3396305 | MD PHL | Maryland Department of Health Laboratories Administration | Eric N. Keller, Jillian Loomis, Kwang Low, Terence L. Moore, Ami Patel, and Robert Myers |
| EPI_ISL_3396306 | Quest Diagnostics Incorporated | Centers for Disease Control and Prevention Division of Viral Diseases, Pathogen Discovery | Dakota Howard, Dhwani Batra, Peter W. Cook, Kara Moser, Adrian Paskey, Jason Caravas, Benjamin Rambo-Martin, Shatavia Morrison, Christopher Gulvick, Scott Sammons, Yvette Unoarumhi, Darlene Wagner, Matthew Schmerer, S. H. Rosenthal, A. Gerasimova, R. M. Kagan, B. Anderson, M. Hua, Y. Liu, L.E. Bernstein, K.E. Livingston, A. Perez, I. A. Shlyakhter, R. V. Rolando, R. Owen, P. Tanpaiboon, F. Lacbawan, Clinton R. Paden, Duncan MacCannell |
| EPI_ISL_3396307 | MD PHL | Maryland Department of Health Laboratories Administration | Eric N. Keller, Jillian Loomis, Kwang Low, Terence L. Moore, Ami Patel, and Robert Myers |
| EPI_ISL_3396308 | Quest Diagnostics Incorporated | Centers for Disease Control and Prevention Division of Viral Diseases, Pathogen Discovery | Dakota Howard, Dhwani Batra, Peter W. Cook, Kara Moser, Adrian Paskey, Jason Caravas, Benjamin Rambo-Martin, Shatavia Morrison, Christopher Gulvick, Scott Sammons, Yvette Unoarumhi, Darlene Wagner, Matthew Schmerer, S. H. Rosenthal, A. Gerasimova, R. M. Kagan, B. Anderson, M. Hua, Y. Liu, L.E. Bernstein, K.E. Livingston, A. Perez, I. A. Shlyakhter, R. V. Rolando, R. Owen, P. Tanpaiboon, F. Lacbawan, Clinton R. Paden, Duncan MacCannell |
| EPI_ISL_3396309 | MD PHL | Maryland Department of Health Laboratories Administration | Eric N. Keller, Jillian Loomis, Kwang Low, Terence L. Moore, Ami Patel, and Robert Myers |
| EPI_ISL_3396310 | Quest Diagnostics Incorporated | Centers for Disease Control and Prevention Division of Viral Diseases, Pathogen Discovery | Dakota Howard, Dhwani Batra, Peter W. Cook, Kara Moser, Adrian Paskey, Jason Caravas, Benjamin Rambo-Martin, Shatavia Morrison, Christopher Gulvick, Scott Sammons, Yvette Unoarumhi, Darlene Wagner, Matthew Schmerer, S. H. Rosenthal, A. Gerasimova, R. M. Kagan, B. Anderson, M. Hua, Y. Liu, L.E. Bernstein, K.E. Livingston, A. Perez, I. A. Shlyakhter, R. V. Rolando, R. Owen, P. Tanpaiboon, F. Lacbawan, Clinton R. Paden, Duncan MacCannell |
| EPI_ISL_3396311, EPI_ISL_3396312 | MD PHL | Maryland Department of Health Laboratories Administration | Eric N. Keller, Jillian Loomis, Kwang Low, Terence L. Moore, Ami Patel, and Robert Myers |
| EPI_ISL_3396313 | Quest Diagnostics Incorporated | Centers for Disease Control and Prevention Division of Viral Diseases, Pathogen Discovery | Dakota Howard, Dhwani Batra, Peter W. Cook, Kara Moser, Adrian Paskey, Jason Caravas, Benjamin Rambo-Martin, Shatavia Morrison, Christopher Gulvick, Scott Sammons, Yvette Unoarumhi, Darlene Wagner, Matthew Schmerer, S. H. Rosenthal, A. Gerasimova, R. M. Kagan, B. Anderson, M. Hua, Y. Liu, L.E. Bernstein, K.E. Livingston, A. Perez, I. A. Shlyakhter, R. V. Rolando, R. Owen, P. Tanpaiboon, F. Lacbawan, Clinton R. Paden, Duncan MacCannell |
| EPI_ISL_3396314, EPI_ISL_3396315 | MD PHL | Maryland Department of Health Laboratories Administration | Eric N. Keller, Jillian Loomis, Kwang Low, Terence L. Moore, Ami Patel, and Robert Myers |
| EPI_ISL_3396316 | Quest Diagnostics Incorporated | Centers for Disease Control and Prevention Division of Viral Diseases, Pathogen Discovery | Dakota Howard, Dhwani Batra, Peter W. Cook, Kara Moser, Adrian Paskey, Jason Caravas, Benjamin Rambo-Martin, Shatavia Morrison, Christopher Gulvick, Scott Sammons, Yvette Unoarumhi, Darlene Wagner, Matthew Schmerer, S. H. Rosenthal, A. Gerasimova, R. M. Kagan, B. Anderson, M. Hua, Y. Liu, L.E. Bernstein, K.E. Livingston, A. Perez, I. A. Shlyakhter, R. V. Rolando, R. Owen, P. Tanpaiboon, F. Lacbawan, Clinton R. Paden, Duncan MacCannell |
| EPI_ISL_3396317 | MD PHL | Maryland Department of Health Laboratories Administration | Eric N. Keller, Jillian Loomis, Kwang Low, Terence L. Moore, Ami Patel, and Robert Myers |
| EPI_ISL_3396318 | Quest Diagnostics Incorporated | Centers for Disease Control and Prevention Division of Viral Diseases, Pathogen Discovery | Dakota Howard, Dhwani Batra, Peter W. Cook, Kara Moser, Adrian Paskey, Jason Caravas, Benjamin Rambo-Martin, Shatavia Morrison, Christopher Gulvick, Scott Sammons, Yvette Unoarumhi, Darlene Wagner, Matthew Schmerer, S. H. Rosenthal, A. Gerasimova, R. M. Kagan, B. Anderson, M. Hua, Y. Liu, L.E. Bernstein, K.E. Livingston, A. Perez, I. A. Shlyakhter, R. V. Rolando, R. Owen, P. Tanpaiboon, F. Lacbawan, Clinton R. Paden, Duncan MacCannell |
| EPI_ISL_3396319 | MD PHL | Maryland Department of Health Laboratories Administration | Eric N. Keller, Jillian Loomis, Kwang Low, Terence L. Moore, Ami Patel, and Robert Myers |
| EPI_ISL_3396320 | Quest Diagnostics Incorporated | Centers for Disease Control and Prevention Division of Viral Diseases, Pathogen Discovery | Dakota Howard, Dhwani Batra, Peter W. Cook, Kara Moser, Adrian Paskey, Jason Caravas, Benjamin Rambo-Martin, Shatavia Morrison, Christopher Gulvick, Scott Sammons, Yvette Unoarumhi, Darlene Wagner, Matthew Schmerer, S. H. Rosenthal, A. Gerasimova, R. M. Kagan, B. Anderson, M. Hua, Y. Liu, L.E. Bernstein, K.E. Livingston, A. Perez, I. A. Shlyakhter, R. V. Rolando, R. Owen, P. Tanpaiboon, F. Lacbawan, Clinton R. Paden, Duncan MacCannell |
| EPI_ISL_3396321, EPI_ISL_3396322 | MD PHL | Maryland Department of Health Laboratories Administration | Eric N. Keller, Jillian Loomis, Kwang Low, Terence L. Moore, Ami Patel, and Robert Myers |
| EPI_ISL_3396323 | Quest Diagnostics Incorporated | Centers for Disease Control and Prevention Division of Viral Diseases, Pathogen Discovery | Dakota Howard, Dhwani Batra, Peter W. Cook, Kara Moser, Adrian Paskey, Jason Caravas, Benjamin Rambo-Martin, Shatavia Morrison, Christopher Gulvick, Scott Sammons, Yvette Unoarumhi, Darlene Wagner, Matthew Schmerer, S. H. Rosenthal, A. Gerasimova, R. M. Kagan, B. Anderson, M. Hua, Y. Liu, L.E. Bernstein, K.E. Livingston, A. Perez, I. A. Shlyakhter, R. V. Rolando, R. Owen, P. Tanpaiboon, F. Lacbawan, Clinton R. Paden, Duncan MacCannell |
| EPI_ISL_3396324 | MD PHL | Maryland Department of Health Laboratories Administration | Eric N. Keller, Jillian Loomis, Kwang Low, Terence L. Moore, Ami Patel, and Robert Myers |
| EPI_ISL_3396325 | Quest Diagnostics Incorporated | Centers for Disease Control and Prevention Division of Viral Diseases, Pathogen Discovery | Dakota Howard, Dhwani Batra, Peter W. Cook, Kara Moser, Adrian Paskey, Jason Caravas, Benjamin Rambo-Martin, Shatavia Morrison, Christopher Gulvick, Scott Sammons, Yvette Unoarumhi, Darlene Wagner, Matthew Schmerer, S. H. Rosenthal, A. Gerasimova, R. M. Kagan, B. Anderson, M. Hua, Y. Liu, L.E. Bernstein, K.E. Livingston, A. Perez, I. A. Shlyakhter, R. V. Rolando, R. Owen, P. Tanpaiboon, F. Lacbawan, Clinton R. Paden, Duncan MacCannell |
| EPI_ISL_3396326 | MD PHL | Maryland Department of Health Laboratories Administration | Eric N. Keller, Jillian Loomis, Kwang Low, Terence L. Moore, Ami Patel, and Robert Myers |
| EPI_ISL_3396327 | Quest Diagnostics Incorporated | Centers for Disease Control and Prevention Division of Viral Diseases, Pathogen Discovery | Dakota Howard, Dhwani Batra, Peter W. Cook, Kara Moser, Adrian Paskey, Jason Caravas, Benjamin Rambo-Martin, Shatavia Morrison, Christopher Gulvick, Scott Sammons, Yvette Unoarumhi, Darlene Wagner, Matthew Schmerer, S. H. Rosenthal, A. Gerasimova, R. M. Kagan, B. Anderson, M. Hua, Y. Liu, L.E. Bernstein, K.E. Livingston, A. Perez, I. A. Shlyakhter, R. V. Rolando, R. Owen, P. Tanpaiboon, F. Lacbawan, Clinton R. Paden, Duncan MacCannell |
| EPI_ISL_3396328 | MD PHL | Maryland Department of Health Laboratories Administration | Eric N. Keller, Jillian Loomis, Kwang Low, Terence L. Moore, Ami Patel, and Robert Myers |
| EPI_ISL_3396329 | Quest Diagnostics Incorporated | Centers for Disease Control and Prevention Division of Viral Diseases, Pathogen Discovery | Dakota Howard, Dhwani Batra, Peter W. Cook, Kara Moser, Adrian Paskey, Jason Caravas, Benjamin Rambo-Martin, Shatavia Morrison, Christopher Gulvick, Scott Sammons, Yvette Unoarumhi, Darlene Wagner, Matthew Schmerer, S. H. Rosenthal, A. Gerasimova, R. M. Kagan, B. Anderson, M. Hua, Y. Liu, L.E. Bernstein, K.E. Livingston, A. Perez, I. A. Shlyakhter, R. V. Rolando, R. Owen, P. Tanpaiboon, F. Lacbawan, Clinton R. Paden, Duncan MacCannell |
| EPI_ISL_3396330, EPI_ISL_3396331 | MD PHL | Maryland Department of Health Laboratories Administration | Eric N. Keller, Jillian Loomis, Kwang Low, Terence L. Moore, Ami Patel, and Robert Myers |
| EPI_ISL_3396332 | Quest Diagnostics Incorporated | Centers for Disease Control and Prevention Division of Viral Diseases, Pathogen Discovery | Dakota Howard, Dhwani Batra, Peter W. Cook, Kara Moser, Adrian Paskey, Jason Caravas, Benjamin Rambo-Martin, Shatavia Morrison, Christopher Gulvick, Scott Sammons, Yvette Unoarumhi, Darlene Wagner, Matthew Schmerer, S. H. Rosenthal, A. Gerasimova, R. M. Kagan, B. Anderson, M. Hua, Y. Liu, L.E. Bernstein, K.E. Livingston, A. Perez, I. A. Shlyakhter, R. V. Rolando, R. Owen, P. Tanpaiboon, F. Lacbawan, Clinton R. Paden, Duncan MacCannell |
| EPI_ISL_3396333 | MD PHL | Maryland Department of Health Laboratories Administration | Eric N. Keller, Jillian Loomis, Kwang Low, Terence L. Moore, Ami Patel, and Robert Myers |
| EPI_ISL_3396334 | Quest Diagnostics Incorporated | Centers for Disease Control and Prevention Division of Viral Diseases, Pathogen Discovery | Dakota Howard, Dhwani Batra, Peter W. Cook, Kara Moser, Adrian Paskey, Jason Caravas, Benjamin Rambo-Martin, Shatavia Morrison, Christopher Gulvick, Scott Sammons, Yvette Unoarumhi, Darlene Wagner, Matthew Schmerer, S. H. Rosenthal, A. Gerasimova, R. M. Kagan, B. Anderson, M. Hua, Y. Liu, L.E. Bernstein, K.E. Livingston, A. Perez, I. A. Shlyakhter, R. V. Rolando, R. Owen, P. Tanpaiboon, F. Lacbawan, Clinton R. Paden, Duncan MacCannell |
| EPI_ISL_3396335, EPI_ISL_3396336 | MD PHL | Maryland Department of Health Laboratories Administration | Eric N. Keller, Jillian Loomis, Kwang Low, Terence L. Moore, Ami Patel, and Robert Myers |
| EPI_ISL_3396337 | Quest Diagnostics Incorporated | Centers for Disease Control and Prevention Division of Viral Diseases, Pathogen Discovery | Dakota Howard, Dhwani Batra, Peter W. Cook, Kara Moser, Adrian Paskey, Jason Caravas, Benjamin Rambo-Martin, Shatavia Morrison, Christopher Gulvick, Scott Sammons, Yvette Unoarumhi, Darlene Wagner, Matthew Schmerer, S. H. Rosenthal, A. Gerasimova, R. M. Kagan, B. Anderson, M. Hua, Y. Liu, L.E. Bernstein, K.E. Livingston, A. Perez, I. A. Shlyakhter, R. V. Rolando, R. Owen, P. Tanpaiboon, F. Lacbawan, Clinton R. Paden, Duncan MacCannell |
| EPI_ISL_3396338 | MD PHL | Maryland Department of Health Laboratories Administration | Eric N. Keller, Jillian Loomis, Kwang Low, Terence L. Moore, Ami Patel, and Robert Myers |
| EPI_ISL_3396339 | Quest Diagnostics Incorporated | Centers for Disease Control and Prevention Division of Viral Diseases, Pathogen Discovery | Dakota Howard, Dhwani Batra, Peter W. Cook, Kara Moser, Adrian Paskey, Jason Caravas, Benjamin Rambo-Martin, Shatavia Morrison, Christopher Gulvick, Scott Sammons, Yvette Unoarumhi, Darlene Wagner, Matthew Schmerer, S. H. Rosenthal, A. Gerasimova, R. M. Kagan, B. Anderson, M. Hua, Y. Liu, L.E. Bernstein, K.E. Livingston, A. Perez, I. A. Shlyakhter, R. V. Rolando, R. Owen, P. Tanpaiboon, F. Lacbawan, Clinton R. Paden, Duncan MacCannell |
| EPI_ISL_3396340 | MD PHL | Maryland Department of Health Laboratories Administration | Eric N. Keller, Jillian Loomis, Kwang Low, Terence L. Moore, Ami Patel, and Robert Myers |
| EPI_ISL_3396341 | Quest Diagnostics Incorporated | Centers for Disease Control and Prevention Division of Viral Diseases, Pathogen Discovery | Dakota Howard, Dhwani Batra, Peter W. Cook, Kara Moser, Adrian Paskey, Jason Caravas, Benjamin Rambo-Martin, Shatavia Morrison, Christopher Gulvick, Scott Sammons, Yvette Unoarumhi, Darlene Wagner, Matthew Schmerer, S. H. Rosenthal, A. Gerasimova, R. M. Kagan, B. Anderson, M. Hua, Y. Liu, L.E. Bernstein, K.E. Livingston, A. Perez, I. A. Shlyakhter, R. V. Rolando, R. Owen, P. Tanpaiboon, F. Lacbawan, Clinton R. Paden, Duncan MacCannell |
| EPI_ISL_3396342, EPI_ISL_3396343 | MD PHL | Maryland Department of Health Laboratories Administration | Eric N. Keller, Jillian Loomis, Kwang Low, Terence L. Moore, Ami Patel, and Robert Myers |
| EPI_ISL_3396344 | Quest Diagnostics Incorporated | Centers for Disease Control and Prevention Division of Viral Diseases, Pathogen Discovery | Dakota Howard, Dhwani Batra, Peter W. Cook, Kara Moser, Adrian Paskey, Jason Caravas, Benjamin Rambo-Martin, Shatavia Morrison, Christopher Gulvick, Scott Sammons, Yvette Unoarumhi, Darlene Wagner, Matthew Schmerer, S. H. Rosenthal, A. Gerasimova, R. M. Kagan, B. Anderson, M. Hua, Y. Liu, L.E. Bernstein, K.E. Livingston, A. Perez, I. A. Shlyakhter, R. V. Rolando, R. Owen, P. Tanpaiboon, F. Lacbawan, Clinton R. Paden, Duncan MacCannell |
| EPI_ISL_3396345 | MD PHL | Maryland Department of Health Laboratories Administration | Eric N. Keller, Jillian Loomis, Kwang Low, Terence L. Moore, Ami Patel, and Robert Myers |
| EPI_ISL_3396346 | Quest Diagnostics Incorporated | Centers for Disease Control and Prevention Division of Viral Diseases, Pathogen Discovery | Dakota Howard, Dhwani Batra, Peter W. Cook, Kara Moser, Adrian Paskey, Jason Caravas, Benjamin Rambo-Martin, Shatavia Morrison, Christopher Gulvick, Scott Sammons, Yvette Unoarumhi, Darlene Wagner, Matthew Schmerer, S. H. Rosenthal, A. Gerasimova, R. M. Kagan, B. Anderson, M. Hua, Y. Liu, L.E. Bernstein, K.E. Livingston, A. Perez, I. A. Shlyakhter, R. V. Rolando, R. Owen, P. Tanpaiboon, F. Lacbawan, Clinton R. Paden, Duncan MacCannell |
| EPI_ISL_3396347 | MD PHL | Maryland Department of Health Laboratories Administration | Eric N. Keller, Jillian Loomis, Kwang Low, Terence L. Moore, Ami Patel, and Robert Myers |
| EPI_ISL_3396348, EPI_ISL_3396349, EPI_ISL_3396350, EPI_ISL_3396351, EPI_ISL_3396352, EPI_ISL_3396353, EPI_ISL_3396354, EPI_ISL_3396356, EPI_ISL_3396357, EPI_ISL_3396358, EPI_ISL_3396359, EPI_ISL_3396360, EPI_ISL_3396361, EPI_ISL_3396362, EPI_ISL_3396363, EPI_ISL_3396364, EPI_ISL_3396365, EPI_ISL_3396366, EPI_ISL_3396367, EPI_ISL_3396368, EPI_ISL_3396370, EPI_ISL_3396371, EPI_ISL_3396373, EPI_ISL_3396374, EPI_ISL_3396375, EPI_ISL_3396376, EPI_ISL_3396377, EPI_ISL_3396378, EPI_ISL_3396379, EPI_ISL_3396380, EPI_ISL_3396381, EPI_ISL_3396382, EPI_ISL_3396384, EPI_ISL_3396385, EPI_ISL_3396386, EPI_ISL_3396387, EPI_ISL_3396388, EPI_ISL_3396389, EPI_ISL_3396390, EPI_ISL_3396391, EPI_ISL_3396392, EPI_ISL_3396393, EPI_ISL_3396394, EPI_ISL_3396395, | | | |

EPI_ISL_3396396, EPI_ISL_3396397, EPI_ISL_3396398, EPI_ISL_3396400, EPI_ISL_3396401, EPI_ISL_3396402, EPI_ISL_3396403, EPI_ISL_3396405, EPI_ISL_3396406, EPI_ISL_3396407, EPI_ISL_3396408, EPI_ISL_3396409, EPI_ISL_3396410, EPI_ISL_3396411, EPI_ISL_3396412, EPI_ISL_3396413, EPI_ISL_3396414, EPI_ISL_3396415, EPI_ISL_3396416, EPI_ISL_3396417, EPI_ISL_3396418, EPI_ISL_3396419, EPI_ISL_3396420, EPI_ISL_3396421, EPI_ISL_3396422, EPI_ISL_3396423, EPI_ISL_3396424, EPI_ISL_3396425, EPI_ISL_3396426, EPI_ISL_3396427, EPI_ISL_3396428, EPI_ISL_3396429, EPI_ISL_3396430, EPI_ISL_3396431, EPI_ISL_3396432, EPI_ISL_3396433, EPI_ISL_3396434, EPI_ISL_3396435, EPI_ISL_3396436

| | | | |
|---|---|---|---|
| see above | Quest Diagnostics Incorporated | Centers for Disease Control and Prevention Division of Viral Diseases, Pathogen Discovery | Dakota Howard, Dhwani Batra, Peter W. Cook, Kara Moser, Adrian Paskey, Jason Caravas, Benjamin Rambo-Martin, Shatavia Morrison, Christopher Gulvick, Scott Sammons, Yvette Unoarumhi, Darlene Wagner, Matthew Schmerer, S. H. Rosenthal, A. Gerasimova, R. M. Kagan, B. Anderson, M. Hua, Y. Liu, L.E. Bernstein, K.E. Livingston, A. Perez, I. A. Shlyakhter, R. V. Rolando, R. Owen, P. Tanpaiboon, F. Lacbawan, Clinton R. Paden, Duncan MacCannell |
| EPI_ISL_3396437 | University Hospital Brno, OKMI | University Hospital Brno, CMBG | Matej Bezdicek, Jan Svaton, Martina Lengerova, Kristyna Dufkova, Pavlina Volfova |
| EPI_ISL_3396438 | Quest Diagnostics Incorporated | Centers for Disease Control and Prevention Division of Viral Diseases, Pathogen Discovery | Dakota Howard, Dhwani Batra, Peter W. Cook, Kara Moser, Adrian Paskey, Jason Caravas, Benjamin Rambo-Martin, Shatavia Morrison, Christopher Gulvick, Scott Sammons, Yvette Unoarumhi, Darlene Wagner, Matthew Schmerer, S. H. Rosenthal, A. Gerasimova, R. M. Kagan, B. Anderson, M. Hua, Y. Liu, L.E. Bernstein, K.E. Livingston, A. Perez, I. A. Shlyakhter, R. V. Rolando, R. Owen, P. Tanpaiboon, F. Lacbawan, Clinton R. Paden, Duncan MacCannell |
| EPI_ISL_3396439 | Nemocnice Trebic | University Hospital Brno, CMBG | Matej Bezdicek, Jan Svaton, Martina Lengerova, Kristyna Dufkova, Pavlina Volfova |
| EPI_ISL_3396440, EPI_ISL_3396441 | Quest Diagnostics Incorporated | Centers for Disease Control and Prevention Division of Viral Diseases, Pathogen Discovery | Dakota Howard, Dhwani Batra, Peter W. Cook, Kara Moser, Adrian Paskey, Jason Caravas, Benjamin Rambo-Martin, Shatavia Morrison, Christopher Gulvick, Scott Sammons, Yvette Unoarumhi, Darlene Wagner, Matthew Schmerer, S. H. Rosenthal, A. Gerasimova, R. M. Kagan, B. Anderson, M. Hua, Y. Liu, L.E. Bernstein, K.E. Livingston, A. Perez, I. A. Shlyakhter, R. V. Rolando, R. Owen, P. Tanpaiboon, F. Lacbawan, Clinton R. Paden, Duncan MacCannell |
| EPI_ISL_3396442 | IFCOR Brno | University Hospital Brno, CMBG | Matej Bezdicek, Jan Svaton, Martina Lengerova, Kristyna Dufkova, Pavlina Volfova |
| EPI_ISL_3396443 | Quest Diagnostics Incorporated | Centers for Disease Control and Prevention Division of Viral Diseases, Pathogen Discovery | Dakota Howard, Dhwani Batra, Peter W. Cook, Kara Moser, Adrian Paskey, Jason Caravas, Benjamin Rambo-Martin, Shatavia Morrison, Christopher Gulvick, Scott Sammons, Yvette Unoarumhi, Darlene Wagner, Matthew Schmerer, S. H. Rosenthal, A. Gerasimova, R. M. Kagan, B. Anderson, M. Hua, Y. Liu, L.E. Bernstein, K.E. Livingston, A. Perez, I. A. Shlyakhter, R. V. Rolando, R. Owen, P. Tanpaiboon, F. Lacbawan, Clinton R. Paden, Duncan MacCannell |
| EPI_ISL_3396444 | IFCOR Brno | University Hospital Brno, CMBG | Matej Bezdicek, Jan Svaton, Martina Lengerova, Kristyna Dufkova, Pavlina Volfova |
| EPI_ISL_3396445, EPI_ISL_3396446 | Quest Diagnostics Incorporated | Centers for Disease Control and Prevention Division of Viral Diseases, Pathogen Discovery | Dakota Howard, Dhwani Batra, Peter W. Cook, Kara Moser, Adrian Paskey, Jason Caravas, Benjamin Rambo-Martin, Shatavia Morrison, Christopher Gulvick, Scott Sammons, Yvette Unoarumhi, Darlene Wagner, Matthew Schmerer, S. H. Rosenthal, A. Gerasimova, R. M. Kagan, B. Anderson, M. Hua, Y. Liu, L.E. Bernstein, K.E. Livingston, A. Perez, I. A. Shlyakhter, R. V. Rolando, R. Owen, P. Tanpaiboon, F. Lacbawan, Clinton R. Paden, Duncan MacCannell |
| EPI_ISL_3396447 | University Hospital Brno, OKMI | University Hospital Brno, CMBG | Matej Bezdicek, Jan Svaton, Martina Lengerova, Kristyna Dufkova, Pavlina Volfova |
| EPI_ISL_3396448 | Quest Diagnostics Incorporated | Centers for Disease Control and Prevention Division of Viral Diseases, Pathogen Discovery | Dakota Howard, Dhwani Batra, Peter W. Cook, Kara Moser, Adrian Paskey, Jason Caravas, Benjamin Rambo-Martin, Shatavia Morrison, Christopher Gulvick, Scott Sammons, Yvette Unoarumhi, Darlene Wagner, Matthew Schmerer, S. H. Rosenthal, A. Gerasimova, R. M. Kagan, B. Anderson, M. Hua, Y. Liu, L.E. Bernstein, K.E. Livingston, A. Perez, I. A. Shlyakhter, R. V. Rolando, R. Owen, P. Tanpaiboon, F. Lacbawan, Clinton R. Paden, Duncan MacCannell |
| EPI_ISL_3396449 | University Hospital Brno, OKMI | University Hospital Brno, CMBG | Matej Bezdicek, Jan Svaton, Martina Lengerova, Kristyna Dufkova, Pavlina Volfova |
| EPI_ISL_3396450, EPI_ISL_3396451 | Quest Diagnostics Incorporated | Centers for Disease Control and Prevention Division of Viral Diseases, Pathogen Discovery | Dakota Howard, Dhwani Batra, Peter W. Cook, Kara Moser, Adrian Paskey, Jason Caravas, Benjamin Rambo-Martin, Shatavia Morrison, Christopher Gulvick, Scott Sammons, Yvette Unoarumhi, Darlene Wagner, Matthew Schmerer, S. H. Rosenthal, A. Gerasimova, R. M. Kagan, B. Anderson, M. Hua, Y. Liu, L.E. Bernstein, K.E. Livingston, A. Perez, I. A. Shlyakhter, R. V. Rolando, R. Owen, P. Tanpaiboon, F. Lacbawan, Clinton R. Paden, Duncan MacCannell |
| EPI_ISL_3396452 | University Hospital Brno, OKMI | University Hospital Brno, CMBG | Matej Bezdicek, Jan Svaton, Martina Lengerova, Kristyna Dufkova, Pavlina Volfova |
| EPI_ISL_3396453, EPI_ISL_3396454 | Quest Diagnostics Incorporated | Centers for Disease Control and Prevention Division of Viral Diseases, Pathogen Discovery | Dakota Howard, Dhwani Batra, Peter W. Cook, Kara Moser, Adrian Paskey, Jason Caravas, Benjamin Rambo-Martin, Shatavia Morrison, Christopher Gulvick, Scott Sammons, Yvette Unoarumhi, Darlene Wagner, Matthew Schmerer, S. H. Rosenthal, A. Gerasimova, R. M. Kagan, B. Anderson, M. Hua, Y. Liu, L.E. Bernstein, K.E. Livingston, A. Perez, I. A. Shlyakhter, R. V. Rolando, R. Owen, P. Tanpaiboon, F. Lacbawan, Clinton R. Paden, Duncan MacCannell |
| EPI_ISL_3396455 | University Hospital Brno, OKMI | University Hospital Brno, CMBG | Matej Bezdicek, Jan Svaton, Martina Lengerova, Kristyna Dufkova, Pavlina Volfova |
| EPI_ISL_3396456, EPI_ISL_3396457 | Quest Diagnostics Incorporated | Centers for Disease Control and Prevention Division of Viral Diseases, Pathogen Discovery | Dakota Howard, Dhwani Batra, Peter W. Cook, Kara Moser, Adrian Paskey, Jason Caravas, Benjamin Rambo-Martin, Shatavia Morrison, Christopher Gulvick, Scott Sammons, Yvette Unoarumhi, Darlene Wagner, Matthew Schmerer, S. H. Rosenthal, A. Gerasimova, R. M. Kagan, B. Anderson, M. Hua, Y. Liu, L.E. Bernstein, K.E. Livingston, A. Perez, I. A. Shlyakhter, R. V. Rolando, R. Owen, P. Tanpaiboon, F. Lacbawan, Clinton R. Paden, Duncan MacCannell |
| EPI_ISL_3396458 | University Hospital Brno, OKMI | University Hospital Brno, CMBG | Matej Bezdicek, Jan Svaton, Martina Lengerova, Kristyna Dufkova, Pavlina Volfova |
| EPI_ISL_3396459 | Quest Diagnostics Incorporated | Centers for Disease Control and Prevention Division of Viral Diseases, Pathogen Discovery | Dakota Howard, Dhwani Batra, Peter W. Cook, Kara Moser, Adrian Paskey, Jason Caravas, Benjamin Rambo-Martin, Shatavia Morrison, Christopher Gulvick, Scott Sammons, Yvette Unoarumhi, Darlene Wagner, Matthew Schmerer, S. H. Rosenthal, A. Gerasimova, R. M. Kagan, B. Anderson, M. Hua, Y. Liu, L.E. Bernstein, K.E. Livingston, A. Perez, I. A. Shlyakhter, R. V. Rolando, R. Owen, P. Tanpaiboon, F. Lacbawan, Clinton R. Paden, Duncan MacCannell |
| EPI_ISL_3396460 | Elisabeth Pharmacon | University Hospital Brno, CMBG | Matej Bezdicek, Jan Svaton, Martina Lengerova, Kristyna Dufkova, Pavlina Volfova |
| EPI_ISL_3396461, EPI_ISL_3396462 | Quest Diagnostics Incorporated | Centers for Disease Control and Prevention Division of Viral Diseases, Pathogen Discovery | Dakota Howard, Dhwani Batra, Peter W. Cook, Kara Moser, Adrian Paskey, Jason Caravas, Benjamin Rambo-Martin, Shatavia Morrison, Christopher Gulvick, Scott Sammons, Yvette Unoarumhi, Darlene Wagner, Matthew Schmerer, S. H. Rosenthal, A. Gerasimova, R. M. Kagan, B. Anderson, M. Hua, Y. Liu, L.E. Bernstein, K.E. Livingston, A. Perez, I. A. Shlyakhter, R. V. Rolando, R. Owen, P. Tanpaiboon, F. Lacbawan, Clinton R. Paden, Duncan MacCannell |
| EPI_ISL_3396463 | Elisabeth Pharmacon | University Hospital Brno, CMBG | Matej Bezdicek, Jan Svaton, Martina Lengerova, Kristyna Dufkova, Pavlina Volfova |
| EPI_ISL_3396464 | Quest Diagnostics Incorporated | Centers for Disease Control and Prevention Division of Viral Diseases, Pathogen Discovery | Dakota Howard, Dhwani Batra, Peter W. Cook, Kara Moser, Adrian Paskey, Jason Caravas, Benjamin Rambo-Martin, Shatavia Morrison, Christopher Gulvick, Scott Sammons, Yvette Unoarumhi, Darlene Wagner, Matthew Schmerer, S. H. Rosenthal, A. Gerasimova, R. M. Kagan, B. Anderson, M. Hua, Y. Liu, L.E. Bernstein, K.E. Livingston, A. Perez, I. A. Shlyakhter, R. V. Rolando, R. Owen, P. Tanpaiboon, F. Lacbawan, Clinton R. Paden, Duncan MacCannell |
| EPI_ISL_3396465 | Elisabeth Pharmacon | University Hospital Brno, CMBG | Matej Bezdicek, Jan Svaton, Martina Lengerova, Kristyna Dufkova, Pavlina Volfova |
| EPI_ISL_3396466 | Quest Diagnostics Incorporated | Centers for Disease Control and Prevention Division of Viral Diseases, Pathogen Discovery | Dakota Howard, Dhwani Batra, Peter W. Cook, Kara Moser, Adrian Paskey, Jason Caravas, Benjamin Rambo-Martin, Shatavia Morrison, Christopher Gulvick, Scott Sammons, Yvette Unoarumhi, Darlene Wagner, Matthew Schmerer, S. H. Rosenthal, A. Gerasimova, R. M. Kagan, B. Anderson, M. Hua, Y. Liu, L.E. Bernstein, K.E. Livingston, A. Perez, I. A. Shlyakhter, R. V. Rolando, R. Owen, P. Tanpaiboon, F. Lacbawan, Clinton R. Paden, Duncan MacCannell |
| EPI_ISL_3396467 | University Hospital Brno, OKMI | University Hospital Brno, CMBG | Matej Bezdicek, Jan Svaton, Martina Lengerova, Kristyna Dufkova, Pavlina Volfova |
| EPI_ISL_3396468 | Quest Diagnostics Incorporated | Centers for Disease Control and Prevention Division of Viral Diseases, Pathogen Discovery | Dakota Howard, Dhwani Batra, Peter W. Cook, Kara Moser, Adrian Paskey, Jason Caravas, Benjamin Rambo-Martin, Shatavia Morrison, Christopher Gulvick, Scott Sammons, Yvette Unoarumhi, Darlene Wagner, Matthew Schmerer, S. H. Rosenthal, A. Gerasimova, R. M. Kagan, B. Anderson, M. Hua, Y. Liu, L.E. Bernstein, K.E. Livingston, A. Perez, I. A. Shlyakhter, R. V. Rolando, R. Owen, P. Tanpaiboon, F. Lacbawan, Clinton R. Paden, Duncan MacCannell |
| EPI_ISL_3396469 | Elisabeth Pharmacon | University Hospital Brno, CMBG | Matej Bezdicek, Jan Svaton, Martina Lengerova, Kristyna Dufkova, Pavlina Volfova |
| EPI_ISL_3396470, EPI_ISL_3396471 | Quest Diagnostics Incorporated | Centers for Disease Control and Prevention Division of Viral Diseases, Pathogen Discovery | Dakota Howard, Dhwani Batra, Peter W. Cook, Kara Moser, Adrian Paskey, Jason Caravas, Benjamin Rambo-Martin, Shatavia Morrison, Christopher Gulvick, Scott Sammons, Yvette Unoarumhi, Darlene Wagner, Matthew Schmerer, S. H. Rosenthal, A. Gerasimova, R. M. Kagan, B. Anderson, M. Hua, Y. Liu, L.E. Bernstein, K.E. Livingston, A. Perez, I. A. Shlyakhter, R. V. Rolando, R. Owen, P. Tanpaiboon, F. Lacbawan, Clinton R. Paden, Duncan MacCannell |
| EPI_ISL_3396472 | St. Anne´s University Hospital Brno | University Hospital Brno, CMBG | Matej Bezdicek, Jan Svaton, Martina Lengerova, Kristyna Dufkova, Pavlina Volfova |
| EPI_ISL_3396473, EPI_ISL_3396474 | Quest Diagnostics Incorporated | Centers for Disease Control and Prevention Division of Viral Diseases, Pathogen Discovery | Dakota Howard, Dhwani Batra, Peter W. Cook, Kara Moser, Adrian Paskey, Jason Caravas, Benjamin Rambo-Martin, Shatavia Morrison, Christopher Gulvick, Scott Sammons, Yvette Unoarumhi, Darlene Wagner, Matthew Schmerer, S. H. Rosenthal, A. Gerasimova, R. M. Kagan, B. Anderson, M. Hua, Y. Liu, L.E. Bernstein, K.E. Livingston, A. Perez, I. A. Shlyakhter, R. V. Rolando, R. Owen, P. Tanpaiboon, F. Lacbawan, Clinton R. Paden, Duncan MacCannell |
| EPI_ISL_3396475 | St. Anne´s University Hospital Brno | University Hospital Brno, CMBG | Matej Bezdicek, Jan Svaton, Martina Lengerova, Kristyna Dufkova, Pavlina Volfova |
| EPI_ISL_3396476 | Quest Diagnostics Incorporated | Centers for Disease Control and Prevention Division of Viral Diseases, Pathogen Discovery | Dakota Howard, Dhwani Batra, Peter W. Cook, Kara Moser, Adrian Paskey, Jason Caravas, Benjamin Rambo-Martin, Shatavia Morrison, Christopher Gulvick, Scott Sammons, Yvette Unoarumhi, Darlene Wagner, Matthew Schmerer, S. H. Rosenthal, A. Gerasimova, R. M. Kagan, B. Anderson, M. Hua, Y. Liu, L.E. Bernstein, K.E. Livingston, A. Perez, I. A. Shlyakhter, R. V. Rolando, R. Owen, P. Tanpaiboon, F. Lacbawan, Clinton R. Paden, Duncan MacCannell |
| EPI_ISL_3396477 | IFCOR Brno | University Hospital Brno, CMBG | Matej Bezdicek, Jan Svaton, Martina Lengerova, Kristyna Dufkova, Pavlina Volfova |
| EPI_ISL_3396478, EPI_ISL_3396479 | Quest Diagnostics Incorporated | Centers for Disease Control and Prevention Division of Viral Diseases, Pathogen Discovery | Dakota Howard, Dhwani Batra, Peter W. Cook, Kara Moser, Adrian Paskey, Jason Caravas, Benjamin Rambo-Martin, Shatavia Morrison, Christopher Gulvick, Scott Sammons, Yvette Unoarumhi, Darlene Wagner, Matthew Schmerer, S. H. Rosenthal, A. Gerasimova, R. M. Kagan, B. Anderson, M. Hua, Y. Liu, L.E. Bernstein, K.E. Livingston, A. Perez, I. A. Shlyakhter, R. V. Rolando, R. Owen, P. Tanpaiboon, F. Lacbawan, Clinton R. Paden, Duncan MacCannell |
| EPI_ISL_3396480 | University Hospital Brno, OKMI | University Hospital Brno, CMBG | Matej Bezdicek, Jan Svaton, Martina Lengerova, Kristyna Dufkova, Pavlina Volfova |
| EPI_ISL_3396481 | Quest Diagnostics Incorporated | Centers for Disease Control and Prevention Division of Viral Diseases, Pathogen Discovery | Dakota Howard, Dhwani Batra, Peter W. Cook, Kara Moser, Adrian Paskey, Jason Caravas, Benjamin Rambo-Martin, Shatavia Morrison, Christopher Gulvick, Scott Sammons, Yvette Unoarumhi, Darlene Wagner, Matthew Schmerer, S. H. Rosenthal, A. Gerasimova, R. M. Kagan, B. Anderson, M. Hua, Y. Liu, L.E. Bernstein, K.E. Livingston, A. Perez, I. A. Shlyakhter, R. V. Rolando, R. Owen, P. Tanpaiboon, F. Lacbawan, Clinton R. Paden, Duncan MacCannell |
| EPI_ISL_3396482 | University Hospital Brno, OKMI | University Hospital Brno, CMBG | Matej Bezdicek, Jan Svaton, Martina Lengerova, Kristyna Dufkova, Pavlina Volfova |
| EPI_ISL_3396483 | Quest Diagnostics Incorporated | Centers for Disease Control and Prevention Division of Viral Diseases, Pathogen Discovery | Dakota Howard, Dhwani Batra, Peter W. Cook, Kara Moser, Adrian Paskey, Jason Caravas, Benjamin Rambo-Martin, Shatavia Morrison, Christopher Gulvick, Scott Sammons, Yvette Unoarumhi, Darlene Wagner, Matthew Schmerer, S. H. Rosenthal, A. Gerasimova, R. M. Kagan, B. Anderson, M. Hua, Y. Liu, L.E. Bernstein, K.E. Livingston, A. Perez, I. A. Shlyakhter, R. V. Rolando, R. Owen, P. Tanpaiboon, F. Lacbawan, Clinton R. Paden, Duncan MacCannell |
| EPI_ISL_3396484 | University Hospital Brno, OKMI | University Hospital Brno, CMBG | Matej Bezdicek, Jan Svaton, Martina Lengerova, Kristyna Dufkova, Pavlina Volfova |
| EPI_ISL_3396485, EPI_ISL_3396486 | Quest Diagnostics Incorporated | Centers for Disease Control and Prevention Division of Viral Diseases, Pathogen Discovery | Dakota Howard, Dhwani Batra, Peter W. Cook, Kara Moser, Adrian Paskey, Jason Caravas, Benjamin Rambo-Martin, Shatavia Morrison, Christopher Gulvick, Scott Sammons, Yvette Unoarumhi, Darlene Wagner, Matthew Schmerer, S. H. Rosenthal, A. Gerasimova, R. M. Kagan, B. Anderson, M. Hua, Y. Liu, L.E. Bernstein, K.E. Livingston, A. Perez, I. A. Shlyakhter, R. V. Rolando, R. Owen, P. Tanpaiboon, F. Lacbawan, Clinton R. Paden, Duncan MacCannell |
| EPI_ISL_3396487 | University Hospital Brno, OKMI | University Hospital Brno, CMBG | Matej Bezdicek, Jan Svaton, Martina Lengerova, Kristyna Dufkova, Pavlina Volfova |
| EPI_ISL_3396488 | Quest Diagnostics Incorporated | Centers for Disease Control and Prevention Division of Viral Diseases, Pathogen Discovery | Dakota Howard, Dhwani Batra, Peter W. Cook, Kara Moser, Adrian Paskey, Jason Caravas, Benjamin Rambo-Martin, Shatavia Morrison, Christopher Gulvick, Scott Sammons, Yvette Unoarumhi, Darlene Wagner, Matthew Schmerer, S. H. Rosenthal, A. Gerasimova, R. M. Kagan, B. Anderson, M. Hua, Y. Liu, L.E. Bernstein, K.E. Livingston, A. Perez, I. A. Shlyakhter, R. V. Rolando, R. Owen, P. Tanpaiboon, F. Lacbawan, Clinton R. Paden, Duncan MacCannell |
| EPI_ISL_3396489 | University Hospital Brno, OKMI | University Hospital Brno, CMBG | Matej Bezdicek, Jan Svaton, Martina Lengerova, Kristyna Dufkova, Pavlina Volfova |
| EPI_ISL_3396490 | Quest Diagnostics Incorporated | Centers for Disease Control and Prevention Division of Viral Diseases, Pathogen Discovery | Dakota Howard, Dhwani Batra, Peter W. Cook, Kara Moser, Adrian Paskey, Jason Caravas, Benjamin Rambo-Martin, Shatavia Morrison, Christopher Gulvick, Scott Sammons, Yvette Unoarumhi, Darlene Wagner, Matthew Schmerer, S. H. Rosenthal, A. Gerasimova, R. M. Kagan, B. Anderson, M. Hua, Y. Liu, L.E. Bernstein, K.E. Livingston, A. Perez, I. A. Shlyakhter, R. V. Rolando, R. Owen, P. Tanpaiboon, F. Lacbawan, Clinton R. Paden, Duncan MacCannell |
| EPI_ISL_3396492 | University Hospital Brno, OKMI | University Hospital Brno, CMBG | Matej Bezdicek, Jan Svaton, Martina Lengerova, Kristyna Dufkova, Pavlina Volfova |
| EPI_ISL_3396493, EPI_ISL_3396494 | Quest Diagnostics Incorporated | Centers for Disease Control and Prevention Division of Viral Diseases, Pathogen Discovery | Dakota Howard, Dhwani Batra, Peter W. Cook, Kara Moser, Adrian Paskey, Jason Caravas, Benjamin Rambo-Martin, Shatavia Morrison, Christopher Gulvick, Scott Sammons, Yvette Unoarumhi, Darlene Wagner, Matthew Schmerer, S. H. Rosenthal, A. Gerasimova, R. M. Kagan, B. Anderson, M. Hua, Y. Liu, L.E. Bernstein, K.E. Livingston, A. Perez, I. A. Shlyakhter, R. V. Rolando, R. Owen, P. Tanpaiboon, F. Lacbawan, Clinton R. Paden, Duncan MacCannell |
| EPI_ISL_3396495 | University Hospital Brno, OKMI | University Hospital Brno, CMBG | Matej Bezdicek, Jan Svaton, Martina Lengerova, Kristyna Dufkova, Pavlina Volfova |
| EPI_ISL_3396496 | Quest Diagnostics Incorporated | Centers for Disease Control and Prevention Division of Viral Diseases, Pathogen Discovery | Dakota Howard, Dhwani Batra, Peter W. Cook, Kara Moser, Adrian Paskey, Jason Caravas, Benjamin Rambo-Martin, Shatavia Morrison, Christopher Gulvick, Scott Sammons, Yvette Unoarumhi, Darlene Wagner, Matthew Schmerer, S. H. Rosenthal, A. Gerasimova, R. M. Kagan, B. Anderson, M. Hua, Y. Liu, L.E. Bernstein, K.E. Livingston, A. Perez, I. A. Shlyakhter, R. V. Rolando, R. Owen, P. Tanpaiboon, F. Lacbawan, Clinton R. Paden, Duncan MacCannell |
| EPI_ISL_3396497 | University Hospital Brno, OKMI | University Hospital Brno, CMBG | Matej Bezdicek, Jan Svaton, Martina Lengerova, Kristyna Dufkova, Pavlina Volfova |
| EPI_ISL_3396498, EPI_ISL_3396499 | Quest Diagnostics Incorporated | Centers for Disease Control and Prevention Division of Viral Diseases, Pathogen Discovery | Dakota Howard, Dhwani Batra, Peter W. Cook, Kara Moser, Adrian Paskey, Jason Caravas, Benjamin Rambo-Martin, Shatavia Morrison, Christopher Gulvick, Scott Sammons, Yvette Unoarumhi, Darlene Wagner, Matthew Schmerer, S. H. Rosenthal, A. Gerasimova, R. M. Kagan, B. Anderson, M. Hua, Y. Liu, L.E. Bernstein, K.E. Livingston, A. Perez, I. A. Shlyakhter, R. V. Rolando, R. Owen, P. Tanpaiboon, F. Lacbawan, Clinton R. Paden, Duncan MacCannell |
| EPI_ISL_3396500 | University Hospital Brno, OKMI | University Hospital Brno, CMBG | Matej Bezdicek, Jan Svaton, Martina Lengerova, Kristyna Dufkova, Pavlina Volfova |
| EPI_ISL_3396501, EPI_ISL_3396502 | Quest Diagnostics Incorporated | Centers for Disease Control and Prevention Division of Viral Diseases, Pathogen Discovery | Dakota Howard, Dhwani Batra, Peter W. Cook, Kara Moser, Adrian Paskey, Jason Caravas, Benjamin Rambo-Martin, Shatavia Morrison, Christopher Gulvick, Scott Sammons, Yvette Unoarumhi, Darlene Wagner, Matthew Schmerer, S. H. Rosenthal, A. Gerasimova, R. M. Kagan, B. Anderson, M. Hua, Y. Liu, L.E. Bernstein, K.E. Livingston, A. Perez, I. A. Shlyakhter, R. V. Rolando, R. Owen, P. Tanpaiboon, F. Lacbawan, Clinton R. Paden, Duncan MacCannell |
| EPI_ISL_3396503 | University Hospital Brno, OKMI | University Hospital Brno, CMBG | Matej Bezdicek, Jan Svaton, Martina Lengerova, Kristyna Dufkova, Pavlina Volfova |
| EPI_ISL_3396504 | Quest Diagnostics Incorporated | Centers for Disease Control and Prevention Division of Viral Diseases, Pathogen Discovery | Dakota Howard, Dhwani Batra, Peter W. Cook, Kara Moser, Adrian Paskey, Jason Caravas, Benjamin Rambo-Martin, Shatavia Morrison, Christopher Gulvick, Scott Sammons, Yvette Unoarumhi, Darlene Wagner, Matthew Schmerer, S. H. Rosenthal, A. Gerasimova, R. M. Kagan, B. Anderson, M. Hua, Y. Liu, L.E. Bernstein, K.E. Livingston, A. Perez, I. A. Shlyakhter, R. V. Rolando, R. Owen, P. Tanpaiboon, F. Lacbawan, Clinton R. Paden, Duncan MacCannell |

| | | | |
|---|---|---|---|
| EPI_ISL_3396505 | University Hospital Brno, OKMI | University Hospital Brno, CMBG | Matej Bezdicek, Jan Svaton, Martina Lengerova, Kristyna Dufkova, Pavlina Volfova |
| EPI_ISL_3396506, EPI_ISL_3396507 | Quest Diagnostics Incorporated | Centers for Disease Control and Prevention Division of Viral Diseases, Pathogen Discovery | Dakota Howard, Dhwani Batra, Peter W. Cook, Kara Moser, Adrian Paskey, Jason Caravas, Benjamin Rambo-Martin, Shatavia Morrison, Christopher Gulvick, Scott Sammons, Yvette Unoarumhi, Darlene Wagner, Matthew Schmerer, S. H. Rosenthal, A. Gerasimova, R. M. Kagan, B. Anderson, M. Hua, Y. Liu, L.E. Bernstein, K.E. Livingston, A. Perez, I. A. Shlyakhter, R. V. Rolando, R. Owen, P. Tanpaiboon, F. Lacbawan, Clinton R. Paden, Duncan MacCannell |
| EPI_ISL_3396508 | University Hospital Brno, OKMI | University Hospital Brno, CMBG | Matej Bezdicek, Jan Svaton, Martina Lengerova, Kristyna Dufkova, Pavlina Volfova |
| EPI_ISL_3396509 | Quest Diagnostics Incorporated | Centers for Disease Control and Prevention Division of Viral Diseases, Pathogen Discovery | Dakota Howard, Dhwani Batra, Peter W. Cook, Kara Moser, Adrian Paskey, Jason Caravas, Benjamin Rambo-Martin, Shatavia Morrison, Christopher Gulvick, Scott Sammons, Yvette Unoarumhi, Darlene Wagner, Matthew Schmerer, S. H. Rosenthal, A. Gerasimova, R. M. Kagan, B. Anderson, M. Hua, Y. Liu, L.E. Bernstein, K.E. Livingston, A. Perez, I. A. Shlyakhter, R. V. Rolando, R. Owen, P. Tanpaiboon, F. Lacbawan, Clinton R. Paden, Duncan MacCannell |
| EPI_ISL_3396510 | University Hospital Brno, OKMI | University Hospital Brno, CMBG | Matej Bezdicek, Jan Svaton, Martina Lengerova, Kristyna Dufkova, Pavlina Volfova |
| EPI_ISL_3396511, EPI_ISL_3396512 | Quest Diagnostics Incorporated | Centers for Disease Control and Prevention Division of Viral Diseases, Pathogen Discovery | Dakota Howard, Dhwani Batra, Peter W. Cook, Kara Moser, Adrian Paskey, Jason Caravas, Benjamin Rambo-Martin, Shatavia Morrison, Christopher Gulvick, Scott Sammons, Yvette Unoarumhi, Darlene Wagner, Matthew Schmerer, S. H. Rosenthal, A. Gerasimova, R. M. Kagan, B. Anderson, M. Hua, Y. Liu, L.E. Bernstein, K.E. Livingston, A. Perez, I. A. Shlyakhter, R. V. Rolando, R. Owen, P. Tanpaiboon, F. Lacbawan, Clinton R. Paden, Duncan MacCannell |
| EPI_ISL_3396513 | University Hospital Brno, OKMI | University Hospital Brno, CMBG | Matej Bezdicek, Jan Svaton, Martina Lengerova, Kristyna Dufkova, Pavlina Volfova |
| EPI_ISL_3396514, EPI_ISL_3396515 | Quest Diagnostics Incorporated | Centers for Disease Control and Prevention Division of Viral Diseases, Pathogen Discovery | Dakota Howard, Dhwani Batra, Peter W. Cook, Kara Moser, Adrian Paskey, Jason Caravas, Benjamin Rambo-Martin, Shatavia Morrison, Christopher Gulvick, Scott Sammons, Yvette Unoarumhi, Darlene Wagner, Matthew Schmerer, S. H. Rosenthal, A. Gerasimova, R. M. Kagan, B. Anderson, M. Hua, Y. Liu, L.E. Bernstein, K.E. Livingston, A. Perez, I. A. Shlyakhter, R. V. Rolando, R. Owen, P. Tanpaiboon, F. Lacbawan, Clinton R. Paden, Duncan MacCannell |
| EPI_ISL_3396516 | University Hospital Brno, OKMI | University Hospital Brno, CMBG | Matej Bezdicek, Jan Svaton, Martina Lengerova, Kristyna Dufkova, Pavlina Volfova |
| EPI_ISL_3396517 | Quest Diagnostics Incorporated | Centers for Disease Control and Prevention Division of Viral Diseases, Pathogen Discovery | Dakota Howard, Dhwani Batra, Peter W. Cook, Kara Moser, Adrian Paskey, Jason Caravas, Benjamin Rambo-Martin, Shatavia Morrison, Christopher Gulvick, Scott Sammons, Yvette Unoarumhi, Darlene Wagner, Matthew Schmerer, S. H. Rosenthal, A. Gerasimova, R. M. Kagan, B. Anderson, M. Hua, Y. Liu, L.E. Bernstein, K.E. Livingston, A. Perez, I. A. Shlyakhter, R. V. Rolando, R. Owen, P. Tanpaiboon, F. Lacbawan, Clinton R. Paden, Duncan MacCannell |
| EPI_ISL_3396518 | University Hospital Brno, OKMI | University Hospital Brno, CMBG | Matej Bezdicek, Jan Svaton, Martina Lengerova, Kristyna Dufkova, Pavlina Volfova |
| EPI_ISL_3396519, EPI_ISL_3396520 | Quest Diagnostics Incorporated | Centers for Disease Control and Prevention Division of Viral Diseases, Pathogen Discovery | Dakota Howard, Dhwani Batra, Peter W. Cook, Kara Moser, Adrian Paskey, Jason Caravas, Benjamin Rambo-Martin, Shatavia Morrison, Christopher Gulvick, Scott Sammons, Yvette Unoarumhi, Darlene Wagner, Matthew Schmerer, S. H. Rosenthal, A. Gerasimova, R. M. Kagan, B. Anderson, M. Hua, Y. Liu, L.E. Bernstein, K.E. Livingston, A. Perez, I. A. Shlyakhter, R. V. Rolando, R. Owen, P. Tanpaiboon, F. Lacbawan, Clinton R. Paden, Duncan MacCannell |
| EPI_ISL_3396521 | University Hospital Brno, OKMI | University Hospital Brno, CMBG | Matej Bezdicek, Jan Svaton, Martina Lengerova, Kristyna Dufkova, Pavlina Volfova |
| EPI_ISL_3396522 | Quest Diagnostics Incorporated | Centers for Disease Control and Prevention Division of Viral Diseases, Pathogen Discovery | Dakota Howard, Dhwani Batra, Peter W. Cook, Kara Moser, Adrian Paskey, Jason Caravas, Benjamin Rambo-Martin, Shatavia Morrison, Christopher Gulvick, Scott Sammons, Yvette Unoarumhi, Darlene Wagner, Matthew Schmerer, S. H. Rosenthal, A. Gerasimova, R. M. Kagan, B. Anderson, M. Hua, Y. Liu, L.E. Bernstein, K.E. Livingston, A. Perez, I. A. Shlyakhter, R. V. Rolando, R. Owen, P. Tanpaiboon, F. Lacbawan, Clinton R. Paden, Duncan MacCannell |
| EPI_ISL_3396523 | University Hospital Brno, OKMI | University Hospital Brno, CMBG | Matej Bezdicek, Jan Svaton, Martina Lengerova, Kristyna Dufkova, Pavlina Volfova |
| EPI_ISL_3396524, EPI_ISL_3396525 | Quest Diagnostics Incorporated | Centers for Disease Control and Prevention Division of Viral Diseases, Pathogen Discovery | Dakota Howard, Dhwani Batra, Peter W. Cook, Kara Moser, Adrian Paskey, Jason Caravas, Benjamin Rambo-Martin, Shatavia Morrison, Christopher Gulvick, Scott Sammons, Yvette Unoarumhi, Darlene Wagner, Matthew Schmerer, S. H. Rosenthal, A. Gerasimova, R. M. Kagan, B. Anderson, M. Hua, Y. Liu, L.E. Bernstein, K.E. Livingston, A. Perez, I. A. Shlyakhter, R. V. Rolando, R. Owen, P. Tanpaiboon, F. Lacbawan, Clinton R. Paden, Duncan MacCannell |
| EPI_ISL_3396526 | University Hospital Brno, OKMI | University Hospital Brno, CMBG | Matej Bezdicek, Jan Svaton, Martina Lengerova, Kristyna Dufkova, Pavlina Volfova |
| EPI_ISL_3396527 | Quest Diagnostics Incorporated | Centers for Disease Control and Prevention Division of Viral Diseases, Pathogen Discovery | Dakota Howard, Dhwani Batra, Peter W. Cook, Kara Moser, Adrian Paskey, Jason Caravas, Benjamin Rambo-Martin, Shatavia Morrison, Christopher Gulvick, Scott Sammons, Yvette Unoarumhi, Darlene Wagner, Matthew Schmerer, S. H. Rosenthal, A. Gerasimova, R. M. Kagan, B. Anderson, M. Hua, Y. Liu, L.E. Bernstein, K.E. Livingston, A. Perez, I. A. Shlyakhter, R. V. Rolando, R. Owen, P. Tanpaiboon, F. Lacbawan, Clinton R. Paden, Duncan MacCannell |
| EPI_ISL_3396528 | Elisabeth Pharmacon | University Hospital Brno, CMBG | Matej Bezdicek, Jan Svaton, Martina Lengerova, Kristyna Dufkova, Pavlina Volfova |
| EPI_ISL_3396529, EPI_ISL_3396530 | Quest Diagnostics Incorporated | Centers for Disease Control and Prevention Division of Viral Diseases, Pathogen Discovery | Dakota Howard, Dhwani Batra, Peter W. Cook, Kara Moser, Adrian Paskey, Jason Caravas, Benjamin Rambo-Martin, Shatavia Morrison, Christopher Gulvick, Scott Sammons, Yvette Unoarumhi, Darlene Wagner, Matthew Schmerer, S. H. Rosenthal, A. Gerasimova, R. M. Kagan, B. Anderson, M. Hua, Y. Liu, L.E. Bernstein, K.E. Livingston, A. Perez, I. A. Shlyakhter, R. V. Rolando, R. Owen, P. Tanpaiboon, F. Lacbawan, Clinton R. Paden, Duncan MacCannell |
| EPI_ISL_3396531 | Elisabeth Pharmacon | University Hospital Brno, CMBG | Matej Bezdicek, Jan Svaton, Martina Lengerova, Kristyna Dufkova, Pavlina Volfova |
| EPI_ISL_3396532 | Quest Diagnostics Incorporated | Centers for Disease Control and Prevention Division of Viral Diseases, Pathogen Discovery | Dakota Howard, Dhwani Batra, Peter W. Cook, Kara Moser, Adrian Paskey, Jason Caravas, Benjamin Rambo-Martin, Shatavia Morrison, Christopher Gulvick, Scott Sammons, Yvette Unoarumhi, Darlene Wagner, Matthew Schmerer, S. H. Rosenthal, A. Gerasimova, R. M. Kagan, B. Anderson, M. Hua, Y. Liu, L.E. Bernstein, K.E. Livingston, A. Perez, I. A. Shlyakhter, R. V. Rolando, R. Owen, P. Tanpaiboon, F. Lacbawan, Clinton R. Paden, Duncan MacCannell |
| EPI_ISL_3396533 | Elisabeth Pharmacon | University Hospital Brno, CMBG | Matej Bezdicek, Jan Svaton, Martina Lengerova, Kristyna Dufkova, Pavlina Volfova |
| EPI_ISL_3396535 | Quest Diagnostics Incorporated | Centers for Disease Control and Prevention Division of Viral Diseases, Pathogen Discovery | Dakota Howard, Dhwani Batra, Peter W. Cook, Kara Moser, Adrian Paskey, Jason Caravas, Benjamin Rambo-Martin, Shatavia Morrison, Christopher Gulvick, Scott Sammons, Yvette Unoarumhi, Darlene Wagner, Matthew Schmerer, S. H. Rosenthal, A. Gerasimova, R. M. Kagan, B. Anderson, M. Hua, Y. Liu, L.E. Bernstein, K.E. Livingston, A. Perez, I. A. Shlyakhter, R. V. Rolando, R. Owen, P. Tanpaiboon, F. Lacbawan, Clinton R. Paden, Duncan MacCannell |
| EPI_ISL_3396536 | Elisabeth Pharmacon | University Hospital Brno, CMBG | Matej Bezdicek, Jan Svaton, Martina Lengerova, Kristyna Dufkova, Pavlina Volfova |
| EPI_ISL_3396537 | Quest Diagnostics Incorporated | Centers for Disease Control and Prevention Division of Viral Diseases, Pathogen Discovery | Dakota Howard, Dhwani Batra, Peter W. Cook, Kara Moser, Adrian Paskey, Jason Caravas, Benjamin Rambo-Martin, Shatavia Morrison, Christopher Gulvick, Scott Sammons, Yvette Unoarumhi, Darlene Wagner, Matthew Schmerer, S. H. Rosenthal, A. Gerasimova, R. M. Kagan, B. Anderson, M. Hua, Y. Liu, L.E. Bernstein, K.E. Livingston, A. Perez, I. A. Shlyakhter, R. V. Rolando, R. Owen, P. Tanpaiboon, F. Lacbawan, Clinton R. Paden, Duncan MacCannell |
| EPI_ISL_3396538 | Elisabeth Pharmacon | University Hospital Brno, CMBG | Matej Bezdicek, Jan Svaton, Martina Lengerova, Kristyna Dufkova, Pavlina Volfova |
| EPI_ISL_3396539, EPI_ISL_3396540 | Quest Diagnostics Incorporated | Centers for Disease Control and Prevention Division of Viral Diseases, Pathogen Discovery | Dakota Howard, Dhwani Batra, Peter W. Cook, Kara Moser, Adrian Paskey, Jason Caravas, Benjamin Rambo-Martin, Shatavia Morrison, Christopher Gulvick, Scott Sammons, Yvette Unoarumhi, Darlene Wagner, Matthew Schmerer, S. H. Rosenthal, A. Gerasimova, R. M. Kagan, B. Anderson, M. Hua, Y. Liu, L.E. Bernstein, K.E. Livingston, A. Perez, I. A. Shlyakhter, R. V. Rolando, R. Owen, P. Tanpaiboon, F. Lacbawan, Clinton R. Paden, Duncan MacCannell |
| EPI_ISL_3396541 | Elisabeth Pharmacon | University Hospital Brno, CMBG | Matej Bezdicek, Jan Svaton, Martina Lengerova, Kristyna Dufkova, Pavlina Volfova |
| EPI_ISL_3396542 | Quest Diagnostics Incorporated | Centers for Disease Control and Prevention Division of Viral Diseases, Pathogen Discovery | Dakota Howard, Dhwani Batra, Peter W. Cook, Kara Moser, Adrian Paskey, Jason Caravas, Benjamin Rambo-Martin, Shatavia Morrison, Christopher Gulvick, Scott Sammons, Yvette Unoarumhi, Darlene Wagner, Matthew Schmerer, S. H. Rosenthal, A. Gerasimova, R. M. Kagan, B. Anderson, M. Hua, Y. Liu, L.E. Bernstein, K.E. Livingston, A. Perez, I. A. Shlyakhter, R. V. Rolando, R. Owen, P. Tanpaiboon, F. Lacbawan, Clinton R. Paden, Duncan MacCannell |
| EPI_ISL_3396543 | Elisabeth Pharmacon | University Hospital Brno, CMBG | Matej Bezdicek, Jan Svaton, Martina Lengerova, Kristyna Dufkova, Pavlina Volfova |
| EPI_ISL_3396544, EPI_ISL_3396545 | Quest Diagnostics Incorporated | Centers for Disease Control and Prevention Division of Viral Diseases, Pathogen Discovery | Dakota Howard, Dhwani Batra, Peter W. Cook, Kara Moser, Adrian Paskey, Jason Caravas, Benjamin Rambo-Martin, Shatavia Morrison, Christopher Gulvick, Scott Sammons, Yvette Unoarumhi, Darlene Wagner, Matthew Schmerer, S. H. Rosenthal, A. Gerasimova, R. M. Kagan, B. Anderson, M. Hua, Y. Liu, L.E. Bernstein, K.E. Livingston, A. Perez, I. A. Shlyakhter, R. V. Rolando, R. Owen, P. Tanpaiboon, F. Lacbawan, Clinton R. Paden, Duncan MacCannell |
| EPI_ISL_3396546 | Elisabeth Pharmacon | University Hospital Brno, CMBG | Matej Bezdicek, Jan Svaton, Martina Lengerova, Kristyna Dufkova, Pavlina Volfova |
| EPI_ISL_3396547, EPI_ISL_3396548 | Quest Diagnostics Incorporated | Centers for Disease Control and Prevention Division of Viral Diseases, Pathogen Discovery | Dakota Howard, Dhwani Batra, Peter W. Cook, Kara Moser, Adrian Paskey, Jason Caravas, Benjamin Rambo-Martin, Shatavia Morrison, Christopher Gulvick, Scott Sammons, Yvette Unoarumhi, Darlene Wagner, Matthew Schmerer, S. H. Rosenthal, A. Gerasimova, R. M. Kagan, B. Anderson, M. Hua, Y. Liu, L.E. Bernstein, K.E. Livingston, A. Perez, I. A. Shlyakhter, R. V. Rolando, R. Owen, P. Tanpaiboon, F. Lacbawan, Clinton R. Paden, Duncan MacCannell |
| EPI_ISL_3396549 | Elisabeth Pharmacon | University Hospital Brno, CMBG | Matej Bezdicek, Jan Svaton, Martina Lengerova, Kristyna Dufkova, Pavlina Volfova |
| EPI_ISL_3396550 | Quest Diagnostics Incorporated | Centers for Disease Control and Prevention Division of Viral Diseases, Pathogen Discovery | Dakota Howard, Dhwani Batra, Peter W. Cook, Kara Moser, Adrian Paskey, Jason Caravas, Benjamin Rambo-Martin, Shatavia Morrison, Christopher Gulvick, Scott Sammons, Yvette Unoarumhi, Darlene Wagner, Matthew Schmerer, S. H. Rosenthal, A. Gerasimova, R. M. Kagan, B. Anderson, M. Hua, Y. Liu, L.E. Bernstein, K.E. Livingston, A. Perez, I. A. Shlyakhter, R. V. Rolando, R. Owen, P. Tanpaiboon, F. Lacbawan, Clinton R. Paden, Duncan MacCannell |
| EPI_ISL_3396551 | Elisabeth Pharmacon | University Hospital Brno, CMBG | Matej Bezdicek, Jan Svaton, Martina Lengerova, Kristyna Dufkova, Pavlina Volfova |
| EPI_ISL_3396552, EPI_ISL_3396553 | Quest Diagnostics Incorporated | Centers for Disease Control and Prevention Division of Viral Diseases, Pathogen Discovery | Dakota Howard, Dhwani Batra, Peter W. Cook, Kara Moser, Adrian Paskey, Jason Caravas, Benjamin Rambo-Martin, Shatavia Morrison, Christopher Gulvick, Scott Sammons, Yvette Unoarumhi, Darlene Wagner, Matthew Schmerer, S. H. Rosenthal, A. Gerasimova, R. M. Kagan, B. Anderson, M. Hua, Y. Liu, L.E. Bernstein, K.E. Livingston, A. Perez, I. A. Shlyakhter, R. V. Rolando, R. Owen, P. Tanpaiboon, F. Lacbawan, Clinton R. Paden, Duncan MacCannell |
| EPI_ISL_3396554 | Elisabeth Pharmacon | University Hospital Brno, CMBG | Matej Bezdicek, Jan Svaton, Martina Lengerova, Kristyna Dufkova, Pavlina Volfova |
| EPI_ISL_3396555 | Quest Diagnostics Incorporated | Centers for Disease Control and Prevention Division of Viral Diseases, Pathogen Discovery | Dakota Howard, Dhwani Batra, Peter W. Cook, Kara Moser, Adrian Paskey, Jason Caravas, Benjamin Rambo-Martin, Shatavia Morrison, Christopher Gulvick, Scott Sammons, Yvette Unoarumhi, Darlene Wagner, Matthew Schmerer, S. H. Rosenthal, A. Gerasimova, R. M. Kagan, B. Anderson, M. Hua, Y. Liu, L.E. Bernstein, K.E. Livingston, A. Perez, I. A. Shlyakhter, R. V. Rolando, R. Owen, P. Tanpaiboon, F. Lacbawan, Clinton R. Paden, Duncan MacCannell |
| EPI_ISL_3396556 | Elisabeth Pharmacon | University Hospital Brno, CMBG | Matej Bezdicek, Jan Svaton, Martina Lengerova, Kristyna Dufkova, Pavlina Volfova |
| EPI_ISL_3396557, EPI_ISL_3396558 | Quest Diagnostics Incorporated | Centers for Disease Control and Prevention Division of Viral Diseases, Pathogen Discovery | Dakota Howard, Dhwani Batra, Peter W. Cook, Kara Moser, Adrian Paskey, Jason Caravas, Benjamin Rambo-Martin, Shatavia Morrison, Christopher Gulvick, Scott Sammons, Yvette Unoarumhi, Darlene Wagner, Matthew Schmerer, S. H. Rosenthal, A. Gerasimova, R. M. Kagan, B. Anderson, M. Hua, Y. Liu, L.E. Bernstein, K.E. Livingston, A. Perez, I. A. Shlyakhter, R. V. Rolando, R. Owen, P. Tanpaiboon, F. Lacbawan, Clinton R. Paden, Duncan MacCannell |
| EPI_ISL_3396559 | Elisabeth Pharmacon | University Hospital Brno, CMBG | Matej Bezdicek, Jan Svaton, Martina Lengerova, Kristyna Dufkova, Pavlina Volfova |
| EPI_ISL_3396561, EPI_ISL_3396562, EPI_ISL_3396563, EPI_ISL_3396564, EPI_ISL_3396565, EPI_ISL_3396567, EPI_ISL_3396569, EPI_ISL_3396570, EPI_ISL_3396571, EPI_ISL_3396572, EPI_ISL_3396573, EPI_ISL_3396574, EPI_ISL_3396575, EPI_ISL_3396576, EPI_ISL_3396577, EPI_ISL_3396578, EPI_ISL_3396579, EPI_ISL_3396580, EPI_ISL_3396581, EPI_ISL_3396582, EPI_ISL_3396583, EPI_ISL_3396584, EPI_ISL_3396585, EPI_ISL_3396587, EPI_ISL_3396588 | | | |
| see above | Quest Diagnostics Incorporated | Centers for Disease Control and Prevention Division of Viral Diseases, Pathogen Discovery | Dakota Howard, Dhwani Batra, Peter W. Cook, Kara Moser, Adrian Paskey, Jason Caravas, Benjamin Rambo-Martin, Shatavia Morrison, Christopher Gulvick, Scott Sammons, Yvette Unoarumhi, Darlene Wagner, Matthew Schmerer, S. H. Rosenthal, A. Gerasimova, R. M. Kagan, B. Anderson, M. Hua, Y. Liu, L.E. Bernstein, K.E. Livingston, A. Perez, I. A. Shlyakhter, R. V. Rolando, R. Owen, P. Tanpaiboon, F. Lacbawan, Clinton R. Paden, Duncan MacCannell |
| EPI_ISL_3396589 | Kansas Health and Environmental Lab | Kansas Health and Environmental Lab | Mike Grose, Jonathan Barnell, Ben Olsen, and Phil Adam |
| EPI_ISL_3396590, EPI_ISL_3396591 | Quest Diagnostics Incorporated | Centers for Disease Control and Prevention Division of Viral Diseases, Pathogen Discovery | Dakota Howard, Dhwani Batra, Peter W. Cook, Kara Moser, Adrian Paskey, Jason Caravas, Benjamin Rambo-Martin, Shatavia Morrison, Christopher Gulvick, Scott Sammons, Yvette Unoarumhi, Darlene Wagner, Matthew Schmerer, S. H. Rosenthal, A. Gerasimova, R. M. Kagan, B. Anderson, M. Hua, Y. Liu, L.E. Bernstein, K.E. Livingston, A. Perez, I. A. Shlyakhter, R. V. Rolando, R. Owen, P. Tanpaiboon, F. Lacbawan, Clinton R. Paden, Duncan MacCannell |
| EPI_ISL_3396592 | Kansas Health and Environmental Lab | Kansas Health and Environmental Lab | Mike Grose, Jonathan Barnell, Ben Olsen, and Phil Adam |
| EPI_ISL_3396593, EPI_ISL_3396594 | Quest Diagnostics Incorporated | Centers for Disease Control and Prevention Division of Viral Diseases, Pathogen Discovery | Dakota Howard, Dhwani Batra, Peter W. Cook, Kara Moser, Adrian Paskey, Jason Caravas, Benjamin Rambo-Martin, Shatavia Morrison, Christopher Gulvick, Scott Sammons, Yvette Unoarumhi, Darlene Wagner, Matthew Schmerer, S. H. Rosenthal, A. Gerasimova, R. M. Kagan, B. Anderson, M. Hua, Y. Liu, L.E. Bernstein, K.E. Livingston, A. Perez, I. A. Shlyakhter, R. V. Rolando, R. Owen, P. Tanpaiboon, F. Lacbawan, Clinton R. Paden, Duncan MacCannell |
| EPI_ISL_3396595 | Kansas Health and Environmental Lab | Kansas Health and Environmental Lab | Mike Grose, Jonathan Barnell, Ben Olsen, and Phil Adam |
| EPI_ISL_3396596, EPI_ISL_3396597 | Quest Diagnostics Incorporated | Centers for Disease Control and Prevention Division of Viral Diseases, Pathogen Discovery | Dakota Howard, Dhwani Batra, Peter W. Cook, Kara Moser, Adrian Paskey, Jason Caravas, Benjamin Rambo-Martin, Shatavia Morrison, Christopher Gulvick, Scott Sammons, Yvette Unoarumhi, Darlene Wagner, Matthew Schmerer, S. H. Rosenthal, A. Gerasimova, R. M. Kagan, B. Anderson, M. Hua, Y. Liu, L.E. Bernstein, K.E. Livingston, A. Perez, I. A. Shlyakhter, R. V. Rolando, R. Owen, P. Tanpaiboon, F. Lacbawan, Clinton R. Paden, Duncan MacCannell |
| EPI_ISL_3396598, EPI_ISL_3396600 | Kansas Health and Environmental Lab | Kansas Health and Environmental Lab | Mike Grose, Jonathan Barnell, Ben Olsen, and Phil Adam |
| EPI_ISL_3396601, EPI_ISL_3396602 | Quest Diagnostics Incorporated | Centers for Disease Control and Prevention Division of Viral Diseases, Pathogen Discovery | Dakota Howard, Dhwani Batra, Peter W. Cook, Kara Moser, Adrian Paskey, Jason Caravas, Benjamin Rambo-Martin, Shatavia Morrison, Christopher Gulvick, Scott Sammons, Yvette Unoarumhi, Darlene Wagner, Matthew Schmerer, S. H. Rosenthal, A. Gerasimova, R. M. Kagan, B. Anderson, M. Hua, Y. Liu, L.E. Bernstein, K.E. Livingston, A. Perez, I. A. Shlyakhter, R. V. Rolando, R. Owen, P. Tanpaiboon, F. Lacbawan, Clinton R. Paden, Duncan MacCannell |
| EPI_ISL_3396603 | Kansas Health and Environmental Lab | Kansas Health and Environmental Lab | Mike Grose, Jonathan Barnell, Ben Olsen, and Phil Adam |
| EPI_ISL_3396604, EPI_ISL_3396605 | Quest Diagnostics Incorporated | Centers for Disease Control and Prevention Division of Viral Diseases, Pathogen Discovery | Dakota Howard, Dhwani Batra, Peter W. Cook, Kara Moser, Adrian Paskey, Jason Caravas, Benjamin Rambo-Martin, Shatavia Morrison, Christopher Gulvick, Scott Sammons, Yvette Unoarumhi, Darlene Wagner, Matthew Schmerer, S. H. Rosenthal, A. Gerasimova, R. M. Kagan, B. Anderson, M. Hua, Y. Liu, L.E. Bernstein, K.E. Livingston, A. Perez, I. A. Shlyakhter, R. V. Rolando, R. Owen, P. Tanpaiboon, F. Lacbawan, Clinton R. Paden, Duncan MacCannell |
| EPI_ISL_3396606 | Kansas Health and Environmental Lab | Kansas Health and Environmental Lab | Mike Grose, Jonathan Barnell, Ben Olsen, and Phil Adam |

| | | | |
|---|---|---|---|
| EPI_ISL_3396607 | Quest Diagnostics Incorporated | Centers for Disease Control and Prevention Division of Viral Diseases, Pathogen Discovery | Dakota Howard, Dhwani Batra, Peter W. Cook, Kara Moser, Adrian Paskey, Jason Caravas, Benjamin Rambo-Martin, Shatavia Morrison, Christopher Gulvick, Scott Sammons, Yvette Unoarumhi, Darlene Wagner, Matthew Schmerer, S. H. Rosenthal, A. Gerasimova, R. M. Kagan, B. Anderson, M. Hua, Y. Liu, L.E. Bernstein, K.E. Livingston, A. Perez, I. A. Shlyakhter, R. V. Rolando, R. Owen, P. Tanpaiboon, F. Lacbawan, Clinton R. Paden, Duncan MacCannell |
| EPI_ISL_3396608 | Kansas Health and Environmental Lab | Kansas Health and Environmental Lab | Mike Grose, Jonathan Barnell, Ben Olsen, and Phil Adam |
| EPI_ISL_3396609, EPI_ISL_3396610 | Quest Diagnostics Incorporated | Centers for Disease Control and Prevention Division of Viral Diseases, Pathogen Discovery | Dakota Howard, Dhwani Batra, Peter W. Cook, Kara Moser, Adrian Paskey, Jason Caravas, Benjamin Rambo-Martin, Shatavia Morrison, Christopher Gulvick, Scott Sammons, Yvette Unoarumhi, Darlene Wagner, Matthew Schmerer, S. H. Rosenthal, A. Gerasimova, R. M. Kagan, B. Anderson, M. Hua, Y. Liu, L.E. Bernstein, K.E. Livingston, A. Perez, I. A. Shlyakhter, R. V. Rolando, R. Owen, P. Tanpaiboon, F. Lacbawan, Clinton R. Paden, Duncan MacCannell |
| EPI_ISL_3396611 | Kansas Health and Environmental Lab | Kansas Health and Environmental Lab | Mike Grose, Jonathan Barnell, Ben Olsen, and Phil Adam |
| EPI_ISL_3396612, EPI_ISL_3396613, EPI_ISL_3396615, EPI_ISL_3396616 | Quest Diagnostics Incorporated | Centers for Disease Control and Prevention Division of Viral Diseases, Pathogen Discovery | Dakota Howard, Dhwani Batra, Peter W. Cook, Kara Moser, Adrian Paskey, Jason Caravas, Benjamin Rambo-Martin, Shatavia Morrison, Christopher Gulvick, Scott Sammons, Yvette Unoarumhi, Darlene Wagner, Matthew Schmerer, S. H. Rosenthal, A. Gerasimova, R. M. Kagan, B. Anderson, M. Hua, Y. Liu, L.E. Bernstein, K.E. Livingston, A. Perez, I. A. Shlyakhter, R. V. Rolando, R. Owen, P. Tanpaiboon, F. Lacbawan, Clinton R. Paden, Duncan MacCannell |
| EPI_ISL_3396617 | Kansas Health and Environmental Lab | Kansas Health and Environmental Lab | Mike Grose, Jonathan Barnell, Ben Olsen, and Phil Adam |
| EPI_ISL_3396618 | Quest Diagnostics Incorporated | Centers for Disease Control and Prevention Division of Viral Diseases, Pathogen Discovery | Dakota Howard, Dhwani Batra, Peter W. Cook, Kara Moser, Adrian Paskey, Jason Caravas, Benjamin Rambo-Martin, Shatavia Morrison, Christopher Gulvick, Scott Sammons, Yvette Unoarumhi, Darlene Wagner, Matthew Schmerer, S. H. Rosenthal, A. Gerasimova, R. M. Kagan, B. Anderson, M. Hua, Y. Liu, L.E. Bernstein, K.E. Livingston, A. Perez, I. A. Shlyakhter, R. V. Rolando, R. Owen, P. Tanpaiboon, F. Lacbawan, Clinton R. Paden, Duncan MacCannell |
| EPI_ISL_3396619 | Kansas Health and Environmental Lab | Kansas Health and Environmental Lab | Mike Grose, Jonathan Barnell, Ben Olsen, and Phil Adam |
| EPI_ISL_3396620, EPI_ISL_3396621 | Quest Diagnostics Incorporated | Centers for Disease Control and Prevention Division of Viral Diseases, Pathogen Discovery | Dakota Howard, Dhwani Batra, Peter W. Cook, Kara Moser, Adrian Paskey, Jason Caravas, Benjamin Rambo-Martin, Shatavia Morrison, Christopher Gulvick, Scott Sammons, Yvette Unoarumhi, Darlene Wagner, Matthew Schmerer, S. H. Rosenthal, A. Gerasimova, R. M. Kagan, B. Anderson, M. Hua, Y. Liu, L.E. Bernstein, K.E. Livingston, A. Perez, I. A. Shlyakhter, R. V. Rolando, R. Owen, P. Tanpaiboon, F. Lacbawan, Clinton R. Paden, Duncan MacCannell |
| EPI_ISL_3396622 | Kansas Health and Environmental Lab | Kansas Health and Environmental Lab | Mike Grose, Jonathan Barnell, Ben Olsen, and Phil Adam |
| EPI_ISL_3396623, EPI_ISL_3396624 | Quest Diagnostics Incorporated | Centers for Disease Control and Prevention Division of Viral Diseases, Pathogen Discovery | Dakota Howard, Dhwani Batra, Peter W. Cook, Kara Moser, Adrian Paskey, Jason Caravas, Benjamin Rambo-Martin, Shatavia Morrison, Christopher Gulvick, Scott Sammons, Yvette Unoarumhi, Darlene Wagner, Matthew Schmerer, S. H. Rosenthal, A. Gerasimova, R. M. Kagan, B. Anderson, M. Hua, Y. Liu, L.E. Bernstein, K.E. Livingston, A. Perez, I. A. Shlyakhter, R. V. Rolando, R. Owen, P. Tanpaiboon, F. Lacbawan, Clinton R. Paden, Duncan MacCannell |
| EPI_ISL_3396625 | Kansas Health and Environmental Lab | Kansas Health and Environmental Lab | Mike Grose, Jonathan Barnell, Ben Olsen, and Phil Adam |
| EPI_ISL_3396626, EPI_ISL_3396627 | Quest Diagnostics Incorporated | Centers for Disease Control and Prevention Division of Viral Diseases, Pathogen Discovery | Dakota Howard, Dhwani Batra, Peter W. Cook, Kara Moser, Adrian Paskey, Jason Caravas, Benjamin Rambo-Martin, Shatavia Morrison, Christopher Gulvick, Scott Sammons, Yvette Unoarumhi, Darlene Wagner, Matthew Schmerer, S. H. Rosenthal, A. Gerasimova, R. M. Kagan, B. Anderson, M. Hua, Y. Liu, L.E. Bernstein, K.E. Livingston, A. Perez, I. A. Shlyakhter, R. V. Rolando, R. Owen, P. Tanpaiboon, F. Lacbawan, Clinton R. Paden, Duncan MacCannell |
| EPI_ISL_3396628 | Kansas Health and Environmental Lab | Kansas Health and Environmental Lab | Mike Grose, Jonathan Barnell, Ben Olsen, and Phil Adam |
| EPI_ISL_3396629, EPI_ISL_3396630 | Quest Diagnostics Incorporated | Centers for Disease Control and Prevention Division of Viral Diseases, Pathogen Discovery | Dakota Howard, Dhwani Batra, Peter W. Cook, Kara Moser, Adrian Paskey, Jason Caravas, Benjamin Rambo-Martin, Shatavia Morrison, Christopher Gulvick, Scott Sammons, Yvette Unoarumhi, Darlene Wagner, Matthew Schmerer, S. H. Rosenthal, A. Gerasimova, R. M. Kagan, B. Anderson, M. Hua, Y. Liu, L.E. Bernstein, K.E. Livingston, A. Perez, I. A. Shlyakhter, R. V. Rolando, R. Owen, P. Tanpaiboon, F. Lacbawan, Clinton R. Paden, Duncan MacCannell |
| EPI_ISL_3396631 | Kansas Health and Environmental Lab | Kansas Health and Environmental Lab | Mike Grose, Jonathan Barnell, Ben Olsen, and Phil Adam |
| EPI_ISL_3396632 | Quest Diagnostics Incorporated | Centers for Disease Control and Prevention Division of Viral Diseases, Pathogen Discovery | Dakota Howard, Dhwani Batra, Peter W. Cook, Kara Moser, Adrian Paskey, Jason Caravas, Benjamin Rambo-Martin, Shatavia Morrison, Christopher Gulvick, Scott Sammons, Yvette Unoarumhi, Darlene Wagner, Matthew Schmerer, S. H. Rosenthal, A. Gerasimova, R. M. Kagan, B. Anderson, M. Hua, Y. Liu, L.E. Bernstein, K.E. Livingston, A. Perez, I. A. Shlyakhter, R. V. Rolando, R. Owen, P. Tanpaiboon, F. Lacbawan, Clinton R. Paden, Duncan MacCannell |
| EPI_ISL_3396633 | Kansas Health and Environmental Lab | Kansas Health and Environmental Lab | Mike Grose, Jonathan Barnell, Ben Olsen, and Phil Adam |
| EPI_ISL_3396634, EPI_ISL_3396635 | Quest Diagnostics Incorporated | Centers for Disease Control and Prevention Division of Viral Diseases, Pathogen Discovery | Dakota Howard, Dhwani Batra, Peter W. Cook, Kara Moser, Adrian Paskey, Jason Caravas, Benjamin Rambo-Martin, Shatavia Morrison, Christopher Gulvick, Scott Sammons, Yvette Unoarumhi, Darlene Wagner, Matthew Schmerer, S. H. Rosenthal, A. Gerasimova, R. M. Kagan, B. Anderson, M. Hua, Y. Liu, L.E. Bernstein, K.E. Livingston, A. Perez, I. A. Shlyakhter, R. V. Rolando, R. Owen, P. Tanpaiboon, F. Lacbawan, Clinton R. Paden, Duncan MacCannell |
| EPI_ISL_3396636 | Kansas Health and Environmental Lab | Kansas Health and Environmental Lab | Mike Grose, Jonathan Barnell, Ben Olsen, and Phil Adam |
| EPI_ISL_3396638 | Quest Diagnostics Incorporated | Centers for Disease Control and Prevention Division of Viral Diseases, Pathogen Discovery | Dakota Howard, Dhwani Batra, Peter W. Cook, Kara Moser, Adrian Paskey, Jason Caravas, Benjamin Rambo-Martin, Shatavia Morrison, Christopher Gulvick, Scott Sammons, Yvette Unoarumhi, Darlene Wagner, Matthew Schmerer, S. H. Rosenthal, A. Gerasimova, R. M. Kagan, B. Anderson, M. Hua, Y. Liu, L.E. Bernstein, K.E. Livingston, A. Perez, I. A. Shlyakhter, R. V. Rolando, R. Owen, P. Tanpaiboon, F. Lacbawan, Clinton R. Paden, Duncan MacCannell |
| EPI_ISL_3396639 | Kansas Health and Environmental Lab | Kansas Health and Environmental Lab | Mike Grose, Jonathan Barnell, Ben Olsen, and Phil Adam |
| EPI_ISL_3396640 | Quest Diagnostics Incorporated | Centers for Disease Control and Prevention Division of Viral Diseases, Pathogen Discovery | Dakota Howard, Dhwani Batra, Peter W. Cook, Kara Moser, Adrian Paskey, Jason Caravas, Benjamin Rambo-Martin, Shatavia Morrison, Christopher Gulvick, Scott Sammons, Yvette Unoarumhi, Darlene Wagner, Matthew Schmerer, S. H. Rosenthal, A. Gerasimova, R. M. Kagan, B. Anderson, M. Hua, Y. Liu, L.E. Bernstein, K.E. Livingston, A. Perez, I. A. Shlyakhter, R. V. Rolando, R. Owen, P. Tanpaiboon, F. Lacbawan, Clinton R. Paden, Duncan MacCannell |
| EPI_ISL_3396641 | Kansas Health and Environmental Lab | Kansas Health and Environmental Lab | Mike Grose, Jonathan Barnell, Ben Olsen, and Phil Adam |
| EPI_ISL_3396643 | Quest Diagnostics Incorporated | Centers for Disease Control and Prevention Division of Viral Diseases, Pathogen Discovery | Dakota Howard, Dhwani Batra, Peter W. Cook, Kara Moser, Adrian Paskey, Jason Caravas, Benjamin Rambo-Martin, Shatavia Morrison, Christopher Gulvick, Scott Sammons, Yvette Unoarumhi, Darlene Wagner, Matthew Schmerer, S. H. Rosenthal, A. Gerasimova, R. M. Kagan, B. Anderson, M. Hua, Y. Liu, L.E. Bernstein, K.E. Livingston, A. Perez, I. A. Shlyakhter, R. V. Rolando, R. Owen, P. Tanpaiboon, F. Lacbawan, Clinton R. Paden, Duncan MacCannell |
| EPI_ISL_3396644 | Kansas Health and Environmental Lab | Kansas Health and Environmental Lab | Mike Grose, Jonathan Barnell, Ben Olsen, and Phil Adam |
| EPI_ISL_3396645 | Quest Diagnostics Incorporated | Centers for Disease Control and Prevention Division of Viral Diseases, Pathogen Discovery | Dakota Howard, Dhwani Batra, Peter W. Cook, Kara Moser, Adrian Paskey, Jason Caravas, Benjamin Rambo-Martin, Shatavia Morrison, Christopher Gulvick, Scott Sammons, Yvette Unoarumhi, Darlene Wagner, Matthew Schmerer, S. H. Rosenthal, A. Gerasimova, R. M. Kagan, B. Anderson, M. Hua, Y. Liu, L.E. Bernstein, K.E. Livingston, A. Perez, I. A. Shlyakhter, R. V. Rolando, R. Owen, P. Tanpaiboon, F. Lacbawan, Clinton R. Paden, Duncan MacCannell |
| EPI_ISL_3396646 | Kansas Health and Environmental Lab | Kansas Health and Environmental Lab | Mike Grose, Jonathan Barnell, Ben Olsen, and Phil Adam |
| EPI_ISL_3396647, EPI_ISL_3396648 | Quest Diagnostics Incorporated | Centers for Disease Control and Prevention Division of Viral Diseases, Pathogen Discovery | Dakota Howard, Dhwani Batra, Peter W. Cook, Kara Moser, Adrian Paskey, Jason Caravas, Benjamin Rambo-Martin, Shatavia Morrison, Christopher Gulvick, Scott Sammons, Yvette Unoarumhi, Darlene Wagner, Matthew Schmerer, S. H. Rosenthal, A. Gerasimova, R. M. Kagan, B. Anderson, M. Hua, Y. Liu, L.E. Bernstein, K.E. Livingston, A. Perez, I. A. Shlyakhter, R. V. Rolando, R. Owen, P. Tanpaiboon, F. Lacbawan, Clinton R. Paden, Duncan MacCannell |
| EPI_ISL_3396649 | Kansas Health and Environmental Lab | Kansas Health and Environmental Lab | Mike Grose, Jonathan Barnell, Ben Olsen, and Phil Adam |
| EPI_ISL_3396650 | Quest Diagnostics Incorporated | Centers for Disease Control and Prevention Division of Viral Diseases, Pathogen Discovery | Dakota Howard, Dhwani Batra, Peter W. Cook, Kara Moser, Adrian Paskey, Jason Caravas, Benjamin Rambo-Martin, Shatavia Morrison, Christopher Gulvick, Scott Sammons, Yvette Unoarumhi, Darlene Wagner, Matthew Schmerer, S. H. Rosenthal, A. Gerasimova, R. M. Kagan, B. Anderson, M. Hua, Y. Liu, L.E. Bernstein, K.E. Livingston, A. Perez, I. A. Shlyakhter, R. V. Rolando, R. Owen, P. Tanpaiboon, F. Lacbawan, Clinton R. Paden, Duncan MacCannell |
| EPI_ISL_3396651 | Kansas Health and Environmental Lab | Kansas Health and Environmental Lab | Mike Grose, Jonathan Barnell, Ben Olsen, and Phil Adam |
| EPI_ISL_3396652, EPI_ISL_3396653 | Quest Diagnostics Incorporated | Centers for Disease Control and Prevention Division of Viral Diseases, Pathogen Discovery | Dakota Howard, Dhwani Batra, Peter W. Cook, Kara Moser, Adrian Paskey, Jason Caravas, Benjamin Rambo-Martin, Shatavia Morrison, Christopher Gulvick, Scott Sammons, Yvette Unoarumhi, Darlene Wagner, Matthew Schmerer, S. H. Rosenthal, A. Gerasimova, R. M. Kagan, B. Anderson, M. Hua, Y. Liu, L.E. Bernstein, K.E. Livingston, A. Perez, I. A. Shlyakhter, R. V. Rolando, R. Owen, P. Tanpaiboon, F. Lacbawan, Clinton R. Paden, Duncan MacCannell |
| EPI_ISL_3396654 | Kansas Health and Environmental Lab | Kansas Health and Environmental Lab | Mike Grose, Jonathan Barnell, Ben Olsen, and Phil Adam |
| EPI_ISL_3396655, EPI_ISL_3396656 | Quest Diagnostics Incorporated | Centers for Disease Control and Prevention Division of Viral Diseases, Pathogen Discovery | Dakota Howard, Dhwani Batra, Peter W. Cook, Kara Moser, Adrian Paskey, Jason Caravas, Benjamin Rambo-Martin, Shatavia Morrison, Christopher Gulvick, Scott Sammons, Yvette Unoarumhi, Darlene Wagner, Matthew Schmerer, S. H. Rosenthal, A. Gerasimova, R. M. Kagan, B. Anderson, M. Hua, Y. Liu, L.E. Bernstein, K.E. Livingston, A. Perez, I. A. Shlyakhter, R. V. Rolando, R. Owen, P. Tanpaiboon, F. Lacbawan, Clinton R. Paden, Duncan MacCannell |
| EPI_ISL_3396657 | Kansas Health and Environmental Lab | Kansas Health and Environmental Lab | Mike Grose, Jonathan Barnell, Ben Olsen, and Phil Adam |
| EPI_ISL_3396658 | Quest Diagnostics Incorporated | Centers for Disease Control and Prevention Division of Viral Diseases, Pathogen Discovery | Dakota Howard, Dhwani Batra, Peter W. Cook, Kara Moser, Adrian Paskey, Jason Caravas, Benjamin Rambo-Martin, Shatavia Morrison, Christopher Gulvick, Scott Sammons, Yvette Unoarumhi, Darlene Wagner, Matthew Schmerer, S. H. Rosenthal, A. Gerasimova, R. M. Kagan, B. Anderson, M. Hua, Y. Liu, L.E. Bernstein, K.E. Livingston, A. Perez, I. A. Shlyakhter, R. V. Rolando, R. Owen, P. Tanpaiboon, F. Lacbawan, Clinton R. Paden, Duncan MacCannell |
| EPI_ISL_3396659 | Kansas Health and Environmental Lab | Kansas Health and Environmental Lab | Mike Grose, Jonathan Barnell, Ben Olsen, and Phil Adam |
| EPI_ISL_3396660, EPI_ISL_3396661 | Quest Diagnostics Incorporated | Centers for Disease Control and Prevention Division of Viral Diseases, Pathogen Discovery | Dakota Howard, Dhwani Batra, Peter W. Cook, Kara Moser, Adrian Paskey, Jason Caravas, Benjamin Rambo-Martin, Shatavia Morrison, Christopher Gulvick, Scott Sammons, Yvette Unoarumhi, Darlene Wagner, Matthew Schmerer, S. H. Rosenthal, A. Gerasimova, R. M. Kagan, B. Anderson, M. Hua, Y. Liu, L.E. Bernstein, K.E. Livingston, A. Perez, I. A. Shlyakhter, R. V. Rolando, R. Owen, P. Tanpaiboon, F. Lacbawan, Clinton R. Paden, Duncan MacCannell |
| EPI_ISL_3396662 | Kansas Health and Environmental Lab | Kansas Health and Environmental Lab | Mike Grose, Jonathan Barnell, Ben Olsen, and Phil Adam |
| EPI_ISL_3396663, EPI_ISL_3396664, EPI_ISL_3396665, EPI_ISL_3396666, EPI_ISL_3396667, EPI_ISL_3396668, EPI_ISL_3396669, EPI_ISL_3396670, EPI_ISL_3396671, EPI_ISL_3396672, EPI_ISL_3396673, EPI_ISL_3396674, EPI_ISL_3396675, EPI_ISL_3396677, EPI_ISL_3396678, EPI_ISL_3396679, EPI_ISL_3396680, EPI_ISL_3396681, EPI_ISL_3396682, EPI_ISL_3396683, EPI_ISL_3396684, EPI_ISL_3396685, EPI_ISL_3396686, EPI_ISL_3396688, EPI_ISL_3396689, EPI_ISL_3396690 | | | |
| see above | Quest Diagnostics Incorporated | Centers for Disease Control and Prevention Division of Viral Diseases, Pathogen Discovery | Dakota Howard, Dhwani Batra, Peter W. Cook, Kara Moser, Adrian Paskey, Jason Caravas, Benjamin Rambo-Martin, Shatavia Morrison, Christopher Gulvick, Scott Sammons, Yvette Unoarumhi, Darlene Wagner, Matthew Schmerer, S. H. Rosenthal, A. Gerasimova, R. M. Kagan, B. Anderson, M. Hua, Y. Liu, L.E. Bernstein, K.E. Livingston, A. Perez, I. A. Shlyakhter, R. V. Rolando, R. Owen, P. Tanpaiboon, F. Lacbawan, Clinton R. Paden, Duncan MacCannell |
| EPI_ISL_3396691 | Kansas Health and Environmental Lab | Kansas Health and Environmental Lab | Mike Grose, Jonathan Barnell, Amanda Bradley, and Phil Adam |
| EPI_ISL_3396692, EPI_ISL_3396693, EPI_ISL_3396694 | Quest Diagnostics Incorporated | Centers for Disease Control and Prevention Division of Viral Diseases, Pathogen Discovery | Dakota Howard, Dhwani Batra, Peter W. Cook, Kara Moser, Adrian Paskey, Jason Caravas, Benjamin Rambo-Martin, Shatavia Morrison, Christopher Gulvick, Scott Sammons, Yvette Unoarumhi, Darlene Wagner, Matthew Schmerer, S. H. Rosenthal, A. Gerasimova, R. M. Kagan, B. Anderson, M. Hua, Y. Liu, L.E. Bernstein, K.E. Livingston, A. Perez, I. A. Shlyakhter, R. V. Rolando, R. Owen, P. Tanpaiboon, F. Lacbawan, Clinton R. Paden, Duncan MacCannell |
| EPI_ISL_3396695 | Kansas Health and Environmental Lab | Kansas Health and Environmental Lab | Mike Grose, Jonathan Barnell, Amanda Bradley, and Phil Adam |
| EPI_ISL_3396696 | Quest Diagnostics Incorporated | Centers for Disease Control and Prevention Division of Viral Diseases, Pathogen Discovery | Dakota Howard, Dhwani Batra, Peter W. Cook, Kara Moser, Adrian Paskey, Jason Caravas, Benjamin Rambo-Martin, Shatavia Morrison, Christopher Gulvick, Scott Sammons, Yvette Unoarumhi, Darlene Wagner, Matthew Schmerer, S. H. Rosenthal, A. Gerasimova, R. M. Kagan, B. Anderson, M. Hua, Y. Liu, L.E. Bernstein, K.E. Livingston, A. Perez, I. A. Shlyakhter, R. V. Rolando, R. Owen, P. Tanpaiboon, F. Lacbawan, Clinton R. Paden, Duncan MacCannell |
| EPI_ISL_3396697 | Kansas Health and Environmental Lab | Kansas Health and Environmental Lab | Mike Grose, Jonathan Barnell, Amanda Bradley, and Phil Adam |
| EPI_ISL_3396698, EPI_ISL_3396699 | Quest Diagnostics Incorporated | Centers for Disease Control and Prevention Division of Viral Diseases, Pathogen Discovery | Dakota Howard, Dhwani Batra, Peter W. Cook, Kara Moser, Adrian Paskey, Jason Caravas, Benjamin Rambo-Martin, Shatavia Morrison, Christopher Gulvick, Scott Sammons, Yvette Unoarumhi, Darlene Wagner, Matthew Schmerer, S. H. Rosenthal, A. Gerasimova, R. M. Kagan, B. Anderson, M. Hua, Y. Liu, L.E. Bernstein, K.E. Livingston, A. Perez, I. A. Shlyakhter, R. V. Rolando, R. Owen, P. Tanpaiboon, F. Lacbawan, Clinton R. Paden, Duncan MacCannell |
| EPI_ISL_3396700 | Kansas Health and Environmental Lab | Kansas Health and Environmental Lab | Mike Grose, Jonathan Barnell, Amanda Bradley, and Phil Adam |
| EPI_ISL_3396701, EPI_ISL_3396702 | Quest Diagnostics Incorporated | Centers for Disease Control and Prevention Division of Viral Diseases, Pathogen Discovery | Dakota Howard, Dhwani Batra, Peter W. Cook, Kara Moser, Adrian Paskey, Jason Caravas, Benjamin Rambo-Martin, Shatavia Morrison, Christopher Gulvick, Scott Sammons, Yvette Unoarumhi, Darlene Wagner, Matthew Schmerer, S. H. Rosenthal, A. Gerasimova, R. M. Kagan, B. Anderson, M. Hua, Y. Liu, L.E. Bernstein, K.E. Livingston, A. Perez, I. A. Shlyakhter, R. V. Rolando, R. Owen, P. Tanpaiboon, F. Lacbawan, Clinton R. Paden, Duncan MacCannell |
| EPI_ISL_3396703 | Kansas Health and Environmental Lab | Kansas Health and Environmental Lab | Mike Grose, Jonathan Barnell, Amanda Bradley, and Phil Adam |
| EPI_ISL_3396704, EPI_ISL_3396705 | Quest Diagnostics Incorporated | Centers for Disease Control and Prevention Division of Viral Diseases, Pathogen Discovery | Dakota Howard, Dhwani Batra, Peter W. Cook, Kara Moser, Adrian Paskey, Jason Caravas, Benjamin Rambo-Martin, Shatavia Morrison, Christopher Gulvick, Scott Sammons, Yvette Unoarumhi, Darlene Wagner, Matthew Schmerer, S. H. Rosenthal, A. Gerasimova, R. M. Kagan, B. Anderson, M. Hua, Y. Liu, L.E. Bernstein, K.E. Livingston, A. Perez, I. A. Shlyakhter, R. V. Rolando, R. Owen, P. Tanpaiboon, F. Lacbawan, Clinton R. Paden, Duncan MacCannell |
| EPI_ISL_3396706 | Kansas Health and Environmental Lab | Kansas Health and Environmental Lab | Mike Grose, Jonathan Barnell, Amanda Bradley, and Phil Adam |
| EPI_ISL_3396707 | Quest Diagnostics Incorporated | Centers for Disease Control and Prevention Division of Viral Diseases, Pathogen Discovery | Dakota Howard, Dhwani Batra, Peter W. Cook, Kara Moser, Adrian Paskey, Jason Caravas, Benjamin Rambo-Martin, Shatavia Morrison, Christopher Gulvick, Scott Sammons, Yvette Unoarumhi, Darlene Wagner, Matthew Schmerer, S. H. Rosenthal, A. Gerasimova, R. M. Kagan, B. Anderson, M. Hua, Y. Liu, L.E. Bernstein, K.E. Livingston, A. Perez, I. A. Shlyakhter, R. V. Rolando, R. Owen, P. Tanpaiboon, F. Lacbawan, Clinton R. Paden, Duncan MacCannell |
| EPI_ISL_3396708 | Kansas Health and Environmental Lab | Kansas Health and Environmental Lab | Mike Grose, Jonathan Barnell, Amanda Bradley, and Phil Adam |

| | | | |
|---|---|---|---|
| EPI_ISL_3396709, EPI_ISL_3396710 | Quest Diagnostics Incorporated | Centers for Disease Control and Prevention Division of Viral Diseases, Pathogen Discovery | Dakota Howard, Dhwani Batra, Peter W. Cook, Kara Moser, Adrian Paskey, Jason Caravas, Benjamin Rambo-Martin, Shatavia Morrison, Christopher Gulvick, Scott Sammons, Yvette Unoarumhi, Darlene Wagner, Matthew Schmerer, S. H. Rosenthal, A. Gerasimova, R. M. Kagan, B. Anderson, M. Hua, Y. Liu, L.E. Bernstein, K.E. Livingston, A. Perez, I. A. Shlyakhter, R. V. Rolando, R. Owen, P. Tanpaiboon, F. Lacbawan, Clinton R. Paden, Duncan MacCannell |
| EPI_ISL_3396711 | Kansas Health and Environmental Lab | Kansas Health and Environmental Lab | Mike Grose, Jonathan Barnell, Amanda Bradley, and Phil Adam |
| EPI_ISL_3396712 | Quest Diagnostics Incorporated | Centers for Disease Control and Prevention Division of Viral Diseases, Pathogen Discovery | Dakota Howard, Dhwani Batra, Peter W. Cook, Kara Moser, Adrian Paskey, Jason Caravas, Benjamin Rambo-Martin, Shatavia Morrison, Christopher Gulvick, Scott Sammons, Yvette Unoarumhi, Darlene Wagner, Matthew Schmerer, S. H. Rosenthal, A. Gerasimova, R. M. Kagan, B. Anderson, M. Hua, Y. Liu, L.E. Bernstein, K.E. Livingston, A. Perez, I. A. Shlyakhter, R. V. Rolando, R. Owen, P. Tanpaiboon, F. Lacbawan, Clinton R. Paden, Duncan MacCannell |
| EPI_ISL_3396713 | Kansas Health and Environmental Lab | Kansas Health and Environmental Lab | Mike Grose, Jonathan Barnell, Amanda Bradley, and Phil Adam |
| EPI_ISL_3396714, EPI_ISL_3396715 | Quest Diagnostics Incorporated | Centers for Disease Control and Prevention Division of Viral Diseases, Pathogen Discovery | Dakota Howard, Dhwani Batra, Peter W. Cook, Kara Moser, Adrian Paskey, Jason Caravas, Benjamin Rambo-Martin, Shatavia Morrison, Christopher Gulvick, Scott Sammons, Yvette Unoarumhi, Darlene Wagner, Matthew Schmerer, S. H. Rosenthal, A. Gerasimova, R. M. Kagan, B. Anderson, M. Hua, Y. Liu, L.E. Bernstein, K.E. Livingston, A. Perez, I. A. Shlyakhter, R. V. Rolando, R. Owen, P. Tanpaiboon, F. Lacbawan, Clinton R. Paden, Duncan MacCannell |
| EPI_ISL_3396716 | Kansas Health and Environmental Lab | Kansas Health and Environmental Lab | Mike Grose, Jonathan Barnell, Amanda Bradley, and Phil Adam |
| EPI_ISL_3396717 | Quest Diagnostics Incorporated | Centers for Disease Control and Prevention Division of Viral Diseases, Pathogen Discovery | Dakota Howard, Dhwani Batra, Peter W. Cook, Kara Moser, Adrian Paskey, Jason Caravas, Benjamin Rambo-Martin, Shatavia Morrison, Christopher Gulvick, Scott Sammons, Yvette Unoarumhi, Darlene Wagner, Matthew Schmerer, S. H. Rosenthal, A. Gerasimova, R. M. Kagan, B. Anderson, M. Hua, Y. Liu, L.E. Bernstein, K.E. Livingston, A. Perez, I. A. Shlyakhter, R. V. Rolando, R. Owen, P. Tanpaiboon, F. Lacbawan, Clinton R. Paden, Duncan MacCannell |
| EPI_ISL_3396718 | Kansas Health and Environmental Lab | Kansas Health and Environmental Lab | Mike Grose, Jonathan Barnell, Amanda Bradley, and Phil Adam |
| EPI_ISL_3396719, EPI_ISL_3396720 | Quest Diagnostics Incorporated | Centers for Disease Control and Prevention Division of Viral Diseases, Pathogen Discovery | Dakota Howard, Dhwani Batra, Peter W. Cook, Kara Moser, Adrian Paskey, Jason Caravas, Benjamin Rambo-Martin, Shatavia Morrison, Christopher Gulvick, Scott Sammons, Yvette Unoarumhi, Darlene Wagner, Matthew Schmerer, S. H. Rosenthal, A. Gerasimova, R. M. Kagan, B. Anderson, M. Hua, Y. Liu, L.E. Bernstein, K.E. Livingston, A. Perez, I. A. Shlyakhter, R. V. Rolando, R. Owen, P. Tanpaiboon, F. Lacbawan, Clinton R. Paden, Duncan MacCannell |
| EPI_ISL_3396721 | Kansas Health and Environmental Lab | Kansas Health and Environmental Lab | Mike Grose, Jonathan Barnell, Amanda Bradley, and Phil Adam |
| EPI_ISL_3396722 | Quest Diagnostics Incorporated | Centers for Disease Control and Prevention Division of Viral Diseases, Pathogen Discovery | Dakota Howard, Dhwani Batra, Peter W. Cook, Kara Moser, Adrian Paskey, Jason Caravas, Benjamin Rambo-Martin, Shatavia Morrison, Christopher Gulvick, Scott Sammons, Yvette Unoarumhi, Darlene Wagner, Matthew Schmerer, S. H. Rosenthal, A. Gerasimova, R. M. Kagan, B. Anderson, M. Hua, Y. Liu, L.E. Bernstein, K.E. Livingston, A. Perez, I. A. Shlyakhter, R. V. Rolando, R. Owen, P. Tanpaiboon, F. Lacbawan, Clinton R. Paden, Duncan MacCannell |
| EPI_ISL_3396723 | Kansas Health and Environmental Lab | Kansas Health and Environmental Lab | Mike Grose, Jonathan Barnell, Amanda Bradley, and Phil Adam |
| EPI_ISL_3396724, EPI_ISL_3396725 | Quest Diagnostics Incorporated | Centers for Disease Control and Prevention Division of Viral Diseases, Pathogen Discovery | Dakota Howard, Dhwani Batra, Peter W. Cook, Kara Moser, Adrian Paskey, Jason Caravas, Benjamin Rambo-Martin, Shatavia Morrison, Christopher Gulvick, Scott Sammons, Yvette Unoarumhi, Darlene Wagner, Matthew Schmerer, S. H. Rosenthal, A. Gerasimova, R. M. Kagan, B. Anderson, M. Hua, Y. Liu, L.E. Bernstein, K.E. Livingston, A. Perez, I. A. Shlyakhter, R. V. Rolando, R. Owen, P. Tanpaiboon, F. Lacbawan, Clinton R. Paden, Duncan MacCannell |
| EPI_ISL_3396726 | Kansas Health and Environmental Lab | Kansas Health and Environmental Lab | Mike Grose, Jonathan Barnell, Amanda Bradley, and Phil Adam |
| EPI_ISL_3396727 | Quest Diagnostics Incorporated | Centers for Disease Control and Prevention Division of Viral Diseases, Pathogen Discovery | Dakota Howard, Dhwani Batra, Peter W. Cook, Kara Moser, Adrian Paskey, Jason Caravas, Benjamin Rambo-Martin, Shatavia Morrison, Christopher Gulvick, Scott Sammons, Yvette Unoarumhi, Darlene Wagner, Matthew Schmerer, S. H. Rosenthal, A. Gerasimova, R. M. Kagan, B. Anderson, M. Hua, Y. Liu, L.E. Bernstein, K.E. Livingston, A. Perez, I. A. Shlyakhter, R. V. Rolando, R. Owen, P. Tanpaiboon, F. Lacbawan, Clinton R. Paden, Duncan MacCannell |
| EPI_ISL_3396728 | Kansas Health and Environmental Lab | Kansas Health and Environmental Lab | Mike Grose, Jonathan Barnell, Amanda Bradley, and Phil Adam |
| EPI_ISL_3396729, EPI_ISL_3396730 | Quest Diagnostics Incorporated | Centers for Disease Control and Prevention Division of Viral Diseases, Pathogen Discovery | Dakota Howard, Dhwani Batra, Peter W. Cook, Kara Moser, Adrian Paskey, Jason Caravas, Benjamin Rambo-Martin, Shatavia Morrison, Christopher Gulvick, Scott Sammons, Yvette Unoarumhi, Darlene Wagner, Matthew Schmerer, S. H. Rosenthal, A. Gerasimova, R. M. Kagan, B. Anderson, M. Hua, Y. Liu, L.E. Bernstein, K.E. Livingston, A. Perez, I. A. Shlyakhter, R. V. Rolando, R. Owen, P. Tanpaiboon, F. Lacbawan, Clinton R. Paden, Duncan MacCannell |
| EPI_ISL_3396731 | Kansas Health and Environmental Lab | Kansas Health and Environmental Lab | Mike Grose, Jonathan Barnell, Amanda Bradley, and Phil Adam |
| EPI_ISL_3396732 | Quest Diagnostics Incorporated | Centers for Disease Control and Prevention Division of Viral Diseases, Pathogen Discovery | Dakota Howard, Dhwani Batra, Peter W. Cook, Kara Moser, Adrian Paskey, Jason Caravas, Benjamin Rambo-Martin, Shatavia Morrison, Christopher Gulvick, Scott Sammons, Yvette Unoarumhi, Darlene Wagner, Matthew Schmerer, S. H. Rosenthal, A. Gerasimova, R. M. Kagan, B. Anderson, M. Hua, Y. Liu, L.E. Bernstein, K.E. Livingston, A. Perez, I. A. Shlyakhter, R. V. Rolando, R. Owen, P. Tanpaiboon, F. Lacbawan, Clinton R. Paden, Duncan MacCannell |
| EPI_ISL_3396733 | Kansas Health and Environmental Lab | Kansas Health and Environmental Lab | Mike Grose, Jonathan Barnell, Amanda Bradley, and Phil Adam |
| EPI_ISL_3396734, EPI_ISL_3396735 | Quest Diagnostics Incorporated | Centers for Disease Control and Prevention Division of Viral Diseases, Pathogen Discovery | Dakota Howard, Dhwani Batra, Peter W. Cook, Kara Moser, Adrian Paskey, Jason Caravas, Benjamin Rambo-Martin, Shatavia Morrison, Christopher Gulvick, Scott Sammons, Yvette Unoarumhi, Darlene Wagner, Matthew Schmerer, S. H. Rosenthal, A. Gerasimova, R. M. Kagan, B. Anderson, M. Hua, Y. Liu, L.E. Bernstein, K.E. Livingston, A. Perez, I. A. Shlyakhter, R. V. Rolando, R. Owen, P. Tanpaiboon, F. Lacbawan, Clinton R. Paden, Duncan MacCannell |
| EPI_ISL_3396736 | Kansas Health and Environmental Lab | Kansas Health and Environmental Lab | Mike Grose, Jonathan Barnell, Amanda Bradley, and Phil Adam |
| EPI_ISL_3396737 | Quest Diagnostics Incorporated | Centers for Disease Control and Prevention Division of Viral Diseases, Pathogen Discovery | Dakota Howard, Dhwani Batra, Peter W. Cook, Kara Moser, Adrian Paskey, Jason Caravas, Benjamin Rambo-Martin, Shatavia Morrison, Christopher Gulvick, Scott Sammons, Yvette Unoarumhi, Darlene Wagner, Matthew Schmerer, S. H. Rosenthal, A. Gerasimova, R. M. Kagan, B. Anderson, M. Hua, Y. Liu, L.E. Bernstein, K.E. Livingston, A. Perez, I. A. Shlyakhter, R. V. Rolando, R. Owen, P. Tanpaiboon, F. Lacbawan, Clinton R. Paden, Duncan MacCannell |
| EPI_ISL_3396738 | Kansas Health and Environmental Lab | Kansas Health and Environmental Lab | Mike Grose, Jonathan Barnell, Amanda Bradley, and Phil Adam |
| EPI_ISL_3396740 | Quest Diagnostics Incorporated | Centers for Disease Control and Prevention Division of Viral Diseases, Pathogen Discovery | Dakota Howard, Dhwani Batra, Peter W. Cook, Kara Moser, Adrian Paskey, Jason Caravas, Benjamin Rambo-Martin, Shatavia Morrison, Christopher Gulvick, Scott Sammons, Yvette Unoarumhi, Darlene Wagner, Matthew Schmerer, S. H. Rosenthal, A. Gerasimova, R. M. Kagan, B. Anderson, M. Hua, Y. Liu, L.E. Bernstein, K.E. Livingston, A. Perez, I. A. Shlyakhter, R. V. Rolando, R. Owen, P. Tanpaiboon, F. Lacbawan, Clinton R. Paden, Duncan MacCannell |
| EPI_ISL_3396741 | Kansas Health and Environmental Lab | Kansas Health and Environmental Lab | Mike Grose, Jonathan Barnell, Amanda Bradley, and Phil Adam |
| EPI_ISL_3396742 | Quest Diagnostics Incorporated | Centers for Disease Control and Prevention Division of Viral Diseases, Pathogen Discovery | Dakota Howard, Dhwani Batra, Peter W. Cook, Kara Moser, Adrian Paskey, Jason Caravas, Benjamin Rambo-Martin, Shatavia Morrison, Christopher Gulvick, Scott Sammons, Yvette Unoarumhi, Darlene Wagner, Matthew Schmerer, S. H. Rosenthal, A. Gerasimova, R. M. Kagan, B. Anderson, M. Hua, Y. Liu, L.E. Bernstein, K.E. Livingston, A. Perez, I. A. Shlyakhter, R. V. Rolando, R. Owen, P. Tanpaiboon, F. Lacbawan, Clinton R. Paden, Duncan MacCannell |
| EPI_ISL_3396743, EPI_ISL_3396744, EPI_ISL_3396745 | Kansas Health and Environmental Lab | Kansas Health and Environmental Lab | Mike Grose, Jonathan Barnell, Amanda Bradley, and Phil Adam |
| EPI_ISL_3396746, EPI_ISL_3396747 | Quest Diagnostics Incorporated | Centers for Disease Control and Prevention Division of Viral Diseases, Pathogen Discovery | Dakota Howard, Dhwani Batra, Peter W. Cook, Kara Moser, Adrian Paskey, Jason Caravas, Benjamin Rambo-Martin, Shatavia Morrison, Christopher Gulvick, Scott Sammons, Yvette Unoarumhi, Darlene Wagner, Matthew Schmerer, S. H. Rosenthal, A. Gerasimova, R. M. Kagan, B. Anderson, M. Hua, Y. Liu, L.E. Bernstein, K.E. Livingston, A. Perez, I. A. Shlyakhter, R. V. Rolando, R. Owen, P. Tanpaiboon, F. Lacbawan, Clinton R. Paden, Duncan MacCannell |
| EPI_ISL_3396748 | Kansas Health and Environmental Lab | Kansas Health and Environmental Lab | Mike Grose, Jonathan Barnell, Amanda Bradley, and Phil Adam |
| EPI_ISL_3396749, EPI_ISL_3396750 | Quest Diagnostics Incorporated | Centers for Disease Control and Prevention Division of Viral Diseases, Pathogen Discovery | Dakota Howard, Dhwani Batra, Peter W. Cook, Kara Moser, Adrian Paskey, Jason Caravas, Benjamin Rambo-Martin, Shatavia Morrison, Christopher Gulvick, Scott Sammons, Yvette Unoarumhi, Darlene Wagner, Matthew Schmerer, S. H. Rosenthal, A. Gerasimova, R. M. Kagan, B. Anderson, M. Hua, Y. Liu, L.E. Bernstein, K.E. Livingston, A. Perez, I. A. Shlyakhter, R. V. Rolando, R. Owen, P. Tanpaiboon, F. Lacbawan, Clinton R. Paden, Duncan MacCannell |
| EPI_ISL_3396751 | Kansas Health and Environmental Lab | Kansas Health and Environmental Lab | Mike Grose, Jonathan Barnell, Amanda Bradley, and Phil Adam |
| EPI_ISL_3396752 | Quest Diagnostics Incorporated | Centers for Disease Control and Prevention Division of Viral Diseases, Pathogen Discovery | Dakota Howard, Dhwani Batra, Peter W. Cook, Kara Moser, Adrian Paskey, Jason Caravas, Benjamin Rambo-Martin, Shatavia Morrison, Christopher Gulvick, Scott Sammons, Yvette Unoarumhi, Darlene Wagner, Matthew Schmerer, S. H. Rosenthal, A. Gerasimova, R. M. Kagan, B. Anderson, M. Hua, Y. Liu, L.E. Bernstein, K.E. Livingston, A. Perez, I. A. Shlyakhter, R. V. Rolando, R. Owen, P. Tanpaiboon, F. Lacbawan, Clinton R. Paden, Duncan MacCannell |
| EPI_ISL_3396753 | Kansas Health and Environmental Lab | Kansas Health and Environmental Lab | Mike Grose, Jonathan Barnell, Amanda Bradley, and Phil Adam |
| EPI_ISL_3396754 | Quest Diagnostics Incorporated | Centers for Disease Control and Prevention Division of Viral Diseases, Pathogen Discovery | Dakota Howard, Dhwani Batra, Peter W. Cook, Kara Moser, Adrian Paskey, Jason Caravas, Benjamin Rambo-Martin, Shatavia Morrison, Christopher Gulvick, Scott Sammons, Yvette Unoarumhi, Darlene Wagner, Matthew Schmerer, S. H. Rosenthal, A. Gerasimova, R. M. Kagan, B. Anderson, M. Hua, Y. Liu, L.E. Bernstein, K.E. Livingston, A. Perez, I. A. Shlyakhter, R. V. Rolando, R. Owen, P. Tanpaiboon, F. Lacbawan, Clinton R. Paden, Duncan MacCannell |
| EPI_ISL_3396755 | Kansas Health and Environmental Lab | Kansas Health and Environmental Lab | Mike Grose, Jonathan Barnell, Amanda Bradley, and Phil Adam |
| EPI_ISL_3396756, EPI_ISL_3396757 | Quest Diagnostics Incorporated | Centers for Disease Control and Prevention Division of Viral Diseases, Pathogen Discovery | Dakota Howard, Dhwani Batra, Peter W. Cook, Kara Moser, Adrian Paskey, Jason Caravas, Benjamin Rambo-Martin, Shatavia Morrison, Christopher Gulvick, Scott Sammons, Yvette Unoarumhi, Darlene Wagner, Matthew Schmerer, S. H. Rosenthal, A. Gerasimova, R. M. Kagan, B. Anderson, M. Hua, Y. Liu, L.E. Bernstein, K.E. Livingston, A. Perez, I. A. Shlyakhter, R. V. Rolando, R. Owen, P. Tanpaiboon, F. Lacbawan, Clinton R. Paden, Duncan MacCannell |
| EPI_ISL_3396758 | Kansas Health and Environmental Lab | Kansas Health and Environmental Lab | Mike Grose, Jonathan Barnell, Amanda Bradley, and Phil Adam |

EPI_ISL_3396759, EPI_ISL_3396760, EPI_ISL_3396762, EPI_ISL_3396763, EPI_ISL_3396764, EPI_ISL_3396765, EPI_ISL_3396766, EPI_ISL_3396767, EPI_ISL_3396768, EPI_ISL_3396769, EPI_ISL_3396770, EPI_ISL_3396771, EPI_ISL_3396772, EPI_ISL_3396773, EPI_ISL_3396774, EPI_ISL_3396775, EPI_ISL_3396777, EPI_ISL_3396778, EPI_ISL_3396779, EPI_ISL_3396780, EPI_ISL_3396781, EPI_ISL_3396782, EPI_ISL_3396783, EPI_ISL_3396784, EPI_ISL_3396785, EPI_ISL_3396786, EPI_ISL_3396787, EPI_ISL_3396788, EPI_ISL_3396789, EPI_ISL_3396790, EPI_ISL_3396791, EPI_ISL_3396792, EPI_ISL_3396793, EPI_ISL_3396794, EPI_ISL_3396795, EPI_ISL_3396796, EPI_ISL_3396797, EPI_ISL_3396798, EPI_ISL_3396799, EPI_ISL_3396800, EPI_ISL_3396801, EPI_ISL_3396802, EPI_ISL_3396803, EPI_ISL_3396804, EPI_ISL_3396805, EPI_ISL_3396806, EPI_ISL_3396807, EPI_ISL_3396808, EPI_ISL_3396809, EPI_ISL_3396810, EPI_ISL_3396811, EPI_ISL_3396812, EPI_ISL_3396813, EPI_ISL_3396814, EPI_ISL_3396815, EPI_ISL_3396816, EPI_ISL_3396817, EPI_ISL_3396818, EPI_ISL_3396819, EPI_ISL_3396820, EPI_ISL_3396822, EPI_ISL_3396823, EPI_ISL_3396825, EPI_ISL_3396826, EPI_ISL_3396827, EPI_ISL_3396828, EPI_ISL_3396829, EPI_ISL_3396830, EPI_ISL_3396831, EPI_ISL_3396832, EPI_ISL_3396833, EPI_ISL_3396834, EPI_ISL_3396835, EPI_ISL_3396836, EPI_ISL_3396837, EPI_ISL_3396838, EPI_ISL_3396839, EPI_ISL_3396840, EPI_ISL_3396841, EPI_ISL_3396842, EPI_ISL_3396843, EPI_ISL_3396844, EPI_ISL_3396845, EPI_ISL_3396846, EPI_ISL_3396847, EPI_ISL_3396848, EPI_ISL_3396849, EPI_ISL_3396850, EPI_ISL_3396851, EPI_ISL_3396853, EPI_ISL_3396854, EPI_ISL_3396855, EPI_ISL_3396856, EPI_ISL_3396857, EPI_ISL_3396858, EPI_ISL_3396859, EPI_ISL_3396860, EPI_ISL_3396861, EPI_ISL_3396863, EPI_ISL_3396864, EPI_ISL_3396865, EPI_ISL_3396866, EPI_ISL_3396867, EPI_ISL_3396868, EPI_ISL_3396869, EPI_ISL_3396870, EPI_ISL_3396871, EPI_ISL_3396872, EPI_ISL_3396873, EPI_ISL_3396874, EPI_ISL_3396875, EPI_ISL_3396876, EPI_ISL_3396878, EPI_ISL_3396879, EPI_ISL_3396880, EPI_ISL_3396881, EPI_ISL_3396882, EPI_ISL_3396883, EPI_ISL_3396884, EPI_ISL_3396885, EPI_ISL_3396886, EPI_ISL_3396887, EPI_ISL_3396888, EPI_ISL_3396889, EPI_ISL_3396890, EPI_ISL_3396891, EPI_ISL_3396892, EPI_ISL_3396893, EPI_ISL_3396895, EPI_ISL_3396896, EPI_ISL_3396897, EPI_ISL_3396898, EPI_ISL_3396899, EPI_ISL_3396900, EPI_ISL_3396901, EPI_ISL_3396902, EPI_ISL_3396903, EPI_ISL_3396904, EPI_ISL_3396905, EPI_ISL_3396906, EPI_ISL_3396907, EPI_ISL_3396908, EPI_ISL_3396909, EPI_ISL_3396910, EPI_ISL_3396911, EPI_ISL_3396912, EPI_ISL_3396913, EPI_ISL_3396914, EPI_ISL_3396915, EPI_ISL_3396916, EPI_ISL_3396917, EPI_ISL_3396918, EPI_ISL_3396920, EPI_ISL_3396921, EPI_ISL_3396922, EPI_ISL_3396923, EPI_ISL_3396924, EPI_ISL_3396925, EPI_ISL_3396926, EPI_ISL_3396927, EPI_ISL_3396928, EPI_ISL_3396929, EPI_ISL_3396930, EPI_ISL_3396931, EPI_ISL_3396932, EPI_ISL_3396933, EPI_ISL_3396934, EPI_ISL_3396935, EPI_ISL_3396936, EPI_ISL_3396937, EPI_ISL_3396938, EPI_ISL_3396939, EPI_ISL_3396940, EPI_ISL_3396942, EPI_ISL_3396943, EPI_ISL_3396944, EPI_ISL_3396946, EPI_ISL_3396947, EPI_ISL_3396948, EPI_ISL_3396949, EPI_ISL_3396950, EPI_ISL_3396951, EPI_ISL_3396952, EPI_ISL_3396953, EPI_ISL_3396954, EPI_ISL_3396955, EPI_ISL_3396956, EPI_ISL_3396957, EPI_ISL_3396958, EPI_ISL_3396959, EPI_ISL_3396960, EPI_ISL_3396961, EPI_ISL_3396962, EPI_ISL_3396963, EPI_ISL_3396964, EPI_ISL_3396965, EPI_ISL_3396966, EPI_ISL_3396967, EPI_ISL_3396968, EPI_ISL_3396969, EPI_ISL_3396970, EPI_ISL_3396971, EPI_ISL_3396972, EPI_ISL_3396973, EPI_ISL_3396974, EPI_ISL_3396976, EPI_ISL_3396977, EPI_ISL_3396978, EPI_ISL_3396979, EPI_ISL_3396980, EPI_ISL_3396981, EPI_ISL_3396982, EPI_ISL_3396983, EPI_ISL_3396984, EPI_ISL_3396985, EPI_ISL_3396986, EPI_ISL_3396987, EPI_ISL_3396988, EPI_ISL_3396989, EPI_ISL_3396990, EPI_ISL_3396991, EPI_ISL_3396992, EPI_ISL_3396993, EPI_ISL_3396994, EPI_ISL_3396995, EPI_ISL_3396996, EPI_ISL_3396997, EPI_ISL_3396998, EPI_ISL_3396999, EPI_ISL_3397000, EPI_ISL_3397001, EPI_ISL_3397002, EPI_ISL_3397003, EPI_ISL_3397004, EPI_ISL_3397005, EPI_ISL_3397007, EPI_ISL_3397008, EPI_ISL_3397009, EPI_ISL_3397010, EPI_ISL_3397011, EPI_ISL_3397012, EPI_ISL_3397013, EPI_ISL_3397014, EPI_ISL_3397015, EPI_ISL_3397016, EPI_ISL_3397017, EPI_ISL_3397018, EPI_ISL_3397019, EPI_ISL_3397020, EPI_ISL_3397021, EPI_ISL_3397022, EPI_ISL_3397023, EPI_ISL_3397024, EPI_ISL_3397025, EPI_ISL_3397026, EPI_ISL_3397027, EPI_ISL_3397028, EPI_ISL_3397029, EPI_ISL_3397030, EPI_ISL_3397031, EPI_ISL_3397032, EPI_ISL_3397034, EPI_ISL_3397035, EPI_ISL_3397036, EPI_ISL_3397037, EPI_ISL_3397038, EPI_ISL_3397039, EPI_ISL_3397041, EPI_ISL_3397042, EPI_ISL_3397043, EPI_ISL_3397044, EPI_ISL_3397045, EPI_ISL_3397046, EPI_ISL_3397047, EPI_ISL_3397048, EPI_ISL_3397049, EPI_ISL_3397050, EPI_ISL_3397051, EPI_ISL_3397052, EPI_ISL_3397053, EPI_ISL_3397054, EPI_ISL_3397055, EPI_ISL_3397056, EPI_ISL_3397057, EPI_ISL_3397058, EPI_ISL_3397059, EPI_ISL_3397060, EPI_ISL_3397061, EPI_ISL_3397062, EPI_ISL_3397064, EPI_ISL_3397065, EPI_ISL_3397066, EPI_ISL_3397067, EPI_ISL_3397068, EPI_ISL_3397069, EPI_ISL_3397070, EPI_ISL_3397072, EPI_ISL_3397073, EPI_ISL_3397074, EPI_ISL_3397075, EPI_ISL_3397076, EPI_ISL_3397077, EPI_ISL_3397078, EPI_ISL_3397079, EPI_ISL_3397080, EPI_ISL_3397081, EPI_ISL_3397082, EPI_ISL_3397083, EPI_ISL_3397084, EPI_ISL_3397085, EPI_ISL_3397086, EPI_ISL_3397087, EPI_ISL_3397088, EPI_ISL_3397089, EPI_ISL_3397090, EPI_ISL_3397091, EPI_ISL_3397092, EPI_ISL_3397093, EPI_ISL_3397094, EPI_ISL_3397095, EPI_ISL_3397096, EPI_ISL_3397097, EPI_ISL_3397098, EPI_ISL_3397099, EPI_ISL_3397100, EPI_ISL_3397101, EPI_ISL_3397102, EPI_ISL_3397103, EPI_ISL_3397104, EPI_ISL_3397105, EPI_ISL_3397106, EPI_ISL_3397107, EPI_ISL_3397108, EPI_ISL_3397109, EPI_ISL_3397110, EPI_ISL_3397111, EPI_ISL_3397112, EPI_ISL_3397113

| | | | |
|---|---|---|---|
| see above | Quest Diagnostics Incorporated | Centers for Disease Control and Prevention Division of Viral Diseases, Pathogen Discovery | Dakota Howard, Dhwani Batra, Peter W. Cook, Kara Moser, Adrian Paskey, Jason Caravas, Benjamin Rambo-Martin, Shatavia Morrison, Christopher Gulvick, Scott Sammons, Yvette Unoarumhi, Darlene Wagner, Matthew Schmerer, S. H. Rosenthal, A. Gerasimova, R. M. Kagan, B. Anderson, M. Hua, Y. Liu, L.E. Bernstein, K.E. Livingston, A. Perez, I. A. Shlyakhter, R. V. Rolando, R. Owen, P. Tanpaiboon, F. Lacbawan, Clinton R. Paden, Duncan MacCannell |

EPI_ISL_3397114, EPI_ISL_3397115, EPI_ISL_3397116, EPI_ISL_3397117, EPI_ISL_3397118, EPI_ISL_3397119, EPI_ISL_3397120, EPI_ISL_3397121, EPI_ISL_3397122, EPI_ISL_3397123, EPI_ISL_3397124, EPI_ISL_3397125, EPI_ISL_3397126, EPI_ISL_3397127, EPI_ISL_3397130, EPI_ISL_3397131, EPI_ISL_3397132, EPI_ISL_3397133, EPI_ISL_3397134, EPI_ISL_3397135, EPI_ISL_3397136, EPI_ISL_3397137, EPI_ISL_3397138, EPI_ISL_3397139, EPI_ISL_3397140, EPI_ISL_3397141, EPI_ISL_3397142, EPI_ISL_3397143, EPI_ISL_3397144, EPI_ISL_3397145, EPI_ISL_3397146, EPI_ISL_3397147, EPI_ISL_3397148, EPI_ISL_3397149, EPI_ISL_3397150, EPI_ISL_3397151, EPI_ISL_3397152, EPI_ISL_3397153, EPI_ISL_3397154, EPI_ISL_3397155, EPI_ISL_3397156, EPI_ISL_3397157, EPI_ISL_3397158, EPI_ISL_3397159, EPI_ISL_3397160, EPI_ISL_3397161, EPI_ISL_3397162, EPI_ISL_3397163, EPI_ISL_3397164, EPI_ISL_3397165, EPI_ISL_3397166, EPI_ISL_3397167, EPI_ISL_3397168, EPI_ISL_3397169, EPI_ISL_3397170, EPI_ISL_3397171, EPI_ISL_3397172, EPI_ISL_3397173, EPI_ISL_3397174, EPI_ISL_3397175, EPI_ISL_3397176, EPI_ISL_3397177, EPI_ISL_3397178, EPI_ISL_3397179, EPI_ISL_3397180, EPI_ISL_3397181, EPI_ISL_3397182, EPI_ISL_3397183, EPI_ISL_3397184, EPI_ISL_3397185, EPI_ISL_3397186, EPI_ISL_3397187, EPI_ISL_3397188, EPI_ISL_3397189, EPI_ISL_3397190, EPI_ISL_3397191, EPI_ISL_3397192, EPI_ISL_3397193, EPI_ISL_3397194, EPI_ISL_3397195, EPI_ISL_3397196, EPI_ISL_3397197, EPI_ISL_3397198, EPI_ISL_3397199, EPI_ISL_3397200, EPI_ISL_3397201, EPI_ISL_3397202, EPI_ISL_3397203, EPI_ISL_3397204, EPI_ISL_3397205, EPI_ISL_3397206, EPI_ISL_3397207, EPI_ISL_3397208, EPI_ISL_3397209, EPI_ISL_3397210, EPI_ISL_3397211, EPI_ISL_3397212, EPI_ISL_3397213, EPI_ISL_3397214, EPI_ISL_3397215, EPI_ISL_3397216, EPI_ISL_3397217, EPI_ISL_3397218, EPI_ISL_3397219, EPI_ISL_3397220, EPI_ISL_3397221, EPI_ISL_3397222, EPI_ISL_3397223, EPI_ISL_3397224, EPI_ISL_3397225, EPI_ISL_3397226, EPI_ISL_3397227, EPI_ISL_3397228, EPI_ISL_3397229, EPI_ISL_3397230, EPI_ISL_3397231, EPI_ISL_3397232, EPI_ISL_3397233, EPI_ISL_3397234, EPI_ISL_3397235, EPI_ISL_3397236, EPI_ISL_3397237, EPI_ISL_3397238, EPI_ISL_3397239, EPI_ISL_3397240, EPI_ISL_3397241, EPI_ISL_3397242, EPI_ISL_3397243, EPI_ISL_3397244, EPI_ISL_3397245, EPI_ISL_3397246, EPI_ISL_3397248, EPI_ISL_3397249, EPI_ISL_3397250, EPI_ISL_3397251, EPI_ISL_3397252, EPI_ISL_3397253, EPI_ISL_3397254, EPI_ISL_3397255, EPI_ISL_3397256, EPI_ISL_3397257, EPI_ISL_3397258, EPI_ISL_3397259, EPI_ISL_3397260, EPI_ISL_3397261, EPI_ISL_3397262, EPI_ISL_3397263, EPI_ISL_3397264, EPI_ISL_3397265, EPI_ISL_3397266, EPI_ISL_3397267, EPI_ISL_3397268

| | | | |
|---|---|---|---|
| see above | Fulgent Genetics | Centers for Disease Control and Prevention Division of Viral Diseases, Pathogen Discovery | Dakota Howard, Dhwani Batra, Peter W. Cook, Kara Moser, Adrian Paskey, Jason Caravas, Benjamin Rambo-Martin, Shatavia Morrison, Christopher Gulvick, Scott Sammons, Yvette Unoarumhi, Darlene Wagner, Matthew Schmerer, Harry Gao, Mickey Li, John Gao, Joseph Fierro, Benafsh Sapra, Becky Tsai, Yan Meng, Doreen Ng, James Xie, Clinton R. Paden, Duncan MacCannell |

EPI_ISL_3397269, EPI_ISL_3397270, EPI_ISL_3397271, EPI_ISL_3397272, EPI_ISL_3397273, EPI_ISL_3397274, EPI_ISL_3397275, EPI_ISL_3397276, EPI_ISL_3397277, EPI_ISL_3397278, EPI_ISL_3397279, EPI_ISL_3397280, EPI_ISL_3397281, EPI_ISL_3397282, EPI_ISL_3397283, EPI_ISL_3397284, EPI_ISL_3397285, EPI_ISL_3397286, EPI_ISL_3397287, EPI_ISL_3397288, EPI_ISL_3397289, EPI_ISL_3397290, EPI_ISL_3397291, EPI_ISL_3397293, EPI_ISL_3397294, EPI_ISL_3397295, EPI_ISL_3397296, EPI_ISL_3397297, EPI_ISL_3397298, EPI_ISL_3397299, EPI_ISL_3397300, EPI_ISL_3397301, EPI_ISL_3397302, EPI_ISL_3397303, EPI_ISL_3397304, EPI_ISL_3397305, EPI_ISL_3397306, EPI_ISL_3397307, EPI_ISL_3397308, EPI_ISL_3397309, EPI_ISL_3397310, EPI_ISL_3397311, EPI_ISL_3397312, EPI_ISL_3397313, EPI_ISL_3397314, EPI_ISL_3397315, EPI_ISL_3397316, EPI_ISL_3397317, EPI_ISL_3397318, EPI_ISL_3397319, EPI_ISL_3397320, EPI_ISL_3397321, EPI_ISL_3397322, EPI_ISL_3397323, EPI_ISL_3397324, EPI_ISL_3397325, EPI_ISL_3397326, EPI_ISL_3397327, EPI_ISL_3397328, EPI_ISL_3397329, EPI_ISL_3397330, EPI_ISL_3397331, EPI_ISL_3397332, EPI_ISL_3397333, EPI_ISL_3397334, EPI_ISL_3397335, EPI_ISL_3397336, EPI_ISL_3397337, EPI_ISL_3397338, EPI_ISL_3397339, EPI_ISL_3397340, EPI_ISL_3397341, EPI_ISL_3397342, EPI_ISL_3397343, EPI_ISL_3397344, EPI_ISL_3397345, EPI_ISL_3397346, EPI_ISL_3397347, EPI_ISL_3397348, EPI_ISL_3397349, EPI_ISL_3397350, EPI_ISL_3397351, EPI_ISL_3397352, EPI_ISL_3397353, EPI_ISL_3397354, EPI_ISL_3397355, EPI_ISL_3397356, EPI_ISL_3397357, EPI_ISL_3397358, EPI_ISL_3397359, EPI_ISL_3397360, EPI_ISL_3397361, EPI_ISL_3397362, EPI_ISL_3397363, EPI_ISL_3397364, EPI_ISL_3397365, EPI_ISL_3397366, EPI_ISL_3397367, EPI_ISL_3397368, EPI_ISL_3397369, EPI_ISL_3397370, EPI_ISL_3397371, EPI_ISL_3397372, EPI_ISL_3397373, EPI_ISL_3397374, EPI_ISL_3397375, EPI_ISL_3397376, EPI_ISL_3397377, EPI_ISL_3397378, EPI_ISL_3397379,

| | | | |
|---|---|---|---|
| EPI_ISL_3397380, EPI_ISL_3397381, EPI_ISL_3397382, EPI_ISL_3397384, EPI_ISL_3397385, EPI_ISL_3397386, EPI_ISL_3397387, EPI_ISL_3397388, EPI_ISL_3397389, EPI_ISL_3397390, EPI_ISL_3397391, EPI_ISL_3397392, EPI_ISL_3397393, EPI_ISL_3397394, EPI_ISL_3397395, EPI_ISL_3397396, EPI_ISL_3397397, EPI_ISL_3397398, EPI_ISL_3397399, EPI_ISL_3397400, EPI_ISL_3397401, EPI_ISL_3397402, EPI_ISL_3397403, EPI_ISL_3397404, EPI_ISL_3397405, EPI_ISL_3397406, EPI_ISL_3397407, EPI_ISL_3397408, EPI_ISL_3397409, EPI_ISL_3397410, EPI_ISL_3397411, EPI_ISL_3397412, EPI_ISL_3397413, EPI_ISL_3397414, EPI_ISL_3397415, EPI_ISL_3397416, EPI_ISL_3397417, EPI_ISL_3397418, EPI_ISL_3397419, EPI_ISL_3397420, EPI_ISL_3397421, EPI_ISL_3397422, EPI_ISL_3397423, EPI_ISL_3397424, EPI_ISL_3397425, EPI_ISL_3397426, EPI_ISL_3397427, EPI_ISL_3397428, EPI_ISL_3397429, EPI_ISL_3397430, EPI_ISL_3397431, EPI_ISL_3397432, EPI_ISL_3397433, EPI_ISL_3397434, EPI_ISL_3397435, EPI_ISL_3397436, EPI_ISL_3397437, EPI_ISL_3397438, EPI_ISL_3397439, EPI_ISL_3397440, EPI_ISL_3397441, EPI_ISL_3397442, EPI_ISL_3397443, EPI_ISL_3397444, EPI_ISL_3397445, EPI_ISL_3397446, EPI_ISL_3397447, EPI_ISL_3397448, EPI_ISL_3397449, EPI_ISL_3397450, EPI_ISL_3397451, EPI_ISL_3397452, EPI_ISL_3397453, EPI_ISL_3397454, EPI_ISL_3397455, EPI_ISL_3397457, EPI_ISL_3397458, EPI_ISL_3397459, EPI_ISL_3397460, EPI_ISL_3397462, EPI_ISL_3397463, EPI_ISL_3397464, EPI_ISL_3397465, EPI_ISL_3397466, EPI_ISL_3397467, EPI_ISL_3397468, EPI_ISL_3397469, EPI_ISL_3397470, EPI_ISL_3397471, EPI_ISL_3397472, EPI_ISL_3397473, EPI_ISL_3397474, EPI_ISL_3397475, EPI_ISL_3397476, EPI_ISL_3397478, EPI_ISL_3397479, EPI_ISL_3397480, EPI_ISL_3397481, EPI_ISL_3397482 | | | |
| see above | Infinity Biologix | Centers for Disease Control and Prevention Division of Viral Diseases, Pathogen Discovery | Dakota Howard, Dhwani Batra, Peter W. Cook, Kara Moser, Adrian Paskey, Jason Caravas, Benjamin Rambo-Martin, Shatavia Morrison, Christopher Gulvick, Scott Sammons, Yvette Unoarumhi, Darlene Wagner, Matthew Schmerer, Christian Bixby, Yihe Wang, Jonathan Schultz, Chirayu Goswami, Russ Hager, Robin Grimwood, Clinton R. Paden, Duncan MacCannell |
| EPI_ISL_3397483, EPI_ISL_3397484, EPI_ISL_3397485, EPI_ISL_3397486, EPI_ISL_3397487, EPI_ISL_3397488, EPI_ISL_3397489, EPI_ISL_3397490, EPI_ISL_3397492, EPI_ISL_3397493, EPI_ISL_3397494, EPI_ISL_3397495, EPI_ISL_3397496, EPI_ISL_3397497, EPI_ISL_3397498, EPI_ISL_3397499, EPI_ISL_3397500, EPI_ISL_3397501, EPI_ISL_3397502, EPI_ISL_3397503, EPI_ISL_3397504, EPI_ISL_3397505, EPI_ISL_3397506, EPI_ISL_3397507, EPI_ISL_3397508, EPI_ISL_3397509, EPI_ISL_3397510, EPI_ISL_3397511, EPI_ISL_3397512, EPI_ISL_3397513, EPI_ISL_3397514, EPI_ISL_3397515, EPI_ISL_3397516, EPI_ISL_3397517, EPI_ISL_3397518, EPI_ISL_3397519, EPI_ISL_3397520, EPI_ISL_3397521, EPI_ISL_3397522, EPI_ISL_3397523, EPI_ISL_3397524, EPI_ISL_3397526, EPI_ISL_3397527, EPI_ISL_3397528, EPI_ISL_3397529, EPI_ISL_3397530, EPI_ISL_3397531, EPI_ISL_3397532, EPI_ISL_3397533, EPI_ISL_3397534, EPI_ISL_3397535, EPI_ISL_3397536, EPI_ISL_3397537, EPI_ISL_3397538, EPI_ISL_3397539, EPI_ISL_3397540, EPI_ISL_3397541, EPI_ISL_3397542, EPI_ISL_3397543, EPI_ISL_3397544, EPI_ISL_3397545, EPI_ISL_3397546, EPI_ISL_3397547, EPI_ISL_3397548, EPI_ISL_3397549, EPI_ISL_3397550, EPI_ISL_3397551, EPI_ISL_3397552, EPI_ISL_3397553, EPI_ISL_3397554, EPI_ISL_3397555, EPI_ISL_3397556, EPI_ISL_3397557, EPI_ISL_3397558, EPI_ISL_3397559, EPI_ISL_3397560, EPI_ISL_3397561, EPI_ISL_3397562, EPI_ISL_3397563, EPI_ISL_3397565, EPI_ISL_3397566, EPI_ISL_3397567, EPI_ISL_3397568, EPI_ISL_3397569, EPI_ISL_3397570, EPI_ISL_3397571, EPI_ISL_3397572, EPI_ISL_3397573, EPI_ISL_3397574, EPI_ISL_3397575, EPI_ISL_3397576, EPI_ISL_3397577, EPI_ISL_3397578, EPI_ISL_3397579, EPI_ISL_3397580, EPI_ISL_3397581, EPI_ISL_3397582, EPI_ISL_3397583, EPI_ISL_3397584, EPI_ISL_3397585, EPI_ISL_3397586, EPI_ISL_3397587, EPI_ISL_3397588, EPI_ISL_3397589, EPI_ISL_3397590, EPI_ISL_3397591, EPI_ISL_3397592, EPI_ISL_3397593, EPI_ISL_3397594, EPI_ISL_3397595, EPI_ISL_3397596, EPI_ISL_3397597, EPI_ISL_3397598, EPI_ISL_3397599, EPI_ISL_3397600, EPI_ISL_3397601, EPI_ISL_3397602, EPI_ISL_3397603, EPI_ISL_3397604, EPI_ISL_3397605, EPI_ISL_3397606, EPI_ISL_3397607, EPI_ISL_3397608, EPI_ISL_3397609, EPI_ISL_3397610, EPI_ISL_3397611, EPI_ISL_3397612, EPI_ISL_3397613, EPI_ISL_3397614, EPI_ISL_3397615, EPI_ISL_3397616, EPI_ISL_3397617, EPI_ISL_3397618, EPI_ISL_3397619, EPI_ISL_3397620, EPI_ISL_3397621, EPI_ISL_3397622, EPI_ISL_3397623, EPI_ISL_3397624, EPI_ISL_3397625, EPI_ISL_3397626, EPI_ISL_3397627, EPI_ISL_3397628, EPI_ISL_3397629, EPI_ISL_3397630, EPI_ISL_3397631, EPI_ISL_3397632, EPI_ISL_3397633, EPI_ISL_3397634, EPI_ISL_3397635, EPI_ISL_3397636, EPI_ISL_3397637, EPI_ISL_3397638, EPI_ISL_3397639, EPI_ISL_3397640, EPI_ISL_3397641, EPI_ISL_3397642, EPI_ISL_3397643, EPI_ISL_3397644, EPI_ISL_3397645, EPI_ISL_3397646, EPI_ISL_3397647, EPI_ISL_3397648, EPI_ISL_3397649, EPI_ISL_3397650, EPI_ISL_3397651, EPI_ISL_3397652, EPI_ISL_3397653, EPI_ISL_3397654, EPI_ISL_3397655, EPI_ISL_3397656, EPI_ISL_3397657, EPI_ISL_3397658, EPI_ISL_3397659, EPI_ISL_3397660, EPI_ISL_3397661, EPI_ISL_3397662, EPI_ISL_3397663, EPI_ISL_3397664, EPI_ISL_3397665, EPI_ISL_3397666, EPI_ISL_3397668, EPI_ISL_3397669, EPI_ISL_3397670, EPI_ISL_3397671, EPI_ISL_3397672, EPI_ISL_3397673, EPI_ISL_3397674, EPI_ISL_3397675, EPI_ISL_3397676, EPI_ISL_3397677, EPI_ISL_3397678, EPI_ISL_3397679, EPI_ISL_3397680, EPI_ISL_3397681, EPI_ISL_3397682, EPI_ISL_3397683, EPI_ISL_3397684, EPI_ISL_3397685, EPI_ISL_3397686, EPI_ISL_3397687, EPI_ISL_3397688, EPI_ISL_3397689, EPI_ISL_3397690, EPI_ISL_3397691, EPI_ISL_3397692, EPI_ISL_3397693, EPI_ISL_3397694, EPI_ISL_3397695, EPI_ISL_3397696, EPI_ISL_3397697, EPI_ISL_3397698, EPI_ISL_3397699, EPI_ISL_3397700, EPI_ISL_3397701, EPI_ISL_3397702, EPI_ISL_3397703, EPI_ISL_3397704, EPI_ISL_3397705, EPI_ISL_3397706, EPI_ISL_3397707, EPI_ISL_3397708, EPI_ISL_3397709, EPI_ISL_3397710, EPI_ISL_3397711, EPI_ISL_3397712, EPI_ISL_3397713, EPI_ISL_3397714, EPI_ISL_3397715, EPI_ISL_3397716, EPI_ISL_3397717, EPI_ISL_3397718, EPI_ISL_3397719, EPI_ISL_3397720, EPI_ISL_3397721, EPI_ISL_3397722, EPI_ISL_3397723, EPI_ISL_3397724, EPI_ISL_3397726, EPI_ISL_3397727, EPI_ISL_3397728, EPI_ISL_3397729, EPI_ISL_3397730, EPI_ISL_3397731, EPI_ISL_3397732, EPI_ISL_3397733, EPI_ISL_3397734, EPI_ISL_3397735, EPI_ISL_3397736, EPI_ISL_3397737, EPI_ISL_3397738, EPI_ISL_3397739, EPI_ISL_3397740, EPI_ISL_3397741, EPI_ISL_3397742, EPI_ISL_3397743, EPI_ISL_3397744, EPI_ISL_3397745, EPI_ISL_3397746, EPI_ISL_3397747, EPI_ISL_3397748, EPI_ISL_3397749, EPI_ISL_3397750, EPI_ISL_3397751, EPI_ISL_3397752, EPI_ISL_3397753, EPI_ISL_3397754, EPI_ISL_3397755, EPI_ISL_3397756, EPI_ISL_3397757, EPI_ISL_3397758, EPI_ISL_3397759, EPI_ISL_3397760, EPI_ISL_3397761, EPI_ISL_3397762, EPI_ISL_3397763, EPI_ISL_3397764, EPI_ISL_3397765, EPI_ISL_3397766, EPI_ISL_3397767, EPI_ISL_3397768, EPI_ISL_3397769, EPI_ISL_3397770, EPI_ISL_3397771, EPI_ISL_3397772, EPI_ISL_3397773, EPI_ISL_3397774, EPI_ISL_3397775, EPI_ISL_3397776, EPI_ISL_3397777, EPI_ISL_3397778, EPI_ISL_3397779, EPI_ISL_3397780, EPI_ISL_3397781, EPI_ISL_3397782, EPI_ISL_3397783, EPI_ISL_3397784, EPI_ISL_3397785, EPI_ISL_3397786, EPI_ISL_3397787, EPI_ISL_3397788, EPI_ISL_3397789, EPI_ISL_3397790, EPI_ISL_3397791, EPI_ISL_3397792, EPI_ISL_3397793, EPI_ISL_3397794, EPI_ISL_3397795, EPI_ISL_3397796, EPI_ISL_3397797, EPI_ISL_3397798, EPI_ISL_3397799, EPI_ISL_3397800, EPI_ISL_3397801, EPI_ISL_3397802, EPI_ISL_3397803, EPI_ISL_3397804, EPI_ISL_3397805, EPI_ISL_3397806, EPI_ISL_3397807, EPI_ISL_3397808, EPI_ISL_3397809, EPI_ISL_3397810, EPI_ISL_3397811, EPI_ISL_3397812, EPI_ISL_3397813, EPI_ISL_3397814, EPI_ISL_3397815, EPI_ISL_3397816, EPI_ISL_3397817, EPI_ISL_3397818, EPI_ISL_3397819, EPI_ISL_3397820, EPI_ISL_3397821, EPI_ISL_3397822, EPI_ISL_3397823, EPI_ISL_3397824, EPI_ISL_3397825, EPI_ISL_3397826, EPI_ISL_3397827, EPI_ISL_3397828, EPI_ISL_3397829, EPI_ISL_3397830, EPI_ISL_3397831, EPI_ISL_3397832, EPI_ISL_3397833, EPI_ISL_3397834, EPI_ISL_3397835, EPI_ISL_3397836, EPI_ISL_3397837, EPI_ISL_3397838, EPI_ISL_3397839, EPI_ISL_3397840, EPI_ISL_3397841, EPI_ISL_3397842, EPI_ISL_3397843, EPI_ISL_3397844, EPI_ISL_3397845, EPI_ISL_3397846, EPI_ISL_3397847, EPI_ISL_3397848, EPI_ISL_3397849, EPI_ISL_3397850, EPI_ISL_3397851, EPI_ISL_3397852, EPI_ISL_3397853, EPI_ISL_3397854, EPI_ISL_3397855, EPI_ISL_3397856, EPI_ISL_3397857, EPI_ISL_3397858, EPI_ISL_3397859, EPI_ISL_3397860, EPI_ISL_3397861, EPI_ISL_3397862, EPI_ISL_3397863, EPI_ISL_3397864, EPI_ISL_3397865, EPI_ISL_3397866, EPI_ISL_3397867, EPI_ISL_3397868, EPI_ISL_3397869, EPI_ISL_3397870, EPI_ISL_3397871, EPI_ISL_3397872, EPI_ISL_3397873, EPI_ISL_3397874, EPI_ISL_3397875, EPI_ISL_3397876, EPI_ISL_3397877, EPI_ISL_3397878, EPI_ISL_3397879, EPI_ISL_3397880, EPI_ISL_3397881, EPI_ISL_3397882, EPI_ISL_3397883, EPI_ISL_3397884 | | | |
| see above | Fulgent Genetics | Centers for Disease Control and Prevention Division of Viral Diseases, Pathogen Discovery | Dakota Howard, Dhwani Batra, Peter W. Cook, Kara Moser, Adrian Paskey, Jason Caravas, Benjamin Rambo-Martin, Shatavia Morrison, Christopher Gulvick, Scott Sammons, Yvette Unoarumhi, Darlene Wagner, Matthew Schmerer, Harry Gao, Mickey Li, John Gao, Joseph Fierro, Benafsh Sapra, Becky Tsai, Yan Meng, Doreen Ng, James Xie, Clinton R. Paden, Duncan MacCannell |
| EPI_ISL_3397885 | CLINIQUE N-D DE GRACE | Institut de Pathologie et Genetique (IPG) | Pascale Hilbert, Jérémie Gras |
| EPI_ISL_3397886 | Fulgent Genetics | Centers for Disease Control and Prevention Division of Viral Diseases, Pathogen Discovery | Dakota Howard, Dhwani Batra, Peter W. Cook, Kara Moser, Adrian Paskey, Jason Caravas, Benjamin Rambo-Martin, Shatavia Morrison, Christopher Gulvick, Scott Sammons, Yvette Unoarumhi, Darlene Wagner, Matthew Schmerer, Harry Gao, Mickey Li, John Gao, Joseph Fierro, Benafsh Sapra, Becky Tsai, Yan Meng, Doreen Ng, James Xie, Clinton R. Paden, Duncan MacCannell |
| EPI_ISL_3397887, EPI_ISL_3397888 | CLINIQUE N-D DE GRACE | Institut de Pathologie et Genetique (IPG) | Pascale Hilbert, Jérémie Gras |
| EPI_ISL_3397889 | Fulgent Genetics | Centers for Disease Control and Prevention Division of Viral Diseases, Pathogen Discovery | Dakota Howard, Dhwani Batra, Peter W. Cook, Kara Moser, Adrian Paskey, Jason Caravas, Benjamin Rambo-Martin, Shatavia Morrison, Christopher Gulvick, Scott Sammons, Yvette Unoarumhi, Darlene Wagner, Matthew Schmerer, Harry Gao, Mickey Li, John Gao, Joseph Fierro, Benafsh Sapra, Becky Tsai, Yan Meng, Doreen Ng, James Xie, Clinton R. Paden, Duncan MacCannell |
| EPI_ISL_3397890 | CLINIQUE N-D DE GRACE | Institut de Pathologie et Genetique (IPG) | Pascale Hilbert, Jérémie Gras |
| EPI_ISL_3397891 | Fulgent Genetics | Centers for Disease Control and Prevention Division of Viral Diseases, Pathogen Discovery | Dakota Howard, Dhwani Batra, Peter W. Cook, Kara Moser, Adrian Paskey, Jason Caravas, Benjamin Rambo-Martin, Shatavia Morrison, Christopher Gulvick, Scott Sammons, Yvette Unoarumhi, Darlene Wagner, Matthew Schmerer, Harry Gao, Mickey Li, John Gao, Joseph Fierro, Benafsh Sapra, Becky Tsai, Yan Meng, Doreen Ng, James Xie, Clinton R. Paden, Duncan MacCannell |
| EPI_ISL_3397892, EPI_ISL_3397893 | CLINIQUE N-D DE GRACE | Institut de Pathologie et Genetique (IPG) | Pascale Hilbert, Jérémie Gras |
| EPI_ISL_3397894 | Fulgent Genetics | Centers for Disease Control and Prevention Division of Viral Diseases, Pathogen Discovery | Dakota Howard, Dhwani Batra, Peter W. Cook, Kara Moser, Adrian Paskey, Jason Caravas, Benjamin Rambo-Martin, Shatavia Morrison, Christopher Gulvick, Scott Sammons, Yvette Unoarumhi, Darlene Wagner, Matthew Schmerer, Harry Gao, Mickey Li, John Gao, Joseph Fierro, Benafsh Sapra, Becky Tsai, Yan Meng, Doreen Ng, James Xie, Clinton R. Paden, Duncan MacCannell |
| EPI_ISL_3397895 | CLINIQUE N-D DE GRACE | Institut de Pathologie et Genetique (IPG) | Pascale Hilbert, Jérémie Gras |
| EPI_ISL_3397896 | Fulgent Genetics | Centers for Disease Control and Prevention Division of Viral Diseases, Pathogen Discovery | Dakota Howard, Dhwani Batra, Peter W. Cook, Kara Moser, Adrian Paskey, Jason Caravas, Benjamin Rambo-Martin, Shatavia Morrison, Christopher Gulvick, Scott Sammons, Yvette Unoarumhi, Darlene Wagner, Matthew Schmerer, Harry Gao, Mickey Li, John Gao, Joseph Fierro, Benafsh Sapra, Becky Tsai, Yan Meng, Doreen Ng, James Xie, Clinton R. Paden, Duncan MacCannell |
| EPI_ISL_3397897 | CLINIQUE N-D DE GRACE | Institut de Pathologie et Genetique (IPG) | Pascale Hilbert, Jérémie Gras |
| EPI_ISL_3397898 | Fulgent Genetics | Centers for Disease Control and Prevention Division of Viral Diseases, Pathogen Discovery | Dakota Howard, Dhwani Batra, Peter W. Cook, Kara Moser, Adrian Paskey, Jason Caravas, Benjamin Rambo-Martin, Shatavia Morrison, Christopher Gulvick, Scott Sammons, Yvette Unoarumhi, Darlene Wagner, Matthew Schmerer, Harry Gao, Mickey Li, John Gao, Joseph Fierro, Benafsh Sapra, Becky Tsai, Yan Meng, Doreen Ng, James Xie, Clinton R. Paden, Duncan MacCannell |
| EPI_ISL_3397899, EPI_ISL_3397900 | CLINIQUE N-D DE GRACE | Institut de Pathologie et Genetique (IPG) | Pascale Hilbert, Jérémie Gras |
| EPI_ISL_3397901 | Fulgent Genetics | Centers for Disease Control and Prevention Division of Viral Diseases, Pathogen Discovery | Dakota Howard, Dhwani Batra, Peter W. Cook, Kara Moser, Adrian Paskey, Jason Caravas, Benjamin Rambo-Martin, Shatavia Morrison, Christopher Gulvick, Scott Sammons, Yvette Unoarumhi, Darlene Wagner, Matthew Schmerer, Harry Gao, Mickey Li, John Gao, Joseph Fierro, Benafsh Sapra, Becky Tsai, Yan Meng, Doreen Ng, James Xie, Clinton R. Paden, Duncan MacCannell |
| EPI_ISL_3397902 | CLINIQUE N-D DE GRACE | Institut de Pathologie et Genetique (IPG) | Pascale Hilbert, Jérémie Gras |
| EPI_ISL_3397903 | Fulgent Genetics | Centers for Disease Control and Prevention Division of Viral Diseases, Pathogen Discovery | Dakota Howard, Dhwani Batra, Peter W. Cook, Kara Moser, Adrian Paskey, Jason Caravas, Benjamin Rambo-Martin, Shatavia Morrison, Christopher Gulvick, Scott Sammons, Yvette Unoarumhi, Darlene Wagner, Matthew Schmerer, Harry Gao, Mickey Li, John Gao, Joseph Fierro, Benafsh Sapra, Becky Tsai, Yan Meng, Doreen Ng, James Xie, Clinton R. Paden, Duncan MacCannell |
| EPI_ISL_3397904, EPI_ISL_3397905 | CLINIQUE N-D DE GRACE | Institut de Pathologie et Genetique (IPG) | Pascale Hilbert, Jérémie Gras |
| EPI_ISL_3397906 | Fulgent Genetics | Centers for Disease Control and Prevention Division of Viral Diseases, Pathogen Discovery | Dakota Howard, Dhwani Batra, Peter W. Cook, Kara Moser, Adrian Paskey, Jason Caravas, Benjamin Rambo-Martin, Shatavia Morrison, Christopher Gulvick, Scott Sammons, Yvette Unoarumhi, Darlene Wagner, Matthew Schmerer, Harry Gao, Mickey Li, John Gao, Joseph Fierro, Benafsh Sapra, Becky Tsai, Yan Meng, Doreen Ng, James Xie, Clinton R. Paden, Duncan MacCannell |
| EPI_ISL_3397907 | CLINIQUE N-D DE GRACE | Institut de Pathologie et Genetique (IPG) | Pascale Hilbert, Jérémie Gras |
| EPI_ISL_3397908 | Fulgent Genetics | Centers for Disease Control and Prevention Division of Viral Diseases, Pathogen Discovery | Dakota Howard, Dhwani Batra, Peter W. Cook, Kara Moser, Adrian Paskey, Jason Caravas, Benjamin Rambo-Martin, Shatavia Morrison, Christopher Gulvick, Scott Sammons, Yvette Unoarumhi, Darlene Wagner, Matthew Schmerer, Harry Gao, Mickey Li, John Gao, Joseph Fierro, Benafsh Sapra, Becky Tsai, Yan Meng, Doreen Ng, James Xie, Clinton R. Paden, Duncan MacCannell |
| EPI_ISL_3397909, EPI_ISL_3397910 | CLINIQUE N-D DE GRACE | Institut de Pathologie et Genetique (IPG) | Pascale Hilbert, Jérémie Gras |
| EPI_ISL_3397911 | Fulgent Genetics | Centers for Disease Control and Prevention Division of Viral Diseases, Pathogen Discovery | Dakota Howard, Dhwani Batra, Peter W. Cook, Kara Moser, Adrian Paskey, Jason Caravas, Benjamin Rambo-Martin, Shatavia Morrison, Christopher Gulvick, Scott Sammons, Yvette Unoarumhi, Darlene Wagner, Matthew Schmerer, Harry Gao, Mickey Li, John Gao, Joseph Fierro, Benafsh Sapra, Becky Tsai, Yan Meng, Doreen Ng, James Xie, Clinton R. Paden, Duncan MacCannell |
| EPI_ISL_3397912 | Cabinet Priv | Institut de Pathologie et Genetique (IPG) | Pascale Hilbert, Jérémie Gras |
| EPI_ISL_3397913 | Fulgent Genetics | Centers for Disease Control and Prevention Division of Viral Diseases, Pathogen Discovery | Dakota Howard, Dhwani Batra, Peter W. Cook, Kara Moser, Adrian Paskey, Jason Caravas, Benjamin Rambo-Martin, Shatavia Morrison, Christopher Gulvick, Scott Sammons, Yvette Unoarumhi, Darlene Wagner, Matthew Schmerer, Harry Gao, Mickey Li, John Gao, Joseph Fierro, Benafsh Sapra, Becky Tsai, Yan Meng, Doreen Ng, James Xie, Clinton R. Paden, Duncan MacCannell |
| EPI_ISL_3397914 | Cabinet Priv | Institut de Pathologie et Genetique (IPG) | Pascale Hilbert, Jérémie Gras |
| EPI_ISL_3397915 | Fulgent Genetics | Centers for Disease Control and Prevention Division of Viral Diseases, Pathogen Discovery | Dakota Howard, Dhwani Batra, Peter W. Cook, Kara Moser, Adrian Paskey, Jason Caravas, Benjamin Rambo-Martin, Shatavia Morrison, Christopher Gulvick, Scott Sammons, Yvette Unoarumhi, Darlene Wagner, Matthew Schmerer, Harry Gao, Mickey Li, John Gao, Joseph Fierro, Benafsh Sapra, Becky Tsai, Yan Meng, Doreen Ng, James Xie, Clinton R. Paden, Duncan MacCannell |
| EPI_ISL_3397916, EPI_ISL_3397917 | CHWAPI - SITE NOTRE DAME | Institut de Pathologie et Genetique (IPG) | Pascale Hilbert, Jérémie Gras |
| EPI_ISL_3397918 | Fulgent Genetics | Centers for Disease Control and Prevention Division of Viral Diseases, Pathogen Discovery | Dakota Howard, Dhwani Batra, Peter W. Cook, Kara Moser, Adrian Paskey, Jason Caravas, Benjamin Rambo-Martin, Shatavia Morrison, Christopher Gulvick, Scott Sammons, Yvette Unoarumhi, Darlene Wagner, Matthew Schmerer, Harry Gao, Mickey Li, John Gao, Joseph Fierro, Benafsh Sapra, Becky Tsai, Yan Meng, Doreen Ng, James Xie, Clinton R. Paden, Duncan MacCannell |
| EPI_ISL_3397919 | CHWAPI - SITE NOTRE DAME | Institut de Pathologie et Genetique (IPG) | Pascale Hilbert, Jérémie Gras |
| EPI_ISL_3397920 | Fulgent Genetics | Centers for Disease Control and Prevention Division of Viral Diseases, Pathogen Discovery | Dakota Howard, Dhwani Batra, Peter W. Cook, Kara Moser, Adrian Paskey, Jason Caravas, Benjamin Rambo-Martin, Shatavia Morrison, Christopher Gulvick, Scott Sammons, Yvette Unoarumhi, Darlene Wagner, Matthew Schmerer, Harry Gao, Mickey Li, John Gao, Joseph Fierro, Benafsh Sapra, Becky Tsai, Yan Meng, Doreen Ng, James Xie, Clinton R. Paden, Duncan MacCannell |
| EPI_ISL_3397921, EPI_ISL_3397922 | CHWAPI - SITE NOTRE DAME | Institut de Pathologie et Genetique (IPG) | Pascale Hilbert, Jérémie Gras |
| EPI_ISL_3397923, EPI_ISL_3397924, EPI_ISL_3397925, EPI_ISL_3397926, EPI_ISL_3397927, EPI_ISL_3397928, EPI_ISL_3397929, EPI_ISL_3397930, EPI_ISL_3397931, EPI_ISL_3397932, EPI_ISL_3397933, EPI_ISL_3397934, EPI_ISL_3397935, EPI_ISL_3397936, EPI_ISL_3397937, EPI_ISL_3397938, EPI_ISL_3397939, EPI_ISL_3397940, EPI_ISL_3397941, EPI_ISL_3397942, EPI_ISL_3397943, EPI_ISL_3397944, EPI_ISL_3397945, EPI_ISL_3397946, EPI_ISL_3397947, EPI_ISL_3397948, EPI_ISL_3397949, EPI_ISL_3397950, EPI_ISL_3397951, EPI_ISL_3397952, EPI_ISL_3397953, EPI_ISL_3397954, EPI_ISL_3397955, EPI_ISL_3397956, EPI_ISL_3397957, EPI_ISL_3397958, EPI_ISL_3397959, EPI_ISL_3397960, EPI_ISL_3397961, EPI_ISL_3397962, EPI_ISL_3397963, EPI_ISL_3397964, EPI_ISL_3397965, EPI_ISL_3397966, EPI_ISL_3397967, EPI_ISL_3397968, EPI_ISL_3397969, EPI_ISL_3397970, EPI_ISL_3397971, EPI_ISL_3397972, EPI_ISL_3397973, EPI_ISL_3397974, EPI_ISL_3397975, EPI_ISL_3397976, EPI_ISL_3397977, EPI_ISL_3397978, EPI_ISL_3397979, EPI_ISL_3397980, EPI_ISL_3397981, EPI_ISL_3397982, EPI_ISL_3397983, EPI_ISL_3397984, EPI_ISL_3397985, EPI_ISL_3397986, EPI_ISL_3397987, EPI_ISL_3397988, EPI_ISL_3397989 | | | |
| see above | Fulgent Genetics | Centers for Disease Control and Prevention Division of Viral Diseases, Pathogen Discovery | Dakota Howard, Dhwani Batra, Peter W. Cook, Kara Moser, Adrian Paskey, Jason Caravas, Benjamin Rambo-Martin, Shatavia Morrison, Christopher Gulvick, Scott Sammons, Yvette Unoarumhi, Darlene Wagner, Matthew Schmerer, Harry Gao, Mickey Li, John Gao, Joseph Fierro, Benafsh Sapra, Becky Tsai, Yan Meng, Doreen Ng, James Xie, Clinton R. Paden, Duncan MacCannell |
| EPI_ISL_3397997, EPI_ISL_3397998, EPI_ISL_3397999, EPI_ISL_3398000, EPI_ISL_3398001, EPI_ISL_3398002, EPI_ISL_3398003, EPI_ISL_3398004, EPI_ISL_3398005, EPI_ISL_3398007, EPI_ISL_3398008, EPI_ISL_3398009, EPI_ISL_3398010, EPI_ISL_3398011, EPI_ISL_3398012 | | | |
| see above | DC Public Health Lab/ Dept. of Forensic Sciences | DC Public Health Lab/ Dept. of Forensic Sciences | Janis Doss, Scott Nguyen, Elizabeth Zelaya, Sarah Scott, Connie Maza, Monica Mann, Brittany Hamilton, David Payne, Jocelyn Hauser |
| EPI_ISL_3398013, EPI_ISL_3398014, EPI_ISL_3398015, EPI_ISL_3398016, EPI_ISL_3398017, EPI_ISL_3398018, EPI_ISL_3398019, EPI_ISL_3398020, EPI_ISL_3398021, EPI_ISL_3398022, EPI_ISL_3398023, EPI_ISL_3398024, EPI_ISL_3398025, EPI_ISL_3398026, EPI_ISL_3398027, EPI_ISL_3398028, EPI_ISL_3398029, EPI_ISL_3398030, EPI_ISL_3398031, EPI_ISL_3398032, EPI_ISL_3398033, EPI_ISL_3398034, EPI_ISL_3398035, EPI_ISL_3398036, EPI_ISL_3398037, EPI_ISL_3398038, EPI_ISL_3398039, EPI_ISL_3398040, EPI_ISL_3398041, EPI_ISL_3398042, EPI_ISL_3398043, EPI_ISL_3398044, EPI_ISL_3398045, EPI_ISL_3398046 | | | |
| see above | State Hygienic Laboratory at the University of Iowa | State Hygienic Laboratory at the University of Iowa | Valerie Reeb, Erik Twait, Anna Yakos, Mariah Knutson, Kris Eveland, Kristen Zanon, Jeff, Benfer, Wes Hottel, Alankar Kampoowale |

| | | | |
|---|---|---|---|
| EPI_ISL_3398047, EPI_ISL_3398048, EPI_ISL_3398049, EPI_ISL_3398050, EPI_ISL_3398051, EPI_ISL_3398052, EPI_ISL_3398053, EPI_ISL_3398054 | Test Iowa | State Hygienic Laboratory at the University of Iowa | Valerie Reeb, Erik Twait, Anna Yakos, Mariah Knutson, Kris Eveland, Kristen Zanon, Jeff, Benfer, Wes Hottel, Alankar Kampoowale |
| EPI_ISL_3398057, EPI_ISL_3398058, EPI_ISL_3398059, EPI_ISL_3398060 | University of Iowa Hospitals and Clinics | State Hygienic Laboratory at the University of Iowa | Valerie Reeb, Erik Twait, Anna Yakos, Mariah Knutson, Kris Eveland, Kristen Zanon, Jeff, Benfer, Wes Hottel, Alankar Kampoowale |
| EPI_ISL_3398061, EPI_ISL_3398062, EPI_ISL_3398063, EPI_ISL_3398064, EPI_ISL_3398065, EPI_ISL_3398066, EPI_ISL_3398067, EPI_ISL_3398068 | State Hygienic Laboratory at the University of Iowa | State Hygienic Laboratory at the University of Iowa | Valerie Reeb, Erik Twait, Anna Yakos, Mariah Knutson, Kris Eveland, Kristen Zanon, Jeff, Benfer, Wes Hottel, Alankar Kampoowale |
| EPI_ISL_3398069, EPI_ISL_3398070, EPI_ISL_3398071, EPI_ISL_3398072, EPI_ISL_3398073, EPI_ISL_3398074, EPI_ISL_3398075, EPI_ISL_3398076, EPI_ISL_3398077, EPI_ISL_3398078, EPI_ISL_3398079, EPI_ISL_3398080, EPI_ISL_3398081, EPI_ISL_3398082, EPI_ISL_3398083, EPI_ISL_3398084, EPI_ISL_3398085, EPI_ISL_3398086, EPI_ISL_3398087, EPI_ISL_3398088, EPI_ISL_3398089, EPI_ISL_3398091, EPI_ISL_3398092, EPI_ISL_3398093, EPI_ISL_3398094 | | | |
| see above | NJDOH, Public Health and Environmental Laboratories | NJ_PHEL | Lindsey Bodnar, Shiv K. Verma, Ryan Pachucki, Chelsea San Filippo, Jacquelyn Deverell, Dana Woell, Allison Roder, Byeong Jeong |
| EPI_ISL_3398095, EPI_ISL_3398096, EPI_ISL_3398097 | CNR Institut Pasteur de la Guyane | Institut Pasteur de la Guyane | Anne Lavergne, Dominique Rousset, Antoine Enfissi, Arielle Salmier |
| EPI_ISL_3398098, EPI_ISL_3398099 | CH de l'Ouest Guyanais Frank Joly | Institut Pasteur de la Guyane | Anne Lavergne, Dominique Rousset, Antoine Enfissi, Arielle Salmier |
| EPI_ISL_3398101, EPI_ISL_3398102, EPI_ISL_3398103 | CHC Andrée Rosemon | Institut Pasteur de la Guyane | Anne Lavergne, Dominique Rousset, Antoine Enfissi, Arielle Salmier |
| EPI_ISL_3398105, EPI_ISL_3398106, EPI_ISL_3398109, EPI_ISL_3398110 | CNR Institut Pasteur de la Guyane | Institut Pasteur de la Guyane | Anne Lavergne, Dominique Rousset, Antoine Enfissi, Arielle Salmier |
| EPI_ISL_3398111, EPI_ISL_3398112, EPI_ISL_3398113, EPI_ISL_3398114, EPI_ISL_3398117 | CHC Andrée Rosemon | Institut Pasteur de la Guyane | Anne Lavergne, Dominique Rousset, Antoine Enfissi, Arielle Salmier |
| EPI_ISL_3398120, EPI_ISL_3398121, EPI_ISL_3398122, EPI_ISL_3398123, EPI_ISL_3398124, EPI_ISL_3398125, EPI_ISL_3398126, EPI_ISL_3398127, EPI_ISL_3398128, EPI_ISL_3398130, EPI_ISL_3398131, EPI_ISL_3398132, EPI_ISL_3398133, EPI_ISL_3398134 | | | |
| see above | Ochsner LSU Health | LSUHS Emerging Viral Threat Laboratory | Jeremy P. Kamil, Jennifer L. Carroll, Maarten Van Diest, Rona S. Scott, Gregory L. Ware, Alexander Mijalis, Andrew D. Yurochko, Christopher G. Kevil, John A. Vanchiere |
| EPI_ISL_3398135 | The Health Hut | LSUHS Emerging Viral Threat Laboratory | Jamie J. Newman, Paul Kim, Jacquelyn K. White |
| EPI_ISL_3398136, EPI_ISL_3398137, EPI_ISL_3398138, EPI_ISL_3398139, EPI_ISL_3398140, EPI_ISL_3398141, EPI_ISL_3398142, EPI_ISL_3398143, EPI_ISL_3398144, EPI_ISL_3398145, EPI_ISL_3398146, EPI_ISL_3398147, EPI_ISL_3398148, EPI_ISL_3398149, EPI_ISL_3398150, EPI_ISL_3398151, EPI_ISL_3398152, EPI_ISL_3398153, EPI_ISL_3398154, EPI_ISL_3398155, EPI_ISL_3398156, EPI_ISL_3398157, EPI_ISL_3398158, EPI_ISL_3398159, EPI_ISL_3398160, EPI_ISL_3398161, EPI_ISL_3398162, EPI_ISL_3398163, EPI_ISL_3398164, EPI_ISL_3398165, EPI_ISL_3398166, EPI_ISL_3398167, EPI_ISL_3398168, EPI_ISL_3398169, EPI_ISL_3398170, EPI_ISL_3398171, EPI_ISL_3398172, EPI_ISL_3398173, EPI_ISL_3398174, EPI_ISL_3398175, EPI_ISL_3398176, EPI_ISL_3398177, EPI_ISL_3398179, EPI_ISL_3398180, EPI_ISL_3398181, EPI_ISL_3398182, EPI_ISL_3398183, EPI_ISL_3398184, EPI_ISL_3398185, EPI_ISL_3398186, EPI_ISL_3398187, EPI_ISL_3398188, EPI_ISL_3398189, EPI_ISL_3398190, EPI_ISL_3398191, EPI_ISL_3398192, EPI_ISL_3398193, EPI_ISL_3398194, EPI_ISL_3398195, EPI_ISL_3398196, EPI_ISL_3398197, EPI_ISL_3398198, EPI_ISL_3398199, EPI_ISL_3398200, EPI_ISL_3398201, EPI_ISL_3398202, EPI_ISL_3398203, EPI_ISL_3398204, EPI_ISL_3398205, EPI_ISL_3398206, EPI_ISL_3398207, EPI_ISL_3398208, EPI_ISL_3398209, EPI_ISL_3398210, EPI_ISL_3398211, EPI_ISL_3398212, EPI_ISL_3398213, EPI_ISL_3398214, EPI_ISL_3398215, EPI_ISL_3398216, EPI_ISL_3398217, EPI_ISL_3398218, EPI_ISL_3398219, EPI_ISL_3398220, EPI_ISL_3398221, EPI_ISL_3398222 | | | |
| see above | LSUHS Emerging Viral Threat Lab | LSUHS Emerging Viral Threat Laboratory | Gregory L. Ware, Alexander Mijalis, Jeremy P. Kamil, Jennifer L. Carroll, Maarten Van Diest, Rona S. Scott, Andrew D. Yurochko, Christopher G. Kevil, John A. Vanchiere |
| EPI_ISL_3398223, EPI_ISL_3398224, EPI_ISL_3398225, EPI_ISL_3398226, EPI_ISL_3398227, EPI_ISL_3398228, EPI_ISL_3398229, EPI_ISL_3398230, EPI_ISL_3398231, EPI_ISL_3398232, EPI_ISL_3398233, EPI_ISL_3398234, EPI_ISL_3398235, EPI_ISL_3398236, EPI_ISL_3398237, EPI_ISL_3398238, EPI_ISL_3398239, EPI_ISL_3398240, EPI_ISL_3398241, EPI_ISL_3398242, EPI_ISL_3398243, EPI_ISL_3398244, EPI_ISL_3398245, EPI_ISL_3398246, EPI_ISL_3398247, EPI_ISL_3398248, EPI_ISL_3398249, EPI_ISL_3398250, EPI_ISL_3398251, EPI_ISL_3398252, EPI_ISL_3398253, EPI_ISL_3398255, EPI_ISL_3398256, EPI_ISL_3398257, EPI_ISL_3398258, EPI_ISL_3398259, EPI_ISL_3398260, EPI_ISL_3398261, EPI_ISL_3398262, EPI_ISL_3398263, EPI_ISL_3398264, EPI_ISL_3398265, EPI_ISL_3398266, EPI_ISL_3398267, EPI_ISL_3398268, EPI_ISL_3398269, EPI_ISL_3398270, EPI_ISL_3398271, EPI_ISL_3398272, EPI_ISL_3398273, EPI_ISL_3398274, EPI_ISL_3398275, EPI_ISL_3398276, EPI_ISL_3398277, EPI_ISL_3398278, EPI_ISL_3398279, EPI_ISL_3398280, EPI_ISL_3398281, EPI_ISL_3398282, EPI_ISL_3398283, EPI_ISL_3398284, EPI_ISL_3398285, EPI_ISL_3398286, EPI_ISL_3398287, EPI_ISL_3398288, EPI_ISL_3398289, EPI_ISL_3398290, EPI_ISL_3398291, EPI_ISL_3398292, EPI_ISL_3398293, EPI_ISL_3398294, EPI_ISL_3398295, EPI_ISL_3398296, EPI_ISL_3398297, EPI_ISL_3398298, EPI_ISL_3398299, EPI_ISL_3398300, EPI_ISL_3398301, EPI_ISL_3398302, EPI_ISL_3398303, EPI_ISL_3398304, EPI_ISL_3398305, EPI_ISL_3398306, EPI_ISL_3398307, EPI_ISL_3398308 | | | |
| see above | Willis-Knighton Medical Center Hospital Laboratory | LSUHS Emerging Viral Threat Laboratory | Lorie M. Atkins, April N. Johnson, Gregory L. Ware, Alexander Mijalis, Jeremy P. Kamil, Jennifer L. Carroll, Maarten Van Diest, Rona S. Scott, Andrew D. Yurochko, Christopher G. Kevil, John A. Vanchiere, Joseph A. Bocchini |
| EPI_ISL_3398309, EPI_ISL_3398310, EPI_ISL_3398311, EPI_ISL_3398312, EPI_ISL_3398313, EPI_ISL_3398314, EPI_ISL_3398315, EPI_ISL_3398316, EPI_ISL_3398317, EPI_ISL_3398318, EPI_ISL_3398319, EPI_ISL_3398320, EPI_ISL_3398321, EPI_ISL_3398322, EPI_ISL_3398323, EPI_ISL_3398324, EPI_ISL_3398325, EPI_ISL_3398326, EPI_ISL_3398327, EPI_ISL_3398328, EPI_ISL_3398329, EPI_ISL_3398330, EPI_ISL_3398331, EPI_ISL_3398332, EPI_ISL_3398333, EPI_ISL_3398334, EPI_ISL_3398335, EPI_ISL_3398336, EPI_ISL_3398337, EPI_ISL_3398338, EPI_ISL_3398339, EPI_ISL_3398340, EPI_ISL_3398341, EPI_ISL_3398342, EPI_ISL_3398343, EPI_ISL_3398344, EPI_ISL_3398345, EPI_ISL_3398346, EPI_ISL_3398347, EPI_ISL_3398348, EPI_ISL_3398349, EPI_ISL_3398350, EPI_ISL_3398351, EPI_ISL_3398352, EPI_ISL_3398353, EPI_ISL_3398354, EPI_ISL_3398355, EPI_ISL_3398356, EPI_ISL_3398357, EPI_ISL_3398358, EPI_ISL_3398359, EPI_ISL_3398360 | | | |
| see above | Louisiana Office of Public Heath Laboratory | LSUHS Emerging Viral Threat Laboratory | Rona S. Scott, Gregory L. Ware, Alexander Mijalis, Jeremy P. Kamil, Jennifer L. Carroll, Maarten Van Diest, Andrew D. Yurochko, Christopher G. Kevil, John A. Vanchiere |
| EPI_ISL_3398374 | Laverty Pathology | NSW Health Pathology - Institute of Clinical Pathology and Medical Research; Westmead Hospital; University of Sydney | Arnott A., Draper J., Gall M., Martinez E., Rockett R., Sintchenko V., on behalf of ICPMR |
| EPI_ISL_3398375 | Douglass Hanly Moir Pathology | NSW Health Pathology - Institute of Clinical Pathology and Medical Research; Westmead Hospital; University of Sydney | Arnott A., Draper J., Gall M., Martinez E., Rockett R., Sintchenko V., on behalf of ICPMR |
| EPI_ISL_3398376 | Laverty Pathology | NSW Health Pathology - Institute of Clinical Pathology and Medical Research; Westmead Hospital; University of Sydney | Arnott A., Draper J., Gall M., Martinez E., Rockett R., Sintchenko V., on behalf of ICPMR |
| EPI_ISL_3398377, EPI_ISL_3398378, EPI_ISL_3398379, EPI_ISL_3398380 | Histopath | NSW Health Pathology - Institute of Clinical Pathology and Medical Research; Westmead Hospital; University of Sydney | Arnott A., Draper J., Gall M., Martinez E., Rockett R., Sintchenko V., on behalf of ICPMR |
| EPI_ISL_3398381, EPI_ISL_3398382, EPI_ISL_3398383 | Sydney South West Pathology Service (SSWPS) - Royal Prince Alfred Hospital - NSW Health Pathology | NSW Health Pathology - Institute of Clinical Pathology and Medical Research; Westmead Hospital; University of Sydney | Arnott A., Draper J., Gall M., Martinez E., Rockett R., Sintchenko V., on behalf of ICPMR |
| EPI_ISL_3398384, EPI_ISL_3398385 | Laverty Pathology | NSW Health Pathology - Institute of Clinical Pathology and Medical Research; Westmead Hospital; University of Sydney | Arnott A., Draper J., Gall M., Martinez E., Rockett R., Sintchenko V., on behalf of ICPMR |
| EPI_ISL_3398386 | Histopath | NSW Health Pathology - Institute of Clinical Pathology and Medical Research; Westmead Hospital; University of Sydney | Arnott A., Draper J., Gall M., Martinez E., Rockett R., Sintchenko V., on behalf of ICPMR |
| EPI_ISL_3398387 | Laverty | NSW Health Pathology - Institute of Clinical Pathology and Medical Research; Westmead Hospital; University of Sydney | Arnott A., Draper J., Gall M., Martinez E., Rockett R., Sintchenko V., on behalf of ICPMR |
| EPI_ISL_3398388 | St Vincent's Pathology (SydPath) | NSW Health Pathology - Institute of Clinical Pathology and Medical Research; Westmead Hospital; University of Sydney | Arnott A., Draper J., Gall M., Martinez E., Rockett R., Sintchenko V., on behalf of ICPMR |
| EPI_ISL_3398389, EPI_ISL_3398390, EPI_ISL_3398391 | Laverty | NSW Health Pathology - Institute of Clinical Pathology and Medical Research; Westmead Hospital; University of Sydney | Arnott A., Draper J., Gall M., Martinez E., Rockett R., Sintchenko V., on behalf of ICPMR |
| EPI_ISL_3398392, EPI_ISL_3398393 | Histopath | NSW Health Pathology - Institute of Clinical Pathology and Medical Research; Westmead Hospital; University of Sydney | Arnott A., Draper J., Gall M., Martinez E., Rockett R., Sintchenko V., on behalf of ICPMR |
| EPI_ISL_3398394, EPI_ISL_3398395, EPI_ISL_3398396, EPI_ISL_3398397 | Sydney South West Pathology Service (SSWPS) - Liverpool Hospital - NSW Health Pathology | NSW Health Pathology - Institute of Clinical Pathology and Medical Research; Westmead Hospital; University of Sydney | Arnott A., Draper J., Gall M., Martinez E., Rockett R., Sintchenko V., on behalf of ICPMR |
| EPI_ISL_3398398, EPI_ISL_3398399, EPI_ISL_3398400, EPI_ISL_3398401 | St Vincent's Pathology (SydPath) | NSW Health Pathology - Institute of Clinical Pathology and Medical Research; Westmead Hospital; University of Sydney | Arnott A., Draper J., Gall M., Martinez E., Rockett R., Sintchenko V., on behalf of ICPMR |
| EPI_ISL_3398402 | 4Cyte Pathology | NSW Health Pathology - Institute of Clinical Pathology and Medical Research; Westmead Hospital; University of Sydney | Arnott A., Draper J., Gall M., Martinez E., Rockett R., Sintchenko V., on behalf of ICPMR |
| EPI_ISL_3398403 | Australian Clinical Labs (formerly Healthscope Pathology) | NSW Health Pathology - Institute of Clinical Pathology and Medical Research; Westmead Hospital; University of Sydney | Arnott A., Draper J., Gall M., Martinez E., Rockett R., Sintchenko V., on behalf of ICPMR |

| | | | |
|---|---|---|---|
| EPI_ISL_3398404 | Pathology West - NSW Health Pathology | NSW Health Pathology - Institute of Clinical Pathology and Medical Research; Westmead Hospital; University of Sydney | Arnott A., Draper J., Gall M., Martinez E., Rockett R., Sintchenko V., on behalf of ICPMR |
| EPI_ISL_3398405 | Sydney South West Pathology Service (SSWPS) - Concord Repatriation General Hospital - NSW Health Pathology | NSW Health Pathology - Institute of Clinical Pathology and Medical Research; Westmead Hospital; University of Sydney | Arnott A., Draper J., Gall M., Martinez E., Rockett R., Sintchenko V., on behalf of ICPMR |
| EPI_ISL_3398406 | St Vincent's Pathology (SydPath) | NSW Health Pathology - Institute of Clinical Pathology and Medical Research; Westmead Hospital; University of Sydney | Arnott A., Draper J., Gall M., Martinez E., Rockett R., Sintchenko V., on behalf of ICPMR |
| EPI_ISL_3398407 | Sydney South West Pathology Service (SSWPS) - Liverpool Hospital - NSW Health Pathology | NSW Health Pathology - Institute of Clinical Pathology and Medical Research; Westmead Hospital; University of Sydney | Arnott A., Draper J., Gall M., Martinez E., Rockett R., Sintchenko V., on behalf of ICPMR |
| EPI_ISL_3398408 | Histopath | NSW Health Pathology - Institute of Clinical Pathology and Medical Research; Westmead Hospital; University of Sydney | Arnott A., Draper J., Gall M., Martinez E., Rockett R., Sintchenko V., on behalf of ICPMR |
| EPI_ISL_3398409 | 4Cyte Pathology | NSW Health Pathology - Institute of Clinical Pathology and Medical Research; Westmead Hospital; University of Sydney | Arnott A., Draper J., Gall M., Martinez E., Rockett R., Sintchenko V., on behalf of ICPMR |
| EPI_ISL_3398410, EPI_ISL_3398411 | St Vincent's Pathology (SydPath) | NSW Health Pathology - Institute of Clinical Pathology and Medical Research; Westmead Hospital; University of Sydney | Arnott A., Draper J., Gall M., Martinez E., Rockett R., Sintchenko V., on behalf of ICPMR |
| EPI_ISL_3398412, EPI_ISL_3398413, EPI_ISL_3398414 | Laverty Pathology | NSW Health Pathology - Institute of Clinical Pathology and Medical Research; Westmead Hospital; University of Sydney | Arnott A., Draper J., Gall M., Martinez E., Rockett R., Sintchenko V., on behalf of ICPMR |
| EPI_ISL_3398415 | 4Cyte Pathology | NSW Health Pathology - Institute of Clinical Pathology and Medical Research; Westmead Hospital; University of Sydney | Arnott A., Draper J., Gall M., Martinez E., Rockett R., Sintchenko V., on behalf of ICPMR |
| EPI_ISL_3398416, EPI_ISL_3398417, EPI_ISL_3398418, EPI_ISL_3398419, EPI_ISL_3398420 | Histopath | NSW Health Pathology - Institute of Clinical Pathology and Medical Research; Westmead Hospital; University of Sydney | Arnott A., Draper J., Gall M., Martinez E., Rockett R., Sintchenko V., on behalf of ICPMR |
| EPI_ISL_3398421, EPI_ISL_3398422, EPI_ISL_3398423 | 4Cyte Pathology | NSW Health Pathology - Institute of Clinical Pathology and Medical Research; Westmead Hospital; University of Sydney | Arnott A., Draper J., Gall M., Martinez E., Rockett R., Sintchenko V., on behalf of ICPMR |
| EPI_ISL_3398424, EPI_ISL_3398425, EPI_ISL_3398426 | Australian Clinical Labs (formerly Healthscope Pathology) | NSW Health Pathology - Institute of Clinical Pathology and Medical Research; Westmead Hospital; University of Sydney | Arnott A., Draper J., Gall M., Martinez E., Rockett R., Sintchenko V., on behalf of ICPMR |
| EPI_ISL_3398427, EPI_ISL_3398428, EPI_ISL_3398429 | Sydney South West Pathology Service (SSWPS) - Liverpool Hospital - NSW Health Pathology | NSW Health Pathology - Institute of Clinical Pathology and Medical Research; Westmead Hospital; University of Sydney | Arnott A., Draper J., Gall M., Martinez E., Rockett R., Sintchenko V., on behalf of ICPMR |
| EPI_ISL_3398430, EPI_ISL_3398431, EPI_ISL_3398432, EPI_ISL_3398433, EPI_ISL_3398434, EPI_ISL_3398435, EPI_ISL_3398436 | Histopath | NSW Health Pathology - Institute of Clinical Pathology and Medical Research; Westmead Hospital; University of Sydney | Arnott A., Draper J., Gall M., Martinez E., Rockett R., Sintchenko V., on behalf of ICPMR |
| EPI_ISL_3398437 | 4Cyte Pathology | NSW Health Pathology - Institute of Clinical Pathology and Medical Research; Westmead Hospital; University of Sydney | Arnott A., Draper J., Gall M., Martinez E., Rockett R., Sintchenko V., on behalf of ICPMR |
| EPI_ISL_3398438 | Douglass Hanly Moir Pathology | NSW Health Pathology - Institute of Clinical Pathology and Medical Research; Westmead Hospital; University of Sydney | Arnott A., Draper J., Gall M., Martinez E., Rockett R., Sintchenko V., on behalf of ICPMR |
| EPI_ISL_3398439 | Pathology West - NSW Health Pathology | NSW Health Pathology - Institute of Clinical Pathology and Medical Research; Westmead Hospital; University of Sydney | Arnott A., Draper J., Gall M., Martinez E., Rockett R., Sintchenko V., on behalf of ICPMR |
| EPI_ISL_3398440, EPI_ISL_3398441, EPI_ISL_3398442, EPI_ISL_3398443, EPI_ISL_3398444, EPI_ISL_3398445, EPI_ISL_3398446, EPI_ISL_3398447, EPI_ISL_3398448, EPI_ISL_3398449, EPI_ISL_3398450 | | | |
| see above | St Vincent's Pathology (SydPath) | NSW Health Pathology - Institute of Clinical Pathology and Medical Research; Westmead Hospital; University of Sydney | Arnott A., Draper J., Gall M., Martinez E., Rockett R., Sintchenko V., on behalf of ICPMR |
| EPI_ISL_3398451, EPI_ISL_3398452, EPI_ISL_3398453 | Histopath | NSW Health Pathology - Institute of Clinical Pathology and Medical Research; Westmead Hospital; University of Sydney | Arnott A., Draper J., Gall M., Martinez E., Rockett R., Sintchenko V., on behalf of ICPMR |
| EPI_ISL_3398454, EPI_ISL_3398455 | 4Cyte Pathology | NSW Health Pathology - Institute of Clinical Pathology and Medical Research; Westmead Hospital; University of Sydney | Arnott A., Draper J., Gall M., Martinez E., Rockett R., Sintchenko V., on behalf of ICPMR |
| EPI_ISL_3398456, EPI_ISL_3398457, EPI_ISL_3398458 | Douglass Hanly Moir Pathology | NSW Health Pathology - Institute of Clinical Pathology and Medical Research; Westmead Hospital; University of Sydney | Arnott A., Draper J., Gall M., Martinez E., Rockett R., Sintchenko V., on behalf of ICPMR |
| EPI_ISL_3398459, EPI_ISL_3398460, EPI_ISL_3398461 | Sydney South West Pathology Service (SSWPS) - Liverpool Hospital - NSW Health Pathology | NSW Health Pathology - Institute of Clinical Pathology and Medical Research; Westmead Hospital; University of Sydney | Arnott A., Draper J., Gall M., Martinez E., Rockett R., Sintchenko V., on behalf of ICPMR |
| EPI_ISL_3398462, EPI_ISL_3398463, EPI_ISL_3398464, EPI_ISL_3398465, EPI_ISL_3398466, EPI_ISL_3398467, EPI_ISL_3398468, EPI_ISL_3398469, EPI_ISL_3398470, EPI_ISL_3398471, EPI_ISL_3398472 | | | |
| see above | St Vincent's Pathology (SydPath) | NSW Health Pathology - Institute of Clinical Pathology and Medical Research; Westmead Hospital; University of Sydney | Arnott A., Draper J., Gall M., Martinez E., Rockett R., Sintchenko V., on behalf of ICPMR |
| EPI_ISL_3398473, EPI_ISL_3398474, EPI_ISL_3398475, EPI_ISL_3398476, EPI_ISL_3398477, EPI_ISL_3398478, EPI_ISL_3398479 | Histopath | NSW Health Pathology - Institute of Clinical Pathology and Medical Research; Westmead Hospital; University of Sydney | Arnott A., Draper J., Gall M., Martinez E., Rockett R., Sintchenko V., on behalf of ICPMR |
| EPI_ISL_3398480, EPI_ISL_3398481 | Medlab Pathology | NSW Health Pathology - Institute of Clinical Pathology and Medical Research; Westmead Hospital; University of Sydney | Arnott A., Draper J., Gall M., Martinez E., Rockett R., Sintchenko V., on behalf of ICPMR |
| EPI_ISL_3398482 | Pathology West - NSW Health Pathology | NSW Health Pathology - Institute of Clinical Pathology and Medical Research; Westmead Hospital; University of Sydney | Arnott A., Draper J., Gall M., Martinez E., Rockett R., Sintchenko V., on behalf of ICPMR |
| EPI_ISL_3398483 | 4Cyte Pathology | NSW Health Pathology - Institute of Clinical Pathology and Medical Research; Westmead Hospital; University of Sydney | Arnott A., Draper J., Gall M., Martinez E., Rockett R., Sintchenko V., on behalf of ICPMR |
| EPI_ISL_3398484, EPI_ISL_3398485 | Laverty Pathology | NSW Health Pathology - Institute of Clinical Pathology and Medical Research; Westmead Hospital; University of Sydney | Arnott A., Draper J., Gall M., Martinez E., Rockett R., Sintchenko V., on behalf of ICPMR |
| EPI_ISL_3398486, EPI_ISL_3398487, EPI_ISL_3398488 | St Vincent's Pathology (SydPath) | NSW Health Pathology - Institute of Clinical Pathology and Medical Research; Westmead Hospital; University of Sydney | Arnott A., Draper J., Gall M., Martinez E., Rockett R., Sintchenko V., on behalf of ICPMR |
| EPI_ISL_3398489, EPI_ISL_3398490, EPI_ISL_3398491 | Histopath | NSW Health Pathology - Institute of Clinical Pathology and Medical Research; Westmead Hospital; University of Sydney | Arnott A., Draper J., Gall M., Martinez E., Rockett R., Sintchenko V., on behalf of ICPMR |
| EPI_ISL_3398492 | Medlab Pathology | NSW Health Pathology - Institute of Clinical Pathology and Medical Research; Westmead Hospital; University of Sydney | Arnott A., Draper J., Gall M., Martinez E., Rockett R., Sintchenko V., on behalf of ICPMR |
| EPI_ISL_3398493, EPI_ISL_3398494, EPI_ISL_3398495 | Laverty Pathology | NSW Health Pathology - Institute of Clinical Pathology and Medical Research; Westmead Hospital; University of Sydney | Arnott A., Draper J., Gall M., Martinez E., Rockett R., Sintchenko V., on behalf of ICPMR |
| EPI_ISL_3398496, EPI_ISL_3398497, EPI_ISL_3398498, EPI_ISL_3398499, EPI_ISL_3398500 | Sydney South West Pathology Service (SSWPS) - Liverpool Hospital - NSW Health Pathology | NSW Health Pathology - Institute of Clinical Pathology and Medical Research; Westmead Hospital; University of Sydney | Arnott A., Draper J., Gall M., Martinez E., Rockett R., Sintchenko V., on behalf of ICPMR |
| EPI_ISL_3398501 | St Vincent's Pathology (SydPath) | NSW Health Pathology - Institute of Clinical Pathology and Medical | Arnott A., Draper J., Gall M., Martinez E., Rockett R., Sintchenko V., on behalf of ICPMR |

| | | | |
|---|---|---|---|
| | | Research; Westmead Hospital; University of Sydney | |
| EPI_ISL_3398502 | Douglass Hanly Moir Pathology | NSW Health Pathology - Institute of Clinical Pathology and Medical Research; Westmead Hospital; University of Sydney | Arnott A., Draper J., Gall M., Martinez E., Rockett R., Sintchenko V., on behalf of ICPMR |
| EPI_ISL_3398503 | Medlab Pathology | NSW Health Pathology - Institute of Clinical Pathology and Medical Research; Westmead Hospital; University of Sydney | Arnott A., Draper J., Gall M., Martinez E., Rockett R., Sintchenko V., on behalf of ICPMR |
| EPI_ISL_3398504, EPI_ISL_3398505, EPI_ISL_3398506, EPI_ISL_3398507 | Histopath | NSW Health Pathology - Institute of Clinical Pathology and Medical Research; Westmead Hospital; University of Sydney | Arnott A., Draper J., Gall M., Martinez E., Rockett R., Sintchenko V., on behalf of ICPMR |
| EPI_ISL_3398508, EPI_ISL_3398509 | 4Cyte Pathology | NSW Health Pathology - Institute of Clinical Pathology and Medical Research; Westmead Hospital; University of Sydney | Arnott A., Draper J., Gall M., Martinez E., Rockett R., Sintchenko V., on behalf of ICPMR |
| EPI_ISL_3398510, EPI_ISL_3398511, EPI_ISL_3398512, EPI_ISL_3398513, EPI_ISL_3398514 | Histopath | NSW Health Pathology - Institute of Clinical Pathology and Medical Research; Westmead Hospital; University of Sydney | Arnott A., Draper J., Gall M., Martinez E., Rockett R., Sintchenko V., on behalf of ICPMR |
| EPI_ISL_3398515, EPI_ISL_3398516, EPI_ISL_3398517 | Laverty Pathology | NSW Health Pathology - Institute of Clinical Pathology and Medical Research; Westmead Hospital; University of Sydney | Arnott A., Draper J., Gall M., Martinez E., Rockett R., Sintchenko V., on behalf of ICPMR |
| EPI_ISL_3398518 | 4Cyte Pathology | NSW Health Pathology - Institute of Clinical Pathology and Medical Research; Westmead Hospital; University of Sydney | Arnott A., Draper J., Gall M., Martinez E., Rockett R., Sintchenko V., on behalf of ICPMR |
| EPI_ISL_3398519, EPI_ISL_3398520 | Pathology West - NSW Health Pathology | NSW Health Pathology - Institute of Clinical Pathology and Medical Research; Westmead Hospital; University of Sydney | Arnott A., Draper J., Gall M., Martinez E., Rockett R., Sintchenko V., on behalf of ICPMR |
| EPI_ISL_3398521, EPI_ISL_3398522, EPI_ISL_3398523, EPI_ISL_3398524, EPI_ISL_3398525, EPI_ISL_3398526, EPI_ISL_3398527, EPI_ISL_3398528, EPI_ISL_3398529, EPI_ISL_3398530, EPI_ISL_3398531, EPI_ISL_3398532 | | | |
| see above | Histopath | NSW Health Pathology - Institute of Clinical Pathology and Medical Research; Westmead Hospital; University of Sydney | Arnott A., Draper J., Gall M., Martinez E., Rockett R., Sintchenko V., on behalf of ICPMR |
| EPI_ISL_3398533 | 4Cyte Pathology | NSW Health Pathology - Institute of Clinical Pathology and Medical Research; Westmead Hospital; University of Sydney | Arnott A., Draper J., Gall M., Martinez E., Rockett R., Sintchenko V., on behalf of ICPMR |
| EPI_ISL_3398534, EPI_ISL_3398535 | The Children's Hospital at Westmead | NSW Health Pathology - Institute of Clinical Pathology and Medical Research; Westmead Hospital; University of Sydney | Arnott A., Draper J., Gall M., Martinez E., Rockett R., Sintchenko V., on behalf of ICPMR |
| EPI_ISL_3398536, EPI_ISL_3398537 | Laverty Pathology | NSW Health Pathology - Institute of Clinical Pathology and Medical Research; Westmead Hospital; University of Sydney | Arnott A., Draper J., Gall M., Martinez E., Rockett R., Sintchenko V., on behalf of ICPMR |
| EPI_ISL_3398538, EPI_ISL_3398539, EPI_ISL_3398540, EPI_ISL_3398541 | Sydney South West Pathology Service (SSWPS) - Liverpool Hospital - NSW Health Pathology | NSW Health Pathology - Institute of Clinical Pathology and Medical Research; Westmead Hospital; University of Sydney | Arnott A., Draper J., Gall M., Martinez E., Rockett R., Sintchenko V., on behalf of ICPMR |
| EPI_ISL_3398542 | St Vincent's Pathology (SydPath) | NSW Health Pathology - Institute of Clinical Pathology and Medical Research; Westmead Hospital; University of Sydney | Arnott A., Draper J., Gall M., Martinez E., Rockett R., Sintchenko V., on behalf of ICPMR |
| EPI_ISL_3398543, EPI_ISL_3398544, EPI_ISL_3398545, EPI_ISL_3398546 | 4Cyte Pathology | NSW Health Pathology - Institute of Clinical Pathology and Medical Research; Westmead Hospital; University of Sydney | Arnott A., Draper J., Gall M., Martinez E., Rockett R., Sintchenko V., on behalf of ICPMR |
| EPI_ISL_3398547 | Laverty Pathology | NSW Health Pathology - Institute of Clinical Pathology and Medical Research; Westmead Hospital; University of Sydney | Arnott A., Draper J., Gall M., Martinez E., Rockett R., Sintchenko V., on behalf of ICPMR |
| EPI_ISL_3398548 | St Vincent's Pathology (SydPath) | NSW Health Pathology - Institute of Clinical Pathology and Medical Research; Westmead Hospital; University of Sydney | Arnott A., Draper J., Gall M., Martinez E., Rockett R., Sintchenko V., on behalf of ICPMR |
| EPI_ISL_3398549, EPI_ISL_3398550, EPI_ISL_3398551 | Histopath | NSW Health Pathology - Institute of Clinical Pathology and Medical Research; Westmead Hospital; University of Sydney | Arnott A., Draper J., Gall M., Martinez E., Rockett R., Sintchenko V., on behalf of ICPMR |
| EPI_ISL_3398552, EPI_ISL_3398553 | 4Cyte Pathology | NSW Health Pathology - Institute of Clinical Pathology and Medical Research; Westmead Hospital; University of Sydney | Arnott A., Draper J., Gall M., Martinez E., Rockett R., Sintchenko V., on behalf of ICPMR |
| EPI_ISL_3398554, EPI_ISL_3398555 | Medlab Pathology | NSW Health Pathology - Institute of Clinical Pathology and Medical Research; Westmead Hospital; University of Sydney | Arnott A., Draper J., Gall M., Martinez E., Rockett R., Sintchenko V., on behalf of ICPMR |
| EPI_ISL_3398556 | St Vincent's Pathology (SydPath) | NSW Health Pathology - Institute of Clinical Pathology and Medical Research; Westmead Hospital; University of Sydney | Arnott A., Draper J., Gall M., Martinez E., Rockett R., Sintchenko V., on behalf of ICPMR |
| EPI_ISL_3398557 | The Children's Hospital at Westmead | NSW Health Pathology - Institute of Clinical Pathology and Medical Research; Westmead Hospital; University of Sydney | Arnott A., Draper J., Gall M., Martinez E., Rockett R., Sintchenko V., on behalf of ICPMR |
| EPI_ISL_3398558 | Sydney South West Pathology Service (SSWPS) - Liverpool Hospital - NSW Health Pathology | NSW Health Pathology - Institute of Clinical Pathology and Medical Research; Westmead Hospital; University of Sydney | Arnott A., Draper J., Gall M., Martinez E., Rockett R., Sintchenko V., on behalf of ICPMR |
| EPI_ISL_3398559, EPI_ISL_3398560 | Histopath | NSW Health Pathology - Institute of Clinical Pathology and Medical Research; Westmead Hospital; University of Sydney | Arnott A., Draper J., Gall M., Martinez E., Rockett R., Sintchenko V., on behalf of ICPMR |
| EPI_ISL_3398561, EPI_ISL_3398562 | Laverty Pathology | NSW Health Pathology - Institute of Clinical Pathology and Medical Research; Westmead Hospital; University of Sydney | Arnott A., Draper J., Gall M., Martinez E., Rockett R., Sintchenko V., on behalf of ICPMR |
| EPI_ISL_3398563, EPI_ISL_3398564 | Sydney South West Pathology Service (SSWPS) - Liverpool Hospital - NSW Health Pathology | NSW Health Pathology - Institute of Clinical Pathology and Medical Research; Westmead Hospital; University of Sydney | Arnott A., Draper J., Gall M., Martinez E., Rockett R., Sintchenko V., on behalf of ICPMR |
| EPI_ISL_3398565 | St Vincent's Pathology (SydPath) | NSW Health Pathology - Institute of Clinical Pathology and Medical Research; Westmead Hospital; University of Sydney | Arnott A., Draper J., Gall M., Martinez E., Rockett R., Sintchenko V., on behalf of ICPMR |
| EPI_ISL_3398566, EPI_ISL_3398567, EPI_ISL_3398568 | 4Cyte Pathology | NSW Health Pathology - Institute of Clinical Pathology and Medical Research; Westmead Hospital; University of Sydney | Arnott A., Draper J., Gall M., Martinez E., Rockett R., Sintchenko V., on behalf of ICPMR |
| EPI_ISL_3398569, EPI_ISL_3398570 | Douglass Hanly Moir Pathology | NSW Health Pathology - Institute of Clinical Pathology and Medical Research; Westmead Hospital; University of Sydney | Arnott A., Draper J., Gall M., Martinez E., Rockett R., Sintchenko V., on behalf of ICPMR |
| EPI_ISL_3398571, EPI_ISL_3398572, EPI_ISL_3398573 | Laverty Pathology | NSW Health Pathology - Institute of Clinical Pathology and Medical Research; Westmead Hospital; University of Sydney | Arnott A., Draper J., Gall M., Martinez E., Rockett R., Sintchenko V., on behalf of ICPMR |
| EPI_ISL_3398574, EPI_ISL_3398575, EPI_ISL_3398576, EPI_ISL_3398577, EPI_ISL_3398578, EPI_ISL_3398579, EPI_ISL_3398580 | Histopath | NSW Health Pathology - Institute of Clinical Pathology and Medical Research; Westmead Hospital; University of Sydney | Arnott A., Draper J., Gall M., Martinez E., Rockett R., Sintchenko V., on behalf of ICPMR |
| EPI_ISL_3398581, EPI_ISL_3398582, EPI_ISL_3398583, EPI_ISL_3398584, EPI_ISL_3398585 | 4Cyte Pathology | NSW Health Pathology - Institute of Clinical Pathology and Medical Research; Westmead Hospital; University of Sydney | Arnott A., Draper J., Gall M., Martinez E., Rockett R., Sintchenko V., on behalf of ICPMR |
| EPI_ISL_3398586 | Australian Clinical Labs (formerly Healthscope Pathology) | NSW Health Pathology - Institute of Clinical Pathology and Medical Research; Westmead Hospital; University of Sydney | Arnott A., Draper J., Gall M., Martinez E., Rockett R., Sintchenko V., on behalf of ICPMR |
| EPI_ISL_3398587 | Austech Medical Laboratories | NSW Health Pathology - Institute of Clinical Pathology and Medical Research; Westmead Hospital; University of Sydney | Arnott A., Draper J., Gall M., Martinez E., Rockett R., Sintchenko V., on behalf of ICPMR |
| EPI_ISL_3398588 | Laverty Pathology | NSW Health Pathology - Institute of Clinical Pathology and Medical Research; Westmead Hospital; University of Sydney | Arnott A., Draper J., Gall M., Martinez E., Rockett R., Sintchenko V., on behalf of ICPMR |
| EPI_ISL_3398589 | Pathology North - Royal North Shore Hospital - NSW Health | NSW Health Pathology - Institute of Clinical Pathology and Medical | Arnott A., Draper J., Gall M., Martinez E., Rockett R., Sintchenko V., on behalf of ICPMR |

| | | | |
|---|---|---|---|
| | Pathology | Research; Westmead Hospital; University of Sydney | |
| EPI_ISL_3398590, EPI_ISL_3398591, EPI_ISL_3398592 | Histopath | NSW Health Pathology - Institute of Clinical Pathology and Medical Research; Westmead Hospital; University of Sydney | Arnott A., Draper J., Gall M., Martinez E., Rockett R., Sintchenko V., on behalf of ICPMR |
| EPI_ISL_3398593, EPI_ISL_3398594, EPI_ISL_3398595 | 4Cyte Pathology | NSW Health Pathology - Institute of Clinical Pathology and Medical Research; Westmead Hospital; University of Sydney | Arnott A., Draper J., Gall M., Martinez E., Rockett R., Sintchenko V., on behalf of ICPMR |
| EPI_ISL_3398596 | Australian Clinical Labs (formerly Healthscope Pathology) | NSW Health Pathology - Institute of Clinical Pathology and Medical Research; Westmead Hospital; University of Sydney | Arnott A., Draper J., Gall M., Martinez E., Rockett R., Sintchenko V., on behalf of ICPMR |
| EPI_ISL_3398597, EPI_ISL_3398598 | Sydney South West Pathology Service (SSWPS) - Liverpool Hospital - NSW Health Pathology | NSW Health Pathology - Institute of Clinical Pathology and Medical Research; Westmead Hospital; University of Sydney | Arnott A., Draper J., Gall M., Martinez E., Rockett R., Sintchenko V., on behalf of ICPMR |
| EPI_ISL_3398599, EPI_ISL_3398600, EPI_ISL_3398601 | St Vincent's Pathology (SydPath) | NSW Health Pathology - Institute of Clinical Pathology and Medical Research; Westmead Hospital; University of Sydney | Arnott A., Draper J., Gall M., Martinez E., Rockett R., Sintchenko V., on behalf of ICPMR |
| EPI_ISL_3398602, EPI_ISL_3398603, EPI_ISL_3398604, EPI_ISL_3398605, EPI_ISL_3398606, EPI_ISL_3398607, EPI_ISL_3398608 | Histopath | NSW Health Pathology - Institute of Clinical Pathology and Medical Research; Westmead Hospital; University of Sydney | Arnott A., Draper J., Gall M., Martinez E., Rockett R., Sintchenko V., on behalf of ICPMR |
| EPI_ISL_3398609, EPI_ISL_3398610, EPI_ISL_3398611 | 4Cyte Pathology | NSW Health Pathology - Institute of Clinical Pathology and Medical Research; Westmead Hospital; University of Sydney | Arnott A., Draper J., Gall M., Martinez E., Rockett R., Sintchenko V., on behalf of ICPMR |
| EPI_ISL_3398612 | Australian Clinical Labs (formerly Healthscope Pathology) | NSW Health Pathology - Institute of Clinical Pathology and Medical Research; Westmead Hospital; University of Sydney | Arnott A., Draper J., Gall M., Martinez E., Rockett R., Sintchenko V., on behalf of ICPMR |
| EPI_ISL_3398613, EPI_ISL_3398614, EPI_ISL_3398615, EPI_ISL_3398616 | Laverty Pathology | NSW Health Pathology - Institute of Clinical Pathology and Medical Research; Westmead Hospital; University of Sydney | Arnott A., Draper J., Gall M., Martinez E., Rockett R., Sintchenko V., on behalf of ICPMR |
| EPI_ISL_3398617 | Douglass Hanly Moir Pathology | NSW Health Pathology - Institute of Clinical Pathology and Medical Research; Westmead Hospital; University of Sydney | Arnott A., Draper J., Gall M., Martinez E., Rockett R., Sintchenko V., on behalf of ICPMR |
| EPI_ISL_3398618, EPI_ISL_3398619, EPI_ISL_3398620, EPI_ISL_3398621 | St Vincent's Pathology (SydPath) | NSW Health Pathology - Institute of Clinical Pathology and Medical Research; Westmead Hospital; University of Sydney | Arnott A., Draper J., Gall M., Martinez E., Rockett R., Sintchenko V., on behalf of ICPMR |
| EPI_ISL_3398622, EPI_ISL_3398623, EPI_ISL_3398624, EPI_ISL_3398625, EPI_ISL_3398626, EPI_ISL_3398627, EPI_ISL_3398628, EPI_ISL_3398629, EPI_ISL_3398630, EPI_ISL_3398631, EPI_ISL_3398632, EPI_ISL_3398633, EPI_ISL_3398634, EPI_ISL_3398635, EPI_ISL_3398636, EPI_ISL_3398637, EPI_ISL_3398638, EPI_ISL_3398639, EPI_ISL_3398640, EPI_ISL_3398641, EPI_ISL_3398642, EPI_ISL_3398643, EPI_ISL_3398644, EPI_ISL_3398645 | | | |
| see above | Histopath | NSW Health Pathology - Institute of Clinical Pathology and Medical Research; Westmead Hospital; University of Sydney | Arnott A., Draper J., Gall M., Martinez E., Rockett R., Sintchenko V., on behalf of ICPMR |
| EPI_ISL_3398646, EPI_ISL_3398647 | 4Cyte Pathology | NSW Health Pathology - Institute of Clinical Pathology and Medical Research; Westmead Hospital; University of Sydney | Arnott A., Draper J., Gall M., Martinez E., Rockett R., Sintchenko V., on behalf of ICPMR |
| EPI_ISL_3398648 | Medlab Pathology | NSW Health Pathology - Institute of Clinical Pathology and Medical Research; Westmead Hospital; University of Sydney | Arnott A., Draper J., Gall M., Martinez E., Rockett R., Sintchenko V., on behalf of ICPMR |
| EPI_ISL_3398649, EPI_ISL_3398650, EPI_ISL_3398651 | Sydney South West Pathology Service (SSWPS) - Liverpool Hospital - NSW Health Pathology | NSW Health Pathology - Institute of Clinical Pathology and Medical Research; Westmead Hospital; University of Sydney | Arnott A., Draper J., Gall M., Martinez E., Rockett R., Sintchenko V., on behalf of ICPMR |
| EPI_ISL_3398652, EPI_ISL_3398653 | Laverty Pathology | NSW Health Pathology - Institute of Clinical Pathology and Medical Research; Westmead Hospital; University of Sydney | Arnott A., Draper J., Gall M., Martinez E., Rockett R., Sintchenko V., on behalf of ICPMR |
| EPI_ISL_3398654, EPI_ISL_3398655 | Douglass Hanly Moir Pathology | NSW Health Pathology - Institute of Clinical Pathology and Medical Research; Westmead Hospital; University of Sydney | Arnott A., Draper J., Gall M., Martinez E., Rockett R., Sintchenko V., on behalf of ICPMR |
| EPI_ISL_3398656 | Australian Clinical Labs (formerly Healthscope Pathology) | NSW Health Pathology - Institute of Clinical Pathology and Medical Research; Westmead Hospital; University of Sydney | Arnott A., Draper J., Gall M., Martinez E., Rockett R., Sintchenko V., on behalf of ICPMR |
| EPI_ISL_3398657, EPI_ISL_3398658 | Laverty Pathology | NSW Health Pathology - Institute of Clinical Pathology and Medical Research; Westmead Hospital; University of Sydney | Arnott A., Draper J., Gall M., Martinez E., Rockett R., Sintchenko V., on behalf of ICPMR |
| EPI_ISL_3398659 | The Children's Hospital at Westmead | NSW Health Pathology - Institute of Clinical Pathology and Medical Research; Westmead Hospital; University of Sydney | Arnott A., Draper J., Gall M., Martinez E., Rockett R., Sintchenko V., on behalf of ICPMR |
| EPI_ISL_3398660, EPI_ISL_3398661, EPI_ISL_3398662, EPI_ISL_3398663, EPI_ISL_3398664, EPI_ISL_3398665, EPI_ISL_3398666, EPI_ISL_3398667, EPI_ISL_3398668, EPI_ISL_3398669, EPI_ISL_3398670, EPI_ISL_3398671, EPI_ISL_3398672, EPI_ISL_3398673, EPI_ISL_3398674, EPI_ISL_3398675, EPI_ISL_3398676, EPI_ISL_3398677 | | | |
| see above | 4Cyte Pathology | NSW Health Pathology - Institute of Clinical Pathology and Medical Research; Westmead Hospital; University of Sydney | Arnott A., Draper J., Gall M., Martinez E., Rockett R., Sintchenko V., on behalf of ICPMR |
| EPI_ISL_3398678, EPI_ISL_3398679, EPI_ISL_3398680, EPI_ISL_3398681, EPI_ISL_3398682, EPI_ISL_3398683, EPI_ISL_3398684 | Australian Clinical Labs (formerly Healthscope Pathology) | NSW Health Pathology - Institute of Clinical Pathology and Medical Research; Westmead Hospital; University of Sydney | Arnott A., Draper J., Gall M., Martinez E., Rockett R., Sintchenko V., on behalf of ICPMR |
| EPI_ISL_3398685, EPI_ISL_3398686, EPI_ISL_3398687, EPI_ISL_3398688, EPI_ISL_3398689, EPI_ISL_3398690 | Douglass Hanly Moir Pathology | NSW Health Pathology - Institute of Clinical Pathology and Medical Research; Westmead Hospital; University of Sydney | Arnott A., Draper J., Gall M., Martinez E., Rockett R., Sintchenko V., on behalf of ICPMR |
| EPI_ISL_3398691, EPI_ISL_3398692, EPI_ISL_3398693, EPI_ISL_3398694, EPI_ISL_3398695, EPI_ISL_3398696, EPI_ISL_3398697, EPI_ISL_3398698, EPI_ISL_3398699, EPI_ISL_3398700, EPI_ISL_3398701 | | | |
| see above | Laverty Pathology | NSW Health Pathology - Institute of Clinical Pathology and Medical Research; Westmead Hospital; University of Sydney | Arnott A., Draper J., Gall M., Martinez E., Rockett R., Sintchenko V., on behalf of ICPMR |
| EPI_ISL_3398702 | Australian Clinical Labs (formerly Healthscope Pathology) | NSW Health Pathology - Institute of Clinical Pathology and Medical Research; Westmead Hospital; University of Sydney | Arnott A., Draper J., Gall M., Martinez E., Rockett R., Sintchenko V., on behalf of ICPMR |
| EPI_ISL_3398703 | Medlab Pathology | NSW Health Pathology - Institute of Clinical Pathology and Medical Research; Westmead Hospital; University of Sydney | Arnott A., Draper J., Gall M., Martinez E., Rockett R., Sintchenko V., on behalf of ICPMR |
| EPI_ISL_3398704, EPI_ISL_3398705, EPI_ISL_3398706 | Australian Clinical Labs (formerly Healthscope Pathology) | NSW Health Pathology - Institute of Clinical Pathology and Medical Research; Westmead Hospital; University of Sydney | Arnott A., Draper J., Gall M., Martinez E., Rockett R., Sintchenko V., on behalf of ICPMR |
| EPI_ISL_3398707 | Medlab Pathology | NSW Health Pathology - Institute of Clinical Pathology and Medical Research; Westmead Hospital; University of Sydney | Arnott A., Draper J., Gall M., Martinez E., Rockett R., Sintchenko V., on behalf of ICPMR |
| EPI_ISL_3398708 | Australian Clinical Labs (formerly Healthscope Pathology) | NSW Health Pathology - Institute of Clinical Pathology and Medical Research; Westmead Hospital; University of Sydney | Arnott A., Draper J., Gall M., Martinez E., Rockett R., Sintchenko V., on behalf of ICPMR |
| EPI_ISL_3398709, EPI_ISL_3398710, EPI_ISL_3398711, EPI_ISL_3398712, EPI_ISL_3398713 | Medlab Pathology | NSW Health Pathology - Institute of Clinical Pathology and Medical Research; Westmead Hospital; University of Sydney | Arnott A., Draper J., Gall M., Martinez E., Rockett R., Sintchenko V., on behalf of ICPMR |
| EPI_ISL_3398714 | Pathology North - Royal North Shore Hospital - NSW Health Pathology | NSW Health Pathology - Institute of Clinical Pathology and Medical Research; Westmead Hospital; University of Sydney | Arnott A., Draper J., Gall M., Martinez E., Rockett R., Sintchenko V., on behalf of ICPMR |
| EPI_ISL_3398715, | Southern. IML Pathology | NSW Health Pathology - Institute of Clinical Pathology and Medical | Arnott A., Draper J., Gall M., Martinez E., Rockett R., Sintchenko V., on behalf of ICPMR |

| | | | |
|---|---|---|---|
| EPI_ISL_3398716 | | Research; Westmead Hospital; University of Sydney | |
| EPI_ISL_3398717,<br>EPI_ISL_3398718,<br>EPI_ISL_3398719,<br>EPI_ISL_3398720,<br>EPI_ISL_3398721,<br>EPI_ISL_3398722,<br>EPI_ISL_3398723,<br>EPI_ISL_3398724,<br>EPI_ISL_3398725,<br>EPI_ISL_3398726 | St Vincent's Pathology (SydPath) | NSW Health Pathology - Institute of Clinical Pathology and Medical Research; Westmead Hospital; University of Sydney | Arnott A., Draper J., Gall M., Martinez E., Rockett R., Sintchenko V., on behalf of ICPMR |
| EPI_ISL_3398727, EPI_ISL_3398728, EPI_ISL_3398729, EPI_ISL_3398730, EPI_ISL_3398731, EPI_ISL_3398732, EPI_ISL_3398733, EPI_ISL_3398734, EPI_ISL_3398735, EPI_ISL_3398736, EPI_ISL_3398737, EPI_ISL_3398738, EPI_ISL_3398739, EPI_ISL_3398740, EPI_ISL_3398741, EPI_ISL_3398742, EPI_ISL_3398743, EPI_ISL_3398744, EPI_ISL_3398745, EPI_ISL_3398746, EPI_ISL_3398747, EPI_ISL_3398748, EPI_ISL_3398749, EPI_ISL_3398750, EPI_ISL_3398751, EPI_ISL_3398752, EPI_ISL_3398753, EPI_ISL_3398754, EPI_ISL_3398755, EPI_ISL_3398756 | | | |
| see above | Histopath | NSW Health Pathology - Institute of Clinical Pathology and Medical Research; Westmead Hospital; University of Sydney | Arnott A., Draper J., Gall M., Martinez E., Rockett R., Sintchenko V., on behalf of ICPMR |
| EPI_ISL_3398757 | Arizona State University | Arizona State University | Ajeet Bains, LaRinda A. Holland, Matthew F. Smith, Peter T. Skidmore, Rabia Maqsood, Nicholas J. Mellor, Nathaniel Johnson, Joshua LaBaer, Vel Murugan, Efrem S. Lim |
| EPI_ISL_3398759, EPI_ISL_3398760, EPI_ISL_3398761, EPI_ISL_3398762, EPI_ISL_3398763, EPI_ISL_3398764, EPI_ISL_3398766, EPI_ISL_3398767, EPI_ISL_3398770, EPI_ISL_3398771, EPI_ISL_3398773, EPI_ISL_3398777, EPI_ISL_3398778, EPI_ISL_3398779, EPI_ISL_3398781, EPI_ISL_3398782, EPI_ISL_3398783, EPI_ISL_3398784, EPI_ISL_3398785, EPI_ISL_3398786, EPI_ISL_3398787, EPI_ISL_3398789, EPI_ISL_3398790 | | | |
| see above | Pathology, UAB | Pathology, UAB | Moates,D., Hendrickson,R.C., Lefkowitz,E.J., Leal,S.M. |
| EPI_ISL_3398791,<br>EPI_ISL_3398792,<br>EPI_ISL_3398793,<br>EPI_ISL_3398795,<br>EPI_ISL_3398798,<br>EPI_ISL_3398799,<br>EPI_ISL_3398800,<br>EPI_ISL_3398801,<br>EPI_ISL_3398802 | Communicable Disease Laboratory, Public Health Directorate | Communicable Disease Laboratory, Public Health Directorate | Alwasti,H., AlHujairi,Z., AlAbbas,Z., Marhoon,A., Almoamen,G. |
| EPI_ISL_3398824, EPI_ISL_3398825, EPI_ISL_3398826, EPI_ISL_3398827, EPI_ISL_3398828, EPI_ISL_3398829, EPI_ISL_3398830, EPI_ISL_3398831, EPI_ISL_3398832, EPI_ISL_3398833, EPI_ISL_3398834, EPI_ISL_3398835, EPI_ISL_3398836, EPI_ISL_3398837, EPI_ISL_3398838, EPI_ISL_3398839, EPI_ISL_3398840 | | | |
| see above | Laboratoire National de Biologie Clinique et de Santé Publique, Bangui, République Centrafricaine | Pathogen Genomics Lab, National Institute for Biomedical Research (INRB) | Placide Mbala-Kingebeni, Clotaire Donatien RAFAÏ , Edith Nkwembe, Eddy Kinganda-Lusamaki, Amuri Aziza, Francisca Muyembe Mawete, Emmanuel Lokilo Lofiko, Jean Claude Makangara, Raphael Lumembe, Gabriel Kabamba, Catherine Pratt, Matthias Pauthner, Josh Quick, Trevor Bedford, Ian Goodfellow, Andrew Rambaut, Nick Loman, Michael Wiley, Steve Ahuka-Mundeke, Jean-Jacques Muyembe Tamfum |
| EPI_ISL_3398842,<br>EPI_ISL_3398843,<br>EPI_ISL_3398844 | S.S.D. Microbiologia, Stabilimento ospedaliero di Sanremo, ASL 1 Liguria | U.O. Igiene, Ospedale Policlinico San Martino | Bruzzone Bianca, Caligiuri Patrizia, De Pace Vanessa, Domnich Alexander, Dusi Pier Andrea, Orsi Andrea, Ricucci Valentina, Icardi Giancarlo on behalf of SARS-CoV-2 ITALIAN RESEARCH ENTERPRISE-(SCIRE) Collaborative Group |
| EPI_ISL_3398845 | Laboratorio di Patologia Clinica, Ospedale San Paolo in Valloria, ASL 2 Liguria | U.O. Igiene, Ospedale Policlinico San Martino | Bruzzone Bianca, Caligiuri Patrizia, De Pace Vanessa, Domnich Alexander, Lillo Flavia, Orsi Andrea, Ricucci Valentina, Icardi Giancarlo on behalf of SARS-CoV-2 ITALIAN RESEARCH ENTERPRISE-(SCIRE) Collaborative Group |
| EPI_ISL_3398846,<br>EPI_ISL_3398847,<br>EPI_ISL_3398848,<br>EPI_ISL_3398849,<br>EPI_ISL_3398850 | S.C. Laboratorio Analisi, ASL 3 Liguria | U.O. Igiene, Ospedale Policlinico San Martino | Bruzzone Bianca, Caligiuri Patrizia, De Pace Vanessa, Domnich Alexander, Orsi Andrea, Ricucci Valentina, Spitaleri Antonino, Icardi Giancarlo on behalf of SARS-CoV-2 ITALIAN RESEARCH ENTERPRISE-(SCIRE) Collaborative Group |
| EPI_ISL_3398851,<br>EPI_ISL_3398852 | S.C. Laboratorio Analisi, Ospedale di Lavagna, ASL 4 Liguria | U.O. Igiene, Ospedale Policlinico San Martino | Bruzzone Bianca, Bandettini Roberto, Caligiuri Patrizia, De Pace Vanessa, Domnich Alexander, Orsi Andrea, Ricucci Valentina, Icardi Giancarlo on behalf of SARS-CoV-2 ITALIAN RESEARCH ENTERPRISE-(SCIRE) Collaborative Group |
| EPI_ISL_3398853,<br>EPI_ISL_3398854,<br>EPI_ISL_3398855 | S.C. Patologia Clinica, Ospedale Sant'Andrea, ASL 5 | U.O. Igiene, Ospedale Policlinico San Martino | Bruzzone Bianca, Battolla Enrico, Caligiuri Patrizia, De Pace Vanessa, Domnich Alexander, Orsi Andrea, Ricucci Valentina, Icardi Giancarlo on behalf of SARS-CoV-2 ITALIAN RESEARCH ENTERPRISE-(SCIRE) Collaborative Group |
| EPI_ISL_3398856 | S.C. Laboratorio Analisi, ASL 3 Liguria | U.O. Igiene, Ospedale Policlinico San Martino | Bruzzone Bianca, Caligiuri Patrizia, De Pace Vanessa, Domnich Alexander, Orsi Andrea, Ricucci Valentina, Spitaleri Antonino, Icardi Giancarlo on behalf of SARS-CoV-2 ITALIAN RESEARCH ENTERPRISE-(SCIRE) Collaborative Group |
| EPI_ISL_3398857 | S.C. Patologia Clinica, Ospedale Sant'Andrea, ASL 5 | U.O. Igiene, Ospedale Policlinico San Martino | Bruzzone Bianca, Battolla Enrico, Caligiuri Patrizia, De Pace Vanessa, Domnich Alexander, Orsi Andrea, Ricucci Valentina, Icardi Giancarlo on behalf of SARS-CoV-2 ITALIAN RESEARCH ENTERPRISE-(SCIRE) Collaborative Group |
| EPI_ISL_3398858,<br>EPI_ISL_3398860,<br>EPI_ISL_3398861,<br>EPI_ISL_3398862,<br>EPI_ISL_3398863 | S.C. Laboratorio Analisi, ASL 3 Liguria | U.O. Igiene, Ospedale Policlinico San Martino | Bruzzone Bianca, Caligiuri Patrizia, De Pace Vanessa, Domnich Alexander, Orsi Andrea, Ricucci Valentina, Spitaleri Antonino, Icardi Giancarlo on behalf of SARS-CoV-2 ITALIAN RESEARCH ENTERPRISE-(SCIRE) Collaborative Group |
| EPI_ISL_3398864 | S.C. Laboratorio di analisi, Ente ospedaliero "Ospedali Galliera" Genova | U.O. Igiene, Ospedale Policlinico San Martino | Bruzzone Bianca, Caligiuri Patrizia, De Pace Vanessa, Domnich Alexander, Innocenti Lionello, Orsi Andrea, Ricucci Valentina, Icardi Giancarlo on behalf of SARS-CoV-2 ITALIAN RESEARCH ENTERPRISE-(SCIRE) Collaborative Group |
| EPI_ISL_3398865,<br>EPI_ISL_3398866,<br>EPI_ISL_3398867 | S.C. Laboratorio Analisi, ASL 3 Liguria | U.O. Igiene, Ospedale Policlinico San Martino | Bruzzone Bianca, Caligiuri Patrizia, De Pace Vanessa, Domnich Alexander, Orsi Andrea, Ricucci Valentina, Spitaleri Antonino, Icardi Giancarlo on behalf of SARS-CoV-2 ITALIAN RESEARCH ENTERPRISE-(SCIRE) Collaborative Group |
| EPI_ISL_3398868, EPI_ISL_3398869, EPI_ISL_3398870, EPI_ISL_3398871, EPI_ISL_3398874, EPI_ISL_3398875, EPI_ISL_3398876, EPI_ISL_3398877, EPI_ISL_3398878, EPI_ISL_3398879, EPI_ISL_3398880, EPI_ISL_3398881, EPI_ISL_3398882, EPI_ISL_3398883, EPI_ISL_3398884, EPI_ISL_3398885, EPI_ISL_3398886, EPI_ISL_3398887, EPI_ISL_3398888, EPI_ISL_3398889, EPI_ISL_3398890, EPI_ISL_3398891, EPI_ISL_3398892, EPI_ISL_3398893, EPI_ISL_3398894, EPI_ISL_3398895, EPI_ISL_3398896, EPI_ISL_3398897, EPI_ISL_3398898, EPI_ISL_3398899, EPI_ISL_3398900, EPI_ISL_3398901, EPI_ISL_3398902, EPI_ISL_3398903, EPI_ISL_3398904, EPI_ISL_3398906, EPI_ISL_3398907, EPI_ISL_3398908, EPI_ISL_3398909, EPI_ISL_3398910, EPI_ISL_3398911, EPI_ISL_3398912, EPI_ISL_3398913, EPI_ISL_3398914 | | | |
| see above | Rhode Island State Health Laboratory | Rhode Island State Health Laboratory | Ewa King, Richard C. Huard, Kristin Carpenter-Azevedo |
| EPI_ISL_3398915, EPI_ISL_3398916, EPI_ISL_3398917, EPI_ISL_3398918, EPI_ISL_3398919, EPI_ISL_3398920, EPI_ISL_3398921, EPI_ISL_3398923, EPI_ISL_3398924, EPI_ISL_3398925, EPI_ISL_3398926, EPI_ISL_3398927, EPI_ISL_3398928, EPI_ISL_3398929, EPI_ISL_3398930, EPI_ISL_3398931, EPI_ISL_3398932, EPI_ISL_3398933, EPI_ISL_3398934, EPI_ISL_3398935, EPI_ISL_3398936, EPI_ISL_3398937, EPI_ISL_3398938, EPI_ISL_3398939, EPI_ISL_3398940, EPI_ISL_3398941, EPI_ISL_3398942, EPI_ISL_3398943, EPI_ISL_3398944, EPI_ISL_3398945, EPI_ISL_3398947, EPI_ISL_3398948, EPI_ISL_3398949, EPI_ISL_3398950, EPI_ISL_3398951, EPI_ISL_3398952, EPI_ISL_3398953, EPI_ISL_3398954, EPI_ISL_3398955, EPI_ISL_3398957, EPI_ISL_3398958, EPI_ISL_3398959, EPI_ISL_3398960, EPI_ISL_3398961, EPI_ISL_3398962, EPI_ISL_3398963, EPI_ISL_3398964, EPI_ISL_3398965, EPI_ISL_3398966, EPI_ISL_3398967 | | | |
| see above | Houston Health Dept. | Houston Health Dept. | Ryker Penn, Pamela Brown, Adolpho Lara, Yanlai Lai |
| EPI_ISL_3398968, EPI_ISL_3398969, EPI_ISL_3398970, EPI_ISL_3398971, EPI_ISL_3398972, EPI_ISL_3398973, EPI_ISL_3398974, EPI_ISL_3398975, EPI_ISL_3398976, EPI_ISL_3398977, EPI_ISL_3398979, EPI_ISL_3398980, EPI_ISL_3398981, EPI_ISL_3398982, EPI_ISL_3398984, EPI_ISL_3398985, EPI_ISL_3398987, EPI_ISL_3398988, EPI_ISL_3398989, EPI_ISL_3398990, EPI_ISL_3398991, EPI_ISL_3398992, EPI_ISL_3398993, EPI_ISL_3398994, EPI_ISL_3398995, EPI_ISL_3398996, EPI_ISL_3398997, EPI_ISL_3398998, EPI_ISL_3398999, EPI_ISL_3399000, EPI_ISL_3399001, EPI_ISL_3399002, EPI_ISL_3399003, EPI_ISL_3399004, EPI_ISL_3399005, EPI_ISL_3399006, EPI_ISL_3399007, EPI_ISL_3399008, EPI_ISL_3399009, EPI_ISL_3399010, EPI_ISL_3399011, EPI_ISL_3399012 | | | |
| see above | Rhode Island State Health Laboratory | Rhode Island State Health Laboratory | Ewa King, Richard C. Huard, Kristin Carpenter-Azevedo |
| EPI_ISL_3399013, EPI_ISL_3399015, EPI_ISL_3399016, EPI_ISL_3399017, EPI_ISL_3399018, EPI_ISL_3399019, EPI_ISL_3399020, EPI_ISL_3399021, EPI_ISL_3399022, EPI_ISL_3399023, EPI_ISL_3399024, EPI_ISL_3399025, EPI_ISL_3399026, EPI_ISL_3399028, EPI_ISL_3399029, EPI_ISL_3399030, EPI_ISL_3399031, EPI_ISL_3399032, EPI_ISL_3399033, EPI_ISL_3399034, EPI_ISL_3399035, EPI_ISL_3399038, EPI_ISL_3399039, EPI_ISL_3399040, EPI_ISL_3399041, EPI_ISL_3399042, EPI_ISL_3399043, EPI_ISL_3399044, EPI_ISL_3399045, EPI_ISL_3399046, EPI_ISL_3399047, EPI_ISL_3399048, EPI_ISL_3399049, EPI_ISL_3399050, EPI_ISL_3399051, EPI_ISL_3399052, EPI_ISL_3399053, EPI_ISL_3399054, EPI_ISL_3399055, EPI_ISL_3399056, EPI_ISL_3399058, EPI_ISL_3399059, EPI_ISL_3399060, EPI_ISL_3399061, EPI_ISL_3399062, EPI_ISL_3399063, EPI_ISL_3399064, EPI_ISL_3399065, EPI_ISL_3399066, EPI_ISL_3399067, EPI_ISL_3399068, EPI_ISL_3399069, EPI_ISL_3399070, EPI_ISL_3399071, EPI_ISL_3399072, EPI_ISL_3399073, EPI_ISL_3399074, EPI_ISL_3399075, EPI_ISL_3399076, EPI_ISL_3399077, EPI_ISL_3399078 | | | |
| see above | Texas Department of State Health Services (TXDSHS) | Texas Department of State Health Services (TXDSHS) | Rashmi Tuladhar, Bonnie Oh, Jenny Zhang, Maliha Rahman, Mayela Pedrueza, Anita Pokharel, Karen Bobier, Lorraine Rodriguez, Myong Koag, Chun Wang, Rachel Lee, Grace Kubin |
| EPI_ISL_3399079, EPI_ISL_3399080, EPI_ISL_3399081, EPI_ISL_3399082, EPI_ISL_3399083, EPI_ISL_3399084, EPI_ISL_3399085, EPI_ISL_3399086, EPI_ISL_3399087, EPI_ISL_3399088, EPI_ISL_3399089, EPI_ISL_3399090, EPI_ISL_3399091, EPI_ISL_3399092, EPI_ISL_3399093 | | | |
| see above | Presidio Ospedaliero "Madonna delle Grazie" di Matera | ASM - P.O. Madonna delle Grazie - Matera | Massimo Dell'Edera |
| EPI_ISL_3399094, EPI_ISL_3399095, EPI_ISL_3399096, EPI_ISL_3399097, EPI_ISL_3399098, EPI_ISL_3399099, EPI_ISL_3399100, EPI_ISL_3399101, EPI_ISL_3399102, EPI_ISL_3399103, EPI_ISL_3399104, EPI_ISL_3399105, EPI_ISL_3399106, EPI_ISL_3399107, EPI_ISL_3399108, EPI_ISL_3399109, EPI_ISL_3399110, EPI_ISL_3399111, EPI_ISL_3399112, EPI_ISL_3399113, EPI_ISL_3399114, EPI_ISL_3399115, EPI_ISL_3399116, EPI_ISL_3399117, EPI_ISL_3399118, EPI_ISL_3399119, EPI_ISL_3399120, EPI_ISL_3399121, EPI_ISL_3399122, EPI_ISL_3399123, EPI_ISL_3399124 | | | |
| see above | Nebraska Public Health Laboratory | NPHL COVID-19 Response Team | NPHL COVID-19 Response Team |
| EPI_ISL_3399125, EPI_ISL_3399126, EPI_ISL_3399127, EPI_ISL_3399128, EPI_ISL_3399129, EPI_ISL_3399130, EPI_ISL_3399131, EPI_ISL_3399132, EPI_ISL_3399133, EPI_ISL_3399134, EPI_ISL_3399135, EPI_ISL_3399136, EPI_ISL_3399137, EPI_ISL_3399138, EPI_ISL_3399139, EPI_ISL_3399140, EPI_ISL_3399141, EPI_ISL_3399142, EPI_ISL_3399143, EPI_ISL_3399144, EPI_ISL_3399145, EPI_ISL_3399146, EPI_ISL_3399147, EPI_ISL_3399148, EPI_ISL_3399149, EPI_ISL_3399150, EPI_ISL_3399151, EPI_ISL_3399152, EPI_ISL_3399153, EPI_ISL_3399154, EPI_ISL_3399155, EPI_ISL_3399156, EPI_ISL_3399157, EPI_ISL_3399158, EPI_ISL_3399159, EPI_ISL_3399160, EPI_ISL_3399161, EPI_ISL_3399162, EPI_ISL_3399163, EPI_ISL_3399164, EPI_ISL_3399165, EPI_ISL_3399166, EPI_ISL_3399167, EPI_ISL_3399168, EPI_ISL_3399169, EPI_ISL_3399170, EPI_ISL_3399171, EPI_ISL_3399172, EPI_ISL_3399173, EPI_ISL_3399174, EPI_ISL_3399175, EPI_ISL_3399176, EPI_ISL_3399177, EPI_ISL_3399178, EPI_ISL_3399179, EPI_ISL_3399180, EPI_ISL_3399181, EPI_ISL_3399182, EPI_ISL_3399183, EPI_ISL_3399184, EPI_ISL_3399185, EPI_ISL_3399186, EPI_ISL_3399187, EPI_ISL_3399188, EPI_ISL_3399189, EPI_ISL_3399190, EPI_ISL_3399191, EPI_ISL_3399192, EPI_ISL_3399193, EPI_ISL_3399194, EPI_ISL_3399195, EPI_ISL_3399196, EPI_ISL_3399197, EPI_ISL_3399198, EPI_ISL_3399199, EPI_ISL_3399200, EPI_ISL_3399201, EPI_ISL_3399202, EPI_ISL_3399203, EPI_ISL_3399204, EPI_ISL_3399205, EPI_ISL_3399206, EPI_ISL_3399207, EPI_ISL_3399208, EPI_ISL_3399209, EPI_ISL_3399210, EPI_ISL_3399211, EPI_ISL_3399212, EPI_ISL_3399213, EPI_ISL_3399214, EPI_ISL_3399215, EPI_ISL_3399216, EPI_ISL_3399217, EPI_ISL_3399218, EPI_ISL_3399219, EPI_ISL_3399220, EPI_ISL_3399221, EPI_ISL_3399222, EPI_ISL_3399223, EPI_ISL_3399224, EPI_ISL_3399225, EPI_ISL_3399226, EPI_ISL_3399227, EPI_ISL_3399228, EPI_ISL_3399229, EPI_ISL_3399230, EPI_ISL_3399231, EPI_ISL_3399232, EPI_ISL_3399233, EPI_ISL_3399234, EPI_ISL_3399235, EPI_ISL_3399236, EPI_ISL_3399237, EPI_ISL_3399238, EPI_ISL_3399239, EPI_ISL_3399240, EPI_ISL_3399241, EPI_ISL_3399242, EPI_ISL_3399243, EPI_ISL_3399244, EPI_ISL_3399245, EPI_ISL_3399246, EPI_ISL_3399247, EPI_ISL_3399248, EPI_ISL_3399249, EPI_ISL_3399250, EPI_ISL_3399251, EPI_ISL_3399252, EPI_ISL_3399253, EPI_ISL_3399254, EPI_ISL_3399255, EPI_ISL_3399256, EPI_ISL_3399257, EPI_ISL_3399258, EPI_ISL_3399259, EPI_ISL_3399260, EPI_ISL_3399261, EPI_ISL_3399262, EPI_ISL_3399263, EPI_ISL_3399264, EPI_ISL_3399265, EPI_ISL_3399266, EPI_ISL_3399267, EPI_ISL_3399268, EPI_ISL_3399269, EPI_ISL_3399270, EPI_ISL_3399271, EPI_ISL_3399272, EPI_ISL_3399273, EPI_ISL_3399274, EPI_ISL_3399275, EPI_ISL_3399276, EPI_ISL_3399277, EPI_ISL_3399278, EPI_ISL_3399279, EPI_ISL_3399280, EPI_ISL_3399281, EPI_ISL_3399282, EPI_ISL_3399283, EPI_ISL_3399284, EPI_ISL_3399285, EPI_ISL_3399286, EPI_ISL_3399287, EPI_ISL_3399288, EPI_ISL_3399289, EPI_ISL_3399290, EPI_ISL_3399291, EPI_ISL_3399292, EPI_ISL_3399293, EPI_ISL_3399294, EPI_ISL_3399295, EPI_ISL_3399296, EPI_ISL_3399297, EPI_ISL_3399298, EPI_ISL_3399299, EPI_ISL_3399300, EPI_ISL_3399301, EPI_ISL_3399302, EPI_ISL_3399303, EPI_ISL_3399304, EPI_ISL_3399305, EPI_ISL_3399306, EPI_ISL_3399307, EPI_ISL_3399308, EPI_ISL_3399309, EPI_ISL_3399310, EPI_ISL_3399311, EPI_ISL_3399312, EPI_ISL_3399313, EPI_ISL_3399314, EPI_ISL_3399315, EPI_ISL_3399316, EPI_ISL_3399317, EPI_ISL_3399318, EPI_ISL_3399319, EPI_ISL_3399320, EPI_ISL_3399321, EPI_ISL_3399322, EPI_ISL_3399323, EPI_ISL_3399324, EPI_ISL_3399325, EPI_ISL_3399326, EPI_ISL_3399327, EPI_ISL_3399328, EPI_ISL_3399329, EPI_ISL_3399330, EPI_ISL_3399331, EPI_ISL_3399332, EPI_ISL_3399333, EPI_ISL_3399334, EPI_ISL_3399335, EPI_ISL_3399336, EPI_ISL_3399337, EPI_ISL_3399338, EPI_ISL_3399339, EPI_ISL_3399340, EPI_ISL_3399341, EPI_ISL_3399342, EPI_ISL_3399343, EPI_ISL_3399344, EPI_ISL_3399345, EPI_ISL_3399346, EPI_ISL_3399347, EPI_ISL_3399348, EPI_ISL_3399349, EPI_ISL_3399350, EPI_ISL_3399351, EPI_ISL_3399352, EPI_ISL_3399353, EPI_ISL_3399354, EPI_ISL_3399355, EPI_ISL_3399356, EPI_ISL_3399357, EPI_ISL_3399358, EPI_ISL_3399359, EPI_ISL_3399360, EPI_ISL_3399361, EPI_ISL_3399362, EPI_ISL_3399363, EPI_ISL_3399364, EPI_ISL_3399365, EPI_ISL_3399366, EPI_ISL_3399367, EPI_ISL_3399368, EPI_ISL_3399369, EPI_ISL_3399370, EPI_ISL_3399371, EPI_ISL_3399372, EPI_ISL_3399373, EPI_ISL_3399374, EPI_ISL_3399375, EPI_ISL_3399376, EPI_ISL_3399377, EPI_ISL_3399378, EPI_ISL_3399379, EPI_ISL_3399380, EPI_ISL_3399381, EPI_ISL_3399382, EPI_ISL_3399383, EPI_ISL_3399384, EPI_ISL_3399385, EPI_ISL_3399386, EPI_ISL_3399387, EPI_ISL_3399388, EPI_ISL_3399389, EPI_ISL_3399390, EPI_ISL_3399391, EPI_ISL_3399392, EPI_ISL_3399393, EPI_ISL_3399394, EPI_ISL_3399395, EPI_ISL_3399396, EPI_ISL_3399397, EPI_ISL_3399398, EPI_ISL_3399399, EPI_ISL_3399400, EPI_ISL_3399401, EPI_ISL_3399402, EPI_ISL_3399403, EPI_ISL_3399404, EPI_ISL_3399405, EPI_ISL_3399406, EPI_ISL_3399407, EPI_ISL_3399408, EPI_ISL_3399409, EPI_ISL_3399410, EPI_ISL_3399411, EPI_ISL_3399412, EPI_ISL_3399413, EPI_ISL_3399414, EPI_ISL_3399415, EPI_ISL_3399416, EPI_ISL_3399417, EPI_ISL_3399418, EPI_ISL_3399419, EPI_ISL_3399420, EPI_ISL_3399421, EPI_ISL_3399422, EPI_ISL_3399423, EPI_ISL_3399424, EPI_ISL_3399425, EPI_ISL_3399426, EPI_ISL_3399427, EPI_ISL_3399428, EPI_ISL_3399429, EPI_ISL_3399430, EPI_ISL_3399431, EPI_ISL_3399432, EPI_ISL_3399433, EPI_ISL_3399434, EPI_ISL_3399435, EPI_ISL_3399436, EPI_ISL_3399437, EPI_ISL_3399438, EPI_ISL_3399439, EPI_ISL_3399440, EPI_ISL_3399441, EPI_ISL_3399442, EPI_ISL_3399443, EPI_ISL_3399444, EPI_ISL_3399445, EPI_ISL_3399446, EPI_ISL_3399447, EPI_ISL_3399448, EPI_ISL_3399449, EPI_ISL_3399450, EPI_ISL_3399451, EPI_ISL_3399452, EPI_ISL_3399453, EPI_ISL_3399454, EPI_ISL_3399455, EPI_ISL_3399456, EPI_ISL_3399457, EPI_ISL_3399458, EPI_ISL_3399459, EPI_ISL_3399460, EPI_ISL_3399461, EPI_ISL_3399462, EPI_ISL_3399463, EPI_ISL_3399464, EPI_ISL_3399465, EPI_ISL_3399466, EPI_ISL_3399467, EPI_ISL_3399468, EPI_ISL_3399469, EPI_ISL_3399470, EPI_ISL_3399471, EPI_ISL_3399472, EPI_ISL_3399473, EPI_ISL_3399474, EPI_ISL_3399475, EPI_ISL_3399476, EPI_ISL_3399477, EPI_ISL_3399478, EPI_ISL_3399479, EPI_ISL_3399480, EPI_ISL_3399481, EPI_ISL_3399482, EPI_ISL_3399483, EPI_ISL_3399484, EPI_ISL_3399485, EPI_ISL_3399486, EPI_ISL_3399487, EPI_ISL_3399488, EPI_ISL_3399489, EPI_ISL_3399492, EPI_ISL_3399493, EPI_ISL_3399494, EPI_ISL_3399495, EPI_ISL_3399496, EPI_ISL_3399497, EPI_ISL_3399498, EPI_ISL_3399499, EPI_ISL_3399500, EPI_ISL_3399501, EPI_ISL_3399502, EPI_ISL_3399503, EPI_ISL_3399504, EPI_ISL_3399505, EPI_ISL_3399506, EPI_ISL_3399507, EPI_ISL_3399508, EPI_ISL_3399509, EPI_ISL_3399510, EPI_ISL_3399511, EPI_ISL_3399512, EPI_ISL_3399513, EPI_ISL_3399514, EPI_ISL_3399515, EPI_ISL_3399516, EPI_ISL_3399517, EPI_ISL_3399518, EPI_ISL_3399519, EPI_ISL_3399520, EPI_ISL_3399521, EPI_ISL_3399522, EPI_ISL_3399523, EPI_ISL_3399524, EPI_ISL_3399525, EPI_ISL_3399526, EPI_ISL_3399527, EPI_ISL_3399528, EPI_ISL_3399529, EPI_ISL_3399530, EPI_ISL_3399531, EPI_ISL_3399532, EPI_ISL_3399533, EPI_ISL_3399535, EPI_ISL_3399536, EPI_ISL_3399537, EPI_ISL_3399538, EPI_ISL_3399539, EPI_ISL_3399540, EPI_ISL_3399541, EPI_ISL_3399542, EPI_ISL_3399543, EPI_ISL_3399544, EPI_ISL_3399545, | | | |

| | | | |
|---|---|---|---|
| EPI_ISL_3399546, EPI_ISL_3399547, EPI_ISL_3399548, EPI_ISL_3399549, EPI_ISL_3399550, EPI_ISL_3399551, EPI_ISL_3399552, EPI_ISL_3399553, EPI_ISL_3399554, EPI_ISL_3399556, EPI_ISL_3399557, EPI_ISL_3399558, EPI_ISL_3399559, EPI_ISL_3399560, EPI_ISL_3399561, EPI_ISL_3399562, EPI_ISL_3399563, EPI_ISL_3399564, EPI_ISL_3399565, EPI_ISL_3399566, EPI_ISL_3399567, EPI_ISL_3399568, EPI_ISL_3399569, EPI_ISL_3399570, EPI_ISL_3399571, EPI_ISL_3399572, EPI_ISL_3399573, EPI_ISL_3399574, EPI_ISL_3399575, EPI_ISL_3399576, EPI_ISL_3399577, EPI_ISL_3399578, EPI_ISL_3399579, EPI_ISL_3399580, EPI_ISL_3399581, EPI_ISL_3399582, EPI_ISL_3399583, EPI_ISL_3399584, EPI_ISL_3399585, EPI_ISL_3399586, EPI_ISL_3399587, EPI_ISL_3399588, EPI_ISL_3399589, EPI_ISL_3399590, EPI_ISL_3399591, EPI_ISL_3399592, EPI_ISL_3399593, EPI_ISL_3399594, EPI_ISL_3399595, EPI_ISL_3399596, EPI_ISL_3399597, EPI_ISL_3399598, EPI_ISL_3399599, EPI_ISL_3399600, EPI_ISL_3399601, EPI_ISL_3399602, EPI_ISL_3399603, EPI_ISL_3399604, EPI_ISL_3399605, EPI_ISL_3399606, EPI_ISL_3399607, EPI_ISL_3399608, EPI_ISL_3399609, EPI_ISL_3399610, EPI_ISL_3399611, EPI_ISL_3399612, EPI_ISL_3399613, EPI_ISL_3399614, EPI_ISL_3399615, EPI_ISL_3399616, EPI_ISL_3399617, EPI_ISL_3399618, EPI_ISL_3399619, EPI_ISL_3399620, EPI_ISL_3399621, EPI_ISL_3399622, EPI_ISL_3399623, EPI_ISL_3399624, EPI_ISL_3399625, EPI_ISL_3399626, EPI_ISL_3399627, EPI_ISL_3399628, EPI_ISL_3399629, EPI_ISL_3399630, EPI_ISL_3399631, EPI_ISL_3399632, EPI_ISL_3399633, EPI_ISL_3399634, EPI_ISL_3399635, EPI_ISL_3399636, EPI_ISL_3399637, EPI_ISL_3399638, EPI_ISL_3399639, EPI_ISL_3399640, EPI_ISL_3399641, EPI_ISL_3399642, EPI_ISL_3399643, EPI_ISL_3399644, EPI_ISL_3399645, EPI_ISL_3399646, EPI_ISL_3399647, EPI_ISL_3399648, EPI_ISL_3399649, EPI_ISL_3399650, EPI_ISL_3399651, EPI_ISL_3399652, EPI_ISL_3399653, EPI_ISL_3399654, EPI_ISL_3399655, EPI_ISL_3399656, EPI_ISL_3399657, EPI_ISL_3399658, EPI_ISL_3399659, EPI_ISL_3399660, EPI_ISL_3399661, EPI_ISL_3399662, EPI_ISL_3399663, EPI_ISL_3399664, EPI_ISL_3399665, EPI_ISL_3399666, EPI_ISL_3399667, EPI_ISL_3399668, EPI_ISL_3399669, EPI_ISL_3399670, EPI_ISL_3399671, EPI_ISL_3399672 | | | |
| see above | Broad Institute Clinical Research Sequencing Platform | Infectious Disease Program, Broad Institute of Harvard and MIT | Siddle,K.J., Adams,G., Pearlman,L., Gladden-Young,A., Vicente,G., Blumenstiel,B., DeFelice,M., Lee,M., McGovern,S., Lagerborg,K., Rudy,M., DeRuff,K., Carter,A., Normandin,E., Bauer,M., Reilly,S., Tomkins-Tinch,C., Loreth,C., Chaluvadi,S., Meldrim,J., Granger,B., Lemieux,J.E., Birren,B.W., Sabeti,P.C., Larkin,K., Dodge,S., Lennon,N., Madoff,L., Brown,C., Gallagher,G., Smole,S., Park,D.J., Gabriel,S., and MacInnis,B.L. |
| EPI_ISL_3399673, EPI_ISL_3399674, EPI_ISL_3399675, EPI_ISL_3399676, EPI_ISL_3399677, EPI_ISL_3399678, EPI_ISL_3399679, EPI_ISL_3399680, EPI_ISL_3399681, EPI_ISL_3399682, EPI_ISL_3399683, EPI_ISL_3399684, EPI_ISL_3399685, EPI_ISL_3399686, EPI_ISL_3399687, EPI_ISL_3399688, EPI_ISL_3399689, EPI_ISL_3399692, EPI_ISL_3399693, EPI_ISL_3399694, EPI_ISL_3399695, EPI_ISL_3399696, EPI_ISL_3399697, EPI_ISL_3399698, EPI_ISL_3399699 | | | |
| see above | University of Rochester Medical Center (URMC) LABS | University of Rochester Medical Center (URMC) LABS | Ghinwa Dumyati, Dwight Hardy, Andrew Cameron, Stepan Voljehn, Kelly DeLary |
| EPI_ISL_3399724, EPI_ISL_3399725, EPI_ISL_3399726, EPI_ISL_3399729, EPI_ISL_3399730, EPI_ISL_3399736, EPI_ISL_3399737, EPI_ISL_3399738, EPI_ISL_3399740, EPI_ISL_3399741, EPI_ISL_3399742, EPI_ISL_3399743, EPI_ISL_3399744, EPI_ISL_3399746, EPI_ISL_3399747, EPI_ISL_3399748, EPI_ISL_3399756, EPI_ISL_3399757, EPI_ISL_3399758, EPI_ISL_3399759 | | | |
| see above | NS-QEII Health Sciences Centre | National Microbiology Laboratory (NML) | Anna Majer, Shari Tyson, Grace Seo, Philip Mabon, Elsie Grudeski, Rhiannon Huzarewich, Russell Mandes, Anneliese Landgraff, Jennifer Tanner, Natalie Knox, Morag Graham, Gary Van Domselaar, Todd Hatchette, Jason LeBlanc, Janice Pettipas, Dan Gaston, Nathal |
| EPI_ISL_3399768, EPI_ISL_3399770, EPI_ISL_3399771, EPI_ISL_3399772, EPI_ISL_3399773, EPI_ISL_3399774, EPI_ISL_3399775, EPI_ISL_3399776, EPI_ISL_3399777, EPI_ISL_3399778, EPI_ISL_3399779, EPI_ISL_3399780, EPI_ISL_3399781, EPI_ISL_3399782, EPI_ISL_3399783, EPI_ISL_3399784, EPI_ISL_3399785, EPI_ISL_3399786, EPI_ISL_3399787, EPI_ISL_3399788, EPI_ISL_3399789, EPI_ISL_3399790, EPI_ISL_3399791, EPI_ISL_3399792, EPI_ISL_3399793, EPI_ISL_3399794, EPI_ISL_3399795, EPI_ISL_3399796, EPI_ISL_3399797, EPI_ISL_3399798, EPI_ISL_3399799, EPI_ISL_3399800, EPI_ISL_3399801, EPI_ISL_3399802, EPI_ISL_3399803, EPI_ISL_3399804, EPI_ISL_3399805, EPI_ISL_3399806, EPI_ISL_3399807, EPI_ISL_3399808, EPI_ISL_3399809, EPI_ISL_3399810, EPI_ISL_3399811 | | | |
| see above | Tampa General Hospital Esoteric Lab | Tampa General Hospital Esoteric Research & Development Lab | Jon Faughn, Amorce Lima, Deanna Becker, Dominic Uy, Jorge Mecias-Frias, Katie Hurless, Vicki Healer, Suzane Silbert |
| EPI_ISL_3399815, EPI_ISL_3399816, EPI_ISL_3399817, EPI_ISL_3399818, EPI_ISL_3399819, EPI_ISL_3399820, EPI_ISL_3399821, EPI_ISL_3399822, EPI_ISL_3399823, EPI_ISL_3399824, EPI_ISL_3399825, EPI_ISL_3399826, EPI_ISL_3399827, EPI_ISL_3399828, EPI_ISL_3399829, EPI_ISL_3399830, EPI_ISL_3399831, EPI_ISL_3399832, EPI_ISL_3399833, EPI_ISL_3399834, EPI_ISL_3399835, EPI_ISL_3399836, EPI_ISL_3399837, EPI_ISL_3399838, EPI_ISL_3399839, EPI_ISL_3399840, EPI_ISL_3399841, EPI_ISL_3399842, EPI_ISL_3399843, EPI_ISL_3399844, EPI_ISL_3399845, EPI_ISL_3399846, EPI_ISL_3399847, EPI_ISL_3399848, EPI_ISL_3399849, EPI_ISL_3399850, EPI_ISL_3399851, EPI_ISL_3399852, EPI_ISL_3399853, EPI_ISL_3399854, EPI_ISL_3399855, EPI_ISL_3399856, EPI_ISL_3399857, EPI_ISL_3399858, EPI_ISL_3399859, EPI_ISL_3399860, EPI_ISL_3399861, EPI_ISL_3399863, EPI_ISL_3399864, EPI_ISL_3399865, EPI_ISL_3399866, EPI_ISL_3399867, EPI_ISL_3399868, EPI_ISL_3399869, EPI_ISL_3399870, EPI_ISL_3399871, EPI_ISL_3399872, EPI_ISL_3399873, EPI_ISL_3399874, EPI_ISL_3399875, EPI_ISL_3399876, EPI_ISL_3399877, EPI_ISL_3399878, EPI_ISL_3399879, EPI_ISL_3399880, EPI_ISL_3399881, EPI_ISL_3399882, EPI_ISL_3399887, EPI_ISL_3399888, EPI_ISL_3399891, EPI_ISL_3399895, EPI_ISL_3399896, EPI_ISL_3399897, EPI_ISL_3399898, EPI_ISL_3399899, EPI_ISL_3399900, EPI_ISL_3399901, EPI_ISL_3399902, EPI_ISL_3399903, EPI_ISL_3399904, EPI_ISL_3399905, EPI_ISL_3399906, EPI_ISL_3399907, EPI_ISL_3399908, EPI_ISL_3399909, EPI_ISL_3399910, EPI_ISL_3399911, EPI_ISL_3399912, EPI_ISL_3399913, EPI_ISL_3399914, EPI_ISL_3399915, EPI_ISL_3399916, EPI_ISL_3399917, EPI_ISL_3399918, EPI_ISL_3399919, EPI_ISL_3399920, EPI_ISL_3399921, EPI_ISL_3399922, EPI_ISL_3399923, EPI_ISL_3399924, EPI_ISL_3399925, EPI_ISL_3399926, EPI_ISL_3399927, EPI_ISL_3399928, EPI_ISL_3399929, EPI_ISL_3399930, EPI_ISL_3399931, EPI_ISL_3399932, EPI_ISL_3399933, EPI_ISL_3399934, EPI_ISL_3399935, EPI_ISL_3399936, EPI_ISL_3399937, EPI_ISL_3399938, EPI_ISL_3399939, EPI_ISL_3399940, EPI_ISL_3399941, EPI_ISL_3399942, EPI_ISL_3399943, EPI_ISL_3399944, EPI_ISL_3399945, EPI_ISL_3399947, EPI_ISL_3399948, EPI_ISL_3399949, EPI_ISL_3399950, EPI_ISL_3399951, EPI_ISL_3399952, EPI_ISL_3399953, EPI_ISL_3399954, EPI_ISL_3399955, EPI_ISL_3399956, EPI_ISL_3399957, EPI_ISL_3399958, EPI_ISL_3399959, EPI_ISL_3399960, EPI_ISL_3399961, EPI_ISL_3399962, EPI_ISL_3399963, EPI_ISL_3399964, EPI_ISL_3399965, EPI_ISL_3399966, EPI_ISL_3399967, EPI_ISL_3399968, EPI_ISL_3399969, EPI_ISL_3399970, EPI_ISL_3399971, EPI_ISL_3399972, EPI_ISL_3399973, EPI_ISL_3399974, EPI_ISL_3399975, EPI_ISL_3399976, EPI_ISL_3399977, EPI_ISL_3399978, EPI_ISL_3399979, EPI_ISL_3399980, EPI_ISL_3399981, EPI_ISL_3399982, EPI_ISL_3399983, EPI_ISL_3399984, EPI_ISL_3399985, EPI_ISL_3399986, EPI_ISL_3399987, EPI_ISL_3399988, EPI_ISL_3399989, EPI_ISL_3399990, EPI_ISL_3399991, EPI_ISL_3399992, EPI_ISL_3399993, EPI_ISL_3399994, EPI_ISL_3399995, EPI_ISL_3399996, EPI_ISL_3399997, EPI_ISL_3399998, EPI_ISL_3399999, EPI_ISL_3400000, EPI_ISL_3400001, EPI_ISL_3400002, EPI_ISL_3400003, EPI_ISL_3400004, EPI_ISL_3400005, EPI_ISL_3400006, EPI_ISL_3400008, EPI_ISL_3400009, EPI_ISL_3400010, EPI_ISL_3400011, EPI_ISL_3400012, EPI_ISL_3400013, EPI_ISL_3400014, EPI_ISL_3400015, EPI_ISL_3400016, EPI_ISL_3400017, EPI_ISL_3400018, EPI_ISL_3400019, EPI_ISL_3400020, EPI_ISL_3400021, EPI_ISL_3400022, EPI_ISL_3400023, EPI_ISL_3400024, EPI_ISL_3400025, EPI_ISL_3400026, EPI_ISL_3400027, EPI_ISL_3400028, EPI_ISL_3400029, EPI_ISL_3400030, EPI_ISL_3400031, EPI_ISL_3400032, EPI_ISL_3400033, EPI_ISL_3400034, EPI_ISL_3400035, EPI_ISL_3400036, EPI_ISL_3400037, EPI_ISL_3400038, EPI_ISL_3400039, EPI_ISL_3400040, EPI_ISL_3400041, EPI_ISL_3400042, EPI_ISL_3400043, EPI_ISL_3400044, EPI_ISL_3400045, EPI_ISL_3400046, EPI_ISL_3400047, EPI_ISL_3400048, EPI_ISL_3400049, EPI_ISL_3400050, EPI_ISL_3400051, EPI_ISL_3400052, EPI_ISL_3400053, EPI_ISL_3400054, EPI_ISL_3400055, EPI_ISL_3400056, EPI_ISL_3400057, EPI_ISL_3400058, EPI_ISL_3400059, EPI_ISL_3400060, EPI_ISL_3400061, EPI_ISL_3400062, EPI_ISL_3400063, EPI_ISL_3400064, EPI_ISL_3400065, EPI_ISL_3400066, EPI_ISL_3400067, EPI_ISL_3400069, EPI_ISL_3400070, EPI_ISL_3400071, EPI_ISL_3400072, EPI_ISL_3400073, EPI_ISL_3400074, EPI_ISL_3400075, EPI_ISL_3400076, EPI_ISL_3400077, EPI_ISL_3400078, EPI_ISL_3400079, EPI_ISL_3400080, EPI_ISL_3400081, EPI_ISL_3400082, EPI_ISL_3400083, EPI_ISL_3400084, EPI_ISL_3400085, EPI_ISL_3400086, EPI_ISL_3400087, EPI_ISL_3400088, EPI_ISL_3400089, EPI_ISL_3400090, EPI_ISL_3400091, EPI_ISL_3400092, EPI_ISL_3400093, EPI_ISL_3400094, EPI_ISL_3400095, EPI_ISL_3400096, EPI_ISL_3400097, EPI_ISL_3400098, EPI_ISL_3400099, EPI_ISL_3400100, EPI_ISL_3400101, EPI_ISL_3400102, EPI_ISL_3400103, EPI_ISL_3400104, EPI_ISL_3400105, EPI_ISL_3400106, EPI_ISL_3400107, EPI_ISL_3400108, EPI_ISL_3400109, EPI_ISL_3400110, EPI_ISL_3400111, EPI_ISL_3400112, EPI_ISL_3400113, EPI_ISL_3400114, EPI_ISL_3400115, EPI_ISL_3400116, EPI_ISL_3400117, EPI_ISL_3400118, EPI_ISL_3400119, EPI_ISL_3400120, EPI_ISL_3400121, EPI_ISL_3400122, EPI_ISL_3400123, EPI_ISL_3400124, EPI_ISL_3400125, EPI_ISL_3400126, EPI_ISL_3400127, EPI_ISL_3400128, EPI_ISL_3400130, EPI_ISL_3400131, EPI_ISL_3400132, EPI_ISL_3400133, EPI_ISL_3400134, EPI_ISL_3400135, EPI_ISL_3400136, EPI_ISL_3400137, EPI_ISL_3400138, EPI_ISL_3400139, EPI_ISL_3400140, EPI_ISL_3400141, EPI_ISL_3400142, EPI_ISL_3400143, EPI_ISL_3400144, EPI_ISL_3400145, EPI_ISL_3400146, EPI_ISL_3400147, EPI_ISL_3400148, EPI_ISL_3400149, EPI_ISL_3400150, EPI_ISL_3400151, EPI_ISL_3400152, EPI_ISL_3400153, EPI_ISL_3400154, EPI_ISL_3400155, EPI_ISL_3400156, EPI_ISL_3400157, EPI_ISL_3400158, EPI_ISL_3400159, EPI_ISL_3400160, EPI_ISL_3400161, EPI_ISL_3400162, EPI_ISL_3400163, EPI_ISL_3400164, EPI_ISL_3400165, EPI_ISL_3400166, EPI_ISL_3400167, EPI_ISL_3400168, EPI_ISL_3400169, EPI_ISL_3400170, EPI_ISL_3400171, EPI_ISL_3400172, EPI_ISL_3400173, EPI_ISL_3400174, EPI_ISL_3400175, EPI_ISL_3400176, EPI_ISL_3400177, EPI_ISL_3400178, EPI_ISL_3400179, EPI_ISL_3400180, EPI_ISL_3400181, EPI_ISL_3400182, EPI_ISL_3400183, EPI_ISL_3400184, EPI_ISL_3400185, EPI_ISL_3400186, EPI_ISL_3400187, EPI_ISL_3400188, EPI_ISL_3400189, EPI_ISL_3400191, EPI_ISL_3400192, EPI_ISL_3400193, EPI_ISL_3400194, EPI_ISL_3400195, EPI_ISL_3400196, EPI_ISL_3400197, EPI_ISL_3400198, EPI_ISL_3400199, EPI_ISL_3400200, EPI_ISL_3400201, EPI_ISL_3400202, EPI_ISL_3400203, EPI_ISL_3400204, EPI_ISL_3400205, EPI_ISL_3400206, EPI_ISL_3400207, EPI_ISL_3400208, EPI_ISL_3400209, EPI_ISL_3400210, EPI_ISL_3400211, EPI_ISL_3400212, EPI_ISL_3400213, EPI_ISL_3400214, EPI_ISL_3400215, EPI_ISL_3400216, EPI_ISL_3400217, EPI_ISL_3400218, EPI_ISL_3400219, EPI_ISL_3400220, EPI_ISL_3400221, EPI_ISL_3400222, EPI_ISL_3400223, EPI_ISL_3400224, EPI_ISL_3400225, EPI_ISL_3400226, EPI_ISL_3400227, EPI_ISL_3400228, EPI_ISL_3400229, EPI_ISL_3400230, EPI_ISL_3400231, EPI_ISL_3400232, EPI_ISL_3400233, EPI_ISL_3400234, EPI_ISL_3400235, EPI_ISL_3400236, EPI_ISL_3400237, EPI_ISL_3400238, EPI_ISL_3400239, EPI_ISL_3400240, EPI_ISL_3400241, EPI_ISL_3400242, EPI_ISL_3400243, EPI_ISL_3400244, EPI_ISL_3400245, EPI_ISL_3400246, EPI_ISL_3400247, EPI_ISL_3400248, EPI_ISL_3400249, EPI_ISL_3400250, EPI_ISL_3400252, EPI_ISL_3400253, EPI_ISL_3400254, EPI_ISL_3400255, EPI_ISL_3400256, EPI_ISL_3400257, EPI_ISL_3400258, EPI_ISL_3400259, EPI_ISL_3400260, EPI_ISL_3400261, EPI_ISL_3400262, EPI_ISL_3400263, EPI_ISL_3400264, EPI_ISL_3400265, EPI_ISL_3400266, EPI_ISL_3400267, EPI_ISL_3400268, EPI_ISL_3400269, EPI_ISL_3400270, EPI_ISL_3400271, EPI_ISL_3400272, EPI_ISL_3400273, EPI_ISL_3400274, EPI_ISL_3400275, EPI_ISL_3400276, EPI_ISL_3400277, EPI_ISL_3400278, EPI_ISL_3400279, EPI_ISL_3400280, EPI_ISL_3400281, EPI_ISL_3400282, EPI_ISL_3400283, EPI_ISL_3400284, EPI_ISL_3400285, EPI_ISL_3400286, EPI_ISL_3400287, EPI_ISL_3400288, EPI_ISL_3400289, EPI_ISL_3400290, EPI_ISL_3400291, EPI_ISL_3400292, EPI_ISL_3400293, EPI_ISL_3400294, EPI_ISL_3400295, EPI_ISL_3400296, EPI_ISL_3400297, EPI_ISL_3400298, EPI_ISL_3400299, EPI_ISL_3400300, EPI_ISL_3400301, EPI_ISL_3400302, EPI_ISL_3400303, EPI_ISL_3400304, EPI_ISL_3400305, EPI_ISL_3400306, EPI_ISL_3400307, EPI_ISL_3400308, EPI_ISL_3400309, EPI_ISL_3400310, EPI_ISL_3400311, EPI_ISL_3400313, EPI_ISL_3400314, EPI_ISL_3400315, EPI_ISL_3400316, EPI_ISL_3400317, EPI_ISL_3400318, EPI_ISL_3400319, EPI_ISL_3400320, EPI_ISL_3400321, EPI_ISL_3400322, EPI_ISL_3400323, EPI_ISL_3400324, EPI_ISL_3400325, EPI_ISL_3400326, EPI_ISL_3400327, EPI_ISL_3400328, EPI_ISL_3400329, EPI_ISL_3400330, EPI_ISL_3400331, EPI_ISL_3400332, EPI_ISL_3400333, EPI_ISL_3400334, EPI_ISL_3400335, EPI_ISL_3400336, EPI_ISL_3400337, EPI_ISL_3400338, EPI_ISL_3400339, EPI_ISL_3400340, EPI_ISL_3400341, EPI_ISL_3400342, EPI_ISL_3400343, EPI_ISL_3400344, EPI_ISL_3400345, EPI_ISL_3400346, EPI_ISL_3400347, EPI_ISL_3400348, EPI_ISL_3400349, EPI_ISL_3400350, EPI_ISL_3400351, EPI_ISL_3400352, EPI_ISL_3400353, EPI_ISL_3400354, EPI_ISL_3400355, EPI_ISL_3400356, EPI_ISL_3400357, EPI_ISL_3400358, EPI_ISL_3400359, EPI_ISL_3400360, EPI_ISL_3400361, EPI_ISL_3400362, EPI_ISL_3400363, EPI_ISL_3400364, EPI_ISL_3400365, EPI_ISL_3400366, EPI_ISL_3400367, EPI_ISL_3400368, EPI_ISL_3400369, EPI_ISL_3400370, EPI_ISL_3400371, EPI_ISL_3400372, EPI_ISL_3400374, EPI_ISL_3400375, EPI_ISL_3400376, EPI_ISL_3400377, EPI_ISL_3400378, EPI_ISL_3400379, EPI_ISL_3400380, EPI_ISL_3400381, EPI_ISL_3400382, EPI_ISL_3400383, EPI_ISL_3400384, EPI_ISL_3400385, EPI_ISL_3400386, EPI_ISL_3400387, EPI_ISL_3400388, EPI_ISL_3400389, EPI_ISL_3400390, EPI_ISL_3400391, EPI_ISL_3400392, EPI_ISL_3400393, EPI_ISL_3400394, EPI_ISL_3400395, EPI_ISL_3400396, EPI_ISL_3400397, EPI_ISL_3400398, EPI_ISL_3400399, EPI_ISL_3400400, EPI_ISL_3400401, EPI_ISL_3400402, EPI_ISL_3400403, EPI_ISL_3400404, EPI_ISL_3400405, EPI_ISL_3400406, EPI_ISL_3400407, EPI_ISL_3400408, EPI_ISL_3400409, EPI_ISL_3400410, EPI_ISL_3400411, EPI_ISL_3400412, EPI_ISL_3400413, EPI_ISL_3400414, EPI_ISL_3400415, EPI_ISL_3400416, EPI_ISL_3400417, EPI_ISL_3400418, EPI_ISL_3400419, EPI_ISL_3400420, EPI_ISL_3400421, EPI_ISL_3400422, EPI_ISL_3400423, EPI_ISL_3400424, EPI_ISL_3400425, EPI_ISL_3400426, EPI_ISL_3400427, EPI_ISL_3400428, EPI_ISL_3400429, EPI_ISL_3400430, EPI_ISL_3400431, EPI_ISL_3400432, EPI_ISL_3400433, EPI_ISL_3400435, EPI_ISL_3400436, EPI_ISL_3400437, EPI_ISL_3400438, EPI_ISL_3400439, EPI_ISL_3400440, EPI_ISL_3400441, EPI_ISL_3400442, EPI_ISL_3400443, EPI_ISL_3400444, EPI_ISL_3400445, EPI_ISL_3400446, EPI_ISL_3400447, EPI_ISL_3400448, EPI_ISL_3400449, EPI_ISL_3400450, EPI_ISL_3400451, EPI_ISL_3400453, EPI_ISL_3400454, EPI_ISL_3400455, EPI_ISL_3400456, EPI_ISL_3400457, EPI_ISL_3400458, EPI_ISL_3400459, EPI_ISL_3400460, EPI_ISL_3400461, EPI_ISL_3400462, EPI_ISL_3400463, EPI_ISL_3400464, EPI_ISL_3400465, EPI_ISL_3400466, EPI_ISL_3400467, EPI_ISL_3400468, EPI_ISL_3400469, EPI_ISL_3400470, EPI_ISL_3400471, EPI_ISL_3400472, EPI_ISL_3400473, EPI_ISL_3400474, EPI_ISL_3400475, EPI_ISL_3400476, EPI_ISL_3400477, EPI_ISL_3400478, EPI_ISL_3400479, EPI_ISL_3400480, EPI_ISL_3400481, EPI_ISL_3400482, EPI_ISL_3400483, EPI_ISL_3400484, EPI_ISL_3400485, EPI_ISL_3400486, EPI_ISL_3400487, EPI_ISL_3400488, EPI_ISL_3400489, EPI_ISL_3400490, EPI_ISL_3400491, EPI_ISL_3400492, EPI_ISL_3400493, EPI_ISL_3400494, EPI_ISL_3400495, EPI_ISL_3400497, EPI_ISL_3400498, EPI_ISL_3400499, EPI_ISL_3400500, EPI_ISL_3400501, EPI_ISL_3400502, EPI_ISL_3400503, EPI_ISL_3400504, EPI_ISL_3400505, EPI_ISL_3400506, EPI_ISL_3400507, EPI_ISL_3400508, EPI_ISL_3400509, EPI_ISL_3400510, EPI_ISL_3400511, EPI_ISL_3400512, EPI_ISL_3400513, EPI_ISL_3400514, EPI_ISL_3400515, EPI_ISL_3400516, EPI_ISL_3400517, EPI_ISL_3400518, EPI_ISL_3400519, EPI_ISL_3400520, EPI_ISL_3400521, EPI_ISL_3400522, EPI_ISL_3400523, EPI_ISL_3400524, EPI_ISL_3400525, EPI_ISL_3400526, EPI_ISL_3400527, EPI_ISL_3400528, EPI_ISL_3400529, EPI_ISL_3400530, EPI_ISL_3400531, EPI_ISL_3400532, EPI_ISL_3400533, EPI_ISL_3400534, EPI_ISL_3400535, EPI_ISL_3400536, EPI_ISL_3400537, EPI_ISL_3400538, EPI_ISL_3400539, EPI_ISL_3400540, EPI_ISL_3400541, EPI_ISL_3400542, EPI_ISL_3400543, EPI_ISL_3400544, EPI_ISL_3400545, EPI_ISL_3400546, EPI_ISL_3400547, EPI_ISL_3400548, EPI_ISL_3400549, EPI_ISL_3400550, EPI_ISL_3400551, EPI_ISL_3400552, EPI_ISL_3400553, EPI_ISL_3400554, EPI_ISL_3400555, EPI_ISL_3400556, EPI_ISL_3400558, EPI_ISL_3400559, EPI_ISL_3400560, EPI_ISL_3400561, EPI_ISL_3400562, EPI_ISL_3400563, EPI_ISL_3400564, EPI_ISL_3400565, EPI_ISL_3400566, EPI_ISL_3400567, EPI_ISL_3400568, EPI_ISL_3400569, EPI_ISL_3400570, EPI_ISL_3400571, EPI_ISL_3400572, EPI_ISL_3400573, EPI_ISL_3400574, EPI_ISL_3400575, EPI_ISL_3400576, EPI_ISL_3400577, EPI_ISL_3400578, EPI_ISL_3400579, EPI_ISL_3400580, EPI_ISL_3400581, EPI_ISL_3400582, EPI_ISL_3400583, EPI_ISL_3400584, EPI_ISL_3400585, EPI_ISL_3400586, EPI_ISL_3400587, EPI_ISL_3400588, EPI_ISL_3400589, EPI_ISL_3400590, EPI_ISL_3400591, EPI_ISL_3400592, EPI_ISL_3400593, EPI_ISL_3400594, EPI_ISL_3400595, EPI_ISL_3400596, EPI_ISL_3400597, EPI_ISL_3400598, EPI_ISL_3400599, EPI_ISL_3400600, EPI_ISL_3400601, EPI_ISL_3400602, EPI_ISL_3400603, EPI_ISL_3400604, EPI_ISL_3400605, EPI_ISL_3400606, EPI_ISL_3400607, EPI_ISL_3400608, EPI_ISL_3400609, EPI_ISL_3400610, EPI_ISL_3400611, EPI_ISL_3400612, EPI_ISL_3400613, EPI_ISL_3400614, EPI_ISL_3400615, EPI_ISL_3400616, EPI_ISL_3400617, EPI_ISL_3400619, EPI_ISL_3400620, EPI_ISL_3400621, EPI_ISL_3400622, EPI_ISL_3400623, EPI_ISL_3400624, EPI_ISL_3400625, EPI_ISL_3400626, EPI_ISL_3400627, EPI_ISL_3400628, EPI_ISL_3400629, EPI_ISL_3400630, EPI_ISL_3400631, EPI_ISL_3400632, EPI_ISL_3400633, EPI_ISL_3400634, EPI_ISL_3400635, EPI_ISL_3400636, EPI_ISL_3400637, EPI_ISL_3400638, EPI_ISL_3400639, EPI_ISL_3400640, EPI_ISL_3400641, EPI_ISL_3400642, EPI_ISL_3400643, EPI_ISL_3400644, EPI_ISL_3400645, EPI_ISL_3400646, EPI_ISL_3400647, EPI_ISL_3400648, EPI_ISL_3400649, EPI_ISL_3400650, EPI_ISL_3400651, EPI_ISL_3400652, EPI_ISL_3400653, EPI_ISL_3400654, EPI_ISL_3400655, EPI_ISL_3400656, EPI_ISL_3400657, EPI_ISL_3400658, EPI_ISL_3400659, EPI_ISL_3400660, EPI_ISL_3400661, EPI_ISL_3400662, EPI_ISL_3400663, EPI_ISL_3400664, EPI_ISL_3400665, EPI_ISL_3400666, EPI_ISL_3400667, EPI_ISL_3400668, EPI_ISL_3400669, EPI_ISL_3400670, EPI_ISL_3400671, EPI_ISL_3400672, EPI_ISL_3400673, EPI_ISL_3400674, EPI_ISL_3400675, EPI_ISL_3400676, EPI_ISL_3400677, EPI_ISL_3400678, EPI_ISL_3400680, EPI_ISL_3400681, EPI_ISL_3400682, EPI_ISL_3400683, EPI_ISL_3400684, EPI_ISL_3400685, EPI_ISL_3400686, EPI_ISL_3400687, EPI_ISL_3400688, EPI_ISL_3400689, EPI_ISL_3400690, EPI_ISL_3400691, EPI_ISL_3400692, EPI_ISL_3400693, EPI_ISL_3400694, EPI_ISL_3400695, EPI_ISL_3400696, EPI_ISL_3400697, EPI_ISL_3400698, EPI_ISL_3400699, EPI_ISL_3400700, EPI_ISL_3400701, EPI_ISL_3400702, EPI_ISL_3400703, EPI_ISL_3400704, EPI_ISL_3400705, EPI_ISL_3400706, EPI_ISL_3400707, EPI_ISL_3400708, EPI_ISL_3400709, EPI_ISL_3400710, EPI_ISL_3400711, EPI_ISL_3400712, EPI_ISL_3400713, EPI_ISL_3400714, EPI_ISL_3400715, EPI_ISL_3400716, EPI_ISL_3400717, EPI_ISL_3400718, EPI_ISL_3400719, EPI_ISL_3400720, EPI_ISL_3400721, EPI_ISL_3400722, EPI_ISL_3400723, EPI_ISL_3400724, EPI_ISL_3400725, EPI_ISL_3400726, EPI_ISL_3400727, EPI_ISL_3400728, EPI_ISL_3400729, EPI_ISL_3400730, EPI_ISL_3400731, EPI_ISL_3400732, EPI_ISL_3400733, EPI_ISL_3400734, EPI_ISL_3400735, EPI_ISL_3400736, EPI_ISL_3400737, EPI_ISL_3400738, EPI_ISL_3400739, EPI_ISL_3400741, EPI_ISL_3400742, EPI_ISL_3400743, EPI_ISL_3400744, EPI_ISL_3400745, EPI_ISL_3400746, EPI_ISL_3400747, EPI_ISL_3400748, EPI_ISL_3400749, EPI_ISL_3400750, EPI_ISL_3400751, EPI_ISL_3400752, EPI_ISL_3400753, EPI_ISL_3400754, EPI_ISL_3400755, EPI_ISL_3400756, EPI_ISL_3400757, EPI_ISL_3400758, EPI_ISL_3400759, EPI_ISL_3400760, EPI_ISL_3400761, EPI_ISL_3400762, EPI_ISL_3400763, EPI_ISL_3400764, EPI_ISL_3400765, EPI_ISL_3400766, EPI_ISL_3400767, EPI_ISL_3400768, EPI_ISL_3400769, EPI_ISL_3400770, EPI_ISL_3400771, EPI_ISL_3400772, EPI_ISL_3400773, EPI_ISL_3400774, EPI_ISL_3400775, EPI_ISL_3400776, EPI_ISL_3400777, EPI_ISL_3400778, EPI_ISL_3400779, EPI_ISL_3400780, EPI_ISL_3400781, EPI_ISL_3400782, EPI_ISL_3400783, EPI_ISL_3400784, EPI_ISL_3400785, EPI_ISL_3400786, EPI_ISL_3400787, EPI_ISL_3400788, EPI_ISL_3400789, EPI_ISL_3400790, EPI_ISL_3400791, EPI_ISL_3400792, EPI_ISL_3400793, EPI_ISL_3400794, EPI_ISL_3400795, EPI_ISL_3400796, EPI_ISL_3400797, EPI_ISL_3400798, EPI_ISL_3400799, EPI_ISL_3400800, EPI_ISL_3400802, EPI_ISL_3400803, EPI_ISL_3400804, EPI_ISL_3400805, EPI_ISL_3400806, EPI_ISL_3400807, EPI_ISL_3400808, EPI_ISL_3400809, EPI_ISL_3400810, EPI_ISL_3400811, EPI_ISL_3400812, EPI_ISL_3400813, EPI_ISL_3400814, EPI_ISL_3400815, EPI_ISL_3400816, EPI_ISL_3400817, EPI_ISL_3400818, EPI_ISL_3400855, EPI_ISL_3400856, EPI_ISL_3400857, EPI_ISL_3400858, EPI_ISL_3400859, EPI_ISL_3400860, EPI_ISL_3400861, EPI_ISL_3400862, EPI_ISL_3400863, EPI_ISL_3400864, EPI_ISL_3400865, EPI_ISL_3400866, EPI_ISL_3400867, EPI_ISL_3400868, EPI_ISL_3400869, EPI_ISL_3400870, EPI_ISL_3400871, EPI_ISL_3400872, EPI_ISL_3400873, EPI_ISL_3400874, EPI_ISL_3400875, EPI_ISL_3400876, EPI_ISL_3400877, EPI_ISL_3400878, EPI_ISL_3400879, EPI_ISL_3400880, EPI_ISL_3400881, EPI_ISL_3400882, EPI_ISL_3400883, EPI_ISL_3400884, EPI_ISL_3400885, EPI_ISL_3400886, EPI_ISL_3400887, EPI_ISL_3400888, EPI_ISL_3400889, EPI_ISL_3400890, EPI_ISL_3400891, EPI_ISL_3400892, EPI_ISL_3400893, EPI_ISL_3400894, EPI_ISL_3400895, EPI_ISL_3400896, EPI_ISL_3400897, EPI_ISL_3400899, EPI_ISL_3400900, EPI_ISL_3400901, EPI_ISL_3400902, EPI_ISL_3400903, EPI_ISL_3400904, EPI_ISL_3400905, EPI_ISL_3400906, EPI_ISL_3400907, EPI_ISL_3400908, EPI_ISL_3400909, EPI_ISL_3400910, EPI_ISL_3400911, EPI_ISL_3400912, EPI_ISL_3400913, EPI_ISL_3400914, EPI_ISL_3400915, EPI_ISL_3400916, EPI_ISL_3400917, EPI_ISL_3400918, EPI_ISL_3400919, EPI_ISL_3400920, EPI_ISL_3400921, EPI_ISL_3400922, EPI_ISL_3400923, EPI_ISL_3400924, EPI_ISL_3400925, EPI_ISL_3400926, EPI_ISL_3400927, EPI_ISL_3400928, EPI_ISL_3400929, EPI_ISL_3400930, EPI_ISL_3400931, EPI_ISL_3400932, EPI_ISL_3400933, EPI_ISL_3400934, EPI_ISL_3400935, EPI_ISL_3400936, EPI_ISL_3400937, EPI_ISL_3400938, EPI_ISL_3400939, EPI_ISL_3400940, EPI_ISL_3400941, EPI_ISL_3400942, EPI_ISL_3400943, EPI_ISL_3400944, EPI_ISL_3400945, EPI_ISL_3400946, EPI_ISL_3400947, EPI_ISL_3400948, EPI_ISL_3400949, EPI_ISL_3400950, EPI_ISL_3400951, EPI_ISL_3400952, EPI_ISL_3400953, EPI_ISL_3400954, EPI_ISL_3400955, EPI_ISL_3400956, EPI_ISL_3400957, EPI_ISL_3400958, EPI_ISL_3400960, EPI_ISL_3400961, EPI_ISL_3400962, EPI_ISL_3400963, EPI_ISL_3400964, EPI_ISL_3400965, EPI_ISL_3400966, EPI_ISL_3400967, EPI_ISL_3400968, EPI_ISL_3400969, EPI_ISL_3400970, EPI_ISL_3400971, EPI_ISL_3400972, EPI_ISL_3400973, EPI_ISL_3400974, EPI_ISL_3400975, EPI_ISL_3400976, EPI_ISL_3400977, EPI_ISL_3400978, EPI_ISL_3400979, EPI_ISL_3400980, EPI_ISL_3400981, EPI_ISL_3400982, EPI_ISL_3400983, EPI_ISL_3400984, EPI_ISL_3400985, EPI_ISL_3400986, EPI_ISL_3400987, EPI_ISL_3400988, EPI_ISL_3400989, EPI_ISL_3400990, EPI_ISL_3400991, EPI_ISL_3400992, EPI_ISL_3400993, EPI_ISL_3400994, EPI_ISL_3400995, EPI_ISL_3400996, EPI_ISL_3400997, EPI_ISL_3400998, EPI_ISL_3400999, EPI_ISL_3401000, EPI_ISL_3401001, EPI_ISL_3401002, EPI_ISL_3401003, EPI_ISL_3401004, EPI_ISL_3401005, EPI_ISL_3401006, EPI_ISL_3401007, EPI_ISL_3401008, EPI_ISL_3401009, EPI_ISL_3401010, EPI_ISL_3401011, EPI_ISL_3401012, EPI_ISL_3401013, EPI_ISL_3401014, EPI_ISL_3401015, EPI_ISL_3401016, EPI_ISL_3401017, EPI_ISL_3401018, EPI_ISL_3401019, EPI_ISL_3401021, EPI_ISL_3401022, EPI_ISL_3401023, EPI_ISL_3401024, EPI_ISL_3401025, EPI_ISL_3401026, EPI_ISL_3401027, EPI_ISL_3401028, EPI_ISL_3401029, EPI_ISL_3401030, EPI_ISL_3401031, EPI_ISL_3401032, EPI_ISL_3401033, EPI_ISL_3401034, EPI_ISL_3401035, EPI_ISL_3401036, EPI_ISL_3401037, EPI_ISL_3401038, EPI_ISL_3401039, EPI_ISL_3401040, EPI_ISL_3401041, EPI_ISL_3401042, EPI_ISL_3401043, EPI_ISL_3401044, EPI_ISL_3401045, EPI_ISL_3401046, EPI_ISL_3401047, EPI_ISL_3401048, EPI_ISL_3401049, EPI_ISL_3401050, EPI_ISL_3401051, EPI_ISL_3401052, EPI_ISL_3401053, EPI_ISL_3401054, EPI_ISL_3401055, EPI_ISL_3401056, EPI_ISL_3401057, EPI_ISL_3401058, EPI_ISL_3401059, EPI_ISL_3401060, EPI_ISL_3401061, EPI_ISL_3401062, EPI_ISL_3401063, EPI_ISL_3401064, EPI_ISL_3401065, EPI_ISL_3401066, EPI_ISL_3401067, EPI_ISL_3401068, EPI_ISL_3401069, EPI_ISL_3401070, EPI_ISL_3401071, EPI_ISL_3401072, EPI_ISL_3401073, EPI_ISL_3401074, EPI_ISL_3401075, EPI_ISL_3401076, EPI_ISL_3401077, EPI_ISL_3401078, EPI_ISL_3401079, EPI_ISL_3401080, EPI_ISL_3401082, EPI_ISL_3401083, EPI_ISL_3401084, EPI_ISL_3401085, EPI_ISL_3401086, EPI_ISL_3401087, EPI_ISL_3401088, EPI_ISL_3401089, EPI_ISL_3401090, EPI_ISL_3401091, EPI_ISL_3401092, EPI_ISL_3401093, EPI_ISL_3401094, EPI_ISL_3401095, EPI_ISL_3401096, EPI_ISL_3401097, EPI_ISL_3401098, EPI_ISL_3401099, EPI_ISL_3401100, EPI_ISL_3401101, EPI_ISL_3401102, EPI_ISL_3401103, EPI_ISL_3401104, EPI_ISL_3401105, EPI_ISL_3401106, EPI_ISL_3401107, EPI_ISL_3401108, EPI_ISL_3401109, EPI_ISL_3401110, EPI_ISL_3401111, EPI_ISL_3401112, EPI_ISL_3401113, EPI_ISL_3401114, EPI_ISL_3401115, EPI_ISL_3401116, EPI_ISL_3401117, EPI_ISL_3401118, EPI_ISL_3401119, EPI_ISL_3401120, EPI_ISL_3401121, EPI_ISL_3401122, EPI_ISL_3401123, EPI_ISL_3401124, EPI_ISL_3401125, EPI_ISL_3401126, EPI_ISL_3401127, EPI_ISL_3401128, EPI_ISL_3401129, EPI_ISL_3401130, EPI_ISL_3401131, EPI_ISL_3401132, EPI_ISL_3401133, EPI_ISL_3401134, EPI_ISL_3401135, EPI_ISL_3401136, EPI_ISL_3401137, EPI_ISL_3401138, EPI_ISL_3401139, EPI_ISL_3401140, EPI_ISL_3401141, EPI_ISL_3401143, EPI_ISL_3401144, EPI_ISL_3401145, EPI_ISL_3401146, EPI_ISL_3401147, EPI_ISL_3401148, EPI_ISL_3401149, EPI_ISL_3401150, EPI_ISL_3401151, EPI_ISL_3401152, EPI_ISL_3401153, EPI_ISL_3401154, EPI_ISL_3401155, EPI_ISL_3401156, EPI_ISL_3401157, EPI_ISL_3401158, EPI_ISL_3401159, EPI_ISL_3401160, EPI_ISL_3401161, EPI_ISL_3401162, EPI_ISL_3401163, EPI_ISL_3401164, EPI_ISL_3401165, EPI_ISL_3401166, EPI_ISL_3401167, EPI_ISL_3401168, EPI_ISL_3401169, EPI_ISL_3401170, EPI_ISL_3401171, EPI_ISL_3401172, EPI_ISL_3401173, EPI_ISL_3401174, EPI_ISL_3401175, EPI_ISL_3401176, EPI_ISL_3401177, EPI_ISL_3401178, EPI_ISL_3401179, EPI_ISL_3401180, EPI_ISL_3401181, EPI_ISL_3401182, EPI_ISL_3401183, EPI_ISL_3401184, EPI_ISL_3401185, EPI_ISL_3401186, EPI_ISL_3401187, EPI_ISL_3401188, EPI_ISL_3401189, EPI_ISL_3401190, EPI_ISL_3401191, EPI_ISL_3401192, EPI_ISL_3401193, EPI_ISL_3401194, EPI_ISL_3401195, EPI_ISL_3401196, EPI_ISL_3401197, EPI_ISL_3401198, EPI_ISL_3401199, EPI_ISL_3401200, EPI_ISL_3401201, EPI_ISL_3401202, EPI_ISL_3401204, EPI_ISL_3401205, EPI_ISL_3401206, EPI_ISL_3401207, EPI_ISL_3401208, EPI_ISL_3401209, EPI_ISL_3401210, EPI_ISL_3401211, EPI_ISL_3401212, EPI_ISL_3401213, EPI_ISL_3401214, EPI_ISL_3401215, EPI_ISL_3401216, EPI_ISL_3401217, EPI_ISL_3401218, EPI_ISL_3401219, EPI_ISL_3401220, EPI_ISL_3401221, EPI_ISL_3401222, EPI_ISL_3401223, EPI_ISL_3401224, EPI_ISL_3401225, EPI_ISL_3401226, EPI_ISL_3401227, EPI_ISL_3401228, EPI_ISL_3401229, EPI_ISL_3401230, EPI_ISL_3401231, EPI_ISL_3401232, EPI_ISL_3401233, EPI_ISL_3401234, EPI_ISL_3401235, EPI_ISL_3401236, EPI_ISL_3401237, EPI_ISL_3401238, EPI_ISL_3401239, EPI_ISL_3401240, EPI_ISL_3401241, EPI_ISL_3401242, EPI_ISL_3401243, EPI_ISL_3401244, EPI_ISL_3401245, EPI_ISL_3401246, EPI_ISL_3401247, EPI_ISL_3401248, EPI_ISL_3401249, EPI_ISL_3401250, EPI_ISL_3401251, EPI_ISL_3401252, EPI_ISL_3401253, EPI_ISL_3401254, EPI_ISL_3401255, EPI_ISL_3401256, EPI_ISL_3401257, EPI_ISL_3401258, EPI_ISL_3401259, EPI_ISL_3401260, EPI_ISL_3401261, EPI_ISL_3401262, EPI_ISL_3401263, EPI_ISL_3401265, EPI_ISL_3401266, EPI_ISL_3401267, EPI_ISL_3401268, EPI_ISL_3401269, EPI_ISL_3401270, EPI_ISL_3401271, EPI_ISL_3401272, EPI_ISL_3401273, EPI_ISL_3401274, EPI_ISL_3401275, EPI_ISL_3401276, EPI_ISL_3401277, EPI_ISL_3401278, EPI_ISL_3401279, EPI_ISL_3401280, EPI_ISL_3401281, EPI_ISL_3401282, EPI_ISL_3401283, EPI_ISL_3401284, EPI_ISL_3401285, EPI_ISL_3401286, EPI_ISL_3401287, EPI_ISL_3401288, EPI_ISL_3401289, EPI_ISL_3401290, EPI_ISL_3401291, EPI_ISL_3401292, EPI_ISL_3401293, EPI_ISL_3401294, EPI_ISL_3401295, EPI_ISL_3401296, EPI_ISL_3401297, EPI_ISL_3401298, EPI_ISL_3401299, EPI_ISL_3401300, EPI_ISL_3401301, EPI_ISL_3401302, EPI_ISL_3401303, EPI_ISL_3401304, EPI_ISL_3401305, EPI_ISL_3401306, EPI_ISL_3401307, EPI_ISL_3401308, EPI_ISL_3401309, EPI_ISL_3401310, EPI_ISL_3401311, EPI_ISL_3401312, EPI_ISL_3401313, EPI_ISL_3401314, EPI_ISL_3401315, EPI_ISL_3401316, EPI_ISL_3401317, EPI_ISL_3401318, EPI_ISL_3401319, EPI_ISL_3401320, EPI_ISL_3401321, EPI_ISL_3401322, EPI_ISL_3401323, EPI_ISL_3401324, EPI_ISL_3401326, EPI_ISL_3401327, EPI_ISL_3401328, EPI_ISL_3401329, EPI_ISL_3401330, EPI_ISL_3401331, EPI_ISL_3401332, EPI_ISL_3401333, EPI_ISL_3401334, EPI_ISL_3401335, EPI_ISL_3401336, EPI_ISL_3401337, EPI_ISL_3401338, EPI_ISL_3401339, EPI_ISL_3401340, EPI_ISL_3401341, EPI_ISL_3401342, EPI_ISL_3401343, EPI_ISL_3401344, EPI_ISL_3401345, EPI_ISL_3401346, EPI_ISL_3401347, EPI_ISL_3401348, EPI_ISL_3401349, EPI_ISL_3401350, EPI_ISL_3401351, EPI_ISL_3401352, EPI_ISL_3401353, EPI_ISL_3401354, EPI_ISL_3401355, EPI_ISL_3401356, EPI_ISL_3401357, EPI_ISL_3401358, EPI_ISL_3401359, EPI_ISL_3401360, EPI_ISL_3401361, EPI_ISL_3401362, EPI_ISL_3401363, EPI_ISL_3401364, EPI_ISL_3401365, EPI_ISL_3401366, EPI_ISL_3401367, EPI_ISL_3401368, EPI_ISL_3401369, EPI_ISL_3401370, EPI_ISL_3401371, EPI_ISL_3401372, EPI_ISL_3401373, EPI_ISL_3401374, EPI_ISL_3401375, EPI_ISL_3401376, EPI_ISL_3401377, EPI_ISL_3401378, EPI_ISL_3401379, EPI_ISL_3401380, EPI_ISL_3401381, EPI_ISL_3401382, EPI_ISL_3401383, EPI_ISL_3401384, EPI_ISL_3401385, EPI_ISL_3401387, EPI_ISL_3401388, EPI_ISL_3401389, EPI_ISL_3401390, EPI_ISL_3401391, EPI_ISL_3401392, EPI_ISL_3401393, EPI_ISL_3401394, EPI_ISL_3401395 | | | |
| see above | Ministry of Health Turkey | Ministry of Health Turkey | Fatma Bayrakdar, Yasemin Cosgun, Suleyman Yalcin, Gulay Korukluoglu |
| EPI_ISL_3401396, EPI_ISL_3401397, EPI_ISL_3401398, EPI_ISL_3401399, EPI_ISL_3401400, EPI_ISL_3401401, EPI_ISL_3401402, EPI_ISL_3401403, EPI_ISL_3401404, EPI_ISL_3401405, EPI_ISL_3401407, EPI_ISL_3401408, EPI_ISL_3401409, EPI_ISL_3401410, EPI_ISL_3401411, EPI_ISL_3401412, EPI_ISL_3401413, EPI_ISL_3401414 | | | |
| see above | Medical Laboratories Duesseldorf | Center of Medical Microbiology, Virology, and Hospital Hygiene, University of Duesseldorf | Maximilian Damagnez;Alexander Dilthey;Angelika Helmer;Torsten Houwaart;Lisanna Hülse;Christian Lange;Malte Kohns Vasconcelos;Nadine Lübke;Jessica Nicolai;Klaus Pfeffer;Daniel Strelow;Jörg Timm;Andreas Walker;Tobias Wienemann |
| EPI_ISL_3401418, EPI_ISL_3401419 | University Campus Bio-Medico of Rome (UCBM) | University Campus Bio-Medico of Rome (UCBM) | Fogolari M., Riva E., De Florio L., Francesconi M., Lintas C., Veralli R., Angeletti S. |
| EPI_ISL_3401425 | Laboratorio de Referencia Nacional de Virus Respiratorios. Centro Nacional de Salud Publica. Instituto Nacional de Salud Peru. | Laboratorio de Referencia Nacional de Virus Respiratorios. Centro Nacional de Salud Publica. Instituto Nacional de Salud Peru. | Carlos Padilla Rojas, Veronica Hurtado Vela, Orson Mestanza Millones, Iris Silva Molina, Luis Barcena Flores, Sandra Morales Ruiz, Steve Acedo Lazo, Wendy Lizarraga Olivares, Victor Jimenez Vasquez, Sara Gordillo Vilchez, Nieves Sevilla Castañeda, Princesa Medrano Alhuay, Omar Caceres Rey, Marco Galarza Perez, Henri Bailon Calderon, Nancy Rojas Serrano, Priscila Lope Pari, Lely Solari Zerpa, Joseph Huayra Niquen |
| EPI_ISL_3401619 | University Campus Bio-Medico of Rome (UCBM) | University Campus Bio-Medico of Rome (UCBM) | Fogolari M., Riva E., De Florio L., Francesconi M., Lintas C., Veralli R., Angeletti S. |
| EPI_ISL_3401626, EPI_ISL_3401628, EPI_ISL_3401629, EPI_ISL_3401630, EPI_ISL_3401632, EPI_ISL_3401634, EPI_ISL_3401637, EPI_ISL_3401638, EPI_ISL_3401640, EPI_ISL_3401641, EPI_ISL_3401642, EPI_ISL_3401644, EPI_ISL_3401645 | | | |
| see above | Wyoming Public Health Laboratory | Wyoming Public Health Laboratory | Jim Mildenberger, Taylor Fearing, Channing Weber, Ashley Norberg, Chayse Rowley, Marley Goetz, Brian Dominguez, Elliot Thomasson, Sam Britz, Cari Sloma, Robert Petit, and Rob Christensen |
| EPI_ISL_3401648, EPI_ISL_3401649, EPI_ISL_3401650, EPI_ISL_3401652, EPI_ISL_3401653, EPI_ISL_3401654, EPI_ISL_3401655, EPI_ISL_3401656, EPI_ISL_3401657, EPI_ISL_3401658, EPI_ISL_3401659, EPI_ISL_3401660, EPI_ISL_3401661, EPI_ISL_3401662, EPI_ISL_3401663, EPI_ISL_3401664, EPI_ISL_3401665, EPI_ISL_3401666, EPI_ISL_3401667, EPI_ISL_3401668, EPI_ISL_3401669 | | | |
| see above | TXDSHS | TXDSHS | Rashmi Tuladhar, Bonnie Oh, Jenny Zhang, Maliha Rahman, Mayela Pedrueza, Anita Pokharel, Karen Bobier, Lorraine Rodriguez, Myong Koag, Chun Wang, Rachel Lee, Grace Kubin |
| EPI_ISL_3401670, EPI_ISL_3401671, EPI_ISL_3401672, EPI_ISL_3401673, EPI_ISL_3401674, EPI_ISL_3401675, EPI_ISL_3401676, EPI_ISL_3401677, EPI_ISL_3401678, EPI_ISL_3401679, EPI_ISL_3401680, EPI_ISL_3401681, EPI_ISL_3401682, EPI_ISL_3401683, EPI_ISL_3401684 | | | |

| | | | |
|---|---|---|---|
| see above | Histopath | NSW Health Pathology - Institute of Clinical Pathology and Medical Research; Westmead Hospital; University of Sydney | Arnott A., Draper J., Gall M., Martinez E., Rockett R., Sintchenko V., on behalf of ICPMR |
| EPI_ISL_3401685 | Pathology North - Hunter - NSW Health Pathology | NSW Health Pathology - Institute of Clinical Pathology and Medical Research; Westmead Hospital; University of Sydney | Arnott A., Draper J., Gall M., Martinez E., Rockett R., Sintchenko V., on behalf of ICPMR |
| EPI_ISL_3401686, EPI_ISL_3401687, EPI_ISL_3401688, EPI_ISL_3401689, EPI_ISL_3401690, EPI_ISL_3401691, EPI_ISL_3401692, EPI_ISL_3401693, EPI_ISL_3401694, EPI_ISL_3401695, EPI_ISL_3401696, EPI_ISL_3401697, EPI_ISL_3401698, EPI_ISL_3401699, EPI_ISL_3401700, EPI_ISL_3401701, EPI_ISL_3401702, EPI_ISL_3401703, EPI_ISL_3401704, EPI_ISL_3401705, EPI_ISL_3401706, EPI_ISL_3401707, EPI_ISL_3401708, EPI_ISL_3401709, EPI_ISL_3401710, EPI_ISL_3401711, EPI_ISL_3401712, EPI_ISL_3401713, EPI_ISL_3401714, EPI_ISL_3401715, EPI_ISL_3401716 | | | |
| see above | Laverty Pathology | NSW Health Pathology - Institute of Clinical Pathology and Medical Research; Westmead Hospital; University of Sydney | Arnott A., Draper J., Gall M., Martinez E., Rockett R., Sintchenko V., on behalf of ICPMR |
| EPI_ISL_3401717, EPI_ISL_3401718, EPI_ISL_3401719, EPI_ISL_3401720, EPI_ISL_3401721, EPI_ISL_3401722, EPI_ISL_3401723, EPI_ISL_3401724, EPI_ISL_3401725, EPI_ISL_3401726, EPI_ISL_3401727, EPI_ISL_3401728, EPI_ISL_3401729, EPI_ISL_3401730, EPI_ISL_3401731, EPI_ISL_3401732, EPI_ISL_3401733, EPI_ISL_3401734, EPI_ISL_3401735, EPI_ISL_3401736, EPI_ISL_3401737, EPI_ISL_3401738, EPI_ISL_3401739 | | | |
| see above | Sydney South West Pathology Service (SSWPS) - Liverpool Hospital - NSW Health Pathology | NSW Health Pathology - Institute of Clinical Pathology and Medical Research; Westmead Hospital; University of Sydney | Arnott A., Draper J., Gall M., Martinez E., Rockett R., Sintchenko V., on behalf of ICPMR |
| EPI_ISL_3401740, EPI_ISL_3401741, EPI_ISL_3401742, EPI_ISL_3401743, EPI_ISL_3401744, EPI_ISL_3401745, EPI_ISL_3401746, EPI_ISL_3401747, EPI_ISL_3401748, EPI_ISL_3401749, EPI_ISL_3401750 | | | |
| see above | Medlab Pathology | NSW Health Pathology - Institute of Clinical Pathology and Medical Research; Westmead Hospital; University of Sydney | Arnott A., Draper J., Gall M., Martinez E., Rockett R., Sintchenko V., on behalf of ICPMR |
| EPI_ISL_3401751, EPI_ISL_3401752 | Sydney South West Pathology Service (SSWPS) - Liverpool Hospital - NSW Health Pathology | NSW Health Pathology - Institute of Clinical Pathology and Medical Research; Westmead Hospital; University of Sydney | Arnott A., Draper J., Gall M., Martinez E., Rockett R., Sintchenko V., on behalf of ICPMR |
| EPI_ISL_3401753, EPI_ISL_3401754, EPI_ISL_3401755, EPI_ISL_3401756, EPI_ISL_3401757, EPI_ISL_3401758, EPI_ISL_3401759, EPI_ISL_3401760, EPI_ISL_3401761, EPI_ISL_3401762, EPI_ISL_3401763, EPI_ISL_3401764, EPI_ISL_3401765, EPI_ISL_3401766, EPI_ISL_3401767 | | | |
| see above | St Vincent's Pathology (SydPath) | NSW Health Pathology - Institute of Clinical Pathology and Medical Research; Westmead Hospital; University of Sydney | Arnott A., Draper J., Gall M., Martinez E., Rockett R., Sintchenko V., on behalf of ICPMR |
| EPI_ISL_3401768, EPI_ISL_3401769, EPI_ISL_3401770, EPI_ISL_3401771, EPI_ISL_3401772, EPI_ISL_3401773, EPI_ISL_3401774, EPI_ISL_3401775, EPI_ISL_3401776, EPI_ISL_3401777, EPI_ISL_3401778, EPI_ISL_3401779, EPI_ISL_3401780, EPI_ISL_3401781, EPI_ISL_3401782 | | | |
| see above | 4Cyte Pathology | NSW Health Pathology - Institute of Clinical Pathology and Medical Research; Westmead Hospital; University of Sydney | Arnott A., Draper J., Gall M., Martinez E., Rockett R., Sintchenko V., on behalf of ICPMR |
| EPI_ISL_3401783 | Sydney South West Pathology Service (SSWPS) - Liverpool Hospital - NSW Health Pathology | NSW Health Pathology - Institute of Clinical Pathology and Medical Research; Westmead Hospital; University of Sydney | Arnott A., Draper J., Gall M., Martinez E., Rockett R., Sintchenko V., on behalf of ICPMR |
| EPI_ISL_3401784, EPI_ISL_3401785 | Pathology North - Hunter - NSW Health Pathology | NSW Health Pathology - Institute of Clinical Pathology and Medical Research; Westmead Hospital; University of Sydney | Arnott A., Draper J., Gall M., Martinez E., Rockett R., Sintchenko V., on behalf of ICPMR |
| EPI_ISL_3401786 | Sydney South West Pathology Service (SSWPS) - Liverpool Hospital - NSW Health Pathology | NSW Health Pathology - Institute of Clinical Pathology and Medical Research; Westmead Hospital; University of Sydney | Arnott A., Draper J., Gall M., Martinez E., Rockett R., Sintchenko V., on behalf of ICPMR |
| EPI_ISL_3401787, EPI_ISL_3401788, EPI_ISL_3401789 | Pathology North - Hunter - NSW Health Pathology | NSW Health Pathology - Institute of Clinical Pathology and Medical Research; Westmead Hospital; University of Sydney | Arnott A., Draper J., Gall M., Martinez E., Rockett R., Sintchenko V., on behalf of ICPMR |
| EPI_ISL_3401790, EPI_ISL_3401791, EPI_ISL_3401792 | Sydney South West Pathology Service (SSWPS) - Liverpool Hospital - NSW Health Pathology | NSW Health Pathology - Institute of Clinical Pathology and Medical Research; Westmead Hospital; University of Sydney | Arnott A., Draper J., Gall M., Martinez E., Rockett R., Sintchenko V., on behalf of ICPMR |
| EPI_ISL_3401793, EPI_ISL_3401794, EPI_ISL_3401795, EPI_ISL_3401796, EPI_ISL_3401797, EPI_ISL_3401798, EPI_ISL_3401799, EPI_ISL_3401800, EPI_ISL_3401801, EPI_ISL_3401802, EPI_ISL_3401803, EPI_ISL_3401804, EPI_ISL_3401805, EPI_ISL_3401806, EPI_ISL_3401807, EPI_ISL_3401808, EPI_ISL_3401809, EPI_ISL_3401810, EPI_ISL_3401811, EPI_ISL_3401812, EPI_ISL_3401813, EPI_ISL_3401814, EPI_ISL_3401815, EPI_ISL_3401816, EPI_ISL_3401817, EPI_ISL_3401818, EPI_ISL_3401819, EPI_ISL_3401820, EPI_ISL_3401821, EPI_ISL_3401822, EPI_ISL_3401823, EPI_ISL_3401824, EPI_ISL_3401825, EPI_ISL_3401826, EPI_ISL_3401827, EPI_ISL_3401828, EPI_ISL_3401829, EPI_ISL_3401830, EPI_ISL_3401831, EPI_ISL_3401832, EPI_ISL_3401833, EPI_ISL_3401834, EPI_ISL_3401835, EPI_ISL_3401836, EPI_ISL_3401837, EPI_ISL_3401838, EPI_ISL_3401839, EPI_ISL_3401840, EPI_ISL_3401841 | | | |
| see above | Histopath | NSW Health Pathology - Institute of Clinical Pathology and Medical Research; Westmead Hospital; University of Sydney | Arnott A., Draper J., Gall M., Martinez E., Rockett R., Sintchenko V., on behalf of ICPMR |
| EPI_ISL_3401842, EPI_ISL_3401843, EPI_ISL_3401844, EPI_ISL_3401845, EPI_ISL_3401846, EPI_ISL_3401847 | Pathology North - Hunter - NSW Health Pathology | NSW Health Pathology - Institute of Clinical Pathology and Medical Research; Westmead Hospital; University of Sydney | Arnott A., Draper J., Gall M., Martinez E., Rockett R., Sintchenko V., on behalf of ICPMR |
| EPI_ISL_3401848 | Pathology North - Royal North Shore Hospital - NSW Health Pathology | NSW Health Pathology - Institute of Clinical Pathology and Medical Research; Westmead Hospital; University of Sydney | Arnott A., Draper J., Gall M., Martinez E., Rockett R., Sintchenko V., on behalf of ICPMR |
| EPI_ISL_3401849 | Pathology West - NSW Health Pathology | NSW Health Pathology - Institute of Clinical Pathology and Medical Research; Westmead Hospital; University of Sydney | Arnott A., Draper J., Gall M., Martinez E., Rockett R., Sintchenko V., on behalf of ICPMR |
| EPI_ISL_3401850 | Genomik Solidaritas Indonesia Laboratorium | Genomik Solidaritas Indonesia Laboratorium | Carissa Sintca Wijaya, Himawan Masyhuri, Vania Gavrila Wikasa, Dhahlia Agustina Cahyono, Annisa Muthiah Sukirman, Gracia Felias Enos Korompis, Anuraj Shankar, Louisa Markus, Meutia Ayuputeri Kumaheri, Ariel Pradipta |
| EPI_ISL_3401851 | RSPI Prof DR. Sulianti Saroso, Rs | Genomik Solidaritas Indonesia Laboratorium | Himawan Masyhuri, Carissa Sintca Wijaya, Dwi Oktavia, Tiranti Vindha, Ngabila Salama, Vania Gavrila Wikasa, Dhahlia Agustina Cahyono, Annisa Muthiah Sukirman, Gracia Felias Enos Korompis, Anuraj Shankar, Louisa Markus, Meutia Ayuputeri Kumaheri, Ariel Pradipta |
| EPI_ISL_3401852, EPI_ISL_3401853 | Labkesda | Genomik Solidaritas Indonesia Laboratorium | Himawan Masyhuri, Carissa Sintca Wijaya, Dwi Oktavia, Tiranti Vindha, Ngabila Salama, Vania Gavrila Wikasa, Dhahlia Agustina Cahyono, Annisa Muthiah Sukirman, Gracia Felias Enos Korompis, Anuraj Shankar, Louisa Markus, Meutia Ayuputeri Kumaheri, Ariel Pradipta |
| EPI_ISL_3401854 | Genomik Solidaritas Indonesia Laboratorium | Genomik Solidaritas Indonesia Laboratorium | Vania Gavrila Wikasa, Carissa Sintca Wijaya, Himawan Masyhuri, Dhahlia Agustina Cahyono, Annisa Muthiah Sukirman, Gracia Felias Enos Korompis, Anuraj Shankar, Louisa Markus, Meutia Ayuputeri Kumaheri, Ariel Pradipta |
| EPI_ISL_3401855 | Puskesmas Kecamatan Tanjung Priok | Genomik Solidaritas Indonesia Laboratorium | Vania Gavrila Wikasa, Carissa Sintca Wijaya, Ahrahayati Wildany, Dwi Oktavia, Tiranti Vindha, Ngabila Salama, Himawan Masyhuri, Dhahlia Agustina Cahyono, Annisa Muthiah Sukirman, Gracia Felias Enos Korompis, Anuraj Shankar, Louisa Markus, Meutia Ayuputeri Kumaheri, Ariel Pradipta |
| EPI_ISL_3401856 | Genomik Solidaritas Indonesia Laboratorium | Genomik Solidaritas Indonesia Laboratorium | Vania Gavrila Wikasa, Carissa Sintca Wijaya, Himawan Masyhuri, Dhahlia Agustina Cahyono, Annisa Muthiah Sukirman, Gracia Felias Enos Korompis, Anuraj Shankar, Louisa Markus, Meutia Ayuputeri Kumaheri, Ariel Pradipta |
| EPI_ISL_3401857 | Genomik Solidaritas Indonesia Laboratorium | Genomik Solidaritas Indonesia Laboratorium | Carissa Sintca Wijaya, Himawan Masyhuri, Vania Gavrila Wikasa, Dhahlia Agustina Cahyono, Annisa Muthiah Sukirman, Gracia Felias Enos Korompis, Anuraj Shankar, Louisa Markus, Meutia Ayuputeri Kumaheri, Ariel Pradipta |
| EPI_ISL_3401858 | Labkesda | Genomik Solidaritas Indonesia Laboratorium | Himawan Masyhuri, Carissa Sintca Wijaya, Dwi Oktavia, Tiranti Vindha, Ngabila Salama, Vania Gavrila Wikasa, Dhahlia Agustina Cahyono, Annisa Muthiah Sukirman, Gracia Felias Enos Korompis, Anuraj Shankar, Louisa Markus, Meutia Ayuputeri Kumaheri, Ariel Pradipta |
| EPI_ISL_3401859 | Laboratorium RSUP Fatmawati | Genomik Solidaritas Indonesia Laboratorium | Himawan Masyhuri, Carissa Sintca Wijaya, Siska Situmorang, Lidya Utami, Loli Simajuntak, Dwi Oktavia, Tiranti Vindha, Ngabila Salama, Vania Gavrila Wikasa, Dhahlia Agustina Cahyono, Annisa Muthiah Sukirman, Gracia Felias Enos Korompis, Anuraj Shankar, Louisa Markus, Meutia Ayuputeri Kumaheri, Ariel Pradipta |
| EPI_ISL_3401860 | Genomik Solidaritas Indonesia Laboratorium | Genomik Solidaritas Indonesia Laboratorium | Carissa Sintca Wijaya, Himawan Masyhuri, Vania Gavrila Wikasa, Dhahlia Agustina Cahyono, Annisa Muthiah Sukirman, Gracia Felias Enos Korompis, Anuraj Shankar, Louisa Markus, Meutia Ayuputeri Kumaheri, Ariel Pradipta |
| EPI_ISL_3401861 | Labkesda | Genomik Solidaritas Indonesia Laboratorium | Himawan Masyhuri, Carissa Sintca Wijaya, Dwi Oktavia, Tiranti Vindha, Ngabila Salama, Vania Gavrila Wikasa, Dhahlia Agustina Cahyono, Annisa Muthiah Sukirman, Gracia Felias Enos Korompis, Anuraj Shankar, Louisa Markus, Meutia Ayuputeri Kumaheri, Ariel Pradipta |
| EPI_ISL_3401863 | RSPI Prof DR. Sulianti Saroso, Rs | Genomik Solidaritas Indonesia Laboratorium | Himawan Masyhuri, Carissa Sintca Wijaya, Dwi Oktavia, Tiranti Vindha, Ngabila Salama, Vania Gavrila Wikasa, Dhahlia Agustina Cahyono, Annisa Muthiah Sukirman, Gracia Felias Enos Korompis, Anuraj Shankar, Louisa Markus, Meutia Ayuputeri Kumaheri, Ariel Pradipta |
| EPI_ISL_3401866 | Genomik Solidaritas Indonesia Laboratorium | Genomik Solidaritas Indonesia Laboratorium | Vania Gavrila Wikasa, Carissa Sintca Wijaya, Himawan Masyhuri, Dhahlia Agustina Cahyono, Annisa Muthiah Sukirman, Gracia Felias Enos Korompis, Anuraj Shankar, Louisa Markus, Meutia Ayuputeri Kumaheri, Ariel Pradipta |
| EPI_ISL_3401867 | RS Atma Jaya | Genomik Solidaritas Indonesia Laboratorium | Vania Gavrila Wikasa, Carissa Sintca Wijaya, Dwi Oktavia, Tiranti Vindha, Ngabila Salama, Himawan Masyhuri, Dhahlia Agustina Cahyono, Annisa Muthiah Sukirman, Gracia Felias Enos Korompis, Anuraj Shankar, Louisa Markus, Meutia Ayuputeri Kumaheri, Ariel Pradipta |
| EPI_ISL_3401868 | Laboratorium RSUP Fatmawati | Genomik Solidaritas Indonesia Laboratorium | Himawan Masyhuri, Carissa Sintca Wijaya, Jerry, Lidya Utami, Loli Simajuntak, Dwi Oktavia, Tiranti Vindha, Ngabila Salama, Vania Gavrila Wikasa, Dhahlia Agustina Cahyono, Annisa Muthiah Sukirman, Gracia Felias Enos Korompis, Anuraj Shankar, Louisa Markus, Meutia Ayuputeri Kumaheri, Ariel Pradipta |
| EPI_ISL_3401869, EPI_ISL_3401870 | Rsia Budhi Jaya | Genomik Solidaritas Indonesia Laboratorium | Himawan Masyhuri, Carissa Sintca Wijaya, Lulu Nursyifa, Dwi Oktavia, Tiranti Vindha, Ngabila Salama, Vania Gavrila Wikasa, Dhahlia Agustina Cahyono, Annisa Muthiah Sukirman, Gracia Felias Enos Korompis, Anuraj Shankar, Louisa Markus, Meutia Ayuputeri Kumaheri, Ariel Pradipta |
| EPI_ISL_3401871 | Rsud Kebayoran Baru | Genomik Solidaritas Indonesia Laboratorium | Himawan Masyhuri, Carissa Sintca Wijaya, Ridho Aulia Rahman, Dwi Oktavia, Tiranti Vindha, Ngabila Salama, Vania Gavrila Wikasa, Dhahlia Agustina Cahyono, Annisa Muthiah Sukirman, Gracia Felias Enos Korompis, Anuraj Shankar, Louisa Markus, Meutia Ayuputeri Kumaheri, Ariel Pradipta |
| EPI_ISL_3401872 | Laboratorium RSUP Fatmawati | Genomik Solidaritas Indonesia Laboratorium | Himawan Masyhuri, Carissa Sintca Wijaya, Triana Puspita Dewi, Lidya Utami, Loli Simajuntak, Dwi Oktavia, Tiranti Vindha, Ngabila Salama, Vania Gavrila Wikasa, Dhahlia Agustina Cahyono, Annisa Muthiah Sukirman, Gracia Felias Enos Korompis, Anuraj Shankar, Louisa Markus, Meutia Ayuputeri Kumaheri, Ariel Pradipta |
| EPI_ISL_3401873 | Laboratorium RSUP Fatmawati | Genomik Solidaritas Indonesia Laboratorium | Himawan Masyhuri, Carissa Sintca Wijaya, Devi, Lidya Utami, Loli Simajuntak, Dwi Oktavia, Tiranti Vindha, Ngabila Salama, Vania Gavrila Wikasa, Dhahlia Agustina Cahyono, Annisa Muthiah Sukirman, Gracia Felias Enos Korompis, Anuraj Shankar, Louisa Markus, Meutia Ayuputeri Kumaheri, Ariel Pradipta |
| EPI_ISL_3401874, EPI_ISL_3401875, EPI_ISL_3401876, EPI_ISL_3401877, EPI_ISL_3401878 | RSPI Prof DR. Sulianti Saroso, Rs | Genomik Solidaritas Indonesia Laboratorium | Himawan Masyhuri, Carissa Sintca Wijaya, Dwi Oktavia, Tiranti Vindha, Ngabila Salama, Vania Gavrila Wikasa, Dhahlia Agustina Cahyono, Annisa Muthiah Sukirman, Gracia Felias Enos Korompis, Anuraj Shankar, Louisa Markus, Meutia Ayuputeri Kumaheri, Ariel Pradipta |
| EPI_ISL_3401879, EPI_ISL_3401880, EPI_ISL_3401881, EPI_ISL_3401882 | Puskesmas Kecamatan Tanjung Priok | Genomik Solidaritas Indonesia Laboratorium | Himawan Masyhuri, Carissa Sintca Wijaya, Ahrahayati Wildany, Dwi Oktavia, Tiranti Vindha, Ngabila Salama, Vania Gavrila Wikasa, Dhahlia Agustina Cahyono, Annisa Muthiah Sukirman, Gracia Felias Enos Korompis, Anuraj Shankar, Louisa Markus, Meutia Ayuputeri Kumaheri, Ariel Pradipta |

| | | | |
|---|---|---|---|
| EPI_ISL_3401883, EPI_ISL_3401884, EPI_ISL_3401885, EPI_ISL_3401886, EPI_ISL_3401887, EPI_ISL_3401888, EPI_ISL_3401889, EPI_ISL_3401890 | Puskesmas Kecamatan Tanjung Priok | Genomik Solidaritas Indonesia Laboratorium | Vania Gavrila Wikasa, Carissa Sintca Wijaya, Ahrahayati Wildany, Dwi Oktavia, Tiranti Vindha, Ngabila Salama, Himawan Masyhuri, Dhahlia Agustina Cahyono, Annisa Muthiah Sukirman, Gracia Felias Enos Korompis, Anuraj Shankar, Louisa Markus, Meutia Ayuputeri Kumaheri, Ariel Pradipta |
| EPI_ISL_3401891, EPI_ISL_3401892 | PKM Kelapa Gading | Genomik Solidaritas Indonesia Laboratorium | Vania Gavrila Wikasa, Carissa Sintca Wijaya, Dwi Oktavia, Tiranti Vindha, Ngabila Salama, Himawan Masyhuri, Dhahlia Agustina Cahyono, Annisa Muthiah Sukirman, Gracia Felias Enos Korompis, Anuraj Shankar, Louisa Markus, Meutia Ayuputeri Kumaheri, Ariel Pradipta |
| EPI_ISL_3401893 | PKM Tanjung Priok | Genomik Solidaritas Indonesia Laboratorium | Vania Gavrila Wikasa, Carissa Sintca Wijaya, Wahyudi, Dwi Oktavia, Tiranti Vindha, Ngabila Salama, Himawan Masyhuri, Dhahlia Agustina Cahyono, Annisa Muthiah Sukirman, Gracia Felias Enos Korompis, Anuraj Shankar, Louisa Markus, Meutia Ayuputeri Kumaheri, Ariel Pradipta |
| EPI_ISL_3401894 | PKM Kelapa Gading | Genomik Solidaritas Indonesia Laboratorium | Vania Gavrila Wikasa, Carissa Sintca Wijaya, Dwi Oktavia, Tiranti Vindha, Ngabila Salama, Himawan Masyhuri, Dhahlia Agustina Cahyono, Annisa Muthiah Sukirman, Gracia Felias Enos Korompis, Anuraj Shankar, Louisa Markus, Meutia Ayuputeri Kumaheri, Ariel Pradipta |
| EPI_ISL_3401895 | RS Atma Jaya | Genomik Solidaritas Indonesia Laboratorium | Vania Gavrila Wikasa, Carissa Sintca Wijaya, Dwi Oktavia, Tiranti Vindha, Ngabila Salama, Himawan Masyhuri, Dhahlia Agustina Cahyono, Annisa Muthiah Sukirman, Gracia Felias Enos Korompis, Anuraj Shankar, Louisa Markus, Meutia Ayuputeri Kumaheri, Ariel Pradipta |
| EPI_ISL_3401896, EPI_ISL_3401897 | Genomik Solidaritas Indonesia Laboratorium | Genomik Solidaritas Indonesia Laboratorium | Vania Gavrila Wikasa, Carissa Sintca Wijaya, Himawan Masyhuri, Dhahlia Agustina Cahyono, Annisa Muthiah Sukirman, Gracia Felias Enos Korompis, Anuraj Shankar, Louisa Markus, Meutia Ayuputeri Kumaheri, Ariel Pradipta |
| EPI_ISL_3401898, EPI_ISL_3401899 | RS Atma Jaya | Genomik Solidaritas Indonesia Laboratorium | Vania Gavrila Wikasa, Carissa Sintca Wijaya, Dwi Oktavia, Tiranti Vindha, Ngabila Salama, Himawan Masyhuri, Dhahlia Agustina Cahyono, Annisa Muthiah Sukirman, Gracia Felias Enos Korompis, Anuraj Shankar, Louisa Markus, Meutia Ayuputeri Kumaheri, Ariel Pradipta |
| EPI_ISL_3401900, EPI_ISL_3401901, EPI_ISL_3401902, EPI_ISL_3401903 | Puskesmas Kecamatan Pandemangan | Genomik Solidaritas Indonesia Laboratorium | Vania Gavrila Wikasa, Carissa Sintca Wijaya, Dwi Oktavia, Tiranti Vindha, Ngabila Salama, Himawan Masyhuri, Dhahlia Agustina Cahyono, Annisa Muthiah Sukirman, Gracia Felias Enos Korompis, Anuraj Shankar, Louisa Markus, Meutia Ayuputeri Kumaheri, Ariel Pradipta |
| EPI_ISL_3401904, EPI_ISL_3401905, EPI_ISL_3401906 | Genomik Solidaritas Indonesia Laboratorium | Genomik Solidaritas Indonesia Laboratorium | Vania Gavrila Wikasa, Carissa Sintca Wijaya, Himawan Masyhuri, Dhahlia Agustina Cahyono, Annisa Muthiah Sukirman, Gracia Felias Enos Korompis, Anuraj Shankar, Louisa Markus, Meutia Ayuputeri Kumaheri, Ariel Pradipta |
| EPI_ISL_3401907, EPI_ISL_3401908, EPI_ISL_3401909, EPI_ISL_3401910, EPI_ISL_3401911, EPI_ISL_3401912, EPI_ISL_3401913, EPI_ISL_3401914, EPI_ISL_3401915, EPI_ISL_3401916, EPI_ISL_3401917, EPI_ISL_3401918, EPI_ISL_3401919, EPI_ISL_3401920, EPI_ISL_3401921, EPI_ISL_3401922, EPI_ISL_3401923 | | | |
| see above | Genomik Solidaritas Indonesia Laboratorium | Genomik Solidaritas Indonesia Laboratorium | Carissa Sintca Wijaya, Himawan Masyhuri, Vania Gavrila Wikasa, Dhahlia Agustina Cahyono, Annisa Muthiah Sukirman, Gracia Felias Enos Korompis, Anuraj Shankar, Louisa Markus, Meutia Ayuputeri Kumaheri, Ariel Pradipta |
| EPI_ISL_3401931, EPI_ISL_3401932, EPI_ISL_3401933, EPI_ISL_3401934 | Hrvatski zavod za javno zdravstvo | Hrvatski zavod za javno zdravstvo | Irena Tabain, Ivana Ferenčak |
| EPI_ISL_3401935 | Nastavni zavod za javno zdravstvo Splitsko- Dalmatinske Županije | Hrvatski zavod za javno zdravstvo | Irena Tabain, Ivana Ferenčak |
| EPI_ISL_3401936 | Hrvatski zavod za javno zdravstvo | Hrvatski zavod za javno zdravstvo | Irena Tabain, Ivana Ferenčak |
| EPI_ISL_3401937 | Zavod za javno zdravstvo Međimurske Županije | Hrvatski zavod za javno zdravstvo | Irena Tabain, Ivana Ferenčak |
| EPI_ISL_3401938 | Hrvatski zavod za javno zdravstvo | Hrvatski zavod za javno zdravstvo | Irena Tabain, Ivana Ferenčak |
| EPI_ISL_3401939 | Zavod za javno zdravstvo Međimurske Županije | Hrvatski zavod za javno zdravstvo | Irena Tabain, Ivana Ferenčak |
| EPI_ISL_3401940, EPI_ISL_3401941 | Nastavni zavod za javno zdravstvo Primorsko- Goranske Županije | Hrvatski zavod za javno zdravstvo | Irena Tabain, Ivana Ferenčak |
| EPI_ISL_3401942, EPI_ISL_3401943 | Zavod za javno zdravstvo Dubrovačko-Neretvanske Županije | Hrvatski zavod za javno zdravstvo | Irena Tabain, Ivana Ferenčak |
| EPI_ISL_3401944 | Nastavni zavod za javno zdravstvo Primorsko- Goranske Županije | Hrvatski zavod za javno zdravstvo | Irena Tabain, Ivana Ferenčak |
| EPI_ISL_3401945, EPI_ISL_3401946 | Hrvatski zavod za javno zdravstvo | Hrvatski zavod za javno zdravstvo | Irena Tabain, Ivana Ferenčak |
| EPI_ISL_3401947 | Nastavni zavod za javno zdravstvo Primorsko- Goranske Županije | Hrvatski zavod za javno zdravstvo | Irena Tabain, Ivana Ferenčak |
| EPI_ISL_3401949 | Zavod za javno zdravstvo Međimurske Županije | Hrvatski zavod za javno zdravstvo | Irena Tabain, Ivana Ferenčak |
| EPI_ISL_3401950 | Zavod za javno zdravstvo Dubrovačko-Neretvanske Županije | Hrvatski zavod za javno zdravstvo | Irena Tabain, Ivana Ferenčak |
| EPI_ISL_3401951, EPI_ISL_3401952, EPI_ISL_3401953 | Hrvatski zavod za javno zdravstvo | Hrvatski zavod za javno zdravstvo | Irena Tabain, Ivana Ferenčak |
| EPI_ISL_3401954 | Nastavni zavod za javno zdravstvo Primorsko- Goranske Županije | Hrvatski zavod za javno zdravstvo | Irena Tabain, Ivana Ferenčak |
| EPI_ISL_3401962, EPI_ISL_3401963 | Zavod za javno zdravstvo Dubrovačko-Neretvanske Županije | Hrvatski zavod za javno zdravstvo | Irena Tabain, Ivana Ferenčak |
| EPI_ISL_3401964 | Nastavni zavod za javno zdravstvo Primorsko- Goranske Županije | Hrvatski zavod za javno zdravstvo | Irena Tabain, Ivana Ferenčak |
| EPI_ISL_3401967, EPI_ISL_3401968 | Hrvatski zavod za javno zdravstvo | Hrvatski zavod za javno zdravstvo | Irena Tabain, Ivana Ferenčak |
| EPI_ISL_3401969 | Nastavni zavod za javno zdravstvo Primorsko- Goranske Županije | Hrvatski zavod za javno zdravstvo | Irena Tabain, Ivana Ferenčak |
| EPI_ISL_3401970 | Hrvatski zavod za javno zdravstvo | Hrvatski zavod za javno zdravstvo | Irena Tabain, Ivana Ferenčak |
| EPI_ISL_3401971 | Zavod za javno zdravstvo Dubrovačko-Neretvanske Županije | Hrvatski zavod za javno zdravstvo | Irena Tabain, Ivana Ferenčak |
| EPI_ISL_3401973 | Klinički bolnički centar Zagreb | Hrvatski zavod za javno zdravstvo | Irena Tabain, Ivana Ferenčak |
| EPI_ISL_3401974, EPI_ISL_3401975, EPI_ISL_3401976 | Zavod za javno zdravstvo Dubrovačko-Neretvanske Županije | Hrvatski zavod za javno zdravstvo | Irena Tabain, Ivana Ferenčak |
| EPI_ISL_3401977, EPI_ISL_3401978 | Klinički bolnički centar Zagreb | Hrvatski zavod za javno zdravstvo | Irena Tabain, Ivana Ferenčak |
| EPI_ISL_3401979 | Zavod za javno zdravstvo Međimurske Županije | Hrvatski zavod za javno zdravstvo | Irena Tabain, Ivana Ferenčak |
| EPI_ISL_3401980, EPI_ISL_3401981, EPI_ISL_3401982, EPI_ISL_3401983, EPI_ISL_3401984, EPI_ISL_3401985, EPI_ISL_3401986, EPI_ISL_3401987, EPI_ISL_3401988, EPI_ISL_3401989, EPI_ISL_3401990, EPI_ISL_3401991, EPI_ISL_3401992, EPI_ISL_3401993, EPI_ISL_3401994, EPI_ISL_3401995 | | | |
| see above | Nastavni zavod za javno zdravstvo Primorsko- Goranske Županije | Hrvatski zavod za javno zdravstvo | Irena Tabain, Ivana Ferenčak |
| EPI_ISL_3401996, EPI_ISL_3401997, EPI_ISL_3401998, EPI_ISL_3401999, EPI_ISL_3402000, EPI_ISL_3402001, EPI_ISL_3402002, EPI_ISL_3402003, EPI_ISL_3402004, EPI_ISL_3402005, EPI_ISL_3402006, EPI_ISL_3402007, EPI_ISL_3402008, EPI_ISL_3402009, EPI_ISL_3402010, EPI_ISL_3402011 | | | |
| see above | Hrvatski zavod za javno zdravstvo | Hrvatski zavod za javno zdravstvo | Irena Tabain, Ivana Ferenčak |
| EPI_ISL_3402012 | Nastavni zavod za javno zdravstvo Splitsko- Dalmatinske Županije | Hrvatski zavod za javno zdravstvo | Irena Tabain, Ivana Ferenčak |
| EPI_ISL_3402015, EPI_ISL_3402016 | Regional Medical Sciences Center 1 Chiang Mai | National Institute of Health, Department of Medical Sciences, Ministry of Public Health, Thailand | Pilailuk Okada; Siripaporn Phuygun; Nuttida Thongpramul; Sittiporn Parnmen; Ratana Tacharoenmuang; Pakorn Piromtong; Natchaya Khiadsang; Thanutsapa Thanadachakul; Warawan Wongboot; sirikanda wimol; Sunthareeya Waicharoen; |
| EPI_ISL_3402018 | Regional Medical Sciences Center 6 Chonburi | National Institute of Health, Department of Medical Sciences, Ministry of Public Health, Thailand | Pilailuk Okada; Siripaporn Phuygun; Nuttida Thongpramul; Sittiporn Parnmen; Ratana Tacharoenmuang; Pakorn Piromtong; Natchaya Khiadsang; Thanutsapa Thanadachakul; Warawan Wongboot; sirikanda wimol; Sunthareeya Waicharoen; |
| EPI_ISL_3402020, EPI_ISL_3402021, EPI_ISL_3402022 | Regional Medical Sciences Center 8 Udonthani | National Institute of Health, Department of Medical Sciences, Ministry of Public Health, Thailand | Pilailuk Okada; Siripaporn Phuygun; Nuttida Thongpramul; Sittiporn Parnmen; Ratana Tacharoenmuang; Pakorn Piromtong; Natchaya Khiadsang; Thanutsapa Thanadachakul; Warawan Wongboot; sirikanda wimol; Sunthareeya Waicharoen; |
| EPI_ISL_3402023, EPI_ISL_3402024, EPI_ISL_3402025, EPI_ISL_3402026, EPI_ISL_3402027, EPI_ISL_3402028, EPI_ISL_3402029, EPI_ISL_3402030, EPI_ISL_3402031, EPI_ISL_3402032, EPI_ISL_3402033, EPI_ISL_3402034, EPI_ISL_3402035, EPI_ISL_3402036, EPI_ISL_3402037, EPI_ISL_3402038, EPI_ISL_3402039, EPI_ISL_3402040, EPI_ISL_3402041, EPI_ISL_3402042, EPI_ISL_3402043, EPI_ISL_3402044, EPI_ISL_3402045, EPI_ISL_3402046, EPI_ISL_3402047, EPI_ISL_3402048, EPI_ISL_3402049, EPI_ISL_3402050, EPI_ISL_3402051, EPI_ISL_3402052, EPI_ISL_3402053, EPI_ISL_3402054, EPI_ISL_3402055, EPI_ISL_3402056, EPI_ISL_3402057, EPI_ISL_3402058, EPI_ISL_3402059, EPI_ISL_3402060, EPI_ISL_3402061, EPI_ISL_3402062, EPI_ISL_3402063, EPI_ISL_3402064, EPI_ISL_3402065, EPI_ISL_3402066, EPI_ISL_3402067, EPI_ISL_3402068, EPI_ISL_3402069, EPI_ISL_3402070, EPI_ISL_3402071, EPI_ISL_3402072, EPI_ISL_3402073, EPI_ISL_3402074, EPI_ISL_3402075, EPI_ISL_3402076, EPI_ISL_3402077, EPI_ISL_3402078, EPI_ISL_3402079, EPI_ISL_3402080, EPI_ISL_3402081, EPI_ISL_3402082, EPI_ISL_3402083, EPI_ISL_3402084, EPI_ISL_3402085, EPI_ISL_3402086, EPI_ISL_3402087, EPI_ISL_3402088, | | | |

| | | | |
|---|---|---|---|
| EPI_ISL_3402089, EPI_ISL_3402090, EPI_ISL_3402091, EPI_ISL_3402092, EPI_ISL_3402093, EPI_ISL_3402094, EPI_ISL_3402095, EPI_ISL_3402096, EPI_ISL_3402097, EPI_ISL_3402098, EPI_ISL_3402099, EPI_ISL_3402100, EPI_ISL_3402101, EPI_ISL_3402102, EPI_ISL_3402103, EPI_ISL_3402104, EPI_ISL_3402105, EPI_ISL_3402106, EPI_ISL_3402107, EPI_ISL_3402108, EPI_ISL_3402109, EPI_ISL_3402110, EPI_ISL_3402111, EPI_ISL_3402112, EPI_ISL_3402113, EPI_ISL_3402114, EPI_ISL_3402115, EPI_ISL_3402116, EPI_ISL_3402117, EPI_ISL_3402118, EPI_ISL_3402119, EPI_ISL_3402120, EPI_ISL_3402121, EPI_ISL_3402122, EPI_ISL_3402123, EPI_ISL_3402124, EPI_ISL_3402125, EPI_ISL_3402126, EPI_ISL_3402127, EPI_ISL_3402129, EPI_ISL_3402130, EPI_ISL_3402131, EPI_ISL_3402132, EPI_ISL_3402133, EPI_ISL_3402134, EPI_ISL_3402135, EPI_ISL_3402136, EPI_ISL_3402137, EPI_ISL_3402138, EPI_ISL_3402139, EPI_ISL_3402140, EPI_ISL_3402141, EPI_ISL_3402142, EPI_ISL_3402143, EPI_ISL_3402144, EPI_ISL_3402145, EPI_ISL_3402146, EPI_ISL_3402147, EPI_ISL_3402148, EPI_ISL_3402149, EPI_ISL_3402150, EPI_ISL_3402151, EPI_ISL_3402152, EPI_ISL_3402153, EPI_ISL_3402154, EPI_ISL_3402155, EPI_ISL_3402156, EPI_ISL_3402157, EPI_ISL_3402158, EPI_ISL_3402159, EPI_ISL_3402160, EPI_ISL_3402161, EPI_ISL_3402162, EPI_ISL_3402163, EPI_ISL_3402164, EPI_ISL_3402165, EPI_ISL_3402166, EPI_ISL_3402167, EPI_ISL_3402168, EPI_ISL_3402169, EPI_ISL_3402170, EPI_ISL_3402171, EPI_ISL_3402172, EPI_ISL_3402173, EPI_ISL_3402174, EPI_ISL_3402175, EPI_ISL_3402176, EPI_ISL_3402177, EPI_ISL_3402178, EPI_ISL_3402179, EPI_ISL_3402180, EPI_ISL_3402181, EPI_ISL_3402182, EPI_ISL_3402183, EPI_ISL_3402184, EPI_ISL_3402185, EPI_ISL_3402186, EPI_ISL_3402187, EPI_ISL_3402188, EPI_ISL_3402189, EPI_ISL_3402190, EPI_ISL_3402191, EPI_ISL_3402192, EPI_ISL_3402193, EPI_ISL_3402194, EPI_ISL_3402195, EPI_ISL_3402196, EPI_ISL_3402197, EPI_ISL_3402198, EPI_ISL_3402199, EPI_ISL_3402200, EPI_ISL_3402201, EPI_ISL_3402202, EPI_ISL_3402203, EPI_ISL_3402204, EPI_ISL_3402206, EPI_ISL_3402207, EPI_ISL_3402208, EPI_ISL_3402209, EPI_ISL_3402210, EPI_ISL_3402211, EPI_ISL_3402212, EPI_ISL_3402213, EPI_ISL_3402214, EPI_ISL_3402215, EPI_ISL_3402216, EPI_ISL_3402217, EPI_ISL_3402218, EPI_ISL_3402219, EPI_ISL_3402220, EPI_ISL_3402221, EPI_ISL_3402222, EPI_ISL_3402223, EPI_ISL_3402224, EPI_ISL_3402225, EPI_ISL_3402226, EPI_ISL_3402227, EPI_ISL_3402228, EPI_ISL_3402229, EPI_ISL_3402230, EPI_ISL_3402231, EPI_ISL_3402232, EPI_ISL_3402233, EPI_ISL_3402234, EPI_ISL_3402235 | | | |
| see above | Clinical Microbiology, Infection Prevention and Control | Section for Molecular Diagnostics | Björn Hallström, Jonas Björkman |
| EPI_ISL_3402244, EPI_ISL_3402245, EPI_ISL_3402246, EPI_ISL_3402247, EPI_ISL_3402249, EPI_ISL_3402250, EPI_ISL_3402253, EPI_ISL_3402254, EPI_ISL_3402257, EPI_ISL_3402258, EPI_ISL_3402259, EPI_ISL_3402265, EPI_ISL_3402266, EPI_ISL_3402268 | | | |
| see above | Wyoming Public Health Laboratory | Wyoming Public Health Laboratory | Jim Mildenberger, Taylor Fearing, Channing Weber, Ashley Norberg, Chayse Rowley, Marley Goetz, Brian Dominguez, Elliot Thomasson, Sam Britz, Cari Sloma, Robert Petit, and Rob Christensen |
| EPI_ISL_3402272, EPI_ISL_3402273, EPI_ISL_3402274, EPI_ISL_3402275, EPI_ISL_3402276, EPI_ISL_3402277, EPI_ISL_3402278, EPI_ISL_3402279, EPI_ISL_3402280, EPI_ISL_3402281, EPI_ISL_3402282, EPI_ISL_3402283, EPI_ISL_3402284, EPI_ISL_3402286, EPI_ISL_3402287, EPI_ISL_3402288, EPI_ISL_3402289, EPI_ISL_3402290, EPI_ISL_3402291, EPI_ISL_3402292, EPI_ISL_3402293, EPI_ISL_3402294, EPI_ISL_3402295, EPI_ISL_3402296, EPI_ISL_3402297, EPI_ISL_3402298, EPI_ISL_3402299, EPI_ISL_3402300, EPI_ISL_3402301, EPI_ISL_3402302, EPI_ISL_3402303, EPI_ISL_3402304, EPI_ISL_3402305, EPI_ISL_3402306, EPI_ISL_3402307, EPI_ISL_3402308, EPI_ISL_3402309, EPI_ISL_3402310, EPI_ISL_3402311, EPI_ISL_3402312, EPI_ISL_3402313, EPI_ISL_3402314, EPI_ISL_3402315, EPI_ISL_3402316, EPI_ISL_3402318, EPI_ISL_3402319, EPI_ISL_3402320, EPI_ISL_3402321, EPI_ISL_3402322, EPI_ISL_3402323, EPI_ISL_3402324, EPI_ISL_3402325, EPI_ISL_3402326, EPI_ISL_3402327, EPI_ISL_3402328, EPI_ISL_3402329, EPI_ISL_3402330, EPI_ISL_3402331, EPI_ISL_3402332, EPI_ISL_3402333, EPI_ISL_3402334, EPI_ISL_3402335, EPI_ISL_3402337, EPI_ISL_3402338, EPI_ISL_3402339, EPI_ISL_3402340, EPI_ISL_3402341, EPI_ISL_3402342, EPI_ISL_3402343, EPI_ISL_3402344, EPI_ISL_3402345, EPI_ISL_3402346, EPI_ISL_3402347, EPI_ISL_3402348, EPI_ISL_3402349, EPI_ISL_3402350, EPI_ISL_3402351, EPI_ISL_3402352, EPI_ISL_3402353, EPI_ISL_3402354, EPI_ISL_3402355 | | | |
| see above | Virginia Division of Consolidated Laboratory Services | Virginia Division of Consolidated Laboratory Services | Virginia Division of Consolidated Laboratory Services |
| EPI_ISL_3402356, EPI_ISL_3402357, EPI_ISL_3402360, EPI_ISL_3402361, EPI_ISL_3402362, EPI_ISL_3402363, EPI_ISL_3402364, EPI_ISL_3402365, EPI_ISL_3402366, EPI_ISL_3402367, EPI_ISL_3402386, EPI_ISL_3402387, EPI_ISL_3402411, EPI_ISL_3402425, EPI_ISL_3402428, EPI_ISL_3402443, EPI_ISL_3402444, EPI_ISL_3402445, EPI_ISL_3402447, EPI_ISL_3402449, EPI_ISL_3402450, EPI_ISL_3402452, EPI_ISL_3402454, EPI_ISL_3402456, EPI_ISL_3402459, EPI_ISL_3402460, EPI_ISL_3402461, EPI_ISL_3402485, EPI_ISL_3402499, EPI_ISL_3402502, EPI_ISL_3402517, EPI_ISL_3402518, EPI_ISL_3402519, EPI_ISL_3402521, EPI_ISL_3402523, EPI_ISL_3402524, EPI_ISL_3402526, EPI_ISL_3402528, EPI_ISL_3402530, EPI_ISL_3402533, EPI_ISL_3402534, EPI_ISL_3402535, EPI_ISL_3402538, EPI_ISL_3402539, EPI_ISL_3402540, EPI_ISL_3402541, EPI_ISL_3402543, EPI_ISL_3402544, EPI_ISL_3402545, EPI_ISL_3402546, EPI_ISL_3402547, EPI_ISL_3402548, EPI_ISL_3402550, EPI_ISL_3402551, EPI_ISL_3402552, EPI_ISL_3402554, EPI_ISL_3402555, EPI_ISL_3402556, EPI_ISL_3402557, EPI_ISL_3402558, EPI_ISL_3402559, EPI_ISL_3402560, EPI_ISL_3402561, EPI_ISL_3402562, EPI_ISL_3402563, EPI_ISL_3402564, EPI_ISL_3402565, EPI_ISL_3402566, EPI_ISL_3402567, EPI_ISL_3402568, EPI_ISL_3402569, EPI_ISL_3402570, EPI_ISL_3402571, EPI_ISL_3402572, EPI_ISL_3402573, EPI_ISL_3402574, EPI_ISL_3402576, EPI_ISL_3402577, EPI_ISL_3402578, EPI_ISL_3402579, EPI_ISL_3402580, EPI_ISL_3402581, EPI_ISL_3402583, EPI_ISL_3402584, EPI_ISL_3402585, EPI_ISL_3402587, EPI_ISL_3402588, EPI_ISL_3402589, EPI_ISL_3402590, EPI_ISL_3402591, EPI_ISL_3402592, EPI_ISL_3402593, EPI_ISL_3402594, EPI_ISL_3402595, EPI_ISL_3402596, EPI_ISL_3402597, EPI_ISL_3402598, EPI_ISL_3402599, EPI_ISL_3402600, EPI_ISL_3402601, EPI_ISL_3402602, EPI_ISL_3402603, EPI_ISL_3402608, EPI_ISL_3402609, EPI_ISL_3402610, EPI_ISL_3402611, EPI_ISL_3402612, EPI_ISL_3402616, EPI_ISL_3402621, EPI_ISL_3402622, EPI_ISL_3402623, EPI_ISL_3402625, EPI_ISL_3402626, EPI_ISL_3402627, EPI_ISL_3402628, EPI_ISL_3402629, EPI_ISL_3402630, EPI_ISL_3402631, EPI_ISL_3402632, EPI_ISL_3402633, EPI_ISL_3402634, EPI_ISL_3402635, EPI_ISL_3402636, EPI_ISL_3402637, EPI_ISL_3402638, EPI_ISL_3402639, EPI_ISL_3402640, EPI_ISL_3402641, EPI_ISL_3402642, EPI_ISL_3402643, EPI_ISL_3402644, EPI_ISL_3402645, EPI_ISL_3402646, EPI_ISL_3402647, EPI_ISL_3402648, EPI_ISL_3402649, EPI_ISL_3402650, EPI_ISL_3402651, EPI_ISL_3402652, EPI_ISL_3402653, EPI_ISL_3402654, EPI_ISL_3402655, EPI_ISL_3402656, EPI_ISL_3402657, EPI_ISL_3402658, EPI_ISL_3402659, EPI_ISL_3402660, EPI_ISL_3402662, EPI_ISL_3402663, EPI_ISL_3402664, EPI_ISL_3402665, EPI_ISL_3402666, EPI_ISL_3402667, EPI_ISL_3402668, EPI_ISL_3402669, EPI_ISL_3402670, EPI_ISL_3402671, EPI_ISL_3402673, EPI_ISL_3402674, EPI_ISL_3402675, EPI_ISL_3402677, EPI_ISL_3402678, EPI_ISL_3402679, EPI_ISL_3402680, EPI_ISL_3402681, EPI_ISL_3402682, EPI_ISL_3402683, EPI_ISL_3402684, EPI_ISL_3402685, EPI_ISL_3402686, EPI_ISL_3402687, EPI_ISL_3402688, EPI_ISL_3402689, EPI_ISL_3402690, EPI_ISL_3402691, EPI_ISL_3402692, EPI_ISL_3402693, EPI_ISL_3402694, EPI_ISL_3402695, EPI_ISL_3402696, EPI_ISL_3402697, EPI_ISL_3402698, EPI_ISL_3402699, EPI_ISL_3402700, EPI_ISL_3402701, EPI_ISL_3402702, EPI_ISL_3402703, EPI_ISL_3402704, EPI_ISL_3402705, EPI_ISL_3402706, EPI_ISL_3402707, EPI_ISL_3402708, EPI_ISL_3402709, EPI_ISL_3402710, EPI_ISL_3402711, EPI_ISL_3402712, EPI_ISL_3402714, EPI_ISL_3402715, EPI_ISL_3402716, EPI_ISL_3402717, EPI_ISL_3402718, EPI_ISL_3402719, EPI_ISL_3402720, EPI_ISL_3402721, EPI_ISL_3402722, EPI_ISL_3402723, EPI_ISL_3402724, EPI_ISL_3402725, EPI_ISL_3402726, EPI_ISL_3402727, EPI_ISL_3402728, EPI_ISL_3402729, EPI_ISL_3402730, EPI_ISL_3402731, EPI_ISL_3402732, EPI_ISL_3402733, EPI_ISL_3402734, EPI_ISL_3402735, EPI_ISL_3402736, EPI_ISL_3402737, EPI_ISL_3402738, EPI_ISL_3402739, EPI_ISL_3402740, EPI_ISL_3402741, EPI_ISL_3402742, EPI_ISL_3402743, EPI_ISL_3402744, EPI_ISL_3402745, EPI_ISL_3402746, EPI_ISL_3402747, EPI_ISL_3402748, EPI_ISL_3402749, EPI_ISL_3402750, EPI_ISL_3402751, EPI_ISL_3402752, EPI_ISL_3402753, EPI_ISL_3402754, EPI_ISL_3402755, EPI_ISL_3402756, EPI_ISL_3402757, EPI_ISL_3402758, EPI_ISL_3402759, EPI_ISL_3402760, EPI_ISL_3402761, EPI_ISL_3402762, EPI_ISL_3402763, EPI_ISL_3402764, EPI_ISL_3402765, EPI_ISL_3402766, EPI_ISL_3402767, EPI_ISL_3402768, EPI_ISL_3402769, EPI_ISL_3402770, EPI_ISL_3402771, EPI_ISL_3402772, EPI_ISL_3402773, EPI_ISL_3402774, EPI_ISL_3402775, EPI_ISL_3402776, EPI_ISL_3402777, EPI_ISL_3402778, EPI_ISL_3402779, EPI_ISL_3402780, EPI_ISL_3402781, EPI_ISL_3402782, EPI_ISL_3402783, EPI_ISL_3402784, EPI_ISL_3402785, EPI_ISL_3402786, EPI_ISL_3402787, EPI_ISL_3402788, EPI_ISL_3402789, EPI_ISL_3402790, EPI_ISL_3402791, EPI_ISL_3402792, EPI_ISL_3402793, EPI_ISL_3402794, EPI_ISL_3402795, EPI_ISL_3402796, EPI_ISL_3402797, EPI_ISL_3402798, EPI_ISL_3402799, EPI_ISL_3402800, EPI_ISL_3402801, EPI_ISL_3402802, EPI_ISL_3402803, EPI_ISL_3402804, EPI_ISL_3402805, EPI_ISL_3402806, EPI_ISL_3402807, EPI_ISL_3402808, EPI_ISL_3402809, EPI_ISL_3402810, EPI_ISL_3402811, EPI_ISL_3402812, EPI_ISL_3402813, EPI_ISL_3402814, EPI_ISL_3402815, EPI_ISL_3402816, EPI_ISL_3402817, EPI_ISL_3402818, EPI_ISL_3402819, EPI_ISL_3402820, EPI_ISL_3402821, EPI_ISL_3402822, EPI_ISL_3402823, EPI_ISL_3402824, EPI_ISL_3402825, EPI_ISL_3402826, EPI_ISL_3402827, EPI_ISL_3402828, EPI_ISL_3402829, EPI_ISL_3402830, EPI_ISL_3402831, EPI_ISL_3402833, EPI_ISL_3402834, EPI_ISL_3402835, EPI_ISL_3402836, EPI_ISL_3402837, EPI_ISL_3402838, EPI_ISL_3402840, EPI_ISL_3402841, EPI_ISL_3402842, EPI_ISL_3402843, EPI_ISL_3402844, EPI_ISL_3402845, EPI_ISL_3402846, EPI_ISL_3402847, EPI_ISL_3402848, EPI_ISL_3402849, EPI_ISL_3402850, EPI_ISL_3402851, EPI_ISL_3402852, EPI_ISL_3402853, EPI_ISL_3402854, EPI_ISL_3402855, EPI_ISL_3402856, EPI_ISL_3402857, EPI_ISL_3402858, EPI_ISL_3402859, EPI_ISL_3402860, EPI_ISL_3402861, EPI_ISL_3402862, EPI_ISL_3402863, EPI_ISL_3402864, EPI_ISL_3402865, EPI_ISL_3402866, EPI_ISL_3402867, EPI_ISL_3402868, EPI_ISL_3402869, EPI_ISL_3402870, EPI_ISL_3402871, EPI_ISL_3402872, EPI_ISL_3402873, EPI_ISL_3402874, EPI_ISL_3402875, EPI_ISL_3402876, EPI_ISL_3402877, EPI_ISL_3402878, EPI_ISL_3402879, EPI_ISL_3402880, EPI_ISL_3402881, EPI_ISL_3402882, EPI_ISL_3402883, EPI_ISL_3402884, EPI_ISL_3402885, EPI_ISL_3402887, EPI_ISL_3402888, EPI_ISL_3402889, EPI_ISL_3402890, EPI_ISL_3402891, EPI_ISL_3402893, EPI_ISL_3402894, EPI_ISL_3402895, EPI_ISL_3402896, EPI_ISL_3402897, EPI_ISL_3402898, EPI_ISL_3402899, EPI_ISL_3402900, EPI_ISL_3402901, EPI_ISL_3402902, EPI_ISL_3402903, EPI_ISL_3402904, EPI_ISL_3402905, EPI_ISL_3402906, EPI_ISL_3402907, EPI_ISL_3402908, EPI_ISL_3402909, EPI_ISL_3402910, EPI_ISL_3402911, EPI_ISL_3402912, EPI_ISL_3402913, EPI_ISL_3402914, EPI_ISL_3402915, EPI_ISL_3402916, EPI_ISL_3402917, EPI_ISL_3402918, EPI_ISL_3402919, EPI_ISL_3402920, EPI_ISL_3402921, EPI_ISL_3402922, EPI_ISL_3402923, EPI_ISL_3402924, EPI_ISL_3402925, EPI_ISL_3402926, EPI_ISL_3402927, EPI_ISL_3402928, EPI_ISL_3402929, EPI_ISL_3402930, EPI_ISL_3402931, EPI_ISL_3402932, EPI_ISL_3402933, EPI_ISL_3402934, EPI_ISL_3402935, EPI_ISL_3402936, EPI_ISL_3402937, EPI_ISL_3402938, EPI_ISL_3402939, EPI_ISL_3402940, EPI_ISL_3402941, EPI_ISL_3402942, EPI_ISL_3402943, EPI_ISL_3402944, EPI_ISL_3402945, EPI_ISL_3402946, EPI_ISL_3402947, EPI_ISL_3402948, EPI_ISL_3402949, EPI_ISL_3402950, EPI_ISL_3402951, EPI_ISL_3402952, EPI_ISL_3402953, EPI_ISL_3402954, EPI_ISL_3402955, EPI_ISL_3402956, EPI_ISL_3402957, EPI_ISL_3402958, EPI_ISL_3402959, EPI_ISL_3402960, EPI_ISL_3402961, EPI_ISL_3402962, EPI_ISL_3402963, EPI_ISL_3402964, EPI_ISL_3402965, EPI_ISL_3402966, EPI_ISL_3402967, EPI_ISL_3402968, EPI_ISL_3402969, EPI_ISL_3402970, EPI_ISL_3402971, EPI_ISL_3402972, EPI_ISL_3402973, EPI_ISL_3402974, EPI_ISL_3402975, EPI_ISL_3402976, EPI_ISL_3402977, EPI_ISL_3402978, EPI_ISL_3402979, EPI_ISL_3402980, EPI_ISL_3402981, EPI_ISL_3402982, EPI_ISL_3402983, EPI_ISL_3402984, EPI_ISL_3402985, EPI_ISL_3402986, EPI_ISL_3402987, EPI_ISL_3402988, EPI_ISL_3402989, EPI_ISL_3402990, EPI_ISL_3402991, EPI_ISL_3402992, EPI_ISL_3402993, EPI_ISL_3402994, EPI_ISL_3402995, EPI_ISL_3402996, EPI_ISL_3402997, EPI_ISL_3402998, EPI_ISL_3402999, EPI_ISL_3403000, EPI_ISL_3403001, EPI_ISL_3403002, EPI_ISL_3403003, EPI_ISL_3403004, EPI_ISL_3403005, EPI_ISL_3403006, EPI_ISL_3403007, EPI_ISL_3403008, EPI_ISL_3403009, EPI_ISL_3403010, EPI_ISL_3403011, EPI_ISL_3403012, EPI_ISL_3403013, EPI_ISL_3403014, EPI_ISL_3403015, EPI_ISL_3403016, EPI_ISL_3403017, EPI_ISL_3403018, EPI_ISL_3403019, EPI_ISL_3403020, EPI_ISL_3403021, EPI_ISL_3403022, EPI_ISL_3403023, EPI_ISL_3403024, EPI_ISL_3403025, EPI_ISL_3403026, EPI_ISL_3403027, EPI_ISL_3403028, EPI_ISL_3403029, EPI_ISL_3403030, EPI_ISL_3403031, EPI_ISL_3403032, EPI_ISL_3403033, EPI_ISL_3403034, EPI_ISL_3403035, EPI_ISL_3403036, EPI_ISL_3403037, EPI_ISL_3403038, EPI_ISL_3403039, EPI_ISL_3403040, EPI_ISL_3403041, EPI_ISL_3403042, EPI_ISL_3403043, EPI_ISL_3403044, EPI_ISL_3403045, EPI_ISL_3403046, EPI_ISL_3403047, EPI_ISL_3403048, EPI_ISL_3403049, EPI_ISL_3403050, EPI_ISL_3403051, EPI_ISL_3403052, EPI_ISL_3403053, EPI_ISL_3403054, EPI_ISL_3403055, EPI_ISL_3403056, EPI_ISL_3403057, EPI_ISL_3403058, EPI_ISL_3403059, EPI_ISL_3403060, EPI_ISL_3403061, EPI_ISL_3403062, EPI_ISL_3403063, EPI_ISL_3403064, EPI_ISL_3403065, EPI_ISL_3403066, EPI_ISL_3403067, EPI_ISL_3403068, EPI_ISL_3403069, EPI_ISL_3403070, EPI_ISL_3403071, EPI_ISL_3403072, EPI_ISL_3403073, EPI_ISL_3403074, EPI_ISL_3403075, EPI_ISL_3403076, EPI_ISL_3403077, EPI_ISL_3403079, EPI_ISL_3403080, EPI_ISL_3403081, EPI_ISL_3403082, EPI_ISL_3403083, EPI_ISL_3403084, EPI_ISL_3403086, EPI_ISL_3403087, EPI_ISL_3403088, EPI_ISL_3403089, EPI_ISL_3403090, EPI_ISL_3403091, EPI_ISL_3403092, EPI_ISL_3403093, EPI_ISL_3403094, EPI_ISL_3403095, EPI_ISL_3403096, EPI_ISL_3403097, EPI_ISL_3403098, EPI_ISL_3403099, EPI_ISL_3403100, EPI_ISL_3403101, EPI_ISL_3403102, EPI_ISL_3403103, EPI_ISL_3403104, EPI_ISL_3403105, EPI_ISL_3403106, EPI_ISL_3403107, EPI_ISL_3403108, EPI_ISL_3403109, EPI_ISL_3403110, EPI_ISL_3403111, EPI_ISL_3403112, EPI_ISL_3403113, EPI_ISL_3403114, EPI_ISL_3403115, EPI_ISL_3403116, EPI_ISL_3403117, EPI_ISL_3403118, EPI_ISL_3403119, EPI_ISL_3403120, EPI_ISL_3403121, EPI_ISL_3403122, EPI_ISL_3403123, EPI_ISL_3403124, EPI_ISL_3403125, EPI_ISL_3403126, EPI_ISL_3403127, EPI_ISL_3403128, EPI_ISL_3403129, EPI_ISL_3403130, EPI_ISL_3403131, EPI_ISL_3403133, EPI_ISL_3403134, EPI_ISL_3403135, EPI_ISL_3403136, EPI_ISL_3403137, EPI_ISL_3403139, EPI_ISL_3403140, EPI_ISL_3403141, EPI_ISL_3403142, EPI_ISL_3403143, EPI_ISL_3403144, EPI_ISL_3403145, EPI_ISL_3403146, EPI_ISL_3403147, EPI_ISL_3403148, EPI_ISL_3403149, EPI_ISL_3403150, EPI_ISL_3403151, EPI_ISL_3403152, EPI_ISL_3403153, EPI_ISL_3403154, EPI_ISL_3403155, EPI_ISL_3403156, EPI_ISL_3403157, EPI_ISL_3403158, EPI_ISL_3403159, EPI_ISL_3403160, EPI_ISL_3403161, EPI_ISL_3403162, EPI_ISL_3403163, EPI_ISL_3403164, EPI_ISL_3403165, EPI_ISL_3403166, EPI_ISL_3403167, EPI_ISL_3403168, EPI_ISL_3403169, EPI_ISL_3403170, EPI_ISL_3403171, EPI_ISL_3403172, EPI_ISL_3403173, EPI_ISL_3403174, EPI_ISL_3403175, EPI_ISL_3403176, EPI_ISL_3403177, EPI_ISL_3403178, EPI_ISL_3403179, EPI_ISL_3403180, EPI_ISL_3403181, EPI_ISL_3403182, EPI_ISL_3403183, EPI_ISL_3403184, EPI_ISL_3403185, EPI_ISL_3403186, EPI_ISL_3403187, EPI_ISL_3403188, EPI_ISL_3403189, EPI_ISL_3403190, EPI_ISL_3403191, EPI_ISL_3403192, EPI_ISL_3403193, EPI_ISL_3403194, EPI_ISL_3403195, EPI_ISL_3403196, EPI_ISL_3403197, EPI_ISL_3403198, EPI_ISL_3403199, EPI_ISL_3403200, EPI_ISL_3403201, EPI_ISL_3403202, EPI_ISL_3403203, EPI_ISL_3403204, EPI_ISL_3403205, EPI_ISL_3403206, EPI_ISL_3403207, EPI_ISL_3403208, EPI_ISL_3403209, EPI_ISL_3403210, EPI_ISL_3403211, EPI_ISL_3403212, EPI_ISL_3403213, EPI_ISL_3403214, EPI_ISL_3403215, EPI_ISL_3403216, EPI_ISL_3403217, EPI_ISL_3403218, EPI_ISL_3403219, EPI_ISL_3403221, EPI_ISL_3403228, EPI_ISL_3403229, EPI_ISL_3403232, EPI_ISL_3403234, EPI_ISL_3403237, EPI_ISL_3403238, EPI_ISL_3403239, EPI_ISL_3403264, EPI_ISL_3403273, EPI_ISL_3403275, EPI_ISL_3403277, EPI_ISL_3403279, EPI_ISL_3403280, EPI_ISL_3403282, EPI_ISL_3403289, EPI_ISL_3403290, EPI_ISL_3403293, EPI_ISL_3403295, EPI_ISL_3403298, EPI_ISL_3403299, EPI_ISL_3403300, EPI_ISL_3403325, EPI_ISL_3403334, EPI_ISL_3403336, EPI_ISL_3403338, EPI_ISL_3403340, EPI_ISL_3403341 | | | |
| see above | Ministry of Health Turkey | Ministry of Health Turkey | Fatma Bayrakdar, Yasemin Cosgun, Suleyman Yalcin, Gulay Korukluoglu |
| EPI_ISL_3403342 | CH VALENCIENNES | CHU Lille - Laboratoire de Virologie | AIT YAHYA Emilie, ALIDJINOU Enagnon Kazali, BOCKET Laurence, CREPIN Michel, DEMAY Christophe, ENGELMANN Ilka, GEFFROY Sandrine, GUIGON Aurélie, LAMBERT Valérie, LAZREK Mouna, NOBILLIAUX Florian, PREVOST Brigitte, THUILLIER Caroline, TINEZ Claire |
| EPI_ISL_3403343 | CH DUNKERQUE | CHU Lille - Laboratoire de Virologie | AIT YAHYA Emilie, ALIDJINOU Enagnon Kazali, BOCKET Laurence, CREPIN Michel, DEMAY Christophe, ENGELMANN Ilka, GEFFROY Sandrine, GUIGON Aurélie, LAMBERT Valérie, LAZREK Mouna, NOBILLIAUX Florian, PREVOST Brigitte, THUILLIER Caroline, TINEZ Claire |
| EPI_ISL_3403345 | National Institute of Health, Department of Medical Sciences, Ministry of Public Health, Thailand | National Institute of Health, Department of Medical Sciences, Ministry of Public Health, Thailand | Pilailuk Okada; Siripaporn Phuygun; Nuttida Thongpramul; Sittiporn Parnmen; Ratana Tacharoenmuang; Pakorn Piromtong; Natchaya Khiadsang; Thanutsapa Thanadachakul; Warawan Wongboot; sirikanda wimol; Suntharееya Waicharoen; |
| EPI_ISL_3403346, EPI_ISL_3403347, EPI_ISL_3403348, EPI_ISL_3403349 | Nonthaburi Public Health Office | National Institute of Health, Department of Medical Sciences, Ministry of Public Health, Thailand | Pilailuk Okada; Siripaporn Phuygun; Nuttida Thongpramul; Sittiporn Parnmen; Ratana Tacharoenmuang; Pakorn Piromtong; Natchaya Khiadsang; Thanutsapa Thanadachakul; Warawan Wongboot; sirikanda wimol; Sunthareeya Waicharoen; |
| EPI_ISL_3403356, EPI_ISL_3403357 | CHU LILLE | CHU Lille - Laboratoire de Virologie | AIT YAHYA Emilie, ALIDJINOU Enagnon Kazali, BOCKET Laurence, CREPIN Michel, DEMAY Christophe, ENGELMANN Ilka, GEFFROY Sandrine, GUIGON Aurélie, LAMBERT Valérie, LAZREK Mouna, NOBILLIAUX Florian, PREVOST Brigitte, THUILLIER Caroline, TINEZ Claire ,TCHANTCHOU NJOSSE YANICK |
| EPI_ISL_3403358 | CH TOURCOING | CHU Lille - Laboratoire de Virologie | AIT YAHYA Emilie, ALIDJINOU Enagnon Kazali, BOCKET Laurence, CREPIN Michel, DEMAY Christophe, ENGELMANN Ilka, GEFFROY Sandrine, GUIGON Aurélie, LAMBERT Valérie, LAZREK Mouna, NOBILLIAUX Florian, PREVOST Brigitte, THUILLIER Caroline, TINEZ Claire ,TCHANTCHOU NJOSSE YANICK |
| EPI_ISL_3403359 | CH ROUBAIX | CHU Lille - Laboratoire de Virologie | AIT YAHYA Emilie, ALIDJINOU Enagnon Kazali, BOCKET Laurence, CREPIN Michel, DEMAY Christophe, ENGELMANN Ilka, GEFFROY Sandrine, GUIGON Aurélie, LAMBERT Valérie, LAZREK Mouna, NOBILLIAUX Florian, PREVOST Brigitte, THUILLIER Caroline, TINEZ Claire ,TCHANTCHOU NJOSSE YANICK |
| EPI_ISL_3403360 | CH BOULOGNE-SUR-MER | CHU Lille - Laboratoire de Virologie | AIT YAHYA Emilie, ALIDJINOU Enagnon Kazali, BOCKET Laurence, CREPIN Michel, DEMAY Christophe, ENGELMANN Ilka, GEFFROY Sandrine, GUIGON Aurélie, LAMBERT Valérie, LAZREK Mouna, NOBILLIAUX Florian, PREVOST Brigitte, THUILLIER Caroline, TINEZ Claire ,TCHANTCHOU NJOSSE YANICK |
| EPI_ISL_3403362 | CH BETHUNE | CHU Lille - Laboratoire de Virologie | AIT YAHYA Emilie, ALIDJINOU Enagnon Kazali, BOCKET Laurence, CREPIN Michel, DEMAY Christophe, ENGELMANN Ilka, GEFFROY Sandrine, GUIGON Aurélie, LAMBERT Valérie, LAZREK Mouna, NOBILLIAUX Florian, PREVOST Brigitte, THUILLIER Caroline, TINEZ Claire ,TCHANTCHOU NJOSSE YANICK |
| EPI_ISL_3403363 | CH VALENCIENNES | CHU Lille - Laboratoire de Virologie | AIT YAHYA Emilie, ALIDJINOU Enagnon Kazali, BOCKET Laurence, CREPIN Michel, DEMAY Christophe, ENGELMANN Ilka, GEFFROY Sandrine, GUIGON Aurélie, LAMBERT Valérie, LAZREK Mouna, NOBILLIAUX Florian, PREVOST Brigitte, THUILLIER Caroline, TINEZ Claire ,TCHANTCHOU NJOSSE YANICK |
| EPI_ISL_3403364 | CH ARMENTIERES | CHU Lille - Laboratoire de Virologie | AIT YAHYA Emilie, ALIDJINOU Enagnon Kazali, BOCKET Laurence, CREPIN Michel, DEMAY Christophe, ENGELMANN Ilka, GEFFROY Sandrine, GUIGON Aurélie, LAMBERT Valérie, LAZREK Mouna, NOBILLIAUX Florian, PREVOST Brigitte, THUILLIER Caroline, TINEZ Claire ,TCHANTCHOU NJOSSE YANICK |
| EPI_ISL_3403365 | CHU LILLE | CHU Lille - Laboratoire de Virologie | AIT YAHYA Emilie, ALIDJINOU Enagnon Kazali, BOCKET Laurence, CREPIN Michel, DEMAY Christophe, ENGELMANN Ilka, GEFFROY Sandrine, GUIGON Aurélie, LAMBERT Valérie, LAZREK Mouna, NOBILLIAUX Florian, PREVOST Brigitte, THUILLIER Caroline, TINEZ Claire ,TCHANTCHOU NJOSSE YANICK |
| EPI_ISL_3403366 | CH VALENCIENNES | CHU Lille - Laboratoire de Virologie | AIT YAHYA Emilie, ALIDJINOU Enagnon Kazali, BOCKET Laurence, CREPIN Michel, DEMAY Christophe, ENGELMANN Ilka, GEFFROY Sandrine, GUIGON Aurélie, LAMBERT Valérie, LAZREK Mouna, NOBILLIAUX Florian, PREVOST Brigitte, THUILLIER Caroline, TINEZ Claire ,TCHANTCHOU NJOSSE YANICK |
| EPI_ISL_3403367 | CH SAINT-PHILIBERT | CHU Lille - Laboratoire de Virologie | AIT YAHYA Emilie, ALIDJINOU Enagnon Kazali, BOCKET Laurence, CREPIN Michel, DEMAY Christophe, ENGELMANN Ilka, GEFFROY Sandrine, GUIGON Aurélie, LAMBERT Valérie, LAZREK Mouna, NOBILLIAUX Florian, PREVOST Brigitte, THUILLIER Caroline, TINEZ Claire ,TCHANTCHOU NJOSSE YANICK |
| EPI_ISL_3403368 | CHU LILLE | CHU Lille - Laboratoire de Virologie | AIT YAHYA Emilie, ALIDJINOU Enagnon Kazali, BOCKET Laurence, CREPIN Michel, DEMAY Christophe, ENGELMANN Ilka, GEFFROY Sandrine, GUIGON Aurélie, LAMBERT Valérie, LAZREK Mouna, NOBILLIAUX Florian, PREVOST Brigitte, THUILLIER Caroline, TINEZ Claire ,TCHANTCHOU NJOSSE YANICK |
| EPI_ISL_3403369 | CH BETHUNE | CHU Lille - Laboratoire de Virologie | AIT YAHYA Emilie, ALIDJINOU Enagnon Kazali, BOCKET Laurence, CREPIN Michel, DEMAY Christophe, ENGELMANN Ilka, GEFFROY Sandrine, GUIGON Aurélie, LAMBERT Valérie, LAZREK Mouna, NOBILLIAUX Florian, PREVOST Brigitte, THUILLIER Caroline, TINEZ Claire ,TCHANTCHOU NJOSSE YANICK |
| EPI_ISL_3403370 | CH BOULOGNE-SUR-MER | CHU Lille - Laboratoire de Virologie | AIT YAHYA Emilie, ALIDJINOU Enagnon Kazali, BOCKET Laurence, CREPIN Michel, DEMAY Christophe, ENGELMANN Ilka, GEFFROY Sandrine, GUIGON Aurélie, LAMBERT Valérie, LAZREK Mouna, NOBILLIAUX Florian, PREVOST Brigitte, THUILLIER Caroline, TINEZ Claire ,TCHANTCHOU NJOSSE YANICK |
| EPI_ISL_3403371, EPI_ISL_3403372, EPI_ISL_3403373 | CH VALENCIENNES | CHU Lille - Laboratoire de Virologie | AIT YAHYA Emilie, ALIDJINOU Enagnon Kazali, BOCKET Laurence, CREPIN Michel, DEMAY Christophe, ENGELMANN Ilka, GEFFROY Sandrine, GUIGON Aurélie, LAMBERT Valérie, LAZREK Mouna, NOBILLIAUX Florian, PREVOST Brigitte, THUILLIER Caroline, TINEZ Claire ,TCHANTCHOU NJOSSE YANICK |
| EPI_ISL_3403374 | CHU LILLE | CHU Lille - Laboratoire de Virologie | AIT YAHYA Emilie, ALIDJINOU Enagnon Kazali, BOCKET Laurence, CREPIN Michel, DEMAY Christophe, ENGELMANN Ilka, GEFFROY Sandrine, GUIGON Aurélie, LAMBERT Valérie, LAZREK Mouna, NOBILLIAUX Florian, PREVOST Brigitte, THUILLIER Caroline, TINEZ Claire ,TCHANTCHOU NJOSSE YANICK |
| EPI_ISL_3403375, EPI_ISL_3403376, EPI_ISL_3403377, EPI_ISL_3403378, EPI_ISL_3403379 | CH VALENCIENNES | CHU Lille - Laboratoire de Virologie | AIT YAHYA Emilie, ALIDJINOU Enagnon Kazali, BOCKET Laurence, CREPIN Michel, DEMAY Christophe, ENGELMANN Ilka, GEFFROY Sandrine, GUIGON Aurélie, LAMBERT Valérie, LAZREK Mouna, NOBILLIAUX Florian, PREVOST Brigitte, THUILLIER Caroline, TINEZ Claire ,TCHANTCHOU NJOSSE YANICK |
| EPI_ISL_3403380, EPI_ISL_3403382, EPI_ISL_3403383 | CHU LILLE | CHU Lille - Laboratoire de Virologie | AIT YAHYA Emilie, ALIDJINOU Enagnon Kazali, BOCKET Laurence, CREPIN Michel, DEMAY Christophe, ENGELMANN Ilka, GEFFROY Sandrine, GUIGON Aurélie, LAMBERT Valérie, LAZREK Mouna, NOBILLIAUX Florian, PREVOST Brigitte, THUILLIER Caroline, TINEZ Claire ,TCHANTCHOU NJOSSE YANICK |

| | | | |
|---|---|---|---|
| EPI_ISL_3403384 | CH DUNKERQUE | CHU Lille - Laboratoire de Virologie | AIT YAHYA Emilie, ALIDJINOU Enagnon Kazali, BOCKET Laurence, CREPIN Michel, DEMAY Christophe, ENGELMANN Ilka, GEFFROY Sandrine, GUIGON Aurélie, LAMBERT Valérie, LAZREK Mouna, NOBILLIAUX Florian, PREVOST Brigitte, THUILLIER Caroline, TINEZ Claire ,TCHANTCHOU NJOSSE YANICK |
| EPI_ISL_3403385, EPI_ISL_3403386, EPI_ISL_3403387 | CHU LILLE | CHU Lille - Laboratoire de Virologie | AIT YAHYA Emilie, ALIDJINOU Enagnon Kazali, BOCKET Laurence, CREPIN Michel, DEMAY Christophe, ENGELMANN Ilka, GEFFROY Sandrine, GUIGON Aurélie, LAMBERT Valérie, LAZREK Mouna, NOBILLIAUX Florian, PREVOST Brigitte, THUILLIER Caroline, TINEZ Claire ,TCHANTCHOU NJOSSE YANICK |
| EPI_ISL_3403388 | CH VALENCIENNES | CHU Lille - Laboratoire de Virologie | AIT YAHYA Emilie, ALIDJINOU Enagnon Kazali, BOCKET Laurence, CREPIN Michel, DEMAY Christophe, ENGELMANN Ilka, GEFFROY Sandrine, GUIGON Aurélie, LAMBERT Valérie, LAZREK Mouna, NOBILLIAUX Florian, PREVOST Brigitte, THUILLIER Caroline, TINEZ Claire ,TCHANTCHOU NJOSSE YANICK |
| EPI_ISL_3403389, EPI_ISL_3403390, EPI_ISL_3403391 | CHU LILLE | CHU Lille - Laboratoire de Virologie | AIT YAHYA Emilie, ALIDJINOU Enagnon Kazali, BOCKET Laurence, CREPIN Michel, DEMAY Christophe, ENGELMANN Ilka, GEFFROY Sandrine, GUIGON Aurélie, LAMBERT Valérie, LAZREK Mouna, NOBILLIAUX Florian, PREVOST Brigitte, THUILLIER Caroline, TINEZ Claire ,TCHANTCHOU NJOSSE YANICK |
| EPI_ISL_3403392, EPI_ISL_3403393 | CH ROUBAIX | CHU Lille - Laboratoire de Virologie | AIT YAHYA Emilie, ALIDJINOU Enagnon Kazali, BOCKET Laurence, CREPIN Michel, DEMAY Christophe, ENGELMANN Ilka, GEFFROY Sandrine, GUIGON Aurélie, LAMBERT Valérie, LAZREK Mouna, NOBILLIAUX Florian, PREVOST Brigitte, THUILLIER Caroline, TINEZ Claire ,TCHANTCHOU NJOSSE YANICK |
| EPI_ISL_3403394 | CHU LILLE | CHU Lille - Laboratoire de Virologie | AIT YAHYA Emilie, ALIDJINOU Enagnon Kazali, BOCKET Laurence, CREPIN Michel, DEMAY Christophe, ENGELMANN Ilka, GEFFROY Sandrine, GUIGON Aurélie, LAMBERT Valérie, LAZREK Mouna, NOBILLIAUX Florian, PREVOST Brigitte, THUILLIER Caroline, TINEZ Claire ,TCHANTCHOU NJOSSE YANICK |
| EPI_ISL_3403395, EPI_ISL_3403396, EPI_ISL_3403397 | CH ROUBAIX | CHU Lille - Laboratoire de Virologie | AIT YAHYA Emilie, ALIDJINOU Enagnon Kazali, BOCKET Laurence, CREPIN Michel, DEMAY Christophe, ENGELMANN Ilka, GEFFROY Sandrine, GUIGON Aurélie, LAMBERT Valérie, LAZREK Mouna, NOBILLIAUX Florian, PREVOST Brigitte, THUILLIER Caroline, TINEZ Claire ,TCHANTCHOU NJOSSE YANICK |
| EPI_ISL_3403398, EPI_ISL_3403399, EPI_ISL_3403400 | CH LENS | CHU Lille - Laboratoire de Virologie | AIT YAHYA Emilie, ALIDJINOU Enagnon Kazali, BOCKET Laurence, CREPIN Michel, DEMAY Christophe, ENGELMANN Ilka, GEFFROY Sandrine, GUIGON Aurélie, LAMBERT Valérie, LAZREK Mouna, NOBILLIAUX Florian, PREVOST Brigitte, THUILLIER Caroline, TINEZ Claire ,TCHANTCHOU NJOSSE YANICK |
| EPI_ISL_3403401, EPI_ISL_3403402, EPI_ISL_3403403, EPI_ISL_3403404 | CH TOURCOING | CHU Lille - Laboratoire de Virologie | AIT YAHYA Emilie, ALIDJINOU Enagnon Kazali, BOCKET Laurence, CREPIN Michel, DEMAY Christophe, ENGELMANN Ilka, GEFFROY Sandrine, GUIGON Aurélie, LAMBERT Valérie, LAZREK Mouna, NOBILLIAUX Florian, PREVOST Brigitte, THUILLIER Caroline, TINEZ Claire ,TCHANTCHOU NJOSSE YANICK |
| EPI_ISL_3403405 | CH BETHUNE | CHU Lille - Laboratoire de Virologie | AIT YAHYA Emilie, ALIDJINOU Enagnon Kazali, BOCKET Laurence, CREPIN Michel, DEMAY Christophe, ENGELMANN Ilka, GEFFROY Sandrine, GUIGON Aurélie, LAMBERT Valérie, LAZREK Mouna, NOBILLIAUX Florian, PREVOST Brigitte, THUILLIER Caroline, TINEZ Claire ,TCHANTCHOU NJOSSE YANICK |
| EPI_ISL_3403406 | CH ARMENTIERES | CHU Lille - Laboratoire de Virologie | AIT YAHYA Emilie, ALIDJINOU Enagnon Kazali, BOCKET Laurence, CREPIN Michel, DEMAY Christophe, ENGELMANN Ilka, GEFFROY Sandrine, GUIGON Aurélie, LAMBERT Valérie, LAZREK Mouna, NOBILLIAUX Florian, PREVOST Brigitte, THUILLIER Caroline, TINEZ Claire ,TCHANTCHOU NJOSSE YANICK |
| EPI_ISL_3403407, EPI_ISL_3403408 | CH BETHUNE | CHU Lille - Laboratoire de Virologie | AIT YAHYA Emilie, ALIDJINOU Enagnon Kazali, BOCKET Laurence, CREPIN Michel, DEMAY Christophe, ENGELMANN Ilka, GEFFROY Sandrine, GUIGON Aurélie, LAMBERT Valérie, LAZREK Mouna, NOBILLIAUX Florian, PREVOST Brigitte, THUILLIER Caroline, TINEZ Claire ,TCHANTCHOU NJOSSE YANICK |
| EPI_ISL_3403409 | CH BOULOGNE-SUR-MER | CHU Lille - Laboratoire de Virologie | AIT YAHYA Emilie, ALIDJINOU Enagnon Kazali, BOCKET Laurence, CREPIN Michel, DEMAY Christophe, ENGELMANN Ilka, GEFFROY Sandrine, GUIGON Aurélie, LAMBERT Valérie, LAZREK Mouna, NOBILLIAUX Florian, PREVOST Brigitte, THUILLIER Caroline, TINEZ Claire ,TCHANTCHOU NJOSSE YANICK |
| EPI_ISL_3403410, EPI_ISL_3403411, EPI_ISL_3403413, EPI_ISL_3403414, EPI_ISL_3403415, EPI_ISL_3403417, EPI_ISL_3403418, EPI_ISL_3403419 | CH VALENCIENNES | CHU Lille - Laboratoire de Virologie | AIT YAHYA Emilie, ALIDJINOU Enagnon Kazali, BOCKET Laurence, CREPIN Michel, DEMAY Christophe, ENGELMANN Ilka, GEFFROY Sandrine, GUIGON Aurélie, LAMBERT Valérie, LAZREK Mouna, NOBILLIAUX Florian, PREVOST Brigitte, THUILLIER Caroline, TINEZ Claire ,TCHANTCHOU NJOSSE YANICK |
| EPI_ISL_3403420 | CHU LILLE | CHU Lille - Laboratoire de Virologie | AIT YAHYA Emilie, ALIDJINOU Enagnon Kazali, BOCKET Laurence, CREPIN Michel, DEMAY Christophe, ENGELMANN Ilka, GEFFROY Sandrine, GUIGON Aurélie, LAMBERT Valérie, LAZREK Mouna, NOBILLIAUX Florian, PREVOST Brigitte, THUILLIER Caroline, TINEZ Claire ,TCHANTCHOU NJOSSE YANICK |
| EPI_ISL_3403421, EPI_ISL_3403422, EPI_ISL_3403423, EPI_ISL_3403424, EPI_ISL_3403425, EPI_ISL_3403426, EPI_ISL_3403427, EPI_ISL_3403428, EPI_ISL_3403429, EPI_ISL_3403430, EPI_ISL_3403431, EPI_ISL_3403432, EPI_ISL_3403433, EPI_ISL_3403434 | | | |
| see above | CH VALENCIENNES | CHU Lille - Laboratoire de Virologie | AIT YAHYA Emilie, ALIDJINOU Enagnon Kazali, BOCKET Laurence, CREPIN Michel, DEMAY Christophe, ENGELMANN Ilka, GEFFROY Sandrine, GUIGON Aurélie, LAMBERT Valérie, LAZREK Mouna, NOBILLIAUX Florian, PREVOST Brigitte, THUILLIER Caroline, TINEZ Claire ,TCHANTCHOU NJOSSE YANICK |
| EPI_ISL_3403435, EPI_ISL_3403436, EPI_ISL_3403437, EPI_ISL_3403438, EPI_ISL_3403439, EPI_ISL_3403440, EPI_ISL_3403441, EPI_ISL_3403442 | CHU LILLE | CHU Lille - Laboratoire de Virologie | AIT YAHYA Emilie, ALIDJINOU Enagnon Kazali, BOCKET Laurence, CREPIN Michel, DEMAY Christophe, ENGELMANN Ilka, GEFFROY Sandrine, GUIGON Aurélie, LAMBERT Valérie, LAZREK Mouna, NOBILLIAUX Florian, PREVOST Brigitte, THUILLIER Caroline, TINEZ Claire ,TCHANTCHOU NJOSSE YANICK |
| EPI_ISL_3403444 | Sweden/Sodermanland/Unilabs | Unilabs/Eskilstuna/Sweden | Emma Arvidsson |
| EPI_ISL_3403445 | Sweden/Vastragotaland/Unilabs | Unilabs/Eskilstuna/Sweden | Emma Arvidsson |
| EPI_ISL_3403446, EPI_ISL_3403447, EPI_ISL_3403448, EPI_ISL_3403449, EPI_ISL_3403450, EPI_ISL_3403451, EPI_ISL_3403452, EPI_ISL_3403453, EPI_ISL_3403454, EPI_ISL_3403455, EPI_ISL_3403458, EPI_ISL_3403459, EPI_ISL_3403460, EPI_ISL_3403461, EPI_ISL_3403462, EPI_ISL_3403463, EPI_ISL_3403464, EPI_ISL_3403465, EPI_ISL_3403466, EPI_ISL_3403468, EPI_ISL_3403469, EPI_ISL_3403470, EPI_ISL_3403471, EPI_ISL_3403472, EPI_ISL_3403473, EPI_ISL_3403474, EPI_ISL_3403476, EPI_ISL_3403478, EPI_ISL_3403479, EPI_ISL_3403480, EPI_ISL_3403481, EPI_ISL_3403482, EPI_ISL_3403483, EPI_ISL_3403484, EPI_ISL_3403485, EPI_ISL_3403486, EPI_ISL_3403487, EPI_ISL_3403488, EPI_ISL_3403489, EPI_ISL_3403491, EPI_ISL_3403493, EPI_ISL_3403494, EPI_ISL_3403495, EPI_ISL_3403496, EPI_ISL_3403497, EPI_ISL_3403499, EPI_ISL_3403500, EPI_ISL_3403501, EPI_ISL_3403502, EPI_ISL_3403503 | | | |
| see above | Sweden/Sodermanland/Unilabs | Unilabs/Eskilstuna/Sweden | Emma Arvidsson |
| EPI_ISL_3403504, EPI_ISL_3403507, EPI_ISL_3403508, EPI_ISL_3403509, EPI_ISL_3403510, EPI_ISL_3403511, EPI_ISL_3403512, EPI_ISL_3403513, EPI_ISL_3403514, EPI_ISL_3403515, EPI_ISL_3403516, EPI_ISL_3403517, EPI_ISL_3403518, EPI_ISL_3403519, EPI_ISL_3403520, EPI_ISL_3403521, EPI_ISL_3403522, EPI_ISL_3403523, EPI_ISL_3403524, EPI_ISL_3403525, EPI_ISL_3403526, EPI_ISL_3403527, EPI_ISL_3403528, EPI_ISL_3403529, EPI_ISL_3403530, EPI_ISL_3403531, EPI_ISL_3403532, EPI_ISL_3403533, EPI_ISL_3403534, EPI_ISL_3403535, EPI_ISL_3403536, EPI_ISL_3403537, EPI_ISL_3403538, EPI_ISL_3403539, EPI_ISL_3403540, EPI_ISL_3403541, EPI_ISL_3403542, EPI_ISL_3403543, EPI_ISL_3403544, EPI_ISL_3403545, EPI_ISL_3403546, EPI_ISL_3403547, EPI_ISL_3403548, EPI_ISL_3403549, EPI_ISL_3403550, EPI_ISL_3403551, EPI_ISL_3403552, EPI_ISL_3403553, EPI_ISL_3403554, EPI_ISL_3403555, EPI_ISL_3403556, EPI_ISL_3403557, EPI_ISL_3403558, EPI_ISL_3403559, EPI_ISL_3403560, EPI_ISL_3403561, EPI_ISL_3403562, EPI_ISL_3403563, EPI_ISL_3403564, EPI_ISL_3403565, EPI_ISL_3403566, EPI_ISL_3403567, EPI_ISL_3403568, EPI_ISL_3403569, EPI_ISL_3403571, EPI_ISL_3403572, EPI_ISL_3403573, EPI_ISL_3403574, EPI_ISL_3403575, EPI_ISL_3403576, EPI_ISL_3403577, EPI_ISL_3403578, EPI_ISL_3403579, EPI_ISL_3403580, EPI_ISL_3403581, EPI_ISL_3403582, EPI_ISL_3403583, EPI_ISL_3403584, EPI_ISL_3403585, EPI_ISL_3403586, EPI_ISL_3403587, EPI_ISL_3403588, EPI_ISL_3403589, EPI_ISL_3403590, EPI_ISL_3403591, EPI_ISL_3403592, EPI_ISL_3403593, EPI_ISL_3403594, EPI_ISL_3403595, EPI_ISL_3403596, EPI_ISL_3403597, EPI_ISL_3403598, EPI_ISL_3403599, EPI_ISL_3403601, EPI_ISL_3403602, EPI_ISL_3403603, EPI_ISL_3403604, EPI_ISL_3403605, EPI_ISL_3403606, EPI_ISL_3403607, EPI_ISL_3403608, EPI_ISL_3403609, EPI_ISL_3403611, EPI_ISL_3403612, EPI_ISL_3403613, EPI_ISL_3403614, EPI_ISL_3403615, EPI_ISL_3403617, EPI_ISL_3403618, EPI_ISL_3403619, EPI_ISL_3403620, EPI_ISL_3403622, EPI_ISL_3403623, EPI_ISL_3403624, EPI_ISL_3403625, EPI_ISL_3403626, EPI_ISL_3403627, EPI_ISL_3403628, EPI_ISL_3403629, EPI_ISL_3403630, EPI_ISL_3403631, EPI_ISL_3403632, EPI_ISL_3403633, EPI_ISL_3403634, EPI_ISL_3403635, EPI_ISL_3403636, EPI_ISL_3403637, EPI_ISL_3403638, EPI_ISL_3403639, EPI_ISL_3403640, EPI_ISL_3403641, EPI_ISL_3403642, EPI_ISL_3403643, EPI_ISL_3403644, EPI_ISL_3403645, EPI_ISL_3403646, EPI_ISL_3403647, EPI_ISL_3403648, EPI_ISL_3403649, EPI_ISL_3403650, EPI_ISL_3403651, EPI_ISL_3403652, EPI_ISL_3403653, EPI_ISL_3403654, EPI_ISL_3403655, EPI_ISL_3403656, EPI_ISL_3403657, EPI_ISL_3403658, EPI_ISL_3403659, EPI_ISL_3403660, EPI_ISL_3403661, EPI_ISL_3403662, EPI_ISL_3403663, EPI_ISL_3403664, EPI_ISL_3403665, EPI_ISL_3403666, EPI_ISL_3403667, EPI_ISL_3403668, EPI_ISL_3403669, EPI_ISL_3403670, EPI_ISL_3403671, EPI_ISL_3403672, EPI_ISL_3403674, EPI_ISL_3403675, EPI_ISL_3403676, EPI_ISL_3403677, EPI_ISL_3403678, EPI_ISL_3403679, EPI_ISL_3403680, EPI_ISL_3403681, EPI_ISL_3403682, EPI_ISL_3403683, EPI_ISL_3403684, EPI_ISL_3403685, EPI_ISL_3403686, EPI_ISL_3403687, EPI_ISL_3403688, EPI_ISL_3403689, EPI_ISL_3403690, EPI_ISL_3403691, EPI_ISL_3403692, EPI_ISL_3403693, EPI_ISL_3403694, EPI_ISL_3403695, EPI_ISL_3403696, EPI_ISL_3403697, EPI_ISL_3403698, EPI_ISL_3403699, EPI_ISL_3403700, EPI_ISL_3403701, EPI_ISL_3403704, EPI_ISL_3403706, EPI_ISL_3403707, EPI_ISL_3403708, EPI_ISL_3403709, EPI_ISL_3403710, EPI_ISL_3403712, EPI_ISL_3403714, EPI_ISL_3403715, EPI_ISL_3403716, EPI_ISL_3403717, EPI_ISL_3403719, EPI_ISL_3403720, EPI_ISL_3403722, EPI_ISL_3403723, EPI_ISL_3403724, EPI_ISL_3403725, EPI_ISL_3403726, EPI_ISL_3403727, EPI_ISL_3403728, EPI_ISL_3403729, EPI_ISL_3403730, EPI_ISL_3403731, EPI_ISL_3403732, EPI_ISL_3403733, EPI_ISL_3403734, EPI_ISL_3403735, EPI_ISL_3403736, EPI_ISL_3403738, EPI_ISL_3403739, EPI_ISL_3403740, EPI_ISL_3403741, EPI_ISL_3403742, EPI_ISL_3403743, EPI_ISL_3403744, EPI_ISL_3403745, EPI_ISL_3403746, EPI_ISL_3403749 | | | |
| see above | Sweden/Vastragotaland/Unilabs | Unilabs/Eskilstuna/Sweden | Emma Arvidsson |
| EPI_ISL_3403789, EPI_ISL_3403790, EPI_ISL_3403791, EPI_ISL_3403792, EPI_ISL_3403793, EPI_ISL_3403794, EPI_ISL_3403795, EPI_ISL_3403796, EPI_ISL_3403797, EPI_ISL_3403798, EPI_ISL_3403799, EPI_ISL_3403800, EPI_ISL_3403801, EPI_ISL_3403802, EPI_ISL_3403803, EPI_ISL_3403804, EPI_ISL_3403864, EPI_ISL_3403865, EPI_ISL_3403866, EPI_ISL_3403867, EPI_ISL_3403868, EPI_ISL_3403869, EPI_ISL_3403870, EPI_ISL_3403871, EPI_ISL_3403872, EPI_ISL_3403873, EPI_ISL_3403874, EPI_ISL_3403875, EPI_ISL_3403876, EPI_ISL_3403877, EPI_ISL_3403878, EPI_ISL_3403879, EPI_ISL_3403906, EPI_ISL_3403907, EPI_ISL_3403908, EPI_ISL_3403909, EPI_ISL_3403910, EPI_ISL_3403911, EPI_ISL_3403912, EPI_ISL_3403913, EPI_ISL_3403914, EPI_ISL_3403915, EPI_ISL_3403916, EPI_ISL_3403917, EPI_ISL_3403918, EPI_ISL_3403919, EPI_ISL_3403920, EPI_ISL_3403921, EPI_ISL_3403922, EPI_ISL_3403923, EPI_ISL_3403924, EPI_ISL_3403925, EPI_ISL_3403926, EPI_ISL_3403927, EPI_ISL_3403928, EPI_ISL_3403929, EPI_ISL_3403930, EPI_ISL_3403931, EPI_ISL_3403932, EPI_ISL_3403933, EPI_ISL_3403934, EPI_ISL_3403935, EPI_ISL_3403936, EPI_ISL_3403937, EPI_ISL_3403938, EPI_ISL_3403939, EPI_ISL_3403940, EPI_ISL_3403941, EPI_ISL_3403942, EPI_ISL_3403943, EPI_ISL_3403944, EPI_ISL_3403945, EPI_ISL_3403962, EPI_ISL_3403963, EPI_ISL_3403964, EPI_ISL_3403965, EPI_ISL_3403966, EPI_ISL_3403967, EPI_ISL_3403968, EPI_ISL_3403969, EPI_ISL_3403970, EPI_ISL_3403971, EPI_ISL_3403972, EPI_ISL_3403973, EPI_ISL_3403974, EPI_ISL_3403975, EPI_ISL_3403976, EPI_ISL_3403977, EPI_ISL_3403978, EPI_ISL_3403979, EPI_ISL_3403980, EPI_ISL_3403981, EPI_ISL_3403982, EPI_ISL_3403983, EPI_ISL_3403984, EPI_ISL_3403985, EPI_ISL_3403986, EPI_ISL_3403987, EPI_ISL_3403988, EPI_ISL_3403989, EPI_ISL_3403990, EPI_ISL_3403991, EPI_ISL_3403992, EPI_ISL_3403993, EPI_ISL_3403994, EPI_ISL_3403995, EPI_ISL_3403996, EPI_ISL_3403997, EPI_ISL_3403998, EPI_ISL_3403999, EPI_ISL_3404000, EPI_ISL_3404001, EPI_ISL_3404002, EPI_ISL_3404003, EPI_ISL_3404004, EPI_ISL_3404005, EPI_ISL_3404006, EPI_ISL_3404007, EPI_ISL_3404008, EPI_ISL_3404009, EPI_ISL_3404010, EPI_ISL_3404011, EPI_ISL_3404012, EPI_ISL_3404013, EPI_ISL_3404014, EPI_ISL_3404015, EPI_ISL_3404016, EPI_ISL_3404017, EPI_ISL_3404018, EPI_ISL_3404019, EPI_ISL_3404020, EPI_ISL_3404021, EPI_ISL_3404022, EPI_ISL_3404023, EPI_ISL_3404024, EPI_ISL_3404025, EPI_ISL_3404026, EPI_ISL_3404027, EPI_ISL_3404028, EPI_ISL_3404029, EPI_ISL_3404030, EPI_ISL_3404031, EPI_ISL_3404032, EPI_ISL_3404033, EPI_ISL_3404082, EPI_ISL_3404083, EPI_ISL_3404084, EPI_ISL_3404085, EPI_ISL_3404086, EPI_ISL_3404087, EPI_ISL_3404088, EPI_ISL_3404089, EPI_ISL_3404090, EPI_ISL_3404091, EPI_ISL_3404092, EPI_ISL_3404093, EPI_ISL_3404094, EPI_ISL_3404095, EPI_ISL_3404096, EPI_ISL_3404097, EPI_ISL_3404098, EPI_ISL_3404099, EPI_ISL_3404100, EPI_ISL_3404101, EPI_ISL_3404102, EPI_ISL_3404103, EPI_ISL_3404104, EPI_ISL_3404105, EPI_ISL_3404106, EPI_ISL_3404107, EPI_ISL_3404108, EPI_ISL_3404109, EPI_ISL_3404110, EPI_ISL_3404111, EPI_ISL_3404112, EPI_ISL_3404113, EPI_ISL_3404114, EPI_ISL_3404115, EPI_ISL_3404116, EPI_ISL_3404117, EPI_ISL_3404142, EPI_ISL_3404143, EPI_ISL_3404144, EPI_ISL_3404145, EPI_ISL_3404146, EPI_ISL_3404147, EPI_ISL_3404148, EPI_ISL_3404149, EPI_ISL_3404150, EPI_ISL_3404151, EPI_ISL_3404152, EPI_ISL_3404153, EPI_ISL_3404154, EPI_ISL_3404155, EPI_ISL_3404156, EPI_ISL_3404157, EPI_ISL_3404158, EPI_ISL_3404159, EPI_ISL_3404160, EPI_ISL_3404161, EPI_ISL_3404162, EPI_ISL_3404163, EPI_ISL_3404164, EPI_ISL_3404165, EPI_ISL_3404166, EPI_ISL_3404167, EPI_ISL_3404168, EPI_ISL_3404169, EPI_ISL_3404170, EPI_ISL_3404171, EPI_ISL_3404172, EPI_ISL_3404173, EPI_ISL_3404174, EPI_ISL_3404175, EPI_ISL_3404176, EPI_ISL_3404177, EPI_ISL_3404178, EPI_ISL_3404179, EPI_ISL_3404214, EPI_ISL_3404215, EPI_ISL_3404216, EPI_ISL_3404217, EPI_ISL_3404218, EPI_ISL_3404219, EPI_ISL_3404220, EPI_ISL_3404221, EPI_ISL_3404222, EPI_ISL_3404223, EPI_ISL_3404224, EPI_ISL_3404225, EPI_ISL_3404226, EPI_ISL_3404227, EPI_ISL_3404228, EPI_ISL_3404229, EPI_ISL_3404230, EPI_ISL_3404231, EPI_ISL_3404232, EPI_ISL_3404233, EPI_ISL_3404234, EPI_ISL_3404235, EPI_ISL_3404236, EPI_ISL_3404237, EPI_ISL_3404238, EPI_ISL_3404239, EPI_ISL_3404240, EPI_ISL_3404241, EPI_ISL_3404242, EPI_ISL_3404243, EPI_ISL_3404244, EPI_ISL_3404245, EPI_ISL_3404246, EPI_ISL_3404247, EPI_ISL_3404248, EPI_ISL_3404249, EPI_ISL_3404250, EPI_ISL_3404251, EPI_ISL_3404252, EPI_ISL_3404253, EPI_ISL_3404254, EPI_ISL_3404255, EPI_ISL_3404256, EPI_ISL_3404257, EPI_ISL_3404258, EPI_ISL_3404259, EPI_ISL_3404260, EPI_ISL_3404261, EPI_ISL_3404262, EPI_ISL_3404263, EPI_ISL_3404275, EPI_ISL_3404324, EPI_ISL_3404325, EPI_ISL_3404326, EPI_ISL_3404327, EPI_ISL_3404328, EPI_ISL_3404329, EPI_ISL_3404330, EPI_ISL_3404331, EPI_ISL_3404332, EPI_ISL_3404333, EPI_ISL_3404334, EPI_ISL_3404335, EPI_ISL_3404336, EPI_ISL_3404337, EPI_ISL_3404338, EPI_ISL_3404339, EPI_ISL_3404340, EPI_ISL_3404341, EPI_ISL_3404342, EPI_ISL_3404343, EPI_ISL_3404344, EPI_ISL_3404345, EPI_ISL_3404346, EPI_ISL_3404347 | | | |
| see above | Colorado Department of Public Health and Environment | Colorado Department of Public Health and Environment | Laura Bankers, Molly C. Hetherington-Rauth, Diana Ir, Alexandria Rossheim, Michael Martin, Mandy Waters, Shannon R. Matzinger, Sarah Elizabeth Totten, Emily A. Travanty |
| EPI_ISL_3404356, EPI_ISL_3404357, EPI_ISL_3404358, EPI_ISL_3404359, EPI_ISL_3404360, EPI_ISL_3404361, EPI_ISL_3404362, EPI_ISL_3404363, EPI_ISL_3404364, EPI_ISL_3404365 | Laboratorium Khusus Speedlab Jakarta | Genomik Solidaritas Indonesia Laboratorium | Neng Hilma Hamidah, Carissa Sintca Wijaya, Vania Gavrila Wikasa, Dhahlia Agustina Cahyono, Gracia Felias Enos Korompis, Annisa Muthiah Sukirman, Anuraj Shankar, Louisa Markus, Meutia Ayuputeri Kumaheri, Ariel Pradipta |
| EPI_ISL_3404366, EPI_ISL_3404367 | Laboratorium Khusus Speedlab Jakarta | Genomik Solidaritas Indonesia Laboratorium | Carissa Sintca Wijaya, Vania Gavrila Wikasa, Neng Hilma Hamidah, Dhahlia Agustina Cahyono, Gracia Felias Enos Korompis, Annisa Muthiah Sukirman, Anuraj Shankar, Louisa Markus, Meutia Ayuputeri Kumaheri, Ariel Pradipta |

| | | | |
|---|---|---|---|
| EPI_ISL_3404368, EPI_ISL_3404369 | RSUD dr. H. Marsidi Judono Belitung | Genomik Solidaritas Indonesia Laboratorium | Carissa Sintca Wijaya, Vania Gavrila Wikasa, Titik Handayani Tanujaya, Teguh A.Maulana Gultom, Neng Hilma Hamidah, Dhahlia Agustina Cahyono, Gracia Felias Enos Korompis, Annisa Muthiah Sukirman, Anuraj Shankar, Louisa Markus, Meutia Ayuputeri Kumaheri, Ariel Pradipta |
| EPI_ISL_3404370 | RSUD dr. H. Marsidi Judono, Belitung | Genomik Solidaritas Indonesia Laboratorium | Carissa Sintca Wijaya, Vania Gavrila Wikasa, Titik Handayani Tanujaya, Teguh A.Maulana Gultom, Neng Hilma Hamidah, Dhahlia Agustina Cahyono, Gracia Felias Enos Korompis, Annisa Muthiah Sukirman, Anuraj Shankar, Louisa Markus, Meutia Ayuputeri Kumaheri, Ariel Pradipta |
| EPI_ISL_3404371, EPI_ISL_3404372 | RSUD dr. H. Marsidi Judono Belitung | Genomik Solidaritas Indonesia Laboratorium | Carissa Sintca Wijaya, Vania Gavrila Wikasa, Titik Handayani Tanujaya, Teguh A.Maulana Gultom, Neng Hilma Hamidah, Dhahlia Agustina Cahyono, Gracia Felias Enos Korompis, Annisa Muthiah Sukirman, Anuraj Shankar, Louisa Markus, Meutia Ayuputeri Kumaheri, Ariel Pradipta |
| EPI_ISL_3404373, EPI_ISL_3404374 | Genomik Solidaritas Indonesia Laboratorium | Genomik Solidaritas Indonesia Laboratorium | Carissa Sintca Wijaya, Vania Gavrila Wikasa, Neng Hilma Hamidah, Dhahlia Agustina Cahyono, Gracia Felias Enos Korompis, Annisa Muthiah Sukirman, Anuraj Shankar, Louisa Markus, Meutia Ayuputeri Kumaheri, Ariel Pradipta |
| EPI_ISL_3404375, EPI_ISL_3404376, EPI_ISL_3404377, EPI_ISL_3404378, EPI_ISL_3404379, EPI_ISL_3404380, EPI_ISL_3404381, EPI_ISL_3404382 | Genomik Solidaritas Indonesia Laboratorium | Genomik Solidaritas Indonesia Laboratorium | Vania Gavrila Wikasa, Neng Hilma Hamidah, Carissa Sintca Wijaya, Dhahlia Agustina Cahyono, Annisa Muthiah Sukirman, Gracia Felias Enos Korompis, Anuraj Shankar, Louisa Markus, Meutia Ayuputeri Kumaheri, Ariel Pradipta |
| EPI_ISL_3404383, EPI_ISL_3404384, EPI_ISL_3404385, EPI_ISL_3404387, EPI_ISL_3404388, EPI_ISL_3404389, EPI_ISL_3404390, EPI_ISL_3404391, EPI_ISL_3404392, EPI_ISL_3404393, EPI_ISL_3404394, EPI_ISL_3404395, EPI_ISL_3404396, EPI_ISL_3404397, EPI_ISL_3404398, EPI_ISL_3404399, EPI_ISL_3404400, EPI_ISL_3404401, EPI_ISL_3404402, EPI_ISL_3404403, EPI_ISL_3404404, EPI_ISL_3404405, EPI_ISL_3404406, EPI_ISL_3404407, EPI_ISL_3404408, EPI_ISL_3404409, EPI_ISL_3404410, EPI_ISL_3404411, EPI_ISL_3404412, EPI_ISL_3404413, EPI_ISL_3404414, EPI_ISL_3404415, EPI_ISL_3404416, EPI_ISL_3404417, EPI_ISL_3404418 | | | |
| see above | Oregon State Public Health Laboratory | Oregon State Public Health Laboratory | Rafia Razzaque, Eugene Yeboah, Vanda Makris, Laura Tsaknaridis, John Fontana and Shane Sevey |
| EPI_ISL_3404422, EPI_ISL_3404423, EPI_ISL_3404424, EPI_ISL_3404425, EPI_ISL_3404426, EPI_ISL_3404427, EPI_ISL_3404428, EPI_ISL_3404430, EPI_ISL_3404431, EPI_ISL_3404432, EPI_ISL_3404433, EPI_ISL_3404434, EPI_ISL_3404435, EPI_ISL_3404436, EPI_ISL_3404437, EPI_ISL_3404438, EPI_ISL_3404439, EPI_ISL_3404440, EPI_ISL_3404441, EPI_ISL_3404442, EPI_ISL_3404443, EPI_ISL_3404444, EPI_ISL_3404445, EPI_ISL_3404446, EPI_ISL_3404447, EPI_ISL_3404452, EPI_ISL_3404459, EPI_ISL_3404473, EPI_ISL_3404474, EPI_ISL_3404475, EPI_ISL_3404476, EPI_ISL_3404478, EPI_ISL_3404480, EPI_ISL_3404481, EPI_ISL_3404482, EPI_ISL_3404483, EPI_ISL_3404484, EPI_ISL_3404485, EPI_ISL_3404486, EPI_ISL_3404487, EPI_ISL_3404488, EPI_ISL_3404490, EPI_ISL_3404491, EPI_ISL_3404493, EPI_ISL_3404516, EPI_ISL_3404527, EPI_ISL_3404529, EPI_ISL_3404530, EPI_ISL_3404543 | | | |
| see above | Ministry of Health Turkey | Ministry of Health Turkey | Fatma Bayrakdar, Yasemin Cosgun, Suleyman Yalcin, Gulay Korukluoglu |
| EPI_ISL_3404549, EPI_ISL_3404609 | University Campus Bio-Medico of Rome (UCBM) | University Campus Bio-Medico of Rome (UCBM) | Fogolari M., Riva E., De Florio L., Francesconi M., Lintas C., Veralli R., Angeletti S. |
| EPI_ISL_3404610, EPI_ISL_3404611, EPI_ISL_3404612, EPI_ISL_3404613, EPI_ISL_3404614, EPI_ISL_3404615, EPI_ISL_3404616, EPI_ISL_3404617, EPI_ISL_3404618, EPI_ISL_3404619, EPI_ISL_3404620, EPI_ISL_3404621, EPI_ISL_3404622, EPI_ISL_3404623, EPI_ISL_3404624, EPI_ISL_3404625, EPI_ISL_3404626, EPI_ISL_3404627, EPI_ISL_3404628, EPI_ISL_3404629, EPI_ISL_3404633, EPI_ISL_3404634, EPI_ISL_3404635, EPI_ISL_3404636, EPI_ISL_3404637, EPI_ISL_3404638, EPI_ISL_3404639, EPI_ISL_3404640, EPI_ISL_3404641, EPI_ISL_3404642, EPI_ISL_3404643, EPI_ISL_3404644, EPI_ISL_3404646, EPI_ISL_3404647, EPI_ISL_3404648, EPI_ISL_3404649, EPI_ISL_3404650, EPI_ISL_3404651, EPI_ISL_3404652, EPI_ISL_3404653, EPI_ISL_3404654, EPI_ISL_3404655, EPI_ISL_3404656, EPI_ISL_3404657, EPI_ISL_3404658, EPI_ISL_3404659, EPI_ISL_3404660, EPI_ISL_3404661, EPI_ISL_3404662, EPI_ISL_3404663, EPI_ISL_3404664, EPI_ISL_3404665, EPI_ISL_3404666, EPI_ISL_3404667, EPI_ISL_3404668, EPI_ISL_3404670, EPI_ISL_3404672, EPI_ISL_3404674, EPI_ISL_3404675, EPI_ISL_3404676, EPI_ISL_3404677, EPI_ISL_3404678, EPI_ISL_3404679, EPI_ISL_3404680, EPI_ISL_3404681, EPI_ISL_3404682, EPI_ISL_3404683, EPI_ISL_3404684, EPI_ISL_3404685, EPI_ISL_3404686, EPI_ISL_3404687, EPI_ISL_3404688, EPI_ISL_3404689, EPI_ISL_3404690, EPI_ISL_3404691, EPI_ISL_3404692, EPI_ISL_3404693, EPI_ISL_3404694, EPI_ISL_3404695, EPI_ISL_3404696, EPI_ISL_3404697, EPI_ISL_3404698, EPI_ISL_3404699, EPI_ISL_3404700, EPI_ISL_3404701, EPI_ISL_3404702, EPI_ISL_3404703, EPI_ISL_3404704, EPI_ISL_3404706, EPI_ISL_3404707, EPI_ISL_3404708, EPI_ISL_3404709, EPI_ISL_3404710, EPI_ISL_3404711, EPI_ISL_3404712, EPI_ISL_3404713, EPI_ISL_3404714, EPI_ISL_3404715, EPI_ISL_3404716, EPI_ISL_3404717, EPI_ISL_3404718, EPI_ISL_3404719, EPI_ISL_3404720, EPI_ISL_3404721, EPI_ISL_3404722, EPI_ISL_3404723, EPI_ISL_3404724, EPI_ISL_3404725, EPI_ISL_3404726, EPI_ISL_3404727, EPI_ISL_3404728, EPI_ISL_3404729, EPI_ISL_3404730 | | | |
| see above | Idaho Bureau of Laboratories | Boise VA Medical Center, PALMS | Matthew Burns, Aimee Ceniseros, Christian Loera, Dr. Christopher Ball, Ying Pei, Cheri Lamb McFarlane, James Razor, Emily Bartlett |
| EPI_ISL_3404731, EPI_ISL_3404732 | University Campus Bio-Medico of Rome (UCBM) | University Campus Bio-Medico of Rome (UCBM) | Fogolari M., Riva E., De Florio L., Francesconi M., Lintas C., Veralli R., Angeletti S. |
| EPI_ISL_3404733, EPI_ISL_3404734, EPI_ISL_3404735, EPI_ISL_3404736, EPI_ISL_3404738, EPI_ISL_3404739, EPI_ISL_3404740, EPI_ISL_3404741, EPI_ISL_3404742, EPI_ISL_3404743, EPI_ISL_3404744, EPI_ISL_3404745, EPI_ISL_3404746, EPI_ISL_3404747, EPI_ISL_3404748, EPI_ISL_3404749, EPI_ISL_3404750, EPI_ISL_3404751 | | | |
| see above | Dianalabs SA | Genesupport | Tanguy ARAUD, Geraldine Jost, Katia Jaton, Nadia Liassine |
| EPI_ISL_3404752, EPI_ISL_3404754, EPI_ISL_3404755, EPI_ISL_3404756, EPI_ISL_3404757, EPI_ISL_3404758, EPI_ISL_3404759, EPI_ISL_3404760, EPI_ISL_3404761, EPI_ISL_3404763, EPI_ISL_3404764, EPI_ISL_3404765, EPI_ISL_3404766, EPI_ISL_3404767, EPI_ISL_3404768, EPI_ISL_3404769, EPI_ISL_3404770, EPI_ISL_3404771, EPI_ISL_3404772, EPI_ISL_3404773, EPI_ISL_3404774, EPI_ISL_3404775, EPI_ISL_3404776, EPI_ISL_3404777, EPI_ISL_3404778, EPI_ISL_3404779, EPI_ISL_3404780, EPI_ISL_3404781, EPI_ISL_3404782, EPI_ISL_3404783, EPI_ISL_3404784, EPI_ISL_3404785, EPI_ISL_3404786, EPI_ISL_3404787, EPI_ISL_3404788, EPI_ISL_3404789, EPI_ISL_3404790, EPI_ISL_3404791, EPI_ISL_3404792, EPI_ISL_3404793, EPI_ISL_3404794, EPI_ISL_3404795, EPI_ISL_3404796, EPI_ISL_3404797, EPI_ISL_3404798 | | | |
| see above | Bioanalytica SA | Genesupport | Tanguy ARAUD, Geraldine Jost, Katia Jaton, Nadia Liassine |
| EPI_ISL_3404799 | Polyanalytic SA | Genesupport | Tanguy ARAUD, Geraldine Jost, Katia Jaton, Nadia Liassine |
| EPI_ISL_3404800, EPI_ISL_3404801, EPI_ISL_3404802, EPI_ISL_3404803, EPI_ISL_3404804, EPI_ISL_3404805, EPI_ISL_3404806 | Dianalabs-Valais SA | Genesupport | Tanguy ARAUD, Geraldine Jost, Katia Jaton, Nadia Liassine |
| EPI_ISL_3404807, EPI_ISL_3404808, EPI_ISL_3404809, EPI_ISL_3404811, EPI_ISL_3404812, EPI_ISL_3404813, EPI_ISL_3404814, EPI_ISL_3404815, EPI_ISL_3404816, EPI_ISL_3404817, EPI_ISL_3404818 | | | |
| see above | Dianalabs SA | Genesupport | Tanguy ARAUD, Geraldine Jost, Katia Jaton, Nadia Liassine |
| EPI_ISL_3404819, EPI_ISL_3404820 | Proxilab SA | Genesupport | Tanguy ARAUD, Geraldine Jost, Katia Jaton, Nadia Liassine |
| EPI_ISL_3404886, EPI_ISL_3404887, EPI_ISL_3404888 | University Campus Bio-Medico of Rome (UCBM) | University Campus Bio-Medico of Rome (UCBM) | Fogolari M., Riva E., De Florio L., Francesconi M., Lintas C., Veralli R., Angeletti S. |
| EPI_ISL_3404889, EPI_ISL_3404890, EPI_ISL_3404891, EPI_ISL_3404892, EPI_ISL_3404893, EPI_ISL_3404894, EPI_ISL_3404895, EPI_ISL_3404896, EPI_ISL_3404897, EPI_ISL_3404898, EPI_ISL_3404899, EPI_ISL_3404900, EPI_ISL_3404901, EPI_ISL_3404902, EPI_ISL_3404903, EPI_ISL_3404904, EPI_ISL_3404905, EPI_ISL_3404907, EPI_ISL_3404908, EPI_ISL_3404909, EPI_ISL_3404910, EPI_ISL_3404911, EPI_ISL_3404912, EPI_ISL_3404913, EPI_ISL_3404914, EPI_ISL_3404915, EPI_ISL_3404916, EPI_ISL_3404917, EPI_ISL_3404918, EPI_ISL_3404919, EPI_ISL_3404920, EPI_ISL_3404921, EPI_ISL_3404922, EPI_ISL_3404923, EPI_ISL_3404925, EPI_ISL_3404926, EPI_ISL_3404927, EPI_ISL_3404928, EPI_ISL_3404929, EPI_ISL_3404931, EPI_ISL_3404932, EPI_ISL_3404933, EPI_ISL_3404934, EPI_ISL_3404935, EPI_ISL_3404936, EPI_ISL_3404937, EPI_ISL_3404938, EPI_ISL_3404939, EPI_ISL_3404940, EPI_ISL_3404941, EPI_ISL_3404942, EPI_ISL_3404943, EPI_ISL_3404944, EPI_ISL_3404945, EPI_ISL_3404946, EPI_ISL_3404947, EPI_ISL_3404948, EPI_ISL_3404949, EPI_ISL_3404951, EPI_ISL_3404952, EPI_ISL_3404953, EPI_ISL_3404955, EPI_ISL_3404956, EPI_ISL_3404957, EPI_ISL_3404958, EPI_ISL_3404959, EPI_ISL_3404960, EPI_ISL_3404961, EPI_ISL_3404962, EPI_ISL_3404963, EPI_ISL_3404964, EPI_ISL_3404966, EPI_ISL_3404967, EPI_ISL_3404968, EPI_ISL_3404969, EPI_ISL_3404970, EPI_ISL_3404971, EPI_ISL_3404972, EPI_ISL_3404973, EPI_ISL_3404974, EPI_ISL_3404975, EPI_ISL_3404976, EPI_ISL_3404977, EPI_ISL_3404978, EPI_ISL_3404979, EPI_ISL_3404980, EPI_ISL_3404981, EPI_ISL_3404983, EPI_ISL_3404984, EPI_ISL_3404985, EPI_ISL_3404986, EPI_ISL_3404987, EPI_ISL_3404988, EPI_ISL_3404989, EPI_ISL_3404990, EPI_ISL_3404992, EPI_ISL_3404993 | | | |
| see above | Ochsner Health | BioInfoExperts | Amy Feehan, Judy Crabtree, Rebecca Rose, Ben Lain, Chris Huston, Samual Moot, David J. Nolan, Tessa LaFleur, Julia-Garcia-Diaz, Lucio Miele, Susanna L. Lamers |
| EPI_ISL_3404995 | ACT Pathology | Schwessinger Lab | Ashley Jones, Benjamin Schwessinger, Emma Crean, Robyn N Hall, Craig Kennedy, Karina Kennedy |
| EPI_ISL_3405008, EPI_ISL_3405009, EPI_ISL_3405010, EPI_ISL_3405011, EPI_ISL_3405012, EPI_ISL_3405014, EPI_ISL_3405015, EPI_ISL_3405016, EPI_ISL_3405017, EPI_ISL_3405018, EPI_ISL_3405019, EPI_ISL_3405020, EPI_ISL_3405021, EPI_ISL_3405022, EPI_ISL_3405023, EPI_ISL_3405024, EPI_ISL_3405025, EPI_ISL_3405026, EPI_ISL_3405027, EPI_ISL_3405028, EPI_ISL_3405029, EPI_ISL_3405030, EPI_ISL_3405031, EPI_ISL_3405032, EPI_ISL_3405033, EPI_ISL_3405035, EPI_ISL_3405036, EPI_ISL_3405037, EPI_ISL_3405038, EPI_ISL_3405039, EPI_ISL_3405040, EPI_ISL_3405041, EPI_ISL_3405042, EPI_ISL_3405043, EPI_ISL_3405044, EPI_ISL_3405045, EPI_ISL_3405046, EPI_ISL_3405047, EPI_ISL_3405048, EPI_ISL_3405049, EPI_ISL_3405050, EPI_ISL_3405051, EPI_ISL_3405052, EPI_ISL_3405053, EPI_ISL_3405054, EPI_ISL_3405055, EPI_ISL_3405056, EPI_ISL_3405057, EPI_ISL_3405058, EPI_ISL_3405060, EPI_ISL_3405061, EPI_ISL_3405062, EPI_ISL_3405063, EPI_ISL_3405066, EPI_ISL_3405067, EPI_ISL_3405068, EPI_ISL_3405069, EPI_ISL_3405070, EPI_ISL_3405071, EPI_ISL_3405072, EPI_ISL_3405073, EPI_ISL_3405074, EPI_ISL_3405075, EPI_ISL_3405076, EPI_ISL_3405077, EPI_ISL_3405078, EPI_ISL_3405079, EPI_ISL_3405080, EPI_ISL_3405081, EPI_ISL_3405082, EPI_ISL_3405083, EPI_ISL_3405084, EPI_ISL_3405085, EPI_ISL_3405086, EPI_ISL_3405087, EPI_ISL_3405088, EPI_ISL_3405089, EPI_ISL_3405090, EPI_ISL_3405091, EPI_ISL_3405092, EPI_ISL_3405093, EPI_ISL_3405094, EPI_ISL_3405095, EPI_ISL_3405096, EPI_ISL_3405097, EPI_ISL_3405098, EPI_ISL_3405099, EPI_ISL_3405100, EPI_ISL_3405101, EPI_ISL_3405102, EPI_ISL_3405103, EPI_ISL_3405104, EPI_ISL_3405105, EPI_ISL_3405106, EPI_ISL_3405107, EPI_ISL_3405108, EPI_ISL_3405109, EPI_ISL_3405110, EPI_ISL_3405111, EPI_ISL_3405112, EPI_ISL_3405113, EPI_ISL_3405114, EPI_ISL_3405115, EPI_ISL_3405116, EPI_ISL_3405117, EPI_ISL_3405118, EPI_ISL_3405119, EPI_ISL_3405120, EPI_ISL_3405121, EPI_ISL_3405122, EPI_ISL_3405123, EPI_ISL_3405124, EPI_ISL_3405125, EPI_ISL_3405126, EPI_ISL_3405127, EPI_ISL_3405128, EPI_ISL_3405129, EPI_ISL_3405130, EPI_ISL_3405131, EPI_ISL_3405133, EPI_ISL_3405134, EPI_ISL_3405135, EPI_ISL_3405136, EPI_ISL_3405137, EPI_ISL_3405138, EPI_ISL_3405139, EPI_ISL_3405140, EPI_ISL_3405141, EPI_ISL_3405142, EPI_ISL_3405143, EPI_ISL_3405144, EPI_ISL_3405145, EPI_ISL_3405146, EPI_ISL_3405147, EPI_ISL_3405148, EPI_ISL_3405149, EPI_ISL_3405150, EPI_ISL_3405152, EPI_ISL_3405153, EPI_ISL_3405155, EPI_ISL_3405156, EPI_ISL_3405157, EPI_ISL_3405158, EPI_ISL_3405159, EPI_ISL_3405161, EPI_ISL_3405162, EPI_ISL_3405163, EPI_ISL_3405164, EPI_ISL_3405165, EPI_ISL_3405166, EPI_ISL_3405167, EPI_ISL_3405168, EPI_ISL_3405169, EPI_ISL_3405170, EPI_ISL_3405171, EPI_ISL_3405172, EPI_ISL_3405173, EPI_ISL_3405174, EPI_ISL_3405175, EPI_ISL_3405176, EPI_ISL_3405177, EPI_ISL_3405178, EPI_ISL_3405179, EPI_ISL_3405180, EPI_ISL_3405182, EPI_ISL_3405183, EPI_ISL_3405184, EPI_ISL_3405185, EPI_ISL_3405186, EPI_ISL_3405187, EPI_ISL_3405188, EPI_ISL_3405189, EPI_ISL_3405190, EPI_ISL_3405191, EPI_ISL_3405192, EPI_ISL_3405193, EPI_ISL_3405194, EPI_ISL_3405195, EPI_ISL_3405196, EPI_ISL_3405197, EPI_ISL_3405198, EPI_ISL_3405199, EPI_ISL_3405200, EPI_ISL_3405201, EPI_ISL_3405202, EPI_ISL_3405203, EPI_ISL_3405204, EPI_ISL_3405205, EPI_ISL_3405207, EPI_ISL_3405208, EPI_ISL_3405209, EPI_ISL_3405210, EPI_ISL_3405213, EPI_ISL_3405214, EPI_ISL_3405215, EPI_ISL_3405216, EPI_ISL_3405217, EPI_ISL_3405218, EPI_ISL_3405219, EPI_ISL_3405220, EPI_ISL_3405221, EPI_ISL_3405222, EPI_ISL_3405223, EPI_ISL_3405224, EPI_ISL_3405225, EPI_ISL_3405226, EPI_ISL_3405227, EPI_ISL_3405228, EPI_ISL_3405229, EPI_ISL_3405230, EPI_ISL_3405231, EPI_ISL_3405232, EPI_ISL_3405233, EPI_ISL_3405234, EPI_ISL_3405235, EPI_ISL_3405236, EPI_ISL_3405237, EPI_ISL_3405238, EPI_ISL_3405239, EPI_ISL_3405240, EPI_ISL_3405241, EPI_ISL_3405242, EPI_ISL_3405243, EPI_ISL_3405244, EPI_ISL_3405245, EPI_ISL_3405246, EPI_ISL_3405247, EPI_ISL_3405248, EPI_ISL_3405249, EPI_ISL_3405250, EPI_ISL_3405251, EPI_ISL_3405252, EPI_ISL_3405253, EPI_ISL_3405254, EPI_ISL_3405255, EPI_ISL_3405256, EPI_ISL_3405257, EPI_ISL_3405258, EPI_ISL_3405259, EPI_ISL_3405260, EPI_ISL_3405261, EPI_ISL_3405262, EPI_ISL_3405263, EPI_ISL_3405264, EPI_ISL_3405265, EPI_ISL_3405266, EPI_ISL_3405267, EPI_ISL_3405268, EPI_ISL_3405269, EPI_ISL_3405270, EPI_ISL_3405271, EPI_ISL_3405272, EPI_ISL_3405273, EPI_ISL_3405274, EPI_ISL_3405275, EPI_ISL_3405276, EPI_ISL_3405277, EPI_ISL_3405278, EPI_ISL_3405279, EPI_ISL_3405280, EPI_ISL_3405282, EPI_ISL_3405283, EPI_ISL_3405284, EPI_ISL_3405285, EPI_ISL_3405286, EPI_ISL_3405287, EPI_ISL_3405288, EPI_ISL_3405289, EPI_ISL_3405290, EPI_ISL_3405291, EPI_ISL_3405292, EPI_ISL_3405293, EPI_ISL_3405294, EPI_ISL_3405295, EPI_ISL_3405296, EPI_ISL_3405297, EPI_ISL_3405298, EPI_ISL_3405299, EPI_ISL_3405301, EPI_ISL_3405302, EPI_ISL_3405303, EPI_ISL_3405305, EPI_ISL_3405306, EPI_ISL_3405307, EPI_ISL_3405308, EPI_ISL_3405309, EPI_ISL_3405311, EPI_ISL_3405312, EPI_ISL_3405313, EPI_ISL_3405314, EPI_ISL_3405315, EPI_ISL_3405316, EPI_ISL_3405317, EPI_ISL_3405318, EPI_ISL_3405319, EPI_ISL_3405320, EPI_ISL_3405321, EPI_ISL_3405322, EPI_ISL_3405323, EPI_ISL_3405324, EPI_ISL_3405325, EPI_ISL_3405326, EPI_ISL_3405327, EPI_ISL_3405328, EPI_ISL_3405329, EPI_ISL_3405330, EPI_ISL_3405332, EPI_ISL_3405333, EPI_ISL_3405334, EPI_ISL_3405335, EPI_ISL_3405336, EPI_ISL_3405337, EPI_ISL_3405338, EPI_ISL_3405339, EPI_ISL_3405340, EPI_ISL_3405341, EPI_ISL_3405342, EPI_ISL_3405343, EPI_ISL_3405344, EPI_ISL_3405345, EPI_ISL_3405346, EPI_ISL_3405347, EPI_ISL_3405348, EPI_ISL_3405349, EPI_ISL_3405350, EPI_ISL_3405351, EPI_ISL_3405352, EPI_ISL_3405353, EPI_ISL_3405354, EPI_ISL_3405355, EPI_ISL_3405357, EPI_ISL_3405358, EPI_ISL_3405359, EPI_ISL_3405360, EPI_ISL_3405363, EPI_ISL_3405364, EPI_ISL_3405365, EPI_ISL_3405366, EPI_ISL_3405367, EPI_ISL_3405368, EPI_ISL_3405369, EPI_ISL_3405370, EPI_ISL_3405371, EPI_ISL_3405372, EPI_ISL_3405373, EPI_ISL_3405374, EPI_ISL_3405375, EPI_ISL_3405376, EPI_ISL_3405377, EPI_ISL_3405378, EPI_ISL_3405379, EPI_ISL_3405380, EPI_ISL_3405381, EPI_ISL_3405382, EPI_ISL_3405383, EPI_ISL_3405384, EPI_ISL_3405385, EPI_ISL_3405386, EPI_ISL_3405387, EPI_ISL_3405388, EPI_ISL_3405389, EPI_ISL_3405390, EPI_ISL_3405391, EPI_ISL_3405392, EPI_ISL_3405393, EPI_ISL_3405394, EPI_ISL_3405395, EPI_ISL_3405396, EPI_ISL_3405397, EPI_ISL_3405398, EPI_ISL_3405399, EPI_ISL_3405400, EPI_ISL_3405401, EPI_ISL_3405402, EPI_ISL_3405403, EPI_ISL_3405404, EPI_ISL_3405405, EPI_ISL_3405406, EPI_ISL_3405407, EPI_ISL_3405408, EPI_ISL_3405409, EPI_ISL_3405410, EPI_ISL_3405411, EPI_ISL_3405412, EPI_ISL_3405413, EPI_ISL_3405414, EPI_ISL_3405415, EPI_ISL_3405416, EPI_ISL_3405417, EPI_ISL_3405418, EPI_ISL_3405419, EPI_ISL_3405420, EPI_ISL_3405421, EPI_ISL_3405422, EPI_ISL_3405423, EPI_ISL_3405424, EPI_ISL_3405425, EPI_ISL_3405426, EPI_ISL_3405427, EPI_ISL_3405428, EPI_ISL_3405429, EPI_ISL_3405430, EPI_ISL_3405432, EPI_ISL_3405433, EPI_ISL_3405434, EPI_ISL_3405435, EPI_ISL_3405436, EPI_ISL_3405437, EPI_ISL_3405438, EPI_ISL_3405439, EPI_ISL_3405440, EPI_ISL_3405441, EPI_ISL_3405442, EPI_ISL_3405443, EPI_ISL_3405444, EPI_ISL_3405445, EPI_ISL_3405446, EPI_ISL_3405447, EPI_ISL_3405448, EPI_ISL_3405449, EPI_ISL_3405451, EPI_ISL_3405452, EPI_ISL_3405453, EPI_ISL_3405455, EPI_ISL_3405456, EPI_ISL_3405457, EPI_ISL_3405458, EPI_ISL_3405459, EPI_ISL_3405461, EPI_ISL_3405462, EPI_ISL_3405463, EPI_ISL_3405464, EPI_ISL_3405465, EPI_ISL_3405466, EPI_ISL_3405467, EPI_ISL_3405468, EPI_ISL_3405469, EPI_ISL_3405470, EPI_ISL_3405471, EPI_ISL_3405472, EPI_ISL_3405473, EPI_ISL_3405474, EPI_ISL_3405475, EPI_ISL_3405476, EPI_ISL_3405477, EPI_ISL_3405478, EPI_ISL_3405479, EPI_ISL_3405480, EPI_ISL_3405482, EPI_ISL_3405483, EPI_ISL_3405484, EPI_ISL_3405485, EPI_ISL_3405486, EPI_ISL_3405487, EPI_ISL_3405488, EPI_ISL_3405489, EPI_ISL_3405490, EPI_ISL_3405491, EPI_ISL_3405492, EPI_ISL_3405493, EPI_ISL_3405494, EPI_ISL_3405495, EPI_ISL_3405496, EPI_ISL_3405497, EPI_ISL_3405498, EPI_ISL_3405499, EPI_ISL_3405500, EPI_ISL_3405501, EPI_ISL_3405502, EPI_ISL_3405503, EPI_ISL_3405504, EPI_ISL_3405505, EPI_ISL_3405507, EPI_ISL_3405508, EPI_ISL_3405509, EPI_ISL_3405510, EPI_ISL_3405513, EPI_ISL_3405514, EPI_ISL_3405515, EPI_ISL_3405516, EPI_ISL_3405517, EPI_ISL_3405518, EPI_ISL_3405519, EPI_ISL_3405520, EPI_ISL_3405521, EPI_ISL_3405522, EPI_ISL_3405523, EPI_ISL_3405524, EPI_ISL_3405525, EPI_ISL_3405526, EPI_ISL_3405527, EPI_ISL_3405528, EPI_ISL_3405529, EPI_ISL_3405530, EPI_ISL_3405531, EPI_ISL_3405532, EPI_ISL_3405533, EPI_ISL_3405534, EPI_ISL_3405535, EPI_ISL_3405536, EPI_ISL_3405537, EPI_ISL_3405538, EPI_ISL_3405539, EPI_ISL_3405540, EPI_ISL_3405541, EPI_ISL_3405542, EPI_ISL_3405543, EPI_ISL_3405544, EPI_ISL_3405545, EPI_ISL_3405546, EPI_ISL_3405547, EPI_ISL_3405548, EPI_ISL_3405549, EPI_ISL_3405550, EPI_ISL_3405551, EPI_ISL_3405552, EPI_ISL_3405553, EPI_ISL_3405554, EPI_ISL_3405555, EPI_ISL_3405556, EPI_ISL_3405557, EPI_ISL_3405558, EPI_ISL_3405559, EPI_ISL_3405560, EPI_ISL_3405561, EPI_ISL_3405562, EPI_ISL_3405563, EPI_ISL_3405564, EPI_ISL_3405565, EPI_ISL_3405566, EPI_ISL_3405567, EPI_ISL_3405568, EPI_ISL_3405569, EPI_ISL_3405570, EPI_ISL_3405571, EPI_ISL_3405572, EPI_ISL_3405573, EPI_ISL_3405574, EPI_ISL_3405575, EPI_ISL_3405577, EPI_ISL_3405578, EPI_ISL_3405579, EPI_ISL_3405580, EPI_ISL_3405581, EPI_ISL_3405582, EPI_ISL_3405583, EPI_ISL_3405584, EPI_ISL_3405585, EPI_ISL_3405586, EPI_ISL_3405587, EPI_ISL_3405588, EPI_ISL_3405589, EPI_ISL_3405590, EPI_ISL_3405591, EPI_ISL_3405592, EPI_ISL_3405594, EPI_ISL_3405595, EPI_ISL_3405596, EPI_ISL_3405598, EPI_ISL_3405625, EPI_ISL_3405696, EPI_ISL_3405725, EPI_ISL_3405796, EPI_ISL_3405825, EPI_ISL_3405896, EPI_ISL_3405925, EPI_ISL_3406021, EPI_ISL_3406028, EPI_ISL_3406041, EPI_ISL_3406048, EPI_ISL_3406057, EPI_ISL_3406058, EPI_ISL_3406059, EPI_ISL_3406060, EPI_ISL_3406061, EPI_ISL_3406062, EPI_ISL_3406063, EPI_ISL_3406064, EPI_ISL_3406065, EPI_ISL_3406066, EPI_ISL_3406067, EPI_ISL_3406068, EPI_ISL_3406069, EPI_ISL_3406070, EPI_ISL_3406071, EPI_ISL_3406072, EPI_ISL_3406073, EPI_ISL_3406074, EPI_ISL_3406075, EPI_ISL_3406076, EPI_ISL_3406077, EPI_ISL_3406078, EPI_ISL_3406079, EPI_ISL_3406080, EPI_ISL_3406081, EPI_ISL_3406082, EPI_ISL_3406083, EPI_ISL_3406084, EPI_ISL_3406085, EPI_ISL_3406086, EPI_ISL_3406087, EPI_ISL_3406088, EPI_ISL_3406089, EPI_ISL_3406090, EPI_ISL_3406091, EPI_ISL_3406092, EPI_ISL_3406093, EPI_ISL_3406094, EPI_ISL_3406095, EPI_ISL_3406096, EPI_ISL_3406097, EPI_ISL_3406098, EPI_ISL_3406099, EPI_ISL_3406101, EPI_ISL_3406102, EPI_ISL_3406103, EPI_ISL_3406104, EPI_ISL_3406105, EPI_ISL_3406106, EPI_ISL_3406107, EPI_ISL_3406108, EPI_ISL_3406109, EPI_ISL_3406110, EPI_ISL_3406111, EPI_ISL_3406112, EPI_ISL_3406113, EPI_ISL_3406114, EPI_ISL_3406115, EPI_ISL_3406116, EPI_ISL_3406117, EPI_ISL_3406118, EPI_ISL_3406119, EPI_ISL_3406120, EPI_ISL_3406121, EPI_ISL_3406122, EPI_ISL_3406123, EPI_ISL_3406124, EPI_ISL_3406125, EPI_ISL_3406126, EPI_ISL_3406127, EPI_ISL_3406128, EPI_ISL_3406129, EPI_ISL_3406130, EPI_ISL_3406131, EPI_ISL_3406132, EPI_ISL_3406133, EPI_ISL_3406134, EPI_ISL_3406135, EPI_ISL_3406136, EPI_ISL_3406137, EPI_ISL_3406138, EPI_ISL_3406139, EPI_ISL_3406140, EPI_ISL_3406141, EPI_ISL_3406142, EPI_ISL_3406143, EPI_ISL_3406144, EPI_ISL_3406145, EPI_ISL_3406146, EPI_ISL_3406147, EPI_ISL_3406148, EPI_ISL_3406149, EPI_ISL_3406150, EPI_ISL_3406151, EPI_ISL_3406152, EPI_ISL_3406153, EPI_ISL_3406154, EPI_ISL_3406155, EPI_ISL_3406156, EPI_ISL_3406157, EPI_ISL_3406158, EPI_ISL_3406159, EPI_ISL_3406160, EPI_ISL_3406161, EPI_ISL_3406162, EPI_ISL_3406163, EPI_ISL_3406164, EPI_ISL_3406165, EPI_ISL_3406166, EPI_ISL_3406167, EPI_ISL_3406168, EPI_ISL_3406169, EPI_ISL_3406170, EPI_ISL_3406171, EPI_ISL_3406172, EPI_ISL_3406173, EPI_ISL_3406174, EPI_ISL_3406175, EPI_ISL_3406176, EPI_ISL_3406177, EPI_ISL_3406179, EPI_ISL_3406180, EPI_ISL_3406181, EPI_ISL_3406182, EPI_ISL_3406183, EPI_ISL_3406184, EPI_ISL_3406185, EPI_ISL_3406186, EPI_ISL_3406187, EPI_ISL_3406188, EPI_ISL_3406189, EPI_ISL_3406190, EPI_ISL_3406191, EPI_ISL_3406192, EPI_ISL_3406193, EPI_ISL_3406194, EPI_ISL_3406195, EPI_ISL_3406196, EPI_ISL_3406197, EPI_ISL_3406198, EPI_ISL_3406199, EPI_ISL_3406200, EPI_ISL_3406201, EPI_ISL_3406202, EPI_ISL_3406203, EPI_ISL_3406204, EPI_ISL_3406205, EPI_ISL_3406206, EPI_ISL_3406207, EPI_ISL_3406208, EPI_ISL_3406209, EPI_ISL_3406210, EPI_ISL_3406211, EPI_ISL_3406212, EPI_ISL_3406213, EPI_ISL_3406214, EPI_ISL_3406215, EPI_ISL_3406216, EPI_ISL_3406217, EPI_ISL_3406218, EPI_ISL_3406219, EPI_ISL_3406220, EPI_ISL_3406221, EPI_ISL_3406222, EPI_ISL_3406223, EPI_ISL_3406224, EPI_ISL_3406225, EPI_ISL_3406226, EPI_ISL_3406227, EPI_ISL_3406228, EPI_ISL_3406229, EPI_ISL_3406230, EPI_ISL_3406231, EPI_ISL_3406232, EPI_ISL_3406233, EPI_ISL_3406234, EPI_ISL_3406235, EPI_ISL_3406236, EPI_ISL_3406237, EPI_ISL_3406238, EPI_ISL_3406239, EPI_ISL_3406240, EPI_ISL_3406241, EPI_ISL_3406242, EPI_ISL_3406243, EPI_ISL_3406244, EPI_ISL_3406245, EPI_ISL_3406246, EPI_ISL_3406247, EPI_ISL_3406248, EPI_ISL_3406249, EPI_ISL_3406250, EPI_ISL_3406251, EPI_ISL_3406252, EPI_ISL_3406253, EPI_ISL_3406254, EPI_ISL_3406255, EPI_ISL_3406257, EPI_ISL_3406258, EPI_ISL_3406259, EPI_ISL_3406260, EPI_ISL_3406261, EPI_ISL_3406262, EPI_ISL_3406263, EPI_ISL_3406264, EPI_ISL_3406265, EPI_ISL_3406266, EPI_ISL_3406267, EPI_ISL_3406268, EPI_ISL_3406269, EPI_ISL_3406270, EPI_ISL_3406271, EPI_ISL_3406272, EPI_ISL_3406273, EPI_ISL_3406446, EPI_ISL_3406505, EPI_ISL_3406689, EPI_ISL_3406748, EPI_ISL_3406932 | | | |

| | | | |
|---|---|---|---|
| see above | Ministry of Health Turkey | Ministry of Health Turkey | Fatma Bayrakdar, Yasemin Cosgun, Suleyman Yalcin, Gulay Korukluoglu |
| EPI_ISL_3406993 | Regional Medical Sciences Center 7 Khon Kaen | National Institute of Health, Department of Medical Sciences, Ministry of Public Health, Thailand | Pilailuk Okada; Siripaporn Phuygun; Nuttida Thongpramul; Sittiporn Parnmen; Ratana Tacharoenmuang; Pakorn Piromtong; Natchaya Khiadsang; Thanutsapa Thanadachakul; Warawan Wongboot; sirikanda wimol; Sunthareeya Waicharoen; |
| EPI_ISL_3406994 | Regional Medical Sciences Center 6 Chonburi | National Institute of Health, Department of Medical Sciences, Ministry of Public Health, Thailand | Pilailuk Okada; Siripaporn Phuygun; Nuttida Thongpramul; Sittiporn Parnmen; Ratana Tacharoenmuang; Pakorn Piromtong; Natchaya Khiadsang; Thanutsapa Thanadachakul; Warawan Wongboot; sirikanda wimol; Sunthareeya Waicharoen; |
| EPI_ISL_3406995 | Regional Medical Sciences Center 2 Phitsanulok | National Institute of Health, Department of Medical Sciences, Ministry of Public Health, Thailand | Pilailuk Okada; Siripaporn Phuygun; Nuttida Thongpramul; Sittiporn Parnmen; Ratana Tacharoenmuang; Pakorn Piromtong; Natchaya Khiadsang; Thanutsapa Thanadachakul; Warawan Wongboot; sirikanda wimol; Sunthareeya Waicharoen; |
| EPI_ISL_3406996 | Regional Medical Sciences Center 7 Khon Kaen | National Institute of Health, Department of Medical Sciences, Ministry of Public Health, Thailand | Pilailuk Okada; Siripaporn Phuygun; Nuttida Thongpramul; Sittiporn Parnmen; Ratana Tacharoenmuang; Pakorn Piromtong; Natchaya Khiadsang; Thanutsapa Thanadachakul; Warawan Wongboot; sirikanda wimol; Sunthareeya Waicharoen; |
| EPI_ISL_3406997, EPI_ISL_3406998 | Phra Nang Klao Hospital | National Institute of Health, Department of Medical Sciences, Ministry of Public Health, Thailand | Pilailuk Okada; Siripaporn Phuygun; Nuttida Thongpramul; Sittiporn Parnmen; Ratana Tacharoenmuang; Pakorn Piromtong; Natchaya Khiadsang; Thanutsapa Thanadachakul; Warawan Wongboot; sirikanda wimol; Sunthareeya Waicharoen; |
| EPI_ISL_3406999, EPI_ISL_3407000 | Regional Medical Sciences Center 5 Samut Songkhram | National Institute of Health, Department of Medical Sciences, Ministry of Public Health, Thailand | Pilailuk Okada; Siripaporn Phuygun; Nuttida Thongpramul; Sittiporn Parnmen; Ratana Tacharoenmuang; Pakorn Piromtong; Natchaya Khiadsang; Thanutsapa Thanadachakul; Warawan Wongboot; sirikanda wimol; Sunthareeya Waicharoen; |
| EPI_ISL_3407001 | Chaophya Hospital | National Institute of Health, Department of Medical Sciences, Ministry of Public Health, Thailand | Pilailuk Okada; Siripaporn Phuygun; Nuttida Thongpramul; Sittiporn Parnmen; Ratana Tacharoenmuang; Pakorn Piromtong; Natchaya Khiadsang; Thanutsapa Thanadachakul; Warawan Wongboot; sirikanda wimol; Sunthareeya Waicharoen; |
| EPI_ISL_3407002, EPI_ISL_3407003 | Regional Medical Sciences Center 7 Khon Kaen | National Institute of Health, Department of Medical Sciences, Ministry of Public Health, Thailand | Pilailuk Okada; Siripaporn Phuygun; Nuttida Thongpramul; Sittiporn Parnmen; Ratana Tacharoenmuang; Pakorn Piromtong; Natchaya Khiadsang; Thanutsapa Thanadachakul; Warawan Wongboot; sirikanda wimol; Sunthareeya Waicharoen; |
| EPI_ISL_3407004 | Phyathai 3 Hospital | National Institute of Health, Department of Medical Sciences, Ministry of Public Health, Thailand | Pilailuk Okada; Siripaporn Phuygun; Nuttida Thongpramul; Sittiporn Parnmen; Ratana Tacharoenmuang; Pakorn Piromtong; Natchaya Khiadsang; Thanutsapa Thanadachakul; Warawan Wongboot; sirikanda wimol; Sunthareeya Waicharoen; |

EPI_ISL_3407005, EPI_ISL_3407006, EPI_ISL_3407007, EPI_ISL_3407008, EPI_ISL_3407009, EPI_ISL_3407010, EPI_ISL_3407011, EPI_ISL_3407012, EPI_ISL_3407013, EPI_ISL_3407014, EPI_ISL_3407015, EPI_ISL_3407016, EPI_ISL_3407017, EPI_ISL_3407018, EPI_ISL_3407019, EPI_ISL_3407020, EPI_ISL_3407021, EPI_ISL_3407022, EPI_ISL_3407023, EPI_ISL_3407024, EPI_ISL_3407025, EPI_ISL_3407026, EPI_ISL_3407027, EPI_ISL_3407028, EPI_ISL_3407029, EPI_ISL_3407030, EPI_ISL_3407031, EPI_ISL_3407032, EPI_ISL_3407033, EPI_ISL_3407034, EPI_ISL_3407035, EPI_ISL_3407036, EPI_ISL_3407037, EPI_ISL_3407038, EPI_ISL_3407039, EPI_ISL_3407040, EPI_ISL_3407041, EPI_ISL_3407042, EPI_ISL_3407043, EPI_ISL_3407045, EPI_ISL_3407046, EPI_ISL_3407047, EPI_ISL_3407048, EPI_ISL_3407049, EPI_ISL_3407050, EPI_ISL_3407051, EPI_ISL_3407052, EPI_ISL_3407053, EPI_ISL_3407054, EPI_ISL_3407055, EPI_ISL_3407056, EPI_ISL_3407057, EPI_ISL_3407058, EPI_ISL_3407059, EPI_ISL_3407060, EPI_ISL_3407061, EPI_ISL_3407062, EPI_ISL_3407063, EPI_ISL_3407064, EPI_ISL_3407065, EPI_ISL_3407066, EPI_ISL_3407067, EPI_ISL_3407069, EPI_ISL_3407070, EPI_ISL_3407071, EPI_ISL_3407072, EPI_ISL_3407073, EPI_ISL_3407074, EPI_ISL_3407075, EPI_ISL_3407076, EPI_ISL_3407077, EPI_ISL_3407078, EPI_ISL_3407079, EPI_ISL_3407080, EPI_ISL_3407081, EPI_ISL_3407082, EPI_ISL_3407083, EPI_ISL_3407084, EPI_ISL_3407085, EPI_ISL_3407086, EPI_ISL_3407087, EPI_ISL_3407088, EPI_ISL_3407089, EPI_ISL_3407090, EPI_ISL_3407091, EPI_ISL_3407092, EPI_ISL_3407093, EPI_ISL_3407094, EPI_ISL_3407095, EPI_ISL_3407096, EPI_ISL_3407097, EPI_ISL_3407098, EPI_ISL_3407099, EPI_ISL_3407100, EPI_ISL_3407101, EPI_ISL_3407102, EPI_ISL_3407103, EPI_ISL_3407104, EPI_ISL_3407105, EPI_ISL_3407106, EPI_ISL_3407107, EPI_ISL_3407108, EPI_ISL_3407109, EPI_ISL_3407110, EPI_ISL_3407111, EPI_ISL_3407112, EPI_ISL_3407113, EPI_ISL_3407114, EPI_ISL_3407115, EPI_ISL_3407116, EPI_ISL_3407117, EPI_ISL_3407118, EPI_ISL_3407119, EPI_ISL_3407120, EPI_ISL_3407121, EPI_ISL_3407122, EPI_ISL_3407123, EPI_ISL_3407124, EPI_ISL_3407125, EPI_ISL_3407127, EPI_ISL_3407128, EPI_ISL_3407129, EPI_ISL_3407130, EPI_ISL_3407131, EPI_ISL_3407132, EPI_ISL_3407133, EPI_ISL_3407134, EPI_ISL_3407135, EPI_ISL_3407136, EPI_ISL_3407137, EPI_ISL_3407138, EPI_ISL_3407139, EPI_ISL_3407140, EPI_ISL_3407141, EPI_ISL_3407142, EPI_ISL_3407143, EPI_ISL_3407144, EPI_ISL_3407145, EPI_ISL_3407146, EPI_ISL_3407147, EPI_ISL_3407148, EPI_ISL_3407149, EPI_ISL_3407150, EPI_ISL_3407151, EPI_ISL_3407152, EPI_ISL_3407153, EPI_ISL_3407154, EPI_ISL_3407155, EPI_ISL_3407156, EPI_ISL_3407157, EPI_ISL_3407158, EPI_ISL_3407159, EPI_ISL_3407160, EPI_ISL_3407161, EPI_ISL_3407162, EPI_ISL_3407163, EPI_ISL_3407164, EPI_ISL_3407165, EPI_ISL_3407166, EPI_ISL_3407167, EPI_ISL_3407168, EPI_ISL_3407169, EPI_ISL_3407170, EPI_ISL_3407171, EPI_ISL_3407172, EPI_ISL_3407173, EPI_ISL_3407174, EPI_ISL_3407175, EPI_ISL_3407176, EPI_ISL_3407177, EPI_ISL_3407178, EPI_ISL_3407179, EPI_ISL_3407180, EPI_ISL_3407181, EPI_ISL_3407182, EPI_ISL_3407183, EPI_ISL_3407184, EPI_ISL_3407185, EPI_ISL_3407186, EPI_ISL_3407187, EPI_ISL_3407188, EPI_ISL_3407189, EPI_ISL_3407191, EPI_ISL_3407192, EPI_ISL_3407193, EPI_ISL_3407194, EPI_ISL_3407195, EPI_ISL_3407197, EPI_ISL_3407198, EPI_ISL_3407199, EPI_ISL_3407200, EPI_ISL_3407201, EPI_ISL_3407202, EPI_ISL_3407203, EPI_ISL_3407204, EPI_ISL_3407205, EPI_ISL_3407206, EPI_ISL_3407207, EPI_ISL_3407208, EPI_ISL_3407209, EPI_ISL_3407210, EPI_ISL_3407211, EPI_ISL_3407212, EPI_ISL_3407213, EPI_ISL_3407214, EPI_ISL_3407215, EPI_ISL_3407216, EPI_ISL_3407217, EPI_ISL_3407218, EPI_ISL_3407219, EPI_ISL_3407220, EPI_ISL_3407221, EPI_ISL_3407222, EPI_ISL_3407223, EPI_ISL_3407224, EPI_ISL_3407225, EPI_ISL_3407226, EPI_ISL_3407227, EPI_ISL_3407228, EPI_ISL_3407229, EPI_ISL_3407230, EPI_ISL_3407231, EPI_ISL_3407232, EPI_ISL_3407233, EPI_ISL_3407234, EPI_ISL_3407235, EPI_ISL_3407236, EPI_ISL_3407237, EPI_ISL_3407238, EPI_ISL_3407239, EPI_ISL_3407240, EPI_ISL_3407241, EPI_ISL_3407242, EPI_ISL_3407243, EPI_ISL_3407244, EPI_ISL_3407245, EPI_ISL_3407246, EPI_ISL_3407247, EPI_ISL_3407248, EPI_ISL_3407249, EPI_ISL_3407250, EPI_ISL_3407251, EPI_ISL_3407252, EPI_ISL_3407253, EPI_ISL_3407254, EPI_ISL_3407256, EPI_ISL_3407257, EPI_ISL_3407258, EPI_ISL_3407260, EPI_ISL_3407261, EPI_ISL_3407262, EPI_ISL_3407263, EPI_ISL_3407264, EPI_ISL_3407265, EPI_ISL_3407266, EPI_ISL_3407267, EPI_ISL_3407268, EPI_ISL_3407269, EPI_ISL_3407270, EPI_ISL_3407271, EPI_ISL_3407272, EPI_ISL_3407273, EPI_ISL_3407274, EPI_ISL_3407275, EPI_ISL_3407276, EPI_ISL_3407277, EPI_ISL_3407278, EPI_ISL_3407279, EPI_ISL_3407280, EPI_ISL_3407281, EPI_ISL_3407282, EPI_ISL_3407283, EPI_ISL_3407284, EPI_ISL_3407285, EPI_ISL_3407286, EPI_ISL_3407287, EPI_ISL_3407288, EPI_ISL_3407289, EPI_ISL_3407290, EPI_ISL_3407291, EPI_ISL_3407292, EPI_ISL_3407293, EPI_ISL_3407294, EPI_ISL_3407295, EPI_ISL_3407296, EPI_ISL_3407297, EPI_ISL_3407298, EPI_ISL_3407299, EPI_ISL_3407300, EPI_ISL_3407301, EPI_ISL_3407302, EPI_ISL_3407303, EPI_ISL_3407304, EPI_ISL_3407305, EPI_ISL_3407306, EPI_ISL_3407307, EPI_ISL_3407308, EPI_ISL_3407309, EPI_ISL_3407310, EPI_ISL_3407311, EPI_ISL_3407312, EPI_ISL_3407313, EPI_ISL_3407314, EPI_ISL_3407315, EPI_ISL_3407316, EPI_ISL_3407317, EPI_ISL_3407318, EPI_ISL_3407319, EPI_ISL_3407320, EPI_ISL_3407321, EPI_ISL_3407322, EPI_ISL_3407323, EPI_ISL_3407324, EPI_ISL_3407325, EPI_ISL_3407326, EPI_ISL_3407327, EPI_ISL_3407328, EPI_ISL_3407329, EPI_ISL_3407330, EPI_ISL_3407331, EPI_ISL_3407332, EPI_ISL_3407333, EPI_ISL_3407334, EPI_ISL_3407335, EPI_ISL_3407336, EPI_ISL_3407337, EPI_ISL_3407338, EPI_ISL_3407339, EPI_ISL_3407340, EPI_ISL_3407341, EPI_ISL_3407342, EPI_ISL_3407343, EPI_ISL_3407344, EPI_ISL_3407346, EPI_ISL_3407347, EPI_ISL_3407348, EPI_ISL_3407349, EPI_ISL_3407350, EPI_ISL_3407351, EPI_ISL_3407352, EPI_ISL_3407353, EPI_ISL_3407354, EPI_ISL_3407355, EPI_ISL_3407356, EPI_ISL_3407357, EPI_ISL_3407358, EPI_ISL_3407359, EPI_ISL_3407360, EPI_ISL_3407361, EPI_ISL_3407362, EPI_ISL_3407363, EPI_ISL_3407364, EPI_ISL_3407365, EPI_ISL_3407366, EPI_ISL_3407367, EPI_ISL_3407368, EPI_ISL_3407369, EPI_ISL_3407370, EPI_ISL_3407371, EPI_ISL_3407372, EPI_ISL_3407373, EPI_ISL_3407374, EPI_ISL_3407375, EPI_ISL_3407376, EPI_ISL_3407377, EPI_ISL_3407378, EPI_ISL_3407379, EPI_ISL_3407380, EPI_ISL_3407381, EPI_ISL_3407382, EPI_ISL_3407383, EPI_ISL_3407384, EPI_ISL_3407385, EPI_ISL_3407386, EPI_ISL_3407387, EPI_ISL_3407388, EPI_ISL_3407389, EPI_ISL_3407390, EPI_ISL_3407391, EPI_ISL_3407392, EPI_ISL_3407393, EPI_ISL_3407394, EPI_ISL_3407395, EPI_ISL_3407396, EPI_ISL_3407397, EPI_ISL_3407398, EPI_ISL_3407399, EPI_ISL_3407400, EPI_ISL_3407401, EPI_ISL_3407402, EPI_ISL_3407403, EPI_ISL_3407404, EPI_ISL_3407405, EPI_ISL_3407406, EPI_ISL_3407407, EPI_ISL_3407408, EPI_ISL_3407409, EPI_ISL_3407410, EPI_ISL_3407411, EPI_ISL_3407412, EPI_ISL_3407413, EPI_ISL_3407414, EPI_ISL_3407415, EPI_ISL_3407416, EPI_ISL_3407417, EPI_ISL_3407418, EPI_ISL_3407419, EPI_ISL_3407420, EPI_ISL_3407421, EPI_ISL_3407422, EPI_ISL_3407423, EPI_ISL_3407424, EPI_ISL_3407425, EPI_ISL_3407426, EPI_ISL_3407427, EPI_ISL_3407428, EPI_ISL_3407429, EPI_ISL_3407430, EPI_ISL_3407431, EPI_ISL_3407432, EPI_ISL_3407433, EPI_ISL_3407434, EPI_ISL_3407435, EPI_ISL_3407436, EPI_ISL_3407437, EPI_ISL_3407438, EPI_ISL_3407439, EPI_ISL_3407440, EPI_ISL_3407441, EPI_ISL_3407442, EPI_ISL_3407443, EPI_ISL_3407444, EPI_ISL_3407445, EPI_ISL_3407446, EPI_ISL_3407447, EPI_ISL_3407449, EPI_ISL_3407450, EPI_ISL_3407451, EPI_ISL_3407452, EPI_ISL_3407453, EPI_ISL_3407454, EPI_ISL_3407455, EPI_ISL_3407456, EPI_ISL_3407457, EPI_ISL_3407458, EPI_ISL_3407459, EPI_ISL_3407460, EPI_ISL_3407461, EPI_ISL_3407462, EPI_ISL_3407463, EPI_ISL_3407464, EPI_ISL_3407465, EPI_ISL_3407466, EPI_ISL_3407467, EPI_ISL_3407468, EPI_ISL_3407469, EPI_ISL_3407470, EPI_ISL_3407471, EPI_ISL_3407472, EPI_ISL_3407473, EPI_ISL_3407474, EPI_ISL_3407475, EPI_ISL_3407476, EPI_ISL_3407477, EPI_ISL_3407478, EPI_ISL_3407479, EPI_ISL_3407480, EPI_ISL_3407481, EPI_ISL_3407482, EPI_ISL_3407483, EPI_ISL_3407484, EPI_ISL_3407485, EPI_ISL_3407486, EPI_ISL_3407487, EPI_ISL_3407488, EPI_ISL_3407489, EPI_ISL_3407490, EPI_ISL_3407491, EPI_ISL_3407492, EPI_ISL_3407493, EPI_ISL_3407494, EPI_ISL_3407495, EPI_ISL_3407496, EPI_ISL_3407497, EPI_ISL_3407498, EPI_ISL_3407499, EPI_ISL_3407500, EPI_ISL_3407501, EPI_ISL_3407502, EPI_ISL_3407503, EPI_ISL_3407504, EPI_ISL_3407505, EPI_ISL_3407506, EPI_ISL_3407507, EPI_ISL_3407508, EPI_ISL_3407509, EPI_ISL_3407510, EPI_ISL_3407511, EPI_ISL_3407512, EPI_ISL_3407513, EPI_ISL_3407514, EPI_ISL_3407515, EPI_ISL_3407516, EPI_ISL_3407517, EPI_ISL_3407518, EPI_ISL_3407519, EPI_ISL_3407520, EPI_ISL_3407521, EPI_ISL_3407522, EPI_ISL_3407523, EPI_ISL_3407524, EPI_ISL_3407525, EPI_ISL_3407526, EPI_ISL_3407527, EPI_ISL_3407528, EPI_ISL_3407529, EPI_ISL_3407530, EPI_ISL_3407531, EPI_ISL_3407532, EPI_ISL_3407533, EPI_ISL_3407534, EPI_ISL_3407535, EPI_ISL_3407536, EPI_ISL_3407537, EPI_ISL_3407538, EPI_ISL_3407539, EPI_ISL_3407540, EPI_ISL_3407541, EPI_ISL_3407542, EPI_ISL_3407543, EPI_ISL_3407544, EPI_ISL_3407545, EPI_ISL_3407546, EPI_ISL_3407547, EPI_ISL_3407548, EPI_ISL_3407549, EPI_ISL_3407550, EPI_ISL_3407551, EPI_ISL_3407552, EPI_ISL_3407553, EPI_ISL_3407554, EPI_ISL_3407555, EPI_ISL_3407556, EPI_ISL_3407557, EPI_ISL_3407558, EPI_ISL_3407559, EPI_ISL_3407560, EPI_ISL_3407561, EPI_ISL_3407562, EPI_ISL_3407563, EPI_ISL_3407564, EPI_ISL_3407565, EPI_ISL_3407566, EPI_ISL_3407567, EPI_ISL_3407568, EPI_ISL_3407569, EPI_ISL_3407570, EPI_ISL_3407571, EPI_ISL_3407572, EPI_ISL_3407573, EPI_ISL_3407574, EPI_ISL_3407575, EPI_ISL_3407576, EPI_ISL_3407577, EPI_ISL_3407578, EPI_ISL_3407579, EPI_ISL_3407580, EPI_ISL_3407581, EPI_ISL_3407582, EPI_ISL_3407583, EPI_ISL_3407584, EPI_ISL_3407585, EPI_ISL_3407586, EPI_ISL_3407587, EPI_ISL_3407588, EPI_ISL_3407589, EPI_ISL_3407590, EPI_ISL_3407591, EPI_ISL_3407592, EPI_ISL_3407593, EPI_ISL_3407594, EPI_ISL_3407595, EPI_ISL_3407596, EPI_ISL_3407597, EPI_ISL_3407598, EPI_ISL_3407599, EPI_ISL_3407600, EPI_ISL_3407601, EPI_ISL_3407602, EPI_ISL_3407603, EPI_ISL_3407604, EPI_ISL_3407605, EPI_ISL_3407606, EPI_ISL_3407607, EPI_ISL_3407608, EPI_ISL_3407609, EPI_ISL_3407610, EPI_ISL_3407611, EPI_ISL_3407612, EPI_ISL_3407613, EPI_ISL_3407614, EPI_ISL_3407615, EPI_ISL_3407616, EPI_ISL_3407617, EPI_ISL_3407618, EPI_ISL_3407619, EPI_ISL_3407620, EPI_ISL_3407621, EPI_ISL_3407622, EPI_ISL_3407623, EPI_ISL_3407624, EPI_ISL_3407625, EPI_ISL_3407626, EPI_ISL_3407627, EPI_ISL_3407628, EPI_ISL_3407629, EPI_ISL_3407630, EPI_ISL_3407631, EPI_ISL_3407632, EPI_ISL_3407633, EPI_ISL_3407634, EPI_ISL_3407635, EPI_ISL_3407636, EPI_ISL_3407637, EPI_ISL_3407638, EPI_ISL_3407639, EPI_ISL_3407640, EPI_ISL_3407641, EPI_ISL_3407642, EPI_ISL_3407643, EPI_ISL_3407644, EPI_ISL_3407645, EPI_ISL_3407646, EPI_ISL_3407647, EPI_ISL_3407648, EPI_ISL_3407649, EPI_ISL_3407650, EPI_ISL_3407651, EPI_ISL_3407652, EPI_ISL_3407653, EPI_ISL_3407654, EPI_ISL_3407655, EPI_ISL_3407656, EPI_ISL_3407657, EPI_ISL_3407658, EPI_ISL_3407659, EPI_ISL_3407660, EPI_ISL_3407661, EPI_ISL_3407662, EPI_ISL_3407663, EPI_ISL_3407664, EPI_ISL_3407665, EPI_ISL_3407666, EPI_ISL_3407667, EPI_ISL_3407668, EPI_ISL_3407669, EPI_ISL_3407670, EPI_ISL_3407671, EPI_ISL_3407672, EPI_ISL_3407673, EPI_ISL_3407674, EPI_ISL_3407675, EPI_ISL_3407676, EPI_ISL_3407677, EPI_ISL_3407678, EPI_ISL_3407679, EPI_ISL_3407680, EPI_ISL_3407681, EPI_ISL_3407682, EPI_ISL_3407683, EPI_ISL_3407684, EPI_ISL_3407685, EPI_ISL_3407686, EPI_ISL_3407687, EPI_ISL_3407688, EPI_ISL_3407689, EPI_ISL_3407690, EPI_ISL_3407691, EPI_ISL_3407692, EPI_ISL_3407693, EPI_ISL_3407694, EPI_ISL_3407695, EPI_ISL_3407696, EPI_ISL_3407697, EPI_ISL_3407698, EPI_ISL_3407699, EPI_ISL_3407700, EPI_ISL_3407701, EPI_ISL_3407702, EPI_ISL_3407703, EPI_ISL_3407704, EPI_ISL_3407705, EPI_ISL_3407706, EPI_ISL_3407707, EPI_ISL_3407708, EPI_ISL_3407709, EPI_ISL_3407710, EPI_ISL_3407711, EPI_ISL_3407712, EPI_ISL_3407713, EPI_ISL_3407714, EPI_ISL_3407715, EPI_ISL_3407717, EPI_ISL_3407718, EPI_ISL_3407719, EPI_ISL_3407720, EPI_ISL_3407721, EPI_ISL_3407722, EPI_ISL_3407723, EPI_ISL_3407724, EPI_ISL_3407725, EPI_ISL_3407726, EPI_ISL_3407727, EPI_ISL_3407728, EPI_ISL_3407729, EPI_ISL_3407730, EPI_ISL_3407731, EPI_ISL_3407732, EPI_ISL_3407733, EPI_ISL_3407734, EPI_ISL_3407735, EPI_ISL_3407736, EPI_ISL_3407737, EPI_ISL_3407738, EPI_ISL_3407739, EPI_ISL_3407740, EPI_ISL_3407741, EPI_ISL_3407742, EPI_ISL_3407743, EPI_ISL_3407744, EPI_ISL_3407745, EPI_ISL_3407746, EPI_ISL_3407747, EPI_ISL_3407748, EPI_ISL_3407749, EPI_ISL_3407750, EPI_ISL_3407751, EPI_ISL_3407752, EPI_ISL_3407753, EPI_ISL_3407754, EPI_ISL_3407755, EPI_ISL_3407756, EPI_ISL_3407757, EPI_ISL_3407762, EPI_ISL_3407763, EPI_ISL_3407764, EPI_ISL_3407765, EPI_ISL_3407766, EPI_ISL_3407767, EPI_ISL_3407768, EPI_ISL_3407769, EPI_ISL_3407770, EPI_ISL_3407771, EPI_ISL_3407772, EPI_ISL_3407773, EPI_ISL_3407774, EPI_ISL_3407775, EPI_ISL_3407776, EPI_ISL_3407777, EPI_ISL_3407778, EPI_ISL_3407779, EPI_ISL_3407780, EPI_ISL_3407781, EPI_ISL_3407782, EPI_ISL_3407783, EPI_ISL_3407784, EPI_ISL_3407785, EPI_ISL_3407786, EPI_ISL_3407787, EPI_ISL_3407788, EPI_ISL_3407789, EPI_ISL_3407790, EPI_ISL_3407791, EPI_ISL_3407792, EPI_ISL_3407793, EPI_ISL_3407794, EPI_ISL_3407795, EPI_ISL_3407796, EPI_ISL_3407797, EPI_ISL_3407798, EPI_ISL_3407799, EPI_ISL_3407800, EPI_ISL_3407801, EPI_ISL_3407802, EPI_ISL_3407803, EPI_ISL_3407804, EPI_ISL_3407805, EPI_ISL_3407806, EPI_ISL_3407807, EPI_ISL_3407808, EPI_ISL_3407809, EPI_ISL_3407810, EPI_ISL_3407811, EPI_ISL_3407812, EPI_ISL_3407813, EPI_ISL_3407814

| | | | |
|---|---|---|---|
| see above | Broad Institute Clinical Research Sequencing Platform | Infectious Disease Program, Broad Institute of Harvard and MIT | Siddle,K.J., Adams,G., Pearlman,L., Gladden-Young,A., Vicente,G., Blumenstiel,B., DeFelice,M., Lee,M., McGovern,S., Lagerborg,K., Rudy,M., DeRuff,K., Carter,A., Normandin,E., Bauer,M., Reilly,S., Tomkins-Tinch,C., Loreth,C., Chaluvadi,S., Meldrim,J., Granger,B., Lemieux,J.E., Birren,B.W., Sabeti,P.C., Larkin,K., Dodge,S., Lennon,N., Madoff,L., Brown,C., Gallagher,G., Smole,S., Park,D.J., Gabriel,S., and MacInnis,B.L. |
| EPI_ISL_3407815 | Regional Medical Sciences Center 1/1 Chiang Rai | National Institute of Health, Department of Medical Sciences, Ministry of Public Health, Thailand | Pilailuk Okada; Siripaporn Phuygun; Nuttida Thongpramul; Sittiporn Parnmen; Ratana Tacharoenmuang; Pakorn Piromtong; Natchaya Khiadsang; Thanutsapa Thanadachakul; Warawan Wongboot; sirikanda wimol; Sunthareeya Waicharoen; |
| EPI_ISL_3407816 | Regional Medical Sciences Center 12/1 Trang | National Institute of Health, Department of Medical Sciences, Ministry of Public Health, Thailand | Pilailuk Okada; Siripaporn Phuygun; Nuttida Thongpramul; Sittiporn Parnmen; Ratana Tacharoenmuang; Pakorn Piromtong; Natchaya Khiadsang; Thanutsapa Thanadachakul; Warawan Wongboot; sirikanda wimol; Sunthareeya Waicharoen; |
| EPI_ISL_3407817 | Nonthaburi Public Health Office | National Institute of Health, Department of Medical Sciences, Ministry of Public Health, Thailand | Pilailuk Okada; Siripaporn Phuygun; Nuttida Thongpramul; Sittiporn Parnmen; Ratana Tacharoenmuang; Pakorn Piromtong; Natchaya Khiadsang; Thanutsapa Thanadachakul; Warawan Wongboot; sirikanda wimol; Sunthareeya Waicharoen; |
| EPI_ISL_3407818 | Regional Medical Sciences Center 2 Phitsanulok | National Institute of Health, Department of Medical Sciences, Ministry of Public Health, Thailand | Pilailuk Okada; Siripaporn Phuygun; Nuttida Thongpramul; Sittiporn Parnmen; Ratana Tacharoenmuang; Pakorn Piromtong; Natchaya Khiadsang; Thanutsapa Thanadachakul; Warawan Wongboot; sirikanda wimol; Sunthareeya Waicharoen; |
| EPI_ISL_3407819 | Regional Medical Sciences Center 3 Nakhon Sawan | National Institute of Health, Department of Medical Sciences, Ministry of Public Health, Thailand | Pilailuk Okada; Siripaporn Phuygun; Nuttida Thongpramul; Sittiporn Parnmen; Ratana Tacharoenmuang; Pakorn Piromtong; Natchaya Khiadsang; Thanutsapa Thanadachakul; Warawan Wongboot; sirikanda wimol; Sunthareeya Waicharoen; |
| EPI_ISL_3407820 | Chaiyaphum Hospital | National Institute of Health, Department of Medical Sciences, Ministry of Public Health, Thailand | Pilailuk Okada; Siripaporn Phuygun; Nuttida Thongpramul; Sittiporn Parnmen; Ratana Tacharoenmuang; Pakorn Piromtong; Natchaya Khiadsang; Thanutsapa Thanadachakul; Warawan Wongboot; sirikanda wimol; Sunthareeya Waicharoen; |
| EPI_ISL_3407821 | Regional Medical Sciences Center 11/1 Phuket | National Institute of Health, Department of Medical Sciences, Ministry of Public Health, Thailand | Pilailuk Okada; Siripaporn Phuygun; Nuttida Thongpramul; Sittiporn Parnmen; Ratana Tacharoenmuang; Pakorn Piromtong; Natchaya Khiadsang; Thanutsapa Thanadachakul; Warawan Wongboot; sirikanda wimol; Sunthareeya Waicharoen; |
| EPI_ISL_3407822 | Regional Medical Sciences Center 3 Nakhon Sawan | National Institute of Health, Department of Medical Sciences, Ministry of Public Health, Thailand | Pilailuk Okada; Siripaporn Phuygun; Nuttida Thongpramul; Sittiporn Parnmen; Ratana Tacharoenmuang; Pakorn Piromtong; Natchaya Khiadsang; Thanutsapa Thanadachakul; Warawan Wongboot; sirikanda wimol; Sunthareeya Waicharoen; |
| EPI_ISL_3407823, EPI_ISL_3407824, EPI_ISL_3407825 | Regional Medical Sciences Center 1 Chiang Mai | National Institute of Health, Department of Medical Sciences, Ministry of Public Health, Thailand | Pilailuk Okada; Siripaporn Phuygun; Nuttida Thongpramul; Sittiporn Parnmen; Ratana Tacharoenmuang; Pakorn Piromtong; Natchaya Khiadsang; Thanutsapa Thanadachakul; Warawan Wongboot; sirikanda wimol; Sunthareeya Waicharoen; |
| EPI_ISL_3407826 | Nonthaburi Public Health Office | National Institute of Health, Department of Medical Sciences, Ministry of Public Health, Thailand | Pilailuk Okada; Siripaporn Phuygun; Nuttida Thongpramul; Sittiporn Parnmen; Ratana Tacharoenmuang; Pakorn Piromtong; Natchaya Khiadsang; Thanutsapa Thanadachakul; Warawan Wongboot; sirikanda wimol; Sunthareeya Waicharoen; |
| EPI_ISL_3407827 | Regional Medical Sciences Center 1 Chiang Mai | National Institute of Health, Department of Medical Sciences, Ministry of Public Health, Thailand | Pilailuk Okada; Siripaporn Phuygun; Nuttida Thongpramul; Sittiporn Parnmen; Ratana Tacharoenmuang; Pakorn Piromtong; Natchaya Khiadsang; Thanutsapa Thanadachakul; Warawan Wongboot; sirikanda wimol; Sunthareeya Waicharoen; |
| EPI_ISL_3407828, EPI_ISL_3407829 | Regional Medical Sciences Center 11 Surat Thani | National Institute of Health, Department of Medical Sciences, Ministry of Public Health, Thailand | Pilailuk Okada; Siripaporn Phuygun; Nuttida Thongpramul; Sittiporn Parnmen; Ratana Tacharoenmuang; Pakorn Piromtong; Natchaya Khiadsang; Thanutsapa Thanadachakul; Warawan Wongboot; sirikanda wimol; Sunthareeya Waicharoen; |
| EPI_ISL_3407830 | Regional Medical Sciences Center 6 Chonburi | National Institute of Health, Department of Medical Sciences, Ministry of Public Health, Thailand | Pilailuk Okada; Siripaporn Phuygun; Nuttida Thongpramul; Sittiporn Parnmen; Ratana Tacharoenmuang; Pakorn Piromtong; Natchaya Khiadsang; Thanutsapa Thanadachakul; Warawan Wongboot; sirikanda wimol; Sunthareeya Waicharoen; |
| EPI_ISL_3407831 | Regional Medical Sciences Center 3 Nakhon Sawan | National Institute of Health, Department of Medical Sciences, Ministry of Public Health, Thailand | Pilailuk Okada; Siripaporn Phuygun; Nuttida Thongpramul; Sittiporn Parnmen; Ratana Tacharoenmuang; Pakorn Piromtong; Natchaya Khiadsang; Thanutsapa Thanadachakul; Warawan Wongboot; sirikanda wimol; Sunthareeya Waicharoen; |
| EPI_ISL_3407832 | Regional Medical Sciences Center 2 Phitsanulok | National Institute of Health, Department of Medical Sciences, Ministry of Public Health, Thailand | Pilailuk Okada; Siripaporn Phuygun; Nuttida Thongpramul; Sittiporn Parnmen; Ratana Tacharoenmuang; Pakorn Piromtong; Natchaya Khiadsang; Thanutsapa Thanadachakul; Warawan Wongboot; sirikanda wimol; Sunthareeya Waicharoen; |
| EPI_ISL_3407833 | Regional Medical Sciences Center 5 Samut Songkhram | National Institute of Health, Department of Medical Sciences, Ministry of Public Health, Thailand | Pilailuk Okada; Siripaporn Phuygun; Nuttida Thongpramul; Sittiporn Parnmen; Ratana Tacharoenmuang; Pakorn Piromtong; Natchaya Khiadsang; Thanutsapa Thanadachakul; Warawan Wongboot; sirikanda wimol; Sunthareeya Waicharoen; |
| EPI_ISL_3407834 | Regional Medical Sciences Center 6 Chonburi | National Institute of Health, Department of Medical Sciences, Ministry of Public Health, Thailand | Pilailuk Okada; Siripaporn Phuygun; Nuttida Thongpramul; Sittiporn Parnmen; Ratana Tacharoenmuang; Pakorn Piromtong; Natchaya Khiadsang; Thanutsapa Thanadachakul; Warawan Wongboot; sirikanda wimol; Sunthareeya Waicharoen; |
| EPI_ISL_3407835, EPI_ISL_3407836 | Regional Medical Sciences Center 2 Phitsanulok | National Institute of Health, Department of Medical Sciences, Ministry of Public Health, Thailand | Pilailuk Okada; Siripaporn Phuygun; Nuttida Thongpramul; Sittiporn Parnmen; Ratana Tacharoenmuang; Pakorn Piromtong; Natchaya Khiadsang; Thanutsapa Thanadachakul; Warawan Wongboot; sirikanda wimol; Sunthareeya Waicharoen; |
| EPI_ISL_3407837 | Regional Medical Sciences Center 11 Surat Thani | National Institute of Health, Department of Medical Sciences, Ministry of Public Health, Thailand | Pilailuk Okada; Siripaporn Phuygun; Nuttida Thongpramul; Sittiporn Parnmen; Ratana Tacharoenmuang; Pakorn Piromtong; Natchaya Khiadsang; Thanutsapa Thanadachakul; Warawan Wongboot; sirikanda wimol; Sunthareeya Waicharoen; |

| | | | |
|---|---|---|---|
| EPI_ISL_3407838 | Nonthaburi Public Health Office | National Institute of Health, Department of Medical Sciences, Ministry of Public Health, Thailand | Pilailuk Okada; Siripaporn Phuygun; Nuttida Thongpramul; Sittiporn Parnmen; Ratana Tacharoenmuang; Pakorn Piromtong; Natchaya Khiadsang; Thanutsapa Thanadachakul; Warawan Wongboot; sirikanda wimol; Sunthareeya Waicharoen; |
| EPI_ISL_3407839, EPI_ISL_3407840 | Regional Medical Sciences Center 11 Surat Thani | National Institute of Health, Department of Medical Sciences, Ministry of Public Health, Thailand | Pilailuk Okada; Siripaporn Phuygun; Nuttida Thongpramul; Sittiporn Parnmen; Ratana Tacharoenmuang; Pakorn Piromtong; Natchaya Khiadsang; Thanutsapa Thanadachakul; Warawan Wongboot; sirikanda wimol; Sunthareeya Waicharoen; |
| EPI_ISL_3407841 | Regional Medical Sciences Center 6 Chonburi | National Institute of Health, Department of Medical Sciences, Ministry of Public Health, Thailand | Pilailuk Okada; Siripaporn Phuygun; Nuttida Thongpramul; Sittiporn Parnmen; Ratana Tacharoenmuang; Pakorn Piromtong; Natchaya Khiadsang; Thanutsapa Thanadachakul; Warawan Wongboot; sirikanda wimol; Sunthareeya Waicharoen; |
| EPI_ISL_3407842 | Regional Medical Sciences Center 7 Khon Kaen | National Institute of Health, Department of Medical Sciences, Ministry of Public Health, Thailand | Pilailuk Okada; Siripaporn Phuygun; Nuttida Thongpramul; Sittiporn Parnmen; Ratana Tacharoenmuang; Pakorn Piromtong; Natchaya Khiadsang; Thanutsapa Thanadachakul; Warawan Wongboot; sirikanda wimol; Sunthareeya Waicharoen; |
| EPI_ISL_3407843 | Chulabhorn Hospital | National Institute of Health, Department of Medical Sciences, Ministry of Public Health, Thailand | Pilailuk Okada; Siripaporn Phuygun; Nuttida Thongpramul; Sittiporn Parnmen; Ratana Tacharoenmuang; Pakorn Piromtong; Natchaya Khiadsang; Thanutsapa Thanadachakul; Warawan Wongboot; sirikanda wimol; Sunthareeya Waicharoen; |
| EPI_ISL_3407853 | Regional Medical Sciences Center 12 Songkhla | National Institute of Health, Department of Medical Sciences, Ministry of Public Health, Thailand | Pilailuk Okada; Siripaporn Phuygun; Nuttida Thongpramul; Sittiporn Parnmen; Ratana Tacharoenmuang; Pakorn Piromtong; Natchaya Khiadsang; Thanutsapa Thanadachakul; Warawan Wongboot; sirikanda wimol; Sunthareeya Waicharoen; |
| EPI_ISL_3407854 | Regional Medical Sciences Center 1 Chiang Mai | National Institute of Health, Department of Medical Sciences, Ministry of Public Health, Thailand | Pilailuk Okada; Siripaporn Phuygun; Nuttida Thongpramul; Sittiporn Parnmen; Ratana Tacharoenmuang; Pakorn Piromtong; Natchaya Khiadsang; Thanutsapa Thanadachakul; Warawan Wongboot; sirikanda wimol; Sunthareeya Waicharoen; |
| EPI_ISL_3407855, EPI_ISL_3407856 | Department of Disease Control | National Institute of Health, Department of Medical Sciences, Ministry of Public Health, Thailand | Pilailuk Okada; Siripaporn Phuygun; Nuttida Thongpramul; Sittiporn Parnmen; Ratana Tacharoenmuang; Pakorn Piromtong; Natchaya Khiadsang; Thanutsapa Thanadachakul; Warawan Wongboot; sirikanda wimol; Sunthareeya Waicharoen; |
| EPI_ISL_3407857 | Regional Medical Sciences Center 9 Nakhon Ratchasima | National Institute of Health, Department of Medical Sciences, Ministry of Public Health, Thailand | Pilailuk Okada; Siripaporn Phuygun; Nuttida Thongpramul; Sittiporn Parnmen; Ratana Tacharoenmuang; Pakorn Piromtong; Natchaya Khiadsang; Thanutsapa Thanadachakul; Warawan Wongboot; sirikanda wimol; Sunthareeya Waicharoen; |
| EPI_ISL_3407858, EPI_ISL_3407859 | Regional Medical Sciences Center 11 Surat Thani | National Institute of Health, Department of Medical Sciences, Ministry of Public Health, Thailand | Pilailuk Okada; Siripaporn Phuygun; Nuttida Thongpramul; Sittiporn Parnmen; Ratana Tacharoenmuang; Pakorn Piromtong; Natchaya Khiadsang; Thanutsapa Thanadachakul; Warawan Wongboot; sirikanda wimol; Sunthareeya Waicharoen; |
| EPI_ISL_3407861 | Regional Medical Sciences Center 11/1 Phuket | National Institute of Health, Department of Medical Sciences, Ministry of Public Health, Thailand | Pilailuk Okada; Siripaporn Phuygun; Nuttida Thongpramul; Sittiporn Parnmen; Ratana Tacharoenmuang; Pakorn Piromtong; Natchaya Khiadsang; Thanutsapa Thanadachakul; Warawan Wongboot; sirikanda wimol; Sunthareeya Waicharoen; |
| EPI_ISL_3407862 | Regional Medical Sciences Center 10 Ubonratchathani | National Institute of Health, Department of Medical Sciences, Ministry of Public Health, Thailand | Pilailuk Okada; Siripaporn Phuygun; Nuttida Thongpramul; Sittiporn Parnmen; Ratana Tacharoenmuang; Pakorn Piromtong; Natchaya Khiadsang; Thanutsapa Thanadachakul; Warawan Wongboot; sirikanda wimol; Sunthareeya Waicharoen; |
| EPI_ISL_3407863 | Regional Medical Sciences Center 3 Nakhon Sawan | National Institute of Health, Department of Medical Sciences, Ministry of Public Health, Thailand | Pilailuk Okada; Siripaporn Phuygun; Nuttida Thongpramul; Sittiporn Parnmen; Ratana Tacharoenmuang; Pakorn Piromtong; Natchaya Khiadsang; Thanutsapa Thanadachakul; Warawan Wongboot; sirikanda wimol; Sunthareeya Waicharoen; |
| EPI_ISL_3407864 | Regional Medical Sciences Center 9 Nakhon Ratchasima | National Institute of Health, Department of Medical Sciences, Ministry of Public Health, Thailand | Pilailuk Okada; Siripaporn Phuygun; Nuttida Thongpramul; Sittiporn Parnmen; Ratana Tacharoenmuang; Pakorn Piromtong; Natchaya Khiadsang; Thanutsapa Thanadachakul; Warawan Wongboot; sirikanda wimol; Sunthareeya Waicharoen; |
| EPI_ISL_3407871 | Regional Medical Sciences Center 2 Phitsanulok | National Institute of Health, Department of Medical Sciences, Ministry of Public Health, Thailand | Pilailuk Okada; Siripaporn Phuygun; Nuttida Thongpramul; Sittiporn Parnmen; Ratana Tacharoenmuang; Pakorn Piromtong; Natchaya Khiadsang; Thanutsapa Thanadachakul; Warawan Wongboot; sirikanda wimol; Sunthareeya Waicharoen; |
| EPI_ISL_3407872 | Regional Medical Sciences Center 5 Samut Songkhram | National Institute of Health, Department of Medical Sciences, Ministry of Public Health, Thailand | Pilailuk Okada; Siripaporn Phuygun; Nuttida Thongpramul; Sittiporn Parnmen; Ratana Tacharoenmuang; Pakorn Piromtong; Natchaya Khiadsang; Thanutsapa Thanadachakul; Warawan Wongboot; sirikanda wimol; Sunthareeya Waicharoen; |
| EPI_ISL_3407873 | Regional Medical Sciences Center 7 Khon Kaen | National Institute of Health, Department of Medical Sciences, Ministry of Public Health, Thailand | Pilailuk Okada; Siripaporn Phuygun; Nuttida Thongpramul; Sittiporn Parnmen; Ratana Tacharoenmuang; Pakorn Piromtong; Natchaya Khiadsang; Thanutsapa Thanadachakul; Warawan Wongboot; sirikanda wimol; Sunthareeya Waicharoen; |
| EPI_ISL_3407874 | Regional Medical Sciences Center 2 Phitsanulok | National Institute of Health, Department of Medical Sciences, Ministry of Public Health, Thailand | Pilailuk Okada; Siripaporn Phuygun; Nuttida Thongpramul; Sittiporn Parnmen; Ratana Tacharoenmuang; Pakorn Piromtong; Natchaya Khiadsang; Thanutsapa Thanadachakul; Warawan Wongboot; sirikanda wimol; Sunthareeya Waicharoen; |
| EPI_ISL_3407876, EPI_ISL_3407877, EPI_ISL_3407878, EPI_ISL_3407880, EPI_ISL_3407893, EPI_ISL_3407894, EPI_ISL_3407895, EPI_ISL_3407896, EPI_ISL_3407897, EPI_ISL_3407898, EPI_ISL_3407899, EPI_ISL_3407900, EPI_ISL_3407901, EPI_ISL_3407902, EPI_ISL_3407903, EPI_ISL_3407904, EPI_ISL_3407905, EPI_ISL_3407906, EPI_ISL_3407907, EPI_ISL_3407908, EPI_ISL_3407909, EPI_ISL_3407910, EPI_ISL_3407911, EPI_ISL_3407912, EPI_ISL_3407913, EPI_ISL_3407914, EPI_ISL_3407915, EPI_ISL_3407916, EPI_ISL_3407917, EPI_ISL_3407918, EPI_ISL_3407919, EPI_ISL_3407920, EPI_ISL_3407921, EPI_ISL_3407922, EPI_ISL_3407923, EPI_ISL_3407924, EPI_ISL_3407925, EPI_ISL_3407926, EPI_ISL_3407927, EPI_ISL_3407928, EPI_ISL_3407929, EPI_ISL_3407930, EPI_ISL_3407931, EPI_ISL_3407932, EPI_ISL_3407933, EPI_ISL_3407934, EPI_ISL_3407935, EPI_ISL_3407936, EPI_ISL_3407937, EPI_ISL_3407938, EPI_ISL_3407939, EPI_ISL_3407940, EPI_ISL_3407941, EPI_ISL_3407942, EPI_ISL_3407943, EPI_ISL_3407944, EPI_ISL_3407945, EPI_ISL_3407946, EPI_ISL_3407947, EPI_ISL_3407948, EPI_ISL_3407949, EPI_ISL_3407950, EPI_ISL_3407951, EPI_ISL_3407952, EPI_ISL_3407953, EPI_ISL_3407954, EPI_ISL_3407955, EPI_ISL_3407956, EPI_ISL_3407957, EPI_ISL_3407958, EPI_ISL_3407959, EPI_ISL_3407960, EPI_ISL_3407961, EPI_ISL_3407962, EPI_ISL_3407963, EPI_ISL_3407964, EPI_ISL_3407965, EPI_ISL_3407966, EPI_ISL_3407967, EPI_ISL_3407968, EPI_ISL_3407969, EPI_ISL_3407970, EPI_ISL_3407971, EPI_ISL_3407972, EPI_ISL_3407973, EPI_ISL_3407974, EPI_ISL_3407975, EPI_ISL_3407976, EPI_ISL_3407977, EPI_ISL_3407978, EPI_ISL_3407979, EPI_ISL_3407980, EPI_ISL_3407981, EPI_ISL_3407982, EPI_ISL_3407983, EPI_ISL_3407984, EPI_ISL_3407985, EPI_ISL_3407986, EPI_ISL_3407987, EPI_ISL_3407988, EPI_ISL_3407989, EPI_ISL_3407990, EPI_ISL_3407991, EPI_ISL_3407992, EPI_ISL_3407993, EPI_ISL_3407994, EPI_ISL_3407995, EPI_ISL_3407996, EPI_ISL_3407997, EPI_ISL_3407998, EPI_ISL_3407999, EPI_ISL_3408000, EPI_ISL_3408001, EPI_ISL_3408002, EPI_ISL_3408003, EPI_ISL_3408004, EPI_ISL_3408005, EPI_ISL_3408006, EPI_ISL_3408007, EPI_ISL_3408008, EPI_ISL_3408009, EPI_ISL_3408010, EPI_ISL_3408011, EPI_ISL_3408012, EPI_ISL_3408013, EPI_ISL_3408014, EPI_ISL_3408015, EPI_ISL_3408016, EPI_ISL_3408017, EPI_ISL_3408018, EPI_ISL_3408019, EPI_ISL_3408020, EPI_ISL_3408021, EPI_ISL_3408022, EPI_ISL_3408023, EPI_ISL_3408024, EPI_ISL_3408025, EPI_ISL_3408026, EPI_ISL_3408027, EPI_ISL_3408028, EPI_ISL_3408029, EPI_ISL_3408030, EPI_ISL_3408031, EPI_ISL_3408032, EPI_ISL_3408033, EPI_ISL_3408034, EPI_ISL_3408035, EPI_ISL_3408036, EPI_ISL_3408037, EPI_ISL_3408038, EPI_ISL_3408039, EPI_ISL_3408040, EPI_ISL_3408041, EPI_ISL_3408042, EPI_ISL_3408043, EPI_ISL_3408044, EPI_ISL_3408045, EPI_ISL_3408046, EPI_ISL_3408047, EPI_ISL_3408048, EPI_ISL_3408049, EPI_ISL_3408050, EPI_ISL_3408051, EPI_ISL_3408052, EPI_ISL_3408053, EPI_ISL_3408054, EPI_ISL_3408055, EPI_ISL_3408056, EPI_ISL_3408057, EPI_ISL_3408058, EPI_ISL_3408059, EPI_ISL_3408060, EPI_ISL_3408061, EPI_ISL_3408062, EPI_ISL_3408063, EPI_ISL_3408064, EPI_ISL_3408065, EPI_ISL_3408066, EPI_ISL_3408067, EPI_ISL_3408068, EPI_ISL_3408069, EPI_ISL_3408070, EPI_ISL_3408071, EPI_ISL_3408072, EPI_ISL_3408073, EPI_ISL_3408074, EPI_ISL_3408075, EPI_ISL_3408076, EPI_ISL_3408077, EPI_ISL_3408078, EPI_ISL_3408079, EPI_ISL_3408080, EPI_ISL_3408081, EPI_ISL_3408082, EPI_ISL_3408083, EPI_ISL_3408084, EPI_ISL_3408085, EPI_ISL_3408086, EPI_ISL_3408087, EPI_ISL_3408088, EPI_ISL_3408089, EPI_ISL_3408090, EPI_ISL_3408091, EPI_ISL_3408092, EPI_ISL_3408093, EPI_ISL_3408094, EPI_ISL_3408095, EPI_ISL_3408096, EPI_ISL_3408097, EPI_ISL_3408098, EPI_ISL_3408099, EPI_ISL_3408100, EPI_ISL_3408101, EPI_ISL_3408102, EPI_ISL_3408103, EPI_ISL_3408104, EPI_ISL_3408105, EPI_ISL_3408106, EPI_ISL_3408107, EPI_ISL_3408108, EPI_ISL_3408109, EPI_ISL_3408110, EPI_ISL_3408111, EPI_ISL_3408112, EPI_ISL_3408113, EPI_ISL_3408114, EPI_ISL_3408115, EPI_ISL_3408116, EPI_ISL_3408118, EPI_ISL_3408119, EPI_ISL_3408120, EPI_ISL_3408121, EPI_ISL_3408122, EPI_ISL_3408123, EPI_ISL_3408124, EPI_ISL_3408125, EPI_ISL_3408126, EPI_ISL_3408127, EPI_ISL_3408128, EPI_ISL_3408129, EPI_ISL_3408130, EPI_ISL_3408131, EPI_ISL_3408132, EPI_ISL_3408133, EPI_ISL_3408134, EPI_ISL_3408135, EPI_ISL_3408136, EPI_ISL_3408137, EPI_ISL_3408138, EPI_ISL_3408139, EPI_ISL_3408140, EPI_ISL_3408141, EPI_ISL_3408142, EPI_ISL_3408143, EPI_ISL_3408144, EPI_ISL_3408145, EPI_ISL_3408147, EPI_ISL_3408148, EPI_ISL_3408149, EPI_ISL_3408150, EPI_ISL_3408151, EPI_ISL_3408152, EPI_ISL_3408153, EPI_ISL_3408154, EPI_ISL_3408155, EPI_ISL_3408156, EPI_ISL_3408157, EPI_ISL_3408158, EPI_ISL_3408159, EPI_ISL_3408160, EPI_ISL_3408161, EPI_ISL_3408162, EPI_ISL_3408163, EPI_ISL_3408164, EPI_ISL_3408165, EPI_ISL_3408166, EPI_ISL_3408167, EPI_ISL_3408168, EPI_ISL_3408169, EPI_ISL_3408170, EPI_ISL_3408171, EPI_ISL_3408172, EPI_ISL_3408173, EPI_ISL_3408174, EPI_ISL_3408175, EPI_ISL_3408176, EPI_ISL_3408177, EPI_ISL_3408178, EPI_ISL_3408179, EPI_ISL_3408180, EPI_ISL_3408181, EPI_ISL_3408182, EPI_ISL_3408183, EPI_ISL_3408184, EPI_ISL_3408185, EPI_ISL_3408186, EPI_ISL_3408187, EPI_ISL_3408188, EPI_ISL_3408189, EPI_ISL_3408190, EPI_ISL_3408191, EPI_ISL_3408192, EPI_ISL_3408193, EPI_ISL_3408194, EPI_ISL_3408195, EPI_ISL_3408196, EPI_ISL_3408197, EPI_ISL_3408198, EPI_ISL_3408199, EPI_ISL_3408200, EPI_ISL_3408201, EPI_ISL_3408202, EPI_ISL_3408203, EPI_ISL_3408204, EPI_ISL_3408205, EPI_ISL_3408206, EPI_ISL_3408207, EPI_ISL_3408208, EPI_ISL_3408209, EPI_ISL_3408210, EPI_ISL_3408211, EPI_ISL_3408212, EPI_ISL_3408213, EPI_ISL_3408214, EPI_ISL_3408215, EPI_ISL_3408216, EPI_ISL_3408217, EPI_ISL_3408218, EPI_ISL_3408219, EPI_ISL_3408220, EPI_ISL_3408221, EPI_ISL_3408222, EPI_ISL_3408223, EPI_ISL_3408224, EPI_ISL_3408225, EPI_ISL_3408226, EPI_ISL_3408227, EPI_ISL_3408228, EPI_ISL_3408229, EPI_ISL_3408230, EPI_ISL_3408231, EPI_ISL_3408232, EPI_ISL_3408233, EPI_ISL_3408234, EPI_ISL_3408235, EPI_ISL_3408236, EPI_ISL_3408237, EPI_ISL_3408238, EPI_ISL_3408239, EPI_ISL_3408240, EPI_ISL_3408241, EPI_ISL_3408242, EPI_ISL_3408243, EPI_ISL_3408244, EPI_ISL_3408245, EPI_ISL_3408246, EPI_ISL_3408247, EPI_ISL_3408248, EPI_ISL_3408249, EPI_ISL_3408250, EPI_ISL_3408251, EPI_ISL_3408252, EPI_ISL_3408253, EPI_ISL_3408254, EPI_ISL_3408255, EPI_ISL_3408256, EPI_ISL_3408257, EPI_ISL_3408258, EPI_ISL_3408259, EPI_ISL_3408260, EPI_ISL_3408261, EPI_ISL_3408262, EPI_ISL_3408263, EPI_ISL_3408264, EPI_ISL_3408265, EPI_ISL_3408266, EPI_ISL_3408267, EPI_ISL_3408268, EPI_ISL_3408269, EPI_ISL_3408270, EPI_ISL_3408271, EPI_ISL_3408272, EPI_ISL_3408273, EPI_ISL_3408274, EPI_ISL_3408275, EPI_ISL_3408276, EPI_ISL_3408277, EPI_ISL_3408278, EPI_ISL_3408279, EPI_ISL_3408280, EPI_ISL_3408281, EPI_ISL_3408282, EPI_ISL_3408284, EPI_ISL_3408285, EPI_ISL_3408286, EPI_ISL_3408287, EPI_ISL_3408288, EPI_ISL_3408289, EPI_ISL_3408290, EPI_ISL_3408291, EPI_ISL_3408292, EPI_ISL_3408293, EPI_ISL_3408294, EPI_ISL_3408295, EPI_ISL_3408296, EPI_ISL_3408297, EPI_ISL_3408298, EPI_ISL_3408299, EPI_ISL_3408300, EPI_ISL_3408301, EPI_ISL_3408302, EPI_ISL_3408303, EPI_ISL_3408304, EPI_ISL_3408305, EPI_ISL_3408306, EPI_ISL_3408307, EPI_ISL_3408308, EPI_ISL_3408309, EPI_ISL_3408310, EPI_ISL_3408312, EPI_ISL_3408313, EPI_ISL_3408314, EPI_ISL_3408315, EPI_ISL_3408316, EPI_ISL_3408317, EPI_ISL_3408318, EPI_ISL_3408319, EPI_ISL_3408320, EPI_ISL_3408321, EPI_ISL_3408322, EPI_ISL_3408323, EPI_ISL_3408324, EPI_ISL_3408325, EPI_ISL_3408326, EPI_ISL_3408327, EPI_ISL_3408329, EPI_ISL_3408330, EPI_ISL_3408331, EPI_ISL_3408332, EPI_ISL_3408333, EPI_ISL_3408334, EPI_ISL_3408335, EPI_ISL_3408338, EPI_ISL_3408339, EPI_ISL_3408340, EPI_ISL_3408341, EPI_ISL_3408342, EPI_ISL_3408343, EPI_ISL_3408348, EPI_ISL_3408349, EPI_ISL_3408350, EPI_ISL_3408351, EPI_ISL_3408352, EPI_ISL_3408353, EPI_ISL_3408354, EPI_ISL_3408355, EPI_ISL_3408356, EPI_ISL_3408357, EPI_ISL_3408358, EPI_ISL_3408359, EPI_ISL_3408360, EPI_ISL_3408361, EPI_ISL_3408362, EPI_ISL_3408363, EPI_ISL_3408364, EPI_ISL_3408365, EPI_ISL_3408366, EPI_ISL_3408367, EPI_ISL_3408368, EPI_ISL_3408369, EPI_ISL_3408370, EPI_ISL_3408371, EPI_ISL_3408372, EPI_ISL_3408373, EPI_ISL_3408374, EPI_ISL_3408375, EPI_ISL_3408376, EPI_ISL_3408377, EPI_ISL_3408378, EPI_ISL_3408379, EPI_ISL_3408380, EPI_ISL_3408381, EPI_ISL_3408382, EPI_ISL_3408383, EPI_ISL_3408384, EPI_ISL_3408385, EPI_ISL_3408386, EPI_ISL_3408387, EPI_ISL_3408388, EPI_ISL_3408389, EPI_ISL_3408390, EPI_ISL_3408391, EPI_ISL_3408392, EPI_ISL_3408393, EPI_ISL_3408394, EPI_ISL_3408395, EPI_ISL_3408396, EPI_ISL_3408397, EPI_ISL_3408398, EPI_ISL_3408399, EPI_ISL_3408400, EPI_ISL_3408401, EPI_ISL_3408402, EPI_ISL_3408403, EPI_ISL_3408404, EPI_ISL_3408405, EPI_ISL_3408406, EPI_ISL_3408407, EPI_ISL_3408408, EPI_ISL_3408409, EPI_ISL_3408410, EPI_ISL_3408411, EPI_ISL_3408412, EPI_ISL_3408413, EPI_ISL_3408414, EPI_ISL_3408415, EPI_ISL_3408416, EPI_ISL_3408417, EPI_ISL_3408418, EPI_ISL_3408419, EPI_ISL_3408420, EPI_ISL_3408421, EPI_ISL_3408422, EPI_ISL_3408423, EPI_ISL_3408424, EPI_ISL_3408425, EPI_ISL_3408426, EPI_ISL_3408427, EPI_ISL_3408428, EPI_ISL_3408429, EPI_ISL_3408430, EPI_ISL_3408431, EPI_ISL_3408432, EPI_ISL_3408433, EPI_ISL_3408434, EPI_ISL_3408435, EPI_ISL_3408436, EPI_ISL_3408437, EPI_ISL_3408438, EPI_ISL_3408439, EPI_ISL_3408440, EPI_ISL_3408441, EPI_ISL_3408442, EPI_ISL_3408443, EPI_ISL_3408444, EPI_ISL_3408445, EPI_ISL_3408446, EPI_ISL_3408447, EPI_ISL_3408448, EPI_ISL_3408449, EPI_ISL_3408450, EPI_ISL_3408451, EPI_ISL_3408452, EPI_ISL_3408453, EPI_ISL_3408454, EPI_ISL_3408455, EPI_ISL_3408456, EPI_ISL_3408457, EPI_ISL_3408458, EPI_ISL_3408459, EPI_ISL_3408460, EPI_ISL_3408461, EPI_ISL_3408462, EPI_ISL_3408463, EPI_ISL_3408464, EPI_ISL_3408465, EPI_ISL_3408466, EPI_ISL_3408467, EPI_ISL_3408468, EPI_ISL_3408469, EPI_ISL_3408470, EPI_ISL_3408471, EPI_ISL_3408472, EPI_ISL_3408473, EPI_ISL_3408474, EPI_ISL_3408475, EPI_ISL_3408476 | | | |
| see above | Broad Institute Clinical Research Sequencing Platform | Infectious Disease Program, Broad Institute of Harvard and MIT | Siddle,K.J., Adams,G., Pearlman,L., Gladden-Young,A., Vicente,G., Blumenstiel,B., DeFelice,M., Lee,M., McGovern,S., Lagerborg,K., Rudy,M., DeRuff,K., Carter,A., Normandin,E., Bauer,M., Reilly,S., Tomkins-Tinch,C., Loreth,C., Chaluvadi,S., Meldrim,J., Granger,B., Lemieux,J.E., Birren,B.W., Sabeti,P.C., Larkin,K., Dodge,S., Lennon,N., Madoff,L., Brown,C., Gallagher,G., Smole,S., Park,D.J., Gabriel,S., and MacInnis,B.L. |
| EPI_ISL_3408477 | Rhode Island Department of Health | Infectious Disease Program, Broad Institute of Harvard and MIT | Siddle,K.J., Adams,G., Pearlman,L., Gladden-Young,A., Vicente,G., Blumenstiel,B., DeFelice,M., Lee,M., McGovern,S., Lagerborg,K., Rudy,M., DeRuff,K., Carter,A., Normandin,E., Bauer,M., Reilly,S., Tomkins-Tinch,C., Loreth,C., Chaluvadi,S., Meldrim,J., Granger,B., Lemieux,J.E., Birren,B.W., Sabeti,P.C., Larkin,K., Dodge,S., Lennon,N., Madoff,L., Brown,C., Gallagher,G., Smole,S., Park,D.J., Gabriel,S., and MacInnis,B.L. |
| EPI_ISL_3408478, EPI_ISL_3408479, EPI_ISL_3408480, EPI_ISL_3408481 | Broad Institute Clinical Research Sequencing Platform | Infectious Disease Program, Broad Institute of Harvard and MIT | Siddle,K.J., Adams,G., Pearlman,L., Gladden-Young,A., Vicente,G., Blumenstiel,B., DeFelice,M., Lee,M., McGovern,S., Lagerborg,K., Rudy,M., DeRuff,K., Carter,A., Normandin,E., Bauer,M., Reilly,S., Tomkins-Tinch,C., Loreth,C., Chaluvadi,S., Meldrim,J., Granger,B., Lemieux,J.E., Birren,B.W., Sabeti,P.C., Larkin,K., Dodge,S., Lennon,N., Madoff,L., Brown,C., Gallagher,G., Smole,S., Park,D.J., Gabriel,S., and MacInnis,B.L. |
| EPI_ISL_3408482 | Rhode Island Department of Health | Infectious Disease Program, Broad Institute of Harvard and MIT | Siddle,K.J., Adams,G., Pearlman,L., Gladden-Young,A., Vicente,G., Blumenstiel,B., DeFelice,M., Lee,M., McGovern,S., Lagerborg,K., Rudy,M., DeRuff,K., Carter,A., Normandin,E., Bauer,M., Reilly,S., Tomkins-Tinch,C., Loreth,C., Chaluvadi,S., Meldrim,J., Granger,B., Lemieux,J.E., Birren,B.W., Sabeti,P.C., Larkin,K., Dodge,S., Lennon,N., Madoff,L., Brown,C., Gallagher,G., Smole,S., Park,D.J., Gabriel,S., and MacInnis,B.L. |
| EPI_ISL_3408483 | Broad Institute Clinical Research Sequencing Platform | Infectious Disease Program, Broad Institute of Harvard and MIT | Siddle,K.J., Adams,G., Pearlman,L., Gladden-Young,A., Vicente,G., Blumenstiel,B., DeFelice,M., Lee,M., McGovern,S., Lagerborg,K., Rudy,M., DeRuff,K., Carter,A., Normandin,E., Bauer,M., Reilly,S., Tomkins-Tinch,C., Loreth,C., Chaluvadi,S., Meldrim,J., Granger,B., Lemieux,J.E., Birren,B.W., Sabeti,P.C., Larkin,K., Dodge,S., Lennon,N., Madoff,L., Brown,C., Gallagher,G., Smole,S., Park,D.J., Gabriel,S., and MacInnis,B.L. |
| EPI_ISL_3408484 | Rhode Island Department of Health | Infectious Disease Program, Broad Institute of Harvard and MIT | Siddle,K.J., Adams,G., Pearlman,L., Gladden-Young,A., Vicente,G., Blumenstiel,B., DeFelice,M., Lee,M., McGovern,S., Lagerborg,K., Rudy,M., DeRuff,K., Carter,A., Normandin,E., Bauer,M., Reilly,S., Tomkins-Tinch,C., Loreth,C., Chaluvadi,S., Meldrim,J., Granger,B., Lemieux,J.E., Birren,B.W., Sabeti,P.C., Larkin,K., Dodge,S., Lennon,N., Madoff,L., Brown,C., Gallagher,G., Smole,S., Park,D.J., Gabriel,S., and MacInnis,B.L. |
| EPI_ISL_3408485, EPI_ISL_3408486, EPI_ISL_3408487, EPI_ISL_3408488, EPI_ISL_3408489, EPI_ISL_3408490, EPI_ISL_3408491, EPI_ISL_3408492 | Broad Institute Clinical Research Sequencing Platform | Infectious Disease Program, Broad Institute of Harvard and MIT | Siddle,K.J., Adams,G., Pearlman,L., Gladden-Young,A., Vicente,G., Blumenstiel,B., DeFelice,M., Lee,M., McGovern,S., Lagerborg,K., Rudy,M., DeRuff,K., Carter,A., Normandin,E., Bauer,M., Reilly,S., Tomkins-Tinch,C., Loreth,C., Chaluvadi,S., Meldrim,J., Granger,B., Lemieux,J.E., Birren,B.W., Sabeti,P.C., Larkin,K., Dodge,S., Lennon,N., Madoff,L., Brown,C., Gallagher,G., Smole,S., Park,D.J., Gabriel,S., and MacInnis,B.L. |
| EPI_ISL_3408493 | Rhode Island Department of Health | Infectious Disease Program, Broad Institute of Harvard and MIT | Siddle,K.J., Adams,G., Pearlman,L., Gladden-Young,A., Vicente,G., Blumenstiel,B., DeFelice,M., Lee,M., McGovern,S., Lagerborg,K., Rudy,M., DeRuff,K., Carter,A., Normandin,E., Bauer,M., Reilly,S., Tomkins-Tinch,C., Loreth,C., Chaluvadi,S., Meldrim,J., Granger,B., Lemieux,J.E., Birren,B.W., Sabeti,P.C., Larkin,K., Dodge,S., Lennon,N., Madoff,L., Brown,C., Gallagher,G., Smole,S., Park,D.J., Gabriel,S., and MacInnis,B.L. |
| EPI_ISL_3408495, EPI_ISL_3408496, EPI_ISL_3408497, EPI_ISL_3408498, EPI_ISL_3408499, EPI_ISL_3408500 | Broad Institute Clinical Research Sequencing Platform | Infectious Disease Program, Broad Institute of Harvard and MIT | Siddle,K.J., Adams,G., Pearlman,L., Gladden-Young,A., Vicente,G., Blumenstiel,B., DeFelice,M., Lee,M., McGovern,S., Lagerborg,K., Rudy,M., DeRuff,K., Carter,A., Normandin,E., Bauer,M., Reilly,S., Tomkins-Tinch,C., Loreth,C., Chaluvadi,S., Meldrim,J., Granger,B., Lemieux,J.E., Birren,B.W., Sabeti,P.C., Larkin,K., Dodge,S., Lennon,N., Madoff,L., Brown,C., Gallagher,G., Smole,S., Park,D.J., Gabriel,S., and MacInnis,B.L. |
| EPI_ISL_3408501 | Rhode Island Department of Health | Infectious Disease Program, Broad Institute of Harvard and MIT | Siddle,K.J., Adams,G., Pearlman,L., Gladden-Young,A., Vicente,G., Blumenstiel,B., DeFelice,M., Lee,M., McGovern,S., Lagerborg,K., Rudy,M., DeRuff,K., Carter,A., Normandin,E., Bauer,M., Reilly,S., Tomkins-Tinch,C., Loreth,C., Chaluvadi,S., Meldrim,J., Granger,B., Lemieux,J.E., Birren,B.W., Sabeti,P.C., Larkin,K., Dodge,S., Lennon,N., Madoff,L., Brown,C., Gallagher,G., Smole,S., Park,D.J., Gabriel,S., and MacInnis,B.L. |
| EPI_ISL_3408502, | Broad Institute Clinical Research Sequencing Platform | Infectious Disease Program, Broad Institute of Harvard and MIT | Siddle,K.J., Adams,G., Pearlman,L., Gladden-Young,A., Vicente,G., Blumenstiel,B., DeFelice,M., Lee,M., McGovern,S., Lagerborg,K., Rudy,M., DeRuff,K., Carter,A., Normandin,E., Bauer,M., Reilly,S., Tomkins-Tinch,C., Loreth,C., Chaluvadi,S., |

| | | | |
|---|---|---|---|
| EPI_ISL_3408503, EPI_ISL_3408504 | | | Meldrim,J., Granger,B., Lemieux,J.E., Birren,B.W., Sabeti,P.C., Larkin,K., Dodge,S., Lennon,N., Madoff,L., Brown,C., Gallagher,G., Smole,S., Park,D.J., Gabriel,S., and MacInnis,B.L. |
| EPI_ISL_3408505, EPI_ISL_3408506 | Rhode Island Department of Health | Infectious Disease Program, Broad Institute of Harvard and MIT | Siddle,K.J., Adams,G., Pearlman,L., Gladden-Young,A., Vicente,G., Blumenstiel,B., DeFelice,M., Lee,M., McGovern,S., Lagerborg,K., Rudy,M., DeRuff,K., Carter,A., Normandin,E., Bauer,M., Reilly,S., Tomkins-Tinch,C., Loreth,C., Chaluvadi,S., Meldrim,J., Granger,B., Lemieux,J.E., Birren,B.W., Sabeti,P.C., Larkin,K., Dodge,S., Lennon,N., Madoff,L., Brown,C., Gallagher,G., Smole,S., Park,D.J., Gabriel,S., and MacInnis,B.L. |
| EPI_ISL_3408507, EPI_ISL_3408508, EPI_ISL_3408509, EPI_ISL_3408510, EPI_ISL_3408511, EPI_ISL_3408512, EPI_ISL_3408513, EPI_ISL_3408514, EPI_ISL_3408515, EPI_ISL_3408516, EPI_ISL_3408517, EPI_ISL_3408518, EPI_ISL_3408519, EPI_ISL_3408520, EPI_ISL_3408521, EPI_ISL_3408522, EPI_ISL_3408523, EPI_ISL_3408524, EPI_ISL_3408525, EPI_ISL_3408526, EPI_ISL_3408527, EPI_ISL_3408528, EPI_ISL_3408529, EPI_ISL_3408530, EPI_ISL_3408531, EPI_ISL_3408532, EPI_ISL_3408533, EPI_ISL_3408534, EPI_ISL_3408535, EPI_ISL_3408536, EPI_ISL_3408537, EPI_ISL_3408538 | | | |
| see above | Broad Institute Clinical Research Sequencing Platform | Infectious Disease Program, Broad Institute of Harvard and MIT | Siddle,K.J., Adams,G., Pearlman,L., Gladden-Young,A., Vicente,G., Blumenstiel,B., DeFelice,M., Lee,M., McGovern,S., Lagerborg,K., Rudy,M., DeRuff,K., Carter,A., Normandin,E., Bauer,M., Reilly,S., Tomkins-Tinch,C., Loreth,C., Chaluvadi,S., Meldrim,J., Granger,B., Lemieux,J.E., Birren,B.W., Sabeti,P.C., Larkin,K., Dodge,S., Lennon,N., Madoff,L., Brown,C., Gallagher,G., Smole,S., Park,D.J., Gabriel,S., and MacInnis,B.L. |
| EPI_ISL_3408539 | Rhode Island Department of Health | Infectious Disease Program, Broad Institute of Harvard and MIT | Siddle,K.J., Adams,G., Pearlman,L., Gladden-Young,A., Vicente,G., Blumenstiel,B., DeFelice,M., Lee,M., McGovern,S., Lagerborg,K., Rudy,M., DeRuff,K., Carter,A., Normandin,E., Bauer,M., Reilly,S., Tomkins-Tinch,C., Loreth,C., Chaluvadi,S., Meldrim,J., Granger,B., Lemieux,J.E., Birren,B.W., Sabeti,P.C., Larkin,K., Dodge,S., Lennon,N., Madoff,L., Brown,C., Gallagher,G., Smole,S., Park,D.J., Gabriel,S., and MacInnis,B.L. |
| EPI_ISL_3408540, EPI_ISL_3408541, EPI_ISL_3408542, EPI_ISL_3408543 | Broad Institute Clinical Research Sequencing Platform | Infectious Disease Program, Broad Institute of Harvard and MIT | Siddle,K.J., Adams,G., Pearlman,L., Gladden-Young,A., Vicente,G., Blumenstiel,B., DeFelice,M., Lee,M., McGovern,S., Lagerborg,K., Rudy,M., DeRuff,K., Carter,A., Normandin,E., Bauer,M., Reilly,S., Tomkins-Tinch,C., Loreth,C., Chaluvadi,S., Meldrim,J., Granger,B., Lemieux,J.E., Birren,B.W., Sabeti,P.C., Larkin,K., Dodge,S., Lennon,N., Madoff,L., Brown,C., Gallagher,G., Smole,S., Park,D.J., Gabriel,S., and MacInnis,B.L. |
| EPI_ISL_3408544 | Rhode Island Department of Health | Infectious Disease Program, Broad Institute of Harvard and MIT | Siddle,K.J., Adams,G., Pearlman,L., Gladden-Young,A., Vicente,G., Blumenstiel,B., DeFelice,M., Lee,M., McGovern,S., Lagerborg,K., Rudy,M., DeRuff,K., Carter,A., Normandin,E., Bauer,M., Reilly,S., Tomkins-Tinch,C., Loreth,C., Chaluvadi,S., Meldrim,J., Granger,B., Lemieux,J.E., Birren,B.W., Sabeti,P.C., Larkin,K., Dodge,S., Lennon,N., Madoff,L., Brown,C., Gallagher,G., Smole,S., Park,D.J., Gabriel,S., and MacInnis,B.L. |
| EPI_ISL_3408545, EPI_ISL_3408546 | Broad Institute Clinical Research Sequencing Platform | Infectious Disease Program, Broad Institute of Harvard and MIT | Siddle,K.J., Adams,G., Pearlman,L., Gladden-Young,A., Vicente,G., Blumenstiel,B., DeFelice,M., Lee,M., McGovern,S., Lagerborg,K., Rudy,M., DeRuff,K., Carter,A., Normandin,E., Bauer,M., Reilly,S., Tomkins-Tinch,C., Loreth,C., Chaluvadi,S., Meldrim,J., Granger,B., Lemieux,J.E., Birren,B.W., Sabeti,P.C., Larkin,K., Dodge,S., Lennon,N., Madoff,L., Brown,C., Gallagher,G., Smole,S., Park,D.J., Gabriel,S., and MacInnis,B.L. |
| EPI_ISL_3408547 | Rhode Island Department of Health | Infectious Disease Program, Broad Institute of Harvard and MIT | Siddle,K.J., Adams,G., Pearlman,L., Gladden-Young,A., Vicente,G., Blumenstiel,B., DeFelice,M., Lee,M., McGovern,S., Lagerborg,K., Rudy,M., DeRuff,K., Carter,A., Normandin,E., Bauer,M., Reilly,S., Tomkins-Tinch,C., Loreth,C., Chaluvadi,S., Meldrim,J., Granger,B., Lemieux,J.E., Birren,B.W., Sabeti,P.C., Larkin,K., Dodge,S., Lennon,N., Madoff,L., Brown,C., Gallagher,G., Smole,S., Park,D.J., Gabriel,S., and MacInnis,B.L. |
| EPI_ISL_3408548, EPI_ISL_3408549, EPI_ISL_3408550, EPI_ISL_3408551, EPI_ISL_3408552, EPI_ISL_3408553, EPI_ISL_3408554, EPI_ISL_3408555, EPI_ISL_3408556, EPI_ISL_3408557, EPI_ISL_3408558, EPI_ISL_3408559, EPI_ISL_3408560, EPI_ISL_3408561, EPI_ISL_3408562, EPI_ISL_3408563, EPI_ISL_3408564, EPI_ISL_3408565, EPI_ISL_3408566 | | | |
| see above | Broad Institute Clinical Research Sequencing Platform | Infectious Disease Program, Broad Institute of Harvard and MIT | Siddle,K.J., Adams,G., Pearlman,L., Gladden-Young,A., Vicente,G., Blumenstiel,B., DeFelice,M., Lee,M., McGovern,S., Lagerborg,K., Rudy,M., DeRuff,K., Carter,A., Normandin,E., Bauer,M., Reilly,S., Tomkins-Tinch,C., Loreth,C., Chaluvadi,S., Meldrim,J., Granger,B., Lemieux,J.E., Birren,B.W., Sabeti,P.C., Larkin,K., Dodge,S., Lennon,N., Madoff,L., Brown,C., Gallagher,G., Smole,S., Park,D.J., Gabriel,S., and MacInnis,B.L. |
| EPI_ISL_3408567, EPI_ISL_3408568 | Rhode Island Department of Health | Infectious Disease Program, Broad Institute of Harvard and MIT | Siddle,K.J., Adams,G., Pearlman,L., Gladden-Young,A., Vicente,G., Blumenstiel,B., DeFelice,M., Lee,M., McGovern,S., Lagerborg,K., Rudy,M., DeRuff,K., Carter,A., Normandin,E., Bauer,M., Reilly,S., Tomkins-Tinch,C., Loreth,C., Chaluvadi,S., Meldrim,J., Granger,B., Lemieux,J.E., Birren,B.W., Sabeti,P.C., Larkin,K., Dodge,S., Lennon,N., Madoff,L., Brown,C., Gallagher,G., Smole,S., Park,D.J., Gabriel,S., and MacInnis,B.L. |
| EPI_ISL_3408569, EPI_ISL_3408570, EPI_ISL_3408571, EPI_ISL_3408572, EPI_ISL_3408573, EPI_ISL_3408575, EPI_ISL_3408576, EPI_ISL_3408577, EPI_ISL_3408578 | Broad Institute Clinical Research Sequencing Platform | Infectious Disease Program, Broad Institute of Harvard and MIT | Siddle,K.J., Adams,G., Pearlman,L., Gladden-Young,A., Vicente,G., Blumenstiel,B., DeFelice,M., Lee,M., McGovern,S., Lagerborg,K., Rudy,M., DeRuff,K., Carter,A., Normandin,E., Bauer,M., Reilly,S., Tomkins-Tinch,C., Loreth,C., Chaluvadi,S., Meldrim,J., Granger,B., Lemieux,J.E., Birren,B.W., Sabeti,P.C., Larkin,K., Dodge,S., Lennon,N., Madoff,L., Brown,C., Gallagher,G., Smole,S., Park,D.J., Gabriel,S., and MacInnis,B.L. |
| EPI_ISL_3408579 | Rhode Island Department of Health | Infectious Disease Program, Broad Institute of Harvard and MIT | Siddle,K.J., Adams,G., Pearlman,L., Gladden-Young,A., Vicente,G., Blumenstiel,B., DeFelice,M., Lee,M., McGovern,S., Lagerborg,K., Rudy,M., DeRuff,K., Carter,A., Normandin,E., Bauer,M., Reilly,S., Tomkins-Tinch,C., Loreth,C., Chaluvadi,S., Meldrim,J., Granger,B., Lemieux,J.E., Birren,B.W., Sabeti,P.C., Larkin,K., Dodge,S., Lennon,N., Madoff,L., Brown,C., Gallagher,G., Smole,S., Park,D.J., Gabriel,S., and MacInnis,B.L. |
| EPI_ISL_3408580, EPI_ISL_3408581, EPI_ISL_3408582, EPI_ISL_3408583, EPI_ISL_3408584, EPI_ISL_3408585, EPI_ISL_3408586, EPI_ISL_3408587, EPI_ISL_3408588, EPI_ISL_3408589, EPI_ISL_3408590, EPI_ISL_3408591, EPI_ISL_3408592, EPI_ISL_3408593 | | | |
| see above | Broad Institute Clinical Research Sequencing Platform | Infectious Disease Program, Broad Institute of Harvard and MIT | Siddle,K.J., Adams,G., Pearlman,L., Gladden-Young,A., Vicente,G., Blumenstiel,B., DeFelice,M., Lee,M., McGovern,S., Lagerborg,K., Rudy,M., DeRuff,K., Carter,A., Normandin,E., Bauer,M., Reilly,S., Tomkins-Tinch,C., Loreth,C., Chaluvadi,S., Meldrim,J., Granger,B., Lemieux,J.E., Birren,B.W., Sabeti,P.C., Larkin,K., Dodge,S., Lennon,N., Madoff,L., Brown,C., Gallagher,G., Smole,S., Park,D.J., Gabriel,S., and MacInnis,B.L. |
| EPI_ISL_3408594 | Rhode Island Department of Health | Infectious Disease Program, Broad Institute of Harvard and MIT | Siddle,K.J., Adams,G., Pearlman,L., Gladden-Young,A., Vicente,G., Blumenstiel,B., DeFelice,M., Lee,M., McGovern,S., Lagerborg,K., Rudy,M., DeRuff,K., Carter,A., Normandin,E., Bauer,M., Reilly,S., Tomkins-Tinch,C., Loreth,C., Chaluvadi,S., Meldrim,J., Granger,B., Lemieux,J.E., Birren,B.W., Sabeti,P.C., Larkin,K., Dodge,S., Lennon,N., Madoff,L., Brown,C., Gallagher,G., Smole,S., Park,D.J., Gabriel,S., and MacInnis,B.L. |
| EPI_ISL_3408595, EPI_ISL_3408596, EPI_ISL_3408597 | Broad Institute Clinical Research Sequencing Platform | Infectious Disease Program, Broad Institute of Harvard and MIT | Siddle,K.J., Adams,G., Pearlman,L., Gladden-Young,A., Vicente,G., Blumenstiel,B., DeFelice,M., Lee,M., McGovern,S., Lagerborg,K., Rudy,M., DeRuff,K., Carter,A., Normandin,E., Bauer,M., Reilly,S., Tomkins-Tinch,C., Loreth,C., Chaluvadi,S., Meldrim,J., Granger,B., Lemieux,J.E., Birren,B.W., Sabeti,P.C., Larkin,K., Dodge,S., Lennon,N., Madoff,L., Brown,C., Gallagher,G., Smole,S., Park,D.J., Gabriel,S., and MacInnis,B.L. |
| EPI_ISL_3408598 | Rhode Island Department of Health | Infectious Disease Program, Broad Institute of Harvard and MIT | Siddle,K.J., Adams,G., Pearlman,L., Gladden-Young,A., Vicente,G., Blumenstiel,B., DeFelice,M., Lee,M., McGovern,S., Lagerborg,K., Rudy,M., DeRuff,K., Carter,A., Normandin,E., Bauer,M., Reilly,S., Tomkins-Tinch,C., Loreth,C., Chaluvadi,S., Meldrim,J., Granger,B., Lemieux,J.E., Birren,B.W., Sabeti,P.C., Larkin,K., Dodge,S., Lennon,N., Madoff,L., Brown,C., Gallagher,G., Smole,S., Park,D.J., Gabriel,S., and MacInnis,B.L. |
| EPI_ISL_3408599 | Broad Institute Clinical Research Sequencing Platform | Infectious Disease Program, Broad Institute of Harvard and MIT | Siddle,K.J., Adams,G., Pearlman,L., Gladden-Young,A., Vicente,G., Blumenstiel,B., DeFelice,M., Lee,M., McGovern,S., Lagerborg,K., Rudy,M., DeRuff,K., Carter,A., Normandin,E., Bauer,M., Reilly,S., Tomkins-Tinch,C., Loreth,C., Chaluvadi,S., Meldrim,J., Granger,B., Lemieux,J.E., Birren,B.W., Sabeti,P.C., Larkin,K., Dodge,S., Lennon,N., Madoff,L., Brown,C., Gallagher,G., Smole,S., Park,D.J., Gabriel,S., and MacInnis,B.L. |
| EPI_ISL_3408600 | Rhode Island Department of Health | Infectious Disease Program, Broad Institute of Harvard and MIT | Siddle,K.J., Adams,G., Pearlman,L., Gladden-Young,A., Vicente,G., Blumenstiel,B., DeFelice,M., Lee,M., McGovern,S., Lagerborg,K., Rudy,M., DeRuff,K., Carter,A., Normandin,E., Bauer,M., Reilly,S., Tomkins-Tinch,C., Loreth,C., Chaluvadi,S., Meldrim,J., Granger,B., Lemieux,J.E., Birren,B.W., Sabeti,P.C., Larkin,K., Dodge,S., Lennon,N., Madoff,L., Brown,C., Gallagher,G., Smole,S., Park,D.J., Gabriel,S., and MacInnis,B.L. |
| EPI_ISL_3408601, EPI_ISL_3408602, EPI_ISL_3408603, EPI_ISL_3408604, EPI_ISL_3408605, EPI_ISL_3408606, EPI_ISL_3408607, EPI_ISL_3408608 | Broad Institute Clinical Research Sequencing Platform | Infectious Disease Program, Broad Institute of Harvard and MIT | Siddle,K.J., Adams,G., Pearlman,L., Gladden-Young,A., Vicente,G., Blumenstiel,B., DeFelice,M., Lee,M., McGovern,S., Lagerborg,K., Rudy,M., DeRuff,K., Carter,A., Normandin,E., Bauer,M., Reilly,S., Tomkins-Tinch,C., Loreth,C., Chaluvadi,S., Meldrim,J., Granger,B., Lemieux,J.E., Birren,B.W., Sabeti,P.C., Larkin,K., Dodge,S., Lennon,N., Madoff,L., Brown,C., Gallagher,G., Smole,S., Park,D.J., Gabriel,S., and MacInnis,B.L. |
| EPI_ISL_3408609 | Rhode Island Department of Health | Infectious Disease Program, Broad Institute of Harvard and MIT | Siddle,K.J., Adams,G., Pearlman,L., Gladden-Young,A., Vicente,G., Blumenstiel,B., DeFelice,M., Lee,M., McGovern,S., Lagerborg,K., Rudy,M., DeRuff,K., Carter,A., Normandin,E., Bauer,M., Reilly,S., Tomkins-Tinch,C., Loreth,C., Chaluvadi,S., Meldrim,J., Granger,B., Lemieux,J.E., Birren,B.W., Sabeti,P.C., Larkin,K., Dodge,S., Lennon,N., Madoff,L., Brown,C., Gallagher,G., Smole,S., Park,D.J., Gabriel,S., and MacInnis,B.L. |
| EPI_ISL_3408610, EPI_ISL_3408611, EPI_ISL_3408612 | Broad Institute Clinical Research Sequencing Platform | Infectious Disease Program, Broad Institute of Harvard and MIT | Siddle,K.J., Adams,G., Pearlman,L., Gladden-Young,A., Vicente,G., Blumenstiel,B., DeFelice,M., Lee,M., McGovern,S., Lagerborg,K., Rudy,M., DeRuff,K., Carter,A., Normandin,E., Bauer,M., Reilly,S., Tomkins-Tinch,C., Loreth,C., Chaluvadi,S., Meldrim,J., Granger,B., Lemieux,J.E., Birren,B.W., Sabeti,P.C., Larkin,K., Dodge,S., Lennon,N., Madoff,L., Brown,C., Gallagher,G., Smole,S., Park,D.J., Gabriel,S., and MacInnis,B.L. |
| EPI_ISL_3408613 | Rhode Island Department of Health | Infectious Disease Program, Broad Institute of Harvard and MIT | Siddle,K.J., Adams,G., Pearlman,L., Gladden-Young,A., Vicente,G., Blumenstiel,B., DeFelice,M., Lee,M., McGovern,S., Lagerborg,K., Rudy,M., DeRuff,K., Carter,A., Normandin,E., Bauer,M., Reilly,S., Tomkins-Tinch,C., Loreth,C., Chaluvadi,S., Meldrim,J., Granger,B., Lemieux,J.E., Birren,B.W., Sabeti,P.C., Larkin,K., Dodge,S., Lennon,N., Madoff,L., Brown,C., Gallagher,G., Smole,S., Park,D.J., Gabriel,S., and MacInnis,B.L. |
| EPI_ISL_3408614, EPI_ISL_3408615, EPI_ISL_3408616, EPI_ISL_3408617, EPI_ISL_3408618, EPI_ISL_3408619, EPI_ISL_3408620 | Broad Institute Clinical Research Sequencing Platform | Infectious Disease Program, Broad Institute of Harvard and MIT | Siddle,K.J., Adams,G., Pearlman,L., Gladden-Young,A., Vicente,G., Blumenstiel,B., DeFelice,M., Lee,M., McGovern,S., Lagerborg,K., Rudy,M., DeRuff,K., Carter,A., Normandin,E., Bauer,M., Reilly,S., Tomkins-Tinch,C., Loreth,C., Chaluvadi,S., Meldrim,J., Granger,B., Lemieux,J.E., Birren,B.W., Sabeti,P.C., Larkin,K., Dodge,S., Lennon,N., Madoff,L., Brown,C., Gallagher,G., Smole,S., Park,D.J., Gabriel,S., and MacInnis,B.L. |
| EPI_ISL_3408621 | Rhode Island Department of Health | Infectious Disease Program, Broad Institute of Harvard and MIT | Siddle,K.J., Adams,G., Pearlman,L., Gladden-Young,A., Vicente,G., Blumenstiel,B., DeFelice,M., Lee,M., McGovern,S., Lagerborg,K., Rudy,M., DeRuff,K., Carter,A., Normandin,E., Bauer,M., Reilly,S., Tomkins-Tinch,C., Loreth,C., Chaluvadi,S., Meldrim,J., Granger,B., Lemieux,J.E., Birren,B.W., Sabeti,P.C., Larkin,K., Dodge,S., Lennon,N., Madoff,L., Brown,C., Gallagher,G., Smole,S., Park,D.J., Gabriel,S., and MacInnis,B.L. |
| EPI_ISL_3408622, EPI_ISL_3408623, EPI_ISL_3408624, EPI_ISL_3408625 | Broad Institute Clinical Research Sequencing Platform | Infectious Disease Program, Broad Institute of Harvard and MIT | Siddle,K.J., Adams,G., Pearlman,L., Gladden-Young,A., Vicente,G., Blumenstiel,B., DeFelice,M., Lee,M., McGovern,S., Lagerborg,K., Rudy,M., DeRuff,K., Carter,A., Normandin,E., Bauer,M., Reilly,S., Tomkins-Tinch,C., Loreth,C., Chaluvadi,S., Meldrim,J., Granger,B., Lemieux,J.E., Birren,B.W., Sabeti,P.C., Larkin,K., Dodge,S., Lennon,N., Madoff,L., Brown,C., Gallagher,G., Smole,S., Park,D.J., Gabriel,S., and MacInnis,B.L. |
| EPI_ISL_3408626 | Rhode Island Department of Health | Infectious Disease Program, Broad Institute of Harvard and MIT | Siddle,K.J., Adams,G., Pearlman,L., Gladden-Young,A., Vicente,G., Blumenstiel,B., DeFelice,M., Lee,M., McGovern,S., Lagerborg,K., Rudy,M., DeRuff,K., Carter,A., Normandin,E., Bauer,M., Reilly,S., Tomkins-Tinch,C., Loreth,C., Chaluvadi,S., Meldrim,J., Granger,B., Lemieux,J.E., Birren,B.W., Sabeti,P.C., Larkin,K., Dodge,S., Lennon,N., Madoff,L., Brown,C., Gallagher,G., Smole,S., Park,D.J., Gabriel,S., and MacInnis,B.L. |
| EPI_ISL_3408627, EPI_ISL_3408628, EPI_ISL_3408629, EPI_ISL_3408630, EPI_ISL_3408631, EPI_ISL_3408632, EPI_ISL_3408633, EPI_ISL_3408634, EPI_ISL_3408635, EPI_ISL_3408636, EPI_ISL_3408637, EPI_ISL_3408638, EPI_ISL_3408639, EPI_ISL_3408640, EPI_ISL_3408641 | | | |
| see above | Broad Institute Clinical Research Sequencing Platform | Infectious Disease Program, Broad Institute of Harvard and MIT | Siddle,K.J., Adams,G., Pearlman,L., Gladden-Young,A., Vicente,G., Blumenstiel,B., DeFelice,M., Lee,M., McGovern,S., Lagerborg,K., Rudy,M., DeRuff,K., Carter,A., Normandin,E., Bauer,M., Reilly,S., Tomkins-Tinch,C., Loreth,C., Chaluvadi,S., Meldrim,J., Granger,B., Lemieux,J.E., Birren,B.W., Sabeti,P.C., Larkin,K., Dodge,S., Lennon,N., Madoff,L., Brown,C., Gallagher,G., Smole,S., Park,D.J., Gabriel,S., and MacInnis,B.L. |
| EPI_ISL_3408642 | Rhode Island Department of Health | Infectious Disease Program, Broad Institute of Harvard and MIT | Siddle,K.J., Adams,G., Pearlman,L., Gladden-Young,A., Vicente,G., Blumenstiel,B., DeFelice,M., Lee,M., McGovern,S., Lagerborg,K., Rudy,M., DeRuff,K., Carter,A., Normandin,E., Bauer,M., Reilly,S., Tomkins-Tinch,C., Loreth,C., Chaluvadi,S., Meldrim,J., Granger,B., Lemieux,J.E., Birren,B.W., Sabeti,P.C., Larkin,K., Dodge,S., Lennon,N., Madoff,L., Brown,C., Gallagher,G., Smole,S., Park,D.J., Gabriel,S., and MacInnis,B.L. |
| EPI_ISL_3408643 | Broad Institute Clinical Research Sequencing Platform | Infectious Disease Program, Broad Institute of Harvard and MIT | Siddle,K.J., Adams,G., Pearlman,L., Gladden-Young,A., Vicente,G., Blumenstiel,B., DeFelice,M., Lee,M., McGovern,S., Lagerborg,K., Rudy,M., DeRuff,K., Carter,A., Normandin,E., Bauer,M., Reilly,S., Tomkins-Tinch,C., Loreth,C., Chaluvadi,S., Meldrim,J., Granger,B., Lemieux,J.E., Birren,B.W., Sabeti,P.C., Larkin,K., Dodge,S., Lennon,N., Madoff,L., Brown,C., Gallagher,G., Smole,S., Park,D.J., Gabriel,S., and MacInnis,B.L. |
| EPI_ISL_3408644, EPI_ISL_3408645, EPI_ISL_3408646 | Rhode Island Department of Health | Infectious Disease Program, Broad Institute of Harvard and MIT | Siddle,K.J., Adams,G., Pearlman,L., Gladden-Young,A., Vicente,G., Blumenstiel,B., DeFelice,M., Lee,M., McGovern,S., Lagerborg,K., Rudy,M., DeRuff,K., Carter,A., Normandin,E., Bauer,M., Reilly,S., Tomkins-Tinch,C., Loreth,C., Chaluvadi,S., Meldrim,J., Granger,B., Lemieux,J.E., Birren,B.W., Sabeti,P.C., Larkin,K., Dodge,S., Lennon,N., Madoff,L., Brown,C., Gallagher,G., Smole,S., Park,D.J., Gabriel,S., and MacInnis,B.L. |
| EPI_ISL_3408647, EPI_ISL_3408648 | Broad Institute Clinical Research Sequencing Platform | Infectious Disease Program, Broad Institute of Harvard and MIT | Siddle,K.J., Adams,G., Pearlman,L., Gladden-Young,A., Vicente,G., Blumenstiel,B., DeFelice,M., Lee,M., McGovern,S., Lagerborg,K., Rudy,M., DeRuff,K., Carter,A., Normandin,E., Bauer,M., Reilly,S., Tomkins-Tinch,C., Loreth,C., Chaluvadi,S., Meldrim,J., Granger,B., Lemieux,J.E., Birren,B.W., Sabeti,P.C., Larkin,K., Dodge,S., Lennon,N., Madoff,L., Brown,C., Gallagher,G., Smole,S., Park,D.J., Gabriel,S., and MacInnis,B.L. |
| EPI_ISL_3408649, EPI_ISL_3408650, | Rhode Island Department of Health | Infectious Disease Program, Broad Institute of Harvard and MIT | Siddle,K.J., Adams,G., Pearlman,L., Gladden-Young,A., Vicente,G., Blumenstiel,B., DeFelice,M., Lee,M., McGovern,S., Lagerborg,K., Rudy,M., DeRuff,K., Carter,A., Normandin,E., Bauer,M., Reilly,S., Tomkins-Tinch,C., Loreth,C., Chaluvadi,S., Meldrim,J., Granger,B., Lemieux,J.E., Birren,B.W., Sabeti,P.C., Larkin,K., Dodge,S., Lennon,N., Madoff,L., Brown,C., Gallagher,G., Smole,S., Park,D.J., Gabriel,S., and MacInnis,B.L. |

| | | | |
|---|---|---|---|
| EPI_ISL_3408651 | | | |
| EPI_ISL_3408652, EPI_ISL_3408653, EPI_ISL_3408654 | Broad Institute Clinical Research Sequencing Platform | Infectious Disease Program, Broad Institute of Harvard and MIT | Siddle,K.J., Adams,G., Pearlman,L., Gladden-Young,A., Vicente,G., Blumenstiel,B., DeFelice,M., Lee,M., McGovern,S., Lagerborg,K., Rudy,M., DeRuff,K., Carter,A., Normandin,E., Bauer,M., Reilly,S., Tomkins-Tinch,C., Loreth,C., Chaluvadi,S., Meldrim,J., Granger,B., Lemieux,J.E., Birren,B.W., Sabeti,P.C., Larkin,K., Dodge,S., Lennon,N., Madoff,L., Brown,C., Gallagher,G., Smole,S., Park,D.J., Gabriel,S., and MacInnis,B.L. |
| EPI_ISL_3408655, EPI_ISL_3408656, EPI_ISL_3408657 | Rhode Island Department of Health | Infectious Disease Program, Broad Institute of Harvard and MIT | Siddle,K.J., Adams,G., Pearlman,L., Gladden-Young,A., Vicente,G., Blumenstiel,B., DeFelice,M., Lee,M., McGovern,S., Lagerborg,K., Rudy,M., DeRuff,K., Carter,A., Normandin,E., Bauer,M., Reilly,S., Tomkins-Tinch,C., Loreth,C., Chaluvadi,S., Meldrim,J., Granger,B., Lemieux,J.E., Birren,B.W., Sabeti,P.C., Larkin,K., Dodge,S., Lennon,N., Madoff,L., Brown,C., Gallagher,G., Smole,S., Park,D.J., Gabriel,S., and MacInnis,B.L. |
| EPI_ISL_3408658, EPI_ISL_3408659, EPI_ISL_3408660 | Broad Institute Clinical Research Sequencing Platform | Infectious Disease Program, Broad Institute of Harvard and MIT | Siddle,K.J., Adams,G., Pearlman,L., Gladden-Young,A., Vicente,G., Blumenstiel,B., DeFelice,M., Lee,M., McGovern,S., Lagerborg,K., Rudy,M., DeRuff,K., Carter,A., Normandin,E., Bauer,M., Reilly,S., Tomkins-Tinch,C., Loreth,C., Chaluvadi,S., Meldrim,J., Granger,B., Lemieux,J.E., Birren,B.W., Sabeti,P.C., Larkin,K., Dodge,S., Lennon,N., Madoff,L., Brown,C., Gallagher,G., Smole,S., Park,D.J., Gabriel,S., and MacInnis,B.L. |
| EPI_ISL_3408661 | Rhode Island Department of Health | Infectious Disease Program, Broad Institute of Harvard and MIT | Siddle,K.J., Adams,G., Pearlman,L., Gladden-Young,A., Vicente,G., Blumenstiel,B., DeFelice,M., Lee,M., McGovern,S., Lagerborg,K., Rudy,M., DeRuff,K., Carter,A., Normandin,E., Bauer,M., Reilly,S., Tomkins-Tinch,C., Loreth,C., Chaluvadi,S., Meldrim,J., Granger,B., Lemieux,J.E., Birren,B.W., Sabeti,P.C., Larkin,K., Dodge,S., Lennon,N., Madoff,L., Brown,C., Gallagher,G., Smole,S., Park,D.J., Gabriel,S., and MacInnis,B.L. |
| EPI_ISL_3408662 | Broad Institute Clinical Research Sequencing Platform | Infectious Disease Program, Broad Institute of Harvard and MIT | Siddle,K.J., Adams,G., Pearlman,L., Gladden-Young,A., Vicente,G., Blumenstiel,B., DeFelice,M., Lee,M., McGovern,S., Lagerborg,K., Rudy,M., DeRuff,K., Carter,A., Normandin,E., Bauer,M., Reilly,S., Tomkins-Tinch,C., Loreth,C., Chaluvadi,S., Meldrim,J., Granger,B., Lemieux,J.E., Birren,B.W., Sabeti,P.C., Larkin,K., Dodge,S., Lennon,N., Madoff,L., Brown,C., Gallagher,G., Smole,S., Park,D.J., Gabriel,S., and MacInnis,B.L. |
| EPI_ISL_3408663, EPI_ISL_3408664, EPI_ISL_3408665 | Rhode Island Department of Health | Infectious Disease Program, Broad Institute of Harvard and MIT | Siddle,K.J., Adams,G., Pearlman,L., Gladden-Young,A., Vicente,G., Blumenstiel,B., DeFelice,M., Lee,M., McGovern,S., Lagerborg,K., Rudy,M., DeRuff,K., Carter,A., Normandin,E., Bauer,M., Reilly,S., Tomkins-Tinch,C., Loreth,C., Chaluvadi,S., Meldrim,J., Granger,B., Lemieux,J.E., Birren,B.W., Sabeti,P.C., Larkin,K., Dodge,S., Lennon,N., Madoff,L., Brown,C., Gallagher,G., Smole,S., Park,D.J., Gabriel,S., and MacInnis,B.L. |
| EPI_ISL_3408666, EPI_ISL_3408667, EPI_ISL_3408668, EPI_ISL_3408669, EPI_ISL_3408670 | Broad Institute Clinical Research Sequencing Platform | Infectious Disease Program, Broad Institute of Harvard and MIT | Siddle,K.J., Adams,G., Pearlman,L., Gladden-Young,A., Vicente,G., Blumenstiel,B., DeFelice,M., Lee,M., McGovern,S., Lagerborg,K., Rudy,M., DeRuff,K., Carter,A., Normandin,E., Bauer,M., Reilly,S., Tomkins-Tinch,C., Loreth,C., Chaluvadi,S., Meldrim,J., Granger,B., Lemieux,J.E., Birren,B.W., Sabeti,P.C., Larkin,K., Dodge,S., Lennon,N., Madoff,L., Brown,C., Gallagher,G., Smole,S., Park,D.J., Gabriel,S., and MacInnis,B.L. |
| EPI_ISL_3408671 | Rhode Island Department of Health | Infectious Disease Program, Broad Institute of Harvard and MIT | Siddle,K.J., Adams,G., Pearlman,L., Gladden-Young,A., Vicente,G., Blumenstiel,B., DeFelice,M., Lee,M., McGovern,S., Lagerborg,K., Rudy,M., DeRuff,K., Carter,A., Normandin,E., Bauer,M., Reilly,S., Tomkins-Tinch,C., Loreth,C., Chaluvadi,S., Meldrim,J., Granger,B., Lemieux,J.E., Birren,B.W., Sabeti,P.C., Larkin,K., Dodge,S., Lennon,N., Madoff,L., Brown,C., Gallagher,G., Smole,S., Park,D.J., Gabriel,S., and MacInnis,B.L. |
| EPI_ISL_3408672 | Broad Institute Clinical Research Sequencing Platform | Infectious Disease Program, Broad Institute of Harvard and MIT | Siddle,K.J., Adams,G., Pearlman,L., Gladden-Young,A., Vicente,G., Blumenstiel,B., DeFelice,M., Lee,M., McGovern,S., Lagerborg,K., Rudy,M., DeRuff,K., Carter,A., Normandin,E., Bauer,M., Reilly,S., Tomkins-Tinch,C., Loreth,C., Chaluvadi,S., Meldrim,J., Granger,B., Lemieux,J.E., Birren,B.W., Sabeti,P.C., Larkin,K., Dodge,S., Lennon,N., Madoff,L., Brown,C., Gallagher,G., Smole,S., Park,D.J., Gabriel,S., and MacInnis,B.L. |
| EPI_ISL_3408673 | Rhode Island Department of Health | Infectious Disease Program, Broad Institute of Harvard and MIT | Siddle,K.J., Adams,G., Pearlman,L., Gladden-Young,A., Vicente,G., Blumenstiel,B., DeFelice,M., Lee,M., McGovern,S., Lagerborg,K., Rudy,M., DeRuff,K., Carter,A., Normandin,E., Bauer,M., Reilly,S., Tomkins-Tinch,C., Loreth,C., Chaluvadi,S., Meldrim,J., Granger,B., Lemieux,J.E., Birren,B.W., Sabeti,P.C., Larkin,K., Dodge,S., Lennon,N., Madoff,L., Brown,C., Gallagher,G., Smole,S., Park,D.J., Gabriel,S., and MacInnis,B.L. |
| EPI_ISL_3408674, EPI_ISL_3408675 | Broad Institute Clinical Research Sequencing Platform | Infectious Disease Program, Broad Institute of Harvard and MIT | Siddle,K.J., Adams,G., Pearlman,L., Gladden-Young,A., Vicente,G., Blumenstiel,B., DeFelice,M., Lee,M., McGovern,S., Lagerborg,K., Rudy,M., DeRuff,K., Carter,A., Normandin,E., Bauer,M., Reilly,S., Tomkins-Tinch,C., Loreth,C., Chaluvadi,S., Meldrim,J., Granger,B., Lemieux,J.E., Birren,B.W., Sabeti,P.C., Larkin,K., Dodge,S., Lennon,N., Madoff,L., Brown,C., Gallagher,G., Smole,S., Park,D.J., Gabriel,S., and MacInnis,B.L. |
| EPI_ISL_3408676, EPI_ISL_3408677, EPI_ISL_3408678, EPI_ISL_3408679, EPI_ISL_3408680 | Rhode Island Department of Health | Infectious Disease Program, Broad Institute of Harvard and MIT | Siddle,K.J., Adams,G., Pearlman,L., Gladden-Young,A., Vicente,G., Blumenstiel,B., DeFelice,M., Lee,M., McGovern,S., Lagerborg,K., Rudy,M., DeRuff,K., Carter,A., Normandin,E., Bauer,M., Reilly,S., Tomkins-Tinch,C., Loreth,C., Chaluvadi,S., Meldrim,J., Granger,B., Lemieux,J.E., Birren,B.W., Sabeti,P.C., Larkin,K., Dodge,S., Lennon,N., Madoff,L., Brown,C., Gallagher,G., Smole,S., Park,D.J., Gabriel,S., and MacInnis,B.L. |
| EPI_ISL_3408681, EPI_ISL_3408682, EPI_ISL_3408683, EPI_ISL_3408684, EPI_ISL_3408685, EPI_ISL_3408686, EPI_ISL_3408687, EPI_ISL_3408688, EPI_ISL_3408689, EPI_ISL_3408690, EPI_ISL_3408691, EPI_ISL_3408692, EPI_ISL_3408693 | | | |
| see above | Broad Institute Clinical Research Sequencing Platform | Infectious Disease Program, Broad Institute of Harvard and MIT | Siddle,K.J., Adams,G., Pearlman,L., Gladden-Young,A., Vicente,G., Blumenstiel,B., DeFelice,M., Lee,M., McGovern,S., Lagerborg,K., Rudy,M., DeRuff,K., Carter,A., Normandin,E., Bauer,M., Reilly,S., Tomkins-Tinch,C., Loreth,C., Chaluvadi,S., Meldrim,J., Granger,B., Lemieux,J.E., Birren,B.W., Sabeti,P.C., Larkin,K., Dodge,S., Lennon,N., Madoff,L., Brown,C., Gallagher,G., Smole,S., Park,D.J., Gabriel,S., and MacInnis,B.L. |
| EPI_ISL_3408694 | Rhode Island Department of Health | Infectious Disease Program, Broad Institute of Harvard and MIT | Siddle,K.J., Adams,G., Pearlman,L., Gladden-Young,A., Vicente,G., Blumenstiel,B., DeFelice,M., Lee,M., McGovern,S., Lagerborg,K., Rudy,M., DeRuff,K., Carter,A., Normandin,E., Bauer,M., Reilly,S., Tomkins-Tinch,C., Loreth,C., Chaluvadi,S., Meldrim,J., Granger,B., Lemieux,J.E., Birren,B.W., Sabeti,P.C., Larkin,K., Dodge,S., Lennon,N., Madoff,L., Brown,C., Gallagher,G., Smole,S., Park,D.J., Gabriel,S., and MacInnis,B.L. |
| EPI_ISL_3408695, EPI_ISL_3408696, EPI_ISL_3408698, EPI_ISL_3408699, EPI_ISL_3408700, EPI_ISL_3408701, EPI_ISL_3408702, EPI_ISL_3408703, EPI_ISL_3408704, EPI_ISL_3408705, EPI_ISL_3408706, EPI_ISL_3408707 | | | |
| see above | Broad Institute Clinical Research Sequencing Platform | Infectious Disease Program, Broad Institute of Harvard and MIT | Siddle,K.J., Adams,G., Pearlman,L., Gladden-Young,A., Vicente,G., Blumenstiel,B., DeFelice,M., Lee,M., McGovern,S., Lagerborg,K., Rudy,M., DeRuff,K., Carter,A., Normandin,E., Bauer,M., Reilly,S., Tomkins-Tinch,C., Loreth,C., Chaluvadi,S., Meldrim,J., Granger,B., Lemieux,J.E., Birren,B.W., Sabeti,P.C., Larkin,K., Dodge,S., Lennon,N., Madoff,L., Brown,C., Gallagher,G., Smole,S., Park,D.J., Gabriel,S., and MacInnis,B.L. |
| EPI_ISL_3408708 | Rhode Island Department of Health | Infectious Disease Program, Broad Institute of Harvard and MIT | Siddle,K.J., Adams,G., Pearlman,L., Gladden-Young,A., Vicente,G., Blumenstiel,B., DeFelice,M., Lee,M., McGovern,S., Lagerborg,K., Rudy,M., DeRuff,K., Carter,A., Normandin,E., Bauer,M., Reilly,S., Tomkins-Tinch,C., Loreth,C., Chaluvadi,S., Meldrim,J., Granger,B., Lemieux,J.E., Birren,B.W., Sabeti,P.C., Larkin,K., Dodge,S., Lennon,N., Madoff,L., Brown,C., Gallagher,G., Smole,S., Park,D.J., Gabriel,S., and MacInnis,B.L. |
| EPI_ISL_3408709, EPI_ISL_3408710, EPI_ISL_3408711, EPI_ISL_3408712, EPI_ISL_3408713, EPI_ISL_3408714, EPI_ISL_3408715, EPI_ISL_3408716, EPI_ISL_3408717, EPI_ISL_3408718, EPI_ISL_3408719, EPI_ISL_3408720, EPI_ISL_3408721, EPI_ISL_3408722, EPI_ISL_3408723, EPI_ISL_3408724, EPI_ISL_3408726, EPI_ISL_3408727, EPI_ISL_3408728, EPI_ISL_3408729, EPI_ISL_3408730, EPI_ISL_3408731 | | | |
| see above | Broad Institute Clinical Research Sequencing Platform | Infectious Disease Program, Broad Institute of Harvard and MIT | Siddle,K.J., Adams,G., Pearlman,L., Gladden-Young,A., Vicente,G., Blumenstiel,B., DeFelice,M., Lee,M., McGovern,S., Lagerborg,K., Rudy,M., DeRuff,K., Carter,A., Normandin,E., Bauer,M., Reilly,S., Tomkins-Tinch,C., Loreth,C., Chaluvadi,S., Meldrim,J., Granger,B., Lemieux,J.E., Birren,B.W., Sabeti,P.C., Larkin,K., Dodge,S., Lennon,N., Madoff,L., Brown,C., Gallagher,G., Smole,S., Park,D.J., Gabriel,S., and MacInnis,B.L. |
| EPI_ISL_3408732 | Rhode Island Department of Health | Infectious Disease Program, Broad Institute of Harvard and MIT | Siddle,K.J., Adams,G., Pearlman,L., Gladden-Young,A., Vicente,G., Blumenstiel,B., DeFelice,M., Lee,M., McGovern,S., Lagerborg,K., Rudy,M., DeRuff,K., Carter,A., Normandin,E., Bauer,M., Reilly,S., Tomkins-Tinch,C., Loreth,C., Chaluvadi,S., Meldrim,J., Granger,B., Lemieux,J.E., Birren,B.W., Sabeti,P.C., Larkin,K., Dodge,S., Lennon,N., Madoff,L., Brown,C., Gallagher,G., Smole,S., Park,D.J., Gabriel,S., and MacInnis,B.L. |
| EPI_ISL_3408733, EPI_ISL_3408734, EPI_ISL_3408735, EPI_ISL_3408736, EPI_ISL_3408737, EPI_ISL_3408738, EPI_ISL_3408739, EPI_ISL_3408740, EPI_ISL_3408741, EPI_ISL_3408742, EPI_ISL_3408743, EPI_ISL_3408744, EPI_ISL_3408745, EPI_ISL_3408746, EPI_ISL_3408747, EPI_ISL_3408748, EPI_ISL_3408749 | | | |
| see above | Broad Institute Clinical Research Sequencing Platform | Infectious Disease Program, Broad Institute of Harvard and MIT | Siddle,K.J., Adams,G., Pearlman,L., Gladden-Young,A., Vicente,G., Blumenstiel,B., DeFelice,M., Lee,M., McGovern,S., Lagerborg,K., Rudy,M., DeRuff,K., Carter,A., Normandin,E., Bauer,M., Reilly,S., Tomkins-Tinch,C., Loreth,C., Chaluvadi,S., Meldrim,J., Granger,B., Lemieux,J.E., Birren,B.W., Sabeti,P.C., Larkin,K., Dodge,S., Lennon,N., Madoff,L., Brown,C., Gallagher,G., Smole,S., Park,D.J., Gabriel,S., and MacInnis,B.L. |
| EPI_ISL_3408750 | Rhode Island Department of Health | Infectious Disease Program, Broad Institute of Harvard and MIT | Siddle,K.J., Adams,G., Pearlman,L., Gladden-Young,A., Vicente,G., Blumenstiel,B., DeFelice,M., Lee,M., McGovern,S., Lagerborg,K., Rudy,M., DeRuff,K., Carter,A., Normandin,E., Bauer,M., Reilly,S., Tomkins-Tinch,C., Loreth,C., Chaluvadi,S., Meldrim,J., Granger,B., Lemieux,J.E., Birren,B.W., Sabeti,P.C., Larkin,K., Dodge,S., Lennon,N., Madoff,L., Brown,C., Gallagher,G., Smole,S., Park,D.J., Gabriel,S., and MacInnis,B.L. |
| EPI_ISL_3408751, EPI_ISL_3408752, EPI_ISL_3408753, EPI_ISL_3408754, EPI_ISL_3408755, EPI_ISL_3408756, EPI_ISL_3408757, EPI_ISL_3408758, EPI_ISL_3408759, EPI_ISL_3408760, EPI_ISL_3408761, EPI_ISL_3408762, EPI_ISL_3408763, EPI_ISL_3408764, EPI_ISL_3408765, EPI_ISL_3408766, EPI_ISL_3408767, EPI_ISL_3408768, EPI_ISL_3408769, EPI_ISL_3408770, EPI_ISL_3408771, EPI_ISL_3408772, EPI_ISL_3408773, EPI_ISL_3408774, EPI_ISL_3408775, EPI_ISL_3408776, EPI_ISL_3408777, EPI_ISL_3408778, EPI_ISL_3408779, EPI_ISL_3408780, EPI_ISL_3408781, EPI_ISL_3408782, EPI_ISL_3408783, EPI_ISL_3408784 | | | |
| see above | Broad Institute Clinical Research Sequencing Platform | Infectious Disease Program, Broad Institute of Harvard and MIT | Siddle,K.J., Adams,G., Pearlman,L., Gladden-Young,A., Vicente,G., Blumenstiel,B., DeFelice,M., Lee,M., McGovern,S., Lagerborg,K., Rudy,M., DeRuff,K., Carter,A., Normandin,E., Bauer,M., Reilly,S., Tomkins-Tinch,C., Loreth,C., Chaluvadi,S., Meldrim,J., Granger,B., Lemieux,J.E., Birren,B.W., Sabeti,P.C., Larkin,K., Dodge,S., Lennon,N., Madoff,L., Brown,C., Gallagher,G., Smole,S., Park,D.J., Gabriel,S., and MacInnis,B.L. |
| EPI_ISL_3408785, EPI_ISL_3408786, EPI_ISL_3408787 | Rhode Island Department of Health | Infectious Disease Program, Broad Institute of Harvard and MIT | Siddle,K.J., Adams,G., Pearlman,L., Gladden-Young,A., Vicente,G., Blumenstiel,B., DeFelice,M., Lee,M., McGovern,S., Lagerborg,K., Rudy,M., DeRuff,K., Carter,A., Normandin,E., Bauer,M., Reilly,S., Tomkins-Tinch,C., Loreth,C., Chaluvadi,S., Meldrim,J., Granger,B., Lemieux,J.E., Birren,B.W., Sabeti,P.C., Larkin,K., Dodge,S., Lennon,N., Madoff,L., Brown,C., Gallagher,G., Smole,S., Park,D.J., Gabriel,S., and MacInnis,B.L. |
| EPI_ISL_3408788, EPI_ISL_3408789, EPI_ISL_3408790, EPI_ISL_3408791, EPI_ISL_3408792 | Broad Institute Clinical Research Sequencing Platform | Infectious Disease Program, Broad Institute of Harvard and MIT | Siddle,K.J., Adams,G., Pearlman,L., Gladden-Young,A., Vicente,G., Blumenstiel,B., DeFelice,M., Lee,M., McGovern,S., Lagerborg,K., Rudy,M., DeRuff,K., Carter,A., Normandin,E., Bauer,M., Reilly,S., Tomkins-Tinch,C., Loreth,C., Chaluvadi,S., Meldrim,J., Granger,B., Lemieux,J.E., Birren,B.W., Sabeti,P.C., Larkin,K., Dodge,S., Lennon,N., Madoff,L., Brown,C., Gallagher,G., Smole,S., Park,D.J., Gabriel,S., and MacInnis,B.L. |
| EPI_ISL_3408793, EPI_ISL_3408794 | Rhode Island Department of Health | Infectious Disease Program, Broad Institute of Harvard and MIT | Siddle,K.J., Adams,G., Pearlman,L., Gladden-Young,A., Vicente,G., Blumenstiel,B., DeFelice,M., Lee,M., McGovern,S., Lagerborg,K., Rudy,M., DeRuff,K., Carter,A., Normandin,E., Bauer,M., Reilly,S., Tomkins-Tinch,C., Loreth,C., Chaluvadi,S., Meldrim,J., Granger,B., Lemieux,J.E., Birren,B.W., Sabeti,P.C., Larkin,K., Dodge,S., Lennon,N., Madoff,L., Brown,C., Gallagher,G., Smole,S., Park,D.J., Gabriel,S., and MacInnis,B.L. |
| EPI_ISL_3408795, EPI_ISL_3408796, EPI_ISL_3408797, EPI_ISL_3408798 | Broad Institute Clinical Research Sequencing Platform | Infectious Disease Program, Broad Institute of Harvard and MIT | Siddle,K.J., Adams,G., Pearlman,L., Gladden-Young,A., Vicente,G., Blumenstiel,B., DeFelice,M., Lee,M., McGovern,S., Lagerborg,K., Rudy,M., DeRuff,K., Carter,A., Normandin,E., Bauer,M., Reilly,S., Tomkins-Tinch,C., Loreth,C., Chaluvadi,S., Meldrim,J., Granger,B., Lemieux,J.E., Birren,B.W., Sabeti,P.C., Larkin,K., Dodge,S., Lennon,N., Madoff,L., Brown,C., Gallagher,G., Smole,S., Park,D.J., Gabriel,S., and MacInnis,B.L. |
| EPI_ISL_3408799, EPI_ISL_3408800, EPI_ISL_3408801 | Rhode Island Department of Health | Infectious Disease Program, Broad Institute of Harvard and MIT | Siddle,K.J., Adams,G., Pearlman,L., Gladden-Young,A., Vicente,G., Blumenstiel,B., DeFelice,M., Lee,M., McGovern,S., Lagerborg,K., Rudy,M., DeRuff,K., Carter,A., Normandin,E., Bauer,M., Reilly,S., Tomkins-Tinch,C., Loreth,C., Chaluvadi,S., Meldrim,J., Granger,B., Lemieux,J.E., Birren,B.W., Sabeti,P.C., Larkin,K., Dodge,S., Lennon,N., Madoff,L., Brown,C., Gallagher,G., Smole,S., Park,D.J., Gabriel,S., and MacInnis,B.L. |
| EPI_ISL_3408802, EPI_ISL_3408803, EPI_ISL_3408804 | Broad Institute Clinical Research Sequencing Platform | Infectious Disease Program, Broad Institute of Harvard and MIT | Siddle,K.J., Adams,G., Pearlman,L., Gladden-Young,A., Vicente,G., Blumenstiel,B., DeFelice,M., Lee,M., McGovern,S., Lagerborg,K., Rudy,M., DeRuff,K., Carter,A., Normandin,E., Bauer,M., Reilly,S., Tomkins-Tinch,C., Loreth,C., Chaluvadi,S., Meldrim,J., Granger,B., Lemieux,J.E., Birren,B.W., Sabeti,P.C., Larkin,K., Dodge,S., Lennon,N., Madoff,L., Brown,C., Gallagher,G., Smole,S., Park,D.J., Gabriel,S., and MacInnis,B.L. |
| EPI_ISL_3408805 | Rhode Island Department of Health | Infectious Disease Program, Broad Institute of Harvard and MIT | Siddle,K.J., Adams,G., Pearlman,L., Gladden-Young,A., Vicente,G., Blumenstiel,B., DeFelice,M., Lee,M., McGovern,S., Lagerborg,K., Rudy,M., DeRuff,K., Carter,A., Normandin,E., Bauer,M., Reilly,S., Tomkins-Tinch,C., Loreth,C., Chaluvadi,S., Meldrim,J., Granger,B., Lemieux,J.E., Birren,B.W., Sabeti,P.C., Larkin,K., Dodge,S., Lennon,N., Madoff,L., Brown,C., Gallagher,G., Smole,S., Park,D.J., Gabriel,S., and MacInnis,B.L. |
| EPI_ISL_3408806, EPI_ISL_3408807, EPI_ISL_3408808, EPI_ISL_3408809, EPI_ISL_3408810, EPI_ISL_3408811, EPI_ISL_3408812, EPI_ISL_3408813, EPI_ISL_3408814, EPI_ISL_3408815, EPI_ISL_3408816, EPI_ISL_3408817, EPI_ISL_3408818, EPI_ISL_3408819, EPI_ISL_3408820, EPI_ISL_3408821, EPI_ISL_3408822, EPI_ISL_3408823, EPI_ISL_3408824, EPI_ISL_3408826, EPI_ISL_3408827, EPI_ISL_3408828, EPI_ISL_3408829, EPI_ISL_3408830, EPI_ISL_3408831, EPI_ISL_3408832, EPI_ISL_3408833, EPI_ISL_3408834, EPI_ISL_3408835, EPI_ISL_3408836, EPI_ISL_3408837, EPI_ISL_3408838, EPI_ISL_3408839, EPI_ISL_3408840, EPI_ISL_3408841, EPI_ISL_3408842, EPI_ISL_3408843, EPI_ISL_3408844, EPI_ISL_3408845 | | | |
| see above | Broad Institute Clinical Research Sequencing Platform | Infectious Disease Program, Broad Institute of Harvard and MIT | Siddle,K.J., Adams,G., Pearlman,L., Gladden-Young,A., Vicente,G., Blumenstiel,B., DeFelice,M., Lee,M., McGovern,S., Lagerborg,K., Rudy,M., DeRuff,K., Carter,A., Normandin,E., Bauer,M., Reilly,S., Tomkins-Tinch,C., Loreth,C., Chaluvadi,S., Meldrim,J., Granger,B., Lemieux,J.E., Birren,B.W., Sabeti,P.C., Larkin,K., Dodge,S., Lennon,N., Madoff,L., Brown,C., Gallagher,G., Smole,S., Park,D.J., Gabriel,S., and MacInnis,B.L. |
| EPI_ISL_3408846, EPI_ISL_3408847, EPI_ISL_3408848 | Rhode Island Department of Health | Infectious Disease Program, Broad Institute of Harvard and MIT | Siddle,K.J., Adams,G., Pearlman,L., Gladden-Young,A., Vicente,G., Blumenstiel,B., DeFelice,M., Lee,M., McGovern,S., Lagerborg,K., Rudy,M., DeRuff,K., Carter,A., Normandin,E., Bauer,M., Reilly,S., Tomkins-Tinch,C., Loreth,C., Chaluvadi,S., Meldrim,J., Granger,B., Lemieux,J.E., Birren,B.W., Sabeti,P.C., Larkin,K., Dodge,S., Lennon,N., Madoff,L., Brown,C., Gallagher,G., Smole,S., Park,D.J., Gabriel,S., and MacInnis,B.L. |
| EPI_ISL_3408849, EPI_ISL_3408850, EPI_ISL_3408851, EPI_ISL_3408852, EPI_ISL_3408853, | Broad Institute Clinical Research Sequencing Platform | Infectious Disease Program, Broad Institute of Harvard and MIT | Siddle,K.J., Adams,G., Pearlman,L., Gladden-Young,A., Vicente,G., Blumenstiel,B., DeFelice,M., Lee,M., McGovern,S., Lagerborg,K., Rudy,M., DeRuff,K., Carter,A., Normandin,E., Bauer,M., Reilly,S., Tomkins-Tinch,C., Loreth,C., Chaluvadi,S., Meldrim,J., Granger,B., Lemieux,J.E., Birren,B.W., Sabeti,P.C., Larkin,K., Dodge,S., Lennon,N., Madoff,L., Brown,C., Gallagher,G., Smole,S., Park,D.J., Gabriel,S., and MacInnis,B.L. |

| | | | |
|---|---|---|---|
| EPI_ISL_3408854, EPI_ISL_3408855, EPI_ISL_3408856, EPI_ISL_3408857 | | | |
| EPI_ISL_3408858 | Rhode Island Department of Health | Infectious Disease Program, Broad Institute of Harvard and MIT | Siddle,K.J., Adams,G., Pearlman,L., Gladden-Young,A., Vicente,G., Blumenstiel,B., DeFelice,M., Lee,M., McGovern,S., Lagerborg,K., Rudy,M., DeRuff,K., Carter,A., Normandin,E., Bauer,M., Reilly,S., Tomkins-Tinch,C., Loreth,C., Chaluvadi,S., Meldrim,J., Granger,B., Lemieux,J.E., Birren,B.W., Sabeti,P.C., Larkin,K., Dodge,S., Lennon,N., Madoff,L., Brown,C., Gallagher,G., Smole,S., Park,D.J., Gabriel,S., and MacInnis,B.L. |
| EPI_ISL_3408859, EPI_ISL_3408860, EPI_ISL_3408861, EPI_ISL_3408862, EPI_ISL_3408863, EPI_ISL_3408864 | Broad Institute Clinical Research Sequencing Platform | Infectious Disease Program, Broad Institute of Harvard and MIT | Siddle,K.J., Adams,G., Pearlman,L., Gladden-Young,A., Vicente,G., Blumenstiel,B., DeFelice,M., Lee,M., McGovern,S., Lagerborg,K., Rudy,M., DeRuff,K., Carter,A., Normandin,E., Bauer,M., Reilly,S., Tomkins-Tinch,C., Loreth,C., Chaluvadi,S., Meldrim,J., Granger,B., Lemieux,J.E., Birren,B.W., Sabeti,P.C., Larkin,K., Dodge,S., Lennon,N., Madoff,L., Brown,C., Gallagher,G., Smole,S., Park,D.J., Gabriel,S., and MacInnis,B.L. |
| EPI_ISL_3408865 | Rhode Island Department of Health | Infectious Disease Program, Broad Institute of Harvard and MIT | Siddle,K.J., Adams,G., Pearlman,L., Gladden-Young,A., Vicente,G., Blumenstiel,B., DeFelice,M., Lee,M., McGovern,S., Lagerborg,K., Rudy,M., DeRuff,K., Carter,A., Normandin,E., Bauer,M., Reilly,S., Tomkins-Tinch,C., Loreth,C., Chaluvadi,S., Meldrim,J., Granger,B., Lemieux,J.E., Birren,B.W., Sabeti,P.C., Larkin,K., Dodge,S., Lennon,N., Madoff,L., Brown,C., Gallagher,G., Smole,S., Park,D.J., Gabriel,S., and MacInnis,B.L. |
| EPI_ISL_3408866, EPI_ISL_3408867, EPI_ISL_3408868 | Broad Institute Clinical Research Sequencing Platform | Infectious Disease Program, Broad Institute of Harvard and MIT | Siddle,K.J., Adams,G., Pearlman,L., Gladden-Young,A., Vicente,G., Blumenstiel,B., DeFelice,M., Lee,M., McGovern,S., Lagerborg,K., Rudy,M., DeRuff,K., Carter,A., Normandin,E., Bauer,M., Reilly,S., Tomkins-Tinch,C., Loreth,C., Chaluvadi,S., Meldrim,J., Granger,B., Lemieux,J.E., Birren,B.W., Sabeti,P.C., Larkin,K., Dodge,S., Lennon,N., Madoff,L., Brown,C., Gallagher,G., Smole,S., Park,D.J., Gabriel,S., and MacInnis,B.L. |
| EPI_ISL_3408869 | Rhode Island Department of Health | Infectious Disease Program, Broad Institute of Harvard and MIT | Siddle,K.J., Adams,G., Pearlman,L., Gladden-Young,A., Vicente,G., Blumenstiel,B., DeFelice,M., Lee,M., McGovern,S., Lagerborg,K., Rudy,M., DeRuff,K., Carter,A., Normandin,E., Bauer,M., Reilly,S., Tomkins-Tinch,C., Loreth,C., Chaluvadi,S., Meldrim,J., Granger,B., Lemieux,J.E., Birren,B.W., Sabeti,P.C., Larkin,K., Dodge,S., Lennon,N., Madoff,L., Brown,C., Gallagher,G., Smole,S., Park,D.J., Gabriel,S., and MacInnis,B.L. |
| EPI_ISL_3408870, EPI_ISL_3408871, EPI_ISL_3408872, EPI_ISL_3408873, EPI_ISL_3408874, EPI_ISL_3408875, EPI_ISL_3408876, EPI_ISL_3408877, EPI_ISL_3408878, EPI_ISL_3408879, EPI_ISL_3408880, EPI_ISL_3408881 | | | |
| see above | Broad Institute Clinical Research Sequencing Platform | Infectious Disease Program, Broad Institute of Harvard and MIT | Siddle,K.J., Adams,G., Pearlman,L., Gladden-Young,A., Vicente,G., Blumenstiel,B., DeFelice,M., Lee,M., McGovern,S., Lagerborg,K., Rudy,M., DeRuff,K., Carter,A., Normandin,E., Bauer,M., Reilly,S., Tomkins-Tinch,C., Loreth,C., Chaluvadi,S., Meldrim,J., Granger,B., Lemieux,J.E., Birren,B.W., Sabeti,P.C., Larkin,K., Dodge,S., Lennon,N., Madoff,L., Brown,C., Gallagher,G., Smole,S., Park,D.J., Gabriel,S., and MacInnis,B.L. |
| EPI_ISL_3408882, EPI_ISL_3408883 | Rhode Island Department of Health | Infectious Disease Program, Broad Institute of Harvard and MIT | Siddle,K.J., Adams,G., Pearlman,L., Gladden-Young,A., Vicente,G., Blumenstiel,B., DeFelice,M., Lee,M., McGovern,S., Lagerborg,K., Rudy,M., DeRuff,K., Carter,A., Normandin,E., Bauer,M., Reilly,S., Tomkins-Tinch,C., Loreth,C., Chaluvadi,S., Meldrim,J., Granger,B., Lemieux,J.E., Birren,B.W., Sabeti,P.C., Larkin,K., Dodge,S., Lennon,N., Madoff,L., Brown,C., Gallagher,G., Smole,S., Park,D.J., Gabriel,S., and MacInnis,B.L. |
| EPI_ISL_3408884, EPI_ISL_3408885, EPI_ISL_3408886 | Broad Institute Clinical Research Sequencing Platform | Infectious Disease Program, Broad Institute of Harvard and MIT | Siddle,K.J., Adams,G., Pearlman,L., Gladden-Young,A., Vicente,G., Blumenstiel,B., DeFelice,M., Lee,M., McGovern,S., Lagerborg,K., Rudy,M., DeRuff,K., Carter,A., Normandin,E., Bauer,M., Reilly,S., Tomkins-Tinch,C., Loreth,C., Chaluvadi,S., Meldrim,J., Granger,B., Lemieux,J.E., Birren,B.W., Sabeti,P.C., Larkin,K., Dodge,S., Lennon,N., Madoff,L., Brown,C., Gallagher,G., Smole,S., Park,D.J., Gabriel,S., and MacInnis,B.L. |
| EPI_ISL_3408887, EPI_ISL_3408888 | Rhode Island Department of Health | Infectious Disease Program, Broad Institute of Harvard and MIT | Siddle,K.J., Adams,G., Pearlman,L., Gladden-Young,A., Vicente,G., Blumenstiel,B., DeFelice,M., Lee,M., McGovern,S., Lagerborg,K., Rudy,M., DeRuff,K., Carter,A., Normandin,E., Bauer,M., Reilly,S., Tomkins-Tinch,C., Loreth,C., Chaluvadi,S., Meldrim,J., Granger,B., Lemieux,J.E., Birren,B.W., Sabeti,P.C., Larkin,K., Dodge,S., Lennon,N., Madoff,L., Brown,C., Gallagher,G., Smole,S., Park,D.J., Gabriel,S., and MacInnis,B.L. |
| EPI_ISL_3408889, EPI_ISL_3408890, EPI_ISL_3408891 | Broad Institute Clinical Research Sequencing Platform | Infectious Disease Program, Broad Institute of Harvard and MIT | Siddle,K.J., Adams,G., Pearlman,L., Gladden-Young,A., Vicente,G., Blumenstiel,B., DeFelice,M., Lee,M., McGovern,S., Lagerborg,K., Rudy,M., DeRuff,K., Carter,A., Normandin,E., Bauer,M., Reilly,S., Tomkins-Tinch,C., Loreth,C., Chaluvadi,S., Meldrim,J., Granger,B., Lemieux,J.E., Birren,B.W., Sabeti,P.C., Larkin,K., Dodge,S., Lennon,N., Madoff,L., Brown,C., Gallagher,G., Smole,S., Park,D.J., Gabriel,S., and MacInnis,B.L. |
| EPI_ISL_3408892 | Rhode Island Department of Health | Infectious Disease Program, Broad Institute of Harvard and MIT | Siddle,K.J., Adams,G., Pearlman,L., Gladden-Young,A., Vicente,G., Blumenstiel,B., DeFelice,M., Lee,M., McGovern,S., Lagerborg,K., Rudy,M., DeRuff,K., Carter,A., Normandin,E., Bauer,M., Reilly,S., Tomkins-Tinch,C., Loreth,C., Chaluvadi,S., Meldrim,J., Granger,B., Lemieux,J.E., Birren,B.W., Sabeti,P.C., Larkin,K., Dodge,S., Lennon,N., Madoff,L., Brown,C., Gallagher,G., Smole,S., Park,D.J., Gabriel,S., and MacInnis,B.L. |
| EPI_ISL_3408893, EPI_ISL_3408894, EPI_ISL_3408895, EPI_ISL_3408896, EPI_ISL_3408897, EPI_ISL_3408898, EPI_ISL_3408899, EPI_ISL_3408900, EPI_ISL_3408901, EPI_ISL_3408902, EPI_ISL_3408903 | | | |
| see above | Broad Institute Clinical Research Sequencing Platform | Infectious Disease Program, Broad Institute of Harvard and MIT | Siddle,K.J., Adams,G., Pearlman,L., Gladden-Young,A., Vicente,G., Blumenstiel,B., DeFelice,M., Lee,M., McGovern,S., Lagerborg,K., Rudy,M., DeRuff,K., Carter,A., Normandin,E., Bauer,M., Reilly,S., Tomkins-Tinch,C., Loreth,C., Chaluvadi,S., Meldrim,J., Granger,B., Lemieux,J.E., Birren,B.W., Sabeti,P.C., Larkin,K., Dodge,S., Lennon,N., Madoff,L., Brown,C., Gallagher,G., Smole,S., Park,D.J., Gabriel,S., and MacInnis,B.L. |
| EPI_ISL_3408904 | Rhode Island Department of Health | Infectious Disease Program, Broad Institute of Harvard and MIT | Siddle,K.J., Adams,G., Pearlman,L., Gladden-Young,A., Vicente,G., Blumenstiel,B., DeFelice,M., Lee,M., McGovern,S., Lagerborg,K., Rudy,M., DeRuff,K., Carter,A., Normandin,E., Bauer,M., Reilly,S., Tomkins-Tinch,C., Loreth,C., Chaluvadi,S., Meldrim,J., Granger,B., Lemieux,J.E., Birren,B.W., Sabeti,P.C., Larkin,K., Dodge,S., Lennon,N., Madoff,L., Brown,C., Gallagher,G., Smole,S., Park,D.J., Gabriel,S., and MacInnis,B.L. |
| EPI_ISL_3408905, EPI_ISL_3408906, EPI_ISL_3408907, EPI_ISL_3408908, EPI_ISL_3408909, EPI_ISL_3408910 | Broad Institute Clinical Research Sequencing Platform | Infectious Disease Program, Broad Institute of Harvard and MIT | Siddle,K.J., Adams,G., Pearlman,L., Gladden-Young,A., Vicente,G., Blumenstiel,B., DeFelice,M., Lee,M., McGovern,S., Lagerborg,K., Rudy,M., DeRuff,K., Carter,A., Normandin,E., Bauer,M., Reilly,S., Tomkins-Tinch,C., Loreth,C., Chaluvadi,S., Meldrim,J., Granger,B., Lemieux,J.E., Birren,B.W., Sabeti,P.C., Larkin,K., Dodge,S., Lennon,N., Madoff,L., Brown,C., Gallagher,G., Smole,S., Park,D.J., Gabriel,S., and MacInnis,B.L. |
| EPI_ISL_3408911 | Rhode Island Department of Health | Infectious Disease Program, Broad Institute of Harvard and MIT | Siddle,K.J., Adams,G., Pearlman,L., Gladden-Young,A., Vicente,G., Blumenstiel,B., DeFelice,M., Lee,M., McGovern,S., Lagerborg,K., Rudy,M., DeRuff,K., Carter,A., Normandin,E., Bauer,M., Reilly,S., Tomkins-Tinch,C., Loreth,C., Chaluvadi,S., Meldrim,J., Granger,B., Lemieux,J.E., Birren,B.W., Sabeti,P.C., Larkin,K., Dodge,S., Lennon,N., Madoff,L., Brown,C., Gallagher,G., Smole,S., Park,D.J., Gabriel,S., and MacInnis,B.L. |
| EPI_ISL_3408912, EPI_ISL_3408913, EPI_ISL_3408914, EPI_ISL_3408915, EPI_ISL_3408916, EPI_ISL_3408917, EPI_ISL_3408918, EPI_ISL_3408919, EPI_ISL_3408920, EPI_ISL_3408921, EPI_ISL_3408922, EPI_ISL_3408923 | | | |
| see above | Broad Institute Clinical Research Sequencing Platform | Infectious Disease Program, Broad Institute of Harvard and MIT | Siddle,K.J., Adams,G., Pearlman,L., Gladden-Young,A., Vicente,G., Blumenstiel,B., DeFelice,M., Lee,M., McGovern,S., Lagerborg,K., Rudy,M., DeRuff,K., Carter,A., Normandin,E., Bauer,M., Reilly,S., Tomkins-Tinch,C., Loreth,C., Chaluvadi,S., Meldrim,J., Granger,B., Lemieux,J.E., Birren,B.W., Sabeti,P.C., Larkin,K., Dodge,S., Lennon,N., Madoff,L., Brown,C., Gallagher,G., Smole,S., Park,D.J., Gabriel,S., and MacInnis,B.L. |
| EPI_ISL_3408924, EPI_ISL_3408925 | Rhode Island Department of Health | Infectious Disease Program, Broad Institute of Harvard and MIT | Siddle,K.J., Adams,G., Pearlman,L., Gladden-Young,A., Vicente,G., Blumenstiel,B., DeFelice,M., Lee,M., McGovern,S., Lagerborg,K., Rudy,M., DeRuff,K., Carter,A., Normandin,E., Bauer,M., Reilly,S., Tomkins-Tinch,C., Loreth,C., Chaluvadi,S., Meldrim,J., Granger,B., Lemieux,J.E., Birren,B.W., Sabeti,P.C., Larkin,K., Dodge,S., Lennon,N., Madoff,L., Brown,C., Gallagher,G., Smole,S., Park,D.J., Gabriel,S., and MacInnis,B.L. |
| EPI_ISL_3408926, EPI_ISL_3408927, EPI_ISL_3408929 | Broad Institute Clinical Research Sequencing Platform | Infectious Disease Program, Broad Institute of Harvard and MIT | Siddle,K.J., Adams,G., Pearlman,L., Gladden-Young,A., Vicente,G., Blumenstiel,B., DeFelice,M., Lee,M., McGovern,S., Lagerborg,K., Rudy,M., DeRuff,K., Carter,A., Normandin,E., Bauer,M., Reilly,S., Tomkins-Tinch,C., Loreth,C., Chaluvadi,S., Meldrim,J., Granger,B., Lemieux,J.E., Birren,B.W., Sabeti,P.C., Larkin,K., Dodge,S., Lennon,N., Madoff,L., Brown,C., Gallagher,G., Smole,S., Park,D.J., Gabriel,S., and MacInnis,B.L. |
| EPI_ISL_3408930, EPI_ISL_3408931 | Rhode Island Department of Health | Infectious Disease Program, Broad Institute of Harvard and MIT | Siddle,K.J., Adams,G., Pearlman,L., Gladden-Young,A., Vicente,G., Blumenstiel,B., DeFelice,M., Lee,M., McGovern,S., Lagerborg,K., Rudy,M., DeRuff,K., Carter,A., Normandin,E., Bauer,M., Reilly,S., Tomkins-Tinch,C., Loreth,C., Chaluvadi,S., Meldrim,J., Granger,B., Lemieux,J.E., Birren,B.W., Sabeti,P.C., Larkin,K., Dodge,S., Lennon,N., Madoff,L., Brown,C., Gallagher,G., Smole,S., Park,D.J., Gabriel,S., and MacInnis,B.L. |
| EPI_ISL_3408932, EPI_ISL_3408933, EPI_ISL_3408934, EPI_ISL_3408935, EPI_ISL_3408936, EPI_ISL_3408937, EPI_ISL_3408938, EPI_ISL_3408939, EPI_ISL_3408940, EPI_ISL_3408941, EPI_ISL_3408942 | | | |
| see above | Broad Institute Clinical Research Sequencing Platform | Infectious Disease Program, Broad Institute of Harvard and MIT | Siddle,K.J., Adams,G., Pearlman,L., Gladden-Young,A., Vicente,G., Blumenstiel,B., DeFelice,M., Lee,M., McGovern,S., Lagerborg,K., Rudy,M., DeRuff,K., Carter,A., Normandin,E., Bauer,M., Reilly,S., Tomkins-Tinch,C., Loreth,C., Chaluvadi,S., Meldrim,J., Granger,B., Lemieux,J.E., Birren,B.W., Sabeti,P.C., Larkin,K., Dodge,S., Lennon,N., Madoff,L., Brown,C., Gallagher,G., Smole,S., Park,D.J., Gabriel,S., and MacInnis,B.L. |
| EPI_ISL_3408943 | Rhode Island Department of Health | Infectious Disease Program, Broad Institute of Harvard and MIT | Siddle,K.J., Adams,G., Pearlman,L., Gladden-Young,A., Vicente,G., Blumenstiel,B., DeFelice,M., Lee,M., McGovern,S., Lagerborg,K., Rudy,M., DeRuff,K., Carter,A., Normandin,E., Bauer,M., Reilly,S., Tomkins-Tinch,C., Loreth,C., Chaluvadi,S., Meldrim,J., Granger,B., Lemieux,J.E., Birren,B.W., Sabeti,P.C., Larkin,K., Dodge,S., Lennon,N., Madoff,L., Brown,C., Gallagher,G., Smole,S., Park,D.J., Gabriel,S., and MacInnis,B.L. |
| EPI_ISL_3408944, EPI_ISL_3408945, EPI_ISL_3408946, EPI_ISL_3408947, EPI_ISL_3408948, EPI_ISL_3408951, EPI_ISL_3408952 | Broad Institute Clinical Research Sequencing Platform | Infectious Disease Program, Broad Institute of Harvard and MIT | Siddle,K.J., Adams,G., Pearlman,L., Gladden-Young,A., Vicente,G., Blumenstiel,B., DeFelice,M., Lee,M., McGovern,S., Lagerborg,K., Rudy,M., DeRuff,K., Carter,A., Normandin,E., Bauer,M., Reilly,S., Tomkins-Tinch,C., Loreth,C., Chaluvadi,S., Meldrim,J., Granger,B., Lemieux,J.E., Birren,B.W., Sabeti,P.C., Larkin,K., Dodge,S., Lennon,N., Madoff,L., Brown,C., Gallagher,G., Smole,S., Park,D.J., Gabriel,S., and MacInnis,B.L. |
| EPI_ISL_3408953 | Rhode Island Department of Health | Infectious Disease Program, Broad Institute of Harvard and MIT | Siddle,K.J., Adams,G., Pearlman,L., Gladden-Young,A., Vicente,G., Blumenstiel,B., DeFelice,M., Lee,M., McGovern,S., Lagerborg,K., Rudy,M., DeRuff,K., Carter,A., Normandin,E., Bauer,M., Reilly,S., Tomkins-Tinch,C., Loreth,C., Chaluvadi,S., Meldrim,J., Granger,B., Lemieux,J.E., Birren,B.W., Sabeti,P.C., Larkin,K., Dodge,S., Lennon,N., Madoff,L., Brown,C., Gallagher,G., Smole,S., Park,D.J., Gabriel,S., and MacInnis,B.L. |
| EPI_ISL_3408954 | Broad Institute Clinical Research Sequencing Platform | Infectious Disease Program, Broad Institute of Harvard and MIT | Siddle,K.J., Adams,G., Pearlman,L., Gladden-Young,A., Vicente,G., Blumenstiel,B., DeFelice,M., Lee,M., McGovern,S., Lagerborg,K., Rudy,M., DeRuff,K., Carter,A., Normandin,E., Bauer,M., Reilly,S., Tomkins-Tinch,C., Loreth,C., Chaluvadi,S., Meldrim,J., Granger,B., Lemieux,J.E., Birren,B.W., Sabeti,P.C., Larkin,K., Dodge,S., Lennon,N., Madoff,L., Brown,C., Gallagher,G., Smole,S., Park,D.J., Gabriel,S., and MacInnis,B.L. |
| EPI_ISL_3408955 | Rhode Island Department of Health | Infectious Disease Program, Broad Institute of Harvard and MIT | Siddle,K.J., Adams,G., Pearlman,L., Gladden-Young,A., Vicente,G., Blumenstiel,B., DeFelice,M., Lee,M., McGovern,S., Lagerborg,K., Rudy,M., DeRuff,K., Carter,A., Normandin,E., Bauer,M., Reilly,S., Tomkins-Tinch,C., Loreth,C., Chaluvadi,S., Meldrim,J., Granger,B., Lemieux,J.E., Birren,B.W., Sabeti,P.C., Larkin,K., Dodge,S., Lennon,N., Madoff,L., Brown,C., Gallagher,G., Smole,S., Park,D.J., Gabriel,S., and MacInnis,B.L. |
| EPI_ISL_3408956, EPI_ISL_3408957, EPI_ISL_3408958, EPI_ISL_3408959, EPI_ISL_3408960, EPI_ISL_3408961 | Broad Institute Clinical Research Sequencing Platform | Infectious Disease Program, Broad Institute of Harvard and MIT | Siddle,K.J., Adams,G., Pearlman,L., Gladden-Young,A., Vicente,G., Blumenstiel,B., DeFelice,M., Lee,M., McGovern,S., Lagerborg,K., Rudy,M., DeRuff,K., Carter,A., Normandin,E., Bauer,M., Reilly,S., Tomkins-Tinch,C., Loreth,C., Chaluvadi,S., Meldrim,J., Granger,B., Lemieux,J.E., Birren,B.W., Sabeti,P.C., Larkin,K., Dodge,S., Lennon,N., Madoff,L., Brown,C., Gallagher,G., Smole,S., Park,D.J., Gabriel,S., and MacInnis,B.L. |
| EPI_ISL_3408962 | Rhode Island Department of Health | Infectious Disease Program, Broad Institute of Harvard and MIT | Siddle,K.J., Adams,G., Pearlman,L., Gladden-Young,A., Vicente,G., Blumenstiel,B., DeFelice,M., Lee,M., McGovern,S., Lagerborg,K., Rudy,M., DeRuff,K., Carter,A., Normandin,E., Bauer,M., Reilly,S., Tomkins-Tinch,C., Loreth,C., Chaluvadi,S., Meldrim,J., Granger,B., Lemieux,J.E., Birren,B.W., Sabeti,P.C., Larkin,K., Dodge,S., Lennon,N., Madoff,L., Brown,C., Gallagher,G., Smole,S., Park,D.J., Gabriel,S., and MacInnis,B.L. |
| EPI_ISL_3408964, EPI_ISL_3408965, EPI_ISL_3408966, EPI_ISL_3408967, EPI_ISL_3408968, EPI_ISL_3408969, EPI_ISL_3408970, EPI_ISL_3408971, EPI_ISL_3408972, EPI_ISL_3408973, EPI_ISL_3408974, EPI_ISL_3408976 | | | |
| see above | Broad Institute Clinical Research Sequencing Platform | Infectious Disease Program, Broad Institute of Harvard and MIT | Siddle,K.J., Adams,G., Pearlman,L., Gladden-Young,A., Vicente,G., Blumenstiel,B., DeFelice,M., Lee,M., McGovern,S., Lagerborg,K., Rudy,M., DeRuff,K., Carter,A., Normandin,E., Bauer,M., Reilly,S., Tomkins-Tinch,C., Loreth,C., Chaluvadi,S., Meldrim,J., Granger,B., Lemieux,J.E., Birren,B.W., Sabeti,P.C., Larkin,K., Dodge,S., Lennon,N., Madoff,L., Brown,C., Gallagher,G., Smole,S., Park,D.J., Gabriel,S., and MacInnis,B.L. |
| EPI_ISL_3408977 | Rhode Island Department of Health | Infectious Disease Program, Broad Institute of Harvard and MIT | Siddle,K.J., Adams,G., Pearlman,L., Gladden-Young,A., Vicente,G., Blumenstiel,B., DeFelice,M., Lee,M., McGovern,S., Lagerborg,K., Rudy,M., DeRuff,K., Carter,A., Normandin,E., Bauer,M., Reilly,S., Tomkins-Tinch,C., Loreth,C., Chaluvadi,S., Meldrim,J., Granger,B., Lemieux,J.E., Birren,B.W., Sabeti,P.C., Larkin,K., Dodge,S., Lennon,N., Madoff,L., Brown,C., Gallagher,G., Smole,S., Park,D.J., Gabriel,S., and MacInnis,B.L. |
| EPI_ISL_3408979, EPI_ISL_3408980, EPI_ISL_3408981, EPI_ISL_3408982, EPI_ISL_3408983, EPI_ISL_3408984, EPI_ISL_3408986, EPI_ISL_3408987, EPI_ISL_3408988, EPI_ISL_3408989, EPI_ISL_3408990, EPI_ISL_3408991, EPI_ISL_3408992 | | | |
| see above | Broad Institute Clinical Research Sequencing Platform | Infectious Disease Program, Broad Institute of Harvard and MIT | Siddle,K.J., Adams,G., Pearlman,L., Gladden-Young,A., Vicente,G., Blumenstiel,B., DeFelice,M., Lee,M., McGovern,S., Lagerborg,K., Rudy,M., DeRuff,K., Carter,A., Normandin,E., Bauer,M., Reilly,S., Tomkins-Tinch,C., Loreth,C., Chaluvadi,S., Meldrim,J., Granger,B., Lemieux,J.E., Birren,B.W., Sabeti,P.C., Larkin,K., Dodge,S., Lennon,N., Madoff,L., Brown,C., Gallagher,G., Smole,S., Park,D.J., Gabriel,S., and MacInnis,B.L. |
| EPI_ISL_3408993 | Rhode Island Department of Health | Infectious Disease Program, Broad Institute of Harvard and MIT | Siddle,K.J., Adams,G., Pearlman,L., Gladden-Young,A., Vicente,G., Blumenstiel,B., DeFelice,M., Lee,M., McGovern,S., Lagerborg,K., Rudy,M., DeRuff,K., Carter,A., Normandin,E., Bauer,M., Reilly,S., Tomkins-Tinch,C., Loreth,C., Chaluvadi,S., Meldrim,J., Granger,B., Lemieux,J.E., Birren,B.W., Sabeti,P.C., Larkin,K., Dodge,S., Lennon,N., Madoff,L., Brown,C., Gallagher,G., Smole,S., Park,D.J., Gabriel,S., and MacInnis,B.L. |

| | | | |
|---|---|---|---|
| EPI_ISL_3408994, EPI_ISL_3408995, EPI_ISL_3408996, EPI_ISL_3408997, EPI_ISL_3408998, EPI_ISL_3408999, EPI_ISL_3409000, EPI_ISL_3409001, EPI_ISL_3409002, EPI_ISL_3409003, EPI_ISL_3409004 | | | |
| see above | Broad Institute Clinical Research Sequencing Platform | Infectious Disease Program, Broad Institute of Harvard and MIT | Siddle,K.J., Adams,G., Pearlman,L., Gladden-Young,A., Vicente,G., Blumenstiel,B., DeFelice,M., Lee,M., McGovern,S., Lagerborg,K., Rudy,M., DeRuff,K., Carter,A., Normandin,E., Bauer,M., Reilly,S., Tomkins-Tinch,C., Loreth,C., Chaluvadi,S., Meldrim,J., Granger,B., Lemieux,J.E., Birren,B.W., Sabeti,P.C., Larkin,K., Dodge,S., Lennon,N., Madoff,L., Brown,C., Gallagher,G., Smole,S., Park,D.J., Gabriel,S., and MacInnis,B.L. |
| EPI_ISL_3409005 | Rhode Island Department of Health | Infectious Disease Program, Broad Institute of Harvard and MIT | Siddle,K.J., Adams,G., Pearlman,L., Gladden-Young,A., Vicente,G., Blumenstiel,B., DeFelice,M., Lee,M., McGovern,S., Lagerborg,K., Rudy,M., DeRuff,K., Carter,A., Normandin,E., Bauer,M., Reilly,S., Tomkins-Tinch,C., Loreth,C., Chaluvadi,S., Meldrim,J., Granger,B., Lemieux,J.E., Birren,B.W., Sabeti,P.C., Larkin,K., Dodge,S., Lennon,N., Madoff,L., Brown,C., Gallagher,G., Smole,S., Park,D.J., Gabriel,S., and MacInnis,B.L. |
| EPI_ISL_3409006 | Broad Institute Clinical Research Sequencing Platform | Infectious Disease Program, Broad Institute of Harvard and MIT | Siddle,K.J., Adams,G., Pearlman,L., Gladden-Young,A., Vicente,G., Blumenstiel,B., DeFelice,M., Lee,M., McGovern,S., Lagerborg,K., Rudy,M., DeRuff,K., Carter,A., Normandin,E., Bauer,M., Reilly,S., Tomkins-Tinch,C., Loreth,C., Chaluvadi,S., Meldrim,J., Granger,B., Lemieux,J.E., Birren,B.W., Sabeti,P.C., Larkin,K., Dodge,S., Lennon,N., Madoff,L., Brown,C., Gallagher,G., Smole,S., Park,D.J., Gabriel,S., and MacInnis,B.L. |
| EPI_ISL_3409007 | Rhode Island Department of Health | Infectious Disease Program, Broad Institute of Harvard and MIT | Siddle,K.J., Adams,G., Pearlman,L., Gladden-Young,A., Vicente,G., Blumenstiel,B., DeFelice,M., Lee,M., McGovern,S., Lagerborg,K., Rudy,M., DeRuff,K., Carter,A., Normandin,E., Bauer,M., Reilly,S., Tomkins-Tinch,C., Loreth,C., Chaluvadi,S., Meldrim,J., Granger,B., Lemieux,J.E., Birren,B.W., Sabeti,P.C., Larkin,K., Dodge,S., Lennon,N., Madoff,L., Brown,C., Gallagher,G., Smole,S., Park,D.J., Gabriel,S., and MacInnis,B.L. |
| EPI_ISL_3409008, EPI_ISL_3409009, EPI_ISL_3409010, EPI_ISL_3409011, EPI_ISL_3409012 | Broad Institute Clinical Research Sequencing Platform | Infectious Disease Program, Broad Institute of Harvard and MIT | Siddle,K.J., Adams,G., Pearlman,L., Gladden-Young,A., Vicente,G., Blumenstiel,B., DeFelice,M., Lee,M., McGovern,S., Lagerborg,K., Rudy,M., DeRuff,K., Carter,A., Normandin,E., Bauer,M., Reilly,S., Tomkins-Tinch,C., Loreth,C., Chaluvadi,S., Meldrim,J., Granger,B., Lemieux,J.E., Birren,B.W., Sabeti,P.C., Larkin,K., Dodge,S., Lennon,N., Madoff,L., Brown,C., Gallagher,G., Smole,S., Park,D.J., Gabriel,S., and MacInnis,B.L. |
| EPI_ISL_3409013 | Rhode Island Department of Health | Infectious Disease Program, Broad Institute of Harvard and MIT | Siddle,K.J., Adams,G., Pearlman,L., Gladden-Young,A., Vicente,G., Blumenstiel,B., DeFelice,M., Lee,M., McGovern,S., Lagerborg,K., Rudy,M., DeRuff,K., Carter,A., Normandin,E., Bauer,M., Reilly,S., Tomkins-Tinch,C., Loreth,C., Chaluvadi,S., Meldrim,J., Granger,B., Lemieux,J.E., Birren,B.W., Sabeti,P.C., Larkin,K., Dodge,S., Lennon,N., Madoff,L., Brown,C., Gallagher,G., Smole,S., Park,D.J., Gabriel,S., and MacInnis,B.L. |
| EPI_ISL_3409014, EPI_ISL_3409015, EPI_ISL_3409016, EPI_ISL_3409017, EPI_ISL_3409018 | Broad Institute Clinical Research Sequencing Platform | Infectious Disease Program, Broad Institute of Harvard and MIT | Siddle,K.J., Adams,G., Pearlman,L., Gladden-Young,A., Vicente,G., Blumenstiel,B., DeFelice,M., Lee,M., McGovern,S., Lagerborg,K., Rudy,M., DeRuff,K., Carter,A., Normandin,E., Bauer,M., Reilly,S., Tomkins-Tinch,C., Loreth,C., Chaluvadi,S., Meldrim,J., Granger,B., Lemieux,J.E., Birren,B.W., Sabeti,P.C., Larkin,K., Dodge,S., Lennon,N., Madoff,L., Brown,C., Gallagher,G., Smole,S., Park,D.J., Gabriel,S., and MacInnis,B.L. |
| EPI_ISL_3409019, EPI_ISL_3409020 | Rhode Island Department of Health | Infectious Disease Program, Broad Institute of Harvard and MIT | Siddle,K.J., Adams,G., Pearlman,L., Gladden-Young,A., Vicente,G., Blumenstiel,B., DeFelice,M., Lee,M., McGovern,S., Lagerborg,K., Rudy,M., DeRuff,K., Carter,A., Normandin,E., Bauer,M., Reilly,S., Tomkins-Tinch,C., Loreth,C., Chaluvadi,S., Meldrim,J., Granger,B., Lemieux,J.E., Birren,B.W., Sabeti,P.C., Larkin,K., Dodge,S., Lennon,N., Madoff,L., Brown,C., Gallagher,G., Smole,S., Park,D.J., Gabriel,S., and MacInnis,B.L. |
| EPI_ISL_3409021, EPI_ISL_3409022, EPI_ISL_3409023, EPI_ISL_3409024, EPI_ISL_3409025, EPI_ISL_3409026, EPI_ISL_3409027, EPI_ISL_3409028 | Broad Institute Clinical Research Sequencing Platform | Infectious Disease Program, Broad Institute of Harvard and MIT | Siddle,K.J., Adams,G., Pearlman,L., Gladden-Young,A., Vicente,G., Blumenstiel,B., DeFelice,M., Lee,M., McGovern,S., Lagerborg,K., Rudy,M., DeRuff,K., Carter,A., Normandin,E., Bauer,M., Reilly,S., Tomkins-Tinch,C., Loreth,C., Chaluvadi,S., Meldrim,J., Granger,B., Lemieux,J.E., Birren,B.W., Sabeti,P.C., Larkin,K., Dodge,S., Lennon,N., Madoff,L., Brown,C., Gallagher,G., Smole,S., Park,D.J., Gabriel,S., and MacInnis,B.L. |
| EPI_ISL_3409029 | Rhode Island Department of Health | Infectious Disease Program, Broad Institute of Harvard and MIT | Siddle,K.J., Adams,G., Pearlman,L., Gladden-Young,A., Vicente,G., Blumenstiel,B., DeFelice,M., Lee,M., McGovern,S., Lagerborg,K., Rudy,M., DeRuff,K., Carter,A., Normandin,E., Bauer,M., Reilly,S., Tomkins-Tinch,C., Loreth,C., Chaluvadi,S., Meldrim,J., Granger,B., Lemieux,J.E., Birren,B.W., Sabeti,P.C., Larkin,K., Dodge,S., Lennon,N., Madoff,L., Brown,C., Gallagher,G., Smole,S., Park,D.J., Gabriel,S., and MacInnis,B.L. |
| EPI_ISL_3409030, EPI_ISL_3409031, EPI_ISL_3409032, EPI_ISL_3409033, EPI_ISL_3409034, EPI_ISL_3409035, EPI_ISL_3409036, EPI_ISL_3409038, EPI_ISL_3409039, EPI_ISL_3409040, EPI_ISL_3409041, EPI_ISL_3409042, EPI_ISL_3409043, EPI_ISL_3409044, EPI_ISL_3409045, EPI_ISL_3409046 | | | |
| see above | Broad Institute Clinical Research Sequencing Platform | Infectious Disease Program, Broad Institute of Harvard and MIT | Siddle,K.J., Adams,G., Pearlman,L., Gladden-Young,A., Vicente,G., Blumenstiel,B., DeFelice,M., Lee,M., McGovern,S., Lagerborg,K., Rudy,M., DeRuff,K., Carter,A., Normandin,E., Bauer,M., Reilly,S., Tomkins-Tinch,C., Loreth,C., Chaluvadi,S., Meldrim,J., Granger,B., Lemieux,J.E., Birren,B.W., Sabeti,P.C., Larkin,K., Dodge,S., Lennon,N., Madoff,L., Brown,C., Gallagher,G., Smole,S., Park,D.J., Gabriel,S., and MacInnis,B.L. |
| EPI_ISL_3409249, EPI_ISL_3409267, EPI_ISL_3409279, EPI_ISL_3409382, EPI_ISL_3409820, EPI_ISL_3409822, EPI_ISL_3410108, EPI_ISL_3410109, EPI_ISL_3410203, EPI_ISL_3410206, EPI_ISL_3410214, EPI_ISL_3410215, EPI_ISL_3410216, EPI_ISL_3410217, EPI_ISL_3410218, EPI_ISL_3410219, EPI_ISL_3410220, EPI_ISL_3410221, EPI_ISL_3410929, EPI_ISL_3410976 | | | |
| see above | UMass Memorial Medical Center | Center for Microbiome Research | Richard T. Ellison III, Karl J. Simon, Doyle V. Ward |
| EPI_ISL_3410991, EPI_ISL_3410992, EPI_ISL_3410993, EPI_ISL_3410994, EPI_ISL_3410995, EPI_ISL_3410996, EPI_ISL_3410997, EPI_ISL_3410998, EPI_ISL_3411001, EPI_ISL_3411002, EPI_ISL_3411003, EPI_ISL_3411004 | | | |
| see above | Ministry of Health Turkey | Ministry of Health Turkey | Fatma Bayrakdar, Yasemin Cosgun, Suleyman Yalcin, Gulay Korukluoglu |
| EPI_ISL_3411005, EPI_ISL_3411006, EPI_ISL_3411007, EPI_ISL_3411008, EPI_ISL_3411009, EPI_ISL_3411010, EPI_ISL_3411011, EPI_ISL_3411012, EPI_ISL_3411013, EPI_ISL_3411014, EPI_ISL_3411015, EPI_ISL_3411016, EPI_ISL_3411017, EPI_ISL_3411018, EPI_ISL_3411019, EPI_ISL_3411020, EPI_ISL_3411021, EPI_ISL_3411022, EPI_ISL_3411024, EPI_ISL_3411025, EPI_ISL_3411026, EPI_ISL_3411027, EPI_ISL_3411028, EPI_ISL_3411029, EPI_ISL_3411030, EPI_ISL_3411031, EPI_ISL_3411032, EPI_ISL_3411033, EPI_ISL_3411034, EPI_ISL_3411035, EPI_ISL_3411036, EPI_ISL_3411037, EPI_ISL_3411038, EPI_ISL_3411039, EPI_ISL_3411040, EPI_ISL_3411041, EPI_ISL_3411042, EPI_ISL_3411043, EPI_ISL_3411044, EPI_ISL_3411045, EPI_ISL_3411046, EPI_ISL_3411047, EPI_ISL_3411048, EPI_ISL_3411049, EPI_ISL_3411050, EPI_ISL_3411051, EPI_ISL_3411052, EPI_ISL_3411053, EPI_ISL_3411054, EPI_ISL_3411055, EPI_ISL_3411056, EPI_ISL_3411057, EPI_ISL_3411058, EPI_ISL_3411059, EPI_ISL_3411060, EPI_ISL_3411061, EPI_ISL_3411062, EPI_ISL_3411063, EPI_ISL_3411064, EPI_ISL_3411065, EPI_ISL_3411066, EPI_ISL_3411067, EPI_ISL_3411068, EPI_ISL_3411069, EPI_ISL_3411070, EPI_ISL_3411071, EPI_ISL_3411072, EPI_ISL_3411073, EPI_ISL_3411074, EPI_ISL_3411075, EPI_ISL_3411076, EPI_ISL_3411077, EPI_ISL_3411078, EPI_ISL_3411079, EPI_ISL_3411080, EPI_ISL_3411081, EPI_ISL_3411082, EPI_ISL_3411083, EPI_ISL_3411084, EPI_ISL_3411085, EPI_ISL_3411086, EPI_ISL_3411087, EPI_ISL_3411088, EPI_ISL_3411089, EPI_ISL_3411090, EPI_ISL_3411091, EPI_ISL_3411092, EPI_ISL_3411093, EPI_ISL_3411094, EPI_ISL_3411095, EPI_ISL_3411096, EPI_ISL_3411097, EPI_ISL_3411098, EPI_ISL_3411099, EPI_ISL_3411100, EPI_ISL_3411101, EPI_ISL_3411102, EPI_ISL_3411103, EPI_ISL_3411104, EPI_ISL_3411105, EPI_ISL_3411106, EPI_ISL_3411107, EPI_ISL_3411108, EPI_ISL_3411109, EPI_ISL_3411110, EPI_ISL_3411111, EPI_ISL_3411112, EPI_ISL_3411113, EPI_ISL_3411114, EPI_ISL_3411115, EPI_ISL_3411116, EPI_ISL_3411117, EPI_ISL_3411118, EPI_ISL_3411119, EPI_ISL_3411120, EPI_ISL_3411121, EPI_ISL_3411122, EPI_ISL_3411123, EPI_ISL_3411124, EPI_ISL_3411125, EPI_ISL_3411126, EPI_ISL_3411127, EPI_ISL_3411128, EPI_ISL_3411129, EPI_ISL_3411130, EPI_ISL_3411131, EPI_ISL_3411132, EPI_ISL_3411133, EPI_ISL_3411134, EPI_ISL_3411135, EPI_ISL_3411136, EPI_ISL_3411137, EPI_ISL_3411138, EPI_ISL_3411139, EPI_ISL_3411140, EPI_ISL_3411141, EPI_ISL_3411142, EPI_ISL_3411143, EPI_ISL_3411144, EPI_ISL_3411146, EPI_ISL_3411148, EPI_ISL_3411149, EPI_ISL_3411150, EPI_ISL_3411151, EPI_ISL_3411152, EPI_ISL_3411153, EPI_ISL_3411154, EPI_ISL_3411155, EPI_ISL_3411156, EPI_ISL_3411157, EPI_ISL_3411158, EPI_ISL_3411159, EPI_ISL_3411160, EPI_ISL_3411161, EPI_ISL_3411162, EPI_ISL_3411163, EPI_ISL_3411164, EPI_ISL_3411165, EPI_ISL_3411166, EPI_ISL_3411167, EPI_ISL_3411168, EPI_ISL_3411169, EPI_ISL_3411170, EPI_ISL_3411171, EPI_ISL_3411172, EPI_ISL_3411173, EPI_ISL_3411174, EPI_ISL_3411175, EPI_ISL_3411176, EPI_ISL_3411177, EPI_ISL_3411178, EPI_ISL_3411179, EPI_ISL_3411180, EPI_ISL_3411181, EPI_ISL_3411182, EPI_ISL_3411183, EPI_ISL_3411184, EPI_ISL_3411185, EPI_ISL_3411186, EPI_ISL_3411187, EPI_ISL_3411188, EPI_ISL_3411189, EPI_ISL_3411190, EPI_ISL_3411191, EPI_ISL_3411192, EPI_ISL_3411193, EPI_ISL_3411194, EPI_ISL_3411195, EPI_ISL_3411196, EPI_ISL_3411197, EPI_ISL_3411198, EPI_ISL_3411199, EPI_ISL_3411200, EPI_ISL_3411201, EPI_ISL_3411202, EPI_ISL_3411203, EPI_ISL_3411204, EPI_ISL_3411205, EPI_ISL_3411206, EPI_ISL_3411207, EPI_ISL_3411208, EPI_ISL_3411209, EPI_ISL_3411210, EPI_ISL_3411211, EPI_ISL_3411212, EPI_ISL_3411213, EPI_ISL_3411214, EPI_ISL_3411215, EPI_ISL_3411216, EPI_ISL_3411217, EPI_ISL_3411218, EPI_ISL_3411219, EPI_ISL_3411220, EPI_ISL_3411221, EPI_ISL_3411222, EPI_ISL_3411223, EPI_ISL_3411224, EPI_ISL_3411225, EPI_ISL_3411226, EPI_ISL_3411227, EPI_ISL_3411228, EPI_ISL_3411229, EPI_ISL_3411230, EPI_ISL_3411231, EPI_ISL_3411232, EPI_ISL_3411233, EPI_ISL_3411234, EPI_ISL_3411235, EPI_ISL_3411236, EPI_ISL_3411237, EPI_ISL_3411238, EPI_ISL_3411239, EPI_ISL_3411241, EPI_ISL_3411242, EPI_ISL_3411243, EPI_ISL_3411244, EPI_ISL_3411245, EPI_ISL_3411246, EPI_ISL_3411247, EPI_ISL_3411248, EPI_ISL_3411249, EPI_ISL_3411250, EPI_ISL_3411251, EPI_ISL_3411252, EPI_ISL_3411253, EPI_ISL_3411254, EPI_ISL_3411255, EPI_ISL_3411256, EPI_ISL_3411257, EPI_ISL_3411258, EPI_ISL_3411259, EPI_ISL_3411260, EPI_ISL_3411261, EPI_ISL_3411262, EPI_ISL_3411263, EPI_ISL_3411264, EPI_ISL_3411265, EPI_ISL_3411266, EPI_ISL_3411267, EPI_ISL_3411268, EPI_ISL_3411269, EPI_ISL_3411270, EPI_ISL_3411271, EPI_ISL_3411272, EPI_ISL_3411273, EPI_ISL_3411274, EPI_ISL_3411275, EPI_ISL_3411276, EPI_ISL_3411277, EPI_ISL_3411278, EPI_ISL_3411279, EPI_ISL_3411280, EPI_ISL_3411281, EPI_ISL_3411282, EPI_ISL_3411283, EPI_ISL_3411284, EPI_ISL_3411285, EPI_ISL_3411286, EPI_ISL_3411287, EPI_ISL_3411288, EPI_ISL_3411289, EPI_ISL_3411290, EPI_ISL_3411291, EPI_ISL_3411292, EPI_ISL_3411293, EPI_ISL_3411294, EPI_ISL_3411295, EPI_ISL_3411296, EPI_ISL_3411297, EPI_ISL_3411298, EPI_ISL_3411299, EPI_ISL_3411300, EPI_ISL_3411301, EPI_ISL_3411302, EPI_ISL_3411303, EPI_ISL_3411304, EPI_ISL_3411305, EPI_ISL_3411306, EPI_ISL_3411307, EPI_ISL_3411308, EPI_ISL_3411309, EPI_ISL_3411310, EPI_ISL_3411311, EPI_ISL_3411312, EPI_ISL_3411313, EPI_ISL_3411314, EPI_ISL_3411315, EPI_ISL_3411316, EPI_ISL_3411317, EPI_ISL_3411318, EPI_ISL_3411319, EPI_ISL_3411320, EPI_ISL_3411321, EPI_ISL_3411322, EPI_ISL_3411323, EPI_ISL_3411324, EPI_ISL_3411325, EPI_ISL_3411326, EPI_ISL_3411327, EPI_ISL_3411328, EPI_ISL_3411329, EPI_ISL_3411330, EPI_ISL_3411331, EPI_ISL_3411332, EPI_ISL_3411334, EPI_ISL_3411335, EPI_ISL_3411336, EPI_ISL_3411337, EPI_ISL_3411338, EPI_ISL_3411340, EPI_ISL_3411341, EPI_ISL_3411342, EPI_ISL_3411343, EPI_ISL_3411344, EPI_ISL_3411345, EPI_ISL_3411346, EPI_ISL_3411347, EPI_ISL_3411348, EPI_ISL_3411349, EPI_ISL_3411350, EPI_ISL_3411351, EPI_ISL_3411352, EPI_ISL_3411353, EPI_ISL_3411354, EPI_ISL_3411356, EPI_ISL_3411357, EPI_ISL_3411358, EPI_ISL_3411359, EPI_ISL_3411360, EPI_ISL_3411361, EPI_ISL_3411362, EPI_ISL_3411363 | | | |
| see above | UW Virology Lab | UW Virology Lab | Pavitra Roychoudhury, Shah Mohamed Bakhash, Ricardo Perez, Hong Xie, Tien V. Nguyen, Maria Lukes, Sean Ellis, Nathan Breit, Robert J. Livingston, Meei-Li Huang, Keith R Jerome, Patrick Mathias, Alexander Greninger |
| EPI_ISL_3411364, EPI_ISL_3411365, EPI_ISL_3411366, EPI_ISL_3411367, EPI_ISL_3411368, EPI_ISL_3411369, EPI_ISL_3411370, EPI_ISL_3411371, EPI_ISL_3411372, EPI_ISL_3411373, EPI_ISL_3411374, EPI_ISL_3411375, EPI_ISL_3411376, EPI_ISL_3411377, EPI_ISL_3411378, EPI_ISL_3411379, EPI_ISL_3411380, EPI_ISL_3411381, EPI_ISL_3411382, EPI_ISL_3411383, EPI_ISL_3411384, EPI_ISL_3411385, EPI_ISL_3411386, EPI_ISL_3411387, EPI_ISL_3411388, EPI_ISL_3411389, EPI_ISL_3411390, EPI_ISL_3411391, EPI_ISL_3411392, EPI_ISL_3411393, EPI_ISL_3411394, EPI_ISL_3411395, EPI_ISL_3411396, EPI_ISL_3411397, EPI_ISL_3411398, EPI_ISL_3411399, EPI_ISL_3411400, EPI_ISL_3411401, EPI_ISL_3411402, EPI_ISL_3411403, EPI_ISL_3411404, EPI_ISL_3411405, EPI_ISL_3411406, EPI_ISL_3411407, EPI_ISL_3411408, EPI_ISL_3411409, EPI_ISL_3411410, EPI_ISL_3411411, EPI_ISL_3411412, EPI_ISL_3411413, EPI_ISL_3411414, EPI_ISL_3411415, EPI_ISL_3411416, EPI_ISL_3411417, EPI_ISL_3411418, EPI_ISL_3411419, EPI_ISL_3411420, EPI_ISL_3411421, EPI_ISL_3411422, EPI_ISL_3411423, EPI_ISL_3411424, EPI_ISL_3411425, EPI_ISL_3411426, EPI_ISL_3411427, EPI_ISL_3411428, EPI_ISL_3411429, EPI_ISL_3411430, EPI_ISL_3411431, EPI_ISL_3411432, EPI_ISL_3411434, EPI_ISL_3411435, EPI_ISL_3411436, EPI_ISL_3411437, EPI_ISL_3411438, EPI_ISL_3411439, EPI_ISL_3411440, EPI_ISL_3411441, EPI_ISL_3411442, EPI_ISL_3411443, EPI_ISL_3411444, EPI_ISL_3411445, EPI_ISL_3411446, EPI_ISL_3411447, EPI_ISL_3411454, EPI_ISL_3411455 | | | |
| see above | UW Virology Lab | UW Virology Lab | Pavitra Roychoudhury, Shah Mohamed Bakhash, Ricardo Perez, Maria Lukes, Hong Xie, Tien V. Nguyen, Sean Ellis, Meei-Li Huang, Keith R Jerome, Alexander Greninger |
| EPI_ISL_3411456 | National Institute for Communicable Diseases of the National Health Laboratory Service | National Institute for Communicable Diseases of the National Health Laboratory Service | Amoako DG, Everatt J, Scheepers C, Mohale T, Ntuli N, Mahlangu B, Mnguni A, Ismail A, Bhiman JN |
| EPI_ISL_3411461 | POLOKWANE LABORATORY | National Institute for Communicable Diseases of the National Health Laboratory Service | Amoako DG, Everatt J, Scheepers C, Mohale T, Ntuli N, Mahlangu B, Mnguni A, Ismail A, Bhiman JN |
| EPI_ISL_3411462, EPI_ISL_3411464, EPI_ISL_3411465 | TSHEPONG LABORATORY | National Institute for Communicable Diseases of the National Health Laboratory Service | Amoako DG, Everatt J, Scheepers C, Mohale T, Ntuli N, Mahlangu B, Mnguni A, Ismail A, Bhiman JN |
| EPI_ISL_3411466 | KOPANONG LABORATORY | National Institute for Communicable Diseases of the National Health Laboratory Service | Amoako DG, Everatt J, Scheepers C, Mohale T, Ntuli N, Mahlangu B, Mnguni A, Ismail A, Bhiman JN |
| EPI_ISL_3411468 | TAMBO MEMORIAL LABORATORY | National Institute for Communicable Diseases of the National Health Laboratory Service | Amoako DG, Everatt J, Scheepers C, Mohale T, Ntuli N, Mahlangu B, Mnguni A, Ismail A, Bhiman JN |
| EPI_ISL_3411473, EPI_ISL_3411474, EPI_ISL_3411475, EPI_ISL_3411476, EPI_ISL_3411477, EPI_ISL_3411478, EPI_ISL_3411479, EPI_ISL_3411480, EPI_ISL_3411481, EPI_ISL_3411482, EPI_ISL_3411483, EPI_ISL_3411484, EPI_ISL_3411485, EPI_ISL_3411486, EPI_ISL_3411487, EPI_ISL_3411488, EPI_ISL_3411489, EPI_ISL_3411490, EPI_ISL_3411491, EPI_ISL_3411492, EPI_ISL_3411493, EPI_ISL_3411494 | | | |
| see above | University Campus Bio-Medico of Rome (UCBM) | University Campus Bio-Medico of Rome (UCBM) | Fogolari M., Riva E., De Florio L., Francesconi M., Lintas C., Veralli R., Angeletti S. |
| EPI_ISL_3411495, EPI_ISL_3411496, EPI_ISL_3411497, EPI_ISL_3411498, EPI_ISL_3411499, EPI_ISL_3411500, EPI_ISL_3411501, EPI_ISL_3411502, EPI_ISL_3411503, EPI_ISL_3411504, EPI_ISL_3411505, EPI_ISL_3411506, EPI_ISL_3411507, EPI_ISL_3411508, EPI_ISL_3411509, EPI_ISL_3411510, EPI_ISL_3411511, EPI_ISL_3411512, EPI_ISL_3411513, EPI_ISL_3411514, EPI_ISL_3411515, EPI_ISL_3411516, EPI_ISL_3411517, EPI_ISL_3411518, EPI_ISL_3411519, EPI_ISL_3411520, EPI_ISL_3411521, EPI_ISL_3411522, EPI_ISL_3411523, EPI_ISL_3411524, EPI_ISL_3411525, EPI_ISL_3411526, EPI_ISL_3411527, EPI_ISL_3411528, EPI_ISL_3411529, EPI_ISL_3411530, EPI_ISL_3411531, EPI_ISL_3411532, EPI_ISL_3411533, EPI_ISL_3411534, EPI_ISL_3411535, EPI_ISL_3411536, EPI_ISL_3411537, EPI_ISL_3411538, EPI_ISL_3411539, EPI_ISL_3411540, EPI_ISL_3411541, EPI_ISL_3411542, EPI_ISL_3411543, EPI_ISL_3411544, EPI_ISL_3411545, EPI_ISL_3411546 | | | |
| see above | Servicio Microbiología Hospital La Paz | Servicio Microbiología Hospital La Paz | Fernando Lázaro, Rubén Cáceres, Iván Bloise, Jesús Mingorance Cruz, Elie Dahdouh |
| EPI_ISL_3411556, EPI_ISL_3411557, EPI_ISL_3411558, EPI_ISL_3411559, EPI_ISL_3411560, EPI_ISL_3411563, EPI_ISL_3411565, EPI_ISL_3411566, EPI_ISL_3411567, EPI_ISL_3411568, EPI_ISL_3411569, EPI_ISL_3411570 | | | |
| see above | Pathcare Vaal | National Institute for Communicable Diseases of the National Health Laboratory Service | Amoako DG, Everatt J, Scheepers C, Mohale T, Ntuli N, Mahlangu B, Mnguni A, Ismail A, Bhiman JN |
| EPI_ISL_3411572, EPI_ISL_3411573, EPI_ISL_3411574, EPI_ISL_3411575 | POLOKWANE LABORATORY | National Institute for Communicable Diseases of the National Health Laboratory Service | Amoako DG, Everatt J, Scheepers C, Mohale T, Ntuli N, Mahlangu B, Mnguni A, Ismail A, Bhiman JN |
| EPI_ISL_3411576, EPI_ISL_3411577, EPI_ISL_3411578, EPI_ISL_3411579 | Pathcare Vaal | National Institute for Communicable Diseases of the National Health Laboratory Service | Amoako DG, Everatt J, Scheepers C, Mohale T, Ntuli N, Mahlangu B, Mnguni A, Ismail A, Bhiman JN |
| EPI_ISL_3411580 | PORT ELIZABETH LABORATORY | National Institute for Communicable Diseases of the National Health Laboratory Service | Amoako DG, Everatt J, Scheepers C, Mohale T, Ntuli N, Mahlangu B, Mnguni A, Ismail A, Bhiman JN |
| EPI_ISL_3411581, EPI_ISL_3411582, EPI_ISL_3411583 | Pathcare Vaal | National Institute for Communicable Diseases of the National Health Laboratory Service | Amoako DG, Everatt J, Scheepers C, Mohale T, Ntuli N, Mahlangu B, Mnguni A, Ismail A, Bhiman JN |

| | | | |
|---|---|---|---|
| EPI_ISL_3411584, EPI_ISL_3411585, EPI_ISL_3411586 | KIMBERLEY LABORATORY | National Institute for Communicable Diseases of the National Health Laboratory Service | Amoako DG, Everatt J, Scheepers C, Mohale T, Ntuli N, Mahlangu B, Mnguni A, Ismail A, Bhiman JN |
| EPI_ISL_3411587, EPI_ISL_3411588, EPI_ISL_3411590, EPI_ISL_3411591, EPI_ISL_3411592, EPI_ISL_3411593, EPI_ISL_3411594, EPI_ISL_3411595, EPI_ISL_3411596, EPI_ISL_3411597, EPI_ISL_3411598, EPI_ISL_3411599, EPI_ISL_3411600, EPI_ISL_3411601, EPI_ISL_3411602, EPI_ISL_3411603, EPI_ISL_3411604, EPI_ISL_3411605, EPI_ISL_3411606 | | | |
| see above | University Pretoria | National Institute for Communicable Diseases of the National Health Laboratory Service | Amoako DG, Everatt J, Scheepers C, Mohale T, Ntuli N, Mahlangu B, Mnguni A, Ismail A, Bhiman JN |
| EPI_ISL_3411609, EPI_ISL_3411610, EPI_ISL_3411611, EPI_ISL_3411612, EPI_ISL_3411613, EPI_ISL_3411615, EPI_ISL_3411616, EPI_ISL_3411617, EPI_ISL_3411618, EPI_ISL_3411619, EPI_ISL_3411620, EPI_ISL_3411621, EPI_ISL_3411622, EPI_ISL_3411623, EPI_ISL_3411624, EPI_ISL_3411625, EPI_ISL_3411626, EPI_ISL_3411627, EPI_ISL_3411628, EPI_ISL_3411629, EPI_ISL_3411630, EPI_ISL_3411631, EPI_ISL_3411632, EPI_ISL_3411633, EPI_ISL_3411634, EPI_ISL_3411635, EPI_ISL_3411636, EPI_ISL_3411637, EPI_ISL_3411638, EPI_ISL_3411639, EPI_ISL_3411640, EPI_ISL_3411641, EPI_ISL_3411642, EPI_ISL_3411643, EPI_ISL_3411645, EPI_ISL_3411646, EPI_ISL_3411647, EPI_ISL_3411648, EPI_ISL_3411649, EPI_ISL_3411650, EPI_ISL_3411651, EPI_ISL_3411652, EPI_ISL_3411653, EPI_ISL_3411654, EPI_ISL_3411655, EPI_ISL_3411656, EPI_ISL_3411657, EPI_ISL_3411658, EPI_ISL_3411659, EPI_ISL_3411660, EPI_ISL_3411661, EPI_ISL_3411662, EPI_ISL_3411663, EPI_ISL_3411664, EPI_ISL_3411665, EPI_ISL_3411666, EPI_ISL_3411667, EPI_ISL_3411668, EPI_ISL_3411669, EPI_ISL_3411670, EPI_ISL_3411671, EPI_ISL_3411672, EPI_ISL_3411673, EPI_ISL_3411674, EPI_ISL_3411675, EPI_ISL_3411676, EPI_ISL_3411677, EPI_ISL_3411678, EPI_ISL_3411679, EPI_ISL_3411680, EPI_ISL_3411682, EPI_ISL_3411683, EPI_ISL_3411684, EPI_ISL_3411685, EPI_ISL_3411686, EPI_ISL_3411687, EPI_ISL_3411688, EPI_ISL_3411689, EPI_ISL_3411690, EPI_ISL_3411691, EPI_ISL_3411692, EPI_ISL_3411693, EPI_ISL_3411694, EPI_ISL_3411695, EPI_ISL_3411696 | | | |
| see above | PORT ELIZABETH LABORATORY | National Institute for Communicable Diseases of the National Health Laboratory Service | Amoako DG, Everatt J, Scheepers C, Mohale T, Ntuli N, Mahlangu B, Mnguni A, Ismail A, Bhiman JN |
| EPI_ISL_3411697 | MANKWENG LABOARTORY | National Institute for Communicable Diseases of the National Health Laboratory Service | Amoako DG, Everatt J, Scheepers C, Mohale T, Ntuli N, Mahlangu B, Mnguni A, Ismail A, Bhiman JN |
| EPI_ISL_3411698, EPI_ISL_3411699, EPI_ISL_3411700, EPI_ISL_3411701 | PORT ELIZABETH LABORATORY | National Institute for Communicable Diseases of the National Health Laboratory Service | Amoako DG, Everatt J, Scheepers C, Mohale T, Ntuli N, Mahlangu B, Mnguni A, Ismail A, Bhiman JN |
| EPI_ISL_3411702, EPI_ISL_3411703, EPI_ISL_3411704, EPI_ISL_3411705, EPI_ISL_3411706, EPI_ISL_3411707, EPI_ISL_3411708, EPI_ISL_3411709, EPI_ISL_3411710, EPI_ISL_3411711, EPI_ISL_3411713 | | | |
| see above | MAFIKENG LABORATORY | National Institute for Communicable Diseases of the National Health Laboratory Service | Amoako DG, Everatt J, Scheepers C, Mohale T, Ntuli N, Mahlangu B, Mnguni A, Ismail A, Bhiman JN |
| EPI_ISL_3411714 | POLOKWANE LABORATORY | National Institute for Communicable Diseases of the National Health Laboratory Service | Amoako DG, Everatt J, Scheepers C, Mohale T, Ntuli N, Mahlangu B, Mnguni A, Ismail A, Bhiman JN |
| EPI_ISL_3411715, EPI_ISL_3411716, EPI_ISL_3411718, EPI_ISL_3411720 | MANKWENG LABOARTORY | National Institute for Communicable Diseases of the National Health Laboratory Service | Amoako DG, Everatt J, Scheepers C, Mohale T, Ntuli N, Mahlangu B, Mnguni A, Ismail A, Bhiman JN |
| EPI_ISL_3411817 | National Institute for Communicable Diseases of the National Health Laboratory Service | National Institute for Communicable Diseases of the National Health Laboratory Service | Amoako DG, Everatt J, Scheepers C, Mohale T, Ntuli N, Mahlangu B, Mnguni A, Ismail A, Bhiman JN |
| EPI_ISL_3411818 | Hospital Virgen De Altagracia | Hospital General Universitario de Ciudad Real | José Martínez-Alarcón, Soledad Illescas Fernández-Bermejo, Marta Torres-Narbona, Cristina Colmenarejo, Lidia García-Agudo |
| EPI_ISL_3411819, EPI_ISL_3411820, EPI_ISL_3411821 | Hospital General Universitario de Ciudad Real | Hospital General Universitario de Ciudad Real | José Martínez-Alarcón, Soledad Illescas Fernández-Bermejo, Marta Torres-Narbona, Cristina Colmenarejo, Lidia García-Agudo |
| EPI_ISL_3411822, EPI_ISL_3411823 | Hospital General de Valdepeñas | Hospital General Universitario de Ciudad Real | José Martínez-Alarcón, Soledad Illescas Fernández-Bermejo, Marta Torres-Narbona, Cristina Colmenarejo, Lidia García-Agudo |
| EPI_ISL_3411824, EPI_ISL_3411825, EPI_ISL_3411826, EPI_ISL_3411827, EPI_ISL_3411828, EPI_ISL_3411829, EPI_ISL_3411830, EPI_ISL_3411831, EPI_ISL_3411832, EPI_ISL_3411833 | Hospital General Universitario de Ciudad Real | Hospital General Universitario de Ciudad Real | José Martínez-Alarcón, Soledad Illescas Fernández-Bermejo, Marta Torres-Narbona, Cristina Colmenarejo, Lidia García-Agudo |
| EPI_ISL_3411836 | Division of Infectious Diseases, Department of Internal Medicine, Yonsei University College of Medicine | Department of Biotechnology, Yonsei University | Minwoo Kim, Youn-Jung Lee, Jung Ho Kim, Jun Yong Choi*, and Jong-Won Oh* |
| EPI_ISL_3411920, EPI_ISL_3411925, EPI_ISL_3411930, EPI_ISL_3411934, EPI_ISL_3411935, EPI_ISL_3411936, EPI_ISL_3411937, EPI_ISL_3411939, EPI_ISL_3411940, EPI_ISL_3411941, EPI_ISL_3411942, EPI_ISL_3411943, EPI_ISL_3411946, EPI_ISL_3411947, EPI_ISL_3411949, EPI_ISL_3411950, EPI_ISL_3411951, EPI_ISL_3411952, EPI_ISL_3411953, EPI_ISL_3411954, EPI_ISL_3411955, EPI_ISL_3411957, EPI_ISL_3411958, EPI_ISL_3411960, EPI_ISL_3411961, EPI_ISL_3411962, EPI_ISL_3411963, EPI_ISL_3411964, EPI_ISL_3411965, EPI_ISL_3411966, EPI_ISL_3411967, EPI_ISL_3411968, EPI_ISL_3411969, EPI_ISL_3411970, EPI_ISL_3411971, EPI_ISL_3411972, EPI_ISL_3411973, EPI_ISL_3411974, EPI_ISL_3411976, EPI_ISL_3411977, EPI_ISL_3411978, EPI_ISL_3411979, EPI_ISL_3411981, EPI_ISL_3411982, EPI_ISL_3411984, EPI_ISL_3411985, EPI_ISL_3411986, EPI_ISL_3411987, EPI_ISL_3411988, EPI_ISL_3411989, EPI_ISL_3411990, EPI_ISL_3411991, EPI_ISL_3411992, EPI_ISL_3411993, EPI_ISL_3411994, EPI_ISL_3411995, EPI_ISL_3411996, EPI_ISL_3411997, EPI_ISL_3411998, EPI_ISL_3411999, EPI_ISL_3412000, EPI_ISL_3412001, EPI_ISL_3412002, EPI_ISL_3412003, EPI_ISL_3412004, EPI_ISL_3412005, EPI_ISL_3412006, EPI_ISL_3412007, EPI_ISL_3412008, EPI_ISL_3412009, EPI_ISL_3412010, EPI_ISL_3412012, EPI_ISL_3412013, EPI_ISL_3412014, EPI_ISL_3412015, EPI_ISL_3412016, EPI_ISL_3412017, EPI_ISL_3412018, EPI_ISL_3412019, EPI_ISL_3412020, EPI_ISL_3412021, EPI_ISL_3412022, EPI_ISL_3412023, EPI_ISL_3412024, EPI_ISL_3412025, EPI_ISL_3412026, EPI_ISL_3412027, EPI_ISL_3412028 | | | |
| see above | Hospital of the University of Pennsylvania Molecular Pathology Lab | Bushman | John K. Everett, Kyle Rodino, Andrew D. Marques, Shantan Reddy, Aoife M. Roche, Young Hwang, Scott Sherrill-Mix, Samantha A. Whiteside, Jevon Graham-Wooten, Layla A. Khatib, Ayannah S. Fitzgerald, Arupa Ganguly, Mike Feldman, Brendan Kelly, Ronald G. Collman and Frederic Bushan |
| EPI_ISL_3412047, EPI_ISL_3412048, EPI_ISL_3412049 | UW Virology Lab | UW Virology Lab | Pavitra Roychoudhury, Shah Mohamed Bakhash, Ricardo Perez, Maria Lukes, Hong Xie, Tien V. Nguyen, Sean Ellis, Meei-Li Huang, Keith R Jerome, Alexander Greninger |
| EPI_ISL_3412050, EPI_ISL_3412051 | UW Virology Lab | UW Virology Lab | Pavitra Roychoudhury, Shah Mohamed Bakhash, Ricardo Perez, Hong Xie, Tien V. Nguyen, Maria Lukes, Sean Ellis, Nathan Breit, Robert J. Livingston, Meei-Li Huang, Keith R Jerome, Patrick Mathias, Alexander Greninger |
| EPI_ISL_3412052, EPI_ISL_3412053, EPI_ISL_3412055, EPI_ISL_3412056, EPI_ISL_3412059, EPI_ISL_3412060, EPI_ISL_3412061, EPI_ISL_3412062, EPI_ISL_3412063, EPI_ISL_3412064, EPI_ISL_3412065, EPI_ISL_3412066, EPI_ISL_3412067, EPI_ISL_3412068, EPI_ISL_3412069, EPI_ISL_3412070, EPI_ISL_3412071, EPI_ISL_3412072, EPI_ISL_3412074, EPI_ISL_3412075, EPI_ISL_3412076, EPI_ISL_3412077, EPI_ISL_3412078, EPI_ISL_3412079, EPI_ISL_3412080, EPI_ISL_3412081, EPI_ISL_3412082, EPI_ISL_3412083, EPI_ISL_3412084, EPI_ISL_3412085, EPI_ISL_3412086, EPI_ISL_3412087, EPI_ISL_3412088, EPI_ISL_3412089, EPI_ISL_3412090, EPI_ISL_3412091, EPI_ISL_3412093, EPI_ISL_3412094, EPI_ISL_3412095, EPI_ISL_3412096, EPI_ISL_3412097, EPI_ISL_3412098, EPI_ISL_3412099, EPI_ISL_3412100, EPI_ISL_3412101, EPI_ISL_3412102, EPI_ISL_3412103, EPI_ISL_3412104, EPI_ISL_3412105, EPI_ISL_3412106, EPI_ISL_3412107, EPI_ISL_3412109, EPI_ISL_3412110, EPI_ISL_3412111, EPI_ISL_3412112, EPI_ISL_3412113, EPI_ISL_3412114, EPI_ISL_3412115, EPI_ISL_3412116, EPI_ISL_3412117, EPI_ISL_3412118, EPI_ISL_3412119, EPI_ISL_3412120, EPI_ISL_3412121, EPI_ISL_3412122, EPI_ISL_3412123, EPI_ISL_3412124, EPI_ISL_3412125, EPI_ISL_3412126, EPI_ISL_3412127, EPI_ISL_3412128, EPI_ISL_3412129, EPI_ISL_3412130, EPI_ISL_3412131, EPI_ISL_3412132, EPI_ISL_3412133, EPI_ISL_3412134, EPI_ISL_3412135, EPI_ISL_3412136, EPI_ISL_3412137, EPI_ISL_3412138, EPI_ISL_3412139, EPI_ISL_3412140, EPI_ISL_3412141, EPI_ISL_3412142, EPI_ISL_3412143, EPI_ISL_3412144, EPI_ISL_3412145, EPI_ISL_3412146, EPI_ISL_3412147, EPI_ISL_3412148, EPI_ISL_3412149, EPI_ISL_3412150, EPI_ISL_3412151, EPI_ISL_3412152, EPI_ISL_3412153, EPI_ISL_3412154, EPI_ISL_3412155, EPI_ISL_3412156, EPI_ISL_3412157, EPI_ISL_3412158, EPI_ISL_3412159, EPI_ISL_3412160, EPI_ISL_3412161, EPI_ISL_3412162, EPI_ISL_3412163, EPI_ISL_3412164, EPI_ISL_3412166, EPI_ISL_3412167, EPI_ISL_3412168, EPI_ISL_3412169, EPI_ISL_3412170, EPI_ISL_3412171, EPI_ISL_3412172, EPI_ISL_3412173, EPI_ISL_3412174, EPI_ISL_3412175, EPI_ISL_3412176, EPI_ISL_3412177, EPI_ISL_3412178, EPI_ISL_3412179, EPI_ISL_3412182, EPI_ISL_3412183, EPI_ISL_3412184, EPI_ISL_3412185, EPI_ISL_3412186, EPI_ISL_3412187, EPI_ISL_3412188, EPI_ISL_3412189, EPI_ISL_3412190, EPI_ISL_3412191, EPI_ISL_3412192, EPI_ISL_3412193, EPI_ISL_3412194, EPI_ISL_3412196, EPI_ISL_3412197, EPI_ISL_3412198, EPI_ISL_3412199, EPI_ISL_3412200, EPI_ISL_3412201, EPI_ISL_3412202, EPI_ISL_3412203, EPI_ISL_3412204, EPI_ISL_3412205, EPI_ISL_3412206, EPI_ISL_3412207, EPI_ISL_3412208, EPI_ISL_3412209, EPI_ISL_3412210, EPI_ISL_3412211, EPI_ISL_3412212, EPI_ISL_3412213, EPI_ISL_3412214, EPI_ISL_3412215, EPI_ISL_3412216, EPI_ISL_3412217, EPI_ISL_3412218, EPI_ISL_3412219, EPI_ISL_3412220, EPI_ISL_3412221, EPI_ISL_3412222, EPI_ISL_3412223, EPI_ISL_3412224, EPI_ISL_3412225, EPI_ISL_3412226, EPI_ISL_3412227, EPI_ISL_3412228, EPI_ISL_3412229, EPI_ISL_3412230, EPI_ISL_3412231, EPI_ISL_3412232, EPI_ISL_3412233, EPI_ISL_3412234, EPI_ISL_3412235, EPI_ISL_3412236, EPI_ISL_3412237, EPI_ISL_3412238, EPI_ISL_3412239, EPI_ISL_3412240, EPI_ISL_3412241, EPI_ISL_3412242, EPI_ISL_3412243, EPI_ISL_3412249, EPI_ISL_3412250, EPI_ISL_3412251, EPI_ISL_3412252, EPI_ISL_3412253, EPI_ISL_3412254, EPI_ISL_3412255, EPI_ISL_3412256, EPI_ISL_3412257, EPI_ISL_3412258, EPI_ISL_3412259, EPI_ISL_3412260, EPI_ISL_3412261, EPI_ISL_3412262, EPI_ISL_3412263, EPI_ISL_3412264, EPI_ISL_3412265, EPI_ISL_3412266, EPI_ISL_3412267, EPI_ISL_3412268, EPI_ISL_3412269, EPI_ISL_3412270, EPI_ISL_3412271, EPI_ISL_3412272, EPI_ISL_3412273, EPI_ISL_3412274, EPI_ISL_3412275, EPI_ISL_3412276, EPI_ISL_3412277, EPI_ISL_3412278, EPI_ISL_3412279, EPI_ISL_3412280, EPI_ISL_3412281, EPI_ISL_3412282, EPI_ISL_3412283, EPI_ISL_3412284, EPI_ISL_3412285, EPI_ISL_3412286, EPI_ISL_3412287, EPI_ISL_3412288, EPI_ISL_3412289, EPI_ISL_3412290, EPI_ISL_3412291, EPI_ISL_3412292, EPI_ISL_3412293, EPI_ISL_3412294, EPI_ISL_3412295, EPI_ISL_3412296, EPI_ISL_3412297, EPI_ISL_3412298, EPI_ISL_3412299, EPI_ISL_3412300, EPI_ISL_3412301, EPI_ISL_3412302, EPI_ISL_3412303, EPI_ISL_3412304, EPI_ISL_3412305, EPI_ISL_3412306, EPI_ISL_3412307, EPI_ISL_3412308, EPI_ISL_3412309, EPI_ISL_3412310, EPI_ISL_3412311, EPI_ISL_3412312, EPI_ISL_3412313, EPI_ISL_3412314, EPI_ISL_3412315, EPI_ISL_3412316, EPI_ISL_3412317, EPI_ISL_3412318, EPI_ISL_3412319, EPI_ISL_3412320, EPI_ISL_3412321, EPI_ISL_3412322, EPI_ISL_3412323, EPI_ISL_3412324, EPI_ISL_3412325, EPI_ISL_3412326, EPI_ISL_3412327, EPI_ISL_3412328, EPI_ISL_3412329, EPI_ISL_3412330, EPI_ISL_3412331, EPI_ISL_3412332, EPI_ISL_3412333, EPI_ISL_3412334, EPI_ISL_3412335, EPI_ISL_3412336, EPI_ISL_3412337, EPI_ISL_3412338, EPI_ISL_3412339, EPI_ISL_3412340, EPI_ISL_3412341, EPI_ISL_3412342, EPI_ISL_3412343, EPI_ISL_3412344, EPI_ISL_3412345, EPI_ISL_3412346, EPI_ISL_3412347, EPI_ISL_3412348, EPI_ISL_3412349, EPI_ISL_3412350, EPI_ISL_3412351, EPI_ISL_3412352, EPI_ISL_3412353, EPI_ISL_3412354, EPI_ISL_3412355, EPI_ISL_3412356, EPI_ISL_3412357, EPI_ISL_3412358, EPI_ISL_3412359, EPI_ISL_3412360, EPI_ISL_3412361, EPI_ISL_3412362, EPI_ISL_3412363, EPI_ISL_3412364, EPI_ISL_3412365, EPI_ISL_3412366, EPI_ISL_3412367, EPI_ISL_3412368, EPI_ISL_3412369, EPI_ISL_3412370, EPI_ISL_3412371, EPI_ISL_3412372, EPI_ISL_3412373, EPI_ISL_3412374, EPI_ISL_3412375, EPI_ISL_3412376, EPI_ISL_3412377, EPI_ISL_3412378, EPI_ISL_3412379, EPI_ISL_3412380, EPI_ISL_3412381, EPI_ISL_3412382, EPI_ISL_3412383, EPI_ISL_3412384, EPI_ISL_3412385, EPI_ISL_3412386, EPI_ISL_3412387, EPI_ISL_3412388, EPI_ISL_3412389, EPI_ISL_3412390, EPI_ISL_3412391, EPI_ISL_3412392, EPI_ISL_3412393, EPI_ISL_3412394, EPI_ISL_3412395, EPI_ISL_3412396, EPI_ISL_3412397, EPI_ISL_3412398, EPI_ISL_3412399, EPI_ISL_3412400, EPI_ISL_3412401, EPI_ISL_3412402, EPI_ISL_3412403, EPI_ISL_3412404, EPI_ISL_3412405, EPI_ISL_3412406, EPI_ISL_3412407, EPI_ISL_3412408, EPI_ISL_3412409 | | | |
| see above | UW Virology Lab | UW Virology Lab | Pavitra Roychoudhury, Shah Mohamed Bakhash, Ricardo Perez, Maria Lukes, Hong Xie, Tien V. Nguyen, Sean Ellis, Meei-Li Huang, Keith R Jerome, Alexander Greninger |
| EPI_ISL_3412464, EPI_ISL_3412488, EPI_ISL_3412497, EPI_ISL_3412521, EPI_ISL_3412971, EPI_ISL_3413721, EPI_ISL_3414339, EPI_ISL_3414398, EPI_ISL_3414613, EPI_ISL_3414672, EPI_ISL_3414685, EPI_ISL_3414720, EPI_ISL_3414805, EPI_ISL_3414807, EPI_ISL_3414842, EPI_ISL_3414927, EPI_ISL_3415022, EPI_ISL_3415057, EPI_ISL_3415142, EPI_ISL_3415144, EPI_ISL_3415179, EPI_ISL_3415264, EPI_ISL_3415457, EPI_ISL_3415481, EPI_ISL_3415490, EPI_ISL_3415514, EPI_ISL_3415929, EPI_ISL_3417364, EPI_ISL_3417365, EPI_ISL_3417366, EPI_ISL_3417367, EPI_ISL_3417368, EPI_ISL_3417369, EPI_ISL_3417370, EPI_ISL_3417371, EPI_ISL_3417372, EPI_ISL_3417373, EPI_ISL_3417374 | | | |
| see above | Ministry of Health Turkey | Ministry of Health Turkey | Fatma Bayrakdar, Yasemin Cosgun, Suleyman Yalcin, Gulay Korukluoglu |
| EPI_ISL_3417410, EPI_ISL_3417411, EPI_ISL_3417412, EPI_ISL_3417413, EPI_ISL_3417414, EPI_ISL_3417415, EPI_ISL_3417416, EPI_ISL_3417417, EPI_ISL_3417418, EPI_ISL_3417419, EPI_ISL_3417420, EPI_ISL_3417421, EPI_ISL_3417422, EPI_ISL_3417423, EPI_ISL_3417424, EPI_ISL_3417425, EPI_ISL_3417426, EPI_ISL_3417427, EPI_ISL_3417429, EPI_ISL_3417430, EPI_ISL_3417431, EPI_ISL_3417432, EPI_ISL_3417433, EPI_ISL_3417434, EPI_ISL_3417435, EPI_ISL_3417436, EPI_ISL_3417437, EPI_ISL_3417438, EPI_ISL_3417439, EPI_ISL_3417440, EPI_ISL_3417441, EPI_ISL_3417442, EPI_ISL_3417443, EPI_ISL_3417444, EPI_ISL_3417445, EPI_ISL_3417446, EPI_ISL_3417447, EPI_ISL_3417448, EPI_ISL_3417449, EPI_ISL_3417450, EPI_ISL_3417451, EPI_ISL_3417452, EPI_ISL_3417453, EPI_ISL_3417454, EPI_ISL_3417455, EPI_ISL_3417456, EPI_ISL_3417457, EPI_ISL_3417458, EPI_ISL_3417459, EPI_ISL_3417460, EPI_ISL_3417461, EPI_ISL_3417462, EPI_ISL_3417463, EPI_ISL_3417464, EPI_ISL_3417465, EPI_ISL_3417466, EPI_ISL_3417467, EPI_ISL_3417468, EPI_ISL_3417469, EPI_ISL_3417470, EPI_ISL_3417471, EPI_ISL_3417472, EPI_ISL_3417473, EPI_ISL_3417474, EPI_ISL_3417475, EPI_ISL_3417476, EPI_ISL_3417477, EPI_ISL_3417478, EPI_ISL_3417479, EPI_ISL_3417480, EPI_ISL_3417481, EPI_ISL_3417482, EPI_ISL_3417483, EPI_ISL_3417484, EPI_ISL_3417485, EPI_ISL_3417486, EPI_ISL_3417487, EPI_ISL_3417488 | | | |
| see above | New South Wales Health Pathology Royal Prince Alfred Hospital | Microbiology RPAH | Foster, C.; Au, J.; Ruiz Silva, M.; Deveson, I.; Bull, R.; Van Hal, S.; Rawlinson, W. |
| EPI_ISL_3417490, EPI_ISL_3417491, EPI_ISL_3417492, EPI_ISL_3417494, EPI_ISL_3417495, EPI_ISL_3417496, EPI_ISL_3417497, EPI_ISL_3417498, EPI_ISL_3417499, EPI_ISL_3417500, EPI_ISL_3417501 | | | |
| see above | Pathology, UAB | Pathology, UAB | Moates,D., Hendrickson,R.C., Lefkowitz,E.J., Leal,S.M. |
| EPI_ISL_3417503, EPI_ISL_3417504, EPI_ISL_3417505, EPI_ISL_3417506, EPI_ISL_3417513, EPI_ISL_3417514, EPI_ISL_3417515, EPI_ISL_3417516, EPI_ISL_3417517, EPI_ISL_3417518, EPI_ISL_3417519 | | | |
| see above | Kawakita General Hospital | Tokyo Medical University Department of Microbiology | Tatsuya Inukai, Takahiro Okai, Kenshirou Kroki, Maya Isogai, Hirofumi Miyake, Takushi Fujiwara, Mami Nagashima, Jun Suzuki, Kenji Sadamasu, Shigeki Nakamura |
| EPI_ISL_3417525, EPI_ISL_3417526, EPI_ISL_3417527, EPI_ISL_3417528, EPI_ISL_3417529, EPI_ISL_3417530, EPI_ISL_3417531, EPI_ISL_3417532 | unknown | PHV-FSS | Chenwei Wang on behalf of Q-PHIRE Genomics |

| | | | |
|---|---|---|---|
| EPI_ISL_3417533 | Queensland Health Forensic and Scientific Services | Queensland Health Forensic and Scientific Services | Chenwei Wang on behalf of Q-PHIRE Genomics |
| EPI_ISL_3417534, EPI_ISL_3417535, EPI_ISL_3417536, EPI_ISL_3417537 | Department of Bacteria, Parasites and Fungi, Statens Serum Institut, Copenhagen, Denmark | Statens Serum Institut Bioinformatics and Microbial Genomics | Danish Covid-19 Genome Consortium |
| EPI_ISL_3417538 | Basurto University Hospital: Clinical Microbiology Laboratory | Biocruces Bizkaia | Mikel Urrutikoetxea-Gutiérrez, Mª Carmen Nieto Toboso, Estibaliz Ugalde Zarraga, Mikel Gallego Rodrigo, Ana de la Hoz, José Luis Díaz de Tuesta del Arco |
| EPI_ISL_3417539 | Department of Bacteria, Parasites and Fungi, Statens Serum Institut, Copenhagen, Denmark | Statens Serum Institut Bioinformatics and Microbial Genomics | Danish Covid-19 Genome Consortium |
| EPI_ISL_3417540 | Molekylær Medicinsk Afdeling, Aarhus University Hospital, Aarhus, Denmark | Statens Serum Institut Bioinformatics and Microbial Genomics | Danish Covid-19 Genome Consortium |
| EPI_ISL_3417541, EPI_ISL_3417542, EPI_ISL_3417543, EPI_ISL_3417544, EPI_ISL_3417545, EPI_ISL_3417546, EPI_ISL_3417547, EPI_ISL_3417548, EPI_ISL_3417549, EPI_ISL_3417550, EPI_ISL_3417551, EPI_ISL_3417552, EPI_ISL_3417553, EPI_ISL_3417554, EPI_ISL_3417555, EPI_ISL_3417556, EPI_ISL_3417557, EPI_ISL_3417558, EPI_ISL_3417559, EPI_ISL_3417560, EPI_ISL_3417561, EPI_ISL_3417562 | | | |
| see above | Department of Bacteria, Parasites and Fungi, Statens Serum Institut, Copenhagen, Denmark | Statens Serum Institut Bioinformatics and Microbial Genomics | Danish Covid-19 Genome Consortium |
| EPI_ISL_3417563 | Molekylær Medicinsk Afdeling, Aarhus University Hospital, Aarhus, Denmark | Statens Serum Institut Bioinformatics and Microbial Genomics | Danish Covid-19 Genome Consortium |
| EPI_ISL_3417564, EPI_ISL_3417565, EPI_ISL_3417566, EPI_ISL_3417567, EPI_ISL_3417568, EPI_ISL_3417569 | Department of Bacteria, Parasites and Fungi, Statens Serum Institut, Copenhagen, Denmark | Statens Serum Institut Bioinformatics and Microbial Genomics | Danish Covid-19 Genome Consortium |
| EPI_ISL_3417570 | Molekylær Medicinsk Afdeling, Aarhus University Hospital, Aarhus, Denmark | Statens Serum Institut Bioinformatics and Microbial Genomics | Danish Covid-19 Genome Consortium |
| EPI_ISL_3417571, EPI_ISL_3417572, EPI_ISL_3417573, EPI_ISL_3417574, EPI_ISL_3417575, EPI_ISL_3417576, EPI_ISL_3417577, EPI_ISL_3417578, EPI_ISL_3417579, EPI_ISL_3417580, EPI_ISL_3417581, EPI_ISL_3417582, EPI_ISL_3417583, EPI_ISL_3417584, EPI_ISL_3417585, EPI_ISL_3417586, EPI_ISL_3417587 | | | |
| see above | Department of Bacteria, Parasites and Fungi, Statens Serum Institut, Copenhagen, Denmark | Statens Serum Institut Bioinformatics and Microbial Genomics | Danish Covid-19 Genome Consortium |
| EPI_ISL_3417588 | Molekylær Medicinsk Afdeling, Aarhus University Hospital, Aarhus, Denmark | Statens Serum Institut Bioinformatics and Microbial Genomics | Danish Covid-19 Genome Consortium |
| EPI_ISL_3417589, EPI_ISL_3417590, EPI_ISL_3417591, EPI_ISL_3417592, EPI_ISL_3417593, EPI_ISL_3417594, EPI_ISL_3417595 | Department of Bacteria, Parasites and Fungi, Statens Serum Institut, Copenhagen, Denmark | Statens Serum Institut Bioinformatics and Microbial Genomics | Danish Covid-19 Genome Consortium |
| EPI_ISL_3417596 | Molekylær Medicinsk Afdeling, Aarhus University Hospital, Aarhus, Denmark | Statens Serum Institut Bioinformatics and Microbial Genomics | Danish Covid-19 Genome Consortium |
| EPI_ISL_3417597, EPI_ISL_3417598, EPI_ISL_3417599, EPI_ISL_3417600, EPI_ISL_3417601, EPI_ISL_3417602, EPI_ISL_3417603, EPI_ISL_3417604, EPI_ISL_3417605, EPI_ISL_3417606, EPI_ISL_3417607, EPI_ISL_3417608, EPI_ISL_3417609, EPI_ISL_3417610 | | | |
| see above | Department of Bacteria, Parasites and Fungi, Statens Serum Institut, Copenhagen, Denmark | Statens Serum Institut Bioinformatics and Microbial Genomics | Danish Covid-19 Genome Consortium |
| EPI_ISL_3417611 | Molekylær Medicinsk Afdeling, Aarhus University Hospital, Aarhus, Denmark | Statens Serum Institut Bioinformatics and Microbial Genomics | Danish Covid-19 Genome Consortium |
| EPI_ISL_3417612, EPI_ISL_3417613, EPI_ISL_3417614, EPI_ISL_3417615, EPI_ISL_3417616, EPI_ISL_3417617, EPI_ISL_3417618, EPI_ISL_3417619, EPI_ISL_3417621, EPI_ISL_3417622, EPI_ISL_3417623, EPI_ISL_3417624, EPI_ISL_3417625, EPI_ISL_3417626, EPI_ISL_3417627, EPI_ISL_3417628, EPI_ISL_3417629, EPI_ISL_3417630, EPI_ISL_3417631, EPI_ISL_3417632, EPI_ISL_3417634, EPI_ISL_3417635, EPI_ISL_3417636, EPI_ISL_3417637, EPI_ISL_3417638, EPI_ISL_3417639, EPI_ISL_3417640, EPI_ISL_3417641, EPI_ISL_3417642, EPI_ISL_3417643, EPI_ISL_3417644, EPI_ISL_3417645, EPI_ISL_3417646, EPI_ISL_3417647, EPI_ISL_3417648, EPI_ISL_3417649, EPI_ISL_3417650 | | | |
| see above | Department of Bacteria, Parasites and Fungi, Statens Serum Institut, Copenhagen, Denmark | Statens Serum Institut Bioinformatics and Microbial Genomics | Danish Covid-19 Genome Consortium |
| EPI_ISL_3417651 | Molekylær Medicinsk Afdeling, Aarhus University Hospital, Aarhus, Denmark | Statens Serum Institut Bioinformatics and Microbial Genomics | Danish Covid-19 Genome Consortium |
| EPI_ISL_3417652, EPI_ISL_3417653, EPI_ISL_3417654, EPI_ISL_3417655, EPI_ISL_3417656, EPI_ISL_3417657, EPI_ISL_3417658, EPI_ISL_3417659, EPI_ISL_3417660, EPI_ISL_3417661, EPI_ISL_3417662, EPI_ISL_3417663, EPI_ISL_3417664, EPI_ISL_3417665, EPI_ISL_3417666, EPI_ISL_3417667, EPI_ISL_3417668, EPI_ISL_3417669, EPI_ISL_3417670, EPI_ISL_3417671, EPI_ISL_3417672, EPI_ISL_3417673, EPI_ISL_3417674, EPI_ISL_3417675, EPI_ISL_3417676, EPI_ISL_3417677, EPI_ISL_3417678, EPI_ISL_3417679, EPI_ISL_3417680, EPI_ISL_3417681, EPI_ISL_3417682, EPI_ISL_3417683, EPI_ISL_3417684, EPI_ISL_3417685, EPI_ISL_3417686 | | | |
| see above | Department of Bacteria, Parasites and Fungi, Statens Serum Institut, Copenhagen, Denmark | Statens Serum Institut Bioinformatics and Microbial Genomics | Danish Covid-19 Genome Consortium |
| EPI_ISL_3417687 | Molekylær Medicinsk Afdeling, Aarhus University Hospital, Aarhus, Denmark | Statens Serum Institut Bioinformatics and Microbial Genomics | Danish Covid-19 Genome Consortium |
| EPI_ISL_3417688, EPI_ISL_3417689, EPI_ISL_3417690, EPI_ISL_3417691, EPI_ISL_3417692, EPI_ISL_3417693, EPI_ISL_3417695, EPI_ISL_3417696, EPI_ISL_3417697, EPI_ISL_3417698, EPI_ISL_3417699, EPI_ISL_3417700, EPI_ISL_3417701, EPI_ISL_3417702, EPI_ISL_3417703, EPI_ISL_3417704, EPI_ISL_3417705, EPI_ISL_3417706, EPI_ISL_3417707, EPI_ISL_3417708, EPI_ISL_3417709, EPI_ISL_3417710, EPI_ISL_3417711, EPI_ISL_3417712, EPI_ISL_3417713, EPI_ISL_3417714, EPI_ISL_3417715, EPI_ISL_3417716, EPI_ISL_3417717, EPI_ISL_3417718, EPI_ISL_3417719, EPI_ISL_3417720, EPI_ISL_3417721, EPI_ISL_3417722, EPI_ISL_3417723, EPI_ISL_3417724, EPI_ISL_3417725, EPI_ISL_3417726, EPI_ISL_3417727, EPI_ISL_3417728, EPI_ISL_3417729, EPI_ISL_3417730, EPI_ISL_3417731, EPI_ISL_3417732, EPI_ISL_3417733, EPI_ISL_3417734, EPI_ISL_3417735, EPI_ISL_3417736, EPI_ISL_3417737, EPI_ISL_3417738, EPI_ISL_3417739, EPI_ISL_3417740, EPI_ISL_3417741, EPI_ISL_3417742, EPI_ISL_3417743, EPI_ISL_3417744, EPI_ISL_3417745, EPI_ISL_3417746, EPI_ISL_3417747, EPI_ISL_3417748, EPI_ISL_3417749, EPI_ISL_3417750, EPI_ISL_3417751, EPI_ISL_3417752 | | | |
| see above | Department of Bacteria, Parasites and Fungi, Statens Serum Institut, Copenhagen, Denmark | Statens Serum Institut Bioinformatics and Microbial Genomics | Danish Covid-19 Genome Consortium |
| EPI_ISL_3417753 | Molekylær Medicinsk Afdeling, Aarhus University Hospital, Aarhus, Denmark | Statens Serum Institut Bioinformatics and Microbial Genomics | Danish Covid-19 Genome Consortium |
| EPI_ISL_3417754, EPI_ISL_3417755, EPI_ISL_3417756, EPI_ISL_3417757, EPI_ISL_3417758, EPI_ISL_3417759, EPI_ISL_3417760, EPI_ISL_3417761, EPI_ISL_3417762, EPI_ISL_3417763, EPI_ISL_3417764, EPI_ISL_3417765, EPI_ISL_3417766, EPI_ISL_3417767, EPI_ISL_3417768, EPI_ISL_3417769, EPI_ISL_3417770, EPI_ISL_3417771, EPI_ISL_3417772, EPI_ISL_3417773, EPI_ISL_3417774, EPI_ISL_3417775, EPI_ISL_3417776 | | | |
| see above | Department of Bacteria, Parasites and Fungi, Statens Serum Institut, Copenhagen, Denmark | Statens Serum Institut Bioinformatics and Microbial Genomics | Danish Covid-19 Genome Consortium |
| EPI_ISL_3417777 | Molekylær Medicinsk Afdeling, Aarhus University Hospital, Aarhus, Denmark | Statens Serum Institut Bioinformatics and Microbial Genomics | Danish Covid-19 Genome Consortium |
| EPI_ISL_3417778, EPI_ISL_3417779, EPI_ISL_3417780, EPI_ISL_3417781, EPI_ISL_3417782, EPI_ISL_3417783, EPI_ISL_3417784, EPI_ISL_3417785, EPI_ISL_3417786, EPI_ISL_3417787, EPI_ISL_3417788, EPI_ISL_3417789, EPI_ISL_3417790, EPI_ISL_3417791, EPI_ISL_3417792, EPI_ISL_3417793, EPI_ISL_3417794, EPI_ISL_3417795, EPI_ISL_3417796, EPI_ISL_3417797, EPI_ISL_3417798, EPI_ISL_3417799, EPI_ISL_3417800, EPI_ISL_3417801, EPI_ISL_3417802, EPI_ISL_3417803, EPI_ISL_3417804, EPI_ISL_3417805, EPI_ISL_3417806, EPI_ISL_3417807, EPI_ISL_3417808, EPI_ISL_3417809, EPI_ISL_3417810, EPI_ISL_3417811, EPI_ISL_3417812, EPI_ISL_3417813, EPI_ISL_3417814, EPI_ISL_3417815, EPI_ISL_3417816, EPI_ISL_3417817, EPI_ISL_3417818, EPI_ISL_3417819, EPI_ISL_3417820, EPI_ISL_3417821, EPI_ISL_3417822, EPI_ISL_3417823, EPI_ISL_3417824, EPI_ISL_3417825, EPI_ISL_3417826, EPI_ISL_3417827, EPI_ISL_3417828, EPI_ISL_3417829, EPI_ISL_3417830, EPI_ISL_3417831, EPI_ISL_3417832, EPI_ISL_3417833, EPI_ISL_3417834, EPI_ISL_3417835, EPI_ISL_3417836, EPI_ISL_3417837, EPI_ISL_3417838, EPI_ISL_3417839, EPI_ISL_3417840, EPI_ISL_3417841, EPI_ISL_3417842, EPI_ISL_3417843, EPI_ISL_3417844, EPI_ISL_3417845, EPI_ISL_3417846, EPI_ISL_3417847, EPI_ISL_3417848, EPI_ISL_3417849, EPI_ISL_3417850, EPI_ISL_3417851, EPI_ISL_3417852, EPI_ISL_3417853, EPI_ISL_3417854, EPI_ISL_3417855 | | | |
| see above | Department of Bacteria, Parasites and Fungi, Statens Serum Institut, Copenhagen, Denmark | Statens Serum Institut Bioinformatics and Microbial Genomics | Danish Covid-19 Genome Consortium |
| EPI_ISL_3417856 | Basurto University Hospital: Clinical Microbiology Laboratory | Biocruces Bizkaia | Mikel Urrutikoetxea-Gutiérrez, Mª Carmen Nieto Toboso, Estibaliz Ugalde Zarraga, Mikel Gallego Rodrigo, Ana de la Hoz, José Luis Díaz de Tuesta del Arco |
| EPI_ISL_3417857, EPI_ISL_3417858, EPI_ISL_3417859, EPI_ISL_3417860, EPI_ISL_3417861, EPI_ISL_3417862, EPI_ISL_3417863, EPI_ISL_3417864, EPI_ISL_3417865, EPI_ISL_3417866, EPI_ISL_3417867, EPI_ISL_3417868, EPI_ISL_3417869, EPI_ISL_3417870, EPI_ISL_3417871, EPI_ISL_3417872, EPI_ISL_3417873, EPI_ISL_3417874, EPI_ISL_3417875, EPI_ISL_3417876, EPI_ISL_3417877, EPI_ISL_3417878, EPI_ISL_3417879, EPI_ISL_3417880, EPI_ISL_3417881, EPI_ISL_3417882, EPI_ISL_3417883, EPI_ISL_3417884, EPI_ISL_3417885, EPI_ISL_3417886, EPI_ISL_3417887, EPI_ISL_3417888, EPI_ISL_3417889, EPI_ISL_3417890, EPI_ISL_3417891, EPI_ISL_3417892, EPI_ISL_3417893, EPI_ISL_3417894, EPI_ISL_3417895, EPI_ISL_3417896, EPI_ISL_3417897, EPI_ISL_3417898, EPI_ISL_3417899, EPI_ISL_3417900, EPI_ISL_3417901, EPI_ISL_3417902, EPI_ISL_3417903, EPI_ISL_3417904, EPI_ISL_3417905, EPI_ISL_3417906, EPI_ISL_3417907, EPI_ISL_3417908, EPI_ISL_3417909, EPI_ISL_3417910, EPI_ISL_3417911, EPI_ISL_3417912, EPI_ISL_3417913, EPI_ISL_3417914, EPI_ISL_3417915, EPI_ISL_3417916, EPI_ISL_3417917, EPI_ISL_3417918, EPI_ISL_3417919, EPI_ISL_3417920, EPI_ISL_3417921, EPI_ISL_3417922, EPI_ISL_3417923, EPI_ISL_3417924, EPI_ISL_3417925, EPI_ISL_3417926, EPI_ISL_3417927, EPI_ISL_3417928, EPI_ISL_3417929, EPI_ISL_3417930, EPI_ISL_3417931, EPI_ISL_3417932, EPI_ISL_3417933, EPI_ISL_3417934, EPI_ISL_3417935, EPI_ISL_3417936, EPI_ISL_3417937, EPI_ISL_3417938, EPI_ISL_3417939, EPI_ISL_3417940, EPI_ISL_3417941, EPI_ISL_3417942, EPI_ISL_3417943, EPI_ISL_3417944, EPI_ISL_3417945, EPI_ISL_3417946, EPI_ISL_3417947, EPI_ISL_3417948, EPI_ISL_3417949, EPI_ISL_3417950, EPI_ISL_3417951, EPI_ISL_3417952, EPI_ISL_3417953, EPI_ISL_3417954, EPI_ISL_3417955, EPI_ISL_3417956, EPI_ISL_3417957, EPI_ISL_3417958, EPI_ISL_3417959, EPI_ISL_3417960, EPI_ISL_3417961, EPI_ISL_3417962, EPI_ISL_3417963, EPI_ISL_3417964, EPI_ISL_3417965, EPI_ISL_3417966, EPI_ISL_3417967, EPI_ISL_3417968, EPI_ISL_3417970, EPI_ISL_3417971, EPI_ISL_3417973, EPI_ISL_3417974, EPI_ISL_3417975, EPI_ISL_3417976, EPI_ISL_3417977, EPI_ISL_3417978, EPI_ISL_3417979, EPI_ISL_3417980, EPI_ISL_3417981, EPI_ISL_3417982, EPI_ISL_3417983, EPI_ISL_3417984, EPI_ISL_3417985, EPI_ISL_3417986, EPI_ISL_3417987, EPI_ISL_3417988, EPI_ISL_3417989, EPI_ISL_3417990, EPI_ISL_3417991, EPI_ISL_3417992, EPI_ISL_3417993 | | | |
| see above | Department of Bacteria, Parasites and Fungi, Statens Serum Institut, Copenhagen, Denmark | Statens Serum Institut Bioinformatics and Microbial Genomics | Danish Covid-19 Genome Consortium |
| EPI_ISL_3417994 | Molekylær Medicinsk Afdeling, Aarhus University Hospital, Aarhus, Denmark | Statens Serum Institut Bioinformatics and Microbial Genomics | Danish Covid-19 Genome Consortium |
| EPI_ISL_3417995, EPI_ISL_3417996, EPI_ISL_3417997, EPI_ISL_3417998, EPI_ISL_3417999, EPI_ISL_3418000, EPI_ISL_3418001, EPI_ISL_3418002, EPI_ISL_3418003, EPI_ISL_3418004, EPI_ISL_3418005, EPI_ISL_3418006, EPI_ISL_3418007, EPI_ISL_3418008, EPI_ISL_3418009, EPI_ISL_3418010, EPI_ISL_3418011, EPI_ISL_3418012, EPI_ISL_3418013, EPI_ISL_3418014, EPI_ISL_3418015, EPI_ISL_3418016, EPI_ISL_3418017, EPI_ISL_3418018, EPI_ISL_3418019, EPI_ISL_3418020, EPI_ISL_3418021, EPI_ISL_3418022, EPI_ISL_3418023, EPI_ISL_3418024, EPI_ISL_3418025, EPI_ISL_3418026, EPI_ISL_3418027, EPI_ISL_3418028, EPI_ISL_3418029, EPI_ISL_3418030, EPI_ISL_3418031, EPI_ISL_3418032, EPI_ISL_3418033, EPI_ISL_3418034, EPI_ISL_3418035, EPI_ISL_3418036, EPI_ISL_3418037 | | | |
| see above | Department of Bacteria, Parasites and Fungi, Statens Serum Institut, Copenhagen, Denmark | Statens Serum Institut Bioinformatics and Microbial Genomics | Danish Covid-19 Genome Consortium |
| EPI_ISL_3418038 | Molekylær Medicinsk Afdeling, Aarhus University Hospital, Aarhus, Denmark | Statens Serum Institut Bioinformatics and Microbial Genomics | Danish Covid-19 Genome Consortium |
| EPI_ISL_3418039, EPI_ISL_3418040, | Department of Bacteria, Parasites and Fungi, Statens Serum Institut, Copenhagen, Denmark | Statens Serum Institut Bioinformatics and Microbial Genomics | Danish Covid-19 Genome Consortium |

| | | | |
|---|---|---|---|
| EPI_ISL_3418041, EPI_ISL_3418042, EPI_ISL_3418043, EPI_ISL_3418044, EPI_ISL_3418045, EPI_ISL_3418046 | | | |
| EPI_ISL_3418047 | Molekylær Medicinsk Afdeling, Aarhus University Hospital, Aarhus, Denmark | Statens Serum Institut Bioinformatics and Microbial Genomics | Danish Covid-19 Genome Consortium |
| EPI_ISL_3418048, EPI_ISL_3418049 | Department of Bacteria, Parasites and Fungi, Statens Serum Institut, Copenhagen, Denmark | Statens Serum Institut Bioinformatics and Microbial Genomics | Danish Covid-19 Genome Consortium |
| EPI_ISL_3418050 | Molekylær Medicinsk Afdeling, Aarhus University Hospital, Aarhus, Denmark | Statens Serum Institut Bioinformatics and Microbial Genomics | Danish Covid-19 Genome Consortium |
| EPI_ISL_3418051, EPI_ISL_3418052, EPI_ISL_3418053 | Department of Bacteria, Parasites and Fungi, Statens Serum Institut, Copenhagen, Denmark | Statens Serum Institut Bioinformatics and Microbial Genomics | Danish Covid-19 Genome Consortium |
| EPI_ISL_3418054 | Molekylær Medicinsk Afdeling, Aarhus University Hospital, Aarhus, Denmark | Statens Serum Institut Bioinformatics and Microbial Genomics | Danish Covid-19 Genome Consortium |
| EPI_ISL_3418055, EPI_ISL_3418056, EPI_ISL_3418057, EPI_ISL_3418058, EPI_ISL_3418059, EPI_ISL_3418060, EPI_ISL_3418061, EPI_ISL_3418062, EPI_ISL_3418063, EPI_ISL_3418064, EPI_ISL_3418065, EPI_ISL_3418066, EPI_ISL_3418067, EPI_ISL_3418068, EPI_ISL_3418069, EPI_ISL_3418070, EPI_ISL_3418071, EPI_ISL_3418072, EPI_ISL_3418073, EPI_ISL_3418074, EPI_ISL_3418075, EPI_ISL_3418076, EPI_ISL_3418077, EPI_ISL_3418078 | | | |
| see above | Department of Bacteria, Parasites and Fungi, Statens Serum Institut, Copenhagen, Denmark | Statens Serum Institut Bioinformatics and Microbial Genomics | Danish Covid-19 Genome Consortium |
| EPI_ISL_3418079 | Molekylær Medicinsk Afdeling, Aarhus University Hospital, Aarhus, Denmark | Statens Serum Institut Bioinformatics and Microbial Genomics | Danish Covid-19 Genome Consortium |
| EPI_ISL_3418080, EPI_ISL_3418081, EPI_ISL_3418082, EPI_ISL_3418083, EPI_ISL_3418084, EPI_ISL_3418085, EPI_ISL_3418086, EPI_ISL_3418087, EPI_ISL_3418089, EPI_ISL_3418090, EPI_ISL_3418091, EPI_ISL_3418092, EPI_ISL_3418093, EPI_ISL_3418094, EPI_ISL_3418095, EPI_ISL_3418096, EPI_ISL_3418097, EPI_ISL_3418098, EPI_ISL_3418099, EPI_ISL_3418100, EPI_ISL_3418101, EPI_ISL_3418102, EPI_ISL_3418103, EPI_ISL_3418104, EPI_ISL_3418105, EPI_ISL_3418106, EPI_ISL_3418107, EPI_ISL_3418108, EPI_ISL_3418109, EPI_ISL_3418110, EPI_ISL_3418111, EPI_ISL_3418112, EPI_ISL_3418113, EPI_ISL_3418114, EPI_ISL_3418115, EPI_ISL_3418116, EPI_ISL_3418117, EPI_ISL_3418118, EPI_ISL_3418119, EPI_ISL_3418120, EPI_ISL_3418121, EPI_ISL_3418122, EPI_ISL_3418123, EPI_ISL_3418124, EPI_ISL_3418125, EPI_ISL_3418126, EPI_ISL_3418127, EPI_ISL_3418128 | | | |
| see above | Department of Bacteria, Parasites and Fungi, Statens Serum Institut, Copenhagen, Denmark | Statens Serum Institut Bioinformatics and Microbial Genomics | Danish Covid-19 Genome Consortium |
| EPI_ISL_3418129 | Molekylær Medicinsk Afdeling, Aarhus University Hospital, Aarhus, Denmark | Statens Serum Institut Bioinformatics and Microbial Genomics | Danish Covid-19 Genome Consortium |
| EPI_ISL_3418130, EPI_ISL_3418131, EPI_ISL_3418132, EPI_ISL_3418133, EPI_ISL_3418134, EPI_ISL_3418135, EPI_ISL_3418136, EPI_ISL_3418137, EPI_ISL_3418138, EPI_ISL_3418139, EPI_ISL_3418140, EPI_ISL_3418141, EPI_ISL_3418142, EPI_ISL_3418143, EPI_ISL_3418144, EPI_ISL_3418145, EPI_ISL_3418146, EPI_ISL_3418147, EPI_ISL_3418148, EPI_ISL_3418149, EPI_ISL_3418150, EPI_ISL_3418151, EPI_ISL_3418152, EPI_ISL_3418153, EPI_ISL_3418154, EPI_ISL_3418155, EPI_ISL_3418156, EPI_ISL_3418157, EPI_ISL_3418158, EPI_ISL_3418159, EPI_ISL_3418160, EPI_ISL_3418161, EPI_ISL_3418162, EPI_ISL_3418163, EPI_ISL_3418165, EPI_ISL_3418166 | | | |
| see above | Department of Bacteria, Parasites and Fungi, Statens Serum Institut, Copenhagen, Denmark | Statens Serum Institut Bioinformatics and Microbial Genomics | Danish Covid-19 Genome Consortium |
| EPI_ISL_3418167 | Molekylær Medicinsk Afdeling, Aarhus University Hospital, Aarhus, Denmark | Statens Serum Institut Bioinformatics and Microbial Genomics | Danish Covid-19 Genome Consortium |
| EPI_ISL_3418168, EPI_ISL_3418169, EPI_ISL_3418170, EPI_ISL_3418171, EPI_ISL_3418172, EPI_ISL_3418173, EPI_ISL_3418174, EPI_ISL_3418175, EPI_ISL_3418176, EPI_ISL_3418177, EPI_ISL_3418178, EPI_ISL_3418179, EPI_ISL_3418180, EPI_ISL_3418181, EPI_ISL_3418183, EPI_ISL_3418184, EPI_ISL_3418185, EPI_ISL_3418186, EPI_ISL_3418187, EPI_ISL_3418188, EPI_ISL_3418189, EPI_ISL_3418190, EPI_ISL_3418191, EPI_ISL_3418192, EPI_ISL_3418193, EPI_ISL_3418194, EPI_ISL_3418195, EPI_ISL_3418196, EPI_ISL_3418197, EPI_ISL_3418198, EPI_ISL_3418199, EPI_ISL_3418200, EPI_ISL_3418201, EPI_ISL_3418202, EPI_ISL_3418203, EPI_ISL_3418204, EPI_ISL_3418205, EPI_ISL_3418206, EPI_ISL_3418207, EPI_ISL_3418208, EPI_ISL_3418209, EPI_ISL_3418210, EPI_ISL_3418211 | | | |
| see above | Department of Bacteria, Parasites and Fungi, Statens Serum Institut, Copenhagen, Denmark | Statens Serum Institut Bioinformatics and Microbial Genomics | Danish Covid-19 Genome Consortium |
| EPI_ISL_3418212 | Molekylær Medicinsk Afdeling, Aarhus University Hospital, Aarhus, Denmark | Statens Serum Institut Bioinformatics and Microbial Genomics | Danish Covid-19 Genome Consortium |
| EPI_ISL_3418213, EPI_ISL_3418214, EPI_ISL_3418215, EPI_ISL_3418216, EPI_ISL_3418217, EPI_ISL_3418218, EPI_ISL_3418219, EPI_ISL_3418220, EPI_ISL_3418221, EPI_ISL_3418222, EPI_ISL_3418223, EPI_ISL_3418224, EPI_ISL_3418225, EPI_ISL_3418226, EPI_ISL_3418227, EPI_ISL_3418228, EPI_ISL_3418229, EPI_ISL_3418230, EPI_ISL_3418231, EPI_ISL_3418232, EPI_ISL_3418233, EPI_ISL_3418234, EPI_ISL_3418235, EPI_ISL_3418236, EPI_ISL_3418237, EPI_ISL_3418238, EPI_ISL_3418239, EPI_ISL_3418240, EPI_ISL_3418241, EPI_ISL_3418242, EPI_ISL_3418243, EPI_ISL_3418244, EPI_ISL_3418245, EPI_ISL_3418246, EPI_ISL_3418247, EPI_ISL_3418248, EPI_ISL_3418249, EPI_ISL_3418250, EPI_ISL_3418251, EPI_ISL_3418252, EPI_ISL_3418253, EPI_ISL_3418254, EPI_ISL_3418255, EPI_ISL_3418256 | | | |
| see above | Department of Bacteria, Parasites and Fungi, Statens Serum Institut, Copenhagen, Denmark | Statens Serum Institut Bioinformatics and Microbial Genomics | Danish Covid-19 Genome Consortium |
| EPI_ISL_3418257 | Molekylær Medicinsk Afdeling, Aarhus University Hospital, Aarhus, Denmark | Statens Serum Institut Bioinformatics and Microbial Genomics | Danish Covid-19 Genome Consortium |
| EPI_ISL_3418258, EPI_ISL_3418259, EPI_ISL_3418260, EPI_ISL_3418261, EPI_ISL_3418262, EPI_ISL_3418263, EPI_ISL_3418264, EPI_ISL_3418265, EPI_ISL_3418266, EPI_ISL_3418267, EPI_ISL_3418268, EPI_ISL_3418269, EPI_ISL_3418270, EPI_ISL_3418271, EPI_ISL_3418273, EPI_ISL_3418274, EPI_ISL_3418275, EPI_ISL_3418276, EPI_ISL_3418277, EPI_ISL_3418278, EPI_ISL_3418279, EPI_ISL_3418280, EPI_ISL_3418281, EPI_ISL_3418282, EPI_ISL_3418283, EPI_ISL_3418284, EPI_ISL_3418285, EPI_ISL_3418286, EPI_ISL_3418287, EPI_ISL_3418288, EPI_ISL_3418289, EPI_ISL_3418290, EPI_ISL_3418291, EPI_ISL_3418292, EPI_ISL_3418293, EPI_ISL_3418294, EPI_ISL_3418295, EPI_ISL_3418296, EPI_ISL_3418297, EPI_ISL_3418298, EPI_ISL_3418299, EPI_ISL_3418300, EPI_ISL_3418301, EPI_ISL_3418302, EPI_ISL_3418303, EPI_ISL_3418304, EPI_ISL_3418305, EPI_ISL_3418306, EPI_ISL_3418307 | | | |
| see above | Department of Bacteria, Parasites and Fungi, Statens Serum Institut, Copenhagen, Denmark | Statens Serum Institut Bioinformatics and Microbial Genomics | Danish Covid-19 Genome Consortium |
| EPI_ISL_3418308 | Molekylær Medicinsk Afdeling, Aarhus University Hospital, Aarhus, Denmark | Statens Serum Institut Bioinformatics and Microbial Genomics | Danish Covid-19 Genome Consortium |
| EPI_ISL_3418309, EPI_ISL_3418310, EPI_ISL_3418311, EPI_ISL_3418312 | Department of Bacteria, Parasites and Fungi, Statens Serum Institut, Copenhagen, Denmark | Statens Serum Institut Bioinformatics and Microbial Genomics | Danish Covid-19 Genome Consortium |
| EPI_ISL_3418313 | Molekylær Medicinsk Afdeling, Aarhus University Hospital, Aarhus, Denmark | Statens Serum Institut Bioinformatics and Microbial Genomics | Danish Covid-19 Genome Consortium |
| EPI_ISL_3418314, EPI_ISL_3418315, EPI_ISL_3418316, EPI_ISL_3418317, EPI_ISL_3418318, EPI_ISL_3418319, EPI_ISL_3418320, EPI_ISL_3418321, EPI_ISL_3418322, EPI_ISL_3418323, EPI_ISL_3418324, EPI_ISL_3418325, EPI_ISL_3418326, EPI_ISL_3418327, EPI_ISL_3418328, EPI_ISL_3418329, EPI_ISL_3418330, EPI_ISL_3418331, EPI_ISL_3418332, EPI_ISL_3418333, EPI_ISL_3418334, EPI_ISL_3418335, EPI_ISL_3418336, EPI_ISL_3418337, EPI_ISL_3418338, EPI_ISL_3418339, EPI_ISL_3418340, EPI_ISL_3418341, EPI_ISL_3418342, EPI_ISL_3418343, EPI_ISL_3418344, EPI_ISL_3418345, EPI_ISL_3418346, EPI_ISL_3418347, EPI_ISL_3418348, EPI_ISL_3418349, EPI_ISL_3418350, EPI_ISL_3418351, EPI_ISL_3418352, EPI_ISL_3418353 | | | |
| see above | Department of Bacteria, Parasites and Fungi, Statens Serum Institut, Copenhagen, Denmark | Statens Serum Institut Bioinformatics and Microbial Genomics | Danish Covid-19 Genome Consortium |
| EPI_ISL_3418354 | Molekylær Medicinsk Afdeling, Aarhus University Hospital, Aarhus, Denmark | Statens Serum Institut Bioinformatics and Microbial Genomics | Danish Covid-19 Genome Consortium |
| EPI_ISL_3418355, EPI_ISL_3418356, EPI_ISL_3418357, EPI_ISL_3418358, EPI_ISL_3418359, EPI_ISL_3418360, EPI_ISL_3418361, EPI_ISL_3418362, EPI_ISL_3418363, EPI_ISL_3418364, EPI_ISL_3418365, EPI_ISL_3418366, EPI_ISL_3418367, EPI_ISL_3418368, EPI_ISL_3418369, EPI_ISL_3418370, EPI_ISL_3418371, EPI_ISL_3418372, EPI_ISL_3418373, EPI_ISL_3418374, EPI_ISL_3418375, EPI_ISL_3418376, EPI_ISL_3418377, EPI_ISL_3418378, EPI_ISL_3418379, EPI_ISL_3418380, EPI_ISL_3418381, EPI_ISL_3418382, EPI_ISL_3418383, EPI_ISL_3418384, EPI_ISL_3418385, EPI_ISL_3418386, EPI_ISL_3418387, EPI_ISL_3418388, EPI_ISL_3418389, EPI_ISL_3418390, EPI_ISL_3418391, EPI_ISL_3418392, EPI_ISL_3418393, EPI_ISL_3418394, EPI_ISL_3418395, EPI_ISL_3418396, EPI_ISL_3418397, EPI_ISL_3418398, EPI_ISL_3418399, EPI_ISL_3418400, EPI_ISL_3418401, EPI_ISL_3418402, EPI_ISL_3418403, EPI_ISL_3418404, EPI_ISL_3418405 | | | |
| see above | Department of Bacteria, Parasites and Fungi, Statens Serum Institut, Copenhagen, Denmark | Statens Serum Institut Bioinformatics and Microbial Genomics | Danish Covid-19 Genome Consortium |
| EPI_ISL_3418406 | Molekylær Medicinsk Afdeling, Aarhus University Hospital, Aarhus, Denmark | Statens Serum Institut Bioinformatics and Microbial Genomics | Danish Covid-19 Genome Consortium |
| EPI_ISL_3418407, EPI_ISL_3418408, EPI_ISL_3418409, EPI_ISL_3418410, EPI_ISL_3418411, EPI_ISL_3418412, EPI_ISL_3418413, EPI_ISL_3418414, EPI_ISL_3418415, EPI_ISL_3418417, EPI_ISL_3418418, EPI_ISL_3418419, EPI_ISL_3418420, EPI_ISL_3418421, EPI_ISL_3418422, EPI_ISL_3418423, EPI_ISL_3418424, EPI_ISL_3418425, EPI_ISL_3418427, EPI_ISL_3418428, EPI_ISL_3418429, EPI_ISL_3418430, EPI_ISL_3418431, EPI_ISL_3418432, EPI_ISL_3418433, EPI_ISL_3418434, EPI_ISL_3418435, EPI_ISL_3418436, EPI_ISL_3418437, EPI_ISL_3418438, EPI_ISL_3418439, EPI_ISL_3418440, EPI_ISL_3418441, EPI_ISL_3418442, EPI_ISL_3418443, EPI_ISL_3418444, EPI_ISL_3418445, EPI_ISL_3418446, EPI_ISL_3418447, EPI_ISL_3418448, EPI_ISL_3418449, EPI_ISL_3418450, EPI_ISL_3418451, EPI_ISL_3418452, EPI_ISL_3418453, EPI_ISL_3418454, EPI_ISL_3418455 | | | |
| see above | Department of Bacteria, Parasites and Fungi, Statens Serum Institut, Copenhagen, Denmark | Statens Serum Institut Bioinformatics and Microbial Genomics | Danish Covid-19 Genome Consortium |
| EPI_ISL_3418457 | Molekylær Medicinsk Afdeling, Aarhus University Hospital, Aarhus, Denmark | Statens Serum Institut Bioinformatics and Microbial Genomics | Danish Covid-19 Genome Consortium |
| EPI_ISL_3418458, EPI_ISL_3418459, EPI_ISL_3418460, EPI_ISL_3418461, EPI_ISL_3418462, EPI_ISL_3418463, EPI_ISL_3418464, EPI_ISL_3418466, EPI_ISL_3418467, EPI_ISL_3418468, EPI_ISL_3418469, EPI_ISL_3418470, EPI_ISL_3418471, EPI_ISL_3418472, EPI_ISL_3418473, EPI_ISL_3418474, EPI_ISL_3418475, EPI_ISL_3418476, EPI_ISL_3418477, EPI_ISL_3418478, EPI_ISL_3418479, EPI_ISL_3418480, EPI_ISL_3418481, EPI_ISL_3418482 | | | |
| see above | Department of Bacteria, Parasites and Fungi, Statens Serum Institut, Copenhagen, Denmark | Statens Serum Institut Bioinformatics and Microbial Genomics | Danish Covid-19 Genome Consortium |
| EPI_ISL_3418483 | Molekylær Medicinsk Afdeling, Aarhus University Hospital, Aarhus, Denmark | Statens Serum Institut Bioinformatics and Microbial Genomics | Danish Covid-19 Genome Consortium |
| EPI_ISL_3418484, EPI_ISL_3418485, EPI_ISL_3418486, EPI_ISL_3418487, EPI_ISL_3418488, EPI_ISL_3418489, | Department of Bacteria, Parasites and Fungi, Statens Serum Institut, Copenhagen, Denmark | Statens Serum Institut Bioinformatics and Microbial Genomics | Danish Covid-19 Genome Consortium |

| | | | |
|---|---|---|---|
| EPI_ISL_3418490 | | | |
| EPI_ISL_3418491, EPI_ISL_3418492 | Molekylær Medicinsk Afdeling, Aarhus University Hospital, Aarhus, Denmark | Statens Serum Institut Bioinformatics and Microbial Genomics | Danish Covid-19 Genome Consortium |
| EPI_ISL_3418493, EPI_ISL_3418494, EPI_ISL_3418495, EPI_ISL_3418496, EPI_ISL_3418497, EPI_ISL_3418498, EPI_ISL_3418499, EPI_ISL_3418500, EPI_ISL_3418501, EPI_ISL_3418502, EPI_ISL_3418503, EPI_ISL_3418504, EPI_ISL_3418505, EPI_ISL_3418506, EPI_ISL_3418507, EPI_ISL_3418508, EPI_ISL_3418509, EPI_ISL_3418510, EPI_ISL_3418511, EPI_ISL_3418512, EPI_ISL_3418513, EPI_ISL_3418514, EPI_ISL_3418515, EPI_ISL_3418516 | | | |
| see above | Department of Bacteria, Parasites and Fungi, Statens Serum Institut, Copenhagen, Denmark | Statens Serum Institut Bioinformatics and Microbial Genomics | Danish Covid-19 Genome Consortium |
| EPI_ISL_3418517 | Molekylær Medicinsk Afdeling, Aarhus University Hospital, Aarhus, Denmark | Statens Serum Institut Bioinformatics and Microbial Genomics | Danish Covid-19 Genome Consortium |
| EPI_ISL_3418518, EPI_ISL_3418519, EPI_ISL_3418520, EPI_ISL_3418521, EPI_ISL_3418522, EPI_ISL_3418523, EPI_ISL_3418524, EPI_ISL_3418525, EPI_ISL_3418526, EPI_ISL_3418527, EPI_ISL_3418528, EPI_ISL_3418529, EPI_ISL_3418530, EPI_ISL_3418531, EPI_ISL_3418532, EPI_ISL_3418533, EPI_ISL_3418534, EPI_ISL_3418535, EPI_ISL_3418536, EPI_ISL_3418537, EPI_ISL_3418538, EPI_ISL_3418539, EPI_ISL_3418540, EPI_ISL_3418541, EPI_ISL_3418542, EPI_ISL_3418543, EPI_ISL_3418544 | | | |
| see above | Department of Bacteria, Parasites and Fungi, Statens Serum Institut, Copenhagen, Denmark | Statens Serum Institut Bioinformatics and Microbial Genomics | Danish Covid-19 Genome Consortium |
| EPI_ISL_3418545 | Molekylær Medicinsk Afdeling, Aarhus University Hospital, Aarhus, Denmark | Statens Serum Institut Bioinformatics and Microbial Genomics | Danish Covid-19 Genome Consortium |
| EPI_ISL_3418546, EPI_ISL_3418547, EPI_ISL_3418548, EPI_ISL_3418549, EPI_ISL_3418550, EPI_ISL_3418551, EPI_ISL_3418552, EPI_ISL_3418553, EPI_ISL_3418554, EPI_ISL_3418555, EPI_ISL_3418556, EPI_ISL_3418557, EPI_ISL_3418558, EPI_ISL_3418559, EPI_ISL_3418560, EPI_ISL_3418561, EPI_ISL_3418562, EPI_ISL_3418563, EPI_ISL_3418564, EPI_ISL_3418565, EPI_ISL_3418566, EPI_ISL_3418567, EPI_ISL_3418568, EPI_ISL_3418569, EPI_ISL_3418570, EPI_ISL_3418571, EPI_ISL_3418572, EPI_ISL_3418573, EPI_ISL_3418574, EPI_ISL_3418575, EPI_ISL_3418576, EPI_ISL_3418577, EPI_ISL_3418578 | | | |
| see above | Department of Bacteria, Parasites and Fungi, Statens Serum Institut, Copenhagen, Denmark | Statens Serum Institut Bioinformatics and Microbial Genomics | Danish Covid-19 Genome Consortium |
| EPI_ISL_3418579 | Molekylær Medicinsk Afdeling, Aarhus University Hospital, Aarhus, Denmark | Statens Serum Institut Bioinformatics and Microbial Genomics | Danish Covid-19 Genome Consortium |
| EPI_ISL_3418580, EPI_ISL_3418581, EPI_ISL_3418582, EPI_ISL_3418583 | Department of Bacteria, Parasites and Fungi, Statens Serum Institut, Copenhagen, Denmark | Statens Serum Institut Bioinformatics and Microbial Genomics | Danish Covid-19 Genome Consortium |
| EPI_ISL_3418584 | Molekylær Medicinsk Afdeling, Aarhus University Hospital, Aarhus, Denmark | Statens Serum Institut Bioinformatics and Microbial Genomics | Danish Covid-19 Genome Consortium |
| EPI_ISL_3418585, EPI_ISL_3418586, EPI_ISL_3418587, EPI_ISL_3418588, EPI_ISL_3418589, EPI_ISL_3418590, EPI_ISL_3418591 | Department of Bacteria, Parasites and Fungi, Statens Serum Institut, Copenhagen, Denmark | Statens Serum Institut Bioinformatics and Microbial Genomics | Danish Covid-19 Genome Consortium |
| EPI_ISL_3418592 | Molekylær Medicinsk Afdeling, Aarhus University Hospital, Aarhus, Denmark | Statens Serum Institut Bioinformatics and Microbial Genomics | Danish Covid-19 Genome Consortium |
| EPI_ISL_3418593, EPI_ISL_3418594, EPI_ISL_3418595, EPI_ISL_3418596, EPI_ISL_3418597, EPI_ISL_3418598, EPI_ISL_3418599, EPI_ISL_3418600, EPI_ISL_3418601, EPI_ISL_3418602, EPI_ISL_3418603, EPI_ISL_3418604, EPI_ISL_3418605, EPI_ISL_3418606, EPI_ISL_3418607, EPI_ISL_3418608, EPI_ISL_3418609, EPI_ISL_3418610, EPI_ISL_3418611, EPI_ISL_3418612, EPI_ISL_3418613, EPI_ISL_3418614, EPI_ISL_3418615, EPI_ISL_3418616, EPI_ISL_3418617, EPI_ISL_3418618, EPI_ISL_3418619, EPI_ISL_3418620, EPI_ISL_3418621, EPI_ISL_3418622, EPI_ISL_3418623, EPI_ISL_3418624, EPI_ISL_3418625, EPI_ISL_3418626, EPI_ISL_3418627, EPI_ISL_3418628, EPI_ISL_3418629, EPI_ISL_3418630, EPI_ISL_3418631, EPI_ISL_3418632, EPI_ISL_3418633, EPI_ISL_3418634, EPI_ISL_3418635 | | | |
| see above | Department of Bacteria, Parasites and Fungi, Statens Serum Institut, Copenhagen, Denmark | Statens Serum Institut Bioinformatics and Microbial Genomics | Danish Covid-19 Genome Consortium |
| EPI_ISL_3418636 | Molekylær Medicinsk Afdeling, Aarhus University Hospital, Aarhus, Denmark | Statens Serum Institut Bioinformatics and Microbial Genomics | Danish Covid-19 Genome Consortium |
| EPI_ISL_3418637, EPI_ISL_3418638, EPI_ISL_3418639, EPI_ISL_3418640, EPI_ISL_3418641, EPI_ISL_3418642, EPI_ISL_3418643, EPI_ISL_3418644, EPI_ISL_3418645, EPI_ISL_3418646, EPI_ISL_3418647, EPI_ISL_3418648, EPI_ISL_3418649, EPI_ISL_3418650, EPI_ISL_3418651, EPI_ISL_3418652, EPI_ISL_3418653, EPI_ISL_3418654, EPI_ISL_3418655, EPI_ISL_3418656, EPI_ISL_3418657, EPI_ISL_3418658, EPI_ISL_3418659, EPI_ISL_3418660, EPI_ISL_3418661 | | | |
| see above | Department of Bacteria, Parasites and Fungi, Statens Serum Institut, Copenhagen, Denmark | Statens Serum Institut Bioinformatics and Microbial Genomics | Danish Covid-19 Genome Consortium |
| EPI_ISL_3418662 | Molekylær Medicinsk Afdeling, Aarhus University Hospital, Aarhus, Denmark | Statens Serum Institut Bioinformatics and Microbial Genomics | Danish Covid-19 Genome Consortium |
| EPI_ISL_3418663, EPI_ISL_3418664, EPI_ISL_3418665, EPI_ISL_3418666, EPI_ISL_3418667, EPI_ISL_3418668 | Department of Bacteria, Parasites and Fungi, Statens Serum Institut, Copenhagen, Denmark | Statens Serum Institut Bioinformatics and Microbial Genomics | Danish Covid-19 Genome Consortium |
| EPI_ISL_3418685 | LAB MIKROBIOLOGI FKUI / DINAS KESEHATAN DKI JAKARTA | Genomik Solidaritas Indonesia Laboratorium | Anna Christina Brazia, Reinhart Gabriel, Vania Gavrila Wikasa, Carissa Sintca Wijaya, Dhahlia Agustina Cahyono, Alfin Mohammad Abdillah, Normastuti Adhini Tantyo, Annisa Muthiah Sukirman, Gracia Felias Enos Korompis, Yulia Rosa, Fera Ibrahim, Pratiwi Sudarmono, Widyastuti, Dwi Oktavia, Tiranti Vindha, Ngabila Salama, Anuraj Shankar, Meutia Ayuputeri Kumaheri, Ariel Pradipta |
| EPI_ISL_3418686, EPI_ISL_3418687, EPI_ISL_3418688, EPI_ISL_3418689, EPI_ISL_3418690, EPI_ISL_3418691, EPI_ISL_3418692, EPI_ISL_3418693, EPI_ISL_3418694, EPI_ISL_3418695 | Area of Virology, Serology and Virology Division (SAViD), New South Wales Health Pathology Randwick | Virology Research Laboratory; Area of Virology, Serology and Virology Division (SAViD), New South Wales Health Pathology Randwick | Foster, C.; Au, J.; Ruiz Silva, M.; Deveson, I.; Bull, R.; Van Hal, S.; Rawlinson, W. |
| EPI_ISL_3418696 | RSUD KEBAYORAN BARU / DINAS KESEHATAN DKI JAKARTA | Genomik Solidaritas Indonesia Laboratorium | Carissa Sintca Wijaya, Vania Gavrila Wikasa, Anna Christina Brazia, Reinhart Gabriel, Dhahlia Agustina Cahyono, Alfin Mohammad Abdillah, Normastuti Adhini Tantyo, Annisa Muthiah Sukirman, Gracia Felias Enos Korompis, Widyastuti, Dwi Oktavia, Tiranti Vindha, Ngabila Salama, Anuraj Shankar, Meutia Ayuputeri Kumaheri, Ariel Pradipta Submitter information |
| EPI_ISL_3418698 | Virology Laboratory, IDD, icddr,b | Virology Laboratory, IDD, icddr,b | Muntasir Alam, M Ishrat Jahan, Shafina Jahan, Mohammad Fakhruddin, Afsana Rashed, Emily S. Gurley, Shams El Arifeen, Mustafizur Rahman |
| EPI_ISL_3418699 | RSUP Dr. Hasan Sadikin Bandung | Genomik Solidaritas Indonesia Laboratorium | Anna Christina Brazia, Reinhart Gabriel, Vania Gavrila Wikasa, Carissa Sintca Wijaya, Dhahlia Agustina Cahyono, Alfin Mohammad Abdillah, Normastuti Adhini Tantyo, Annisa Muthiah Sukirman, Gracia Felias Enos Korompis, Louisa Markus, Basti Andriyoko, Delita Prihatni, Dewi Kartika Turbawati, Vivi Setyawati, Anuraj Shankar, Meutia Ayuputeri Kumaheri, Ariel Pradipta |
| EPI_ISL_3418706, EPI_ISL_3418707, EPI_ISL_3418708, EPI_ISL_3418709, EPI_ISL_3418710, EPI_ISL_3418711, EPI_ISL_3418712, EPI_ISL_3418713, EPI_ISL_3418714, EPI_ISL_3418715, EPI_ISL_3418716, EPI_ISL_3418717, EPI_ISL_3418718, EPI_ISL_3418719, EPI_ISL_3418720, EPI_ISL_3418721, EPI_ISL_3418722, EPI_ISL_3418723, EPI_ISL_3418724, EPI_ISL_3418725, EPI_ISL_3418726, EPI_ISL_3418727, EPI_ISL_3418728, EPI_ISL_3418729, EPI_ISL_3418730, EPI_ISL_3418731, EPI_ISL_3418732, EPI_ISL_3418733, EPI_ISL_3418734, EPI_ISL_3418735, EPI_ISL_3418736, EPI_ISL_3418737, EPI_ISL_3418738, EPI_ISL_3418739, EPI_ISL_3418740, EPI_ISL_3418741, EPI_ISL_3418742, EPI_ISL_3418743, EPI_ISL_3418744, EPI_ISL_3418745, EPI_ISL_3418747, EPI_ISL_3418748, EPI_ISL_3418749, EPI_ISL_3418750, EPI_ISL_3418751, EPI_ISL_3418752, EPI_ISL_3418753, EPI_ISL_3418754, EPI_ISL_3418755, EPI_ISL_3418756, EPI_ISL_3418757, EPI_ISL_3418758, EPI_ISL_3418759, EPI_ISL_3418760, EPI_ISL_3418761, EPI_ISL_3418762, EPI_ISL_3418763, EPI_ISL_3418764, EPI_ISL_3418765, EPI_ISL_3418766, EPI_ISL_3418767, EPI_ISL_3418768, EPI_ISL_3418769, EPI_ISL_3418770, EPI_ISL_3418771, EPI_ISL_3418772, EPI_ISL_3418773, EPI_ISL_3418774, EPI_ISL_3418775, EPI_ISL_3418776, EPI_ISL_3418777, EPI_ISL_3418778, EPI_ISL_3418779, EPI_ISL_3418780, EPI_ISL_3418781, EPI_ISL_3418782, EPI_ISL_3418783, EPI_ISL_3418784, EPI_ISL_3418785, EPI_ISL_3418786 | | | |
| see above | Baylor Scott & White-Temple | Baylor Scott & White-Temple | Ari Rao, Linden Morales, Kimberly Walker, Marcus Volz, Shelby Johnson |
| EPI_ISL_3418787 | Dr. Yoshitaka Tamura Department of Clinical Laboratory, Osaka Habikino Medical Center | Department of Developmental Medicine, Research Institute, Osaka Women's and Children's Hospital | Itaru Yanagihara, Yukiko Nakura |
| EPI_ISL_3418788, EPI_ISL_3418789, EPI_ISL_3418790, EPI_ISL_3418793, EPI_ISL_3418795, EPI_ISL_3418796, EPI_ISL_3418797, EPI_ISL_3418799, EPI_ISL_3418802, EPI_ISL_3418803, EPI_ISL_3418804, EPI_ISL_3418805, EPI_ISL_3418806, EPI_ISL_3418808, EPI_ISL_3418809, EPI_ISL_3418810, EPI_ISL_3418811, EPI_ISL_3418812, EPI_ISL_3418813, EPI_ISL_3418814, EPI_ISL_3418815, EPI_ISL_3418816, EPI_ISL_3418817, EPI_ISL_3418818, EPI_ISL_3418819, EPI_ISL_3418820, EPI_ISL_3418821, EPI_ISL_3418822, EPI_ISL_3418823, EPI_ISL_3418824, EPI_ISL_3418825, EPI_ISL_3418826, EPI_ISL_3418829, EPI_ISL_3418831, EPI_ISL_3418832, EPI_ISL_3418833, EPI_ISL_3418834, EPI_ISL_3418835, EPI_ISL_3418836, EPI_ISL_3418837, EPI_ISL_3418838, EPI_ISL_3418839, EPI_ISL_3418840, EPI_ISL_3418843, EPI_ISL_3418844, EPI_ISL_3418845, EPI_ISL_3418846, EPI_ISL_3418847, EPI_ISL_3418848, EPI_ISL_3418849, EPI_ISL_3418850, EPI_ISL_3418852, EPI_ISL_3418853, EPI_ISL_3418854, EPI_ISL_3418856, EPI_ISL_3418857, EPI_ISL_3418858, EPI_ISL_3418859, EPI_ISL_3418860, EPI_ISL_3418861, EPI_ISL_3418862, EPI_ISL_3418863 | | | |
| see above | Baylor Scott & White-Temple | Baylor Scott & White-Temple | Ari Rao, Linden Morales, Kimberly Walker, Marcus Volz, Shelby Johnson |
| EPI_ISL_3418864 | Dr. Yoshitaka Tamura Department of Clinical Laboratory, Osaka Habikino Medical Center | Department of Developmental Medicine, Research Institute, Osaka Women's and Children's Hospital | Itaru Yanagihara, Yukiko Nakura |
| EPI_ISL_3418866 | RSUP Dr. Hasan Sadikin Bandung | Genomik Solidaritas Indonesia Laboratorium | Anna Christina Brazia, Reinhart Gabriel, Vania Gavrila Wikasa, Carissa Sintca Wijaya, Dhahlia Agustina Cahyono, Alfin Mohammad Abdillah, Normastuti Adhini Tantyo, Annisa Muthiah Sukirman, Gracia Felias Enos Korompis, Louisa Markus, Basti Andriyoko, Delita Prihatni, Dewi Kartika Turbawati, Vivi Setyawati, Anuraj Shankar, Meutia Ayuputeri Kumaheri, Ariel Pradipta |
| EPI_ISL_3418867, EPI_ISL_3418868, EPI_ISL_3418869, EPI_ISL_3418870, EPI_ISL_3418871, EPI_ISL_3418984, EPI_ISL_3418985, EPI_ISL_3418986, EPI_ISL_3418992, EPI_ISL_3418993, EPI_ISL_3418994, EPI_ISL_3418996, EPI_ISL_3418997, EPI_ISL_3418998, EPI_ISL_3419002, EPI_ISL_3419006, EPI_ISL_3419007, EPI_ISL_3419008, EPI_ISL_3419009, EPI_ISL_3419011, EPI_ISL_3419012, EPI_ISL_3419014, EPI_ISL_3419015, EPI_ISL_3419016, EPI_ISL_3419017, EPI_ISL_3419019, EPI_ISL_3419029, EPI_ISL_3419030, EPI_ISL_3419032, EPI_ISL_3419033, EPI_ISL_3419034, EPI_ISL_3419035, EPI_ISL_3419037, EPI_ISL_3419038, EPI_ISL_3419039, EPI_ISL_3419040, EPI_ISL_3419041, EPI_ISL_3419042, EPI_ISL_3419043, EPI_ISL_3419044, EPI_ISL_3419045, EPI_ISL_3419046, EPI_ISL_3419047, EPI_ISL_3419048, EPI_ISL_3419049, EPI_ISL_3419050, EPI_ISL_3419052, EPI_ISL_3419053, EPI_ISL_3419054, EPI_ISL_3419055, EPI_ISL_3419056 | | | |
| see above | StarMed Healthcare | UNC Charlotte Environmental Monitoring Laboratory | Jessica Schlueter, Jannatul Ferdous, Erin Stiers, Adam France, Sam Kunkleman, Kevin Lambirth, Cynthia Gibas |
| EPI_ISL_3419057 | A.S.L. CUNEO 1 | Fondazione del Piemonte per l'Oncologia IRCCS | Antonino Sottile, Silvia Brossa, Paola Marino, Giorgio Giardina |
| EPI_ISL_3419058 | StarMed Healthcare | UNC Charlotte Environmental Monitoring Laboratory | Jessica Schlueter, Jannatul Ferdous, Erin Stiers, Adam France, Sam Kunkleman, Kevin Lambirth, Cynthia Gibas |
| EPI_ISL_3419059 | A.S.L. NOVARA | Fondazione del Piemonte per l'Oncologia IRCCS | Antonino Sottile, Silvia Brossa, Paola Marino, Giorgio Giardina |

| | | | |
|---|---|---|---|
| EPI_ISL_3419060 | StarMed Healthcare | UNC Charlotte Environmental Monitoring Laboratory | Jessica Schlueter, Jannatul Ferdous, Erin Stiers, Adam France, Sam Kunkleman, Kevin Lambirth, Cynthia Gibas |
| EPI_ISL_3419061 | A.S.L. CUNEO 1 | Fondazione del Piemonte per l'Oncologia IRCCS | Antonino Sottile, Silvia Brossa, Paola Marino, Giorgio Giardina |
| EPI_ISL_3419062 | StarMed Healthcare | UNC Charlotte Environmental Monitoring Laboratory | Jessica Schlueter, Jannatul Ferdous, Erin Stiers, Adam France, Sam Kunkleman, Kevin Lambirth, Cynthia Gibas |
| EPI_ISL_3419063, EPI_ISL_3419064 | A.S.L. CUNEO 1 | Fondazione del Piemonte per l'Oncologia IRCCS | Antonino Sottile, Silvia Brossa, Paola Marino, Giorgio Giardina |
| EPI_ISL_3419065, EPI_ISL_3419066 | StarMed Healthcare | UNC Charlotte Environmental Monitoring Laboratory | Jessica Schlueter, Jannatul Ferdous, Erin Stiers, Adam France, Sam Kunkleman, Kevin Lambirth, Cynthia Gibas |
| EPI_ISL_3419068, EPI_ISL_3419070 | A.S.L. CUNEO 1 | Fondazione del Piemonte per l'Oncologia IRCCS | Antonino Sottile, Silvia Brossa, Paola Marino, Giorgio Giardina |
| EPI_ISL_3419071 | StarMed Healthcare | UNC Charlotte Environmental Monitoring Laboratory | Jessica Schlueter, Jannatul Ferdous, Erin Stiers, Adam France, Sam Kunkleman, Kevin Lambirth, Cynthia Gibas |
| EPI_ISL_3419072 | A.S.L. CUNEO 1 | Fondazione del Piemonte per l'Oncologia IRCCS | Antonino Sottile, Silvia Brossa, Paola Marino, Giorgio Giardina |
| EPI_ISL_3419073 | StarMed Healthcare | UNC Charlotte Environmental Monitoring Laboratory | Jessica Schlueter, Jannatul Ferdous, Erin Stiers, Adam France, Sam Kunkleman, Kevin Lambirth, Cynthia Gibas |
| EPI_ISL_3419074 | A.S.L. CUNEO 1 | Fondazione del Piemonte per l'Oncologia IRCCS | Antonino Sottile, Silvia Brossa, Paola Marino, Giorgio Giardina |
| EPI_ISL_3419075 | StarMed Healthcare | UNC Charlotte Environmental Monitoring Laboratory | Jessica Schlueter, Jannatul Ferdous, Erin Stiers, Adam France, Sam Kunkleman, Kevin Lambirth, Cynthia Gibas |
| EPI_ISL_3419076 | A.S.L. TO4 | Fondazione del Piemonte per l'Oncologia IRCCS | Antonino Sottile, Silvia Brossa, Paola Marino, Giorgio Giardina |
| EPI_ISL_3419077 | StarMed Healthcare | UNC Charlotte Environmental Monitoring Laboratory | Jessica Schlueter, Jannatul Ferdous, Erin Stiers, Adam France, Sam Kunkleman, Kevin Lambirth, Cynthia Gibas |
| EPI_ISL_3419078 | A.S.L. TO4 | Fondazione del Piemonte per l'Oncologia IRCCS | Antonino Sottile, Silvia Brossa, Paola Marino, Giorgio Giardina |
| EPI_ISL_3419079 | StarMed Healthcare | UNC Charlotte Environmental Monitoring Laboratory | Jessica Schlueter, Jannatul Ferdous, Erin Stiers, Adam France, Sam Kunkleman, Kevin Lambirth, Cynthia Gibas |
| EPI_ISL_3419080 | A.S.L. CUNEO 1 | Fondazione del Piemonte per l'Oncologia IRCCS | Antonino Sottile, Silvia Brossa, Paola Marino, Giorgio Giardina |
| EPI_ISL_3419081 | StarMed Healthcare | UNC Charlotte Environmental Monitoring Laboratory | Jessica Schlueter, Jannatul Ferdous, Erin Stiers, Adam France, Sam Kunkleman, Kevin Lambirth, Cynthia Gibas |
| EPI_ISL_3419082 | A.S.L. TO5 | Fondazione del Piemonte per l'Oncologia IRCCS | Antonino Sottile, Silvia Brossa, Paola Marino, Giorgio Giardina |
| EPI_ISL_3419083 | StarMed Healthcare | UNC Charlotte Environmental Monitoring Laboratory | Jessica Schlueter, Jannatul Ferdous, Erin Stiers, Adam France, Sam Kunkleman, Kevin Lambirth, Cynthia Gibas |
| EPI_ISL_3419084 | A.S.L. TO4 | Fondazione del Piemonte per l'Oncologia IRCCS | Antonino Sottile, Silvia Brossa, Paola Marino, Giorgio Giardina |
| EPI_ISL_3419085 | StarMed Healthcare | UNC Charlotte Environmental Monitoring Laboratory | Jessica Schlueter, Jannatul Ferdous, Erin Stiers, Adam France, Sam Kunkleman, Kevin Lambirth, Cynthia Gibas |
| EPI_ISL_3419086, EPI_ISL_3419087 | A.S.L. CITTA' DI TORINO - OSPEDALE AMEDEO DI SAVOIA | Fondazione del Piemonte per l'Oncologia IRCCS | Antonino Sottile, Silvia Brossa, Paola Marino, Giorgio Giardina |
| EPI_ISL_3419088 | StarMed Healthcare | UNC Charlotte Environmental Monitoring Laboratory | Jessica Schlueter, Jannatul Ferdous, Erin Stiers, Adam France, Sam Kunkleman, Kevin Lambirth, Cynthia Gibas |
| EPI_ISL_3419089 | A.S.L. NOVARA | Fondazione del Piemonte per l'Oncologia IRCCS | Antonino Sottile, Silvia Brossa, Paola Marino, Giorgio Giardina |
| EPI_ISL_3419090 | StarMed Healthcare | UNC Charlotte Environmental Monitoring Laboratory | Jessica Schlueter, Jannatul Ferdous, Erin Stiers, Adam France, Sam Kunkleman, Kevin Lambirth, Cynthia Gibas |
| EPI_ISL_3419091 | A.S.L. NOVARA | Fondazione del Piemonte per l'Oncologia IRCCS | Antonino Sottile, Silvia Brossa, Paola Marino, Giorgio Giardina |
| EPI_ISL_3419092 | StarMed Healthcare | UNC Charlotte Environmental Monitoring Laboratory | Jessica Schlueter, Jannatul Ferdous, Erin Stiers, Adam France, Sam Kunkleman, Kevin Lambirth, Cynthia Gibas |
| EPI_ISL_3419093 | A.S.L. CITTA' DI TORINO - OSPEDALE MAURIZIANO | Fondazione del Piemonte per l'Oncologia IRCCS | Antonino Sottile, Silvia Brossa, Paola Marino, Giorgio Giardina |
| EPI_ISL_3419095, EPI_ISL_3419097 | A.S.L. NOVARA | Fondazione del Piemonte per l'Oncologia IRCCS | Antonino Sottile, Silvia Brossa, Paola Marino, Giorgio Giardina |
| EPI_ISL_3419098 | StarMed Healthcare | UNC Charlotte Environmental Monitoring Laboratory | Jessica Schlueter, Jannatul Ferdous, Erin Stiers, Adam France, Sam Kunkleman, Kevin Lambirth, Cynthia Gibas |
| EPI_ISL_3419099 | A.S.L. NOVARA | Fondazione del Piemonte per l'Oncologia IRCCS | Antonino Sottile, Silvia Brossa, Paola Marino, Giorgio Giardina |
| EPI_ISL_3419100 | StarMed Healthcare | UNC Charlotte Environmental Monitoring Laboratory | Jessica Schlueter, Jannatul Ferdous, Erin Stiers, Adam France, Sam Kunkleman, Kevin Lambirth, Cynthia Gibas |
| EPI_ISL_3419101 | A.S.L. NOVARA | Fondazione del Piemonte per l'Oncologia IRCCS | Antonino Sottile, Silvia Brossa, Paola Marino, Giorgio Giardina |
| EPI_ISL_3419102 | StarMed Healthcare | UNC Charlotte Environmental Monitoring Laboratory | Jessica Schlueter, Jannatul Ferdous, Erin Stiers, Adam France, Sam Kunkleman, Kevin Lambirth, Cynthia Gibas |
| EPI_ISL_3419103 | A.S.L. NOVARA | Fondazione del Piemonte per l'Oncologia IRCCS | Antonino Sottile, Silvia Brossa, Paola Marino, Giorgio Giardina |
| EPI_ISL_3419104 | StarMed Healthcare | UNC Charlotte Environmental Monitoring Laboratory | Jessica Schlueter, Jannatul Ferdous, Erin Stiers, Adam France, Sam Kunkleman, Kevin Lambirth, Cynthia Gibas |
| EPI_ISL_3419105, EPI_ISL_3419106 | A.S.L. NOVARA | Fondazione del Piemonte per l'Oncologia IRCCS | Antonino Sottile, Silvia Brossa, Paola Marino, Giorgio Giardina |
| EPI_ISL_3419107 | StarMed Healthcare | UNC Charlotte Environmental Monitoring Laboratory | Jessica Schlueter, Jannatul Ferdous, Erin Stiers, Adam France, Sam Kunkleman, Kevin Lambirth, Cynthia Gibas |
| EPI_ISL_3419108, EPI_ISL_3419110 | A.S.L. NOVARA | Fondazione del Piemonte per l'Oncologia IRCCS | Antonino Sottile, Silvia Brossa, Paola Marino, Giorgio Giardina |
| EPI_ISL_3419111 | StarMed Healthcare | UNC Charlotte Environmental Monitoring Laboratory | Jessica Schlueter, Jannatul Ferdous, Erin Stiers, Adam France, Sam Kunkleman, Kevin Lambirth, Cynthia Gibas |
| EPI_ISL_3419112 | A.S.L. NOVARA | Fondazione del Piemonte per l'Oncologia IRCCS | Antonino Sottile, Silvia Brossa, Paola Marino, Giorgio Giardina |
| EPI_ISL_3419113 | StarMed Healthcare | UNC Charlotte Environmental Monitoring Laboratory | Jessica Schlueter, Jannatul Ferdous, Erin Stiers, Adam France, Sam Kunkleman, Kevin Lambirth, Cynthia Gibas |
| EPI_ISL_3419114 | A.S.L. NOVARA | Fondazione del Piemonte per l'Oncologia IRCCS | Antonino Sottile, Silvia Brossa, Paola Marino, Giorgio Giardina |
| EPI_ISL_3419115 | StarMed Healthcare | UNC Charlotte Environmental Monitoring Laboratory | Jessica Schlueter, Jannatul Ferdous, Erin Stiers, Adam France, Sam Kunkleman, Kevin Lambirth, Cynthia Gibas |
| EPI_ISL_3419116 | A.S.L. NOVARA | Fondazione del Piemonte per l'Oncologia IRCCS | Antonino Sottile, Silvia Brossa, Paola Marino, Giorgio Giardina |
| EPI_ISL_3419117 | StarMed Healthcare | UNC Charlotte Environmental Monitoring Laboratory | Jessica Schlueter, Jannatul Ferdous, Erin Stiers, Adam France, Sam Kunkleman, Kevin Lambirth, Cynthia Gibas |
| EPI_ISL_3419118 | A.S.L. NOVARA | Fondazione del Piemonte per l'Oncologia IRCCS | Antonino Sottile, Silvia Brossa, Paola Marino, Giorgio Giardina |
| EPI_ISL_3419119 | StarMed Healthcare | UNC Charlotte Environmental Monitoring Laboratory | Jessica Schlueter, Jannatul Ferdous, Erin Stiers, Adam France, Sam Kunkleman, Kevin Lambirth, Cynthia Gibas |
| EPI_ISL_3419120 | A.S.L. NOVARA | Fondazione del Piemonte per l'Oncologia IRCCS | Antonino Sottile, Silvia Brossa, Paola Marino, Giorgio Giardina |
| EPI_ISL_3419121 | StarMed Healthcare | UNC Charlotte Environmental Monitoring Laboratory | Jessica Schlueter, Jannatul Ferdous, Erin Stiers, Adam France, Sam Kunkleman, Kevin Lambirth, Cynthia Gibas |
| EPI_ISL_3419122 | A.S.L. NOVARA | Fondazione del Piemonte per l'Oncologia IRCCS | Antonino Sottile, Silvia Brossa, Paola Marino, Giorgio Giardina |
| EPI_ISL_3419123 | StarMed Healthcare | UNC Charlotte Environmental Monitoring Laboratory | Jessica Schlueter, Jannatul Ferdous, Erin Stiers, Adam France, Sam Kunkleman, Kevin Lambirth, Cynthia Gibas |
| EPI_ISL_3419124 | A.S.L. NOVARA | Fondazione del Piemonte per l'Oncologia IRCCS | Antonino Sottile, Silvia Brossa, Paola Marino, Giorgio Giardina |
| EPI_ISL_3419125 | StarMed Healthcare | UNC Charlotte Environmental Monitoring Laboratory | Jessica Schlueter, Jannatul Ferdous, Erin Stiers, Adam France, Sam Kunkleman, Kevin Lambirth, Cynthia Gibas |
| EPI_ISL_3419126 | A.S.L. NOVARA | Fondazione del Piemonte per l'Oncologia IRCCS | Antonino Sottile, Silvia Brossa, Paola Marino, Giorgio Giardina |
| EPI_ISL_3419127, EPI_ISL_3419129 | StarMed Healthcare | UNC Charlotte Environmental Monitoring Laboratory | Jessica Schlueter, Jannatul Ferdous, Erin Stiers, Adam France, Sam Kunkleman, Kevin Lambirth, Cynthia Gibas |
| EPI_ISL_3419130, EPI_ISL_3419132 | A.S.L. NOVARA | Fondazione del Piemonte per l'Oncologia IRCCS | Antonino Sottile, Silvia Brossa, Paola Marino, Giorgio Giardina |
| EPI_ISL_3419133 | StarMed Healthcare | UNC Charlotte Environmental Monitoring Laboratory | Jessica Schlueter, Jannatul Ferdous, Erin Stiers, Adam France, Sam Kunkleman, Kevin Lambirth, Cynthia Gibas |
| EPI_ISL_3419134 | A.S.L. NOVARA | Fondazione del Piemonte per l'Oncologia IRCCS | Antonino Sottile, Silvia Brossa, Paola Marino, Giorgio Giardina |
| EPI_ISL_3419135 | StarMed Healthcare | UNC Charlotte Environmental Monitoring Laboratory | Jessica Schlueter, Jannatul Ferdous, Erin Stiers, Adam France, Sam Kunkleman, Kevin Lambirth, Cynthia Gibas |
| EPI_ISL_3419136, EPI_ISL_3419137 | A.S.L. NOVARA | Fondazione del Piemonte per l'Oncologia IRCCS | Antonino Sottile, Silvia Brossa, Paola Marino, Giorgio Giardina |
| EPI_ISL_3419138 | StarMed Healthcare | UNC Charlotte Environmental Monitoring Laboratory | Jessica Schlueter, Jannatul Ferdous, Erin Stiers, Adam France, Sam Kunkleman, Kevin Lambirth, Cynthia Gibas |
| EPI_ISL_3419139 | A.S.L. NOVARA | Fondazione del Piemonte per l'Oncologia IRCCS | Antonino Sottile, Silvia Brossa, Paola Marino, Giorgio Giardina |
| EPI_ISL_3419140, EPI_ISL_3419141 | StarMed Healthcare | UNC Charlotte Environmental Monitoring Laboratory | Jessica Schlueter, Jannatul Ferdous, Erin Stiers, Adam France, Sam Kunkleman, Kevin Lambirth, Cynthia Gibas |
| EPI_ISL_3419142 | A.S.L. NOVARA | Fondazione del Piemonte per l'Oncologia IRCCS | Antonino Sottile, Silvia Brossa, Paola Marino, Giorgio Giardina |
| EPI_ISL_3419143 | StarMed Healthcare | UNC Charlotte Environmental Monitoring Laboratory | Jessica Schlueter, Jannatul Ferdous, Erin Stiers, Adam France, Sam Kunkleman, Kevin Lambirth, Cynthia Gibas |
| EPI_ISL_3419144, EPI_ISL_3419145 | A.S.L. NOVARA | Fondazione del Piemonte per l'Oncologia IRCCS | Antonino Sottile, Silvia Brossa, Paola Marino, Giorgio Giardina |
| EPI_ISL_3419146 | StarMed Healthcare | UNC Charlotte Environmental Monitoring Laboratory | Jessica Schlueter, Jannatul Ferdous, Erin Stiers, Adam France, Sam Kunkleman, Kevin Lambirth, Cynthia Gibas |

| | | | |
|---|---|---|---|
| EPI_ISL_3419147 | A.S.L. NOVARA | Fondazione del Piemonte per l'Oncologia IRCCS | Antonino Sottile, Silvia Brossa, Paola Marino, Giorgio Giardina |
| EPI_ISL_3419148 | StarMed Healthcare | UNC Charlotte Environmental Monitoring Laboratory | Jessica Schlueter, Jannatul Ferdous, Erin Stiers, Adam France, Sam Kunkleman, Kevin Lambirth, Cynthia Gibas |
| EPI_ISL_3419149, EPI_ISL_3419151 | A.S.L. NOVARA | Fondazione del Piemonte per l'Oncologia IRCCS | Antonino Sottile, Silvia Brossa, Paola Marino, Giorgio Giardina |
| EPI_ISL_3419152 | StarMed Healthcare | UNC Charlotte Environmental Monitoring Laboratory | Jessica Schlueter, Jannatul Ferdous, Erin Stiers, Adam France, Sam Kunkleman, Kevin Lambirth, Cynthia Gibas |
| EPI_ISL_3419153 | A.S.L. NOVARA | Fondazione del Piemonte per l'Oncologia IRCCS | Antonino Sottile, Silvia Brossa, Paola Marino, Giorgio Giardina |
| EPI_ISL_3419154 | StarMed Healthcare | UNC Charlotte Environmental Monitoring Laboratory | Jessica Schlueter, Jannatul Ferdous, Erin Stiers, Adam France, Sam Kunkleman, Kevin Lambirth, Cynthia Gibas |
| EPI_ISL_3419155 | A.S.L. NOVARA | Fondazione del Piemonte per l'Oncologia IRCCS | Antonino Sottile, Silvia Brossa, Paola Marino, Giorgio Giardina |
| EPI_ISL_3419156 | StarMed Healthcare | UNC Charlotte Environmental Monitoring Laboratory | Jessica Schlueter, Jannatul Ferdous, Erin Stiers, Adam France, Sam Kunkleman, Kevin Lambirth, Cynthia Gibas |
| EPI_ISL_3419157, EPI_ISL_3419158 | A.S.L. NOVARA | Fondazione del Piemonte per l'Oncologia IRCCS | Antonino Sottile, Silvia Brossa, Paola Marino, Giorgio Giardina |
| EPI_ISL_3419159 | StarMed Healthcare | UNC Charlotte Environmental Monitoring Laboratory | Jessica Schlueter, Jannatul Ferdous, Erin Stiers, Adam France, Sam Kunkleman, Kevin Lambirth, Cynthia Gibas |
| EPI_ISL_3419160 | A.S.L. NOVARA | Fondazione del Piemonte per l'Oncologia IRCCS | Antonino Sottile, Silvia Brossa, Paola Marino, Giorgio Giardina |
| EPI_ISL_3419161 | StarMed Healthcare | UNC Charlotte Environmental Monitoring Laboratory | Jessica Schlueter, Jannatul Ferdous, Erin Stiers, Adam France, Sam Kunkleman, Kevin Lambirth, Cynthia Gibas |
| EPI_ISL_3419162 | A.S.L. NOVARA | Fondazione del Piemonte per l'Oncologia IRCCS | Antonino Sottile, Silvia Brossa, Paola Marino, Giorgio Giardina |
| EPI_ISL_3419163 | StarMed Healthcare | UNC Charlotte Environmental Monitoring Laboratory | Jessica Schlueter, Jannatul Ferdous, Erin Stiers, Adam France, Sam Kunkleman, Kevin Lambirth, Cynthia Gibas |
| EPI_ISL_3419164 | A.S.L. NOVARA | Fondazione del Piemonte per l'Oncologia IRCCS | Antonino Sottile, Silvia Brossa, Paola Marino, Giorgio Giardina |
| EPI_ISL_3419165 | StarMed Healthcare | UNC Charlotte Environmental Monitoring Laboratory | Jessica Schlueter, Jannatul Ferdous, Erin Stiers, Adam France, Sam Kunkleman, Kevin Lambirth, Cynthia Gibas |
| EPI_ISL_3419166 | A.S.L. NOVARA | Fondazione del Piemonte per l'Oncologia IRCCS | Antonino Sottile, Silvia Brossa, Paola Marino, Giorgio Giardina |
| EPI_ISL_3419167 | StarMed Healthcare | UNC Charlotte Environmental Monitoring Laboratory | Jessica Schlueter, Jannatul Ferdous, Erin Stiers, Adam France, Sam Kunkleman, Kevin Lambirth, Cynthia Gibas |
| EPI_ISL_3419168 | A.S.L. NOVARA | Fondazione del Piemonte per l'Oncologia IRCCS | Antonino Sottile, Silvia Brossa, Paola Marino, Giorgio Giardina |
| EPI_ISL_3419169 | StarMed Healthcare | UNC Charlotte Environmental Monitoring Laboratory | Jessica Schlueter, Jannatul Ferdous, Erin Stiers, Adam France, Sam Kunkleman, Kevin Lambirth, Cynthia Gibas |
| EPI_ISL_3419170 | A.S.L. NOVARA | Fondazione del Piemonte per l'Oncologia IRCCS | Antonino Sottile, Silvia Brossa, Paola Marino, Giorgio Giardina |
| EPI_ISL_3419171 | StarMed Healthcare | UNC Charlotte Environmental Monitoring Laboratory | Jessica Schlueter, Jannatul Ferdous, Erin Stiers, Adam France, Sam Kunkleman, Kevin Lambirth, Cynthia Gibas |
| EPI_ISL_3419173 | A.S.L. NOVARA | Fondazione del Piemonte per l'Oncologia IRCCS | Antonino Sottile, Silvia Brossa, Paola Marino, Giorgio Giardina |
| EPI_ISL_3419174 | StarMed Healthcare | UNC Charlotte Environmental Monitoring Laboratory | Jessica Schlueter, Jannatul Ferdous, Erin Stiers, Adam France, Sam Kunkleman, Kevin Lambirth, Cynthia Gibas |
| EPI_ISL_3419175 | A.S.L. NOVARA | Fondazione del Piemonte per l'Oncologia IRCCS | Antonino Sottile, Silvia Brossa, Paola Marino, Giorgio Giardina |
| EPI_ISL_3419176 | StarMed Healthcare | UNC Charlotte Environmental Monitoring Laboratory | Jessica Schlueter, Jannatul Ferdous, Erin Stiers, Adam France, Sam Kunkleman, Kevin Lambirth, Cynthia Gibas |
| EPI_ISL_3419177 | A.S.L. NOVARA | Fondazione del Piemonte per l'Oncologia IRCCS | Antonino Sottile, Silvia Brossa, Paola Marino, Giorgio Giardina |
| EPI_ISL_3419178 | StarMed Healthcare | UNC Charlotte Environmental Monitoring Laboratory | Jessica Schlueter, Jannatul Ferdous, Erin Stiers, Adam France, Sam Kunkleman, Kevin Lambirth, Cynthia Gibas |
| EPI_ISL_3419179 | A.S.L. NOVARA | Fondazione del Piemonte per l'Oncologia IRCCS | Antonino Sottile, Silvia Brossa, Paola Marino, Giorgio Giardina |
| EPI_ISL_3419180 | StarMed Healthcare | UNC Charlotte Environmental Monitoring Laboratory | Jessica Schlueter, Jannatul Ferdous, Erin Stiers, Adam France, Sam Kunkleman, Kevin Lambirth, Cynthia Gibas |
| EPI_ISL_3419181 | A.S.L. NOVARA | Fondazione del Piemonte per l'Oncologia IRCCS | Antonino Sottile, Silvia Brossa, Paola Marino, Giorgio Giardina |
| EPI_ISL_3419182 | StarMed Healthcare | UNC Charlotte Environmental Monitoring Laboratory | Jessica Schlueter, Jannatul Ferdous, Erin Stiers, Adam France, Sam Kunkleman, Kevin Lambirth, Cynthia Gibas |
| EPI_ISL_3419183, EPI_ISL_3419184 | A.S.L. NOVARA | Fondazione del Piemonte per l'Oncologia IRCCS | Antonino Sottile, Silvia Brossa, Paola Marino, Giorgio Giardina |
| EPI_ISL_3419185 | StarMed Healthcare | UNC Charlotte Environmental Monitoring Laboratory | Jessica Schlueter, Jannatul Ferdous, Erin Stiers, Adam France, Sam Kunkleman, Kevin Lambirth, Cynthia Gibas |
| EPI_ISL_3419186 | A.S.L. NOVARA | Fondazione del Piemonte per l'Oncologia IRCCS | Antonino Sottile, Silvia Brossa, Paola Marino, Giorgio Giardina |
| EPI_ISL_3419187 | StarMed Healthcare | UNC Charlotte Environmental Monitoring Laboratory | Jessica Schlueter, Jannatul Ferdous, Erin Stiers, Adam France, Sam Kunkleman, Kevin Lambirth, Cynthia Gibas |
| EPI_ISL_3419188 | A.S.L. NOVARA | Fondazione del Piemonte per l'Oncologia IRCCS | Antonino Sottile, Silvia Brossa, Paola Marino, Giorgio Giardina |
| EPI_ISL_3419189 | StarMed Healthcare | UNC Charlotte Environmental Monitoring Laboratory | Jessica Schlueter, Jannatul Ferdous, Erin Stiers, Adam France, Sam Kunkleman, Kevin Lambirth, Cynthia Gibas |
| EPI_ISL_3419190 | A.S.L. NOVARA | Fondazione del Piemonte per l'Oncologia IRCCS | Antonino Sottile, Silvia Brossa, Paola Marino, Giorgio Giardina |
| EPI_ISL_3419191 | StarMed Healthcare | UNC Charlotte Environmental Monitoring Laboratory | Jessica Schlueter, Jannatul Ferdous, Erin Stiers, Adam France, Sam Kunkleman, Kevin Lambirth, Cynthia Gibas |
| EPI_ISL_3419192 | A.S.L. NOVARA | Fondazione del Piemonte per l'Oncologia IRCCS | Antonino Sottile, Silvia Brossa, Paola Marino, Giorgio Giardina |
| EPI_ISL_3419193 | StarMed Healthcare | UNC Charlotte Environmental Monitoring Laboratory | Jessica Schlueter, Jannatul Ferdous, Erin Stiers, Adam France, Sam Kunkleman, Kevin Lambirth, Cynthia Gibas |
| EPI_ISL_3419194 | A.S.L. NOVARA | Fondazione del Piemonte per l'Oncologia IRCCS | Antonino Sottile, Silvia Brossa, Paola Marino, Giorgio Giardina |
| EPI_ISL_3419195 | StarMed Healthcare | UNC Charlotte Environmental Monitoring Laboratory | Jessica Schlueter, Jannatul Ferdous, Erin Stiers, Adam France, Sam Kunkleman, Kevin Lambirth, Cynthia Gibas |
| EPI_ISL_3419196 | A.S.L. NOVARA | Fondazione del Piemonte per l'Oncologia IRCCS | Antonino Sottile, Silvia Brossa, Paola Marino, Giorgio Giardina |
| EPI_ISL_3419197 | StarMed Healthcare | UNC Charlotte Environmental Monitoring Laboratory | Jessica Schlueter, Jannatul Ferdous, Erin Stiers, Adam France, Sam Kunkleman, Kevin Lambirth, Cynthia Gibas |
| EPI_ISL_3419198 | A.S.L. NOVARA | Fondazione del Piemonte per l'Oncologia IRCCS | Antonino Sottile, Silvia Brossa, Paola Marino, Giorgio Giardina |
| EPI_ISL_3419199 | StarMed Healthcare | UNC Charlotte Environmental Monitoring Laboratory | Jessica Schlueter, Jannatul Ferdous, Erin Stiers, Adam France, Sam Kunkleman, Kevin Lambirth, Cynthia Gibas |
| EPI_ISL_3419200 | A.S.L. NOVARA | Fondazione del Piemonte per l'Oncologia IRCCS | Antonino Sottile, Silvia Brossa, Paola Marino, Giorgio Giardina |
| EPI_ISL_3419201 | StarMed Healthcare | UNC Charlotte Environmental Monitoring Laboratory | Jessica Schlueter, Jannatul Ferdous, Erin Stiers, Adam France, Sam Kunkleman, Kevin Lambirth, Cynthia Gibas |
| EPI_ISL_3419202 | A.S.L. NOVARA | Fondazione del Piemonte per l'Oncologia IRCCS | Antonino Sottile, Silvia Brossa, Paola Marino, Giorgio Giardina |
| EPI_ISL_3419203 | StarMed Healthcare | UNC Charlotte Environmental Monitoring Laboratory | Jessica Schlueter, Jannatul Ferdous, Erin Stiers, Adam France, Sam Kunkleman, Kevin Lambirth, Cynthia Gibas |
| EPI_ISL_3419204 | A.S.L. NOVARA | Fondazione del Piemonte per l'Oncologia IRCCS | Antonino Sottile, Silvia Brossa, Paola Marino, Giorgio Giardina |
| EPI_ISL_3419205 | StarMed Healthcare | UNC Charlotte Environmental Monitoring Laboratory | Jessica Schlueter, Jannatul Ferdous, Erin Stiers, Adam France, Sam Kunkleman, Kevin Lambirth, Cynthia Gibas |
| EPI_ISL_3419206 | A.S.L. NOVARA | Fondazione del Piemonte per l'Oncologia IRCCS | Antonino Sottile, Silvia Brossa, Paola Marino, Giorgio Giardina |
| EPI_ISL_3419207 | StarMed Healthcare | UNC Charlotte Environmental Monitoring Laboratory | Jessica Schlueter, Jannatul Ferdous, Erin Stiers, Adam France, Sam Kunkleman, Kevin Lambirth, Cynthia Gibas |
| EPI_ISL_3419208, EPI_ISL_3419209 | A.S.L. NOVARA | Fondazione del Piemonte per l'Oncologia IRCCS | Antonino Sottile, Silvia Brossa, Paola Marino, Giorgio Giardina |
| EPI_ISL_3419210 | StarMed Healthcare | UNC Charlotte Environmental Monitoring Laboratory | Jessica Schlueter, Jannatul Ferdous, Erin Stiers, Adam France, Sam Kunkleman, Kevin Lambirth, Cynthia Gibas |
| EPI_ISL_3419211 | A.S.L.NOVARA-BORGOMANERO | Fondazione del Piemonte per l'Oncologia IRCCS | Antonino Sottile, Silvia Brossa, Paola Marino, Giorgio Giardina |
| EPI_ISL_3419212 | StarMed Healthcare | UNC Charlotte Environmental Monitoring Laboratory | Jessica Schlueter, Jannatul Ferdous, Erin Stiers, Adam France, Sam Kunkleman, Kevin Lambirth, Cynthia Gibas |
| EPI_ISL_3419213 | A.S.L. CUNEO 1 | Fondazione del Piemonte per l'Oncologia IRCCS | Antonino Sottile, Silvia Brossa, Paola Marino, Giorgio Giardina |
| EPI_ISL_3419214, EPI_ISL_3419215 | StarMed Healthcare | UNC Charlotte Environmental Monitoring Laboratory | Jessica Schlueter, Jannatul Ferdous, Erin Stiers, Adam France, Sam Kunkleman, Kevin Lambirth, Cynthia Gibas |
| EPI_ISL_3419216 | A.S.L. VERCELLI | Fondazione del Piemonte per l'Oncologia IRCCS | Antonino Sottile, Silvia Brossa, Paola Marino, Giorgio Giardina |
| EPI_ISL_3419217 | StarMed Healthcare | UNC Charlotte Environmental Monitoring Laboratory | Jessica Schlueter, Jannatul Ferdous, Erin Stiers, Adam France, Sam Kunkleman, Kevin Lambirth, Cynthia Gibas |
| EPI_ISL_3419218 | A.S.L. VERCELLI | Fondazione del Piemonte per l'Oncologia IRCCS | Antonino Sottile, Silvia Brossa, Paola Marino, Giorgio Giardina |
| EPI_ISL_3419219 | StarMed Healthcare | UNC Charlotte Environmental Monitoring Laboratory | Jessica Schlueter, Jannatul Ferdous, Erin Stiers, Adam France, Sam Kunkleman, Kevin Lambirth, Cynthia Gibas |
| EPI_ISL_3419220 | A.S.L. VERCELLI | Fondazione del Piemonte per l'Oncologia IRCCS | Antonino Sottile, Silvia Brossa, Paola Marino, Giorgio Giardina |
| EPI_ISL_3419222, EPI_ISL_3419223 | A.S.L. NOVARA | Fondazione del Piemonte per l'Oncologia IRCCS | Antonino Sottile, Silvia Brossa, Paola Marino, Giorgio Giardina |
| EPI_ISL_3419224 | StarMed Healthcare | UNC Charlotte Environmental Monitoring Laboratory | Jessica Schlueter, Jannatul Ferdous, Erin Stiers, Adam France, Sam Kunkleman, Kevin Lambirth, Cynthia Gibas |
| EPI_ISL_3419225 | A.S.L. CUNEO 1 | Fondazione del Piemonte per l'Oncologia IRCCS | Antonino Sottile, Silvia Brossa, Paola Marino, Giorgio Giardina |
| EPI_ISL_3419226 | StarMed Healthcare | UNC Charlotte Environmental Monitoring Laboratory | Jessica Schlueter, Jannatul Ferdous, Erin Stiers, Adam France, Sam Kunkleman, Kevin Lambirth, Cynthia Gibas |
| EPI_ISL_3419227 | A.S.L. NOVARA | Fondazione del Piemonte per l'Oncologia IRCCS | Antonino Sottile, Silvia Brossa, Paola Marino, Giorgio Giardina |

| | | | |
|---|---|---|---|
| EPI_ISL_3419228 | StarMed Healthcare | UNC Charlotte Environmental Monitoring Laboratory | Jessica Schlueter, Jannatul Ferdous, Erin Stiers, Adam France, Sam Kunkleman, Kevin Lambirth, Cynthia Gibas |
| EPI_ISL_3419229 | A.S.L. NOVARA | Fondazione del Piemonte per l'Oncologia IRCCS | Antonino Sottile, Silvia Brossa, Paola Marino, Giorgio Giardina |
| EPI_ISL_3419230, EPI_ISL_3419231, EPI_ISL_3419232, EPI_ISL_3419233, EPI_ISL_3419234, EPI_ISL_3419235, EPI_ISL_3419236, EPI_ISL_3419237, EPI_ISL_3419238, EPI_ISL_3419239, EPI_ISL_3419240, EPI_ISL_3419242, EPI_ISL_3419243, EPI_ISL_3419244, EPI_ISL_3419245, EPI_ISL_3419246, EPI_ISL_3419247, EPI_ISL_3419248, EPI_ISL_3419249, EPI_ISL_3419250, EPI_ISL_3419251, EPI_ISL_3419252, EPI_ISL_3419253, EPI_ISL_3419254, EPI_ISL_3419255, EPI_ISL_3419256, EPI_ISL_3419257, EPI_ISL_3419258, EPI_ISL_3419259, EPI_ISL_3419260, EPI_ISL_3419261, EPI_ISL_3419262, EPI_ISL_3419263, EPI_ISL_3419264, EPI_ISL_3419265, EPI_ISL_3419266, EPI_ISL_3419267, EPI_ISL_3419268, EPI_ISL_3419269, EPI_ISL_3419270, EPI_ISL_3419271, EPI_ISL_3419272, EPI_ISL_3419273, EPI_ISL_3419274, EPI_ISL_3419275, EPI_ISL_3419276, EPI_ISL_3419277, EPI_ISL_3419278, EPI_ISL_3419279, EPI_ISL_3419280, EPI_ISL_3419281, EPI_ISL_3419282, EPI_ISL_3419283, EPI_ISL_3419284, EPI_ISL_3419285, EPI_ISL_3419286, EPI_ISL_3419288, EPI_ISL_3419289, EPI_ISL_3419290, EPI_ISL_3419291, EPI_ISL_3419292, EPI_ISL_3419293, EPI_ISL_3419294, EPI_ISL_3419295, EPI_ISL_3419296, EPI_ISL_3419297, EPI_ISL_3419298, EPI_ISL_3419299, EPI_ISL_3419300, EPI_ISL_3419301, EPI_ISL_3419302, EPI_ISL_3419303, EPI_ISL_3419304, EPI_ISL_3419305, EPI_ISL_3419306, EPI_ISL_3419307, EPI_ISL_3419308, EPI_ISL_3419309, EPI_ISL_3419310, EPI_ISL_3419311, EPI_ISL_3419312, EPI_ISL_3419313, EPI_ISL_3419314, EPI_ISL_3419315 | | | |
| see above | StarMed Healthcare | UNC Charlotte Environmental Monitoring Laboratory | Jessica Schlueter, Jannatul Ferdous, Erin Stiers, Adam France, Sam Kunkleman, Kevin Lambirth, Cynthia Gibas |
| EPI_ISL_3419316 | Regional Medical Sciences Center 6 Chonburi | National Institute of Health, Department of Medical Sciences, Ministry of Public Health, Thailand | Pilailuk Okada; Siripaporn Phuygun; Nuttida Thongpramul; Sittiporn Parnmen; Ratana Tacharoenmuang; Pakorn Piromtong; Natchaya Khiadsang; Thanutsapa Thanadachakul; Warawan Wongboot; sirikanda wimol; Sunthareeya Waicharoen; |
| EPI_ISL_3419317 | Regional Medical Sciences Center 9 Nakhon Ratchasima | National Institute of Health, Department of Medical Sciences, Ministry of Public Health, Thailand | Pilailuk Okada; Siripaporn Phuygun; Nuttida Thongpramul; Sittiporn Parnmen; Ratana Tacharoenmuang; Pakorn Piromtong; Natchaya Khiadsang; Thanutsapa Thanadachakul; Warawan Wongboot; sirikanda wimol; Sunthareeya Waicharoen; |
| EPI_ISL_3419318 | Regional Medical Sciences Center 5 Samut Songkhram | National Institute of Health, Department of Medical Sciences, Ministry of Public Health, Thailand | Pilailuk Okada; Siripaporn Phuygun; Nuttida Thongpramul; Sittiporn Parnmen; Ratana Tacharoenmuang; Pakorn Piromtong; Natchaya Khiadsang; Thanutsapa Thanadachakul; Warawan Wongboot; sirikanda wimol; Sunthareeya Waicharoen; |
| EPI_ISL_3419319 | Microbiology Department - University Hospital Brussel | Microbiology Department - University Hospital Brussel | Oriane Soetens, Thomas Demuyser |
| EPI_ISL_3419331, EPI_ISL_3419333 | Dr. Yoshitaka Tamura Department of Clinical Laboratory, Osaka Habikino Medical Center | Department of Developmental Medicine, Research Institute, Osaka Women's and Children's Hospital | Itaru Yanagihara, Yukiko Nakura |
| EPI_ISL_3419335 | MCL Med. Laboratorien AG | Institute for Infectious Diseases | Stefan Neuenschwander, Christian Baumann, Miguel A Terrazos Miani, Cora SÃ¤gesser, Pascal Bittel, Peter Keller, Franziska Suter-Riniker, Stephen L Leib, Alban Ramette |
| EPI_ISL_3419336, EPI_ISL_3419337 | Institute for Infectious Diseases | Institute for Infectious Diseases | Stefan Neuenschwander, Christian Baumann, Miguel A Terrazos Miani, Cora SÃ¤gesser, Pascal Bittel, Peter Keller, Franziska Suter-Riniker, Stephen L Leib, Alban Ramette |
| EPI_ISL_3419339, EPI_ISL_3419340, EPI_ISL_3419341 | Bakt. Institut Olten Bio AG | Institute for Infectious Diseases | Stefan Neuenschwander, Christian Baumann, Miguel A Terrazos Miani, Cora SÃ¤gesser, Pascal Bittel, Peter Keller, Franziska Suter-Riniker, Stephen L Leib, Alban Ramette |
| EPI_ISL_3419342 | Institute for Infectious Diseases | Institute for Infectious Diseases | Stefan Neuenschwander, Christian Baumann, Miguel A Terrazos Miani, Cora SÃ¤gesser, Pascal Bittel, Peter Keller, Franziska Suter-Riniker, Stephen L Leib, Alban Ramette |
| EPI_ISL_3419343 | Bio Analytica AG | Institute for Infectious Diseases | Stefan Neuenschwander, Christian Baumann, Miguel A Terrazos Miani, Cora SÃ¤gesser, Pascal Bittel, Peter Keller, Franziska Suter-Riniker, Stephen L Leib, Alban Ramette |
| EPI_ISL_3419344 | Luzerner Kantonsspital | Institute for Infectious Diseases | Stefan Neuenschwander, Christian Baumann, Miguel A Terrazos Miani, Cora SÃ¤gesser, Pascal Bittel, Peter Keller, Franziska Suter-Riniker, Stephen L Leib, Alban Ramette |
| EPI_ISL_3419345 | MCL Med. Laboratorien AG | Institute for Infectious Diseases | Stefan Neuenschwander, Christian Baumann, Miguel A Terrazos Miani, Cora SÃ¤gesser, Pascal Bittel, Peter Keller, Franziska Suter-Riniker, Stephen L Leib, Alban Ramette |
| EPI_ISL_3419346, EPI_ISL_3419347 | Institute for Infectious Diseases | Institute for Infectious Diseases | Stefan Neuenschwander, Christian Baumann, Miguel A Terrazos Miani, Cora SÃ¤gesser, Pascal Bittel, Peter Keller, Franziska Suter-Riniker, Stephen L Leib, Alban Ramette |
| EPI_ISL_3419348, EPI_ISL_3419349 | Bakt. Institut Olten Bio AG | Institute for Infectious Diseases | Stefan Neuenschwander, Christian Baumann, Miguel A Terrazos Miani, Cora SÃ¤gesser, Pascal Bittel, Peter Keller, Franziska Suter-Riniker, Stephen L Leib, Alban Ramette |
| EPI_ISL_3419350 | Bio Analytica AG | Institute for Infectious Diseases | Stefan Neuenschwander, Christian Baumann, Miguel A Terrazos Miani, Cora SÃ¤gesser, Pascal Bittel, Peter Keller, Franziska Suter-Riniker, Stephen L Leib, Alban Ramette |
| EPI_ISL_3419351, EPI_ISL_3419352 | Luzerner Kantonsspital | Institute for Infectious Diseases | Stefan Neuenschwander, Christian Baumann, Miguel A Terrazos Miani, Cora SÃ¤gesser, Pascal Bittel, Peter Keller, Franziska Suter-Riniker, Stephen L Leib, Alban Ramette |
| EPI_ISL_3419353, EPI_ISL_3419354 | MCL Med. Laboratorien AG | Institute for Infectious Diseases | Stefan Neuenschwander, Christian Baumann, Miguel A Terrazos Miani, Cora SÃ¤gesser, Pascal Bittel, Peter Keller, Franziska Suter-Riniker, Stephen L Leib, Alban Ramette |
| EPI_ISL_3419355, EPI_ISL_3419356, EPI_ISL_3419357, EPI_ISL_3419358, EPI_ISL_3419359 | Bakt. Institut Olten Bio AG | Institute for Infectious Diseases | Stefan Neuenschwander, Christian Baumann, Miguel A Terrazos Miani, Cora SÃ¤gesser, Pascal Bittel, Peter Keller, Franziska Suter-Riniker, Stephen L Leib, Alban Ramette |
| EPI_ISL_3419360, EPI_ISL_3419361, EPI_ISL_3419362 | Institute for Infectious Diseases | Institute for Infectious Diseases | Stefan Neuenschwander, Christian Baumann, Miguel A Terrazos Miani, Cora SÃ¤gesser, Pascal Bittel, Peter Keller, Franziska Suter-Riniker, Stephen L Leib, Alban Ramette |
| EPI_ISL_3419363 | RSUP Dr. Hasan Sadikin Bandung | Genomik Solidaritas Indonesia Laboratorium | Anna Christina Brazia, Reinhart Gabriel, Vania Gavrila Wikasa, Carissa Sintca Wijaya, Dhahlia Agustina Cahyono, Alfin Mohammad Abdillah, Normastuti Adhini Tantyo, Annisa Muthiah Sukirman, Gracia Felias Enos Korompis, Louisa Markus, Basti Andriyoko, Delita Prihatni, Dewi Kartika Turbawati, Vivi Setyawati, Anuraj Shankar, Meutia Ayuputeri Kumaheri, Ariel Pradipta |
| EPI_ISL_3419364 | Dr. Yoshitaka Tamura Department of Clinical Laboratory, Osaka Habikino Medical Center | Department of Developmental Medicine, Research Institute, Osaka Women's and Children's Hospital | Itaru Yanagihara, Yukiko Nakura |
| EPI_ISL_3419365 | Microbiology Department - University Hospital Brussel | Microbiology Department - University Hospital Brussel | Oriane Soetens, Thomas Demuyser |
| EPI_ISL_3419369 | CAP El Remei | Banc de Sang i Teixits | Francisco Vidal, Irene Corrales, Nina Borràs, Carlos Hobeich, Natália Comes, Lorena Ramírez, Noemí Gonzalez, Silvia Sauleda, Maria Glòria Soria |
| EPI_ISL_3419370 | Microbiology Department - University Hospital Brussel | Microbiology Department - University Hospital Brussel | Oriane Soetens, Thomas Demuyser |
| EPI_ISL_3419372, EPI_ISL_3419373, EPI_ISL_3419374, EPI_ISL_3419375, EPI_ISL_3419376, EPI_ISL_3419377, EPI_ISL_3419378, EPI_ISL_3419379, EPI_ISL_3419380, EPI_ISL_3419381, EPI_ISL_3419382, EPI_ISL_3419383, EPI_ISL_3419384, EPI_ISL_3419385, EPI_ISL_3419386, EPI_ISL_3419387, EPI_ISL_3419388, EPI_ISL_3419389 | | | |
| see above | Institute for Urban Disease Control and Prevention | Division of Genomic Medicine and Innovation support,Department of Medical Sciences, Ministry of Public Health, Thailand | Surakameth Mahasirimongkol,Nuanjun Wichukchinda,Archawin Rojanawiwat,Pilailuk Akkapaiboon Okada ,Waritta Sawaengdee,Penpitcha Thawong,Pundharika Piboonsiri,Jirapha Pakdee,Natthakul Bunneang |
| EPI_ISL_3419391, EPI_ISL_3419392 | Regional Medical Sciences Center 11 Surat Thani | National Institute of Health, Department of Medical Sciences, Ministry of Public Health, Thailand | Pilailuk Okada; Siripaporn Phuygun; Nuttida Thongpramul; Sittiporn Parnmen; Ratana Tacharoenmuang; Pakorn Piromtong; Natchaya Khiadsang; Thanutsapa Thanadachakul; Warawan Wongboot; sirikanda wimol; Sunthareeya Waicharoen; |
| EPI_ISL_3419393 | Chaiyaphum Hospital | National Institute of Health, Department of Medical Sciences, Ministry of Public Health, Thailand | Pilailuk Okada; Siripaporn Phuygun; Nuttida Thongpramul; Sittiporn Parnmen; Ratana Tacharoenmuang; Pakorn Piromtong; Natchaya Khiadsang; Thanutsapa Thanadachakul; Warawan Wongboot; sirikanda wimol; Sunthareeya Waicharoen; |
| EPI_ISL_3419395, EPI_ISL_3419397 | Regional Medical Sciences Center 6 Chonburi | National Institute of Health, Department of Medical Sciences, Ministry of Public Health, Thailand | Pilailuk Okada; Siripaporn Phuygun; Nuttida Thongpramul; Sittiporn Parnmen; Ratana Tacharoenmuang; Pakorn Piromtong; Natchaya Khiadsang; Thanutsapa Thanadachakul; Warawan Wongboot; sirikanda wimol; Sunthareeya Waicharoen; |
| EPI_ISL_3419398, EPI_ISL_3419399, EPI_ISL_3419400, EPI_ISL_3419401, EPI_ISL_3419402, EPI_ISL_3419403, EPI_ISL_3419404, EPI_ISL_3419405, EPI_ISL_3419406, EPI_ISL_3419407, EPI_ISL_3419408, EPI_ISL_3419409, EPI_ISL_3419410, EPI_ISL_3419411, EPI_ISL_3419412 | | | |
| see above | Microbiology Department, Laboratori Clínic Metropolitana Nord. Hospital Universitari Germans Trias i Pujol | Can Ruti SARS-CoV-2 Sequencing Hub (HUGTiP/IrsiCaixa/IGTP) | Marc Noguera-Julian, Pilar Armengol, Ignacio Blanco, Antoni E Bordoy, Francesc Catala-Moll, Pere-Joan Cardona, Maria Casadellà, Cristina Casañ, Gemma Clara, Bonaventura Clotet, Montserrat Giménez, Mercedes Guerrero, Anna Not, David Panisello, Roger Paredes, Mariona Parera, Alexia París, Verónica Saludes, Alba Sánchez, Laia Soler, Lauro Sumoy, and Elisa Martró on behalf of the Can Ruti SARS-CoV-2 Sequencing Hub |
| EPI_ISL_3419413 | Microbiologia CATLAB | Can Ruti SARS-CoV-2 Sequencing Hub (HUGTiP/IrsiCaixa/IGTP) | Marc Noguera-Julian, Pilar Armengol, Ignacio Blanco, Antoni E Bordoy, Francesc Catala-Moll, Pere-Joan Cardona, Maria Casadellà, Cristina Casañ, Gemma Clara, Bonaventura Clotet, Montserrat Giménez, Mercedes Guerrero, Anna Not, David Panisello, Roger Paredes, Mariona Parera, Alexia París, Verónica Saludes, Alba Sánchez, Laia Soler, Lauro Sumoy, and Elisa Martró on behalf of the Can Ruti SARS-CoV-2 Sequencing Hub |
| EPI_ISL_3419414, EPI_ISL_3419415, EPI_ISL_3419416, EPI_ISL_3419417, EPI_ISL_3419418, EPI_ISL_3419419, EPI_ISL_3419420, EPI_ISL_3419421, EPI_ISL_3419422, EPI_ISL_3419423, EPI_ISL_3419424, EPI_ISL_3419425, EPI_ISL_3419426, EPI_ISL_3419427, EPI_ISL_3419428, EPI_ISL_3419429, EPI_ISL_3419430, EPI_ISL_3419431, EPI_ISL_3419432, EPI_ISL_3419433, EPI_ISL_3419434, EPI_ISL_3419435, EPI_ISL_3419436, EPI_ISL_3419437, EPI_ISL_3419438, EPI_ISL_3419439, EPI_ISL_3419440, EPI_ISL_3419441, EPI_ISL_3419442, EPI_ISL_3419443, EPI_ISL_3419444, EPI_ISL_3419446, EPI_ISL_3419447, EPI_ISL_3419448, EPI_ISL_3419449, EPI_ISL_3419450, EPI_ISL_3419451, EPI_ISL_3419452, EPI_ISL_3419454, EPI_ISL_3419455, EPI_ISL_3419456, EPI_ISL_3419457, EPI_ISL_3419458, EPI_ISL_3419459, EPI_ISL_3419460 | | | |
| see above | Microbiology Department, Laboratori Clínic Metropolitana Nord. Hospital Universitari Germans Trias i Pujol | Can Ruti SARS-CoV-2 Sequencing Hub (HUGTiP/IrsiCaixa/IGTP) | Marc Noguera-Julian, Pilar Armengol, Ignacio Blanco, Antoni E Bordoy, Francesc Catala-Moll, Pere-Joan Cardona, Maria Casadellà, Cristina Casañ, Gemma Clara, Bonaventura Clotet, Montserrat Giménez, Mercedes Guerrero, Anna Not, David Panisello, Roger Paredes, Mariona Parera, Alexia París, Verónica Saludes, Alba Sánchez, Laia Soler, Lauro Sumoy, and Elisa Martró on behalf of the Can Ruti SARS-CoV-2 Sequencing Hub |
| EPI_ISL_3419461 | Microbiologia CATLAB | Can Ruti SARS-CoV-2 Sequencing Hub (HUGTiP/IrsiCaixa/IGTP) | Marc Noguera-Julian, Pilar Armengol, Ignacio Blanco, Antoni E Bordoy, Francesc Catala-Moll, Pere-Joan Cardona, Maria Casadellà, Cristina Casañ, Gemma Clara, Bonaventura Clotet, Montserrat Giménez, Mercedes Guerrero, Anna Not, David Panisello, Roger Paredes, Mariona Parera, Alexia París, Verónica Saludes, Alba Sánchez, Laia Soler, Lauro Sumoy, and Elisa Martró on behalf of the Can Ruti SARS-CoV-2 Sequencing Hub |
| EPI_ISL_3419462, EPI_ISL_3419463, EPI_ISL_3419464, EPI_ISL_3419465, EPI_ISL_3419466, EPI_ISL_3419467, EPI_ISL_3419468, EPI_ISL_3419469, EPI_ISL_3419470, EPI_ISL_3419471, EPI_ISL_3419472, EPI_ISL_3419473, EPI_ISL_3419474, EPI_ISL_3419475, EPI_ISL_3419476, EPI_ISL_3419477, EPI_ISL_3419478, EPI_ISL_3419479, EPI_ISL_3419481, EPI_ISL_3419482, EPI_ISL_3419483, EPI_ISL_3419484, EPI_ISL_3419485, EPI_ISL_3419486, EPI_ISL_3419487, EPI_ISL_3419488, EPI_ISL_3419489, EPI_ISL_3419490, EPI_ISL_3419491 | | | |
| see above | Microbiology Department, Laboratori Clínic Metropolitana Nord. Hospital Universitari Germans Trias i Pujol | Can Ruti SARS-CoV-2 Sequencing Hub (HUGTiP/IrsiCaixa/IGTP) | Marc Noguera-Julian, Pilar Armengol, Ignacio Blanco, Antoni E Bordoy, Francesc Catala-Moll, Pere-Joan Cardona, Maria Casadellà, Cristina Casañ, Gemma Clara, Bonaventura Clotet, Montserrat Giménez, Mercedes Guerrero, Anna Not, David Panisello, Roger Paredes, Mariona Parera, Alexia París, Verónica Saludes, Alba Sánchez, Laia Soler, Lauro Sumoy, and Elisa Martró on behalf of the Can Ruti SARS-CoV-2 Sequencing Hub |
| EPI_ISL_3419577, EPI_ISL_3419595, EPI_ISL_3419596, EPI_ISL_3419598, EPI_ISL_3419599, EPI_ISL_3419608, EPI_ISL_3419615, EPI_ISL_3419632, EPI_ISL_3419633, EPI_ISL_3419634, EPI_ISL_3419635, EPI_ISL_3419638, EPI_ISL_3419640, EPI_ISL_3419645, EPI_ISL_3419647, EPI_ISL_3419648, EPI_ISL_3419649, EPI_ISL_3419652, EPI_ISL_3419653, EPI_ISL_3419654, EPI_ISL_3419656, EPI_ISL_3419657, EPI_ISL_3419658, EPI_ISL_3419659, EPI_ISL_3419660, EPI_ISL_3419661, EPI_ISL_3419662, EPI_ISL_3419664, EPI_ISL_3419666, EPI_ISL_3419667, EPI_ISL_3419668, EPI_ISL_3419669, EPI_ISL_3419671, EPI_ISL_3419672, EPI_ISL_3419673, EPI_ISL_3419675, EPI_ISL_3419676, EPI_ISL_3419677, EPI_ISL_3419678, EPI_ISL_3419679, EPI_ISL_3419680, EPI_ISL_3419681, EPI_ISL_3419682, EPI_ISL_3419685, EPI_ISL_3419686, EPI_ISL_3419687, EPI_ISL_3419688, EPI_ISL_3419689, EPI_ISL_3419691, EPI_ISL_3419692, EPI_ISL_3419693, EPI_ISL_3419694, EPI_ISL_3419695, EPI_ISL_3419696, EPI_ISL_3419697, EPI_ISL_3419698, EPI_ISL_3419700, EPI_ISL_3419701, EPI_ISL_3419702 | | | |
| see above | UC Davis Genome Center and Healthy Davis Together | UC Davis Genome Center | UC Davis |
| EPI_ISL_3419704, EPI_ISL_3419706 | Osaka General Medical Center | Department of Developmental Medicine, Research Institute, Osaka Women's and Children's Hospital | Itaru Yanagihara, Yukiko Nakura |
| EPI_ISL_3419708 | Department of clinical lab., Osaka Women's and Children's Hospital | Department of Developmental Medicine, Research Institute, Osaka Women's and Children's Hospital | Itaru Yanagihara, Yukiko Nakura, Satoshi Yanagihara |
| EPI_ISL_3419710 | Johor Bahru Public Health Laboratory | Institute for Medical Research, Infectious Disease Research Centre, National Institutes of Health, Ministry of Health Malaysia | Suppiah J, Kamel K, Mohd Zawawi Z, Azizan MA, Ramly N, Robert F, Anasir MI, Thayan R |
| EPI_ISL_3419711, EPI_ISL_3419712 | Dr. Yoshitaka Tamura Department of Clinical Laboratory, Osaka Habikino Medical Center | Department of Developmental Medicine, Research Institute, Osaka Women's and Children's Hospital | Itaru Yanagihara, Yukiko Nakura |
| EPI_ISL_3419713 | IRCCS San Gallicano Dermatological Institute | IRCCS Regina Elena National Cancer Institute | Sabrina Strano, Sara Donzelli, Matteo Pallocca, Grazia Prignano, Fabrizio Ensoli, Fulvia Pimpinelli, Maurizio Fanciulli, Giovanni Blandino, Aldo Morrone, Gennaro Ciliberto |
| EPI_ISL_3419714 | IRCCS San Gallicano Dermatological Institute | IRCCS Regina Elena National Cancer Institute | Francesca De Nicola, Giulia Orlandi, Eleonora Sperandio, Elisabetta Trento, Fabrizio Ensoli, Fulvia Pimpinelli, Maurizio Fanciulli, Giovanni Blandino, Aldo Morrone, Gennaro Ciliberto |

| | | | |
|---|---|---|---|
| EPI_ISL_3419715 | IRCCS San Gallicano Dermatological Institute | IRCCS Regina Elena National Cancer Institute | Ludovica Ciuffreda, Sara Donzelli, Alice Massacci, Arianna Mastrofrancesco, Fabrizio Ensoli, Fulvia Pimpinelli, Maurizio Fanciulli, Giovanni Blandino, Aldo Morrone, Gennaro Ciliberto |
| EPI_ISL_3419717 | IRCCS San Gallicano Dermatological Institute | IRCCS Regina Elena National Cancer Institute | Sabrina Strano, Sara Donzelli, Eleonora Sperandio, Carla Mottini, Fabrizio Ensoli, Fulvia Pimpinelli, Maurizio Fanciulli, Giovanni Blandino, Aldo Morrone, Gennaro Ciliberto |
| EPI_ISL_3419718 | IRCCS San Gallicano Dermatological Institute | IRCCS Regina Elena National Cancer Institute | Francesca De Nicola, Giulia Orlandi, Alice Massacci, Martina Pontone, Fabrizio Ensoli, Fulvia Pimpinelli, Maurizio Fanciulli, Giovanni Blandino, Aldo Morrone, Gennaro Ciliberto |
| EPI_ISL_3419719 | IRCCS San Gallicano Dermatological Institute | IRCCS Regina Elena National Cancer Institute | Ludovica Ciuffreda, Sara Donzelli, Matteo Pallocca, di Domenico Enea Gino, Fabrizio Ensoli, Fulvia Pimpinelli, Maurizio Fanciulli, Giovanni Blandino, Aldo Morrone, Gennaro Ciliberto |
| EPI_ISL_3419720 | IRCCS San Gallicano Dermatological Institute | IRCCS Regina Elena National Cancer Institute | Frauke Goeman, Giulia Orlandi, Eleonora Sperandio, Ilaria Cavallo, Fabrizio Ensoli, Fulvia Pimpinelli, Maurizio Fanciulli, Giovanni Blandino, Aldo Morrone, Gennaro Ciliberto |
| EPI_ISL_3419721 | IRCCS San Gallicano Dermatological Institute | IRCCS Regina Elena National Cancer Institute | Sabrina Strano, Sara Donzelli, Alice Massacci, Francesca Sivori, Fabrizio Ensoli, Fulvia Pimpinelli, Maurizio Fanciulli, Giovanni Blandino, Aldo Morrone, Gennaro Ciliberto |
| EPI_ISL_3419722 | IRCCS San Gallicano Dermatological Institute | IRCCS Regina Elena National Cancer Institute | Francesca De Nicola, Giulia Orlandi, Matteo Pallocca, Andrea Cazzani, Fabrizio Ensoli, Fulvia Pimpinelli, Maurizio Fanciulli, Giovanni Blandino, Aldo Morrone, Gennaro Ciliberto |
| EPI_ISL_3419723 | IRCCS San Gallicano Dermatological Institute | IRCCS Regina Elena National Cancer Institute | Ludovica Ciuffreda, Sara Donzelli, Eleonora Sperandio, Ilaria Celesti, Fabrizio Ensoli, Fulvia Pimpinelli, Maurizio Fanciulli, Giovanni Blandino, Aldo Morrone, Gennaro Ciliberto |
| EPI_ISL_3419724 | IRCCS San Gallicano Dermatological Institute | IRCCS Regina Elena National Cancer Institute | Frauke Goeman, Giulia Orlandi, Alice Massacci, Martina Diano, Fabrizio Ensoli, Fulvia Pimpinelli, Maurizio Fanciulli, Giovanni Blandino, Aldo Morrone, Gennaro Ciliberto |
| EPI_ISL_3419725 | IRCCS San Gallicano Dermatological Institute | IRCCS Regina Elena National Cancer Institute | Sabrina Strano, Sara Donzelli, Matteo Pallocca, Antonio Federico, Fabrizio Ensoli, Fulvia Pimpinelli, Maurizio Fanciulli, Giovanni Blandino, Aldo Morrone, Gennaro Ciliberto |
| EPI_ISL_3419726 | IRCCS San Gallicano Dermatological Institute | IRCCS Regina Elena National Cancer Institute | Francesca De Nicola, Giulia Orlandi, Eleonora Sperandio, Fulvia Fraticelli, Fabrizio Ensoli, Fulvia Pimpinelli, Maurizio Fanciulli, Giovanni Blandino, Aldo Morrone, Gennaro Ciliberto |
| EPI_ISL_3419727 | IRCCS San Gallicano Dermatological Institute | IRCCS Regina Elena National Cancer Institute | Ludovica Ciuffreda, Sara Donzelli, Alice Massacci, Lorenzo Furzi, Fabrizio Ensoli, Fulvia Pimpinelli, Maurizio Fanciulli, Giovanni Blandino, Aldo Morrone, Gennaro Ciliberto |
| EPI_ISL_3419728 | IRCCS San Gallicano Dermatological Institute | IRCCS Regina Elena National Cancer Institute | Frauke Goeman, Giulia Orlandi, Matteo Pallocca, Alessia Lauretti, Fabrizio Ensoli, Fulvia Pimpinelli, Maurizio Fanciulli, Giovanni Blandino, Aldo Morrone, Gennaro Ciliberto |
| EPI_ISL_3419729 | IRCCS San Gallicano Dermatological Institute | IRCCS Regina Elena National Cancer Institute | Sabrina Strano, Sara Donzelli, Eleonora Sperandio, Francesca Maione, Fabrizio Ensoli, Fulvia Pimpinelli, Maurizio Fanciulli, Giovanni Blandino, Aldo Morrone, Gennaro Ciliberto |
| EPI_ISL_3419730 | IRCCS San Gallicano Dermatological Institute | IRCCS Regina Elena National Cancer Institute | Francesca De Nicola, Giulia Orlandi, Alice Massacci, Francisco Obregon, Fabrizio Ensoli, Fulvia Pimpinelli, Maurizio Fanciulli, Giovanni Blandino, Aldo Morrone, Gennaro Ciliberto |
| EPI_ISL_3419731 | IRCCS San Gallicano Dermatological Institute | IRCCS Regina Elena National Cancer Institute | Ludovica Ciuffreda, Sara Donzelli, Matteo Pallocca, Silvia Paluzzi, Fabrizio Ensoli, Fulvia Pimpinelli, Maurizio Fanciulli, Giovanni Blandino, Aldo Morrone, Gennaro Ciliberto |
| EPI_ISL_3419732 | IRCCS San Gallicano Dermatological Institute | IRCCS Regina Elena National Cancer Institute | Frauke Goeman, Giulia Orlandi, Eleonora Sperandio, Sara Petrolo, Fabrizio Ensoli, Fulvia Pimpinelli, Maurizio Fanciulli, Giovanni Blandino, Aldo Morrone, Gennaro Ciliberto |
| EPI_ISL_3419733 | IRCCS San Gallicano Dermatological Institute | IRCCS Regina Elena National Cancer Institute | Sabrina Strano, Sara Donzelli, Alice Massacci, Valentina Ricca, Fabrizio Ensoli, Fulvia Pimpinelli, Maurizio Fanciulli, Giovanni Blandino, Aldo Morrone, Gennaro Ciliberto |
| EPI_ISL_3419734 | IRCCS San Gallicano Dermatological Institute | IRCCS Regina Elena National Cancer Institute | Francesca De Nicola, Giulia Orlandi, Matteo Pallocca, Serena Salvo, Fabrizio Ensoli, Fulvia Pimpinelli, Maurizio Fanciulli, Giovanni Blandino, Aldo Morrone, Gennaro Ciliberto |
| EPI_ISL_3419735 | IRCCS San Gallicano Dermatological Institute | IRCCS Regina Elena National Cancer Institute | Ludovica Ciuffreda, Sara Donzelli, Eleonora Sperandio, Grazia Prignano, Fabrizio Ensoli, Fulvia Pimpinelli, Maurizio Fanciulli, Giovanni Blandino, Aldo Morrone, Gennaro Ciliberto |
| EPI_ISL_3419736 | IRCCS San Gallicano Dermatological Institute | IRCCS Regina Elena National Cancer Institute | Frauke Goeman, Giulia Orlandi, Alice Massacci, Elisabetta Trento, Fabrizio Ensoli, Fulvia Pimpinelli, Maurizio Fanciulli, Giovanni Blandino, Aldo Morrone, Gennaro Ciliberto |
| EPI_ISL_3419737 | IRCCS San Gallicano Dermatological Institute | IRCCS Regina Elena National Cancer Institute | Sabrina Strano, Sara Donzelli, Matteo Pallocca, Arianna Mastrofrancesco, Fabrizio Ensoli, Fulvia Pimpinelli, Maurizio Fanciulli, Giovanni Blandino, Aldo Morrone, Gennaro Ciliberto |
| EPI_ISL_3419738 | IRCCS San Gallicano Dermatological Institute | IRCCS Regina Elena National Cancer Institute | Francesca De Nicola, Giulia Orlandi, Eleonora Sperandio, Giovanna D’agosto, Fabrizio Ensoli, Fulvia Pimpinelli, Maurizio Fanciulli, Giovanni Blandino, Aldo Morrone, Gennaro Ciliberto |
| EPI_ISL_3419739 | IRCCS San Gallicano Dermatological Institute | IRCCS Regina Elena National Cancer Institute | Ludovica Ciuffreda, Sara Donzelli, Alice Massacci, Carla Mottini, Fabrizio Ensoli, Fulvia Pimpinelli, Maurizio Fanciulli, Giovanni Blandino, Aldo Morrone, Gennaro Ciliberto |
| EPI_ISL_3419741 | IRCCS San Gallicano Dermatological Institute | IRCCS Regina Elena National Cancer Institute | Sabrina Strano, Sara Donzelli, Eleonora Sperandio, di Domenico Enea Gino, Fabrizio Ensoli, Fulvia Pimpinelli, Maurizio Fanciulli, Giovanni Blandino, Aldo Morrone, Gennaro Ciliberto |
| EPI_ISL_3419742 | IRCCS San Gallicano Dermatological Institute | IRCCS Regina Elena National Cancer Institute | Francesca De Nicola, Giulia Orlandi, Alice Massacci, Ilaria Cavallo, Fabrizio Ensoli, Fulvia Pimpinelli, Maurizio Fanciulli, Giovanni Blandino, Aldo Morrone, Gennaro Ciliberto |
| EPI_ISL_3419743 | IRCCS San Gallicano Dermatological Institute | IRCCS Regina Elena National Cancer Institute | Ludovica Ciuffreda, Sara Donzelli, Matteo Pallocca, Francesca Sivori, Fabrizio Ensoli, Fulvia Pimpinelli, Maurizio Fanciulli, Giovanni Blandino, Aldo Morrone, Gennaro Ciliberto |
| EPI_ISL_3419744 | IRCCS San Gallicano Dermatological Institute | IRCCS Regina Elena National Cancer Institute | Frauke Goeman, Giulia Orlandi, Eleonora Sperandio, Andrea Cazzani, Fabrizio Ensoli, Fulvia Pimpinelli, Maurizio Fanciulli, Giovanni Blandino, Aldo Morrone, Gennaro Ciliberto |
| EPI_ISL_3419745 | IRCCS San Gallicano Dermatological Institute | IRCCS Regina Elena National Cancer Institute | Sabrina Strano, Sara Donzelli, Alice Massacci, Ilaria Celesti, Fabrizio Ensoli, Fulvia Pimpinelli, Maurizio Fanciulli, Giovanni Blandino, Aldo Morrone, Gennaro Ciliberto |
| EPI_ISL_3419746 | IRCCS San Gallicano Dermatological Institute | IRCCS Regina Elena National Cancer Institute | Francesca De Nicola, Giulia Orlandi, Matteo Pallocca, Martina Diano, Fabrizio Ensoli, Fulvia Pimpinelli, Maurizio Fanciulli, Giovanni Blandino, Aldo Morrone, Gennaro Ciliberto |
| EPI_ISL_3419747 | IRCCS San Gallicano Dermatological Institute | IRCCS Regina Elena National Cancer Institute | Ludovica Ciuffreda, Sara Donzelli, Eleonora Sperandio, Antonio Federico, Fabrizio Ensoli, Fulvia Pimpinelli, Maurizio Fanciulli, Giovanni Blandino, Aldo Morrone, Gennaro Ciliberto |
| EPI_ISL_3419748 | IRCCS San Gallicano Dermatological Institute | IRCCS Regina Elena National Cancer Institute | Frauke Goeman, Giulia Orlandi, Alice Massacci, Fulvia Fraticelli, Fabrizio Ensoli, Fulvia Pimpinelli, Maurizio Fanciulli, Giovanni Blandino, Aldo Morrone, Gennaro Ciliberto |
| EPI_ISL_3419749 | IRCCS San Gallicano Dermatological Institute | IRCCS Regina Elena National Cancer Institute | Sabrina Strano, Sara Donzelli, Matteo Pallocca, Lorenzo Furzi, Fabrizio Ensoli, Fulvia Pimpinelli, Maurizio Fanciulli, Giovanni Blandino, Aldo Morrone, Gennaro Ciliberto |
| EPI_ISL_3419750 | IRCCS San Gallicano Dermatological Institute | IRCCS Regina Elena National Cancer Institute | Francesca De Nicola, Giulia Orlandi, Eleonora Sperandio, Alessia Lauretti, Fabrizio Ensoli, Fulvia Pimpinelli, Maurizio Fanciulli, Giovanni Blandino, Aldo Morrone, Gennaro Ciliberto |
| EPI_ISL_3419751 | IRCCS San Gallicano Dermatological Institute | IRCCS Regina Elena National Cancer Institute | Ludovica Ciuffreda, Sara Donzelli, Alice Massacci, Francesca Maione, Fabrizio Ensoli, Fulvia Pimpinelli, Maurizio Fanciulli, Giovanni Blandino, Aldo Morrone, Gennaro Ciliberto |
| EPI_ISL_3419752 | IRCCS San Gallicano Dermatological Institute | IRCCS Regina Elena National Cancer Institute | Frauke Goeman, Giulia Orlandi, Matteo Pallocca, Francisco Obregon, Fabrizio Ensoli, Fulvia Pimpinelli, Maurizio Fanciulli, Giovanni Blandino, Aldo Morrone, Gennaro Ciliberto |
| EPI_ISL_3419753 | IRCCS San Gallicano Dermatological Institute | IRCCS Regina Elena National Cancer Institute | Sabrina Strano, Sara Donzelli, Eleonora Sperandio, Silvia Paluzzi, Fabrizio Ensoli, Fulvia Pimpinelli, Maurizio Fanciulli, Giovanni Blandino, Aldo Morrone, Gennaro Ciliberto |
| EPI_ISL_3419754 | IRCCS San Gallicano Dermatological Institute | IRCCS Regina Elena National Cancer Institute | Francesca De Nicola, Giulia Orlandi, Alice Massacci, Sara Petrolo, Fabrizio Ensoli, Fulvia Pimpinelli, Maurizio Fanciulli, Giovanni Blandino, Aldo Morrone, Gennaro Ciliberto |
| EPI_ISL_3419756 | IRCCS San Gallicano Dermatological Institute | IRCCS Regina Elena National Cancer Institute | Frauke Goeman, Giulia Orlandi, Eleonora Sperandio, Serena Salvo, Fabrizio Ensoli, Fulvia Pimpinelli, Maurizio Fanciulli, Giovanni Blandino, Aldo Morrone, Gennaro Ciliberto |
| EPI_ISL_3419757 | IRCCS San Gallicano Dermatological Institute | IRCCS Regina Elena National Cancer Institute | Sabrina Strano, Sara Donzelli, Alice Massacci, Grazia Prignano, Fabrizio Ensoli, Fulvia Pimpinelli, Maurizio Fanciulli, Giovanni Blandino, Aldo Morrone, Gennaro Ciliberto |
| EPI_ISL_3419758 | IRCCS San Gallicano Dermatological Institute | IRCCS Regina Elena National Cancer Institute | Francesca De Nicola, Giulia Orlandi, Matteo Pallocca, Elisabetta Trento, Fabrizio Ensoli, Fulvia Pimpinelli, Maurizio Fanciulli, Giovanni Blandino, Aldo Morrone, Gennaro Ciliberto |
| EPI_ISL_3419759 | IRCCS San Gallicano Dermatological Institute | IRCCS Regina Elena National Cancer Institute | Ludovica Ciuffreda, Sara Donzelli, Eleonora Sperandio, Arianna Mastrofrancesco, Fabrizio Ensoli, Fulvia Pimpinelli, Maurizio Fanciulli, Giovanni Blandino, Aldo Morrone, Gennaro Ciliberto |
| EPI_ISL_3419760 | IRCCS San Gallicano Dermatological Institute | IRCCS Regina Elena National Cancer Institute | Frauke Goeman, Giulia Orlandi, Alice Massacci, Giovanna D’agosto, Fabrizio Ensoli, Fulvia Pimpinelli, Maurizio Fanciulli, Giovanni Blandino, Aldo Morrone, Gennaro Ciliberto |
| EPI_ISL_3419761 | IRCCS San Gallicano Dermatological Institute | IRCCS Regina Elena National Cancer Institute | Sabrina Strano, Sara Donzelli, Matteo Pallocca, Carla Mottini, Fabrizio Ensoli, Fulvia Pimpinelli, Maurizio Fanciulli, Giovanni Blandino, Aldo Morrone, Gennaro Ciliberto |
| EPI_ISL_3419762 | IRCCS San Gallicano Dermatological Institute | IRCCS Regina Elena National Cancer Institute | Francesca De Nicola, Giulia Orlandi, Eleonora Sperandio, Martina Pontone, Fabrizio Ensoli, Fulvia Pimpinelli, Maurizio Fanciulli, Giovanni Blandino, Aldo Morrone, Gennaro Ciliberto |
| EPI_ISL_3419763 | IRCCS San Gallicano Dermatological Institute | IRCCS Regina Elena National Cancer Institute | Ludovica Ciuffreda, Sara Donzelli, Alice Massacci, di Domenico Enea Gino, Fabrizio Ensoli, Fulvia Pimpinelli, Maurizio Fanciulli, Giovanni Blandino, Aldo Morrone, Gennaro Ciliberto |
| EPI_ISL_3419764 | IRCCS San Gallicano Dermatological Institute | IRCCS Regina Elena National Cancer Institute | Frauke Goeman, Giulia Orlandi, Matteo Pallocca, Ilaria Cavallo, Fabrizio Ensoli, Fulvia Pimpinelli, Maurizio Fanciulli, Giovanni Blandino, Aldo Morrone, Gennaro Ciliberto |
| EPI_ISL_3419765 | IRCCS San Gallicano Dermatological Institute | IRCCS Regina Elena National Cancer Institute | Sabrina Strano, Sara Donzelli, Eleonora Sperandio, Francesca Sivori, Fabrizio Ensoli, Fulvia Pimpinelli, Maurizio Fanciulli, Giovanni Blandino, Aldo Morrone, Gennaro Ciliberto |
| EPI_ISL_3419766 | IRCCS San Gallicano Dermatological Institute | IRCCS Regina Elena National Cancer Institute | Francesca De Nicola, Giulia Orlandi, Alice Massacci, Andrea Cazzani, Fabrizio Ensoli, Fulvia Pimpinelli, Maurizio Fanciulli, Giovanni Blandino, Aldo Morrone, Gennaro Ciliberto |
| EPI_ISL_3419767 | IRCCS San Gallicano Dermatological Institute | IRCCS Regina Elena National Cancer Institute | Ludovica Ciuffreda, Sara Donzelli, Matteo Pallocca, Ilaria Celesti, Fabrizio Ensoli, Fulvia Pimpinelli, Maurizio Fanciulli, Giovanni Blandino, Aldo Morrone, Gennaro Ciliberto |
| EPI_ISL_3419768 | IRCCS San Gallicano Dermatological Institute | IRCCS Regina Elena National Cancer Institute | Frauke Goeman, Giulia Orlandi, Eleonora Sperandio, Martina Diano, Fabrizio Ensoli, Fulvia Pimpinelli, Maurizio Fanciulli, Giovanni Blandino, Aldo Morrone, Gennaro Ciliberto |
| EPI_ISL_3419769 | IRCCS San Gallicano Dermatological Institute | IRCCS Regina Elena National Cancer Institute | Sabrina Strano, Sara Donzelli, Alice Massacci, Antonio Federico, Fabrizio Ensoli, Fulvia Pimpinelli, Maurizio Fanciulli, Giovanni Blandino, Aldo Morrone, Gennaro Ciliberto |
| EPI_ISL_3419770 | IRCCS San Gallicano Dermatological Institute | IRCCS Regina Elena National Cancer Institute | Francesca De Nicola, Giulia Orlandi, Matteo Pallocca, Fulvia Fraticelli, Fabrizio Ensoli, Fulvia Pimpinelli, Maurizio Fanciulli, Giovanni Blandino, Aldo Morrone, Gennaro Ciliberto |
| EPI_ISL_3419771 | IRCCS San Gallicano Dermatological Institute | IRCCS Regina Elena National Cancer Institute | Ludovica Ciuffreda, Sara Donzelli, Eleonora Sperandio, Lorenzo Furzi, Fabrizio Ensoli, Fulvia Pimpinelli, Maurizio Fanciulli, Giovanni Blandino, Aldo Morrone, Gennaro Ciliberto |
| EPI_ISL_3419772 | IRCCS San Gallicano Dermatological Institute | IRCCS Regina Elena National Cancer Institute | Frauke Goeman, Giulia Orlandi, Alice Massacci, Alessia Lauretti, Fabrizio Ensoli, Fulvia Pimpinelli, Maurizio Fanciulli, Giovanni Blandino, Aldo Morrone, Gennaro Ciliberto |
| EPI_ISL_3419773 | IRCCS San Gallicano Dermatological Institute | IRCCS Regina Elena National Cancer Institute | Sabrina Strano, Sara Donzelli, Matteo Pallocca, Francesca Maione, Fabrizio Ensoli, Fulvia Pimpinelli, Maurizio Fanciulli, Giovanni Blandino, Aldo Morrone, Gennaro Ciliberto |
| EPI_ISL_3419774 | IRCCS San Gallicano Dermatological Institute | IRCCS Regina Elena National Cancer Institute | Francesca De Nicola, Giulia Orlandi, Eleonora Sperandio, Francisco Obregon, Fabrizio Ensoli, Fulvia Pimpinelli, Maurizio Fanciulli, Giovanni Blandino, Aldo Morrone, Gennaro Ciliberto |
| EPI_ISL_3419775 | IRCCS San Gallicano Dermatological Institute | IRCCS Regina Elena National Cancer Institute | Ludovica Ciuffreda, Sara Donzelli, Alice Massacci, Silvia Paluzzi, Fabrizio Ensoli, Fulvia Pimpinelli, Maurizio Fanciulli, Giovanni Blandino, Aldo Morrone, Gennaro Ciliberto |
| EPI_ISL_3419776 | IRCCS San Gallicano Dermatological Institute | IRCCS Regina Elena National Cancer Institute | Frauke Goeman, Giulia Orlandi, Matteo Pallocca, Sara Petrolo, Fabrizio Ensoli, Fulvia Pimpinelli, Maurizio Fanciulli, Giovanni Blandino, Aldo Morrone, Gennaro Ciliberto |
| EPI_ISL_3419777 | IRCCS San Gallicano Dermatological Institute | IRCCS Regina Elena National Cancer Institute | Sabrina Strano, Sara Donzelli, Eleonora Sperandio, Valentina Ricca, Fabrizio Ensoli, Fulvia Pimpinelli, Maurizio Fanciulli, Giovanni Blandino, Aldo Morrone, Gennaro Ciliberto |
| EPI_ISL_3419778 | IRCCS San Gallicano Dermatological Institute | IRCCS Regina Elena National Cancer Institute | Francesca De Nicola, Giulia Orlandi, Alice Massacci, Serena Salvo, Fabrizio Ensoli, Fulvia Pimpinelli, Maurizio Fanciulli, Giovanni Blandino, Aldo Morrone, Gennaro Ciliberto |
| EPI_ISL_3419779 | IRCCS San Gallicano Dermatological Institute | IRCCS Regina Elena National Cancer Institute | Ludovica Ciuffreda, Sara Donzelli, Matteo Pallocca, Grazia Prignano, Fabrizio Ensoli, Fulvia Pimpinelli, Maurizio Fanciulli, Giovanni Blandino, Aldo Morrone, Gennaro Ciliberto |
| EPI_ISL_3419780 | IRCCS San Gallicano Dermatological Institute | IRCCS Regina Elena National Cancer Institute | Frauke Goeman, Giulia Orlandi, Eleonora Sperandio, Elisabetta Trento, Fabrizio Ensoli, Fulvia Pimpinelli, Maurizio Fanciulli, Giovanni Blandino, Aldo Morrone, Gennaro Ciliberto |
| EPI_ISL_3419781 | IRCCS San Gallicano Dermatological Institute | IRCCS Regina Elena National Cancer Institute | Sabrina Strano, Sara Donzelli, Alice Massacci, Arianna Mastrofrancesco, Fabrizio Ensoli, Fulvia Pimpinelli, Maurizio Fanciulli, Giovanni Blandino, Aldo Morrone, Gennaro Ciliberto |
| EPI_ISL_3419782 | IRCCS San Gallicano Dermatological Institute | IRCCS Regina Elena National Cancer Institute | Francesca De Nicola, Giulia Orlandi, Matteo Pallocca, Giovanna D’agosto, Fabrizio Ensoli, Fulvia Pimpinelli, Maurizio Fanciulli, Giovanni Blandino, Aldo Morrone, Gennaro Ciliberto |
| EPI_ISL_3419783 | IRCCS San Gallicano Dermatological Institute | IRCCS Regina Elena National Cancer Institute | Ludovica Ciuffreda, Sara Donzelli, Eleonora Sperandio, Carla Mottini, Fabrizio Ensoli, Fulvia Pimpinelli, Maurizio Fanciulli, Giovanni Blandino, Aldo Morrone, Gennaro Ciliberto |
| EPI_ISL_3419784 | IRCCS San Gallicano Dermatological Institute | IRCCS Regina Elena National Cancer Institute | Frauke Goeman, Giulia Orlandi, Alice Massacci, Martina Pontone, Fabrizio Ensoli, Fulvia Pimpinelli, Maurizio Fanciulli, Giovanni Blandino, Aldo Morrone, Gennaro Ciliberto |
| EPI_ISL_3419785 | IRCCS San Gallicano Dermatological Institute | IRCCS Regina Elena National Cancer Institute | Sabrina Strano, Sara Donzelli, Matteo Pallocca, di Domenico Enea Gino, Fabrizio Ensoli, Fulvia Pimpinelli, Maurizio Fanciulli, Giovanni Blandino, Aldo Morrone, Gennaro Ciliberto |
| EPI_ISL_3419786 | IRCCS San Gallicano Dermatological Institute | IRCCS Regina Elena National Cancer Institute | Francesca De Nicola, Giulia Orlandi, Eleonora Sperandio, Ilaria Cavallo, Fabrizio Ensoli, Fulvia Pimpinelli, Maurizio Fanciulli, Giovanni Blandino, Aldo Morrone, Gennaro Ciliberto |
| EPI_ISL_3419787 | IRCCS San Gallicano Dermatological Institute | IRCCS Regina Elena National Cancer Institute | Ludovica Ciuffreda, Sara Donzelli, Alice Massacci, Francesca Sivori, Fabrizio Ensoli, Fulvia Pimpinelli, Maurizio Fanciulli, Giovanni Blandino, Aldo Morrone, Gennaro Ciliberto |
| EPI_ISL_3419788 | IRCCS San Gallicano Dermatological Institute | IRCCS Regina Elena National Cancer Institute | Frauke Goeman, Giulia Orlandi, Matteo Pallocca, Andrea Cazzani, Fabrizio Ensoli, Fulvia Pimpinelli, Maurizio Fanciulli, Giovanni Blandino, Aldo Morrone, Gennaro Ciliberto |
| EPI_ISL_3419789 | IRCCS San Gallicano Dermatological Institute | IRCCS Regina Elena National Cancer Institute | Sabrina Strano, Sara Donzelli, Eleonora Sperandio, Ilaria Celesti, Fabrizio Ensoli, Fulvia Pimpinelli, Maurizio Fanciulli, Giovanni Blandino, Aldo Morrone, Gennaro Ciliberto |
| EPI_ISL_3419790 | IRCCS San Gallicano Dermatological Institute | IRCCS Regina Elena National Cancer Institute | Francesca De Nicola, Giulia Orlandi, Alice Massacci, Martina Diano, Fabrizio Ensoli, Fulvia Pimpinelli, Maurizio Fanciulli, Giovanni Blandino, Aldo Morrone, Gennaro Ciliberto |
| EPI_ISL_3419791 | IRCCS San Gallicano Dermatological Institute | IRCCS Regina Elena National Cancer Institute | Ludovica Ciuffreda, Sara Donzelli, Matteo Pallocca, Antonio Federico, Fabrizio Ensoli, Fulvia Pimpinelli, Maurizio Fanciulli, Giovanni Blandino, Aldo Morrone, Gennaro Ciliberto |
| EPI_ISL_3419792 | IRCCS San Gallicano Dermatological Institute | IRCCS Regina Elena National Cancer Institute | Frauke Goeman, Giulia Orlandi, Eleonora Sperandio, Fulvia Fraticelli, Fabrizio Ensoli, Fulvia Pimpinelli, Maurizio Fanciulli, Giovanni Blandino, Aldo Morrone, Gennaro Ciliberto |
| EPI_ISL_3419793 | IRCCS San Gallicano Dermatological Institute | IRCCS Regina Elena National Cancer Institute | Sabrina Strano, Sara Donzelli, Alice Massacci, Lorenzo Furzi, Fabrizio Ensoli, Fulvia Pimpinelli, Maurizio Fanciulli, Giovanni Blandino, Aldo Morrone, Gennaro Ciliberto |
| EPI_ISL_3419794 | IRCCS San Gallicano Dermatological Institute | IRCCS Regina Elena National Cancer Institute | Francesca De Nicola, Giulia Orlandi, Matteo Pallocca, Alessia Lauretti, Fabrizio Ensoli, Fulvia Pimpinelli, Maurizio Fanciulli, Giovanni Blandino, Aldo Morrone, Gennaro Ciliberto |

| | | | |
|---|---|---|---|
| EPI_ISL_3419795 | IRCCS San Gallicano Dermatological Institute | IRCCS Regina Elena National Cancer Institute | Ludovica Ciuffreda, Sara Donzelli, Eleonora Sperandio, Francesca Maione, Fabrizio Ensoli, Fulvia Pimpinelli, Maurizio Fanciulli, Giovanni Blandino, Aldo Morrone, Gennaro Ciliberto |
| EPI_ISL_3419796 | IRCCS San Gallicano Dermatological Institute | IRCCS Regina Elena National Cancer Institute | Frauke Goeman, Giulia Orlandi, Alice Massacci, Francisco Obregon, Fabrizio Ensoli, Fulvia Pimpinelli, Maurizio Fanciulli, Giovanni Blandino, Aldo Morrone, Gennaro Ciliberto |
| EPI_ISL_3419797 | IRCCS San Gallicano Dermatological Institute | IRCCS Regina Elena National Cancer Institute | Sabrina Strano, Sara Donzelli, Matteo Pallocca, Silvia Paluzzi, Fabrizio Ensoli, Fulvia Pimpinelli, Maurizio Fanciulli, Giovanni Blandino, Aldo Morrone, Gennaro Ciliberto |
| EPI_ISL_3419798 | IRCCS San Gallicano Dermatological Institute | IRCCS Regina Elena National Cancer Institute | Francesca De Nicola, Giulia Orlandi, Eleonora Sperandio, Sara Petrolo, Fabrizio Ensoli, Fulvia Pimpinelli, Maurizio Fanciulli, Giovanni Blandino, Aldo Morrone, Gennaro Ciliberto |
| EPI_ISL_3419799 | IRCCS San Gallicano Dermatological Institute | IRCCS Regina Elena National Cancer Institute | Ludovica Ciuffreda, Sara Donzelli, Alice Massacci, Valentina Ricca, Fabrizio Ensoli, Fulvia Pimpinelli, Maurizio Fanciulli, Giovanni Blandino, Aldo Morrone, Gennaro Ciliberto |
| EPI_ISL_3419800 | IRCCS San Gallicano Dermatological Institute | IRCCS Regina Elena National Cancer Institute | Frauke Goeman, Giulia Orlandi, Matteo Pallocca, Serena Salvo, Fabrizio Ensoli, Fulvia Pimpinelli, Maurizio Fanciulli, Giovanni Blandino, Aldo Morrone, Gennaro Ciliberto |
| EPI_ISL_3419801 | IRCCS San Gallicano Dermatological Institute | IRCCS Regina Elena National Cancer Institute | Sabrina Strano, Sara Donzelli, Eleonora Sperandio, Grazia Prignano, Fabrizio Ensoli, Fulvia Pimpinelli, Maurizio Fanciulli, Giovanni Blandino, Aldo Morrone, Gennaro Ciliberto |
| EPI_ISL_3419802 | IRCCS San Gallicano Dermatological Institute | IRCCS Regina Elena National Cancer Institute | Ludovica Ciuffreda, Sara Donzelli, Matteo Pallocca, Arianna Mastrofrancesco, Fabrizio Ensoli, Fulvia Pimpinelli, Maurizio Fanciulli, Giovanni Blandino, Aldo Morrone, Gennaro Ciliberto |
| EPI_ISL_3419803 | IRCCS San Gallicano Dermatological Institute | IRCCS Regina Elena National Cancer Institute | Frauke Goeman, Giulia Orlandi, Eleonora Sperandio, Giovanna D'agosto, Fabrizio Ensoli, Fulvia Pimpinelli, Maurizio Fanciulli, Giovanni Blandino, Aldo Morrone, Gennaro Ciliberto |
| EPI_ISL_3419804 | IRCCS San Gallicano Dermatological Institute | IRCCS Regina Elena National Cancer Institute | Sabrina Strano, Sara Donzelli, Alice Massacci, Carla Mottini, Fabrizio Ensoli, Fulvia Pimpinelli, Maurizio Fanciulli, Giovanni Blandino, Aldo Morrone, Gennaro Ciliberto |
| EPI_ISL_3419805 | IRCCS San Gallicano Dermatological Institute | IRCCS Regina Elena National Cancer Institute | Francesca De Nicola, Giulia Orlandi, Matteo Pallocca, Martina Pontone, Fabrizio Ensoli, Fulvia Pimpinelli, Maurizio Fanciulli, Giovanni Blandino, Aldo Morrone, Gennaro Ciliberto |
| EPI_ISL_3419806 | IRCCS San Gallicano Dermatological Institute | IRCCS Regina Elena National Cancer Institute | Ludovica Ciuffreda, Sara Donzelli, Eleonora Sperandio, di Domenico Enea Gino, Fabrizio Ensoli, Fulvia Pimpinelli, Maurizio Fanciulli, Giovanni Blandino, Aldo Morrone, Gennaro Ciliberto |
| EPI_ISL_3419807 | IRCCS San Gallicano Dermatological Institute | IRCCS Regina Elena National Cancer Institute | Frauke Goeman, Giulia Orlandi, Alice Massacci, Ilaria Cavallo, Fabrizio Ensoli, Fulvia Pimpinelli, Maurizio Fanciulli, Giovanni Blandino, Aldo Morrone, Gennaro Ciliberto |
| EPI_ISL_3419808 | IRCCS San Gallicano Dermatological Institute | IRCCS Regina Elena National Cancer Institute | Sabrina Strano, Sara Donzelli, Matteo Pallocca, Francesca Sivori, Fabrizio Ensoli, Fulvia Pimpinelli, Maurizio Fanciulli, Giovanni Blandino, Aldo Morrone, Gennaro Ciliberto |
| EPI_ISL_3419809 | IRCCS San Gallicano Dermatological Institute | IRCCS Regina Elena National Cancer Institute | Francesca De Nicola, Giulia Orlandi, Eleonora Sperandio, Andrea Cazzani, Fabrizio Ensoli, Fulvia Pimpinelli, Maurizio Fanciulli, Giovanni Blandino, Aldo Morrone, Gennaro Ciliberto |
| EPI_ISL_3419810 | IRCCS San Gallicano Dermatological Institute | IRCCS Regina Elena National Cancer Institute | Ludovica Ciuffreda, Sara Donzelli, Alice Massacci, Ilaria Celesti, Fabrizio Ensoli, Fulvia Pimpinelli, Maurizio Fanciulli, Giovanni Blandino, Aldo Morrone, Gennaro Ciliberto |
| EPI_ISL_3419811 | IRCCS San Gallicano Dermatological Institute | IRCCS Regina Elena National Cancer Institute | Frauke Goeman, Giulia Orlandi, Matteo Pallocca, Martina Diano, Fabrizio Ensoli, Fulvia Pimpinelli, Maurizio Fanciulli, Giovanni Blandino, Aldo Morrone, Gennaro Ciliberto |
| EPI_ISL_3419812 | IRCCS San Gallicano Dermatological Institute | IRCCS Regina Elena National Cancer Institute | Sabrina Strano, Sara Donzelli, Eleonora Sperandio, Antonio Federico, Fabrizio Ensoli, Fulvia Pimpinelli, Maurizio Fanciulli, Giovanni Blandino, Aldo Morrone, Gennaro Ciliberto |
| EPI_ISL_3419813 | IRCCS San Gallicano Dermatological Institute | IRCCS Regina Elena National Cancer Institute | Francesca De Nicola, Giulia Orlandi, Alice Massacci, Fulvia Fraticelli, Fabrizio Ensoli, Fulvia Pimpinelli, Maurizio Fanciulli, Giovanni Blandino, Aldo Morrone, Gennaro Ciliberto |
| EPI_ISL_3419814 | IRCCS San Gallicano Dermatological Institute | IRCCS Regina Elena National Cancer Institute | Ludovica Ciuffreda, Sara Donzelli, Matteo Pallocca, Lorenzo Furzi, Fabrizio Ensoli, Fulvia Pimpinelli, Maurizio Fanciulli, Giovanni Blandino, Aldo Morrone, Gennaro Ciliberto |
| EPI_ISL_3419815 | IRCCS San Gallicano Dermatological Institute | IRCCS Regina Elena National Cancer Institute | Frauke Goeman, Giulia Orlandi, Eleonora Sperandio, Alessia Lauretti, Fabrizio Ensoli, Fulvia Pimpinelli, Maurizio Fanciulli, Giovanni Blandino, Aldo Morrone, Gennaro Ciliberto |
| EPI_ISL_3419816 | IRCCS San Gallicano Dermatological Institute | IRCCS Regina Elena National Cancer Institute | Sabrina Strano, Sara Donzelli, Alice Massacci, Francesca Maione, Fabrizio Ensoli, Fulvia Pimpinelli, Maurizio Fanciulli, Giovanni Blandino, Aldo Morrone, Gennaro Ciliberto |
| EPI_ISL_3419817 | IRCCS San Gallicano Dermatological Institute | IRCCS Regina Elena National Cancer Institute | Francesca De Nicola, Giulia Orlandi, Matteo Pallocca, Francisco Obregon, Fabrizio Ensoli, Fulvia Pimpinelli, Maurizio Fanciulli, Giovanni Blandino, Aldo Morrone, Gennaro Ciliberto |
| EPI_ISL_3419818 | IRCCS San Gallicano Dermatological Institute | IRCCS Regina Elena National Cancer Institute | Ludovica Ciuffreda, Sara Donzelli, Eleonora Sperandio, Silvia Paluzzi, Fabrizio Ensoli, Fulvia Pimpinelli, Maurizio Fanciulli, Giovanni Blandino, Aldo Morrone, Gennaro Ciliberto |
| EPI_ISL_3419819 | IRCCS San Gallicano Dermatological Institute | IRCCS Regina Elena National Cancer Institute | Frauke Goeman, Giulia Orlandi, Alice Massacci, Sara Petrolo, Fabrizio Ensoli, Fulvia Pimpinelli, Maurizio Fanciulli, Giovanni Blandino, Aldo Morrone, Gennaro Ciliberto |
| EPI_ISL_3419821 | IRCCS San Gallicano Dermatological Institute | IRCCS Regina Elena National Cancer Institute | Francesca De Nicola, Giulia Orlandi, Eleonora Sperandio, Serena Salvo, Fabrizio Ensoli, Fulvia Pimpinelli, Maurizio Fanciulli, Giovanni Blandino, Aldo Morrone, Gennaro Ciliberto |
| EPI_ISL_3419822 | IRCCS San Gallicano Dermatological Institute | IRCCS Regina Elena National Cancer Institute | Ludovica Ciuffreda, Sara Donzelli, Alice Massacci, Grazia Prignano, Fabrizio Ensoli, Fulvia Pimpinelli, Maurizio Fanciulli, Giovanni Blandino, Aldo Morrone, Gennaro Ciliberto |
| EPI_ISL_3419823 | IRCCS San Gallicano Dermatological Institute | IRCCS Regina Elena National Cancer Institute | Frauke Goeman, Giulia Orlandi, Matteo Pallocca, Elisabetta Trento, Fabrizio Ensoli, Fulvia Pimpinelli, Maurizio Fanciulli, Giovanni Blandino, Aldo Morrone, Gennaro Ciliberto |
| EPI_ISL_3419824 | IRCCS San Gallicano Dermatological Institute | IRCCS Regina Elena National Cancer Institute | Sabrina Strano, Sara Donzelli, Eleonora Sperandio, Arianna Mastrofrancesco, Fabrizio Ensoli, Fulvia Pimpinelli, Maurizio Fanciulli, Giovanni Blandino, Aldo Morrone, Gennaro Ciliberto |
| EPI_ISL_3419825 | IRCCS San Gallicano Dermatological Institute | IRCCS Regina Elena National Cancer Institute | Francesca De Nicola, Giulia Orlandi, Alice Massacci, Giovanna D'agosto, Fabrizio Ensoli, Fulvia Pimpinelli, Maurizio Fanciulli, Giovanni Blandino, Aldo Morrone, Gennaro Ciliberto |
| EPI_ISL_3419826 | IRCCS San Gallicano Dermatological Institute | IRCCS Regina Elena National Cancer Institute | Ludovica Ciuffreda, Sara Donzelli, Matteo Pallocca, Carla Mottini, Fabrizio Ensoli, Fulvia Pimpinelli, Maurizio Fanciulli, Giovanni Blandino, Aldo Morrone, Gennaro Ciliberto |
| EPI_ISL_3419827 | IRCCS San Gallicano Dermatological Institute | IRCCS Regina Elena National Cancer Institute | Frauke Goeman, Giulia Orlandi, Eleonora Sperandio, Martina Pontone, Fabrizio Ensoli, Fulvia Pimpinelli, Maurizio Fanciulli, Giovanni Blandino, Aldo Morrone, Gennaro Ciliberto |
| EPI_ISL_3419828 | IRCCS San Gallicano Dermatological Institute | IRCCS Regina Elena National Cancer Institute | Sabrina Strano, Sara Donzelli, Alice Massacci, di Domenico Enea Gino, Fabrizio Ensoli, Fulvia Pimpinelli, Maurizio Fanciulli, Giovanni Blandino, Aldo Morrone, Gennaro Ciliberto |
| EPI_ISL_3419829 | IRCCS San Gallicano Dermatological Institute | IRCCS Regina Elena National Cancer Institute | Francesca De Nicola, Giulia Orlandi, Matteo Pallocca, Ilaria Cavallo, Fabrizio Ensoli, Fulvia Pimpinelli, Maurizio Fanciulli, Giovanni Blandino, Aldo Morrone, Gennaro Ciliberto |
| EPI_ISL_3419830 | IRCCS San Gallicano Dermatological Institute | IRCCS Regina Elena National Cancer Institute | Ludovica Ciuffreda, Sara Donzelli, Eleonora Sperandio, Francesca Sivori, Fabrizio Ensoli, Fulvia Pimpinelli, Maurizio Fanciulli, Giovanni Blandino, Aldo Morrone, Gennaro Ciliberto |
| EPI_ISL_3419831 | IRCCS San Gallicano Dermatological Institute | IRCCS Regina Elena National Cancer Institute | Frauke Goeman, Giulia Orlandi, Alice Massacci, Andrea Cazzani, Fabrizio Ensoli, Fulvia Pimpinelli, Maurizio Fanciulli, Giovanni Blandino, Aldo Morrone, Gennaro Ciliberto |
| EPI_ISL_3419832 | IRCCS San Gallicano Dermatological Institute | IRCCS Regina Elena National Cancer Institute | Sabrina Strano, Sara Donzelli, Matteo Pallocca, Ilaria Celesti, Fabrizio Ensoli, Fulvia Pimpinelli, Maurizio Fanciulli, Giovanni Blandino, Aldo Morrone, Gennaro Ciliberto |
| EPI_ISL_3419833 | IRCCS San Gallicano Dermatological Institute | IRCCS Regina Elena National Cancer Institute | Francesca De Nicola, Giulia Orlandi, Eleonora Sperandio, Martina Diano, Fabrizio Ensoli, Fulvia Pimpinelli, Maurizio Fanciulli, Giovanni Blandino, Aldo Morrone, Gennaro Ciliberto |
| EPI_ISL_3419834 | IRCCS San Gallicano Dermatological Institute | IRCCS Regina Elena National Cancer Institute | Ludovica Ciuffreda, Sara Donzelli, Alice Massacci, Antonio Federico, Fabrizio Ensoli, Fulvia Pimpinelli, Maurizio Fanciulli, Giovanni Blandino, Aldo Morrone, Gennaro Ciliberto |
| EPI_ISL_3419835 | IRCCS San Gallicano Dermatological Institute | IRCCS Regina Elena National Cancer Institute | Frauke Goeman, Giulia Orlandi, Matteo Pallocca, Fulvia Fraticelli, Fabrizio Ensoli, Fulvia Pimpinelli, Maurizio Fanciulli, Giovanni Blandino, Aldo Morrone, Gennaro Ciliberto |
| EPI_ISL_3419836 | IRCCS San Gallicano Dermatological Institute | IRCCS Regina Elena National Cancer Institute | Sabrina Strano, Sara Donzelli, Eleonora Sperandio, Lorenzo Furzi, Fabrizio Ensoli, Fulvia Pimpinelli, Maurizio Fanciulli, Giovanni Blandino, Aldo Morrone, Gennaro Ciliberto |
| EPI_ISL_3419837 | IRCCS San Gallicano Dermatological Institute | IRCCS Regina Elena National Cancer Institute | Francesca De Nicola, Giulia Orlandi, Alice Massacci, Alessia Lauretti, Fabrizio Ensoli, Fulvia Pimpinelli, Maurizio Fanciulli, Giovanni Blandino, Aldo Morrone, Gennaro Ciliberto |
| EPI_ISL_3419838 | IRCCS San Gallicano Dermatological Institute | IRCCS Regina Elena National Cancer Institute | Ludovica Ciuffreda, Sara Donzelli, Matteo Pallocca, Francesca Maione, Fabrizio Ensoli, Fulvia Pimpinelli, Maurizio Fanciulli, Giovanni Blandino, Aldo Morrone, Gennaro Ciliberto |
| EPI_ISL_3419839 | IRCCS San Gallicano Dermatological Institute | IRCCS Regina Elena National Cancer Institute | Frauke Goeman, Giulia Orlandi, Eleonora Sperandio, Francisco Obregon, Fabrizio Ensoli, Fulvia Pimpinelli, Maurizio Fanciulli, Giovanni Blandino, Aldo Morrone, Gennaro Ciliberto |
| EPI_ISL_3419841 | IRCCS San Gallicano Dermatological Institute | IRCCS Regina Elena National Cancer Institute | Francesca De Nicola, Giulia Orlandi, Matteo Pallocca, Sara Petrolo, Fabrizio Ensoli, Fulvia Pimpinelli, Maurizio Fanciulli, Giovanni Blandino, Aldo Morrone, Gennaro Ciliberto |
| EPI_ISL_3419842 | IRCCS San Gallicano Dermatological Institute | IRCCS Regina Elena National Cancer Institute | Ludovica Ciuffreda, Sara Donzelli, Eleonora Sperandio, Valentina Ricca, Fabrizio Ensoli, Fulvia Pimpinelli, Maurizio Fanciulli, Giovanni Blandino, Aldo Morrone, Gennaro Ciliberto |
| EPI_ISL_3419843 | IRCCS San Gallicano Dermatological Institute | IRCCS Regina Elena National Cancer Institute | Frauke Goeman, Giulia Orlandi, Alice Massacci, Serena Salvo, Fabrizio Ensoli, Fulvia Pimpinelli, Maurizio Fanciulli, Giovanni Blandino, Aldo Morrone, Gennaro Ciliberto |
| EPI_ISL_3419844 | IRCCS San Gallicano Dermatological Institute | IRCCS Regina Elena National Cancer Institute | Sabrina Strano, Sara Donzelli, Matteo Pallocca, Grazia Prignano, Fabrizio Ensoli, Fulvia Pimpinelli, Maurizio Fanciulli, Giovanni Blandino, Aldo Morrone, Gennaro Ciliberto |
| EPI_ISL_3419845 | IRCCS San Gallicano Dermatological Institute | IRCCS Regina Elena National Cancer Institute | Francesca De Nicola, Giulia Orlandi, Eleonora Sperandio, Elisabetta Trento, Fabrizio Ensoli, Fulvia Pimpinelli, Maurizio Fanciulli, Giovanni Blandino, Aldo Morrone, Gennaro Ciliberto |
| EPI_ISL_3419846 | IRCCS San Gallicano Dermatological Institute | IRCCS Regina Elena National Cancer Institute | Ludovica Ciuffreda, Sara Donzelli, Alice Massacci, Arianna Mastrofrancesco, Fabrizio Ensoli, Fulvia Pimpinelli, Maurizio Fanciulli, Giovanni Blandino, Aldo Morrone, Gennaro Ciliberto |
| EPI_ISL_3419847 | IRCCS San Gallicano Dermatological Institute | IRCCS Regina Elena National Cancer Institute | Frauke Goeman, Giulia Orlandi, Matteo Pallocca, Giovanna D'agosto, Fabrizio Ensoli, Fulvia Pimpinelli, Maurizio Fanciulli, Giovanni Blandino, Aldo Morrone, Gennaro Ciliberto |
| EPI_ISL_3419848 | IRCCS San Gallicano Dermatological Institute | IRCCS Regina Elena National Cancer Institute | Sabrina Strano, Sara Donzelli, Eleonora Sperandio, Carla Mottini, Fabrizio Ensoli, Fulvia Pimpinelli, Maurizio Fanciulli, Giovanni Blandino, Aldo Morrone, Gennaro Ciliberto |
| EPI_ISL_3419849 | IRCCS San Gallicano Dermatological Institute | IRCCS Regina Elena National Cancer Institute | Francesca De Nicola, Giulia Orlandi, Alice Massacci, Martina Pontone, Fabrizio Ensoli, Fulvia Pimpinelli, Maurizio Fanciulli, Giovanni Blandino, Aldo Morrone, Gennaro Ciliberto |
| EPI_ISL_3419850 | IRCCS San Gallicano Dermatological Institute | IRCCS Regina Elena National Cancer Institute | Ludovica Ciuffreda, Sara Donzelli, Matteo Pallocca, di Domenico Enea Gino, Fabrizio Ensoli, Fulvia Pimpinelli, Maurizio Fanciulli, Giovanni Blandino, Aldo Morrone, Gennaro Ciliberto |
| EPI_ISL_3419851 | IRCCS San Gallicano Dermatological Institute | IRCCS Regina Elena National Cancer Institute | Frauke Goeman, Giulia Orlandi, Eleonora Sperandio, Ilaria Cavallo, Fabrizio Ensoli, Fulvia Pimpinelli, Maurizio Fanciulli, Giovanni Blandino, Aldo Morrone, Gennaro Ciliberto |
| EPI_ISL_3419852 | IRCCS San Gallicano Dermatological Institute | IRCCS Regina Elena National Cancer Institute | Sabrina Strano, Sara Donzelli, Alice Massacci, Francesca Sivori, Fabrizio Ensoli, Fulvia Pimpinelli, Maurizio Fanciulli, Giovanni Blandino, Aldo Morrone, Gennaro Ciliberto |
| EPI_ISL_3419853 | IRCCS San Gallicano Dermatological Institute | IRCCS Regina Elena National Cancer Institute | Francesca De Nicola, Giulia Orlandi, Matteo Pallocca, Andrea Cazzani, Fabrizio Ensoli, Fulvia Pimpinelli, Maurizio Fanciulli, Giovanni Blandino, Aldo Morrone, Gennaro Ciliberto |
| EPI_ISL_3419854 | IRCCS San Gallicano Dermatological Institute | IRCCS Regina Elena National Cancer Institute | Ludovica Ciuffreda, Sara Donzelli, Eleonora Sperandio, Ilaria Celesti, Fabrizio Ensoli, Fulvia Pimpinelli, Maurizio Fanciulli, Giovanni Blandino, Aldo Morrone, Gennaro Ciliberto |
| EPI_ISL_3419855 | IRCCS San Gallicano Dermatological Institute | IRCCS Regina Elena National Cancer Institute | Frauke Goeman, Giulia Orlandi, Alice Massacci, Martina Diano, Fabrizio Ensoli, Fulvia Pimpinelli, Maurizio Fanciulli, Giovanni Blandino, Aldo Morrone, Gennaro Ciliberto |
| EPI_ISL_3419856 | IRCCS San Gallicano Dermatological Institute | IRCCS Regina Elena National Cancer Institute | Sabrina Strano, Sara Donzelli, Matteo Pallocca, Antonio Federico, Fabrizio Ensoli, Fulvia Pimpinelli, Maurizio Fanciulli, Giovanni Blandino, Aldo Morrone, Gennaro Ciliberto |
| EPI_ISL_3419857 | IRCCS San Gallicano Dermatological Institute | IRCCS Regina Elena National Cancer Institute | Francesca De Nicola, Giulia Orlandi, Eleonora Sperandio, Fulvia Fraticelli, Fabrizio Ensoli, Fulvia Pimpinelli, Maurizio Fanciulli, Giovanni Blandino, Aldo Morrone, Gennaro Ciliberto |
| EPI_ISL_3419858 | IRCCS San Gallicano Dermatological Institute | IRCCS Regina Elena National Cancer Institute | Ludovica Ciuffreda, Sara Donzelli, Alice Massacci, Lorenzo Furzi, Fabrizio Ensoli, Fulvia Pimpinelli, Maurizio Fanciulli, Giovanni Blandino, Aldo Morrone, Gennaro Ciliberto |
| EPI_ISL_3419859 | IRCCS San Gallicano Dermatological Institute | IRCCS Regina Elena National Cancer Institute | Frauke Goeman, Giulia Orlandi, Matteo Pallocca, Alessia Lauretti, Fabrizio Ensoli, Fulvia Pimpinelli, Maurizio Fanciulli, Giovanni Blandino, Aldo Morrone, Gennaro Ciliberto |
| EPI_ISL_3419860 | IRCCS San Gallicano Dermatological Institute | IRCCS Regina Elena National Cancer Institute | Francesca De Nicola, Giulia Orlandi, Alice Massacci, Francisco Obregon, Fabrizio Ensoli, Fulvia Pimpinelli, Maurizio Fanciulli, Giovanni Blandino, Aldo Morrone, Gennaro Ciliberto |
| EPI_ISL_3419861 | IRCCS San Gallicano Dermatological Institute | IRCCS Regina Elena National Cancer Institute | Ludovica Ciuffreda, Sara Donzelli, Matteo Pallocca, Silvia Paluzzi, Fabrizio Ensoli, Fulvia Pimpinelli, Maurizio Fanciulli, Giovanni Blandino, Aldo Morrone, Gennaro Ciliberto |
| EPI_ISL_3419862 | IRCCS San Gallicano Dermatological Institute | IRCCS Regina Elena National Cancer Institute | Frauke Goeman, Giulia Orlandi, Eleonora Sperandio, Sara Petrolo, Fabrizio Ensoli, Fulvia Pimpinelli, Maurizio Fanciulli, Giovanni Blandino, Aldo Morrone, Gennaro Ciliberto |
| EPI_ISL_3419864 | IRCCS San Gallicano Dermatological Institute | IRCCS Regina Elena National Cancer Institute | Francesca De Nicola, Giulia Orlandi, Matteo Pallocca, Serena Salvo, Fabrizio Ensoli, Fulvia Pimpinelli, Maurizio Fanciulli, Giovanni Blandino, Aldo Morrone, Gennaro Ciliberto |
| EPI_ISL_3419865 | IRCCS San Gallicano Dermatological Institute | IRCCS Regina Elena National Cancer Institute | Ludovica Ciuffreda, Sara Donzelli, Eleonora Sperandio, Grazia Prignano, Fabrizio Ensoli, Fulvia Pimpinelli, Maurizio Fanciulli, Giovanni Blandino, Aldo Morrone, Gennaro Ciliberto |
| EPI_ISL_3419866 | IRCCS San Gallicano Dermatological Institute | IRCCS Regina Elena National Cancer Institute | Frauke Goeman, Giulia Orlandi, Alice Massacci, Elisabetta Trento, Fabrizio Ensoli, Fulvia Pimpinelli, Maurizio Fanciulli, Giovanni Blandino, Aldo Morrone, Gennaro Ciliberto |
| EPI_ISL_3419867 | IRCCS San Gallicano Dermatological Institute | IRCCS Regina Elena National Cancer Institute | Sabrina Strano, Sara Donzelli, Matteo Pallocca, Arianna Mastrofrancesco, Fabrizio Ensoli, Fulvia Pimpinelli, Maurizio Fanciulli, Giovanni Blandino, Aldo Morrone, Gennaro Ciliberto |
| EPI_ISL_3419869 | IRCCS San Gallicano Dermatological Institute | IRCCS Regina Elena National Cancer Institute | Ludovica Ciuffreda, Sara Donzelli, Alice Massacci, Carla Mottini, Fabrizio Ensoli, Fulvia Pimpinelli, Maurizio Fanciulli, Giovanni Blandino, Aldo Morrone, Gennaro Ciliberto |
| EPI_ISL_3419871 | IRCCS San Gallicano Dermatological Institute | IRCCS Regina Elena National Cancer Institute | Sabrina Strano, Sara Donzelli, Eleonora Sperandio, di Domenico Enea Gino, Fabrizio Ensoli, Fulvia Pimpinelli, Maurizio Fanciulli, Giovanni Blandino, Aldo Morrone, Gennaro Ciliberto |
| EPI_ISL_3419872 | IRCCS San Gallicano Dermatological Institute | IRCCS Regina Elena National Cancer Institute | Francesca De Nicola, Giulia Orlandi, Alice Massacci, Ilaria Cavallo, Fabrizio Ensoli, Fulvia Pimpinelli, Maurizio Fanciulli, Giovanni Blandino, Aldo Morrone, Gennaro Ciliberto |
| EPI_ISL_3419873 | IRCCS San Gallicano Dermatological Institute | IRCCS Regina Elena National Cancer Institute | Ludovica Ciuffreda, Sara Donzelli, Matteo Pallocca, Francesca Sivori, Fabrizio Ensoli, Fulvia Pimpinelli, Maurizio Fanciulli, Giovanni Blandino, Aldo Morrone, Gennaro Ciliberto |
| EPI_ISL_3419874 | IRCCS San Gallicano Dermatological Institute | IRCCS Regina Elena National Cancer Institute | Frauke Goeman, Giulia Orlandi, Eleonora Sperandio, Andrea Cazzani, Fabrizio Ensoli, Fulvia Pimpinelli, Maurizio Fanciulli, Giovanni Blandino, Aldo Morrone, Gennaro Ciliberto |
| EPI_ISL_3419875 | IRCCS San Gallicano Dermatological Institute | IRCCS Regina Elena National Cancer Institute | Sabrina Strano, Sara Donzelli, Alice Massacci, Ilaria Celesti, Fabrizio Ensoli, Fulvia Pimpinelli, Maurizio Fanciulli, Giovanni Blandino, Aldo Morrone, Gennaro Ciliberto |

| | | | |
|---|---|---|---|
| EPI_ISL_3419876 | IRCCS San Gallicano Dermatological Institute | IRCCS Regina Elena National Cancer Institute | Francesca De Nicola, Giulia Orlandi, Matteo Pallocca, Martina Diano, Fabrizio Ensoli, Fulvia Pimpinelli, Maurizio Fanciulli, Giovanni Blandino, Aldo Morrone, Gennaro Ciliberto |
| EPI_ISL_3419877 | IRCCS San Gallicano Dermatological Institute | IRCCS Regina Elena National Cancer Institute | Ludovica Ciuffreda, Sara Donzelli, Eleonora Sperandio, Antonio Federico, Fabrizio Ensoli, Fulvia Pimpinelli, Maurizio Fanciulli, Giovanni Blandino, Aldo Morrone, Gennaro Ciliberto |
| EPI_ISL_3419878 | IRCCS San Gallicano Dermatological Institute | IRCCS Regina Elena National Cancer Institute | Frauke Goeman, Giulia Orlandi, Alice Massacci, Fulvia Fraticelli, Fabrizio Ensoli, Fulvia Pimpinelli, Maurizio Fanciulli, Giovanni Blandino, Aldo Morrone, Gennaro Ciliberto |
| EPI_ISL_3419879 | IRCCS San Gallicano Dermatological Institute | IRCCS Regina Elena National Cancer Institute | Francesca De Nicola, Giulia Orlandi, Eleonora Sperandio, Alessia Lauretti, Fabrizio Ensoli, Fulvia Pimpinelli, Maurizio Fanciulli, Giovanni Blandino, Aldo Morrone, Gennaro Ciliberto |
| EPI_ISL_3419880 | IRCCS San Gallicano Dermatological Institute | IRCCS Regina Elena National Cancer Institute | Ludovica Ciuffreda, Sara Donzelli, Alice Massacci, Francesca Maione, Fabrizio Ensoli, Fulvia Pimpinelli, Maurizio Fanciulli, Giovanni Blandino, Aldo Morrone, Gennaro Ciliberto |
| EPI_ISL_3419881 | IRCCS San Gallicano Dermatological Institute | IRCCS Regina Elena National Cancer Institute | Sabrina Strano, Sara Donzelli, Eleonora Sperandio, Silvia Paluzzi, Fabrizio Ensoli, Fulvia Pimpinelli, Maurizio Fanciulli, Giovanni Blandino, Aldo Morrone, Gennaro Ciliberto |
| EPI_ISL_3419882 | IRCCS San Gallicano Dermatological Institute | IRCCS Regina Elena National Cancer Institute | Francesca De Nicola, Giulia Orlandi, Alice Massacci, Sara Petrolo, Fabrizio Ensoli, Fulvia Pimpinelli, Maurizio Fanciulli, Giovanni Blandino, Aldo Morrone, Gennaro Ciliberto |
| EPI_ISL_3419883 | IRCCS San Gallicano Dermatological Institute | IRCCS Regina Elena National Cancer Institute | Ludovica Ciuffreda, Sara Donzelli, Matteo Pallocca, Valentina Ricca, Fabrizio Ensoli, Fulvia Pimpinelli, Maurizio Fanciulli, Giovanni Blandino, Aldo Morrone, Gennaro Ciliberto |
| EPI_ISL_3419884 | IRCCS San Gallicano Dermatological Institute | IRCCS Regina Elena National Cancer Institute | Frauke Goeman, Giulia Orlandi, Eleonora Sperandio, Serena Salvo, Fabrizio Ensoli, Fulvia Pimpinelli, Maurizio Fanciulli, Giovanni Blandino, Aldo Morrone, Gennaro Ciliberto |
| EPI_ISL_3419885 | IRCCS San Gallicano Dermatological Institute | IRCCS Regina Elena National Cancer Institute | Sabrina Strano, Sara Donzelli, Alice Massacci, Grazia Prignano, Fabrizio Ensoli, Fulvia Pimpinelli, Maurizio Fanciulli, Giovanni Blandino, Aldo Morrone, Gennaro Ciliberto |
| EPI_ISL_3419886 | IRCCS San Gallicano Dermatological Institute | IRCCS Regina Elena National Cancer Institute | Francesca De Nicola, Giulia Orlandi, Matteo Pallocca, Elisabetta Trento, Fabrizio Ensoli, Fulvia Pimpinelli, Maurizio Fanciulli, Giovanni Blandino, Aldo Morrone, Gennaro Ciliberto |
| EPI_ISL_3419887 | IRCCS San Gallicano Dermatological Institute | IRCCS Regina Elena National Cancer Institute | Ludovica Ciuffreda, Sara Donzelli, Eleonora Sperandio, Arianna Mastrofrancesco, Fabrizio Ensoli, Fulvia Pimpinelli, Maurizio Fanciulli, Giovanni Blandino, Aldo Morrone, Gennaro Ciliberto |
| EPI_ISL_3419888 | IRCCS San Gallicano Dermatological Institute | IRCCS Regina Elena National Cancer Institute | Frauke Goeman, Giulia Orlandi, Alice Massacci, Giovanna D'agosto, Fabrizio Ensoli, Fulvia Pimpinelli, Maurizio Fanciulli, Giovanni Blandino, Aldo Morrone, Gennaro Ciliberto |
| EPI_ISL_3419889 | IRCCS San Gallicano Dermatological Institute | IRCCS Regina Elena National Cancer Institute | Sabrina Strano, Sara Donzelli, Matteo Pallocca, Carla Mottini, Fabrizio Ensoli, Fulvia Pimpinelli, Maurizio Fanciulli, Giovanni Blandino, Aldo Morrone, Gennaro Ciliberto |
| EPI_ISL_3419890 | IRCCS San Gallicano Dermatological Institute | IRCCS Regina Elena National Cancer Institute | Francesca De Nicola, Giulia Orlandi, Eleonora Sperandio, Martina Pontone, Fabrizio Ensoli, Fulvia Pimpinelli, Maurizio Fanciulli, Giovanni Blandino, Aldo Morrone, Gennaro Ciliberto |
| EPI_ISL_3419891 | IRCCS San Gallicano Dermatological Institute | IRCCS Regina Elena National Cancer Institute | Ludovica Ciuffreda, Sara Donzelli, Alice Massacci, di Domenico Enea Gino, Fabrizio Ensoli, Fulvia Pimpinelli, Maurizio Fanciulli, Giovanni Blandino, Aldo Morrone, Gennaro Ciliberto |
| EPI_ISL_3419892 | IRCCS San Gallicano Dermatological Institute | IRCCS Regina Elena National Cancer Institute | Frauke Goeman, Giulia Orlandi, Matteo Pallocca, Ilaria Cavallo, Fabrizio Ensoli, Fulvia Pimpinelli, Maurizio Fanciulli, Giovanni Blandino, Aldo Morrone, Gennaro Ciliberto |
| EPI_ISL_3419893 | IRCCS San Gallicano Dermatological Institute | IRCCS Regina Elena National Cancer Institute | Sabrina Strano, Sara Donzelli, Eleonora Sperandio, Francesca Sivori, Fabrizio Ensoli, Fulvia Pimpinelli, Maurizio Fanciulli, Giovanni Blandino, Aldo Morrone, Gennaro Ciliberto |
| EPI_ISL_3419894 | IRCCS San Gallicano Dermatological Institute | IRCCS Regina Elena National Cancer Institute | Francesca De Nicola, Giulia Orlandi, Alice Massacci, Andrea Cazzani, Fabrizio Ensoli, Fulvia Pimpinelli, Maurizio Fanciulli, Giovanni Blandino, Aldo Morrone, Gennaro Ciliberto |
| EPI_ISL_3419896 | IRCCS San Gallicano Dermatological Institute | IRCCS Regina Elena National Cancer Institute | Frauke Goeman, Giulia Orlandi, Eleonora Sperandio, Martina Diano, Fabrizio Ensoli, Fulvia Pimpinelli, Maurizio Fanciulli, Giovanni Blandino, Aldo Morrone, Gennaro Ciliberto |
| EPI_ISL_3419897 | IRCCS San Gallicano Dermatological Institute | IRCCS Regina Elena National Cancer Institute | Sabrina Strano, Sara Donzelli, Alice Massacci, Antonio Federico, Fabrizio Ensoli, Fulvia Pimpinelli, Maurizio Fanciulli, Giovanni Blandino, Aldo Morrone, Gennaro Ciliberto |
| EPI_ISL_3419898 | IRCCS San Gallicano Dermatological Institute | IRCCS Regina Elena National Cancer Institute | Francesca De Nicola, Giulia Orlandi, Matteo Pallocca, Fulvia Fraticelli, Fabrizio Ensoli, Fulvia Pimpinelli, Maurizio Fanciulli, Giovanni Blandino, Aldo Morrone, Gennaro Ciliberto |
| EPI_ISL_3419899 | IRCCS San Gallicano Dermatological Institute | IRCCS Regina Elena National Cancer Institute | Ludovica Ciuffreda, Sara Donzelli, Eleonora Sperandio, Lorenzo Furzi, Fabrizio Ensoli, Fulvia Pimpinelli, Maurizio Fanciulli, Giovanni Blandino, Aldo Morrone, Gennaro Ciliberto |
| EPI_ISL_3419900 | IRCCS San Gallicano Dermatological Institute | IRCCS Regina Elena National Cancer Institute | Frauke Goeman, Giulia Orlandi, Alice Massacci, Alessia Lauretti, Fabrizio Ensoli, Fulvia Pimpinelli, Maurizio Fanciulli, Giovanni Blandino, Aldo Morrone, Gennaro Ciliberto |
| EPI_ISL_3419901 | IRCCS San Gallicano Dermatological Institute | IRCCS Regina Elena National Cancer Institute | Sabrina Strano, Sara Donzelli, Matteo Pallocca, Francesca Maione, Fabrizio Ensoli, Fulvia Pimpinelli, Maurizio Fanciulli, Giovanni Blandino, Aldo Morrone, Gennaro Ciliberto |
| EPI_ISL_3419902 | IRCCS San Gallicano Dermatological Institute | IRCCS Regina Elena National Cancer Institute | Francesca De Nicola, Giulia Orlandi, Eleonora Sperandio, Francisco Obregon, Fabrizio Ensoli, Fulvia Pimpinelli, Maurizio Fanciulli, Giovanni Blandino, Aldo Morrone, Gennaro Ciliberto |
| EPI_ISL_3419903 | IRCCS San Gallicano Dermatological Institute | IRCCS Regina Elena National Cancer Institute | Ludovica Ciuffreda, Sara Donzelli, Alice Massacci, Silvia Paluzzi, Fabrizio Ensoli, Fulvia Pimpinelli, Maurizio Fanciulli, Giovanni Blandino, Aldo Morrone, Gennaro Ciliberto |
| EPI_ISL_3419904 | IRCCS San Gallicano Dermatological Institute | IRCCS Regina Elena National Cancer Institute | Frauke Goeman, Giulia Orlandi, Matteo Pallocca, Sara Petrolo, Fabrizio Ensoli, Fulvia Pimpinelli, Maurizio Fanciulli, Giovanni Blandino, Aldo Morrone, Gennaro Ciliberto |
| EPI_ISL_3419905 | IRCCS San Gallicano Dermatological Institute | IRCCS Regina Elena National Cancer Institute | Sabrina Strano, Sara Donzelli, Eleonora Sperandio, Valentina Ricca, Fabrizio Ensoli, Fulvia Pimpinelli, Maurizio Fanciulli, Giovanni Blandino, Aldo Morrone, Gennaro Ciliberto |
| EPI_ISL_3419906 | IRCCS San Gallicano Dermatological Institute | IRCCS Regina Elena National Cancer Institute | Francesca De Nicola, Giulia Orlandi, Alice Massacci, Serena Salvo, Fabrizio Ensoli, Fulvia Pimpinelli, Maurizio Fanciulli, Giovanni Blandino, Aldo Morrone, Gennaro Ciliberto |
| EPI_ISL_3419907 | IRCCS San Gallicano Dermatological Institute | IRCCS Regina Elena National Cancer Institute | Ludovica Ciuffreda, Sara Donzelli, Matteo Pallocca, Grazia Prignano, Fabrizio Ensoli, Fulvia Pimpinelli, Maurizio Fanciulli, Giovanni Blandino, Aldo Morrone, Gennaro Ciliberto |
| EPI_ISL_3419908 | IRCCS San Gallicano Dermatological Institute | IRCCS Regina Elena National Cancer Institute | Frauke Goeman, Giulia Orlandi, Eleonora Sperandio, Elisabetta Trento, Fabrizio Ensoli, Fulvia Pimpinelli, Maurizio Fanciulli, Giovanni Blandino, Aldo Morrone, Gennaro Ciliberto |
| EPI_ISL_3419909 | IRCCS San Gallicano Dermatological Institute | IRCCS Regina Elena National Cancer Institute | Sabrina Strano, Sara Donzelli, Alice Massacci, Arianna Mastrofrancesco, Fabrizio Ensoli, Fulvia Pimpinelli, Maurizio Fanciulli, Giovanni Blandino, Aldo Morrone, Gennaro Ciliberto |
| EPI_ISL_3419910 | IRCCS San Gallicano Dermatological Institute | IRCCS Regina Elena National Cancer Institute | Francesca De Nicola, Giulia Orlandi, Matteo Pallocca, Giovanna D'agosto, Fabrizio Ensoli, Fulvia Pimpinelli, Maurizio Fanciulli, Giovanni Blandino, Aldo Morrone, Gennaro Ciliberto |
| EPI_ISL_3419911 | IRCCS San Gallicano Dermatological Institute | IRCCS Regina Elena National Cancer Institute | Ludovica Ciuffreda, Sara Donzelli, Eleonora Sperandio, Carla Mottini, Fabrizio Ensoli, Fulvia Pimpinelli, Maurizio Fanciulli, Giovanni Blandino, Aldo Morrone, Gennaro Ciliberto |
| EPI_ISL_3419912 | IRCCS San Gallicano Dermatological Institute | IRCCS Regina Elena National Cancer Institute | Frauke Goeman, Giulia Orlandi, Alice Massacci, Martina Pontone, Fabrizio Ensoli, Fulvia Pimpinelli, Maurizio Fanciulli, Giovanni Blandino, Aldo Morrone, Gennaro Ciliberto |
| EPI_ISL_3419913 | IRCCS San Gallicano Dermatological Institute | IRCCS Regina Elena National Cancer Institute | Sabrina Strano, Sara Donzelli, Matteo Pallocca, di Domenico Enea Gino, Fabrizio Ensoli, Fulvia Pimpinelli, Maurizio Fanciulli, Giovanni Blandino, Aldo Morrone, Gennaro Ciliberto |
| EPI_ISL_3419914 | IRCCS San Gallicano Dermatological Institute | IRCCS Regina Elena National Cancer Institute | Francesca De Nicola, Giulia Orlandi, Eleonora Sperandio, Ilaria Cavallo, Fabrizio Ensoli, Fulvia Pimpinelli, Maurizio Fanciulli, Giovanni Blandino, Aldo Morrone, Gennaro Ciliberto |
| EPI_ISL_3419915 | IRCCS San Gallicano Dermatological Institute | IRCCS Regina Elena National Cancer Institute | Ludovica Ciuffreda, Sara Donzelli, Alice Massacci, Francesca Sivori, Fabrizio Ensoli, Fulvia Pimpinelli, Maurizio Fanciulli, Giovanni Blandino, Aldo Morrone, Gennaro Ciliberto |
| EPI_ISL_3419916 | IRCCS San Gallicano Dermatological Institute | IRCCS Regina Elena National Cancer Institute | Frauke Goeman, Giulia Orlandi, Matteo Pallocca, Andrea Cazzani, Fabrizio Ensoli, Fulvia Pimpinelli, Maurizio Fanciulli, Giovanni Blandino, Aldo Morrone, Gennaro Ciliberto |
| EPI_ISL_3419917 | IRCCS San Gallicano Dermatological Institute | IRCCS Regina Elena National Cancer Institute | Sabrina Strano, Sara Donzelli, Eleonora Sperandio, Ilaria Celesti, Fabrizio Ensoli, Fulvia Pimpinelli, Maurizio Fanciulli, Giovanni Blandino, Aldo Morrone, Gennaro Ciliberto |
| EPI_ISL_3419918 | IRCCS San Gallicano Dermatological Institute | IRCCS Regina Elena National Cancer Institute | Francesca De Nicola, Giulia Orlandi, Alice Massacci, Martina Diano, Fabrizio Ensoli, Fulvia Pimpinelli, Maurizio Fanciulli, Giovanni Blandino, Aldo Morrone, Gennaro Ciliberto |
| EPI_ISL_3419919 | IRCCS San Gallicano Dermatological Institute | IRCCS Regina Elena National Cancer Institute | Ludovica Ciuffreda, Sara Donzelli, Matteo Pallocca, Antonio Federico, Fabrizio Ensoli, Fulvia Pimpinelli, Maurizio Fanciulli, Giovanni Blandino, Aldo Morrone, Gennaro Ciliberto |
| EPI_ISL_3419920 | IRCCS San Gallicano Dermatological Institute | IRCCS Regina Elena National Cancer Institute | Frauke Goeman, Giulia Orlandi, Eleonora Sperandio, Fulvia Fraticelli, Fabrizio Ensoli, Fulvia Pimpinelli, Maurizio Fanciulli, Giovanni Blandino, Aldo Morrone, Gennaro Ciliberto |
| EPI_ISL_3419921 | IRCCS San Gallicano Dermatological Institute | IRCCS Regina Elena National Cancer Institute | Sabrina Strano, Sara Donzelli, Alice Massacci, Lorenzo Furzi, Fabrizio Ensoli, Fulvia Pimpinelli, Maurizio Fanciulli, Giovanni Blandino, Aldo Morrone, Gennaro Ciliberto |
| EPI_ISL_3419922 | IRCCS San Gallicano Dermatological Institute | IRCCS Regina Elena National Cancer Institute | Francesca De Nicola, Giulia Orlandi, Matteo Pallocca, Alessia Lauretti, Fabrizio Ensoli, Fulvia Pimpinelli, Maurizio Fanciulli, Giovanni Blandino, Aldo Morrone, Gennaro Ciliberto |
| EPI_ISL_3419923 | IRCCS San Gallicano Dermatological Institute | IRCCS Regina Elena National Cancer Institute | Ludovica Ciuffreda, Sara Donzelli, Eleonora Sperandio, Francesca Maione, Fabrizio Ensoli, Fulvia Pimpinelli, Maurizio Fanciulli, Giovanni Blandino, Aldo Morrone, Gennaro Ciliberto |
| EPI_ISL_3419924 | IRCCS San Gallicano Dermatological Institute | IRCCS Regina Elena National Cancer Institute | Frauke Goeman, Giulia Orlandi, Alice Massacci, Francisco Obregon, Fabrizio Ensoli, Fulvia Pimpinelli, Maurizio Fanciulli, Giovanni Blandino, Aldo Morrone, Gennaro Ciliberto |
| EPI_ISL_3419925 | IRCCS San Gallicano Dermatological Institute | IRCCS Regina Elena National Cancer Institute | Sabrina Strano, Sara Donzelli, Matteo Pallocca, Silvia Paluzzi, Fabrizio Ensoli, Fulvia Pimpinelli, Maurizio Fanciulli, Giovanni Blandino, Aldo Morrone, Gennaro Ciliberto |
| EPI_ISL_3419926 | IRCCS San Gallicano Dermatological Institute | IRCCS Regina Elena National Cancer Institute | Francesca De Nicola, Giulia Orlandi, Eleonora Sperandio, Sara Petrolo, Fabrizio Ensoli, Fulvia Pimpinelli, Maurizio Fanciulli, Giovanni Blandino, Aldo Morrone, Gennaro Ciliberto |
| EPI_ISL_3419927 | IRCCS San Gallicano Dermatological Institute | IRCCS Regina Elena National Cancer Institute | Ludovica Ciuffreda, Sara Donzelli, Alice Massacci, Valentina Ricca, Fabrizio Ensoli, Fulvia Pimpinelli, Maurizio Fanciulli, Giovanni Blandino, Aldo Morrone, Gennaro Ciliberto |
| EPI_ISL_3419928 | IRCCS San Gallicano Dermatological Institute | IRCCS Regina Elena National Cancer Institute | Frauke Goeman, Giulia Orlandi, Matteo Pallocca, Serena Salvo, Fabrizio Ensoli, Fulvia Pimpinelli, Maurizio Fanciulli, Giovanni Blandino, Aldo Morrone, Gennaro Ciliberto |
| EPI_ISL_3419929 | IRCCS San Gallicano Dermatological Institute | IRCCS Regina Elena National Cancer Institute | Sabrina Strano, Sara Donzelli, Eleonora Sperandio, Grazia Prignano, Fabrizio Ensoli, Fulvia Pimpinelli, Maurizio Fanciulli, Giovanni Blandino, Aldo Morrone, Gennaro Ciliberto |
| EPI_ISL_3419930 | IRCCS San Gallicano Dermatological Institute | IRCCS Regina Elena National Cancer Institute | Francesca De Nicola, Giulia Orlandi, Alice Massacci, Elisabetta Trento, Fabrizio Ensoli, Fulvia Pimpinelli, Maurizio Fanciulli, Giovanni Blandino, Aldo Morrone, Gennaro Ciliberto |
| EPI_ISL_3419931 | IRCCS San Gallicano Dermatological Institute | IRCCS Regina Elena National Cancer Institute | Ludovica Ciuffreda, Sara Donzelli, Matteo Pallocca, Arianna Mastrofrancesco, Fabrizio Ensoli, Fulvia Pimpinelli, Maurizio Fanciulli, Giovanni Blandino, Aldo Morrone, Gennaro Ciliberto |
| EPI_ISL_3419932 | IRCCS San Gallicano Dermatological Institute | IRCCS Regina Elena National Cancer Institute | Frauke Goeman, Giulia Orlandi, Eleonora Sperandio, Giovanna D'agosto, Fabrizio Ensoli, Fulvia Pimpinelli, Maurizio Fanciulli, Giovanni Blandino, Aldo Morrone, Gennaro Ciliberto |
| EPI_ISL_3419933 | IRCCS San Gallicano Dermatological Institute | IRCCS Regina Elena National Cancer Institute | Sabrina Strano, Sara Donzelli, Alice Massacci, Carla Mottini, Fabrizio Ensoli, Fulvia Pimpinelli, Maurizio Fanciulli, Giovanni Blandino, Aldo Morrone, Gennaro Ciliberto |
| EPI_ISL_3419934 | IRCCS San Gallicano Dermatological Institute | IRCCS Regina Elena National Cancer Institute | Francesca De Nicola, Giulia Orlandi, Matteo Pallocca, Martina Pontone, Fabrizio Ensoli, Fulvia Pimpinelli, Maurizio Fanciulli, Giovanni Blandino, Aldo Morrone, Gennaro Ciliberto |
| EPI_ISL_3419935 | IRCCS San Gallicano Dermatological Institute | IRCCS Regina Elena National Cancer Institute | Ludovica Ciuffreda, Sara Donzelli, Eleonora Sperandio, di Domenico Enea Gino, Fabrizio Ensoli, Fulvia Pimpinelli, Maurizio Fanciulli, Giovanni Blandino, Aldo Morrone, Gennaro Ciliberto |
| EPI_ISL_3419936 | IRCCS San Gallicano Dermatological Institute | IRCCS Regina Elena National Cancer Institute | Frauke Goeman, Giulia Orlandi, Alice Massacci, Ilaria Cavallo, Fabrizio Ensoli, Fulvia Pimpinelli, Maurizio Fanciulli, Giovanni Blandino, Aldo Morrone, Gennaro Ciliberto |
| EPI_ISL_3419937 | IRCCS San Gallicano Dermatological Institute | IRCCS Regina Elena National Cancer Institute | Sabrina Strano, Sara Donzelli, Matteo Pallocca, Francesca Sivori, Fabrizio Ensoli, Fulvia Pimpinelli, Maurizio Fanciulli, Giovanni Blandino, Aldo Morrone, Gennaro Ciliberto |
| EPI_ISL_3419938 | IRCCS San Gallicano Dermatological Institute | IRCCS Regina Elena National Cancer Institute | Francesca De Nicola, Giulia Orlandi, Eleonora Sperandio, Andrea Cazzani, Fabrizio Ensoli, Fulvia Pimpinelli, Maurizio Fanciulli, Giovanni Blandino, Aldo Morrone, Gennaro Ciliberto |
| EPI_ISL_3419939 | IRCCS San Gallicano Dermatological Institute | IRCCS Regina Elena National Cancer Institute | Ludovica Ciuffreda, Sara Donzelli, Alice Massacci, Ilaria Celesti, Fabrizio Ensoli, Fulvia Pimpinelli, Maurizio Fanciulli, Giovanni Blandino, Aldo Morrone, Gennaro Ciliberto |
| EPI_ISL_3419942 | IRCCS San Gallicano Dermatological Institute | IRCCS Regina Elena National Cancer Institute | Francesca De Nicola, Giulia Orlandi, Alice Massacci, Fulvia Fraticelli, Fabrizio Ensoli, Fulvia Pimpinelli, Maurizio Fanciulli, Giovanni Blandino, Aldo Morrone, Gennaro Ciliberto |
| EPI_ISL_3419943 | IRCCS San Gallicano Dermatological Institute | IRCCS Regina Elena National Cancer Institute | Ludovica Ciuffreda, Sara Donzelli, Matteo Pallocca, Lorenzo Furzi, Fabrizio Ensoli, Fulvia Pimpinelli, Maurizio Fanciulli, Giovanni Blandino, Aldo Morrone, Gennaro Ciliberto |
| EPI_ISL_3419944 | IRCCS San Gallicano Dermatological Institute | IRCCS Regina Elena National Cancer Institute | Frauke Goeman, Giulia Orlandi, Eleonora Sperandio, Alessia Lauretti, Fabrizio Ensoli, Fulvia Pimpinelli, Maurizio Fanciulli, Giovanni Blandino, Aldo Morrone, Gennaro Ciliberto |
| EPI_ISL_3419945 | IRCCS San Gallicano Dermatological Institute | IRCCS Regina Elena National Cancer Institute | Sabrina Strano, Sara Donzelli, Alice Massacci, Francesca Maione, Fabrizio Ensoli, Fulvia Pimpinelli, Maurizio Fanciulli, Giovanni Blandino, Aldo Morrone, Gennaro Ciliberto |
| EPI_ISL_3419946 | IRCCS San Gallicano Dermatological Institute | IRCCS Regina Elena National Cancer Institute | Francesca De Nicola, Giulia Orlandi, Matteo Pallocca, Francisco Obregon, Fabrizio Ensoli, Fulvia Pimpinelli, Maurizio Fanciulli, Giovanni Blandino, Aldo Morrone, Gennaro Ciliberto |
| EPI_ISL_3419947 | IRCCS San Gallicano Dermatological Institute | IRCCS Regina Elena National Cancer Institute | Ludovica Ciuffreda, Sara Donzelli, Eleonora Sperandio, Silvia Paluzzi, Fabrizio Ensoli, Fulvia Pimpinelli, Maurizio Fanciulli, Giovanni Blandino, Aldo Morrone, Gennaro Ciliberto |
| EPI_ISL_3419949 | IRCCS San Gallicano Dermatological Institute | IRCCS Regina Elena National Cancer Institute | Sabrina Strano, Sara Donzelli, Matteo Pallocca, Valentina Ricca, Fabrizio Ensoli, Fulvia Pimpinelli, Maurizio Fanciulli, Giovanni Blandino, Aldo Morrone, Gennaro Ciliberto |
| EPI_ISL_3419950 | IRCCS San Gallicano Dermatological Institute | IRCCS Regina Elena National Cancer Institute | Francesca De Nicola, Giulia Orlandi, Eleonora Sperandio, Serena Salvo, Fabrizio Ensoli, Fulvia Pimpinelli, Maurizio Fanciulli, Giovanni Blandino, Aldo Morrone, Gennaro Ciliberto |
| EPI_ISL_3419951 | IRCCS San Gallicano Dermatological Institute | IRCCS Regina Elena National Cancer Institute | Ludovica Ciuffreda, Sara Donzelli, Alice Massacci, Grazia Prignano, Fabrizio Ensoli, Fulvia Pimpinelli, Maurizio Fanciulli, Giovanni Blandino, Aldo Morrone, Gennaro Ciliberto |
| EPI_ISL_3419952 | IRCCS San Gallicano Dermatological Institute | IRCCS Regina Elena National Cancer Institute | Frauke Goeman, Giulia Orlandi, Matteo Pallocca, Elisabetta Trento, Fabrizio Ensoli, Fulvia Pimpinelli, Maurizio Fanciulli, Giovanni Blandino, Aldo Morrone, Gennaro Ciliberto |
| EPI_ISL_3419953 | IRCCS San Gallicano Dermatological Institute | IRCCS Regina Elena National Cancer Institute | Sabrina Strano, Sara Donzelli, Eleonora Sperandio, Arianna Mastrofrancesco, Fabrizio Ensoli, Fulvia Pimpinelli, Maurizio Fanciulli, Giovanni Blandino, Aldo Morrone, Gennaro Ciliberto |
| EPI_ISL_3419954 | IRCCS San Gallicano Dermatological Institute | IRCCS Regina Elena National Cancer Institute | Francesca De Nicola, Giulia Orlandi, Alice Massacci, Giovanna D'agosto, Fabrizio Ensoli, Fulvia Pimpinelli, Maurizio Fanciulli, Giovanni Blandino, Aldo Morrone, Gennaro Ciliberto |
| EPI_ISL_3419955 | IRCCS San Gallicano Dermatological Institute | IRCCS Regina Elena National Cancer Institute | Ludovica Ciuffreda, Sara Donzelli, Matteo Pallocca, Carla Mottini, Fabrizio Ensoli, Fulvia Pimpinelli, Maurizio Fanciulli, Giovanni Blandino, Aldo Morrone, Gennaro Ciliberto |

| | | | |
|---|---|---|---|
| EPI_ISL_3419956 | IRCCS San Gallicano Dermatological Institute | IRCCS Regina Elena National Cancer Institute | Frauke Goeman, Giulia Orlandi, Eleonora Sperandio, Martina Pontone, Fabrizio Ensoli, Fulvia Pimpinelli, Maurizio Fanciulli, Giovanni Blandino, Aldo Morrone, Gennaro Ciliberto |
| EPI_ISL_3419957 | IRCCS San Gallicano Dermatological Institute | IRCCS Regina Elena National Cancer Institute | Sabrina Strano, Sara Donzelli, Alice Massacci, di Domenico Enea Gino, Fabrizio Ensoli, Fulvia Pimpinelli, Maurizio Fanciulli, Giovanni Blandino, Aldo Morrone, Gennaro Ciliberto |
| EPI_ISL_3419958 | IRCCS San Gallicano Dermatological Institute | IRCCS Regina Elena National Cancer Institute | Francesca De Nicola, Giulia Orlandi, Matteo Pallocca, Ilaria Cavallo, Fabrizio Ensoli, Fulvia Pimpinelli, Maurizio Fanciulli, Giovanni Blandino, Aldo Morrone, Gennaro Ciliberto |
| EPI_ISL_3419959 | IRCCS San Gallicano Dermatological Institute | IRCCS Regina Elena National Cancer Institute | Ludovica Ciuffreda, Sara Donzelli, Eleonora Sperandio, Francesca Sivori, Fabrizio Ensoli, Fulvia Pimpinelli, Maurizio Fanciulli, Giovanni Blandino, Aldo Morrone, Gennaro Ciliberto |
| EPI_ISL_3419960 | IRCCS San Gallicano Dermatological Institute | IRCCS Regina Elena National Cancer Institute | Frauke Goeman, Giulia Orlandi, Alice Massacci, Andrea Cazzani, Fabrizio Ensoli, Fulvia Pimpinelli, Maurizio Fanciulli, Giovanni Blandino, Aldo Morrone, Gennaro Ciliberto |
| EPI_ISL_3419961 | IRCCS San Gallicano Dermatological Institute | IRCCS Regina Elena National Cancer Institute | Sabrina Strano, Sara Donzelli, Matteo Pallocca, Ilaria Celesti, Fabrizio Ensoli, Fulvia Pimpinelli, Maurizio Fanciulli, Giovanni Blandino, Aldo Morrone, Gennaro Ciliberto |
| EPI_ISL_3419962 | IRCCS San Gallicano Dermatological Institute | IRCCS Regina Elena National Cancer Institute | Francesca De Nicola, Giulia Orlandi, Eleonora Sperandio, Martina Diano, Fabrizio Ensoli, Fulvia Pimpinelli, Maurizio Fanciulli, Giovanni Blandino, Aldo Morrone, Gennaro Ciliberto |
| EPI_ISL_3419963 | IRCCS San Gallicano Dermatological Institute | IRCCS Regina Elena National Cancer Institute | Ludovica Ciuffreda, Sara Donzelli, Alice Massacci, Antonio Federico, Fabrizio Ensoli, Fulvia Pimpinelli, Maurizio Fanciulli, Giovanni Blandino, Aldo Morrone, Gennaro Ciliberto |
| EPI_ISL_3419964 | IRCCS San Gallicano Dermatological Institute | IRCCS Regina Elena National Cancer Institute | Frauke Goeman, Giulia Orlandi, Matteo Pallocca, Fulvia Fraticelli, Fabrizio Ensoli, Fulvia Pimpinelli, Maurizio Fanciulli, Giovanni Blandino, Aldo Morrone, Gennaro Ciliberto |
| EPI_ISL_3419965 | IRCCS San Gallicano Dermatological Institute | IRCCS Regina Elena National Cancer Institute | Sabrina Strano, Sara Donzelli, Eleonora Sperandio, Lorenzo Furzi, Fabrizio Ensoli, Fulvia Pimpinelli, Maurizio Fanciulli, Giovanni Blandino, Aldo Morrone, Gennaro Ciliberto |
| EPI_ISL_3419966 | IRCCS San Gallicano Dermatological Institute | IRCCS Regina Elena National Cancer Institute | Francesca De Nicola, Giulia Orlandi, Alice Massacci, Alessia Lauretti, Fabrizio Ensoli, Fulvia Pimpinelli, Maurizio Fanciulli, Giovanni Blandino, Aldo Morrone, Gennaro Ciliberto |
| EPI_ISL_3419967 | IRCCS San Gallicano Dermatological Institute | IRCCS Regina Elena National Cancer Institute | Ludovica Ciuffreda, Sara Donzelli, Matteo Pallocca, Francesca Maione, Fabrizio Ensoli, Fulvia Pimpinelli, Maurizio Fanciulli, Giovanni Blandino, Aldo Morrone, Gennaro Ciliberto |
| EPI_ISL_3419968 | IRCCS San Gallicano Dermatological Institute | IRCCS Regina Elena National Cancer Institute | Frauke Goeman, Giulia Orlandi, Eleonora Sperandio, Francisco Obregon, Fabrizio Ensoli, Fulvia Pimpinelli, Maurizio Fanciulli, Giovanni Blandino, Aldo Morrone, Gennaro Ciliberto |
| EPI_ISL_3419969 | IRCCS San Gallicano Dermatological Institute | IRCCS Regina Elena National Cancer Institute | Sabrina Strano, Sara Donzelli, Alice Massacci, Silvia Paluzzi, Fabrizio Ensoli, Fulvia Pimpinelli, Maurizio Fanciulli, Giovanni Blandino, Aldo Morrone, Gennaro Ciliberto |
| EPI_ISL_3419970 | IRCCS San Gallicano Dermatological Institute | IRCCS Regina Elena National Cancer Institute | Francesca De Nicola, Giulia Orlandi, Matteo Pallocca, Sara Petrolo, Fabrizio Ensoli, Fulvia Pimpinelli, Maurizio Fanciulli, Giovanni Blandino, Aldo Morrone, Gennaro Ciliberto |
| EPI_ISL_3419971 | IRCCS San Gallicano Dermatological Institute | IRCCS Regina Elena National Cancer Institute | Ludovica Ciuffreda, Sara Donzelli, Eleonora Sperandio, Valentina Ricca, Fabrizio Ensoli, Fulvia Pimpinelli, Maurizio Fanciulli, Giovanni Blandino, Aldo Morrone, Gennaro Ciliberto |
| EPI_ISL_3419972 | IRCCS San Gallicano Dermatological Institute | IRCCS Regina Elena National Cancer Institute | Frauke Goeman, Giulia Orlandi, Alice Massacci, Serena Salvo, Fabrizio Ensoli, Fulvia Pimpinelli, Maurizio Fanciulli, Giovanni Blandino, Aldo Morrone, Gennaro Ciliberto |
| EPI_ISL_3419973 | IRCCS San Gallicano Dermatological Institute | IRCCS Regina Elena National Cancer Institute | Sabrina Strano, Sara Donzelli, Matteo Pallocca, Grazia Prignano, Fabrizio Ensoli, Fulvia Pimpinelli, Maurizio Fanciulli, Giovanni Blandino, Aldo Morrone, Gennaro Ciliberto |
| EPI_ISL_3419974 | IRCCS San Gallicano Dermatological Institute | IRCCS Regina Elena National Cancer Institute | Francesca De Nicola, Giulia Orlandi, Eleonora Sperandio, Elisabetta Trento, Fabrizio Ensoli, Fulvia Pimpinelli, Maurizio Fanciulli, Giovanni Blandino, Aldo Morrone, Gennaro Ciliberto |
| EPI_ISL_3419975 | IRCCS San Gallicano Dermatological Institute | IRCCS Regina Elena National Cancer Institute | Ludovica Ciuffreda, Sara Donzelli, Alice Massacci, Arianna Mastrofrancesco, Fabrizio Ensoli, Fulvia Pimpinelli, Maurizio Fanciulli, Giovanni Blandino, Aldo Morrone, Gennaro Ciliberto |
| EPI_ISL_3419976 | IRCCS San Gallicano Dermatological Institute | IRCCS Regina Elena National Cancer Institute | Frauke Goeman, Giulia Orlandi, Matteo Pallocca, Giovanna D’agosto, Fabrizio Ensoli, Fulvia Pimpinelli, Maurizio Fanciulli, Giovanni Blandino, Aldo Morrone, Gennaro Ciliberto |
| EPI_ISL_3419977 | IRCCS San Gallicano Dermatological Institute | IRCCS Regina Elena National Cancer Institute | Sabrina Strano, Sara Donzelli, Eleonora Sperandio, Carla Mottini, Fabrizio Ensoli, Fulvia Pimpinelli, Maurizio Fanciulli, Giovanni Blandino, Aldo Morrone, Gennaro Ciliberto |
| EPI_ISL_3419978 | IRCCS San Gallicano Dermatological Institute | IRCCS Regina Elena National Cancer Institute | Francesca De Nicola, Giulia Orlandi, Alice Massacci, Martina Pontone, Fabrizio Ensoli, Fulvia Pimpinelli, Maurizio Fanciulli, Giovanni Blandino, Aldo Morrone, Gennaro Ciliberto |
| EPI_ISL_3419979 | IRCCS San Gallicano Dermatological Institute | IRCCS Regina Elena National Cancer Institute | Ludovica Ciuffreda, Sara Donzelli, Matteo Pallocca, di Domenico Enea Gino, Fabrizio Ensoli, Fulvia Pimpinelli, Maurizio Fanciulli, Giovanni Blandino, Aldo Morrone, Gennaro Ciliberto |
| EPI_ISL_3419980 | IRCCS San Gallicano Dermatological Institute | IRCCS Regina Elena National Cancer Institute | Frauke Goeman, Giulia Orlandi, Eleonora Sperandio, Ilaria Cavallo, Fabrizio Ensoli, Fulvia Pimpinelli, Maurizio Fanciulli, Giovanni Blandino, Aldo Morrone, Gennaro Ciliberto |
| EPI_ISL_3419981 | IRCCS San Gallicano Dermatological Institute | IRCCS Regina Elena National Cancer Institute | Sabrina Strano, Sara Donzelli, Alice Massacci, Francesca Sivori, Fabrizio Ensoli, Fulvia Pimpinelli, Maurizio Fanciulli, Giovanni Blandino, Aldo Morrone, Gennaro Ciliberto |
| EPI_ISL_3419982 | IRCCS San Gallicano Dermatological Institute | IRCCS Regina Elena National Cancer Institute | Francesca De Nicola, Giulia Orlandi, Matteo Pallocca, Andrea Cazzani, Fabrizio Ensoli, Fulvia Pimpinelli, Maurizio Fanciulli, Giovanni Blandino, Aldo Morrone, Gennaro Ciliberto |
| EPI_ISL_3419983 | IRCCS San Gallicano Dermatological Institute | IRCCS Regina Elena National Cancer Institute | Ludovica Ciuffreda, Sara Donzelli, Eleonora Sperandio, Ilaria Celesti, Fabrizio Ensoli, Fulvia Pimpinelli, Maurizio Fanciulli, Giovanni Blandino, Aldo Morrone, Gennaro Ciliberto |
| EPI_ISL_3419984 | IRCCS San Gallicano Dermatological Institute | IRCCS Regina Elena National Cancer Institute | Frauke Goeman, Giulia Orlandi, Alice Massacci, Martina Diano, Fabrizio Ensoli, Fulvia Pimpinelli, Maurizio Fanciulli, Giovanni Blandino, Aldo Morrone, Gennaro Ciliberto |
| EPI_ISL_3419985 | IRCCS San Gallicano Dermatological Institute | IRCCS Regina Elena National Cancer Institute | Sabrina Strano, Sara Donzelli, Matteo Pallocca, Antonio Federico, Fabrizio Ensoli, Fulvia Pimpinelli, Maurizio Fanciulli, Giovanni Blandino, Aldo Morrone, Gennaro Ciliberto |
| EPI_ISL_3419986 | IRCCS San Gallicano Dermatological Institute | IRCCS Regina Elena National Cancer Institute | Francesca De Nicola, Giulia Orlandi, Eleonora Sperandio, Fulvia Fraticelli, Fabrizio Ensoli, Fulvia Pimpinelli, Maurizio Fanciulli, Giovanni Blandino, Aldo Morrone, Gennaro Ciliberto |
| EPI_ISL_3419987 | IRCCS San Gallicano Dermatological Institute | IRCCS Regina Elena National Cancer Institute | Ludovica Ciuffreda, Sara Donzelli, Alice Massacci, Lorenzo Furzi, Fabrizio Ensoli, Fulvia Pimpinelli, Maurizio Fanciulli, Giovanni Blandino, Aldo Morrone, Gennaro Ciliberto |
| EPI_ISL_3419988 | IRCCS San Gallicano Dermatological Institute | IRCCS Regina Elena National Cancer Institute | Frauke Goeman, Giulia Orlandi, Matteo Pallocca, Alessia Lauretti, Fabrizio Ensoli, Fulvia Pimpinelli, Maurizio Fanciulli, Giovanni Blandino, Aldo Morrone, Gennaro Ciliberto |
| EPI_ISL_3419989 | IRCCS San Gallicano Dermatological Institute | IRCCS Regina Elena National Cancer Institute | Sabrina Strano, Sara Donzelli, Eleonora Sperandio, Francesca Maione, Fabrizio Ensoli, Fulvia Pimpinelli, Maurizio Fanciulli, Giovanni Blandino, Aldo Morrone, Gennaro Ciliberto |
| EPI_ISL_3419990 | IRCCS San Gallicano Dermatological Institute | IRCCS Regina Elena National Cancer Institute | Francesca De Nicola, Giulia Orlandi, Alice Massacci, Francisco Obregon, Fabrizio Ensoli, Fulvia Pimpinelli, Maurizio Fanciulli, Giovanni Blandino, Aldo Morrone, Gennaro Ciliberto |
| EPI_ISL_3419991 | IRCCS San Gallicano Dermatological Institute | IRCCS Regina Elena National Cancer Institute | Ludovica Ciuffreda, Sara Donzelli, Matteo Pallocca, Silvia Paluzzi, Fabrizio Ensoli, Fulvia Pimpinelli, Maurizio Fanciulli, Giovanni Blandino, Aldo Morrone, Gennaro Ciliberto |
| EPI_ISL_3419992 | IRCCS San Gallicano Dermatological Institute | IRCCS Regina Elena National Cancer Institute | Frauke Goeman, Giulia Orlandi, Eleonora Sperandio, Sara Petrolo, Fabrizio Ensoli, Fulvia Pimpinelli, Maurizio Fanciulli, Giovanni Blandino, Aldo Morrone, Gennaro Ciliberto |
| EPI_ISL_3419993 | IRCCS San Gallicano Dermatological Institute | IRCCS Regina Elena National Cancer Institute | Sabrina Strano, Sara Donzelli, Alice Massacci, Valentina Ricca, Fabrizio Ensoli, Fulvia Pimpinelli, Maurizio Fanciulli, Giovanni Blandino, Aldo Morrone, Gennaro Ciliberto |
| EPI_ISL_3419994 | IRCCS San Gallicano Dermatological Institute | IRCCS Regina Elena National Cancer Institute | Francesca De Nicola, Giulia Orlandi, Matteo Pallocca, Serena Salvo, Fabrizio Ensoli, Fulvia Pimpinelli, Maurizio Fanciulli, Giovanni Blandino, Aldo Morrone, Gennaro Ciliberto |
| EPI_ISL_3419995 | IRCCS San Gallicano Dermatological Institute | IRCCS Regina Elena National Cancer Institute | Ludovica Ciuffreda, Sara Donzelli, Eleonora Sperandio, Grazia Prignano, Fabrizio Ensoli, Fulvia Pimpinelli, Maurizio Fanciulli, Giovanni Blandino, Aldo Morrone, Gennaro Ciliberto |
| EPI_ISL_3419996 | IRCCS San Gallicano Dermatological Institute | IRCCS Regina Elena National Cancer Institute | Frauke Goeman, Giulia Orlandi, Alice Massacci, Elisabetta Trento, Fabrizio Ensoli, Fulvia Pimpinelli, Maurizio Fanciulli, Giovanni Blandino, Aldo Morrone, Gennaro Ciliberto |
| EPI_ISL_3419997 | IRCCS San Gallicano Dermatological Institute | IRCCS Regina Elena National Cancer Institute | Sabrina Strano, Sara Donzelli, Matteo Pallocca, Arianna Mastrofrancesco, Fabrizio Ensoli, Fulvia Pimpinelli, Maurizio Fanciulli, Giovanni Blandino, Aldo Morrone, Gennaro Ciliberto |
| EPI_ISL_3419998 | IRCCS San Gallicano Dermatological Institute | IRCCS Regina Elena National Cancer Institute | Francesca De Nicola, Giulia Orlandi, Eleonora Sperandio, Giovanna D’agosto, Fabrizio Ensoli, Fulvia Pimpinelli, Maurizio Fanciulli, Giovanni Blandino, Aldo Morrone, Gennaro Ciliberto |
| EPI_ISL_3419999 | IRCCS San Gallicano Dermatological Institute | IRCCS Regina Elena National Cancer Institute | Ludovica Ciuffreda, Sara Donzelli, Alice Massacci, Carla Mottini, Fabrizio Ensoli, Fulvia Pimpinelli, Maurizio Fanciulli, Giovanni Blandino, Aldo Morrone, Gennaro Ciliberto |
| EPI_ISL_3420000 | IRCCS San Gallicano Dermatological Institute | IRCCS Regina Elena National Cancer Institute | Frauke Goeman, Giulia Orlandi, Matteo Pallocca, Martina Pontone, Fabrizio Ensoli, Fulvia Pimpinelli, Maurizio Fanciulli, Giovanni Blandino, Aldo Morrone, Gennaro Ciliberto |
| EPI_ISL_3420001 | IRCCS San Gallicano Dermatological Institute | IRCCS Regina Elena National Cancer Institute | Sabrina Strano, Sara Donzelli, Eleonora Sperandio, di Domenico Enea Gino, Fabrizio Ensoli, Fulvia Pimpinelli, Maurizio Fanciulli, Giovanni Blandino, Aldo Morrone, Gennaro Ciliberto |
| EPI_ISL_3420002 | IRCCS San Gallicano Dermatological Institute | IRCCS Regina Elena National Cancer Institute | Francesca De Nicola, Giulia Orlandi, Alice Massacci, Ilaria Cavallo, Fabrizio Ensoli, Fulvia Pimpinelli, Maurizio Fanciulli, Giovanni Blandino, Aldo Morrone, Gennaro Ciliberto |
| EPI_ISL_3420003 | IRCCS San Gallicano Dermatological Institute | IRCCS Regina Elena National Cancer Institute | Ludovica Ciuffreda, Sara Donzelli, Matteo Pallocca, Francesca Sivori, Fabrizio Ensoli, Fulvia Pimpinelli, Maurizio Fanciulli, Giovanni Blandino, Aldo Morrone, Gennaro Ciliberto |
| EPI_ISL_3420004 | IRCCS San Gallicano Dermatological Institute | IRCCS Regina Elena National Cancer Institute | Frauke Goeman, Giulia Orlandi, Eleonora Sperandio, Andrea Cazzani, Fabrizio Ensoli, Fulvia Pimpinelli, Maurizio Fanciulli, Giovanni Blandino, Aldo Morrone, Gennaro Ciliberto |
| EPI_ISL_3420005 | IRCCS San Gallicano Dermatological Institute | IRCCS Regina Elena National Cancer Institute | Sabrina Strano, Sara Donzelli, Alice Massacci, Ilaria Celesti, Fabrizio Ensoli, Fulvia Pimpinelli, Maurizio Fanciulli, Giovanni Blandino, Aldo Morrone, Gennaro Ciliberto |
| EPI_ISL_3420006 | IRCCS San Gallicano Dermatological Institute | IRCCS Regina Elena National Cancer Institute | Francesca De Nicola, Giulia Orlandi, Matteo Pallocca, Martina Diano, Fabrizio Ensoli, Fulvia Pimpinelli, Maurizio Fanciulli, Giovanni Blandino, Aldo Morrone, Gennaro Ciliberto |
| EPI_ISL_3420007 | IRCCS San Gallicano Dermatological Institute | IRCCS Regina Elena National Cancer Institute | Ludovica Ciuffreda, Sara Donzelli, Eleonora Sperandio, Antonio Federico, Fabrizio Ensoli, Fulvia Pimpinelli, Maurizio Fanciulli, Giovanni Blandino, Aldo Morrone, Gennaro Ciliberto |
| EPI_ISL_3420008 | IRCCS San Gallicano Dermatological Institute | IRCCS Regina Elena National Cancer Institute | Frauke Goeman, Giulia Orlandi, Alice Massacci, Fulvia Fraticelli, Fabrizio Ensoli, Fulvia Pimpinelli, Maurizio Fanciulli, Giovanni Blandino, Aldo Morrone, Gennaro Ciliberto |
| EPI_ISL_3420009 | IRCCS San Gallicano Dermatological Institute | IRCCS Regina Elena National Cancer Institute | Sabrina Strano, Sara Donzelli, Matteo Pallocca, Lorenzo Furzi, Fabrizio Ensoli, Fulvia Pimpinelli, Maurizio Fanciulli, Giovanni Blandino, Aldo Morrone, Gennaro Ciliberto |
| EPI_ISL_3420010 | IRCCS San Gallicano Dermatological Institute | IRCCS Regina Elena National Cancer Institute | Francesca De Nicola, Giulia Orlandi, Eleonora Sperandio, Alessia Lauretti, Fabrizio Ensoli, Fulvia Pimpinelli, Maurizio Fanciulli, Giovanni Blandino, Aldo Morrone, Gennaro Ciliberto |
| EPI_ISL_3420011 | IRCCS San Gallicano Dermatological Institute | IRCCS Regina Elena National Cancer Institute | Ludovica Ciuffreda, Sara Donzelli, Alice Massacci, Francesca Maione, Fabrizio Ensoli, Fulvia Pimpinelli, Maurizio Fanciulli, Giovanni Blandino, Aldo Morrone, Gennaro Ciliberto |
| EPI_ISL_3420012 | IRCCS San Gallicano Dermatological Institute | IRCCS Regina Elena National Cancer Institute | Frauke Goeman, Giulia Orlandi, Matteo Pallocca, Francisco Obregon, Fabrizio Ensoli, Fulvia Pimpinelli, Maurizio Fanciulli, Giovanni Blandino, Aldo Morrone, Gennaro Ciliberto |
| EPI_ISL_3420013 | IRCCS San Gallicano Dermatological Institute | IRCCS Regina Elena National Cancer Institute | Sabrina Strano, Sara Donzelli, Eleonora Sperandio, Silvia Paluzzi, Fabrizio Ensoli, Fulvia Pimpinelli, Maurizio Fanciulli, Giovanni Blandino, Aldo Morrone, Gennaro Ciliberto |
| EPI_ISL_3420014 | IRCCS San Gallicano Dermatological Institute | IRCCS Regina Elena National Cancer Institute | Francesca De Nicola, Giulia Orlandi, Alice Massacci, Sara Petrolo, Fabrizio Ensoli, Fulvia Pimpinelli, Maurizio Fanciulli, Giovanni Blandino, Aldo Morrone, Gennaro Ciliberto |
| EPI_ISL_3420015 | IRCCS San Gallicano Dermatological Institute | IRCCS Regina Elena National Cancer Institute | Ludovica Ciuffreda, Sara Donzelli, Matteo Pallocca, Valentina Ricca, Fabrizio Ensoli, Fulvia Pimpinelli, Maurizio Fanciulli, Giovanni Blandino, Aldo Morrone, Gennaro Ciliberto |
| EPI_ISL_3420016 | IRCCS San Gallicano Dermatological Institute | IRCCS Regina Elena National Cancer Institute | Frauke Goeman, Giulia Orlandi, Eleonora Sperandio, Serena Salvo, Fabrizio Ensoli, Fulvia Pimpinelli, Maurizio Fanciulli, Giovanni Blandino, Aldo Morrone, Gennaro Ciliberto |
| EPI_ISL_3420017 | IRCCS San Gallicano Dermatological Institute | IRCCS Regina Elena National Cancer Institute | Sabrina Strano, Sara Donzelli, Alice Massacci, Grazia Prignano, Fabrizio Ensoli, Fulvia Pimpinelli, Maurizio Fanciulli, Giovanni Blandino, Aldo Morrone, Gennaro Ciliberto |
| EPI_ISL_3420018 | IRCCS San Gallicano Dermatological Institute | IRCCS Regina Elena National Cancer Institute | Francesca De Nicola, Giulia Orlandi, Matteo Pallocca, Elisabetta Trento, Fabrizio Ensoli, Fulvia Pimpinelli, Maurizio Fanciulli, Giovanni Blandino, Aldo Morrone, Gennaro Ciliberto |
| EPI_ISL_3420019 | IRCCS San Gallicano Dermatological Institute | IRCCS Regina Elena National Cancer Institute | Ludovica Ciuffreda, Sara Donzelli, Eleonora Sperandio, Arianna Mastrofrancesco, Fabrizio Ensoli, Fulvia Pimpinelli, Maurizio Fanciulli, Giovanni Blandino, Aldo Morrone, Gennaro Ciliberto |
| EPI_ISL_3420020 | IRCCS San Gallicano Dermatological Institute | IRCCS Regina Elena National Cancer Institute | Frauke Goeman, Giulia Orlandi, Alice Massacci, Giovanna D’agosto, Fabrizio Ensoli, Fulvia Pimpinelli, Maurizio Fanciulli, Giovanni Blandino, Aldo Morrone, Gennaro Ciliberto |
| EPI_ISL_3420021 | IRCCS San Gallicano Dermatological Institute | IRCCS Regina Elena National Cancer Institute | Sabrina Strano, Sara Donzelli, Matteo Pallocca, Carla Mottini, Fabrizio Ensoli, Fulvia Pimpinelli, Maurizio Fanciulli, Giovanni Blandino, Aldo Morrone, Gennaro Ciliberto |
| EPI_ISL_3420022 | IRCCS San Gallicano Dermatological Institute | IRCCS Regina Elena National Cancer Institute | Francesca De Nicola, Giulia Orlandi, Eleonora Sperandio, Martina Pontone, Fabrizio Ensoli, Fulvia Pimpinelli, Maurizio Fanciulli, Giovanni Blandino, Aldo Morrone, Gennaro Ciliberto |

EPI_ISL_3420024, EPI_ISL_3420025, EPI_ISL_3420026, EPI_ISL_3420027, EPI_ISL_3420028, EPI_ISL_3420029, EPI_ISL_3420030, EPI_ISL_3420031, EPI_ISL_3420032, EPI_ISL_3420033, EPI_ISL_3420034, EPI_ISL_3420035, EPI_ISL_3420036, EPI_ISL_3420037, EPI_ISL_3420038, EPI_ISL_3420039, EPI_ISL_3420040, EPI_ISL_3420041, EPI_ISL_3420042, EPI_ISL_3420043, EPI_ISL_3420044, EPI_ISL_3420045, EPI_ISL_3420046, EPI_ISL_3420047, EPI_ISL_3420048, EPI_ISL_3420049, EPI_ISL_3420050, EPI_ISL_3420051, EPI_ISL_3420052, EPI_ISL_3420053, EPI_ISL_3420054, EPI_ISL_3420055, EPI_ISL_3420056, EPI_ISL_3420057, EPI_ISL_3420058, EPI_ISL_3420059, EPI_ISL_3420060, EPI_ISL_3420061, EPI_ISL_3420062, EPI_ISL_3420063, EPI_ISL_3420064, EPI_ISL_3420065, EPI_ISL_3420066, EPI_ISL_3420067, EPI_ISL_3420068, EPI_ISL_3420069, EPI_ISL_3420070, EPI_ISL_3420071, EPI_ISL_3420072, EPI_ISL_3420073, EPI_ISL_3420074, EPI_ISL_3420075, EPI_ISL_3420076, EPI_ISL_3420077, EPI_ISL_3420078, EPI_ISL_3420079, EPI_ISL_3420080, EPI_ISL_3420081, EPI_ISL_3420082, EPI_ISL_3420083, EPI_ISL_3420084, EPI_ISL_3420085, EPI_ISL_3420086, EPI_ISL_3420087, EPI_ISL_3420088, EPI_ISL_3420089, EPI_ISL_3420090, EPI_ISL_3420091, EPI_ISL_3420092, EPI_ISL_3420093, EPI_ISL_3420094, EPI_ISL_3420095, EPI_ISL_3420096, EPI_ISL_3420097, EPI_ISL_3420098, EPI_ISL_3420099, EPI_ISL_3420100, EPI_ISL_3420101, EPI_ISL_3420102, EPI_ISL_3420103, EPI_ISL_3420104, EPI_ISL_3420105, EPI_ISL_3420106, EPI_ISL_3420107, EPI_ISL_3420108, EPI_ISL_3420109, EPI_ISL_3420110, EPI_ISL_3420111, EPI_ISL_3420112, EPI_ISL_3420113, EPI_ISL_3420114, EPI_ISL_3420115, EPI_ISL_3420116, EPI_ISL_3420117, EPI_ISL_3420118, EPI_ISL_3420119, EPI_ISL_3420120, EPI_ISL_3420121, EPI_ISL_3420122, EPI_ISL_3420123, EPI_ISL_3420124, EPI_ISL_3420125, EPI_ISL_3420126, EPI_ISL_3420127, EPI_ISL_3420128, EPI_ISL_3420129, EPI_ISL_3420130, EPI_ISL_3420131, EPI_ISL_3420132, EPI_ISL_3420133, EPI_ISL_3420134, EPI_ISL_3420135, EPI_ISL_3420136, EPI_ISL_3420137, EPI_ISL_3420138, EPI_ISL_3420139, EPI_ISL_3420140, EPI_ISL_3420141, EPI_ISL_3420142, EPI_ISL_3420143, EPI_ISL_3420144, EPI_ISL_3420145, EPI_ISL_3420146, EPI_ISL_3420147, EPI_ISL_3420148, EPI_ISL_3420149, EPI_ISL_3420150, EPI_ISL_3420151, EPI_ISL_3420152, EPI_ISL_3420153, EPI_ISL_3420154, EPI_ISL_3420155, EPI_ISL_3420156, EPI_ISL_3420157, EPI_ISL_3420158, EPI_ISL_3420159, EPI_ISL_3420160, EPI_ISL_3420161, EPI_ISL_3420162, EPI_ISL_3420163, EPI_ISL_3420164, EPI_ISL_3420165, EPI_ISL_3420166, EPI_ISL_3420167, EPI_ISL_3420168, EPI_ISL_3420169, EPI_ISL_3420170, EPI_ISL_3420171, EPI_ISL_3420172, EPI_ISL_3420173, EPI_ISL_3420174, EPI_ISL_3420175, EPI_ISL_3420176, EPI_ISL_3420177, EPI_ISL_3420178, EPI_ISL_3420179, EPI_ISL_3420180, EPI_ISL_3420181, EPI_ISL_3420182, EPI_ISL_3420183, EPI_ISL_3420184, EPI_ISL_3420185, EPI_ISL_3420186, EPI_ISL_3420187, EPI_ISL_3420188, EPI_ISL_3420189, EPI_ISL_3420190, EPI_ISL_3420191, EPI_ISL_3420192, EPI_ISL_3420193, EPI_ISL_3420194, EPI_ISL_3420195, EPI_ISL_3420196, EPI_ISL_3420197, EPI_ISL_3420198, EPI_ISL_3420199, EPI_ISL_3420200, EPI_ISL_3420201, EPI_ISL_3420202, EPI_ISL_3420203, EPI_ISL_3420204, EPI_ISL_3420205, EPI_ISL_3420206, EPI_ISL_3420207, EPI_ISL_3420208, EPI_ISL_3420209, EPI_ISL_3420210, EPI_ISL_3420211, EPI_ISL_3420212, EPI_ISL_3420213, EPI_ISL_3420214, EPI_ISL_3420215, EPI_ISL_3420216, EPI_ISL_3420217, EPI_ISL_3420218, EPI_ISL_3420219, EPI_ISL_3420220, EPI_ISL_3420221, EPI_ISL_3420222, EPI_ISL_3420223, EPI_ISL_3420224, EPI_ISL_3420225, EPI_ISL_3420226, EPI_ISL_3420227, EPI_ISL_3420228, EPI_ISL_3420229, EPI_ISL_3420230, EPI_ISL_3420231, EPI_ISL_3420232, EPI_ISL_3420233, EPI_ISL_3420234, EPI_ISL_3420235, EPI_ISL_3420236, EPI_ISL_3420237, EPI_ISL_3420238, EPI_ISL_3420239, EPI_ISL_3420240, EPI_ISL_3420241, EPI_ISL_3420242, EPI_ISL_3420243, EPI_ISL_3420244, EPI_ISL_3420245, EPI_ISL_3420246, EPI_ISL_3420247, EPI_ISL_3420248, EPI_ISL_3420249, EPI_ISL_3420250, EPI_ISL_3420251, EPI_ISL_3420252, EPI_ISL_3420253, EPI_ISL_3420254, EPI_ISL_3420255, EPI_ISL_3420256, EPI_ISL_3420257, EPI_ISL_3420258, EPI_ISL_3420259, EPI_ISL_3420260, EPI_ISL_3420261, EPI_ISL_3420262, EPI_ISL_3420263, EPI_ISL_3420264, EPI_ISL_3420265, EPI_ISL_3420266, EPI_ISL_3420267, EPI_ISL_3420268, EPI_ISL_3420269, EPI_ISL_3420270, EPI_ISL_3420271, EPI_ISL_3420272, EPI_ISL_3420273, EPI_ISL_3420274, EPI_ISL_3420275, EPI_ISL_3420276, EPI_ISL_3420277, EPI_ISL_3420278, EPI_ISL_3420279, EPI_ISL_3420280, EPI_ISL_3420281, EPI_ISL_3420282, EPI_ISL_3420283, EPI_ISL_3420284, EPI_ISL_3420285, EPI_ISL_3420286, EPI_ISL_3420287, EPI_ISL_3420288, EPI_ISL_3420289, EPI_ISL_3420290, EPI_ISL_3420291, EPI_ISL_3420292, EPI_ISL_3420293, EPI_ISL_3420294, EPI_ISL_3420295, EPI_ISL_3420296, EPI_ISL_3420297, EPI_ISL_3420298, EPI_ISL_3420299, EPI_ISL_3420300, EPI_ISL_3420301, EPI_ISL_3420302, EPI_ISL_3420303, EPI_ISL_3420304, EPI_ISL_3420305, EPI_ISL_3420306, EPI_ISL_3420307, EPI_ISL_3420308, EPI_ISL_3420309, EPI_ISL_3420310, EPI_ISL_3420311, EPI_ISL_3420312, EPI_ISL_3420313, EPI_ISL_3420314, EPI_ISL_3420315, EPI_ISL_3420316, EPI_ISL_3420317, EPI_ISL_3420318, EPI_ISL_3420319, EPI_ISL_3420320, EPI_ISL_3420321, EPI_ISL_3420322, EPI_ISL_3420323, EPI_ISL_3420324, EPI_ISL_3420325, EPI_ISL_3420326, EPI_ISL_3420327, EPI_ISL_3420328, EPI_ISL_3420329, EPI_ISL_3420330, EPI_ISL_3420331,

EPI_ISL_3420332, EPI_ISL_3420333, EPI_ISL_3420334, EPI_ISL_3420335, EPI_ISL_3420336, EPI_ISL_3420337, EPI_ISL_3420338, EPI_ISL_3420339, EPI_ISL_3420340, EPI_ISL_3420341, EPI_ISL_3420342, EPI_ISL_3420343, EPI_ISL_3420344, EPI_ISL_3420345, EPI_ISL_3420346, EPI_ISL_3420347, EPI_ISL_3420348, EPI_ISL_3420349, EPI_ISL_3420350

see above | West of Scotland Specialist Virology Centre, NHSGGC / MRC-University of Glasgow Centre for Virus Research | COVID-19 Genomics UK (COG-UK) Consortium | Ana da Silva Filipe, Natasha Johnson, Kathy Smollett, Daniel Mair, Stephen Carmichael, Alice Broos, Lily Tong, Jenna Nichols, Kyriaki Nomikou; Sarah McDonald; Ioulia Tsatsani, Richard Orton, Joseph Hughes, Sreenu Vattipally, David L Robertson; Alasdair MacLean, Rory Gunson; Sharif Shaaban, Matthew Holden; Rachel Blacow, Guy Mollett, Kathy Li, James Shepherd, Antonia Ho, Emma Thomson

EPI_ISL_3420351, EPI_ISL_3420352, EPI_ISL_3420353, EPI_ISL_3420354, EPI_ISL_3420355, EPI_ISL_3420356, EPI_ISL_3420357, EPI_ISL_3420358, EPI_ISL_3420359, EPI_ISL_3420360, EPI_ISL_3420361, EPI_ISL_3420362, EPI_ISL_3420363, EPI_ISL_3420364, EPI_ISL_3420365, EPI_ISL_3420366, EPI_ISL_3420367, EPI_ISL_3420368, EPI_ISL_3420369, EPI_ISL_3420370, EPI_ISL_3420371, EPI_ISL_3420372, EPI_ISL_3420373, EPI_ISL_3420374, EPI_ISL_3420375, EPI_ISL_3420376, EPI_ISL_3420377, EPI_ISL_3420378, EPI_ISL_3420379, EPI_ISL_3420380, EPI_ISL_3420381, EPI_ISL_3420382, EPI_ISL_3420383, EPI_ISL_3420384, EPI_ISL_3420385, EPI_ISL_3420386, EPI_ISL_3420387, EPI_ISL_3420388, EPI_ISL_3420389, EPI_ISL_3420390, EPI_ISL_3420391, EPI_ISL_3420392, EPI_ISL_3420393, EPI_ISL_3420394, EPI_ISL_3420395, EPI_ISL_3420396, EPI_ISL_3420397, EPI_ISL_3420398, EPI_ISL_3420399, EPI_ISL_3420400, EPI_ISL_3420401, EPI_ISL_3420402, EPI_ISL_3420403, EPI_ISL_3420404, EPI_ISL_3420405, EPI_ISL_3420406, EPI_ISL_3420407, EPI_ISL_3420408, EPI_ISL_3420409, EPI_ISL_3420410, EPI_ISL_3420411, EPI_ISL_3420412, EPI_ISL_3420413, EPI_ISL_3420414, EPI_ISL_3420415, EPI_ISL_3420416, EPI_ISL_3420417, EPI_ISL_3420418, EPI_ISL_3420419, EPI_ISL_3420420, EPI_ISL_3420421, EPI_ISL_3420422, EPI_ISL_3420423, EPI_ISL_3420424, EPI_ISL_3420425, EPI_ISL_3420426, EPI_ISL_3420427, EPI_ISL_3420428, EPI_ISL_3420429, EPI_ISL_3420430, EPI_ISL_3420431, EPI_ISL_3420432, EPI_ISL_3420433, EPI_ISL_3420434, EPI_ISL_3420435, EPI_ISL_3420436, EPI_ISL_3420437, EPI_ISL_3420438, EPI_ISL_3420439, EPI_ISL_3420440, EPI_ISL_3420441, EPI_ISL_3420442, EPI_ISL_3420443, EPI_ISL_3420444, EPI_ISL_3420445, EPI_ISL_3420446, EPI_ISL_3420447, EPI_ISL_3420448, EPI_ISL_3420449, EPI_ISL_3420450, EPI_ISL_3420451, EPI_ISL_3420452, EPI_ISL_3420453, EPI_ISL_3420454, EPI_ISL_3420455, EPI_ISL_3420456, EPI_ISL_3420457, EPI_ISL_3420458, EPI_ISL_3420459, EPI_ISL_3420460, EPI_ISL_3420461, EPI_ISL_3420462, EPI_ISL_3420463, EPI_ISL_3420464, EPI_ISL_3420465, EPI_ISL_3420466, EPI_ISL_3420467, EPI_ISL_3420468, EPI_ISL_3420469, EPI_ISL_3420470, EPI_ISL_3420471, EPI_ISL_3420472, EPI_ISL_3420473, EPI_ISL_3420474, EPI_ISL_3420475, EPI_ISL_3420476, EPI_ISL_3420477, EPI_ISL_3420478, EPI_ISL_3420479, EPI_ISL_3420480, EPI_ISL_3420481, EPI_ISL_3420482, EPI_ISL_3420483, EPI_ISL_3420484, EPI_ISL_3420485, EPI_ISL_3420486, EPI_ISL_3420487, EPI_ISL_3420488, EPI_ISL_3420489, EPI_ISL_3420490, EPI_ISL_3420491, EPI_ISL_3420492, EPI_ISL_3420493, EPI_ISL_3420494, EPI_ISL_3420495, EPI_ISL_3420496, EPI_ISL_3420497, EPI_ISL_3420498, EPI_ISL_3420499, EPI_ISL_3420500, EPI_ISL_3420501, EPI_ISL_3420502, EPI_ISL_3420503, EPI_ISL_3420504, EPI_ISL_3420505, EPI_ISL_3420506, EPI_ISL_3420507, EPI_ISL_3420508, EPI_ISL_3420509, EPI_ISL_3420510, EPI_ISL_3420511, EPI_ISL_3420512, EPI_ISL_3420513, EPI_ISL_3420514, EPI_ISL_3420515, EPI_ISL_3420516, EPI_ISL_3420517

see above | Virology Department, Royal Infirmary of Edinburgh, NHS Lothian / School of Biological Sciences, University of Edinburgh | COVID-19 Genomics UK (COG-UK) Consortium | McHugh M, Dewar R, Cotton S, Gallagher A, Maloney D, Fernandez G, O'Toole Á, Scher E, Hill V, McCrone JT, Colquhoun R, Yu X, Jackson B, Rambaut A, Templeton K

EPI_ISL_3420521, EPI_ISL_3420522, EPI_ISL_3420523, EPI_ISL_3420524, EPI_ISL_3420525, EPI_ISL_3420526, EPI_ISL_3420527, EPI_ISL_3420528, EPI_ISL_3420529, EPI_ISL_3420530, EPI_ISL_3420531, EPI_ISL_3420532, EPI_ISL_3420533, EPI_ISL_3420534, EPI_ISL_3420535, EPI_ISL_3420536, EPI_ISL_3420537, EPI_ISL_3420538

see above | Barts Health NHS Trust | COVID-19 Genomics UK (COG-UK) Consortium | CUTINO-MOGUEL, Maria-Teresa; HARRINGTON, David; OWOYEMI, Dola; KULASEGARAN-SHYLINI, Raghavendran; BROAD, Claire; KELE, Beatrix

EPI_ISL_3420539, EPI_ISL_3420540, EPI_ISL_3420541, EPI_ISL_3420542, EPI_ISL_3420543, EPI_ISL_3420544, EPI_ISL_3420545, EPI_ISL_3420546, EPI_ISL_3420547, EPI_ISL_3420548, EPI_ISL_3420549, EPI_ISL_3420550, EPI_ISL_3420551, EPI_ISL_3420552, EPI_ISL_3420553, EPI_ISL_3420554, EPI_ISL_3420555, EPI_ISL_3420556, EPI_ISL_3420557, EPI_ISL_3420558, EPI_ISL_3420559, EPI_ISL_3420560, EPI_ISL_3420561, EPI_ISL_3420562, EPI_ISL_3420563, EPI_ISL_3420564, EPI_ISL_3420565, EPI_ISL_3420566, EPI_ISL_3420567, EPI_ISL_3420568, EPI_ISL_3420569, EPI_ISL_3420570, EPI_ISL_3420571, EPI_ISL_3420572, EPI_ISL_3420573, EPI_ISL_3420574, EPI_ISL_3420575, EPI_ISL_3420576, EPI_ISL_3420577, EPI_ISL_3420578, EPI_ISL_3420579, EPI_ISL_3420580, EPI_ISL_3420581, EPI_ISL_3420582, EPI_ISL_3420583, EPI_ISL_3420584, EPI_ISL_3420585, EPI_ISL_3420586, EPI_ISL_3420587, EPI_ISL_3420588, EPI_ISL_3420589, EPI_ISL_3420590, EPI_ISL_3420591, EPI_ISL_3420592, EPI_ISL_3420593, EPI_ISL_3420594, EPI_ISL_3420595, EPI_ISL_3420596, EPI_ISL_3420597, EPI_ISL_3420598, EPI_ISL_3420599, EPI_ISL_3420600, EPI_ISL_3420601, EPI_ISL_3420602, EPI_ISL_3420603, EPI_ISL_3420604, EPI_ISL_3420605, EPI_ISL_3420606, EPI_ISL_3420607, EPI_ISL_3420608, EPI_ISL_3420609, EPI_ISL_3420610, EPI_ISL_3420611, EPI_ISL_3420612, EPI_ISL_3420613, EPI_ISL_3420614, EPI_ISL_3420615, EPI_ISL_3420616, EPI_ISL_3420617, EPI_ISL_3420618, EPI_ISL_3420619, EPI_ISL_3420620, EPI_ISL_3420621, EPI_ISL_3420622, EPI_ISL_3420623, EPI_ISL_3420624, EPI_ISL_3420625, EPI_ISL_3420626, EPI_ISL_3420627, EPI_ISL_3420628, EPI_ISL_3420629, EPI_ISL_3420630, EPI_ISL_3420631, EPI_ISL_3420632, EPI_ISL_3420633, EPI_ISL_3420634, EPI_ISL_3420635, EPI_ISL_3420636, EPI_ISL_3420637, EPI_ISL_3420638, EPI_ISL_3420639, EPI_ISL_3420640, EPI_ISL_3420641, EPI_ISL_3420642, EPI_ISL_3420643, EPI_ISL_3420644, EPI_ISL_3420645, EPI_ISL_3420646, EPI_ISL_3420647, EPI_ISL_3420648, EPI_ISL_3420649, EPI_ISL_3420650, EPI_ISL_3420651, EPI_ISL_3420652, EPI_ISL_3420653, EPI_ISL_3420654, EPI_ISL_3420655, EPI_ISL_3420656, EPI_ISL_3420657, EPI_ISL_3420658, EPI_ISL_3420659, EPI_ISL_3420660, EPI_ISL_3420661, EPI_ISL_3420662, EPI_ISL_3420663, EPI_ISL_3420664, EPI_ISL_3420665, EPI_ISL_3420666, EPI_ISL_3420667, EPI_ISL_3420668, EPI_ISL_3420669, EPI_ISL_3420670, EPI_ISL_3420671, EPI_ISL_3420672, EPI_ISL_3420673, EPI_ISL_3420674, EPI_ISL_3420675, EPI_ISL_3420676, EPI_ISL_3420677, EPI_ISL_3420678, EPI_ISL_3420679, EPI_ISL_3420680, EPI_ISL_3420681

see above | University College London, Great Ormond Street Hospital for Children NHS Foundation Trust, Imperial College Healthcare NHS Trust | COVID-19 Genomics UK (COG-UK) Consortium | Sergi Castellano, Rachel Williams, Mark Kristiansen, Sunando Roy, Charlotte Williams, Nadua Bayzid, Marius Cotic, Helena Tutill, Patricia Dyal, Judith Breuer

EPI_ISL_3420682, EPI_ISL_3420683, EPI_ISL_3420684, EPI_ISL_3420685, EPI_ISL_3420686, EPI_ISL_3420687, EPI_ISL_3420688, EPI_ISL_3420689, EPI_ISL_3420690, EPI_ISL_3420691, EPI_ISL_3420692, EPI_ISL_3420693, EPI_ISL_3420694, EPI_ISL_3420695, EPI_ISL_3420696, EPI_ISL_3420697, EPI_ISL_3420698, EPI_ISL_3420699, EPI_ISL_3420700, EPI_ISL_3420701, EPI_ISL_3420702, EPI_ISL_3420703, EPI_ISL_3420704, EPI_ISL_3420705, EPI_ISL_3420706, EPI_ISL_3420707, EPI_ISL_3420708, EPI_ISL_3420709, EPI_ISL_3420710, EPI_ISL_3420711, EPI_ISL_3420712, EPI_ISL_3420713, EPI_ISL_3420714, EPI_ISL_3420715, EPI_ISL_3420716, EPI_ISL_3420717, EPI_ISL_3420718, EPI_ISL_3420719, EPI_ISL_3420720, EPI_ISL_3420721, EPI_ISL_3420722, EPI_ISL_3420723, EPI_ISL_3420724, EPI_ISL_3420725, EPI_ISL_3420726, EPI_ISL_3420727, EPI_ISL_3420728, EPI_ISL_3420729, EPI_ISL_3420730, EPI_ISL_3420731, EPI_ISL_3420732, EPI_ISL_3420733, EPI_ISL_3420734, EPI_ISL_3420735, EPI_ISL_3420736, EPI_ISL_3420737, EPI_ISL_3420738, EPI_ISL_3420739, EPI_ISL_3420740, EPI_ISL_3420741, EPI_ISL_3420742, EPI_ISL_3420743, EPI_ISL_3420744, EPI_ISL_3420745, EPI_ISL_3420746, EPI_ISL_3420747, EPI_ISL_3420748, EPI_ISL_3420749, EPI_ISL_3420750, EPI_ISL_3420751, EPI_ISL_3420752, EPI_ISL_3420753, EPI_ISL_3420754, EPI_ISL_3420755, EPI_ISL_3420756, EPI_ISL_3420757, EPI_ISL_3420758, EPI_ISL_3420759, EPI_ISL_3420760, EPI_ISL_3420761, EPI_ISL_3420762, EPI_ISL_3420763, EPI_ISL_3420764, EPI_ISL_3420765, EPI_ISL_3420766, EPI_ISL_3420767, EPI_ISL_3420768, EPI_ISL_3420769, EPI_ISL_3420770, EPI_ISL_3420771, EPI_ISL_3420772, EPI_ISL_3420773, EPI_ISL_3420774, EPI_ISL_3420775, EPI_ISL_3420776, EPI_ISL_3420777, EPI_ISL_3420778, EPI_ISL_3420779, EPI_ISL_3420780, EPI_ISL_3420781, EPI_ISL_3420782, EPI_ISL_3420783, EPI_ISL_3420784, EPI_ISL_3420785, EPI_ISL_3420786, EPI_ISL_3420787, EPI_ISL_3420788, EPI_ISL_3420789, EPI_ISL_3420790, EPI_ISL_3420791, EPI_ISL_3420792, EPI_ISL_3420793, EPI_ISL_3420794, EPI_ISL_3420795, EPI_ISL_3420796, EPI_ISL_3420797, EPI_ISL_3420798, EPI_ISL_3420799, EPI_ISL_3420800, EPI_ISL_3420801, EPI_ISL_3420802, EPI_ISL_3420803, EPI_ISL_3420804, EPI_ISL_3420805, EPI_ISL_3420806, EPI_ISL_3420807, EPI_ISL_3420808, EPI_ISL_3420809, EPI_ISL_3420810, EPI_ISL_3420811, EPI_ISL_3420812, EPI_ISL_3420813, EPI_ISL_3420814, EPI_ISL_3420815, EPI_ISL_3420816, EPI_ISL_3420817, EPI_ISL_3420818, EPI_ISL_3420819, EPI_ISL_3420820, EPI_ISL_3420821, EPI_ISL_3420822, EPI_ISL_3420823, EPI_ISL_3420824, EPI_ISL_3420825, EPI_ISL_3420826, EPI_ISL_3420827, EPI_ISL_3420828, EPI_ISL_3420829, EPI_ISL_3420830, EPI_ISL_3420831, EPI_ISL_3420832, EPI_ISL_3420833, EPI_ISL_3420834, EPI_ISL_3420835, EPI_ISL_3420836, EPI_ISL_3420837, EPI_ISL_3420838, EPI_ISL_3420839, EPI_ISL_3420840, EPI_ISL_3420841, EPI_ISL_3420842, EPI_ISL_3420843, EPI_ISL_3420844, EPI_ISL_3420845, EPI_ISL_3420846, EPI_ISL_3420847, EPI_ISL_3420848, EPI_ISL_3420849, EPI_ISL_3420850, EPI_ISL_3420851, EPI_ISL_3420852, EPI_ISL_3420853, EPI_ISL_3420854, EPI_ISL_3420855, EPI_ISL_3420856, EPI_ISL_3420857, EPI_ISL_3420858, EPI_ISL_3420859, EPI_ISL_3420860, EPI_ISL_3420861, EPI_ISL_3420862, EPI_ISL_3420863, EPI_ISL_3420864, EPI_ISL_3420865, EPI_ISL_3420866, EPI_ISL_3420867, EPI_ISL_3420868, EPI_ISL_3420869, EPI_ISL_3420870, EPI_ISL_3420871, EPI_ISL_3420872, EPI_ISL_3420873, EPI_ISL_3420874, EPI_ISL_3420875, EPI_ISL_3420876, EPI_ISL_3420877, EPI_ISL_3420878, EPI_ISL_3420879, EPI_ISL_3420880, EPI_ISL_3420881, EPI_ISL_3420882, EPI_ISL_3420883, EPI_ISL_3420884, EPI_ISL_3420885, EPI_ISL_3420886, EPI_ISL_3420887, EPI_ISL_3420888, EPI_ISL_3420889, EPI_ISL_3420890, EPI_ISL_3420891, EPI_ISL_3420892, EPI_ISL_3420893, EPI_ISL_3420894, EPI_ISL_3420895, EPI_ISL_3420896, EPI_ISL_3420897, EPI_ISL_3420898, EPI_ISL_3420899, EPI_ISL_3420900, EPI_ISL_3420901, EPI_ISL_3420902, EPI_ISL_3420903, EPI_ISL_3420904, EPI_ISL_3420905, EPI_ISL_3420906, EPI_ISL_3420907, EPI_ISL_3420908, EPI_ISL_3420909, EPI_ISL_3420910, EPI_ISL_3420911, EPI_ISL_3420912, EPI_ISL_3420913, EPI_ISL_3420914, EPI_ISL_3420915, EPI_ISL_3420916, EPI_ISL_3420917, EPI_ISL_3420918, EPI_ISL_3420919, EPI_ISL_3420920, EPI_ISL_3420921, EPI_ISL_3420922, EPI_ISL_3420923, EPI_ISL_3420924, EPI_ISL_3420925, EPI_ISL_3420926, EPI_ISL_3420927, EPI_ISL_3420928, EPI_ISL_3420929, EPI_ISL_3420930, EPI_ISL_3420931, EPI_ISL_3420932, EPI_ISL_3420933, EPI_ISL_3420934, EPI_ISL_3420935, EPI_ISL_3420936, EPI_ISL_3420937, EPI_ISL_3420938, EPI_ISL_3420939, EPI_ISL_3420940, EPI_ISL_3420941, EPI_ISL_3420942, EPI_ISL_3420943, EPI_ISL_3420944, EPI_ISL_3420945, EPI_ISL_3420946, EPI_ISL_3420947, EPI_ISL_3420948, EPI_ISL_3420949, EPI_ISL_3420950, EPI_ISL_3420951, EPI_ISL_3420952, EPI_ISL_3420953, EPI_ISL_3420954, EPI_ISL_3420955, EPI_ISL_3420956, EPI_ISL_3420957, EPI_ISL_3420958, EPI_ISL_3420959, EPI_ISL_3420960, EPI_ISL_3420961, EPI_ISL_3420962, EPI_ISL_3420963, EPI_ISL_3420964, EPI_ISL_3420965, EPI_ISL_3420966, EPI_ISL_3420967, EPI_ISL_3420968, EPI_ISL_3420969, EPI_ISL_3420970, EPI_ISL_3420971, EPI_ISL_3420972, EPI_ISL_3420973, EPI_ISL_3420974, EPI_ISL_3420975, EPI_ISL_3420976, EPI_ISL_3420977, EPI_ISL_3420978, EPI_ISL_3420979, EPI_ISL_3420980, EPI_ISL_3420981, EPI_ISL_3420982, EPI_ISL_3420983, EPI_ISL_3420984, EPI_ISL_3420985, EPI_ISL_3420986, EPI_ISL_3420987, EPI_ISL_3420988, EPI_ISL_3420989, EPI_ISL_3420990, EPI_ISL_3420991, EPI_ISL_3420992, EPI_ISL_3420993, EPI_ISL_3420994, EPI_ISL_3420995, EPI_ISL_3420996, EPI_ISL_3420997, EPI_ISL_3420998, EPI_ISL_3420999, EPI_ISL_3421000, EPI_ISL_3421001, EPI_ISL_3421002, EPI_ISL_3421003, EPI_ISL_3421004, EPI_ISL_3421005, EPI_ISL_3421006, EPI_ISL_3421007, EPI_ISL_3421008, EPI_ISL_3421009, EPI_ISL_3421010, EPI_ISL_3421011, EPI_ISL_3421012, EPI_ISL_3421013, EPI_ISL_3421014, EPI_ISL_3421015, EPI_ISL_3421016, EPI_ISL_3421017, EPI_ISL_3421018, EPI_ISL_3421019, EPI_ISL_3421020, EPI_ISL_3421021, EPI_ISL_3421022, EPI_ISL_3421023, EPI_ISL_3421024, EPI_ISL_3421025, EPI_ISL_3421026, EPI_ISL_3421027, EPI_ISL_3421028, EPI_ISL_3421029, EPI_ISL_3421030, EPI_ISL_3421031, EPI_ISL_3421032, EPI_ISL_3421033, EPI_ISL_3421034, EPI_ISL_3421035, EPI_ISL_3421036, EPI_ISL_3421037, EPI_ISL_3421038, EPI_ISL_3421039, EPI_ISL_3421040, EPI_ISL_3421041, EPI_ISL_3421042, EPI_ISL_3421043, EPI_ISL_3421044, EPI_ISL_3421045, EPI_ISL_3421046, EPI_ISL_3421047, EPI_ISL_3421048, EPI_ISL_3421049, EPI_ISL_3421050, EPI_ISL_3421051, EPI_ISL_3421052, EPI_ISL_3421053, EPI_ISL_3421054, EPI_ISL_3421055, EPI_ISL_3421056, EPI_ISL_3421057, EPI_ISL_3421058, EPI_ISL_3421059, EPI_ISL_3421060, EPI_ISL_3421061, EPI_ISL_3421062, EPI_ISL_3421063, EPI_ISL_3421064, EPI_ISL_3421065, EPI_ISL_3421066, EPI_ISL_3421067, EPI_ISL_3421068, EPI_ISL_3421069, EPI_ISL_3421070, EPI_ISL_3421071, EPI_ISL_3421072, EPI_ISL_3421073, EPI_ISL_3421074, EPI_ISL_3421075, EPI_ISL_3421076, EPI_ISL_3421077, EPI_ISL_3421078, EPI_ISL_3421079, EPI_ISL_3421080, EPI_ISL_3421081, EPI_ISL_3421082, EPI_ISL_3421083, EPI_ISL_3421084, EPI_ISL_3421085, EPI_ISL_3421086, EPI_ISL_3421087, EPI_ISL_3421088, EPI_ISL_3421089, EPI_ISL_3421090, EPI_ISL_3421091, EPI_ISL_3421092, EPI_ISL_3421093, EPI_ISL_3421094, EPI_ISL_3421095, EPI_ISL_3421096, EPI_ISL_3421097, EPI_ISL_3421098, EPI_ISL_3421099, EPI_ISL_3421100, EPI_ISL_3421101, EPI_ISL_3421102, EPI_ISL_3421103, EPI_ISL_3421104, EPI_ISL_3421105, EPI_ISL_3421106, EPI_ISL_3421107, EPI_ISL_3421108, EPI_ISL_3421109, EPI_ISL_3421110, EPI_ISL_3421111, EPI_ISL_3421112, EPI_ISL_3421113, EPI_ISL_3421114, EPI_ISL_3421115, EPI_ISL_3421116, EPI_ISL_3421117, EPI_ISL_3421118, EPI_ISL_3421119, EPI_ISL_3421120, EPI_ISL_3421121, EPI_ISL_3421122, EPI_ISL_3421123, EPI_ISL_3421124, EPI_ISL_3421125, EPI_ISL_3421126, EPI_ISL_3421127, EPI_ISL_3421128, EPI_ISL_3421129, EPI_ISL_3421130, EPI_ISL_3421131, EPI_ISL_3421132, EPI_ISL_3421133, EPI_ISL_3421134, EPI_ISL_3421135, EPI_ISL_3421136, EPI_ISL_3421137, EPI_ISL_3421138, EPI_ISL_3421139, EPI_ISL_3421140, EPI_ISL_3421141, EPI_ISL_3421142, EPI_ISL_3421143, EPI_ISL_3421144, EPI_ISL_3421145, EPI_ISL_3421146, EPI_ISL_3421147, EPI_ISL_3421148, EPI_ISL_3421149, EPI_ISL_3421150, EPI_ISL_3421151, EPI_ISL_3421152, EPI_ISL_3421153, EPI_ISL_3421154, EPI_ISL_3421155, EPI_ISL_3421156, EPI_ISL_3421157, EPI_ISL_3421158, EPI_ISL_3421159, EPI_ISL_3421160, EPI_ISL_3421161, EPI_ISL_3421162, EPI_ISL_3421163, EPI_ISL_3421164, EPI_ISL_3421165, EPI_ISL_3421166, EPI_ISL_3421167, EPI_ISL_3421168, EPI_ISL_3421169, EPI_ISL_3421170, EPI_ISL_3421171, EPI_ISL_3421172, EPI_ISL_3421173, EPI_ISL_3421174, EPI_ISL_3421175, EPI_ISL_3421176, EPI_ISL_3421177, EPI_ISL_3421178, EPI_ISL_3421179, EPI_ISL_3421180, EPI_ISL_3421181, EPI_ISL_3421182, EPI_ISL_3421183, EPI_ISL_3421184, EPI_ISL_3421185, EPI_ISL_3421186, EPI_ISL_3421187, EPI_ISL_3421188, EPI_ISL_3421189, EPI_ISL_3421190, EPI_ISL_3421191, EPI_ISL_3421192, EPI_ISL_3421193, EPI_ISL_3421194, EPI_ISL_3421195, EPI_ISL_3421196, EPI_ISL_3421197, EPI_ISL_3421198, EPI_ISL_3421199, EPI_ISL_3421200, EPI_ISL_3421201, EPI_ISL_3421202, EPI_ISL_3421203, EPI_ISL_3421204, EPI_ISL_3421205, EPI_ISL_3421206, EPI_ISL_3421207, EPI_ISL_3421208, EPI_ISL_3421209, EPI_ISL_3421210, EPI_ISL_3421211, EPI_ISL_3421212, EPI_ISL_3421213, EPI_ISL_3421214, EPI_ISL_3421215, EPI_ISL_3421216, EPI_ISL_3421217, EPI_ISL_3421218, EPI_ISL_3421219, EPI_ISL_3421220, EPI_ISL_3421221, EPI_ISL_3421222, EPI_ISL_3421223, EPI_ISL_3421224, EPI_ISL_3421225, EPI_ISL_3421226, EPI_ISL_3421227, EPI_ISL_3421228, EPI_ISL_3421229, EPI_ISL_3421230, EPI_ISL_3421231, EPI_ISL_3421232, EPI_ISL_3421233, EPI_ISL_3421234, EPI_ISL_3421235, EPI_ISL_3421236, EPI_ISL_3421237, EPI_ISL_3421238, EPI_ISL_3421239, EPI_ISL_3421240, EPI_ISL_3421241, EPI_ISL_3421242, EPI_ISL_3421243, EPI_ISL_3421244, EPI_ISL_3421245, EPI_ISL_3421246, EPI_ISL_3421247, EPI_ISL_3421248, EPI_ISL_3421249, EPI_ISL_3421250, EPI_ISL_3421251, EPI_ISL_3421252, EPI_ISL_3421253, EPI_ISL_3421254, EPI_ISL_3421255, EPI_ISL_3421256, EPI_ISL_3421257, EPI_ISL_3421258, EPI_ISL_3421259, EPI_ISL_3421260, EPI_ISL_3421261, EPI_ISL_3421262, EPI_ISL_3421263, EPI_ISL_3421264, EPI_ISL_3421265, EPI_ISL_3421266, EPI_ISL_3421267, EPI_ISL_3421268, EPI_ISL_3421269, EPI_ISL_3421270, EPI_ISL_3421271, EPI_ISL_3421272, EPI_ISL_3421273, EPI_ISL_3421274, EPI_ISL_3421275, EPI_ISL_3421276, EPI_ISL_3421277, EPI_ISL_3421278, EPI_ISL_3421279, EPI_ISL_3421280, EPI_ISL_3421281, EPI_ISL_3421282, EPI_ISL_3421283, EPI_ISL_3421284, EPI_ISL_3421285, EPI_ISL_3421286, EPI_ISL_3421287, EPI_ISL_3421288, EPI_ISL_3421289, EPI_ISL_3421290, EPI_ISL_3421291, EPI_ISL_3421292, EPI_ISL_3421293, EPI_ISL_3421294, EPI_ISL_3421295, EPI_ISL_3421296, EPI_ISL_3421297, EPI_ISL_3421298, EPI_ISL_3421299, EPI_ISL_3421300, EPI_ISL_3421301, EPI_ISL_3421302, EPI_ISL_3421303, EPI_ISL_3421304, EPI_ISL_3421305, EPI_ISL_3421306, EPI_ISL_3421307, EPI_ISL_3421308, EPI_ISL_3421309, EPI_ISL_3421310, EPI_ISL_3421311, EPI_ISL_3421312, EPI_ISL_3421313, EPI_ISL_3421314, EPI_ISL_3421315, EPI_ISL_3421316, EPI_ISL_3421317, EPI_ISL_3421318, EPI_ISL_3421319, EPI_ISL_3421320, EPI_ISL_3421321, EPI_ISL_3421322, EPI_ISL_3421323, EPI_ISL_3421324, EPI_ISL_3421325, EPI_ISL_3421326, EPI_ISL_3421327, EPI_ISL_3421328, EPI_ISL_3421329, EPI_ISL_3421330, EPI_ISL_3421331, EPI_ISL_3421332, EPI_ISL_3421333, EPI_ISL_3421334, EPI_ISL_3421335, EPI_ISL_3421336, EPI_ISL_3421337, EPI_ISL_3421338, EPI_ISL_3421339, EPI_ISL_3421340, EPI_ISL_3421341, EPI_ISL_3421342, EPI_ISL_3421343, EPI_ISL_3421344, EPI_ISL_3421345, EPI_ISL_3421346, EPI_ISL_3421347, EPI_ISL_3421348, EPI_ISL_3421349, EPI_ISL_3421350, EPI_ISL_3421351, EPI_ISL_3421352, EPI_ISL_3421353, EPI_ISL_3421354, EPI_ISL_3421355, EPI_ISL_3421356, EPI_ISL_3421357, EPI_ISL_3421358, EPI_ISL_3421359, EPI_ISL_3421360, EPI_ISL_3421361, EPI_ISL_3421362, EPI_ISL_3421363, EPI_ISL_3421364, EPI_ISL_3421365, EPI_ISL_3421366, EPI_ISL_3421367, EPI_ISL_3421368, EPI_ISL_3421369, EPI_ISL_3421370, EPI_ISL_3421371, EPI_ISL_3421372, EPI_ISL_3421373, EPI_ISL_3421374, EPI_ISL_3421375, EPI_ISL_3421376, EPI_ISL_3421377, EPI_ISL_3421378, EPI_ISL_3421379, EPI_ISL_3421380, EPI_ISL_3421381, EPI_ISL_3421382, EPI_ISL_3421383, EPI_ISL_3421384, EPI_ISL_3421385, EPI_ISL_3421386, EPI_ISL_3421387, EPI_ISL_3421388, EPI_ISL_3421389, EPI_ISL_3421390, EPI_ISL_3421391, EPI_ISL_3421392, EPI_ISL_3421393, EPI_ISL_3421394, EPI_ISL_3421395, EPI_ISL_3421396, EPI_ISL_3421397, EPI_ISL_3421398, EPI_ISL_3421399, EPI_ISL_3421400, EPI_ISL_3421401, EPI_ISL_3421402, EPI_ISL_3421403, EPI_ISL_3421404, EPI_ISL_3421405, EPI_ISL_3421406, EPI_ISL_3421407, EPI_ISL_3421408, EPI_ISL_3421409, EPI_ISL_3421410, EPI_ISL_3421411, EPI_ISL_3421412, EPI_ISL_3421413, EPI_ISL_3421414, EPI_ISL_3421415, EPI_ISL_3421416, EPI_ISL_3421417, EPI_ISL_3421418, EPI_ISL_3421419, EPI_ISL_3421420, EPI_ISL_3421421, EPI_ISL_3421422, EPI_ISL_3421423, EPI_ISL_3421424, EPI_ISL_3421425, EPI_ISL_3421426, EPI_ISL_3421427, EPI_ISL_3421428, EPI_ISL_3421429, EPI_ISL_3421430, EPI_ISL_3421431, EPI_ISL_3421432, EPI_ISL_3421433, EPI_ISL_3421434, EPI_ISL_3421435, EPI_ISL_3421436, EPI_ISL_3421437, EPI_ISL_3421438, EPI_ISL_3421439, EPI_ISL_3421440, EPI_ISL_3421441, EPI_ISL_3421442, EPI_ISL_3421443, EPI_ISL_3421444, EPI_ISL_3421445, EPI_ISL_3421446, EPI_ISL_3421447, EPI_ISL_3421448, EPI_ISL_3421449, EPI_ISL_3421450, EPI_ISL_3421451, EPI_ISL_3421452, EPI_ISL_3421453, EPI_ISL_3421454, EPI_ISL_3421455, EPI_ISL_3421456, EPI_ISL_3421457, EPI_ISL_3421458, EPI_ISL_3421459, EPI_ISL_3421460, EPI_ISL_3421461, EPI_ISL_3421462, EPI_ISL_3421463, EPI_ISL_3421464, EPI_ISL_3421465, EPI_ISL_3421466, EPI_ISL_3421467, EPI_ISL_3421468, EPI_ISL_3421469, EPI_ISL_3421470, EPI_ISL_3421471, EPI_ISL_3421472, EPI_ISL_3421473, EPI_ISL_3421474, EPI_ISL_3421475, EPI_ISL_3421476, EPI_ISL_3421477, EPI_ISL_3421478, EPI_ISL_3421479, EPI_ISL_3421480, EPI_ISL_3421481, EPI_ISL_3421482, EPI_ISL_3421483, EPI_ISL_3421484, EPI_ISL_3421485, EPI_ISL_3421486, EPI_ISL_3421487, EPI_ISL_3421488, EPI_ISL_3421489, EPI_ISL_3421490, EPI_ISL_3421491, EPI_ISL_3421492, EPI_ISL_3421493, EPI_ISL_3421494, EPI_ISL_3421495, EPI_ISL_3421496, EPI_ISL_3421497, EPI_ISL_3421498, EPI_ISL_3421499, EPI_ISL_3421500, EPI_ISL_3421501, EPI_ISL_3421502, EPI_ISL_3421503, EPI_ISL_3421504, EPI_ISL_3421505, EPI_ISL_3421506, EPI_ISL_3421507, EPI_ISL_3421508, EPI_ISL_3421509, EPI_ISL_3421510, EPI_ISL_3421511, EPI_ISL_3421512, EPI_ISL_3421513, EPI_ISL_3421514, EPI_ISL_3421515, EPI_ISL_3421516, EPI_ISL_3421517, EPI_ISL_3421518, EPI_ISL_3421519, EPI_ISL_3421520, EPI_ISL_3421521, EPI_ISL_3421522, EPI_ISL_3421523, EPI_ISL_3421524, EPI_ISL_3421525, EPI_ISL_3421526, EPI_ISL_3421527, EPI_ISL_3421528, EPI_ISL_3421529, EPI_ISL_3421530, EPI_ISL_3421531, EPI_ISL_3421532, EPI_ISL_3421533, EPI_ISL_3421534, EPI_ISL_3421535, EPI_ISL_3421536, EPI_ISL_3421537, EPI_ISL_3421538, EPI_ISL_3421539, EPI_ISL_3421540, EPI_ISL_3421541, EPI_ISL_3421542, EPI_ISL_3421543, EPI_ISL_3421544, EPI_ISL_3421545, EPI_ISL_3421546, EPI_ISL_3421547, EPI_ISL_3421548, EPI_ISL_3421549, EPI_ISL_3421550, EPI_ISL_3421551, EPI_ISL_3421552, EPI_ISL_3421553, EPI_ISL_3421554, EPI_ISL_3421555, EPI_ISL_3421556, EPI_ISL_3421557, EPI_ISL_3421558, EPI_ISL_3421559, EPI_ISL_3421560, EPI_ISL_3421561, EPI_ISL_3421562, EPI_ISL_3421563, EPI_ISL_3421564, EPI_ISL_3421565, EPI_ISL_3421566, EPI_ISL_3421567, EPI_ISL_3421568, EPI_ISL_3421569, EPI_ISL_3421570, EPI_ISL_3421571, EPI_ISL_3421572, EPI_ISL_3421573, EPI_ISL_3421574, EPI_ISL_3421575, EPI_ISL_3421576, EPI_ISL_3421577, EPI_ISL_3421578, EPI_ISL_3421579, EPI_ISL_3421580, EPI_ISL_3421581, EPI_ISL_3421582, EPI_ISL_3421583, EPI_ISL_3421584, EPI_ISL_3421585, EPI_ISL_3421586, EPI_ISL_3421587, EPI_ISL_3421588, EPI_ISL_3421589, EPI_ISL_3421590, EPI_ISL_3421591, EPI_ISL_3421592, EPI_ISL_3421593, EPI_ISL_3421594, EPI_ISL_3421595, EPI_ISL_3421596, EPI_ISL_3421597, EPI_ISL_3421598, EPI_ISL_3421599, EPI_ISL_3421600, EPI_ISL_3421601, EPI_ISL_3421602, EPI_ISL_3421603, EPI_ISL_3421604, EPI_ISL_3421605, EPI_ISL_3421606, EPI_ISL_3421607, EPI_ISL_3421608, EPI_ISL_3421609, EPI_ISL_3421610, EPI_ISL_3421611, EPI_ISL_3421612, EPI_ISL_3421613, EPI_ISL_3421614, EPI_ISL_3421615, EPI_ISL_3421616, EPI_ISL_3421617, EPI_ISL_3421618, EPI_ISL_3421619, EPI_ISL_3421620, EPI_ISL_3421621, EPI_ISL_3421622, EPI_ISL_3421623, EPI_ISL_3421624, EPI_ISL_3421625, EPI_ISL_3421626, EPI_ISL_3421627, EPI_ISL_3421628, EPI_ISL_3421629, EPI_ISL_3421630, EPI_ISL_3421631, EPI_ISL_3421632

see above | Northumbria University / South Tees Hospitals NHS Foundation Trust / North Cumbria Integrated Care NHS Foundation Trust / North Tees and Hartlepool NHS Foundation Trust / Newcastle Hospitals NHS Foundation Trust | COVID-19 Genomics UK (COG-UK) Consortium | Darren L Smith,Andrew Nelson,Matthew Bashton,Greg R Young,Joshua Loh,John Allan,Mohammad A Tariq,Giles S Holt,Gary Black,Wen C Yew,Lynn Dover,Paul Baker,Steve Liggett,Sarah Essex,Jane Greenaway,Debra Padgett,Clive Graham,Garren Scott,Edward Barton,Emma Swindells,Brendan Payne,Jennifer Collins,Yusri Taha,Gary Eltringham

EPI_ISL_3421633, EPI_ISL_3421634, EPI_ISL_3421635, EPI_ISL_3421636, EPI_ISL_3421637, EPI_ISL_3421638, EPI_ISL_3421639, EPI_ISL_3421640, EPI_ISL_3421641, EPI_ISL_3421642, EPI_ISL_3421643, EPI_ISL_3421644, EPI_ISL_3421645, EPI_ISL_3421646, EPI_ISL_3421647, EPI_ISL_3421648, EPI_ISL_3421649, EPI_ISL_3421650, EPI_ISL_3421651, EPI_ISL_3421652, EPI_ISL_3421653, EPI_ISL_3421654, EPI_ISL_3421655, EPI_ISL_3421656, EPI_ISL_3421657, EPI_ISL_3421658, EPI_ISL_3421659, EPI_ISL_3421660, EPI_ISL_3421661, EPI_ISL_3421662, EPI_ISL_3421663, EPI_ISL_3421664, EPI_ISL_3421665, EPI_ISL_3421666, EPI_ISL_3421667, EPI_ISL_3421668, EPI_ISL_3421669, EPI_ISL_3421670, EPI_ISL_3421671, EPI_ISL_3421672, EPI_ISL_3421673, EPI_ISL_3421674, EPI_ISL_3421675, EPI_ISL_3421676, EPI_ISL_3421677, EPI_ISL_3421678, EPI_ISL_3421679

see above | Quadram Institute Bioscience | COVID-19 Genomics UK (COG-UK) Consortium | Dave J. Baker, Gemma L. Kay, Alp Aydin, Thanh Le-Viet, Steven Rudder, Ana P. Tedim, Anastasia Kolyva, Maria Diaz, Leonardo de Oliveira Martins, Nabil-Fareed Alikhan, Lizzie Meadows, Rachael Stanley, Rose K Davidson, Ngozi Elumogo, Muhammed Yasir, Sophie J. Prosolek, Ebenezer Foster-Nyarko, Martin Lott, Christopher Jeanes, Lewis G. Spurgin, Nicholas M. Thomson, Alexander J Trotter, Rachel Gilroy, Luke Bedford, Samuel Bloomfield, Claire Stuart, Andrew Bell, Reenesh Prakash, Lindsay Coupland, Samir Dervisevic, Alison E. Mather, John Wain, Mark Webber, Andrew J. Page, Justin O'Grady

EPI_ISL_3421854, EPI_ISL_3421855, EPI_ISL_3421856, EPI_ISL_3421857, EPI_ISL_3421858, EPI_ISL_3421859, EPI_ISL_3421860, EPI_ISL_3421861, EPI_ISL_3421863, EPI_ISL_3421864, EPI_ISL_3421865, EPI_ISL_3421866, EPI_ISL_3421867, EPI_ISL_3421868, EPI_ISL_3421869, EPI_ISL_3421870, EPI_ISL_3421871, EPI_ISL_3421872, EPI_ISL_3421873, EPI_ISL_3421874, EPI_ISL_3421875, EPI_ISL_3421876, EPI_ISL_3421879, EPI_ISL_3421880, EPI_ISL_3421882, EPI_ISL_3421883, EPI_ISL_3421884, EPI_ISL_3421885, EPI_ISL_3421886, EPI_ISL_3421887, EPI_ISL_3421888, EPI_ISL_3421889, EPI_ISL_3421891, EPI_ISL_3421892, EPI_ISL_3421893, EPI_ISL_3421894, EPI_ISL_3421895, EPI_ISL_3421896, EPI_ISL_3421897, EPI_ISL_3421898, EPI_ISL_3421899, EPI_ISL_3421900, EPI_ISL_3421901, EPI_ISL_3421902, EPI_ISL_3421904, EPI_ISL_3421906, EPI_ISL_3421907, EPI_ISL_3421908, EPI_ISL_3421910, EPI_ISL_3421912, EPI_ISL_3421913, EPI_ISL_3421914, EPI_ISL_3421915, EPI_ISL_3421916, EPI_ISL_3421917, EPI_ISL_3421918, EPI_ISL_3421919, EPI_ISL_3421920, EPI_ISL_3421921, EPI_ISL_3421922, EPI_ISL_3421923, EPI_ISL_3421924, EPI_ISL_3421925, EPI_ISL_3421926, EPI_ISL_3421927, EPI_ISL_3421928, EPI_ISL_3421930, EPI_ISL_3421931, EPI_ISL_3421932, EPI_ISL_3421934, EPI_ISL_3421935, EPI_ISL_3421936, EPI_ISL_3421937, EPI_ISL_3421938, EPI_ISL_3421939, EPI_ISL_3421940, EPI_ISL_3421941, EPI_ISL_3421942, EPI_ISL_3421943, EPI_ISL_3421944, EPI_ISL_3421946, EPI_ISL_3421948, EPI_ISL_3421949, EPI_ISL_3421950, EPI_ISL_3421952

see above | Oxford Viromics, NDM, University of Oxford; Oxford University Hospitals; Basingstoke and North Hampshire Hospital | COVID-19 Genomics UK (COG-UK) Consortium | Tanya Golubchik, David Bonsall, George Macintyre, Amy Trebes, Mariateresa de Cesare, Catrin Moore, Alex Mobbs, Anita Justice, Robert Shaw, Monique Andersson, Timothy Peto, Emma Wise, Nathan Moore, Jessica Lynch, Nick Cortes, Matilde Mori, Stephen Kidd, David Buck, John Todd, Christophe Fraser

EPI_ISL_3421953, EPI_ISL_3421954, EPI_ISL_3421955, EPI_ISL_3421956, EPI_ISL_3421957, EPI_ISL_3421958, EPI_ISL_3421959, EPI_ISL_3421960, EPI_ISL_3421961, EPI_ISL_3421962, EPI_ISL_3421963, EPI_ISL_3421964, EPI_ISL_3421965, EPI_ISL_3421966, EPI_ISL_3421967, EPI_ISL_3421968, EPI_ISL_3421969, EPI_ISL_3421970, EPI_ISL_3421971, EPI_ISL_3421972, EPI_ISL_3421973, EPI_ISL_3421974, EPI_ISL_3421975, EPI_ISL_3421976, EPI_ISL_3421977, EPI_ISL_3421978, EPI_ISL_3421979, EPI_ISL_3421980, EPI_ISL_3421981, EPI_ISL_3421982, EPI_ISL_3421983, EPI_ISL_3421984, EPI_ISL_3421985, EPI_ISL_3421986, EPI_ISL_3421987, EPI_ISL_3421988, EPI_ISL_3421989, EPI_ISL_3421990, EPI_ISL_3421992, EPI_ISL_3421993, EPI_ISL_3421994, EPI_ISL_3421995, EPI_ISL_3421996, EPI_ISL_3421997, EPI_ISL_3421998, EPI_ISL_3421999, EPI_ISL_3422000, EPI_ISL_3422001, EPI_ISL_3422002, EPI_ISL_3422003, EPI_ISL_3422004, EPI_ISL_3422005, EPI_ISL_3422006, EPI_ISL_3422007, EPI_ISL_3422008, EPI_ISL_3422009, EPI_ISL_3422010, EPI_ISL_3422011, EPI_ISL_3422012, EPI_ISL_3422013, EPI_ISL_3422014, EPI_ISL_3422015, EPI_ISL_3422016, EPI_ISL_3422017, EPI_ISL_3422018, EPI_ISL_3422019, EPI_ISL_3422020, EPI_ISL_3422021, EPI_ISL_3422022, EPI_ISL_3422023, EPI_ISL_3422024, EPI_ISL_3422025, EPI_ISL_3422026, EPI_ISL_3422027, EPI_ISL_3422028, EPI_ISL_3422029, EPI_ISL_3422030, EPI_ISL_3422031, EPI_ISL_3422032, EPI_ISL_3422033, EPI_ISL_3422034, EPI_ISL_3422035, EPI_ISL_3422036, EPI_ISL_3422037, EPI_ISL_3422038, EPI_ISL_3422039, EPI_ISL_3422040, EPI_ISL_3422041, EPI_ISL_3422042, EPI_ISL_3422043, EPI_ISL_3422044, EPI_ISL_3422045, EPI_ISL_3422046, EPI_ISL_3422047, EPI_ISL_3422048, EPI_ISL_3422049, EPI_ISL_3422050, EPI_ISL_3422051, EPI_ISL_3422052, EPI_ISL_3422053, EPI_ISL_3422054, EPI_ISL_3422055, EPI_ISL_3422056, EPI_ISL_3422057, EPI_ISL_3422058, EPI_ISL_3422059, EPI_ISL_3422060, EPI_ISL_3422061, EPI_ISL_3422062, EPI_ISL_3422063, EPI_ISL_3422064, EPI_ISL_3422065, EPI_ISL_3422066, EPI_ISL_3422067, EPI_ISL_3422068, EPI_ISL_3422069, EPI_ISL_3422070, EPI_ISL_3422071, EPI_ISL_3422072, EPI_ISL_3422073, EPI_ISL_3422074, EPI_ISL_3422075, EPI_ISL_3422076, EPI_ISL_3422077, EPI_ISL_3422078, EPI_ISL_3422079, EPI_ISL_3422080, EPI_ISL_3422081, EPI_ISL_3422082, EPI_ISL_3422083, EPI_ISL_3422084, EPI_ISL_3422085, EPI_ISL_3422086, EPI_ISL_3422087, EPI_ISL_3422088, EPI_ISL_3422089, EPI_ISL_3422090, EPI_ISL_3422091, EPI_ISL_3422092, EPI_ISL_3422093, EPI_ISL_3422094, EPI_ISL_3422095, EPI_ISL_3422096, EPI_ISL_3422097, EPI_ISL_3422098, EPI_ISL_3422099, EPI_ISL_3422100, EPI_ISL_3422101, EPI_ISL_3422102, EPI_ISL_3422103, EPI_ISL_3422104, EPI_ISL_3422105, EPI_ISL_3422106, EPI_ISL_3422107, EPI_ISL_3422108, EPI_ISL_3422109, EPI_ISL_3422110, EPI_ISL_3422111, EPI_ISL_3422112, EPI_ISL_3422113, EPI_ISL_3422115, EPI_ISL_3422116, EPI_ISL_3422117, EPI_ISL_3422118, EPI_ISL_3422119, EPI_ISL_3422120, EPI_ISL_3422121, EPI_ISL_3422122, EPI_ISL_3422123, EPI_ISL_3422124, EPI_ISL_3422125, EPI_ISL_3422126, EPI_ISL_3422127, EPI_ISL_3422128, EPI_ISL_3422129, EPI_ISL_3422130, EPI_ISL_3422131, EPI_ISL_3422132, EPI_ISL_3422133, EPI_ISL_3422134, EPI_ISL_3422135, EPI_ISL_3422136, EPI_ISL_3422137, EPI_ISL_3422138, EPI_ISL_3422139, EPI_ISL_3422140, EPI_ISL_3422141, EPI_ISL_3422142, EPI_ISL_3422143, EPI_ISL_3422144, EPI_ISL_3422145, EPI_ISL_3422146, EPI_ISL_3422147, EPI_ISL_3422148, EPI_ISL_3422149, EPI_ISL_3422150, EPI_ISL_3422151, EPI_ISL_3422152, EPI_ISL_3422153, EPI_ISL_3422154, EPI_ISL_3422155, EPI_ISL_3422156, EPI_ISL_3422157, EPI_ISL_3422158, EPI_ISL_3422159, EPI_ISL_3422160, EPI_ISL_3422161, EPI_ISL_3422162, EPI_ISL_3422163, EPI_ISL_3422164, EPI_ISL_3422165, EPI_ISL_3422166, EPI_ISL_3422167, EPI_ISL_3422168, EPI_ISL_3422169, EPI_ISL_3422170, EPI_ISL_3422171, EPI_ISL_3422172, EPI_ISL_3422173, EPI_ISL_3422174, EPI_ISL_3422175, EPI_ISL_3422176, EPI_ISL_3422177, EPI_ISL_3422178, EPI_ISL_3422179, EPI_ISL_3422180, EPI_ISL_3422181, EPI_ISL_3422182, EPI_ISL_3422183, EPI_ISL_3422184, EPI_ISL_3422185, EPI_ISL_3422186, EPI_ISL_3422188, EPI_ISL_3422189, EPI_ISL_3422190, EPI_ISL_3422191, EPI_ISL_3422192, EPI_ISL_3422193, EPI_ISL_3422194, EPI_ISL_3422195, EPI_ISL_3422196, EPI_ISL_3422197, EPI_ISL_3422198, EPI_ISL_3422199, EPI_ISL_3422200, EPI_ISL_3422201, EPI_ISL_3422202, EPI_ISL_3422203, EPI_ISL_3422204, EPI_ISL_3422205, EPI_ISL_3422206, EPI_ISL_3422207, EPI_ISL_3422208, EPI_ISL_3422209, EPI_ISL_3422210, EPI_ISL_3422211, EPI_ISL_3422212, EPI_ISL_3422213, EPI_ISL_3422214, EPI_ISL_3422215, EPI_ISL_3422216, EPI_ISL_3422217, EPI_ISL_3422218, EPI_ISL_3422219, EPI_ISL_3422220, EPI_ISL_3422221, EPI_ISL_3422222, EPI_ISL_3422223, EPI_ISL_3422224, EPI_ISL_3422225, EPI_ISL_3422226, EPI_ISL_3422227, EPI_ISL_3422228, EPI_ISL_3422229, EPI_ISL_3422230, EPI_ISL_3422231, EPI_ISL_3422232, EPI_ISL_3422233, EPI_ISL_3422234, EPI_ISL_3422235, EPI_ISL_3422236, EPI_ISL_3422237, EPI_ISL_3422238, EPI_ISL_3422239, EPI_ISL_3422240, EPI_ISL_3422241, EPI_ISL_3422242, EPI_ISL_3422243, EPI_ISL_3422244, EPI_ISL_3422245, EPI_ISL_3422246, EPI_ISL_3422247, EPI_ISL_3422248, EPI_ISL_3422249, EPI_ISL_3422250, EPI_ISL_3422251, EPI_ISL_3422252, EPI_ISL_3422253, EPI_ISL_3422254, EPI_ISL_3422255, EPI_ISL_3422256, EPI_ISL_3422257, EPI_ISL_3422258, EPI_ISL_3422259, EPI_ISL_3422260, EPI_ISL_3422261, EPI_ISL_3422262, EPI_ISL_3422263, EPI_ISL_3422264, EPI_ISL_3422265, EPI_ISL_3422266, EPI_ISL_3422267, EPI_ISL_3422268, EPI_ISL_3422269, EPI_ISL_3422270, EPI_ISL_3422271, EPI_ISL_3422272, EPI_ISL_3422273, EPI_ISL_3422274, EPI_ISL_3422275, EPI_ISL_3422276, EPI_ISL_3422277, EPI_ISL_3422278, EPI_ISL_3422279, EPI_ISL_3422280, EPI_ISL_3422281, EPI_ISL_3422282, EPI_ISL_3422283, EPI_ISL_3422284, EPI_ISL_3422285, EPI_ISL_3422286, EPI_ISL_3422287, EPI_ISL_3422288, EPI_ISL_3422289, EPI_ISL_3422290, EPI_ISL_3422291, EPI_ISL_3422292, EPI_ISL_3422293, EPI_ISL_3422295, EPI_ISL_3422296, EPI_ISL_3422297, EPI_ISL_3422298, EPI_ISL_3422299, EPI_ISL_3422300, EPI_ISL_3422301, EPI_ISL_3422302, EPI_ISL_3422303, EPI_ISL_3422304, EPI_ISL_3422305, EPI_ISL_3422306, EPI_ISL_3422307, EPI_ISL_3422308, EPI_ISL_3422309, EPI_ISL_3422310, EPI_ISL_3422311, EPI_ISL_3422312, EPI_ISL_3422313, EPI_ISL_3422314, EPI_ISL_3422315, EPI_ISL_3422316, EPI_ISL_3422317, EPI_ISL_3422318, EPI_ISL_3422319, EPI_ISL_3422320, EPI_ISL_3422321, EPI_ISL_3422322, EPI_ISL_3422323, EPI_ISL_3422324, EPI_ISL_3422325, EPI_ISL_3422326, EPI_ISL_3422327, EPI_ISL_3422328, EPI_ISL_3422329, EPI_ISL_3422330, EPI_ISL_3422331, EPI_ISL_3422332, EPI_ISL_3422333, EPI_ISL_3422334, EPI_ISL_3422335, EPI_ISL_3422336, EPI_ISL_3422337, EPI_ISL_3422338, EPI_ISL_3422339

see above | Respiratory Virus Unit, Microbiology Services Colindale, Public | COVID-19 Genomics UK (COG-UK) Consortium | PHE Covid Sequencing Team

Health England

EPI_ISL_3422340, EPI_ISL_3422341, EPI_ISL_3422342, EPI_ISL_3422343, EPI_ISL_3422344, EPI_ISL_3422345, EPI_ISL_3422346, EPI_ISL_3422347, EPI_ISL_3422348, EPI_ISL_3422349, EPI_ISL_3422350, EPI_ISL_3422351, EPI_ISL_3422352, EPI_ISL_3422353, EPI_ISL_3422354, EPI_ISL_3422355, EPI_ISL_3422356, EPI_ISL_3422357, EPI_ISL_3422358, EPI_ISL_3422359, EPI_ISL_3422360, EPI_ISL_3422361,
EPI_ISL_3422362, EPI_ISL_3422363, EPI_ISL_3422364, EPI_ISL_3422365, EPI_ISL_3422366, EPI_ISL_3422367, EPI_ISL_3422368, EPI_ISL_3422369, EPI_ISL_3422370, EPI_ISL_3422371, EPI_ISL_3422372, EPI_ISL_3422373, EPI_ISL_3422374, EPI_ISL_3422375, EPI_ISL_3422376, EPI_ISL_3422377, EPI_ISL_3422378, EPI_ISL_3422379, EPI_ISL_3422380, EPI_ISL_3422381, EPI_ISL_3422382, EPI_ISL_3422383,
EPI_ISL_3422384, EPI_ISL_3422385, EPI_ISL_3422386, EPI_ISL_3422387, EPI_ISL_3422388, EPI_ISL_3422389, EPI_ISL_3422390, EPI_ISL_3422391, EPI_ISL_3422392, EPI_ISL_3422393, EPI_ISL_3422394, EPI_ISL_3422395, EPI_ISL_3422396, EPI_ISL_3422397, EPI_ISL_3422398, EPI_ISL_3422399, EPI_ISL_3422400, EPI_ISL_3422401, EPI_ISL_3422402, EPI_ISL_3422403, EPI_ISL_3422404, EPI_ISL_3422405,
EPI_ISL_3422406, EPI_ISL_3422407, EPI_ISL_3422408, EPI_ISL_3422409, EPI_ISL_3422410, EPI_ISL_3422411, EPI_ISL_3422412, EPI_ISL_3422413, EPI_ISL_3422414, EPI_ISL_3422415, EPI_ISL_3422416, EPI_ISL_3422417, EPI_ISL_3422418, EPI_ISL_3422419, EPI_ISL_3422420, EPI_ISL_3422421, EPI_ISL_3422422, EPI_ISL_3422423, EPI_ISL_3422424, EPI_ISL_3422425, EPI_ISL_3422426, EPI_ISL_3422427,
EPI_ISL_3422428, EPI_ISL_3422429, EPI_ISL_3422430, EPI_ISL_3422431, EPI_ISL_3422432, EPI_ISL_3422433, EPI_ISL_3422434, EPI_ISL_3422435, EPI_ISL_3422436, EPI_ISL_3422437, EPI_ISL_3422438, EPI_ISL_3422439, EPI_ISL_3422440, EPI_ISL_3422441, EPI_ISL_3422442, EPI_ISL_3422443, EPI_ISL_3422444, EPI_ISL_3422445, EPI_ISL_3422446, EPI_ISL_3422447, EPI_ISL_3422448, EPI_ISL_3422449,
EPI_ISL_3422450, EPI_ISL_3422451, EPI_ISL_3422452, EPI_ISL_3422453, EPI_ISL_3422454, EPI_ISL_3422455, EPI_ISL_3422456, EPI_ISL_3422457, EPI_ISL_3422458, EPI_ISL_3422459, EPI_ISL_3422460, EPI_ISL_3422461, EPI_ISL_3422462, EPI_ISL_3422463, EPI_ISL_3422464, EPI_ISL_3422465, EPI_ISL_3422466, EPI_ISL_3422467, EPI_ISL_3422468, EPI_ISL_3422469, EPI_ISL_3422470, EPI_ISL_3422471,
EPI_ISL_3422472, EPI_ISL_3422473, EPI_ISL_3422474, EPI_ISL_3422475, EPI_ISL_3422476, EPI_ISL_3422477, EPI_ISL_3422478, EPI_ISL_3422479, EPI_ISL_3422480, EPI_ISL_3422481, EPI_ISL_3422482, EPI_ISL_3422483, EPI_ISL_3422484, EPI_ISL_3422485, EPI_ISL_3422486, EPI_ISL_3422487, EPI_ISL_3422488, EPI_ISL_3422489, EPI_ISL_3422490, EPI_ISL_3422491, EPI_ISL_3422492, EPI_ISL_3422493,
EPI_ISL_3422494, EPI_ISL_3422495, EPI_ISL_3422496, EPI_ISL_3422497, EPI_ISL_3422498, EPI_ISL_3422499, EPI_ISL_3422500, EPI_ISL_3422501, EPI_ISL_3422502, EPI_ISL_3422503, EPI_ISL_3422504, EPI_ISL_3422505, EPI_ISL_3422506, EPI_ISL_3422507, EPI_ISL_3422508, EPI_ISL_3422509, EPI_ISL_3422510, EPI_ISL_3422511, EPI_ISL_3422512, EPI_ISL_3422513, EPI_ISL_3422514, EPI_ISL_3422515,
EPI_ISL_3422516, EPI_ISL_3422517, EPI_ISL_3422518, EPI_ISL_3422519, EPI_ISL_3422520, EPI_ISL_3422521, EPI_ISL_3422522, EPI_ISL_3422523, EPI_ISL_3422524, EPI_ISL_3422525, EPI_ISL_3422526, EPI_ISL_3422527, EPI_ISL_3422528, EPI_ISL_3422529, EPI_ISL_3422530, EPI_ISL_3422531, EPI_ISL_3422532, EPI_ISL_3422533, EPI_ISL_3422534, EPI_ISL_3422535, EPI_ISL_3422536, EPI_ISL_3422537,
EPI_ISL_3422538, EPI_ISL_3422539, EPI_ISL_3422540, EPI_ISL_3422541, EPI_ISL_3422542, EPI_ISL_3422543, EPI_ISL_3422544, EPI_ISL_3422545, EPI_ISL_3422546, EPI_ISL_3422547, EPI_ISL_3422548, EPI_ISL_3422549, EPI_ISL_3422550, EPI_ISL_3422551, EPI_ISL_3422552, EPI_ISL_3422553, EPI_ISL_3422554, EPI_ISL_3422555, EPI_ISL_3422556, EPI_ISL_3422557, EPI_ISL_3422558, EPI_ISL_3422559,
EPI_ISL_3422560, EPI_ISL_3422561, EPI_ISL_3422562, EPI_ISL_3422563, EPI_ISL_3422564, EPI_ISL_3422565, EPI_ISL_3422566, EPI_ISL_3422567, EPI_ISL_3422568, EPI_ISL_3422569, EPI_ISL_3422570, EPI_ISL_3422571, EPI_ISL_3422572, EPI_ISL_3422573, EPI_ISL_3422574, EPI_ISL_3422575, EPI_ISL_3422576, EPI_ISL_3422577, EPI_ISL_3422578, EPI_ISL_3422579, EPI_ISL_3422580, EPI_ISL_3422581,
EPI_ISL_3422582, EPI_ISL_3422583, EPI_ISL_3422584, EPI_ISL_3422585, EPI_ISL_3422586, EPI_ISL_3422587, EPI_ISL_3422588, EPI_ISL_3422589, EPI_ISL_3422590, EPI_ISL_3422591, EPI_ISL_3422592, EPI_ISL_3422593, EPI_ISL_3422594, EPI_ISL_3422595, EPI_ISL_3422596, EPI_ISL_3422597, EPI_ISL_3422598, EPI_ISL_3422599, EPI_ISL_3422600, EPI_ISL_3422601, EPI_ISL_3422602, EPI_ISL_3422603,
EPI_ISL_3422604, EPI_ISL_3422605, EPI_ISL_3422606, EPI_ISL_3422607, EPI_ISL_3422608, EPI_ISL_3422609, EPI_ISL_3422610, EPI_ISL_3422611, EPI_ISL_3422612, EPI_ISL_3422613, EPI_ISL_3422614, EPI_ISL_3422615, EPI_ISL_3422616, EPI_ISL_3422617, EPI_ISL_3422618, EPI_ISL_3422619, EPI_ISL_3422620, EPI_ISL_3422621, EPI_ISL_3422622, EPI_ISL_3422623, EPI_ISL_3422624, EPI_ISL_3422625,
EPI_ISL_3422626, EPI_ISL_3422627, EPI_ISL_3422628, EPI_ISL_3422629, EPI_ISL_3422630, EPI_ISL_3422631, EPI_ISL_3422632, EPI_ISL_3422633, EPI_ISL_3422634, EPI_ISL_3422635, EPI_ISL_3422636, EPI_ISL_3422637, EPI_ISL_3422638, EPI_ISL_3422639, EPI_ISL_3422640, EPI_ISL_3422641, EPI_ISL_3422642, EPI_ISL_3422643, EPI_ISL_3422644, EPI_ISL_3422645, EPI_ISL_3422646, EPI_ISL_3422647,
EPI_ISL_3422648, EPI_ISL_3422649, EPI_ISL_3422650, EPI_ISL_3422651, EPI_ISL_3422652, EPI_ISL_3422653, EPI_ISL_3422654, EPI_ISL_3422655, EPI_ISL_3422656, EPI_ISL_3422657, EPI_ISL_3422658, EPI_ISL_3422659, EPI_ISL_3422660, EPI_ISL_3422661, EPI_ISL_3422662, EPI_ISL_3422663, EPI_ISL_3422664, EPI_ISL_3422665, EPI_ISL_3422666, EPI_ISL_3422667, EPI_ISL_3422668, EPI_ISL_3422669,
EPI_ISL_3422670, EPI_ISL_3422671, EPI_ISL_3422672, EPI_ISL_3422673, EPI_ISL_3422674, EPI_ISL_3422675, EPI_ISL_3422676, EPI_ISL_3422677, EPI_ISL_3422678, EPI_ISL_3422679, EPI_ISL_3422680, EPI_ISL_3422681, EPI_ISL_3422682, EPI_ISL_3422683, EPI_ISL_3422684, EPI_ISL_3422685, EPI_ISL_3422686, EPI_ISL_3422687, EPI_ISL_3422688, EPI_ISL_3422689, EPI_ISL_3422690, EPI_ISL_3422691,
EPI_ISL_3422692, EPI_ISL_3422693, EPI_ISL_3422694, EPI_ISL_3422695, EPI_ISL_3422696, EPI_ISL_3422697, EPI_ISL_3422698, EPI_ISL_3422699, EPI_ISL_3422700, EPI_ISL_3422701, EPI_ISL_3422702, EPI_ISL_3422703, EPI_ISL_3422704, EPI_ISL_3422705, EPI_ISL_3422706, EPI_ISL_3422707, EPI_ISL_3422708, EPI_ISL_3422709, EPI_ISL_3422710, EPI_ISL_3422711, EPI_ISL_3422712, EPI_ISL_3422713,
EPI_ISL_3422714, EPI_ISL_3422715, EPI_ISL_3422716, EPI_ISL_3422717, EPI_ISL_3422718, EPI_ISL_3422719, EPI_ISL_3422720, EPI_ISL_3422721, EPI_ISL_3422722, EPI_ISL_3422723, EPI_ISL_3422724, EPI_ISL_3422725, EPI_ISL_3422726, EPI_ISL_3422727, EPI_ISL_3422728, EPI_ISL_3422729, EPI_ISL_3422730, EPI_ISL_3422731, EPI_ISL_3422732, EPI_ISL_3422733, EPI_ISL_3422734, EPI_ISL_3422735,
EPI_ISL_3422736, EPI_ISL_3422737, EPI_ISL_3422738, EPI_ISL_3422739, EPI_ISL_3422740, EPI_ISL_3422741, EPI_ISL_3422742, EPI_ISL_3422743, EPI_ISL_3422744, EPI_ISL_3422745, EPI_ISL_3422746, EPI_ISL_3422747, EPI_ISL_3422748, EPI_ISL_3422749, EPI_ISL_3422750, EPI_ISL_3422751, EPI_ISL_3422752, EPI_ISL_3422753, EPI_ISL_3422754, EPI_ISL_3422755, EPI_ISL_3422756, EPI_ISL_3422757,
EPI_ISL_3422758, EPI_ISL_3422759, EPI_ISL_3422760, EPI_ISL_3422761, EPI_ISL_3422762, EPI_ISL_3422763, EPI_ISL_3422764, EPI_ISL_3422765, EPI_ISL_3422766, EPI_ISL_3422767, EPI_ISL_3422768, EPI_ISL_3422769, EPI_ISL_3422770, EPI_ISL_3422771, EPI_ISL_3422772, EPI_ISL_3422773, EPI_ISL_3422774, EPI_ISL_3422775, EPI_ISL_3422776, EPI_ISL_3422777, EPI_ISL_3422778, EPI_ISL_3422779,
EPI_ISL_3422780, EPI_ISL_3422781, EPI_ISL_3422782, EPI_ISL_3422783, EPI_ISL_3422784, EPI_ISL_3422785, EPI_ISL_3422786, EPI_ISL_3422787, EPI_ISL_3422788, EPI_ISL_3422789, EPI_ISL_3422790, EPI_ISL_3422791, EPI_ISL_3422792, EPI_ISL_3422793, EPI_ISL_3422794, EPI_ISL_3422795, EPI_ISL_3422796, EPI_ISL_3422797, EPI_ISL_3422798, EPI_ISL_3422799, EPI_ISL_3422800, EPI_ISL_3422801,
EPI_ISL_3422802, EPI_ISL_3422803, EPI_ISL_3422804, EPI_ISL_3422805, EPI_ISL_3422806, EPI_ISL_3422807, EPI_ISL_3422808, EPI_ISL_3422809, EPI_ISL_3422810, EPI_ISL_3422811, EPI_ISL_3422812, EPI_ISL_3422813, EPI_ISL_3422814, EPI_ISL_3422815, EPI_ISL_3422816, EPI_ISL_3422817, EPI_ISL_3422818, EPI_ISL_3422819, EPI_ISL_3422820, EPI_ISL_3422821, EPI_ISL_3422822, EPI_ISL_3422823,
EPI_ISL_3422824, EPI_ISL_3422825, EPI_ISL_3422826, EPI_ISL_3422827, EPI_ISL_3422828, EPI_ISL_3422829, EPI_ISL_3422830, EPI_ISL_3422831, EPI_ISL_3422832, EPI_ISL_3422833, EPI_ISL_3422834, EPI_ISL_3422835, EPI_ISL_3422836, EPI_ISL_3422837, EPI_ISL_3422838, EPI_ISL_3422839, EPI_ISL_3422840, EPI_ISL_3422841, EPI_ISL_3422842, EPI_ISL_3422843, EPI_ISL_3422844, EPI_ISL_3422845,
EPI_ISL_3422846, EPI_ISL_3422847, EPI_ISL_3422848, EPI_ISL_3422849, EPI_ISL_3422850, EPI_ISL_3422851, EPI_ISL_3422852, EPI_ISL_3422853, EPI_ISL_3422854, EPI_ISL_3422855, EPI_ISL_3422856, EPI_ISL_3422857, EPI_ISL_3422858, EPI_ISL_3422859, EPI_ISL_3422860, EPI_ISL_3422861, EPI_ISL_3422862, EPI_ISL_3422863, EPI_ISL_3422864, EPI_ISL_3422865, EPI_ISL_3422866, EPI_ISL_3422867,
EPI_ISL_3422868, EPI_ISL_3422869, EPI_ISL_3422870, EPI_ISL_3422871, EPI_ISL_3422872, EPI_ISL_3422873, EPI_ISL_3422874, EPI_ISL_3422875, EPI_ISL_3422876, EPI_ISL_3422877, EPI_ISL_3422878, EPI_ISL_3422879, EPI_ISL_3422880, EPI_ISL_3422881, EPI_ISL_3422882, EPI_ISL_3422883, EPI_ISL_3422884, EPI_ISL_3422885, EPI_ISL_3422886, EPI_ISL_3422887, EPI_ISL_3422888, EPI_ISL_3422889,
EPI_ISL_3422890, EPI_ISL_3422891, EPI_ISL_3422892, EPI_ISL_3422893, EPI_ISL_3422894, EPI_ISL_3422895, EPI_ISL_3422896, EPI_ISL_3422897, EPI_ISL_3422898, EPI_ISL_3422899, EPI_ISL_3422900, EPI_ISL_3422901, EPI_ISL_3422902, EPI_ISL_3422903, EPI_ISL_3422904, EPI_ISL_3422905, EPI_ISL_3422906, EPI_ISL_3422907, EPI_ISL_3422908, EPI_ISL_3422909, EPI_ISL_3422910, EPI_ISL_3422911,
EPI_ISL_3422912, EPI_ISL_3422913, EPI_ISL_3422914, EPI_ISL_3422915, EPI_ISL_3422916, EPI_ISL_3422917, EPI_ISL_3422918, EPI_ISL_3422919, EPI_ISL_3422920, EPI_ISL_3422921, EPI_ISL_3422922, EPI_ISL_3422923, EPI_ISL_3422924, EPI_ISL_3422925, EPI_ISL_3422926, EPI_ISL_3422927, EPI_ISL_3422928, EPI_ISL_3422929, EPI_ISL_3422930, EPI_ISL_3422931, EPI_ISL_3422932, EPI_ISL_3422933,
EPI_ISL_3422934, EPI_ISL_3422935, EPI_ISL_3422936, EPI_ISL_3422937, EPI_ISL_3422938, EPI_ISL_3422939, EPI_ISL_3422940, EPI_ISL_3422941, EPI_ISL_3422942, EPI_ISL_3422943, EPI_ISL_3422944, EPI_ISL_3422945, EPI_ISL_3422946, EPI_ISL_3422947, EPI_ISL_3422948, EPI_ISL_3422949, EPI_ISL_3422950, EPI_ISL_3422951, EPI_ISL_3422952, EPI_ISL_3422953, EPI_ISL_3422954, EPI_ISL_3422955,
EPI_ISL_3422956, EPI_ISL_3422957, EPI_ISL_3422958, EPI_ISL_3422959, EPI_ISL_3422960, EPI_ISL_3422961, EPI_ISL_3422962, EPI_ISL_3422963, EPI_ISL_3422964, EPI_ISL_3422965, EPI_ISL_3422966, EPI_ISL_3422967, EPI_ISL_3422968, EPI_ISL_3422969, EPI_ISL_3422970, EPI_ISL_3422971, EPI_ISL_3422972, EPI_ISL_3422973, EPI_ISL_3422974, EPI_ISL_3422975, EPI_ISL_3422976, EPI_ISL_3422977,
EPI_ISL_3422978, EPI_ISL_3422979, EPI_ISL_3422980, EPI_ISL_3422981, EPI_ISL_3422982, EPI_ISL_3422983, EPI_ISL_3422984, EPI_ISL_3422985, EPI_ISL_3422986, EPI_ISL_3422987, EPI_ISL_3422988, EPI_ISL_3422989, EPI_ISL_3422990, EPI_ISL_3422991, EPI_ISL_3422992, EPI_ISL_3422993, EPI_ISL_3422994, EPI_ISL_3422995, EPI_ISL_3422996, EPI_ISL_3422997, EPI_ISL_3422998, EPI_ISL_3422999,
EPI_ISL_3423000, EPI_ISL_3423001, EPI_ISL_3423002, EPI_ISL_3423003, EPI_ISL_3423004, EPI_ISL_3423005, EPI_ISL_3423006, EPI_ISL_3423007, EPI_ISL_3423008, EPI_ISL_3423009, EPI_ISL_3423010, EPI_ISL_3423011, EPI_ISL_3423012, EPI_ISL_3423013, EPI_ISL_3423014, EPI_ISL_3423015, EPI_ISL_3423016, EPI_ISL_3423017, EPI_ISL_3423018, EPI_ISL_3423019, EPI_ISL_3423020, EPI_ISL_3423021,
EPI_ISL_3423022, EPI_ISL_3423023, EPI_ISL_3423024, EPI_ISL_3423025, EPI_ISL_3423026, EPI_ISL_3423027, EPI_ISL_3423028, EPI_ISL_3423029, EPI_ISL_3423030, EPI_ISL_3423031, EPI_ISL_3423032, EPI_ISL_3423033, EPI_ISL_3423034, EPI_ISL_3423035, EPI_ISL_3423036, EPI_ISL_3423037, EPI_ISL_3423038, EPI_ISL_3423039, EPI_ISL_3423040, EPI_ISL_3423041, EPI_ISL_3423042, EPI_ISL_3423043,
EPI_ISL_3423044, EPI_ISL_3423045, EPI_ISL_3423046, EPI_ISL_3423047, EPI_ISL_3423048, EPI_ISL_3423049, EPI_ISL_3423050, EPI_ISL_3423051, EPI_ISL_3423052, EPI_ISL_3423053, EPI_ISL_3423054, EPI_ISL_3423055, EPI_ISL_3423056, EPI_ISL_3423057, EPI_ISL_3423058, EPI_ISL_3423059, EPI_ISL_3423060, EPI_ISL_3423061, EPI_ISL_3423062, EPI_ISL_3423063, EPI_ISL_3423064, EPI_ISL_3423065,
EPI_ISL_3423066, EPI_ISL_3423067, EPI_ISL_3423068, EPI_ISL_3423069, EPI_ISL_3423070, EPI_ISL_3423071, EPI_ISL_3423072, EPI_ISL_3423073, EPI_ISL_3423074, EPI_ISL_3423075, EPI_ISL_3423076, EPI_ISL_3423077, EPI_ISL_3423078, EPI_ISL_3423079, EPI_ISL_3423080, EPI_ISL_3423081, EPI_ISL_3423082, EPI_ISL_3423083, EPI_ISL_3423084, EPI_ISL_3423085, EPI_ISL_3423086, EPI_ISL_3423087,
EPI_ISL_3423088, EPI_ISL_3423089, EPI_ISL_3423090, EPI_ISL_3423091, EPI_ISL_3423092, EPI_ISL_3423093, EPI_ISL_3423094, EPI_ISL_3423095, EPI_ISL_3423096, EPI_ISL_3423097, EPI_ISL_3423098, EPI_ISL_3423099, EPI_ISL_3423100, EPI_ISL_3423101, EPI_ISL_3423102, EPI_ISL_3423103, EPI_ISL_3423104, EPI_ISL_3423105, EPI_ISL_3423106, EPI_ISL_3423107, EPI_ISL_3423108, EPI_ISL_3423109,
EPI_ISL_3423110, EPI_ISL_3423111, EPI_ISL_3423112, EPI_ISL_3423113, EPI_ISL_3423114, EPI_ISL_3423115, EPI_ISL_3423116, EPI_ISL_3423117, EPI_ISL_3423118, EPI_ISL_3423119, EPI_ISL_3423120, EPI_ISL_3423121, EPI_ISL_3423122, EPI_ISL_3423123, EPI_ISL_3423124, EPI_ISL_3423125, EPI_ISL_3423126, EPI_ISL_3423127, EPI_ISL_3423128, EPI_ISL_3423129, EPI_ISL_3423130, EPI_ISL_3423131,
EPI_ISL_3423132, EPI_ISL_3423133, EPI_ISL_3423134, EPI_ISL_3423135, EPI_ISL_3423136, EPI_ISL_3423137, EPI_ISL_3423138, EPI_ISL_3423139, EPI_ISL_3423140, EPI_ISL_3423141, EPI_ISL_3423142, EPI_ISL_3423143, EPI_ISL_3423144, EPI_ISL_3423145, EPI_ISL_3423146, EPI_ISL_3423147, EPI_ISL_3423148, EPI_ISL_3423149, EPI_ISL_3423150, EPI_ISL_3423151, EPI_ISL_3423152, EPI_ISL_3423153,
EPI_ISL_3423154, EPI_ISL_3423155, EPI_ISL_3423156, EPI_ISL_3423157, EPI_ISL_3423158, EPI_ISL_3423159, EPI_ISL_3423160, EPI_ISL_3423161, EPI_ISL_3423162, EPI_ISL_3423163, EPI_ISL_3423164, EPI_ISL_3423165, EPI_ISL_3423166, EPI_ISL_3423167, EPI_ISL_3423168, EPI_ISL_3423169, EPI_ISL_3423170, EPI_ISL_3423171, EPI_ISL_3423172, EPI_ISL_3423173, EPI_ISL_3423174, EPI_ISL_3423175,
EPI_ISL_3423176, EPI_ISL_3423177, EPI_ISL_3423178, EPI_ISL_3423179, EPI_ISL_3423180, EPI_ISL_3423181, EPI_ISL_3423182, EPI_ISL_3423183, EPI_ISL_3423184, EPI_ISL_3423185, EPI_ISL_3423186, EPI_ISL_3423187, EPI_ISL_3423188, EPI_ISL_3423189, EPI_ISL_3423190, EPI_ISL_3423191, EPI_ISL_3423192, EPI_ISL_3423193, EPI_ISL_3423194, EPI_ISL_3423195, EPI_ISL_3423196, EPI_ISL_3423197,
EPI_ISL_3423198, EPI_ISL_3423199, EPI_ISL_3423200, EPI_ISL_3423201, EPI_ISL_3423202, EPI_ISL_3423203, EPI_ISL_3423204, EPI_ISL_3423205, EPI_ISL_3423206, EPI_ISL_3423207, EPI_ISL_3423208, EPI_ISL_3423209, EPI_ISL_3423210, EPI_ISL_3423211, EPI_ISL_3423212, EPI_ISL_3423213, EPI_ISL_3423214, EPI_ISL_3423215, EPI_ISL_3423216, EPI_ISL_3423217, EPI_ISL_3423218, EPI_ISL_3423219,
EPI_ISL_3423220, EPI_ISL_3423221, EPI_ISL_3423222, EPI_ISL_3423223, EPI_ISL_3423224, EPI_ISL_3423225, EPI_ISL_3423226, EPI_ISL_3423227, EPI_ISL_3423228, EPI_ISL_3423229, EPI_ISL_3423230, EPI_ISL_3423231, EPI_ISL_3423232, EPI_ISL_3423233, EPI_ISL_3423234, EPI_ISL_3423235, EPI_ISL_3423236, EPI_ISL_3423237, EPI_ISL_3423238, EPI_ISL_3423239, EPI_ISL_3423240, EPI_ISL_3423241,
EPI_ISL_3423242, EPI_ISL_3423243, EPI_ISL_3423244, EPI_ISL_3423245, EPI_ISL_3423246, EPI_ISL_3423247, EPI_ISL_3423248, EPI_ISL_3423249, EPI_ISL_3423250, EPI_ISL_3423251, EPI_ISL_3423252, EPI_ISL_3423253, EPI_ISL_3423254, EPI_ISL_3423255, EPI_ISL_3423256, EPI_ISL_3423257, EPI_ISL_3423258, EPI_ISL_3423259, EPI_ISL_3423260, EPI_ISL_3423261, EPI_ISL_3423262, EPI_ISL_3423263,
EPI_ISL_3423264, EPI_ISL_3423265, EPI_ISL_3423266, EPI_ISL_3423267, EPI_ISL_3423268, EPI_ISL_3423269, EPI_ISL_3423270, EPI_ISL_3423271, EPI_ISL_3423272, EPI_ISL_3423273, EPI_ISL_3423274, EPI_ISL_3423275, EPI_ISL_3423276, EPI_ISL_3423277, EPI_ISL_3423278, EPI_ISL_3423279, EPI_ISL_3423280, EPI_ISL_3423281, EPI_ISL_3423282, EPI_ISL_3423283, EPI_ISL_3423284, EPI_ISL_3423285,
EPI_ISL_3423286, EPI_ISL_3423287, EPI_ISL_3423288, EPI_ISL_3423289, EPI_ISL_3423290, EPI_ISL_3423291, EPI_ISL_3423292, EPI_ISL_3423293, EPI_ISL_3423294, EPI_ISL_3423295, EPI_ISL_3423296, EPI_ISL_3423297, EPI_ISL_3423298, EPI_ISL_3423299, EPI_ISL_3423300, EPI_ISL_3423301, EPI_ISL_3423302, EPI_ISL_3423303, EPI_ISL_3423304, EPI_ISL_3423305, EPI_ISL_3423306, EPI_ISL_3423307,
EPI_ISL_3423308, EPI_ISL_3423309, EPI_ISL_3423310, EPI_ISL_3423311, EPI_ISL_3423312, EPI_ISL_3423313, EPI_ISL_3423314, EPI_ISL_3423315, EPI_ISL_3423316, EPI_ISL_3423317, EPI_ISL_3423318, EPI_ISL_3423319, EPI_ISL_3423320, EPI_ISL_3423321, EPI_ISL_3423322, EPI_ISL_3423323, EPI_ISL_3423324, EPI_ISL_3423325, EPI_ISL_3423326, EPI_ISL_3423327, EPI_ISL_3423328, EPI_ISL_3423329,
EPI_ISL_3423330, EPI_ISL_3423331, EPI_ISL_3423332, EPI_ISL_3423333, EPI_ISL_3423334, EPI_ISL_3423335, EPI_ISL_3423336, EPI_ISL_3423337, EPI_ISL_3423338, EPI_ISL_3423339, EPI_ISL_3423340, EPI_ISL_3423341, EPI_ISL_3423342, EPI_ISL_3423343, EPI_ISL_3423344, EPI_ISL_3423345, EPI_ISL_3423346, EPI_ISL_3423347, EPI_ISL_3423348, EPI_ISL_3423349, EPI_ISL_3423350, EPI_ISL_3423351,
EPI_ISL_3423352, EPI_ISL_3423353, EPI_ISL_3423354, EPI_ISL_3423355, EPI_ISL_3423356, EPI_ISL_3423357, EPI_ISL_3423358, EPI_ISL_3423359, EPI_ISL_3423360, EPI_ISL_3423361, EPI_ISL_3423362, EPI_ISL_3423363, EPI_ISL_3423364, EPI_ISL_3423365, EPI_ISL_3423366, EPI_ISL_3423367, EPI_ISL_3423368, EPI_ISL_3423369, EPI_ISL_3423370, EPI_ISL_3423371, EPI_ISL_3423372, EPI_ISL_3423373,
EPI_ISL_3423374, EPI_ISL_3423375, EPI_ISL_3423376, EPI_ISL_3423377, EPI_ISL_3423378, EPI_ISL_3423379, EPI_ISL_3423380, EPI_ISL_3423381, EPI_ISL_3423382, EPI_ISL_3423383, EPI_ISL_3423384, EPI_ISL_3423385, EPI_ISL_3423386, EPI_ISL_3423387, EPI_ISL_3423388, EPI_ISL_3423389, EPI_ISL_3423390, EPI_ISL_3423391, EPI_ISL_3423392, EPI_ISL_3423393, EPI_ISL_3423394, EPI_ISL_3423395, EPI_ISL_3423396,
EPI_ISL_3423397, EPI_ISL_3423398, EPI_ISL_3423399, EPI_ISL_3423400, EPI_ISL_3423401, EPI_ISL_3423402, EPI_ISL_3423403, EPI_ISL_3423404, EPI_ISL_3423405, EPI_ISL_3423406, EPI_ISL_3423407, EPI_ISL_3423408, EPI_ISL_3423409, EPI_ISL_3423410, EPI_ISL_3423411, EPI_ISL_3423412, EPI_ISL_3423413, EPI_ISL_3423414, EPI_ISL_3423415, EPI_ISL_3423416, EPI_ISL_3423417, EPI_ISL_3423418,
EPI_ISL_3423419, EPI_ISL_3423420, EPI_ISL_3423421, EPI_ISL_3423422, EPI_ISL_3423423, EPI_ISL_3423424, EPI_ISL_3423425, EPI_ISL_3423426, EPI_ISL_3423427, EPI_ISL_3423428, EPI_ISL_3423429, EPI_ISL_3423430, EPI_ISL_3423431, EPI_ISL_3423432, EPI_ISL_3423433, EPI_ISL_3423434, EPI_ISL_3423435, EPI_ISL_3423436, EPI_ISL_3423437, EPI_ISL_3423438, EPI_ISL_3423439, EPI_ISL_3423440,
EPI_ISL_3423441, EPI_ISL_3423442, EPI_ISL_3423443, EPI_ISL_3423444, EPI_ISL_3423445, EPI_ISL_3423446, EPI_ISL_3423447, EPI_ISL_3423448, EPI_ISL_3423449, EPI_ISL_3423450, EPI_ISL_3423451, EPI_ISL_3423452, EPI_ISL_3423453, EPI_ISL_3423454, EPI_ISL_3423455, EPI_ISL_3423456, EPI_ISL_3423457, EPI_ISL_3423458, EPI_ISL_3423459, EPI_ISL_3423460, EPI_ISL_3423461, EPI_ISL_3423462,
EPI_ISL_3423463, EPI_ISL_3423464, EPI_ISL_3423465, EPI_ISL_3423466, EPI_ISL_3423467, EPI_ISL_3423468, EPI_ISL_3423469, EPI_ISL_3423470, EPI_ISL_3423471, EPI_ISL_3423472, EPI_ISL_3423473, EPI_ISL_3423474, EPI_ISL_3423475, EPI_ISL_3423476, EPI_ISL_3423477, EPI_ISL_3423478, EPI_ISL_3423479, EPI_ISL_3423480, EPI_ISL_3423481, EPI_ISL_3423482, EPI_ISL_3423483, EPI_ISL_3423484,
EPI_ISL_3423485, EPI_ISL_3423486, EPI_ISL_3423487, EPI_ISL_3423488, EPI_ISL_3423489, EPI_ISL_3423490, EPI_ISL_3423491, EPI_ISL_3423492, EPI_ISL_3423493, EPI_ISL_3423494, EPI_ISL_3423495, EPI_ISL_3423496, EPI_ISL_3423497, EPI_ISL_3423498, EPI_ISL_3423499, EPI_ISL_3423500, EPI_ISL_3423501, EPI_ISL_3423502, EPI_ISL_3423503, EPI_ISL_3423504, EPI_ISL_3423505, EPI_ISL_3423506,
EPI_ISL_3423507, EPI_ISL_3423508, EPI_ISL_3423509, EPI_ISL_3423510, EPI_ISL_3423511, EPI_ISL_3423512, EPI_ISL_3423513, EPI_ISL_3423514, EPI_ISL_3423515, EPI_ISL_3423516, EPI_ISL_3423517, EPI_ISL_3423518, EPI_ISL_3423519, EPI_ISL_3423520, EPI_ISL_3423521, EPI_ISL_3423522, EPI_ISL_3423523, EPI_ISL_3423524, EPI_ISL_3423525, EPI_ISL_3423526, EPI_ISL_3423527, EPI_ISL_3423528,
EPI_ISL_3423529, EPI_ISL_3423530, EPI_ISL_3423531, EPI_ISL_3423532, EPI_ISL_3423533, EPI_ISL_3423534, EPI_ISL_3423535, EPI_ISL_3423536, EPI_ISL_3423537, EPI_ISL_3423538, EPI_ISL_3423539, EPI_ISL_3423540, EPI_ISL_3423541, EPI_ISL_3423542, EPI_ISL_3423543, EPI_ISL_3423544, EPI_ISL_3423545, EPI_ISL_3423546, EPI_ISL_3423547, EPI_ISL_3423548, EPI_ISL_3423549, EPI_ISL_3423550,
EPI_ISL_3423551, EPI_ISL_3423552, EPI_ISL_3423553, EPI_ISL_3423554, EPI_ISL_3423555, EPI_ISL_3423556, EPI_ISL_3423557, EPI_ISL_3423558, EPI_ISL_3423559, EPI_ISL_3423560, EPI_ISL_3423561, EPI_ISL_3423562, EPI_ISL_3423563, EPI_ISL_3423564, EPI_ISL_3423565, EPI_ISL_3423566, EPI_ISL_3423567, EPI_ISL_3423568, EPI_ISL_3423569, EPI_ISL_3423570, EPI_ISL_3423571, EPI_ISL_3423572,
EPI_ISL_3423573, EPI_ISL_3423574, EPI_ISL_3423575, EPI_ISL_3423576, EPI_ISL_3423577, EPI_ISL_3423578, EPI_ISL_3423579, EPI_ISL_3423580, EPI_ISL_3423581, EPI_ISL_3423582, EPI_ISL_3423583, EPI_ISL_3423584, EPI_ISL_3423585, EPI_ISL_3423586, EPI_ISL_3423587, EPI_ISL_3423588, EPI_ISL_3423589, EPI_ISL_3423590, EPI_ISL_3423591, EPI_ISL_3423592, EPI_ISL_3423593, EPI_ISL_3423594,
EPI_ISL_3423595, EPI_ISL_3423596, EPI_ISL_3423597, EPI_ISL_3423598, EPI_ISL_3423599, EPI_ISL_3423600, EPI_ISL_3423601, EPI_ISL_3423602, EPI_ISL_3423603, EPI_ISL_3423604, EPI_ISL_3423605, EPI_ISL_3423606, EPI_ISL_3423607, EPI_ISL_3423608, EPI_ISL_3423609, EPI_ISL_3423610, EPI_ISL_3423611, EPI_ISL_3423612, EPI_ISL_3423613, EPI_ISL_3423614, EPI_ISL_3423615, EPI_ISL_3423616,
EPI_ISL_3423617, EPI_ISL_3423618, EPI_ISL_3423619, EPI_ISL_3423620, EPI_ISL_3423621, EPI_ISL_3423622, EPI_ISL_3423623, EPI_ISL_3423624, EPI_ISL_3423625, EPI_ISL_3423626, EPI_ISL_3423627, EPI_ISL_3423628, EPI_ISL_3423629, EPI_ISL_3423630, EPI_ISL_3423631, EPI_ISL_3423632, EPI_ISL_3423633, EPI_ISL_3423634, EPI_ISL_3423635, EPI_ISL_3423636, EPI_ISL_3423637, EPI_ISL_3423638,
EPI_ISL_3423639, EPI_ISL_3423640, EPI_ISL_3423641, EPI_ISL_3423642, EPI_ISL_3423643, EPI_ISL_3423644, EPI_ISL_3423645, EPI_ISL_3423646, EPI_ISL_3423647, EPI_ISL_3423648, EPI_ISL_3423649, EPI_ISL_3423650, EPI_ISL_3423651, EPI_ISL_3423652, EPI_ISL_3423653, EPI_ISL_3423654, EPI_ISL_3423655, EPI_ISL_3423656, EPI_ISL_3423657, EPI_ISL_3423658, EPI_ISL_3423659, EPI_ISL_3423660,
EPI_ISL_3423661, EPI_ISL_3423662, EPI_ISL_3423663, EPI_ISL_3423664, EPI_ISL_3423665, EPI_ISL_3423666, EPI_ISL_3423667, EPI_ISL_3423668, EPI_ISL_3423669, EPI_ISL_3423670, EPI_ISL_3423671, EPI_ISL_3423672, EPI_ISL_3423673, EPI_ISL_3423674, EPI_ISL_3423675, EPI_ISL_3423676, EPI_ISL_3423677, EPI_ISL_3423678, EPI_ISL_3423679, EPI_ISL_3423680, EPI_ISL_3423681, EPI_ISL_3423682,
EPI_ISL_3423683, EPI_ISL_3423684, EPI_ISL_3423685, EPI_ISL_3423686, EPI_ISL_3423687, EPI_ISL_3423688, EPI_ISL_3423689, EPI_ISL_3423690, EPI_ISL_3423691, EPI_ISL_3423692, EPI_ISL_3423693, EPI_ISL_3423694, EPI_ISL_3423695, EPI_ISL_3423696, EPI_ISL_3423697, EPI_ISL_3423698, EPI_ISL_3423699, EPI_ISL_3423700, EPI_ISL_3423701, EPI_ISL_3423702, EPI_ISL_3423703, EPI_ISL_3423704,
EPI_ISL_3423705, EPI_ISL_3423706, EPI_ISL_3423707, EPI_ISL_3423708, EPI_ISL_3423709, EPI_ISL_3423710, EPI_ISL_3423711, EPI_ISL_3423712, EPI_ISL_3423713, EPI_ISL_3423714, EPI_ISL_3423715, EPI_ISL_3423716, EPI_ISL_3423717, EPI_ISL_3423718, EPI_ISL_3423719, EPI_ISL_3423720, EPI_ISL_3423721, EPI_ISL_3423722, EPI_ISL_3423723, EPI_ISL_3423724, EPI_ISL_3423725, EPI_ISL_3423726,
EPI_ISL_3423727, EPI_ISL_3423728, EPI_ISL_3423729, EPI_ISL_3423730, EPI_ISL_3423731, EPI_ISL_3423732, EPI_ISL_3423733, EPI_ISL_3423734, EPI_ISL_3423735, EPI_ISL_3423736, EPI_ISL_3423737, EPI_ISL_3423738, EPI_ISL_3423739, EPI_ISL_3423740, EPI_ISL_3423741, EPI_ISL_3423742, EPI_ISL_3423743, EPI_ISL_3423744, EPI_ISL_3423745, EPI_ISL_3423746, EPI_ISL_3423747, EPI_ISL_3423748,
EPI_ISL_3423749, EPI_ISL_3423750, EPI_ISL_3423751, EPI_ISL_3423752, EPI_ISL_3423753, EPI_ISL_3423754, EPI_ISL_3423755, EPI_ISL_3423756, EPI_ISL_3423757, EPI_ISL_3423758, EPI_ISL_3423759, EPI_ISL_3423760, EPI_ISL_3423761, EPI_ISL_3423762, EPI_ISL_3423763, EPI_ISL_3423764, EPI_ISL_3423765, EPI_ISL_3423766, EPI_ISL_3423767, EPI_ISL_3423768, EPI_ISL_3423769, EPI_ISL_3423770,
EPI_ISL_3423771, EPI_ISL_3423772, EPI_ISL_3423773, EPI_ISL_3423774, EPI_ISL_3423775, EPI_ISL_3423776, EPI_ISL_3423777, EPI_ISL_3423778, EPI_ISL_3423779, EPI_ISL_3423780, EPI_ISL_3423781, EPI_ISL_3423782, EPI_ISL_3423783, EPI_ISL_3423784, EPI_ISL_3423785, EPI_ISL_3423786, EPI_ISL_3423787, EPI_ISL_3423788, EPI_ISL_3423789, EPI_ISL_3423790, EPI_ISL_3423791, EPI_ISL_3423792,
EPI_ISL_3423793, EPI_ISL_3423794, EPI_ISL_3423795, EPI_ISL_3423796, EPI_ISL_3423797, EPI_ISL_3423798, EPI_ISL_3423799, EPI_ISL_3423800, EPI_ISL_3423801, EPI_ISL_3423802, EPI_ISL_3423803, EPI_ISL_3423804, EPI_ISL_3423805, EPI_ISL_3423806, EPI_ISL_3423807, EPI_ISL_3423808, EPI_ISL_3423809, EPI_ISL_3423810, EPI_ISL_3423811, EPI_ISL_3423812, EPI_ISL_3423813, EPI_ISL_3423814,
EPI_ISL_3423815, EPI_ISL_3423816, EPI_ISL_3423817, EPI_ISL_3423818, EPI_ISL_3423819, EPI_ISL_3423820, EPI_ISL_3423821, EPI_ISL_3423822, EPI_ISL_3423823, EPI_ISL_3423824, EPI_ISL_3423825, EPI_ISL_3423826, EPI_ISL_3423827, EPI_ISL_3423828, EPI_ISL_3423829, EPI_ISL_3423830, EPI_ISL_3423831, EPI_ISL_3423832, EPI_ISL_3423833, EPI_ISL_3423834, EPI_ISL_3423835, EPI_ISL_3423836,
EPI_ISL_3423837, EPI_ISL_3423838, EPI_ISL_3423839, EPI_ISL_3423840, EPI_ISL_3423841, EPI_ISL_3423842, EPI_ISL_3423843, EPI_ISL_3423844, EPI_ISL_3423845, EPI_ISL_3423846, EPI_ISL_3423847, EPI_ISL_3423848, EPI_ISL_3423849, EPI_ISL_3423850, EPI_ISL_3423851, EPI_ISL_3423852, EPI_ISL_3423853, EPI_ISL_3423854, EPI_ISL_3423855, EPI_ISL_3423856, EPI_ISL_3423857, EPI_ISL_3423858,
EPI_ISL_3423859, EPI_ISL_3423860, EPI_ISL_3423861, EPI_ISL_3423862, EPI_ISL_3423863, EPI_ISL_3423864, EPI_ISL_3423865, EPI_ISL_3423866, EPI_ISL_3423867, EPI_ISL_3423868, EPI_ISL_3423869, EPI_ISL_3423870, EPI_ISL_3423871, EPI_ISL_3423872, EPI_ISL_3423873, EPI_ISL_3423874, EPI_ISL_3423875, EPI_ISL_3423876, EPI_ISL_3423877, EPI_ISL_3423878, EPI_ISL_3423879, EPI_ISL_3423880,
EPI_ISL_3423881, EPI_ISL_3423882, EPI_ISL_3423883, EPI_ISL_3423884, EPI_ISL_3423885, EPI_ISL_3423886, EPI_ISL_3423887, EPI_ISL_3423888, EPI_ISL_3423889, EPI_ISL_3423890, EPI_ISL_3423891, EPI_ISL_3423892, EPI_ISL_3423893, EPI_ISL_3423894, EPI_ISL_3423895, EPI_ISL_3423896, EPI_ISL_3423897, EPI_ISL_3423898, EPI_ISL_3423899, EPI_ISL_3423900, EPI_ISL_3423901, EPI_ISL_3423902,
EPI_ISL_3423903, EPI_ISL_3423904, EPI_ISL_3423905, EPI_ISL_3423906, EPI_ISL_3423907, EPI_ISL_3423908, EPI_ISL_3423909, EPI_ISL_3423910, EPI_ISL_3423911, EPI_ISL_3423912, EPI_ISL_3423913, EPI_ISL_3423914, EPI_ISL_3423915, EPI_ISL_3423916, EPI_ISL_3423917, EPI_ISL_3423918, EPI_ISL_3423919, EPI_ISL_3423920, EPI_ISL_3423921, EPI_ISL_3423922, EPI_ISL_3423923, EPI_ISL_3423924,
EPI_ISL_3423925, EPI_ISL_3423926, EPI_ISL_3423927, EPI_ISL_3423928, EPI_ISL_3423929, EPI_ISL_3423930, EPI_ISL_3423931, EPI_ISL_3423932, EPI_ISL_3423933, EPI_ISL_3423934, EPI_ISL_3423935, EPI_ISL_3423936, EPI_ISL_3423937, EPI_ISL_3423938, EPI_ISL_3423939, EPI_ISL_3423940, EPI_ISL_3423941, EPI_ISL_3423942, EPI_ISL_3423943, EPI_ISL_3423944, EPI_ISL_3423945, EPI_ISL_3423946,
EPI_ISL_3423947, EPI_ISL_3423948, EPI_ISL_3423949, EPI_ISL_3423950, EPI_ISL_3423951, EPI_ISL_3423952, EPI_ISL_3423953, EPI_ISL_3423954, EPI_ISL_3423955, EPI_ISL_3423956, EPI_ISL_3423957, EPI_ISL_3423958, EPI_ISL_3423959, EPI_ISL_3423960, EPI_ISL_3423961, EPI_ISL_3423962, EPI_ISL_3423963, EPI_ISL_3423964, EPI_ISL_3423965, EPI_ISL_3423966, EPI_ISL_3423967, EPI_ISL_3423968,
EPI_ISL_3423969, EPI_ISL_3423970, EPI_ISL_3423971, EPI_ISL_3423972, EPI_ISL_3423973, EPI_ISL_3423974, EPI_ISL_3423975, EPI_ISL_3423976, EPI_ISL_3423977, EPI_ISL_3423978, EPI_ISL_3423979, EPI_ISL_3423980, EPI_ISL_3423981, EPI_ISL_3423982, EPI_ISL_3423983, EPI_ISL_3423984, EPI_ISL_3423985, EPI_ISL_3423986, EPI_ISL_3423987, EPI_ISL_3423988, EPI_ISL_3423989, EPI_ISL_3423990,
EPI_ISL_3423991, EPI_ISL_3423992, EPI_ISL_3423993, EPI_ISL_3423994, EPI_ISL_3423995, EPI_ISL_3423996, EPI_ISL_3423997, EPI_ISL_3423998, EPI_ISL_3423999, EPI_ISL_3424000, EPI_ISL_3424001, EPI_ISL_3424002, EPI_ISL_3424003, EPI_ISL_3424004, EPI_ISL_3424005, EPI_ISL_3424006, EPI_ISL_3424007, EPI_ISL_3424008, EPI_ISL_3424009, EPI_ISL_3424010, EPI_ISL_3424011, EPI_ISL_3424012,
EPI_ISL_3424013, EPI_ISL_3424014, EPI_ISL_3424015, EPI_ISL_3424016, EPI_ISL_3424017, EPI_ISL_3424018, EPI_ISL_3424019, EPI_ISL_3424020, EPI_ISL_3424021, EPI_ISL_3424022, EPI_ISL_3424023, EPI_ISL_3424024, EPI_ISL_3424025, EPI_ISL_3424026, EPI_ISL_3424027, EPI_ISL_3424028, EPI_ISL_3424029, EPI_ISL_3424030, EPI_ISL_3424031, EPI_ISL_3424032, EPI_ISL_3424033, EPI_ISL_3424034,
EPI_ISL_3424035, EPI_ISL_3424036, EPI_ISL_3424037, EPI_ISL_3424038, EPI_ISL_3424039, EPI_ISL_3424040, EPI_ISL_3424041, EPI_ISL_3424042, EPI_ISL_3424043, EPI_ISL_3424044, EPI_ISL_3424045, EPI_ISL_3424046, EPI_ISL_3424047, EPI_ISL_3424048, EPI_ISL_3424049, EPI_ISL_3424050, EPI_ISL_3424051, EPI_ISL_3424052, EPI_ISL_3424053, EPI_ISL_3424054, EPI_ISL_3424055, EPI_ISL_3424056,
EPI_ISL_3424057, EPI_ISL_3424058, EPI_ISL_3424059, EPI_ISL_3424060, EPI_ISL_3424061, EPI_ISL_3424062, EPI_ISL_3424063, EPI_ISL_3424064, EPI_ISL_3424065, EPI_ISL_3424066, EPI_ISL_3424067, EPI_ISL_3424068, EPI_ISL_3424069, EPI_ISL_3424070, EPI_ISL_3424071, EPI_ISL_3424072, EPI_ISL_3424073, EPI_ISL_3424074, EPI_ISL_3424075, EPI_ISL_3424076, EPI_ISL_3424077, EPI_ISL_3424078,
EPI_ISL_3424079, EPI_ISL_3424080, EPI_ISL_3424081, EPI_ISL_3424082, EPI_ISL_3424083, EPI_ISL_3424084, EPI_ISL_3424085, EPI_ISL_3424086, EPI_ISL_3424087, EPI_ISL_3424088, EPI_ISL_3424089, EPI_ISL_3424090, EPI_ISL_3424091, EPI_ISL_3424092, EPI_ISL_3424093, EPI_ISL_3424094, EPI_ISL_3424095, EPI_ISL_3424096, EPI_ISL_3424097, EPI_ISL_3424098, EPI_ISL_3424099, EPI_ISL_3424100,
EPI_ISL_3424101, EPI_ISL_3424102, EPI_ISL_3424103, EPI_ISL_3424104, EPI_ISL_3424105, EPI_ISL_3424106, EPI_ISL_3424107, EPI_ISL_3424108, EPI_ISL_3424109, EPI_ISL_3424110, EPI_ISL_3424111, EPI_ISL_3424112, EPI_ISL_3424113, EPI_ISL_3424114, EPI_ISL_3424115, EPI_ISL_3424116, EPI_ISL_3424117, EPI_ISL_3424118, EPI_ISL_3424119, EPI_ISL_3424120, EPI_ISL_3424121, EPI_ISL_3424122,
EPI_ISL_3424123, EPI_ISL_3424124, EPI_ISL_3424125, EPI_ISL_3424126, EPI_ISL_3424127, EPI_ISL_3424128, EPI_ISL_3424129, EPI_ISL_3424130, EPI_ISL_3424131, EPI_ISL_3424132, EPI_ISL_3424133, EPI_ISL_3424134, EPI_ISL_3424135, EPI_ISL_3424136, EPI_ISL_3424137, EPI_ISL_3424138, EPI_ISL_3424139, EPI_ISL_3424140, EPI_ISL_3424141, EPI_ISL_3424142, EPI_ISL_3424143, EPI_ISL_3424144,
EPI_ISL_3424145, EPI_ISL_3424146, EPI_ISL_3424147, EPI_ISL_3424148, EPI_ISL_3424149, EPI_ISL_3424150, EPI_ISL_3424151, EPI_ISL_3424152, EPI_ISL_3424153, EPI_ISL_3424154, EPI_ISL_3424155, EPI_ISL_3424156, EPI_ISL_3424157, EPI_ISL_3424158, EPI_ISL_3424159, EPI_ISL_3424160, EPI_ISL_3424161, EPI_ISL_3424162, EPI_ISL_3424163, EPI_ISL_3424164, EPI_ISL_3424165, EPI_ISL_3424166,
EPI_ISL_3424167, EPI_ISL_3424168, EPI_ISL_3424169, EPI_ISL_3424170, EPI_ISL_3424171, EPI_ISL_3424172, EPI_ISL_3424173, EPI_ISL_3424174, EPI_ISL_3424175, EPI_ISL_3424176, EPI_ISL_3424177, EPI_ISL_3424178, EPI_ISL_3424179, EPI_ISL_3424180, EPI_ISL_3424181, EPI_ISL_3424182, EPI_ISL_3424183, EPI_ISL_3424184, EPI_ISL_3424185, EPI_ISL_3424186, EPI_ISL_3424187, EPI_ISL_3424188,
EPI_ISL_3424189, EPI_ISL_3424190, EPI_ISL_3424191, EPI_ISL_3424192, EPI_ISL_3424193, EPI_ISL_3424194, EPI_ISL_3424195, EPI_ISL_3424196, EPI_ISL_3424197, EPI_ISL_3424198, EPI_ISL_3424199, EPI_ISL_3424200, EPI_ISL_3424201, EPI_ISL_3424202, EPI_ISL_3424203, EPI_ISL_3424204, EPI_ISL_3424205, EPI_ISL_3424206, EPI_ISL_3424207, EPI_ISL_3424208, EPI_ISL_3424209, EPI_ISL_3424210,
EPI_ISL_3424211, EPI_ISL_3424212, EPI_ISL_3424213, EPI_ISL_3424214, EPI_ISL_3424215, EPI_ISL_3424216, EPI_ISL_3424217, EPI_ISL_3424218, EPI_ISL_3424219, EPI_ISL_3424220, EPI_ISL_3424221, EPI_ISL_3424222, EPI_ISL_3424223, EPI_ISL_3424224, EPI_ISL_3424225, EPI_ISL_3424226, EPI_ISL_3424227, EPI_ISL_3424228, EPI_ISL_3424229, EPI_ISL_3424230, EPI_ISL_3424231, EPI_ISL_3424232,
EPI_ISL_3424233, EPI_ISL_3424234, EPI_ISL_3424235, EPI_ISL_3424236, EPI_ISL_3424237, EPI_ISL_3424238, EPI_ISL_3424239, EPI_ISL_3424240, EPI_ISL_3424241, EPI_ISL_3424242, EPI_ISL_3424243, EPI_ISL_3424244, EPI_ISL_3424245, EPI_ISL_3424246, EPI_ISL_3424247, EPI_ISL_3424248, EPI_ISL_3424249, EPI_ISL_3424250, EPI_ISL_3424251, EPI_ISL_3424252, EPI_ISL_3424253, EPI_ISL_3424254,
EPI_ISL_3424255, EPI_ISL_3424256, EPI_ISL_3424257, EPI_ISL_3424258, EPI_ISL_3424259, EPI_ISL_3424260, EPI_ISL_3424261, EPI_ISL_3424262, EPI_ISL_3424263, EPI_ISL_3424264, EPI_ISL_3424265, EPI_ISL_3424266, EPI_ISL_3424267, EPI_ISL_3424268, EPI_ISL_3424269, EPI_ISL_3424270, EPI_ISL_3424271, EPI_ISL_3424272, EPI_ISL_3424273, EPI_ISL_3424274, EPI_ISL_3424275, EPI_ISL_3424276,
EPI_ISL_3424277, EPI_ISL_3424278, EPI_ISL_3424279, EPI_ISL_3424280, EPI_ISL_3424281, EPI_ISL_3424282, EPI_ISL_3424283, EPI_ISL_3424284, EPI_ISL_3424285, EPI_ISL_3424286, EPI_ISL_3424287, EPI_ISL_3424288, EPI_ISL_3424289, EPI_ISL_3424290, EPI_ISL_3424291, EPI_ISL_3424292, EPI_ISL_3424293, EPI_ISL_3424294, EPI_ISL_3424295, EPI_ISL_3424296, EPI_ISL_3424297, EPI_ISL_3424298,
EPI_ISL_3424299, EPI_ISL_3424300, EPI_ISL_3424301, EPI_ISL_3424302, EPI_ISL_3424303, EPI_ISL_3424304, EPI_ISL_3424305, EPI_ISL_3424306, EPI_ISL_3424307, EPI_ISL_3424308, EPI_ISL_3424309, EPI_ISL_3424310, EPI_ISL_3424311, EPI_ISL_3424312, EPI_ISL_3424313, EPI_ISL_3424314, EPI_ISL_3424315, EPI_ISL_3424316, EPI_ISL_3424317, EPI_ISL_3424318, EPI_ISL_3424319, EPI_ISL_3424320,
EPI_ISL_3424321, EPI_ISL_3424322, EPI_ISL_3424323, EPI_ISL_3424324, EPI_ISL_3424325, EPI_ISL_3424326, EPI_ISL_3424327, EPI_ISL_3424328, EPI_ISL_3424329, EPI_ISL_3424330, EPI_ISL_3424331, EPI_ISL_3424332, EPI_ISL_3424333, EPI_ISL_3424334, EPI_ISL_3424335, EPI_ISL_3424336, EPI_ISL_3424337, EPI_ISL_3424338, EPI_ISL_3424339, EPI_ISL_3424340, EPI_ISL_3424341, EPI_ISL_3424342,
EPI_ISL_3424343, EPI_ISL_3424344, EPI_ISL_3424345, EPI_ISL_3424346, EPI_ISL_3424347, EPI_ISL_3424348, EPI_ISL_3424349, EPI_ISL_3424350, EPI_ISL_3424351, EPI_ISL_3424352, EPI_ISL_3424353, EPI_ISL_3424354, EPI_ISL_3424355, EPI_ISL_3424356, EPI_ISL_3424357, EPI_ISL_3424358, EPI_ISL_3424359, EPI_ISL_3424360, EPI_ISL_3424361, EPI_ISL_3424362, EPI_ISL_3424363, EPI_ISL_3424364,
EPI_ISL_3424365, EPI_ISL_3424366, EPI_ISL_3424367, EPI_ISL_3424368, EPI_ISL_3424369, EPI_ISL_3424370, EPI_ISL_3424371, EPI_ISL_3424372, EPI_ISL_3424373, EPI_ISL_3424374, EPI_ISL_3424375, EPI_ISL_3424376, EPI_ISL_3424377, EPI_ISL_3424378, EPI_ISL_3424379, EPI_ISL_3424380, EPI_ISL_3424381, EPI_ISL_3424382, EPI_ISL_3424383, EPI_ISL_3424384, EPI_ISL_3424385, EPI_ISL_3424386,
EPI_ISL_3424387, EPI_ISL_3424388, EPI_ISL_3424389, EPI_ISL_3424390, EPI_ISL_3424391, EPI_ISL_3424392, EPI_ISL_3424393, EPI_ISL_3424394, EPI_ISL_3424395, EPI_ISL_3424396, EPI_ISL_3424397, EPI_ISL_3424398, EPI_ISL_3424399, EPI_ISL_3424400, EPI_ISL_3424401, EPI_ISL_3424402, EPI_ISL_3424403, EPI_ISL_3424404, EPI_ISL_3424405, EPI_ISL_3424406, EPI_ISL_3424407, EPI_ISL_3424408,
EPI_ISL_3424409, EPI_ISL_3424410, EPI_ISL_3424411, EPI_ISL_3424412, EPI_ISL_3424413, EPI_ISL_3424414, EPI_ISL_3424415, EPI_ISL_3424416, EPI_ISL_3424417, EPI_ISL_3424418, EPI_ISL_3424419, EPI_ISL_3424420, EPI_ISL_3424421, EPI_ISL_3424422, EPI_ISL_3424423, EPI_ISL_3424424, EPI_ISL_3424425, EPI_ISL_3424426, EPI_ISL_3424427, EPI_ISL_3424428, EPI_ISL_3424429, EPI_ISL_3424430,
EPI_ISL_3424431, EPI_ISL_3424432, EPI_ISL_3424433, EPI_ISL_3424434, EPI_ISL_3424435, EPI_ISL_3424436, EPI_ISL_3424437, EPI_ISL_3424438, EPI_ISL_3424439, EPI_ISL_3424440, EPI_ISL_3424441, EPI_ISL_3424442, EPI_ISL_3424443, EPI_ISL_3424444, EPI_ISL_3424445, EPI_ISL_3424446, EPI_ISL_3424447, EPI_ISL_3424448, EPI_ISL_3424449, EPI_ISL_3424450, EPI_ISL_3424451, EPI_ISL_3424452,
EPI_ISL_3424453, EPI_ISL_3424454, EPI_ISL_3424455, EPI_ISL_3424456, EPI_ISL_3424457, EPI_ISL_3424458, EPI_ISL_3424459, EPI_ISL_3424460, EPI_ISL_3424461, EPI_ISL_3424462, EPI_ISL_3424463, EPI_ISL_3424464, EPI_ISL_3424465, EPI_ISL_3424466, EPI_ISL_3424467, EPI_ISL_3424468, EPI_ISL_3424469, EPI_ISL_3424470, EPI_ISL_3424471, EPI_ISL_3424472, EPI_ISL_3424473, EPI_ISL_3424474,
EPI_ISL_3424475, EPI_ISL_3424476, EPI_ISL_3424477, EPI_ISL_3424478, EPI_ISL_3424479, EPI_ISL_3424480, EPI_ISL_3424481, EPI_ISL_3424482, EPI_ISL_3424483, EPI_ISL_3424484, EPI_ISL_3424485, EPI_ISL_3424486, EPI_ISL_3424487, EPI_ISL_3424488, EPI_ISL_3424489, EPI_ISL_3424490, EPI_ISL_3424491, EPI_ISL_3424492, EPI_ISL_3424493, EPI_ISL_3424494, EPI_ISL_3424495, EPI_ISL_3424496,
EPI_ISL_3424497, EPI_ISL_3424498, EPI_ISL_3424499, EPI_ISL_3424500, EPI_ISL_3424501, EPI_ISL_3424502, EPI_ISL_3424503, EPI_ISL_3424504, EPI_ISL_3424505, EPI_ISL_3424506, EPI_ISL_3424507, EPI_ISL_3424508, EPI_ISL_3424509, EPI_ISL_3424510, EPI_ISL_3424511, EPI_ISL_3424512, EPI_ISL_3424513, EPI_ISL_3424514, EPI_ISL_3424515, EPI_ISL_3424516, EPI_ISL_3424517, EPI_ISL_3424518,
EPI_ISL_3424519, EPI_ISL_3424520, EPI_ISL_3424521, EPI_ISL_3424522, EPI_ISL_3424523, EPI_ISL_3424524, EPI_ISL_3424525, EPI_ISL_3424526, EPI_ISL_3424527, EPI_ISL_3424528, EPI_ISL_3424529, EPI_ISL_3424530, EPI_ISL_3424531, EPI_ISL_3424532, EPI_ISL_3424533, EPI_ISL_3424534, EPI_ISL_3424535, EPI_ISL_3424536, EPI_ISL_3424537, EPI_ISL_3424538, EPI_ISL_3424539, EPI_ISL_3424540,
EPI_ISL_3424541, EPI_ISL_3424542, EPI_ISL_3424543, EPI_ISL_3424544, EPI_ISL_3424545, EPI_ISL_3424546, EPI_ISL_3424547, EPI_ISL_3424548, EPI_ISL_3424549, EPI_ISL_3424550, EPI_ISL_3424551, EPI_ISL_3424552, EPI_ISL_3424553, EPI_ISL_3424554, EPI_ISL_3424555, EPI_ISL_3424556, EPI_ISL_3424557, EPI_ISL_3424558, EPI_ISL_3424559, EPI_ISL_3424560, EPI_ISL_3424561, EPI_ISL_3424562, EPI_ISL_3424563,
EPI_ISL_3424564, EPI_ISL_3424565, EPI_ISL_3424566, EPI_ISL_3424567, EPI_ISL_3424568, EPI_ISL_3424569, EPI_ISL_3424570, EPI_ISL_3424571, EPI_ISL_3424572, EPI_ISL_3424573, EPI_ISL_3424574, EPI_ISL_3424575, EPI_ISL_3424576, EPI_ISL_3424577, EPI_ISL_3424578, EPI_ISL_3424579, EPI_ISL_3424580, EPI_ISL_3424581, EPI_ISL_3424582, EPI_ISL_3424583, EPI_ISL_3424584, EPI_ISL_3424585,
EPI_ISL_3424586, EPI_ISL_3424587, EPI_ISL_3424588, EPI_ISL_3424589, EPI_ISL_3424590, EPI_ISL_3424591, EPI_ISL_3424592, EPI_ISL_3424593, EPI_ISL_3424594, EPI_ISL_3424595, EPI_ISL_3424596, EPI_ISL_3424597, EPI_ISL_3424598, EPI_ISL_3424599, EPI_ISL_3424600, EPI_ISL_3424601, EPI_ISL_3424602, EPI_ISL_3424603, EPI_ISL_3424604, EPI_ISL_3424605

| | | | |
|---|---|---|---|
| see above | Originating lab: Wales Specialist Virology Centre Sequencing lab: Pathogen Genomics Unit | Public Health Wales Microbiology Cardiff Wales Specialist Virology Centre | Catherine Moore, Johnathan Evans, Laura Gifford, Malorie Perry, Simon Cottrell, Angela Marchbank, Alec Birchley, Alexander Adams, Amy Gaskin, Bree Gatica-Wilcox, Jason Coombes, Joel Southgate, Lauren Gilbert, Lee Graham, Nicole Pacchiarini, Sara Kumziene-Summerhayes, Sarah Taylor, Sophie Jones, Sara Rey, Matthew Bull, Joanne Watkins, Sally Corden, Tom Connor |

EPI_ISL_3424606, EPI_ISL_3424607, EPI_ISL_3424608, EPI_ISL_3424609, EPI_ISL_3424610, EPI_ISL_3424611, EPI_ISL_3424612, EPI_ISL_3424613, EPI_ISL_3424614, EPI_ISL_3424615, EPI_ISL_3424616, EPI_ISL_3424617, EPI_ISL_3424618, EPI_ISL_3424619, EPI_ISL_3424620, EPI_ISL_3424621, EPI_ISL_3424622, EPI_ISL_3424623, EPI_ISL_3424624, EPI_ISL_3424625, EPI_ISL_3424626, EPI_ISL_3424627,
EPI_ISL_3424628, EPI_ISL_3424629, EPI_ISL_3424630, EPI_ISL_3424631, EPI_ISL_3424632, EPI_ISL_3424633, EPI_ISL_3424634, EPI_ISL_3424635, EPI_ISL_3424636, EPI_ISL_3424637, EPI_ISL_3424638, EPI_ISL_3424639, EPI_ISL_3424640, EPI_ISL_3424641, EPI_ISL_3424642, EPI_ISL_3424643, EPI_ISL_3424644, EPI_ISL_3424645, EPI_ISL_3424646, EPI_ISL_3424647, EPI_ISL_3424648, EPI_ISL_3424649,

| | | | |
|---|---|---|---|
| EPI_ISL_3424650, EPI_ISL_3424651, EPI_ISL_3424652, EPI_ISL_3424653, EPI_ISL_3424654, EPI_ISL_3424655, EPI_ISL_3424656, EPI_ISL_3424657, EPI_ISL_3424658, EPI_ISL_3424659, EPI_ISL_3424660, EPI_ISL_3424661, EPI_ISL_3424662, EPI_ISL_3424663, EPI_ISL_3424664, EPI_ISL_3424665, EPI_ISL_3424666, EPI_ISL_3424667, EPI_ISL_3424668, EPI_ISL_3424669, EPI_ISL_3424670, EPI_ISL_3424671, EPI_ISL_3424672, EPI_ISL_3424673, EPI_ISL_3424674, EPI_ISL_3424675, EPI_ISL_3424676, EPI_ISL_3424677, EPI_ISL_3424678, EPI_ISL_3424679, EPI_ISL_3424680, EPI_ISL_3424681, EPI_ISL_3424682, EPI_ISL_3424683, EPI_ISL_3424684, EPI_ISL_3424685, EPI_ISL_3424686, EPI_ISL_3424687, EPI_ISL_3424688, EPI_ISL_3424689, EPI_ISL_3424690, EPI_ISL_3424691, EPI_ISL_3424692, EPI_ISL_3424693, EPI_ISL_3424694, EPI_ISL_3424695, EPI_ISL_3424696, EPI_ISL_3424697, EPI_ISL_3424698, EPI_ISL_3424699, EPI_ISL_3424700, EPI_ISL_3424701, EPI_ISL_3424702, EPI_ISL_3424703, EPI_ISL_3424704, EPI_ISL_3424705, EPI_ISL_3424706, EPI_ISL_3424707, EPI_ISL_3424708, EPI_ISL_3424709, EPI_ISL_3424710, EPI_ISL_3424711, EPI_ISL_3424712, EPI_ISL_3424713, EPI_ISL_3424714, EPI_ISL_3424715, EPI_ISL_3424716, EPI_ISL_3424717, EPI_ISL_3424718, EPI_ISL_3424719, EPI_ISL_3424720, EPI_ISL_3424721, EPI_ISL_3424722, EPI_ISL_3424723, EPI_ISL_3424724, EPI_ISL_3424725, EPI_ISL_3424726, EPI_ISL_3424727, EPI_ISL_3424728, EPI_ISL_3424729, EPI_ISL_3424730, EPI_ISL_3424731, EPI_ISL_3424732, EPI_ISL_3424733, EPI_ISL_3424734, EPI_ISL_3424735, EPI_ISL_3424736, EPI_ISL_3424737, EPI_ISL_3424738, EPI_ISL_3424739, EPI_ISL_3424740, EPI_ISL_3424741, EPI_ISL_3424742, EPI_ISL_3424743, EPI_ISL_3424744, EPI_ISL_3424745, EPI_ISL_3424746, EPI_ISL_3424747, EPI_ISL_3424748, EPI_ISL_3424749, EPI_ISL_3424750, EPI_ISL_3424751, EPI_ISL_3424752, EPI_ISL_3424753, EPI_ISL_3424754, EPI_ISL_3424755, EPI_ISL_3424756, EPI_ISL_3424757, EPI_ISL_3424758, EPI_ISL_3424759, EPI_ISL_3424760, EPI_ISL_3424761, EPI_ISL_3424762, EPI_ISL_3424763, EPI_ISL_3424764, EPI_ISL_3424765, EPI_ISL_3424766, EPI_ISL_3424767, EPI_ISL_3424768, EPI_ISL_3424769, EPI_ISL_3424770, EPI_ISL_3424771, EPI_ISL_3424772, EPI_ISL_3424773, EPI_ISL_3424774, EPI_ISL_3424775, EPI_ISL_3424776, EPI_ISL_3424777, EPI_ISL_3424778, EPI_ISL_3424779, EPI_ISL_3424780, EPI_ISL_3424781, EPI_ISL_3424782, EPI_ISL_3424783, EPI_ISL_3424784, EPI_ISL_3424785, EPI_ISL_3424786, EPI_ISL_3424787, EPI_ISL_3424788, EPI_ISL_3424789, EPI_ISL_3424790, EPI_ISL_3424791, EPI_ISL_3424792, EPI_ISL_3424793, EPI_ISL_3424794, EPI_ISL_3424795, EPI_ISL_3424796, EPI_ISL_3424797, EPI_ISL_3424798, EPI_ISL_3424799, EPI_ISL_3424800, EPI_ISL_3424801, EPI_ISL_3424802, EPI_ISL_3424803, EPI_ISL_3424804, EPI_ISL_3424805, EPI_ISL_3424806, EPI_ISL_3424807, EPI_ISL_3424808, EPI_ISL_3424809, EPI_ISL_3424810, EPI_ISL_3424811, EPI_ISL_3424812, EPI_ISL_3424813, EPI_ISL_3424814, EPI_ISL_3424815, EPI_ISL_3424816, EPI_ISL_3424817, EPI_ISL_3424818, EPI_ISL_3424819, EPI_ISL_3424820, EPI_ISL_3424821, EPI_ISL_3424822, EPI_ISL_3424823, EPI_ISL_3424824, EPI_ISL_3424825, EPI_ISL_3424826, EPI_ISL_3424827, EPI_ISL_3424828, EPI_ISL_3424829, EPI_ISL_3424830, EPI_ISL_3424831, EPI_ISL_3424832, EPI_ISL_3424833, EPI_ISL_3424834, EPI_ISL_3424835, EPI_ISL_3424836, EPI_ISL_3424837, EPI_ISL_3424838, EPI_ISL_3424839, EPI_ISL_3424840, EPI_ISL_3424841, EPI_ISL_3424842, EPI_ISL_3424843, EPI_ISL_3424844, EPI_ISL_3424845, EPI_ISL_3424846, EPI_ISL_3424847, EPI_ISL_3424848, EPI_ISL_3424849, EPI_ISL_3424850, EPI_ISL_3424851, EPI_ISL_3424852, EPI_ISL_3424853, EPI_ISL_3424854, EPI_ISL_3424855, EPI_ISL_3424856, EPI_ISL_3424857, EPI_ISL_3424858, EPI_ISL_3424859, EPI_ISL_3424860, EPI_ISL_3424861, EPI_ISL_3424862, EPI_ISL_3424863, EPI_ISL_3424864, EPI_ISL_3424865, EPI_ISL_3424866, EPI_ISL_3424867, EPI_ISL_3424868, EPI_ISL_3424869, EPI_ISL_3424870, EPI_ISL_3424871, EPI_ISL_3424872, EPI_ISL_3424873, EPI_ISL_3424874, EPI_ISL_3424875, EPI_ISL_3424876, EPI_ISL_3424877, EPI_ISL_3424878, EPI_ISL_3424879, EPI_ISL_3424880, EPI_ISL_3424881, EPI_ISL_3424882, EPI_ISL_3424883, EPI_ISL_3424884, EPI_ISL_3424885, EPI_ISL_3424886, EPI_ISL_3424887, EPI_ISL_3424888, EPI_ISL_3424889, EPI_ISL_3424890, EPI_ISL_3424891, EPI_ISL_3424892, EPI_ISL_3424893, EPI_ISL_3424894, EPI_ISL_3424895, EPI_ISL_3424896, EPI_ISL_3424897, EPI_ISL_3424898, EPI_ISL_3424899, EPI_ISL_3424900, EPI_ISL_3424901, EPI_ISL_3424902, EPI_ISL_3424903, EPI_ISL_3424904, EPI_ISL_3424905, EPI_ISL_3424906, EPI_ISL_3424907, EPI_ISL_3424908, EPI_ISL_3424909, EPI_ISL_3424910, EPI_ISL_3424911, EPI_ISL_3424912, EPI_ISL_3424913, EPI_ISL_3424914, EPI_ISL_3424915, EPI_ISL_3424916, EPI_ISL_3424917, EPI_ISL_3424918, EPI_ISL_3424919, EPI_ISL_3424920, EPI_ISL_3424921, EPI_ISL_3424922, EPI_ISL_3424923, EPI_ISL_3424924, EPI_ISL_3424925, EPI_ISL_3424926, EPI_ISL_3424927, EPI_ISL_3424928, EPI_ISL_3424929, EPI_ISL_3424930, EPI_ISL_3424931, EPI_ISL_3424932, EPI_ISL_3424933, EPI_ISL_3424934, EPI_ISL_3424935, EPI_ISL_3424936, EPI_ISL_3424937, EPI_ISL_3424938, EPI_ISL_3424939, EPI_ISL_3424940, EPI_ISL_3424941, EPI_ISL_3424942, EPI_ISL_3424943, EPI_ISL_3424944, EPI_ISL_3424945, EPI_ISL_3424946, EPI_ISL_3424947, EPI_ISL_3424948, EPI_ISL_3424949, EPI_ISL_3424950, EPI_ISL_3424951, EPI_ISL_3424952, EPI_ISL_3424953, EPI_ISL_3424954, EPI_ISL_3424955, EPI_ISL_3424956, EPI_ISL_3424957, EPI_ISL_3424958, EPI_ISL_3424959, EPI_ISL_3424960, EPI_ISL_3424961, EPI_ISL_3424962, EPI_ISL_3424963, EPI_ISL_3424964, EPI_ISL_3424965, EPI_ISL_3424966, EPI_ISL_3424967, EPI_ISL_3424968, EPI_ISL_3424969, EPI_ISL_3424970, EPI_ISL_3424971, EPI_ISL_3424972, EPI_ISL_3424973, EPI_ISL_3424974, EPI_ISL_3424975, EPI_ISL_3424976, EPI_ISL_3424977, EPI_ISL_3424978, EPI_ISL_3424979, EPI_ISL_3424980, EPI_ISL_3424981, EPI_ISL_3424982, EPI_ISL_3424983, EPI_ISL_3424984, EPI_ISL_3424985, EPI_ISL_3424986, EPI_ISL_3424987, EPI_ISL_3424988, EPI_ISL_3424989, EPI_ISL_3424990, EPI_ISL_3424991, EPI_ISL_3424992, EPI_ISL_3424993, EPI_ISL_3424994, EPI_ISL_3424995, EPI_ISL_3424996, EPI_ISL_3424997, EPI_ISL_3424998, EPI_ISL_3424999, EPI_ISL_3425000, EPI_ISL_3425001, EPI_ISL_3425002, EPI_ISL_3425003, EPI_ISL_3425004, EPI_ISL_3425005, EPI_ISL_3425006, EPI_ISL_3425007, EPI_ISL_3425008, EPI_ISL_3425009, EPI_ISL_3425010, EPI_ISL_3425011, EPI_ISL_3425012, EPI_ISL_3425013, EPI_ISL_3425014, EPI_ISL_3425015, EPI_ISL_3425016, EPI_ISL_3425017, EPI_ISL_3425018, EPI_ISL_3425019, EPI_ISL_3425020, EPI_ISL_3425021, EPI_ISL_3425022, EPI_ISL_3425023, EPI_ISL_3425024, EPI_ISL_3425025, EPI_ISL_3425026, EPI_ISL_3425027, EPI_ISL_3425028, EPI_ISL_3425029, EPI_ISL_3425030, EPI_ISL_3425031, EPI_ISL_3425032, EPI_ISL_3425033, EPI_ISL_3425034, EPI_ISL_3425035, EPI_ISL_3425036, EPI_ISL_3425037, EPI_ISL_3425038, EPI_ISL_3425039, EPI_ISL_3425040, EPI_ISL_3425041, EPI_ISL_3425042, EPI_ISL_3425043, EPI_ISL_3425044, EPI_ISL_3425045, EPI_ISL_3425046, EPI_ISL_3425047, EPI_ISL_3425048, EPI_ISL_3425049, EPI_ISL_3425050, EPI_ISL_3425051, EPI_ISL_3425052, EPI_ISL_3425053, EPI_ISL_3425054, EPI_ISL_3425055, EPI_ISL_3425056, EPI_ISL_3425057, EPI_ISL_3425058, EPI_ISL_3425059, EPI_ISL_3425060, EPI_ISL_3425061, EPI_ISL_3425062, EPI_ISL_3425063, EPI_ISL_3425064, EPI_ISL_3425065, EPI_ISL_3425066, EPI_ISL_3425067, EPI_ISL_3425068, EPI_ISL_3425069, EPI_ISL_3425070, EPI_ISL_3425071, EPI_ISL_3425072, EPI_ISL_3425073, EPI_ISL_3425074, EPI_ISL_3425075, EPI_ISL_3425076, EPI_ISL_3425077, EPI_ISL_3425078, EPI_ISL_3425079, EPI_ISL_3425080, EPI_ISL_3425081, EPI_ISL_3425082, EPI_ISL_3425083, EPI_ISL_3425084, EPI_ISL_3425085, EPI_ISL_3425086, EPI_ISL_3425087, EPI_ISL_3425088, EPI_ISL_3425089, EPI_ISL_3425090, EPI_ISL_3425091, EPI_ISL_3425092, EPI_ISL_3425093, EPI_ISL_3425094, EPI_ISL_3425095, EPI_ISL_3425096, EPI_ISL_3425097, EPI_ISL_3425098, EPI_ISL_3425099, EPI_ISL_3425100, EPI_ISL_3425101, EPI_ISL_3425102 | | | |
| see above | Wales Specialist Virology Centre Sequencing lab: Pathogen Genomics Unit | Public Health Wales Microbiology Cardiff Wales Specialist Virology Centre | Catherine Moore, Johnathan Evans, Laura Gifford, Malorie Perry, Simon Cottrell, Angela Marchbank, Alec Birchley, Alexander Adams, Amy Gaskin, Bree Gatica-Wilcox, Jason Coombes, Joel Southgate, Lauren Gilbert, Lee Graham, Nicole Pacchiarini, Sara Kumziene-Summerhayes, Sarah Taylor, Sophie Jones, Sara Rey, Matthew Bull, Joanne Watkins, Sally Corden, Tom Connor |
| EPI_ISL_3425103, EPI_ISL_3425104, EPI_ISL_3425105, EPI_ISL_3425106, EPI_ISL_3425107, EPI_ISL_3425108, EPI_ISL_3425109, EPI_ISL_3425110, EPI_ISL_3425111, EPI_ISL_3425112, EPI_ISL_3425113, EPI_ISL_3425114, EPI_ISL_3425115, EPI_ISL_3425116, EPI_ISL_3425117, EPI_ISL_3425118, EPI_ISL_3425119, EPI_ISL_3425120, EPI_ISL_3425121, EPI_ISL_3425122, EPI_ISL_3425123, EPI_ISL_3425124, EPI_ISL_3425125, EPI_ISL_3425126, EPI_ISL_3425127, EPI_ISL_3425128, EPI_ISL_3425129, EPI_ISL_3425130, EPI_ISL_3425131, EPI_ISL_3425132, EPI_ISL_3425133, EPI_ISL_3425134, EPI_ISL_3425135, EPI_ISL_3425136, EPI_ISL_3425137, EPI_ISL_3425138, EPI_ISL_3425139, EPI_ISL_3425140, EPI_ISL_3425141, EPI_ISL_3425142, EPI_ISL_3425143, EPI_ISL_3425144, EPI_ISL_3425145, EPI_ISL_3425146, EPI_ISL_3425147, EPI_ISL_3425148, EPI_ISL_3425149, EPI_ISL_3425150, EPI_ISL_3425151, EPI_ISL_3425152, EPI_ISL_3425153, EPI_ISL_3425154, EPI_ISL_3425155, EPI_ISL_3425156, EPI_ISL_3425157, EPI_ISL_3425158, EPI_ISL_3425159, EPI_ISL_3425160, EPI_ISL_3425161, EPI_ISL_3425162, EPI_ISL_3425163, EPI_ISL_3425164, EPI_ISL_3425165, EPI_ISL_3425166, EPI_ISL_3425167, EPI_ISL_3425168, EPI_ISL_3425169, EPI_ISL_3425170, EPI_ISL_3425171, EPI_ISL_3425172, EPI_ISL_3425173, EPI_ISL_3425174, EPI_ISL_3425175, EPI_ISL_3425176, EPI_ISL_3425177, EPI_ISL_3425178, EPI_ISL_3425179, EPI_ISL_3425180, EPI_ISL_3425181, EPI_ISL_3425182, EPI_ISL_3425183, EPI_ISL_3425184, EPI_ISL_3425185, EPI_ISL_3425186, EPI_ISL_3425187, EPI_ISL_3425188, EPI_ISL_3425189, EPI_ISL_3425190, EPI_ISL_3425191, EPI_ISL_3425192, EPI_ISL_3425193, EPI_ISL_3425194, EPI_ISL_3425195, EPI_ISL_3425196, EPI_ISL_3425197 | | | |
| see above | Originating lab: Wales Specialist Virology Centre Sequencing lab: Pathogen Genomics Unit | Public Health Wales Microbiology Cardiff Wales Specialist Virology Centre | Catherine Moore, Johnathan Evans, Laura Gifford, Malorie Perry, Simon Cottrell, Angela Marchbank, Alec Birchley, Alexander Adams, Amy Gaskin, Bree Gatica-Wilcox, Jason Coombes, Joel Southgate, Lauren Gilbert, Lee Graham, Nicole Pacchiarini, Sara Kumziene-Summerhayes, Sarah Taylor, Sophie Jones, Sara Rey, Matthew Bull, Joanne Watkins, Sally Corden, Tom Connor |
| EPI_ISL_3425198, EPI_ISL_3425199, EPI_ISL_3425200, EPI_ISL_3425201, EPI_ISL_3425202, EPI_ISL_3425203, EPI_ISL_3425204, EPI_ISL_3425205, EPI_ISL_3425206, EPI_ISL_3425207, EPI_ISL_3425208, EPI_ISL_3425209, EPI_ISL_3425210, EPI_ISL_3425211, EPI_ISL_3425212, EPI_ISL_3425213, EPI_ISL_3425214, EPI_ISL_3425215, EPI_ISL_3425216, EPI_ISL_3425217, EPI_ISL_3425218, EPI_ISL_3425219, EPI_ISL_3425220, EPI_ISL_3425221, EPI_ISL_3425222, EPI_ISL_3425223, EPI_ISL_3425224, EPI_ISL_3425225, EPI_ISL_3425226, EPI_ISL_3425227, EPI_ISL_3425228, EPI_ISL_3425229, EPI_ISL_3425230, EPI_ISL_3425231, EPI_ISL_3425232, EPI_ISL_3425233, EPI_ISL_3425234, EPI_ISL_3425235, EPI_ISL_3425236, EPI_ISL_3425237, EPI_ISL_3425238, EPI_ISL_3425239, EPI_ISL_3425240, EPI_ISL_3425241, EPI_ISL_3425242, EPI_ISL_3425243, EPI_ISL_3425244, EPI_ISL_3425245, EPI_ISL_3425246, EPI_ISL_3425247, EPI_ISL_3425248, EPI_ISL_3425249, EPI_ISL_3425250, EPI_ISL_3425251, EPI_ISL_3425252, EPI_ISL_3425253, EPI_ISL_3425254, EPI_ISL_3425255, EPI_ISL_3425256, EPI_ISL_3425257, EPI_ISL_3425258, EPI_ISL_3425259, EPI_ISL_3425260, EPI_ISL_3425261, EPI_ISL_3425262, EPI_ISL_3425263, EPI_ISL_3425264, EPI_ISL_3425265, EPI_ISL_3425266, EPI_ISL_3425267, EPI_ISL_3425268, EPI_ISL_3425269, EPI_ISL_3425270, EPI_ISL_3425271, EPI_ISL_3425272, EPI_ISL_3425273 | | | |
| see above | Centre for Enzyme Innovation, University of Portsmouth / Translational Research Laboratory, Portsmouth Hospitals NHS Trust / Hampshire Hospitals NHS Foundation Trust / Bournemouth University / University Hospitals Dorset NHS Foundation Trust / Brighton and Sussex University Hospitals NHS Trust / Dartford and Gravesham NHS Trust / Isle of Wight NHS Trust / Maidstone and Tunbridge Wells NHS Trust / University Hospital Southampton NHS Foundation Trust | COVID-19 Genomics UK (COG-UK) Consortium | Samuel Robson, Angela Beckett, Salman Goudarzi, Christopher Fearn, Kate Cook, Katie Loveson, Sharon Glaysher, Scott Elliott, Kelly Bicknell, Sarah Wyllie, Allyson Lloyd, Robert Impey, Anoop Chauhan, Stephen Kidd, Nathan Moore, Nick Cortes, Claire Thomas, Anna Mantzaouratou, Sarah Buchan, Magdalena Barrow, Andrew Butt, Liz Sheridan, Jonnie Seymour, Dorian Crudgington, Ben Macklin, Mohammed Hassan-Ibrahim, Cassandra Malone, Benjamin Cogger, Kevin Tucker, Samirakhon Raupova, Rachael Jeremiah, Anibolina Castigador, Emily Macnaughton, Karen Withell, Kordo Saeed, Jacqui Prieto, Adhyana Mahanama, Buddhini Samaraweera, Siona Silvieri, Emanuela Pelosi, Eleri Wilson-Davies, Sarah Jeremiah, Helen Wheeler, Matthew Harvey, Thea Sass, Helen Umpleby, Stephen Aplin |
| EPI_ISL_3425282, EPI_ISL_3425283, EPI_ISL_3425284, EPI_ISL_3425285, EPI_ISL_3425286, EPI_ISL_3425287, EPI_ISL_3425290, EPI_ISL_3425291, EPI_ISL_3425292, EPI_ISL_3425293, EPI_ISL_3425295 | | | |
| see above | Department of Microbiology, Yamanashi Institute for Public Health and Environment | Genome Analysis Center, Yamanashi Central Hospital | Yosuke Hirotsu |
| EPI_ISL_3425296, EPI_ISL_3425297, EPI_ISL_3425298, EPI_ISL_3425299, EPI_ISL_3425300, EPI_ISL_3425301, EPI_ISL_3425302, EPI_ISL_3425303, EPI_ISL_3425304, EPI_ISL_3425305, EPI_ISL_3425306, EPI_ISL_3425307, EPI_ISL_3425308, EPI_ISL_3425309, EPI_ISL_3425311 | | | |
| see above | Genome Analysis Center, Yamanashi Central Hospital | Genome Analysis Center, Yamanashi Central Hospital | Yosuke Hirotsu |
| EPI_ISL_3425312 | Kuala Lumpur International Airport (KLIA) Health Office | Institute for Medical Research, Infectious Disease Research Centre, National Institutes of Health, Ministry of Health Malaysia | Suppiah J, Kamel K, Mohd Zawawi Z, Azizan MA, Ramly N, Robert F, Anasir MI, Thayan R |
| EPI_ISL_3425313, EPI_ISL_3425314, EPI_ISL_3425315 | Arlon | Plateforme de testing Namuroise | Lesly Nyinkeu Kemamen; Nicolas Gilliard; Demars Aurore ; Maschietto Céline ; Nobis Chloé ; Otto Gaetan ; Denis Olivier ; Degosserie Jonathan ; Mullier François |
| EPI_ISL_3425316, EPI_ISL_3425317 | Plateforme de testing Namuroise | Plateforme de testing Namuroise | Lesly Nyinkeu Kemamen; Nicolas Gilliard; Demars Aurore ; Maschietto Céline ; Nobis Chloé ; Otto Gaetan ; Denis Olivier ; Degosserie Jonathan ; Mullier François |
| EPI_ISL_3425318, EPI_ISL_3425319 | Arlon | Plateforme de testing Namuroise | Lesly Nyinkeu Kemamen; Nicolas Gilliard; Demars Aurore ; Maschietto Céline ; Nobis Chloé ; Otto Gaetan ; Denis Olivier ; Degosserie Jonathan ; Mullier François |
| EPI_ISL_3425320 | Plateforme de testing Namuroise | Plateforme de testing Namuroise | Lesly Nyinkeu Kemamen; Nicolas Gilliard; Demars Aurore ; Maschietto Céline ; Nobis Chloé ; Otto Gaetan ; Denis Olivier ; Degosserie Jonathan ; Mullier François |
| EPI_ISL_3425321 | Marche en Famenne | Plateforme de testing Namuroise | Lesly Nyinkeu Kemamen; Nicolas Gilliard; Demars Aurore ; Maschietto Céline ; Nobis Chloé ; Otto Gaetan ; Denis Olivier ; Degosserie Jonathan ; Mullier François |
| EPI_ISL_3425322, EPI_ISL_3425323 | Arlon | Plateforme de testing Namuroise | Lesly Nyinkeu Kemamen; Nicolas Gilliard; Demars Aurore ; Maschietto Céline ; Nobis Chloé ; Otto Gaetan ; Denis Olivier ; Degosserie Jonathan ; Mullier François |
| EPI_ISL_3425324 | Marche en Famenne | Plateforme de testing Namuroise | Lesly Nyinkeu Kemamen; Nicolas Gilliard; Demars Aurore ; Maschietto Céline ; Nobis Chloé ; Otto Gaetan ; Denis Olivier ; Degosserie Jonathan ; Mullier François |
| EPI_ISL_3425325 | Arlon | Plateforme de testing Namuroise | Lesly Nyinkeu Kemamen; Nicolas Gilliard; Demars Aurore ; Maschietto Céline ; Nobis Chloé ; Otto Gaetan ; Denis Olivier ; Degosserie Jonathan ; Mullier François |
| EPI_ISL_3425326, EPI_ISL_3425327 | Plateforme de testing Namuroise | Plateforme de testing Namuroise | Lesly Nyinkeu Kemamen; Nicolas Gilliard; Demars Aurore ; Maschietto Céline ; Nobis Chloé ; Otto Gaetan ; Denis Olivier ; Degosserie Jonathan ; Mullier François |
| EPI_ISL_3425328 | Marche en Famenne | Plateforme de testing Namuroise | Lesly Nyinkeu Kemamen; Nicolas Gilliard; Demars Aurore ; Maschietto Céline ; Nobis Chloé ; Otto Gaetan ; Denis Olivier ; Degosserie Jonathan ; Mullier François |
| EPI_ISL_3425329 | Arlon | Plateforme de testing Namuroise | Lesly Nyinkeu Kemamen; Nicolas Gilliard; Demars Aurore ; Maschietto Céline ; Nobis Chloé ; Otto Gaetan ; Denis Olivier ; Degosserie Jonathan ; Mullier François |
| EPI_ISL_3425331, EPI_ISL_3425332 | Plateforme de testing Namuroise | Plateforme de testing Namuroise | Lesly Nyinkeu Kemamen; Nicolas Gilliard; Demars Aurore ; Maschietto Céline ; Nobis Chloé ; Otto Gaetan ; Denis Olivier ; Degosserie Jonathan ; Mullier François |
| EPI_ISL_3425401, EPI_ISL_3425402, EPI_ISL_3425403, EPI_ISL_3425404, EPI_ISL_3425405, EPI_ISL_3425407, EPI_ISL_3425408, EPI_ISL_3425409, EPI_ISL_3425410, EPI_ISL_3425411, EPI_ISL_3425412, EPI_ISL_3425413, EPI_ISL_3425414, EPI_ISL_3425415, EPI_ISL_3425416, EPI_ISL_3425417, EPI_ISL_3425418, EPI_ISL_3425419, EPI_ISL_3425420, EPI_ISL_3425421, EPI_ISL_3425422, EPI_ISL_3425423, EPI_ISL_3425424, EPI_ISL_3425425, EPI_ISL_3425426, EPI_ISL_3425427, EPI_ISL_3425428, EPI_ISL_3425429, EPI_ISL_3425430, EPI_ISL_3425431, EPI_ISL_3425432, EPI_ISL_3425433, EPI_ISL_3425434, EPI_ISL_3425435, EPI_ISL_3425436, EPI_ISL_3425437, EPI_ISL_3425438, EPI_ISL_3425439, EPI_ISL_3425440, EPI_ISL_3425441, EPI_ISL_3425442, EPI_ISL_3425443, EPI_ISL_3425444, EPI_ISL_3425445, EPI_ISL_3425446, EPI_ISL_3425447, EPI_ISL_3425448, EPI_ISL_3425449, EPI_ISL_3425450, EPI_ISL_3425451, EPI_ISL_3425452, EPI_ISL_3425453, EPI_ISL_3425454, EPI_ISL_3425455, EPI_ISL_3425456, EPI_ISL_3425457, EPI_ISL_3425459, EPI_ISL_3425460, EPI_ISL_3425461 | | | |
| see above | Maine Health and Environmental Testing Laboratory | Tewhey Lab, The Jackson Laboratory | Matluk,N., Dewey,H., Iosue,F., Barter,M., Lynch,R., Munger,H. and Tewhey,R. |
| EPI_ISL_3425462 | Kuala Lumpur International Airport (KLIA) Health Office | Institute for Medical Research, Infectious Disease Research Centre, National Institutes of Health, Ministry of Health Malaysia | Suppiah J, Kamel K, Mohd Zawawi Z, Azizan MA, Ramly N, Robert F, Anasir MI, Thayan R |
| EPI_ISL_3425463, EPI_ISL_3425464, EPI_ISL_3425466, EPI_ISL_3425467, EPI_ISL_3425468, EPI_ISL_3425469, EPI_ISL_3425470, EPI_ISL_3425471, EPI_ISL_3425472, EPI_ISL_3425473, EPI_ISL_3425475, EPI_ISL_3425478, EPI_ISL_3425479, EPI_ISL_3425480, EPI_ISL_3425481, EPI_ISL_3425482, EPI_ISL_3425484, EPI_ISL_3425485, EPI_ISL_3425488, EPI_ISL_3425489, EPI_ISL_3425490, EPI_ISL_3425491, EPI_ISL_3425492, EPI_ISL_3425493, EPI_ISL_3425494, EPI_ISL_3425495, EPI_ISL_3425496, EPI_ISL_3425497, EPI_ISL_3425498, EPI_ISL_3425499, EPI_ISL_3425500, EPI_ISL_3425501, EPI_ISL_3425502, EPI_ISL_3425503, EPI_ISL_3425504, EPI_ISL_3425505, EPI_ISL_3425507, EPI_ISL_3425508, EPI_ISL_3425509, EPI_ISL_3425510, EPI_ISL_3425512, EPI_ISL_3425513, EPI_ISL_3425514, EPI_ISL_3425515, EPI_ISL_3425516, EPI_ISL_3425517, EPI_ISL_3425518, EPI_ISL_3425519, EPI_ISL_3425520, EPI_ISL_3425521, EPI_ISL_3425522, EPI_ISL_3425523, EPI_ISL_3425524, EPI_ISL_3425525, EPI_ISL_3425526, EPI_ISL_3425527, EPI_ISL_3425528, EPI_ISL_3425529, EPI_ISL_3425530, EPI_ISL_3425531, EPI_ISL_3425532, EPI_ISL_3425533, EPI_ISL_3425534, EPI_ISL_3425535, EPI_ISL_3425536, EPI_ISL_3425537, EPI_ISL_3425538, EPI_ISL_3425539, EPI_ISL_3425540, EPI_ISL_3425541, EPI_ISL_3425542, EPI_ISL_3425543, EPI_ISL_3425544, EPI_ISL_3425545, EPI_ISL_3425546, EPI_ISL_3425547, EPI_ISL_3425548, EPI_ISL_3425549, EPI_ISL_3425550, EPI_ISL_3425551, EPI_ISL_3425552, EPI_ISL_3425553 | | | |
| see above | Maine Health and Environmental Testing Laboratory | Tewhey Lab, The Jackson Laboratory | Matluk,N., Dewey,H., Iosue,F., Barter,M., Lynch,R., Munger,H. and Tewhey,R. |
| EPI_ISL_3425554, EPI_ISL_3425555, EPI_ISL_3425556, EPI_ISL_3425557, EPI_ISL_3425558, EPI_ISL_3425559, EPI_ISL_3425560, EPI_ISL_3425561, EPI_ISL_3425562, EPI_ISL_3425563, EPI_ISL_3425564, EPI_ISL_3425565, EPI_ISL_3425566, EPI_ISL_3425567, EPI_ISL_3425568, EPI_ISL_3425569, EPI_ISL_3425570, EPI_ISL_3425571, EPI_ISL_3425572, EPI_ISL_3425573, EPI_ISL_3425574, EPI_ISL_3425575, EPI_ISL_3425576, EPI_ISL_3425577, EPI_ISL_3425578, EPI_ISL_3425579, EPI_ISL_3425580, EPI_ISL_3425581, EPI_ISL_3425582, EPI_ISL_3425583, EPI_ISL_3425584, EPI_ISL_3425585, EPI_ISL_3425586, EPI_ISL_3425587, EPI_ISL_3425588, EPI_ISL_3425589, EPI_ISL_3425590, EPI_ISL_3425591, EPI_ISL_3425592, EPI_ISL_3425593, EPI_ISL_3425594, EPI_ISL_3425595, EPI_ISL_3425596, EPI_ISL_3425597, EPI_ISL_3425598, EPI_ISL_3425599, EPI_ISL_3425601, EPI_ISL_3425602, EPI_ISL_3425603, EPI_ISL_3425604, EPI_ISL_3425605, EPI_ISL_3425606, EPI_ISL_3425607, EPI_ISL_3425608, EPI_ISL_3425609, EPI_ISL_3425610, EPI_ISL_3425611, EPI_ISL_3425612, EPI_ISL_3425613, EPI_ISL_3425614, EPI_ISL_3425615, EPI_ISL_3425616, EPI_ISL_3425617, EPI_ISL_3425618, EPI_ISL_3425619, EPI_ISL_3425620, EPI_ISL_3425621, EPI_ISL_3425623, EPI_ISL_3425624, EPI_ISL_3425625, EPI_ISL_3425626, EPI_ISL_3425627, EPI_ISL_3425628, EPI_ISL_3425629, EPI_ISL_3425630, EPI_ISL_3425631, EPI_ISL_3425632, EPI_ISL_3425633, EPI_ISL_3425634, EPI_ISL_3425635, EPI_ISL_3425636 | | | |
| see above | Michigan Department of Health and Human Services, Bureau of Laboratories | Michigan Department of Health and Human Services, Bureau of Laboratories | Blankenship HM, Riner D, Soehnlen MK |
| EPI_ISL_3425637 | Hospital De Vichy | CHU Clermont-Ferrand, service de virologie | Bisseux Maxime, Mirand Audrey, Combes Patricia, Henquell Cécile |
| EPI_ISL_3425638 | Michigan Department of Health and Human Services, Bureau of Laboratories | Michigan Department of Health and Human Services, Bureau of Laboratories | Blankenship HM, Riner D, Soehnlen MK |
| EPI_ISL_3425639 | Hospital De Vichy | CHU Clermont-Ferrand, service de virologie | Bisseux Maxime, Mirand Audrey, Combes Patricia, Henquell Cécile |
| EPI_ISL_3425640 | Michigan Department of Health and Human Services, Bureau of Laboratories | Michigan Department of Health and Human Services, Bureau of Laboratories | Blankenship HM, Riner D, Soehnlen MK |
| EPI_ISL_3425641 | Hospital De Vichy | CHU Clermont-Ferrand, service de virologie | Bisseux Maxime, Mirand Audrey, Combes Patricia, Henquell Cécile |
| EPI_ISL_3425642 | Michigan Department of Health and Human Services, Bureau of Laboratories | Michigan Department of Health and Human Services, Bureau of Laboratories | Blankenship HM, Riner D, Soehnlen MK |

| | | | |
|---|---|---|---|
| EPI_ISL_3425643 | Centre hospitalier D'aurillac | CHU Clermont-Ferrand, service de virologie | Bisseux Maxime, Mirand Audrey, Combes Patricia, Henquell Cécile |
| EPI_ISL_3425644 | Hospital De Vichy | CHU Clermont-Ferrand, service de virologie | Bisseux Maxime, Mirand Audrey, Combes Patricia, Henquell Cécile |
| EPI_ISL_3425645 | Michigan Department of Health and Human Services, Bureau of Laboratories | Michigan Department of Health and Human Services, Bureau of Laboratories | Blankenship HM, Riner D, Soehnlen MK |
| EPI_ISL_3425646 | Hospital De Vichy | CHU Clermont-Ferrand, service de virologie | Bisseux Maxime, Mirand Audrey, Combes Patricia, Henquell Cécile |
| EPI_ISL_3425647 | Michigan Department of Health and Human Services, Bureau of Laboratories | Michigan Department of Health and Human Services, Bureau of Laboratories | Blankenship HM, Riner D, Soehnlen MK |
| EPI_ISL_3425648 | Centre hospitalier D'aurillac | CHU Clermont-Ferrand, service de virologie | Bisseux Maxime, Mirand Audrey, Combes Patricia, Henquell Cécile |
| EPI_ISL_3425649 | Michigan Department of Health and Human Services, Bureau of Laboratories | Michigan Department of Health and Human Services, Bureau of Laboratories | Blankenship HM, Riner D, Soehnlen MK |
| EPI_ISL_3425650 | Hospital De Vichy | CHU Clermont-Ferrand, service de virologie | Bisseux Maxime, Mirand Audrey, Combes Patricia, Henquell Cécile |
| EPI_ISL_3425651 | Michigan Department of Health and Human Services, Bureau of Laboratories | Michigan Department of Health and Human Services, Bureau of Laboratories | Blankenship HM, Riner D, Soehnlen MK |
| EPI_ISL_3425652, EPI_ISL_3425653 | Centre Hospitalier Universitaire Clermont-Ferrand | CHU Clermont-Ferrand, service de virologie | Bisseux Maxime, Mirand Audrey, Combes Patricia, Henquell Cécile |
| EPI_ISL_3425654 | Michigan Department of Health and Human Services, Bureau of Laboratories | Michigan Department of Health and Human Services, Bureau of Laboratories | Blankenship HM, Riner D, Soehnlen MK |
| EPI_ISL_3425655, EPI_ISL_3425657, EPI_ISL_3425658 | Centre Hospitalier Universitaire Clermont-Ferrand | CHU Clermont-Ferrand, service de virologie | Bisseux Maxime, Mirand Audrey, Combes Patricia, Henquell Cécile |
| EPI_ISL_3425659, EPI_ISL_3425660, EPI_ISL_3425661 | Michigan Department of Health and Human Services, Bureau of Laboratories | Michigan Department of Health and Human Services, Bureau of Laboratories | Blankenship HM, Riner D, Soehnlen MK |
| EPI_ISL_3425662 | Centre Hospitalier Universitaire Clermont-Ferrand | CHU Clermont-Ferrand, service de virologie | Bisseux Maxime, Mirand Audrey, Combes Patricia, Henquell Cécile |
| EPI_ISL_3425663 | Michigan Department of Health and Human Services, Bureau of Laboratories | Michigan Department of Health and Human Services, Bureau of Laboratories | Blankenship HM, Riner D, Soehnlen MK |
| EPI_ISL_3425664 | Centre Hospitalier Universitaire Clermont-Ferrand | CHU Clermont-Ferrand, service de virologie | Bisseux Maxime, Mirand Audrey, Combes Patricia, Henquell Cécile |
| EPI_ISL_3425665 | Michigan Department of Health and Human Services, Bureau of Laboratories | Michigan Department of Health and Human Services, Bureau of Laboratories | Blankenship HM, Riner D, Soehnlen MK |
| EPI_ISL_3425666 | Centre Hospitalier Universitaire Clermont-Ferrand | CHU Clermont-Ferrand, service de virologie | Bisseux Maxime, Mirand Audrey, Combes Patricia, Henquell Cécile |
| EPI_ISL_3425667 | Michigan Department of Health and Human Services, Bureau of Laboratories | Michigan Department of Health and Human Services, Bureau of Laboratories | Blankenship HM, Riner D, Soehnlen MK |
| EPI_ISL_3425668, EPI_ISL_3425669 | Centre Hospitalier Universitaire Clermont-Ferrand | CHU Clermont-Ferrand, service de virologie | Bisseux Maxime, Mirand Audrey, Combes Patricia, Henquell Cécile |
| EPI_ISL_3425670 | Michigan Department of Health and Human Services, Bureau of Laboratories | Michigan Department of Health and Human Services, Bureau of Laboratories | Blankenship HM, Riner D, Soehnlen MK |
| EPI_ISL_3425671 | Centre Hospitalier Universitaire Clermont-Ferrand | CHU Clermont-Ferrand, service de virologie | Bisseux Maxime, Mirand Audrey, Combes Patricia, Henquell Cécile |
| EPI_ISL_3425672 | Michigan Department of Health and Human Services, Bureau of Laboratories | Michigan Department of Health and Human Services, Bureau of Laboratories | Blankenship HM, Riner D, Soehnlen MK |
| EPI_ISL_3425673 | Hospital Du Mont Dore | CHU Clermont-Ferrand, service de virologie | Bisseux Maxime, Mirand Audrey, Combes Patricia, Henquell Cécile |
| EPI_ISL_3425674 | Michigan Department of Health and Human Services, Bureau of Laboratories | Michigan Department of Health and Human Services, Bureau of Laboratories | Blankenship HM, Riner D, Soehnlen MK |
| EPI_ISL_3425675 | Centre Hospitalier Universitaire Clermont-Ferrand | CHU Clermont-Ferrand, service de virologie | Bisseux Maxime, Mirand Audrey, Combes Patricia, Henquell Cécile |
| EPI_ISL_3425676 | Michigan Department of Health and Human Services, Bureau of Laboratories | Michigan Department of Health and Human Services, Bureau of Laboratories | Blankenship HM, Riner D, Soehnlen MK |
| EPI_ISL_3425677 | Centre Hospitalier Universitaire Clermont-Ferrand | CHU Clermont-Ferrand, service de virologie | Bisseux Maxime, Mirand Audrey, Combes Patricia, Henquell Cécile |
| EPI_ISL_3425678 | Hospital Du Mont Dore | CHU Clermont-Ferrand, service de virologie | Bisseux Maxime, Mirand Audrey, Combes Patricia, Henquell Cécile |
| EPI_ISL_3425679 | Michigan Department of Health and Human Services, Bureau of Laboratories | Michigan Department of Health and Human Services, Bureau of Laboratories | Blankenship HM, Riner D, Soehnlen MK |
| EPI_ISL_3425680 | Hospital Du Mont Dore | CHU Clermont-Ferrand, service de virologie | Bisseux Maxime, Mirand Audrey, Combes Patricia, Henquell Cécile |
| EPI_ISL_3425681 | Michigan Department of Health and Human Services, Bureau of Laboratories | Michigan Department of Health and Human Services, Bureau of Laboratories | Blankenship HM, Riner D, Soehnlen MK |
| EPI_ISL_3425682 | Hospital Du Mont Dore | CHU Clermont-Ferrand, service de virologie | Bisseux Maxime, Mirand Audrey, Combes Patricia, Henquell Cécile |
| EPI_ISL_3425683 | Michigan Department of Health and Human Services, Bureau of Laboratories | Michigan Department of Health and Human Services, Bureau of Laboratories | Blankenship HM, Riner D, Soehnlen MK |
| EPI_ISL_3425684 | Centre Hospitalier Universitaire Clermont-Ferrand | CHU Clermont-Ferrand, service de virologie | Bisseux Maxime, Mirand Audrey, Combes Patricia, Henquell Cécile |
| EPI_ISL_3425685 | Hospital Du Mont Dore | CHU Clermont-Ferrand, service de virologie | Bisseux Maxime, Mirand Audrey, Combes Patricia, Henquell Cécile |
| EPI_ISL_3425686 | Michigan Department of Health and Human Services, Bureau of Laboratories | Michigan Department of Health and Human Services, Bureau of Laboratories | Blankenship HM, Riner D, Soehnlen MK |
| EPI_ISL_3425687, EPI_ISL_3425688, EPI_ISL_3425689 | Hospital Du Mont Dore | CHU Clermont-Ferrand, service de virologie | Bisseux Maxime, Mirand Audrey, Combes Patricia, Henquell Cécile |
| EPI_ISL_3425690 | Centre Hospitalier Universitaire Clermont-Ferrand | CHU Clermont-Ferrand, service de virologie | Bisseux Maxime, Mirand Audrey, Combes Patricia, Henquell Cécile |
| EPI_ISL_3425691, EPI_ISL_3425692 | Centre De Prelevement COVID RIOM | CHU Clermont-Ferrand, service de virologie | Bisseux Maxime, Mirand Audrey, Combes Patricia, Henquell Cécile |
| EPI_ISL_3425693 | Centre Hospitalier Universitaire Clermont-Ferrand | CHU Clermont-Ferrand, service de virologie | Bisseux Maxime, Mirand Audrey, Combes Patricia, Henquell Cécile |
| EPI_ISL_3425694 | Hospital De Vichy | CHU Clermont-Ferrand, service de virologie | Bisseux Maxime, Mirand Audrey, Combes Patricia, Henquell Cécile |
| EPI_ISL_3425695 | Centre Hospitalier Universitaire Clermont-Ferrand | CHU Clermont-Ferrand, service de virologie | Bisseux Maxime, Mirand Audrey, Combes Patricia, Henquell Cécile |
| EPI_ISL_3425696 | Centre De Prelevement COVID RIOM | CHU Clermont-Ferrand, service de virologie | Bisseux Maxime, Mirand Audrey, Combes Patricia, Henquell Cécile |
| EPI_ISL_3425697 | Hospital Du Mont Dore | CHU Clermont-Ferrand, service de virologie | Bisseux Maxime, Mirand Audrey, Combes Patricia, Henquell Cécile |
| EPI_ISL_3425698, EPI_ISL_3425699, EPI_ISL_3425700 | Centre De Prelevement COVID RIOM | CHU Clermont-Ferrand, service de virologie | Bisseux Maxime, Mirand Audrey, Combes Patricia, Henquell Cécile |
| EPI_ISL_3425701 | Centre Hospitalier Universitaire Clermont-Ferrand | CHU Clermont-Ferrand, service de virologie | Bisseux Maxime, Mirand Audrey, Combes Patricia, Henquell Cécile |
| EPI_ISL_3425702, EPI_ISL_3425703, EPI_ISL_3425704 | Hospital De Vichy | CHU Clermont-Ferrand, service de virologie | Bisseux Maxime, Mirand Audrey, Combes Patricia, Henquell Cécile |
| EPI_ISL_3425705 | Centre Hospitalier Universitaire Clermont-Ferrand | CHU Clermont-Ferrand, service de virologie | Bisseux Maxime, Mirand Audrey, Combes Patricia, Henquell Cécile |
| EPI_ISL_3425706, EPI_ISL_3425707 | Sungai Buloh Hospital | Institute for Medical Research, Infectious Disease Research Centre, National Institutes of Health, Ministry of Health Malaysia | Suppiah J, Kamel K, Mohd Zawawi Z, Azizan MA, Ramly N, Robert F, Anasir MI, Thayan R |
| EPI_ISL_3425708, EPI_ISL_3425709, EPI_ISL_3425710, EPI_ISL_3425711, EPI_ISL_3425712, EPI_ISL_3425713, EPI_ISL_3425714, EPI_ISL_3425715, EPI_ISL_3425716, EPI_ISL_3425717, EPI_ISL_3425718, EPI_ISL_3425720, EPI_ISL_3425721, EPI_ISL_3425722, EPI_ISL_3425723, EPI_ISL_3425724, EPI_ISL_3425725, EPI_ISL_3425726, EPI_ISL_3425727, EPI_ISL_3425728, EPI_ISL_3425729, EPI_ISL_3425732, EPI_ISL_3425733, EPI_ISL_3425734, EPI_ISL_3425735, EPI_ISL_3425736, EPI_ISL_3425737, EPI_ISL_3425738, EPI_ISL_3425739, EPI_ISL_3425740, EPI_ISL_3425741, EPI_ISL_3425742, EPI_ISL_3425743, EPI_ISL_3425744, EPI_ISL_3425745, EPI_ISL_3425746, EPI_ISL_3425747, EPI_ISL_3425748, EPI_ISL_3425749, EPI_ISL_3425750, EPI_ISL_3425751, EPI_ISL_3425752, EPI_ISL_3425753, EPI_ISL_3425754, EPI_ISL_3425755, EPI_ISL_3425756, EPI_ISL_3425757, EPI_ISL_3425758, EPI_ISL_3425759, EPI_ISL_3425760, EPI_ISL_3425761, EPI_ISL_3425762, EPI_ISL_3425763, EPI_ISL_3425764, EPI_ISL_3425765, EPI_ISL_3425766, EPI_ISL_3425767, EPI_ISL_3425768, EPI_ISL_3425769, EPI_ISL_3425770, EPI_ISL_3425771, EPI_ISL_3425772, EPI_ISL_3425773, EPI_ISL_3425774, EPI_ISL_3425775, EPI_ISL_3425776, EPI_ISL_3425777, EPI_ISL_3425778, EPI_ISL_3425779, EPI_ISL_3425780, EPI_ISL_3425781, EPI_ISL_3425782 | | | |
| see above | National Institute of Public Health | National Institute of Public Health | Helena Jirincova, Jaromira Vecerova, Timotej Suri, Dusan Trnka, Alexander Nagy |
| EPI_ISL_3425788 | Synlab Suisse SA | Clinical Bacteriology | Tim Roloff, Fanny Wegner, Helena MB Seth-Smith, Alfredo Mari, Karoline Leuzinger, Julia Bielicki, Manuel Battegay, Hans Hirsch, Adrian Egli |

| | | | |
|---|---|---|---|
| EPI_ISL_3425789 | Sungai Buloh Hospital | Institute for Medical Research, Infectious Disease Research Centre, National Institutes of Health, Ministry of Health Malaysia | Suppiah J, Kamel K, Mohd Zawawi Z, Azizan MA, Ramly N, Robert F, Anasir MI, Thayan R |
| EPI_ISL_3425792, EPI_ISL_3425793 | Synlab Suisse SA | Clinical Bacteriology | Tim Roloff, Fanny Wegner, Helena MB Seth-Smith, Alfredo Mari, Karoline Leuzinger, Julia Bielicki, Manuel Battegay, Hans Hirsch, Adrian Egli |
| EPI_ISL_3425795, EPI_ISL_3425796, EPI_ISL_3425797, EPI_ISL_3425798, EPI_ISL_3425799, EPI_ISL_3425800, EPI_ISL_3425801, EPI_ISL_3425802, EPI_ISL_3425803, EPI_ISL_3425804 | Medics Labor AG | Clinical Bacteriology | Tim Roloff, Fanny Wegner, Helena MB Seth-Smith, Alfredo Mari, Karoline Leuzinger, Julia Bielicki, Manuel Battegay, Hans Hirsch, Adrian Egli |
| EPI_ISL_3425805, EPI_ISL_3425806 | Rothen Medizinische Laboratorien AG | Clinical Bacteriology | Tim Roloff, Fanny Wegner, Helena MB Seth-Smith, Alfredo Mari, Karoline Leuzinger, Julia Bielicki, Ingrid Steffen, Manuel Battegay, Hans Hirsch, Adrian Egli |
| EPI_ISL_3425807, EPI_ISL_3425808, EPI_ISL_3425809, EPI_ISL_3425810, EPI_ISL_3425811, EPI_ISL_3425812, EPI_ISL_3425813, EPI_ISL_3425814, EPI_ISL_3425815 | Synlab Suisse SA | Clinical Bacteriology | Tim Roloff, Fanny Wegner, Helena MB Seth-Smith, Alfredo Mari, Karoline Leuzinger, Julia Bielicki, Manuel Battegay, Hans Hirsch, Adrian Egli |
| EPI_ISL_3425816, EPI_ISL_3425817, EPI_ISL_3425818, EPI_ISL_3425819, EPI_ISL_3425820, EPI_ISL_3425821 | Hopital du Jura: Delemont | Clinical Bacteriology | Tim Roloff, Fanny Wegner, Helena MB Seth-Smith, Alfredo Mari, Karoline Leuzinger, Julia Bielicki, Manuel Battegay, Hans Hirsch, Adrian Egli |
| EPI_ISL_3425822, EPI_ISL_3425823, EPI_ISL_3425824, EPI_ISL_3425825 | UniversitätsSpital Zürich | Institute of Medical Virology | Daniel Ehrsam, Isabel Stürmer, Catharine Aquino, Joel Wirz, Weihong Qi, Hubert Rehrauer, Verena Kufner, Gabriela Ziltener, Maryam Zaheri, Stefan Schmutz, Annette Audigé, Maria Grünberg, Kevin Steiner, Jon Huder, Cyril Shah, Riccarda Capaul, Guido Bloemberg, Jürg Böni, Michael Huber, Alexandra Trkola |
| EPI_ISL_3425826 | Universitäts-Kinderspital Zürich | Institute of Medical Virology | Daniel Ehrsam, Isabel Stürmer, Catharine Aquino, Joel Wirz, Weihong Qi, Hubert Rehrauer, Verena Kufner, Gabriela Ziltener, Maryam Zaheri, Stefan Schmutz, Annette Audigé, Maria Grünberg, Kevin Steiner, Jon Huder, Cyril Shah, Riccarda Capaul, Guido Bloemberg, Jürg Böni, Michael Huber, Alexandra Trkola |
| EPI_ISL_3425827, EPI_ISL_3425828 | Universität Zürich | Institute of Medical Virology | Daniel Ehrsam, Isabel Stürmer, Catharine Aquino, Joel Wirz, Weihong Qi, Hubert Rehrauer, Verena Kufner, Gabriela Ziltener, Maryam Zaheri, Stefan Schmutz, Annette Audigé, Maria Grünberg, Kevin Steiner, Jon Huder, Cyril Shah, Riccarda Capaul, Guido Bloemberg, Jürg Böni, Michael Huber, Alexandra Trkola |
| EPI_ISL_3425829, EPI_ISL_3425830, EPI_ISL_3425831, EPI_ISL_3425832, EPI_ISL_3425833, EPI_ISL_3425834, EPI_ISL_3425835, EPI_ISL_3425836, EPI_ISL_3425837, EPI_ISL_3425838, EPI_ISL_3425839, EPI_ISL_3425840, EPI_ISL_3425841, EPI_ISL_3425842, EPI_ISL_3425843, EPI_ISL_3425844, EPI_ISL_3425845, EPI_ISL_3425846, EPI_ISL_3425847, EPI_ISL_3425848, EPI_ISL_3425849, EPI_ISL_3425850, EPI_ISL_3425851, EPI_ISL_3425852 | | | |
| see above | Stadtspital Triemli | Institute of Medical Virology | Daniel Ehrsam, Isabel Stürmer, Catharine Aquino, Joel Wirz, Weihong Qi, Hubert Rehrauer, Verena Kufner, Gabriela Ziltener, Maryam Zaheri, Stefan Schmutz, Annette Audigé, Maria Grünberg, Kevin Steiner, Jon Huder, Cyril Shah, Riccarda Capaul, Guido Bloemberg, Jürg Böni, Michael Huber, Alexandra Trkola |
| EPI_ISL_3425853, EPI_ISL_3425854, EPI_ISL_3425855, EPI_ISL_3425856 | UniversitätsSpital Zürich | Institute of Medical Virology | Daniel Ehrsam, Isabel Stürmer, Catharine Aquino, Joel Wirz, Weihong Qi, Hubert Rehrauer, Verena Kufner, Gabriela Ziltener, Maryam Zaheri, Stefan Schmutz, Annette Audigé, Maria Grünberg, Kevin Steiner, Jon Huder, Cyril Shah, Riccarda Capaul, Guido Bloemberg, Jürg Böni, Michael Huber, Alexandra Trkola |
| EPI_ISL_3425857 | Stadtspital Waid | Institute of Medical Virology | Daniel Ehrsam, Isabel Stürmer, Catharine Aquino, Joel Wirz, Weihong Qi, Hubert Rehrauer, Verena Kufner, Gabriela Ziltener, Maryam Zaheri, Stefan Schmutz, Annette Audigé, Maria Grünberg, Kevin Steiner, Jon Huder, Cyril Shah, Riccarda Capaul, Guido Bloemberg, Jürg Böni, Michael Huber, Alexandra Trkola |
| EPI_ISL_3425858 | Universitäts-Kinderspital Zürich | Institute of Medical Virology | Daniel Ehrsam, Isabel Stürmer, Catharine Aquino, Joel Wirz, Weihong Qi, Hubert Rehrauer, Verena Kufner, Gabriela Ziltener, Maryam Zaheri, Stefan Schmutz, Annette Audigé, Maria Grünberg, Kevin Steiner, Jon Huder, Cyril Shah, Riccarda Capaul, Guido Bloemberg, Jürg Böni, Michael Huber, Alexandra Trkola |
| EPI_ISL_3425859, EPI_ISL_3425860 | UniversitätsSpital Zürich | Institute of Medical Virology | Daniel Ehrsam, Isabel Stürmer, Catharine Aquino, Joel Wirz, Weihong Qi, Hubert Rehrauer, Verena Kufner, Gabriela Ziltener, Maryam Zaheri, Stefan Schmutz, Annette Audigé, Maria Grünberg, Kevin Steiner, Jon Huder, Cyril Shah, Riccarda Capaul, Guido Bloemberg, Jürg Böni, Michael Huber, Alexandra Trkola |
| EPI_ISL_3425861, EPI_ISL_3425862, EPI_ISL_3425863, EPI_ISL_3425864, EPI_ISL_3425865, EPI_ISL_3425866, EPI_ISL_3425867, EPI_ISL_3425868, EPI_ISL_3425869, EPI_ISL_3425870, EPI_ISL_3425871, EPI_ISL_3425872, EPI_ISL_3425873 | | | |
| see above | Stadtspital Triemli | Institute of Medical Virology | Daniel Ehrsam, Isabel Stürmer, Catharine Aquino, Joel Wirz, Weihong Qi, Hubert Rehrauer, Verena Kufner, Gabriela Ziltener, Maryam Zaheri, Stefan Schmutz, Annette Audigé, Maria Grünberg, Kevin Steiner, Jon Huder, Cyril Shah, Riccarda Capaul, Guido Bloemberg, Jürg Böni, Michael Huber, Alexandra Trkola |
| EPI_ISL_3425874, EPI_ISL_3425875 | Universität Zürich | Institute of Medical Virology | Daniel Ehrsam, Isabel Stürmer, Catharine Aquino, Joel Wirz, Weihong Qi, Hubert Rehrauer, Verena Kufner, Gabriela Ziltener, Maryam Zaheri, Stefan Schmutz, Annette Audigé, Maria Grünberg, Kevin Steiner, Jon Huder, Cyril Shah, Riccarda Capaul, Guido Bloemberg, Jürg Böni, Michael Huber, Alexandra Trkola |
| EPI_ISL_3425876 | UniversitätsSpital Zürich 044 | Institute of Medical Virology | Daniel Ehrsam, Isabel Stürmer, Catharine Aquino, Joel Wirz, Weihong Qi, Hubert Rehrauer, Verena Kufner, Gabriela Ziltener, Maryam Zaheri, Stefan Schmutz, Annette Audigé, Maria Grünberg, Kevin Steiner, Jon Huder, Cyril Shah, Riccarda Capaul, Guido Bloemberg, Jürg Böni, Michael Huber, Alexandra Trkola |
| EPI_ISL_3425877 | Kinderspital Zürich | Institute of Medical Virology | Daniel Ehrsam, Isabel Stürmer, Catharine Aquino, Joel Wirz, Weihong Qi, Hubert Rehrauer, Verena Kufner, Gabriela Ziltener, Maryam Zaheri, Stefan Schmutz, Annette Audigé, Maria Grünberg, Kevin Steiner, Jon Huder, Cyril Shah, Riccarda Capaul, Guido Bloemberg, Jürg Böni, Michael Huber, Alexandra Trkola |
| EPI_ISL_3425878 | Universität Zürich | Institute of Medical Virology | Daniel Ehrsam, Isabel Stürmer, Catharine Aquino, Joel Wirz, Weihong Qi, Hubert Rehrauer, Verena Kufner, Gabriela Ziltener, Maryam Zaheri, Stefan Schmutz, Annette Audigé, Maria Grünberg, Kevin Steiner, Jon Huder, Cyril Shah, Riccarda Capaul, Guido Bloemberg, Jürg Böni, Michael Huber, Alexandra Trkola |
| EPI_ISL_3425879, EPI_ISL_3425880, EPI_ISL_3425881, EPI_ISL_3425882, EPI_ISL_3425883 | UniversitätsSpital Zürich | Institute of Medical Virology | Daniel Ehrsam, Isabel Stürmer, Catharine Aquino, Joel Wirz, Weihong Qi, Hubert Rehrauer, Verena Kufner, Gabriela Ziltener, Maryam Zaheri, Stefan Schmutz, Annette Audigé, Maria Grünberg, Kevin Steiner, Jon Huder, Cyril Shah, Riccarda Capaul, Guido Bloemberg, Jürg Böni, Michael Huber, Alexandra Trkola |
| EPI_ISL_3425884 | Ampang Hospital | Institute for Medical Research, Infectious Disease Research Centre, National Institutes of Health, Ministry of Health Malaysia | Suppiah J, Kamel K, Mohd Zawawi Z, Azizan MA, Ramly N, Robert F, Anasir MI, Thayan R |
| EPI_ISL_3425885 | UniversitätsSpital Zürich | Institute of Medical Virology | Verena Kufner, Gabriela Ziltener, Maryam Zaheri, Stefan Schmutz, Annette Audigé, Maria Grünberg, Kevin Steiner, Jon Huder, Cyril Shah, Riccarda Capaul, Guido Bloemberg, Jürg Böni, Michael Huber, Alexandra Trkola |
| EPI_ISL_3425886 | Kantonsspital Aarau | Institute of Medical Virology | Verena Kufner, Gabriela Ziltener, Maryam Zaheri, Stefan Schmutz, Annette Audigé, Maria Grünberg, Kevin Steiner, Jon Huder, Cyril Shah, Riccarda Capaul, Guido Bloemberg, Jürg Böni, Michael Huber, Alexandra Trkola |
| EPI_ISL_3425887, EPI_ISL_3425888 | Zentrum für Labormedizin | Institute of Medical Virology | Verena Kufner, Gabriela Ziltener, Maryam Zaheri, Stefan Schmutz, Annette Audigé, Maria Grünberg, Kevin Steiner, Jon Huder, Cyril Shah, Riccarda Capaul, Guido Bloemberg, Jürg Böni, Michael Huber, Alexandra Trkola |
| EPI_ISL_3425889 | Universität Zürich | Institute of Medical Virology | Verena Kufner, Gabriela Ziltener, Maryam Zaheri, Stefan Schmutz, Annette Audigé, Maria Grünberg, Kevin Steiner, Jon Huder, Cyril Shah, Riccarda Capaul, Guido Bloemberg, Jürg Böni, Michael Huber, Alexandra Trkola |
| EPI_ISL_3425890 | Kantonsspital Baden AG | Institute of Medical Virology | Verena Kufner, Gabriela Ziltener, Maryam Zaheri, Stefan Schmutz, Annette Audigé, Maria Grünberg, Kevin Steiner, Jon Huder, Cyril Shah, Riccarda Capaul, Guido Bloemberg, Jürg Böni, Michael Huber, Alexandra Trkola |
| EPI_ISL_3425891, EPI_ISL_3425892 | Stadtspital Waid | Institute of Medical Virology | Verena Kufner, Gabriela Ziltener, Maryam Zaheri, Stefan Schmutz, Annette Audigé, Maria Grünberg, Kevin Steiner, Jon Huder, Cyril Shah, Riccarda Capaul, Guido Bloemberg, Jürg Böni, Michael Huber, Alexandra Trkola |
| EPI_ISL_3425893 | Invenimus AG | Institute of Medical Virology | Verena Kufner, Gabriela Ziltener, Maryam Zaheri, Stefan Schmutz, Annette Audigé, Maria Grünberg, Kevin Steiner, Jon Huder, Cyril Shah, Riccarda Capaul, Guido Bloemberg, Jürg Böni, Michael Huber, Alexandra Trkola |
| EPI_ISL_3425895, EPI_ISL_3425896, EPI_ISL_3425897, EPI_ISL_3425898, EPI_ISL_3425899 | Stadtspital Triemli | Institute of Medical Virology | Verena Kufner, Gabriela Ziltener, Maryam Zaheri, Stefan Schmutz, Annette Audigé, Maria Grünberg, Kevin Steiner, Jon Huder, Cyril Shah, Riccarda Capaul, Guido Bloemberg, Jürg Böni, Michael Huber, Alexandra Trkola |
| EPI_ISL_3425900 | Unilabs | Institute of Medical Virology | Verena Kufner, Gabriela Ziltener, Maryam Zaheri, Stefan Schmutz, Annette Audigé, Maria Grünberg, Kevin Steiner, Jon Huder, Cyril Shah, Riccarda Capaul, Guido Bloemberg, Jürg Böni, Michael Huber, Alexandra Trkola |
| EPI_ISL_3425901, EPI_ISL_3425902, EPI_ISL_3425903, EPI_ISL_3425904, EPI_ISL_3425905 | Medica | Institute of Medical Virology | Verena Kufner, Gabriela Ziltener, Maryam Zaheri, Stefan Schmutz, Annette Audigé, Maria Grünberg, Kevin Steiner, Jon Huder, Cyril Shah, Riccarda Capaul, Guido Bloemberg, Jürg Böni, Michael Huber, Alexandra Trkola |
| EPI_ISL_3425906, EPI_ISL_3425907, EPI_ISL_3425908 | Analytica | Institute of Medical Virology | Verena Kufner, Gabriela Ziltener, Maryam Zaheri, Stefan Schmutz, Annette Audigé, Maria Grünberg, Kevin Steiner, Jon Huder, Cyril Shah, Riccarda Capaul, Guido Bloemberg, Jürg Böni, Michael Huber, Alexandra Trkola |
| EPI_ISL_3425909, | Gesundheitsamt SO | Institute of Medical Virology | Verena Kufner, Gabriela Ziltener, Maryam Zaheri, Stefan Schmutz, Annette Audigé, Maria Grünberg, Kevin Steiner, Jon Huder, Cyril Shah, Riccarda Capaul, Guido Bloemberg, Jürg Böni, Michael Huber, Alexandra Trkola |

| | | | |
|---|---|---|---|
| EPI_ISL_3425910, EPI_ISL_3425911 | | | |
| EPI_ISL_3425912 | UniversitätsSpital Zürich | Institute of Medical Virology | Verena Kufner, Gabriela Ziltener, Maryam Zaheri, Stefan Schmutz, Annette Audigé, Maria Grünberg, Kevin Steiner, Jon Huder, Cyril Shah, Riccarda Capaul, Guido Bloemberg, Jürg Böni, Michael Huber, Alexandra Trkola |
| EPI_ISL_3425913 | Zentrum für Labormedizin | Institute of Medical Virology | Verena Kufner, Gabriela Ziltener, Maryam Zaheri, Stefan Schmutz, Annette Audigé, Maria Grünberg, Kevin Steiner, Jon Huder, Cyril Shah, Riccarda Capaul, Guido Bloemberg, Jürg Böni, Michael Huber, Alexandra Trkola |
| EPI_ISL_3425914, EPI_ISL_3425915 | Diamisanto GmbH | Institute of Medical Virology | Verena Kufner, Gabriela Ziltener, Maryam Zaheri, Stefan Schmutz, Annette Audigé, Maria Grünberg, Kevin Steiner, Jon Huder, Cyril Shah, Riccarda Capaul, Guido Bloemberg, Jürg Böni, Michael Huber, Alexandra Trkola |
| EPI_ISL_3425916, EPI_ISL_3425917, EPI_ISL_3425918 | Kantonsspital Aarau | Institute of Medical Virology | Verena Kufner, Gabriela Ziltener, Maryam Zaheri, Stefan Schmutz, Annette Audigé, Maria Grünberg, Kevin Steiner, Jon Huder, Cyril Shah, Riccarda Capaul, Guido Bloemberg, Jürg Böni, Michael Huber, Alexandra Trkola |
| EPI_ISL_3425919, EPI_ISL_3425920, EPI_ISL_3425921, EPI_ISL_3425922 | Universität Zürich | Institute of Medical Virology | Verena Kufner, Gabriela Ziltener, Maryam Zaheri, Stefan Schmutz, Annette Audigé, Maria Grünberg, Kevin Steiner, Jon Huder, Cyril Shah, Riccarda Capaul, Guido Bloemberg, Jürg Böni, Michael Huber, Alexandra Trkola |
| EPI_ISL_3425923 | nan | Institute of Medical Virology | Verena Kufner, Gabriela Ziltener, Maryam Zaheri, Stefan Schmutz, Annette Audigé, Maria Grünberg, Kevin Steiner, Jon Huder, Cyril Shah, Riccarda Capaul, Guido Bloemberg, Jürg Böni, Michael Huber, Alexandra Trkola |
| EPI_ISL_3425924, EPI_ISL_3425925, EPI_ISL_3425926 | Universität Zürich | Institute of Medical Virology | Verena Kufner, Gabriela Ziltener, Maryam Zaheri, Stefan Schmutz, Annette Audigé, Maria Grünberg, Kevin Steiner, Jon Huder, Cyril Shah, Riccarda Capaul, Guido Bloemberg, Jürg Böni, Michael Huber, Alexandra Trkola |
| EPI_ISL_3425927 | Medlab Pathology | NSW Health Pathology - Institute of Clinical Pathology and Medical Research; Westmead Hospital; University of Sydney | Arnott A., Draper J., Gall M., Martinez E., Rockett R., Sintchenko V., on behalf of ICPMR |
| EPI_ISL_3425928, EPI_ISL_3425929 | St Vincent's Pathology (SydPath) | NSW Health Pathology - Institute of Clinical Pathology and Medical Research; Westmead Hospital; University of Sydney | Arnott A., Draper J., Gall M., Martinez E., Rockett R., Sintchenko V., on behalf of ICPMR |
| EPI_ISL_3425930, EPI_ISL_3425931 | Medlab Pathology | NSW Health Pathology - Institute of Clinical Pathology and Medical Research; Westmead Hospital; University of Sydney | Arnott A., Draper J., Gall M., Martinez E., Rockett R., Sintchenko V., on behalf of ICPMR |
| EPI_ISL_3425932 | Laverty Pathology | NSW Health Pathology - Institute of Clinical Pathology and Medical Research; Westmead Hospital; University of Sydney | Arnott A., Draper J., Gall M., Martinez E., Rockett R., Sintchenko V., on behalf of ICPMR |
| EPI_ISL_3425933 | Pathology North - Hunter - NSW Health Pathology | NSW Health Pathology - Institute of Clinical Pathology and Medical Research; Westmead Hospital; University of Sydney | Arnott A., Draper J., Gall M., Martinez E., Rockett R., Sintchenko V., on behalf of ICPMR |
| EPI_ISL_3425934, EPI_ISL_3425935 | Histopath | NSW Health Pathology - Institute of Clinical Pathology and Medical Research; Westmead Hospital; University of Sydney | Arnott A., Draper J., Gall M., Martinez E., Rockett R., Sintchenko V., on behalf of ICPMR |
| EPI_ISL_3425936 | Pathology North - Hunter - NSW Health Pathology | NSW Health Pathology - Institute of Clinical Pathology and Medical Research; Westmead Hospital; University of Sydney | Arnott A., Draper J., Gall M., Martinez E., Rockett R., Sintchenko V., on behalf of ICPMR |
| EPI_ISL_3425937, EPI_ISL_3425938 | 4Cyte Pathology | NSW Health Pathology - Institute of Clinical Pathology and Medical Research; Westmead Hospital; University of Sydney | Arnott A., Draper J., Gall M., Martinez E., Rockett R., Sintchenko V., on behalf of ICPMR |
| EPI_ISL_3425939 | Medlab Pathology | NSW Health Pathology - Institute of Clinical Pathology and Medical Research; Westmead Hospital; University of Sydney | Arnott A., Draper J., Gall M., Martinez E., Rockett R., Sintchenko V., on behalf of ICPMR |
| EPI_ISL_3425940 | Capital Pathology | NSW Health Pathology - Institute of Clinical Pathology and Medical Research; Westmead Hospital; University of Sydney | Arnott A., Draper J., Gall M., Martinez E., Rockett R., Sintchenko V., on behalf of ICPMR |
| EPI_ISL_3425941 | Histopath | NSW Health Pathology - Institute of Clinical Pathology and Medical Research; Westmead Hospital; University of Sydney | Arnott A., Draper J., Gall M., Martinez E., Rockett R., Sintchenko V., on behalf of ICPMR |
| EPI_ISL_3425942, EPI_ISL_3425943 | St Vincent's Pathology (SydPath) | NSW Health Pathology - Institute of Clinical Pathology and Medical Research; Westmead Hospital; University of Sydney | Arnott A., Draper J., Gall M., Martinez E., Rockett R., Sintchenko V., on behalf of ICPMR |
| EPI_ISL_3425944 | Histopath | NSW Health Pathology - Institute of Clinical Pathology and Medical Research; Westmead Hospital; University of Sydney | Arnott A., Draper J., Gall M., Martinez E., Rockett R., Sintchenko V., on behalf of ICPMR |
| EPI_ISL_3425945, EPI_ISL_3425946 | 4Cyte Pathology | NSW Health Pathology - Institute of Clinical Pathology and Medical Research; Westmead Hospital; University of Sydney | Arnott A., Draper J., Gall M., Martinez E., Rockett R., Sintchenko V., on behalf of ICPMR |
| EPI_ISL_3425947, EPI_ISL_3425948 | Histopath | NSW Health Pathology - Institute of Clinical Pathology and Medical Research; Westmead Hospital; University of Sydney | Arnott A., Draper J., Gall M., Martinez E., Rockett R., Sintchenko V., on behalf of ICPMR |
| EPI_ISL_3425949 | St Vincent's Pathology (SydPath) | NSW Health Pathology - Institute of Clinical Pathology and Medical Research; Westmead Hospital; University of Sydney | Arnott A., Draper J., Gall M., Martinez E., Rockett R., Sintchenko V., on behalf of ICPMR |
| EPI_ISL_3425950 | Histopath | NSW Health Pathology - Institute of Clinical Pathology and Medical Research; Westmead Hospital; University of Sydney | Arnott A., Draper J., Gall M., Martinez E., Rockett R., Sintchenko V., on behalf of ICPMR |
| EPI_ISL_3425951, EPI_ISL_3425952 | Medlab Pathology | NSW Health Pathology - Institute of Clinical Pathology and Medical Research; Westmead Hospital; University of Sydney | Arnott A., Draper J., Gall M., Martinez E., Rockett R., Sintchenko V., on behalf of ICPMR |
| EPI_ISL_3425953 | St Vincent's Pathology (SydPath) | NSW Health Pathology - Institute of Clinical Pathology and Medical Research; Westmead Hospital; University of Sydney | Arnott A., Draper J., Gall M., Martinez E., Rockett R., Sintchenko V., on behalf of ICPMR |
| EPI_ISL_3425954 | Histopath | NSW Health Pathology - Institute of Clinical Pathology and Medical Research; Westmead Hospital; University of Sydney | Arnott A., Draper J., Gall M., Martinez E., Rockett R., Sintchenko V., on behalf of ICPMR |
| EPI_ISL_3425955 | St Vincent's Pathology (SydPath) | NSW Health Pathology - Institute of Clinical Pathology and Medical Research; Westmead Hospital; University of Sydney | Arnott A., Draper J., Gall M., Martinez E., Rockett R., Sintchenko V., on behalf of ICPMR |
| EPI_ISL_3425956 | Histopath | NSW Health Pathology - Institute of Clinical Pathology and Medical Research; Westmead Hospital; University of Sydney | Arnott A., Draper J., Gall M., Martinez E., Rockett R., Sintchenko V., on behalf of ICPMR |
| EPI_ISL_3425957 | 4Cyte Pathology | NSW Health Pathology - Institute of Clinical Pathology and Medical Research; Westmead Hospital; University of Sydney | Arnott A., Draper J., Gall M., Martinez E., Rockett R., Sintchenko V., on behalf of ICPMR |
| EPI_ISL_3425958 | Medlab Pathology | NSW Health Pathology - Institute of Clinical Pathology and Medical Research; Westmead Hospital; University of Sydney | Arnott A., Draper J., Gall M., Martinez E., Rockett R., Sintchenko V., on behalf of ICPMR |
| EPI_ISL_3425959 | Laverty Pathology | NSW Health Pathology - Institute of Clinical Pathology and Medical Research; Westmead Hospital; University of Sydney | Arnott A., Draper J., Gall M., Martinez E., Rockett R., Sintchenko V., on behalf of ICPMR |
| EPI_ISL_3425960 | Histopath | NSW Health Pathology - Institute of Clinical Pathology and Medical Research; Westmead Hospital; University of Sydney | Arnott A., Draper J., Gall M., Martinez E., Rockett R., Sintchenko V., on behalf of ICPMR |
| EPI_ISL_3425961 | 4Cyte Pathology | NSW Health Pathology - Institute of Clinical Pathology and Medical Research; Westmead Hospital; University of Sydney | Arnott A., Draper J., Gall M., Martinez E., Rockett R., Sintchenko V., on behalf of ICPMR |
| EPI_ISL_3425962 | Medlab Pathology | NSW Health Pathology - Institute of Clinical Pathology and Medical Research; Westmead Hospital; University of Sydney | Arnott A., Draper J., Gall M., Martinez E., Rockett R., Sintchenko V., on behalf of ICPMR |
| EPI_ISL_3425963 | Laverty Pathology | NSW Health Pathology - Institute of Clinical Pathology and Medical Research; Westmead Hospital; University of Sydney | Arnott A., Draper J., Gall M., Martinez E., Rockett R., Sintchenko V., on behalf of ICPMR |
| EPI_ISL_3425964 | 4Cyte Pathology | NSW Health Pathology - Institute of Clinical Pathology and Medical Research; Westmead Hospital; University of Sydney | Arnott A., Draper J., Gall M., Martinez E., Rockett R., Sintchenko V., on behalf of ICPMR |
| EPI_ISL_3425965 | Pathology North - Royal North Shore Hospital - NSW Health Pathology | NSW Health Pathology - Institute of Clinical Pathology and Medical Research; Westmead Hospital; University of Sydney | Arnott A., Draper J., Gall M., Martinez E., Rockett R., Sintchenko V., on behalf of ICPMR |
| EPI_ISL_3425966 | Medlab Pathology | NSW Health Pathology - Institute of Clinical Pathology and Medical Research; Westmead Hospital; University of Sydney | Arnott A., Draper J., Gall M., Martinez E., Rockett R., Sintchenko V., on behalf of ICPMR |
| EPI_ISL_3425967 | 4Cyte Pathology | NSW Health Pathology - Institute of Clinical Pathology and Medical Research; Westmead Hospital; University of Sydney | Arnott A., Draper J., Gall M., Martinez E., Rockett R., Sintchenko V., on behalf of ICPMR |
| EPI_ISL_3425968 | The Children's Hospital at Westmead | NSW Health Pathology - Institute of Clinical Pathology and Medical Research; Westmead Hospital; University of Sydney | Arnott A., Draper J., Gall M., Martinez E., Rockett R., Sintchenko V., on behalf of ICPMR |
| EPI_ISL_3425969 | 4Cyte Pathology | NSW Health Pathology - Institute of Clinical Pathology and Medical Research; Westmead Hospital; University of Sydney | Arnott A., Draper J., Gall M., Martinez E., Rockett R., Sintchenko V., on behalf of ICPMR |
| EPI_ISL_3425970 | Austech Medical Laboratories | NSW Health Pathology - Institute of Clinical Pathology and Medical Research; Westmead Hospital; University of Sydney | Arnott A., Draper J., Gall M., Martinez E., Rockett R., Sintchenko V., on behalf of ICPMR |
| EPI_ISL_3425971 | Pathology West - NSW Health Pathology | NSW Health Pathology - Institute of Clinical Pathology and Medical Research; Westmead Hospital; University of Sydney | Arnott A., Draper J., Gall M., Martinez E., Rockett R., Sintchenko V., on behalf of ICPMR |

| | | | |
|---|---|---|---|
| EPI_ISL_3425972 | Medlab Pathology | NSW Health Pathology - Institute of Clinical Pathology and Medical Research; Westmead Hospital; University of Sydney | Arnott A., Draper J., Gall M., Martinez E., Rockett R., Sintchenko V., on behalf of ICPMR |
| EPI_ISL_3425973 | 4Cyte Pathology | NSW Health Pathology - Institute of Clinical Pathology and Medical Research; Westmead Hospital; University of Sydney | Arnott A., Draper J., Gall M., Martinez E., Rockett R., Sintchenko V., on behalf of ICPMR |
| EPI_ISL_3425974 | Sydney South West Pathology Service (SSWPS) - Concord Repatriation General Hospital - NSW Health Pathology | NSW Health Pathology - Institute of Clinical Pathology and Medical Research; Westmead Hospital; University of Sydney | Arnott A., Draper J., Gall M., Martinez E., Rockett R., Sintchenko V., on behalf of ICPMR |
| EPI_ISL_3425975 | Sydney South West Pathology Service (SSWPS) - Liverpool Hospital - NSW Health Pathology | NSW Health Pathology - Institute of Clinical Pathology and Medical Research; Westmead Hospital; University of Sydney | Arnott A., Draper J., Gall M., Martinez E., Rockett R., Sintchenko V., on behalf of ICPMR |
| EPI_ISL_3425976 | Histopath | NSW Health Pathology - Institute of Clinical Pathology and Medical Research; Westmead Hospital; University of Sydney | Arnott A., Draper J., Gall M., Martinez E., Rockett R., Sintchenko V., on behalf of ICPMR |
| EPI_ISL_3425977 | 4Cyte Pathology | NSW Health Pathology - Institute of Clinical Pathology and Medical Research; Westmead Hospital; University of Sydney | Arnott A., Draper J., Gall M., Martinez E., Rockett R., Sintchenko V., on behalf of ICPMR |
| EPI_ISL_3425978 | Medlab Pathology | NSW Health Pathology - Institute of Clinical Pathology and Medical Research; Westmead Hospital; University of Sydney | Arnott A., Draper J., Gall M., Martinez E., Rockett R., Sintchenko V., on behalf of ICPMR |
| EPI_ISL_3425979, EPI_ISL_3425980 | Histopath | NSW Health Pathology - Institute of Clinical Pathology and Medical Research; Westmead Hospital; University of Sydney | Arnott A., Draper J., Gall M., Martinez E., Rockett R., Sintchenko V., on behalf of ICPMR |
| EPI_ISL_3425981 | Medlab Pathology | NSW Health Pathology - Institute of Clinical Pathology and Medical Research; Westmead Hospital; University of Sydney | Arnott A., Draper J., Gall M., Martinez E., Rockett R., Sintchenko V., on behalf of ICPMR |
| EPI_ISL_3425982 | St Vincent's Pathology (SydPath) | NSW Health Pathology - Institute of Clinical Pathology and Medical Research; Westmead Hospital; University of Sydney | Arnott A., Draper J., Gall M., Martinez E., Rockett R., Sintchenko V., on behalf of ICPMR |
| EPI_ISL_3425983 | Australian Clinical Labs (formerly Healthscope Pathology) | NSW Health Pathology - Institute of Clinical Pathology and Medical Research; Westmead Hospital; University of Sydney | Arnott A., Draper J., Gall M., Martinez E., Rockett R., Sintchenko V., on behalf of ICPMR |
| EPI_ISL_3425984 | Austech Medical Laboratories | NSW Health Pathology - Institute of Clinical Pathology and Medical Research; Westmead Hospital; University of Sydney | Arnott A., Draper J., Gall M., Martinez E., Rockett R., Sintchenko V., on behalf of ICPMR |
| EPI_ISL_3425985 | Laverty Pathology | NSW Health Pathology - Institute of Clinical Pathology and Medical Research; Westmead Hospital; University of Sydney | Arnott A., Draper J., Gall M., Martinez E., Rockett R., Sintchenko V., on behalf of ICPMR |
| EPI_ISL_3425986, EPI_ISL_3425987 | Histopath | NSW Health Pathology - Institute of Clinical Pathology and Medical Research; Westmead Hospital; University of Sydney | Arnott A., Draper J., Gall M., Martinez E., Rockett R., Sintchenko V., on behalf of ICPMR |
| EPI_ISL_3425988 | The Children's Hospital at Westmead | NSW Health Pathology - Institute of Clinical Pathology and Medical Research; Westmead Hospital; University of Sydney | Arnott A., Draper J., Gall M., Martinez E., Rockett R., Sintchenko V., on behalf of ICPMR |
| EPI_ISL_3425989 | Laverty Pathology | NSW Health Pathology - Institute of Clinical Pathology and Medical Research; Westmead Hospital; University of Sydney | Arnott A., Draper J., Gall M., Martinez E., Rockett R., Sintchenko V., on behalf of ICPMR |
| EPI_ISL_3425990 | 4Cyte Pathology | NSW Health Pathology - Institute of Clinical Pathology and Medical Research; Westmead Hospital; University of Sydney | Arnott A., Draper J., Gall M., Martinez E., Rockett R., Sintchenko V., on behalf of ICPMR |
| EPI_ISL_3425991 | Medlab Pathology | NSW Health Pathology - Institute of Clinical Pathology and Medical Research; Westmead Hospital; University of Sydney | Arnott A., Draper J., Gall M., Martinez E., Rockett R., Sintchenko V., on behalf of ICPMR |
| EPI_ISL_3425992 | Australian Clinical Labs (formerly Healthscope Pathology) | NSW Health Pathology - Institute of Clinical Pathology and Medical Research; Westmead Hospital; University of Sydney | Arnott A., Draper J., Gall M., Martinez E., Rockett R., Sintchenko V., on behalf of ICPMR |
| EPI_ISL_3425993 | 4Cyte Pathology | NSW Health Pathology - Institute of Clinical Pathology and Medical Research; Westmead Hospital; University of Sydney | Arnott A., Draper J., Gall M., Martinez E., Rockett R., Sintchenko V., on behalf of ICPMR |
| EPI_ISL_3425994 | Australian Clinical Labs (formerly Healthscope Pathology) | NSW Health Pathology - Institute of Clinical Pathology and Medical Research; Westmead Hospital; University of Sydney | Arnott A., Draper J., Gall M., Martinez E., Rockett R., Sintchenko V., on behalf of ICPMR |
| EPI_ISL_3425995, EPI_ISL_3425996 | Histopath | NSW Health Pathology - Institute of Clinical Pathology and Medical Research; Westmead Hospital; University of Sydney | Arnott A., Draper J., Gall M., Martinez E., Rockett R., Sintchenko V., on behalf of ICPMR |
| EPI_ISL_3425997 | Medlab Pathology | NSW Health Pathology - Institute of Clinical Pathology and Medical Research; Westmead Hospital; University of Sydney | Arnott A., Draper J., Gall M., Martinez E., Rockett R., Sintchenko V., on behalf of ICPMR |
| EPI_ISL_3425998, EPI_ISL_3425999 | St Vincent's Pathology (SydPath) | NSW Health Pathology - Institute of Clinical Pathology and Medical Research; Westmead Hospital; University of Sydney | Arnott A., Draper J., Gall M., Martinez E., Rockett R., Sintchenko V., on behalf of ICPMR |
| EPI_ISL_3426000 | Medlab Pathology | NSW Health Pathology - Institute of Clinical Pathology and Medical Research; Westmead Hospital; University of Sydney | Arnott A., Draper J., Gall M., Martinez E., Rockett R., Sintchenko V., on behalf of ICPMR |
| EPI_ISL_3426001, EPI_ISL_3426002, EPI_ISL_3426003 | Histopath | NSW Health Pathology - Institute of Clinical Pathology and Medical Research; Westmead Hospital; University of Sydney | Arnott A., Draper J., Gall M., Martinez E., Rockett R., Sintchenko V., on behalf of ICPMR |
| EPI_ISL_3426004 | Pathology West - NSW Health Pathology | NSW Health Pathology - Institute of Clinical Pathology and Medical Research; Westmead Hospital; University of Sydney | Arnott A., Draper J., Gall M., Martinez E., Rockett R., Sintchenko V., on behalf of ICPMR |
| EPI_ISL_3426005 | Histopath | NSW Health Pathology - Institute of Clinical Pathology and Medical Research; Westmead Hospital; University of Sydney | Arnott A., Draper J., Gall M., Martinez E., Rockett R., Sintchenko V., on behalf of ICPMR |
| EPI_ISL_3426006 | 4Cyte Pathology | NSW Health Pathology - Institute of Clinical Pathology and Medical Research; Westmead Hospital; University of Sydney | Arnott A., Draper J., Gall M., Martinez E., Rockett R., Sintchenko V., on behalf of ICPMR |
| EPI_ISL_3426007, EPI_ISL_3426008 | St Vincent's Pathology (SydPath) | NSW Health Pathology - Institute of Clinical Pathology and Medical Research; Westmead Hospital; University of Sydney | Arnott A., Draper J., Gall M., Martinez E., Rockett R., Sintchenko V., on behalf of ICPMR |
| EPI_ISL_3426009 | Pathology West - NSW Health Pathology | NSW Health Pathology - Institute of Clinical Pathology and Medical Research; Westmead Hospital; University of Sydney | Arnott A., Draper J., Gall M., Martinez E., Rockett R., Sintchenko V., on behalf of ICPMR |
| EPI_ISL_3426010 | Histopath | NSW Health Pathology - Institute of Clinical Pathology and Medical Research; Westmead Hospital; University of Sydney | Arnott A., Draper J., Gall M., Martinez E., Rockett R., Sintchenko V., on behalf of ICPMR |
| EPI_ISL_3426011 | Pathology West - NSW Health Pathology | NSW Health Pathology - Institute of Clinical Pathology and Medical Research; Westmead Hospital; University of Sydney | Arnott A., Draper J., Gall M., Martinez E., Rockett R., Sintchenko V., on behalf of ICPMR |
| EPI_ISL_3426012 | Histopath | NSW Health Pathology - Institute of Clinical Pathology and Medical Research; Westmead Hospital; University of Sydney | Arnott A., Draper J., Gall M., Martinez E., Rockett R., Sintchenko V., on behalf of ICPMR |
| EPI_ISL_3426013, EPI_ISL_3426014 | Laverty Pathology | NSW Health Pathology - Institute of Clinical Pathology and Medical Research; Westmead Hospital; University of Sydney | Arnott A., Draper J., Gall M., Martinez E., Rockett R., Sintchenko V., on behalf of ICPMR |
| EPI_ISL_3426015 | Medlab Pathology | NSW Health Pathology - Institute of Clinical Pathology and Medical Research; Westmead Hospital; University of Sydney | Arnott A., Draper J., Gall M., Martinez E., Rockett R., Sintchenko V., on behalf of ICPMR |
| EPI_ISL_3426016 | Laverty Pathology | NSW Health Pathology - Institute of Clinical Pathology and Medical Research; Westmead Hospital; University of Sydney | Arnott A., Draper J., Gall M., Martinez E., Rockett R., Sintchenko V., on behalf of ICPMR |
| EPI_ISL_3426017 | Histopath | NSW Health Pathology - Institute of Clinical Pathology and Medical Research; Westmead Hospital; University of Sydney | Arnott A., Draper J., Gall M., Martinez E., Rockett R., Sintchenko V., on behalf of ICPMR |
| EPI_ISL_3426018 | 4Cyte Pathology | NSW Health Pathology - Institute of Clinical Pathology and Medical Research; Westmead Hospital; University of Sydney | Arnott A., Draper J., Gall M., Martinez E., Rockett R., Sintchenko V., on behalf of ICPMR |
| EPI_ISL_3426019 | Histopath | NSW Health Pathology - Institute of Clinical Pathology and Medical Research; Westmead Hospital; University of Sydney | Arnott A., Draper J., Gall M., Martinez E., Rockett R., Sintchenko V., on behalf of ICPMR |
| EPI_ISL_3426020 | Laverty Pathology | NSW Health Pathology - Institute of Clinical Pathology and Medical Research; Westmead Hospital; University of Sydney | Arnott A., Draper J., Gall M., Martinez E., Rockett R., Sintchenko V., on behalf of ICPMR |
| EPI_ISL_3426021 | Histopath | NSW Health Pathology - Institute of Clinical Pathology and Medical Research; Westmead Hospital; University of Sydney | Arnott A., Draper J., Gall M., Martinez E., Rockett R., Sintchenko V., on behalf of ICPMR |
| EPI_ISL_3426022 | Pathology West - NSW Health Pathology | NSW Health Pathology - Institute of Clinical Pathology and Medical Research; Westmead Hospital; University of Sydney | Arnott A., Draper J., Gall M., Martinez E., Rockett R., Sintchenko V., on behalf of ICPMR |
| EPI_ISL_3426023 | Histopath | NSW Health Pathology - Institute of Clinical Pathology and Medical Research; Westmead Hospital; University of Sydney | Arnott A., Draper J., Gall M., Martinez E., Rockett R., Sintchenko V., on behalf of ICPMR |
| EPI_ISL_3426024 | St Vincent's Pathology (SydPath) | NSW Health Pathology - Institute of Clinical Pathology and Medical Research; Westmead Hospital; University of Sydney | Arnott A., Draper J., Gall M., Martinez E., Rockett R., Sintchenko V., on behalf of ICPMR |

| | | | |
|---|---|---|---|
| EPI_ISL_3426025 | Histopath | NSW Health Pathology - Institute of Clinical Pathology and Medical Research; Westmead Hospital; University of Sydney | Arnott A., Draper J., Gall M., Martinez E., Rockett R., Sintchenko V., on behalf of ICPMR |
| EPI_ISL_3426026 | Laverty Pathology | NSW Health Pathology - Institute of Clinical Pathology and Medical Research; Westmead Hospital; University of Sydney | Arnott A., Draper J., Gall M., Martinez E., Rockett R., Sintchenko V., on behalf of ICPMR |
| EPI_ISL_3426027 | 4Cyte Pathology | NSW Health Pathology - Institute of Clinical Pathology and Medical Research; Westmead Hospital; University of Sydney | Arnott A., Draper J., Gall M., Martinez E., Rockett R., Sintchenko V., on behalf of ICPMR |
| EPI_ISL_3426028 | Medlab Pathology | NSW Health Pathology - Institute of Clinical Pathology and Medical Research; Westmead Hospital; University of Sydney | Arnott A., Draper J., Gall M., Martinez E., Rockett R., Sintchenko V., on behalf of ICPMR |
| EPI_ISL_3426029 | Histopath | NSW Health Pathology - Institute of Clinical Pathology and Medical Research; Westmead Hospital; University of Sydney | Arnott A., Draper J., Gall M., Martinez E., Rockett R., Sintchenko V., on behalf of ICPMR |
| EPI_ISL_3426030 | 4Cyte Pathology | NSW Health Pathology - Institute of Clinical Pathology and Medical Research; Westmead Hospital; University of Sydney | Arnott A., Draper J., Gall M., Martinez E., Rockett R., Sintchenko V., on behalf of ICPMR |
| EPI_ISL_3426031 | The Children's Hospital at Westmead | NSW Health Pathology - Institute of Clinical Pathology and Medical Research; Westmead Hospital; University of Sydney | Arnott A., Draper J., Gall M., Martinez E., Rockett R., Sintchenko V., on behalf of ICPMR |
| EPI_ISL_3426032, EPI_ISL_3426033 | Laverty Pathology | NSW Health Pathology - Institute of Clinical Pathology and Medical Research; Westmead Hospital; University of Sydney | Arnott A., Draper J., Gall M., Martinez E., Rockett R., Sintchenko V., on behalf of ICPMR |
| EPI_ISL_3426034 | Austech Medical Laboratories | NSW Health Pathology - Institute of Clinical Pathology and Medical Research; Westmead Hospital; University of Sydney | Arnott A., Draper J., Gall M., Martinez E., Rockett R., Sintchenko V., on behalf of ICPMR |
| EPI_ISL_3426035 | 4Cyte Pathology | NSW Health Pathology - Institute of Clinical Pathology and Medical Research; Westmead Hospital; University of Sydney | Arnott A., Draper J., Gall M., Martinez E., Rockett R., Sintchenko V., on behalf of ICPMR |
| EPI_ISL_3426036 | The Children's Hospital at Westmead | NSW Health Pathology - Institute of Clinical Pathology and Medical Research; Westmead Hospital; University of Sydney | Arnott A., Draper J., Gall M., Martinez E., Rockett R., Sintchenko V., on behalf of ICPMR |
| EPI_ISL_3426037 | St Vincent's Pathology (SydPath) | NSW Health Pathology - Institute of Clinical Pathology and Medical Research; Westmead Hospital; University of Sydney | Arnott A., Draper J., Gall M., Martinez E., Rockett R., Sintchenko V., on behalf of ICPMR |
| EPI_ISL_3426038 | Pathology West - NSW Health Pathology | NSW Health Pathology - Institute of Clinical Pathology and Medical Research; Westmead Hospital; University of Sydney | Arnott A., Draper J., Gall M., Martinez E., Rockett R., Sintchenko V., on behalf of ICPMR |
| EPI_ISL_3426039 | Histopath | NSW Health Pathology - Institute of Clinical Pathology and Medical Research; Westmead Hospital; University of Sydney | Arnott A., Draper J., Gall M., Martinez E., Rockett R., Sintchenko V., on behalf of ICPMR |
| EPI_ISL_3426040 | Laverty Pathology | NSW Health Pathology - Institute of Clinical Pathology and Medical Research; Westmead Hospital; University of Sydney | Arnott A., Draper J., Gall M., Martinez E., Rockett R., Sintchenko V., on behalf of ICPMR |
| EPI_ISL_3426041, EPI_ISL_3426042 | Histopath | NSW Health Pathology - Institute of Clinical Pathology and Medical Research; Westmead Hospital; University of Sydney | Arnott A., Draper J., Gall M., Martinez E., Rockett R., Sintchenko V., on behalf of ICPMR |
| EPI_ISL_3426043 | Pathology North - Royal North Shore Hospital - NSW Health Pathology | NSW Health Pathology - Institute of Clinical Pathology and Medical Research; Westmead Hospital; University of Sydney | Arnott A., Draper J., Gall M., Martinez E., Rockett R., Sintchenko V., on behalf of ICPMR |
| EPI_ISL_3426044 | Capital Pathology | NSW Health Pathology - Institute of Clinical Pathology and Medical Research; Westmead Hospital; University of Sydney | Arnott A., Draper J., Gall M., Martinez E., Rockett R., Sintchenko V., on behalf of ICPMR |
| EPI_ISL_3426045 | Pathology West - NSW Health Pathology | NSW Health Pathology - Institute of Clinical Pathology and Medical Research; Westmead Hospital; University of Sydney | Arnott A., Draper J., Gall M., Martinez E., Rockett R., Sintchenko V., on behalf of ICPMR |
| EPI_ISL_3426046 | Australian Clinical Labs (formerly Healthscope Pathology) | NSW Health Pathology - Institute of Clinical Pathology and Medical Research; Westmead Hospital; University of Sydney | Arnott A., Draper J., Gall M., Martinez E., Rockett R., Sintchenko V., on behalf of ICPMR |
| EPI_ISL_3426047 | Austech Medical Laboratories | NSW Health Pathology - Institute of Clinical Pathology and Medical Research; Westmead Hospital; University of Sydney | Arnott A., Draper J., Gall M., Martinez E., Rockett R., Sintchenko V., on behalf of ICPMR |
| EPI_ISL_3426048, EPI_ISL_3426049, EPI_ISL_3426050, EPI_ISL_3426051 | Histopath | NSW Health Pathology - Institute of Clinical Pathology and Medical Research; Westmead Hospital; University of Sydney | Arnott A., Draper J., Gall M., Martinez E., Rockett R., Sintchenko V., on behalf of ICPMR |
| EPI_ISL_3426052 | Australian Clinical Labs (formerly Healthscope Pathology) | NSW Health Pathology - Institute of Clinical Pathology and Medical Research; Westmead Hospital; University of Sydney | Arnott A., Draper J., Gall M., Martinez E., Rockett R., Sintchenko V., on behalf of ICPMR |
| EPI_ISL_3426053 | St Vincent's Pathology (SydPath) | NSW Health Pathology - Institute of Clinical Pathology and Medical Research; Westmead Hospital; University of Sydney | Arnott A., Draper J., Gall M., Martinez E., Rockett R., Sintchenko V., on behalf of ICPMR |
| EPI_ISL_3426054, EPI_ISL_3426055 | Histopath | NSW Health Pathology - Institute of Clinical Pathology and Medical Research; Westmead Hospital; University of Sydney | Arnott A., Draper J., Gall M., Martinez E., Rockett R., Sintchenko V., on behalf of ICPMR |
| EPI_ISL_3426056 | The Children's Hospital at Westmead | NSW Health Pathology - Institute of Clinical Pathology and Medical Research; Westmead Hospital; University of Sydney | Arnott A., Draper J., Gall M., Martinez E., Rockett R., Sintchenko V., on behalf of ICPMR |
| EPI_ISL_3426057, EPI_ISL_3426058, EPI_ISL_3426059 | Histopath | NSW Health Pathology - Institute of Clinical Pathology and Medical Research; Westmead Hospital; University of Sydney | Arnott A., Draper J., Gall M., Martinez E., Rockett R., Sintchenko V., on behalf of ICPMR |
| EPI_ISL_3426060, EPI_ISL_3426061 | Pathology West - NSW Health Pathology | NSW Health Pathology - Institute of Clinical Pathology and Medical Research; Westmead Hospital; University of Sydney | Arnott A., Draper J., Gall M., Martinez E., Rockett R., Sintchenko V., on behalf of ICPMR |
| EPI_ISL_3426062, EPI_ISL_3426063, EPI_ISL_3426064 | Australian Clinical Labs (formerly Healthscope Pathology) | NSW Health Pathology - Institute of Clinical Pathology and Medical Research; Westmead Hospital; University of Sydney | Arnott A., Draper J., Gall M., Martinez E., Rockett R., Sintchenko V., on behalf of ICPMR |
| EPI_ISL_3426065 | Laverty Pathology | NSW Health Pathology - Institute of Clinical Pathology and Medical Research; Westmead Hospital; University of Sydney | Arnott A., Draper J., Gall M., Martinez E., Rockett R., Sintchenko V., on behalf of ICPMR |
| EPI_ISL_3426066 | 4Cyte Pathology | NSW Health Pathology - Institute of Clinical Pathology and Medical Research; Westmead Hospital; University of Sydney | Arnott A., Draper J., Gall M., Martinez E., Rockett R., Sintchenko V., on behalf of ICPMR |
| EPI_ISL_3426067, EPI_ISL_3426068 | Histopath | NSW Health Pathology - Institute of Clinical Pathology and Medical Research; Westmead Hospital; University of Sydney | Arnott A., Draper J., Gall M., Martinez E., Rockett R., Sintchenko V., on behalf of ICPMR |
| EPI_ISL_3426069 | Laverty Pathology | NSW Health Pathology - Institute of Clinical Pathology and Medical Research; Westmead Hospital; University of Sydney | Arnott A., Draper J., Gall M., Martinez E., Rockett R., Sintchenko V., on behalf of ICPMR |
| EPI_ISL_3426070 | Histopath | NSW Health Pathology - Institute of Clinical Pathology and Medical Research; Westmead Hospital; University of Sydney | Arnott A., Draper J., Gall M., Martinez E., Rockett R., Sintchenko V., on behalf of ICPMR |
| EPI_ISL_3426071 | Pathology North - Hunter - NSW Health Pathology | NSW Health Pathology - Institute of Clinical Pathology and Medical Research; Westmead Hospital; University of Sydney | Arnott A., Draper J., Gall M., Martinez E., Rockett R., Sintchenko V., on behalf of ICPMR |
| EPI_ISL_3426072, EPI_ISL_3426073, EPI_ISL_3426074 | Histopath | NSW Health Pathology - Institute of Clinical Pathology and Medical Research; Westmead Hospital; University of Sydney | Arnott A., Draper J., Gall M., Martinez E., Rockett R., Sintchenko V., on behalf of ICPMR |
| EPI_ISL_3426075 | Laverty Pathology | NSW Health Pathology - Institute of Clinical Pathology and Medical Research; Westmead Hospital; University of Sydney | Arnott A., Draper J., Gall M., Martinez E., Rockett R., Sintchenko V., on behalf of ICPMR |
| EPI_ISL_3426076 | 4Cyte Pathology | NSW Health Pathology - Institute of Clinical Pathology and Medical Research; Westmead Hospital; University of Sydney | Arnott A., Draper J., Gall M., Martinez E., Rockett R., Sintchenko V., on behalf of ICPMR |
| EPI_ISL_3426077, EPI_ISL_3426078, EPI_ISL_3426079 | Histopath | NSW Health Pathology - Institute of Clinical Pathology and Medical Research; Westmead Hospital; University of Sydney | Arnott A., Draper J., Gall M., Martinez E., Rockett R., Sintchenko V., on behalf of ICPMR |
| EPI_ISL_3426080 | Australian Clinical Labs (formerly Healthscope Pathology) | NSW Health Pathology - Institute of Clinical Pathology and Medical Research; Westmead Hospital; University of Sydney | Arnott A., Draper J., Gall M., Martinez E., Rockett R., Sintchenko V., on behalf of ICPMR |
| EPI_ISL_3426081, EPI_ISL_3426082, EPI_ISL_3426083 | Histopath | NSW Health Pathology - Institute of Clinical Pathology and Medical Research; Westmead Hospital; University of Sydney | Arnott A., Draper J., Gall M., Martinez E., Rockett R., Sintchenko V., on behalf of ICPMR |
| EPI_ISL_3426084, EPI_ISL_3426085 | 4Cyte Pathology | NSW Health Pathology - Institute of Clinical Pathology and Medical Research; Westmead Hospital; University of Sydney | Arnott A., Draper J., Gall M., Martinez E., Rockett R., Sintchenko V., on behalf of ICPMR |

| | | | |
|---|---|---|---|
| EPI_ISL_3426086, EPI_ISL_3426087 | Histopath | NSW Health Pathology - Institute of Clinical Pathology and Medical Research; Westmead Hospital; University of Sydney | Arnott A., Draper J., Gall M., Martinez E., Rockett R., Sintchenko V., on behalf of ICPMR |
| EPI_ISL_3426088 | 4Cyte Pathology | NSW Health Pathology - Institute of Clinical Pathology and Medical Research; Westmead Hospital; University of Sydney | Arnott A., Draper J., Gall M., Martinez E., Rockett R., Sintchenko V., on behalf of ICPMR |
| EPI_ISL_3426089, EPI_ISL_3426090 | Histopath | NSW Health Pathology - Institute of Clinical Pathology and Medical Research; Westmead Hospital; University of Sydney | Arnott A., Draper J., Gall M., Martinez E., Rockett R., Sintchenko V., on behalf of ICPMR |
| EPI_ISL_3426091, EPI_ISL_3426092, EPI_ISL_3426093, EPI_ISL_3426094 | Laverty Pathology | NSW Health Pathology - Institute of Clinical Pathology and Medical Research; Westmead Hospital; University of Sydney | Arnott A., Draper J., Gall M., Martinez E., Rockett R., Sintchenko V., on behalf of ICPMR |
| EPI_ISL_3426095 | 4Cyte Pathology | NSW Health Pathology - Institute of Clinical Pathology and Medical Research; Westmead Hospital; University of Sydney | Arnott A., Draper J., Gall M., Martinez E., Rockett R., Sintchenko V., on behalf of ICPMR |
| EPI_ISL_3426096, EPI_ISL_3426097, EPI_ISL_3426098 | Histopath | NSW Health Pathology - Institute of Clinical Pathology and Medical Research; Westmead Hospital; University of Sydney | Arnott A., Draper J., Gall M., Martinez E., Rockett R., Sintchenko V., on behalf of ICPMR |
| EPI_ISL_3426099 | 4Cyte Pathology | NSW Health Pathology - Institute of Clinical Pathology and Medical Research; Westmead Hospital; University of Sydney | Arnott A., Draper J., Gall M., Martinez E., Rockett R., Sintchenko V., on behalf of ICPMR |
| EPI_ISL_3426100, EPI_ISL_3426101, EPI_ISL_3426102 | Histopath | NSW Health Pathology - Institute of Clinical Pathology and Medical Research; Westmead Hospital; University of Sydney | Arnott A., Draper J., Gall M., Martinez E., Rockett R., Sintchenko V., on behalf of ICPMR |
| EPI_ISL_3426103 | St Vincent's Pathology (SydPath) | NSW Health Pathology - Institute of Clinical Pathology and Medical Research; Westmead Hospital; University of Sydney | Arnott A., Draper J., Gall M., Martinez E., Rockett R., Sintchenko V., on behalf of ICPMR |
| EPI_ISL_3426104 | Sydney South West Pathology Service (SSWPS) - Liverpool Hospital - NSW Health Pathology | NSW Health Pathology - Institute of Clinical Pathology and Medical Research; Westmead Hospital; University of Sydney | Arnott A., Draper J., Gall M., Martinez E., Rockett R., Sintchenko V., on behalf of ICPMR |
| EPI_ISL_3426105 | Laverty Pathology | NSW Health Pathology - Institute of Clinical Pathology and Medical Research; Westmead Hospital; University of Sydney | Arnott A., Draper J., Gall M., Martinez E., Rockett R., Sintchenko V., on behalf of ICPMR |
| EPI_ISL_3426106 | 4Cyte Pathology | NSW Health Pathology - Institute of Clinical Pathology and Medical Research; Westmead Hospital; University of Sydney | Arnott A., Draper J., Gall M., Martinez E., Rockett R., Sintchenko V., on behalf of ICPMR |
| EPI_ISL_3426107 | Histopath | NSW Health Pathology - Institute of Clinical Pathology and Medical Research; Westmead Hospital; University of Sydney | Arnott A., Draper J., Gall M., Martinez E., Rockett R., Sintchenko V., on behalf of ICPMR |
| EPI_ISL_3426108 | 4Cyte Pathology | NSW Health Pathology - Institute of Clinical Pathology and Medical Research; Westmead Hospital; University of Sydney | Arnott A., Draper J., Gall M., Martinez E., Rockett R., Sintchenko V., on behalf of ICPMR |
| EPI_ISL_3426109 | Histopath | NSW Health Pathology - Institute of Clinical Pathology and Medical Research; Westmead Hospital; University of Sydney | Arnott A., Draper J., Gall M., Martinez E., Rockett R., Sintchenko V., on behalf of ICPMR |
| EPI_ISL_3426110 | Capital Pathology | NSW Health Pathology - Institute of Clinical Pathology and Medical Research; Westmead Hospital; University of Sydney | Arnott A., Draper J., Gall M., Martinez E., Rockett R., Sintchenko V., on behalf of ICPMR |
| EPI_ISL_3426111 | Histopath | NSW Health Pathology - Institute of Clinical Pathology and Medical Research; Westmead Hospital; University of Sydney | Arnott A., Draper J., Gall M., Martinez E., Rockett R., Sintchenko V., on behalf of ICPMR |
| EPI_ISL_3426112 | Laverty Pathology | NSW Health Pathology - Institute of Clinical Pathology and Medical Research; Westmead Hospital; University of Sydney | Arnott A., Draper J., Gall M., Martinez E., Rockett R., Sintchenko V., on behalf of ICPMR |
| EPI_ISL_3426113, EPI_ISL_3426115, EPI_ISL_3426116 | Histopath | NSW Health Pathology - Institute of Clinical Pathology and Medical Research; Westmead Hospital; University of Sydney | Arnott A., Draper J., Gall M., Martinez E., Rockett R., Sintchenko V., on behalf of ICPMR |
| EPI_ISL_3426117 | 4Cyte Pathology | NSW Health Pathology - Institute of Clinical Pathology and Medical Research; Westmead Hospital; University of Sydney | Arnott A., Draper J., Gall M., Martinez E., Rockett R., Sintchenko V., on behalf of ICPMR |
| EPI_ISL_3426118 | Histopath | NSW Health Pathology - Institute of Clinical Pathology and Medical Research; Westmead Hospital; University of Sydney | Arnott A., Draper J., Gall M., Martinez E., Rockett R., Sintchenko V., on behalf of ICPMR |
| EPI_ISL_3426119 | St Vincent's Pathology (SydPath) | NSW Health Pathology - Institute of Clinical Pathology and Medical Research; Westmead Hospital; University of Sydney | Arnott A., Draper J., Gall M., Martinez E., Rockett R., Sintchenko V., on behalf of ICPMR |
| EPI_ISL_3426120 | Medlab Pathology | NSW Health Pathology - Institute of Clinical Pathology and Medical Research; Westmead Hospital; University of Sydney | Arnott A., Draper J., Gall M., Martinez E., Rockett R., Sintchenko V., on behalf of ICPMR |
| EPI_ISL_3426121 | Australian Clinical Labs (formerly Healthscope Pathology) | NSW Health Pathology - Institute of Clinical Pathology and Medical Research; Westmead Hospital; University of Sydney | Arnott A., Draper J., Gall M., Martinez E., Rockett R., Sintchenko V., on behalf of ICPMR |
| EPI_ISL_3426122 | 4Cyte Pathology | NSW Health Pathology - Institute of Clinical Pathology and Medical Research; Westmead Hospital; University of Sydney | Arnott A., Draper J., Gall M., Martinez E., Rockett R., Sintchenko V., on behalf of ICPMR |
| EPI_ISL_3426123 | Medlab Pathology | NSW Health Pathology - Institute of Clinical Pathology and Medical Research; Westmead Hospital; University of Sydney | Arnott A., Draper J., Gall M., Martinez E., Rockett R., Sintchenko V., on behalf of ICPMR |
| EPI_ISL_3426124 | 4Cyte Pathology | NSW Health Pathology - Institute of Clinical Pathology and Medical Research; Westmead Hospital; University of Sydney | Arnott A., Draper J., Gall M., Martinez E., Rockett R., Sintchenko V., on behalf of ICPMR |
| EPI_ISL_3426125 | Medlab Pathology | NSW Health Pathology - Institute of Clinical Pathology and Medical Research; Westmead Hospital; University of Sydney | Arnott A., Draper J., Gall M., Martinez E., Rockett R., Sintchenko V., on behalf of ICPMR |
| EPI_ISL_3426126 | Australian Clinical Labs (formerly Healthscope Pathology) | NSW Health Pathology - Institute of Clinical Pathology and Medical Research; Westmead Hospital; University of Sydney | Arnott A., Draper J., Gall M., Martinez E., Rockett R., Sintchenko V., on behalf of ICPMR |
| EPI_ISL_3426127 | 4Cyte Pathology | NSW Health Pathology - Institute of Clinical Pathology and Medical Research; Westmead Hospital; University of Sydney | Arnott A., Draper J., Gall M., Martinez E., Rockett R., Sintchenko V., on behalf of ICPMR |
| EPI_ISL_3426128 | Medlab Pathology | NSW Health Pathology - Institute of Clinical Pathology and Medical Research; Westmead Hospital; University of Sydney | Arnott A., Draper J., Gall M., Martinez E., Rockett R., Sintchenko V., on behalf of ICPMR |
| EPI_ISL_3426129 | St Vincent's Pathology (SydPath) | NSW Health Pathology - Institute of Clinical Pathology and Medical Research; Westmead Hospital; University of Sydney | Arnott A., Draper J., Gall M., Martinez E., Rockett R., Sintchenko V., on behalf of ICPMR |
| EPI_ISL_3426130, EPI_ISL_3426131 | Histopath | NSW Health Pathology - Institute of Clinical Pathology and Medical Research; Westmead Hospital; University of Sydney | Arnott A., Draper J., Gall M., Martinez E., Rockett R., Sintchenko V., on behalf of ICPMR |
| EPI_ISL_3426132 | Laverty Pathology | NSW Health Pathology - Institute of Clinical Pathology and Medical Research; Westmead Hospital; University of Sydney | Arnott A., Draper J., Gall M., Martinez E., Rockett R., Sintchenko V., on behalf of ICPMR |
| EPI_ISL_3426133 | Pathology West - NSW Health Pathology | NSW Health Pathology - Institute of Clinical Pathology and Medical Research; Westmead Hospital; University of Sydney | Arnott A., Draper J., Gall M., Martinez E., Rockett R., Sintchenko V., on behalf of ICPMR |
| EPI_ISL_3426134 | The Children's Hospital at Westmead | NSW Health Pathology - Institute of Clinical Pathology and Medical Research; Westmead Hospital; University of Sydney | Arnott A., Draper J., Gall M., Martinez E., Rockett R., Sintchenko V., on behalf of ICPMR |
| EPI_ISL_3426135 | St Vincent's Pathology (SydPath) | NSW Health Pathology - Institute of Clinical Pathology and Medical Research; Westmead Hospital; University of Sydney | Arnott A., Draper J., Gall M., Martinez E., Rockett R., Sintchenko V., on behalf of ICPMR |
| EPI_ISL_3426136, EPI_ISL_3426137 | Histopath | NSW Health Pathology - Institute of Clinical Pathology and Medical Research; Westmead Hospital; University of Sydney | Arnott A., Draper J., Gall M., Martinez E., Rockett R., Sintchenko V., on behalf of ICPMR |
| EPI_ISL_3426138 | Australian Clinical Labs (formerly Healthscope Pathology) | NSW Health Pathology - Institute of Clinical Pathology and Medical Research; Westmead Hospital; University of Sydney | Arnott A., Draper J., Gall M., Martinez E., Rockett R., Sintchenko V., on behalf of ICPMR |
| EPI_ISL_3426139 | Laverty Pathology | NSW Health Pathology - Institute of Clinical Pathology and Medical Research; Westmead Hospital; University of Sydney | Arnott A., Draper J., Gall M., Martinez E., Rockett R., Sintchenko V., on behalf of ICPMR |
| EPI_ISL_3426140 | St Vincent's Pathology (SydPath) | NSW Health Pathology - Institute of Clinical Pathology and Medical Research; Westmead Hospital; University of Sydney | Arnott A., Draper J., Gall M., Martinez E., Rockett R., Sintchenko V., on behalf of ICPMR |
| EPI_ISL_3426141 | 4Cyte Pathology | NSW Health Pathology - Institute of Clinical Pathology and Medical Research; Westmead Hospital; University of Sydney | Arnott A., Draper J., Gall M., Martinez E., Rockett R., Sintchenko V., on behalf of ICPMR |
| EPI_ISL_3426142 | Pathology West - NSW Health Pathology | NSW Health Pathology - Institute of Clinical Pathology and Medical Research; Westmead Hospital; University of Sydney | Arnott A., Draper J., Gall M., Martinez E., Rockett R., Sintchenko V., on behalf of ICPMR |

| | | | |
|---|---|---|---|
| EPI_ISL_3426143 | Australian Clinical Labs (formerly Healthscope Pathology) | NSW Health Pathology - Institute of Clinical Pathology and Medical Research; Westmead Hospital; University of Sydney | Arnott A., Draper J., Gall M., Martinez E., Rockett R., Sintchenko V., on behalf of ICPMR |
| EPI_ISL_3426144 | Pathology North - Hunter - NSW Health Pathology | NSW Health Pathology - Institute of Clinical Pathology and Medical Research; Westmead Hospital; University of Sydney | Arnott A., Draper J., Gall M., Martinez E., Rockett R., Sintchenko V., on behalf of ICPMR |
| EPI_ISL_3426145 | Pathology West - NSW Health Pathology | NSW Health Pathology - Institute of Clinical Pathology and Medical Research; Westmead Hospital; University of Sydney | Arnott A., Draper J., Gall M., Martinez E., Rockett R., Sintchenko V., on behalf of ICPMR |
| EPI_ISL_3426146 | 4Cyte Pathology | NSW Health Pathology - Institute of Clinical Pathology and Medical Research; Westmead Hospital; University of Sydney | Arnott A., Draper J., Gall M., Martinez E., Rockett R., Sintchenko V., on behalf of ICPMR |
| EPI_ISL_3426147 | Histopath | NSW Health Pathology - Institute of Clinical Pathology and Medical Research; Westmead Hospital; University of Sydney | Arnott A., Draper J., Gall M., Martinez E., Rockett R., Sintchenko V., on behalf of ICPMR |
| EPI_ISL_3426148 | Laverty Pathology | NSW Health Pathology - Institute of Clinical Pathology and Medical Research; Westmead Hospital; University of Sydney | Arnott A., Draper J., Gall M., Martinez E., Rockett R., Sintchenko V., on behalf of ICPMR |
| EPI_ISL_3426149, EPI_ISL_3426150 | The Children's Hospital at Westmead | NSW Health Pathology - Institute of Clinical Pathology and Medical Research; Westmead Hospital; University of Sydney | Arnott A., Draper J., Gall M., Martinez E., Rockett R., Sintchenko V., on behalf of ICPMR |
| EPI_ISL_3426151 | 4Cyte Pathology | NSW Health Pathology - Institute of Clinical Pathology and Medical Research; Westmead Hospital; University of Sydney | Arnott A., Draper J., Gall M., Martinez E., Rockett R., Sintchenko V., on behalf of ICPMR |
| EPI_ISL_3426152 | The Children's Hospital at Westmead | NSW Health Pathology - Institute of Clinical Pathology and Medical Research; Westmead Hospital; University of Sydney | Arnott A., Draper J., Gall M., Martinez E., Rockett R., Sintchenko V., on behalf of ICPMR |
| EPI_ISL_3426153 | Histopath | NSW Health Pathology - Institute of Clinical Pathology and Medical Research; Westmead Hospital; University of Sydney | Arnott A., Draper J., Gall M., Martinez E., Rockett R., Sintchenko V., on behalf of ICPMR |
| EPI_ISL_3426154 | Laverty Pathology | NSW Health Pathology - Institute of Clinical Pathology and Medical Research; Westmead Hospital; University of Sydney | Arnott A., Draper J., Gall M., Martinez E., Rockett R., Sintchenko V., on behalf of ICPMR |
| EPI_ISL_3426155, EPI_ISL_3426156 | St Vincent's Pathology (SydPath) | NSW Health Pathology - Institute of Clinical Pathology and Medical Research; Westmead Hospital; University of Sydney | Arnott A., Draper J., Gall M., Martinez E., Rockett R., Sintchenko V., on behalf of ICPMR |
| EPI_ISL_3426157 | Histopath | NSW Health Pathology - Institute of Clinical Pathology and Medical Research; Westmead Hospital; University of Sydney | Arnott A., Draper J., Gall M., Martinez E., Rockett R., Sintchenko V., on behalf of ICPMR |
| EPI_ISL_3426158 | Pathology West - NSW Health Pathology | NSW Health Pathology - Institute of Clinical Pathology and Medical Research; Westmead Hospital; University of Sydney | Arnott A., Draper J., Gall M., Martinez E., Rockett R., Sintchenko V., on behalf of ICPMR |
| EPI_ISL_3426159 | 4Cyte Pathology | NSW Health Pathology - Institute of Clinical Pathology and Medical Research; Westmead Hospital; University of Sydney | Arnott A., Draper J., Gall M., Martinez E., Rockett R., Sintchenko V., on behalf of ICPMR |
| EPI_ISL_3426160 | The Children's Hospital at Westmead | NSW Health Pathology - Institute of Clinical Pathology and Medical Research; Westmead Hospital; University of Sydney | Arnott A., Draper J., Gall M., Martinez E., Rockett R., Sintchenko V., on behalf of ICPMR |
| EPI_ISL_3426161, EPI_ISL_3426162 | Histopath | NSW Health Pathology - Institute of Clinical Pathology and Medical Research; Westmead Hospital; University of Sydney | Arnott A., Draper J., Gall M., Martinez E., Rockett R., Sintchenko V., on behalf of ICPMR |
| EPI_ISL_3426163 | Australian Clinical Labs (formerly Healthscope Pathology) | NSW Health Pathology - Institute of Clinical Pathology and Medical Research; Westmead Hospital; University of Sydney | Arnott A., Draper J., Gall M., Martinez E., Rockett R., Sintchenko V., on behalf of ICPMR |
| EPI_ISL_3426164 | Laverty Pathology | NSW Health Pathology - Institute of Clinical Pathology and Medical Research; Westmead Hospital; University of Sydney | Arnott A., Draper J., Gall M., Martinez E., Rockett R., Sintchenko V., on behalf of ICPMR |
| EPI_ISL_3426165 | Histopath | NSW Health Pathology - Institute of Clinical Pathology and Medical Research; Westmead Hospital; University of Sydney | Arnott A., Draper J., Gall M., Martinez E., Rockett R., Sintchenko V., on behalf of ICPMR |
| EPI_ISL_3426166 | Pathology West - NSW Health Pathology | NSW Health Pathology - Institute of Clinical Pathology and Medical Research; Westmead Hospital; University of Sydney | Arnott A., Draper J., Gall M., Martinez E., Rockett R., Sintchenko V., on behalf of ICPMR |
| EPI_ISL_3426167 | St Vincent's Pathology (SydPath) | NSW Health Pathology - Institute of Clinical Pathology and Medical Research; Westmead Hospital; University of Sydney | Arnott A., Draper J., Gall M., Martinez E., Rockett R., Sintchenko V., on behalf of ICPMR |
| EPI_ISL_3426168 | Laverty Pathology | NSW Health Pathology - Institute of Clinical Pathology and Medical Research; Westmead Hospital; University of Sydney | Arnott A., Draper J., Gall M., Martinez E., Rockett R., Sintchenko V., on behalf of ICPMR |
| EPI_ISL_3426169 | Capital Pathology | NSW Health Pathology - Institute of Clinical Pathology and Medical Research; Westmead Hospital; University of Sydney | Arnott A., Draper J., Gall M., Martinez E., Rockett R., Sintchenko V., on behalf of ICPMR |
| EPI_ISL_3426170, EPI_ISL_3426171, EPI_ISL_3426172 | Histopath | NSW Health Pathology - Institute of Clinical Pathology and Medical Research; Westmead Hospital; University of Sydney | Arnott A., Draper J., Gall M., Martinez E., Rockett R., Sintchenko V., on behalf of ICPMR |
| EPI_ISL_3426173 | 4Cyte Pathology | NSW Health Pathology - Institute of Clinical Pathology and Medical Research; Westmead Hospital; University of Sydney | Arnott A., Draper J., Gall M., Martinez E., Rockett R., Sintchenko V., on behalf of ICPMR |
| EPI_ISL_3426174 | Histopath | NSW Health Pathology - Institute of Clinical Pathology and Medical Research; Westmead Hospital; University of Sydney | Arnott A., Draper J., Gall M., Martinez E., Rockett R., Sintchenko V., on behalf of ICPMR |
| EPI_ISL_3426175 | The Children's Hospital at Westmead | NSW Health Pathology - Institute of Clinical Pathology and Medical Research; Westmead Hospital; University of Sydney | Arnott A., Draper J., Gall M., Martinez E., Rockett R., Sintchenko V., on behalf of ICPMR |
| EPI_ISL_3426176 | St Vincent's Pathology (SydPath) | NSW Health Pathology - Institute of Clinical Pathology and Medical Research; Westmead Hospital; University of Sydney | Arnott A., Draper J., Gall M., Martinez E., Rockett R., Sintchenko V., on behalf of ICPMR |
| EPI_ISL_3426177 | Australian Clinical Labs (formerly Healthscope Pathology) | NSW Health Pathology - Institute of Clinical Pathology and Medical Research; Westmead Hospital; University of Sydney | Arnott A., Draper J., Gall M., Martinez E., Rockett R., Sintchenko V., on behalf of ICPMR |
| EPI_ISL_3426178, EPI_ISL_3426179, EPI_ISL_3426180 | Histopath | NSW Health Pathology - Institute of Clinical Pathology and Medical Research; Westmead Hospital; University of Sydney | Arnott A., Draper J., Gall M., Martinez E., Rockett R., Sintchenko V., on behalf of ICPMR |
| EPI_ISL_3426181 | St Vincent's Pathology (SydPath) | NSW Health Pathology - Institute of Clinical Pathology and Medical Research; Westmead Hospital; University of Sydney | Arnott A., Draper J., Gall M., Martinez E., Rockett R., Sintchenko V., on behalf of ICPMR |
| EPI_ISL_3426182 | Medlab Pathology | NSW Health Pathology - Institute of Clinical Pathology and Medical Research; Westmead Hospital; University of Sydney | Arnott A., Draper J., Gall M., Martinez E., Rockett R., Sintchenko V., on behalf of ICPMR |
| EPI_ISL_3426183, EPI_ISL_3426184 | Histopath | NSW Health Pathology - Institute of Clinical Pathology and Medical Research; Westmead Hospital; University of Sydney | Arnott A., Draper J., Gall M., Martinez E., Rockett R., Sintchenko V., on behalf of ICPMR |
| EPI_ISL_3426185 | Capital Pathology | NSW Health Pathology - Institute of Clinical Pathology and Medical Research; Westmead Hospital; University of Sydney | Arnott A., Draper J., Gall M., Martinez E., Rockett R., Sintchenko V., on behalf of ICPMR |
| EPI_ISL_3426186 | 4Cyte Pathology | NSW Health Pathology - Institute of Clinical Pathology and Medical Research; Westmead Hospital; University of Sydney | Arnott A., Draper J., Gall M., Martinez E., Rockett R., Sintchenko V., on behalf of ICPMR |
| EPI_ISL_3426187 | St Vincent's Pathology (SydPath) | NSW Health Pathology - Institute of Clinical Pathology and Medical Research; Westmead Hospital; University of Sydney | Arnott A., Draper J., Gall M., Martinez E., Rockett R., Sintchenko V., on behalf of ICPMR |
| EPI_ISL_3426188 | 4Cyte Pathology | NSW Health Pathology - Institute of Clinical Pathology and Medical Research; Westmead Hospital; University of Sydney | Arnott A., Draper J., Gall M., Martinez E., Rockett R., Sintchenko V., on behalf of ICPMR |
| EPI_ISL_3426189 | Pathology North - Hunter - NSW Health Pathology | NSW Health Pathology - Institute of Clinical Pathology and Medical Research; Westmead Hospital; University of Sydney | Arnott A., Draper J., Gall M., Martinez E., Rockett R., Sintchenko V., on behalf of ICPMR |
| EPI_ISL_3426190, EPI_ISL_3426191, EPI_ISL_3426192, EPI_ISL_3426193, EPI_ISL_3426194, EPI_ISL_3426195, EPI_ISL_3426196, EPI_ISL_3426197, EPI_ISL_3426198, EPI_ISL_3426199, EPI_ISL_3426200, EPI_ISL_3426201, EPI_ISL_3426202, EPI_ISL_3426203, EPI_ISL_3426204, EPI_ISL_3426205, EPI_ISL_3426206, EPI_ISL_3426207, EPI_ISL_3426208, EPI_ISL_3426209, EPI_ISL_3426210, EPI_ISL_3426211, EPI_ISL_3426212, EPI_ISL_3426213, EPI_ISL_3426214, EPI_ISL_3426215 | | | |
| see above | 4Cyte Pathology | NSW Health Pathology - Institute of Clinical Pathology and Medical Research; Westmead Hospital; University of Sydney | Arnott A., Draper J., Gall M., Martinez E., Rockett R., Sintchenko V., on behalf of ICPMR |
| EPI_ISL_3426216, EPI_ISL_3426217, EPI_ISL_3426218, EPI_ISL_3426219, EPI_ISL_3426220, EPI_ISL_3426221 | Sydney South West Pathology Service (SSWPS) - Liverpool Hospital - NSW Health Pathology | NSW Health Pathology - Institute of Clinical Pathology and Medical Research; Westmead Hospital; University of Sydney | Arnott A., Draper J., Gall M., Martinez E., Rockett R., Sintchenko V., on behalf of ICPMR |
| EPI_ISL_3426222 | Capital Pathology | NSW Health Pathology - Institute of Clinical Pathology and Medical Research; Westmead Hospital; University of Sydney | Arnott A., Draper J., Gall M., Martinez E., Rockett R., Sintchenko V., on behalf of ICPMR |

| | | | |
|---|---|---|---|
| EPI_ISL_3426223, EPI_ISL_3426224, EPI_ISL_3426225, EPI_ISL_3426226, EPI_ISL_3426227, EPI_ISL_3426228, EPI_ISL_3426229, EPI_ISL_3426230, EPI_ISL_3426231, EPI_ISL_3426232, EPI_ISL_3426233, EPI_ISL_3426234, EPI_ISL_3426235, EPI_ISL_3426236, EPI_ISL_3426237, EPI_ISL_3426238, EPI_ISL_3426239, EPI_ISL_3426240, EPI_ISL_3426241 | | | |
| see above | Histopath | NSW Health Pathology - Institute of Clinical Pathology and Medical Research; Westmead Hospital; University of Sydney | Arnott A., Draper J., Gall M., Martinez E., Rockett R., Sintchenko V., on behalf of ICPMR |
| EPI_ISL_3426242, EPI_ISL_3426243, EPI_ISL_3426244, EPI_ISL_3426245, EPI_ISL_3426246, EPI_ISL_3426247, EPI_ISL_3426248 | 4Cyte Pathology | NSW Health Pathology - Institute of Clinical Pathology and Medical Research; Westmead Hospital; University of Sydney | Arnott A., Draper J., Gall M., Martinez E., Rockett R., Sintchenko V., on behalf of ICPMR |
| EPI_ISL_3426249, EPI_ISL_3426250, EPI_ISL_3426251, EPI_ISL_3426252, EPI_ISL_3426253, EPI_ISL_3426254 | Australian Clinical Labs (formerly Healthscope Pathology) | NSW Health Pathology - Institute of Clinical Pathology and Medical Research; Westmead Hospital; University of Sydney | Arnott A., Draper J., Gall M., Martinez E., Rockett R., Sintchenko V., on behalf of ICPMR |
| EPI_ISL_3426255 | Pathology North - Gosford Hospital - NSW Health Pathology | NSW Health Pathology - Institute of Clinical Pathology and Medical Research; Westmead Hospital; University of Sydney | Arnott A., Draper J., Gall M., Martinez E., Rockett R., Sintchenko V., on behalf of ICPMR |
| EPI_ISL_3426256, EPI_ISL_3426257, EPI_ISL_3426258, EPI_ISL_3426259, EPI_ISL_3426260, EPI_ISL_3426261, EPI_ISL_3426262, EPI_ISL_3426263, EPI_ISL_3426264, EPI_ISL_3426265, EPI_ISL_3426266, EPI_ISL_3426267, EPI_ISL_3426268, EPI_ISL_3426269, EPI_ISL_3426270, EPI_ISL_3426271, EPI_ISL_3426272 | | | |
| see above | Histopath | NSW Health Pathology - Institute of Clinical Pathology and Medical Research; Westmead Hospital; University of Sydney | Arnott A., Draper J., Gall M., Martinez E., Rockett R., Sintchenko V., on behalf of ICPMR |
| EPI_ISL_3426273, EPI_ISL_3426274 | Pathology North - Hunter - NSW Health Pathology | NSW Health Pathology - Institute of Clinical Pathology and Medical Research; Westmead Hospital; University of Sydney | Arnott A., Draper J., Gall M., Martinez E., Rockett R., Sintchenko V., on behalf of ICPMR |
| EPI_ISL_3426275, EPI_ISL_3426276, EPI_ISL_3426277, EPI_ISL_3426278, EPI_ISL_3426279, EPI_ISL_3426280, EPI_ISL_3426281, EPI_ISL_3426282, EPI_ISL_3426283, EPI_ISL_3426284, EPI_ISL_3426285, EPI_ISL_3426286, EPI_ISL_3426287, EPI_ISL_3426288 | | | |
| see above | Laverty Pathology | NSW Health Pathology - Institute of Clinical Pathology and Medical Research; Westmead Hospital; University of Sydney | Arnott A., Draper J., Gall M., Martinez E., Rockett R., Sintchenko V., on behalf of ICPMR |
| EPI_ISL_3426289, EPI_ISL_3426290, EPI_ISL_3426291, EPI_ISL_3426292 | Medlab Pathology | NSW Health Pathology - Institute of Clinical Pathology and Medical Research; Westmead Hospital; University of Sydney | Arnott A., Draper J., Gall M., Martinez E., Rockett R., Sintchenko V., on behalf of ICPMR |
| EPI_ISL_3426293 | Pathology North - Royal North Shore Hospital - NSW Health Pathology | NSW Health Pathology - Institute of Clinical Pathology and Medical Research; Westmead Hospital; University of Sydney | Arnott A., Draper J., Gall M., Martinez E., Rockett R., Sintchenko V., on behalf of ICPMR |
| EPI_ISL_3426294, EPI_ISL_3426295, EPI_ISL_3426296, EPI_ISL_3426297, EPI_ISL_3426298, EPI_ISL_3426299 | The Children's Hospital at Westmead | NSW Health Pathology - Institute of Clinical Pathology and Medical Research; Westmead Hospital; University of Sydney | Arnott A., Draper J., Gall M., Martinez E., Rockett R., Sintchenko V., on behalf of ICPMR |
| EPI_ISL_3426300, EPI_ISL_3426301, EPI_ISL_3426302, EPI_ISL_3426303, EPI_ISL_3426304, EPI_ISL_3426305, EPI_ISL_3426306 | St Vincent's Pathology (SydPath) | NSW Health Pathology - Institute of Clinical Pathology and Medical Research; Westmead Hospital; University of Sydney | Arnott A., Draper J., Gall M., Martinez E., Rockett R., Sintchenko V., on behalf of ICPMR |
| EPI_ISL_3426307, EPI_ISL_3426308, EPI_ISL_3426309, EPI_ISL_3426310, EPI_ISL_3426311, EPI_ISL_3426312, EPI_ISL_3426313, EPI_ISL_3426314 | Pathology West - NSW Health Pathology | NSW Health Pathology - Institute of Clinical Pathology and Medical Research; Westmead Hospital; University of Sydney | Arnott A., Draper J., Gall M., Martinez E., Rockett R., Sintchenko V., on behalf of ICPMR |
| EPI_ISL_3426315, EPI_ISL_3426316, EPI_ISL_3426317, EPI_ISL_3426318, EPI_ISL_3426319, EPI_ISL_3426320, EPI_ISL_3426321, EPI_ISL_3426322, EPI_ISL_3426323, EPI_ISL_3426324, EPI_ISL_3426325, EPI_ISL_3426326, EPI_ISL_3426327, EPI_ISL_3426328, EPI_ISL_3426329, EPI_ISL_3426330, EPI_ISL_3426331, EPI_ISL_3426332 | | | |
| see above | 4Cyte Pathology | NSW Health Pathology - Institute of Clinical Pathology and Medical Research; Westmead Hospital; University of Sydney | Arnott A., Draper J., Gall M., Martinez E., Rockett R., Sintchenko V., on behalf of ICPMR |
| EPI_ISL_3426333, EPI_ISL_3426334, EPI_ISL_3426335, EPI_ISL_3426336, EPI_ISL_3426337, EPI_ISL_3426338, EPI_ISL_3426339, EPI_ISL_3426340, EPI_ISL_3426341 | Australian Clinical Labs (formerly Healthscope Pathology) | NSW Health Pathology - Institute of Clinical Pathology and Medical Research; Westmead Hospital; University of Sydney | Arnott A., Draper J., Gall M., Martinez E., Rockett R., Sintchenko V., on behalf of ICPMR |
| EPI_ISL_3426342, EPI_ISL_3426343, EPI_ISL_3426344, EPI_ISL_3426345, EPI_ISL_3426346, EPI_ISL_3426347, EPI_ISL_3426348, EPI_ISL_3426349, EPI_ISL_3426350, EPI_ISL_3426351, EPI_ISL_3426352, EPI_ISL_3426353, EPI_ISL_3426354, EPI_ISL_3426355, EPI_ISL_3426356, EPI_ISL_3426357, EPI_ISL_3426358, EPI_ISL_3426359, EPI_ISL_3426360, EPI_ISL_3426361, EPI_ISL_3426362, EPI_ISL_3426363, EPI_ISL_3426364, EPI_ISL_3426365, EPI_ISL_3426366, EPI_ISL_3426367, EPI_ISL_3426368, EPI_ISL_3426369, EPI_ISL_3426370, EPI_ISL_3426371 | | | |
| see above | Histopath | NSW Health Pathology - Institute of Clinical Pathology and Medical Research; Westmead Hospital; University of Sydney | Arnott A., Draper J., Gall M., Martinez E., Rockett R., Sintchenko V., on behalf of ICPMR |
| EPI_ISL_3426372, EPI_ISL_3426373, EPI_ISL_3426374, EPI_ISL_3426375, EPI_ISL_3426376, EPI_ISL_3426377 | Pathology North - Hunter - NSW Health Pathology | NSW Health Pathology - Institute of Clinical Pathology and Medical Research; Westmead Hospital; University of Sydney | Arnott A., Draper J., Gall M., Martinez E., Rockett R., Sintchenko V., on behalf of ICPMR |
| EPI_ISL_3426378 | Laverty Pathology | NSW Health Pathology - Institute of Clinical Pathology and Medical Research; Westmead Hospital; University of Sydney | Arnott A., Draper J., Gall M., Martinez E., Rockett R., Sintchenko V., on behalf of ICPMR |
| EPI_ISL_3426379, EPI_ISL_3426380 | St Vincent's Pathology (SydPath) | NSW Health Pathology - Institute of Clinical Pathology and Medical Research; Westmead Hospital; University of Sydney | Arnott A., Draper J., Gall M., Martinez E., Rockett R., Sintchenko V., on behalf of ICPMR |
| EPI_ISL_3426381 | 4Cyte Pathology | NSW Health Pathology - Institute of Clinical Pathology and Medical Research; Westmead Hospital; University of Sydney | Arnott A., Draper J., Gall M., Martinez E., Rockett R., Sintchenko V., on behalf of ICPMR |
| EPI_ISL_3426382 | Laverty Pathology | NSW Health Pathology - Institute of Clinical Pathology and Medical Research; Westmead Hospital; University of Sydney | Arnott A., Draper J., Gall M., Martinez E., Rockett R., Sintchenko V., on behalf of ICPMR |
| EPI_ISL_3426383, EPI_ISL_3426384, EPI_ISL_3426385, EPI_ISL_3426386, EPI_ISL_3426387, EPI_ISL_3426388 | 4Cyte Pathology | NSW Health Pathology - Institute of Clinical Pathology and Medical Research; Westmead Hospital; University of Sydney | Arnott A., Draper J., Gall M., Martinez E., Rockett R., Sintchenko V., on behalf of ICPMR |
| EPI_ISL_3426389, EPI_ISL_3426390 | Australian Clinical Labs (formerly Healthscope Pathology) | NSW Health Pathology - Institute of Clinical Pathology and Medical Research; Westmead Hospital; University of Sydney | Arnott A., Draper J., Gall M., Martinez E., Rockett R., Sintchenko V., on behalf of ICPMR |
| EPI_ISL_3426391 | Austech Medical Laboratories | NSW Health Pathology - Institute of Clinical Pathology and Medical Research; Westmead Hospital; University of Sydney | Arnott A., Draper J., Gall M., Martinez E., Rockett R., Sintchenko V., on behalf of ICPMR |
| EPI_ISL_3426392 | Sydney South West Pathology Service (SSWPS) - Liverpool Hospital - NSW Health Pathology | NSW Health Pathology - Institute of Clinical Pathology and Medical Research; Westmead Hospital; University of Sydney | Arnott A., Draper J., Gall M., Martinez E., Rockett R., Sintchenko V., on behalf of ICPMR |
| EPI_ISL_3426393, EPI_ISL_3426394, | Histopath | NSW Health Pathology - Institute of Clinical Pathology and Medical Research; Westmead Hospital; University of Sydney | Arnott A., Draper J., Gall M., Martinez E., Rockett R., Sintchenko V., on behalf of ICPMR |

| | | | |
|---|---|---|---|
| EPI_ISL_3426395, EPI_ISL_3426396, EPI_ISL_3426397, EPI_ISL_3426398, EPI_ISL_3426399, EPI_ISL_3426400 | | | |
| EPI_ISL_3426401, EPI_ISL_3426402 | 4Cyte Pathology | NSW Health Pathology - Institute of Clinical Pathology and Medical Research; Westmead Hospital; University of Sydney | Arnott A., Draper J., Gall M., Martinez E., Rockett R., Sintchenko V., on behalf of ICPMR |
| EPI_ISL_3426403, EPI_ISL_3426404 | Capital Pathology | NSW Health Pathology - Institute of Clinical Pathology and Medical Research; Westmead Hospital; University of Sydney | Arnott A., Draper J., Gall M., Martinez E., Rockett R., Sintchenko V., on behalf of ICPMR |
| EPI_ISL_3426405 | Sydney South West Pathology Service (SSWPS) - Concord Repatriation General Hospital - NSW Health Pathology | NSW Health Pathology - Institute of Clinical Pathology and Medical Research; Westmead Hospital; University of Sydney | Arnott A., Draper J., Gall M., Martinez E., Rockett R., Sintchenko V., on behalf of ICPMR |
| EPI_ISL_3426406 | Histopath | NSW Health Pathology - Institute of Clinical Pathology and Medical Research; Westmead Hospital; University of Sydney | Arnott A., Draper J., Gall M., Martinez E., Rockett R., Sintchenko V., on behalf of ICPMR |
| EPI_ISL_3426407 | Australian Clinical Labs (formerly Healthscope Pathology) | NSW Health Pathology - Institute of Clinical Pathology and Medical Research; Westmead Hospital; University of Sydney | Arnott A., Draper J., Gall M., Martinez E., Rockett R., Sintchenko V., on behalf of ICPMR |
| EPI_ISL_3426408, EPI_ISL_3426409 | Sydney South West Pathology Service (SSWPS) - Concord Repatriation General Hospital - NSW Health Pathology | NSW Health Pathology - Institute of Clinical Pathology and Medical Research; Westmead Hospital; University of Sydney | Arnott A., Draper J., Gall M., Martinez E., Rockett R., Sintchenko V., on behalf of ICPMR |
| EPI_ISL_3426410, EPI_ISL_3426411, EPI_ISL_3426412 | Histopath | NSW Health Pathology - Institute of Clinical Pathology and Medical Research; Westmead Hospital; University of Sydney | Arnott A., Draper J., Gall M., Martinez E., Rockett R., Sintchenko V., on behalf of ICPMR |
| EPI_ISL_3426413 | The Children's Hospital at Westmead | NSW Health Pathology - Institute of Clinical Pathology and Medical Research; Westmead Hospital; University of Sydney | Arnott A., Draper J., Gall M., Martinez E., Rockett R., Sintchenko V., on behalf of ICPMR |
| EPI_ISL_3426414, EPI_ISL_3426415 | Pathology West - NSW Health Pathology | NSW Health Pathology - Institute of Clinical Pathology and Medical Research; Westmead Hospital; University of Sydney | Arnott A., Draper J., Gall M., Martinez E., Rockett R., Sintchenko V., on behalf of ICPMR |
| EPI_ISL_3426416, EPI_ISL_3426417 | 4Cyte Pathology | NSW Health Pathology - Institute of Clinical Pathology and Medical Research; Westmead Hospital; University of Sydney | Arnott A., Draper J., Gall M., Martinez E., Rockett R., Sintchenko V., on behalf of ICPMR |
| EPI_ISL_3426418, EPI_ISL_3426419, EPI_ISL_3426420 | Australian Clinical Labs (formerly Healthscope Pathology) | NSW Health Pathology - Institute of Clinical Pathology and Medical Research; Westmead Hospital; University of Sydney | Arnott A., Draper J., Gall M., Martinez E., Rockett R., Sintchenko V., on behalf of ICPMR |
| EPI_ISL_3426421, EPI_ISL_3426422, EPI_ISL_3426423, EPI_ISL_3426424, EPI_ISL_3426425, EPI_ISL_3426426, EPI_ISL_3426427, EPI_ISL_3426428, EPI_ISL_3426429, EPI_ISL_3426430 | Histopath | NSW Health Pathology - Institute of Clinical Pathology and Medical Research; Westmead Hospital; University of Sydney | Arnott A., Draper J., Gall M., Martinez E., Rockett R., Sintchenko V., on behalf of ICPMR |
| EPI_ISL_3426431 | Laverty Pathology | NSW Health Pathology - Institute of Clinical Pathology and Medical Research; Westmead Hospital; University of Sydney | Arnott A., Draper J., Gall M., Martinez E., Rockett R., Sintchenko V., on behalf of ICPMR |
| EPI_ISL_3426432 | Medlab Pathology | NSW Health Pathology - Institute of Clinical Pathology and Medical Research; Westmead Hospital; University of Sydney | Arnott A., Draper J., Gall M., Martinez E., Rockett R., Sintchenko V., on behalf of ICPMR |
| EPI_ISL_3426433, EPI_ISL_3426434 | Pathology West - NSW Health Pathology | NSW Health Pathology - Institute of Clinical Pathology and Medical Research; Westmead Hospital; University of Sydney | Arnott A., Draper J., Gall M., Martinez E., Rockett R., Sintchenko V., on behalf of ICPMR |
| EPI_ISL_3426435 | 4Cyte Pathology | NSW Health Pathology - Institute of Clinical Pathology and Medical Research; Westmead Hospital; University of Sydney | Arnott A., Draper J., Gall M., Martinez E., Rockett R., Sintchenko V., on behalf of ICPMR |
| EPI_ISL_3426436 | Sydney South West Pathology Service (SSWPS) - Liverpool Hospital - NSW Health Pathology | NSW Health Pathology - Institute of Clinical Pathology and Medical Research; Westmead Hospital; University of Sydney | Arnott A., Draper J., Gall M., Martinez E., Rockett R., Sintchenko V., on behalf of ICPMR |
| EPI_ISL_3426437 | Histopath | NSW Health Pathology - Institute of Clinical Pathology and Medical Research; Westmead Hospital; University of Sydney | Arnott A., Draper J., Gall M., Martinez E., Rockett R., Sintchenko V., on behalf of ICPMR |
| EPI_ISL_3426440, EPI_ISL_3426446, EPI_ISL_3426448, EPI_ISL_3426470, EPI_ISL_3426471, EPI_ISL_3426472, EPI_ISL_3426499, EPI_ISL_3426500, EPI_ISL_3426501, EPI_ISL_3426502, EPI_ISL_3426503, EPI_ISL_3426504, EPI_ISL_3426505, EPI_ISL_3426506, EPI_ISL_3426507, EPI_ISL_3426508, EPI_ISL_3426509, EPI_ISL_3426510, EPI_ISL_3426511, EPI_ISL_3426512, EPI_ISL_3426513, EPI_ISL_3426514, EPI_ISL_3426515, EPI_ISL_3426516, EPI_ISL_3426517, EPI_ISL_3426518, EPI_ISL_3426519, EPI_ISL_3426520, EPI_ISL_3426521, EPI_ISL_3426522, EPI_ISL_3426523, EPI_ISL_3426524, EPI_ISL_3426525, EPI_ISL_3426526, EPI_ISL_3426527, EPI_ISL_3426528, EPI_ISL_3426529, EPI_ISL_3426530, EPI_ISL_3426531, EPI_ISL_3426532, EPI_ISL_3426533, EPI_ISL_3426534, EPI_ISL_3426535, EPI_ISL_3426536, EPI_ISL_3426537, EPI_ISL_3426538, EPI_ISL_3426539, EPI_ISL_3426540, EPI_ISL_3426541, EPI_ISL_3426542, EPI_ISL_3426543, EPI_ISL_3426544 | | | |
| see above | Institute of Microbiology and Immunology, Faculty of Medicine, University of Ljubljana | Institute of Microbiology and Immunology, Faculty of Medicine, University of Ljubljana | Alen Suljič, Samo Zakotnik, Tomaž Mark Zorec, Matic Brvar, Doroteja Vlaj, Andraž Celar, Dominika Šturm, Patricija Pozvek, Špela Pleh, Miša Korva, Mario Poljak, Tatjana Avšič – Županc |
| EPI_ISL_3426550, EPI_ISL_3426551, EPI_ISL_3426552, EPI_ISL_3426553, EPI_ISL_3426554, EPI_ISL_3426555, EPI_ISL_3426556, EPI_ISL_3426557, EPI_ISL_3426558, EPI_ISL_3426559, EPI_ISL_3426560, EPI_ISL_3426561, EPI_ISL_3426562, EPI_ISL_3426563, EPI_ISL_3426564, EPI_ISL_3426565, EPI_ISL_3426566, EPI_ISL_3426568, EPI_ISL_3426570, EPI_ISL_3426571, EPI_ISL_3426572, EPI_ISL_3426573, EPI_ISL_3426574, EPI_ISL_3426575, EPI_ISL_3426576, EPI_ISL_3426577, EPI_ISL_3426578, EPI_ISL_3426579, EPI_ISL_3426580, EPI_ISL_3426581, EPI_ISL_3426582, EPI_ISL_3426583, EPI_ISL_3426584, EPI_ISL_3426585, EPI_ISL_3426586, EPI_ISL_3426587, EPI_ISL_3426588, EPI_ISL_3426589, EPI_ISL_3426590, EPI_ISL_3426591, EPI_ISL_3426592, EPI_ISL_3426593, EPI_ISL_3426594, EPI_ISL_3426595, EPI_ISL_3426596, EPI_ISL_3426597, EPI_ISL_3426598, EPI_ISL_3426599, EPI_ISL_3426600, EPI_ISL_3426601, EPI_ISL_3426602, EPI_ISL_3426603, EPI_ISL_3426604, EPI_ISL_3426605, EPI_ISL_3426606, EPI_ISL_3426607, EPI_ISL_3426608, EPI_ISL_3426609, EPI_ISL_3426610, EPI_ISL_3426620, EPI_ISL_3426621, EPI_ISL_3426622, EPI_ISL_3426623, EPI_ISL_3426624, EPI_ISL_3426625, EPI_ISL_3426626, EPI_ISL_3426627, EPI_ISL_3426628, EPI_ISL_3426629, EPI_ISL_3426630, EPI_ISL_3426631, EPI_ISL_3426632, EPI_ISL_3426633, EPI_ISL_3426634, EPI_ISL_3426635, EPI_ISL_3426636, EPI_ISL_3426637, EPI_ISL_3426638, EPI_ISL_3426639, EPI_ISL_3426640, EPI_ISL_3426641, EPI_ISL_3426642, EPI_ISL_3426643, EPI_ISL_3426644, EPI_ISL_3426645, EPI_ISL_3426646, EPI_ISL_3426647, EPI_ISL_3426648, EPI_ISL_3426649, EPI_ISL_3426650, EPI_ISL_3426651, EPI_ISL_3426652, EPI_ISL_3426653, EPI_ISL_3426654, EPI_ISL_3426655, EPI_ISL_3426656, EPI_ISL_3426657, EPI_ISL_3426658, EPI_ISL_3426659, EPI_ISL_3426660, EPI_ISL_3426661, EPI_ISL_3426662, EPI_ISL_3426663, EPI_ISL_3426664, EPI_ISL_3426665, EPI_ISL_3426666, EPI_ISL_3426667, EPI_ISL_3426668, EPI_ISL_3426669, EPI_ISL_3426670, EPI_ISL_3426671, EPI_ISL_3426672, EPI_ISL_3426674, EPI_ISL_3426675, EPI_ISL_3426676, EPI_ISL_3426677, EPI_ISL_3426678, EPI_ISL_3426679, EPI_ISL_3426680, EPI_ISL_3426681, EPI_ISL_3426682, EPI_ISL_3426683, EPI_ISL_3426684, EPI_ISL_3426685, EPI_ISL_3426686, EPI_ISL_3426687, EPI_ISL_3426688, EPI_ISL_3426689, EPI_ISL_3426690, EPI_ISL_3426691, EPI_ISL_3426692, EPI_ISL_3426693, EPI_ISL_3426694, EPI_ISL_3426695, EPI_ISL_3426696, EPI_ISL_3426697, EPI_ISL_3426698, EPI_ISL_3426699, EPI_ISL_3426700, EPI_ISL_3426701, EPI_ISL_3426702, EPI_ISL_3426703, EPI_ISL_3426704, EPI_ISL_3426705, EPI_ISL_3426707, EPI_ISL_3426708, EPI_ISL_3426709, EPI_ISL_3426710, EPI_ISL_3426711, EPI_ISL_3426712, EPI_ISL_3426713, EPI_ISL_3426714, EPI_ISL_3426715, EPI_ISL_3426716, EPI_ISL_3426717, EPI_ISL_3426718, EPI_ISL_3426719, EPI_ISL_3426720, EPI_ISL_3426721, EPI_ISL_3426722, EPI_ISL_3426723, EPI_ISL_3426724, EPI_ISL_3426725, EPI_ISL_3426726, EPI_ISL_3426727, EPI_ISL_3426729, EPI_ISL_3426730, EPI_ISL_3426731, EPI_ISL_3426732, EPI_ISL_3426733, EPI_ISL_3426734, EPI_ISL_3426735, EPI_ISL_3426736, EPI_ISL_3426737, EPI_ISL_3426738, EPI_ISL_3426739, EPI_ISL_3426740, EPI_ISL_3426741, EPI_ISL_3426742, EPI_ISL_3426743, EPI_ISL_3426744, EPI_ISL_3426745, EPI_ISL_3426746, EPI_ISL_3426747, EPI_ISL_3426748, EPI_ISL_3426749, EPI_ISL_3426750, EPI_ISL_3426751, EPI_ISL_3426752, EPI_ISL_3426753, EPI_ISL_3426754, EPI_ISL_3426755, EPI_ISL_3426756, EPI_ISL_3426757, EPI_ISL_3426758, EPI_ISL_3426759, EPI_ISL_3426760, EPI_ISL_3426761, EPI_ISL_3426762, EPI_ISL_3426763, EPI_ISL_3426764, EPI_ISL_3426765, EPI_ISL_3426766, EPI_ISL_3426767 | | | |
| see above | Microvida | Microvida | Suzan D. Pas, Jaco J. Verweij, Joep J. J. M. Stohr |
| EPI_ISL_3426768, EPI_ISL_3426769 | Innlandet Hospital Trust, Division Lillehammer, Department for Medical Microbiology | Norwegian Institute of Public Health, Department of Virology | Kathrine Stene-Johansen, Kamilla Heddeland Instefjord, Hilde Elshaug, Garcia Llorente Ignacio, Jon Bråte, Engebretsen Serina Beate,Pedersen Benedikte Nevjen, Line Victoria Moen, Debech Nadia, Atiya R Ali,Marie Paulsen Madsen, Rasmus Riis Kopperud, Hilde Vollan, Karoline Bragstad, Olav Hungnes |
| EPI_ISL_3426770 | Medical Microbiology Unit, Department for Laboratory Medicine, Drammen Hospital, Vestre Viken Health Trust, | Norwegian Institute of Public Health, Department of Virology | Kathrine Stene-Johansen, Kamilla Heddeland Instefjord, Hilde Elshaug, Garcia Llorente Ignacio, Jon Bråte, Engebretsen Serina Beate,Pedersen Benedikte Nevjen, Line Victoria Moen, Debech Nadia, Atiya R Ali,Marie Paulsen Madsen, Rasmus Riis Kopperud, Hilde Vollan, Karoline Bragstad, Olav Hungnes |
| EPI_ISL_3426771 | Furst Medical Laboratory | Norwegian Institute of Public Health, Department of Virology | Kathrine Stene-Johansen, Kamilla Heddeland Instefjord, Hilde Elshaug, Garcia Llorente Ignacio, Jon Bråte, Engebretsen Serina Beate,Pedersen Benedikte Nevjen, Line Victoria Moen, Debech Nadia, Atiya R Ali,Marie Paulsen Madsen, Rasmus Riis Kopperud, Hilde Vollan, Karoline Bragstad, Olav Hungnes |
| EPI_ISL_3426772 | Vestfold Hospital, Toensberg Department of Microbiology | Norwegian Institute of Public Health, Department of Virology | Kathrine Stene-Johansen, Kamilla Heddeland Instefjord, Hilde Elshaug, Garcia Llorente Ignacio, Jon Bråte, Engebretsen Serina Beate,Pedersen Benedikte Nevjen, Line Victoria Moen, Debech Nadia, Atiya R Ali,Marie Paulsen Madsen, Rasmus Riis Kopperud, Hilde Vollan, Karoline Bragstad, Olav Hungnes |
| EPI_ISL_3426773 | Furst Medical Laboratory | Norwegian Institute of Public Health, Department of Virology | Kathrine Stene-Johansen, Kamilla Heddeland Instefjord, Hilde Elshaug, Garcia Llorente Ignacio, Jon Bråte, Engebretsen Serina Beate,Pedersen Benedikte Nevjen, Line Victoria Moen, Debech Nadia, Atiya R Ali,Marie Paulsen Madsen, Rasmus Riis Kopperud, Hilde Vollan, Karoline Bragstad, Olav Hungnes |
| EPI_ISL_3426775 | Nordland Hospital - Bodo, Laboratory Department, Molecular Biology Unit | Norwegian Institute of Public Health, Department of Virology | Kathrine Stene-Johansen, Kamilla Heddeland Instefjord, Hilde Elshaug, Garcia Llorente Ignacio, Jon Bråte, Engebretsen Serina Beate,Pedersen Benedikte Nevjen, Line Victoria Moen, Debech Nadia, Atiya R Ali,Marie Paulsen Madsen, Rasmus Riis Kopperud, Hilde Vollan, Karoline Bragstad, Olav Hungnes |
| EPI_ISL_3426776 | Medical Microbiology Unit, Department for Laboratory Medicine, Drammen Hospital, Vestre Viken Health Trust, | Norwegian Institute of Public Health, Department of Virology | Kathrine Stene-Johansen, Kamilla Heddeland Instefjord, Hilde Elshaug, Garcia Llorente Ignacio, Jon Bråte, Engebretsen Serina Beate,Pedersen Benedikte Nevjen, Line Victoria Moen, Debech Nadia, Atiya R Ali,Marie Paulsen Madsen, Rasmus Riis Kopperud, Hilde Vollan, Karoline Bragstad, Olav Hungnes |
| EPI_ISL_3426777 | Nordland Hospital - Bodo, Laboratory Department, Molecular Biology Unit | Norwegian Institute of Public Health, Department of Virology | Kathrine Stene-Johansen, Kamilla Heddeland Instefjord, Hilde Elshaug, Garcia Llorente Ignacio, Jon Bråte, Engebretsen Serina Beate,Pedersen Benedikte Nevjen, Line Victoria Moen, Debech Nadia, Atiya R Ali,Marie Paulsen Madsen, Rasmus Riis Kopperud, Hilde Vollan, Karoline Bragstad, Olav Hungnes |
| EPI_ISL_3426779 | Unilabs Laboratory Medicine | Norwegian Institute of Public Health, Department of Virology | Kathrine Stene-Johansen, Kamilla Heddeland Instefjord, Hilde Elshaug, Garcia Llorente Ignacio, Jon Bråte, Engebretsen Serina Beate,Pedersen Benedikte Nevjen, Line Victoria Moen, Debech Nadia, Atiya R Ali,Marie Paulsen Madsen, Rasmus Riis Kopperud, Hilde Vollan, Karoline Bragstad, Olav Hungnes |
| EPI_ISL_3426780, EPI_ISL_3426781 | Vestfold Hospital, Toensberg Department of Microbiology | Norwegian Institute of Public Health, Department of Virology | Kathrine Stene-Johansen, Kamilla Heddeland Instefjord, Hilde Elshaug, Garcia Llorente Ignacio, Jon Bråte, Engebretsen Serina Beate,Pedersen Benedikte Nevjen, Line Victoria Moen, Debech Nadia, Atiya R Ali,Marie Paulsen Madsen, Rasmus Riis Kopperud, Hilde Vollan, Karoline Bragstad, Olav Hungnes |
| EPI_ISL_3426782, EPI_ISL_3426784 | Unilabs Laboratory Medicine | Norwegian Institute of Public Health, Department of Virology | Kathrine Stene-Johansen, Kamilla Heddeland Instefjord, Hilde Elshaug, Garcia Llorente Ignacio, Jon Bråte, Engebretsen Serina Beate,Pedersen Benedikte Nevjen, Line Victoria Moen, Debech Nadia, Atiya R Ali,Marie Paulsen Madsen, Rasmus Riis Kopperud, Hilde Vollan, Karoline Bragstad, Olav Hungnes |
| EPI_ISL_3426785 | Furst Medical Laboratory | Norwegian Institute of Public Health, Department of Virology | Kathrine Stene-Johansen, Kamilla Heddeland Instefjord, Hilde Elshaug, Garcia Llorente Ignacio, Jon Bråte, Engebretsen Serina Beate,Pedersen Benedikte Nevjen, Line Victoria Moen, Debech Nadia, Atiya R Ali,Marie Paulsen Madsen, Rasmus Riis Kopperud, Hilde Vollan, Karoline Bragstad, Olav Hungnes |
| EPI_ISL_3426786 | Unilabs Laboratory Medicine | Norwegian Institute of Public Health, Department of Virology | Kathrine Stene-Johansen, Kamilla Heddeland Instefjord, Hilde Elshaug, Garcia Llorente Ignacio, Jon Bråte, Engebretsen Serina Beate,Pedersen Benedikte Nevjen, Line Victoria Moen, Debech Nadia, Atiya R Ali,Marie Paulsen Madsen, Rasmus Riis Kopperud, Hilde Vollan, Karoline Bragstad, Olav Hungnes |

| | | | |
|---|---|---|---|
| EPI_ISL_3426787 | Medical Microbiology Unit, Department for Laboratory Medicine, Drammen Hospital, Vestre Viken Health Trust, | Norwegian Institute of Public Health, Department of Virology | Kathrine Stene-Johansen, Kamilla Heddeland Instefjord, Hilde Elshaug, Garcia Llorente Ignacio, Jon Bråte, Engebretsen Serina Beate,Pedersen Benedikte Nevjen, Line Victoria Moen, Debech Nadia, Atiya R Ali,Marie Paulsen Madsen, Rasmus Riis Kopperud, Hilde Vollan, Karoline Bragstad, Olav Hungnes |
| EPI_ISL_3426788 | Furst Medical Laboratory | Norwegian Institute of Public Health, Department of Virology | Kathrine Stene-Johansen, Kamilla Heddeland Instefjord, Hilde Elshaug, Garcia Llorente Ignacio, Jon Bråte, Engebretsen Serina Beate,Pedersen Benedikte Nevjen, Line Victoria Moen, Debech Nadia, Atiya R Ali,Marie Paulsen Madsen, Rasmus Riis Kopperud, Hilde Vollan, Karoline Bragstad, Olav Hungnes |
| EPI_ISL_3426789 | Medical Microbiology Unit, Department for Laboratory Medicine, Drammen Hospital, Vestre Viken Health Trust, | Norwegian Institute of Public Health, Department of Virology | Kathrine Stene-Johansen, Kamilla Heddeland Instefjord, Hilde Elshaug, Garcia Llorente Ignacio, Jon Bråte, Engebretsen Serina Beate,Pedersen Benedikte Nevjen, Line Victoria Moen, Debech Nadia, Atiya R Ali,Marie Paulsen Madsen, Rasmus Riis Kopperud, Hilde Vollan, Karoline Bragstad, Olav Hungnes |
| EPI_ISL_3426790 | Furst Medical Laboratory | Norwegian Institute of Public Health, Department of Virology | Kathrine Stene-Johansen, Kamilla Heddeland Instefjord, Hilde Elshaug, Garcia Llorente Ignacio, Jon Bråte, Engebretsen Serina Beate,Pedersen Benedikte Nevjen, Line Victoria Moen, Debech Nadia, Atiya R Ali,Marie Paulsen Madsen, Rasmus Riis Kopperud, Hilde Vollan, Karoline Bragstad, Olav Hungnes |
| EPI_ISL_3426791, EPI_ISL_3426792 | Department of Medical Microbiology - section Molde, Molde Hospital | Norwegian Institute of Public Health, Department of Virology | Kathrine Stene-Johansen, Kamilla Heddeland Instefjord, Hilde Elshaug, Garcia Llorente Ignacio, Jon Bråte, Engebretsen Serina Beate,Pedersen Benedikte Nevjen, Line Victoria Moen, Debech Nadia, Atiya R Ali,Marie Paulsen Madsen, Rasmus Riis Kopperud, Hilde Vollan, Karoline Bragstad, Olav Hungnes |
| EPI_ISL_3426793 | Haukeland University Hospital, Dept. of Microbiology | Norwegian Institute of Public Health, Department of Virology | Kathrine Stene-Johansen, Kamilla Heddeland Instefjord, Hilde Elshaug, Garcia Llorente Ignacio, Jon Bråte, Engebretsen Serina Beate,Pedersen Benedikte Nevjen, Line Victoria Moen, Debech Nadia, Atiya R Ali,Marie Paulsen Madsen, Rasmus Riis Kopperud, Hilde Vollan, Karoline Bragstad, Olav Hungnes |
| EPI_ISL_3426794 | Department of Medical Microbiology - section Molde, Molde Hospital | Norwegian Institute of Public Health, Department of Virology | Kathrine Stene-Johansen, Kamilla Heddeland Instefjord, Hilde Elshaug, Garcia Llorente Ignacio, Jon Bråte, Engebretsen Serina Beate,Pedersen Benedikte Nevjen, Line Victoria Moen, Debech Nadia, Atiya R Ali,Marie Paulsen Madsen, Rasmus Riis Kopperud, Hilde Vollan, Karoline Bragstad, Olav Hungnes |
| EPI_ISL_3426795 | Haukeland University Hospital, Dept. of Microbiology | Norwegian Institute of Public Health, Department of Virology | Kathrine Stene-Johansen, Kamilla Heddeland Instefjord, Hilde Elshaug, Garcia Llorente Ignacio, Jon Bråte, Engebretsen Serina Beate,Pedersen Benedikte Nevjen, Line Victoria Moen, Debech Nadia, Atiya R Ali,Marie Paulsen Madsen, Rasmus Riis Kopperud, Hilde Vollan, Karoline Bragstad, Olav Hungnes |
| EPI_ISL_3426796, EPI_ISL_3426797, EPI_ISL_3426798 | Department of Medical Microbiology - section Molde, Molde Hospital | Norwegian Institute of Public Health, Department of Virology | Kathrine Stene-Johansen, Kamilla Heddeland Instefjord, Hilde Elshaug, Garcia Llorente Ignacio, Jon Bråte, Engebretsen Serina Beate,Pedersen Benedikte Nevjen, Line Victoria Moen, Debech Nadia, Atiya R Ali,Marie Paulsen Madsen, Rasmus Riis Kopperud, Hilde Vollan, Karoline Bragstad, Olav Hungnes |
| EPI_ISL_3426799 | Haukeland University Hospital, Dept. of Microbiology | Norwegian Institute of Public Health, Department of Virology | Kathrine Stene-Johansen, Kamilla Heddeland Instefjord, Hilde Elshaug, Garcia Llorente Ignacio, Jon Bråte, Engebretsen Serina Beate,Pedersen Benedikte Nevjen, Line Victoria Moen, Debech Nadia, Atiya R Ali,Marie Paulsen Madsen, Rasmus Riis Kopperud, Hilde Vollan, Karoline Bragstad, Olav Hungnes |
| EPI_ISL_3426800, EPI_ISL_3426801 | Department of Medical Microbiology - section Molde, Molde Hospital | Norwegian Institute of Public Health, Department of Virology | Kathrine Stene-Johansen, Kamilla Heddeland Instefjord, Hilde Elshaug, Garcia Llorente Ignacio, Jon Bråte, Engebretsen Serina Beate,Pedersen Benedikte Nevjen, Line Victoria Moen, Debech Nadia, Atiya R Ali,Marie Paulsen Madsen, Rasmus Riis Kopperud, Hilde Vollan, Karoline Bragstad, Olav Hungnes |
| EPI_ISL_3426802 | Haukeland University Hospital, Dept. of Microbiology | Norwegian Institute of Public Health, Department of Virology | Kathrine Stene-Johansen, Kamilla Heddeland Instefjord, Hilde Elshaug, Garcia Llorente Ignacio, Jon Bråte, Engebretsen Serina Beate,Pedersen Benedikte Nevjen, Line Victoria Moen, Debech Nadia, Atiya R Ali,Marie Paulsen Madsen, Rasmus Riis Kopperud, Hilde Vollan, Karoline Bragstad, Olav Hungnes |
| EPI_ISL_3426803 | Department of Medical Microbiology - section Molde, Molde Hospital | Norwegian Institute of Public Health, Department of Virology | Kathrine Stene-Johansen, Kamilla Heddeland Instefjord, Hilde Elshaug, Garcia Llorente Ignacio, Jon Bråte, Engebretsen Serina Beate,Pedersen Benedikte Nevjen, Line Victoria Moen, Debech Nadia, Atiya R Ali,Marie Paulsen Madsen, Rasmus Riis Kopperud, Hilde Vollan, Karoline Bragstad, Olav Hungnes |
| EPI_ISL_3426804 | Haukeland University Hospital, Dept. of Microbiology | Norwegian Institute of Public Health, Department of Virology | Kathrine Stene-Johansen, Kamilla Heddeland Instefjord, Hilde Elshaug, Garcia Llorente Ignacio, Jon Bråte, Engebretsen Serina Beate,Pedersen Benedikte Nevjen, Line Victoria Moen, Debech Nadia, Atiya R Ali,Marie Paulsen Madsen, Rasmus Riis Kopperud, Hilde Vollan, Karoline Bragstad, Olav Hungnes |
| EPI_ISL_3426805, EPI_ISL_3426806 | Department of Medical Microbiology - section Molde, Molde Hospital | Norwegian Institute of Public Health, Department of Virology | Kathrine Stene-Johansen, Kamilla Heddeland Instefjord, Hilde Elshaug, Garcia Llorente Ignacio, Jon Bråte, Engebretsen Serina Beate,Pedersen Benedikte Nevjen, Line Victoria Moen, Debech Nadia, Atiya R Ali,Marie Paulsen Madsen, Rasmus Riis Kopperud, Hilde Vollan, Karoline Bragstad, Olav Hungnes |
| EPI_ISL_3426807, EPI_ISL_3426808 | Nordland Hospital - Bodo, Laboratory Department, Molecular Biology Unit | Norwegian Institute of Public Health, Department of Virology | Kathrine Stene-Johansen, Kamilla Heddeland Instefjord, Hilde Elshaug, Garcia Llorente Ignacio, Jon Bråte, Engebretsen Serina Beate,Pedersen Benedikte Nevjen, Line Victoria Moen, Debech Nadia, Atiya R Ali,Marie Paulsen Madsen, Rasmus Riis Kopperud, Hilde Vollan, Karoline Bragstad, Olav Hungnes |
| EPI_ISL_3426809 | Unilabs Laboratory Medicine | Norwegian Institute of Public Health, Department of Virology | Kathrine Stene-Johansen, Kamilla Heddeland Instefjord, Hilde Elshaug, Garcia Llorente Ignacio, Jon Bråte, Engebretsen Serina Beate,Pedersen Benedikte Nevjen, Line Victoria Moen, Debech Nadia, Atiya R Ali,Marie Paulsen Madsen, Rasmus Riis Kopperud, Hilde Vollan, Karoline Bragstad, Olav Hungnes |
| EPI_ISL_3426810 | Nordland Hospital - Bodo, Laboratory Department, Molecular Biology Unit | Norwegian Institute of Public Health, Department of Virology | Kathrine Stene-Johansen, Kamilla Heddeland Instefjord, Hilde Elshaug, Garcia Llorente Ignacio, Jon Bråte, Engebretsen Serina Beate,Pedersen Benedikte Nevjen, Line Victoria Moen, Debech Nadia, Atiya R Ali,Marie Paulsen Madsen, Rasmus Riis Kopperud, Hilde Vollan, Karoline Bragstad, Olav Hungnes |
| EPI_ISL_3426811 | Vestfold Hospital, Toensberg Department of Microbiology | Norwegian Institute of Public Health, Department of Virology | Kathrine Stene-Johansen, Kamilla Heddeland Instefjord, Hilde Elshaug, Garcia Llorente Ignacio, Jon Bråte, Engebretsen Serina Beate,Pedersen Benedikte Nevjen, Line Victoria Moen, Debech Nadia, Atiya R Ali,Marie Paulsen Madsen, Rasmus Riis Kopperud, Hilde Vollan, Karoline Bragstad, Olav Hungnes |
| EPI_ISL_3426812 | Medical Microbiology Unit, Department for Laboratory Medicine, Drammen Hospital, Vestre Viken Health Trust, | Norwegian Institute of Public Health, Department of Virology | Kathrine Stene-Johansen, Kamilla Heddeland Instefjord, Hilde Elshaug, Garcia Llorente Ignacio, Jon Bråte, Engebretsen Serina Beate,Pedersen Benedikte Nevjen, Line Victoria Moen, Debech Nadia, Atiya R Ali,Marie Paulsen Madsen, Rasmus Riis Kopperud, Hilde Vollan, Karoline Bragstad, Olav Hungnes |
| EPI_ISL_3426813 | Unilabs Laboratory Medicine | Norwegian Institute of Public Health, Department of Virology | Kathrine Stene-Johansen, Kamilla Heddeland Instefjord, Hilde Elshaug, Garcia Llorente Ignacio, Jon Bråte, Engebretsen Serina Beate,Pedersen Benedikte Nevjen, Line Victoria Moen, Debech Nadia, Atiya R Ali,Marie Paulsen Madsen, Rasmus Riis Kopperud, Hilde Vollan, Karoline Bragstad, Olav Hungnes |
| EPI_ISL_3426814 | Nordland Hospital - Bodo, Laboratory Department, Molecular Biology Unit | Norwegian Institute of Public Health, Department of Virology | Kathrine Stene-Johansen, Kamilla Heddeland Instefjord, Hilde Elshaug, Garcia Llorente Ignacio, Jon Bråte, Engebretsen Serina Beate,Pedersen Benedikte Nevjen, Line Victoria Moen, Debech Nadia, Atiya R Ali,Marie Paulsen Madsen, Rasmus Riis Kopperud, Hilde Vollan, Karoline Bragstad, Olav Hungnes |
| EPI_ISL_3426815 | Vestfold Hospital, Toensberg Department of Microbiology | Norwegian Institute of Public Health, Department of Virology | Kathrine Stene-Johansen, Kamilla Heddeland Instefjord, Hilde Elshaug, Garcia Llorente Ignacio, Jon Bråte, Engebretsen Serina Beate,Pedersen Benedikte Nevjen, Line Victoria Moen, Debech Nadia, Atiya R Ali,Marie Paulsen Madsen, Rasmus Riis Kopperud, Hilde Vollan, Karoline Bragstad, Olav Hungnes |
| EPI_ISL_3426816 | Medical Microbiology Unit, Department for Laboratory Medicine, Drammen Hospital, Vestre Viken Health Trust, | Norwegian Institute of Public Health, Department of Virology | Kathrine Stene-Johansen, Kamilla Heddeland Instefjord, Hilde Elshaug, Garcia Llorente Ignacio, Jon Bråte, Engebretsen Serina Beate,Pedersen Benedikte Nevjen, Line Victoria Moen, Debech Nadia, Atiya R Ali,Marie Paulsen Madsen, Rasmus Riis Kopperud, Hilde Vollan, Karoline Bragstad, Olav Hungnes |
| EPI_ISL_3426817 | Furst Medical Laboratory | Norwegian Institute of Public Health, Department of Virology | Kathrine Stene-Johansen, Kamilla Heddeland Instefjord, Hilde Elshaug, Garcia Llorente Ignacio, Jon Bråte, Engebretsen Serina Beate,Pedersen Benedikte Nevjen, Line Victoria Moen, Debech Nadia, Atiya R Ali,Marie Paulsen Madsen, Rasmus Riis Kopperud, Hilde Vollan, Karoline Bragstad, Olav Hungnes |
| EPI_ISL_3426818 | Department of Medical Microbiology - section Molde, Molde Hospital | Norwegian Institute of Public Health, Department of Virology | Kathrine Stene-Johansen, Kamilla Heddeland Instefjord, Hilde Elshaug, Garcia Llorente Ignacio, Jon Bråte, Engebretsen Serina Beate,Pedersen Benedikte Nevjen, Line Victoria Moen, Debech Nadia, Atiya R Ali,Marie Paulsen Madsen, Rasmus Riis Kopperud, Hilde Vollan, Karoline Bragstad, Olav Hungnes |
| EPI_ISL_3426819 | Vestfold Hospital, Toensberg Department of Microbiology | Norwegian Institute of Public Health, Department of Virology | Kathrine Stene-Johansen, Kamilla Heddeland Instefjord, Hilde Elshaug, Garcia Llorente Ignacio, Jon Bråte, Engebretsen Serina Beate,Pedersen Benedikte Nevjen, Line Victoria Moen, Debech Nadia, Atiya R Ali,Marie Paulsen Madsen, Rasmus Riis Kopperud, Hilde Vollan, Karoline Bragstad, Olav Hungnes |
| EPI_ISL_3426820 | Department of Medical Microbiology - section Molde, Molde Hospital | Norwegian Institute of Public Health, Department of Virology | Kathrine Stene-Johansen, Kamilla Heddeland Instefjord, Hilde Elshaug, Garcia Llorente Ignacio, Jon Bråte, Engebretsen Serina Beate,Pedersen Benedikte Nevjen, Line Victoria Moen, Debech Nadia, Atiya R Ali,Marie Paulsen Madsen, Rasmus Riis Kopperud, Hilde Vollan, Karoline Bragstad, Olav Hungnes |
| EPI_ISL_3426821 | Vestfold Hospital, Toensberg Department of Microbiology | Norwegian Institute of Public Health, Department of Virology | Kathrine Stene-Johansen, Kamilla Heddeland Instefjord, Hilde Elshaug, Garcia Llorente Ignacio, Jon Bråte, Engebretsen Serina Beate,Pedersen Benedikte Nevjen, Line Victoria Moen, Debech Nadia, Atiya R Ali,Marie Paulsen Madsen, Rasmus Riis Kopperud, Hilde Vollan, Karoline Bragstad, Olav Hungnes |
| EPI_ISL_3426822 | Department of Medical Microbiology - section Molde, Molde Hospital | Norwegian Institute of Public Health, Department of Virology | Kathrine Stene-Johansen, Kamilla Heddeland Instefjord, Hilde Elshaug, Garcia Llorente Ignacio, Jon Bråte, Engebretsen Serina Beate,Pedersen Benedikte Nevjen, Line Victoria Moen, Debech Nadia, Atiya R Ali,Marie Paulsen Madsen, Rasmus Riis Kopperud, Hilde Vollan, Karoline Bragstad, Olav Hungnes |
| EPI_ISL_3426823 | Vestfold Hospital, Toensberg Department of Microbiology | Norwegian Institute of Public Health, Department of Virology | Kathrine Stene-Johansen, Kamilla Heddeland Instefjord, Hilde Elshaug, Garcia Llorente Ignacio, Jon Bråte, Engebretsen Serina Beate,Pedersen Benedikte Nevjen, Line Victoria Moen, Debech Nadia, Atiya R Ali,Marie Paulsen Madsen, Rasmus Riis Kopperud, Hilde Vollan, Karoline Bragstad, Olav Hungnes |
| EPI_ISL_3426824 | Medical Microbiology Unit, Department for Laboratory Medicine, Drammen Hospital, Vestre Viken Health Trust, | Norwegian Institute of Public Health, Department of Virology | Kathrine Stene-Johansen, Kamilla Heddeland Instefjord, Hilde Elshaug, Garcia Llorente Ignacio, Jon Bråte, Engebretsen Serina Beate,Pedersen Benedikte Nevjen, Line Victoria Moen, Debech Nadia, Atiya R Ali,Marie Paulsen Madsen, Rasmus Riis Kopperud, Hilde Vollan, Karoline Bragstad, Olav Hungnes |
| EPI_ISL_3426825 | Department of Medical Microbiology - section Molde, Molde Hospital | Norwegian Institute of Public Health, Department of Virology | Kathrine Stene-Johansen, Kamilla Heddeland Instefjord, Hilde Elshaug, Garcia Llorente Ignacio, Jon Bråte, Engebretsen Serina Beate,Pedersen Benedikte Nevjen, Line Victoria Moen, Debech Nadia, Atiya R Ali,Marie Paulsen Madsen, Rasmus Riis Kopperud, Hilde Vollan, Karoline Bragstad, Olav Hungnes |
| EPI_ISL_3426826 | Vestfold Hospital, Toensberg Department of Microbiology | Norwegian Institute of Public Health, Department of Virology | Kathrine Stene-Johansen, Kamilla Heddeland Instefjord, Hilde Elshaug, Garcia Llorente Ignacio, Jon Bråte, Engebretsen Serina Beate,Pedersen Benedikte Nevjen, Line Victoria Moen, Debech Nadia, Atiya R Ali,Marie Paulsen Madsen, Rasmus Riis Kopperud, Hilde Vollan, Karoline Bragstad, Olav Hungnes |
| EPI_ISL_3426827 | Unilabs Laboratory Medicine | Norwegian Institute of Public Health, Department of Virology | Kathrine Stene-Johansen, Kamilla Heddeland Instefjord, Hilde Elshaug, Garcia Llorente Ignacio, Jon Bråte, Engebretsen Serina Beate,Pedersen Benedikte Nevjen, Line Victoria Moen, Debech Nadia, Atiya R Ali,Marie Paulsen Madsen, Rasmus Riis Kopperud, Hilde Vollan, Karoline Bragstad, Olav Hungnes |
| EPI_ISL_3426828 | Department of Medical Microbiology - section Molde, Molde Hospital | Norwegian Institute of Public Health, Department of Virology | Kathrine Stene-Johansen, Kamilla Heddeland Instefjord, Hilde Elshaug, Garcia Llorente Ignacio, Jon Bråte, Engebretsen Serina Beate,Pedersen Benedikte Nevjen, Line Victoria Moen, Debech Nadia, Atiya R Ali,Marie Paulsen Madsen, Rasmus Riis Kopperud, Hilde Vollan, Karoline Bragstad, Olav Hungnes |
| EPI_ISL_3426830 | Vestfold Hospital, Toensberg Department of Microbiology | Norwegian Institute of Public Health, Department of Virology | Kathrine Stene-Johansen, Kamilla Heddeland Instefjord, Hilde Elshaug, Garcia Llorente Ignacio, Jon Bråte, Engebretsen Serina Beate,Pedersen Benedikte Nevjen, Line Victoria Moen, Debech Nadia, Atiya R Ali,Marie Paulsen Madsen, Rasmus Riis Kopperud, Hilde Vollan, Karoline Bragstad, Olav Hungnes |
| EPI_ISL_3426831, EPI_ISL_3426832, EPI_ISL_3426833, EPI_ISL_3426834, EPI_ISL_3426835, EPI_ISL_3426836, EPI_ISL_3426837 | Department of Medical Microbiology - section Molde, Molde Hospital | Norwegian Institute of Public Health, Department of Virology | Kathrine Stene-Johansen, Kamilla Heddeland Instefjord, Hilde Elshaug, Garcia Llorente Ignacio, Jon Bråte, Engebretsen Serina Beate,Pedersen Benedikte Nevjen, Line Victoria Moen, Debech Nadia, Atiya R Ali,Marie Paulsen Madsen, Rasmus Riis Kopperud, Hilde Vollan, Karoline Bragstad, Olav Hungnes |
| EPI_ISL_3426838 | Haukeland University Hospital, Dept. of Microbiology | Norwegian Institute of Public Health, Department of Virology | Kathrine Stene-Johansen, Kamilla Heddeland Instefjord, Hilde Elshaug, Garcia Llorente Ignacio, Jon Bråte, Engebretsen Serina Beate,Pedersen Benedikte Nevjen, Line Victoria Moen, Debech Nadia, Atiya R Ali,Marie Paulsen Madsen, Rasmus Riis Kopperud, Hilde Vollan, Karoline Bragstad, Olav Hungnes |
| EPI_ISL_3426839 | Department of Medical Microbiology - section Molde, Molde Hospital | Norwegian Institute of Public Health, Department of Virology | Kathrine Stene-Johansen, Kamilla Heddeland Instefjord, Hilde Elshaug, Garcia Llorente Ignacio, Jon Bråte, Engebretsen Serina Beate,Pedersen Benedikte Nevjen, Line Victoria Moen, Debech Nadia, Atiya R Ali,Marie Paulsen Madsen, Rasmus Riis Kopperud, Hilde Vollan, Karoline Bragstad, Olav Hungnes |
| EPI_ISL_3426840, EPI_ISL_3426841, EPI_ISL_3426842 | Medical Microbiology Unit, Department for Laboratory Medicine, Drammen Hospital, Vestre Viken Health Trust, | Norwegian Institute of Public Health, Department of Virology | Kathrine Stene-Johansen, Kamilla Heddeland Instefjord, Hilde Elshaug, Garcia Llorente Ignacio, Jon Bråte, Engebretsen Serina Beate,Pedersen Benedikte Nevjen, Line Victoria Moen, Debech Nadia, Atiya R Ali,Marie Paulsen Madsen, Rasmus Riis Kopperud, Hilde Vollan, Karoline Bragstad, Olav Hungnes |
| EPI_ISL_3426843 | Department of Medical Microbiology - section Molde, Molde Hospital | Norwegian Institute of Public Health, Department of Virology | Kathrine Stene-Johansen, Kamilla Heddeland Instefjord, Hilde Elshaug, Garcia Llorente Ignacio, Jon Bråte, Engebretsen Serina Beate,Pedersen Benedikte Nevjen, Line Victoria Moen, Debech Nadia, Atiya R Ali,Marie Paulsen Madsen, Rasmus Riis Kopperud, Hilde Vollan, Karoline Bragstad, Olav Hungnes |

| | | | |
|---|---|---|---|
| EPI_ISL_3426844, EPI_ISL_3426845 | Haukeland University Hospital, Dept. of Microbiology | Norwegian Institute of Public Health, Department of Virology | Kathrine Stene-Johansen, Kamilla Heddeland Instefjord, Hilde Elshaug, Garcia Llorente Ignacio, Jon Bråte, Engebretsen Serina Beate,Pedersen Benedikte Nevjen, Line Victoria Moen, Debech Nadia, Atiya R Ali,Marie Paulsen Madsen, Rasmus Riis Kopperud, Hilde Vollan, Karoline Bragstad, Olav Hungnes |
| EPI_ISL_3426846 | Department of Medical Microbiology - section Molde, Molde Hospital | Norwegian Institute of Public Health, Department of Virology | Kathrine Stene-Johansen, Kamilla Heddeland Instefjord, Hilde Elshaug, Garcia Llorente Ignacio, Jon Bråte, Engebretsen Serina Beate,Pedersen Benedikte Nevjen, Line Victoria Moen, Debech Nadia, Atiya R Ali,Marie Paulsen Madsen, Rasmus Riis Kopperud, Hilde Vollan, Karoline Bragstad, Olav Hungnes |
| EPI_ISL_3426847 | Haukeland University Hospital, Dept. of Microbiology | Norwegian Institute of Public Health, Department of Virology | Kathrine Stene-Johansen, Kamilla Heddeland Instefjord, Hilde Elshaug, Garcia Llorente Ignacio, Jon Bråte, Engebretsen Serina Beate,Pedersen Benedikte Nevjen, Line Victoria Moen, Debech Nadia, Atiya R Ali,Marie Paulsen Madsen, Rasmus Riis Kopperud, Hilde Vollan, Karoline Bragstad, Olav Hungnes |
| EPI_ISL_3426848, EPI_ISL_3426849, EPI_ISL_3426850 | Department of Medical Microbiology - section Molde, Molde Hospital | Norwegian Institute of Public Health, Department of Virology | Kathrine Stene-Johansen, Kamilla Heddeland Instefjord, Hilde Elshaug, Garcia Llorente Ignacio, Jon Bråte, Engebretsen Serina Beate,Pedersen Benedikte Nevjen, Line Victoria Moen, Debech Nadia, Atiya R Ali,Marie Paulsen Madsen, Rasmus Riis Kopperud, Hilde Vollan, Karoline Bragstad, Olav Hungnes |
| EPI_ISL_3426851 | Haukeland University Hospital, Dept. of Microbiology | Norwegian Institute of Public Health, Department of Virology | Kathrine Stene-Johansen, Kamilla Heddeland Instefjord, Hilde Elshaug, Garcia Llorente Ignacio, Jon Bråte, Engebretsen Serina Beate,Pedersen Benedikte Nevjen, Line Victoria Moen, Debech Nadia, Atiya R Ali,Marie Paulsen Madsen, Rasmus Riis Kopperud, Hilde Vollan, Karoline Bragstad, Olav Hungnes |
| EPI_ISL_3426852, EPI_ISL_3426853 | Department of Medical Microbiology - section Molde, Molde Hospital | Norwegian Institute of Public Health, Department of Virology | Kathrine Stene-Johansen, Kamilla Heddeland Instefjord, Hilde Elshaug, Garcia Llorente Ignacio, Jon Bråte, Engebretsen Serina Beate,Pedersen Benedikte Nevjen, Line Victoria Moen, Debech Nadia, Atiya R Ali,Marie Paulsen Madsen, Rasmus Riis Kopperud, Hilde Vollan, Karoline Bragstad, Olav Hungnes |
| EPI_ISL_3426854 | Haukeland University Hospital, Dept. of Microbiology | Norwegian Institute of Public Health, Department of Virology | Kathrine Stene-Johansen, Kamilla Heddeland Instefjord, Hilde Elshaug, Garcia Llorente Ignacio, Jon Bråte, Engebretsen Serina Beate,Pedersen Benedikte Nevjen, Line Victoria Moen, Debech Nadia, Atiya R Ali,Marie Paulsen Madsen, Rasmus Riis Kopperud, Hilde Vollan, Karoline Bragstad, Olav Hungnes |
| EPI_ISL_3426855, EPI_ISL_3426856 | Department of Medical Microbiology - section Molde, Molde Hospital | Norwegian Institute of Public Health, Department of Virology | Kathrine Stene-Johansen, Kamilla Heddeland Instefjord, Hilde Elshaug, Garcia Llorente Ignacio, Jon Bråte, Engebretsen Serina Beate,Pedersen Benedikte Nevjen, Line Victoria Moen, Debech Nadia, Atiya R Ali,Marie Paulsen Madsen, Rasmus Riis Kopperud, Hilde Vollan, Karoline Bragstad, Olav Hungnes |
| EPI_ISL_3426857 | Haukeland University Hospital, Dept. of Microbiology | Norwegian Institute of Public Health, Department of Virology | Kathrine Stene-Johansen, Kamilla Heddeland Instefjord, Hilde Elshaug, Garcia Llorente Ignacio, Jon Bråte, Engebretsen Serina Beate,Pedersen Benedikte Nevjen, Line Victoria Moen, Debech Nadia, Atiya R Ali,Marie Paulsen Madsen, Rasmus Riis Kopperud, Hilde Vollan, Karoline Bragstad, Olav Hungnes |
| EPI_ISL_3426858 | Department of Medical Microbiology - section Molde, Molde Hospital | Norwegian Institute of Public Health, Department of Virology | Kathrine Stene-Johansen, Kamilla Heddeland Instefjord, Hilde Elshaug, Garcia Llorente Ignacio, Jon Bråte, Engebretsen Serina Beate,Pedersen Benedikte Nevjen, Line Victoria Moen, Debech Nadia, Atiya R Ali,Marie Paulsen Madsen, Rasmus Riis Kopperud, Hilde Vollan, Karoline Bragstad, Olav Hungnes |
| EPI_ISL_3426859, EPI_ISL_3426860 | University Hospital of Northern Norway, Department for Microbiology and Infectious Disease Control | Norwegian Institute of Public Health, Department of Virology | Kathrine Stene-Johansen, Kamilla Heddeland Instefjord, Hilde Elshaug, Garcia Llorente Ignacio, Jon Bråte, Engebretsen Serina Beate,Pedersen Benedikte Nevjen, Line Victoria Moen, Debech Nadia, Atiya R Ali,Marie Paulsen Madsen, Rasmus Riis Kopperud, Hilde Vollan, Karoline Bragstad, Olav Hungnes |
| EPI_ISL_3426861, EPI_ISL_3426862 | Innlandet Hospital Trust, Division Lillehammer, Department for Medical Microbiology | Norwegian Institute of Public Health, Department of Virology | Kathrine Stene-Johansen, Kamilla Heddeland Instefjord, Hilde Elshaug, Garcia Llorente Ignacio, Jon Bråte, Engebretsen Serina Beate,Pedersen Benedikte Nevjen, Line Victoria Moen, Debech Nadia, Atiya R Ali,Marie Paulsen Madsen, Rasmus Riis Kopperud, Hilde Vollan, Karoline Bragstad, Olav Hungnes |
| EPI_ISL_3426863 | Nordland Hospital - Bodo, Laboratory Department, Molecular Biology Unit | Norwegian Institute of Public Health, Department of Virology | Kathrine Stene-Johansen, Kamilla Heddeland Instefjord, Hilde Elshaug, Garcia Llorente Ignacio, Jon Bråte, Engebretsen Serina Beate,Pedersen Benedikte Nevjen, Line Victoria Moen, Debech Nadia, Atiya R Ali,Marie Paulsen Madsen, Rasmus Riis Kopperud, Hilde Vollan, Karoline Bragstad, Olav Hungnes |
| EPI_ISL_3426864 | Vestfold Hospital, Toensberg Department of Microbiology | Norwegian Institute of Public Health, Department of Virology | Kathrine Stene-Johansen, Kamilla Heddeland Instefjord, Hilde Elshaug, Garcia Llorente Ignacio, Jon Bråte, Engebretsen Serina Beate,Pedersen Benedikte Nevjen, Line Victoria Moen, Debech Nadia, Atiya R Ali,Marie Paulsen Madsen, Rasmus Riis Kopperud, Hilde Vollan, Karoline Bragstad, Olav Hungnes |
| EPI_ISL_3426865 | Unilabs Laboratory Medicine | Norwegian Institute of Public Health, Department of Virology | Kathrine Stene-Johansen, Kamilla Heddeland Instefjord, Hilde Elshaug, Garcia Llorente Ignacio, Jon Bråte, Engebretsen Serina Beate,Pedersen Benedikte Nevjen, Line Victoria Moen, Debech Nadia, Atiya R Ali,Marie Paulsen Madsen, Rasmus Riis Kopperud, Hilde Vollan, Karoline Bragstad, Olav Hungnes |
| EPI_ISL_3426866, EPI_ISL_3426867 | Vestfold Hospital, Toensberg Department of Microbiology | Norwegian Institute of Public Health, Department of Virology | Kathrine Stene-Johansen, Kamilla Heddeland Instefjord, Hilde Elshaug, Garcia Llorente Ignacio, Jon Bråte, Engebretsen Serina Beate,Pedersen Benedikte Nevjen, Line Victoria Moen, Debech Nadia, Atiya R Ali,Marie Paulsen Madsen, Rasmus Riis Kopperud, Hilde Vollan, Karoline Bragstad, Olav Hungnes |
| EPI_ISL_3426868 | Nordland Hospital - Bodo, Laboratory Department, Molecular Biology Unit | Norwegian Institute of Public Health, Department of Virology | Kathrine Stene-Johansen, Kamilla Heddeland Instefjord, Hilde Elshaug, Garcia Llorente Ignacio, Jon Bråte, Engebretsen Serina Beate,Pedersen Benedikte Nevjen, Line Victoria Moen, Debech Nadia, Atiya R Ali,Marie Paulsen Madsen, Rasmus Riis Kopperud, Hilde Vollan, Karoline Bragstad, Olav Hungnes |
| EPI_ISL_3426869 | Medical Microbiology Unit, Department for Laboratory Medicine, Drammen Hospital, Vestre Viken Health Trust, | Norwegian Institute of Public Health, Department of Virology | Kathrine Stene-Johansen, Kamilla Heddeland Instefjord, Hilde Elshaug, Garcia Llorente Ignacio, Jon Bråte, Engebretsen Serina Beate,Pedersen Benedikte Nevjen, Line Victoria Moen, Debech Nadia, Atiya R Ali,Marie Paulsen Madsen, Rasmus Riis Kopperud, Hilde Vollan, Karoline Bragstad, Olav Hungnes |
| EPI_ISL_3426870 | Department of Medical Microbiology - section Molde, Molde Hospital | Norwegian Institute of Public Health, Department of Virology | Kathrine Stene-Johansen, Kamilla Heddeland Instefjord, Hilde Elshaug, Garcia Llorente Ignacio, Jon Bråte, Engebretsen Serina Beate,Pedersen Benedikte Nevjen, Line Victoria Moen, Debech Nadia, Atiya R Ali,Marie Paulsen Madsen, Rasmus Riis Kopperud, Hilde Vollan, Karoline Bragstad, Olav Hungnes |
| EPI_ISL_3426871 | Nordland Hospital - Bodo, Laboratory Department, Molecular Biology Unit | Norwegian Institute of Public Health, Department of Virology | Kathrine Stene-Johansen, Kamilla Heddeland Instefjord, Hilde Elshaug, Garcia Llorente Ignacio, Jon Bråte, Engebretsen Serina Beate,Pedersen Benedikte Nevjen, Line Victoria Moen, Debech Nadia, Atiya R Ali,Marie Paulsen Madsen, Rasmus Riis Kopperud, Hilde Vollan, Karoline Bragstad, Olav Hungnes |
| EPI_ISL_3426872, EPI_ISL_3426873 | Vestfold Hospital, Toensberg Department of Microbiology | Norwegian Institute of Public Health, Department of Virology | Kathrine Stene-Johansen, Kamilla Heddeland Instefjord, Hilde Elshaug, Garcia Llorente Ignacio, Jon Bråte, Engebretsen Serina Beate,Pedersen Benedikte Nevjen, Line Victoria Moen, Debech Nadia, Atiya R Ali,Marie Paulsen Madsen, Rasmus Riis Kopperud, Hilde Vollan, Karoline Bragstad, Olav Hungnes |
| EPI_ISL_3426874, EPI_ISL_3426875 | Medical Microbiology Unit, Department for Laboratory Medicine, Drammen Hospital, Vestre Viken Health Trust, | Norwegian Institute of Public Health, Department of Virology | Kathrine Stene-Johansen, Kamilla Heddeland Instefjord, Hilde Elshaug, Garcia Llorente Ignacio, Jon Bråte, Engebretsen Serina Beate,Pedersen Benedikte Nevjen, Line Victoria Moen, Debech Nadia, Atiya R Ali,Marie Paulsen Madsen, Rasmus Riis Kopperud, Hilde Vollan, Karoline Bragstad, Olav Hungnes |
| EPI_ISL_3426876 | Dept. of Medical Microbiology, Stavanger University Hospital, Helse Stavanger HF | Norwegian Institute of Public Health, Department of Virology | Kathrine Stene-Johansen, Kamilla Heddeland Instefjord, Hilde Elshaug, Garcia Llorente Ignacio, Jon Bråte, Engebretsen Serina Beate,Pedersen Benedikte Nevjen, Line Victoria Moen, Debech Nadia, Atiya R Ali,Marie Paulsen Madsen, Rasmus Riis Kopperud, Hilde Vollan, Karoline Bragstad, Olav Hungnes |
| EPI_ISL_3426877 | Medical Microbiology Unit, Department for Laboratory Medicine, Drammen Hospital, Vestre Viken Health Trust, | Norwegian Institute of Public Health, Department of Virology | Kathrine Stene-Johansen, Kamilla Heddeland Instefjord, Hilde Elshaug, Garcia Llorente Ignacio, Jon Bråte, Engebretsen Serina Beate,Pedersen Benedikte Nevjen, Line Victoria Moen, Debech Nadia, Atiya R Ali,Marie Paulsen Madsen, Rasmus Riis Kopperud, Hilde Vollan, Karoline Bragstad, Olav Hungnes |
| EPI_ISL_3426878 | Dept. of Medical Microbiology, Stavanger University Hospital, Helse Stavanger HF | Norwegian Institute of Public Health, Department of Virology | Kathrine Stene-Johansen, Kamilla Heddeland Instefjord, Hilde Elshaug, Garcia Llorente Ignacio, Jon Bråte, Engebretsen Serina Beate,Pedersen Benedikte Nevjen, Line Victoria Moen, Debech Nadia, Atiya R Ali,Marie Paulsen Madsen, Rasmus Riis Kopperud, Hilde Vollan, Karoline Bragstad, Olav Hungnes |
| EPI_ISL_3426879 | Medical Microbiology Unit, Department for Laboratory Medicine, Drammen Hospital, Vestre Viken Health Trust, | Norwegian Institute of Public Health, Department of Virology | Kathrine Stene-Johansen, Kamilla Heddeland Instefjord, Hilde Elshaug, Garcia Llorente Ignacio, Jon Bråte, Engebretsen Serina Beate,Pedersen Benedikte Nevjen, Line Victoria Moen, Debech Nadia, Atiya R Ali,Marie Paulsen Madsen, Rasmus Riis Kopperud, Hilde Vollan, Karoline Bragstad, Olav Hungnes |
| EPI_ISL_3426880 | Dept. of Medical Microbiology, Stavanger University Hospital, Helse Stavanger HF | Norwegian Institute of Public Health, Department of Virology | Kathrine Stene-Johansen, Kamilla Heddeland Instefjord, Hilde Elshaug, Garcia Llorente Ignacio, Jon Bråte, Engebretsen Serina Beate,Pedersen Benedikte Nevjen, Line Victoria Moen, Debech Nadia, Atiya R Ali,Marie Paulsen Madsen, Rasmus Riis Kopperud, Hilde Vollan, Karoline Bragstad, Olav Hungnes |
| EPI_ISL_3426881 | Medical Microbiology Unit, Department for Laboratory Medicine, Drammen Hospital, Vestre Viken Health Trust, | Norwegian Institute of Public Health, Department of Virology | Kathrine Stene-Johansen, Kamilla Heddeland Instefjord, Hilde Elshaug, Garcia Llorente Ignacio, Jon Bråte, Engebretsen Serina Beate,Pedersen Benedikte Nevjen, Line Victoria Moen, Debech Nadia, Atiya R Ali,Marie Paulsen Madsen, Rasmus Riis Kopperud, Hilde Vollan, Karoline Bragstad, Olav Hungnes |
| EPI_ISL_3426886, EPI_ISL_3426894 | Ostfold Hospital Trust - Kalnes, Centre for Laboratory Medicine, Section for gene technology and infection serology | Norwegian Institute of Public Health, Department of Virology | Kathrine Stene-Johansen, Kamilla Heddeland Instefjord, Hilde Elshaug, Garcia Llorente Ignacio, Jon Bråte, Engebretsen Serina Beate,Pedersen Benedikte Nevjen, Line Victoria Moen, Debech Nadia, Atiya R Ali,Marie Paulsen Madsen, Rasmus Riis Kopperud, Hilde Vollan, Karoline Bragstad, Olav Hungnes |
| EPI_ISL_3426897 | Furst Medical Laboratory | Norwegian Institute of Public Health, Department of Virology | Kathrine Stene-Johansen, Kamilla Heddeland Instefjord, Hilde Elshaug, Garcia Llorente Ignacio, Jon Bråte, Engebretsen Serina Beate,Pedersen Benedikte Nevjen, Line Victoria Moen, Debech Nadia, Atiya R Ali,Marie Paulsen Madsen, Rasmus Riis Kopperud, Hilde Vollan, Karoline Bragstad, Olav Hungnes |
| EPI_ISL_3426901, EPI_ISL_3426902 | Haukeland University Hospital, Dept. of Microbiology | Norwegian Institute of Public Health, Department of Virology | Kathrine Stene-Johansen, Kamilla Heddeland Instefjord, Hilde Elshaug, Garcia Llorente Ignacio, Jon Bråte, Engebretsen Serina Beate,Pedersen Benedikte Nevjen, Line Victoria Moen, Debech Nadia, Atiya R Ali,Marie Paulsen Madsen, Rasmus Riis Kopperud, Hilde Vollan, Karoline Bragstad, Olav Hungnes |
| EPI_ISL_3426905 | Ostfold Hospital Trust - Kalnes, Centre for Laboratory Medicine, Section for gene technology and infection serology | Norwegian Institute of Public Health, Department of Virology | Kathrine Stene-Johansen, Kamilla Heddeland Instefjord, Hilde Elshaug, Garcia Llorente Ignacio, Jon Bråte, Engebretsen Serina Beate,Pedersen Benedikte Nevjen, Line Victoria Moen, Debech Nadia, Atiya R Ali,Marie Paulsen Madsen, Rasmus Riis Kopperud, Hilde Vollan, Karoline Bragstad, Olav Hungnes |
| EPI_ISL_3426907 | Dept. of Medical Microbiology, Stavanger University Hospital, Helse Stavanger HF | Norwegian Institute of Public Health, Department of Virology | Kathrine Stene-Johansen, Kamilla Heddeland Instefjord, Hilde Elshaug, Garcia Llorente Ignacio, Jon Bråte, Engebretsen Serina Beate,Pedersen Benedikte Nevjen, Line Victoria Moen, Debech Nadia, Atiya R Ali,Marie Paulsen Madsen, Rasmus Riis Kopperud, Hilde Vollan, Karoline Bragstad, Olav Hungnes |
| EPI_ISL_3426908, EPI_ISL_3426909 | Haukeland University Hospital, Dept. of Microbiology | Norwegian Institute of Public Health, Department of Virology | Kathrine Stene-Johansen, Kamilla Heddeland Instefjord, Hilde Elshaug, Garcia Llorente Ignacio, Jon Bråte, Engebretsen Serina Beate,Pedersen Benedikte Nevjen, Line Victoria Moen, Debech Nadia, Atiya R Ali,Marie Paulsen Madsen, Rasmus Riis Kopperud, Hilde Vollan, Karoline Bragstad, Olav Hungnes |
| EPI_ISL_3426912 | Ostfold Hospital Trust - Kalnes, Centre for Laboratory Medicine, Section for gene technology and infection serology | Norwegian Institute of Public Health, Department of Virology | Kathrine Stene-Johansen, Kamilla Heddeland Instefjord, Hilde Elshaug, Garcia Llorente Ignacio, Jon Bråte, Engebretsen Serina Beate,Pedersen Benedikte Nevjen, Line Victoria Moen, Debech Nadia, Atiya R Ali,Marie Paulsen Madsen, Rasmus Riis Kopperud, Hilde Vollan, Karoline Bragstad, Olav Hungnes |
| EPI_ISL_3426913 | Medical Microbiology Unit, Department for Laboratory Medicine, Drammen Hospital, Vestre Viken Health Trust, | Norwegian Institute of Public Health, Department of Virology | Kathrine Stene-Johansen, Kamilla Heddeland Instefjord, Hilde Elshaug, Garcia Llorente Ignacio, Jon Bråte, Engebretsen Serina Beate,Pedersen Benedikte Nevjen, Line Victoria Moen, Debech Nadia, Atiya R Ali,Marie Paulsen Madsen, Rasmus Riis Kopperud, Hilde Vollan, Karoline Bragstad, Olav Hungnes |
| EPI_ISL_3426914 | Haukeland University Hospital, Dept. of Microbiology | Norwegian Institute of Public Health, Department of Virology | Kathrine Stene-Johansen, Kamilla Heddeland Instefjord, Hilde Elshaug, Garcia Llorente Ignacio, Jon Bråte, Engebretsen Serina Beate,Pedersen Benedikte Nevjen, Line Victoria Moen, Debech Nadia, Atiya R Ali,Marie Paulsen Madsen, Rasmus Riis Kopperud, Hilde Vollan, Karoline Bragstad, Olav Hungnes |
| EPI_ISL_3426915 | Medical Microbiology Unit, Department for Laboratory Medicine, Drammen Hospital, Vestre Viken Health Trust, | Norwegian Institute of Public Health, Department of Virology | Kathrine Stene-Johansen, Kamilla Heddeland Instefjord, Hilde Elshaug, Garcia Llorente Ignacio, Jon Bråte, Engebretsen Serina Beate,Pedersen Benedikte Nevjen, Line Victoria Moen, Debech Nadia, Atiya R Ali,Marie Paulsen Madsen, Rasmus Riis Kopperud, Hilde Vollan, Karoline Bragstad, Olav Hungnes |
| EPI_ISL_3426916 | Haukeland University Hospital, Dept. of Microbiology | Norwegian Institute of Public Health, Department of Virology | Kathrine Stene-Johansen, Kamilla Heddeland Instefjord, Hilde Elshaug, Garcia Llorente Ignacio, Jon Bråte, Engebretsen Serina Beate,Pedersen Benedikte Nevjen, Line Victoria Moen, Debech Nadia, Atiya R Ali,Marie Paulsen Madsen, Rasmus Riis Kopperud, Hilde Vollan, Karoline Bragstad, Olav Hungnes |
| EPI_ISL_3426918 | Ostfold Hospital Trust - Kalnes, Centre for Laboratory Medicine, Section for gene technology and infection serology | Norwegian Institute of Public Health, Department of Virology | Kathrine Stene-Johansen, Kamilla Heddeland Instefjord, Hilde Elshaug, Garcia Llorente Ignacio, Jon Bråte, Engebretsen Serina Beate,Pedersen Benedikte Nevjen, Line Victoria Moen, Debech Nadia, Atiya R Ali,Marie Paulsen Madsen, Rasmus Riis Kopperud, Hilde Vollan, Karoline Bragstad, Olav Hungnes |
| EPI_ISL_3426919 | Furst Medical Laboratory | Norwegian Institute of Public Health, Department of Virology | Kathrine Stene-Johansen, Kamilla Heddeland Instefjord, Hilde Elshaug, Garcia Llorente Ignacio, Jon Bråte, Engebretsen Serina Beate,Pedersen Benedikte Nevjen, Line Victoria Moen, Debech Nadia, Atiya R Ali,Marie Paulsen Madsen, Rasmus Riis Kopperud, Hilde Vollan, Karoline Bragstad, Olav Hungnes |
| EPI_ISL_3426921 | Ostfold Hospital Trust - Kalnes, Centre for Laboratory Medicine, Section for gene technology and infection serology | Norwegian Institute of Public Health, Department of Virology | Kathrine Stene-Johansen, Kamilla Heddeland Instefjord, Hilde Elshaug, Garcia Llorente Ignacio, Jon Bråte, Engebretsen Serina Beate,Pedersen Benedikte Nevjen, Line Victoria Moen, Debech Nadia, Atiya R Ali,Marie Paulsen Madsen, Rasmus Riis Kopperud, Hilde Vollan, Karoline Bragstad, Olav Hungnes |
| EPI_ISL_3426922, EPI_ISL_3426924 | Furst Medical Laboratory | Norwegian Institute of Public Health, Department of Virology | Kathrine Stene-Johansen, Kamilla Heddeland Instefjord, Hilde Elshaug, Garcia Llorente Ignacio, Jon Bråte, Engebretsen Serina Beate,Pedersen Benedikte Nevjen, Line Victoria Moen, Debech Nadia, Atiya R Ali,Marie Paulsen Madsen, Rasmus Riis Kopperud, Hilde Vollan, Karoline Bragstad, Olav Hungnes |
| EPI_ISL_3426925 | Dept. of Medical Microbiology, Stavanger University Hospital, Helse Stavanger HF | Norwegian Institute of Public Health, Department of Virology | Kathrine Stene-Johansen, Kamilla Heddeland Instefjord, Hilde Elshaug, Garcia Llorente Ignacio, Jon Bråte, Engebretsen Serina Beate,Pedersen Benedikte Nevjen, Line Victoria Moen, Debech Nadia, Atiya R Ali,Marie Paulsen Madsen, Rasmus Riis Kopperud, Hilde Vollan, Karoline Bragstad, Olav Hungnes |
| EPI_ISL_3426926 | Unilabs Laboratory Medicine | Norwegian Institute of Public Health, Department of Virology | Kathrine Stene-Johansen, Kamilla Heddeland Instefjord, Hilde Elshaug, Garcia Llorente Ignacio, Jon Bråte, Engebretsen Serina Beate,Pedersen Benedikte Nevjen, Line Victoria Moen, Debech Nadia, Atiya R Ali,Marie Paulsen Madsen, Rasmus Riis Kopperud, Hilde Vollan, Karoline Bragstad, Olav Hungnes |

| | | | |
|---|---|---|---|
| EPI_ISL_3426927, EPI_ISL_3426928, EPI_ISL_3426929, EPI_ISL_3426930, EPI_ISL_3426931, EPI_ISL_3426932, EPI_ISL_3426933, EPI_ISL_3426934, EPI_ISL_3426935, EPI_ISL_3426936, EPI_ISL_3426937, EPI_ISL_3426938, EPI_ISL_3426939, EPI_ISL_3426940, EPI_ISL_3426941, EPI_ISL_3426942, EPI_ISL_3426943, EPI_ISL_3426944, EPI_ISL_3426945, EPI_ISL_3426946, EPI_ISL_3426947, EPI_ISL_3426948, EPI_ISL_3426949, EPI_ISL_3426950, EPI_ISL_3426951, EPI_ISL_3426952, EPI_ISL_3426953, EPI_ISL_3426954, EPI_ISL_3426955, EPI_ISL_3426956, EPI_ISL_3426957, EPI_ISL_3426958, EPI_ISL_3426959, EPI_ISL_3426960, EPI_ISL_3426961, EPI_ISL_3426962, EPI_ISL_3426963, EPI_ISL_3426964, EPI_ISL_3426965, EPI_ISL_3426966, EPI_ISL_3426967, EPI_ISL_3426968, EPI_ISL_3426969, EPI_ISL_3426970, EPI_ISL_3426971, EPI_ISL_3426972, EPI_ISL_3426973, EPI_ISL_3426974, EPI_ISL_3426975, EPI_ISL_3426976, EPI_ISL_3426977, EPI_ISL_3426978, EPI_ISL_3426979, EPI_ISL_3426980, EPI_ISL_3426981, EPI_ISL_3426982, EPI_ISL_3426983, EPI_ISL_3426984, EPI_ISL_3426985, EPI_ISL_3426986, EPI_ISL_3426987, EPI_ISL_3426988, EPI_ISL_3426989, EPI_ISL_3426990, EPI_ISL_3426991, EPI_ISL_3426992, EPI_ISL_3426993, EPI_ISL_3426994, EPI_ISL_3426995, EPI_ISL_3426996, EPI_ISL_3426997, EPI_ISL_3426998, EPI_ISL_3426999, EPI_ISL_3427000, EPI_ISL_3427001, EPI_ISL_3427002, EPI_ISL_3427003, EPI_ISL_3427004, EPI_ISL_3427005, EPI_ISL_3427006, EPI_ISL_3427007, EPI_ISL_3427008, EPI_ISL_3427009, EPI_ISL_3427010, EPI_ISL_3427011, EPI_ISL_3427012, EPI_ISL_3427013, EPI_ISL_3427014, EPI_ISL_3427015, EPI_ISL_3427016, EPI_ISL_3427017, EPI_ISL_3427018, EPI_ISL_3427019, EPI_ISL_3427020, EPI_ISL_3427021, EPI_ISL_3427022, EPI_ISL_3427023, EPI_ISL_3427024, EPI_ISL_3427025, EPI_ISL_3427026, EPI_ISL_3427027, EPI_ISL_3427028, EPI_ISL_3427029, EPI_ISL_3427030, EPI_ISL_3427031, EPI_ISL_3427032, EPI_ISL_3427033, EPI_ISL_3427034, EPI_ISL_3427035, EPI_ISL_3427036, EPI_ISL_3427037, EPI_ISL_3427038, EPI_ISL_3427039, EPI_ISL_3427040, EPI_ISL_3427041, EPI_ISL_3427042, EPI_ISL_3427043, EPI_ISL_3427044, EPI_ISL_3427045, EPI_ISL_3427046, EPI_ISL_3427047, EPI_ISL_3427048, EPI_ISL_3427049, EPI_ISL_3427050, EPI_ISL_3427051, EPI_ISL_3427052, EPI_ISL_3427053, EPI_ISL_3427054, EPI_ISL_3427055, EPI_ISL_3427056, EPI_ISL_3427057, EPI_ISL_3427058, EPI_ISL_3427059, EPI_ISL_3427060, EPI_ISL_3427061, EPI_ISL_3427062, EPI_ISL_3427063, EPI_ISL_3427064, EPI_ISL_3427065, EPI_ISL_3427066, EPI_ISL_3427067, EPI_ISL_3427068, EPI_ISL_3427069, EPI_ISL_3427070, EPI_ISL_3427071, EPI_ISL_3427072, EPI_ISL_3427073, EPI_ISL_3427074, EPI_ISL_3427075, EPI_ISL_3427076, EPI_ISL_3427077, EPI_ISL_3427078, EPI_ISL_3427079, EPI_ISL_3427080, EPI_ISL_3427081, EPI_ISL_3427082, EPI_ISL_3427083, EPI_ISL_3427084, EPI_ISL_3427085, EPI_ISL_3427086, EPI_ISL_3427087, EPI_ISL_3427088, EPI_ISL_3427089, EPI_ISL_3427090, EPI_ISL_3427091, EPI_ISL_3427092, EPI_ISL_3427093, EPI_ISL_3427094, EPI_ISL_3427095, EPI_ISL_3427096, EPI_ISL_3427097, EPI_ISL_3427098, EPI_ISL_3427099, EPI_ISL_3427100, EPI_ISL_3427101, EPI_ISL_3427102, EPI_ISL_3427103, EPI_ISL_3427104, EPI_ISL_3427105, EPI_ISL_3427106, EPI_ISL_3427107, EPI_ISL_3427108, EPI_ISL_3427109, EPI_ISL_3427110, EPI_ISL_3427111, EPI_ISL_3427112 | | | |
| see above | National Public Health Laboratory, National Centre for Infectious Diseases | National Public Health Laboratory, National Centre for Infectious Diseases | Katherine Ching, Zhenyang Zhou, Royce Ang, Lin Cui, Raymond Tzer Pin Lin |
| EPI_ISL_3427113, EPI_ISL_3427114, EPI_ISL_3427115, EPI_ISL_3427116, EPI_ISL_3427117, EPI_ISL_3427119, EPI_ISL_3427120, EPI_ISL_3427121, EPI_ISL_3427122 | Università degli Studi di Perugia | Istituto Zooprofilattico Sperimentale dell'Abruzzo e Molise "G. Caporale" | Mencacci A, Camilloni B, Biagetti M, Giammarioli M, Lorusso A, Marcacci M, Di Domenico M, Ancora M, Curini V, Mangone I, Rinaldi A, Delli Compagni E, Scialabba S, Di Pasquale A, Cammà C, Puglia I, Calistri P, Savini G |
| EPI_ISL_3427123 | Laboratorio Analisi Osp. Città di Castello - Azienda USL Umbria1 | Istituto Zooprofilattico Sperimentale dell'Abruzzo e Molise "G. Caporale" | Malagigi V, Taccani P,Lorusso A, Marcacci M, Di Domenico M, Ancora M, Curini V, Mangone I, Rinaldi A, Delli Compagni E, Scialabba S, Di Pasquale A, Cammà C, Puglia I, Calistri P, Savini G |
| EPI_ISL_3427124, EPI_ISL_3427126, EPI_ISL_3427127, EPI_ISL_3427128, EPI_ISL_3427129, EPI_ISL_3427130 | Università degli Studi di Perugia | Istituto Zooprofilattico Sperimentale dell'Abruzzo e Molise "G. Caporale" | Mencacci A, Camilloni B, Biagetti M, Giammarioli M, Lorusso A, Marcacci M, Di Domenico M, Ancora M, Curini V, Mangone I, Rinaldi A, Delli Compagni E, Scialabba S, Di Pasquale A, Cammà C, Puglia I, Calistri P, Savini G |
| EPI_ISL_3427133 | Azienda USL Umbria 2 | Istituto Zooprofilattico Sperimentale dell'Abruzzo e Molise "G. Caporale" | Proietti A, Pistoni E, Lorusso A, Marcacci M, Di Domenico M, Ancora M, Curini V, Mangone I, Rinaldi A, Delli Compagni E, Scialabba S, Di Pasquale A, Cammà C, Puglia I, Calistri P, Savini G |
| EPI_ISL_3427134 | Laboratorio Analisi Osp. Città di Castello - Azienda USL Umbria1 | Istituto Zooprofilattico Sperimentale dell'Abruzzo e Molise "G. Caporale" | Malagigi V, Taccani P,Lorusso A, Marcacci M, Di Domenico M, Ancora M, Curini V, Mangone I, Rinaldi A, Delli Compagni E, Scialabba S, Di Pasquale A, Cammà C, Puglia I, Calistri P, Savini G |
| EPI_ISL_3427135, EPI_ISL_3427136, EPI_ISL_3427137, EPI_ISL_3427138 | Azienda USL Umbria 2 | Istituto Zooprofilattico Sperimentale dell'Abruzzo e Molise "G. Caporale" | Proietti A, Pistoni E, Lorusso A, Marcacci M, Di Domenico M, Ancora M, Curini V, Mangone I, Rinaldi A, Delli Compagni E, Scialabba S, Di Pasquale A, Cammà C, Puglia I, Calistri P, Savini G |
| EPI_ISL_3427139, EPI_ISL_3427140, EPI_ISL_3427141, EPI_ISL_3427142, EPI_ISL_3427143, EPI_ISL_3427144 | Azienda Ospedaliera Terni | Istituto Zooprofilattico Sperimentale dell'Abruzzo e Molise "G. Caporale" | Palumbo M, Scaccetti A, Lorusso A, Marcacci M, Di Domenico M, Ancora M, Curini V, Mangone I, Rinaldi A, Delli Compagni E, Scialabba S, Di Pasquale A, Cammà C, Puglia I, Calistri P, Savini G |
| EPI_ISL_3427145 | Laboratorio Analisi Osp. Città di Castello - Azienda USL Umbria1 | Istituto Zooprofilattico Sperimentale dell'Abruzzo e Molise "G. Caporale" | Malagigi V, Taccani P,Lorusso A, Marcacci M, Di Domenico M, Ancora M, Curini V, Mangone I, Rinaldi A, Delli Compagni E, Scialabba S, Di Pasquale A, Cammà C, Puglia I, Calistri P, Savini G |
| EPI_ISL_3427146, EPI_ISL_3427147, EPI_ISL_3427148, EPI_ISL_3427149, EPI_ISL_3427150, EPI_ISL_3427151, EPI_ISL_3427152, EPI_ISL_3427153, EPI_ISL_3427154, EPI_ISL_3427155 | Azienda Ospedaliera Terni | Istituto Zooprofilattico Sperimentale dell'Abruzzo e Molise "G. Caporale" | Palumbo M, Scaccetti A, Lorusso A, Marcacci M, Di Domenico M, Ancora M, Curini V, Mangone I, Rinaldi A, Delli Compagni E, Scialabba S, Di Pasquale A, Cammà C, Puglia I, Calistri P, Savini G |
| EPI_ISL_3427156, EPI_ISL_3427157 | Laboratorio Analisi Osp. Città di Castello - Azienda USL Umbria1 | Istituto Zooprofilattico Sperimentale dell'Abruzzo e Molise "G. Caporale" | Malagigi V, Taccani P,Lorusso A, Marcacci M, Di Domenico M, Ancora M, Curini V, Mangone I, Rinaldi A, Delli Compagni E, Scialabba S, Di Pasquale A, Cammà C, Puglia I, Calistri P, Savini G |
| EPI_ISL_3427158, EPI_ISL_3427159, EPI_ISL_3427160 | Università degli Studi di Perugia | Istituto Zooprofilattico Sperimentale dell'Abruzzo e Molise "G. Caporale" | Mencacci A, Camilloni B, Biagetti M, Giammarioli M, Lorusso A, Marcacci M, Di Domenico M, Ancora M, Curini V, Mangone I, Rinaldi A, Delli Compagni E, Scialabba S, Di Pasquale A, Cammà C, Puglia I, Calistri P, Savini G |
| EPI_ISL_3427161 | Laboratorio Analisi Osp. Città di Castello - Azienda USL Umbria1 | Istituto Zooprofilattico Sperimentale dell'Abruzzo e Molise "G. Caporale" | Malagigi V, Taccani P,Lorusso A, Marcacci M, Di Domenico M, Ancora M, Curini V, Mangone I, Rinaldi A, Delli Compagni E, Scialabba S, Di Pasquale A, Cammà C, Puglia I, Calistri P, Savini G |
| EPI_ISL_3427162, EPI_ISL_3427163, EPI_ISL_3427164, EPI_ISL_3427165, EPI_ISL_3427166, EPI_ISL_3427167, EPI_ISL_3427168, EPI_ISL_3427169, EPI_ISL_3427170, EPI_ISL_3427171, EPI_ISL_3427172, EPI_ISL_3427173, EPI_ISL_3427174, EPI_ISL_3427175, EPI_ISL_3427176, EPI_ISL_3427177, EPI_ISL_3427180, EPI_ISL_3427181, EPI_ISL_3427183, EPI_ISL_3427184, EPI_ISL_3427185, EPI_ISL_3427186, EPI_ISL_3427187, EPI_ISL_3427188, EPI_ISL_3427189, EPI_ISL_3427190, EPI_ISL_3427191, EPI_ISL_3427192, EPI_ISL_3427193, EPI_ISL_3427194, EPI_ISL_3427195, EPI_ISL_3427196, EPI_ISL_3427197, EPI_ISL_3427198, EPI_ISL_3427199, EPI_ISL_3427200, EPI_ISL_3427201, EPI_ISL_3427202, EPI_ISL_3427203, EPI_ISL_3427204, EPI_ISL_3427205, EPI_ISL_3427206, EPI_ISL_3427208, EPI_ISL_3427209, EPI_ISL_3427210, EPI_ISL_3427211, EPI_ISL_3427212, EPI_ISL_3427213, EPI_ISL_3427214, EPI_ISL_3427215, EPI_ISL_3427216, EPI_ISL_3427217, EPI_ISL_3427218, EPI_ISL_3427219, EPI_ISL_3427220, EPI_ISL_3427221, EPI_ISL_3427222, EPI_ISL_3427223, EPI_ISL_3427224, EPI_ISL_3427225, EPI_ISL_3427226, EPI_ISL_3427227, EPI_ISL_3427230, EPI_ISL_3427231, EPI_ISL_3427233, EPI_ISL_3427234, EPI_ISL_3427236, EPI_ISL_3427237, EPI_ISL_3427238, EPI_ISL_3427239, EPI_ISL_3427240, EPI_ISL_3427241, EPI_ISL_3427242, EPI_ISL_3427243, EPI_ISL_3427244, EPI_ISL_3427245, EPI_ISL_3427246, EPI_ISL_3427247, EPI_ISL_3427248, EPI_ISL_3427249, EPI_ISL_3427250, EPI_ISL_3427252, EPI_ISL_3427253, EPI_ISL_3427254, EPI_ISL_3427255, EPI_ISL_3427256, EPI_ISL_3427257, EPI_ISL_3427258, EPI_ISL_3427259, EPI_ISL_3427260, EPI_ISL_3427261, EPI_ISL_3427262, EPI_ISL_3427263, EPI_ISL_3427264, EPI_ISL_3427265, EPI_ISL_3427266, EPI_ISL_3427268, EPI_ISL_3427269, EPI_ISL_3427270, EPI_ISL_3427271, EPI_ISL_3427272, EPI_ISL_3427273, EPI_ISL_3427274, EPI_ISL_3427275, EPI_ISL_3427276, EPI_ISL_3427277, EPI_ISL_3427278, EPI_ISL_3427280, EPI_ISL_3427281, EPI_ISL_3427283, EPI_ISL_3427284, EPI_ISL_3427285, EPI_ISL_3427287, EPI_ISL_3427288, EPI_ISL_3427289, EPI_ISL_3427290, EPI_ISL_3427291, EPI_ISL_3427292, EPI_ISL_3427294, EPI_ISL_3427295, EPI_ISL_3427296, EPI_ISL_3427298, EPI_ISL_3427299, EPI_ISL_3427300, EPI_ISL_3427301, EPI_ISL_3427302, EPI_ISL_3427303, EPI_ISL_3427304, EPI_ISL_3427305, EPI_ISL_3427306, EPI_ISL_3427307, EPI_ISL_3427308, EPI_ISL_3427309, EPI_ISL_3427310, EPI_ISL_3427311, EPI_ISL_3427312, EPI_ISL_3427313, EPI_ISL_3427314 | | | |
| see above | Mako Medical | Centers for Disease Control and Prevention Division of Viral Diseases, Pathogen Discovery | Dakota Howard, Dhwani Batra, Peter W. Cook, Kara Moser, Adrian Paskey, Jason Caravas, Benjamin Rambo-Martin, Shatavia Morrison, Christopher Gulvick, Scott Sammons, Yvette Unoarumhi, Darlene Wagner, Matthew Schmerer, Matthew Tugwell, Lauren Moon, Clinton R. Paden, Duncan MacCannell |
| EPI_ISL_3427315 | OSPEDALE CIVILE TERAMO - CENTRO TRASFUSIONALE | Istituto Zooprofilattico Sperimentale dell'Abruzzo e Molise "G. Caporale" | Lorusso A, Marcacci M, Di Domenico M, Ancora M, Curini V, Di Lollo Valeria, Mangone I, Rinaldi A, Delli Compagni E, Scialabba S, Caporale M, Di Pasquale A, Cammà C, Puglia I, Calistri P, Savini G |
| EPI_ISL_3427316 | Mako Medical | Centers for Disease Control and Prevention Division of Viral Diseases, Pathogen Discovery | Dakota Howard, Dhwani Batra, Peter W. Cook, Kara Moser, Adrian Paskey, Jason Caravas, Benjamin Rambo-Martin, Shatavia Morrison, Christopher Gulvick, Scott Sammons, Yvette Unoarumhi, Darlene Wagner, Matthew Schmerer, Matthew Tugwell, Lauren Moon, Clinton R. Paden, Duncan MacCannell |
| EPI_ISL_3427317, EPI_ISL_3427318 | OSPEDALE CIVILE TERAMO - CENTRO TRASFUSIONALE | Istituto Zooprofilattico Sperimentale dell'Abruzzo e Molise "G. Caporale" | Lorusso A, Marcacci M, Di Domenico M, Ancora M, Curini V, Di Lollo Valeria, Mangone I, Rinaldi A, Delli Compagni E, Scialabba S, Caporale M, Di Pasquale A, Cammà C, Puglia I, Calistri P, Savini G |
| EPI_ISL_3427319 | Mako Medical | Centers for Disease Control and Prevention Division of Viral Diseases, Pathogen Discovery | Dakota Howard, Dhwani Batra, Peter W. Cook, Kara Moser, Adrian Paskey, Jason Caravas, Benjamin Rambo-Martin, Shatavia Morrison, Christopher Gulvick, Scott Sammons, Yvette Unoarumhi, Darlene Wagner, Matthew Schmerer, Matthew Tugwell, Lauren Moon, Clinton R. Paden, Duncan MacCannell |
| EPI_ISL_3427320 | OSPEDALE CIVILE TERAMO - CENTRO TRASFUSIONALE | Istituto Zooprofilattico Sperimentale dell'Abruzzo e Molise "G. Caporale" | Lorusso A, Marcacci M, Di Domenico M, Ancora M, Curini V, Di Lollo Valeria, Mangone I, Rinaldi A, Delli Compagni E, Scialabba S, Caporale M, Di Pasquale A, Cammà C, Puglia I, Calistri P, Savini G |
| EPI_ISL_3427321 | Mako Medical | Centers for Disease Control and Prevention Division of Viral Diseases, Pathogen Discovery | Dakota Howard, Dhwani Batra, Peter W. Cook, Kara Moser, Adrian Paskey, Jason Caravas, Benjamin Rambo-Martin, Shatavia Morrison, Christopher Gulvick, Scott Sammons, Yvette Unoarumhi, Darlene Wagner, Matthew Schmerer, Matthew Tugwell, Lauren Moon, Clinton R. Paden, Duncan MacCannell |
| EPI_ISL_3427322 | OSPEDALE CIVILE TERAMO - CENTRO TRASFUSIONALE | Istituto Zooprofilattico Sperimentale dell'Abruzzo e Molise "G. Caporale" | Lorusso A, Marcacci M, Di Domenico M, Ancora M, Curini V, Di Lollo Valeria, Mangone I, Rinaldi A, Delli Compagni E, Scialabba S, Caporale M, Di Pasquale A, Cammà C, Puglia I, Calistri P, Savini G |
| EPI_ISL_3427323 | Mako Medical | Centers for Disease Control and Prevention Division of Viral Diseases, Pathogen Discovery | Dakota Howard, Dhwani Batra, Peter W. Cook, Kara Moser, Adrian Paskey, Jason Caravas, Benjamin Rambo-Martin, Shatavia Morrison, Christopher Gulvick, Scott Sammons, Yvette Unoarumhi, Darlene Wagner, Matthew Schmerer, Matthew Tugwell, Lauren Moon, Clinton R. Paden, Duncan MacCannell |
| EPI_ISL_3427324, EPI_ISL_3427325 | OSPEDALE CIVILE TERAMO - CENTRO TRASFUSIONALE | Istituto Zooprofilattico Sperimentale dell'Abruzzo e Molise "G. Caporale" | Lorusso A, Marcacci M, Di Domenico M, Ancora M, Curini V, Di Lollo Valeria, Mangone I, Rinaldi A, Delli Compagni E, Scialabba S, Caporale M, Di Pasquale A, Cammà C, Puglia I, Calistri P, Savini G |
| EPI_ISL_3427326 | Mako Medical | Centers for Disease Control and Prevention Division of Viral Diseases, Pathogen Discovery | Dakota Howard, Dhwani Batra, Peter W. Cook, Kara Moser, Adrian Paskey, Jason Caravas, Benjamin Rambo-Martin, Shatavia Morrison, Christopher Gulvick, Scott Sammons, Yvette Unoarumhi, Darlene Wagner, Matthew Schmerer, Matthew Tugwell, Lauren Moon, Clinton R. Paden, Duncan MacCannell |
| EPI_ISL_3427327 | OSPEDALE CIVILE TERAMO - CENTRO TRASFUSIONALE | Istituto Zooprofilattico Sperimentale dell'Abruzzo e Molise "G. Caporale" | Lorusso A, Marcacci M, Di Domenico M, Ancora M, Curini V, Di Lollo Valeria, Mangone I, Rinaldi A, Delli Compagni E, Scialabba S, Caporale M, Di Pasquale A, Cammà C, Puglia I, Calistri P, Savini G |
| EPI_ISL_3427328 | Mako Medical | Centers for Disease Control and Prevention Division of Viral Diseases, Pathogen Discovery | Dakota Howard, Dhwani Batra, Peter W. Cook, Kara Moser, Adrian Paskey, Jason Caravas, Benjamin Rambo-Martin, Shatavia Morrison, Christopher Gulvick, Scott Sammons, Yvette Unoarumhi, Darlene Wagner, Matthew Schmerer, Matthew Tugwell, Lauren Moon, Clinton R. Paden, Duncan MacCannell |
| EPI_ISL_3427329, EPI_ISL_3427330 | OSPEDALE CIVILE TERAMO - CENTRO TRASFUSIONALE | Istituto Zooprofilattico Sperimentale dell'Abruzzo e Molise "G. Caporale" | Lorusso A, Marcacci M, Di Domenico M, Ancora M, Curini V, Di Lollo Valeria, Mangone I, Rinaldi A, Delli Compagni E, Scialabba S, Caporale M, Di Pasquale A, Cammà C, Puglia I, Calistri P, Savini G |

| | | | |
|---|---|---|---|
| EPI_ISL_3427331 | Mako Medical | Centers for Disease Control and Prevention Division of Viral Diseases, Pathogen Discovery | Dakota Howard, Dhwani Batra, Peter W. Cook, Kara Moser, Adrian Paskey, Jason Caravas, Benjamin Rambo-Martin, Shatavia Morrison, Christopher Gulvick, Scott Sammons, Yvette Unoarumhi, Darlene Wagner, Matthew Schmerer, Matthew Tugwell, Lauren Moon, Clinton R. Paden, Duncan MacCannell |
| EPI_ISL_3427332 | OSPEDALE CIVILE TERAMO - CENTRO TRASFUSIONALE | Istituto Zooprofilattico Sperimentale dell'Abruzzo e Molise "G. Caporale" | Lorusso A, Marcacci M, Di Domenico M, Ancora M, Curini V, Di Lollo Valeria, Mangone I, Rinaldi A, Delli Compagni E, Scialabba S, Caporale M, Di Pasquale A, Cammà C, Puglia I, Calistri P, Savini G |
| EPI_ISL_3427333 | Mako Medical | Centers for Disease Control and Prevention Division of Viral Diseases, Pathogen Discovery | Dakota Howard, Dhwani Batra, Peter W. Cook, Kara Moser, Adrian Paskey, Jason Caravas, Benjamin Rambo-Martin, Shatavia Morrison, Christopher Gulvick, Scott Sammons, Yvette Unoarumhi, Darlene Wagner, Matthew Schmerer, Matthew Tugwell, Lauren Moon, Clinton R. Paden, Duncan MacCannell |
| EPI_ISL_3427334, EPI_ISL_3427335 | OSPEDALE CIVILE TERAMO - CENTRO TRASFUSIONALE | Istituto Zooprofilattico Sperimentale dell'Abruzzo e Molise "G. Caporale" | Lorusso A, Marcacci M, Di Domenico M, Ancora M, Curini V, Di Lollo Valeria, Mangone I, Rinaldi A, Delli Compagni E, Scialabba S, Caporale M, Di Pasquale A, Cammà C, Puglia I, Calistri P, Savini G |
| EPI_ISL_3427336 | Mako Medical | Centers for Disease Control and Prevention Division of Viral Diseases, Pathogen Discovery | Dakota Howard, Dhwani Batra, Peter W. Cook, Kara Moser, Adrian Paskey, Jason Caravas, Benjamin Rambo-Martin, Shatavia Morrison, Christopher Gulvick, Scott Sammons, Yvette Unoarumhi, Darlene Wagner, Matthew Schmerer, Matthew Tugwell, Lauren Moon, Clinton R. Paden, Duncan MacCannell |
| EPI_ISL_3427337, EPI_ISL_3427339, EPI_ISL_3427341, EPI_ISL_3427342 | OSPEDALE CIVILE TERAMO - CENTRO TRASFUSIONALE | Istituto Zooprofilattico Sperimentale dell'Abruzzo e Molise "G. Caporale" | Lorusso A, Marcacci M, Di Domenico M, Ancora M, Curini V, Di Lollo Valeria, Mangone I, Rinaldi A, Delli Compagni E, Scialabba S, Caporale M, Di Pasquale A, Cammà C, Puglia I, Calistri P, Savini G |
| EPI_ISL_3427343 | Mako Medical | Centers for Disease Control and Prevention Division of Viral Diseases, Pathogen Discovery | Dakota Howard, Dhwani Batra, Peter W. Cook, Kara Moser, Adrian Paskey, Jason Caravas, Benjamin Rambo-Martin, Shatavia Morrison, Christopher Gulvick, Scott Sammons, Yvette Unoarumhi, Darlene Wagner, Matthew Schmerer, Matthew Tugwell, Lauren Moon, Clinton R. Paden, Duncan MacCannell |
| EPI_ISL_3427344 | OSPEDALE CIVILE TERAMO - CENTRO TRASFUSIONALE | Istituto Zooprofilattico Sperimentale dell'Abruzzo e Molise "G. Caporale" | Lorusso A, Marcacci M, Di Domenico M, Ancora M, Curini V, Di Lollo Valeria, Mangone I, Rinaldi A, Delli Compagni E, Scialabba S, Caporale M, Di Pasquale A, Cammà C, Puglia I, Calistri P, Savini G |
| EPI_ISL_3427345 | Mako Medical | Centers for Disease Control and Prevention Division of Viral Diseases, Pathogen Discovery | Dakota Howard, Dhwani Batra, Peter W. Cook, Kara Moser, Adrian Paskey, Jason Caravas, Benjamin Rambo-Martin, Shatavia Morrison, Christopher Gulvick, Scott Sammons, Yvette Unoarumhi, Darlene Wagner, Matthew Schmerer, Matthew Tugwell, Lauren Moon, Clinton R. Paden, Duncan MacCannell |
| EPI_ISL_3427346, EPI_ISL_3427347 | OSPEDALE CIVILE TERAMO - CENTRO TRASFUSIONALE | Istituto Zooprofilattico Sperimentale dell'Abruzzo e Molise "G. Caporale" | Lorusso A, Marcacci M, Di Domenico M, Ancora M, Curini V, Di Lollo Valeria, Mangone I, Rinaldi A, Delli Compagni E, Scialabba S, Caporale M, Di Pasquale A, Cammà C, Puglia I, Calistri P, Savini G |
| EPI_ISL_3427348 | Mako Medical | Centers for Disease Control and Prevention Division of Viral Diseases, Pathogen Discovery | Dakota Howard, Dhwani Batra, Peter W. Cook, Kara Moser, Adrian Paskey, Jason Caravas, Benjamin Rambo-Martin, Shatavia Morrison, Christopher Gulvick, Scott Sammons, Yvette Unoarumhi, Darlene Wagner, Matthew Schmerer, Matthew Tugwell, Lauren Moon, Clinton R. Paden, Duncan MacCannell |
| EPI_ISL_3427349 | OSPEDALE CIVILE TERAMO - CENTRO TRASFUSIONALE | Istituto Zooprofilattico Sperimentale dell'Abruzzo e Molise "G. Caporale" | Lorusso A, Marcacci M, Di Domenico M, Ancora M, Curini V, Di Lollo Valeria, Mangone I, Rinaldi A, Delli Compagni E, Scialabba S, Caporale M, Di Pasquale A, Cammà C, Puglia I, Calistri P, Savini G |
| EPI_ISL_3427350 | Mako Medical | Centers for Disease Control and Prevention Division of Viral Diseases, Pathogen Discovery | Dakota Howard, Dhwani Batra, Peter W. Cook, Kara Moser, Adrian Paskey, Jason Caravas, Benjamin Rambo-Martin, Shatavia Morrison, Christopher Gulvick, Scott Sammons, Yvette Unoarumhi, Darlene Wagner, Matthew Schmerer, Matthew Tugwell, Lauren Moon, Clinton R. Paden, Duncan MacCannell |
| EPI_ISL_3427351 | OSPEDALE CIVILE TERAMO - CENTRO TRASFUSIONALE | Istituto Zooprofilattico Sperimentale dell'Abruzzo e Molise "G. Caporale" | Lorusso A, Marcacci M, Di Domenico M, Ancora M, Curini V, Di Lollo Valeria, Mangone I, Rinaldi A, Delli Compagni E, Scialabba S, Caporale M, Di Pasquale A, Cammà C, Puglia I, Calistri P, Savini G |
| EPI_ISL_3427352 | Mako Medical | Centers for Disease Control and Prevention Division of Viral Diseases, Pathogen Discovery | Dakota Howard, Dhwani Batra, Peter W. Cook, Kara Moser, Adrian Paskey, Jason Caravas, Benjamin Rambo-Martin, Shatavia Morrison, Christopher Gulvick, Scott Sammons, Yvette Unoarumhi, Darlene Wagner, Matthew Schmerer, Matthew Tugwell, Lauren Moon, Clinton R. Paden, Duncan MacCannell |
| EPI_ISL_3427353, EPI_ISL_3427354 | OSPEDALE CIVILE TERAMO - CENTRO TRASFUSIONALE | Istituto Zooprofilattico Sperimentale dell'Abruzzo e Molise "G. Caporale" | Lorusso A, Marcacci M, Di Domenico M, Ancora M, Curini V, Di Lollo Valeria, Mangone I, Rinaldi A, Delli Compagni E, Scialabba S, Caporale M, Di Pasquale A, Cammà C, Puglia I, Calistri P, Savini G |
| EPI_ISL_3427355 | Mako Medical | Centers for Disease Control and Prevention Division of Viral Diseases, Pathogen Discovery | Dakota Howard, Dhwani Batra, Peter W. Cook, Kara Moser, Adrian Paskey, Jason Caravas, Benjamin Rambo-Martin, Shatavia Morrison, Christopher Gulvick, Scott Sammons, Yvette Unoarumhi, Darlene Wagner, Matthew Schmerer, Matthew Tugwell, Lauren Moon, Clinton R. Paden, Duncan MacCannell |
| EPI_ISL_3427356, EPI_ISL_3427357 | OSPEDALE CIVILE TERAMO - CENTRO TRASFUSIONALE | Istituto Zooprofilattico Sperimentale dell'Abruzzo e Molise "G. Caporale" | Lorusso A, Marcacci M, Di Domenico M, Ancora M, Curini V, Di Lollo Valeria, Mangone I, Rinaldi A, Delli Compagni E, Scialabba S, Caporale M, Di Pasquale A, Cammà C, Puglia I, Calistri P, Savini G |
| EPI_ISL_3427358 | Mako Medical | Centers for Disease Control and Prevention Division of Viral Diseases, Pathogen Discovery | Dakota Howard, Dhwani Batra, Peter W. Cook, Kara Moser, Adrian Paskey, Jason Caravas, Benjamin Rambo-Martin, Shatavia Morrison, Christopher Gulvick, Scott Sammons, Yvette Unoarumhi, Darlene Wagner, Matthew Schmerer, Matthew Tugwell, Lauren Moon, Clinton R. Paden, Duncan MacCannell |
| EPI_ISL_3427359 | OSPEDALE CIVILE TERAMO - CENTRO TRASFUSIONALE | Istituto Zooprofilattico Sperimentale dell'Abruzzo e Molise "G. Caporale" | Lorusso A, Marcacci M, Di Domenico M, Ancora M, Curini V, Di Lollo Valeria, Mangone I, Rinaldi A, Delli Compagni E, Scialabba S, Caporale M, Di Pasquale A, Cammà C, Puglia I, Calistri P, Savini G |
| EPI_ISL_3427360 | Mako Medical | Centers for Disease Control and Prevention Division of Viral Diseases, Pathogen Discovery | Dakota Howard, Dhwani Batra, Peter W. Cook, Kara Moser, Adrian Paskey, Jason Caravas, Benjamin Rambo-Martin, Shatavia Morrison, Christopher Gulvick, Scott Sammons, Yvette Unoarumhi, Darlene Wagner, Matthew Schmerer, Matthew Tugwell, Lauren Moon, Clinton R. Paden, Duncan MacCannell |
| EPI_ISL_3427361 | OSPEDALE CIVILE TERAMO - CENTRO TRASFUSIONALE | Istituto Zooprofilattico Sperimentale dell'Abruzzo e Molise "G. Caporale" | Lorusso A, Marcacci M, Di Domenico M, Ancora M, Curini V, Di Lollo Valeria, Mangone I, Rinaldi A, Delli Compagni E, Scialabba S, Caporale M, Di Pasquale A, Cammà C, Puglia I, Calistri P, Savini G |
| EPI_ISL_3427362 | Mako Medical | Centers for Disease Control and Prevention Division of Viral Diseases, Pathogen Discovery | Dakota Howard, Dhwani Batra, Peter W. Cook, Kara Moser, Adrian Paskey, Jason Caravas, Benjamin Rambo-Martin, Shatavia Morrison, Christopher Gulvick, Scott Sammons, Yvette Unoarumhi, Darlene Wagner, Matthew Schmerer, Matthew Tugwell, Lauren Moon, Clinton R. Paden, Duncan MacCannell |
| EPI_ISL_3427363 | OSPEDALE CIVILE TERAMO - CENTRO TRASFUSIONALE | Istituto Zooprofilattico Sperimentale dell'Abruzzo e Molise "G. Caporale" | Lorusso A, Marcacci M, Di Domenico M, Ancora M, Curini V, Di Lollo Valeria, Mangone I, Rinaldi A, Delli Compagni E, Scialabba S, Caporale M, Di Pasquale A, Cammà C, Puglia I, Calistri P, Savini G |
| EPI_ISL_3427364 | Mako Medical | Centers for Disease Control and Prevention Division of Viral Diseases, Pathogen Discovery | Dakota Howard, Dhwani Batra, Peter W. Cook, Kara Moser, Adrian Paskey, Jason Caravas, Benjamin Rambo-Martin, Shatavia Morrison, Christopher Gulvick, Scott Sammons, Yvette Unoarumhi, Darlene Wagner, Matthew Schmerer, Matthew Tugwell, Lauren Moon, Clinton R. Paden, Duncan MacCannell |
| EPI_ISL_3427365, EPI_ISL_3427366 | OSPEDALE SAN SALVATORE - MEDICINA DI LABORATORIO | Istituto Zooprofilattico Sperimentale dell'Abruzzo e Molise "G. Caporale" | Lorusso A, Marcacci M, Di Domenico M, Ancora M, Curini V, Di Lollo Valeria, Mangone I, Rinaldi A, Delli Compagni E, Scialabba S, Caporale M, Di Pasquale A, Cammà C, Puglia I, Calistri P, Savini G |
| EPI_ISL_3427367 | Mako Medical | Centers for Disease Control and Prevention Division of Viral Diseases, Pathogen Discovery | Dakota Howard, Dhwani Batra, Peter W. Cook, Kara Moser, Adrian Paskey, Jason Caravas, Benjamin Rambo-Martin, Shatavia Morrison, Christopher Gulvick, Scott Sammons, Yvette Unoarumhi, Darlene Wagner, Matthew Schmerer, Matthew Tugwell, Lauren Moon, Clinton R. Paden, Duncan MacCannell |
| EPI_ISL_3427368 | OSPEDALE SAN SALVATORE - MEDICINA DI LABORATORIO | Istituto Zooprofilattico Sperimentale dell'Abruzzo e Molise "G. Caporale" | Lorusso A, Marcacci M, Di Domenico M, Ancora M, Curini V, Di Lollo Valeria, Mangone I, Rinaldi A, Delli Compagni E, Scialabba S, Caporale M, Di Pasquale A, Cammà C, Puglia I, Calistri P, Savini G |
| EPI_ISL_3427369 | Mako Medical | Centers for Disease Control and Prevention Division of Viral Diseases, Pathogen Discovery | Dakota Howard, Dhwani Batra, Peter W. Cook, Kara Moser, Adrian Paskey, Jason Caravas, Benjamin Rambo-Martin, Shatavia Morrison, Christopher Gulvick, Scott Sammons, Yvette Unoarumhi, Darlene Wagner, Matthew Schmerer, Matthew Tugwell, Lauren Moon, Clinton R. Paden, Duncan MacCannell |
| EPI_ISL_3427370, EPI_ISL_3427371 | OSPEDALE SAN SALVATORE - MEDICINA DI LABORATORIO | Istituto Zooprofilattico Sperimentale dell'Abruzzo e Molise "G. Caporale" | Lorusso A, Marcacci M, Di Domenico M, Ancora M, Curini V, Di Lollo Valeria, Mangone I, Rinaldi A, Delli Compagni E, Scialabba S, Caporale M, Di Pasquale A, Cammà C, Puglia I, Calistri P, Savini G |
| EPI_ISL_3427372 | Mako Medical | Centers for Disease Control and Prevention Division of Viral Diseases, Pathogen Discovery | Dakota Howard, Dhwani Batra, Peter W. Cook, Kara Moser, Adrian Paskey, Jason Caravas, Benjamin Rambo-Martin, Shatavia Morrison, Christopher Gulvick, Scott Sammons, Yvette Unoarumhi, Darlene Wagner, Matthew Schmerer, Matthew Tugwell, Lauren Moon, Clinton R. Paden, Duncan MacCannell |
| EPI_ISL_3427373 | OSPEDALE SAN SALVATORE - MEDICINA DI LABORATORIO | Istituto Zooprofilattico Sperimentale dell'Abruzzo e Molise "G. Caporale" | Lorusso A, Marcacci M, Di Domenico M, Ancora M, Curini V, Di Lollo Valeria, Mangone I, Rinaldi A, Delli Compagni E, Scialabba S, Caporale M, Di Pasquale A, Cammà C, Puglia I, Calistri P, Savini G |
| EPI_ISL_3427374 | Mako Medical | Centers for Disease Control and Prevention Division of Viral Diseases, Pathogen Discovery | Dakota Howard, Dhwani Batra, Peter W. Cook, Kara Moser, Adrian Paskey, Jason Caravas, Benjamin Rambo-Martin, Shatavia Morrison, Christopher Gulvick, Scott Sammons, Yvette Unoarumhi, Darlene Wagner, Matthew Schmerer, Matthew Tugwell, Lauren Moon, Clinton R. Paden, Duncan MacCannell |
| EPI_ISL_3427375, EPI_ISL_3427376 | OSPEDALE SAN SALVATORE - MEDICINA DI LABORATORIO | Istituto Zooprofilattico Sperimentale dell'Abruzzo e Molise "G. Caporale" | Lorusso A, Marcacci M, Di Domenico M, Ancora M, Curini V, Di Lollo Valeria, Mangone I, Rinaldi A, Delli Compagni E, Scialabba S, Caporale M, Di Pasquale A, Cammà C, Puglia I, Calistri P, Savini G |
| EPI_ISL_3427377 | Mako Medical | Centers for Disease Control and Prevention Division of Viral Diseases, Pathogen Discovery | Dakota Howard, Dhwani Batra, Peter W. Cook, Kara Moser, Adrian Paskey, Jason Caravas, Benjamin Rambo-Martin, Shatavia Morrison, Christopher Gulvick, Scott Sammons, Yvette Unoarumhi, Darlene Wagner, Matthew Schmerer, Matthew Tugwell, Lauren Moon, Clinton R. Paden, Duncan MacCannell |
| EPI_ISL_3427378 | OSPEDALE SAN SALVATORE - MEDICINA DI LABORATORIO | Istituto Zooprofilattico Sperimentale dell'Abruzzo e Molise "G. Caporale" | Lorusso A, Marcacci M, Di Domenico M, Ancora M, Curini V, Di Lollo Valeria, Mangone I, Rinaldi A, Delli Compagni E, Scialabba S, Caporale M, Di Pasquale A, Cammà C, Puglia I, Calistri P, Savini G |
| EPI_ISL_3427379 | Mako Medical | Centers for Disease Control and Prevention Division of Viral Diseases, Pathogen Discovery | Dakota Howard, Dhwani Batra, Peter W. Cook, Kara Moser, Adrian Paskey, Jason Caravas, Benjamin Rambo-Martin, Shatavia Morrison, Christopher Gulvick, Scott Sammons, Yvette Unoarumhi, Darlene Wagner, Matthew Schmerer, Matthew Tugwell, Lauren Moon, Clinton R. Paden, Duncan MacCannell |
| EPI_ISL_3427380, EPI_ISL_3427381 | OSPEDALE SAN SALVATORE - MEDICINA DI LABORATORIO | Istituto Zooprofilattico Sperimentale dell'Abruzzo e Molise "G. Caporale" | Lorusso A, Marcacci M, Di Domenico M, Ancora M, Curini V, Di Lollo Valeria, Mangone I, Rinaldi A, Delli Compagni E, Scialabba S, Caporale M, Di Pasquale A, Cammà C, Puglia I, Calistri P, Savini G |
| EPI_ISL_3427382 | Mako Medical | Centers for Disease Control and Prevention Division of Viral Diseases, Pathogen Discovery | Dakota Howard, Dhwani Batra, Peter W. Cook, Kara Moser, Adrian Paskey, Jason Caravas, Benjamin Rambo-Martin, Shatavia Morrison, Christopher Gulvick, Scott Sammons, Yvette Unoarumhi, Darlene Wagner, Matthew Schmerer, Matthew Tugwell, Lauren Moon, Clinton R. Paden, Duncan MacCannell |
| EPI_ISL_3427383 | OSPEDALE SAN SALVATORE - MEDICINA DI LABORATORIO | Istituto Zooprofilattico Sperimentale dell'Abruzzo e Molise "G. Caporale" | Lorusso A, Marcacci M, Di Domenico M, Ancora M, Curini V, Di Lollo Valeria, Mangone I, Rinaldi A, Delli Compagni E, Scialabba S, Caporale M, Di Pasquale A, Cammà C, Puglia I, Calistri P, Savini G |
| EPI_ISL_3427384 | Mako Medical | Centers for Disease Control and Prevention Division of Viral Diseases, Pathogen Discovery | Dakota Howard, Dhwani Batra, Peter W. Cook, Kara Moser, Adrian Paskey, Jason Caravas, Benjamin Rambo-Martin, Shatavia Morrison, Christopher Gulvick, Scott Sammons, Yvette Unoarumhi, Darlene Wagner, Matthew Schmerer, Matthew Tugwell, Lauren Moon, Clinton R. Paden, Duncan MacCannell |
| EPI_ISL_3427385, EPI_ISL_3427386 | OSPEDALE CIVILE TERAMO - CENTRO TRASFUSIONALE | Istituto Zooprofilattico Sperimentale dell'Abruzzo e Molise "G. Caporale" | Lorusso A, Marcacci M, Di Domenico M, Ancora M, Curini V, Di Lollo Valeria, Mangone I, Rinaldi A, Delli Compagni E, Scialabba S, Caporale M, Di Pasquale A, Cammà C, Puglia I, Calistri P, Savini G |
| EPI_ISL_3427387 | Mako Medical | Centers for Disease Control and Prevention Division of Viral Diseases, Pathogen Discovery | Dakota Howard, Dhwani Batra, Peter W. Cook, Kara Moser, Adrian Paskey, Jason Caravas, Benjamin Rambo-Martin, Shatavia Morrison, Christopher Gulvick, Scott Sammons, Yvette Unoarumhi, Darlene Wagner, Matthew Schmerer, Matthew Tugwell, Lauren Moon, Clinton R. Paden, Duncan MacCannell |
| EPI_ISL_3427388 | OSPEDALE CIVILE TERAMO - CENTRO TRASFUSIONALE | Istituto Zooprofilattico Sperimentale dell'Abruzzo e Molise "G. Caporale" | Lorusso A, Marcacci M, Di Domenico M, Ancora M, Curini V, Di Lollo Valeria, Mangone I, Rinaldi A, Delli Compagni E, Scialabba S, Caporale M, Di Pasquale A, Cammà C, Puglia I, Calistri P, Savini G |

| | | | |
|---|---|---|---|
| EPI_ISL_3427389 | Mako Medical | Centers for Disease Control and Prevention Division of Viral Diseases, Pathogen Discovery | Dakota Howard, Dhwani Batra, Peter W. Cook, Kara Moser, Adrian Paskey, Jason Caravas, Benjamin Rambo-Martin, Shatavia Morrison, Christopher Gulvick, Scott Sammons, Yvette Unoarumhi, Darlene Wagner, Matthew Schmerer, Matthew Tugwell, Lauren Moon, Clinton R. Paden, Duncan MacCannell |
| EPI_ISL_3427390 | OSPEDALE CIVILE TERAMO - CENTRO TRASFUSIONALE | Istituto Zooprofilattico Sperimentale dell’Abruzzo e Molise “G. Caporale” | Lorusso A, Marcacci M, Di Domenico M, Ancora M, Curini V, Di Lollo Valeria, Mangone I, Rinaldi A, Delli Compagni E, Scialabba S, Caporale M, Di Pasquale A, Cammà C, Puglia I, Calistri P, Savini G |
| EPI_ISL_3427391, EPI_ISL_3427392, EPI_ISL_3427393, EPI_ISL_3427394, EPI_ISL_3427395, EPI_ISL_3427396, EPI_ISL_3427397, EPI_ISL_3427398, EPI_ISL_3427399, EPI_ISL_3427400, EPI_ISL_3427401, EPI_ISL_3427402, EPI_ISL_3427404, EPI_ISL_3427405, EPI_ISL_3427406, EPI_ISL_3427408, EPI_ISL_3427409, EPI_ISL_3427410, EPI_ISL_3427411, EPI_ISL_3427412, EPI_ISL_3427413, EPI_ISL_3427414, EPI_ISL_3427415, EPI_ISL_3427416, EPI_ISL_3427418, EPI_ISL_3427419, EPI_ISL_3427421, EPI_ISL_3427422, EPI_ISL_3427423, EPI_ISL_3427424, EPI_ISL_3427425, EPI_ISL_3427426, EPI_ISL_3427427, EPI_ISL_3427428, EPI_ISL_3427429, EPI_ISL_3427430, EPI_ISL_3427431, EPI_ISL_3427432, EPI_ISL_3427433, EPI_ISL_3427434, EPI_ISL_3427435, EPI_ISL_3427436, EPI_ISL_3427437, EPI_ISL_3427438, EPI_ISL_3427439, EPI_ISL_3427440, EPI_ISL_3427441, EPI_ISL_3427442, EPI_ISL_3427443, EPI_ISL_3427444, EPI_ISL_3427445, EPI_ISL_3427446, EPI_ISL_3427447, EPI_ISL_3427448, EPI_ISL_3427449, EPI_ISL_3427450, EPI_ISL_3427451, EPI_ISL_3427452, EPI_ISL_3427453, EPI_ISL_3427454, EPI_ISL_3427456, EPI_ISL_3427458, EPI_ISL_3427459, EPI_ISL_3427460, EPI_ISL_3427461, EPI_ISL_3427462, EPI_ISL_3427463, EPI_ISL_3427464, EPI_ISL_3427465, EPI_ISL_3427466, EPI_ISL_3427467, EPI_ISL_3427468, EPI_ISL_3427469, EPI_ISL_3427470, EPI_ISL_3427471, EPI_ISL_3427472, EPI_ISL_3427473, EPI_ISL_3427474, EPI_ISL_3427475, EPI_ISL_3427476, EPI_ISL_3427477, EPI_ISL_3427478, EPI_ISL_3427479, EPI_ISL_3427480, EPI_ISL_3427481, EPI_ISL_3427482, EPI_ISL_3427483, EPI_ISL_3427484, EPI_ISL_3427485, EPI_ISL_3427486, EPI_ISL_3427487, EPI_ISL_3427488, EPI_ISL_3427489, EPI_ISL_3427490, EPI_ISL_3427491, EPI_ISL_3427492, EPI_ISL_3427493, EPI_ISL_3427494, EPI_ISL_3427495, EPI_ISL_3427496, EPI_ISL_3427497, EPI_ISL_3427498, EPI_ISL_3427499, EPI_ISL_3427503, EPI_ISL_3427504, EPI_ISL_3427505, EPI_ISL_3427506, EPI_ISL_3427507, EPI_ISL_3427508, EPI_ISL_3427509, EPI_ISL_3427510, EPI_ISL_3427511, EPI_ISL_3427512, EPI_ISL_3427514, EPI_ISL_3427515, EPI_ISL_3427516, EPI_ISL_3427517, EPI_ISL_3427518, EPI_ISL_3427519, EPI_ISL_3427520, EPI_ISL_3427521, EPI_ISL_3427522, EPI_ISL_3427523, EPI_ISL_3427524, EPI_ISL_3427525, EPI_ISL_3427526, EPI_ISL_3427527, EPI_ISL_3427528, EPI_ISL_3427529, EPI_ISL_3427531, EPI_ISL_3427532, EPI_ISL_3427533, EPI_ISL_3427534, EPI_ISL_3427535, EPI_ISL_3427536, EPI_ISL_3427537, EPI_ISL_3427538, EPI_ISL_3427539, EPI_ISL_3427540, EPI_ISL_3427541, EPI_ISL_3427542, EPI_ISL_3427543, EPI_ISL_3427544, EPI_ISL_3427545, EPI_ISL_3427546, EPI_ISL_3427547, EPI_ISL_3427548, EPI_ISL_3427550, EPI_ISL_3427551, EPI_ISL_3427552, EPI_ISL_3427553, EPI_ISL_3427554, EPI_ISL_3427555, EPI_ISL_3427556, EPI_ISL_3427557, EPI_ISL_3427558, EPI_ISL_3427559, EPI_ISL_3427560, EPI_ISL_3427561, EPI_ISL_3427562, EPI_ISL_3427563, EPI_ISL_3427564, EPI_ISL_3427565, EPI_ISL_3427566, EPI_ISL_3427567, EPI_ISL_3427568, EPI_ISL_3427569, EPI_ISL_3427571, EPI_ISL_3427572, EPI_ISL_3427573, EPI_ISL_3427574, EPI_ISL_3427575, EPI_ISL_3427576, EPI_ISL_3427577, EPI_ISL_3427578, EPI_ISL_3427579, EPI_ISL_3427580, EPI_ISL_3427582, EPI_ISL_3427583, EPI_ISL_3427584, EPI_ISL_3427585, EPI_ISL_3427586, EPI_ISL_3427587, EPI_ISL_3427588, EPI_ISL_3427589, EPI_ISL_3427591, EPI_ISL_3427592, EPI_ISL_3427594, EPI_ISL_3427596, EPI_ISL_3427597, EPI_ISL_3427598, EPI_ISL_3427600, EPI_ISL_3427601, EPI_ISL_3427602, EPI_ISL_3427605, EPI_ISL_3427606, EPI_ISL_3427607, EPI_ISL_3427608, EPI_ISL_3427609, EPI_ISL_3427610, EPI_ISL_3427611, EPI_ISL_3427612, EPI_ISL_3427613, EPI_ISL_3427614, EPI_ISL_3427616, EPI_ISL_3427617, EPI_ISL_3427619, EPI_ISL_3427621, EPI_ISL_3427622, EPI_ISL_3427623, EPI_ISL_3427624, EPI_ISL_3427625, EPI_ISL_3427626, EPI_ISL_3427627, EPI_ISL_3427628, EPI_ISL_3427629, EPI_ISL_3427630, EPI_ISL_3427631, EPI_ISL_3427632, EPI_ISL_3427633, EPI_ISL_3427634, EPI_ISL_3427635, EPI_ISL_3427636, EPI_ISL_3427637, EPI_ISL_3427638, EPI_ISL_3427639, EPI_ISL_3427640, EPI_ISL_3427641, EPI_ISL_3427642, EPI_ISL_3427643, EPI_ISL_3427644, EPI_ISL_3427645, EPI_ISL_3427646, EPI_ISL_3427647, EPI_ISL_3427648, EPI_ISL_3427649, EPI_ISL_3427650, EPI_ISL_3427651, EPI_ISL_3427652, EPI_ISL_3427653, EPI_ISL_3427654, EPI_ISL_3427655, EPI_ISL_3427656, EPI_ISL_3427657, EPI_ISL_3427659, EPI_ISL_3427660, EPI_ISL_3427661, EPI_ISL_3427662, EPI_ISL_3427663, EPI_ISL_3427664, EPI_ISL_3427665, EPI_ISL_3427666, EPI_ISL_3427667, EPI_ISL_3427669, EPI_ISL_3427670, EPI_ISL_3427671, EPI_ISL_3427672, EPI_ISL_3427673, EPI_ISL_3427674, EPI_ISL_3427675, EPI_ISL_3427676, EPI_ISL_3427677, EPI_ISL_3427678, EPI_ISL_3427679, EPI_ISL_3427680, EPI_ISL_3427681, EPI_ISL_3427682, EPI_ISL_3427683, EPI_ISL_3427684, EPI_ISL_3427685, EPI_ISL_3427686, EPI_ISL_3427688, EPI_ISL_3427689, EPI_ISL_3427690, EPI_ISL_3427691, EPI_ISL_3427692, EPI_ISL_3427693, EPI_ISL_3427694, EPI_ISL_3427695, EPI_ISL_3427696, EPI_ISL_3427697, EPI_ISL_3427699, EPI_ISL_3427700, EPI_ISL_3427701, EPI_ISL_3427702, EPI_ISL_3427703, EPI_ISL_3427704, EPI_ISL_3427705, EPI_ISL_3427706, EPI_ISL_3427707, EPI_ISL_3427708, EPI_ISL_3427709, EPI_ISL_3427710, EPI_ISL_3427712, EPI_ISL_3427713, EPI_ISL_3427714, EPI_ISL_3427715, EPI_ISL_3427716, EPI_ISL_3427717, EPI_ISL_3427718, EPI_ISL_3427719, EPI_ISL_3427720, EPI_ISL_3427721, EPI_ISL_3427722, EPI_ISL_3427723, EPI_ISL_3427724, EPI_ISL_3427725, EPI_ISL_3427726, EPI_ISL_3427728, EPI_ISL_3427729, EPI_ISL_3427731, EPI_ISL_3427732, EPI_ISL_3427733, EPI_ISL_3427734, EPI_ISL_3427735, EPI_ISL_3427736, EPI_ISL_3427737, EPI_ISL_3427738, EPI_ISL_3427739, EPI_ISL_3427740, EPI_ISL_3427742, EPI_ISL_3427743, EPI_ISL_3427744, EPI_ISL_3427745, EPI_ISL_3427746, EPI_ISL_3427747, EPI_ISL_3427748, EPI_ISL_3427749, EPI_ISL_3427750, EPI_ISL_3427751, EPI_ISL_3427752, EPI_ISL_3427753, EPI_ISL_3427754, EPI_ISL_3427755, EPI_ISL_3427756, EPI_ISL_3427757, EPI_ISL_3427758, EPI_ISL_3427759, EPI_ISL_3427760, EPI_ISL_3427761, EPI_ISL_3427762, EPI_ISL_3427763, EPI_ISL_3427764, EPI_ISL_3427765, EPI_ISL_3427766, EPI_ISL_3427767, EPI_ISL_3427768, EPI_ISL_3427769, EPI_ISL_3427770, EPI_ISL_3427771, EPI_ISL_3427772, EPI_ISL_3427773, EPI_ISL_3427774, EPI_ISL_3427775, EPI_ISL_3427776, EPI_ISL_3427777, EPI_ISL_3427778, EPI_ISL_3427779, EPI_ISL_3427780, EPI_ISL_3427781, EPI_ISL_3427782, EPI_ISL_3427783, EPI_ISL_3427784, EPI_ISL_3427785, EPI_ISL_3427786, EPI_ISL_3427787, EPI_ISL_3427788, EPI_ISL_3427789, EPI_ISL_3427790, EPI_ISL_3427791, EPI_ISL_3427792, EPI_ISL_3427793, EPI_ISL_3427794, EPI_ISL_3427795, EPI_ISL_3427796, EPI_ISL_3427797, EPI_ISL_3427798, EPI_ISL_3427800, EPI_ISL_3427801, EPI_ISL_3427802, EPI_ISL_3427803, EPI_ISL_3427804, EPI_ISL_3427805, EPI_ISL_3427806, EPI_ISL_3427808, EPI_ISL_3427809, EPI_ISL_3427810, EPI_ISL_3427811, EPI_ISL_3427812, EPI_ISL_3427813, EPI_ISL_3427814, EPI_ISL_3427815, EPI_ISL_3427816, EPI_ISL_3427817, EPI_ISL_3427818, EPI_ISL_3427819, EPI_ISL_3427820, EPI_ISL_3427821, EPI_ISL_3427822, EPI_ISL_3427823, EPI_ISL_3427824, EPI_ISL_3427825, EPI_ISL_3427826, EPI_ISL_3427827, EPI_ISL_3427828, EPI_ISL_3427829, EPI_ISL_3427830, EPI_ISL_3427831, EPI_ISL_3427832, EPI_ISL_3427833, EPI_ISL_3427834, EPI_ISL_3427835, EPI_ISL_3427836, EPI_ISL_3427837, EPI_ISL_3427838, EPI_ISL_3427839, EPI_ISL_3427840, EPI_ISL_3427841, EPI_ISL_3427842, EPI_ISL_3427843, EPI_ISL_3427844, EPI_ISL_3427845, EPI_ISL_3427846, EPI_ISL_3427847, EPI_ISL_3427848, EPI_ISL_3427849, EPI_ISL_3427850, EPI_ISL_3427851, EPI_ISL_3427853, EPI_ISL_3427854, EPI_ISL_3427855, EPI_ISL_3427856, EPI_ISL_3427857, EPI_ISL_3427858, EPI_ISL_3427859, EPI_ISL_3427860, EPI_ISL_3427861, EPI_ISL_3427862, EPI_ISL_3427863, EPI_ISL_3427864, EPI_ISL_3427866, EPI_ISL_3427867, EPI_ISL_3427868, EPI_ISL_3427870, EPI_ISL_3427871, EPI_ISL_3427872, EPI_ISL_3427873, EPI_ISL_3427874, EPI_ISL_3427875, EPI_ISL_3427876, EPI_ISL_3427877, EPI_ISL_3427878, EPI_ISL_3427879, EPI_ISL_3427880, EPI_ISL_3427882, EPI_ISL_3427883, EPI_ISL_3427884, EPI_ISL_3427885, EPI_ISL_3427886, EPI_ISL_3427887, EPI_ISL_3427888, EPI_ISL_3427889, EPI_ISL_3427890, EPI_ISL_3427891, EPI_ISL_3427892, EPI_ISL_3427893, EPI_ISL_3427894, EPI_ISL_3427895, EPI_ISL_3427896, EPI_ISL_3427897, EPI_ISL_3427898, EPI_ISL_3427899, EPI_ISL_3427900, EPI_ISL_3427901, EPI_ISL_3427903, EPI_ISL_3427904, EPI_ISL_3427905, EPI_ISL_3427906, EPI_ISL_3427907, EPI_ISL_3427908, EPI_ISL_3427909, EPI_ISL_3427910, EPI_ISL_3427911, EPI_ISL_3427912, EPI_ISL_3427914, EPI_ISL_3427915, EPI_ISL_3427916, EPI_ISL_3427918, EPI_ISL_3427919, EPI_ISL_3427920, EPI_ISL_3427921, EPI_ISL_3427922, EPI_ISL_3427923, EPI_ISL_3427924, EPI_ISL_3427925, EPI_ISL_3427926, EPI_ISL_3427927, EPI_ISL_3427928, EPI_ISL_3427929, EPI_ISL_3427930, EPI_ISL_3427931, EPI_ISL_3427932, EPI_ISL_3427933, EPI_ISL_3427935, EPI_ISL_3427936, EPI_ISL_3427937, EPI_ISL_3427938, EPI_ISL_3427939, EPI_ISL_3427940, EPI_ISL_3427941, EPI_ISL_3427942, EPI_ISL_3427943, EPI_ISL_3427944, EPI_ISL_3427945, EPI_ISL_3427946, EPI_ISL_3427947, EPI_ISL_3427948, EPI_ISL_3427949, EPI_ISL_3427950, EPI_ISL_3427951, EPI_ISL_3427952, EPI_ISL_3427953, EPI_ISL_3427954, EPI_ISL_3427955, EPI_ISL_3427956 | | | |
| see above | Mako Medical | Centers for Disease Control and Prevention Division of Viral Diseases, Pathogen Discovery | Dakota Howard, Dhwani Batra, Peter W. Cook, Kara Moser, Adrian Paskey, Jason Caravas, Benjamin Rambo-Martin, Shatavia Morrison, Christopher Gulvick, Scott Sammons, Yvette Unoarumhi, Darlene Wagner, Matthew Schmerer, Matthew Tugwell, Lauren Moon, Clinton R. Paden, Duncan MacCannell |
| EPI_ISL_3427957 | UMC Utrecht | UMC Utrecht | Rob Schuurman, Anniek Tanja, Annemarie Wensing |
| EPI_ISL_3427958 | Mako Medical | Centers for Disease Control and Prevention Division of Viral Diseases, Pathogen Discovery | Dakota Howard, Dhwani Batra, Peter W. Cook, Kara Moser, Adrian Paskey, Jason Caravas, Benjamin Rambo-Martin, Shatavia Morrison, Christopher Gulvick, Scott Sammons, Yvette Unoarumhi, Darlene Wagner, Matthew Schmerer, Matthew Tugwell, Lauren Moon, Clinton R. Paden, Duncan MacCannell |
| EPI_ISL_3427959 | UMC Utrecht | UMC Utrecht | Rob Schuurman, Anniek Tanja, Annemarie Wensing |
| EPI_ISL_3427960 | Mako Medical | Centers for Disease Control and Prevention Division of Viral Diseases, Pathogen Discovery | Dakota Howard, Dhwani Batra, Peter W. Cook, Kara Moser, Adrian Paskey, Jason Caravas, Benjamin Rambo-Martin, Shatavia Morrison, Christopher Gulvick, Scott Sammons, Yvette Unoarumhi, Darlene Wagner, Matthew Schmerer, Matthew Tugwell, Lauren Moon, Clinton R. Paden, Duncan MacCannell |
| EPI_ISL_3427961 | UMC Utrecht | UMC Utrecht | Rob Schuurman, Anniek Tanja, Annemarie Wensing |
| EPI_ISL_3427962 | Mako Medical | Centers for Disease Control and Prevention Division of Viral Diseases, Pathogen Discovery | Dakota Howard, Dhwani Batra, Peter W. Cook, Kara Moser, Adrian Paskey, Jason Caravas, Benjamin Rambo-Martin, Shatavia Morrison, Christopher Gulvick, Scott Sammons, Yvette Unoarumhi, Darlene Wagner, Matthew Schmerer, Matthew Tugwell, Lauren Moon, Clinton R. Paden, Duncan MacCannell |
| EPI_ISL_3427963 | UMC Utrecht | UMC Utrecht | Rob Schuurman, Anniek Tanja, Annemarie Wensing |
| EPI_ISL_3427964 | Mako Medical | Centers for Disease Control and Prevention Division of Viral Diseases, Pathogen Discovery | Dakota Howard, Dhwani Batra, Peter W. Cook, Kara Moser, Adrian Paskey, Jason Caravas, Benjamin Rambo-Martin, Shatavia Morrison, Christopher Gulvick, Scott Sammons, Yvette Unoarumhi, Darlene Wagner, Matthew Schmerer, Matthew Tugwell, Lauren Moon, Clinton R. Paden, Duncan MacCannell |
| EPI_ISL_3427965 | UMC Utrecht | UMC Utrecht | Rob Schuurman, Anniek Tanja, Annemarie Wensing |
| EPI_ISL_3427966 | Mako Medical | Centers for Disease Control and Prevention Division of Viral Diseases, Pathogen Discovery | Dakota Howard, Dhwani Batra, Peter W. Cook, Kara Moser, Adrian Paskey, Jason Caravas, Benjamin Rambo-Martin, Shatavia Morrison, Christopher Gulvick, Scott Sammons, Yvette Unoarumhi, Darlene Wagner, Matthew Schmerer, Matthew Tugwell, Lauren Moon, Clinton R. Paden, Duncan MacCannell |
| EPI_ISL_3427967 | UMC Utrecht | UMC Utrecht | Rob Schuurman, Anniek Tanja, Annemarie Wensing |
| EPI_ISL_3427968 | Mako Medical | Centers for Disease Control and Prevention Division of Viral Diseases, Pathogen Discovery | Dakota Howard, Dhwani Batra, Peter W. Cook, Kara Moser, Adrian Paskey, Jason Caravas, Benjamin Rambo-Martin, Shatavia Morrison, Christopher Gulvick, Scott Sammons, Yvette Unoarumhi, Darlene Wagner, Matthew Schmerer, Matthew Tugwell, Lauren Moon, Clinton R. Paden, Duncan MacCannell |
| EPI_ISL_3427969 | UMC Utrecht | UMC Utrecht | Rob Schuurman, Anniek Tanja, Annemarie Wensing |
| EPI_ISL_3427970 | Mako Medical | Centers for Disease Control and Prevention Division of Viral Diseases, Pathogen Discovery | Dakota Howard, Dhwani Batra, Peter W. Cook, Kara Moser, Adrian Paskey, Jason Caravas, Benjamin Rambo-Martin, Shatavia Morrison, Christopher Gulvick, Scott Sammons, Yvette Unoarumhi, Darlene Wagner, Matthew Schmerer, Matthew Tugwell, Lauren Moon, Clinton R. Paden, Duncan MacCannell |
| EPI_ISL_3427971 | UMC Utrecht | UMC Utrecht | Rob Schuurman, Anniek Tanja, Annemarie Wensing |
| EPI_ISL_3427972 | Mako Medical | Centers for Disease Control and Prevention Division of Viral Diseases, Pathogen Discovery | Dakota Howard, Dhwani Batra, Peter W. Cook, Kara Moser, Adrian Paskey, Jason Caravas, Benjamin Rambo-Martin, Shatavia Morrison, Christopher Gulvick, Scott Sammons, Yvette Unoarumhi, Darlene Wagner, Matthew Schmerer, Matthew Tugwell, Lauren Moon, Clinton R. Paden, Duncan MacCannell |
| EPI_ISL_3427973 | UMC Utrecht | UMC Utrecht | Rob Schuurman, Anniek Tanja, Annemarie Wensing |
| EPI_ISL_3427974 | Mako Medical | Centers for Disease Control and Prevention Division of Viral Diseases, Pathogen Discovery | Dakota Howard, Dhwani Batra, Peter W. Cook, Kara Moser, Adrian Paskey, Jason Caravas, Benjamin Rambo-Martin, Shatavia Morrison, Christopher Gulvick, Scott Sammons, Yvette Unoarumhi, Darlene Wagner, Matthew Schmerer, Matthew Tugwell, Lauren Moon, Clinton R. Paden, Duncan MacCannell |
| EPI_ISL_3427975 | UMC Utrecht | UMC Utrecht | Rob Schuurman, Anniek Tanja, Annemarie Wensing |
| EPI_ISL_3427976 | Mako Medical | Centers for Disease Control and Prevention Division of Viral Diseases, Pathogen Discovery | Dakota Howard, Dhwani Batra, Peter W. Cook, Kara Moser, Adrian Paskey, Jason Caravas, Benjamin Rambo-Martin, Shatavia Morrison, Christopher Gulvick, Scott Sammons, Yvette Unoarumhi, Darlene Wagner, Matthew Schmerer, Matthew Tugwell, Lauren Moon, Clinton R. Paden, Duncan MacCannell |
| EPI_ISL_3427977 | UMC Utrecht | UMC Utrecht | Rob Schuurman, Anniek Tanja, Annemarie Wensing |
| EPI_ISL_3427978 | Mako Medical | Centers for Disease Control and Prevention Division of Viral Diseases, Pathogen Discovery | Dakota Howard, Dhwani Batra, Peter W. Cook, Kara Moser, Adrian Paskey, Jason Caravas, Benjamin Rambo-Martin, Shatavia Morrison, Christopher Gulvick, Scott Sammons, Yvette Unoarumhi, Darlene Wagner, Matthew Schmerer, Matthew Tugwell, Lauren Moon, Clinton R. Paden, Duncan MacCannell |
| EPI_ISL_3427979 | UMC Utrecht | UMC Utrecht | Rob Schuurman, Anniek Tanja, Annemarie Wensing |
| EPI_ISL_3427980 | Mako Medical | Centers for Disease Control and Prevention Division of Viral Diseases, Pathogen Discovery | Dakota Howard, Dhwani Batra, Peter W. Cook, Kara Moser, Adrian Paskey, Jason Caravas, Benjamin Rambo-Martin, Shatavia Morrison, Christopher Gulvick, Scott Sammons, Yvette Unoarumhi, Darlene Wagner, Matthew Schmerer, Matthew Tugwell, Lauren Moon, Clinton R. Paden, Duncan MacCannell |
| EPI_ISL_3427981 | UMC Utrecht | UMC Utrecht | Rob Schuurman, Anniek Tanja, Annemarie Wensing |
| EPI_ISL_3427982 | Mako Medical | Centers for Disease Control and Prevention Division of Viral Diseases, Pathogen Discovery | Dakota Howard, Dhwani Batra, Peter W. Cook, Kara Moser, Adrian Paskey, Jason Caravas, Benjamin Rambo-Martin, Shatavia Morrison, Christopher Gulvick, Scott Sammons, Yvette Unoarumhi, Darlene Wagner, Matthew Schmerer, Matthew Tugwell, Lauren Moon, Clinton R. Paden, Duncan MacCannell |
| EPI_ISL_3427983 | UMC Utrecht | UMC Utrecht | Rob Schuurman, Anniek Tanja, Annemarie Wensing |
| EPI_ISL_3427984 | Mako Medical | Centers for Disease Control and Prevention Division of Viral Diseases, Pathogen Discovery | Dakota Howard, Dhwani Batra, Peter W. Cook, Kara Moser, Adrian Paskey, Jason Caravas, Benjamin Rambo-Martin, Shatavia Morrison, Christopher Gulvick, Scott Sammons, Yvette Unoarumhi, Darlene Wagner, Matthew Schmerer, Matthew Tugwell, Lauren Moon, Clinton R. Paden, Duncan MacCannell |
| EPI_ISL_3427985 | UMC Utrecht | UMC Utrecht | Rob Schuurman, Anniek Tanja, Annemarie Wensing |
| EPI_ISL_3427986 | Mako Medical | Centers for Disease Control and Prevention Division of Viral Diseases, Pathogen Discovery | Dakota Howard, Dhwani Batra, Peter W. Cook, Kara Moser, Adrian Paskey, Jason Caravas, Benjamin Rambo-Martin, Shatavia Morrison, Christopher Gulvick, Scott Sammons, Yvette Unoarumhi, Darlene Wagner, Matthew Schmerer, Matthew Tugwell, Lauren Moon, Clinton R. Paden, Duncan MacCannell |
| EPI_ISL_3427987 | UMC Utrecht | UMC Utrecht | Rob Schuurman, Anniek Tanja, Annemarie Wensing |
| EPI_ISL_3427988 | Mako Medical | Centers for Disease Control and Prevention Division of Viral Diseases, Pathogen Discovery | Dakota Howard, Dhwani Batra, Peter W. Cook, Kara Moser, Adrian Paskey, Jason Caravas, Benjamin Rambo-Martin, Shatavia Morrison, Christopher Gulvick, Scott Sammons, Yvette Unoarumhi, Darlene Wagner, Matthew Schmerer, Matthew Tugwell, Lauren Moon, Clinton R. Paden, Duncan MacCannell |
| EPI_ISL_3427989 | UMC Utrecht | UMC Utrecht | Rob Schuurman, Anniek Tanja, Annemarie Wensing |
| EPI_ISL_3427990 | Mako Medical | Centers for Disease Control and Prevention Division of Viral Diseases, Pathogen Discovery | Dakota Howard, Dhwani Batra, Peter W. Cook, Kara Moser, Adrian Paskey, Jason Caravas, Benjamin Rambo-Martin, Shatavia Morrison, Christopher Gulvick, Scott Sammons, Yvette Unoarumhi, Darlene Wagner, Matthew Schmerer, Matthew Tugwell, Lauren Moon, Clinton R. Paden, Duncan MacCannell |
| EPI_ISL_3427991 | UMC Utrecht | UMC Utrecht | Rob Schuurman, Anniek Tanja, Annemarie Wensing |
| EPI_ISL_3427992 | Mako Medical | Centers for Disease Control and Prevention Division of Viral Diseases, Pathogen Discovery | Dakota Howard, Dhwani Batra, Peter W. Cook, Kara Moser, Adrian Paskey, Jason Caravas, Benjamin Rambo-Martin, Shatavia Morrison, Christopher Gulvick, Scott Sammons, Yvette Unoarumhi, Darlene Wagner, Matthew Schmerer, Matthew Tugwell, Lauren Moon, Clinton R. Paden, Duncan MacCannell |
| EPI_ISL_3427993 | UMC Utrecht | UMC Utrecht | Rob Schuurman, Anniek Tanja, Annemarie Wensing |
| EPI_ISL_3427994 | Mako Medical | Centers for Disease Control and Prevention Division of Viral Diseases, Pathogen Discovery | Dakota Howard, Dhwani Batra, Peter W. Cook, Kara Moser, Adrian Paskey, Jason Caravas, Benjamin Rambo-Martin, Shatavia Morrison, Christopher Gulvick, Scott Sammons, Yvette Unoarumhi, Darlene Wagner, Matthew Schmerer, Matthew Tugwell, Lauren Moon, Clinton R. Paden, Duncan MacCannell |
| EPI_ISL_3427995 | UMC Utrecht | UMC Utrecht | Rob Schuurman, Anniek Tanja, Annemarie Wensing |
| EPI_ISL_3427996 | Mako Medical | Centers for Disease Control and Prevention Division of Viral Diseases, Pathogen Discovery | Dakota Howard, Dhwani Batra, Peter W. Cook, Kara Moser, Adrian Paskey, Jason Caravas, Benjamin Rambo-Martin, Shatavia Morrison, Christopher Gulvick, Scott Sammons, Yvette Unoarumhi, Darlene Wagner, Matthew Schmerer, Matthew Tugwell, Lauren Moon, Clinton R. Paden, Duncan MacCannell |

| | | | |
|---|---|---|---|
| EPI_ISL_3427997 | UMC Utrecht | UMC Utrecht | Rob Schuurman, Anniek Tanja, Annemarie Wensing |
| EPI_ISL_3427998 | Mako Medical | Centers for Disease Control and Prevention Division of Viral Diseases, Pathogen Discovery | Dakota Howard, Dhwani Batra, Peter W. Cook, Kara Moser, Adrian Paskey, Jason Caravas, Benjamin Rambo-Martin, Shatavia Morrison, Christopher Gulvick, Scott Sammons, Yvette Unoarumhi, Darlene Wagner, Matthew Schmerer, Matthew Tugwell, Lauren Moon, Clinton R. Paden, Duncan MacCannell |
| EPI_ISL_3427999 | UMC Utrecht | UMC Utrecht | Rob Schuurman, Anniek Tanja, Annemarie Wensing |
| EPI_ISL_3428000 | Mako Medical | Centers for Disease Control and Prevention Division of Viral Diseases, Pathogen Discovery | Dakota Howard, Dhwani Batra, Peter W. Cook, Kara Moser, Adrian Paskey, Jason Caravas, Benjamin Rambo-Martin, Shatavia Morrison, Christopher Gulvick, Scott Sammons, Yvette Unoarumhi, Darlene Wagner, Matthew Schmerer, Matthew Tugwell, Lauren Moon, Clinton R. Paden, Duncan MacCannell |
| EPI_ISL_3428001 | UMC Utrecht | UMC Utrecht | Rob Schuurman, Anniek Tanja, Annemarie Wensing |
| EPI_ISL_3428002 | Mako Medical | Centers for Disease Control and Prevention Division of Viral Diseases, Pathogen Discovery | Dakota Howard, Dhwani Batra, Peter W. Cook, Kara Moser, Adrian Paskey, Jason Caravas, Benjamin Rambo-Martin, Shatavia Morrison, Christopher Gulvick, Scott Sammons, Yvette Unoarumhi, Darlene Wagner, Matthew Schmerer, Matthew Tugwell, Lauren Moon, Clinton R. Paden, Duncan MacCannell |
| EPI_ISL_3428003 | UMC Utrecht | UMC Utrecht | Rob Schuurman, Anniek Tanja, Annemarie Wensing |
| EPI_ISL_3428004, EPI_ISL_3428006 | Mako Medical | Centers for Disease Control and Prevention Division of Viral Diseases, Pathogen Discovery | Dakota Howard, Dhwani Batra, Peter W. Cook, Kara Moser, Adrian Paskey, Jason Caravas, Benjamin Rambo-Martin, Shatavia Morrison, Christopher Gulvick, Scott Sammons, Yvette Unoarumhi, Darlene Wagner, Matthew Schmerer, Matthew Tugwell, Lauren Moon, Clinton R. Paden, Duncan MacCannell |
| EPI_ISL_3428007 | UMC Utrecht | UMC Utrecht | Rob Schuurman, Anniek Tanja, Annemarie Wensing |
| EPI_ISL_3428008 | Mako Medical | Centers for Disease Control and Prevention Division of Viral Diseases, Pathogen Discovery | Dakota Howard, Dhwani Batra, Peter W. Cook, Kara Moser, Adrian Paskey, Jason Caravas, Benjamin Rambo-Martin, Shatavia Morrison, Christopher Gulvick, Scott Sammons, Yvette Unoarumhi, Darlene Wagner, Matthew Schmerer, Matthew Tugwell, Lauren Moon, Clinton R. Paden, Duncan MacCannell |
| EPI_ISL_3428009 | UMC Utrecht | UMC Utrecht | Rob Schuurman, Anniek Tanja, Annemarie Wensing |
| EPI_ISL_3428010 | Mako Medical | Centers for Disease Control and Prevention Division of Viral Diseases, Pathogen Discovery | Dakota Howard, Dhwani Batra, Peter W. Cook, Kara Moser, Adrian Paskey, Jason Caravas, Benjamin Rambo-Martin, Shatavia Morrison, Christopher Gulvick, Scott Sammons, Yvette Unoarumhi, Darlene Wagner, Matthew Schmerer, Matthew Tugwell, Lauren Moon, Clinton R. Paden, Duncan MacCannell |
| EPI_ISL_3428011 | UMC Utrecht | UMC Utrecht | Rob Schuurman, Anniek Tanja, Annemarie Wensing |
| EPI_ISL_3428012 | Mako Medical | Centers for Disease Control and Prevention Division of Viral Diseases, Pathogen Discovery | Dakota Howard, Dhwani Batra, Peter W. Cook, Kara Moser, Adrian Paskey, Jason Caravas, Benjamin Rambo-Martin, Shatavia Morrison, Christopher Gulvick, Scott Sammons, Yvette Unoarumhi, Darlene Wagner, Matthew Schmerer, Matthew Tugwell, Lauren Moon, Clinton R. Paden, Duncan MacCannell |
| EPI_ISL_3428013 | UMC Utrecht | UMC Utrecht | Rob Schuurman, Anniek Tanja, Annemarie Wensing |
| EPI_ISL_3428014 | Mako Medical | Centers for Disease Control and Prevention Division of Viral Diseases, Pathogen Discovery | Dakota Howard, Dhwani Batra, Peter W. Cook, Kara Moser, Adrian Paskey, Jason Caravas, Benjamin Rambo-Martin, Shatavia Morrison, Christopher Gulvick, Scott Sammons, Yvette Unoarumhi, Darlene Wagner, Matthew Schmerer, Matthew Tugwell, Lauren Moon, Clinton R. Paden, Duncan MacCannell |
| EPI_ISL_3428015 | UMC Utrecht | UMC Utrecht | Rob Schuurman, Anniek Tanja, Annemarie Wensing |
| EPI_ISL_3428016 | Mako Medical | Centers for Disease Control and Prevention Division of Viral Diseases, Pathogen Discovery | Dakota Howard, Dhwani Batra, Peter W. Cook, Kara Moser, Adrian Paskey, Jason Caravas, Benjamin Rambo-Martin, Shatavia Morrison, Christopher Gulvick, Scott Sammons, Yvette Unoarumhi, Darlene Wagner, Matthew Schmerer, Matthew Tugwell, Lauren Moon, Clinton R. Paden, Duncan MacCannell |
| EPI_ISL_3428017 | UMC Utrecht | UMC Utrecht | Rob Schuurman, Anniek Tanja, Annemarie Wensing |
| EPI_ISL_3428018 | Mako Medical | Centers for Disease Control and Prevention Division of Viral Diseases, Pathogen Discovery | Dakota Howard, Dhwani Batra, Peter W. Cook, Kara Moser, Adrian Paskey, Jason Caravas, Benjamin Rambo-Martin, Shatavia Morrison, Christopher Gulvick, Scott Sammons, Yvette Unoarumhi, Darlene Wagner, Matthew Schmerer, Matthew Tugwell, Lauren Moon, Clinton R. Paden, Duncan MacCannell |
| EPI_ISL_3428019 | UMC Utrecht | UMC Utrecht | Rob Schuurman, Anniek Tanja, Annemarie Wensing |
| EPI_ISL_3428020 | Mako Medical | Centers for Disease Control and Prevention Division of Viral Diseases, Pathogen Discovery | Dakota Howard, Dhwani Batra, Peter W. Cook, Kara Moser, Adrian Paskey, Jason Caravas, Benjamin Rambo-Martin, Shatavia Morrison, Christopher Gulvick, Scott Sammons, Yvette Unoarumhi, Darlene Wagner, Matthew Schmerer, Matthew Tugwell, Lauren Moon, Clinton R. Paden, Duncan MacCannell |
| EPI_ISL_3428021 | UMC Utrecht | UMC Utrecht | Rob Schuurman, Anniek Tanja, Annemarie Wensing |
| EPI_ISL_3428022 | Mako Medical | Centers for Disease Control and Prevention Division of Viral Diseases, Pathogen Discovery | Dakota Howard, Dhwani Batra, Peter W. Cook, Kara Moser, Adrian Paskey, Jason Caravas, Benjamin Rambo-Martin, Shatavia Morrison, Christopher Gulvick, Scott Sammons, Yvette Unoarumhi, Darlene Wagner, Matthew Schmerer, Matthew Tugwell, Lauren Moon, Clinton R. Paden, Duncan MacCannell |
| EPI_ISL_3428023 | UMC Utrecht | UMC Utrecht | Rob Schuurman, Anniek Tanja, Annemarie Wensing |
| EPI_ISL_3428024 | Mako Medical | Centers for Disease Control and Prevention Division of Viral Diseases, Pathogen Discovery | Dakota Howard, Dhwani Batra, Peter W. Cook, Kara Moser, Adrian Paskey, Jason Caravas, Benjamin Rambo-Martin, Shatavia Morrison, Christopher Gulvick, Scott Sammons, Yvette Unoarumhi, Darlene Wagner, Matthew Schmerer, Matthew Tugwell, Lauren Moon, Clinton R. Paden, Duncan MacCannell |
| EPI_ISL_3428025 | UMC Utrecht | UMC Utrecht | Rob Schuurman, Anniek Tanja, Annemarie Wensing |
| EPI_ISL_3428026 | Mako Medical | Centers for Disease Control and Prevention Division of Viral Diseases, Pathogen Discovery | Dakota Howard, Dhwani Batra, Peter W. Cook, Kara Moser, Adrian Paskey, Jason Caravas, Benjamin Rambo-Martin, Shatavia Morrison, Christopher Gulvick, Scott Sammons, Yvette Unoarumhi, Darlene Wagner, Matthew Schmerer, Matthew Tugwell, Lauren Moon, Clinton R. Paden, Duncan MacCannell |
| EPI_ISL_3428027 | UMC Utrecht | UMC Utrecht | Rob Schuurman, Anniek Tanja, Annemarie Wensing |
| EPI_ISL_3428028 | Mako Medical | Centers for Disease Control and Prevention Division of Viral Diseases, Pathogen Discovery | Dakota Howard, Dhwani Batra, Peter W. Cook, Kara Moser, Adrian Paskey, Jason Caravas, Benjamin Rambo-Martin, Shatavia Morrison, Christopher Gulvick, Scott Sammons, Yvette Unoarumhi, Darlene Wagner, Matthew Schmerer, Matthew Tugwell, Lauren Moon, Clinton R. Paden, Duncan MacCannell |
| EPI_ISL_3428029 | UMC Utrecht | UMC Utrecht | Rob Schuurman, Anniek Tanja, Annemarie Wensing |
| EPI_ISL_3428030, EPI_ISL_3428031, EPI_ISL_3428032, EPI_ISL_3428033, EPI_ISL_3428034, EPI_ISL_3428035, EPI_ISL_3428036, EPI_ISL_3428037, EPI_ISL_3428038, EPI_ISL_3428039, EPI_ISL_3428040, EPI_ISL_3428041, EPI_ISL_3428042, EPI_ISL_3428043, EPI_ISL_3428044, EPI_ISL_3428045, EPI_ISL_3428046, EPI_ISL_3428047, EPI_ISL_3428048, EPI_ISL_3428049, EPI_ISL_3428050, EPI_ISL_3428051, EPI_ISL_3428052, EPI_ISL_3428053, EPI_ISL_3428054, EPI_ISL_3428055, EPI_ISL_3428056, EPI_ISL_3428057, EPI_ISL_3428058, EPI_ISL_3428059, EPI_ISL_3428060, EPI_ISL_3428061, EPI_ISL_3428062, EPI_ISL_3428063, EPI_ISL_3428064, EPI_ISL_3428065, EPI_ISL_3428066, EPI_ISL_3428067, EPI_ISL_3428068, EPI_ISL_3428069, EPI_ISL_3428070, EPI_ISL_3428071, EPI_ISL_3428072, EPI_ISL_3428073, EPI_ISL_3428074, EPI_ISL_3428075, EPI_ISL_3428076, EPI_ISL_3428077, EPI_ISL_3428078, EPI_ISL_3428079, EPI_ISL_3428080, EPI_ISL_3428081, EPI_ISL_3428082, EPI_ISL_3428083, EPI_ISL_3428084, EPI_ISL_3428085, EPI_ISL_3428086, EPI_ISL_3428087, EPI_ISL_3428088, EPI_ISL_3428089, EPI_ISL_3428090, EPI_ISL_3428091, EPI_ISL_3428092, EPI_ISL_3428093, EPI_ISL_3428095, EPI_ISL_3428096, EPI_ISL_3428097, EPI_ISL_3428098, EPI_ISL_3428099, EPI_ISL_3428100, EPI_ISL_3428101, EPI_ISL_3428102, EPI_ISL_3428103, EPI_ISL_3428104, EPI_ISL_3428105, EPI_ISL_3428106, EPI_ISL_3428107, EPI_ISL_3428108, EPI_ISL_3428109, EPI_ISL_3428110, EPI_ISL_3428111, EPI_ISL_3428112, EPI_ISL_3428113, EPI_ISL_3428114, EPI_ISL_3428115, EPI_ISL_3428116, EPI_ISL_3428117, EPI_ISL_3428118, EPI_ISL_3428119, EPI_ISL_3428120, EPI_ISL_3428121, EPI_ISL_3428122, EPI_ISL_3428124, EPI_ISL_3428125, EPI_ISL_3428126, EPI_ISL_3428127, EPI_ISL_3428128, EPI_ISL_3428129, EPI_ISL_3428130, EPI_ISL_3428131, EPI_ISL_3428132, EPI_ISL_3428133, EPI_ISL_3428134, EPI_ISL_3428135, EPI_ISL_3428136, EPI_ISL_3428137, EPI_ISL_3428138, EPI_ISL_3428139, EPI_ISL_3428140, EPI_ISL_3428141, EPI_ISL_3428142, EPI_ISL_3428143, EPI_ISL_3428144, EPI_ISL_3428145, EPI_ISL_3428146, EPI_ISL_3428147, EPI_ISL_3428148, EPI_ISL_3428149, EPI_ISL_3428150, EPI_ISL_3428151, EPI_ISL_3428152, EPI_ISL_3428153, EPI_ISL_3428154, EPI_ISL_3428155, EPI_ISL_3428156, EPI_ISL_3428157, EPI_ISL_3428158, EPI_ISL_3428159, EPI_ISL_3428160, EPI_ISL_3428161, EPI_ISL_3428162, EPI_ISL_3428163, EPI_ISL_3428164, EPI_ISL_3428165, EPI_ISL_3428166, EPI_ISL_3428167, EPI_ISL_3428168, EPI_ISL_3428169, EPI_ISL_3428170, EPI_ISL_3428171, EPI_ISL_3428172, EPI_ISL_3428173, EPI_ISL_3428174, EPI_ISL_3428175, EPI_ISL_3428176, EPI_ISL_3428177, EPI_ISL_3428178, EPI_ISL_3428179, EPI_ISL_3428180, EPI_ISL_3428181, EPI_ISL_3428182, EPI_ISL_3428183, EPI_ISL_3428184, EPI_ISL_3428185, EPI_ISL_3428186, EPI_ISL_3428187, EPI_ISL_3428188, EPI_ISL_3428189, EPI_ISL_3428190, EPI_ISL_3428191, EPI_ISL_3428192, EPI_ISL_3428193, EPI_ISL_3428194, EPI_ISL_3428195, EPI_ISL_3428196, EPI_ISL_3428197, EPI_ISL_3428198, EPI_ISL_3428199, EPI_ISL_3428200, EPI_ISL_3428201, EPI_ISL_3428202, EPI_ISL_3428203, EPI_ISL_3428204, EPI_ISL_3428205, EPI_ISL_3428206, EPI_ISL_3428207, EPI_ISL_3428208, EPI_ISL_3428209, EPI_ISL_3428210, EPI_ISL_3428211, EPI_ISL_3428212, EPI_ISL_3428213, EPI_ISL_3428214, EPI_ISL_3428215, EPI_ISL_3428216, EPI_ISL_3428217, EPI_ISL_3428219, EPI_ISL_3428220, EPI_ISL_3428222, EPI_ISL_3428223, EPI_ISL_3428224, EPI_ISL_3428225, EPI_ISL_3428226, EPI_ISL_3428227, EPI_ISL_3428228, EPI_ISL_3428229, EPI_ISL_3428230, EPI_ISL_3428231, EPI_ISL_3428232, EPI_ISL_3428233, EPI_ISL_3428234, EPI_ISL_3428235, EPI_ISL_3428236, EPI_ISL_3428237, EPI_ISL_3428238, EPI_ISL_3428239, EPI_ISL_3428240, EPI_ISL_3428241, EPI_ISL_3428242, EPI_ISL_3428243, EPI_ISL_3428244, EPI_ISL_3428245, EPI_ISL_3428246, EPI_ISL_3428247, EPI_ISL_3428248, EPI_ISL_3428249, EPI_ISL_3428250, EPI_ISL_3428251, EPI_ISL_3428253, EPI_ISL_3428254, EPI_ISL_3428255, EPI_ISL_3428256, EPI_ISL_3428257, EPI_ISL_3428258, EPI_ISL_3428259, EPI_ISL_3428260, EPI_ISL_3428261, EPI_ISL_3428262, EPI_ISL_3428263, EPI_ISL_3428264, EPI_ISL_3428265, EPI_ISL_3428266, EPI_ISL_3428268, EPI_ISL_3428269, EPI_ISL_3428270, EPI_ISL_3428271, EPI_ISL_3428272, EPI_ISL_3428273, EPI_ISL_3428274, EPI_ISL_3428275, EPI_ISL_3428276, EPI_ISL_3428277, EPI_ISL_3428278, EPI_ISL_3428279, EPI_ISL_3428280, EPI_ISL_3428281, EPI_ISL_3428282, EPI_ISL_3428283, EPI_ISL_3428284, EPI_ISL_3428285, EPI_ISL_3428286, EPI_ISL_3428287, EPI_ISL_3428288, EPI_ISL_3428289, EPI_ISL_3428290, EPI_ISL_3428291, EPI_ISL_3428292, EPI_ISL_3428293, EPI_ISL_3428294, EPI_ISL_3428295, EPI_ISL_3428296, EPI_ISL_3428297, EPI_ISL_3428298, EPI_ISL_3428299, EPI_ISL_3428301, EPI_ISL_3428302, EPI_ISL_3428304, EPI_ISL_3428305, EPI_ISL_3428306, EPI_ISL_3428307, EPI_ISL_3428308, EPI_ISL_3428309, EPI_ISL_3428310, EPI_ISL_3428311, EPI_ISL_3428312, EPI_ISL_3428313, EPI_ISL_3428314, EPI_ISL_3428315, EPI_ISL_3428316, EPI_ISL_3428317, EPI_ISL_3428318, EPI_ISL_3428319, EPI_ISL_3428320, EPI_ISL_3428321, EPI_ISL_3428322, EPI_ISL_3428323, EPI_ISL_3428324, EPI_ISL_3428325, EPI_ISL_3428326, EPI_ISL_3428327, EPI_ISL_3428328, EPI_ISL_3428329, EPI_ISL_3428330, EPI_ISL_3428331, EPI_ISL_3428333, EPI_ISL_3428335, EPI_ISL_3428336, EPI_ISL_3428337, EPI_ISL_3428338, EPI_ISL_3428339, EPI_ISL_3428340, EPI_ISL_3428341, EPI_ISL_3428342, EPI_ISL_3428343, EPI_ISL_3428344, EPI_ISL_3428345, EPI_ISL_3428346, EPI_ISL_3428347, EPI_ISL_3428348, EPI_ISL_3428349, EPI_ISL_3428350, EPI_ISL_3428351, EPI_ISL_3428352, EPI_ISL_3428353, EPI_ISL_3428354, EPI_ISL_3428355, EPI_ISL_3428356, EPI_ISL_3428357, EPI_ISL_3428358, EPI_ISL_3428359, EPI_ISL_3428360, EPI_ISL_3428361, EPI_ISL_3428362, EPI_ISL_3428363, EPI_ISL_3428364, EPI_ISL_3428365, EPI_ISL_3428366, EPI_ISL_3428367, EPI_ISL_3428369 | | | |
| see above | Mako Medical | Centers for Disease Control and Prevention Division of Viral Diseases, Pathogen Discovery | Dakota Howard, Dhwani Batra, Peter W. Cook, Kara Moser, Adrian Paskey, Jason Caravas, Benjamin Rambo-Martin, Shatavia Morrison, Christopher Gulvick, Scott Sammons, Yvette Unoarumhi, Darlene Wagner, Matthew Schmerer, Matthew Tugwell, Lauren Moon, Clinton R. Paden, Duncan MacCannell |
| EPI_ISL_3428370 | UMC Utrecht | UMC Utrecht | Rob Schuurman, Anniek Tanja, Annemarie Wensing |
| EPI_ISL_3428371 | Mako Medical | Centers for Disease Control and Prevention Division of Viral Diseases, Pathogen Discovery | Dakota Howard, Dhwani Batra, Peter W. Cook, Kara Moser, Adrian Paskey, Jason Caravas, Benjamin Rambo-Martin, Shatavia Morrison, Christopher Gulvick, Scott Sammons, Yvette Unoarumhi, Darlene Wagner, Matthew Schmerer, Matthew Tugwell, Lauren Moon, Clinton R. Paden, Duncan MacCannell |
| EPI_ISL_3428372 | UMC Utrecht | UMC Utrecht | Rob Schuurman, Anniek Tanja, Annemarie Wensing |
| EPI_ISL_3428373 | Mako Medical | Centers for Disease Control and Prevention Division of Viral Diseases, Pathogen Discovery | Dakota Howard, Dhwani Batra, Peter W. Cook, Kara Moser, Adrian Paskey, Jason Caravas, Benjamin Rambo-Martin, Shatavia Morrison, Christopher Gulvick, Scott Sammons, Yvette Unoarumhi, Darlene Wagner, Matthew Schmerer, Matthew Tugwell, Lauren Moon, Clinton R. Paden, Duncan MacCannell |
| EPI_ISL_3428374 | UMC Utrecht | UMC Utrecht | Rob Schuurman, Anniek Tanja, Annemarie Wensing |
| EPI_ISL_3428375 | Mako Medical | Centers for Disease Control and Prevention Division of Viral Diseases, Pathogen Discovery | Dakota Howard, Dhwani Batra, Peter W. Cook, Kara Moser, Adrian Paskey, Jason Caravas, Benjamin Rambo-Martin, Shatavia Morrison, Christopher Gulvick, Scott Sammons, Yvette Unoarumhi, Darlene Wagner, Matthew Schmerer, Matthew Tugwell, Lauren Moon, Clinton R. Paden, Duncan MacCannell |
| EPI_ISL_3428376 | UMC Utrecht | UMC Utrecht | Rob Schuurman, Anniek Tanja, Annemarie Wensing |
| EPI_ISL_3428377 | Mako Medical | Centers for Disease Control and Prevention Division of Viral Diseases, Pathogen Discovery | Dakota Howard, Dhwani Batra, Peter W. Cook, Kara Moser, Adrian Paskey, Jason Caravas, Benjamin Rambo-Martin, Shatavia Morrison, Christopher Gulvick, Scott Sammons, Yvette Unoarumhi, Darlene Wagner, Matthew Schmerer, Matthew Tugwell, Lauren Moon, Clinton R. Paden, Duncan MacCannell |
| EPI_ISL_3428378, EPI_ISL_3428379 | UMC Utrecht | UMC Utrecht | Rob Schuurman, Anniek Tanja, Annemarie Wensing |
| EPI_ISL_3428380, EPI_ISL_3428381 | Mako Medical | Centers for Disease Control and Prevention Division of Viral Diseases, Pathogen Discovery | Dakota Howard, Dhwani Batra, Peter W. Cook, Kara Moser, Adrian Paskey, Jason Caravas, Benjamin Rambo-Martin, Shatavia Morrison, Christopher Gulvick, Scott Sammons, Yvette Unoarumhi, Darlene Wagner, Matthew Schmerer, Matthew Tugwell, Lauren Moon, Clinton R. Paden, Duncan MacCannell |
| EPI_ISL_3428382, EPI_ISL_3428383 | UMC Utrecht | UMC Utrecht | Rob Schuurman, Anniek Tanja, Annemarie Wensing |
| EPI_ISL_3428384 | Mako Medical | Centers for Disease Control and Prevention Division of Viral Diseases, Pathogen Discovery | Dakota Howard, Dhwani Batra, Peter W. Cook, Kara Moser, Adrian Paskey, Jason Caravas, Benjamin Rambo-Martin, Shatavia Morrison, Christopher Gulvick, Scott Sammons, Yvette Unoarumhi, Darlene Wagner, Matthew Schmerer, Matthew Tugwell, Lauren Moon, Clinton R. Paden, Duncan MacCannell |
| EPI_ISL_3428385 | UMC Utrecht | UMC Utrecht | Rob Schuurman, Anniek Tanja, Annemarie Wensing |
| EPI_ISL_3428386 | Mako Medical | Centers for Disease Control and Prevention Division of Viral Diseases, Pathogen Discovery | Dakota Howard, Dhwani Batra, Peter W. Cook, Kara Moser, Adrian Paskey, Jason Caravas, Benjamin Rambo-Martin, Shatavia Morrison, Christopher Gulvick, Scott Sammons, Yvette Unoarumhi, Darlene Wagner, Matthew Schmerer, Matthew Tugwell, Lauren Moon, Clinton R. Paden, Duncan MacCannell |
| EPI_ISL_3428387 | UMC Utrecht | UMC Utrecht | Rob Schuurman, Anniek Tanja, Annemarie Wensing |
| EPI_ISL_3428388 | Mako Medical | Centers for Disease Control and Prevention Division of Viral Diseases, Pathogen Discovery | Dakota Howard, Dhwani Batra, Peter W. Cook, Kara Moser, Adrian Paskey, Jason Caravas, Benjamin Rambo-Martin, Shatavia Morrison, Christopher Gulvick, Scott Sammons, Yvette Unoarumhi, Darlene Wagner, Matthew Schmerer, Matthew Tugwell, Lauren Moon, Clinton R. Paden, Duncan MacCannell |

| | | | |
|---|---|---|---|
| EPI_ISL_3428389 | UMC Utrecht | UMC Utrecht | Rob Schuurman, Anniek Tanja, Annemarie Wensing |
| EPI_ISL_3428390 | Mako Medical | Centers for Disease Control and Prevention Division of Viral Diseases, Pathogen Discovery | Dakota Howard, Dhwani Batra, Peter W. Cook, Kara Moser, Adrian Paskey, Jason Caravas, Benjamin Rambo-Martin, Shatavia Morrison, Christopher Gulvick, Scott Sammons, Yvette Unoarumhi, Darlene Wagner, Matthew Schmerer, Matthew Tugwell, Lauren Moon, Clinton R. Paden, Duncan MacCannell |
| EPI_ISL_3428391, EPI_ISL_3428392 | UMC Utrecht | UMC Utrecht | Rob Schuurman, Anniek Tanja, Annemarie Wensing |
| EPI_ISL_3428393 | Mako Medical | Centers for Disease Control and Prevention Division of Viral Diseases, Pathogen Discovery | Dakota Howard, Dhwani Batra, Peter W. Cook, Kara Moser, Adrian Paskey, Jason Caravas, Benjamin Rambo-Martin, Shatavia Morrison, Christopher Gulvick, Scott Sammons, Yvette Unoarumhi, Darlene Wagner, Matthew Schmerer, Matthew Tugwell, Lauren Moon, Clinton R. Paden, Duncan MacCannell |
| EPI_ISL_3428394 | UMC Utrecht | UMC Utrecht | Rob Schuurman, Anniek Tanja, Annemarie Wensing |
| EPI_ISL_3428395 | Mako Medical | Centers for Disease Control and Prevention Division of Viral Diseases, Pathogen Discovery | Dakota Howard, Dhwani Batra, Peter W. Cook, Kara Moser, Adrian Paskey, Jason Caravas, Benjamin Rambo-Martin, Shatavia Morrison, Christopher Gulvick, Scott Sammons, Yvette Unoarumhi, Darlene Wagner, Matthew Schmerer, Matthew Tugwell, Lauren Moon, Clinton R. Paden, Duncan MacCannell |
| EPI_ISL_3428396 | UMC Utrecht | UMC Utrecht | Rob Schuurman, Anniek Tanja, Annemarie Wensing |
| EPI_ISL_3428397 | Mako Medical | Centers for Disease Control and Prevention Division of Viral Diseases, Pathogen Discovery | Dakota Howard, Dhwani Batra, Peter W. Cook, Kara Moser, Adrian Paskey, Jason Caravas, Benjamin Rambo-Martin, Shatavia Morrison, Christopher Gulvick, Scott Sammons, Yvette Unoarumhi, Darlene Wagner, Matthew Schmerer, Matthew Tugwell, Lauren Moon, Clinton R. Paden, Duncan MacCannell |
| EPI_ISL_3428398 | UMC Utrecht | UMC Utrecht | Rob Schuurman, Anniek Tanja, Annemarie Wensing |
| EPI_ISL_3428399 | Mako Medical | Centers for Disease Control and Prevention Division of Viral Diseases, Pathogen Discovery | Dakota Howard, Dhwani Batra, Peter W. Cook, Kara Moser, Adrian Paskey, Jason Caravas, Benjamin Rambo-Martin, Shatavia Morrison, Christopher Gulvick, Scott Sammons, Yvette Unoarumhi, Darlene Wagner, Matthew Schmerer, Matthew Tugwell, Lauren Moon, Clinton R. Paden, Duncan MacCannell |
| EPI_ISL_3428400, EPI_ISL_3428401 | UMC Utrecht | UMC Utrecht | Rob Schuurman, Anniek Tanja, Annemarie Wensing |
| EPI_ISL_3428402 | Mako Medical | Centers for Disease Control and Prevention Division of Viral Diseases, Pathogen Discovery | Dakota Howard, Dhwani Batra, Peter W. Cook, Kara Moser, Adrian Paskey, Jason Caravas, Benjamin Rambo-Martin, Shatavia Morrison, Christopher Gulvick, Scott Sammons, Yvette Unoarumhi, Darlene Wagner, Matthew Schmerer, Matthew Tugwell, Lauren Moon, Clinton R. Paden, Duncan MacCannell |
| EPI_ISL_3428403 | UMC Utrecht | UMC Utrecht | Rob Schuurman, Anniek Tanja, Annemarie Wensing |
| EPI_ISL_3428404 | Mako Medical | Centers for Disease Control and Prevention Division of Viral Diseases, Pathogen Discovery | Dakota Howard, Dhwani Batra, Peter W. Cook, Kara Moser, Adrian Paskey, Jason Caravas, Benjamin Rambo-Martin, Shatavia Morrison, Christopher Gulvick, Scott Sammons, Yvette Unoarumhi, Darlene Wagner, Matthew Schmerer, Matthew Tugwell, Lauren Moon, Clinton R. Paden, Duncan MacCannell |
| EPI_ISL_3428405 | UMC Utrecht | UMC Utrecht | Rob Schuurman, Anniek Tanja, Annemarie Wensing |
| EPI_ISL_3428406 | Mako Medical | Centers for Disease Control and Prevention Division of Viral Diseases, Pathogen Discovery | Dakota Howard, Dhwani Batra, Peter W. Cook, Kara Moser, Adrian Paskey, Jason Caravas, Benjamin Rambo-Martin, Shatavia Morrison, Christopher Gulvick, Scott Sammons, Yvette Unoarumhi, Darlene Wagner, Matthew Schmerer, Matthew Tugwell, Lauren Moon, Clinton R. Paden, Duncan MacCannell |
| EPI_ISL_3428407 | UMC Utrecht | UMC Utrecht | Rob Schuurman, Anniek Tanja, Annemarie Wensing |
| EPI_ISL_3428408 | Mako Medical | Centers for Disease Control and Prevention Division of Viral Diseases, Pathogen Discovery | Dakota Howard, Dhwani Batra, Peter W. Cook, Kara Moser, Adrian Paskey, Jason Caravas, Benjamin Rambo-Martin, Shatavia Morrison, Christopher Gulvick, Scott Sammons, Yvette Unoarumhi, Darlene Wagner, Matthew Schmerer, Matthew Tugwell, Lauren Moon, Clinton R. Paden, Duncan MacCannell |
| EPI_ISL_3428409, EPI_ISL_3428410 | UMC Utrecht | UMC Utrecht | Rob Schuurman, Anniek Tanja, Annemarie Wensing |
| EPI_ISL_3428411 | Mako Medical | Centers for Disease Control and Prevention Division of Viral Diseases, Pathogen Discovery | Dakota Howard, Dhwani Batra, Peter W. Cook, Kara Moser, Adrian Paskey, Jason Caravas, Benjamin Rambo-Martin, Shatavia Morrison, Christopher Gulvick, Scott Sammons, Yvette Unoarumhi, Darlene Wagner, Matthew Schmerer, Matthew Tugwell, Lauren Moon, Clinton R. Paden, Duncan MacCannell |
| EPI_ISL_3428412 | UMC Utrecht | UMC Utrecht | Rob Schuurman, Anniek Tanja, Annemarie Wensing |
| EPI_ISL_3428413 | Mako Medical | Centers for Disease Control and Prevention Division of Viral Diseases, Pathogen Discovery | Dakota Howard, Dhwani Batra, Peter W. Cook, Kara Moser, Adrian Paskey, Jason Caravas, Benjamin Rambo-Martin, Shatavia Morrison, Christopher Gulvick, Scott Sammons, Yvette Unoarumhi, Darlene Wagner, Matthew Schmerer, Matthew Tugwell, Lauren Moon, Clinton R. Paden, Duncan MacCannell |
| EPI_ISL_3428414 | UMC Utrecht | UMC Utrecht | Rob Schuurman, Anniek Tanja, Annemarie Wensing |
| EPI_ISL_3428415 | Mako Medical | Centers for Disease Control and Prevention Division of Viral Diseases, Pathogen Discovery | Dakota Howard, Dhwani Batra, Peter W. Cook, Kara Moser, Adrian Paskey, Jason Caravas, Benjamin Rambo-Martin, Shatavia Morrison, Christopher Gulvick, Scott Sammons, Yvette Unoarumhi, Darlene Wagner, Matthew Schmerer, Matthew Tugwell, Lauren Moon, Clinton R. Paden, Duncan MacCannell |
| EPI_ISL_3428416 | UMC Utrecht | UMC Utrecht | Rob Schuurman, Anniek Tanja, Annemarie Wensing |
| EPI_ISL_3428417 | Mako Medical | Centers for Disease Control and Prevention Division of Viral Diseases, Pathogen Discovery | Dakota Howard, Dhwani Batra, Peter W. Cook, Kara Moser, Adrian Paskey, Jason Caravas, Benjamin Rambo-Martin, Shatavia Morrison, Christopher Gulvick, Scott Sammons, Yvette Unoarumhi, Darlene Wagner, Matthew Schmerer, Matthew Tugwell, Lauren Moon, Clinton R. Paden, Duncan MacCannell |
| EPI_ISL_3428418, EPI_ISL_3428419 | UMC Utrecht | UMC Utrecht | Rob Schuurman, Anniek Tanja, Annemarie Wensing |
| EPI_ISL_3428420 | Mako Medical | Centers for Disease Control and Prevention Division of Viral Diseases, Pathogen Discovery | Dakota Howard, Dhwani Batra, Peter W. Cook, Kara Moser, Adrian Paskey, Jason Caravas, Benjamin Rambo-Martin, Shatavia Morrison, Christopher Gulvick, Scott Sammons, Yvette Unoarumhi, Darlene Wagner, Matthew Schmerer, Matthew Tugwell, Lauren Moon, Clinton R. Paden, Duncan MacCannell |
| EPI_ISL_3428421 | UMC Utrecht | UMC Utrecht | Rob Schuurman, Anniek Tanja, Annemarie Wensing |
| EPI_ISL_3428422 | Mako Medical | Centers for Disease Control and Prevention Division of Viral Diseases, Pathogen Discovery | Dakota Howard, Dhwani Batra, Peter W. Cook, Kara Moser, Adrian Paskey, Jason Caravas, Benjamin Rambo-Martin, Shatavia Morrison, Christopher Gulvick, Scott Sammons, Yvette Unoarumhi, Darlene Wagner, Matthew Schmerer, Matthew Tugwell, Lauren Moon, Clinton R. Paden, Duncan MacCannell |
| EPI_ISL_3428423 | UMC Utrecht | UMC Utrecht | Rob Schuurman, Anniek Tanja, Annemarie Wensing |
| EPI_ISL_3428424 | Mako Medical | Centers for Disease Control and Prevention Division of Viral Diseases, Pathogen Discovery | Dakota Howard, Dhwani Batra, Peter W. Cook, Kara Moser, Adrian Paskey, Jason Caravas, Benjamin Rambo-Martin, Shatavia Morrison, Christopher Gulvick, Scott Sammons, Yvette Unoarumhi, Darlene Wagner, Matthew Schmerer, Matthew Tugwell, Lauren Moon, Clinton R. Paden, Duncan MacCannell |
| EPI_ISL_3428425 | UMC Utrecht | UMC Utrecht | Rob Schuurman, Anniek Tanja, Annemarie Wensing |
| EPI_ISL_3428426 | Mako Medical | Centers for Disease Control and Prevention Division of Viral Diseases, Pathogen Discovery | Dakota Howard, Dhwani Batra, Peter W. Cook, Kara Moser, Adrian Paskey, Jason Caravas, Benjamin Rambo-Martin, Shatavia Morrison, Christopher Gulvick, Scott Sammons, Yvette Unoarumhi, Darlene Wagner, Matthew Schmerer, Matthew Tugwell, Lauren Moon, Clinton R. Paden, Duncan MacCannell |
| EPI_ISL_3428427, EPI_ISL_3428428 | UMC Utrecht | UMC Utrecht | Rob Schuurman, Anniek Tanja, Annemarie Wensing |
| EPI_ISL_3428429 | Mako Medical | Centers for Disease Control and Prevention Division of Viral Diseases, Pathogen Discovery | Dakota Howard, Dhwani Batra, Peter W. Cook, Kara Moser, Adrian Paskey, Jason Caravas, Benjamin Rambo-Martin, Shatavia Morrison, Christopher Gulvick, Scott Sammons, Yvette Unoarumhi, Darlene Wagner, Matthew Schmerer, Matthew Tugwell, Lauren Moon, Clinton R. Paden, Duncan MacCannell |
| EPI_ISL_3428430 | UMC Utrecht | UMC Utrecht | Rob Schuurman, Anniek Tanja, Annemarie Wensing |
| EPI_ISL_3428431 | Mako Medical | Centers for Disease Control and Prevention Division of Viral Diseases, Pathogen Discovery | Dakota Howard, Dhwani Batra, Peter W. Cook, Kara Moser, Adrian Paskey, Jason Caravas, Benjamin Rambo-Martin, Shatavia Morrison, Christopher Gulvick, Scott Sammons, Yvette Unoarumhi, Darlene Wagner, Matthew Schmerer, Matthew Tugwell, Lauren Moon, Clinton R. Paden, Duncan MacCannell |
| EPI_ISL_3428432, EPI_ISL_3428434 | UMC Utrecht | UMC Utrecht | Rob Schuurman, Anniek Tanja, Annemarie Wensing |
| EPI_ISL_3428435 | Mako Medical | Centers for Disease Control and Prevention Division of Viral Diseases, Pathogen Discovery | Dakota Howard, Dhwani Batra, Peter W. Cook, Kara Moser, Adrian Paskey, Jason Caravas, Benjamin Rambo-Martin, Shatavia Morrison, Christopher Gulvick, Scott Sammons, Yvette Unoarumhi, Darlene Wagner, Matthew Schmerer, Matthew Tugwell, Lauren Moon, Clinton R. Paden, Duncan MacCannell |
| EPI_ISL_3428436, EPI_ISL_3428437 | UMC Utrecht | UMC Utrecht | Rob Schuurman, Anniek Tanja, Annemarie Wensing |
| EPI_ISL_3428438 | Mako Medical | Centers for Disease Control and Prevention Division of Viral Diseases, Pathogen Discovery | Dakota Howard, Dhwani Batra, Peter W. Cook, Kara Moser, Adrian Paskey, Jason Caravas, Benjamin Rambo-Martin, Shatavia Morrison, Christopher Gulvick, Scott Sammons, Yvette Unoarumhi, Darlene Wagner, Matthew Schmerer, Matthew Tugwell, Lauren Moon, Clinton R. Paden, Duncan MacCannell |
| EPI_ISL_3428439 | UMC Utrecht | UMC Utrecht | Rob Schuurman, Anniek Tanja, Annemarie Wensing |
| EPI_ISL_3428440 | Mako Medical | Centers for Disease Control and Prevention Division of Viral Diseases, Pathogen Discovery | Dakota Howard, Dhwani Batra, Peter W. Cook, Kara Moser, Adrian Paskey, Jason Caravas, Benjamin Rambo-Martin, Shatavia Morrison, Christopher Gulvick, Scott Sammons, Yvette Unoarumhi, Darlene Wagner, Matthew Schmerer, Matthew Tugwell, Lauren Moon, Clinton R. Paden, Duncan MacCannell |
| EPI_ISL_3428441 | UMC Utrecht | UMC Utrecht | Rob Schuurman, Anniek Tanja, Annemarie Wensing |
| EPI_ISL_3428442 | Mako Medical | Centers for Disease Control and Prevention Division of Viral Diseases, Pathogen Discovery | Dakota Howard, Dhwani Batra, Peter W. Cook, Kara Moser, Adrian Paskey, Jason Caravas, Benjamin Rambo-Martin, Shatavia Morrison, Christopher Gulvick, Scott Sammons, Yvette Unoarumhi, Darlene Wagner, Matthew Schmerer, Matthew Tugwell, Lauren Moon, Clinton R. Paden, Duncan MacCannell |
| EPI_ISL_3428443 | UMC Utrecht | UMC Utrecht | Rob Schuurman, Anniek Tanja, Annemarie Wensing |
| EPI_ISL_3428444 | Mako Medical | Centers for Disease Control and Prevention Division of Viral Diseases, Pathogen Discovery | Dakota Howard, Dhwani Batra, Peter W. Cook, Kara Moser, Adrian Paskey, Jason Caravas, Benjamin Rambo-Martin, Shatavia Morrison, Christopher Gulvick, Scott Sammons, Yvette Unoarumhi, Darlene Wagner, Matthew Schmerer, Matthew Tugwell, Lauren Moon, Clinton R. Paden, Duncan MacCannell |
| EPI_ISL_3428445 | UMC Utrecht | UMC Utrecht | Rob Schuurman, Anniek Tanja, Annemarie Wensing |
| EPI_ISL_3428446 | Mako Medical | Centers for Disease Control and Prevention Division of Viral Diseases, Pathogen Discovery | Dakota Howard, Dhwani Batra, Peter W. Cook, Kara Moser, Adrian Paskey, Jason Caravas, Benjamin Rambo-Martin, Shatavia Morrison, Christopher Gulvick, Scott Sammons, Yvette Unoarumhi, Darlene Wagner, Matthew Schmerer, Matthew Tugwell, Lauren Moon, Clinton R. Paden, Duncan MacCannell |
| EPI_ISL_3428447 | UMC Utrecht | UMC Utrecht | Rob Schuurman, Anniek Tanja, Annemarie Wensing |
| EPI_ISL_3428448 | Mako Medical | Centers for Disease Control and Prevention Division of Viral Diseases, Pathogen Discovery | Dakota Howard, Dhwani Batra, Peter W. Cook, Kara Moser, Adrian Paskey, Jason Caravas, Benjamin Rambo-Martin, Shatavia Morrison, Christopher Gulvick, Scott Sammons, Yvette Unoarumhi, Darlene Wagner, Matthew Schmerer, Matthew Tugwell, Lauren Moon, Clinton R. Paden, Duncan MacCannell |
| EPI_ISL_3428449, EPI_ISL_3428450, | UMC Utrecht | UMC Utrecht | Rob Schuurman, Anniek Tanja, Annemarie Wensing |

| | | | |
|---|---|---|---|
| EPI_ISL_3428452 | | | |
| EPI_ISL_3428453, EPI_ISL_3428454, EPI_ISL_3428455, EPI_ISL_3428456, EPI_ISL_3428457, EPI_ISL_3428458, EPI_ISL_3428459, EPI_ISL_3428460, EPI_ISL_3428461, EPI_ISL_3428462, EPI_ISL_3428463, EPI_ISL_3428464, EPI_ISL_3428465, EPI_ISL_3428466, EPI_ISL_3428467, EPI_ISL_3428468, EPI_ISL_3428469, EPI_ISL_3428470, EPI_ISL_3428471, EPI_ISL_3428472, EPI_ISL_3428473, EPI_ISL_3428474, EPI_ISL_3428475, EPI_ISL_3428476, EPI_ISL_3428477, EPI_ISL_3428478, EPI_ISL_3428479, EPI_ISL_3428480, EPI_ISL_3428481, EPI_ISL_3428482, EPI_ISL_3428483, EPI_ISL_3428484, EPI_ISL_3428485, EPI_ISL_3428486, EPI_ISL_3428487, EPI_ISL_3428488, EPI_ISL_3428490, EPI_ISL_3428492, EPI_ISL_3428493, EPI_ISL_3428494, EPI_ISL_3428495, EPI_ISL_3428496, EPI_ISL_3428497, EPI_ISL_3428498, EPI_ISL_3428499, EPI_ISL_3428500, EPI_ISL_3428501, EPI_ISL_3428502, EPI_ISL_3428503, EPI_ISL_3428504, EPI_ISL_3428505, EPI_ISL_3428506, EPI_ISL_3428507, EPI_ISL_3428508, EPI_ISL_3428509, EPI_ISL_3428513, EPI_ISL_3428514, EPI_ISL_3428515, EPI_ISL_3428516, EPI_ISL_3428517, EPI_ISL_3428518, EPI_ISL_3428519, EPI_ISL_3428520, EPI_ISL_3428521, EPI_ISL_3428522, EPI_ISL_3428523, EPI_ISL_3428524, EPI_ISL_3428525, EPI_ISL_3428526, EPI_ISL_3428527, EPI_ISL_3428528, EPI_ISL_3428529, EPI_ISL_3428530, EPI_ISL_3428531, EPI_ISL_3428532, EPI_ISL_3428534, EPI_ISL_3428535, EPI_ISL_3428536, EPI_ISL_3428537, EPI_ISL_3428538, EPI_ISL_3428539, EPI_ISL_3428540, EPI_ISL_3428541, EPI_ISL_3428542, EPI_ISL_3428543, EPI_ISL_3428544, EPI_ISL_3428545, EPI_ISL_3428546, EPI_ISL_3428547, EPI_ISL_3428548, EPI_ISL_3428549, EPI_ISL_3428550, EPI_ISL_3428551, EPI_ISL_3428552, EPI_ISL_3428553, EPI_ISL_3428554, EPI_ISL_3428555, EPI_ISL_3428556, EPI_ISL_3428557, EPI_ISL_3428558, EPI_ISL_3428559, EPI_ISL_3428560, EPI_ISL_3428561, EPI_ISL_3428562, EPI_ISL_3428563, EPI_ISL_3428566, EPI_ISL_3428567, EPI_ISL_3428568, EPI_ISL_3428569, EPI_ISL_3428570, EPI_ISL_3428572, EPI_ISL_3428575, EPI_ISL_3428576, EPI_ISL_3428577, EPI_ISL_3428578, EPI_ISL_3428579, EPI_ISL_3428580, EPI_ISL_3428581, EPI_ISL_3428582, EPI_ISL_3428583, EPI_ISL_3428584, EPI_ISL_3428585, EPI_ISL_3428586, EPI_ISL_3428587, EPI_ISL_3428588, EPI_ISL_3428589, EPI_ISL_3428590, EPI_ISL_3428591, EPI_ISL_3428592, EPI_ISL_3428593, EPI_ISL_3428595, EPI_ISL_3428596, EPI_ISL_3428597, EPI_ISL_3428598, EPI_ISL_3428599, EPI_ISL_3428600, EPI_ISL_3428601, EPI_ISL_3428602, EPI_ISL_3428603, EPI_ISL_3428604, EPI_ISL_3428605, EPI_ISL_3428606, EPI_ISL_3428607, EPI_ISL_3428608, EPI_ISL_3428609, EPI_ISL_3428610, EPI_ISL_3428612, EPI_ISL_3428613, EPI_ISL_3428614, EPI_ISL_3428615, EPI_ISL_3428616, EPI_ISL_3428617, EPI_ISL_3428618, EPI_ISL_3428619, EPI_ISL_3428620, EPI_ISL_3428621, EPI_ISL_3428622, EPI_ISL_3428623, EPI_ISL_3428624, EPI_ISL_3428625, EPI_ISL_3428626, EPI_ISL_3428627, EPI_ISL_3428628, EPI_ISL_3428629, EPI_ISL_3428630, EPI_ISL_3428631, EPI_ISL_3428632, EPI_ISL_3428633, EPI_ISL_3428634, EPI_ISL_3428635, EPI_ISL_3428636, EPI_ISL_3428637, EPI_ISL_3428638, EPI_ISL_3428639, EPI_ISL_3428640, EPI_ISL_3428641, EPI_ISL_3428642, EPI_ISL_3428643, EPI_ISL_3428644, EPI_ISL_3428646, EPI_ISL_3428647, EPI_ISL_3428648, EPI_ISL_3428649, EPI_ISL_3428650, EPI_ISL_3428651, EPI_ISL_3428652, EPI_ISL_3428653, EPI_ISL_3428654, EPI_ISL_3428655, EPI_ISL_3428656, EPI_ISL_3428658, EPI_ISL_3428660, EPI_ISL_3428661, EPI_ISL_3428662, EPI_ISL_3428663, EPI_ISL_3428664, EPI_ISL_3428665, EPI_ISL_3428666, EPI_ISL_3428667, EPI_ISL_3428669, EPI_ISL_3428670, EPI_ISL_3428671, EPI_ISL_3428672, EPI_ISL_3428673, EPI_ISL_3428674, EPI_ISL_3428675, EPI_ISL_3428676, EPI_ISL_3428677, EPI_ISL_3428678, EPI_ISL_3428679, EPI_ISL_3428680, EPI_ISL_3428681, EPI_ISL_3428682, EPI_ISL_3428683, EPI_ISL_3428684, EPI_ISL_3428685, EPI_ISL_3428686, EPI_ISL_3428687, EPI_ISL_3428688, EPI_ISL_3428689, EPI_ISL_3428690, EPI_ISL_3428691, EPI_ISL_3428692, EPI_ISL_3428693, EPI_ISL_3428694 | | | |
| see above | Mako Medical | Centers for Disease Control and Prevention Division of Viral Diseases, Pathogen Discovery | Dakota Howard, Dhwani Batra, Peter W. Cook, Kara Moser, Adrian Paskey, Jason Caravas, Benjamin Rambo-Martin, Shatavia Morrison, Christopher Gulvick, Scott Sammons, Yvette Unoarumhi, Darlene Wagner, Matthew Schmerer, Matthew Tugwell, Lauren Moon, Clinton R. Paden, Duncan MacCannell |
| EPI_ISL_3428695 | UMC Utrecht | UMC Utrecht | Rob Schuurman, Anniek Tanja, Annemarie Wensing |
| EPI_ISL_3428696 | Mako Medical | Centers for Disease Control and Prevention Division of Viral Diseases, Pathogen Discovery | Dakota Howard, Dhwani Batra, Peter W. Cook, Kara Moser, Adrian Paskey, Jason Caravas, Benjamin Rambo-Martin, Shatavia Morrison, Christopher Gulvick, Scott Sammons, Yvette Unoarumhi, Darlene Wagner, Matthew Schmerer, Matthew Tugwell, Lauren Moon, Clinton R. Paden, Duncan MacCannell |
| EPI_ISL_3428697 | UMC Utrecht | UMC Utrecht | Rob Schuurman, Anniek Tanja, Annemarie Wensing |
| EPI_ISL_3428698 | Mako Medical | Centers for Disease Control and Prevention Division of Viral Diseases, Pathogen Discovery | Dakota Howard, Dhwani Batra, Peter W. Cook, Kara Moser, Adrian Paskey, Jason Caravas, Benjamin Rambo-Martin, Shatavia Morrison, Christopher Gulvick, Scott Sammons, Yvette Unoarumhi, Darlene Wagner, Matthew Schmerer, Matthew Tugwell, Lauren Moon, Clinton R. Paden, Duncan MacCannell |
| EPI_ISL_3428699 | UMC Utrecht | UMC Utrecht | Rob Schuurman, Anniek Tanja, Annemarie Wensing |
| EPI_ISL_3428700 | Mako Medical | Centers for Disease Control and Prevention Division of Viral Diseases, Pathogen Discovery | Dakota Howard, Dhwani Batra, Peter W. Cook, Kara Moser, Adrian Paskey, Jason Caravas, Benjamin Rambo-Martin, Shatavia Morrison, Christopher Gulvick, Scott Sammons, Yvette Unoarumhi, Darlene Wagner, Matthew Schmerer, Matthew Tugwell, Lauren Moon, Clinton R. Paden, Duncan MacCannell |
| EPI_ISL_3428701 | UMC Utrecht | UMC Utrecht | Rob Schuurman, Anniek Tanja, Annemarie Wensing |
| EPI_ISL_3428702, EPI_ISL_3428703 | Mako Medical | Centers for Disease Control and Prevention Division of Viral Diseases, Pathogen Discovery | Dakota Howard, Dhwani Batra, Peter W. Cook, Kara Moser, Adrian Paskey, Jason Caravas, Benjamin Rambo-Martin, Shatavia Morrison, Christopher Gulvick, Scott Sammons, Yvette Unoarumhi, Darlene Wagner, Matthew Schmerer, Matthew Tugwell, Lauren Moon, Clinton R. Paden, Duncan MacCannell |
| EPI_ISL_3428704 | UMC Utrecht | UMC Utrecht | Rob Schuurman, Anniek Tanja, Annemarie Wensing |
| EPI_ISL_3428705, EPI_ISL_3428707 | Mako Medical | Centers for Disease Control and Prevention Division of Viral Diseases, Pathogen Discovery | Dakota Howard, Dhwani Batra, Peter W. Cook, Kara Moser, Adrian Paskey, Jason Caravas, Benjamin Rambo-Martin, Shatavia Morrison, Christopher Gulvick, Scott Sammons, Yvette Unoarumhi, Darlene Wagner, Matthew Schmerer, Matthew Tugwell, Lauren Moon, Clinton R. Paden, Duncan MacCannell |
| EPI_ISL_3428708 | UMC Utrecht | UMC Utrecht | Rob Schuurman, Anniek Tanja, Annemarie Wensing |
| EPI_ISL_3428709, EPI_ISL_3428710 | Mako Medical | Centers for Disease Control and Prevention Division of Viral Diseases, Pathogen Discovery | Dakota Howard, Dhwani Batra, Peter W. Cook, Kara Moser, Adrian Paskey, Jason Caravas, Benjamin Rambo-Martin, Shatavia Morrison, Christopher Gulvick, Scott Sammons, Yvette Unoarumhi, Darlene Wagner, Matthew Schmerer, Matthew Tugwell, Lauren Moon, Clinton R. Paden, Duncan MacCannell |
| EPI_ISL_3428711 | UMC Utrecht | UMC Utrecht | Rob Schuurman, Anniek Tanja, Annemarie Wensing |
| EPI_ISL_3428712 | Mako Medical | Centers for Disease Control and Prevention Division of Viral Diseases, Pathogen Discovery | Dakota Howard, Dhwani Batra, Peter W. Cook, Kara Moser, Adrian Paskey, Jason Caravas, Benjamin Rambo-Martin, Shatavia Morrison, Christopher Gulvick, Scott Sammons, Yvette Unoarumhi, Darlene Wagner, Matthew Schmerer, Matthew Tugwell, Lauren Moon, Clinton R. Paden, Duncan MacCannell |
| EPI_ISL_3428713 | UMC Utrecht | UMC Utrecht | Rob Schuurman, Anniek Tanja, Annemarie Wensing |
| EPI_ISL_3428714, EPI_ISL_3428715 | Mako Medical | Centers for Disease Control and Prevention Division of Viral Diseases, Pathogen Discovery | Dakota Howard, Dhwani Batra, Peter W. Cook, Kara Moser, Adrian Paskey, Jason Caravas, Benjamin Rambo-Martin, Shatavia Morrison, Christopher Gulvick, Scott Sammons, Yvette Unoarumhi, Darlene Wagner, Matthew Schmerer, Matthew Tugwell, Lauren Moon, Clinton R. Paden, Duncan MacCannell |
| EPI_ISL_3428716 | UMC Utrecht | UMC Utrecht | Rob Schuurman, Anniek Tanja, Annemarie Wensing |
| EPI_ISL_3428717, EPI_ISL_3428718 | Mako Medical | Centers for Disease Control and Prevention Division of Viral Diseases, Pathogen Discovery | Dakota Howard, Dhwani Batra, Peter W. Cook, Kara Moser, Adrian Paskey, Jason Caravas, Benjamin Rambo-Martin, Shatavia Morrison, Christopher Gulvick, Scott Sammons, Yvette Unoarumhi, Darlene Wagner, Matthew Schmerer, Matthew Tugwell, Lauren Moon, Clinton R. Paden, Duncan MacCannell |
| EPI_ISL_3428719, EPI_ISL_3428721, EPI_ISL_3428722 | UMC Utrecht | UMC Utrecht | Rob Schuurman, Anniek Tanja, Annemarie Wensing |
| EPI_ISL_3428723, EPI_ISL_3428724 | Mako Medical | Centers for Disease Control and Prevention Division of Viral Diseases, Pathogen Discovery | Dakota Howard, Dhwani Batra, Peter W. Cook, Kara Moser, Adrian Paskey, Jason Caravas, Benjamin Rambo-Martin, Shatavia Morrison, Christopher Gulvick, Scott Sammons, Yvette Unoarumhi, Darlene Wagner, Matthew Schmerer, Matthew Tugwell, Lauren Moon, Clinton R. Paden, Duncan MacCannell |
| EPI_ISL_3428725 | UMC Utrecht | UMC Utrecht | Rob Schuurman, Anniek Tanja, Annemarie Wensing |
| EPI_ISL_3428726, EPI_ISL_3428727 | Mako Medical | Centers for Disease Control and Prevention Division of Viral Diseases, Pathogen Discovery | Dakota Howard, Dhwani Batra, Peter W. Cook, Kara Moser, Adrian Paskey, Jason Caravas, Benjamin Rambo-Martin, Shatavia Morrison, Christopher Gulvick, Scott Sammons, Yvette Unoarumhi, Darlene Wagner, Matthew Schmerer, Matthew Tugwell, Lauren Moon, Clinton R. Paden, Duncan MacCannell |
| EPI_ISL_3428728 | UMC Utrecht | UMC Utrecht | Rob Schuurman, Anniek Tanja, Annemarie Wensing |
| EPI_ISL_3428729, EPI_ISL_3428730 | Mako Medical | Centers for Disease Control and Prevention Division of Viral Diseases, Pathogen Discovery | Dakota Howard, Dhwani Batra, Peter W. Cook, Kara Moser, Adrian Paskey, Jason Caravas, Benjamin Rambo-Martin, Shatavia Morrison, Christopher Gulvick, Scott Sammons, Yvette Unoarumhi, Darlene Wagner, Matthew Schmerer, Matthew Tugwell, Lauren Moon, Clinton R. Paden, Duncan MacCannell |
| EPI_ISL_3428731 | UMC Utrecht | UMC Utrecht | Rob Schuurman, Anniek Tanja, Annemarie Wensing |
| EPI_ISL_3428732 | Mako Medical | Centers for Disease Control and Prevention Division of Viral Diseases, Pathogen Discovery | Dakota Howard, Dhwani Batra, Peter W. Cook, Kara Moser, Adrian Paskey, Jason Caravas, Benjamin Rambo-Martin, Shatavia Morrison, Christopher Gulvick, Scott Sammons, Yvette Unoarumhi, Darlene Wagner, Matthew Schmerer, Matthew Tugwell, Lauren Moon, Clinton R. Paden, Duncan MacCannell |
| EPI_ISL_3428733, EPI_ISL_3428734 | UMC Utrecht | UMC Utrecht | Rob Schuurman, Anniek Tanja, Annemarie Wensing |
| EPI_ISL_3428735, EPI_ISL_3428736 | Mako Medical | Centers for Disease Control and Prevention Division of Viral Diseases, Pathogen Discovery | Dakota Howard, Dhwani Batra, Peter W. Cook, Kara Moser, Adrian Paskey, Jason Caravas, Benjamin Rambo-Martin, Shatavia Morrison, Christopher Gulvick, Scott Sammons, Yvette Unoarumhi, Darlene Wagner, Matthew Schmerer, Matthew Tugwell, Lauren Moon, Clinton R. Paden, Duncan MacCannell |
| EPI_ISL_3428737 | UMC Utrecht | UMC Utrecht | Rob Schuurman, Anniek Tanja, Annemarie Wensing |
| EPI_ISL_3428738 | Mako Medical | Centers for Disease Control and Prevention Division of Viral Diseases, Pathogen Discovery | Dakota Howard, Dhwani Batra, Peter W. Cook, Kara Moser, Adrian Paskey, Jason Caravas, Benjamin Rambo-Martin, Shatavia Morrison, Christopher Gulvick, Scott Sammons, Yvette Unoarumhi, Darlene Wagner, Matthew Schmerer, Matthew Tugwell, Lauren Moon, Clinton R. Paden, Duncan MacCannell |
| EPI_ISL_3428739 | UMC Utrecht | UMC Utrecht | Rob Schuurman, Anniek Tanja, Annemarie Wensing |
| EPI_ISL_3428740, EPI_ISL_3428741 | Mako Medical | Centers for Disease Control and Prevention Division of Viral Diseases, Pathogen Discovery | Dakota Howard, Dhwani Batra, Peter W. Cook, Kara Moser, Adrian Paskey, Jason Caravas, Benjamin Rambo-Martin, Shatavia Morrison, Christopher Gulvick, Scott Sammons, Yvette Unoarumhi, Darlene Wagner, Matthew Schmerer, Matthew Tugwell, Lauren Moon, Clinton R. Paden, Duncan MacCannell |
| EPI_ISL_3428742 | UMC Utrecht | UMC Utrecht | Rob Schuurman, Anniek Tanja, Annemarie Wensing |
| EPI_ISL_3428743, EPI_ISL_3428744 | Mako Medical | Centers for Disease Control and Prevention Division of Viral Diseases, Pathogen Discovery | Dakota Howard, Dhwani Batra, Peter W. Cook, Kara Moser, Adrian Paskey, Jason Caravas, Benjamin Rambo-Martin, Shatavia Morrison, Christopher Gulvick, Scott Sammons, Yvette Unoarumhi, Darlene Wagner, Matthew Schmerer, Matthew Tugwell, Lauren Moon, Clinton R. Paden, Duncan MacCannell |
| EPI_ISL_3428745 | UMC Utrecht | UMC Utrecht | Rob Schuurman, Anniek Tanja, Annemarie Wensing |
| EPI_ISL_3428746, EPI_ISL_3428747 | Mako Medical | Centers for Disease Control and Prevention Division of Viral Diseases, Pathogen Discovery | Dakota Howard, Dhwani Batra, Peter W. Cook, Kara Moser, Adrian Paskey, Jason Caravas, Benjamin Rambo-Martin, Shatavia Morrison, Christopher Gulvick, Scott Sammons, Yvette Unoarumhi, Darlene Wagner, Matthew Schmerer, Matthew Tugwell, Lauren Moon, Clinton R. Paden, Duncan MacCannell |
| EPI_ISL_3428748 | UMC Utrecht | UMC Utrecht | Rob Schuurman, Anniek Tanja, Annemarie Wensing |
| EPI_ISL_3428749, EPI_ISL_3428750 | Mako Medical | Centers for Disease Control and Prevention Division of Viral Diseases, Pathogen Discovery | Dakota Howard, Dhwani Batra, Peter W. Cook, Kara Moser, Adrian Paskey, Jason Caravas, Benjamin Rambo-Martin, Shatavia Morrison, Christopher Gulvick, Scott Sammons, Yvette Unoarumhi, Darlene Wagner, Matthew Schmerer, Matthew Tugwell, Lauren Moon, Clinton R. Paden, Duncan MacCannell |
| EPI_ISL_3428751 | UMC Utrecht | UMC Utrecht | Rob Schuurman, Anniek Tanja, Annemarie Wensing |
| EPI_ISL_3428752 | Mako Medical | Centers for Disease Control and Prevention Division of Viral Diseases, Pathogen Discovery | Dakota Howard, Dhwani Batra, Peter W. Cook, Kara Moser, Adrian Paskey, Jason Caravas, Benjamin Rambo-Martin, Shatavia Morrison, Christopher Gulvick, Scott Sammons, Yvette Unoarumhi, Darlene Wagner, Matthew Schmerer, Matthew Tugwell, Lauren Moon, Clinton R. Paden, Duncan MacCannell |
| EPI_ISL_3428753, EPI_ISL_3428754 | UMC Utrecht | UMC Utrecht | Rob Schuurman, Anniek Tanja, Annemarie Wensing |
| EPI_ISL_3428755, EPI_ISL_3428756 | Mako Medical | Centers for Disease Control and Prevention Division of Viral Diseases, Pathogen Discovery | Dakota Howard, Dhwani Batra, Peter W. Cook, Kara Moser, Adrian Paskey, Jason Caravas, Benjamin Rambo-Martin, Shatavia Morrison, Christopher Gulvick, Scott Sammons, Yvette Unoarumhi, Darlene Wagner, Matthew Schmerer, Matthew Tugwell, Lauren Moon, Clinton R. Paden, Duncan MacCannell |
| EPI_ISL_3428757 | UMC Utrecht | UMC Utrecht | Rob Schuurman, Anniek Tanja, Annemarie Wensing |
| EPI_ISL_3428758 | Mako Medical | Centers for Disease Control and Prevention Division of Viral Diseases, Pathogen Discovery | Dakota Howard, Dhwani Batra, Peter W. Cook, Kara Moser, Adrian Paskey, Jason Caravas, Benjamin Rambo-Martin, Shatavia Morrison, Christopher Gulvick, Scott Sammons, Yvette Unoarumhi, Darlene Wagner, Matthew Schmerer, Matthew Tugwell, Lauren Moon, Clinton R. Paden, Duncan MacCannell |
| EPI_ISL_3428759, EPI_ISL_3428760 | UMC Utrecht | UMC Utrecht | Rob Schuurman, Anniek Tanja, Annemarie Wensing |
| EPI_ISL_3428761 | Mako Medical | Centers for Disease Control and Prevention Division of Viral Diseases, Pathogen Discovery | Dakota Howard, Dhwani Batra, Peter W. Cook, Kara Moser, Adrian Paskey, Jason Caravas, Benjamin Rambo-Martin, Shatavia Morrison, Christopher Gulvick, Scott Sammons, Yvette Unoarumhi, Darlene Wagner, Matthew Schmerer, Matthew Tugwell, Lauren Moon, Clinton R. Paden, Duncan MacCannell |
| EPI_ISL_3428762 | UMC Utrecht | UMC Utrecht | Rob Schuurman, Anniek Tanja, Annemarie Wensing |

| | | | |
|---|---|---|---|
| EPI_ISL_3428763 | Mako Medical | Centers for Disease Control and Prevention Division of Viral Diseases, Pathogen Discovery | Dakota Howard, Dhwani Batra, Peter W. Cook, Kara Moser, Adrian Paskey, Jason Caravas, Benjamin Rambo-Martin, Shatavia Morrison, Christopher Gulvick, Scott Sammons, Yvette Unoarumhi, Darlene Wagner, Matthew Schmerer, Matthew Tugwell, Lauren Moon, Clinton R. Paden, Duncan MacCannell |
| EPI_ISL_3428764 | UMC Utrecht | UMC Utrecht | Rob Schuurman, Anniek Tanja, Annemarie Wensing |
| EPI_ISL_3428765 | Mako Medical | Centers for Disease Control and Prevention Division of Viral Diseases, Pathogen Discovery | Dakota Howard, Dhwani Batra, Peter W. Cook, Kara Moser, Adrian Paskey, Jason Caravas, Benjamin Rambo-Martin, Shatavia Morrison, Christopher Gulvick, Scott Sammons, Yvette Unoarumhi, Darlene Wagner, Matthew Schmerer, Matthew Tugwell, Lauren Moon, Clinton R. Paden, Duncan MacCannell |
| EPI_ISL_3428766, EPI_ISL_3428768 | UMC Utrecht | UMC Utrecht | Rob Schuurman, Anniek Tanja, Annemarie Wensing |
| EPI_ISL_3428769 | Mako Medical | Centers for Disease Control and Prevention Division of Viral Diseases, Pathogen Discovery | Dakota Howard, Dhwani Batra, Peter W. Cook, Kara Moser, Adrian Paskey, Jason Caravas, Benjamin Rambo-Martin, Shatavia Morrison, Christopher Gulvick, Scott Sammons, Yvette Unoarumhi, Darlene Wagner, Matthew Schmerer, Matthew Tugwell, Lauren Moon, Clinton R. Paden, Duncan MacCannell |
| EPI_ISL_3428770 | UMC Utrecht | UMC Utrecht | Rob Schuurman, Anniek Tanja, Annemarie Wensing |
| EPI_ISL_3428771 | Mako Medical | Centers for Disease Control and Prevention Division of Viral Diseases, Pathogen Discovery | Dakota Howard, Dhwani Batra, Peter W. Cook, Kara Moser, Adrian Paskey, Jason Caravas, Benjamin Rambo-Martin, Shatavia Morrison, Christopher Gulvick, Scott Sammons, Yvette Unoarumhi, Darlene Wagner, Matthew Schmerer, Matthew Tugwell, Lauren Moon, Clinton R. Paden, Duncan MacCannell |
| EPI_ISL_3428772, EPI_ISL_3428773 | UMC Utrecht | UMC Utrecht | Rob Schuurman, Anniek Tanja, Annemarie Wensing |
| EPI_ISL_3428774 | Mako Medical | Centers for Disease Control and Prevention Division of Viral Diseases, Pathogen Discovery | Dakota Howard, Dhwani Batra, Peter W. Cook, Kara Moser, Adrian Paskey, Jason Caravas, Benjamin Rambo-Martin, Shatavia Morrison, Christopher Gulvick, Scott Sammons, Yvette Unoarumhi, Darlene Wagner, Matthew Schmerer, Matthew Tugwell, Lauren Moon, Clinton R. Paden, Duncan MacCannell |
| EPI_ISL_3428775 | UMC Utrecht | UMC Utrecht | Rob Schuurman, Anniek Tanja, Annemarie Wensing |
| EPI_ISL_3428776 | Mako Medical | Centers for Disease Control and Prevention Division of Viral Diseases, Pathogen Discovery | Dakota Howard, Dhwani Batra, Peter W. Cook, Kara Moser, Adrian Paskey, Jason Caravas, Benjamin Rambo-Martin, Shatavia Morrison, Christopher Gulvick, Scott Sammons, Yvette Unoarumhi, Darlene Wagner, Matthew Schmerer, Matthew Tugwell, Lauren Moon, Clinton R. Paden, Duncan MacCannell |
| EPI_ISL_3428777 | UMC Utrecht | UMC Utrecht | Rob Schuurman, Anniek Tanja, Annemarie Wensing |
| EPI_ISL_3428778 | Mako Medical | Centers for Disease Control and Prevention Division of Viral Diseases, Pathogen Discovery | Dakota Howard, Dhwani Batra, Peter W. Cook, Kara Moser, Adrian Paskey, Jason Caravas, Benjamin Rambo-Martin, Shatavia Morrison, Christopher Gulvick, Scott Sammons, Yvette Unoarumhi, Darlene Wagner, Matthew Schmerer, Matthew Tugwell, Lauren Moon, Clinton R. Paden, Duncan MacCannell |
| EPI_ISL_3428779, EPI_ISL_3428780 | UMC Utrecht | UMC Utrecht | Rob Schuurman, Anniek Tanja, Annemarie Wensing |
| EPI_ISL_3428781 | Mako Medical | Centers for Disease Control and Prevention Division of Viral Diseases, Pathogen Discovery | Dakota Howard, Dhwani Batra, Peter W. Cook, Kara Moser, Adrian Paskey, Jason Caravas, Benjamin Rambo-Martin, Shatavia Morrison, Christopher Gulvick, Scott Sammons, Yvette Unoarumhi, Darlene Wagner, Matthew Schmerer, Matthew Tugwell, Lauren Moon, Clinton R. Paden, Duncan MacCannell |
| EPI_ISL_3428782 | UMC Utrecht | UMC Utrecht | Rob Schuurman, Anniek Tanja, Annemarie Wensing |
| EPI_ISL_3428783 | Mako Medical | Centers for Disease Control and Prevention Division of Viral Diseases, Pathogen Discovery | Dakota Howard, Dhwani Batra, Peter W. Cook, Kara Moser, Adrian Paskey, Jason Caravas, Benjamin Rambo-Martin, Shatavia Morrison, Christopher Gulvick, Scott Sammons, Yvette Unoarumhi, Darlene Wagner, Matthew Schmerer, Matthew Tugwell, Lauren Moon, Clinton R. Paden, Duncan MacCannell |
| EPI_ISL_3428784 | UMC Utrecht | UMC Utrecht | Rob Schuurman, Anniek Tanja, Annemarie Wensing |
| EPI_ISL_3428785 | Mako Medical | Centers for Disease Control and Prevention Division of Viral Diseases, Pathogen Discovery | Dakota Howard, Dhwani Batra, Peter W. Cook, Kara Moser, Adrian Paskey, Jason Caravas, Benjamin Rambo-Martin, Shatavia Morrison, Christopher Gulvick, Scott Sammons, Yvette Unoarumhi, Darlene Wagner, Matthew Schmerer, Matthew Tugwell, Lauren Moon, Clinton R. Paden, Duncan MacCannell |
| EPI_ISL_3428786 | UMC Utrecht | UMC Utrecht | Rob Schuurman, Anniek Tanja, Annemarie Wensing |
| EPI_ISL_3428787 | Mako Medical | Centers for Disease Control and Prevention Division of Viral Diseases, Pathogen Discovery | Dakota Howard, Dhwani Batra, Peter W. Cook, Kara Moser, Adrian Paskey, Jason Caravas, Benjamin Rambo-Martin, Shatavia Morrison, Christopher Gulvick, Scott Sammons, Yvette Unoarumhi, Darlene Wagner, Matthew Schmerer, Matthew Tugwell, Lauren Moon, Clinton R. Paden, Duncan MacCannell |
| EPI_ISL_3428788, EPI_ISL_3428789 | UMC Utrecht | UMC Utrecht | Rob Schuurman, Anniek Tanja, Annemarie Wensing |
| EPI_ISL_3428790 | Mako Medical | Centers for Disease Control and Prevention Division of Viral Diseases, Pathogen Discovery | Dakota Howard, Dhwani Batra, Peter W. Cook, Kara Moser, Adrian Paskey, Jason Caravas, Benjamin Rambo-Martin, Shatavia Morrison, Christopher Gulvick, Scott Sammons, Yvette Unoarumhi, Darlene Wagner, Matthew Schmerer, Matthew Tugwell, Lauren Moon, Clinton R. Paden, Duncan MacCannell |
| EPI_ISL_3428791 | UMC Utrecht | UMC Utrecht | Rob Schuurman, Anniek Tanja, Annemarie Wensing |
| EPI_ISL_3428792 | Mako Medical | Centers for Disease Control and Prevention Division of Viral Diseases, Pathogen Discovery | Dakota Howard, Dhwani Batra, Peter W. Cook, Kara Moser, Adrian Paskey, Jason Caravas, Benjamin Rambo-Martin, Shatavia Morrison, Christopher Gulvick, Scott Sammons, Yvette Unoarumhi, Darlene Wagner, Matthew Schmerer, Matthew Tugwell, Lauren Moon, Clinton R. Paden, Duncan MacCannell |
| EPI_ISL_3428793 | UMC Utrecht | UMC Utrecht | Rob Schuurman, Anniek Tanja, Annemarie Wensing |
| EPI_ISL_3428795, EPI_ISL_3428796, EPI_ISL_3428798, EPI_ISL_3428799, EPI_ISL_3428800, EPI_ISL_3428801, EPI_ISL_3428802, EPI_ISL_3428803, EPI_ISL_3428804, EPI_ISL_3428805, EPI_ISL_3428806, EPI_ISL_3428807, EPI_ISL_3428808, EPI_ISL_3428809, EPI_ISL_3428810, EPI_ISL_3428812, EPI_ISL_3428813, EPI_ISL_3428814, EPI_ISL_3428815, EPI_ISL_3428816, EPI_ISL_3428817, EPI_ISL_3428818, EPI_ISL_3428819, EPI_ISL_3428820, EPI_ISL_3428821, EPI_ISL_3428822, EPI_ISL_3428823, EPI_ISL_3428824, EPI_ISL_3428825, EPI_ISL_3428826, EPI_ISL_3428827, EPI_ISL_3428828, EPI_ISL_3428829, EPI_ISL_3428830, EPI_ISL_3428832, EPI_ISL_3428833, EPI_ISL_3428834, EPI_ISL_3428835, EPI_ISL_3428836, EPI_ISL_3428837, EPI_ISL_3428838, EPI_ISL_3428839, EPI_ISL_3428840, EPI_ISL_3428841, EPI_ISL_3428842, EPI_ISL_3428843, EPI_ISL_3428844, EPI_ISL_3428845, EPI_ISL_3428846, EPI_ISL_3428847, EPI_ISL_3428848, EPI_ISL_3428849, EPI_ISL_3428850, EPI_ISL_3428851, EPI_ISL_3428852, EPI_ISL_3428853, EPI_ISL_3428855, EPI_ISL_3428856, EPI_ISL_3428857, EPI_ISL_3428858, EPI_ISL_3428859, EPI_ISL_3428860, EPI_ISL_3428861, EPI_ISL_3428862, EPI_ISL_3428863, EPI_ISL_3428864, EPI_ISL_3428865, EPI_ISL_3428866, EPI_ISL_3428867, EPI_ISL_3428868, EPI_ISL_3428869, EPI_ISL_3428871, EPI_ISL_3428872, EPI_ISL_3428873, EPI_ISL_3428874, EPI_ISL_3428875, EPI_ISL_3428876, EPI_ISL_3428877, EPI_ISL_3428878, EPI_ISL_3428879, EPI_ISL_3428880, EPI_ISL_3428881, EPI_ISL_3428882, EPI_ISL_3428883, EPI_ISL_3428884, EPI_ISL_3428885, EPI_ISL_3428886, EPI_ISL_3428887, EPI_ISL_3428888, EPI_ISL_3428889, EPI_ISL_3428890, EPI_ISL_3428891, EPI_ISL_3428892, EPI_ISL_3428893, EPI_ISL_3428894, EPI_ISL_3428895, EPI_ISL_3428897, EPI_ISL_3428898, EPI_ISL_3428899, EPI_ISL_3428900, EPI_ISL_3428901, EPI_ISL_3428903, EPI_ISL_3428904, EPI_ISL_3428905, EPI_ISL_3428906, EPI_ISL_3428907, EPI_ISL_3428908, EPI_ISL_3428909, EPI_ISL_3428910, EPI_ISL_3428911, EPI_ISL_3428912, EPI_ISL_3428913, EPI_ISL_3428914, EPI_ISL_3428915, EPI_ISL_3428916, EPI_ISL_3428917, EPI_ISL_3428918, EPI_ISL_3428919, EPI_ISL_3428920, EPI_ISL_3428921, EPI_ISL_3428922, EPI_ISL_3428923, EPI_ISL_3428924, EPI_ISL_3428925, EPI_ISL_3428926, EPI_ISL_3428927, EPI_ISL_3428928, EPI_ISL_3428929, EPI_ISL_3428930, EPI_ISL_3428931, EPI_ISL_3428932, EPI_ISL_3428933, EPI_ISL_3428934, EPI_ISL_3428935, EPI_ISL_3428936, EPI_ISL_3428937, EPI_ISL_3428938, EPI_ISL_3428939, EPI_ISL_3428940, EPI_ISL_3428941, EPI_ISL_3428942, EPI_ISL_3428943, EPI_ISL_3428944, EPI_ISL_3428945, EPI_ISL_3428946, EPI_ISL_3428947, EPI_ISL_3428948, EPI_ISL_3428949, EPI_ISL_3428950, EPI_ISL_3428951, EPI_ISL_3428953, EPI_ISL_3428954, EPI_ISL_3428955, EPI_ISL_3428956, EPI_ISL_3428957, EPI_ISL_3428958, EPI_ISL_3428960, EPI_ISL_3428961, EPI_ISL_3428962, EPI_ISL_3428963, EPI_ISL_3428964, EPI_ISL_3428965, EPI_ISL_3428966, EPI_ISL_3428967, EPI_ISL_3428968, EPI_ISL_3428969, EPI_ISL_3428970, EPI_ISL_3428971, EPI_ISL_3428972, EPI_ISL_3428973, EPI_ISL_3428974, EPI_ISL_3428975, EPI_ISL_3428976, EPI_ISL_3428977, EPI_ISL_3428978, EPI_ISL_3428979, EPI_ISL_3428980, EPI_ISL_3428981, EPI_ISL_3428982, EPI_ISL_3428983, EPI_ISL_3428984, EPI_ISL_3428985, EPI_ISL_3428986, EPI_ISL_3428987, EPI_ISL_3428988, EPI_ISL_3428989, EPI_ISL_3428990, EPI_ISL_3428991, EPI_ISL_3428992, EPI_ISL_3428993, EPI_ISL_3428994, EPI_ISL_3428995 | | | |
| see above | Mako Medical | Centers for Disease Control and Prevention Division of Viral Diseases, Pathogen Discovery | Dakota Howard, Dhwani Batra, Peter W. Cook, Kara Moser, Adrian Paskey, Jason Caravas, Benjamin Rambo-Martin, Shatavia Morrison, Christopher Gulvick, Scott Sammons, Yvette Unoarumhi, Darlene Wagner, Matthew Schmerer, Matthew Tugwell, Lauren Moon, Clinton R. Paden, Duncan MacCannell |
| EPI_ISL_3428996, EPI_ISL_3428997, EPI_ISL_3428998, EPI_ISL_3428999, EPI_ISL_3429000, EPI_ISL_3429001, EPI_ISL_3429002, EPI_ISL_3429003, EPI_ISL_3429004, EPI_ISL_3429005, EPI_ISL_3429006, EPI_ISL_3429007, EPI_ISL_3429008, EPI_ISL_3429009, EPI_ISL_3429010, EPI_ISL_3429011, EPI_ISL_3429012, EPI_ISL_3429013, EPI_ISL_3429014, EPI_ISL_3429015, EPI_ISL_3429016, EPI_ISL_3429017, EPI_ISL_3429018, EPI_ISL_3429019, EPI_ISL_3429020, EPI_ISL_3429021, EPI_ISL_3429022, EPI_ISL_3429023, EPI_ISL_3429024, EPI_ISL_3429025, EPI_ISL_3429026, EPI_ISL_3429027, EPI_ISL_3429028, EPI_ISL_3429029, EPI_ISL_3429030, EPI_ISL_3429031, EPI_ISL_3429032, EPI_ISL_3429033, EPI_ISL_3429034, EPI_ISL_3429035, EPI_ISL_3429036, EPI_ISL_3429037, EPI_ISL_3429038, EPI_ISL_3429039, EPI_ISL_3429040, EPI_ISL_3429041, EPI_ISL_3429042, EPI_ISL_3429043, EPI_ISL_3429044, EPI_ISL_3429045, EPI_ISL_3429046, EPI_ISL_3429047, EPI_ISL_3429048, EPI_ISL_3429049, EPI_ISL_3429050, EPI_ISL_3429051, EPI_ISL_3429052, EPI_ISL_3429053, EPI_ISL_3429054, EPI_ISL_3429055, EPI_ISL_3429056, EPI_ISL_3429057, EPI_ISL_3429058, EPI_ISL_3429059, EPI_ISL_3429060, EPI_ISL_3429061, EPI_ISL_3429062, EPI_ISL_3429063, EPI_ISL_3429064, EPI_ISL_3429065, EPI_ISL_3429066, EPI_ISL_3429067, EPI_ISL_3429068, EPI_ISL_3429069, EPI_ISL_3429070, EPI_ISL_3429071, EPI_ISL_3429072, EPI_ISL_3429073, EPI_ISL_3429074, EPI_ISL_3429075, EPI_ISL_3429076, EPI_ISL_3429077, EPI_ISL_3429078, EPI_ISL_3429079, EPI_ISL_3429080, EPI_ISL_3429081, EPI_ISL_3429082, EPI_ISL_3429083, EPI_ISL_3429085, EPI_ISL_3429086, EPI_ISL_3429087, EPI_ISL_3429088, EPI_ISL_3429089, EPI_ISL_3429090, EPI_ISL_3429091, EPI_ISL_3429092, EPI_ISL_3429093, EPI_ISL_3429094, EPI_ISL_3429095, EPI_ISL_3429096, EPI_ISL_3429097, EPI_ISL_3429098, EPI_ISL_3429099, EPI_ISL_3429100, EPI_ISL_3429101, EPI_ISL_3429102, EPI_ISL_3429103, EPI_ISL_3429104, EPI_ISL_3429105, EPI_ISL_3429106, EPI_ISL_3429107, EPI_ISL_3429108, EPI_ISL_3429109, EPI_ISL_3429110, EPI_ISL_3429111, EPI_ISL_3429112, EPI_ISL_3429113, EPI_ISL_3429114, EPI_ISL_3429115, EPI_ISL_3429116, EPI_ISL_3429117, EPI_ISL_3429118, EPI_ISL_3429120, EPI_ISL_3429121, EPI_ISL_3429122, EPI_ISL_3429123, EPI_ISL_3429125, EPI_ISL_3429126, EPI_ISL_3429127, EPI_ISL_3429128, EPI_ISL_3429129, EPI_ISL_3429130, EPI_ISL_3429131, EPI_ISL_3429132, EPI_ISL_3429133, EPI_ISL_3429134, EPI_ISL_3429135, EPI_ISL_3429136, EPI_ISL_3429137, EPI_ISL_3429138, EPI_ISL_3429139, EPI_ISL_3429140, EPI_ISL_3429141, EPI_ISL_3429142, EPI_ISL_3429143, EPI_ISL_3429144, EPI_ISL_3429145, EPI_ISL_3429146, EPI_ISL_3429147, EPI_ISL_3429148, EPI_ISL_3429149, EPI_ISL_3429150, EPI_ISL_3429151, EPI_ISL_3429152, EPI_ISL_3429153, EPI_ISL_3429154, EPI_ISL_3429155, EPI_ISL_3429156, EPI_ISL_3429157, EPI_ISL_3429158, EPI_ISL_3429159, EPI_ISL_3429160, EPI_ISL_3429161, EPI_ISL_3429162, EPI_ISL_3429163, EPI_ISL_3429164, EPI_ISL_3429165, EPI_ISL_3429166, EPI_ISL_3429167, EPI_ISL_3429168, EPI_ISL_3429169, EPI_ISL_3429170, EPI_ISL_3429171, EPI_ISL_3429172, EPI_ISL_3429173, EPI_ISL_3429174, EPI_ISL_3429175, EPI_ISL_3429176, EPI_ISL_3429177, EPI_ISL_3429178, EPI_ISL_3429179, EPI_ISL_3429180, EPI_ISL_3429181, EPI_ISL_3429182, EPI_ISL_3429183, EPI_ISL_3429184, EPI_ISL_3429185, EPI_ISL_3429186, EPI_ISL_3429188, EPI_ISL_3429189, EPI_ISL_3429190, EPI_ISL_3429191, EPI_ISL_3429192, EPI_ISL_3429193, EPI_ISL_3429194, EPI_ISL_3429195, EPI_ISL_3429196 | | | |
| see above | Fulgent Genetics | Centers for Disease Control and Prevention Division of Viral Diseases, Pathogen Discovery | Dakota Howard, Dhwani Batra, Peter W. Cook, Kara Moser, Adrian Paskey, Jason Caravas, Benjamin Rambo-Martin, Shatavia Morrison, Christopher Gulvick, Scott Sammons, Yvette Unoarumhi, Darlene Wagner, Matthew Schmerer, Harry Gao, Mickey Li, John Gao, Joseph Fierro, Benafsh Sapra, Becky Tsai, Yan Meng, Doreen Ng, James Xie, Clinton R. Paden, Duncan MacCannell |
| EPI_ISL_3429197 | UMC Utrecht | UMC Utrecht | Rob Schuurman, Anniek Tanja, Annemarie Wensing |
| EPI_ISL_3429198, EPI_ISL_3429199 | Fulgent Genetics | Centers for Disease Control and Prevention Division of Viral Diseases, Pathogen Discovery | Dakota Howard, Dhwani Batra, Peter W. Cook, Kara Moser, Adrian Paskey, Jason Caravas, Benjamin Rambo-Martin, Shatavia Morrison, Christopher Gulvick, Scott Sammons, Yvette Unoarumhi, Darlene Wagner, Matthew Schmerer, Harry Gao, Mickey Li, John Gao, Joseph Fierro, Benafsh Sapra, Becky Tsai, Yan Meng, Doreen Ng, James Xie, Clinton R. Paden, Duncan MacCannell |
| EPI_ISL_3429200 | UMC Utrecht | UMC Utrecht | Rob Schuurman, Anniek Tanja, Annemarie Wensing |
| EPI_ISL_3429201 | Fulgent Genetics | Centers for Disease Control and Prevention Division of Viral Diseases, Pathogen Discovery | Dakota Howard, Dhwani Batra, Peter W. Cook, Kara Moser, Adrian Paskey, Jason Caravas, Benjamin Rambo-Martin, Shatavia Morrison, Christopher Gulvick, Scott Sammons, Yvette Unoarumhi, Darlene Wagner, Matthew Schmerer, Harry Gao, Mickey Li, John Gao, Joseph Fierro, Benafsh Sapra, Becky Tsai, Yan Meng, Doreen Ng, James Xie, Clinton R. Paden, Duncan MacCannell |
| EPI_ISL_3429202 | UMC Utrecht | UMC Utrecht | Rob Schuurman, Anniek Tanja, Annemarie Wensing |
| EPI_ISL_3429203 | Fulgent Genetics | Centers for Disease Control and Prevention Division of Viral Diseases, Pathogen Discovery | Dakota Howard, Dhwani Batra, Peter W. Cook, Kara Moser, Adrian Paskey, Jason Caravas, Benjamin Rambo-Martin, Shatavia Morrison, Christopher Gulvick, Scott Sammons, Yvette Unoarumhi, Darlene Wagner, Matthew Schmerer, Harry Gao, Mickey Li, John Gao, Joseph Fierro, Benafsh Sapra, Becky Tsai, Yan Meng, Doreen Ng, James Xie, Clinton R. Paden, Duncan MacCannell |
| EPI_ISL_3429204 | UMC Utrecht | UMC Utrecht | Rob Schuurman, Anniek Tanja, Annemarie Wensing |
| EPI_ISL_3429205, EPI_ISL_3429206 | Fulgent Genetics | Centers for Disease Control and Prevention Division of Viral Diseases, Pathogen Discovery | Dakota Howard, Dhwani Batra, Peter W. Cook, Kara Moser, Adrian Paskey, Jason Caravas, Benjamin Rambo-Martin, Shatavia Morrison, Christopher Gulvick, Scott Sammons, Yvette Unoarumhi, Darlene Wagner, Matthew Schmerer, Harry Gao, Mickey Li, John Gao, Joseph Fierro, Benafsh Sapra, Becky Tsai, Yan Meng, Doreen Ng, James Xie, Clinton R. Paden, Duncan MacCannell |
| EPI_ISL_3429207 | UMC Utrecht | UMC Utrecht | Rob Schuurman, Anniek Tanja, Annemarie Wensing |
| EPI_ISL_3429208, EPI_ISL_3429209 | Fulgent Genetics | Centers for Disease Control and Prevention Division of Viral Diseases, Pathogen Discovery | Dakota Howard, Dhwani Batra, Peter W. Cook, Kara Moser, Adrian Paskey, Jason Caravas, Benjamin Rambo-Martin, Shatavia Morrison, Christopher Gulvick, Scott Sammons, Yvette Unoarumhi, Darlene Wagner, Matthew Schmerer, Harry Gao, Mickey Li, John Gao, Joseph Fierro, Benafsh Sapra, Becky Tsai, Yan Meng, Doreen Ng, James Xie, Clinton R. Paden, Duncan MacCannell |
| EPI_ISL_3429210 | UMC Utrecht | UMC Utrecht | Rob Schuurman, Anniek Tanja, Annemarie Wensing |
| EPI_ISL_3429211 | Fulgent Genetics | Centers for Disease Control and Prevention Division of Viral Diseases, Pathogen Discovery | Dakota Howard, Dhwani Batra, Peter W. Cook, Kara Moser, Adrian Paskey, Jason Caravas, Benjamin Rambo-Martin, Shatavia Morrison, Christopher Gulvick, Scott Sammons, Yvette Unoarumhi, Darlene Wagner, Matthew Schmerer, Harry Gao, Mickey Li, John Gao, Joseph Fierro, Benafsh Sapra, Becky Tsai, Yan Meng, Doreen Ng, James Xie, Clinton R. Paden, Duncan MacCannell |
| EPI_ISL_3429212 | UMC Utrecht | UMC Utrecht | Rob Schuurman, Anniek Tanja, Annemarie Wensing |
| EPI_ISL_3429213 | Fulgent Genetics | Centers for Disease Control and Prevention Division of Viral Diseases, Pathogen Discovery | Dakota Howard, Dhwani Batra, Peter W. Cook, Kara Moser, Adrian Paskey, Jason Caravas, Benjamin Rambo-Martin, Shatavia Morrison, Christopher Gulvick, Scott Sammons, Yvette Unoarumhi, Darlene Wagner, Matthew Schmerer, Harry Gao, Mickey Li, John Gao, Joseph Fierro, Benafsh Sapra, Becky Tsai, Yan Meng, Doreen Ng, James Xie, Clinton R. Paden, Duncan MacCannell |
| EPI_ISL_3429214 | UMC Utrecht | UMC Utrecht | Rob Schuurman, Anniek Tanja, Annemarie Wensing |
| EPI_ISL_3429215, EPI_ISL_3429216 | Fulgent Genetics | Centers for Disease Control and Prevention Division of Viral Diseases, Pathogen Discovery | Dakota Howard, Dhwani Batra, Peter W. Cook, Kara Moser, Adrian Paskey, Jason Caravas, Benjamin Rambo-Martin, Shatavia Morrison, Christopher Gulvick, Scott Sammons, Yvette Unoarumhi, Darlene Wagner, Matthew Schmerer, Harry Gao, Mickey Li, John Gao, Joseph Fierro, Benafsh Sapra, Becky Tsai, Yan Meng, Doreen Ng, James Xie, Clinton R. Paden, Duncan MacCannell |

| | | | |
|---|---|---|---|
| EPI_ISL_3429217 | UMC Utrecht | UMC Utrecht | Rob Schuurman, Anniek Tanja, Annemarie Wensing |
| EPI_ISL_3429218 | Fulgent Genetics | Centers for Disease Control and Prevention Division of Viral Diseases, Pathogen Discovery | Dakota Howard, Dhwani Batra, Peter W. Cook, Kara Moser, Adrian Paskey, Jason Caravas, Benjamin Rambo-Martin, Shatavia Morrison, Christopher Gulvick, Scott Sammons, Yvette Unoarumhi, Darlene Wagner, Matthew Schmerer, Harry Gao, Mickey Li, John Gao, Joseph Fierro, Benafsh Sapra, Becky Tsai, Yan Meng, Doreen Ng, James Xie, Clinton R. Paden, Duncan MacCannell |
| EPI_ISL_3429219 | UMC Utrecht | UMC Utrecht | Rob Schuurman, Anniek Tanja, Annemarie Wensing |
| EPI_ISL_3429220, EPI_ISL_3429221 | Fulgent Genetics | Centers for Disease Control and Prevention Division of Viral Diseases, Pathogen Discovery | Dakota Howard, Dhwani Batra, Peter W. Cook, Kara Moser, Adrian Paskey, Jason Caravas, Benjamin Rambo-Martin, Shatavia Morrison, Christopher Gulvick, Scott Sammons, Yvette Unoarumhi, Darlene Wagner, Matthew Schmerer, Harry Gao, Mickey Li, John Gao, Joseph Fierro, Benafsh Sapra, Becky Tsai, Yan Meng, Doreen Ng, James Xie, Clinton R. Paden, Duncan MacCannell |
| EPI_ISL_3429222 | UMC Utrecht | UMC Utrecht | Rob Schuurman, Anniek Tanja, Annemarie Wensing |
| EPI_ISL_3429223, EPI_ISL_3429224 | Fulgent Genetics | Centers for Disease Control and Prevention Division of Viral Diseases, Pathogen Discovery | Dakota Howard, Dhwani Batra, Peter W. Cook, Kara Moser, Adrian Paskey, Jason Caravas, Benjamin Rambo-Martin, Shatavia Morrison, Christopher Gulvick, Scott Sammons, Yvette Unoarumhi, Darlene Wagner, Matthew Schmerer, Harry Gao, Mickey Li, John Gao, Joseph Fierro, Benafsh Sapra, Becky Tsai, Yan Meng, Doreen Ng, James Xie, Clinton R. Paden, Duncan MacCannell |
| EPI_ISL_3429225 | UMC Utrecht | UMC Utrecht | Rob Schuurman, Anniek Tanja, Annemarie Wensing |
| EPI_ISL_3429226 | Fulgent Genetics | Centers for Disease Control and Prevention Division of Viral Diseases, Pathogen Discovery | Dakota Howard, Dhwani Batra, Peter W. Cook, Kara Moser, Adrian Paskey, Jason Caravas, Benjamin Rambo-Martin, Shatavia Morrison, Christopher Gulvick, Scott Sammons, Yvette Unoarumhi, Darlene Wagner, Matthew Schmerer, Harry Gao, Mickey Li, John Gao, Joseph Fierro, Benafsh Sapra, Becky Tsai, Yan Meng, Doreen Ng, James Xie, Clinton R. Paden, Duncan MacCannell |
| EPI_ISL_3429227 | UMC Utrecht | UMC Utrecht | Rob Schuurman, Anniek Tanja, Annemarie Wensing |
| EPI_ISL_3429228 | Fulgent Genetics | Centers for Disease Control and Prevention Division of Viral Diseases, Pathogen Discovery | Dakota Howard, Dhwani Batra, Peter W. Cook, Kara Moser, Adrian Paskey, Jason Caravas, Benjamin Rambo-Martin, Shatavia Morrison, Christopher Gulvick, Scott Sammons, Yvette Unoarumhi, Darlene Wagner, Matthew Schmerer, Harry Gao, Mickey Li, John Gao, Joseph Fierro, Benafsh Sapra, Becky Tsai, Yan Meng, Doreen Ng, James Xie, Clinton R. Paden, Duncan MacCannell |
| EPI_ISL_3429229 | UMC Utrecht | UMC Utrecht | Rob Schuurman, Anniek Tanja, Annemarie Wensing |
| EPI_ISL_3429230 | Fulgent Genetics | Centers for Disease Control and Prevention Division of Viral Diseases, Pathogen Discovery | Dakota Howard, Dhwani Batra, Peter W. Cook, Kara Moser, Adrian Paskey, Jason Caravas, Benjamin Rambo-Martin, Shatavia Morrison, Christopher Gulvick, Scott Sammons, Yvette Unoarumhi, Darlene Wagner, Matthew Schmerer, Harry Gao, Mickey Li, John Gao, Joseph Fierro, Benafsh Sapra, Becky Tsai, Yan Meng, Doreen Ng, James Xie, Clinton R. Paden, Duncan MacCannell |
| EPI_ISL_3429231 | UMC Utrecht | UMC Utrecht | Rob Schuurman, Anniek Tanja, Annemarie Wensing |
| EPI_ISL_3429232 | Fulgent Genetics | Centers for Disease Control and Prevention Division of Viral Diseases, Pathogen Discovery | Dakota Howard, Dhwani Batra, Peter W. Cook, Kara Moser, Adrian Paskey, Jason Caravas, Benjamin Rambo-Martin, Shatavia Morrison, Christopher Gulvick, Scott Sammons, Yvette Unoarumhi, Darlene Wagner, Matthew Schmerer, Harry Gao, Mickey Li, John Gao, Joseph Fierro, Benafsh Sapra, Becky Tsai, Yan Meng, Doreen Ng, James Xie, Clinton R. Paden, Duncan MacCannell |
| EPI_ISL_3429233 | UMC Utrecht | UMC Utrecht | Rob Schuurman, Anniek Tanja, Annemarie Wensing |
| EPI_ISL_3429234, EPI_ISL_3429235 | Fulgent Genetics | Centers for Disease Control and Prevention Division of Viral Diseases, Pathogen Discovery | Dakota Howard, Dhwani Batra, Peter W. Cook, Kara Moser, Adrian Paskey, Jason Caravas, Benjamin Rambo-Martin, Shatavia Morrison, Christopher Gulvick, Scott Sammons, Yvette Unoarumhi, Darlene Wagner, Matthew Schmerer, Harry Gao, Mickey Li, John Gao, Joseph Fierro, Benafsh Sapra, Becky Tsai, Yan Meng, Doreen Ng, James Xie, Clinton R. Paden, Duncan MacCannell |
| EPI_ISL_3429236, EPI_ISL_3429237 | UMC Utrecht | UMC Utrecht | Rob Schuurman, Anniek Tanja, Annemarie Wensing |
| EPI_ISL_3429238 | Fulgent Genetics | Centers for Disease Control and Prevention Division of Viral Diseases, Pathogen Discovery | Dakota Howard, Dhwani Batra, Peter W. Cook, Kara Moser, Adrian Paskey, Jason Caravas, Benjamin Rambo-Martin, Shatavia Morrison, Christopher Gulvick, Scott Sammons, Yvette Unoarumhi, Darlene Wagner, Matthew Schmerer, Harry Gao, Mickey Li, John Gao, Joseph Fierro, Benafsh Sapra, Becky Tsai, Yan Meng, Doreen Ng, James Xie, Clinton R. Paden, Duncan MacCannell |
| EPI_ISL_3429239 | UMC Utrecht | UMC Utrecht | Rob Schuurman, Anniek Tanja, Annemarie Wensing |
| EPI_ISL_3429240 | Fulgent Genetics | Centers for Disease Control and Prevention Division of Viral Diseases, Pathogen Discovery | Dakota Howard, Dhwani Batra, Peter W. Cook, Kara Moser, Adrian Paskey, Jason Caravas, Benjamin Rambo-Martin, Shatavia Morrison, Christopher Gulvick, Scott Sammons, Yvette Unoarumhi, Darlene Wagner, Matthew Schmerer, Harry Gao, Mickey Li, John Gao, Joseph Fierro, Benafsh Sapra, Becky Tsai, Yan Meng, Doreen Ng, James Xie, Clinton R. Paden, Duncan MacCannell |
| EPI_ISL_3429241 | UMC Utrecht | UMC Utrecht | Rob Schuurman, Anniek Tanja, Annemarie Wensing |
| EPI_ISL_3429242 | Fulgent Genetics | Centers for Disease Control and Prevention Division of Viral Diseases, Pathogen Discovery | Dakota Howard, Dhwani Batra, Peter W. Cook, Kara Moser, Adrian Paskey, Jason Caravas, Benjamin Rambo-Martin, Shatavia Morrison, Christopher Gulvick, Scott Sammons, Yvette Unoarumhi, Darlene Wagner, Matthew Schmerer, Harry Gao, Mickey Li, John Gao, Joseph Fierro, Benafsh Sapra, Becky Tsai, Yan Meng, Doreen Ng, James Xie, Clinton R. Paden, Duncan MacCannell |
| EPI_ISL_3429243 | UMC Utrecht | UMC Utrecht | Rob Schuurman, Anniek Tanja, Annemarie Wensing |
| EPI_ISL_3429244 | Fulgent Genetics | Centers for Disease Control and Prevention Division of Viral Diseases, Pathogen Discovery | Dakota Howard, Dhwani Batra, Peter W. Cook, Kara Moser, Adrian Paskey, Jason Caravas, Benjamin Rambo-Martin, Shatavia Morrison, Christopher Gulvick, Scott Sammons, Yvette Unoarumhi, Darlene Wagner, Matthew Schmerer, Harry Gao, Mickey Li, John Gao, Joseph Fierro, Benafsh Sapra, Becky Tsai, Yan Meng, Doreen Ng, James Xie, Clinton R. Paden, Duncan MacCannell |
| EPI_ISL_3429245 | UMC Utrecht | UMC Utrecht | Rob Schuurman, Anniek Tanja, Annemarie Wensing |
| EPI_ISL_3429246 | Fulgent Genetics | Centers for Disease Control and Prevention Division of Viral Diseases, Pathogen Discovery | Dakota Howard, Dhwani Batra, Peter W. Cook, Kara Moser, Adrian Paskey, Jason Caravas, Benjamin Rambo-Martin, Shatavia Morrison, Christopher Gulvick, Scott Sammons, Yvette Unoarumhi, Darlene Wagner, Matthew Schmerer, Harry Gao, Mickey Li, John Gao, Joseph Fierro, Benafsh Sapra, Becky Tsai, Yan Meng, Doreen Ng, James Xie, Clinton R. Paden, Duncan MacCannell |
| EPI_ISL_3429247 | UMC Utrecht | UMC Utrecht | Rob Schuurman, Anniek Tanja, Annemarie Wensing |
| EPI_ISL_3429248 | Fulgent Genetics | Centers for Disease Control and Prevention Division of Viral Diseases, Pathogen Discovery | Dakota Howard, Dhwani Batra, Peter W. Cook, Kara Moser, Adrian Paskey, Jason Caravas, Benjamin Rambo-Martin, Shatavia Morrison, Christopher Gulvick, Scott Sammons, Yvette Unoarumhi, Darlene Wagner, Matthew Schmerer, Harry Gao, Mickey Li, John Gao, Joseph Fierro, Benafsh Sapra, Becky Tsai, Yan Meng, Doreen Ng, James Xie, Clinton R. Paden, Duncan MacCannell |
| EPI_ISL_3429249 | UMC Utrecht | UMC Utrecht | Rob Schuurman, Anniek Tanja, Annemarie Wensing |
| EPI_ISL_3429250 | Fulgent Genetics | Centers for Disease Control and Prevention Division of Viral Diseases, Pathogen Discovery | Dakota Howard, Dhwani Batra, Peter W. Cook, Kara Moser, Adrian Paskey, Jason Caravas, Benjamin Rambo-Martin, Shatavia Morrison, Christopher Gulvick, Scott Sammons, Yvette Unoarumhi, Darlene Wagner, Matthew Schmerer, Harry Gao, Mickey Li, John Gao, Joseph Fierro, Benafsh Sapra, Becky Tsai, Yan Meng, Doreen Ng, James Xie, Clinton R. Paden, Duncan MacCannell |
| EPI_ISL_3429251 | UMC Utrecht | UMC Utrecht | Rob Schuurman, Anniek Tanja, Annemarie Wensing |
| EPI_ISL_3429252 | Fulgent Genetics | Centers for Disease Control and Prevention Division of Viral Diseases, Pathogen Discovery | Dakota Howard, Dhwani Batra, Peter W. Cook, Kara Moser, Adrian Paskey, Jason Caravas, Benjamin Rambo-Martin, Shatavia Morrison, Christopher Gulvick, Scott Sammons, Yvette Unoarumhi, Darlene Wagner, Matthew Schmerer, Harry Gao, Mickey Li, John Gao, Joseph Fierro, Benafsh Sapra, Becky Tsai, Yan Meng, Doreen Ng, James Xie, Clinton R. Paden, Duncan MacCannell |
| EPI_ISL_3429253 | UMC Utrecht | UMC Utrecht | Rob Schuurman, Anniek Tanja, Annemarie Wensing |
| EPI_ISL_3429254 | Fulgent Genetics | Centers for Disease Control and Prevention Division of Viral Diseases, Pathogen Discovery | Dakota Howard, Dhwani Batra, Peter W. Cook, Kara Moser, Adrian Paskey, Jason Caravas, Benjamin Rambo-Martin, Shatavia Morrison, Christopher Gulvick, Scott Sammons, Yvette Unoarumhi, Darlene Wagner, Matthew Schmerer, Harry Gao, Mickey Li, John Gao, Joseph Fierro, Benafsh Sapra, Becky Tsai, Yan Meng, Doreen Ng, James Xie, Clinton R. Paden, Duncan MacCannell |
| EPI_ISL_3429255 | UMC Utrecht | UMC Utrecht | Rob Schuurman, Anniek Tanja, Annemarie Wensing |
| EPI_ISL_3429256 | Fulgent Genetics | Centers for Disease Control and Prevention Division of Viral Diseases, Pathogen Discovery | Dakota Howard, Dhwani Batra, Peter W. Cook, Kara Moser, Adrian Paskey, Jason Caravas, Benjamin Rambo-Martin, Shatavia Morrison, Christopher Gulvick, Scott Sammons, Yvette Unoarumhi, Darlene Wagner, Matthew Schmerer, Harry Gao, Mickey Li, John Gao, Joseph Fierro, Benafsh Sapra, Becky Tsai, Yan Meng, Doreen Ng, James Xie, Clinton R. Paden, Duncan MacCannell |
| EPI_ISL_3429257 | UMC Utrecht | UMC Utrecht | Rob Schuurman, Anniek Tanja, Annemarie Wensing |
| EPI_ISL_3429258, EPI_ISL_3429259 | Fulgent Genetics | Centers for Disease Control and Prevention Division of Viral Diseases, Pathogen Discovery | Dakota Howard, Dhwani Batra, Peter W. Cook, Kara Moser, Adrian Paskey, Jason Caravas, Benjamin Rambo-Martin, Shatavia Morrison, Christopher Gulvick, Scott Sammons, Yvette Unoarumhi, Darlene Wagner, Matthew Schmerer, Harry Gao, Mickey Li, John Gao, Joseph Fierro, Benafsh Sapra, Becky Tsai, Yan Meng, Doreen Ng, James Xie, Clinton R. Paden, Duncan MacCannell |
| EPI_ISL_3429260 | UMC Utrecht | UMC Utrecht | Rob Schuurman, Anniek Tanja, Annemarie Wensing |
| EPI_ISL_3429261, EPI_ISL_3429262, EPI_ISL_3429263, EPI_ISL_3429264, EPI_ISL_3429265, EPI_ISL_3429266, EPI_ISL_3429267, EPI_ISL_3429268, EPI_ISL_3429269, EPI_ISL_3429270, EPI_ISL_3429271, EPI_ISL_3429272, EPI_ISL_3429273, EPI_ISL_3429274, EPI_ISL_3429275, EPI_ISL_3429276, EPI_ISL_3429277, EPI_ISL_3429279, EPI_ISL_3429280, EPI_ISL_3429281, EPI_ISL_3429282, EPI_ISL_3429283, EPI_ISL_3429284, EPI_ISL_3429285, EPI_ISL_3429286, EPI_ISL_3429287, EPI_ISL_3429288, EPI_ISL_3429289, EPI_ISL_3429290, EPI_ISL_3429291, EPI_ISL_3429292, EPI_ISL_3429293, EPI_ISL_3429294, EPI_ISL_3429295, EPI_ISL_3429296, EPI_ISL_3429297, EPI_ISL_3429298, EPI_ISL_3429299, EPI_ISL_3429300, EPI_ISL_3429301, EPI_ISL_3429302, EPI_ISL_3429303, EPI_ISL_3429304, EPI_ISL_3429305, EPI_ISL_3429306, EPI_ISL_3429307, EPI_ISL_3429308, EPI_ISL_3429309, EPI_ISL_3429310, EPI_ISL_3429311, EPI_ISL_3429312, EPI_ISL_3429313, EPI_ISL_3429314, EPI_ISL_3429315, EPI_ISL_3429316, EPI_ISL_3429317, EPI_ISL_3429318, EPI_ISL_3429319, EPI_ISL_3429320, EPI_ISL_3429321, EPI_ISL_3429323, EPI_ISL_3429324, EPI_ISL_3429325, EPI_ISL_3429326, EPI_ISL_3429327, EPI_ISL_3429328, EPI_ISL_3429329, EPI_ISL_3429330, EPI_ISL_3429331, EPI_ISL_3429332, EPI_ISL_3429333, EPI_ISL_3429335, EPI_ISL_3429336, EPI_ISL_3429337, EPI_ISL_3429338, EPI_ISL_3429339, EPI_ISL_3429340, EPI_ISL_3429341, EPI_ISL_3429342, EPI_ISL_3429343, EPI_ISL_3429344, EPI_ISL_3429345, EPI_ISL_3429346, EPI_ISL_3429347, EPI_ISL_3429348, EPI_ISL_3429349, EPI_ISL_3429350, EPI_ISL_3429351, EPI_ISL_3429352, EPI_ISL_3429353, EPI_ISL_3429354, EPI_ISL_3429355, EPI_ISL_3429356, EPI_ISL_3429357, EPI_ISL_3429358, EPI_ISL_3429359, EPI_ISL_3429360, EPI_ISL_3429361, EPI_ISL_3429362, EPI_ISL_3429363, EPI_ISL_3429364, EPI_ISL_3429365, EPI_ISL_3429366, EPI_ISL_3429367, EPI_ISL_3429368, EPI_ISL_3429369, EPI_ISL_3429370, EPI_ISL_3429371, EPI_ISL_3429372, EPI_ISL_3429373, EPI_ISL_3429374, EPI_ISL_3429375, EPI_ISL_3429376, EPI_ISL_3429377, EPI_ISL_3429378, EPI_ISL_3429379, EPI_ISL_3429380, EPI_ISL_3429381, EPI_ISL_3429382, EPI_ISL_3429383, EPI_ISL_3429384, EPI_ISL_3429385, EPI_ISL_3429386, EPI_ISL_3429387, EPI_ISL_3429388, EPI_ISL_3429389, EPI_ISL_3429390, EPI_ISL_3429391, EPI_ISL_3429392, EPI_ISL_3429393, EPI_ISL_3429394, EPI_ISL_3429395, EPI_ISL_3429396, EPI_ISL_3429397, EPI_ISL_3429398, EPI_ISL_3429399, EPI_ISL_3429400, EPI_ISL_3429401, EPI_ISL_3429402, EPI_ISL_3429403, EPI_ISL_3429404, EPI_ISL_3429405, EPI_ISL_3429406, EPI_ISL_3429407, EPI_ISL_3429408, EPI_ISL_3429409, EPI_ISL_3429410, EPI_ISL_3429411, EPI_ISL_3429412, EPI_ISL_3429413, EPI_ISL_3429414, EPI_ISL_3429415, EPI_ISL_3429416, EPI_ISL_3429417, EPI_ISL_3429418, EPI_ISL_3429419, EPI_ISL_3429420, EPI_ISL_3429421, EPI_ISL_3429422, EPI_ISL_3429423, EPI_ISL_3429424, EPI_ISL_3429425, EPI_ISL_3429426, EPI_ISL_3429427, EPI_ISL_3429428, EPI_ISL_3429429, EPI_ISL_3429430, EPI_ISL_3429431, EPI_ISL_3429432, EPI_ISL_3429433, EPI_ISL_3429434, EPI_ISL_3429435, EPI_ISL_3429437, EPI_ISL_3429438, EPI_ISL_3429439, EPI_ISL_3429440, EPI_ISL_3429441 | | | |
| see above | Fulgent Genetics | Centers for Disease Control and Prevention Division of Viral Diseases, Pathogen Discovery | Dakota Howard, Dhwani Batra, Peter W. Cook, Kara Moser, Adrian Paskey, Jason Caravas, Benjamin Rambo-Martin, Shatavia Morrison, Christopher Gulvick, Scott Sammons, Yvette Unoarumhi, Darlene Wagner, Matthew Schmerer, Harry Gao, Mickey Li, John Gao, Joseph Fierro, Benafsh Sapra, Becky Tsai, Yan Meng, Doreen Ng, James Xie, Clinton R. Paden, Duncan MacCannell |
| EPI_ISL_3429442 | UMC Utrecht | UMC Utrecht | Rob Schuurman, Anniek Tanja, Annemarie Wensing |
| EPI_ISL_3429443, EPI_ISL_3429445 | Fulgent Genetics | Centers for Disease Control and Prevention Division of Viral Diseases, Pathogen Discovery | Dakota Howard, Dhwani Batra, Peter W. Cook, Kara Moser, Adrian Paskey, Jason Caravas, Benjamin Rambo-Martin, Shatavia Morrison, Christopher Gulvick, Scott Sammons, Yvette Unoarumhi, Darlene Wagner, Matthew Schmerer, Harry Gao, Mickey Li, John Gao, Joseph Fierro, Benafsh Sapra, Becky Tsai, Yan Meng, Doreen Ng, James Xie, Clinton R. Paden, Duncan MacCannell |
| EPI_ISL_3429446 | UMC Utrecht | UMC Utrecht | Rob Schuurman, Anniek Tanja, Annemarie Wensing |
| EPI_ISL_3429447 | Fulgent Genetics | Centers for Disease Control and Prevention Division of Viral Diseases, Pathogen Discovery | Dakota Howard, Dhwani Batra, Peter W. Cook, Kara Moser, Adrian Paskey, Jason Caravas, Benjamin Rambo-Martin, Shatavia Morrison, Christopher Gulvick, Scott Sammons, Yvette Unoarumhi, Darlene Wagner, Matthew Schmerer, Harry Gao, Mickey Li, John Gao, Joseph Fierro, Benafsh Sapra, Becky Tsai, Yan Meng, Doreen Ng, James Xie, Clinton R. Paden, Duncan MacCannell |
| EPI_ISL_3429448 | UMC Utrecht | UMC Utrecht | Rob Schuurman, Anniek Tanja, Annemarie Wensing |
| EPI_ISL_3429449 | Fulgent Genetics | Centers for Disease Control and Prevention Division of Viral Diseases, Pathogen Discovery | Dakota Howard, Dhwani Batra, Peter W. Cook, Kara Moser, Adrian Paskey, Jason Caravas, Benjamin Rambo-Martin, Shatavia Morrison, Christopher Gulvick, Scott Sammons, Yvette Unoarumhi, Darlene Wagner, Matthew Schmerer, Harry Gao, Mickey Li, John Gao, Joseph Fierro, Benafsh Sapra, Becky Tsai, Yan Meng, Doreen Ng, James Xie, Clinton R. Paden, Duncan MacCannell |
| EPI_ISL_3429450 | UMC Utrecht | UMC Utrecht | Rob Schuurman, Anniek Tanja, Annemarie Wensing |
| EPI_ISL_3429451, EPI_ISL_3429452 | Fulgent Genetics | Centers for Disease Control and Prevention Division of Viral Diseases, Pathogen Discovery | Dakota Howard, Dhwani Batra, Peter W. Cook, Kara Moser, Adrian Paskey, Jason Caravas, Benjamin Rambo-Martin, Shatavia Morrison, Christopher Gulvick, Scott Sammons, Yvette Unoarumhi, Darlene Wagner, Matthew Schmerer, Harry Gao, Mickey Li, John Gao, Joseph Fierro, Benafsh Sapra, Becky Tsai, Yan Meng, Doreen Ng, James Xie, Clinton R. Paden, Duncan MacCannell |
| EPI_ISL_3429453 | UMC Utrecht | UMC Utrecht | Rob Schuurman, Anniek Tanja, Annemarie Wensing |
| EPI_ISL_3429454, EPI_ISL_3429455, EPI_ISL_3429456 | Fulgent Genetics | Centers for Disease Control and Prevention Division of Viral Diseases, Pathogen Discovery | Dakota Howard, Dhwani Batra, Peter W. Cook, Kara Moser, Adrian Paskey, Jason Caravas, Benjamin Rambo-Martin, Shatavia Morrison, Christopher Gulvick, Scott Sammons, Yvette Unoarumhi, Darlene Wagner, Matthew Schmerer, Harry Gao, Mickey Li, John Gao, Joseph Fierro, Benafsh Sapra, Becky Tsai, Yan Meng, Doreen Ng, James Xie, Clinton R. Paden, Duncan MacCannell |

| | | | |
|---|---|---|---|
| EPI_ISL_3429457, EPI_ISL_3429458 | UMC Utrecht | UMC Utrecht | Rob Schuurman, Anniek Tanja, Annemarie Wensing |
| EPI_ISL_3429459 | Fulgent Genetics | Centers for Disease Control and Prevention Division of Viral Diseases, Pathogen Discovery | Dakota Howard, Dhwani Batra, Peter W. Cook, Kara Moser, Adrian Paskey, Jason Caravas, Benjamin Rambo-Martin, Shatavia Morrison, Christopher Gulvick, Scott Sammons, Yvette Unoarumhi, Darlene Wagner, Matthew Schmerer, Harry Gao, Mickey Li, John Gao, Joseph Fierro, Benafsh Sapra, Becky Tsai, Yan Meng, Doreen Ng, James Xie, Clinton R. Paden, Duncan MacCannell |
| EPI_ISL_3429460 | UMC Utrecht | UMC Utrecht | Rob Schuurman, Anniek Tanja, Annemarie Wensing |
| EPI_ISL_3429461, EPI_ISL_3429462 | Fulgent Genetics | Centers for Disease Control and Prevention Division of Viral Diseases, Pathogen Discovery | Dakota Howard, Dhwani Batra, Peter W. Cook, Kara Moser, Adrian Paskey, Jason Caravas, Benjamin Rambo-Martin, Shatavia Morrison, Christopher Gulvick, Scott Sammons, Yvette Unoarumhi, Darlene Wagner, Matthew Schmerer, Harry Gao, Mickey Li, John Gao, Joseph Fierro, Benafsh Sapra, Becky Tsai, Yan Meng, Doreen Ng, James Xie, Clinton R. Paden, Duncan MacCannell |
| EPI_ISL_3429463 | UMC Utrecht | UMC Utrecht | Rob Schuurman, Anniek Tanja, Annemarie Wensing |
| EPI_ISL_3429464, EPI_ISL_3429466 | Fulgent Genetics | Centers for Disease Control and Prevention Division of Viral Diseases, Pathogen Discovery | Dakota Howard, Dhwani Batra, Peter W. Cook, Kara Moser, Adrian Paskey, Jason Caravas, Benjamin Rambo-Martin, Shatavia Morrison, Christopher Gulvick, Scott Sammons, Yvette Unoarumhi, Darlene Wagner, Matthew Schmerer, Harry Gao, Mickey Li, John Gao, Joseph Fierro, Benafsh Sapra, Becky Tsai, Yan Meng, Doreen Ng, James Xie, Clinton R. Paden, Duncan MacCannell |
| EPI_ISL_3429467 | UMC Utrecht | UMC Utrecht | Rob Schuurman, Anniek Tanja, Annemarie Wensing |
| EPI_ISL_3429468, EPI_ISL_3429470 | Fulgent Genetics | Centers for Disease Control and Prevention Division of Viral Diseases, Pathogen Discovery | Dakota Howard, Dhwani Batra, Peter W. Cook, Kara Moser, Adrian Paskey, Jason Caravas, Benjamin Rambo-Martin, Shatavia Morrison, Christopher Gulvick, Scott Sammons, Yvette Unoarumhi, Darlene Wagner, Matthew Schmerer, Harry Gao, Mickey Li, John Gao, Joseph Fierro, Benafsh Sapra, Becky Tsai, Yan Meng, Doreen Ng, James Xie, Clinton R. Paden, Duncan MacCannell |
| EPI_ISL_3429471 | UMC Utrecht | UMC Utrecht | Rob Schuurman, Anniek Tanja, Annemarie Wensing |
| EPI_ISL_3429472 | Fulgent Genetics | Centers for Disease Control and Prevention Division of Viral Diseases, Pathogen Discovery | Dakota Howard, Dhwani Batra, Peter W. Cook, Kara Moser, Adrian Paskey, Jason Caravas, Benjamin Rambo-Martin, Shatavia Morrison, Christopher Gulvick, Scott Sammons, Yvette Unoarumhi, Darlene Wagner, Matthew Schmerer, Harry Gao, Mickey Li, John Gao, Joseph Fierro, Benafsh Sapra, Becky Tsai, Yan Meng, Doreen Ng, James Xie, Clinton R. Paden, Duncan MacCannell |
| EPI_ISL_3429473 | UMC Utrecht | UMC Utrecht | Rob Schuurman, Anniek Tanja, Annemarie Wensing |
| EPI_ISL_3429474 | Fulgent Genetics | Centers for Disease Control and Prevention Division of Viral Diseases, Pathogen Discovery | Dakota Howard, Dhwani Batra, Peter W. Cook, Kara Moser, Adrian Paskey, Jason Caravas, Benjamin Rambo-Martin, Shatavia Morrison, Christopher Gulvick, Scott Sammons, Yvette Unoarumhi, Darlene Wagner, Matthew Schmerer, Harry Gao, Mickey Li, John Gao, Joseph Fierro, Benafsh Sapra, Becky Tsai, Yan Meng, Doreen Ng, James Xie, Clinton R. Paden, Duncan MacCannell |
| EPI_ISL_3429475 | UMC Utrecht | UMC Utrecht | Rob Schuurman, Anniek Tanja, Annemarie Wensing |
| EPI_ISL_3429476, EPI_ISL_3429477 | Fulgent Genetics | Centers for Disease Control and Prevention Division of Viral Diseases, Pathogen Discovery | Dakota Howard, Dhwani Batra, Peter W. Cook, Kara Moser, Adrian Paskey, Jason Caravas, Benjamin Rambo-Martin, Shatavia Morrison, Christopher Gulvick, Scott Sammons, Yvette Unoarumhi, Darlene Wagner, Matthew Schmerer, Harry Gao, Mickey Li, John Gao, Joseph Fierro, Benafsh Sapra, Becky Tsai, Yan Meng, Doreen Ng, James Xie, Clinton R. Paden, Duncan MacCannell |
| EPI_ISL_3429478 | UMC Utrecht | UMC Utrecht | Rob Schuurman, Anniek Tanja, Annemarie Wensing |
| EPI_ISL_3429479 | Fulgent Genetics | Centers for Disease Control and Prevention Division of Viral Diseases, Pathogen Discovery | Dakota Howard, Dhwani Batra, Peter W. Cook, Kara Moser, Adrian Paskey, Jason Caravas, Benjamin Rambo-Martin, Shatavia Morrison, Christopher Gulvick, Scott Sammons, Yvette Unoarumhi, Darlene Wagner, Matthew Schmerer, Harry Gao, Mickey Li, John Gao, Joseph Fierro, Benafsh Sapra, Becky Tsai, Yan Meng, Doreen Ng, James Xie, Clinton R. Paden, Duncan MacCannell |
| EPI_ISL_3429480 | UMC Utrecht | UMC Utrecht | Rob Schuurman, Anniek Tanja, Annemarie Wensing |
| EPI_ISL_3429481 | Fulgent Genetics | Centers for Disease Control and Prevention Division of Viral Diseases, Pathogen Discovery | Dakota Howard, Dhwani Batra, Peter W. Cook, Kara Moser, Adrian Paskey, Jason Caravas, Benjamin Rambo-Martin, Shatavia Morrison, Christopher Gulvick, Scott Sammons, Yvette Unoarumhi, Darlene Wagner, Matthew Schmerer, Harry Gao, Mickey Li, John Gao, Joseph Fierro, Benafsh Sapra, Becky Tsai, Yan Meng, Doreen Ng, James Xie, Clinton R. Paden, Duncan MacCannell |
| EPI_ISL_3429482 | UMC Utrecht | UMC Utrecht | Rob Schuurman, Anniek Tanja, Annemarie Wensing |
| EPI_ISL_3429483, EPI_ISL_3429484, EPI_ISL_3429485 | Fulgent Genetics | Centers for Disease Control and Prevention Division of Viral Diseases, Pathogen Discovery | Dakota Howard, Dhwani Batra, Peter W. Cook, Kara Moser, Adrian Paskey, Jason Caravas, Benjamin Rambo-Martin, Shatavia Morrison, Christopher Gulvick, Scott Sammons, Yvette Unoarumhi, Darlene Wagner, Matthew Schmerer, Harry Gao, Mickey Li, John Gao, Joseph Fierro, Benafsh Sapra, Becky Tsai, Yan Meng, Doreen Ng, James Xie, Clinton R. Paden, Duncan MacCannell |
| EPI_ISL_3429486 | UMC Utrecht | UMC Utrecht | Rob Schuurman, Anniek Tanja, Annemarie Wensing |
| EPI_ISL_3429487, EPI_ISL_3429488, EPI_ISL_3429490, EPI_ISL_3429491, EPI_ISL_3429493 | Fulgent Genetics | Centers for Disease Control and Prevention Division of Viral Diseases, Pathogen Discovery | Dakota Howard, Dhwani Batra, Peter W. Cook, Kara Moser, Adrian Paskey, Jason Caravas, Benjamin Rambo-Martin, Shatavia Morrison, Christopher Gulvick, Scott Sammons, Yvette Unoarumhi, Darlene Wagner, Matthew Schmerer, Harry Gao, Mickey Li, John Gao, Joseph Fierro, Benafsh Sapra, Becky Tsai, Yan Meng, Doreen Ng, James Xie, Clinton R. Paden, Duncan MacCannell |
| EPI_ISL_3429494 | UMC Utrecht | UMC Utrecht | Rob Schuurman, Anniek Tanja, Annemarie Wensing |
| EPI_ISL_3429495 | Fulgent Genetics | Centers for Disease Control and Prevention Division of Viral Diseases, Pathogen Discovery | Dakota Howard, Dhwani Batra, Peter W. Cook, Kara Moser, Adrian Paskey, Jason Caravas, Benjamin Rambo-Martin, Shatavia Morrison, Christopher Gulvick, Scott Sammons, Yvette Unoarumhi, Darlene Wagner, Matthew Schmerer, Harry Gao, Mickey Li, John Gao, Joseph Fierro, Benafsh Sapra, Becky Tsai, Yan Meng, Doreen Ng, James Xie, Clinton R. Paden, Duncan MacCannell |
| EPI_ISL_3429496, EPI_ISL_3429497 | UMC Utrecht | UMC Utrecht | Rob Schuurman, Anniek Tanja, Annemarie Wensing |
| EPI_ISL_3429498 | Fulgent Genetics | Centers for Disease Control and Prevention Division of Viral Diseases, Pathogen Discovery | Dakota Howard, Dhwani Batra, Peter W. Cook, Kara Moser, Adrian Paskey, Jason Caravas, Benjamin Rambo-Martin, Shatavia Morrison, Christopher Gulvick, Scott Sammons, Yvette Unoarumhi, Darlene Wagner, Matthew Schmerer, Harry Gao, Mickey Li, John Gao, Joseph Fierro, Benafsh Sapra, Becky Tsai, Yan Meng, Doreen Ng, James Xie, Clinton R. Paden, Duncan MacCannell |
| EPI_ISL_3429499 | UMC Utrecht | UMC Utrecht | Rob Schuurman, Anniek Tanja, Annemarie Wensing |
| EPI_ISL_3429500, EPI_ISL_3429502 | Fulgent Genetics | Centers for Disease Control and Prevention Division of Viral Diseases, Pathogen Discovery | Dakota Howard, Dhwani Batra, Peter W. Cook, Kara Moser, Adrian Paskey, Jason Caravas, Benjamin Rambo-Martin, Shatavia Morrison, Christopher Gulvick, Scott Sammons, Yvette Unoarumhi, Darlene Wagner, Matthew Schmerer, Harry Gao, Mickey Li, John Gao, Joseph Fierro, Benafsh Sapra, Becky Tsai, Yan Meng, Doreen Ng, James Xie, Clinton R. Paden, Duncan MacCannell |
| EPI_ISL_3429503 | UMC Utrecht | UMC Utrecht | Rob Schuurman, Anniek Tanja, Annemarie Wensing |
| EPI_ISL_3429504, EPI_ISL_3429505 | Fulgent Genetics | Centers for Disease Control and Prevention Division of Viral Diseases, Pathogen Discovery | Dakota Howard, Dhwani Batra, Peter W. Cook, Kara Moser, Adrian Paskey, Jason Caravas, Benjamin Rambo-Martin, Shatavia Morrison, Christopher Gulvick, Scott Sammons, Yvette Unoarumhi, Darlene Wagner, Matthew Schmerer, Harry Gao, Mickey Li, John Gao, Joseph Fierro, Benafsh Sapra, Becky Tsai, Yan Meng, Doreen Ng, James Xie, Clinton R. Paden, Duncan MacCannell |
| EPI_ISL_3429506 | UMC Utrecht | UMC Utrecht | Rob Schuurman, Anniek Tanja, Annemarie Wensing |

EPI_ISL_3429507, EPI_ISL_3429509, EPI_ISL_3429510, EPI_ISL_3429511, EPI_ISL_3429512, EPI_ISL_3429513, EPI_ISL_3429514, EPI_ISL_3429515, EPI_ISL_3429516, EPI_ISL_3429517, EPI_ISL_3429518, EPI_ISL_3429519, EPI_ISL_3429520, EPI_ISL_3429521, EPI_ISL_3429522, EPI_ISL_3429523, EPI_ISL_3429524, EPI_ISL_3429525, EPI_ISL_3429526, EPI_ISL_3429527, EPI_ISL_3429528, EPI_ISL_3429529, EPI_ISL_3429530, EPI_ISL_3429531, EPI_ISL_3429532, EPI_ISL_3429533, EPI_ISL_3429534, EPI_ISL_3429535, EPI_ISL_3429536, EPI_ISL_3429537, EPI_ISL_3429538, EPI_ISL_3429539, EPI_ISL_3429540, EPI_ISL_3429541, EPI_ISL_3429542, EPI_ISL_3429543, EPI_ISL_3429544, EPI_ISL_3429545, EPI_ISL_3429546, EPI_ISL_3429547, EPI_ISL_3429548, EPI_ISL_3429549, EPI_ISL_3429550, EPI_ISL_3429551, EPI_ISL_3429553, EPI_ISL_3429554, EPI_ISL_3429555, EPI_ISL_3429556, EPI_ISL_3429557, EPI_ISL_3429558, EPI_ISL_3429559, EPI_ISL_3429560, EPI_ISL_3429561, EPI_ISL_3429562, EPI_ISL_3429563, EPI_ISL_3429564, EPI_ISL_3429565, EPI_ISL_3429566, EPI_ISL_3429567, EPI_ISL_3429568, EPI_ISL_3429569, EPI_ISL_3429570, EPI_ISL_3429571, EPI_ISL_3429572, EPI_ISL_3429573, EPI_ISL_3429574, EPI_ISL_3429575, EPI_ISL_3429576, EPI_ISL_3429577, EPI_ISL_3429578, EPI_ISL_3429579, EPI_ISL_3429580, EPI_ISL_3429581, EPI_ISL_3429582, EPI_ISL_3429583, EPI_ISL_3429584, EPI_ISL_3429585, EPI_ISL_3429586, EPI_ISL_3429587, EPI_ISL_3429588, EPI_ISL_3429589, EPI_ISL_3429590, EPI_ISL_3429591, EPI_ISL_3429592, EPI_ISL_3429593, EPI_ISL_3429594, EPI_ISL_3429595, EPI_ISL_3429596, EPI_ISL_3429598, EPI_ISL_3429599, EPI_ISL_3429601, EPI_ISL_3429602, EPI_ISL_3429603, EPI_ISL_3429604, EPI_ISL_3429605, EPI_ISL_3429607, EPI_ISL_3429608, EPI_ISL_3429609, EPI_ISL_3429610, EPI_ISL_3429611, EPI_ISL_3429612, EPI_ISL_3429613, EPI_ISL_3429614, EPI_ISL_3429615, EPI_ISL_3429616, EPI_ISL_3429617, EPI_ISL_3429618, EPI_ISL_3429619, EPI_ISL_3429620, EPI_ISL_3429621, EPI_ISL_3429622, EPI_ISL_3429623, EPI_ISL_3429624, EPI_ISL_3429625, EPI_ISL_3429626, EPI_ISL_3429627, EPI_ISL_3429628, EPI_ISL_3429629, EPI_ISL_3429630, EPI_ISL_3429631, EPI_ISL_3429632, EPI_ISL_3429633, EPI_ISL_3429634, EPI_ISL_3429635, EPI_ISL_3429636, EPI_ISL_3429637, EPI_ISL_3429638, EPI_ISL_3429639, EPI_ISL_3429640, EPI_ISL_3429641, EPI_ISL_3429642, EPI_ISL_3429643, EPI_ISL_3429644, EPI_ISL_3429645, EPI_ISL_3429646, EPI_ISL_3429647, EPI_ISL_3429648, EPI_ISL_3429649, EPI_ISL_3429650, EPI_ISL_3429651, EPI_ISL_3429653, EPI_ISL_3429654, EPI_ISL_3429655, EPI_ISL_3429656, EPI_ISL_3429657, EPI_ISL_3429658, EPI_ISL_3429659, EPI_ISL_3429660, EPI_ISL_3429661, EPI_ISL_3429662, EPI_ISL_3429663, EPI_ISL_3429664, EPI_ISL_3429666, EPI_ISL_3429667, EPI_ISL_3429668, EPI_ISL_3429670, EPI_ISL_3429671, EPI_ISL_3429672, EPI_ISL_3429673, EPI_ISL_3429674, EPI_ISL_3429675, EPI_ISL_3429676, EPI_ISL_3429677, EPI_ISL_3429678, EPI_ISL_3429679, EPI_ISL_3429680, EPI_ISL_3429681, EPI_ISL_3429682, EPI_ISL_3429683, EPI_ISL_3429684, EPI_ISL_3429685, EPI_ISL_3429686, EPI_ISL_3429687, EPI_ISL_3429688, EPI_ISL_3429689, EPI_ISL_3429690, EPI_ISL_3429691, EPI_ISL_3429692, EPI_ISL_3429693, EPI_ISL_3429694, EPI_ISL_3429695, EPI_ISL_3429696, EPI_ISL_3429697, EPI_ISL_3429698, EPI_ISL_3429699, EPI_ISL_3429700, EPI_ISL_3429701, EPI_ISL_3429702, EPI_ISL_3429703, EPI_ISL_3429704, EPI_ISL_3429705, EPI_ISL_3429706, EPI_ISL_3429707, EPI_ISL_3429708, EPI_ISL_3429709, EPI_ISL_3429710, EPI_ISL_3429712, EPI_ISL_3429713, EPI_ISL_3429714, EPI_ISL_3429715, EPI_ISL_3429716, EPI_ISL_3429717, EPI_ISL_3429718, EPI_ISL_3429719, EPI_ISL_3429720, EPI_ISL_3429721

| | | | |
|---|---|---|---|
| see above | Fulgent Genetics | Centers for Disease Control and Prevention Division of Viral Diseases, Pathogen Discovery | Dakota Howard, Dhwani Batra, Peter W. Cook, Kara Moser, Adrian Paskey, Jason Caravas, Benjamin Rambo-Martin, Shatavia Morrison, Christopher Gulvick, Scott Sammons, Yvette Unoarumhi, Darlene Wagner, Matthew Schmerer, Harry Gao, Mickey Li, John Gao, Joseph Fierro, Benafsh Sapra, Becky Tsai, Yan Meng, Doreen Ng, James Xie, Clinton R. Paden, Duncan MacCannell |
| EPI_ISL_3429722 | UMC Utrecht | UMC Utrecht | Rob Schuurman, Anniek Tanja, Annemarie Wensing |
| EPI_ISL_3429723 | Fulgent Genetics | Centers for Disease Control and Prevention Division of Viral Diseases, Pathogen Discovery | Dakota Howard, Dhwani Batra, Peter W. Cook, Kara Moser, Adrian Paskey, Jason Caravas, Benjamin Rambo-Martin, Shatavia Morrison, Christopher Gulvick, Scott Sammons, Yvette Unoarumhi, Darlene Wagner, Matthew Schmerer, Harry Gao, Mickey Li, John Gao, Joseph Fierro, Benafsh Sapra, Becky Tsai, Yan Meng, Doreen Ng, James Xie, Clinton R. Paden, Duncan MacCannell |
| EPI_ISL_3429724 | UMC Utrecht | UMC Utrecht | Rob Schuurman, Anniek Tanja, Annemarie Wensing |
| EPI_ISL_3429725 | Fulgent Genetics | Centers for Disease Control and Prevention Division of Viral Diseases, Pathogen Discovery | Dakota Howard, Dhwani Batra, Peter W. Cook, Kara Moser, Adrian Paskey, Jason Caravas, Benjamin Rambo-Martin, Shatavia Morrison, Christopher Gulvick, Scott Sammons, Yvette Unoarumhi, Darlene Wagner, Matthew Schmerer, Harry Gao, Mickey Li, John Gao, Joseph Fierro, Benafsh Sapra, Becky Tsai, Yan Meng, Doreen Ng, James Xie, Clinton R. Paden, Duncan MacCannell |
| EPI_ISL_3429726, EPI_ISL_3429727 | UMC Utrecht | UMC Utrecht | Rob Schuurman, Anniek Tanja, Annemarie Wensing |
| EPI_ISL_3429728 | Fulgent Genetics | Centers for Disease Control and Prevention Division of Viral Diseases, Pathogen Discovery | Dakota Howard, Dhwani Batra, Peter W. Cook, Kara Moser, Adrian Paskey, Jason Caravas, Benjamin Rambo-Martin, Shatavia Morrison, Christopher Gulvick, Scott Sammons, Yvette Unoarumhi, Darlene Wagner, Matthew Schmerer, Harry Gao, Mickey Li, John Gao, Joseph Fierro, Benafsh Sapra, Becky Tsai, Yan Meng, Doreen Ng, James Xie, Clinton R. Paden, Duncan MacCannell |
| EPI_ISL_3429729 | UMC Utrecht | UMC Utrecht | Rob Schuurman, Anniek Tanja, Annemarie Wensing |
| EPI_ISL_3429730, EPI_ISL_3429731, EPI_ISL_3429732 | Fulgent Genetics | Centers for Disease Control and Prevention Division of Viral Diseases, Pathogen Discovery | Dakota Howard, Dhwani Batra, Peter W. Cook, Kara Moser, Adrian Paskey, Jason Caravas, Benjamin Rambo-Martin, Shatavia Morrison, Christopher Gulvick, Scott Sammons, Yvette Unoarumhi, Darlene Wagner, Matthew Schmerer, Harry Gao, Mickey Li, John Gao, Joseph Fierro, Benafsh Sapra, Becky Tsai, Yan Meng, Doreen Ng, James Xie, Clinton R. Paden, Duncan MacCannell |
| EPI_ISL_3429733 | UMC Utrecht | UMC Utrecht | Rob Schuurman, Anniek Tanja, Annemarie Wensing |
| EPI_ISL_3429734 | Fulgent Genetics | Centers for Disease Control and Prevention Division of Viral Diseases, Pathogen Discovery | Dakota Howard, Dhwani Batra, Peter W. Cook, Kara Moser, Adrian Paskey, Jason Caravas, Benjamin Rambo-Martin, Shatavia Morrison, Christopher Gulvick, Scott Sammons, Yvette Unoarumhi, Darlene Wagner, Matthew Schmerer, Harry Gao, Mickey Li, John Gao, Joseph Fierro, Benafsh Sapra, Becky Tsai, Yan Meng, Doreen Ng, James Xie, Clinton R. Paden, Duncan MacCannell |
| EPI_ISL_3429735 | UMC Utrecht | UMC Utrecht | Rob Schuurman, Anniek Tanja, Annemarie Wensing |
| EPI_ISL_3429736, EPI_ISL_3429737, EPI_ISL_3429738, EPI_ISL_3429739 | Fulgent Genetics | Centers for Disease Control and Prevention Division of Viral Diseases, Pathogen Discovery | Dakota Howard, Dhwani Batra, Peter W. Cook, Kara Moser, Adrian Paskey, Jason Caravas, Benjamin Rambo-Martin, Shatavia Morrison, Christopher Gulvick, Scott Sammons, Yvette Unoarumhi, Darlene Wagner, Matthew Schmerer, Harry Gao, Mickey Li, John Gao, Joseph Fierro, Benafsh Sapra, Becky Tsai, Yan Meng, Doreen Ng, James Xie, Clinton R. Paden, Duncan MacCannell |
| EPI_ISL_3429740 | UMC Utrecht | UMC Utrecht | Rob Schuurman, Anniek Tanja, Annemarie Wensing |
| EPI_ISL_3429741, EPI_ISL_3429742 | Fulgent Genetics | Centers for Disease Control and Prevention Division of Viral Diseases, Pathogen Discovery | Dakota Howard, Dhwani Batra, Peter W. Cook, Kara Moser, Adrian Paskey, Jason Caravas, Benjamin Rambo-Martin, Shatavia Morrison, Christopher Gulvick, Scott Sammons, Yvette Unoarumhi, Darlene Wagner, Matthew Schmerer, Harry Gao, Mickey Li, John Gao, Joseph Fierro, Benafsh Sapra, Becky Tsai, Yan Meng, Doreen Ng, James Xie, Clinton R. Paden, Duncan MacCannell |

| | | | |
|---|---|---|---|
| EPI_ISL_3429743 | UMC Utrecht | UMC Utrecht | Rob Schuurman, Anniek Tanja, Annemarie Wensing |
| EPI_ISL_3429744 | Fulgent Genetics | Centers for Disease Control and Prevention Division of Viral Diseases, Pathogen Discovery | Dakota Howard, Dhwani Batra, Peter W. Cook, Kara Moser, Adrian Paskey, Jason Caravas, Benjamin Rambo-Martin, Shatavia Morrison, Christopher Gulvick, Scott Sammons, Yvette Unoarumhi, Darlene Wagner, Matthew Schmerer, Harry Gao, Mickey Li, John Gao, Joseph Fierro, Benafsh Sapra, Becky Tsai, Yan Meng, Doreen Ng, James Xie, Clinton R. Paden, Duncan MacCannell |
| EPI_ISL_3429745 | UMC Utrecht | UMC Utrecht | Rob Schuurman, Anniek Tanja, Annemarie Wensing |
| EPI_ISL_3429746, EPI_ISL_3429747 | Fulgent Genetics | Centers for Disease Control and Prevention Division of Viral Diseases, Pathogen Discovery | Dakota Howard, Dhwani Batra, Peter W. Cook, Kara Moser, Adrian Paskey, Jason Caravas, Benjamin Rambo-Martin, Shatavia Morrison, Christopher Gulvick, Scott Sammons, Yvette Unoarumhi, Darlene Wagner, Matthew Schmerer, Harry Gao, Mickey Li, John Gao, Joseph Fierro, Benafsh Sapra, Becky Tsai, Yan Meng, Doreen Ng, James Xie, Clinton R. Paden, Duncan MacCannell |
| EPI_ISL_3429748, EPI_ISL_3429749 | UMC Utrecht | UMC Utrecht | Rob Schuurman, Anniek Tanja, Annemarie Wensing |
| EPI_ISL_3429750, EPI_ISL_3429751 | Fulgent Genetics | Centers for Disease Control and Prevention Division of Viral Diseases, Pathogen Discovery | Dakota Howard, Dhwani Batra, Peter W. Cook, Kara Moser, Adrian Paskey, Jason Caravas, Benjamin Rambo-Martin, Shatavia Morrison, Christopher Gulvick, Scott Sammons, Yvette Unoarumhi, Darlene Wagner, Matthew Schmerer, Harry Gao, Mickey Li, John Gao, Joseph Fierro, Benafsh Sapra, Becky Tsai, Yan Meng, Doreen Ng, James Xie, Clinton R. Paden, Duncan MacCannell |
| EPI_ISL_3429752 | UMC Utrecht | UMC Utrecht | Rob Schuurman, Anniek Tanja, Annemarie Wensing |
| EPI_ISL_3429753 | Fulgent Genetics | Centers for Disease Control and Prevention Division of Viral Diseases, Pathogen Discovery | Dakota Howard, Dhwani Batra, Peter W. Cook, Kara Moser, Adrian Paskey, Jason Caravas, Benjamin Rambo-Martin, Shatavia Morrison, Christopher Gulvick, Scott Sammons, Yvette Unoarumhi, Darlene Wagner, Matthew Schmerer, Harry Gao, Mickey Li, John Gao, Joseph Fierro, Benafsh Sapra, Becky Tsai, Yan Meng, Doreen Ng, James Xie, Clinton R. Paden, Duncan MacCannell |
| EPI_ISL_3429754 | UMC Utrecht | UMC Utrecht | Rob Schuurman, Anniek Tanja, Annemarie Wensing |
| EPI_ISL_3429755 | Fulgent Genetics | Centers for Disease Control and Prevention Division of Viral Diseases, Pathogen Discovery | Dakota Howard, Dhwani Batra, Peter W. Cook, Kara Moser, Adrian Paskey, Jason Caravas, Benjamin Rambo-Martin, Shatavia Morrison, Christopher Gulvick, Scott Sammons, Yvette Unoarumhi, Darlene Wagner, Matthew Schmerer, Harry Gao, Mickey Li, John Gao, Joseph Fierro, Benafsh Sapra, Becky Tsai, Yan Meng, Doreen Ng, James Xie, Clinton R. Paden, Duncan MacCannell |
| EPI_ISL_3429756 | UMC Utrecht | UMC Utrecht | Rob Schuurman, Anniek Tanja, Annemarie Wensing |
| EPI_ISL_3429757, EPI_ISL_3429758 | Fulgent Genetics | Centers for Disease Control and Prevention Division of Viral Diseases, Pathogen Discovery | Dakota Howard, Dhwani Batra, Peter W. Cook, Kara Moser, Adrian Paskey, Jason Caravas, Benjamin Rambo-Martin, Shatavia Morrison, Christopher Gulvick, Scott Sammons, Yvette Unoarumhi, Darlene Wagner, Matthew Schmerer, Harry Gao, Mickey Li, John Gao, Joseph Fierro, Benafsh Sapra, Becky Tsai, Yan Meng, Doreen Ng, James Xie, Clinton R. Paden, Duncan MacCannell |
| EPI_ISL_3429759 | UMC Utrecht | UMC Utrecht | Rob Schuurman, Anniek Tanja, Annemarie Wensing |
| EPI_ISL_3429760, EPI_ISL_3429762, EPI_ISL_3429763 | Fulgent Genetics | Centers for Disease Control and Prevention Division of Viral Diseases, Pathogen Discovery | Dakota Howard, Dhwani Batra, Peter W. Cook, Kara Moser, Adrian Paskey, Jason Caravas, Benjamin Rambo-Martin, Shatavia Morrison, Christopher Gulvick, Scott Sammons, Yvette Unoarumhi, Darlene Wagner, Matthew Schmerer, Harry Gao, Mickey Li, John Gao, Joseph Fierro, Benafsh Sapra, Becky Tsai, Yan Meng, Doreen Ng, James Xie, Clinton R. Paden, Duncan MacCannell |
| EPI_ISL_3429764 | UMC Utrecht | UMC Utrecht | Rob Schuurman, Anniek Tanja, Annemarie Wensing |
| EPI_ISL_3429765 | Fulgent Genetics | Centers for Disease Control and Prevention Division of Viral Diseases, Pathogen Discovery | Dakota Howard, Dhwani Batra, Peter W. Cook, Kara Moser, Adrian Paskey, Jason Caravas, Benjamin Rambo-Martin, Shatavia Morrison, Christopher Gulvick, Scott Sammons, Yvette Unoarumhi, Darlene Wagner, Matthew Schmerer, Harry Gao, Mickey Li, John Gao, Joseph Fierro, Benafsh Sapra, Becky Tsai, Yan Meng, Doreen Ng, James Xie, Clinton R. Paden, Duncan MacCannell |
| EPI_ISL_3429766 | UMC Utrecht | UMC Utrecht | Rob Schuurman, Anniek Tanja, Annemarie Wensing |
| EPI_ISL_3429767, EPI_ISL_3429768 | Fulgent Genetics | Centers for Disease Control and Prevention Division of Viral Diseases, Pathogen Discovery | Dakota Howard, Dhwani Batra, Peter W. Cook, Kara Moser, Adrian Paskey, Jason Caravas, Benjamin Rambo-Martin, Shatavia Morrison, Christopher Gulvick, Scott Sammons, Yvette Unoarumhi, Darlene Wagner, Matthew Schmerer, Harry Gao, Mickey Li, John Gao, Joseph Fierro, Benafsh Sapra, Becky Tsai, Yan Meng, Doreen Ng, James Xie, Clinton R. Paden, Duncan MacCannell |
| EPI_ISL_3429769 | UMC Utrecht | UMC Utrecht | Rob Schuurman, Anniek Tanja, Annemarie Wensing |
| EPI_ISL_3429770 | Fulgent Genetics | Centers for Disease Control and Prevention Division of Viral Diseases, Pathogen Discovery | Dakota Howard, Dhwani Batra, Peter W. Cook, Kara Moser, Adrian Paskey, Jason Caravas, Benjamin Rambo-Martin, Shatavia Morrison, Christopher Gulvick, Scott Sammons, Yvette Unoarumhi, Darlene Wagner, Matthew Schmerer, Harry Gao, Mickey Li, John Gao, Joseph Fierro, Benafsh Sapra, Becky Tsai, Yan Meng, Doreen Ng, James Xie, Clinton R. Paden, Duncan MacCannell |
| EPI_ISL_3429771 | UMC Utrecht | UMC Utrecht | Rob Schuurman, Anniek Tanja, Annemarie Wensing |
| EPI_ISL_3429772, EPI_ISL_3429773 | Fulgent Genetics | Centers for Disease Control and Prevention Division of Viral Diseases, Pathogen Discovery | Dakota Howard, Dhwani Batra, Peter W. Cook, Kara Moser, Adrian Paskey, Jason Caravas, Benjamin Rambo-Martin, Shatavia Morrison, Christopher Gulvick, Scott Sammons, Yvette Unoarumhi, Darlene Wagner, Matthew Schmerer, Harry Gao, Mickey Li, John Gao, Joseph Fierro, Benafsh Sapra, Becky Tsai, Yan Meng, Doreen Ng, James Xie, Clinton R. Paden, Duncan MacCannell |
| EPI_ISL_3429774 | UMC Utrecht | UMC Utrecht | Rob Schuurman, Anniek Tanja, Annemarie Wensing |
| EPI_ISL_3429775 | Fulgent Genetics | Centers for Disease Control and Prevention Division of Viral Diseases, Pathogen Discovery | Dakota Howard, Dhwani Batra, Peter W. Cook, Kara Moser, Adrian Paskey, Jason Caravas, Benjamin Rambo-Martin, Shatavia Morrison, Christopher Gulvick, Scott Sammons, Yvette Unoarumhi, Darlene Wagner, Matthew Schmerer, Harry Gao, Mickey Li, John Gao, Joseph Fierro, Benafsh Sapra, Becky Tsai, Yan Meng, Doreen Ng, James Xie, Clinton R. Paden, Duncan MacCannell |
| EPI_ISL_3429776 | UMC Utrecht | UMC Utrecht | Rob Schuurman, Anniek Tanja, Annemarie Wensing |
| EPI_ISL_3429777 | Fulgent Genetics | Centers for Disease Control and Prevention Division of Viral Diseases, Pathogen Discovery | Dakota Howard, Dhwani Batra, Peter W. Cook, Kara Moser, Adrian Paskey, Jason Caravas, Benjamin Rambo-Martin, Shatavia Morrison, Christopher Gulvick, Scott Sammons, Yvette Unoarumhi, Darlene Wagner, Matthew Schmerer, Harry Gao, Mickey Li, John Gao, Joseph Fierro, Benafsh Sapra, Becky Tsai, Yan Meng, Doreen Ng, James Xie, Clinton R. Paden, Duncan MacCannell |
| EPI_ISL_3429778 | UMC Utrecht | UMC Utrecht | Rob Schuurman, Anniek Tanja, Annemarie Wensing |
| EPI_ISL_3429779, EPI_ISL_3429780 | Fulgent Genetics | Centers for Disease Control and Prevention Division of Viral Diseases, Pathogen Discovery | Dakota Howard, Dhwani Batra, Peter W. Cook, Kara Moser, Adrian Paskey, Jason Caravas, Benjamin Rambo-Martin, Shatavia Morrison, Christopher Gulvick, Scott Sammons, Yvette Unoarumhi, Darlene Wagner, Matthew Schmerer, Harry Gao, Mickey Li, John Gao, Joseph Fierro, Benafsh Sapra, Becky Tsai, Yan Meng, Doreen Ng, James Xie, Clinton R. Paden, Duncan MacCannell |
| EPI_ISL_3429781 | UMC Utrecht | UMC Utrecht | Rob Schuurman, Anniek Tanja, Annemarie Wensing |
| EPI_ISL_3429782, EPI_ISL_3429783 | Fulgent Genetics | Centers for Disease Control and Prevention Division of Viral Diseases, Pathogen Discovery | Dakota Howard, Dhwani Batra, Peter W. Cook, Kara Moser, Adrian Paskey, Jason Caravas, Benjamin Rambo-Martin, Shatavia Morrison, Christopher Gulvick, Scott Sammons, Yvette Unoarumhi, Darlene Wagner, Matthew Schmerer, Harry Gao, Mickey Li, John Gao, Joseph Fierro, Benafsh Sapra, Becky Tsai, Yan Meng, Doreen Ng, James Xie, Clinton R. Paden, Duncan MacCannell |
| EPI_ISL_3429784 | UMC Utrecht | UMC Utrecht | Rob Schuurman, Anniek Tanja, Annemarie Wensing |
| EPI_ISL_3429785 | Fulgent Genetics | Centers for Disease Control and Prevention Division of Viral Diseases, Pathogen Discovery | Dakota Howard, Dhwani Batra, Peter W. Cook, Kara Moser, Adrian Paskey, Jason Caravas, Benjamin Rambo-Martin, Shatavia Morrison, Christopher Gulvick, Scott Sammons, Yvette Unoarumhi, Darlene Wagner, Matthew Schmerer, Harry Gao, Mickey Li, John Gao, Joseph Fierro, Benafsh Sapra, Becky Tsai, Yan Meng, Doreen Ng, James Xie, Clinton R. Paden, Duncan MacCannell |
| EPI_ISL_3429786 | UMC Utrecht | UMC Utrecht | Rob Schuurman, Anniek Tanja, Annemarie Wensing |
| EPI_ISL_3429787, EPI_ISL_3429788 | Fulgent Genetics | Centers for Disease Control and Prevention Division of Viral Diseases, Pathogen Discovery | Dakota Howard, Dhwani Batra, Peter W. Cook, Kara Moser, Adrian Paskey, Jason Caravas, Benjamin Rambo-Martin, Shatavia Morrison, Christopher Gulvick, Scott Sammons, Yvette Unoarumhi, Darlene Wagner, Matthew Schmerer, Harry Gao, Mickey Li, John Gao, Joseph Fierro, Benafsh Sapra, Becky Tsai, Yan Meng, Doreen Ng, James Xie, Clinton R. Paden, Duncan MacCannell |
| EPI_ISL_3429789 | UMC Utrecht | UMC Utrecht | Rob Schuurman, Anniek Tanja, Annemarie Wensing |
| EPI_ISL_3429790 | Fulgent Genetics | Centers for Disease Control and Prevention Division of Viral Diseases, Pathogen Discovery | Dakota Howard, Dhwani Batra, Peter W. Cook, Kara Moser, Adrian Paskey, Jason Caravas, Benjamin Rambo-Martin, Shatavia Morrison, Christopher Gulvick, Scott Sammons, Yvette Unoarumhi, Darlene Wagner, Matthew Schmerer, Harry Gao, Mickey Li, John Gao, Joseph Fierro, Benafsh Sapra, Becky Tsai, Yan Meng, Doreen Ng, James Xie, Clinton R. Paden, Duncan MacCannell |
| EPI_ISL_3429791 | UMC Utrecht | UMC Utrecht | Rob Schuurman, Anniek Tanja, Annemarie Wensing |
| EPI_ISL_3429792, EPI_ISL_3429793 | Fulgent Genetics | Centers for Disease Control and Prevention Division of Viral Diseases, Pathogen Discovery | Dakota Howard, Dhwani Batra, Peter W. Cook, Kara Moser, Adrian Paskey, Jason Caravas, Benjamin Rambo-Martin, Shatavia Morrison, Christopher Gulvick, Scott Sammons, Yvette Unoarumhi, Darlene Wagner, Matthew Schmerer, Harry Gao, Mickey Li, John Gao, Joseph Fierro, Benafsh Sapra, Becky Tsai, Yan Meng, Doreen Ng, James Xie, Clinton R. Paden, Duncan MacCannell |
| EPI_ISL_3429794 | UMC Utrecht | UMC Utrecht | Rob Schuurman, Anniek Tanja, Annemarie Wensing |
| EPI_ISL_3429795 | Fulgent Genetics | Centers for Disease Control and Prevention Division of Viral Diseases, Pathogen Discovery | Dakota Howard, Dhwani Batra, Peter W. Cook, Kara Moser, Adrian Paskey, Jason Caravas, Benjamin Rambo-Martin, Shatavia Morrison, Christopher Gulvick, Scott Sammons, Yvette Unoarumhi, Darlene Wagner, Matthew Schmerer, Harry Gao, Mickey Li, John Gao, Joseph Fierro, Benafsh Sapra, Becky Tsai, Yan Meng, Doreen Ng, James Xie, Clinton R. Paden, Duncan MacCannell |
| EPI_ISL_3429796 | UMC Utrecht | UMC Utrecht | Rob Schuurman, Anniek Tanja, Annemarie Wensing |
| EPI_ISL_3429797, EPI_ISL_3429798 | Fulgent Genetics | Centers for Disease Control and Prevention Division of Viral Diseases, Pathogen Discovery | Dakota Howard, Dhwani Batra, Peter W. Cook, Kara Moser, Adrian Paskey, Jason Caravas, Benjamin Rambo-Martin, Shatavia Morrison, Christopher Gulvick, Scott Sammons, Yvette Unoarumhi, Darlene Wagner, Matthew Schmerer, Harry Gao, Mickey Li, John Gao, Joseph Fierro, Benafsh Sapra, Becky Tsai, Yan Meng, Doreen Ng, James Xie, Clinton R. Paden, Duncan MacCannell |
| EPI_ISL_3429799 | UMC Utrecht | UMC Utrecht | Rob Schuurman, Anniek Tanja, Annemarie Wensing |
| EPI_ISL_3429800, EPI_ISL_3429802, EPI_ISL_3429803 | Fulgent Genetics | Centers for Disease Control and Prevention Division of Viral Diseases, Pathogen Discovery | Dakota Howard, Dhwani Batra, Peter W. Cook, Kara Moser, Adrian Paskey, Jason Caravas, Benjamin Rambo-Martin, Shatavia Morrison, Christopher Gulvick, Scott Sammons, Yvette Unoarumhi, Darlene Wagner, Matthew Schmerer, Harry Gao, Mickey Li, John Gao, Joseph Fierro, Benafsh Sapra, Becky Tsai, Yan Meng, Doreen Ng, James Xie, Clinton R. Paden, Duncan MacCannell |
| EPI_ISL_3429804 | UMC Utrecht | UMC Utrecht | Rob Schuurman, Anniek Tanja, Annemarie Wensing |
| EPI_ISL_3429805, EPI_ISL_3429806 | Fulgent Genetics | Centers for Disease Control and Prevention Division of Viral Diseases, Pathogen Discovery | Dakota Howard, Dhwani Batra, Peter W. Cook, Kara Moser, Adrian Paskey, Jason Caravas, Benjamin Rambo-Martin, Shatavia Morrison, Christopher Gulvick, Scott Sammons, Yvette Unoarumhi, Darlene Wagner, Matthew Schmerer, Harry Gao, Mickey Li, John Gao, Joseph Fierro, Benafsh Sapra, Becky Tsai, Yan Meng, Doreen Ng, James Xie, Clinton R. Paden, Duncan MacCannell |
| EPI_ISL_3429807 | UMC Utrecht | UMC Utrecht | Rob Schuurman, Anniek Tanja, Annemarie Wensing |
| EPI_ISL_3429808 | Fulgent Genetics | Centers for Disease Control and Prevention Division of Viral Diseases, Pathogen Discovery | Dakota Howard, Dhwani Batra, Peter W. Cook, Kara Moser, Adrian Paskey, Jason Caravas, Benjamin Rambo-Martin, Shatavia Morrison, Christopher Gulvick, Scott Sammons, Yvette Unoarumhi, Darlene Wagner, Matthew Schmerer, Harry Gao, Mickey Li, John Gao, Joseph Fierro, Benafsh Sapra, Becky Tsai, Yan Meng, Doreen Ng, James Xie, Clinton R. Paden, Duncan MacCannell |
| EPI_ISL_3429809 | UMC Utrecht | UMC Utrecht | Rob Schuurman, Anniek Tanja, Annemarie Wensing |
| EPI_ISL_3429810, EPI_ISL_3429811 | Fulgent Genetics | Centers for Disease Control and Prevention Division of Viral Diseases, Pathogen Discovery | Dakota Howard, Dhwani Batra, Peter W. Cook, Kara Moser, Adrian Paskey, Jason Caravas, Benjamin Rambo-Martin, Shatavia Morrison, Christopher Gulvick, Scott Sammons, Yvette Unoarumhi, Darlene Wagner, Matthew Schmerer, Harry Gao, Mickey Li, John Gao, Joseph Fierro, Benafsh Sapra, Becky Tsai, Yan Meng, Doreen Ng, James Xie, Clinton R. Paden, Duncan MacCannell |
| EPI_ISL_3429812 | UMC Utrecht | UMC Utrecht | Rob Schuurman, Anniek Tanja, Annemarie Wensing |
| EPI_ISL_3429813, EPI_ISL_3429814 | Fulgent Genetics | Centers for Disease Control and Prevention Division of Viral Diseases, Pathogen Discovery | Dakota Howard, Dhwani Batra, Peter W. Cook, Kara Moser, Adrian Paskey, Jason Caravas, Benjamin Rambo-Martin, Shatavia Morrison, Christopher Gulvick, Scott Sammons, Yvette Unoarumhi, Darlene Wagner, Matthew Schmerer, Harry Gao, Mickey Li, John Gao, Joseph Fierro, Benafsh Sapra, Becky Tsai, Yan Meng, Doreen Ng, James Xie, Clinton R. Paden, Duncan MacCannell |
| EPI_ISL_3429815 | UMC Utrecht | UMC Utrecht | Rob Schuurman, Anniek Tanja, Annemarie Wensing |
| EPI_ISL_3429816 | Fulgent Genetics | Centers for Disease Control and Prevention Division of Viral Diseases, Pathogen Discovery | Dakota Howard, Dhwani Batra, Peter W. Cook, Kara Moser, Adrian Paskey, Jason Caravas, Benjamin Rambo-Martin, Shatavia Morrison, Christopher Gulvick, Scott Sammons, Yvette Unoarumhi, Darlene Wagner, Matthew Schmerer, Harry Gao, Mickey Li, John Gao, Joseph Fierro, Benafsh Sapra, Becky Tsai, Yan Meng, Doreen Ng, James Xie, Clinton R. Paden, Duncan MacCannell |
| EPI_ISL_3429817 | UMC Utrecht | UMC Utrecht | Rob Schuurman, Anniek Tanja, Annemarie Wensing |

EPI_ISL_3429818, EPI_ISL_3429819, EPI_ISL_3429820, EPI_ISL_3429821, EPI_ISL_3429822, EPI_ISL_3429823, EPI_ISL_3429824, EPI_ISL_3429825, EPI_ISL_3429826, EPI_ISL_3429827, EPI_ISL_3429828, EPI_ISL_3429829, EPI_ISL_3429830, EPI_ISL_3429831, EPI_ISL_3429832, EPI_ISL_3429833, EPI_ISL_3429834, EPI_ISL_3429835, EPI_ISL_3429836, EPI_ISL_3429837, EPI_ISL_3429838, EPI_ISL_3429839, EPI_ISL_3429840, EPI_ISL_3429841, EPI_ISL_3429842, EPI_ISL_3429843, EPI_ISL_3429844, EPI_ISL_3429845, EPI_ISL_3429846, EPI_ISL_3429847, EPI_ISL_3429848, EPI_ISL_3429849, EPI_ISL_3429850, EPI_ISL_3429851, EPI_ISL_3429852, EPI_ISL_3429853, EPI_ISL_3429854, EPI_ISL_3429855, EPI_ISL_3429856, EPI_ISL_3429857, EPI_ISL_3429858, EPI_ISL_3429859, EPI_ISL_3429860, EPI_ISL_3429861, EPI_ISL_3429862, EPI_ISL_3429863, EPI_ISL_3429864, EPI_ISL_3429865, EPI_ISL_3429866, EPI_ISL_3429867, EPI_ISL_3429868, EPI_ISL_3429869, EPI_ISL_3429870, EPI_ISL_3429871, EPI_ISL_3429872, EPI_ISL_3429873, EPI_ISL_3429874, EPI_ISL_3429875, EPI_ISL_3429876, EPI_ISL_3429877, EPI_ISL_3429878, EPI_ISL_3429879, EPI_ISL_3429880, EPI_ISL_3429881, EPI_ISL_3429882, EPI_ISL_3429883, EPI_ISL_3429884, EPI_ISL_3429885, EPI_ISL_3429886, EPI_ISL_3429887, EPI_ISL_3429889, EPI_ISL_3429890, EPI_ISL_3429891, EPI_ISL_3429892, EPI_ISL_3429893, EPI_ISL_3429894, EPI_ISL_3429895, EPI_ISL_3429896, EPI_ISL_3429897, EPI_ISL_3429898, EPI_ISL_3429899, EPI_ISL_3429900, EPI_ISL_3429901, EPI_ISL_3429902, EPI_ISL_3429903, EPI_ISL_3429904, EPI_ISL_3429905, EPI_ISL_3429906, EPI_ISL_3429907, EPI_ISL_3429908, EPI_ISL_3429909, EPI_ISL_3429910, EPI_ISL_3429912, EPI_ISL_3429913, EPI_ISL_3429914, EPI_ISL_3429915, EPI_ISL_3429916, EPI_ISL_3429917, EPI_ISL_3429918, EPI_ISL_3429919, EPI_ISL_3429920, EPI_ISL_3429921, EPI_ISL_3429922, EPI_ISL_3429923, EPI_ISL_3429927, EPI_ISL_3429929, EPI_ISL_3429930, EPI_ISL_3429931, EPI_ISL_3429932, EPI_ISL_3429933, EPI_ISL_3429934, EPI_ISL_3429935, EPI_ISL_3429936, EPI_ISL_3429937, EPI_ISL_3429938, EPI_ISL_3429939, EPI_ISL_3429940, EPI_ISL_3429941, EPI_ISL_3429942, EPI_ISL_3429943, EPI_ISL_3429944, EPI_ISL_3429945, EPI_ISL_3429946, EPI_ISL_3429947, EPI_ISL_3429948, EPI_ISL_3429949, EPI_ISL_3429951, EPI_ISL_3429953, EPI_ISL_3429955, EPI_ISL_3429957, EPI_ISL_3429959, EPI_ISL_3429961, EPI_ISL_3429963, EPI_ISL_3429965

| | | | |
|---|---|---|---|
| see above | Fulgent Genetics | Centers for Disease Control and Prevention Division of Viral Diseases, Pathogen Discovery | Dakota Howard, Dhwani Batra, Peter W. Cook, Kara Moser, Adrian Paskey, Jason Caravas, Benjamin Rambo-Martin, Shatavia Morrison, Christopher Gulvick, Scott Sammons, Yvette Unoarumhi, Darlene Wagner, Matthew Schmerer, Harry Gao, Mickey Li, John Gao, Joseph Fierro, Benafsh Sapra, Becky Tsai, Yan Meng, Doreen Ng, James Xie, Clinton R. Paden, Duncan MacCannell |
| EPI_ISL_3429966 | Aegis Sciences Corporation | Centers for Disease Control and Prevention Division of Viral Diseases, Pathogen Discovery | Dakota Howard, Dhwani Batra, Peter W. Cook, Kara Moser, Adrian Paskey, Jason Caravas, Benjamin Rambo-Martin, Shatavia Morrison, Christopher Gulvick, Scott Sammons, Yvette Unoarumhi, Darlene Wagner, Matthew Schmerer, Cyndi Clark, Patrick Campbell, Rob Case, Vikramsinha Ghorpade, Holly Houdeshell, Ola Kvalvaag, Dillon Nall, Ethan Sanders, Alec Vest, Shaun Westlund, Matthew Hardison, Clinton R. Paden, Duncan MacCannell |
| EPI_ISL_3429967 | Fulgent Genetics | Centers for Disease Control and Prevention Division of Viral Diseases, Pathogen Discovery | Dakota Howard, Dhwani Batra, Peter W. Cook, Kara Moser, Adrian Paskey, Jason Caravas, Benjamin Rambo-Martin, Shatavia Morrison, Christopher Gulvick, Scott Sammons, Yvette Unoarumhi, Darlene Wagner, Matthew Schmerer, Harry Gao, Mickey Li, John Gao, Joseph Fierro, Benafsh Sapra, Becky Tsai, Yan Meng, Doreen Ng, James Xie, Clinton R. Paden, Duncan MacCannell |
| EPI_ISL_3429968 | Aegis Sciences Corporation | Centers for Disease Control and Prevention Division of Viral Diseases, Pathogen Discovery | Dakota Howard, Dhwani Batra, Peter W. Cook, Kara Moser, Adrian Paskey, Jason Caravas, Benjamin Rambo-Martin, Shatavia Morrison, Christopher Gulvick, Scott Sammons, Yvette Unoarumhi, Darlene Wagner, Matthew Schmerer, Cyndi Clark, Patrick Campbell, Rob Case, Vikramsinha Ghorpade, Holly Houdeshell, Ola Kvalvaag, Dillon Nall, Ethan Sanders, Alec Vest, Shaun Westlund, Matthew Hardison, Clinton R. Paden, Duncan MacCannell |
| EPI_ISL_3429969 | Fulgent Genetics | Centers for Disease Control and Prevention Division of Viral Diseases, Pathogen Discovery | Dakota Howard, Dhwani Batra, Peter W. Cook, Kara Moser, Adrian Paskey, Jason Caravas, Benjamin Rambo-Martin, Shatavia Morrison, Christopher Gulvick, Scott Sammons, Yvette Unoarumhi, Darlene Wagner, Matthew Schmerer, Harry Gao, Mickey Li, John Gao, Joseph Fierro, Benafsh Sapra, Becky Tsai, Yan Meng, Doreen Ng, James Xie, Clinton R. Paden, Duncan MacCannell |
| EPI_ISL_3429970 | Aegis Sciences Corporation | Centers for Disease Control and Prevention Division of Viral Diseases, Pathogen Discovery | Dakota Howard, Dhwani Batra, Peter W. Cook, Kara Moser, Adrian Paskey, Jason Caravas, Benjamin Rambo-Martin, Shatavia Morrison, Christopher Gulvick, Scott Sammons, Yvette Unoarumhi, Darlene Wagner, Matthew Schmerer, Cyndi Clark, Patrick Campbell, Rob Case, Vikramsinha Ghorpade, Holly Houdeshell, Ola Kvalvaag, Dillon Nall, Ethan Sanders, Alec Vest, Shaun Westlund, Matthew Hardison, Clinton R. Paden, Duncan MacCannell |
| EPI_ISL_3429971 | Fulgent Genetics | Centers for Disease Control and Prevention Division of Viral Diseases, Pathogen Discovery | Dakota Howard, Dhwani Batra, Peter W. Cook, Kara Moser, Adrian Paskey, Jason Caravas, Benjamin Rambo-Martin, Shatavia Morrison, Christopher Gulvick, Scott Sammons, Yvette Unoarumhi, Darlene Wagner, Matthew Schmerer, Harry Gao, Mickey Li, John Gao, Joseph Fierro, Benafsh Sapra, Becky Tsai, Yan Meng, Doreen Ng, James Xie, Clinton R. Paden, Duncan MacCannell |
| EPI_ISL_3429972 | Aegis Sciences Corporation | Centers for Disease Control and Prevention Division of Viral Diseases, Pathogen Discovery | Dakota Howard, Dhwani Batra, Peter W. Cook, Kara Moser, Adrian Paskey, Jason Caravas, Benjamin Rambo-Martin, Shatavia Morrison, Christopher Gulvick, Scott Sammons, Yvette Unoarumhi, Darlene Wagner, Matthew Schmerer, Cyndi Clark, Patrick Campbell, Rob Case, Vikramsinha Ghorpade, Holly Houdeshell, Ola Kvalvaag, Dillon Nall, Ethan Sanders, Alec Vest, Shaun Westlund, Matthew Hardison, Clinton R. Paden, Duncan MacCannell |
| EPI_ISL_3429973, EPI_ISL_3429975, EPI_ISL_3429977 | Fulgent Genetics | Centers for Disease Control and Prevention Division of Viral Diseases, Pathogen Discovery | Dakota Howard, Dhwani Batra, Peter W. Cook, Kara Moser, Adrian Paskey, Jason Caravas, Benjamin Rambo-Martin, Shatavia Morrison, Christopher Gulvick, Scott Sammons, Yvette Unoarumhi, Darlene Wagner, Matthew Schmerer, Harry Gao, Mickey Li, John Gao, Joseph Fierro, Benafsh Sapra, Becky Tsai, Yan Meng, Doreen Ng, James Xie, Clinton R. Paden, Duncan MacCannell |
| EPI_ISL_3429978 | Aegis Sciences Corporation | Centers for Disease Control and Prevention Division of Viral Diseases, Pathogen Discovery | Dakota Howard, Dhwani Batra, Peter W. Cook, Kara Moser, Adrian Paskey, Jason Caravas, Benjamin Rambo-Martin, Shatavia Morrison, Christopher Gulvick, Scott Sammons, Yvette Unoarumhi, Darlene Wagner, Matthew Schmerer, Cyndi Clark, Patrick Campbell, Rob Case, Vikramsinha Ghorpade, Holly Houdeshell, Ola Kvalvaag, Dillon Nall, Ethan Sanders, Alec Vest, Shaun Westlund, Matthew Hardison, Clinton R. Paden, Duncan MacCannell |
| EPI_ISL_3429979 | Fulgent Genetics | Centers for Disease Control and Prevention Division of Viral Diseases, Pathogen Discovery | Dakota Howard, Dhwani Batra, Peter W. Cook, Kara Moser, Adrian Paskey, Jason Caravas, Benjamin Rambo-Martin, Shatavia Morrison, Christopher Gulvick, Scott Sammons, Yvette Unoarumhi, Darlene Wagner, Matthew Schmerer, Harry Gao, Mickey Li, John Gao, Joseph Fierro, Benafsh Sapra, Becky Tsai, Yan Meng, Doreen Ng, James Xie, Clinton R. Paden, Duncan MacCannell |
| EPI_ISL_3429981 | Aegis Sciences Corporation | Centers for Disease Control and Prevention Division of Viral Diseases, Pathogen Discovery | Dakota Howard, Dhwani Batra, Peter W. Cook, Kara Moser, Adrian Paskey, Jason Caravas, Benjamin Rambo-Martin, Shatavia Morrison, Christopher Gulvick, Scott Sammons, Yvette Unoarumhi, Darlene Wagner, Matthew Schmerer, Cyndi Clark, Patrick Campbell, Rob Case, Vikramsinha Ghorpade, Holly Houdeshell, Ola Kvalvaag, Dillon Nall, Ethan Sanders, Alec Vest, Shaun Westlund, Matthew Hardison, Clinton R. Paden, Duncan MacCannell |
| EPI_ISL_3429982 | Fulgent Genetics | Centers for Disease Control and Prevention Division of Viral Diseases, Pathogen Discovery | Dakota Howard, Dhwani Batra, Peter W. Cook, Kara Moser, Adrian Paskey, Jason Caravas, Benjamin Rambo-Martin, Shatavia Morrison, Christopher Gulvick, Scott Sammons, Yvette Unoarumhi, Darlene Wagner, Matthew Schmerer, Harry Gao, Mickey Li, John Gao, Joseph Fierro, Benafsh Sapra, Becky Tsai, Yan Meng, Doreen Ng, James Xie, Clinton R. Paden, Duncan MacCannell |
| EPI_ISL_3429983 | Aegis Sciences Corporation | Centers for Disease Control and Prevention Division of Viral Diseases, Pathogen Discovery | Dakota Howard, Dhwani Batra, Peter W. Cook, Kara Moser, Adrian Paskey, Jason Caravas, Benjamin Rambo-Martin, Shatavia Morrison, Christopher Gulvick, Scott Sammons, Yvette Unoarumhi, Darlene Wagner, Matthew Schmerer, Cyndi Clark, Patrick Campbell, Rob Case, Vikramsinha Ghorpade, Holly Houdeshell, Ola Kvalvaag, Dillon Nall, Ethan Sanders, Alec Vest, Shaun Westlund, Matthew Hardison, Clinton R. Paden, Duncan MacCannell |
| EPI_ISL_3429984, EPI_ISL_3429985, EPI_ISL_3429987, EPI_ISL_3429989, EPI_ISL_3429991, EPI_ISL_3429993 | Fulgent Genetics | Centers for Disease Control and Prevention Division of Viral Diseases, Pathogen Discovery | Dakota Howard, Dhwani Batra, Peter W. Cook, Kara Moser, Adrian Paskey, Jason Caravas, Benjamin Rambo-Martin, Shatavia Morrison, Christopher Gulvick, Scott Sammons, Yvette Unoarumhi, Darlene Wagner, Matthew Schmerer, Harry Gao, Mickey Li, John Gao, Joseph Fierro, Benafsh Sapra, Becky Tsai, Yan Meng, Doreen Ng, James Xie, Clinton R. Paden, Duncan MacCannell |
| EPI_ISL_3429994 | Aegis Sciences Corporation | Centers for Disease Control and Prevention Division of Viral Diseases, Pathogen Discovery | Dakota Howard, Dhwani Batra, Peter W. Cook, Kara Moser, Adrian Paskey, Jason Caravas, Benjamin Rambo-Martin, Shatavia Morrison, Christopher Gulvick, Scott Sammons, Yvette Unoarumhi, Darlene Wagner, Matthew Schmerer, Cyndi Clark, Patrick Campbell, Rob Case, Vikramsinha Ghorpade, Holly Houdeshell, Ola Kvalvaag, Dillon Nall, Ethan Sanders, Alec Vest, Shaun Westlund, Matthew Hardison, Clinton R. Paden, Duncan MacCannell |
| EPI_ISL_3429995 | Fulgent Genetics | Centers for Disease Control and Prevention Division of Viral Diseases, Pathogen Discovery | Dakota Howard, Dhwani Batra, Peter W. Cook, Kara Moser, Adrian Paskey, Jason Caravas, Benjamin Rambo-Martin, Shatavia Morrison, Christopher Gulvick, Scott Sammons, Yvette Unoarumhi, Darlene Wagner, Matthew Schmerer, Harry Gao, Mickey Li, John Gao, Joseph Fierro, Benafsh Sapra, Becky Tsai, Yan Meng, Doreen Ng, James Xie, Clinton R. Paden, Duncan MacCannell |
| EPI_ISL_3429996 | Aegis Sciences Corporation | Centers for Disease Control and Prevention Division of Viral Diseases, Pathogen Discovery | Dakota Howard, Dhwani Batra, Peter W. Cook, Kara Moser, Adrian Paskey, Jason Caravas, Benjamin Rambo-Martin, Shatavia Morrison, Christopher Gulvick, Scott Sammons, Yvette Unoarumhi, Darlene Wagner, Matthew Schmerer, Cyndi Clark, Patrick Campbell, Rob Case, Vikramsinha Ghorpade, Holly Houdeshell, Ola Kvalvaag, Dillon Nall, Ethan Sanders, Alec Vest, Shaun Westlund, Matthew Hardison, Clinton R. Paden, Duncan MacCannell |
| EPI_ISL_3429997, EPI_ISL_3430000, EPI_ISL_3430002, EPI_ISL_3430004 | Fulgent Genetics | Centers for Disease Control and Prevention Division of Viral Diseases, Pathogen Discovery | Dakota Howard, Dhwani Batra, Peter W. Cook, Kara Moser, Adrian Paskey, Jason Caravas, Benjamin Rambo-Martin, Shatavia Morrison, Christopher Gulvick, Scott Sammons, Yvette Unoarumhi, Darlene Wagner, Matthew Schmerer, Harry Gao, Mickey Li, John Gao, Joseph Fierro, Benafsh Sapra, Becky Tsai, Yan Meng, Doreen Ng, James Xie, Clinton R. Paden, Duncan MacCannell |
| EPI_ISL_3430005 | Aegis Sciences Corporation | Centers for Disease Control and Prevention Division of Viral Diseases, Pathogen Discovery | Dakota Howard, Dhwani Batra, Peter W. Cook, Kara Moser, Adrian Paskey, Jason Caravas, Benjamin Rambo-Martin, Shatavia Morrison, Christopher Gulvick, Scott Sammons, Yvette Unoarumhi, Darlene Wagner, Matthew Schmerer, Cyndi Clark, Patrick Campbell, Rob Case, Vikramsinha Ghorpade, Holly Houdeshell, Ola Kvalvaag, Dillon Nall, Ethan Sanders, Alec Vest, Shaun Westlund, Matthew Hardison, Clinton R. Paden, Duncan MacCannell |
| EPI_ISL_3430006 | Fulgent Genetics | Centers for Disease Control and Prevention Division of Viral Diseases, Pathogen Discovery | Dakota Howard, Dhwani Batra, Peter W. Cook, Kara Moser, Adrian Paskey, Jason Caravas, Benjamin Rambo-Martin, Shatavia Morrison, Christopher Gulvick, Scott Sammons, Yvette Unoarumhi, Darlene Wagner, Matthew Schmerer, Harry Gao, Mickey Li, John Gao, Joseph Fierro, Benafsh Sapra, Becky Tsai, Yan Meng, Doreen Ng, James Xie, Clinton R. Paden, Duncan MacCannell |
| EPI_ISL_3430007 | Aegis Sciences Corporation | Centers for Disease Control and Prevention Division of Viral Diseases, Pathogen Discovery | Dakota Howard, Dhwani Batra, Peter W. Cook, Kara Moser, Adrian Paskey, Jason Caravas, Benjamin Rambo-Martin, Shatavia Morrison, Christopher Gulvick, Scott Sammons, Yvette Unoarumhi, Darlene Wagner, Matthew Schmerer, Cyndi Clark, Patrick Campbell, Rob Case, Vikramsinha Ghorpade, Holly Houdeshell, Ola Kvalvaag, Dillon Nall, Ethan Sanders, Alec Vest, Shaun Westlund, Matthew Hardison, Clinton R. Paden, Duncan MacCannell |
| EPI_ISL_3430008 | Fulgent Genetics | Centers for Disease Control and Prevention Division of Viral Diseases, Pathogen Discovery | Dakota Howard, Dhwani Batra, Peter W. Cook, Kara Moser, Adrian Paskey, Jason Caravas, Benjamin Rambo-Martin, Shatavia Morrison, Christopher Gulvick, Scott Sammons, Yvette Unoarumhi, Darlene Wagner, Matthew Schmerer, Harry Gao, Mickey Li, John Gao, Joseph Fierro, Benafsh Sapra, Becky Tsai, Yan Meng, Doreen Ng, James Xie, Clinton R. Paden, Duncan MacCannell |
| EPI_ISL_3430009 | Aegis Sciences Corporation | Centers for Disease Control and Prevention Division of Viral Diseases, Pathogen Discovery | Dakota Howard, Dhwani Batra, Peter W. Cook, Kara Moser, Adrian Paskey, Jason Caravas, Benjamin Rambo-Martin, Shatavia Morrison, Christopher Gulvick, Scott Sammons, Yvette Unoarumhi, Darlene Wagner, Matthew Schmerer, Cyndi Clark, Patrick Campbell, Rob Case, Vikramsinha Ghorpade, Holly Houdeshell, Ola Kvalvaag, Dillon Nall, Ethan Sanders, Alec Vest, Shaun Westlund, Matthew Hardison, Clinton R. Paden, Duncan MacCannell |
| EPI_ISL_3430010, EPI_ISL_3430011 | Fulgent Genetics | Centers for Disease Control and Prevention Division of Viral Diseases, Pathogen Discovery | Dakota Howard, Dhwani Batra, Peter W. Cook, Kara Moser, Adrian Paskey, Jason Caravas, Benjamin Rambo-Martin, Shatavia Morrison, Christopher Gulvick, Scott Sammons, Yvette Unoarumhi, Darlene Wagner, Matthew Schmerer, Harry Gao, Mickey Li, John Gao, Joseph Fierro, Benafsh Sapra, Becky Tsai, Yan Meng, Doreen Ng, James Xie, Clinton R. Paden, Duncan MacCannell |
| EPI_ISL_3430012 | Aegis Sciences Corporation | Centers for Disease Control and Prevention Division of Viral Diseases, Pathogen Discovery | Dakota Howard, Dhwani Batra, Peter W. Cook, Kara Moser, Adrian Paskey, Jason Caravas, Benjamin Rambo-Martin, Shatavia Morrison, Christopher Gulvick, Scott Sammons, Yvette Unoarumhi, Darlene Wagner, Matthew Schmerer, Cyndi Clark, Patrick Campbell, Rob Case, Vikramsinha Ghorpade, Holly Houdeshell, Ola Kvalvaag, Dillon Nall, Ethan Sanders, Alec Vest, Shaun Westlund, Matthew Hardison, Clinton R. Paden, Duncan MacCannell |
| EPI_ISL_3430013 | Fulgent Genetics | Centers for Disease Control and Prevention Division of Viral Diseases, Pathogen Discovery | Dakota Howard, Dhwani Batra, Peter W. Cook, Kara Moser, Adrian Paskey, Jason Caravas, Benjamin Rambo-Martin, Shatavia Morrison, Christopher Gulvick, Scott Sammons, Yvette Unoarumhi, Darlene Wagner, Matthew Schmerer, Harry Gao, Mickey Li, John Gao, Joseph Fierro, Benafsh Sapra, Becky Tsai, Yan Meng, Doreen Ng, James Xie, Clinton R. Paden, Duncan MacCannell |
| EPI_ISL_3430014 | Aegis Sciences Corporation | Centers for Disease Control and Prevention Division of Viral Diseases, Pathogen Discovery | Dakota Howard, Dhwani Batra, Peter W. Cook, Kara Moser, Adrian Paskey, Jason Caravas, Benjamin Rambo-Martin, Shatavia Morrison, Christopher Gulvick, Scott Sammons, Yvette Unoarumhi, Darlene Wagner, Matthew Schmerer, Cyndi Clark, Patrick Campbell, Rob Case, Vikramsinha Ghorpade, Holly Houdeshell, Ola Kvalvaag, Dillon Nall, Ethan Sanders, Alec Vest, Shaun Westlund, Matthew Hardison, Clinton R. Paden, Duncan MacCannell |
| EPI_ISL_3430015, EPI_ISL_3430017 | Fulgent Genetics | Centers for Disease Control and Prevention Division of Viral Diseases, Pathogen Discovery | Dakota Howard, Dhwani Batra, Peter W. Cook, Kara Moser, Adrian Paskey, Jason Caravas, Benjamin Rambo-Martin, Shatavia Morrison, Christopher Gulvick, Scott Sammons, Yvette Unoarumhi, Darlene Wagner, Matthew Schmerer, Harry Gao, Mickey Li, John Gao, Joseph Fierro, Benafsh Sapra, Becky Tsai, Yan Meng, Doreen Ng, James Xie, Clinton R. Paden, Duncan MacCannell |
| EPI_ISL_3430019 | Aegis Sciences Corporation | Centers for Disease Control and Prevention Division of Viral Diseases, Pathogen Discovery | Dakota Howard, Dhwani Batra, Peter W. Cook, Kara Moser, Adrian Paskey, Jason Caravas, Benjamin Rambo-Martin, Shatavia Morrison, Christopher Gulvick, Scott Sammons, Yvette Unoarumhi, Darlene Wagner, Matthew Schmerer, Cyndi Clark, Patrick Campbell, Rob Case, Vikramsinha Ghorpade, Holly Houdeshell, Ola Kvalvaag, Dillon Nall, Ethan Sanders, Alec Vest, Shaun Westlund, Matthew Hardison, Clinton R. Paden, Duncan MacCannell |
| EPI_ISL_3430020 | Fulgent Genetics | Centers for Disease Control and Prevention Division of Viral Diseases, Pathogen Discovery | Dakota Howard, Dhwani Batra, Peter W. Cook, Kara Moser, Adrian Paskey, Jason Caravas, Benjamin Rambo-Martin, Shatavia Morrison, Christopher Gulvick, Scott Sammons, Yvette Unoarumhi, Darlene Wagner, Matthew Schmerer, Harry Gao, Mickey Li, John Gao, Joseph Fierro, Benafsh Sapra, Becky Tsai, Yan Meng, Doreen Ng, James Xie, Clinton R. Paden, Duncan MacCannell |
| EPI_ISL_3430021 | Aegis Sciences Corporation | Centers for Disease Control and Prevention Division of Viral Diseases, Pathogen Discovery | Dakota Howard, Dhwani Batra, Peter W. Cook, Kara Moser, Adrian Paskey, Jason Caravas, Benjamin Rambo-Martin, Shatavia Morrison, Christopher Gulvick, Scott Sammons, Yvette Unoarumhi, Darlene Wagner, Matthew Schmerer, Cyndi Clark, Patrick Campbell, Rob Case, Vikramsinha Ghorpade, Holly Houdeshell, Ola Kvalvaag, Dillon Nall, Ethan Sanders, Alec Vest, Shaun Westlund, Matthew Hardison, Clinton R. Paden, Duncan MacCannell |
| EPI_ISL_3430022 | Fulgent Genetics | Centers for Disease Control and Prevention Division of Viral Diseases, Pathogen Discovery | Dakota Howard, Dhwani Batra, Peter W. Cook, Kara Moser, Adrian Paskey, Jason Caravas, Benjamin Rambo-Martin, Shatavia Morrison, Christopher Gulvick, Scott Sammons, Yvette Unoarumhi, Darlene Wagner, Matthew Schmerer, Harry Gao, Mickey Li, John Gao, Joseph Fierro, Benafsh Sapra, Becky Tsai, Yan Meng, Doreen Ng, James Xie, Clinton R. Paden, Duncan MacCannell |
| EPI_ISL_3430023 | Aegis Sciences Corporation | Centers for Disease Control and Prevention Division of Viral Diseases, Pathogen Discovery | Dakota Howard, Dhwani Batra, Peter W. Cook, Kara Moser, Adrian Paskey, Jason Caravas, Benjamin Rambo-Martin, Shatavia Morrison, Christopher Gulvick, Scott Sammons, Yvette Unoarumhi, Darlene Wagner, Matthew Schmerer, Cyndi Clark, Patrick Campbell, Rob Case, Vikramsinha Ghorpade, Holly Houdeshell, Ola Kvalvaag, Dillon Nall, Ethan Sanders, Alec Vest, Shaun Westlund, Matthew Hardison, Clinton R. Paden, Duncan MacCannell |
| EPI_ISL_3430024 | Fulgent Genetics | Centers for Disease Control and Prevention Division of Viral Diseases, Pathogen Discovery | Dakota Howard, Dhwani Batra, Peter W. Cook, Kara Moser, Adrian Paskey, Jason Caravas, Benjamin Rambo-Martin, Shatavia Morrison, Christopher Gulvick, Scott Sammons, Yvette Unoarumhi, Darlene Wagner, Matthew Schmerer, Harry Gao, Mickey Li, John Gao, Joseph Fierro, Benafsh Sapra, Becky Tsai, Yan Meng, Doreen Ng, James Xie, Clinton R. Paden, Duncan MacCannell |
| EPI_ISL_3430025 | Aegis Sciences Corporation | Centers for Disease Control and Prevention Division of Viral Diseases, Pathogen Discovery | Dakota Howard, Dhwani Batra, Peter W. Cook, Kara Moser, Adrian Paskey, Jason Caravas, Benjamin Rambo-Martin, Shatavia Morrison, Christopher Gulvick, Scott Sammons, Yvette Unoarumhi, Darlene Wagner, Matthew Schmerer, Cyndi Clark, Patrick Campbell, Rob Case, Vikramsinha Ghorpade, Holly Houdeshell, Ola Kvalvaag, Dillon Nall, Ethan Sanders, Alec Vest, Shaun Westlund, Matthew Hardison, Clinton R. Paden, Duncan MacCannell |
| EPI_ISL_3430026 | Fulgent Genetics | Centers for Disease Control and Prevention Division of Viral Diseases, Pathogen Discovery | Dakota Howard, Dhwani Batra, Peter W. Cook, Kara Moser, Adrian Paskey, Jason Caravas, Benjamin Rambo-Martin, Shatavia Morrison, Christopher Gulvick, Scott Sammons, Yvette Unoarumhi, Darlene Wagner, Matthew Schmerer, Harry Gao, Mickey Li, John Gao, Joseph Fierro, Benafsh Sapra, Becky Tsai, Yan Meng, Doreen Ng, James Xie, Clinton R. Paden, Duncan MacCannell |
| EPI_ISL_3430027 | Aegis Sciences Corporation | Centers for Disease Control and Prevention Division of Viral Diseases, Pathogen Discovery | Dakota Howard, Dhwani Batra, Peter W. Cook, Kara Moser, Adrian Paskey, Jason Caravas, Benjamin Rambo-Martin, Shatavia Morrison, Christopher Gulvick, Scott Sammons, Yvette Unoarumhi, Darlene Wagner, Matthew Schmerer, Cyndi Clark, Patrick Campbell, Rob Case, Vikramsinha Ghorpade, Holly Houdeshell, Ola Kvalvaag, Dillon Nall, Ethan Sanders, Alec Vest, Shaun Westlund, Matthew Hardison, Clinton R. Paden, Duncan MacCannell |
| EPI_ISL_3430028 | Fulgent Genetics | Centers for Disease Control and Prevention Division of Viral Diseases, Pathogen Discovery | Dakota Howard, Dhwani Batra, Peter W. Cook, Kara Moser, Adrian Paskey, Jason Caravas, Benjamin Rambo-Martin, Shatavia Morrison, Christopher Gulvick, Scott Sammons, Yvette Unoarumhi, Darlene Wagner, Matthew Schmerer, Harry Gao, Mickey Li, John Gao, Joseph Fierro, Benafsh Sapra, Becky Tsai, Yan Meng, Doreen Ng, James Xie, Clinton R. Paden, Duncan MacCannell |
| EPI_ISL_3430029 | Aegis Sciences Corporation | Centers for Disease Control and Prevention Division of Viral | Dakota Howard, Dhwani Batra, Peter W. Cook, Kara Moser, Adrian Paskey, Jason Caravas, Benjamin Rambo-Martin, Shatavia Morrison, Christopher Gulvick, Scott Sammons, Yvette Unoarumhi, Darlene Wagner, Matthew Schmerer, Cyndi Clark, |

| | | | |
|---|---|---|---|
| | | Diseases, Pathogen Discovery | Patrick Campbell, Rob Case, Vikramsinha Ghorpade, Holly Houdeshell, Ola Kvalvaag, Dillon Nall, Ethan Sanders, Alec Vest, Shaun Westlund, Matthew Hardison, Clinton R. Paden, Duncan MacCannell |
| EPI_ISL_3430030 | Fulgent Genetics | Centers for Disease Control and Prevention Division of Viral Diseases, Pathogen Discovery | Dakota Howard, Dhwani Batra, Peter W. Cook, Kara Moser, Adrian Paskey, Jason Caravas, Benjamin Rambo-Martin, Shatavia Morrison, Christopher Gulvick, Scott Sammons, Yvette Unoarumhi, Darlene Wagner, Matthew Schmerer, Harry Gao, Mickey Li, John Gao, Joseph Fierro, Benafsh Sapra, Becky Tsai, Yan Meng, Doreen Ng, James Xie, Clinton R. Paden, Duncan MacCannell |
| EPI_ISL_3430031 | Aegis Sciences Corporation | Centers for Disease Control and Prevention Division of Viral Diseases, Pathogen Discovery | Dakota Howard, Dhwani Batra, Peter W. Cook, Kara Moser, Adrian Paskey, Jason Caravas, Benjamin Rambo-Martin, Shatavia Morrison, Christopher Gulvick, Scott Sammons, Yvette Unoarumhi, Darlene Wagner, Matthew Schmerer, Cyndi Clark, Patrick Campbell, Rob Case, Vikramsinha Ghorpade, Holly Houdeshell, Ola Kvalvaag, Dillon Nall, Ethan Sanders, Alec Vest, Shaun Westlund, Matthew Hardison, Clinton R. Paden, Duncan MacCannell |
| EPI_ISL_3430032, EPI_ISL_3430034 | Fulgent Genetics | Centers for Disease Control and Prevention Division of Viral Diseases, Pathogen Discovery | Dakota Howard, Dhwani Batra, Peter W. Cook, Kara Moser, Adrian Paskey, Jason Caravas, Benjamin Rambo-Martin, Shatavia Morrison, Christopher Gulvick, Scott Sammons, Yvette Unoarumhi, Darlene Wagner, Matthew Schmerer, Harry Gao, Mickey Li, John Gao, Joseph Fierro, Benafsh Sapra, Becky Tsai, Yan Meng, Doreen Ng, James Xie, Clinton R. Paden, Duncan MacCannell |
| EPI_ISL_3430035 | Aegis Sciences Corporation | Centers for Disease Control and Prevention Division of Viral Diseases, Pathogen Discovery | Dakota Howard, Dhwani Batra, Peter W. Cook, Kara Moser, Adrian Paskey, Jason Caravas, Benjamin Rambo-Martin, Shatavia Morrison, Christopher Gulvick, Scott Sammons, Yvette Unoarumhi, Darlene Wagner, Matthew Schmerer, Cyndi Clark, Patrick Campbell, Rob Case, Vikramsinha Ghorpade, Holly Houdeshell, Ola Kvalvaag, Dillon Nall, Ethan Sanders, Alec Vest, Shaun Westlund, Matthew Hardison, Clinton R. Paden, Duncan MacCannell |
| EPI_ISL_3430036 | Fulgent Genetics | Centers for Disease Control and Prevention Division of Viral Diseases, Pathogen Discovery | Dakota Howard, Dhwani Batra, Peter W. Cook, Kara Moser, Adrian Paskey, Jason Caravas, Benjamin Rambo-Martin, Shatavia Morrison, Christopher Gulvick, Scott Sammons, Yvette Unoarumhi, Darlene Wagner, Matthew Schmerer, Harry Gao, Mickey Li, John Gao, Joseph Fierro, Benafsh Sapra, Becky Tsai, Yan Meng, Doreen Ng, James Xie, Clinton R. Paden, Duncan MacCannell |
| EPI_ISL_3430037 | Aegis Sciences Corporation | Centers for Disease Control and Prevention Division of Viral Diseases, Pathogen Discovery | Dakota Howard, Dhwani Batra, Peter W. Cook, Kara Moser, Adrian Paskey, Jason Caravas, Benjamin Rambo-Martin, Shatavia Morrison, Christopher Gulvick, Scott Sammons, Yvette Unoarumhi, Darlene Wagner, Matthew Schmerer, Cyndi Clark, Patrick Campbell, Rob Case, Vikramsinha Ghorpade, Holly Houdeshell, Ola Kvalvaag, Dillon Nall, Ethan Sanders, Alec Vest, Shaun Westlund, Matthew Hardison, Clinton R. Paden, Duncan MacCannell |
| EPI_ISL_3430038 | Fulgent Genetics | Centers for Disease Control and Prevention Division of Viral Diseases, Pathogen Discovery | Dakota Howard, Dhwani Batra, Peter W. Cook, Kara Moser, Adrian Paskey, Jason Caravas, Benjamin Rambo-Martin, Shatavia Morrison, Christopher Gulvick, Scott Sammons, Yvette Unoarumhi, Darlene Wagner, Matthew Schmerer, Harry Gao, Mickey Li, John Gao, Joseph Fierro, Benafsh Sapra, Becky Tsai, Yan Meng, Doreen Ng, James Xie, Clinton R. Paden, Duncan MacCannell |
| EPI_ISL_3430039 | Aegis Sciences Corporation | Centers for Disease Control and Prevention Division of Viral Diseases, Pathogen Discovery | Dakota Howard, Dhwani Batra, Peter W. Cook, Kara Moser, Adrian Paskey, Jason Caravas, Benjamin Rambo-Martin, Shatavia Morrison, Christopher Gulvick, Scott Sammons, Yvette Unoarumhi, Darlene Wagner, Matthew Schmerer, Cyndi Clark, Patrick Campbell, Rob Case, Vikramsinha Ghorpade, Holly Houdeshell, Ola Kvalvaag, Dillon Nall, Ethan Sanders, Alec Vest, Shaun Westlund, Matthew Hardison, Clinton R. Paden, Duncan MacCannell |
| EPI_ISL_3430040, EPI_ISL_3430042, EPI_ISL_3430043 | Fulgent Genetics | Centers for Disease Control and Prevention Division of Viral Diseases, Pathogen Discovery | Dakota Howard, Dhwani Batra, Peter W. Cook, Kara Moser, Adrian Paskey, Jason Caravas, Benjamin Rambo-Martin, Shatavia Morrison, Christopher Gulvick, Scott Sammons, Yvette Unoarumhi, Darlene Wagner, Matthew Schmerer, Harry Gao, Mickey Li, John Gao, Joseph Fierro, Benafsh Sapra, Becky Tsai, Yan Meng, Doreen Ng, James Xie, Clinton R. Paden, Duncan MacCannell |
| EPI_ISL_3430044 | Aegis Sciences Corporation | Centers for Disease Control and Prevention Division of Viral Diseases, Pathogen Discovery | Dakota Howard, Dhwani Batra, Peter W. Cook, Kara Moser, Adrian Paskey, Jason Caravas, Benjamin Rambo-Martin, Shatavia Morrison, Christopher Gulvick, Scott Sammons, Yvette Unoarumhi, Darlene Wagner, Matthew Schmerer, Cyndi Clark, Patrick Campbell, Rob Case, Vikramsinha Ghorpade, Holly Houdeshell, Ola Kvalvaag, Dillon Nall, Ethan Sanders, Alec Vest, Shaun Westlund, Matthew Hardison, Clinton R. Paden, Duncan MacCannell |
| EPI_ISL_3430045, EPI_ISL_3430047, EPI_ISL_3430051, EPI_ISL_3430053 | Fulgent Genetics | Centers for Disease Control and Prevention Division of Viral Diseases, Pathogen Discovery | Dakota Howard, Dhwani Batra, Peter W. Cook, Kara Moser, Adrian Paskey, Jason Caravas, Benjamin Rambo-Martin, Shatavia Morrison, Christopher Gulvick, Scott Sammons, Yvette Unoarumhi, Darlene Wagner, Matthew Schmerer, Harry Gao, Mickey Li, John Gao, Joseph Fierro, Benafsh Sapra, Becky Tsai, Yan Meng, Doreen Ng, James Xie, Clinton R. Paden, Duncan MacCannell |
| EPI_ISL_3430054 | Aegis Sciences Corporation | Centers for Disease Control and Prevention Division of Viral Diseases, Pathogen Discovery | Dakota Howard, Dhwani Batra, Peter W. Cook, Kara Moser, Adrian Paskey, Jason Caravas, Benjamin Rambo-Martin, Shatavia Morrison, Christopher Gulvick, Scott Sammons, Yvette Unoarumhi, Darlene Wagner, Matthew Schmerer, Cyndi Clark, Patrick Campbell, Rob Case, Vikramsinha Ghorpade, Holly Houdeshell, Ola Kvalvaag, Dillon Nall, Ethan Sanders, Alec Vest, Shaun Westlund, Matthew Hardison, Clinton R. Paden, Duncan MacCannell |
| EPI_ISL_3430055 | Fulgent Genetics | Centers for Disease Control and Prevention Division of Viral Diseases, Pathogen Discovery | Dakota Howard, Dhwani Batra, Peter W. Cook, Kara Moser, Adrian Paskey, Jason Caravas, Benjamin Rambo-Martin, Shatavia Morrison, Christopher Gulvick, Scott Sammons, Yvette Unoarumhi, Darlene Wagner, Matthew Schmerer, Harry Gao, Mickey Li, John Gao, Joseph Fierro, Benafsh Sapra, Becky Tsai, Yan Meng, Doreen Ng, James Xie, Clinton R. Paden, Duncan MacCannell |
| EPI_ISL_3430056 | Aegis Sciences Corporation | Centers for Disease Control and Prevention Division of Viral Diseases, Pathogen Discovery | Dakota Howard, Dhwani Batra, Peter W. Cook, Kara Moser, Adrian Paskey, Jason Caravas, Benjamin Rambo-Martin, Shatavia Morrison, Christopher Gulvick, Scott Sammons, Yvette Unoarumhi, Darlene Wagner, Matthew Schmerer, Cyndi Clark, Patrick Campbell, Rob Case, Vikramsinha Ghorpade, Holly Houdeshell, Ola Kvalvaag, Dillon Nall, Ethan Sanders, Alec Vest, Shaun Westlund, Matthew Hardison, Clinton R. Paden, Duncan MacCannell |
| EPI_ISL_3430057, EPI_ISL_3430059 | Fulgent Genetics | Centers for Disease Control and Prevention Division of Viral Diseases, Pathogen Discovery | Dakota Howard, Dhwani Batra, Peter W. Cook, Kara Moser, Adrian Paskey, Jason Caravas, Benjamin Rambo-Martin, Shatavia Morrison, Christopher Gulvick, Scott Sammons, Yvette Unoarumhi, Darlene Wagner, Matthew Schmerer, Harry Gao, Mickey Li, John Gao, Joseph Fierro, Benafsh Sapra, Becky Tsai, Yan Meng, Doreen Ng, James Xie, Clinton R. Paden, Duncan MacCannell |
| EPI_ISL_3430060 | Aegis Sciences Corporation | Centers for Disease Control and Prevention Division of Viral Diseases, Pathogen Discovery | Dakota Howard, Dhwani Batra, Peter W. Cook, Kara Moser, Adrian Paskey, Jason Caravas, Benjamin Rambo-Martin, Shatavia Morrison, Christopher Gulvick, Scott Sammons, Yvette Unoarumhi, Darlene Wagner, Matthew Schmerer, Cyndi Clark, Patrick Campbell, Rob Case, Vikramsinha Ghorpade, Holly Houdeshell, Ola Kvalvaag, Dillon Nall, Ethan Sanders, Alec Vest, Shaun Westlund, Matthew Hardison, Clinton R. Paden, Duncan MacCannell |
| EPI_ISL_3430061 | Fulgent Genetics | Centers for Disease Control and Prevention Division of Viral Diseases, Pathogen Discovery | Dakota Howard, Dhwani Batra, Peter W. Cook, Kara Moser, Adrian Paskey, Jason Caravas, Benjamin Rambo-Martin, Shatavia Morrison, Christopher Gulvick, Scott Sammons, Yvette Unoarumhi, Darlene Wagner, Matthew Schmerer, Harry Gao, Mickey Li, John Gao, Joseph Fierro, Benafsh Sapra, Becky Tsai, Yan Meng, Doreen Ng, James Xie, Clinton R. Paden, Duncan MacCannell |
| EPI_ISL_3430062 | Aegis Sciences Corporation | Centers for Disease Control and Prevention Division of Viral Diseases, Pathogen Discovery | Dakota Howard, Dhwani Batra, Peter W. Cook, Kara Moser, Adrian Paskey, Jason Caravas, Benjamin Rambo-Martin, Shatavia Morrison, Christopher Gulvick, Scott Sammons, Yvette Unoarumhi, Darlene Wagner, Matthew Schmerer, Cyndi Clark, Patrick Campbell, Rob Case, Vikramsinha Ghorpade, Holly Houdeshell, Ola Kvalvaag, Dillon Nall, Ethan Sanders, Alec Vest, Shaun Westlund, Matthew Hardison, Clinton R. Paden, Duncan MacCannell |
| EPI_ISL_3430063, EPI_ISL_3430065 | Fulgent Genetics | Centers for Disease Control and Prevention Division of Viral Diseases, Pathogen Discovery | Dakota Howard, Dhwani Batra, Peter W. Cook, Kara Moser, Adrian Paskey, Jason Caravas, Benjamin Rambo-Martin, Shatavia Morrison, Christopher Gulvick, Scott Sammons, Yvette Unoarumhi, Darlene Wagner, Matthew Schmerer, Harry Gao, Mickey Li, John Gao, Joseph Fierro, Benafsh Sapra, Becky Tsai, Yan Meng, Doreen Ng, James Xie, Clinton R. Paden, Duncan MacCannell |
| EPI_ISL_3430066 | Aegis Sciences Corporation | Centers for Disease Control and Prevention Division of Viral Diseases, Pathogen Discovery | Dakota Howard, Dhwani Batra, Peter W. Cook, Kara Moser, Adrian Paskey, Jason Caravas, Benjamin Rambo-Martin, Shatavia Morrison, Christopher Gulvick, Scott Sammons, Yvette Unoarumhi, Darlene Wagner, Matthew Schmerer, Cyndi Clark, Patrick Campbell, Rob Case, Vikramsinha Ghorpade, Holly Houdeshell, Ola Kvalvaag, Dillon Nall, Ethan Sanders, Alec Vest, Shaun Westlund, Matthew Hardison, Clinton R. Paden, Duncan MacCannell |
| EPI_ISL_3430067, EPI_ISL_3430069 | Fulgent Genetics | Centers for Disease Control and Prevention Division of Viral Diseases, Pathogen Discovery | Dakota Howard, Dhwani Batra, Peter W. Cook, Kara Moser, Adrian Paskey, Jason Caravas, Benjamin Rambo-Martin, Shatavia Morrison, Christopher Gulvick, Scott Sammons, Yvette Unoarumhi, Darlene Wagner, Matthew Schmerer, Harry Gao, Mickey Li, John Gao, Joseph Fierro, Benafsh Sapra, Becky Tsai, Yan Meng, Doreen Ng, James Xie, Clinton R. Paden, Duncan MacCannell |
| EPI_ISL_3430070 | Aegis Sciences Corporation | Centers for Disease Control and Prevention Division of Viral Diseases, Pathogen Discovery | Dakota Howard, Dhwani Batra, Peter W. Cook, Kara Moser, Adrian Paskey, Jason Caravas, Benjamin Rambo-Martin, Shatavia Morrison, Christopher Gulvick, Scott Sammons, Yvette Unoarumhi, Darlene Wagner, Matthew Schmerer, Cyndi Clark, Patrick Campbell, Rob Case, Vikramsinha Ghorpade, Holly Houdeshell, Ola Kvalvaag, Dillon Nall, Ethan Sanders, Alec Vest, Shaun Westlund, Matthew Hardison, Clinton R. Paden, Duncan MacCannell |
| EPI_ISL_3430071, EPI_ISL_3430073 | Fulgent Genetics | Centers for Disease Control and Prevention Division of Viral Diseases, Pathogen Discovery | Dakota Howard, Dhwani Batra, Peter W. Cook, Kara Moser, Adrian Paskey, Jason Caravas, Benjamin Rambo-Martin, Shatavia Morrison, Christopher Gulvick, Scott Sammons, Yvette Unoarumhi, Darlene Wagner, Matthew Schmerer, Harry Gao, Mickey Li, John Gao, Joseph Fierro, Benafsh Sapra, Becky Tsai, Yan Meng, Doreen Ng, James Xie, Clinton R. Paden, Duncan MacCannell |
| EPI_ISL_3430074, EPI_ISL_3430075 | Aegis Sciences Corporation | Centers for Disease Control and Prevention Division of Viral Diseases, Pathogen Discovery | Dakota Howard, Dhwani Batra, Peter W. Cook, Kara Moser, Adrian Paskey, Jason Caravas, Benjamin Rambo-Martin, Shatavia Morrison, Christopher Gulvick, Scott Sammons, Yvette Unoarumhi, Darlene Wagner, Matthew Schmerer, Cyndi Clark, Patrick Campbell, Rob Case, Vikramsinha Ghorpade, Holly Houdeshell, Ola Kvalvaag, Dillon Nall, Ethan Sanders, Alec Vest, Shaun Westlund, Matthew Hardison, Clinton R. Paden, Duncan MacCannell |
| EPI_ISL_3430076 | Fulgent Genetics | Centers for Disease Control and Prevention Division of Viral Diseases, Pathogen Discovery | Dakota Howard, Dhwani Batra, Peter W. Cook, Kara Moser, Adrian Paskey, Jason Caravas, Benjamin Rambo-Martin, Shatavia Morrison, Christopher Gulvick, Scott Sammons, Yvette Unoarumhi, Darlene Wagner, Matthew Schmerer, Harry Gao, Mickey Li, John Gao, Joseph Fierro, Benafsh Sapra, Becky Tsai, Yan Meng, Doreen Ng, James Xie, Clinton R. Paden, Duncan MacCannell |
| EPI_ISL_3430077 | Aegis Sciences Corporation | Centers for Disease Control and Prevention Division of Viral Diseases, Pathogen Discovery | Dakota Howard, Dhwani Batra, Peter W. Cook, Kara Moser, Adrian Paskey, Jason Caravas, Benjamin Rambo-Martin, Shatavia Morrison, Christopher Gulvick, Scott Sammons, Yvette Unoarumhi, Darlene Wagner, Matthew Schmerer, Cyndi Clark, Patrick Campbell, Rob Case, Vikramsinha Ghorpade, Holly Houdeshell, Ola Kvalvaag, Dillon Nall, Ethan Sanders, Alec Vest, Shaun Westlund, Matthew Hardison, Clinton R. Paden, Duncan MacCannell |
| EPI_ISL_3430078, EPI_ISL_3430080 | Fulgent Genetics | Centers for Disease Control and Prevention Division of Viral Diseases, Pathogen Discovery | Dakota Howard, Dhwani Batra, Peter W. Cook, Kara Moser, Adrian Paskey, Jason Caravas, Benjamin Rambo-Martin, Shatavia Morrison, Christopher Gulvick, Scott Sammons, Yvette Unoarumhi, Darlene Wagner, Matthew Schmerer, Harry Gao, Mickey Li, John Gao, Joseph Fierro, Benafsh Sapra, Becky Tsai, Yan Meng, Doreen Ng, James Xie, Clinton R. Paden, Duncan MacCannell |
| EPI_ISL_3430081 | Aegis Sciences Corporation | Centers for Disease Control and Prevention Division of Viral Diseases, Pathogen Discovery | Dakota Howard, Dhwani Batra, Peter W. Cook, Kara Moser, Adrian Paskey, Jason Caravas, Benjamin Rambo-Martin, Shatavia Morrison, Christopher Gulvick, Scott Sammons, Yvette Unoarumhi, Darlene Wagner, Matthew Schmerer, Cyndi Clark, Patrick Campbell, Rob Case, Vikramsinha Ghorpade, Holly Houdeshell, Ola Kvalvaag, Dillon Nall, Ethan Sanders, Alec Vest, Shaun Westlund, Matthew Hardison, Clinton R. Paden, Duncan MacCannell |
| EPI_ISL_3430082, EPI_ISL_3430084 | Fulgent Genetics | Centers for Disease Control and Prevention Division of Viral Diseases, Pathogen Discovery | Dakota Howard, Dhwani Batra, Peter W. Cook, Kara Moser, Adrian Paskey, Jason Caravas, Benjamin Rambo-Martin, Shatavia Morrison, Christopher Gulvick, Scott Sammons, Yvette Unoarumhi, Darlene Wagner, Matthew Schmerer, Harry Gao, Mickey Li, John Gao, Joseph Fierro, Benafsh Sapra, Becky Tsai, Yan Meng, Doreen Ng, James Xie, Clinton R. Paden, Duncan MacCannell |
| EPI_ISL_3430085 | Aegis Sciences Corporation | Centers for Disease Control and Prevention Division of Viral Diseases, Pathogen Discovery | Dakota Howard, Dhwani Batra, Peter W. Cook, Kara Moser, Adrian Paskey, Jason Caravas, Benjamin Rambo-Martin, Shatavia Morrison, Christopher Gulvick, Scott Sammons, Yvette Unoarumhi, Darlene Wagner, Matthew Schmerer, Cyndi Clark, Patrick Campbell, Rob Case, Vikramsinha Ghorpade, Holly Houdeshell, Ola Kvalvaag, Dillon Nall, Ethan Sanders, Alec Vest, Shaun Westlund, Matthew Hardison, Clinton R. Paden, Duncan MacCannell |
| EPI_ISL_3430086, EPI_ISL_3430088 | Fulgent Genetics | Centers for Disease Control and Prevention Division of Viral Diseases, Pathogen Discovery | Dakota Howard, Dhwani Batra, Peter W. Cook, Kara Moser, Adrian Paskey, Jason Caravas, Benjamin Rambo-Martin, Shatavia Morrison, Christopher Gulvick, Scott Sammons, Yvette Unoarumhi, Darlene Wagner, Matthew Schmerer, Harry Gao, Mickey Li, John Gao, Joseph Fierro, Benafsh Sapra, Becky Tsai, Yan Meng, Doreen Ng, James Xie, Clinton R. Paden, Duncan MacCannell |
| EPI_ISL_3430089 | Aegis Sciences Corporation | Centers for Disease Control and Prevention Division of Viral Diseases, Pathogen Discovery | Dakota Howard, Dhwani Batra, Peter W. Cook, Kara Moser, Adrian Paskey, Jason Caravas, Benjamin Rambo-Martin, Shatavia Morrison, Christopher Gulvick, Scott Sammons, Yvette Unoarumhi, Darlene Wagner, Matthew Schmerer, Cyndi Clark, Patrick Campbell, Rob Case, Vikramsinha Ghorpade, Holly Houdeshell, Ola Kvalvaag, Dillon Nall, Ethan Sanders, Alec Vest, Shaun Westlund, Matthew Hardison, Clinton R. Paden, Duncan MacCannell |
| EPI_ISL_3430090 | Fulgent Genetics | Centers for Disease Control and Prevention Division of Viral Diseases, Pathogen Discovery | Dakota Howard, Dhwani Batra, Peter W. Cook, Kara Moser, Adrian Paskey, Jason Caravas, Benjamin Rambo-Martin, Shatavia Morrison, Christopher Gulvick, Scott Sammons, Yvette Unoarumhi, Darlene Wagner, Matthew Schmerer, Harry Gao, Mickey Li, John Gao, Joseph Fierro, Benafsh Sapra, Becky Tsai, Yan Meng, Doreen Ng, James Xie, Clinton R. Paden, Duncan MacCannell |
| EPI_ISL_3430091 | Aegis Sciences Corporation | Centers for Disease Control and Prevention Division of Viral Diseases, Pathogen Discovery | Dakota Howard, Dhwani Batra, Peter W. Cook, Kara Moser, Adrian Paskey, Jason Caravas, Benjamin Rambo-Martin, Shatavia Morrison, Christopher Gulvick, Scott Sammons, Yvette Unoarumhi, Darlene Wagner, Matthew Schmerer, Cyndi Clark, Patrick Campbell, Rob Case, Vikramsinha Ghorpade, Holly Houdeshell, Ola Kvalvaag, Dillon Nall, Ethan Sanders, Alec Vest, Shaun Westlund, Matthew Hardison, Clinton R. Paden, Duncan MacCannell |
| EPI_ISL_3430092, EPI_ISL_3430094 | Fulgent Genetics | Centers for Disease Control and Prevention Division of Viral Diseases, Pathogen Discovery | Dakota Howard, Dhwani Batra, Peter W. Cook, Kara Moser, Adrian Paskey, Jason Caravas, Benjamin Rambo-Martin, Shatavia Morrison, Christopher Gulvick, Scott Sammons, Yvette Unoarumhi, Darlene Wagner, Matthew Schmerer, Harry Gao, Mickey Li, John Gao, Joseph Fierro, Benafsh Sapra, Becky Tsai, Yan Meng, Doreen Ng, James Xie, Clinton R. Paden, Duncan MacCannell |
| EPI_ISL_3430095 | Aegis Sciences Corporation | Centers for Disease Control and Prevention Division of Viral Diseases, Pathogen Discovery | Dakota Howard, Dhwani Batra, Peter W. Cook, Kara Moser, Adrian Paskey, Jason Caravas, Benjamin Rambo-Martin, Shatavia Morrison, Christopher Gulvick, Scott Sammons, Yvette Unoarumhi, Darlene Wagner, Matthew Schmerer, Cyndi Clark, Patrick Campbell, Rob Case, Vikramsinha Ghorpade, Holly Houdeshell, Ola Kvalvaag, Dillon Nall, Ethan Sanders, Alec Vest, Shaun Westlund, Matthew Hardison, Clinton R. Paden, Duncan MacCannell |
| EPI_ISL_3430096, EPI_ISL_3430098 | Fulgent Genetics | Centers for Disease Control and Prevention Division of Viral Diseases, Pathogen Discovery | Dakota Howard, Dhwani Batra, Peter W. Cook, Kara Moser, Adrian Paskey, Jason Caravas, Benjamin Rambo-Martin, Shatavia Morrison, Christopher Gulvick, Scott Sammons, Yvette Unoarumhi, Darlene Wagner, Matthew Schmerer, Harry Gao, Mickey Li, John Gao, Joseph Fierro, Benafsh Sapra, Becky Tsai, Yan Meng, Doreen Ng, James Xie, Clinton R. Paden, Duncan MacCannell |
| EPI_ISL_3430099 | Aegis Sciences Corporation | Centers for Disease Control and Prevention Division of Viral Diseases, Pathogen Discovery | Dakota Howard, Dhwani Batra, Peter W. Cook, Kara Moser, Adrian Paskey, Jason Caravas, Benjamin Rambo-Martin, Shatavia Morrison, Christopher Gulvick, Scott Sammons, Yvette Unoarumhi, Darlene Wagner, Matthew Schmerer, Cyndi Clark, Patrick Campbell, Rob Case, Vikramsinha Ghorpade, Holly Houdeshell, Ola Kvalvaag, Dillon Nall, Ethan Sanders, Alec Vest, Shaun Westlund, Matthew Hardison, Clinton R. Paden, Duncan MacCannell |
| EPI_ISL_3430100 | Fulgent Genetics | Centers for Disease Control and Prevention Division of Viral Diseases, Pathogen Discovery | Dakota Howard, Dhwani Batra, Peter W. Cook, Kara Moser, Adrian Paskey, Jason Caravas, Benjamin Rambo-Martin, Shatavia Morrison, Christopher Gulvick, Scott Sammons, Yvette Unoarumhi, Darlene Wagner, Matthew Schmerer, Harry Gao, Mickey Li, John Gao, Joseph Fierro, Benafsh Sapra, Becky Tsai, Yan Meng, Doreen Ng, James Xie, Clinton R. Paden, Duncan MacCannell |
| EPI_ISL_3430101 | Aegis Sciences Corporation | Centers for Disease Control and Prevention Division of Viral Diseases, Pathogen Discovery | Dakota Howard, Dhwani Batra, Peter W. Cook, Kara Moser, Adrian Paskey, Jason Caravas, Benjamin Rambo-Martin, Shatavia Morrison, Christopher Gulvick, Scott Sammons, Yvette Unoarumhi, Darlene Wagner, Matthew Schmerer, Cyndi Clark, Patrick Campbell, Rob Case, Vikramsinha Ghorpade, Holly Houdeshell, Ola Kvalvaag, Dillon Nall, Ethan Sanders, Alec Vest, Shaun Westlund, Matthew Hardison, Clinton R. Paden, Duncan MacCannell |
| EPI_ISL_3430102, EPI_ISL_3430104, EPI_ISL_3430106, EPI_ISL_3430108 | Fulgent Genetics | Centers for Disease Control and Prevention Division of Viral Diseases, Pathogen Discovery | Dakota Howard, Dhwani Batra, Peter W. Cook, Kara Moser, Adrian Paskey, Jason Caravas, Benjamin Rambo-Martin, Shatavia Morrison, Christopher Gulvick, Scott Sammons, Yvette Unoarumhi, Darlene Wagner, Matthew Schmerer, Harry Gao, Mickey Li, John Gao, Joseph Fierro, Benafsh Sapra, Becky Tsai, Yan Meng, Doreen Ng, James Xie, Clinton R. Paden, Duncan MacCannell |
| EPI_ISL_3430109 | Aegis Sciences Corporation | Centers for Disease Control and Prevention Division of Viral Diseases, Pathogen Discovery | Dakota Howard, Dhwani Batra, Peter W. Cook, Kara Moser, Adrian Paskey, Jason Caravas, Benjamin Rambo-Martin, Shatavia Morrison, Christopher Gulvick, Scott Sammons, Yvette Unoarumhi, Darlene Wagner, Matthew Schmerer, Cyndi Clark, Patrick Campbell, Rob Case, Vikramsinha Ghorpade, Holly Houdeshell, Ola Kvalvaag, Dillon Nall, Ethan Sanders, Alec Vest, Shaun Westlund, Matthew Hardison, Clinton R. Paden, Duncan MacCannell |
| EPI_ISL_3430110, | Fulgent Genetics | Centers for Disease Control and Prevention Division of Viral | Dakota Howard, Dhwani Batra, Peter W. Cook, Kara Moser, Adrian Paskey, Jason Caravas, Benjamin Rambo-Martin, Shatavia Morrison, Christopher Gulvick, Scott Sammons, Yvette Unoarumhi, Darlene Wagner, Matthew Schmerer, Harry Gao, |

| | | | |
|---|---|---|---|
| EPI_ISL_3430112, EPI_ISL_3430114, EPI_ISL_3430116, EPI_ISL_3430118 | | Diseases, Pathogen Discovery | Mickey Li, John Gao, Joseph Fierro, Benafsh Sapra, Becky Tsai, Yan Meng, Doreen Ng, James Xie, Clinton R. Paden, Duncan MacCannell |
| EPI_ISL_3430119 | Aegis Sciences Corporation | Centers for Disease Control and Prevention Division of Viral Diseases, Pathogen Discovery | Dakota Howard, Dhwani Batra, Peter W. Cook, Kara Moser, Adrian Paskey, Jason Caravas, Benjamin Rambo-Martin, Shatavia Morrison, Christopher Gulvick, Scott Sammons, Yvette Unoarumhi, Darlene Wagner, Matthew Schmerer, Cyndi Clark, Patrick Campbell, Rob Case, Vikramsinha Ghorpade, Holly Houdeshell, Ola Kvalvaag, Dillon Nall, Ethan Sanders, Alec Vest, Shaun Westlund, Matthew Hardison, Clinton R. Paden, Duncan MacCannell |
| EPI_ISL_3430120, EPI_ISL_3430122 | Fulgent Genetics | Centers for Disease Control and Prevention Division of Viral Diseases, Pathogen Discovery | Dakota Howard, Dhwani Batra, Peter W. Cook, Kara Moser, Adrian Paskey, Jason Caravas, Benjamin Rambo-Martin, Shatavia Morrison, Christopher Gulvick, Scott Sammons, Yvette Unoarumhi, Darlene Wagner, Matthew Schmerer, Harry Gao, Mickey Li, John Gao, Joseph Fierro, Benafsh Sapra, Becky Tsai, Yan Meng, Doreen Ng, James Xie, Clinton R. Paden, Duncan MacCannell |
| EPI_ISL_3430123 | Aegis Sciences Corporation | Centers for Disease Control and Prevention Division of Viral Diseases, Pathogen Discovery | Dakota Howard, Dhwani Batra, Peter W. Cook, Kara Moser, Adrian Paskey, Jason Caravas, Benjamin Rambo-Martin, Shatavia Morrison, Christopher Gulvick, Scott Sammons, Yvette Unoarumhi, Darlene Wagner, Matthew Schmerer, Cyndi Clark, Patrick Campbell, Rob Case, Vikramsinha Ghorpade, Holly Houdeshell, Ola Kvalvaag, Dillon Nall, Ethan Sanders, Alec Vest, Shaun Westlund, Matthew Hardison, Clinton R. Paden, Duncan MacCannell |
| EPI_ISL_3430124, EPI_ISL_3430126, EPI_ISL_3430128 | Fulgent Genetics | Centers for Disease Control and Prevention Division of Viral Diseases, Pathogen Discovery | Dakota Howard, Dhwani Batra, Peter W. Cook, Kara Moser, Adrian Paskey, Jason Caravas, Benjamin Rambo-Martin, Shatavia Morrison, Christopher Gulvick, Scott Sammons, Yvette Unoarumhi, Darlene Wagner, Matthew Schmerer, Harry Gao, Mickey Li, John Gao, Joseph Fierro, Benafsh Sapra, Becky Tsai, Yan Meng, Doreen Ng, James Xie, Clinton R. Paden, Duncan MacCannell |
| EPI_ISL_3430129 | Aegis Sciences Corporation | Centers for Disease Control and Prevention Division of Viral Diseases, Pathogen Discovery | Dakota Howard, Dhwani Batra, Peter W. Cook, Kara Moser, Adrian Paskey, Jason Caravas, Benjamin Rambo-Martin, Shatavia Morrison, Christopher Gulvick, Scott Sammons, Yvette Unoarumhi, Darlene Wagner, Matthew Schmerer, Cyndi Clark, Patrick Campbell, Rob Case, Vikramsinha Ghorpade, Holly Houdeshell, Ola Kvalvaag, Dillon Nall, Ethan Sanders, Alec Vest, Shaun Westlund, Matthew Hardison, Clinton R. Paden, Duncan MacCannell |
| EPI_ISL_3430130 | Fulgent Genetics | Centers for Disease Control and Prevention Division of Viral Diseases, Pathogen Discovery | Dakota Howard, Dhwani Batra, Peter W. Cook, Kara Moser, Adrian Paskey, Jason Caravas, Benjamin Rambo-Martin, Shatavia Morrison, Christopher Gulvick, Scott Sammons, Yvette Unoarumhi, Darlene Wagner, Matthew Schmerer, Harry Gao, Mickey Li, John Gao, Joseph Fierro, Benafsh Sapra, Becky Tsai, Yan Meng, Doreen Ng, James Xie, Clinton R. Paden, Duncan MacCannell |
| EPI_ISL_3430131 | Aegis Sciences Corporation | Centers for Disease Control and Prevention Division of Viral Diseases, Pathogen Discovery | Dakota Howard, Dhwani Batra, Peter W. Cook, Kara Moser, Adrian Paskey, Jason Caravas, Benjamin Rambo-Martin, Shatavia Morrison, Christopher Gulvick, Scott Sammons, Yvette Unoarumhi, Darlene Wagner, Matthew Schmerer, Cyndi Clark, Patrick Campbell, Rob Case, Vikramsinha Ghorpade, Holly Houdeshell, Ola Kvalvaag, Dillon Nall, Ethan Sanders, Alec Vest, Shaun Westlund, Matthew Hardison, Clinton R. Paden, Duncan MacCannell |
| EPI_ISL_3430132, EPI_ISL_3430134 | Fulgent Genetics | Centers for Disease Control and Prevention Division of Viral Diseases, Pathogen Discovery | Dakota Howard, Dhwani Batra, Peter W. Cook, Kara Moser, Adrian Paskey, Jason Caravas, Benjamin Rambo-Martin, Shatavia Morrison, Christopher Gulvick, Scott Sammons, Yvette Unoarumhi, Darlene Wagner, Matthew Schmerer, Harry Gao, Mickey Li, John Gao, Joseph Fierro, Benafsh Sapra, Becky Tsai, Yan Meng, Doreen Ng, James Xie, Clinton R. Paden, Duncan MacCannell |
| EPI_ISL_3430135 | Aegis Sciences Corporation | Centers for Disease Control and Prevention Division of Viral Diseases, Pathogen Discovery | Dakota Howard, Dhwani Batra, Peter W. Cook, Kara Moser, Adrian Paskey, Jason Caravas, Benjamin Rambo-Martin, Shatavia Morrison, Christopher Gulvick, Scott Sammons, Yvette Unoarumhi, Darlene Wagner, Matthew Schmerer, Cyndi Clark, Patrick Campbell, Rob Case, Vikramsinha Ghorpade, Holly Houdeshell, Ola Kvalvaag, Dillon Nall, Ethan Sanders, Alec Vest, Shaun Westlund, Matthew Hardison, Clinton R. Paden, Duncan MacCannell |
| EPI_ISL_3430136 | Fulgent Genetics | Centers for Disease Control and Prevention Division of Viral Diseases, Pathogen Discovery | Dakota Howard, Dhwani Batra, Peter W. Cook, Kara Moser, Adrian Paskey, Jason Caravas, Benjamin Rambo-Martin, Shatavia Morrison, Christopher Gulvick, Scott Sammons, Yvette Unoarumhi, Darlene Wagner, Matthew Schmerer, Harry Gao, Mickey Li, John Gao, Joseph Fierro, Benafsh Sapra, Becky Tsai, Yan Meng, Doreen Ng, James Xie, Clinton R. Paden, Duncan MacCannell |
| EPI_ISL_3430137 | Aegis Sciences Corporation | Centers for Disease Control and Prevention Division of Viral Diseases, Pathogen Discovery | Dakota Howard, Dhwani Batra, Peter W. Cook, Kara Moser, Adrian Paskey, Jason Caravas, Benjamin Rambo-Martin, Shatavia Morrison, Christopher Gulvick, Scott Sammons, Yvette Unoarumhi, Darlene Wagner, Matthew Schmerer, Cyndi Clark, Patrick Campbell, Rob Case, Vikramsinha Ghorpade, Holly Houdeshell, Ola Kvalvaag, Dillon Nall, Ethan Sanders, Alec Vest, Shaun Westlund, Matthew Hardison, Clinton R. Paden, Duncan MacCannell |
| EPI_ISL_3430138 | Fulgent Genetics | Centers for Disease Control and Prevention Division of Viral Diseases, Pathogen Discovery | Dakota Howard, Dhwani Batra, Peter W. Cook, Kara Moser, Adrian Paskey, Jason Caravas, Benjamin Rambo-Martin, Shatavia Morrison, Christopher Gulvick, Scott Sammons, Yvette Unoarumhi, Darlene Wagner, Matthew Schmerer, Harry Gao, Mickey Li, John Gao, Joseph Fierro, Benafsh Sapra, Becky Tsai, Yan Meng, Doreen Ng, James Xie, Clinton R. Paden, Duncan MacCannell |
| EPI_ISL_3430139 | Aegis Sciences Corporation | Centers for Disease Control and Prevention Division of Viral Diseases, Pathogen Discovery | Dakota Howard, Dhwani Batra, Peter W. Cook, Kara Moser, Adrian Paskey, Jason Caravas, Benjamin Rambo-Martin, Shatavia Morrison, Christopher Gulvick, Scott Sammons, Yvette Unoarumhi, Darlene Wagner, Matthew Schmerer, Cyndi Clark, Patrick Campbell, Rob Case, Vikramsinha Ghorpade, Holly Houdeshell, Ola Kvalvaag, Dillon Nall, Ethan Sanders, Alec Vest, Shaun Westlund, Matthew Hardison, Clinton R. Paden, Duncan MacCannell |
| EPI_ISL_3430140, EPI_ISL_3430143 | Fulgent Genetics | Centers for Disease Control and Prevention Division of Viral Diseases, Pathogen Discovery | Dakota Howard, Dhwani Batra, Peter W. Cook, Kara Moser, Adrian Paskey, Jason Caravas, Benjamin Rambo-Martin, Shatavia Morrison, Christopher Gulvick, Scott Sammons, Yvette Unoarumhi, Darlene Wagner, Matthew Schmerer, Harry Gao, Mickey Li, John Gao, Joseph Fierro, Benafsh Sapra, Becky Tsai, Yan Meng, Doreen Ng, James Xie, Clinton R. Paden, Duncan MacCannell |
| EPI_ISL_3430144 | Aegis Sciences Corporation | Centers for Disease Control and Prevention Division of Viral Diseases, Pathogen Discovery | Dakota Howard, Dhwani Batra, Peter W. Cook, Kara Moser, Adrian Paskey, Jason Caravas, Benjamin Rambo-Martin, Shatavia Morrison, Christopher Gulvick, Scott Sammons, Yvette Unoarumhi, Darlene Wagner, Matthew Schmerer, Cyndi Clark, Patrick Campbell, Rob Case, Vikramsinha Ghorpade, Holly Houdeshell, Ola Kvalvaag, Dillon Nall, Ethan Sanders, Alec Vest, Shaun Westlund, Matthew Hardison, Clinton R. Paden, Duncan MacCannell |
| EPI_ISL_3430145, EPI_ISL_3430146, EPI_ISL_3430148 | Fulgent Genetics | Centers for Disease Control and Prevention Division of Viral Diseases, Pathogen Discovery | Dakota Howard, Dhwani Batra, Peter W. Cook, Kara Moser, Adrian Paskey, Jason Caravas, Benjamin Rambo-Martin, Shatavia Morrison, Christopher Gulvick, Scott Sammons, Yvette Unoarumhi, Darlene Wagner, Matthew Schmerer, Harry Gao, Mickey Li, John Gao, Joseph Fierro, Benafsh Sapra, Becky Tsai, Yan Meng, Doreen Ng, James Xie, Clinton R. Paden, Duncan MacCannell |
| EPI_ISL_3430149, EPI_ISL_3430150 | Aegis Sciences Corporation | Centers for Disease Control and Prevention Division of Viral Diseases, Pathogen Discovery | Dakota Howard, Dhwani Batra, Peter W. Cook, Kara Moser, Adrian Paskey, Jason Caravas, Benjamin Rambo-Martin, Shatavia Morrison, Christopher Gulvick, Scott Sammons, Yvette Unoarumhi, Darlene Wagner, Matthew Schmerer, Cyndi Clark, Patrick Campbell, Rob Case, Vikramsinha Ghorpade, Holly Houdeshell, Ola Kvalvaag, Dillon Nall, Ethan Sanders, Alec Vest, Shaun Westlund, Matthew Hardison, Clinton R. Paden, Duncan MacCannell |
| EPI_ISL_3430151, EPI_ISL_3430153, EPI_ISL_3430154 | Fulgent Genetics | Centers for Disease Control and Prevention Division of Viral Diseases, Pathogen Discovery | Dakota Howard, Dhwani Batra, Peter W. Cook, Kara Moser, Adrian Paskey, Jason Caravas, Benjamin Rambo-Martin, Shatavia Morrison, Christopher Gulvick, Scott Sammons, Yvette Unoarumhi, Darlene Wagner, Matthew Schmerer, Harry Gao, Mickey Li, John Gao, Joseph Fierro, Benafsh Sapra, Becky Tsai, Yan Meng, Doreen Ng, James Xie, Clinton R. Paden, Duncan MacCannell |
| EPI_ISL_3430155 | Aegis Sciences Corporation | Centers for Disease Control and Prevention Division of Viral Diseases, Pathogen Discovery | Dakota Howard, Dhwani Batra, Peter W. Cook, Kara Moser, Adrian Paskey, Jason Caravas, Benjamin Rambo-Martin, Shatavia Morrison, Christopher Gulvick, Scott Sammons, Yvette Unoarumhi, Darlene Wagner, Matthew Schmerer, Cyndi Clark, Patrick Campbell, Rob Case, Vikramsinha Ghorpade, Holly Houdeshell, Ola Kvalvaag, Dillon Nall, Ethan Sanders, Alec Vest, Shaun Westlund, Matthew Hardison, Clinton R. Paden, Duncan MacCannell |
| EPI_ISL_3430157, EPI_ISL_3430158, EPI_ISL_3430160, EPI_ISL_3430161, EPI_ISL_3430162, EPI_ISL_3430163, EPI_ISL_3430164, EPI_ISL_3430165, EPI_ISL_3430166, EPI_ISL_3430168, EPI_ISL_3430169 | | | |
| see above | Fulgent Genetics | Centers for Disease Control and Prevention Division of Viral Diseases, Pathogen Discovery | Dakota Howard, Dhwani Batra, Peter W. Cook, Kara Moser, Adrian Paskey, Jason Caravas, Benjamin Rambo-Martin, Shatavia Morrison, Christopher Gulvick, Scott Sammons, Yvette Unoarumhi, Darlene Wagner, Matthew Schmerer, Harry Gao, Mickey Li, John Gao, Joseph Fierro, Benafsh Sapra, Becky Tsai, Yan Meng, Doreen Ng, James Xie, Clinton R. Paden, Duncan MacCannell |
| EPI_ISL_3430170 | Aegis Sciences Corporation | Centers for Disease Control and Prevention Division of Viral Diseases, Pathogen Discovery | Dakota Howard, Dhwani Batra, Peter W. Cook, Kara Moser, Adrian Paskey, Jason Caravas, Benjamin Rambo-Martin, Shatavia Morrison, Christopher Gulvick, Scott Sammons, Yvette Unoarumhi, Darlene Wagner, Matthew Schmerer, Cyndi Clark, Patrick Campbell, Rob Case, Vikramsinha Ghorpade, Holly Houdeshell, Ola Kvalvaag, Dillon Nall, Ethan Sanders, Alec Vest, Shaun Westlund, Matthew Hardison, Clinton R. Paden, Duncan MacCannell |
| EPI_ISL_3430172 | Fulgent Genetics | Centers for Disease Control and Prevention Division of Viral Diseases, Pathogen Discovery | Dakota Howard, Dhwani Batra, Peter W. Cook, Kara Moser, Adrian Paskey, Jason Caravas, Benjamin Rambo-Martin, Shatavia Morrison, Christopher Gulvick, Scott Sammons, Yvette Unoarumhi, Darlene Wagner, Matthew Schmerer, Harry Gao, Mickey Li, John Gao, Joseph Fierro, Benafsh Sapra, Becky Tsai, Yan Meng, Doreen Ng, James Xie, Clinton R. Paden, Duncan MacCannell |
| EPI_ISL_3430173, EPI_ISL_3430176, EPI_ISL_3430178, EPI_ISL_3430179, EPI_ISL_3430180, EPI_ISL_3430183, EPI_ISL_3430184, EPI_ISL_3430186, EPI_ISL_3430187, EPI_ISL_3430188, EPI_ISL_3430189, EPI_ISL_3430192, EPI_ISL_3430194, EPI_ISL_3430195, EPI_ISL_3430197, EPI_ISL_3430198, EPI_ISL_3430199, EPI_ISL_3430201, EPI_ISL_3430202, EPI_ISL_3430203, EPI_ISL_3430204, EPI_ISL_3430205, EPI_ISL_3430206, EPI_ISL_3430207, EPI_ISL_3430209, EPI_ISL_3430210, EPI_ISL_3430212, EPI_ISL_3430214, EPI_ISL_3430215, EPI_ISL_3430216, EPI_ISL_3430217, EPI_ISL_3430219, EPI_ISL_3430220, EPI_ISL_3430223, EPI_ISL_3430228, EPI_ISL_3430229, EPI_ISL_3430231, EPI_ISL_3430232, EPI_ISL_3430234, EPI_ISL_3430235, EPI_ISL_3430236, EPI_ISL_3430237, EPI_ISL_3430240, EPI_ISL_3430241, EPI_ISL_3430243, EPI_ISL_3430244, EPI_ISL_3430247, EPI_ISL_3430249, EPI_ISL_3430250, EPI_ISL_3430251, EPI_ISL_3430252, EPI_ISL_3430253, EPI_ISL_3430254, EPI_ISL_3430255, EPI_ISL_3430256, EPI_ISL_3430258, EPI_ISL_3430259, EPI_ISL_3430261, EPI_ISL_3430262, EPI_ISL_3430263, EPI_ISL_3430264, EPI_ISL_3430265, EPI_ISL_3430267, EPI_ISL_3430268, EPI_ISL_3430269, EPI_ISL_3430271, EPI_ISL_3430273, EPI_ISL_3430274, EPI_ISL_3430278, EPI_ISL_3430279, EPI_ISL_3430280, EPI_ISL_3430281, EPI_ISL_3430282, EPI_ISL_3430283, EPI_ISL_3430284, EPI_ISL_3430285, EPI_ISL_3430286, EPI_ISL_3430287, EPI_ISL_3430288, EPI_ISL_3430289, EPI_ISL_3430290, EPI_ISL_3430293, EPI_ISL_3430294 | | | |
| see above | Aegis Sciences Corporation | Centers for Disease Control and Prevention Division of Viral Diseases, Pathogen Discovery | Dakota Howard, Dhwani Batra, Peter W. Cook, Kara Moser, Adrian Paskey, Jason Caravas, Benjamin Rambo-Martin, Shatavia Morrison, Christopher Gulvick, Scott Sammons, Yvette Unoarumhi, Darlene Wagner, Matthew Schmerer, Cyndi Clark, Patrick Campbell, Rob Case, Vikramsinha Ghorpade, Holly Houdeshell, Ola Kvalvaag, Dillon Nall, Ethan Sanders, Alec Vest, Shaun Westlund, Matthew Hardison, Clinton R. Paden, Duncan MacCannell |
| EPI_ISL_3430295, EPI_ISL_3430297 | Molecular Diagnostics Laboratory | National Reference Laboratory, Nigeria Centre for Disease Control | Dr Ndodo Nnaemeka, Olusola Anuoluwapo Akanbi, Chimaobi Chukwu, Dr Omoare Adesuyi, Celestina Obiekea, Anthony Ahumibe, Catherine Okoi, Nwando Mba, Dr Chikwe Ihekweazu |
| EPI_ISL_3430298 | Aegis Sciences Corporation | Centers for Disease Control and Prevention Division of Viral Diseases, Pathogen Discovery | Dakota Howard, Dhwani Batra, Peter W. Cook, Kara Moser, Adrian Paskey, Jason Caravas, Benjamin Rambo-Martin, Shatavia Morrison, Christopher Gulvick, Scott Sammons, Yvette Unoarumhi, Darlene Wagner, Matthew Schmerer, Cyndi Clark, Patrick Campbell, Rob Case, Vikramsinha Ghorpade, Holly Houdeshell, Ola Kvalvaag, Dillon Nall, Ethan Sanders, Alec Vest, Shaun Westlund, Matthew Hardison, Clinton R. Paden, Duncan MacCannell |
| EPI_ISL_3430299, EPI_ISL_3430301, EPI_ISL_3430303 | Molecular Diagnostics Laboratory | National Reference Laboratory, Nigeria Centre for Disease Control | Dr Ndodo Nnaemeka, Olusola Anuoluwapo Akanbi, Chimaobi Chukwu, Dr Omoare Adesuyi, Celestina Obiekea, Anthony Ahumibe, Catherine Okoi, Nwando Mba, Dr Chikwe Ihekweazu |
| EPI_ISL_3430304 | Aegis Sciences Corporation | Centers for Disease Control and Prevention Division of Viral Diseases, Pathogen Discovery | Dakota Howard, Dhwani Batra, Peter W. Cook, Kara Moser, Adrian Paskey, Jason Caravas, Benjamin Rambo-Martin, Shatavia Morrison, Christopher Gulvick, Scott Sammons, Yvette Unoarumhi, Darlene Wagner, Matthew Schmerer, Cyndi Clark, Patrick Campbell, Rob Case, Vikramsinha Ghorpade, Holly Houdeshell, Ola Kvalvaag, Dillon Nall, Ethan Sanders, Alec Vest, Shaun Westlund, Matthew Hardison, Clinton R. Paden, Duncan MacCannell |
| EPI_ISL_3430307 | Molecular Diagnostics Laboratory | National Reference Laboratory, Nigeria Centre for Disease Control | Dr Ndodo Nnaemeka, Olusola Anuoluwapo Akanbi, Chimaobi Chukwu, Dr Omoare Adesuyi, Celestina Obiekea, Anthony Ahumibe, Catherine Okoi, Nwando Mba, Dr Chikwe Ihekweazu |
| EPI_ISL_3430308 | Aegis Sciences Corporation | Centers for Disease Control and Prevention Division of Viral Diseases, Pathogen Discovery | Dakota Howard, Dhwani Batra, Peter W. Cook, Kara Moser, Adrian Paskey, Jason Caravas, Benjamin Rambo-Martin, Shatavia Morrison, Christopher Gulvick, Scott Sammons, Yvette Unoarumhi, Darlene Wagner, Matthew Schmerer, Cyndi Clark, Patrick Campbell, Rob Case, Vikramsinha Ghorpade, Holly Houdeshell, Ola Kvalvaag, Dillon Nall, Ethan Sanders, Alec Vest, Shaun Westlund, Matthew Hardison, Clinton R. Paden, Duncan MacCannell |
| EPI_ISL_3430310 | DemyHealth Laboratory | National Reference Laboratory, Nigeria Centre for Disease Control | Dr Ndodo Nnaemeka, Olusola Anuoluwapo Akanbi, Chimaobi Chukwu, Dr Omoare Adesuyi, Celestina Obiekea, Anthony Ahumibe, Catherine Okoi, Nwando Mba, Dr Chikwe Ihekweazu |
| EPI_ISL_3430312 | Bayelsa | National Reference Laboratory, Nigeria Centre for Disease Control | Dr Ndodo Nnaemeka, Olusola Anuoluwapo Akanbi, Chimaobi Chukwu, Dr Omoare Adesuyi, Celestina Obiekea, Anthony Ahumibe, Catherine Okoi, Nwando Mba, Dr Chikwe Ihekweazu |
| EPI_ISL_3430313 | Aegis Sciences Corporation | Centers for Disease Control and Prevention Division of Viral Diseases, Pathogen Discovery | Dakota Howard, Dhwani Batra, Peter W. Cook, Kara Moser, Adrian Paskey, Jason Caravas, Benjamin Rambo-Martin, Shatavia Morrison, Christopher Gulvick, Scott Sammons, Yvette Unoarumhi, Darlene Wagner, Matthew Schmerer, Cyndi Clark, Patrick Campbell, Rob Case, Vikramsinha Ghorpade, Holly Houdeshell, Ola Kvalvaag, Dillon Nall, Ethan Sanders, Alec Vest, Shaun Westlund, Matthew Hardison, Clinton R. Paden, Duncan MacCannell |
| EPI_ISL_3430314 | Bayelsa | National Reference Laboratory, Nigeria Centre for Disease Control | Dr Ndodo Nnaemeka, Olusola Anuoluwapo Akanbi, Chimaobi Chukwu, Dr Omoare Adesuyi, Celestina Obiekea, Anthony Ahumibe, Catherine Okoi, Nwando Mba, Dr Chikwe Ihekweazu |
| EPI_ISL_3430315 | Aegis Sciences Corporation | Centers for Disease Control and Prevention Division of Viral Diseases, Pathogen Discovery | Dakota Howard, Dhwani Batra, Peter W. Cook, Kara Moser, Adrian Paskey, Jason Caravas, Benjamin Rambo-Martin, Shatavia Morrison, Christopher Gulvick, Scott Sammons, Yvette Unoarumhi, Darlene Wagner, Matthew Schmerer, Cyndi Clark, Patrick Campbell, Rob Case, Vikramsinha Ghorpade, Holly Houdeshell, Ola Kvalvaag, Dillon Nall, Ethan Sanders, Alec Vest, Shaun Westlund, Matthew Hardison, Clinton R. Paden, Duncan MacCannell |
| EPI_ISL_3430316 | Kaduna | National Reference Laboratory, Nigeria Centre for Disease Control | Dr Ndodo Nnaemeka, Olusola Anuoluwapo Akanbi, Chimaobi Chukwu, Dr Omoare Adesuyi, Celestina Obiekea, Anthony Ahumibe, Catherine Okoi, Nwando Mba, Dr Chikwe Ihekweazu |
| EPI_ISL_3430317 | Aegis Sciences Corporation | Centers for Disease Control and Prevention Division of Viral Diseases, Pathogen Discovery | Dakota Howard, Dhwani Batra, Peter W. Cook, Kara Moser, Adrian Paskey, Jason Caravas, Benjamin Rambo-Martin, Shatavia Morrison, Christopher Gulvick, Scott Sammons, Yvette Unoarumhi, Darlene Wagner, Matthew Schmerer, Cyndi Clark, Patrick Campbell, Rob Case, Vikramsinha Ghorpade, Holly Houdeshell, Ola Kvalvaag, Dillon Nall, Ethan Sanders, Alec Vest, Shaun Westlund, Matthew Hardison, Clinton R. Paden, Duncan MacCannell |
| EPI_ISL_3430318, EPI_ISL_3430320 | Kaduna | National Reference Laboratory, Nigeria Centre for Disease Control | Dr Ndodo Nnaemeka, Olusola Anuoluwapo Akanbi, Chimaobi Chukwu, Dr Omoare Adesuyi, Celestina Obiekea, Anthony Ahumibe, Catherine Okoi, Nwando Mba, Dr Chikwe Ihekweazu |
| EPI_ISL_3430321 | Aegis Sciences Corporation | Centers for Disease Control and Prevention Division of Viral Diseases, Pathogen Discovery | Dakota Howard, Dhwani Batra, Peter W. Cook, Kara Moser, Adrian Paskey, Jason Caravas, Benjamin Rambo-Martin, Shatavia Morrison, Christopher Gulvick, Scott Sammons, Yvette Unoarumhi, Darlene Wagner, Matthew Schmerer, Cyndi Clark, Patrick Campbell, Rob Case, Vikramsinha Ghorpade, Holly Houdeshell, Ola Kvalvaag, Dillon Nall, Ethan Sanders, Alec Vest, Shaun Westlund, Matthew Hardison, Clinton R. Paden, Duncan MacCannell |
| EPI_ISL_3430322, EPI_ISL_3430323 | Cedarcrest Molecular laboratory | National Reference Laboratory, Nigeria Centre for Disease Control | Dr Ndodo Nnaemeka, Olusola Anuoluwapo Akanbi, Chimaobi Chukwu, Dr Omoare Adesuyi, Celestina Obiekea, Anthony Ahumibe, Catherine Okoi, Nwando Mba, Dr Chikwe Ihekweazu |
| EPI_ISL_3430325, EPI_ISL_3430326, EPI_ISL_3430327, EPI_ISL_3430328, EPI_ISL_3430331, EPI_ISL_3430332, EPI_ISL_3430335, EPI_ISL_3430336, EPI_ISL_3430337, EPI_ISL_3430342, EPI_ISL_3430343, EPI_ISL_3430345, EPI_ISL_3430346, EPI_ISL_3430347, EPI_ISL_3430348, EPI_ISL_3430349, EPI_ISL_3430353, EPI_ISL_3430354, EPI_ISL_3430355, EPI_ISL_3430356, EPI_ISL_3430359, EPI_ISL_3430360, EPI_ISL_3430362, EPI_ISL_3430366, EPI_ISL_3430368, EPI_ISL_3430369, EPI_ISL_3430371, EPI_ISL_3430372, EPI_ISL_3430375, EPI_ISL_3430376, EPI_ISL_3430377, EPI_ISL_3430379, EPI_ISL_3430380, EPI_ISL_3430382, EPI_ISL_3430384, EPI_ISL_3430385, EPI_ISL_3430386, EPI_ISL_3430387, EPI_ISL_3430389, EPI_ISL_3430391, EPI_ISL_3430393, EPI_ISL_3430394, EPI_ISL_3430395, EPI_ISL_3430399, EPI_ISL_3430400, EPI_ISL_3430401, EPI_ISL_3430404, EPI_ISL_3430405, EPI_ISL_3430406, EPI_ISL_3430407, EPI_ISL_3430408, EPI_ISL_3430409, EPI_ISL_3430411, EPI_ISL_3430412, EPI_ISL_3430413, EPI_ISL_3430415, EPI_ISL_3430417, EPI_ISL_3430418, EPI_ISL_3430419, EPI_ISL_3430421, EPI_ISL_3430423, EPI_ISL_3430424, EPI_ISL_3430425, EPI_ISL_3430426, EPI_ISL_3430428, EPI_ISL_3430430, EPI_ISL_3430431, EPI_ISL_3430432, EPI_ISL_3430436, EPI_ISL_3430439, EPI_ISL_3430440, EPI_ISL_3430442, EPI_ISL_3430443, EPI_ISL_3430444, EPI_ISL_3430446, EPI_ISL_3430447, EPI_ISL_3430448, EPI_ISL_3430449, EPI_ISL_3430450, EPI_ISL_3430451, EPI_ISL_3430453, EPI_ISL_3430457, EPI_ISL_3430458, EPI_ISL_3430459, EPI_ISL_3430460, EPI_ISL_3430461, EPI_ISL_3430463, EPI_ISL_3430464, EPI_ISL_3430465, EPI_ISL_3430467, EPI_ISL_3430468, EPI_ISL_3430469, EPI_ISL_3430471, EPI_ISL_3430472, EPI_ISL_3430473, EPI_ISL_3430474, EPI_ISL_3430475, EPI_ISL_3430477, EPI_ISL_3430478, EPI_ISL_3430479, EPI_ISL_3430480, EPI_ISL_3430482, EPI_ISL_3430485, EPI_ISL_3430486, EPI_ISL_3430487, EPI_ISL_3430488, EPI_ISL_3430489, EPI_ISL_3430490, EPI_ISL_3430491, EPI_ISL_3430492, | | | |

| | | | |
|---|---|---|---|
| EPI_ISL_3430494, EPI_ISL_3430495, EPI_ISL_3430496, EPI_ISL_3430497, EPI_ISL_3430498, EPI_ISL_3430500, EPI_ISL_3430501, EPI_ISL_3430505, EPI_ISL_3430507, EPI_ISL_3430508, EPI_ISL_3430511, EPI_ISL_3430516, EPI_ISL_3430517, EPI_ISL_3430519, EPI_ISL_3430520, EPI_ISL_3430521, EPI_ISL_3430522, EPI_ISL_3430523, EPI_ISL_3430525, EPI_ISL_3430526, EPI_ISL_3430527, EPI_ISL_3430528, EPI_ISL_3430529, EPI_ISL_3430533, EPI_ISL_3430535, EPI_ISL_3430536, EPI_ISL_3430537, EPI_ISL_3430539, EPI_ISL_3430540, EPI_ISL_3430542, EPI_ISL_3430544, EPI_ISL_3430548, EPI_ISL_3430549, EPI_ISL_3430550, EPI_ISL_3430551, EPI_ISL_3430555, EPI_ISL_3430557, EPI_ISL_3430558, EPI_ISL_3430560, EPI_ISL_3430561, EPI_ISL_3430562, EPI_ISL_3430563, EPI_ISL_3430564, EPI_ISL_3430565, EPI_ISL_3430566, EPI_ISL_3430567, EPI_ISL_3430568, EPI_ISL_3430569, EPI_ISL_3430570, EPI_ISL_3430571, EPI_ISL_3430573, EPI_ISL_3430574, EPI_ISL_3430575, EPI_ISL_3430576, EPI_ISL_3430577, EPI_ISL_3430578, EPI_ISL_3430580, EPI_ISL_3430582, EPI_ISL_3430583, EPI_ISL_3430584, EPI_ISL_3430585, EPI_ISL_3430587, EPI_ISL_3430588, EPI_ISL_3430589, EPI_ISL_3430590, EPI_ISL_3430591, EPI_ISL_3430592, EPI_ISL_3430593, EPI_ISL_3430594, EPI_ISL_3430595, EPI_ISL_3430596, EPI_ISL_3430597, EPI_ISL_3430599, EPI_ISL_3430600, EPI_ISL_3430601, EPI_ISL_3430602, EPI_ISL_3430603, EPI_ISL_3430604, EPI_ISL_3430605, EPI_ISL_3430606, EPI_ISL_3430609, EPI_ISL_3430610, EPI_ISL_3430611, EPI_ISL_3430613, EPI_ISL_3430614, EPI_ISL_3430615, EPI_ISL_3430616, EPI_ISL_3430617, EPI_ISL_3430618, EPI_ISL_3430619, EPI_ISL_3430620, EPI_ISL_3430621, EPI_ISL_3430622, EPI_ISL_3430624, EPI_ISL_3430625, EPI_ISL_3430626, EPI_ISL_3430628, EPI_ISL_3430629, EPI_ISL_3430630, EPI_ISL_3430631, EPI_ISL_3430632, EPI_ISL_3430633, EPI_ISL_3430634, EPI_ISL_3430635, EPI_ISL_3430637, EPI_ISL_3430638, EPI_ISL_3430639, EPI_ISL_3430641, EPI_ISL_3430642, EPI_ISL_3430643, EPI_ISL_3430644, EPI_ISL_3430645, EPI_ISL_3430647, EPI_ISL_3430648, EPI_ISL_3430650, EPI_ISL_3430651, EPI_ISL_3430652, EPI_ISL_3430653, EPI_ISL_3430654, EPI_ISL_3430655, EPI_ISL_3430656, EPI_ISL_3430657, EPI_ISL_3430659, EPI_ISL_3430660, EPI_ISL_3430661, EPI_ISL_3430662, EPI_ISL_3430663, EPI_ISL_3430664, EPI_ISL_3430665, EPI_ISL_3430666, EPI_ISL_3430669, EPI_ISL_3430670, EPI_ISL_3430671, EPI_ISL_3430673, EPI_ISL_3430674, EPI_ISL_3430675, EPI_ISL_3430676, EPI_ISL_3430677, EPI_ISL_3430679, EPI_ISL_3430681, EPI_ISL_3430682, EPI_ISL_3430683, EPI_ISL_3430685, EPI_ISL_3430688, EPI_ISL_3430689, EPI_ISL_3430690, EPI_ISL_3430691, EPI_ISL_3430693, EPI_ISL_3430695, EPI_ISL_3430697, EPI_ISL_3430698, EPI_ISL_3430699, EPI_ISL_3430704, EPI_ISL_3430706 | | | |
| see above | Aegis Sciences Corporation | Centers for Disease Control and Prevention Division of Viral Diseases, Pathogen Discovery | Dakota Howard, Dhwani Batra, Peter W. Cook, Kara Moser, Adrian Paskey, Jason Caravas, Benjamin Rambo-Martin, Shatavia Morrison, Christopher Gulvick, Scott Sammons, Yvette Unoarumhi, Darlene Wagner, Matthew Schmerer, Cyndi Clark, Patrick Campbell, Rob Case, Vikramsinha Ghorpade, Holly Houdeshell, Ola Kvalvaag, Dillon Nall, Ethan Sanders, Alec Vest, Shaun Westlund, Matthew Hardison, Clinton R. Paden, Duncan MacCannell |
| EPI_ISL_3430711, EPI_ISL_3430715, EPI_ISL_3430717, EPI_ISL_3430719, EPI_ISL_3430722, EPI_ISL_3430723, EPI_ISL_3430728, EPI_ISL_3430729, EPI_ISL_3430735, EPI_ISL_3430736, EPI_ISL_3430738, EPI_ISL_3430742, EPI_ISL_3430744, EPI_ISL_3430745, EPI_ISL_3430747, EPI_ISL_3430749, EPI_ISL_3430751, EPI_ISL_3430760, EPI_ISL_3430766, EPI_ISL_3430767, EPI_ISL_3430768, EPI_ISL_3430770, EPI_ISL_3430772, EPI_ISL_3430773, EPI_ISL_3430784, EPI_ISL_3430785, EPI_ISL_3430789, EPI_ISL_3430794, EPI_ISL_3430798, EPI_ISL_3430799, EPI_ISL_3430801, EPI_ISL_3430802, EPI_ISL_3430803, EPI_ISL_3430805, EPI_ISL_3430806, EPI_ISL_3430814, EPI_ISL_3430816, EPI_ISL_3430817, EPI_ISL_3430818, EPI_ISL_3430819, EPI_ISL_3430824, EPI_ISL_3430829, EPI_ISL_3430835, EPI_ISL_3430842, EPI_ISL_3430843, EPI_ISL_3430845, EPI_ISL_3430847, EPI_ISL_3430852, EPI_ISL_3430866, EPI_ISL_3430872, EPI_ISL_3430875, EPI_ISL_3430877, EPI_ISL_3430879, EPI_ISL_3430881, EPI_ISL_3430883, EPI_ISL_3430887, EPI_ISL_3430889, EPI_ISL_3430891, EPI_ISL_3430893, EPI_ISL_3430897, EPI_ISL_3430899, EPI_ISL_3430912, EPI_ISL_3430914, EPI_ISL_3430917, EPI_ISL_3430920, EPI_ISL_3430922, EPI_ISL_3430924, EPI_ISL_3430926, EPI_ISL_3430927, EPI_ISL_3430929, EPI_ISL_3430930, EPI_ISL_3430935, EPI_ISL_3430937, EPI_ISL_3430938, EPI_ISL_3430940, EPI_ISL_3430945, EPI_ISL_3430946, EPI_ISL_3430948, EPI_ISL_3430949, EPI_ISL_3430951, EPI_ISL_3430952, EPI_ISL_3430954, EPI_ISL_3430956, EPI_ISL_3430958, EPI_ISL_3430961, EPI_ISL_3430962, EPI_ISL_3430963, EPI_ISL_3430967, EPI_ISL_3430971, EPI_ISL_3430972, EPI_ISL_3430975, EPI_ISL_3430976, EPI_ISL_3430977, EPI_ISL_3430978, EPI_ISL_3430979, EPI_ISL_3430980, EPI_ISL_3430981, EPI_ISL_3430983, EPI_ISL_3430987, EPI_ISL_3430988, EPI_ISL_3430990, EPI_ISL_3430991, EPI_ISL_3430992, EPI_ISL_3430993, EPI_ISL_3430994, EPI_ISL_3430995, EPI_ISL_3430996, EPI_ISL_3431000, EPI_ISL_3431001, EPI_ISL_3431003, EPI_ISL_3431005, EPI_ISL_3431008, EPI_ISL_3431009, EPI_ISL_3431011, EPI_ISL_3431012, EPI_ISL_3431013, EPI_ISL_3431016, EPI_ISL_3431020, EPI_ISL_3431022, EPI_ISL_3431025, EPI_ISL_3431026, EPI_ISL_3431027, EPI_ISL_3431028, EPI_ISL_3431032, EPI_ISL_3431037, EPI_ISL_3431038, EPI_ISL_3431040, EPI_ISL_3431041, EPI_ISL_3431044, EPI_ISL_3431049, EPI_ISL_3431051, EPI_ISL_3431053, EPI_ISL_3431054, EPI_ISL_3431057 | | | |
| see above | Laboratory Corporation of America | Centers for Disease Control and Prevention Division of Viral Diseases, Pathogen Discovery | Dakota Howard, Dhwani Batra, Peter W. Cook, Kara Moser, Adrian Paskey, Jason Caravas, Benjamin Rambo-Martin, Shatavia Morrison, Christopher Gulvick, Scott Sammons, Yvette Unoarumhi, Darlene Wagner, Matthew Schmerer, Minoo Agarwal, Eyad Almasri, Debbie Boles, Ayla Burns, Nuthawin Charoensri, Oren Cohen, Susan Countryman, Mary Ann Cristobal, Bobbi Croy, Suzanne Dale, Hrushikesh Deshmukh, Amanda Douglas, Vincent Drouillon, Marcia Eisenberg, Howard Engler, Rama Ghatti, Prashant Gupta, Susan Hicks, Jake Humphrey, Lax Iyer, Lisa Pfefferle, Manoj Jain, Matthew Robinson, Mohan Kolli, Brian Krueger, Tim Kuphal, Stanley Letovsky, Michael Levandoski, Craig Lukasik, Jonathan Meltzer, Brian Norvell, Mindy Nye, Scott Parker, Christos Petropoulos, John Pruitt, Steven Ragan, Scott Ryan, Mike Sapeta, Jana Schroth, Suresh Babu Selvaraju, Goran Stevovic, Amanda Suchanek, Andrea Throop, Lyndon Tilson, Thomas Urban, Joe Voshell, Kimberly Wagner, Jonathan Williams, Mary Williamson, Qian Zeng, Tricia Zwiefelhofer, Clinton R. Paden, Duncan MacCannell |
| EPI_ISL_3431068, EPI_ISL_3431069, EPI_ISL_3431070, EPI_ISL_3431072, EPI_ISL_3431077, EPI_ISL_3431081, EPI_ISL_3431082, EPI_ISL_3431084, EPI_ISL_3431086, EPI_ISL_3431088, EPI_ISL_3431089, EPI_ISL_3431091, EPI_ISL_3431092, EPI_ISL_3431093 | | | |
| see above | Quest Diagnostics Incorporated | Centers for Disease Control and Prevention Division of Viral Diseases, Pathogen Discovery | Dakota Howard, Dhwani Batra, Peter W. Cook, Kara Moser, Adrian Paskey, Jason Caravas, Benjamin Rambo-Martin, Shatavia Morrison, Christopher Gulvick, Scott Sammons, Yvette Unoarumhi, Darlene Wagner, Matthew Schmerer, S. H. Rosenthal, A. Gerasimova, R. M. Kagan, B. Anderson, M. Hua, Y. Liu, L.E. Bernstein, K.E. Livingston, A. Perez, I. A. Shlyakhter, R. V. Rolando, R. Owen, P. Tanpaiboon, F. Lacbawan, Clinton R. Paden, Duncan MacCannell |
| EPI_ISL_3431094, EPI_ISL_3431095, EPI_ISL_3431096, EPI_ISL_3431097, EPI_ISL_3431098, EPI_ISL_3431099, EPI_ISL_3431100, EPI_ISL_3431102, EPI_ISL_3431103, EPI_ISL_3431105, EPI_ISL_3431108 | | | |
| see above | Fulgent Genetics | Centers for Disease Control and Prevention Division of Viral Diseases, Pathogen Discovery | Dakota Howard, Dhwani Batra, Peter W. Cook, Kara Moser, Adrian Paskey, Jason Caravas, Benjamin Rambo-Martin, Shatavia Morrison, Christopher Gulvick, Scott Sammons, Yvette Unoarumhi, Darlene Wagner, Matthew Schmerer, Harry Gao, Mickey Li, John Gao, Joseph Fierro, Benafsh Sapra, Becky Tsai, Yan Meng, Doreen Ng, James Xie, Clinton R. Paden, Duncan MacCannell |
| EPI_ISL_3431111, EPI_ISL_3431112, EPI_ISL_3431113, EPI_ISL_3431115, EPI_ISL_3431116, EPI_ISL_3431117, EPI_ISL_3431118, EPI_ISL_3431119, EPI_ISL_3431120, EPI_ISL_3431121, EPI_ISL_3431122, EPI_ISL_3431123, EPI_ISL_3431124, EPI_ISL_3431125, EPI_ISL_3431126, EPI_ISL_3431127, EPI_ISL_3431128, EPI_ISL_3431129, EPI_ISL_3431130, EPI_ISL_3431131, EPI_ISL_3431132, EPI_ISL_3431133, EPI_ISL_3431134, EPI_ISL_3431135, EPI_ISL_3431136, EPI_ISL_3431137, EPI_ISL_3431138, EPI_ISL_3431139, EPI_ISL_3431140, EPI_ISL_3431141, EPI_ISL_3431142, EPI_ISL_3431143, EPI_ISL_3431144, EPI_ISL_3431145, EPI_ISL_3431146, EPI_ISL_3431147, EPI_ISL_3431148, EPI_ISL_3431149, EPI_ISL_3431150, EPI_ISL_3431151, EPI_ISL_3431152, EPI_ISL_3431153, EPI_ISL_3431154, EPI_ISL_3431155, EPI_ISL_3431156, EPI_ISL_3431157, EPI_ISL_3431158, EPI_ISL_3431159, EPI_ISL_3431160, EPI_ISL_3431161, EPI_ISL_3431162, EPI_ISL_3431163, EPI_ISL_3431164, EPI_ISL_3431165, EPI_ISL_3431166, EPI_ISL_3431167, EPI_ISL_3431168, EPI_ISL_3431169, EPI_ISL_3431170, EPI_ISL_3431171, EPI_ISL_3431172, EPI_ISL_3431173, EPI_ISL_3431174, EPI_ISL_3431175, EPI_ISL_3431176, EPI_ISL_3431177, EPI_ISL_3431178, EPI_ISL_3431179, EPI_ISL_3431180, EPI_ISL_3431181, EPI_ISL_3431182, EPI_ISL_3431183, EPI_ISL_3431184, EPI_ISL_3431185, EPI_ISL_3431186, EPI_ISL_3431187, EPI_ISL_3431188, EPI_ISL_3431189, EPI_ISL_3431190, EPI_ISL_3431191, EPI_ISL_3431192, EPI_ISL_3431193, EPI_ISL_3431194, EPI_ISL_3431195, EPI_ISL_3431196, EPI_ISL_3431197, EPI_ISL_3431198, EPI_ISL_3431199, EPI_ISL_3431200, EPI_ISL_3431201, EPI_ISL_3431202, EPI_ISL_3431203, EPI_ISL_3431204, EPI_ISL_3431205, EPI_ISL_3431206, EPI_ISL_3431207, EPI_ISL_3431208, EPI_ISL_3431209, EPI_ISL_3431210, EPI_ISL_3431211, EPI_ISL_3431212, EPI_ISL_3431213, EPI_ISL_3431214, EPI_ISL_3431215, EPI_ISL_3431216, EPI_ISL_3431217, EPI_ISL_3431218, EPI_ISL_3431219, EPI_ISL_3431220, EPI_ISL_3431221, EPI_ISL_3431222, EPI_ISL_3431223, EPI_ISL_3431224, EPI_ISL_3431225, EPI_ISL_3431227, EPI_ISL_3431228, EPI_ISL_3431229, EPI_ISL_3431230, EPI_ISL_3431231, EPI_ISL_3431232, EPI_ISL_3431233, EPI_ISL_3431234, EPI_ISL_3431235, EPI_ISL_3431236, EPI_ISL_3431237, EPI_ISL_3431238, EPI_ISL_3431239, EPI_ISL_3431240, EPI_ISL_3431241, EPI_ISL_3431243, EPI_ISL_3431244, EPI_ISL_3431245, EPI_ISL_3431246, EPI_ISL_3431247, EPI_ISL_3431249, EPI_ISL_3431250, EPI_ISL_3431251, EPI_ISL_3431252, EPI_ISL_3431253, EPI_ISL_3431254, EPI_ISL_3431255, EPI_ISL_3431256, EPI_ISL_3431257, EPI_ISL_3431258, EPI_ISL_3431259, EPI_ISL_3431260, EPI_ISL_3431261, EPI_ISL_3431262, EPI_ISL_3431263, EPI_ISL_3431264, EPI_ISL_3431265, EPI_ISL_3431266, EPI_ISL_3431267, EPI_ISL_3431268, EPI_ISL_3431269, EPI_ISL_3431270, EPI_ISL_3431271, EPI_ISL_3431272, EPI_ISL_3431273, EPI_ISL_3431274, EPI_ISL_3431275, EPI_ISL_3431276, EPI_ISL_3431277, EPI_ISL_3431278, EPI_ISL_3431279, EPI_ISL_3431280, EPI_ISL_3431281, EPI_ISL_3431282, EPI_ISL_3431283, EPI_ISL_3431284, EPI_ISL_3431285, EPI_ISL_3431286, EPI_ISL_3431287, EPI_ISL_3431289, EPI_ISL_3431290, EPI_ISL_3431291, EPI_ISL_3431292, EPI_ISL_3431293, EPI_ISL_3431294, EPI_ISL_3431295, EPI_ISL_3431296, EPI_ISL_3431297, EPI_ISL_3431298, EPI_ISL_3431299, EPI_ISL_3431300, EPI_ISL_3431301, EPI_ISL_3431302, EPI_ISL_3431303, EPI_ISL_3431304, EPI_ISL_3431305, EPI_ISL_3431306, EPI_ISL_3431307, EPI_ISL_3431308, EPI_ISL_3431309, EPI_ISL_3431310, EPI_ISL_3431311, EPI_ISL_3431312, EPI_ISL_3431313, EPI_ISL_3431314, EPI_ISL_3431315, EPI_ISL_3431316, EPI_ISL_3431317, EPI_ISL_3431318, EPI_ISL_3431319, EPI_ISL_3431320, EPI_ISL_3431321, EPI_ISL_3431322, EPI_ISL_3431323, EPI_ISL_3431324, EPI_ISL_3431325, EPI_ISL_3431326, EPI_ISL_3431327, EPI_ISL_3431328, EPI_ISL_3431329, EPI_ISL_3431330, EPI_ISL_3431331, EPI_ISL_3431332, EPI_ISL_3431333, EPI_ISL_3431334, EPI_ISL_3431335, EPI_ISL_3431336, EPI_ISL_3431337, EPI_ISL_3431338, EPI_ISL_3431339, EPI_ISL_3431340, EPI_ISL_3431341, EPI_ISL_3431342, EPI_ISL_3431343, EPI_ISL_3431344, EPI_ISL_3431345, EPI_ISL_3431346, EPI_ISL_3431347, EPI_ISL_3431348, EPI_ISL_3431349, EPI_ISL_3431351, EPI_ISL_3431352, EPI_ISL_3431353, EPI_ISL_3431354, EPI_ISL_3431355, EPI_ISL_3431356, EPI_ISL_3431357, EPI_ISL_3431358, EPI_ISL_3431359, EPI_ISL_3431360, EPI_ISL_3431361, EPI_ISL_3431362, EPI_ISL_3431363, EPI_ISL_3431364, EPI_ISL_3431365, EPI_ISL_3431366, EPI_ISL_3431367, EPI_ISL_3431368, EPI_ISL_3431369, EPI_ISL_3431370, EPI_ISL_3431371, EPI_ISL_3431372, EPI_ISL_3431373, EPI_ISL_3431374, EPI_ISL_3431375, EPI_ISL_3431376, EPI_ISL_3431377, EPI_ISL_3431378, EPI_ISL_3431379, EPI_ISL_3431380, EPI_ISL_3431381, EPI_ISL_3431382, EPI_ISL_3431383, EPI_ISL_3431384, EPI_ISL_3431385, EPI_ISL_3431386, EPI_ISL_3431387, EPI_ISL_3431388, EPI_ISL_3431389, EPI_ISL_3431390, EPI_ISL_3431391, EPI_ISL_3431393, EPI_ISL_3431394, EPI_ISL_3431395, EPI_ISL_3431396, EPI_ISL_3431397, EPI_ISL_3431398, EPI_ISL_3431399, EPI_ISL_3431400, EPI_ISL_3431401, EPI_ISL_3431402, EPI_ISL_3431403, EPI_ISL_3431404, EPI_ISL_3431405, EPI_ISL_3431406, EPI_ISL_3431407, EPI_ISL_3431408, EPI_ISL_3431409, EPI_ISL_3431410, EPI_ISL_3431411, EPI_ISL_3431412, EPI_ISL_3431413, EPI_ISL_3431414, EPI_ISL_3431415, EPI_ISL_3431416, EPI_ISL_3431417, EPI_ISL_3431418, EPI_ISL_3431419, EPI_ISL_3431420, EPI_ISL_3431421, EPI_ISL_3431422, EPI_ISL_3431423, EPI_ISL_3431424, EPI_ISL_3431425, EPI_ISL_3431426, EPI_ISL_3431427, EPI_ISL_3431428, EPI_ISL_3431429, EPI_ISL_3431430, EPI_ISL_3431431, EPI_ISL_3431432, EPI_ISL_3431433, EPI_ISL_3431434, EPI_ISL_3431435, EPI_ISL_3431436, EPI_ISL_3431437, EPI_ISL_3431438, EPI_ISL_3431439, EPI_ISL_3431440, EPI_ISL_3431441, EPI_ISL_3431442, EPI_ISL_3431443, EPI_ISL_3431444, EPI_ISL_3431445, EPI_ISL_3431446, EPI_ISL_3431447, EPI_ISL_3431448, EPI_ISL_3431449, EPI_ISL_3431450, EPI_ISL_3431451, EPI_ISL_3431452, EPI_ISL_3431453, EPI_ISL_3431454, EPI_ISL_3431455, EPI_ISL_3431456, EPI_ISL_3431457, EPI_ISL_3431458, EPI_ISL_3431459, EPI_ISL_3431460, EPI_ISL_3431461, EPI_ISL_3431462, EPI_ISL_3431463, EPI_ISL_3431464, EPI_ISL_3431465, EPI_ISL_3431466, EPI_ISL_3431467, EPI_ISL_3431468, EPI_ISL_3431469, EPI_ISL_3431470, EPI_ISL_3431471, EPI_ISL_3431472, EPI_ISL_3431473, EPI_ISL_3431474, EPI_ISL_3431475, EPI_ISL_3431476, EPI_ISL_3431477, EPI_ISL_3431478, EPI_ISL_3431479, EPI_ISL_3431480, EPI_ISL_3431481, EPI_ISL_3431482, EPI_ISL_3431483, EPI_ISL_3431484, EPI_ISL_3431485, EPI_ISL_3431486, EPI_ISL_3431488, EPI_ISL_3431489, EPI_ISL_3431490, EPI_ISL_3431491, EPI_ISL_3431492, EPI_ISL_3431493, EPI_ISL_3431494, EPI_ISL_3431496, EPI_ISL_3431497, EPI_ISL_3431498, EPI_ISL_3431499, EPI_ISL_3431500, EPI_ISL_3431501, EPI_ISL_3431502, EPI_ISL_3431503, EPI_ISL_3431504, EPI_ISL_3431505, EPI_ISL_3431506, EPI_ISL_3431507, EPI_ISL_3431508, EPI_ISL_3431509, EPI_ISL_3431510, EPI_ISL_3431511, EPI_ISL_3431512, EPI_ISL_3431513, EPI_ISL_3431514, EPI_ISL_3431515, EPI_ISL_3431516, EPI_ISL_3431517, EPI_ISL_3431518, EPI_ISL_3431519, EPI_ISL_3431520, EPI_ISL_3431521, EPI_ISL_3431523, EPI_ISL_3431524, EPI_ISL_3431525, EPI_ISL_3431526, EPI_ISL_3431527, EPI_ISL_3431528, EPI_ISL_3431529, EPI_ISL_3431530, EPI_ISL_3431531, EPI_ISL_3431532, EPI_ISL_3431533, EPI_ISL_3431534, EPI_ISL_3431535, EPI_ISL_3431536, EPI_ISL_3431537, EPI_ISL_3431538, EPI_ISL_3431539, EPI_ISL_3431540, EPI_ISL_3431541, EPI_ISL_3431542, EPI_ISL_3431543, EPI_ISL_3431544, EPI_ISL_3431545, EPI_ISL_3431546, EPI_ISL_3431547, EPI_ISL_3431548, EPI_ISL_3431549, EPI_ISL_3431550, EPI_ISL_3431551, EPI_ISL_3431552, EPI_ISL_3431553, EPI_ISL_3431554, EPI_ISL_3431555, EPI_ISL_3431556, EPI_ISL_3431557, EPI_ISL_3431558, EPI_ISL_3431559, EPI_ISL_3431560, EPI_ISL_3431561, EPI_ISL_3431562, EPI_ISL_3431563, EPI_ISL_3431564, EPI_ISL_3431565, EPI_ISL_3431566, EPI_ISL_3431567, EPI_ISL_3431568, EPI_ISL_3431569, EPI_ISL_3431570, EPI_ISL_3431571, EPI_ISL_3431572, EPI_ISL_3431573, EPI_ISL_3431574, EPI_ISL_3431575, EPI_ISL_3431576, EPI_ISL_3431577, EPI_ISL_3431578, EPI_ISL_3431579, EPI_ISL_3431580, EPI_ISL_3431581, EPI_ISL_3431582, EPI_ISL_3431583, EPI_ISL_3431584, EPI_ISL_3431585, EPI_ISL_3431586, EPI_ISL_3431587, EPI_ISL_3431588, EPI_ISL_3431589, EPI_ISL_3431590, EPI_ISL_3431591, EPI_ISL_3431592, EPI_ISL_3431593, EPI_ISL_3431594, EPI_ISL_3431595, EPI_ISL_3431596, EPI_ISL_3431597, EPI_ISL_3431598, EPI_ISL_3431599, EPI_ISL_3431600, EPI_ISL_3431601, EPI_ISL_3431602, EPI_ISL_3431603, EPI_ISL_3431604, EPI_ISL_3431605, EPI_ISL_3431606, EPI_ISL_3431607, EPI_ISL_3431608, EPI_ISL_3431609, EPI_ISL_3431610, EPI_ISL_3431611, EPI_ISL_3431612, EPI_ISL_3431613, EPI_ISL_3431614, EPI_ISL_3431615, EPI_ISL_3431616, EPI_ISL_3431617, EPI_ISL_3431618, EPI_ISL_3431619, EPI_ISL_3431620, EPI_ISL_3431621, EPI_ISL_3431622, EPI_ISL_3431623, EPI_ISL_3431624, EPI_ISL_3431625, EPI_ISL_3431626, EPI_ISL_3431627, EPI_ISL_3431628, EPI_ISL_3431629, EPI_ISL_3431630, EPI_ISL_3431631, EPI_ISL_3431632, EPI_ISL_3431633, EPI_ISL_3431634, EPI_ISL_3431635, EPI_ISL_3431636, EPI_ISL_3431637, EPI_ISL_3431638, EPI_ISL_3431639, EPI_ISL_3431640, EPI_ISL_3431641, EPI_ISL_3431642, EPI_ISL_3431643, EPI_ISL_3431644, EPI_ISL_3431645, EPI_ISL_3431646, EPI_ISL_3431647, EPI_ISL_3431648, EPI_ISL_3431649, EPI_ISL_3431650, EPI_ISL_3431651, EPI_ISL_3431652, EPI_ISL_3431653, EPI_ISL_3431654, EPI_ISL_3431655, EPI_ISL_3431656, EPI_ISL_3431657, EPI_ISL_3431658, EPI_ISL_3431659, EPI_ISL_3431660, EPI_ISL_3431661, EPI_ISL_3431662, EPI_ISL_3431663, EPI_ISL_3431664, EPI_ISL_3431665, EPI_ISL_3431666, EPI_ISL_3431667, EPI_ISL_3431668, EPI_ISL_3431669, EPI_ISL_3431670, EPI_ISL_3431671, EPI_ISL_3431672, EPI_ISL_3431673, EPI_ISL_3431674, EPI_ISL_3431675, EPI_ISL_3431676, EPI_ISL_3431677, EPI_ISL_3431678, EPI_ISL_3431679, EPI_ISL_3431680, EPI_ISL_3431681 | | | |
| see above | Broad Institute Clinical Research Sequencing Platform | Infectious Disease Program, Broad Institute of Harvard and MIT | Siddle,K.J., Adams,G., Pearlman,L., Gladden-Young,A., Vicente,G., Blumenstiel,B., DeFelice,M., Lee,M., McGovern,S., Lagerborg,K., Rudy,M., DeRuff,K., Carter,A., Normandin,E., Bauer,M., Reilly,S., Tomkins-Tinch,C., Loreth,C., Chaluvadi,S., Meldrim,J., Granger,B., Lemieux,J.E., Birren,B.W., Sabeti,P.C., Larkin,K., Dodge,S., Lennon,N., Madoff,L., Brown,C., Gallagher,G., Smole,S., Park,D.J., Gabriel,S., and MacInnis,B.L. |
| EPI_ISL_3431682, EPI_ISL_3431683, EPI_ISL_3431684, EPI_ISL_3431685, EPI_ISL_3431686, EPI_ISL_3431687, EPI_ISL_3431688, EPI_ISL_3431689, EPI_ISL_3431690, EPI_ISL_3431691, EPI_ISL_3431692, EPI_ISL_3431693, EPI_ISL_3431694, EPI_ISL_3431695, EPI_ISL_3431696, EPI_ISL_3431697, EPI_ISL_3431698, EPI_ISL_3431699, EPI_ISL_3431700, EPI_ISL_3431701, EPI_ISL_3431702, EPI_ISL_3431703, EPI_ISL_3431704, EPI_ISL_3431705, EPI_ISL_3431706, EPI_ISL_3431707, EPI_ISL_3431708, EPI_ISL_3431709, EPI_ISL_3431710, EPI_ISL_3431711, EPI_ISL_3431712, EPI_ISL_3431713, EPI_ISL_3431714, EPI_ISL_3431715, EPI_ISL_3431716, EPI_ISL_3431717, EPI_ISL_3431718, EPI_ISL_3431719, EPI_ISL_3431720, EPI_ISL_3431721, EPI_ISL_3431722, EPI_ISL_3431723, EPI_ISL_3431724, EPI_ISL_3431725, EPI_ISL_3431726, EPI_ISL_3431727 | | | |
| see above | Orebro University Hospital, Dept Laboratory Medicine, Clinical Microbiology | Orebro University Hospital | Sundqvist M et al |
| EPI_ISL_3431731, EPI_ISL_3431732 | HOSPITAL CAN MISSES | HOSPITAL UNIVERSITARIO SON ESPASES | Dr. Antonio Oliver, Dr. Carla López-Causapé, Dr. Gabriel Cabot, on behalf of Servicio de Microbiología, Hospital Universitario Son Espases |
| EPI_ISL_3431733 | HOSPITAL UNIVERSITARIO SON LLATZER | HOSPITAL UNIVERSITARIO SON ESPASES | Dr. Antonio Oliver, Dr. Carla López-Causapé, Dr. Gabriel Cabot, on behalf of Servicio de Microbiología, Hospital Universitario Son Espases |
| EPI_ISL_3431734 | HOSPITAL MATEU ORFILA | HOSPITAL UNIVERSITARIO SON ESPASES | Dr. Antonio Oliver, Dr. Carla López-Causapé, Dr. Gabriel Cabot, on behalf of Servicio de Microbiología, Hospital Universitario Son Espases |
| EPI_ISL_3431735 | HOSPITAL MANACOR | HOSPITAL UNIVERSITARIO SON ESPASES | Dr. Antonio Oliver, Dr. Carla López-Causapé, Dr. Gabriel Cabot, on behalf of Servicio de Microbiología, Hospital Universitario Son Espases |
| EPI_ISL_3431736, EPI_ISL_3431737 | HOSPITAL UNIVERSITARIO SON ESPASES | HOSPITAL UNIVERSITARIO SON ESPASES | Dr. Antonio Oliver, Dr. Carla López-Causapé, Dr. Gabriel Cabot, on behalf of Servicio de Microbiología, Hospital Universitario Son Espases |
| EPI_ISL_3431738 | HOSPITAL CAN MISSES | HOSPITAL UNIVERSITARIO SON ESPASES | Dr. Antonio Oliver, Dr. Carla López-Causapé, Dr. Gabriel Cabot, on behalf of Servicio de Microbiología, Hospital Universitario Son Espases |
| EPI_ISL_3431739 | HOSPITAL UNIVERSITARIO SON ESPASES | HOSPITAL UNIVERSITARIO SON ESPASES | Dr. Antonio Oliver, Dr. Carla López-Causapé, Dr. Gabriel Cabot, on behalf of Servicio de Microbiología, Hospital Universitario Son Espases |
| EPI_ISL_3431740 | HOSPITAL CAN MISSES | HOSPITAL UNIVERSITARIO SON ESPASES | Dr. Antonio Oliver, Dr. Carla López-Causapé, Dr. Gabriel Cabot, on behalf of Servicio de Microbiología, Hospital Universitario Son Espases |
| EPI_ISL_3431741, EPI_ISL_3431742 | HOSPITAL UNIVERSITARIO SON ESPASES | HOSPITAL UNIVERSITARIO SON ESPASES | Dr. Antonio Oliver, Dr. Carla López-Causapé, Dr. Gabriel Cabot, on behalf of Servicio de Microbiología, Hospital Universitario Son Espases |
| EPI_ISL_3431743 | HOSPITAL CAN MISSES | HOSPITAL UNIVERSITARIO SON ESPASES | Dr. Antonio Oliver, Dr. Carla López-Causapé, Dr. Gabriel Cabot, on behalf of Servicio de Microbiología, Hospital Universitario Son Espases |
| EPI_ISL_3431744 | HOSPITAL UNIVERSITARIO SON ESPASES | HOSPITAL UNIVERSITARIO SON ESPASES | Dr. Antonio Oliver, Dr. Carla López-Causapé, Dr. Gabriel Cabot, on behalf of Servicio de Microbiología, Hospital Universitario Son Espases |
| EPI_ISL_3431745, EPI_ISL_3431746, EPI_ISL_3431747 | HOSPITAL CAN MISSES | HOSPITAL UNIVERSITARIO SON ESPASES | Dr. Antonio Oliver, Dr. Carla López-Causapé, Dr. Gabriel Cabot, on behalf of Servicio de Microbiología, Hospital Universitario Son Espases |
| EPI_ISL_3431748, EPI_ISL_3431749 | HOSPITAL UNIVERSITARIO SON ESPASES | HOSPITAL UNIVERSITARIO SON ESPASES | Dr. Antonio Oliver, Dr. Carla López-Causapé, Dr. Gabriel Cabot, on behalf of Servicio de Microbiología, Hospital Universitario Son Espases |
| EPI_ISL_3431750 | HOSPITAL CAN MISSES | HOSPITAL UNIVERSITARIO SON ESPASES | Dr. Antonio Oliver, Dr. Carla López-Causapé, Dr. Gabriel Cabot, on behalf of Servicio de Microbiología, Hospital Universitario Son Espases |
| EPI_ISL_3431751 | HOSPITAL MANACOR | HOSPITAL UNIVERSITARIO SON ESPASES | Dr. Antonio Oliver, Dr. Carla López-Causapé, Dr. Gabriel Cabot, on behalf of Servicio de Microbiología, Hospital Universitario Son Espases |
| EPI_ISL_3431752, EPI_ISL_3431753, EPI_ISL_3431754 | HOSPITAL UNIVERSITARIO SON ESPASES | HOSPITAL UNIVERSITARIO SON ESPASES | Dr. Antonio Oliver, Dr. Carla López-Causapé, Dr. Gabriel Cabot, on behalf of Servicio de Microbiología, Hospital Universitario Son Espases |
| EPI_ISL_3431755 | HOSPITAL MANACOR | HOSPITAL UNIVERSITARIO SON ESPASES | Dr. Antonio Oliver, Dr. Carla López-Causapé, Dr. Gabriel Cabot, on behalf of Servicio de Microbiología, Hospital Universitario Son Espases |
| EPI_ISL_3431756, EPI_ISL_3431757, EPI_ISL_3431758, EPI_ISL_3431759, EPI_ISL_3431760, EPI_ISL_3431761, EPI_ISL_3431762 | HOSPITAL CAN MISSES | HOSPITAL UNIVERSITARIO SON ESPASES | Dr. Antonio Oliver, Dr. Carla López-Causapé, Dr. Gabriel Cabot, on behalf of Servicio de Microbiología, Hospital Universitario Son Espases |
| EPI_ISL_3431763 | HOSPITAL MATEU ORFILA | HOSPITAL UNIVERSITARIO SON ESPASES | Dr. Antonio Oliver, Dr. Carla López-Causapé, Dr. Gabriel Cabot, on behalf of Servicio de Microbiología, Hospital Universitario Son Espases |
| EPI_ISL_3431764, EPI_ISL_3431765, EPI_ISL_3431766 | HOSPITAL UNIVERSITARIO SON ESPASES | HOSPITAL UNIVERSITARIO SON ESPASES | Dr. Antonio Oliver, Dr. Carla López-Causapé, Dr. Gabriel Cabot, on behalf of Servicio de Microbiología, Hospital Universitario Son Espases |

| | | | |
|---|---|---|---|
| EPI_ISL_3431767 | HOSPITAL UNIVERSITARIO SON LLATZER | HOSPITAL UNIVERSITARIO SON ESPASES | Dr. Antonio Oliver, Dr. Carla López-Causapé, Dr. Gabriel Cabot, on behalf of Servicio de Microbiología, Hospital Universitario Son Espases |
| EPI_ISL_3431768 | HOSPITAL MANACOR | HOSPITAL UNIVERSITARIO SON ESPASES | Dr. Antonio Oliver, Dr. Carla López-Causapé, Dr. Gabriel Cabot, on behalf of Servicio de Microbiología, Hospital Universitario Son Espases |
| EPI_ISL_3431769, EPI_ISL_3431770 | HOSPITAL UNIVERSITARIO SON ESPASES | HOSPITAL UNIVERSITARIO SON ESPASES | Dr. Antonio Oliver, Dr. Carla López-Causapé, Dr. Gabriel Cabot, on behalf of Servicio de Microbiología, Hospital Universitario Son Espases |
| EPI_ISL_3431771 | HOSPITAL MATEU ORFILA | HOSPITAL UNIVERSITARIO SON ESPASES | Dr. Antonio Oliver, Dr. Carla López-Causapé, Dr. Gabriel Cabot, on behalf of Servicio de Microbiología, Hospital Universitario Son Espases |
| EPI_ISL_3431772 | HOSPITAL UNIVERSITARIO SON ESPASES | HOSPITAL UNIVERSITARIO SON ESPASES | Dr. Antonio Oliver, Dr. Carla López-Causapé, Dr. Gabriel Cabot, on behalf of Servicio de Microbiología, Hospital Universitario Son Espases |
| EPI_ISL_3431773, EPI_ISL_3431774 | HOSPITAL MANACOR | HOSPITAL UNIVERSITARIO SON ESPASES | Dr. Antonio Oliver, Dr. Carla López-Causapé, Dr. Gabriel Cabot, on behalf of Servicio de Microbiología, Hospital Universitario Son Espases |
| EPI_ISL_3431775 | HOSPITAL UNIVERSITARIO SON ESPASES | HOSPITAL UNIVERSITARIO SON ESPASES | Dr. Antonio Oliver, Dr. Carla López-Causapé, Dr. Gabriel Cabot, on behalf of Servicio de Microbiología, Hospital Universitario Son Espases |
| EPI_ISL_3431776, EPI_ISL_3431777 | HOSPITAL CAN MISSES | HOSPITAL UNIVERSITARIO SON ESPASES | Dr. Antonio Oliver, Dr. Carla López-Causapé, Dr. Gabriel Cabot, on behalf of Servicio de Microbiología, Hospital Universitario Son Espases |
| EPI_ISL_3431778, EPI_ISL_3431779, EPI_ISL_3431780 | HOSPITAL UNIVERSITARIO SON ESPASES | HOSPITAL UNIVERSITARIO SON ESPASES | Dr. Antonio Oliver, Dr. Carla López-Causapé, Dr. Gabriel Cabot, on behalf of Servicio de Microbiología, Hospital Universitario Son Espases |
| EPI_ISL_3431781 | HOSPITAL CAN MISSES | HOSPITAL UNIVERSITARIO SON ESPASES | Dr. Antonio Oliver, Dr. Carla López-Causapé, Dr. Gabriel Cabot, on behalf of Servicio de Microbiología, Hospital Universitario Son Espases |
| EPI_ISL_3431782 | HOSPITAL MATEU ORFILA | HOSPITAL UNIVERSITARIO SON ESPASES | Dr. Antonio Oliver, Dr. Carla López-Causapé, Dr. Gabriel Cabot, on behalf of Servicio de Microbiología, Hospital Universitario Son Espases |
| EPI_ISL_3431783, EPI_ISL_3431784 | HOSPITAL UNIVERSITARIO SON ESPASES | HOSPITAL UNIVERSITARIO SON ESPASES | Dr. Antonio Oliver, Dr. Carla López-Causapé, Dr. Gabriel Cabot, on behalf of Servicio de Microbiología, Hospital Universitario Son Espases |
| EPI_ISL_3431785 | HOSPITAL CAN MISSES | HOSPITAL UNIVERSITARIO SON ESPASES | Dr. Antonio Oliver, Dr. Carla López-Causapé, Dr. Gabriel Cabot, on behalf of Servicio de Microbiología, Hospital Universitario Son Espases |
| EPI_ISL_3431786 | HOSPITAL MANACOR | HOSPITAL UNIVERSITARIO SON ESPASES | Dr. Antonio Oliver, Dr. Carla López-Causapé, Dr. Gabriel Cabot, on behalf of Servicio de Microbiología, Hospital Universitario Son Espases |
| EPI_ISL_3431787 | HOSPITAL UNIVERSITARIO SON ESPASES | HOSPITAL UNIVERSITARIO SON ESPASES | Dr. Antonio Oliver, Dr. Carla López-Causapé, Dr. Gabriel Cabot, on behalf of Servicio de Microbiología, Hospital Universitario Son Espases |
| EPI_ISL_3431788, EPI_ISL_3431789 | HOSPITAL CAN MISSES | HOSPITAL UNIVERSITARIO SON ESPASES | Dr. Antonio Oliver, Dr. Carla López-Causapé, Dr. Gabriel Cabot, on behalf of Servicio de Microbiología, Hospital Universitario Son Espases |
| EPI_ISL_3431790 | HOSPITAL UNIVERSITARIO SON ESPASES | HOSPITAL UNIVERSITARIO SON ESPASES | Dr. Antonio Oliver, Dr. Carla López-Causapé, Dr. Gabriel Cabot, on behalf of Servicio de Microbiología, Hospital Universitario Son Espases |
| EPI_ISL_3431791, EPI_ISL_3431792 | HOSPITAL MANACOR | HOSPITAL UNIVERSITARIO SON ESPASES | Dr. Antonio Oliver, Dr. Carla López-Causapé, Dr. Gabriel Cabot, on behalf of Servicio de Microbiología, Hospital Universitario Son Espases |
| EPI_ISL_3431793 | HOSPITAL MATEU ORFILA | HOSPITAL UNIVERSITARIO SON ESPASES | Dr. Antonio Oliver, Dr. Carla López-Causapé, Dr. Gabriel Cabot, on behalf of Servicio de Microbiología, Hospital Universitario Son Espases |
| EPI_ISL_3431794 | HOSPITAL UNIVERSITARIO SON ESPASES | HOSPITAL UNIVERSITARIO SON ESPASES | Dr. Antonio Oliver, Dr. Carla López-Causapé, Dr. Gabriel Cabot, on behalf of Servicio de Microbiología, Hospital Universitario Son Espases |
| EPI_ISL_3431795 | HOSPITAL MANACOR | HOSPITAL UNIVERSITARIO SON ESPASES | Dr. Antonio Oliver, Dr. Carla López-Causapé, Dr. Gabriel Cabot, on behalf of Servicio de Microbiología, Hospital Universitario Son Espases |
| EPI_ISL_3431796 | HOSPITAL UNIVERSITARIO SON ESPASES | HOSPITAL UNIVERSITARIO SON ESPASES | Dr. Antonio Oliver, Dr. Carla López-Causapé, Dr. Gabriel Cabot, on behalf of Servicio de Microbiología, Hospital Universitario Son Espases |
| EPI_ISL_3431799 | HOSPITAL MANACOR | HOSPITAL UNIVERSITARIO SON ESPASES | Dr. Antonio Oliver, Dr. Carla López-Causapé, Dr. Gabriel Cabot, on behalf of Servicio de Microbiología, Hospital Universitario Son Espases |
| EPI_ISL_3431800 | HOSPITAL CAN MISSES | HOSPITAL UNIVERSITARIO SON ESPASES | Dr. Antonio Oliver, Dr. Carla López-Causapé, Dr. Gabriel Cabot, on behalf of Servicio de Microbiología, Hospital Universitario Son Espases |
| EPI_ISL_3431801 | HOSPITAL UNIVERSITARIO SON ESPASES | HOSPITAL UNIVERSITARIO SON ESPASES | Dr. Antonio Oliver, Dr. Carla López-Causapé, Dr. Gabriel Cabot, on behalf of Servicio de Microbiología, Hospital Universitario Son Espases |
| EPI_ISL_3431802 | HOSPITAL MANACOR | HOSPITAL UNIVERSITARIO SON ESPASES | Dr. Antonio Oliver, Dr. Carla López-Causapé, Dr. Gabriel Cabot, on behalf of Servicio de Microbiología, Hospital Universitario Son Espases |
| EPI_ISL_3431803 | HOSPITAL UNIVERSITARIO SON LLATZER | HOSPITAL UNIVERSITARIO SON ESPASES | Dr. Antonio Oliver, Dr. Carla López-Causapé, Dr. Gabriel Cabot, on behalf of Servicio de Microbiología, Hospital Universitario Son Espases |
| EPI_ISL_3431804, EPI_ISL_3431806 | HOSPITAL UNIVERSITARIO SON ESPASES | HOSPITAL UNIVERSITARIO SON ESPASES | Dr. Antonio Oliver, Dr. Carla López-Causapé, Dr. Gabriel Cabot, on behalf of Servicio de Microbiología, Hospital Universitario Son Espases |
| EPI_ISL_3431807 | HOSPITAL CAN MISSES | HOSPITAL UNIVERSITARIO SON ESPASES | Dr. Antonio Oliver, Dr. Carla López-Causapé, Dr. Gabriel Cabot, on behalf of Servicio de Microbiología, Hospital Universitario Son Espases |
| EPI_ISL_3431808 | HOSPITAL MATEU ORFILA | HOSPITAL UNIVERSITARIO SON ESPASES | Dr. Antonio Oliver, Dr. Carla López-Causapé, Dr. Gabriel Cabot, on behalf of Servicio de Microbiología, Hospital Universitario Son Espases |
| EPI_ISL_3431809, EPI_ISL_3431810 | HOSPITAL UNIVERSITARIO SON ESPASES | HOSPITAL UNIVERSITARIO SON ESPASES | Dr. Antonio Oliver, Dr. Carla López-Causapé, Dr. Gabriel Cabot, on behalf of Servicio de Microbiología, Hospital Universitario Son Espases |
| EPI_ISL_3431811 | HOSPITAL UNIVERSITARIO SON LLATZER | HOSPITAL UNIVERSITARIO SON ESPASES | Dr. Antonio Oliver, Dr. Carla López-Causapé, Dr. Gabriel Cabot, on behalf of Servicio de Microbiología, Hospital Universitario Son Espases |
| EPI_ISL_3431812 | Cliniques universitaires Saint-Luc | UCLouvain/IREC/MBLG | Jean Ruelle, Bertrand Bearzatto, Benoit Kabamba Mukadi |
| EPI_ISL_3431813 | Department of Virology, AP-HP Avicenne hospital | Department of Virology, Pitié-Salpêtrière hospital | Stéphane Marot, Valentin Leducq, Karen Zafilaza, Vincent Calvez, Anne-Geneviève Marcelin, Aude Jary |
| EPI_ISL_3431832, EPI_ISL_3431833, EPI_ISL_3431834, EPI_ISL_3431835, EPI_ISL_3431836, EPI_ISL_3431837 | ULS Norte Alentejano | Instituto Nacional de Saude (INSA) | Borges et al |
| EPI_ISL_3431838, EPI_ISL_3431839, EPI_ISL_3431840, EPI_ISL_3431841, EPI_ISL_3431842, EPI_ISL_3431843, EPI_ISL_3431844, EPI_ISL_3431845, EPI_ISL_3431846, EPI_ISL_3431847, EPI_ISL_3431848, EPI_ISL_3431849, EPI_ISL_3431850, EPI_ISL_3431851, EPI_ISL_3431852, EPI_ISL_3431853, EPI_ISL_3431854, EPI_ISL_3431855, EPI_ISL_3431856, EPI_ISL_3431857, EPI_ISL_3431858, EPI_ISL_3431859 | | | |
| see above | SYNLAB | Instituto Nacional de Saude (INSA) | Borges et al |
| EPI_ISL_3431860, EPI_ISL_3431861, EPI_ISL_3431862, EPI_ISL_3431863, EPI_ISL_3431864, EPI_ISL_3431865, EPI_ISL_3431866, EPI_ISL_3431867, EPI_ISL_3431868, EPI_ISL_3431869, EPI_ISL_3431870, EPI_ISL_3431871, EPI_ISL_3431872, EPI_ISL_3431873, EPI_ISL_3431874, EPI_ISL_3431875, EPI_ISL_3431876, EPI_ISL_3431877 | | | |
| see above | H Santarem | Instituto Nacional de Saude (INSA) | Borges et al |
| EPI_ISL_3431878, EPI_ISL_3431879, EPI_ISL_3431880, EPI_ISL_3431881, EPI_ISL_3431882, EPI_ISL_3431883, EPI_ISL_3431884, EPI_ISL_3431885, EPI_ISL_3431886, EPI_ISL_3431887, EPI_ISL_3431888, EPI_ISL_3431889, EPI_ISL_3431890, EPI_ISL_3431891, EPI_ISL_3431892 | | | |
| see above | H Evora | Instituto Nacional de Saude (INSA) | Borges et al |
| EPI_ISL_3431893, EPI_ISL_3431894, EPI_ISL_3431895, EPI_ISL_3431896, EPI_ISL_3431897, EPI_ISL_3431898, EPI_ISL_3431899, EPI_ISL_3431900, EPI_ISL_3431901, EPI_ISL_3431902, EPI_ISL_3431903, EPI_ISL_3431904, EPI_ISL_3431905, EPI_ISL_3431906, EPI_ISL_3431907, EPI_ISL_3431908, EPI_ISL_3431909, EPI_ISL_3431910, EPI_ISL_3431911 | | | |
| see above | ULS Castelo Branco | Instituto Nacional de Saude (INSA) | Borges et al |
| EPI_ISL_3431912, EPI_ISL_3431913, EPI_ISL_3431914, EPI_ISL_3431915, EPI_ISL_3431916, EPI_ISL_3431917, EPI_ISL_3431918, EPI_ISL_3431919, EPI_ISL_3431920 | H Vila Franca Xira | Instituto Nacional de Saude (INSA) | Borges et al |
| EPI_ISL_3431921, EPI_ISL_3431922, EPI_ISL_3431923, EPI_ISL_3431924, EPI_ISL_3431925, EPI_ISL_3431926, EPI_ISL_3431927, EPI_ISL_3431928, EPI_ISL_3431929, EPI_ISL_3431930, EPI_ISL_3431931, EPI_ISL_3431932, EPI_ISL_3431933, EPI_ISL_3431934, EPI_ISL_3431935, EPI_ISL_3431936 | | | |
| see above | CHLC | Instituto Nacional de Saude (INSA) | Borges et al |
| EPI_ISL_3431937, EPI_ISL_3431938, EPI_ISL_3431939, EPI_ISL_3431940, EPI_ISL_3431941, EPI_ISL_3431942, EPI_ISL_3431943, EPI_ISL_3431944, EPI_ISL_3431945, EPI_ISL_3431946, EPI_ISL_3431947, EPI_ISL_3431948, EPI_ISL_3431949, EPI_ISL_3431950 | | | |
| see above | ULS Litoral Alentejano | Instituto Nacional de Saude (INSA) | Borges et al |
| EPI_ISL_3431951, EPI_ISL_3431952, EPI_ISL_3431953, EPI_ISL_3431954, EPI_ISL_3431955, EPI_ISL_3431956, EPI_ISL_3431957, EPI_ISL_3431958, EPI_ISL_3431959, EPI_ISL_3431960, EPI_ISL_3431961 | | | |
| see above | H Fernando Fonseca | Instituto Nacional de Saude (INSA) | Borges et al |
| EPI_ISL_3431962, EPI_ISL_3431963 | CH Setubal | Instituto Nacional de Saude (INSA) | Borges et al |
| EPI_ISL_3431964, EPI_ISL_3431965, EPI_ISL_3431966, EPI_ISL_3431967, EPI_ISL_3431968, EPI_ISL_3431969, EPI_ISL_3431970, EPI_ISL_3431971, EPI_ISL_3431972, EPI_ISL_3431973 | ARS Algarve - Laboratorio Laura Ayres | Instituto Nacional de Saude (INSA) | Borges et al |
| EPI_ISL_3431974 | Department of Virology, Pitié-Salpêtrière hospital | Department of Virology, Pitié-Salpêtrière hospital | Stéphane Marot, Valentin Leducq, Karen Zafilaza, Vincent Calvez, Anne-Geneviève Marcelin, Aude Jary |
| EPI_ISL_3431975, EPI_ISL_3431976, EPI_ISL_3431977, EPI_ISL_3431978, EPI_ISL_3431979, EPI_ISL_3431980, EPI_ISL_3431981, EPI_ISL_3431982, EPI_ISL_3431983, EPI_ISL_3431984, EPI_ISL_3431985, EPI_ISL_3431986, EPI_ISL_3431987, EPI_ISL_3431988, EPI_ISL_3431989, EPI_ISL_3431990, EPI_ISL_3431991, EPI_ISL_3431992, EPI_ISL_3431993, EPI_ISL_3431994, EPI_ISL_3431995, EPI_ISL_3431996, | | | |

| | | | |
|---|---|---|---|
| EPI_ISL_3431997, EPI_ISL_3431998, EPI_ISL_3431999, EPI_ISL_3432000, EPI_ISL_3432001, EPI_ISL_3432002, EPI_ISL_3432003, EPI_ISL_3432004, EPI_ISL_3432005, EPI_ISL_3432006 | | | |
| see above | ARS Algarve - Laboratorio Laura Ayres | Instituto Nacional de Saude (INSA) | Borges et al |
| EPI_ISL_3432007, EPI_ISL_3432008, EPI_ISL_3432009, EPI_ISL_3432010, EPI_ISL_3432011, EPI_ISL_3432012, EPI_ISL_3432013, EPI_ISL_3432014, EPI_ISL_3432015, EPI_ISL_3432016, EPI_ISL_3432017 | | | |
| see above | CH Tondela Viseu | Instituto Nacional de Saude (INSA) | Borges et al |
| EPI_ISL_3432018, EPI_ISL_3432019, EPI_ISL_3432020, EPI_ISL_3432021, EPI_ISL_3432022, EPI_ISL_3432023, EPI_ISL_3432024, EPI_ISL_3432025, EPI_ISL_3432026, EPI_ISL_3432027, EPI_ISL_3432028, EPI_ISL_3432029 | | | |
| see above | CH Porto - H Sto Antonio | Instituto Nacional de Saude (INSA) | Borges et al |
| EPI_ISL_3432030 | CHEDV | Instituto Nacional de Saude (INSA) | Borges et al |
| EPI_ISL_3432031 | CH Porto - H Sto Antonio | Instituto Nacional de Saude (INSA) | Borges et al |
| EPI_ISL_3432032, EPI_ISL_3432033, EPI_ISL_3432034, EPI_ISL_3432035, EPI_ISL_3432036, EPI_ISL_3432037, EPI_ISL_3432038, EPI_ISL_3432039, EPI_ISL_3432040, EPI_ISL_3432041 | CHEDV | Instituto Nacional de Saude (INSA) | Borges et al |
| EPI_ISL_3432042, EPI_ISL_3432043, EPI_ISL_3432044, EPI_ISL_3432045, EPI_ISL_3432046, EPI_ISL_3432047, EPI_ISL_3432048, EPI_ISL_3432049, EPI_ISL_3432050, EPI_ISL_3432051, EPI_ISL_3432052, EPI_ISL_3432053, EPI_ISL_3432054, EPI_ISL_3432055, EPI_ISL_3432056 | | | |
| see above | CHU Sao Joao, Porto | Instituto Nacional de Saude (INSA) | Borges et al |
| EPI_ISL_3432057, EPI_ISL_3432058 | ARS Alentejo | Instituto Nacional de Saude (INSA) | Borges et al |
| EPI_ISL_3432059, EPI_ISL_3432060, EPI_ISL_3432061, EPI_ISL_3432062, EPI_ISL_3432063, EPI_ISL_3432064, EPI_ISL_3432065, EPI_ISL_3432066, EPI_ISL_3432067, EPI_ISL_3432068, EPI_ISL_3432069, EPI_ISL_3432070, EPI_ISL_3432071, EPI_ISL_3432072 | | | |
| see above | Labeto - CAB - Leiria | Instituto Nacional de Saude (INSA) | Borges et al |
| EPI_ISL_3432073, EPI_ISL_3432074, EPI_ISL_3432075, EPI_ISL_3432076, EPI_ISL_3432077, EPI_ISL_3432078, EPI_ISL_3432079, EPI_ISL_3432080, EPI_ISL_3432081, EPI_ISL_3432082, EPI_ISL_3432083, EPI_ISL_3432084, EPI_ISL_3432085, EPI_ISL_3432086, EPI_ISL_3432087, EPI_ISL_3432088, EPI_ISL_3432089, EPI_ISL_3432090, EPI_ISL_3432091, EPI_ISL_3432092, EPI_ISL_3432093, EPI_ISL_3432094, EPI_ISL_3432095, EPI_ISL_3432096, EPI_ISL_3432097 | | | |
| see above | H Divino Espirito Santo - Ponta Delgada | Instituto Nacional de Saude (INSA) | Borges et al |
| EPI_ISL_3432098, EPI_ISL_3432099, EPI_ISL_3432100, EPI_ISL_3432101, EPI_ISL_3432102, EPI_ISL_3432103, EPI_ISL_3432104, EPI_ISL_3432105, EPI_ISL_3432106, EPI_ISL_3432107, EPI_ISL_3432108, EPI_ISL_3432109, EPI_ISL_3432110, EPI_ISL_3432111, EPI_ISL_3432112, EPI_ISL_3432113, EPI_ISL_3432114, EPI_ISL_3432115, EPI_ISL_3432116, EPI_ISL_3432117, EPI_ISL_3432118, EPI_ISL_3432119, EPI_ISL_3432120, EPI_ISL_3432121, EPI_ISL_3432122 | | | |
| see above | SEEBMO | Instituto Nacional de Saude (INSA) | Borges et al |
| EPI_ISL_3432123, EPI_ISL_3432124, EPI_ISL_3432125, EPI_ISL_3432126, EPI_ISL_3432127, EPI_ISL_3432128, EPI_ISL_3432129, EPI_ISL_3432130, EPI_ISL_3432131, EPI_ISL_3432132, EPI_ISL_3432134, EPI_ISL_3432135, EPI_ISL_3432136, EPI_ISL_3432137 | | | |
| see above | CHTMAD | Instituto Nacional de Saude (INSA) | Borges et al |
| EPI_ISL_3432138, EPI_ISL_3432139, EPI_ISL_3432140, EPI_ISL_3432141, EPI_ISL_3432142 | H Guimaraes | Instituto Nacional de Saude (INSA) | Borges et al |
| EPI_ISL_3432143, EPI_ISL_3432144, EPI_ISL_3432145, EPI_ISL_3432146, EPI_ISL_3432148, EPI_ISL_3432149, EPI_ISL_3432150, EPI_ISL_3432151, EPI_ISL_3432152, EPI_ISL_3432153, EPI_ISL_3432154, EPI_ISL_3432155, EPI_ISL_3432156 | | | |
| see above | H Braga | Instituto Nacional de Saude (INSA) | Borges et al |
| EPI_ISL_3432157, EPI_ISL_3432158, EPI_ISL_3432159, EPI_ISL_3432160, EPI_ISL_3432161, EPI_ISL_3432162, EPI_ISL_3432163, EPI_ISL_3432164, EPI_ISL_3432165, EPI_ISL_3432166, EPI_ISL_3432167, EPI_ISL_3432168, EPI_ISL_3432169, EPI_ISL_3432170, EPI_ISL_3432171 | | | |
| see above | CHULN - H Santa Maria | Instituto Nacional de Saude (INSA) | Borges et al |
| EPI_ISL_3432172, EPI_ISL_3432173, EPI_ISL_3432174, EPI_ISL_3432175, EPI_ISL_3432176, EPI_ISL_3432177, EPI_ISL_3432178, EPI_ISL_3432179, EPI_ISL_3432180, EPI_ISL_3432181, EPI_ISL_3432182, EPI_ISL_3432183, EPI_ISL_3432184, EPI_ISL_3432185, EPI_ISL_3432186, EPI_ISL_3432187, EPI_ISL_3432188, EPI_ISL_3432189, EPI_ISL_3432190, EPI_ISL_3432191, EPI_ISL_3432192, EPI_ISL_3432193, EPI_ISL_3432194, EPI_ISL_3432195, EPI_ISL_3432196, EPI_ISL_3432197, EPI_ISL_3432198, EPI_ISL_3432199, EPI_ISL_3432200, EPI_ISL_3432201, EPI_ISL_3432202, EPI_ISL_3432203, EPI_ISL_3432204, EPI_ISL_3432205, EPI_ISL_3432206 | | | |
| see above | SESARAM | Instituto Nacional de Saude (INSA) | Borges et al |
| EPI_ISL_3432207 | Hospital Universitari Joan XXIII | Hospital Universitari Joan XXIII | Gemma Recio, Cristina Gutiérrez, Clara Benavent, Carla Martín, Margarida Terrón, Ester Picó, Natalia Bastón |
| EPI_ISL_3432208, EPI_ISL_3432209 | Laboratori de Referencia de Catalunya | Laboratori de Referencia de Catalunya | Ramirez A., Bellosillo B., Padilla E., Vilas A., Hernandez JJ., Canal M. |
| EPI_ISL_3432210, EPI_ISL_3432211, EPI_ISL_3432212, EPI_ISL_3432213, EPI_ISL_3432214, EPI_ISL_3432217, EPI_ISL_3432218, EPI_ISL_3432219, EPI_ISL_3432220, EPI_ISL_3432221, EPI_ISL_3432222, EPI_ISL_3432223, EPI_ISL_3432224, EPI_ISL_3432225, EPI_ISL_3432226, EPI_ISL_3432227, EPI_ISL_3432228, EPI_ISL_3432229, EPI_ISL_3432231, EPI_ISL_3432232, EPI_ISL_3432233, EPI_ISL_3432234, EPI_ISL_3432235, EPI_ISL_3432236, EPI_ISL_3432237, EPI_ISL_3432238, EPI_ISL_3432239, EPI_ISL_3432240, EPI_ISL_3432241, EPI_ISL_3432242, EPI_ISL_3432243, EPI_ISL_3432244, EPI_ISL_3432245, EPI_ISL_3432246, EPI_ISL_3432247, EPI_ISL_3432248, EPI_ISL_3432249, EPI_ISL_3432252, EPI_ISL_3432253, EPI_ISL_3432254, EPI_ISL_3432255, EPI_ISL_3432256, EPI_ISL_3432257, EPI_ISL_3432259, EPI_ISL_3432260, EPI_ISL_3432261, EPI_ISL_3432262, EPI_ISL_3432263, EPI_ISL_3432264, EPI_ISL_3432265, EPI_ISL_3432266, EPI_ISL_3432267, EPI_ISL_3432268, EPI_ISL_3432269, EPI_ISL_3432270, EPI_ISL_3432271, EPI_ISL_3432272, EPI_ISL_3432273, EPI_ISL_3432274, EPI_ISL_3432275, EPI_ISL_3432276, EPI_ISL_3432277, EPI_ISL_3432278, EPI_ISL_3432279, EPI_ISL_3432280, EPI_ISL_3432281, EPI_ISL_3432282, EPI_ISL_3432283, EPI_ISL_3432284, EPI_ISL_3432285, EPI_ISL_3432286 | | | |
| see above | Indian Council of Medical Research-National Institute of Virology, Microbial Containment Complex | Indian Council of Medical Research-National Institute of Virology, Microbial Containment Complex | Pragya D Yadav |
| EPI_ISL_3432291, EPI_ISL_3432292, EPI_ISL_3432293, EPI_ISL_3432294, EPI_ISL_3432295, EPI_ISL_3432296, EPI_ISL_3432297, EPI_ISL_3432298, EPI_ISL_3432299, EPI_ISL_3432300, EPI_ISL_3432301, EPI_ISL_3432302, EPI_ISL_3432303, EPI_ISL_3432304, EPI_ISL_3432305, EPI_ISL_3432306, EPI_ISL_3432307, EPI_ISL_3432308, EPI_ISL_3432309, EPI_ISL_3432310, EPI_ISL_3432311, EPI_ISL_3432312, EPI_ISL_3432313, EPI_ISL_3432314 | | | |
| see above | Kansas Health and Environmental Lab | Kansas Health and Environmental Lab | Mike Grose, Jonathan Barnell, Amanda Bradley, and Phil Adam |
| EPI_ISL_3432316, EPI_ISL_3432317, EPI_ISL_3432318, EPI_ISL_3432319, EPI_ISL_3432320, EPI_ISL_3432321, EPI_ISL_3432322, EPI_ISL_3432323, EPI_ISL_3432324, EPI_ISL_3432325, EPI_ISL_3432326, EPI_ISL_3432327, EPI_ISL_3432328, EPI_ISL_3432329 | | | |
| see above | Plateforme de testing Namuroise | Plateforme de testing Namuroise | Lesly Nyinkeu Kemamen: Nicolas Gilliard; Demars Aurore ; Maschietto Céline ; Nobis Chloé ; Otto Gaetan ; Denis Olivier ; Degosserie Jonathan ; Mullier François |
| EPI_ISL_3432330 | Centre Hospitalier Universitaire Clermont-Ferrand | CHU Clermont-Ferrand, service de virologie | Bisseux Maxime, Mirand Audrey, Combes Patricia, Henquell Cécile |
| EPI_ISL_3432334 | HOPITAL DE VICHY | CHU Clermont-Ferrand, service de virologie | Bisseux Maxime, Mirand Audrey, Combes Patricia, Henquell Cécile |
| EPI_ISL_3432335 | Centre hospitalier D'AURILLAC | CHU Clermont-Ferrand, service de virologie | Bisseux Maxime, Mirand Audrey, Combes Patricia, Henquell Cécile |
| EPI_ISL_3432336, EPI_ISL_3432337 | HOPITAL DE VICHY | CHU Clermont-Ferrand, service de virologie | Bisseux Maxime, Mirand Audrey, Combes Patricia, Henquell Cécile |
| EPI_ISL_3432338, EPI_ISL_3432339 | Centre Hospitalier Universitaire Clermont-Ferrand | CHU Clermont-Ferrand, service de virologie | Bisseux Maxime, Mirand Audrey, Combes Patricia, Henquell Cécile |
| EPI_ISL_3432340 | HOPITAL DE MONTLUCON | CHU Clermont-Ferrand, service de virologie | Bisseux Maxime, Mirand Audrey, Combes Patricia, Henquell Cécile |
| EPI_ISL_3432341 | HOPITAL DE VICHY | CHU Clermont-Ferrand, service de virologie | Bisseux Maxime, Mirand Audrey, Combes Patricia, Henquell Cécile |
| EPI_ISL_3432342, EPI_ISL_3432343 | Centre Hospitalier Universitaire Clermont-Ferrand | CHU Clermont-Ferrand, service de virologie | Bisseux Maxime, Mirand Audrey, Combes Patricia, Henquell Cécile |
| EPI_ISL_3432344 | Centre Hospitalier Paul Ardier | CHU Clermont-Ferrand, service de virologie | Bisseux Maxime, Mirand Audrey, Combes Patricia, Henquell Cécile |
| EPI_ISL_3432345 | Centre Hospitalier Universitaire Clermont-Ferrand | CHU Clermont-Ferrand, service de virologie | Bisseux Maxime, Mirand Audrey, Combes Patricia, Henquell Cécile |
| EPI_ISL_3432346 | HOPITAL DE MONTLUCON | CHU Clermont-Ferrand, service de virologie | Bisseux Maxime, Mirand Audrey, Combes Patricia, Henquell Cécile |
| EPI_ISL_3432347, EPI_ISL_3432348, EPI_ISL_3432349 | HOPITAL DE VICHY | CHU Clermont-Ferrand, service de virologie | Bisseux Maxime, Mirand Audrey, Combes Patricia, Henquell Cécile |
| EPI_ISL_3432350 | Centre Hospitalier Universitaire Clermont-Ferrand | CHU Clermont-Ferrand, service de virologie | Bisseux Maxime, Mirand Audrey, Combes Patricia, Henquell Cécile |
| EPI_ISL_3432351 | HOPITAL DE VICHY | CHU Clermont-Ferrand, service de virologie | Bisseux Maxime, Mirand Audrey, Combes Patricia, Henquell Cécile |
| EPI_ISL_3432352, EPI_ISL_3432353, EPI_ISL_3432354, EPI_ISL_3432355, EPI_ISL_3432356, EPI_ISL_3432357 | Centre Hospitalier Universitaire Clermont-Ferrand | CHU Clermont-Ferrand, service de virologie | Bisseux Maxime, Mirand Audrey, Combes Patricia, Henquell Cécile |
| EPI_ISL_3432358, EPI_ISL_3432359 | HOPITAL DU MONT DORE | CHU Clermont-Ferrand, service de virologie | Bisseux Maxime, Mirand Audrey, Combes Patricia, Henquell Cécile |
| EPI_ISL_3432360 | Centre De Prelevement COVID RIOM | CHU Clermont-Ferrand, service de virologie | Bisseux Maxime, Mirand Audrey, Combes Patricia, Henquell Cécile |
| EPI_ISL_3432361 | CDP COVID MONTLUCON | CHU Clermont-Ferrand, service de virologie | Bisseux Maxime, Mirand Audrey, Combes Patricia, Henquell Cécile |

EPI_ISL_3432362, EPI_ISL_3432363 Centre Hospitalier Universitaire Clermont-Ferrand CHU Clermont-Ferrand, service de virologie Bisseux Maxime, Mirand Audrey, Combes Patricia, Henquell Cécile

EPI_ISL_3432365, EPI_ISL_3432366, EPI_ISL_3432367, EPI_ISL_3432368, EPI_ISL_3432369, EPI_ISL_3432370, EPI_ISL_3432371, EPI_ISL_3432372, EPI_ISL_3432373, EPI_ISL_3432374, EPI_ISL_3432375, EPI_ISL_3432377, EPI_ISL_3432379, EPI_ISL_3432380, EPI_ISL_3432383, EPI_ISL_3432384, EPI_ISL_3432386, EPI_ISL_3432387, EPI_ISL_3432388, EPI_ISL_3432389, EPI_ISL_3432390, EPI_ISL_3432391, EPI_ISL_3432392, EPI_ISL_3432393, EPI_ISL_3432394, EPI_ISL_3432395, EPI_ISL_3432396, EPI_ISL_3432397, EPI_ISL_3432398, EPI_ISL_3432399, EPI_ISL_3432400, EPI_ISL_3432401, EPI_ISL_3432402, EPI_ISL_3432403, EPI_ISL_3432404, EPI_ISL_3432405, EPI_ISL_3432406, EPI_ISL_3432407, EPI_ISL_3432408, EPI_ISL_3432411, EPI_ISL_3432412

see above Servicio de Microbiologia Hospital Ramon y Cajal Servicio de Microbiologia Hospital Ramon y Cajal Ponce M, Galan JC, Martinez L. Abreu M, y Gonzalez-Alba JM

EPI_ISL_3432413, EPI_ISL_3432414, EPI_ISL_3432415, EPI_ISL_3432416, EPI_ISL_3432417, EPI_ISL_3432418, EPI_ISL_3432419, EPI_ISL_3432420, EPI_ISL_3432421, EPI_ISL_3432422, EPI_ISL_3432423, EPI_ISL_3432424, EPI_ISL_3432425, EPI_ISL_3432426, EPI_ISL_3432427, EPI_ISL_3432428, EPI_ISL_3432429, EPI_ISL_3432430, EPI_ISL_3432431, EPI_ISL_3432432, EPI_ISL_3432433, EPI_ISL_3432434, EPI_ISL_3432435, EPI_ISL_3432436, EPI_ISL_3432437, EPI_ISL_3432438, EPI_ISL_3432439, EPI_ISL_3432440, EPI_ISL_3432445, EPI_ISL_3432446, EPI_ISL_3432447, EPI_ISL_3432448, EPI_ISL_3432449, EPI_ISL_3432450, EPI_ISL_3432451, EPI_ISL_3432452, EPI_ISL_3432453, EPI_ISL_3432454, EPI_ISL_3432455, EPI_ISL_3432456, EPI_ISL_3432457, EPI_ISL_3432458, EPI_ISL_3432459, EPI_ISL_3432460, EPI_ISL_3432461, EPI_ISL_3432462, EPI_ISL_3432463, EPI_ISL_3432464, EPI_ISL_3432465, EPI_ISL_3432466, EPI_ISL_3432467, EPI_ISL_3432468, EPI_ISL_3432469, EPI_ISL_3432470, EPI_ISL_3432471, EPI_ISL_3432472, EPI_ISL_3432473, EPI_ISL_3432474, EPI_ISL_3432475, EPI_ISL_3432476, EPI_ISL_3432477, EPI_ISL_3432478, EPI_ISL_3432479, EPI_ISL_3432480, EPI_ISL_3432481, EPI_ISL_3432482, EPI_ISL_3432483, EPI_ISL_3432484

see above ADPH ADPH Adrianna Walker

EPI_ISL_3432485, EPI_ISL_3432487 Laboratori d'anàlisis clíniques, Hospital Nostra Senyora de Meritxell LABORATORI NACIONAL EPIDEMIOLOGIA C, Lobaco et Al

EPI_ISL_3432488, EPI_ISL_3432492, EPI_ISL_3432493, EPI_ISL_3432494, EPI_ISL_3432496, EPI_ISL_3432498, EPI_ISL_3432499, EPI_ISL_3432502, EPI_ISL_3432503, EPI_ISL_3432509, EPI_ISL_3432511, EPI_ISL_3432514, EPI_ISL_3432516, EPI_ISL_3432518, EPI_ISL_3432520, EPI_ISL_3432521, EPI_ISL_3432522, EPI_ISL_3432524, EPI_ISL_3432525, EPI_ISL_3432528, EPI_ISL_3432529, EPI_ISL_3432530, EPI_ISL_3432531, EPI_ISL_3432532, EPI_ISL_3432534, EPI_ISL_3432535, EPI_ISL_3432536, EPI_ISL_3432537, EPI_ISL_3432538, EPI_ISL_3432539, EPI_ISL_3432540, EPI_ISL_3432541, EPI_ISL_3432545, EPI_ISL_3432547, EPI_ISL_3432549, EPI_ISL_3432550, EPI_ISL_3432551, EPI_ISL_3432554, EPI_ISL_3432555, EPI_ISL_3432556, EPI_ISL_3432558, EPI_ISL_3432559, EPI_ISL_3432560, EPI_ISL_3432561, EPI_ISL_3432562, EPI_ISL_3432563, EPI_ISL_3432565, EPI_ISL_3432567, EPI_ISL_3432568, EPI_ISL_3432569, EPI_ISL_3432570, EPI_ISL_3432571, EPI_ISL_3432573, EPI_ISL_3432575, EPI_ISL_3432577, EPI_ISL_3432578, EPI_ISL_3432579, EPI_ISL_3432580, EPI_ISL_3432581, EPI_ISL_3432582, EPI_ISL_3432583, EPI_ISL_3432584, EPI_ISL_3432586, EPI_ISL_3432588, EPI_ISL_3432589, EPI_ISL_3432590, EPI_ISL_3432592, EPI_ISL_3432594, EPI_ISL_3432595, EPI_ISL_3432596, EPI_ISL_3432597, EPI_ISL_3432598, EPI_ISL_3432601, EPI_ISL_3432603, EPI_ISL_3432604, EPI_ISL_3432605, EPI_ISL_3432606, EPI_ISL_3432607, EPI_ISL_3432608, EPI_ISL_3432609, EPI_ISL_3432610, EPI_ISL_3432612, EPI_ISL_3432613, EPI_ISL_3432614, EPI_ISL_3432615, EPI_ISL_3432616, EPI_ISL_3432617, EPI_ISL_3432618, EPI_ISL_3432619, EPI_ISL_3432620, EPI_ISL_3432621, EPI_ISL_3432622, EPI_ISL_3432623, EPI_ISL_3432625, EPI_ISL_3432626, EPI_ISL_3432627, EPI_ISL_3432628, EPI_ISL_3432629, EPI_ISL_3432630, EPI_ISL_3432631, EPI_ISL_3432632, EPI_ISL_3432633, EPI_ISL_3432634, EPI_ISL_3432635, EPI_ISL_3432636, EPI_ISL_3432637, EPI_ISL_3432638, EPI_ISL_3432639, EPI_ISL_3432640, EPI_ISL_3432641, EPI_ISL_3432642, EPI_ISL_3432643, EPI_ISL_3432644, EPI_ISL_3432645, EPI_ISL_3432646, EPI_ISL_3432647, EPI_ISL_3432648, EPI_ISL_3432649, EPI_ISL_3432650, EPI_ISL_3432651, EPI_ISL_3432652, EPI_ISL_3432653, EPI_ISL_3432654, EPI_ISL_3432655, EPI_ISL_3432656, EPI_ISL_3432658, EPI_ISL_3432659, EPI_ISL_3432660, EPI_ISL_3432661, EPI_ISL_3432662, EPI_ISL_3432663, EPI_ISL_3432664, EPI_ISL_3432665, EPI_ISL_3432666, EPI_ISL_3432667, EPI_ISL_3432668, EPI_ISL_3432669, EPI_ISL_3432671, EPI_ISL_3432672, EPI_ISL_3432673, EPI_ISL_3432674, EPI_ISL_3432675, EPI_ISL_3432676, EPI_ISL_3432677, EPI_ISL_3432679, EPI_ISL_3432681, EPI_ISL_3432682, EPI_ISL_3432683, EPI_ISL_3432684, EPI_ISL_3432685, EPI_ISL_3432686, EPI_ISL_3432688, EPI_ISL_3432689, EPI_ISL_3432690, EPI_ISL_3432691, EPI_ISL_3432692, EPI_ISL_3432693, EPI_ISL_3432696, EPI_ISL_3432697, EPI_ISL_3432698, EPI_ISL_3432699, EPI_ISL_3432700, EPI_ISL_3432701, EPI_ISL_3432702, EPI_ISL_3432703, EPI_ISL_3432704, EPI_ISL_3432705, EPI_ISL_3432706, EPI_ISL_3432707, EPI_ISL_3432708, EPI_ISL_3432709, EPI_ISL_3432710, EPI_ISL_3432711, EPI_ISL_3432715, EPI_ISL_3432716, EPI_ISL_3432717, EPI_ISL_3432718, EPI_ISL_3432720, EPI_ISL_3432721, EPI_ISL_3432722, EPI_ISL_3432723, EPI_ISL_3432724, EPI_ISL_3432725, EPI_ISL_3432726, EPI_ISL_3432727, EPI_ISL_3432728, EPI_ISL_3432729, EPI_ISL_3432730, EPI_ISL_3432731, EPI_ISL_3432732, EPI_ISL_3432733, EPI_ISL_3432734, EPI_ISL_3432735, EPI_ISL_3432736, EPI_ISL_3432737, EPI_ISL_3432738, EPI_ISL_3432739, EPI_ISL_3432740, EPI_ISL_3432741, EPI_ISL_3432742, EPI_ISL_3432743, EPI_ISL_3432744, EPI_ISL_3432745, EPI_ISL_3432746, EPI_ISL_3432747, EPI_ISL_3432748, EPI_ISL_3432749, EPI_ISL_3432750, EPI_ISL_3432751, EPI_ISL_3432753, EPI_ISL_3432754, EPI_ISL_3432755, EPI_ISL_3432756, EPI_ISL_3432757, EPI_ISL_3432758, EPI_ISL_3432759, EPI_ISL_3432760, EPI_ISL_3432761, EPI_ISL_3432762, EPI_ISL_3432763, EPI_ISL_3432764, EPI_ISL_3432765, EPI_ISL_3432766, EPI_ISL_3432767, EPI_ISL_3432768, EPI_ISL_3432769, EPI_ISL_3432770, EPI_ISL_3432771, EPI_ISL_3432772, EPI_ISL_3432773, EPI_ISL_3432791, EPI_ISL_3432792, EPI_ISL_3432793, EPI_ISL_3432813, EPI_ISL_3432814, EPI_ISL_3432815, EPI_ISL_3432816, EPI_ISL_3432817, EPI_ISL_3432818, EPI_ISL_3432819, EPI_ISL_3432825, EPI_ISL_3432827, EPI_ISL_3432829, EPI_ISL_3432830, EPI_ISL_3432831, EPI_ISL_3432832, EPI_ISL_3432833, EPI_ISL_3432837, EPI_ISL_3432840, EPI_ISL_3432846, EPI_ISL_3432849, EPI_ISL_3432850, EPI_ISL_3432855, EPI_ISL_3432856, EPI_ISL_3432858, EPI_ISL_3432859, EPI_ISL_3432861, EPI_ISL_3432862, EPI_ISL_3432865, EPI_ISL_3432866, EPI_ISL_3432867, EPI_ISL_3432868, EPI_ISL_3432877, EPI_ISL_3432878, EPI_ISL_3432881, EPI_ISL_3432882, EPI_ISL_3432884, EPI_ISL_3432885, EPI_ISL_3432886, EPI_ISL_3432888, EPI_ISL_3432890, EPI_ISL_3432892, EPI_ISL_3432893, EPI_ISL_3432894, EPI_ISL_3432895, EPI_ISL_3432898, EPI_ISL_3432899, EPI_ISL_3432900, EPI_ISL_3432908, EPI_ISL_3432910, EPI_ISL_3432913, EPI_ISL_3432914, EPI_ISL_3432916, EPI_ISL_3432917, EPI_ISL_3432918, EPI_ISL_3432919, EPI_ISL_3432920, EPI_ISL_3432921, EPI_ISL_3432922, EPI_ISL_3432923, EPI_ISL_3432925, EPI_ISL_3432928, EPI_ISL_3432930, EPI_ISL_3432931, EPI_ISL_3432932, EPI_ISL_3432934, EPI_ISL_3432952, EPI_ISL_3432953, EPI_ISL_3432954, EPI_ISL_3432955, EPI_ISL_3432956, EPI_ISL_3432957, EPI_ISL_3432958, EPI_ISL_3432959, EPI_ISL_3432960, EPI_ISL_3432961, EPI_ISL_3432962, EPI_ISL_3432963, EPI_ISL_3432964, EPI_ISL_3432965, EPI_ISL_3432966, EPI_ISL_3432967, EPI_ISL_3432968, EPI_ISL_3432970, EPI_ISL_3432971, EPI_ISL_3432974, EPI_ISL_3432976, EPI_ISL_3432977, EPI_ISL_3432978, EPI_ISL_3432979, EPI_ISL_3432980, EPI_ISL_3432981, EPI_ISL_3432982, EPI_ISL_3432983, EPI_ISL_3432984, EPI_ISL_3432985, EPI_ISL_3432986, EPI_ISL_3432987, EPI_ISL_3432988, EPI_ISL_3432990, EPI_ISL_3432991, EPI_ISL_3432997, EPI_ISL_3432998, EPI_ISL_3432999, EPI_ISL_3433000, EPI_ISL_3433001, EPI_ISL_3433002, EPI_ISL_3433003, EPI_ISL_3433004, EPI_ISL_3433005, EPI_ISL_3433006, EPI_ISL_3433007, EPI_ISL_3433008, EPI_ISL_3433009, EPI_ISL_3433010, EPI_ISL_3433011, EPI_ISL_3433012, EPI_ISL_3433013, EPI_ISL_3433014, EPI_ISL_3433015, EPI_ISL_3433016, EPI_ISL_3433017, EPI_ISL_3433018, EPI_ISL_3433019, EPI_ISL_3433020, EPI_ISL_3433021, EPI_ISL_3433022, EPI_ISL_3433023, EPI_ISL_3433024, EPI_ISL_3433025, EPI_ISL_3433026, EPI_ISL_3433027, EPI_ISL_3433028, EPI_ISL_3433029, EPI_ISL_3433030, EPI_ISL_3433031, EPI_ISL_3433032, EPI_ISL_3433033, EPI_ISL_3433034, EPI_ISL_3433035, EPI_ISL_3433036, EPI_ISL_3433037, EPI_ISL_3433038, EPI_ISL_3433039, EPI_ISL_3433040, EPI_ISL_3433041, EPI_ISL_3433042, EPI_ISL_3433043, EPI_ISL_3433044, EPI_ISL_3433045, EPI_ISL_3433046, EPI_ISL_3433047, EPI_ISL_3433048, EPI_ISL_3433049, EPI_ISL_3433050, EPI_ISL_3433051, EPI_ISL_3433052, EPI_ISL_3433053, EPI_ISL_3433054, EPI_ISL_3433055, EPI_ISL_3433056, EPI_ISL_3433057, EPI_ISL_3433058, EPI_ISL_3433059, EPI_ISL_3433060, EPI_ISL_3433061, EPI_ISL_3433062, EPI_ISL_3433063, EPI_ISL_3433064, EPI_ISL_3433065, EPI_ISL_3433066, EPI_ISL_3433067, EPI_ISL_3433068, EPI_ISL_3433069, EPI_ISL_3433070, EPI_ISL_3433071, EPI_ISL_3433072, EPI_ISL_3433073, EPI_ISL_3433074, EPI_ISL_3433075, EPI_ISL_3433076, EPI_ISL_3433077, EPI_ISL_3433078, EPI_ISL_3433079, EPI_ISL_3433080, EPI_ISL_3433081, EPI_ISL_3433082, EPI_ISL_3433083, EPI_ISL_3433084, EPI_ISL_3433085, EPI_ISL_3433086, EPI_ISL_3433087, EPI_ISL_3433088, EPI_ISL_3433089, EPI_ISL_3433090, EPI_ISL_3433091, EPI_ISL_3433092, EPI_ISL_3433093, EPI_ISL_3433094, EPI_ISL_3433095, EPI_ISL_3433096, EPI_ISL_3433097, EPI_ISL_3433098, EPI_ISL_3433099, EPI_ISL_3433100, EPI_ISL_3433101, EPI_ISL_3433102, EPI_ISL_3433103, EPI_ISL_3433104, EPI_ISL_3433105, EPI_ISL_3433106, EPI_ISL_3433107, EPI_ISL_3433108, EPI_ISL_3433109, EPI_ISL_3433110, EPI_ISL_3433111, EPI_ISL_3433112, EPI_ISL_3433113, EPI_ISL_3433114, EPI_ISL_3433115, EPI_ISL_3433116, EPI_ISL_3433117, EPI_ISL_3433118, EPI_ISL_3433119, EPI_ISL_3433120, EPI_ISL_3433121, EPI_ISL_3433122, EPI_ISL_3433123, EPI_ISL_3433124, EPI_ISL_3433125, EPI_ISL_3433126, EPI_ISL_3433127, EPI_ISL_3433128, EPI_ISL_3433129, EPI_ISL_3433130, EPI_ISL_3433131, EPI_ISL_3433132, EPI_ISL_3433133, EPI_ISL_3433134, EPI_ISL_3433135, EPI_ISL_3433136, EPI_ISL_3433137, EPI_ISL_3433138, EPI_ISL_3433139, EPI_ISL_3433140, EPI_ISL_3433141, EPI_ISL_3433142, EPI_ISL_3433143, EPI_ISL_3433144, EPI_ISL_3433145, EPI_ISL_3433146, EPI_ISL_3433147, EPI_ISL_3433148, EPI_ISL_3433149, EPI_ISL_3433150, EPI_ISL_3433151, EPI_ISL_3433152, EPI_ISL_3433153, EPI_ISL_3433154, EPI_ISL_3433155, EPI_ISL_3433156, EPI_ISL_3433157, EPI_ISL_3433158, EPI_ISL_3433159, EPI_ISL_3433160, EPI_ISL_3433161, EPI_ISL_3433162, EPI_ISL_3433163, EPI_ISL_3433164, EPI_ISL_3433165, EPI_ISL_3433166, EPI_ISL_3433167, EPI_ISL_3433168, EPI_ISL_3433169, EPI_ISL_3433170, EPI_ISL_3433171, EPI_ISL_3433172, EPI_ISL_3433173, EPI_ISL_3433174, EPI_ISL_3433175, EPI_ISL_3433176, EPI_ISL_3433177, EPI_ISL_3433178, EPI_ISL_3433179, EPI_ISL_3433180, EPI_ISL_3433181, EPI_ISL_3433182, EPI_ISL_3433183, EPI_ISL_3433184, EPI_ISL_3433185, EPI_ISL_3433186, EPI_ISL_3433187, EPI_ISL_3433188, EPI_ISL_3433189, EPI_ISL_3433190, EPI_ISL_3433191, EPI_ISL_3433192, EPI_ISL_3433193, EPI_ISL_3433194, EPI_ISL_3433195, EPI_ISL_3433196, EPI_ISL_3433197, EPI_ISL_3433198, EPI_ISL_3433199, EPI_ISL_3433200, EPI_ISL_3433201, EPI_ISL_3433202, EPI_ISL_3433203, EPI_ISL_3433204, EPI_ISL_3433205, EPI_ISL_3433206, EPI_ISL_3433207, EPI_ISL_3433208, EPI_ISL_3433209, EPI_ISL_3433210, EPI_ISL_3433211, EPI_ISL_3433212, EPI_ISL_3433213, EPI_ISL_3433214, EPI_ISL_3433215, EPI_ISL_3433216, EPI_ISL_3433217, EPI_ISL_3433218, EPI_ISL_3433219, EPI_ISL_3433220, EPI_ISL_3433221, EPI_ISL_3433222, EPI_ISL_3433223, EPI_ISL_3433224, EPI_ISL_3433225, EPI_ISL_3433226, EPI_ISL_3433227, EPI_ISL_3433228, EPI_ISL_3433229, EPI_ISL_3433230, EPI_ISL_3433231, EPI_ISL_3433232, EPI_ISL_3433233, EPI_ISL_3433234, EPI_ISL_3433235, EPI_ISL_3433236, EPI_ISL_3433237, EPI_ISL_3433238, EPI_ISL_3433239, EPI_ISL_3433240, EPI_ISL_3433241, EPI_ISL_3433242, EPI_ISL_3433243, EPI_ISL_3433244, EPI_ISL_3433245, EPI_ISL_3433246, EPI_ISL_3433247, EPI_ISL_3433248, EPI_ISL_3433249, EPI_ISL_3433250, EPI_ISL_3433251, EPI_ISL_3433252, EPI_ISL_3433253, EPI_ISL_3433254, EPI_ISL_3433255, EPI_ISL_3433256, EPI_ISL_3433257, EPI_ISL_3433258, EPI_ISL_3433259, EPI_ISL_3433260, EPI_ISL_3433261, EPI_ISL_3433262, EPI_ISL_3433263, EPI_ISL_3433264, EPI_ISL_3433265, EPI_ISL_3433266, EPI_ISL_3433267, EPI_ISL_3433268, EPI_ISL_3433269, EPI_ISL_3433270, EPI_ISL_3433271, EPI_ISL_3433272, EPI_ISL_3433273, EPI_ISL_3433274, EPI_ISL_3433275, EPI_ISL_3433276, EPI_ISL_3433277, EPI_ISL_3433278, EPI_ISL_3433279, EPI_ISL_3433280, EPI_ISL_3433281, EPI_ISL_3433282, EPI_ISL_3433283, EPI_ISL_3433284, EPI_ISL_3433285, EPI_ISL_3433286, EPI_ISL_3433287, EPI_ISL_3433288, EPI_ISL_3433289, EPI_ISL_3433290, EPI_ISL_3433291, EPI_ISL_3433292, EPI_ISL_3433293, EPI_ISL_3433294, EPI_ISL_3433295, EPI_ISL_3433296, EPI_ISL_3433297, EPI_ISL_3433298, EPI_ISL_3433299, EPI_ISL_3433300, EPI_ISL_3433301, EPI_ISL_3433302, EPI_ISL_3433303, EPI_ISL_3433304, EPI_ISL_3433305, EPI_ISL_3433306, EPI_ISL_3433307, EPI_ISL_3433308, EPI_ISL_3433309, EPI_ISL_3433310, EPI_ISL_3433311, EPI_ISL_3433312, EPI_ISL_3433313, EPI_ISL_3433314, EPI_ISL_3433315, EPI_ISL_3433316, EPI_ISL_3433317, EPI_ISL_3433318, EPI_ISL_3433319, EPI_ISL_3433320, EPI_ISL_3433321, EPI_ISL_3433322, EPI_ISL_3433323, EPI_ISL_3433324, EPI_ISL_3433325, EPI_ISL_3433326, EPI_ISL_3433327, EPI_ISL_3433328, EPI_ISL_3433329, EPI_ISL_3433330, EPI_ISL_3433331, EPI_ISL_3433332, EPI_ISL_3433333, EPI_ISL_3433334, EPI_ISL_3433335, EPI_ISL_3433336, EPI_ISL_3433337, EPI_ISL_3433338, EPI_ISL_3433339, EPI_ISL_3433340, EPI_ISL_3433341, EPI_ISL_3433342, EPI_ISL_3433343, EPI_ISL_3433344, EPI_ISL_3433345, EPI_ISL_3433346, EPI_ISL_3433347, EPI_ISL_3433348, EPI_ISL_3433349, EPI_ISL_3433350, EPI_ISL_3433351, EPI_ISL_3433352, EPI_ISL_3433353, EPI_ISL_3433354, EPI_ISL_3433355, EPI_ISL_3433356, EPI_ISL_3433357, EPI_ISL_3433358, EPI_ISL_3433359, EPI_ISL_3433360, EPI_ISL_3433361, EPI_ISL_3433362, EPI_ISL_3433363, EPI_ISL_3433364, EPI_ISL_3433365, EPI_ISL_3433366, EPI_ISL_3433367, EPI_ISL_3433368, EPI_ISL_3433369, EPI_ISL_3433370, EPI_ISL_3433371, EPI_ISL_3433372, EPI_ISL_3433373, EPI_ISL_3433374, EPI_ISL_3433375, EPI_ISL_3433376, EPI_ISL_3433377, EPI_ISL_3433378, EPI_ISL_3433379, EPI_ISL_3433380, EPI_ISL_3433381, EPI_ISL_3433382, EPI_ISL_3433383, EPI_ISL_3433384, EPI_ISL_3433385, EPI_ISL_3433386, EPI_ISL_3433387, EPI_ISL_3433388, EPI_ISL_3433389, EPI_ISL_3433390, EPI_ISL_3433391, EPI_ISL_3433392, EPI_ISL_3433393, EPI_ISL_3433394, EPI_ISL_3433395, EPI_ISL_3433396, EPI_ISL_3433397, EPI_ISL_3433398, EPI_ISL_3433399, EPI_ISL_3433400, EPI_ISL_3433401, EPI_ISL_3433402, EPI_ISL_3433403, EPI_ISL_3433404, EPI_ISL_3433405, EPI_ISL_3433406, EPI_ISL_3433407, EPI_ISL_3433408, EPI_ISL_3433409, EPI_ISL_3433410, EPI_ISL_3433411, EPI_ISL_3433412, EPI_ISL_3433413, EPI_ISL_3433414, EPI_ISL_3433415, EPI_ISL_3433416, EPI_ISL_3433417, EPI_ISL_3433418, EPI_ISL_3433419, EPI_ISL_3433420, EPI_ISL_3433421, EPI_ISL_3433422, EPI_ISL_3433423, EPI_ISL_3433424, EPI_ISL_3433425, EPI_ISL_3433426, EPI_ISL_3433427, EPI_ISL_3433428, EPI_ISL_3433429, EPI_ISL_3433430, EPI_ISL_3433431, EPI_ISL_3433432, EPI_ISL_3433433, EPI_ISL_3433434, EPI_ISL_3433435, EPI_ISL_3433436, EPI_ISL_3433437, EPI_ISL_3433438, EPI_ISL_3433439, EPI_ISL_3433440, EPI_ISL_3433441, EPI_ISL_3433442, EPI_ISL_3433443, EPI_ISL_3433444, EPI_ISL_3433445, EPI_ISL_3433446, EPI_ISL_3433447, EPI_ISL_3433448, EPI_ISL_3433449, EPI_ISL_3433450, EPI_ISL_3433451, EPI_ISL_3433452, EPI_ISL_3433453, EPI_ISL_3433454, EPI_ISL_3433455, EPI_ISL_3433456, EPI_ISL_3433457, EPI_ISL_3433458, EPI_ISL_3433459, EPI_ISL_3433460, EPI_ISL_3433461, EPI_ISL_3433462, EPI_ISL_3433463, EPI_ISL_3433464, EPI_ISL_3433465, EPI_ISL_3433466, EPI_ISL_3433467, EPI_ISL_3433468, EPI_ISL_3433469, EPI_ISL_3433470, EPI_ISL_3433471, EPI_ISL_3433472, EPI_ISL_3433473, EPI_ISL_3433474, EPI_ISL_3433475, EPI_ISL_3433476, EPI_ISL_3433477, EPI_ISL_3433478, EPI_ISL_3433479, EPI_ISL_3433480, EPI_ISL_3433481, EPI_ISL_3433482, EPI_ISL_3433483, EPI_ISL_3433484, EPI_ISL_3433485, EPI_ISL_3433486, EPI_ISL_3433487, EPI_ISL_3433488, EPI_ISL_3433489, EPI_ISL_3433490, EPI_ISL_3433491, EPI_ISL_3433492, EPI_ISL_3433493, EPI_ISL_3433494, EPI_ISL_3433495, EPI_ISL_3433496, EPI_ISL_3433497, EPI_ISL_3433498, EPI_ISL_3433499, EPI_ISL_3433500, EPI_ISL_3433501, EPI_ISL_3433502, EPI_ISL_3433503, EPI_ISL_3433504, EPI_ISL_3433505, EPI_ISL_3433506, EPI_ISL_3433507, EPI_ISL_3433508, EPI_ISL_3433509, EPI_ISL_3433510, EPI_ISL_3433511, EPI_ISL_3433512, EPI_ISL_3433513, EPI_ISL_3433514, EPI_ISL_3433515, EPI_ISL_3433516, EPI_ISL_3433517, EPI_ISL_3433518, EPI_ISL_3433519, EPI_ISL_3433520, EPI_ISL_3433521, EPI_ISL_3433522, EPI_ISL_3433523, EPI_ISL_3433524, EPI_ISL_3433525, EPI_ISL_3433526, EPI_ISL_3433527, EPI_ISL_3433528, EPI_ISL_3433529, EPI_ISL_3433530, EPI_ISL_3433531, EPI_ISL_3433532, EPI_ISL_3433533, EPI_ISL_3433534, EPI_ISL_3433535, EPI_ISL_3433536, EPI_ISL_3433537, EPI_ISL_3433538, EPI_ISL_3433539, EPI_ISL_3433540, EPI_ISL_3433541, EPI_ISL_3433542, EPI_ISL_3433543, EPI_ISL_3433544, EPI_ISL_3433545, EPI_ISL_3433546, EPI_ISL_3433547, EPI_ISL_3433548, EPI_ISL_3433549, EPI_ISL_3433550, EPI_ISL_3433551, EPI_ISL_3433552, EPI_ISL_3433553, EPI_ISL_3433554, EPI_ISL_3433555, EPI_ISL_3433556, EPI_ISL_3433557, EPI_ISL_3433558, EPI_ISL_3433559, EPI_ISL_3433560, EPI_ISL_3433561, EPI_ISL_3433562, EPI_ISL_3433563, EPI_ISL_3433564, EPI_ISL_3433565, EPI_ISL_3433566, EPI_ISL_3433567, EPI_ISL_3433568, EPI_ISL_3433569, EPI_ISL_3433570, EPI_ISL_3433571, EPI_ISL_3433572, EPI_ISL_3433573, EPI_ISL_3433574, EPI_ISL_3433575, EPI_ISL_3433576, EPI_ISL_3433577, EPI_ISL_3433578, EPI_ISL_3433579, EPI_ISL_3433580, EPI_ISL_3433581, EPI_ISL_3433582, EPI_ISL_3433583, EPI_ISL_3433584, EPI_ISL_3433585, EPI_ISL_3433586, EPI_ISL_3433587, EPI_ISL_3433588, EPI_ISL_3433589, EPI_ISL_3433590, EPI_ISL_3433591, EPI_ISL_3433592, EPI_ISL_3433593, EPI_ISL_3433594, EPI_ISL_3433595, EPI_ISL_3433596, EPI_ISL_3433597, EPI_ISL_3433598, EPI_ISL_3433599, EPI_ISL_3433600, EPI_ISL_3433601, EPI_ISL_3433602, EPI_ISL_3433603, EPI_ISL_3433604, EPI_ISL_3433605, EPI_ISL_3433606, EPI_ISL_3433607, EPI_ISL_3433608, EPI_ISL_3433609, EPI_ISL_3433610, EPI_ISL_3433611, EPI_ISL_3433612, EPI_ISL_3433613, EPI_ISL_3433614, EPI_ISL_3433615, EPI_ISL_3433616, EPI_ISL_3433617, EPI_ISL_3433618, EPI_ISL_3433619, EPI_ISL_3433620, EPI_ISL_3433621, EPI_ISL_3433622, EPI_ISL_3433623, EPI_ISL_3433624, EPI_ISL_3433625, EPI_ISL_3433626, EPI_ISL_3433627, EPI_ISL_3433628, EPI_ISL_3433629, EPI_ISL_3433630, EPI_ISL_3433631, EPI_ISL_3433632, EPI_ISL_3433633, EPI_ISL_3433634, EPI_ISL_3433635, EPI_ISL_3433636, EPI_ISL_3433637, EPI_ISL_3433638, EPI_ISL_3433639, EPI_ISL_3433640, EPI_ISL_3433641, EPI_ISL_3433642, EPI_ISL_3433643, EPI_ISL_3433644, EPI_ISL_3433645, EPI_ISL_3433646, EPI_ISL_3433647, EPI_ISL_3433648, EPI_ISL_3433649, EPI_ISL_3433650, EPI_ISL_3433651, EPI_ISL_3433652, EPI_ISL_3433653, EPI_ISL_3433654, EPI_ISL_3433655, EPI_ISL_3433656, EPI_ISL_3433657, EPI_ISL_3433658, EPI_ISL_3433659, EPI_ISL_3433660, EPI_ISL_3433661, EPI_ISL_3433662, EPI_ISL_3433663, EPI_ISL_3433664, EPI_ISL_3433665, EPI_ISL_3433666, EPI_ISL_3433667, EPI_ISL_3433668, EPI_ISL_3433669, EPI_ISL_3433670, EPI_ISL_3433671, EPI_ISL_3433672, EPI_ISL_3433673, EPI_ISL_3433674, EPI_ISL_3433675, EPI_ISL_3433676, EPI_ISL_3433677, EPI_ISL_3433678, EPI_ISL_3433679, EPI_ISL_3433680, EPI_ISL_3433681, EPI_ISL_3433682, EPI_ISL_3433683, EPI_ISL_3433684, EPI_ISL_3433685, EPI_ISL_3433686, EPI_ISL_3433687, EPI_ISL_3433688, EPI_ISL_3433689, EPI_ISL_3433690, EPI_ISL_3433691, EPI_ISL_3433692, EPI_ISL_3433693, EPI_ISL_3433694, EPI_ISL_3433695, EPI_ISL_3433696, EPI_ISL_3433697, EPI_ISL_3433698, EPI_ISL_3433699, EPI_ISL_3433700, EPI_ISL_3433701, EPI_ISL_3433702, EPI_ISL_3433703, EPI_ISL_3433704, EPI_ISL_3433705, EPI_ISL_3433706, EPI_ISL_3433707, EPI_ISL_3433708, EPI_ISL_3433709, EPI_ISL_3433710, EPI_ISL_3433711, EPI_ISL_3433712, EPI_ISL_3433713, EPI_ISL_3433714, EPI_ISL_3433715, EPI_ISL_3433716, EPI_ISL_3433717, EPI_ISL_3433718, EPI_ISL_3433719, EPI_ISL_3433720, EPI_ISL_3433721, EPI_ISL_3433722, EPI_ISL_3433723, EPI_ISL_3433724, EPI_ISL_3433725, EPI_ISL_3433726, EPI_ISL_3433727, EPI_ISL_3433728, EPI_ISL_3433729, EPI_ISL_3433730, EPI_ISL_3433731, EPI_ISL_3433732, EPI_ISL_3433733, EPI_ISL_3433734, EPI_ISL_3433735, EPI_ISL_3433736, EPI_ISL_3433737, EPI_ISL_3433738, EPI_ISL_3433739, EPI_ISL_3433740, EPI_ISL_3433741, EPI_ISL_3433742, EPI_ISL_3433743, EPI_ISL_3433744, EPI_ISL_3433745, EPI_ISL_3433746, EPI_ISL_3433747, EPI_ISL_3433748, EPI_ISL_3433749, EPI_ISL_3433750, EPI_ISL_3433751, EPI_ISL_3433752, EPI_ISL_3433753, EPI_ISL_3433754, EPI_ISL_3433755, EPI_ISL_3433756, EPI_ISL_3433757, EPI_ISL_3433758, EPI_ISL_3433759, EPI_ISL_3433760, EPI_ISL_3433761, EPI_ISL_3433762, EPI_ISL_3433763, EPI_ISL_3433764, EPI_ISL_3433765, EPI_ISL_3433766, EPI_ISL_3433767, EPI_ISL_3433768, EPI_ISL_3433769, EPI_ISL_3433770, EPI_ISL_3433771, EPI_ISL_3433772, EPI_ISL_3433773, EPI_ISL_3433774, EPI_ISL_3433775, EPI_ISL_3433776, EPI_ISL_3433777, EPI_ISL_3433778, EPI_ISL_3433779, EPI_ISL_3433780, EPI_ISL_3433781, EPI_ISL_3433782, EPI_ISL_3433783, EPI_ISL_3433784, EPI_ISL_3433785, EPI_ISL_3433786, EPI_ISL_3433787, EPI_ISL_3433788, EPI_ISL_3433789, EPI_ISL_3433790, EPI_ISL_3433791, EPI_ISL_3433792, EPI_ISL_3433793, EPI_ISL_3433794, EPI_ISL_3433795, EPI_ISL_3433796, EPI_ISL_3433797, EPI_ISL_3433798, EPI_ISL_3433799, EPI_ISL_3433800, EPI_ISL_3433801, EPI_ISL_3433802, EPI_ISL_3433803, EPI_ISL_3433804, EPI_ISL_3433805, EPI_ISL_3433806, EPI_ISL_3433807, EPI_ISL_3433808, EPI_ISL_3433809, EPI_ISL_3433810, EPI_ISL_3433811, EPI_ISL_3433812, EPI_ISL_3433813, EPI_ISL_3433814, EPI_ISL_3433815, EPI_ISL_3433816, EPI_ISL_3433817, EPI_ISL_3433818, EPI_ISL_3433819, EPI_ISL_3433820, EPI_ISL_3433821, EPI_ISL_3433822, EPI_ISL_3433823, EPI_ISL_3433824, EPI_ISL_3433825, EPI_ISL_3433826, EPI_ISL_3433827, EPI_ISL_3433828, EPI_ISL_3433829, EPI_ISL_3433830, EPI_ISL_3433831, EPI_ISL_3433832, EPI_ISL_3433833, EPI_ISL_3433834, EPI_ISL_3433835, EPI_ISL_3433836, EPI_ISL_3433837, EPI_ISL_3433838, EPI_ISL_3433839, EPI_ISL_3433840, EPI_ISL_3433841, EPI_ISL_3433842, EPI_ISL_3433843, EPI_ISL_3433844, EPI_ISL_3433845, EPI_ISL_3433846, EPI_ISL_3433847, EPI_ISL_3433848, EPI_ISL_3433849, EPI_ISL_3433850, EPI_ISL_3433851, EPI_ISL_3433852, EPI_ISL_3433853, EPI_ISL_3433854, EPI_ISL_3433855, EPI_ISL_3433856, EPI_ISL_3433857, EPI_ISL_3433858, EPI_ISL_3433859, EPI_ISL_3433860, EPI_ISL_3433861, EPI_ISL_3433862, EPI_ISL_3433863, EPI_ISL_3433864, EPI_ISL_3433865, EPI_ISL_3433866, EPI_ISL_3433867, EPI_ISL_3433868, EPI_ISL_3433869, EPI_ISL_3433870, EPI_ISL_3433871, EPI_ISL_3433872, EPI_ISL_3433873, EPI_ISL_3433874, EPI_ISL_3433875, EPI_ISL_3433876, EPI_ISL_3433877, EPI_ISL_3433878, EPI_ISL_3433879, EPI_ISL_3433880, EPI_ISL_3433881, EPI_ISL_3433882, EPI_ISL_3433883, EPI_ISL_3433884, EPI_ISL_3433885, EPI_ISL_3433886, EPI_ISL_3433887, EPI_ISL_3433888, EPI_ISL_3433889, EPI_ISL_3433890, EPI_ISL_3433891, EPI_ISL_3433892, EPI_ISL_3433893, EPI_ISL_3433894, EPI_ISL_3433895, EPI_ISL_3433896, EPI_ISL_3433897, EPI_ISL_3433898, EPI_ISL_3433899, EPI_ISL_3433900, EPI_ISL_3433901, EPI_ISL_3433902, EPI_ISL_3433903, EPI_ISL_3433904, EPI_ISL_3433905, EPI_ISL_3433906, EPI_ISL_3433907, EPI_ISL_3433908, EPI_ISL_3433909, EPI_ISL_3433910, EPI_ISL_3433911, EPI_ISL_3433912, EPI_ISL_3433913, EPI_ISL_3433914, EPI_ISL_3433915, EPI_ISL_3433916, EPI_ISL_3433917, EPI_ISL_3433918, EPI_ISL_3433919, EPI_ISL_3433920, EPI_ISL_3433921, EPI_ISL_3433922, EPI_ISL_3433923, EPI_ISL_3433924, EPI_ISL_3433925, EPI_ISL_3433926, EPI_ISL_3433927, EPI_ISL_3433928, EPI_ISL_3433929, EPI_ISL_3433930, EPI_ISL_3433931, EPI_ISL_3433932, EPI_ISL_3433933, EPI_ISL_3433934, EPI_ISL_3433935, EPI_ISL_3433936, EPI_ISL_3433937, EPI_ISL_3433938, EPI_ISL_3433939, EPI_ISL_3433940, EPI_ISL_3433941, EPI_ISL_3433942, EPI_ISL_3433943, EPI_ISL_3433944, EPI_ISL_3433945, EPI_ISL_3433946, EPI_ISL_3433947, EPI_ISL_3433948, EPI_ISL_3433949, EPI_ISL_3433950, EPI_ISL_3433951, EPI_ISL_3433952, EPI_ISL_3433953, EPI_ISL_3433954, EPI_ISL_3433955, EPI_ISL_3433956, EPI_ISL_3433957, EPI_ISL_3433958, EPI_ISL_3433959, EPI_ISL_3433960, EPI_ISL_3433961, EPI_ISL_3433962, EPI_ISL_3433963, EPI_ISL_3433964, EPI_ISL_3433965, EPI_ISL_3433966, EPI_ISL_3433967, EPI_ISL_3433968, EPI_ISL_3433969, EPI_ISL_3433970, EPI_ISL_3433971, EPI_ISL_3433972, EPI_ISL_3433973, EPI_ISL_3433974, EPI_ISL_3433975, EPI_ISL_3433976, EPI_ISL_3433977, EPI_ISL_3433978, EPI_ISL_3433979, EPI_ISL_3433980, EPI_ISL_3433981, EPI_ISL_3433982, EPI_ISL_3433983, EPI_ISL_3433984, EPI_ISL_3433985, EPI_ISL_3433986, EPI_ISL_3433987, EPI_ISL_3433988, EPI_ISL_3433989, EPI_ISL_3433990, EPI_ISL_3433991, EPI_ISL_3433992, EPI_ISL_3433993, EPI_ISL_3433994, EPI_ISL_3433995, EPI_ISL_3433996, EPI_ISL_3433997, EPI_ISL_3433998, EPI_ISL_3433999, EPI_ISL_3434000, EPI_ISL_3434001, EPI_ISL_3434002, EPI_ISL_3434003, EPI_ISL_3434004, EPI_ISL_3434005, EPI_ISL_3434006, EPI_ISL_3434007, EPI_ISL_3434008, EPI_ISL_3434009, EPI_ISL_3434010, EPI_ISL_3434011, EPI_ISL_3434012, EPI_ISL_3434013, EPI_ISL_3434014, EPI_ISL_3434015, EPI_ISL_3434016, EPI_ISL_3434017, EPI_ISL_3434018, EPI_ISL_3434019, EPI_ISL_3434020, EPI_ISL_3434021, EPI_ISL_3434022, EPI_ISL_3434023, EPI_ISL_3434024, EPI_ISL_3434025, EPI_ISL_3434026, EPI_ISL_3434027, EPI_ISL_3434028, EPI_ISL_3434029, EPI_ISL_3434030, EPI_ISL_3434031, EPI_ISL_3434032, EPI_ISL_3434033, EPI_ISL_3434034, EPI_ISL_3434035, EPI_ISL_3434036, EPI_ISL_3434037, EPI_ISL_3434038, EPI_ISL_3434039, EPI_ISL_3434040, EPI_ISL_3434041, EPI_ISL_3434042, EPI_ISL_3434043, EPI_ISL_3434044, EPI_ISL_3434045, EPI_ISL_3434046, EPI_ISL_3434047, EPI_ISL_3434048, EPI_ISL_3434049, EPI_ISL_3434050, EPI_ISL_3434051, EPI_ISL_3434052, EPI_ISL_3434053, EPI_ISL_3434054, EPI_ISL_3434055, EPI_ISL_3434056, EPI_ISL_3434057, EPI_ISL_3434058, EPI_ISL_3434059, EPI_ISL_3434060, EPI_ISL_3434061, EPI_ISL_3434062, EPI_ISL_3434063, EPI_ISL_3434064, EPI_ISL_3434065, EPI_ISL_3434066, EPI_ISL_3434067, EPI_ISL_3434068, EPI_ISL_3434069, EPI_ISL_3434070, EPI_ISL_3434071, EPI_ISL_3434072, EPI_ISL_3434073, EPI_ISL_3434074, EPI_ISL_3434075, EPI_ISL_3434076, EPI_ISL_3434077, EPI_ISL_3434078, EPI_ISL_3434079, EPI_ISL_3434080, EPI_ISL_3434081, EPI_ISL_3434082, EPI_ISL_3434083, EPI_ISL_3434084, EPI_ISL_3434085, EPI_ISL_3434086, EPI_ISL_3434087, EPI_ISL_3434088, EPI_ISL_3434089, EPI_ISL_3434090, EPI_ISL_3434091, EPI_ISL_3434092, EPI_ISL_3434093, EPI_ISL_3434094, EPI_ISL_3434095, EPI_ISL_3434096, EPI_ISL_3434097, EPI_ISL_3434098, EPI_ISL_3434099, EPI_ISL_3434100, EPI_ISL_3434101, EPI_ISL_3434102, EPI_ISL_3434103, EPI_ISL_3434104, EPI_ISL_3434105, EPI_ISL_3434106, EPI_ISL_3434107, EPI_ISL_3434108, EPI_ISL_3434109, EPI_ISL_3434110, EPI_ISL_3434111, EPI_ISL_3434112, EPI_ISL_3434113, EPI_ISL_3434114, EPI_ISL_3434115, EPI_ISL_3434116, EPI_ISL_3434117, EPI_ISL_3434118, EPI_ISL_3434119, EPI_ISL_3434120, EPI_ISL_3434121, EPI_ISL_3434122, EPI_ISL_3434123, EPI_ISL_3434124, EPI_ISL_3434125, EPI_ISL_3434126, EPI_ISL_3434127, EPI_ISL_3434128, EPI_ISL_3434129, EPI_ISL_3434130, EPI_ISL_3434131, EPI_ISL_3434132, EPI_ISL_3434133, EPI_ISL_3434134, EPI_ISL_3434135, EPI_ISL_3434136, EPI_ISL_3434137, EPI_ISL_3434138, EPI_ISL_3434139, EPI_ISL_3434140, EPI_ISL_3434141, EPI_ISL_3434142, EPI_ISL_3434143, EPI_ISL_3434144, EPI_ISL_3434145, EPI_ISL_3434146, EPI_ISL_3434147, EPI_ISL_3434148, EPI_ISL_3434149, EPI_ISL_3434150, EPI_ISL_3434151, EPI_ISL_3434152, EPI_ISL_3434153, EPI_ISL_3434154, EPI_ISL_3434155, EPI_ISL_3434156, EPI_ISL_3434157, EPI_ISL_3434158, EPI_ISL_3434159, EPI_ISL_3434160, EPI_ISL_3434161, EPI_ISL_3434162, EPI_ISL_3434163, EPI_ISL_3434164, EPI_ISL_3434165, EPI_ISL_3434166, EPI_ISL_3434167, EPI_ISL_3434168, EPI_ISL_3434169, EPI_ISL_3434170, EPI_ISL_3434171, EPI_ISL_3434172, EPI_ISL_3434173, EPI_ISL_3434174, EPI_ISL_3434175, EPI_ISL_3434176, EPI_ISL_3434177, EPI_ISL_3434178, EPI_ISL_3434179, EPI_ISL_3434180, EPI_ISL_3434181, EPI_ISL_3434182, EPI_ISL_3434183, EPI_ISL_3434184, EPI_ISL_3434185, EPI_ISL_3434186, EPI_ISL_3434187, EPI_ISL_3434188, EPI_ISL_3434189, EPI_ISL_3434190, EPI_ISL_3434191, EPI_ISL_3434192, EPI_ISL_3434193, EPI_ISL_3434194, EPI_ISL_3434195, EPI_ISL_3434196, EPI_ISL_3434197, EPI_ISL_3434198, EPI_ISL_3434199, EPI_ISL_3434200, EPI_ISL_3434201, EPI_ISL_3434202, EPI_ISL_3434203, EPI_ISL_3434204, EPI_ISL_3434205, EPI_ISL_3434206, EPI_ISL_3434207, EPI_ISL_3434208, EPI_ISL_3434209, EPI_ISL_3434210, EPI_ISL_3434211, EPI_ISL_3434212, EPI_ISL_3434213, EPI_ISL_3434214, EPI_ISL_3434215, EPI_ISL_3434216, EPI_ISL_3434217, EPI_ISL_3434218, EPI_ISL_3434219, EPI_ISL_3434220, EPI_ISL_3434221, EPI_ISL_3434222, EPI_ISL_3434223, EPI_ISL_3434224, EPI_ISL_3434225, EPI_ISL_3434226, EPI_ISL_3434227, EPI_ISL_3434228, EPI_ISL_3434229, EPI_ISL_3434230, EPI_ISL_3434231, EPI_ISL_3434232, EPI_ISL_3434233, EPI_ISL_3434234, EPI_ISL_3434235, EPI_ISL_3434236, EPI_ISL_3434237, EPI_ISL_3434238, EPI_ISL_3434239, EPI_ISL_3434240, EPI_ISL_3434241, EPI_ISL_3434242, EPI_ISL_3434243, EPI_ISL_3434244, EPI_ISL_3434245, EPI_ISL_3434246, EPI_ISL_3434247, EPI_ISL_3434248, EPI_ISL_3434249, EPI_ISL_3434250, EPI_ISL_3434251, EPI_ISL_3434252, EPI_ISL_3434253, EPI_ISL_3434254, EPI_ISL_3434255, EPI_ISL_3434256, EPI_ISL_3434257, EPI_ISL_3434258, EPI_ISL_3434259, EPI_ISL_3434260, EPI_ISL_3434261, EPI_ISL_3434262, EPI_ISL_3434263, EPI_ISL_3434264, EPI_ISL_3434265, EPI_ISL_3434266, EPI_ISL_3434267, EPI_ISL_3434268, EPI_ISL_3434269, EPI_ISL_3434270, EPI_ISL_3434271, EPI_ISL_3434272, EPI_ISL_3434273, EPI_ISL_3434274, EPI_ISL_3434275, EPI_ISL_3434276, EPI_ISL_3434277, EPI_ISL_3434278, EPI_ISL_3434279, EPI_ISL_3434280, EPI_ISL_3434281, EPI_ISL_3434282, EPI_ISL_3434283, EPI_ISL_3434284, EPI_ISL_3434285, EPI_ISL_3434286, EPI_ISL_3434287, EPI_ISL_3434288, EPI_ISL_3434289, EPI_ISL_3434290, EPI_ISL_3434291, EPI_ISL_3434292, EPI_ISL_3434293, EPI_ISL_3434294, EPI_ISL_3434295, EPI_ISL_3434296, EPI_ISL_3434297, EPI_ISL_3434298, EPI_ISL_3434299, EPI_ISL_3434300, EPI_ISL_3434301, EPI_ISL_3434302, EPI_ISL_3434303, EPI_ISL_3434304, EPI_ISL_3434305, EPI_ISL_3434306, EPI_ISL_3434307, EPI_ISL_3434308, EPI_ISL_3434309, EPI_ISL_3434310, EPI_ISL_3434311, EPI_ISL_3434312, EPI_ISL_3434313, EPI_ISL_3434314, EPI_ISL_3434315, EPI_ISL_3434316, EPI_ISL_3434317, EPI_ISL_3434318, EPI_ISL_3434319, EPI_ISL_3434320, EPI_ISL_3434321, EPI_ISL_3434322, EPI_ISL_3434323, EPI_ISL_3434324, EPI_ISL_3434325, EPI_ISL_3434326, EPI_ISL_3434327, EPI_ISL_3434328, EPI_ISL_3434329, EPI_ISL_3434330, EPI_ISL_3434331, EPI_ISL_3434332, EPI_ISL_3434333, EPI_ISL_3434334, EPI_ISL_3434335, EPI_ISL_3434336, EPI_ISL_3434337, EPI_ISL_3434338, EPI_ISL_3434339, EPI_ISL_3434340, EPI_ISL_3434341, EPI_ISL_3434342, EPI_ISL_3434343, EPI_ISL_3434344, EPI_ISL_3434345, EPI_ISL_3434346, EPI_ISL_3434347, EPI_ISL_3434348, EPI_ISL_3434349, EPI_ISL_3434350, EPI_ISL_3434351, EPI_ISL_3434352, EPI_ISL_3434353, EPI_ISL_3434354, EPI_ISL_3434355, EPI_ISL_3434356, EPI_ISL_3434357, EPI_ISL_3434358, EPI_ISL_3434359, EPI_ISL_3434360, EPI_ISL_3434361, EPI_ISL_3434362, EPI_ISL_3434363, EPI_ISL_3434364, EPI_ISL_3434365, EPI_ISL_3434366, EPI_ISL_3434367, EPI_ISL_3434368, EPI_ISL_3434369, EPI_ISL_3434370, EPI_ISL_3434371, EPI_ISL_3434372, EPI_ISL_3434373, EPI_ISL_3434374, EPI_ISL_3434375, EPI_ISL_3434376, EPI_ISL_3434377, EPI_ISL_3434378, EPI_ISL_3434379, EPI_ISL_3434380, EPI_ISL_3434381, EPI_ISL_3434382, EPI_ISL_3434383, EPI_ISL_3434384, EPI_ISL_3434385, EPI_ISL_3434386, EPI_ISL_3434387, EPI_ISL_3434388, EPI_ISL_3434389, EPI_ISL_3434390, EPI_ISL_3434391, EPI_ISL_3434392, EPI_ISL_3434393, EPI_ISL_3434394, EPI_ISL_3434395, EPI_ISL_3434396, EPI_ISL_3434397, EPI_ISL_3434398, EPI_ISL_3434399, EPI_ISL_3434400, EPI_ISL_3434401, EPI_ISL_3434402, EPI_ISL_3434403, EPI_ISL_3434404, EPI_ISL_3434405, EPI_ISL_3434406, EPI_ISL_3434407, EPI_ISL_3434408, EPI_ISL_3434409, EPI_ISL_3434410, EPI_ISL_3434411, EPI_ISL_3434412, EPI_ISL_3434413, EPI_ISL_3434414, EPI_ISL_3434415, EPI_ISL_3434416, EPI_ISL_3434417, EPI_ISL_3434418, EPI_ISL_3434419, EPI_ISL_3434420, EPI_ISL_3434421, EPI_ISL_3434422, EPI_ISL_3434423, EPI_ISL_3434424, EPI_ISL_3434425, EPI_ISL_3434426, EPI_ISL_3434427, EPI_ISL_3434428, EPI_ISL_3434429, EPI_ISL_3434430, EPI_ISL_3434431, EPI_ISL_3434432, EPI_ISL_3434433, EPI_ISL_3434434, EPI_ISL_3434435, EPI_ISL_3434436, EPI_ISL_3434437, EPI_ISL_3434438, EPI_ISL_3434439, EPI_ISL_3434440, EPI_ISL_3434441, EPI_ISL_3434442, EPI_ISL_3434443, EPI_ISL_3434444, EPI_ISL_3434445, EPI_ISL_3434446, EPI_ISL_3434447, EPI_ISL_3434448, EPI_ISL_3434449, EPI_ISL_3434450, EPI_ISL_3434451, EPI_ISL_3434452, EPI_ISL_3434453, EPI_ISL_3434454, EPI_ISL_3434455, EPI_ISL_3434456, EPI_ISL_3434457, EPI_ISL_3434458, EPI_ISL_3434459, EPI_ISL_3434460, EPI_ISL_3434461, EPI_ISL_3434462, EPI_ISL_3434463, EPI_ISL_3434464, EPI_ISL_3434465, EPI_ISL_3434466, EPI_ISL_3434467, EPI_ISL_3434468, EPI_ISL_3434469, EPI_ISL_3434470, EPI_ISL_3434471, EPI_ISL_3434472, EPI_ISL_3434473, EPI_ISL_3434474, EPI_ISL_3434475, EPI_ISL_3434476, EPI_ISL_3434477, EPI_ISL_3434478, EPI_ISL_3434479, EPI_ISL_3434480, EPI_ISL_3434481, EPI_ISL_3434482, EPI_ISL_3434483, EPI_ISL_3434484, EPI_ISL_3434485, EPI_ISL_3434486, EPI_ISL_3434487, EPI_ISL_3434488, EPI_ISL_3434489, EPI_ISL_3434490, EPI_ISL_3434491, EPI_ISL_3434492, EPI_ISL_3434493, EPI_ISL_3434494, EPI_ISL_3434495, EPI_ISL_3434496, EPI_ISL_3434497, EPI_ISL_3434498, EPI_ISL_3434499, EPI_ISL_3434500, EPI_ISL_3434501, EPI_ISL_3434502, EPI_ISL_3434503, EPI_ISL_3434504, EPI_ISL_3434505, EPI_ISL_3434506, EPI_ISL_3434507, EPI_ISL_3434508, EPI_ISL_3434509, EPI_ISL_3434510, EPI_ISL_3434511, EPI_ISL_3434512, EPI_ISL_3434513, EPI_ISL_3434514, EPI_ISL_3434515, EPI_ISL_3434516, EPI_ISL_3434517, EPI_ISL_3434518, EPI_ISL_3434519, EPI_ISL_3434520, EPI_ISL_3434521, EPI_ISL_3434522, EPI_ISL_3434523, EPI_ISL_3434524, EPI_ISL_3434525, EPI_ISL_3434526, EPI_ISL_3434527, EPI_ISL_3434528, EPI_ISL_3434529, EPI_ISL_3434530, EPI_ISL_3434531, EPI_ISL_3434532, EPI_ISL_3434533, EPI_ISL_3434534, EPI_ISL_3434535, EPI_ISL_3434536, EPI_ISL_3434537, EPI_ISL_3434538, EPI_ISL_3434539, EPI_ISL_3434540, EPI_ISL_3434541, EPI_ISL_3434542, EPI_ISL_3434543, EPI_ISL_3434544, EPI_ISL_3434545, EPI_ISL_3434546, EPI_ISL_3434547, EPI_ISL_3434548, EPI_ISL_3434549, EPI_ISL_3434550, EPI_ISL_3434551, EPI_ISL_3434552, EPI_ISL_3434553, EPI_ISL_3434554, EPI_ISL_3434555, EPI_ISL_3434556, EPI_ISL_3434557, EPI_ISL_3434558, EPI_ISL_3434559, EPI_ISL_3434560, EPI_ISL_3434561, EPI_ISL_3434562, EPI_ISL_3434563, EPI_ISL_3434564, EPI_ISL_3434565, EPI_ISL_3434566, EPI_ISL_3434567, EPI_ISL_3434568, EPI_ISL_3434569, EPI_ISL_3434570, EPI_ISL_3434571, EPI_ISL_3434572, EPI_ISL_3434573, EPI_ISL_3434574, EPI_ISL_3434575, EPI_ISL_3434576, EPI_ISL_3434577, EPI_ISL_3434578, EPI_ISL_3434579, EPI_ISL_3434580, EPI_ISL_3434581, EPI_ISL_3434582, EPI_ISL_3434583, EPI_ISL_3434584, EPI_ISL_3434585, EPI_ISL_3434586, EPI_ISL_3434587, EPI_ISL_3434588, EPI_ISL_3434589, EPI_ISL_3434590, EPI_ISL_3434591, EPI_ISL_3434592, EPI_ISL_3434593, EPI_ISL_3434594, EPI_ISL_3434595, EPI_ISL_3434596, EPI_ISL_3434597, EPI_ISL_3434598, EPI_ISL_3434599, EPI_ISL_3434600, EPI_ISL_3434601, EPI_ISL_3434602, EPI_ISL_3434603, EPI_ISL_3434604, EPI_ISL_3434605, EPI_ISL_3434606, EPI_ISL_3434607, EPI_ISL_3434608, EPI_ISL_3434609, EPI_ISL_3434610, EPI_ISL_3434611, EPI_ISL_3434614

see above MEPHI, Aix Marseille University MEPHI, Aix Marseille University Anthony LEVASSEUR

EPI_ISL_3434658, EPI_ISL_3434659, EPI_ISL_3434660, EPI_ISL_3434661, EPI_ISL_3434662, EPI_ISL_3434663, EPI_ISL_3434664, EPI_ISL_3434665, EPI_ISL_3434666, EPI_ISL_3434667, EPI_ISL_3434668, EPI_ISL_3434669, EPI_ISL_3434670, EPI_ISL_3434671, EPI_ISL_3434672, EPI_ISL_3434673, EPI_ISL_3434674, EPI_ISL_3434675, EPI_ISL_3434676, EPI_ISL_3434677, EPI_ISL_3434678, EPI_ISL_3434679, EPI_ISL_3434680, EPI_ISL_3434681, EPI_ISL_3434682, EPI_ISL_3434684, EPI_ISL_3434685, EPI_ISL_3434686, EPI_ISL_3434687, EPI_ISL_3434688

see above IN State Department of Health Laboratory Services IN State Department of Health Laboratory Services Cassandra Campion, Jamie Yeadon, Brian Pope, Lixia Liu, Kyle Brownlee, Melissa Hindenlang, Mark Glazier

EPI_ISL_3434690, EPI_ISL_3434691, EPI_ISL_3434692, EPI_ISL_3434693, EPI_ISL_3434694, EPI_ISL_3434695, EPI_ISL_3434696, EPI_ISL_3434697, EPI_ISL_3434698, EPI_ISL_3434699, EPI_ISL_3434700, EPI_ISL_3434701, EPI_ISL_3434702, EPI_ISL_3434703, EPI_ISL_3434704, EPI_ISL_3434705, EPI_ISL_3434706, EPI_ISL_3434707, EPI_ISL_3434708, EPI_ISL_3434709, EPI_ISL_3434710, EPI_ISL_3434711,

| | | | |
|---|---|---|---|
| EPI_ISL_3434712, EPI_ISL_3434713, EPI_ISL_3434714, EPI_ISL_3434715, EPI_ISL_3434716, EPI_ISL_3434717 | | | |
| see above | Kansas Health and Environmental Lab | Kansas Health and Environmental Lab | Mike Grose, Jonathan Barnell, Amanda Bradley, and Phil Adam |
| EPI_ISL_3434720 | Azarian Lab at Burnett School of Biomedical Sciences | Azarian Lab at Burnett School of Biomedical Sciences | Catherine Johnston, Eleonora Cella, Taj Azarian |
| EPI_ISL_3435076, EPI_ISL_3435080, EPI_ISL_3435081, EPI_ISL_3435082, EPI_ISL_3435083, EPI_ISL_3435084, EPI_ISL_3435085, EPI_ISL_3435086, EPI_ISL_3435087, EPI_ISL_3435088, EPI_ISL_3435089, EPI_ISL_3435090, EPI_ISL_3435091, EPI_ISL_3435092, EPI_ISL_3435093, EPI_ISL_3435094, EPI_ISL_3435095, EPI_ISL_3435096, EPI_ISL_3435097, EPI_ISL_3435098, EPI_ISL_3435099, EPI_ISL_3435100, EPI_ISL_3435101, EPI_ISL_3435102, EPI_ISL_3435103, EPI_ISL_3435104, EPI_ISL_3435105, EPI_ISL_3435106, EPI_ISL_3435107, EPI_ISL_3435108, EPI_ISL_3435109, EPI_ISL_3435110, EPI_ISL_3435111, EPI_ISL_3435112, EPI_ISL_3435113, EPI_ISL_3435114, EPI_ISL_3435115, EPI_ISL_3435116, EPI_ISL_3435117, EPI_ISL_3435118, EPI_ISL_3435119, EPI_ISL_3435120, EPI_ISL_3435121, EPI_ISL_3435122, EPI_ISL_3435123, EPI_ISL_3435124, EPI_ISL_3435125, EPI_ISL_3435126, EPI_ISL_3435127, EPI_ISL_3435128, EPI_ISL_3435129, EPI_ISL_3435130, EPI_ISL_3435131, EPI_ISL_3435132, EPI_ISL_3435133, EPI_ISL_3435134, EPI_ISL_3435135, EPI_ISL_3435136, EPI_ISL_3435137, EPI_ISL_3435138, EPI_ISL_3435139, EPI_ISL_3435140, EPI_ISL_3435141, EPI_ISL_3435142, EPI_ISL_3435143, EPI_ISL_3435144, EPI_ISL_3435145, EPI_ISL_3435146, EPI_ISL_3435147, EPI_ISL_3435148, EPI_ISL_3435149, EPI_ISL_3435150, EPI_ISL_3435151, EPI_ISL_3435152, EPI_ISL_3435153, EPI_ISL_3435154, EPI_ISL_3435155, EPI_ISL_3435156, EPI_ISL_3435157, EPI_ISL_3435158, EPI_ISL_3435159, EPI_ISL_3435160, EPI_ISL_3435161, EPI_ISL_3435162, EPI_ISL_3435163, EPI_ISL_3435164, EPI_ISL_3435165, EPI_ISL_3435166, EPI_ISL_3435167, EPI_ISL_3435168, EPI_ISL_3435169, EPI_ISL_3435170, EPI_ISL_3435171, EPI_ISL_3435172, EPI_ISL_3435173, EPI_ISL_3435174, EPI_ISL_3435175, EPI_ISL_3435176, EPI_ISL_3435177, EPI_ISL_3435178, EPI_ISL_3435179, EPI_ISL_3435180, EPI_ISL_3435181, EPI_ISL_3435182, EPI_ISL_3435183, EPI_ISL_3435184, EPI_ISL_3435185, EPI_ISL_3435186, EPI_ISL_3435187, EPI_ISL_3435188, EPI_ISL_3435189, EPI_ISL_3435190, EPI_ISL_3435191, EPI_ISL_3435192, EPI_ISL_3435193, EPI_ISL_3435194, EPI_ISL_3435195, EPI_ISL_3435196, EPI_ISL_3435197, EPI_ISL_3435198, EPI_ISL_3435199, EPI_ISL_3435200, EPI_ISL_3435201, EPI_ISL_3435202, EPI_ISL_3435203, EPI_ISL_3435204, EPI_ISL_3435205, EPI_ISL_3435206, EPI_ISL_3435207, EPI_ISL_3435208, EPI_ISL_3435209, EPI_ISL_3435210, EPI_ISL_3435211, EPI_ISL_3435212, EPI_ISL_3435213, EPI_ISL_3435214, EPI_ISL_3435215, EPI_ISL_3435216, EPI_ISL_3435217, EPI_ISL_3435218, EPI_ISL_3435219, EPI_ISL_3435220, EPI_ISL_3435221, EPI_ISL_3435222, EPI_ISL_3435223, EPI_ISL_3435224, EPI_ISL_3435225, EPI_ISL_3435226, EPI_ISL_3435227, EPI_ISL_3435228, EPI_ISL_3435229, EPI_ISL_3435230, EPI_ISL_3435231, EPI_ISL_3435232, EPI_ISL_3435233, EPI_ISL_3435234, EPI_ISL_3435235, EPI_ISL_3435236, EPI_ISL_3435237, EPI_ISL_3435238, EPI_ISL_3435239, EPI_ISL_3435240, EPI_ISL_3435241, EPI_ISL_3435242, EPI_ISL_3435243, EPI_ISL_3435244, EPI_ISL_3435245, EPI_ISL_3435246, EPI_ISL_3435247, EPI_ISL_3435248, EPI_ISL_3435249, EPI_ISL_3435250, EPI_ISL_3435251, EPI_ISL_3435252, EPI_ISL_3435253, EPI_ISL_3435254, EPI_ISL_3435255, EPI_ISL_3435256, EPI_ISL_3435257, EPI_ISL_3435258, EPI_ISL_3435259, EPI_ISL_3435260, EPI_ISL_3435261, EPI_ISL_3435262, EPI_ISL_3435263, EPI_ISL_3435264, EPI_ISL_3435265, EPI_ISL_3435266, EPI_ISL_3435267, EPI_ISL_3435268, EPI_ISL_3435269, EPI_ISL_3435270, EPI_ISL_3435271, EPI_ISL_3435272, EPI_ISL_3435273, EPI_ISL_3435274, EPI_ISL_3435275, EPI_ISL_3435276, EPI_ISL_3435277, EPI_ISL_3435278 | | | |
| see above | GA Department of Public Health | GA Department of Public Health | Stacy Reeves, Sharmila Talekar, Jonathan Edwards, Cynthia Dixey, Tonia Parrott, Aliyah Fields, Taylor Smith |
| EPI_ISL_3435279, EPI_ISL_3435280, EPI_ISL_3435281, EPI_ISL_3435282, EPI_ISL_3435283, EPI_ISL_3435284, EPI_ISL_3435286, EPI_ISL_3435287, EPI_ISL_3435288, EPI_ISL_3435290, EPI_ISL_3435291, EPI_ISL_3435292, EPI_ISL_3435293, EPI_ISL_3435294, EPI_ISL_3435295, EPI_ISL_3435296, EPI_ISL_3435297, EPI_ISL_3435299, EPI_ISL_3435305, EPI_ISL_3435307, EPI_ISL_3435308, EPI_ISL_3435315, EPI_ISL_3435344, EPI_ISL_3435345, EPI_ISL_3435346, EPI_ISL_3435349, EPI_ISL_3435350, EPI_ISL_3435351, EPI_ISL_3435353, EPI_ISL_3435354, EPI_ISL_3435360, EPI_ISL_3435361, EPI_ISL_3435367, EPI_ISL_3435368, EPI_ISL_3435369, EPI_ISL_3435378, EPI_ISL_3435379, EPI_ISL_3435380, EPI_ISL_3435381, EPI_ISL_3435382, EPI_ISL_3435383, EPI_ISL_3435384, EPI_ISL_3435385, EPI_ISL_3435386, EPI_ISL_3435387, EPI_ISL_3435388, EPI_ISL_3435389, EPI_ISL_3435390, EPI_ISL_3435391, EPI_ISL_3435392, EPI_ISL_3435393, EPI_ISL_3435394, EPI_ISL_3435395, EPI_ISL_3435396, EPI_ISL_3435397, EPI_ISL_3435398, EPI_ISL_3435399, EPI_ISL_3435400, EPI_ISL_3435401, EPI_ISL_3435402, EPI_ISL_3435403, EPI_ISL_3435404, EPI_ISL_3435405, EPI_ISL_3435406, EPI_ISL_3435407, EPI_ISL_3435408, EPI_ISL_3435409, EPI_ISL_3435410, EPI_ISL_3435411, EPI_ISL_3435412, EPI_ISL_3435413, EPI_ISL_3435414, EPI_ISL_3435415, EPI_ISL_3435416, EPI_ISL_3435417, EPI_ISL_3435418, EPI_ISL_3435419, EPI_ISL_3435420, EPI_ISL_3435421, EPI_ISL_3435422, EPI_ISL_3435423, EPI_ISL_3435424, EPI_ISL_3435425, EPI_ISL_3435426, EPI_ISL_3435427, EPI_ISL_3435428, EPI_ISL_3435429, EPI_ISL_3435430, EPI_ISL_3435431, EPI_ISL_3435432, EPI_ISL_3435433, EPI_ISL_3435434, EPI_ISL_3435435, EPI_ISL_3435436, EPI_ISL_3435437, EPI_ISL_3435438, EPI_ISL_3435439, EPI_ISL_3435440, EPI_ISL_3435441, EPI_ISL_3435442, EPI_ISL_3435443, EPI_ISL_3435444, EPI_ISL_3435445, EPI_ISL_3435446, EPI_ISL_3435447, EPI_ISL_3435448, EPI_ISL_3435449, EPI_ISL_3435450, EPI_ISL_3435451, EPI_ISL_3435452, EPI_ISL_3435454, EPI_ISL_3435455 | | | |
| see above | Maryland Genomics, Institute for Genome Sciences, University of Maryland School of Medicine | Maryland Genomics, Institute for Genome Sciences, University of Maryland School of Medicine | Tallon, Luke J; Sadzewicz, Lisa D; Humphrys, Mike; Ott, Sandra; Roussey, Holly; Santana-Cruz, Ivette; Vavikolanu, Kranthi; Lim, Jonathan;Hazen, Tracy;Fraser, Claire M; Ravel, Jacques |
| EPI_ISL_3435456 | Lighthouse Lab in Alderley Park | Wellcome Sanger Institute for the COVID-19 Genomics UK (COG-UK) Consortium | Jacquelyn Wynn, Mairead Hyland, The Lighthouse Lab in Alderley Park and Alex Alderton, Roberto Amato, Jeffrey Barrett, Sonia Goncalves, Ewan Harrison, David K. Jackson, Ian Johnston, Dominic Kwiatkowski, Cordelia Langford, John Sillitoe on behalf of the Wellcome Sanger Institute COVID-19 Surveillance Team |
| EPI_ISL_3435457 | Lighthouse Lab in Milton Keynes | Wellcome Sanger Institute for the COVID-19 Genomics UK (COG-UK) Consortium | The Lighthouse Lab in Milton Keynes and Alex Alderton, Roberto Amato, Jeffrey Barrett, Sonia Goncalves, Ewan Harrison, David K. Jackson, Ian Johnston, Dominic Kwiatkowski, Cordelia Langford, John Sillitoe on behalf of the Wellcome Sanger Institute COVID-19 Surveillance Team |
| EPI_ISL_3435458 | Lighthouse Lab in Glasgow | Wellcome Sanger Institute for the COVID-19 Genomics UK (COG-UK) Consortium | Harper VanSteenhouse, Yumi Kasai, David Gray, Carol Clugston, Anna Dominiczak and Alex Alderton, Roberto Amato, Jeffrey Barrett, Sonia Goncalves, Ewan Harrison, David K. Jackson, Ian Johnston, Dominic Kwiatkowski, Cordelia Langford, John Sillitoe on behalf of the Wellcome Sanger Institute COVID-19 Surveillance Team |
| EPI_ISL_3435459, EPI_ISL_3435460, EPI_ISL_3435461, EPI_ISL_3435462 | Lighthouse Lab in Alderley Park | Wellcome Sanger Institute for the COVID-19 Genomics UK (COG-UK) Consortium | Jacquelyn Wynn, Mairead Hyland, The Lighthouse Lab in Alderley Park and Alex Alderton, Roberto Amato, Jeffrey Barrett, Sonia Goncalves, Ewan Harrison, David K. Jackson, Ian Johnston, Dominic Kwiatkowski, Cordelia Langford, John Sillitoe on behalf of the Wellcome Sanger Institute COVID-19 Surveillance Team |
| EPI_ISL_3435463 | Lighthouse Lab in Glasgow | Wellcome Sanger Institute for the COVID-19 Genomics UK (COG-UK) Consortium | Harper VanSteenhouse, Yumi Kasai, David Gray, Carol Clugston, Anna Dominiczak and Alex Alderton, Roberto Amato, Jeffrey Barrett, Sonia Goncalves, Ewan Harrison, David K. Jackson, Ian Johnston, Dominic Kwiatkowski, Cordelia Langford, John Sillitoe on behalf of the Wellcome Sanger Institute COVID-19 Surveillance Team |
| EPI_ISL_3435464, EPI_ISL_3435465, EPI_ISL_3435466 | Lighthouse Lab in Milton Keynes | Wellcome Sanger Institute for the COVID-19 Genomics UK (COG-UK) Consortium | The Lighthouse Lab in Milton Keynes and Alex Alderton, Roberto Amato, Jeffrey Barrett, Sonia Goncalves, Ewan Harrison, David K. Jackson, Ian Johnston, Dominic Kwiatkowski, Cordelia Langford, John Sillitoe on behalf of the Wellcome Sanger Institute COVID-19 Surveillance Team |
| EPI_ISL_3435467 | Lighthouse Lab in Alderley Park | Wellcome Sanger Institute for the COVID-19 Genomics UK (COG-UK) Consortium | Jacquelyn Wynn, Mairead Hyland, The Lighthouse Lab in Alderley Park and Alex Alderton, Roberto Amato, Jeffrey Barrett, Sonia Goncalves, Ewan Harrison, David K. Jackson, Ian Johnston, Dominic Kwiatkowski, Cordelia Langford, John Sillitoe on behalf of the Wellcome Sanger Institute COVID-19 Surveillance Team |
| EPI_ISL_3435468 | Lighthouse Lab in Glasgow | Wellcome Sanger Institute for the COVID-19 Genomics UK (COG-UK) Consortium | Harper VanSteenhouse, Yumi Kasai, David Gray, Carol Clugston, Anna Dominiczak and Alex Alderton, Roberto Amato, Jeffrey Barrett, Sonia Goncalves, Ewan Harrison, David K. Jackson, Ian Johnston, Dominic Kwiatkowski, Cordelia Langford, John Sillitoe on behalf of the Wellcome Sanger Institute COVID-19 Surveillance Team |
| EPI_ISL_3435469 | Lighthouse Lab in Alderley Park | Wellcome Sanger Institute for the COVID-19 Genomics UK (COG-UK) Consortium | Jacquelyn Wynn, Mairead Hyland, The Lighthouse Lab in Alderley Park and Alex Alderton, Roberto Amato, Jeffrey Barrett, Sonia Goncalves, Ewan Harrison, David K. Jackson, Ian Johnston, Dominic Kwiatkowski, Cordelia Langford, John Sillitoe on behalf of the Wellcome Sanger Institute COVID-19 Surveillance Team |
| EPI_ISL_3435470 | Lighthouse Lab in Glasgow | Wellcome Sanger Institute for the COVID-19 Genomics UK (COG-UK) Consortium | Harper VanSteenhouse, Yumi Kasai, David Gray, Carol Clugston, Anna Dominiczak and Alex Alderton, Roberto Amato, Jeffrey Barrett, Sonia Goncalves, Ewan Harrison, David K. Jackson, Ian Johnston, Dominic Kwiatkowski, Cordelia Langford, John Sillitoe on behalf of the Wellcome Sanger Institute COVID-19 Surveillance Team |
| EPI_ISL_3435471 | Lighthouse Lab in Milton Keynes | Wellcome Sanger Institute for the COVID-19 Genomics UK (COG-UK) Consortium | The Lighthouse Lab in Milton Keynes and Alex Alderton, Roberto Amato, Jeffrey Barrett, Sonia Goncalves, Ewan Harrison, David K. Jackson, Ian Johnston, Dominic Kwiatkowski, Cordelia Langford, John Sillitoe on behalf of the Wellcome Sanger Institute COVID-19 Surveillance Team |
| EPI_ISL_3435472, EPI_ISL_3435473 | Lighthouse Lab in Glasgow | Wellcome Sanger Institute for the COVID-19 Genomics UK (COG-UK) Consortium | Harper VanSteenhouse, Yumi Kasai, David Gray, Carol Clugston, Anna Dominiczak and Alex Alderton, Roberto Amato, Jeffrey Barrett, Sonia Goncalves, Ewan Harrison, David K. Jackson, Ian Johnston, Dominic Kwiatkowski, Cordelia Langford, John Sillitoe on behalf of the Wellcome Sanger Institute COVID-19 Surveillance Team |
| EPI_ISL_3435474 | Lighthouse Lab in Alderley Park | Wellcome Sanger Institute for the COVID-19 Genomics UK (COG-UK) Consortium | Jacquelyn Wynn, Mairead Hyland, The Lighthouse Lab in Alderley Park and Alex Alderton, Roberto Amato, Jeffrey Barrett, Sonia Goncalves, Ewan Harrison, David K. Jackson, Ian Johnston, Dominic Kwiatkowski, Cordelia Langford, John Sillitoe on behalf of the Wellcome Sanger Institute COVID-19 Surveillance Team |
| EPI_ISL_3435475 | Lighthouse Lab in Milton Keynes | Wellcome Sanger Institute for the COVID-19 Genomics UK (COG-UK) Consortium | The Lighthouse Lab in Milton Keynes and Alex Alderton, Roberto Amato, Jeffrey Barrett, Sonia Goncalves, Ewan Harrison, David K. Jackson, Ian Johnston, Dominic Kwiatkowski, Cordelia Langford, John Sillitoe on behalf of the Wellcome Sanger Institute COVID-19 Surveillance Team |
| EPI_ISL_3435476, EPI_ISL_3435477 | Lighthouse Lab in Glasgow | Wellcome Sanger Institute for the COVID-19 Genomics UK (COG-UK) Consortium | Harper VanSteenhouse, Yumi Kasai, David Gray, Carol Clugston, Anna Dominiczak and Alex Alderton, Roberto Amato, Jeffrey Barrett, Sonia Goncalves, Ewan Harrison, David K. Jackson, Ian Johnston, Dominic Kwiatkowski, Cordelia Langford, John Sillitoe on behalf of the Wellcome Sanger Institute COVID-19 Surveillance Team |
| EPI_ISL_3435478 | Lighthouse Lab in Alderley Park | Wellcome Sanger Institute for the COVID-19 Genomics UK (COG-UK) Consortium | Jacquelyn Wynn, Mairead Hyland, The Lighthouse Lab in Alderley Park and Alex Alderton, Roberto Amato, Jeffrey Barrett, Sonia Goncalves, Ewan Harrison, David K. Jackson, Ian Johnston, Dominic Kwiatkowski, Cordelia Langford, John Sillitoe on behalf of the Wellcome Sanger Institute COVID-19 Surveillance Team |
| EPI_ISL_3435479 | Lighthouse Lab in Glasgow | Wellcome Sanger Institute for the COVID-19 Genomics UK (COG-UK) Consortium | Harper VanSteenhouse, Yumi Kasai, David Gray, Carol Clugston, Anna Dominiczak and Alex Alderton, Roberto Amato, Jeffrey Barrett, Sonia Goncalves, Ewan Harrison, David K. Jackson, Ian Johnston, Dominic Kwiatkowski, Cordelia Langford, John Sillitoe on behalf of the Wellcome Sanger Institute COVID-19 Surveillance Team |
| EPI_ISL_3435480, EPI_ISL_3435481 | Lighthouse Lab in Alderley Park | Wellcome Sanger Institute for the COVID-19 Genomics UK (COG-UK) Consortium | Jacquelyn Wynn, Mairead Hyland, The Lighthouse Lab in Alderley Park and Alex Alderton, Roberto Amato, Jeffrey Barrett, Sonia Goncalves, Ewan Harrison, David K. Jackson, Ian Johnston, Dominic Kwiatkowski, Cordelia Langford, John Sillitoe on behalf of the Wellcome Sanger Institute COVID-19 Surveillance Team |
| EPI_ISL_3435482, EPI_ISL_3435483, EPI_ISL_3435484 | Lighthouse Lab in Milton Keynes | Wellcome Sanger Institute for the COVID-19 Genomics UK (COG-UK) Consortium | The Lighthouse Lab in Milton Keynes and Alex Alderton, Roberto Amato, Jeffrey Barrett, Sonia Goncalves, Ewan Harrison, David K. Jackson, Ian Johnston, Dominic Kwiatkowski, Cordelia Langford, John Sillitoe on behalf of the Wellcome Sanger Institute COVID-19 Surveillance Team |
| EPI_ISL_3435485, EPI_ISL_3435486 | Lighthouse Lab in Glasgow | Wellcome Sanger Institute for the COVID-19 Genomics UK (COG-UK) Consortium | Harper VanSteenhouse, Yumi Kasai, David Gray, Carol Clugston, Anna Dominiczak and Alex Alderton, Roberto Amato, Jeffrey Barrett, Sonia Goncalves, Ewan Harrison, David K. Jackson, Ian Johnston, Dominic Kwiatkowski, Cordelia Langford, John Sillitoe on behalf of the Wellcome Sanger Institute COVID-19 Surveillance Team |
| EPI_ISL_3435487, EPI_ISL_3435488 | Lighthouse Lab in Alderley Park | Wellcome Sanger Institute for the COVID-19 Genomics UK (COG-UK) Consortium | Jacquelyn Wynn, Mairead Hyland, The Lighthouse Lab in Alderley Park and Alex Alderton, Roberto Amato, Jeffrey Barrett, Sonia Goncalves, Ewan Harrison, David K. Jackson, Ian Johnston, Dominic Kwiatkowski, Cordelia Langford, John Sillitoe on behalf of the Wellcome Sanger Institute COVID-19 Surveillance Team |
| EPI_ISL_3435489, EPI_ISL_3435490, EPI_ISL_3435491, EPI_ISL_3435492 | Lighthouse Lab in Milton Keynes | Wellcome Sanger Institute for the COVID-19 Genomics UK (COG-UK) Consortium | The Lighthouse Lab in Milton Keynes and Alex Alderton, Roberto Amato, Jeffrey Barrett, Sonia Goncalves, Ewan Harrison, David K. Jackson, Ian Johnston, Dominic Kwiatkowski, Cordelia Langford, John Sillitoe on behalf of the Wellcome Sanger Institute COVID-19 Surveillance Team |
| EPI_ISL_3435493, EPI_ISL_3435494, EPI_ISL_3435495 | Lighthouse Lab in Glasgow | Wellcome Sanger Institute for the COVID-19 Genomics UK (COG-UK) Consortium | Harper VanSteenhouse, Yumi Kasai, David Gray, Carol Clugston, Anna Dominiczak and Alex Alderton, Roberto Amato, Jeffrey Barrett, Sonia Goncalves, Ewan Harrison, David K. Jackson, Ian Johnston, Dominic Kwiatkowski, Cordelia Langford, John Sillitoe on behalf of the Wellcome Sanger Institute COVID-19 Surveillance Team |
| EPI_ISL_3435496 | Lighthouse Lab in Milton Keynes | Wellcome Sanger Institute for the COVID-19 Genomics UK (COG-UK) Consortium | The Lighthouse Lab in Milton Keynes and Alex Alderton, Roberto Amato, Jeffrey Barrett, Sonia Goncalves, Ewan Harrison, David K. Jackson, Ian Johnston, Dominic Kwiatkowski, Cordelia Langford, John Sillitoe on behalf of the Wellcome Sanger Institute COVID-19 Surveillance Team |
| EPI_ISL_3435497, EPI_ISL_3435498 | Lighthouse Lab in Alderley Park | Wellcome Sanger Institute for the COVID-19 Genomics UK (COG-UK) Consortium | Jacquelyn Wynn, Mairead Hyland, The Lighthouse Lab in Alderley Park and Alex Alderton, Roberto Amato, Jeffrey Barrett, Sonia Goncalves, Ewan Harrison, David K. Jackson, Ian Johnston, Dominic Kwiatkowski, Cordelia Langford, John Sillitoe on behalf of the Wellcome Sanger Institute COVID-19 Surveillance Team |
| EPI_ISL_3435499 | Lighthouse Lab in Glasgow | Wellcome Sanger Institute for the COVID-19 Genomics UK (COG-UK) Consortium | Harper VanSteenhouse, Yumi Kasai, David Gray, Carol Clugston, Anna Dominiczak and Alex Alderton, Roberto Amato, Jeffrey Barrett, Sonia Goncalves, Ewan Harrison, David K. Jackson, Ian Johnston, Dominic Kwiatkowski, Cordelia Langford, John Sillitoe on behalf of the Wellcome Sanger Institute COVID-19 Surveillance Team |
| EPI_ISL_3435500 | Lighthouse Lab in Alderley Park | Wellcome Sanger Institute for the COVID-19 Genomics UK (COG-UK) Consortium | Jacquelyn Wynn, Mairead Hyland, The Lighthouse Lab in Alderley Park and Alex Alderton, Roberto Amato, Jeffrey Barrett, Sonia Goncalves, Ewan Harrison, David K. Jackson, Ian Johnston, Dominic Kwiatkowski, Cordelia Langford, John Sillitoe on behalf of the Wellcome Sanger Institute COVID-19 Surveillance Team |
| EPI_ISL_3435501, EPI_ISL_3435502, EPI_ISL_3435503, EPI_ISL_3435504 | Lighthouse Lab in Milton Keynes | Wellcome Sanger Institute for the COVID-19 Genomics UK (COG-UK) Consortium | The Lighthouse Lab in Milton Keynes and Alex Alderton, Roberto Amato, Jeffrey Barrett, Sonia Goncalves, Ewan Harrison, David K. Jackson, Ian Johnston, Dominic Kwiatkowski, Cordelia Langford, John Sillitoe on behalf of the Wellcome Sanger Institute COVID-19 Surveillance Team |
| EPI_ISL_3435505 | Lighthouse Lab in Glasgow | Wellcome Sanger Institute for the COVID-19 Genomics UK (COG-UK) Consortium | Harper VanSteenhouse, Yumi Kasai, David Gray, Carol Clugston, Anna Dominiczak and Alex Alderton, Roberto Amato, Jeffrey Barrett, Sonia Goncalves, Ewan Harrison, David K. Jackson, Ian Johnston, Dominic Kwiatkowski, Cordelia Langford, John Sillitoe on behalf of the Wellcome Sanger Institute COVID-19 Surveillance Team |
| EPI_ISL_3435506 | Lighthouse Lab in Milton Keynes | Wellcome Sanger Institute for the COVID-19 Genomics UK (COG-UK) Consortium | The Lighthouse Lab in Milton Keynes and Alex Alderton, Roberto Amato, Jeffrey Barrett, Sonia Goncalves, Ewan Harrison, David K. Jackson, Ian Johnston, Dominic Kwiatkowski, Cordelia Langford, John Sillitoe on behalf of the Wellcome Sanger Institute COVID-19 Surveillance Team |
| EPI_ISL_3435507, EPI_ISL_3435508, EPI_ISL_3435509 | Lighthouse Lab in Glasgow | Wellcome Sanger Institute for the COVID-19 Genomics UK (COG-UK) Consortium | Harper VanSteenhouse, Yumi Kasai, David Gray, Carol Clugston, Anna Dominiczak and Alex Alderton, Roberto Amato, Jeffrey Barrett, Sonia Goncalves, Ewan Harrison, David K. Jackson, Ian Johnston, Dominic Kwiatkowski, Cordelia Langford, John Sillitoe on behalf of the Wellcome Sanger Institute COVID-19 Surveillance Team |

| | | | |
|---|---|---|---|
| EPI_ISL_3435510 | Lighthouse Lab in Alderley Park | Wellcome Sanger Institute for the COVID-19 Genomics UK (COG-UK) Consortium | Jacquelyn Wynn, Mairead Hyland, The Lighthouse Lab in Alderley Park and Alex Alderton, Roberto Amato, Jeffrey Barrett, Sonia Goncalves, Ewan Harrison, David K. Jackson, Ian Johnston, Dominic Kwiatkowski, Cordelia Langford, John Sillitoe on behalf of the Wellcome Sanger Institute COVID-19 Surveillance Team |
| EPI_ISL_3435511, EPI_ISL_3435512 | Lighthouse Lab in Milton Keynes | Wellcome Sanger Institute for the COVID-19 Genomics UK (COG-UK) Consortium | The Lighthouse Lab in Milton Keynes and Alex Alderton, Roberto Amato, Jeffrey Barrett, Sonia Goncalves, Ewan Harrison, David K. Jackson, Ian Johnston, Dominic Kwiatkowski, Cordelia Langford, John Sillitoe on behalf of the Wellcome Sanger Institute COVID-19 Surveillance Team |
| EPI_ISL_3435513, EPI_ISL_3435514 | Lighthouse Lab in Glasgow | Wellcome Sanger Institute for the COVID-19 Genomics UK (COG-UK) Consortium | Harper VanSteenhouse, Yumi Kasai, David Gray, Carol Clugston, Anna Dominiczak and Alex Alderton, Roberto Amato, Jeffrey Barrett, Sonia Goncalves, Ewan Harrison, David K. Jackson, Ian Johnston, Dominic Kwiatkowski, Cordelia Langford, John Sillitoe on behalf of the Wellcome Sanger Institute COVID-19 Surveillance Team |
| EPI_ISL_3435515 | Lighthouse Lab in Milton Keynes | Wellcome Sanger Institute for the COVID-19 Genomics UK (COG-UK) Consortium | The Lighthouse Lab in Milton Keynes and Alex Alderton, Roberto Amato, Jeffrey Barrett, Sonia Goncalves, Ewan Harrison, David K. Jackson, Ian Johnston, Dominic Kwiatkowski, Cordelia Langford, John Sillitoe on behalf of the Wellcome Sanger Institute COVID-19 Surveillance Team |
| EPI_ISL_3435516 | Lighthouse Lab in Alderley Park | Wellcome Sanger Institute for the COVID-19 Genomics UK (COG-UK) Consortium | Jacquelyn Wynn, Mairead Hyland, The Lighthouse Lab in Alderley Park and Alex Alderton, Roberto Amato, Jeffrey Barrett, Sonia Goncalves, Ewan Harrison, David K. Jackson, Ian Johnston, Dominic Kwiatkowski, Cordelia Langford, John Sillitoe on behalf of the Wellcome Sanger Institute COVID-19 Surveillance Team |
| EPI_ISL_3435517, EPI_ISL_3435518 | Lighthouse Lab in Milton Keynes | Wellcome Sanger Institute for the COVID-19 Genomics UK (COG-UK) Consortium | The Lighthouse Lab in Milton Keynes and Alex Alderton, Roberto Amato, Jeffrey Barrett, Sonia Goncalves, Ewan Harrison, David K. Jackson, Ian Johnston, Dominic Kwiatkowski, Cordelia Langford, John Sillitoe on behalf of the Wellcome Sanger Institute COVID-19 Surveillance Team |
| EPI_ISL_3435519 | Lighthouse Lab in Alderley Park | Wellcome Sanger Institute for the COVID-19 Genomics UK (COG-UK) Consortium | Jacquelyn Wynn, Mairead Hyland, The Lighthouse Lab in Alderley Park and Alex Alderton, Roberto Amato, Jeffrey Barrett, Sonia Goncalves, Ewan Harrison, David K. Jackson, Ian Johnston, Dominic Kwiatkowski, Cordelia Langford, John Sillitoe on behalf of the Wellcome Sanger Institute COVID-19 Surveillance Team |
| EPI_ISL_3435520 | Lighthouse Lab in Glasgow | Wellcome Sanger Institute for the COVID-19 Genomics UK (COG-UK) Consortium | Harper VanSteenhouse, Yumi Kasai, David Gray, Carol Clugston, Anna Dominiczak and Alex Alderton, Roberto Amato, Jeffrey Barrett, Sonia Goncalves, Ewan Harrison, David K. Jackson, Ian Johnston, Dominic Kwiatkowski, Cordelia Langford, John Sillitoe on behalf of the Wellcome Sanger Institute COVID-19 Surveillance Team |
| EPI_ISL_3435521 | Lighthouse Lab in Milton Keynes | Wellcome Sanger Institute for the COVID-19 Genomics UK (COG-UK) Consortium | The Lighthouse Lab in Milton Keynes and Alex Alderton, Roberto Amato, Jeffrey Barrett, Sonia Goncalves, Ewan Harrison, David K. Jackson, Ian Johnston, Dominic Kwiatkowski, Cordelia Langford, John Sillitoe on behalf of the Wellcome Sanger Institute COVID-19 Surveillance Team |
| EPI_ISL_3435522 | Lighthouse Lab in Glasgow | Wellcome Sanger Institute for the COVID-19 Genomics UK (COG-UK) Consortium | Harper VanSteenhouse, Yumi Kasai, David Gray, Carol Clugston, Anna Dominiczak and Alex Alderton, Roberto Amato, Jeffrey Barrett, Sonia Goncalves, Ewan Harrison, David K. Jackson, Ian Johnston, Dominic Kwiatkowski, Cordelia Langford, John Sillitoe on behalf of the Wellcome Sanger Institute COVID-19 Surveillance Team |
| EPI_ISL_3435523 | Lighthouse Lab in Milton Keynes | Wellcome Sanger Institute for the COVID-19 Genomics UK (COG-UK) Consortium | The Lighthouse Lab in Milton Keynes and Alex Alderton, Roberto Amato, Jeffrey Barrett, Sonia Goncalves, Ewan Harrison, David K. Jackson, Ian Johnston, Dominic Kwiatkowski, Cordelia Langford, John Sillitoe on behalf of the Wellcome Sanger Institute COVID-19 Surveillance Team |
| EPI_ISL_3435524, EPI_ISL_3435525 | Lighthouse Lab in Glasgow | Wellcome Sanger Institute for the COVID-19 Genomics UK (COG-UK) Consortium | Harper VanSteenhouse, Yumi Kasai, David Gray, Carol Clugston, Anna Dominiczak and Alex Alderton, Roberto Amato, Jeffrey Barrett, Sonia Goncalves, Ewan Harrison, David K. Jackson, Ian Johnston, Dominic Kwiatkowski, Cordelia Langford, John Sillitoe on behalf of the Wellcome Sanger Institute COVID-19 Surveillance Team |
| EPI_ISL_3435526, EPI_ISL_3435527 | Lighthouse Lab in Milton Keynes | Wellcome Sanger Institute for the COVID-19 Genomics UK (COG-UK) Consortium | The Lighthouse Lab in Milton Keynes and Alex Alderton, Roberto Amato, Jeffrey Barrett, Sonia Goncalves, Ewan Harrison, David K. Jackson, Ian Johnston, Dominic Kwiatkowski, Cordelia Langford, John Sillitoe on behalf of the Wellcome Sanger Institute COVID-19 Surveillance Team |
| EPI_ISL_3435528 | Lighthouse Lab in Alderley Park | Wellcome Sanger Institute for the COVID-19 Genomics UK (COG-UK) Consortium | Jacquelyn Wynn, Mairead Hyland, The Lighthouse Lab in Alderley Park and Alex Alderton, Roberto Amato, Jeffrey Barrett, Sonia Goncalves, Ewan Harrison, David K. Jackson, Ian Johnston, Dominic Kwiatkowski, Cordelia Langford, John Sillitoe on behalf of the Wellcome Sanger Institute COVID-19 Surveillance Team |
| EPI_ISL_3435529 | Lighthouse Lab in Milton Keynes | Wellcome Sanger Institute for the COVID-19 Genomics UK (COG-UK) Consortium | The Lighthouse Lab in Milton Keynes and Alex Alderton, Roberto Amato, Jeffrey Barrett, Sonia Goncalves, Ewan Harrison, David K. Jackson, Ian Johnston, Dominic Kwiatkowski, Cordelia Langford, John Sillitoe on behalf of the Wellcome Sanger Institute COVID-19 Surveillance Team |
| EPI_ISL_3435530 | Lighthouse Lab in Glasgow | Wellcome Sanger Institute for the COVID-19 Genomics UK (COG-UK) Consortium | Harper VanSteenhouse, Yumi Kasai, David Gray, Carol Clugston, Anna Dominiczak and Alex Alderton, Roberto Amato, Jeffrey Barrett, Sonia Goncalves, Ewan Harrison, David K. Jackson, Ian Johnston, Dominic Kwiatkowski, Cordelia Langford, John Sillitoe on behalf of the Wellcome Sanger Institute COVID-19 Surveillance Team |
| EPI_ISL_3435531 | Lighthouse Lab in Milton Keynes | Wellcome Sanger Institute for the COVID-19 Genomics UK (COG-UK) Consortium | The Lighthouse Lab in Milton Keynes and Alex Alderton, Roberto Amato, Jeffrey Barrett, Sonia Goncalves, Ewan Harrison, David K. Jackson, Ian Johnston, Dominic Kwiatkowski, Cordelia Langford, John Sillitoe on behalf of the Wellcome Sanger Institute COVID-19 Surveillance Team |
| EPI_ISL_3435532 | Lighthouse Lab in Alderley Park | Wellcome Sanger Institute for the COVID-19 Genomics UK (COG-UK) Consortium | Jacquelyn Wynn, Mairead Hyland, The Lighthouse Lab in Alderley Park and Alex Alderton, Roberto Amato, Jeffrey Barrett, Sonia Goncalves, Ewan Harrison, David K. Jackson, Ian Johnston, Dominic Kwiatkowski, Cordelia Langford, John Sillitoe on behalf of the Wellcome Sanger Institute COVID-19 Surveillance Team |
| EPI_ISL_3435533, EPI_ISL_3435534 | Lighthouse Lab in Glasgow | Wellcome Sanger Institute for the COVID-19 Genomics UK (COG-UK) Consortium | Harper VanSteenhouse, Yumi Kasai, David Gray, Carol Clugston, Anna Dominiczak and Alex Alderton, Roberto Amato, Jeffrey Barrett, Sonia Goncalves, Ewan Harrison, David K. Jackson, Ian Johnston, Dominic Kwiatkowski, Cordelia Langford, John Sillitoe on behalf of the Wellcome Sanger Institute COVID-19 Surveillance Team |
| EPI_ISL_3435535 | Lighthouse Lab in Milton Keynes | Wellcome Sanger Institute for the COVID-19 Genomics UK (COG-UK) Consortium | The Lighthouse Lab in Milton Keynes and Alex Alderton, Roberto Amato, Jeffrey Barrett, Sonia Goncalves, Ewan Harrison, David K. Jackson, Ian Johnston, Dominic Kwiatkowski, Cordelia Langford, John Sillitoe on behalf of the Wellcome Sanger Institute COVID-19 Surveillance Team |
| EPI_ISL_3435536, EPI_ISL_3435537, EPI_ISL_3435538 | Lighthouse Lab in Glasgow | Wellcome Sanger Institute for the COVID-19 Genomics UK (COG-UK) Consortium | Harper VanSteenhouse, Yumi Kasai, David Gray, Carol Clugston, Anna Dominiczak and Alex Alderton, Roberto Amato, Jeffrey Barrett, Sonia Goncalves, Ewan Harrison, David K. Jackson, Ian Johnston, Dominic Kwiatkowski, Cordelia Langford, John Sillitoe on behalf of the Wellcome Sanger Institute COVID-19 Surveillance Team |
| EPI_ISL_3435539 | Lighthouse Lab in Alderley Park | Wellcome Sanger Institute for the COVID-19 Genomics UK (COG-UK) Consortium | Jacquelyn Wynn, Mairead Hyland, The Lighthouse Lab in Alderley Park and Alex Alderton, Roberto Amato, Jeffrey Barrett, Sonia Goncalves, Ewan Harrison, David K. Jackson, Ian Johnston, Dominic Kwiatkowski, Cordelia Langford, John Sillitoe on behalf of the Wellcome Sanger Institute COVID-19 Surveillance Team |
| EPI_ISL_3435540 | Lighthouse Lab in Glasgow | Wellcome Sanger Institute for the COVID-19 Genomics UK (COG-UK) Consortium | Harper VanSteenhouse, Yumi Kasai, David Gray, Carol Clugston, Anna Dominiczak and Alex Alderton, Roberto Amato, Jeffrey Barrett, Sonia Goncalves, Ewan Harrison, David K. Jackson, Ian Johnston, Dominic Kwiatkowski, Cordelia Langford, John Sillitoe on behalf of the Wellcome Sanger Institute COVID-19 Surveillance Team |
| EPI_ISL_3435541, EPI_ISL_3435542 | Lighthouse Lab in Milton Keynes | Wellcome Sanger Institute for the COVID-19 Genomics UK (COG-UK) Consortium | The Lighthouse Lab in Milton Keynes and Alex Alderton, Roberto Amato, Jeffrey Barrett, Sonia Goncalves, Ewan Harrison, David K. Jackson, Ian Johnston, Dominic Kwiatkowski, Cordelia Langford, John Sillitoe on behalf of the Wellcome Sanger Institute COVID-19 Surveillance Team |
| EPI_ISL_3435543 | Lighthouse Lab in Alderley Park | Wellcome Sanger Institute for the COVID-19 Genomics UK (COG-UK) Consortium | Jacquelyn Wynn, Mairead Hyland, The Lighthouse Lab in Alderley Park and Alex Alderton, Roberto Amato, Jeffrey Barrett, Sonia Goncalves, Ewan Harrison, David K. Jackson, Ian Johnston, Dominic Kwiatkowski, Cordelia Langford, John Sillitoe on behalf of the Wellcome Sanger Institute COVID-19 Surveillance Team |
| EPI_ISL_3435544 | Lighthouse Lab in Glasgow | Wellcome Sanger Institute for the COVID-19 Genomics UK (COG-UK) Consortium | Harper VanSteenhouse, Yumi Kasai, David Gray, Carol Clugston, Anna Dominiczak and Alex Alderton, Roberto Amato, Jeffrey Barrett, Sonia Goncalves, Ewan Harrison, David K. Jackson, Ian Johnston, Dominic Kwiatkowski, Cordelia Langford, John Sillitoe on behalf of the Wellcome Sanger Institute COVID-19 Surveillance Team |
| EPI_ISL_3435545 | Lighthouse Lab in Milton Keynes | Wellcome Sanger Institute for the COVID-19 Genomics UK (COG-UK) Consortium | The Lighthouse Lab in Milton Keynes and Alex Alderton, Roberto Amato, Jeffrey Barrett, Sonia Goncalves, Ewan Harrison, David K. Jackson, Ian Johnston, Dominic Kwiatkowski, Cordelia Langford, John Sillitoe on behalf of the Wellcome Sanger Institute COVID-19 Surveillance Team |
| EPI_ISL_3435546 | Lighthouse Lab in Glasgow | Wellcome Sanger Institute for the COVID-19 Genomics UK (COG-UK) Consortium | Harper VanSteenhouse, Yumi Kasai, David Gray, Carol Clugston, Anna Dominiczak and Alex Alderton, Roberto Amato, Jeffrey Barrett, Sonia Goncalves, Ewan Harrison, David K. Jackson, Ian Johnston, Dominic Kwiatkowski, Cordelia Langford, John Sillitoe on behalf of the Wellcome Sanger Institute COVID-19 Surveillance Team |
| EPI_ISL_3435547 | Lighthouse Lab in Alderley Park | Wellcome Sanger Institute for the COVID-19 Genomics UK (COG-UK) Consortium | Jacquelyn Wynn, Mairead Hyland, The Lighthouse Lab in Alderley Park and Alex Alderton, Roberto Amato, Jeffrey Barrett, Sonia Goncalves, Ewan Harrison, David K. Jackson, Ian Johnston, Dominic Kwiatkowski, Cordelia Langford, John Sillitoe on behalf of the Wellcome Sanger Institute COVID-19 Surveillance Team |
| EPI_ISL_3435548 | Lighthouse Lab in Glasgow | Wellcome Sanger Institute for the COVID-19 Genomics UK (COG-UK) Consortium | Harper VanSteenhouse, Yumi Kasai, David Gray, Carol Clugston, Anna Dominiczak and Alex Alderton, Roberto Amato, Jeffrey Barrett, Sonia Goncalves, Ewan Harrison, David K. Jackson, Ian Johnston, Dominic Kwiatkowski, Cordelia Langford, John Sillitoe on behalf of the Wellcome Sanger Institute COVID-19 Surveillance Team |
| EPI_ISL_3435549, EPI_ISL_3435550 | Lighthouse Lab in Alderley Park | Wellcome Sanger Institute for the COVID-19 Genomics UK (COG-UK) Consortium | Jacquelyn Wynn, Mairead Hyland, The Lighthouse Lab in Alderley Park and Alex Alderton, Roberto Amato, Jeffrey Barrett, Sonia Goncalves, Ewan Harrison, David K. Jackson, Ian Johnston, Dominic Kwiatkowski, Cordelia Langford, John Sillitoe on behalf of the Wellcome Sanger Institute COVID-19 Surveillance Team |
| EPI_ISL_3435551, EPI_ISL_3435552, EPI_ISL_3435553 | Lighthouse Lab in Milton Keynes | Wellcome Sanger Institute for the COVID-19 Genomics UK (COG-UK) Consortium | The Lighthouse Lab in Milton Keynes and Alex Alderton, Roberto Amato, Jeffrey Barrett, Sonia Goncalves, Ewan Harrison, David K. Jackson, Ian Johnston, Dominic Kwiatkowski, Cordelia Langford, John Sillitoe on behalf of the Wellcome Sanger Institute COVID-19 Surveillance Team |
| EPI_ISL_3435554, EPI_ISL_3435555 | Lighthouse Lab in Alderley Park | Wellcome Sanger Institute for the COVID-19 Genomics UK (COG-UK) Consortium | Jacquelyn Wynn, Mairead Hyland, The Lighthouse Lab in Alderley Park and Alex Alderton, Roberto Amato, Jeffrey Barrett, Sonia Goncalves, Ewan Harrison, David K. Jackson, Ian Johnston, Dominic Kwiatkowski, Cordelia Langford, John Sillitoe on behalf of the Wellcome Sanger Institute COVID-19 Surveillance Team |
| EPI_ISL_3435556 | Lighthouse Lab in Glasgow | Wellcome Sanger Institute for the COVID-19 Genomics UK (COG-UK) Consortium | Harper VanSteenhouse, Yumi Kasai, David Gray, Carol Clugston, Anna Dominiczak and Alex Alderton, Roberto Amato, Jeffrey Barrett, Sonia Goncalves, Ewan Harrison, David K. Jackson, Ian Johnston, Dominic Kwiatkowski, Cordelia Langford, John Sillitoe on behalf of the Wellcome Sanger Institute COVID-19 Surveillance Team |
| EPI_ISL_3435557, EPI_ISL_3435558, EPI_ISL_3435559 | Lighthouse Lab in Milton Keynes | Wellcome Sanger Institute for the COVID-19 Genomics UK (COG-UK) Consortium | The Lighthouse Lab in Milton Keynes and Alex Alderton, Roberto Amato, Jeffrey Barrett, Sonia Goncalves, Ewan Harrison, David K. Jackson, Ian Johnston, Dominic Kwiatkowski, Cordelia Langford, John Sillitoe on behalf of the Wellcome Sanger Institute COVID-19 Surveillance Team |
| EPI_ISL_3435560 | Lighthouse Lab in Glasgow | Wellcome Sanger Institute for the COVID-19 Genomics UK (COG-UK) Consortium | Harper VanSteenhouse, Yumi Kasai, David Gray, Carol Clugston, Anna Dominiczak and Alex Alderton, Roberto Amato, Jeffrey Barrett, Sonia Goncalves, Ewan Harrison, David K. Jackson, Ian Johnston, Dominic Kwiatkowski, Cordelia Langford, John Sillitoe on behalf of the Wellcome Sanger Institute COVID-19 Surveillance Team |
| EPI_ISL_3435561, EPI_ISL_3435562 | Lighthouse Lab in Alderley Park | Wellcome Sanger Institute for the COVID-19 Genomics UK (COG-UK) Consortium | Jacquelyn Wynn, Mairead Hyland, The Lighthouse Lab in Alderley Park and Alex Alderton, Roberto Amato, Jeffrey Barrett, Sonia Goncalves, Ewan Harrison, David K. Jackson, Ian Johnston, Dominic Kwiatkowski, Cordelia Langford, John Sillitoe on behalf of the Wellcome Sanger Institute COVID-19 Surveillance Team |
| EPI_ISL_3435563 | Lighthouse Lab in Milton Keynes | Wellcome Sanger Institute for the COVID-19 Genomics UK (COG-UK) Consortium | The Lighthouse Lab in Milton Keynes and Alex Alderton, Roberto Amato, Jeffrey Barrett, Sonia Goncalves, Ewan Harrison, David K. Jackson, Ian Johnston, Dominic Kwiatkowski, Cordelia Langford, John Sillitoe on behalf of the Wellcome Sanger Institute COVID-19 Surveillance Team |
| EPI_ISL_3435564 | Lighthouse Lab in Alderley Park | Wellcome Sanger Institute for the COVID-19 Genomics UK (COG-UK) Consortium | Jacquelyn Wynn, Mairead Hyland, The Lighthouse Lab in Alderley Park and Alex Alderton, Roberto Amato, Jeffrey Barrett, Sonia Goncalves, Ewan Harrison, David K. Jackson, Ian Johnston, Dominic Kwiatkowski, Cordelia Langford, John Sillitoe on behalf of the Wellcome Sanger Institute COVID-19 Surveillance Team |
| EPI_ISL_3435565, EPI_ISL_3435566, EPI_ISL_3435567 | Lighthouse Lab in Milton Keynes | Wellcome Sanger Institute for the COVID-19 Genomics UK (COG-UK) Consortium | The Lighthouse Lab in Milton Keynes and Alex Alderton, Roberto Amato, Jeffrey Barrett, Sonia Goncalves, Ewan Harrison, David K. Jackson, Ian Johnston, Dominic Kwiatkowski, Cordelia Langford, John Sillitoe on behalf of the Wellcome Sanger Institute COVID-19 Surveillance Team |
| EPI_ISL_3435568, EPI_ISL_3435569 | Lighthouse Lab in Glasgow | Wellcome Sanger Institute for the COVID-19 Genomics UK (COG-UK) Consortium | Harper VanSteenhouse, Yumi Kasai, David Gray, Carol Clugston, Anna Dominiczak and Alex Alderton, Roberto Amato, Jeffrey Barrett, Sonia Goncalves, Ewan Harrison, David K. Jackson, Ian Johnston, Dominic Kwiatkowski, Cordelia Langford, John Sillitoe on behalf of the Wellcome Sanger Institute COVID-19 Surveillance Team |
| EPI_ISL_3435570 | Lighthouse Lab in Milton Keynes | Wellcome Sanger Institute for the COVID-19 Genomics UK (COG-UK) Consortium | The Lighthouse Lab in Milton Keynes and Alex Alderton, Roberto Amato, Jeffrey Barrett, Sonia Goncalves, Ewan Harrison, David K. Jackson, Ian Johnston, Dominic Kwiatkowski, Cordelia Langford, John Sillitoe on behalf of the Wellcome Sanger Institute COVID-19 Surveillance Team |
| EPI_ISL_3435571 | Lighthouse Lab in Alderley Park | Wellcome Sanger Institute for the COVID-19 Genomics UK (COG-UK) Consortium | Jacquelyn Wynn, Mairead Hyland, The Lighthouse Lab in Alderley Park and Alex Alderton, Roberto Amato, Jeffrey Barrett, Sonia Goncalves, Ewan Harrison, David K. Jackson, Ian Johnston, Dominic Kwiatkowski, Cordelia Langford, John Sillitoe on behalf of the Wellcome Sanger Institute COVID-19 Surveillance Team |
| EPI_ISL_3435572, | Lighthouse Lab in Milton Keynes | Wellcome Sanger Institute for the COVID-19 Genomics UK (COG-UK) | The Lighthouse Lab in Milton Keynes and Alex Alderton, Roberto Amato, Jeffrey Barrett, Sonia Goncalves, Ewan Harrison, David K. Jackson, Ian Johnston, Dominic Kwiatkowski, Cordelia Langford, John Sillitoe on behalf of the Wellcome Sanger |

| | | | |
|---|---|---|---|
| EPI_ISL_3435573 | | Consortium | Institute COVID-19 Surveillance Team |
| EPI_ISL_3435574, EPI_ISL_3435575 | Lighthouse Lab in Glasgow | Wellcome Sanger Institute for the COVID-19 Genomics UK (COG-UK) Consortium | Harper VanSteenhouse, Yumi Kasai, David Gray, Carol Clugston, Anna Dominiczak and Alex Alderton, Roberto Amato, Jeffrey Barrett, Sonia Goncalves, Ewan Harrison, David K. Jackson, Ian Johnston, Dominic Kwiatkowski, Cordelia Langford, John Sillitoe on behalf of the Wellcome Sanger Institute COVID-19 Surveillance Team |
| EPI_ISL_3435576 | Lighthouse Lab in Milton Keynes | Wellcome Sanger Institute for the COVID-19 Genomics UK (COG-UK) Consortium | The Lighthouse Lab in Milton Keynes and Alex Alderton, Roberto Amato, Jeffrey Barrett, Sonia Goncalves, Ewan Harrison, David K. Jackson, Ian Johnston, Dominic Kwiatkowski, Cordelia Langford, John Sillitoe on behalf of the Wellcome Sanger Institute COVID-19 Surveillance Team |
| EPI_ISL_3435577, EPI_ISL_3435578 | Lighthouse Lab in Alderley Park | Wellcome Sanger Institute for the COVID-19 Genomics UK (COG-UK) Consortium | Jacquelyn Wynn, Mairead Hyland, The Lighthouse Lab in Alderley Park and Alex Alderton, Roberto Amato, Jeffrey Barrett, Sonia Goncalves, Ewan Harrison, David K. Jackson, Ian Johnston, Dominic Kwiatkowski, Cordelia Langford, John Sillitoe on behalf of the Wellcome Sanger Institute COVID-19 Surveillance Team |
| EPI_ISL_3435579 | Lighthouse Lab in Milton Keynes | Wellcome Sanger Institute for the COVID-19 Genomics UK (COG-UK) Consortium | The Lighthouse Lab in Milton Keynes and Alex Alderton, Roberto Amato, Jeffrey Barrett, Sonia Goncalves, Ewan Harrison, David K. Jackson, Ian Johnston, Dominic Kwiatkowski, Cordelia Langford, John Sillitoe on behalf of the Wellcome Sanger Institute COVID-19 Surveillance Team |
| EPI_ISL_3435580 | Lighthouse Lab in Alderley Park | Wellcome Sanger Institute for the COVID-19 Genomics UK (COG-UK) Consortium | Jacquelyn Wynn, Mairead Hyland, The Lighthouse Lab in Alderley Park and Alex Alderton, Roberto Amato, Jeffrey Barrett, Sonia Goncalves, Ewan Harrison, David K. Jackson, Ian Johnston, Dominic Kwiatkowski, Cordelia Langford, John Sillitoe on behalf of the Wellcome Sanger Institute COVID-19 Surveillance Team |
| EPI_ISL_3435581 | Lighthouse Lab in Glasgow | Wellcome Sanger Institute for the COVID-19 Genomics UK (COG-UK) Consortium | Harper VanSteenhouse, Yumi Kasai, David Gray, Carol Clugston, Anna Dominiczak and Alex Alderton, Roberto Amato, Jeffrey Barrett, Sonia Goncalves, Ewan Harrison, David K. Jackson, Ian Johnston, Dominic Kwiatkowski, Cordelia Langford, John Sillitoe on behalf of the Wellcome Sanger Institute COVID-19 Surveillance Team |
| EPI_ISL_3435582 | Lighthouse Lab in Milton Keynes | Wellcome Sanger Institute for the COVID-19 Genomics UK (COG-UK) Consortium | The Lighthouse Lab in Milton Keynes and Alex Alderton, Roberto Amato, Jeffrey Barrett, Sonia Goncalves, Ewan Harrison, David K. Jackson, Ian Johnston, Dominic Kwiatkowski, Cordelia Langford, John Sillitoe on behalf of the Wellcome Sanger Institute COVID-19 Surveillance Team |
| EPI_ISL_3435583 | Lighthouse Lab in Alderley Park | Wellcome Sanger Institute for the COVID-19 Genomics UK (COG-UK) Consortium | Jacquelyn Wynn, Mairead Hyland, The Lighthouse Lab in Alderley Park and Alex Alderton, Roberto Amato, Jeffrey Barrett, Sonia Goncalves, Ewan Harrison, David K. Jackson, Ian Johnston, Dominic Kwiatkowski, Cordelia Langford, John Sillitoe on behalf of the Wellcome Sanger Institute COVID-19 Surveillance Team |
| EPI_ISL_3435584, EPI_ISL_3435585, EPI_ISL_3435586, EPI_ISL_3435587 | Lighthouse Lab in Milton Keynes | Wellcome Sanger Institute for the COVID-19 Genomics UK (COG-UK) Consortium | The Lighthouse Lab in Milton Keynes and Alex Alderton, Roberto Amato, Jeffrey Barrett, Sonia Goncalves, Ewan Harrison, David K. Jackson, Ian Johnston, Dominic Kwiatkowski, Cordelia Langford, John Sillitoe on behalf of the Wellcome Sanger Institute COVID-19 Surveillance Team |
| EPI_ISL_3435588, EPI_ISL_3435589, EPI_ISL_3435590, EPI_ISL_3435591 | Lighthouse Lab in Glasgow | Wellcome Sanger Institute for the COVID-19 Genomics UK (COG-UK) Consortium | Harper VanSteenhouse, Yumi Kasai, David Gray, Carol Clugston, Anna Dominiczak and Alex Alderton, Roberto Amato, Jeffrey Barrett, Sonia Goncalves, Ewan Harrison, David K. Jackson, Ian Johnston, Dominic Kwiatkowski, Cordelia Langford, John Sillitoe on behalf of the Wellcome Sanger Institute COVID-19 Surveillance Team |
| EPI_ISL_3435592 | Lighthouse Lab in Alderley Park | Wellcome Sanger Institute for the COVID-19 Genomics UK (COG-UK) Consortium | Jacquelyn Wynn, Mairead Hyland, The Lighthouse Lab in Alderley Park and Alex Alderton, Roberto Amato, Jeffrey Barrett, Sonia Goncalves, Ewan Harrison, David K. Jackson, Ian Johnston, Dominic Kwiatkowski, Cordelia Langford, John Sillitoe on behalf of the Wellcome Sanger Institute COVID-19 Surveillance Team |
| EPI_ISL_3435593, EPI_ISL_3435594 | Lighthouse Lab in Milton Keynes | Wellcome Sanger Institute for the COVID-19 Genomics UK (COG-UK) Consortium | The Lighthouse Lab in Milton Keynes and Alex Alderton, Roberto Amato, Jeffrey Barrett, Sonia Goncalves, Ewan Harrison, David K. Jackson, Ian Johnston, Dominic Kwiatkowski, Cordelia Langford, John Sillitoe on behalf of the Wellcome Sanger Institute COVID-19 Surveillance Team |
| EPI_ISL_3435595 | Lighthouse Lab in Glasgow | Wellcome Sanger Institute for the COVID-19 Genomics UK (COG-UK) Consortium | Harper VanSteenhouse, Yumi Kasai, David Gray, Carol Clugston, Anna Dominiczak and Alex Alderton, Roberto Amato, Jeffrey Barrett, Sonia Goncalves, Ewan Harrison, David K. Jackson, Ian Johnston, Dominic Kwiatkowski, Cordelia Langford, John Sillitoe on behalf of the Wellcome Sanger Institute COVID-19 Surveillance Team |
| EPI_ISL_3435596, EPI_ISL_3435597, EPI_ISL_3435598, EPI_ISL_3435599 | Lighthouse Lab in Milton Keynes | Wellcome Sanger Institute for the COVID-19 Genomics UK (COG-UK) Consortium | The Lighthouse Lab in Milton Keynes and Alex Alderton, Roberto Amato, Jeffrey Barrett, Sonia Goncalves, Ewan Harrison, David K. Jackson, Ian Johnston, Dominic Kwiatkowski, Cordelia Langford, John Sillitoe on behalf of the Wellcome Sanger Institute COVID-19 Surveillance Team |
| EPI_ISL_3435600 | Lighthouse Lab in Alderley Park | Wellcome Sanger Institute for the COVID-19 Genomics UK (COG-UK) Consortium | Jacquelyn Wynn, Mairead Hyland, The Lighthouse Lab in Alderley Park and Alex Alderton, Roberto Amato, Jeffrey Barrett, Sonia Goncalves, Ewan Harrison, David K. Jackson, Ian Johnston, Dominic Kwiatkowski, Cordelia Langford, John Sillitoe on behalf of the Wellcome Sanger Institute COVID-19 Surveillance Team |
| EPI_ISL_3435601, EPI_ISL_3435602 | Lighthouse Lab in Milton Keynes | Wellcome Sanger Institute for the COVID-19 Genomics UK (COG-UK) Consortium | The Lighthouse Lab in Milton Keynes and Alex Alderton, Roberto Amato, Jeffrey Barrett, Sonia Goncalves, Ewan Harrison, David K. Jackson, Ian Johnston, Dominic Kwiatkowski, Cordelia Langford, John Sillitoe on behalf of the Wellcome Sanger Institute COVID-19 Surveillance Team |
| EPI_ISL_3435603 | Lighthouse Lab in Alderley Park | Wellcome Sanger Institute for the COVID-19 Genomics UK (COG-UK) Consortium | Jacquelyn Wynn, Mairead Hyland, The Lighthouse Lab in Alderley Park and Alex Alderton, Roberto Amato, Jeffrey Barrett, Sonia Goncalves, Ewan Harrison, David K. Jackson, Ian Johnston, Dominic Kwiatkowski, Cordelia Langford, John Sillitoe on behalf of the Wellcome Sanger Institute COVID-19 Surveillance Team |
| EPI_ISL_3435604, EPI_ISL_3435605 | Lighthouse Lab in Milton Keynes | Wellcome Sanger Institute for the COVID-19 Genomics UK (COG-UK) Consortium | The Lighthouse Lab in Milton Keynes and Alex Alderton, Roberto Amato, Jeffrey Barrett, Sonia Goncalves, Ewan Harrison, David K. Jackson, Ian Johnston, Dominic Kwiatkowski, Cordelia Langford, John Sillitoe on behalf of the Wellcome Sanger Institute COVID-19 Surveillance Team |
| EPI_ISL_3435606 | Lighthouse Lab in Glasgow | Wellcome Sanger Institute for the COVID-19 Genomics UK (COG-UK) Consortium | Harper VanSteenhouse, Yumi Kasai, David Gray, Carol Clugston, Anna Dominiczak and Alex Alderton, Roberto Amato, Jeffrey Barrett, Sonia Goncalves, Ewan Harrison, David K. Jackson, Ian Johnston, Dominic Kwiatkowski, Cordelia Langford, John Sillitoe on behalf of the Wellcome Sanger Institute COVID-19 Surveillance Team |
| EPI_ISL_3435607, EPI_ISL_3435608, EPI_ISL_3435609 | Lighthouse Lab in Alderley Park | Wellcome Sanger Institute for the COVID-19 Genomics UK (COG-UK) Consortium | Jacquelyn Wynn, Mairead Hyland, The Lighthouse Lab in Alderley Park and Alex Alderton, Roberto Amato, Jeffrey Barrett, Sonia Goncalves, Ewan Harrison, David K. Jackson, Ian Johnston, Dominic Kwiatkowski, Cordelia Langford, John Sillitoe on behalf of the Wellcome Sanger Institute COVID-19 Surveillance Team |
| EPI_ISL_3435610, EPI_ISL_3435611 | Lighthouse Lab in Glasgow | Wellcome Sanger Institute for the COVID-19 Genomics UK (COG-UK) Consortium | Harper VanSteenhouse, Yumi Kasai, David Gray, Carol Clugston, Anna Dominiczak and Alex Alderton, Roberto Amato, Jeffrey Barrett, Sonia Goncalves, Ewan Harrison, David K. Jackson, Ian Johnston, Dominic Kwiatkowski, Cordelia Langford, John Sillitoe on behalf of the Wellcome Sanger Institute COVID-19 Surveillance Team |
| EPI_ISL_3435612, EPI_ISL_3435613 | Lighthouse Lab in Milton Keynes | Wellcome Sanger Institute for the COVID-19 Genomics UK (COG-UK) Consortium | The Lighthouse Lab in Milton Keynes and Alex Alderton, Roberto Amato, Jeffrey Barrett, Sonia Goncalves, Ewan Harrison, David K. Jackson, Ian Johnston, Dominic Kwiatkowski, Cordelia Langford, John Sillitoe on behalf of the Wellcome Sanger Institute COVID-19 Surveillance Team |
| EPI_ISL_3435614 | Lighthouse Lab in Glasgow | Wellcome Sanger Institute for the COVID-19 Genomics UK (COG-UK) Consortium | Harper VanSteenhouse, Yumi Kasai, David Gray, Carol Clugston, Anna Dominiczak and Alex Alderton, Roberto Amato, Jeffrey Barrett, Sonia Goncalves, Ewan Harrison, David K. Jackson, Ian Johnston, Dominic Kwiatkowski, Cordelia Langford, John Sillitoe on behalf of the Wellcome Sanger Institute COVID-19 Surveillance Team |
| EPI_ISL_3435615 | Lighthouse Lab in Alderley Park | Wellcome Sanger Institute for the COVID-19 Genomics UK (COG-UK) Consortium | Jacquelyn Wynn, Mairead Hyland, The Lighthouse Lab in Alderley Park and Alex Alderton, Roberto Amato, Jeffrey Barrett, Sonia Goncalves, Ewan Harrison, David K. Jackson, Ian Johnston, Dominic Kwiatkowski, Cordelia Langford, John Sillitoe on behalf of the Wellcome Sanger Institute COVID-19 Surveillance Team |
| EPI_ISL_3435616 | Lighthouse Lab in Milton Keynes | Wellcome Sanger Institute for the COVID-19 Genomics UK (COG-UK) Consortium | The Lighthouse Lab in Milton Keynes and Alex Alderton, Roberto Amato, Jeffrey Barrett, Sonia Goncalves, Ewan Harrison, David K. Jackson, Ian Johnston, Dominic Kwiatkowski, Cordelia Langford, John Sillitoe on behalf of the Wellcome Sanger Institute COVID-19 Surveillance Team |
| EPI_ISL_3435617 | Lighthouse Lab in Glasgow | Wellcome Sanger Institute for the COVID-19 Genomics UK (COG-UK) Consortium | Harper VanSteenhouse, Yumi Kasai, David Gray, Carol Clugston, Anna Dominiczak and Alex Alderton, Roberto Amato, Jeffrey Barrett, Sonia Goncalves, Ewan Harrison, David K. Jackson, Ian Johnston, Dominic Kwiatkowski, Cordelia Langford, John Sillitoe on behalf of the Wellcome Sanger Institute COVID-19 Surveillance Team |
| EPI_ISL_3435618 | Lighthouse Lab in Milton Keynes | Wellcome Sanger Institute for the COVID-19 Genomics UK (COG-UK) Consortium | The Lighthouse Lab in Milton Keynes and Alex Alderton, Roberto Amato, Jeffrey Barrett, Sonia Goncalves, Ewan Harrison, David K. Jackson, Ian Johnston, Dominic Kwiatkowski, Cordelia Langford, John Sillitoe on behalf of the Wellcome Sanger Institute COVID-19 Surveillance Team |
| EPI_ISL_3435619 | Lighthouse Lab in Alderley Park | Wellcome Sanger Institute for the COVID-19 Genomics UK (COG-UK) Consortium | Jacquelyn Wynn, Mairead Hyland, The Lighthouse Lab in Alderley Park and Alex Alderton, Roberto Amato, Jeffrey Barrett, Sonia Goncalves, Ewan Harrison, David K. Jackson, Ian Johnston, Dominic Kwiatkowski, Cordelia Langford, John Sillitoe on behalf of the Wellcome Sanger Institute COVID-19 Surveillance Team |
| EPI_ISL_3435620 | Lighthouse Lab in Milton Keynes | Wellcome Sanger Institute for the COVID-19 Genomics UK (COG-UK) Consortium | The Lighthouse Lab in Milton Keynes and Alex Alderton, Roberto Amato, Jeffrey Barrett, Sonia Goncalves, Ewan Harrison, David K. Jackson, Ian Johnston, Dominic Kwiatkowski, Cordelia Langford, John Sillitoe on behalf of the Wellcome Sanger Institute COVID-19 Surveillance Team |
| EPI_ISL_3435621 | Lighthouse Lab in Glasgow | Wellcome Sanger Institute for the COVID-19 Genomics UK (COG-UK) Consortium | Harper VanSteenhouse, Yumi Kasai, David Gray, Carol Clugston, Anna Dominiczak and Alex Alderton, Roberto Amato, Jeffrey Barrett, Sonia Goncalves, Ewan Harrison, David K. Jackson, Ian Johnston, Dominic Kwiatkowski, Cordelia Langford, John Sillitoe on behalf of the Wellcome Sanger Institute COVID-19 Surveillance Team |
| EPI_ISL_3435622 | Lighthouse Lab in Alderley Park | Wellcome Sanger Institute for the COVID-19 Genomics UK (COG-UK) Consortium | Jacquelyn Wynn, Mairead Hyland, The Lighthouse Lab in Alderley Park and Alex Alderton, Roberto Amato, Jeffrey Barrett, Sonia Goncalves, Ewan Harrison, David K. Jackson, Ian Johnston, Dominic Kwiatkowski, Cordelia Langford, John Sillitoe on behalf of the Wellcome Sanger Institute COVID-19 Surveillance Team |
| EPI_ISL_3435623 | Lighthouse Lab in Glasgow | Wellcome Sanger Institute for the COVID-19 Genomics UK (COG-UK) Consortium | Harper VanSteenhouse, Yumi Kasai, David Gray, Carol Clugston, Anna Dominiczak and Alex Alderton, Roberto Amato, Jeffrey Barrett, Sonia Goncalves, Ewan Harrison, David K. Jackson, Ian Johnston, Dominic Kwiatkowski, Cordelia Langford, John Sillitoe on behalf of the Wellcome Sanger Institute COVID-19 Surveillance Team |
| EPI_ISL_3435624 | Lighthouse Lab in Alderley Park | Wellcome Sanger Institute for the COVID-19 Genomics UK (COG-UK) Consortium | Jacquelyn Wynn, Mairead Hyland, The Lighthouse Lab in Alderley Park and Alex Alderton, Roberto Amato, Jeffrey Barrett, Sonia Goncalves, Ewan Harrison, David K. Jackson, Ian Johnston, Dominic Kwiatkowski, Cordelia Langford, John Sillitoe on behalf of the Wellcome Sanger Institute COVID-19 Surveillance Team |
| EPI_ISL_3435625 | Lighthouse Lab in Milton Keynes | Wellcome Sanger Institute for the COVID-19 Genomics UK (COG-UK) Consortium | The Lighthouse Lab in Milton Keynes and Alex Alderton, Roberto Amato, Jeffrey Barrett, Sonia Goncalves, Ewan Harrison, David K. Jackson, Ian Johnston, Dominic Kwiatkowski, Cordelia Langford, John Sillitoe on behalf of the Wellcome Sanger Institute COVID-19 Surveillance Team |
| EPI_ISL_3435626, EPI_ISL_3435627 | Lighthouse Lab in Alderley Park | Wellcome Sanger Institute for the COVID-19 Genomics UK (COG-UK) Consortium | Jacquelyn Wynn, Mairead Hyland, The Lighthouse Lab in Alderley Park and Alex Alderton, Roberto Amato, Jeffrey Barrett, Sonia Goncalves, Ewan Harrison, David K. Jackson, Ian Johnston, Dominic Kwiatkowski, Cordelia Langford, John Sillitoe on behalf of the Wellcome Sanger Institute COVID-19 Surveillance Team |
| EPI_ISL_3435628, EPI_ISL_3435629 | Lighthouse Lab in Milton Keynes | Wellcome Sanger Institute for the COVID-19 Genomics UK (COG-UK) Consortium | The Lighthouse Lab in Milton Keynes and Alex Alderton, Roberto Amato, Jeffrey Barrett, Sonia Goncalves, Ewan Harrison, David K. Jackson, Ian Johnston, Dominic Kwiatkowski, Cordelia Langford, John Sillitoe on behalf of the Wellcome Sanger Institute COVID-19 Surveillance Team |
| EPI_ISL_3435630 | Lighthouse Lab in Alderley Park | Wellcome Sanger Institute for the COVID-19 Genomics UK (COG-UK) Consortium | Jacquelyn Wynn, Mairead Hyland, The Lighthouse Lab in Alderley Park and Alex Alderton, Roberto Amato, Jeffrey Barrett, Sonia Goncalves, Ewan Harrison, David K. Jackson, Ian Johnston, Dominic Kwiatkowski, Cordelia Langford, John Sillitoe on behalf of the Wellcome Sanger Institute COVID-19 Surveillance Team |
| EPI_ISL_3435631 | Lighthouse Lab in Glasgow | Wellcome Sanger Institute for the COVID-19 Genomics UK (COG-UK) Consortium | Harper VanSteenhouse, Yumi Kasai, David Gray, Carol Clugston, Anna Dominiczak and Alex Alderton, Roberto Amato, Jeffrey Barrett, Sonia Goncalves, Ewan Harrison, David K. Jackson, Ian Johnston, Dominic Kwiatkowski, Cordelia Langford, John Sillitoe on behalf of the Wellcome Sanger Institute COVID-19 Surveillance Team |
| EPI_ISL_3435632, EPI_ISL_3435633 | Lighthouse Lab in Alderley Park | Wellcome Sanger Institute for the COVID-19 Genomics UK (COG-UK) Consortium | Jacquelyn Wynn, Mairead Hyland, The Lighthouse Lab in Alderley Park and Alex Alderton, Roberto Amato, Jeffrey Barrett, Sonia Goncalves, Ewan Harrison, David K. Jackson, Ian Johnston, Dominic Kwiatkowski, Cordelia Langford, John Sillitoe on behalf of the Wellcome Sanger Institute COVID-19 Surveillance Team |
| EPI_ISL_3435634 | Lighthouse Lab in Glasgow | Wellcome Sanger Institute for the COVID-19 Genomics UK (COG-UK) Consortium | Harper VanSteenhouse, Yumi Kasai, David Gray, Carol Clugston, Anna Dominiczak and Alex Alderton, Roberto Amato, Jeffrey Barrett, Sonia Goncalves, Ewan Harrison, David K. Jackson, Ian Johnston, Dominic Kwiatkowski, Cordelia Langford, John Sillitoe on behalf of the Wellcome Sanger Institute COVID-19 Surveillance Team |
| EPI_ISL_3435635 | Lighthouse Lab in Milton Keynes | Wellcome Sanger Institute for the COVID-19 Genomics UK (COG-UK) Consortium | The Lighthouse Lab in Milton Keynes and Alex Alderton, Roberto Amato, Jeffrey Barrett, Sonia Goncalves, Ewan Harrison, David K. Jackson, Ian Johnston, Dominic Kwiatkowski, Cordelia Langford, John Sillitoe on behalf of the Wellcome Sanger Institute COVID-19 Surveillance Team |
| EPI_ISL_3435636 | Lighthouse Lab in Alderley Park | Wellcome Sanger Institute for the COVID-19 Genomics UK (COG-UK) Consortium | Jacquelyn Wynn, Mairead Hyland, The Lighthouse Lab in Alderley Park and Alex Alderton, Roberto Amato, Jeffrey Barrett, Sonia Goncalves, Ewan Harrison, David K. Jackson, Ian Johnston, Dominic Kwiatkowski, Cordelia Langford, John Sillitoe on behalf of the Wellcome Sanger Institute COVID-19 Surveillance Team |

| | | | |
|---|---|---|---|
| EPI_ISL_3435637 | Lighthouse Lab in Glasgow | Wellcome Sanger Institute for the COVID-19 Genomics UK (COG-UK) Consortium | Harper VanSteenhouse, Yumi Kasai, David Gray, Carol Clugston, Anna Dominiczak and Alex Alderton, Roberto Amato, Jeffrey Barrett, Sonia Goncalves, Ewan Harrison, David K. Jackson, Ian Johnston, Dominic Kwiatkowski, Cordelia Langford, John Sillitoe on behalf of the Wellcome Sanger Institute COVID-19 Surveillance Team |
| EPI_ISL_3435638, EPI_ISL_3435639, EPI_ISL_3435640, EPI_ISL_3435641, EPI_ISL_3435642, EPI_ISL_3435643, EPI_ISL_3435644, EPI_ISL_3435645, EPI_ISL_3435646 | Lighthouse Lab in Alderley Park | Wellcome Sanger Institute for the COVID-19 Genomics UK (COG-UK) Consortium | Jacquelyn Wynn, Mairead Hyland, The Lighthouse Lab in Alderley Park and Alex Alderton, Roberto Amato, Jeffrey Barrett, Sonia Goncalves, Ewan Harrison, David K. Jackson, Ian Johnston, Dominic Kwiatkowski, Cordelia Langford, John Sillitoe on behalf of the Wellcome Sanger Institute COVID-19 Surveillance Team |
| EPI_ISL_3435647 | Lighthouse Lab in Milton Keynes | Wellcome Sanger Institute for the COVID-19 Genomics UK (COG-UK) Consortium | The Lighthouse Lab in Milton Keynes and Alex Alderton, Roberto Amato, Jeffrey Barrett, Sonia Goncalves, Ewan Harrison, David K. Jackson, Ian Johnston, Dominic Kwiatkowski, Cordelia Langford, John Sillitoe on behalf of the Wellcome Sanger Institute COVID-19 Surveillance Team |
| EPI_ISL_3435648, EPI_ISL_3435649 | Lighthouse Lab in Alderley Park | Wellcome Sanger Institute for the COVID-19 Genomics UK (COG-UK) Consortium | Jacquelyn Wynn, Mairead Hyland, The Lighthouse Lab in Alderley Park and Alex Alderton, Roberto Amato, Jeffrey Barrett, Sonia Goncalves, Ewan Harrison, David K. Jackson, Ian Johnston, Dominic Kwiatkowski, Cordelia Langford, John Sillitoe on behalf of the Wellcome Sanger Institute COVID-19 Surveillance Team |
| EPI_ISL_3435650, EPI_ISL_3435651 | Lighthouse Lab in Milton Keynes | Wellcome Sanger Institute for the COVID-19 Genomics UK (COG-UK) Consortium | The Lighthouse Lab in Milton Keynes and Alex Alderton, Roberto Amato, Jeffrey Barrett, Sonia Goncalves, Ewan Harrison, David K. Jackson, Ian Johnston, Dominic Kwiatkowski, Cordelia Langford, John Sillitoe on behalf of the Wellcome Sanger Institute COVID-19 Surveillance Team |
| EPI_ISL_3435652 | Lighthouse Lab in Alderley Park | Wellcome Sanger Institute for the COVID-19 Genomics UK (COG-UK) Consortium | Jacquelyn Wynn, Mairead Hyland, The Lighthouse Lab in Alderley Park and Alex Alderton, Roberto Amato, Jeffrey Barrett, Sonia Goncalves, Ewan Harrison, David K. Jackson, Ian Johnston, Dominic Kwiatkowski, Cordelia Langford, John Sillitoe on behalf of the Wellcome Sanger Institute COVID-19 Surveillance Team |
| EPI_ISL_3435653, EPI_ISL_3435654, EPI_ISL_3435655, EPI_ISL_3435656 | Lighthouse Lab in Milton Keynes | Wellcome Sanger Institute for the COVID-19 Genomics UK (COG-UK) Consortium | The Lighthouse Lab in Milton Keynes and Alex Alderton, Roberto Amato, Jeffrey Barrett, Sonia Goncalves, Ewan Harrison, David K. Jackson, Ian Johnston, Dominic Kwiatkowski, Cordelia Langford, John Sillitoe on behalf of the Wellcome Sanger Institute COVID-19 Surveillance Team |
| EPI_ISL_3435657 | Lighthouse Lab in Glasgow | Wellcome Sanger Institute for the COVID-19 Genomics UK (COG-UK) Consortium | Harper VanSteenhouse, Yumi Kasai, David Gray, Carol Clugston, Anna Dominiczak and Alex Alderton, Roberto Amato, Jeffrey Barrett, Sonia Goncalves, Ewan Harrison, David K. Jackson, Ian Johnston, Dominic Kwiatkowski, Cordelia Langford, John Sillitoe on behalf of the Wellcome Sanger Institute COVID-19 Surveillance Team |
| EPI_ISL_3435658, EPI_ISL_3435659 | Lighthouse Lab in Milton Keynes | Wellcome Sanger Institute for the COVID-19 Genomics UK (COG-UK) Consortium | The Lighthouse Lab in Milton Keynes and Alex Alderton, Roberto Amato, Jeffrey Barrett, Sonia Goncalves, Ewan Harrison, David K. Jackson, Ian Johnston, Dominic Kwiatkowski, Cordelia Langford, John Sillitoe on behalf of the Wellcome Sanger Institute COVID-19 Surveillance Team |
| EPI_ISL_3435660 | Lighthouse Lab in Glasgow | Wellcome Sanger Institute for the COVID-19 Genomics UK (COG-UK) Consortium | Harper VanSteenhouse, Yumi Kasai, David Gray, Carol Clugston, Anna Dominiczak and Alex Alderton, Roberto Amato, Jeffrey Barrett, Sonia Goncalves, Ewan Harrison, David K. Jackson, Ian Johnston, Dominic Kwiatkowski, Cordelia Langford, John Sillitoe on behalf of the Wellcome Sanger Institute COVID-19 Surveillance Team |
| EPI_ISL_3435661 | Lighthouse Lab in Milton Keynes | Wellcome Sanger Institute for the COVID-19 Genomics UK (COG-UK) Consortium | The Lighthouse Lab in Milton Keynes and Alex Alderton, Roberto Amato, Jeffrey Barrett, Sonia Goncalves, Ewan Harrison, David K. Jackson, Ian Johnston, Dominic Kwiatkowski, Cordelia Langford, John Sillitoe on behalf of the Wellcome Sanger Institute COVID-19 Surveillance Team |
| EPI_ISL_3435662, EPI_ISL_3435663, EPI_ISL_3435664 | Lighthouse Lab in Alderley Park | Wellcome Sanger Institute for the COVID-19 Genomics UK (COG-UK) Consortium | Jacquelyn Wynn, Mairead Hyland, The Lighthouse Lab in Alderley Park and Alex Alderton, Roberto Amato, Jeffrey Barrett, Sonia Goncalves, Ewan Harrison, David K. Jackson, Ian Johnston, Dominic Kwiatkowski, Cordelia Langford, John Sillitoe on behalf of the Wellcome Sanger Institute COVID-19 Surveillance Team |
| EPI_ISL_3435665, EPI_ISL_3435666 | Lighthouse Lab in Milton Keynes | Wellcome Sanger Institute for the COVID-19 Genomics UK (COG-UK) Consortium | The Lighthouse Lab in Milton Keynes and Alex Alderton, Roberto Amato, Jeffrey Barrett, Sonia Goncalves, Ewan Harrison, David K. Jackson, Ian Johnston, Dominic Kwiatkowski, Cordelia Langford, John Sillitoe on behalf of the Wellcome Sanger Institute COVID-19 Surveillance Team |
| EPI_ISL_3435667 | Lighthouse Lab in Alderley Park | Wellcome Sanger Institute for the COVID-19 Genomics UK (COG-UK) Consortium | Jacquelyn Wynn, Mairead Hyland, The Lighthouse Lab in Alderley Park and Alex Alderton, Roberto Amato, Jeffrey Barrett, Sonia Goncalves, Ewan Harrison, David K. Jackson, Ian Johnston, Dominic Kwiatkowski, Cordelia Langford, John Sillitoe on behalf of the Wellcome Sanger Institute COVID-19 Surveillance Team |
| EPI_ISL_3435668, EPI_ISL_3435669 | Lighthouse Lab in Glasgow | Wellcome Sanger Institute for the COVID-19 Genomics UK (COG-UK) Consortium | Harper VanSteenhouse, Yumi Kasai, David Gray, Carol Clugston, Anna Dominiczak and Alex Alderton, Roberto Amato, Jeffrey Barrett, Sonia Goncalves, Ewan Harrison, David K. Jackson, Ian Johnston, Dominic Kwiatkowski, Cordelia Langford, John Sillitoe on behalf of the Wellcome Sanger Institute COVID-19 Surveillance Team |
| EPI_ISL_3435670 | Lighthouse Lab in Alderley Park | Wellcome Sanger Institute for the COVID-19 Genomics UK (COG-UK) Consortium | Jacquelyn Wynn, Mairead Hyland, The Lighthouse Lab in Alderley Park and Alex Alderton, Roberto Amato, Jeffrey Barrett, Sonia Goncalves, Ewan Harrison, David K. Jackson, Ian Johnston, Dominic Kwiatkowski, Cordelia Langford, John Sillitoe on behalf of the Wellcome Sanger Institute COVID-19 Surveillance Team |
| EPI_ISL_3435671 | Lighthouse Lab in Milton Keynes | Wellcome Sanger Institute for the COVID-19 Genomics UK (COG-UK) Consortium | The Lighthouse Lab in Milton Keynes and Alex Alderton, Roberto Amato, Jeffrey Barrett, Sonia Goncalves, Ewan Harrison, David K. Jackson, Ian Johnston, Dominic Kwiatkowski, Cordelia Langford, John Sillitoe on behalf of the Wellcome Sanger Institute COVID-19 Surveillance Team |
| EPI_ISL_3435672 | Lighthouse Lab in Alderley Park | Wellcome Sanger Institute for the COVID-19 Genomics UK (COG-UK) Consortium | Jacquelyn Wynn, Mairead Hyland, The Lighthouse Lab in Alderley Park and Alex Alderton, Roberto Amato, Jeffrey Barrett, Sonia Goncalves, Ewan Harrison, David K. Jackson, Ian Johnston, Dominic Kwiatkowski, Cordelia Langford, John Sillitoe on behalf of the Wellcome Sanger Institute COVID-19 Surveillance Team |
| EPI_ISL_3435673 | Lighthouse Lab in Milton Keynes | Wellcome Sanger Institute for the COVID-19 Genomics UK (COG-UK) Consortium | The Lighthouse Lab in Milton Keynes and Alex Alderton, Roberto Amato, Jeffrey Barrett, Sonia Goncalves, Ewan Harrison, David K. Jackson, Ian Johnston, Dominic Kwiatkowski, Cordelia Langford, John Sillitoe on behalf of the Wellcome Sanger Institute COVID-19 Surveillance Team |
| EPI_ISL_3435674 | Berkshire and Surrey Pathology Services Lighthouse Laboratory | Wellcome Sanger Institute for the COVID-19 Genomics UK (COG-UK) Consortium | Berkshire and Surrey Pathology Services Lighthouse Laboratory and Alex Alderton, Roberto Amato, Jeffrey Barrett, Sonia Goncalves, Ewan Harrison, David K. Jackson, Ian Johnston, Dominic Kwiatkowski, Cordelia Langford, John Sillitoe on behalf of the Wellcome Sanger Institute COVID-19 Surveillance Team |
| EPI_ISL_3435675, EPI_ISL_3435676, EPI_ISL_3435677, EPI_ISL_3435678, EPI_ISL_3435679, EPI_ISL_3435680 | Lighthouse Lab in Milton Keynes | Wellcome Sanger Institute for the COVID-19 Genomics UK (COG-UK) Consortium | The Lighthouse Lab in Milton Keynes and Alex Alderton, Roberto Amato, Jeffrey Barrett, Sonia Goncalves, Ewan Harrison, David K. Jackson, Ian Johnston, Dominic Kwiatkowski, Cordelia Langford, John Sillitoe on behalf of the Wellcome Sanger Institute COVID-19 Surveillance Team |
| EPI_ISL_3435681 | Berkshire and Surrey Pathology Services Lighthouse Laboratory | Wellcome Sanger Institute for the COVID-19 Genomics UK (COG-UK) Consortium | Berkshire and Surrey Pathology Services Lighthouse Laboratory and Alex Alderton, Roberto Amato, Jeffrey Barrett, Sonia Goncalves, Ewan Harrison, David K. Jackson, Ian Johnston, Dominic Kwiatkowski, Cordelia Langford, John Sillitoe on behalf of the Wellcome Sanger Institute COVID-19 Surveillance Team |
| EPI_ISL_3435682, EPI_ISL_3435683 | Lighthouse Lab in Milton Keynes | Wellcome Sanger Institute for the COVID-19 Genomics UK (COG-UK) Consortium | The Lighthouse Lab in Milton Keynes and Alex Alderton, Roberto Amato, Jeffrey Barrett, Sonia Goncalves, Ewan Harrison, David K. Jackson, Ian Johnston, Dominic Kwiatkowski, Cordelia Langford, John Sillitoe on behalf of the Wellcome Sanger Institute COVID-19 Surveillance Team |
| EPI_ISL_3435684, EPI_ISL_3435685 | Berkshire and Surrey Pathology Services Lighthouse Laboratory | Wellcome Sanger Institute for the COVID-19 Genomics UK (COG-UK) Consortium | Berkshire and Surrey Pathology Services Lighthouse Laboratory and Alex Alderton, Roberto Amato, Jeffrey Barrett, Sonia Goncalves, Ewan Harrison, David K. Jackson, Ian Johnston, Dominic Kwiatkowski, Cordelia Langford, John Sillitoe on behalf of the Wellcome Sanger Institute COVID-19 Surveillance Team |
| EPI_ISL_3435686, EPI_ISL_3435687, EPI_ISL_3435688 | Lighthouse Lab in Milton Keynes | Wellcome Sanger Institute for the COVID-19 Genomics UK (COG-UK) Consortium | The Lighthouse Lab in Milton Keynes and Alex Alderton, Roberto Amato, Jeffrey Barrett, Sonia Goncalves, Ewan Harrison, David K. Jackson, Ian Johnston, Dominic Kwiatkowski, Cordelia Langford, John Sillitoe on behalf of the Wellcome Sanger Institute COVID-19 Surveillance Team |
| EPI_ISL_3435689 | Berkshire and Surrey Pathology Services Lighthouse Laboratory | Wellcome Sanger Institute for the COVID-19 Genomics UK (COG-UK) Consortium | Berkshire and Surrey Pathology Services Lighthouse Laboratory and Alex Alderton, Roberto Amato, Jeffrey Barrett, Sonia Goncalves, Ewan Harrison, David K. Jackson, Ian Johnston, Dominic Kwiatkowski, Cordelia Langford, John Sillitoe on behalf of the Wellcome Sanger Institute COVID-19 Surveillance Team |
| EPI_ISL_3435690, EPI_ISL_3435691, EPI_ISL_3435692 | Lighthouse Lab in Milton Keynes | Wellcome Sanger Institute for the COVID-19 Genomics UK (COG-UK) Consortium | The Lighthouse Lab in Milton Keynes and Alex Alderton, Roberto Amato, Jeffrey Barrett, Sonia Goncalves, Ewan Harrison, David K. Jackson, Ian Johnston, Dominic Kwiatkowski, Cordelia Langford, John Sillitoe on behalf of the Wellcome Sanger Institute COVID-19 Surveillance Team |
| EPI_ISL_3435693, EPI_ISL_3435694, EPI_ISL_3435695 | Berkshire and Surrey Pathology Services Lighthouse Laboratory | Wellcome Sanger Institute for the COVID-19 Genomics UK (COG-UK) Consortium | Berkshire and Surrey Pathology Services Lighthouse Laboratory and Alex Alderton, Roberto Amato, Jeffrey Barrett, Sonia Goncalves, Ewan Harrison, David K. Jackson, Ian Johnston, Dominic Kwiatkowski, Cordelia Langford, John Sillitoe on behalf of the Wellcome Sanger Institute COVID-19 Surveillance Team |
| EPI_ISL_3435696 | Lighthouse Lab in Milton Keynes | Wellcome Sanger Institute for the COVID-19 Genomics UK (COG-UK) Consortium | The Lighthouse Lab in Milton Keynes and Alex Alderton, Roberto Amato, Jeffrey Barrett, Sonia Goncalves, Ewan Harrison, David K. Jackson, Ian Johnston, Dominic Kwiatkowski, Cordelia Langford, John Sillitoe on behalf of the Wellcome Sanger Institute COVID-19 Surveillance Team |
| EPI_ISL_3435697 | Berkshire and Surrey Pathology Services Lighthouse Laboratory | Wellcome Sanger Institute for the COVID-19 Genomics UK (COG-UK) Consortium | Berkshire and Surrey Pathology Services Lighthouse Laboratory and Alex Alderton, Roberto Amato, Jeffrey Barrett, Sonia Goncalves, Ewan Harrison, David K. Jackson, Ian Johnston, Dominic Kwiatkowski, Cordelia Langford, John Sillitoe on behalf of the Wellcome Sanger Institute COVID-19 Surveillance Team |
| EPI_ISL_3435698 | Lighthouse Lab in Milton Keynes | Wellcome Sanger Institute for the COVID-19 Genomics UK (COG-UK) Consortium | The Lighthouse Lab in Milton Keynes and Alex Alderton, Roberto Amato, Jeffrey Barrett, Sonia Goncalves, Ewan Harrison, David K. Jackson, Ian Johnston, Dominic Kwiatkowski, Cordelia Langford, John Sillitoe on behalf of the Wellcome Sanger Institute COVID-19 Surveillance Team |
| EPI_ISL_3435699 | Berkshire and Surrey Pathology Services Lighthouse Laboratory | Wellcome Sanger Institute for the COVID-19 Genomics UK (COG-UK) Consortium | Berkshire and Surrey Pathology Services Lighthouse Laboratory and Alex Alderton, Roberto Amato, Jeffrey Barrett, Sonia Goncalves, Ewan Harrison, David K. Jackson, Ian Johnston, Dominic Kwiatkowski, Cordelia Langford, John Sillitoe on behalf of the Wellcome Sanger Institute COVID-19 Surveillance Team |
| EPI_ISL_3435700, EPI_ISL_3435701, EPI_ISL_3435702, EPI_ISL_3435703, EPI_ISL_3435704 | Lighthouse Lab in Milton Keynes | Wellcome Sanger Institute for the COVID-19 Genomics UK (COG-UK) Consortium | The Lighthouse Lab in Milton Keynes and Alex Alderton, Roberto Amato, Jeffrey Barrett, Sonia Goncalves, Ewan Harrison, David K. Jackson, Ian Johnston, Dominic Kwiatkowski, Cordelia Langford, John Sillitoe on behalf of the Wellcome Sanger Institute COVID-19 Surveillance Team |
| EPI_ISL_3435705, EPI_ISL_3435706 | Berkshire and Surrey Pathology Services Lighthouse Laboratory | Wellcome Sanger Institute for the COVID-19 Genomics UK (COG-UK) Consortium | Berkshire and Surrey Pathology Services Lighthouse Laboratory and Alex Alderton, Roberto Amato, Jeffrey Barrett, Sonia Goncalves, Ewan Harrison, David K. Jackson, Ian Johnston, Dominic Kwiatkowski, Cordelia Langford, John Sillitoe on behalf of the Wellcome Sanger Institute COVID-19 Surveillance Team |
| EPI_ISL_3435707 | Lighthouse Lab in Milton Keynes | Wellcome Sanger Institute for the COVID-19 Genomics UK (COG-UK) Consortium | The Lighthouse Lab in Milton Keynes and Alex Alderton, Roberto Amato, Jeffrey Barrett, Sonia Goncalves, Ewan Harrison, David K. Jackson, Ian Johnston, Dominic Kwiatkowski, Cordelia Langford, John Sillitoe on behalf of the Wellcome Sanger Institute COVID-19 Surveillance Team |
| EPI_ISL_3435708, EPI_ISL_3435709 | Berkshire and Surrey Pathology Services Lighthouse Laboratory | Wellcome Sanger Institute for the COVID-19 Genomics UK (COG-UK) Consortium | Berkshire and Surrey Pathology Services Lighthouse Laboratory and Alex Alderton, Roberto Amato, Jeffrey Barrett, Sonia Goncalves, Ewan Harrison, David K. Jackson, Ian Johnston, Dominic Kwiatkowski, Cordelia Langford, John Sillitoe on behalf of the Wellcome Sanger Institute COVID-19 Surveillance Team |

| | | | |
|---|---|---|---|
| EPI_ISL_3435710, EPI_ISL_3435711 | Lighthouse Lab in Milton Keynes | Wellcome Sanger Institute for the COVID-19 Genomics UK (COG-UK) Consortium | The Lighthouse Lab in Milton Keynes and Alex Alderton, Roberto Amato, Jeffrey Barrett, Sonia Goncalves, Ewan Harrison, David K. Jackson, Ian Johnston, Dominic Kwiatkowski, Cordelia Langford, John Sillitoe on behalf of the Wellcome Sanger Institute COVID-19 Surveillance Team |
| EPI_ISL_3435712, EPI_ISL_3435713, EPI_ISL_3435714, EPI_ISL_3435715 | Berkshire and Surrey Pathology Services Lighthouse Laboratory | Wellcome Sanger Institute for the COVID-19 Genomics UK (COG-UK) Consortium | Berkshire and Surrey Pathology Services Lighthouse Laboratory and Alex Alderton, Roberto Amato, Jeffrey Barrett, Sonia Goncalves, Ewan Harrison, David K. Jackson, Ian Johnston, Dominic Kwiatkowski, Cordelia Langford, John Sillitoe on behalf of the Wellcome Sanger Institute COVID-19 Surveillance Team |
| EPI_ISL_3435716 | Lighthouse Lab in Milton Keynes | Wellcome Sanger Institute for the COVID-19 Genomics UK (COG-UK) Consortium | The Lighthouse Lab in Milton Keynes and Alex Alderton, Roberto Amato, Jeffrey Barrett, Sonia Goncalves, Ewan Harrison, David K. Jackson, Ian Johnston, Dominic Kwiatkowski, Cordelia Langford, John Sillitoe on behalf of the Wellcome Sanger Institute COVID-19 Surveillance Team |
| EPI_ISL_3435717 | Berkshire and Surrey Pathology Services Lighthouse Laboratory | Wellcome Sanger Institute for the COVID-19 Genomics UK (COG-UK) Consortium | Berkshire and Surrey Pathology Services Lighthouse Laboratory and Alex Alderton, Roberto Amato, Jeffrey Barrett, Sonia Goncalves, Ewan Harrison, David K. Jackson, Ian Johnston, Dominic Kwiatkowski, Cordelia Langford, John Sillitoe on behalf of the Wellcome Sanger Institute COVID-19 Surveillance Team |
| EPI_ISL_3435718 | Lighthouse Lab in Milton Keynes | Wellcome Sanger Institute for the COVID-19 Genomics UK (COG-UK) Consortium | The Lighthouse Lab in Milton Keynes and Alex Alderton, Roberto Amato, Jeffrey Barrett, Sonia Goncalves, Ewan Harrison, David K. Jackson, Ian Johnston, Dominic Kwiatkowski, Cordelia Langford, John Sillitoe on behalf of the Wellcome Sanger Institute COVID-19 Surveillance Team |
| EPI_ISL_3435719, EPI_ISL_3435720, EPI_ISL_3435721, EPI_ISL_3435722 | Berkshire and Surrey Pathology Services Lighthouse Laboratory | Wellcome Sanger Institute for the COVID-19 Genomics UK (COG-UK) Consortium | Berkshire and Surrey Pathology Services Lighthouse Laboratory and Alex Alderton, Roberto Amato, Jeffrey Barrett, Sonia Goncalves, Ewan Harrison, David K. Jackson, Ian Johnston, Dominic Kwiatkowski, Cordelia Langford, John Sillitoe on behalf of the Wellcome Sanger Institute COVID-19 Surveillance Team |
| EPI_ISL_3435723 | Lighthouse Lab in Milton Keynes | Wellcome Sanger Institute for the COVID-19 Genomics UK (COG-UK) Consortium | The Lighthouse Lab in Milton Keynes and Alex Alderton, Roberto Amato, Jeffrey Barrett, Sonia Goncalves, Ewan Harrison, David K. Jackson, Ian Johnston, Dominic Kwiatkowski, Cordelia Langford, John Sillitoe on behalf of the Wellcome Sanger Institute COVID-19 Surveillance Team |
| EPI_ISL_3435724, EPI_ISL_3435725, EPI_ISL_3435726 | Berkshire and Surrey Pathology Services Lighthouse Laboratory | Wellcome Sanger Institute for the COVID-19 Genomics UK (COG-UK) Consortium | Berkshire and Surrey Pathology Services Lighthouse Laboratory and Alex Alderton, Roberto Amato, Jeffrey Barrett, Sonia Goncalves, Ewan Harrison, David K. Jackson, Ian Johnston, Dominic Kwiatkowski, Cordelia Langford, John Sillitoe on behalf of the Wellcome Sanger Institute COVID-19 Surveillance Team |
| EPI_ISL_3435727, EPI_ISL_3435728 | Lighthouse Lab in Milton Keynes | Wellcome Sanger Institute for the COVID-19 Genomics UK (COG-UK) Consortium | The Lighthouse Lab in Milton Keynes and Alex Alderton, Roberto Amato, Jeffrey Barrett, Sonia Goncalves, Ewan Harrison, David K. Jackson, Ian Johnston, Dominic Kwiatkowski, Cordelia Langford, John Sillitoe on behalf of the Wellcome Sanger Institute COVID-19 Surveillance Team |
| EPI_ISL_3435729 | Berkshire and Surrey Pathology Services Lighthouse Laboratory | Wellcome Sanger Institute for the COVID-19 Genomics UK (COG-UK) Consortium | Berkshire and Surrey Pathology Services Lighthouse Laboratory and Alex Alderton, Roberto Amato, Jeffrey Barrett, Sonia Goncalves, Ewan Harrison, David K. Jackson, Ian Johnston, Dominic Kwiatkowski, Cordelia Langford, John Sillitoe on behalf of the Wellcome Sanger Institute COVID-19 Surveillance Team |
| EPI_ISL_3435730, EPI_ISL_3435731, EPI_ISL_3435732, EPI_ISL_3435733, EPI_ISL_3435734 | Lighthouse Lab in Milton Keynes | Wellcome Sanger Institute for the COVID-19 Genomics UK (COG-UK) Consortium | The Lighthouse Lab in Milton Keynes and Alex Alderton, Roberto Amato, Jeffrey Barrett, Sonia Goncalves, Ewan Harrison, David K. Jackson, Ian Johnston, Dominic Kwiatkowski, Cordelia Langford, John Sillitoe on behalf of the Wellcome Sanger Institute COVID-19 Surveillance Team |
| EPI_ISL_3435735 | Berkshire and Surrey Pathology Services Lighthouse Laboratory | Wellcome Sanger Institute for the COVID-19 Genomics UK (COG-UK) Consortium | Berkshire and Surrey Pathology Services Lighthouse Laboratory and Alex Alderton, Roberto Amato, Jeffrey Barrett, Sonia Goncalves, Ewan Harrison, David K. Jackson, Ian Johnston, Dominic Kwiatkowski, Cordelia Langford, John Sillitoe on behalf of the Wellcome Sanger Institute COVID-19 Surveillance Team |
| EPI_ISL_3435736 | Lighthouse Lab in Milton Keynes | Wellcome Sanger Institute for the COVID-19 Genomics UK (COG-UK) Consortium | The Lighthouse Lab in Milton Keynes and Alex Alderton, Roberto Amato, Jeffrey Barrett, Sonia Goncalves, Ewan Harrison, David K. Jackson, Ian Johnston, Dominic Kwiatkowski, Cordelia Langford, John Sillitoe on behalf of the Wellcome Sanger Institute COVID-19 Surveillance Team |
| EPI_ISL_3435737, EPI_ISL_3435738 | Berkshire and Surrey Pathology Services Lighthouse Laboratory | Wellcome Sanger Institute for the COVID-19 Genomics UK (COG-UK) Consortium | Berkshire and Surrey Pathology Services Lighthouse Laboratory and Alex Alderton, Roberto Amato, Jeffrey Barrett, Sonia Goncalves, Ewan Harrison, David K. Jackson, Ian Johnston, Dominic Kwiatkowski, Cordelia Langford, John Sillitoe on behalf of the Wellcome Sanger Institute COVID-19 Surveillance Team |
| EPI_ISL_3435739, EPI_ISL_3435740, EPI_ISL_3435741 | Lighthouse Lab in Milton Keynes | Wellcome Sanger Institute for the COVID-19 Genomics UK (COG-UK) Consortium | The Lighthouse Lab in Milton Keynes and Alex Alderton, Roberto Amato, Jeffrey Barrett, Sonia Goncalves, Ewan Harrison, David K. Jackson, Ian Johnston, Dominic Kwiatkowski, Cordelia Langford, John Sillitoe on behalf of the Wellcome Sanger Institute COVID-19 Surveillance Team |
| EPI_ISL_3435742, EPI_ISL_3435743 | Berkshire and Surrey Pathology Services Lighthouse Laboratory | Wellcome Sanger Institute for the COVID-19 Genomics UK (COG-UK) Consortium | Berkshire and Surrey Pathology Services Lighthouse Laboratory and Alex Alderton, Roberto Amato, Jeffrey Barrett, Sonia Goncalves, Ewan Harrison, David K. Jackson, Ian Johnston, Dominic Kwiatkowski, Cordelia Langford, John Sillitoe on behalf of the Wellcome Sanger Institute COVID-19 Surveillance Team |
| EPI_ISL_3435744 | Lighthouse Lab in Milton Keynes | Wellcome Sanger Institute for the COVID-19 Genomics UK (COG-UK) Consortium | The Lighthouse Lab in Milton Keynes and Alex Alderton, Roberto Amato, Jeffrey Barrett, Sonia Goncalves, Ewan Harrison, David K. Jackson, Ian Johnston, Dominic Kwiatkowski, Cordelia Langford, John Sillitoe on behalf of the Wellcome Sanger Institute COVID-19 Surveillance Team |
| EPI_ISL_3435745, EPI_ISL_3435746 | Berkshire and Surrey Pathology Services Lighthouse Laboratory | Wellcome Sanger Institute for the COVID-19 Genomics UK (COG-UK) Consortium | Berkshire and Surrey Pathology Services Lighthouse Laboratory and Alex Alderton, Roberto Amato, Jeffrey Barrett, Sonia Goncalves, Ewan Harrison, David K. Jackson, Ian Johnston, Dominic Kwiatkowski, Cordelia Langford, John Sillitoe on behalf of the Wellcome Sanger Institute COVID-19 Surveillance Team |
| EPI_ISL_3435747, EPI_ISL_3435748, EPI_ISL_3435749 | Lighthouse Lab in Milton Keynes | Wellcome Sanger Institute for the COVID-19 Genomics UK (COG-UK) Consortium | The Lighthouse Lab in Milton Keynes and Alex Alderton, Roberto Amato, Jeffrey Barrett, Sonia Goncalves, Ewan Harrison, David K. Jackson, Ian Johnston, Dominic Kwiatkowski, Cordelia Langford, John Sillitoe on behalf of the Wellcome Sanger Institute COVID-19 Surveillance Team |
| EPI_ISL_3435750 | Berkshire and Surrey Pathology Services Lighthouse Laboratory | Wellcome Sanger Institute for the COVID-19 Genomics UK (COG-UK) Consortium | Berkshire and Surrey Pathology Services Lighthouse Laboratory and Alex Alderton, Roberto Amato, Jeffrey Barrett, Sonia Goncalves, Ewan Harrison, David K. Jackson, Ian Johnston, Dominic Kwiatkowski, Cordelia Langford, John Sillitoe on behalf of the Wellcome Sanger Institute COVID-19 Surveillance Team |
| EPI_ISL_3435751 | Lighthouse Lab in Milton Keynes | Wellcome Sanger Institute for the COVID-19 Genomics UK (COG-UK) Consortium | The Lighthouse Lab in Milton Keynes and Alex Alderton, Roberto Amato, Jeffrey Barrett, Sonia Goncalves, Ewan Harrison, David K. Jackson, Ian Johnston, Dominic Kwiatkowski, Cordelia Langford, John Sillitoe on behalf of the Wellcome Sanger Institute COVID-19 Surveillance Team |
| EPI_ISL_3435752, EPI_ISL_3435753, EPI_ISL_3435754 | Berkshire and Surrey Pathology Services Lighthouse Laboratory | Wellcome Sanger Institute for the COVID-19 Genomics UK (COG-UK) Consortium | Berkshire and Surrey Pathology Services Lighthouse Laboratory and Alex Alderton, Roberto Amato, Jeffrey Barrett, Sonia Goncalves, Ewan Harrison, David K. Jackson, Ian Johnston, Dominic Kwiatkowski, Cordelia Langford, John Sillitoe on behalf of the Wellcome Sanger Institute COVID-19 Surveillance Team |
| EPI_ISL_3435755, EPI_ISL_3435756, EPI_ISL_3435757 | Lighthouse Lab in Milton Keynes | Wellcome Sanger Institute for the COVID-19 Genomics UK (COG-UK) Consortium | The Lighthouse Lab in Milton Keynes and Alex Alderton, Roberto Amato, Jeffrey Barrett, Sonia Goncalves, Ewan Harrison, David K. Jackson, Ian Johnston, Dominic Kwiatkowski, Cordelia Langford, John Sillitoe on behalf of the Wellcome Sanger Institute COVID-19 Surveillance Team |
| EPI_ISL_3435758, EPI_ISL_3435759 | Berkshire and Surrey Pathology Services Lighthouse Laboratory | Wellcome Sanger Institute for the COVID-19 Genomics UK (COG-UK) Consortium | Berkshire and Surrey Pathology Services Lighthouse Laboratory and Alex Alderton, Roberto Amato, Jeffrey Barrett, Sonia Goncalves, Ewan Harrison, David K. Jackson, Ian Johnston, Dominic Kwiatkowski, Cordelia Langford, John Sillitoe on behalf of the Wellcome Sanger Institute COVID-19 Surveillance Team |
| EPI_ISL_3435760 | Lighthouse Lab in Milton Keynes | Wellcome Sanger Institute for the COVID-19 Genomics UK (COG-UK) Consortium | The Lighthouse Lab in Milton Keynes and Alex Alderton, Roberto Amato, Jeffrey Barrett, Sonia Goncalves, Ewan Harrison, David K. Jackson, Ian Johnston, Dominic Kwiatkowski, Cordelia Langford, John Sillitoe on behalf of the Wellcome Sanger Institute COVID-19 Surveillance Team |
| EPI_ISL_3435761 | Berkshire and Surrey Pathology Services Lighthouse Laboratory | Wellcome Sanger Institute for the COVID-19 Genomics UK (COG-UK) Consortium | Berkshire and Surrey Pathology Services Lighthouse Laboratory and Alex Alderton, Roberto Amato, Jeffrey Barrett, Sonia Goncalves, Ewan Harrison, David K. Jackson, Ian Johnston, Dominic Kwiatkowski, Cordelia Langford, John Sillitoe on behalf of the Wellcome Sanger Institute COVID-19 Surveillance Team |
| EPI_ISL_3435762, EPI_ISL_3435763, EPI_ISL_3435764, EPI_ISL_3435765 | Lighthouse Lab in Milton Keynes | Wellcome Sanger Institute for the COVID-19 Genomics UK (COG-UK) Consortium | The Lighthouse Lab in Milton Keynes and Alex Alderton, Roberto Amato, Jeffrey Barrett, Sonia Goncalves, Ewan Harrison, David K. Jackson, Ian Johnston, Dominic Kwiatkowski, Cordelia Langford, John Sillitoe on behalf of the Wellcome Sanger Institute COVID-19 Surveillance Team |
| EPI_ISL_3435766 | Berkshire and Surrey Pathology Services Lighthouse Laboratory | Wellcome Sanger Institute for the COVID-19 Genomics UK (COG-UK) Consortium | Berkshire and Surrey Pathology Services Lighthouse Laboratory and Alex Alderton, Roberto Amato, Jeffrey Barrett, Sonia Goncalves, Ewan Harrison, David K. Jackson, Ian Johnston, Dominic Kwiatkowski, Cordelia Langford, John Sillitoe on behalf of the Wellcome Sanger Institute COVID-19 Surveillance Team |
| EPI_ISL_3435767 | Lighthouse Lab in Milton Keynes | Wellcome Sanger Institute for the COVID-19 Genomics UK (COG-UK) Consortium | The Lighthouse Lab in Milton Keynes and Alex Alderton, Roberto Amato, Jeffrey Barrett, Sonia Goncalves, Ewan Harrison, David K. Jackson, Ian Johnston, Dominic Kwiatkowski, Cordelia Langford, John Sillitoe on behalf of the Wellcome Sanger Institute COVID-19 Surveillance Team |
| EPI_ISL_3435768, EPI_ISL_3435769, EPI_ISL_3435770 | Berkshire and Surrey Pathology Services Lighthouse Laboratory | Wellcome Sanger Institute for the COVID-19 Genomics UK (COG-UK) Consortium | Berkshire and Surrey Pathology Services Lighthouse Laboratory and Alex Alderton, Roberto Amato, Jeffrey Barrett, Sonia Goncalves, Ewan Harrison, David K. Jackson, Ian Johnston, Dominic Kwiatkowski, Cordelia Langford, John Sillitoe on behalf of the Wellcome Sanger Institute COVID-19 Surveillance Team |
| EPI_ISL_3435771 | Lighthouse Lab in Milton Keynes | Wellcome Sanger Institute for the COVID-19 Genomics UK (COG-UK) Consortium | The Lighthouse Lab in Milton Keynes and Alex Alderton, Roberto Amato, Jeffrey Barrett, Sonia Goncalves, Ewan Harrison, David K. Jackson, Ian Johnston, Dominic Kwiatkowski, Cordelia Langford, John Sillitoe on behalf of the Wellcome Sanger Institute COVID-19 Surveillance Team |
| EPI_ISL_3435772, EPI_ISL_3435773 | Berkshire and Surrey Pathology Services Lighthouse Laboratory | Wellcome Sanger Institute for the COVID-19 Genomics UK (COG-UK) Consortium | Berkshire and Surrey Pathology Services Lighthouse Laboratory and Alex Alderton, Roberto Amato, Jeffrey Barrett, Sonia Goncalves, Ewan Harrison, David K. Jackson, Ian Johnston, Dominic Kwiatkowski, Cordelia Langford, John Sillitoe on behalf of the Wellcome Sanger Institute COVID-19 Surveillance Team |
| EPI_ISL_3435774, EPI_ISL_3435775 | Lighthouse Lab in Milton Keynes | Wellcome Sanger Institute for the COVID-19 Genomics UK (COG-UK) Consortium | The Lighthouse Lab in Milton Keynes and Alex Alderton, Roberto Amato, Jeffrey Barrett, Sonia Goncalves, Ewan Harrison, David K. Jackson, Ian Johnston, Dominic Kwiatkowski, Cordelia Langford, John Sillitoe on behalf of the Wellcome Sanger Institute COVID-19 Surveillance Team |
| EPI_ISL_3435776, EPI_ISL_3435777, EPI_ISL_3435778, EPI_ISL_3435779, EPI_ISL_3435780, EPI_ISL_3435781 | Berkshire and Surrey Pathology Services Lighthouse Laboratory | Wellcome Sanger Institute for the COVID-19 Genomics UK (COG-UK) Consortium | Berkshire and Surrey Pathology Services Lighthouse Laboratory and Alex Alderton, Roberto Amato, Jeffrey Barrett, Sonia Goncalves, Ewan Harrison, David K. Jackson, Ian Johnston, Dominic Kwiatkowski, Cordelia Langford, John Sillitoe on behalf of the Wellcome Sanger Institute COVID-19 Surveillance Team |
| EPI_ISL_3435782, EPI_ISL_3435783, EPI_ISL_3435784, EPI_ISL_3435785, EPI_ISL_3435786 | Lighthouse Lab in Milton Keynes | Wellcome Sanger Institute for the COVID-19 Genomics UK (COG-UK) Consortium | The Lighthouse Lab in Milton Keynes and Alex Alderton, Roberto Amato, Jeffrey Barrett, Sonia Goncalves, Ewan Harrison, David K. Jackson, Ian Johnston, Dominic Kwiatkowski, Cordelia Langford, John Sillitoe on behalf of the Wellcome Sanger Institute COVID-19 Surveillance Team |
| EPI_ISL_3435787 | Berkshire and Surrey Pathology Services Lighthouse Laboratory | Wellcome Sanger Institute for the COVID-19 Genomics UK (COG-UK) Consortium | Berkshire and Surrey Pathology Services Lighthouse Laboratory and Alex Alderton, Roberto Amato, Jeffrey Barrett, Sonia Goncalves, Ewan Harrison, David K. Jackson, Ian Johnston, Dominic Kwiatkowski, Cordelia Langford, John Sillitoe on behalf of the Wellcome Sanger Institute COVID-19 Surveillance Team |

| | | | |
|---|---|---|---|
| EPI_ISL_3435788 | Lighthouse Lab in Milton Keynes | Wellcome Sanger Institute for the COVID-19 Genomics UK (COG-UK) Consortium | The Lighthouse Lab in Milton Keynes and Alex Alderton, Roberto Amato, Jeffrey Barrett, Sonia Goncalves, Ewan Harrison, David K. Jackson, Ian Johnston, Dominic Kwiatkowski, Cordelia Langford, John Sillitoe on behalf of the Wellcome Sanger Institute COVID-19 Surveillance Team |
| EPI_ISL_3435789 | Berkshire and Surrey Pathology Services Lighthouse Laboratory | Wellcome Sanger Institute for the COVID-19 Genomics UK (COG-UK) Consortium | Berkshire and Surrey Pathology Services Lighthouse Laboratory and Alex Alderton, Roberto Amato, Jeffrey Barrett, Sonia Goncalves, Ewan Harrison, David K. Jackson, Ian Johnston, Dominic Kwiatkowski, Cordelia Langford, John Sillitoe on behalf of the Wellcome Sanger Institute COVID-19 Surveillance Team |
| EPI_ISL_3435790, EPI_ISL_3435791 | Lighthouse Lab in Milton Keynes | Wellcome Sanger Institute for the COVID-19 Genomics UK (COG-UK) Consortium | The Lighthouse Lab in Milton Keynes and Alex Alderton, Roberto Amato, Jeffrey Barrett, Sonia Goncalves, Ewan Harrison, David K. Jackson, Ian Johnston, Dominic Kwiatkowski, Cordelia Langford, John Sillitoe on behalf of the Wellcome Sanger Institute COVID-19 Surveillance Team |
| EPI_ISL_3435792, EPI_ISL_3435793, EPI_ISL_3435794, EPI_ISL_3435795 | Berkshire and Surrey Pathology Services Lighthouse Laboratory | Wellcome Sanger Institute for the COVID-19 Genomics UK (COG-UK) Consortium | Berkshire and Surrey Pathology Services Lighthouse Laboratory and Alex Alderton, Roberto Amato, Jeffrey Barrett, Sonia Goncalves, Ewan Harrison, David K. Jackson, Ian Johnston, Dominic Kwiatkowski, Cordelia Langford, John Sillitoe on behalf of the Wellcome Sanger Institute COVID-19 Surveillance Team |
| EPI_ISL_3435796, EPI_ISL_3435797 | Lighthouse Lab in Milton Keynes | Wellcome Sanger Institute for the COVID-19 Genomics UK (COG-UK) Consortium | The Lighthouse Lab in Milton Keynes and Alex Alderton, Roberto Amato, Jeffrey Barrett, Sonia Goncalves, Ewan Harrison, David K. Jackson, Ian Johnston, Dominic Kwiatkowski, Cordelia Langford, John Sillitoe on behalf of the Wellcome Sanger Institute COVID-19 Surveillance Team |
| EPI_ISL_3435798 | Berkshire and Surrey Pathology Services Lighthouse Laboratory | Wellcome Sanger Institute for the COVID-19 Genomics UK (COG-UK) Consortium | Berkshire and Surrey Pathology Services Lighthouse Laboratory and Alex Alderton, Roberto Amato, Jeffrey Barrett, Sonia Goncalves, Ewan Harrison, David K. Jackson, Ian Johnston, Dominic Kwiatkowski, Cordelia Langford, John Sillitoe on behalf of the Wellcome Sanger Institute COVID-19 Surveillance Team |
| EPI_ISL_3435799, EPI_ISL_3435800 | Lighthouse Lab in Milton Keynes | Wellcome Sanger Institute for the COVID-19 Genomics UK (COG-UK) Consortium | The Lighthouse Lab in Milton Keynes and Alex Alderton, Roberto Amato, Jeffrey Barrett, Sonia Goncalves, Ewan Harrison, David K. Jackson, Ian Johnston, Dominic Kwiatkowski, Cordelia Langford, John Sillitoe on behalf of the Wellcome Sanger Institute COVID-19 Surveillance Team |
| EPI_ISL_3435801, EPI_ISL_3435802, EPI_ISL_3435803, EPI_ISL_3435804 | Berkshire and Surrey Pathology Services Lighthouse Laboratory | Wellcome Sanger Institute for the COVID-19 Genomics UK (COG-UK) Consortium | Berkshire and Surrey Pathology Services Lighthouse Laboratory and Alex Alderton, Roberto Amato, Jeffrey Barrett, Sonia Goncalves, Ewan Harrison, David K. Jackson, Ian Johnston, Dominic Kwiatkowski, Cordelia Langford, John Sillitoe on behalf of the Wellcome Sanger Institute COVID-19 Surveillance Team |
| EPI_ISL_3435805, EPI_ISL_3435806, EPI_ISL_3435807 | Lighthouse Lab in Milton Keynes | Wellcome Sanger Institute for the COVID-19 Genomics UK (COG-UK) Consortium | The Lighthouse Lab in Milton Keynes and Alex Alderton, Roberto Amato, Jeffrey Barrett, Sonia Goncalves, Ewan Harrison, David K. Jackson, Ian Johnston, Dominic Kwiatkowski, Cordelia Langford, John Sillitoe on behalf of the Wellcome Sanger Institute COVID-19 Surveillance Team |
| EPI_ISL_3435808, EPI_ISL_3435809, EPI_ISL_3435810 | Berkshire and Surrey Pathology Services Lighthouse Laboratory | Wellcome Sanger Institute for the COVID-19 Genomics UK (COG-UK) Consortium | Berkshire and Surrey Pathology Services Lighthouse Laboratory and Alex Alderton, Roberto Amato, Jeffrey Barrett, Sonia Goncalves, Ewan Harrison, David K. Jackson, Ian Johnston, Dominic Kwiatkowski, Cordelia Langford, John Sillitoe on behalf of the Wellcome Sanger Institute COVID-19 Surveillance Team |
| EPI_ISL_3435811, EPI_ISL_3435812, EPI_ISL_3435813, EPI_ISL_3435814, EPI_ISL_3435815, EPI_ISL_3435816, EPI_ISL_3435817, EPI_ISL_3435818, EPI_ISL_3435819 | Lighthouse Lab in Milton Keynes | Wellcome Sanger Institute for the COVID-19 Genomics UK (COG-UK) Consortium | The Lighthouse Lab in Milton Keynes and Alex Alderton, Roberto Amato, Jeffrey Barrett, Sonia Goncalves, Ewan Harrison, David K. Jackson, Ian Johnston, Dominic Kwiatkowski, Cordelia Langford, John Sillitoe on behalf of the Wellcome Sanger Institute COVID-19 Surveillance Team |
| EPI_ISL_3435820, EPI_ISL_3435821, EPI_ISL_3435822, EPI_ISL_3435823, EPI_ISL_3435824, EPI_ISL_3435825, EPI_ISL_3435826 | Berkshire and Surrey Pathology Services Lighthouse Laboratory | Wellcome Sanger Institute for the COVID-19 Genomics UK (COG-UK) Consortium | Berkshire and Surrey Pathology Services Lighthouse Laboratory and Alex Alderton, Roberto Amato, Jeffrey Barrett, Sonia Goncalves, Ewan Harrison, David K. Jackson, Ian Johnston, Dominic Kwiatkowski, Cordelia Langford, John Sillitoe on behalf of the Wellcome Sanger Institute COVID-19 Surveillance Team |
| EPI_ISL_3435827 | Lighthouse Lab in Milton Keynes | Wellcome Sanger Institute for the COVID-19 Genomics UK (COG-UK) Consortium | The Lighthouse Lab in Milton Keynes and Alex Alderton, Roberto Amato, Jeffrey Barrett, Sonia Goncalves, Ewan Harrison, David K. Jackson, Ian Johnston, Dominic Kwiatkowski, Cordelia Langford, John Sillitoe on behalf of the Wellcome Sanger Institute COVID-19 Surveillance Team |
| EPI_ISL_3435828 | Berkshire and Surrey Pathology Services Lighthouse Laboratory | Wellcome Sanger Institute for the COVID-19 Genomics UK (COG-UK) Consortium | Berkshire and Surrey Pathology Services Lighthouse Laboratory and Alex Alderton, Roberto Amato, Jeffrey Barrett, Sonia Goncalves, Ewan Harrison, David K. Jackson, Ian Johnston, Dominic Kwiatkowski, Cordelia Langford, John Sillitoe on behalf of the Wellcome Sanger Institute COVID-19 Surveillance Team |
| EPI_ISL_3435829, EPI_ISL_3435830 | Lighthouse Lab in Milton Keynes | Wellcome Sanger Institute for the COVID-19 Genomics UK (COG-UK) Consortium | The Lighthouse Lab in Milton Keynes and Alex Alderton, Roberto Amato, Jeffrey Barrett, Sonia Goncalves, Ewan Harrison, David K. Jackson, Ian Johnston, Dominic Kwiatkowski, Cordelia Langford, John Sillitoe on behalf of the Wellcome Sanger Institute COVID-19 Surveillance Team |
| EPI_ISL_3435831 | Berkshire and Surrey Pathology Services Lighthouse Laboratory | Wellcome Sanger Institute for the COVID-19 Genomics UK (COG-UK) Consortium | Berkshire and Surrey Pathology Services Lighthouse Laboratory and Alex Alderton, Roberto Amato, Jeffrey Barrett, Sonia Goncalves, Ewan Harrison, David K. Jackson, Ian Johnston, Dominic Kwiatkowski, Cordelia Langford, John Sillitoe on behalf of the Wellcome Sanger Institute COVID-19 Surveillance Team |
| EPI_ISL_3435832, EPI_ISL_3435833, EPI_ISL_3435834 | Lighthouse Lab in Milton Keynes | Wellcome Sanger Institute for the COVID-19 Genomics UK (COG-UK) Consortium | The Lighthouse Lab in Milton Keynes and Alex Alderton, Roberto Amato, Jeffrey Barrett, Sonia Goncalves, Ewan Harrison, David K. Jackson, Ian Johnston, Dominic Kwiatkowski, Cordelia Langford, John Sillitoe on behalf of the Wellcome Sanger Institute COVID-19 Surveillance Team |
| EPI_ISL_3435835, EPI_ISL_3435836, EPI_ISL_3435837 | Berkshire and Surrey Pathology Services Lighthouse Laboratory | Wellcome Sanger Institute for the COVID-19 Genomics UK (COG-UK) Consortium | Berkshire and Surrey Pathology Services Lighthouse Laboratory and Alex Alderton, Roberto Amato, Jeffrey Barrett, Sonia Goncalves, Ewan Harrison, David K. Jackson, Ian Johnston, Dominic Kwiatkowski, Cordelia Langford, John Sillitoe on behalf of the Wellcome Sanger Institute COVID-19 Surveillance Team |
| EPI_ISL_3435838 | Lighthouse Lab in Milton Keynes | Wellcome Sanger Institute for the COVID-19 Genomics UK (COG-UK) Consortium | The Lighthouse Lab in Milton Keynes and Alex Alderton, Roberto Amato, Jeffrey Barrett, Sonia Goncalves, Ewan Harrison, David K. Jackson, Ian Johnston, Dominic Kwiatkowski, Cordelia Langford, John Sillitoe on behalf of the Wellcome Sanger Institute COVID-19 Surveillance Team |
| EPI_ISL_3435839, EPI_ISL_3435840 | Berkshire and Surrey Pathology Services Lighthouse Laboratory | Wellcome Sanger Institute for the COVID-19 Genomics UK (COG-UK) Consortium | Berkshire and Surrey Pathology Services Lighthouse Laboratory and Alex Alderton, Roberto Amato, Jeffrey Barrett, Sonia Goncalves, Ewan Harrison, David K. Jackson, Ian Johnston, Dominic Kwiatkowski, Cordelia Langford, John Sillitoe on behalf of the Wellcome Sanger Institute COVID-19 Surveillance Team |
| EPI_ISL_3435841, EPI_ISL_3435842 | Lighthouse Lab in Milton Keynes | Wellcome Sanger Institute for the COVID-19 Genomics UK (COG-UK) Consortium | The Lighthouse Lab in Milton Keynes and Alex Alderton, Roberto Amato, Jeffrey Barrett, Sonia Goncalves, Ewan Harrison, David K. Jackson, Ian Johnston, Dominic Kwiatkowski, Cordelia Langford, John Sillitoe on behalf of the Wellcome Sanger Institute COVID-19 Surveillance Team |
| EPI_ISL_3435843, EPI_ISL_3435844 | Berkshire and Surrey Pathology Services Lighthouse Laboratory | Wellcome Sanger Institute for the COVID-19 Genomics UK (COG-UK) Consortium | Berkshire and Surrey Pathology Services Lighthouse Laboratory and Alex Alderton, Roberto Amato, Jeffrey Barrett, Sonia Goncalves, Ewan Harrison, David K. Jackson, Ian Johnston, Dominic Kwiatkowski, Cordelia Langford, John Sillitoe on behalf of the Wellcome Sanger Institute COVID-19 Surveillance Team |
| EPI_ISL_3435845, EPI_ISL_3435846 | Lighthouse Lab in Milton Keynes | Wellcome Sanger Institute for the COVID-19 Genomics UK (COG-UK) Consortium | The Lighthouse Lab in Milton Keynes and Alex Alderton, Roberto Amato, Jeffrey Barrett, Sonia Goncalves, Ewan Harrison, David K. Jackson, Ian Johnston, Dominic Kwiatkowski, Cordelia Langford, John Sillitoe on behalf of the Wellcome Sanger Institute COVID-19 Surveillance Team |
| EPI_ISL_3435847, EPI_ISL_3435848, EPI_ISL_3435849 | Berkshire and Surrey Pathology Services Lighthouse Laboratory | Wellcome Sanger Institute for the COVID-19 Genomics UK (COG-UK) Consortium | Berkshire and Surrey Pathology Services Lighthouse Laboratory and Alex Alderton, Roberto Amato, Jeffrey Barrett, Sonia Goncalves, Ewan Harrison, David K. Jackson, Ian Johnston, Dominic Kwiatkowski, Cordelia Langford, John Sillitoe on behalf of the Wellcome Sanger Institute COVID-19 Surveillance Team |
| EPI_ISL_3435850 | Lighthouse Lab in Milton Keynes | Wellcome Sanger Institute for the COVID-19 Genomics UK (COG-UK) Consortium | The Lighthouse Lab in Milton Keynes and Alex Alderton, Roberto Amato, Jeffrey Barrett, Sonia Goncalves, Ewan Harrison, David K. Jackson, Ian Johnston, Dominic Kwiatkowski, Cordelia Langford, John Sillitoe on behalf of the Wellcome Sanger Institute COVID-19 Surveillance Team |
| EPI_ISL_3435851 | Berkshire and Surrey Pathology Services Lighthouse Laboratory | Wellcome Sanger Institute for the COVID-19 Genomics UK (COG-UK) Consortium | Berkshire and Surrey Pathology Services Lighthouse Laboratory and Alex Alderton, Roberto Amato, Jeffrey Barrett, Sonia Goncalves, Ewan Harrison, David K. Jackson, Ian Johnston, Dominic Kwiatkowski, Cordelia Langford, John Sillitoe on behalf of the Wellcome Sanger Institute COVID-19 Surveillance Team |
| EPI_ISL_3435852 | Lighthouse Lab in Milton Keynes | Wellcome Sanger Institute for the COVID-19 Genomics UK (COG-UK) Consortium | The Lighthouse Lab in Milton Keynes and Alex Alderton, Roberto Amato, Jeffrey Barrett, Sonia Goncalves, Ewan Harrison, David K. Jackson, Ian Johnston, Dominic Kwiatkowski, Cordelia Langford, John Sillitoe on behalf of the Wellcome Sanger Institute COVID-19 Surveillance Team |
| EPI_ISL_3435853 | Berkshire and Surrey Pathology Services Lighthouse Laboratory | Wellcome Sanger Institute for the COVID-19 Genomics UK (COG-UK) Consortium | Berkshire and Surrey Pathology Services Lighthouse Laboratory and Alex Alderton, Roberto Amato, Jeffrey Barrett, Sonia Goncalves, Ewan Harrison, David K. Jackson, Ian Johnston, Dominic Kwiatkowski, Cordelia Langford, John Sillitoe on behalf of the Wellcome Sanger Institute COVID-19 Surveillance Team |
| EPI_ISL_3435854, EPI_ISL_3435855, EPI_ISL_3435856 | Lighthouse Lab in Milton Keynes | Wellcome Sanger Institute for the COVID-19 Genomics UK (COG-UK) Consortium | The Lighthouse Lab in Milton Keynes and Alex Alderton, Roberto Amato, Jeffrey Barrett, Sonia Goncalves, Ewan Harrison, David K. Jackson, Ian Johnston, Dominic Kwiatkowski, Cordelia Langford, John Sillitoe on behalf of the Wellcome Sanger Institute COVID-19 Surveillance Team |
| EPI_ISL_3435857, EPI_ISL_3435858 | Berkshire and Surrey Pathology Services Lighthouse Laboratory | Wellcome Sanger Institute for the COVID-19 Genomics UK (COG-UK) Consortium | Berkshire and Surrey Pathology Services Lighthouse Laboratory and Alex Alderton, Roberto Amato, Jeffrey Barrett, Sonia Goncalves, Ewan Harrison, David K. Jackson, Ian Johnston, Dominic Kwiatkowski, Cordelia Langford, John Sillitoe on behalf of the Wellcome Sanger Institute COVID-19 Surveillance Team |
| EPI_ISL_3435859 | Lighthouse Lab in Milton Keynes | Wellcome Sanger Institute for the COVID-19 Genomics UK (COG-UK) Consortium | The Lighthouse Lab in Milton Keynes and Alex Alderton, Roberto Amato, Jeffrey Barrett, Sonia Goncalves, Ewan Harrison, David K. Jackson, Ian Johnston, Dominic Kwiatkowski, Cordelia Langford, John Sillitoe on behalf of the Wellcome Sanger Institute COVID-19 Surveillance Team |
| EPI_ISL_3435860, EPI_ISL_3435861 | Berkshire and Surrey Pathology Services Lighthouse Laboratory | Wellcome Sanger Institute for the COVID-19 Genomics UK (COG-UK) Consortium | Berkshire and Surrey Pathology Services Lighthouse Laboratory and Alex Alderton, Roberto Amato, Jeffrey Barrett, Sonia Goncalves, Ewan Harrison, David K. Jackson, Ian Johnston, Dominic Kwiatkowski, Cordelia Langford, John Sillitoe on behalf of the Wellcome Sanger Institute COVID-19 Surveillance Team |
| EPI_ISL_3435862 | Lighthouse Lab in Milton Keynes | Wellcome Sanger Institute for the COVID-19 Genomics UK (COG-UK) Consortium | The Lighthouse Lab in Milton Keynes and Alex Alderton, Roberto Amato, Jeffrey Barrett, Sonia Goncalves, Ewan Harrison, David K. Jackson, Ian Johnston, Dominic Kwiatkowski, Cordelia Langford, John Sillitoe on behalf of the Wellcome Sanger Institute COVID-19 Surveillance Team |
| EPI_ISL_3435863 | Berkshire and Surrey Pathology Services Lighthouse Laboratory | Wellcome Sanger Institute for the COVID-19 Genomics UK (COG-UK) Consortium | Berkshire and Surrey Pathology Services Lighthouse Laboratory and Alex Alderton, Roberto Amato, Jeffrey Barrett, Sonia Goncalves, Ewan Harrison, David K. Jackson, Ian Johnston, Dominic Kwiatkowski, Cordelia Langford, John Sillitoe on behalf of the Wellcome Sanger Institute COVID-19 Surveillance Team |
| EPI_ISL_3435864 | Lighthouse Lab in Milton Keynes | Wellcome Sanger Institute for the COVID-19 Genomics UK (COG-UK) Consortium | The Lighthouse Lab in Milton Keynes and Alex Alderton, Roberto Amato, Jeffrey Barrett, Sonia Goncalves, Ewan Harrison, David K. Jackson, Ian Johnston, Dominic Kwiatkowski, Cordelia Langford, John Sillitoe on behalf of the Wellcome Sanger Institute COVID-19 Surveillance Team |
| EPI_ISL_3435865 | Berkshire and Surrey Pathology Services Lighthouse Laboratory | Wellcome Sanger Institute for the COVID-19 Genomics UK (COG-UK) Consortium | Berkshire and Surrey Pathology Services Lighthouse Laboratory and Alex Alderton, Roberto Amato, Jeffrey Barrett, Sonia Goncalves, Ewan Harrison, David K. Jackson, Ian Johnston, Dominic Kwiatkowski, Cordelia Langford, John Sillitoe on behalf of the Wellcome Sanger Institute COVID-19 Surveillance Team |

| | | | |
|---|---|---|---|
| EPI_ISL_3435866 | Lighthouse Lab in Milton Keynes | Wellcome Sanger Institute for the COVID-19 Genomics UK (COG-UK) Consortium | The Lighthouse Lab in Milton Keynes and Alex Alderton, Roberto Amato, Jeffrey Barrett, Sonia Goncalves, Ewan Harrison, David K. Jackson, Ian Johnston, Dominic Kwiatkowski, Cordelia Langford, John Sillitoe on behalf of the Wellcome Sanger Institute COVID-19 Surveillance Team |
| EPI_ISL_3435867 | Berkshire and Surrey Pathology Services Lighthouse Laboratory | Wellcome Sanger Institute for the COVID-19 Genomics UK (COG-UK) Consortium | Berkshire and Surrey Pathology Services Lighthouse Laboratory and Alex Alderton, Roberto Amato, Jeffrey Barrett, Sonia Goncalves, Ewan Harrison, David K. Jackson, Ian Johnston, Dominic Kwiatkowski, Cordelia Langford, John Sillitoe on behalf of the Wellcome Sanger Institute COVID-19 Surveillance Team |
| EPI_ISL_3435868 | Lighthouse Lab in Milton Keynes | Wellcome Sanger Institute for the COVID-19 Genomics UK (COG-UK) Consortium | The Lighthouse Lab in Milton Keynes and Alex Alderton, Roberto Amato, Jeffrey Barrett, Sonia Goncalves, Ewan Harrison, David K. Jackson, Ian Johnston, Dominic Kwiatkowski, Cordelia Langford, John Sillitoe on behalf of the Wellcome Sanger Institute COVID-19 Surveillance Team |
| EPI_ISL_3435869, EPI_ISL_3435870 | Berkshire and Surrey Pathology Services Lighthouse Laboratory | Wellcome Sanger Institute for the COVID-19 Genomics UK (COG-UK) Consortium | Berkshire and Surrey Pathology Services Lighthouse Laboratory and Alex Alderton, Roberto Amato, Jeffrey Barrett, Sonia Goncalves, Ewan Harrison, David K. Jackson, Ian Johnston, Dominic Kwiatkowski, Cordelia Langford, John Sillitoe on behalf of the Wellcome Sanger Institute COVID-19 Surveillance Team |
| EPI_ISL_3435871, EPI_ISL_3435872 | Lighthouse Lab in Milton Keynes | Wellcome Sanger Institute for the COVID-19 Genomics UK (COG-UK) Consortium | The Lighthouse Lab in Milton Keynes and Alex Alderton, Roberto Amato, Jeffrey Barrett, Sonia Goncalves, Ewan Harrison, David K. Jackson, Ian Johnston, Dominic Kwiatkowski, Cordelia Langford, John Sillitoe on behalf of the Wellcome Sanger Institute COVID-19 Surveillance Team |
| EPI_ISL_3435873, EPI_ISL_3435874 | Berkshire and Surrey Pathology Services Lighthouse Laboratory | Wellcome Sanger Institute for the COVID-19 Genomics UK (COG-UK) Consortium | Berkshire and Surrey Pathology Services Lighthouse Laboratory and Alex Alderton, Roberto Amato, Jeffrey Barrett, Sonia Goncalves, Ewan Harrison, David K. Jackson, Ian Johnston, Dominic Kwiatkowski, Cordelia Langford, John Sillitoe on behalf of the Wellcome Sanger Institute COVID-19 Surveillance Team |
| EPI_ISL_3435875, EPI_ISL_3435876, EPI_ISL_3435877 | Lighthouse Lab in Milton Keynes | Wellcome Sanger Institute for the COVID-19 Genomics UK (COG-UK) Consortium | The Lighthouse Lab in Milton Keynes and Alex Alderton, Roberto Amato, Jeffrey Barrett, Sonia Goncalves, Ewan Harrison, David K. Jackson, Ian Johnston, Dominic Kwiatkowski, Cordelia Langford, John Sillitoe on behalf of the Wellcome Sanger Institute COVID-19 Surveillance Team |
| EPI_ISL_3435878, EPI_ISL_3435879 | Berkshire and Surrey Pathology Services Lighthouse Laboratory | Wellcome Sanger Institute for the COVID-19 Genomics UK (COG-UK) Consortium | Berkshire and Surrey Pathology Services Lighthouse Laboratory and Alex Alderton, Roberto Amato, Jeffrey Barrett, Sonia Goncalves, Ewan Harrison, David K. Jackson, Ian Johnston, Dominic Kwiatkowski, Cordelia Langford, John Sillitoe on behalf of the Wellcome Sanger Institute COVID-19 Surveillance Team |
| EPI_ISL_3435880, EPI_ISL_3435881, EPI_ISL_3435882 | Lighthouse Lab in Milton Keynes | Wellcome Sanger Institute for the COVID-19 Genomics UK (COG-UK) Consortium | The Lighthouse Lab in Milton Keynes and Alex Alderton, Roberto Amato, Jeffrey Barrett, Sonia Goncalves, Ewan Harrison, David K. Jackson, Ian Johnston, Dominic Kwiatkowski, Cordelia Langford, John Sillitoe on behalf of the Wellcome Sanger Institute COVID-19 Surveillance Team |
| EPI_ISL_3435883, EPI_ISL_3435884 | Berkshire and Surrey Pathology Services Lighthouse Laboratory | Wellcome Sanger Institute for the COVID-19 Genomics UK (COG-UK) Consortium | Berkshire and Surrey Pathology Services Lighthouse Laboratory and Alex Alderton, Roberto Amato, Jeffrey Barrett, Sonia Goncalves, Ewan Harrison, David K. Jackson, Ian Johnston, Dominic Kwiatkowski, Cordelia Langford, John Sillitoe on behalf of the Wellcome Sanger Institute COVID-19 Surveillance Team |
| EPI_ISL_3435885 | Lighthouse Lab in Milton Keynes | Wellcome Sanger Institute for the COVID-19 Genomics UK (COG-UK) Consortium | The Lighthouse Lab in Milton Keynes and Alex Alderton, Roberto Amato, Jeffrey Barrett, Sonia Goncalves, Ewan Harrison, David K. Jackson, Ian Johnston, Dominic Kwiatkowski, Cordelia Langford, John Sillitoe on behalf of the Wellcome Sanger Institute COVID-19 Surveillance Team |
| EPI_ISL_3435886 | Berkshire and Surrey Pathology Services Lighthouse Laboratory | Wellcome Sanger Institute for the COVID-19 Genomics UK (COG-UK) Consortium | Berkshire and Surrey Pathology Services Lighthouse Laboratory and Alex Alderton, Roberto Amato, Jeffrey Barrett, Sonia Goncalves, Ewan Harrison, David K. Jackson, Ian Johnston, Dominic Kwiatkowski, Cordelia Langford, John Sillitoe on behalf of the Wellcome Sanger Institute COVID-19 Surveillance Team |
| EPI_ISL_3435887, EPI_ISL_3435888, EPI_ISL_3435889 | Lighthouse Lab in Milton Keynes | Wellcome Sanger Institute for the COVID-19 Genomics UK (COG-UK) Consortium | The Lighthouse Lab in Milton Keynes and Alex Alderton, Roberto Amato, Jeffrey Barrett, Sonia Goncalves, Ewan Harrison, David K. Jackson, Ian Johnston, Dominic Kwiatkowski, Cordelia Langford, John Sillitoe on behalf of the Wellcome Sanger Institute COVID-19 Surveillance Team |
| EPI_ISL_3435890 | Berkshire and Surrey Pathology Services Lighthouse Laboratory | Wellcome Sanger Institute for the COVID-19 Genomics UK (COG-UK) Consortium | Berkshire and Surrey Pathology Services Lighthouse Laboratory and Alex Alderton, Roberto Amato, Jeffrey Barrett, Sonia Goncalves, Ewan Harrison, David K. Jackson, Ian Johnston, Dominic Kwiatkowski, Cordelia Langford, John Sillitoe on behalf of the Wellcome Sanger Institute COVID-19 Surveillance Team |
| EPI_ISL_3435891 | Lighthouse Lab in Milton Keynes | Wellcome Sanger Institute for the COVID-19 Genomics UK (COG-UK) Consortium | The Lighthouse Lab in Milton Keynes and Alex Alderton, Roberto Amato, Jeffrey Barrett, Sonia Goncalves, Ewan Harrison, David K. Jackson, Ian Johnston, Dominic Kwiatkowski, Cordelia Langford, John Sillitoe on behalf of the Wellcome Sanger Institute COVID-19 Surveillance Team |
| EPI_ISL_3435892, EPI_ISL_3435893, EPI_ISL_3435894 | Berkshire and Surrey Pathology Services Lighthouse Laboratory | Wellcome Sanger Institute for the COVID-19 Genomics UK (COG-UK) Consortium | Berkshire and Surrey Pathology Services Lighthouse Laboratory and Alex Alderton, Roberto Amato, Jeffrey Barrett, Sonia Goncalves, Ewan Harrison, David K. Jackson, Ian Johnston, Dominic Kwiatkowski, Cordelia Langford, John Sillitoe on behalf of the Wellcome Sanger Institute COVID-19 Surveillance Team |
| EPI_ISL_3435895, EPI_ISL_3435896, EPI_ISL_3435897, EPI_ISL_3435898, EPI_ISL_3435899 | Lighthouse Lab in Milton Keynes | Wellcome Sanger Institute for the COVID-19 Genomics UK (COG-UK) Consortium | The Lighthouse Lab in Milton Keynes and Alex Alderton, Roberto Amato, Jeffrey Barrett, Sonia Goncalves, Ewan Harrison, David K. Jackson, Ian Johnston, Dominic Kwiatkowski, Cordelia Langford, John Sillitoe on behalf of the Wellcome Sanger Institute COVID-19 Surveillance Team |
| EPI_ISL_3435900 | Berkshire and Surrey Pathology Services Lighthouse Laboratory | Wellcome Sanger Institute for the COVID-19 Genomics UK (COG-UK) Consortium | Berkshire and Surrey Pathology Services Lighthouse Laboratory and Alex Alderton, Roberto Amato, Jeffrey Barrett, Sonia Goncalves, Ewan Harrison, David K. Jackson, Ian Johnston, Dominic Kwiatkowski, Cordelia Langford, John Sillitoe on behalf of the Wellcome Sanger Institute COVID-19 Surveillance Team |
| EPI_ISL_3435901, EPI_ISL_3435902 | Lighthouse Lab in Milton Keynes | Wellcome Sanger Institute for the COVID-19 Genomics UK (COG-UK) Consortium | The Lighthouse Lab in Milton Keynes and Alex Alderton, Roberto Amato, Jeffrey Barrett, Sonia Goncalves, Ewan Harrison, David K. Jackson, Ian Johnston, Dominic Kwiatkowski, Cordelia Langford, John Sillitoe on behalf of the Wellcome Sanger Institute COVID-19 Surveillance Team |
| EPI_ISL_3435903 | Berkshire and Surrey Pathology Services Lighthouse Laboratory | Wellcome Sanger Institute for the COVID-19 Genomics UK (COG-UK) Consortium | Berkshire and Surrey Pathology Services Lighthouse Laboratory and Alex Alderton, Roberto Amato, Jeffrey Barrett, Sonia Goncalves, Ewan Harrison, David K. Jackson, Ian Johnston, Dominic Kwiatkowski, Cordelia Langford, John Sillitoe on behalf of the Wellcome Sanger Institute COVID-19 Surveillance Team |
| EPI_ISL_3435904, EPI_ISL_3435905, EPI_ISL_3435906, EPI_ISL_3435907 | Lighthouse Lab in Milton Keynes | Wellcome Sanger Institute for the COVID-19 Genomics UK (COG-UK) Consortium | The Lighthouse Lab in Milton Keynes and Alex Alderton, Roberto Amato, Jeffrey Barrett, Sonia Goncalves, Ewan Harrison, David K. Jackson, Ian Johnston, Dominic Kwiatkowski, Cordelia Langford, John Sillitoe on behalf of the Wellcome Sanger Institute COVID-19 Surveillance Team |
| EPI_ISL_3435908 | Berkshire and Surrey Pathology Services Lighthouse Laboratory | Wellcome Sanger Institute for the COVID-19 Genomics UK (COG-UK) Consortium | Berkshire and Surrey Pathology Services Lighthouse Laboratory and Alex Alderton, Roberto Amato, Jeffrey Barrett, Sonia Goncalves, Ewan Harrison, David K. Jackson, Ian Johnston, Dominic Kwiatkowski, Cordelia Langford, John Sillitoe on behalf of the Wellcome Sanger Institute COVID-19 Surveillance Team |
| EPI_ISL_3435909 | Lighthouse Lab in Milton Keynes | Wellcome Sanger Institute for the COVID-19 Genomics UK (COG-UK) Consortium | The Lighthouse Lab in Milton Keynes and Alex Alderton, Roberto Amato, Jeffrey Barrett, Sonia Goncalves, Ewan Harrison, David K. Jackson, Ian Johnston, Dominic Kwiatkowski, Cordelia Langford, John Sillitoe on behalf of the Wellcome Sanger Institute COVID-19 Surveillance Team |
| EPI_ISL_3435910, EPI_ISL_3435911 | Berkshire and Surrey Pathology Services Lighthouse Laboratory | Wellcome Sanger Institute for the COVID-19 Genomics UK (COG-UK) Consortium | Berkshire and Surrey Pathology Services Lighthouse Laboratory and Alex Alderton, Roberto Amato, Jeffrey Barrett, Sonia Goncalves, Ewan Harrison, David K. Jackson, Ian Johnston, Dominic Kwiatkowski, Cordelia Langford, John Sillitoe on behalf of the Wellcome Sanger Institute COVID-19 Surveillance Team |
| EPI_ISL_3435912, EPI_ISL_3435913 | Lighthouse Lab in Milton Keynes | Wellcome Sanger Institute for the COVID-19 Genomics UK (COG-UK) Consortium | The Lighthouse Lab in Milton Keynes and Alex Alderton, Roberto Amato, Jeffrey Barrett, Sonia Goncalves, Ewan Harrison, David K. Jackson, Ian Johnston, Dominic Kwiatkowski, Cordelia Langford, John Sillitoe on behalf of the Wellcome Sanger Institute COVID-19 Surveillance Team |
| EPI_ISL_3435914 | Berkshire and Surrey Pathology Services Lighthouse Laboratory | Wellcome Sanger Institute for the COVID-19 Genomics UK (COG-UK) Consortium | Berkshire and Surrey Pathology Services Lighthouse Laboratory and Alex Alderton, Roberto Amato, Jeffrey Barrett, Sonia Goncalves, Ewan Harrison, David K. Jackson, Ian Johnston, Dominic Kwiatkowski, Cordelia Langford, John Sillitoe on behalf of the Wellcome Sanger Institute COVID-19 Surveillance Team |
| EPI_ISL_3435915, EPI_ISL_3435916 | Lighthouse Lab in Milton Keynes | Wellcome Sanger Institute for the COVID-19 Genomics UK (COG-UK) Consortium | The Lighthouse Lab in Milton Keynes and Alex Alderton, Roberto Amato, Jeffrey Barrett, Sonia Goncalves, Ewan Harrison, David K. Jackson, Ian Johnston, Dominic Kwiatkowski, Cordelia Langford, John Sillitoe on behalf of the Wellcome Sanger Institute COVID-19 Surveillance Team |
| EPI_ISL_3435917 | Berkshire and Surrey Pathology Services Lighthouse Laboratory | Wellcome Sanger Institute for the COVID-19 Genomics UK (COG-UK) Consortium | Berkshire and Surrey Pathology Services Lighthouse Laboratory and Alex Alderton, Roberto Amato, Jeffrey Barrett, Sonia Goncalves, Ewan Harrison, David K. Jackson, Ian Johnston, Dominic Kwiatkowski, Cordelia Langford, John Sillitoe on behalf of the Wellcome Sanger Institute COVID-19 Surveillance Team |
| EPI_ISL_3435918, EPI_ISL_3435919, EPI_ISL_3435920, EPI_ISL_3435921, EPI_ISL_3435922, EPI_ISL_3435923 | Lighthouse Lab in Milton Keynes | Wellcome Sanger Institute for the COVID-19 Genomics UK (COG-UK) Consortium | The Lighthouse Lab in Milton Keynes and Alex Alderton, Roberto Amato, Jeffrey Barrett, Sonia Goncalves, Ewan Harrison, David K. Jackson, Ian Johnston, Dominic Kwiatkowski, Cordelia Langford, John Sillitoe on behalf of the Wellcome Sanger Institute COVID-19 Surveillance Team |
| EPI_ISL_3435924 | Berkshire and Surrey Pathology Services Lighthouse Laboratory | Wellcome Sanger Institute for the COVID-19 Genomics UK (COG-UK) Consortium | Berkshire and Surrey Pathology Services Lighthouse Laboratory and Alex Alderton, Roberto Amato, Jeffrey Barrett, Sonia Goncalves, Ewan Harrison, David K. Jackson, Ian Johnston, Dominic Kwiatkowski, Cordelia Langford, John Sillitoe on behalf of the Wellcome Sanger Institute COVID-19 Surveillance Team |
| EPI_ISL_3435925 | Lighthouse Lab in Milton Keynes | Wellcome Sanger Institute for the COVID-19 Genomics UK (COG-UK) Consortium | The Lighthouse Lab in Milton Keynes and Alex Alderton, Roberto Amato, Jeffrey Barrett, Sonia Goncalves, Ewan Harrison, David K. Jackson, Ian Johnston, Dominic Kwiatkowski, Cordelia Langford, John Sillitoe on behalf of the Wellcome Sanger Institute COVID-19 Surveillance Team |
| EPI_ISL_3435926, EPI_ISL_3435927, EPI_ISL_3435928, EPI_ISL_3435929 | Berkshire and Surrey Pathology Services Lighthouse Laboratory | Wellcome Sanger Institute for the COVID-19 Genomics UK (COG-UK) Consortium | Berkshire and Surrey Pathology Services Lighthouse Laboratory and Alex Alderton, Roberto Amato, Jeffrey Barrett, Sonia Goncalves, Ewan Harrison, David K. Jackson, Ian Johnston, Dominic Kwiatkowski, Cordelia Langford, John Sillitoe on behalf of the Wellcome Sanger Institute COVID-19 Surveillance Team |
| EPI_ISL_3435930, EPI_ISL_3435931 | Lighthouse Lab in Milton Keynes | Wellcome Sanger Institute for the COVID-19 Genomics UK (COG-UK) Consortium | The Lighthouse Lab in Milton Keynes and Alex Alderton, Roberto Amato, Jeffrey Barrett, Sonia Goncalves, Ewan Harrison, David K. Jackson, Ian Johnston, Dominic Kwiatkowski, Cordelia Langford, John Sillitoe on behalf of the Wellcome Sanger Institute COVID-19 Surveillance Team |
| EPI_ISL_3435932, EPI_ISL_3435933, EPI_ISL_3435934 | Berkshire and Surrey Pathology Services Lighthouse Laboratory | Wellcome Sanger Institute for the COVID-19 Genomics UK (COG-UK) Consortium | Berkshire and Surrey Pathology Services Lighthouse Laboratory and Alex Alderton, Roberto Amato, Jeffrey Barrett, Sonia Goncalves, Ewan Harrison, David K. Jackson, Ian Johnston, Dominic Kwiatkowski, Cordelia Langford, John Sillitoe on behalf of the Wellcome Sanger Institute COVID-19 Surveillance Team |
| EPI_ISL_3435935, EPI_ISL_3435936 | Lighthouse Lab in Milton Keynes | Wellcome Sanger Institute for the COVID-19 Genomics UK (COG-UK) Consortium | The Lighthouse Lab in Milton Keynes and Alex Alderton, Roberto Amato, Jeffrey Barrett, Sonia Goncalves, Ewan Harrison, David K. Jackson, Ian Johnston, Dominic Kwiatkowski, Cordelia Langford, John Sillitoe on behalf of the Wellcome Sanger Institute COVID-19 Surveillance Team |
| EPI_ISL_3435937 | Berkshire and Surrey Pathology Services Lighthouse Laboratory | Wellcome Sanger Institute for the COVID-19 Genomics UK (COG-UK) Consortium | Berkshire and Surrey Pathology Services Lighthouse Laboratory and Alex Alderton, Roberto Amato, Jeffrey Barrett, Sonia Goncalves, Ewan Harrison, David K. Jackson, Ian Johnston, Dominic Kwiatkowski, Cordelia Langford, John Sillitoe on behalf of the Wellcome Sanger Institute COVID-19 Surveillance Team |
| EPI_ISL_3435938 | Lighthouse Lab in Milton Keynes | Wellcome Sanger Institute for the COVID-19 Genomics UK (COG-UK) Consortium | The Lighthouse Lab in Milton Keynes and Alex Alderton, Roberto Amato, Jeffrey Barrett, Sonia Goncalves, Ewan Harrison, David K. Jackson, Ian Johnston, Dominic Kwiatkowski, Cordelia Langford, John Sillitoe on behalf of the Wellcome Sanger Institute COVID-19 Surveillance Team |
| EPI_ISL_3435939 | Berkshire and Surrey Pathology Services Lighthouse Laboratory | Wellcome Sanger Institute for the COVID-19 Genomics UK (COG-UK) Consortium | Berkshire and Surrey Pathology Services Lighthouse Laboratory and Alex Alderton, Roberto Amato, Jeffrey Barrett, Sonia Goncalves, Ewan Harrison, David K. Jackson, Ian Johnston, Dominic Kwiatkowski, Cordelia Langford, John Sillitoe on behalf of the Wellcome Sanger Institute COVID-19 Surveillance Team |
| EPI_ISL_3435940, | Lighthouse Lab in Milton Keynes | Wellcome Sanger Institute for the COVID-19 Genomics UK (COG-UK) | The Lighthouse Lab in Milton Keynes and Alex Alderton, Roberto Amato, Jeffrey Barrett, Sonia Goncalves, Ewan Harrison, David K. Jackson, Ian Johnston, Dominic Kwiatkowski, Cordelia Langford, John Sillitoe on behalf of the Wellcome Sanger |

| | | | |
|---|---|---|---|
| EPI_ISL_3435941, EPI_ISL_3435942 | | Consortium | Institute COVID-19 Surveillance Team |
| EPI_ISL_3435943 | Berkshire and Surrey Pathology Services Lighthouse Laboratory | Wellcome Sanger Institute for the COVID-19 Genomics UK (COG-UK) Consortium | Berkshire and Surrey Pathology Services Lighthouse Laboratory and Alex Alderton, Roberto Amato, Jeffrey Barrett, Sonia Goncalves, Ewan Harrison, David K. Jackson, Ian Johnston, Dominic Kwiatkowski, Cordelia Langford, John Sillitoe on behalf of the Wellcome Sanger Institute COVID-19 Surveillance Team |
| EPI_ISL_3435944, EPI_ISL_3435945, EPI_ISL_3435946, EPI_ISL_3435947, EPI_ISL_3435948, EPI_ISL_3435949 | Lighthouse Lab in Milton Keynes | Wellcome Sanger Institute for the COVID-19 Genomics UK (COG-UK) Consortium | The Lighthouse Lab in Milton Keynes and Alex Alderton, Roberto Amato, Jeffrey Barrett, Sonia Goncalves, Ewan Harrison, David K. Jackson, Ian Johnston, Dominic Kwiatkowski, Cordelia Langford, John Sillitoe on behalf of the Wellcome Sanger Institute COVID-19 Surveillance Team |
| EPI_ISL_3435950, EPI_ISL_3435951 | Berkshire and Surrey Pathology Services Lighthouse Laboratory | Wellcome Sanger Institute for the COVID-19 Genomics UK (COG-UK) Consortium | Berkshire and Surrey Pathology Services Lighthouse Laboratory and Alex Alderton, Roberto Amato, Jeffrey Barrett, Sonia Goncalves, Ewan Harrison, David K. Jackson, Ian Johnston, Dominic Kwiatkowski, Cordelia Langford, John Sillitoe on behalf of the Wellcome Sanger Institute COVID-19 Surveillance Team |
| EPI_ISL_3435952, EPI_ISL_3435953, EPI_ISL_3435954, EPI_ISL_3435955 | Lighthouse Lab in Milton Keynes | Wellcome Sanger Institute for the COVID-19 Genomics UK (COG-UK) Consortium | The Lighthouse Lab in Milton Keynes and Alex Alderton, Roberto Amato, Jeffrey Barrett, Sonia Goncalves, Ewan Harrison, David K. Jackson, Ian Johnston, Dominic Kwiatkowski, Cordelia Langford, John Sillitoe on behalf of the Wellcome Sanger Institute COVID-19 Surveillance Team |
| EPI_ISL_3435956, EPI_ISL_3435957 | Berkshire and Surrey Pathology Services Lighthouse Laboratory | Wellcome Sanger Institute for the COVID-19 Genomics UK (COG-UK) Consortium | Berkshire and Surrey Pathology Services Lighthouse Laboratory and Alex Alderton, Roberto Amato, Jeffrey Barrett, Sonia Goncalves, Ewan Harrison, David K. Jackson, Ian Johnston, Dominic Kwiatkowski, Cordelia Langford, John Sillitoe on behalf of the Wellcome Sanger Institute COVID-19 Surveillance Team |
| EPI_ISL_3435958, EPI_ISL_3435959 | Lighthouse Lab in Milton Keynes | Wellcome Sanger Institute for the COVID-19 Genomics UK (COG-UK) Consortium | The Lighthouse Lab in Milton Keynes and Alex Alderton, Roberto Amato, Jeffrey Barrett, Sonia Goncalves, Ewan Harrison, David K. Jackson, Ian Johnston, Dominic Kwiatkowski, Cordelia Langford, John Sillitoe on behalf of the Wellcome Sanger Institute COVID-19 Surveillance Team |
| EPI_ISL_3435960 | Berkshire and Surrey Pathology Services Lighthouse Laboratory | Wellcome Sanger Institute for the COVID-19 Genomics UK (COG-UK) Consortium | Berkshire and Surrey Pathology Services Lighthouse Laboratory and Alex Alderton, Roberto Amato, Jeffrey Barrett, Sonia Goncalves, Ewan Harrison, David K. Jackson, Ian Johnston, Dominic Kwiatkowski, Cordelia Langford, John Sillitoe on behalf of the Wellcome Sanger Institute COVID-19 Surveillance Team |
| EPI_ISL_3435961 | Lighthouse Lab in Milton Keynes | Wellcome Sanger Institute for the COVID-19 Genomics UK (COG-UK) Consortium | The Lighthouse Lab in Milton Keynes and Alex Alderton, Roberto Amato, Jeffrey Barrett, Sonia Goncalves, Ewan Harrison, David K. Jackson, Ian Johnston, Dominic Kwiatkowski, Cordelia Langford, John Sillitoe on behalf of the Wellcome Sanger Institute COVID-19 Surveillance Team |
| EPI_ISL_3435962, EPI_ISL_3435963, EPI_ISL_3435964, EPI_ISL_3435965, EPI_ISL_3435966 | Berkshire and Surrey Pathology Services Lighthouse Laboratory | Wellcome Sanger Institute for the COVID-19 Genomics UK (COG-UK) Consortium | Berkshire and Surrey Pathology Services Lighthouse Laboratory and Alex Alderton, Roberto Amato, Jeffrey Barrett, Sonia Goncalves, Ewan Harrison, David K. Jackson, Ian Johnston, Dominic Kwiatkowski, Cordelia Langford, John Sillitoe on behalf of the Wellcome Sanger Institute COVID-19 Surveillance Team |
| EPI_ISL_3435967, EPI_ISL_3435968, EPI_ISL_3435969, EPI_ISL_3435970, EPI_ISL_3435971, EPI_ISL_3435972, EPI_ISL_3435973 | Lighthouse Lab in Milton Keynes | Wellcome Sanger Institute for the COVID-19 Genomics UK (COG-UK) Consortium | The Lighthouse Lab in Milton Keynes and Alex Alderton, Roberto Amato, Jeffrey Barrett, Sonia Goncalves, Ewan Harrison, David K. Jackson, Ian Johnston, Dominic Kwiatkowski, Cordelia Langford, John Sillitoe on behalf of the Wellcome Sanger Institute COVID-19 Surveillance Team |
| EPI_ISL_3435974 | Lighthouse Lab in Glasgow | Wellcome Sanger Institute for the COVID-19 Genomics UK (COG-UK) Consortium | Harper VanSteenhouse, Yumi Kasai, David Gray, Carol Clugston, Anna Dominiczak and Alex Alderton, Roberto Amato, Jeffrey Barrett, Sonia Goncalves, Ewan Harrison, David K. Jackson, Ian Johnston, Dominic Kwiatkowski, Cordelia Langford, John Sillitoe on behalf of the Wellcome Sanger Institute COVID-19 Surveillance Team |
| EPI_ISL_3435975, EPI_ISL_3435976 | Lighthouse Lab in Milton Keynes | Wellcome Sanger Institute for the COVID-19 Genomics UK (COG-UK) Consortium | The Lighthouse Lab in Milton Keynes and Alex Alderton, Roberto Amato, Jeffrey Barrett, Sonia Goncalves, Ewan Harrison, David K. Jackson, Ian Johnston, Dominic Kwiatkowski, Cordelia Langford, John Sillitoe on behalf of the Wellcome Sanger Institute COVID-19 Surveillance Team |
| EPI_ISL_3435977 | Lighthouse Lab in Glasgow | Wellcome Sanger Institute for the COVID-19 Genomics UK (COG-UK) Consortium | Harper VanSteenhouse, Yumi Kasai, David Gray, Carol Clugston, Anna Dominiczak and Alex Alderton, Roberto Amato, Jeffrey Barrett, Sonia Goncalves, Ewan Harrison, David K. Jackson, Ian Johnston, Dominic Kwiatkowski, Cordelia Langford, John Sillitoe on behalf of the Wellcome Sanger Institute COVID-19 Surveillance Team |
| EPI_ISL_3435978, EPI_ISL_3435979, EPI_ISL_3435980, EPI_ISL_3435981, EPI_ISL_3435982, EPI_ISL_3435983, EPI_ISL_3435984, EPI_ISL_3435985, EPI_ISL_3435986 | Lighthouse Lab in Milton Keynes | Wellcome Sanger Institute for the COVID-19 Genomics UK (COG-UK) Consortium | The Lighthouse Lab in Milton Keynes and Alex Alderton, Roberto Amato, Jeffrey Barrett, Sonia Goncalves, Ewan Harrison, David K. Jackson, Ian Johnston, Dominic Kwiatkowski, Cordelia Langford, John Sillitoe on behalf of the Wellcome Sanger Institute COVID-19 Surveillance Team |
| EPI_ISL_3435987 | Lighthouse Lab in Glasgow | Wellcome Sanger Institute for the COVID-19 Genomics UK (COG-UK) Consortium | Harper VanSteenhouse, Yumi Kasai, David Gray, Carol Clugston, Anna Dominiczak and Alex Alderton, Roberto Amato, Jeffrey Barrett, Sonia Goncalves, Ewan Harrison, David K. Jackson, Ian Johnston, Dominic Kwiatkowski, Cordelia Langford, John Sillitoe on behalf of the Wellcome Sanger Institute COVID-19 Surveillance Team |
| EPI_ISL_3435988, EPI_ISL_3435989, EPI_ISL_3435990, EPI_ISL_3435991, EPI_ISL_3435992, EPI_ISL_3435993, EPI_ISL_3435994, EPI_ISL_3435995, EPI_ISL_3435996, EPI_ISL_3435997 | Lighthouse Lab in Milton Keynes | Wellcome Sanger Institute for the COVID-19 Genomics UK (COG-UK) Consortium | The Lighthouse Lab in Milton Keynes and Alex Alderton, Roberto Amato, Jeffrey Barrett, Sonia Goncalves, Ewan Harrison, David K. Jackson, Ian Johnston, Dominic Kwiatkowski, Cordelia Langford, John Sillitoe on behalf of the Wellcome Sanger Institute COVID-19 Surveillance Team |
| EPI_ISL_3435998 | Lighthouse Lab in Glasgow | Wellcome Sanger Institute for the COVID-19 Genomics UK (COG-UK) Consortium | Harper VanSteenhouse, Yumi Kasai, David Gray, Carol Clugston, Anna Dominiczak and Alex Alderton, Roberto Amato, Jeffrey Barrett, Sonia Goncalves, Ewan Harrison, David K. Jackson, Ian Johnston, Dominic Kwiatkowski, Cordelia Langford, John Sillitoe on behalf of the Wellcome Sanger Institute COVID-19 Surveillance Team |
| EPI_ISL_3435999, EPI_ISL_3436000, EPI_ISL_3436001, EPI_ISL_3436002, EPI_ISL_3436003, EPI_ISL_3436004, EPI_ISL_3436005, EPI_ISL_3436006 | Lighthouse Lab in Milton Keynes | Wellcome Sanger Institute for the COVID-19 Genomics UK (COG-UK) Consortium | The Lighthouse Lab in Milton Keynes and Alex Alderton, Roberto Amato, Jeffrey Barrett, Sonia Goncalves, Ewan Harrison, David K. Jackson, Ian Johnston, Dominic Kwiatkowski, Cordelia Langford, John Sillitoe on behalf of the Wellcome Sanger Institute COVID-19 Surveillance Team |
| EPI_ISL_3436007 | Lighthouse Lab in Glasgow | Wellcome Sanger Institute for the COVID-19 Genomics UK (COG-UK) Consortium | Harper VanSteenhouse, Yumi Kasai, David Gray, Carol Clugston, Anna Dominiczak and Alex Alderton, Roberto Amato, Jeffrey Barrett, Sonia Goncalves, Ewan Harrison, David K. Jackson, Ian Johnston, Dominic Kwiatkowski, Cordelia Langford, John Sillitoe on behalf of the Wellcome Sanger Institute COVID-19 Surveillance Team |
| EPI_ISL_3436008, EPI_ISL_3436009, EPI_ISL_3436010, EPI_ISL_3436011, EPI_ISL_3436012 | Lighthouse Lab in Milton Keynes | Wellcome Sanger Institute for the COVID-19 Genomics UK (COG-UK) Consortium | The Lighthouse Lab in Milton Keynes and Alex Alderton, Roberto Amato, Jeffrey Barrett, Sonia Goncalves, Ewan Harrison, David K. Jackson, Ian Johnston, Dominic Kwiatkowski, Cordelia Langford, John Sillitoe on behalf of the Wellcome Sanger Institute COVID-19 Surveillance Team |
| EPI_ISL_3436013, EPI_ISL_3436014 | Lighthouse Lab in Glasgow | Wellcome Sanger Institute for the COVID-19 Genomics UK (COG-UK) Consortium | Harper VanSteenhouse, Yumi Kasai, David Gray, Carol Clugston, Anna Dominiczak and Alex Alderton, Roberto Amato, Jeffrey Barrett, Sonia Goncalves, Ewan Harrison, David K. Jackson, Ian Johnston, Dominic Kwiatkowski, Cordelia Langford, John Sillitoe on behalf of the Wellcome Sanger Institute COVID-19 Surveillance Team |
| EPI_ISL_3436015, EPI_ISL_3436016, EPI_ISL_3436017, EPI_ISL_3436018, EPI_ISL_3436019, EPI_ISL_3436020, EPI_ISL_3436021, EPI_ISL_3436022, EPI_ISL_3436023, EPI_ISL_3436024, EPI_ISL_3436025, EPI_ISL_3436026, EPI_ISL_3436027, EPI_ISL_3436028, EPI_ISL_3436029, EPI_ISL_3436030, EPI_ISL_3436031, EPI_ISL_3436032, EPI_ISL_3436033, EPI_ISL_3436034, EPI_ISL_3436035, EPI_ISL_3436036, EPI_ISL_3436037, EPI_ISL_3436038, EPI_ISL_3436039, EPI_ISL_3436040, EPI_ISL_3436041, EPI_ISL_3436042, EPI_ISL_3436043, EPI_ISL_3436044, EPI_ISL_3436045, EPI_ISL_3436046, EPI_ISL_3436047, EPI_ISL_3436048, EPI_ISL_3436049, EPI_ISL_3436050, EPI_ISL_3436051, EPI_ISL_3436052, EPI_ISL_3436053, EPI_ISL_3436054, EPI_ISL_3436055, EPI_ISL_3436056, EPI_ISL_3436057, EPI_ISL_3436058, EPI_ISL_3436059, EPI_ISL_3436060, EPI_ISL_3436061, EPI_ISL_3436062, EPI_ISL_3436063, EPI_ISL_3436064, EPI_ISL_3436065, EPI_ISL_3436066, EPI_ISL_3436067, EPI_ISL_3436068, EPI_ISL_3436069, EPI_ISL_3436070, EPI_ISL_3436071, EPI_ISL_3436072, EPI_ISL_3436073, EPI_ISL_3436074, EPI_ISL_3436075 | | | |
| see above | Lighthouse Lab in Milton Keynes | Wellcome Sanger Institute for the COVID-19 Genomics UK (COG-UK) Consortium | The Lighthouse Lab in Milton Keynes and Alex Alderton, Roberto Amato, Jeffrey Barrett, Sonia Goncalves, Ewan Harrison, David K. Jackson, Ian Johnston, Dominic Kwiatkowski, Cordelia Langford, John Sillitoe on behalf of the Wellcome Sanger Institute COVID-19 Surveillance Team |
| EPI_ISL_3436076 | Lighthouse Lab in Glasgow | Wellcome Sanger Institute for the COVID-19 Genomics UK (COG-UK) Consortium | Harper VanSteenhouse, Yumi Kasai, David Gray, Carol Clugston, Anna Dominiczak and Alex Alderton, Roberto Amato, Jeffrey Barrett, Sonia Goncalves, Ewan Harrison, David K. Jackson, Ian Johnston, Dominic Kwiatkowski, Cordelia Langford, John Sillitoe on behalf of the Wellcome Sanger Institute COVID-19 Surveillance Team |
| EPI_ISL_3436077, EPI_ISL_3436078, EPI_ISL_3436079 | Lighthouse Lab in Milton Keynes | Wellcome Sanger Institute for the COVID-19 Genomics UK (COG-UK) Consortium | The Lighthouse Lab in Milton Keynes and Alex Alderton, Roberto Amato, Jeffrey Barrett, Sonia Goncalves, Ewan Harrison, David K. Jackson, Ian Johnston, Dominic Kwiatkowski, Cordelia Langford, John Sillitoe on behalf of the Wellcome Sanger Institute COVID-19 Surveillance Team |
| EPI_ISL_3436080 | Lighthouse Lab in Glasgow | Wellcome Sanger Institute for the COVID-19 Genomics UK (COG-UK) Consortium | Harper VanSteenhouse, Yumi Kasai, David Gray, Carol Clugston, Anna Dominiczak and Alex Alderton, Roberto Amato, Jeffrey Barrett, Sonia Goncalves, Ewan Harrison, David K. Jackson, Ian Johnston, Dominic Kwiatkowski, Cordelia Langford, John Sillitoe on behalf of the Wellcome Sanger Institute COVID-19 Surveillance Team |
| EPI_ISL_3436081, EPI_ISL_3436082, EPI_ISL_3436083, EPI_ISL_3436084, EPI_ISL_3436085, EPI_ISL_3436086, EPI_ISL_3436087, EPI_ISL_3436088, EPI_ISL_3436089, EPI_ISL_3436090, EPI_ISL_3436091, EPI_ISL_3436092, EPI_ISL_3436093 | | | |
| see above | Lighthouse Lab in Milton Keynes | Wellcome Sanger Institute for the COVID-19 Genomics UK (COG-UK) Consortium | The Lighthouse Lab in Milton Keynes and Alex Alderton, Roberto Amato, Jeffrey Barrett, Sonia Goncalves, Ewan Harrison, David K. Jackson, Ian Johnston, Dominic Kwiatkowski, Cordelia Langford, John Sillitoe on behalf of the Wellcome Sanger Institute COVID-19 Surveillance Team |

| | | | |
|---|---|---|---|
| EPI_ISL_3436094 | Lighthouse Lab in Glasgow | Wellcome Sanger Institute for the COVID-19 Genomics UK (COG-UK) Consortium | Harper VanSteenhouse, Yumi Kasai, David Gray, Carol Clugston, Anna Dominiczak and Alex Alderton, Roberto Amato, Jeffrey Barrett, Sonia Goncalves, Ewan Harrison, David K. Jackson, Ian Johnston, Dominic Kwiatkowski, Cordelia Langford, John Sillitoe on behalf of the Wellcome Sanger Institute COVID-19 Surveillance Team |
| EPI_ISL_3436095, EPI_ISL_3436096 | Lighthouse Lab in Milton Keynes | Wellcome Sanger Institute for the COVID-19 Genomics UK (COG-UK) Consortium | The Lighthouse Lab in Milton Keynes and Alex Alderton, Roberto Amato, Jeffrey Barrett, Sonia Goncalves, Ewan Harrison, David K. Jackson, Ian Johnston, Dominic Kwiatkowski, Cordelia Langford, John Sillitoe on behalf of the Wellcome Sanger Institute COVID-19 Surveillance Team |
| EPI_ISL_3436097 | Lighthouse Lab in Glasgow | Wellcome Sanger Institute for the COVID-19 Genomics UK (COG-UK) Consortium | Harper VanSteenhouse, Yumi Kasai, David Gray, Carol Clugston, Anna Dominiczak and Alex Alderton, Roberto Amato, Jeffrey Barrett, Sonia Goncalves, Ewan Harrison, David K. Jackson, Ian Johnston, Dominic Kwiatkowski, Cordelia Langford, John Sillitoe on behalf of the Wellcome Sanger Institute COVID-19 Surveillance Team |
| EPI_ISL_3436098, EPI_ISL_3436099, EPI_ISL_3436100, EPI_ISL_3436101, EPI_ISL_3436102, EPI_ISL_3436103, EPI_ISL_3436104, EPI_ISL_3436105, EPI_ISL_3436106, EPI_ISL_3436107, EPI_ISL_3436108, EPI_ISL_3436109, EPI_ISL_3436110, EPI_ISL_3436111, EPI_ISL_3436112 | | | |
| see above | Lighthouse Lab in Milton Keynes | Wellcome Sanger Institute for the COVID-19 Genomics UK (COG-UK) Consortium | The Lighthouse Lab in Milton Keynes and Alex Alderton, Roberto Amato, Jeffrey Barrett, Sonia Goncalves, Ewan Harrison, David K. Jackson, Ian Johnston, Dominic Kwiatkowski, Cordelia Langford, John Sillitoe on behalf of the Wellcome Sanger Institute COVID-19 Surveillance Team |
| EPI_ISL_3436113 | Lighthouse Lab in Glasgow | Wellcome Sanger Institute for the COVID-19 Genomics UK (COG-UK) Consortium | Harper VanSteenhouse, Yumi Kasai, David Gray, Carol Clugston, Anna Dominiczak and Alex Alderton, Roberto Amato, Jeffrey Barrett, Sonia Goncalves, Ewan Harrison, David K. Jackson, Ian Johnston, Dominic Kwiatkowski, Cordelia Langford, John Sillitoe on behalf of the Wellcome Sanger Institute COVID-19 Surveillance Team |
| EPI_ISL_3436114, EPI_ISL_3436115, EPI_ISL_3436116 | Lighthouse Lab in Milton Keynes | Wellcome Sanger Institute for the COVID-19 Genomics UK (COG-UK) Consortium | The Lighthouse Lab in Milton Keynes and Alex Alderton, Roberto Amato, Jeffrey Barrett, Sonia Goncalves, Ewan Harrison, David K. Jackson, Ian Johnston, Dominic Kwiatkowski, Cordelia Langford, John Sillitoe on behalf of the Wellcome Sanger Institute COVID-19 Surveillance Team |
| EPI_ISL_3436117 | Lighthouse Lab in Glasgow | Wellcome Sanger Institute for the COVID-19 Genomics UK (COG-UK) Consortium | Harper VanSteenhouse, Yumi Kasai, David Gray, Carol Clugston, Anna Dominiczak and Alex Alderton, Roberto Amato, Jeffrey Barrett, Sonia Goncalves, Ewan Harrison, David K. Jackson, Ian Johnston, Dominic Kwiatkowski, Cordelia Langford, John Sillitoe on behalf of the Wellcome Sanger Institute COVID-19 Surveillance Team |
| EPI_ISL_3436118, EPI_ISL_3436119, EPI_ISL_3436120, EPI_ISL_3436121, EPI_ISL_3436122, EPI_ISL_3436123, EPI_ISL_3436124, EPI_ISL_3436125, EPI_ISL_3436126, EPI_ISL_3436127, EPI_ISL_3436128, EPI_ISL_3436129, EPI_ISL_3436130, EPI_ISL_3436131, EPI_ISL_3436132, EPI_ISL_3436133 | | | |
| see above | Lighthouse Lab in Milton Keynes | Wellcome Sanger Institute for the COVID-19 Genomics UK (COG-UK) Consortium | The Lighthouse Lab in Milton Keynes and Alex Alderton, Roberto Amato, Jeffrey Barrett, Sonia Goncalves, Ewan Harrison, David K. Jackson, Ian Johnston, Dominic Kwiatkowski, Cordelia Langford, John Sillitoe on behalf of the Wellcome Sanger Institute COVID-19 Surveillance Team |
| EPI_ISL_3436134 | Lighthouse Lab in Glasgow | Wellcome Sanger Institute for the COVID-19 Genomics UK (COG-UK) Consortium | Harper VanSteenhouse, Yumi Kasai, David Gray, Carol Clugston, Anna Dominiczak and Alex Alderton, Roberto Amato, Jeffrey Barrett, Sonia Goncalves, Ewan Harrison, David K. Jackson, Ian Johnston, Dominic Kwiatkowski, Cordelia Langford, John Sillitoe on behalf of the Wellcome Sanger Institute COVID-19 Surveillance Team |
| EPI_ISL_3436135 | Lighthouse Lab in Milton Keynes | Wellcome Sanger Institute for the COVID-19 Genomics UK (COG-UK) Consortium | The Lighthouse Lab in Milton Keynes and Alex Alderton, Roberto Amato, Jeffrey Barrett, Sonia Goncalves, Ewan Harrison, David K. Jackson, Ian Johnston, Dominic Kwiatkowski, Cordelia Langford, John Sillitoe on behalf of the Wellcome Sanger Institute COVID-19 Surveillance Team |
| EPI_ISL_3436136, EPI_ISL_3436137 | Lighthouse Lab in Glasgow | Wellcome Sanger Institute for the COVID-19 Genomics UK (COG-UK) Consortium | Harper VanSteenhouse, Yumi Kasai, David Gray, Carol Clugston, Anna Dominiczak and Alex Alderton, Roberto Amato, Jeffrey Barrett, Sonia Goncalves, Ewan Harrison, David K. Jackson, Ian Johnston, Dominic Kwiatkowski, Cordelia Langford, John Sillitoe on behalf of the Wellcome Sanger Institute COVID-19 Surveillance Team |
| EPI_ISL_3436138, EPI_ISL_3436139, EPI_ISL_3436140, EPI_ISL_3436141 | Lighthouse Lab in Milton Keynes | Wellcome Sanger Institute for the COVID-19 Genomics UK (COG-UK) Consortium | The Lighthouse Lab in Milton Keynes and Alex Alderton, Roberto Amato, Jeffrey Barrett, Sonia Goncalves, Ewan Harrison, David K. Jackson, Ian Johnston, Dominic Kwiatkowski, Cordelia Langford, John Sillitoe on behalf of the Wellcome Sanger Institute COVID-19 Surveillance Team |
| EPI_ISL_3436142 | Lighthouse Lab in Glasgow | Wellcome Sanger Institute for the COVID-19 Genomics UK (COG-UK) Consortium | Harper VanSteenhouse, Yumi Kasai, David Gray, Carol Clugston, Anna Dominiczak and Alex Alderton, Roberto Amato, Jeffrey Barrett, Sonia Goncalves, Ewan Harrison, David K. Jackson, Ian Johnston, Dominic Kwiatkowski, Cordelia Langford, John Sillitoe on behalf of the Wellcome Sanger Institute COVID-19 Surveillance Team |
| EPI_ISL_3436143, EPI_ISL_3436144 | Lighthouse Lab in Milton Keynes | Wellcome Sanger Institute for the COVID-19 Genomics UK (COG-UK) Consortium | The Lighthouse Lab in Milton Keynes and Alex Alderton, Roberto Amato, Jeffrey Barrett, Sonia Goncalves, Ewan Harrison, David K. Jackson, Ian Johnston, Dominic Kwiatkowski, Cordelia Langford, John Sillitoe on behalf of the Wellcome Sanger Institute COVID-19 Surveillance Team |
| EPI_ISL_3436145 | Lighthouse Lab in Glasgow | Wellcome Sanger Institute for the COVID-19 Genomics UK (COG-UK) Consortium | Harper VanSteenhouse, Yumi Kasai, David Gray, Carol Clugston, Anna Dominiczak and Alex Alderton, Roberto Amato, Jeffrey Barrett, Sonia Goncalves, Ewan Harrison, David K. Jackson, Ian Johnston, Dominic Kwiatkowski, Cordelia Langford, John Sillitoe on behalf of the Wellcome Sanger Institute COVID-19 Surveillance Team |
| EPI_ISL_3436146, EPI_ISL_3436147, EPI_ISL_3436148, EPI_ISL_3436149, EPI_ISL_3436150 | Lighthouse Lab in Milton Keynes | Wellcome Sanger Institute for the COVID-19 Genomics UK (COG-UK) Consortium | The Lighthouse Lab in Milton Keynes and Alex Alderton, Roberto Amato, Jeffrey Barrett, Sonia Goncalves, Ewan Harrison, David K. Jackson, Ian Johnston, Dominic Kwiatkowski, Cordelia Langford, John Sillitoe on behalf of the Wellcome Sanger Institute COVID-19 Surveillance Team |
| EPI_ISL_3436151 | Lighthouse Lab in Glasgow | Wellcome Sanger Institute for the COVID-19 Genomics UK (COG-UK) Consortium | Harper VanSteenhouse, Yumi Kasai, David Gray, Carol Clugston, Anna Dominiczak and Alex Alderton, Roberto Amato, Jeffrey Barrett, Sonia Goncalves, Ewan Harrison, David K. Jackson, Ian Johnston, Dominic Kwiatkowski, Cordelia Langford, John Sillitoe on behalf of the Wellcome Sanger Institute COVID-19 Surveillance Team |
| EPI_ISL_3436152, EPI_ISL_3436153, EPI_ISL_3436154, EPI_ISL_3436155, EPI_ISL_3436156, EPI_ISL_3436157, EPI_ISL_3436158, EPI_ISL_3436159 | Lighthouse Lab in Milton Keynes | Wellcome Sanger Institute for the COVID-19 Genomics UK (COG-UK) Consortium | The Lighthouse Lab in Milton Keynes and Alex Alderton, Roberto Amato, Jeffrey Barrett, Sonia Goncalves, Ewan Harrison, David K. Jackson, Ian Johnston, Dominic Kwiatkowski, Cordelia Langford, John Sillitoe on behalf of the Wellcome Sanger Institute COVID-19 Surveillance Team |
| EPI_ISL_3436160 | Lighthouse Lab in Glasgow | Wellcome Sanger Institute for the COVID-19 Genomics UK (COG-UK) Consortium | Harper VanSteenhouse, Yumi Kasai, David Gray, Carol Clugston, Anna Dominiczak and Alex Alderton, Roberto Amato, Jeffrey Barrett, Sonia Goncalves, Ewan Harrison, David K. Jackson, Ian Johnston, Dominic Kwiatkowski, Cordelia Langford, John Sillitoe on behalf of the Wellcome Sanger Institute COVID-19 Surveillance Team |
| EPI_ISL_3436161, EPI_ISL_3436162, EPI_ISL_3436163, EPI_ISL_3436164, EPI_ISL_3436165, EPI_ISL_3436166, EPI_ISL_3436167 | Lighthouse Lab in Milton Keynes | Wellcome Sanger Institute for the COVID-19 Genomics UK (COG-UK) Consortium | The Lighthouse Lab in Milton Keynes and Alex Alderton, Roberto Amato, Jeffrey Barrett, Sonia Goncalves, Ewan Harrison, David K. Jackson, Ian Johnston, Dominic Kwiatkowski, Cordelia Langford, John Sillitoe on behalf of the Wellcome Sanger Institute COVID-19 Surveillance Team |
| EPI_ISL_3436168 | Lighthouse Lab in Glasgow | Wellcome Sanger Institute for the COVID-19 Genomics UK (COG-UK) Consortium | Harper VanSteenhouse, Yumi Kasai, David Gray, Carol Clugston, Anna Dominiczak and Alex Alderton, Roberto Amato, Jeffrey Barrett, Sonia Goncalves, Ewan Harrison, David K. Jackson, Ian Johnston, Dominic Kwiatkowski, Cordelia Langford, John Sillitoe on behalf of the Wellcome Sanger Institute COVID-19 Surveillance Team |
| EPI_ISL_3436169, EPI_ISL_3436170, EPI_ISL_3436171, EPI_ISL_3436172, EPI_ISL_3436173, EPI_ISL_3436174, EPI_ISL_3436175, EPI_ISL_3436176, EPI_ISL_3436177, EPI_ISL_3436178, EPI_ISL_3436179, EPI_ISL_3436180, EPI_ISL_3436181, EPI_ISL_3436182, EPI_ISL_3436183, EPI_ISL_3436184, EPI_ISL_3436185, EPI_ISL_3436186, EPI_ISL_3436187, EPI_ISL_3436188, EPI_ISL_3436189, EPI_ISL_3436190, EPI_ISL_3436191, EPI_ISL_3436192, EPI_ISL_3436193, EPI_ISL_3436194 | | | |
| see above | Lighthouse Lab in Milton Keynes | Wellcome Sanger Institute for the COVID-19 Genomics UK (COG-UK) Consortium | The Lighthouse Lab in Milton Keynes and Alex Alderton, Roberto Amato, Jeffrey Barrett, Sonia Goncalves, Ewan Harrison, David K. Jackson, Ian Johnston, Dominic Kwiatkowski, Cordelia Langford, John Sillitoe on behalf of the Wellcome Sanger Institute COVID-19 Surveillance Team |
| EPI_ISL_3436195 | Lighthouse Lab in Glasgow | Wellcome Sanger Institute for the COVID-19 Genomics UK (COG-UK) Consortium | Harper VanSteenhouse, Yumi Kasai, David Gray, Carol Clugston, Anna Dominiczak and Alex Alderton, Roberto Amato, Jeffrey Barrett, Sonia Goncalves, Ewan Harrison, David K. Jackson, Ian Johnston, Dominic Kwiatkowski, Cordelia Langford, John Sillitoe on behalf of the Wellcome Sanger Institute COVID-19 Surveillance Team |
| EPI_ISL_3436196, EPI_ISL_3436197, EPI_ISL_3436198, EPI_ISL_3436199, EPI_ISL_3436200, EPI_ISL_3436201, EPI_ISL_3436202, EPI_ISL_3436203, EPI_ISL_3436204 | Lighthouse Lab in Milton Keynes | Wellcome Sanger Institute for the COVID-19 Genomics UK (COG-UK) Consortium | The Lighthouse Lab in Milton Keynes and Alex Alderton, Roberto Amato, Jeffrey Barrett, Sonia Goncalves, Ewan Harrison, David K. Jackson, Ian Johnston, Dominic Kwiatkowski, Cordelia Langford, John Sillitoe on behalf of the Wellcome Sanger Institute COVID-19 Surveillance Team |
| EPI_ISL_3436205 | Lighthouse Lab in Glasgow | Wellcome Sanger Institute for the COVID-19 Genomics UK (COG-UK) Consortium | Harper VanSteenhouse, Yumi Kasai, David Gray, Carol Clugston, Anna Dominiczak and Alex Alderton, Roberto Amato, Jeffrey Barrett, Sonia Goncalves, Ewan Harrison, David K. Jackson, Ian Johnston, Dominic Kwiatkowski, Cordelia Langford, John Sillitoe on behalf of the Wellcome Sanger Institute COVID-19 Surveillance Team |
| EPI_ISL_3436206, EPI_ISL_3436207 | Lighthouse Lab in Milton Keynes | Wellcome Sanger Institute for the COVID-19 Genomics UK (COG-UK) Consortium | The Lighthouse Lab in Milton Keynes and Alex Alderton, Roberto Amato, Jeffrey Barrett, Sonia Goncalves, Ewan Harrison, David K. Jackson, Ian Johnston, Dominic Kwiatkowski, Cordelia Langford, John Sillitoe on behalf of the Wellcome Sanger Institute COVID-19 Surveillance Team |
| EPI_ISL_3436208 | Lighthouse Lab in Glasgow | Wellcome Sanger Institute for the COVID-19 Genomics UK (COG-UK) Consortium | Harper VanSteenhouse, Yumi Kasai, David Gray, Carol Clugston, Anna Dominiczak and Alex Alderton, Roberto Amato, Jeffrey Barrett, Sonia Goncalves, Ewan Harrison, David K. Jackson, Ian Johnston, Dominic Kwiatkowski, Cordelia Langford, John Sillitoe on behalf of the Wellcome Sanger Institute COVID-19 Surveillance Team |
| EPI_ISL_3436209, EPI_ISL_3436210 | Lighthouse Lab in Milton Keynes | Wellcome Sanger Institute for the COVID-19 Genomics UK (COG-UK) Consortium | The Lighthouse Lab in Milton Keynes and Alex Alderton, Roberto Amato, Jeffrey Barrett, Sonia Goncalves, Ewan Harrison, David K. Jackson, Ian Johnston, Dominic Kwiatkowski, Cordelia Langford, John Sillitoe on behalf of the Wellcome Sanger Institute COVID-19 Surveillance Team |
| EPI_ISL_3436211 | Lighthouse Lab in Glasgow | Wellcome Sanger Institute for the COVID-19 Genomics UK (COG-UK) Consortium | Harper VanSteenhouse, Yumi Kasai, David Gray, Carol Clugston, Anna Dominiczak and Alex Alderton, Roberto Amato, Jeffrey Barrett, Sonia Goncalves, Ewan Harrison, David K. Jackson, Ian Johnston, Dominic Kwiatkowski, Cordelia Langford, John Sillitoe on behalf of the Wellcome Sanger Institute COVID-19 Surveillance Team |
| EPI_ISL_3436212, EPI_ISL_3436213 | Lighthouse Lab in Milton Keynes | Wellcome Sanger Institute for the COVID-19 Genomics UK (COG-UK) Consortium | The Lighthouse Lab in Milton Keynes and Alex Alderton, Roberto Amato, Jeffrey Barrett, Sonia Goncalves, Ewan Harrison, David K. Jackson, Ian Johnston, Dominic Kwiatkowski, Cordelia Langford, John Sillitoe on behalf of the Wellcome Sanger Institute COVID-19 Surveillance Team |
| EPI_ISL_3436214, EPI_ISL_3436215 | Lighthouse Lab in Glasgow | Wellcome Sanger Institute for the COVID-19 Genomics UK (COG-UK) Consortium | Harper VanSteenhouse, Yumi Kasai, David Gray, Carol Clugston, Anna Dominiczak and Alex Alderton, Roberto Amato, Jeffrey Barrett, Sonia Goncalves, Ewan Harrison, David K. Jackson, Ian Johnston, Dominic Kwiatkowski, Cordelia Langford, John Sillitoe on behalf of the Wellcome Sanger Institute COVID-19 Surveillance Team |
| EPI_ISL_3436216, EPI_ISL_3436217, EPI_ISL_3436218, EPI_ISL_3436219, EPI_ISL_3436220, EPI_ISL_3436221, EPI_ISL_3436222, EPI_ISL_3436223, EPI_ISL_3436224, EPI_ISL_3436225, EPI_ISL_3436226, EPI_ISL_3436227 | | | |
| see above | Lighthouse Lab in Milton Keynes | Wellcome Sanger Institute for the COVID-19 Genomics UK (COG-UK) Consortium | The Lighthouse Lab in Milton Keynes and Alex Alderton, Roberto Amato, Jeffrey Barrett, Sonia Goncalves, Ewan Harrison, David K. Jackson, Ian Johnston, Dominic Kwiatkowski, Cordelia Langford, John Sillitoe on behalf of the Wellcome Sanger Institute COVID-19 Surveillance Team |
| EPI_ISL_3436228 | Lighthouse Lab in Glasgow | Wellcome Sanger Institute for the COVID-19 Genomics UK (COG-UK) | Harper VanSteenhouse, Yumi Kasai, David Gray, Carol Clugston, Anna Dominiczak and Alex Alderton, Roberto Amato, Jeffrey Barrett, Sonia Goncalves, Ewan Harrison, David K. Jackson, Ian Johnston, Dominic Kwiatkowski, Cordelia Langford, |

| | | | |
|---|---|---|---|
| | | Consortium | John Sillitoe on behalf of the Wellcome Sanger Institute COVID-19 Surveillance Team |
| EPI_ISL_3436229, EPI_ISL_3436230, EPI_ISL_3436231, EPI_ISL_3436232, EPI_ISL_3436233 | Lighthouse Lab in Milton Keynes | Wellcome Sanger Institute for the COVID-19 Genomics UK (COG-UK) Consortium | The Lighthouse Lab in Milton Keynes and Alex Alderton, Roberto Amato, Jeffrey Barrett, Sonia Goncalves, Ewan Harrison, David K. Jackson, Ian Johnston, Dominic Kwiatkowski, Cordelia Langford, John Sillitoe on behalf of the Wellcome Sanger Institute COVID-19 Surveillance Team |
| EPI_ISL_3436234, EPI_ISL_3436235, EPI_ISL_3436236 | Lighthouse Lab in Glasgow | Wellcome Sanger Institute for the COVID-19 Genomics UK (COG-UK) Consortium | Harper VanSteenhouse, Yumi Kasai, David Gray, Carol Clugston, Anna Dominiczak and Alex Alderton, Roberto Amato, Jeffrey Barrett, Sonia Goncalves, Ewan Harrison, David K. Jackson, Ian Johnston, Dominic Kwiatkowski, Cordelia Langford, John Sillitoe on behalf of the Wellcome Sanger Institute COVID-19 Surveillance Team |
| EPI_ISL_3436237, EPI_ISL_3436238 | Lighthouse Lab in Milton Keynes | Wellcome Sanger Institute for the COVID-19 Genomics UK (COG-UK) Consortium | The Lighthouse Lab in Milton Keynes and Alex Alderton, Roberto Amato, Jeffrey Barrett, Sonia Goncalves, Ewan Harrison, David K. Jackson, Ian Johnston, Dominic Kwiatkowski, Cordelia Langford, John Sillitoe on behalf of the Wellcome Sanger Institute COVID-19 Surveillance Team |
| EPI_ISL_3436239, EPI_ISL_3436240, EPI_ISL_3436241 | Lighthouse Lab in Glasgow | Wellcome Sanger Institute for the COVID-19 Genomics UK (COG-UK) Consortium | Harper VanSteenhouse, Yumi Kasai, David Gray, Carol Clugston, Anna Dominiczak and Alex Alderton, Roberto Amato, Jeffrey Barrett, Sonia Goncalves, Ewan Harrison, David K. Jackson, Ian Johnston, Dominic Kwiatkowski, Cordelia Langford, John Sillitoe on behalf of the Wellcome Sanger Institute COVID-19 Surveillance Team |
| EPI_ISL_3436242, EPI_ISL_3436243, EPI_ISL_3436244, EPI_ISL_3436245, EPI_ISL_3436246, EPI_ISL_3436247, EPI_ISL_3436248, EPI_ISL_3436249, EPI_ISL_3436250, EPI_ISL_3436251, EPI_ISL_3436252, EPI_ISL_3436253 | | | |
| see above | Lighthouse Lab in Milton Keynes | Wellcome Sanger Institute for the COVID-19 Genomics UK (COG-UK) Consortium | The Lighthouse Lab in Milton Keynes and Alex Alderton, Roberto Amato, Jeffrey Barrett, Sonia Goncalves, Ewan Harrison, David K. Jackson, Ian Johnston, Dominic Kwiatkowski, Cordelia Langford, John Sillitoe on behalf of the Wellcome Sanger Institute COVID-19 Surveillance Team |
| EPI_ISL_3436254 | Lighthouse Lab in Glasgow | Wellcome Sanger Institute for the COVID-19 Genomics UK (COG-UK) Consortium | Harper VanSteenhouse, Yumi Kasai, David Gray, Carol Clugston, Anna Dominiczak and Alex Alderton, Roberto Amato, Jeffrey Barrett, Sonia Goncalves, Ewan Harrison, David K. Jackson, Ian Johnston, Dominic Kwiatkowski, Cordelia Langford, John Sillitoe on behalf of the Wellcome Sanger Institute COVID-19 Surveillance Team |
| EPI_ISL_3436255, EPI_ISL_3436256, EPI_ISL_3436257, EPI_ISL_3436258, EPI_ISL_3436259, EPI_ISL_3436260, EPI_ISL_3436261, EPI_ISL_3436262, EPI_ISL_3436263, EPI_ISL_3436264, EPI_ISL_3436265 | | | |
| see above | Lighthouse Lab in Milton Keynes | Wellcome Sanger Institute for the COVID-19 Genomics UK (COG-UK) Consortium | The Lighthouse Lab in Milton Keynes and Alex Alderton, Roberto Amato, Jeffrey Barrett, Sonia Goncalves, Ewan Harrison, David K. Jackson, Ian Johnston, Dominic Kwiatkowski, Cordelia Langford, John Sillitoe on behalf of the Wellcome Sanger Institute COVID-19 Surveillance Team |
| EPI_ISL_3436266 | Lighthouse Lab in Alderley Park | Wellcome Sanger Institute for the COVID-19 Genomics UK (COG-UK) Consortium | Jacquelyn Wynn, Mairead Hyland, The Lighthouse Lab in Alderley Park and Alex Alderton, Roberto Amato, Jeffrey Barrett, Sonia Goncalves, Ewan Harrison, David K. Jackson, Ian Johnston, Dominic Kwiatkowski, Cordelia Langford, John Sillitoe on behalf of the Wellcome Sanger Institute COVID-19 Surveillance Team |
| EPI_ISL_3436267, EPI_ISL_3436268, EPI_ISL_3436269 | Lighthouse Lab in Milton Keynes | Wellcome Sanger Institute for the COVID-19 Genomics UK (COG-UK) Consortium | The Lighthouse Lab in Milton Keynes and Alex Alderton, Roberto Amato, Jeffrey Barrett, Sonia Goncalves, Ewan Harrison, David K. Jackson, Ian Johnston, Dominic Kwiatkowski, Cordelia Langford, John Sillitoe on behalf of the Wellcome Sanger Institute COVID-19 Surveillance Team |
| EPI_ISL_3436270 | Lighthouse Lab in Alderley Park | Wellcome Sanger Institute for the COVID-19 Genomics UK (COG-UK) Consortium | Jacquelyn Wynn, Mairead Hyland, The Lighthouse Lab in Alderley Park and Alex Alderton, Roberto Amato, Jeffrey Barrett, Sonia Goncalves, Ewan Harrison, David K. Jackson, Ian Johnston, Dominic Kwiatkowski, Cordelia Langford, John Sillitoe on behalf of the Wellcome Sanger Institute COVID-19 Surveillance Team |
| EPI_ISL_3436271, EPI_ISL_3436272, EPI_ISL_3436273, EPI_ISL_3436274 | Lighthouse Lab in Milton Keynes | Wellcome Sanger Institute for the COVID-19 Genomics UK (COG-UK) Consortium | The Lighthouse Lab in Milton Keynes and Alex Alderton, Roberto Amato, Jeffrey Barrett, Sonia Goncalves, Ewan Harrison, David K. Jackson, Ian Johnston, Dominic Kwiatkowski, Cordelia Langford, John Sillitoe on behalf of the Wellcome Sanger Institute COVID-19 Surveillance Team |
| EPI_ISL_3436275 | Lighthouse Lab in Glasgow | Wellcome Sanger Institute for the COVID-19 Genomics UK (COG-UK) Consortium | Harper VanSteenhouse, Yumi Kasai, David Gray, Carol Clugston, Anna Dominiczak and Alex Alderton, Roberto Amato, Jeffrey Barrett, Sonia Goncalves, Ewan Harrison, David K. Jackson, Ian Johnston, Dominic Kwiatkowski, Cordelia Langford, John Sillitoe on behalf of the Wellcome Sanger Institute COVID-19 Surveillance Team |
| EPI_ISL_3436276, EPI_ISL_3436277 | Lighthouse Lab in Milton Keynes | Wellcome Sanger Institute for the COVID-19 Genomics UK (COG-UK) Consortium | The Lighthouse Lab in Milton Keynes and Alex Alderton, Roberto Amato, Jeffrey Barrett, Sonia Goncalves, Ewan Harrison, David K. Jackson, Ian Johnston, Dominic Kwiatkowski, Cordelia Langford, John Sillitoe on behalf of the Wellcome Sanger Institute COVID-19 Surveillance Team |
| EPI_ISL_3436278, EPI_ISL_3436279 | Lighthouse Lab in Glasgow | Wellcome Sanger Institute for the COVID-19 Genomics UK (COG-UK) Consortium | Harper VanSteenhouse, Yumi Kasai, David Gray, Carol Clugston, Anna Dominiczak and Alex Alderton, Roberto Amato, Jeffrey Barrett, Sonia Goncalves, Ewan Harrison, David K. Jackson, Ian Johnston, Dominic Kwiatkowski, Cordelia Langford, John Sillitoe on behalf of the Wellcome Sanger Institute COVID-19 Surveillance Team |
| EPI_ISL_3436280, EPI_ISL_3436281, EPI_ISL_3436282, EPI_ISL_3436283 | Lighthouse Lab in Milton Keynes | Wellcome Sanger Institute for the COVID-19 Genomics UK (COG-UK) Consortium | The Lighthouse Lab in Milton Keynes and Alex Alderton, Roberto Amato, Jeffrey Barrett, Sonia Goncalves, Ewan Harrison, David K. Jackson, Ian Johnston, Dominic Kwiatkowski, Cordelia Langford, John Sillitoe on behalf of the Wellcome Sanger Institute COVID-19 Surveillance Team |
| EPI_ISL_3436284, EPI_ISL_3436285, EPI_ISL_3436286 | Lighthouse Lab in Alderley Park | Wellcome Sanger Institute for the COVID-19 Genomics UK (COG-UK) Consortium | Jacquelyn Wynn, Mairead Hyland, The Lighthouse Lab in Alderley Park and Alex Alderton, Roberto Amato, Jeffrey Barrett, Sonia Goncalves, Ewan Harrison, David K. Jackson, Ian Johnston, Dominic Kwiatkowski, Cordelia Langford, John Sillitoe on behalf of the Wellcome Sanger Institute COVID-19 Surveillance Team |
| EPI_ISL_3436287, EPI_ISL_3436288, EPI_ISL_3436289 | Lighthouse Lab in Milton Keynes | Wellcome Sanger Institute for the COVID-19 Genomics UK (COG-UK) Consortium | The Lighthouse Lab in Milton Keynes and Alex Alderton, Roberto Amato, Jeffrey Barrett, Sonia Goncalves, Ewan Harrison, David K. Jackson, Ian Johnston, Dominic Kwiatkowski, Cordelia Langford, John Sillitoe on behalf of the Wellcome Sanger Institute COVID-19 Surveillance Team |
| EPI_ISL_3436290 | Lighthouse Lab in Alderley Park | Wellcome Sanger Institute for the COVID-19 Genomics UK (COG-UK) Consortium | Jacquelyn Wynn, Mairead Hyland, The Lighthouse Lab in Alderley Park and Alex Alderton, Roberto Amato, Jeffrey Barrett, Sonia Goncalves, Ewan Harrison, David K. Jackson, Ian Johnston, Dominic Kwiatkowski, Cordelia Langford, John Sillitoe on behalf of the Wellcome Sanger Institute COVID-19 Surveillance Team |
| EPI_ISL_3436291, EPI_ISL_3436292, EPI_ISL_3436293 | Lighthouse Lab in Milton Keynes | Wellcome Sanger Institute for the COVID-19 Genomics UK (COG-UK) Consortium | The Lighthouse Lab in Milton Keynes and Alex Alderton, Roberto Amato, Jeffrey Barrett, Sonia Goncalves, Ewan Harrison, David K. Jackson, Ian Johnston, Dominic Kwiatkowski, Cordelia Langford, John Sillitoe on behalf of the Wellcome Sanger Institute COVID-19 Surveillance Team |
| EPI_ISL_3436294, EPI_ISL_3436295 | Lighthouse Lab in Glasgow | Wellcome Sanger Institute for the COVID-19 Genomics UK (COG-UK) Consortium | Harper VanSteenhouse, Yumi Kasai, David Gray, Carol Clugston, Anna Dominiczak and Alex Alderton, Roberto Amato, Jeffrey Barrett, Sonia Goncalves, Ewan Harrison, David K. Jackson, Ian Johnston, Dominic Kwiatkowski, Cordelia Langford, John Sillitoe on behalf of the Wellcome Sanger Institute COVID-19 Surveillance Team |
| EPI_ISL_3436296 | Lighthouse Lab in Milton Keynes | Wellcome Sanger Institute for the COVID-19 Genomics UK (COG-UK) Consortium | The Lighthouse Lab in Milton Keynes and Alex Alderton, Roberto Amato, Jeffrey Barrett, Sonia Goncalves, Ewan Harrison, David K. Jackson, Ian Johnston, Dominic Kwiatkowski, Cordelia Langford, John Sillitoe on behalf of the Wellcome Sanger Institute COVID-19 Surveillance Team |
| EPI_ISL_3436297 | Lighthouse Lab in Alderley Park | Wellcome Sanger Institute for the COVID-19 Genomics UK (COG-UK) Consortium | Jacquelyn Wynn, Mairead Hyland, The Lighthouse Lab in Alderley Park and Alex Alderton, Roberto Amato, Jeffrey Barrett, Sonia Goncalves, Ewan Harrison, David K. Jackson, Ian Johnston, Dominic Kwiatkowski, Cordelia Langford, John Sillitoe on behalf of the Wellcome Sanger Institute COVID-19 Surveillance Team |
| EPI_ISL_3436298, EPI_ISL_3436299, EPI_ISL_3436300, EPI_ISL_3436301, EPI_ISL_3436302 | Lighthouse Lab in Milton Keynes | Wellcome Sanger Institute for the COVID-19 Genomics UK (COG-UK) Consortium | The Lighthouse Lab in Milton Keynes and Alex Alderton, Roberto Amato, Jeffrey Barrett, Sonia Goncalves, Ewan Harrison, David K. Jackson, Ian Johnston, Dominic Kwiatkowski, Cordelia Langford, John Sillitoe on behalf of the Wellcome Sanger Institute COVID-19 Surveillance Team |
| EPI_ISL_3436303 | Lighthouse Lab in Alderley Park | Wellcome Sanger Institute for the COVID-19 Genomics UK (COG-UK) Consortium | Jacquelyn Wynn, Mairead Hyland, The Lighthouse Lab in Alderley Park and Alex Alderton, Roberto Amato, Jeffrey Barrett, Sonia Goncalves, Ewan Harrison, David K. Jackson, Ian Johnston, Dominic Kwiatkowski, Cordelia Langford, John Sillitoe on behalf of the Wellcome Sanger Institute COVID-19 Surveillance Team |
| EPI_ISL_3436304, EPI_ISL_3436305, EPI_ISL_3436306 | Lighthouse Lab in Milton Keynes | Wellcome Sanger Institute for the COVID-19 Genomics UK (COG-UK) Consortium | The Lighthouse Lab in Milton Keynes and Alex Alderton, Roberto Amato, Jeffrey Barrett, Sonia Goncalves, Ewan Harrison, David K. Jackson, Ian Johnston, Dominic Kwiatkowski, Cordelia Langford, John Sillitoe on behalf of the Wellcome Sanger Institute COVID-19 Surveillance Team |
| EPI_ISL_3436307 | Lighthouse Lab in Glasgow | Wellcome Sanger Institute for the COVID-19 Genomics UK (COG-UK) Consortium | Harper VanSteenhouse, Yumi Kasai, David Gray, Carol Clugston, Anna Dominiczak and Alex Alderton, Roberto Amato, Jeffrey Barrett, Sonia Goncalves, Ewan Harrison, David K. Jackson, Ian Johnston, Dominic Kwiatkowski, Cordelia Langford, John Sillitoe on behalf of the Wellcome Sanger Institute COVID-19 Surveillance Team |
| EPI_ISL_3436308 | Lighthouse Lab in Milton Keynes | Wellcome Sanger Institute for the COVID-19 Genomics UK (COG-UK) Consortium | The Lighthouse Lab in Milton Keynes and Alex Alderton, Roberto Amato, Jeffrey Barrett, Sonia Goncalves, Ewan Harrison, David K. Jackson, Ian Johnston, Dominic Kwiatkowski, Cordelia Langford, John Sillitoe on behalf of the Wellcome Sanger Institute COVID-19 Surveillance Team |
| EPI_ISL_3436309 | Lighthouse Lab in Glasgow | Wellcome Sanger Institute for the COVID-19 Genomics UK (COG-UK) Consortium | Harper VanSteenhouse, Yumi Kasai, David Gray, Carol Clugston, Anna Dominiczak and Alex Alderton, Roberto Amato, Jeffrey Barrett, Sonia Goncalves, Ewan Harrison, David K. Jackson, Ian Johnston, Dominic Kwiatkowski, Cordelia Langford, John Sillitoe on behalf of the Wellcome Sanger Institute COVID-19 Surveillance Team |
| EPI_ISL_3436310 | Lighthouse Lab in Alderley Park | Wellcome Sanger Institute for the COVID-19 Genomics UK (COG-UK) Consortium | Jacquelyn Wynn, Mairead Hyland, The Lighthouse Lab in Alderley Park and Alex Alderton, Roberto Amato, Jeffrey Barrett, Sonia Goncalves, Ewan Harrison, David K. Jackson, Ian Johnston, Dominic Kwiatkowski, Cordelia Langford, John Sillitoe on behalf of the Wellcome Sanger Institute COVID-19 Surveillance Team |
| EPI_ISL_3436311, EPI_ISL_3436312, EPI_ISL_3436313, EPI_ISL_3436314, EPI_ISL_3436315, EPI_ISL_3436316, EPI_ISL_3436317 | Lighthouse Lab in Milton Keynes | Wellcome Sanger Institute for the COVID-19 Genomics UK (COG-UK) Consortium | The Lighthouse Lab in Milton Keynes and Alex Alderton, Roberto Amato, Jeffrey Barrett, Sonia Goncalves, Ewan Harrison, David K. Jackson, Ian Johnston, Dominic Kwiatkowski, Cordelia Langford, John Sillitoe on behalf of the Wellcome Sanger Institute COVID-19 Surveillance Team |
| EPI_ISL_3436318 | Lighthouse Lab in Glasgow | Wellcome Sanger Institute for the COVID-19 Genomics UK (COG-UK) Consortium | Harper VanSteenhouse, Yumi Kasai, David Gray, Carol Clugston, Anna Dominiczak and Alex Alderton, Roberto Amato, Jeffrey Barrett, Sonia Goncalves, Ewan Harrison, David K. Jackson, Ian Johnston, Dominic Kwiatkowski, Cordelia Langford, John Sillitoe on behalf of the Wellcome Sanger Institute COVID-19 Surveillance Team |
| EPI_ISL_3436319 | Lighthouse Lab in Milton Keynes | Wellcome Sanger Institute for the COVID-19 Genomics UK (COG-UK) Consortium | The Lighthouse Lab in Milton Keynes and Alex Alderton, Roberto Amato, Jeffrey Barrett, Sonia Goncalves, Ewan Harrison, David K. Jackson, Ian Johnston, Dominic Kwiatkowski, Cordelia Langford, John Sillitoe on behalf of the Wellcome Sanger Institute COVID-19 Surveillance Team |
| EPI_ISL_3436320 | Lighthouse Lab in Alderley Park | Wellcome Sanger Institute for the COVID-19 Genomics UK (COG-UK) Consortium | Jacquelyn Wynn, Mairead Hyland, The Lighthouse Lab in Alderley Park and Alex Alderton, Roberto Amato, Jeffrey Barrett, Sonia Goncalves, Ewan Harrison, David K. Jackson, Ian Johnston, Dominic Kwiatkowski, Cordelia Langford, John Sillitoe on behalf of the Wellcome Sanger Institute COVID-19 Surveillance Team |
| EPI_ISL_3436321, EPI_ISL_3436322, EPI_ISL_3436323, | Lighthouse Lab in Milton Keynes | Wellcome Sanger Institute for the COVID-19 Genomics UK (COG-UK) Consortium | The Lighthouse Lab in Milton Keynes and Alex Alderton, Roberto Amato, Jeffrey Barrett, Sonia Goncalves, Ewan Harrison, David K. Jackson, Ian Johnston, Dominic Kwiatkowski, Cordelia Langford, John Sillitoe on behalf of the Wellcome Sanger Institute COVID-19 Surveillance Team |

| | | | |
|---|---|---|---|
| EPI_ISL_3436324, EPI_ISL_3436325, EPI_ISL_3436326, EPI_ISL_3436327, EPI_ISL_3436328 | | | |
| EPI_ISL_3436329 | Lighthouse Lab in Glasgow | Wellcome Sanger Institute for the COVID-19 Genomics UK (COG-UK) Consortium | Harper VanSteenhouse, Yumi Kasai, David Gray, Carol Clugston, Anna Dominiczak and Alex Alderton, Roberto Amato, Jeffrey Barrett, Sonia Goncalves, Ewan Harrison, David K. Jackson, Ian Johnston, Dominic Kwiatkowski, Cordelia Langford, John Sillitoe on behalf of the Wellcome Sanger Institute COVID-19 Surveillance Team |
| EPI_ISL_3436330, EPI_ISL_3436331, EPI_ISL_3436332 | Lighthouse Lab in Alderley Park | Wellcome Sanger Institute for the COVID-19 Genomics UK (COG-UK) Consortium | Jacquelyn Wynn, Mairead Hyland, The Lighthouse Lab in Alderley Park and Alex Alderton, Roberto Amato, Jeffrey Barrett, Sonia Goncalves, Ewan Harrison, David K. Jackson, Ian Johnston, Dominic Kwiatkowski, Cordelia Langford, John Sillitoe on behalf of the Wellcome Sanger Institute COVID-19 Surveillance Team |
| EPI_ISL_3436333, EPI_ISL_3436334, EPI_ISL_3436335, EPI_ISL_3436336, EPI_ISL_3436337, EPI_ISL_3436338 | Lighthouse Lab in Milton Keynes | Wellcome Sanger Institute for the COVID-19 Genomics UK (COG-UK) Consortium | The Lighthouse Lab in Milton Keynes and Alex Alderton, Roberto Amato, Jeffrey Barrett, Sonia Goncalves, Ewan Harrison, David K. Jackson, Ian Johnston, Dominic Kwiatkowski, Cordelia Langford, John Sillitoe on behalf of the Wellcome Sanger Institute COVID-19 Surveillance Team |
| EPI_ISL_3436339 | Lighthouse Lab in Glasgow | Wellcome Sanger Institute for the COVID-19 Genomics UK (COG-UK) Consortium | Harper VanSteenhouse, Yumi Kasai, David Gray, Carol Clugston, Anna Dominiczak and Alex Alderton, Roberto Amato, Jeffrey Barrett, Sonia Goncalves, Ewan Harrison, David K. Jackson, Ian Johnston, Dominic Kwiatkowski, Cordelia Langford, John Sillitoe on behalf of the Wellcome Sanger Institute COVID-19 Surveillance Team |
| EPI_ISL_3436340, EPI_ISL_3436341, EPI_ISL_3436342, EPI_ISL_3436343, EPI_ISL_3436344 | Lighthouse Lab in Milton Keynes | Wellcome Sanger Institute for the COVID-19 Genomics UK (COG-UK) Consortium | The Lighthouse Lab in Milton Keynes and Alex Alderton, Roberto Amato, Jeffrey Barrett, Sonia Goncalves, Ewan Harrison, David K. Jackson, Ian Johnston, Dominic Kwiatkowski, Cordelia Langford, John Sillitoe on behalf of the Wellcome Sanger Institute COVID-19 Surveillance Team |
| EPI_ISL_3436345 | Lighthouse Lab in Glasgow | Wellcome Sanger Institute for the COVID-19 Genomics UK (COG-UK) Consortium | Harper VanSteenhouse, Yumi Kasai, David Gray, Carol Clugston, Anna Dominiczak and Alex Alderton, Roberto Amato, Jeffrey Barrett, Sonia Goncalves, Ewan Harrison, David K. Jackson, Ian Johnston, Dominic Kwiatkowski, Cordelia Langford, John Sillitoe on behalf of the Wellcome Sanger Institute COVID-19 Surveillance Team |
| EPI_ISL_3436346 | Lighthouse Lab in Alderley Park | Wellcome Sanger Institute for the COVID-19 Genomics UK (COG-UK) Consortium | Jacquelyn Wynn, Mairead Hyland, The Lighthouse Lab in Alderley Park and Alex Alderton, Roberto Amato, Jeffrey Barrett, Sonia Goncalves, Ewan Harrison, David K. Jackson, Ian Johnston, Dominic Kwiatkowski, Cordelia Langford, John Sillitoe on behalf of the Wellcome Sanger Institute COVID-19 Surveillance Team |
| EPI_ISL_3436347 | Lighthouse Lab in Glasgow | Wellcome Sanger Institute for the COVID-19 Genomics UK (COG-UK) Consortium | Harper VanSteenhouse, Yumi Kasai, David Gray, Carol Clugston, Anna Dominiczak and Alex Alderton, Roberto Amato, Jeffrey Barrett, Sonia Goncalves, Ewan Harrison, David K. Jackson, Ian Johnston, Dominic Kwiatkowski, Cordelia Langford, John Sillitoe on behalf of the Wellcome Sanger Institute COVID-19 Surveillance Team |
| EPI_ISL_3436348, EPI_ISL_3436349, EPI_ISL_3436350, EPI_ISL_3436351, EPI_ISL_3436352 | Lighthouse Lab in Milton Keynes | Wellcome Sanger Institute for the COVID-19 Genomics UK (COG-UK) Consortium | The Lighthouse Lab in Milton Keynes and Alex Alderton, Roberto Amato, Jeffrey Barrett, Sonia Goncalves, Ewan Harrison, David K. Jackson, Ian Johnston, Dominic Kwiatkowski, Cordelia Langford, John Sillitoe on behalf of the Wellcome Sanger Institute COVID-19 Surveillance Team |
| EPI_ISL_3436353 | Lighthouse Lab in Glasgow | Wellcome Sanger Institute for the COVID-19 Genomics UK (COG-UK) Consortium | Harper VanSteenhouse, Yumi Kasai, David Gray, Carol Clugston, Anna Dominiczak and Alex Alderton, Roberto Amato, Jeffrey Barrett, Sonia Goncalves, Ewan Harrison, David K. Jackson, Ian Johnston, Dominic Kwiatkowski, Cordelia Langford, John Sillitoe on behalf of the Wellcome Sanger Institute COVID-19 Surveillance Team |
| EPI_ISL_3436354, EPI_ISL_3436355 | Lighthouse Lab in Milton Keynes | Wellcome Sanger Institute for the COVID-19 Genomics UK (COG-UK) Consortium | The Lighthouse Lab in Milton Keynes and Alex Alderton, Roberto Amato, Jeffrey Barrett, Sonia Goncalves, Ewan Harrison, David K. Jackson, Ian Johnston, Dominic Kwiatkowski, Cordelia Langford, John Sillitoe on behalf of the Wellcome Sanger Institute COVID-19 Surveillance Team |
| EPI_ISL_3436356, EPI_ISL_3436357 | Lighthouse Lab in Glasgow | Wellcome Sanger Institute for the COVID-19 Genomics UK (COG-UK) Consortium | Harper VanSteenhouse, Yumi Kasai, David Gray, Carol Clugston, Anna Dominiczak and Alex Alderton, Roberto Amato, Jeffrey Barrett, Sonia Goncalves, Ewan Harrison, David K. Jackson, Ian Johnston, Dominic Kwiatkowski, Cordelia Langford, John Sillitoe on behalf of the Wellcome Sanger Institute COVID-19 Surveillance Team |
| EPI_ISL_3436358 | Lighthouse Lab in Milton Keynes | Wellcome Sanger Institute for the COVID-19 Genomics UK (COG-UK) Consortium | The Lighthouse Lab in Milton Keynes and Alex Alderton, Roberto Amato, Jeffrey Barrett, Sonia Goncalves, Ewan Harrison, David K. Jackson, Ian Johnston, Dominic Kwiatkowski, Cordelia Langford, John Sillitoe on behalf of the Wellcome Sanger Institute COVID-19 Surveillance Team |
| EPI_ISL_3436359 | Lighthouse Lab in Glasgow | Wellcome Sanger Institute for the COVID-19 Genomics UK (COG-UK) Consortium | Harper VanSteenhouse, Yumi Kasai, David Gray, Carol Clugston, Anna Dominiczak and Alex Alderton, Roberto Amato, Jeffrey Barrett, Sonia Goncalves, Ewan Harrison, David K. Jackson, Ian Johnston, Dominic Kwiatkowski, Cordelia Langford, John Sillitoe on behalf of the Wellcome Sanger Institute COVID-19 Surveillance Team |
| EPI_ISL_3436360 | Lighthouse Lab in Milton Keynes | Wellcome Sanger Institute for the COVID-19 Genomics UK (COG-UK) Consortium | The Lighthouse Lab in Milton Keynes and Alex Alderton, Roberto Amato, Jeffrey Barrett, Sonia Goncalves, Ewan Harrison, David K. Jackson, Ian Johnston, Dominic Kwiatkowski, Cordelia Langford, John Sillitoe on behalf of the Wellcome Sanger Institute COVID-19 Surveillance Team |
| EPI_ISL_3436361 | Lighthouse Lab in Glasgow | Wellcome Sanger Institute for the COVID-19 Genomics UK (COG-UK) Consortium | Harper VanSteenhouse, Yumi Kasai, David Gray, Carol Clugston, Anna Dominiczak and Alex Alderton, Roberto Amato, Jeffrey Barrett, Sonia Goncalves, Ewan Harrison, David K. Jackson, Ian Johnston, Dominic Kwiatkowski, Cordelia Langford, John Sillitoe on behalf of the Wellcome Sanger Institute COVID-19 Surveillance Team |
| EPI_ISL_3436362 | Lighthouse Lab in Milton Keynes | Wellcome Sanger Institute for the COVID-19 Genomics UK (COG-UK) Consortium | The Lighthouse Lab in Milton Keynes and Alex Alderton, Roberto Amato, Jeffrey Barrett, Sonia Goncalves, Ewan Harrison, David K. Jackson, Ian Johnston, Dominic Kwiatkowski, Cordelia Langford, John Sillitoe on behalf of the Wellcome Sanger Institute COVID-19 Surveillance Team |
| EPI_ISL_3436363 | Lighthouse Lab in Alderley Park | Wellcome Sanger Institute for the COVID-19 Genomics UK (COG-UK) Consortium | Jacquelyn Wynn, Mairead Hyland, The Lighthouse Lab in Alderley Park and Alex Alderton, Roberto Amato, Jeffrey Barrett, Sonia Goncalves, Ewan Harrison, David K. Jackson, Ian Johnston, Dominic Kwiatkowski, Cordelia Langford, John Sillitoe on behalf of the Wellcome Sanger Institute COVID-19 Surveillance Team |
| EPI_ISL_3436364 | Lighthouse Lab in Glasgow | Wellcome Sanger Institute for the COVID-19 Genomics UK (COG-UK) Consortium | Harper VanSteenhouse, Yumi Kasai, David Gray, Carol Clugston, Anna Dominiczak and Alex Alderton, Roberto Amato, Jeffrey Barrett, Sonia Goncalves, Ewan Harrison, David K. Jackson, Ian Johnston, Dominic Kwiatkowski, Cordelia Langford, John Sillitoe on behalf of the Wellcome Sanger Institute COVID-19 Surveillance Team |
| EPI_ISL_3436365, EPI_ISL_3436366, EPI_ISL_3436367 | Lighthouse Lab in Milton Keynes | Wellcome Sanger Institute for the COVID-19 Genomics UK (COG-UK) Consortium | The Lighthouse Lab in Milton Keynes and Alex Alderton, Roberto Amato, Jeffrey Barrett, Sonia Goncalves, Ewan Harrison, David K. Jackson, Ian Johnston, Dominic Kwiatkowski, Cordelia Langford, John Sillitoe on behalf of the Wellcome Sanger Institute COVID-19 Surveillance Team |
| EPI_ISL_3436368 | Lighthouse Lab in Glasgow | Wellcome Sanger Institute for the COVID-19 Genomics UK (COG-UK) Consortium | Harper VanSteenhouse, Yumi Kasai, David Gray, Carol Clugston, Anna Dominiczak and Alex Alderton, Roberto Amato, Jeffrey Barrett, Sonia Goncalves, Ewan Harrison, David K. Jackson, Ian Johnston, Dominic Kwiatkowski, Cordelia Langford, John Sillitoe on behalf of the Wellcome Sanger Institute COVID-19 Surveillance Team |
| EPI_ISL_3436369 | Lighthouse Lab in Milton Keynes | Wellcome Sanger Institute for the COVID-19 Genomics UK (COG-UK) Consortium | The Lighthouse Lab in Milton Keynes and Alex Alderton, Roberto Amato, Jeffrey Barrett, Sonia Goncalves, Ewan Harrison, David K. Jackson, Ian Johnston, Dominic Kwiatkowski, Cordelia Langford, John Sillitoe on behalf of the Wellcome Sanger Institute COVID-19 Surveillance Team |
| EPI_ISL_3436370 | Lighthouse Lab in Glasgow | Wellcome Sanger Institute for the COVID-19 Genomics UK (COG-UK) Consortium | Harper VanSteenhouse, Yumi Kasai, David Gray, Carol Clugston, Anna Dominiczak and Alex Alderton, Roberto Amato, Jeffrey Barrett, Sonia Goncalves, Ewan Harrison, David K. Jackson, Ian Johnston, Dominic Kwiatkowski, Cordelia Langford, John Sillitoe on behalf of the Wellcome Sanger Institute COVID-19 Surveillance Team |
| EPI_ISL_3436371 | Lighthouse Lab in Milton Keynes | Wellcome Sanger Institute for the COVID-19 Genomics UK (COG-UK) Consortium | The Lighthouse Lab in Milton Keynes and Alex Alderton, Roberto Amato, Jeffrey Barrett, Sonia Goncalves, Ewan Harrison, David K. Jackson, Ian Johnston, Dominic Kwiatkowski, Cordelia Langford, John Sillitoe on behalf of the Wellcome Sanger Institute COVID-19 Surveillance Team |
| EPI_ISL_3436372 | Lighthouse Lab in Glasgow | Wellcome Sanger Institute for the COVID-19 Genomics UK (COG-UK) Consortium | Harper VanSteenhouse, Yumi Kasai, David Gray, Carol Clugston, Anna Dominiczak and Alex Alderton, Roberto Amato, Jeffrey Barrett, Sonia Goncalves, Ewan Harrison, David K. Jackson, Ian Johnston, Dominic Kwiatkowski, Cordelia Langford, John Sillitoe on behalf of the Wellcome Sanger Institute COVID-19 Surveillance Team |
| EPI_ISL_3436373 | Lighthouse Lab in Alderley Park | Wellcome Sanger Institute for the COVID-19 Genomics UK (COG-UK) Consortium | Jacquelyn Wynn, Mairead Hyland, The Lighthouse Lab in Alderley Park and Alex Alderton, Roberto Amato, Jeffrey Barrett, Sonia Goncalves, Ewan Harrison, David K. Jackson, Ian Johnston, Dominic Kwiatkowski, Cordelia Langford, John Sillitoe on behalf of the Wellcome Sanger Institute COVID-19 Surveillance Team |
| EPI_ISL_3436374 | Lighthouse Lab in Milton Keynes | Wellcome Sanger Institute for the COVID-19 Genomics UK (COG-UK) Consortium | The Lighthouse Lab in Milton Keynes and Alex Alderton, Roberto Amato, Jeffrey Barrett, Sonia Goncalves, Ewan Harrison, David K. Jackson, Ian Johnston, Dominic Kwiatkowski, Cordelia Langford, John Sillitoe on behalf of the Wellcome Sanger Institute COVID-19 Surveillance Team |
| EPI_ISL_3436375 | Lighthouse Lab in Glasgow | Wellcome Sanger Institute for the COVID-19 Genomics UK (COG-UK) Consortium | Harper VanSteenhouse, Yumi Kasai, David Gray, Carol Clugston, Anna Dominiczak and Alex Alderton, Roberto Amato, Jeffrey Barrett, Sonia Goncalves, Ewan Harrison, David K. Jackson, Ian Johnston, Dominic Kwiatkowski, Cordelia Langford, John Sillitoe on behalf of the Wellcome Sanger Institute COVID-19 Surveillance Team |
| EPI_ISL_3436376, EPI_ISL_3436377, EPI_ISL_3436378, EPI_ISL_3436379, EPI_ISL_3436380, EPI_ISL_3436381, EPI_ISL_3436382 | Lighthouse Lab in Milton Keynes | Wellcome Sanger Institute for the COVID-19 Genomics UK (COG-UK) Consortium | The Lighthouse Lab in Milton Keynes and Alex Alderton, Roberto Amato, Jeffrey Barrett, Sonia Goncalves, Ewan Harrison, David K. Jackson, Ian Johnston, Dominic Kwiatkowski, Cordelia Langford, John Sillitoe on behalf of the Wellcome Sanger Institute COVID-19 Surveillance Team |
| EPI_ISL_3436383 | Lighthouse Lab in Alderley Park | Wellcome Sanger Institute for the COVID-19 Genomics UK (COG-UK) Consortium | Jacquelyn Wynn, Mairead Hyland, The Lighthouse Lab in Alderley Park and Alex Alderton, Roberto Amato, Jeffrey Barrett, Sonia Goncalves, Ewan Harrison, David K. Jackson, Ian Johnston, Dominic Kwiatkowski, Cordelia Langford, John Sillitoe on behalf of the Wellcome Sanger Institute COVID-19 Surveillance Team |
| EPI_ISL_3436384 | Lighthouse Lab in Glasgow | Wellcome Sanger Institute for the COVID-19 Genomics UK (COG-UK) Consortium | Harper VanSteenhouse, Yumi Kasai, David Gray, Carol Clugston, Anna Dominiczak and Alex Alderton, Roberto Amato, Jeffrey Barrett, Sonia Goncalves, Ewan Harrison, David K. Jackson, Ian Johnston, Dominic Kwiatkowski, Cordelia Langford, John Sillitoe on behalf of the Wellcome Sanger Institute COVID-19 Surveillance Team |
| EPI_ISL_3436385, EPI_ISL_3436386, EPI_ISL_3436387 | Lighthouse Lab in Milton Keynes | Wellcome Sanger Institute for the COVID-19 Genomics UK (COG-UK) Consortium | The Lighthouse Lab in Milton Keynes and Alex Alderton, Roberto Amato, Jeffrey Barrett, Sonia Goncalves, Ewan Harrison, David K. Jackson, Ian Johnston, Dominic Kwiatkowski, Cordelia Langford, John Sillitoe on behalf of the Wellcome Sanger Institute COVID-19 Surveillance Team |
| EPI_ISL_3436388 | Lighthouse Lab in Alderley Park | Wellcome Sanger Institute for the COVID-19 Genomics UK (COG-UK) Consortium | Jacquelyn Wynn, Mairead Hyland, The Lighthouse Lab in Alderley Park and Alex Alderton, Roberto Amato, Jeffrey Barrett, Sonia Goncalves, Ewan Harrison, David K. Jackson, Ian Johnston, Dominic Kwiatkowski, Cordelia Langford, John Sillitoe on behalf of the Wellcome Sanger Institute COVID-19 Surveillance Team |
| EPI_ISL_3436389 | Lighthouse Lab in Milton Keynes | Wellcome Sanger Institute for the COVID-19 Genomics UK (COG-UK) Consortium | The Lighthouse Lab in Milton Keynes and Alex Alderton, Roberto Amato, Jeffrey Barrett, Sonia Goncalves, Ewan Harrison, David K. Jackson, Ian Johnston, Dominic Kwiatkowski, Cordelia Langford, John Sillitoe on behalf of the Wellcome Sanger Institute COVID-19 Surveillance Team |
| EPI_ISL_3436390 | Lighthouse Lab in Alderley Park | Wellcome Sanger Institute for the COVID-19 Genomics UK (COG-UK) Consortium | Jacquelyn Wynn, Mairead Hyland, The Lighthouse Lab in Alderley Park and Alex Alderton, Roberto Amato, Jeffrey Barrett, Sonia Goncalves, Ewan Harrison, David K. Jackson, Ian Johnston, Dominic Kwiatkowski, Cordelia Langford, John Sillitoe on behalf of the Wellcome Sanger Institute COVID-19 Surveillance Team |
| EPI_ISL_3436391 | Lighthouse Lab in Milton Keynes | Wellcome Sanger Institute for the COVID-19 Genomics UK (COG-UK) | The Lighthouse Lab in Milton Keynes and Alex Alderton, Roberto Amato, Jeffrey Barrett, Sonia Goncalves, Ewan Harrison, David K. Jackson, Ian Johnston, Dominic Kwiatkowski, Cordelia Langford, John Sillitoe on behalf of the Wellcome Sanger |

| | | | |
|---|---|---|---|
| | | Consortium | Institute COVID-19 Surveillance Team |
| EPI_ISL_3436392 | Lighthouse Lab in Glasgow | Wellcome Sanger Institute for the COVID-19 Genomics UK (COG-UK) Consortium | Harper VanSteenhouse, Yumi Kasai, David Gray, Carol Clugston, Anna Dominiczak and Alex Alderton, Roberto Amato, Jeffrey Barrett, Sonia Goncalves, Ewan Harrison, David K. Jackson, Ian Johnston, Dominic Kwiatkowski, Cordelia Langford, John Sillitoe on behalf of the Wellcome Sanger Institute COVID-19 Surveillance Team |
| EPI_ISL_3436393, EPI_ISL_3436394, EPI_ISL_3436395, EPI_ISL_3436396, EPI_ISL_3436397 | Lighthouse Lab in Milton Keynes | Wellcome Sanger Institute for the COVID-19 Genomics UK (COG-UK) Consortium | The Lighthouse Lab in Milton Keynes and Alex Alderton, Roberto Amato, Jeffrey Barrett, Sonia Goncalves, Ewan Harrison, David K. Jackson, Ian Johnston, Dominic Kwiatkowski, Cordelia Langford, John Sillitoe on behalf of the Wellcome Sanger Institute COVID-19 Surveillance Team |
| EPI_ISL_3436398 | Lighthouse Lab in Alderley Park | Wellcome Sanger Institute for the COVID-19 Genomics UK (COG-UK) Consortium | Jacquelyn Wynn, Mairead Hyland, The Lighthouse Lab in Alderley Park and Alex Alderton, Roberto Amato, Jeffrey Barrett, Sonia Goncalves, Ewan Harrison, David K. Jackson, Ian Johnston, Dominic Kwiatkowski, Cordelia Langford, John Sillitoe on behalf of the Wellcome Sanger Institute COVID-19 Surveillance Team |
| EPI_ISL_3436399 | Lighthouse Lab in Glasgow | Wellcome Sanger Institute for the COVID-19 Genomics UK (COG-UK) Consortium | Harper VanSteenhouse, Yumi Kasai, David Gray, Carol Clugston, Anna Dominiczak and Alex Alderton, Roberto Amato, Jeffrey Barrett, Sonia Goncalves, Ewan Harrison, David K. Jackson, Ian Johnston, Dominic Kwiatkowski, Cordelia Langford, John Sillitoe on behalf of the Wellcome Sanger Institute COVID-19 Surveillance Team |
| EPI_ISL_3436400, EPI_ISL_3436401, EPI_ISL_3436402, EPI_ISL_3436403 | Lighthouse Lab in Milton Keynes | Wellcome Sanger Institute for the COVID-19 Genomics UK (COG-UK) Consortium | The Lighthouse Lab in Milton Keynes and Alex Alderton, Roberto Amato, Jeffrey Barrett, Sonia Goncalves, Ewan Harrison, David K. Jackson, Ian Johnston, Dominic Kwiatkowski, Cordelia Langford, John Sillitoe on behalf of the Wellcome Sanger Institute COVID-19 Surveillance Team |
| EPI_ISL_3436404 | Lighthouse Lab in Alderley Park | Wellcome Sanger Institute for the COVID-19 Genomics UK (COG-UK) Consortium | Jacquelyn Wynn, Mairead Hyland, The Lighthouse Lab in Alderley Park and Alex Alderton, Roberto Amato, Jeffrey Barrett, Sonia Goncalves, Ewan Harrison, David K. Jackson, Ian Johnston, Dominic Kwiatkowski, Cordelia Langford, John Sillitoe on behalf of the Wellcome Sanger Institute COVID-19 Surveillance Team |
| EPI_ISL_3436405 | Lighthouse Lab in Milton Keynes | Wellcome Sanger Institute for the COVID-19 Genomics UK (COG-UK) Consortium | The Lighthouse Lab in Milton Keynes and Alex Alderton, Roberto Amato, Jeffrey Barrett, Sonia Goncalves, Ewan Harrison, David K. Jackson, Ian Johnston, Dominic Kwiatkowski, Cordelia Langford, John Sillitoe on behalf of the Wellcome Sanger Institute COVID-19 Surveillance Team |
| EPI_ISL_3436406 | Lighthouse Lab in Alderley Park | Wellcome Sanger Institute for the COVID-19 Genomics UK (COG-UK) Consortium | Jacquelyn Wynn, Mairead Hyland, The Lighthouse Lab in Alderley Park and Alex Alderton, Roberto Amato, Jeffrey Barrett, Sonia Goncalves, Ewan Harrison, David K. Jackson, Ian Johnston, Dominic Kwiatkowski, Cordelia Langford, John Sillitoe on behalf of the Wellcome Sanger Institute COVID-19 Surveillance Team |
| EPI_ISL_3436407, EPI_ISL_3436408 | Lighthouse Lab in Milton Keynes | Wellcome Sanger Institute for the COVID-19 Genomics UK (COG-UK) Consortium | The Lighthouse Lab in Milton Keynes and Alex Alderton, Roberto Amato, Jeffrey Barrett, Sonia Goncalves, Ewan Harrison, David K. Jackson, Ian Johnston, Dominic Kwiatkowski, Cordelia Langford, John Sillitoe on behalf of the Wellcome Sanger Institute COVID-19 Surveillance Team |
| EPI_ISL_3436409, EPI_ISL_3436410 | Lighthouse Lab in Glasgow | Wellcome Sanger Institute for the COVID-19 Genomics UK (COG-UK) Consortium | Harper VanSteenhouse, Yumi Kasai, David Gray, Carol Clugston, Anna Dominiczak and Alex Alderton, Roberto Amato, Jeffrey Barrett, Sonia Goncalves, Ewan Harrison, David K. Jackson, Ian Johnston, Dominic Kwiatkowski, Cordelia Langford, John Sillitoe on behalf of the Wellcome Sanger Institute COVID-19 Surveillance Team |
| EPI_ISL_3436411 | Lighthouse Lab in Alderley Park | Wellcome Sanger Institute for the COVID-19 Genomics UK (COG-UK) Consortium | Jacquelyn Wynn, Mairead Hyland, The Lighthouse Lab in Alderley Park and Alex Alderton, Roberto Amato, Jeffrey Barrett, Sonia Goncalves, Ewan Harrison, David K. Jackson, Ian Johnston, Dominic Kwiatkowski, Cordelia Langford, John Sillitoe on behalf of the Wellcome Sanger Institute COVID-19 Surveillance Team |
| EPI_ISL_3436412, EPI_ISL_3436413 | Lighthouse Lab in Milton Keynes | Wellcome Sanger Institute for the COVID-19 Genomics UK (COG-UK) Consortium | The Lighthouse Lab in Milton Keynes and Alex Alderton, Roberto Amato, Jeffrey Barrett, Sonia Goncalves, Ewan Harrison, David K. Jackson, Ian Johnston, Dominic Kwiatkowski, Cordelia Langford, John Sillitoe on behalf of the Wellcome Sanger Institute COVID-19 Surveillance Team |
| EPI_ISL_3436414 | Lighthouse Lab in Alderley Park | Wellcome Sanger Institute for the COVID-19 Genomics UK (COG-UK) Consortium | Jacquelyn Wynn, Mairead Hyland, The Lighthouse Lab in Alderley Park and Alex Alderton, Roberto Amato, Jeffrey Barrett, Sonia Goncalves, Ewan Harrison, David K. Jackson, Ian Johnston, Dominic Kwiatkowski, Cordelia Langford, John Sillitoe on behalf of the Wellcome Sanger Institute COVID-19 Surveillance Team |
| EPI_ISL_3436415 | Lighthouse Lab in Milton Keynes | Wellcome Sanger Institute for the COVID-19 Genomics UK (COG-UK) Consortium | The Lighthouse Lab in Milton Keynes and Alex Alderton, Roberto Amato, Jeffrey Barrett, Sonia Goncalves, Ewan Harrison, David K. Jackson, Ian Johnston, Dominic Kwiatkowski, Cordelia Langford, John Sillitoe on behalf of the Wellcome Sanger Institute COVID-19 Surveillance Team |
| EPI_ISL_3436416 | Lighthouse Lab in Glasgow | Wellcome Sanger Institute for the COVID-19 Genomics UK (COG-UK) Consortium | Harper VanSteenhouse, Yumi Kasai, David Gray, Carol Clugston, Anna Dominiczak and Alex Alderton, Roberto Amato, Jeffrey Barrett, Sonia Goncalves, Ewan Harrison, David K. Jackson, Ian Johnston, Dominic Kwiatkowski, Cordelia Langford, John Sillitoe on behalf of the Wellcome Sanger Institute COVID-19 Surveillance Team |
| EPI_ISL_3436417, EPI_ISL_3436418 | Lighthouse Lab in Milton Keynes | Wellcome Sanger Institute for the COVID-19 Genomics UK (COG-UK) Consortium | The Lighthouse Lab in Milton Keynes and Alex Alderton, Roberto Amato, Jeffrey Barrett, Sonia Goncalves, Ewan Harrison, David K. Jackson, Ian Johnston, Dominic Kwiatkowski, Cordelia Langford, John Sillitoe on behalf of the Wellcome Sanger Institute COVID-19 Surveillance Team |
| EPI_ISL_3436419 | Lighthouse Lab in Alderley Park | Wellcome Sanger Institute for the COVID-19 Genomics UK (COG-UK) Consortium | Jacquelyn Wynn, Mairead Hyland, The Lighthouse Lab in Alderley Park and Alex Alderton, Roberto Amato, Jeffrey Barrett, Sonia Goncalves, Ewan Harrison, David K. Jackson, Ian Johnston, Dominic Kwiatkowski, Cordelia Langford, John Sillitoe on behalf of the Wellcome Sanger Institute COVID-19 Surveillance Team |
| EPI_ISL_3436420 | Lighthouse Lab in Glasgow | Wellcome Sanger Institute for the COVID-19 Genomics UK (COG-UK) Consortium | Harper VanSteenhouse, Yumi Kasai, David Gray, Carol Clugston, Anna Dominiczak and Alex Alderton, Roberto Amato, Jeffrey Barrett, Sonia Goncalves, Ewan Harrison, David K. Jackson, Ian Johnston, Dominic Kwiatkowski, Cordelia Langford, John Sillitoe on behalf of the Wellcome Sanger Institute COVID-19 Surveillance Team |
| EPI_ISL_3436421 | Lighthouse Lab in Alderley Park | Wellcome Sanger Institute for the COVID-19 Genomics UK (COG-UK) Consortium | Jacquelyn Wynn, Mairead Hyland, The Lighthouse Lab in Alderley Park and Alex Alderton, Roberto Amato, Jeffrey Barrett, Sonia Goncalves, Ewan Harrison, David K. Jackson, Ian Johnston, Dominic Kwiatkowski, Cordelia Langford, John Sillitoe on behalf of the Wellcome Sanger Institute COVID-19 Surveillance Team |
| EPI_ISL_3436422, EPI_ISL_3436423, EPI_ISL_3436424, EPI_ISL_3436425 | Lighthouse Lab in Milton Keynes | Wellcome Sanger Institute for the COVID-19 Genomics UK (COG-UK) Consortium | The Lighthouse Lab in Milton Keynes and Alex Alderton, Roberto Amato, Jeffrey Barrett, Sonia Goncalves, Ewan Harrison, David K. Jackson, Ian Johnston, Dominic Kwiatkowski, Cordelia Langford, John Sillitoe on behalf of the Wellcome Sanger Institute COVID-19 Surveillance Team |
| EPI_ISL_3436426 | Lighthouse Lab in Glasgow | Wellcome Sanger Institute for the COVID-19 Genomics UK (COG-UK) Consortium | Harper VanSteenhouse, Yumi Kasai, David Gray, Carol Clugston, Anna Dominiczak and Alex Alderton, Roberto Amato, Jeffrey Barrett, Sonia Goncalves, Ewan Harrison, David K. Jackson, Ian Johnston, Dominic Kwiatkowski, Cordelia Langford, John Sillitoe on behalf of the Wellcome Sanger Institute COVID-19 Surveillance Team |
| EPI_ISL_3436427 | Lighthouse Lab in Alderley Park | Wellcome Sanger Institute for the COVID-19 Genomics UK (COG-UK) Consortium | Jacquelyn Wynn, Mairead Hyland, The Lighthouse Lab in Alderley Park and Alex Alderton, Roberto Amato, Jeffrey Barrett, Sonia Goncalves, Ewan Harrison, David K. Jackson, Ian Johnston, Dominic Kwiatkowski, Cordelia Langford, John Sillitoe on behalf of the Wellcome Sanger Institute COVID-19 Surveillance Team |
| EPI_ISL_3436428, EPI_ISL_3436429, EPI_ISL_3436430, EPI_ISL_3436431 | Lighthouse Lab in Milton Keynes | Wellcome Sanger Institute for the COVID-19 Genomics UK (COG-UK) Consortium | The Lighthouse Lab in Milton Keynes and Alex Alderton, Roberto Amato, Jeffrey Barrett, Sonia Goncalves, Ewan Harrison, David K. Jackson, Ian Johnston, Dominic Kwiatkowski, Cordelia Langford, John Sillitoe on behalf of the Wellcome Sanger Institute COVID-19 Surveillance Team |
| EPI_ISL_3436432, EPI_ISL_3436433 | Lighthouse Lab in Alderley Park | Wellcome Sanger Institute for the COVID-19 Genomics UK (COG-UK) Consortium | Jacquelyn Wynn, Mairead Hyland, The Lighthouse Lab in Alderley Park and Alex Alderton, Roberto Amato, Jeffrey Barrett, Sonia Goncalves, Ewan Harrison, David K. Jackson, Ian Johnston, Dominic Kwiatkowski, Cordelia Langford, John Sillitoe on behalf of the Wellcome Sanger Institute COVID-19 Surveillance Team |
| EPI_ISL_3436434, EPI_ISL_3436435, EPI_ISL_3436436, EPI_ISL_3436437 | Lighthouse Lab in Milton Keynes | Wellcome Sanger Institute for the COVID-19 Genomics UK (COG-UK) Consortium | The Lighthouse Lab in Milton Keynes and Alex Alderton, Roberto Amato, Jeffrey Barrett, Sonia Goncalves, Ewan Harrison, David K. Jackson, Ian Johnston, Dominic Kwiatkowski, Cordelia Langford, John Sillitoe on behalf of the Wellcome Sanger Institute COVID-19 Surveillance Team |
| EPI_ISL_3436438 | Lighthouse Lab in Alderley Park | Wellcome Sanger Institute for the COVID-19 Genomics UK (COG-UK) Consortium | Jacquelyn Wynn, Mairead Hyland, The Lighthouse Lab in Alderley Park and Alex Alderton, Roberto Amato, Jeffrey Barrett, Sonia Goncalves, Ewan Harrison, David K. Jackson, Ian Johnston, Dominic Kwiatkowski, Cordelia Langford, John Sillitoe on behalf of the Wellcome Sanger Institute COVID-19 Surveillance Team |
| EPI_ISL_3436439, EPI_ISL_3436440 | Lighthouse Lab in Milton Keynes | Wellcome Sanger Institute for the COVID-19 Genomics UK (COG-UK) Consortium | The Lighthouse Lab in Milton Keynes and Alex Alderton, Roberto Amato, Jeffrey Barrett, Sonia Goncalves, Ewan Harrison, David K. Jackson, Ian Johnston, Dominic Kwiatkowski, Cordelia Langford, John Sillitoe on behalf of the Wellcome Sanger Institute COVID-19 Surveillance Team |
| EPI_ISL_3436441 | Lighthouse Lab in Glasgow | Wellcome Sanger Institute for the COVID-19 Genomics UK (COG-UK) Consortium | Harper VanSteenhouse, Yumi Kasai, David Gray, Carol Clugston, Anna Dominiczak and Alex Alderton, Roberto Amato, Jeffrey Barrett, Sonia Goncalves, Ewan Harrison, David K. Jackson, Ian Johnston, Dominic Kwiatkowski, Cordelia Langford, John Sillitoe on behalf of the Wellcome Sanger Institute COVID-19 Surveillance Team |
| EPI_ISL_3436442 | Lighthouse Lab in Milton Keynes | Wellcome Sanger Institute for the COVID-19 Genomics UK (COG-UK) Consortium | The Lighthouse Lab in Milton Keynes and Alex Alderton, Roberto Amato, Jeffrey Barrett, Sonia Goncalves, Ewan Harrison, David K. Jackson, Ian Johnston, Dominic Kwiatkowski, Cordelia Langford, John Sillitoe on behalf of the Wellcome Sanger Institute COVID-19 Surveillance Team |
| EPI_ISL_3436443 | Lighthouse Lab in Glasgow | Wellcome Sanger Institute for the COVID-19 Genomics UK (COG-UK) Consortium | Harper VanSteenhouse, Yumi Kasai, David Gray, Carol Clugston, Anna Dominiczak and Alex Alderton, Roberto Amato, Jeffrey Barrett, Sonia Goncalves, Ewan Harrison, David K. Jackson, Ian Johnston, Dominic Kwiatkowski, Cordelia Langford, John Sillitoe on behalf of the Wellcome Sanger Institute COVID-19 Surveillance Team |
| EPI_ISL_3436444 | Lighthouse Lab in Milton Keynes | Wellcome Sanger Institute for the COVID-19 Genomics UK (COG-UK) Consortium | The Lighthouse Lab in Milton Keynes and Alex Alderton, Roberto Amato, Jeffrey Barrett, Sonia Goncalves, Ewan Harrison, David K. Jackson, Ian Johnston, Dominic Kwiatkowski, Cordelia Langford, John Sillitoe on behalf of the Wellcome Sanger Institute COVID-19 Surveillance Team |
| EPI_ISL_3436445 | Lighthouse Lab in Glasgow | Wellcome Sanger Institute for the COVID-19 Genomics UK (COG-UK) Consortium | Harper VanSteenhouse, Yumi Kasai, David Gray, Carol Clugston, Anna Dominiczak and Alex Alderton, Roberto Amato, Jeffrey Barrett, Sonia Goncalves, Ewan Harrison, David K. Jackson, Ian Johnston, Dominic Kwiatkowski, Cordelia Langford, John Sillitoe on behalf of the Wellcome Sanger Institute COVID-19 Surveillance Team |
| EPI_ISL_3436446, EPI_ISL_3436447, EPI_ISL_3436448 | Lighthouse Lab in Milton Keynes | Wellcome Sanger Institute for the COVID-19 Genomics UK (COG-UK) Consortium | The Lighthouse Lab in Milton Keynes and Alex Alderton, Roberto Amato, Jeffrey Barrett, Sonia Goncalves, Ewan Harrison, David K. Jackson, Ian Johnston, Dominic Kwiatkowski, Cordelia Langford, John Sillitoe on behalf of the Wellcome Sanger Institute COVID-19 Surveillance Team |
| EPI_ISL_3436449 | Lighthouse Lab in Alderley Park | Wellcome Sanger Institute for the COVID-19 Genomics UK (COG-UK) Consortium | Jacquelyn Wynn, Mairead Hyland, The Lighthouse Lab in Alderley Park and Alex Alderton, Roberto Amato, Jeffrey Barrett, Sonia Goncalves, Ewan Harrison, David K. Jackson, Ian Johnston, Dominic Kwiatkowski, Cordelia Langford, John Sillitoe on behalf of the Wellcome Sanger Institute COVID-19 Surveillance Team |
| EPI_ISL_3436450 | Lighthouse Lab in Milton Keynes | Wellcome Sanger Institute for the COVID-19 Genomics UK (COG-UK) Consortium | The Lighthouse Lab in Milton Keynes and Alex Alderton, Roberto Amato, Jeffrey Barrett, Sonia Goncalves, Ewan Harrison, David K. Jackson, Ian Johnston, Dominic Kwiatkowski, Cordelia Langford, John Sillitoe on behalf of the Wellcome Sanger Institute COVID-19 Surveillance Team |
| EPI_ISL_3436451 | Lighthouse Lab in Glasgow | Wellcome Sanger Institute for the COVID-19 Genomics UK (COG-UK) Consortium | Harper VanSteenhouse, Yumi Kasai, David Gray, Carol Clugston, Anna Dominiczak and Alex Alderton, Roberto Amato, Jeffrey Barrett, Sonia Goncalves, Ewan Harrison, David K. Jackson, Ian Johnston, Dominic Kwiatkowski, Cordelia Langford, John Sillitoe on behalf of the Wellcome Sanger Institute COVID-19 Surveillance Team |
| EPI_ISL_3436452, EPI_ISL_3436453 | Lighthouse Lab in Milton Keynes | Wellcome Sanger Institute for the COVID-19 Genomics UK (COG-UK) Consortium | The Lighthouse Lab in Milton Keynes and Alex Alderton, Roberto Amato, Jeffrey Barrett, Sonia Goncalves, Ewan Harrison, David K. Jackson, Ian Johnston, Dominic Kwiatkowski, Cordelia Langford, John Sillitoe on behalf of the Wellcome Sanger Institute COVID-19 Surveillance Team |
| EPI_ISL_3436454 | Lighthouse Lab in Alderley Park | Wellcome Sanger Institute for the COVID-19 Genomics UK (COG-UK) Consortium | Jacquelyn Wynn, Mairead Hyland, The Lighthouse Lab in Alderley Park and Alex Alderton, Roberto Amato, Jeffrey Barrett, Sonia Goncalves, Ewan Harrison, David K. Jackson, Ian Johnston, Dominic Kwiatkowski, Cordelia Langford, John Sillitoe on behalf of the Wellcome Sanger Institute COVID-19 Surveillance Team |
| EPI_ISL_3436455, EPI_ISL_3436456 | Lighthouse Lab in Milton Keynes | Wellcome Sanger Institute for the COVID-19 Genomics UK (COG-UK) Consortium | The Lighthouse Lab in Milton Keynes and Alex Alderton, Roberto Amato, Jeffrey Barrett, Sonia Goncalves, Ewan Harrison, David K. Jackson, Ian Johnston, Dominic Kwiatkowski, Cordelia Langford, John Sillitoe on behalf of the Wellcome Sanger Institute COVID-19 Surveillance Team |
| EPI_ISL_3436457, | Lighthouse Lab in Glasgow | Wellcome Sanger Institute for the COVID-19 Genomics UK (COG-UK) | Harper VanSteenhouse, Yumi Kasai, David Gray, Carol Clugston, Anna Dominiczak and Alex Alderton, Roberto Amato, Jeffrey Barrett, Sonia Goncalves, Ewan Harrison, David K. Jackson, Ian Johnston, Dominic Kwiatkowski, Cordelia Langford, |

| | | | |
|---|---|---|---|
| EPI_ISL_3436458 | | Consortium | John Sillitoe on behalf of the Wellcome Sanger Institute COVID-19 Surveillance Team |
| EPI_ISL_3436459 | Lighthouse Lab in Milton Keynes | Wellcome Sanger Institute for the COVID-19 Genomics UK (COG-UK) Consortium | The Lighthouse Lab in Milton Keynes and Alex Alderton, Roberto Amato, Jeffrey Barrett, Sonia Goncalves, Ewan Harrison, David K. Jackson, Ian Johnston, Dominic Kwiatkowski, Cordelia Langford, John Sillitoe on behalf of the Wellcome Sanger Institute COVID-19 Surveillance Team |
| EPI_ISL_3436460 | Lighthouse Lab in Alderley Park | Wellcome Sanger Institute for the COVID-19 Genomics UK (COG-UK) Consortium | Jacquelyn Wynn, Mairead Hyland, The Lighthouse Lab in Alderley Park and Alex Alderton, Roberto Amato, Jeffrey Barrett, Sonia Goncalves, Ewan Harrison, David K. Jackson, Ian Johnston, Dominic Kwiatkowski, Cordelia Langford, John Sillitoe on behalf of the Wellcome Sanger Institute COVID-19 Surveillance Team |
| EPI_ISL_3436461, EPI_ISL_3436462, EPI_ISL_3436463 | Lighthouse Lab in Milton Keynes | Wellcome Sanger Institute for the COVID-19 Genomics UK (COG-UK) Consortium | The Lighthouse Lab in Milton Keynes and Alex Alderton, Roberto Amato, Jeffrey Barrett, Sonia Goncalves, Ewan Harrison, David K. Jackson, Ian Johnston, Dominic Kwiatkowski, Cordelia Langford, John Sillitoe on behalf of the Wellcome Sanger Institute COVID-19 Surveillance Team |
| EPI_ISL_3436464, EPI_ISL_3436465, EPI_ISL_3436466 | Lighthouse Lab in Alderley Park | Wellcome Sanger Institute for the COVID-19 Genomics UK (COG-UK) Consortium | Jacquelyn Wynn, Mairead Hyland, The Lighthouse Lab in Alderley Park and Alex Alderton, Roberto Amato, Jeffrey Barrett, Sonia Goncalves, Ewan Harrison, David K. Jackson, Ian Johnston, Dominic Kwiatkowski, Cordelia Langford, John Sillitoe on behalf of the Wellcome Sanger Institute COVID-19 Surveillance Team |
| EPI_ISL_3436467 | Lighthouse Lab in Milton Keynes | Wellcome Sanger Institute for the COVID-19 Genomics UK (COG-UK) Consortium | The Lighthouse Lab in Milton Keynes and Alex Alderton, Roberto Amato, Jeffrey Barrett, Sonia Goncalves, Ewan Harrison, David K. Jackson, Ian Johnston, Dominic Kwiatkowski, Cordelia Langford, John Sillitoe on behalf of the Wellcome Sanger Institute COVID-19 Surveillance Team |
| EPI_ISL_3436468 | Lighthouse Lab in Glasgow | Wellcome Sanger Institute for the COVID-19 Genomics UK (COG-UK) Consortium | Harper VanSteenhouse, Yumi Kasai, David Gray, Carol Clugston, Anna Dominiczak and Alex Alderton, Roberto Amato, Jeffrey Barrett, Sonia Goncalves, Ewan Harrison, David K. Jackson, Ian Johnston, Dominic Kwiatkowski, Cordelia Langford, John Sillitoe on behalf of the Wellcome Sanger Institute COVID-19 Surveillance Team |
| EPI_ISL_3436469, EPI_ISL_3436470 | Lighthouse Lab in Milton Keynes | Wellcome Sanger Institute for the COVID-19 Genomics UK (COG-UK) Consortium | The Lighthouse Lab in Milton Keynes and Alex Alderton, Roberto Amato, Jeffrey Barrett, Sonia Goncalves, Ewan Harrison, David K. Jackson, Ian Johnston, Dominic Kwiatkowski, Cordelia Langford, John Sillitoe on behalf of the Wellcome Sanger Institute COVID-19 Surveillance Team |
| EPI_ISL_3436471, EPI_ISL_3436472 | Lighthouse Lab in Alderley Park | Wellcome Sanger Institute for the COVID-19 Genomics UK (COG-UK) Consortium | Jacquelyn Wynn, Mairead Hyland, The Lighthouse Lab in Alderley Park and Alex Alderton, Roberto Amato, Jeffrey Barrett, Sonia Goncalves, Ewan Harrison, David K. Jackson, Ian Johnston, Dominic Kwiatkowski, Cordelia Langford, John Sillitoe on behalf of the Wellcome Sanger Institute COVID-19 Surveillance Team |
| EPI_ISL_3436473, EPI_ISL_3436474, EPI_ISL_3436475, EPI_ISL_3436476 | Lighthouse Lab in Milton Keynes | Wellcome Sanger Institute for the COVID-19 Genomics UK (COG-UK) Consortium | The Lighthouse Lab in Milton Keynes and Alex Alderton, Roberto Amato, Jeffrey Barrett, Sonia Goncalves, Ewan Harrison, David K. Jackson, Ian Johnston, Dominic Kwiatkowski, Cordelia Langford, John Sillitoe on behalf of the Wellcome Sanger Institute COVID-19 Surveillance Team |
| EPI_ISL_3436477 | Lighthouse Lab in Glasgow | Wellcome Sanger Institute for the COVID-19 Genomics UK (COG-UK) Consortium | Harper VanSteenhouse, Yumi Kasai, David Gray, Carol Clugston, Anna Dominiczak and Alex Alderton, Roberto Amato, Jeffrey Barrett, Sonia Goncalves, Ewan Harrison, David K. Jackson, Ian Johnston, Dominic Kwiatkowski, Cordelia Langford, John Sillitoe on behalf of the Wellcome Sanger Institute COVID-19 Surveillance Team |
| EPI_ISL_3436478, EPI_ISL_3436479 | Lighthouse Lab in Milton Keynes | Wellcome Sanger Institute for the COVID-19 Genomics UK (COG-UK) Consortium | The Lighthouse Lab in Milton Keynes and Alex Alderton, Roberto Amato, Jeffrey Barrett, Sonia Goncalves, Ewan Harrison, David K. Jackson, Ian Johnston, Dominic Kwiatkowski, Cordelia Langford, John Sillitoe on behalf of the Wellcome Sanger Institute COVID-19 Surveillance Team |
| EPI_ISL_3436480 | Lighthouse Lab in Glasgow | Wellcome Sanger Institute for the COVID-19 Genomics UK (COG-UK) Consortium | Harper VanSteenhouse, Yumi Kasai, David Gray, Carol Clugston, Anna Dominiczak and Alex Alderton, Roberto Amato, Jeffrey Barrett, Sonia Goncalves, Ewan Harrison, David K. Jackson, Ian Johnston, Dominic Kwiatkowski, Cordelia Langford, John Sillitoe on behalf of the Wellcome Sanger Institute COVID-19 Surveillance Team |
| EPI_ISL_3436481 | Lighthouse Lab in Alderley Park | Wellcome Sanger Institute for the COVID-19 Genomics UK (COG-UK) Consortium | Jacquelyn Wynn, Mairead Hyland, The Lighthouse Lab in Alderley Park and Alex Alderton, Roberto Amato, Jeffrey Barrett, Sonia Goncalves, Ewan Harrison, David K. Jackson, Ian Johnston, Dominic Kwiatkowski, Cordelia Langford, John Sillitoe on behalf of the Wellcome Sanger Institute COVID-19 Surveillance Team |
| EPI_ISL_3436482, EPI_ISL_3436483, EPI_ISL_3436484, EPI_ISL_3436485, EPI_ISL_3436486, EPI_ISL_3436487 | Lighthouse Lab in Milton Keynes | Wellcome Sanger Institute for the COVID-19 Genomics UK (COG-UK) Consortium | The Lighthouse Lab in Milton Keynes and Alex Alderton, Roberto Amato, Jeffrey Barrett, Sonia Goncalves, Ewan Harrison, David K. Jackson, Ian Johnston, Dominic Kwiatkowski, Cordelia Langford, John Sillitoe on behalf of the Wellcome Sanger Institute COVID-19 Surveillance Team |
| EPI_ISL_3436488 | Lighthouse Lab in Glasgow | Wellcome Sanger Institute for the COVID-19 Genomics UK (COG-UK) Consortium | Harper VanSteenhouse, Yumi Kasai, David Gray, Carol Clugston, Anna Dominiczak and Alex Alderton, Roberto Amato, Jeffrey Barrett, Sonia Goncalves, Ewan Harrison, David K. Jackson, Ian Johnston, Dominic Kwiatkowski, Cordelia Langford, John Sillitoe on behalf of the Wellcome Sanger Institute COVID-19 Surveillance Team |
| EPI_ISL_3436489, EPI_ISL_3436490 | Lighthouse Lab in Milton Keynes | Wellcome Sanger Institute for the COVID-19 Genomics UK (COG-UK) Consortium | The Lighthouse Lab in Milton Keynes and Alex Alderton, Roberto Amato, Jeffrey Barrett, Sonia Goncalves, Ewan Harrison, David K. Jackson, Ian Johnston, Dominic Kwiatkowski, Cordelia Langford, John Sillitoe on behalf of the Wellcome Sanger Institute COVID-19 Surveillance Team |
| EPI_ISL_3436491 | Lighthouse Lab in Glasgow | Wellcome Sanger Institute for the COVID-19 Genomics UK (COG-UK) Consortium | Harper VanSteenhouse, Yumi Kasai, David Gray, Carol Clugston, Anna Dominiczak and Alex Alderton, Roberto Amato, Jeffrey Barrett, Sonia Goncalves, Ewan Harrison, David K. Jackson, Ian Johnston, Dominic Kwiatkowski, Cordelia Langford, John Sillitoe on behalf of the Wellcome Sanger Institute COVID-19 Surveillance Team |
| EPI_ISL_3436492, EPI_ISL_3436493 | Lighthouse Lab in Milton Keynes | Wellcome Sanger Institute for the COVID-19 Genomics UK (COG-UK) Consortium | The Lighthouse Lab in Milton Keynes and Alex Alderton, Roberto Amato, Jeffrey Barrett, Sonia Goncalves, Ewan Harrison, David K. Jackson, Ian Johnston, Dominic Kwiatkowski, Cordelia Langford, John Sillitoe on behalf of the Wellcome Sanger Institute COVID-19 Surveillance Team |
| EPI_ISL_3436494 | Lighthouse Lab in Alderley Park | Wellcome Sanger Institute for the COVID-19 Genomics UK (COG-UK) Consortium | Jacquelyn Wynn, Mairead Hyland, The Lighthouse Lab in Alderley Park and Alex Alderton, Roberto Amato, Jeffrey Barrett, Sonia Goncalves, Ewan Harrison, David K. Jackson, Ian Johnston, Dominic Kwiatkowski, Cordelia Langford, John Sillitoe on behalf of the Wellcome Sanger Institute COVID-19 Surveillance Team |
| EPI_ISL_3436495 | Lighthouse Lab in Glasgow | Wellcome Sanger Institute for the COVID-19 Genomics UK (COG-UK) Consortium | Harper VanSteenhouse, Yumi Kasai, David Gray, Carol Clugston, Anna Dominiczak and Alex Alderton, Roberto Amato, Jeffrey Barrett, Sonia Goncalves, Ewan Harrison, David K. Jackson, Ian Johnston, Dominic Kwiatkowski, Cordelia Langford, John Sillitoe on behalf of the Wellcome Sanger Institute COVID-19 Surveillance Team |
| EPI_ISL_3436496, EPI_ISL_3436497 | Lighthouse Lab in Milton Keynes | Wellcome Sanger Institute for the COVID-19 Genomics UK (COG-UK) Consortium | The Lighthouse Lab in Milton Keynes and Alex Alderton, Roberto Amato, Jeffrey Barrett, Sonia Goncalves, Ewan Harrison, David K. Jackson, Ian Johnston, Dominic Kwiatkowski, Cordelia Langford, John Sillitoe on behalf of the Wellcome Sanger Institute COVID-19 Surveillance Team |
| EPI_ISL_3436498 | Lighthouse Lab in Alderley Park | Wellcome Sanger Institute for the COVID-19 Genomics UK (COG-UK) Consortium | Jacquelyn Wynn, Mairead Hyland, The Lighthouse Lab in Alderley Park and Alex Alderton, Roberto Amato, Jeffrey Barrett, Sonia Goncalves, Ewan Harrison, David K. Jackson, Ian Johnston, Dominic Kwiatkowski, Cordelia Langford, John Sillitoe on behalf of the Wellcome Sanger Institute COVID-19 Surveillance Team |
| EPI_ISL_3436499, EPI_ISL_3436500, EPI_ISL_3436501 | Lighthouse Lab in Glasgow | Wellcome Sanger Institute for the COVID-19 Genomics UK (COG-UK) Consortium | Harper VanSteenhouse, Yumi Kasai, David Gray, Carol Clugston, Anna Dominiczak and Alex Alderton, Roberto Amato, Jeffrey Barrett, Sonia Goncalves, Ewan Harrison, David K. Jackson, Ian Johnston, Dominic Kwiatkowski, Cordelia Langford, John Sillitoe on behalf of the Wellcome Sanger Institute COVID-19 Surveillance Team |
| EPI_ISL_3436502, EPI_ISL_3436503 | Lighthouse Lab in Milton Keynes | Wellcome Sanger Institute for the COVID-19 Genomics UK (COG-UK) Consortium | The Lighthouse Lab in Milton Keynes and Alex Alderton, Roberto Amato, Jeffrey Barrett, Sonia Goncalves, Ewan Harrison, David K. Jackson, Ian Johnston, Dominic Kwiatkowski, Cordelia Langford, John Sillitoe on behalf of the Wellcome Sanger Institute COVID-19 Surveillance Team |
| EPI_ISL_3436504 | Lighthouse Lab in Alderley Park | Wellcome Sanger Institute for the COVID-19 Genomics UK (COG-UK) Consortium | Jacquelyn Wynn, Mairead Hyland, The Lighthouse Lab in Alderley Park and Alex Alderton, Roberto Amato, Jeffrey Barrett, Sonia Goncalves, Ewan Harrison, David K. Jackson, Ian Johnston, Dominic Kwiatkowski, Cordelia Langford, John Sillitoe on behalf of the Wellcome Sanger Institute COVID-19 Surveillance Team |
| EPI_ISL_3436505, EPI_ISL_3436506 | Lighthouse Lab in Milton Keynes | Wellcome Sanger Institute for the COVID-19 Genomics UK (COG-UK) Consortium | The Lighthouse Lab in Milton Keynes and Alex Alderton, Roberto Amato, Jeffrey Barrett, Sonia Goncalves, Ewan Harrison, David K. Jackson, Ian Johnston, Dominic Kwiatkowski, Cordelia Langford, John Sillitoe on behalf of the Wellcome Sanger Institute COVID-19 Surveillance Team |
| EPI_ISL_3436507, EPI_ISL_3436508 | Lighthouse Lab in Alderley Park | Wellcome Sanger Institute for the COVID-19 Genomics UK (COG-UK) Consortium | Jacquelyn Wynn, Mairead Hyland, The Lighthouse Lab in Alderley Park and Alex Alderton, Roberto Amato, Jeffrey Barrett, Sonia Goncalves, Ewan Harrison, David K. Jackson, Ian Johnston, Dominic Kwiatkowski, Cordelia Langford, John Sillitoe on behalf of the Wellcome Sanger Institute COVID-19 Surveillance Team |
| EPI_ISL_3436509 | Lighthouse Lab in Glasgow | Wellcome Sanger Institute for the COVID-19 Genomics UK (COG-UK) Consortium | Harper VanSteenhouse, Yumi Kasai, David Gray, Carol Clugston, Anna Dominiczak and Alex Alderton, Roberto Amato, Jeffrey Barrett, Sonia Goncalves, Ewan Harrison, David K. Jackson, Ian Johnston, Dominic Kwiatkowski, Cordelia Langford, John Sillitoe on behalf of the Wellcome Sanger Institute COVID-19 Surveillance Team |
| EPI_ISL_3436510 | Lighthouse Lab in Alderley Park | Wellcome Sanger Institute for the COVID-19 Genomics UK (COG-UK) Consortium | Jacquelyn Wynn, Mairead Hyland, The Lighthouse Lab in Alderley Park and Alex Alderton, Roberto Amato, Jeffrey Barrett, Sonia Goncalves, Ewan Harrison, David K. Jackson, Ian Johnston, Dominic Kwiatkowski, Cordelia Langford, John Sillitoe on behalf of the Wellcome Sanger Institute COVID-19 Surveillance Team |
| EPI_ISL_3436511 | Lighthouse Lab in Milton Keynes | Wellcome Sanger Institute for the COVID-19 Genomics UK (COG-UK) Consortium | The Lighthouse Lab in Milton Keynes and Alex Alderton, Roberto Amato, Jeffrey Barrett, Sonia Goncalves, Ewan Harrison, David K. Jackson, Ian Johnston, Dominic Kwiatkowski, Cordelia Langford, John Sillitoe on behalf of the Wellcome Sanger Institute COVID-19 Surveillance Team |
| EPI_ISL_3436512, EPI_ISL_3436513 | Lighthouse Lab in Alderley Park | Wellcome Sanger Institute for the COVID-19 Genomics UK (COG-UK) Consortium | Jacquelyn Wynn, Mairead Hyland, The Lighthouse Lab in Alderley Park and Alex Alderton, Roberto Amato, Jeffrey Barrett, Sonia Goncalves, Ewan Harrison, David K. Jackson, Ian Johnston, Dominic Kwiatkowski, Cordelia Langford, John Sillitoe on behalf of the Wellcome Sanger Institute COVID-19 Surveillance Team |
| EPI_ISL_3436514, EPI_ISL_3436515 | Lighthouse Lab in Glasgow | Wellcome Sanger Institute for the COVID-19 Genomics UK (COG-UK) Consortium | Harper VanSteenhouse, Yumi Kasai, David Gray, Carol Clugston, Anna Dominiczak and Alex Alderton, Roberto Amato, Jeffrey Barrett, Sonia Goncalves, Ewan Harrison, David K. Jackson, Ian Johnston, Dominic Kwiatkowski, Cordelia Langford, John Sillitoe on behalf of the Wellcome Sanger Institute COVID-19 Surveillance Team |
| EPI_ISL_3436516 | Berkshire and Surrey Pathology Services Lighthouse Laboratory | Wellcome Sanger Institute for the COVID-19 Genomics UK (COG-UK) Consortium | Berkshire and Surrey Pathology Services Lighthouse Laboratory and Alex Alderton, Roberto Amato, Jeffrey Barrett, Sonia Goncalves, Ewan Harrison, David K. Jackson, Ian Johnston, Dominic Kwiatkowski, Cordelia Langford, John Sillitoe on behalf of the Wellcome Sanger Institute COVID-19 Surveillance Team |
| EPI_ISL_3436517, EPI_ISL_3436518 | Lighthouse Lab in Alderley Park | Wellcome Sanger Institute for the COVID-19 Genomics UK (COG-UK) Consortium | Jacquelyn Wynn, Mairead Hyland, The Lighthouse Lab in Alderley Park and Alex Alderton, Roberto Amato, Jeffrey Barrett, Sonia Goncalves, Ewan Harrison, David K. Jackson, Ian Johnston, Dominic Kwiatkowski, Cordelia Langford, John Sillitoe on behalf of the Wellcome Sanger Institute COVID-19 Surveillance Team |
| EPI_ISL_3436519 | Berkshire and Surrey Pathology Services Lighthouse Laboratory | Wellcome Sanger Institute for the COVID-19 Genomics UK (COG-UK) Consortium | Berkshire and Surrey Pathology Services Lighthouse Laboratory and Alex Alderton, Roberto Amato, Jeffrey Barrett, Sonia Goncalves, Ewan Harrison, David K. Jackson, Ian Johnston, Dominic Kwiatkowski, Cordelia Langford, John Sillitoe on behalf of the Wellcome Sanger Institute COVID-19 Surveillance Team |
| EPI_ISL_3436520 | Lighthouse Lab in Alderley Park | Wellcome Sanger Institute for the COVID-19 Genomics UK (COG-UK) Consortium | Jacquelyn Wynn, Mairead Hyland, The Lighthouse Lab in Alderley Park and Alex Alderton, Roberto Amato, Jeffrey Barrett, Sonia Goncalves, Ewan Harrison, David K. Jackson, Ian Johnston, Dominic Kwiatkowski, Cordelia Langford, John Sillitoe on behalf of the Wellcome Sanger Institute COVID-19 Surveillance Team |
| EPI_ISL_3436521 | Berkshire and Surrey Pathology Services Lighthouse Laboratory | Wellcome Sanger Institute for the COVID-19 Genomics UK (COG-UK) Consortium | Berkshire and Surrey Pathology Services Lighthouse Laboratory and Alex Alderton, Roberto Amato, Jeffrey Barrett, Sonia Goncalves, Ewan Harrison, David K. Jackson, Ian Johnston, Dominic Kwiatkowski, Cordelia Langford, John Sillitoe on behalf of the Wellcome Sanger Institute COVID-19 Surveillance Team |
| EPI_ISL_3436522, EPI_ISL_3436523, EPI_ISL_3436524 | Lighthouse Lab in Alderley Park | Wellcome Sanger Institute for the COVID-19 Genomics UK (COG-UK) Consortium | Jacquelyn Wynn, Mairead Hyland, The Lighthouse Lab in Alderley Park and Alex Alderton, Roberto Amato, Jeffrey Barrett, Sonia Goncalves, Ewan Harrison, David K. Jackson, Ian Johnston, Dominic Kwiatkowski, Cordelia Langford, John Sillitoe on behalf of the Wellcome Sanger Institute COVID-19 Surveillance Team |
| EPI_ISL_3436525 | Lighthouse Lab in Glasgow | Wellcome Sanger Institute for the COVID-19 Genomics UK (COG-UK) Consortium | Harper VanSteenhouse, Yumi Kasai, David Gray, Carol Clugston, Anna Dominiczak and Alex Alderton, Roberto Amato, Jeffrey Barrett, Sonia Goncalves, Ewan Harrison, David K. Jackson, Ian Johnston, Dominic Kwiatkowski, Cordelia Langford, John Sillitoe on behalf of the Wellcome Sanger Institute COVID-19 Surveillance Team |
| EPI_ISL_3436526, EPI_ISL_3436527, EPI_ISL_3436528 | Berkshire and Surrey Pathology Services Lighthouse Laboratory | Wellcome Sanger Institute for the COVID-19 Genomics UK (COG-UK) Consortium | Berkshire and Surrey Pathology Services Lighthouse Laboratory and Alex Alderton, Roberto Amato, Jeffrey Barrett, Sonia Goncalves, Ewan Harrison, David K. Jackson, Ian Johnston, Dominic Kwiatkowski, Cordelia Langford, John Sillitoe on behalf of the Wellcome Sanger Institute COVID-19 Surveillance Team |

| | | | |
|---|---|---|---|
| EPI_ISL_3436529 | Lighthouse Lab in Alderley Park | Wellcome Sanger Institute for the COVID-19 Genomics UK (COG-UK) Consortium | Jacquelyn Wynn, Mairead Hyland, The Lighthouse Lab in Alderley Park and Alex Alderton, Roberto Amato, Jeffrey Barrett, Sonia Goncalves, Ewan Harrison, David K. Jackson, Ian Johnston, Dominic Kwiatkowski, Cordelia Langford, John Sillitoe on behalf of the Wellcome Sanger Institute COVID-19 Surveillance Team |
| EPI_ISL_3436530, EPI_ISL_3436531 | Berkshire and Surrey Pathology Services Lighthouse Laboratory | Wellcome Sanger Institute for the COVID-19 Genomics UK (COG-UK) Consortium | Berkshire and Surrey Pathology Services Lighthouse Laboratory and Alex Alderton, Roberto Amato, Jeffrey Barrett, Sonia Goncalves, Ewan Harrison, David K. Jackson, Ian Johnston, Dominic Kwiatkowski, Cordelia Langford, John Sillitoe on behalf of the Wellcome Sanger Institute COVID-19 Surveillance Team |
| EPI_ISL_3436532 | Lighthouse Lab in Alderley Park | Wellcome Sanger Institute for the COVID-19 Genomics UK (COG-UK) Consortium | Jacquelyn Wynn, Mairead Hyland, The Lighthouse Lab in Alderley Park and Alex Alderton, Roberto Amato, Jeffrey Barrett, Sonia Goncalves, Ewan Harrison, David K. Jackson, Ian Johnston, Dominic Kwiatkowski, Cordelia Langford, John Sillitoe on behalf of the Wellcome Sanger Institute COVID-19 Surveillance Team |
| EPI_ISL_3436533 | Lighthouse Lab in Glasgow | Wellcome Sanger Institute for the COVID-19 Genomics UK (COG-UK) Consortium | Harper VanSteenhouse, Yumi Kasai, David Gray, Carol Clugston, Anna Dominiczak and Alex Alderton, Roberto Amato, Jeffrey Barrett, Sonia Goncalves, Ewan Harrison, David K. Jackson, Ian Johnston, Dominic Kwiatkowski, Cordelia Langford, John Sillitoe on behalf of the Wellcome Sanger Institute COVID-19 Surveillance Team |
| EPI_ISL_3436534 | Berkshire and Surrey Pathology Services Lighthouse Laboratory | Wellcome Sanger Institute for the COVID-19 Genomics UK (COG-UK) Consortium | Berkshire and Surrey Pathology Services Lighthouse Laboratory and Alex Alderton, Roberto Amato, Jeffrey Barrett, Sonia Goncalves, Ewan Harrison, David K. Jackson, Ian Johnston, Dominic Kwiatkowski, Cordelia Langford, John Sillitoe on behalf of the Wellcome Sanger Institute COVID-19 Surveillance Team |
| EPI_ISL_3436535 | Lighthouse Lab in Alderley Park | Wellcome Sanger Institute for the COVID-19 Genomics UK (COG-UK) Consortium | Jacquelyn Wynn, Mairead Hyland, The Lighthouse Lab in Alderley Park and Alex Alderton, Roberto Amato, Jeffrey Barrett, Sonia Goncalves, Ewan Harrison, David K. Jackson, Ian Johnston, Dominic Kwiatkowski, Cordelia Langford, John Sillitoe on behalf of the Wellcome Sanger Institute COVID-19 Surveillance Team |
| EPI_ISL_3436536 | Berkshire and Surrey Pathology Services Lighthouse Laboratory | Wellcome Sanger Institute for the COVID-19 Genomics UK (COG-UK) Consortium | Berkshire and Surrey Pathology Services Lighthouse Laboratory and Alex Alderton, Roberto Amato, Jeffrey Barrett, Sonia Goncalves, Ewan Harrison, David K. Jackson, Ian Johnston, Dominic Kwiatkowski, Cordelia Langford, John Sillitoe on behalf of the Wellcome Sanger Institute COVID-19 Surveillance Team |
| EPI_ISL_3436537 | Lighthouse Lab in Glasgow | Wellcome Sanger Institute for the COVID-19 Genomics UK (COG-UK) Consortium | Harper VanSteenhouse, Yumi Kasai, David Gray, Carol Clugston, Anna Dominiczak and Alex Alderton, Roberto Amato, Jeffrey Barrett, Sonia Goncalves, Ewan Harrison, David K. Jackson, Ian Johnston, Dominic Kwiatkowski, Cordelia Langford, John Sillitoe on behalf of the Wellcome Sanger Institute COVID-19 Surveillance Team |
| EPI_ISL_3436538, EPI_ISL_3436539, EPI_ISL_3436540 | Berkshire and Surrey Pathology Services Lighthouse Laboratory | Wellcome Sanger Institute for the COVID-19 Genomics UK (COG-UK) Consortium | Berkshire and Surrey Pathology Services Lighthouse Laboratory and Alex Alderton, Roberto Amato, Jeffrey Barrett, Sonia Goncalves, Ewan Harrison, David K. Jackson, Ian Johnston, Dominic Kwiatkowski, Cordelia Langford, John Sillitoe on behalf of the Wellcome Sanger Institute COVID-19 Surveillance Team |
| EPI_ISL_3436541, EPI_ISL_3436542 | Lighthouse Lab in Glasgow | Wellcome Sanger Institute for the COVID-19 Genomics UK (COG-UK) Consortium | Harper VanSteenhouse, Yumi Kasai, David Gray, Carol Clugston, Anna Dominiczak and Alex Alderton, Roberto Amato, Jeffrey Barrett, Sonia Goncalves, Ewan Harrison, David K. Jackson, Ian Johnston, Dominic Kwiatkowski, Cordelia Langford, John Sillitoe on behalf of the Wellcome Sanger Institute COVID-19 Surveillance Team |
| EPI_ISL_3436543, EPI_ISL_3436544, EPI_ISL_3436545, EPI_ISL_3436546, EPI_ISL_3436547, EPI_ISL_3436548, EPI_ISL_3436549, EPI_ISL_3436550 | Berkshire and Surrey Pathology Services Lighthouse Laboratory | Wellcome Sanger Institute for the COVID-19 Genomics UK (COG-UK) Consortium | Berkshire and Surrey Pathology Services Lighthouse Laboratory and Alex Alderton, Roberto Amato, Jeffrey Barrett, Sonia Goncalves, Ewan Harrison, David K. Jackson, Ian Johnston, Dominic Kwiatkowski, Cordelia Langford, John Sillitoe on behalf of the Wellcome Sanger Institute COVID-19 Surveillance Team |
| EPI_ISL_3436551 | Lighthouse Lab in Alderley Park | Wellcome Sanger Institute for the COVID-19 Genomics UK (COG-UK) Consortium | Jacquelyn Wynn, Mairead Hyland, The Lighthouse Lab in Alderley Park and Alex Alderton, Roberto Amato, Jeffrey Barrett, Sonia Goncalves, Ewan Harrison, David K. Jackson, Ian Johnston, Dominic Kwiatkowski, Cordelia Langford, John Sillitoe on behalf of the Wellcome Sanger Institute COVID-19 Surveillance Team |
| EPI_ISL_3436552, EPI_ISL_3436553 | Berkshire and Surrey Pathology Services Lighthouse Laboratory | Wellcome Sanger Institute for the COVID-19 Genomics UK (COG-UK) Consortium | Berkshire and Surrey Pathology Services Lighthouse Laboratory and Alex Alderton, Roberto Amato, Jeffrey Barrett, Sonia Goncalves, Ewan Harrison, David K. Jackson, Ian Johnston, Dominic Kwiatkowski, Cordelia Langford, John Sillitoe on behalf of the Wellcome Sanger Institute COVID-19 Surveillance Team |
| EPI_ISL_3436554 | Lighthouse Lab in Glasgow | Wellcome Sanger Institute for the COVID-19 Genomics UK (COG-UK) Consortium | Harper VanSteenhouse, Yumi Kasai, David Gray, Carol Clugston, Anna Dominiczak and Alex Alderton, Roberto Amato, Jeffrey Barrett, Sonia Goncalves, Ewan Harrison, David K. Jackson, Ian Johnston, Dominic Kwiatkowski, Cordelia Langford, John Sillitoe on behalf of the Wellcome Sanger Institute COVID-19 Surveillance Team |
| EPI_ISL_3436555, EPI_ISL_3436556, EPI_ISL_3436557 | Berkshire and Surrey Pathology Services Lighthouse Laboratory | Wellcome Sanger Institute for the COVID-19 Genomics UK (COG-UK) Consortium | Berkshire and Surrey Pathology Services Lighthouse Laboratory and Alex Alderton, Roberto Amato, Jeffrey Barrett, Sonia Goncalves, Ewan Harrison, David K. Jackson, Ian Johnston, Dominic Kwiatkowski, Cordelia Langford, John Sillitoe on behalf of the Wellcome Sanger Institute COVID-19 Surveillance Team |
| EPI_ISL_3436558 | Lighthouse Lab in Alderley Park | Wellcome Sanger Institute for the COVID-19 Genomics UK (COG-UK) Consortium | Jacquelyn Wynn, Mairead Hyland, The Lighthouse Lab in Alderley Park and Alex Alderton, Roberto Amato, Jeffrey Barrett, Sonia Goncalves, Ewan Harrison, David K. Jackson, Ian Johnston, Dominic Kwiatkowski, Cordelia Langford, John Sillitoe on behalf of the Wellcome Sanger Institute COVID-19 Surveillance Team |
| EPI_ISL_3436559, EPI_ISL_3436560 | Berkshire and Surrey Pathology Services Lighthouse Laboratory | Wellcome Sanger Institute for the COVID-19 Genomics UK (COG-UK) Consortium | Berkshire and Surrey Pathology Services Lighthouse Laboratory and Alex Alderton, Roberto Amato, Jeffrey Barrett, Sonia Goncalves, Ewan Harrison, David K. Jackson, Ian Johnston, Dominic Kwiatkowski, Cordelia Langford, John Sillitoe on behalf of the Wellcome Sanger Institute COVID-19 Surveillance Team |
| EPI_ISL_3436561 | Lighthouse Lab in Alderley Park | Wellcome Sanger Institute for the COVID-19 Genomics UK (COG-UK) Consortium | Jacquelyn Wynn, Mairead Hyland, The Lighthouse Lab in Alderley Park and Alex Alderton, Roberto Amato, Jeffrey Barrett, Sonia Goncalves, Ewan Harrison, David K. Jackson, Ian Johnston, Dominic Kwiatkowski, Cordelia Langford, John Sillitoe on behalf of the Wellcome Sanger Institute COVID-19 Surveillance Team |
| EPI_ISL_3436562, EPI_ISL_3436563 | Berkshire and Surrey Pathology Services Lighthouse Laboratory | Wellcome Sanger Institute for the COVID-19 Genomics UK (COG-UK) Consortium | Berkshire and Surrey Pathology Services Lighthouse Laboratory and Alex Alderton, Roberto Amato, Jeffrey Barrett, Sonia Goncalves, Ewan Harrison, David K. Jackson, Ian Johnston, Dominic Kwiatkowski, Cordelia Langford, John Sillitoe on behalf of the Wellcome Sanger Institute COVID-19 Surveillance Team |
| EPI_ISL_3436564 | Lighthouse Lab in Alderley Park | Wellcome Sanger Institute for the COVID-19 Genomics UK (COG-UK) Consortium | Jacquelyn Wynn, Mairead Hyland, The Lighthouse Lab in Alderley Park and Alex Alderton, Roberto Amato, Jeffrey Barrett, Sonia Goncalves, Ewan Harrison, David K. Jackson, Ian Johnston, Dominic Kwiatkowski, Cordelia Langford, John Sillitoe on behalf of the Wellcome Sanger Institute COVID-19 Surveillance Team |
| EPI_ISL_3436565 | Lighthouse Lab in Glasgow | Wellcome Sanger Institute for the COVID-19 Genomics UK (COG-UK) Consortium | Harper VanSteenhouse, Yumi Kasai, David Gray, Carol Clugston, Anna Dominiczak and Alex Alderton, Roberto Amato, Jeffrey Barrett, Sonia Goncalves, Ewan Harrison, David K. Jackson, Ian Johnston, Dominic Kwiatkowski, Cordelia Langford, John Sillitoe on behalf of the Wellcome Sanger Institute COVID-19 Surveillance Team |
| EPI_ISL_3436566, EPI_ISL_3436567 | Berkshire and Surrey Pathology Services Lighthouse Laboratory | Wellcome Sanger Institute for the COVID-19 Genomics UK (COG-UK) Consortium | Berkshire and Surrey Pathology Services Lighthouse Laboratory and Alex Alderton, Roberto Amato, Jeffrey Barrett, Sonia Goncalves, Ewan Harrison, David K. Jackson, Ian Johnston, Dominic Kwiatkowski, Cordelia Langford, John Sillitoe on behalf of the Wellcome Sanger Institute COVID-19 Surveillance Team |
| EPI_ISL_3436568 | Lighthouse Lab in Alderley Park | Wellcome Sanger Institute for the COVID-19 Genomics UK (COG-UK) Consortium | Jacquelyn Wynn, Mairead Hyland, The Lighthouse Lab in Alderley Park and Alex Alderton, Roberto Amato, Jeffrey Barrett, Sonia Goncalves, Ewan Harrison, David K. Jackson, Ian Johnston, Dominic Kwiatkowski, Cordelia Langford, John Sillitoe on behalf of the Wellcome Sanger Institute COVID-19 Surveillance Team |
| EPI_ISL_3436569, EPI_ISL_3436570, EPI_ISL_3436571 | Berkshire and Surrey Pathology Services Lighthouse Laboratory | Wellcome Sanger Institute for the COVID-19 Genomics UK (COG-UK) Consortium | Berkshire and Surrey Pathology Services Lighthouse Laboratory and Alex Alderton, Roberto Amato, Jeffrey Barrett, Sonia Goncalves, Ewan Harrison, David K. Jackson, Ian Johnston, Dominic Kwiatkowski, Cordelia Langford, John Sillitoe on behalf of the Wellcome Sanger Institute COVID-19 Surveillance Team |
| EPI_ISL_3436572, EPI_ISL_3436573 | Lighthouse Lab in Alderley Park | Wellcome Sanger Institute for the COVID-19 Genomics UK (COG-UK) Consortium | Jacquelyn Wynn, Mairead Hyland, The Lighthouse Lab in Alderley Park and Alex Alderton, Roberto Amato, Jeffrey Barrett, Sonia Goncalves, Ewan Harrison, David K. Jackson, Ian Johnston, Dominic Kwiatkowski, Cordelia Langford, John Sillitoe on behalf of the Wellcome Sanger Institute COVID-19 Surveillance Team |
| EPI_ISL_3436574 | Berkshire and Surrey Pathology Services Lighthouse Laboratory | Wellcome Sanger Institute for the COVID-19 Genomics UK (COG-UK) Consortium | Berkshire and Surrey Pathology Services Lighthouse Laboratory and Alex Alderton, Roberto Amato, Jeffrey Barrett, Sonia Goncalves, Ewan Harrison, David K. Jackson, Ian Johnston, Dominic Kwiatkowski, Cordelia Langford, John Sillitoe on behalf of the Wellcome Sanger Institute COVID-19 Surveillance Team |
| EPI_ISL_3436575 | Lighthouse Lab in Glasgow | Wellcome Sanger Institute for the COVID-19 Genomics UK (COG-UK) Consortium | Harper VanSteenhouse, Yumi Kasai, David Gray, Carol Clugston, Anna Dominiczak and Alex Alderton, Roberto Amato, Jeffrey Barrett, Sonia Goncalves, Ewan Harrison, David K. Jackson, Ian Johnston, Dominic Kwiatkowski, Cordelia Langford, John Sillitoe on behalf of the Wellcome Sanger Institute COVID-19 Surveillance Team |
| EPI_ISL_3436576 | Berkshire and Surrey Pathology Services Lighthouse Laboratory | Wellcome Sanger Institute for the COVID-19 Genomics UK (COG-UK) Consortium | Berkshire and Surrey Pathology Services Lighthouse Laboratory and Alex Alderton, Roberto Amato, Jeffrey Barrett, Sonia Goncalves, Ewan Harrison, David K. Jackson, Ian Johnston, Dominic Kwiatkowski, Cordelia Langford, John Sillitoe on behalf of the Wellcome Sanger Institute COVID-19 Surveillance Team |
| EPI_ISL_3436577 | Lighthouse Lab in Alderley Park | Wellcome Sanger Institute for the COVID-19 Genomics UK (COG-UK) Consortium | Jacquelyn Wynn, Mairead Hyland, The Lighthouse Lab in Alderley Park and Alex Alderton, Roberto Amato, Jeffrey Barrett, Sonia Goncalves, Ewan Harrison, David K. Jackson, Ian Johnston, Dominic Kwiatkowski, Cordelia Langford, John Sillitoe on behalf of the Wellcome Sanger Institute COVID-19 Surveillance Team |
| EPI_ISL_3436578, EPI_ISL_3436579 | Lighthouse Lab in Glasgow | Wellcome Sanger Institute for the COVID-19 Genomics UK (COG-UK) Consortium | Harper VanSteenhouse, Yumi Kasai, David Gray, Carol Clugston, Anna Dominiczak and Alex Alderton, Roberto Amato, Jeffrey Barrett, Sonia Goncalves, Ewan Harrison, David K. Jackson, Ian Johnston, Dominic Kwiatkowski, Cordelia Langford, John Sillitoe on behalf of the Wellcome Sanger Institute COVID-19 Surveillance Team |
| EPI_ISL_3436580, EPI_ISL_3436581 | Berkshire and Surrey Pathology Services Lighthouse Laboratory | Wellcome Sanger Institute for the COVID-19 Genomics UK (COG-UK) Consortium | Berkshire and Surrey Pathology Services Lighthouse Laboratory and Alex Alderton, Roberto Amato, Jeffrey Barrett, Sonia Goncalves, Ewan Harrison, David K. Jackson, Ian Johnston, Dominic Kwiatkowski, Cordelia Langford, John Sillitoe on behalf of the Wellcome Sanger Institute COVID-19 Surveillance Team |
| EPI_ISL_3436582 | Lighthouse Lab in Alderley Park | Wellcome Sanger Institute for the COVID-19 Genomics UK (COG-UK) Consortium | Jacquelyn Wynn, Mairead Hyland, The Lighthouse Lab in Alderley Park and Alex Alderton, Roberto Amato, Jeffrey Barrett, Sonia Goncalves, Ewan Harrison, David K. Jackson, Ian Johnston, Dominic Kwiatkowski, Cordelia Langford, John Sillitoe on behalf of the Wellcome Sanger Institute COVID-19 Surveillance Team |
| EPI_ISL_3436583, EPI_ISL_3436584 | Berkshire and Surrey Pathology Services Lighthouse Laboratory | Wellcome Sanger Institute for the COVID-19 Genomics UK (COG-UK) Consortium | Berkshire and Surrey Pathology Services Lighthouse Laboratory and Alex Alderton, Roberto Amato, Jeffrey Barrett, Sonia Goncalves, Ewan Harrison, David K. Jackson, Ian Johnston, Dominic Kwiatkowski, Cordelia Langford, John Sillitoe on behalf of the Wellcome Sanger Institute COVID-19 Surveillance Team |
| EPI_ISL_3436585 | Lighthouse Lab in Alderley Park | Wellcome Sanger Institute for the COVID-19 Genomics UK (COG-UK) Consortium | Jacquelyn Wynn, Mairead Hyland, The Lighthouse Lab in Alderley Park and Alex Alderton, Roberto Amato, Jeffrey Barrett, Sonia Goncalves, Ewan Harrison, David K. Jackson, Ian Johnston, Dominic Kwiatkowski, Cordelia Langford, John Sillitoe on behalf of the Wellcome Sanger Institute COVID-19 Surveillance Team |
| EPI_ISL_3436586, EPI_ISL_3436587 | Berkshire and Surrey Pathology Services Lighthouse Laboratory | Wellcome Sanger Institute for the COVID-19 Genomics UK (COG-UK) Consortium | Berkshire and Surrey Pathology Services Lighthouse Laboratory and Alex Alderton, Roberto Amato, Jeffrey Barrett, Sonia Goncalves, Ewan Harrison, David K. Jackson, Ian Johnston, Dominic Kwiatkowski, Cordelia Langford, John Sillitoe on behalf of the Wellcome Sanger Institute COVID-19 Surveillance Team |
| EPI_ISL_3436588, EPI_ISL_3436589 | Lighthouse Lab in Alderley Park | Wellcome Sanger Institute for the COVID-19 Genomics UK (COG-UK) Consortium | Jacquelyn Wynn, Mairead Hyland, The Lighthouse Lab in Alderley Park and Alex Alderton, Roberto Amato, Jeffrey Barrett, Sonia Goncalves, Ewan Harrison, David K. Jackson, Ian Johnston, Dominic Kwiatkowski, Cordelia Langford, John Sillitoe on behalf of the Wellcome Sanger Institute COVID-19 Surveillance Team |
| EPI_ISL_3436590, EPI_ISL_3436591 | Berkshire and Surrey Pathology Services Lighthouse Laboratory | Wellcome Sanger Institute for the COVID-19 Genomics UK (COG-UK) Consortium | Berkshire and Surrey Pathology Services Lighthouse Laboratory and Alex Alderton, Roberto Amato, Jeffrey Barrett, Sonia Goncalves, Ewan Harrison, David K. Jackson, Ian Johnston, Dominic Kwiatkowski, Cordelia Langford, John Sillitoe on behalf of the Wellcome Sanger Institute COVID-19 Surveillance Team |
| EPI_ISL_3436592 | Lighthouse Lab in Glasgow | Wellcome Sanger Institute for the COVID-19 Genomics UK (COG-UK) Consortium | Harper VanSteenhouse, Yumi Kasai, David Gray, Carol Clugston, Anna Dominiczak and Alex Alderton, Roberto Amato, Jeffrey Barrett, Sonia Goncalves, Ewan Harrison, David K. Jackson, Ian Johnston, Dominic Kwiatkowski, Cordelia Langford, John Sillitoe on behalf of the Wellcome Sanger Institute COVID-19 Surveillance Team |
| EPI_ISL_3436593 | Berkshire and Surrey Pathology Services Lighthouse Laboratory | Wellcome Sanger Institute for the COVID-19 Genomics UK (COG-UK) Consortium | Berkshire and Surrey Pathology Services Lighthouse Laboratory and Alex Alderton, Roberto Amato, Jeffrey Barrett, Sonia Goncalves, Ewan Harrison, David K. Jackson, Ian Johnston, Dominic Kwiatkowski, Cordelia Langford, John Sillitoe on behalf of the Wellcome Sanger Institute COVID-19 Surveillance Team |
| EPI_ISL_3436594 | Lighthouse Lab in Alderley Park | Wellcome Sanger Institute for the COVID-19 Genomics UK (COG-UK) Consortium | Jacquelyn Wynn, Mairead Hyland, The Lighthouse Lab in Alderley Park and Alex Alderton, Roberto Amato, Jeffrey Barrett, Sonia Goncalves, Ewan Harrison, David K. Jackson, Ian Johnston, Dominic Kwiatkowski, Cordelia Langford, John Sillitoe on behalf of the Wellcome Sanger Institute COVID-19 Surveillance Team |
| EPI_ISL_3436595, EPI_ISL_3436596 | Berkshire and Surrey Pathology Services Lighthouse Laboratory | Wellcome Sanger Institute for the COVID-19 Genomics UK (COG-UK) Consortium | Berkshire and Surrey Pathology Services Lighthouse Laboratory and Alex Alderton, Roberto Amato, Jeffrey Barrett, Sonia Goncalves, Ewan Harrison, David K. Jackson, Ian Johnston, Dominic Kwiatkowski, Cordelia Langford, John Sillitoe on behalf of the Wellcome Sanger Institute COVID-19 Surveillance Team |
| EPI_ISL_3436597 | Lighthouse Lab in Alderley Park | Wellcome Sanger Institute for the COVID-19 Genomics UK (COG-UK) | Jacquelyn Wynn, Mairead Hyland, The Lighthouse Lab in Alderley Park and Alex Alderton, Roberto Amato, Jeffrey Barrett, Sonia Goncalves, Ewan Harrison, David K. Jackson, Ian Johnston, Dominic Kwiatkowski, Cordelia Langford, John Sillitoe on |

| | | | |
|---|---|---|---|
| | | Consortium | behalf of the Wellcome Sanger Institute COVID-19 Surveillance Team |
| EPI_ISL_3436598, EPI_ISL_3436599 | Berkshire and Surrey Pathology Services Lighthouse Laboratory | Wellcome Sanger Institute for the COVID-19 Genomics UK (COG-UK) Consortium | Berkshire and Surrey Pathology Services Lighthouse Laboratory and Alex Alderton, Roberto Amato, Jeffrey Barrett, Sonia Goncalves, Ewan Harrison, David K. Jackson, Ian Johnston, Dominic Kwiatkowski, Cordelia Langford, John Sillitoe on behalf of the Wellcome Sanger Institute COVID-19 Surveillance Team |
| EPI_ISL_3436600 | Lighthouse Lab in Alderley Park | Wellcome Sanger Institute for the COVID-19 Genomics UK (COG-UK) Consortium | Jacquelyn Wynn, Mairead Hyland, The Lighthouse Lab in Alderley Park and Alex Alderton, Roberto Amato, Jeffrey Barrett, Sonia Goncalves, Ewan Harrison, David K. Jackson, Ian Johnston, Dominic Kwiatkowski, Cordelia Langford, John Sillitoe on behalf of the Wellcome Sanger Institute COVID-19 Surveillance Team |
| EPI_ISL_3436601, EPI_ISL_3436602 | Berkshire and Surrey Pathology Services Lighthouse Laboratory | Wellcome Sanger Institute for the COVID-19 Genomics UK (COG-UK) Consortium | Berkshire and Surrey Pathology Services Lighthouse Laboratory and Alex Alderton, Roberto Amato, Jeffrey Barrett, Sonia Goncalves, Ewan Harrison, David K. Jackson, Ian Johnston, Dominic Kwiatkowski, Cordelia Langford, John Sillitoe on behalf of the Wellcome Sanger Institute COVID-19 Surveillance Team |
| EPI_ISL_3436603 | Lighthouse Lab in Alderley Park | Wellcome Sanger Institute for the COVID-19 Genomics UK (COG-UK) Consortium | Jacquelyn Wynn, Mairead Hyland, The Lighthouse Lab in Alderley Park and Alex Alderton, Roberto Amato, Jeffrey Barrett, Sonia Goncalves, Ewan Harrison, David K. Jackson, Ian Johnston, Dominic Kwiatkowski, Cordelia Langford, John Sillitoe on behalf of the Wellcome Sanger Institute COVID-19 Surveillance Team |
| EPI_ISL_3436604 | Berkshire and Surrey Pathology Services Lighthouse Laboratory | Wellcome Sanger Institute for the COVID-19 Genomics UK (COG-UK) Consortium | Berkshire and Surrey Pathology Services Lighthouse Laboratory and Alex Alderton, Roberto Amato, Jeffrey Barrett, Sonia Goncalves, Ewan Harrison, David K. Jackson, Ian Johnston, Dominic Kwiatkowski, Cordelia Langford, John Sillitoe on behalf of the Wellcome Sanger Institute COVID-19 Surveillance Team |
| EPI_ISL_3436605, EPI_ISL_3436606 | Lighthouse Lab in Alderley Park | Wellcome Sanger Institute for the COVID-19 Genomics UK (COG-UK) Consortium | Jacquelyn Wynn, Mairead Hyland, The Lighthouse Lab in Alderley Park and Alex Alderton, Roberto Amato, Jeffrey Barrett, Sonia Goncalves, Ewan Harrison, David K. Jackson, Ian Johnston, Dominic Kwiatkowski, Cordelia Langford, John Sillitoe on behalf of the Wellcome Sanger Institute COVID-19 Surveillance Team |
| EPI_ISL_3436607, EPI_ISL_3436608, EPI_ISL_3436609 | Berkshire and Surrey Pathology Services Lighthouse Laboratory | Wellcome Sanger Institute for the COVID-19 Genomics UK (COG-UK) Consortium | Berkshire and Surrey Pathology Services Lighthouse Laboratory and Alex Alderton, Roberto Amato, Jeffrey Barrett, Sonia Goncalves, Ewan Harrison, David K. Jackson, Ian Johnston, Dominic Kwiatkowski, Cordelia Langford, John Sillitoe on behalf of the Wellcome Sanger Institute COVID-19 Surveillance Team |
| EPI_ISL_3436610 | Lighthouse Lab in Glasgow | Wellcome Sanger Institute for the COVID-19 Genomics UK (COG-UK) Consortium | Harper VanSteenhouse, Yumi Kasai, David Gray, Carol Clugston, Anna Dominiczak and Alex Alderton, Roberto Amato, Jeffrey Barrett, Sonia Goncalves, Ewan Harrison, David K. Jackson, Ian Johnston, Dominic Kwiatkowski, Cordelia Langford, John Sillitoe on behalf of the Wellcome Sanger Institute COVID-19 Surveillance Team |
| EPI_ISL_3436611 | Berkshire and Surrey Pathology Services Lighthouse Laboratory | Wellcome Sanger Institute for the COVID-19 Genomics UK (COG-UK) Consortium | Berkshire and Surrey Pathology Services Lighthouse Laboratory and Alex Alderton, Roberto Amato, Jeffrey Barrett, Sonia Goncalves, Ewan Harrison, David K. Jackson, Ian Johnston, Dominic Kwiatkowski, Cordelia Langford, John Sillitoe on behalf of the Wellcome Sanger Institute COVID-19 Surveillance Team |
| EPI_ISL_3436612 | Lighthouse Lab in Glasgow | Wellcome Sanger Institute for the COVID-19 Genomics UK (COG-UK) Consortium | Harper VanSteenhouse, Yumi Kasai, David Gray, Carol Clugston, Anna Dominiczak and Alex Alderton, Roberto Amato, Jeffrey Barrett, Sonia Goncalves, Ewan Harrison, David K. Jackson, Ian Johnston, Dominic Kwiatkowski, Cordelia Langford, John Sillitoe on behalf of the Wellcome Sanger Institute COVID-19 Surveillance Team |
| EPI_ISL_3436613 | Berkshire and Surrey Pathology Services Lighthouse Laboratory | Wellcome Sanger Institute for the COVID-19 Genomics UK (COG-UK) Consortium | Berkshire and Surrey Pathology Services Lighthouse Laboratory and Alex Alderton, Roberto Amato, Jeffrey Barrett, Sonia Goncalves, Ewan Harrison, David K. Jackson, Ian Johnston, Dominic Kwiatkowski, Cordelia Langford, John Sillitoe on behalf of the Wellcome Sanger Institute COVID-19 Surveillance Team |
| EPI_ISL_3436614 | Lighthouse Lab in Alderley Park | Wellcome Sanger Institute for the COVID-19 Genomics UK (COG-UK) Consortium | Jacquelyn Wynn, Mairead Hyland, The Lighthouse Lab in Alderley Park and Alex Alderton, Roberto Amato, Jeffrey Barrett, Sonia Goncalves, Ewan Harrison, David K. Jackson, Ian Johnston, Dominic Kwiatkowski, Cordelia Langford, John Sillitoe on behalf of the Wellcome Sanger Institute COVID-19 Surveillance Team |
| EPI_ISL_3436615 | Berkshire and Surrey Pathology Services Lighthouse Laboratory | Wellcome Sanger Institute for the COVID-19 Genomics UK (COG-UK) Consortium | Berkshire and Surrey Pathology Services Lighthouse Laboratory and Alex Alderton, Roberto Amato, Jeffrey Barrett, Sonia Goncalves, Ewan Harrison, David K. Jackson, Ian Johnston, Dominic Kwiatkowski, Cordelia Langford, John Sillitoe on behalf of the Wellcome Sanger Institute COVID-19 Surveillance Team |
| EPI_ISL_3436616 | Lighthouse Lab in Alderley Park | Wellcome Sanger Institute for the COVID-19 Genomics UK (COG-UK) Consortium | Jacquelyn Wynn, Mairead Hyland, The Lighthouse Lab in Alderley Park and Alex Alderton, Roberto Amato, Jeffrey Barrett, Sonia Goncalves, Ewan Harrison, David K. Jackson, Ian Johnston, Dominic Kwiatkowski, Cordelia Langford, John Sillitoe on behalf of the Wellcome Sanger Institute COVID-19 Surveillance Team |
| EPI_ISL_3436617, EPI_ISL_3436618 | Berkshire and Surrey Pathology Services Lighthouse Laboratory | Wellcome Sanger Institute for the COVID-19 Genomics UK (COG-UK) Consortium | Berkshire and Surrey Pathology Services Lighthouse Laboratory and Alex Alderton, Roberto Amato, Jeffrey Barrett, Sonia Goncalves, Ewan Harrison, David K. Jackson, Ian Johnston, Dominic Kwiatkowski, Cordelia Langford, John Sillitoe on behalf of the Wellcome Sanger Institute COVID-19 Surveillance Team |
| EPI_ISL_3436619 | Lighthouse Lab in Alderley Park | Wellcome Sanger Institute for the COVID-19 Genomics UK (COG-UK) Consortium | Jacquelyn Wynn, Mairead Hyland, The Lighthouse Lab in Alderley Park and Alex Alderton, Roberto Amato, Jeffrey Barrett, Sonia Goncalves, Ewan Harrison, David K. Jackson, Ian Johnston, Dominic Kwiatkowski, Cordelia Langford, John Sillitoe on behalf of the Wellcome Sanger Institute COVID-19 Surveillance Team |
| EPI_ISL_3436620 | Berkshire and Surrey Pathology Services Lighthouse Laboratory | Wellcome Sanger Institute for the COVID-19 Genomics UK (COG-UK) Consortium | Berkshire and Surrey Pathology Services Lighthouse Laboratory and Alex Alderton, Roberto Amato, Jeffrey Barrett, Sonia Goncalves, Ewan Harrison, David K. Jackson, Ian Johnston, Dominic Kwiatkowski, Cordelia Langford, John Sillitoe on behalf of the Wellcome Sanger Institute COVID-19 Surveillance Team |
| EPI_ISL_3436621 | Lighthouse Lab in Glasgow | Wellcome Sanger Institute for the COVID-19 Genomics UK (COG-UK) Consortium | Harper VanSteenhouse, Yumi Kasai, David Gray, Carol Clugston, Anna Dominiczak and Alex Alderton, Roberto Amato, Jeffrey Barrett, Sonia Goncalves, Ewan Harrison, David K. Jackson, Ian Johnston, Dominic Kwiatkowski, Cordelia Langford, John Sillitoe on behalf of the Wellcome Sanger Institute COVID-19 Surveillance Team |
| EPI_ISL_3436622 | Berkshire and Surrey Pathology Services Lighthouse Laboratory | Wellcome Sanger Institute for the COVID-19 Genomics UK (COG-UK) Consortium | Berkshire and Surrey Pathology Services Lighthouse Laboratory and Alex Alderton, Roberto Amato, Jeffrey Barrett, Sonia Goncalves, Ewan Harrison, David K. Jackson, Ian Johnston, Dominic Kwiatkowski, Cordelia Langford, John Sillitoe on behalf of the Wellcome Sanger Institute COVID-19 Surveillance Team |
| EPI_ISL_3436623, EPI_ISL_3436624 | Lighthouse Lab in Glasgow | Wellcome Sanger Institute for the COVID-19 Genomics UK (COG-UK) Consortium | Harper VanSteenhouse, Yumi Kasai, David Gray, Carol Clugston, Anna Dominiczak and Alex Alderton, Roberto Amato, Jeffrey Barrett, Sonia Goncalves, Ewan Harrison, David K. Jackson, Ian Johnston, Dominic Kwiatkowski, Cordelia Langford, John Sillitoe on behalf of the Wellcome Sanger Institute COVID-19 Surveillance Team |
| EPI_ISL_3436625 | Berkshire and Surrey Pathology Services Lighthouse Laboratory | Wellcome Sanger Institute for the COVID-19 Genomics UK (COG-UK) Consortium | Berkshire and Surrey Pathology Services Lighthouse Laboratory and Alex Alderton, Roberto Amato, Jeffrey Barrett, Sonia Goncalves, Ewan Harrison, David K. Jackson, Ian Johnston, Dominic Kwiatkowski, Cordelia Langford, John Sillitoe on behalf of the Wellcome Sanger Institute COVID-19 Surveillance Team |
| EPI_ISL_3436626 | Lighthouse Lab in Alderley Park | Wellcome Sanger Institute for the COVID-19 Genomics UK (COG-UK) Consortium | Jacquelyn Wynn, Mairead Hyland, The Lighthouse Lab in Alderley Park and Alex Alderton, Roberto Amato, Jeffrey Barrett, Sonia Goncalves, Ewan Harrison, David K. Jackson, Ian Johnston, Dominic Kwiatkowski, Cordelia Langford, John Sillitoe on behalf of the Wellcome Sanger Institute COVID-19 Surveillance Team |
| EPI_ISL_3436627 | Berkshire and Surrey Pathology Services Lighthouse Laboratory | Wellcome Sanger Institute for the COVID-19 Genomics UK (COG-UK) Consortium | Berkshire and Surrey Pathology Services Lighthouse Laboratory and Alex Alderton, Roberto Amato, Jeffrey Barrett, Sonia Goncalves, Ewan Harrison, David K. Jackson, Ian Johnston, Dominic Kwiatkowski, Cordelia Langford, John Sillitoe on behalf of the Wellcome Sanger Institute COVID-19 Surveillance Team |
| EPI_ISL_3436628 | Lighthouse Lab in Glasgow | Wellcome Sanger Institute for the COVID-19 Genomics UK (COG-UK) Consortium | Harper VanSteenhouse, Yumi Kasai, David Gray, Carol Clugston, Anna Dominiczak and Alex Alderton, Roberto Amato, Jeffrey Barrett, Sonia Goncalves, Ewan Harrison, David K. Jackson, Ian Johnston, Dominic Kwiatkowski, Cordelia Langford, John Sillitoe on behalf of the Wellcome Sanger Institute COVID-19 Surveillance Team |
| EPI_ISL_3436629, EPI_ISL_3436630 | Berkshire and Surrey Pathology Services Lighthouse Laboratory | Wellcome Sanger Institute for the COVID-19 Genomics UK (COG-UK) Consortium | Berkshire and Surrey Pathology Services Lighthouse Laboratory and Alex Alderton, Roberto Amato, Jeffrey Barrett, Sonia Goncalves, Ewan Harrison, David K. Jackson, Ian Johnston, Dominic Kwiatkowski, Cordelia Langford, John Sillitoe on behalf of the Wellcome Sanger Institute COVID-19 Surveillance Team |
| EPI_ISL_3436631 | Lighthouse Lab in Alderley Park | Wellcome Sanger Institute for the COVID-19 Genomics UK (COG-UK) Consortium | Jacquelyn Wynn, Mairead Hyland, The Lighthouse Lab in Alderley Park and Alex Alderton, Roberto Amato, Jeffrey Barrett, Sonia Goncalves, Ewan Harrison, David K. Jackson, Ian Johnston, Dominic Kwiatkowski, Cordelia Langford, John Sillitoe on behalf of the Wellcome Sanger Institute COVID-19 Surveillance Team |
| EPI_ISL_3436632 | Berkshire and Surrey Pathology Services Lighthouse Laboratory | Wellcome Sanger Institute for the COVID-19 Genomics UK (COG-UK) Consortium | Berkshire and Surrey Pathology Services Lighthouse Laboratory and Alex Alderton, Roberto Amato, Jeffrey Barrett, Sonia Goncalves, Ewan Harrison, David K. Jackson, Ian Johnston, Dominic Kwiatkowski, Cordelia Langford, John Sillitoe on behalf of the Wellcome Sanger Institute COVID-19 Surveillance Team |
| EPI_ISL_3436633 | Lighthouse Lab in Glasgow | Wellcome Sanger Institute for the COVID-19 Genomics UK (COG-UK) Consortium | Harper VanSteenhouse, Yumi Kasai, David Gray, Carol Clugston, Anna Dominiczak and Alex Alderton, Roberto Amato, Jeffrey Barrett, Sonia Goncalves, Ewan Harrison, David K. Jackson, Ian Johnston, Dominic Kwiatkowski, Cordelia Langford, John Sillitoe on behalf of the Wellcome Sanger Institute COVID-19 Surveillance Team |
| EPI_ISL_3436634 | Lighthouse Lab in Alderley Park | Wellcome Sanger Institute for the COVID-19 Genomics UK (COG-UK) Consortium | Jacquelyn Wynn, Mairead Hyland, The Lighthouse Lab in Alderley Park and Alex Alderton, Roberto Amato, Jeffrey Barrett, Sonia Goncalves, Ewan Harrison, David K. Jackson, Ian Johnston, Dominic Kwiatkowski, Cordelia Langford, John Sillitoe on behalf of the Wellcome Sanger Institute COVID-19 Surveillance Team |
| EPI_ISL_3436635 | Berkshire and Surrey Pathology Services Lighthouse Laboratory | Wellcome Sanger Institute for the COVID-19 Genomics UK (COG-UK) Consortium | Berkshire and Surrey Pathology Services Lighthouse Laboratory and Alex Alderton, Roberto Amato, Jeffrey Barrett, Sonia Goncalves, Ewan Harrison, David K. Jackson, Ian Johnston, Dominic Kwiatkowski, Cordelia Langford, John Sillitoe on behalf of the Wellcome Sanger Institute COVID-19 Surveillance Team |
| EPI_ISL_3436636 | Lighthouse Lab in Alderley Park | Wellcome Sanger Institute for the COVID-19 Genomics UK (COG-UK) Consortium | Jacquelyn Wynn, Mairead Hyland, The Lighthouse Lab in Alderley Park and Alex Alderton, Roberto Amato, Jeffrey Barrett, Sonia Goncalves, Ewan Harrison, David K. Jackson, Ian Johnston, Dominic Kwiatkowski, Cordelia Langford, John Sillitoe on behalf of the Wellcome Sanger Institute COVID-19 Surveillance Team |
| EPI_ISL_3436637 | Lighthouse Lab in Glasgow | Wellcome Sanger Institute for the COVID-19 Genomics UK (COG-UK) Consortium | Harper VanSteenhouse, Yumi Kasai, David Gray, Carol Clugston, Anna Dominiczak and Alex Alderton, Roberto Amato, Jeffrey Barrett, Sonia Goncalves, Ewan Harrison, David K. Jackson, Ian Johnston, Dominic Kwiatkowski, Cordelia Langford, John Sillitoe on behalf of the Wellcome Sanger Institute COVID-19 Surveillance Team |
| EPI_ISL_3436638 | Berkshire and Surrey Pathology Services Lighthouse Laboratory | Wellcome Sanger Institute for the COVID-19 Genomics UK (COG-UK) Consortium | Berkshire and Surrey Pathology Services Lighthouse Laboratory and Alex Alderton, Roberto Amato, Jeffrey Barrett, Sonia Goncalves, Ewan Harrison, David K. Jackson, Ian Johnston, Dominic Kwiatkowski, Cordelia Langford, John Sillitoe on behalf of the Wellcome Sanger Institute COVID-19 Surveillance Team |
| EPI_ISL_3436639 | Lighthouse Lab in Alderley Park | Wellcome Sanger Institute for the COVID-19 Genomics UK (COG-UK) Consortium | Jacquelyn Wynn, Mairead Hyland, The Lighthouse Lab in Alderley Park and Alex Alderton, Roberto Amato, Jeffrey Barrett, Sonia Goncalves, Ewan Harrison, David K. Jackson, Ian Johnston, Dominic Kwiatkowski, Cordelia Langford, John Sillitoe on behalf of the Wellcome Sanger Institute COVID-19 Surveillance Team |
| EPI_ISL_3436640, EPI_ISL_3436641 | Berkshire and Surrey Pathology Services Lighthouse Laboratory | Wellcome Sanger Institute for the COVID-19 Genomics UK (COG-UK) Consortium | Berkshire and Surrey Pathology Services Lighthouse Laboratory and Alex Alderton, Roberto Amato, Jeffrey Barrett, Sonia Goncalves, Ewan Harrison, David K. Jackson, Ian Johnston, Dominic Kwiatkowski, Cordelia Langford, John Sillitoe on behalf of the Wellcome Sanger Institute COVID-19 Surveillance Team |
| EPI_ISL_3436642, EPI_ISL_3436643 | Lighthouse Lab in Alderley Park | Wellcome Sanger Institute for the COVID-19 Genomics UK (COG-UK) Consortium | Jacquelyn Wynn, Mairead Hyland, The Lighthouse Lab in Alderley Park and Alex Alderton, Roberto Amato, Jeffrey Barrett, Sonia Goncalves, Ewan Harrison, David K. Jackson, Ian Johnston, Dominic Kwiatkowski, Cordelia Langford, John Sillitoe on behalf of the Wellcome Sanger Institute COVID-19 Surveillance Team |
| EPI_ISL_3436644, EPI_ISL_3436645, EPI_ISL_3436646 | Berkshire and Surrey Pathology Services Lighthouse Laboratory | Wellcome Sanger Institute for the COVID-19 Genomics UK (COG-UK) Consortium | Berkshire and Surrey Pathology Services Lighthouse Laboratory and Alex Alderton, Roberto Amato, Jeffrey Barrett, Sonia Goncalves, Ewan Harrison, David K. Jackson, Ian Johnston, Dominic Kwiatkowski, Cordelia Langford, John Sillitoe on behalf of the Wellcome Sanger Institute COVID-19 Surveillance Team |
| EPI_ISL_3436647, EPI_ISL_3436648, EPI_ISL_3436649 | Lighthouse Lab in Alderley Park | Wellcome Sanger Institute for the COVID-19 Genomics UK (COG-UK) Consortium | Jacquelyn Wynn, Mairead Hyland, The Lighthouse Lab in Alderley Park and Alex Alderton, Roberto Amato, Jeffrey Barrett, Sonia Goncalves, Ewan Harrison, David K. Jackson, Ian Johnston, Dominic Kwiatkowski, Cordelia Langford, John Sillitoe on behalf of the Wellcome Sanger Institute COVID-19 Surveillance Team |
| EPI_ISL_3436650 | Lighthouse Lab in Glasgow | Wellcome Sanger Institute for the COVID-19 Genomics UK (COG-UK) Consortium | Harper VanSteenhouse, Yumi Kasai, David Gray, Carol Clugston, Anna Dominiczak and Alex Alderton, Roberto Amato, Jeffrey Barrett, Sonia Goncalves, Ewan Harrison, David K. Jackson, Ian Johnston, Dominic Kwiatkowski, Cordelia Langford, John Sillitoe on behalf of the Wellcome Sanger Institute COVID-19 Surveillance Team |
| EPI_ISL_3436651, EPI_ISL_3436652 | Berkshire and Surrey Pathology Services Lighthouse Laboratory | Wellcome Sanger Institute for the COVID-19 Genomics UK (COG-UK) Consortium | Berkshire and Surrey Pathology Services Lighthouse Laboratory and Alex Alderton, Roberto Amato, Jeffrey Barrett, Sonia Goncalves, Ewan Harrison, David K. Jackson, Ian Johnston, Dominic Kwiatkowski, Cordelia Langford, John Sillitoe on behalf of the Wellcome Sanger Institute COVID-19 Surveillance Team |
| EPI_ISL_3436653 | Lighthouse Lab in Glasgow | Wellcome Sanger Institute for the COVID-19 Genomics UK (COG-UK) Consortium | Harper VanSteenhouse, Yumi Kasai, David Gray, Carol Clugston, Anna Dominiczak and Alex Alderton, Roberto Amato, Jeffrey Barrett, Sonia Goncalves, Ewan Harrison, David K. Jackson, Ian Johnston, Dominic Kwiatkowski, Cordelia Langford, John Sillitoe on behalf of the Wellcome Sanger Institute COVID-19 Surveillance Team |
| EPI_ISL_3436654, EPI_ISL_3436655 | Lighthouse Lab in Alderley Park | Wellcome Sanger Institute for the COVID-19 Genomics UK (COG-UK) Consortium | Jacquelyn Wynn, Mairead Hyland, The Lighthouse Lab in Alderley Park and Alex Alderton, Roberto Amato, Jeffrey Barrett, Sonia Goncalves, Ewan Harrison, David K. Jackson, Ian Johnston, Dominic Kwiatkowski, Cordelia Langford, John Sillitoe on behalf of the Wellcome Sanger Institute COVID-19 Surveillance Team |
| EPI_ISL_3436656, EPI_ISL_3436657, EPI_ISL_3436658 | Berkshire and Surrey Pathology Services Lighthouse Laboratory | Wellcome Sanger Institute for the COVID-19 Genomics UK (COG-UK) Consortium | Berkshire and Surrey Pathology Services Lighthouse Laboratory and Alex Alderton, Roberto Amato, Jeffrey Barrett, Sonia Goncalves, Ewan Harrison, David K. Jackson, Ian Johnston, Dominic Kwiatkowski, Cordelia Langford, John Sillitoe on behalf of the Wellcome Sanger Institute COVID-19 Surveillance Team |

| | | | |
|---|---|---|---|
| EPI_ISL_3436659 | Lighthouse Lab in Glasgow | Wellcome Sanger Institute for the COVID-19 Genomics UK (COG-UK) Consortium | Harper VanSteenhouse, Yumi Kasai, David Gray, Carol Clugston, Anna Dominiczak and Alex Alderton, Roberto Amato, Jeffrey Barrett, Sonia Goncalves, Ewan Harrison, David K. Jackson, Ian Johnston, Dominic Kwiatkowski, Cordelia Langford, John Sillitoe on behalf of the Wellcome Sanger Institute COVID-19 Surveillance Team |
| EPI_ISL_3436660 | Lighthouse Lab in Alderley Park | Wellcome Sanger Institute for the COVID-19 Genomics UK (COG-UK) Consortium | Jacquelyn Wynn, Mairead Hyland, The Lighthouse Lab in Alderley Park and Alex Alderton, Roberto Amato, Jeffrey Barrett, Sonia Goncalves, Ewan Harrison, David K. Jackson, Ian Johnston, Dominic Kwiatkowski, Cordelia Langford, John Sillitoe on behalf of the Wellcome Sanger Institute COVID-19 Surveillance Team |
| EPI_ISL_3436661, EPI_ISL_3436662 | Berkshire and Surrey Pathology Services Lighthouse Laboratory | Wellcome Sanger Institute for the COVID-19 Genomics UK (COG-UK) Consortium | Berkshire and Surrey Pathology Services Lighthouse Laboratory and Alex Alderton, Roberto Amato, Jeffrey Barrett, Sonia Goncalves, Ewan Harrison, David K. Jackson, Ian Johnston, Dominic Kwiatkowski, Cordelia Langford, John Sillitoe on behalf of the Wellcome Sanger Institute COVID-19 Surveillance Team |
| EPI_ISL_3436663 | Lighthouse Lab in Alderley Park | Wellcome Sanger Institute for the COVID-19 Genomics UK (COG-UK) Consortium | Jacquelyn Wynn, Mairead Hyland, The Lighthouse Lab in Alderley Park and Alex Alderton, Roberto Amato, Jeffrey Barrett, Sonia Goncalves, Ewan Harrison, David K. Jackson, Ian Johnston, Dominic Kwiatkowski, Cordelia Langford, John Sillitoe on behalf of the Wellcome Sanger Institute COVID-19 Surveillance Team |
| EPI_ISL_3436664, EPI_ISL_3436665 | Lighthouse Lab in Glasgow | Wellcome Sanger Institute for the COVID-19 Genomics UK (COG-UK) Consortium | Harper VanSteenhouse, Yumi Kasai, David Gray, Carol Clugston, Anna Dominiczak and Alex Alderton, Roberto Amato, Jeffrey Barrett, Sonia Goncalves, Ewan Harrison, David K. Jackson, Ian Johnston, Dominic Kwiatkowski, Cordelia Langford, John Sillitoe on behalf of the Wellcome Sanger Institute COVID-19 Surveillance Team |
| EPI_ISL_3436666, EPI_ISL_3436667, EPI_ISL_3436668, EPI_ISL_3436669 | Berkshire and Surrey Pathology Services Lighthouse Laboratory | Wellcome Sanger Institute for the COVID-19 Genomics UK (COG-UK) Consortium | Berkshire and Surrey Pathology Services Lighthouse Laboratory and Alex Alderton, Roberto Amato, Jeffrey Barrett, Sonia Goncalves, Ewan Harrison, David K. Jackson, Ian Johnston, Dominic Kwiatkowski, Cordelia Langford, John Sillitoe on behalf of the Wellcome Sanger Institute COVID-19 Surveillance Team |
| EPI_ISL_3436670, EPI_ISL_3436671 | Lighthouse Lab in Glasgow | Wellcome Sanger Institute for the COVID-19 Genomics UK (COG-UK) Consortium | Harper VanSteenhouse, Yumi Kasai, David Gray, Carol Clugston, Anna Dominiczak and Alex Alderton, Roberto Amato, Jeffrey Barrett, Sonia Goncalves, Ewan Harrison, David K. Jackson, Ian Johnston, Dominic Kwiatkowski, Cordelia Langford, John Sillitoe on behalf of the Wellcome Sanger Institute COVID-19 Surveillance Team |
| EPI_ISL_3436672 | Lighthouse Lab in Alderley Park | Wellcome Sanger Institute for the COVID-19 Genomics UK (COG-UK) Consortium | Jacquelyn Wynn, Mairead Hyland, The Lighthouse Lab in Alderley Park and Alex Alderton, Roberto Amato, Jeffrey Barrett, Sonia Goncalves, Ewan Harrison, David K. Jackson, Ian Johnston, Dominic Kwiatkowski, Cordelia Langford, John Sillitoe on behalf of the Wellcome Sanger Institute COVID-19 Surveillance Team |
| EPI_ISL_3436673 | Berkshire and Surrey Pathology Services Lighthouse Laboratory | Wellcome Sanger Institute for the COVID-19 Genomics UK (COG-UK) Consortium | Berkshire and Surrey Pathology Services Lighthouse Laboratory and Alex Alderton, Roberto Amato, Jeffrey Barrett, Sonia Goncalves, Ewan Harrison, David K. Jackson, Ian Johnston, Dominic Kwiatkowski, Cordelia Langford, John Sillitoe on behalf of the Wellcome Sanger Institute COVID-19 Surveillance Team |
| EPI_ISL_3436674 | Lighthouse Lab in Alderley Park | Wellcome Sanger Institute for the COVID-19 Genomics UK (COG-UK) Consortium | Jacquelyn Wynn, Mairead Hyland, The Lighthouse Lab in Alderley Park and Alex Alderton, Roberto Amato, Jeffrey Barrett, Sonia Goncalves, Ewan Harrison, David K. Jackson, Ian Johnston, Dominic Kwiatkowski, Cordelia Langford, John Sillitoe on behalf of the Wellcome Sanger Institute COVID-19 Surveillance Team |
| EPI_ISL_3436675, EPI_ISL_3436676 | Berkshire and Surrey Pathology Services Lighthouse Laboratory | Wellcome Sanger Institute for the COVID-19 Genomics UK (COG-UK) Consortium | Berkshire and Surrey Pathology Services Lighthouse Laboratory and Alex Alderton, Roberto Amato, Jeffrey Barrett, Sonia Goncalves, Ewan Harrison, David K. Jackson, Ian Johnston, Dominic Kwiatkowski, Cordelia Langford, John Sillitoe on behalf of the Wellcome Sanger Institute COVID-19 Surveillance Team |
| EPI_ISL_3436677 | Lighthouse Lab in Glasgow | Wellcome Sanger Institute for the COVID-19 Genomics UK (COG-UK) Consortium | Harper VanSteenhouse, Yumi Kasai, David Gray, Carol Clugston, Anna Dominiczak and Alex Alderton, Roberto Amato, Jeffrey Barrett, Sonia Goncalves, Ewan Harrison, David K. Jackson, Ian Johnston, Dominic Kwiatkowski, Cordelia Langford, John Sillitoe on behalf of the Wellcome Sanger Institute COVID-19 Surveillance Team |
| EPI_ISL_3436678 | Berkshire and Surrey Pathology Services Lighthouse Laboratory | Wellcome Sanger Institute for the COVID-19 Genomics UK (COG-UK) Consortium | Berkshire and Surrey Pathology Services Lighthouse Laboratory and Alex Alderton, Roberto Amato, Jeffrey Barrett, Sonia Goncalves, Ewan Harrison, David K. Jackson, Ian Johnston, Dominic Kwiatkowski, Cordelia Langford, John Sillitoe on behalf of the Wellcome Sanger Institute COVID-19 Surveillance Team |
| EPI_ISL_3436679, EPI_ISL_3436680 | Lighthouse Lab in Alderley Park | Wellcome Sanger Institute for the COVID-19 Genomics UK (COG-UK) Consortium | Jacquelyn Wynn, Mairead Hyland, The Lighthouse Lab in Alderley Park and Alex Alderton, Roberto Amato, Jeffrey Barrett, Sonia Goncalves, Ewan Harrison, David K. Jackson, Ian Johnston, Dominic Kwiatkowski, Cordelia Langford, John Sillitoe on behalf of the Wellcome Sanger Institute COVID-19 Surveillance Team |
| EPI_ISL_3436681, EPI_ISL_3436682 | Berkshire and Surrey Pathology Services Lighthouse Laboratory | Wellcome Sanger Institute for the COVID-19 Genomics UK (COG-UK) Consortium | Berkshire and Surrey Pathology Services Lighthouse Laboratory and Alex Alderton, Roberto Amato, Jeffrey Barrett, Sonia Goncalves, Ewan Harrison, David K. Jackson, Ian Johnston, Dominic Kwiatkowski, Cordelia Langford, John Sillitoe on behalf of the Wellcome Sanger Institute COVID-19 Surveillance Team |
| EPI_ISL_3436683, EPI_ISL_3436684 | Lighthouse Lab in Alderley Park | Wellcome Sanger Institute for the COVID-19 Genomics UK (COG-UK) Consortium | Jacquelyn Wynn, Mairead Hyland, The Lighthouse Lab in Alderley Park and Alex Alderton, Roberto Amato, Jeffrey Barrett, Sonia Goncalves, Ewan Harrison, David K. Jackson, Ian Johnston, Dominic Kwiatkowski, Cordelia Langford, John Sillitoe on behalf of the Wellcome Sanger Institute COVID-19 Surveillance Team |
| EPI_ISL_3436685 | Lighthouse Lab in Glasgow | Wellcome Sanger Institute for the COVID-19 Genomics UK (COG-UK) Consortium | Harper VanSteenhouse, Yumi Kasai, David Gray, Carol Clugston, Anna Dominiczak and Alex Alderton, Roberto Amato, Jeffrey Barrett, Sonia Goncalves, Ewan Harrison, David K. Jackson, Ian Johnston, Dominic Kwiatkowski, Cordelia Langford, John Sillitoe on behalf of the Wellcome Sanger Institute COVID-19 Surveillance Team |
| EPI_ISL_3436686 | Berkshire and Surrey Pathology Services Lighthouse Laboratory | Wellcome Sanger Institute for the COVID-19 Genomics UK (COG-UK) Consortium | Berkshire and Surrey Pathology Services Lighthouse Laboratory and Alex Alderton, Roberto Amato, Jeffrey Barrett, Sonia Goncalves, Ewan Harrison, David K. Jackson, Ian Johnston, Dominic Kwiatkowski, Cordelia Langford, John Sillitoe on behalf of the Wellcome Sanger Institute COVID-19 Surveillance Team |
| EPI_ISL_3436687 | Lighthouse Lab in Alderley Park | Wellcome Sanger Institute for the COVID-19 Genomics UK (COG-UK) Consortium | Jacquelyn Wynn, Mairead Hyland, The Lighthouse Lab in Alderley Park and Alex Alderton, Roberto Amato, Jeffrey Barrett, Sonia Goncalves, Ewan Harrison, David K. Jackson, Ian Johnston, Dominic Kwiatkowski, Cordelia Langford, John Sillitoe on behalf of the Wellcome Sanger Institute COVID-19 Surveillance Team |
| EPI_ISL_3436688 | Berkshire and Surrey Pathology Services Lighthouse Laboratory | Wellcome Sanger Institute for the COVID-19 Genomics UK (COG-UK) Consortium | Berkshire and Surrey Pathology Services Lighthouse Laboratory and Alex Alderton, Roberto Amato, Jeffrey Barrett, Sonia Goncalves, Ewan Harrison, David K. Jackson, Ian Johnston, Dominic Kwiatkowski, Cordelia Langford, John Sillitoe on behalf of the Wellcome Sanger Institute COVID-19 Surveillance Team |
| EPI_ISL_3436689 | Lighthouse Lab in Alderley Park | Wellcome Sanger Institute for the COVID-19 Genomics UK (COG-UK) Consortium | Jacquelyn Wynn, Mairead Hyland, The Lighthouse Lab in Alderley Park and Alex Alderton, Roberto Amato, Jeffrey Barrett, Sonia Goncalves, Ewan Harrison, David K. Jackson, Ian Johnston, Dominic Kwiatkowski, Cordelia Langford, John Sillitoe on behalf of the Wellcome Sanger Institute COVID-19 Surveillance Team |
| EPI_ISL_3436690 | Lighthouse Lab in Glasgow | Wellcome Sanger Institute for the COVID-19 Genomics UK (COG-UK) Consortium | Harper VanSteenhouse, Yumi Kasai, David Gray, Carol Clugston, Anna Dominiczak and Alex Alderton, Roberto Amato, Jeffrey Barrett, Sonia Goncalves, Ewan Harrison, David K. Jackson, Ian Johnston, Dominic Kwiatkowski, Cordelia Langford, John Sillitoe on behalf of the Wellcome Sanger Institute COVID-19 Surveillance Team |
| EPI_ISL_3436691 | Berkshire and Surrey Pathology Services Lighthouse Laboratory | Wellcome Sanger Institute for the COVID-19 Genomics UK (COG-UK) Consortium | Berkshire and Surrey Pathology Services Lighthouse Laboratory and Alex Alderton, Roberto Amato, Jeffrey Barrett, Sonia Goncalves, Ewan Harrison, David K. Jackson, Ian Johnston, Dominic Kwiatkowski, Cordelia Langford, John Sillitoe on behalf of the Wellcome Sanger Institute COVID-19 Surveillance Team |
| EPI_ISL_3436692, EPI_ISL_3436693 | Lighthouse Lab in Glasgow | Wellcome Sanger Institute for the COVID-19 Genomics UK (COG-UK) Consortium | Harper VanSteenhouse, Yumi Kasai, David Gray, Carol Clugston, Anna Dominiczak and Alex Alderton, Roberto Amato, Jeffrey Barrett, Sonia Goncalves, Ewan Harrison, David K. Jackson, Ian Johnston, Dominic Kwiatkowski, Cordelia Langford, John Sillitoe on behalf of the Wellcome Sanger Institute COVID-19 Surveillance Team |
| EPI_ISL_3436694, EPI_ISL_3436695 | Berkshire and Surrey Pathology Services Lighthouse Laboratory | Wellcome Sanger Institute for the COVID-19 Genomics UK (COG-UK) Consortium | Berkshire and Surrey Pathology Services Lighthouse Laboratory and Alex Alderton, Roberto Amato, Jeffrey Barrett, Sonia Goncalves, Ewan Harrison, David K. Jackson, Ian Johnston, Dominic Kwiatkowski, Cordelia Langford, John Sillitoe on behalf of the Wellcome Sanger Institute COVID-19 Surveillance Team |
| EPI_ISL_3436696 | Lighthouse Lab in Glasgow | Wellcome Sanger Institute for the COVID-19 Genomics UK (COG-UK) Consortium | Harper VanSteenhouse, Yumi Kasai, David Gray, Carol Clugston, Anna Dominiczak and Alex Alderton, Roberto Amato, Jeffrey Barrett, Sonia Goncalves, Ewan Harrison, David K. Jackson, Ian Johnston, Dominic Kwiatkowski, Cordelia Langford, John Sillitoe on behalf of the Wellcome Sanger Institute COVID-19 Surveillance Team |
| EPI_ISL_3436697 | Berkshire and Surrey Pathology Services Lighthouse Laboratory | Wellcome Sanger Institute for the COVID-19 Genomics UK (COG-UK) Consortium | Berkshire and Surrey Pathology Services Lighthouse Laboratory and Alex Alderton, Roberto Amato, Jeffrey Barrett, Sonia Goncalves, Ewan Harrison, David K. Jackson, Ian Johnston, Dominic Kwiatkowski, Cordelia Langford, John Sillitoe on behalf of the Wellcome Sanger Institute COVID-19 Surveillance Team |
| EPI_ISL_3436698 | Lighthouse Lab in Alderley Park | Wellcome Sanger Institute for the COVID-19 Genomics UK (COG-UK) Consortium | Jacquelyn Wynn, Mairead Hyland, The Lighthouse Lab in Alderley Park and Alex Alderton, Roberto Amato, Jeffrey Barrett, Sonia Goncalves, Ewan Harrison, David K. Jackson, Ian Johnston, Dominic Kwiatkowski, Cordelia Langford, John Sillitoe on behalf of the Wellcome Sanger Institute COVID-19 Surveillance Team |
| EPI_ISL_3436699 | Berkshire and Surrey Pathology Services Lighthouse Laboratory | Wellcome Sanger Institute for the COVID-19 Genomics UK (COG-UK) Consortium | Berkshire and Surrey Pathology Services Lighthouse Laboratory and Alex Alderton, Roberto Amato, Jeffrey Barrett, Sonia Goncalves, Ewan Harrison, David K. Jackson, Ian Johnston, Dominic Kwiatkowski, Cordelia Langford, John Sillitoe on behalf of the Wellcome Sanger Institute COVID-19 Surveillance Team |
| EPI_ISL_3436700 | Lighthouse Lab in Alderley Park | Wellcome Sanger Institute for the COVID-19 Genomics UK (COG-UK) Consortium | Jacquelyn Wynn, Mairead Hyland, The Lighthouse Lab in Alderley Park and Alex Alderton, Roberto Amato, Jeffrey Barrett, Sonia Goncalves, Ewan Harrison, David K. Jackson, Ian Johnston, Dominic Kwiatkowski, Cordelia Langford, John Sillitoe on behalf of the Wellcome Sanger Institute COVID-19 Surveillance Team |
| EPI_ISL_3436701, EPI_ISL_3436702 | Berkshire and Surrey Pathology Services Lighthouse Laboratory | Wellcome Sanger Institute for the COVID-19 Genomics UK (COG-UK) Consortium | Berkshire and Surrey Pathology Services Lighthouse Laboratory and Alex Alderton, Roberto Amato, Jeffrey Barrett, Sonia Goncalves, Ewan Harrison, David K. Jackson, Ian Johnston, Dominic Kwiatkowski, Cordelia Langford, John Sillitoe on behalf of the Wellcome Sanger Institute COVID-19 Surveillance Team |
| EPI_ISL_3436703, EPI_ISL_3436704 | Lighthouse Lab in Alderley Park | Wellcome Sanger Institute for the COVID-19 Genomics UK (COG-UK) Consortium | Jacquelyn Wynn, Mairead Hyland, The Lighthouse Lab in Alderley Park and Alex Alderton, Roberto Amato, Jeffrey Barrett, Sonia Goncalves, Ewan Harrison, David K. Jackson, Ian Johnston, Dominic Kwiatkowski, Cordelia Langford, John Sillitoe on behalf of the Wellcome Sanger Institute COVID-19 Surveillance Team |
| EPI_ISL_3436705, EPI_ISL_3436706, EPI_ISL_3436707 | Berkshire and Surrey Pathology Services Lighthouse Laboratory | Wellcome Sanger Institute for the COVID-19 Genomics UK (COG-UK) Consortium | Berkshire and Surrey Pathology Services Lighthouse Laboratory and Alex Alderton, Roberto Amato, Jeffrey Barrett, Sonia Goncalves, Ewan Harrison, David K. Jackson, Ian Johnston, Dominic Kwiatkowski, Cordelia Langford, John Sillitoe on behalf of the Wellcome Sanger Institute COVID-19 Surveillance Team |
| EPI_ISL_3436708, EPI_ISL_3436709 | Lighthouse Lab in Glasgow | Wellcome Sanger Institute for the COVID-19 Genomics UK (COG-UK) Consortium | Harper VanSteenhouse, Yumi Kasai, David Gray, Carol Clugston, Anna Dominiczak and Alex Alderton, Roberto Amato, Jeffrey Barrett, Sonia Goncalves, Ewan Harrison, David K. Jackson, Ian Johnston, Dominic Kwiatkowski, Cordelia Langford, John Sillitoe on behalf of the Wellcome Sanger Institute COVID-19 Surveillance Team |
| EPI_ISL_3436710 | Lighthouse Lab in Alderley Park | Wellcome Sanger Institute for the COVID-19 Genomics UK (COG-UK) Consortium | Jacquelyn Wynn, Mairead Hyland, The Lighthouse Lab in Alderley Park and Alex Alderton, Roberto Amato, Jeffrey Barrett, Sonia Goncalves, Ewan Harrison, David K. Jackson, Ian Johnston, Dominic Kwiatkowski, Cordelia Langford, John Sillitoe on behalf of the Wellcome Sanger Institute COVID-19 Surveillance Team |
| EPI_ISL_3436711 | Lighthouse Lab in Glasgow | Wellcome Sanger Institute for the COVID-19 Genomics UK (COG-UK) Consortium | Harper VanSteenhouse, Yumi Kasai, David Gray, Carol Clugston, Anna Dominiczak and Alex Alderton, Roberto Amato, Jeffrey Barrett, Sonia Goncalves, Ewan Harrison, David K. Jackson, Ian Johnston, Dominic Kwiatkowski, Cordelia Langford, John Sillitoe on behalf of the Wellcome Sanger Institute COVID-19 Surveillance Team |
| EPI_ISL_3436712 | Berkshire and Surrey Pathology Services Lighthouse Laboratory | Wellcome Sanger Institute for the COVID-19 Genomics UK (COG-UK) Consortium | Berkshire and Surrey Pathology Services Lighthouse Laboratory and Alex Alderton, Roberto Amato, Jeffrey Barrett, Sonia Goncalves, Ewan Harrison, David K. Jackson, Ian Johnston, Dominic Kwiatkowski, Cordelia Langford, John Sillitoe on behalf of the Wellcome Sanger Institute COVID-19 Surveillance Team |
| EPI_ISL_3436713 | Lighthouse Lab in Alderley Park | Wellcome Sanger Institute for the COVID-19 Genomics UK (COG-UK) Consortium | Jacquelyn Wynn, Mairead Hyland, The Lighthouse Lab in Alderley Park and Alex Alderton, Roberto Amato, Jeffrey Barrett, Sonia Goncalves, Ewan Harrison, David K. Jackson, Ian Johnston, Dominic Kwiatkowski, Cordelia Langford, John Sillitoe on behalf of the Wellcome Sanger Institute COVID-19 Surveillance Team |
| EPI_ISL_3436714 | Berkshire and Surrey Pathology Services Lighthouse Laboratory | Wellcome Sanger Institute for the COVID-19 Genomics UK (COG-UK) Consortium | Berkshire and Surrey Pathology Services Lighthouse Laboratory and Alex Alderton, Roberto Amato, Jeffrey Barrett, Sonia Goncalves, Ewan Harrison, David K. Jackson, Ian Johnston, Dominic Kwiatkowski, Cordelia Langford, John Sillitoe on behalf of the Wellcome Sanger Institute COVID-19 Surveillance Team |
| EPI_ISL_3436715 | Lighthouse Lab in Alderley Park | Wellcome Sanger Institute for the COVID-19 Genomics UK (COG-UK) Consortium | Jacquelyn Wynn, Mairead Hyland, The Lighthouse Lab in Alderley Park and Alex Alderton, Roberto Amato, Jeffrey Barrett, Sonia Goncalves, Ewan Harrison, David K. Jackson, Ian Johnston, Dominic Kwiatkowski, Cordelia Langford, John Sillitoe on behalf of the Wellcome Sanger Institute COVID-19 Surveillance Team |
| EPI_ISL_3436716 | Berkshire and Surrey Pathology Services Lighthouse Laboratory | Wellcome Sanger Institute for the COVID-19 Genomics UK (COG-UK) Consortium | Berkshire and Surrey Pathology Services Lighthouse Laboratory and Alex Alderton, Roberto Amato, Jeffrey Barrett, Sonia Goncalves, Ewan Harrison, David K. Jackson, Ian Johnston, Dominic Kwiatkowski, Cordelia Langford, John Sillitoe on behalf of the Wellcome Sanger Institute COVID-19 Surveillance Team |
| EPI_ISL_3436717 | Lighthouse Lab in Alderley Park | Wellcome Sanger Institute for the COVID-19 Genomics UK (COG-UK) Consortium | Jacquelyn Wynn, Mairead Hyland, The Lighthouse Lab in Alderley Park and Alex Alderton, Roberto Amato, Jeffrey Barrett, Sonia Goncalves, Ewan Harrison, David K. Jackson, Ian Johnston, Dominic Kwiatkowski, Cordelia Langford, John Sillitoe on behalf of the Wellcome Sanger Institute COVID-19 Surveillance Team |
| EPI_ISL_3436718, EPI_ISL_3436719 | Berkshire and Surrey Pathology Services Lighthouse Laboratory | Wellcome Sanger Institute for the COVID-19 Genomics UK (COG-UK) Consortium | Berkshire and Surrey Pathology Services Lighthouse Laboratory and Alex Alderton, Roberto Amato, Jeffrey Barrett, Sonia Goncalves, Ewan Harrison, David K. Jackson, Ian Johnston, Dominic Kwiatkowski, Cordelia Langford, John Sillitoe on behalf of the Wellcome Sanger Institute COVID-19 Surveillance Team |
| EPI_ISL_3436720 | Lighthouse Lab in Glasgow | Wellcome Sanger Institute for the COVID-19 Genomics UK (COG-UK) | Harper VanSteenhouse, Yumi Kasai, David Gray, Carol Clugston, Anna Dominiczak and Alex Alderton, Roberto Amato, Jeffrey Barrett, Sonia Goncalves, Ewan Harrison, David K. Jackson, Ian Johnston, Dominic Kwiatkowski, Cordelia Langford, |

| | | | |
|---|---|---|---|
| | | Consortium | John Sillitoe on behalf of the Wellcome Sanger Institute COVID-19 Surveillance Team |
| EPI_ISL_3436721, EPI_ISL_3436722 | Berkshire and Surrey Pathology Services Lighthouse Laboratory | Wellcome Sanger Institute for the COVID-19 Genomics UK (COG-UK) Consortium | Berkshire and Surrey Pathology Services Lighthouse Laboratory and Alex Alderton, Roberto Amato, Jeffrey Barrett, Sonia Goncalves, Ewan Harrison, David K. Jackson, Ian Johnston, Dominic Kwiatkowski, Cordelia Langford, John Sillitoe on behalf of the Wellcome Sanger Institute COVID-19 Surveillance Team |
| EPI_ISL_3436723 | Lighthouse Lab in Alderley Park | Wellcome Sanger Institute for the COVID-19 Genomics UK (COG-UK) Consortium | Jacquelyn Wynn, Mairead Hyland, The Lighthouse Lab in Alderley Park and Alex Alderton, Roberto Amato, Jeffrey Barrett, Sonia Goncalves, Ewan Harrison, David K. Jackson, Ian Johnston, Dominic Kwiatkowski, Cordelia Langford, John Sillitoe on behalf of the Wellcome Sanger Institute COVID-19 Surveillance Team |
| EPI_ISL_3436724, EPI_ISL_3436725 | Berkshire and Surrey Pathology Services Lighthouse Laboratory | Wellcome Sanger Institute for the COVID-19 Genomics UK (COG-UK) Consortium | Berkshire and Surrey Pathology Services Lighthouse Laboratory and Alex Alderton, Roberto Amato, Jeffrey Barrett, Sonia Goncalves, Ewan Harrison, David K. Jackson, Ian Johnston, Dominic Kwiatkowski, Cordelia Langford, John Sillitoe on behalf of the Wellcome Sanger Institute COVID-19 Surveillance Team |
| EPI_ISL_3436726 | Lighthouse Lab in Alderley Park | Wellcome Sanger Institute for the COVID-19 Genomics UK (COG-UK) Consortium | Jacquelyn Wynn, Mairead Hyland, The Lighthouse Lab in Alderley Park and Alex Alderton, Roberto Amato, Jeffrey Barrett, Sonia Goncalves, Ewan Harrison, David K. Jackson, Ian Johnston, Dominic Kwiatkowski, Cordelia Langford, John Sillitoe on behalf of the Wellcome Sanger Institute COVID-19 Surveillance Team |
| EPI_ISL_3436727 | Berkshire and Surrey Pathology Services Lighthouse Laboratory | Wellcome Sanger Institute for the COVID-19 Genomics UK (COG-UK) Consortium | Berkshire and Surrey Pathology Services Lighthouse Laboratory and Alex Alderton, Roberto Amato, Jeffrey Barrett, Sonia Goncalves, Ewan Harrison, David K. Jackson, Ian Johnston, Dominic Kwiatkowski, Cordelia Langford, John Sillitoe on behalf of the Wellcome Sanger Institute COVID-19 Surveillance Team |
| EPI_ISL_3436728, EPI_ISL_3436729, EPI_ISL_3436730 | Lighthouse Lab in Alderley Park | Wellcome Sanger Institute for the COVID-19 Genomics UK (COG-UK) Consortium | Jacquelyn Wynn, Mairead Hyland, The Lighthouse Lab in Alderley Park and Alex Alderton, Roberto Amato, Jeffrey Barrett, Sonia Goncalves, Ewan Harrison, David K. Jackson, Ian Johnston, Dominic Kwiatkowski, Cordelia Langford, John Sillitoe on behalf of the Wellcome Sanger Institute COVID-19 Surveillance Team |
| EPI_ISL_3436731, EPI_ISL_3436732, EPI_ISL_3436733 | Berkshire and Surrey Pathology Services Lighthouse Laboratory | Wellcome Sanger Institute for the COVID-19 Genomics UK (COG-UK) Consortium | Berkshire and Surrey Pathology Services Lighthouse Laboratory and Alex Alderton, Roberto Amato, Jeffrey Barrett, Sonia Goncalves, Ewan Harrison, David K. Jackson, Ian Johnston, Dominic Kwiatkowski, Cordelia Langford, John Sillitoe on behalf of the Wellcome Sanger Institute COVID-19 Surveillance Team |
| EPI_ISL_3436734 | Lighthouse Lab in Glasgow | Wellcome Sanger Institute for the COVID-19 Genomics UK (COG-UK) Consortium | Harper VanSteenhouse, Yumi Kasai, David Gray, Carol Clugston, Anna Dominiczak and Alex Alderton, Roberto Amato, Jeffrey Barrett, Sonia Goncalves, Ewan Harrison, David K. Jackson, Ian Johnston, Dominic Kwiatkowski, Cordelia Langford, John Sillitoe on behalf of the Wellcome Sanger Institute COVID-19 Surveillance Team |
| EPI_ISL_3436735 | Berkshire and Surrey Pathology Services Lighthouse Laboratory | Wellcome Sanger Institute for the COVID-19 Genomics UK (COG-UK) Consortium | Berkshire and Surrey Pathology Services Lighthouse Laboratory and Alex Alderton, Roberto Amato, Jeffrey Barrett, Sonia Goncalves, Ewan Harrison, David K. Jackson, Ian Johnston, Dominic Kwiatkowski, Cordelia Langford, John Sillitoe on behalf of the Wellcome Sanger Institute COVID-19 Surveillance Team |
| EPI_ISL_3436736, EPI_ISL_3436737, EPI_ISL_3436738, EPI_ISL_3436739 | Lighthouse Lab in Alderley Park | Wellcome Sanger Institute for the COVID-19 Genomics UK (COG-UK) Consortium | Jacquelyn Wynn, Mairead Hyland, The Lighthouse Lab in Alderley Park and Alex Alderton, Roberto Amato, Jeffrey Barrett, Sonia Goncalves, Ewan Harrison, David K. Jackson, Ian Johnston, Dominic Kwiatkowski, Cordelia Langford, John Sillitoe on behalf of the Wellcome Sanger Institute COVID-19 Surveillance Team |
| EPI_ISL_3436740 | Berkshire and Surrey Pathology Services Lighthouse Laboratory | Wellcome Sanger Institute for the COVID-19 Genomics UK (COG-UK) Consortium | Berkshire and Surrey Pathology Services Lighthouse Laboratory and Alex Alderton, Roberto Amato, Jeffrey Barrett, Sonia Goncalves, Ewan Harrison, David K. Jackson, Ian Johnston, Dominic Kwiatkowski, Cordelia Langford, John Sillitoe on behalf of the Wellcome Sanger Institute COVID-19 Surveillance Team |
| EPI_ISL_3436741 | Lighthouse Lab in Alderley Park | Wellcome Sanger Institute for the COVID-19 Genomics UK (COG-UK) Consortium | Jacquelyn Wynn, Mairead Hyland, The Lighthouse Lab in Alderley Park and Alex Alderton, Roberto Amato, Jeffrey Barrett, Sonia Goncalves, Ewan Harrison, David K. Jackson, Ian Johnston, Dominic Kwiatkowski, Cordelia Langford, John Sillitoe on behalf of the Wellcome Sanger Institute COVID-19 Surveillance Team |
| EPI_ISL_3436742 | Berkshire and Surrey Pathology Services Lighthouse Laboratory | Wellcome Sanger Institute for the COVID-19 Genomics UK (COG-UK) Consortium | Berkshire and Surrey Pathology Services Lighthouse Laboratory and Alex Alderton, Roberto Amato, Jeffrey Barrett, Sonia Goncalves, Ewan Harrison, David K. Jackson, Ian Johnston, Dominic Kwiatkowski, Cordelia Langford, John Sillitoe on behalf of the Wellcome Sanger Institute COVID-19 Surveillance Team |
| EPI_ISL_3436743 | Lighthouse Lab in Alderley Park | Wellcome Sanger Institute for the COVID-19 Genomics UK (COG-UK) Consortium | Jacquelyn Wynn, Mairead Hyland, The Lighthouse Lab in Alderley Park and Alex Alderton, Roberto Amato, Jeffrey Barrett, Sonia Goncalves, Ewan Harrison, David K. Jackson, Ian Johnston, Dominic Kwiatkowski, Cordelia Langford, John Sillitoe on behalf of the Wellcome Sanger Institute COVID-19 Surveillance Team |
| EPI_ISL_3436744 | Berkshire and Surrey Pathology Services Lighthouse Laboratory | Wellcome Sanger Institute for the COVID-19 Genomics UK (COG-UK) Consortium | Berkshire and Surrey Pathology Services Lighthouse Laboratory and Alex Alderton, Roberto Amato, Jeffrey Barrett, Sonia Goncalves, Ewan Harrison, David K. Jackson, Ian Johnston, Dominic Kwiatkowski, Cordelia Langford, John Sillitoe on behalf of the Wellcome Sanger Institute COVID-19 Surveillance Team |
| EPI_ISL_3436745, EPI_ISL_3436746 | Lighthouse Lab in Alderley Park | Wellcome Sanger Institute for the COVID-19 Genomics UK (COG-UK) Consortium | Jacquelyn Wynn, Mairead Hyland, The Lighthouse Lab in Alderley Park and Alex Alderton, Roberto Amato, Jeffrey Barrett, Sonia Goncalves, Ewan Harrison, David K. Jackson, Ian Johnston, Dominic Kwiatkowski, Cordelia Langford, John Sillitoe on behalf of the Wellcome Sanger Institute COVID-19 Surveillance Team |
| EPI_ISL_3436747, EPI_ISL_3436748 | Berkshire and Surrey Pathology Services Lighthouse Laboratory | Wellcome Sanger Institute for the COVID-19 Genomics UK (COG-UK) Consortium | Berkshire and Surrey Pathology Services Lighthouse Laboratory and Alex Alderton, Roberto Amato, Jeffrey Barrett, Sonia Goncalves, Ewan Harrison, David K. Jackson, Ian Johnston, Dominic Kwiatkowski, Cordelia Langford, John Sillitoe on behalf of the Wellcome Sanger Institute COVID-19 Surveillance Team |
| EPI_ISL_3436749 | Lighthouse Lab in Alderley Park | Wellcome Sanger Institute for the COVID-19 Genomics UK (COG-UK) Consortium | Jacquelyn Wynn, Mairead Hyland, The Lighthouse Lab in Alderley Park and Alex Alderton, Roberto Amato, Jeffrey Barrett, Sonia Goncalves, Ewan Harrison, David K. Jackson, Ian Johnston, Dominic Kwiatkowski, Cordelia Langford, John Sillitoe on behalf of the Wellcome Sanger Institute COVID-19 Surveillance Team |
| EPI_ISL_3436750 | Berkshire and Surrey Pathology Services Lighthouse Laboratory | Wellcome Sanger Institute for the COVID-19 Genomics UK (COG-UK) Consortium | Berkshire and Surrey Pathology Services Lighthouse Laboratory and Alex Alderton, Roberto Amato, Jeffrey Barrett, Sonia Goncalves, Ewan Harrison, David K. Jackson, Ian Johnston, Dominic Kwiatkowski, Cordelia Langford, John Sillitoe on behalf of the Wellcome Sanger Institute COVID-19 Surveillance Team |
| EPI_ISL_3436751 | Lighthouse Lab in Alderley Park | Wellcome Sanger Institute for the COVID-19 Genomics UK (COG-UK) Consortium | Jacquelyn Wynn, Mairead Hyland, The Lighthouse Lab in Alderley Park and Alex Alderton, Roberto Amato, Jeffrey Barrett, Sonia Goncalves, Ewan Harrison, David K. Jackson, Ian Johnston, Dominic Kwiatkowski, Cordelia Langford, John Sillitoe on behalf of the Wellcome Sanger Institute COVID-19 Surveillance Team |
| EPI_ISL_3436752, EPI_ISL_3436753 | Berkshire and Surrey Pathology Services Lighthouse Laboratory | Wellcome Sanger Institute for the COVID-19 Genomics UK (COG-UK) Consortium | Berkshire and Surrey Pathology Services Lighthouse Laboratory and Alex Alderton, Roberto Amato, Jeffrey Barrett, Sonia Goncalves, Ewan Harrison, David K. Jackson, Ian Johnston, Dominic Kwiatkowski, Cordelia Langford, John Sillitoe on behalf of the Wellcome Sanger Institute COVID-19 Surveillance Team |
| EPI_ISL_3436754 | Lighthouse Lab in Alderley Park | Wellcome Sanger Institute for the COVID-19 Genomics UK (COG-UK) Consortium | Jacquelyn Wynn, Mairead Hyland, The Lighthouse Lab in Alderley Park and Alex Alderton, Roberto Amato, Jeffrey Barrett, Sonia Goncalves, Ewan Harrison, David K. Jackson, Ian Johnston, Dominic Kwiatkowski, Cordelia Langford, John Sillitoe on behalf of the Wellcome Sanger Institute COVID-19 Surveillance Team |
| EPI_ISL_3436755, EPI_ISL_3436756, EPI_ISL_3436757 | Berkshire and Surrey Pathology Services Lighthouse Laboratory | Wellcome Sanger Institute for the COVID-19 Genomics UK (COG-UK) Consortium | Berkshire and Surrey Pathology Services Lighthouse Laboratory and Alex Alderton, Roberto Amato, Jeffrey Barrett, Sonia Goncalves, Ewan Harrison, David K. Jackson, Ian Johnston, Dominic Kwiatkowski, Cordelia Langford, John Sillitoe on behalf of the Wellcome Sanger Institute COVID-19 Surveillance Team |
| EPI_ISL_3436758 | Lighthouse Lab in Alderley Park | Wellcome Sanger Institute for the COVID-19 Genomics UK (COG-UK) Consortium | Jacquelyn Wynn, Mairead Hyland, The Lighthouse Lab in Alderley Park and Alex Alderton, Roberto Amato, Jeffrey Barrett, Sonia Goncalves, Ewan Harrison, David K. Jackson, Ian Johnston, Dominic Kwiatkowski, Cordelia Langford, John Sillitoe on behalf of the Wellcome Sanger Institute COVID-19 Surveillance Team |
| EPI_ISL_3436759, EPI_ISL_3436760 | Berkshire and Surrey Pathology Services Lighthouse Laboratory | Wellcome Sanger Institute for the COVID-19 Genomics UK (COG-UK) Consortium | Berkshire and Surrey Pathology Services Lighthouse Laboratory and Alex Alderton, Roberto Amato, Jeffrey Barrett, Sonia Goncalves, Ewan Harrison, David K. Jackson, Ian Johnston, Dominic Kwiatkowski, Cordelia Langford, John Sillitoe on behalf of the Wellcome Sanger Institute COVID-19 Surveillance Team |
| EPI_ISL_3436761 | Lighthouse Lab in Glasgow | Wellcome Sanger Institute for the COVID-19 Genomics UK (COG-UK) Consortium | Harper VanSteenhouse, Yumi Kasai, David Gray, Carol Clugston, Anna Dominiczak and Alex Alderton, Roberto Amato, Jeffrey Barrett, Sonia Goncalves, Ewan Harrison, David K. Jackson, Ian Johnston, Dominic Kwiatkowski, Cordelia Langford, John Sillitoe on behalf of the Wellcome Sanger Institute COVID-19 Surveillance Team |
| EPI_ISL_3436762, EPI_ISL_3436763 | Berkshire and Surrey Pathology Services Lighthouse Laboratory | Wellcome Sanger Institute for the COVID-19 Genomics UK (COG-UK) Consortium | Berkshire and Surrey Pathology Services Lighthouse Laboratory and Alex Alderton, Roberto Amato, Jeffrey Barrett, Sonia Goncalves, Ewan Harrison, David K. Jackson, Ian Johnston, Dominic Kwiatkowski, Cordelia Langford, John Sillitoe on behalf of the Wellcome Sanger Institute COVID-19 Surveillance Team |
| EPI_ISL_3436764, EPI_ISL_3436765, EPI_ISL_3436766 | Lighthouse Lab in Alderley Park | Wellcome Sanger Institute for the COVID-19 Genomics UK (COG-UK) Consortium | Jacquelyn Wynn, Mairead Hyland, The Lighthouse Lab in Alderley Park and Alex Alderton, Roberto Amato, Jeffrey Barrett, Sonia Goncalves, Ewan Harrison, David K. Jackson, Ian Johnston, Dominic Kwiatkowski, Cordelia Langford, John Sillitoe on behalf of the Wellcome Sanger Institute COVID-19 Surveillance Team |
| EPI_ISL_3436767, EPI_ISL_3436768, EPI_ISL_3436769 | Berkshire and Surrey Pathology Services Lighthouse Laboratory | Wellcome Sanger Institute for the COVID-19 Genomics UK (COG-UK) Consortium | Berkshire and Surrey Pathology Services Lighthouse Laboratory and Alex Alderton, Roberto Amato, Jeffrey Barrett, Sonia Goncalves, Ewan Harrison, David K. Jackson, Ian Johnston, Dominic Kwiatkowski, Cordelia Langford, John Sillitoe on behalf of the Wellcome Sanger Institute COVID-19 Surveillance Team |
| EPI_ISL_3436770 | Lighthouse Lab in Alderley Park | Wellcome Sanger Institute for the COVID-19 Genomics UK (COG-UK) Consortium | Jacquelyn Wynn, Mairead Hyland, The Lighthouse Lab in Alderley Park and Alex Alderton, Roberto Amato, Jeffrey Barrett, Sonia Goncalves, Ewan Harrison, David K. Jackson, Ian Johnston, Dominic Kwiatkowski, Cordelia Langford, John Sillitoe on behalf of the Wellcome Sanger Institute COVID-19 Surveillance Team |
| EPI_ISL_3436771 | Berkshire and Surrey Pathology Services Lighthouse Laboratory | Wellcome Sanger Institute for the COVID-19 Genomics UK (COG-UK) Consortium | Berkshire and Surrey Pathology Services Lighthouse Laboratory and Alex Alderton, Roberto Amato, Jeffrey Barrett, Sonia Goncalves, Ewan Harrison, David K. Jackson, Ian Johnston, Dominic Kwiatkowski, Cordelia Langford, John Sillitoe on behalf of the Wellcome Sanger Institute COVID-19 Surveillance Team |
| EPI_ISL_3436772 | Lighthouse Lab in Glasgow | Wellcome Sanger Institute for the COVID-19 Genomics UK (COG-UK) Consortium | Harper VanSteenhouse, Yumi Kasai, David Gray, Carol Clugston, Anna Dominiczak and Alex Alderton, Roberto Amato, Jeffrey Barrett, Sonia Goncalves, Ewan Harrison, David K. Jackson, Ian Johnston, Dominic Kwiatkowski, Cordelia Langford, John Sillitoe on behalf of the Wellcome Sanger Institute COVID-19 Surveillance Team |
| EPI_ISL_3436773 | Berkshire and Surrey Pathology Services Lighthouse Laboratory | Wellcome Sanger Institute for the COVID-19 Genomics UK (COG-UK) Consortium | Berkshire and Surrey Pathology Services Lighthouse Laboratory and Alex Alderton, Roberto Amato, Jeffrey Barrett, Sonia Goncalves, Ewan Harrison, David K. Jackson, Ian Johnston, Dominic Kwiatkowski, Cordelia Langford, John Sillitoe on behalf of the Wellcome Sanger Institute COVID-19 Surveillance Team |
| EPI_ISL_3436774 | Lighthouse Lab in Alderley Park | Wellcome Sanger Institute for the COVID-19 Genomics UK (COG-UK) Consortium | Jacquelyn Wynn, Mairead Hyland, The Lighthouse Lab in Alderley Park and Alex Alderton, Roberto Amato, Jeffrey Barrett, Sonia Goncalves, Ewan Harrison, David K. Jackson, Ian Johnston, Dominic Kwiatkowski, Cordelia Langford, John Sillitoe on behalf of the Wellcome Sanger Institute COVID-19 Surveillance Team |
| EPI_ISL_3436775 | Lighthouse Lab in Glasgow | Wellcome Sanger Institute for the COVID-19 Genomics UK (COG-UK) Consortium | Harper VanSteenhouse, Yumi Kasai, David Gray, Carol Clugston, Anna Dominiczak and Alex Alderton, Roberto Amato, Jeffrey Barrett, Sonia Goncalves, Ewan Harrison, David K. Jackson, Ian Johnston, Dominic Kwiatkowski, Cordelia Langford, John Sillitoe on behalf of the Wellcome Sanger Institute COVID-19 Surveillance Team |
| EPI_ISL_3436776, EPI_ISL_3436777 | Berkshire and Surrey Pathology Services Lighthouse Laboratory | Wellcome Sanger Institute for the COVID-19 Genomics UK (COG-UK) Consortium | Berkshire and Surrey Pathology Services Lighthouse Laboratory and Alex Alderton, Roberto Amato, Jeffrey Barrett, Sonia Goncalves, Ewan Harrison, David K. Jackson, Ian Johnston, Dominic Kwiatkowski, Cordelia Langford, John Sillitoe on behalf of the Wellcome Sanger Institute COVID-19 Surveillance Team |
| EPI_ISL_3436778 | Lighthouse Lab in Alderley Park | Wellcome Sanger Institute for the COVID-19 Genomics UK (COG-UK) Consortium | Jacquelyn Wynn, Mairead Hyland, The Lighthouse Lab in Alderley Park and Alex Alderton, Roberto Amato, Jeffrey Barrett, Sonia Goncalves, Ewan Harrison, David K. Jackson, Ian Johnston, Dominic Kwiatkowski, Cordelia Langford, John Sillitoe on behalf of the Wellcome Sanger Institute COVID-19 Surveillance Team |
| EPI_ISL_3436779, EPI_ISL_3436780, EPI_ISL_3436781, EPI_ISL_3436782 | Berkshire and Surrey Pathology Services Lighthouse Laboratory | Wellcome Sanger Institute for the COVID-19 Genomics UK (COG-UK) Consortium | Berkshire and Surrey Pathology Services Lighthouse Laboratory and Alex Alderton, Roberto Amato, Jeffrey Barrett, Sonia Goncalves, Ewan Harrison, David K. Jackson, Ian Johnston, Dominic Kwiatkowski, Cordelia Langford, John Sillitoe on behalf of the Wellcome Sanger Institute COVID-19 Surveillance Team |
| EPI_ISL_3436783 | Lighthouse Lab in Glasgow | Wellcome Sanger Institute for the COVID-19 Genomics UK (COG-UK) Consortium | Harper VanSteenhouse, Yumi Kasai, David Gray, Carol Clugston, Anna Dominiczak and Alex Alderton, Roberto Amato, Jeffrey Barrett, Sonia Goncalves, Ewan Harrison, David K. Jackson, Ian Johnston, Dominic Kwiatkowski, Cordelia Langford, John Sillitoe on behalf of the Wellcome Sanger Institute COVID-19 Surveillance Team |
| EPI_ISL_3436784 | Berkshire and Surrey Pathology Services Lighthouse Laboratory | Wellcome Sanger Institute for the COVID-19 Genomics UK (COG-UK) Consortium | Berkshire and Surrey Pathology Services Lighthouse Laboratory and Alex Alderton, Roberto Amato, Jeffrey Barrett, Sonia Goncalves, Ewan Harrison, David K. Jackson, Ian Johnston, Dominic Kwiatkowski, Cordelia Langford, John Sillitoe on behalf of the Wellcome Sanger Institute COVID-19 Surveillance Team |
| EPI_ISL_3436785, EPI_ISL_3436786 | Lighthouse Lab in Glasgow | Wellcome Sanger Institute for the COVID-19 Genomics UK (COG-UK) Consortium | Harper VanSteenhouse, Yumi Kasai, David Gray, Carol Clugston, Anna Dominiczak and Alex Alderton, Roberto Amato, Jeffrey Barrett, Sonia Goncalves, Ewan Harrison, David K. Jackson, Ian Johnston, Dominic Kwiatkowski, Cordelia Langford, John Sillitoe on behalf of the Wellcome Sanger Institute COVID-19 Surveillance Team |

| | | | |
|---|---|---|---|
| EPI_ISL_3436787 | Lighthouse Lab in Alderley Park | Wellcome Sanger Institute for the COVID-19 Genomics UK (COG-UK) Consortium | Jacquelyn Wynn, Mairead Hyland, The Lighthouse Lab in Alderley Park and Alex Alderton, Roberto Amato, Jeffrey Barrett, Sonia Goncalves, Ewan Harrison, David K. Jackson, Ian Johnston, Dominic Kwiatkowski, Cordelia Langford, John Sillitoe on behalf of the Wellcome Sanger Institute COVID-19 Surveillance Team |
| EPI_ISL_3436788 | Lighthouse Lab in Glasgow | Wellcome Sanger Institute for the COVID-19 Genomics UK (COG-UK) Consortium | Harper VanSteenhouse, Yumi Kasai, David Gray, Carol Clugston, Anna Dominiczak and Alex Alderton, Roberto Amato, Jeffrey Barrett, Sonia Goncalves, Ewan Harrison, David K. Jackson, Ian Johnston, Dominic Kwiatkowski, Cordelia Langford, John Sillitoe on behalf of the Wellcome Sanger Institute COVID-19 Surveillance Team |
| EPI_ISL_3436789, EPI_ISL_3436790, EPI_ISL_3436791 | Berkshire and Surrey Pathology Services Lighthouse Laboratory | Wellcome Sanger Institute for the COVID-19 Genomics UK (COG-UK) Consortium | Berkshire and Surrey Pathology Services Lighthouse Laboratory and Alex Alderton, Roberto Amato, Jeffrey Barrett, Sonia Goncalves, Ewan Harrison, David K. Jackson, Ian Johnston, Dominic Kwiatkowski, Cordelia Langford, John Sillitoe on behalf of the Wellcome Sanger Institute COVID-19 Surveillance Team |
| EPI_ISL_3436792 | Lighthouse Lab in Alderley Park | Wellcome Sanger Institute for the COVID-19 Genomics UK (COG-UK) Consortium | Jacquelyn Wynn, Mairead Hyland, The Lighthouse Lab in Alderley Park and Alex Alderton, Roberto Amato, Jeffrey Barrett, Sonia Goncalves, Ewan Harrison, David K. Jackson, Ian Johnston, Dominic Kwiatkowski, Cordelia Langford, John Sillitoe on behalf of the Wellcome Sanger Institute COVID-19 Surveillance Team |
| EPI_ISL_3436793 | Berkshire and Surrey Pathology Services Lighthouse Laboratory | Wellcome Sanger Institute for the COVID-19 Genomics UK (COG-UK) Consortium | Berkshire and Surrey Pathology Services Lighthouse Laboratory and Alex Alderton, Roberto Amato, Jeffrey Barrett, Sonia Goncalves, Ewan Harrison, David K. Jackson, Ian Johnston, Dominic Kwiatkowski, Cordelia Langford, John Sillitoe on behalf of the Wellcome Sanger Institute COVID-19 Surveillance Team |
| EPI_ISL_3436794 | Lighthouse Lab in Glasgow | Wellcome Sanger Institute for the COVID-19 Genomics UK (COG-UK) Consortium | Harper VanSteenhouse, Yumi Kasai, David Gray, Carol Clugston, Anna Dominiczak and Alex Alderton, Roberto Amato, Jeffrey Barrett, Sonia Goncalves, Ewan Harrison, David K. Jackson, Ian Johnston, Dominic Kwiatkowski, Cordelia Langford, John Sillitoe on behalf of the Wellcome Sanger Institute COVID-19 Surveillance Team |
| EPI_ISL_3436795 | Lighthouse Lab in Alderley Park | Wellcome Sanger Institute for the COVID-19 Genomics UK (COG-UK) Consortium | Jacquelyn Wynn, Mairead Hyland, The Lighthouse Lab in Alderley Park and Alex Alderton, Roberto Amato, Jeffrey Barrett, Sonia Goncalves, Ewan Harrison, David K. Jackson, Ian Johnston, Dominic Kwiatkowski, Cordelia Langford, John Sillitoe on behalf of the Wellcome Sanger Institute COVID-19 Surveillance Team |
| EPI_ISL_3436796 | Lighthouse Lab in Glasgow | Wellcome Sanger Institute for the COVID-19 Genomics UK (COG-UK) Consortium | Harper VanSteenhouse, Yumi Kasai, David Gray, Carol Clugston, Anna Dominiczak and Alex Alderton, Roberto Amato, Jeffrey Barrett, Sonia Goncalves, Ewan Harrison, David K. Jackson, Ian Johnston, Dominic Kwiatkowski, Cordelia Langford, John Sillitoe on behalf of the Wellcome Sanger Institute COVID-19 Surveillance Team |
| EPI_ISL_3436797, EPI_ISL_3436798, EPI_ISL_3436799, EPI_ISL_3436800 | Berkshire and Surrey Pathology Services Lighthouse Laboratory | Wellcome Sanger Institute for the COVID-19 Genomics UK (COG-UK) Consortium | Berkshire and Surrey Pathology Services Lighthouse Laboratory and Alex Alderton, Roberto Amato, Jeffrey Barrett, Sonia Goncalves, Ewan Harrison, David K. Jackson, Ian Johnston, Dominic Kwiatkowski, Cordelia Langford, John Sillitoe on behalf of the Wellcome Sanger Institute COVID-19 Surveillance Team |
| EPI_ISL_3436801 | Lighthouse Lab in Alderley Park | Wellcome Sanger Institute for the COVID-19 Genomics UK (COG-UK) Consortium | Jacquelyn Wynn, Mairead Hyland, The Lighthouse Lab in Alderley Park and Alex Alderton, Roberto Amato, Jeffrey Barrett, Sonia Goncalves, Ewan Harrison, David K. Jackson, Ian Johnston, Dominic Kwiatkowski, Cordelia Langford, John Sillitoe on behalf of the Wellcome Sanger Institute COVID-19 Surveillance Team |
| EPI_ISL_3436802, EPI_ISL_3436803 | Berkshire and Surrey Pathology Services Lighthouse Laboratory | Wellcome Sanger Institute for the COVID-19 Genomics UK (COG-UK) Consortium | Berkshire and Surrey Pathology Services Lighthouse Laboratory and Alex Alderton, Roberto Amato, Jeffrey Barrett, Sonia Goncalves, Ewan Harrison, David K. Jackson, Ian Johnston, Dominic Kwiatkowski, Cordelia Langford, John Sillitoe on behalf of the Wellcome Sanger Institute COVID-19 Surveillance Team |
| EPI_ISL_3436804 | Lighthouse Lab in Alderley Park | Wellcome Sanger Institute for the COVID-19 Genomics UK (COG-UK) Consortium | Jacquelyn Wynn, Mairead Hyland, The Lighthouse Lab in Alderley Park and Alex Alderton, Roberto Amato, Jeffrey Barrett, Sonia Goncalves, Ewan Harrison, David K. Jackson, Ian Johnston, Dominic Kwiatkowski, Cordelia Langford, John Sillitoe on behalf of the Wellcome Sanger Institute COVID-19 Surveillance Team |
| EPI_ISL_3436805 | Berkshire and Surrey Pathology Services Lighthouse Laboratory | Wellcome Sanger Institute for the COVID-19 Genomics UK (COG-UK) Consortium | Berkshire and Surrey Pathology Services Lighthouse Laboratory and Alex Alderton, Roberto Amato, Jeffrey Barrett, Sonia Goncalves, Ewan Harrison, David K. Jackson, Ian Johnston, Dominic Kwiatkowski, Cordelia Langford, John Sillitoe on behalf of the Wellcome Sanger Institute COVID-19 Surveillance Team |
| EPI_ISL_3436806 | Lighthouse Lab in Glasgow | Wellcome Sanger Institute for the COVID-19 Genomics UK (COG-UK) Consortium | Harper VanSteenhouse, Yumi Kasai, David Gray, Carol Clugston, Anna Dominiczak and Alex Alderton, Roberto Amato, Jeffrey Barrett, Sonia Goncalves, Ewan Harrison, David K. Jackson, Ian Johnston, Dominic Kwiatkowski, Cordelia Langford, John Sillitoe on behalf of the Wellcome Sanger Institute COVID-19 Surveillance Team |
| EPI_ISL_3436807, EPI_ISL_3436808 | Lighthouse Lab in Alderley Park | Wellcome Sanger Institute for the COVID-19 Genomics UK (COG-UK) Consortium | Jacquelyn Wynn, Mairead Hyland, The Lighthouse Lab in Alderley Park and Alex Alderton, Roberto Amato, Jeffrey Barrett, Sonia Goncalves, Ewan Harrison, David K. Jackson, Ian Johnston, Dominic Kwiatkowski, Cordelia Langford, John Sillitoe on behalf of the Wellcome Sanger Institute COVID-19 Surveillance Team |
| EPI_ISL_3436809, EPI_ISL_3436810 | Lighthouse Lab in Glasgow | Wellcome Sanger Institute for the COVID-19 Genomics UK (COG-UK) Consortium | Harper VanSteenhouse, Yumi Kasai, David Gray, Carol Clugston, Anna Dominiczak and Alex Alderton, Roberto Amato, Jeffrey Barrett, Sonia Goncalves, Ewan Harrison, David K. Jackson, Ian Johnston, Dominic Kwiatkowski, Cordelia Langford, John Sillitoe on behalf of the Wellcome Sanger Institute COVID-19 Surveillance Team |
| EPI_ISL_3436811 | Berkshire and Surrey Pathology Services Lighthouse Laboratory | Wellcome Sanger Institute for the COVID-19 Genomics UK (COG-UK) Consortium | Berkshire and Surrey Pathology Services Lighthouse Laboratory and Alex Alderton, Roberto Amato, Jeffrey Barrett, Sonia Goncalves, Ewan Harrison, David K. Jackson, Ian Johnston, Dominic Kwiatkowski, Cordelia Langford, John Sillitoe on behalf of the Wellcome Sanger Institute COVID-19 Surveillance Team |
| EPI_ISL_3436812 | Lighthouse Lab in Alderley Park | Wellcome Sanger Institute for the COVID-19 Genomics UK (COG-UK) Consortium | Jacquelyn Wynn, Mairead Hyland, The Lighthouse Lab in Alderley Park and Alex Alderton, Roberto Amato, Jeffrey Barrett, Sonia Goncalves, Ewan Harrison, David K. Jackson, Ian Johnston, Dominic Kwiatkowski, Cordelia Langford, John Sillitoe on behalf of the Wellcome Sanger Institute COVID-19 Surveillance Team |
| EPI_ISL_3436813, EPI_ISL_3436814 | Berkshire and Surrey Pathology Services Lighthouse Laboratory | Wellcome Sanger Institute for the COVID-19 Genomics UK (COG-UK) Consortium | Berkshire and Surrey Pathology Services Lighthouse Laboratory and Alex Alderton, Roberto Amato, Jeffrey Barrett, Sonia Goncalves, Ewan Harrison, David K. Jackson, Ian Johnston, Dominic Kwiatkowski, Cordelia Langford, John Sillitoe on behalf of the Wellcome Sanger Institute COVID-19 Surveillance Team |
| EPI_ISL_3436815, EPI_ISL_3436816, EPI_ISL_3436817 | Lighthouse Lab in Alderley Park | Wellcome Sanger Institute for the COVID-19 Genomics UK (COG-UK) Consortium | Jacquelyn Wynn, Mairead Hyland, The Lighthouse Lab in Alderley Park and Alex Alderton, Roberto Amato, Jeffrey Barrett, Sonia Goncalves, Ewan Harrison, David K. Jackson, Ian Johnston, Dominic Kwiatkowski, Cordelia Langford, John Sillitoe on behalf of the Wellcome Sanger Institute COVID-19 Surveillance Team |
| EPI_ISL_3436818, EPI_ISL_3436819, EPI_ISL_3436820, EPI_ISL_3436821, EPI_ISL_3436822, EPI_ISL_3436823, EPI_ISL_3436824, EPI_ISL_3436825 | Berkshire and Surrey Pathology Services Lighthouse Laboratory | Wellcome Sanger Institute for the COVID-19 Genomics UK (COG-UK) Consortium | Berkshire and Surrey Pathology Services Lighthouse Laboratory and Alex Alderton, Roberto Amato, Jeffrey Barrett, Sonia Goncalves, Ewan Harrison, David K. Jackson, Ian Johnston, Dominic Kwiatkowski, Cordelia Langford, John Sillitoe on behalf of the Wellcome Sanger Institute COVID-19 Surveillance Team |
| EPI_ISL_3436826, EPI_ISL_3436827 | Lighthouse Lab in Alderley Park | Wellcome Sanger Institute for the COVID-19 Genomics UK (COG-UK) Consortium | Jacquelyn Wynn, Mairead Hyland, The Lighthouse Lab in Alderley Park and Alex Alderton, Roberto Amato, Jeffrey Barrett, Sonia Goncalves, Ewan Harrison, David K. Jackson, Ian Johnston, Dominic Kwiatkowski, Cordelia Langford, John Sillitoe on behalf of the Wellcome Sanger Institute COVID-19 Surveillance Team |
| EPI_ISL_3436828, EPI_ISL_3436829 | Berkshire and Surrey Pathology Services Lighthouse Laboratory | Wellcome Sanger Institute for the COVID-19 Genomics UK (COG-UK) Consortium | Berkshire and Surrey Pathology Services Lighthouse Laboratory and Alex Alderton, Roberto Amato, Jeffrey Barrett, Sonia Goncalves, Ewan Harrison, David K. Jackson, Ian Johnston, Dominic Kwiatkowski, Cordelia Langford, John Sillitoe on behalf of the Wellcome Sanger Institute COVID-19 Surveillance Team |
| EPI_ISL_3436830 | Lighthouse Lab in Alderley Park | Wellcome Sanger Institute for the COVID-19 Genomics UK (COG-UK) Consortium | Jacquelyn Wynn, Mairead Hyland, The Lighthouse Lab in Alderley Park and Alex Alderton, Roberto Amato, Jeffrey Barrett, Sonia Goncalves, Ewan Harrison, David K. Jackson, Ian Johnston, Dominic Kwiatkowski, Cordelia Langford, John Sillitoe on behalf of the Wellcome Sanger Institute COVID-19 Surveillance Team |
| EPI_ISL_3436831, EPI_ISL_3436832 | Berkshire and Surrey Pathology Services Lighthouse Laboratory | Wellcome Sanger Institute for the COVID-19 Genomics UK (COG-UK) Consortium | Berkshire and Surrey Pathology Services Lighthouse Laboratory and Alex Alderton, Roberto Amato, Jeffrey Barrett, Sonia Goncalves, Ewan Harrison, David K. Jackson, Ian Johnston, Dominic Kwiatkowski, Cordelia Langford, John Sillitoe on behalf of the Wellcome Sanger Institute COVID-19 Surveillance Team |
| EPI_ISL_3436833 | Lighthouse Lab in Alderley Park | Wellcome Sanger Institute for the COVID-19 Genomics UK (COG-UK) Consortium | Jacquelyn Wynn, Mairead Hyland, The Lighthouse Lab in Alderley Park and Alex Alderton, Roberto Amato, Jeffrey Barrett, Sonia Goncalves, Ewan Harrison, David K. Jackson, Ian Johnston, Dominic Kwiatkowski, Cordelia Langford, John Sillitoe on behalf of the Wellcome Sanger Institute COVID-19 Surveillance Team |
| EPI_ISL_3436834, EPI_ISL_3436835, EPI_ISL_3436836 | Berkshire and Surrey Pathology Services Lighthouse Laboratory | Wellcome Sanger Institute for the COVID-19 Genomics UK (COG-UK) Consortium | Berkshire and Surrey Pathology Services Lighthouse Laboratory and Alex Alderton, Roberto Amato, Jeffrey Barrett, Sonia Goncalves, Ewan Harrison, David K. Jackson, Ian Johnston, Dominic Kwiatkowski, Cordelia Langford, John Sillitoe on behalf of the Wellcome Sanger Institute COVID-19 Surveillance Team |
| EPI_ISL_3436837, EPI_ISL_3436838 | Lighthouse Lab in Alderley Park | Wellcome Sanger Institute for the COVID-19 Genomics UK (COG-UK) Consortium | Jacquelyn Wynn, Mairead Hyland, The Lighthouse Lab in Alderley Park and Alex Alderton, Roberto Amato, Jeffrey Barrett, Sonia Goncalves, Ewan Harrison, David K. Jackson, Ian Johnston, Dominic Kwiatkowski, Cordelia Langford, John Sillitoe on behalf of the Wellcome Sanger Institute COVID-19 Surveillance Team |
| EPI_ISL_3436839, EPI_ISL_3436840, EPI_ISL_3436841, EPI_ISL_3436842 | Berkshire and Surrey Pathology Services Lighthouse Laboratory | Wellcome Sanger Institute for the COVID-19 Genomics UK (COG-UK) Consortium | Berkshire and Surrey Pathology Services Lighthouse Laboratory and Alex Alderton, Roberto Amato, Jeffrey Barrett, Sonia Goncalves, Ewan Harrison, David K. Jackson, Ian Johnston, Dominic Kwiatkowski, Cordelia Langford, John Sillitoe on behalf of the Wellcome Sanger Institute COVID-19 Surveillance Team |
| EPI_ISL_3436843 | Lighthouse Lab in Alderley Park | Wellcome Sanger Institute for the COVID-19 Genomics UK (COG-UK) Consortium | Jacquelyn Wynn, Mairead Hyland, The Lighthouse Lab in Alderley Park and Alex Alderton, Roberto Amato, Jeffrey Barrett, Sonia Goncalves, Ewan Harrison, David K. Jackson, Ian Johnston, Dominic Kwiatkowski, Cordelia Langford, John Sillitoe on behalf of the Wellcome Sanger Institute COVID-19 Surveillance Team |
| EPI_ISL_3436844 | Berkshire and Surrey Pathology Services Lighthouse Laboratory | Wellcome Sanger Institute for the COVID-19 Genomics UK (COG-UK) Consortium | Berkshire and Surrey Pathology Services Lighthouse Laboratory and Alex Alderton, Roberto Amato, Jeffrey Barrett, Sonia Goncalves, Ewan Harrison, David K. Jackson, Ian Johnston, Dominic Kwiatkowski, Cordelia Langford, John Sillitoe on behalf of the Wellcome Sanger Institute COVID-19 Surveillance Team |
| EPI_ISL_3436845, EPI_ISL_3436846 | Lighthouse Lab in Alderley Park | Wellcome Sanger Institute for the COVID-19 Genomics UK (COG-UK) Consortium | Jacquelyn Wynn, Mairead Hyland, The Lighthouse Lab in Alderley Park and Alex Alderton, Roberto Amato, Jeffrey Barrett, Sonia Goncalves, Ewan Harrison, David K. Jackson, Ian Johnston, Dominic Kwiatkowski, Cordelia Langford, John Sillitoe on behalf of the Wellcome Sanger Institute COVID-19 Surveillance Team |
| EPI_ISL_3436847 | Berkshire and Surrey Pathology Services Lighthouse Laboratory | Wellcome Sanger Institute for the COVID-19 Genomics UK (COG-UK) Consortium | Berkshire and Surrey Pathology Services Lighthouse Laboratory and Alex Alderton, Roberto Amato, Jeffrey Barrett, Sonia Goncalves, Ewan Harrison, David K. Jackson, Ian Johnston, Dominic Kwiatkowski, Cordelia Langford, John Sillitoe on behalf of the Wellcome Sanger Institute COVID-19 Surveillance Team |
| EPI_ISL_3436848 | Lighthouse Lab in Glasgow | Wellcome Sanger Institute for the COVID-19 Genomics UK (COG-UK) Consortium | Harper VanSteenhouse, Yumi Kasai, David Gray, Carol Clugston, Anna Dominiczak and Alex Alderton, Roberto Amato, Jeffrey Barrett, Sonia Goncalves, Ewan Harrison, David K. Jackson, Ian Johnston, Dominic Kwiatkowski, Cordelia Langford, John Sillitoe on behalf of the Wellcome Sanger Institute COVID-19 Surveillance Team |
| EPI_ISL_3436849 | Berkshire and Surrey Pathology Services Lighthouse Laboratory | Wellcome Sanger Institute for the COVID-19 Genomics UK (COG-UK) Consortium | Berkshire and Surrey Pathology Services Lighthouse Laboratory and Alex Alderton, Roberto Amato, Jeffrey Barrett, Sonia Goncalves, Ewan Harrison, David K. Jackson, Ian Johnston, Dominic Kwiatkowski, Cordelia Langford, John Sillitoe on behalf of the Wellcome Sanger Institute COVID-19 Surveillance Team |
| EPI_ISL_3436850 | Lighthouse Lab in Alderley Park | Wellcome Sanger Institute for the COVID-19 Genomics UK (COG-UK) Consortium | Jacquelyn Wynn, Mairead Hyland, The Lighthouse Lab in Alderley Park and Alex Alderton, Roberto Amato, Jeffrey Barrett, Sonia Goncalves, Ewan Harrison, David K. Jackson, Ian Johnston, Dominic Kwiatkowski, Cordelia Langford, John Sillitoe on behalf of the Wellcome Sanger Institute COVID-19 Surveillance Team |
| EPI_ISL_3436851 | Lighthouse Lab in Glasgow | Wellcome Sanger Institute for the COVID-19 Genomics UK (COG-UK) Consortium | Harper VanSteenhouse, Yumi Kasai, David Gray, Carol Clugston, Anna Dominiczak and Alex Alderton, Roberto Amato, Jeffrey Barrett, Sonia Goncalves, Ewan Harrison, David K. Jackson, Ian Johnston, Dominic Kwiatkowski, Cordelia Langford, John Sillitoe on behalf of the Wellcome Sanger Institute COVID-19 Surveillance Team |
| EPI_ISL_3436852 | Lighthouse Lab in Alderley Park | Wellcome Sanger Institute for the COVID-19 Genomics UK (COG-UK) Consortium | Jacquelyn Wynn, Mairead Hyland, The Lighthouse Lab in Alderley Park and Alex Alderton, Roberto Amato, Jeffrey Barrett, Sonia Goncalves, Ewan Harrison, David K. Jackson, Ian Johnston, Dominic Kwiatkowski, Cordelia Langford, John Sillitoe on behalf of the Wellcome Sanger Institute COVID-19 Surveillance Team |
| EPI_ISL_3436853 | Berkshire and Surrey Pathology Services Lighthouse Laboratory | Wellcome Sanger Institute for the COVID-19 Genomics UK (COG-UK) Consortium | Berkshire and Surrey Pathology Services Lighthouse Laboratory and Alex Alderton, Roberto Amato, Jeffrey Barrett, Sonia Goncalves, Ewan Harrison, David K. Jackson, Ian Johnston, Dominic Kwiatkowski, Cordelia Langford, John Sillitoe on behalf of the Wellcome Sanger Institute COVID-19 Surveillance Team |
| EPI_ISL_3436854 | Lighthouse Lab in Alderley Park | Wellcome Sanger Institute for the COVID-19 Genomics UK (COG-UK) | Jacquelyn Wynn, Mairead Hyland, The Lighthouse Lab in Alderley Park and Alex Alderton, Roberto Amato, Jeffrey Barrett, Sonia Goncalves, Ewan Harrison, David K. Jackson, Ian Johnston, Dominic Kwiatkowski, Cordelia Langford, John Sillitoe on |

| | | | |
|---|---|---|---|
| | | Consortium | behalf of the Wellcome Sanger Institute COVID-19 Surveillance Team |
| EPI_ISL_3436855 | Berkshire and Surrey Pathology Services Lighthouse Laboratory | Wellcome Sanger Institute for the COVID-19 Genomics UK (COG-UK) Consortium | Berkshire and Surrey Pathology Services Lighthouse Laboratory and Alex Alderton, Roberto Amato, Jeffrey Barrett, Sonia Goncalves, Ewan Harrison, David K. Jackson, Ian Johnston, Dominic Kwiatkowski, Cordelia Langford, John Sillitoe on behalf of the Wellcome Sanger Institute COVID-19 Surveillance Team |
| EPI_ISL_3436856 | Lighthouse Lab in Alderley Park | Wellcome Sanger Institute for the COVID-19 Genomics UK (COG-UK) Consortium | Jacquelyn Wynn, Mairead Hyland, The Lighthouse Lab in Alderley Park and Alex Alderton, Roberto Amato, Jeffrey Barrett, Sonia Goncalves, Ewan Harrison, David K. Jackson, Ian Johnston, Dominic Kwiatkowski, Cordelia Langford, John Sillitoe on behalf of the Wellcome Sanger Institute COVID-19 Surveillance Team |
| EPI_ISL_3436857, EPI_ISL_3436858 | Berkshire and Surrey Pathology Services Lighthouse Laboratory | Wellcome Sanger Institute for the COVID-19 Genomics UK (COG-UK) Consortium | Berkshire and Surrey Pathology Services Lighthouse Laboratory and Alex Alderton, Roberto Amato, Jeffrey Barrett, Sonia Goncalves, Ewan Harrison, David K. Jackson, Ian Johnston, Dominic Kwiatkowski, Cordelia Langford, John Sillitoe on behalf of the Wellcome Sanger Institute COVID-19 Surveillance Team |
| EPI_ISL_3436859 | Lighthouse Lab in Alderley Park | Wellcome Sanger Institute for the COVID-19 Genomics UK (COG-UK) Consortium | Jacquelyn Wynn, Mairead Hyland, The Lighthouse Lab in Alderley Park and Alex Alderton, Roberto Amato, Jeffrey Barrett, Sonia Goncalves, Ewan Harrison, David K. Jackson, Ian Johnston, Dominic Kwiatkowski, Cordelia Langford, John Sillitoe on behalf of the Wellcome Sanger Institute COVID-19 Surveillance Team |
| EPI_ISL_3436860 | Berkshire and Surrey Pathology Services Lighthouse Laboratory | Wellcome Sanger Institute for the COVID-19 Genomics UK (COG-UK) Consortium | Berkshire and Surrey Pathology Services Lighthouse Laboratory and Alex Alderton, Roberto Amato, Jeffrey Barrett, Sonia Goncalves, Ewan Harrison, David K. Jackson, Ian Johnston, Dominic Kwiatkowski, Cordelia Langford, John Sillitoe on behalf of the Wellcome Sanger Institute COVID-19 Surveillance Team |
| EPI_ISL_3436861, EPI_ISL_3436862 | Lighthouse Lab in Alderley Park | Wellcome Sanger Institute for the COVID-19 Genomics UK (COG-UK) Consortium | Jacquelyn Wynn, Mairead Hyland, The Lighthouse Lab in Alderley Park and Alex Alderton, Roberto Amato, Jeffrey Barrett, Sonia Goncalves, Ewan Harrison, David K. Jackson, Ian Johnston, Dominic Kwiatkowski, Cordelia Langford, John Sillitoe on behalf of the Wellcome Sanger Institute COVID-19 Surveillance Team |
| EPI_ISL_3436863, EPI_ISL_3436864 | Berkshire and Surrey Pathology Services Lighthouse Laboratory | Wellcome Sanger Institute for the COVID-19 Genomics UK (COG-UK) Consortium | Berkshire and Surrey Pathology Services Lighthouse Laboratory and Alex Alderton, Roberto Amato, Jeffrey Barrett, Sonia Goncalves, Ewan Harrison, David K. Jackson, Ian Johnston, Dominic Kwiatkowski, Cordelia Langford, John Sillitoe on behalf of the Wellcome Sanger Institute COVID-19 Surveillance Team |
| EPI_ISL_3436865, EPI_ISL_3436866, EPI_ISL_3436867 | Lighthouse Lab in Alderley Park | Wellcome Sanger Institute for the COVID-19 Genomics UK (COG-UK) Consortium | Jacquelyn Wynn, Mairead Hyland, The Lighthouse Lab in Alderley Park and Alex Alderton, Roberto Amato, Jeffrey Barrett, Sonia Goncalves, Ewan Harrison, David K. Jackson, Ian Johnston, Dominic Kwiatkowski, Cordelia Langford, John Sillitoe on behalf of the Wellcome Sanger Institute COVID-19 Surveillance Team |
| EPI_ISL_3436868, EPI_ISL_3436869, EPI_ISL_3436870, EPI_ISL_3436871, EPI_ISL_3436872, EPI_ISL_3436873, EPI_ISL_3436874, EPI_ISL_3436875, EPI_ISL_3436876, EPI_ISL_3436877, EPI_ISL_3436878 | | | |
| see above | Berkshire and Surrey Pathology Services Lighthouse Laboratory | Wellcome Sanger Institute for the COVID-19 Genomics UK (COG-UK) Consortium | Berkshire and Surrey Pathology Services Lighthouse Laboratory and Alex Alderton, Roberto Amato, Jeffrey Barrett, Sonia Goncalves, Ewan Harrison, David K. Jackson, Ian Johnston, Dominic Kwiatkowski, Cordelia Langford, John Sillitoe on behalf of the Wellcome Sanger Institute COVID-19 Surveillance Team |
| EPI_ISL_3436879 | Lighthouse Lab in Alderley Park | Wellcome Sanger Institute for the COVID-19 Genomics UK (COG-UK) Consortium | Jacquelyn Wynn, Mairead Hyland, The Lighthouse Lab in Alderley Park and Alex Alderton, Roberto Amato, Jeffrey Barrett, Sonia Goncalves, Ewan Harrison, David K. Jackson, Ian Johnston, Dominic Kwiatkowski, Cordelia Langford, John Sillitoe on behalf of the Wellcome Sanger Institute COVID-19 Surveillance Team |
| EPI_ISL_3436880 | Berkshire and Surrey Pathology Services Lighthouse Laboratory | Wellcome Sanger Institute for the COVID-19 Genomics UK (COG-UK) Consortium | Berkshire and Surrey Pathology Services Lighthouse Laboratory and Alex Alderton, Roberto Amato, Jeffrey Barrett, Sonia Goncalves, Ewan Harrison, David K. Jackson, Ian Johnston, Dominic Kwiatkowski, Cordelia Langford, John Sillitoe on behalf of the Wellcome Sanger Institute COVID-19 Surveillance Team |
| EPI_ISL_3436881 | Lighthouse Lab in Alderley Park | Wellcome Sanger Institute for the COVID-19 Genomics UK (COG-UK) Consortium | Jacquelyn Wynn, Mairead Hyland, The Lighthouse Lab in Alderley Park and Alex Alderton, Roberto Amato, Jeffrey Barrett, Sonia Goncalves, Ewan Harrison, David K. Jackson, Ian Johnston, Dominic Kwiatkowski, Cordelia Langford, John Sillitoe on behalf of the Wellcome Sanger Institute COVID-19 Surveillance Team |
| EPI_ISL_3436882, EPI_ISL_3436883 | Berkshire and Surrey Pathology Services Lighthouse Laboratory | Wellcome Sanger Institute for the COVID-19 Genomics UK (COG-UK) Consortium | Berkshire and Surrey Pathology Services Lighthouse Laboratory and Alex Alderton, Roberto Amato, Jeffrey Barrett, Sonia Goncalves, Ewan Harrison, David K. Jackson, Ian Johnston, Dominic Kwiatkowski, Cordelia Langford, John Sillitoe on behalf of the Wellcome Sanger Institute COVID-19 Surveillance Team |
| EPI_ISL_3436884 | Lighthouse Lab in Glasgow | Wellcome Sanger Institute for the COVID-19 Genomics UK (COG-UK) Consortium | Harper VanSteenhouse, Yumi Kasai, David Gray, Carol Clugston, Anna Dominiczak and Alex Alderton, Roberto Amato, Jeffrey Barrett, Sonia Goncalves, Ewan Harrison, David K. Jackson, Ian Johnston, Dominic Kwiatkowski, Cordelia Langford, John Sillitoe on behalf of the Wellcome Sanger Institute COVID-19 Surveillance Team |
| EPI_ISL_3436885 | Lighthouse Lab in Alderley Park | Wellcome Sanger Institute for the COVID-19 Genomics UK (COG-UK) Consortium | Jacquelyn Wynn, Mairead Hyland, The Lighthouse Lab in Alderley Park and Alex Alderton, Roberto Amato, Jeffrey Barrett, Sonia Goncalves, Ewan Harrison, David K. Jackson, Ian Johnston, Dominic Kwiatkowski, Cordelia Langford, John Sillitoe on behalf of the Wellcome Sanger Institute COVID-19 Surveillance Team |
| EPI_ISL_3436886 | Berkshire and Surrey Pathology Services Lighthouse Laboratory | Wellcome Sanger Institute for the COVID-19 Genomics UK (COG-UK) Consortium | Berkshire and Surrey Pathology Services Lighthouse Laboratory and Alex Alderton, Roberto Amato, Jeffrey Barrett, Sonia Goncalves, Ewan Harrison, David K. Jackson, Ian Johnston, Dominic Kwiatkowski, Cordelia Langford, John Sillitoe on behalf of the Wellcome Sanger Institute COVID-19 Surveillance Team |
| EPI_ISL_3436887 | Lighthouse Lab in Alderley Park | Wellcome Sanger Institute for the COVID-19 Genomics UK (COG-UK) Consortium | Jacquelyn Wynn, Mairead Hyland, The Lighthouse Lab in Alderley Park and Alex Alderton, Roberto Amato, Jeffrey Barrett, Sonia Goncalves, Ewan Harrison, David K. Jackson, Ian Johnston, Dominic Kwiatkowski, Cordelia Langford, John Sillitoe on behalf of the Wellcome Sanger Institute COVID-19 Surveillance Team |
| EPI_ISL_3436888 | Berkshire and Surrey Pathology Services Lighthouse Laboratory | Wellcome Sanger Institute for the COVID-19 Genomics UK (COG-UK) Consortium | Berkshire and Surrey Pathology Services Lighthouse Laboratory and Alex Alderton, Roberto Amato, Jeffrey Barrett, Sonia Goncalves, Ewan Harrison, David K. Jackson, Ian Johnston, Dominic Kwiatkowski, Cordelia Langford, John Sillitoe on behalf of the Wellcome Sanger Institute COVID-19 Surveillance Team |
| EPI_ISL_3436889 | Lighthouse Lab in Alderley Park | Wellcome Sanger Institute for the COVID-19 Genomics UK (COG-UK) Consortium | Jacquelyn Wynn, Mairead Hyland, The Lighthouse Lab in Alderley Park and Alex Alderton, Roberto Amato, Jeffrey Barrett, Sonia Goncalves, Ewan Harrison, David K. Jackson, Ian Johnston, Dominic Kwiatkowski, Cordelia Langford, John Sillitoe on behalf of the Wellcome Sanger Institute COVID-19 Surveillance Team |
| EPI_ISL_3436890, EPI_ISL_3436891, EPI_ISL_3436892 | Berkshire and Surrey Pathology Services Lighthouse Laboratory | Wellcome Sanger Institute for the COVID-19 Genomics UK (COG-UK) Consortium | Berkshire and Surrey Pathology Services Lighthouse Laboratory and Alex Alderton, Roberto Amato, Jeffrey Barrett, Sonia Goncalves, Ewan Harrison, David K. Jackson, Ian Johnston, Dominic Kwiatkowski, Cordelia Langford, John Sillitoe on behalf of the Wellcome Sanger Institute COVID-19 Surveillance Team |
| EPI_ISL_3436893 | Lighthouse Lab in Alderley Park | Wellcome Sanger Institute for the COVID-19 Genomics UK (COG-UK) Consortium | Jacquelyn Wynn, Mairead Hyland, The Lighthouse Lab in Alderley Park and Alex Alderton, Roberto Amato, Jeffrey Barrett, Sonia Goncalves, Ewan Harrison, David K. Jackson, Ian Johnston, Dominic Kwiatkowski, Cordelia Langford, John Sillitoe on behalf of the Wellcome Sanger Institute COVID-19 Surveillance Team |
| EPI_ISL_3436894 | Berkshire and Surrey Pathology Services Lighthouse Laboratory | Wellcome Sanger Institute for the COVID-19 Genomics UK (COG-UK) Consortium | Berkshire and Surrey Pathology Services Lighthouse Laboratory and Alex Alderton, Roberto Amato, Jeffrey Barrett, Sonia Goncalves, Ewan Harrison, David K. Jackson, Ian Johnston, Dominic Kwiatkowski, Cordelia Langford, John Sillitoe on behalf of the Wellcome Sanger Institute COVID-19 Surveillance Team |
| EPI_ISL_3436895 | Lighthouse Lab in Glasgow | Wellcome Sanger Institute for the COVID-19 Genomics UK (COG-UK) Consortium | Harper VanSteenhouse, Yumi Kasai, David Gray, Carol Clugston, Anna Dominiczak and Alex Alderton, Roberto Amato, Jeffrey Barrett, Sonia Goncalves, Ewan Harrison, David K. Jackson, Ian Johnston, Dominic Kwiatkowski, Cordelia Langford, John Sillitoe on behalf of the Wellcome Sanger Institute COVID-19 Surveillance Team |
| EPI_ISL_3436896 | Lighthouse Lab in Alderley Park | Wellcome Sanger Institute for the COVID-19 Genomics UK (COG-UK) Consortium | Jacquelyn Wynn, Mairead Hyland, The Lighthouse Lab in Alderley Park and Alex Alderton, Roberto Amato, Jeffrey Barrett, Sonia Goncalves, Ewan Harrison, David K. Jackson, Ian Johnston, Dominic Kwiatkowski, Cordelia Langford, John Sillitoe on behalf of the Wellcome Sanger Institute COVID-19 Surveillance Team |
| EPI_ISL_3436897, EPI_ISL_3436898, EPI_ISL_3436899, EPI_ISL_3436900, EPI_ISL_3436901, EPI_ISL_3436902, EPI_ISL_3436903 | Berkshire and Surrey Pathology Services Lighthouse Laboratory | Wellcome Sanger Institute for the COVID-19 Genomics UK (COG-UK) Consortium | Berkshire and Surrey Pathology Services Lighthouse Laboratory and Alex Alderton, Roberto Amato, Jeffrey Barrett, Sonia Goncalves, Ewan Harrison, David K. Jackson, Ian Johnston, Dominic Kwiatkowski, Cordelia Langford, John Sillitoe on behalf of the Wellcome Sanger Institute COVID-19 Surveillance Team |
| EPI_ISL_3436904, EPI_ISL_3436905, EPI_ISL_3436906 | Lighthouse Lab in Alderley Park | Wellcome Sanger Institute for the COVID-19 Genomics UK (COG-UK) Consortium | Jacquelyn Wynn, Mairead Hyland, The Lighthouse Lab in Alderley Park and Alex Alderton, Roberto Amato, Jeffrey Barrett, Sonia Goncalves, Ewan Harrison, David K. Jackson, Ian Johnston, Dominic Kwiatkowski, Cordelia Langford, John Sillitoe on behalf of the Wellcome Sanger Institute COVID-19 Surveillance Team |
| EPI_ISL_3436907, EPI_ISL_3436908 | Lighthouse Lab in Glasgow | Wellcome Sanger Institute for the COVID-19 Genomics UK (COG-UK) Consortium | Harper VanSteenhouse, Yumi Kasai, David Gray, Carol Clugston, Anna Dominiczak and Alex Alderton, Roberto Amato, Jeffrey Barrett, Sonia Goncalves, Ewan Harrison, David K. Jackson, Ian Johnston, Dominic Kwiatkowski, Cordelia Langford, John Sillitoe on behalf of the Wellcome Sanger Institute COVID-19 Surveillance Team |
| EPI_ISL_3436909 | Lighthouse Lab in Alderley Park | Wellcome Sanger Institute for the COVID-19 Genomics UK (COG-UK) Consortium | Jacquelyn Wynn, Mairead Hyland, The Lighthouse Lab in Alderley Park and Alex Alderton, Roberto Amato, Jeffrey Barrett, Sonia Goncalves, Ewan Harrison, David K. Jackson, Ian Johnston, Dominic Kwiatkowski, Cordelia Langford, John Sillitoe on behalf of the Wellcome Sanger Institute COVID-19 Surveillance Team |
| EPI_ISL_3436910 | Berkshire and Surrey Pathology Services Lighthouse Laboratory | Wellcome Sanger Institute for the COVID-19 Genomics UK (COG-UK) Consortium | Berkshire and Surrey Pathology Services Lighthouse Laboratory and Alex Alderton, Roberto Amato, Jeffrey Barrett, Sonia Goncalves, Ewan Harrison, David K. Jackson, Ian Johnston, Dominic Kwiatkowski, Cordelia Langford, John Sillitoe on behalf of the Wellcome Sanger Institute COVID-19 Surveillance Team |
| EPI_ISL_3436911 | Lighthouse Lab in Glasgow | Wellcome Sanger Institute for the COVID-19 Genomics UK (COG-UK) Consortium | Harper VanSteenhouse, Yumi Kasai, David Gray, Carol Clugston, Anna Dominiczak and Alex Alderton, Roberto Amato, Jeffrey Barrett, Sonia Goncalves, Ewan Harrison, David K. Jackson, Ian Johnston, Dominic Kwiatkowski, Cordelia Langford, John Sillitoe on behalf of the Wellcome Sanger Institute COVID-19 Surveillance Team |
| EPI_ISL_3436912 | Lighthouse Lab in Alderley Park | Wellcome Sanger Institute for the COVID-19 Genomics UK (COG-UK) Consortium | Jacquelyn Wynn, Mairead Hyland, The Lighthouse Lab in Alderley Park and Alex Alderton, Roberto Amato, Jeffrey Barrett, Sonia Goncalves, Ewan Harrison, David K. Jackson, Ian Johnston, Dominic Kwiatkowski, Cordelia Langford, John Sillitoe on behalf of the Wellcome Sanger Institute COVID-19 Surveillance Team |
| EPI_ISL_3436913 | Lighthouse Lab in Glasgow | Wellcome Sanger Institute for the COVID-19 Genomics UK (COG-UK) Consortium | Harper VanSteenhouse, Yumi Kasai, David Gray, Carol Clugston, Anna Dominiczak and Alex Alderton, Roberto Amato, Jeffrey Barrett, Sonia Goncalves, Ewan Harrison, David K. Jackson, Ian Johnston, Dominic Kwiatkowski, Cordelia Langford, John Sillitoe on behalf of the Wellcome Sanger Institute COVID-19 Surveillance Team |
| EPI_ISL_3436914 | Berkshire and Surrey Pathology Services Lighthouse Laboratory | Wellcome Sanger Institute for the COVID-19 Genomics UK (COG-UK) Consortium | Berkshire and Surrey Pathology Services Lighthouse Laboratory and Alex Alderton, Roberto Amato, Jeffrey Barrett, Sonia Goncalves, Ewan Harrison, David K. Jackson, Ian Johnston, Dominic Kwiatkowski, Cordelia Langford, John Sillitoe on behalf of the Wellcome Sanger Institute COVID-19 Surveillance Team |
| EPI_ISL_3436915 | Lighthouse Lab in Alderley Park | Wellcome Sanger Institute for the COVID-19 Genomics UK (COG-UK) Consortium | Jacquelyn Wynn, Mairead Hyland, The Lighthouse Lab in Alderley Park and Alex Alderton, Roberto Amato, Jeffrey Barrett, Sonia Goncalves, Ewan Harrison, David K. Jackson, Ian Johnston, Dominic Kwiatkowski, Cordelia Langford, John Sillitoe on behalf of the Wellcome Sanger Institute COVID-19 Surveillance Team |
| EPI_ISL_3436916 | Lighthouse Lab in Glasgow | Wellcome Sanger Institute for the COVID-19 Genomics UK (COG-UK) Consortium | Harper VanSteenhouse, Yumi Kasai, David Gray, Carol Clugston, Anna Dominiczak and Alex Alderton, Roberto Amato, Jeffrey Barrett, Sonia Goncalves, Ewan Harrison, David K. Jackson, Ian Johnston, Dominic Kwiatkowski, Cordelia Langford, John Sillitoe on behalf of the Wellcome Sanger Institute COVID-19 Surveillance Team |
| EPI_ISL_3436917 | Lighthouse Lab in Alderley Park | Wellcome Sanger Institute for the COVID-19 Genomics UK (COG-UK) Consortium | Jacquelyn Wynn, Mairead Hyland, The Lighthouse Lab in Alderley Park and Alex Alderton, Roberto Amato, Jeffrey Barrett, Sonia Goncalves, Ewan Harrison, David K. Jackson, Ian Johnston, Dominic Kwiatkowski, Cordelia Langford, John Sillitoe on behalf of the Wellcome Sanger Institute COVID-19 Surveillance Team |
| EPI_ISL_3436918 | Berkshire and Surrey Pathology Services Lighthouse Laboratory | Wellcome Sanger Institute for the COVID-19 Genomics UK (COG-UK) Consortium | Berkshire and Surrey Pathology Services Lighthouse Laboratory and Alex Alderton, Roberto Amato, Jeffrey Barrett, Sonia Goncalves, Ewan Harrison, David K. Jackson, Ian Johnston, Dominic Kwiatkowski, Cordelia Langford, John Sillitoe on behalf of the Wellcome Sanger Institute COVID-19 Surveillance Team |
| EPI_ISL_3436919 | Lighthouse Lab in Glasgow | Wellcome Sanger Institute for the COVID-19 Genomics UK (COG-UK) Consortium | Harper VanSteenhouse, Yumi Kasai, David Gray, Carol Clugston, Anna Dominiczak and Alex Alderton, Roberto Amato, Jeffrey Barrett, Sonia Goncalves, Ewan Harrison, David K. Jackson, Ian Johnston, Dominic Kwiatkowski, Cordelia Langford, John Sillitoe on behalf of the Wellcome Sanger Institute COVID-19 Surveillance Team |
| EPI_ISL_3436920 | Lighthouse Lab in Alderley Park | Wellcome Sanger Institute for the COVID-19 Genomics UK (COG-UK) Consortium | Jacquelyn Wynn, Mairead Hyland, The Lighthouse Lab in Alderley Park and Alex Alderton, Roberto Amato, Jeffrey Barrett, Sonia Goncalves, Ewan Harrison, David K. Jackson, Ian Johnston, Dominic Kwiatkowski, Cordelia Langford, John Sillitoe on behalf of the Wellcome Sanger Institute COVID-19 Surveillance Team |
| EPI_ISL_3436921, EPI_ISL_3436922 | Lighthouse Lab in Glasgow | Wellcome Sanger Institute for the COVID-19 Genomics UK (COG-UK) Consortium | Harper VanSteenhouse, Yumi Kasai, David Gray, Carol Clugston, Anna Dominiczak and Alex Alderton, Roberto Amato, Jeffrey Barrett, Sonia Goncalves, Ewan Harrison, David K. Jackson, Ian Johnston, Dominic Kwiatkowski, Cordelia Langford, John Sillitoe on behalf of the Wellcome Sanger Institute COVID-19 Surveillance Team |
| EPI_ISL_3436923, | Lighthouse Lab in Alderley Park | Wellcome Sanger Institute for the COVID-19 Genomics UK (COG-UK) | Jacquelyn Wynn, Mairead Hyland, The Lighthouse Lab in Alderley Park and Alex Alderton, Roberto Amato, Jeffrey Barrett, Sonia Goncalves, Ewan Harrison, David K. Jackson, Ian Johnston, Dominic Kwiatkowski, Cordelia Langford, John Sillitoe on |

| | | | |
|---|---|---|---|
| EPI_ISL_3436924 | | Consortium | behalf of the Wellcome Sanger Institute COVID-19 Surveillance Team |
| EPI_ISL_3436925, EPI_ISL_3436926, EPI_ISL_3436927 | Berkshire and Surrey Pathology Services Lighthouse Laboratory | Wellcome Sanger Institute for the COVID-19 Genomics UK (COG-UK) Consortium | Berkshire and Surrey Pathology Services Lighthouse Laboratory and Alex Alderton, Roberto Amato, Jeffrey Barrett, Sonia Goncalves, Ewan Harrison, David K. Jackson, Ian Johnston, Dominic Kwiatkowski, Cordelia Langford, John Sillitoe on behalf of the Wellcome Sanger Institute COVID-19 Surveillance Team |
| EPI_ISL_3436928 | Lighthouse Lab in Glasgow | Wellcome Sanger Institute for the COVID-19 Genomics UK (COG-UK) Consortium | Harper VanSteenhouse, Yumi Kasai, David Gray, Carol Clugston, Anna Dominiczak and Alex Alderton, Roberto Amato, Jeffrey Barrett, Sonia Goncalves, Ewan Harrison, David K. Jackson, Ian Johnston, Dominic Kwiatkowski, Cordelia Langford, John Sillitoe on behalf of the Wellcome Sanger Institute COVID-19 Surveillance Team |
| EPI_ISL_3436929, EPI_ISL_3436930 | Berkshire and Surrey Pathology Services Lighthouse Laboratory | Wellcome Sanger Institute for the COVID-19 Genomics UK (COG-UK) Consortium | Berkshire and Surrey Pathology Services Lighthouse Laboratory and Alex Alderton, Roberto Amato, Jeffrey Barrett, Sonia Goncalves, Ewan Harrison, David K. Jackson, Ian Johnston, Dominic Kwiatkowski, Cordelia Langford, John Sillitoe on behalf of the Wellcome Sanger Institute COVID-19 Surveillance Team |
| EPI_ISL_3436931 | Lighthouse Lab in Alderley Park | Wellcome Sanger Institute for the COVID-19 Genomics UK (COG-UK) Consortium | Jacquelyn Wynn, Mairead Hyland, The Lighthouse Lab in Alderley Park and Alex Alderton, Roberto Amato, Jeffrey Barrett, Sonia Goncalves, Ewan Harrison, David K. Jackson, Ian Johnston, Dominic Kwiatkowski, Cordelia Langford, John Sillitoe on behalf of the Wellcome Sanger Institute COVID-19 Surveillance Team |
| EPI_ISL_3436932, EPI_ISL_3436933, EPI_ISL_3436934 | Berkshire and Surrey Pathology Services Lighthouse Laboratory | Wellcome Sanger Institute for the COVID-19 Genomics UK (COG-UK) Consortium | Berkshire and Surrey Pathology Services Lighthouse Laboratory and Alex Alderton, Roberto Amato, Jeffrey Barrett, Sonia Goncalves, Ewan Harrison, David K. Jackson, Ian Johnston, Dominic Kwiatkowski, Cordelia Langford, John Sillitoe on behalf of the Wellcome Sanger Institute COVID-19 Surveillance Team |
| EPI_ISL_3436935 | Lighthouse Lab in Alderley Park | Wellcome Sanger Institute for the COVID-19 Genomics UK (COG-UK) Consortium | Jacquelyn Wynn, Mairead Hyland, The Lighthouse Lab in Alderley Park and Alex Alderton, Roberto Amato, Jeffrey Barrett, Sonia Goncalves, Ewan Harrison, David K. Jackson, Ian Johnston, Dominic Kwiatkowski, Cordelia Langford, John Sillitoe on behalf of the Wellcome Sanger Institute COVID-19 Surveillance Team |
| EPI_ISL_3436936 | Berkshire and Surrey Pathology Services Lighthouse Laboratory | Wellcome Sanger Institute for the COVID-19 Genomics UK (COG-UK) Consortium | Berkshire and Surrey Pathology Services Lighthouse Laboratory and Alex Alderton, Roberto Amato, Jeffrey Barrett, Sonia Goncalves, Ewan Harrison, David K. Jackson, Ian Johnston, Dominic Kwiatkowski, Cordelia Langford, John Sillitoe on behalf of the Wellcome Sanger Institute COVID-19 Surveillance Team |
| EPI_ISL_3436937 | Lighthouse Lab in Glasgow | Wellcome Sanger Institute for the COVID-19 Genomics UK (COG-UK) Consortium | Harper VanSteenhouse, Yumi Kasai, David Gray, Carol Clugston, Anna Dominiczak and Alex Alderton, Roberto Amato, Jeffrey Barrett, Sonia Goncalves, Ewan Harrison, David K. Jackson, Ian Johnston, Dominic Kwiatkowski, Cordelia Langford, John Sillitoe on behalf of the Wellcome Sanger Institute COVID-19 Surveillance Team |
| EPI_ISL_3436938 | Lighthouse Lab in Alderley Park | Wellcome Sanger Institute for the COVID-19 Genomics UK (COG-UK) Consortium | Jacquelyn Wynn, Mairead Hyland, The Lighthouse Lab in Alderley Park and Alex Alderton, Roberto Amato, Jeffrey Barrett, Sonia Goncalves, Ewan Harrison, David K. Jackson, Ian Johnston, Dominic Kwiatkowski, Cordelia Langford, John Sillitoe on behalf of the Wellcome Sanger Institute COVID-19 Surveillance Team |
| EPI_ISL_3436939, EPI_ISL_3436940 | Lighthouse Lab in Glasgow | Wellcome Sanger Institute for the COVID-19 Genomics UK (COG-UK) Consortium | Harper VanSteenhouse, Yumi Kasai, David Gray, Carol Clugston, Anna Dominiczak and Alex Alderton, Roberto Amato, Jeffrey Barrett, Sonia Goncalves, Ewan Harrison, David K. Jackson, Ian Johnston, Dominic Kwiatkowski, Cordelia Langford, John Sillitoe on behalf of the Wellcome Sanger Institute COVID-19 Surveillance Team |
| EPI_ISL_3436941 | Berkshire and Surrey Pathology Services Lighthouse Laboratory | Wellcome Sanger Institute for the COVID-19 Genomics UK (COG-UK) Consortium | Berkshire and Surrey Pathology Services Lighthouse Laboratory and Alex Alderton, Roberto Amato, Jeffrey Barrett, Sonia Goncalves, Ewan Harrison, David K. Jackson, Ian Johnston, Dominic Kwiatkowski, Cordelia Langford, John Sillitoe on behalf of the Wellcome Sanger Institute COVID-19 Surveillance Team |
| EPI_ISL_3436942 | Lighthouse Lab in Glasgow | Wellcome Sanger Institute for the COVID-19 Genomics UK (COG-UK) Consortium | Harper VanSteenhouse, Yumi Kasai, David Gray, Carol Clugston, Anna Dominiczak and Alex Alderton, Roberto Amato, Jeffrey Barrett, Sonia Goncalves, Ewan Harrison, David K. Jackson, Ian Johnston, Dominic Kwiatkowski, Cordelia Langford, John Sillitoe on behalf of the Wellcome Sanger Institute COVID-19 Surveillance Team |
| EPI_ISL_3436943 | Berkshire and Surrey Pathology Services Lighthouse Laboratory | Wellcome Sanger Institute for the COVID-19 Genomics UK (COG-UK) Consortium | Berkshire and Surrey Pathology Services Lighthouse Laboratory and Alex Alderton, Roberto Amato, Jeffrey Barrett, Sonia Goncalves, Ewan Harrison, David K. Jackson, Ian Johnston, Dominic Kwiatkowski, Cordelia Langford, John Sillitoe on behalf of the Wellcome Sanger Institute COVID-19 Surveillance Team |
| EPI_ISL_3436944 | Lighthouse Lab in Glasgow | Wellcome Sanger Institute for the COVID-19 Genomics UK (COG-UK) Consortium | Harper VanSteenhouse, Yumi Kasai, David Gray, Carol Clugston, Anna Dominiczak and Alex Alderton, Roberto Amato, Jeffrey Barrett, Sonia Goncalves, Ewan Harrison, David K. Jackson, Ian Johnston, Dominic Kwiatkowski, Cordelia Langford, John Sillitoe on behalf of the Wellcome Sanger Institute COVID-19 Surveillance Team |
| EPI_ISL_3436945, EPI_ISL_3436946 | Lighthouse Lab in Alderley Park | Wellcome Sanger Institute for the COVID-19 Genomics UK (COG-UK) Consortium | Jacquelyn Wynn, Mairead Hyland, The Lighthouse Lab in Alderley Park and Alex Alderton, Roberto Amato, Jeffrey Barrett, Sonia Goncalves, Ewan Harrison, David K. Jackson, Ian Johnston, Dominic Kwiatkowski, Cordelia Langford, John Sillitoe on behalf of the Wellcome Sanger Institute COVID-19 Surveillance Team |
| EPI_ISL_3436947, EPI_ISL_3436948, EPI_ISL_3436949 | Berkshire and Surrey Pathology Services Lighthouse Laboratory | Wellcome Sanger Institute for the COVID-19 Genomics UK (COG-UK) Consortium | Berkshire and Surrey Pathology Services Lighthouse Laboratory and Alex Alderton, Roberto Amato, Jeffrey Barrett, Sonia Goncalves, Ewan Harrison, David K. Jackson, Ian Johnston, Dominic Kwiatkowski, Cordelia Langford, John Sillitoe on behalf of the Wellcome Sanger Institute COVID-19 Surveillance Team |
| EPI_ISL_3436950 | Lighthouse Lab in Glasgow | Wellcome Sanger Institute for the COVID-19 Genomics UK (COG-UK) Consortium | Harper VanSteenhouse, Yumi Kasai, David Gray, Carol Clugston, Anna Dominiczak and Alex Alderton, Roberto Amato, Jeffrey Barrett, Sonia Goncalves, Ewan Harrison, David K. Jackson, Ian Johnston, Dominic Kwiatkowski, Cordelia Langford, John Sillitoe on behalf of the Wellcome Sanger Institute COVID-19 Surveillance Team |
| EPI_ISL_3436951 | Berkshire and Surrey Pathology Services Lighthouse Laboratory | Wellcome Sanger Institute for the COVID-19 Genomics UK (COG-UK) Consortium | Berkshire and Surrey Pathology Services Lighthouse Laboratory and Alex Alderton, Roberto Amato, Jeffrey Barrett, Sonia Goncalves, Ewan Harrison, David K. Jackson, Ian Johnston, Dominic Kwiatkowski, Cordelia Langford, John Sillitoe on behalf of the Wellcome Sanger Institute COVID-19 Surveillance Team |
| EPI_ISL_3436952 | Lighthouse Lab in Alderley Park | Wellcome Sanger Institute for the COVID-19 Genomics UK (COG-UK) Consortium | Jacquelyn Wynn, Mairead Hyland, The Lighthouse Lab in Alderley Park and Alex Alderton, Roberto Amato, Jeffrey Barrett, Sonia Goncalves, Ewan Harrison, David K. Jackson, Ian Johnston, Dominic Kwiatkowski, Cordelia Langford, John Sillitoe on behalf of the Wellcome Sanger Institute COVID-19 Surveillance Team |
| EPI_ISL_3436953, EPI_ISL_3436954, EPI_ISL_3436955, EPI_ISL_3436956 | Berkshire and Surrey Pathology Services Lighthouse Laboratory | Wellcome Sanger Institute for the COVID-19 Genomics UK (COG-UK) Consortium | Berkshire and Surrey Pathology Services Lighthouse Laboratory and Alex Alderton, Roberto Amato, Jeffrey Barrett, Sonia Goncalves, Ewan Harrison, David K. Jackson, Ian Johnston, Dominic Kwiatkowski, Cordelia Langford, John Sillitoe on behalf of the Wellcome Sanger Institute COVID-19 Surveillance Team |
| EPI_ISL_3436957 | Lighthouse Lab in Glasgow | Wellcome Sanger Institute for the COVID-19 Genomics UK (COG-UK) Consortium | Harper VanSteenhouse, Yumi Kasai, David Gray, Carol Clugston, Anna Dominiczak and Alex Alderton, Roberto Amato, Jeffrey Barrett, Sonia Goncalves, Ewan Harrison, David K. Jackson, Ian Johnston, Dominic Kwiatkowski, Cordelia Langford, John Sillitoe on behalf of the Wellcome Sanger Institute COVID-19 Surveillance Team |
| EPI_ISL_3436958 | Berkshire and Surrey Pathology Services Lighthouse Laboratory | Wellcome Sanger Institute for the COVID-19 Genomics UK (COG-UK) Consortium | Berkshire and Surrey Pathology Services Lighthouse Laboratory and Alex Alderton, Roberto Amato, Jeffrey Barrett, Sonia Goncalves, Ewan Harrison, David K. Jackson, Ian Johnston, Dominic Kwiatkowski, Cordelia Langford, John Sillitoe on behalf of the Wellcome Sanger Institute COVID-19 Surveillance Team |
| EPI_ISL_3436959 | Lighthouse Lab in Alderley Park | Wellcome Sanger Institute for the COVID-19 Genomics UK (COG-UK) Consortium | Jacquelyn Wynn, Mairead Hyland, The Lighthouse Lab in Alderley Park and Alex Alderton, Roberto Amato, Jeffrey Barrett, Sonia Goncalves, Ewan Harrison, David K. Jackson, Ian Johnston, Dominic Kwiatkowski, Cordelia Langford, John Sillitoe on behalf of the Wellcome Sanger Institute COVID-19 Surveillance Team |
| EPI_ISL_3436960, EPI_ISL_3436961, EPI_ISL_3436962, EPI_ISL_3436963 | Berkshire and Surrey Pathology Services Lighthouse Laboratory | Wellcome Sanger Institute for the COVID-19 Genomics UK (COG-UK) Consortium | Berkshire and Surrey Pathology Services Lighthouse Laboratory and Alex Alderton, Roberto Amato, Jeffrey Barrett, Sonia Goncalves, Ewan Harrison, David K. Jackson, Ian Johnston, Dominic Kwiatkowski, Cordelia Langford, John Sillitoe on behalf of the Wellcome Sanger Institute COVID-19 Surveillance Team |
| EPI_ISL_3436964 | Lighthouse Lab in Glasgow | Wellcome Sanger Institute for the COVID-19 Genomics UK (COG-UK) Consortium | Harper VanSteenhouse, Yumi Kasai, David Gray, Carol Clugston, Anna Dominiczak and Alex Alderton, Roberto Amato, Jeffrey Barrett, Sonia Goncalves, Ewan Harrison, David K. Jackson, Ian Johnston, Dominic Kwiatkowski, Cordelia Langford, John Sillitoe on behalf of the Wellcome Sanger Institute COVID-19 Surveillance Team |
| EPI_ISL_3436965 | Berkshire and Surrey Pathology Services Lighthouse Laboratory | Wellcome Sanger Institute for the COVID-19 Genomics UK (COG-UK) Consortium | Berkshire and Surrey Pathology Services Lighthouse Laboratory and Alex Alderton, Roberto Amato, Jeffrey Barrett, Sonia Goncalves, Ewan Harrison, David K. Jackson, Ian Johnston, Dominic Kwiatkowski, Cordelia Langford, John Sillitoe on behalf of the Wellcome Sanger Institute COVID-19 Surveillance Team |
| EPI_ISL_3436966 | Lighthouse Lab in Alderley Park | Wellcome Sanger Institute for the COVID-19 Genomics UK (COG-UK) Consortium | Jacquelyn Wynn, Mairead Hyland, The Lighthouse Lab in Alderley Park and Alex Alderton, Roberto Amato, Jeffrey Barrett, Sonia Goncalves, Ewan Harrison, David K. Jackson, Ian Johnston, Dominic Kwiatkowski, Cordelia Langford, John Sillitoe on behalf of the Wellcome Sanger Institute COVID-19 Surveillance Team |
| EPI_ISL_3436967, EPI_ISL_3436968 | Berkshire and Surrey Pathology Services Lighthouse Laboratory | Wellcome Sanger Institute for the COVID-19 Genomics UK (COG-UK) Consortium | Berkshire and Surrey Pathology Services Lighthouse Laboratory and Alex Alderton, Roberto Amato, Jeffrey Barrett, Sonia Goncalves, Ewan Harrison, David K. Jackson, Ian Johnston, Dominic Kwiatkowski, Cordelia Langford, John Sillitoe on behalf of the Wellcome Sanger Institute COVID-19 Surveillance Team |
| EPI_ISL_3436969 | Lighthouse Lab in Glasgow | Wellcome Sanger Institute for the COVID-19 Genomics UK (COG-UK) Consortium | Harper VanSteenhouse, Yumi Kasai, David Gray, Carol Clugston, Anna Dominiczak and Alex Alderton, Roberto Amato, Jeffrey Barrett, Sonia Goncalves, Ewan Harrison, David K. Jackson, Ian Johnston, Dominic Kwiatkowski, Cordelia Langford, John Sillitoe on behalf of the Wellcome Sanger Institute COVID-19 Surveillance Team |
| EPI_ISL_3436970, EPI_ISL_3436971 | Berkshire and Surrey Pathology Services Lighthouse Laboratory | Wellcome Sanger Institute for the COVID-19 Genomics UK (COG-UK) Consortium | Berkshire and Surrey Pathology Services Lighthouse Laboratory and Alex Alderton, Roberto Amato, Jeffrey Barrett, Sonia Goncalves, Ewan Harrison, David K. Jackson, Ian Johnston, Dominic Kwiatkowski, Cordelia Langford, John Sillitoe on behalf of the Wellcome Sanger Institute COVID-19 Surveillance Team |
| EPI_ISL_3436972 | Lighthouse Lab in Glasgow | Wellcome Sanger Institute for the COVID-19 Genomics UK (COG-UK) Consortium | Harper VanSteenhouse, Yumi Kasai, David Gray, Carol Clugston, Anna Dominiczak and Alex Alderton, Roberto Amato, Jeffrey Barrett, Sonia Goncalves, Ewan Harrison, David K. Jackson, Ian Johnston, Dominic Kwiatkowski, Cordelia Langford, John Sillitoe on behalf of the Wellcome Sanger Institute COVID-19 Surveillance Team |
| EPI_ISL_3436973 | Berkshire and Surrey Pathology Services Lighthouse Laboratory | Wellcome Sanger Institute for the COVID-19 Genomics UK (COG-UK) Consortium | Berkshire and Surrey Pathology Services Lighthouse Laboratory and Alex Alderton, Roberto Amato, Jeffrey Barrett, Sonia Goncalves, Ewan Harrison, David K. Jackson, Ian Johnston, Dominic Kwiatkowski, Cordelia Langford, John Sillitoe on behalf of the Wellcome Sanger Institute COVID-19 Surveillance Team |
| EPI_ISL_3436974 | Lighthouse Lab in Alderley Park | Wellcome Sanger Institute for the COVID-19 Genomics UK (COG-UK) Consortium | Jacquelyn Wynn, Mairead Hyland, The Lighthouse Lab in Alderley Park and Alex Alderton, Roberto Amato, Jeffrey Barrett, Sonia Goncalves, Ewan Harrison, David K. Jackson, Ian Johnston, Dominic Kwiatkowski, Cordelia Langford, John Sillitoe on behalf of the Wellcome Sanger Institute COVID-19 Surveillance Team |
| EPI_ISL_3436975 | Lighthouse Lab in Glasgow | Wellcome Sanger Institute for the COVID-19 Genomics UK (COG-UK) Consortium | Harper VanSteenhouse, Yumi Kasai, David Gray, Carol Clugston, Anna Dominiczak and Alex Alderton, Roberto Amato, Jeffrey Barrett, Sonia Goncalves, Ewan Harrison, David K. Jackson, Ian Johnston, Dominic Kwiatkowski, Cordelia Langford, John Sillitoe on behalf of the Wellcome Sanger Institute COVID-19 Surveillance Team |
| EPI_ISL_3436976, EPI_ISL_3436977 | Berkshire and Surrey Pathology Services Lighthouse Laboratory | Wellcome Sanger Institute for the COVID-19 Genomics UK (COG-UK) Consortium | Berkshire and Surrey Pathology Services Lighthouse Laboratory and Alex Alderton, Roberto Amato, Jeffrey Barrett, Sonia Goncalves, Ewan Harrison, David K. Jackson, Ian Johnston, Dominic Kwiatkowski, Cordelia Langford, John Sillitoe on behalf of the Wellcome Sanger Institute COVID-19 Surveillance Team |
| EPI_ISL_3436978, EPI_ISL_3436979 | Lighthouse Lab in Alderley Park | Wellcome Sanger Institute for the COVID-19 Genomics UK (COG-UK) Consortium | Jacquelyn Wynn, Mairead Hyland, The Lighthouse Lab in Alderley Park and Alex Alderton, Roberto Amato, Jeffrey Barrett, Sonia Goncalves, Ewan Harrison, David K. Jackson, Ian Johnston, Dominic Kwiatkowski, Cordelia Langford, John Sillitoe on behalf of the Wellcome Sanger Institute COVID-19 Surveillance Team |
| EPI_ISL_3436980, EPI_ISL_3436981 | Berkshire and Surrey Pathology Services Lighthouse Laboratory | Wellcome Sanger Institute for the COVID-19 Genomics UK (COG-UK) Consortium | Berkshire and Surrey Pathology Services Lighthouse Laboratory and Alex Alderton, Roberto Amato, Jeffrey Barrett, Sonia Goncalves, Ewan Harrison, David K. Jackson, Ian Johnston, Dominic Kwiatkowski, Cordelia Langford, John Sillitoe on behalf of the Wellcome Sanger Institute COVID-19 Surveillance Team |
| EPI_ISL_3436982, EPI_ISL_3436983 | Lighthouse Lab in Alderley Park | Wellcome Sanger Institute for the COVID-19 Genomics UK (COG-UK) Consortium | Jacquelyn Wynn, Mairead Hyland, The Lighthouse Lab in Alderley Park and Alex Alderton, Roberto Amato, Jeffrey Barrett, Sonia Goncalves, Ewan Harrison, David K. Jackson, Ian Johnston, Dominic Kwiatkowski, Cordelia Langford, John Sillitoe on behalf of the Wellcome Sanger Institute COVID-19 Surveillance Team |
| EPI_ISL_3436984 | Berkshire and Surrey Pathology Services Lighthouse Laboratory | Wellcome Sanger Institute for the COVID-19 Genomics UK (COG-UK) Consortium | Berkshire and Surrey Pathology Services Lighthouse Laboratory and Alex Alderton, Roberto Amato, Jeffrey Barrett, Sonia Goncalves, Ewan Harrison, David K. Jackson, Ian Johnston, Dominic Kwiatkowski, Cordelia Langford, John Sillitoe on behalf of the Wellcome Sanger Institute COVID-19 Surveillance Team |
| EPI_ISL_3436985 | Lighthouse Lab in Alderley Park | Wellcome Sanger Institute for the COVID-19 Genomics UK (COG-UK) Consortium | Jacquelyn Wynn, Mairead Hyland, The Lighthouse Lab in Alderley Park and Alex Alderton, Roberto Amato, Jeffrey Barrett, Sonia Goncalves, Ewan Harrison, David K. Jackson, Ian Johnston, Dominic Kwiatkowski, Cordelia Langford, John Sillitoe on behalf of the Wellcome Sanger Institute COVID-19 Surveillance Team |
| EPI_ISL_3436986 | Lighthouse Lab in Glasgow | Wellcome Sanger Institute for the COVID-19 Genomics UK (COG-UK) Consortium | Harper VanSteenhouse, Yumi Kasai, David Gray, Carol Clugston, Anna Dominiczak and Alex Alderton, Roberto Amato, Jeffrey Barrett, Sonia Goncalves, Ewan Harrison, David K. Jackson, Ian Johnston, Dominic Kwiatkowski, Cordelia Langford, John Sillitoe on behalf of the Wellcome Sanger Institute COVID-19 Surveillance Team |
| EPI_ISL_3436987, | Berkshire and Surrey Pathology Services Lighthouse Laboratory | Wellcome Sanger Institute for the COVID-19 Genomics UK (COG-UK) | Berkshire and Surrey Pathology Services Lighthouse Laboratory and Alex Alderton, Roberto Amato, Jeffrey Barrett, Sonia Goncalves, Ewan Harrison, David K. Jackson, Ian Johnston, Dominic Kwiatkowski, Cordelia Langford, John Sillitoe on behalf |

| | | | |
|---|---|---|---|
| EPI_ISL_3436988, EPI_ISL_3436989 | | Consortium | of the Wellcome Sanger Institute COVID-19 Surveillance Team |
| EPI_ISL_3436990 | Lighthouse Lab in Alderley Park | Wellcome Sanger Institute for the COVID-19 Genomics UK (COG-UK) Consortium | Jacquelyn Wynn, Mairead Hyland, The Lighthouse Lab in Alderley Park and Alex Alderton, Roberto Amato, Jeffrey Barrett, Sonia Goncalves, Ewan Harrison, David K. Jackson, Ian Johnston, Dominic Kwiatkowski, Cordelia Langford, John Sillitoe on behalf of the Wellcome Sanger Institute COVID-19 Surveillance Team |
| EPI_ISL_3436991, EPI_ISL_3436992 | Berkshire and Surrey Pathology Services Lighthouse Laboratory | Wellcome Sanger Institute for the COVID-19 Genomics UK (COG-UK) Consortium | Berkshire and Surrey Pathology Services Lighthouse Laboratory and Alex Alderton, Roberto Amato, Jeffrey Barrett, Sonia Goncalves, Ewan Harrison, David K. Jackson, Ian Johnston, Dominic Kwiatkowski, Cordelia Langford, John Sillitoe on behalf of the Wellcome Sanger Institute COVID-19 Surveillance Team |
| EPI_ISL_3436993 | Lighthouse Lab in Alderley Park | Wellcome Sanger Institute for the COVID-19 Genomics UK (COG-UK) Consortium | Jacquelyn Wynn, Mairead Hyland, The Lighthouse Lab in Alderley Park and Alex Alderton, Roberto Amato, Jeffrey Barrett, Sonia Goncalves, Ewan Harrison, David K. Jackson, Ian Johnston, Dominic Kwiatkowski, Cordelia Langford, John Sillitoe on behalf of the Wellcome Sanger Institute COVID-19 Surveillance Team |
| EPI_ISL_3436994 | Lighthouse Lab in Glasgow | Wellcome Sanger Institute for the COVID-19 Genomics UK (COG-UK) Consortium | Harper VanSteenhouse, Yumi Kasai, David Gray, Carol Clugston, Anna Dominiczak and Alex Alderton, Roberto Amato, Jeffrey Barrett, Sonia Goncalves, Ewan Harrison, David K. Jackson, Ian Johnston, Dominic Kwiatkowski, Cordelia Langford, John Sillitoe on behalf of the Wellcome Sanger Institute COVID-19 Surveillance Team |
| EPI_ISL_3436995, EPI_ISL_3436996 | Berkshire and Surrey Pathology Services Lighthouse Laboratory | Wellcome Sanger Institute for the COVID-19 Genomics UK (COG-UK) Consortium | Berkshire and Surrey Pathology Services Lighthouse Laboratory and Alex Alderton, Roberto Amato, Jeffrey Barrett, Sonia Goncalves, Ewan Harrison, David K. Jackson, Ian Johnston, Dominic Kwiatkowski, Cordelia Langford, John Sillitoe on behalf of the Wellcome Sanger Institute COVID-19 Surveillance Team |
| EPI_ISL_3436997 | Lighthouse Lab in Alderley Park | Wellcome Sanger Institute for the COVID-19 Genomics UK (COG-UK) Consortium | Jacquelyn Wynn, Mairead Hyland, The Lighthouse Lab in Alderley Park and Alex Alderton, Roberto Amato, Jeffrey Barrett, Sonia Goncalves, Ewan Harrison, David K. Jackson, Ian Johnston, Dominic Kwiatkowski, Cordelia Langford, John Sillitoe on behalf of the Wellcome Sanger Institute COVID-19 Surveillance Team |
| EPI_ISL_3436998, EPI_ISL_3436999, EPI_ISL_3437000 | Berkshire and Surrey Pathology Services Lighthouse Laboratory | Wellcome Sanger Institute for the COVID-19 Genomics UK (COG-UK) Consortium | Berkshire and Surrey Pathology Services Lighthouse Laboratory and Alex Alderton, Roberto Amato, Jeffrey Barrett, Sonia Goncalves, Ewan Harrison, David K. Jackson, Ian Johnston, Dominic Kwiatkowski, Cordelia Langford, John Sillitoe on behalf of the Wellcome Sanger Institute COVID-19 Surveillance Team |
| EPI_ISL_3437001 | Lighthouse Lab in Alderley Park | Wellcome Sanger Institute for the COVID-19 Genomics UK (COG-UK) Consortium | Jacquelyn Wynn, Mairead Hyland, The Lighthouse Lab in Alderley Park and Alex Alderton, Roberto Amato, Jeffrey Barrett, Sonia Goncalves, Ewan Harrison, David K. Jackson, Ian Johnston, Dominic Kwiatkowski, Cordelia Langford, John Sillitoe on behalf of the Wellcome Sanger Institute COVID-19 Surveillance Team |
| EPI_ISL_3437002 | Berkshire and Surrey Pathology Services Lighthouse Laboratory | Wellcome Sanger Institute for the COVID-19 Genomics UK (COG-UK) Consortium | Berkshire and Surrey Pathology Services Lighthouse Laboratory and Alex Alderton, Roberto Amato, Jeffrey Barrett, Sonia Goncalves, Ewan Harrison, David K. Jackson, Ian Johnston, Dominic Kwiatkowski, Cordelia Langford, John Sillitoe on behalf of the Wellcome Sanger Institute COVID-19 Surveillance Team |
| EPI_ISL_3437003 | Lighthouse Lab in Glasgow | Wellcome Sanger Institute for the COVID-19 Genomics UK (COG-UK) Consortium | Harper VanSteenhouse, Yumi Kasai, David Gray, Carol Clugston, Anna Dominiczak and Alex Alderton, Roberto Amato, Jeffrey Barrett, Sonia Goncalves, Ewan Harrison, David K. Jackson, Ian Johnston, Dominic Kwiatkowski, Cordelia Langford, John Sillitoe on behalf of the Wellcome Sanger Institute COVID-19 Surveillance Team |
| EPI_ISL_3437004, EPI_ISL_3437005 | Berkshire and Surrey Pathology Services Lighthouse Laboratory | Wellcome Sanger Institute for the COVID-19 Genomics UK (COG-UK) Consortium | Berkshire and Surrey Pathology Services Lighthouse Laboratory and Alex Alderton, Roberto Amato, Jeffrey Barrett, Sonia Goncalves, Ewan Harrison, David K. Jackson, Ian Johnston, Dominic Kwiatkowski, Cordelia Langford, John Sillitoe on behalf of the Wellcome Sanger Institute COVID-19 Surveillance Team |
| EPI_ISL_3437006, EPI_ISL_3437007, EPI_ISL_3437008, EPI_ISL_3437009 | Lighthouse Lab in Alderley Park | Wellcome Sanger Institute for the COVID-19 Genomics UK (COG-UK) Consortium | Jacquelyn Wynn, Mairead Hyland, The Lighthouse Lab in Alderley Park and Alex Alderton, Roberto Amato, Jeffrey Barrett, Sonia Goncalves, Ewan Harrison, David K. Jackson, Ian Johnston, Dominic Kwiatkowski, Cordelia Langford, John Sillitoe on behalf of the Wellcome Sanger Institute COVID-19 Surveillance Team |
| EPI_ISL_3437010, EPI_ISL_3437011 | Berkshire and Surrey Pathology Services Lighthouse Laboratory | Wellcome Sanger Institute for the COVID-19 Genomics UK (COG-UK) Consortium | Berkshire and Surrey Pathology Services Lighthouse Laboratory and Alex Alderton, Roberto Amato, Jeffrey Barrett, Sonia Goncalves, Ewan Harrison, David K. Jackson, Ian Johnston, Dominic Kwiatkowski, Cordelia Langford, John Sillitoe on behalf of the Wellcome Sanger Institute COVID-19 Surveillance Team |
| EPI_ISL_3437012 | Lighthouse Lab in Alderley Park | Wellcome Sanger Institute for the COVID-19 Genomics UK (COG-UK) Consortium | Jacquelyn Wynn, Mairead Hyland, The Lighthouse Lab in Alderley Park and Alex Alderton, Roberto Amato, Jeffrey Barrett, Sonia Goncalves, Ewan Harrison, David K. Jackson, Ian Johnston, Dominic Kwiatkowski, Cordelia Langford, John Sillitoe on behalf of the Wellcome Sanger Institute COVID-19 Surveillance Team |
| EPI_ISL_3437013 | Berkshire and Surrey Pathology Services Lighthouse Laboratory | Wellcome Sanger Institute for the COVID-19 Genomics UK (COG-UK) Consortium | Berkshire and Surrey Pathology Services Lighthouse Laboratory and Alex Alderton, Roberto Amato, Jeffrey Barrett, Sonia Goncalves, Ewan Harrison, David K. Jackson, Ian Johnston, Dominic Kwiatkowski, Cordelia Langford, John Sillitoe on behalf of the Wellcome Sanger Institute COVID-19 Surveillance Team |
| EPI_ISL_3437014 | Lighthouse Lab in Glasgow | Wellcome Sanger Institute for the COVID-19 Genomics UK (COG-UK) Consortium | Harper VanSteenhouse, Yumi Kasai, David Gray, Carol Clugston, Anna Dominiczak and Alex Alderton, Roberto Amato, Jeffrey Barrett, Sonia Goncalves, Ewan Harrison, David K. Jackson, Ian Johnston, Dominic Kwiatkowski, Cordelia Langford, John Sillitoe on behalf of the Wellcome Sanger Institute COVID-19 Surveillance Team |
| EPI_ISL_3437015, EPI_ISL_3437016, EPI_ISL_3437017 | Berkshire and Surrey Pathology Services Lighthouse Laboratory | Wellcome Sanger Institute for the COVID-19 Genomics UK (COG-UK) Consortium | Berkshire and Surrey Pathology Services Lighthouse Laboratory and Alex Alderton, Roberto Amato, Jeffrey Barrett, Sonia Goncalves, Ewan Harrison, David K. Jackson, Ian Johnston, Dominic Kwiatkowski, Cordelia Langford, John Sillitoe on behalf of the Wellcome Sanger Institute COVID-19 Surveillance Team |
| EPI_ISL_3437018 | Lighthouse Lab in Alderley Park | Wellcome Sanger Institute for the COVID-19 Genomics UK (COG-UK) Consortium | Jacquelyn Wynn, Mairead Hyland, The Lighthouse Lab in Alderley Park and Alex Alderton, Roberto Amato, Jeffrey Barrett, Sonia Goncalves, Ewan Harrison, David K. Jackson, Ian Johnston, Dominic Kwiatkowski, Cordelia Langford, John Sillitoe on behalf of the Wellcome Sanger Institute COVID-19 Surveillance Team |
| EPI_ISL_3437019, EPI_ISL_3437020 | Berkshire and Surrey Pathology Services Lighthouse Laboratory | Wellcome Sanger Institute for the COVID-19 Genomics UK (COG-UK) Consortium | Berkshire and Surrey Pathology Services Lighthouse Laboratory and Alex Alderton, Roberto Amato, Jeffrey Barrett, Sonia Goncalves, Ewan Harrison, David K. Jackson, Ian Johnston, Dominic Kwiatkowski, Cordelia Langford, John Sillitoe on behalf of the Wellcome Sanger Institute COVID-19 Surveillance Team |
| EPI_ISL_3437021 | Lighthouse Lab in Alderley Park | Wellcome Sanger Institute for the COVID-19 Genomics UK (COG-UK) Consortium | Jacquelyn Wynn, Mairead Hyland, The Lighthouse Lab in Alderley Park and Alex Alderton, Roberto Amato, Jeffrey Barrett, Sonia Goncalves, Ewan Harrison, David K. Jackson, Ian Johnston, Dominic Kwiatkowski, Cordelia Langford, John Sillitoe on behalf of the Wellcome Sanger Institute COVID-19 Surveillance Team |
| EPI_ISL_3437022 | Lighthouse Lab in Glasgow | Wellcome Sanger Institute for the COVID-19 Genomics UK (COG-UK) Consortium | Harper VanSteenhouse, Yumi Kasai, David Gray, Carol Clugston, Anna Dominiczak and Alex Alderton, Roberto Amato, Jeffrey Barrett, Sonia Goncalves, Ewan Harrison, David K. Jackson, Ian Johnston, Dominic Kwiatkowski, Cordelia Langford, John Sillitoe on behalf of the Wellcome Sanger Institute COVID-19 Surveillance Team |
| EPI_ISL_3437023 | Lighthouse Lab in Alderley Park | Wellcome Sanger Institute for the COVID-19 Genomics UK (COG-UK) Consortium | Jacquelyn Wynn, Mairead Hyland, The Lighthouse Lab in Alderley Park and Alex Alderton, Roberto Amato, Jeffrey Barrett, Sonia Goncalves, Ewan Harrison, David K. Jackson, Ian Johnston, Dominic Kwiatkowski, Cordelia Langford, John Sillitoe on behalf of the Wellcome Sanger Institute COVID-19 Surveillance Team |
| EPI_ISL_3437024 | Berkshire and Surrey Pathology Services Lighthouse Laboratory | Wellcome Sanger Institute for the COVID-19 Genomics UK (COG-UK) Consortium | Berkshire and Surrey Pathology Services Lighthouse Laboratory and Alex Alderton, Roberto Amato, Jeffrey Barrett, Sonia Goncalves, Ewan Harrison, David K. Jackson, Ian Johnston, Dominic Kwiatkowski, Cordelia Langford, John Sillitoe on behalf of the Wellcome Sanger Institute COVID-19 Surveillance Team |
| EPI_ISL_3437025 | Lighthouse Lab in Glasgow | Wellcome Sanger Institute for the COVID-19 Genomics UK (COG-UK) Consortium | Harper VanSteenhouse, Yumi Kasai, David Gray, Carol Clugston, Anna Dominiczak and Alex Alderton, Roberto Amato, Jeffrey Barrett, Sonia Goncalves, Ewan Harrison, David K. Jackson, Ian Johnston, Dominic Kwiatkowski, Cordelia Langford, John Sillitoe on behalf of the Wellcome Sanger Institute COVID-19 Surveillance Team |
| EPI_ISL_3437026, EPI_ISL_3437027, EPI_ISL_3437028 | Berkshire and Surrey Pathology Services Lighthouse Laboratory | Wellcome Sanger Institute for the COVID-19 Genomics UK (COG-UK) Consortium | Berkshire and Surrey Pathology Services Lighthouse Laboratory and Alex Alderton, Roberto Amato, Jeffrey Barrett, Sonia Goncalves, Ewan Harrison, David K. Jackson, Ian Johnston, Dominic Kwiatkowski, Cordelia Langford, John Sillitoe on behalf of the Wellcome Sanger Institute COVID-19 Surveillance Team |
| EPI_ISL_3437029 | Lighthouse Lab in Alderley Park | Wellcome Sanger Institute for the COVID-19 Genomics UK (COG-UK) Consortium | Jacquelyn Wynn, Mairead Hyland, The Lighthouse Lab in Alderley Park and Alex Alderton, Roberto Amato, Jeffrey Barrett, Sonia Goncalves, Ewan Harrison, David K. Jackson, Ian Johnston, Dominic Kwiatkowski, Cordelia Langford, John Sillitoe on behalf of the Wellcome Sanger Institute COVID-19 Surveillance Team |
| EPI_ISL_3437030 | Lighthouse Lab in Glasgow | Wellcome Sanger Institute for the COVID-19 Genomics UK (COG-UK) Consortium | Harper VanSteenhouse, Yumi Kasai, David Gray, Carol Clugston, Anna Dominiczak and Alex Alderton, Roberto Amato, Jeffrey Barrett, Sonia Goncalves, Ewan Harrison, David K. Jackson, Ian Johnston, Dominic Kwiatkowski, Cordelia Langford, John Sillitoe on behalf of the Wellcome Sanger Institute COVID-19 Surveillance Team |
| EPI_ISL_3437031, EPI_ISL_3437032 | Lighthouse Lab in Alderley Park | Wellcome Sanger Institute for the COVID-19 Genomics UK (COG-UK) Consortium | Jacquelyn Wynn, Mairead Hyland, The Lighthouse Lab in Alderley Park and Alex Alderton, Roberto Amato, Jeffrey Barrett, Sonia Goncalves, Ewan Harrison, David K. Jackson, Ian Johnston, Dominic Kwiatkowski, Cordelia Langford, John Sillitoe on behalf of the Wellcome Sanger Institute COVID-19 Surveillance Team |
| EPI_ISL_3437033 | Berkshire and Surrey Pathology Services Lighthouse Laboratory | Wellcome Sanger Institute for the COVID-19 Genomics UK (COG-UK) Consortium | Berkshire and Surrey Pathology Services Lighthouse Laboratory and Alex Alderton, Roberto Amato, Jeffrey Barrett, Sonia Goncalves, Ewan Harrison, David K. Jackson, Ian Johnston, Dominic Kwiatkowski, Cordelia Langford, John Sillitoe on behalf of the Wellcome Sanger Institute COVID-19 Surveillance Team |
| EPI_ISL_3437034 | Lighthouse Lab in Glasgow | Wellcome Sanger Institute for the COVID-19 Genomics UK (COG-UK) Consortium | Harper VanSteenhouse, Yumi Kasai, David Gray, Carol Clugston, Anna Dominiczak and Alex Alderton, Roberto Amato, Jeffrey Barrett, Sonia Goncalves, Ewan Harrison, David K. Jackson, Ian Johnston, Dominic Kwiatkowski, Cordelia Langford, John Sillitoe on behalf of the Wellcome Sanger Institute COVID-19 Surveillance Team |
| EPI_ISL_3437035 | Berkshire and Surrey Pathology Services Lighthouse Laboratory | Wellcome Sanger Institute for the COVID-19 Genomics UK (COG-UK) Consortium | Berkshire and Surrey Pathology Services Lighthouse Laboratory and Alex Alderton, Roberto Amato, Jeffrey Barrett, Sonia Goncalves, Ewan Harrison, David K. Jackson, Ian Johnston, Dominic Kwiatkowski, Cordelia Langford, John Sillitoe on behalf of the Wellcome Sanger Institute COVID-19 Surveillance Team |
| EPI_ISL_3437036, EPI_ISL_3437037, EPI_ISL_3437038 | Lighthouse Lab in Alderley Park | Wellcome Sanger Institute for the COVID-19 Genomics UK (COG-UK) Consortium | Jacquelyn Wynn, Mairead Hyland, The Lighthouse Lab in Alderley Park and Alex Alderton, Roberto Amato, Jeffrey Barrett, Sonia Goncalves, Ewan Harrison, David K. Jackson, Ian Johnston, Dominic Kwiatkowski, Cordelia Langford, John Sillitoe on behalf of the Wellcome Sanger Institute COVID-19 Surveillance Team |
| EPI_ISL_3437039 | Berkshire and Surrey Pathology Services Lighthouse Laboratory | Wellcome Sanger Institute for the COVID-19 Genomics UK (COG-UK) Consortium | Berkshire and Surrey Pathology Services Lighthouse Laboratory and Alex Alderton, Roberto Amato, Jeffrey Barrett, Sonia Goncalves, Ewan Harrison, David K. Jackson, Ian Johnston, Dominic Kwiatkowski, Cordelia Langford, John Sillitoe on behalf of the Wellcome Sanger Institute COVID-19 Surveillance Team |
| EPI_ISL_3437040 | Lighthouse Lab in Alderley Park | Wellcome Sanger Institute for the COVID-19 Genomics UK (COG-UK) Consortium | Jacquelyn Wynn, Mairead Hyland, The Lighthouse Lab in Alderley Park and Alex Alderton, Roberto Amato, Jeffrey Barrett, Sonia Goncalves, Ewan Harrison, David K. Jackson, Ian Johnston, Dominic Kwiatkowski, Cordelia Langford, John Sillitoe on behalf of the Wellcome Sanger Institute COVID-19 Surveillance Team |
| EPI_ISL_3437041 | Lighthouse Lab in Glasgow | Wellcome Sanger Institute for the COVID-19 Genomics UK (COG-UK) Consortium | Harper VanSteenhouse, Yumi Kasai, David Gray, Carol Clugston, Anna Dominiczak and Alex Alderton, Roberto Amato, Jeffrey Barrett, Sonia Goncalves, Ewan Harrison, David K. Jackson, Ian Johnston, Dominic Kwiatkowski, Cordelia Langford, John Sillitoe on behalf of the Wellcome Sanger Institute COVID-19 Surveillance Team |
| EPI_ISL_3437042 | Berkshire and Surrey Pathology Services Lighthouse Laboratory | Wellcome Sanger Institute for the COVID-19 Genomics UK (COG-UK) Consortium | Berkshire and Surrey Pathology Services Lighthouse Laboratory and Alex Alderton, Roberto Amato, Jeffrey Barrett, Sonia Goncalves, Ewan Harrison, David K. Jackson, Ian Johnston, Dominic Kwiatkowski, Cordelia Langford, John Sillitoe on behalf of the Wellcome Sanger Institute COVID-19 Surveillance Team |
| EPI_ISL_3437043 | Lighthouse Lab in Alderley Park | Wellcome Sanger Institute for the COVID-19 Genomics UK (COG-UK) Consortium | Jacquelyn Wynn, Mairead Hyland, The Lighthouse Lab in Alderley Park and Alex Alderton, Roberto Amato, Jeffrey Barrett, Sonia Goncalves, Ewan Harrison, David K. Jackson, Ian Johnston, Dominic Kwiatkowski, Cordelia Langford, John Sillitoe on behalf of the Wellcome Sanger Institute COVID-19 Surveillance Team |
| EPI_ISL_3437044 | Lighthouse Laboratory Plymouth | Wellcome Sanger Institute for the COVID-19 Genomics UK (COG-UK) Consortium | Lighthouse Laboratory Plymouth and Alex Alderton, Roberto Amato, Jeffrey Barrett, Sonia Goncalves, Ewan Harrison, David K. Jackson, Ian Johnston, Dominic Kwiatkowski, Cordelia Langford, John Sillitoe on behalf of the Wellcome Sanger Institute COVID-19 Surveillance Team |
| EPI_ISL_3437045 | Berkshire and Surrey Pathology Services Lighthouse Laboratory | Wellcome Sanger Institute for the COVID-19 Genomics UK (COG-UK) Consortium | Berkshire and Surrey Pathology Services Lighthouse Laboratory and Alex Alderton, Roberto Amato, Jeffrey Barrett, Sonia Goncalves, Ewan Harrison, David K. Jackson, Ian Johnston, Dominic Kwiatkowski, Cordelia Langford, John Sillitoe on behalf of the Wellcome Sanger Institute COVID-19 Surveillance Team |
| EPI_ISL_3437046 | Lighthouse Laboratory Plymouth | Wellcome Sanger Institute for the COVID-19 Genomics UK (COG-UK) Consortium | Lighthouse Laboratory Plymouth and Alex Alderton, Roberto Amato, Jeffrey Barrett, Sonia Goncalves, Ewan Harrison, David K. Jackson, Ian Johnston, Dominic Kwiatkowski, Cordelia Langford, John Sillitoe on behalf of the Wellcome Sanger Institute COVID-19 Surveillance Team |
| EPI_ISL_3437047 | Berkshire and Surrey Pathology Services Lighthouse Laboratory | Wellcome Sanger Institute for the COVID-19 Genomics UK (COG-UK) Consortium | Berkshire and Surrey Pathology Services Lighthouse Laboratory and Alex Alderton, Roberto Amato, Jeffrey Barrett, Sonia Goncalves, Ewan Harrison, David K. Jackson, Ian Johnston, Dominic Kwiatkowski, Cordelia Langford, John Sillitoe on behalf of the Wellcome Sanger Institute COVID-19 Surveillance Team |
| EPI_ISL_3437048 | Lighthouse Lab in Alderley Park | Wellcome Sanger Institute for the COVID-19 Genomics UK (COG-UK) Consortium | Jacquelyn Wynn, Mairead Hyland, The Lighthouse Lab in Alderley Park and Alex Alderton, Roberto Amato, Jeffrey Barrett, Sonia Goncalves, Ewan Harrison, David K. Jackson, Ian Johnston, Dominic Kwiatkowski, Cordelia Langford, John Sillitoe on behalf of the Wellcome Sanger Institute COVID-19 Surveillance Team |

| | | | |
|---|---|---|---|
| EPI_ISL_3437049 | Health Services Laboratories | Wellcome Sanger Institute for the COVID-19 Genomics UK (COG-UK) Consortium | Health Services Laboratories and Alex Alderton, Roberto Amato, Jeffrey Barrett, Sonia Goncalves, Ewan Harrison, David K. Jackson, Ian Johnston, Dominic Kwiatkowski, Cordelia Langford, John Sillitoe on behalf of the Wellcome Sanger Institute COVID-19 Surveillance Team |
| EPI_ISL_3437050 | Lighthouse Lab in Alderley Park | Wellcome Sanger Institute for the COVID-19 Genomics UK (COG-UK) Consortium | Jacquelyn Wynn, Mairead Hyland, The Lighthouse Lab in Alderley Park and Alex Alderton, Roberto Amato, Jeffrey Barrett, Sonia Goncalves, Ewan Harrison, David K. Jackson, Ian Johnston, Dominic Kwiatkowski, Cordelia Langford, John Sillitoe on behalf of the Wellcome Sanger Institute COVID-19 Surveillance Team |
| EPI_ISL_3437051 | Lighthouse Laboratory Plymouth | Wellcome Sanger Institute for the COVID-19 Genomics UK (COG-UK) Consortium | Lighthouse Laboratory Plymouth and Alex Alderton, Roberto Amato, Jeffrey Barrett, Sonia Goncalves, Ewan Harrison, David K. Jackson, Ian Johnston, Dominic Kwiatkowski, Cordelia Langford, John Sillitoe on behalf of the Wellcome Sanger Institute COVID-19 Surveillance Team |
| EPI_ISL_3437052 | Berkshire and Surrey Pathology Services Lighthouse Laboratory | Wellcome Sanger Institute for the COVID-19 Genomics UK (COG-UK) Consortium | Berkshire and Surrey Pathology Services Lighthouse Laboratory and Alex Alderton, Roberto Amato, Jeffrey Barrett, Sonia Goncalves, Ewan Harrison, David K. Jackson, Ian Johnston, Dominic Kwiatkowski, Cordelia Langford, John Sillitoe on behalf of the Wellcome Sanger Institute COVID-19 Surveillance Team |
| EPI_ISL_3437053 | Health Services Laboratories | Wellcome Sanger Institute for the COVID-19 Genomics UK (COG-UK) Consortium | Health Services Laboratories and Alex Alderton, Roberto Amato, Jeffrey Barrett, Sonia Goncalves, Ewan Harrison, David K. Jackson, Ian Johnston, Dominic Kwiatkowski, Cordelia Langford, John Sillitoe on behalf of the Wellcome Sanger Institute COVID-19 Surveillance Team |
| EPI_ISL_3437054 | Berkshire and Surrey Pathology Services Lighthouse Laboratory | Wellcome Sanger Institute for the COVID-19 Genomics UK (COG-UK) Consortium | Berkshire and Surrey Pathology Services Lighthouse Laboratory and Alex Alderton, Roberto Amato, Jeffrey Barrett, Sonia Goncalves, Ewan Harrison, David K. Jackson, Ian Johnston, Dominic Kwiatkowski, Cordelia Langford, John Sillitoe on behalf of the Wellcome Sanger Institute COVID-19 Surveillance Team |
| EPI_ISL_3437055 | Lighthouse Lab in Alderley Park | Wellcome Sanger Institute for the COVID-19 Genomics UK (COG-UK) Consortium | Jacquelyn Wynn, Mairead Hyland, The Lighthouse Lab in Alderley Park and Alex Alderton, Roberto Amato, Jeffrey Barrett, Sonia Goncalves, Ewan Harrison, David K. Jackson, Ian Johnston, Dominic Kwiatkowski, Cordelia Langford, John Sillitoe on behalf of the Wellcome Sanger Institute COVID-19 Surveillance Team |
| EPI_ISL_3437056 | Berkshire and Surrey Pathology Services Lighthouse Laboratory | Wellcome Sanger Institute for the COVID-19 Genomics UK (COG-UK) Consortium | Berkshire and Surrey Pathology Services Lighthouse Laboratory and Alex Alderton, Roberto Amato, Jeffrey Barrett, Sonia Goncalves, Ewan Harrison, David K. Jackson, Ian Johnston, Dominic Kwiatkowski, Cordelia Langford, John Sillitoe on behalf of the Wellcome Sanger Institute COVID-19 Surveillance Team |
| EPI_ISL_3437057 | Lighthouse Lab in Alderley Park | Wellcome Sanger Institute for the COVID-19 Genomics UK (COG-UK) Consortium | Jacquelyn Wynn, Mairead Hyland, The Lighthouse Lab in Alderley Park and Alex Alderton, Roberto Amato, Jeffrey Barrett, Sonia Goncalves, Ewan Harrison, David K. Jackson, Ian Johnston, Dominic Kwiatkowski, Cordelia Langford, John Sillitoe on behalf of the Wellcome Sanger Institute COVID-19 Surveillance Team |
| EPI_ISL_3437058 | Lighthouse Laboratory Plymouth | Wellcome Sanger Institute for the COVID-19 Genomics UK (COG-UK) Consortium | Lighthouse Laboratory Plymouth and Alex Alderton, Roberto Amato, Jeffrey Barrett, Sonia Goncalves, Ewan Harrison, David K. Jackson, Ian Johnston, Dominic Kwiatkowski, Cordelia Langford, John Sillitoe on behalf of the Wellcome Sanger Institute COVID-19 Surveillance Team |
| EPI_ISL_3437059 | Health Services Laboratories | Wellcome Sanger Institute for the COVID-19 Genomics UK (COG-UK) Consortium | Health Services Laboratories and Alex Alderton, Roberto Amato, Jeffrey Barrett, Sonia Goncalves, Ewan Harrison, David K. Jackson, Ian Johnston, Dominic Kwiatkowski, Cordelia Langford, John Sillitoe on behalf of the Wellcome Sanger Institute COVID-19 Surveillance Team |
| EPI_ISL_3437060 | Lighthouse Lab in Alderley Park | Wellcome Sanger Institute for the COVID-19 Genomics UK (COG-UK) Consortium | Jacquelyn Wynn, Mairead Hyland, The Lighthouse Lab in Alderley Park and Alex Alderton, Roberto Amato, Jeffrey Barrett, Sonia Goncalves, Ewan Harrison, David K. Jackson, Ian Johnston, Dominic Kwiatkowski, Cordelia Langford, John Sillitoe on behalf of the Wellcome Sanger Institute COVID-19 Surveillance Team |
| EPI_ISL_3437061 | Berkshire and Surrey Pathology Services Lighthouse Laboratory | Wellcome Sanger Institute for the COVID-19 Genomics UK (COG-UK) Consortium | Berkshire and Surrey Pathology Services Lighthouse Laboratory and Alex Alderton, Roberto Amato, Jeffrey Barrett, Sonia Goncalves, Ewan Harrison, David K. Jackson, Ian Johnston, Dominic Kwiatkowski, Cordelia Langford, John Sillitoe on behalf of the Wellcome Sanger Institute COVID-19 Surveillance Team |
| EPI_ISL_3437062 | Lighthouse Lab in Alderley Park | Wellcome Sanger Institute for the COVID-19 Genomics UK (COG-UK) Consortium | Jacquelyn Wynn, Mairead Hyland, The Lighthouse Lab in Alderley Park and Alex Alderton, Roberto Amato, Jeffrey Barrett, Sonia Goncalves, Ewan Harrison, David K. Jackson, Ian Johnston, Dominic Kwiatkowski, Cordelia Langford, John Sillitoe on behalf of the Wellcome Sanger Institute COVID-19 Surveillance Team |
| EPI_ISL_3437063, EPI_ISL_3437064 | Health Services Laboratories | Wellcome Sanger Institute for the COVID-19 Genomics UK (COG-UK) Consortium | Health Services Laboratories and Alex Alderton, Roberto Amato, Jeffrey Barrett, Sonia Goncalves, Ewan Harrison, David K. Jackson, Ian Johnston, Dominic Kwiatkowski, Cordelia Langford, John Sillitoe on behalf of the Wellcome Sanger Institute COVID-19 Surveillance Team |
| EPI_ISL_3437065 | Lighthouse Lab in Alderley Park | Wellcome Sanger Institute for the COVID-19 Genomics UK (COG-UK) Consortium | Jacquelyn Wynn, Mairead Hyland, The Lighthouse Lab in Alderley Park and Alex Alderton, Roberto Amato, Jeffrey Barrett, Sonia Goncalves, Ewan Harrison, David K. Jackson, Ian Johnston, Dominic Kwiatkowski, Cordelia Langford, John Sillitoe on behalf of the Wellcome Sanger Institute COVID-19 Surveillance Team |
| EPI_ISL_3437066, EPI_ISL_3437067 | Berkshire and Surrey Pathology Services Lighthouse Laboratory | Wellcome Sanger Institute for the COVID-19 Genomics UK (COG-UK) Consortium | Berkshire and Surrey Pathology Services Lighthouse Laboratory and Alex Alderton, Roberto Amato, Jeffrey Barrett, Sonia Goncalves, Ewan Harrison, David K. Jackson, Ian Johnston, Dominic Kwiatkowski, Cordelia Langford, John Sillitoe on behalf of the Wellcome Sanger Institute COVID-19 Surveillance Team |
| EPI_ISL_3437068 | Health Services Laboratories | Wellcome Sanger Institute for the COVID-19 Genomics UK (COG-UK) Consortium | Health Services Laboratories and Alex Alderton, Roberto Amato, Jeffrey Barrett, Sonia Goncalves, Ewan Harrison, David K. Jackson, Ian Johnston, Dominic Kwiatkowski, Cordelia Langford, John Sillitoe on behalf of the Wellcome Sanger Institute COVID-19 Surveillance Team |
| EPI_ISL_3437069 | Lighthouse Laboratory Plymouth | Wellcome Sanger Institute for the COVID-19 Genomics UK (COG-UK) Consortium | Lighthouse Laboratory Plymouth and Alex Alderton, Roberto Amato, Jeffrey Barrett, Sonia Goncalves, Ewan Harrison, David K. Jackson, Ian Johnston, Dominic Kwiatkowski, Cordelia Langford, John Sillitoe on behalf of the Wellcome Sanger Institute COVID-19 Surveillance Team |
| EPI_ISL_3437070, EPI_ISL_3437071, EPI_ISL_3437072 | Lighthouse Lab in Alderley Park | Wellcome Sanger Institute for the COVID-19 Genomics UK (COG-UK) Consortium | Jacquelyn Wynn, Mairead Hyland, The Lighthouse Lab in Alderley Park and Alex Alderton, Roberto Amato, Jeffrey Barrett, Sonia Goncalves, Ewan Harrison, David K. Jackson, Ian Johnston, Dominic Kwiatkowski, Cordelia Langford, John Sillitoe on behalf of the Wellcome Sanger Institute COVID-19 Surveillance Team |
| EPI_ISL_3437073 | Health Services Laboratories | Wellcome Sanger Institute for the COVID-19 Genomics UK (COG-UK) Consortium | Health Services Laboratories and Alex Alderton, Roberto Amato, Jeffrey Barrett, Sonia Goncalves, Ewan Harrison, David K. Jackson, Ian Johnston, Dominic Kwiatkowski, Cordelia Langford, John Sillitoe on behalf of the Wellcome Sanger Institute COVID-19 Surveillance Team |
| EPI_ISL_3437074, EPI_ISL_3437075 | Berkshire and Surrey Pathology Services Lighthouse Laboratory | Wellcome Sanger Institute for the COVID-19 Genomics UK (COG-UK) Consortium | Berkshire and Surrey Pathology Services Lighthouse Laboratory and Alex Alderton, Roberto Amato, Jeffrey Barrett, Sonia Goncalves, Ewan Harrison, David K. Jackson, Ian Johnston, Dominic Kwiatkowski, Cordelia Langford, John Sillitoe on behalf of the Wellcome Sanger Institute COVID-19 Surveillance Team |
| EPI_ISL_3437076 | Lighthouse Lab in Alderley Park | Wellcome Sanger Institute for the COVID-19 Genomics UK (COG-UK) Consortium | Jacquelyn Wynn, Mairead Hyland, The Lighthouse Lab in Alderley Park and Alex Alderton, Roberto Amato, Jeffrey Barrett, Sonia Goncalves, Ewan Harrison, David K. Jackson, Ian Johnston, Dominic Kwiatkowski, Cordelia Langford, John Sillitoe on behalf of the Wellcome Sanger Institute COVID-19 Surveillance Team |
| EPI_ISL_3437077 | Lighthouse Laboratory Plymouth | Wellcome Sanger Institute for the COVID-19 Genomics UK (COG-UK) Consortium | Lighthouse Laboratory Plymouth and Alex Alderton, Roberto Amato, Jeffrey Barrett, Sonia Goncalves, Ewan Harrison, David K. Jackson, Ian Johnston, Dominic Kwiatkowski, Cordelia Langford, John Sillitoe on behalf of the Wellcome Sanger Institute COVID-19 Surveillance Team |
| EPI_ISL_3437078 | Health Services Laboratories | Wellcome Sanger Institute for the COVID-19 Genomics UK (COG-UK) Consortium | Health Services Laboratories and Alex Alderton, Roberto Amato, Jeffrey Barrett, Sonia Goncalves, Ewan Harrison, David K. Jackson, Ian Johnston, Dominic Kwiatkowski, Cordelia Langford, John Sillitoe on behalf of the Wellcome Sanger Institute COVID-19 Surveillance Team |
| EPI_ISL_3437079, EPI_ISL_3437080, EPI_ISL_3437081 | Berkshire and Surrey Pathology Services Lighthouse Laboratory | Wellcome Sanger Institute for the COVID-19 Genomics UK (COG-UK) Consortium | Berkshire and Surrey Pathology Services Lighthouse Laboratory and Alex Alderton, Roberto Amato, Jeffrey Barrett, Sonia Goncalves, Ewan Harrison, David K. Jackson, Ian Johnston, Dominic Kwiatkowski, Cordelia Langford, John Sillitoe on behalf of the Wellcome Sanger Institute COVID-19 Surveillance Team |
| EPI_ISL_3437082 | Lighthouse Laboratory Plymouth | Wellcome Sanger Institute for the COVID-19 Genomics UK (COG-UK) Consortium | Lighthouse Laboratory Plymouth and Alex Alderton, Roberto Amato, Jeffrey Barrett, Sonia Goncalves, Ewan Harrison, David K. Jackson, Ian Johnston, Dominic Kwiatkowski, Cordelia Langford, John Sillitoe on behalf of the Wellcome Sanger Institute COVID-19 Surveillance Team |
| EPI_ISL_3437083, EPI_ISL_3437084 | Lighthouse Lab in Alderley Park | Wellcome Sanger Institute for the COVID-19 Genomics UK (COG-UK) Consortium | Jacquelyn Wynn, Mairead Hyland, The Lighthouse Lab in Alderley Park and Alex Alderton, Roberto Amato, Jeffrey Barrett, Sonia Goncalves, Ewan Harrison, David K. Jackson, Ian Johnston, Dominic Kwiatkowski, Cordelia Langford, John Sillitoe on behalf of the Wellcome Sanger Institute COVID-19 Surveillance Team |
| EPI_ISL_3437085, EPI_ISL_3437086 | Berkshire and Surrey Pathology Services Lighthouse Laboratory | Wellcome Sanger Institute for the COVID-19 Genomics UK (COG-UK) Consortium | Berkshire and Surrey Pathology Services Lighthouse Laboratory and Alex Alderton, Roberto Amato, Jeffrey Barrett, Sonia Goncalves, Ewan Harrison, David K. Jackson, Ian Johnston, Dominic Kwiatkowski, Cordelia Langford, John Sillitoe on behalf of the Wellcome Sanger Institute COVID-19 Surveillance Team |
| EPI_ISL_3437087 | Lighthouse Lab in Alderley Park | Wellcome Sanger Institute for the COVID-19 Genomics UK (COG-UK) Consortium | Jacquelyn Wynn, Mairead Hyland, The Lighthouse Lab in Alderley Park and Alex Alderton, Roberto Amato, Jeffrey Barrett, Sonia Goncalves, Ewan Harrison, David K. Jackson, Ian Johnston, Dominic Kwiatkowski, Cordelia Langford, John Sillitoe on behalf of the Wellcome Sanger Institute COVID-19 Surveillance Team |
| EPI_ISL_3437088, EPI_ISL_3437089, EPI_ISL_3437090 | Berkshire and Surrey Pathology Services Lighthouse Laboratory | Wellcome Sanger Institute for the COVID-19 Genomics UK (COG-UK) Consortium | Berkshire and Surrey Pathology Services Lighthouse Laboratory and Alex Alderton, Roberto Amato, Jeffrey Barrett, Sonia Goncalves, Ewan Harrison, David K. Jackson, Ian Johnston, Dominic Kwiatkowski, Cordelia Langford, John Sillitoe on behalf of the Wellcome Sanger Institute COVID-19 Surveillance Team |
| EPI_ISL_3437091, EPI_ISL_3437092 | Lighthouse Lab in Alderley Park | Wellcome Sanger Institute for the COVID-19 Genomics UK (COG-UK) Consortium | Jacquelyn Wynn, Mairead Hyland, The Lighthouse Lab in Alderley Park and Alex Alderton, Roberto Amato, Jeffrey Barrett, Sonia Goncalves, Ewan Harrison, David K. Jackson, Ian Johnston, Dominic Kwiatkowski, Cordelia Langford, John Sillitoe on behalf of the Wellcome Sanger Institute COVID-19 Surveillance Team |
| EPI_ISL_3437093 | Lighthouse Laboratory Plymouth | Wellcome Sanger Institute for the COVID-19 Genomics UK (COG-UK) Consortium | Lighthouse Laboratory Plymouth and Alex Alderton, Roberto Amato, Jeffrey Barrett, Sonia Goncalves, Ewan Harrison, David K. Jackson, Ian Johnston, Dominic Kwiatkowski, Cordelia Langford, John Sillitoe on behalf of the Wellcome Sanger Institute COVID-19 Surveillance Team |
| EPI_ISL_3437094 | Health Services Laboratories | Wellcome Sanger Institute for the COVID-19 Genomics UK (COG-UK) Consortium | Health Services Laboratories and Alex Alderton, Roberto Amato, Jeffrey Barrett, Sonia Goncalves, Ewan Harrison, David K. Jackson, Ian Johnston, Dominic Kwiatkowski, Cordelia Langford, John Sillitoe on behalf of the Wellcome Sanger Institute COVID-19 Surveillance Team |
| EPI_ISL_3437095, EPI_ISL_3437096 | Lighthouse Lab in Alderley Park | Wellcome Sanger Institute for the COVID-19 Genomics UK (COG-UK) Consortium | Jacquelyn Wynn, Mairead Hyland, The Lighthouse Lab in Alderley Park and Alex Alderton, Roberto Amato, Jeffrey Barrett, Sonia Goncalves, Ewan Harrison, David K. Jackson, Ian Johnston, Dominic Kwiatkowski, Cordelia Langford, John Sillitoe on behalf of the Wellcome Sanger Institute COVID-19 Surveillance Team |
| EPI_ISL_3437097 | Lighthouse Laboratory Plymouth | Wellcome Sanger Institute for the COVID-19 Genomics UK (COG-UK) Consortium | Lighthouse Laboratory Plymouth and Alex Alderton, Roberto Amato, Jeffrey Barrett, Sonia Goncalves, Ewan Harrison, David K. Jackson, Ian Johnston, Dominic Kwiatkowski, Cordelia Langford, John Sillitoe on behalf of the Wellcome Sanger Institute COVID-19 Surveillance Team |
| EPI_ISL_3437098 | Berkshire and Surrey Pathology Services Lighthouse Laboratory | Wellcome Sanger Institute for the COVID-19 Genomics UK (COG-UK) Consortium | Berkshire and Surrey Pathology Services Lighthouse Laboratory and Alex Alderton, Roberto Amato, Jeffrey Barrett, Sonia Goncalves, Ewan Harrison, David K. Jackson, Ian Johnston, Dominic Kwiatkowski, Cordelia Langford, John Sillitoe on behalf of the Wellcome Sanger Institute COVID-19 Surveillance Team |
| EPI_ISL_3437099 | Lighthouse Laboratory Plymouth | Wellcome Sanger Institute for the COVID-19 Genomics UK (COG-UK) Consortium | Lighthouse Laboratory Plymouth and Alex Alderton, Roberto Amato, Jeffrey Barrett, Sonia Goncalves, Ewan Harrison, David K. Jackson, Ian Johnston, Dominic Kwiatkowski, Cordelia Langford, John Sillitoe on behalf of the Wellcome Sanger Institute COVID-19 Surveillance Team |
| EPI_ISL_3437100 | Berkshire and Surrey Pathology Services Lighthouse Laboratory | Wellcome Sanger Institute for the COVID-19 Genomics UK (COG-UK) Consortium | Berkshire and Surrey Pathology Services Lighthouse Laboratory and Alex Alderton, Roberto Amato, Jeffrey Barrett, Sonia Goncalves, Ewan Harrison, David K. Jackson, Ian Johnston, Dominic Kwiatkowski, Cordelia Langford, John Sillitoe on behalf of the Wellcome Sanger Institute COVID-19 Surveillance Team |
| EPI_ISL_3437101 | Lighthouse Laboratory Plymouth | Wellcome Sanger Institute for the COVID-19 Genomics UK (COG-UK) Consortium | Lighthouse Laboratory Plymouth and Alex Alderton, Roberto Amato, Jeffrey Barrett, Sonia Goncalves, Ewan Harrison, David K. Jackson, Ian Johnston, Dominic Kwiatkowski, Cordelia Langford, John Sillitoe on behalf of the Wellcome Sanger Institute COVID-19 Surveillance Team |
| EPI_ISL_3437102 | Lighthouse Lab in Alderley Park | Wellcome Sanger Institute for the COVID-19 Genomics UK (COG-UK) Consortium | Jacquelyn Wynn, Mairead Hyland, The Lighthouse Lab in Alderley Park and Alex Alderton, Roberto Amato, Jeffrey Barrett, Sonia Goncalves, Ewan Harrison, David K. Jackson, Ian Johnston, Dominic Kwiatkowski, Cordelia Langford, John Sillitoe on behalf of the Wellcome Sanger Institute COVID-19 Surveillance Team |
| EPI_ISL_3437103 | Health Services Laboratories | Wellcome Sanger Institute for the COVID-19 Genomics UK (COG-UK) Consortium | Health Services Laboratories and Alex Alderton, Roberto Amato, Jeffrey Barrett, Sonia Goncalves, Ewan Harrison, David K. Jackson, Ian Johnston, Dominic Kwiatkowski, Cordelia Langford, John Sillitoe on behalf of the Wellcome Sanger Institute COVID-19 Surveillance Team |
| EPI_ISL_3437104, EPI_ISL_3437105 | Lighthouse Laboratory Plymouth | Wellcome Sanger Institute for the COVID-19 Genomics UK (COG-UK) Consortium | Lighthouse Laboratory Plymouth and Alex Alderton, Roberto Amato, Jeffrey Barrett, Sonia Goncalves, Ewan Harrison, David K. Jackson, Ian Johnston, Dominic Kwiatkowski, Cordelia Langford, John Sillitoe on behalf of the Wellcome Sanger Institute COVID-19 Surveillance Team |
| EPI_ISL_3437106 | Berkshire and Surrey Pathology Services Lighthouse Laboratory | Wellcome Sanger Institute for the COVID-19 Genomics UK (COG-UK) Consortium | Berkshire and Surrey Pathology Services Lighthouse Laboratory and Alex Alderton, Roberto Amato, Jeffrey Barrett, Sonia Goncalves, Ewan Harrison, David K. Jackson, Ian Johnston, Dominic Kwiatkowski, Cordelia Langford, John Sillitoe on behalf of the Wellcome Sanger Institute COVID-19 Surveillance Team |

| | | | |
|---|---|---|---|
| EPI_ISL_3437107 | Lighthouse Lab in Alderley Park | Wellcome Sanger Institute for the COVID-19 Genomics UK (COG-UK) Consortium | Jacquelyn Wynn, Mairead Hyland, The Lighthouse Lab in Alderley Park and Alex Alderton, Roberto Amato, Jeffrey Barrett, Sonia Goncalves, Ewan Harrison, David K. Jackson, Ian Johnston, Dominic Kwiatkowski, Cordelia Langford, John Sillitoe on behalf of the Wellcome Sanger Institute COVID-19 Surveillance Team |
| EPI_ISL_3437108, EPI_ISL_3437109 | Berkshire and Surrey Pathology Services Lighthouse Laboratory | Wellcome Sanger Institute for the COVID-19 Genomics UK (COG-UK) Consortium | Berkshire and Surrey Pathology Services Lighthouse Laboratory and Alex Alderton, Roberto Amato, Jeffrey Barrett, Sonia Goncalves, Ewan Harrison, David K. Jackson, Ian Johnston, Dominic Kwiatkowski, Cordelia Langford, John Sillitoe on behalf of the Wellcome Sanger Institute COVID-19 Surveillance Team |
| EPI_ISL_3437110 | Lighthouse Laboratory Plymouth | Wellcome Sanger Institute for the COVID-19 Genomics UK (COG-UK) Consortium | Lighthouse Laboratory Plymouth and Alex Alderton, Roberto Amato, Jeffrey Barrett, Sonia Goncalves, Ewan Harrison, David K. Jackson, Ian Johnston, Dominic Kwiatkowski, Cordelia Langford, John Sillitoe on behalf of the Wellcome Sanger Institute COVID-19 Surveillance Team |
| EPI_ISL_3437111, EPI_ISL_3437112 | Lighthouse Lab in Alderley Park | Wellcome Sanger Institute for the COVID-19 Genomics UK (COG-UK) Consortium | Jacquelyn Wynn, Mairead Hyland, The Lighthouse Lab in Alderley Park and Alex Alderton, Roberto Amato, Jeffrey Barrett, Sonia Goncalves, Ewan Harrison, David K. Jackson, Ian Johnston, Dominic Kwiatkowski, Cordelia Langford, John Sillitoe on behalf of the Wellcome Sanger Institute COVID-19 Surveillance Team |
| EPI_ISL_3437113 | Health Services Laboratories | Wellcome Sanger Institute for the COVID-19 Genomics UK (COG-UK) Consortium | Health Services Laboratories and Alex Alderton, Roberto Amato, Jeffrey Barrett, Sonia Goncalves, Ewan Harrison, David K. Jackson, Ian Johnston, Dominic Kwiatkowski, Cordelia Langford, John Sillitoe on behalf of the Wellcome Sanger Institute COVID-19 Surveillance Team |
| EPI_ISL_3437114 | Lighthouse Laboratory Plymouth | Wellcome Sanger Institute for the COVID-19 Genomics UK (COG-UK) Consortium | Lighthouse Laboratory Plymouth and Alex Alderton, Roberto Amato, Jeffrey Barrett, Sonia Goncalves, Ewan Harrison, David K. Jackson, Ian Johnston, Dominic Kwiatkowski, Cordelia Langford, John Sillitoe on behalf of the Wellcome Sanger Institute COVID-19 Surveillance Team |
| EPI_ISL_3437115 | Health Services Laboratories | Wellcome Sanger Institute for the COVID-19 Genomics UK (COG-UK) Consortium | Health Services Laboratories and Alex Alderton, Roberto Amato, Jeffrey Barrett, Sonia Goncalves, Ewan Harrison, David K. Jackson, Ian Johnston, Dominic Kwiatkowski, Cordelia Langford, John Sillitoe on behalf of the Wellcome Sanger Institute COVID-19 Surveillance Team |
| EPI_ISL_3437116, EPI_ISL_3437117, EPI_ISL_3437118 | Berkshire and Surrey Pathology Services Lighthouse Laboratory | Wellcome Sanger Institute for the COVID-19 Genomics UK (COG-UK) Consortium | Berkshire and Surrey Pathology Services Lighthouse Laboratory and Alex Alderton, Roberto Amato, Jeffrey Barrett, Sonia Goncalves, Ewan Harrison, David K. Jackson, Ian Johnston, Dominic Kwiatkowski, Cordelia Langford, John Sillitoe on behalf of the Wellcome Sanger Institute COVID-19 Surveillance Team |
| EPI_ISL_3437119 | Health Services Laboratories | Wellcome Sanger Institute for the COVID-19 Genomics UK (COG-UK) Consortium | Health Services Laboratories and Alex Alderton, Roberto Amato, Jeffrey Barrett, Sonia Goncalves, Ewan Harrison, David K. Jackson, Ian Johnston, Dominic Kwiatkowski, Cordelia Langford, John Sillitoe on behalf of the Wellcome Sanger Institute COVID-19 Surveillance Team |
| EPI_ISL_3437120 | Berkshire and Surrey Pathology Services Lighthouse Laboratory | Wellcome Sanger Institute for the COVID-19 Genomics UK (COG-UK) Consortium | Berkshire and Surrey Pathology Services Lighthouse Laboratory and Alex Alderton, Roberto Amato, Jeffrey Barrett, Sonia Goncalves, Ewan Harrison, David K. Jackson, Ian Johnston, Dominic Kwiatkowski, Cordelia Langford, John Sillitoe on behalf of the Wellcome Sanger Institute COVID-19 Surveillance Team |
| EPI_ISL_3437121 | Lighthouse Lab in Alderley Park | Wellcome Sanger Institute for the COVID-19 Genomics UK (COG-UK) Consortium | Jacquelyn Wynn, Mairead Hyland, The Lighthouse Lab in Alderley Park and Alex Alderton, Roberto Amato, Jeffrey Barrett, Sonia Goncalves, Ewan Harrison, David K. Jackson, Ian Johnston, Dominic Kwiatkowski, Cordelia Langford, John Sillitoe on behalf of the Wellcome Sanger Institute COVID-19 Surveillance Team |
| EPI_ISL_3437122 | Lighthouse Laboratory Plymouth | Wellcome Sanger Institute for the COVID-19 Genomics UK (COG-UK) Consortium | Lighthouse Laboratory Plymouth and Alex Alderton, Roberto Amato, Jeffrey Barrett, Sonia Goncalves, Ewan Harrison, David K. Jackson, Ian Johnston, Dominic Kwiatkowski, Cordelia Langford, John Sillitoe on behalf of the Wellcome Sanger Institute COVID-19 Surveillance Team |
| EPI_ISL_3437123 | Lighthouse Lab in Alderley Park | Wellcome Sanger Institute for the COVID-19 Genomics UK (COG-UK) Consortium | Jacquelyn Wynn, Mairead Hyland, The Lighthouse Lab in Alderley Park and Alex Alderton, Roberto Amato, Jeffrey Barrett, Sonia Goncalves, Ewan Harrison, David K. Jackson, Ian Johnston, Dominic Kwiatkowski, Cordelia Langford, John Sillitoe on behalf of the Wellcome Sanger Institute COVID-19 Surveillance Team |
| EPI_ISL_3437124 | Health Services Laboratories | Wellcome Sanger Institute for the COVID-19 Genomics UK (COG-UK) Consortium | Health Services Laboratories and Alex Alderton, Roberto Amato, Jeffrey Barrett, Sonia Goncalves, Ewan Harrison, David K. Jackson, Ian Johnston, Dominic Kwiatkowski, Cordelia Langford, John Sillitoe on behalf of the Wellcome Sanger Institute COVID-19 Surveillance Team |
| EPI_ISL_3437125 | Lighthouse Lab in Alderley Park | Wellcome Sanger Institute for the COVID-19 Genomics UK (COG-UK) Consortium | Jacquelyn Wynn, Mairead Hyland, The Lighthouse Lab in Alderley Park and Alex Alderton, Roberto Amato, Jeffrey Barrett, Sonia Goncalves, Ewan Harrison, David K. Jackson, Ian Johnston, Dominic Kwiatkowski, Cordelia Langford, John Sillitoe on behalf of the Wellcome Sanger Institute COVID-19 Surveillance Team |
| EPI_ISL_3437126 | Berkshire and Surrey Pathology Services Lighthouse Laboratory | Wellcome Sanger Institute for the COVID-19 Genomics UK (COG-UK) Consortium | Berkshire and Surrey Pathology Services Lighthouse Laboratory and Alex Alderton, Roberto Amato, Jeffrey Barrett, Sonia Goncalves, Ewan Harrison, David K. Jackson, Ian Johnston, Dominic Kwiatkowski, Cordelia Langford, John Sillitoe on behalf of the Wellcome Sanger Institute COVID-19 Surveillance Team |
| EPI_ISL_3437127, EPI_ISL_3437128 | Lighthouse Laboratory Plymouth | Wellcome Sanger Institute for the COVID-19 Genomics UK (COG-UK) Consortium | Lighthouse Laboratory Plymouth and Alex Alderton, Roberto Amato, Jeffrey Barrett, Sonia Goncalves, Ewan Harrison, David K. Jackson, Ian Johnston, Dominic Kwiatkowski, Cordelia Langford, John Sillitoe on behalf of the Wellcome Sanger Institute COVID-19 Surveillance Team |
| EPI_ISL_3437129 | Lighthouse Lab in Alderley Park | Wellcome Sanger Institute for the COVID-19 Genomics UK (COG-UK) Consortium | Jacquelyn Wynn, Mairead Hyland, The Lighthouse Lab in Alderley Park and Alex Alderton, Roberto Amato, Jeffrey Barrett, Sonia Goncalves, Ewan Harrison, David K. Jackson, Ian Johnston, Dominic Kwiatkowski, Cordelia Langford, John Sillitoe on behalf of the Wellcome Sanger Institute COVID-19 Surveillance Team |
| EPI_ISL_3437130 | Lighthouse Laboratory Plymouth | Wellcome Sanger Institute for the COVID-19 Genomics UK (COG-UK) Consortium | Lighthouse Laboratory Plymouth and Alex Alderton, Roberto Amato, Jeffrey Barrett, Sonia Goncalves, Ewan Harrison, David K. Jackson, Ian Johnston, Dominic Kwiatkowski, Cordelia Langford, John Sillitoe on behalf of the Wellcome Sanger Institute COVID-19 Surveillance Team |
| EPI_ISL_3437131 | Berkshire and Surrey Pathology Services Lighthouse Laboratory | Wellcome Sanger Institute for the COVID-19 Genomics UK (COG-UK) Consortium | Berkshire and Surrey Pathology Services Lighthouse Laboratory and Alex Alderton, Roberto Amato, Jeffrey Barrett, Sonia Goncalves, Ewan Harrison, David K. Jackson, Ian Johnston, Dominic Kwiatkowski, Cordelia Langford, John Sillitoe on behalf of the Wellcome Sanger Institute COVID-19 Surveillance Team |
| EPI_ISL_3437132 | Lighthouse Laboratory Plymouth | Wellcome Sanger Institute for the COVID-19 Genomics UK (COG-UK) Consortium | Lighthouse Laboratory Plymouth and Alex Alderton, Roberto Amato, Jeffrey Barrett, Sonia Goncalves, Ewan Harrison, David K. Jackson, Ian Johnston, Dominic Kwiatkowski, Cordelia Langford, John Sillitoe on behalf of the Wellcome Sanger Institute COVID-19 Surveillance Team |
| EPI_ISL_3437133 | Berkshire and Surrey Pathology Services Lighthouse Laboratory | Wellcome Sanger Institute for the COVID-19 Genomics UK (COG-UK) Consortium | Berkshire and Surrey Pathology Services Lighthouse Laboratory and Alex Alderton, Roberto Amato, Jeffrey Barrett, Sonia Goncalves, Ewan Harrison, David K. Jackson, Ian Johnston, Dominic Kwiatkowski, Cordelia Langford, John Sillitoe on behalf of the Wellcome Sanger Institute COVID-19 Surveillance Team |
| EPI_ISL_3437134, EPI_ISL_3437135 | Lighthouse Laboratory Plymouth | Wellcome Sanger Institute for the COVID-19 Genomics UK (COG-UK) Consortium | Lighthouse Laboratory Plymouth and Alex Alderton, Roberto Amato, Jeffrey Barrett, Sonia Goncalves, Ewan Harrison, David K. Jackson, Ian Johnston, Dominic Kwiatkowski, Cordelia Langford, John Sillitoe on behalf of the Wellcome Sanger Institute COVID-19 Surveillance Team |
| EPI_ISL_3437136 | Lighthouse Lab in Alderley Park | Wellcome Sanger Institute for the COVID-19 Genomics UK (COG-UK) Consortium | Jacquelyn Wynn, Mairead Hyland, The Lighthouse Lab in Alderley Park and Alex Alderton, Roberto Amato, Jeffrey Barrett, Sonia Goncalves, Ewan Harrison, David K. Jackson, Ian Johnston, Dominic Kwiatkowski, Cordelia Langford, John Sillitoe on behalf of the Wellcome Sanger Institute COVID-19 Surveillance Team |
| EPI_ISL_3437137 | Berkshire and Surrey Pathology Services Lighthouse Laboratory | Wellcome Sanger Institute for the COVID-19 Genomics UK (COG-UK) Consortium | Berkshire and Surrey Pathology Services Lighthouse Laboratory and Alex Alderton, Roberto Amato, Jeffrey Barrett, Sonia Goncalves, Ewan Harrison, David K. Jackson, Ian Johnston, Dominic Kwiatkowski, Cordelia Langford, John Sillitoe on behalf of the Wellcome Sanger Institute COVID-19 Surveillance Team |
| EPI_ISL_3437138, EPI_ISL_3437139 | Lighthouse Laboratory Plymouth | Wellcome Sanger Institute for the COVID-19 Genomics UK (COG-UK) Consortium | Lighthouse Laboratory Plymouth and Alex Alderton, Roberto Amato, Jeffrey Barrett, Sonia Goncalves, Ewan Harrison, David K. Jackson, Ian Johnston, Dominic Kwiatkowski, Cordelia Langford, John Sillitoe on behalf of the Wellcome Sanger Institute COVID-19 Surveillance Team |
| EPI_ISL_3437140 | Lighthouse Lab in Alderley Park | Wellcome Sanger Institute for the COVID-19 Genomics UK (COG-UK) Consortium | Jacquelyn Wynn, Mairead Hyland, The Lighthouse Lab in Alderley Park and Alex Alderton, Roberto Amato, Jeffrey Barrett, Sonia Goncalves, Ewan Harrison, David K. Jackson, Ian Johnston, Dominic Kwiatkowski, Cordelia Langford, John Sillitoe on behalf of the Wellcome Sanger Institute COVID-19 Surveillance Team |
| EPI_ISL_3437141 | Health Services Laboratories | Wellcome Sanger Institute for the COVID-19 Genomics UK (COG-UK) Consortium | Health Services Laboratories and Alex Alderton, Roberto Amato, Jeffrey Barrett, Sonia Goncalves, Ewan Harrison, David K. Jackson, Ian Johnston, Dominic Kwiatkowski, Cordelia Langford, John Sillitoe on behalf of the Wellcome Sanger Institute COVID-19 Surveillance Team |
| EPI_ISL_3437142, EPI_ISL_3437143 | Lighthouse Laboratory Plymouth | Wellcome Sanger Institute for the COVID-19 Genomics UK (COG-UK) Consortium | Lighthouse Laboratory Plymouth and Alex Alderton, Roberto Amato, Jeffrey Barrett, Sonia Goncalves, Ewan Harrison, David K. Jackson, Ian Johnston, Dominic Kwiatkowski, Cordelia Langford, John Sillitoe on behalf of the Wellcome Sanger Institute COVID-19 Surveillance Team |
| EPI_ISL_3437144 | Health Services Laboratories | Wellcome Sanger Institute for the COVID-19 Genomics UK (COG-UK) Consortium | Health Services Laboratories and Alex Alderton, Roberto Amato, Jeffrey Barrett, Sonia Goncalves, Ewan Harrison, David K. Jackson, Ian Johnston, Dominic Kwiatkowski, Cordelia Langford, John Sillitoe on behalf of the Wellcome Sanger Institute COVID-19 Surveillance Team |
| EPI_ISL_3437145, EPI_ISL_3437146 | Lighthouse Laboratory Plymouth | Wellcome Sanger Institute for the COVID-19 Genomics UK (COG-UK) Consortium | Lighthouse Laboratory Plymouth and Alex Alderton, Roberto Amato, Jeffrey Barrett, Sonia Goncalves, Ewan Harrison, David K. Jackson, Ian Johnston, Dominic Kwiatkowski, Cordelia Langford, John Sillitoe on behalf of the Wellcome Sanger Institute COVID-19 Surveillance Team |
| EPI_ISL_3437147 | Lighthouse Lab in Alderley Park | Wellcome Sanger Institute for the COVID-19 Genomics UK (COG-UK) Consortium | Jacquelyn Wynn, Mairead Hyland, The Lighthouse Lab in Alderley Park and Alex Alderton, Roberto Amato, Jeffrey Barrett, Sonia Goncalves, Ewan Harrison, David K. Jackson, Ian Johnston, Dominic Kwiatkowski, Cordelia Langford, John Sillitoe on behalf of the Wellcome Sanger Institute COVID-19 Surveillance Team |
| EPI_ISL_3437148 | Lighthouse Laboratory Plymouth | Wellcome Sanger Institute for the COVID-19 Genomics UK (COG-UK) Consortium | Lighthouse Laboratory Plymouth and Alex Alderton, Roberto Amato, Jeffrey Barrett, Sonia Goncalves, Ewan Harrison, David K. Jackson, Ian Johnston, Dominic Kwiatkowski, Cordelia Langford, John Sillitoe on behalf of the Wellcome Sanger Institute COVID-19 Surveillance Team |
| EPI_ISL_3437149 | Health Services Laboratories | Wellcome Sanger Institute for the COVID-19 Genomics UK (COG-UK) Consortium | Health Services Laboratories and Alex Alderton, Roberto Amato, Jeffrey Barrett, Sonia Goncalves, Ewan Harrison, David K. Jackson, Ian Johnston, Dominic Kwiatkowski, Cordelia Langford, John Sillitoe on behalf of the Wellcome Sanger Institute COVID-19 Surveillance Team |
| EPI_ISL_3437150 | Berkshire and Surrey Pathology Services Lighthouse Laboratory | Wellcome Sanger Institute for the COVID-19 Genomics UK (COG-UK) Consortium | Berkshire and Surrey Pathology Services Lighthouse Laboratory and Alex Alderton, Roberto Amato, Jeffrey Barrett, Sonia Goncalves, Ewan Harrison, David K. Jackson, Ian Johnston, Dominic Kwiatkowski, Cordelia Langford, John Sillitoe on behalf of the Wellcome Sanger Institute COVID-19 Surveillance Team |
| EPI_ISL_3437151 | Lighthouse Lab in Alderley Park | Wellcome Sanger Institute for the COVID-19 Genomics UK (COG-UK) Consortium | Jacquelyn Wynn, Mairead Hyland, The Lighthouse Lab in Alderley Park and Alex Alderton, Roberto Amato, Jeffrey Barrett, Sonia Goncalves, Ewan Harrison, David K. Jackson, Ian Johnston, Dominic Kwiatkowski, Cordelia Langford, John Sillitoe on behalf of the Wellcome Sanger Institute COVID-19 Surveillance Team |
| EPI_ISL_3437152 | Berkshire and Surrey Pathology Services Lighthouse Laboratory | Wellcome Sanger Institute for the COVID-19 Genomics UK (COG-UK) Consortium | Berkshire and Surrey Pathology Services Lighthouse Laboratory and Alex Alderton, Roberto Amato, Jeffrey Barrett, Sonia Goncalves, Ewan Harrison, David K. Jackson, Ian Johnston, Dominic Kwiatkowski, Cordelia Langford, John Sillitoe on behalf of the Wellcome Sanger Institute COVID-19 Surveillance Team |
| EPI_ISL_3437153 | Lighthouse Lab in Alderley Park | Wellcome Sanger Institute for the COVID-19 Genomics UK (COG-UK) Consortium | Jacquelyn Wynn, Mairead Hyland, The Lighthouse Lab in Alderley Park and Alex Alderton, Roberto Amato, Jeffrey Barrett, Sonia Goncalves, Ewan Harrison, David K. Jackson, Ian Johnston, Dominic Kwiatkowski, Cordelia Langford, John Sillitoe on behalf of the Wellcome Sanger Institute COVID-19 Surveillance Team |
| EPI_ISL_3437154 | Lighthouse Laboratory Plymouth | Wellcome Sanger Institute for the COVID-19 Genomics UK (COG-UK) Consortium | Lighthouse Laboratory Plymouth and Alex Alderton, Roberto Amato, Jeffrey Barrett, Sonia Goncalves, Ewan Harrison, David K. Jackson, Ian Johnston, Dominic Kwiatkowski, Cordelia Langford, John Sillitoe on behalf of the Wellcome Sanger Institute COVID-19 Surveillance Team |
| EPI_ISL_3437155 | Lighthouse Lab in Alderley Park | Wellcome Sanger Institute for the COVID-19 Genomics UK (COG-UK) Consortium | Jacquelyn Wynn, Mairead Hyland, The Lighthouse Lab in Alderley Park and Alex Alderton, Roberto Amato, Jeffrey Barrett, Sonia Goncalves, Ewan Harrison, David K. Jackson, Ian Johnston, Dominic Kwiatkowski, Cordelia Langford, John Sillitoe on behalf of the Wellcome Sanger Institute COVID-19 Surveillance Team |
| EPI_ISL_3437156 | Berkshire and Surrey Pathology Services Lighthouse Laboratory | Wellcome Sanger Institute for the COVID-19 Genomics UK (COG-UK) Consortium | Berkshire and Surrey Pathology Services Lighthouse Laboratory and Alex Alderton, Roberto Amato, Jeffrey Barrett, Sonia Goncalves, Ewan Harrison, David K. Jackson, Ian Johnston, Dominic Kwiatkowski, Cordelia Langford, John Sillitoe on behalf of the Wellcome Sanger Institute COVID-19 Surveillance Team |
| EPI_ISL_3437157, EPI_ISL_3437158, EPI_ISL_3437159 | Lighthouse Laboratory Plymouth | Wellcome Sanger Institute for the COVID-19 Genomics UK (COG-UK) Consortium | Lighthouse Laboratory Plymouth and Alex Alderton, Roberto Amato, Jeffrey Barrett, Sonia Goncalves, Ewan Harrison, David K. Jackson, Ian Johnston, Dominic Kwiatkowski, Cordelia Langford, John Sillitoe on behalf of the Wellcome Sanger Institute COVID-19 Surveillance Team |
| EPI_ISL_3437160 | Lighthouse Lab in Alderley Park | Wellcome Sanger Institute for the COVID-19 Genomics UK (COG-UK) Consortium | Jacquelyn Wynn, Mairead Hyland, The Lighthouse Lab in Alderley Park and Alex Alderton, Roberto Amato, Jeffrey Barrett, Sonia Goncalves, Ewan Harrison, David K. Jackson, Ian Johnston, Dominic Kwiatkowski, Cordelia Langford, John Sillitoe on behalf of the Wellcome Sanger Institute COVID-19 Surveillance Team |
| EPI_ISL_3437161, EPI_ISL_3437162 | Lighthouse Laboratory Plymouth | Wellcome Sanger Institute for the COVID-19 Genomics UK (COG-UK) Consortium | Lighthouse Laboratory Plymouth and Alex Alderton, Roberto Amato, Jeffrey Barrett, Sonia Goncalves, Ewan Harrison, David K. Jackson, Ian Johnston, Dominic Kwiatkowski, Cordelia Langford, John Sillitoe on behalf of the Wellcome Sanger Institute COVID-19 Surveillance Team |

| | | | |
|---|---|---|---|
| EPI_ISL_3437163 | Lighthouse Lab in Alderley Park | Wellcome Sanger Institute for the COVID-19 Genomics UK (COG-UK) Consortium | Jacquelyn Wynn, Mairead Hyland, The Lighthouse Lab in Alderley Park and Alex Alderton, Roberto Amato, Jeffrey Barrett, Sonia Goncalves, Ewan Harrison, David K. Jackson, Ian Johnston, Dominic Kwiatkowski, Cordelia Langford, John Sillitoe on behalf of the Wellcome Sanger Institute COVID-19 Surveillance Team |
| EPI_ISL_3437164 | Lighthouse Laboratory Plymouth | Wellcome Sanger Institute for the COVID-19 Genomics UK (COG-UK) Consortium | Lighthouse Laboratory Plymouth and Alex Alderton, Roberto Amato, Jeffrey Barrett, Sonia Goncalves, Ewan Harrison, David K. Jackson, Ian Johnston, Dominic Kwiatkowski, Cordelia Langford, John Sillitoe on behalf of the Wellcome Sanger Institute COVID-19 Surveillance Team |
| EPI_ISL_3437165, EPI_ISL_3437166 | Lighthouse Lab in Alderley Park | Wellcome Sanger Institute for the COVID-19 Genomics UK (COG-UK) Consortium | Jacquelyn Wynn, Mairead Hyland, The Lighthouse Lab in Alderley Park and Alex Alderton, Roberto Amato, Jeffrey Barrett, Sonia Goncalves, Ewan Harrison, David K. Jackson, Ian Johnston, Dominic Kwiatkowski, Cordelia Langford, John Sillitoe on behalf of the Wellcome Sanger Institute COVID-19 Surveillance Team |
| EPI_ISL_3437167 | Berkshire and Surrey Pathology Services Lighthouse Laboratory | Wellcome Sanger Institute for the COVID-19 Genomics UK (COG-UK) Consortium | Berkshire and Surrey Pathology Services Lighthouse Laboratory and Alex Alderton, Roberto Amato, Jeffrey Barrett, Sonia Goncalves, Ewan Harrison, David K. Jackson, Ian Johnston, Dominic Kwiatkowski, Cordelia Langford, John Sillitoe on behalf of the Wellcome Sanger Institute COVID-19 Surveillance Team |
| EPI_ISL_3437168 | Lighthouse Lab in Alderley Park | Wellcome Sanger Institute for the COVID-19 Genomics UK (COG-UK) Consortium | Jacquelyn Wynn, Mairead Hyland, The Lighthouse Lab in Alderley Park and Alex Alderton, Roberto Amato, Jeffrey Barrett, Sonia Goncalves, Ewan Harrison, David K. Jackson, Ian Johnston, Dominic Kwiatkowski, Cordelia Langford, John Sillitoe on behalf of the Wellcome Sanger Institute COVID-19 Surveillance Team |
| EPI_ISL_3437169 | Health Services Laboratories | Wellcome Sanger Institute for the COVID-19 Genomics UK (COG-UK) Consortium | Health Services Laboratories and Alex Alderton, Roberto Amato, Jeffrey Barrett, Sonia Goncalves, Ewan Harrison, David K. Jackson, Ian Johnston, Dominic Kwiatkowski, Cordelia Langford, John Sillitoe on behalf of the Wellcome Sanger Institute COVID-19 Surveillance Team |
| EPI_ISL_3437170, EPI_ISL_3437171 | Lighthouse Laboratory Plymouth | Wellcome Sanger Institute for the COVID-19 Genomics UK (COG-UK) Consortium | Lighthouse Laboratory Plymouth and Alex Alderton, Roberto Amato, Jeffrey Barrett, Sonia Goncalves, Ewan Harrison, David K. Jackson, Ian Johnston, Dominic Kwiatkowski, Cordelia Langford, John Sillitoe on behalf of the Wellcome Sanger Institute COVID-19 Surveillance Team |
| EPI_ISL_3437172 | Lighthouse Lab in Alderley Park | Wellcome Sanger Institute for the COVID-19 Genomics UK (COG-UK) Consortium | Jacquelyn Wynn, Mairead Hyland, The Lighthouse Lab in Alderley Park and Alex Alderton, Roberto Amato, Jeffrey Barrett, Sonia Goncalves, Ewan Harrison, David K. Jackson, Ian Johnston, Dominic Kwiatkowski, Cordelia Langford, John Sillitoe on behalf of the Wellcome Sanger Institute COVID-19 Surveillance Team |
| EPI_ISL_3437173 | Lighthouse Laboratory Plymouth | Wellcome Sanger Institute for the COVID-19 Genomics UK (COG-UK) Consortium | Lighthouse Laboratory Plymouth and Alex Alderton, Roberto Amato, Jeffrey Barrett, Sonia Goncalves, Ewan Harrison, David K. Jackson, Ian Johnston, Dominic Kwiatkowski, Cordelia Langford, John Sillitoe on behalf of the Wellcome Sanger Institute COVID-19 Surveillance Team |
| EPI_ISL_3437174 | Lighthouse Lab in Alderley Park | Wellcome Sanger Institute for the COVID-19 Genomics UK (COG-UK) Consortium | Jacquelyn Wynn, Mairead Hyland, The Lighthouse Lab in Alderley Park and Alex Alderton, Roberto Amato, Jeffrey Barrett, Sonia Goncalves, Ewan Harrison, David K. Jackson, Ian Johnston, Dominic Kwiatkowski, Cordelia Langford, John Sillitoe on behalf of the Wellcome Sanger Institute COVID-19 Surveillance Team |
| EPI_ISL_3437175 | Berkshire and Surrey Pathology Services Lighthouse Laboratory | Wellcome Sanger Institute for the COVID-19 Genomics UK (COG-UK) Consortium | Berkshire and Surrey Pathology Services Lighthouse Laboratory and Alex Alderton, Roberto Amato, Jeffrey Barrett, Sonia Goncalves, Ewan Harrison, David K. Jackson, Ian Johnston, Dominic Kwiatkowski, Cordelia Langford, John Sillitoe on behalf of the Wellcome Sanger Institute COVID-19 Surveillance Team |
| EPI_ISL_3437176 | Lighthouse Laboratory Plymouth | Wellcome Sanger Institute for the COVID-19 Genomics UK (COG-UK) Consortium | Lighthouse Laboratory Plymouth and Alex Alderton, Roberto Amato, Jeffrey Barrett, Sonia Goncalves, Ewan Harrison, David K. Jackson, Ian Johnston, Dominic Kwiatkowski, Cordelia Langford, John Sillitoe on behalf of the Wellcome Sanger Institute COVID-19 Surveillance Team |
| EPI_ISL_3437177 | Lighthouse Lab in Alderley Park | Wellcome Sanger Institute for the COVID-19 Genomics UK (COG-UK) Consortium | Jacquelyn Wynn, Mairead Hyland, The Lighthouse Lab in Alderley Park and Alex Alderton, Roberto Amato, Jeffrey Barrett, Sonia Goncalves, Ewan Harrison, David K. Jackson, Ian Johnston, Dominic Kwiatkowski, Cordelia Langford, John Sillitoe on behalf of the Wellcome Sanger Institute COVID-19 Surveillance Team |
| EPI_ISL_3437178 | Lighthouse Laboratory Plymouth | Wellcome Sanger Institute for the COVID-19 Genomics UK (COG-UK) Consortium | Lighthouse Laboratory Plymouth and Alex Alderton, Roberto Amato, Jeffrey Barrett, Sonia Goncalves, Ewan Harrison, David K. Jackson, Ian Johnston, Dominic Kwiatkowski, Cordelia Langford, John Sillitoe on behalf of the Wellcome Sanger Institute COVID-19 Surveillance Team |
| EPI_ISL_3437179 | Health Services Laboratories | Wellcome Sanger Institute for the COVID-19 Genomics UK (COG-UK) Consortium | Health Services Laboratories and Alex Alderton, Roberto Amato, Jeffrey Barrett, Sonia Goncalves, Ewan Harrison, David K. Jackson, Ian Johnston, Dominic Kwiatkowski, Cordelia Langford, John Sillitoe on behalf of the Wellcome Sanger Institute COVID-19 Surveillance Team |
| EPI_ISL_3437180, EPI_ISL_3437181 | Lighthouse Lab in Alderley Park | Wellcome Sanger Institute for the COVID-19 Genomics UK (COG-UK) Consortium | Jacquelyn Wynn, Mairead Hyland, The Lighthouse Lab in Alderley Park and Alex Alderton, Roberto Amato, Jeffrey Barrett, Sonia Goncalves, Ewan Harrison, David K. Jackson, Ian Johnston, Dominic Kwiatkowski, Cordelia Langford, John Sillitoe on behalf of the Wellcome Sanger Institute COVID-19 Surveillance Team |
| EPI_ISL_3437182 | Berkshire and Surrey Pathology Services Lighthouse Laboratory | Wellcome Sanger Institute for the COVID-19 Genomics UK (COG-UK) Consortium | Berkshire and Surrey Pathology Services Lighthouse Laboratory and Alex Alderton, Roberto Amato, Jeffrey Barrett, Sonia Goncalves, Ewan Harrison, David K. Jackson, Ian Johnston, Dominic Kwiatkowski, Cordelia Langford, John Sillitoe on behalf of the Wellcome Sanger Institute COVID-19 Surveillance Team |
| EPI_ISL_3437183 | Lighthouse Laboratory Plymouth | Wellcome Sanger Institute for the COVID-19 Genomics UK (COG-UK) Consortium | Lighthouse Laboratory Plymouth and Alex Alderton, Roberto Amato, Jeffrey Barrett, Sonia Goncalves, Ewan Harrison, David K. Jackson, Ian Johnston, Dominic Kwiatkowski, Cordelia Langford, John Sillitoe on behalf of the Wellcome Sanger Institute COVID-19 Surveillance Team |
| EPI_ISL_3437184 | Lighthouse Lab in Alderley Park | Wellcome Sanger Institute for the COVID-19 Genomics UK (COG-UK) Consortium | Jacquelyn Wynn, Mairead Hyland, The Lighthouse Lab in Alderley Park and Alex Alderton, Roberto Amato, Jeffrey Barrett, Sonia Goncalves, Ewan Harrison, David K. Jackson, Ian Johnston, Dominic Kwiatkowski, Cordelia Langford, John Sillitoe on behalf of the Wellcome Sanger Institute COVID-19 Surveillance Team |
| EPI_ISL_3437185 | Berkshire and Surrey Pathology Services Lighthouse Laboratory | Wellcome Sanger Institute for the COVID-19 Genomics UK (COG-UK) Consortium | Berkshire and Surrey Pathology Services Lighthouse Laboratory and Alex Alderton, Roberto Amato, Jeffrey Barrett, Sonia Goncalves, Ewan Harrison, David K. Jackson, Ian Johnston, Dominic Kwiatkowski, Cordelia Langford, John Sillitoe on behalf of the Wellcome Sanger Institute COVID-19 Surveillance Team |
| EPI_ISL_3437186 | Lighthouse Laboratory Plymouth | Wellcome Sanger Institute for the COVID-19 Genomics UK (COG-UK) Consortium | Lighthouse Laboratory Plymouth and Alex Alderton, Roberto Amato, Jeffrey Barrett, Sonia Goncalves, Ewan Harrison, David K. Jackson, Ian Johnston, Dominic Kwiatkowski, Cordelia Langford, John Sillitoe on behalf of the Wellcome Sanger Institute COVID-19 Surveillance Team |
| EPI_ISL_3437187, EPI_ISL_3437188 | Lighthouse Lab in Alderley Park | Wellcome Sanger Institute for the COVID-19 Genomics UK (COG-UK) Consortium | Jacquelyn Wynn, Mairead Hyland, The Lighthouse Lab in Alderley Park and Alex Alderton, Roberto Amato, Jeffrey Barrett, Sonia Goncalves, Ewan Harrison, David K. Jackson, Ian Johnston, Dominic Kwiatkowski, Cordelia Langford, John Sillitoe on behalf of the Wellcome Sanger Institute COVID-19 Surveillance Team |
| EPI_ISL_3437189 | Berkshire and Surrey Pathology Services Lighthouse Laboratory | Wellcome Sanger Institute for the COVID-19 Genomics UK (COG-UK) Consortium | Berkshire and Surrey Pathology Services Lighthouse Laboratory and Alex Alderton, Roberto Amato, Jeffrey Barrett, Sonia Goncalves, Ewan Harrison, David K. Jackson, Ian Johnston, Dominic Kwiatkowski, Cordelia Langford, John Sillitoe on behalf of the Wellcome Sanger Institute COVID-19 Surveillance Team |
| EPI_ISL_3437190 | Lighthouse Lab in Alderley Park | Wellcome Sanger Institute for the COVID-19 Genomics UK (COG-UK) Consortium | Jacquelyn Wynn, Mairead Hyland, The Lighthouse Lab in Alderley Park and Alex Alderton, Roberto Amato, Jeffrey Barrett, Sonia Goncalves, Ewan Harrison, David K. Jackson, Ian Johnston, Dominic Kwiatkowski, Cordelia Langford, John Sillitoe on behalf of the Wellcome Sanger Institute COVID-19 Surveillance Team |
| EPI_ISL_3437191 | Berkshire and Surrey Pathology Services Lighthouse Laboratory | Wellcome Sanger Institute for the COVID-19 Genomics UK (COG-UK) Consortium | Berkshire and Surrey Pathology Services Lighthouse Laboratory and Alex Alderton, Roberto Amato, Jeffrey Barrett, Sonia Goncalves, Ewan Harrison, David K. Jackson, Ian Johnston, Dominic Kwiatkowski, Cordelia Langford, John Sillitoe on behalf of the Wellcome Sanger Institute COVID-19 Surveillance Team |
| EPI_ISL_3437192, EPI_ISL_3437193 | Lighthouse Laboratory Plymouth | Wellcome Sanger Institute for the COVID-19 Genomics UK (COG-UK) Consortium | Lighthouse Laboratory Plymouth and Alex Alderton, Roberto Amato, Jeffrey Barrett, Sonia Goncalves, Ewan Harrison, David K. Jackson, Ian Johnston, Dominic Kwiatkowski, Cordelia Langford, John Sillitoe on behalf of the Wellcome Sanger Institute COVID-19 Surveillance Team |
| EPI_ISL_3437194 | Lighthouse Lab in Alderley Park | Wellcome Sanger Institute for the COVID-19 Genomics UK (COG-UK) Consortium | Jacquelyn Wynn, Mairead Hyland, The Lighthouse Lab in Alderley Park and Alex Alderton, Roberto Amato, Jeffrey Barrett, Sonia Goncalves, Ewan Harrison, David K. Jackson, Ian Johnston, Dominic Kwiatkowski, Cordelia Langford, John Sillitoe on behalf of the Wellcome Sanger Institute COVID-19 Surveillance Team |
| EPI_ISL_3437195 | Berkshire and Surrey Pathology Services Lighthouse Laboratory | Wellcome Sanger Institute for the COVID-19 Genomics UK (COG-UK) Consortium | Berkshire and Surrey Pathology Services Lighthouse Laboratory and Alex Alderton, Roberto Amato, Jeffrey Barrett, Sonia Goncalves, Ewan Harrison, David K. Jackson, Ian Johnston, Dominic Kwiatkowski, Cordelia Langford, John Sillitoe on behalf of the Wellcome Sanger Institute COVID-19 Surveillance Team |
| EPI_ISL_3437196, EPI_ISL_3437197 | Lighthouse Lab in Alderley Park | Wellcome Sanger Institute for the COVID-19 Genomics UK (COG-UK) Consortium | Jacquelyn Wynn, Mairead Hyland, The Lighthouse Lab in Alderley Park and Alex Alderton, Roberto Amato, Jeffrey Barrett, Sonia Goncalves, Ewan Harrison, David K. Jackson, Ian Johnston, Dominic Kwiatkowski, Cordelia Langford, John Sillitoe on behalf of the Wellcome Sanger Institute COVID-19 Surveillance Team |
| EPI_ISL_3437198, EPI_ISL_3437199 | Lighthouse Laboratory Plymouth | Wellcome Sanger Institute for the COVID-19 Genomics UK (COG-UK) Consortium | Lighthouse Laboratory Plymouth and Alex Alderton, Roberto Amato, Jeffrey Barrett, Sonia Goncalves, Ewan Harrison, David K. Jackson, Ian Johnston, Dominic Kwiatkowski, Cordelia Langford, John Sillitoe on behalf of the Wellcome Sanger Institute COVID-19 Surveillance Team |
| EPI_ISL_3437200 | Health Services Laboratories | Wellcome Sanger Institute for the COVID-19 Genomics UK (COG-UK) Consortium | Health Services Laboratories and Alex Alderton, Roberto Amato, Jeffrey Barrett, Sonia Goncalves, Ewan Harrison, David K. Jackson, Ian Johnston, Dominic Kwiatkowski, Cordelia Langford, John Sillitoe on behalf of the Wellcome Sanger Institute COVID-19 Surveillance Team |
| EPI_ISL_3437201 | Lighthouse Lab in Alderley Park | Wellcome Sanger Institute for the COVID-19 Genomics UK (COG-UK) Consortium | Jacquelyn Wynn, Mairead Hyland, The Lighthouse Lab in Alderley Park and Alex Alderton, Roberto Amato, Jeffrey Barrett, Sonia Goncalves, Ewan Harrison, David K. Jackson, Ian Johnston, Dominic Kwiatkowski, Cordelia Langford, John Sillitoe on behalf of the Wellcome Sanger Institute COVID-19 Surveillance Team |
| EPI_ISL_3437202, EPI_ISL_3437203, EPI_ISL_3437204 | Health Services Laboratories | Wellcome Sanger Institute for the COVID-19 Genomics UK (COG-UK) Consortium | Health Services Laboratories and Alex Alderton, Roberto Amato, Jeffrey Barrett, Sonia Goncalves, Ewan Harrison, David K. Jackson, Ian Johnston, Dominic Kwiatkowski, Cordelia Langford, John Sillitoe on behalf of the Wellcome Sanger Institute COVID-19 Surveillance Team |
| EPI_ISL_3437205, EPI_ISL_3437206, EPI_ISL_3437207 | Lighthouse Laboratory Plymouth | Wellcome Sanger Institute for the COVID-19 Genomics UK (COG-UK) Consortium | Lighthouse Laboratory Plymouth and Alex Alderton, Roberto Amato, Jeffrey Barrett, Sonia Goncalves, Ewan Harrison, David K. Jackson, Ian Johnston, Dominic Kwiatkowski, Cordelia Langford, John Sillitoe on behalf of the Wellcome Sanger Institute COVID-19 Surveillance Team |
| EPI_ISL_3437208 | Berkshire and Surrey Pathology Services Lighthouse Laboratory | Wellcome Sanger Institute for the COVID-19 Genomics UK (COG-UK) Consortium | Berkshire and Surrey Pathology Services Lighthouse Laboratory and Alex Alderton, Roberto Amato, Jeffrey Barrett, Sonia Goncalves, Ewan Harrison, David K. Jackson, Ian Johnston, Dominic Kwiatkowski, Cordelia Langford, John Sillitoe on behalf of the Wellcome Sanger Institute COVID-19 Surveillance Team |
| EPI_ISL_3437209 | Health Services Laboratories | Wellcome Sanger Institute for the COVID-19 Genomics UK (COG-UK) Consortium | Health Services Laboratories and Alex Alderton, Roberto Amato, Jeffrey Barrett, Sonia Goncalves, Ewan Harrison, David K. Jackson, Ian Johnston, Dominic Kwiatkowski, Cordelia Langford, John Sillitoe on behalf of the Wellcome Sanger Institute COVID-19 Surveillance Team |
| EPI_ISL_3437210 | Lighthouse Laboratory Plymouth | Wellcome Sanger Institute for the COVID-19 Genomics UK (COG-UK) Consortium | Lighthouse Laboratory Plymouth and Alex Alderton, Roberto Amato, Jeffrey Barrett, Sonia Goncalves, Ewan Harrison, David K. Jackson, Ian Johnston, Dominic Kwiatkowski, Cordelia Langford, John Sillitoe on behalf of the Wellcome Sanger Institute COVID-19 Surveillance Team |
| EPI_ISL_3437211, EPI_ISL_3437212 | Health Services Laboratories | Wellcome Sanger Institute for the COVID-19 Genomics UK (COG-UK) Consortium | Health Services Laboratories and Alex Alderton, Roberto Amato, Jeffrey Barrett, Sonia Goncalves, Ewan Harrison, David K. Jackson, Ian Johnston, Dominic Kwiatkowski, Cordelia Langford, John Sillitoe on behalf of the Wellcome Sanger Institute COVID-19 Surveillance Team |
| EPI_ISL_3437213 | Lighthouse Laboratory Plymouth | Wellcome Sanger Institute for the COVID-19 Genomics UK (COG-UK) Consortium | Lighthouse Laboratory Plymouth and Alex Alderton, Roberto Amato, Jeffrey Barrett, Sonia Goncalves, Ewan Harrison, David K. Jackson, Ian Johnston, Dominic Kwiatkowski, Cordelia Langford, John Sillitoe on behalf of the Wellcome Sanger Institute COVID-19 Surveillance Team |
| EPI_ISL_3437214 | Lighthouse Lab in Alderley Park | Wellcome Sanger Institute for the COVID-19 Genomics UK (COG-UK) Consortium | Jacquelyn Wynn, Mairead Hyland, The Lighthouse Lab in Alderley Park and Alex Alderton, Roberto Amato, Jeffrey Barrett, Sonia Goncalves, Ewan Harrison, David K. Jackson, Ian Johnston, Dominic Kwiatkowski, Cordelia Langford, John Sillitoe on behalf of the Wellcome Sanger Institute COVID-19 Surveillance Team |
| EPI_ISL_3437215 | Berkshire and Surrey Pathology Services Lighthouse Laboratory | Wellcome Sanger Institute for the COVID-19 Genomics UK (COG-UK) Consortium | Berkshire and Surrey Pathology Services Lighthouse Laboratory and Alex Alderton, Roberto Amato, Jeffrey Barrett, Sonia Goncalves, Ewan Harrison, David K. Jackson, Ian Johnston, Dominic Kwiatkowski, Cordelia Langford, John Sillitoe on behalf of the Wellcome Sanger Institute COVID-19 Surveillance Team |
| EPI_ISL_3437216 | Lighthouse Lab in Alderley Park | Wellcome Sanger Institute for the COVID-19 Genomics UK (COG-UK) Consortium | Jacquelyn Wynn, Mairead Hyland, The Lighthouse Lab in Alderley Park and Alex Alderton, Roberto Amato, Jeffrey Barrett, Sonia Goncalves, Ewan Harrison, David K. Jackson, Ian Johnston, Dominic Kwiatkowski, Cordelia Langford, John Sillitoe on behalf of the Wellcome Sanger Institute COVID-19 Surveillance Team |
| EPI_ISL_3437217 | Berkshire and Surrey Pathology Services Lighthouse Laboratory | Wellcome Sanger Institute for the COVID-19 Genomics UK (COG-UK) Consortium | Berkshire and Surrey Pathology Services Lighthouse Laboratory and Alex Alderton, Roberto Amato, Jeffrey Barrett, Sonia Goncalves, Ewan Harrison, David K. Jackson, Ian Johnston, Dominic Kwiatkowski, Cordelia Langford, John Sillitoe on behalf of the Wellcome Sanger Institute COVID-19 Surveillance Team |
| EPI_ISL_3437218 | Lighthouse Lab in Alderley Park | Wellcome Sanger Institute for the COVID-19 Genomics UK (COG-UK) Consortium | Jacquelyn Wynn, Mairead Hyland, The Lighthouse Lab in Alderley Park and Alex Alderton, Roberto Amato, Jeffrey Barrett, Sonia Goncalves, Ewan Harrison, David K. Jackson, Ian Johnston, Dominic Kwiatkowski, Cordelia Langford, John Sillitoe on behalf of the Wellcome Sanger Institute COVID-19 Surveillance Team |
| EPI_ISL_3437219 | Health Services Laboratories | Wellcome Sanger Institute for the COVID-19 Genomics UK (COG-UK) | Health Services Laboratories and Alex Alderton, Roberto Amato, Jeffrey Barrett, Sonia Goncalves, Ewan Harrison, David K. Jackson, Ian Johnston, Dominic Kwiatkowski, Cordelia Langford, John Sillitoe on behalf of the Wellcome Sanger Institute |

| | | | |
|---|---|---|---|
| | | Consortium | COVID-19 Surveillance Team |
| EPI_ISL_3437220 | Berkshire and Surrey Pathology Services Lighthouse Laboratory | Wellcome Sanger Institute for the COVID-19 Genomics UK (COG-UK) Consortium | Berkshire and Surrey Pathology Services Lighthouse Laboratory and Alex Alderton, Roberto Amato, Jeffrey Barrett, Sonia Goncalves, Ewan Harrison, David K. Jackson, Ian Johnston, Dominic Kwiatkowski, Cordelia Langford, John Sillitoe on behalf of the Wellcome Sanger Institute COVID-19 Surveillance Team |
| EPI_ISL_3437221 | Health Services Laboratories | Wellcome Sanger Institute for the COVID-19 Genomics UK (COG-UK) Consortium | Health Services Laboratories and Alex Alderton, Roberto Amato, Jeffrey Barrett, Sonia Goncalves, Ewan Harrison, David K. Jackson, Ian Johnston, Dominic Kwiatkowski, Cordelia Langford, John Sillitoe on behalf of the Wellcome Sanger Institute COVID-19 Surveillance Team |
| EPI_ISL_3437222 | Lighthouse Lab in Alderley Park | Wellcome Sanger Institute for the COVID-19 Genomics UK (COG-UK) Consortium | Jacquelyn Wynn, Mairead Hyland, The Lighthouse Lab in Alderley Park and Alex Alderton, Roberto Amato, Jeffrey Barrett, Sonia Goncalves, Ewan Harrison, David K. Jackson, Ian Johnston, Dominic Kwiatkowski, Cordelia Langford, John Sillitoe on behalf of the Wellcome Sanger Institute COVID-19 Surveillance Team |
| EPI_ISL_3437223, EPI_ISL_3437224 | Health Services Laboratories | Wellcome Sanger Institute for the COVID-19 Genomics UK (COG-UK) Consortium | Health Services Laboratories and Alex Alderton, Roberto Amato, Jeffrey Barrett, Sonia Goncalves, Ewan Harrison, David K. Jackson, Ian Johnston, Dominic Kwiatkowski, Cordelia Langford, John Sillitoe on behalf of the Wellcome Sanger Institute COVID-19 Surveillance Team |
| EPI_ISL_3437225 | Lighthouse Lab in Alderley Park | Wellcome Sanger Institute for the COVID-19 Genomics UK (COG-UK) Consortium | Jacquelyn Wynn, Mairead Hyland, The Lighthouse Lab in Alderley Park and Alex Alderton, Roberto Amato, Jeffrey Barrett, Sonia Goncalves, Ewan Harrison, David K. Jackson, Ian Johnston, Dominic Kwiatkowski, Cordelia Langford, John Sillitoe on behalf of the Wellcome Sanger Institute COVID-19 Surveillance Team |
| EPI_ISL_3437226, EPI_ISL_3437227, EPI_ISL_3437228, EPI_ISL_3437229 | Lighthouse Laboratory Plymouth | Wellcome Sanger Institute for the COVID-19 Genomics UK (COG-UK) Consortium | Lighthouse Laboratory Plymouth and Alex Alderton, Roberto Amato, Jeffrey Barrett, Sonia Goncalves, Ewan Harrison, David K. Jackson, Ian Johnston, Dominic Kwiatkowski, Cordelia Langford, John Sillitoe on behalf of the Wellcome Sanger Institute COVID-19 Surveillance Team |
| EPI_ISL_3437230 | Health Services Laboratories | Wellcome Sanger Institute for the COVID-19 Genomics UK (COG-UK) Consortium | Health Services Laboratories and Alex Alderton, Roberto Amato, Jeffrey Barrett, Sonia Goncalves, Ewan Harrison, David K. Jackson, Ian Johnston, Dominic Kwiatkowski, Cordelia Langford, John Sillitoe on behalf of the Wellcome Sanger Institute COVID-19 Surveillance Team |
| EPI_ISL_3437231 | Lighthouse Laboratory Plymouth | Wellcome Sanger Institute for the COVID-19 Genomics UK (COG-UK) Consortium | Lighthouse Laboratory Plymouth and Alex Alderton, Roberto Amato, Jeffrey Barrett, Sonia Goncalves, Ewan Harrison, David K. Jackson, Ian Johnston, Dominic Kwiatkowski, Cordelia Langford, John Sillitoe on behalf of the Wellcome Sanger Institute COVID-19 Surveillance Team |
| EPI_ISL_3437232 | Berkshire and Surrey Pathology Services Lighthouse Laboratory | Wellcome Sanger Institute for the COVID-19 Genomics UK (COG-UK) Consortium | Berkshire and Surrey Pathology Services Lighthouse Laboratory and Alex Alderton, Roberto Amato, Jeffrey Barrett, Sonia Goncalves, Ewan Harrison, David K. Jackson, Ian Johnston, Dominic Kwiatkowski, Cordelia Langford, John Sillitoe on behalf of the Wellcome Sanger Institute COVID-19 Surveillance Team |
| EPI_ISL_3437233 | Lighthouse Laboratory Plymouth | Wellcome Sanger Institute for the COVID-19 Genomics UK (COG-UK) Consortium | Lighthouse Laboratory Plymouth and Alex Alderton, Roberto Amato, Jeffrey Barrett, Sonia Goncalves, Ewan Harrison, David K. Jackson, Ian Johnston, Dominic Kwiatkowski, Cordelia Langford, John Sillitoe on behalf of the Wellcome Sanger Institute COVID-19 Surveillance Team |
| EPI_ISL_3437234 | Berkshire and Surrey Pathology Services Lighthouse Laboratory | Wellcome Sanger Institute for the COVID-19 Genomics UK (COG-UK) Consortium | Berkshire and Surrey Pathology Services Lighthouse Laboratory and Alex Alderton, Roberto Amato, Jeffrey Barrett, Sonia Goncalves, Ewan Harrison, David K. Jackson, Ian Johnston, Dominic Kwiatkowski, Cordelia Langford, John Sillitoe on behalf of the Wellcome Sanger Institute COVID-19 Surveillance Team |
| EPI_ISL_3437235, EPI_ISL_3437236 | Lighthouse Laboratory Plymouth | Wellcome Sanger Institute for the COVID-19 Genomics UK (COG-UK) Consortium | Lighthouse Laboratory Plymouth and Alex Alderton, Roberto Amato, Jeffrey Barrett, Sonia Goncalves, Ewan Harrison, David K. Jackson, Ian Johnston, Dominic Kwiatkowski, Cordelia Langford, John Sillitoe on behalf of the Wellcome Sanger Institute COVID-19 Surveillance Team |
| EPI_ISL_3437237 | Lighthouse Lab in Alderley Park | Wellcome Sanger Institute for the COVID-19 Genomics UK (COG-UK) Consortium | Jacquelyn Wynn, Mairead Hyland, The Lighthouse Lab in Alderley Park and Alex Alderton, Roberto Amato, Jeffrey Barrett, Sonia Goncalves, Ewan Harrison, David K. Jackson, Ian Johnston, Dominic Kwiatkowski, Cordelia Langford, John Sillitoe on behalf of the Wellcome Sanger Institute COVID-19 Surveillance Team |
| EPI_ISL_3437238, EPI_ISL_3437239, EPI_ISL_3437240 | Berkshire and Surrey Pathology Services Lighthouse Laboratory | Wellcome Sanger Institute for the COVID-19 Genomics UK (COG-UK) Consortium | Berkshire and Surrey Pathology Services Lighthouse Laboratory and Alex Alderton, Roberto Amato, Jeffrey Barrett, Sonia Goncalves, Ewan Harrison, David K. Jackson, Ian Johnston, Dominic Kwiatkowski, Cordelia Langford, John Sillitoe on behalf of the Wellcome Sanger Institute COVID-19 Surveillance Team |
| EPI_ISL_3437241 | Health Services Laboratories | Wellcome Sanger Institute for the COVID-19 Genomics UK (COG-UK) Consortium | Health Services Laboratories and Alex Alderton, Roberto Amato, Jeffrey Barrett, Sonia Goncalves, Ewan Harrison, David K. Jackson, Ian Johnston, Dominic Kwiatkowski, Cordelia Langford, John Sillitoe on behalf of the Wellcome Sanger Institute COVID-19 Surveillance Team |
| EPI_ISL_3437242 | Berkshire and Surrey Pathology Services Lighthouse Laboratory | Wellcome Sanger Institute for the COVID-19 Genomics UK (COG-UK) Consortium | Berkshire and Surrey Pathology Services Lighthouse Laboratory and Alex Alderton, Roberto Amato, Jeffrey Barrett, Sonia Goncalves, Ewan Harrison, David K. Jackson, Ian Johnston, Dominic Kwiatkowski, Cordelia Langford, John Sillitoe on behalf of the Wellcome Sanger Institute COVID-19 Surveillance Team |
| EPI_ISL_3437243, EPI_ISL_3437244, EPI_ISL_3437245 | Health Services Laboratories | Wellcome Sanger Institute for the COVID-19 Genomics UK (COG-UK) Consortium | Health Services Laboratories and Alex Alderton, Roberto Amato, Jeffrey Barrett, Sonia Goncalves, Ewan Harrison, David K. Jackson, Ian Johnston, Dominic Kwiatkowski, Cordelia Langford, John Sillitoe on behalf of the Wellcome Sanger Institute COVID-19 Surveillance Team |
| EPI_ISL_3437246 | Berkshire and Surrey Pathology Services Lighthouse Laboratory | Wellcome Sanger Institute for the COVID-19 Genomics UK (COG-UK) Consortium | Berkshire and Surrey Pathology Services Lighthouse Laboratory and Alex Alderton, Roberto Amato, Jeffrey Barrett, Sonia Goncalves, Ewan Harrison, David K. Jackson, Ian Johnston, Dominic Kwiatkowski, Cordelia Langford, John Sillitoe on behalf of the Wellcome Sanger Institute COVID-19 Surveillance Team |
| EPI_ISL_3437247 | Lighthouse Laboratory Plymouth | Wellcome Sanger Institute for the COVID-19 Genomics UK (COG-UK) Consortium | Lighthouse Laboratory Plymouth and Alex Alderton, Roberto Amato, Jeffrey Barrett, Sonia Goncalves, Ewan Harrison, David K. Jackson, Ian Johnston, Dominic Kwiatkowski, Cordelia Langford, John Sillitoe on behalf of the Wellcome Sanger Institute COVID-19 Surveillance Team |
| EPI_ISL_3437248 | Lighthouse Lab in Alderley Park | Wellcome Sanger Institute for the COVID-19 Genomics UK (COG-UK) Consortium | Jacquelyn Wynn, Mairead Hyland, The Lighthouse Lab in Alderley Park and Alex Alderton, Roberto Amato, Jeffrey Barrett, Sonia Goncalves, Ewan Harrison, David K. Jackson, Ian Johnston, Dominic Kwiatkowski, Cordelia Langford, John Sillitoe on behalf of the Wellcome Sanger Institute COVID-19 Surveillance Team |
| EPI_ISL_3437249 | Berkshire and Surrey Pathology Services Lighthouse Laboratory | Wellcome Sanger Institute for the COVID-19 Genomics UK (COG-UK) Consortium | Berkshire and Surrey Pathology Services Lighthouse Laboratory and Alex Alderton, Roberto Amato, Jeffrey Barrett, Sonia Goncalves, Ewan Harrison, David K. Jackson, Ian Johnston, Dominic Kwiatkowski, Cordelia Langford, John Sillitoe on behalf of the Wellcome Sanger Institute COVID-19 Surveillance Team |
| EPI_ISL_3437250 | Health Services Laboratories | Wellcome Sanger Institute for the COVID-19 Genomics UK (COG-UK) Consortium | Health Services Laboratories and Alex Alderton, Roberto Amato, Jeffrey Barrett, Sonia Goncalves, Ewan Harrison, David K. Jackson, Ian Johnston, Dominic Kwiatkowski, Cordelia Langford, John Sillitoe on behalf of the Wellcome Sanger Institute COVID-19 Surveillance Team |
| EPI_ISL_3437251 | Lighthouse Laboratory Plymouth | Wellcome Sanger Institute for the COVID-19 Genomics UK (COG-UK) Consortium | Lighthouse Laboratory Plymouth and Alex Alderton, Roberto Amato, Jeffrey Barrett, Sonia Goncalves, Ewan Harrison, David K. Jackson, Ian Johnston, Dominic Kwiatkowski, Cordelia Langford, John Sillitoe on behalf of the Wellcome Sanger Institute COVID-19 Surveillance Team |
| EPI_ISL_3437252 | Berkshire and Surrey Pathology Services Lighthouse Laboratory | Wellcome Sanger Institute for the COVID-19 Genomics UK (COG-UK) Consortium | Berkshire and Surrey Pathology Services Lighthouse Laboratory and Alex Alderton, Roberto Amato, Jeffrey Barrett, Sonia Goncalves, Ewan Harrison, David K. Jackson, Ian Johnston, Dominic Kwiatkowski, Cordelia Langford, John Sillitoe on behalf of the Wellcome Sanger Institute COVID-19 Surveillance Team |
| EPI_ISL_3437253 | Lighthouse Laboratory Plymouth | Wellcome Sanger Institute for the COVID-19 Genomics UK (COG-UK) Consortium | Lighthouse Laboratory Plymouth and Alex Alderton, Roberto Amato, Jeffrey Barrett, Sonia Goncalves, Ewan Harrison, David K. Jackson, Ian Johnston, Dominic Kwiatkowski, Cordelia Langford, John Sillitoe on behalf of the Wellcome Sanger Institute COVID-19 Surveillance Team |
| EPI_ISL_3437254, EPI_ISL_3437255 | Lighthouse Lab in Alderley Park | Wellcome Sanger Institute for the COVID-19 Genomics UK (COG-UK) Consortium | Jacquelyn Wynn, Mairead Hyland, The Lighthouse Lab in Alderley Park and Alex Alderton, Roberto Amato, Jeffrey Barrett, Sonia Goncalves, Ewan Harrison, David K. Jackson, Ian Johnston, Dominic Kwiatkowski, Cordelia Langford, John Sillitoe on behalf of the Wellcome Sanger Institute COVID-19 Surveillance Team |
| EPI_ISL_3437256 | Berkshire and Surrey Pathology Services Lighthouse Laboratory | Wellcome Sanger Institute for the COVID-19 Genomics UK (COG-UK) Consortium | Berkshire and Surrey Pathology Services Lighthouse Laboratory and Alex Alderton, Roberto Amato, Jeffrey Barrett, Sonia Goncalves, Ewan Harrison, David K. Jackson, Ian Johnston, Dominic Kwiatkowski, Cordelia Langford, John Sillitoe on behalf of the Wellcome Sanger Institute COVID-19 Surveillance Team |
| EPI_ISL_3437257 | Lighthouse Lab in Alderley Park | Wellcome Sanger Institute for the COVID-19 Genomics UK (COG-UK) Consortium | Jacquelyn Wynn, Mairead Hyland, The Lighthouse Lab in Alderley Park and Alex Alderton, Roberto Amato, Jeffrey Barrett, Sonia Goncalves, Ewan Harrison, David K. Jackson, Ian Johnston, Dominic Kwiatkowski, Cordelia Langford, John Sillitoe on behalf of the Wellcome Sanger Institute COVID-19 Surveillance Team |
| EPI_ISL_3437258 | Lighthouse Laboratory Plymouth | Wellcome Sanger Institute for the COVID-19 Genomics UK (COG-UK) Consortium | Lighthouse Laboratory Plymouth and Alex Alderton, Roberto Amato, Jeffrey Barrett, Sonia Goncalves, Ewan Harrison, David K. Jackson, Ian Johnston, Dominic Kwiatkowski, Cordelia Langford, John Sillitoe on behalf of the Wellcome Sanger Institute COVID-19 Surveillance Team |
| EPI_ISL_3437259 | Health Services Laboratories | Wellcome Sanger Institute for the COVID-19 Genomics UK (COG-UK) Consortium | Health Services Laboratories and Alex Alderton, Roberto Amato, Jeffrey Barrett, Sonia Goncalves, Ewan Harrison, David K. Jackson, Ian Johnston, Dominic Kwiatkowski, Cordelia Langford, John Sillitoe on behalf of the Wellcome Sanger Institute COVID-19 Surveillance Team |
| EPI_ISL_3437260 | Lighthouse Lab in Alderley Park | Wellcome Sanger Institute for the COVID-19 Genomics UK (COG-UK) Consortium | Jacquelyn Wynn, Mairead Hyland, The Lighthouse Lab in Alderley Park and Alex Alderton, Roberto Amato, Jeffrey Barrett, Sonia Goncalves, Ewan Harrison, David K. Jackson, Ian Johnston, Dominic Kwiatkowski, Cordelia Langford, John Sillitoe on behalf of the Wellcome Sanger Institute COVID-19 Surveillance Team |
| EPI_ISL_3437261, EPI_ISL_3437262 | Lighthouse Laboratory Plymouth | Wellcome Sanger Institute for the COVID-19 Genomics UK (COG-UK) Consortium | Lighthouse Laboratory Plymouth and Alex Alderton, Roberto Amato, Jeffrey Barrett, Sonia Goncalves, Ewan Harrison, David K. Jackson, Ian Johnston, Dominic Kwiatkowski, Cordelia Langford, John Sillitoe on behalf of the Wellcome Sanger Institute COVID-19 Surveillance Team |
| EPI_ISL_3437263 | Health Services Laboratories | Wellcome Sanger Institute for the COVID-19 Genomics UK (COG-UK) Consortium | Health Services Laboratories and Alex Alderton, Roberto Amato, Jeffrey Barrett, Sonia Goncalves, Ewan Harrison, David K. Jackson, Ian Johnston, Dominic Kwiatkowski, Cordelia Langford, John Sillitoe on behalf of the Wellcome Sanger Institute COVID-19 Surveillance Team |
| EPI_ISL_3437264, EPI_ISL_3437265 | Lighthouse Lab in Alderley Park | Wellcome Sanger Institute for the COVID-19 Genomics UK (COG-UK) Consortium | Jacquelyn Wynn, Mairead Hyland, The Lighthouse Lab in Alderley Park and Alex Alderton, Roberto Amato, Jeffrey Barrett, Sonia Goncalves, Ewan Harrison, David K. Jackson, Ian Johnston, Dominic Kwiatkowski, Cordelia Langford, John Sillitoe on behalf of the Wellcome Sanger Institute COVID-19 Surveillance Team |
| EPI_ISL_3437266, EPI_ISL_3437267 | Berkshire and Surrey Pathology Services Lighthouse Laboratory | Wellcome Sanger Institute for the COVID-19 Genomics UK (COG-UK) Consortium | Berkshire and Surrey Pathology Services Lighthouse Laboratory and Alex Alderton, Roberto Amato, Jeffrey Barrett, Sonia Goncalves, Ewan Harrison, David K. Jackson, Ian Johnston, Dominic Kwiatkowski, Cordelia Langford, John Sillitoe on behalf of the Wellcome Sanger Institute COVID-19 Surveillance Team |
| EPI_ISL_3437268 | Lighthouse Laboratory Plymouth | Wellcome Sanger Institute for the COVID-19 Genomics UK (COG-UK) Consortium | Lighthouse Laboratory Plymouth and Alex Alderton, Roberto Amato, Jeffrey Barrett, Sonia Goncalves, Ewan Harrison, David K. Jackson, Ian Johnston, Dominic Kwiatkowski, Cordelia Langford, John Sillitoe on behalf of the Wellcome Sanger Institute COVID-19 Surveillance Team |
| EPI_ISL_3437269 | Berkshire and Surrey Pathology Services Lighthouse Laboratory | Wellcome Sanger Institute for the COVID-19 Genomics UK (COG-UK) Consortium | Berkshire and Surrey Pathology Services Lighthouse Laboratory and Alex Alderton, Roberto Amato, Jeffrey Barrett, Sonia Goncalves, Ewan Harrison, David K. Jackson, Ian Johnston, Dominic Kwiatkowski, Cordelia Langford, John Sillitoe on behalf of the Wellcome Sanger Institute COVID-19 Surveillance Team |
| EPI_ISL_3437270, EPI_ISL_3437271 | Lighthouse Lab in Alderley Park | Wellcome Sanger Institute for the COVID-19 Genomics UK (COG-UK) Consortium | Jacquelyn Wynn, Mairead Hyland, The Lighthouse Lab in Alderley Park and Alex Alderton, Roberto Amato, Jeffrey Barrett, Sonia Goncalves, Ewan Harrison, David K. Jackson, Ian Johnston, Dominic Kwiatkowski, Cordelia Langford, John Sillitoe on behalf of the Wellcome Sanger Institute COVID-19 Surveillance Team |
| EPI_ISL_3437272 | Berkshire and Surrey Pathology Services Lighthouse Laboratory | Wellcome Sanger Institute for the COVID-19 Genomics UK (COG-UK) Consortium | Berkshire and Surrey Pathology Services Lighthouse Laboratory and Alex Alderton, Roberto Amato, Jeffrey Barrett, Sonia Goncalves, Ewan Harrison, David K. Jackson, Ian Johnston, Dominic Kwiatkowski, Cordelia Langford, John Sillitoe on behalf of the Wellcome Sanger Institute COVID-19 Surveillance Team |
| EPI_ISL_3437273 | Lighthouse Laboratory Plymouth | Wellcome Sanger Institute for the COVID-19 Genomics UK (COG-UK) Consortium | Lighthouse Laboratory Plymouth and Alex Alderton, Roberto Amato, Jeffrey Barrett, Sonia Goncalves, Ewan Harrison, David K. Jackson, Ian Johnston, Dominic Kwiatkowski, Cordelia Langford, John Sillitoe on behalf of the Wellcome Sanger Institute COVID-19 Surveillance Team |
| EPI_ISL_3437274 | Lighthouse Lab in Alderley Park | Wellcome Sanger Institute for the COVID-19 Genomics UK (COG-UK) Consortium | Jacquelyn Wynn, Mairead Hyland, The Lighthouse Lab in Alderley Park and Alex Alderton, Roberto Amato, Jeffrey Barrett, Sonia Goncalves, Ewan Harrison, David K. Jackson, Ian Johnston, Dominic Kwiatkowski, Cordelia Langford, John Sillitoe on behalf of the Wellcome Sanger Institute COVID-19 Surveillance Team |
| EPI_ISL_3437275 | Lighthouse Laboratory Plymouth | Wellcome Sanger Institute for the COVID-19 Genomics UK (COG-UK) Consortium | Lighthouse Laboratory Plymouth and Alex Alderton, Roberto Amato, Jeffrey Barrett, Sonia Goncalves, Ewan Harrison, David K. Jackson, Ian Johnston, Dominic Kwiatkowski, Cordelia Langford, John Sillitoe on behalf of the Wellcome Sanger Institute COVID-19 Surveillance Team |
| EPI_ISL_3437276 | Berkshire and Surrey Pathology Services Lighthouse Laboratory | Wellcome Sanger Institute for the COVID-19 Genomics UK (COG-UK) | Berkshire and Surrey Pathology Services Lighthouse Laboratory and Alex Alderton, Roberto Amato, Jeffrey Barrett, Sonia Goncalves, Ewan Harrison, David K. Jackson, Ian Johnston, Dominic Kwiatkowski, Cordelia Langford, John Sillitoe on behalf |

| | | | |
|---|---|---|---|
| | | Consortium | of the Wellcome Sanger Institute COVID-19 Surveillance Team |
| EPI_ISL_3437277 | Lighthouse Laboratory Plymouth | Wellcome Sanger Institute for the COVID-19 Genomics UK (COG-UK) Consortium | Lighthouse Laboratory Plymouth and Alex Alderton, Roberto Amato, Jeffrey Barrett, Sonia Goncalves, Ewan Harrison, David K. Jackson, Ian Johnston, Dominic Kwiatkowski, Cordelia Langford, John Sillitoe on behalf of the Wellcome Sanger Institute COVID-19 Surveillance Team |
| EPI_ISL_3437278 | Berkshire and Surrey Pathology Services Lighthouse Laboratory | Wellcome Sanger Institute for the COVID-19 Genomics UK (COG-UK) Consortium | Berkshire and Surrey Pathology Services Lighthouse Laboratory and Alex Alderton, Roberto Amato, Jeffrey Barrett, Sonia Goncalves, Ewan Harrison, David K. Jackson, Ian Johnston, Dominic Kwiatkowski, Cordelia Langford, John Sillitoe on behalf of the Wellcome Sanger Institute COVID-19 Surveillance Team |
| EPI_ISL_3437279 | Lighthouse Laboratory Plymouth | Wellcome Sanger Institute for the COVID-19 Genomics UK (COG-UK) Consortium | Lighthouse Laboratory Plymouth and Alex Alderton, Roberto Amato, Jeffrey Barrett, Sonia Goncalves, Ewan Harrison, David K. Jackson, Ian Johnston, Dominic Kwiatkowski, Cordelia Langford, John Sillitoe on behalf of the Wellcome Sanger Institute COVID-19 Surveillance Team |
| EPI_ISL_3437280, EPI_ISL_3437281 | Lighthouse Lab in Alderley Park | Wellcome Sanger Institute for the COVID-19 Genomics UK (COG-UK) Consortium | Jacquelyn Wynn, Mairead Hyland, The Lighthouse Lab in Alderley Park and Alex Alderton, Roberto Amato, Jeffrey Barrett, Sonia Goncalves, Ewan Harrison, David K. Jackson, Ian Johnston, Dominic Kwiatkowski, Cordelia Langford, John Sillitoe on behalf of the Wellcome Sanger Institute COVID-19 Surveillance Team |
| EPI_ISL_3437282 | Berkshire and Surrey Pathology Services Lighthouse Laboratory | Wellcome Sanger Institute for the COVID-19 Genomics UK (COG-UK) Consortium | Berkshire and Surrey Pathology Services Lighthouse Laboratory and Alex Alderton, Roberto Amato, Jeffrey Barrett, Sonia Goncalves, Ewan Harrison, David K. Jackson, Ian Johnston, Dominic Kwiatkowski, Cordelia Langford, John Sillitoe on behalf of the Wellcome Sanger Institute COVID-19 Surveillance Team |
| EPI_ISL_3437283, EPI_ISL_3437284, EPI_ISL_3437285 | Lighthouse Laboratory Plymouth | Wellcome Sanger Institute for the COVID-19 Genomics UK (COG-UK) Consortium | Lighthouse Laboratory Plymouth and Alex Alderton, Roberto Amato, Jeffrey Barrett, Sonia Goncalves, Ewan Harrison, David K. Jackson, Ian Johnston, Dominic Kwiatkowski, Cordelia Langford, John Sillitoe on behalf of the Wellcome Sanger Institute COVID-19 Surveillance Team |
| EPI_ISL_3437286, EPI_ISL_3437287 | Lighthouse Lab in Alderley Park | Wellcome Sanger Institute for the COVID-19 Genomics UK (COG-UK) Consortium | Jacquelyn Wynn, Mairead Hyland, The Lighthouse Lab in Alderley Park and Alex Alderton, Roberto Amato, Jeffrey Barrett, Sonia Goncalves, Ewan Harrison, David K. Jackson, Ian Johnston, Dominic Kwiatkowski, Cordelia Langford, John Sillitoe on behalf of the Wellcome Sanger Institute COVID-19 Surveillance Team |
| EPI_ISL_3437288, EPI_ISL_3437289 | Berkshire and Surrey Pathology Services Lighthouse Laboratory | Wellcome Sanger Institute for the COVID-19 Genomics UK (COG-UK) Consortium | Berkshire and Surrey Pathology Services Lighthouse Laboratory and Alex Alderton, Roberto Amato, Jeffrey Barrett, Sonia Goncalves, Ewan Harrison, David K. Jackson, Ian Johnston, Dominic Kwiatkowski, Cordelia Langford, John Sillitoe on behalf of the Wellcome Sanger Institute COVID-19 Surveillance Team |
| EPI_ISL_3437290 | Lighthouse Lab in Alderley Park | Wellcome Sanger Institute for the COVID-19 Genomics UK (COG-UK) Consortium | Jacquelyn Wynn, Mairead Hyland, The Lighthouse Lab in Alderley Park and Alex Alderton, Roberto Amato, Jeffrey Barrett, Sonia Goncalves, Ewan Harrison, David K. Jackson, Ian Johnston, Dominic Kwiatkowski, Cordelia Langford, John Sillitoe on behalf of the Wellcome Sanger Institute COVID-19 Surveillance Team |
| EPI_ISL_3437291, EPI_ISL_3437292 | Berkshire and Surrey Pathology Services Lighthouse Laboratory | Wellcome Sanger Institute for the COVID-19 Genomics UK (COG-UK) Consortium | Berkshire and Surrey Pathology Services Lighthouse Laboratory and Alex Alderton, Roberto Amato, Jeffrey Barrett, Sonia Goncalves, Ewan Harrison, David K. Jackson, Ian Johnston, Dominic Kwiatkowski, Cordelia Langford, John Sillitoe on behalf of the Wellcome Sanger Institute COVID-19 Surveillance Team |
| EPI_ISL_3437293 | Lighthouse Lab in Alderley Park | Wellcome Sanger Institute for the COVID-19 Genomics UK (COG-UK) Consortium | Jacquelyn Wynn, Mairead Hyland, The Lighthouse Lab in Alderley Park and Alex Alderton, Roberto Amato, Jeffrey Barrett, Sonia Goncalves, Ewan Harrison, David K. Jackson, Ian Johnston, Dominic Kwiatkowski, Cordelia Langford, John Sillitoe on behalf of the Wellcome Sanger Institute COVID-19 Surveillance Team |
| EPI_ISL_3437294, EPI_ISL_3437295, EPI_ISL_3437296, EPI_ISL_3437297, EPI_ISL_3437298, EPI_ISL_3437299 | Berkshire and Surrey Pathology Services Lighthouse Laboratory | Wellcome Sanger Institute for the COVID-19 Genomics UK (COG-UK) Consortium | Berkshire and Surrey Pathology Services Lighthouse Laboratory and Alex Alderton, Roberto Amato, Jeffrey Barrett, Sonia Goncalves, Ewan Harrison, David K. Jackson, Ian Johnston, Dominic Kwiatkowski, Cordelia Langford, John Sillitoe on behalf of the Wellcome Sanger Institute COVID-19 Surveillance Team |
| EPI_ISL_3437300 | Lighthouse Lab in Alderley Park | Wellcome Sanger Institute for the COVID-19 Genomics UK (COG-UK) Consortium | Jacquelyn Wynn, Mairead Hyland, The Lighthouse Lab in Alderley Park and Alex Alderton, Roberto Amato, Jeffrey Barrett, Sonia Goncalves, Ewan Harrison, David K. Jackson, Ian Johnston, Dominic Kwiatkowski, Cordelia Langford, John Sillitoe on behalf of the Wellcome Sanger Institute COVID-19 Surveillance Team |
| EPI_ISL_3437301 | Berkshire and Surrey Pathology Services Lighthouse Laboratory | Wellcome Sanger Institute for the COVID-19 Genomics UK (COG-UK) Consortium | Berkshire and Surrey Pathology Services Lighthouse Laboratory and Alex Alderton, Roberto Amato, Jeffrey Barrett, Sonia Goncalves, Ewan Harrison, David K. Jackson, Ian Johnston, Dominic Kwiatkowski, Cordelia Langford, John Sillitoe on behalf of the Wellcome Sanger Institute COVID-19 Surveillance Team |
| EPI_ISL_3437302, EPI_ISL_3437303, EPI_ISL_3437304 | Lighthouse Lab in Alderley Park | Wellcome Sanger Institute for the COVID-19 Genomics UK (COG-UK) Consortium | Jacquelyn Wynn, Mairead Hyland, The Lighthouse Lab in Alderley Park and Alex Alderton, Roberto Amato, Jeffrey Barrett, Sonia Goncalves, Ewan Harrison, David K. Jackson, Ian Johnston, Dominic Kwiatkowski, Cordelia Langford, John Sillitoe on behalf of the Wellcome Sanger Institute COVID-19 Surveillance Team |
| EPI_ISL_3437305 | Berkshire and Surrey Pathology Services Lighthouse Laboratory | Wellcome Sanger Institute for the COVID-19 Genomics UK (COG-UK) Consortium | Berkshire and Surrey Pathology Services Lighthouse Laboratory and Alex Alderton, Roberto Amato, Jeffrey Barrett, Sonia Goncalves, Ewan Harrison, David K. Jackson, Ian Johnston, Dominic Kwiatkowski, Cordelia Langford, John Sillitoe on behalf of the Wellcome Sanger Institute COVID-19 Surveillance Team |
| EPI_ISL_3437306 | Lighthouse Lab in Alderley Park | Wellcome Sanger Institute for the COVID-19 Genomics UK (COG-UK) Consortium | Jacquelyn Wynn, Mairead Hyland, The Lighthouse Lab in Alderley Park and Alex Alderton, Roberto Amato, Jeffrey Barrett, Sonia Goncalves, Ewan Harrison, David K. Jackson, Ian Johnston, Dominic Kwiatkowski, Cordelia Langford, John Sillitoe on behalf of the Wellcome Sanger Institute COVID-19 Surveillance Team |
| EPI_ISL_3437307 | Berkshire and Surrey Pathology Services Lighthouse Laboratory | Wellcome Sanger Institute for the COVID-19 Genomics UK (COG-UK) Consortium | Berkshire and Surrey Pathology Services Lighthouse Laboratory and Alex Alderton, Roberto Amato, Jeffrey Barrett, Sonia Goncalves, Ewan Harrison, David K. Jackson, Ian Johnston, Dominic Kwiatkowski, Cordelia Langford, John Sillitoe on behalf of the Wellcome Sanger Institute COVID-19 Surveillance Team |
| EPI_ISL_3437308, EPI_ISL_3437309, EPI_ISL_3437310, EPI_ISL_3437311 | Lighthouse Lab in Alderley Park | Wellcome Sanger Institute for the COVID-19 Genomics UK (COG-UK) Consortium | Jacquelyn Wynn, Mairead Hyland, The Lighthouse Lab in Alderley Park and Alex Alderton, Roberto Amato, Jeffrey Barrett, Sonia Goncalves, Ewan Harrison, David K. Jackson, Ian Johnston, Dominic Kwiatkowski, Cordelia Langford, John Sillitoe on behalf of the Wellcome Sanger Institute COVID-19 Surveillance Team |
| EPI_ISL_3437312 | Berkshire and Surrey Pathology Services Lighthouse Laboratory | Wellcome Sanger Institute for the COVID-19 Genomics UK (COG-UK) Consortium | Berkshire and Surrey Pathology Services Lighthouse Laboratory and Alex Alderton, Roberto Amato, Jeffrey Barrett, Sonia Goncalves, Ewan Harrison, David K. Jackson, Ian Johnston, Dominic Kwiatkowski, Cordelia Langford, John Sillitoe on behalf of the Wellcome Sanger Institute COVID-19 Surveillance Team |
| EPI_ISL_3437313, EPI_ISL_3437314 | Lighthouse Lab in Alderley Park | Wellcome Sanger Institute for the COVID-19 Genomics UK (COG-UK) Consortium | Jacquelyn Wynn, Mairead Hyland, The Lighthouse Lab in Alderley Park and Alex Alderton, Roberto Amato, Jeffrey Barrett, Sonia Goncalves, Ewan Harrison, David K. Jackson, Ian Johnston, Dominic Kwiatkowski, Cordelia Langford, John Sillitoe on behalf of the Wellcome Sanger Institute COVID-19 Surveillance Team |
| EPI_ISL_3437315 | Berkshire and Surrey Pathology Services Lighthouse Laboratory | Wellcome Sanger Institute for the COVID-19 Genomics UK (COG-UK) Consortium | Berkshire and Surrey Pathology Services Lighthouse Laboratory and Alex Alderton, Roberto Amato, Jeffrey Barrett, Sonia Goncalves, Ewan Harrison, David K. Jackson, Ian Johnston, Dominic Kwiatkowski, Cordelia Langford, John Sillitoe on behalf of the Wellcome Sanger Institute COVID-19 Surveillance Team |
| EPI_ISL_3437316 | Lighthouse Lab in Alderley Park | Wellcome Sanger Institute for the COVID-19 Genomics UK (COG-UK) Consortium | Jacquelyn Wynn, Mairead Hyland, The Lighthouse Lab in Alderley Park and Alex Alderton, Roberto Amato, Jeffrey Barrett, Sonia Goncalves, Ewan Harrison, David K. Jackson, Ian Johnston, Dominic Kwiatkowski, Cordelia Langford, John Sillitoe on behalf of the Wellcome Sanger Institute COVID-19 Surveillance Team |
| EPI_ISL_3437317, EPI_ISL_3437318, EPI_ISL_3437319 | Berkshire and Surrey Pathology Services Lighthouse Laboratory | Wellcome Sanger Institute for the COVID-19 Genomics UK (COG-UK) Consortium | Berkshire and Surrey Pathology Services Lighthouse Laboratory and Alex Alderton, Roberto Amato, Jeffrey Barrett, Sonia Goncalves, Ewan Harrison, David K. Jackson, Ian Johnston, Dominic Kwiatkowski, Cordelia Langford, John Sillitoe on behalf of the Wellcome Sanger Institute COVID-19 Surveillance Team |
| EPI_ISL_3437320 | Lighthouse Lab in Alderley Park | Wellcome Sanger Institute for the COVID-19 Genomics UK (COG-UK) Consortium | Jacquelyn Wynn, Mairead Hyland, The Lighthouse Lab in Alderley Park and Alex Alderton, Roberto Amato, Jeffrey Barrett, Sonia Goncalves, Ewan Harrison, David K. Jackson, Ian Johnston, Dominic Kwiatkowski, Cordelia Langford, John Sillitoe on behalf of the Wellcome Sanger Institute COVID-19 Surveillance Team |
| EPI_ISL_3437321, EPI_ISL_3437322, EPI_ISL_3437323, EPI_ISL_3437324, EPI_ISL_3437325, EPI_ISL_3437326, EPI_ISL_3437327, EPI_ISL_3437328, EPI_ISL_3437329, EPI_ISL_3437330 | Berkshire and Surrey Pathology Services Lighthouse Laboratory | Wellcome Sanger Institute for the COVID-19 Genomics UK (COG-UK) Consortium | Berkshire and Surrey Pathology Services Lighthouse Laboratory and Alex Alderton, Roberto Amato, Jeffrey Barrett, Sonia Goncalves, Ewan Harrison, David K. Jackson, Ian Johnston, Dominic Kwiatkowski, Cordelia Langford, John Sillitoe on behalf of the Wellcome Sanger Institute COVID-19 Surveillance Team |
| EPI_ISL_3437331 | Lighthouse Lab in Alderley Park | Wellcome Sanger Institute for the COVID-19 Genomics UK (COG-UK) Consortium | Jacquelyn Wynn, Mairead Hyland, The Lighthouse Lab in Alderley Park and Alex Alderton, Roberto Amato, Jeffrey Barrett, Sonia Goncalves, Ewan Harrison, David K. Jackson, Ian Johnston, Dominic Kwiatkowski, Cordelia Langford, John Sillitoe on behalf of the Wellcome Sanger Institute COVID-19 Surveillance Team |
| EPI_ISL_3437332 | Berkshire and Surrey Pathology Services Lighthouse Laboratory | Wellcome Sanger Institute for the COVID-19 Genomics UK (COG-UK) Consortium | Berkshire and Surrey Pathology Services Lighthouse Laboratory and Alex Alderton, Roberto Amato, Jeffrey Barrett, Sonia Goncalves, Ewan Harrison, David K. Jackson, Ian Johnston, Dominic Kwiatkowski, Cordelia Langford, John Sillitoe on behalf of the Wellcome Sanger Institute COVID-19 Surveillance Team |
| EPI_ISL_3437333 | Lighthouse Lab in Alderley Park | Wellcome Sanger Institute for the COVID-19 Genomics UK (COG-UK) Consortium | Jacquelyn Wynn, Mairead Hyland, The Lighthouse Lab in Alderley Park and Alex Alderton, Roberto Amato, Jeffrey Barrett, Sonia Goncalves, Ewan Harrison, David K. Jackson, Ian Johnston, Dominic Kwiatkowski, Cordelia Langford, John Sillitoe on behalf of the Wellcome Sanger Institute COVID-19 Surveillance Team |
| EPI_ISL_3437334 | Berkshire and Surrey Pathology Services Lighthouse Laboratory | Wellcome Sanger Institute for the COVID-19 Genomics UK (COG-UK) Consortium | Berkshire and Surrey Pathology Services Lighthouse Laboratory and Alex Alderton, Roberto Amato, Jeffrey Barrett, Sonia Goncalves, Ewan Harrison, David K. Jackson, Ian Johnston, Dominic Kwiatkowski, Cordelia Langford, John Sillitoe on behalf of the Wellcome Sanger Institute COVID-19 Surveillance Team |
| EPI_ISL_3437335 | Lighthouse Lab in Alderley Park | Wellcome Sanger Institute for the COVID-19 Genomics UK (COG-UK) Consortium | Jacquelyn Wynn, Mairead Hyland, The Lighthouse Lab in Alderley Park and Alex Alderton, Roberto Amato, Jeffrey Barrett, Sonia Goncalves, Ewan Harrison, David K. Jackson, Ian Johnston, Dominic Kwiatkowski, Cordelia Langford, John Sillitoe on behalf of the Wellcome Sanger Institute COVID-19 Surveillance Team |
| EPI_ISL_3437336 | Berkshire and Surrey Pathology Services Lighthouse Laboratory | Wellcome Sanger Institute for the COVID-19 Genomics UK (COG-UK) Consortium | Berkshire and Surrey Pathology Services Lighthouse Laboratory and Alex Alderton, Roberto Amato, Jeffrey Barrett, Sonia Goncalves, Ewan Harrison, David K. Jackson, Ian Johnston, Dominic Kwiatkowski, Cordelia Langford, John Sillitoe on behalf of the Wellcome Sanger Institute COVID-19 Surveillance Team |
| EPI_ISL_3437337 | Lighthouse Lab in Alderley Park | Wellcome Sanger Institute for the COVID-19 Genomics UK (COG-UK) Consortium | Jacquelyn Wynn, Mairead Hyland, The Lighthouse Lab in Alderley Park and Alex Alderton, Roberto Amato, Jeffrey Barrett, Sonia Goncalves, Ewan Harrison, David K. Jackson, Ian Johnston, Dominic Kwiatkowski, Cordelia Langford, John Sillitoe on behalf of the Wellcome Sanger Institute COVID-19 Surveillance Team |
| EPI_ISL_3437338 | Berkshire and Surrey Pathology Services Lighthouse Laboratory | Wellcome Sanger Institute for the COVID-19 Genomics UK (COG-UK) Consortium | Berkshire and Surrey Pathology Services Lighthouse Laboratory and Alex Alderton, Roberto Amato, Jeffrey Barrett, Sonia Goncalves, Ewan Harrison, David K. Jackson, Ian Johnston, Dominic Kwiatkowski, Cordelia Langford, John Sillitoe on behalf of the Wellcome Sanger Institute COVID-19 Surveillance Team |
| EPI_ISL_3437339, EPI_ISL_3437340, EPI_ISL_3437341 | Lighthouse Lab in Alderley Park | Wellcome Sanger Institute for the COVID-19 Genomics UK (COG-UK) Consortium | Jacquelyn Wynn, Mairead Hyland, The Lighthouse Lab in Alderley Park and Alex Alderton, Roberto Amato, Jeffrey Barrett, Sonia Goncalves, Ewan Harrison, David K. Jackson, Ian Johnston, Dominic Kwiatkowski, Cordelia Langford, John Sillitoe on behalf of the Wellcome Sanger Institute COVID-19 Surveillance Team |
| EPI_ISL_3437342, EPI_ISL_3437343, EPI_ISL_3437344, EPI_ISL_3437345 | Berkshire and Surrey Pathology Services Lighthouse Laboratory | Wellcome Sanger Institute for the COVID-19 Genomics UK (COG-UK) Consortium | Berkshire and Surrey Pathology Services Lighthouse Laboratory and Alex Alderton, Roberto Amato, Jeffrey Barrett, Sonia Goncalves, Ewan Harrison, David K. Jackson, Ian Johnston, Dominic Kwiatkowski, Cordelia Langford, John Sillitoe on behalf of the Wellcome Sanger Institute COVID-19 Surveillance Team |

| | | | |
|---|---|---|---|
| EPI_ISL_3437346 | Lighthouse Lab in Alderley Park | Wellcome Sanger Institute for the COVID-19 Genomics UK (COG-UK) Consortium | Jacquelyn Wynn, Mairead Hyland, The Lighthouse Lab in Alderley Park and Alex Alderton, Roberto Amato, Jeffrey Barrett, Sonia Goncalves, Ewan Harrison, David K. Jackson, Ian Johnston, Dominic Kwiatkowski, Cordelia Langford, John Sillitoe on behalf of the Wellcome Sanger Institute COVID-19 Surveillance Team |
| EPI_ISL_3437347, EPI_ISL_3437348, EPI_ISL_3437349, EPI_ISL_3437350 | Berkshire and Surrey Pathology Services Lighthouse Laboratory | Wellcome Sanger Institute for the COVID-19 Genomics UK (COG-UK) Consortium | Berkshire and Surrey Pathology Services Lighthouse Laboratory and Alex Alderton, Roberto Amato, Jeffrey Barrett, Sonia Goncalves, Ewan Harrison, David K. Jackson, Ian Johnston, Dominic Kwiatkowski, Cordelia Langford, John Sillitoe on behalf of the Wellcome Sanger Institute COVID-19 Surveillance Team |
| EPI_ISL_3437351 | Lighthouse Lab in Alderley Park | Wellcome Sanger Institute for the COVID-19 Genomics UK (COG-UK) Consortium | Jacquelyn Wynn, Mairead Hyland, The Lighthouse Lab in Alderley Park and Alex Alderton, Roberto Amato, Jeffrey Barrett, Sonia Goncalves, Ewan Harrison, David K. Jackson, Ian Johnston, Dominic Kwiatkowski, Cordelia Langford, John Sillitoe on behalf of the Wellcome Sanger Institute COVID-19 Surveillance Team |
| EPI_ISL_3437352 | Berkshire and Surrey Pathology Services Lighthouse Laboratory | Wellcome Sanger Institute for the COVID-19 Genomics UK (COG-UK) Consortium | Berkshire and Surrey Pathology Services Lighthouse Laboratory and Alex Alderton, Roberto Amato, Jeffrey Barrett, Sonia Goncalves, Ewan Harrison, David K. Jackson, Ian Johnston, Dominic Kwiatkowski, Cordelia Langford, John Sillitoe on behalf of the Wellcome Sanger Institute COVID-19 Surveillance Team |
| EPI_ISL_3437353 | Lighthouse Lab in Alderley Park | Wellcome Sanger Institute for the COVID-19 Genomics UK (COG-UK) Consortium | Jacquelyn Wynn, Mairead Hyland, The Lighthouse Lab in Alderley Park and Alex Alderton, Roberto Amato, Jeffrey Barrett, Sonia Goncalves, Ewan Harrison, David K. Jackson, Ian Johnston, Dominic Kwiatkowski, Cordelia Langford, John Sillitoe on behalf of the Wellcome Sanger Institute COVID-19 Surveillance Team |
| EPI_ISL_3437354, EPI_ISL_3437355 | Berkshire and Surrey Pathology Services Lighthouse Laboratory | Wellcome Sanger Institute for the COVID-19 Genomics UK (COG-UK) Consortium | Berkshire and Surrey Pathology Services Lighthouse Laboratory and Alex Alderton, Roberto Amato, Jeffrey Barrett, Sonia Goncalves, Ewan Harrison, David K. Jackson, Ian Johnston, Dominic Kwiatkowski, Cordelia Langford, John Sillitoe on behalf of the Wellcome Sanger Institute COVID-19 Surveillance Team |
| EPI_ISL_3437356 | Lighthouse Lab in Alderley Park | Wellcome Sanger Institute for the COVID-19 Genomics UK (COG-UK) Consortium | Jacquelyn Wynn, Mairead Hyland, The Lighthouse Lab in Alderley Park and Alex Alderton, Roberto Amato, Jeffrey Barrett, Sonia Goncalves, Ewan Harrison, David K. Jackson, Ian Johnston, Dominic Kwiatkowski, Cordelia Langford, John Sillitoe on behalf of the Wellcome Sanger Institute COVID-19 Surveillance Team |
| EPI_ISL_3437357 | Berkshire and Surrey Pathology Services Lighthouse Laboratory | Wellcome Sanger Institute for the COVID-19 Genomics UK (COG-UK) Consortium | Berkshire and Surrey Pathology Services Lighthouse Laboratory and Alex Alderton, Roberto Amato, Jeffrey Barrett, Sonia Goncalves, Ewan Harrison, David K. Jackson, Ian Johnston, Dominic Kwiatkowski, Cordelia Langford, John Sillitoe on behalf of the Wellcome Sanger Institute COVID-19 Surveillance Team |
| EPI_ISL_3437358 | Lighthouse Lab in Alderley Park | Wellcome Sanger Institute for the COVID-19 Genomics UK (COG-UK) Consortium | Jacquelyn Wynn, Mairead Hyland, The Lighthouse Lab in Alderley Park and Alex Alderton, Roberto Amato, Jeffrey Barrett, Sonia Goncalves, Ewan Harrison, David K. Jackson, Ian Johnston, Dominic Kwiatkowski, Cordelia Langford, John Sillitoe on behalf of the Wellcome Sanger Institute COVID-19 Surveillance Team |
| EPI_ISL_3437359, EPI_ISL_3437360, EPI_ISL_3437361, EPI_ISL_3437362, EPI_ISL_3437363 | Berkshire and Surrey Pathology Services Lighthouse Laboratory | Wellcome Sanger Institute for the COVID-19 Genomics UK (COG-UK) Consortium | Berkshire and Surrey Pathology Services Lighthouse Laboratory and Alex Alderton, Roberto Amato, Jeffrey Barrett, Sonia Goncalves, Ewan Harrison, David K. Jackson, Ian Johnston, Dominic Kwiatkowski, Cordelia Langford, John Sillitoe on behalf of the Wellcome Sanger Institute COVID-19 Surveillance Team |
| EPI_ISL_3437364, EPI_ISL_3437365, EPI_ISL_3437366, EPI_ISL_3437367, EPI_ISL_3437368 | Lighthouse Lab in Alderley Park | Wellcome Sanger Institute for the COVID-19 Genomics UK (COG-UK) Consortium | Jacquelyn Wynn, Mairead Hyland, The Lighthouse Lab in Alderley Park and Alex Alderton, Roberto Amato, Jeffrey Barrett, Sonia Goncalves, Ewan Harrison, David K. Jackson, Ian Johnston, Dominic Kwiatkowski, Cordelia Langford, John Sillitoe on behalf of the Wellcome Sanger Institute COVID-19 Surveillance Team |
| EPI_ISL_3437369, EPI_ISL_3437370 | Berkshire and Surrey Pathology Services Lighthouse Laboratory | Wellcome Sanger Institute for the COVID-19 Genomics UK (COG-UK) Consortium | Berkshire and Surrey Pathology Services Lighthouse Laboratory and Alex Alderton, Roberto Amato, Jeffrey Barrett, Sonia Goncalves, Ewan Harrison, David K. Jackson, Ian Johnston, Dominic Kwiatkowski, Cordelia Langford, John Sillitoe on behalf of the Wellcome Sanger Institute COVID-19 Surveillance Team |
| EPI_ISL_3437371 | Lighthouse Lab in Alderley Park | Wellcome Sanger Institute for the COVID-19 Genomics UK (COG-UK) Consortium | Jacquelyn Wynn, Mairead Hyland, The Lighthouse Lab in Alderley Park and Alex Alderton, Roberto Amato, Jeffrey Barrett, Sonia Goncalves, Ewan Harrison, David K. Jackson, Ian Johnston, Dominic Kwiatkowski, Cordelia Langford, John Sillitoe on behalf of the Wellcome Sanger Institute COVID-19 Surveillance Team |
| EPI_ISL_3437372 | Berkshire and Surrey Pathology Services Lighthouse Laboratory | Wellcome Sanger Institute for the COVID-19 Genomics UK (COG-UK) Consortium | Berkshire and Surrey Pathology Services Lighthouse Laboratory and Alex Alderton, Roberto Amato, Jeffrey Barrett, Sonia Goncalves, Ewan Harrison, David K. Jackson, Ian Johnston, Dominic Kwiatkowski, Cordelia Langford, John Sillitoe on behalf of the Wellcome Sanger Institute COVID-19 Surveillance Team |
| EPI_ISL_3437373 | Lighthouse Lab in Alderley Park | Wellcome Sanger Institute for the COVID-19 Genomics UK (COG-UK) Consortium | Jacquelyn Wynn, Mairead Hyland, The Lighthouse Lab in Alderley Park and Alex Alderton, Roberto Amato, Jeffrey Barrett, Sonia Goncalves, Ewan Harrison, David K. Jackson, Ian Johnston, Dominic Kwiatkowski, Cordelia Langford, John Sillitoe on behalf of the Wellcome Sanger Institute COVID-19 Surveillance Team |
| EPI_ISL_3437374 | Berkshire and Surrey Pathology Services Lighthouse Laboratory | Wellcome Sanger Institute for the COVID-19 Genomics UK (COG-UK) Consortium | Berkshire and Surrey Pathology Services Lighthouse Laboratory and Alex Alderton, Roberto Amato, Jeffrey Barrett, Sonia Goncalves, Ewan Harrison, David K. Jackson, Ian Johnston, Dominic Kwiatkowski, Cordelia Langford, John Sillitoe on behalf of the Wellcome Sanger Institute COVID-19 Surveillance Team |
| EPI_ISL_3437375 | Lighthouse Lab in Alderley Park | Wellcome Sanger Institute for the COVID-19 Genomics UK (COG-UK) Consortium | Jacquelyn Wynn, Mairead Hyland, The Lighthouse Lab in Alderley Park and Alex Alderton, Roberto Amato, Jeffrey Barrett, Sonia Goncalves, Ewan Harrison, David K. Jackson, Ian Johnston, Dominic Kwiatkowski, Cordelia Langford, John Sillitoe on behalf of the Wellcome Sanger Institute COVID-19 Surveillance Team |
| EPI_ISL_3437376, EPI_ISL_3437377 | Berkshire and Surrey Pathology Services Lighthouse Laboratory | Wellcome Sanger Institute for the COVID-19 Genomics UK (COG-UK) Consortium | Berkshire and Surrey Pathology Services Lighthouse Laboratory and Alex Alderton, Roberto Amato, Jeffrey Barrett, Sonia Goncalves, Ewan Harrison, David K. Jackson, Ian Johnston, Dominic Kwiatkowski, Cordelia Langford, John Sillitoe on behalf of the Wellcome Sanger Institute COVID-19 Surveillance Team |
| EPI_ISL_3437378 | Lighthouse Lab in Alderley Park | Wellcome Sanger Institute for the COVID-19 Genomics UK (COG-UK) Consortium | Jacquelyn Wynn, Mairead Hyland, The Lighthouse Lab in Alderley Park and Alex Alderton, Roberto Amato, Jeffrey Barrett, Sonia Goncalves, Ewan Harrison, David K. Jackson, Ian Johnston, Dominic Kwiatkowski, Cordelia Langford, John Sillitoe on behalf of the Wellcome Sanger Institute COVID-19 Surveillance Team |
| EPI_ISL_3437379, EPI_ISL_3437380, EPI_ISL_3437381 | Berkshire and Surrey Pathology Services Lighthouse Laboratory | Wellcome Sanger Institute for the COVID-19 Genomics UK (COG-UK) Consortium | Berkshire and Surrey Pathology Services Lighthouse Laboratory and Alex Alderton, Roberto Amato, Jeffrey Barrett, Sonia Goncalves, Ewan Harrison, David K. Jackson, Ian Johnston, Dominic Kwiatkowski, Cordelia Langford, John Sillitoe on behalf of the Wellcome Sanger Institute COVID-19 Surveillance Team |
| EPI_ISL_3437382 | Lighthouse Lab in Alderley Park | Wellcome Sanger Institute for the COVID-19 Genomics UK (COG-UK) Consortium | Jacquelyn Wynn, Mairead Hyland, The Lighthouse Lab in Alderley Park and Alex Alderton, Roberto Amato, Jeffrey Barrett, Sonia Goncalves, Ewan Harrison, David K. Jackson, Ian Johnston, Dominic Kwiatkowski, Cordelia Langford, John Sillitoe on behalf of the Wellcome Sanger Institute COVID-19 Surveillance Team |
| EPI_ISL_3437383, EPI_ISL_3437384 | Berkshire and Surrey Pathology Services Lighthouse Laboratory | Wellcome Sanger Institute for the COVID-19 Genomics UK (COG-UK) Consortium | Berkshire and Surrey Pathology Services Lighthouse Laboratory and Alex Alderton, Roberto Amato, Jeffrey Barrett, Sonia Goncalves, Ewan Harrison, David K. Jackson, Ian Johnston, Dominic Kwiatkowski, Cordelia Langford, John Sillitoe on behalf of the Wellcome Sanger Institute COVID-19 Surveillance Team |
| EPI_ISL_3437385 | Lighthouse Lab in Alderley Park | Wellcome Sanger Institute for the COVID-19 Genomics UK (COG-UK) Consortium | Jacquelyn Wynn, Mairead Hyland, The Lighthouse Lab in Alderley Park and Alex Alderton, Roberto Amato, Jeffrey Barrett, Sonia Goncalves, Ewan Harrison, David K. Jackson, Ian Johnston, Dominic Kwiatkowski, Cordelia Langford, John Sillitoe on behalf of the Wellcome Sanger Institute COVID-19 Surveillance Team |
| EPI_ISL_3437386, EPI_ISL_3437387, EPI_ISL_3437388, EPI_ISL_3437389 | Berkshire and Surrey Pathology Services Lighthouse Laboratory | Wellcome Sanger Institute for the COVID-19 Genomics UK (COG-UK) Consortium | Berkshire and Surrey Pathology Services Lighthouse Laboratory and Alex Alderton, Roberto Amato, Jeffrey Barrett, Sonia Goncalves, Ewan Harrison, David K. Jackson, Ian Johnston, Dominic Kwiatkowski, Cordelia Langford, John Sillitoe on behalf of the Wellcome Sanger Institute COVID-19 Surveillance Team |
| EPI_ISL_3437390 | Lighthouse Lab in Alderley Park | Wellcome Sanger Institute for the COVID-19 Genomics UK (COG-UK) Consortium | Jacquelyn Wynn, Mairead Hyland, The Lighthouse Lab in Alderley Park and Alex Alderton, Roberto Amato, Jeffrey Barrett, Sonia Goncalves, Ewan Harrison, David K. Jackson, Ian Johnston, Dominic Kwiatkowski, Cordelia Langford, John Sillitoe on behalf of the Wellcome Sanger Institute COVID-19 Surveillance Team |
| EPI_ISL_3437391, EPI_ISL_3437392, EPI_ISL_3437393, EPI_ISL_3437394, EPI_ISL_3437395, EPI_ISL_3437396 | Berkshire and Surrey Pathology Services Lighthouse Laboratory | Wellcome Sanger Institute for the COVID-19 Genomics UK (COG-UK) Consortium | Berkshire and Surrey Pathology Services Lighthouse Laboratory and Alex Alderton, Roberto Amato, Jeffrey Barrett, Sonia Goncalves, Ewan Harrison, David K. Jackson, Ian Johnston, Dominic Kwiatkowski, Cordelia Langford, John Sillitoe on behalf of the Wellcome Sanger Institute COVID-19 Surveillance Team |
| EPI_ISL_3437397, EPI_ISL_3437398 | Lighthouse Lab in Alderley Park | Wellcome Sanger Institute for the COVID-19 Genomics UK (COG-UK) Consortium | Jacquelyn Wynn, Mairead Hyland, The Lighthouse Lab in Alderley Park and Alex Alderton, Roberto Amato, Jeffrey Barrett, Sonia Goncalves, Ewan Harrison, David K. Jackson, Ian Johnston, Dominic Kwiatkowski, Cordelia Langford, John Sillitoe on behalf of the Wellcome Sanger Institute COVID-19 Surveillance Team |
| EPI_ISL_3437399 | Berkshire and Surrey Pathology Services Lighthouse Laboratory | Wellcome Sanger Institute for the COVID-19 Genomics UK (COG-UK) Consortium | Berkshire and Surrey Pathology Services Lighthouse Laboratory and Alex Alderton, Roberto Amato, Jeffrey Barrett, Sonia Goncalves, Ewan Harrison, David K. Jackson, Ian Johnston, Dominic Kwiatkowski, Cordelia Langford, John Sillitoe on behalf of the Wellcome Sanger Institute COVID-19 Surveillance Team |
| EPI_ISL_3437400, EPI_ISL_3437401, EPI_ISL_3437402 | Lighthouse Lab in Alderley Park | Wellcome Sanger Institute for the COVID-19 Genomics UK (COG-UK) Consortium | Jacquelyn Wynn, Mairead Hyland, The Lighthouse Lab in Alderley Park and Alex Alderton, Roberto Amato, Jeffrey Barrett, Sonia Goncalves, Ewan Harrison, David K. Jackson, Ian Johnston, Dominic Kwiatkowski, Cordelia Langford, John Sillitoe on behalf of the Wellcome Sanger Institute COVID-19 Surveillance Team |
| EPI_ISL_3437403 | Berkshire and Surrey Pathology Services Lighthouse Laboratory | Wellcome Sanger Institute for the COVID-19 Genomics UK (COG-UK) Consortium | Berkshire and Surrey Pathology Services Lighthouse Laboratory and Alex Alderton, Roberto Amato, Jeffrey Barrett, Sonia Goncalves, Ewan Harrison, David K. Jackson, Ian Johnston, Dominic Kwiatkowski, Cordelia Langford, John Sillitoe on behalf of the Wellcome Sanger Institute COVID-19 Surveillance Team |
| EPI_ISL_3437404 | Lighthouse Lab in Alderley Park | Wellcome Sanger Institute for the COVID-19 Genomics UK (COG-UK) Consortium | Jacquelyn Wynn, Mairead Hyland, The Lighthouse Lab in Alderley Park and Alex Alderton, Roberto Amato, Jeffrey Barrett, Sonia Goncalves, Ewan Harrison, David K. Jackson, Ian Johnston, Dominic Kwiatkowski, Cordelia Langford, John Sillitoe on behalf of the Wellcome Sanger Institute COVID-19 Surveillance Team |
| EPI_ISL_3437405, EPI_ISL_3437406 | Berkshire and Surrey Pathology Services Lighthouse Laboratory | Wellcome Sanger Institute for the COVID-19 Genomics UK (COG-UK) Consortium | Berkshire and Surrey Pathology Services Lighthouse Laboratory and Alex Alderton, Roberto Amato, Jeffrey Barrett, Sonia Goncalves, Ewan Harrison, David K. Jackson, Ian Johnston, Dominic Kwiatkowski, Cordelia Langford, John Sillitoe on behalf of the Wellcome Sanger Institute COVID-19 Surveillance Team |
| EPI_ISL_3437407, EPI_ISL_3437408 | Lighthouse Lab in Alderley Park | Wellcome Sanger Institute for the COVID-19 Genomics UK (COG-UK) Consortium | Jacquelyn Wynn, Mairead Hyland, The Lighthouse Lab in Alderley Park and Alex Alderton, Roberto Amato, Jeffrey Barrett, Sonia Goncalves, Ewan Harrison, David K. Jackson, Ian Johnston, Dominic Kwiatkowski, Cordelia Langford, John Sillitoe on behalf of the Wellcome Sanger Institute COVID-19 Surveillance Team |
| EPI_ISL_3437409 | Berkshire and Surrey Pathology Services Lighthouse Laboratory | Wellcome Sanger Institute for the COVID-19 Genomics UK (COG-UK) Consortium | Berkshire and Surrey Pathology Services Lighthouse Laboratory and Alex Alderton, Roberto Amato, Jeffrey Barrett, Sonia Goncalves, Ewan Harrison, David K. Jackson, Ian Johnston, Dominic Kwiatkowski, Cordelia Langford, John Sillitoe on behalf of the Wellcome Sanger Institute COVID-19 Surveillance Team |
| EPI_ISL_3437410 | Lighthouse Lab in Alderley Park | Wellcome Sanger Institute for the COVID-19 Genomics UK (COG-UK) Consortium | Jacquelyn Wynn, Mairead Hyland, The Lighthouse Lab in Alderley Park and Alex Alderton, Roberto Amato, Jeffrey Barrett, Sonia Goncalves, Ewan Harrison, David K. Jackson, Ian Johnston, Dominic Kwiatkowski, Cordelia Langford, John Sillitoe on behalf of the Wellcome Sanger Institute COVID-19 Surveillance Team |
| EPI_ISL_3437411 | Berkshire and Surrey Pathology Services Lighthouse Laboratory | Wellcome Sanger Institute for the COVID-19 Genomics UK (COG-UK) Consortium | Berkshire and Surrey Pathology Services Lighthouse Laboratory and Alex Alderton, Roberto Amato, Jeffrey Barrett, Sonia Goncalves, Ewan Harrison, David K. Jackson, Ian Johnston, Dominic Kwiatkowski, Cordelia Langford, John Sillitoe on behalf of the Wellcome Sanger Institute COVID-19 Surveillance Team |
| EPI_ISL_3437412, EPI_ISL_3437413 | Lighthouse Lab in Alderley Park | Wellcome Sanger Institute for the COVID-19 Genomics UK (COG-UK) Consortium | Jacquelyn Wynn, Mairead Hyland, The Lighthouse Lab in Alderley Park and Alex Alderton, Roberto Amato, Jeffrey Barrett, Sonia Goncalves, Ewan Harrison, David K. Jackson, Ian Johnston, Dominic Kwiatkowski, Cordelia Langford, John Sillitoe on behalf of the Wellcome Sanger Institute COVID-19 Surveillance Team |
| EPI_ISL_3437414 | Berkshire and Surrey Pathology Services Lighthouse Laboratory | Wellcome Sanger Institute for the COVID-19 Genomics UK (COG-UK) Consortium | Berkshire and Surrey Pathology Services Lighthouse Laboratory and Alex Alderton, Roberto Amato, Jeffrey Barrett, Sonia Goncalves, Ewan Harrison, David K. Jackson, Ian Johnston, Dominic Kwiatkowski, Cordelia Langford, John Sillitoe on behalf of the Wellcome Sanger Institute COVID-19 Surveillance Team |
| EPI_ISL_3437415 | Lighthouse Lab in Alderley Park | Wellcome Sanger Institute for the COVID-19 Genomics UK (COG-UK) | Jacquelyn Wynn, Mairead Hyland, The Lighthouse Lab in Alderley Park and Alex Alderton, Roberto Amato, Jeffrey Barrett, Sonia Goncalves, Ewan Harrison, David K. Jackson, Ian Johnston, Dominic Kwiatkowski, Cordelia Langford, John Sillitoe on |

| | | | |
|---|---|---|---|
| | | Consortium | behalf of the Wellcome Sanger Institute COVID-19 Surveillance Team |
| EPI_ISL_3437416 | Berkshire and Surrey Pathology Services Lighthouse Laboratory | Wellcome Sanger Institute for the COVID-19 Genomics UK (COG-UK) Consortium | Berkshire and Surrey Pathology Services Lighthouse Laboratory and Alex Alderton, Roberto Amato, Jeffrey Barrett, Sonia Goncalves, Ewan Harrison, David K. Jackson, Ian Johnston, Dominic Kwiatkowski, Cordelia Langford, John Sillitoe on behalf of the Wellcome Sanger Institute COVID-19 Surveillance Team |
| EPI_ISL_3437417, EPI_ISL_3437418 | Lighthouse Lab in Alderley Park | Wellcome Sanger Institute for the COVID-19 Genomics UK (COG-UK) Consortium | Jacquelyn Wynn, Mairead Hyland, The Lighthouse Lab in Alderley Park and Alex Alderton, Roberto Amato, Jeffrey Barrett, Sonia Goncalves, Ewan Harrison, David K. Jackson, Ian Johnston, Dominic Kwiatkowski, Cordelia Langford, John Sillitoe on behalf of the Wellcome Sanger Institute COVID-19 Surveillance Team |
| EPI_ISL_3437419 | Berkshire and Surrey Pathology Services Lighthouse Laboratory | Wellcome Sanger Institute for the COVID-19 Genomics UK (COG-UK) Consortium | Berkshire and Surrey Pathology Services Lighthouse Laboratory and Alex Alderton, Roberto Amato, Jeffrey Barrett, Sonia Goncalves, Ewan Harrison, David K. Jackson, Ian Johnston, Dominic Kwiatkowski, Cordelia Langford, John Sillitoe on behalf of the Wellcome Sanger Institute COVID-19 Surveillance Team |
| EPI_ISL_3437420 | Lighthouse Lab in Alderley Park | Wellcome Sanger Institute for the COVID-19 Genomics UK (COG-UK) Consortium | Jacquelyn Wynn, Mairead Hyland, The Lighthouse Lab in Alderley Park and Alex Alderton, Roberto Amato, Jeffrey Barrett, Sonia Goncalves, Ewan Harrison, David K. Jackson, Ian Johnston, Dominic Kwiatkowski, Cordelia Langford, John Sillitoe on behalf of the Wellcome Sanger Institute COVID-19 Surveillance Team |
| EPI_ISL_3437421, EPI_ISL_3437422 | Berkshire and Surrey Pathology Services Lighthouse Laboratory | Wellcome Sanger Institute for the COVID-19 Genomics UK (COG-UK) Consortium | Berkshire and Surrey Pathology Services Lighthouse Laboratory and Alex Alderton, Roberto Amato, Jeffrey Barrett, Sonia Goncalves, Ewan Harrison, David K. Jackson, Ian Johnston, Dominic Kwiatkowski, Cordelia Langford, John Sillitoe on behalf of the Wellcome Sanger Institute COVID-19 Surveillance Team |
| EPI_ISL_3437423 | Lighthouse Lab in Alderley Park | Wellcome Sanger Institute for the COVID-19 Genomics UK (COG-UK) Consortium | Jacquelyn Wynn, Mairead Hyland, The Lighthouse Lab in Alderley Park and Alex Alderton, Roberto Amato, Jeffrey Barrett, Sonia Goncalves, Ewan Harrison, David K. Jackson, Ian Johnston, Dominic Kwiatkowski, Cordelia Langford, John Sillitoe on behalf of the Wellcome Sanger Institute COVID-19 Surveillance Team |
| EPI_ISL_3437424 | Berkshire and Surrey Pathology Services Lighthouse Laboratory | Wellcome Sanger Institute for the COVID-19 Genomics UK (COG-UK) Consortium | Berkshire and Surrey Pathology Services Lighthouse Laboratory and Alex Alderton, Roberto Amato, Jeffrey Barrett, Sonia Goncalves, Ewan Harrison, David K. Jackson, Ian Johnston, Dominic Kwiatkowski, Cordelia Langford, John Sillitoe on behalf of the Wellcome Sanger Institute COVID-19 Surveillance Team |
| EPI_ISL_3437425, EPI_ISL_3437426 | Lighthouse Lab in Alderley Park | Wellcome Sanger Institute for the COVID-19 Genomics UK (COG-UK) Consortium | Jacquelyn Wynn, Mairead Hyland, The Lighthouse Lab in Alderley Park and Alex Alderton, Roberto Amato, Jeffrey Barrett, Sonia Goncalves, Ewan Harrison, David K. Jackson, Ian Johnston, Dominic Kwiatkowski, Cordelia Langford, John Sillitoe on behalf of the Wellcome Sanger Institute COVID-19 Surveillance Team |
| EPI_ISL_3437427, EPI_ISL_3437428, EPI_ISL_3437429 | Berkshire and Surrey Pathology Services Lighthouse Laboratory | Wellcome Sanger Institute for the COVID-19 Genomics UK (COG-UK) Consortium | Berkshire and Surrey Pathology Services Lighthouse Laboratory and Alex Alderton, Roberto Amato, Jeffrey Barrett, Sonia Goncalves, Ewan Harrison, David K. Jackson, Ian Johnston, Dominic Kwiatkowski, Cordelia Langford, John Sillitoe on behalf of the Wellcome Sanger Institute COVID-19 Surveillance Team |
| EPI_ISL_3437430 | Lighthouse Lab in Alderley Park | Wellcome Sanger Institute for the COVID-19 Genomics UK (COG-UK) Consortium | Jacquelyn Wynn, Mairead Hyland, The Lighthouse Lab in Alderley Park and Alex Alderton, Roberto Amato, Jeffrey Barrett, Sonia Goncalves, Ewan Harrison, David K. Jackson, Ian Johnston, Dominic Kwiatkowski, Cordelia Langford, John Sillitoe on behalf of the Wellcome Sanger Institute COVID-19 Surveillance Team |
| EPI_ISL_3437431, EPI_ISL_3437432 | Berkshire and Surrey Pathology Services Lighthouse Laboratory | Wellcome Sanger Institute for the COVID-19 Genomics UK (COG-UK) Consortium | Berkshire and Surrey Pathology Services Lighthouse Laboratory and Alex Alderton, Roberto Amato, Jeffrey Barrett, Sonia Goncalves, Ewan Harrison, David K. Jackson, Ian Johnston, Dominic Kwiatkowski, Cordelia Langford, John Sillitoe on behalf of the Wellcome Sanger Institute COVID-19 Surveillance Team |
| EPI_ISL_3437433 | Lighthouse Lab in Alderley Park | Wellcome Sanger Institute for the COVID-19 Genomics UK (COG-UK) Consortium | Jacquelyn Wynn, Mairead Hyland, The Lighthouse Lab in Alderley Park and Alex Alderton, Roberto Amato, Jeffrey Barrett, Sonia Goncalves, Ewan Harrison, David K. Jackson, Ian Johnston, Dominic Kwiatkowski, Cordelia Langford, John Sillitoe on behalf of the Wellcome Sanger Institute COVID-19 Surveillance Team |
| EPI_ISL_3437434, EPI_ISL_3437435, EPI_ISL_3437436, EPI_ISL_3437437, EPI_ISL_3437438, EPI_ISL_3437439 | Berkshire and Surrey Pathology Services Lighthouse Laboratory | Wellcome Sanger Institute for the COVID-19 Genomics UK (COG-UK) Consortium | Berkshire and Surrey Pathology Services Lighthouse Laboratory and Alex Alderton, Roberto Amato, Jeffrey Barrett, Sonia Goncalves, Ewan Harrison, David K. Jackson, Ian Johnston, Dominic Kwiatkowski, Cordelia Langford, John Sillitoe on behalf of the Wellcome Sanger Institute COVID-19 Surveillance Team |
| EPI_ISL_3437440, EPI_ISL_3437441 | Lighthouse Lab in Alderley Park | Wellcome Sanger Institute for the COVID-19 Genomics UK (COG-UK) Consortium | Jacquelyn Wynn, Mairead Hyland, The Lighthouse Lab in Alderley Park and Alex Alderton, Roberto Amato, Jeffrey Barrett, Sonia Goncalves, Ewan Harrison, David K. Jackson, Ian Johnston, Dominic Kwiatkowski, Cordelia Langford, John Sillitoe on behalf of the Wellcome Sanger Institute COVID-19 Surveillance Team |
| EPI_ISL_3437442, EPI_ISL_3437443, EPI_ISL_3437444, EPI_ISL_3437445 | Berkshire and Surrey Pathology Services Lighthouse Laboratory | Wellcome Sanger Institute for the COVID-19 Genomics UK (COG-UK) Consortium | Berkshire and Surrey Pathology Services Lighthouse Laboratory and Alex Alderton, Roberto Amato, Jeffrey Barrett, Sonia Goncalves, Ewan Harrison, David K. Jackson, Ian Johnston, Dominic Kwiatkowski, Cordelia Langford, John Sillitoe on behalf of the Wellcome Sanger Institute COVID-19 Surveillance Team |
| EPI_ISL_3437446, EPI_ISL_3437447 | Lighthouse Lab in Alderley Park | Wellcome Sanger Institute for the COVID-19 Genomics UK (COG-UK) Consortium | Jacquelyn Wynn, Mairead Hyland, The Lighthouse Lab in Alderley Park and Alex Alderton, Roberto Amato, Jeffrey Barrett, Sonia Goncalves, Ewan Harrison, David K. Jackson, Ian Johnston, Dominic Kwiatkowski, Cordelia Langford, John Sillitoe on behalf of the Wellcome Sanger Institute COVID-19 Surveillance Team |
| EPI_ISL_3437448, EPI_ISL_3437449 | Berkshire and Surrey Pathology Services Lighthouse Laboratory | Wellcome Sanger Institute for the COVID-19 Genomics UK (COG-UK) Consortium | Berkshire and Surrey Pathology Services Lighthouse Laboratory and Alex Alderton, Roberto Amato, Jeffrey Barrett, Sonia Goncalves, Ewan Harrison, David K. Jackson, Ian Johnston, Dominic Kwiatkowski, Cordelia Langford, John Sillitoe on behalf of the Wellcome Sanger Institute COVID-19 Surveillance Team |
| EPI_ISL_3437450, EPI_ISL_3437451 | Lighthouse Lab in Alderley Park | Wellcome Sanger Institute for the COVID-19 Genomics UK (COG-UK) Consortium | Jacquelyn Wynn, Mairead Hyland, The Lighthouse Lab in Alderley Park and Alex Alderton, Roberto Amato, Jeffrey Barrett, Sonia Goncalves, Ewan Harrison, David K. Jackson, Ian Johnston, Dominic Kwiatkowski, Cordelia Langford, John Sillitoe on behalf of the Wellcome Sanger Institute COVID-19 Surveillance Team |
| EPI_ISL_3437452 | Berkshire and Surrey Pathology Services Lighthouse Laboratory | Wellcome Sanger Institute for the COVID-19 Genomics UK (COG-UK) Consortium | Berkshire and Surrey Pathology Services Lighthouse Laboratory and Alex Alderton, Roberto Amato, Jeffrey Barrett, Sonia Goncalves, Ewan Harrison, David K. Jackson, Ian Johnston, Dominic Kwiatkowski, Cordelia Langford, John Sillitoe on behalf of the Wellcome Sanger Institute COVID-19 Surveillance Team |
| EPI_ISL_3437453 | Lighthouse Lab in Alderley Park | Wellcome Sanger Institute for the COVID-19 Genomics UK (COG-UK) Consortium | Jacquelyn Wynn, Mairead Hyland, The Lighthouse Lab in Alderley Park and Alex Alderton, Roberto Amato, Jeffrey Barrett, Sonia Goncalves, Ewan Harrison, David K. Jackson, Ian Johnston, Dominic Kwiatkowski, Cordelia Langford, John Sillitoe on behalf of the Wellcome Sanger Institute COVID-19 Surveillance Team |
| EPI_ISL_3437454, EPI_ISL_3437455 | Berkshire and Surrey Pathology Services Lighthouse Laboratory | Wellcome Sanger Institute for the COVID-19 Genomics UK (COG-UK) Consortium | Berkshire and Surrey Pathology Services Lighthouse Laboratory and Alex Alderton, Roberto Amato, Jeffrey Barrett, Sonia Goncalves, Ewan Harrison, David K. Jackson, Ian Johnston, Dominic Kwiatkowski, Cordelia Langford, John Sillitoe on behalf of the Wellcome Sanger Institute COVID-19 Surveillance Team |
| EPI_ISL_3437456 | Lighthouse Lab in Alderley Park | Wellcome Sanger Institute for the COVID-19 Genomics UK (COG-UK) Consortium | Jacquelyn Wynn, Mairead Hyland, The Lighthouse Lab in Alderley Park and Alex Alderton, Roberto Amato, Jeffrey Barrett, Sonia Goncalves, Ewan Harrison, David K. Jackson, Ian Johnston, Dominic Kwiatkowski, Cordelia Langford, John Sillitoe on behalf of the Wellcome Sanger Institute COVID-19 Surveillance Team |
| EPI_ISL_3437457 | Berkshire and Surrey Pathology Services Lighthouse Laboratory | Wellcome Sanger Institute for the COVID-19 Genomics UK (COG-UK) Consortium | Berkshire and Surrey Pathology Services Lighthouse Laboratory and Alex Alderton, Roberto Amato, Jeffrey Barrett, Sonia Goncalves, Ewan Harrison, David K. Jackson, Ian Johnston, Dominic Kwiatkowski, Cordelia Langford, John Sillitoe on behalf of the Wellcome Sanger Institute COVID-19 Surveillance Team |
| EPI_ISL_3437458 | Lighthouse Lab in Alderley Park | Wellcome Sanger Institute for the COVID-19 Genomics UK (COG-UK) Consortium | Jacquelyn Wynn, Mairead Hyland, The Lighthouse Lab in Alderley Park and Alex Alderton, Roberto Amato, Jeffrey Barrett, Sonia Goncalves, Ewan Harrison, David K. Jackson, Ian Johnston, Dominic Kwiatkowski, Cordelia Langford, John Sillitoe on behalf of the Wellcome Sanger Institute COVID-19 Surveillance Team |
| EPI_ISL_3437459 | Berkshire and Surrey Pathology Services Lighthouse Laboratory | Wellcome Sanger Institute for the COVID-19 Genomics UK (COG-UK) Consortium | Berkshire and Surrey Pathology Services Lighthouse Laboratory and Alex Alderton, Roberto Amato, Jeffrey Barrett, Sonia Goncalves, Ewan Harrison, David K. Jackson, Ian Johnston, Dominic Kwiatkowski, Cordelia Langford, John Sillitoe on behalf of the Wellcome Sanger Institute COVID-19 Surveillance Team |
| EPI_ISL_3437460 | Lighthouse Lab in Alderley Park | Wellcome Sanger Institute for the COVID-19 Genomics UK (COG-UK) Consortium | Jacquelyn Wynn, Mairead Hyland, The Lighthouse Lab in Alderley Park and Alex Alderton, Roberto Amato, Jeffrey Barrett, Sonia Goncalves, Ewan Harrison, David K. Jackson, Ian Johnston, Dominic Kwiatkowski, Cordelia Langford, John Sillitoe on behalf of the Wellcome Sanger Institute COVID-19 Surveillance Team |
| EPI_ISL_3437461, EPI_ISL_3437462, EPI_ISL_3437463 | Berkshire and Surrey Pathology Services Lighthouse Laboratory | Wellcome Sanger Institute for the COVID-19 Genomics UK (COG-UK) Consortium | Berkshire and Surrey Pathology Services Lighthouse Laboratory and Alex Alderton, Roberto Amato, Jeffrey Barrett, Sonia Goncalves, Ewan Harrison, David K. Jackson, Ian Johnston, Dominic Kwiatkowski, Cordelia Langford, John Sillitoe on behalf of the Wellcome Sanger Institute COVID-19 Surveillance Team |
| EPI_ISL_3437464 | Lighthouse Lab in Alderley Park | Wellcome Sanger Institute for the COVID-19 Genomics UK (COG-UK) Consortium | Jacquelyn Wynn, Mairead Hyland, The Lighthouse Lab in Alderley Park and Alex Alderton, Roberto Amato, Jeffrey Barrett, Sonia Goncalves, Ewan Harrison, David K. Jackson, Ian Johnston, Dominic Kwiatkowski, Cordelia Langford, John Sillitoe on behalf of the Wellcome Sanger Institute COVID-19 Surveillance Team |
| EPI_ISL_3437465 | Berkshire and Surrey Pathology Services Lighthouse Laboratory | Wellcome Sanger Institute for the COVID-19 Genomics UK (COG-UK) Consortium | Berkshire and Surrey Pathology Services Lighthouse Laboratory and Alex Alderton, Roberto Amato, Jeffrey Barrett, Sonia Goncalves, Ewan Harrison, David K. Jackson, Ian Johnston, Dominic Kwiatkowski, Cordelia Langford, John Sillitoe on behalf of the Wellcome Sanger Institute COVID-19 Surveillance Team |
| EPI_ISL_3437466, EPI_ISL_3437467 | Lighthouse Lab in Alderley Park | Wellcome Sanger Institute for the COVID-19 Genomics UK (COG-UK) Consortium | Jacquelyn Wynn, Mairead Hyland, The Lighthouse Lab in Alderley Park and Alex Alderton, Roberto Amato, Jeffrey Barrett, Sonia Goncalves, Ewan Harrison, David K. Jackson, Ian Johnston, Dominic Kwiatkowski, Cordelia Langford, John Sillitoe on behalf of the Wellcome Sanger Institute COVID-19 Surveillance Team |
| EPI_ISL_3437468, EPI_ISL_3437469, EPI_ISL_3437470 | Berkshire and Surrey Pathology Services Lighthouse Laboratory | Wellcome Sanger Institute for the COVID-19 Genomics UK (COG-UK) Consortium | Berkshire and Surrey Pathology Services Lighthouse Laboratory and Alex Alderton, Roberto Amato, Jeffrey Barrett, Sonia Goncalves, Ewan Harrison, David K. Jackson, Ian Johnston, Dominic Kwiatkowski, Cordelia Langford, John Sillitoe on behalf of the Wellcome Sanger Institute COVID-19 Surveillance Team |
| EPI_ISL_3437471, EPI_ISL_3437472 | Lighthouse Lab in Alderley Park | Wellcome Sanger Institute for the COVID-19 Genomics UK (COG-UK) Consortium | Jacquelyn Wynn, Mairead Hyland, The Lighthouse Lab in Alderley Park and Alex Alderton, Roberto Amato, Jeffrey Barrett, Sonia Goncalves, Ewan Harrison, David K. Jackson, Ian Johnston, Dominic Kwiatkowski, Cordelia Langford, John Sillitoe on behalf of the Wellcome Sanger Institute COVID-19 Surveillance Team |
| EPI_ISL_3437473, EPI_ISL_3437474 | Berkshire and Surrey Pathology Services Lighthouse Laboratory | Wellcome Sanger Institute for the COVID-19 Genomics UK (COG-UK) Consortium | Berkshire and Surrey Pathology Services Lighthouse Laboratory and Alex Alderton, Roberto Amato, Jeffrey Barrett, Sonia Goncalves, Ewan Harrison, David K. Jackson, Ian Johnston, Dominic Kwiatkowski, Cordelia Langford, John Sillitoe on behalf of the Wellcome Sanger Institute COVID-19 Surveillance Team |
| EPI_ISL_3437475 | Lighthouse Lab in Alderley Park | Wellcome Sanger Institute for the COVID-19 Genomics UK (COG-UK) Consortium | Jacquelyn Wynn, Mairead Hyland, The Lighthouse Lab in Alderley Park and Alex Alderton, Roberto Amato, Jeffrey Barrett, Sonia Goncalves, Ewan Harrison, David K. Jackson, Ian Johnston, Dominic Kwiatkowski, Cordelia Langford, John Sillitoe on behalf of the Wellcome Sanger Institute COVID-19 Surveillance Team |
| EPI_ISL_3437476, EPI_ISL_3437477 | Berkshire and Surrey Pathology Services Lighthouse Laboratory | Wellcome Sanger Institute for the COVID-19 Genomics UK (COG-UK) Consortium | Berkshire and Surrey Pathology Services Lighthouse Laboratory and Alex Alderton, Roberto Amato, Jeffrey Barrett, Sonia Goncalves, Ewan Harrison, David K. Jackson, Ian Johnston, Dominic Kwiatkowski, Cordelia Langford, John Sillitoe on behalf of the Wellcome Sanger Institute COVID-19 Surveillance Team |
| EPI_ISL_3437478 | Lighthouse Lab in Alderley Park | Wellcome Sanger Institute for the COVID-19 Genomics UK (COG-UK) Consortium | Jacquelyn Wynn, Mairead Hyland, The Lighthouse Lab in Alderley Park and Alex Alderton, Roberto Amato, Jeffrey Barrett, Sonia Goncalves, Ewan Harrison, David K. Jackson, Ian Johnston, Dominic Kwiatkowski, Cordelia Langford, John Sillitoe on behalf of the Wellcome Sanger Institute COVID-19 Surveillance Team |
| EPI_ISL_3437479, EPI_ISL_3437480 | Berkshire and Surrey Pathology Services Lighthouse Laboratory | Wellcome Sanger Institute for the COVID-19 Genomics UK (COG-UK) Consortium | Berkshire and Surrey Pathology Services Lighthouse Laboratory and Alex Alderton, Roberto Amato, Jeffrey Barrett, Sonia Goncalves, Ewan Harrison, David K. Jackson, Ian Johnston, Dominic Kwiatkowski, Cordelia Langford, John Sillitoe on behalf of the Wellcome Sanger Institute COVID-19 Surveillance Team |
| EPI_ISL_3437481 | Lighthouse Lab in Alderley Park | Wellcome Sanger Institute for the COVID-19 Genomics UK (COG-UK) Consortium | Jacquelyn Wynn, Mairead Hyland, The Lighthouse Lab in Alderley Park and Alex Alderton, Roberto Amato, Jeffrey Barrett, Sonia Goncalves, Ewan Harrison, David K. Jackson, Ian Johnston, Dominic Kwiatkowski, Cordelia Langford, John Sillitoe on behalf of the Wellcome Sanger Institute COVID-19 Surveillance Team |
| EPI_ISL_3437482, EPI_ISL_3437483, EPI_ISL_3437484 | Berkshire and Surrey Pathology Services Lighthouse Laboratory | Wellcome Sanger Institute for the COVID-19 Genomics UK (COG-UK) Consortium | Berkshire and Surrey Pathology Services Lighthouse Laboratory and Alex Alderton, Roberto Amato, Jeffrey Barrett, Sonia Goncalves, Ewan Harrison, David K. Jackson, Ian Johnston, Dominic Kwiatkowski, Cordelia Langford, John Sillitoe on behalf of the Wellcome Sanger Institute COVID-19 Surveillance Team |
| EPI_ISL_3437485 | Lighthouse Lab in Glasgow | Wellcome Sanger Institute for the COVID-19 Genomics UK (COG-UK) Consortium | Harper VanSteenhouse, Yumi Kasai, David Gray, Carol Clugston, Anna Dominiczak and Alex Alderton, Roberto Amato, Jeffrey Barrett, Sonia Goncalves, Ewan Harrison, David K. Jackson, Ian Johnston, Dominic Kwiatkowski, Cordelia Langford, John Sillitoe on behalf of the Wellcome Sanger Institute COVID-19 Surveillance Team |
| EPI_ISL_3437486, | Lighthouse Lab in Alderley Park | Wellcome Sanger Institute for the COVID-19 Genomics UK (COG-UK) | Jacquelyn Wynn, Mairead Hyland, The Lighthouse Lab in Alderley Park and Alex Alderton, Roberto Amato, Jeffrey Barrett, Sonia Goncalves, Ewan Harrison, David K. Jackson, Ian Johnston, Dominic Kwiatkowski, Cordelia Langford, John Sillitoe on |

| | | | |
|---|---|---|---|
| EPI_ISL_3437487, EPI_ISL_3437488 | | Consortium | behalf of the Wellcome Sanger Institute COVID-19 Surveillance Team |
| EPI_ISL_3437489, EPI_ISL_3437490 | Lighthouse Lab in Milton Keynes | Wellcome Sanger Institute for the COVID-19 Genomics UK (COG-UK) Consortium | The Lighthouse Lab in Milton Keynes and Alex Alderton, Roberto Amato, Jeffrey Barrett, Sonia Goncalves, Ewan Harrison, David K. Jackson, Ian Johnston, Dominic Kwiatkowski, Cordelia Langford, John Sillitoe on behalf of the Wellcome Sanger Institute COVID-19 Surveillance Team |
| EPI_ISL_3437491 | Lighthouse Lab in Alderley Park | Wellcome Sanger Institute for the COVID-19 Genomics UK (COG-UK) Consortium | Jacquelyn Wynn, Mairead Hyland, The Lighthouse Lab in Alderley Park and Alex Alderton, Roberto Amato, Jeffrey Barrett, Sonia Goncalves, Ewan Harrison, David K. Jackson, Ian Johnston, Dominic Kwiatkowski, Cordelia Langford, John Sillitoe on behalf of the Wellcome Sanger Institute COVID-19 Surveillance Team |
| EPI_ISL_3437492, EPI_ISL_3437493, EPI_ISL_3437494, EPI_ISL_3437495 | Lighthouse Lab in Milton Keynes | Wellcome Sanger Institute for the COVID-19 Genomics UK (COG-UK) Consortium | The Lighthouse Lab in Milton Keynes and Alex Alderton, Roberto Amato, Jeffrey Barrett, Sonia Goncalves, Ewan Harrison, David K. Jackson, Ian Johnston, Dominic Kwiatkowski, Cordelia Langford, John Sillitoe on behalf of the Wellcome Sanger Institute COVID-19 Surveillance Team |
| EPI_ISL_3437496, EPI_ISL_3437497 | Lighthouse Lab in Alderley Park | Wellcome Sanger Institute for the COVID-19 Genomics UK (COG-UK) Consortium | Jacquelyn Wynn, Mairead Hyland, The Lighthouse Lab in Alderley Park and Alex Alderton, Roberto Amato, Jeffrey Barrett, Sonia Goncalves, Ewan Harrison, David K. Jackson, Ian Johnston, Dominic Kwiatkowski, Cordelia Langford, John Sillitoe on behalf of the Wellcome Sanger Institute COVID-19 Surveillance Team |
| EPI_ISL_3437498, EPI_ISL_3437499, EPI_ISL_3437500 | Lighthouse Lab in Milton Keynes | Wellcome Sanger Institute for the COVID-19 Genomics UK (COG-UK) Consortium | The Lighthouse Lab in Milton Keynes and Alex Alderton, Roberto Amato, Jeffrey Barrett, Sonia Goncalves, Ewan Harrison, David K. Jackson, Ian Johnston, Dominic Kwiatkowski, Cordelia Langford, John Sillitoe on behalf of the Wellcome Sanger Institute COVID-19 Surveillance Team |
| EPI_ISL_3437501 | Lighthouse Lab in Alderley Park | Wellcome Sanger Institute for the COVID-19 Genomics UK (COG-UK) Consortium | Jacquelyn Wynn, Mairead Hyland, The Lighthouse Lab in Alderley Park and Alex Alderton, Roberto Amato, Jeffrey Barrett, Sonia Goncalves, Ewan Harrison, David K. Jackson, Ian Johnston, Dominic Kwiatkowski, Cordelia Langford, John Sillitoe on behalf of the Wellcome Sanger Institute COVID-19 Surveillance Team |
| EPI_ISL_3437502, EPI_ISL_3437503, EPI_ISL_3437504, EPI_ISL_3437505, EPI_ISL_3437506, EPI_ISL_3437507, EPI_ISL_3437508, EPI_ISL_3437509, EPI_ISL_3437510, EPI_ISL_3437511, EPI_ISL_3437512, EPI_ISL_3437513, EPI_ISL_3437514 | | | |
| see above | Lighthouse Lab in Milton Keynes | Wellcome Sanger Institute for the COVID-19 Genomics UK (COG-UK) Consortium | The Lighthouse Lab in Milton Keynes and Alex Alderton, Roberto Amato, Jeffrey Barrett, Sonia Goncalves, Ewan Harrison, David K. Jackson, Ian Johnston, Dominic Kwiatkowski, Cordelia Langford, John Sillitoe on behalf of the Wellcome Sanger Institute COVID-19 Surveillance Team |
| EPI_ISL_3437515 | Lighthouse Lab in Alderley Park | Wellcome Sanger Institute for the COVID-19 Genomics UK (COG-UK) Consortium | Jacquelyn Wynn, Mairead Hyland, The Lighthouse Lab in Alderley Park and Alex Alderton, Roberto Amato, Jeffrey Barrett, Sonia Goncalves, Ewan Harrison, David K. Jackson, Ian Johnston, Dominic Kwiatkowski, Cordelia Langford, John Sillitoe on behalf of the Wellcome Sanger Institute COVID-19 Surveillance Team |
| EPI_ISL_3437516, EPI_ISL_3437517 | Lighthouse Lab in Milton Keynes | Wellcome Sanger Institute for the COVID-19 Genomics UK (COG-UK) Consortium | The Lighthouse Lab in Milton Keynes and Alex Alderton, Roberto Amato, Jeffrey Barrett, Sonia Goncalves, Ewan Harrison, David K. Jackson, Ian Johnston, Dominic Kwiatkowski, Cordelia Langford, John Sillitoe on behalf of the Wellcome Sanger Institute COVID-19 Surveillance Team |
| EPI_ISL_3437518 | Lighthouse Lab in Alderley Park | Wellcome Sanger Institute for the COVID-19 Genomics UK (COG-UK) Consortium | Jacquelyn Wynn, Mairead Hyland, The Lighthouse Lab in Alderley Park and Alex Alderton, Roberto Amato, Jeffrey Barrett, Sonia Goncalves, Ewan Harrison, David K. Jackson, Ian Johnston, Dominic Kwiatkowski, Cordelia Langford, John Sillitoe on behalf of the Wellcome Sanger Institute COVID-19 Surveillance Team |
| EPI_ISL_3437519, EPI_ISL_3437520, EPI_ISL_3437521 | Lighthouse Lab in Milton Keynes | Wellcome Sanger Institute for the COVID-19 Genomics UK (COG-UK) Consortium | The Lighthouse Lab in Milton Keynes and Alex Alderton, Roberto Amato, Jeffrey Barrett, Sonia Goncalves, Ewan Harrison, David K. Jackson, Ian Johnston, Dominic Kwiatkowski, Cordelia Langford, John Sillitoe on behalf of the Wellcome Sanger Institute COVID-19 Surveillance Team |
| EPI_ISL_3437522, EPI_ISL_3437523 | Lighthouse Lab in Alderley Park | Wellcome Sanger Institute for the COVID-19 Genomics UK (COG-UK) Consortium | Jacquelyn Wynn, Mairead Hyland, The Lighthouse Lab in Alderley Park and Alex Alderton, Roberto Amato, Jeffrey Barrett, Sonia Goncalves, Ewan Harrison, David K. Jackson, Ian Johnston, Dominic Kwiatkowski, Cordelia Langford, John Sillitoe on behalf of the Wellcome Sanger Institute COVID-19 Surveillance Team |
| EPI_ISL_3437524 | Lighthouse Lab in Milton Keynes | Wellcome Sanger Institute for the COVID-19 Genomics UK (COG-UK) Consortium | The Lighthouse Lab in Milton Keynes and Alex Alderton, Roberto Amato, Jeffrey Barrett, Sonia Goncalves, Ewan Harrison, David K. Jackson, Ian Johnston, Dominic Kwiatkowski, Cordelia Langford, John Sillitoe on behalf of the Wellcome Sanger Institute COVID-19 Surveillance Team |
| EPI_ISL_3437525 | Lighthouse Lab in Alderley Park | Wellcome Sanger Institute for the COVID-19 Genomics UK (COG-UK) Consortium | Jacquelyn Wynn, Mairead Hyland, The Lighthouse Lab in Alderley Park and Alex Alderton, Roberto Amato, Jeffrey Barrett, Sonia Goncalves, Ewan Harrison, David K. Jackson, Ian Johnston, Dominic Kwiatkowski, Cordelia Langford, John Sillitoe on behalf of the Wellcome Sanger Institute COVID-19 Surveillance Team |
| EPI_ISL_3437526 | Lighthouse Lab in Milton Keynes | Wellcome Sanger Institute for the COVID-19 Genomics UK (COG-UK) Consortium | The Lighthouse Lab in Milton Keynes and Alex Alderton, Roberto Amato, Jeffrey Barrett, Sonia Goncalves, Ewan Harrison, David K. Jackson, Ian Johnston, Dominic Kwiatkowski, Cordelia Langford, John Sillitoe on behalf of the Wellcome Sanger Institute COVID-19 Surveillance Team |
| EPI_ISL_3437527 | Lighthouse Lab in Alderley Park | Wellcome Sanger Institute for the COVID-19 Genomics UK (COG-UK) Consortium | Jacquelyn Wynn, Mairead Hyland, The Lighthouse Lab in Alderley Park and Alex Alderton, Roberto Amato, Jeffrey Barrett, Sonia Goncalves, Ewan Harrison, David K. Jackson, Ian Johnston, Dominic Kwiatkowski, Cordelia Langford, John Sillitoe on behalf of the Wellcome Sanger Institute COVID-19 Surveillance Team |
| EPI_ISL_3437528, EPI_ISL_3437529 | Lighthouse Lab in Milton Keynes | Wellcome Sanger Institute for the COVID-19 Genomics UK (COG-UK) Consortium | The Lighthouse Lab in Milton Keynes and Alex Alderton, Roberto Amato, Jeffrey Barrett, Sonia Goncalves, Ewan Harrison, David K. Jackson, Ian Johnston, Dominic Kwiatkowski, Cordelia Langford, John Sillitoe on behalf of the Wellcome Sanger Institute COVID-19 Surveillance Team |
| EPI_ISL_3437530 | Lighthouse Lab in Alderley Park | Wellcome Sanger Institute for the COVID-19 Genomics UK (COG-UK) Consortium | Jacquelyn Wynn, Mairead Hyland, The Lighthouse Lab in Alderley Park and Alex Alderton, Roberto Amato, Jeffrey Barrett, Sonia Goncalves, Ewan Harrison, David K. Jackson, Ian Johnston, Dominic Kwiatkowski, Cordelia Langford, John Sillitoe on behalf of the Wellcome Sanger Institute COVID-19 Surveillance Team |
| EPI_ISL_3437531, EPI_ISL_3437532, EPI_ISL_3437533, EPI_ISL_3437534, EPI_ISL_3437535, EPI_ISL_3437536, EPI_ISL_3437537, EPI_ISL_3437538, EPI_ISL_3437539 | Lighthouse Lab in Milton Keynes | Wellcome Sanger Institute for the COVID-19 Genomics UK (COG-UK) Consortium | The Lighthouse Lab in Milton Keynes and Alex Alderton, Roberto Amato, Jeffrey Barrett, Sonia Goncalves, Ewan Harrison, David K. Jackson, Ian Johnston, Dominic Kwiatkowski, Cordelia Langford, John Sillitoe on behalf of the Wellcome Sanger Institute COVID-19 Surveillance Team |
| EPI_ISL_3437540, EPI_ISL_3437541, EPI_ISL_3437542 | Lighthouse Lab in Alderley Park | Wellcome Sanger Institute for the COVID-19 Genomics UK (COG-UK) Consortium | Jacquelyn Wynn, Mairead Hyland, The Lighthouse Lab in Alderley Park and Alex Alderton, Roberto Amato, Jeffrey Barrett, Sonia Goncalves, Ewan Harrison, David K. Jackson, Ian Johnston, Dominic Kwiatkowski, Cordelia Langford, John Sillitoe on behalf of the Wellcome Sanger Institute COVID-19 Surveillance Team |
| EPI_ISL_3437543 | Lighthouse Lab in Milton Keynes | Wellcome Sanger Institute for the COVID-19 Genomics UK (COG-UK) Consortium | The Lighthouse Lab in Milton Keynes and Alex Alderton, Roberto Amato, Jeffrey Barrett, Sonia Goncalves, Ewan Harrison, David K. Jackson, Ian Johnston, Dominic Kwiatkowski, Cordelia Langford, John Sillitoe on behalf of the Wellcome Sanger Institute COVID-19 Surveillance Team |
| EPI_ISL_3437544 | Lighthouse Lab in Alderley Park | Wellcome Sanger Institute for the COVID-19 Genomics UK (COG-UK) Consortium | Jacquelyn Wynn, Mairead Hyland, The Lighthouse Lab in Alderley Park and Alex Alderton, Roberto Amato, Jeffrey Barrett, Sonia Goncalves, Ewan Harrison, David K. Jackson, Ian Johnston, Dominic Kwiatkowski, Cordelia Langford, John Sillitoe on behalf of the Wellcome Sanger Institute COVID-19 Surveillance Team |
| EPI_ISL_3437545 | Lighthouse Lab in Milton Keynes | Wellcome Sanger Institute for the COVID-19 Genomics UK (COG-UK) Consortium | The Lighthouse Lab in Milton Keynes and Alex Alderton, Roberto Amato, Jeffrey Barrett, Sonia Goncalves, Ewan Harrison, David K. Jackson, Ian Johnston, Dominic Kwiatkowski, Cordelia Langford, John Sillitoe on behalf of the Wellcome Sanger Institute COVID-19 Surveillance Team |
| EPI_ISL_3437546, EPI_ISL_3437547 | Lighthouse Lab in Alderley Park | Wellcome Sanger Institute for the COVID-19 Genomics UK (COG-UK) Consortium | Jacquelyn Wynn, Mairead Hyland, The Lighthouse Lab in Alderley Park and Alex Alderton, Roberto Amato, Jeffrey Barrett, Sonia Goncalves, Ewan Harrison, David K. Jackson, Ian Johnston, Dominic Kwiatkowski, Cordelia Langford, John Sillitoe on behalf of the Wellcome Sanger Institute COVID-19 Surveillance Team |
| EPI_ISL_3437548 | Lighthouse Lab in Milton Keynes | Wellcome Sanger Institute for the COVID-19 Genomics UK (COG-UK) Consortium | The Lighthouse Lab in Milton Keynes and Alex Alderton, Roberto Amato, Jeffrey Barrett, Sonia Goncalves, Ewan Harrison, David K. Jackson, Ian Johnston, Dominic Kwiatkowski, Cordelia Langford, John Sillitoe on behalf of the Wellcome Sanger Institute COVID-19 Surveillance Team |
| EPI_ISL_3437549 | Lighthouse Lab in Alderley Park | Wellcome Sanger Institute for the COVID-19 Genomics UK (COG-UK) Consortium | Jacquelyn Wynn, Mairead Hyland, The Lighthouse Lab in Alderley Park and Alex Alderton, Roberto Amato, Jeffrey Barrett, Sonia Goncalves, Ewan Harrison, David K. Jackson, Ian Johnston, Dominic Kwiatkowski, Cordelia Langford, John Sillitoe on behalf of the Wellcome Sanger Institute COVID-19 Surveillance Team |
| EPI_ISL_3437550, EPI_ISL_3437551, EPI_ISL_3437552, EPI_ISL_3437553, EPI_ISL_3437554 | Lighthouse Lab in Milton Keynes | Wellcome Sanger Institute for the COVID-19 Genomics UK (COG-UK) Consortium | The Lighthouse Lab in Milton Keynes and Alex Alderton, Roberto Amato, Jeffrey Barrett, Sonia Goncalves, Ewan Harrison, David K. Jackson, Ian Johnston, Dominic Kwiatkowski, Cordelia Langford, John Sillitoe on behalf of the Wellcome Sanger Institute COVID-19 Surveillance Team |
| EPI_ISL_3437555 | Lighthouse Lab in Alderley Park | Wellcome Sanger Institute for the COVID-19 Genomics UK (COG-UK) Consortium | Jacquelyn Wynn, Mairead Hyland, The Lighthouse Lab in Alderley Park and Alex Alderton, Roberto Amato, Jeffrey Barrett, Sonia Goncalves, Ewan Harrison, David K. Jackson, Ian Johnston, Dominic Kwiatkowski, Cordelia Langford, John Sillitoe on behalf of the Wellcome Sanger Institute COVID-19 Surveillance Team |
| EPI_ISL_3437556, EPI_ISL_3437557, EPI_ISL_3437558 | Lighthouse Lab in Milton Keynes | Wellcome Sanger Institute for the COVID-19 Genomics UK (COG-UK) Consortium | The Lighthouse Lab in Milton Keynes and Alex Alderton, Roberto Amato, Jeffrey Barrett, Sonia Goncalves, Ewan Harrison, David K. Jackson, Ian Johnston, Dominic Kwiatkowski, Cordelia Langford, John Sillitoe on behalf of the Wellcome Sanger Institute COVID-19 Surveillance Team |
| EPI_ISL_3437559, EPI_ISL_3437560 | Lighthouse Lab in Alderley Park | Wellcome Sanger Institute for the COVID-19 Genomics UK (COG-UK) Consortium | Jacquelyn Wynn, Mairead Hyland, The Lighthouse Lab in Alderley Park and Alex Alderton, Roberto Amato, Jeffrey Barrett, Sonia Goncalves, Ewan Harrison, David K. Jackson, Ian Johnston, Dominic Kwiatkowski, Cordelia Langford, John Sillitoe on behalf of the Wellcome Sanger Institute COVID-19 Surveillance Team |
| EPI_ISL_3437561 | Lighthouse Lab in Milton Keynes | Wellcome Sanger Institute for the COVID-19 Genomics UK (COG-UK) Consortium | The Lighthouse Lab in Milton Keynes and Alex Alderton, Roberto Amato, Jeffrey Barrett, Sonia Goncalves, Ewan Harrison, David K. Jackson, Ian Johnston, Dominic Kwiatkowski, Cordelia Langford, John Sillitoe on behalf of the Wellcome Sanger Institute COVID-19 Surveillance Team |
| EPI_ISL_3437562 | Lighthouse Lab in Alderley Park | Wellcome Sanger Institute for the COVID-19 Genomics UK (COG-UK) Consortium | Jacquelyn Wynn, Mairead Hyland, The Lighthouse Lab in Alderley Park and Alex Alderton, Roberto Amato, Jeffrey Barrett, Sonia Goncalves, Ewan Harrison, David K. Jackson, Ian Johnston, Dominic Kwiatkowski, Cordelia Langford, John Sillitoe on behalf of the Wellcome Sanger Institute COVID-19 Surveillance Team |
| EPI_ISL_3437563, EPI_ISL_3437564, EPI_ISL_3437565, EPI_ISL_3437566, EPI_ISL_3437567, EPI_ISL_3437568, EPI_ISL_3437569 | Lighthouse Lab in Milton Keynes | Wellcome Sanger Institute for the COVID-19 Genomics UK (COG-UK) Consortium | The Lighthouse Lab in Milton Keynes and Alex Alderton, Roberto Amato, Jeffrey Barrett, Sonia Goncalves, Ewan Harrison, David K. Jackson, Ian Johnston, Dominic Kwiatkowski, Cordelia Langford, John Sillitoe on behalf of the Wellcome Sanger Institute COVID-19 Surveillance Team |
| EPI_ISL_3437570 | Lighthouse Lab in Alderley Park | Wellcome Sanger Institute for the COVID-19 Genomics UK (COG-UK) Consortium | Jacquelyn Wynn, Mairead Hyland, The Lighthouse Lab in Alderley Park and Alex Alderton, Roberto Amato, Jeffrey Barrett, Sonia Goncalves, Ewan Harrison, David K. Jackson, Ian Johnston, Dominic Kwiatkowski, Cordelia Langford, John Sillitoe on behalf of the Wellcome Sanger Institute COVID-19 Surveillance Team |
| EPI_ISL_3437571, EPI_ISL_3437572, | Lighthouse Lab in Milton Keynes | Wellcome Sanger Institute for the COVID-19 Genomics UK (COG-UK) Consortium | The Lighthouse Lab in Milton Keynes and Alex Alderton, Roberto Amato, Jeffrey Barrett, Sonia Goncalves, Ewan Harrison, David K. Jackson, Ian Johnston, Dominic Kwiatkowski, Cordelia Langford, John Sillitoe on behalf of the Wellcome Sanger Institute COVID-19 Surveillance Team |

| | | | |
|---|---|---|---|
| EPI_ISL_3437573 | | | |
| EPI_ISL_3437574 | Lighthouse Lab in Alderley Park | Wellcome Sanger Institute for the COVID-19 Genomics UK (COG-UK) Consortium | Jacquelyn Wynn, Mairead Hyland, The Lighthouse Lab in Alderley Park and Alex Alderton, Roberto Amato, Jeffrey Barrett, Sonia Goncalves, Ewan Harrison, David K. Jackson, Ian Johnston, Dominic Kwiatkowski, Cordelia Langford, John Sillitoe on behalf of the Wellcome Sanger Institute COVID-19 Surveillance Team |
| EPI_ISL_3437575 | Lighthouse Lab in Milton Keynes | Wellcome Sanger Institute for the COVID-19 Genomics UK (COG-UK) Consortium | The Lighthouse Lab in Milton Keynes and Alex Alderton, Roberto Amato, Jeffrey Barrett, Sonia Goncalves, Ewan Harrison, David K. Jackson, Ian Johnston, Dominic Kwiatkowski, Cordelia Langford, John Sillitoe on behalf of the Wellcome Sanger Institute COVID-19 Surveillance Team |
| EPI_ISL_3437576 | Lighthouse Lab in Alderley Park | Wellcome Sanger Institute for the COVID-19 Genomics UK (COG-UK) Consortium | Jacquelyn Wynn, Mairead Hyland, The Lighthouse Lab in Alderley Park and Alex Alderton, Roberto Amato, Jeffrey Barrett, Sonia Goncalves, Ewan Harrison, David K. Jackson, Ian Johnston, Dominic Kwiatkowski, Cordelia Langford, John Sillitoe on behalf of the Wellcome Sanger Institute COVID-19 Surveillance Team |
| EPI_ISL_3437577, EPI_ISL_3437578, EPI_ISL_3437579, EPI_ISL_3437580, EPI_ISL_3437581, EPI_ISL_3437582, EPI_ISL_3437583 | Lighthouse Lab in Milton Keynes | Wellcome Sanger Institute for the COVID-19 Genomics UK (COG-UK) Consortium | The Lighthouse Lab in Milton Keynes and Alex Alderton, Roberto Amato, Jeffrey Barrett, Sonia Goncalves, Ewan Harrison, David K. Jackson, Ian Johnston, Dominic Kwiatkowski, Cordelia Langford, John Sillitoe on behalf of the Wellcome Sanger Institute COVID-19 Surveillance Team |
| EPI_ISL_3437584 | Lighthouse Lab in Alderley Park | Wellcome Sanger Institute for the COVID-19 Genomics UK (COG-UK) Consortium | Jacquelyn Wynn, Mairead Hyland, The Lighthouse Lab in Alderley Park and Alex Alderton, Roberto Amato, Jeffrey Barrett, Sonia Goncalves, Ewan Harrison, David K. Jackson, Ian Johnston, Dominic Kwiatkowski, Cordelia Langford, John Sillitoe on behalf of the Wellcome Sanger Institute COVID-19 Surveillance Team |
| EPI_ISL_3437585, EPI_ISL_3437586, EPI_ISL_3437587, EPI_ISL_3437588, EPI_ISL_3437589, EPI_ISL_3437590, EPI_ISL_3437591 | Lighthouse Lab in Milton Keynes | Wellcome Sanger Institute for the COVID-19 Genomics UK (COG-UK) Consortium | The Lighthouse Lab in Milton Keynes and Alex Alderton, Roberto Amato, Jeffrey Barrett, Sonia Goncalves, Ewan Harrison, David K. Jackson, Ian Johnston, Dominic Kwiatkowski, Cordelia Langford, John Sillitoe on behalf of the Wellcome Sanger Institute COVID-19 Surveillance Team |
| EPI_ISL_3437592 | Lighthouse Lab in Alderley Park | Wellcome Sanger Institute for the COVID-19 Genomics UK (COG-UK) Consortium | Jacquelyn Wynn, Mairead Hyland, The Lighthouse Lab in Alderley Park and Alex Alderton, Roberto Amato, Jeffrey Barrett, Sonia Goncalves, Ewan Harrison, David K. Jackson, Ian Johnston, Dominic Kwiatkowski, Cordelia Langford, John Sillitoe on behalf of the Wellcome Sanger Institute COVID-19 Surveillance Team |
| EPI_ISL_3437593, EPI_ISL_3437594, EPI_ISL_3437595, EPI_ISL_3437596 | Lighthouse Lab in Milton Keynes | Wellcome Sanger Institute for the COVID-19 Genomics UK (COG-UK) Consortium | The Lighthouse Lab in Milton Keynes and Alex Alderton, Roberto Amato, Jeffrey Barrett, Sonia Goncalves, Ewan Harrison, David K. Jackson, Ian Johnston, Dominic Kwiatkowski, Cordelia Langford, John Sillitoe on behalf of the Wellcome Sanger Institute COVID-19 Surveillance Team |
| EPI_ISL_3437597, EPI_ISL_3437598 | Lighthouse Lab in Alderley Park | Wellcome Sanger Institute for the COVID-19 Genomics UK (COG-UK) Consortium | Jacquelyn Wynn, Mairead Hyland, The Lighthouse Lab in Alderley Park and Alex Alderton, Roberto Amato, Jeffrey Barrett, Sonia Goncalves, Ewan Harrison, David K. Jackson, Ian Johnston, Dominic Kwiatkowski, Cordelia Langford, John Sillitoe on behalf of the Wellcome Sanger Institute COVID-19 Surveillance Team |
| EPI_ISL_3437599, EPI_ISL_3437600, EPI_ISL_3437601, EPI_ISL_3437602, EPI_ISL_3437603, EPI_ISL_3437604, EPI_ISL_3437605, EPI_ISL_3437606 | Lighthouse Lab in Milton Keynes | Wellcome Sanger Institute for the COVID-19 Genomics UK (COG-UK) Consortium | The Lighthouse Lab in Milton Keynes and Alex Alderton, Roberto Amato, Jeffrey Barrett, Sonia Goncalves, Ewan Harrison, David K. Jackson, Ian Johnston, Dominic Kwiatkowski, Cordelia Langford, John Sillitoe on behalf of the Wellcome Sanger Institute COVID-19 Surveillance Team |
| EPI_ISL_3437607, EPI_ISL_3437608 | Lighthouse Lab in Alderley Park | Wellcome Sanger Institute for the COVID-19 Genomics UK (COG-UK) Consortium | Jacquelyn Wynn, Mairead Hyland, The Lighthouse Lab in Alderley Park and Alex Alderton, Roberto Amato, Jeffrey Barrett, Sonia Goncalves, Ewan Harrison, David K. Jackson, Ian Johnston, Dominic Kwiatkowski, Cordelia Langford, John Sillitoe on behalf of the Wellcome Sanger Institute COVID-19 Surveillance Team |
| EPI_ISL_3437609, EPI_ISL_3437610, EPI_ISL_3437611, EPI_ISL_3437612, EPI_ISL_3437613 | Lighthouse Lab in Milton Keynes | Wellcome Sanger Institute for the COVID-19 Genomics UK (COG-UK) Consortium | The Lighthouse Lab in Milton Keynes and Alex Alderton, Roberto Amato, Jeffrey Barrett, Sonia Goncalves, Ewan Harrison, David K. Jackson, Ian Johnston, Dominic Kwiatkowski, Cordelia Langford, John Sillitoe on behalf of the Wellcome Sanger Institute COVID-19 Surveillance Team |
| EPI_ISL_3437614 | Lighthouse Lab in Alderley Park | Wellcome Sanger Institute for the COVID-19 Genomics UK (COG-UK) Consortium | Jacquelyn Wynn, Mairead Hyland, The Lighthouse Lab in Alderley Park and Alex Alderton, Roberto Amato, Jeffrey Barrett, Sonia Goncalves, Ewan Harrison, David K. Jackson, Ian Johnston, Dominic Kwiatkowski, Cordelia Langford, John Sillitoe on behalf of the Wellcome Sanger Institute COVID-19 Surveillance Team |
| EPI_ISL_3437615, EPI_ISL_3437616, EPI_ISL_3437617, EPI_ISL_3437618, EPI_ISL_3437619, EPI_ISL_3437620, EPI_ISL_3437621, EPI_ISL_3437622 | Lighthouse Lab in Milton Keynes | Wellcome Sanger Institute for the COVID-19 Genomics UK (COG-UK) Consortium | The Lighthouse Lab in Milton Keynes and Alex Alderton, Roberto Amato, Jeffrey Barrett, Sonia Goncalves, Ewan Harrison, David K. Jackson, Ian Johnston, Dominic Kwiatkowski, Cordelia Langford, John Sillitoe on behalf of the Wellcome Sanger Institute COVID-19 Surveillance Team |
| EPI_ISL_3437623 | Lighthouse Lab in Alderley Park | Wellcome Sanger Institute for the COVID-19 Genomics UK (COG-UK) Consortium | Jacquelyn Wynn, Mairead Hyland, The Lighthouse Lab in Alderley Park and Alex Alderton, Roberto Amato, Jeffrey Barrett, Sonia Goncalves, Ewan Harrison, David K. Jackson, Ian Johnston, Dominic Kwiatkowski, Cordelia Langford, John Sillitoe on behalf of the Wellcome Sanger Institute COVID-19 Surveillance Team |
| EPI_ISL_3437624, EPI_ISL_3437625, EPI_ISL_3437626, EPI_ISL_3437627, EPI_ISL_3437628 | Lighthouse Lab in Milton Keynes | Wellcome Sanger Institute for the COVID-19 Genomics UK (COG-UK) Consortium | The Lighthouse Lab in Milton Keynes and Alex Alderton, Roberto Amato, Jeffrey Barrett, Sonia Goncalves, Ewan Harrison, David K. Jackson, Ian Johnston, Dominic Kwiatkowski, Cordelia Langford, John Sillitoe on behalf of the Wellcome Sanger Institute COVID-19 Surveillance Team |
| EPI_ISL_3437629, EPI_ISL_3437630, EPI_ISL_3437631 | Lighthouse Lab in Alderley Park | Wellcome Sanger Institute for the COVID-19 Genomics UK (COG-UK) Consortium | Jacquelyn Wynn, Mairead Hyland, The Lighthouse Lab in Alderley Park and Alex Alderton, Roberto Amato, Jeffrey Barrett, Sonia Goncalves, Ewan Harrison, David K. Jackson, Ian Johnston, Dominic Kwiatkowski, Cordelia Langford, John Sillitoe on behalf of the Wellcome Sanger Institute COVID-19 Surveillance Team |
| EPI_ISL_3437632, EPI_ISL_3437633, EPI_ISL_3437634 | Lighthouse Lab in Milton Keynes | Wellcome Sanger Institute for the COVID-19 Genomics UK (COG-UK) Consortium | The Lighthouse Lab in Milton Keynes and Alex Alderton, Roberto Amato, Jeffrey Barrett, Sonia Goncalves, Ewan Harrison, David K. Jackson, Ian Johnston, Dominic Kwiatkowski, Cordelia Langford, John Sillitoe on behalf of the Wellcome Sanger Institute COVID-19 Surveillance Team |
| EPI_ISL_3437635 | Lighthouse Lab in Alderley Park | Wellcome Sanger Institute for the COVID-19 Genomics UK (COG-UK) Consortium | Jacquelyn Wynn, Mairead Hyland, The Lighthouse Lab in Alderley Park and Alex Alderton, Roberto Amato, Jeffrey Barrett, Sonia Goncalves, Ewan Harrison, David K. Jackson, Ian Johnston, Dominic Kwiatkowski, Cordelia Langford, John Sillitoe on behalf of the Wellcome Sanger Institute COVID-19 Surveillance Team |
| EPI_ISL_3437636 | Lighthouse Lab in Milton Keynes | Wellcome Sanger Institute for the COVID-19 Genomics UK (COG-UK) Consortium | The Lighthouse Lab in Milton Keynes and Alex Alderton, Roberto Amato, Jeffrey Barrett, Sonia Goncalves, Ewan Harrison, David K. Jackson, Ian Johnston, Dominic Kwiatkowski, Cordelia Langford, John Sillitoe on behalf of the Wellcome Sanger Institute COVID-19 Surveillance Team |
| EPI_ISL_3437637 | Lighthouse Lab in Alderley Park | Wellcome Sanger Institute for the COVID-19 Genomics UK (COG-UK) Consortium | Jacquelyn Wynn, Mairead Hyland, The Lighthouse Lab in Alderley Park and Alex Alderton, Roberto Amato, Jeffrey Barrett, Sonia Goncalves, Ewan Harrison, David K. Jackson, Ian Johnston, Dominic Kwiatkowski, Cordelia Langford, John Sillitoe on behalf of the Wellcome Sanger Institute COVID-19 Surveillance Team |
| EPI_ISL_3437638, EPI_ISL_3437639, EPI_ISL_3437640, EPI_ISL_3437641, EPI_ISL_3437642 | Lighthouse Lab in Milton Keynes | Wellcome Sanger Institute for the COVID-19 Genomics UK (COG-UK) Consortium | The Lighthouse Lab in Milton Keynes and Alex Alderton, Roberto Amato, Jeffrey Barrett, Sonia Goncalves, Ewan Harrison, David K. Jackson, Ian Johnston, Dominic Kwiatkowski, Cordelia Langford, John Sillitoe on behalf of the Wellcome Sanger Institute COVID-19 Surveillance Team |
| EPI_ISL_3437643 | Lighthouse Lab in Alderley Park | Wellcome Sanger Institute for the COVID-19 Genomics UK (COG-UK) Consortium | Jacquelyn Wynn, Mairead Hyland, The Lighthouse Lab in Alderley Park and Alex Alderton, Roberto Amato, Jeffrey Barrett, Sonia Goncalves, Ewan Harrison, David K. Jackson, Ian Johnston, Dominic Kwiatkowski, Cordelia Langford, John Sillitoe on behalf of the Wellcome Sanger Institute COVID-19 Surveillance Team |
| EPI_ISL_3437644, EPI_ISL_3437645, EPI_ISL_3437646 | Lighthouse Lab in Milton Keynes | Wellcome Sanger Institute for the COVID-19 Genomics UK (COG-UK) Consortium | The Lighthouse Lab in Milton Keynes and Alex Alderton, Roberto Amato, Jeffrey Barrett, Sonia Goncalves, Ewan Harrison, David K. Jackson, Ian Johnston, Dominic Kwiatkowski, Cordelia Langford, John Sillitoe on behalf of the Wellcome Sanger Institute COVID-19 Surveillance Team |
| EPI_ISL_3437647, EPI_ISL_3437648 | Lighthouse Lab in Alderley Park | Wellcome Sanger Institute for the COVID-19 Genomics UK (COG-UK) Consortium | Jacquelyn Wynn, Mairead Hyland, The Lighthouse Lab in Alderley Park and Alex Alderton, Roberto Amato, Jeffrey Barrett, Sonia Goncalves, Ewan Harrison, David K. Jackson, Ian Johnston, Dominic Kwiatkowski, Cordelia Langford, John Sillitoe on behalf of the Wellcome Sanger Institute COVID-19 Surveillance Team |
| EPI_ISL_3437649 | Lighthouse Lab in Milton Keynes | Wellcome Sanger Institute for the COVID-19 Genomics UK (COG-UK) Consortium | The Lighthouse Lab in Milton Keynes and Alex Alderton, Roberto Amato, Jeffrey Barrett, Sonia Goncalves, Ewan Harrison, David K. Jackson, Ian Johnston, Dominic Kwiatkowski, Cordelia Langford, John Sillitoe on behalf of the Wellcome Sanger Institute COVID-19 Surveillance Team |
| EPI_ISL_3437650 | Lighthouse Lab in Alderley Park | Wellcome Sanger Institute for the COVID-19 Genomics UK (COG-UK) Consortium | Jacquelyn Wynn, Mairead Hyland, The Lighthouse Lab in Alderley Park and Alex Alderton, Roberto Amato, Jeffrey Barrett, Sonia Goncalves, Ewan Harrison, David K. Jackson, Ian Johnston, Dominic Kwiatkowski, Cordelia Langford, John Sillitoe on behalf of the Wellcome Sanger Institute COVID-19 Surveillance Team |
| EPI_ISL_3437651, EPI_ISL_3437652, EPI_ISL_3437653, EPI_ISL_3437654 | Lighthouse Lab in Milton Keynes | Wellcome Sanger Institute for the COVID-19 Genomics UK (COG-UK) Consortium | The Lighthouse Lab in Milton Keynes and Alex Alderton, Roberto Amato, Jeffrey Barrett, Sonia Goncalves, Ewan Harrison, David K. Jackson, Ian Johnston, Dominic Kwiatkowski, Cordelia Langford, John Sillitoe on behalf of the Wellcome Sanger Institute COVID-19 Surveillance Team |
| EPI_ISL_3437655 | Lighthouse Lab in Alderley Park | Wellcome Sanger Institute for the COVID-19 Genomics UK (COG-UK) Consortium | Jacquelyn Wynn, Mairead Hyland, The Lighthouse Lab in Alderley Park and Alex Alderton, Roberto Amato, Jeffrey Barrett, Sonia Goncalves, Ewan Harrison, David K. Jackson, Ian Johnston, Dominic Kwiatkowski, Cordelia Langford, John Sillitoe on behalf of the Wellcome Sanger Institute COVID-19 Surveillance Team |

| | | | |
|---|---|---|---|
| EPI_ISL_3437656 | Lighthouse Lab in Milton Keynes | Wellcome Sanger Institute for the COVID-19 Genomics UK (COG-UK) Consortium | The Lighthouse Lab in Milton Keynes and Alex Alderton, Roberto Amato, Jeffrey Barrett, Sonia Goncalves, Ewan Harrison, David K. Jackson, Ian Johnston, Dominic Kwiatkowski, Cordelia Langford, John Sillitoe on behalf of the Wellcome Sanger Institute COVID-19 Surveillance Team |
| EPI_ISL_3437657 | Lighthouse Lab in Alderley Park | Wellcome Sanger Institute for the COVID-19 Genomics UK (COG-UK) Consortium | Jacquelyn Wynn, Mairead Hyland, The Lighthouse Lab in Alderley Park and Alex Alderton, Roberto Amato, Jeffrey Barrett, Sonia Goncalves, Ewan Harrison, David K. Jackson, Ian Johnston, Dominic Kwiatkowski, Cordelia Langford, John Sillitoe on behalf of the Wellcome Sanger Institute COVID-19 Surveillance Team |
| EPI_ISL_3437658, EPI_ISL_3437659, EPI_ISL_3437660, EPI_ISL_3437661, EPI_ISL_3437662, EPI_ISL_3437663, EPI_ISL_3437664, EPI_ISL_3437665, EPI_ISL_3437666, EPI_ISL_3437667, EPI_ISL_3437668, EPI_ISL_3437669, EPI_ISL_3437670, EPI_ISL_3437671 | | | |
| see above | Lighthouse Lab in Milton Keynes | Wellcome Sanger Institute for the COVID-19 Genomics UK (COG-UK) Consortium | The Lighthouse Lab in Milton Keynes and Alex Alderton, Roberto Amato, Jeffrey Barrett, Sonia Goncalves, Ewan Harrison, David K. Jackson, Ian Johnston, Dominic Kwiatkowski, Cordelia Langford, John Sillitoe on behalf of the Wellcome Sanger Institute COVID-19 Surveillance Team |
| EPI_ISL_3437672 | Lighthouse Lab in Alderley Park | Wellcome Sanger Institute for the COVID-19 Genomics UK (COG-UK) Consortium | Jacquelyn Wynn, Mairead Hyland, The Lighthouse Lab in Alderley Park and Alex Alderton, Roberto Amato, Jeffrey Barrett, Sonia Goncalves, Ewan Harrison, David K. Jackson, Ian Johnston, Dominic Kwiatkowski, Cordelia Langford, John Sillitoe on behalf of the Wellcome Sanger Institute COVID-19 Surveillance Team |
| EPI_ISL_3437673, EPI_ISL_3437674 | Lighthouse Lab in Milton Keynes | Wellcome Sanger Institute for the COVID-19 Genomics UK (COG-UK) Consortium | The Lighthouse Lab in Milton Keynes and Alex Alderton, Roberto Amato, Jeffrey Barrett, Sonia Goncalves, Ewan Harrison, David K. Jackson, Ian Johnston, Dominic Kwiatkowski, Cordelia Langford, John Sillitoe on behalf of the Wellcome Sanger Institute COVID-19 Surveillance Team |
| EPI_ISL_3437675 | Lighthouse Lab in Alderley Park | Wellcome Sanger Institute for the COVID-19 Genomics UK (COG-UK) Consortium | Jacquelyn Wynn, Mairead Hyland, The Lighthouse Lab in Alderley Park and Alex Alderton, Roberto Amato, Jeffrey Barrett, Sonia Goncalves, Ewan Harrison, David K. Jackson, Ian Johnston, Dominic Kwiatkowski, Cordelia Langford, John Sillitoe on behalf of the Wellcome Sanger Institute COVID-19 Surveillance Team |
| EPI_ISL_3437676, EPI_ISL_3437677 | Lighthouse Lab in Milton Keynes | Wellcome Sanger Institute for the COVID-19 Genomics UK (COG-UK) Consortium | The Lighthouse Lab in Milton Keynes and Alex Alderton, Roberto Amato, Jeffrey Barrett, Sonia Goncalves, Ewan Harrison, David K. Jackson, Ian Johnston, Dominic Kwiatkowski, Cordelia Langford, John Sillitoe on behalf of the Wellcome Sanger Institute COVID-19 Surveillance Team |
| EPI_ISL_3437678 | Lighthouse Lab in Alderley Park | Wellcome Sanger Institute for the COVID-19 Genomics UK (COG-UK) Consortium | Jacquelyn Wynn, Mairead Hyland, The Lighthouse Lab in Alderley Park and Alex Alderton, Roberto Amato, Jeffrey Barrett, Sonia Goncalves, Ewan Harrison, David K. Jackson, Ian Johnston, Dominic Kwiatkowski, Cordelia Langford, John Sillitoe on behalf of the Wellcome Sanger Institute COVID-19 Surveillance Team |
| EPI_ISL_3437679, EPI_ISL_3437680, EPI_ISL_3437681, EPI_ISL_3437682 | Lighthouse Lab in Milton Keynes | Wellcome Sanger Institute for the COVID-19 Genomics UK (COG-UK) Consortium | The Lighthouse Lab in Milton Keynes and Alex Alderton, Roberto Amato, Jeffrey Barrett, Sonia Goncalves, Ewan Harrison, David K. Jackson, Ian Johnston, Dominic Kwiatkowski, Cordelia Langford, John Sillitoe on behalf of the Wellcome Sanger Institute COVID-19 Surveillance Team |
| EPI_ISL_3437683, EPI_ISL_3437684 | Lighthouse Lab in Alderley Park | Wellcome Sanger Institute for the COVID-19 Genomics UK (COG-UK) Consortium | Jacquelyn Wynn, Mairead Hyland, The Lighthouse Lab in Alderley Park and Alex Alderton, Roberto Amato, Jeffrey Barrett, Sonia Goncalves, Ewan Harrison, David K. Jackson, Ian Johnston, Dominic Kwiatkowski, Cordelia Langford, John Sillitoe on behalf of the Wellcome Sanger Institute COVID-19 Surveillance Team |
| EPI_ISL_3437685, EPI_ISL_3437686, EPI_ISL_3437687, EPI_ISL_3437688 | Lighthouse Lab in Milton Keynes | Wellcome Sanger Institute for the COVID-19 Genomics UK (COG-UK) Consortium | The Lighthouse Lab in Milton Keynes and Alex Alderton, Roberto Amato, Jeffrey Barrett, Sonia Goncalves, Ewan Harrison, David K. Jackson, Ian Johnston, Dominic Kwiatkowski, Cordelia Langford, John Sillitoe on behalf of the Wellcome Sanger Institute COVID-19 Surveillance Team |
| EPI_ISL_3437689 | Lighthouse Lab in Alderley Park | Wellcome Sanger Institute for the COVID-19 Genomics UK (COG-UK) Consortium | Jacquelyn Wynn, Mairead Hyland, The Lighthouse Lab in Alderley Park and Alex Alderton, Roberto Amato, Jeffrey Barrett, Sonia Goncalves, Ewan Harrison, David K. Jackson, Ian Johnston, Dominic Kwiatkowski, Cordelia Langford, John Sillitoe on behalf of the Wellcome Sanger Institute COVID-19 Surveillance Team |
| EPI_ISL_3437690, EPI_ISL_3437691, EPI_ISL_3437692, EPI_ISL_3437693 | Lighthouse Lab in Milton Keynes | Wellcome Sanger Institute for the COVID-19 Genomics UK (COG-UK) Consortium | The Lighthouse Lab in Milton Keynes and Alex Alderton, Roberto Amato, Jeffrey Barrett, Sonia Goncalves, Ewan Harrison, David K. Jackson, Ian Johnston, Dominic Kwiatkowski, Cordelia Langford, John Sillitoe on behalf of the Wellcome Sanger Institute COVID-19 Surveillance Team |
| EPI_ISL_3437694 | Lighthouse Lab in Alderley Park | Wellcome Sanger Institute for the COVID-19 Genomics UK (COG-UK) Consortium | Jacquelyn Wynn, Mairead Hyland, The Lighthouse Lab in Alderley Park and Alex Alderton, Roberto Amato, Jeffrey Barrett, Sonia Goncalves, Ewan Harrison, David K. Jackson, Ian Johnston, Dominic Kwiatkowski, Cordelia Langford, John Sillitoe on behalf of the Wellcome Sanger Institute COVID-19 Surveillance Team |
| EPI_ISL_3437695, EPI_ISL_3437696, EPI_ISL_3437697, EPI_ISL_3437698, EPI_ISL_3437699, EPI_ISL_3437700 | Lighthouse Lab in Milton Keynes | Wellcome Sanger Institute for the COVID-19 Genomics UK (COG-UK) Consortium | The Lighthouse Lab in Milton Keynes and Alex Alderton, Roberto Amato, Jeffrey Barrett, Sonia Goncalves, Ewan Harrison, David K. Jackson, Ian Johnston, Dominic Kwiatkowski, Cordelia Langford, John Sillitoe on behalf of the Wellcome Sanger Institute COVID-19 Surveillance Team |
| EPI_ISL_3437701 | Lighthouse Lab in Alderley Park | Wellcome Sanger Institute for the COVID-19 Genomics UK (COG-UK) Consortium | Jacquelyn Wynn, Mairead Hyland, The Lighthouse Lab in Alderley Park and Alex Alderton, Roberto Amato, Jeffrey Barrett, Sonia Goncalves, Ewan Harrison, David K. Jackson, Ian Johnston, Dominic Kwiatkowski, Cordelia Langford, John Sillitoe on behalf of the Wellcome Sanger Institute COVID-19 Surveillance Team |
| EPI_ISL_3437702, EPI_ISL_3437703, EPI_ISL_3437704 | Lighthouse Lab in Milton Keynes | Wellcome Sanger Institute for the COVID-19 Genomics UK (COG-UK) Consortium | The Lighthouse Lab in Milton Keynes and Alex Alderton, Roberto Amato, Jeffrey Barrett, Sonia Goncalves, Ewan Harrison, David K. Jackson, Ian Johnston, Dominic Kwiatkowski, Cordelia Langford, John Sillitoe on behalf of the Wellcome Sanger Institute COVID-19 Surveillance Team |
| EPI_ISL_3437705, EPI_ISL_3437706 | Lighthouse Lab in Alderley Park | Wellcome Sanger Institute for the COVID-19 Genomics UK (COG-UK) Consortium | Jacquelyn Wynn, Mairead Hyland, The Lighthouse Lab in Alderley Park and Alex Alderton, Roberto Amato, Jeffrey Barrett, Sonia Goncalves, Ewan Harrison, David K. Jackson, Ian Johnston, Dominic Kwiatkowski, Cordelia Langford, John Sillitoe on behalf of the Wellcome Sanger Institute COVID-19 Surveillance Team |
| EPI_ISL_3437707, EPI_ISL_3437708, EPI_ISL_3437709 | Lighthouse Lab in Milton Keynes | Wellcome Sanger Institute for the COVID-19 Genomics UK (COG-UK) Consortium | The Lighthouse Lab in Milton Keynes and Alex Alderton, Roberto Amato, Jeffrey Barrett, Sonia Goncalves, Ewan Harrison, David K. Jackson, Ian Johnston, Dominic Kwiatkowski, Cordelia Langford, John Sillitoe on behalf of the Wellcome Sanger Institute COVID-19 Surveillance Team |
| EPI_ISL_3437710 | Lighthouse Lab in Alderley Park | Wellcome Sanger Institute for the COVID-19 Genomics UK (COG-UK) Consortium | Jacquelyn Wynn, Mairead Hyland, The Lighthouse Lab in Alderley Park and Alex Alderton, Roberto Amato, Jeffrey Barrett, Sonia Goncalves, Ewan Harrison, David K. Jackson, Ian Johnston, Dominic Kwiatkowski, Cordelia Langford, John Sillitoe on behalf of the Wellcome Sanger Institute COVID-19 Surveillance Team |
| EPI_ISL_3437711, EPI_ISL_3437712, EPI_ISL_3437713 | Lighthouse Lab in Milton Keynes | Wellcome Sanger Institute for the COVID-19 Genomics UK (COG-UK) Consortium | The Lighthouse Lab in Milton Keynes and Alex Alderton, Roberto Amato, Jeffrey Barrett, Sonia Goncalves, Ewan Harrison, David K. Jackson, Ian Johnston, Dominic Kwiatkowski, Cordelia Langford, John Sillitoe on behalf of the Wellcome Sanger Institute COVID-19 Surveillance Team |
| EPI_ISL_3437714, EPI_ISL_3437715 | Lighthouse Lab in Alderley Park | Wellcome Sanger Institute for the COVID-19 Genomics UK (COG-UK) Consortium | Jacquelyn Wynn, Mairead Hyland, The Lighthouse Lab in Alderley Park and Alex Alderton, Roberto Amato, Jeffrey Barrett, Sonia Goncalves, Ewan Harrison, David K. Jackson, Ian Johnston, Dominic Kwiatkowski, Cordelia Langford, John Sillitoe on behalf of the Wellcome Sanger Institute COVID-19 Surveillance Team |
| EPI_ISL_3437716, EPI_ISL_3437717 | Lighthouse Lab in Milton Keynes | Wellcome Sanger Institute for the COVID-19 Genomics UK (COG-UK) Consortium | The Lighthouse Lab in Milton Keynes and Alex Alderton, Roberto Amato, Jeffrey Barrett, Sonia Goncalves, Ewan Harrison, David K. Jackson, Ian Johnston, Dominic Kwiatkowski, Cordelia Langford, John Sillitoe on behalf of the Wellcome Sanger Institute COVID-19 Surveillance Team |
| EPI_ISL_3437718 | Lighthouse Lab in Alderley Park | Wellcome Sanger Institute for the COVID-19 Genomics UK (COG-UK) Consortium | Jacquelyn Wynn, Mairead Hyland, The Lighthouse Lab in Alderley Park and Alex Alderton, Roberto Amato, Jeffrey Barrett, Sonia Goncalves, Ewan Harrison, David K. Jackson, Ian Johnston, Dominic Kwiatkowski, Cordelia Langford, John Sillitoe on behalf of the Wellcome Sanger Institute COVID-19 Surveillance Team |
| EPI_ISL_3437719, EPI_ISL_3437720 | Lighthouse Lab in Milton Keynes | Wellcome Sanger Institute for the COVID-19 Genomics UK (COG-UK) Consortium | The Lighthouse Lab in Milton Keynes and Alex Alderton, Roberto Amato, Jeffrey Barrett, Sonia Goncalves, Ewan Harrison, David K. Jackson, Ian Johnston, Dominic Kwiatkowski, Cordelia Langford, John Sillitoe on behalf of the Wellcome Sanger Institute COVID-19 Surveillance Team |
| EPI_ISL_3437721 | Lighthouse Lab in Alderley Park | Wellcome Sanger Institute for the COVID-19 Genomics UK (COG-UK) Consortium | Jacquelyn Wynn, Mairead Hyland, The Lighthouse Lab in Alderley Park and Alex Alderton, Roberto Amato, Jeffrey Barrett, Sonia Goncalves, Ewan Harrison, David K. Jackson, Ian Johnston, Dominic Kwiatkowski, Cordelia Langford, John Sillitoe on behalf of the Wellcome Sanger Institute COVID-19 Surveillance Team |
| EPI_ISL_3437722 | Lighthouse Lab in Milton Keynes | Wellcome Sanger Institute for the COVID-19 Genomics UK (COG-UK) Consortium | The Lighthouse Lab in Milton Keynes and Alex Alderton, Roberto Amato, Jeffrey Barrett, Sonia Goncalves, Ewan Harrison, David K. Jackson, Ian Johnston, Dominic Kwiatkowski, Cordelia Langford, John Sillitoe on behalf of the Wellcome Sanger Institute COVID-19 Surveillance Team |
| EPI_ISL_3437723 | Lighthouse Lab in Alderley Park | Wellcome Sanger Institute for the COVID-19 Genomics UK (COG-UK) Consortium | Jacquelyn Wynn, Mairead Hyland, The Lighthouse Lab in Alderley Park and Alex Alderton, Roberto Amato, Jeffrey Barrett, Sonia Goncalves, Ewan Harrison, David K. Jackson, Ian Johnston, Dominic Kwiatkowski, Cordelia Langford, John Sillitoe on behalf of the Wellcome Sanger Institute COVID-19 Surveillance Team |
| EPI_ISL_3437724, EPI_ISL_3437725, EPI_ISL_3437726, EPI_ISL_3437727, EPI_ISL_3437728, EPI_ISL_3437729 | Lighthouse Lab in Milton Keynes | Wellcome Sanger Institute for the COVID-19 Genomics UK (COG-UK) Consortium | The Lighthouse Lab in Milton Keynes and Alex Alderton, Roberto Amato, Jeffrey Barrett, Sonia Goncalves, Ewan Harrison, David K. Jackson, Ian Johnston, Dominic Kwiatkowski, Cordelia Langford, John Sillitoe on behalf of the Wellcome Sanger Institute COVID-19 Surveillance Team |
| EPI_ISL_3437730 | Lighthouse Lab in Alderley Park | Wellcome Sanger Institute for the COVID-19 Genomics UK (COG-UK) Consortium | Jacquelyn Wynn, Mairead Hyland, The Lighthouse Lab in Alderley Park and Alex Alderton, Roberto Amato, Jeffrey Barrett, Sonia Goncalves, Ewan Harrison, David K. Jackson, Ian Johnston, Dominic Kwiatkowski, Cordelia Langford, John Sillitoe on behalf of the Wellcome Sanger Institute COVID-19 Surveillance Team |
| EPI_ISL_3437731, EPI_ISL_3437732, EPI_ISL_3437733 | Lighthouse Lab in Milton Keynes | Wellcome Sanger Institute for the COVID-19 Genomics UK (COG-UK) Consortium | The Lighthouse Lab in Milton Keynes and Alex Alderton, Roberto Amato, Jeffrey Barrett, Sonia Goncalves, Ewan Harrison, David K. Jackson, Ian Johnston, Dominic Kwiatkowski, Cordelia Langford, John Sillitoe on behalf of the Wellcome Sanger Institute COVID-19 Surveillance Team |
| EPI_ISL_3437734 | Lighthouse Lab in Alderley Park | Wellcome Sanger Institute for the COVID-19 Genomics UK (COG-UK) Consortium | Jacquelyn Wynn, Mairead Hyland, The Lighthouse Lab in Alderley Park and Alex Alderton, Roberto Amato, Jeffrey Barrett, Sonia Goncalves, Ewan Harrison, David K. Jackson, Ian Johnston, Dominic Kwiatkowski, Cordelia Langford, John Sillitoe on behalf of the Wellcome Sanger Institute COVID-19 Surveillance Team |
| EPI_ISL_3437735 | Lighthouse Lab in Milton Keynes | Wellcome Sanger Institute for the COVID-19 Genomics UK (COG-UK) Consortium | The Lighthouse Lab in Milton Keynes and Alex Alderton, Roberto Amato, Jeffrey Barrett, Sonia Goncalves, Ewan Harrison, David K. Jackson, Ian Johnston, Dominic Kwiatkowski, Cordelia Langford, John Sillitoe on behalf of the Wellcome Sanger Institute COVID-19 Surveillance Team |
| EPI_ISL_3437736, EPI_ISL_3437737 | Lighthouse Lab in Alderley Park | Wellcome Sanger Institute for the COVID-19 Genomics UK (COG-UK) Consortium | Jacquelyn Wynn, Mairead Hyland, The Lighthouse Lab in Alderley Park and Alex Alderton, Roberto Amato, Jeffrey Barrett, Sonia Goncalves, Ewan Harrison, David K. Jackson, Ian Johnston, Dominic Kwiatkowski, Cordelia Langford, John Sillitoe on behalf of the Wellcome Sanger Institute COVID-19 Surveillance Team |
| EPI_ISL_3437738, EPI_ISL_3437739 | Lighthouse Lab in Milton Keynes | Wellcome Sanger Institute for the COVID-19 Genomics UK (COG-UK) Consortium | The Lighthouse Lab in Milton Keynes and Alex Alderton, Roberto Amato, Jeffrey Barrett, Sonia Goncalves, Ewan Harrison, David K. Jackson, Ian Johnston, Dominic Kwiatkowski, Cordelia Langford, John Sillitoe on behalf of the Wellcome Sanger Institute COVID-19 Surveillance Team |
| EPI_ISL_3437740 | Lighthouse Lab in Alderley Park | Wellcome Sanger Institute for the COVID-19 Genomics UK (COG-UK) Consortium | Jacquelyn Wynn, Mairead Hyland, The Lighthouse Lab in Alderley Park and Alex Alderton, Roberto Amato, Jeffrey Barrett, Sonia Goncalves, Ewan Harrison, David K. Jackson, Ian Johnston, Dominic Kwiatkowski, Cordelia Langford, John Sillitoe on behalf of the Wellcome Sanger Institute COVID-19 Surveillance Team |
| EPI_ISL_3437741, EPI_ISL_3437742, EPI_ISL_3437743, | Lighthouse Lab in Milton Keynes | Wellcome Sanger Institute for the COVID-19 Genomics UK (COG-UK) Consortium | The Lighthouse Lab in Milton Keynes and Alex Alderton, Roberto Amato, Jeffrey Barrett, Sonia Goncalves, Ewan Harrison, David K. Jackson, Ian Johnston, Dominic Kwiatkowski, Cordelia Langford, John Sillitoe on behalf of the Wellcome Sanger Institute COVID-19 Surveillance Team |

| | | | |
|---|---|---|---|
| EPI_ISL_3437744, EPI_ISL_3437745, EPI_ISL_3437746, EPI_ISL_3437747, EPI_ISL_3437748 | | | |
| EPI_ISL_3437749 | Lighthouse Lab in Alderley Park | Wellcome Sanger Institute for the COVID-19 Genomics UK (COG-UK) Consortium | Jacquelyn Wynn, Mairead Hyland, The Lighthouse Lab in Alderley Park and Alex Alderton, Roberto Amato, Jeffrey Barrett, Sonia Goncalves, Ewan Harrison, David K. Jackson, Ian Johnston, Dominic Kwiatkowski, Cordelia Langford, John Sillitoe on behalf of the Wellcome Sanger Institute COVID-19 Surveillance Team |
| EPI_ISL_3437750 | Lighthouse Lab in Milton Keynes | Wellcome Sanger Institute for the COVID-19 Genomics UK (COG-UK) Consortium | The Lighthouse Lab in Milton Keynes and Alex Alderton, Roberto Amato, Jeffrey Barrett, Sonia Goncalves, Ewan Harrison, David K. Jackson, Ian Johnston, Dominic Kwiatkowski, Cordelia Langford, John Sillitoe on behalf of the Wellcome Sanger Institute COVID-19 Surveillance Team |
| EPI_ISL_3437751, EPI_ISL_3437752 | Lighthouse Lab in Alderley Park | Wellcome Sanger Institute for the COVID-19 Genomics UK (COG-UK) Consortium | Jacquelyn Wynn, Mairead Hyland, The Lighthouse Lab in Alderley Park and Alex Alderton, Roberto Amato, Jeffrey Barrett, Sonia Goncalves, Ewan Harrison, David K. Jackson, Ian Johnston, Dominic Kwiatkowski, Cordelia Langford, John Sillitoe on behalf of the Wellcome Sanger Institute COVID-19 Surveillance Team |
| EPI_ISL_3437753, EPI_ISL_3437754, EPI_ISL_3437755, EPI_ISL_3437756, EPI_ISL_3437757 | Lighthouse Lab in Milton Keynes | Wellcome Sanger Institute for the COVID-19 Genomics UK (COG-UK) Consortium | The Lighthouse Lab in Milton Keynes and Alex Alderton, Roberto Amato, Jeffrey Barrett, Sonia Goncalves, Ewan Harrison, David K. Jackson, Ian Johnston, Dominic Kwiatkowski, Cordelia Langford, John Sillitoe on behalf of the Wellcome Sanger Institute COVID-19 Surveillance Team |
| EPI_ISL_3437758 | Lighthouse Lab in Alderley Park | Wellcome Sanger Institute for the COVID-19 Genomics UK (COG-UK) Consortium | Jacquelyn Wynn, Mairead Hyland, The Lighthouse Lab in Alderley Park and Alex Alderton, Roberto Amato, Jeffrey Barrett, Sonia Goncalves, Ewan Harrison, David K. Jackson, Ian Johnston, Dominic Kwiatkowski, Cordelia Langford, John Sillitoe on behalf of the Wellcome Sanger Institute COVID-19 Surveillance Team |
| EPI_ISL_3437759, EPI_ISL_3437760 | Lighthouse Lab in Milton Keynes | Wellcome Sanger Institute for the COVID-19 Genomics UK (COG-UK) Consortium | The Lighthouse Lab in Milton Keynes and Alex Alderton, Roberto Amato, Jeffrey Barrett, Sonia Goncalves, Ewan Harrison, David K. Jackson, Ian Johnston, Dominic Kwiatkowski, Cordelia Langford, John Sillitoe on behalf of the Wellcome Sanger Institute COVID-19 Surveillance Team |
| EPI_ISL_3437761 | Lighthouse Lab in Alderley Park | Wellcome Sanger Institute for the COVID-19 Genomics UK (COG-UK) Consortium | Jacquelyn Wynn, Mairead Hyland, The Lighthouse Lab in Alderley Park and Alex Alderton, Roberto Amato, Jeffrey Barrett, Sonia Goncalves, Ewan Harrison, David K. Jackson, Ian Johnston, Dominic Kwiatkowski, Cordelia Langford, John Sillitoe on behalf of the Wellcome Sanger Institute COVID-19 Surveillance Team |
| EPI_ISL_3437762, EPI_ISL_3437763, EPI_ISL_3437764, EPI_ISL_3437765, EPI_ISL_3437766, EPI_ISL_3437767, EPI_ISL_3437768, EPI_ISL_3437769 | Lighthouse Lab in Milton Keynes | Wellcome Sanger Institute for the COVID-19 Genomics UK (COG-UK) Consortium | The Lighthouse Lab in Milton Keynes and Alex Alderton, Roberto Amato, Jeffrey Barrett, Sonia Goncalves, Ewan Harrison, David K. Jackson, Ian Johnston, Dominic Kwiatkowski, Cordelia Langford, John Sillitoe on behalf of the Wellcome Sanger Institute COVID-19 Surveillance Team |
| EPI_ISL_3437770 | Lighthouse Lab in Alderley Park | Wellcome Sanger Institute for the COVID-19 Genomics UK (COG-UK) Consortium | Jacquelyn Wynn, Mairead Hyland, The Lighthouse Lab in Alderley Park and Alex Alderton, Roberto Amato, Jeffrey Barrett, Sonia Goncalves, Ewan Harrison, David K. Jackson, Ian Johnston, Dominic Kwiatkowski, Cordelia Langford, John Sillitoe on behalf of the Wellcome Sanger Institute COVID-19 Surveillance Team |
| EPI_ISL_3437771, EPI_ISL_3437772, EPI_ISL_3437773, EPI_ISL_3437774 | Lighthouse Lab in Milton Keynes | Wellcome Sanger Institute for the COVID-19 Genomics UK (COG-UK) Consortium | The Lighthouse Lab in Milton Keynes and Alex Alderton, Roberto Amato, Jeffrey Barrett, Sonia Goncalves, Ewan Harrison, David K. Jackson, Ian Johnston, Dominic Kwiatkowski, Cordelia Langford, John Sillitoe on behalf of the Wellcome Sanger Institute COVID-19 Surveillance Team |
| EPI_ISL_3437775, EPI_ISL_3437776 | Lighthouse Lab in Alderley Park | Wellcome Sanger Institute for the COVID-19 Genomics UK (COG-UK) Consortium | Jacquelyn Wynn, Mairead Hyland, The Lighthouse Lab in Alderley Park and Alex Alderton, Roberto Amato, Jeffrey Barrett, Sonia Goncalves, Ewan Harrison, David K. Jackson, Ian Johnston, Dominic Kwiatkowski, Cordelia Langford, John Sillitoe on behalf of the Wellcome Sanger Institute COVID-19 Surveillance Team |
| EPI_ISL_3437777, EPI_ISL_3437778, EPI_ISL_3437779 | Lighthouse Lab in Milton Keynes | Wellcome Sanger Institute for the COVID-19 Genomics UK (COG-UK) Consortium | The Lighthouse Lab in Milton Keynes and Alex Alderton, Roberto Amato, Jeffrey Barrett, Sonia Goncalves, Ewan Harrison, David K. Jackson, Ian Johnston, Dominic Kwiatkowski, Cordelia Langford, John Sillitoe on behalf of the Wellcome Sanger Institute COVID-19 Surveillance Team |
| EPI_ISL_3437780 | Lighthouse Lab in Alderley Park | Wellcome Sanger Institute for the COVID-19 Genomics UK (COG-UK) Consortium | Jacquelyn Wynn, Mairead Hyland, The Lighthouse Lab in Alderley Park and Alex Alderton, Roberto Amato, Jeffrey Barrett, Sonia Goncalves, Ewan Harrison, David K. Jackson, Ian Johnston, Dominic Kwiatkowski, Cordelia Langford, John Sillitoe on behalf of the Wellcome Sanger Institute COVID-19 Surveillance Team |
| EPI_ISL_3437781, EPI_ISL_3437782, EPI_ISL_3437783, EPI_ISL_3437784, EPI_ISL_3437785, EPI_ISL_3437786, EPI_ISL_3437787 | Lighthouse Lab in Milton Keynes | Wellcome Sanger Institute for the COVID-19 Genomics UK (COG-UK) Consortium | The Lighthouse Lab in Milton Keynes and Alex Alderton, Roberto Amato, Jeffrey Barrett, Sonia Goncalves, Ewan Harrison, David K. Jackson, Ian Johnston, Dominic Kwiatkowski, Cordelia Langford, John Sillitoe on behalf of the Wellcome Sanger Institute COVID-19 Surveillance Team |
| EPI_ISL_3437788 | Lighthouse Lab in Alderley Park | Wellcome Sanger Institute for the COVID-19 Genomics UK (COG-UK) Consortium | Jacquelyn Wynn, Mairead Hyland, The Lighthouse Lab in Alderley Park and Alex Alderton, Roberto Amato, Jeffrey Barrett, Sonia Goncalves, Ewan Harrison, David K. Jackson, Ian Johnston, Dominic Kwiatkowski, Cordelia Langford, John Sillitoe on behalf of the Wellcome Sanger Institute COVID-19 Surveillance Team |
| EPI_ISL_3437789, EPI_ISL_3437790, EPI_ISL_3437791, EPI_ISL_3437792, EPI_ISL_3437793, EPI_ISL_3437794, EPI_ISL_3437795 | Lighthouse Lab in Milton Keynes | Wellcome Sanger Institute for the COVID-19 Genomics UK (COG-UK) Consortium | The Lighthouse Lab in Milton Keynes and Alex Alderton, Roberto Amato, Jeffrey Barrett, Sonia Goncalves, Ewan Harrison, David K. Jackson, Ian Johnston, Dominic Kwiatkowski, Cordelia Langford, John Sillitoe on behalf of the Wellcome Sanger Institute COVID-19 Surveillance Team |
| EPI_ISL_3437796, EPI_ISL_3437797 | Lighthouse Lab in Alderley Park | Wellcome Sanger Institute for the COVID-19 Genomics UK (COG-UK) Consortium | Jacquelyn Wynn, Mairead Hyland, The Lighthouse Lab in Alderley Park and Alex Alderton, Roberto Amato, Jeffrey Barrett, Sonia Goncalves, Ewan Harrison, David K. Jackson, Ian Johnston, Dominic Kwiatkowski, Cordelia Langford, John Sillitoe on behalf of the Wellcome Sanger Institute COVID-19 Surveillance Team |
| EPI_ISL_3437798, EPI_ISL_3437799, EPI_ISL_3437800, EPI_ISL_3437801, EPI_ISL_3437802, EPI_ISL_3437803, EPI_ISL_3437804, EPI_ISL_3437805, EPI_ISL_3437806, EPI_ISL_3437807, EPI_ISL_3437808, EPI_ISL_3437809, EPI_ISL_3437810 | | | |
| see above | Lighthouse Lab in Milton Keynes | Wellcome Sanger Institute for the COVID-19 Genomics UK (COG-UK) Consortium | The Lighthouse Lab in Milton Keynes and Alex Alderton, Roberto Amato, Jeffrey Barrett, Sonia Goncalves, Ewan Harrison, David K. Jackson, Ian Johnston, Dominic Kwiatkowski, Cordelia Langford, John Sillitoe on behalf of the Wellcome Sanger Institute COVID-19 Surveillance Team |
| EPI_ISL_3437811 | Lighthouse Lab in Glasgow | Wellcome Sanger Institute for the COVID-19 Genomics UK (COG-UK) Consortium | Harper VanSteenhouse, Yumi Kasai, David Gray, Carol Clugston, Anna Dominiczak and Alex Alderton, Roberto Amato, Jeffrey Barrett, Sonia Goncalves, Ewan Harrison, David K. Jackson, Ian Johnston, Dominic Kwiatkowski, Cordelia Langford, John Sillitoe on behalf of the Wellcome Sanger Institute COVID-19 Surveillance Team |
| EPI_ISL_3437812, EPI_ISL_3437813, EPI_ISL_3437814, EPI_ISL_3437815, EPI_ISL_3437816, EPI_ISL_3437817, EPI_ISL_3437818 | Lighthouse Lab in Milton Keynes | Wellcome Sanger Institute for the COVID-19 Genomics UK (COG-UK) Consortium | The Lighthouse Lab in Milton Keynes and Alex Alderton, Roberto Amato, Jeffrey Barrett, Sonia Goncalves, Ewan Harrison, David K. Jackson, Ian Johnston, Dominic Kwiatkowski, Cordelia Langford, John Sillitoe on behalf of the Wellcome Sanger Institute COVID-19 Surveillance Team |
| EPI_ISL_3437819 | Lighthouse Lab in Glasgow | Wellcome Sanger Institute for the COVID-19 Genomics UK (COG-UK) Consortium | Harper VanSteenhouse, Yumi Kasai, David Gray, Carol Clugston, Anna Dominiczak and Alex Alderton, Roberto Amato, Jeffrey Barrett, Sonia Goncalves, Ewan Harrison, David K. Jackson, Ian Johnston, Dominic Kwiatkowski, Cordelia Langford, John Sillitoe on behalf of the Wellcome Sanger Institute COVID-19 Surveillance Team |
| EPI_ISL_3437820, EPI_ISL_3437821 | Lighthouse Lab in Milton Keynes | Wellcome Sanger Institute for the COVID-19 Genomics UK (COG-UK) Consortium | The Lighthouse Lab in Milton Keynes and Alex Alderton, Roberto Amato, Jeffrey Barrett, Sonia Goncalves, Ewan Harrison, David K. Jackson, Ian Johnston, Dominic Kwiatkowski, Cordelia Langford, John Sillitoe on behalf of the Wellcome Sanger Institute COVID-19 Surveillance Team |
| EPI_ISL_3437822 | Lighthouse Lab in Glasgow | Wellcome Sanger Institute for the COVID-19 Genomics UK (COG-UK) Consortium | Harper VanSteenhouse, Yumi Kasai, David Gray, Carol Clugston, Anna Dominiczak and Alex Alderton, Roberto Amato, Jeffrey Barrett, Sonia Goncalves, Ewan Harrison, David K. Jackson, Ian Johnston, Dominic Kwiatkowski, Cordelia Langford, John Sillitoe on behalf of the Wellcome Sanger Institute COVID-19 Surveillance Team |
| EPI_ISL_3437823 | Lighthouse Lab in Milton Keynes | Wellcome Sanger Institute for the COVID-19 Genomics UK (COG-UK) Consortium | The Lighthouse Lab in Milton Keynes and Alex Alderton, Roberto Amato, Jeffrey Barrett, Sonia Goncalves, Ewan Harrison, David K. Jackson, Ian Johnston, Dominic Kwiatkowski, Cordelia Langford, John Sillitoe on behalf of the Wellcome Sanger Institute COVID-19 Surveillance Team |
| EPI_ISL_3437824, EPI_ISL_3437825, EPI_ISL_3437826 | Lighthouse Lab in Glasgow | Wellcome Sanger Institute for the COVID-19 Genomics UK (COG-UK) Consortium | Harper VanSteenhouse, Yumi Kasai, David Gray, Carol Clugston, Anna Dominiczak and Alex Alderton, Roberto Amato, Jeffrey Barrett, Sonia Goncalves, Ewan Harrison, David K. Jackson, Ian Johnston, Dominic Kwiatkowski, Cordelia Langford, John Sillitoe on behalf of the Wellcome Sanger Institute COVID-19 Surveillance Team |
| EPI_ISL_3437827, EPI_ISL_3437828, EPI_ISL_3437829, EPI_ISL_3437830, EPI_ISL_3437831, EPI_ISL_3437832, EPI_ISL_3437833, EPI_ISL_3437834, EPI_ISL_3437835, EPI_ISL_3437836, EPI_ISL_3437837, EPI_ISL_3437838 | | | |
| see above | Lighthouse Lab in Milton Keynes | Wellcome Sanger Institute for the COVID-19 Genomics UK (COG-UK) Consortium | The Lighthouse Lab in Milton Keynes and Alex Alderton, Roberto Amato, Jeffrey Barrett, Sonia Goncalves, Ewan Harrison, David K. Jackson, Ian Johnston, Dominic Kwiatkowski, Cordelia Langford, John Sillitoe on behalf of the Wellcome Sanger Institute COVID-19 Surveillance Team |
| EPI_ISL_3437839 | Lighthouse Lab in Glasgow | Wellcome Sanger Institute for the COVID-19 Genomics UK (COG-UK) Consortium | Harper VanSteenhouse, Yumi Kasai, David Gray, Carol Clugston, Anna Dominiczak and Alex Alderton, Roberto Amato, Jeffrey Barrett, Sonia Goncalves, Ewan Harrison, David K. Jackson, Ian Johnston, Dominic Kwiatkowski, Cordelia Langford, John Sillitoe on behalf of the Wellcome Sanger Institute COVID-19 Surveillance Team |
| EPI_ISL_3437840, EPI_ISL_3437841, EPI_ISL_3437842, EPI_ISL_3437843, EPI_ISL_3437844, EPI_ISL_3437845, EPI_ISL_3437846 | Lighthouse Lab in Milton Keynes | Wellcome Sanger Institute for the COVID-19 Genomics UK (COG-UK) Consortium | The Lighthouse Lab in Milton Keynes and Alex Alderton, Roberto Amato, Jeffrey Barrett, Sonia Goncalves, Ewan Harrison, David K. Jackson, Ian Johnston, Dominic Kwiatkowski, Cordelia Langford, John Sillitoe on behalf of the Wellcome Sanger Institute COVID-19 Surveillance Team |
| EPI_ISL_3437847 | Lighthouse Lab in Glasgow | Wellcome Sanger Institute for the COVID-19 Genomics UK (COG-UK) | Harper VanSteenhouse, Yumi Kasai, David Gray, Carol Clugston, Anna Dominiczak and Alex Alderton, Roberto Amato, Jeffrey Barrett, Sonia Goncalves, Ewan Harrison, David K. Jackson, Ian Johnston, Dominic Kwiatkowski, Cordelia Langford, |

| | | | |
|---|---|---|---|
| | | Consortium | John Sillitoe on behalf of the Wellcome Sanger Institute COVID-19 Surveillance Team |
| EPI_ISL_3437848, EPI_ISL_3437849 | Lighthouse Lab in Milton Keynes | Wellcome Sanger Institute for the COVID-19 Genomics UK (COG-UK) Consortium | The Lighthouse Lab in Milton Keynes and Alex Alderton, Roberto Amato, Jeffrey Barrett, Sonia Goncalves, Ewan Harrison, David K. Jackson, Ian Johnston, Dominic Kwiatkowski, Cordelia Langford, John Sillitoe on behalf of the Wellcome Sanger Institute COVID-19 Surveillance Team |
| EPI_ISL_3437850 | Lighthouse Lab in Glasgow | Wellcome Sanger Institute for the COVID-19 Genomics UK (COG-UK) Consortium | Harper VanSteenhouse, Yumi Kasai, David Gray, Carol Clugston, Anna Dominiczak and Alex Alderton, Roberto Amato, Jeffrey Barrett, Sonia Goncalves, Ewan Harrison, David K. Jackson, Ian Johnston, Dominic Kwiatkowski, Cordelia Langford, John Sillitoe on behalf of the Wellcome Sanger Institute COVID-19 Surveillance Team |
| EPI_ISL_3437851 | Lighthouse Lab in Milton Keynes | Wellcome Sanger Institute for the COVID-19 Genomics UK (COG-UK) Consortium | The Lighthouse Lab in Milton Keynes and Alex Alderton, Roberto Amato, Jeffrey Barrett, Sonia Goncalves, Ewan Harrison, David K. Jackson, Ian Johnston, Dominic Kwiatkowski, Cordelia Langford, John Sillitoe on behalf of the Wellcome Sanger Institute COVID-19 Surveillance Team |
| EPI_ISL_3437852 | Lighthouse Lab in Glasgow | Wellcome Sanger Institute for the COVID-19 Genomics UK (COG-UK) Consortium | Harper VanSteenhouse, Yumi Kasai, David Gray, Carol Clugston, Anna Dominiczak and Alex Alderton, Roberto Amato, Jeffrey Barrett, Sonia Goncalves, Ewan Harrison, David K. Jackson, Ian Johnston, Dominic Kwiatkowski, Cordelia Langford, John Sillitoe on behalf of the Wellcome Sanger Institute COVID-19 Surveillance Team |
| EPI_ISL_3437853, EPI_ISL_3437854, EPI_ISL_3437855 | Lighthouse Lab in Milton Keynes | Wellcome Sanger Institute for the COVID-19 Genomics UK (COG-UK) Consortium | The Lighthouse Lab in Milton Keynes and Alex Alderton, Roberto Amato, Jeffrey Barrett, Sonia Goncalves, Ewan Harrison, David K. Jackson, Ian Johnston, Dominic Kwiatkowski, Cordelia Langford, John Sillitoe on behalf of the Wellcome Sanger Institute COVID-19 Surveillance Team |
| EPI_ISL_3437856 | Lighthouse Lab in Glasgow | Wellcome Sanger Institute for the COVID-19 Genomics UK (COG-UK) Consortium | Harper VanSteenhouse, Yumi Kasai, David Gray, Carol Clugston, Anna Dominiczak and Alex Alderton, Roberto Amato, Jeffrey Barrett, Sonia Goncalves, Ewan Harrison, David K. Jackson, Ian Johnston, Dominic Kwiatkowski, Cordelia Langford, John Sillitoe on behalf of the Wellcome Sanger Institute COVID-19 Surveillance Team |
| EPI_ISL_3437857, EPI_ISL_3437858, EPI_ISL_3437859, EPI_ISL_3437860, EPI_ISL_3437861 | Lighthouse Lab in Milton Keynes | Wellcome Sanger Institute for the COVID-19 Genomics UK (COG-UK) Consortium | The Lighthouse Lab in Milton Keynes and Alex Alderton, Roberto Amato, Jeffrey Barrett, Sonia Goncalves, Ewan Harrison, David K. Jackson, Ian Johnston, Dominic Kwiatkowski, Cordelia Langford, John Sillitoe on behalf of the Wellcome Sanger Institute COVID-19 Surveillance Team |
| EPI_ISL_3437862 | Lighthouse Lab in Glasgow | Wellcome Sanger Institute for the COVID-19 Genomics UK (COG-UK) Consortium | Harper VanSteenhouse, Yumi Kasai, David Gray, Carol Clugston, Anna Dominiczak and Alex Alderton, Roberto Amato, Jeffrey Barrett, Sonia Goncalves, Ewan Harrison, David K. Jackson, Ian Johnston, Dominic Kwiatkowski, Cordelia Langford, John Sillitoe on behalf of the Wellcome Sanger Institute COVID-19 Surveillance Team |
| EPI_ISL_3437863, EPI_ISL_3437864 | Lighthouse Lab in Milton Keynes | Wellcome Sanger Institute for the COVID-19 Genomics UK (COG-UK) Consortium | The Lighthouse Lab in Milton Keynes and Alex Alderton, Roberto Amato, Jeffrey Barrett, Sonia Goncalves, Ewan Harrison, David K. Jackson, Ian Johnston, Dominic Kwiatkowski, Cordelia Langford, John Sillitoe on behalf of the Wellcome Sanger Institute COVID-19 Surveillance Team |
| EPI_ISL_3437865 | Lighthouse Lab in Glasgow | Wellcome Sanger Institute for the COVID-19 Genomics UK (COG-UK) Consortium | Harper VanSteenhouse, Yumi Kasai, David Gray, Carol Clugston, Anna Dominiczak and Alex Alderton, Roberto Amato, Jeffrey Barrett, Sonia Goncalves, Ewan Harrison, David K. Jackson, Ian Johnston, Dominic Kwiatkowski, Cordelia Langford, John Sillitoe on behalf of the Wellcome Sanger Institute COVID-19 Surveillance Team |
| EPI_ISL_3437866, EPI_ISL_3437867, EPI_ISL_3437868, EPI_ISL_3437869, EPI_ISL_3437870, EPI_ISL_3437871, EPI_ISL_3437872, EPI_ISL_3437873, EPI_ISL_3437874, EPI_ISL_3437875, EPI_ISL_3437876, EPI_ISL_3437877, EPI_ISL_3437878, EPI_ISL_3437879, EPI_ISL_3437880, EPI_ISL_3437881, EPI_ISL_3437882, EPI_ISL_3437883, EPI_ISL_3437884, EPI_ISL_3437885, EPI_ISL_3437886, EPI_ISL_3437887 | | | |
| see above | Lighthouse Lab in Milton Keynes | Wellcome Sanger Institute for the COVID-19 Genomics UK (COG-UK) Consortium | The Lighthouse Lab in Milton Keynes and Alex Alderton, Roberto Amato, Jeffrey Barrett, Sonia Goncalves, Ewan Harrison, David K. Jackson, Ian Johnston, Dominic Kwiatkowski, Cordelia Langford, John Sillitoe on behalf of the Wellcome Sanger Institute COVID-19 Surveillance Team |
| EPI_ISL_3437888 | Lighthouse Lab in Glasgow | Wellcome Sanger Institute for the COVID-19 Genomics UK (COG-UK) Consortium | Harper VanSteenhouse, Yumi Kasai, David Gray, Carol Clugston, Anna Dominiczak and Alex Alderton, Roberto Amato, Jeffrey Barrett, Sonia Goncalves, Ewan Harrison, David K. Jackson, Ian Johnston, Dominic Kwiatkowski, Cordelia Langford, John Sillitoe on behalf of the Wellcome Sanger Institute COVID-19 Surveillance Team |
| EPI_ISL_3437889, EPI_ISL_3437890, EPI_ISL_3437891 | Lighthouse Lab in Milton Keynes | Wellcome Sanger Institute for the COVID-19 Genomics UK (COG-UK) Consortium | The Lighthouse Lab in Milton Keynes and Alex Alderton, Roberto Amato, Jeffrey Barrett, Sonia Goncalves, Ewan Harrison, David K. Jackson, Ian Johnston, Dominic Kwiatkowski, Cordelia Langford, John Sillitoe on behalf of the Wellcome Sanger Institute COVID-19 Surveillance Team |
| EPI_ISL_3437892 | Lighthouse Lab in Glasgow | Wellcome Sanger Institute for the COVID-19 Genomics UK (COG-UK) Consortium | Harper VanSteenhouse, Yumi Kasai, David Gray, Carol Clugston, Anna Dominiczak and Alex Alderton, Roberto Amato, Jeffrey Barrett, Sonia Goncalves, Ewan Harrison, David K. Jackson, Ian Johnston, Dominic Kwiatkowski, Cordelia Langford, John Sillitoe on behalf of the Wellcome Sanger Institute COVID-19 Surveillance Team |
| EPI_ISL_3437893 | Lighthouse Lab in Milton Keynes | Wellcome Sanger Institute for the COVID-19 Genomics UK (COG-UK) Consortium | The Lighthouse Lab in Milton Keynes and Alex Alderton, Roberto Amato, Jeffrey Barrett, Sonia Goncalves, Ewan Harrison, David K. Jackson, Ian Johnston, Dominic Kwiatkowski, Cordelia Langford, John Sillitoe on behalf of the Wellcome Sanger Institute COVID-19 Surveillance Team |
| EPI_ISL_3437894 | Lighthouse Lab in Glasgow | Wellcome Sanger Institute for the COVID-19 Genomics UK (COG-UK) Consortium | Harper VanSteenhouse, Yumi Kasai, David Gray, Carol Clugston, Anna Dominiczak and Alex Alderton, Roberto Amato, Jeffrey Barrett, Sonia Goncalves, Ewan Harrison, David K. Jackson, Ian Johnston, Dominic Kwiatkowski, Cordelia Langford, John Sillitoe on behalf of the Wellcome Sanger Institute COVID-19 Surveillance Team |
| EPI_ISL_3437895 | Lighthouse Lab in Milton Keynes | Wellcome Sanger Institute for the COVID-19 Genomics UK (COG-UK) Consortium | The Lighthouse Lab in Milton Keynes and Alex Alderton, Roberto Amato, Jeffrey Barrett, Sonia Goncalves, Ewan Harrison, David K. Jackson, Ian Johnston, Dominic Kwiatkowski, Cordelia Langford, John Sillitoe on behalf of the Wellcome Sanger Institute COVID-19 Surveillance Team |
| EPI_ISL_3437896 | Lighthouse Lab in Glasgow | Wellcome Sanger Institute for the COVID-19 Genomics UK (COG-UK) Consortium | Harper VanSteenhouse, Yumi Kasai, David Gray, Carol Clugston, Anna Dominiczak and Alex Alderton, Roberto Amato, Jeffrey Barrett, Sonia Goncalves, Ewan Harrison, David K. Jackson, Ian Johnston, Dominic Kwiatkowski, Cordelia Langford, John Sillitoe on behalf of the Wellcome Sanger Institute COVID-19 Surveillance Team |
| EPI_ISL_3437897, EPI_ISL_3437898, EPI_ISL_3437899, EPI_ISL_3437900, EPI_ISL_3437901, EPI_ISL_3437902 | Lighthouse Lab in Milton Keynes | Wellcome Sanger Institute for the COVID-19 Genomics UK (COG-UK) Consortium | The Lighthouse Lab in Milton Keynes and Alex Alderton, Roberto Amato, Jeffrey Barrett, Sonia Goncalves, Ewan Harrison, David K. Jackson, Ian Johnston, Dominic Kwiatkowski, Cordelia Langford, John Sillitoe on behalf of the Wellcome Sanger Institute COVID-19 Surveillance Team |
| EPI_ISL_3437903 | Lighthouse Lab in Glasgow | Wellcome Sanger Institute for the COVID-19 Genomics UK (COG-UK) Consortium | Harper VanSteenhouse, Yumi Kasai, David Gray, Carol Clugston, Anna Dominiczak and Alex Alderton, Roberto Amato, Jeffrey Barrett, Sonia Goncalves, Ewan Harrison, David K. Jackson, Ian Johnston, Dominic Kwiatkowski, Cordelia Langford, John Sillitoe on behalf of the Wellcome Sanger Institute COVID-19 Surveillance Team |
| EPI_ISL_3437904, EPI_ISL_3437905, EPI_ISL_3437906, EPI_ISL_3437907, EPI_ISL_3437908, EPI_ISL_3437909, EPI_ISL_3437910, EPI_ISL_3437911 | Lighthouse Lab in Milton Keynes | Wellcome Sanger Institute for the COVID-19 Genomics UK (COG-UK) Consortium | The Lighthouse Lab in Milton Keynes and Alex Alderton, Roberto Amato, Jeffrey Barrett, Sonia Goncalves, Ewan Harrison, David K. Jackson, Ian Johnston, Dominic Kwiatkowski, Cordelia Langford, John Sillitoe on behalf of the Wellcome Sanger Institute COVID-19 Surveillance Team |
| EPI_ISL_3437912 | Lighthouse Lab in Glasgow | Wellcome Sanger Institute for the COVID-19 Genomics UK (COG-UK) Consortium | Harper VanSteenhouse, Yumi Kasai, David Gray, Carol Clugston, Anna Dominiczak and Alex Alderton, Roberto Amato, Jeffrey Barrett, Sonia Goncalves, Ewan Harrison, David K. Jackson, Ian Johnston, Dominic Kwiatkowski, Cordelia Langford, John Sillitoe on behalf of the Wellcome Sanger Institute COVID-19 Surveillance Team |
| EPI_ISL_3437913, EPI_ISL_3437914, EPI_ISL_3437915 | Lighthouse Lab in Milton Keynes | Wellcome Sanger Institute for the COVID-19 Genomics UK (COG-UK) Consortium | The Lighthouse Lab in Milton Keynes and Alex Alderton, Roberto Amato, Jeffrey Barrett, Sonia Goncalves, Ewan Harrison, David K. Jackson, Ian Johnston, Dominic Kwiatkowski, Cordelia Langford, John Sillitoe on behalf of the Wellcome Sanger Institute COVID-19 Surveillance Team |
| EPI_ISL_3437916 | Lighthouse Lab in Glasgow | Wellcome Sanger Institute for the COVID-19 Genomics UK (COG-UK) Consortium | Harper VanSteenhouse, Yumi Kasai, David Gray, Carol Clugston, Anna Dominiczak and Alex Alderton, Roberto Amato, Jeffrey Barrett, Sonia Goncalves, Ewan Harrison, David K. Jackson, Ian Johnston, Dominic Kwiatkowski, Cordelia Langford, John Sillitoe on behalf of the Wellcome Sanger Institute COVID-19 Surveillance Team |
| EPI_ISL_3437917, EPI_ISL_3437918, EPI_ISL_3437919, EPI_ISL_3437920, EPI_ISL_3437921 | Lighthouse Lab in Milton Keynes | Wellcome Sanger Institute for the COVID-19 Genomics UK (COG-UK) Consortium | The Lighthouse Lab in Milton Keynes and Alex Alderton, Roberto Amato, Jeffrey Barrett, Sonia Goncalves, Ewan Harrison, David K. Jackson, Ian Johnston, Dominic Kwiatkowski, Cordelia Langford, John Sillitoe on behalf of the Wellcome Sanger Institute COVID-19 Surveillance Team |
| EPI_ISL_3437922 | Lighthouse Lab in Glasgow | Wellcome Sanger Institute for the COVID-19 Genomics UK (COG-UK) Consortium | Harper VanSteenhouse, Yumi Kasai, David Gray, Carol Clugston, Anna Dominiczak and Alex Alderton, Roberto Amato, Jeffrey Barrett, Sonia Goncalves, Ewan Harrison, David K. Jackson, Ian Johnston, Dominic Kwiatkowski, Cordelia Langford, John Sillitoe on behalf of the Wellcome Sanger Institute COVID-19 Surveillance Team |
| EPI_ISL_3437923, EPI_ISL_3437924, EPI_ISL_3437925, EPI_ISL_3437926, EPI_ISL_3437927, EPI_ISL_3437928 | Lighthouse Lab in Milton Keynes | Wellcome Sanger Institute for the COVID-19 Genomics UK (COG-UK) Consortium | The Lighthouse Lab in Milton Keynes and Alex Alderton, Roberto Amato, Jeffrey Barrett, Sonia Goncalves, Ewan Harrison, David K. Jackson, Ian Johnston, Dominic Kwiatkowski, Cordelia Langford, John Sillitoe on behalf of the Wellcome Sanger Institute COVID-19 Surveillance Team |
| EPI_ISL_3437929 | Lighthouse Lab in Glasgow | Wellcome Sanger Institute for the COVID-19 Genomics UK (COG-UK) Consortium | Harper VanSteenhouse, Yumi Kasai, David Gray, Carol Clugston, Anna Dominiczak and Alex Alderton, Roberto Amato, Jeffrey Barrett, Sonia Goncalves, Ewan Harrison, David K. Jackson, Ian Johnston, Dominic Kwiatkowski, Cordelia Langford, John Sillitoe on behalf of the Wellcome Sanger Institute COVID-19 Surveillance Team |
| EPI_ISL_3437930, EPI_ISL_3437931 | Lighthouse Lab in Milton Keynes | Wellcome Sanger Institute for the COVID-19 Genomics UK (COG-UK) Consortium | The Lighthouse Lab in Milton Keynes and Alex Alderton, Roberto Amato, Jeffrey Barrett, Sonia Goncalves, Ewan Harrison, David K. Jackson, Ian Johnston, Dominic Kwiatkowski, Cordelia Langford, John Sillitoe on behalf of the Wellcome Sanger Institute COVID-19 Surveillance Team |
| EPI_ISL_3437932 | Lighthouse Lab in Glasgow | Wellcome Sanger Institute for the COVID-19 Genomics UK (COG-UK) Consortium | Harper VanSteenhouse, Yumi Kasai, David Gray, Carol Clugston, Anna Dominiczak and Alex Alderton, Roberto Amato, Jeffrey Barrett, Sonia Goncalves, Ewan Harrison, David K. Jackson, Ian Johnston, Dominic Kwiatkowski, Cordelia Langford, John Sillitoe on behalf of the Wellcome Sanger Institute COVID-19 Surveillance Team |
| EPI_ISL_3437933, EPI_ISL_3437934, EPI_ISL_3437935, EPI_ISL_3437936 | Lighthouse Lab in Milton Keynes | Wellcome Sanger Institute for the COVID-19 Genomics UK (COG-UK) Consortium | The Lighthouse Lab in Milton Keynes and Alex Alderton, Roberto Amato, Jeffrey Barrett, Sonia Goncalves, Ewan Harrison, David K. Jackson, Ian Johnston, Dominic Kwiatkowski, Cordelia Langford, John Sillitoe on behalf of the Wellcome Sanger Institute COVID-19 Surveillance Team |
| EPI_ISL_3437937, EPI_ISL_3437938, EPI_ISL_3437939, EPI_ISL_3437940 | Lighthouse Lab in Glasgow | Wellcome Sanger Institute for the COVID-19 Genomics UK (COG-UK) Consortium | Harper VanSteenhouse, Yumi Kasai, David Gray, Carol Clugston, Anna Dominiczak and Alex Alderton, Roberto Amato, Jeffrey Barrett, Sonia Goncalves, Ewan Harrison, David K. Jackson, Ian Johnston, Dominic Kwiatkowski, Cordelia Langford, John Sillitoe on behalf of the Wellcome Sanger Institute COVID-19 Surveillance Team |
| EPI_ISL_3437941, | Lighthouse Lab in Milton Keynes | Wellcome Sanger Institute for the COVID-19 Genomics UK (COG-UK) | The Lighthouse Lab in Milton Keynes and Alex Alderton, Roberto Amato, Jeffrey Barrett, Sonia Goncalves, Ewan Harrison, David K. Jackson, Ian Johnston, Dominic Kwiatkowski, Cordelia Langford, John Sillitoe on behalf of the Wellcome Sanger |

| | | | |
|---|---|---|---|
| EPI_ISL_3437942, EPI_ISL_3437943, EPI_ISL_3437944 | | Consortium | Institute COVID-19 Surveillance Team |
| EPI_ISL_3437945 | Lighthouse Lab in Glasgow | Wellcome Sanger Institute for the COVID-19 Genomics UK (COG-UK) Consortium | Harper VanSteenhouse, Yumi Kasai, David Gray, Carol Clugston, Anna Dominiczak and Alex Alderton, Roberto Amato, Jeffrey Barrett, Sonia Goncalves, Ewan Harrison, David K. Jackson, Ian Johnston, Dominic Kwiatkowski, Cordelia Langford, John Sillitoe on behalf of the Wellcome Sanger Institute COVID-19 Surveillance Team |
| EPI_ISL_3437946, EPI_ISL_3437947 | Lighthouse Lab in Milton Keynes | Wellcome Sanger Institute for the COVID-19 Genomics UK (COG-UK) Consortium | The Lighthouse Lab in Milton Keynes and Alex Alderton, Roberto Amato, Jeffrey Barrett, Sonia Goncalves, Ewan Harrison, David K. Jackson, Ian Johnston, Dominic Kwiatkowski, Cordelia Langford, John Sillitoe on behalf of the Wellcome Sanger Institute COVID-19 Surveillance Team |
| EPI_ISL_3437948 | Lighthouse Lab in Glasgow | Wellcome Sanger Institute for the COVID-19 Genomics UK (COG-UK) Consortium | Harper VanSteenhouse, Yumi Kasai, David Gray, Carol Clugston, Anna Dominiczak and Alex Alderton, Roberto Amato, Jeffrey Barrett, Sonia Goncalves, Ewan Harrison, David K. Jackson, Ian Johnston, Dominic Kwiatkowski, Cordelia Langford, John Sillitoe on behalf of the Wellcome Sanger Institute COVID-19 Surveillance Team |
| EPI_ISL_3437949, EPI_ISL_3437950, EPI_ISL_3437951 | Lighthouse Lab in Milton Keynes | Wellcome Sanger Institute for the COVID-19 Genomics UK (COG-UK) Consortium | The Lighthouse Lab in Milton Keynes and Alex Alderton, Roberto Amato, Jeffrey Barrett, Sonia Goncalves, Ewan Harrison, David K. Jackson, Ian Johnston, Dominic Kwiatkowski, Cordelia Langford, John Sillitoe on behalf of the Wellcome Sanger Institute COVID-19 Surveillance Team |
| EPI_ISL_3437952 | Lighthouse Lab in Glasgow | Wellcome Sanger Institute for the COVID-19 Genomics UK (COG-UK) Consortium | Harper VanSteenhouse, Yumi Kasai, David Gray, Carol Clugston, Anna Dominiczak and Alex Alderton, Roberto Amato, Jeffrey Barrett, Sonia Goncalves, Ewan Harrison, David K. Jackson, Ian Johnston, Dominic Kwiatkowski, Cordelia Langford, John Sillitoe on behalf of the Wellcome Sanger Institute COVID-19 Surveillance Team |
| EPI_ISL_3437953, EPI_ISL_3437954, EPI_ISL_3437955, EPI_ISL_3437956 | Lighthouse Lab in Milton Keynes | Wellcome Sanger Institute for the COVID-19 Genomics UK (COG-UK) Consortium | The Lighthouse Lab in Milton Keynes and Alex Alderton, Roberto Amato, Jeffrey Barrett, Sonia Goncalves, Ewan Harrison, David K. Jackson, Ian Johnston, Dominic Kwiatkowski, Cordelia Langford, John Sillitoe on behalf of the Wellcome Sanger Institute COVID-19 Surveillance Team |
| EPI_ISL_3437957 | Lighthouse Lab in Glasgow | Wellcome Sanger Institute for the COVID-19 Genomics UK (COG-UK) Consortium | Harper VanSteenhouse, Yumi Kasai, David Gray, Carol Clugston, Anna Dominiczak and Alex Alderton, Roberto Amato, Jeffrey Barrett, Sonia Goncalves, Ewan Harrison, David K. Jackson, Ian Johnston, Dominic Kwiatkowski, Cordelia Langford, John Sillitoe on behalf of the Wellcome Sanger Institute COVID-19 Surveillance Team |
| EPI_ISL_3437958, EPI_ISL_3437959 | Lighthouse Lab in Milton Keynes | Wellcome Sanger Institute for the COVID-19 Genomics UK (COG-UK) Consortium | The Lighthouse Lab in Milton Keynes and Alex Alderton, Roberto Amato, Jeffrey Barrett, Sonia Goncalves, Ewan Harrison, David K. Jackson, Ian Johnston, Dominic Kwiatkowski, Cordelia Langford, John Sillitoe on behalf of the Wellcome Sanger Institute COVID-19 Surveillance Team |
| EPI_ISL_3437960 | Lighthouse Lab in Glasgow | Wellcome Sanger Institute for the COVID-19 Genomics UK (COG-UK) Consortium | Harper VanSteenhouse, Yumi Kasai, David Gray, Carol Clugston, Anna Dominiczak and Alex Alderton, Roberto Amato, Jeffrey Barrett, Sonia Goncalves, Ewan Harrison, David K. Jackson, Ian Johnston, Dominic Kwiatkowski, Cordelia Langford, John Sillitoe on behalf of the Wellcome Sanger Institute COVID-19 Surveillance Team |
| EPI_ISL_3437961, EPI_ISL_3437962, EPI_ISL_3437963, EPI_ISL_3437964 | Lighthouse Lab in Milton Keynes | Wellcome Sanger Institute for the COVID-19 Genomics UK (COG-UK) Consortium | The Lighthouse Lab in Milton Keynes and Alex Alderton, Roberto Amato, Jeffrey Barrett, Sonia Goncalves, Ewan Harrison, David K. Jackson, Ian Johnston, Dominic Kwiatkowski, Cordelia Langford, John Sillitoe on behalf of the Wellcome Sanger Institute COVID-19 Surveillance Team |
| EPI_ISL_3437965 | Lighthouse Lab in Glasgow | Wellcome Sanger Institute for the COVID-19 Genomics UK (COG-UK) Consortium | Harper VanSteenhouse, Yumi Kasai, David Gray, Carol Clugston, Anna Dominiczak and Alex Alderton, Roberto Amato, Jeffrey Barrett, Sonia Goncalves, Ewan Harrison, David K. Jackson, Ian Johnston, Dominic Kwiatkowski, Cordelia Langford, John Sillitoe on behalf of the Wellcome Sanger Institute COVID-19 Surveillance Team |
| EPI_ISL_3437966, EPI_ISL_3437967, EPI_ISL_3437968, EPI_ISL_3437969, EPI_ISL_3437970, EPI_ISL_3437971, EPI_ISL_3437972, EPI_ISL_3437973, EPI_ISL_3437974, EPI_ISL_3437975, EPI_ISL_3437976, EPI_ISL_3437977, EPI_ISL_3437978, EPI_ISL_3437979, EPI_ISL_3437980 | | | |
| see above | Lighthouse Lab in Milton Keynes | Wellcome Sanger Institute for the COVID-19 Genomics UK (COG-UK) Consortium | The Lighthouse Lab in Milton Keynes and Alex Alderton, Roberto Amato, Jeffrey Barrett, Sonia Goncalves, Ewan Harrison, David K. Jackson, Ian Johnston, Dominic Kwiatkowski, Cordelia Langford, John Sillitoe on behalf of the Wellcome Sanger Institute COVID-19 Surveillance Team |
| EPI_ISL_3437981 | Lighthouse Lab in Glasgow | Wellcome Sanger Institute for the COVID-19 Genomics UK (COG-UK) Consortium | Harper VanSteenhouse, Yumi Kasai, David Gray, Carol Clugston, Anna Dominiczak and Alex Alderton, Roberto Amato, Jeffrey Barrett, Sonia Goncalves, Ewan Harrison, David K. Jackson, Ian Johnston, Dominic Kwiatkowski, Cordelia Langford, John Sillitoe on behalf of the Wellcome Sanger Institute COVID-19 Surveillance Team |
| EPI_ISL_3437982, EPI_ISL_3437983, EPI_ISL_3437984, EPI_ISL_3437985, EPI_ISL_3437986, EPI_ISL_3437987, EPI_ISL_3437988, EPI_ISL_3437989, EPI_ISL_3437990, EPI_ISL_3437991, EPI_ISL_3437992, EPI_ISL_3437993 | | | |
| see above | Lighthouse Lab in Milton Keynes | Wellcome Sanger Institute for the COVID-19 Genomics UK (COG-UK) Consortium | The Lighthouse Lab in Milton Keynes and Alex Alderton, Roberto Amato, Jeffrey Barrett, Sonia Goncalves, Ewan Harrison, David K. Jackson, Ian Johnston, Dominic Kwiatkowski, Cordelia Langford, John Sillitoe on behalf of the Wellcome Sanger Institute COVID-19 Surveillance Team |
| EPI_ISL_3437994 | Lighthouse Lab in Glasgow | Wellcome Sanger Institute for the COVID-19 Genomics UK (COG-UK) Consortium | Harper VanSteenhouse, Yumi Kasai, David Gray, Carol Clugston, Anna Dominiczak and Alex Alderton, Roberto Amato, Jeffrey Barrett, Sonia Goncalves, Ewan Harrison, David K. Jackson, Ian Johnston, Dominic Kwiatkowski, Cordelia Langford, John Sillitoe on behalf of the Wellcome Sanger Institute COVID-19 Surveillance Team |
| EPI_ISL_3437995, EPI_ISL_3437996, EPI_ISL_3437997, EPI_ISL_3437998, EPI_ISL_3437999, EPI_ISL_3438000, EPI_ISL_3438001 | Lighthouse Lab in Milton Keynes | Wellcome Sanger Institute for the COVID-19 Genomics UK (COG-UK) Consortium | The Lighthouse Lab in Milton Keynes and Alex Alderton, Roberto Amato, Jeffrey Barrett, Sonia Goncalves, Ewan Harrison, David K. Jackson, Ian Johnston, Dominic Kwiatkowski, Cordelia Langford, John Sillitoe on behalf of the Wellcome Sanger Institute COVID-19 Surveillance Team |
| EPI_ISL_3438002, EPI_ISL_3438003 | Lighthouse Lab in Glasgow | Wellcome Sanger Institute for the COVID-19 Genomics UK (COG-UK) Consortium | Harper VanSteenhouse, Yumi Kasai, David Gray, Carol Clugston, Anna Dominiczak and Alex Alderton, Roberto Amato, Jeffrey Barrett, Sonia Goncalves, Ewan Harrison, David K. Jackson, Ian Johnston, Dominic Kwiatkowski, Cordelia Langford, John Sillitoe on behalf of the Wellcome Sanger Institute COVID-19 Surveillance Team |
| EPI_ISL_3438004, EPI_ISL_3438005, EPI_ISL_3438006 | Lighthouse Lab in Milton Keynes | Wellcome Sanger Institute for the COVID-19 Genomics UK (COG-UK) Consortium | The Lighthouse Lab in Milton Keynes and Alex Alderton, Roberto Amato, Jeffrey Barrett, Sonia Goncalves, Ewan Harrison, David K. Jackson, Ian Johnston, Dominic Kwiatkowski, Cordelia Langford, John Sillitoe on behalf of the Wellcome Sanger Institute COVID-19 Surveillance Team |
| EPI_ISL_3438007 | Lighthouse Lab in Glasgow | Wellcome Sanger Institute for the COVID-19 Genomics UK (COG-UK) Consortium | Harper VanSteenhouse, Yumi Kasai, David Gray, Carol Clugston, Anna Dominiczak and Alex Alderton, Roberto Amato, Jeffrey Barrett, Sonia Goncalves, Ewan Harrison, David K. Jackson, Ian Johnston, Dominic Kwiatkowski, Cordelia Langford, John Sillitoe on behalf of the Wellcome Sanger Institute COVID-19 Surveillance Team |
| EPI_ISL_3438008, EPI_ISL_3438009, EPI_ISL_3438010 | Lighthouse Lab in Milton Keynes | Wellcome Sanger Institute for the COVID-19 Genomics UK (COG-UK) Consortium | The Lighthouse Lab in Milton Keynes and Alex Alderton, Roberto Amato, Jeffrey Barrett, Sonia Goncalves, Ewan Harrison, David K. Jackson, Ian Johnston, Dominic Kwiatkowski, Cordelia Langford, John Sillitoe on behalf of the Wellcome Sanger Institute COVID-19 Surveillance Team |
| EPI_ISL_3438011 | Lighthouse Lab in Glasgow | Wellcome Sanger Institute for the COVID-19 Genomics UK (COG-UK) Consortium | Harper VanSteenhouse, Yumi Kasai, David Gray, Carol Clugston, Anna Dominiczak and Alex Alderton, Roberto Amato, Jeffrey Barrett, Sonia Goncalves, Ewan Harrison, David K. Jackson, Ian Johnston, Dominic Kwiatkowski, Cordelia Langford, John Sillitoe on behalf of the Wellcome Sanger Institute COVID-19 Surveillance Team |
| EPI_ISL_3438012 | Lighthouse Lab in Milton Keynes | Wellcome Sanger Institute for the COVID-19 Genomics UK (COG-UK) Consortium | The Lighthouse Lab in Milton Keynes and Alex Alderton, Roberto Amato, Jeffrey Barrett, Sonia Goncalves, Ewan Harrison, David K. Jackson, Ian Johnston, Dominic Kwiatkowski, Cordelia Langford, John Sillitoe on behalf of the Wellcome Sanger Institute COVID-19 Surveillance Team |
| EPI_ISL_3438013, EPI_ISL_3438014 | Lighthouse Lab in Glasgow | Wellcome Sanger Institute for the COVID-19 Genomics UK (COG-UK) Consortium | Harper VanSteenhouse, Yumi Kasai, David Gray, Carol Clugston, Anna Dominiczak and Alex Alderton, Roberto Amato, Jeffrey Barrett, Sonia Goncalves, Ewan Harrison, David K. Jackson, Ian Johnston, Dominic Kwiatkowski, Cordelia Langford, John Sillitoe on behalf of the Wellcome Sanger Institute COVID-19 Surveillance Team |
| EPI_ISL_3438015, EPI_ISL_3438016, EPI_ISL_3438017, EPI_ISL_3438018, EPI_ISL_3438019, EPI_ISL_3438020, EPI_ISL_3438021, EPI_ISL_3438022, EPI_ISL_3438023 | Lighthouse Lab in Milton Keynes | Wellcome Sanger Institute for the COVID-19 Genomics UK (COG-UK) Consortium | The Lighthouse Lab in Milton Keynes and Alex Alderton, Roberto Amato, Jeffrey Barrett, Sonia Goncalves, Ewan Harrison, David K. Jackson, Ian Johnston, Dominic Kwiatkowski, Cordelia Langford, John Sillitoe on behalf of the Wellcome Sanger Institute COVID-19 Surveillance Team |
| EPI_ISL_3438024 | Lighthouse Lab in Glasgow | Wellcome Sanger Institute for the COVID-19 Genomics UK (COG-UK) Consortium | Harper VanSteenhouse, Yumi Kasai, David Gray, Carol Clugston, Anna Dominiczak and Alex Alderton, Roberto Amato, Jeffrey Barrett, Sonia Goncalves, Ewan Harrison, David K. Jackson, Ian Johnston, Dominic Kwiatkowski, Cordelia Langford, John Sillitoe on behalf of the Wellcome Sanger Institute COVID-19 Surveillance Team |
| EPI_ISL_3438025 | Lighthouse Lab in Milton Keynes | Wellcome Sanger Institute for the COVID-19 Genomics UK (COG-UK) Consortium | The Lighthouse Lab in Milton Keynes and Alex Alderton, Roberto Amato, Jeffrey Barrett, Sonia Goncalves, Ewan Harrison, David K. Jackson, Ian Johnston, Dominic Kwiatkowski, Cordelia Langford, John Sillitoe on behalf of the Wellcome Sanger Institute COVID-19 Surveillance Team |
| EPI_ISL_3438026 | Lighthouse Lab in Glasgow | Wellcome Sanger Institute for the COVID-19 Genomics UK (COG-UK) Consortium | Harper VanSteenhouse, Yumi Kasai, David Gray, Carol Clugston, Anna Dominiczak and Alex Alderton, Roberto Amato, Jeffrey Barrett, Sonia Goncalves, Ewan Harrison, David K. Jackson, Ian Johnston, Dominic Kwiatkowski, Cordelia Langford, John Sillitoe on behalf of the Wellcome Sanger Institute COVID-19 Surveillance Team |
| EPI_ISL_3438027, EPI_ISL_3438028, EPI_ISL_3438029 | Lighthouse Lab in Milton Keynes | Wellcome Sanger Institute for the COVID-19 Genomics UK (COG-UK) Consortium | The Lighthouse Lab in Milton Keynes and Alex Alderton, Roberto Amato, Jeffrey Barrett, Sonia Goncalves, Ewan Harrison, David K. Jackson, Ian Johnston, Dominic Kwiatkowski, Cordelia Langford, John Sillitoe on behalf of the Wellcome Sanger Institute COVID-19 Surveillance Team |
| EPI_ISL_3438030 | Lighthouse Lab in Glasgow | Wellcome Sanger Institute for the COVID-19 Genomics UK (COG-UK) Consortium | Harper VanSteenhouse, Yumi Kasai, David Gray, Carol Clugston, Anna Dominiczak and Alex Alderton, Roberto Amato, Jeffrey Barrett, Sonia Goncalves, Ewan Harrison, David K. Jackson, Ian Johnston, Dominic Kwiatkowski, Cordelia Langford, John Sillitoe on behalf of the Wellcome Sanger Institute COVID-19 Surveillance Team |
| EPI_ISL_3438031, EPI_ISL_3438032, EPI_ISL_3438033 | Lighthouse Lab in Milton Keynes | Wellcome Sanger Institute for the COVID-19 Genomics UK (COG-UK) Consortium | The Lighthouse Lab in Milton Keynes and Alex Alderton, Roberto Amato, Jeffrey Barrett, Sonia Goncalves, Ewan Harrison, David K. Jackson, Ian Johnston, Dominic Kwiatkowski, Cordelia Langford, John Sillitoe on behalf of the Wellcome Sanger Institute COVID-19 Surveillance Team |
| EPI_ISL_3438034 | Lighthouse Lab in Glasgow | Wellcome Sanger Institute for the COVID-19 Genomics UK (COG-UK) Consortium | Harper VanSteenhouse, Yumi Kasai, David Gray, Carol Clugston, Anna Dominiczak and Alex Alderton, Roberto Amato, Jeffrey Barrett, Sonia Goncalves, Ewan Harrison, David K. Jackson, Ian Johnston, Dominic Kwiatkowski, Cordelia Langford, John Sillitoe on behalf of the Wellcome Sanger Institute COVID-19 Surveillance Team |
| EPI_ISL_3438035, EPI_ISL_3438036, EPI_ISL_3438037, EPI_ISL_3438038, EPI_ISL_3438039, EPI_ISL_3438040, EPI_ISL_3438041, EPI_ISL_3438042, EPI_ISL_3438043, EPI_ISL_3438044, EPI_ISL_3438045, EPI_ISL_3438046, EPI_ISL_3438047, EPI_ISL_3438048, EPI_ISL_3438049, EPI_ISL_3438050, EPI_ISL_3438051 | | | |
| see above | Lighthouse Lab in Milton Keynes | Wellcome Sanger Institute for the COVID-19 Genomics UK (COG-UK) Consortium | The Lighthouse Lab in Milton Keynes and Alex Alderton, Roberto Amato, Jeffrey Barrett, Sonia Goncalves, Ewan Harrison, David K. Jackson, Ian Johnston, Dominic Kwiatkowski, Cordelia Langford, John Sillitoe on behalf of the Wellcome Sanger Institute COVID-19 Surveillance Team |
| EPI_ISL_3438052 | Lighthouse Lab in Glasgow | Wellcome Sanger Institute for the COVID-19 Genomics UK (COG-UK) Consortium | Harper VanSteenhouse, Yumi Kasai, David Gray, Carol Clugston, Anna Dominiczak and Alex Alderton, Roberto Amato, Jeffrey Barrett, Sonia Goncalves, Ewan Harrison, David K. Jackson, Ian Johnston, Dominic Kwiatkowski, Cordelia Langford, John Sillitoe on behalf of the Wellcome Sanger Institute COVID-19 Surveillance Team |
| EPI_ISL_3438053, | Lighthouse Lab in Milton Keynes | Wellcome Sanger Institute for the COVID-19 Genomics UK (COG-UK) | The Lighthouse Lab in Milton Keynes and Alex Alderton, Roberto Amato, Jeffrey Barrett, Sonia Goncalves, Ewan Harrison, David K. Jackson, Ian Johnston, Dominic Kwiatkowski, Cordelia Langford, John Sillitoe on behalf of the Wellcome Sanger |

| | | | |
|---|---|---|---|
| EPI_ISL_3438054, EPI_ISL_3438055, EPI_ISL_3438056, EPI_ISL_3438057 | | Consortium | Institute COVID-19 Surveillance Team |
| EPI_ISL_3438058 | Lighthouse Lab in Glasgow | Wellcome Sanger Institute for the COVID-19 Genomics UK (COG-UK) Consortium | Harper VanSteenhouse, Yumi Kasai, David Gray, Carol Clugston, Anna Dominiczak and Alex Alderton, Roberto Amato, Jeffrey Barrett, Sonia Goncalves, Ewan Harrison, David K. Jackson, Ian Johnston, Dominic Kwiatkowski, Cordelia Langford, John Sillitoe on behalf of the Wellcome Sanger Institute COVID-19 Surveillance Team |
| EPI_ISL_3438059 | Lighthouse Lab in Milton Keynes | Wellcome Sanger Institute for the COVID-19 Genomics UK (COG-UK) Consortium | The Lighthouse Lab in Milton Keynes and Alex Alderton, Roberto Amato, Jeffrey Barrett, Sonia Goncalves, Ewan Harrison, David K. Jackson, Ian Johnston, Dominic Kwiatkowski, Cordelia Langford, John Sillitoe on behalf of the Wellcome Sanger Institute COVID-19 Surveillance Team |
| EPI_ISL_3438060, EPI_ISL_3438061 | Lighthouse Lab in Glasgow | Wellcome Sanger Institute for the COVID-19 Genomics UK (COG-UK) Consortium | Harper VanSteenhouse, Yumi Kasai, David Gray, Carol Clugston, Anna Dominiczak and Alex Alderton, Roberto Amato, Jeffrey Barrett, Sonia Goncalves, Ewan Harrison, David K. Jackson, Ian Johnston, Dominic Kwiatkowski, Cordelia Langford, John Sillitoe on behalf of the Wellcome Sanger Institute COVID-19 Surveillance Team |
| EPI_ISL_3438062, EPI_ISL_3438063, EPI_ISL_3438064 | Lighthouse Lab in Milton Keynes | Wellcome Sanger Institute for the COVID-19 Genomics UK (COG-UK) Consortium | The Lighthouse Lab in Milton Keynes and Alex Alderton, Roberto Amato, Jeffrey Barrett, Sonia Goncalves, Ewan Harrison, David K. Jackson, Ian Johnston, Dominic Kwiatkowski, Cordelia Langford, John Sillitoe on behalf of the Wellcome Sanger Institute COVID-19 Surveillance Team |
| EPI_ISL_3438065, EPI_ISL_3438066 | Lighthouse Lab in Glasgow | Wellcome Sanger Institute for the COVID-19 Genomics UK (COG-UK) Consortium | Harper VanSteenhouse, Yumi Kasai, David Gray, Carol Clugston, Anna Dominiczak and Alex Alderton, Roberto Amato, Jeffrey Barrett, Sonia Goncalves, Ewan Harrison, David K. Jackson, Ian Johnston, Dominic Kwiatkowski, Cordelia Langford, John Sillitoe on behalf of the Wellcome Sanger Institute COVID-19 Surveillance Team |
| EPI_ISL_3438067 | Lighthouse Lab in Milton Keynes | Wellcome Sanger Institute for the COVID-19 Genomics UK (COG-UK) Consortium | The Lighthouse Lab in Milton Keynes and Alex Alderton, Roberto Amato, Jeffrey Barrett, Sonia Goncalves, Ewan Harrison, David K. Jackson, Ian Johnston, Dominic Kwiatkowski, Cordelia Langford, John Sillitoe on behalf of the Wellcome Sanger Institute COVID-19 Surveillance Team |
| EPI_ISL_3438068 | Lighthouse Lab in Glasgow | Wellcome Sanger Institute for the COVID-19 Genomics UK (COG-UK) Consortium | Harper VanSteenhouse, Yumi Kasai, David Gray, Carol Clugston, Anna Dominiczak and Alex Alderton, Roberto Amato, Jeffrey Barrett, Sonia Goncalves, Ewan Harrison, David K. Jackson, Ian Johnston, Dominic Kwiatkowski, Cordelia Langford, John Sillitoe on behalf of the Wellcome Sanger Institute COVID-19 Surveillance Team |
| EPI_ISL_3438069, EPI_ISL_3438070 | Lighthouse Lab in Milton Keynes | Wellcome Sanger Institute for the COVID-19 Genomics UK (COG-UK) Consortium | The Lighthouse Lab in Milton Keynes and Alex Alderton, Roberto Amato, Jeffrey Barrett, Sonia Goncalves, Ewan Harrison, David K. Jackson, Ian Johnston, Dominic Kwiatkowski, Cordelia Langford, John Sillitoe on behalf of the Wellcome Sanger Institute COVID-19 Surveillance Team |
| EPI_ISL_3438071 | Lighthouse Lab in Glasgow | Wellcome Sanger Institute for the COVID-19 Genomics UK (COG-UK) Consortium | Harper VanSteenhouse, Yumi Kasai, David Gray, Carol Clugston, Anna Dominiczak and Alex Alderton, Roberto Amato, Jeffrey Barrett, Sonia Goncalves, Ewan Harrison, David K. Jackson, Ian Johnston, Dominic Kwiatkowski, Cordelia Langford, John Sillitoe on behalf of the Wellcome Sanger Institute COVID-19 Surveillance Team |
| EPI_ISL_3438072, EPI_ISL_3438073, EPI_ISL_3438074, EPI_ISL_3438075 | Lighthouse Lab in Milton Keynes | Wellcome Sanger Institute for the COVID-19 Genomics UK (COG-UK) Consortium | The Lighthouse Lab in Milton Keynes and Alex Alderton, Roberto Amato, Jeffrey Barrett, Sonia Goncalves, Ewan Harrison, David K. Jackson, Ian Johnston, Dominic Kwiatkowski, Cordelia Langford, John Sillitoe on behalf of the Wellcome Sanger Institute COVID-19 Surveillance Team |
| EPI_ISL_3438076 | Lighthouse Lab in Glasgow | Wellcome Sanger Institute for the COVID-19 Genomics UK (COG-UK) Consortium | Harper VanSteenhouse, Yumi Kasai, David Gray, Carol Clugston, Anna Dominiczak and Alex Alderton, Roberto Amato, Jeffrey Barrett, Sonia Goncalves, Ewan Harrison, David K. Jackson, Ian Johnston, Dominic Kwiatkowski, Cordelia Langford, John Sillitoe on behalf of the Wellcome Sanger Institute COVID-19 Surveillance Team |
| EPI_ISL_3438077, EPI_ISL_3438078, EPI_ISL_3438079, EPI_ISL_3438080 | Lighthouse Lab in Milton Keynes | Wellcome Sanger Institute for the COVID-19 Genomics UK (COG-UK) Consortium | The Lighthouse Lab in Milton Keynes and Alex Alderton, Roberto Amato, Jeffrey Barrett, Sonia Goncalves, Ewan Harrison, David K. Jackson, Ian Johnston, Dominic Kwiatkowski, Cordelia Langford, John Sillitoe on behalf of the Wellcome Sanger Institute COVID-19 Surveillance Team |
| EPI_ISL_3438081 | Lighthouse Lab in Glasgow | Wellcome Sanger Institute for the COVID-19 Genomics UK (COG-UK) Consortium | Harper VanSteenhouse, Yumi Kasai, David Gray, Carol Clugston, Anna Dominiczak and Alex Alderton, Roberto Amato, Jeffrey Barrett, Sonia Goncalves, Ewan Harrison, David K. Jackson, Ian Johnston, Dominic Kwiatkowski, Cordelia Langford, John Sillitoe on behalf of the Wellcome Sanger Institute COVID-19 Surveillance Team |
| EPI_ISL_3438082, EPI_ISL_3438083 | Lighthouse Lab in Milton Keynes | Wellcome Sanger Institute for the COVID-19 Genomics UK (COG-UK) Consortium | The Lighthouse Lab in Milton Keynes and Alex Alderton, Roberto Amato, Jeffrey Barrett, Sonia Goncalves, Ewan Harrison, David K. Jackson, Ian Johnston, Dominic Kwiatkowski, Cordelia Langford, John Sillitoe on behalf of the Wellcome Sanger Institute COVID-19 Surveillance Team |
| EPI_ISL_3438084 | Lighthouse Lab in Glasgow | Wellcome Sanger Institute for the COVID-19 Genomics UK (COG-UK) Consortium | Harper VanSteenhouse, Yumi Kasai, David Gray, Carol Clugston, Anna Dominiczak and Alex Alderton, Roberto Amato, Jeffrey Barrett, Sonia Goncalves, Ewan Harrison, David K. Jackson, Ian Johnston, Dominic Kwiatkowski, Cordelia Langford, John Sillitoe on behalf of the Wellcome Sanger Institute COVID-19 Surveillance Team |
| EPI_ISL_3438085, EPI_ISL_3438086, EPI_ISL_3438087 | Lighthouse Lab in Milton Keynes | Wellcome Sanger Institute for the COVID-19 Genomics UK (COG-UK) Consortium | The Lighthouse Lab in Milton Keynes and Alex Alderton, Roberto Amato, Jeffrey Barrett, Sonia Goncalves, Ewan Harrison, David K. Jackson, Ian Johnston, Dominic Kwiatkowski, Cordelia Langford, John Sillitoe on behalf of the Wellcome Sanger Institute COVID-19 Surveillance Team |
| EPI_ISL_3438088 | Lighthouse Lab in Glasgow | Wellcome Sanger Institute for the COVID-19 Genomics UK (COG-UK) Consortium | Harper VanSteenhouse, Yumi Kasai, David Gray, Carol Clugston, Anna Dominiczak and Alex Alderton, Roberto Amato, Jeffrey Barrett, Sonia Goncalves, Ewan Harrison, David K. Jackson, Ian Johnston, Dominic Kwiatkowski, Cordelia Langford, John Sillitoe on behalf of the Wellcome Sanger Institute COVID-19 Surveillance Team |
| EPI_ISL_3438089, EPI_ISL_3438090, EPI_ISL_3438091, EPI_ISL_3438092, EPI_ISL_3438093, EPI_ISL_3438094 | Lighthouse Lab in Milton Keynes | Wellcome Sanger Institute for the COVID-19 Genomics UK (COG-UK) Consortium | The Lighthouse Lab in Milton Keynes and Alex Alderton, Roberto Amato, Jeffrey Barrett, Sonia Goncalves, Ewan Harrison, David K. Jackson, Ian Johnston, Dominic Kwiatkowski, Cordelia Langford, John Sillitoe on behalf of the Wellcome Sanger Institute COVID-19 Surveillance Team |
| EPI_ISL_3438095 | Lighthouse Lab in Glasgow | Wellcome Sanger Institute for the COVID-19 Genomics UK (COG-UK) Consortium | Harper VanSteenhouse, Yumi Kasai, David Gray, Carol Clugston, Anna Dominiczak and Alex Alderton, Roberto Amato, Jeffrey Barrett, Sonia Goncalves, Ewan Harrison, David K. Jackson, Ian Johnston, Dominic Kwiatkowski, Cordelia Langford, John Sillitoe on behalf of the Wellcome Sanger Institute COVID-19 Surveillance Team |
| EPI_ISL_3438096, EPI_ISL_3438097, EPI_ISL_3438098, EPI_ISL_3438099, EPI_ISL_3438100 | Lighthouse Lab in Milton Keynes | Wellcome Sanger Institute for the COVID-19 Genomics UK (COG-UK) Consortium | The Lighthouse Lab in Milton Keynes and Alex Alderton, Roberto Amato, Jeffrey Barrett, Sonia Goncalves, Ewan Harrison, David K. Jackson, Ian Johnston, Dominic Kwiatkowski, Cordelia Langford, John Sillitoe on behalf of the Wellcome Sanger Institute COVID-19 Surveillance Team |
| EPI_ISL_3438101 | Lighthouse Lab in Glasgow | Wellcome Sanger Institute for the COVID-19 Genomics UK (COG-UK) Consortium | Harper VanSteenhouse, Yumi Kasai, David Gray, Carol Clugston, Anna Dominiczak and Alex Alderton, Roberto Amato, Jeffrey Barrett, Sonia Goncalves, Ewan Harrison, David K. Jackson, Ian Johnston, Dominic Kwiatkowski, Cordelia Langford, John Sillitoe on behalf of the Wellcome Sanger Institute COVID-19 Surveillance Team |
| EPI_ISL_3438102, EPI_ISL_3438103, EPI_ISL_3438104, EPI_ISL_3438105 | Lighthouse Lab in Milton Keynes | Wellcome Sanger Institute for the COVID-19 Genomics UK (COG-UK) Consortium | The Lighthouse Lab in Milton Keynes and Alex Alderton, Roberto Amato, Jeffrey Barrett, Sonia Goncalves, Ewan Harrison, David K. Jackson, Ian Johnston, Dominic Kwiatkowski, Cordelia Langford, John Sillitoe on behalf of the Wellcome Sanger Institute COVID-19 Surveillance Team |
| EPI_ISL_3438106 | Lighthouse Lab in Glasgow | Wellcome Sanger Institute for the COVID-19 Genomics UK (COG-UK) Consortium | Harper VanSteenhouse, Yumi Kasai, David Gray, Carol Clugston, Anna Dominiczak and Alex Alderton, Roberto Amato, Jeffrey Barrett, Sonia Goncalves, Ewan Harrison, David K. Jackson, Ian Johnston, Dominic Kwiatkowski, Cordelia Langford, John Sillitoe on behalf of the Wellcome Sanger Institute COVID-19 Surveillance Team |
| EPI_ISL_3438107, EPI_ISL_3438108, EPI_ISL_3438109, EPI_ISL_3438110 | Lighthouse Lab in Milton Keynes | Wellcome Sanger Institute for the COVID-19 Genomics UK (COG-UK) Consortium | The Lighthouse Lab in Milton Keynes and Alex Alderton, Roberto Amato, Jeffrey Barrett, Sonia Goncalves, Ewan Harrison, David K. Jackson, Ian Johnston, Dominic Kwiatkowski, Cordelia Langford, John Sillitoe on behalf of the Wellcome Sanger Institute COVID-19 Surveillance Team |
| EPI_ISL_3438111, EPI_ISL_3438112, EPI_ISL_3438113, EPI_ISL_3438114, EPI_ISL_3438115, EPI_ISL_3438116, EPI_ISL_3438117, EPI_ISL_3438118, EPI_ISL_3438119, EPI_ISL_3438120, EPI_ISL_3438121, EPI_ISL_3438122, EPI_ISL_3438123, EPI_ISL_3438124, EPI_ISL_3438125, EPI_ISL_3438126, EPI_ISL_3438127, EPI_ISL_3438128, EPI_ISL_3438129, EPI_ISL_3438130, EPI_ISL_3438131, EPI_ISL_3438132, EPI_ISL_3438133, EPI_ISL_3438134, EPI_ISL_3438135, EPI_ISL_3438136, EPI_ISL_3438137, EPI_ISL_3438138, EPI_ISL_3438139, EPI_ISL_3438140, EPI_ISL_3438141, EPI_ISL_3438142, EPI_ISL_3438143, EPI_ISL_3438144, EPI_ISL_3438145, EPI_ISL_3438146, EPI_ISL_3438147, EPI_ISL_3438148, EPI_ISL_3438149, EPI_ISL_3438150, EPI_ISL_3438151, EPI_ISL_3438152, EPI_ISL_3438153, EPI_ISL_3438154, EPI_ISL_3438155, EPI_ISL_3438156, EPI_ISL_3438157, EPI_ISL_3438158, EPI_ISL_3438159, EPI_ISL_3438160, EPI_ISL_3438161, EPI_ISL_3438162, EPI_ISL_3438163, EPI_ISL_3438164, EPI_ISL_3438165, EPI_ISL_3438166, EPI_ISL_3438167, EPI_ISL_3438168, EPI_ISL_3438169, EPI_ISL_3438170, EPI_ISL_3438171, EPI_ISL_3438172, EPI_ISL_3438173, EPI_ISL_3438174, EPI_ISL_3438175, EPI_ISL_3438176, EPI_ISL_3438177, EPI_ISL_3438178, EPI_ISL_3438179, EPI_ISL_3438180, EPI_ISL_3438181 | | | |
| see above | Berkshire and Surrey Pathology Services Lighthouse Laboratory | Wellcome Sanger Institute for the COVID-19 Genomics UK (COG-UK) Consortium | Berkshire and Surrey Pathology Services Lighthouse Laboratory and Alex Alderton, Roberto Amato, Jeffrey Barrett, Sonia Goncalves, Ewan Harrison, David K. Jackson, Ian Johnston, Dominic Kwiatkowski, Cordelia Langford, John Sillitoe on behalf of the Wellcome Sanger Institute COVID-19 Surveillance Team |
| EPI_ISL_3438182, EPI_ISL_3438183, EPI_ISL_3438184, EPI_ISL_3438185, EPI_ISL_3438186, EPI_ISL_3438187, EPI_ISL_3438188, EPI_ISL_3438189, EPI_ISL_3438190, EPI_ISL_3438191, EPI_ISL_3438192, EPI_ISL_3438193, EPI_ISL_3438194, EPI_ISL_3438195, EPI_ISL_3438196, EPI_ISL_3438197, EPI_ISL_3438198, EPI_ISL_3438199, EPI_ISL_3438200, EPI_ISL_3438201, EPI_ISL_3438202, EPI_ISL_3438203, EPI_ISL_3438204, EPI_ISL_3438205, EPI_ISL_3438206, EPI_ISL_3438207, EPI_ISL_3438208, EPI_ISL_3438209, EPI_ISL_3438210, EPI_ISL_3438211, EPI_ISL_3438212, EPI_ISL_3438213, EPI_ISL_3438214, EPI_ISL_3438215, EPI_ISL_3438216, EPI_ISL_3438217, EPI_ISL_3438218, EPI_ISL_3438219, EPI_ISL_3438220, EPI_ISL_3438221, EPI_ISL_3438222, EPI_ISL_3438223, EPI_ISL_3438224, EPI_ISL_3438225, EPI_ISL_3438226, EPI_ISL_3438227, EPI_ISL_3438228, EPI_ISL_3438229, EPI_ISL_3438230, EPI_ISL_3438231, EPI_ISL_3438232, EPI_ISL_3438233, EPI_ISL_3438234, EPI_ISL_3438235, EPI_ISL_3438236, EPI_ISL_3438237, EPI_ISL_3438238, EPI_ISL_3438239, EPI_ISL_3438240, EPI_ISL_3438241, EPI_ISL_3438242, EPI_ISL_3438243, EPI_ISL_3438244, EPI_ISL_3438245, EPI_ISL_3438246, EPI_ISL_3438247, EPI_ISL_3438248, EPI_ISL_3438249, EPI_ISL_3438250, EPI_ISL_3438251, EPI_ISL_3438252, EPI_ISL_3438253, EPI_ISL_3438254, EPI_ISL_3438255, EPI_ISL_3438256, EPI_ISL_3438257, EPI_ISL_3438258, EPI_ISL_3438259, EPI_ISL_3438260, EPI_ISL_3438261, EPI_ISL_3438262, EPI_ISL_3438263, EPI_ISL_3438264, EPI_ISL_3438265, EPI_ISL_3438266, EPI_ISL_3438267, EPI_ISL_3438268, EPI_ISL_3438269, EPI_ISL_3438270, EPI_ISL_3438271, EPI_ISL_3438272, EPI_ISL_3438273, EPI_ISL_3438274, EPI_ISL_3438275, EPI_ISL_3438276, EPI_ISL_3438277, EPI_ISL_3438278, EPI_ISL_3438279, EPI_ISL_3438280, EPI_ISL_3438281, EPI_ISL_3438282, EPI_ISL_3438283, EPI_ISL_3438284, EPI_ISL_3438285, EPI_ISL_3438286, EPI_ISL_3438287, EPI_ISL_3438288, EPI_ISL_3438289, EPI_ISL_3438290, EPI_ISL_3438291, EPI_ISL_3438292, EPI_ISL_3438293, EPI_ISL_3438294, EPI_ISL_3438295, EPI_ISL_3438296, EPI_ISL_3438297, EPI_ISL_3438298, EPI_ISL_3438299, EPI_ISL_3438300, EPI_ISL_3438301, EPI_ISL_3438302, EPI_ISL_3438303, EPI_ISL_3438304, EPI_ISL_3438305, EPI_ISL_3438306, EPI_ISL_3438307, EPI_ISL_3438308, EPI_ISL_3438309, EPI_ISL_3438310, EPI_ISL_3438311, EPI_ISL_3438312, EPI_ISL_3438313, EPI_ISL_3438314, EPI_ISL_3438315, EPI_ISL_3438316, EPI_ISL_3438317, EPI_ISL_3438318, EPI_ISL_3438319, EPI_ISL_3438320, EPI_ISL_3438321, EPI_ISL_3438322, EPI_ISL_3438323, EPI_ISL_3438324, EPI_ISL_3438325, EPI_ISL_3438326, EPI_ISL_3438327, EPI_ISL_3438328, EPI_ISL_3438329, EPI_ISL_3438330, EPI_ISL_3438331, EPI_ISL_3438332, EPI_ISL_3438333, EPI_ISL_3438334, EPI_ISL_3438335, EPI_ISL_3438336, EPI_ISL_3438337, EPI_ISL_3438338, EPI_ISL_3438339, EPI_ISL_3438340, EPI_ISL_3438341, EPI_ISL_3438342, EPI_ISL_3438343, EPI_ISL_3438344, EPI_ISL_3438345, EPI_ISL_3438346, EPI_ISL_3438347, EPI_ISL_3438348, EPI_ISL_3438349, EPI_ISL_3438350, EPI_ISL_3438351, EPI_ISL_3438352, EPI_ISL_3438353, EPI_ISL_3438354, EPI_ISL_3438355, EPI_ISL_3438356, EPI_ISL_3438357, EPI_ISL_3438358, EPI_ISL_3438359, EPI_ISL_3438360, EPI_ISL_3438361, EPI_ISL_3438362, EPI_ISL_3438363, EPI_ISL_3438364, EPI_ISL_3438365, EPI_ISL_3438366, EPI_ISL_3438367, EPI_ISL_3438368, EPI_ISL_3438369, EPI_ISL_3438370, EPI_ISL_3438371, EPI_ISL_3438372, EPI_ISL_3438373, EPI_ISL_3438374, EPI_ISL_3438375, EPI_ISL_3438376, EPI_ISL_3438377, EPI_ISL_3438378, EPI_ISL_3438379, EPI_ISL_3438380, EPI_ISL_3438381, EPI_ISL_3438382, EPI_ISL_3438383, EPI_ISL_3438384, EPI_ISL_3438385, EPI_ISL_3438386, EPI_ISL_3438387 | | | |
| see above | Lighthouse Lab in Glasgow | Wellcome Sanger Institute for the COVID-19 Genomics UK (COG-UK) Consortium | Harper VanSteenhouse, Yumi Kasai, David Gray, Carol Clugston, Anna Dominiczak and Alex Alderton, Roberto Amato, Jeffrey Barrett, Sonia Goncalves, Ewan Harrison, David K. Jackson, Ian Johnston, Dominic Kwiatkowski, Cordelia Langford, John Sillitoe on behalf of the Wellcome Sanger Institute COVID-19 Surveillance Team |
| EPI_ISL_3438388 | Lighthouse Lab in Alderley Park | Wellcome Sanger Institute for the COVID-19 Genomics UK (COG-UK) Consortium | Jacquelyn Wynn, Mairead Hyland, The Lighthouse Lab in Alderley Park and Alex Alderton, Roberto Amato, Jeffrey Barrett, Sonia Goncalves, Ewan Harrison, David K. Jackson, Ian Johnston, Dominic Kwiatkowski, Cordelia Langford, John Sillitoe on behalf of the Wellcome Sanger Institute COVID-19 Surveillance Team |
| EPI_ISL_3438389 | Lighthouse Lab in Glasgow | Wellcome Sanger Institute for the COVID-19 Genomics UK (COG-UK) Consortium | Harper VanSteenhouse, Yumi Kasai, David Gray, Carol Clugston, Anna Dominiczak and Alex Alderton, Roberto Amato, Jeffrey Barrett, Sonia Goncalves, Ewan Harrison, David K. Jackson, Ian Johnston, Dominic Kwiatkowski, Cordelia Langford, John Sillitoe on behalf of the Wellcome Sanger Institute COVID-19 Surveillance Team |
| EPI_ISL_3438390 | Lighthouse Lab in Alderley Park | Wellcome Sanger Institute for the COVID-19 Genomics UK (COG-UK) Consortium | Jacquelyn Wynn, Mairead Hyland, The Lighthouse Lab in Alderley Park and Alex Alderton, Roberto Amato, Jeffrey Barrett, Sonia Goncalves, Ewan Harrison, David K. Jackson, Ian Johnston, Dominic Kwiatkowski, Cordelia Langford, John Sillitoe on behalf of the Wellcome Sanger Institute COVID-19 Surveillance Team |
| EPI_ISL_3438391 | Berkshire and Surrey Pathology Services Lighthouse Laboratory | Wellcome Sanger Institute for the COVID-19 Genomics UK (COG-UK) Consortium | Berkshire and Surrey Pathology Services Lighthouse Laboratory and Alex Alderton, Roberto Amato, Jeffrey Barrett, Sonia Goncalves, Ewan Harrison, David K. Jackson, Ian Johnston, Dominic Kwiatkowski, Cordelia Langford, John Sillitoe on behalf of the Wellcome Sanger Institute COVID-19 Surveillance Team |

| | | | |
|---|---|---|---|
| EPI_ISL_3438392 | Lighthouse Lab in Glasgow | Wellcome Sanger Institute for the COVID-19 Genomics UK (COG-UK) Consortium | Harper VanSteenhouse, Yumi Kasai, David Gray, Carol Clugston, Anna Dominiczak and Alex Alderton, Roberto Amato, Jeffrey Barrett, Sonia Goncalves, Ewan Harrison, David K. Jackson, Ian Johnston, Dominic Kwiatkowski, Cordelia Langford, John Sillitoe on behalf of the Wellcome Sanger Institute COVID-19 Surveillance Team |
| EPI_ISL_3438393 | Lighthouse Lab in Alderley Park | Wellcome Sanger Institute for the COVID-19 Genomics UK (COG-UK) Consortium | Jacquelyn Wynn, Mairead Hyland, The Lighthouse Lab in Alderley Park and Alex Alderton, Roberto Amato, Jeffrey Barrett, Sonia Goncalves, Ewan Harrison, David K. Jackson, Ian Johnston, Dominic Kwiatkowski, Cordelia Langford, John Sillitoe on behalf of the Wellcome Sanger Institute COVID-19 Surveillance Team |
| EPI_ISL_3438394 | Lighthouse Lab in Glasgow | Wellcome Sanger Institute for the COVID-19 Genomics UK (COG-UK) Consortium | Harper VanSteenhouse, Yumi Kasai, David Gray, Carol Clugston, Anna Dominiczak and Alex Alderton, Roberto Amato, Jeffrey Barrett, Sonia Goncalves, Ewan Harrison, David K. Jackson, Ian Johnston, Dominic Kwiatkowski, Cordelia Langford, John Sillitoe on behalf of the Wellcome Sanger Institute COVID-19 Surveillance Team |
| EPI_ISL_3438395, EPI_ISL_3438396, EPI_ISL_3438397, EPI_ISL_3438398 | Berkshire and Surrey Pathology Services Lighthouse Laboratory | Wellcome Sanger Institute for the COVID-19 Genomics UK (COG-UK) Consortium | Berkshire and Surrey Pathology Services Lighthouse Laboratory and Alex Alderton, Roberto Amato, Jeffrey Barrett, Sonia Goncalves, Ewan Harrison, David K. Jackson, Ian Johnston, Dominic Kwiatkowski, Cordelia Langford, John Sillitoe on behalf of the Wellcome Sanger Institute COVID-19 Surveillance Team |
| EPI_ISL_3438399, EPI_ISL_3438400, EPI_ISL_3438401 | Lighthouse Lab in Glasgow | Wellcome Sanger Institute for the COVID-19 Genomics UK (COG-UK) Consortium | Harper VanSteenhouse, Yumi Kasai, David Gray, Carol Clugston, Anna Dominiczak and Alex Alderton, Roberto Amato, Jeffrey Barrett, Sonia Goncalves, Ewan Harrison, David K. Jackson, Ian Johnston, Dominic Kwiatkowski, Cordelia Langford, John Sillitoe on behalf of the Wellcome Sanger Institute COVID-19 Surveillance Team |
| EPI_ISL_3438402 | Berkshire and Surrey Pathology Services Lighthouse Laboratory | Wellcome Sanger Institute for the COVID-19 Genomics UK (COG-UK) Consortium | Berkshire and Surrey Pathology Services Lighthouse Laboratory and Alex Alderton, Roberto Amato, Jeffrey Barrett, Sonia Goncalves, Ewan Harrison, David K. Jackson, Ian Johnston, Dominic Kwiatkowski, Cordelia Langford, John Sillitoe on behalf of the Wellcome Sanger Institute COVID-19 Surveillance Team |
| EPI_ISL_3438403 | Lighthouse Lab in Glasgow | Wellcome Sanger Institute for the COVID-19 Genomics UK (COG-UK) Consortium | Harper VanSteenhouse, Yumi Kasai, David Gray, Carol Clugston, Anna Dominiczak and Alex Alderton, Roberto Amato, Jeffrey Barrett, Sonia Goncalves, Ewan Harrison, David K. Jackson, Ian Johnston, Dominic Kwiatkowski, Cordelia Langford, John Sillitoe on behalf of the Wellcome Sanger Institute COVID-19 Surveillance Team |
| EPI_ISL_3438404 | Lighthouse Lab in Alderley Park | Wellcome Sanger Institute for the COVID-19 Genomics UK (COG-UK) Consortium | Jacquelyn Wynn, Mairead Hyland, The Lighthouse Lab in Alderley Park and Alex Alderton, Roberto Amato, Jeffrey Barrett, Sonia Goncalves, Ewan Harrison, David K. Jackson, Ian Johnston, Dominic Kwiatkowski, Cordelia Langford, John Sillitoe on behalf of the Wellcome Sanger Institute COVID-19 Surveillance Team |
| EPI_ISL_3438405 | Berkshire and Surrey Pathology Services Lighthouse Laboratory | Wellcome Sanger Institute for the COVID-19 Genomics UK (COG-UK) Consortium | Berkshire and Surrey Pathology Services Lighthouse Laboratory and Alex Alderton, Roberto Amato, Jeffrey Barrett, Sonia Goncalves, Ewan Harrison, David K. Jackson, Ian Johnston, Dominic Kwiatkowski, Cordelia Langford, John Sillitoe on behalf of the Wellcome Sanger Institute COVID-19 Surveillance Team |
| EPI_ISL_3438406 | Lighthouse Lab in Glasgow | Wellcome Sanger Institute for the COVID-19 Genomics UK (COG-UK) Consortium | Harper VanSteenhouse, Yumi Kasai, David Gray, Carol Clugston, Anna Dominiczak and Alex Alderton, Roberto Amato, Jeffrey Barrett, Sonia Goncalves, Ewan Harrison, David K. Jackson, Ian Johnston, Dominic Kwiatkowski, Cordelia Langford, John Sillitoe on behalf of the Wellcome Sanger Institute COVID-19 Surveillance Team |
| EPI_ISL_3438407, EPI_ISL_3438408 | Berkshire and Surrey Pathology Services Lighthouse Laboratory | Wellcome Sanger Institute for the COVID-19 Genomics UK (COG-UK) Consortium | Berkshire and Surrey Pathology Services Lighthouse Laboratory and Alex Alderton, Roberto Amato, Jeffrey Barrett, Sonia Goncalves, Ewan Harrison, David K. Jackson, Ian Johnston, Dominic Kwiatkowski, Cordelia Langford, John Sillitoe on behalf of the Wellcome Sanger Institute COVID-19 Surveillance Team |
| EPI_ISL_3438409, EPI_ISL_3438410 | Lighthouse Lab in Glasgow | Wellcome Sanger Institute for the COVID-19 Genomics UK (COG-UK) Consortium | Harper VanSteenhouse, Yumi Kasai, David Gray, Carol Clugston, Anna Dominiczak and Alex Alderton, Roberto Amato, Jeffrey Barrett, Sonia Goncalves, Ewan Harrison, David K. Jackson, Ian Johnston, Dominic Kwiatkowski, Cordelia Langford, John Sillitoe on behalf of the Wellcome Sanger Institute COVID-19 Surveillance Team |
| EPI_ISL_3438411 | Lighthouse Lab in Alderley Park | Wellcome Sanger Institute for the COVID-19 Genomics UK (COG-UK) Consortium | Jacquelyn Wynn, Mairead Hyland, The Lighthouse Lab in Alderley Park and Alex Alderton, Roberto Amato, Jeffrey Barrett, Sonia Goncalves, Ewan Harrison, David K. Jackson, Ian Johnston, Dominic Kwiatkowski, Cordelia Langford, John Sillitoe on behalf of the Wellcome Sanger Institute COVID-19 Surveillance Team |
| EPI_ISL_3438412 | Berkshire and Surrey Pathology Services Lighthouse Laboratory | Wellcome Sanger Institute for the COVID-19 Genomics UK (COG-UK) Consortium | Berkshire and Surrey Pathology Services Lighthouse Laboratory and Alex Alderton, Roberto Amato, Jeffrey Barrett, Sonia Goncalves, Ewan Harrison, David K. Jackson, Ian Johnston, Dominic Kwiatkowski, Cordelia Langford, John Sillitoe on behalf of the Wellcome Sanger Institute COVID-19 Surveillance Team |
| EPI_ISL_3438413 | Lighthouse Lab in Alderley Park | Wellcome Sanger Institute for the COVID-19 Genomics UK (COG-UK) Consortium | Jacquelyn Wynn, Mairead Hyland, The Lighthouse Lab in Alderley Park and Alex Alderton, Roberto Amato, Jeffrey Barrett, Sonia Goncalves, Ewan Harrison, David K. Jackson, Ian Johnston, Dominic Kwiatkowski, Cordelia Langford, John Sillitoe on behalf of the Wellcome Sanger Institute COVID-19 Surveillance Team |
| EPI_ISL_3438414 | Berkshire and Surrey Pathology Services Lighthouse Laboratory | Wellcome Sanger Institute for the COVID-19 Genomics UK (COG-UK) Consortium | Berkshire and Surrey Pathology Services Lighthouse Laboratory and Alex Alderton, Roberto Amato, Jeffrey Barrett, Sonia Goncalves, Ewan Harrison, David K. Jackson, Ian Johnston, Dominic Kwiatkowski, Cordelia Langford, John Sillitoe on behalf of the Wellcome Sanger Institute COVID-19 Surveillance Team |
| EPI_ISL_3438415 | Lighthouse Lab in Alderley Park | Wellcome Sanger Institute for the COVID-19 Genomics UK (COG-UK) Consortium | Jacquelyn Wynn, Mairead Hyland, The Lighthouse Lab in Alderley Park and Alex Alderton, Roberto Amato, Jeffrey Barrett, Sonia Goncalves, Ewan Harrison, David K. Jackson, Ian Johnston, Dominic Kwiatkowski, Cordelia Langford, John Sillitoe on behalf of the Wellcome Sanger Institute COVID-19 Surveillance Team |
| EPI_ISL_3438416 | Lighthouse Lab in Glasgow | Wellcome Sanger Institute for the COVID-19 Genomics UK (COG-UK) Consortium | Harper VanSteenhouse, Yumi Kasai, David Gray, Carol Clugston, Anna Dominiczak and Alex Alderton, Roberto Amato, Jeffrey Barrett, Sonia Goncalves, Ewan Harrison, David K. Jackson, Ian Johnston, Dominic Kwiatkowski, Cordelia Langford, John Sillitoe on behalf of the Wellcome Sanger Institute COVID-19 Surveillance Team |
| EPI_ISL_3438417 | Lighthouse Lab in Alderley Park | Wellcome Sanger Institute for the COVID-19 Genomics UK (COG-UK) Consortium | Jacquelyn Wynn, Mairead Hyland, The Lighthouse Lab in Alderley Park and Alex Alderton, Roberto Amato, Jeffrey Barrett, Sonia Goncalves, Ewan Harrison, David K. Jackson, Ian Johnston, Dominic Kwiatkowski, Cordelia Langford, John Sillitoe on behalf of the Wellcome Sanger Institute COVID-19 Surveillance Team |
| EPI_ISL_3438418, EPI_ISL_3438419 | Lighthouse Lab in Glasgow | Wellcome Sanger Institute for the COVID-19 Genomics UK (COG-UK) Consortium | Harper VanSteenhouse, Yumi Kasai, David Gray, Carol Clugston, Anna Dominiczak and Alex Alderton, Roberto Amato, Jeffrey Barrett, Sonia Goncalves, Ewan Harrison, David K. Jackson, Ian Johnston, Dominic Kwiatkowski, Cordelia Langford, John Sillitoe on behalf of the Wellcome Sanger Institute COVID-19 Surveillance Team |
| EPI_ISL_3438420 | Lighthouse Lab in Alderley Park | Wellcome Sanger Institute for the COVID-19 Genomics UK (COG-UK) Consortium | Jacquelyn Wynn, Mairead Hyland, The Lighthouse Lab in Alderley Park and Alex Alderton, Roberto Amato, Jeffrey Barrett, Sonia Goncalves, Ewan Harrison, David K. Jackson, Ian Johnston, Dominic Kwiatkowski, Cordelia Langford, John Sillitoe on behalf of the Wellcome Sanger Institute COVID-19 Surveillance Team |
| EPI_ISL_3438421, EPI_ISL_3438422 | Lighthouse Lab in Glasgow | Wellcome Sanger Institute for the COVID-19 Genomics UK (COG-UK) Consortium | Harper VanSteenhouse, Yumi Kasai, David Gray, Carol Clugston, Anna Dominiczak and Alex Alderton, Roberto Amato, Jeffrey Barrett, Sonia Goncalves, Ewan Harrison, David K. Jackson, Ian Johnston, Dominic Kwiatkowski, Cordelia Langford, John Sillitoe on behalf of the Wellcome Sanger Institute COVID-19 Surveillance Team |
| EPI_ISL_3438423 | Berkshire and Surrey Pathology Services Lighthouse Laboratory | Wellcome Sanger Institute for the COVID-19 Genomics UK (COG-UK) Consortium | Berkshire and Surrey Pathology Services Lighthouse Laboratory and Alex Alderton, Roberto Amato, Jeffrey Barrett, Sonia Goncalves, Ewan Harrison, David K. Jackson, Ian Johnston, Dominic Kwiatkowski, Cordelia Langford, John Sillitoe on behalf of the Wellcome Sanger Institute COVID-19 Surveillance Team |
| EPI_ISL_3438424, EPI_ISL_3438425 | Lighthouse Lab in Glasgow | Wellcome Sanger Institute for the COVID-19 Genomics UK (COG-UK) Consortium | Harper VanSteenhouse, Yumi Kasai, David Gray, Carol Clugston, Anna Dominiczak and Alex Alderton, Roberto Amato, Jeffrey Barrett, Sonia Goncalves, Ewan Harrison, David K. Jackson, Ian Johnston, Dominic Kwiatkowski, Cordelia Langford, John Sillitoe on behalf of the Wellcome Sanger Institute COVID-19 Surveillance Team |
| EPI_ISL_3438426, EPI_ISL_3438427, EPI_ISL_3438428 | Berkshire and Surrey Pathology Services Lighthouse Laboratory | Wellcome Sanger Institute for the COVID-19 Genomics UK (COG-UK) Consortium | Berkshire and Surrey Pathology Services Lighthouse Laboratory and Alex Alderton, Roberto Amato, Jeffrey Barrett, Sonia Goncalves, Ewan Harrison, David K. Jackson, Ian Johnston, Dominic Kwiatkowski, Cordelia Langford, John Sillitoe on behalf of the Wellcome Sanger Institute COVID-19 Surveillance Team |
| EPI_ISL_3438429 | Lighthouse Lab in Glasgow | Wellcome Sanger Institute for the COVID-19 Genomics UK (COG-UK) Consortium | Harper VanSteenhouse, Yumi Kasai, David Gray, Carol Clugston, Anna Dominiczak and Alex Alderton, Roberto Amato, Jeffrey Barrett, Sonia Goncalves, Ewan Harrison, David K. Jackson, Ian Johnston, Dominic Kwiatkowski, Cordelia Langford, John Sillitoe on behalf of the Wellcome Sanger Institute COVID-19 Surveillance Team |
| EPI_ISL_3438430 | Lighthouse Lab in Alderley Park | Wellcome Sanger Institute for the COVID-19 Genomics UK (COG-UK) Consortium | Jacquelyn Wynn, Mairead Hyland, The Lighthouse Lab in Alderley Park and Alex Alderton, Roberto Amato, Jeffrey Barrett, Sonia Goncalves, Ewan Harrison, David K. Jackson, Ian Johnston, Dominic Kwiatkowski, Cordelia Langford, John Sillitoe on behalf of the Wellcome Sanger Institute COVID-19 Surveillance Team |
| EPI_ISL_3438431 | Berkshire and Surrey Pathology Services Lighthouse Laboratory | Wellcome Sanger Institute for the COVID-19 Genomics UK (COG-UK) Consortium | Berkshire and Surrey Pathology Services Lighthouse Laboratory and Alex Alderton, Roberto Amato, Jeffrey Barrett, Sonia Goncalves, Ewan Harrison, David K. Jackson, Ian Johnston, Dominic Kwiatkowski, Cordelia Langford, John Sillitoe on behalf of the Wellcome Sanger Institute COVID-19 Surveillance Team |
| EPI_ISL_3438432 | Lighthouse Lab in Glasgow | Wellcome Sanger Institute for the COVID-19 Genomics UK (COG-UK) Consortium | Harper VanSteenhouse, Yumi Kasai, David Gray, Carol Clugston, Anna Dominiczak and Alex Alderton, Roberto Amato, Jeffrey Barrett, Sonia Goncalves, Ewan Harrison, David K. Jackson, Ian Johnston, Dominic Kwiatkowski, Cordelia Langford, John Sillitoe on behalf of the Wellcome Sanger Institute COVID-19 Surveillance Team |
| EPI_ISL_3438433 | Lighthouse Lab in Alderley Park | Wellcome Sanger Institute for the COVID-19 Genomics UK (COG-UK) Consortium | Jacquelyn Wynn, Mairead Hyland, The Lighthouse Lab in Alderley Park and Alex Alderton, Roberto Amato, Jeffrey Barrett, Sonia Goncalves, Ewan Harrison, David K. Jackson, Ian Johnston, Dominic Kwiatkowski, Cordelia Langford, John Sillitoe on behalf of the Wellcome Sanger Institute COVID-19 Surveillance Team |
| EPI_ISL_3438434 | Lighthouse Lab in Glasgow | Wellcome Sanger Institute for the COVID-19 Genomics UK (COG-UK) Consortium | Harper VanSteenhouse, Yumi Kasai, David Gray, Carol Clugston, Anna Dominiczak and Alex Alderton, Roberto Amato, Jeffrey Barrett, Sonia Goncalves, Ewan Harrison, David K. Jackson, Ian Johnston, Dominic Kwiatkowski, Cordelia Langford, John Sillitoe on behalf of the Wellcome Sanger Institute COVID-19 Surveillance Team |
| EPI_ISL_3438435 | Berkshire and Surrey Pathology Services Lighthouse Laboratory | Wellcome Sanger Institute for the COVID-19 Genomics UK (COG-UK) Consortium | Berkshire and Surrey Pathology Services Lighthouse Laboratory and Alex Alderton, Roberto Amato, Jeffrey Barrett, Sonia Goncalves, Ewan Harrison, David K. Jackson, Ian Johnston, Dominic Kwiatkowski, Cordelia Langford, John Sillitoe on behalf of the Wellcome Sanger Institute COVID-19 Surveillance Team |
| EPI_ISL_3438436 | Lighthouse Lab in Alderley Park | Wellcome Sanger Institute for the COVID-19 Genomics UK (COG-UK) Consortium | Jacquelyn Wynn, Mairead Hyland, The Lighthouse Lab in Alderley Park and Alex Alderton, Roberto Amato, Jeffrey Barrett, Sonia Goncalves, Ewan Harrison, David K. Jackson, Ian Johnston, Dominic Kwiatkowski, Cordelia Langford, John Sillitoe on behalf of the Wellcome Sanger Institute COVID-19 Surveillance Team |
| EPI_ISL_3438437 | Lighthouse Lab in Glasgow | Wellcome Sanger Institute for the COVID-19 Genomics UK (COG-UK) Consortium | Harper VanSteenhouse, Yumi Kasai, David Gray, Carol Clugston, Anna Dominiczak and Alex Alderton, Roberto Amato, Jeffrey Barrett, Sonia Goncalves, Ewan Harrison, David K. Jackson, Ian Johnston, Dominic Kwiatkowski, Cordelia Langford, John Sillitoe on behalf of the Wellcome Sanger Institute COVID-19 Surveillance Team |
| EPI_ISL_3438438, EPI_ISL_3438439 | Lighthouse Lab in Alderley Park | Wellcome Sanger Institute for the COVID-19 Genomics UK (COG-UK) Consortium | Jacquelyn Wynn, Mairead Hyland, The Lighthouse Lab in Alderley Park and Alex Alderton, Roberto Amato, Jeffrey Barrett, Sonia Goncalves, Ewan Harrison, David K. Jackson, Ian Johnston, Dominic Kwiatkowski, Cordelia Langford, John Sillitoe on behalf of the Wellcome Sanger Institute COVID-19 Surveillance Team |
| EPI_ISL_3438440 | Lighthouse Lab in Glasgow | Wellcome Sanger Institute for the COVID-19 Genomics UK (COG-UK) Consortium | Harper VanSteenhouse, Yumi Kasai, David Gray, Carol Clugston, Anna Dominiczak and Alex Alderton, Roberto Amato, Jeffrey Barrett, Sonia Goncalves, Ewan Harrison, David K. Jackson, Ian Johnston, Dominic Kwiatkowski, Cordelia Langford, John Sillitoe on behalf of the Wellcome Sanger Institute COVID-19 Surveillance Team |
| EPI_ISL_3438441 | Berkshire and Surrey Pathology Services Lighthouse Laboratory | Wellcome Sanger Institute for the COVID-19 Genomics UK (COG-UK) Consortium | Berkshire and Surrey Pathology Services Lighthouse Laboratory and Alex Alderton, Roberto Amato, Jeffrey Barrett, Sonia Goncalves, Ewan Harrison, David K. Jackson, Ian Johnston, Dominic Kwiatkowski, Cordelia Langford, John Sillitoe on behalf of the Wellcome Sanger Institute COVID-19 Surveillance Team |
| EPI_ISL_3438442 | Lighthouse Lab in Glasgow | Wellcome Sanger Institute for the COVID-19 Genomics UK (COG-UK) Consortium | Harper VanSteenhouse, Yumi Kasai, David Gray, Carol Clugston, Anna Dominiczak and Alex Alderton, Roberto Amato, Jeffrey Barrett, Sonia Goncalves, Ewan Harrison, David K. Jackson, Ian Johnston, Dominic Kwiatkowski, Cordelia Langford, John Sillitoe on behalf of the Wellcome Sanger Institute COVID-19 Surveillance Team |
| EPI_ISL_3438443 | Lighthouse Lab in Alderley Park | Wellcome Sanger Institute for the COVID-19 Genomics UK (COG-UK) Consortium | Jacquelyn Wynn, Mairead Hyland, The Lighthouse Lab in Alderley Park and Alex Alderton, Roberto Amato, Jeffrey Barrett, Sonia Goncalves, Ewan Harrison, David K. Jackson, Ian Johnston, Dominic Kwiatkowski, Cordelia Langford, John Sillitoe on behalf of the Wellcome Sanger Institute COVID-19 Surveillance Team |
| EPI_ISL_3438444 | Lighthouse Lab in Glasgow | Wellcome Sanger Institute for the COVID-19 Genomics UK (COG-UK) Consortium | Harper VanSteenhouse, Yumi Kasai, David Gray, Carol Clugston, Anna Dominiczak and Alex Alderton, Roberto Amato, Jeffrey Barrett, Sonia Goncalves, Ewan Harrison, David K. Jackson, Ian Johnston, Dominic Kwiatkowski, Cordelia Langford, John Sillitoe on behalf of the Wellcome Sanger Institute COVID-19 Surveillance Team |
| EPI_ISL_3438445 | Lighthouse Lab in Alderley Park | Wellcome Sanger Institute for the COVID-19 Genomics UK (COG-UK) Consortium | Jacquelyn Wynn, Mairead Hyland, The Lighthouse Lab in Alderley Park and Alex Alderton, Roberto Amato, Jeffrey Barrett, Sonia Goncalves, Ewan Harrison, David K. Jackson, Ian Johnston, Dominic Kwiatkowski, Cordelia Langford, John Sillitoe on behalf of the Wellcome Sanger Institute COVID-19 Surveillance Team |
| EPI_ISL_3438446 | Berkshire and Surrey Pathology Services Lighthouse Laboratory | Wellcome Sanger Institute for the COVID-19 Genomics UK (COG-UK) Consortium | Berkshire and Surrey Pathology Services Lighthouse Laboratory and Alex Alderton, Roberto Amato, Jeffrey Barrett, Sonia Goncalves, Ewan Harrison, David K. Jackson, Ian Johnston, Dominic Kwiatkowski, Cordelia Langford, John Sillitoe on behalf of the Wellcome Sanger Institute COVID-19 Surveillance Team |
| EPI_ISL_3438447 | Lighthouse Lab in Glasgow | Wellcome Sanger Institute for the COVID-19 Genomics UK (COG-UK) Consortium | Harper VanSteenhouse, Yumi Kasai, David Gray, Carol Clugston, Anna Dominiczak and Alex Alderton, Roberto Amato, Jeffrey Barrett, Sonia Goncalves, Ewan Harrison, David K. Jackson, Ian Johnston, Dominic Kwiatkowski, Cordelia Langford, John Sillitoe on behalf of the Wellcome Sanger Institute COVID-19 Surveillance Team |
| EPI_ISL_3438448 | Lighthouse Lab in Alderley Park | Wellcome Sanger Institute for the COVID-19 Genomics UK (COG-UK) | Jacquelyn Wynn, Mairead Hyland, The Lighthouse Lab in Alderley Park and Alex Alderton, Roberto Amato, Jeffrey Barrett, Sonia Goncalves, Ewan Harrison, David K. Jackson, Ian Johnston, Dominic Kwiatkowski, Cordelia Langford, John Sillitoe on |

| | | | |
|---|---|---|---|
| | | Consortium | behalf of the Wellcome Sanger Institute COVID-19 Surveillance Team |
| EPI_ISL_3438449, EPI_ISL_3438450, EPI_ISL_3438451 | Lighthouse Lab in Glasgow | Wellcome Sanger Institute for the COVID-19 Genomics UK (COG-UK) Consortium | Harper VanSteenhouse, Yumi Kasai, David Gray, Carol Clugston, Anna Dominiczak and Alex Alderton, Roberto Amato, Jeffrey Barrett, Sonia Goncalves, Ewan Harrison, David K. Jackson, Ian Johnston, Dominic Kwiatkowski, Cordelia Langford, John Sillitoe on behalf of the Wellcome Sanger Institute COVID-19 Surveillance Team |
| EPI_ISL_3438452 | Berkshire and Surrey Pathology Services Lighthouse Laboratory | Wellcome Sanger Institute for the COVID-19 Genomics UK (COG-UK) Consortium | Berkshire and Surrey Pathology Services Lighthouse Laboratory and Alex Alderton, Roberto Amato, Jeffrey Barrett, Sonia Goncalves, Ewan Harrison, David K. Jackson, Ian Johnston, Dominic Kwiatkowski, Cordelia Langford, John Sillitoe on behalf of the Wellcome Sanger Institute COVID-19 Surveillance Team |
| EPI_ISL_3438453, EPI_ISL_3438454, EPI_ISL_3438455, EPI_ISL_3438456, EPI_ISL_3438457 | Lighthouse Lab in Glasgow | Wellcome Sanger Institute for the COVID-19 Genomics UK (COG-UK) Consortium | Harper VanSteenhouse, Yumi Kasai, David Gray, Carol Clugston, Anna Dominiczak and Alex Alderton, Roberto Amato, Jeffrey Barrett, Sonia Goncalves, Ewan Harrison, David K. Jackson, Ian Johnston, Dominic Kwiatkowski, Cordelia Langford, John Sillitoe on behalf of the Wellcome Sanger Institute COVID-19 Surveillance Team |
| EPI_ISL_3438458 | Berkshire and Surrey Pathology Services Lighthouse Laboratory | Wellcome Sanger Institute for the COVID-19 Genomics UK (COG-UK) Consortium | Berkshire and Surrey Pathology Services Lighthouse Laboratory and Alex Alderton, Roberto Amato, Jeffrey Barrett, Sonia Goncalves, Ewan Harrison, David K. Jackson, Ian Johnston, Dominic Kwiatkowski, Cordelia Langford, John Sillitoe on behalf of the Wellcome Sanger Institute COVID-19 Surveillance Team |
| EPI_ISL_3438459 | Lighthouse Lab in Alderley Park | Wellcome Sanger Institute for the COVID-19 Genomics UK (COG-UK) Consortium | Jacquelyn Wynn, Mairead Hyland, The Lighthouse Lab in Alderley Park and Alex Alderton, Roberto Amato, Jeffrey Barrett, Sonia Goncalves, Ewan Harrison, David K. Jackson, Ian Johnston, Dominic Kwiatkowski, Cordelia Langford, John Sillitoe on behalf of the Wellcome Sanger Institute COVID-19 Surveillance Team |
| EPI_ISL_3438460 | Lighthouse Lab in Glasgow | Wellcome Sanger Institute for the COVID-19 Genomics UK (COG-UK) Consortium | Harper VanSteenhouse, Yumi Kasai, David Gray, Carol Clugston, Anna Dominiczak and Alex Alderton, Roberto Amato, Jeffrey Barrett, Sonia Goncalves, Ewan Harrison, David K. Jackson, Ian Johnston, Dominic Kwiatkowski, Cordelia Langford, John Sillitoe on behalf of the Wellcome Sanger Institute COVID-19 Surveillance Team |
| EPI_ISL_3438461 | Berkshire and Surrey Pathology Services Lighthouse Laboratory | Wellcome Sanger Institute for the COVID-19 Genomics UK (COG-UK) Consortium | Berkshire and Surrey Pathology Services Lighthouse Laboratory and Alex Alderton, Roberto Amato, Jeffrey Barrett, Sonia Goncalves, Ewan Harrison, David K. Jackson, Ian Johnston, Dominic Kwiatkowski, Cordelia Langford, John Sillitoe on behalf of the Wellcome Sanger Institute COVID-19 Surveillance Team |
| EPI_ISL_3438462, EPI_ISL_3438463 | Lighthouse Lab in Alderley Park | Wellcome Sanger Institute for the COVID-19 Genomics UK (COG-UK) Consortium | Jacquelyn Wynn, Mairead Hyland, The Lighthouse Lab in Alderley Park and Alex Alderton, Roberto Amato, Jeffrey Barrett, Sonia Goncalves, Ewan Harrison, David K. Jackson, Ian Johnston, Dominic Kwiatkowski, Cordelia Langford, John Sillitoe on behalf of the Wellcome Sanger Institute COVID-19 Surveillance Team |
| EPI_ISL_3438464 | Lighthouse Lab in Glasgow | Wellcome Sanger Institute for the COVID-19 Genomics UK (COG-UK) Consortium | Harper VanSteenhouse, Yumi Kasai, David Gray, Carol Clugston, Anna Dominiczak and Alex Alderton, Roberto Amato, Jeffrey Barrett, Sonia Goncalves, Ewan Harrison, David K. Jackson, Ian Johnston, Dominic Kwiatkowski, Cordelia Langford, John Sillitoe on behalf of the Wellcome Sanger Institute COVID-19 Surveillance Team |
| EPI_ISL_3438465, EPI_ISL_3438466 | Berkshire and Surrey Pathology Services Lighthouse Laboratory | Wellcome Sanger Institute for the COVID-19 Genomics UK (COG-UK) Consortium | Berkshire and Surrey Pathology Services Lighthouse Laboratory and Alex Alderton, Roberto Amato, Jeffrey Barrett, Sonia Goncalves, Ewan Harrison, David K. Jackson, Ian Johnston, Dominic Kwiatkowski, Cordelia Langford, John Sillitoe on behalf of the Wellcome Sanger Institute COVID-19 Surveillance Team |
| EPI_ISL_3438467, EPI_ISL_3438468 | Lighthouse Lab in Alderley Park | Wellcome Sanger Institute for the COVID-19 Genomics UK (COG-UK) Consortium | Jacquelyn Wynn, Mairead Hyland, The Lighthouse Lab in Alderley Park and Alex Alderton, Roberto Amato, Jeffrey Barrett, Sonia Goncalves, Ewan Harrison, David K. Jackson, Ian Johnston, Dominic Kwiatkowski, Cordelia Langford, John Sillitoe on behalf of the Wellcome Sanger Institute COVID-19 Surveillance Team |
| EPI_ISL_3438469, EPI_ISL_3438470, EPI_ISL_3438471 | Lighthouse Lab in Glasgow | Wellcome Sanger Institute for the COVID-19 Genomics UK (COG-UK) Consortium | Harper VanSteenhouse, Yumi Kasai, David Gray, Carol Clugston, Anna Dominiczak and Alex Alderton, Roberto Amato, Jeffrey Barrett, Sonia Goncalves, Ewan Harrison, David K. Jackson, Ian Johnston, Dominic Kwiatkowski, Cordelia Langford, John Sillitoe on behalf of the Wellcome Sanger Institute COVID-19 Surveillance Team |
| EPI_ISL_3438472, EPI_ISL_3438473 | Lighthouse Lab in Alderley Park | Wellcome Sanger Institute for the COVID-19 Genomics UK (COG-UK) Consortium | Jacquelyn Wynn, Mairead Hyland, The Lighthouse Lab in Alderley Park and Alex Alderton, Roberto Amato, Jeffrey Barrett, Sonia Goncalves, Ewan Harrison, David K. Jackson, Ian Johnston, Dominic Kwiatkowski, Cordelia Langford, John Sillitoe on behalf of the Wellcome Sanger Institute COVID-19 Surveillance Team |
| EPI_ISL_3438474, EPI_ISL_3438475, EPI_ISL_3438476, EPI_ISL_3438477 | Lighthouse Lab in Glasgow | Wellcome Sanger Institute for the COVID-19 Genomics UK (COG-UK) Consortium | Harper VanSteenhouse, Yumi Kasai, David Gray, Carol Clugston, Anna Dominiczak and Alex Alderton, Roberto Amato, Jeffrey Barrett, Sonia Goncalves, Ewan Harrison, David K. Jackson, Ian Johnston, Dominic Kwiatkowski, Cordelia Langford, John Sillitoe on behalf of the Wellcome Sanger Institute COVID-19 Surveillance Team |
| EPI_ISL_3438478 | Lighthouse Lab in Alderley Park | Wellcome Sanger Institute for the COVID-19 Genomics UK (COG-UK) Consortium | Jacquelyn Wynn, Mairead Hyland, The Lighthouse Lab in Alderley Park and Alex Alderton, Roberto Amato, Jeffrey Barrett, Sonia Goncalves, Ewan Harrison, David K. Jackson, Ian Johnston, Dominic Kwiatkowski, Cordelia Langford, John Sillitoe on behalf of the Wellcome Sanger Institute COVID-19 Surveillance Team |
| EPI_ISL_3438479, EPI_ISL_3438480 | Lighthouse Lab in Glasgow | Wellcome Sanger Institute for the COVID-19 Genomics UK (COG-UK) Consortium | Harper VanSteenhouse, Yumi Kasai, David Gray, Carol Clugston, Anna Dominiczak and Alex Alderton, Roberto Amato, Jeffrey Barrett, Sonia Goncalves, Ewan Harrison, David K. Jackson, Ian Johnston, Dominic Kwiatkowski, Cordelia Langford, John Sillitoe on behalf of the Wellcome Sanger Institute COVID-19 Surveillance Team |
| EPI_ISL_3438481 | Berkshire and Surrey Pathology Services Lighthouse Laboratory | Wellcome Sanger Institute for the COVID-19 Genomics UK (COG-UK) Consortium | Berkshire and Surrey Pathology Services Lighthouse Laboratory and Alex Alderton, Roberto Amato, Jeffrey Barrett, Sonia Goncalves, Ewan Harrison, David K. Jackson, Ian Johnston, Dominic Kwiatkowski, Cordelia Langford, John Sillitoe on behalf of the Wellcome Sanger Institute COVID-19 Surveillance Team |
| EPI_ISL_3438482 | Lighthouse Lab in Glasgow | Wellcome Sanger Institute for the COVID-19 Genomics UK (COG-UK) Consortium | Harper VanSteenhouse, Yumi Kasai, David Gray, Carol Clugston, Anna Dominiczak and Alex Alderton, Roberto Amato, Jeffrey Barrett, Sonia Goncalves, Ewan Harrison, David K. Jackson, Ian Johnston, Dominic Kwiatkowski, Cordelia Langford, John Sillitoe on behalf of the Wellcome Sanger Institute COVID-19 Surveillance Team |
| EPI_ISL_3438483 | Berkshire and Surrey Pathology Services Lighthouse Laboratory | Wellcome Sanger Institute for the COVID-19 Genomics UK (COG-UK) Consortium | Berkshire and Surrey Pathology Services Lighthouse Laboratory and Alex Alderton, Roberto Amato, Jeffrey Barrett, Sonia Goncalves, Ewan Harrison, David K. Jackson, Ian Johnston, Dominic Kwiatkowski, Cordelia Langford, John Sillitoe on behalf of the Wellcome Sanger Institute COVID-19 Surveillance Team |
| EPI_ISL_3438484 | Lighthouse Lab in Alderley Park | Wellcome Sanger Institute for the COVID-19 Genomics UK (COG-UK) Consortium | Jacquelyn Wynn, Mairead Hyland, The Lighthouse Lab in Alderley Park and Alex Alderton, Roberto Amato, Jeffrey Barrett, Sonia Goncalves, Ewan Harrison, David K. Jackson, Ian Johnston, Dominic Kwiatkowski, Cordelia Langford, John Sillitoe on behalf of the Wellcome Sanger Institute COVID-19 Surveillance Team |
| EPI_ISL_3438485, EPI_ISL_3438486 | Lighthouse Lab in Glasgow | Wellcome Sanger Institute for the COVID-19 Genomics UK (COG-UK) Consortium | Harper VanSteenhouse, Yumi Kasai, David Gray, Carol Clugston, Anna Dominiczak and Alex Alderton, Roberto Amato, Jeffrey Barrett, Sonia Goncalves, Ewan Harrison, David K. Jackson, Ian Johnston, Dominic Kwiatkowski, Cordelia Langford, John Sillitoe on behalf of the Wellcome Sanger Institute COVID-19 Surveillance Team |
| EPI_ISL_3438487 | Lighthouse Lab in Alderley Park | Wellcome Sanger Institute for the COVID-19 Genomics UK (COG-UK) Consortium | Jacquelyn Wynn, Mairead Hyland, The Lighthouse Lab in Alderley Park and Alex Alderton, Roberto Amato, Jeffrey Barrett, Sonia Goncalves, Ewan Harrison, David K. Jackson, Ian Johnston, Dominic Kwiatkowski, Cordelia Langford, John Sillitoe on behalf of the Wellcome Sanger Institute COVID-19 Surveillance Team |
| EPI_ISL_3438488, EPI_ISL_3438489 | Lighthouse Lab in Glasgow | Wellcome Sanger Institute for the COVID-19 Genomics UK (COG-UK) Consortium | Harper VanSteenhouse, Yumi Kasai, David Gray, Carol Clugston, Anna Dominiczak and Alex Alderton, Roberto Amato, Jeffrey Barrett, Sonia Goncalves, Ewan Harrison, David K. Jackson, Ian Johnston, Dominic Kwiatkowski, Cordelia Langford, John Sillitoe on behalf of the Wellcome Sanger Institute COVID-19 Surveillance Team |
| EPI_ISL_3438490 | Lighthouse Lab in Alderley Park | Wellcome Sanger Institute for the COVID-19 Genomics UK (COG-UK) Consortium | Jacquelyn Wynn, Mairead Hyland, The Lighthouse Lab in Alderley Park and Alex Alderton, Roberto Amato, Jeffrey Barrett, Sonia Goncalves, Ewan Harrison, David K. Jackson, Ian Johnston, Dominic Kwiatkowski, Cordelia Langford, John Sillitoe on behalf of the Wellcome Sanger Institute COVID-19 Surveillance Team |
| EPI_ISL_3438491 | Lighthouse Lab in Glasgow | Wellcome Sanger Institute for the COVID-19 Genomics UK (COG-UK) Consortium | Harper VanSteenhouse, Yumi Kasai, David Gray, Carol Clugston, Anna Dominiczak and Alex Alderton, Roberto Amato, Jeffrey Barrett, Sonia Goncalves, Ewan Harrison, David K. Jackson, Ian Johnston, Dominic Kwiatkowski, Cordelia Langford, John Sillitoe on behalf of the Wellcome Sanger Institute COVID-19 Surveillance Team |
| EPI_ISL_3438492 | Berkshire and Surrey Pathology Services Lighthouse Laboratory | Wellcome Sanger Institute for the COVID-19 Genomics UK (COG-UK) Consortium | Berkshire and Surrey Pathology Services Lighthouse Laboratory and Alex Alderton, Roberto Amato, Jeffrey Barrett, Sonia Goncalves, Ewan Harrison, David K. Jackson, Ian Johnston, Dominic Kwiatkowski, Cordelia Langford, John Sillitoe on behalf of the Wellcome Sanger Institute COVID-19 Surveillance Team |
| EPI_ISL_3438493, EPI_ISL_3438494, EPI_ISL_3438495 | Lighthouse Lab in Alderley Park | Wellcome Sanger Institute for the COVID-19 Genomics UK (COG-UK) Consortium | Jacquelyn Wynn, Mairead Hyland, The Lighthouse Lab in Alderley Park and Alex Alderton, Roberto Amato, Jeffrey Barrett, Sonia Goncalves, Ewan Harrison, David K. Jackson, Ian Johnston, Dominic Kwiatkowski, Cordelia Langford, John Sillitoe on behalf of the Wellcome Sanger Institute COVID-19 Surveillance Team |
| EPI_ISL_3438496, EPI_ISL_3438497 | Lighthouse Lab in Glasgow | Wellcome Sanger Institute for the COVID-19 Genomics UK (COG-UK) Consortium | Harper VanSteenhouse, Yumi Kasai, David Gray, Carol Clugston, Anna Dominiczak and Alex Alderton, Roberto Amato, Jeffrey Barrett, Sonia Goncalves, Ewan Harrison, David K. Jackson, Ian Johnston, Dominic Kwiatkowski, Cordelia Langford, John Sillitoe on behalf of the Wellcome Sanger Institute COVID-19 Surveillance Team |
| EPI_ISL_3438498 | Berkshire and Surrey Pathology Services Lighthouse Laboratory | Wellcome Sanger Institute for the COVID-19 Genomics UK (COG-UK) Consortium | Berkshire and Surrey Pathology Services Lighthouse Laboratory and Alex Alderton, Roberto Amato, Jeffrey Barrett, Sonia Goncalves, Ewan Harrison, David K. Jackson, Ian Johnston, Dominic Kwiatkowski, Cordelia Langford, John Sillitoe on behalf of the Wellcome Sanger Institute COVID-19 Surveillance Team |
| EPI_ISL_3438499 | Lighthouse Lab in Glasgow | Wellcome Sanger Institute for the COVID-19 Genomics UK (COG-UK) Consortium | Harper VanSteenhouse, Yumi Kasai, David Gray, Carol Clugston, Anna Dominiczak and Alex Alderton, Roberto Amato, Jeffrey Barrett, Sonia Goncalves, Ewan Harrison, David K. Jackson, Ian Johnston, Dominic Kwiatkowski, Cordelia Langford, John Sillitoe on behalf of the Wellcome Sanger Institute COVID-19 Surveillance Team |
| EPI_ISL_3438500 | Lighthouse Lab in Alderley Park | Wellcome Sanger Institute for the COVID-19 Genomics UK (COG-UK) Consortium | Jacquelyn Wynn, Mairead Hyland, The Lighthouse Lab in Alderley Park and Alex Alderton, Roberto Amato, Jeffrey Barrett, Sonia Goncalves, Ewan Harrison, David K. Jackson, Ian Johnston, Dominic Kwiatkowski, Cordelia Langford, John Sillitoe on behalf of the Wellcome Sanger Institute COVID-19 Surveillance Team |
| EPI_ISL_3438501, EPI_ISL_3438502, EPI_ISL_3438503, EPI_ISL_3438504, EPI_ISL_3438505 | Lighthouse Lab in Glasgow | Wellcome Sanger Institute for the COVID-19 Genomics UK (COG-UK) Consortium | Harper VanSteenhouse, Yumi Kasai, David Gray, Carol Clugston, Anna Dominiczak and Alex Alderton, Roberto Amato, Jeffrey Barrett, Sonia Goncalves, Ewan Harrison, David K. Jackson, Ian Johnston, Dominic Kwiatkowski, Cordelia Langford, John Sillitoe on behalf of the Wellcome Sanger Institute COVID-19 Surveillance Team |
| EPI_ISL_3438506 | Berkshire and Surrey Pathology Services Lighthouse Laboratory | Wellcome Sanger Institute for the COVID-19 Genomics UK (COG-UK) Consortium | Berkshire and Surrey Pathology Services Lighthouse Laboratory and Alex Alderton, Roberto Amato, Jeffrey Barrett, Sonia Goncalves, Ewan Harrison, David K. Jackson, Ian Johnston, Dominic Kwiatkowski, Cordelia Langford, John Sillitoe on behalf of the Wellcome Sanger Institute COVID-19 Surveillance Team |
| EPI_ISL_3438507 | Lighthouse Lab in Alderley Park | Wellcome Sanger Institute for the COVID-19 Genomics UK (COG-UK) Consortium | Jacquelyn Wynn, Mairead Hyland, The Lighthouse Lab in Alderley Park and Alex Alderton, Roberto Amato, Jeffrey Barrett, Sonia Goncalves, Ewan Harrison, David K. Jackson, Ian Johnston, Dominic Kwiatkowski, Cordelia Langford, John Sillitoe on behalf of the Wellcome Sanger Institute COVID-19 Surveillance Team |
| EPI_ISL_3438508 | Lighthouse Lab in Glasgow | Wellcome Sanger Institute for the COVID-19 Genomics UK (COG-UK) Consortium | Harper VanSteenhouse, Yumi Kasai, David Gray, Carol Clugston, Anna Dominiczak and Alex Alderton, Roberto Amato, Jeffrey Barrett, Sonia Goncalves, Ewan Harrison, David K. Jackson, Ian Johnston, Dominic Kwiatkowski, Cordelia Langford, John Sillitoe on behalf of the Wellcome Sanger Institute COVID-19 Surveillance Team |
| EPI_ISL_3438509, EPI_ISL_3438510 | Lighthouse Lab in Alderley Park | Wellcome Sanger Institute for the COVID-19 Genomics UK (COG-UK) Consortium | Jacquelyn Wynn, Mairead Hyland, The Lighthouse Lab in Alderley Park and Alex Alderton, Roberto Amato, Jeffrey Barrett, Sonia Goncalves, Ewan Harrison, David K. Jackson, Ian Johnston, Dominic Kwiatkowski, Cordelia Langford, John Sillitoe on behalf of the Wellcome Sanger Institute COVID-19 Surveillance Team |
| EPI_ISL_3438511, EPI_ISL_3438512 | Berkshire and Surrey Pathology Services Lighthouse Laboratory | Wellcome Sanger Institute for the COVID-19 Genomics UK (COG-UK) Consortium | Berkshire and Surrey Pathology Services Lighthouse Laboratory and Alex Alderton, Roberto Amato, Jeffrey Barrett, Sonia Goncalves, Ewan Harrison, David K. Jackson, Ian Johnston, Dominic Kwiatkowski, Cordelia Langford, John Sillitoe on behalf of the Wellcome Sanger Institute COVID-19 Surveillance Team |
| EPI_ISL_3438513, EPI_ISL_3438514 | Lighthouse Lab in Alderley Park | Wellcome Sanger Institute for the COVID-19 Genomics UK (COG-UK) Consortium | Jacquelyn Wynn, Mairead Hyland, The Lighthouse Lab in Alderley Park and Alex Alderton, Roberto Amato, Jeffrey Barrett, Sonia Goncalves, Ewan Harrison, David K. Jackson, Ian Johnston, Dominic Kwiatkowski, Cordelia Langford, John Sillitoe on behalf of the Wellcome Sanger Institute COVID-19 Surveillance Team |
| EPI_ISL_3438515 | Lighthouse Lab in Glasgow | Wellcome Sanger Institute for the COVID-19 Genomics UK (COG-UK) Consortium | Harper VanSteenhouse, Yumi Kasai, David Gray, Carol Clugston, Anna Dominiczak and Alex Alderton, Roberto Amato, Jeffrey Barrett, Sonia Goncalves, Ewan Harrison, David K. Jackson, Ian Johnston, Dominic Kwiatkowski, Cordelia Langford, John Sillitoe on behalf of the Wellcome Sanger Institute COVID-19 Surveillance Team |
| EPI_ISL_3438516, EPI_ISL_3438517 | Lighthouse Lab in Alderley Park | Wellcome Sanger Institute for the COVID-19 Genomics UK (COG-UK) Consortium | Jacquelyn Wynn, Mairead Hyland, The Lighthouse Lab in Alderley Park and Alex Alderton, Roberto Amato, Jeffrey Barrett, Sonia Goncalves, Ewan Harrison, David K. Jackson, Ian Johnston, Dominic Kwiatkowski, Cordelia Langford, John Sillitoe on behalf of the Wellcome Sanger Institute COVID-19 Surveillance Team |

| | | | |
|---|---|---|---|
| EPI_ISL_3438518 | Berkshire and Surrey Pathology Services Lighthouse Laboratory | Wellcome Sanger Institute for the COVID-19 Genomics UK (COG-UK) Consortium | Berkshire and Surrey Pathology Services Lighthouse Laboratory and Alex Alderton, Roberto Amato, Jeffrey Barrett, Sonia Goncalves, Ewan Harrison, David K. Jackson, Ian Johnston, Dominic Kwiatkowski, Cordelia Langford, John Sillitoe on behalf of the Wellcome Sanger Institute COVID-19 Surveillance Team |
| EPI_ISL_3438519, EPI_ISL_3438520 | Lighthouse Lab in Glasgow | Wellcome Sanger Institute for the COVID-19 Genomics UK (COG-UK) Consortium | Harper VanSteenhouse, Yumi Kasai, David Gray, Carol Clugston, Anna Dominiczak and Alex Alderton, Roberto Amato, Jeffrey Barrett, Sonia Goncalves, Ewan Harrison, David K. Jackson, Ian Johnston, Dominic Kwiatkowski, Cordelia Langford, John Sillitoe on behalf of the Wellcome Sanger Institute COVID-19 Surveillance Team |
| EPI_ISL_3438521 | Lighthouse Lab in Alderley Park | Wellcome Sanger Institute for the COVID-19 Genomics UK (COG-UK) Consortium | Jacquelyn Wynn, Mairead Hyland, The Lighthouse Lab in Alderley Park and Alex Alderton, Roberto Amato, Jeffrey Barrett, Sonia Goncalves, Ewan Harrison, David K. Jackson, Ian Johnston, Dominic Kwiatkowski, Cordelia Langford, John Sillitoe on behalf of the Wellcome Sanger Institute COVID-19 Surveillance Team |
| EPI_ISL_3438522 | Lighthouse Lab in Glasgow | Wellcome Sanger Institute for the COVID-19 Genomics UK (COG-UK) Consortium | Harper VanSteenhouse, Yumi Kasai, David Gray, Carol Clugston, Anna Dominiczak and Alex Alderton, Roberto Amato, Jeffrey Barrett, Sonia Goncalves, Ewan Harrison, David K. Jackson, Ian Johnston, Dominic Kwiatkowski, Cordelia Langford, John Sillitoe on behalf of the Wellcome Sanger Institute COVID-19 Surveillance Team |
| EPI_ISL_3438523, EPI_ISL_3438524, EPI_ISL_3438525 | Berkshire and Surrey Pathology Services Lighthouse Laboratory | Wellcome Sanger Institute for the COVID-19 Genomics UK (COG-UK) Consortium | Berkshire and Surrey Pathology Services Lighthouse Laboratory and Alex Alderton, Roberto Amato, Jeffrey Barrett, Sonia Goncalves, Ewan Harrison, David K. Jackson, Ian Johnston, Dominic Kwiatkowski, Cordelia Langford, John Sillitoe on behalf of the Wellcome Sanger Institute COVID-19 Surveillance Team |
| EPI_ISL_3438526 | Lighthouse Lab in Alderley Park | Wellcome Sanger Institute for the COVID-19 Genomics UK (COG-UK) Consortium | Jacquelyn Wynn, Mairead Hyland, The Lighthouse Lab in Alderley Park and Alex Alderton, Roberto Amato, Jeffrey Barrett, Sonia Goncalves, Ewan Harrison, David K. Jackson, Ian Johnston, Dominic Kwiatkowski, Cordelia Langford, John Sillitoe on behalf of the Wellcome Sanger Institute COVID-19 Surveillance Team |
| EPI_ISL_3438527, EPI_ISL_3438528 | Lighthouse Lab in Glasgow | Wellcome Sanger Institute for the COVID-19 Genomics UK (COG-UK) Consortium | Harper VanSteenhouse, Yumi Kasai, David Gray, Carol Clugston, Anna Dominiczak and Alex Alderton, Roberto Amato, Jeffrey Barrett, Sonia Goncalves, Ewan Harrison, David K. Jackson, Ian Johnston, Dominic Kwiatkowski, Cordelia Langford, John Sillitoe on behalf of the Wellcome Sanger Institute COVID-19 Surveillance Team |
| EPI_ISL_3438529 | Berkshire and Surrey Pathology Services Lighthouse Laboratory | Wellcome Sanger Institute for the COVID-19 Genomics UK (COG-UK) Consortium | Berkshire and Surrey Pathology Services Lighthouse Laboratory and Alex Alderton, Roberto Amato, Jeffrey Barrett, Sonia Goncalves, Ewan Harrison, David K. Jackson, Ian Johnston, Dominic Kwiatkowski, Cordelia Langford, John Sillitoe on behalf of the Wellcome Sanger Institute COVID-19 Surveillance Team |
| EPI_ISL_3438530 | Lighthouse Lab in Glasgow | Wellcome Sanger Institute for the COVID-19 Genomics UK (COG-UK) Consortium | Harper VanSteenhouse, Yumi Kasai, David Gray, Carol Clugston, Anna Dominiczak and Alex Alderton, Roberto Amato, Jeffrey Barrett, Sonia Goncalves, Ewan Harrison, David K. Jackson, Ian Johnston, Dominic Kwiatkowski, Cordelia Langford, John Sillitoe on behalf of the Wellcome Sanger Institute COVID-19 Surveillance Team |
| EPI_ISL_3438531, EPI_ISL_3438532, EPI_ISL_3438533 | Berkshire and Surrey Pathology Services Lighthouse Laboratory | Wellcome Sanger Institute for the COVID-19 Genomics UK (COG-UK) Consortium | Berkshire and Surrey Pathology Services Lighthouse Laboratory and Alex Alderton, Roberto Amato, Jeffrey Barrett, Sonia Goncalves, Ewan Harrison, David K. Jackson, Ian Johnston, Dominic Kwiatkowski, Cordelia Langford, John Sillitoe on behalf of the Wellcome Sanger Institute COVID-19 Surveillance Team |
| EPI_ISL_3438534, EPI_ISL_3438535 | Lighthouse Lab in Glasgow | Wellcome Sanger Institute for the COVID-19 Genomics UK (COG-UK) Consortium | Harper VanSteenhouse, Yumi Kasai, David Gray, Carol Clugston, Anna Dominiczak and Alex Alderton, Roberto Amato, Jeffrey Barrett, Sonia Goncalves, Ewan Harrison, David K. Jackson, Ian Johnston, Dominic Kwiatkowski, Cordelia Langford, John Sillitoe on behalf of the Wellcome Sanger Institute COVID-19 Surveillance Team |
| EPI_ISL_3438536 | Berkshire and Surrey Pathology Services Lighthouse Laboratory | Wellcome Sanger Institute for the COVID-19 Genomics UK (COG-UK) Consortium | Berkshire and Surrey Pathology Services Lighthouse Laboratory and Alex Alderton, Roberto Amato, Jeffrey Barrett, Sonia Goncalves, Ewan Harrison, David K. Jackson, Ian Johnston, Dominic Kwiatkowski, Cordelia Langford, John Sillitoe on behalf of the Wellcome Sanger Institute COVID-19 Surveillance Team |
| EPI_ISL_3438537 | Lighthouse Lab in Glasgow | Wellcome Sanger Institute for the COVID-19 Genomics UK (COG-UK) Consortium | Harper VanSteenhouse, Yumi Kasai, David Gray, Carol Clugston, Anna Dominiczak and Alex Alderton, Roberto Amato, Jeffrey Barrett, Sonia Goncalves, Ewan Harrison, David K. Jackson, Ian Johnston, Dominic Kwiatkowski, Cordelia Langford, John Sillitoe on behalf of the Wellcome Sanger Institute COVID-19 Surveillance Team |
| EPI_ISL_3438538 | Berkshire and Surrey Pathology Services Lighthouse Laboratory | Wellcome Sanger Institute for the COVID-19 Genomics UK (COG-UK) Consortium | Berkshire and Surrey Pathology Services Lighthouse Laboratory and Alex Alderton, Roberto Amato, Jeffrey Barrett, Sonia Goncalves, Ewan Harrison, David K. Jackson, Ian Johnston, Dominic Kwiatkowski, Cordelia Langford, John Sillitoe on behalf of the Wellcome Sanger Institute COVID-19 Surveillance Team |
| EPI_ISL_3438539, EPI_ISL_3438540 | Lighthouse Lab in Alderley Park | Wellcome Sanger Institute for the COVID-19 Genomics UK (COG-UK) Consortium | Jacquelyn Wynn, Mairead Hyland, The Lighthouse Lab in Alderley Park and Alex Alderton, Roberto Amato, Jeffrey Barrett, Sonia Goncalves, Ewan Harrison, David K. Jackson, Ian Johnston, Dominic Kwiatkowski, Cordelia Langford, John Sillitoe on behalf of the Wellcome Sanger Institute COVID-19 Surveillance Team |
| EPI_ISL_3438541 | Berkshire and Surrey Pathology Services Lighthouse Laboratory | Wellcome Sanger Institute for the COVID-19 Genomics UK (COG-UK) Consortium | Berkshire and Surrey Pathology Services Lighthouse Laboratory and Alex Alderton, Roberto Amato, Jeffrey Barrett, Sonia Goncalves, Ewan Harrison, David K. Jackson, Ian Johnston, Dominic Kwiatkowski, Cordelia Langford, John Sillitoe on behalf of the Wellcome Sanger Institute COVID-19 Surveillance Team |
| EPI_ISL_3438542 | Lighthouse Lab in Alderley Park | Wellcome Sanger Institute for the COVID-19 Genomics UK (COG-UK) Consortium | Jacquelyn Wynn, Mairead Hyland, The Lighthouse Lab in Alderley Park and Alex Alderton, Roberto Amato, Jeffrey Barrett, Sonia Goncalves, Ewan Harrison, David K. Jackson, Ian Johnston, Dominic Kwiatkowski, Cordelia Langford, John Sillitoe on behalf of the Wellcome Sanger Institute COVID-19 Surveillance Team |
| EPI_ISL_3438543 | Berkshire and Surrey Pathology Services Lighthouse Laboratory | Wellcome Sanger Institute for the COVID-19 Genomics UK (COG-UK) Consortium | Berkshire and Surrey Pathology Services Lighthouse Laboratory and Alex Alderton, Roberto Amato, Jeffrey Barrett, Sonia Goncalves, Ewan Harrison, David K. Jackson, Ian Johnston, Dominic Kwiatkowski, Cordelia Langford, John Sillitoe on behalf of the Wellcome Sanger Institute COVID-19 Surveillance Team |
| EPI_ISL_3438544, EPI_ISL_3438545 | Lighthouse Lab in Alderley Park | Wellcome Sanger Institute for the COVID-19 Genomics UK (COG-UK) Consortium | Jacquelyn Wynn, Mairead Hyland, The Lighthouse Lab in Alderley Park and Alex Alderton, Roberto Amato, Jeffrey Barrett, Sonia Goncalves, Ewan Harrison, David K. Jackson, Ian Johnston, Dominic Kwiatkowski, Cordelia Langford, John Sillitoe on behalf of the Wellcome Sanger Institute COVID-19 Surveillance Team |
| EPI_ISL_3438546 | Lighthouse Lab in Glasgow | Wellcome Sanger Institute for the COVID-19 Genomics UK (COG-UK) Consortium | Harper VanSteenhouse, Yumi Kasai, David Gray, Carol Clugston, Anna Dominiczak and Alex Alderton, Roberto Amato, Jeffrey Barrett, Sonia Goncalves, Ewan Harrison, David K. Jackson, Ian Johnston, Dominic Kwiatkowski, Cordelia Langford, John Sillitoe on behalf of the Wellcome Sanger Institute COVID-19 Surveillance Team |
| EPI_ISL_3438547, EPI_ISL_3438548 | Lighthouse Lab in Alderley Park | Wellcome Sanger Institute for the COVID-19 Genomics UK (COG-UK) Consortium | Jacquelyn Wynn, Mairead Hyland, The Lighthouse Lab in Alderley Park and Alex Alderton, Roberto Amato, Jeffrey Barrett, Sonia Goncalves, Ewan Harrison, David K. Jackson, Ian Johnston, Dominic Kwiatkowski, Cordelia Langford, John Sillitoe on behalf of the Wellcome Sanger Institute COVID-19 Surveillance Team |
| EPI_ISL_3438549 | Lighthouse Lab in Glasgow | Wellcome Sanger Institute for the COVID-19 Genomics UK (COG-UK) Consortium | Harper VanSteenhouse, Yumi Kasai, David Gray, Carol Clugston, Anna Dominiczak and Alex Alderton, Roberto Amato, Jeffrey Barrett, Sonia Goncalves, Ewan Harrison, David K. Jackson, Ian Johnston, Dominic Kwiatkowski, Cordelia Langford, John Sillitoe on behalf of the Wellcome Sanger Institute COVID-19 Surveillance Team |
| EPI_ISL_3438550, EPI_ISL_3438551, EPI_ISL_3438552 | Lighthouse Lab in Alderley Park | Wellcome Sanger Institute for the COVID-19 Genomics UK (COG-UK) Consortium | Jacquelyn Wynn, Mairead Hyland, The Lighthouse Lab in Alderley Park and Alex Alderton, Roberto Amato, Jeffrey Barrett, Sonia Goncalves, Ewan Harrison, David K. Jackson, Ian Johnston, Dominic Kwiatkowski, Cordelia Langford, John Sillitoe on behalf of the Wellcome Sanger Institute COVID-19 Surveillance Team |
| EPI_ISL_3438553, EPI_ISL_3438554 | Lighthouse Lab in Glasgow | Wellcome Sanger Institute for the COVID-19 Genomics UK (COG-UK) Consortium | Harper VanSteenhouse, Yumi Kasai, David Gray, Carol Clugston, Anna Dominiczak and Alex Alderton, Roberto Amato, Jeffrey Barrett, Sonia Goncalves, Ewan Harrison, David K. Jackson, Ian Johnston, Dominic Kwiatkowski, Cordelia Langford, John Sillitoe on behalf of the Wellcome Sanger Institute COVID-19 Surveillance Team |
| EPI_ISL_3438555 | Lighthouse Lab in Alderley Park | Wellcome Sanger Institute for the COVID-19 Genomics UK (COG-UK) Consortium | Jacquelyn Wynn, Mairead Hyland, The Lighthouse Lab in Alderley Park and Alex Alderton, Roberto Amato, Jeffrey Barrett, Sonia Goncalves, Ewan Harrison, David K. Jackson, Ian Johnston, Dominic Kwiatkowski, Cordelia Langford, John Sillitoe on behalf of the Wellcome Sanger Institute COVID-19 Surveillance Team |
| EPI_ISL_3438556 | Berkshire and Surrey Pathology Services Lighthouse Laboratory | Wellcome Sanger Institute for the COVID-19 Genomics UK (COG-UK) Consortium | Berkshire and Surrey Pathology Services Lighthouse Laboratory and Alex Alderton, Roberto Amato, Jeffrey Barrett, Sonia Goncalves, Ewan Harrison, David K. Jackson, Ian Johnston, Dominic Kwiatkowski, Cordelia Langford, John Sillitoe on behalf of the Wellcome Sanger Institute COVID-19 Surveillance Team |
| EPI_ISL_3438557 | Lighthouse Lab in Alderley Park | Wellcome Sanger Institute for the COVID-19 Genomics UK (COG-UK) Consortium | Jacquelyn Wynn, Mairead Hyland, The Lighthouse Lab in Alderley Park and Alex Alderton, Roberto Amato, Jeffrey Barrett, Sonia Goncalves, Ewan Harrison, David K. Jackson, Ian Johnston, Dominic Kwiatkowski, Cordelia Langford, John Sillitoe on behalf of the Wellcome Sanger Institute COVID-19 Surveillance Team |
| EPI_ISL_3438558, EPI_ISL_3438559, EPI_ISL_3438560 | Berkshire and Surrey Pathology Services Lighthouse Laboratory | Wellcome Sanger Institute for the COVID-19 Genomics UK (COG-UK) Consortium | Berkshire and Surrey Pathology Services Lighthouse Laboratory and Alex Alderton, Roberto Amato, Jeffrey Barrett, Sonia Goncalves, Ewan Harrison, David K. Jackson, Ian Johnston, Dominic Kwiatkowski, Cordelia Langford, John Sillitoe on behalf of the Wellcome Sanger Institute COVID-19 Surveillance Team |
| EPI_ISL_3438561 | Lighthouse Lab in Glasgow | Wellcome Sanger Institute for the COVID-19 Genomics UK (COG-UK) Consortium | Harper VanSteenhouse, Yumi Kasai, David Gray, Carol Clugston, Anna Dominiczak and Alex Alderton, Roberto Amato, Jeffrey Barrett, Sonia Goncalves, Ewan Harrison, David K. Jackson, Ian Johnston, Dominic Kwiatkowski, Cordelia Langford, John Sillitoe on behalf of the Wellcome Sanger Institute COVID-19 Surveillance Team |
| EPI_ISL_3438562 | Berkshire and Surrey Pathology Services Lighthouse Laboratory | Wellcome Sanger Institute for the COVID-19 Genomics UK (COG-UK) Consortium | Berkshire and Surrey Pathology Services Lighthouse Laboratory and Alex Alderton, Roberto Amato, Jeffrey Barrett, Sonia Goncalves, Ewan Harrison, David K. Jackson, Ian Johnston, Dominic Kwiatkowski, Cordelia Langford, John Sillitoe on behalf of the Wellcome Sanger Institute COVID-19 Surveillance Team |
| EPI_ISL_3438563 | Lighthouse Lab in Glasgow | Wellcome Sanger Institute for the COVID-19 Genomics UK (COG-UK) Consortium | Harper VanSteenhouse, Yumi Kasai, David Gray, Carol Clugston, Anna Dominiczak and Alex Alderton, Roberto Amato, Jeffrey Barrett, Sonia Goncalves, Ewan Harrison, David K. Jackson, Ian Johnston, Dominic Kwiatkowski, Cordelia Langford, John Sillitoe on behalf of the Wellcome Sanger Institute COVID-19 Surveillance Team |
| EPI_ISL_3438564 | Lighthouse Lab in Alderley Park | Wellcome Sanger Institute for the COVID-19 Genomics UK (COG-UK) Consortium | Jacquelyn Wynn, Mairead Hyland, The Lighthouse Lab in Alderley Park and Alex Alderton, Roberto Amato, Jeffrey Barrett, Sonia Goncalves, Ewan Harrison, David K. Jackson, Ian Johnston, Dominic Kwiatkowski, Cordelia Langford, John Sillitoe on behalf of the Wellcome Sanger Institute COVID-19 Surveillance Team |
| EPI_ISL_3438565, EPI_ISL_3438566 | Lighthouse Lab in Glasgow | Wellcome Sanger Institute for the COVID-19 Genomics UK (COG-UK) Consortium | Harper VanSteenhouse, Yumi Kasai, David Gray, Carol Clugston, Anna Dominiczak and Alex Alderton, Roberto Amato, Jeffrey Barrett, Sonia Goncalves, Ewan Harrison, David K. Jackson, Ian Johnston, Dominic Kwiatkowski, Cordelia Langford, John Sillitoe on behalf of the Wellcome Sanger Institute COVID-19 Surveillance Team |
| EPI_ISL_3438567, EPI_ISL_3438568, EPI_ISL_3438569, EPI_ISL_3438570 | Lighthouse Lab in Alderley Park | Wellcome Sanger Institute for the COVID-19 Genomics UK (COG-UK) Consortium | Jacquelyn Wynn, Mairead Hyland, The Lighthouse Lab in Alderley Park and Alex Alderton, Roberto Amato, Jeffrey Barrett, Sonia Goncalves, Ewan Harrison, David K. Jackson, Ian Johnston, Dominic Kwiatkowski, Cordelia Langford, John Sillitoe on behalf of the Wellcome Sanger Institute COVID-19 Surveillance Team |
| EPI_ISL_3438571 | Berkshire and Surrey Pathology Services Lighthouse Laboratory | Wellcome Sanger Institute for the COVID-19 Genomics UK (COG-UK) Consortium | Berkshire and Surrey Pathology Services Lighthouse Laboratory and Alex Alderton, Roberto Amato, Jeffrey Barrett, Sonia Goncalves, Ewan Harrison, David K. Jackson, Ian Johnston, Dominic Kwiatkowski, Cordelia Langford, John Sillitoe on behalf of the Wellcome Sanger Institute COVID-19 Surveillance Team |
| EPI_ISL_3438572 | Lighthouse Lab in Glasgow | Wellcome Sanger Institute for the COVID-19 Genomics UK (COG-UK) Consortium | Harper VanSteenhouse, Yumi Kasai, David Gray, Carol Clugston, Anna Dominiczak and Alex Alderton, Roberto Amato, Jeffrey Barrett, Sonia Goncalves, Ewan Harrison, David K. Jackson, Ian Johnston, Dominic Kwiatkowski, Cordelia Langford, John Sillitoe on behalf of the Wellcome Sanger Institute COVID-19 Surveillance Team |
| EPI_ISL_3438573 | Lighthouse Lab in Alderley Park | Wellcome Sanger Institute for the COVID-19 Genomics UK (COG-UK) Consortium | Jacquelyn Wynn, Mairead Hyland, The Lighthouse Lab in Alderley Park and Alex Alderton, Roberto Amato, Jeffrey Barrett, Sonia Goncalves, Ewan Harrison, David K. Jackson, Ian Johnston, Dominic Kwiatkowski, Cordelia Langford, John Sillitoe on behalf of the Wellcome Sanger Institute COVID-19 Surveillance Team |
| EPI_ISL_3438574 | Berkshire and Surrey Pathology Services Lighthouse Laboratory | Wellcome Sanger Institute for the COVID-19 Genomics UK (COG-UK) Consortium | Berkshire and Surrey Pathology Services Lighthouse Laboratory and Alex Alderton, Roberto Amato, Jeffrey Barrett, Sonia Goncalves, Ewan Harrison, David K. Jackson, Ian Johnston, Dominic Kwiatkowski, Cordelia Langford, John Sillitoe on behalf of the Wellcome Sanger Institute COVID-19 Surveillance Team |
| EPI_ISL_3438575 | Lighthouse Lab in Glasgow | Wellcome Sanger Institute for the COVID-19 Genomics UK (COG-UK) Consortium | Harper VanSteenhouse, Yumi Kasai, David Gray, Carol Clugston, Anna Dominiczak and Alex Alderton, Roberto Amato, Jeffrey Barrett, Sonia Goncalves, Ewan Harrison, David K. Jackson, Ian Johnston, Dominic Kwiatkowski, Cordelia Langford, John Sillitoe on behalf of the Wellcome Sanger Institute COVID-19 Surveillance Team |
| EPI_ISL_3438576 | Lighthouse Lab in Alderley Park | Wellcome Sanger Institute for the COVID-19 Genomics UK (COG-UK) Consortium | Jacquelyn Wynn, Mairead Hyland, The Lighthouse Lab in Alderley Park and Alex Alderton, Roberto Amato, Jeffrey Barrett, Sonia Goncalves, Ewan Harrison, David K. Jackson, Ian Johnston, Dominic Kwiatkowski, Cordelia Langford, John Sillitoe on behalf of the Wellcome Sanger Institute COVID-19 Surveillance Team |
| EPI_ISL_3438577, EPI_ISL_3438578 | Berkshire and Surrey Pathology Services Lighthouse Laboratory | Wellcome Sanger Institute for the COVID-19 Genomics UK (COG-UK) Consortium | Berkshire and Surrey Pathology Services Lighthouse Laboratory and Alex Alderton, Roberto Amato, Jeffrey Barrett, Sonia Goncalves, Ewan Harrison, David K. Jackson, Ian Johnston, Dominic Kwiatkowski, Cordelia Langford, John Sillitoe on behalf of the Wellcome Sanger Institute COVID-19 Surveillance Team |
| EPI_ISL_3438579 | Lighthouse Lab in Alderley Park | Wellcome Sanger Institute for the COVID-19 Genomics UK (COG-UK) Consortium | Jacquelyn Wynn, Mairead Hyland, The Lighthouse Lab in Alderley Park and Alex Alderton, Roberto Amato, Jeffrey Barrett, Sonia Goncalves, Ewan Harrison, David K. Jackson, Ian Johnston, Dominic Kwiatkowski, Cordelia Langford, John Sillitoe on behalf of the Wellcome Sanger Institute COVID-19 Surveillance Team |
| EPI_ISL_3438580 | Lighthouse Lab in Glasgow | Wellcome Sanger Institute for the COVID-19 Genomics UK (COG-UK) | Harper VanSteenhouse, Yumi Kasai, David Gray, Carol Clugston, Anna Dominiczak and Alex Alderton, Roberto Amato, Jeffrey Barrett, Sonia Goncalves, Ewan Harrison, David K. Jackson, Ian Johnston, Dominic Kwiatkowski, Cordelia Langford, |

| | | | |
|---|---|---|---|
| | | Consortium | John Sillitoe on behalf of the Wellcome Sanger Institute COVID-19 Surveillance Team |
| EPI_ISL_3438581 | Lighthouse Lab in Alderley Park | Wellcome Sanger Institute for the COVID-19 Genomics UK (COG-UK) Consortium | Jacquelyn Wynn, Mairead Hyland, The Lighthouse Lab in Alderley Park and Alex Alderton, Roberto Amato, Jeffrey Barrett, Sonia Goncalves, Ewan Harrison, David K. Jackson, Ian Johnston, Dominic Kwiatkowski, Cordelia Langford, John Sillitoe on behalf of the Wellcome Sanger Institute COVID-19 Surveillance Team |
| EPI_ISL_3438582 | Berkshire and Surrey Pathology Services Lighthouse Laboratory | Wellcome Sanger Institute for the COVID-19 Genomics UK (COG-UK) Consortium | Berkshire and Surrey Pathology Services Lighthouse Laboratory and Alex Alderton, Roberto Amato, Jeffrey Barrett, Sonia Goncalves, Ewan Harrison, David K. Jackson, Ian Johnston, Dominic Kwiatkowski, Cordelia Langford, John Sillitoe on behalf of the Wellcome Sanger Institute COVID-19 Surveillance Team |
| EPI_ISL_3438583 | Lighthouse Lab in Glasgow | Wellcome Sanger Institute for the COVID-19 Genomics UK (COG-UK) Consortium | Harper VanSteenhouse, Yumi Kasai, David Gray, Carol Clugston, Anna Dominiczak and Alex Alderton, Roberto Amato, Jeffrey Barrett, Sonia Goncalves, Ewan Harrison, David K. Jackson, Ian Johnston, Dominic Kwiatkowski, Cordelia Langford, John Sillitoe on behalf of the Wellcome Sanger Institute COVID-19 Surveillance Team |
| EPI_ISL_3438584 | Berkshire and Surrey Pathology Services Lighthouse Laboratory | Wellcome Sanger Institute for the COVID-19 Genomics UK (COG-UK) Consortium | Berkshire and Surrey Pathology Services Lighthouse Laboratory and Alex Alderton, Roberto Amato, Jeffrey Barrett, Sonia Goncalves, Ewan Harrison, David K. Jackson, Ian Johnston, Dominic Kwiatkowski, Cordelia Langford, John Sillitoe on behalf of the Wellcome Sanger Institute COVID-19 Surveillance Team |
| EPI_ISL_3438585 | Lighthouse Lab in Alderley Park | Wellcome Sanger Institute for the COVID-19 Genomics UK (COG-UK) Consortium | Jacquelyn Wynn, Mairead Hyland, The Lighthouse Lab in Alderley Park and Alex Alderton, Roberto Amato, Jeffrey Barrett, Sonia Goncalves, Ewan Harrison, David K. Jackson, Ian Johnston, Dominic Kwiatkowski, Cordelia Langford, John Sillitoe on behalf of the Wellcome Sanger Institute COVID-19 Surveillance Team |
| EPI_ISL_3438586 | Lighthouse Lab in Glasgow | Wellcome Sanger Institute for the COVID-19 Genomics UK (COG-UK) Consortium | Harper VanSteenhouse, Yumi Kasai, David Gray, Carol Clugston, Anna Dominiczak and Alex Alderton, Roberto Amato, Jeffrey Barrett, Sonia Goncalves, Ewan Harrison, David K. Jackson, Ian Johnston, Dominic Kwiatkowski, Cordelia Langford, John Sillitoe on behalf of the Wellcome Sanger Institute COVID-19 Surveillance Team |
| EPI_ISL_3438587, EPI_ISL_3438588 | Lighthouse Lab in Alderley Park | Wellcome Sanger Institute for the COVID-19 Genomics UK (COG-UK) Consortium | Jacquelyn Wynn, Mairead Hyland, The Lighthouse Lab in Alderley Park and Alex Alderton, Roberto Amato, Jeffrey Barrett, Sonia Goncalves, Ewan Harrison, David K. Jackson, Ian Johnston, Dominic Kwiatkowski, Cordelia Langford, John Sillitoe on behalf of the Wellcome Sanger Institute COVID-19 Surveillance Team |
| EPI_ISL_3438589 | Berkshire and Surrey Pathology Services Lighthouse Laboratory | Wellcome Sanger Institute for the COVID-19 Genomics UK (COG-UK) Consortium | Berkshire and Surrey Pathology Services Lighthouse Laboratory and Alex Alderton, Roberto Amato, Jeffrey Barrett, Sonia Goncalves, Ewan Harrison, David K. Jackson, Ian Johnston, Dominic Kwiatkowski, Cordelia Langford, John Sillitoe on behalf of the Wellcome Sanger Institute COVID-19 Surveillance Team |
| EPI_ISL_3438590 | Lighthouse Lab in Alderley Park | Wellcome Sanger Institute for the COVID-19 Genomics UK (COG-UK) Consortium | Jacquelyn Wynn, Mairead Hyland, The Lighthouse Lab in Alderley Park and Alex Alderton, Roberto Amato, Jeffrey Barrett, Sonia Goncalves, Ewan Harrison, David K. Jackson, Ian Johnston, Dominic Kwiatkowski, Cordelia Langford, John Sillitoe on behalf of the Wellcome Sanger Institute COVID-19 Surveillance Team |
| EPI_ISL_3438591, EPI_ISL_3438592 | Lighthouse Lab in Glasgow | Wellcome Sanger Institute for the COVID-19 Genomics UK (COG-UK) Consortium | Harper VanSteenhouse, Yumi Kasai, David Gray, Carol Clugston, Anna Dominiczak and Alex Alderton, Roberto Amato, Jeffrey Barrett, Sonia Goncalves, Ewan Harrison, David K. Jackson, Ian Johnston, Dominic Kwiatkowski, Cordelia Langford, John Sillitoe on behalf of the Wellcome Sanger Institute COVID-19 Surveillance Team |
| EPI_ISL_3438593 | Berkshire and Surrey Pathology Services Lighthouse Laboratory | Wellcome Sanger Institute for the COVID-19 Genomics UK (COG-UK) Consortium | Berkshire and Surrey Pathology Services Lighthouse Laboratory and Alex Alderton, Roberto Amato, Jeffrey Barrett, Sonia Goncalves, Ewan Harrison, David K. Jackson, Ian Johnston, Dominic Kwiatkowski, Cordelia Langford, John Sillitoe on behalf of the Wellcome Sanger Institute COVID-19 Surveillance Team |
| EPI_ISL_3438594 | Lighthouse Lab in Glasgow | Wellcome Sanger Institute for the COVID-19 Genomics UK (COG-UK) Consortium | Harper VanSteenhouse, Yumi Kasai, David Gray, Carol Clugston, Anna Dominiczak and Alex Alderton, Roberto Amato, Jeffrey Barrett, Sonia Goncalves, Ewan Harrison, David K. Jackson, Ian Johnston, Dominic Kwiatkowski, Cordelia Langford, John Sillitoe on behalf of the Wellcome Sanger Institute COVID-19 Surveillance Team |
| EPI_ISL_3438595 | Lighthouse Lab in Alderley Park | Wellcome Sanger Institute for the COVID-19 Genomics UK (COG-UK) Consortium | Jacquelyn Wynn, Mairead Hyland, The Lighthouse Lab in Alderley Park and Alex Alderton, Roberto Amato, Jeffrey Barrett, Sonia Goncalves, Ewan Harrison, David K. Jackson, Ian Johnston, Dominic Kwiatkowski, Cordelia Langford, John Sillitoe on behalf of the Wellcome Sanger Institute COVID-19 Surveillance Team |
| EPI_ISL_3438596 | Lighthouse Lab in Glasgow | Wellcome Sanger Institute for the COVID-19 Genomics UK (COG-UK) Consortium | Harper VanSteenhouse, Yumi Kasai, David Gray, Carol Clugston, Anna Dominiczak and Alex Alderton, Roberto Amato, Jeffrey Barrett, Sonia Goncalves, Ewan Harrison, David K. Jackson, Ian Johnston, Dominic Kwiatkowski, Cordelia Langford, John Sillitoe on behalf of the Wellcome Sanger Institute COVID-19 Surveillance Team |
| EPI_ISL_3438597 | Berkshire and Surrey Pathology Services Lighthouse Laboratory | Wellcome Sanger Institute for the COVID-19 Genomics UK (COG-UK) Consortium | Berkshire and Surrey Pathology Services Lighthouse Laboratory and Alex Alderton, Roberto Amato, Jeffrey Barrett, Sonia Goncalves, Ewan Harrison, David K. Jackson, Ian Johnston, Dominic Kwiatkowski, Cordelia Langford, John Sillitoe on behalf of the Wellcome Sanger Institute COVID-19 Surveillance Team |
| EPI_ISL_3438598, EPI_ISL_3438599 | Lighthouse Lab in Glasgow | Wellcome Sanger Institute for the COVID-19 Genomics UK (COG-UK) Consortium | Harper VanSteenhouse, Yumi Kasai, David Gray, Carol Clugston, Anna Dominiczak and Alex Alderton, Roberto Amato, Jeffrey Barrett, Sonia Goncalves, Ewan Harrison, David K. Jackson, Ian Johnston, Dominic Kwiatkowski, Cordelia Langford, John Sillitoe on behalf of the Wellcome Sanger Institute COVID-19 Surveillance Team |
| EPI_ISL_3438600 | Berkshire and Surrey Pathology Services Lighthouse Laboratory | Wellcome Sanger Institute for the COVID-19 Genomics UK (COG-UK) Consortium | Berkshire and Surrey Pathology Services Lighthouse Laboratory and Alex Alderton, Roberto Amato, Jeffrey Barrett, Sonia Goncalves, Ewan Harrison, David K. Jackson, Ian Johnston, Dominic Kwiatkowski, Cordelia Langford, John Sillitoe on behalf of the Wellcome Sanger Institute COVID-19 Surveillance Team |
| EPI_ISL_3438601 | Lighthouse Lab in Alderley Park | Wellcome Sanger Institute for the COVID-19 Genomics UK (COG-UK) Consortium | Jacquelyn Wynn, Mairead Hyland, The Lighthouse Lab in Alderley Park and Alex Alderton, Roberto Amato, Jeffrey Barrett, Sonia Goncalves, Ewan Harrison, David K. Jackson, Ian Johnston, Dominic Kwiatkowski, Cordelia Langford, John Sillitoe on behalf of the Wellcome Sanger Institute COVID-19 Surveillance Team |
| EPI_ISL_3438602 | Lighthouse Lab in Glasgow | Wellcome Sanger Institute for the COVID-19 Genomics UK (COG-UK) Consortium | Harper VanSteenhouse, Yumi Kasai, David Gray, Carol Clugston, Anna Dominiczak and Alex Alderton, Roberto Amato, Jeffrey Barrett, Sonia Goncalves, Ewan Harrison, David K. Jackson, Ian Johnston, Dominic Kwiatkowski, Cordelia Langford, John Sillitoe on behalf of the Wellcome Sanger Institute COVID-19 Surveillance Team |
| EPI_ISL_3438603 | Berkshire and Surrey Pathology Services Lighthouse Laboratory | Wellcome Sanger Institute for the COVID-19 Genomics UK (COG-UK) Consortium | Berkshire and Surrey Pathology Services Lighthouse Laboratory and Alex Alderton, Roberto Amato, Jeffrey Barrett, Sonia Goncalves, Ewan Harrison, David K. Jackson, Ian Johnston, Dominic Kwiatkowski, Cordelia Langford, John Sillitoe on behalf of the Wellcome Sanger Institute COVID-19 Surveillance Team |
| EPI_ISL_3438604 | Lighthouse Lab in Alderley Park | Wellcome Sanger Institute for the COVID-19 Genomics UK (COG-UK) Consortium | Jacquelyn Wynn, Mairead Hyland, The Lighthouse Lab in Alderley Park and Alex Alderton, Roberto Amato, Jeffrey Barrett, Sonia Goncalves, Ewan Harrison, David K. Jackson, Ian Johnston, Dominic Kwiatkowski, Cordelia Langford, John Sillitoe on behalf of the Wellcome Sanger Institute COVID-19 Surveillance Team |
| EPI_ISL_3438605, EPI_ISL_3438606 | Lighthouse Lab in Glasgow | Wellcome Sanger Institute for the COVID-19 Genomics UK (COG-UK) Consortium | Harper VanSteenhouse, Yumi Kasai, David Gray, Carol Clugston, Anna Dominiczak and Alex Alderton, Roberto Amato, Jeffrey Barrett, Sonia Goncalves, Ewan Harrison, David K. Jackson, Ian Johnston, Dominic Kwiatkowski, Cordelia Langford, John Sillitoe on behalf of the Wellcome Sanger Institute COVID-19 Surveillance Team |
| EPI_ISL_3438607, EPI_ISL_3438608, EPI_ISL_3438609 | Berkshire and Surrey Pathology Services Lighthouse Laboratory | Wellcome Sanger Institute for the COVID-19 Genomics UK (COG-UK) Consortium | Berkshire and Surrey Pathology Services Lighthouse Laboratory and Alex Alderton, Roberto Amato, Jeffrey Barrett, Sonia Goncalves, Ewan Harrison, David K. Jackson, Ian Johnston, Dominic Kwiatkowski, Cordelia Langford, John Sillitoe on behalf of the Wellcome Sanger Institute COVID-19 Surveillance Team |
| EPI_ISL_3438610 | Lighthouse Lab in Alderley Park | Wellcome Sanger Institute for the COVID-19 Genomics UK (COG-UK) Consortium | Jacquelyn Wynn, Mairead Hyland, The Lighthouse Lab in Alderley Park and Alex Alderton, Roberto Amato, Jeffrey Barrett, Sonia Goncalves, Ewan Harrison, David K. Jackson, Ian Johnston, Dominic Kwiatkowski, Cordelia Langford, John Sillitoe on behalf of the Wellcome Sanger Institute COVID-19 Surveillance Team |
| EPI_ISL_3438611 | Berkshire and Surrey Pathology Services Lighthouse Laboratory | Wellcome Sanger Institute for the COVID-19 Genomics UK (COG-UK) Consortium | Berkshire and Surrey Pathology Services Lighthouse Laboratory and Alex Alderton, Roberto Amato, Jeffrey Barrett, Sonia Goncalves, Ewan Harrison, David K. Jackson, Ian Johnston, Dominic Kwiatkowski, Cordelia Langford, John Sillitoe on behalf of the Wellcome Sanger Institute COVID-19 Surveillance Team |
| EPI_ISL_3438612, EPI_ISL_3438613 | Lighthouse Lab in Alderley Park | Wellcome Sanger Institute for the COVID-19 Genomics UK (COG-UK) Consortium | Jacquelyn Wynn, Mairead Hyland, The Lighthouse Lab in Alderley Park and Alex Alderton, Roberto Amato, Jeffrey Barrett, Sonia Goncalves, Ewan Harrison, David K. Jackson, Ian Johnston, Dominic Kwiatkowski, Cordelia Langford, John Sillitoe on behalf of the Wellcome Sanger Institute COVID-19 Surveillance Team |
| EPI_ISL_3438614, EPI_ISL_3438615 | Berkshire and Surrey Pathology Services Lighthouse Laboratory | Wellcome Sanger Institute for the COVID-19 Genomics UK (COG-UK) Consortium | Berkshire and Surrey Pathology Services Lighthouse Laboratory and Alex Alderton, Roberto Amato, Jeffrey Barrett, Sonia Goncalves, Ewan Harrison, David K. Jackson, Ian Johnston, Dominic Kwiatkowski, Cordelia Langford, John Sillitoe on behalf of the Wellcome Sanger Institute COVID-19 Surveillance Team |
| EPI_ISL_3438616, EPI_ISL_3438617, EPI_ISL_3438618 | Lighthouse Lab in Alderley Park | Wellcome Sanger Institute for the COVID-19 Genomics UK (COG-UK) Consortium | Jacquelyn Wynn, Mairead Hyland, The Lighthouse Lab in Alderley Park and Alex Alderton, Roberto Amato, Jeffrey Barrett, Sonia Goncalves, Ewan Harrison, David K. Jackson, Ian Johnston, Dominic Kwiatkowski, Cordelia Langford, John Sillitoe on behalf of the Wellcome Sanger Institute COVID-19 Surveillance Team |
| EPI_ISL_3438619 | Lighthouse Lab in Glasgow | Wellcome Sanger Institute for the COVID-19 Genomics UK (COG-UK) Consortium | Harper VanSteenhouse, Yumi Kasai, David Gray, Carol Clugston, Anna Dominiczak and Alex Alderton, Roberto Amato, Jeffrey Barrett, Sonia Goncalves, Ewan Harrison, David K. Jackson, Ian Johnston, Dominic Kwiatkowski, Cordelia Langford, John Sillitoe on behalf of the Wellcome Sanger Institute COVID-19 Surveillance Team |
| EPI_ISL_3438620 | Lighthouse Lab in Alderley Park | Wellcome Sanger Institute for the COVID-19 Genomics UK (COG-UK) Consortium | Jacquelyn Wynn, Mairead Hyland, The Lighthouse Lab in Alderley Park and Alex Alderton, Roberto Amato, Jeffrey Barrett, Sonia Goncalves, Ewan Harrison, David K. Jackson, Ian Johnston, Dominic Kwiatkowski, Cordelia Langford, John Sillitoe on behalf of the Wellcome Sanger Institute COVID-19 Surveillance Team |
| EPI_ISL_3438621 | Berkshire and Surrey Pathology Services Lighthouse Laboratory | Wellcome Sanger Institute for the COVID-19 Genomics UK (COG-UK) Consortium | Berkshire and Surrey Pathology Services Lighthouse Laboratory and Alex Alderton, Roberto Amato, Jeffrey Barrett, Sonia Goncalves, Ewan Harrison, David K. Jackson, Ian Johnston, Dominic Kwiatkowski, Cordelia Langford, John Sillitoe on behalf of the Wellcome Sanger Institute COVID-19 Surveillance Team |
| EPI_ISL_3438622 | Lighthouse Lab in Alderley Park | Wellcome Sanger Institute for the COVID-19 Genomics UK (COG-UK) Consortium | Jacquelyn Wynn, Mairead Hyland, The Lighthouse Lab in Alderley Park and Alex Alderton, Roberto Amato, Jeffrey Barrett, Sonia Goncalves, Ewan Harrison, David K. Jackson, Ian Johnston, Dominic Kwiatkowski, Cordelia Langford, John Sillitoe on behalf of the Wellcome Sanger Institute COVID-19 Surveillance Team |
| EPI_ISL_3438623, EPI_ISL_3438624 | Lighthouse Laboratory Plymouth | Wellcome Sanger Institute for the COVID-19 Genomics UK (COG-UK) Consortium | Lighthouse Laboratory Plymouth and Alex Alderton, Roberto Amato, Jeffrey Barrett, Sonia Goncalves, Ewan Harrison, David K. Jackson, Ian Johnston, Dominic Kwiatkowski, Cordelia Langford, John Sillitoe on behalf of the Wellcome Sanger Institute COVID-19 Surveillance Team |
| EPI_ISL_3438625 | Lighthouse Lab in Alderley Park | Wellcome Sanger Institute for the COVID-19 Genomics UK (COG-UK) Consortium | Jacquelyn Wynn, Mairead Hyland, The Lighthouse Lab in Alderley Park and Alex Alderton, Roberto Amato, Jeffrey Barrett, Sonia Goncalves, Ewan Harrison, David K. Jackson, Ian Johnston, Dominic Kwiatkowski, Cordelia Langford, John Sillitoe on behalf of the Wellcome Sanger Institute COVID-19 Surveillance Team |
| EPI_ISL_3438626 | Lighthouse Laboratory Plymouth | Wellcome Sanger Institute for the COVID-19 Genomics UK (COG-UK) Consortium | Lighthouse Laboratory Plymouth and Alex Alderton, Roberto Amato, Jeffrey Barrett, Sonia Goncalves, Ewan Harrison, David K. Jackson, Ian Johnston, Dominic Kwiatkowski, Cordelia Langford, John Sillitoe on behalf of the Wellcome Sanger Institute COVID-19 Surveillance Team |
| EPI_ISL_3438627 | Lighthouse Lab in Alderley Park | Wellcome Sanger Institute for the COVID-19 Genomics UK (COG-UK) Consortium | Jacquelyn Wynn, Mairead Hyland, The Lighthouse Lab in Alderley Park and Alex Alderton, Roberto Amato, Jeffrey Barrett, Sonia Goncalves, Ewan Harrison, David K. Jackson, Ian Johnston, Dominic Kwiatkowski, Cordelia Langford, John Sillitoe on behalf of the Wellcome Sanger Institute COVID-19 Surveillance Team |
| EPI_ISL_3438628 | Lighthouse Laboratory Plymouth | Wellcome Sanger Institute for the COVID-19 Genomics UK (COG-UK) Consortium | Lighthouse Laboratory Plymouth and Alex Alderton, Roberto Amato, Jeffrey Barrett, Sonia Goncalves, Ewan Harrison, David K. Jackson, Ian Johnston, Dominic Kwiatkowski, Cordelia Langford, John Sillitoe on behalf of the Wellcome Sanger Institute COVID-19 Surveillance Team |
| EPI_ISL_3438629 | Lighthouse Lab in Alderley Park | Wellcome Sanger Institute for the COVID-19 Genomics UK (COG-UK) Consortium | Jacquelyn Wynn, Mairead Hyland, The Lighthouse Lab in Alderley Park and Alex Alderton, Roberto Amato, Jeffrey Barrett, Sonia Goncalves, Ewan Harrison, David K. Jackson, Ian Johnston, Dominic Kwiatkowski, Cordelia Langford, John Sillitoe on behalf of the Wellcome Sanger Institute COVID-19 Surveillance Team |
| EPI_ISL_3438630 | Berkshire and Surrey Pathology Services Lighthouse Laboratory | Wellcome Sanger Institute for the COVID-19 Genomics UK (COG-UK) Consortium | Berkshire and Surrey Pathology Services Lighthouse Laboratory and Alex Alderton, Roberto Amato, Jeffrey Barrett, Sonia Goncalves, Ewan Harrison, David K. Jackson, Ian Johnston, Dominic Kwiatkowski, Cordelia Langford, John Sillitoe on behalf of the Wellcome Sanger Institute COVID-19 Surveillance Team |
| EPI_ISL_3438631 | Lighthouse Laboratory Plymouth | Wellcome Sanger Institute for the COVID-19 Genomics UK (COG-UK) Consortium | Lighthouse Laboratory Plymouth and Alex Alderton, Roberto Amato, Jeffrey Barrett, Sonia Goncalves, Ewan Harrison, David K. Jackson, Ian Johnston, Dominic Kwiatkowski, Cordelia Langford, John Sillitoe on behalf of the Wellcome Sanger Institute COVID-19 Surveillance Team |
| EPI_ISL_3438632 | Lighthouse Lab in Alderley Park | Wellcome Sanger Institute for the COVID-19 Genomics UK (COG-UK) Consortium | Jacquelyn Wynn, Mairead Hyland, The Lighthouse Lab in Alderley Park and Alex Alderton, Roberto Amato, Jeffrey Barrett, Sonia Goncalves, Ewan Harrison, David K. Jackson, Ian Johnston, Dominic Kwiatkowski, Cordelia Langford, John Sillitoe on behalf of the Wellcome Sanger Institute COVID-19 Surveillance Team |
| EPI_ISL_3438633 | Lighthouse Laboratory Plymouth | Wellcome Sanger Institute for the COVID-19 Genomics UK (COG-UK) Consortium | Lighthouse Laboratory Plymouth and Alex Alderton, Roberto Amato, Jeffrey Barrett, Sonia Goncalves, Ewan Harrison, David K. Jackson, Ian Johnston, Dominic Kwiatkowski, Cordelia Langford, John Sillitoe on behalf of the Wellcome Sanger Institute COVID-19 Surveillance Team |
| EPI_ISL_3438634 | Lighthouse Lab in Alderley Park | Wellcome Sanger Institute for the COVID-19 Genomics UK (COG-UK) Consortium | Jacquelyn Wynn, Mairead Hyland, The Lighthouse Lab in Alderley Park and Alex Alderton, Roberto Amato, Jeffrey Barrett, Sonia Goncalves, Ewan Harrison, David K. Jackson, Ian Johnston, Dominic Kwiatkowski, Cordelia Langford, John Sillitoe on behalf of the Wellcome Sanger Institute COVID-19 Surveillance Team |
| EPI_ISL_3438635, EPI_ISL_3438636 | Berkshire and Surrey Pathology Services Lighthouse Laboratory | Wellcome Sanger Institute for the COVID-19 Genomics UK (COG-UK) Consortium | Berkshire and Surrey Pathology Services Lighthouse Laboratory and Alex Alderton, Roberto Amato, Jeffrey Barrett, Sonia Goncalves, Ewan Harrison, David K. Jackson, Ian Johnston, Dominic Kwiatkowski, Cordelia Langford, John Sillitoe on behalf of the Wellcome Sanger Institute COVID-19 Surveillance Team |

| | | | |
|---|---|---|---|
| EPI_ISL_3438637, EPI_ISL_3438638, EPI_ISL_3438639 | Lighthouse Laboratory Plymouth | Wellcome Sanger Institute for the COVID-19 Genomics UK (COG-UK) Consortium | Lighthouse Laboratory Plymouth and Alex Alderton, Roberto Amato, Jeffrey Barrett, Sonia Goncalves, Ewan Harrison, David K. Jackson, Ian Johnston, Dominic Kwiatkowski, Cordelia Langford, John Sillitoe on behalf of the Wellcome Sanger Institute COVID-19 Surveillance Team |
| EPI_ISL_3438640 | Lighthouse Lab in Alderley Park | Wellcome Sanger Institute for the COVID-19 Genomics UK (COG-UK) Consortium | Jacquelyn Wynn, Mairead Hyland, The Lighthouse Lab in Alderley Park and Alex Alderton, Roberto Amato, Jeffrey Barrett, Sonia Goncalves, Ewan Harrison, David K. Jackson, Ian Johnston, Dominic Kwiatkowski, Cordelia Langford, John Sillitoe on behalf of the Wellcome Sanger Institute COVID-19 Surveillance Team |
| EPI_ISL_3438641, EPI_ISL_3438642, EPI_ISL_3438643 | Lighthouse Laboratory Plymouth | Wellcome Sanger Institute for the COVID-19 Genomics UK (COG-UK) Consortium | Lighthouse Laboratory Plymouth and Alex Alderton, Roberto Amato, Jeffrey Barrett, Sonia Goncalves, Ewan Harrison, David K. Jackson, Ian Johnston, Dominic Kwiatkowski, Cordelia Langford, John Sillitoe on behalf of the Wellcome Sanger Institute COVID-19 Surveillance Team |
| EPI_ISL_3438644 | Lighthouse Lab in Alderley Park | Wellcome Sanger Institute for the COVID-19 Genomics UK (COG-UK) Consortium | Jacquelyn Wynn, Mairead Hyland, The Lighthouse Lab in Alderley Park and Alex Alderton, Roberto Amato, Jeffrey Barrett, Sonia Goncalves, Ewan Harrison, David K. Jackson, Ian Johnston, Dominic Kwiatkowski, Cordelia Langford, John Sillitoe on behalf of the Wellcome Sanger Institute COVID-19 Surveillance Team |
| EPI_ISL_3438645, EPI_ISL_3438646 | Lighthouse Laboratory Plymouth | Wellcome Sanger Institute for the COVID-19 Genomics UK (COG-UK) Consortium | Lighthouse Laboratory Plymouth and Alex Alderton, Roberto Amato, Jeffrey Barrett, Sonia Goncalves, Ewan Harrison, David K. Jackson, Ian Johnston, Dominic Kwiatkowski, Cordelia Langford, John Sillitoe on behalf of the Wellcome Sanger Institute COVID-19 Surveillance Team |
| EPI_ISL_3438647 | Berkshire and Surrey Pathology Services Lighthouse Laboratory | Wellcome Sanger Institute for the COVID-19 Genomics UK (COG-UK) Consortium | Berkshire and Surrey Pathology Services Lighthouse Laboratory and Alex Alderton, Roberto Amato, Jeffrey Barrett, Sonia Goncalves, Ewan Harrison, David K. Jackson, Ian Johnston, Dominic Kwiatkowski, Cordelia Langford, John Sillitoe on behalf of the Wellcome Sanger Institute COVID-19 Surveillance Team |
| EPI_ISL_3438648 | Lighthouse Lab in Alderley Park | Wellcome Sanger Institute for the COVID-19 Genomics UK (COG-UK) Consortium | Jacquelyn Wynn, Mairead Hyland, The Lighthouse Lab in Alderley Park and Alex Alderton, Roberto Amato, Jeffrey Barrett, Sonia Goncalves, Ewan Harrison, David K. Jackson, Ian Johnston, Dominic Kwiatkowski, Cordelia Langford, John Sillitoe on behalf of the Wellcome Sanger Institute COVID-19 Surveillance Team |
| EPI_ISL_3438649 | Lighthouse Laboratory Plymouth | Wellcome Sanger Institute for the COVID-19 Genomics UK (COG-UK) Consortium | Lighthouse Laboratory Plymouth and Alex Alderton, Roberto Amato, Jeffrey Barrett, Sonia Goncalves, Ewan Harrison, David K. Jackson, Ian Johnston, Dominic Kwiatkowski, Cordelia Langford, John Sillitoe on behalf of the Wellcome Sanger Institute COVID-19 Surveillance Team |
| EPI_ISL_3438650 | Lighthouse Lab in Alderley Park | Wellcome Sanger Institute for the COVID-19 Genomics UK (COG-UK) Consortium | Jacquelyn Wynn, Mairead Hyland, The Lighthouse Lab in Alderley Park and Alex Alderton, Roberto Amato, Jeffrey Barrett, Sonia Goncalves, Ewan Harrison, David K. Jackson, Ian Johnston, Dominic Kwiatkowski, Cordelia Langford, John Sillitoe on behalf of the Wellcome Sanger Institute COVID-19 Surveillance Team |
| EPI_ISL_3438651, EPI_ISL_3438652 | Lighthouse Laboratory Plymouth | Wellcome Sanger Institute for the COVID-19 Genomics UK (COG-UK) Consortium | Lighthouse Laboratory Plymouth and Alex Alderton, Roberto Amato, Jeffrey Barrett, Sonia Goncalves, Ewan Harrison, David K. Jackson, Ian Johnston, Dominic Kwiatkowski, Cordelia Langford, John Sillitoe on behalf of the Wellcome Sanger Institute COVID-19 Surveillance Team |
| EPI_ISL_3438653 | Berkshire and Surrey Pathology Services Lighthouse Laboratory | Wellcome Sanger Institute for the COVID-19 Genomics UK (COG-UK) Consortium | Berkshire and Surrey Pathology Services Lighthouse Laboratory and Alex Alderton, Roberto Amato, Jeffrey Barrett, Sonia Goncalves, Ewan Harrison, David K. Jackson, Ian Johnston, Dominic Kwiatkowski, Cordelia Langford, John Sillitoe on behalf of the Wellcome Sanger Institute COVID-19 Surveillance Team |
| EPI_ISL_3438654 | Lighthouse Lab in Alderley Park | Wellcome Sanger Institute for the COVID-19 Genomics UK (COG-UK) Consortium | Jacquelyn Wynn, Mairead Hyland, The Lighthouse Lab in Alderley Park and Alex Alderton, Roberto Amato, Jeffrey Barrett, Sonia Goncalves, Ewan Harrison, David K. Jackson, Ian Johnston, Dominic Kwiatkowski, Cordelia Langford, John Sillitoe on behalf of the Wellcome Sanger Institute COVID-19 Surveillance Team |
| EPI_ISL_3438655 | Berkshire and Surrey Pathology Services Lighthouse Laboratory | Wellcome Sanger Institute for the COVID-19 Genomics UK (COG-UK) Consortium | Berkshire and Surrey Pathology Services Lighthouse Laboratory and Alex Alderton, Roberto Amato, Jeffrey Barrett, Sonia Goncalves, Ewan Harrison, David K. Jackson, Ian Johnston, Dominic Kwiatkowski, Cordelia Langford, John Sillitoe on behalf of the Wellcome Sanger Institute COVID-19 Surveillance Team |
| EPI_ISL_3438656, EPI_ISL_3438657 | Lighthouse Laboratory Plymouth | Wellcome Sanger Institute for the COVID-19 Genomics UK (COG-UK) Consortium | Lighthouse Laboratory Plymouth and Alex Alderton, Roberto Amato, Jeffrey Barrett, Sonia Goncalves, Ewan Harrison, David K. Jackson, Ian Johnston, Dominic Kwiatkowski, Cordelia Langford, John Sillitoe on behalf of the Wellcome Sanger Institute COVID-19 Surveillance Team |
| EPI_ISL_3438658 | Health Services Laboratories | Wellcome Sanger Institute for the COVID-19 Genomics UK (COG-UK) Consortium | Health Services Laboratories and Alex Alderton, Roberto Amato, Jeffrey Barrett, Sonia Goncalves, Ewan Harrison, David K. Jackson, Ian Johnston, Dominic Kwiatkowski, Cordelia Langford, John Sillitoe on behalf of the Wellcome Sanger Institute COVID-19 Surveillance Team |
| EPI_ISL_3438659 | Lighthouse Laboratory Plymouth | Wellcome Sanger Institute for the COVID-19 Genomics UK (COG-UK) Consortium | Lighthouse Laboratory Plymouth and Alex Alderton, Roberto Amato, Jeffrey Barrett, Sonia Goncalves, Ewan Harrison, David K. Jackson, Ian Johnston, Dominic Kwiatkowski, Cordelia Langford, John Sillitoe on behalf of the Wellcome Sanger Institute COVID-19 Surveillance Team |
| EPI_ISL_3438660 | Lighthouse Lab in Alderley Park | Wellcome Sanger Institute for the COVID-19 Genomics UK (COG-UK) Consortium | Jacquelyn Wynn, Mairead Hyland, The Lighthouse Lab in Alderley Park and Alex Alderton, Roberto Amato, Jeffrey Barrett, Sonia Goncalves, Ewan Harrison, David K. Jackson, Ian Johnston, Dominic Kwiatkowski, Cordelia Langford, John Sillitoe on behalf of the Wellcome Sanger Institute COVID-19 Surveillance Team |
| EPI_ISL_3438661 | Lighthouse Laboratory Plymouth | Wellcome Sanger Institute for the COVID-19 Genomics UK (COG-UK) Consortium | Lighthouse Laboratory Plymouth and Alex Alderton, Roberto Amato, Jeffrey Barrett, Sonia Goncalves, Ewan Harrison, David K. Jackson, Ian Johnston, Dominic Kwiatkowski, Cordelia Langford, John Sillitoe on behalf of the Wellcome Sanger Institute COVID-19 Surveillance Team |
| EPI_ISL_3438662 | Berkshire and Surrey Pathology Services Lighthouse Laboratory | Wellcome Sanger Institute for the COVID-19 Genomics UK (COG-UK) Consortium | Berkshire and Surrey Pathology Services Lighthouse Laboratory and Alex Alderton, Roberto Amato, Jeffrey Barrett, Sonia Goncalves, Ewan Harrison, David K. Jackson, Ian Johnston, Dominic Kwiatkowski, Cordelia Langford, John Sillitoe on behalf of the Wellcome Sanger Institute COVID-19 Surveillance Team |
| EPI_ISL_3438663 | Health Services Laboratories | Wellcome Sanger Institute for the COVID-19 Genomics UK (COG-UK) Consortium | Health Services Laboratories and Alex Alderton, Roberto Amato, Jeffrey Barrett, Sonia Goncalves, Ewan Harrison, David K. Jackson, Ian Johnston, Dominic Kwiatkowski, Cordelia Langford, John Sillitoe on behalf of the Wellcome Sanger Institute COVID-19 Surveillance Team |
| EPI_ISL_3438664 | Lighthouse Lab in Alderley Park | Wellcome Sanger Institute for the COVID-19 Genomics UK (COG-UK) Consortium | Jacquelyn Wynn, Mairead Hyland, The Lighthouse Lab in Alderley Park and Alex Alderton, Roberto Amato, Jeffrey Barrett, Sonia Goncalves, Ewan Harrison, David K. Jackson, Ian Johnston, Dominic Kwiatkowski, Cordelia Langford, John Sillitoe on behalf of the Wellcome Sanger Institute COVID-19 Surveillance Team |
| EPI_ISL_3438665 | Berkshire and Surrey Pathology Services Lighthouse Laboratory | Wellcome Sanger Institute for the COVID-19 Genomics UK (COG-UK) Consortium | Berkshire and Surrey Pathology Services Lighthouse Laboratory and Alex Alderton, Roberto Amato, Jeffrey Barrett, Sonia Goncalves, Ewan Harrison, David K. Jackson, Ian Johnston, Dominic Kwiatkowski, Cordelia Langford, John Sillitoe on behalf of the Wellcome Sanger Institute COVID-19 Surveillance Team |
| EPI_ISL_3438666 | Lighthouse Lab in Alderley Park | Wellcome Sanger Institute for the COVID-19 Genomics UK (COG-UK) Consortium | Jacquelyn Wynn, Mairead Hyland, The Lighthouse Lab in Alderley Park and Alex Alderton, Roberto Amato, Jeffrey Barrett, Sonia Goncalves, Ewan Harrison, David K. Jackson, Ian Johnston, Dominic Kwiatkowski, Cordelia Langford, John Sillitoe on behalf of the Wellcome Sanger Institute COVID-19 Surveillance Team |
| EPI_ISL_3438667 | Lighthouse Laboratory Plymouth | Wellcome Sanger Institute for the COVID-19 Genomics UK (COG-UK) Consortium | Lighthouse Laboratory Plymouth and Alex Alderton, Roberto Amato, Jeffrey Barrett, Sonia Goncalves, Ewan Harrison, David K. Jackson, Ian Johnston, Dominic Kwiatkowski, Cordelia Langford, John Sillitoe on behalf of the Wellcome Sanger Institute COVID-19 Surveillance Team |
| EPI_ISL_3438668 | Lighthouse Lab in Alderley Park | Wellcome Sanger Institute for the COVID-19 Genomics UK (COG-UK) Consortium | Jacquelyn Wynn, Mairead Hyland, The Lighthouse Lab in Alderley Park and Alex Alderton, Roberto Amato, Jeffrey Barrett, Sonia Goncalves, Ewan Harrison, David K. Jackson, Ian Johnston, Dominic Kwiatkowski, Cordelia Langford, John Sillitoe on behalf of the Wellcome Sanger Institute COVID-19 Surveillance Team |
| EPI_ISL_3438669, EPI_ISL_3438670, EPI_ISL_3438671 | Lighthouse Laboratory Plymouth | Wellcome Sanger Institute for the COVID-19 Genomics UK (COG-UK) Consortium | Lighthouse Laboratory Plymouth and Alex Alderton, Roberto Amato, Jeffrey Barrett, Sonia Goncalves, Ewan Harrison, David K. Jackson, Ian Johnston, Dominic Kwiatkowski, Cordelia Langford, John Sillitoe on behalf of the Wellcome Sanger Institute COVID-19 Surveillance Team |
| EPI_ISL_3438672 | Lighthouse Lab in Alderley Park | Wellcome Sanger Institute for the COVID-19 Genomics UK (COG-UK) Consortium | Jacquelyn Wynn, Mairead Hyland, The Lighthouse Lab in Alderley Park and Alex Alderton, Roberto Amato, Jeffrey Barrett, Sonia Goncalves, Ewan Harrison, David K. Jackson, Ian Johnston, Dominic Kwiatkowski, Cordelia Langford, John Sillitoe on behalf of the Wellcome Sanger Institute COVID-19 Surveillance Team |
| EPI_ISL_3438673, EPI_ISL_3438674, EPI_ISL_3438675, EPI_ISL_3438676, EPI_ISL_3438677, EPI_ISL_3438678, EPI_ISL_3438679 | Lighthouse Laboratory Plymouth | Wellcome Sanger Institute for the COVID-19 Genomics UK (COG-UK) Consortium | Lighthouse Laboratory Plymouth and Alex Alderton, Roberto Amato, Jeffrey Barrett, Sonia Goncalves, Ewan Harrison, David K. Jackson, Ian Johnston, Dominic Kwiatkowski, Cordelia Langford, John Sillitoe on behalf of the Wellcome Sanger Institute COVID-19 Surveillance Team |
| EPI_ISL_3438680 | Lighthouse Lab in Alderley Park | Wellcome Sanger Institute for the COVID-19 Genomics UK (COG-UK) Consortium | Jacquelyn Wynn, Mairead Hyland, The Lighthouse Lab in Alderley Park and Alex Alderton, Roberto Amato, Jeffrey Barrett, Sonia Goncalves, Ewan Harrison, David K. Jackson, Ian Johnston, Dominic Kwiatkowski, Cordelia Langford, John Sillitoe on behalf of the Wellcome Sanger Institute COVID-19 Surveillance Team |
| EPI_ISL_3438681, EPI_ISL_3438682, EPI_ISL_3438683 | Lighthouse Laboratory Plymouth | Wellcome Sanger Institute for the COVID-19 Genomics UK (COG-UK) Consortium | Lighthouse Laboratory Plymouth and Alex Alderton, Roberto Amato, Jeffrey Barrett, Sonia Goncalves, Ewan Harrison, David K. Jackson, Ian Johnston, Dominic Kwiatkowski, Cordelia Langford, John Sillitoe on behalf of the Wellcome Sanger Institute COVID-19 Surveillance Team |
| EPI_ISL_3438684 | Lighthouse Lab in Alderley Park | Wellcome Sanger Institute for the COVID-19 Genomics UK (COG-UK) Consortium | Jacquelyn Wynn, Mairead Hyland, The Lighthouse Lab in Alderley Park and Alex Alderton, Roberto Amato, Jeffrey Barrett, Sonia Goncalves, Ewan Harrison, David K. Jackson, Ian Johnston, Dominic Kwiatkowski, Cordelia Langford, John Sillitoe on behalf of the Wellcome Sanger Institute COVID-19 Surveillance Team |
| EPI_ISL_3438685 | Berkshire and Surrey Pathology Services Lighthouse Laboratory | Wellcome Sanger Institute for the COVID-19 Genomics UK (COG-UK) Consortium | Berkshire and Surrey Pathology Services Lighthouse Laboratory and Alex Alderton, Roberto Amato, Jeffrey Barrett, Sonia Goncalves, Ewan Harrison, David K. Jackson, Ian Johnston, Dominic Kwiatkowski, Cordelia Langford, John Sillitoe on behalf of the Wellcome Sanger Institute COVID-19 Surveillance Team |
| EPI_ISL_3438686 | Lighthouse Lab in Alderley Park | Wellcome Sanger Institute for the COVID-19 Genomics UK (COG-UK) Consortium | Jacquelyn Wynn, Mairead Hyland, The Lighthouse Lab in Alderley Park and Alex Alderton, Roberto Amato, Jeffrey Barrett, Sonia Goncalves, Ewan Harrison, David K. Jackson, Ian Johnston, Dominic Kwiatkowski, Cordelia Langford, John Sillitoe on behalf of the Wellcome Sanger Institute COVID-19 Surveillance Team |
| EPI_ISL_3438687 | Lighthouse Laboratory Plymouth | Wellcome Sanger Institute for the COVID-19 Genomics UK (COG-UK) Consortium | Lighthouse Laboratory Plymouth and Alex Alderton, Roberto Amato, Jeffrey Barrett, Sonia Goncalves, Ewan Harrison, David K. Jackson, Ian Johnston, Dominic Kwiatkowski, Cordelia Langford, John Sillitoe on behalf of the Wellcome Sanger Institute COVID-19 Surveillance Team |
| EPI_ISL_3438688, EPI_ISL_3438689 | Lighthouse Lab in Alderley Park | Wellcome Sanger Institute for the COVID-19 Genomics UK (COG-UK) Consortium | Jacquelyn Wynn, Mairead Hyland, The Lighthouse Lab in Alderley Park and Alex Alderton, Roberto Amato, Jeffrey Barrett, Sonia Goncalves, Ewan Harrison, David K. Jackson, Ian Johnston, Dominic Kwiatkowski, Cordelia Langford, John Sillitoe on behalf of the Wellcome Sanger Institute COVID-19 Surveillance Team |
| EPI_ISL_3438690, EPI_ISL_3438691 | Lighthouse Laboratory Plymouth | Wellcome Sanger Institute for the COVID-19 Genomics UK (COG-UK) Consortium | Lighthouse Laboratory Plymouth and Alex Alderton, Roberto Amato, Jeffrey Barrett, Sonia Goncalves, Ewan Harrison, David K. Jackson, Ian Johnston, Dominic Kwiatkowski, Cordelia Langford, John Sillitoe on behalf of the Wellcome Sanger Institute COVID-19 Surveillance Team |
| EPI_ISL_3438692 | Lighthouse Lab in Alderley Park | Wellcome Sanger Institute for the COVID-19 Genomics UK (COG-UK) Consortium | Jacquelyn Wynn, Mairead Hyland, The Lighthouse Lab in Alderley Park and Alex Alderton, Roberto Amato, Jeffrey Barrett, Sonia Goncalves, Ewan Harrison, David K. Jackson, Ian Johnston, Dominic Kwiatkowski, Cordelia Langford, John Sillitoe on behalf of the Wellcome Sanger Institute COVID-19 Surveillance Team |
| EPI_ISL_3438693 | Lighthouse Laboratory Plymouth | Wellcome Sanger Institute for the COVID-19 Genomics UK (COG-UK) Consortium | Lighthouse Laboratory Plymouth and Alex Alderton, Roberto Amato, Jeffrey Barrett, Sonia Goncalves, Ewan Harrison, David K. Jackson, Ian Johnston, Dominic Kwiatkowski, Cordelia Langford, John Sillitoe on behalf of the Wellcome Sanger Institute COVID-19 Surveillance Team |
| EPI_ISL_3438694 | Lighthouse Lab in Alderley Park | Wellcome Sanger Institute for the COVID-19 Genomics UK (COG-UK) Consortium | Jacquelyn Wynn, Mairead Hyland, The Lighthouse Lab in Alderley Park and Alex Alderton, Roberto Amato, Jeffrey Barrett, Sonia Goncalves, Ewan Harrison, David K. Jackson, Ian Johnston, Dominic Kwiatkowski, Cordelia Langford, John Sillitoe on behalf of the Wellcome Sanger Institute COVID-19 Surveillance Team |
| EPI_ISL_3438695 | Lighthouse Laboratory Plymouth | Wellcome Sanger Institute for the COVID-19 Genomics UK (COG-UK) Consortium | Lighthouse Laboratory Plymouth and Alex Alderton, Roberto Amato, Jeffrey Barrett, Sonia Goncalves, Ewan Harrison, David K. Jackson, Ian Johnston, Dominic Kwiatkowski, Cordelia Langford, John Sillitoe on behalf of the Wellcome Sanger Institute COVID-19 Surveillance Team |
| EPI_ISL_3438696 | Berkshire and Surrey Pathology Services Lighthouse Laboratory | Wellcome Sanger Institute for the COVID-19 Genomics UK (COG-UK) Consortium | Berkshire and Surrey Pathology Services Lighthouse Laboratory and Alex Alderton, Roberto Amato, Jeffrey Barrett, Sonia Goncalves, Ewan Harrison, David K. Jackson, Ian Johnston, Dominic Kwiatkowski, Cordelia Langford, John Sillitoe on behalf of the Wellcome Sanger Institute COVID-19 Surveillance Team |

| | | | |
|---|---|---|---|
| EPI_ISL_3438697 | Lighthouse Laboratory Plymouth | Wellcome Sanger Institute for the COVID-19 Genomics UK (COG-UK) Consortium | Lighthouse Laboratory Plymouth and Alex Alderton, Roberto Amato, Jeffrey Barrett, Sonia Goncalves, Ewan Harrison, David K. Jackson, Ian Johnston, Dominic Kwiatkowski, Cordelia Langford, John Sillitoe on behalf of the Wellcome Sanger Institute COVID-19 Surveillance Team |
| EPI_ISL_3438698 | Lighthouse Lab in Alderley Park | Wellcome Sanger Institute for the COVID-19 Genomics UK (COG-UK) Consortium | Jacquelyn Wynn, Mairead Hyland, The Lighthouse Lab in Alderley Park and Alex Alderton, Roberto Amato, Jeffrey Barrett, Sonia Goncalves, Ewan Harrison, David K. Jackson, Ian Johnston, Dominic Kwiatkowski, Cordelia Langford, John Sillitoe on behalf of the Wellcome Sanger Institute COVID-19 Surveillance Team |
| EPI_ISL_3438699, EPI_ISL_3438700, EPI_ISL_3438701, EPI_ISL_3438702, EPI_ISL_3438703 | Lighthouse Laboratory Plymouth | Wellcome Sanger Institute for the COVID-19 Genomics UK (COG-UK) Consortium | Lighthouse Laboratory Plymouth and Alex Alderton, Roberto Amato, Jeffrey Barrett, Sonia Goncalves, Ewan Harrison, David K. Jackson, Ian Johnston, Dominic Kwiatkowski, Cordelia Langford, John Sillitoe on behalf of the Wellcome Sanger Institute COVID-19 Surveillance Team |
| EPI_ISL_3438704 | Lighthouse Lab in Alderley Park | Wellcome Sanger Institute for the COVID-19 Genomics UK (COG-UK) Consortium | Jacquelyn Wynn, Mairead Hyland, The Lighthouse Lab in Alderley Park and Alex Alderton, Roberto Amato, Jeffrey Barrett, Sonia Goncalves, Ewan Harrison, David K. Jackson, Ian Johnston, Dominic Kwiatkowski, Cordelia Langford, John Sillitoe on behalf of the Wellcome Sanger Institute COVID-19 Surveillance Team |
| EPI_ISL_3438705 | Lighthouse Laboratory Plymouth | Wellcome Sanger Institute for the COVID-19 Genomics UK (COG-UK) Consortium | Lighthouse Laboratory Plymouth and Alex Alderton, Roberto Amato, Jeffrey Barrett, Sonia Goncalves, Ewan Harrison, David K. Jackson, Ian Johnston, Dominic Kwiatkowski, Cordelia Langford, John Sillitoe on behalf of the Wellcome Sanger Institute COVID-19 Surveillance Team |
| EPI_ISL_3438706, EPI_ISL_3438707 | Berkshire and Surrey Pathology Services Lighthouse Laboratory | Wellcome Sanger Institute for the COVID-19 Genomics UK (COG-UK) Consortium | Berkshire and Surrey Pathology Services Lighthouse Laboratory and Alex Alderton, Roberto Amato, Jeffrey Barrett, Sonia Goncalves, Ewan Harrison, David K. Jackson, Ian Johnston, Dominic Kwiatkowski, Cordelia Langford, John Sillitoe on behalf of the Wellcome Sanger Institute COVID-19 Surveillance Team |
| EPI_ISL_3438708 | Lighthouse Laboratory Plymouth | Wellcome Sanger Institute for the COVID-19 Genomics UK (COG-UK) Consortium | Lighthouse Laboratory Plymouth and Alex Alderton, Roberto Amato, Jeffrey Barrett, Sonia Goncalves, Ewan Harrison, David K. Jackson, Ian Johnston, Dominic Kwiatkowski, Cordelia Langford, John Sillitoe on behalf of the Wellcome Sanger Institute COVID-19 Surveillance Team |
| EPI_ISL_3438709 | Lighthouse Lab in Alderley Park | Wellcome Sanger Institute for the COVID-19 Genomics UK (COG-UK) Consortium | Jacquelyn Wynn, Mairead Hyland, The Lighthouse Lab in Alderley Park and Alex Alderton, Roberto Amato, Jeffrey Barrett, Sonia Goncalves, Ewan Harrison, David K. Jackson, Ian Johnston, Dominic Kwiatkowski, Cordelia Langford, John Sillitoe on behalf of the Wellcome Sanger Institute COVID-19 Surveillance Team |
| EPI_ISL_3438710, EPI_ISL_3438711, EPI_ISL_3438712, EPI_ISL_3438713, EPI_ISL_3438714, EPI_ISL_3438715, EPI_ISL_3438716, EPI_ISL_3438717 | Lighthouse Laboratory Plymouth | Wellcome Sanger Institute for the COVID-19 Genomics UK (COG-UK) Consortium | Lighthouse Laboratory Plymouth and Alex Alderton, Roberto Amato, Jeffrey Barrett, Sonia Goncalves, Ewan Harrison, David K. Jackson, Ian Johnston, Dominic Kwiatkowski, Cordelia Langford, John Sillitoe on behalf of the Wellcome Sanger Institute COVID-19 Surveillance Team |
| EPI_ISL_3438718 | Lighthouse Lab in Alderley Park | Wellcome Sanger Institute for the COVID-19 Genomics UK (COG-UK) Consortium | Jacquelyn Wynn, Mairead Hyland, The Lighthouse Lab in Alderley Park and Alex Alderton, Roberto Amato, Jeffrey Barrett, Sonia Goncalves, Ewan Harrison, David K. Jackson, Ian Johnston, Dominic Kwiatkowski, Cordelia Langford, John Sillitoe on behalf of the Wellcome Sanger Institute COVID-19 Surveillance Team |
| EPI_ISL_3438719 | Lighthouse Laboratory Plymouth | Wellcome Sanger Institute for the COVID-19 Genomics UK (COG-UK) Consortium | Lighthouse Laboratory Plymouth and Alex Alderton, Roberto Amato, Jeffrey Barrett, Sonia Goncalves, Ewan Harrison, David K. Jackson, Ian Johnston, Dominic Kwiatkowski, Cordelia Langford, John Sillitoe on behalf of the Wellcome Sanger Institute COVID-19 Surveillance Team |
| EPI_ISL_3438720 | Health Services Laboratories | Wellcome Sanger Institute for the COVID-19 Genomics UK (COG-UK) Consortium | Health Services Laboratories and Alex Alderton, Roberto Amato, Jeffrey Barrett, Sonia Goncalves, Ewan Harrison, David K. Jackson, Ian Johnston, Dominic Kwiatkowski, Cordelia Langford, John Sillitoe on behalf of the Wellcome Sanger Institute COVID-19 Surveillance Team |
| EPI_ISL_3438721, EPI_ISL_3438722, EPI_ISL_3438723, EPI_ISL_3438724 | Lighthouse Laboratory Plymouth | Wellcome Sanger Institute for the COVID-19 Genomics UK (COG-UK) Consortium | Lighthouse Laboratory Plymouth and Alex Alderton, Roberto Amato, Jeffrey Barrett, Sonia Goncalves, Ewan Harrison, David K. Jackson, Ian Johnston, Dominic Kwiatkowski, Cordelia Langford, John Sillitoe on behalf of the Wellcome Sanger Institute COVID-19 Surveillance Team |
| EPI_ISL_3438725 | Lighthouse Lab in Alderley Park | Wellcome Sanger Institute for the COVID-19 Genomics UK (COG-UK) Consortium | Jacquelyn Wynn, Mairead Hyland, The Lighthouse Lab in Alderley Park and Alex Alderton, Roberto Amato, Jeffrey Barrett, Sonia Goncalves, Ewan Harrison, David K. Jackson, Ian Johnston, Dominic Kwiatkowski, Cordelia Langford, John Sillitoe on behalf of the Wellcome Sanger Institute COVID-19 Surveillance Team |
| EPI_ISL_3438726, EPI_ISL_3438727, EPI_ISL_3438728, EPI_ISL_3438729, EPI_ISL_3438730 | Lighthouse Laboratory Plymouth | Wellcome Sanger Institute for the COVID-19 Genomics UK (COG-UK) Consortium | Lighthouse Laboratory Plymouth and Alex Alderton, Roberto Amato, Jeffrey Barrett, Sonia Goncalves, Ewan Harrison, David K. Jackson, Ian Johnston, Dominic Kwiatkowski, Cordelia Langford, John Sillitoe on behalf of the Wellcome Sanger Institute COVID-19 Surveillance Team |
| EPI_ISL_3438731 | Lighthouse Lab in Alderley Park | Wellcome Sanger Institute for the COVID-19 Genomics UK (COG-UK) Consortium | Jacquelyn Wynn, Mairead Hyland, The Lighthouse Lab in Alderley Park and Alex Alderton, Roberto Amato, Jeffrey Barrett, Sonia Goncalves, Ewan Harrison, David K. Jackson, Ian Johnston, Dominic Kwiatkowski, Cordelia Langford, John Sillitoe on behalf of the Wellcome Sanger Institute COVID-19 Surveillance Team |
| EPI_ISL_3438732, EPI_ISL_3438733, EPI_ISL_3438734 | Lighthouse Laboratory Plymouth | Wellcome Sanger Institute for the COVID-19 Genomics UK (COG-UK) Consortium | Lighthouse Laboratory Plymouth and Alex Alderton, Roberto Amato, Jeffrey Barrett, Sonia Goncalves, Ewan Harrison, David K. Jackson, Ian Johnston, Dominic Kwiatkowski, Cordelia Langford, John Sillitoe on behalf of the Wellcome Sanger Institute COVID-19 Surveillance Team |
| EPI_ISL_3438735, EPI_ISL_3438736 | Berkshire and Surrey Pathology Services Lighthouse Laboratory | Wellcome Sanger Institute for the COVID-19 Genomics UK (COG-UK) Consortium | Berkshire and Surrey Pathology Services Lighthouse Laboratory and Alex Alderton, Roberto Amato, Jeffrey Barrett, Sonia Goncalves, Ewan Harrison, David K. Jackson, Ian Johnston, Dominic Kwiatkowski, Cordelia Langford, John Sillitoe on behalf of the Wellcome Sanger Institute COVID-19 Surveillance Team |
| EPI_ISL_3438737, EPI_ISL_3438738 | Lighthouse Lab in Alderley Park | Wellcome Sanger Institute for the COVID-19 Genomics UK (COG-UK) Consortium | Jacquelyn Wynn, Mairead Hyland, The Lighthouse Lab in Alderley Park and Alex Alderton, Roberto Amato, Jeffrey Barrett, Sonia Goncalves, Ewan Harrison, David K. Jackson, Ian Johnston, Dominic Kwiatkowski, Cordelia Langford, John Sillitoe on behalf of the Wellcome Sanger Institute COVID-19 Surveillance Team |
| EPI_ISL_3438739 | Lighthouse Laboratory Plymouth | Wellcome Sanger Institute for the COVID-19 Genomics UK (COG-UK) Consortium | Lighthouse Laboratory Plymouth and Alex Alderton, Roberto Amato, Jeffrey Barrett, Sonia Goncalves, Ewan Harrison, David K. Jackson, Ian Johnston, Dominic Kwiatkowski, Cordelia Langford, John Sillitoe on behalf of the Wellcome Sanger Institute COVID-19 Surveillance Team |
| EPI_ISL_3438740 | Lighthouse Lab in Alderley Park | Wellcome Sanger Institute for the COVID-19 Genomics UK (COG-UK) Consortium | Jacquelyn Wynn, Mairead Hyland, The Lighthouse Lab in Alderley Park and Alex Alderton, Roberto Amato, Jeffrey Barrett, Sonia Goncalves, Ewan Harrison, David K. Jackson, Ian Johnston, Dominic Kwiatkowski, Cordelia Langford, John Sillitoe on behalf of the Wellcome Sanger Institute COVID-19 Surveillance Team |
| EPI_ISL_3438741 | Lighthouse Laboratory Plymouth | Wellcome Sanger Institute for the COVID-19 Genomics UK (COG-UK) Consortium | Lighthouse Laboratory Plymouth and Alex Alderton, Roberto Amato, Jeffrey Barrett, Sonia Goncalves, Ewan Harrison, David K. Jackson, Ian Johnston, Dominic Kwiatkowski, Cordelia Langford, John Sillitoe on behalf of the Wellcome Sanger Institute COVID-19 Surveillance Team |
| EPI_ISL_3438742 | Berkshire and Surrey Pathology Services Lighthouse Laboratory | Wellcome Sanger Institute for the COVID-19 Genomics UK (COG-UK) Consortium | Berkshire and Surrey Pathology Services Lighthouse Laboratory and Alex Alderton, Roberto Amato, Jeffrey Barrett, Sonia Goncalves, Ewan Harrison, David K. Jackson, Ian Johnston, Dominic Kwiatkowski, Cordelia Langford, John Sillitoe on behalf of the Wellcome Sanger Institute COVID-19 Surveillance Team |
| EPI_ISL_3438743 | Lighthouse Lab in Alderley Park | Wellcome Sanger Institute for the COVID-19 Genomics UK (COG-UK) Consortium | Jacquelyn Wynn, Mairead Hyland, The Lighthouse Lab in Alderley Park and Alex Alderton, Roberto Amato, Jeffrey Barrett, Sonia Goncalves, Ewan Harrison, David K. Jackson, Ian Johnston, Dominic Kwiatkowski, Cordelia Langford, John Sillitoe on behalf of the Wellcome Sanger Institute COVID-19 Surveillance Team |
| EPI_ISL_3438744, EPI_ISL_3438745, EPI_ISL_3438746, EPI_ISL_3438747 | Lighthouse Laboratory Plymouth | Wellcome Sanger Institute for the COVID-19 Genomics UK (COG-UK) Consortium | Lighthouse Laboratory Plymouth and Alex Alderton, Roberto Amato, Jeffrey Barrett, Sonia Goncalves, Ewan Harrison, David K. Jackson, Ian Johnston, Dominic Kwiatkowski, Cordelia Langford, John Sillitoe on behalf of the Wellcome Sanger Institute COVID-19 Surveillance Team |
| EPI_ISL_3438748 | Berkshire and Surrey Pathology Services Lighthouse Laboratory | Wellcome Sanger Institute for the COVID-19 Genomics UK (COG-UK) Consortium | Berkshire and Surrey Pathology Services Lighthouse Laboratory and Alex Alderton, Roberto Amato, Jeffrey Barrett, Sonia Goncalves, Ewan Harrison, David K. Jackson, Ian Johnston, Dominic Kwiatkowski, Cordelia Langford, John Sillitoe on behalf of the Wellcome Sanger Institute COVID-19 Surveillance Team |
| EPI_ISL_3438749, EPI_ISL_3438750 | Lighthouse Lab in Alderley Park | Wellcome Sanger Institute for the COVID-19 Genomics UK (COG-UK) Consortium | Jacquelyn Wynn, Mairead Hyland, The Lighthouse Lab in Alderley Park and Alex Alderton, Roberto Amato, Jeffrey Barrett, Sonia Goncalves, Ewan Harrison, David K. Jackson, Ian Johnston, Dominic Kwiatkowski, Cordelia Langford, John Sillitoe on behalf of the Wellcome Sanger Institute COVID-19 Surveillance Team |
| EPI_ISL_3438751, EPI_ISL_3438752 | Lighthouse Laboratory Plymouth | Wellcome Sanger Institute for the COVID-19 Genomics UK (COG-UK) Consortium | Lighthouse Laboratory Plymouth and Alex Alderton, Roberto Amato, Jeffrey Barrett, Sonia Goncalves, Ewan Harrison, David K. Jackson, Ian Johnston, Dominic Kwiatkowski, Cordelia Langford, John Sillitoe on behalf of the Wellcome Sanger Institute COVID-19 Surveillance Team |
| EPI_ISL_3438753 | Lighthouse Lab in Alderley Park | Wellcome Sanger Institute for the COVID-19 Genomics UK (COG-UK) Consortium | Jacquelyn Wynn, Mairead Hyland, The Lighthouse Lab in Alderley Park and Alex Alderton, Roberto Amato, Jeffrey Barrett, Sonia Goncalves, Ewan Harrison, David K. Jackson, Ian Johnston, Dominic Kwiatkowski, Cordelia Langford, John Sillitoe on behalf of the Wellcome Sanger Institute COVID-19 Surveillance Team |
| EPI_ISL_3438754, EPI_ISL_3438755, EPI_ISL_3438756, EPI_ISL_3438757, EPI_ISL_3438758, EPI_ISL_3438759, EPI_ISL_3438760 | Lighthouse Laboratory Plymouth | Wellcome Sanger Institute for the COVID-19 Genomics UK (COG-UK) Consortium | Lighthouse Laboratory Plymouth and Alex Alderton, Roberto Amato, Jeffrey Barrett, Sonia Goncalves, Ewan Harrison, David K. Jackson, Ian Johnston, Dominic Kwiatkowski, Cordelia Langford, John Sillitoe on behalf of the Wellcome Sanger Institute COVID-19 Surveillance Team |
| EPI_ISL_3438761 | Lighthouse Lab in Alderley Park | Wellcome Sanger Institute for the COVID-19 Genomics UK (COG-UK) Consortium | Jacquelyn Wynn, Mairead Hyland, The Lighthouse Lab in Alderley Park and Alex Alderton, Roberto Amato, Jeffrey Barrett, Sonia Goncalves, Ewan Harrison, David K. Jackson, Ian Johnston, Dominic Kwiatkowski, Cordelia Langford, John Sillitoe on behalf of the Wellcome Sanger Institute COVID-19 Surveillance Team |
| EPI_ISL_3438762 | Berkshire and Surrey Pathology Services Lighthouse Laboratory | Wellcome Sanger Institute for the COVID-19 Genomics UK (COG-UK) Consortium | Berkshire and Surrey Pathology Services Lighthouse Laboratory and Alex Alderton, Roberto Amato, Jeffrey Barrett, Sonia Goncalves, Ewan Harrison, David K. Jackson, Ian Johnston, Dominic Kwiatkowski, Cordelia Langford, John Sillitoe on behalf of the Wellcome Sanger Institute COVID-19 Surveillance Team |
| EPI_ISL_3438763 | Lighthouse Laboratory Plymouth | Wellcome Sanger Institute for the COVID-19 Genomics UK (COG-UK) Consortium | Lighthouse Laboratory Plymouth and Alex Alderton, Roberto Amato, Jeffrey Barrett, Sonia Goncalves, Ewan Harrison, David K. Jackson, Ian Johnston, Dominic Kwiatkowski, Cordelia Langford, John Sillitoe on behalf of the Wellcome Sanger Institute COVID-19 Surveillance Team |
| EPI_ISL_3438764 | Lighthouse Lab in Alderley Park | Wellcome Sanger Institute for the COVID-19 Genomics UK (COG-UK) Consortium | Jacquelyn Wynn, Mairead Hyland, The Lighthouse Lab in Alderley Park and Alex Alderton, Roberto Amato, Jeffrey Barrett, Sonia Goncalves, Ewan Harrison, David K. Jackson, Ian Johnston, Dominic Kwiatkowski, Cordelia Langford, John Sillitoe on behalf of the Wellcome Sanger Institute COVID-19 Surveillance Team |
| EPI_ISL_3438765, EPI_ISL_3438766, EPI_ISL_3438767 | Lighthouse Laboratory Plymouth | Wellcome Sanger Institute for the COVID-19 Genomics UK (COG-UK) Consortium | Lighthouse Laboratory Plymouth and Alex Alderton, Roberto Amato, Jeffrey Barrett, Sonia Goncalves, Ewan Harrison, David K. Jackson, Ian Johnston, Dominic Kwiatkowski, Cordelia Langford, John Sillitoe on behalf of the Wellcome Sanger Institute COVID-19 Surveillance Team |
| EPI_ISL_3438768 | Health Services Laboratories | Wellcome Sanger Institute for the COVID-19 Genomics UK (COG-UK) | Health Services Laboratories and Alex Alderton, Roberto Amato, Jeffrey Barrett, Sonia Goncalves, Ewan Harrison, David K. Jackson, Ian Johnston, Dominic Kwiatkowski, Cordelia Langford, John Sillitoe on behalf of the Wellcome Sanger Institute |

| | | | |
|---|---|---|---|
| | | Consortium | COVID-19 Surveillance Team |
| EPI_ISL_3438769, EPI_ISL_3438770 | Lighthouse Laboratory Plymouth | Wellcome Sanger Institute for the COVID-19 Genomics UK (COG-UK) Consortium | Lighthouse Laboratory Plymouth and Alex Alderton, Roberto Amato, Jeffrey Barrett, Sonia Goncalves, Ewan Harrison, David K. Jackson, Ian Johnston, Dominic Kwiatkowski, Cordelia Langford, John Sillitoe on behalf of the Wellcome Sanger Institute COVID-19 Surveillance Team |
| EPI_ISL_3438771, EPI_ISL_3438772 | Lighthouse Lab in Alderley Park | Wellcome Sanger Institute for the COVID-19 Genomics UK (COG-UK) Consortium | Jacquelyn Wynn, Mairead Hyland, The Lighthouse Lab in Alderley Park and Alex Alderton, Roberto Amato, Jeffrey Barrett, Sonia Goncalves, Ewan Harrison, David K. Jackson, Ian Johnston, Dominic Kwiatkowski, Cordelia Langford, John Sillitoe on behalf of the Wellcome Sanger Institute COVID-19 Surveillance Team |
| EPI_ISL_3438773 | Lighthouse Laboratory Plymouth | Wellcome Sanger Institute for the COVID-19 Genomics UK (COG-UK) Consortium | Lighthouse Laboratory Plymouth and Alex Alderton, Roberto Amato, Jeffrey Barrett, Sonia Goncalves, Ewan Harrison, David K. Jackson, Ian Johnston, Dominic Kwiatkowski, Cordelia Langford, John Sillitoe on behalf of the Wellcome Sanger Institute COVID-19 Surveillance Team |
| EPI_ISL_3438774 | Berkshire and Surrey Pathology Services Lighthouse Laboratory | Wellcome Sanger Institute for the COVID-19 Genomics UK (COG-UK) Consortium | Berkshire and Surrey Pathology Services Lighthouse Laboratory and Alex Alderton, Roberto Amato, Jeffrey Barrett, Sonia Goncalves, Ewan Harrison, David K. Jackson, Ian Johnston, Dominic Kwiatkowski, Cordelia Langford, John Sillitoe on behalf of the Wellcome Sanger Institute COVID-19 Surveillance Team |
| EPI_ISL_3438775 | Lighthouse Lab in Alderley Park | Wellcome Sanger Institute for the COVID-19 Genomics UK (COG-UK) Consortium | Jacquelyn Wynn, Mairead Hyland, The Lighthouse Lab in Alderley Park and Alex Alderton, Roberto Amato, Jeffrey Barrett, Sonia Goncalves, Ewan Harrison, David K. Jackson, Ian Johnston, Dominic Kwiatkowski, Cordelia Langford, John Sillitoe on behalf of the Wellcome Sanger Institute COVID-19 Surveillance Team |
| EPI_ISL_3438776, EPI_ISL_3438777, EPI_ISL_3438778 | Lighthouse Laboratory Plymouth | Wellcome Sanger Institute for the COVID-19 Genomics UK (COG-UK) Consortium | Lighthouse Laboratory Plymouth and Alex Alderton, Roberto Amato, Jeffrey Barrett, Sonia Goncalves, Ewan Harrison, David K. Jackson, Ian Johnston, Dominic Kwiatkowski, Cordelia Langford, John Sillitoe on behalf of the Wellcome Sanger Institute COVID-19 Surveillance Team |
| EPI_ISL_3438779 | Berkshire and Surrey Pathology Services Lighthouse Laboratory | Wellcome Sanger Institute for the COVID-19 Genomics UK (COG-UK) Consortium | Berkshire and Surrey Pathology Services Lighthouse Laboratory and Alex Alderton, Roberto Amato, Jeffrey Barrett, Sonia Goncalves, Ewan Harrison, David K. Jackson, Ian Johnston, Dominic Kwiatkowski, Cordelia Langford, John Sillitoe on behalf of the Wellcome Sanger Institute COVID-19 Surveillance Team |
| EPI_ISL_3438780 | Lighthouse Laboratory Plymouth | Wellcome Sanger Institute for the COVID-19 Genomics UK (COG-UK) Consortium | Lighthouse Laboratory Plymouth and Alex Alderton, Roberto Amato, Jeffrey Barrett, Sonia Goncalves, Ewan Harrison, David K. Jackson, Ian Johnston, Dominic Kwiatkowski, Cordelia Langford, John Sillitoe on behalf of the Wellcome Sanger Institute COVID-19 Surveillance Team |
| EPI_ISL_3438781 | Lighthouse Lab in Alderley Park | Wellcome Sanger Institute for the COVID-19 Genomics UK (COG-UK) Consortium | Jacquelyn Wynn, Mairead Hyland, The Lighthouse Lab in Alderley Park and Alex Alderton, Roberto Amato, Jeffrey Barrett, Sonia Goncalves, Ewan Harrison, David K. Jackson, Ian Johnston, Dominic Kwiatkowski, Cordelia Langford, John Sillitoe on behalf of the Wellcome Sanger Institute COVID-19 Surveillance Team |
| EPI_ISL_3438782, EPI_ISL_3438783 | Lighthouse Laboratory Plymouth | Wellcome Sanger Institute for the COVID-19 Genomics UK (COG-UK) Consortium | Lighthouse Laboratory Plymouth and Alex Alderton, Roberto Amato, Jeffrey Barrett, Sonia Goncalves, Ewan Harrison, David K. Jackson, Ian Johnston, Dominic Kwiatkowski, Cordelia Langford, John Sillitoe on behalf of the Wellcome Sanger Institute COVID-19 Surveillance Team |
| EPI_ISL_3438784 | Lighthouse Lab in Alderley Park | Wellcome Sanger Institute for the COVID-19 Genomics UK (COG-UK) Consortium | Jacquelyn Wynn, Mairead Hyland, The Lighthouse Lab in Alderley Park and Alex Alderton, Roberto Amato, Jeffrey Barrett, Sonia Goncalves, Ewan Harrison, David K. Jackson, Ian Johnston, Dominic Kwiatkowski, Cordelia Langford, John Sillitoe on behalf of the Wellcome Sanger Institute COVID-19 Surveillance Team |
| EPI_ISL_3438785, EPI_ISL_3438786 | Lighthouse Laboratory Plymouth | Wellcome Sanger Institute for the COVID-19 Genomics UK (COG-UK) Consortium | Lighthouse Laboratory Plymouth and Alex Alderton, Roberto Amato, Jeffrey Barrett, Sonia Goncalves, Ewan Harrison, David K. Jackson, Ian Johnston, Dominic Kwiatkowski, Cordelia Langford, John Sillitoe on behalf of the Wellcome Sanger Institute COVID-19 Surveillance Team |
| EPI_ISL_3438787 | Berkshire and Surrey Pathology Services Lighthouse Laboratory | Wellcome Sanger Institute for the COVID-19 Genomics UK (COG-UK) Consortium | Berkshire and Surrey Pathology Services Lighthouse Laboratory and Alex Alderton, Roberto Amato, Jeffrey Barrett, Sonia Goncalves, Ewan Harrison, David K. Jackson, Ian Johnston, Dominic Kwiatkowski, Cordelia Langford, John Sillitoe on behalf of the Wellcome Sanger Institute COVID-19 Surveillance Team |
| EPI_ISL_3438788, EPI_ISL_3438789, EPI_ISL_3438790 | Lighthouse Laboratory Plymouth | Wellcome Sanger Institute for the COVID-19 Genomics UK (COG-UK) Consortium | Lighthouse Laboratory Plymouth and Alex Alderton, Roberto Amato, Jeffrey Barrett, Sonia Goncalves, Ewan Harrison, David K. Jackson, Ian Johnston, Dominic Kwiatkowski, Cordelia Langford, John Sillitoe on behalf of the Wellcome Sanger Institute COVID-19 Surveillance Team |
| EPI_ISL_3438791, EPI_ISL_3438792 | Berkshire and Surrey Pathology Services Lighthouse Laboratory | Wellcome Sanger Institute for the COVID-19 Genomics UK (COG-UK) Consortium | Berkshire and Surrey Pathology Services Lighthouse Laboratory and Alex Alderton, Roberto Amato, Jeffrey Barrett, Sonia Goncalves, Ewan Harrison, David K. Jackson, Ian Johnston, Dominic Kwiatkowski, Cordelia Langford, John Sillitoe on behalf of the Wellcome Sanger Institute COVID-19 Surveillance Team |
| EPI_ISL_3438793 | Lighthouse Lab in Alderley Park | Wellcome Sanger Institute for the COVID-19 Genomics UK (COG-UK) Consortium | Jacquelyn Wynn, Mairead Hyland, The Lighthouse Lab in Alderley Park and Alex Alderton, Roberto Amato, Jeffrey Barrett, Sonia Goncalves, Ewan Harrison, David K. Jackson, Ian Johnston, Dominic Kwiatkowski, Cordelia Langford, John Sillitoe on behalf of the Wellcome Sanger Institute COVID-19 Surveillance Team |
| EPI_ISL_3438794 | Berkshire and Surrey Pathology Services Lighthouse Laboratory | Wellcome Sanger Institute for the COVID-19 Genomics UK (COG-UK) Consortium | Berkshire and Surrey Pathology Services Lighthouse Laboratory and Alex Alderton, Roberto Amato, Jeffrey Barrett, Sonia Goncalves, Ewan Harrison, David K. Jackson, Ian Johnston, Dominic Kwiatkowski, Cordelia Langford, John Sillitoe on behalf of the Wellcome Sanger Institute COVID-19 Surveillance Team |
| EPI_ISL_3438795, EPI_ISL_3438796 | Lighthouse Laboratory Plymouth | Wellcome Sanger Institute for the COVID-19 Genomics UK (COG-UK) Consortium | Lighthouse Laboratory Plymouth and Alex Alderton, Roberto Amato, Jeffrey Barrett, Sonia Goncalves, Ewan Harrison, David K. Jackson, Ian Johnston, Dominic Kwiatkowski, Cordelia Langford, John Sillitoe on behalf of the Wellcome Sanger Institute COVID-19 Surveillance Team |
| EPI_ISL_3438797 | Lighthouse Lab in Alderley Park | Wellcome Sanger Institute for the COVID-19 Genomics UK (COG-UK) Consortium | Jacquelyn Wynn, Mairead Hyland, The Lighthouse Lab in Alderley Park and Alex Alderton, Roberto Amato, Jeffrey Barrett, Sonia Goncalves, Ewan Harrison, David K. Jackson, Ian Johnston, Dominic Kwiatkowski, Cordelia Langford, John Sillitoe on behalf of the Wellcome Sanger Institute COVID-19 Surveillance Team |
| EPI_ISL_3438798 | Lighthouse Laboratory Plymouth | Wellcome Sanger Institute for the COVID-19 Genomics UK (COG-UK) Consortium | Lighthouse Laboratory Plymouth and Alex Alderton, Roberto Amato, Jeffrey Barrett, Sonia Goncalves, Ewan Harrison, David K. Jackson, Ian Johnston, Dominic Kwiatkowski, Cordelia Langford, John Sillitoe on behalf of the Wellcome Sanger Institute COVID-19 Surveillance Team |
| EPI_ISL_3438799 | Berkshire and Surrey Pathology Services Lighthouse Laboratory | Wellcome Sanger Institute for the COVID-19 Genomics UK (COG-UK) Consortium | Berkshire and Surrey Pathology Services Lighthouse Laboratory and Alex Alderton, Roberto Amato, Jeffrey Barrett, Sonia Goncalves, Ewan Harrison, David K. Jackson, Ian Johnston, Dominic Kwiatkowski, Cordelia Langford, John Sillitoe on behalf of the Wellcome Sanger Institute COVID-19 Surveillance Team |
| EPI_ISL_3438800 | Lighthouse Lab in Alderley Park | Wellcome Sanger Institute for the COVID-19 Genomics UK (COG-UK) Consortium | Jacquelyn Wynn, Mairead Hyland, The Lighthouse Lab in Alderley Park and Alex Alderton, Roberto Amato, Jeffrey Barrett, Sonia Goncalves, Ewan Harrison, David K. Jackson, Ian Johnston, Dominic Kwiatkowski, Cordelia Langford, John Sillitoe on behalf of the Wellcome Sanger Institute COVID-19 Surveillance Team |
| EPI_ISL_3438801 | Lighthouse Laboratory Plymouth | Wellcome Sanger Institute for the COVID-19 Genomics UK (COG-UK) Consortium | Lighthouse Laboratory Plymouth and Alex Alderton, Roberto Amato, Jeffrey Barrett, Sonia Goncalves, Ewan Harrison, David K. Jackson, Ian Johnston, Dominic Kwiatkowski, Cordelia Langford, John Sillitoe on behalf of the Wellcome Sanger Institute COVID-19 Surveillance Team |
| EPI_ISL_3438802 | Lighthouse Lab in Alderley Park | Wellcome Sanger Institute for the COVID-19 Genomics UK (COG-UK) Consortium | Jacquelyn Wynn, Mairead Hyland, The Lighthouse Lab in Alderley Park and Alex Alderton, Roberto Amato, Jeffrey Barrett, Sonia Goncalves, Ewan Harrison, David K. Jackson, Ian Johnston, Dominic Kwiatkowski, Cordelia Langford, John Sillitoe on behalf of the Wellcome Sanger Institute COVID-19 Surveillance Team |
| EPI_ISL_3438803 | Lighthouse Laboratory Plymouth | Wellcome Sanger Institute for the COVID-19 Genomics UK (COG-UK) Consortium | Lighthouse Laboratory Plymouth and Alex Alderton, Roberto Amato, Jeffrey Barrett, Sonia Goncalves, Ewan Harrison, David K. Jackson, Ian Johnston, Dominic Kwiatkowski, Cordelia Langford, John Sillitoe on behalf of the Wellcome Sanger Institute COVID-19 Surveillance Team |
| EPI_ISL_3438804 | Berkshire and Surrey Pathology Services Lighthouse Laboratory | Wellcome Sanger Institute for the COVID-19 Genomics UK (COG-UK) Consortium | Berkshire and Surrey Pathology Services Lighthouse Laboratory and Alex Alderton, Roberto Amato, Jeffrey Barrett, Sonia Goncalves, Ewan Harrison, David K. Jackson, Ian Johnston, Dominic Kwiatkowski, Cordelia Langford, John Sillitoe on behalf of the Wellcome Sanger Institute COVID-19 Surveillance Team |
| EPI_ISL_3438805 | Lighthouse Lab in Alderley Park | Wellcome Sanger Institute for the COVID-19 Genomics UK (COG-UK) Consortium | Jacquelyn Wynn, Mairead Hyland, The Lighthouse Lab in Alderley Park and Alex Alderton, Roberto Amato, Jeffrey Barrett, Sonia Goncalves, Ewan Harrison, David K. Jackson, Ian Johnston, Dominic Kwiatkowski, Cordelia Langford, John Sillitoe on behalf of the Wellcome Sanger Institute COVID-19 Surveillance Team |
| EPI_ISL_3438806 | Lighthouse Laboratory Plymouth | Wellcome Sanger Institute for the COVID-19 Genomics UK (COG-UK) Consortium | Lighthouse Laboratory Plymouth and Alex Alderton, Roberto Amato, Jeffrey Barrett, Sonia Goncalves, Ewan Harrison, David K. Jackson, Ian Johnston, Dominic Kwiatkowski, Cordelia Langford, John Sillitoe on behalf of the Wellcome Sanger Institute COVID-19 Surveillance Team |
| EPI_ISL_3438807 | Lighthouse Lab in Alderley Park | Wellcome Sanger Institute for the COVID-19 Genomics UK (COG-UK) Consortium | Jacquelyn Wynn, Mairead Hyland, The Lighthouse Lab in Alderley Park and Alex Alderton, Roberto Amato, Jeffrey Barrett, Sonia Goncalves, Ewan Harrison, David K. Jackson, Ian Johnston, Dominic Kwiatkowski, Cordelia Langford, John Sillitoe on behalf of the Wellcome Sanger Institute COVID-19 Surveillance Team |
| EPI_ISL_3438808 | Lighthouse Laboratory Plymouth | Wellcome Sanger Institute for the COVID-19 Genomics UK (COG-UK) Consortium | Lighthouse Laboratory Plymouth and Alex Alderton, Roberto Amato, Jeffrey Barrett, Sonia Goncalves, Ewan Harrison, David K. Jackson, Ian Johnston, Dominic Kwiatkowski, Cordelia Langford, John Sillitoe on behalf of the Wellcome Sanger Institute COVID-19 Surveillance Team |
| EPI_ISL_3438809 | Berkshire and Surrey Pathology Services Lighthouse Laboratory | Wellcome Sanger Institute for the COVID-19 Genomics UK (COG-UK) Consortium | Berkshire and Surrey Pathology Services Lighthouse Laboratory and Alex Alderton, Roberto Amato, Jeffrey Barrett, Sonia Goncalves, Ewan Harrison, David K. Jackson, Ian Johnston, Dominic Kwiatkowski, Cordelia Langford, John Sillitoe on behalf of the Wellcome Sanger Institute COVID-19 Surveillance Team |
| EPI_ISL_3438810, EPI_ISL_3438811, EPI_ISL_3438812, EPI_ISL_3438813, EPI_ISL_3438814 | Lighthouse Laboratory Plymouth | Wellcome Sanger Institute for the COVID-19 Genomics UK (COG-UK) Consortium | Lighthouse Laboratory Plymouth and Alex Alderton, Roberto Amato, Jeffrey Barrett, Sonia Goncalves, Ewan Harrison, David K. Jackson, Ian Johnston, Dominic Kwiatkowski, Cordelia Langford, John Sillitoe on behalf of the Wellcome Sanger Institute COVID-19 Surveillance Team |
| EPI_ISL_3438815 | Lighthouse Lab in Alderley Park | Wellcome Sanger Institute for the COVID-19 Genomics UK (COG-UK) Consortium | Jacquelyn Wynn, Mairead Hyland, The Lighthouse Lab in Alderley Park and Alex Alderton, Roberto Amato, Jeffrey Barrett, Sonia Goncalves, Ewan Harrison, David K. Jackson, Ian Johnston, Dominic Kwiatkowski, Cordelia Langford, John Sillitoe on behalf of the Wellcome Sanger Institute COVID-19 Surveillance Team |
| EPI_ISL_3438816, EPI_ISL_3438817, EPI_ISL_3438818, EPI_ISL_3438819 | Lighthouse Laboratory Plymouth | Wellcome Sanger Institute for the COVID-19 Genomics UK (COG-UK) Consortium | Lighthouse Laboratory Plymouth and Alex Alderton, Roberto Amato, Jeffrey Barrett, Sonia Goncalves, Ewan Harrison, David K. Jackson, Ian Johnston, Dominic Kwiatkowski, Cordelia Langford, John Sillitoe on behalf of the Wellcome Sanger Institute COVID-19 Surveillance Team |
| EPI_ISL_3438820 | Berkshire and Surrey Pathology Services Lighthouse Laboratory | Wellcome Sanger Institute for the COVID-19 Genomics UK (COG-UK) Consortium | Berkshire and Surrey Pathology Services Lighthouse Laboratory and Alex Alderton, Roberto Amato, Jeffrey Barrett, Sonia Goncalves, Ewan Harrison, David K. Jackson, Ian Johnston, Dominic Kwiatkowski, Cordelia Langford, John Sillitoe on behalf of the Wellcome Sanger Institute COVID-19 Surveillance Team |
| EPI_ISL_3438821, EPI_ISL_3438822, EPI_ISL_3438823 | Lighthouse Laboratory Plymouth | Wellcome Sanger Institute for the COVID-19 Genomics UK (COG-UK) Consortium | Lighthouse Laboratory Plymouth and Alex Alderton, Roberto Amato, Jeffrey Barrett, Sonia Goncalves, Ewan Harrison, David K. Jackson, Ian Johnston, Dominic Kwiatkowski, Cordelia Langford, John Sillitoe on behalf of the Wellcome Sanger Institute COVID-19 Surveillance Team |
| EPI_ISL_3438824 | Berkshire and Surrey Pathology Services Lighthouse Laboratory | Wellcome Sanger Institute for the COVID-19 Genomics UK (COG-UK) Consortium | Berkshire and Surrey Pathology Services Lighthouse Laboratory and Alex Alderton, Roberto Amato, Jeffrey Barrett, Sonia Goncalves, Ewan Harrison, David K. Jackson, Ian Johnston, Dominic Kwiatkowski, Cordelia Langford, John Sillitoe on behalf of the Wellcome Sanger Institute COVID-19 Surveillance Team |
| EPI_ISL_3438825 | Lighthouse Laboratory Plymouth | Wellcome Sanger Institute for the COVID-19 Genomics UK (COG-UK) Consortium | Lighthouse Laboratory Plymouth and Alex Alderton, Roberto Amato, Jeffrey Barrett, Sonia Goncalves, Ewan Harrison, David K. Jackson, Ian Johnston, Dominic Kwiatkowski, Cordelia Langford, John Sillitoe on behalf of the Wellcome Sanger Institute COVID-19 Surveillance Team |
| EPI_ISL_3438826, EPI_ISL_3438827 | Berkshire and Surrey Pathology Services Lighthouse Laboratory | Wellcome Sanger Institute for the COVID-19 Genomics UK (COG-UK) Consortium | Berkshire and Surrey Pathology Services Lighthouse Laboratory and Alex Alderton, Roberto Amato, Jeffrey Barrett, Sonia Goncalves, Ewan Harrison, David K. Jackson, Ian Johnston, Dominic Kwiatkowski, Cordelia Langford, John Sillitoe on behalf of the Wellcome Sanger Institute COVID-19 Surveillance Team |
| EPI_ISL_3438828, EPI_ISL_3438829 | Lighthouse Laboratory Plymouth | Wellcome Sanger Institute for the COVID-19 Genomics UK (COG-UK) Consortium | Lighthouse Laboratory Plymouth and Alex Alderton, Roberto Amato, Jeffrey Barrett, Sonia Goncalves, Ewan Harrison, David K. Jackson, Ian Johnston, Dominic Kwiatkowski, Cordelia Langford, John Sillitoe on behalf of the Wellcome Sanger Institute COVID-19 Surveillance Team |
| EPI_ISL_3438830 | Lighthouse Lab in Alderley Park | Wellcome Sanger Institute for the COVID-19 Genomics UK (COG-UK) Consortium | Jacquelyn Wynn, Mairead Hyland, The Lighthouse Lab in Alderley Park and Alex Alderton, Roberto Amato, Jeffrey Barrett, Sonia Goncalves, Ewan Harrison, David K. Jackson, Ian Johnston, Dominic Kwiatkowski, Cordelia Langford, John Sillitoe on behalf of the Wellcome Sanger Institute COVID-19 Surveillance Team |

| | | | |
|---|---|---|---|
| EPI_ISL_3438831, EPI_ISL_3438832 | Lighthouse Laboratory Plymouth | Wellcome Sanger Institute for the COVID-19 Genomics UK (COG-UK) Consortium | Lighthouse Laboratory Plymouth and Alex Alderton, Roberto Amato, Jeffrey Barrett, Sonia Goncalves, Ewan Harrison, David K. Jackson, Ian Johnston, Dominic Kwiatkowski, Cordelia Langford, John Sillitoe on behalf of the Wellcome Sanger Institute COVID-19 Surveillance Team |
| EPI_ISL_3438833, EPI_ISL_3438834 | Berkshire and Surrey Pathology Services Lighthouse Laboratory | Wellcome Sanger Institute for the COVID-19 Genomics UK (COG-UK) Consortium | Berkshire and Surrey Pathology Services Lighthouse Laboratory and Alex Alderton, Roberto Amato, Jeffrey Barrett, Sonia Goncalves, Ewan Harrison, David K. Jackson, Ian Johnston, Dominic Kwiatkowski, Cordelia Langford, John Sillitoe on behalf of the Wellcome Sanger Institute COVID-19 Surveillance Team |
| EPI_ISL_3438835 | Lighthouse Laboratory Plymouth | Wellcome Sanger Institute for the COVID-19 Genomics UK (COG-UK) Consortium | Lighthouse Laboratory Plymouth and Alex Alderton, Roberto Amato, Jeffrey Barrett, Sonia Goncalves, Ewan Harrison, David K. Jackson, Ian Johnston, Dominic Kwiatkowski, Cordelia Langford, John Sillitoe on behalf of the Wellcome Sanger Institute COVID-19 Surveillance Team |
| EPI_ISL_3438836, EPI_ISL_3438837 | Berkshire and Surrey Pathology Services Lighthouse Laboratory | Wellcome Sanger Institute for the COVID-19 Genomics UK (COG-UK) Consortium | Berkshire and Surrey Pathology Services Lighthouse Laboratory and Alex Alderton, Roberto Amato, Jeffrey Barrett, Sonia Goncalves, Ewan Harrison, David K. Jackson, Ian Johnston, Dominic Kwiatkowski, Cordelia Langford, John Sillitoe on behalf of the Wellcome Sanger Institute COVID-19 Surveillance Team |
| EPI_ISL_3438838 | Lighthouse Laboratory Plymouth | Wellcome Sanger Institute for the COVID-19 Genomics UK (COG-UK) Consortium | Lighthouse Laboratory Plymouth and Alex Alderton, Roberto Amato, Jeffrey Barrett, Sonia Goncalves, Ewan Harrison, David K. Jackson, Ian Johnston, Dominic Kwiatkowski, Cordelia Langford, John Sillitoe on behalf of the Wellcome Sanger Institute COVID-19 Surveillance Team |
| EPI_ISL_3438839, EPI_ISL_3438840 | Lighthouse Lab in Alderley Park | Wellcome Sanger Institute for the COVID-19 Genomics UK (COG-UK) Consortium | Jacquelyn Wynn, Mairead Hyland, The Lighthouse Lab in Alderley Park and Alex Alderton, Roberto Amato, Jeffrey Barrett, Sonia Goncalves, Ewan Harrison, David K. Jackson, Ian Johnston, Dominic Kwiatkowski, Cordelia Langford, John Sillitoe on behalf of the Wellcome Sanger Institute COVID-19 Surveillance Team |
| EPI_ISL_3438841, EPI_ISL_3438842 | Berkshire and Surrey Pathology Services Lighthouse Laboratory | Wellcome Sanger Institute for the COVID-19 Genomics UK (COG-UK) Consortium | Berkshire and Surrey Pathology Services Lighthouse Laboratory and Alex Alderton, Roberto Amato, Jeffrey Barrett, Sonia Goncalves, Ewan Harrison, David K. Jackson, Ian Johnston, Dominic Kwiatkowski, Cordelia Langford, John Sillitoe on behalf of the Wellcome Sanger Institute COVID-19 Surveillance Team |
| EPI_ISL_3438843 | Lighthouse Laboratory Plymouth | Wellcome Sanger Institute for the COVID-19 Genomics UK (COG-UK) Consortium | Lighthouse Laboratory Plymouth and Alex Alderton, Roberto Amato, Jeffrey Barrett, Sonia Goncalves, Ewan Harrison, David K. Jackson, Ian Johnston, Dominic Kwiatkowski, Cordelia Langford, John Sillitoe on behalf of the Wellcome Sanger Institute COVID-19 Surveillance Team |
| EPI_ISL_3438844 | Lighthouse Lab in Alderley Park | Wellcome Sanger Institute for the COVID-19 Genomics UK (COG-UK) Consortium | Jacquelyn Wynn, Mairead Hyland, The Lighthouse Lab in Alderley Park and Alex Alderton, Roberto Amato, Jeffrey Barrett, Sonia Goncalves, Ewan Harrison, David K. Jackson, Ian Johnston, Dominic Kwiatkowski, Cordelia Langford, John Sillitoe on behalf of the Wellcome Sanger Institute COVID-19 Surveillance Team |
| EPI_ISL_3438845, EPI_ISL_3438846 | Lighthouse Laboratory Plymouth | Wellcome Sanger Institute for the COVID-19 Genomics UK (COG-UK) Consortium | Lighthouse Laboratory Plymouth and Alex Alderton, Roberto Amato, Jeffrey Barrett, Sonia Goncalves, Ewan Harrison, David K. Jackson, Ian Johnston, Dominic Kwiatkowski, Cordelia Langford, John Sillitoe on behalf of the Wellcome Sanger Institute COVID-19 Surveillance Team |
| EPI_ISL_3438847 | Berkshire and Surrey Pathology Services Lighthouse Laboratory | Wellcome Sanger Institute for the COVID-19 Genomics UK (COG-UK) Consortium | Berkshire and Surrey Pathology Services Lighthouse Laboratory and Alex Alderton, Roberto Amato, Jeffrey Barrett, Sonia Goncalves, Ewan Harrison, David K. Jackson, Ian Johnston, Dominic Kwiatkowski, Cordelia Langford, John Sillitoe on behalf of the Wellcome Sanger Institute COVID-19 Surveillance Team |
| EPI_ISL_3438848 | Lighthouse Lab in Alderley Park | Wellcome Sanger Institute for the COVID-19 Genomics UK (COG-UK) Consortium | Jacquelyn Wynn, Mairead Hyland, The Lighthouse Lab in Alderley Park and Alex Alderton, Roberto Amato, Jeffrey Barrett, Sonia Goncalves, Ewan Harrison, David K. Jackson, Ian Johnston, Dominic Kwiatkowski, Cordelia Langford, John Sillitoe on behalf of the Wellcome Sanger Institute COVID-19 Surveillance Team |
| EPI_ISL_3438849, EPI_ISL_3438850 | Lighthouse Laboratory Plymouth | Wellcome Sanger Institute for the COVID-19 Genomics UK (COG-UK) Consortium | Lighthouse Laboratory Plymouth and Alex Alderton, Roberto Amato, Jeffrey Barrett, Sonia Goncalves, Ewan Harrison, David K. Jackson, Ian Johnston, Dominic Kwiatkowski, Cordelia Langford, John Sillitoe on behalf of the Wellcome Sanger Institute COVID-19 Surveillance Team |
| EPI_ISL_3438851 | Berkshire and Surrey Pathology Services Lighthouse Laboratory | Wellcome Sanger Institute for the COVID-19 Genomics UK (COG-UK) Consortium | Berkshire and Surrey Pathology Services Lighthouse Laboratory and Alex Alderton, Roberto Amato, Jeffrey Barrett, Sonia Goncalves, Ewan Harrison, David K. Jackson, Ian Johnston, Dominic Kwiatkowski, Cordelia Langford, John Sillitoe on behalf of the Wellcome Sanger Institute COVID-19 Surveillance Team |
| EPI_ISL_3438852, EPI_ISL_3438853, EPI_ISL_3438854 | Lighthouse Laboratory Plymouth | Wellcome Sanger Institute for the COVID-19 Genomics UK (COG-UK) Consortium | Lighthouse Laboratory Plymouth and Alex Alderton, Roberto Amato, Jeffrey Barrett, Sonia Goncalves, Ewan Harrison, David K. Jackson, Ian Johnston, Dominic Kwiatkowski, Cordelia Langford, John Sillitoe on behalf of the Wellcome Sanger Institute COVID-19 Surveillance Team |
| EPI_ISL_3438855, EPI_ISL_3438856 | Berkshire and Surrey Pathology Services Lighthouse Laboratory | Wellcome Sanger Institute for the COVID-19 Genomics UK (COG-UK) Consortium | Berkshire and Surrey Pathology Services Lighthouse Laboratory and Alex Alderton, Roberto Amato, Jeffrey Barrett, Sonia Goncalves, Ewan Harrison, David K. Jackson, Ian Johnston, Dominic Kwiatkowski, Cordelia Langford, John Sillitoe on behalf of the Wellcome Sanger Institute COVID-19 Surveillance Team |
| EPI_ISL_3438857, EPI_ISL_3438858 | Lighthouse Lab in Alderley Park | Wellcome Sanger Institute for the COVID-19 Genomics UK (COG-UK) Consortium | Jacquelyn Wynn, Mairead Hyland, The Lighthouse Lab in Alderley Park and Alex Alderton, Roberto Amato, Jeffrey Barrett, Sonia Goncalves, Ewan Harrison, David K. Jackson, Ian Johnston, Dominic Kwiatkowski, Cordelia Langford, John Sillitoe on behalf of the Wellcome Sanger Institute COVID-19 Surveillance Team |
| EPI_ISL_3438859, EPI_ISL_3438860, EPI_ISL_3438861 | Lighthouse Laboratory Plymouth | Wellcome Sanger Institute for the COVID-19 Genomics UK (COG-UK) Consortium | Lighthouse Laboratory Plymouth and Alex Alderton, Roberto Amato, Jeffrey Barrett, Sonia Goncalves, Ewan Harrison, David K. Jackson, Ian Johnston, Dominic Kwiatkowski, Cordelia Langford, John Sillitoe on behalf of the Wellcome Sanger Institute COVID-19 Surveillance Team |
| EPI_ISL_3438862 | Berkshire and Surrey Pathology Services Lighthouse Laboratory | Wellcome Sanger Institute for the COVID-19 Genomics UK (COG-UK) Consortium | Berkshire and Surrey Pathology Services Lighthouse Laboratory and Alex Alderton, Roberto Amato, Jeffrey Barrett, Sonia Goncalves, Ewan Harrison, David K. Jackson, Ian Johnston, Dominic Kwiatkowski, Cordelia Langford, John Sillitoe on behalf of the Wellcome Sanger Institute COVID-19 Surveillance Team |
| EPI_ISL_3438863, EPI_ISL_3438864 | Lighthouse Laboratory Plymouth | Wellcome Sanger Institute for the COVID-19 Genomics UK (COG-UK) Consortium | Lighthouse Laboratory Plymouth and Alex Alderton, Roberto Amato, Jeffrey Barrett, Sonia Goncalves, Ewan Harrison, David K. Jackson, Ian Johnston, Dominic Kwiatkowski, Cordelia Langford, John Sillitoe on behalf of the Wellcome Sanger Institute COVID-19 Surveillance Team |
| EPI_ISL_3438865 | Berkshire and Surrey Pathology Services Lighthouse Laboratory | Wellcome Sanger Institute for the COVID-19 Genomics UK (COG-UK) Consortium | Berkshire and Surrey Pathology Services Lighthouse Laboratory and Alex Alderton, Roberto Amato, Jeffrey Barrett, Sonia Goncalves, Ewan Harrison, David K. Jackson, Ian Johnston, Dominic Kwiatkowski, Cordelia Langford, John Sillitoe on behalf of the Wellcome Sanger Institute COVID-19 Surveillance Team |
| EPI_ISL_3438866, EPI_ISL_3438867 | Lighthouse Laboratory Plymouth | Wellcome Sanger Institute for the COVID-19 Genomics UK (COG-UK) Consortium | Lighthouse Laboratory Plymouth and Alex Alderton, Roberto Amato, Jeffrey Barrett, Sonia Goncalves, Ewan Harrison, David K. Jackson, Ian Johnston, Dominic Kwiatkowski, Cordelia Langford, John Sillitoe on behalf of the Wellcome Sanger Institute COVID-19 Surveillance Team |
| EPI_ISL_3438868 | Lighthouse Lab in Alderley Park | Wellcome Sanger Institute for the COVID-19 Genomics UK (COG-UK) Consortium | Jacquelyn Wynn, Mairead Hyland, The Lighthouse Lab in Alderley Park and Alex Alderton, Roberto Amato, Jeffrey Barrett, Sonia Goncalves, Ewan Harrison, David K. Jackson, Ian Johnston, Dominic Kwiatkowski, Cordelia Langford, John Sillitoe on behalf of the Wellcome Sanger Institute COVID-19 Surveillance Team |
| EPI_ISL_3438869, EPI_ISL_3438870 | Lighthouse Laboratory Plymouth | Wellcome Sanger Institute for the COVID-19 Genomics UK (COG-UK) Consortium | Lighthouse Laboratory Plymouth and Alex Alderton, Roberto Amato, Jeffrey Barrett, Sonia Goncalves, Ewan Harrison, David K. Jackson, Ian Johnston, Dominic Kwiatkowski, Cordelia Langford, John Sillitoe on behalf of the Wellcome Sanger Institute COVID-19 Surveillance Team |
| EPI_ISL_3438871 | Lighthouse Lab in Alderley Park | Wellcome Sanger Institute for the COVID-19 Genomics UK (COG-UK) Consortium | Jacquelyn Wynn, Mairead Hyland, The Lighthouse Lab in Alderley Park and Alex Alderton, Roberto Amato, Jeffrey Barrett, Sonia Goncalves, Ewan Harrison, David K. Jackson, Ian Johnston, Dominic Kwiatkowski, Cordelia Langford, John Sillitoe on behalf of the Wellcome Sanger Institute COVID-19 Surveillance Team |
| EPI_ISL_3438872, EPI_ISL_3438873 | Berkshire and Surrey Pathology Services Lighthouse Laboratory | Wellcome Sanger Institute for the COVID-19 Genomics UK (COG-UK) Consortium | Berkshire and Surrey Pathology Services Lighthouse Laboratory and Alex Alderton, Roberto Amato, Jeffrey Barrett, Sonia Goncalves, Ewan Harrison, David K. Jackson, Ian Johnston, Dominic Kwiatkowski, Cordelia Langford, John Sillitoe on behalf of the Wellcome Sanger Institute COVID-19 Surveillance Team |
| EPI_ISL_3438874 | Lighthouse Lab in Alderley Park | Wellcome Sanger Institute for the COVID-19 Genomics UK (COG-UK) Consortium | Jacquelyn Wynn, Mairead Hyland, The Lighthouse Lab in Alderley Park and Alex Alderton, Roberto Amato, Jeffrey Barrett, Sonia Goncalves, Ewan Harrison, David K. Jackson, Ian Johnston, Dominic Kwiatkowski, Cordelia Langford, John Sillitoe on behalf of the Wellcome Sanger Institute COVID-19 Surveillance Team |
| EPI_ISL_3438875 | Lighthouse Laboratory Plymouth | Wellcome Sanger Institute for the COVID-19 Genomics UK (COG-UK) Consortium | Lighthouse Laboratory Plymouth and Alex Alderton, Roberto Amato, Jeffrey Barrett, Sonia Goncalves, Ewan Harrison, David K. Jackson, Ian Johnston, Dominic Kwiatkowski, Cordelia Langford, John Sillitoe on behalf of the Wellcome Sanger Institute COVID-19 Surveillance Team |
| EPI_ISL_3438876, EPI_ISL_3438877 | Lighthouse Lab in Alderley Park | Wellcome Sanger Institute for the COVID-19 Genomics UK (COG-UK) Consortium | Jacquelyn Wynn, Mairead Hyland, The Lighthouse Lab in Alderley Park and Alex Alderton, Roberto Amato, Jeffrey Barrett, Sonia Goncalves, Ewan Harrison, David K. Jackson, Ian Johnston, Dominic Kwiatkowski, Cordelia Langford, John Sillitoe on behalf of the Wellcome Sanger Institute COVID-19 Surveillance Team |
| EPI_ISL_3438878 | Lighthouse Laboratory Plymouth | Wellcome Sanger Institute for the COVID-19 Genomics UK (COG-UK) Consortium | Lighthouse Laboratory Plymouth and Alex Alderton, Roberto Amato, Jeffrey Barrett, Sonia Goncalves, Ewan Harrison, David K. Jackson, Ian Johnston, Dominic Kwiatkowski, Cordelia Langford, John Sillitoe on behalf of the Wellcome Sanger Institute COVID-19 Surveillance Team |
| EPI_ISL_3438879 | Berkshire and Surrey Pathology Services Lighthouse Laboratory | Wellcome Sanger Institute for the COVID-19 Genomics UK (COG-UK) Consortium | Berkshire and Surrey Pathology Services Lighthouse Laboratory and Alex Alderton, Roberto Amato, Jeffrey Barrett, Sonia Goncalves, Ewan Harrison, David K. Jackson, Ian Johnston, Dominic Kwiatkowski, Cordelia Langford, John Sillitoe on behalf of the Wellcome Sanger Institute COVID-19 Surveillance Team |
| EPI_ISL_3438880 | Lighthouse Laboratory Plymouth | Wellcome Sanger Institute for the COVID-19 Genomics UK (COG-UK) Consortium | Lighthouse Laboratory Plymouth and Alex Alderton, Roberto Amato, Jeffrey Barrett, Sonia Goncalves, Ewan Harrison, David K. Jackson, Ian Johnston, Dominic Kwiatkowski, Cordelia Langford, John Sillitoe on behalf of the Wellcome Sanger Institute COVID-19 Surveillance Team |
| EPI_ISL_3438881 | Berkshire and Surrey Pathology Services Lighthouse Laboratory | Wellcome Sanger Institute for the COVID-19 Genomics UK (COG-UK) Consortium | Berkshire and Surrey Pathology Services Lighthouse Laboratory and Alex Alderton, Roberto Amato, Jeffrey Barrett, Sonia Goncalves, Ewan Harrison, David K. Jackson, Ian Johnston, Dominic Kwiatkowski, Cordelia Langford, John Sillitoe on behalf of the Wellcome Sanger Institute COVID-19 Surveillance Team |
| EPI_ISL_3438882 | Lighthouse Laboratory Plymouth | Wellcome Sanger Institute for the COVID-19 Genomics UK (COG-UK) Consortium | Lighthouse Laboratory Plymouth and Alex Alderton, Roberto Amato, Jeffrey Barrett, Sonia Goncalves, Ewan Harrison, David K. Jackson, Ian Johnston, Dominic Kwiatkowski, Cordelia Langford, John Sillitoe on behalf of the Wellcome Sanger Institute COVID-19 Surveillance Team |
| EPI_ISL_3438883 | Health Services Laboratories | Wellcome Sanger Institute for the COVID-19 Genomics UK (COG-UK) Consortium | Health Services Laboratories and Alex Alderton, Roberto Amato, Jeffrey Barrett, Sonia Goncalves, Ewan Harrison, David K. Jackson, Ian Johnston, Dominic Kwiatkowski, Cordelia Langford, John Sillitoe on behalf of the Wellcome Sanger Institute COVID-19 Surveillance Team |
| EPI_ISL_3438884, EPI_ISL_3438885, EPI_ISL_3438886 | Lighthouse Lab in Alderley Park | Wellcome Sanger Institute for the COVID-19 Genomics UK (COG-UK) Consortium | Jacquelyn Wynn, Mairead Hyland, The Lighthouse Lab in Alderley Park and Alex Alderton, Roberto Amato, Jeffrey Barrett, Sonia Goncalves, Ewan Harrison, David K. Jackson, Ian Johnston, Dominic Kwiatkowski, Cordelia Langford, John Sillitoe on behalf of the Wellcome Sanger Institute COVID-19 Surveillance Team |
| EPI_ISL_3438887 | Berkshire and Surrey Pathology Services Lighthouse Laboratory | Wellcome Sanger Institute for the COVID-19 Genomics UK (COG-UK) Consortium | Berkshire and Surrey Pathology Services Lighthouse Laboratory and Alex Alderton, Roberto Amato, Jeffrey Barrett, Sonia Goncalves, Ewan Harrison, David K. Jackson, Ian Johnston, Dominic Kwiatkowski, Cordelia Langford, John Sillitoe on behalf of the Wellcome Sanger Institute COVID-19 Surveillance Team |
| EPI_ISL_3438888 | Lighthouse Laboratory Plymouth | Wellcome Sanger Institute for the COVID-19 Genomics UK (COG-UK) Consortium | Lighthouse Laboratory Plymouth and Alex Alderton, Roberto Amato, Jeffrey Barrett, Sonia Goncalves, Ewan Harrison, David K. Jackson, Ian Johnston, Dominic Kwiatkowski, Cordelia Langford, John Sillitoe on behalf of the Wellcome Sanger Institute COVID-19 Surveillance Team |
| EPI_ISL_3438889 | Lighthouse Lab in Alderley Park | Wellcome Sanger Institute for the COVID-19 Genomics UK (COG-UK) Consortium | Jacquelyn Wynn, Mairead Hyland, The Lighthouse Lab in Alderley Park and Alex Alderton, Roberto Amato, Jeffrey Barrett, Sonia Goncalves, Ewan Harrison, David K. Jackson, Ian Johnston, Dominic Kwiatkowski, Cordelia Langford, John Sillitoe on behalf of the Wellcome Sanger Institute COVID-19 Surveillance Team |
| EPI_ISL_3438890 | Health Services Laboratories | Wellcome Sanger Institute for the COVID-19 Genomics UK (COG-UK) Consortium | Health Services Laboratories and Alex Alderton, Roberto Amato, Jeffrey Barrett, Sonia Goncalves, Ewan Harrison, David K. Jackson, Ian Johnston, Dominic Kwiatkowski, Cordelia Langford, John Sillitoe on behalf of the Wellcome Sanger Institute COVID-19 Surveillance Team |
| EPI_ISL_3438891, EPI_ISL_3438892 | Lighthouse Laboratory Plymouth | Wellcome Sanger Institute for the COVID-19 Genomics UK (COG-UK) Consortium | Lighthouse Laboratory Plymouth and Alex Alderton, Roberto Amato, Jeffrey Barrett, Sonia Goncalves, Ewan Harrison, David K. Jackson, Ian Johnston, Dominic Kwiatkowski, Cordelia Langford, John Sillitoe on behalf of the Wellcome Sanger Institute COVID-19 Surveillance Team |
| EPI_ISL_3438893, EPI_ISL_3438894 | Berkshire and Surrey Pathology Services Lighthouse Laboratory | Wellcome Sanger Institute for the COVID-19 Genomics UK (COG-UK) Consortium | Berkshire and Surrey Pathology Services Lighthouse Laboratory and Alex Alderton, Roberto Amato, Jeffrey Barrett, Sonia Goncalves, Ewan Harrison, David K. Jackson, Ian Johnston, Dominic Kwiatkowski, Cordelia Langford, John Sillitoe on behalf of the Wellcome Sanger Institute COVID-19 Surveillance Team |
| EPI_ISL_3438895 | Lighthouse Lab in Alderley Park | Wellcome Sanger Institute for the COVID-19 Genomics UK (COG-UK) Consortium | Jacquelyn Wynn, Mairead Hyland, The Lighthouse Lab in Alderley Park and Alex Alderton, Roberto Amato, Jeffrey Barrett, Sonia Goncalves, Ewan Harrison, David K. Jackson, Ian Johnston, Dominic Kwiatkowski, Cordelia Langford, John Sillitoe on behalf of the Wellcome Sanger Institute COVID-19 Surveillance Team |
| EPI_ISL_3438896, EPI_ISL_3438897 | Lighthouse Laboratory Plymouth | Wellcome Sanger Institute for the COVID-19 Genomics UK (COG-UK) Consortium | Lighthouse Laboratory Plymouth and Alex Alderton, Roberto Amato, Jeffrey Barrett, Sonia Goncalves, Ewan Harrison, David K. Jackson, Ian Johnston, Dominic Kwiatkowski, Cordelia Langford, John Sillitoe on behalf of the Wellcome Sanger Institute COVID-19 Surveillance Team |

| | | | |
|---|---|---|---|
| EPI_ISL_3438898 | Health Services Laboratories | Wellcome Sanger Institute for the COVID-19 Genomics UK (COG-UK) Consortium | Health Services Laboratories and Alex Alderton, Roberto Amato, Jeffrey Barrett, Sonia Goncalves, Ewan Harrison, David K. Jackson, Ian Johnston, Dominic Kwiatkowski, Cordelia Langford, John Sillitoe on behalf of the Wellcome Sanger Institute COVID-19 Surveillance Team |
| EPI_ISL_3438899 | Lighthouse Lab in Alderley Park | Wellcome Sanger Institute for the COVID-19 Genomics UK (COG-UK) Consortium | Jacquelyn Wynn, Mairead Hyland, The Lighthouse Lab in Alderley Park and Alex Alderton, Roberto Amato, Jeffrey Barrett, Sonia Goncalves, Ewan Harrison, David K. Jackson, Ian Johnston, Dominic Kwiatkowski, Cordelia Langford, John Sillitoe on behalf of the Wellcome Sanger Institute COVID-19 Surveillance Team |
| EPI_ISL_3438900, EPI_ISL_3438901, EPI_ISL_3438902 | Lighthouse Laboratory Plymouth | Wellcome Sanger Institute for the COVID-19 Genomics UK (COG-UK) Consortium | Lighthouse Laboratory Plymouth and Alex Alderton, Roberto Amato, Jeffrey Barrett, Sonia Goncalves, Ewan Harrison, David K. Jackson, Ian Johnston, Dominic Kwiatkowski, Cordelia Langford, John Sillitoe on behalf of the Wellcome Sanger Institute COVID-19 Surveillance Team |
| EPI_ISL_3438903 | Berkshire and Surrey Pathology Services Lighthouse Laboratory | Wellcome Sanger Institute for the COVID-19 Genomics UK (COG-UK) Consortium | Berkshire and Surrey Pathology Services Lighthouse Laboratory and Alex Alderton, Roberto Amato, Jeffrey Barrett, Sonia Goncalves, Ewan Harrison, David K. Jackson, Ian Johnston, Dominic Kwiatkowski, Cordelia Langford, John Sillitoe on behalf of the Wellcome Sanger Institute COVID-19 Surveillance Team |
| EPI_ISL_3438904 | Health Services Laboratories | Wellcome Sanger Institute for the COVID-19 Genomics UK (COG-UK) Consortium | Health Services Laboratories and Alex Alderton, Roberto Amato, Jeffrey Barrett, Sonia Goncalves, Ewan Harrison, David K. Jackson, Ian Johnston, Dominic Kwiatkowski, Cordelia Langford, John Sillitoe on behalf of the Wellcome Sanger Institute COVID-19 Surveillance Team |
| EPI_ISL_3438905 | Lighthouse Lab in Alderley Park | Wellcome Sanger Institute for the COVID-19 Genomics UK (COG-UK) Consortium | Jacquelyn Wynn, Mairead Hyland, The Lighthouse Lab in Alderley Park and Alex Alderton, Roberto Amato, Jeffrey Barrett, Sonia Goncalves, Ewan Harrison, David K. Jackson, Ian Johnston, Dominic Kwiatkowski, Cordelia Langford, John Sillitoe on behalf of the Wellcome Sanger Institute COVID-19 Surveillance Team |
| EPI_ISL_3438906 | Berkshire and Surrey Pathology Services Lighthouse Laboratory | Wellcome Sanger Institute for the COVID-19 Genomics UK (COG-UK) Consortium | Berkshire and Surrey Pathology Services Lighthouse Laboratory and Alex Alderton, Roberto Amato, Jeffrey Barrett, Sonia Goncalves, Ewan Harrison, David K. Jackson, Ian Johnston, Dominic Kwiatkowski, Cordelia Langford, John Sillitoe on behalf of the Wellcome Sanger Institute COVID-19 Surveillance Team |
| EPI_ISL_3438907, EPI_ISL_3438908, EPI_ISL_3438909 | Lighthouse Lab in Alderley Park | Wellcome Sanger Institute for the COVID-19 Genomics UK (COG-UK) Consortium | Jacquelyn Wynn, Mairead Hyland, The Lighthouse Lab in Alderley Park and Alex Alderton, Roberto Amato, Jeffrey Barrett, Sonia Goncalves, Ewan Harrison, David K. Jackson, Ian Johnston, Dominic Kwiatkowski, Cordelia Langford, John Sillitoe on behalf of the Wellcome Sanger Institute COVID-19 Surveillance Team |
| EPI_ISL_3438910, EPI_ISL_3438911 | Berkshire and Surrey Pathology Services Lighthouse Laboratory | Wellcome Sanger Institute for the COVID-19 Genomics UK (COG-UK) Consortium | Berkshire and Surrey Pathology Services Lighthouse Laboratory and Alex Alderton, Roberto Amato, Jeffrey Barrett, Sonia Goncalves, Ewan Harrison, David K. Jackson, Ian Johnston, Dominic Kwiatkowski, Cordelia Langford, John Sillitoe on behalf of the Wellcome Sanger Institute COVID-19 Surveillance Team |
| EPI_ISL_3438912, EPI_ISL_3438913 | Lighthouse Lab in Alderley Park | Wellcome Sanger Institute for the COVID-19 Genomics UK (COG-UK) Consortium | Jacquelyn Wynn, Mairead Hyland, The Lighthouse Lab in Alderley Park and Alex Alderton, Roberto Amato, Jeffrey Barrett, Sonia Goncalves, Ewan Harrison, David K. Jackson, Ian Johnston, Dominic Kwiatkowski, Cordelia Langford, John Sillitoe on behalf of the Wellcome Sanger Institute COVID-19 Surveillance Team |
| EPI_ISL_3438914 | Berkshire and Surrey Pathology Services Lighthouse Laboratory | Wellcome Sanger Institute for the COVID-19 Genomics UK (COG-UK) Consortium | Berkshire and Surrey Pathology Services Lighthouse Laboratory and Alex Alderton, Roberto Amato, Jeffrey Barrett, Sonia Goncalves, Ewan Harrison, David K. Jackson, Ian Johnston, Dominic Kwiatkowski, Cordelia Langford, John Sillitoe on behalf of the Wellcome Sanger Institute COVID-19 Surveillance Team |
| EPI_ISL_3438915 | Lighthouse Lab in Alderley Park | Wellcome Sanger Institute for the COVID-19 Genomics UK (COG-UK) Consortium | Jacquelyn Wynn, Mairead Hyland, The Lighthouse Lab in Alderley Park and Alex Alderton, Roberto Amato, Jeffrey Barrett, Sonia Goncalves, Ewan Harrison, David K. Jackson, Ian Johnston, Dominic Kwiatkowski, Cordelia Langford, John Sillitoe on behalf of the Wellcome Sanger Institute COVID-19 Surveillance Team |
| EPI_ISL_3438916, EPI_ISL_3438917 | Berkshire and Surrey Pathology Services Lighthouse Laboratory | Wellcome Sanger Institute for the COVID-19 Genomics UK (COG-UK) Consortium | Berkshire and Surrey Pathology Services Lighthouse Laboratory and Alex Alderton, Roberto Amato, Jeffrey Barrett, Sonia Goncalves, Ewan Harrison, David K. Jackson, Ian Johnston, Dominic Kwiatkowski, Cordelia Langford, John Sillitoe on behalf of the Wellcome Sanger Institute COVID-19 Surveillance Team |
| EPI_ISL_3438918, EPI_ISL_3438919, EPI_ISL_3438920 | Lighthouse Lab in Alderley Park | Wellcome Sanger Institute for the COVID-19 Genomics UK (COG-UK) Consortium | Jacquelyn Wynn, Mairead Hyland, The Lighthouse Lab in Alderley Park and Alex Alderton, Roberto Amato, Jeffrey Barrett, Sonia Goncalves, Ewan Harrison, David K. Jackson, Ian Johnston, Dominic Kwiatkowski, Cordelia Langford, John Sillitoe on behalf of the Wellcome Sanger Institute COVID-19 Surveillance Team |
| EPI_ISL_3438921 | Berkshire and Surrey Pathology Services Lighthouse Laboratory | Wellcome Sanger Institute for the COVID-19 Genomics UK (COG-UK) Consortium | Berkshire and Surrey Pathology Services Lighthouse Laboratory and Alex Alderton, Roberto Amato, Jeffrey Barrett, Sonia Goncalves, Ewan Harrison, David K. Jackson, Ian Johnston, Dominic Kwiatkowski, Cordelia Langford, John Sillitoe on behalf of the Wellcome Sanger Institute COVID-19 Surveillance Team |
| EPI_ISL_3438922, EPI_ISL_3438923, EPI_ISL_3438924, EPI_ISL_3438925 | Lighthouse Lab in Alderley Park | Wellcome Sanger Institute for the COVID-19 Genomics UK (COG-UK) Consortium | Jacquelyn Wynn, Mairead Hyland, The Lighthouse Lab in Alderley Park and Alex Alderton, Roberto Amato, Jeffrey Barrett, Sonia Goncalves, Ewan Harrison, David K. Jackson, Ian Johnston, Dominic Kwiatkowski, Cordelia Langford, John Sillitoe on behalf of the Wellcome Sanger Institute COVID-19 Surveillance Team |
| EPI_ISL_3438926 | Berkshire and Surrey Pathology Services Lighthouse Laboratory | Wellcome Sanger Institute for the COVID-19 Genomics UK (COG-UK) Consortium | Berkshire and Surrey Pathology Services Lighthouse Laboratory and Alex Alderton, Roberto Amato, Jeffrey Barrett, Sonia Goncalves, Ewan Harrison, David K. Jackson, Ian Johnston, Dominic Kwiatkowski, Cordelia Langford, John Sillitoe on behalf of the Wellcome Sanger Institute COVID-19 Surveillance Team |
| EPI_ISL_3438927, EPI_ISL_3438928, EPI_ISL_3438929, EPI_ISL_3438930 | Lighthouse Lab in Alderley Park | Wellcome Sanger Institute for the COVID-19 Genomics UK (COG-UK) Consortium | Jacquelyn Wynn, Mairead Hyland, The Lighthouse Lab in Alderley Park and Alex Alderton, Roberto Amato, Jeffrey Barrett, Sonia Goncalves, Ewan Harrison, David K. Jackson, Ian Johnston, Dominic Kwiatkowski, Cordelia Langford, John Sillitoe on behalf of the Wellcome Sanger Institute COVID-19 Surveillance Team |
| EPI_ISL_3438931 | Berkshire and Surrey Pathology Services Lighthouse Laboratory | Wellcome Sanger Institute for the COVID-19 Genomics UK (COG-UK) Consortium | Berkshire and Surrey Pathology Services Lighthouse Laboratory and Alex Alderton, Roberto Amato, Jeffrey Barrett, Sonia Goncalves, Ewan Harrison, David K. Jackson, Ian Johnston, Dominic Kwiatkowski, Cordelia Langford, John Sillitoe on behalf of the Wellcome Sanger Institute COVID-19 Surveillance Team |
| EPI_ISL_3438932, EPI_ISL_3438933, EPI_ISL_3438934 | Lighthouse Lab in Alderley Park | Wellcome Sanger Institute for the COVID-19 Genomics UK (COG-UK) Consortium | Jacquelyn Wynn, Mairead Hyland, The Lighthouse Lab in Alderley Park and Alex Alderton, Roberto Amato, Jeffrey Barrett, Sonia Goncalves, Ewan Harrison, David K. Jackson, Ian Johnston, Dominic Kwiatkowski, Cordelia Langford, John Sillitoe on behalf of the Wellcome Sanger Institute COVID-19 Surveillance Team |
| EPI_ISL_3438935 | Berkshire and Surrey Pathology Services Lighthouse Laboratory | Wellcome Sanger Institute for the COVID-19 Genomics UK (COG-UK) Consortium | Berkshire and Surrey Pathology Services Lighthouse Laboratory and Alex Alderton, Roberto Amato, Jeffrey Barrett, Sonia Goncalves, Ewan Harrison, David K. Jackson, Ian Johnston, Dominic Kwiatkowski, Cordelia Langford, John Sillitoe on behalf of the Wellcome Sanger Institute COVID-19 Surveillance Team |
| EPI_ISL_3438936, EPI_ISL_3438937, EPI_ISL_3438938, EPI_ISL_3438939, EPI_ISL_3438940, EPI_ISL_3438941, EPI_ISL_3438942, EPI_ISL_3438943 | Lighthouse Lab in Alderley Park | Wellcome Sanger Institute for the COVID-19 Genomics UK (COG-UK) Consortium | Jacquelyn Wynn, Mairead Hyland, The Lighthouse Lab in Alderley Park and Alex Alderton, Roberto Amato, Jeffrey Barrett, Sonia Goncalves, Ewan Harrison, David K. Jackson, Ian Johnston, Dominic Kwiatkowski, Cordelia Langford, John Sillitoe on behalf of the Wellcome Sanger Institute COVID-19 Surveillance Team |
| EPI_ISL_3438944, EPI_ISL_3438945 | Berkshire and Surrey Pathology Services Lighthouse Laboratory | Wellcome Sanger Institute for the COVID-19 Genomics UK (COG-UK) Consortium | Berkshire and Surrey Pathology Services Lighthouse Laboratory and Alex Alderton, Roberto Amato, Jeffrey Barrett, Sonia Goncalves, Ewan Harrison, David K. Jackson, Ian Johnston, Dominic Kwiatkowski, Cordelia Langford, John Sillitoe on behalf of the Wellcome Sanger Institute COVID-19 Surveillance Team |
| EPI_ISL_3438946 | Lighthouse Lab in Alderley Park | Wellcome Sanger Institute for the COVID-19 Genomics UK (COG-UK) Consortium | Jacquelyn Wynn, Mairead Hyland, The Lighthouse Lab in Alderley Park and Alex Alderton, Roberto Amato, Jeffrey Barrett, Sonia Goncalves, Ewan Harrison, David K. Jackson, Ian Johnston, Dominic Kwiatkowski, Cordelia Langford, John Sillitoe on behalf of the Wellcome Sanger Institute COVID-19 Surveillance Team |
| EPI_ISL_3438947 | Berkshire and Surrey Pathology Services Lighthouse Laboratory | Wellcome Sanger Institute for the COVID-19 Genomics UK (COG-UK) Consortium | Berkshire and Surrey Pathology Services Lighthouse Laboratory and Alex Alderton, Roberto Amato, Jeffrey Barrett, Sonia Goncalves, Ewan Harrison, David K. Jackson, Ian Johnston, Dominic Kwiatkowski, Cordelia Langford, John Sillitoe on behalf of the Wellcome Sanger Institute COVID-19 Surveillance Team |
| EPI_ISL_3438948, EPI_ISL_3438949 | Lighthouse Lab in Alderley Park | Wellcome Sanger Institute for the COVID-19 Genomics UK (COG-UK) Consortium | Jacquelyn Wynn, Mairead Hyland, The Lighthouse Lab in Alderley Park and Alex Alderton, Roberto Amato, Jeffrey Barrett, Sonia Goncalves, Ewan Harrison, David K. Jackson, Ian Johnston, Dominic Kwiatkowski, Cordelia Langford, John Sillitoe on behalf of the Wellcome Sanger Institute COVID-19 Surveillance Team |
| EPI_ISL_3438950 | Berkshire and Surrey Pathology Services Lighthouse Laboratory | Wellcome Sanger Institute for the COVID-19 Genomics UK (COG-UK) Consortium | Berkshire and Surrey Pathology Services Lighthouse Laboratory and Alex Alderton, Roberto Amato, Jeffrey Barrett, Sonia Goncalves, Ewan Harrison, David K. Jackson, Ian Johnston, Dominic Kwiatkowski, Cordelia Langford, John Sillitoe on behalf of the Wellcome Sanger Institute COVID-19 Surveillance Team |
| EPI_ISL_3438951, EPI_ISL_3438952 | Lighthouse Lab in Alderley Park | Wellcome Sanger Institute for the COVID-19 Genomics UK (COG-UK) Consortium | Jacquelyn Wynn, Mairead Hyland, The Lighthouse Lab in Alderley Park and Alex Alderton, Roberto Amato, Jeffrey Barrett, Sonia Goncalves, Ewan Harrison, David K. Jackson, Ian Johnston, Dominic Kwiatkowski, Cordelia Langford, John Sillitoe on behalf of the Wellcome Sanger Institute COVID-19 Surveillance Team |
| EPI_ISL_3438953 | Berkshire and Surrey Pathology Services Lighthouse Laboratory | Wellcome Sanger Institute for the COVID-19 Genomics UK (COG-UK) Consortium | Berkshire and Surrey Pathology Services Lighthouse Laboratory and Alex Alderton, Roberto Amato, Jeffrey Barrett, Sonia Goncalves, Ewan Harrison, David K. Jackson, Ian Johnston, Dominic Kwiatkowski, Cordelia Langford, John Sillitoe on behalf of the Wellcome Sanger Institute COVID-19 Surveillance Team |
| EPI_ISL_3438954, EPI_ISL_3438955, EPI_ISL_3438956, EPI_ISL_3438957 | Lighthouse Lab in Alderley Park | Wellcome Sanger Institute for the COVID-19 Genomics UK (COG-UK) Consortium | Jacquelyn Wynn, Mairead Hyland, The Lighthouse Lab in Alderley Park and Alex Alderton, Roberto Amato, Jeffrey Barrett, Sonia Goncalves, Ewan Harrison, David K. Jackson, Ian Johnston, Dominic Kwiatkowski, Cordelia Langford, John Sillitoe on behalf of the Wellcome Sanger Institute COVID-19 Surveillance Team |
| EPI_ISL_3438958, EPI_ISL_3438959 | Berkshire and Surrey Pathology Services Lighthouse Laboratory | Wellcome Sanger Institute for the COVID-19 Genomics UK (COG-UK) Consortium | Berkshire and Surrey Pathology Services Lighthouse Laboratory and Alex Alderton, Roberto Amato, Jeffrey Barrett, Sonia Goncalves, Ewan Harrison, David K. Jackson, Ian Johnston, Dominic Kwiatkowski, Cordelia Langford, John Sillitoe on behalf of the Wellcome Sanger Institute COVID-19 Surveillance Team |
| EPI_ISL_3438960 | Lighthouse Lab in Alderley Park | Wellcome Sanger Institute for the COVID-19 Genomics UK (COG-UK) Consortium | Jacquelyn Wynn, Mairead Hyland, The Lighthouse Lab in Alderley Park and Alex Alderton, Roberto Amato, Jeffrey Barrett, Sonia Goncalves, Ewan Harrison, David K. Jackson, Ian Johnston, Dominic Kwiatkowski, Cordelia Langford, John Sillitoe on behalf of the Wellcome Sanger Institute COVID-19 Surveillance Team |
| EPI_ISL_3438961 | Berkshire and Surrey Pathology Services Lighthouse Laboratory | Wellcome Sanger Institute for the COVID-19 Genomics UK (COG-UK) Consortium | Berkshire and Surrey Pathology Services Lighthouse Laboratory and Alex Alderton, Roberto Amato, Jeffrey Barrett, Sonia Goncalves, Ewan Harrison, David K. Jackson, Ian Johnston, Dominic Kwiatkowski, Cordelia Langford, John Sillitoe on behalf of the Wellcome Sanger Institute COVID-19 Surveillance Team |
| EPI_ISL_3438962 | Lighthouse Lab in Alderley Park | Wellcome Sanger Institute for the COVID-19 Genomics UK (COG-UK) Consortium | Jacquelyn Wynn, Mairead Hyland, The Lighthouse Lab in Alderley Park and Alex Alderton, Roberto Amato, Jeffrey Barrett, Sonia Goncalves, Ewan Harrison, David K. Jackson, Ian Johnston, Dominic Kwiatkowski, Cordelia Langford, John Sillitoe on behalf of the Wellcome Sanger Institute COVID-19 Surveillance Team |
| EPI_ISL_3438963 | Berkshire and Surrey Pathology Services Lighthouse Laboratory | Wellcome Sanger Institute for the COVID-19 Genomics UK (COG-UK) Consortium | Berkshire and Surrey Pathology Services Lighthouse Laboratory and Alex Alderton, Roberto Amato, Jeffrey Barrett, Sonia Goncalves, Ewan Harrison, David K. Jackson, Ian Johnston, Dominic Kwiatkowski, Cordelia Langford, John Sillitoe on behalf of the Wellcome Sanger Institute COVID-19 Surveillance Team |
| EPI_ISL_3438964, EPI_ISL_3438965 | Lighthouse Lab in Alderley Park | Wellcome Sanger Institute for the COVID-19 Genomics UK (COG-UK) Consortium | Jacquelyn Wynn, Mairead Hyland, The Lighthouse Lab in Alderley Park and Alex Alderton, Roberto Amato, Jeffrey Barrett, Sonia Goncalves, Ewan Harrison, David K. Jackson, Ian Johnston, Dominic Kwiatkowski, Cordelia Langford, John Sillitoe on behalf of the Wellcome Sanger Institute COVID-19 Surveillance Team |
| EPI_ISL_3438966 | Berkshire and Surrey Pathology Services Lighthouse Laboratory | Wellcome Sanger Institute for the COVID-19 Genomics UK (COG-UK) Consortium | Berkshire and Surrey Pathology Services Lighthouse Laboratory and Alex Alderton, Roberto Amato, Jeffrey Barrett, Sonia Goncalves, Ewan Harrison, David K. Jackson, Ian Johnston, Dominic Kwiatkowski, Cordelia Langford, John Sillitoe on behalf of the Wellcome Sanger Institute COVID-19 Surveillance Team |
| EPI_ISL_3438967, EPI_ISL_3438968, EPI_ISL_3438969, EPI_ISL_3438970 | Lighthouse Lab in Alderley Park | Wellcome Sanger Institute for the COVID-19 Genomics UK (COG-UK) Consortium | Jacquelyn Wynn, Mairead Hyland, The Lighthouse Lab in Alderley Park and Alex Alderton, Roberto Amato, Jeffrey Barrett, Sonia Goncalves, Ewan Harrison, David K. Jackson, Ian Johnston, Dominic Kwiatkowski, Cordelia Langford, John Sillitoe on behalf of the Wellcome Sanger Institute COVID-19 Surveillance Team |

| | | | |
|---|---|---|---|
| EPI_ISL_3438971 | Berkshire and Surrey Pathology Services Lighthouse Laboratory | Wellcome Sanger Institute for the COVID-19 Genomics UK (COG-UK) Consortium | Berkshire and Surrey Pathology Services Lighthouse Laboratory and Alex Alderton, Roberto Amato, Jeffrey Barrett, Sonia Goncalves, Ewan Harrison, David K. Jackson, Ian Johnston, Dominic Kwiatkowski, Cordelia Langford, John Sillitoe on behalf of the Wellcome Sanger Institute COVID-19 Surveillance Team |
| EPI_ISL_3438972, EPI_ISL_3438973, EPI_ISL_3438974, EPI_ISL_3438975, EPI_ISL_3438976, EPI_ISL_3438977, EPI_ISL_3438978, EPI_ISL_3438979 | Lighthouse Lab in Alderley Park | Wellcome Sanger Institute for the COVID-19 Genomics UK (COG-UK) Consortium | Jacquelyn Wynn, Mairead Hyland, The Lighthouse Lab in Alderley Park and Alex Alderton, Roberto Amato, Jeffrey Barrett, Sonia Goncalves, Ewan Harrison, David K. Jackson, Ian Johnston, Dominic Kwiatkowski, Cordelia Langford, John Sillitoe on behalf of the Wellcome Sanger Institute COVID-19 Surveillance Team |
| EPI_ISL_3438980 | Berkshire and Surrey Pathology Services Lighthouse Laboratory | Wellcome Sanger Institute for the COVID-19 Genomics UK (COG-UK) Consortium | Berkshire and Surrey Pathology Services Lighthouse Laboratory and Alex Alderton, Roberto Amato, Jeffrey Barrett, Sonia Goncalves, Ewan Harrison, David K. Jackson, Ian Johnston, Dominic Kwiatkowski, Cordelia Langford, John Sillitoe on behalf of the Wellcome Sanger Institute COVID-19 Surveillance Team |
| EPI_ISL_3438981, EPI_ISL_3438982, EPI_ISL_3438983, EPI_ISL_3438984, EPI_ISL_3438985, EPI_ISL_3438986, EPI_ISL_3438987, EPI_ISL_3438988, EPI_ISL_3438989 | Lighthouse Lab in Alderley Park | Wellcome Sanger Institute for the COVID-19 Genomics UK (COG-UK) Consortium | Jacquelyn Wynn, Mairead Hyland, The Lighthouse Lab in Alderley Park and Alex Alderton, Roberto Amato, Jeffrey Barrett, Sonia Goncalves, Ewan Harrison, David K. Jackson, Ian Johnston, Dominic Kwiatkowski, Cordelia Langford, John Sillitoe on behalf of the Wellcome Sanger Institute COVID-19 Surveillance Team |
| EPI_ISL_3438990, EPI_ISL_3438991 | Berkshire and Surrey Pathology Services Lighthouse Laboratory | Wellcome Sanger Institute for the COVID-19 Genomics UK (COG-UK) Consortium | Berkshire and Surrey Pathology Services Lighthouse Laboratory and Alex Alderton, Roberto Amato, Jeffrey Barrett, Sonia Goncalves, Ewan Harrison, David K. Jackson, Ian Johnston, Dominic Kwiatkowski, Cordelia Langford, John Sillitoe on behalf of the Wellcome Sanger Institute COVID-19 Surveillance Team |
| EPI_ISL_3438992, EPI_ISL_3438993 | Lighthouse Lab in Alderley Park | Wellcome Sanger Institute for the COVID-19 Genomics UK (COG-UK) Consortium | Jacquelyn Wynn, Mairead Hyland, The Lighthouse Lab in Alderley Park and Alex Alderton, Roberto Amato, Jeffrey Barrett, Sonia Goncalves, Ewan Harrison, David K. Jackson, Ian Johnston, Dominic Kwiatkowski, Cordelia Langford, John Sillitoe on behalf of the Wellcome Sanger Institute COVID-19 Surveillance Team |
| EPI_ISL_3438994 | Berkshire and Surrey Pathology Services Lighthouse Laboratory | Wellcome Sanger Institute for the COVID-19 Genomics UK (COG-UK) Consortium | Berkshire and Surrey Pathology Services Lighthouse Laboratory and Alex Alderton, Roberto Amato, Jeffrey Barrett, Sonia Goncalves, Ewan Harrison, David K. Jackson, Ian Johnston, Dominic Kwiatkowski, Cordelia Langford, John Sillitoe on behalf of the Wellcome Sanger Institute COVID-19 Surveillance Team |
| EPI_ISL_3438995, EPI_ISL_3438996, EPI_ISL_3438997, EPI_ISL_3438998 | Lighthouse Lab in Alderley Park | Wellcome Sanger Institute for the COVID-19 Genomics UK (COG-UK) Consortium | Jacquelyn Wynn, Mairead Hyland, The Lighthouse Lab in Alderley Park and Alex Alderton, Roberto Amato, Jeffrey Barrett, Sonia Goncalves, Ewan Harrison, David K. Jackson, Ian Johnston, Dominic Kwiatkowski, Cordelia Langford, John Sillitoe on behalf of the Wellcome Sanger Institute COVID-19 Surveillance Team |
| EPI_ISL_3438999 | Berkshire and Surrey Pathology Services Lighthouse Laboratory | Wellcome Sanger Institute for the COVID-19 Genomics UK (COG-UK) Consortium | Berkshire and Surrey Pathology Services Lighthouse Laboratory and Alex Alderton, Roberto Amato, Jeffrey Barrett, Sonia Goncalves, Ewan Harrison, David K. Jackson, Ian Johnston, Dominic Kwiatkowski, Cordelia Langford, John Sillitoe on behalf of the Wellcome Sanger Institute COVID-19 Surveillance Team |
| EPI_ISL_3439000, EPI_ISL_3439001, EPI_ISL_3439002 | Lighthouse Lab in Alderley Park | Wellcome Sanger Institute for the COVID-19 Genomics UK (COG-UK) Consortium | Jacquelyn Wynn, Mairead Hyland, The Lighthouse Lab in Alderley Park and Alex Alderton, Roberto Amato, Jeffrey Barrett, Sonia Goncalves, Ewan Harrison, David K. Jackson, Ian Johnston, Dominic Kwiatkowski, Cordelia Langford, John Sillitoe on behalf of the Wellcome Sanger Institute COVID-19 Surveillance Team |
| EPI_ISL_3439003, EPI_ISL_3439004 | Berkshire and Surrey Pathology Services Lighthouse Laboratory | Wellcome Sanger Institute for the COVID-19 Genomics UK (COG-UK) Consortium | Berkshire and Surrey Pathology Services Lighthouse Laboratory and Alex Alderton, Roberto Amato, Jeffrey Barrett, Sonia Goncalves, Ewan Harrison, David K. Jackson, Ian Johnston, Dominic Kwiatkowski, Cordelia Langford, John Sillitoe on behalf of the Wellcome Sanger Institute COVID-19 Surveillance Team |
| EPI_ISL_3439005, EPI_ISL_3439006 | Lighthouse Lab in Alderley Park | Wellcome Sanger Institute for the COVID-19 Genomics UK (COG-UK) Consortium | Jacquelyn Wynn, Mairead Hyland, The Lighthouse Lab in Alderley Park and Alex Alderton, Roberto Amato, Jeffrey Barrett, Sonia Goncalves, Ewan Harrison, David K. Jackson, Ian Johnston, Dominic Kwiatkowski, Cordelia Langford, John Sillitoe on behalf of the Wellcome Sanger Institute COVID-19 Surveillance Team |
| EPI_ISL_3439007 | Berkshire and Surrey Pathology Services Lighthouse Laboratory | Wellcome Sanger Institute for the COVID-19 Genomics UK (COG-UK) Consortium | Berkshire and Surrey Pathology Services Lighthouse Laboratory and Alex Alderton, Roberto Amato, Jeffrey Barrett, Sonia Goncalves, Ewan Harrison, David K. Jackson, Ian Johnston, Dominic Kwiatkowski, Cordelia Langford, John Sillitoe on behalf of the Wellcome Sanger Institute COVID-19 Surveillance Team |
| EPI_ISL_3439008, EPI_ISL_3439009, EPI_ISL_3439010, EPI_ISL_3439011, EPI_ISL_3439012 | Lighthouse Lab in Alderley Park | Wellcome Sanger Institute for the COVID-19 Genomics UK (COG-UK) Consortium | Jacquelyn Wynn, Mairead Hyland, The Lighthouse Lab in Alderley Park and Alex Alderton, Roberto Amato, Jeffrey Barrett, Sonia Goncalves, Ewan Harrison, David K. Jackson, Ian Johnston, Dominic Kwiatkowski, Cordelia Langford, John Sillitoe on behalf of the Wellcome Sanger Institute COVID-19 Surveillance Team |
| EPI_ISL_3439013, EPI_ISL_3439014 | Berkshire and Surrey Pathology Services Lighthouse Laboratory | Wellcome Sanger Institute for the COVID-19 Genomics UK (COG-UK) Consortium | Berkshire and Surrey Pathology Services Lighthouse Laboratory and Alex Alderton, Roberto Amato, Jeffrey Barrett, Sonia Goncalves, Ewan Harrison, David K. Jackson, Ian Johnston, Dominic Kwiatkowski, Cordelia Langford, John Sillitoe on behalf of the Wellcome Sanger Institute COVID-19 Surveillance Team |
| EPI_ISL_3439015, EPI_ISL_3439016, EPI_ISL_3439017, EPI_ISL_3439018, EPI_ISL_3439019 | Lighthouse Lab in Alderley Park | Wellcome Sanger Institute for the COVID-19 Genomics UK (COG-UK) Consortium | Jacquelyn Wynn, Mairead Hyland, The Lighthouse Lab in Alderley Park and Alex Alderton, Roberto Amato, Jeffrey Barrett, Sonia Goncalves, Ewan Harrison, David K. Jackson, Ian Johnston, Dominic Kwiatkowski, Cordelia Langford, John Sillitoe on behalf of the Wellcome Sanger Institute COVID-19 Surveillance Team |
| EPI_ISL_3439020 | Berkshire and Surrey Pathology Services Lighthouse Laboratory | Wellcome Sanger Institute for the COVID-19 Genomics UK (COG-UK) Consortium | Berkshire and Surrey Pathology Services Lighthouse Laboratory and Alex Alderton, Roberto Amato, Jeffrey Barrett, Sonia Goncalves, Ewan Harrison, David K. Jackson, Ian Johnston, Dominic Kwiatkowski, Cordelia Langford, John Sillitoe on behalf of the Wellcome Sanger Institute COVID-19 Surveillance Team |
| EPI_ISL_3439021, EPI_ISL_3439022, EPI_ISL_3439023, EPI_ISL_3439024, EPI_ISL_3439025, EPI_ISL_3439026 | Lighthouse Lab in Alderley Park | Wellcome Sanger Institute for the COVID-19 Genomics UK (COG-UK) Consortium | Jacquelyn Wynn, Mairead Hyland, The Lighthouse Lab in Alderley Park and Alex Alderton, Roberto Amato, Jeffrey Barrett, Sonia Goncalves, Ewan Harrison, David K. Jackson, Ian Johnston, Dominic Kwiatkowski, Cordelia Langford, John Sillitoe on behalf of the Wellcome Sanger Institute COVID-19 Surveillance Team |
| EPI_ISL_3439027 | Berkshire and Surrey Pathology Services Lighthouse Laboratory | Wellcome Sanger Institute for the COVID-19 Genomics UK (COG-UK) Consortium | Berkshire and Surrey Pathology Services Lighthouse Laboratory and Alex Alderton, Roberto Amato, Jeffrey Barrett, Sonia Goncalves, Ewan Harrison, David K. Jackson, Ian Johnston, Dominic Kwiatkowski, Cordelia Langford, John Sillitoe on behalf of the Wellcome Sanger Institute COVID-19 Surveillance Team |
| EPI_ISL_3439028, EPI_ISL_3439029 | Lighthouse Lab in Alderley Park | Wellcome Sanger Institute for the COVID-19 Genomics UK (COG-UK) Consortium | Jacquelyn Wynn, Mairead Hyland, The Lighthouse Lab in Alderley Park and Alex Alderton, Roberto Amato, Jeffrey Barrett, Sonia Goncalves, Ewan Harrison, David K. Jackson, Ian Johnston, Dominic Kwiatkowski, Cordelia Langford, John Sillitoe on behalf of the Wellcome Sanger Institute COVID-19 Surveillance Team |
| EPI_ISL_3439030 | Berkshire and Surrey Pathology Services Lighthouse Laboratory | Wellcome Sanger Institute for the COVID-19 Genomics UK (COG-UK) Consortium | Berkshire and Surrey Pathology Services Lighthouse Laboratory and Alex Alderton, Roberto Amato, Jeffrey Barrett, Sonia Goncalves, Ewan Harrison, David K. Jackson, Ian Johnston, Dominic Kwiatkowski, Cordelia Langford, John Sillitoe on behalf of the Wellcome Sanger Institute COVID-19 Surveillance Team |
| EPI_ISL_3439031, EPI_ISL_3439032 | Lighthouse Lab in Alderley Park | Wellcome Sanger Institute for the COVID-19 Genomics UK (COG-UK) Consortium | Jacquelyn Wynn, Mairead Hyland, The Lighthouse Lab in Alderley Park and Alex Alderton, Roberto Amato, Jeffrey Barrett, Sonia Goncalves, Ewan Harrison, David K. Jackson, Ian Johnston, Dominic Kwiatkowski, Cordelia Langford, John Sillitoe on behalf of the Wellcome Sanger Institute COVID-19 Surveillance Team |
| EPI_ISL_3439033 | Berkshire and Surrey Pathology Services Lighthouse Laboratory | Wellcome Sanger Institute for the COVID-19 Genomics UK (COG-UK) Consortium | Berkshire and Surrey Pathology Services Lighthouse Laboratory and Alex Alderton, Roberto Amato, Jeffrey Barrett, Sonia Goncalves, Ewan Harrison, David K. Jackson, Ian Johnston, Dominic Kwiatkowski, Cordelia Langford, John Sillitoe on behalf of the Wellcome Sanger Institute COVID-19 Surveillance Team |
| EPI_ISL_3439034, EPI_ISL_3439035, EPI_ISL_3439036, EPI_ISL_3439037, EPI_ISL_3439038, EPI_ISL_3439039 | Lighthouse Lab in Alderley Park | Wellcome Sanger Institute for the COVID-19 Genomics UK (COG-UK) Consortium | Jacquelyn Wynn, Mairead Hyland, The Lighthouse Lab in Alderley Park and Alex Alderton, Roberto Amato, Jeffrey Barrett, Sonia Goncalves, Ewan Harrison, David K. Jackson, Ian Johnston, Dominic Kwiatkowski, Cordelia Langford, John Sillitoe on behalf of the Wellcome Sanger Institute COVID-19 Surveillance Team |
| EPI_ISL_3439040 | Berkshire and Surrey Pathology Services Lighthouse Laboratory | Wellcome Sanger Institute for the COVID-19 Genomics UK (COG-UK) Consortium | Berkshire and Surrey Pathology Services Lighthouse Laboratory and Alex Alderton, Roberto Amato, Jeffrey Barrett, Sonia Goncalves, Ewan Harrison, David K. Jackson, Ian Johnston, Dominic Kwiatkowski, Cordelia Langford, John Sillitoe on behalf of the Wellcome Sanger Institute COVID-19 Surveillance Team |
| EPI_ISL_3439041, EPI_ISL_3439042, EPI_ISL_3439043, EPI_ISL_3439044, EPI_ISL_3439045, EPI_ISL_3439046, EPI_ISL_3439047, EPI_ISL_3439048, EPI_ISL_3439049, EPI_ISL_3439050, EPI_ISL_3439051, EPI_ISL_3439052, EPI_ISL_3439053, EPI_ISL_3439054, EPI_ISL_3439055 | | | |
| see above | Lighthouse Lab in Alderley Park | Wellcome Sanger Institute for the COVID-19 Genomics UK (COG-UK) Consortium | Jacquelyn Wynn, Mairead Hyland, The Lighthouse Lab in Alderley Park and Alex Alderton, Roberto Amato, Jeffrey Barrett, Sonia Goncalves, Ewan Harrison, David K. Jackson, Ian Johnston, Dominic Kwiatkowski, Cordelia Langford, John Sillitoe on behalf of the Wellcome Sanger Institute COVID-19 Surveillance Team |
| EPI_ISL_3439056 | Berkshire and Surrey Pathology Services Lighthouse Laboratory | Wellcome Sanger Institute for the COVID-19 Genomics UK (COG-UK) Consortium | Berkshire and Surrey Pathology Services Lighthouse Laboratory and Alex Alderton, Roberto Amato, Jeffrey Barrett, Sonia Goncalves, Ewan Harrison, David K. Jackson, Ian Johnston, Dominic Kwiatkowski, Cordelia Langford, John Sillitoe on behalf of the Wellcome Sanger Institute COVID-19 Surveillance Team |
| EPI_ISL_3439057 | Lighthouse Lab in Alderley Park | Wellcome Sanger Institute for the COVID-19 Genomics UK (COG-UK) Consortium | Jacquelyn Wynn, Mairead Hyland, The Lighthouse Lab in Alderley Park and Alex Alderton, Roberto Amato, Jeffrey Barrett, Sonia Goncalves, Ewan Harrison, David K. Jackson, Ian Johnston, Dominic Kwiatkowski, Cordelia Langford, John Sillitoe on behalf of the Wellcome Sanger Institute COVID-19 Surveillance Team |
| EPI_ISL_3439058 | Berkshire and Surrey Pathology Services Lighthouse Laboratory | Wellcome Sanger Institute for the COVID-19 Genomics UK (COG-UK) Consortium | Berkshire and Surrey Pathology Services Lighthouse Laboratory and Alex Alderton, Roberto Amato, Jeffrey Barrett, Sonia Goncalves, Ewan Harrison, David K. Jackson, Ian Johnston, Dominic Kwiatkowski, Cordelia Langford, John Sillitoe on behalf of the Wellcome Sanger Institute COVID-19 Surveillance Team |
| EPI_ISL_3439059, EPI_ISL_3439060 | Lighthouse Lab in Alderley Park | Wellcome Sanger Institute for the COVID-19 Genomics UK (COG-UK) Consortium | Jacquelyn Wynn, Mairead Hyland, The Lighthouse Lab in Alderley Park and Alex Alderton, Roberto Amato, Jeffrey Barrett, Sonia Goncalves, Ewan Harrison, David K. Jackson, Ian Johnston, Dominic Kwiatkowski, Cordelia Langford, John Sillitoe on behalf of the Wellcome Sanger Institute COVID-19 Surveillance Team |
| EPI_ISL_3439061 | Berkshire and Surrey Pathology Services Lighthouse Laboratory | Wellcome Sanger Institute for the COVID-19 Genomics UK (COG-UK) Consortium | Berkshire and Surrey Pathology Services Lighthouse Laboratory and Alex Alderton, Roberto Amato, Jeffrey Barrett, Sonia Goncalves, Ewan Harrison, David K. Jackson, Ian Johnston, Dominic Kwiatkowski, Cordelia Langford, John Sillitoe on behalf of the Wellcome Sanger Institute COVID-19 Surveillance Team |
| EPI_ISL_3439062, EPI_ISL_3439063, EPI_ISL_3439064, | Lighthouse Lab in Alderley Park | Wellcome Sanger Institute for the COVID-19 Genomics UK (COG-UK) Consortium | Jacquelyn Wynn, Mairead Hyland, The Lighthouse Lab in Alderley Park and Alex Alderton, Roberto Amato, Jeffrey Barrett, Sonia Goncalves, Ewan Harrison, David K. Jackson, Ian Johnston, Dominic Kwiatkowski, Cordelia Langford, John Sillitoe on behalf of the Wellcome Sanger Institute COVID-19 Surveillance Team |

| | | | |
|---|---|---|---|
| EPI_ISL_3439065, EPI_ISL_3439066, EPI_ISL_3439067 | | | |
| EPI_ISL_3439068 | Berkshire and Surrey Pathology Services Lighthouse Laboratory | Wellcome Sanger Institute for the COVID-19 Genomics UK (COG-UK) Consortium | Berkshire and Surrey Pathology Services Lighthouse Laboratory and Alex Alderton, Roberto Amato, Jeffrey Barrett, Sonia Goncalves, Ewan Harrison, David K. Jackson, Ian Johnston, Dominic Kwiatkowski, Cordelia Langford, John Sillitoe on behalf of the Wellcome Sanger Institute COVID-19 Surveillance Team |
| EPI_ISL_3439069, EPI_ISL_3439070 | Lighthouse Lab in Alderley Park | Wellcome Sanger Institute for the COVID-19 Genomics UK (COG-UK) Consortium | Jacquelyn Wynn, Mairead Hyland, The Lighthouse Lab in Alderley Park and Alex Alderton, Roberto Amato, Jeffrey Barrett, Sonia Goncalves, Ewan Harrison, David K. Jackson, Ian Johnston, Dominic Kwiatkowski, Cordelia Langford, John Sillitoe on behalf of the Wellcome Sanger Institute COVID-19 Surveillance Team |
| EPI_ISL_3439071, EPI_ISL_3439072 | Berkshire and Surrey Pathology Services Lighthouse Laboratory | Wellcome Sanger Institute for the COVID-19 Genomics UK (COG-UK) Consortium | Berkshire and Surrey Pathology Services Lighthouse Laboratory and Alex Alderton, Roberto Amato, Jeffrey Barrett, Sonia Goncalves, Ewan Harrison, David K. Jackson, Ian Johnston, Dominic Kwiatkowski, Cordelia Langford, John Sillitoe on behalf of the Wellcome Sanger Institute COVID-19 Surveillance Team |
| EPI_ISL_3439073, EPI_ISL_3439074 | Lighthouse Lab in Alderley Park | Wellcome Sanger Institute for the COVID-19 Genomics UK (COG-UK) Consortium | Jacquelyn Wynn, Mairead Hyland, The Lighthouse Lab in Alderley Park and Alex Alderton, Roberto Amato, Jeffrey Barrett, Sonia Goncalves, Ewan Harrison, David K. Jackson, Ian Johnston, Dominic Kwiatkowski, Cordelia Langford, John Sillitoe on behalf of the Wellcome Sanger Institute COVID-19 Surveillance Team |
| EPI_ISL_3439075 | Berkshire and Surrey Pathology Services Lighthouse Laboratory | Wellcome Sanger Institute for the COVID-19 Genomics UK (COG-UK) Consortium | Berkshire and Surrey Pathology Services Lighthouse Laboratory and Alex Alderton, Roberto Amato, Jeffrey Barrett, Sonia Goncalves, Ewan Harrison, David K. Jackson, Ian Johnston, Dominic Kwiatkowski, Cordelia Langford, John Sillitoe on behalf of the Wellcome Sanger Institute COVID-19 Surveillance Team |
| EPI_ISL_3439076 | Lighthouse Lab in Alderley Park | Wellcome Sanger Institute for the COVID-19 Genomics UK (COG-UK) Consortium | Jacquelyn Wynn, Mairead Hyland, The Lighthouse Lab in Alderley Park and Alex Alderton, Roberto Amato, Jeffrey Barrett, Sonia Goncalves, Ewan Harrison, David K. Jackson, Ian Johnston, Dominic Kwiatkowski, Cordelia Langford, John Sillitoe on behalf of the Wellcome Sanger Institute COVID-19 Surveillance Team |
| EPI_ISL_3439077 | Berkshire and Surrey Pathology Services Lighthouse Laboratory | Wellcome Sanger Institute for the COVID-19 Genomics UK (COG-UK) Consortium | Berkshire and Surrey Pathology Services Lighthouse Laboratory and Alex Alderton, Roberto Amato, Jeffrey Barrett, Sonia Goncalves, Ewan Harrison, David K. Jackson, Ian Johnston, Dominic Kwiatkowski, Cordelia Langford, John Sillitoe on behalf of the Wellcome Sanger Institute COVID-19 Surveillance Team |
| EPI_ISL_3439078, EPI_ISL_3439079 | Lighthouse Lab in Alderley Park | Wellcome Sanger Institute for the COVID-19 Genomics UK (COG-UK) Consortium | Jacquelyn Wynn, Mairead Hyland, The Lighthouse Lab in Alderley Park and Alex Alderton, Roberto Amato, Jeffrey Barrett, Sonia Goncalves, Ewan Harrison, David K. Jackson, Ian Johnston, Dominic Kwiatkowski, Cordelia Langford, John Sillitoe on behalf of the Wellcome Sanger Institute COVID-19 Surveillance Team |
| EPI_ISL_3439080 | Berkshire and Surrey Pathology Services Lighthouse Laboratory | Wellcome Sanger Institute for the COVID-19 Genomics UK (COG-UK) Consortium | Berkshire and Surrey Pathology Services Lighthouse Laboratory and Alex Alderton, Roberto Amato, Jeffrey Barrett, Sonia Goncalves, Ewan Harrison, David K. Jackson, Ian Johnston, Dominic Kwiatkowski, Cordelia Langford, John Sillitoe on behalf of the Wellcome Sanger Institute COVID-19 Surveillance Team |
| EPI_ISL_3439081, EPI_ISL_3439082 | Lighthouse Lab in Alderley Park | Wellcome Sanger Institute for the COVID-19 Genomics UK (COG-UK) Consortium | Jacquelyn Wynn, Mairead Hyland, The Lighthouse Lab in Alderley Park and Alex Alderton, Roberto Amato, Jeffrey Barrett, Sonia Goncalves, Ewan Harrison, David K. Jackson, Ian Johnston, Dominic Kwiatkowski, Cordelia Langford, John Sillitoe on behalf of the Wellcome Sanger Institute COVID-19 Surveillance Team |
| EPI_ISL_3439083 | Berkshire and Surrey Pathology Services Lighthouse Laboratory | Wellcome Sanger Institute for the COVID-19 Genomics UK (COG-UK) Consortium | Berkshire and Surrey Pathology Services Lighthouse Laboratory and Alex Alderton, Roberto Amato, Jeffrey Barrett, Sonia Goncalves, Ewan Harrison, David K. Jackson, Ian Johnston, Dominic Kwiatkowski, Cordelia Langford, John Sillitoe on behalf of the Wellcome Sanger Institute COVID-19 Surveillance Team |
| EPI_ISL_3439084, EPI_ISL_3439085 | Lighthouse Lab in Alderley Park | Wellcome Sanger Institute for the COVID-19 Genomics UK (COG-UK) Consortium | Jacquelyn Wynn, Mairead Hyland, The Lighthouse Lab in Alderley Park and Alex Alderton, Roberto Amato, Jeffrey Barrett, Sonia Goncalves, Ewan Harrison, David K. Jackson, Ian Johnston, Dominic Kwiatkowski, Cordelia Langford, John Sillitoe on behalf of the Wellcome Sanger Institute COVID-19 Surveillance Team |
| EPI_ISL_3439086 | Berkshire and Surrey Pathology Services Lighthouse Laboratory | Wellcome Sanger Institute for the COVID-19 Genomics UK (COG-UK) Consortium | Berkshire and Surrey Pathology Services Lighthouse Laboratory and Alex Alderton, Roberto Amato, Jeffrey Barrett, Sonia Goncalves, Ewan Harrison, David K. Jackson, Ian Johnston, Dominic Kwiatkowski, Cordelia Langford, John Sillitoe on behalf of the Wellcome Sanger Institute COVID-19 Surveillance Team |
| EPI_ISL_3439087, EPI_ISL_3439088 | Lighthouse Lab in Alderley Park | Wellcome Sanger Institute for the COVID-19 Genomics UK (COG-UK) Consortium | Jacquelyn Wynn, Mairead Hyland, The Lighthouse Lab in Alderley Park and Alex Alderton, Roberto Amato, Jeffrey Barrett, Sonia Goncalves, Ewan Harrison, David K. Jackson, Ian Johnston, Dominic Kwiatkowski, Cordelia Langford, John Sillitoe on behalf of the Wellcome Sanger Institute COVID-19 Surveillance Team |
| EPI_ISL_3439089 | Berkshire and Surrey Pathology Services Lighthouse Laboratory | Wellcome Sanger Institute for the COVID-19 Genomics UK (COG-UK) Consortium | Berkshire and Surrey Pathology Services Lighthouse Laboratory and Alex Alderton, Roberto Amato, Jeffrey Barrett, Sonia Goncalves, Ewan Harrison, David K. Jackson, Ian Johnston, Dominic Kwiatkowski, Cordelia Langford, John Sillitoe on behalf of the Wellcome Sanger Institute COVID-19 Surveillance Team |
| EPI_ISL_3439090 | Lighthouse Lab in Alderley Park | Wellcome Sanger Institute for the COVID-19 Genomics UK (COG-UK) Consortium | Jacquelyn Wynn, Mairead Hyland, The Lighthouse Lab in Alderley Park and Alex Alderton, Roberto Amato, Jeffrey Barrett, Sonia Goncalves, Ewan Harrison, David K. Jackson, Ian Johnston, Dominic Kwiatkowski, Cordelia Langford, John Sillitoe on behalf of the Wellcome Sanger Institute COVID-19 Surveillance Team |
| EPI_ISL_3439091, EPI_ISL_3439092 | Berkshire and Surrey Pathology Services Lighthouse Laboratory | Wellcome Sanger Institute for the COVID-19 Genomics UK (COG-UK) Consortium | Berkshire and Surrey Pathology Services Lighthouse Laboratory and Alex Alderton, Roberto Amato, Jeffrey Barrett, Sonia Goncalves, Ewan Harrison, David K. Jackson, Ian Johnston, Dominic Kwiatkowski, Cordelia Langford, John Sillitoe on behalf of the Wellcome Sanger Institute COVID-19 Surveillance Team |
| EPI_ISL_3439093, EPI_ISL_3439094 | Lighthouse Lab in Alderley Park | Wellcome Sanger Institute for the COVID-19 Genomics UK (COG-UK) Consortium | Jacquelyn Wynn, Mairead Hyland, The Lighthouse Lab in Alderley Park and Alex Alderton, Roberto Amato, Jeffrey Barrett, Sonia Goncalves, Ewan Harrison, David K. Jackson, Ian Johnston, Dominic Kwiatkowski, Cordelia Langford, John Sillitoe on behalf of the Wellcome Sanger Institute COVID-19 Surveillance Team |
| EPI_ISL_3439095 | Berkshire and Surrey Pathology Services Lighthouse Laboratory | Wellcome Sanger Institute for the COVID-19 Genomics UK (COG-UK) Consortium | Berkshire and Surrey Pathology Services Lighthouse Laboratory and Alex Alderton, Roberto Amato, Jeffrey Barrett, Sonia Goncalves, Ewan Harrison, David K. Jackson, Ian Johnston, Dominic Kwiatkowski, Cordelia Langford, John Sillitoe on behalf of the Wellcome Sanger Institute COVID-19 Surveillance Team |
| EPI_ISL_3439096, EPI_ISL_3439097 | Lighthouse Lab in Alderley Park | Wellcome Sanger Institute for the COVID-19 Genomics UK (COG-UK) Consortium | Jacquelyn Wynn, Mairead Hyland, The Lighthouse Lab in Alderley Park and Alex Alderton, Roberto Amato, Jeffrey Barrett, Sonia Goncalves, Ewan Harrison, David K. Jackson, Ian Johnston, Dominic Kwiatkowski, Cordelia Langford, John Sillitoe on behalf of the Wellcome Sanger Institute COVID-19 Surveillance Team |
| EPI_ISL_3439098 | Berkshire and Surrey Pathology Services Lighthouse Laboratory | Wellcome Sanger Institute for the COVID-19 Genomics UK (COG-UK) Consortium | Berkshire and Surrey Pathology Services Lighthouse Laboratory and Alex Alderton, Roberto Amato, Jeffrey Barrett, Sonia Goncalves, Ewan Harrison, David K. Jackson, Ian Johnston, Dominic Kwiatkowski, Cordelia Langford, John Sillitoe on behalf of the Wellcome Sanger Institute COVID-19 Surveillance Team |
| EPI_ISL_3439099 | Lighthouse Lab in Alderley Park | Wellcome Sanger Institute for the COVID-19 Genomics UK (COG-UK) Consortium | Jacquelyn Wynn, Mairead Hyland, The Lighthouse Lab in Alderley Park and Alex Alderton, Roberto Amato, Jeffrey Barrett, Sonia Goncalves, Ewan Harrison, David K. Jackson, Ian Johnston, Dominic Kwiatkowski, Cordelia Langford, John Sillitoe on behalf of the Wellcome Sanger Institute COVID-19 Surveillance Team |
| EPI_ISL_3439100 | Berkshire and Surrey Pathology Services Lighthouse Laboratory | Wellcome Sanger Institute for the COVID-19 Genomics UK (COG-UK) Consortium | Berkshire and Surrey Pathology Services Lighthouse Laboratory and Alex Alderton, Roberto Amato, Jeffrey Barrett, Sonia Goncalves, Ewan Harrison, David K. Jackson, Ian Johnston, Dominic Kwiatkowski, Cordelia Langford, John Sillitoe on behalf of the Wellcome Sanger Institute COVID-19 Surveillance Team |
| EPI_ISL_3439101, EPI_ISL_3439102, EPI_ISL_3439103 | Lighthouse Lab in Alderley Park | Wellcome Sanger Institute for the COVID-19 Genomics UK (COG-UK) Consortium | Jacquelyn Wynn, Mairead Hyland, The Lighthouse Lab in Alderley Park and Alex Alderton, Roberto Amato, Jeffrey Barrett, Sonia Goncalves, Ewan Harrison, David K. Jackson, Ian Johnston, Dominic Kwiatkowski, Cordelia Langford, John Sillitoe on behalf of the Wellcome Sanger Institute COVID-19 Surveillance Team |
| EPI_ISL_3439104 | Berkshire and Surrey Pathology Services Lighthouse Laboratory | Wellcome Sanger Institute for the COVID-19 Genomics UK (COG-UK) Consortium | Berkshire and Surrey Pathology Services Lighthouse Laboratory and Alex Alderton, Roberto Amato, Jeffrey Barrett, Sonia Goncalves, Ewan Harrison, David K. Jackson, Ian Johnston, Dominic Kwiatkowski, Cordelia Langford, John Sillitoe on behalf of the Wellcome Sanger Institute COVID-19 Surveillance Team |
| EPI_ISL_3439105, EPI_ISL_3439106, EPI_ISL_3439107, EPI_ISL_3439108 | Lighthouse Lab in Alderley Park | Wellcome Sanger Institute for the COVID-19 Genomics UK (COG-UK) Consortium | Jacquelyn Wynn, Mairead Hyland, The Lighthouse Lab in Alderley Park and Alex Alderton, Roberto Amato, Jeffrey Barrett, Sonia Goncalves, Ewan Harrison, David K. Jackson, Ian Johnston, Dominic Kwiatkowski, Cordelia Langford, John Sillitoe on behalf of the Wellcome Sanger Institute COVID-19 Surveillance Team |
| EPI_ISL_3439109 | Berkshire and Surrey Pathology Services Lighthouse Laboratory | Wellcome Sanger Institute for the COVID-19 Genomics UK (COG-UK) Consortium | Berkshire and Surrey Pathology Services Lighthouse Laboratory and Alex Alderton, Roberto Amato, Jeffrey Barrett, Sonia Goncalves, Ewan Harrison, David K. Jackson, Ian Johnston, Dominic Kwiatkowski, Cordelia Langford, John Sillitoe on behalf of the Wellcome Sanger Institute COVID-19 Surveillance Team |
| EPI_ISL_3439110, EPI_ISL_3439111, EPI_ISL_3439112, EPI_ISL_3439113, EPI_ISL_3439114, EPI_ISL_3439115, EPI_ISL_3439116 | Lighthouse Lab in Alderley Park | Wellcome Sanger Institute for the COVID-19 Genomics UK (COG-UK) Consortium | Jacquelyn Wynn, Mairead Hyland, The Lighthouse Lab in Alderley Park and Alex Alderton, Roberto Amato, Jeffrey Barrett, Sonia Goncalves, Ewan Harrison, David K. Jackson, Ian Johnston, Dominic Kwiatkowski, Cordelia Langford, John Sillitoe on behalf of the Wellcome Sanger Institute COVID-19 Surveillance Team |
| EPI_ISL_3439117 | Berkshire and Surrey Pathology Services Lighthouse Laboratory | Wellcome Sanger Institute for the COVID-19 Genomics UK (COG-UK) Consortium | Berkshire and Surrey Pathology Services Lighthouse Laboratory and Alex Alderton, Roberto Amato, Jeffrey Barrett, Sonia Goncalves, Ewan Harrison, David K. Jackson, Ian Johnston, Dominic Kwiatkowski, Cordelia Langford, John Sillitoe on behalf of the Wellcome Sanger Institute COVID-19 Surveillance Team |
| EPI_ISL_3439118, EPI_ISL_3439119 | Lighthouse Lab in Alderley Park | Wellcome Sanger Institute for the COVID-19 Genomics UK (COG-UK) Consortium | Jacquelyn Wynn, Mairead Hyland, The Lighthouse Lab in Alderley Park and Alex Alderton, Roberto Amato, Jeffrey Barrett, Sonia Goncalves, Ewan Harrison, David K. Jackson, Ian Johnston, Dominic Kwiatkowski, Cordelia Langford, John Sillitoe on behalf of the Wellcome Sanger Institute COVID-19 Surveillance Team |
| EPI_ISL_3439120 | Berkshire and Surrey Pathology Services Lighthouse Laboratory | Wellcome Sanger Institute for the COVID-19 Genomics UK (COG-UK) Consortium | Berkshire and Surrey Pathology Services Lighthouse Laboratory and Alex Alderton, Roberto Amato, Jeffrey Barrett, Sonia Goncalves, Ewan Harrison, David K. Jackson, Ian Johnston, Dominic Kwiatkowski, Cordelia Langford, John Sillitoe on behalf of the Wellcome Sanger Institute COVID-19 Surveillance Team |
| EPI_ISL_3439121 | Lighthouse Lab in Alderley Park | Wellcome Sanger Institute for the COVID-19 Genomics UK (COG-UK) Consortium | Jacquelyn Wynn, Mairead Hyland, The Lighthouse Lab in Alderley Park and Alex Alderton, Roberto Amato, Jeffrey Barrett, Sonia Goncalves, Ewan Harrison, David K. Jackson, Ian Johnston, Dominic Kwiatkowski, Cordelia Langford, John Sillitoe on behalf of the Wellcome Sanger Institute COVID-19 Surveillance Team |
| EPI_ISL_3439122, EPI_ISL_3439123 | Berkshire and Surrey Pathology Services Lighthouse Laboratory | Wellcome Sanger Institute for the COVID-19 Genomics UK (COG-UK) Consortium | Berkshire and Surrey Pathology Services Lighthouse Laboratory and Alex Alderton, Roberto Amato, Jeffrey Barrett, Sonia Goncalves, Ewan Harrison, David K. Jackson, Ian Johnston, Dominic Kwiatkowski, Cordelia Langford, John Sillitoe on behalf of the Wellcome Sanger Institute COVID-19 Surveillance Team |
| EPI_ISL_3439124, EPI_ISL_3439125 | Lighthouse Lab in Alderley Park | Wellcome Sanger Institute for the COVID-19 Genomics UK (COG-UK) Consortium | Jacquelyn Wynn, Mairead Hyland, The Lighthouse Lab in Alderley Park and Alex Alderton, Roberto Amato, Jeffrey Barrett, Sonia Goncalves, Ewan Harrison, David K. Jackson, Ian Johnston, Dominic Kwiatkowski, Cordelia Langford, John Sillitoe on behalf of the Wellcome Sanger Institute COVID-19 Surveillance Team |
| EPI_ISL_3439126 | Berkshire and Surrey Pathology Services Lighthouse Laboratory | Wellcome Sanger Institute for the COVID-19 Genomics UK (COG-UK) Consortium | Berkshire and Surrey Pathology Services Lighthouse Laboratory and Alex Alderton, Roberto Amato, Jeffrey Barrett, Sonia Goncalves, Ewan Harrison, David K. Jackson, Ian Johnston, Dominic Kwiatkowski, Cordelia Langford, John Sillitoe on behalf of the Wellcome Sanger Institute COVID-19 Surveillance Team |
| EPI_ISL_3439127 | Lighthouse Lab in Alderley Park | Wellcome Sanger Institute for the COVID-19 Genomics UK (COG-UK) Consortium | Jacquelyn Wynn, Mairead Hyland, The Lighthouse Lab in Alderley Park and Alex Alderton, Roberto Amato, Jeffrey Barrett, Sonia Goncalves, Ewan Harrison, David K. Jackson, Ian Johnston, Dominic Kwiatkowski, Cordelia Langford, John Sillitoe on behalf of the Wellcome Sanger Institute COVID-19 Surveillance Team |
| EPI_ISL_3439128 | Berkshire and Surrey Pathology Services Lighthouse Laboratory | Wellcome Sanger Institute for the COVID-19 Genomics UK (COG-UK) Consortium | Berkshire and Surrey Pathology Services Lighthouse Laboratory and Alex Alderton, Roberto Amato, Jeffrey Barrett, Sonia Goncalves, Ewan Harrison, David K. Jackson, Ian Johnston, Dominic Kwiatkowski, Cordelia Langford, John Sillitoe on behalf of the Wellcome Sanger Institute COVID-19 Surveillance Team |
| EPI_ISL_3439129 | Lighthouse Lab in Alderley Park | Wellcome Sanger Institute for the COVID-19 Genomics UK (COG-UK) Consortium | Jacquelyn Wynn, Mairead Hyland, The Lighthouse Lab in Alderley Park and Alex Alderton, Roberto Amato, Jeffrey Barrett, Sonia Goncalves, Ewan Harrison, David K. Jackson, Ian Johnston, Dominic Kwiatkowski, Cordelia Langford, John Sillitoe on behalf of the Wellcome Sanger Institute COVID-19 Surveillance Team |
| EPI_ISL_3439130, EPI_ISL_3439131 | Berkshire and Surrey Pathology Services Lighthouse Laboratory | Wellcome Sanger Institute for the COVID-19 Genomics UK (COG-UK) Consortium | Berkshire and Surrey Pathology Services Lighthouse Laboratory and Alex Alderton, Roberto Amato, Jeffrey Barrett, Sonia Goncalves, Ewan Harrison, David K. Jackson, Ian Johnston, Dominic Kwiatkowski, Cordelia Langford, John Sillitoe on behalf of the Wellcome Sanger Institute COVID-19 Surveillance Team |
| EPI_ISL_3439132, EPI_ISL_3439133, | Lighthouse Lab in Alderley Park | Wellcome Sanger Institute for the COVID-19 Genomics UK (COG-UK) Consortium | Jacquelyn Wynn, Mairead Hyland, The Lighthouse Lab in Alderley Park and Alex Alderton, Roberto Amato, Jeffrey Barrett, Sonia Goncalves, Ewan Harrison, David K. Jackson, Ian Johnston, Dominic Kwiatkowski, Cordelia Langford, John Sillitoe on behalf of the Wellcome Sanger Institute COVID-19 Surveillance Team |

| | | | |
|---|---|---|---|
| EPI_ISL_3439134, EPI_ISL_3439135 | | | |
| EPI_ISL_3439136 | Berkshire and Surrey Pathology Services Lighthouse Laboratory | Wellcome Sanger Institute for the COVID-19 Genomics UK (COG-UK) Consortium | Berkshire and Surrey Pathology Services Lighthouse Laboratory and Alex Alderton, Roberto Amato, Jeffrey Barrett, Sonia Goncalves, Ewan Harrison, David K. Jackson, Ian Johnston, Dominic Kwiatkowski, Cordelia Langford, John Sillitoe on behalf of the Wellcome Sanger Institute COVID-19 Surveillance Team |
| EPI_ISL_3439137 | Lighthouse Lab in Alderley Park | Wellcome Sanger Institute for the COVID-19 Genomics UK (COG-UK) Consortium | Jacquelyn Wynn, Mairead Hyland, The Lighthouse Lab in Alderley Park and Alex Alderton, Roberto Amato, Jeffrey Barrett, Sonia Goncalves, Ewan Harrison, David K. Jackson, Ian Johnston, Dominic Kwiatkowski, Cordelia Langford, John Sillitoe on behalf of the Wellcome Sanger Institute COVID-19 Surveillance Team |
| EPI_ISL_3439138 | Berkshire and Surrey Pathology Services Lighthouse Laboratory | Wellcome Sanger Institute for the COVID-19 Genomics UK (COG-UK) Consortium | Berkshire and Surrey Pathology Services Lighthouse Laboratory and Alex Alderton, Roberto Amato, Jeffrey Barrett, Sonia Goncalves, Ewan Harrison, David K. Jackson, Ian Johnston, Dominic Kwiatkowski, Cordelia Langford, John Sillitoe on behalf of the Wellcome Sanger Institute COVID-19 Surveillance Team |
| EPI_ISL_3439139 | Lighthouse Lab in Alderley Park | Wellcome Sanger Institute for the COVID-19 Genomics UK (COG-UK) Consortium | Jacquelyn Wynn, Mairead Hyland, The Lighthouse Lab in Alderley Park and Alex Alderton, Roberto Amato, Jeffrey Barrett, Sonia Goncalves, Ewan Harrison, David K. Jackson, Ian Johnston, Dominic Kwiatkowski, Cordelia Langford, John Sillitoe on behalf of the Wellcome Sanger Institute COVID-19 Surveillance Team |
| EPI_ISL_3439140, EPI_ISL_3439141, EPI_ISL_3439142, EPI_ISL_3439143 | Berkshire and Surrey Pathology Services Lighthouse Laboratory | Wellcome Sanger Institute for the COVID-19 Genomics UK (COG-UK) Consortium | Berkshire and Surrey Pathology Services Lighthouse Laboratory and Alex Alderton, Roberto Amato, Jeffrey Barrett, Sonia Goncalves, Ewan Harrison, David K. Jackson, Ian Johnston, Dominic Kwiatkowski, Cordelia Langford, John Sillitoe on behalf of the Wellcome Sanger Institute COVID-19 Surveillance Team |
| EPI_ISL_3439144 | Lighthouse Lab in Alderley Park | Wellcome Sanger Institute for the COVID-19 Genomics UK (COG-UK) Consortium | Jacquelyn Wynn, Mairead Hyland, The Lighthouse Lab in Alderley Park and Alex Alderton, Roberto Amato, Jeffrey Barrett, Sonia Goncalves, Ewan Harrison, David K. Jackson, Ian Johnston, Dominic Kwiatkowski, Cordelia Langford, John Sillitoe on behalf of the Wellcome Sanger Institute COVID-19 Surveillance Team |
| EPI_ISL_3439145 | Berkshire and Surrey Pathology Services Lighthouse Laboratory | Wellcome Sanger Institute for the COVID-19 Genomics UK (COG-UK) Consortium | Berkshire and Surrey Pathology Services Lighthouse Laboratory and Alex Alderton, Roberto Amato, Jeffrey Barrett, Sonia Goncalves, Ewan Harrison, David K. Jackson, Ian Johnston, Dominic Kwiatkowski, Cordelia Langford, John Sillitoe on behalf of the Wellcome Sanger Institute COVID-19 Surveillance Team |
| EPI_ISL_3439146, EPI_ISL_3439147 | Lighthouse Lab in Alderley Park | Wellcome Sanger Institute for the COVID-19 Genomics UK (COG-UK) Consortium | Jacquelyn Wynn, Mairead Hyland, The Lighthouse Lab in Alderley Park and Alex Alderton, Roberto Amato, Jeffrey Barrett, Sonia Goncalves, Ewan Harrison, David K. Jackson, Ian Johnston, Dominic Kwiatkowski, Cordelia Langford, John Sillitoe on behalf of the Wellcome Sanger Institute COVID-19 Surveillance Team |
| EPI_ISL_3439148 | Berkshire and Surrey Pathology Services Lighthouse Laboratory | Wellcome Sanger Institute for the COVID-19 Genomics UK (COG-UK) Consortium | Berkshire and Surrey Pathology Services Lighthouse Laboratory and Alex Alderton, Roberto Amato, Jeffrey Barrett, Sonia Goncalves, Ewan Harrison, David K. Jackson, Ian Johnston, Dominic Kwiatkowski, Cordelia Langford, John Sillitoe on behalf of the Wellcome Sanger Institute COVID-19 Surveillance Team |
| EPI_ISL_3439149 | Lighthouse Lab in Alderley Park | Wellcome Sanger Institute for the COVID-19 Genomics UK (COG-UK) Consortium | Jacquelyn Wynn, Mairead Hyland, The Lighthouse Lab in Alderley Park and Alex Alderton, Roberto Amato, Jeffrey Barrett, Sonia Goncalves, Ewan Harrison, David K. Jackson, Ian Johnston, Dominic Kwiatkowski, Cordelia Langford, John Sillitoe on behalf of the Wellcome Sanger Institute COVID-19 Surveillance Team |
| EPI_ISL_3439150 | Berkshire and Surrey Pathology Services Lighthouse Laboratory | Wellcome Sanger Institute for the COVID-19 Genomics UK (COG-UK) Consortium | Berkshire and Surrey Pathology Services Lighthouse Laboratory and Alex Alderton, Roberto Amato, Jeffrey Barrett, Sonia Goncalves, Ewan Harrison, David K. Jackson, Ian Johnston, Dominic Kwiatkowski, Cordelia Langford, John Sillitoe on behalf of the Wellcome Sanger Institute COVID-19 Surveillance Team |
| EPI_ISL_3439151, EPI_ISL_3439152, EPI_ISL_3439153, EPI_ISL_3439154, EPI_ISL_3439155, EPI_ISL_3439156, EPI_ISL_3439157, EPI_ISL_3439158, EPI_ISL_3439159, EPI_ISL_3439160, EPI_ISL_3439161, EPI_ISL_3439162, EPI_ISL_3439163, EPI_ISL_3439164, EPI_ISL_3439165, EPI_ISL_3439166, EPI_ISL_3439167, EPI_ISL_3439168, EPI_ISL_3439169, EPI_ISL_3439170, EPI_ISL_3439171, EPI_ISL_3439172, EPI_ISL_3439173, EPI_ISL_3439174, EPI_ISL_3439175, EPI_ISL_3439176, EPI_ISL_3439177, EPI_ISL_3439178, EPI_ISL_3439179, EPI_ISL_3439180, EPI_ISL_3439181, EPI_ISL_3439182, EPI_ISL_3439183, EPI_ISL_3439184, EPI_ISL_3439185, EPI_ISL_3439186, EPI_ISL_3439187, EPI_ISL_3439188, EPI_ISL_3439189, EPI_ISL_3439190, EPI_ISL_3439191, EPI_ISL_3439192, EPI_ISL_3439193, EPI_ISL_3439194, EPI_ISL_3439195, EPI_ISL_3439196, EPI_ISL_3439197, EPI_ISL_3439198, EPI_ISL_3439199, EPI_ISL_3439200, EPI_ISL_3439201, EPI_ISL_3439202, EPI_ISL_3439203, EPI_ISL_3439204, EPI_ISL_3439205, EPI_ISL_3439206, EPI_ISL_3439207, EPI_ISL_3439208, EPI_ISL_3439209, EPI_ISL_3439210, EPI_ISL_3439211, EPI_ISL_3439212, EPI_ISL_3439213, EPI_ISL_3439214, EPI_ISL_3439215, EPI_ISL_3439216, EPI_ISL_3439217, EPI_ISL_3439218, EPI_ISL_3439219, EPI_ISL_3439220, EPI_ISL_3439221, EPI_ISL_3439222, EPI_ISL_3439223, EPI_ISL_3439224, EPI_ISL_3439225, EPI_ISL_3439226, EPI_ISL_3439227, EPI_ISL_3439228, EPI_ISL_3439229, EPI_ISL_3439230, EPI_ISL_3439231, EPI_ISL_3439232, EPI_ISL_3439233, EPI_ISL_3439234, EPI_ISL_3439235, EPI_ISL_3439236, EPI_ISL_3439237, EPI_ISL_3439238, EPI_ISL_3439239, EPI_ISL_3439240, EPI_ISL_3439241, EPI_ISL_3439242, EPI_ISL_3439243, EPI_ISL_3439244, EPI_ISL_3439245, EPI_ISL_3439246, EPI_ISL_3439247, EPI_ISL_3439248, EPI_ISL_3439249, EPI_ISL_3439250, EPI_ISL_3439251, EPI_ISL_3439252, EPI_ISL_3439253, EPI_ISL_3439254, EPI_ISL_3439255, EPI_ISL_3439256, EPI_ISL_3439257, EPI_ISL_3439258, EPI_ISL_3439259, EPI_ISL_3439260, EPI_ISL_3439261, EPI_ISL_3439262, EPI_ISL_3439263, EPI_ISL_3439264, EPI_ISL_3439265, EPI_ISL_3439266, EPI_ISL_3439267, EPI_ISL_3439268, EPI_ISL_3439269, EPI_ISL_3439270, EPI_ISL_3439271, EPI_ISL_3439272, EPI_ISL_3439273, EPI_ISL_3439274, EPI_ISL_3439275, EPI_ISL_3439276, EPI_ISL_3439277, EPI_ISL_3439278, EPI_ISL_3439279, EPI_ISL_3439280, EPI_ISL_3439281, EPI_ISL_3439282, EPI_ISL_3439283, EPI_ISL_3439284, EPI_ISL_3439285, EPI_ISL_3439286, EPI_ISL_3439287, EPI_ISL_3439288, EPI_ISL_3439289, EPI_ISL_3439290, EPI_ISL_3439291, EPI_ISL_3439292, EPI_ISL_3439293, EPI_ISL_3439294, EPI_ISL_3439295, EPI_ISL_3439296, EPI_ISL_3439297, EPI_ISL_3439298, EPI_ISL_3439299, EPI_ISL_3439300, EPI_ISL_3439301, EPI_ISL_3439302, EPI_ISL_3439303, EPI_ISL_3439304, EPI_ISL_3439305, EPI_ISL_3439306, EPI_ISL_3439307, EPI_ISL_3439308, EPI_ISL_3439309, EPI_ISL_3439310, EPI_ISL_3439311, EPI_ISL_3439312, EPI_ISL_3439313, EPI_ISL_3439314, EPI_ISL_3439315, EPI_ISL_3439316, EPI_ISL_3439317, EPI_ISL_3439318, EPI_ISL_3439319, EPI_ISL_3439320, EPI_ISL_3439321, EPI_ISL_3439322, EPI_ISL_3439323, EPI_ISL_3439324, EPI_ISL_3439325, EPI_ISL_3439326, EPI_ISL_3439327, EPI_ISL_3439328, EPI_ISL_3439329, EPI_ISL_3439330, EPI_ISL_3439331, EPI_ISL_3439332, EPI_ISL_3439333, EPI_ISL_3439334, EPI_ISL_3439335, EPI_ISL_3439336, EPI_ISL_3439337, EPI_ISL_3439338, EPI_ISL_3439339, EPI_ISL_3439340, EPI_ISL_3439341, EPI_ISL_3439342, EPI_ISL_3439343, EPI_ISL_3439344, EPI_ISL_3439345, EPI_ISL_3439346, EPI_ISL_3439347, EPI_ISL_3439348, EPI_ISL_3439349, EPI_ISL_3439350, EPI_ISL_3439351, EPI_ISL_3439352, EPI_ISL_3439353, EPI_ISL_3439354, EPI_ISL_3439355, EPI_ISL_3439356, EPI_ISL_3439357, EPI_ISL_3439358, EPI_ISL_3439359, EPI_ISL_3439360, EPI_ISL_3439361, EPI_ISL_3439362, EPI_ISL_3439363, EPI_ISL_3439364, EPI_ISL_3439365, EPI_ISL_3439366, EPI_ISL_3439367, EPI_ISL_3439368, EPI_ISL_3439369, EPI_ISL_3439370, EPI_ISL_3439371, EPI_ISL_3439372, EPI_ISL_3439373, EPI_ISL_3439374, EPI_ISL_3439375, EPI_ISL_3439376, EPI_ISL_3439377, EPI_ISL_3439378, EPI_ISL_3439379, EPI_ISL_3439380, EPI_ISL_3439381, EPI_ISL_3439382, EPI_ISL_3439383, EPI_ISL_3439384, EPI_ISL_3439385, EPI_ISL_3439386, EPI_ISL_3439387, EPI_ISL_3439388 | | | |
| see above | Lighthouse Laboratory Plymouth | Wellcome Sanger Institute for the COVID-19 Genomics UK (COG-UK) Consortium | Lighthouse Laboratory Plymouth and Alex Alderton, Roberto Amato, Jeffrey Barrett, Sonia Goncalves, Ewan Harrison, David K. Jackson, Ian Johnston, Dominic Kwiatkowski, Cordelia Langford, John Sillitoe on behalf of the Wellcome Sanger Institute COVID-19 Surveillance Team |
| EPI_ISL_3439389 | Lighthouse Lab in Milton Keynes | Wellcome Sanger Institute for the COVID-19 Genomics UK (COG-UK) Consortium | The Lighthouse Lab in Milton Keynes and Alex Alderton, Roberto Amato, Jeffrey Barrett, Sonia Goncalves, Ewan Harrison, David K. Jackson, Ian Johnston, Dominic Kwiatkowski, Cordelia Langford, John Sillitoe on behalf of the Wellcome Sanger Institute COVID-19 Surveillance Team |
| EPI_ISL_3439390 | Lighthouse Lab in Glasgow | Wellcome Sanger Institute for the COVID-19 Genomics UK (COG-UK) Consortium | Harper VanSteenhouse, Yumi Kasai, David Gray, Carol Clugston, Anna Dominiczak and Alex Alderton, Roberto Amato, Jeffrey Barrett, Sonia Goncalves, Ewan Harrison, David K. Jackson, Ian Johnston, Dominic Kwiatkowski, Cordelia Langford, John Sillitoe on behalf of the Wellcome Sanger Institute COVID-19 Surveillance Team |
| EPI_ISL_3439391 | Lighthouse Lab in Milton Keynes | Wellcome Sanger Institute for the COVID-19 Genomics UK (COG-UK) Consortium | The Lighthouse Lab in Milton Keynes and Alex Alderton, Roberto Amato, Jeffrey Barrett, Sonia Goncalves, Ewan Harrison, David K. Jackson, Ian Johnston, Dominic Kwiatkowski, Cordelia Langford, John Sillitoe on behalf of the Wellcome Sanger Institute COVID-19 Surveillance Team |
| EPI_ISL_3439392 | Lighthouse Lab in Glasgow | Wellcome Sanger Institute for the COVID-19 Genomics UK (COG-UK) Consortium | Harper VanSteenhouse, Yumi Kasai, David Gray, Carol Clugston, Anna Dominiczak and Alex Alderton, Roberto Amato, Jeffrey Barrett, Sonia Goncalves, Ewan Harrison, David K. Jackson, Ian Johnston, Dominic Kwiatkowski, Cordelia Langford, John Sillitoe on behalf of the Wellcome Sanger Institute COVID-19 Surveillance Team |
| EPI_ISL_3439393, EPI_ISL_3439394, EPI_ISL_3439395, EPI_ISL_3439396, EPI_ISL_3439397, EPI_ISL_3439398, EPI_ISL_3439399, EPI_ISL_3439400, EPI_ISL_3439401, EPI_ISL_3439402, EPI_ISL_3439403, EPI_ISL_3439404, EPI_ISL_3439405 | | | |
| see above | Lighthouse Lab in Milton Keynes | Wellcome Sanger Institute for the COVID-19 Genomics UK (COG-UK) Consortium | The Lighthouse Lab in Milton Keynes and Alex Alderton, Roberto Amato, Jeffrey Barrett, Sonia Goncalves, Ewan Harrison, David K. Jackson, Ian Johnston, Dominic Kwiatkowski, Cordelia Langford, John Sillitoe on behalf of the Wellcome Sanger Institute COVID-19 Surveillance Team |
| EPI_ISL_3439406 | Lighthouse Lab in Glasgow | Wellcome Sanger Institute for the COVID-19 Genomics UK (COG-UK) Consortium | Harper VanSteenhouse, Yumi Kasai, David Gray, Carol Clugston, Anna Dominiczak and Alex Alderton, Roberto Amato, Jeffrey Barrett, Sonia Goncalves, Ewan Harrison, David K. Jackson, Ian Johnston, Dominic Kwiatkowski, Cordelia Langford, John Sillitoe on behalf of the Wellcome Sanger Institute COVID-19 Surveillance Team |
| EPI_ISL_3439407 | Lighthouse Lab in Milton Keynes | Wellcome Sanger Institute for the COVID-19 Genomics UK (COG-UK) Consortium | The Lighthouse Lab in Milton Keynes and Alex Alderton, Roberto Amato, Jeffrey Barrett, Sonia Goncalves, Ewan Harrison, David K. Jackson, Ian Johnston, Dominic Kwiatkowski, Cordelia Langford, John Sillitoe on behalf of the Wellcome Sanger Institute COVID-19 Surveillance Team |
| EPI_ISL_3439408 | Lighthouse Lab in Glasgow | Wellcome Sanger Institute for the COVID-19 Genomics UK (COG-UK) Consortium | Harper VanSteenhouse, Yumi Kasai, David Gray, Carol Clugston, Anna Dominiczak and Alex Alderton, Roberto Amato, Jeffrey Barrett, Sonia Goncalves, Ewan Harrison, David K. Jackson, Ian Johnston, Dominic Kwiatkowski, Cordelia Langford, John Sillitoe on behalf of the Wellcome Sanger Institute COVID-19 Surveillance Team |
| EPI_ISL_3439409, EPI_ISL_3439410 | Lighthouse Lab in Milton Keynes | Wellcome Sanger Institute for the COVID-19 Genomics UK (COG-UK) Consortium | The Lighthouse Lab in Milton Keynes and Alex Alderton, Roberto Amato, Jeffrey Barrett, Sonia Goncalves, Ewan Harrison, David K. Jackson, Ian Johnston, Dominic Kwiatkowski, Cordelia Langford, John Sillitoe on behalf of the Wellcome Sanger Institute COVID-19 Surveillance Team |
| EPI_ISL_3439411, EPI_ISL_3439412 | Lighthouse Lab in Glasgow | Wellcome Sanger Institute for the COVID-19 Genomics UK (COG-UK) Consortium | Harper VanSteenhouse, Yumi Kasai, David Gray, Carol Clugston, Anna Dominiczak and Alex Alderton, Roberto Amato, Jeffrey Barrett, Sonia Goncalves, Ewan Harrison, David K. Jackson, Ian Johnston, Dominic Kwiatkowski, Cordelia Langford, John Sillitoe on behalf of the Wellcome Sanger Institute COVID-19 Surveillance Team |
| EPI_ISL_3439413, EPI_ISL_3439414, EPI_ISL_3439415, EPI_ISL_3439416, EPI_ISL_3439417, EPI_ISL_3439418 | Lighthouse Lab in Milton Keynes | Wellcome Sanger Institute for the COVID-19 Genomics UK (COG-UK) Consortium | The Lighthouse Lab in Milton Keynes and Alex Alderton, Roberto Amato, Jeffrey Barrett, Sonia Goncalves, Ewan Harrison, David K. Jackson, Ian Johnston, Dominic Kwiatkowski, Cordelia Langford, John Sillitoe on behalf of the Wellcome Sanger Institute COVID-19 Surveillance Team |
| EPI_ISL_3439419 | Lighthouse Lab in Glasgow | Wellcome Sanger Institute for the COVID-19 Genomics UK (COG-UK) Consortium | Harper VanSteenhouse, Yumi Kasai, David Gray, Carol Clugston, Anna Dominiczak and Alex Alderton, Roberto Amato, Jeffrey Barrett, Sonia Goncalves, Ewan Harrison, David K. Jackson, Ian Johnston, Dominic Kwiatkowski, Cordelia Langford, John Sillitoe on behalf of the Wellcome Sanger Institute COVID-19 Surveillance Team |
| EPI_ISL_3439420 | Lighthouse Lab in Milton Keynes | Wellcome Sanger Institute for the COVID-19 Genomics UK (COG-UK) Consortium | The Lighthouse Lab in Milton Keynes and Alex Alderton, Roberto Amato, Jeffrey Barrett, Sonia Goncalves, Ewan Harrison, David K. Jackson, Ian Johnston, Dominic Kwiatkowski, Cordelia Langford, John Sillitoe on behalf of the Wellcome Sanger Institute COVID-19 Surveillance Team |
| EPI_ISL_3439421, EPI_ISL_3439422, EPI_ISL_3439423 | Lighthouse Lab in Glasgow | Wellcome Sanger Institute for the COVID-19 Genomics UK (COG-UK) Consortium | Harper VanSteenhouse, Yumi Kasai, David Gray, Carol Clugston, Anna Dominiczak and Alex Alderton, Roberto Amato, Jeffrey Barrett, Sonia Goncalves, Ewan Harrison, David K. Jackson, Ian Johnston, Dominic Kwiatkowski, Cordelia Langford, John Sillitoe on behalf of the Wellcome Sanger Institute COVID-19 Surveillance Team |
| EPI_ISL_3439424, EPI_ISL_3439425, EPI_ISL_3439426, EPI_ISL_3439427, EPI_ISL_3439428 | Lighthouse Lab in Milton Keynes | Wellcome Sanger Institute for the COVID-19 Genomics UK (COG-UK) Consortium | The Lighthouse Lab in Milton Keynes and Alex Alderton, Roberto Amato, Jeffrey Barrett, Sonia Goncalves, Ewan Harrison, David K. Jackson, Ian Johnston, Dominic Kwiatkowski, Cordelia Langford, John Sillitoe on behalf of the Wellcome Sanger Institute COVID-19 Surveillance Team |
| EPI_ISL_3439429 | Lighthouse Lab in Glasgow | Wellcome Sanger Institute for the COVID-19 Genomics UK (COG-UK) Consortium | Harper VanSteenhouse, Yumi Kasai, David Gray, Carol Clugston, Anna Dominiczak and Alex Alderton, Roberto Amato, Jeffrey Barrett, Sonia Goncalves, Ewan Harrison, David K. Jackson, Ian Johnston, Dominic Kwiatkowski, Cordelia Langford, John Sillitoe on behalf of the Wellcome Sanger Institute COVID-19 Surveillance Team |
| EPI_ISL_3439430, EPI_ISL_3439431, EPI_ISL_3439432 | Lighthouse Lab in Milton Keynes | Wellcome Sanger Institute for the COVID-19 Genomics UK (COG-UK) Consortium | The Lighthouse Lab in Milton Keynes and Alex Alderton, Roberto Amato, Jeffrey Barrett, Sonia Goncalves, Ewan Harrison, David K. Jackson, Ian Johnston, Dominic Kwiatkowski, Cordelia Langford, John Sillitoe on behalf of the Wellcome Sanger Institute COVID-19 Surveillance Team |
| EPI_ISL_3439433 | Lighthouse Lab in Glasgow | Wellcome Sanger Institute for the COVID-19 Genomics UK (COG-UK) Consortium | Harper VanSteenhouse, Yumi Kasai, David Gray, Carol Clugston, Anna Dominiczak and Alex Alderton, Roberto Amato, Jeffrey Barrett, Sonia Goncalves, Ewan Harrison, David K. Jackson, Ian Johnston, Dominic Kwiatkowski, Cordelia Langford, John Sillitoe on behalf of the Wellcome Sanger Institute COVID-19 Surveillance Team |
| EPI_ISL_3439434, EPI_ISL_3439435, EPI_ISL_3439436, EPI_ISL_3439437, EPI_ISL_3439438 | Lighthouse Lab in Milton Keynes | Wellcome Sanger Institute for the COVID-19 Genomics UK (COG-UK) Consortium | The Lighthouse Lab in Milton Keynes and Alex Alderton, Roberto Amato, Jeffrey Barrett, Sonia Goncalves, Ewan Harrison, David K. Jackson, Ian Johnston, Dominic Kwiatkowski, Cordelia Langford, John Sillitoe on behalf of the Wellcome Sanger Institute COVID-19 Surveillance Team |
| EPI_ISL_3439439 | Lighthouse Lab in Glasgow | Wellcome Sanger Institute for the COVID-19 Genomics UK (COG-UK) Consortium | Harper VanSteenhouse, Yumi Kasai, David Gray, Carol Clugston, Anna Dominiczak and Alex Alderton, Roberto Amato, Jeffrey Barrett, Sonia Goncalves, Ewan Harrison, David K. Jackson, Ian Johnston, Dominic Kwiatkowski, Cordelia Langford, John Sillitoe on behalf of the Wellcome Sanger Institute COVID-19 Surveillance Team |
| EPI_ISL_3439440, EPI_ISL_3439441 | Lighthouse Lab in Milton Keynes | Wellcome Sanger Institute for the COVID-19 Genomics UK (COG-UK) Consortium | The Lighthouse Lab in Milton Keynes and Alex Alderton, Roberto Amato, Jeffrey Barrett, Sonia Goncalves, Ewan Harrison, David K. Jackson, Ian Johnston, Dominic Kwiatkowski, Cordelia Langford, John Sillitoe on behalf of the Wellcome Sanger Institute COVID-19 Surveillance Team |

| | | | |
|---|---|---|---|
| EPI_ISL_3439442 | Lighthouse Lab in Glasgow | Wellcome Sanger Institute for the COVID-19 Genomics UK (COG-UK) Consortium | Harper VanSteenhouse, Yumi Kasai, David Gray, Carol Clugston, Anna Dominiczak and Alex Alderton, Roberto Amato, Jeffrey Barrett, Sonia Goncalves, Ewan Harrison, David K. Jackson, Ian Johnston, Dominic Kwiatkowski, Cordelia Langford, John Sillitoe on behalf of the Wellcome Sanger Institute COVID-19 Surveillance Team |
| EPI_ISL_3439443, EPI_ISL_3439444, EPI_ISL_3439445, EPI_ISL_3439446, EPI_ISL_3439447, EPI_ISL_3439448, EPI_ISL_3439449, EPI_ISL_3439450, EPI_ISL_3439451, EPI_ISL_3439452 | Lighthouse Lab in Milton Keynes | Wellcome Sanger Institute for the COVID-19 Genomics UK (COG-UK) Consortium | The Lighthouse Lab in Milton Keynes and Alex Alderton, Roberto Amato, Jeffrey Barrett, Sonia Goncalves, Ewan Harrison, David K. Jackson, Ian Johnston, Dominic Kwiatkowski, Cordelia Langford, John Sillitoe on behalf of the Wellcome Sanger Institute COVID-19 Surveillance Team |
| EPI_ISL_3439453 | Lighthouse Lab in Glasgow | Wellcome Sanger Institute for the COVID-19 Genomics UK (COG-UK) Consortium | Harper VanSteenhouse, Yumi Kasai, David Gray, Carol Clugston, Anna Dominiczak and Alex Alderton, Roberto Amato, Jeffrey Barrett, Sonia Goncalves, Ewan Harrison, David K. Jackson, Ian Johnston, Dominic Kwiatkowski, Cordelia Langford, John Sillitoe on behalf of the Wellcome Sanger Institute COVID-19 Surveillance Team |
| EPI_ISL_3439454, EPI_ISL_3439455, EPI_ISL_3439456, EPI_ISL_3439457 | Lighthouse Lab in Milton Keynes | Wellcome Sanger Institute for the COVID-19 Genomics UK (COG-UK) Consortium | The Lighthouse Lab in Milton Keynes and Alex Alderton, Roberto Amato, Jeffrey Barrett, Sonia Goncalves, Ewan Harrison, David K. Jackson, Ian Johnston, Dominic Kwiatkowski, Cordelia Langford, John Sillitoe on behalf of the Wellcome Sanger Institute COVID-19 Surveillance Team |
| EPI_ISL_3439458, EPI_ISL_3439459 | Lighthouse Lab in Glasgow | Wellcome Sanger Institute for the COVID-19 Genomics UK (COG-UK) Consortium | Harper VanSteenhouse, Yumi Kasai, David Gray, Carol Clugston, Anna Dominiczak and Alex Alderton, Roberto Amato, Jeffrey Barrett, Sonia Goncalves, Ewan Harrison, David K. Jackson, Ian Johnston, Dominic Kwiatkowski, Cordelia Langford, John Sillitoe on behalf of the Wellcome Sanger Institute COVID-19 Surveillance Team |
| EPI_ISL_3439460, EPI_ISL_3439461, EPI_ISL_3439462, EPI_ISL_3439463, EPI_ISL_3439464, EPI_ISL_3439465, EPI_ISL_3439466, EPI_ISL_3439467, EPI_ISL_3439468, EPI_ISL_3439469, EPI_ISL_3439470 | | | |
| see above | Lighthouse Lab in Milton Keynes | Wellcome Sanger Institute for the COVID-19 Genomics UK (COG-UK) Consortium | The Lighthouse Lab in Milton Keynes and Alex Alderton, Roberto Amato, Jeffrey Barrett, Sonia Goncalves, Ewan Harrison, David K. Jackson, Ian Johnston, Dominic Kwiatkowski, Cordelia Langford, John Sillitoe on behalf of the Wellcome Sanger Institute COVID-19 Surveillance Team |
| EPI_ISL_3439471 | Lighthouse Lab in Glasgow | Wellcome Sanger Institute for the COVID-19 Genomics UK (COG-UK) Consortium | Harper VanSteenhouse, Yumi Kasai, David Gray, Carol Clugston, Anna Dominiczak and Alex Alderton, Roberto Amato, Jeffrey Barrett, Sonia Goncalves, Ewan Harrison, David K. Jackson, Ian Johnston, Dominic Kwiatkowski, Cordelia Langford, John Sillitoe on behalf of the Wellcome Sanger Institute COVID-19 Surveillance Team |
| EPI_ISL_3439472, EPI_ISL_3439473 | Lighthouse Lab in Milton Keynes | Wellcome Sanger Institute for the COVID-19 Genomics UK (COG-UK) Consortium | The Lighthouse Lab in Milton Keynes and Alex Alderton, Roberto Amato, Jeffrey Barrett, Sonia Goncalves, Ewan Harrison, David K. Jackson, Ian Johnston, Dominic Kwiatkowski, Cordelia Langford, John Sillitoe on behalf of the Wellcome Sanger Institute COVID-19 Surveillance Team |
| EPI_ISL_3439474 | Lighthouse Lab in Glasgow | Wellcome Sanger Institute for the COVID-19 Genomics UK (COG-UK) Consortium | Harper VanSteenhouse, Yumi Kasai, David Gray, Carol Clugston, Anna Dominiczak and Alex Alderton, Roberto Amato, Jeffrey Barrett, Sonia Goncalves, Ewan Harrison, David K. Jackson, Ian Johnston, Dominic Kwiatkowski, Cordelia Langford, John Sillitoe on behalf of the Wellcome Sanger Institute COVID-19 Surveillance Team |
| EPI_ISL_3439475, EPI_ISL_3439476 | Lighthouse Lab in Milton Keynes | Wellcome Sanger Institute for the COVID-19 Genomics UK (COG-UK) Consortium | The Lighthouse Lab in Milton Keynes and Alex Alderton, Roberto Amato, Jeffrey Barrett, Sonia Goncalves, Ewan Harrison, David K. Jackson, Ian Johnston, Dominic Kwiatkowski, Cordelia Langford, John Sillitoe on behalf of the Wellcome Sanger Institute COVID-19 Surveillance Team |
| EPI_ISL_3439477 | Lighthouse Lab in Glasgow | Wellcome Sanger Institute for the COVID-19 Genomics UK (COG-UK) Consortium | Harper VanSteenhouse, Yumi Kasai, David Gray, Carol Clugston, Anna Dominiczak and Alex Alderton, Roberto Amato, Jeffrey Barrett, Sonia Goncalves, Ewan Harrison, David K. Jackson, Ian Johnston, Dominic Kwiatkowski, Cordelia Langford, John Sillitoe on behalf of the Wellcome Sanger Institute COVID-19 Surveillance Team |
| EPI_ISL_3439478, EPI_ISL_3439479, EPI_ISL_3439480, EPI_ISL_3439481, EPI_ISL_3439482, EPI_ISL_3439483, EPI_ISL_3439484 | Lighthouse Lab in Milton Keynes | Wellcome Sanger Institute for the COVID-19 Genomics UK (COG-UK) Consortium | The Lighthouse Lab in Milton Keynes and Alex Alderton, Roberto Amato, Jeffrey Barrett, Sonia Goncalves, Ewan Harrison, David K. Jackson, Ian Johnston, Dominic Kwiatkowski, Cordelia Langford, John Sillitoe on behalf of the Wellcome Sanger Institute COVID-19 Surveillance Team |
| EPI_ISL_3439485, EPI_ISL_3439486 | Lighthouse Lab in Glasgow | Wellcome Sanger Institute for the COVID-19 Genomics UK (COG-UK) Consortium | Harper VanSteenhouse, Yumi Kasai, David Gray, Carol Clugston, Anna Dominiczak and Alex Alderton, Roberto Amato, Jeffrey Barrett, Sonia Goncalves, Ewan Harrison, David K. Jackson, Ian Johnston, Dominic Kwiatkowski, Cordelia Langford, John Sillitoe on behalf of the Wellcome Sanger Institute COVID-19 Surveillance Team |
| EPI_ISL_3439487, EPI_ISL_3439488, EPI_ISL_3439489, EPI_ISL_3439490, EPI_ISL_3439491 | Lighthouse Lab in Milton Keynes | Wellcome Sanger Institute for the COVID-19 Genomics UK (COG-UK) Consortium | The Lighthouse Lab in Milton Keynes and Alex Alderton, Roberto Amato, Jeffrey Barrett, Sonia Goncalves, Ewan Harrison, David K. Jackson, Ian Johnston, Dominic Kwiatkowski, Cordelia Langford, John Sillitoe on behalf of the Wellcome Sanger Institute COVID-19 Surveillance Team |
| EPI_ISL_3439492 | Lighthouse Lab in Glasgow | Wellcome Sanger Institute for the COVID-19 Genomics UK (COG-UK) Consortium | Harper VanSteenhouse, Yumi Kasai, David Gray, Carol Clugston, Anna Dominiczak and Alex Alderton, Roberto Amato, Jeffrey Barrett, Sonia Goncalves, Ewan Harrison, David K. Jackson, Ian Johnston, Dominic Kwiatkowski, Cordelia Langford, John Sillitoe on behalf of the Wellcome Sanger Institute COVID-19 Surveillance Team |
| EPI_ISL_3439493, EPI_ISL_3439494 | Lighthouse Lab in Milton Keynes | Wellcome Sanger Institute for the COVID-19 Genomics UK (COG-UK) Consortium | The Lighthouse Lab in Milton Keynes and Alex Alderton, Roberto Amato, Jeffrey Barrett, Sonia Goncalves, Ewan Harrison, David K. Jackson, Ian Johnston, Dominic Kwiatkowski, Cordelia Langford, John Sillitoe on behalf of the Wellcome Sanger Institute COVID-19 Surveillance Team |
| EPI_ISL_3439495 | Lighthouse Lab in Glasgow | Wellcome Sanger Institute for the COVID-19 Genomics UK (COG-UK) Consortium | Harper VanSteenhouse, Yumi Kasai, David Gray, Carol Clugston, Anna Dominiczak and Alex Alderton, Roberto Amato, Jeffrey Barrett, Sonia Goncalves, Ewan Harrison, David K. Jackson, Ian Johnston, Dominic Kwiatkowski, Cordelia Langford, John Sillitoe on behalf of the Wellcome Sanger Institute COVID-19 Surveillance Team |
| EPI_ISL_3439496, EPI_ISL_3439497, EPI_ISL_3439498, EPI_ISL_3439499, EPI_ISL_3439500 | Lighthouse Lab in Milton Keynes | Wellcome Sanger Institute for the COVID-19 Genomics UK (COG-UK) Consortium | The Lighthouse Lab in Milton Keynes and Alex Alderton, Roberto Amato, Jeffrey Barrett, Sonia Goncalves, Ewan Harrison, David K. Jackson, Ian Johnston, Dominic Kwiatkowski, Cordelia Langford, John Sillitoe on behalf of the Wellcome Sanger Institute COVID-19 Surveillance Team |
| EPI_ISL_3439501 | Lighthouse Lab in Glasgow | Wellcome Sanger Institute for the COVID-19 Genomics UK (COG-UK) Consortium | Harper VanSteenhouse, Yumi Kasai, David Gray, Carol Clugston, Anna Dominiczak and Alex Alderton, Roberto Amato, Jeffrey Barrett, Sonia Goncalves, Ewan Harrison, David K. Jackson, Ian Johnston, Dominic Kwiatkowski, Cordelia Langford, John Sillitoe on behalf of the Wellcome Sanger Institute COVID-19 Surveillance Team |
| EPI_ISL_3439502, EPI_ISL_3439503, EPI_ISL_3439504, EPI_ISL_3439505, EPI_ISL_3439506, EPI_ISL_3439507 | Lighthouse Lab in Milton Keynes | Wellcome Sanger Institute for the COVID-19 Genomics UK (COG-UK) Consortium | The Lighthouse Lab in Milton Keynes and Alex Alderton, Roberto Amato, Jeffrey Barrett, Sonia Goncalves, Ewan Harrison, David K. Jackson, Ian Johnston, Dominic Kwiatkowski, Cordelia Langford, John Sillitoe on behalf of the Wellcome Sanger Institute COVID-19 Surveillance Team |
| EPI_ISL_3439508 | Lighthouse Lab in Glasgow | Wellcome Sanger Institute for the COVID-19 Genomics UK (COG-UK) Consortium | Harper VanSteenhouse, Yumi Kasai, David Gray, Carol Clugston, Anna Dominiczak and Alex Alderton, Roberto Amato, Jeffrey Barrett, Sonia Goncalves, Ewan Harrison, David K. Jackson, Ian Johnston, Dominic Kwiatkowski, Cordelia Langford, John Sillitoe on behalf of the Wellcome Sanger Institute COVID-19 Surveillance Team |
| EPI_ISL_3439509, EPI_ISL_3439510, EPI_ISL_3439511 | Lighthouse Lab in Milton Keynes | Wellcome Sanger Institute for the COVID-19 Genomics UK (COG-UK) Consortium | The Lighthouse Lab in Milton Keynes and Alex Alderton, Roberto Amato, Jeffrey Barrett, Sonia Goncalves, Ewan Harrison, David K. Jackson, Ian Johnston, Dominic Kwiatkowski, Cordelia Langford, John Sillitoe on behalf of the Wellcome Sanger Institute COVID-19 Surveillance Team |
| EPI_ISL_3439512 | Lighthouse Lab in Glasgow | Wellcome Sanger Institute for the COVID-19 Genomics UK (COG-UK) Consortium | Harper VanSteenhouse, Yumi Kasai, David Gray, Carol Clugston, Anna Dominiczak and Alex Alderton, Roberto Amato, Jeffrey Barrett, Sonia Goncalves, Ewan Harrison, David K. Jackson, Ian Johnston, Dominic Kwiatkowski, Cordelia Langford, John Sillitoe on behalf of the Wellcome Sanger Institute COVID-19 Surveillance Team |
| EPI_ISL_3439513, EPI_ISL_3439514 | Lighthouse Lab in Milton Keynes | Wellcome Sanger Institute for the COVID-19 Genomics UK (COG-UK) Consortium | The Lighthouse Lab in Milton Keynes and Alex Alderton, Roberto Amato, Jeffrey Barrett, Sonia Goncalves, Ewan Harrison, David K. Jackson, Ian Johnston, Dominic Kwiatkowski, Cordelia Langford, John Sillitoe on behalf of the Wellcome Sanger Institute COVID-19 Surveillance Team |
| EPI_ISL_3439515 | Lighthouse Lab in Glasgow | Wellcome Sanger Institute for the COVID-19 Genomics UK (COG-UK) Consortium | Harper VanSteenhouse, Yumi Kasai, David Gray, Carol Clugston, Anna Dominiczak and Alex Alderton, Roberto Amato, Jeffrey Barrett, Sonia Goncalves, Ewan Harrison, David K. Jackson, Ian Johnston, Dominic Kwiatkowski, Cordelia Langford, John Sillitoe on behalf of the Wellcome Sanger Institute COVID-19 Surveillance Team |
| EPI_ISL_3439516, EPI_ISL_3439517, EPI_ISL_3439518 | Lighthouse Lab in Milton Keynes | Wellcome Sanger Institute for the COVID-19 Genomics UK (COG-UK) Consortium | The Lighthouse Lab in Milton Keynes and Alex Alderton, Roberto Amato, Jeffrey Barrett, Sonia Goncalves, Ewan Harrison, David K. Jackson, Ian Johnston, Dominic Kwiatkowski, Cordelia Langford, John Sillitoe on behalf of the Wellcome Sanger Institute COVID-19 Surveillance Team |
| EPI_ISL_3439519 | Lighthouse Lab in Glasgow | Wellcome Sanger Institute for the COVID-19 Genomics UK (COG-UK) Consortium | Harper VanSteenhouse, Yumi Kasai, David Gray, Carol Clugston, Anna Dominiczak and Alex Alderton, Roberto Amato, Jeffrey Barrett, Sonia Goncalves, Ewan Harrison, David K. Jackson, Ian Johnston, Dominic Kwiatkowski, Cordelia Langford, John Sillitoe on behalf of the Wellcome Sanger Institute COVID-19 Surveillance Team |
| EPI_ISL_3439520, EPI_ISL_3439521, EPI_ISL_3439522 | Lighthouse Lab in Milton Keynes | Wellcome Sanger Institute for the COVID-19 Genomics UK (COG-UK) Consortium | The Lighthouse Lab in Milton Keynes and Alex Alderton, Roberto Amato, Jeffrey Barrett, Sonia Goncalves, Ewan Harrison, David K. Jackson, Ian Johnston, Dominic Kwiatkowski, Cordelia Langford, John Sillitoe on behalf of the Wellcome Sanger Institute COVID-19 Surveillance Team |
| EPI_ISL_3439523 | Lighthouse Lab in Glasgow | Wellcome Sanger Institute for the COVID-19 Genomics UK (COG-UK) Consortium | Harper VanSteenhouse, Yumi Kasai, David Gray, Carol Clugston, Anna Dominiczak and Alex Alderton, Roberto Amato, Jeffrey Barrett, Sonia Goncalves, Ewan Harrison, David K. Jackson, Ian Johnston, Dominic Kwiatkowski, Cordelia Langford, John Sillitoe on behalf of the Wellcome Sanger Institute COVID-19 Surveillance Team |
| EPI_ISL_3439524, EPI_ISL_3439525, EPI_ISL_3439526, EPI_ISL_3439527, EPI_ISL_3439528, EPI_ISL_3439529, EPI_ISL_3439530, EPI_ISL_3439531 | Lighthouse Lab in Milton Keynes | Wellcome Sanger Institute for the COVID-19 Genomics UK (COG-UK) Consortium | The Lighthouse Lab in Milton Keynes and Alex Alderton, Roberto Amato, Jeffrey Barrett, Sonia Goncalves, Ewan Harrison, David K. Jackson, Ian Johnston, Dominic Kwiatkowski, Cordelia Langford, John Sillitoe on behalf of the Wellcome Sanger Institute COVID-19 Surveillance Team |
| EPI_ISL_3439532 | Lighthouse Lab in Glasgow | Wellcome Sanger Institute for the COVID-19 Genomics UK (COG-UK) Consortium | Harper VanSteenhouse, Yumi Kasai, David Gray, Carol Clugston, Anna Dominiczak and Alex Alderton, Roberto Amato, Jeffrey Barrett, Sonia Goncalves, Ewan Harrison, David K. Jackson, Ian Johnston, Dominic Kwiatkowski, Cordelia Langford, John Sillitoe on behalf of the Wellcome Sanger Institute COVID-19 Surveillance Team |

| | | | |
|---|---|---|---|
| EPI_ISL_3439533, EPI_ISL_3439534, EPI_ISL_3439535, EPI_ISL_3439536, EPI_ISL_3439537, EPI_ISL_3439538, EPI_ISL_3439539, EPI_ISL_3439540, EPI_ISL_3439541, EPI_ISL_3439542, EPI_ISL_3439543, EPI_ISL_3439544, EPI_ISL_3439545 | | | |
| see above | Lighthouse Lab in Milton Keynes | Wellcome Sanger Institute for the COVID-19 Genomics UK (COG-UK) Consortium | The Lighthouse Lab in Milton Keynes and Alex Alderton, Roberto Amato, Jeffrey Barrett, Sonia Goncalves, Ewan Harrison, David K. Jackson, Ian Johnston, Dominic Kwiatkowski, Cordelia Langford, John Sillitoe on behalf of the Wellcome Sanger Institute COVID-19 Surveillance Team |
| EPI_ISL_3439546 | Lighthouse Lab in Glasgow | Wellcome Sanger Institute for the COVID-19 Genomics UK (COG-UK) Consortium | Harper VanSteenhouse, Yumi Kasai, David Gray, Carol Clugston, Anna Dominiczak and Alex Alderton, Roberto Amato, Jeffrey Barrett, Sonia Goncalves, Ewan Harrison, David K. Jackson, Ian Johnston, Dominic Kwiatkowski, Cordelia Langford, John Sillitoe on behalf of the Wellcome Sanger Institute COVID-19 Surveillance Team |
| EPI_ISL_3439547, EPI_ISL_3439548, EPI_ISL_3439549, EPI_ISL_3439550 | Lighthouse Lab in Milton Keynes | Wellcome Sanger Institute for the COVID-19 Genomics UK (COG-UK) Consortium | The Lighthouse Lab in Milton Keynes and Alex Alderton, Roberto Amato, Jeffrey Barrett, Sonia Goncalves, Ewan Harrison, David K. Jackson, Ian Johnston, Dominic Kwiatkowski, Cordelia Langford, John Sillitoe on behalf of the Wellcome Sanger Institute COVID-19 Surveillance Team |
| EPI_ISL_3439551 | Lighthouse Lab in Glasgow | Wellcome Sanger Institute for the COVID-19 Genomics UK (COG-UK) Consortium | Harper VanSteenhouse, Yumi Kasai, David Gray, Carol Clugston, Anna Dominiczak and Alex Alderton, Roberto Amato, Jeffrey Barrett, Sonia Goncalves, Ewan Harrison, David K. Jackson, Ian Johnston, Dominic Kwiatkowski, Cordelia Langford, John Sillitoe on behalf of the Wellcome Sanger Institute COVID-19 Surveillance Team |
| EPI_ISL_3439552 | Lighthouse Lab in Milton Keynes | Wellcome Sanger Institute for the COVID-19 Genomics UK (COG-UK) Consortium | The Lighthouse Lab in Milton Keynes and Alex Alderton, Roberto Amato, Jeffrey Barrett, Sonia Goncalves, Ewan Harrison, David K. Jackson, Ian Johnston, Dominic Kwiatkowski, Cordelia Langford, John Sillitoe on behalf of the Wellcome Sanger Institute COVID-19 Surveillance Team |
| EPI_ISL_3439553 | Lighthouse Lab in Glasgow | Wellcome Sanger Institute for the COVID-19 Genomics UK (COG-UK) Consortium | Harper VanSteenhouse, Yumi Kasai, David Gray, Carol Clugston, Anna Dominiczak and Alex Alderton, Roberto Amato, Jeffrey Barrett, Sonia Goncalves, Ewan Harrison, David K. Jackson, Ian Johnston, Dominic Kwiatkowski, Cordelia Langford, John Sillitoe on behalf of the Wellcome Sanger Institute COVID-19 Surveillance Team |
| EPI_ISL_3439554, EPI_ISL_3439555, EPI_ISL_3439556, EPI_ISL_3439557, EPI_ISL_3439558, EPI_ISL_3439559 | Lighthouse Lab in Milton Keynes | Wellcome Sanger Institute for the COVID-19 Genomics UK (COG-UK) Consortium | The Lighthouse Lab in Milton Keynes and Alex Alderton, Roberto Amato, Jeffrey Barrett, Sonia Goncalves, Ewan Harrison, David K. Jackson, Ian Johnston, Dominic Kwiatkowski, Cordelia Langford, John Sillitoe on behalf of the Wellcome Sanger Institute COVID-19 Surveillance Team |
| EPI_ISL_3439560 | Lighthouse Lab in Glasgow | Wellcome Sanger Institute for the COVID-19 Genomics UK (COG-UK) Consortium | Harper VanSteenhouse, Yumi Kasai, David Gray, Carol Clugston, Anna Dominiczak and Alex Alderton, Roberto Amato, Jeffrey Barrett, Sonia Goncalves, Ewan Harrison, David K. Jackson, Ian Johnston, Dominic Kwiatkowski, Cordelia Langford, John Sillitoe on behalf of the Wellcome Sanger Institute COVID-19 Surveillance Team |
| EPI_ISL_3439561 | Lighthouse Lab in Milton Keynes | Wellcome Sanger Institute for the COVID-19 Genomics UK (COG-UK) Consortium | The Lighthouse Lab in Milton Keynes and Alex Alderton, Roberto Amato, Jeffrey Barrett, Sonia Goncalves, Ewan Harrison, David K. Jackson, Ian Johnston, Dominic Kwiatkowski, Cordelia Langford, John Sillitoe on behalf of the Wellcome Sanger Institute COVID-19 Surveillance Team |
| EPI_ISL_3439562 | Lighthouse Lab in Glasgow | Wellcome Sanger Institute for the COVID-19 Genomics UK (COG-UK) Consortium | Harper VanSteenhouse, Yumi Kasai, David Gray, Carol Clugston, Anna Dominiczak and Alex Alderton, Roberto Amato, Jeffrey Barrett, Sonia Goncalves, Ewan Harrison, David K. Jackson, Ian Johnston, Dominic Kwiatkowski, Cordelia Langford, John Sillitoe on behalf of the Wellcome Sanger Institute COVID-19 Surveillance Team |
| EPI_ISL_3439563, EPI_ISL_3439564, EPI_ISL_3439565, EPI_ISL_3439566, EPI_ISL_3439567, EPI_ISL_3439568, EPI_ISL_3439569, EPI_ISL_3439570, EPI_ISL_3439571, EPI_ISL_3439572, EPI_ISL_3439573, EPI_ISL_3439574, EPI_ISL_3439575 | | | |
| see above | Lighthouse Lab in Milton Keynes | Wellcome Sanger Institute for the COVID-19 Genomics UK (COG-UK) Consortium | The Lighthouse Lab in Milton Keynes and Alex Alderton, Roberto Amato, Jeffrey Barrett, Sonia Goncalves, Ewan Harrison, David K. Jackson, Ian Johnston, Dominic Kwiatkowski, Cordelia Langford, John Sillitoe on behalf of the Wellcome Sanger Institute COVID-19 Surveillance Team |
| EPI_ISL_3439576 | Lighthouse Lab in Glasgow | Wellcome Sanger Institute for the COVID-19 Genomics UK (COG-UK) Consortium | Harper VanSteenhouse, Yumi Kasai, David Gray, Carol Clugston, Anna Dominiczak and Alex Alderton, Roberto Amato, Jeffrey Barrett, Sonia Goncalves, Ewan Harrison, David K. Jackson, Ian Johnston, Dominic Kwiatkowski, Cordelia Langford, John Sillitoe on behalf of the Wellcome Sanger Institute COVID-19 Surveillance Team |
| EPI_ISL_3439577, EPI_ISL_3439578, EPI_ISL_3439579, EPI_ISL_3439580, EPI_ISL_3439581, EPI_ISL_3439582, EPI_ISL_3439583, EPI_ISL_3439584, EPI_ISL_3439585, EPI_ISL_3439586, EPI_ISL_3439587, EPI_ISL_3439588, EPI_ISL_3439589, EPI_ISL_3439590, EPI_ISL_3439591, EPI_ISL_3439592, EPI_ISL_3439593, EPI_ISL_3439594, EPI_ISL_3439595, EPI_ISL_3439596, EPI_ISL_3439597, EPI_ISL_3439598, EPI_ISL_3439599, EPI_ISL_3439600, EPI_ISL_3439601, EPI_ISL_3439602 | | | |
| see above | Lighthouse Lab in Milton Keynes | Wellcome Sanger Institute for the COVID-19 Genomics UK (COG-UK) Consortium | The Lighthouse Lab in Milton Keynes and Alex Alderton, Roberto Amato, Jeffrey Barrett, Sonia Goncalves, Ewan Harrison, David K. Jackson, Ian Johnston, Dominic Kwiatkowski, Cordelia Langford, John Sillitoe on behalf of the Wellcome Sanger Institute COVID-19 Surveillance Team |
| EPI_ISL_3439603 | Lighthouse Lab in Glasgow | Wellcome Sanger Institute for the COVID-19 Genomics UK (COG-UK) Consortium | Harper VanSteenhouse, Yumi Kasai, David Gray, Carol Clugston, Anna Dominiczak and Alex Alderton, Roberto Amato, Jeffrey Barrett, Sonia Goncalves, Ewan Harrison, David K. Jackson, Ian Johnston, Dominic Kwiatkowski, Cordelia Langford, John Sillitoe on behalf of the Wellcome Sanger Institute COVID-19 Surveillance Team |
| EPI_ISL_3439604, EPI_ISL_3439605, EPI_ISL_3439606 | Lighthouse Lab in Milton Keynes | Wellcome Sanger Institute for the COVID-19 Genomics UK (COG-UK) Consortium | The Lighthouse Lab in Milton Keynes and Alex Alderton, Roberto Amato, Jeffrey Barrett, Sonia Goncalves, Ewan Harrison, David K. Jackson, Ian Johnston, Dominic Kwiatkowski, Cordelia Langford, John Sillitoe on behalf of the Wellcome Sanger Institute COVID-19 Surveillance Team |
| EPI_ISL_3439607 | Lighthouse Lab in Glasgow | Wellcome Sanger Institute for the COVID-19 Genomics UK (COG-UK) Consortium | Harper VanSteenhouse, Yumi Kasai, David Gray, Carol Clugston, Anna Dominiczak and Alex Alderton, Roberto Amato, Jeffrey Barrett, Sonia Goncalves, Ewan Harrison, David K. Jackson, Ian Johnston, Dominic Kwiatkowski, Cordelia Langford, John Sillitoe on behalf of the Wellcome Sanger Institute COVID-19 Surveillance Team |
| EPI_ISL_3439608, EPI_ISL_3439609, EPI_ISL_3439610, EPI_ISL_3439611, EPI_ISL_3439612, EPI_ISL_3439613, EPI_ISL_3439614, EPI_ISL_3439615, EPI_ISL_3439616, EPI_ISL_3439617 | Lighthouse Lab in Milton Keynes | Wellcome Sanger Institute for the COVID-19 Genomics UK (COG-UK) Consortium | The Lighthouse Lab in Milton Keynes and Alex Alderton, Roberto Amato, Jeffrey Barrett, Sonia Goncalves, Ewan Harrison, David K. Jackson, Ian Johnston, Dominic Kwiatkowski, Cordelia Langford, John Sillitoe on behalf of the Wellcome Sanger Institute COVID-19 Surveillance Team |
| EPI_ISL_3439618 | Lighthouse Lab in Glasgow | Wellcome Sanger Institute for the COVID-19 Genomics UK (COG-UK) Consortium | Harper VanSteenhouse, Yumi Kasai, David Gray, Carol Clugston, Anna Dominiczak and Alex Alderton, Roberto Amato, Jeffrey Barrett, Sonia Goncalves, Ewan Harrison, David K. Jackson, Ian Johnston, Dominic Kwiatkowski, Cordelia Langford, John Sillitoe on behalf of the Wellcome Sanger Institute COVID-19 Surveillance Team |
| EPI_ISL_3439619 | Lighthouse Lab in Milton Keynes | Wellcome Sanger Institute for the COVID-19 Genomics UK (COG-UK) Consortium | The Lighthouse Lab in Milton Keynes and Alex Alderton, Roberto Amato, Jeffrey Barrett, Sonia Goncalves, Ewan Harrison, David K. Jackson, Ian Johnston, Dominic Kwiatkowski, Cordelia Langford, John Sillitoe on behalf of the Wellcome Sanger Institute COVID-19 Surveillance Team |
| EPI_ISL_3439620 | Lighthouse Lab in Glasgow | Wellcome Sanger Institute for the COVID-19 Genomics UK (COG-UK) Consortium | Harper VanSteenhouse, Yumi Kasai, David Gray, Carol Clugston, Anna Dominiczak and Alex Alderton, Roberto Amato, Jeffrey Barrett, Sonia Goncalves, Ewan Harrison, David K. Jackson, Ian Johnston, Dominic Kwiatkowski, Cordelia Langford, John Sillitoe on behalf of the Wellcome Sanger Institute COVID-19 Surveillance Team |
| EPI_ISL_3439621 | Lighthouse Lab in Milton Keynes | Wellcome Sanger Institute for the COVID-19 Genomics UK (COG-UK) Consortium | The Lighthouse Lab in Milton Keynes and Alex Alderton, Roberto Amato, Jeffrey Barrett, Sonia Goncalves, Ewan Harrison, David K. Jackson, Ian Johnston, Dominic Kwiatkowski, Cordelia Langford, John Sillitoe on behalf of the Wellcome Sanger Institute COVID-19 Surveillance Team |
| EPI_ISL_3439622 | Lighthouse Lab in Glasgow | Wellcome Sanger Institute for the COVID-19 Genomics UK (COG-UK) Consortium | Harper VanSteenhouse, Yumi Kasai, David Gray, Carol Clugston, Anna Dominiczak and Alex Alderton, Roberto Amato, Jeffrey Barrett, Sonia Goncalves, Ewan Harrison, David K. Jackson, Ian Johnston, Dominic Kwiatkowski, Cordelia Langford, John Sillitoe on behalf of the Wellcome Sanger Institute COVID-19 Surveillance Team |
| EPI_ISL_3439623, EPI_ISL_3439624, EPI_ISL_3439625 | Lighthouse Lab in Milton Keynes | Wellcome Sanger Institute for the COVID-19 Genomics UK (COG-UK) Consortium | The Lighthouse Lab in Milton Keynes and Alex Alderton, Roberto Amato, Jeffrey Barrett, Sonia Goncalves, Ewan Harrison, David K. Jackson, Ian Johnston, Dominic Kwiatkowski, Cordelia Langford, John Sillitoe on behalf of the Wellcome Sanger Institute COVID-19 Surveillance Team |
| EPI_ISL_3439626 | Lighthouse Lab in Glasgow | Wellcome Sanger Institute for the COVID-19 Genomics UK (COG-UK) Consortium | Harper VanSteenhouse, Yumi Kasai, David Gray, Carol Clugston, Anna Dominiczak and Alex Alderton, Roberto Amato, Jeffrey Barrett, Sonia Goncalves, Ewan Harrison, David K. Jackson, Ian Johnston, Dominic Kwiatkowski, Cordelia Langford, John Sillitoe on behalf of the Wellcome Sanger Institute COVID-19 Surveillance Team |
| EPI_ISL_3439627 | Lighthouse Lab in Milton Keynes | Wellcome Sanger Institute for the COVID-19 Genomics UK (COG-UK) Consortium | The Lighthouse Lab in Milton Keynes and Alex Alderton, Roberto Amato, Jeffrey Barrett, Sonia Goncalves, Ewan Harrison, David K. Jackson, Ian Johnston, Dominic Kwiatkowski, Cordelia Langford, John Sillitoe on behalf of the Wellcome Sanger Institute COVID-19 Surveillance Team |
| EPI_ISL_3439628 | Lighthouse Lab in Glasgow | Wellcome Sanger Institute for the COVID-19 Genomics UK (COG-UK) Consortium | Harper VanSteenhouse, Yumi Kasai, David Gray, Carol Clugston, Anna Dominiczak and Alex Alderton, Roberto Amato, Jeffrey Barrett, Sonia Goncalves, Ewan Harrison, David K. Jackson, Ian Johnston, Dominic Kwiatkowski, Cordelia Langford, John Sillitoe on behalf of the Wellcome Sanger Institute COVID-19 Surveillance Team |
| EPI_ISL_3439629, EPI_ISL_3439630, EPI_ISL_3439631, EPI_ISL_3439632 | Lighthouse Lab in Milton Keynes | Wellcome Sanger Institute for the COVID-19 Genomics UK (COG-UK) Consortium | The Lighthouse Lab in Milton Keynes and Alex Alderton, Roberto Amato, Jeffrey Barrett, Sonia Goncalves, Ewan Harrison, David K. Jackson, Ian Johnston, Dominic Kwiatkowski, Cordelia Langford, John Sillitoe on behalf of the Wellcome Sanger Institute COVID-19 Surveillance Team |
| EPI_ISL_3439633, EPI_ISL_3439634 | Lighthouse Lab in Glasgow | Wellcome Sanger Institute for the COVID-19 Genomics UK (COG-UK) Consortium | Harper VanSteenhouse, Yumi Kasai, David Gray, Carol Clugston, Anna Dominiczak and Alex Alderton, Roberto Amato, Jeffrey Barrett, Sonia Goncalves, Ewan Harrison, David K. Jackson, Ian Johnston, Dominic Kwiatkowski, Cordelia Langford, John Sillitoe on behalf of the Wellcome Sanger Institute COVID-19 Surveillance Team |
| EPI_ISL_3439635, EPI_ISL_3439636, EPI_ISL_3439637, EPI_ISL_3439638 | Lighthouse Lab in Milton Keynes | Wellcome Sanger Institute for the COVID-19 Genomics UK (COG-UK) Consortium | The Lighthouse Lab in Milton Keynes and Alex Alderton, Roberto Amato, Jeffrey Barrett, Sonia Goncalves, Ewan Harrison, David K. Jackson, Ian Johnston, Dominic Kwiatkowski, Cordelia Langford, John Sillitoe on behalf of the Wellcome Sanger Institute COVID-19 Surveillance Team |
| EPI_ISL_3439639, EPI_ISL_3439640 | Lighthouse Lab in Glasgow | Wellcome Sanger Institute for the COVID-19 Genomics UK (COG-UK) Consortium | Harper VanSteenhouse, Yumi Kasai, David Gray, Carol Clugston, Anna Dominiczak and Alex Alderton, Roberto Amato, Jeffrey Barrett, Sonia Goncalves, Ewan Harrison, David K. Jackson, Ian Johnston, Dominic Kwiatkowski, Cordelia Langford, John Sillitoe on behalf of the Wellcome Sanger Institute COVID-19 Surveillance Team |
| EPI_ISL_3439641, EPI_ISL_3439642, EPI_ISL_3439643, EPI_ISL_3439644, EPI_ISL_3439645, EPI_ISL_3439646, EPI_ISL_3439647, EPI_ISL_3439648, EPI_ISL_3439649 | Lighthouse Lab in Milton Keynes | Wellcome Sanger Institute for the COVID-19 Genomics UK (COG-UK) Consortium | The Lighthouse Lab in Milton Keynes and Alex Alderton, Roberto Amato, Jeffrey Barrett, Sonia Goncalves, Ewan Harrison, David K. Jackson, Ian Johnston, Dominic Kwiatkowski, Cordelia Langford, John Sillitoe on behalf of the Wellcome Sanger Institute COVID-19 Surveillance Team |
| EPI_ISL_3439650 | Lighthouse Lab in Glasgow | Wellcome Sanger Institute for the COVID-19 Genomics UK (COG-UK) | Harper VanSteenhouse, Yumi Kasai, David Gray, Carol Clugston, Anna Dominiczak and Alex Alderton, Roberto Amato, Jeffrey Barrett, Sonia Goncalves, Ewan Harrison, David K. Jackson, Ian Johnston, Dominic Kwiatkowski, Cordelia Langford, |

| | | | |
|---|---|---|---|
| | | Consortium | John Sillitoe on behalf of the Wellcome Sanger Institute COVID-19 Surveillance Team |
| EPI_ISL_3439651 | Lighthouse Lab in Milton Keynes | Wellcome Sanger Institute for the COVID-19 Genomics UK (COG-UK) Consortium | The Lighthouse Lab in Milton Keynes and Alex Alderton, Roberto Amato, Jeffrey Barrett, Sonia Goncalves, Ewan Harrison, David K. Jackson, Ian Johnston, Dominic Kwiatkowski, Cordelia Langford, John Sillitoe on behalf of the Wellcome Sanger Institute COVID-19 Surveillance Team |
| EPI_ISL_3439652 | Lighthouse Lab in Glasgow | Wellcome Sanger Institute for the COVID-19 Genomics UK (COG-UK) Consortium | Harper VanSteenhouse, Yumi Kasai, David Gray, Carol Clugston, Anna Dominiczak and Alex Alderton, Roberto Amato, Jeffrey Barrett, Sonia Goncalves, Ewan Harrison, David K. Jackson, Ian Johnston, Dominic Kwiatkowski, Cordelia Langford, John Sillitoe on behalf of the Wellcome Sanger Institute COVID-19 Surveillance Team |
| EPI_ISL_3439653 | Lighthouse Lab in Milton Keynes | Wellcome Sanger Institute for the COVID-19 Genomics UK (COG-UK) Consortium | The Lighthouse Lab in Milton Keynes and Alex Alderton, Roberto Amato, Jeffrey Barrett, Sonia Goncalves, Ewan Harrison, David K. Jackson, Ian Johnston, Dominic Kwiatkowski, Cordelia Langford, John Sillitoe on behalf of the Wellcome Sanger Institute COVID-19 Surveillance Team |
| EPI_ISL_3439654 | Lighthouse Lab in Glasgow | Wellcome Sanger Institute for the COVID-19 Genomics UK (COG-UK) Consortium | Harper VanSteenhouse, Yumi Kasai, David Gray, Carol Clugston, Anna Dominiczak and Alex Alderton, Roberto Amato, Jeffrey Barrett, Sonia Goncalves, Ewan Harrison, David K. Jackson, Ian Johnston, Dominic Kwiatkowski, Cordelia Langford, John Sillitoe on behalf of the Wellcome Sanger Institute COVID-19 Surveillance Team |
| EPI_ISL_3439655, EPI_ISL_3439656, EPI_ISL_3439657 | Lighthouse Lab in Milton Keynes | Wellcome Sanger Institute for the COVID-19 Genomics UK (COG-UK) Consortium | The Lighthouse Lab in Milton Keynes and Alex Alderton, Roberto Amato, Jeffrey Barrett, Sonia Goncalves, Ewan Harrison, David K. Jackson, Ian Johnston, Dominic Kwiatkowski, Cordelia Langford, John Sillitoe on behalf of the Wellcome Sanger Institute COVID-19 Surveillance Team |
| EPI_ISL_3439658 | Lighthouse Lab in Glasgow | Wellcome Sanger Institute for the COVID-19 Genomics UK (COG-UK) Consortium | Harper VanSteenhouse, Yumi Kasai, David Gray, Carol Clugston, Anna Dominiczak and Alex Alderton, Roberto Amato, Jeffrey Barrett, Sonia Goncalves, Ewan Harrison, David K. Jackson, Ian Johnston, Dominic Kwiatkowski, Cordelia Langford, John Sillitoe on behalf of the Wellcome Sanger Institute COVID-19 Surveillance Team |
| EPI_ISL_3439659, EPI_ISL_3439660, EPI_ISL_3439661, EPI_ISL_3439662 | Lighthouse Lab in Milton Keynes | Wellcome Sanger Institute for the COVID-19 Genomics UK (COG-UK) Consortium | The Lighthouse Lab in Milton Keynes and Alex Alderton, Roberto Amato, Jeffrey Barrett, Sonia Goncalves, Ewan Harrison, David K. Jackson, Ian Johnston, Dominic Kwiatkowski, Cordelia Langford, John Sillitoe on behalf of the Wellcome Sanger Institute COVID-19 Surveillance Team |
| EPI_ISL_3439663, EPI_ISL_3439664, EPI_ISL_3439665 | Lighthouse Lab in Glasgow | Wellcome Sanger Institute for the COVID-19 Genomics UK (COG-UK) Consortium | Harper VanSteenhouse, Yumi Kasai, David Gray, Carol Clugston, Anna Dominiczak and Alex Alderton, Roberto Amato, Jeffrey Barrett, Sonia Goncalves, Ewan Harrison, David K. Jackson, Ian Johnston, Dominic Kwiatkowski, Cordelia Langford, John Sillitoe on behalf of the Wellcome Sanger Institute COVID-19 Surveillance Team |
| EPI_ISL_3439666, EPI_ISL_3439667, EPI_ISL_3439668 | Lighthouse Lab in Milton Keynes | Wellcome Sanger Institute for the COVID-19 Genomics UK (COG-UK) Consortium | The Lighthouse Lab in Milton Keynes and Alex Alderton, Roberto Amato, Jeffrey Barrett, Sonia Goncalves, Ewan Harrison, David K. Jackson, Ian Johnston, Dominic Kwiatkowski, Cordelia Langford, John Sillitoe on behalf of the Wellcome Sanger Institute COVID-19 Surveillance Team |
| EPI_ISL_3439669, EPI_ISL_3439670 | Lighthouse Lab in Glasgow | Wellcome Sanger Institute for the COVID-19 Genomics UK (COG-UK) Consortium | Harper VanSteenhouse, Yumi Kasai, David Gray, Carol Clugston, Anna Dominiczak and Alex Alderton, Roberto Amato, Jeffrey Barrett, Sonia Goncalves, Ewan Harrison, David K. Jackson, Ian Johnston, Dominic Kwiatkowski, Cordelia Langford, John Sillitoe on behalf of the Wellcome Sanger Institute COVID-19 Surveillance Team |
| EPI_ISL_3439671, EPI_ISL_3439672, EPI_ISL_3439673, EPI_ISL_3439674, EPI_ISL_3439675, EPI_ISL_3439676, EPI_ISL_3439677, EPI_ISL_3439678, EPI_ISL_3439679, EPI_ISL_3439680 | Lighthouse Lab in Milton Keynes | Wellcome Sanger Institute for the COVID-19 Genomics UK (COG-UK) Consortium | The Lighthouse Lab in Milton Keynes and Alex Alderton, Roberto Amato, Jeffrey Barrett, Sonia Goncalves, Ewan Harrison, David K. Jackson, Ian Johnston, Dominic Kwiatkowski, Cordelia Langford, John Sillitoe on behalf of the Wellcome Sanger Institute COVID-19 Surveillance Team |
| EPI_ISL_3439681 | Lighthouse Lab in Glasgow | Wellcome Sanger Institute for the COVID-19 Genomics UK (COG-UK) Consortium | Harper VanSteenhouse, Yumi Kasai, David Gray, Carol Clugston, Anna Dominiczak and Alex Alderton, Roberto Amato, Jeffrey Barrett, Sonia Goncalves, Ewan Harrison, David K. Jackson, Ian Johnston, Dominic Kwiatkowski, Cordelia Langford, John Sillitoe on behalf of the Wellcome Sanger Institute COVID-19 Surveillance Team |
| EPI_ISL_3439682, EPI_ISL_3439683, EPI_ISL_3439684 | Lighthouse Lab in Milton Keynes | Wellcome Sanger Institute for the COVID-19 Genomics UK (COG-UK) Consortium | The Lighthouse Lab in Milton Keynes and Alex Alderton, Roberto Amato, Jeffrey Barrett, Sonia Goncalves, Ewan Harrison, David K. Jackson, Ian Johnston, Dominic Kwiatkowski, Cordelia Langford, John Sillitoe on behalf of the Wellcome Sanger Institute COVID-19 Surveillance Team |
| EPI_ISL_3439685 | Lighthouse Lab in Glasgow | Wellcome Sanger Institute for the COVID-19 Genomics UK (COG-UK) Consortium | Harper VanSteenhouse, Yumi Kasai, David Gray, Carol Clugston, Anna Dominiczak and Alex Alderton, Roberto Amato, Jeffrey Barrett, Sonia Goncalves, Ewan Harrison, David K. Jackson, Ian Johnston, Dominic Kwiatkowski, Cordelia Langford, John Sillitoe on behalf of the Wellcome Sanger Institute COVID-19 Surveillance Team |
| EPI_ISL_3439686, EPI_ISL_3439687, EPI_ISL_3439688, EPI_ISL_3439689, EPI_ISL_3439690, EPI_ISL_3439691, EPI_ISL_3439692, EPI_ISL_3439693 | Lighthouse Lab in Milton Keynes | Wellcome Sanger Institute for the COVID-19 Genomics UK (COG-UK) Consortium | The Lighthouse Lab in Milton Keynes and Alex Alderton, Roberto Amato, Jeffrey Barrett, Sonia Goncalves, Ewan Harrison, David K. Jackson, Ian Johnston, Dominic Kwiatkowski, Cordelia Langford, John Sillitoe on behalf of the Wellcome Sanger Institute COVID-19 Surveillance Team |
| EPI_ISL_3439694 | Lighthouse Lab in Alderley Park | Wellcome Sanger Institute for the COVID-19 Genomics UK (COG-UK) Consortium | Jacquelyn Wynn, Mairead Hyland, The Lighthouse Lab in Alderley Park and Alex Alderton, Roberto Amato, Jeffrey Barrett, Sonia Goncalves, Ewan Harrison, David K. Jackson, Ian Johnston, Dominic Kwiatkowski, Cordelia Langford, John Sillitoe on behalf of the Wellcome Sanger Institute COVID-19 Surveillance Team |
| EPI_ISL_3439695, EPI_ISL_3439696, EPI_ISL_3439697 | Lighthouse Lab in Milton Keynes | Wellcome Sanger Institute for the COVID-19 Genomics UK (COG-UK) Consortium | The Lighthouse Lab in Milton Keynes and Alex Alderton, Roberto Amato, Jeffrey Barrett, Sonia Goncalves, Ewan Harrison, David K. Jackson, Ian Johnston, Dominic Kwiatkowski, Cordelia Langford, John Sillitoe on behalf of the Wellcome Sanger Institute COVID-19 Surveillance Team |
| EPI_ISL_3439698 | Lighthouse Lab in Alderley Park | Wellcome Sanger Institute for the COVID-19 Genomics UK (COG-UK) Consortium | Jacquelyn Wynn, Mairead Hyland, The Lighthouse Lab in Alderley Park and Alex Alderton, Roberto Amato, Jeffrey Barrett, Sonia Goncalves, Ewan Harrison, David K. Jackson, Ian Johnston, Dominic Kwiatkowski, Cordelia Langford, John Sillitoe on behalf of the Wellcome Sanger Institute COVID-19 Surveillance Team |
| EPI_ISL_3439699, EPI_ISL_3439700, EPI_ISL_3439701, EPI_ISL_3439702, EPI_ISL_3439703, EPI_ISL_3439704 | Lighthouse Lab in Milton Keynes | Wellcome Sanger Institute for the COVID-19 Genomics UK (COG-UK) Consortium | The Lighthouse Lab in Milton Keynes and Alex Alderton, Roberto Amato, Jeffrey Barrett, Sonia Goncalves, Ewan Harrison, David K. Jackson, Ian Johnston, Dominic Kwiatkowski, Cordelia Langford, John Sillitoe on behalf of the Wellcome Sanger Institute COVID-19 Surveillance Team |
| EPI_ISL_3439705, EPI_ISL_3439706 | Lighthouse Lab in Alderley Park | Wellcome Sanger Institute for the COVID-19 Genomics UK (COG-UK) Consortium | Jacquelyn Wynn, Mairead Hyland, The Lighthouse Lab in Alderley Park and Alex Alderton, Roberto Amato, Jeffrey Barrett, Sonia Goncalves, Ewan Harrison, David K. Jackson, Ian Johnston, Dominic Kwiatkowski, Cordelia Langford, John Sillitoe on behalf of the Wellcome Sanger Institute COVID-19 Surveillance Team |
| EPI_ISL_3439707, EPI_ISL_3439708 | Lighthouse Lab in Milton Keynes | Wellcome Sanger Institute for the COVID-19 Genomics UK (COG-UK) Consortium | The Lighthouse Lab in Milton Keynes and Alex Alderton, Roberto Amato, Jeffrey Barrett, Sonia Goncalves, Ewan Harrison, David K. Jackson, Ian Johnston, Dominic Kwiatkowski, Cordelia Langford, John Sillitoe on behalf of the Wellcome Sanger Institute COVID-19 Surveillance Team |
| EPI_ISL_3439709 | Lighthouse Lab in Alderley Park | Wellcome Sanger Institute for the COVID-19 Genomics UK (COG-UK) Consortium | Jacquelyn Wynn, Mairead Hyland, The Lighthouse Lab in Alderley Park and Alex Alderton, Roberto Amato, Jeffrey Barrett, Sonia Goncalves, Ewan Harrison, David K. Jackson, Ian Johnston, Dominic Kwiatkowski, Cordelia Langford, John Sillitoe on behalf of the Wellcome Sanger Institute COVID-19 Surveillance Team |
| EPI_ISL_3439710, EPI_ISL_3439711, EPI_ISL_3439712, EPI_ISL_3439713, EPI_ISL_3439714 | Lighthouse Lab in Milton Keynes | Wellcome Sanger Institute for the COVID-19 Genomics UK (COG-UK) Consortium | The Lighthouse Lab in Milton Keynes and Alex Alderton, Roberto Amato, Jeffrey Barrett, Sonia Goncalves, Ewan Harrison, David K. Jackson, Ian Johnston, Dominic Kwiatkowski, Cordelia Langford, John Sillitoe on behalf of the Wellcome Sanger Institute COVID-19 Surveillance Team |
| EPI_ISL_3439715 | Lighthouse Lab in Alderley Park | Wellcome Sanger Institute for the COVID-19 Genomics UK (COG-UK) Consortium | Jacquelyn Wynn, Mairead Hyland, The Lighthouse Lab in Alderley Park and Alex Alderton, Roberto Amato, Jeffrey Barrett, Sonia Goncalves, Ewan Harrison, David K. Jackson, Ian Johnston, Dominic Kwiatkowski, Cordelia Langford, John Sillitoe on behalf of the Wellcome Sanger Institute COVID-19 Surveillance Team |
| EPI_ISL_3439716, EPI_ISL_3439717, EPI_ISL_3439718, EPI_ISL_3439719, EPI_ISL_3439720, EPI_ISL_3439721, EPI_ISL_3439722 | Lighthouse Lab in Milton Keynes | Wellcome Sanger Institute for the COVID-19 Genomics UK (COG-UK) Consortium | The Lighthouse Lab in Milton Keynes and Alex Alderton, Roberto Amato, Jeffrey Barrett, Sonia Goncalves, Ewan Harrison, David K. Jackson, Ian Johnston, Dominic Kwiatkowski, Cordelia Langford, John Sillitoe on behalf of the Wellcome Sanger Institute COVID-19 Surveillance Team |
| EPI_ISL_3439723, EPI_ISL_3439724, EPI_ISL_3439725 | Lighthouse Lab in Alderley Park | Wellcome Sanger Institute for the COVID-19 Genomics UK (COG-UK) Consortium | Jacquelyn Wynn, Mairead Hyland, The Lighthouse Lab in Alderley Park and Alex Alderton, Roberto Amato, Jeffrey Barrett, Sonia Goncalves, Ewan Harrison, David K. Jackson, Ian Johnston, Dominic Kwiatkowski, Cordelia Langford, John Sillitoe on behalf of the Wellcome Sanger Institute COVID-19 Surveillance Team |
| EPI_ISL_3439726, EPI_ISL_3439727, EPI_ISL_3439728, EPI_ISL_3439729, EPI_ISL_3439730, EPI_ISL_3439731, EPI_ISL_3439732, EPI_ISL_3439733, EPI_ISL_3439734, EPI_ISL_3439735, EPI_ISL_3439736, EPI_ISL_3439737, EPI_ISL_3439738 | | | |
| see above | Lighthouse Lab in Milton Keynes | Wellcome Sanger Institute for the COVID-19 Genomics UK (COG-UK) Consortium | The Lighthouse Lab in Milton Keynes and Alex Alderton, Roberto Amato, Jeffrey Barrett, Sonia Goncalves, Ewan Harrison, David K. Jackson, Ian Johnston, Dominic Kwiatkowski, Cordelia Langford, John Sillitoe on behalf of the Wellcome Sanger Institute COVID-19 Surveillance Team |
| EPI_ISL_3439739, EPI_ISL_3439740 | Lighthouse Lab in Alderley Park | Wellcome Sanger Institute for the COVID-19 Genomics UK (COG-UK) Consortium | Jacquelyn Wynn, Mairead Hyland, The Lighthouse Lab in Alderley Park and Alex Alderton, Roberto Amato, Jeffrey Barrett, Sonia Goncalves, Ewan Harrison, David K. Jackson, Ian Johnston, Dominic Kwiatkowski, Cordelia Langford, John Sillitoe on behalf of the Wellcome Sanger Institute COVID-19 Surveillance Team |
| EPI_ISL_3439741, EPI_ISL_3439742, EPI_ISL_3439743, EPI_ISL_3439744, | Lighthouse Lab in Milton Keynes | Wellcome Sanger Institute for the COVID-19 Genomics UK (COG-UK) Consortium | The Lighthouse Lab in Milton Keynes and Alex Alderton, Roberto Amato, Jeffrey Barrett, Sonia Goncalves, Ewan Harrison, David K. Jackson, Ian Johnston, Dominic Kwiatkowski, Cordelia Langford, John Sillitoe on behalf of the Wellcome Sanger Institute COVID-19 Surveillance Team |

| | | | |
|---|---|---|---|
| EPI_ISL_3439745, EPI_ISL_3439746, EPI_ISL_3439747 | | | |
| EPI_ISL_3439748 | Lighthouse Lab in Alderley Park | Wellcome Sanger Institute for the COVID-19 Genomics UK (COG-UK) Consortium | Jacquelyn Wynn, Mairead Hyland, The Lighthouse Lab in Alderley Park and Alex Alderton, Roberto Amato, Jeffrey Barrett, Sonia Goncalves, Ewan Harrison, David K. Jackson, Ian Johnston, Dominic Kwiatkowski, Cordelia Langford, John Sillitoe on behalf of the Wellcome Sanger Institute COVID-19 Surveillance Team |
| EPI_ISL_3439749, EPI_ISL_3439750, EPI_ISL_3439751, EPI_ISL_3439752, EPI_ISL_3439753, EPI_ISL_3439754 | Lighthouse Lab in Milton Keynes | Wellcome Sanger Institute for the COVID-19 Genomics UK (COG-UK) Consortium | The Lighthouse Lab in Milton Keynes and Alex Alderton, Roberto Amato, Jeffrey Barrett, Sonia Goncalves, Ewan Harrison, David K. Jackson, Ian Johnston, Dominic Kwiatkowski, Cordelia Langford, John Sillitoe on behalf of the Wellcome Sanger Institute COVID-19 Surveillance Team |
| EPI_ISL_3439755 | Lighthouse Lab in Alderley Park | Wellcome Sanger Institute for the COVID-19 Genomics UK (COG-UK) Consortium | Jacquelyn Wynn, Mairead Hyland, The Lighthouse Lab in Alderley Park and Alex Alderton, Roberto Amato, Jeffrey Barrett, Sonia Goncalves, Ewan Harrison, David K. Jackson, Ian Johnston, Dominic Kwiatkowski, Cordelia Langford, John Sillitoe on behalf of the Wellcome Sanger Institute COVID-19 Surveillance Team |
| EPI_ISL_3439756 | Lighthouse Lab in Milton Keynes | Wellcome Sanger Institute for the COVID-19 Genomics UK (COG-UK) Consortium | The Lighthouse Lab in Milton Keynes and Alex Alderton, Roberto Amato, Jeffrey Barrett, Sonia Goncalves, Ewan Harrison, David K. Jackson, Ian Johnston, Dominic Kwiatkowski, Cordelia Langford, John Sillitoe on behalf of the Wellcome Sanger Institute COVID-19 Surveillance Team |
| EPI_ISL_3439757, EPI_ISL_3439758, EPI_ISL_3439759 | Lighthouse Lab in Alderley Park | Wellcome Sanger Institute for the COVID-19 Genomics UK (COG-UK) Consortium | Jacquelyn Wynn, Mairead Hyland, The Lighthouse Lab in Alderley Park and Alex Alderton, Roberto Amato, Jeffrey Barrett, Sonia Goncalves, Ewan Harrison, David K. Jackson, Ian Johnston, Dominic Kwiatkowski, Cordelia Langford, John Sillitoe on behalf of the Wellcome Sanger Institute COVID-19 Surveillance Team |
| EPI_ISL_3439760, EPI_ISL_3439761, EPI_ISL_3439762, EPI_ISL_3439763, EPI_ISL_3439764, EPI_ISL_3439765, EPI_ISL_3439766, EPI_ISL_3439767, EPI_ISL_3439768, EPI_ISL_3439769, EPI_ISL_3439770, EPI_ISL_3439771, EPI_ISL_3439772, EPI_ISL_3439773, EPI_ISL_3439774 | | | |
| see above | Lighthouse Lab in Milton Keynes | Wellcome Sanger Institute for the COVID-19 Genomics UK (COG-UK) Consortium | The Lighthouse Lab in Milton Keynes and Alex Alderton, Roberto Amato, Jeffrey Barrett, Sonia Goncalves, Ewan Harrison, David K. Jackson, Ian Johnston, Dominic Kwiatkowski, Cordelia Langford, John Sillitoe on behalf of the Wellcome Sanger Institute COVID-19 Surveillance Team |
| EPI_ISL_3439775 | Lighthouse Lab in Alderley Park | Wellcome Sanger Institute for the COVID-19 Genomics UK (COG-UK) Consortium | Jacquelyn Wynn, Mairead Hyland, The Lighthouse Lab in Alderley Park and Alex Alderton, Roberto Amato, Jeffrey Barrett, Sonia Goncalves, Ewan Harrison, David K. Jackson, Ian Johnston, Dominic Kwiatkowski, Cordelia Langford, John Sillitoe on behalf of the Wellcome Sanger Institute COVID-19 Surveillance Team |
| EPI_ISL_3439776, EPI_ISL_3439777, EPI_ISL_3439778, EPI_ISL_3439779, EPI_ISL_3439780 | Lighthouse Lab in Milton Keynes | Wellcome Sanger Institute for the COVID-19 Genomics UK (COG-UK) Consortium | The Lighthouse Lab in Milton Keynes and Alex Alderton, Roberto Amato, Jeffrey Barrett, Sonia Goncalves, Ewan Harrison, David K. Jackson, Ian Johnston, Dominic Kwiatkowski, Cordelia Langford, John Sillitoe on behalf of the Wellcome Sanger Institute COVID-19 Surveillance Team |
| EPI_ISL_3439781, EPI_ISL_3439782 | Lighthouse Lab in Alderley Park | Wellcome Sanger Institute for the COVID-19 Genomics UK (COG-UK) Consortium | Jacquelyn Wynn, Mairead Hyland, The Lighthouse Lab in Alderley Park and Alex Alderton, Roberto Amato, Jeffrey Barrett, Sonia Goncalves, Ewan Harrison, David K. Jackson, Ian Johnston, Dominic Kwiatkowski, Cordelia Langford, John Sillitoe on behalf of the Wellcome Sanger Institute COVID-19 Surveillance Team |
| EPI_ISL_3439783, EPI_ISL_3439784, EPI_ISL_3439785, EPI_ISL_3439786 | Lighthouse Lab in Milton Keynes | Wellcome Sanger Institute for the COVID-19 Genomics UK (COG-UK) Consortium | The Lighthouse Lab in Milton Keynes and Alex Alderton, Roberto Amato, Jeffrey Barrett, Sonia Goncalves, Ewan Harrison, David K. Jackson, Ian Johnston, Dominic Kwiatkowski, Cordelia Langford, John Sillitoe on behalf of the Wellcome Sanger Institute COVID-19 Surveillance Team |
| EPI_ISL_3439787, EPI_ISL_3439788 | Lighthouse Lab in Alderley Park | Wellcome Sanger Institute for the COVID-19 Genomics UK (COG-UK) Consortium | Jacquelyn Wynn, Mairead Hyland, The Lighthouse Lab in Alderley Park and Alex Alderton, Roberto Amato, Jeffrey Barrett, Sonia Goncalves, Ewan Harrison, David K. Jackson, Ian Johnston, Dominic Kwiatkowski, Cordelia Langford, John Sillitoe on behalf of the Wellcome Sanger Institute COVID-19 Surveillance Team |
| EPI_ISL_3439789, EPI_ISL_3439790, EPI_ISL_3439791 | Lighthouse Lab in Milton Keynes | Wellcome Sanger Institute for the COVID-19 Genomics UK (COG-UK) Consortium | The Lighthouse Lab in Milton Keynes and Alex Alderton, Roberto Amato, Jeffrey Barrett, Sonia Goncalves, Ewan Harrison, David K. Jackson, Ian Johnston, Dominic Kwiatkowski, Cordelia Langford, John Sillitoe on behalf of the Wellcome Sanger Institute COVID-19 Surveillance Team |
| EPI_ISL_3439792 | Lighthouse Lab in Alderley Park | Wellcome Sanger Institute for the COVID-19 Genomics UK (COG-UK) Consortium | Jacquelyn Wynn, Mairead Hyland, The Lighthouse Lab in Alderley Park and Alex Alderton, Roberto Amato, Jeffrey Barrett, Sonia Goncalves, Ewan Harrison, David K. Jackson, Ian Johnston, Dominic Kwiatkowski, Cordelia Langford, John Sillitoe on behalf of the Wellcome Sanger Institute COVID-19 Surveillance Team |
| EPI_ISL_3439793, EPI_ISL_3439794, EPI_ISL_3439795 | Lighthouse Lab in Milton Keynes | Wellcome Sanger Institute for the COVID-19 Genomics UK (COG-UK) Consortium | The Lighthouse Lab in Milton Keynes and Alex Alderton, Roberto Amato, Jeffrey Barrett, Sonia Goncalves, Ewan Harrison, David K. Jackson, Ian Johnston, Dominic Kwiatkowski, Cordelia Langford, John Sillitoe on behalf of the Wellcome Sanger Institute COVID-19 Surveillance Team |
| EPI_ISL_3439796 | Lighthouse Lab in Alderley Park | Wellcome Sanger Institute for the COVID-19 Genomics UK (COG-UK) Consortium | Jacquelyn Wynn, Mairead Hyland, The Lighthouse Lab in Alderley Park and Alex Alderton, Roberto Amato, Jeffrey Barrett, Sonia Goncalves, Ewan Harrison, David K. Jackson, Ian Johnston, Dominic Kwiatkowski, Cordelia Langford, John Sillitoe on behalf of the Wellcome Sanger Institute COVID-19 Surveillance Team |
| EPI_ISL_3439797 | Lighthouse Lab in Milton Keynes | Wellcome Sanger Institute for the COVID-19 Genomics UK (COG-UK) Consortium | The Lighthouse Lab in Milton Keynes and Alex Alderton, Roberto Amato, Jeffrey Barrett, Sonia Goncalves, Ewan Harrison, David K. Jackson, Ian Johnston, Dominic Kwiatkowski, Cordelia Langford, John Sillitoe on behalf of the Wellcome Sanger Institute COVID-19 Surveillance Team |
| EPI_ISL_3439798 | Lighthouse Lab in Alderley Park | Wellcome Sanger Institute for the COVID-19 Genomics UK (COG-UK) Consortium | Jacquelyn Wynn, Mairead Hyland, The Lighthouse Lab in Alderley Park and Alex Alderton, Roberto Amato, Jeffrey Barrett, Sonia Goncalves, Ewan Harrison, David K. Jackson, Ian Johnston, Dominic Kwiatkowski, Cordelia Langford, John Sillitoe on behalf of the Wellcome Sanger Institute COVID-19 Surveillance Team |
| EPI_ISL_3439799, EPI_ISL_3439800, EPI_ISL_3439801, EPI_ISL_3439802 | Lighthouse Lab in Milton Keynes | Wellcome Sanger Institute for the COVID-19 Genomics UK (COG-UK) Consortium | The Lighthouse Lab in Milton Keynes and Alex Alderton, Roberto Amato, Jeffrey Barrett, Sonia Goncalves, Ewan Harrison, David K. Jackson, Ian Johnston, Dominic Kwiatkowski, Cordelia Langford, John Sillitoe on behalf of the Wellcome Sanger Institute COVID-19 Surveillance Team |
| EPI_ISL_3439803 | Lighthouse Lab in Alderley Park | Wellcome Sanger Institute for the COVID-19 Genomics UK (COG-UK) Consortium | Jacquelyn Wynn, Mairead Hyland, The Lighthouse Lab in Alderley Park and Alex Alderton, Roberto Amato, Jeffrey Barrett, Sonia Goncalves, Ewan Harrison, David K. Jackson, Ian Johnston, Dominic Kwiatkowski, Cordelia Langford, John Sillitoe on behalf of the Wellcome Sanger Institute COVID-19 Surveillance Team |
| EPI_ISL_3439804, EPI_ISL_3439805, EPI_ISL_3439806 | Lighthouse Lab in Milton Keynes | Wellcome Sanger Institute for the COVID-19 Genomics UK (COG-UK) Consortium | The Lighthouse Lab in Milton Keynes and Alex Alderton, Roberto Amato, Jeffrey Barrett, Sonia Goncalves, Ewan Harrison, David K. Jackson, Ian Johnston, Dominic Kwiatkowski, Cordelia Langford, John Sillitoe on behalf of the Wellcome Sanger Institute COVID-19 Surveillance Team |
| EPI_ISL_3439807 | Lighthouse Lab in Alderley Park | Wellcome Sanger Institute for the COVID-19 Genomics UK (COG-UK) Consortium | Jacquelyn Wynn, Mairead Hyland, The Lighthouse Lab in Alderley Park and Alex Alderton, Roberto Amato, Jeffrey Barrett, Sonia Goncalves, Ewan Harrison, David K. Jackson, Ian Johnston, Dominic Kwiatkowski, Cordelia Langford, John Sillitoe on behalf of the Wellcome Sanger Institute COVID-19 Surveillance Team |
| EPI_ISL_3439808 | Lighthouse Lab in Milton Keynes | Wellcome Sanger Institute for the COVID-19 Genomics UK (COG-UK) Consortium | The Lighthouse Lab in Milton Keynes and Alex Alderton, Roberto Amato, Jeffrey Barrett, Sonia Goncalves, Ewan Harrison, David K. Jackson, Ian Johnston, Dominic Kwiatkowski, Cordelia Langford, John Sillitoe on behalf of the Wellcome Sanger Institute COVID-19 Surveillance Team |
| EPI_ISL_3439809, EPI_ISL_3439810, EPI_ISL_3439811, EPI_ISL_3439812 | Lighthouse Lab in Alderley Park | Wellcome Sanger Institute for the COVID-19 Genomics UK (COG-UK) Consortium | Jacquelyn Wynn, Mairead Hyland, The Lighthouse Lab in Alderley Park and Alex Alderton, Roberto Amato, Jeffrey Barrett, Sonia Goncalves, Ewan Harrison, David K. Jackson, Ian Johnston, Dominic Kwiatkowski, Cordelia Langford, John Sillitoe on behalf of the Wellcome Sanger Institute COVID-19 Surveillance Team |
| EPI_ISL_3439813, EPI_ISL_3439814, EPI_ISL_3439815, EPI_ISL_3439816, EPI_ISL_3439817, EPI_ISL_3439818, EPI_ISL_3439819 | Lighthouse Lab in Milton Keynes | Wellcome Sanger Institute for the COVID-19 Genomics UK (COG-UK) Consortium | The Lighthouse Lab in Milton Keynes and Alex Alderton, Roberto Amato, Jeffrey Barrett, Sonia Goncalves, Ewan Harrison, David K. Jackson, Ian Johnston, Dominic Kwiatkowski, Cordelia Langford, John Sillitoe on behalf of the Wellcome Sanger Institute COVID-19 Surveillance Team |
| EPI_ISL_3439820, EPI_ISL_3439821 | Lighthouse Lab in Alderley Park | Wellcome Sanger Institute for the COVID-19 Genomics UK (COG-UK) Consortium | Jacquelyn Wynn, Mairead Hyland, The Lighthouse Lab in Alderley Park and Alex Alderton, Roberto Amato, Jeffrey Barrett, Sonia Goncalves, Ewan Harrison, David K. Jackson, Ian Johnston, Dominic Kwiatkowski, Cordelia Langford, John Sillitoe on behalf of the Wellcome Sanger Institute COVID-19 Surveillance Team |
| EPI_ISL_3439822, EPI_ISL_3439823, EPI_ISL_3439824, EPI_ISL_3439825, EPI_ISL_3439826, EPI_ISL_3439827, EPI_ISL_3439828, EPI_ISL_3439829, EPI_ISL_3439830, EPI_ISL_3439831 | Lighthouse Lab in Milton Keynes | Wellcome Sanger Institute for the COVID-19 Genomics UK (COG-UK) Consortium | The Lighthouse Lab in Milton Keynes and Alex Alderton, Roberto Amato, Jeffrey Barrett, Sonia Goncalves, Ewan Harrison, David K. Jackson, Ian Johnston, Dominic Kwiatkowski, Cordelia Langford, John Sillitoe on behalf of the Wellcome Sanger Institute COVID-19 Surveillance Team |
| EPI_ISL_3439832 | Lighthouse Lab in Alderley Park | Wellcome Sanger Institute for the COVID-19 Genomics UK (COG-UK) Consortium | Jacquelyn Wynn, Mairead Hyland, The Lighthouse Lab in Alderley Park and Alex Alderton, Roberto Amato, Jeffrey Barrett, Sonia Goncalves, Ewan Harrison, David K. Jackson, Ian Johnston, Dominic Kwiatkowski, Cordelia Langford, John Sillitoe on behalf of the Wellcome Sanger Institute COVID-19 Surveillance Team |
| EPI_ISL_3439833, EPI_ISL_3439834 | Lighthouse Lab in Milton Keynes | Wellcome Sanger Institute for the COVID-19 Genomics UK (COG-UK) Consortium | The Lighthouse Lab in Milton Keynes and Alex Alderton, Roberto Amato, Jeffrey Barrett, Sonia Goncalves, Ewan Harrison, David K. Jackson, Ian Johnston, Dominic Kwiatkowski, Cordelia Langford, John Sillitoe on behalf of the Wellcome Sanger Institute COVID-19 Surveillance Team |
| EPI_ISL_3439835 | Lighthouse Lab in Alderley Park | Wellcome Sanger Institute for the COVID-19 Genomics UK (COG-UK) Consortium | Jacquelyn Wynn, Mairead Hyland, The Lighthouse Lab in Alderley Park and Alex Alderton, Roberto Amato, Jeffrey Barrett, Sonia Goncalves, Ewan Harrison, David K. Jackson, Ian Johnston, Dominic Kwiatkowski, Cordelia Langford, John Sillitoe on behalf of the Wellcome Sanger Institute COVID-19 Surveillance Team |
| EPI_ISL_3439836 | Lighthouse Lab in Milton Keynes | Wellcome Sanger Institute for the COVID-19 Genomics UK (COG-UK) Consortium | The Lighthouse Lab in Milton Keynes and Alex Alderton, Roberto Amato, Jeffrey Barrett, Sonia Goncalves, Ewan Harrison, David K. Jackson, Ian Johnston, Dominic Kwiatkowski, Cordelia Langford, John Sillitoe on behalf of the Wellcome Sanger Institute COVID-19 Surveillance Team |
| EPI_ISL_3439837 | Lighthouse Lab in Alderley Park | Wellcome Sanger Institute for the COVID-19 Genomics UK (COG-UK) Consortium | Jacquelyn Wynn, Mairead Hyland, The Lighthouse Lab in Alderley Park and Alex Alderton, Roberto Amato, Jeffrey Barrett, Sonia Goncalves, Ewan Harrison, David K. Jackson, Ian Johnston, Dominic Kwiatkowski, Cordelia Langford, John Sillitoe on behalf of the Wellcome Sanger Institute COVID-19 Surveillance Team |

| | | | |
|---|---|---|---|
| EPI_ISL_3439838, EPI_ISL_3439839, EPI_ISL_3439840, EPI_ISL_3439841, EPI_ISL_3439842, EPI_ISL_3439843, EPI_ISL_3439844 | Lighthouse Lab in Milton Keynes | Wellcome Sanger Institute for the COVID-19 Genomics UK (COG-UK) Consortium | The Lighthouse Lab in Milton Keynes and Alex Alderton, Roberto Amato, Jeffrey Barrett, Sonia Goncalves, Ewan Harrison, David K. Jackson, Ian Johnston, Dominic Kwiatkowski, Cordelia Langford, John Sillitoe on behalf of the Wellcome Sanger Institute COVID-19 Surveillance Team |
| EPI_ISL_3439845 | Lighthouse Lab in Alderley Park | Wellcome Sanger Institute for the COVID-19 Genomics UK (COG-UK) Consortium | Jacquelyn Wynn, Mairead Hyland, The Lighthouse Lab in Alderley Park and Alex Alderton, Roberto Amato, Jeffrey Barrett, Sonia Goncalves, Ewan Harrison, David K. Jackson, Ian Johnston, Dominic Kwiatkowski, Cordelia Langford, John Sillitoe on behalf of the Wellcome Sanger Institute COVID-19 Surveillance Team |
| EPI_ISL_3439846 | Lighthouse Lab in Milton Keynes | Wellcome Sanger Institute for the COVID-19 Genomics UK (COG-UK) Consortium | The Lighthouse Lab in Milton Keynes and Alex Alderton, Roberto Amato, Jeffrey Barrett, Sonia Goncalves, Ewan Harrison, David K. Jackson, Ian Johnston, Dominic Kwiatkowski, Cordelia Langford, John Sillitoe on behalf of the Wellcome Sanger Institute COVID-19 Surveillance Team |
| EPI_ISL_3439847, EPI_ISL_3439848 | Lighthouse Lab in Alderley Park | Wellcome Sanger Institute for the COVID-19 Genomics UK (COG-UK) Consortium | Jacquelyn Wynn, Mairead Hyland, The Lighthouse Lab in Alderley Park and Alex Alderton, Roberto Amato, Jeffrey Barrett, Sonia Goncalves, Ewan Harrison, David K. Jackson, Ian Johnston, Dominic Kwiatkowski, Cordelia Langford, John Sillitoe on behalf of the Wellcome Sanger Institute COVID-19 Surveillance Team |
| EPI_ISL_3439849 | Lighthouse Lab in Milton Keynes | Wellcome Sanger Institute for the COVID-19 Genomics UK (COG-UK) Consortium | The Lighthouse Lab in Milton Keynes and Alex Alderton, Roberto Amato, Jeffrey Barrett, Sonia Goncalves, Ewan Harrison, David K. Jackson, Ian Johnston, Dominic Kwiatkowski, Cordelia Langford, John Sillitoe on behalf of the Wellcome Sanger Institute COVID-19 Surveillance Team |
| EPI_ISL_3439850, EPI_ISL_3439851 | Lighthouse Lab in Alderley Park | Wellcome Sanger Institute for the COVID-19 Genomics UK (COG-UK) Consortium | Jacquelyn Wynn, Mairead Hyland, The Lighthouse Lab in Alderley Park and Alex Alderton, Roberto Amato, Jeffrey Barrett, Sonia Goncalves, Ewan Harrison, David K. Jackson, Ian Johnston, Dominic Kwiatkowski, Cordelia Langford, John Sillitoe on behalf of the Wellcome Sanger Institute COVID-19 Surveillance Team |
| EPI_ISL_3439852, EPI_ISL_3439853 | Lighthouse Lab in Milton Keynes | Wellcome Sanger Institute for the COVID-19 Genomics UK (COG-UK) Consortium | The Lighthouse Lab in Milton Keynes and Alex Alderton, Roberto Amato, Jeffrey Barrett, Sonia Goncalves, Ewan Harrison, David K. Jackson, Ian Johnston, Dominic Kwiatkowski, Cordelia Langford, John Sillitoe on behalf of the Wellcome Sanger Institute COVID-19 Surveillance Team |
| EPI_ISL_3439854, EPI_ISL_3439855 | Lighthouse Lab in Alderley Park | Wellcome Sanger Institute for the COVID-19 Genomics UK (COG-UK) Consortium | Jacquelyn Wynn, Mairead Hyland, The Lighthouse Lab in Alderley Park and Alex Alderton, Roberto Amato, Jeffrey Barrett, Sonia Goncalves, Ewan Harrison, David K. Jackson, Ian Johnston, Dominic Kwiatkowski, Cordelia Langford, John Sillitoe on behalf of the Wellcome Sanger Institute COVID-19 Surveillance Team |
| EPI_ISL_3439856 | Lighthouse Lab in Milton Keynes | Wellcome Sanger Institute for the COVID-19 Genomics UK (COG-UK) Consortium | The Lighthouse Lab in Milton Keynes and Alex Alderton, Roberto Amato, Jeffrey Barrett, Sonia Goncalves, Ewan Harrison, David K. Jackson, Ian Johnston, Dominic Kwiatkowski, Cordelia Langford, John Sillitoe on behalf of the Wellcome Sanger Institute COVID-19 Surveillance Team |
| EPI_ISL_3439857 | Lighthouse Lab in Alderley Park | Wellcome Sanger Institute for the COVID-19 Genomics UK (COG-UK) Consortium | Jacquelyn Wynn, Mairead Hyland, The Lighthouse Lab in Alderley Park and Alex Alderton, Roberto Amato, Jeffrey Barrett, Sonia Goncalves, Ewan Harrison, David K. Jackson, Ian Johnston, Dominic Kwiatkowski, Cordelia Langford, John Sillitoe on behalf of the Wellcome Sanger Institute COVID-19 Surveillance Team |
| EPI_ISL_3439858, EPI_ISL_3439859, EPI_ISL_3439860, EPI_ISL_3439861, EPI_ISL_3439862, EPI_ISL_3439863, EPI_ISL_3439864, EPI_ISL_3439865, EPI_ISL_3439866 | Lighthouse Lab in Milton Keynes | Wellcome Sanger Institute for the COVID-19 Genomics UK (COG-UK) Consortium | The Lighthouse Lab in Milton Keynes and Alex Alderton, Roberto Amato, Jeffrey Barrett, Sonia Goncalves, Ewan Harrison, David K. Jackson, Ian Johnston, Dominic Kwiatkowski, Cordelia Langford, John Sillitoe on behalf of the Wellcome Sanger Institute COVID-19 Surveillance Team |
| EPI_ISL_3439867, EPI_ISL_3439868, EPI_ISL_3439869 | Lighthouse Lab in Alderley Park | Wellcome Sanger Institute for the COVID-19 Genomics UK (COG-UK) Consortium | Jacquelyn Wynn, Mairead Hyland, The Lighthouse Lab in Alderley Park and Alex Alderton, Roberto Amato, Jeffrey Barrett, Sonia Goncalves, Ewan Harrison, David K. Jackson, Ian Johnston, Dominic Kwiatkowski, Cordelia Langford, John Sillitoe on behalf of the Wellcome Sanger Institute COVID-19 Surveillance Team |
| EPI_ISL_3439870, EPI_ISL_3439871, EPI_ISL_3439872 | Lighthouse Lab in Milton Keynes | Wellcome Sanger Institute for the COVID-19 Genomics UK (COG-UK) Consortium | The Lighthouse Lab in Milton Keynes and Alex Alderton, Roberto Amato, Jeffrey Barrett, Sonia Goncalves, Ewan Harrison, David K. Jackson, Ian Johnston, Dominic Kwiatkowski, Cordelia Langford, John Sillitoe on behalf of the Wellcome Sanger Institute COVID-19 Surveillance Team |
| EPI_ISL_3439873, EPI_ISL_3439874 | Lighthouse Lab in Alderley Park | Wellcome Sanger Institute for the COVID-19 Genomics UK (COG-UK) Consortium | Jacquelyn Wynn, Mairead Hyland, The Lighthouse Lab in Alderley Park and Alex Alderton, Roberto Amato, Jeffrey Barrett, Sonia Goncalves, Ewan Harrison, David K. Jackson, Ian Johnston, Dominic Kwiatkowski, Cordelia Langford, John Sillitoe on behalf of the Wellcome Sanger Institute COVID-19 Surveillance Team |
| EPI_ISL_3439875, EPI_ISL_3439876, EPI_ISL_3439877, EPI_ISL_3439878, EPI_ISL_3439879, EPI_ISL_3439880, EPI_ISL_3439881, EPI_ISL_3439882, EPI_ISL_3439883, EPI_ISL_3439884, EPI_ISL_3439885, EPI_ISL_3439886, EPI_ISL_3439887, EPI_ISL_3439888 | | | |
| see above | Lighthouse Lab in Milton Keynes | Wellcome Sanger Institute for the COVID-19 Genomics UK (COG-UK) Consortium | The Lighthouse Lab in Milton Keynes and Alex Alderton, Roberto Amato, Jeffrey Barrett, Sonia Goncalves, Ewan Harrison, David K. Jackson, Ian Johnston, Dominic Kwiatkowski, Cordelia Langford, John Sillitoe on behalf of the Wellcome Sanger Institute COVID-19 Surveillance Team |
| EPI_ISL_3439889 | Lighthouse Lab in Alderley Park | Wellcome Sanger Institute for the COVID-19 Genomics UK (COG-UK) Consortium | Jacquelyn Wynn, Mairead Hyland, The Lighthouse Lab in Alderley Park and Alex Alderton, Roberto Amato, Jeffrey Barrett, Sonia Goncalves, Ewan Harrison, David K. Jackson, Ian Johnston, Dominic Kwiatkowski, Cordelia Langford, John Sillitoe on behalf of the Wellcome Sanger Institute COVID-19 Surveillance Team |
| EPI_ISL_3439890, EPI_ISL_3439891 | Lighthouse Lab in Milton Keynes | Wellcome Sanger Institute for the COVID-19 Genomics UK (COG-UK) Consortium | The Lighthouse Lab in Milton Keynes and Alex Alderton, Roberto Amato, Jeffrey Barrett, Sonia Goncalves, Ewan Harrison, David K. Jackson, Ian Johnston, Dominic Kwiatkowski, Cordelia Langford, John Sillitoe on behalf of the Wellcome Sanger Institute COVID-19 Surveillance Team |
| EPI_ISL_3439892, EPI_ISL_3439893 | Lighthouse Lab in Alderley Park | Wellcome Sanger Institute for the COVID-19 Genomics UK (COG-UK) Consortium | Jacquelyn Wynn, Mairead Hyland, The Lighthouse Lab in Alderley Park and Alex Alderton, Roberto Amato, Jeffrey Barrett, Sonia Goncalves, Ewan Harrison, David K. Jackson, Ian Johnston, Dominic Kwiatkowski, Cordelia Langford, John Sillitoe on behalf of the Wellcome Sanger Institute COVID-19 Surveillance Team |
| EPI_ISL_3439894, EPI_ISL_3439895, EPI_ISL_3439896, EPI_ISL_3439897 | Lighthouse Lab in Milton Keynes | Wellcome Sanger Institute for the COVID-19 Genomics UK (COG-UK) Consortium | The Lighthouse Lab in Milton Keynes and Alex Alderton, Roberto Amato, Jeffrey Barrett, Sonia Goncalves, Ewan Harrison, David K. Jackson, Ian Johnston, Dominic Kwiatkowski, Cordelia Langford, John Sillitoe on behalf of the Wellcome Sanger Institute COVID-19 Surveillance Team |
| EPI_ISL_3439898 | Lighthouse Lab in Alderley Park | Wellcome Sanger Institute for the COVID-19 Genomics UK (COG-UK) Consortium | Jacquelyn Wynn, Mairead Hyland, The Lighthouse Lab in Alderley Park and Alex Alderton, Roberto Amato, Jeffrey Barrett, Sonia Goncalves, Ewan Harrison, David K. Jackson, Ian Johnston, Dominic Kwiatkowski, Cordelia Langford, John Sillitoe on behalf of the Wellcome Sanger Institute COVID-19 Surveillance Team |
| EPI_ISL_3439899, EPI_ISL_3439900, EPI_ISL_3439901, EPI_ISL_3439902 | Lighthouse Lab in Milton Keynes | Wellcome Sanger Institute for the COVID-19 Genomics UK (COG-UK) Consortium | The Lighthouse Lab in Milton Keynes and Alex Alderton, Roberto Amato, Jeffrey Barrett, Sonia Goncalves, Ewan Harrison, David K. Jackson, Ian Johnston, Dominic Kwiatkowski, Cordelia Langford, John Sillitoe on behalf of the Wellcome Sanger Institute COVID-19 Surveillance Team |
| EPI_ISL_3439903, EPI_ISL_3439904 | Lighthouse Lab in Alderley Park | Wellcome Sanger Institute for the COVID-19 Genomics UK (COG-UK) Consortium | Jacquelyn Wynn, Mairead Hyland, The Lighthouse Lab in Alderley Park and Alex Alderton, Roberto Amato, Jeffrey Barrett, Sonia Goncalves, Ewan Harrison, David K. Jackson, Ian Johnston, Dominic Kwiatkowski, Cordelia Langford, John Sillitoe on behalf of the Wellcome Sanger Institute COVID-19 Surveillance Team |
| EPI_ISL_3439905, EPI_ISL_3439906, EPI_ISL_3439907 | Lighthouse Lab in Milton Keynes | Wellcome Sanger Institute for the COVID-19 Genomics UK (COG-UK) Consortium | The Lighthouse Lab in Milton Keynes and Alex Alderton, Roberto Amato, Jeffrey Barrett, Sonia Goncalves, Ewan Harrison, David K. Jackson, Ian Johnston, Dominic Kwiatkowski, Cordelia Langford, John Sillitoe on behalf of the Wellcome Sanger Institute COVID-19 Surveillance Team |
| EPI_ISL_3439908 | Lighthouse Lab in Alderley Park | Wellcome Sanger Institute for the COVID-19 Genomics UK (COG-UK) Consortium | Jacquelyn Wynn, Mairead Hyland, The Lighthouse Lab in Alderley Park and Alex Alderton, Roberto Amato, Jeffrey Barrett, Sonia Goncalves, Ewan Harrison, David K. Jackson, Ian Johnston, Dominic Kwiatkowski, Cordelia Langford, John Sillitoe on behalf of the Wellcome Sanger Institute COVID-19 Surveillance Team |
| EPI_ISL_3439909, EPI_ISL_3439910, EPI_ISL_3439911, EPI_ISL_3439912, EPI_ISL_3439913, EPI_ISL_3439914, EPI_ISL_3439915, EPI_ISL_3439916, EPI_ISL_3439917 | Lighthouse Lab in Milton Keynes | Wellcome Sanger Institute for the COVID-19 Genomics UK (COG-UK) Consortium | The Lighthouse Lab in Milton Keynes and Alex Alderton, Roberto Amato, Jeffrey Barrett, Sonia Goncalves, Ewan Harrison, David K. Jackson, Ian Johnston, Dominic Kwiatkowski, Cordelia Langford, John Sillitoe on behalf of the Wellcome Sanger Institute COVID-19 Surveillance Team |
| EPI_ISL_3439918 | Lighthouse Lab in Alderley Park | Wellcome Sanger Institute for the COVID-19 Genomics UK (COG-UK) Consortium | Jacquelyn Wynn, Mairead Hyland, The Lighthouse Lab in Alderley Park and Alex Alderton, Roberto Amato, Jeffrey Barrett, Sonia Goncalves, Ewan Harrison, David K. Jackson, Ian Johnston, Dominic Kwiatkowski, Cordelia Langford, John Sillitoe on behalf of the Wellcome Sanger Institute COVID-19 Surveillance Team |
| EPI_ISL_3439919, EPI_ISL_3439920, EPI_ISL_3439921, EPI_ISL_3439922, EPI_ISL_3439923, EPI_ISL_3439924 | Lighthouse Lab in Milton Keynes | Wellcome Sanger Institute for the COVID-19 Genomics UK (COG-UK) Consortium | The Lighthouse Lab in Milton Keynes and Alex Alderton, Roberto Amato, Jeffrey Barrett, Sonia Goncalves, Ewan Harrison, David K. Jackson, Ian Johnston, Dominic Kwiatkowski, Cordelia Langford, John Sillitoe on behalf of the Wellcome Sanger Institute COVID-19 Surveillance Team |
| EPI_ISL_3439925 | Lighthouse Lab in Alderley Park | Wellcome Sanger Institute for the COVID-19 Genomics UK (COG-UK) Consortium | Jacquelyn Wynn, Mairead Hyland, The Lighthouse Lab in Alderley Park and Alex Alderton, Roberto Amato, Jeffrey Barrett, Sonia Goncalves, Ewan Harrison, David K. Jackson, Ian Johnston, Dominic Kwiatkowski, Cordelia Langford, John Sillitoe on behalf of the Wellcome Sanger Institute COVID-19 Surveillance Team |
| EPI_ISL_3439926, EPI_ISL_3439927, EPI_ISL_3439928, EPI_ISL_3439929, EPI_ISL_3439930, EPI_ISL_3439931, EPI_ISL_3439932, EPI_ISL_3439933, EPI_ISL_3439934 | Lighthouse Lab in Milton Keynes | Wellcome Sanger Institute for the COVID-19 Genomics UK (COG-UK) Consortium | The Lighthouse Lab in Milton Keynes and Alex Alderton, Roberto Amato, Jeffrey Barrett, Sonia Goncalves, Ewan Harrison, David K. Jackson, Ian Johnston, Dominic Kwiatkowski, Cordelia Langford, John Sillitoe on behalf of the Wellcome Sanger Institute COVID-19 Surveillance Team |
| EPI_ISL_3439935 | Lighthouse Lab in Alderley Park | Wellcome Sanger Institute for the COVID-19 Genomics UK (COG-UK) | Jacquelyn Wynn, Mairead Hyland, The Lighthouse Lab in Alderley Park and Alex Alderton, Roberto Amato, Jeffrey Barrett, Sonia Goncalves, Ewan Harrison, David K. Jackson, Ian Johnston, Dominic Kwiatkowski, Cordelia Langford, John Sillitoe on |

| | | | |
|---|---|---|---|
| | | Consortium | behalf of the Wellcome Sanger Institute COVID-19 Surveillance Team |
| EPI_ISL_3439936, EPI_ISL_3439937, EPI_ISL_3439938, EPI_ISL_3439939, EPI_ISL_3439940 | Lighthouse Lab in Milton Keynes | Wellcome Sanger Institute for the COVID-19 Genomics UK (COG-UK) Consortium | The Lighthouse Lab in Milton Keynes and Alex Alderton, Roberto Amato, Jeffrey Barrett, Sonia Goncalves, Ewan Harrison, David K. Jackson, Ian Johnston, Dominic Kwiatkowski, Cordelia Langford, John Sillitoe on behalf of the Wellcome Sanger Institute COVID-19 Surveillance Team |
| EPI_ISL_3439941, EPI_ISL_3439942 | Lighthouse Lab in Alderley Park | Wellcome Sanger Institute for the COVID-19 Genomics UK (COG-UK) Consortium | Jacquelyn Wynn, Mairead Hyland, The Lighthouse Lab in Alderley Park and Alex Alderton, Roberto Amato, Jeffrey Barrett, Sonia Goncalves, Ewan Harrison, David K. Jackson, Ian Johnston, Dominic Kwiatkowski, Cordelia Langford, John Sillitoe on behalf of the Wellcome Sanger Institute COVID-19 Surveillance Team |
| EPI_ISL_3439943, EPI_ISL_3439944, EPI_ISL_3439945 | Lighthouse Lab in Milton Keynes | Wellcome Sanger Institute for the COVID-19 Genomics UK (COG-UK) Consortium | The Lighthouse Lab in Milton Keynes and Alex Alderton, Roberto Amato, Jeffrey Barrett, Sonia Goncalves, Ewan Harrison, David K. Jackson, Ian Johnston, Dominic Kwiatkowski, Cordelia Langford, John Sillitoe on behalf of the Wellcome Sanger Institute COVID-19 Surveillance Team |
| EPI_ISL_3439946 | Lighthouse Lab in Alderley Park | Wellcome Sanger Institute for the COVID-19 Genomics UK (COG-UK) Consortium | Jacquelyn Wynn, Mairead Hyland, The Lighthouse Lab in Alderley Park and Alex Alderton, Roberto Amato, Jeffrey Barrett, Sonia Goncalves, Ewan Harrison, David K. Jackson, Ian Johnston, Dominic Kwiatkowski, Cordelia Langford, John Sillitoe on behalf of the Wellcome Sanger Institute COVID-19 Surveillance Team |
| EPI_ISL_3439947, EPI_ISL_3439948, EPI_ISL_3439949, EPI_ISL_3439950 | Lighthouse Lab in Milton Keynes | Wellcome Sanger Institute for the COVID-19 Genomics UK (COG-UK) Consortium | The Lighthouse Lab in Milton Keynes and Alex Alderton, Roberto Amato, Jeffrey Barrett, Sonia Goncalves, Ewan Harrison, David K. Jackson, Ian Johnston, Dominic Kwiatkowski, Cordelia Langford, John Sillitoe on behalf of the Wellcome Sanger Institute COVID-19 Surveillance Team |
| EPI_ISL_3439951 | Lighthouse Lab in Alderley Park | Wellcome Sanger Institute for the COVID-19 Genomics UK (COG-UK) Consortium | Jacquelyn Wynn, Mairead Hyland, The Lighthouse Lab in Alderley Park and Alex Alderton, Roberto Amato, Jeffrey Barrett, Sonia Goncalves, Ewan Harrison, David K. Jackson, Ian Johnston, Dominic Kwiatkowski, Cordelia Langford, John Sillitoe on behalf of the Wellcome Sanger Institute COVID-19 Surveillance Team |
| EPI_ISL_3439952 | Lighthouse Lab in Milton Keynes | Wellcome Sanger Institute for the COVID-19 Genomics UK (COG-UK) Consortium | The Lighthouse Lab in Milton Keynes and Alex Alderton, Roberto Amato, Jeffrey Barrett, Sonia Goncalves, Ewan Harrison, David K. Jackson, Ian Johnston, Dominic Kwiatkowski, Cordelia Langford, John Sillitoe on behalf of the Wellcome Sanger Institute COVID-19 Surveillance Team |
| EPI_ISL_3439953 | Lighthouse Lab in Alderley Park | Wellcome Sanger Institute for the COVID-19 Genomics UK (COG-UK) Consortium | Jacquelyn Wynn, Mairead Hyland, The Lighthouse Lab in Alderley Park and Alex Alderton, Roberto Amato, Jeffrey Barrett, Sonia Goncalves, Ewan Harrison, David K. Jackson, Ian Johnston, Dominic Kwiatkowski, Cordelia Langford, John Sillitoe on behalf of the Wellcome Sanger Institute COVID-19 Surveillance Team |
| EPI_ISL_3439954, EPI_ISL_3439955, EPI_ISL_3439956, EPI_ISL_3439957, EPI_ISL_3439958, EPI_ISL_3439959, EPI_ISL_3439960 | Lighthouse Lab in Milton Keynes | Wellcome Sanger Institute for the COVID-19 Genomics UK (COG-UK) Consortium | The Lighthouse Lab in Milton Keynes and Alex Alderton, Roberto Amato, Jeffrey Barrett, Sonia Goncalves, Ewan Harrison, David K. Jackson, Ian Johnston, Dominic Kwiatkowski, Cordelia Langford, John Sillitoe on behalf of the Wellcome Sanger Institute COVID-19 Surveillance Team |
| EPI_ISL_3439961 | Lighthouse Lab in Alderley Park | Wellcome Sanger Institute for the COVID-19 Genomics UK (COG-UK) Consortium | Jacquelyn Wynn, Mairead Hyland, The Lighthouse Lab in Alderley Park and Alex Alderton, Roberto Amato, Jeffrey Barrett, Sonia Goncalves, Ewan Harrison, David K. Jackson, Ian Johnston, Dominic Kwiatkowski, Cordelia Langford, John Sillitoe on behalf of the Wellcome Sanger Institute COVID-19 Surveillance Team |
| EPI_ISL_3439962 | Lighthouse Lab in Milton Keynes | Wellcome Sanger Institute for the COVID-19 Genomics UK (COG-UK) Consortium | The Lighthouse Lab in Milton Keynes and Alex Alderton, Roberto Amato, Jeffrey Barrett, Sonia Goncalves, Ewan Harrison, David K. Jackson, Ian Johnston, Dominic Kwiatkowski, Cordelia Langford, John Sillitoe on behalf of the Wellcome Sanger Institute COVID-19 Surveillance Team |
| EPI_ISL_3439963 | Lighthouse Lab in Alderley Park | Wellcome Sanger Institute for the COVID-19 Genomics UK (COG-UK) Consortium | Jacquelyn Wynn, Mairead Hyland, The Lighthouse Lab in Alderley Park and Alex Alderton, Roberto Amato, Jeffrey Barrett, Sonia Goncalves, Ewan Harrison, David K. Jackson, Ian Johnston, Dominic Kwiatkowski, Cordelia Langford, John Sillitoe on behalf of the Wellcome Sanger Institute COVID-19 Surveillance Team |
| EPI_ISL_3439964, EPI_ISL_3439965 | Lighthouse Lab in Milton Keynes | Wellcome Sanger Institute for the COVID-19 Genomics UK (COG-UK) Consortium | The Lighthouse Lab in Milton Keynes and Alex Alderton, Roberto Amato, Jeffrey Barrett, Sonia Goncalves, Ewan Harrison, David K. Jackson, Ian Johnston, Dominic Kwiatkowski, Cordelia Langford, John Sillitoe on behalf of the Wellcome Sanger Institute COVID-19 Surveillance Team |
| EPI_ISL_3439966 | Lighthouse Lab in Alderley Park | Wellcome Sanger Institute for the COVID-19 Genomics UK (COG-UK) Consortium | Jacquelyn Wynn, Mairead Hyland, The Lighthouse Lab in Alderley Park and Alex Alderton, Roberto Amato, Jeffrey Barrett, Sonia Goncalves, Ewan Harrison, David K. Jackson, Ian Johnston, Dominic Kwiatkowski, Cordelia Langford, John Sillitoe on behalf of the Wellcome Sanger Institute COVID-19 Surveillance Team |
| EPI_ISL_3439967, EPI_ISL_3439968 | Lighthouse Lab in Milton Keynes | Wellcome Sanger Institute for the COVID-19 Genomics UK (COG-UK) Consortium | The Lighthouse Lab in Milton Keynes and Alex Alderton, Roberto Amato, Jeffrey Barrett, Sonia Goncalves, Ewan Harrison, David K. Jackson, Ian Johnston, Dominic Kwiatkowski, Cordelia Langford, John Sillitoe on behalf of the Wellcome Sanger Institute COVID-19 Surveillance Team |
| EPI_ISL_3439969 | Lighthouse Lab in Alderley Park | Wellcome Sanger Institute for the COVID-19 Genomics UK (COG-UK) Consortium | Jacquelyn Wynn, Mairead Hyland, The Lighthouse Lab in Alderley Park and Alex Alderton, Roberto Amato, Jeffrey Barrett, Sonia Goncalves, Ewan Harrison, David K. Jackson, Ian Johnston, Dominic Kwiatkowski, Cordelia Langford, John Sillitoe on behalf of the Wellcome Sanger Institute COVID-19 Surveillance Team |
| EPI_ISL_3439970, EPI_ISL_3439971 | Lighthouse Lab in Milton Keynes | Wellcome Sanger Institute for the COVID-19 Genomics UK (COG-UK) Consortium | The Lighthouse Lab in Milton Keynes and Alex Alderton, Roberto Amato, Jeffrey Barrett, Sonia Goncalves, Ewan Harrison, David K. Jackson, Ian Johnston, Dominic Kwiatkowski, Cordelia Langford, John Sillitoe on behalf of the Wellcome Sanger Institute COVID-19 Surveillance Team |
| EPI_ISL_3439972, EPI_ISL_3439973, EPI_ISL_3439974 | Lighthouse Lab in Alderley Park | Wellcome Sanger Institute for the COVID-19 Genomics UK (COG-UK) Consortium | Jacquelyn Wynn, Mairead Hyland, The Lighthouse Lab in Alderley Park and Alex Alderton, Roberto Amato, Jeffrey Barrett, Sonia Goncalves, Ewan Harrison, David K. Jackson, Ian Johnston, Dominic Kwiatkowski, Cordelia Langford, John Sillitoe on behalf of the Wellcome Sanger Institute COVID-19 Surveillance Team |
| EPI_ISL_3439975, EPI_ISL_3439976, EPI_ISL_3439977, EPI_ISL_3439978 | Lighthouse Lab in Milton Keynes | Wellcome Sanger Institute for the COVID-19 Genomics UK (COG-UK) Consortium | The Lighthouse Lab in Milton Keynes and Alex Alderton, Roberto Amato, Jeffrey Barrett, Sonia Goncalves, Ewan Harrison, David K. Jackson, Ian Johnston, Dominic Kwiatkowski, Cordelia Langford, John Sillitoe on behalf of the Wellcome Sanger Institute COVID-19 Surveillance Team |
| EPI_ISL_3439979 | Lighthouse Lab in Alderley Park | Wellcome Sanger Institute for the COVID-19 Genomics UK (COG-UK) Consortium | Jacquelyn Wynn, Mairead Hyland, The Lighthouse Lab in Alderley Park and Alex Alderton, Roberto Amato, Jeffrey Barrett, Sonia Goncalves, Ewan Harrison, David K. Jackson, Ian Johnston, Dominic Kwiatkowski, Cordelia Langford, John Sillitoe on behalf of the Wellcome Sanger Institute COVID-19 Surveillance Team |
| EPI_ISL_3439980, EPI_ISL_3439981, EPI_ISL_3439982, EPI_ISL_3439983, EPI_ISL_3439984, EPI_ISL_3439985, EPI_ISL_3439986 | Lighthouse Lab in Milton Keynes | Wellcome Sanger Institute for the COVID-19 Genomics UK (COG-UK) Consortium | The Lighthouse Lab in Milton Keynes and Alex Alderton, Roberto Amato, Jeffrey Barrett, Sonia Goncalves, Ewan Harrison, David K. Jackson, Ian Johnston, Dominic Kwiatkowski, Cordelia Langford, John Sillitoe on behalf of the Wellcome Sanger Institute COVID-19 Surveillance Team |
| EPI_ISL_3439987, EPI_ISL_3439988, EPI_ISL_3439989 | Lighthouse Lab in Alderley Park | Wellcome Sanger Institute for the COVID-19 Genomics UK (COG-UK) Consortium | Jacquelyn Wynn, Mairead Hyland, The Lighthouse Lab in Alderley Park and Alex Alderton, Roberto Amato, Jeffrey Barrett, Sonia Goncalves, Ewan Harrison, David K. Jackson, Ian Johnston, Dominic Kwiatkowski, Cordelia Langford, John Sillitoe on behalf of the Wellcome Sanger Institute COVID-19 Surveillance Team |
| EPI_ISL_3439990, EPI_ISL_3439991, EPI_ISL_3439992 | Lighthouse Lab in Milton Keynes | Wellcome Sanger Institute for the COVID-19 Genomics UK (COG-UK) Consortium | The Lighthouse Lab in Milton Keynes and Alex Alderton, Roberto Amato, Jeffrey Barrett, Sonia Goncalves, Ewan Harrison, David K. Jackson, Ian Johnston, Dominic Kwiatkowski, Cordelia Langford, John Sillitoe on behalf of the Wellcome Sanger Institute COVID-19 Surveillance Team |
| EPI_ISL_3439993, EPI_ISL_3439994, EPI_ISL_3439995 | Lighthouse Lab in Alderley Park | Wellcome Sanger Institute for the COVID-19 Genomics UK (COG-UK) Consortium | Jacquelyn Wynn, Mairead Hyland, The Lighthouse Lab in Alderley Park and Alex Alderton, Roberto Amato, Jeffrey Barrett, Sonia Goncalves, Ewan Harrison, David K. Jackson, Ian Johnston, Dominic Kwiatkowski, Cordelia Langford, John Sillitoe on behalf of the Wellcome Sanger Institute COVID-19 Surveillance Team |
| EPI_ISL_3439996, EPI_ISL_3439997, EPI_ISL_3439998 | Lighthouse Lab in Milton Keynes | Wellcome Sanger Institute for the COVID-19 Genomics UK (COG-UK) Consortium | The Lighthouse Lab in Milton Keynes and Alex Alderton, Roberto Amato, Jeffrey Barrett, Sonia Goncalves, Ewan Harrison, David K. Jackson, Ian Johnston, Dominic Kwiatkowski, Cordelia Langford, John Sillitoe on behalf of the Wellcome Sanger Institute COVID-19 Surveillance Team |
| EPI_ISL_3439999 | Lighthouse Lab in Alderley Park | Wellcome Sanger Institute for the COVID-19 Genomics UK (COG-UK) Consortium | Jacquelyn Wynn, Mairead Hyland, The Lighthouse Lab in Alderley Park and Alex Alderton, Roberto Amato, Jeffrey Barrett, Sonia Goncalves, Ewan Harrison, David K. Jackson, Ian Johnston, Dominic Kwiatkowski, Cordelia Langford, John Sillitoe on behalf of the Wellcome Sanger Institute COVID-19 Surveillance Team |
| EPI_ISL_3440000, EPI_ISL_3440001, EPI_ISL_3440002, EPI_ISL_3440004, EPI_ISL_3440005, EPI_ISL_3440006, EPI_ISL_3440007, EPI_ISL_3440008, EPI_ISL_3440009, EPI_ISL_3440010, EPI_ISL_3440011 | | | |
| see above | Lighthouse Lab in Milton Keynes | Wellcome Sanger Institute for the COVID-19 Genomics UK (COG-UK) Consortium | The Lighthouse Lab in Milton Keynes and Alex Alderton, Roberto Amato, Jeffrey Barrett, Sonia Goncalves, Ewan Harrison, David K. Jackson, Ian Johnston, Dominic Kwiatkowski, Cordelia Langford, John Sillitoe on behalf of the Wellcome Sanger Institute COVID-19 Surveillance Team |
| EPI_ISL_3440012, EPI_ISL_3440013, EPI_ISL_3440014 | Lighthouse Lab in Alderley Park | Wellcome Sanger Institute for the COVID-19 Genomics UK (COG-UK) Consortium | Jacquelyn Wynn, Mairead Hyland, The Lighthouse Lab in Alderley Park and Alex Alderton, Roberto Amato, Jeffrey Barrett, Sonia Goncalves, Ewan Harrison, David K. Jackson, Ian Johnston, Dominic Kwiatkowski, Cordelia Langford, John Sillitoe on behalf of the Wellcome Sanger Institute COVID-19 Surveillance Team |
| EPI_ISL_3440015, EPI_ISL_3440016 | Lighthouse Lab in Milton Keynes | Wellcome Sanger Institute for the COVID-19 Genomics UK (COG-UK) Consortium | The Lighthouse Lab in Milton Keynes and Alex Alderton, Roberto Amato, Jeffrey Barrett, Sonia Goncalves, Ewan Harrison, David K. Jackson, Ian Johnston, Dominic Kwiatkowski, Cordelia Langford, John Sillitoe on behalf of the Wellcome Sanger Institute COVID-19 Surveillance Team |
| EPI_ISL_3440017 | Lighthouse Lab in Alderley Park | Wellcome Sanger Institute for the COVID-19 Genomics UK (COG-UK) Consortium | Jacquelyn Wynn, Mairead Hyland, The Lighthouse Lab in Alderley Park and Alex Alderton, Roberto Amato, Jeffrey Barrett, Sonia Goncalves, Ewan Harrison, David K. Jackson, Ian Johnston, Dominic Kwiatkowski, Cordelia Langford, John Sillitoe on behalf of the Wellcome Sanger Institute COVID-19 Surveillance Team |
| EPI_ISL_3440018 | Lighthouse Lab in Milton Keynes | Wellcome Sanger Institute for the COVID-19 Genomics UK (COG-UK) Consortium | The Lighthouse Lab in Milton Keynes and Alex Alderton, Roberto Amato, Jeffrey Barrett, Sonia Goncalves, Ewan Harrison, David K. Jackson, Ian Johnston, Dominic Kwiatkowski, Cordelia Langford, John Sillitoe on behalf of the Wellcome Sanger Institute COVID-19 Surveillance Team |
| EPI_ISL_3440019 | Lighthouse Lab in Alderley Park | Wellcome Sanger Institute for the COVID-19 Genomics UK (COG-UK) Consortium | Jacquelyn Wynn, Mairead Hyland, The Lighthouse Lab in Alderley Park and Alex Alderton, Roberto Amato, Jeffrey Barrett, Sonia Goncalves, Ewan Harrison, David K. Jackson, Ian Johnston, Dominic Kwiatkowski, Cordelia Langford, John Sillitoe on behalf of the Wellcome Sanger Institute COVID-19 Surveillance Team |
| EPI_ISL_3440020 | Lighthouse Lab in Glasgow | Wellcome Sanger Institute for the COVID-19 Genomics UK (COG-UK) Consortium | Harper VanSteenhouse, Yumi Kasai, David Gray, Carol Clugston, Anna Dominiczak and Alex Alderton, Roberto Amato, Jeffrey Barrett, Sonia Goncalves, Ewan Harrison, David K. Jackson, Ian Johnston, Dominic Kwiatkowski, Cordelia Langford, John Sillitoe on behalf of the Wellcome Sanger Institute COVID-19 Surveillance Team |
| EPI_ISL_3440021 | Lighthouse Laboratory Plymouth | Wellcome Sanger Institute for the COVID-19 Genomics UK (COG-UK) Consortium | Lighthouse Laboratory Plymouth and Alex Alderton, Roberto Amato, Jeffrey Barrett, Sonia Goncalves, Ewan Harrison, David K. Jackson, Ian Johnston, Dominic Kwiatkowski, Cordelia Langford, John Sillitoe on behalf of the Wellcome Sanger Institute COVID-19 Surveillance Team |

| | | | |
|---|---|---|---|
| EPI_ISL_3440022, EPI_ISL_3440023, EPI_ISL_3440024 | Lighthouse Lab in Glasgow | Wellcome Sanger Institute for the COVID-19 Genomics UK (COG-UK) Consortium | Harper VanSteenhouse, Yumi Kasai, David Gray, Carol Clugston, Anna Dominiczak and Alex Alderton, Roberto Amato, Jeffrey Barrett, Sonia Goncalves, Ewan Harrison, David K. Jackson, Ian Johnston, Dominic Kwiatkowski, Cordelia Langford, John Sillitoe on behalf of the Wellcome Sanger Institute COVID-19 Surveillance Team |
| EPI_ISL_3440025, EPI_ISL_3440026, EPI_ISL_3440027 | Lighthouse Laboratory Plymouth | Wellcome Sanger Institute for the COVID-19 Genomics UK (COG-UK) Consortium | Lighthouse Laboratory Plymouth and Alex Alderton, Roberto Amato, Jeffrey Barrett, Sonia Goncalves, Ewan Harrison, David K. Jackson, Ian Johnston, Dominic Kwiatkowski, Cordelia Langford, John Sillitoe on behalf of the Wellcome Sanger Institute COVID-19 Surveillance Team |
| EPI_ISL_3440028, EPI_ISL_3440029 | Lighthouse Lab in Glasgow | Wellcome Sanger Institute for the COVID-19 Genomics UK (COG-UK) Consortium | Harper VanSteenhouse, Yumi Kasai, David Gray, Carol Clugston, Anna Dominiczak and Alex Alderton, Roberto Amato, Jeffrey Barrett, Sonia Goncalves, Ewan Harrison, David K. Jackson, Ian Johnston, Dominic Kwiatkowski, Cordelia Langford, John Sillitoe on behalf of the Wellcome Sanger Institute COVID-19 Surveillance Team |
| EPI_ISL_3440030 | Lighthouse Laboratory Plymouth | Wellcome Sanger Institute for the COVID-19 Genomics UK (COG-UK) Consortium | Lighthouse Laboratory Plymouth and Alex Alderton, Roberto Amato, Jeffrey Barrett, Sonia Goncalves, Ewan Harrison, David K. Jackson, Ian Johnston, Dominic Kwiatkowski, Cordelia Langford, John Sillitoe on behalf of the Wellcome Sanger Institute COVID-19 Surveillance Team |
| EPI_ISL_3440031 | Lighthouse Lab in Glasgow | Wellcome Sanger Institute for the COVID-19 Genomics UK (COG-UK) Consortium | Harper VanSteenhouse, Yumi Kasai, David Gray, Carol Clugston, Anna Dominiczak and Alex Alderton, Roberto Amato, Jeffrey Barrett, Sonia Goncalves, Ewan Harrison, David K. Jackson, Ian Johnston, Dominic Kwiatkowski, Cordelia Langford, John Sillitoe on behalf of the Wellcome Sanger Institute COVID-19 Surveillance Team |
| EPI_ISL_3440032 | Lighthouse Laboratory Plymouth | Wellcome Sanger Institute for the COVID-19 Genomics UK (COG-UK) Consortium | Lighthouse Laboratory Plymouth and Alex Alderton, Roberto Amato, Jeffrey Barrett, Sonia Goncalves, Ewan Harrison, David K. Jackson, Ian Johnston, Dominic Kwiatkowski, Cordelia Langford, John Sillitoe on behalf of the Wellcome Sanger Institute COVID-19 Surveillance Team |
| EPI_ISL_3440033 | Lighthouse Lab in Glasgow | Wellcome Sanger Institute for the COVID-19 Genomics UK (COG-UK) Consortium | Harper VanSteenhouse, Yumi Kasai, David Gray, Carol Clugston, Anna Dominiczak and Alex Alderton, Roberto Amato, Jeffrey Barrett, Sonia Goncalves, Ewan Harrison, David K. Jackson, Ian Johnston, Dominic Kwiatkowski, Cordelia Langford, John Sillitoe on behalf of the Wellcome Sanger Institute COVID-19 Surveillance Team |
| EPI_ISL_3440034 | Lighthouse Laboratory Plymouth | Wellcome Sanger Institute for the COVID-19 Genomics UK (COG-UK) Consortium | Lighthouse Laboratory Plymouth and Alex Alderton, Roberto Amato, Jeffrey Barrett, Sonia Goncalves, Ewan Harrison, David K. Jackson, Ian Johnston, Dominic Kwiatkowski, Cordelia Langford, John Sillitoe on behalf of the Wellcome Sanger Institute COVID-19 Surveillance Team |
| EPI_ISL_3440035, EPI_ISL_3440036 | Lighthouse Lab in Glasgow | Wellcome Sanger Institute for the COVID-19 Genomics UK (COG-UK) Consortium | Harper VanSteenhouse, Yumi Kasai, David Gray, Carol Clugston, Anna Dominiczak and Alex Alderton, Roberto Amato, Jeffrey Barrett, Sonia Goncalves, Ewan Harrison, David K. Jackson, Ian Johnston, Dominic Kwiatkowski, Cordelia Langford, John Sillitoe on behalf of the Wellcome Sanger Institute COVID-19 Surveillance Team |
| EPI_ISL_3440037, EPI_ISL_3440038 | Lighthouse Laboratory Plymouth | Wellcome Sanger Institute for the COVID-19 Genomics UK (COG-UK) Consortium | Lighthouse Laboratory Plymouth and Alex Alderton, Roberto Amato, Jeffrey Barrett, Sonia Goncalves, Ewan Harrison, David K. Jackson, Ian Johnston, Dominic Kwiatkowski, Cordelia Langford, John Sillitoe on behalf of the Wellcome Sanger Institute COVID-19 Surveillance Team |
| EPI_ISL_3440039 | Lighthouse Lab in Glasgow | Wellcome Sanger Institute for the COVID-19 Genomics UK (COG-UK) Consortium | Harper VanSteenhouse, Yumi Kasai, David Gray, Carol Clugston, Anna Dominiczak and Alex Alderton, Roberto Amato, Jeffrey Barrett, Sonia Goncalves, Ewan Harrison, David K. Jackson, Ian Johnston, Dominic Kwiatkowski, Cordelia Langford, John Sillitoe on behalf of the Wellcome Sanger Institute COVID-19 Surveillance Team |
| EPI_ISL_3440040 | Lighthouse Laboratory Plymouth | Wellcome Sanger Institute for the COVID-19 Genomics UK (COG-UK) Consortium | Lighthouse Laboratory Plymouth and Alex Alderton, Roberto Amato, Jeffrey Barrett, Sonia Goncalves, Ewan Harrison, David K. Jackson, Ian Johnston, Dominic Kwiatkowski, Cordelia Langford, John Sillitoe on behalf of the Wellcome Sanger Institute COVID-19 Surveillance Team |
| EPI_ISL_3440041 | Lighthouse Lab in Glasgow | Wellcome Sanger Institute for the COVID-19 Genomics UK (COG-UK) Consortium | Harper VanSteenhouse, Yumi Kasai, David Gray, Carol Clugston, Anna Dominiczak and Alex Alderton, Roberto Amato, Jeffrey Barrett, Sonia Goncalves, Ewan Harrison, David K. Jackson, Ian Johnston, Dominic Kwiatkowski, Cordelia Langford, John Sillitoe on behalf of the Wellcome Sanger Institute COVID-19 Surveillance Team |
| EPI_ISL_3440042 | Lighthouse Laboratory Plymouth | Wellcome Sanger Institute for the COVID-19 Genomics UK (COG-UK) Consortium | Lighthouse Laboratory Plymouth and Alex Alderton, Roberto Amato, Jeffrey Barrett, Sonia Goncalves, Ewan Harrison, David K. Jackson, Ian Johnston, Dominic Kwiatkowski, Cordelia Langford, John Sillitoe on behalf of the Wellcome Sanger Institute COVID-19 Surveillance Team |
| EPI_ISL_3440043, EPI_ISL_3440044, EPI_ISL_3440045, EPI_ISL_3440046 | Lighthouse Lab in Glasgow | Wellcome Sanger Institute for the COVID-19 Genomics UK (COG-UK) Consortium | Harper VanSteenhouse, Yumi Kasai, David Gray, Carol Clugston, Anna Dominiczak and Alex Alderton, Roberto Amato, Jeffrey Barrett, Sonia Goncalves, Ewan Harrison, David K. Jackson, Ian Johnston, Dominic Kwiatkowski, Cordelia Langford, John Sillitoe on behalf of the Wellcome Sanger Institute COVID-19 Surveillance Team |
| EPI_ISL_3440047, EPI_ISL_3440048, EPI_ISL_3440049 | Lighthouse Laboratory Plymouth | Wellcome Sanger Institute for the COVID-19 Genomics UK (COG-UK) Consortium | Lighthouse Laboratory Plymouth and Alex Alderton, Roberto Amato, Jeffrey Barrett, Sonia Goncalves, Ewan Harrison, David K. Jackson, Ian Johnston, Dominic Kwiatkowski, Cordelia Langford, John Sillitoe on behalf of the Wellcome Sanger Institute COVID-19 Surveillance Team |
| EPI_ISL_3440050, EPI_ISL_3440051, EPI_ISL_3440052, EPI_ISL_3440053, EPI_ISL_3440054, EPI_ISL_3440055 | Lighthouse Lab in Glasgow | Wellcome Sanger Institute for the COVID-19 Genomics UK (COG-UK) Consortium | Harper VanSteenhouse, Yumi Kasai, David Gray, Carol Clugston, Anna Dominiczak and Alex Alderton, Roberto Amato, Jeffrey Barrett, Sonia Goncalves, Ewan Harrison, David K. Jackson, Ian Johnston, Dominic Kwiatkowski, Cordelia Langford, John Sillitoe on behalf of the Wellcome Sanger Institute COVID-19 Surveillance Team |
| EPI_ISL_3440056 | Lighthouse Laboratory Plymouth | Wellcome Sanger Institute for the COVID-19 Genomics UK (COG-UK) Consortium | Lighthouse Laboratory Plymouth and Alex Alderton, Roberto Amato, Jeffrey Barrett, Sonia Goncalves, Ewan Harrison, David K. Jackson, Ian Johnston, Dominic Kwiatkowski, Cordelia Langford, John Sillitoe on behalf of the Wellcome Sanger Institute COVID-19 Surveillance Team |
| EPI_ISL_3440057, EPI_ISL_3440058, EPI_ISL_3440059, EPI_ISL_3440060 | Lighthouse Lab in Glasgow | Wellcome Sanger Institute for the COVID-19 Genomics UK (COG-UK) Consortium | Harper VanSteenhouse, Yumi Kasai, David Gray, Carol Clugston, Anna Dominiczak and Alex Alderton, Roberto Amato, Jeffrey Barrett, Sonia Goncalves, Ewan Harrison, David K. Jackson, Ian Johnston, Dominic Kwiatkowski, Cordelia Langford, John Sillitoe on behalf of the Wellcome Sanger Institute COVID-19 Surveillance Team |
| EPI_ISL_3440061 | Lighthouse Laboratory Plymouth | Wellcome Sanger Institute for the COVID-19 Genomics UK (COG-UK) Consortium | Lighthouse Laboratory Plymouth and Alex Alderton, Roberto Amato, Jeffrey Barrett, Sonia Goncalves, Ewan Harrison, David K. Jackson, Ian Johnston, Dominic Kwiatkowski, Cordelia Langford, John Sillitoe on behalf of the Wellcome Sanger Institute COVID-19 Surveillance Team |
| EPI_ISL_3440062, EPI_ISL_3440063, EPI_ISL_3440064, EPI_ISL_3440065, EPI_ISL_3440066, EPI_ISL_3440067 | Lighthouse Lab in Glasgow | Wellcome Sanger Institute for the COVID-19 Genomics UK (COG-UK) Consortium | Harper VanSteenhouse, Yumi Kasai, David Gray, Carol Clugston, Anna Dominiczak and Alex Alderton, Roberto Amato, Jeffrey Barrett, Sonia Goncalves, Ewan Harrison, David K. Jackson, Ian Johnston, Dominic Kwiatkowski, Cordelia Langford, John Sillitoe on behalf of the Wellcome Sanger Institute COVID-19 Surveillance Team |
| EPI_ISL_3440068 | Lighthouse Laboratory Plymouth | Wellcome Sanger Institute for the COVID-19 Genomics UK (COG-UK) Consortium | Lighthouse Laboratory Plymouth and Alex Alderton, Roberto Amato, Jeffrey Barrett, Sonia Goncalves, Ewan Harrison, David K. Jackson, Ian Johnston, Dominic Kwiatkowski, Cordelia Langford, John Sillitoe on behalf of the Wellcome Sanger Institute COVID-19 Surveillance Team |
| EPI_ISL_3440069, EPI_ISL_3440070, EPI_ISL_3440071, EPI_ISL_3440072 | Lighthouse Lab in Glasgow | Wellcome Sanger Institute for the COVID-19 Genomics UK (COG-UK) Consortium | Harper VanSteenhouse, Yumi Kasai, David Gray, Carol Clugston, Anna Dominiczak and Alex Alderton, Roberto Amato, Jeffrey Barrett, Sonia Goncalves, Ewan Harrison, David K. Jackson, Ian Johnston, Dominic Kwiatkowski, Cordelia Langford, John Sillitoe on behalf of the Wellcome Sanger Institute COVID-19 Surveillance Team |
| EPI_ISL_3440073 | Lighthouse Laboratory Plymouth | Wellcome Sanger Institute for the COVID-19 Genomics UK (COG-UK) Consortium | Lighthouse Laboratory Plymouth and Alex Alderton, Roberto Amato, Jeffrey Barrett, Sonia Goncalves, Ewan Harrison, David K. Jackson, Ian Johnston, Dominic Kwiatkowski, Cordelia Langford, John Sillitoe on behalf of the Wellcome Sanger Institute COVID-19 Surveillance Team |
| EPI_ISL_3440074, EPI_ISL_3440075, EPI_ISL_3440076, EPI_ISL_3440077, EPI_ISL_3440078 | Lighthouse Lab in Glasgow | Wellcome Sanger Institute for the COVID-19 Genomics UK (COG-UK) Consortium | Harper VanSteenhouse, Yumi Kasai, David Gray, Carol Clugston, Anna Dominiczak and Alex Alderton, Roberto Amato, Jeffrey Barrett, Sonia Goncalves, Ewan Harrison, David K. Jackson, Ian Johnston, Dominic Kwiatkowski, Cordelia Langford, John Sillitoe on behalf of the Wellcome Sanger Institute COVID-19 Surveillance Team |
| EPI_ISL_3440079 | Lighthouse Laboratory Plymouth | Wellcome Sanger Institute for the COVID-19 Genomics UK (COG-UK) Consortium | Lighthouse Laboratory Plymouth and Alex Alderton, Roberto Amato, Jeffrey Barrett, Sonia Goncalves, Ewan Harrison, David K. Jackson, Ian Johnston, Dominic Kwiatkowski, Cordelia Langford, John Sillitoe on behalf of the Wellcome Sanger Institute COVID-19 Surveillance Team |
| EPI_ISL_3440080, EPI_ISL_3440081, EPI_ISL_3440082, EPI_ISL_3440083, EPI_ISL_3440084, EPI_ISL_3440085, EPI_ISL_3440086, EPI_ISL_3440087, EPI_ISL_3440088, EPI_ISL_3440089, EPI_ISL_3440090, EPI_ISL_3440091, EPI_ISL_3440092 | | | |
| see above | Lighthouse Lab in Glasgow | Wellcome Sanger Institute for the COVID-19 Genomics UK (COG-UK) Consortium | Harper VanSteenhouse, Yumi Kasai, David Gray, Carol Clugston, Anna Dominiczak and Alex Alderton, Roberto Amato, Jeffrey Barrett, Sonia Goncalves, Ewan Harrison, David K. Jackson, Ian Johnston, Dominic Kwiatkowski, Cordelia Langford, John Sillitoe on behalf of the Wellcome Sanger Institute COVID-19 Surveillance Team |
| EPI_ISL_3440093 | Lighthouse Laboratory Plymouth | Wellcome Sanger Institute for the COVID-19 Genomics UK (COG-UK) Consortium | Lighthouse Laboratory Plymouth and Alex Alderton, Roberto Amato, Jeffrey Barrett, Sonia Goncalves, Ewan Harrison, David K. Jackson, Ian Johnston, Dominic Kwiatkowski, Cordelia Langford, John Sillitoe on behalf of the Wellcome Sanger Institute COVID-19 Surveillance Team |
| EPI_ISL_3440094, EPI_ISL_3440095 | Lighthouse Lab in Glasgow | Wellcome Sanger Institute for the COVID-19 Genomics UK (COG-UK) Consortium | Harper VanSteenhouse, Yumi Kasai, David Gray, Carol Clugston, Anna Dominiczak and Alex Alderton, Roberto Amato, Jeffrey Barrett, Sonia Goncalves, Ewan Harrison, David K. Jackson, Ian Johnston, Dominic Kwiatkowski, Cordelia Langford, John Sillitoe on behalf of the Wellcome Sanger Institute COVID-19 Surveillance Team |
| EPI_ISL_3440096 | Lighthouse Laboratory Plymouth | Wellcome Sanger Institute for the COVID-19 Genomics UK (COG-UK) Consortium | Lighthouse Laboratory Plymouth and Alex Alderton, Roberto Amato, Jeffrey Barrett, Sonia Goncalves, Ewan Harrison, David K. Jackson, Ian Johnston, Dominic Kwiatkowski, Cordelia Langford, John Sillitoe on behalf of the Wellcome Sanger Institute COVID-19 Surveillance Team |
| EPI_ISL_3440097 | Lighthouse Lab in Glasgow | Wellcome Sanger Institute for the COVID-19 Genomics UK (COG-UK) Consortium | Harper VanSteenhouse, Yumi Kasai, David Gray, Carol Clugston, Anna Dominiczak and Alex Alderton, Roberto Amato, Jeffrey Barrett, Sonia Goncalves, Ewan Harrison, David K. Jackson, Ian Johnston, Dominic Kwiatkowski, Cordelia Langford, John Sillitoe on behalf of the Wellcome Sanger Institute COVID-19 Surveillance Team |
| EPI_ISL_3440098 | Lighthouse Laboratory Plymouth | Wellcome Sanger Institute for the COVID-19 Genomics UK (COG-UK) Consortium | Lighthouse Laboratory Plymouth and Alex Alderton, Roberto Amato, Jeffrey Barrett, Sonia Goncalves, Ewan Harrison, David K. Jackson, Ian Johnston, Dominic Kwiatkowski, Cordelia Langford, John Sillitoe on behalf of the Wellcome Sanger Institute COVID-19 Surveillance Team |
| EPI_ISL_3440099 | Lighthouse Lab in Glasgow | Wellcome Sanger Institute for the COVID-19 Genomics UK (COG-UK) Consortium | Harper VanSteenhouse, Yumi Kasai, David Gray, Carol Clugston, Anna Dominiczak and Alex Alderton, Roberto Amato, Jeffrey Barrett, Sonia Goncalves, Ewan Harrison, David K. Jackson, Ian Johnston, Dominic Kwiatkowski, Cordelia Langford, John Sillitoe on behalf of the Wellcome Sanger Institute COVID-19 Surveillance Team |
| EPI_ISL_3440100 | Lighthouse Laboratory Plymouth | Wellcome Sanger Institute for the COVID-19 Genomics UK (COG-UK) Consortium | Lighthouse Laboratory Plymouth and Alex Alderton, Roberto Amato, Jeffrey Barrett, Sonia Goncalves, Ewan Harrison, David K. Jackson, Ian Johnston, Dominic Kwiatkowski, Cordelia Langford, John Sillitoe on behalf of the Wellcome Sanger Institute COVID-19 Surveillance Team |
| EPI_ISL_3440101 | Lighthouse Lab in Glasgow | Wellcome Sanger Institute for the COVID-19 Genomics UK (COG-UK) Consortium | Harper VanSteenhouse, Yumi Kasai, David Gray, Carol Clugston, Anna Dominiczak and Alex Alderton, Roberto Amato, Jeffrey Barrett, Sonia Goncalves, Ewan Harrison, David K. Jackson, Ian Johnston, Dominic Kwiatkowski, Cordelia Langford, John Sillitoe on behalf of the Wellcome Sanger Institute COVID-19 Surveillance Team |
| EPI_ISL_3440102 | Lighthouse Laboratory Plymouth | Wellcome Sanger Institute for the COVID-19 Genomics UK (COG-UK) Consortium | Lighthouse Laboratory Plymouth and Alex Alderton, Roberto Amato, Jeffrey Barrett, Sonia Goncalves, Ewan Harrison, David K. Jackson, Ian Johnston, Dominic Kwiatkowski, Cordelia Langford, John Sillitoe on behalf of the Wellcome Sanger Institute COVID-19 Surveillance Team |

| | | | |
|---|---|---|---|
| EPI_ISL_3440103, EPI_ISL_3440104, EPI_ISL_3440105, EPI_ISL_3440106, EPI_ISL_3440107, EPI_ISL_3440108, EPI_ISL_3440109, EPI_ISL_3440110, EPI_ISL_3440111, EPI_ISL_3440112, EPI_ISL_3440113, EPI_ISL_3440114, EPI_ISL_3440115, EPI_ISL_3440116, EPI_ISL_3440117, EPI_ISL_3440118, EPI_ISL_3440119, EPI_ISL_3440120, EPI_ISL_3440121, EPI_ISL_3440122, EPI_ISL_3440123, EPI_ISL_3440124, EPI_ISL_3440125, EPI_ISL_3440126, EPI_ISL_3440127, EPI_ISL_3440128, EPI_ISL_3440129, EPI_ISL_3440130, EPI_ISL_3440131, EPI_ISL_3440132, EPI_ISL_3440133, EPI_ISL_3440134, EPI_ISL_3440135, EPI_ISL_3440136, EPI_ISL_3440137, EPI_ISL_3440138, EPI_ISL_3440139 | | | |
| see above | Lighthouse Lab in Glasgow | Wellcome Sanger Institute for the COVID-19 Genomics UK (COG-UK) Consortium | Harper VanSteenhouse, Yumi Kasai, David Gray, Carol Clugston, Anna Dominiczak and Alex Alderton, Roberto Amato, Jeffrey Barrett, Sonia Goncalves, Ewan Harrison, David K. Jackson, Ian Johnston, Dominic Kwiatkowski, Cordelia Langford, John Sillitoe on behalf of the Wellcome Sanger Institute COVID-19 Surveillance Team |
| EPI_ISL_3440140 | Lighthouse Laboratory Plymouth | Wellcome Sanger Institute for the COVID-19 Genomics UK (COG-UK) Consortium | Lighthouse Laboratory Plymouth and Alex Alderton, Roberto Amato, Jeffrey Barrett, Sonia Goncalves, Ewan Harrison, David K. Jackson, Ian Johnston, Dominic Kwiatkowski, Cordelia Langford, John Sillitoe on behalf of the Wellcome Sanger Institute COVID-19 Surveillance Team |
| EPI_ISL_3440141, EPI_ISL_3440142 | Lighthouse Lab in Glasgow | Wellcome Sanger Institute for the COVID-19 Genomics UK (COG-UK) Consortium | Harper VanSteenhouse, Yumi Kasai, David Gray, Carol Clugston, Anna Dominiczak and Alex Alderton, Roberto Amato, Jeffrey Barrett, Sonia Goncalves, Ewan Harrison, David K. Jackson, Ian Johnston, Dominic Kwiatkowski, Cordelia Langford, John Sillitoe on behalf of the Wellcome Sanger Institute COVID-19 Surveillance Team |
| EPI_ISL_3440143 | Lighthouse Laboratory Plymouth | Wellcome Sanger Institute for the COVID-19 Genomics UK (COG-UK) Consortium | Lighthouse Laboratory Plymouth and Alex Alderton, Roberto Amato, Jeffrey Barrett, Sonia Goncalves, Ewan Harrison, David K. Jackson, Ian Johnston, Dominic Kwiatkowski, Cordelia Langford, John Sillitoe on behalf of the Wellcome Sanger Institute COVID-19 Surveillance Team |
| EPI_ISL_3440144 | Lighthouse Lab in Glasgow | Wellcome Sanger Institute for the COVID-19 Genomics UK (COG-UK) Consortium | Harper VanSteenhouse, Yumi Kasai, David Gray, Carol Clugston, Anna Dominiczak and Alex Alderton, Roberto Amato, Jeffrey Barrett, Sonia Goncalves, Ewan Harrison, David K. Jackson, Ian Johnston, Dominic Kwiatkowski, Cordelia Langford, John Sillitoe on behalf of the Wellcome Sanger Institute COVID-19 Surveillance Team |
| EPI_ISL_3440145 | Lighthouse Laboratory Plymouth | Wellcome Sanger Institute for the COVID-19 Genomics UK (COG-UK) Consortium | Lighthouse Laboratory Plymouth and Alex Alderton, Roberto Amato, Jeffrey Barrett, Sonia Goncalves, Ewan Harrison, David K. Jackson, Ian Johnston, Dominic Kwiatkowski, Cordelia Langford, John Sillitoe on behalf of the Wellcome Sanger Institute COVID-19 Surveillance Team |
| EPI_ISL_3440146, EPI_ISL_3440147, EPI_ISL_3440148 | Lighthouse Lab in Glasgow | Wellcome Sanger Institute for the COVID-19 Genomics UK (COG-UK) Consortium | Harper VanSteenhouse, Yumi Kasai, David Gray, Carol Clugston, Anna Dominiczak and Alex Alderton, Roberto Amato, Jeffrey Barrett, Sonia Goncalves, Ewan Harrison, David K. Jackson, Ian Johnston, Dominic Kwiatkowski, Cordelia Langford, John Sillitoe on behalf of the Wellcome Sanger Institute COVID-19 Surveillance Team |
| EPI_ISL_3440149, EPI_ISL_3440150 | Lighthouse Laboratory Plymouth | Wellcome Sanger Institute for the COVID-19 Genomics UK (COG-UK) Consortium | Lighthouse Laboratory Plymouth and Alex Alderton, Roberto Amato, Jeffrey Barrett, Sonia Goncalves, Ewan Harrison, David K. Jackson, Ian Johnston, Dominic Kwiatkowski, Cordelia Langford, John Sillitoe on behalf of the Wellcome Sanger Institute COVID-19 Surveillance Team |
| EPI_ISL_3440151, EPI_ISL_3440152, EPI_ISL_3440153, EPI_ISL_3440154, EPI_ISL_3440155 | Lighthouse Lab in Glasgow | Wellcome Sanger Institute for the COVID-19 Genomics UK (COG-UK) Consortium | Harper VanSteenhouse, Yumi Kasai, David Gray, Carol Clugston, Anna Dominiczak and Alex Alderton, Roberto Amato, Jeffrey Barrett, Sonia Goncalves, Ewan Harrison, David K. Jackson, Ian Johnston, Dominic Kwiatkowski, Cordelia Langford, John Sillitoe on behalf of the Wellcome Sanger Institute COVID-19 Surveillance Team |
| EPI_ISL_3440156 | Lighthouse Laboratory Plymouth | Wellcome Sanger Institute for the COVID-19 Genomics UK (COG-UK) Consortium | Lighthouse Laboratory Plymouth and Alex Alderton, Roberto Amato, Jeffrey Barrett, Sonia Goncalves, Ewan Harrison, David K. Jackson, Ian Johnston, Dominic Kwiatkowski, Cordelia Langford, John Sillitoe on behalf of the Wellcome Sanger Institute COVID-19 Surveillance Team |
| EPI_ISL_3440157, EPI_ISL_3440158, EPI_ISL_3440159, EPI_ISL_3440160 | Lighthouse Lab in Glasgow | Wellcome Sanger Institute for the COVID-19 Genomics UK (COG-UK) Consortium | Harper VanSteenhouse, Yumi Kasai, David Gray, Carol Clugston, Anna Dominiczak and Alex Alderton, Roberto Amato, Jeffrey Barrett, Sonia Goncalves, Ewan Harrison, David K. Jackson, Ian Johnston, Dominic Kwiatkowski, Cordelia Langford, John Sillitoe on behalf of the Wellcome Sanger Institute COVID-19 Surveillance Team |
| EPI_ISL_3440161 | Lighthouse Laboratory Plymouth | Wellcome Sanger Institute for the COVID-19 Genomics UK (COG-UK) Consortium | Lighthouse Laboratory Plymouth and Alex Alderton, Roberto Amato, Jeffrey Barrett, Sonia Goncalves, Ewan Harrison, David K. Jackson, Ian Johnston, Dominic Kwiatkowski, Cordelia Langford, John Sillitoe on behalf of the Wellcome Sanger Institute COVID-19 Surveillance Team |
| EPI_ISL_3440162, EPI_ISL_3440163, EPI_ISL_3440164, EPI_ISL_3440165, EPI_ISL_3440166, EPI_ISL_3440167, EPI_ISL_3440168, EPI_ISL_3440169 | Lighthouse Lab in Glasgow | Wellcome Sanger Institute for the COVID-19 Genomics UK (COG-UK) Consortium | Harper VanSteenhouse, Yumi Kasai, David Gray, Carol Clugston, Anna Dominiczak and Alex Alderton, Roberto Amato, Jeffrey Barrett, Sonia Goncalves, Ewan Harrison, David K. Jackson, Ian Johnston, Dominic Kwiatkowski, Cordelia Langford, John Sillitoe on behalf of the Wellcome Sanger Institute COVID-19 Surveillance Team |
| EPI_ISL_3440170 | Lighthouse Laboratory Plymouth | Wellcome Sanger Institute for the COVID-19 Genomics UK (COG-UK) Consortium | Lighthouse Laboratory Plymouth and Alex Alderton, Roberto Amato, Jeffrey Barrett, Sonia Goncalves, Ewan Harrison, David K. Jackson, Ian Johnston, Dominic Kwiatkowski, Cordelia Langford, John Sillitoe on behalf of the Wellcome Sanger Institute COVID-19 Surveillance Team |
| EPI_ISL_3440171, EPI_ISL_3440172, EPI_ISL_3440173, EPI_ISL_3440174 | Lighthouse Lab in Glasgow | Wellcome Sanger Institute for the COVID-19 Genomics UK (COG-UK) Consortium | Harper VanSteenhouse, Yumi Kasai, David Gray, Carol Clugston, Anna Dominiczak and Alex Alderton, Roberto Amato, Jeffrey Barrett, Sonia Goncalves, Ewan Harrison, David K. Jackson, Ian Johnston, Dominic Kwiatkowski, Cordelia Langford, John Sillitoe on behalf of the Wellcome Sanger Institute COVID-19 Surveillance Team |
| EPI_ISL_3440175, EPI_ISL_3440176, EPI_ISL_3440177, EPI_ISL_3440178 | Lighthouse Laboratory Plymouth | Wellcome Sanger Institute for the COVID-19 Genomics UK (COG-UK) Consortium | Lighthouse Laboratory Plymouth and Alex Alderton, Roberto Amato, Jeffrey Barrett, Sonia Goncalves, Ewan Harrison, David K. Jackson, Ian Johnston, Dominic Kwiatkowski, Cordelia Langford, John Sillitoe on behalf of the Wellcome Sanger Institute COVID-19 Surveillance Team |
| EPI_ISL_3440179, EPI_ISL_3440180 | Lighthouse Lab in Glasgow | Wellcome Sanger Institute for the COVID-19 Genomics UK (COG-UK) Consortium | Harper VanSteenhouse, Yumi Kasai, David Gray, Carol Clugston, Anna Dominiczak and Alex Alderton, Roberto Amato, Jeffrey Barrett, Sonia Goncalves, Ewan Harrison, David K. Jackson, Ian Johnston, Dominic Kwiatkowski, Cordelia Langford, John Sillitoe on behalf of the Wellcome Sanger Institute COVID-19 Surveillance Team |
| EPI_ISL_3440181 | Lighthouse Laboratory Plymouth | Wellcome Sanger Institute for the COVID-19 Genomics UK (COG-UK) Consortium | Lighthouse Laboratory Plymouth and Alex Alderton, Roberto Amato, Jeffrey Barrett, Sonia Goncalves, Ewan Harrison, David K. Jackson, Ian Johnston, Dominic Kwiatkowski, Cordelia Langford, John Sillitoe on behalf of the Wellcome Sanger Institute COVID-19 Surveillance Team |
| EPI_ISL_3440182, EPI_ISL_3440183 | Lighthouse Lab in Glasgow | Wellcome Sanger Institute for the COVID-19 Genomics UK (COG-UK) Consortium | Harper VanSteenhouse, Yumi Kasai, David Gray, Carol Clugston, Anna Dominiczak and Alex Alderton, Roberto Amato, Jeffrey Barrett, Sonia Goncalves, Ewan Harrison, David K. Jackson, Ian Johnston, Dominic Kwiatkowski, Cordelia Langford, John Sillitoe on behalf of the Wellcome Sanger Institute COVID-19 Surveillance Team |
| EPI_ISL_3440184 | Lighthouse Laboratory Plymouth | Wellcome Sanger Institute for the COVID-19 Genomics UK (COG-UK) Consortium | Lighthouse Laboratory Plymouth and Alex Alderton, Roberto Amato, Jeffrey Barrett, Sonia Goncalves, Ewan Harrison, David K. Jackson, Ian Johnston, Dominic Kwiatkowski, Cordelia Langford, John Sillitoe on behalf of the Wellcome Sanger Institute COVID-19 Surveillance Team |
| EPI_ISL_3440185, EPI_ISL_3440186, EPI_ISL_3440187, EPI_ISL_3440188, EPI_ISL_3440189, EPI_ISL_3440190, EPI_ISL_3440191, EPI_ISL_3440192 | Lighthouse Lab in Glasgow | Wellcome Sanger Institute for the COVID-19 Genomics UK (COG-UK) Consortium | Harper VanSteenhouse, Yumi Kasai, David Gray, Carol Clugston, Anna Dominiczak and Alex Alderton, Roberto Amato, Jeffrey Barrett, Sonia Goncalves, Ewan Harrison, David K. Jackson, Ian Johnston, Dominic Kwiatkowski, Cordelia Langford, John Sillitoe on behalf of the Wellcome Sanger Institute COVID-19 Surveillance Team |
| EPI_ISL_3440193 | Lighthouse Laboratory Plymouth | Wellcome Sanger Institute for the COVID-19 Genomics UK (COG-UK) Consortium | Lighthouse Laboratory Plymouth and Alex Alderton, Roberto Amato, Jeffrey Barrett, Sonia Goncalves, Ewan Harrison, David K. Jackson, Ian Johnston, Dominic Kwiatkowski, Cordelia Langford, John Sillitoe on behalf of the Wellcome Sanger Institute COVID-19 Surveillance Team |
| EPI_ISL_3440194, EPI_ISL_3440195, EPI_ISL_3440196, EPI_ISL_3440197, EPI_ISL_3440198, EPI_ISL_3440199, EPI_ISL_3440200, EPI_ISL_3440201, EPI_ISL_3440202 | Lighthouse Lab in Glasgow | Wellcome Sanger Institute for the COVID-19 Genomics UK (COG-UK) Consortium | Harper VanSteenhouse, Yumi Kasai, David Gray, Carol Clugston, Anna Dominiczak and Alex Alderton, Roberto Amato, Jeffrey Barrett, Sonia Goncalves, Ewan Harrison, David K. Jackson, Ian Johnston, Dominic Kwiatkowski, Cordelia Langford, John Sillitoe on behalf of the Wellcome Sanger Institute COVID-19 Surveillance Team |
| EPI_ISL_3440203, EPI_ISL_3440204 | Lighthouse Laboratory Plymouth | Wellcome Sanger Institute for the COVID-19 Genomics UK (COG-UK) Consortium | Lighthouse Laboratory Plymouth and Alex Alderton, Roberto Amato, Jeffrey Barrett, Sonia Goncalves, Ewan Harrison, David K. Jackson, Ian Johnston, Dominic Kwiatkowski, Cordelia Langford, John Sillitoe on behalf of the Wellcome Sanger Institute COVID-19 Surveillance Team |
| EPI_ISL_3440205 | Lighthouse Lab in Glasgow | Wellcome Sanger Institute for the COVID-19 Genomics UK (COG-UK) Consortium | Harper VanSteenhouse, Yumi Kasai, David Gray, Carol Clugston, Anna Dominiczak and Alex Alderton, Roberto Amato, Jeffrey Barrett, Sonia Goncalves, Ewan Harrison, David K. Jackson, Ian Johnston, Dominic Kwiatkowski, Cordelia Langford, John Sillitoe on behalf of the Wellcome Sanger Institute COVID-19 Surveillance Team |
| EPI_ISL_3440206, EPI_ISL_3440207 | Lighthouse Laboratory Plymouth | Wellcome Sanger Institute for the COVID-19 Genomics UK (COG-UK) Consortium | Lighthouse Laboratory Plymouth and Alex Alderton, Roberto Amato, Jeffrey Barrett, Sonia Goncalves, Ewan Harrison, David K. Jackson, Ian Johnston, Dominic Kwiatkowski, Cordelia Langford, John Sillitoe on behalf of the Wellcome Sanger Institute COVID-19 Surveillance Team |
| EPI_ISL_3440208, EPI_ISL_3440209, EPI_ISL_3440210, EPI_ISL_3440211 | Lighthouse Lab in Glasgow | Wellcome Sanger Institute for the COVID-19 Genomics UK (COG-UK) Consortium | Harper VanSteenhouse, Yumi Kasai, David Gray, Carol Clugston, Anna Dominiczak and Alex Alderton, Roberto Amato, Jeffrey Barrett, Sonia Goncalves, Ewan Harrison, David K. Jackson, Ian Johnston, Dominic Kwiatkowski, Cordelia Langford, John Sillitoe on behalf of the Wellcome Sanger Institute COVID-19 Surveillance Team |
| EPI_ISL_3440212, EPI_ISL_3440213 | Lighthouse Laboratory Plymouth | Wellcome Sanger Institute for the COVID-19 Genomics UK (COG-UK) Consortium | Lighthouse Laboratory Plymouth and Alex Alderton, Roberto Amato, Jeffrey Barrett, Sonia Goncalves, Ewan Harrison, David K. Jackson, Ian Johnston, Dominic Kwiatkowski, Cordelia Langford, John Sillitoe on behalf of the Wellcome Sanger Institute COVID-19 Surveillance Team |
| EPI_ISL_3440214, EPI_ISL_3440215, EPI_ISL_3440216 | Lighthouse Lab in Glasgow | Wellcome Sanger Institute for the COVID-19 Genomics UK (COG-UK) Consortium | Harper VanSteenhouse, Yumi Kasai, David Gray, Carol Clugston, Anna Dominiczak and Alex Alderton, Roberto Amato, Jeffrey Barrett, Sonia Goncalves, Ewan Harrison, David K. Jackson, Ian Johnston, Dominic Kwiatkowski, Cordelia Langford, John Sillitoe on behalf of the Wellcome Sanger Institute COVID-19 Surveillance Team |
| EPI_ISL_3440217 | Lighthouse Laboratory Plymouth | Wellcome Sanger Institute for the COVID-19 Genomics UK (COG-UK) Consortium | Lighthouse Laboratory Plymouth and Alex Alderton, Roberto Amato, Jeffrey Barrett, Sonia Goncalves, Ewan Harrison, David K. Jackson, Ian Johnston, Dominic Kwiatkowski, Cordelia Langford, John Sillitoe on behalf of the Wellcome Sanger Institute COVID-19 Surveillance Team |
| EPI_ISL_3440218, EPI_ISL_3440219, EPI_ISL_3440220, EPI_ISL_3440221, EPI_ISL_3440222, EPI_ISL_3440223, EPI_ISL_3440224, EPI_ISL_3440225, EPI_ISL_3440226, EPI_ISL_3440227, EPI_ISL_3440228, EPI_ISL_3440229, EPI_ISL_3440230 | | | |
| see above | Lighthouse Lab in Glasgow | Wellcome Sanger Institute for the COVID-19 Genomics UK (COG-UK) | Harper VanSteenhouse, Yumi Kasai, David Gray, Carol Clugston, Anna Dominiczak and Alex Alderton, Roberto Amato, Jeffrey Barrett, Sonia Goncalves, Ewan Harrison, David K. Jackson, Ian Johnston, Dominic Kwiatkowski, Cordelia Langford, |

| | | | |
|---|---|---|---|
| | | Consortium | John Sillitoe on behalf of the Wellcome Sanger Institute COVID-19 Surveillance Team |
| EPI_ISL_3440231 | Lighthouse Laboratory Plymouth | Wellcome Sanger Institute for the COVID-19 Genomics UK (COG-UK) Consortium | Lighthouse Laboratory Plymouth and Alex Alderton, Roberto Amato, Jeffrey Barrett, Sonia Goncalves, Ewan Harrison, David K. Jackson, Ian Johnston, Dominic Kwiatkowski, Cordelia Langford, John Sillitoe on behalf of the Wellcome Sanger Institute COVID-19 Surveillance Team |
| EPI_ISL_3440232, EPI_ISL_3440233, EPI_ISL_3440234, EPI_ISL_3440235 | Lighthouse Lab in Glasgow | Wellcome Sanger Institute for the COVID-19 Genomics UK (COG-UK) Consortium | Harper VanSteenhouse, Yumi Kasai, David Gray, Carol Clugston, Anna Dominiczak and Alex Alderton, Roberto Amato, Jeffrey Barrett, Sonia Goncalves, Ewan Harrison, David K. Jackson, Ian Johnston, Dominic Kwiatkowski, Cordelia Langford, John Sillitoe on behalf of the Wellcome Sanger Institute COVID-19 Surveillance Team |
| EPI_ISL_3440236, EPI_ISL_3440237 | Lighthouse Laboratory Plymouth | Wellcome Sanger Institute for the COVID-19 Genomics UK (COG-UK) Consortium | Lighthouse Laboratory Plymouth and Alex Alderton, Roberto Amato, Jeffrey Barrett, Sonia Goncalves, Ewan Harrison, David K. Jackson, Ian Johnston, Dominic Kwiatkowski, Cordelia Langford, John Sillitoe on behalf of the Wellcome Sanger Institute COVID-19 Surveillance Team |
| EPI_ISL_3440238, EPI_ISL_3440239, EPI_ISL_3440240, EPI_ISL_3440241, EPI_ISL_3440242, EPI_ISL_3440243, EPI_ISL_3440244, EPI_ISL_3440245, EPI_ISL_3440246, EPI_ISL_3440247, EPI_ISL_3440248 | | | |
| see above | Lighthouse Lab in Glasgow | Wellcome Sanger Institute for the COVID-19 Genomics UK (COG-UK) Consortium | Harper VanSteenhouse, Yumi Kasai, David Gray, Carol Clugston, Anna Dominiczak and Alex Alderton, Roberto Amato, Jeffrey Barrett, Sonia Goncalves, Ewan Harrison, David K. Jackson, Ian Johnston, Dominic Kwiatkowski, Cordelia Langford, John Sillitoe on behalf of the Wellcome Sanger Institute COVID-19 Surveillance Team |
| EPI_ISL_3440249 | Lighthouse Laboratory Plymouth | Wellcome Sanger Institute for the COVID-19 Genomics UK (COG-UK) Consortium | Lighthouse Laboratory Plymouth and Alex Alderton, Roberto Amato, Jeffrey Barrett, Sonia Goncalves, Ewan Harrison, David K. Jackson, Ian Johnston, Dominic Kwiatkowski, Cordelia Langford, John Sillitoe on behalf of the Wellcome Sanger Institute COVID-19 Surveillance Team |
| EPI_ISL_3440250, EPI_ISL_3440251 | Lighthouse Lab in Glasgow | Wellcome Sanger Institute for the COVID-19 Genomics UK (COG-UK) Consortium | Harper VanSteenhouse, Yumi Kasai, David Gray, Carol Clugston, Anna Dominiczak and Alex Alderton, Roberto Amato, Jeffrey Barrett, Sonia Goncalves, Ewan Harrison, David K. Jackson, Ian Johnston, Dominic Kwiatkowski, Cordelia Langford, John Sillitoe on behalf of the Wellcome Sanger Institute COVID-19 Surveillance Team |
| EPI_ISL_3440252 | Lighthouse Laboratory Plymouth | Wellcome Sanger Institute for the COVID-19 Genomics UK (COG-UK) Consortium | Lighthouse Laboratory Plymouth and Alex Alderton, Roberto Amato, Jeffrey Barrett, Sonia Goncalves, Ewan Harrison, David K. Jackson, Ian Johnston, Dominic Kwiatkowski, Cordelia Langford, John Sillitoe on behalf of the Wellcome Sanger Institute COVID-19 Surveillance Team |
| EPI_ISL_3440253 | Lighthouse Lab in Glasgow | Wellcome Sanger Institute for the COVID-19 Genomics UK (COG-UK) Consortium | Harper VanSteenhouse, Yumi Kasai, David Gray, Carol Clugston, Anna Dominiczak and Alex Alderton, Roberto Amato, Jeffrey Barrett, Sonia Goncalves, Ewan Harrison, David K. Jackson, Ian Johnston, Dominic Kwiatkowski, Cordelia Langford, John Sillitoe on behalf of the Wellcome Sanger Institute COVID-19 Surveillance Team |
| EPI_ISL_3440254 | Lighthouse Laboratory Plymouth | Wellcome Sanger Institute for the COVID-19 Genomics UK (COG-UK) Consortium | Lighthouse Laboratory Plymouth and Alex Alderton, Roberto Amato, Jeffrey Barrett, Sonia Goncalves, Ewan Harrison, David K. Jackson, Ian Johnston, Dominic Kwiatkowski, Cordelia Langford, John Sillitoe on behalf of the Wellcome Sanger Institute COVID-19 Surveillance Team |
| EPI_ISL_3440255, EPI_ISL_3440256, EPI_ISL_3440257, EPI_ISL_3440258 | Lighthouse Lab in Glasgow | Wellcome Sanger Institute for the COVID-19 Genomics UK (COG-UK) Consortium | Harper VanSteenhouse, Yumi Kasai, David Gray, Carol Clugston, Anna Dominiczak and Alex Alderton, Roberto Amato, Jeffrey Barrett, Sonia Goncalves, Ewan Harrison, David K. Jackson, Ian Johnston, Dominic Kwiatkowski, Cordelia Langford, John Sillitoe on behalf of the Wellcome Sanger Institute COVID-19 Surveillance Team |
| EPI_ISL_3440259 | Lighthouse Laboratory Plymouth | Wellcome Sanger Institute for the COVID-19 Genomics UK (COG-UK) Consortium | Lighthouse Laboratory Plymouth and Alex Alderton, Roberto Amato, Jeffrey Barrett, Sonia Goncalves, Ewan Harrison, David K. Jackson, Ian Johnston, Dominic Kwiatkowski, Cordelia Langford, John Sillitoe on behalf of the Wellcome Sanger Institute COVID-19 Surveillance Team |
| EPI_ISL_3440260, EPI_ISL_3440261, EPI_ISL_3440262, EPI_ISL_3440263, EPI_ISL_3440264, EPI_ISL_3440265, EPI_ISL_3440266, EPI_ISL_3440267, EPI_ISL_3440268, EPI_ISL_3440269 | Lighthouse Lab in Glasgow | Wellcome Sanger Institute for the COVID-19 Genomics UK (COG-UK) Consortium | Harper VanSteenhouse, Yumi Kasai, David Gray, Carol Clugston, Anna Dominiczak and Alex Alderton, Roberto Amato, Jeffrey Barrett, Sonia Goncalves, Ewan Harrison, David K. Jackson, Ian Johnston, Dominic Kwiatkowski, Cordelia Langford, John Sillitoe on behalf of the Wellcome Sanger Institute COVID-19 Surveillance Team |
| EPI_ISL_3440270 | Lighthouse Laboratory Plymouth | Wellcome Sanger Institute for the COVID-19 Genomics UK (COG-UK) Consortium | Lighthouse Laboratory Plymouth and Alex Alderton, Roberto Amato, Jeffrey Barrett, Sonia Goncalves, Ewan Harrison, David K. Jackson, Ian Johnston, Dominic Kwiatkowski, Cordelia Langford, John Sillitoe on behalf of the Wellcome Sanger Institute COVID-19 Surveillance Team |
| EPI_ISL_3440271, EPI_ISL_3440272, EPI_ISL_3440273 | Lighthouse Lab in Glasgow | Wellcome Sanger Institute for the COVID-19 Genomics UK (COG-UK) Consortium | Harper VanSteenhouse, Yumi Kasai, David Gray, Carol Clugston, Anna Dominiczak and Alex Alderton, Roberto Amato, Jeffrey Barrett, Sonia Goncalves, Ewan Harrison, David K. Jackson, Ian Johnston, Dominic Kwiatkowski, Cordelia Langford, John Sillitoe on behalf of the Wellcome Sanger Institute COVID-19 Surveillance Team |
| EPI_ISL_3440274 | Lighthouse Laboratory Plymouth | Wellcome Sanger Institute for the COVID-19 Genomics UK (COG-UK) Consortium | Lighthouse Laboratory Plymouth and Alex Alderton, Roberto Amato, Jeffrey Barrett, Sonia Goncalves, Ewan Harrison, David K. Jackson, Ian Johnston, Dominic Kwiatkowski, Cordelia Langford, John Sillitoe on behalf of the Wellcome Sanger Institute COVID-19 Surveillance Team |
| EPI_ISL_3440275, EPI_ISL_3440276 | Lighthouse Lab in Glasgow | Wellcome Sanger Institute for the COVID-19 Genomics UK (COG-UK) Consortium | Harper VanSteenhouse, Yumi Kasai, David Gray, Carol Clugston, Anna Dominiczak and Alex Alderton, Roberto Amato, Jeffrey Barrett, Sonia Goncalves, Ewan Harrison, David K. Jackson, Ian Johnston, Dominic Kwiatkowski, Cordelia Langford, John Sillitoe on behalf of the Wellcome Sanger Institute COVID-19 Surveillance Team |
| EPI_ISL_3440277 | Lighthouse Laboratory Plymouth | Wellcome Sanger Institute for the COVID-19 Genomics UK (COG-UK) Consortium | Lighthouse Laboratory Plymouth and Alex Alderton, Roberto Amato, Jeffrey Barrett, Sonia Goncalves, Ewan Harrison, David K. Jackson, Ian Johnston, Dominic Kwiatkowski, Cordelia Langford, John Sillitoe on behalf of the Wellcome Sanger Institute COVID-19 Surveillance Team |
| EPI_ISL_3440278 | Lighthouse Lab in Glasgow | Wellcome Sanger Institute for the COVID-19 Genomics UK (COG-UK) Consortium | Harper VanSteenhouse, Yumi Kasai, David Gray, Carol Clugston, Anna Dominiczak and Alex Alderton, Roberto Amato, Jeffrey Barrett, Sonia Goncalves, Ewan Harrison, David K. Jackson, Ian Johnston, Dominic Kwiatkowski, Cordelia Langford, John Sillitoe on behalf of the Wellcome Sanger Institute COVID-19 Surveillance Team |
| EPI_ISL_3440279 | Lighthouse Laboratory Plymouth | Wellcome Sanger Institute for the COVID-19 Genomics UK (COG-UK) Consortium | Lighthouse Laboratory Plymouth and Alex Alderton, Roberto Amato, Jeffrey Barrett, Sonia Goncalves, Ewan Harrison, David K. Jackson, Ian Johnston, Dominic Kwiatkowski, Cordelia Langford, John Sillitoe on behalf of the Wellcome Sanger Institute COVID-19 Surveillance Team |
| EPI_ISL_3440280, EPI_ISL_3440281, EPI_ISL_3440282, EPI_ISL_3440283, EPI_ISL_3440284 | Lighthouse Lab in Glasgow | Wellcome Sanger Institute for the COVID-19 Genomics UK (COG-UK) Consortium | Harper VanSteenhouse, Yumi Kasai, David Gray, Carol Clugston, Anna Dominiczak and Alex Alderton, Roberto Amato, Jeffrey Barrett, Sonia Goncalves, Ewan Harrison, David K. Jackson, Ian Johnston, Dominic Kwiatkowski, Cordelia Langford, John Sillitoe on behalf of the Wellcome Sanger Institute COVID-19 Surveillance Team |
| EPI_ISL_3440285 | Lighthouse Laboratory Plymouth | Wellcome Sanger Institute for the COVID-19 Genomics UK (COG-UK) Consortium | Lighthouse Laboratory Plymouth and Alex Alderton, Roberto Amato, Jeffrey Barrett, Sonia Goncalves, Ewan Harrison, David K. Jackson, Ian Johnston, Dominic Kwiatkowski, Cordelia Langford, John Sillitoe on behalf of the Wellcome Sanger Institute COVID-19 Surveillance Team |
| EPI_ISL_3440286, EPI_ISL_3440287, EPI_ISL_3440288, EPI_ISL_3440289, EPI_ISL_3440290, EPI_ISL_3440291 | Lighthouse Lab in Glasgow | Wellcome Sanger Institute for the COVID-19 Genomics UK (COG-UK) Consortium | Harper VanSteenhouse, Yumi Kasai, David Gray, Carol Clugston, Anna Dominiczak and Alex Alderton, Roberto Amato, Jeffrey Barrett, Sonia Goncalves, Ewan Harrison, David K. Jackson, Ian Johnston, Dominic Kwiatkowski, Cordelia Langford, John Sillitoe on behalf of the Wellcome Sanger Institute COVID-19 Surveillance Team |
| EPI_ISL_3440292, EPI_ISL_3440293 | Lighthouse Laboratory Plymouth | Wellcome Sanger Institute for the COVID-19 Genomics UK (COG-UK) Consortium | Lighthouse Laboratory Plymouth and Alex Alderton, Roberto Amato, Jeffrey Barrett, Sonia Goncalves, Ewan Harrison, David K. Jackson, Ian Johnston, Dominic Kwiatkowski, Cordelia Langford, John Sillitoe on behalf of the Wellcome Sanger Institute COVID-19 Surveillance Team |
| EPI_ISL_3440294, EPI_ISL_3440295, EPI_ISL_3440296, EPI_ISL_3440297, EPI_ISL_3440298, EPI_ISL_3440299 | Lighthouse Lab in Glasgow | Wellcome Sanger Institute for the COVID-19 Genomics UK (COG-UK) Consortium | Harper VanSteenhouse, Yumi Kasai, David Gray, Carol Clugston, Anna Dominiczak and Alex Alderton, Roberto Amato, Jeffrey Barrett, Sonia Goncalves, Ewan Harrison, David K. Jackson, Ian Johnston, Dominic Kwiatkowski, Cordelia Langford, John Sillitoe on behalf of the Wellcome Sanger Institute COVID-19 Surveillance Team |
| EPI_ISL_3440300 | Lighthouse Laboratory Plymouth | Wellcome Sanger Institute for the COVID-19 Genomics UK (COG-UK) Consortium | Lighthouse Laboratory Plymouth and Alex Alderton, Roberto Amato, Jeffrey Barrett, Sonia Goncalves, Ewan Harrison, David K. Jackson, Ian Johnston, Dominic Kwiatkowski, Cordelia Langford, John Sillitoe on behalf of the Wellcome Sanger Institute COVID-19 Surveillance Team |
| EPI_ISL_3440301, EPI_ISL_3440302 | Lighthouse Lab in Glasgow | Wellcome Sanger Institute for the COVID-19 Genomics UK (COG-UK) Consortium | Harper VanSteenhouse, Yumi Kasai, David Gray, Carol Clugston, Anna Dominiczak and Alex Alderton, Roberto Amato, Jeffrey Barrett, Sonia Goncalves, Ewan Harrison, David K. Jackson, Ian Johnston, Dominic Kwiatkowski, Cordelia Langford, John Sillitoe on behalf of the Wellcome Sanger Institute COVID-19 Surveillance Team |
| EPI_ISL_3440303 | Lighthouse Laboratory Plymouth | Wellcome Sanger Institute for the COVID-19 Genomics UK (COG-UK) Consortium | Lighthouse Laboratory Plymouth and Alex Alderton, Roberto Amato, Jeffrey Barrett, Sonia Goncalves, Ewan Harrison, David K. Jackson, Ian Johnston, Dominic Kwiatkowski, Cordelia Langford, John Sillitoe on behalf of the Wellcome Sanger Institute COVID-19 Surveillance Team |
| EPI_ISL_3440304 | Lighthouse Lab in Glasgow | Wellcome Sanger Institute for the COVID-19 Genomics UK (COG-UK) Consortium | Harper VanSteenhouse, Yumi Kasai, David Gray, Carol Clugston, Anna Dominiczak and Alex Alderton, Roberto Amato, Jeffrey Barrett, Sonia Goncalves, Ewan Harrison, David K. Jackson, Ian Johnston, Dominic Kwiatkowski, Cordelia Langford, John Sillitoe on behalf of the Wellcome Sanger Institute COVID-19 Surveillance Team |
| EPI_ISL_3440305 | Lighthouse Laboratory Plymouth | Wellcome Sanger Institute for the COVID-19 Genomics UK (COG-UK) Consortium | Lighthouse Laboratory Plymouth and Alex Alderton, Roberto Amato, Jeffrey Barrett, Sonia Goncalves, Ewan Harrison, David K. Jackson, Ian Johnston, Dominic Kwiatkowski, Cordelia Langford, John Sillitoe on behalf of the Wellcome Sanger Institute COVID-19 Surveillance Team |
| EPI_ISL_3440306 | Lighthouse Lab in Glasgow | Wellcome Sanger Institute for the COVID-19 Genomics UK (COG-UK) Consortium | Harper VanSteenhouse, Yumi Kasai, David Gray, Carol Clugston, Anna Dominiczak and Alex Alderton, Roberto Amato, Jeffrey Barrett, Sonia Goncalves, Ewan Harrison, David K. Jackson, Ian Johnston, Dominic Kwiatkowski, Cordelia Langford, John Sillitoe on behalf of the Wellcome Sanger Institute COVID-19 Surveillance Team |
| EPI_ISL_3440307 | Lighthouse Laboratory Plymouth | Wellcome Sanger Institute for the COVID-19 Genomics UK (COG-UK) Consortium | Lighthouse Laboratory Plymouth and Alex Alderton, Roberto Amato, Jeffrey Barrett, Sonia Goncalves, Ewan Harrison, David K. Jackson, Ian Johnston, Dominic Kwiatkowski, Cordelia Langford, John Sillitoe on behalf of the Wellcome Sanger Institute COVID-19 Surveillance Team |
| EPI_ISL_3440308, EPI_ISL_3440309, EPI_ISL_3440310 | Lighthouse Lab in Glasgow | Wellcome Sanger Institute for the COVID-19 Genomics UK (COG-UK) Consortium | Harper VanSteenhouse, Yumi Kasai, David Gray, Carol Clugston, Anna Dominiczak and Alex Alderton, Roberto Amato, Jeffrey Barrett, Sonia Goncalves, Ewan Harrison, David K. Jackson, Ian Johnston, Dominic Kwiatkowski, Cordelia Langford, John Sillitoe on behalf of the Wellcome Sanger Institute COVID-19 Surveillance Team |
| EPI_ISL_3440311 | Lighthouse Laboratory Plymouth | Wellcome Sanger Institute for the COVID-19 Genomics UK (COG-UK) Consortium | Lighthouse Laboratory Plymouth and Alex Alderton, Roberto Amato, Jeffrey Barrett, Sonia Goncalves, Ewan Harrison, David K. Jackson, Ian Johnston, Dominic Kwiatkowski, Cordelia Langford, John Sillitoe on behalf of the Wellcome Sanger Institute COVID-19 Surveillance Team |
| EPI_ISL_3440312, EPI_ISL_3440313 | Lighthouse Lab in Glasgow | Wellcome Sanger Institute for the COVID-19 Genomics UK (COG-UK) Consortium | Harper VanSteenhouse, Yumi Kasai, David Gray, Carol Clugston, Anna Dominiczak and Alex Alderton, Roberto Amato, Jeffrey Barrett, Sonia Goncalves, Ewan Harrison, David K. Jackson, Ian Johnston, Dominic Kwiatkowski, Cordelia Langford, John Sillitoe on behalf of the Wellcome Sanger Institute COVID-19 Surveillance Team |

| | | | |
|---|---|---|---|
| EPI_ISL_3440314 | Lighthouse Laboratory Plymouth | Wellcome Sanger Institute for the COVID-19 Genomics UK (COG-UK) Consortium | Lighthouse Laboratory Plymouth and Alex Alderton, Roberto Amato, Jeffrey Barrett, Sonia Goncalves, Ewan Harrison, David K. Jackson, Ian Johnston, Dominic Kwiatkowski, Cordelia Langford, John Sillitoe on behalf of the Wellcome Sanger Institute COVID-19 Surveillance Team |
| EPI_ISL_3440315, EPI_ISL_3440316 | Lighthouse Lab in Glasgow | Wellcome Sanger Institute for the COVID-19 Genomics UK (COG-UK) Consortium | Harper VanSteenhouse, Yumi Kasai, David Gray, Carol Clugston, Anna Dominiczak and Alex Alderton, Roberto Amato, Jeffrey Barrett, Sonia Goncalves, Ewan Harrison, David K. Jackson, Ian Johnston, Dominic Kwiatkowski, Cordelia Langford, John Sillitoe on behalf of the Wellcome Sanger Institute COVID-19 Surveillance Team |
| EPI_ISL_3440317 | Lighthouse Laboratory Plymouth | Wellcome Sanger Institute for the COVID-19 Genomics UK (COG-UK) Consortium | Lighthouse Laboratory Plymouth and Alex Alderton, Roberto Amato, Jeffrey Barrett, Sonia Goncalves, Ewan Harrison, David K. Jackson, Ian Johnston, Dominic Kwiatkowski, Cordelia Langford, John Sillitoe on behalf of the Wellcome Sanger Institute COVID-19 Surveillance Team |
| EPI_ISL_3440318, EPI_ISL_3440319, EPI_ISL_3440320, EPI_ISL_3440321, EPI_ISL_3440322 | Lighthouse Lab in Glasgow | Wellcome Sanger Institute for the COVID-19 Genomics UK (COG-UK) Consortium | Harper VanSteenhouse, Yumi Kasai, David Gray, Carol Clugston, Anna Dominiczak and Alex Alderton, Roberto Amato, Jeffrey Barrett, Sonia Goncalves, Ewan Harrison, David K. Jackson, Ian Johnston, Dominic Kwiatkowski, Cordelia Langford, John Sillitoe on behalf of the Wellcome Sanger Institute COVID-19 Surveillance Team |
| EPI_ISL_3440323 | Lighthouse Laboratory Plymouth | Wellcome Sanger Institute for the COVID-19 Genomics UK (COG-UK) Consortium | Lighthouse Laboratory Plymouth and Alex Alderton, Roberto Amato, Jeffrey Barrett, Sonia Goncalves, Ewan Harrison, David K. Jackson, Ian Johnston, Dominic Kwiatkowski, Cordelia Langford, John Sillitoe on behalf of the Wellcome Sanger Institute COVID-19 Surveillance Team |
| EPI_ISL_3440324 | Lighthouse Lab in Glasgow | Wellcome Sanger Institute for the COVID-19 Genomics UK (COG-UK) Consortium | Harper VanSteenhouse, Yumi Kasai, David Gray, Carol Clugston, Anna Dominiczak and Alex Alderton, Roberto Amato, Jeffrey Barrett, Sonia Goncalves, Ewan Harrison, David K. Jackson, Ian Johnston, Dominic Kwiatkowski, Cordelia Langford, John Sillitoe on behalf of the Wellcome Sanger Institute COVID-19 Surveillance Team |
| EPI_ISL_3440325 | Lighthouse Laboratory Plymouth | Wellcome Sanger Institute for the COVID-19 Genomics UK (COG-UK) Consortium | Lighthouse Laboratory Plymouth and Alex Alderton, Roberto Amato, Jeffrey Barrett, Sonia Goncalves, Ewan Harrison, David K. Jackson, Ian Johnston, Dominic Kwiatkowski, Cordelia Langford, John Sillitoe on behalf of the Wellcome Sanger Institute COVID-19 Surveillance Team |
| EPI_ISL_3440326 | Lighthouse Lab in Glasgow | Wellcome Sanger Institute for the COVID-19 Genomics UK (COG-UK) Consortium | Harper VanSteenhouse, Yumi Kasai, David Gray, Carol Clugston, Anna Dominiczak and Alex Alderton, Roberto Amato, Jeffrey Barrett, Sonia Goncalves, Ewan Harrison, David K. Jackson, Ian Johnston, Dominic Kwiatkowski, Cordelia Langford, John Sillitoe on behalf of the Wellcome Sanger Institute COVID-19 Surveillance Team |
| EPI_ISL_3440327 | Lighthouse Laboratory Plymouth | Wellcome Sanger Institute for the COVID-19 Genomics UK (COG-UK) Consortium | Lighthouse Laboratory Plymouth and Alex Alderton, Roberto Amato, Jeffrey Barrett, Sonia Goncalves, Ewan Harrison, David K. Jackson, Ian Johnston, Dominic Kwiatkowski, Cordelia Langford, John Sillitoe on behalf of the Wellcome Sanger Institute COVID-19 Surveillance Team |
| EPI_ISL_3440328, EPI_ISL_3440329 | Lighthouse Lab in Glasgow | Wellcome Sanger Institute for the COVID-19 Genomics UK (COG-UK) Consortium | Harper VanSteenhouse, Yumi Kasai, David Gray, Carol Clugston, Anna Dominiczak and Alex Alderton, Roberto Amato, Jeffrey Barrett, Sonia Goncalves, Ewan Harrison, David K. Jackson, Ian Johnston, Dominic Kwiatkowski, Cordelia Langford, John Sillitoe on behalf of the Wellcome Sanger Institute COVID-19 Surveillance Team |
| EPI_ISL_3440330 | Lighthouse Laboratory Plymouth | Wellcome Sanger Institute for the COVID-19 Genomics UK (COG-UK) Consortium | Lighthouse Laboratory Plymouth and Alex Alderton, Roberto Amato, Jeffrey Barrett, Sonia Goncalves, Ewan Harrison, David K. Jackson, Ian Johnston, Dominic Kwiatkowski, Cordelia Langford, John Sillitoe on behalf of the Wellcome Sanger Institute COVID-19 Surveillance Team |
| EPI_ISL_3440331, EPI_ISL_3440332 | Lighthouse Lab in Glasgow | Wellcome Sanger Institute for the COVID-19 Genomics UK (COG-UK) Consortium | Harper VanSteenhouse, Yumi Kasai, David Gray, Carol Clugston, Anna Dominiczak and Alex Alderton, Roberto Amato, Jeffrey Barrett, Sonia Goncalves, Ewan Harrison, David K. Jackson, Ian Johnston, Dominic Kwiatkowski, Cordelia Langford, John Sillitoe on behalf of the Wellcome Sanger Institute COVID-19 Surveillance Team |
| EPI_ISL_3440333, EPI_ISL_3440334 | Lighthouse Laboratory Plymouth | Wellcome Sanger Institute for the COVID-19 Genomics UK (COG-UK) Consortium | Lighthouse Laboratory Plymouth and Alex Alderton, Roberto Amato, Jeffrey Barrett, Sonia Goncalves, Ewan Harrison, David K. Jackson, Ian Johnston, Dominic Kwiatkowski, Cordelia Langford, John Sillitoe on behalf of the Wellcome Sanger Institute COVID-19 Surveillance Team |
| EPI_ISL_3440335, EPI_ISL_3440336 | Lighthouse Lab in Glasgow | Wellcome Sanger Institute for the COVID-19 Genomics UK (COG-UK) Consortium | Harper VanSteenhouse, Yumi Kasai, David Gray, Carol Clugston, Anna Dominiczak and Alex Alderton, Roberto Amato, Jeffrey Barrett, Sonia Goncalves, Ewan Harrison, David K. Jackson, Ian Johnston, Dominic Kwiatkowski, Cordelia Langford, John Sillitoe on behalf of the Wellcome Sanger Institute COVID-19 Surveillance Team |
| EPI_ISL_3440337, EPI_ISL_3440338 | Lighthouse Laboratory Plymouth | Wellcome Sanger Institute for the COVID-19 Genomics UK (COG-UK) Consortium | Lighthouse Laboratory Plymouth and Alex Alderton, Roberto Amato, Jeffrey Barrett, Sonia Goncalves, Ewan Harrison, David K. Jackson, Ian Johnston, Dominic Kwiatkowski, Cordelia Langford, John Sillitoe on behalf of the Wellcome Sanger Institute COVID-19 Surveillance Team |
| EPI_ISL_3440339, EPI_ISL_3440340, EPI_ISL_3440341, EPI_ISL_3440342, EPI_ISL_3440343 | Lighthouse Lab in Glasgow | Wellcome Sanger Institute for the COVID-19 Genomics UK (COG-UK) Consortium | Harper VanSteenhouse, Yumi Kasai, David Gray, Carol Clugston, Anna Dominiczak and Alex Alderton, Roberto Amato, Jeffrey Barrett, Sonia Goncalves, Ewan Harrison, David K. Jackson, Ian Johnston, Dominic Kwiatkowski, Cordelia Langford, John Sillitoe on behalf of the Wellcome Sanger Institute COVID-19 Surveillance Team |
| EPI_ISL_3440344 | Lighthouse Laboratory Plymouth | Wellcome Sanger Institute for the COVID-19 Genomics UK (COG-UK) Consortium | Lighthouse Laboratory Plymouth and Alex Alderton, Roberto Amato, Jeffrey Barrett, Sonia Goncalves, Ewan Harrison, David K. Jackson, Ian Johnston, Dominic Kwiatkowski, Cordelia Langford, John Sillitoe on behalf of the Wellcome Sanger Institute COVID-19 Surveillance Team |
| EPI_ISL_3440345 | Lighthouse Lab in Glasgow | Wellcome Sanger Institute for the COVID-19 Genomics UK (COG-UK) Consortium | Harper VanSteenhouse, Yumi Kasai, David Gray, Carol Clugston, Anna Dominiczak and Alex Alderton, Roberto Amato, Jeffrey Barrett, Sonia Goncalves, Ewan Harrison, David K. Jackson, Ian Johnston, Dominic Kwiatkowski, Cordelia Langford, John Sillitoe on behalf of the Wellcome Sanger Institute COVID-19 Surveillance Team |
| EPI_ISL_3440346, EPI_ISL_3440347 | Lighthouse Laboratory Plymouth | Wellcome Sanger Institute for the COVID-19 Genomics UK (COG-UK) Consortium | Lighthouse Laboratory Plymouth and Alex Alderton, Roberto Amato, Jeffrey Barrett, Sonia Goncalves, Ewan Harrison, David K. Jackson, Ian Johnston, Dominic Kwiatkowski, Cordelia Langford, John Sillitoe on behalf of the Wellcome Sanger Institute COVID-19 Surveillance Team |
| EPI_ISL_3440348 | Lighthouse Lab in Glasgow | Wellcome Sanger Institute for the COVID-19 Genomics UK (COG-UK) Consortium | Harper VanSteenhouse, Yumi Kasai, David Gray, Carol Clugston, Anna Dominiczak and Alex Alderton, Roberto Amato, Jeffrey Barrett, Sonia Goncalves, Ewan Harrison, David K. Jackson, Ian Johnston, Dominic Kwiatkowski, Cordelia Langford, John Sillitoe on behalf of the Wellcome Sanger Institute COVID-19 Surveillance Team |
| EPI_ISL_3440349, EPI_ISL_3440350 | Lighthouse Laboratory Plymouth | Wellcome Sanger Institute for the COVID-19 Genomics UK (COG-UK) Consortium | Lighthouse Laboratory Plymouth and Alex Alderton, Roberto Amato, Jeffrey Barrett, Sonia Goncalves, Ewan Harrison, David K. Jackson, Ian Johnston, Dominic Kwiatkowski, Cordelia Langford, John Sillitoe on behalf of the Wellcome Sanger Institute COVID-19 Surveillance Team |
| EPI_ISL_3440351, EPI_ISL_3440352, EPI_ISL_3440353 | Lighthouse Lab in Glasgow | Wellcome Sanger Institute for the COVID-19 Genomics UK (COG-UK) Consortium | Harper VanSteenhouse, Yumi Kasai, David Gray, Carol Clugston, Anna Dominiczak and Alex Alderton, Roberto Amato, Jeffrey Barrett, Sonia Goncalves, Ewan Harrison, David K. Jackson, Ian Johnston, Dominic Kwiatkowski, Cordelia Langford, John Sillitoe on behalf of the Wellcome Sanger Institute COVID-19 Surveillance Team |
| EPI_ISL_3440354 | Lighthouse Laboratory Plymouth | Wellcome Sanger Institute for the COVID-19 Genomics UK (COG-UK) Consortium | Lighthouse Laboratory Plymouth and Alex Alderton, Roberto Amato, Jeffrey Barrett, Sonia Goncalves, Ewan Harrison, David K. Jackson, Ian Johnston, Dominic Kwiatkowski, Cordelia Langford, John Sillitoe on behalf of the Wellcome Sanger Institute COVID-19 Surveillance Team |
| EPI_ISL_3440355, EPI_ISL_3440356, EPI_ISL_3440357, EPI_ISL_3440358, EPI_ISL_3440359, EPI_ISL_3440360, EPI_ISL_3440361, EPI_ISL_3440362, EPI_ISL_3440363, EPI_ISL_3440364, EPI_ISL_3440365, EPI_ISL_3440366, EPI_ISL_3440367, EPI_ISL_3440368, EPI_ISL_3440369, EPI_ISL_3440370, EPI_ISL_3440371, EPI_ISL_3440372, EPI_ISL_3440373, EPI_ISL_3440374, EPI_ISL_3440375, EPI_ISL_3440376, EPI_ISL_3440377, EPI_ISL_3440378, EPI_ISL_3440379, EPI_ISL_3440380, EPI_ISL_3440381, EPI_ISL_3440382, EPI_ISL_3440383, EPI_ISL_3440384, EPI_ISL_3440385, EPI_ISL_3440386, EPI_ISL_3440387, EPI_ISL_3440388, EPI_ISL_3440389, EPI_ISL_3440390, EPI_ISL_3440391, EPI_ISL_3440392, EPI_ISL_3440393, EPI_ISL_3440394, EPI_ISL_3440395, EPI_ISL_3440396, EPI_ISL_3440397, EPI_ISL_3440398, EPI_ISL_3440399, EPI_ISL_3440400, EPI_ISL_3440401, EPI_ISL_3440402, EPI_ISL_3440403, EPI_ISL_3440404, EPI_ISL_3440405, EPI_ISL_3440406, EPI_ISL_3440407, EPI_ISL_3440408, EPI_ISL_3440409, EPI_ISL_3440410, EPI_ISL_3440411, EPI_ISL_3440412, EPI_ISL_3440413, EPI_ISL_3440414, EPI_ISL_3440415, EPI_ISL_3440416, EPI_ISL_3440417, EPI_ISL_3440418, EPI_ISL_3440419, EPI_ISL_3440420, EPI_ISL_3440421, EPI_ISL_3440422, EPI_ISL_3440423, EPI_ISL_3440424, EPI_ISL_3440425, EPI_ISL_3440426, EPI_ISL_3440427, EPI_ISL_3440428, EPI_ISL_3440429, EPI_ISL_3440430, EPI_ISL_3440431, EPI_ISL_3440432, EPI_ISL_3440433, EPI_ISL_3440434, EPI_ISL_3440435, EPI_ISL_3440436, EPI_ISL_3440437, EPI_ISL_3440438, EPI_ISL_3440439, EPI_ISL_3440440, EPI_ISL_3440441, EPI_ISL_3440442, EPI_ISL_3440443, EPI_ISL_3440444, EPI_ISL_3440445, EPI_ISL_3440446, EPI_ISL_3440447, EPI_ISL_3440448, EPI_ISL_3440449, EPI_ISL_3440450, EPI_ISL_3440451, EPI_ISL_3440452, EPI_ISL_3440453, EPI_ISL_3440454, EPI_ISL_3440455, EPI_ISL_3440456, EPI_ISL_3440457, EPI_ISL_3440458, EPI_ISL_3440459, EPI_ISL_3440460, EPI_ISL_3440461, EPI_ISL_3440462, EPI_ISL_3440463, EPI_ISL_3440464, EPI_ISL_3440465, EPI_ISL_3440466, EPI_ISL_3440467, EPI_ISL_3440468, EPI_ISL_3440469, EPI_ISL_3440470, EPI_ISL_3440471, EPI_ISL_3440472, EPI_ISL_3440473, EPI_ISL_3440474, EPI_ISL_3440475, EPI_ISL_3440476, EPI_ISL_3440477, EPI_ISL_3440478, EPI_ISL_3440479, EPI_ISL_3440480, EPI_ISL_3440481, EPI_ISL_3440482, EPI_ISL_3440483, EPI_ISL_3440484, EPI_ISL_3440485, EPI_ISL_3440486, EPI_ISL_3440487, EPI_ISL_3440488, EPI_ISL_3440489, EPI_ISL_3440490, EPI_ISL_3440491, EPI_ISL_3440492, EPI_ISL_3440493, EPI_ISL_3440494, EPI_ISL_3440495, EPI_ISL_3440496, EPI_ISL_3440497, EPI_ISL_3440498, EPI_ISL_3440499, EPI_ISL_3440500, EPI_ISL_3440501, EPI_ISL_3440502, EPI_ISL_3440503, EPI_ISL_3440504, EPI_ISL_3440505, EPI_ISL_3440506, EPI_ISL_3440507, EPI_ISL_3440508, EPI_ISL_3440509, EPI_ISL_3440510, EPI_ISL_3440511, EPI_ISL_3440512, EPI_ISL_3440513, EPI_ISL_3440514, EPI_ISL_3440515, EPI_ISL_3440516, EPI_ISL_3440517, EPI_ISL_3440518, EPI_ISL_3440519, EPI_ISL_3440520, EPI_ISL_3440521, EPI_ISL_3440522, EPI_ISL_3440523, EPI_ISL_3440524, EPI_ISL_3440525, EPI_ISL_3440526, EPI_ISL_3440527, EPI_ISL_3440528, EPI_ISL_3440529, EPI_ISL_3440530, EPI_ISL_3440531, EPI_ISL_3440532, EPI_ISL_3440533, EPI_ISL_3440534, EPI_ISL_3440535, EPI_ISL_3440536, EPI_ISL_3440537, EPI_ISL_3440538, EPI_ISL_3440539, EPI_ISL_3440540, EPI_ISL_3440541, EPI_ISL_3440542, EPI_ISL_3440543, EPI_ISL_3440544, EPI_ISL_3440545, EPI_ISL_3440546, EPI_ISL_3440547, EPI_ISL_3440548, EPI_ISL_3440549, EPI_ISL_3440550, EPI_ISL_3440551, EPI_ISL_3440552, EPI_ISL_3440553, EPI_ISL_3440554, EPI_ISL_3440555, EPI_ISL_3440556, EPI_ISL_3440557, EPI_ISL_3440558, EPI_ISL_3440559, EPI_ISL_3440560, EPI_ISL_3440561, EPI_ISL_3440562, EPI_ISL_3440563, EPI_ISL_3440564, EPI_ISL_3440565, EPI_ISL_3440566, EPI_ISL_3440567, EPI_ISL_3440568, EPI_ISL_3440569, EPI_ISL_3440570, EPI_ISL_3440571, EPI_ISL_3440572, EPI_ISL_3440573, EPI_ISL_3440574, EPI_ISL_3440575, EPI_ISL_3440576, EPI_ISL_3440577, EPI_ISL_3440578, EPI_ISL_3440579, EPI_ISL_3440580, EPI_ISL_3440581, EPI_ISL_3440582, EPI_ISL_3440583, EPI_ISL_3440584, EPI_ISL_3440585, EPI_ISL_3440586, EPI_ISL_3440587, EPI_ISL_3440588, EPI_ISL_3440589, EPI_ISL_3440590, EPI_ISL_3440591, EPI_ISL_3440592, EPI_ISL_3440593, EPI_ISL_3440594, EPI_ISL_3440595, EPI_ISL_3440596, EPI_ISL_3440597, EPI_ISL_3440598, EPI_ISL_3440599, EPI_ISL_3440600, EPI_ISL_3440601, EPI_ISL_3440602, EPI_ISL_3440603, EPI_ISL_3440604, EPI_ISL_3440605, EPI_ISL_3440606, EPI_ISL_3440607, EPI_ISL_3440608, EPI_ISL_3440609, EPI_ISL_3440610, EPI_ISL_3440611 | | | |
| see above | Lighthouse Lab in Alderley Park | Wellcome Sanger Institute for the COVID-19 Genomics UK (COG-UK) Consortium | Jacquelyn Wynn, Mairead Hyland, The Lighthouse Lab in Alderley Park and Alex Alderton, Roberto Amato, Jeffrey Barrett, Sonia Goncalves, Ewan Harrison, David K. Jackson, Ian Johnston, Dominic Kwiatkowski, Cordelia Langford, John Sillitoe on behalf of the Wellcome Sanger Institute COVID-19 Surveillance Team |
| EPI_ISL_3440612 | Lighthouse Laboratory Plymouth | Wellcome Sanger Institute for the COVID-19 Genomics UK (COG-UK) Consortium | Lighthouse Laboratory Plymouth and Alex Alderton, Roberto Amato, Jeffrey Barrett, Sonia Goncalves, Ewan Harrison, David K. Jackson, Ian Johnston, Dominic Kwiatkowski, Cordelia Langford, John Sillitoe on behalf of the Wellcome Sanger Institute COVID-19 Surveillance Team |
| EPI_ISL_3440613, EPI_ISL_3440614, EPI_ISL_3440615 | Lighthouse Lab in Milton Keynes | Wellcome Sanger Institute for the COVID-19 Genomics UK (COG-UK) Consortium | The Lighthouse Lab in Milton Keynes and Alex Alderton, Roberto Amato, Jeffrey Barrett, Sonia Goncalves, Ewan Harrison, David K. Jackson, Ian Johnston, Dominic Kwiatkowski, Cordelia Langford, John Sillitoe on behalf of the Wellcome Sanger Institute COVID-19 Surveillance Team |
| EPI_ISL_3440616 | Lighthouse Laboratory Plymouth | Wellcome Sanger Institute for the COVID-19 Genomics UK (COG-UK) Consortium | Lighthouse Laboratory Plymouth and Alex Alderton, Roberto Amato, Jeffrey Barrett, Sonia Goncalves, Ewan Harrison, David K. Jackson, Ian Johnston, Dominic Kwiatkowski, Cordelia Langford, John Sillitoe on behalf of the Wellcome Sanger Institute COVID-19 Surveillance Team |
| EPI_ISL_3440617, EPI_ISL_3440618, EPI_ISL_3440619 | Lighthouse Lab in Milton Keynes | Wellcome Sanger Institute for the COVID-19 Genomics UK (COG-UK) Consortium | The Lighthouse Lab in Milton Keynes and Alex Alderton, Roberto Amato, Jeffrey Barrett, Sonia Goncalves, Ewan Harrison, David K. Jackson, Ian Johnston, Dominic Kwiatkowski, Cordelia Langford, John Sillitoe on behalf of the Wellcome Sanger Institute COVID-19 Surveillance Team |
| EPI_ISL_3440620 | Lighthouse Laboratory Plymouth | Wellcome Sanger Institute for the COVID-19 Genomics UK (COG-UK) Consortium | Lighthouse Laboratory Plymouth and Alex Alderton, Roberto Amato, Jeffrey Barrett, Sonia Goncalves, Ewan Harrison, David K. Jackson, Ian Johnston, Dominic Kwiatkowski, Cordelia Langford, John Sillitoe on behalf of the Wellcome Sanger Institute COVID-19 Surveillance Team |
| EPI_ISL_3440621 | Lighthouse Lab in Milton Keynes | Wellcome Sanger Institute for the COVID-19 Genomics UK (COG-UK) Consortium | The Lighthouse Lab in Milton Keynes and Alex Alderton, Roberto Amato, Jeffrey Barrett, Sonia Goncalves, Ewan Harrison, David K. Jackson, Ian Johnston, Dominic Kwiatkowski, Cordelia Langford, John Sillitoe on behalf of the Wellcome Sanger Institute COVID-19 Surveillance Team |
| EPI_ISL_3440622, EPI_ISL_3440623, EPI_ISL_3440624 | Lighthouse Laboratory Plymouth | Wellcome Sanger Institute for the COVID-19 Genomics UK (COG-UK) Consortium | Lighthouse Laboratory Plymouth and Alex Alderton, Roberto Amato, Jeffrey Barrett, Sonia Goncalves, Ewan Harrison, David K. Jackson, Ian Johnston, Dominic Kwiatkowski, Cordelia Langford, John Sillitoe on behalf of the Wellcome Sanger Institute COVID-19 Surveillance Team |
| EPI_ISL_3440625 | Lighthouse Lab in Glasgow | Wellcome Sanger Institute for the COVID-19 Genomics UK (COG-UK) Consortium | Harper VanSteenhouse, Yumi Kasai, David Gray, Carol Clugston, Anna Dominiczak and Alex Alderton, Roberto Amato, Jeffrey Barrett, Sonia Goncalves, Ewan Harrison, David K. Jackson, Ian Johnston, Dominic Kwiatkowski, Cordelia Langford, John Sillitoe on behalf of the Wellcome Sanger Institute COVID-19 Surveillance Team |
| EPI_ISL_3440626 | Lighthouse Lab in Milton Keynes | Wellcome Sanger Institute for the COVID-19 Genomics UK (COG-UK) Consortium | The Lighthouse Lab in Milton Keynes and Alex Alderton, Roberto Amato, Jeffrey Barrett, Sonia Goncalves, Ewan Harrison, David K. Jackson, Ian Johnston, Dominic Kwiatkowski, Cordelia Langford, John Sillitoe on behalf of the Wellcome Sanger Institute COVID-19 Surveillance Team |
| EPI_ISL_3440627 | Lighthouse Lab in Glasgow | Wellcome Sanger Institute for the COVID-19 Genomics UK (COG-UK) Consortium | Harper VanSteenhouse, Yumi Kasai, David Gray, Carol Clugston, Anna Dominiczak and Alex Alderton, Roberto Amato, Jeffrey Barrett, Sonia Goncalves, Ewan Harrison, David K. Jackson, Ian Johnston, Dominic Kwiatkowski, Cordelia Langford, John Sillitoe on behalf of the Wellcome Sanger Institute COVID-19 Surveillance Team |
| EPI_ISL_3440628 | Lighthouse Lab in Milton Keynes | Wellcome Sanger Institute for the COVID-19 Genomics UK (COG-UK) Consortium | The Lighthouse Lab in Milton Keynes and Alex Alderton, Roberto Amato, Jeffrey Barrett, Sonia Goncalves, Ewan Harrison, David K. Jackson, Ian Johnston, Dominic Kwiatkowski, Cordelia Langford, John Sillitoe on behalf of the Wellcome Sanger Institute COVID-19 Surveillance Team |
| EPI_ISL_3440629 | Lighthouse Laboratory Plymouth | Wellcome Sanger Institute for the COVID-19 Genomics UK (COG-UK) | Lighthouse Laboratory Plymouth and Alex Alderton, Roberto Amato, Jeffrey Barrett, Sonia Goncalves, Ewan Harrison, David K. Jackson, Ian Johnston, Dominic Kwiatkowski, Cordelia Langford, John Sillitoe on behalf of the Wellcome Sanger |

| | | | |
|---|---|---|---|
| | | Consortium | Institute COVID-19 Surveillance Team |
| EPI_ISL_3440630, EPI_ISL_3440631, EPI_ISL_3440632, EPI_ISL_3440633 | Lighthouse Lab in Milton Keynes | Wellcome Sanger Institute for the COVID-19 Genomics UK (COG-UK) Consortium | The Lighthouse Lab in Milton Keynes and Alex Alderton, Roberto Amato, Jeffrey Barrett, Sonia Goncalves, Ewan Harrison, David K. Jackson, Ian Johnston, Dominic Kwiatkowski, Cordelia Langford, John Sillitoe on behalf of the Wellcome Sanger Institute COVID-19 Surveillance Team |
| EPI_ISL_3440634 | Lighthouse Laboratory Plymouth | Wellcome Sanger Institute for the COVID-19 Genomics UK (COG-UK) Consortium | Lighthouse Laboratory Plymouth and Alex Alderton, Roberto Amato, Jeffrey Barrett, Sonia Goncalves, Ewan Harrison, David K. Jackson, Ian Johnston, Dominic Kwiatkowski, Cordelia Langford, John Sillitoe on behalf of the Wellcome Sanger Institute COVID-19 Surveillance Team |
| EPI_ISL_3440635, EPI_ISL_3440636 | Lighthouse Lab in Milton Keynes | Wellcome Sanger Institute for the COVID-19 Genomics UK (COG-UK) Consortium | The Lighthouse Lab in Milton Keynes and Alex Alderton, Roberto Amato, Jeffrey Barrett, Sonia Goncalves, Ewan Harrison, David K. Jackson, Ian Johnston, Dominic Kwiatkowski, Cordelia Langford, John Sillitoe on behalf of the Wellcome Sanger Institute COVID-19 Surveillance Team |
| EPI_ISL_3440637, EPI_ISL_3440638 | Lighthouse Laboratory Plymouth | Wellcome Sanger Institute for the COVID-19 Genomics UK (COG-UK) Consortium | Lighthouse Laboratory Plymouth and Alex Alderton, Roberto Amato, Jeffrey Barrett, Sonia Goncalves, Ewan Harrison, David K. Jackson, Ian Johnston, Dominic Kwiatkowski, Cordelia Langford, John Sillitoe on behalf of the Wellcome Sanger Institute COVID-19 Surveillance Team |
| EPI_ISL_3440639 | Lighthouse Lab in Glasgow | Wellcome Sanger Institute for the COVID-19 Genomics UK (COG-UK) Consortium | Harper VanSteenhouse, Yumi Kasai, David Gray, Carol Clugston, Anna Dominiczak and Alex Alderton, Roberto Amato, Jeffrey Barrett, Sonia Goncalves, Ewan Harrison, David K. Jackson, Ian Johnston, Dominic Kwiatkowski, Cordelia Langford, John Sillitoe on behalf of the Wellcome Sanger Institute COVID-19 Surveillance Team |
| EPI_ISL_3440640, EPI_ISL_3440641, EPI_ISL_3440642 | Lighthouse Lab in Milton Keynes | Wellcome Sanger Institute for the COVID-19 Genomics UK (COG-UK) Consortium | The Lighthouse Lab in Milton Keynes and Alex Alderton, Roberto Amato, Jeffrey Barrett, Sonia Goncalves, Ewan Harrison, David K. Jackson, Ian Johnston, Dominic Kwiatkowski, Cordelia Langford, John Sillitoe on behalf of the Wellcome Sanger Institute COVID-19 Surveillance Team |
| EPI_ISL_3440643, EPI_ISL_3440644, EPI_ISL_3440645 | Lighthouse Laboratory Plymouth | Wellcome Sanger Institute for the COVID-19 Genomics UK (COG-UK) Consortium | Lighthouse Laboratory Plymouth and Alex Alderton, Roberto Amato, Jeffrey Barrett, Sonia Goncalves, Ewan Harrison, David K. Jackson, Ian Johnston, Dominic Kwiatkowski, Cordelia Langford, John Sillitoe on behalf of the Wellcome Sanger Institute COVID-19 Surveillance Team |
| EPI_ISL_3440646 | Lighthouse Lab in Milton Keynes | Wellcome Sanger Institute for the COVID-19 Genomics UK (COG-UK) Consortium | The Lighthouse Lab in Milton Keynes and Alex Alderton, Roberto Amato, Jeffrey Barrett, Sonia Goncalves, Ewan Harrison, David K. Jackson, Ian Johnston, Dominic Kwiatkowski, Cordelia Langford, John Sillitoe on behalf of the Wellcome Sanger Institute COVID-19 Surveillance Team |
| EPI_ISL_3440647 | Lighthouse Laboratory Plymouth | Wellcome Sanger Institute for the COVID-19 Genomics UK (COG-UK) Consortium | Lighthouse Laboratory Plymouth and Alex Alderton, Roberto Amato, Jeffrey Barrett, Sonia Goncalves, Ewan Harrison, David K. Jackson, Ian Johnston, Dominic Kwiatkowski, Cordelia Langford, John Sillitoe on behalf of the Wellcome Sanger Institute COVID-19 Surveillance Team |
| EPI_ISL_3440648 | Lighthouse Lab in Milton Keynes | Wellcome Sanger Institute for the COVID-19 Genomics UK (COG-UK) Consortium | The Lighthouse Lab in Milton Keynes and Alex Alderton, Roberto Amato, Jeffrey Barrett, Sonia Goncalves, Ewan Harrison, David K. Jackson, Ian Johnston, Dominic Kwiatkowski, Cordelia Langford, John Sillitoe on behalf of the Wellcome Sanger Institute COVID-19 Surveillance Team |
| EPI_ISL_3440649, EPI_ISL_3440650 | Lighthouse Lab in Glasgow | Wellcome Sanger Institute for the COVID-19 Genomics UK (COG-UK) Consortium | Harper VanSteenhouse, Yumi Kasai, David Gray, Carol Clugston, Anna Dominiczak and Alex Alderton, Roberto Amato, Jeffrey Barrett, Sonia Goncalves, Ewan Harrison, David K. Jackson, Ian Johnston, Dominic Kwiatkowski, Cordelia Langford, John Sillitoe on behalf of the Wellcome Sanger Institute COVID-19 Surveillance Team |
| EPI_ISL_3440651 | Lighthouse Lab in Milton Keynes | Wellcome Sanger Institute for the COVID-19 Genomics UK (COG-UK) Consortium | The Lighthouse Lab in Milton Keynes and Alex Alderton, Roberto Amato, Jeffrey Barrett, Sonia Goncalves, Ewan Harrison, David K. Jackson, Ian Johnston, Dominic Kwiatkowski, Cordelia Langford, John Sillitoe on behalf of the Wellcome Sanger Institute COVID-19 Surveillance Team |
| EPI_ISL_3440652 | Lighthouse Lab in Glasgow | Wellcome Sanger Institute for the COVID-19 Genomics UK (COG-UK) Consortium | Harper VanSteenhouse, Yumi Kasai, David Gray, Carol Clugston, Anna Dominiczak and Alex Alderton, Roberto Amato, Jeffrey Barrett, Sonia Goncalves, Ewan Harrison, David K. Jackson, Ian Johnston, Dominic Kwiatkowski, Cordelia Langford, John Sillitoe on behalf of the Wellcome Sanger Institute COVID-19 Surveillance Team |
| EPI_ISL_3440653 | Lighthouse Laboratory Plymouth | Wellcome Sanger Institute for the COVID-19 Genomics UK (COG-UK) Consortium | Lighthouse Laboratory Plymouth and Alex Alderton, Roberto Amato, Jeffrey Barrett, Sonia Goncalves, Ewan Harrison, David K. Jackson, Ian Johnston, Dominic Kwiatkowski, Cordelia Langford, John Sillitoe on behalf of the Wellcome Sanger Institute COVID-19 Surveillance Team |
| EPI_ISL_3440654 | Lighthouse Lab in Glasgow | Wellcome Sanger Institute for the COVID-19 Genomics UK (COG-UK) Consortium | Harper VanSteenhouse, Yumi Kasai, David Gray, Carol Clugston, Anna Dominiczak and Alex Alderton, Roberto Amato, Jeffrey Barrett, Sonia Goncalves, Ewan Harrison, David K. Jackson, Ian Johnston, Dominic Kwiatkowski, Cordelia Langford, John Sillitoe on behalf of the Wellcome Sanger Institute COVID-19 Surveillance Team |
| EPI_ISL_3440655 | Lighthouse Lab in Milton Keynes | Wellcome Sanger Institute for the COVID-19 Genomics UK (COG-UK) Consortium | The Lighthouse Lab in Milton Keynes and Alex Alderton, Roberto Amato, Jeffrey Barrett, Sonia Goncalves, Ewan Harrison, David K. Jackson, Ian Johnston, Dominic Kwiatkowski, Cordelia Langford, John Sillitoe on behalf of the Wellcome Sanger Institute COVID-19 Surveillance Team |
| EPI_ISL_3440656 | Lighthouse Laboratory Plymouth | Wellcome Sanger Institute for the COVID-19 Genomics UK (COG-UK) Consortium | Lighthouse Laboratory Plymouth and Alex Alderton, Roberto Amato, Jeffrey Barrett, Sonia Goncalves, Ewan Harrison, David K. Jackson, Ian Johnston, Dominic Kwiatkowski, Cordelia Langford, John Sillitoe on behalf of the Wellcome Sanger Institute COVID-19 Surveillance Team |
| EPI_ISL_3440657 | Lighthouse Lab in Glasgow | Wellcome Sanger Institute for the COVID-19 Genomics UK (COG-UK) Consortium | Harper VanSteenhouse, Yumi Kasai, David Gray, Carol Clugston, Anna Dominiczak and Alex Alderton, Roberto Amato, Jeffrey Barrett, Sonia Goncalves, Ewan Harrison, David K. Jackson, Ian Johnston, Dominic Kwiatkowski, Cordelia Langford, John Sillitoe on behalf of the Wellcome Sanger Institute COVID-19 Surveillance Team |
| EPI_ISL_3440658 | Lighthouse Lab in Milton Keynes | Wellcome Sanger Institute for the COVID-19 Genomics UK (COG-UK) Consortium | The Lighthouse Lab in Milton Keynes and Alex Alderton, Roberto Amato, Jeffrey Barrett, Sonia Goncalves, Ewan Harrison, David K. Jackson, Ian Johnston, Dominic Kwiatkowski, Cordelia Langford, John Sillitoe on behalf of the Wellcome Sanger Institute COVID-19 Surveillance Team |
| EPI_ISL_3440659, EPI_ISL_3440660 | Lighthouse Laboratory Plymouth | Wellcome Sanger Institute for the COVID-19 Genomics UK (COG-UK) Consortium | Lighthouse Laboratory Plymouth and Alex Alderton, Roberto Amato, Jeffrey Barrett, Sonia Goncalves, Ewan Harrison, David K. Jackson, Ian Johnston, Dominic Kwiatkowski, Cordelia Langford, John Sillitoe on behalf of the Wellcome Sanger Institute COVID-19 Surveillance Team |
| EPI_ISL_3440661, EPI_ISL_3440662 | Lighthouse Lab in Milton Keynes | Wellcome Sanger Institute for the COVID-19 Genomics UK (COG-UK) Consortium | The Lighthouse Lab in Milton Keynes and Alex Alderton, Roberto Amato, Jeffrey Barrett, Sonia Goncalves, Ewan Harrison, David K. Jackson, Ian Johnston, Dominic Kwiatkowski, Cordelia Langford, John Sillitoe on behalf of the Wellcome Sanger Institute COVID-19 Surveillance Team |
| EPI_ISL_3440663 | Lighthouse Laboratory Plymouth | Wellcome Sanger Institute for the COVID-19 Genomics UK (COG-UK) Consortium | Lighthouse Laboratory Plymouth and Alex Alderton, Roberto Amato, Jeffrey Barrett, Sonia Goncalves, Ewan Harrison, David K. Jackson, Ian Johnston, Dominic Kwiatkowski, Cordelia Langford, John Sillitoe on behalf of the Wellcome Sanger Institute COVID-19 Surveillance Team |
| EPI_ISL_3440664, EPI_ISL_3440665, EPI_ISL_3440666, EPI_ISL_3440667 | Lighthouse Lab in Milton Keynes | Wellcome Sanger Institute for the COVID-19 Genomics UK (COG-UK) Consortium | The Lighthouse Lab in Milton Keynes and Alex Alderton, Roberto Amato, Jeffrey Barrett, Sonia Goncalves, Ewan Harrison, David K. Jackson, Ian Johnston, Dominic Kwiatkowski, Cordelia Langford, John Sillitoe on behalf of the Wellcome Sanger Institute COVID-19 Surveillance Team |
| EPI_ISL_3440668 | Lighthouse Laboratory Plymouth | Wellcome Sanger Institute for the COVID-19 Genomics UK (COG-UK) Consortium | Lighthouse Laboratory Plymouth and Alex Alderton, Roberto Amato, Jeffrey Barrett, Sonia Goncalves, Ewan Harrison, David K. Jackson, Ian Johnston, Dominic Kwiatkowski, Cordelia Langford, John Sillitoe on behalf of the Wellcome Sanger Institute COVID-19 Surveillance Team |
| EPI_ISL_3440669 | Lighthouse Lab in Milton Keynes | Wellcome Sanger Institute for the COVID-19 Genomics UK (COG-UK) Consortium | The Lighthouse Lab in Milton Keynes and Alex Alderton, Roberto Amato, Jeffrey Barrett, Sonia Goncalves, Ewan Harrison, David K. Jackson, Ian Johnston, Dominic Kwiatkowski, Cordelia Langford, John Sillitoe on behalf of the Wellcome Sanger Institute COVID-19 Surveillance Team |
| EPI_ISL_3440670 | Lighthouse Laboratory Plymouth | Wellcome Sanger Institute for the COVID-19 Genomics UK (COG-UK) Consortium | Lighthouse Laboratory Plymouth and Alex Alderton, Roberto Amato, Jeffrey Barrett, Sonia Goncalves, Ewan Harrison, David K. Jackson, Ian Johnston, Dominic Kwiatkowski, Cordelia Langford, John Sillitoe on behalf of the Wellcome Sanger Institute COVID-19 Surveillance Team |
| EPI_ISL_3440671 | Lighthouse Lab in Milton Keynes | Wellcome Sanger Institute for the COVID-19 Genomics UK (COG-UK) Consortium | The Lighthouse Lab in Milton Keynes and Alex Alderton, Roberto Amato, Jeffrey Barrett, Sonia Goncalves, Ewan Harrison, David K. Jackson, Ian Johnston, Dominic Kwiatkowski, Cordelia Langford, John Sillitoe on behalf of the Wellcome Sanger Institute COVID-19 Surveillance Team |
| EPI_ISL_3440672 | Lighthouse Laboratory Plymouth | Wellcome Sanger Institute for the COVID-19 Genomics UK (COG-UK) Consortium | Lighthouse Laboratory Plymouth and Alex Alderton, Roberto Amato, Jeffrey Barrett, Sonia Goncalves, Ewan Harrison, David K. Jackson, Ian Johnston, Dominic Kwiatkowski, Cordelia Langford, John Sillitoe on behalf of the Wellcome Sanger Institute COVID-19 Surveillance Team |
| EPI_ISL_3440673, EPI_ISL_3440674 | Lighthouse Lab in Glasgow | Wellcome Sanger Institute for the COVID-19 Genomics UK (COG-UK) Consortium | Harper VanSteenhouse, Yumi Kasai, David Gray, Carol Clugston, Anna Dominiczak and Alex Alderton, Roberto Amato, Jeffrey Barrett, Sonia Goncalves, Ewan Harrison, David K. Jackson, Ian Johnston, Dominic Kwiatkowski, Cordelia Langford, John Sillitoe on behalf of the Wellcome Sanger Institute COVID-19 Surveillance Team |
| EPI_ISL_3440675 | Lighthouse Laboratory Plymouth | Wellcome Sanger Institute for the COVID-19 Genomics UK (COG-UK) Consortium | Lighthouse Laboratory Plymouth and Alex Alderton, Roberto Amato, Jeffrey Barrett, Sonia Goncalves, Ewan Harrison, David K. Jackson, Ian Johnston, Dominic Kwiatkowski, Cordelia Langford, John Sillitoe on behalf of the Wellcome Sanger Institute COVID-19 Surveillance Team |
| EPI_ISL_3440676 | Lighthouse Lab in Glasgow | Wellcome Sanger Institute for the COVID-19 Genomics UK (COG-UK) Consortium | Harper VanSteenhouse, Yumi Kasai, David Gray, Carol Clugston, Anna Dominiczak and Alex Alderton, Roberto Amato, Jeffrey Barrett, Sonia Goncalves, Ewan Harrison, David K. Jackson, Ian Johnston, Dominic Kwiatkowski, Cordelia Langford, John Sillitoe on behalf of the Wellcome Sanger Institute COVID-19 Surveillance Team |
| EPI_ISL_3440677, EPI_ISL_3440678 | Lighthouse Lab in Milton Keynes | Wellcome Sanger Institute for the COVID-19 Genomics UK (COG-UK) Consortium | The Lighthouse Lab in Milton Keynes and Alex Alderton, Roberto Amato, Jeffrey Barrett, Sonia Goncalves, Ewan Harrison, David K. Jackson, Ian Johnston, Dominic Kwiatkowski, Cordelia Langford, John Sillitoe on behalf of the Wellcome Sanger Institute COVID-19 Surveillance Team |
| EPI_ISL_3440679 | Lighthouse Lab in Glasgow | Wellcome Sanger Institute for the COVID-19 Genomics UK (COG-UK) Consortium | Harper VanSteenhouse, Yumi Kasai, David Gray, Carol Clugston, Anna Dominiczak and Alex Alderton, Roberto Amato, Jeffrey Barrett, Sonia Goncalves, Ewan Harrison, David K. Jackson, Ian Johnston, Dominic Kwiatkowski, Cordelia Langford, John Sillitoe on behalf of the Wellcome Sanger Institute COVID-19 Surveillance Team |
| EPI_ISL_3440680 | Lighthouse Lab in Milton Keynes | Wellcome Sanger Institute for the COVID-19 Genomics UK (COG-UK) Consortium | The Lighthouse Lab in Milton Keynes and Alex Alderton, Roberto Amato, Jeffrey Barrett, Sonia Goncalves, Ewan Harrison, David K. Jackson, Ian Johnston, Dominic Kwiatkowski, Cordelia Langford, John Sillitoe on behalf of the Wellcome Sanger Institute COVID-19 Surveillance Team |
| EPI_ISL_3440681 | Lighthouse Laboratory Plymouth | Wellcome Sanger Institute for the COVID-19 Genomics UK (COG-UK) Consortium | Lighthouse Laboratory Plymouth and Alex Alderton, Roberto Amato, Jeffrey Barrett, Sonia Goncalves, Ewan Harrison, David K. Jackson, Ian Johnston, Dominic Kwiatkowski, Cordelia Langford, John Sillitoe on behalf of the Wellcome Sanger Institute COVID-19 Surveillance Team |
| EPI_ISL_3440682 | Lighthouse Lab in Milton Keynes | Wellcome Sanger Institute for the COVID-19 Genomics UK (COG-UK) Consortium | The Lighthouse Lab in Milton Keynes and Alex Alderton, Roberto Amato, Jeffrey Barrett, Sonia Goncalves, Ewan Harrison, David K. Jackson, Ian Johnston, Dominic Kwiatkowski, Cordelia Langford, John Sillitoe on behalf of the Wellcome Sanger Institute COVID-19 Surveillance Team |
| EPI_ISL_3440683, EPI_ISL_3440684 | Lighthouse Lab in Glasgow | Wellcome Sanger Institute for the COVID-19 Genomics UK (COG-UK) Consortium | Harper VanSteenhouse, Yumi Kasai, David Gray, Carol Clugston, Anna Dominiczak and Alex Alderton, Roberto Amato, Jeffrey Barrett, Sonia Goncalves, Ewan Harrison, David K. Jackson, Ian Johnston, Dominic Kwiatkowski, Cordelia Langford, John Sillitoe on behalf of the Wellcome Sanger Institute COVID-19 Surveillance Team |
| EPI_ISL_3440685, EPI_ISL_3440686 | Lighthouse Lab in Milton Keynes | Wellcome Sanger Institute for the COVID-19 Genomics UK (COG-UK) Consortium | The Lighthouse Lab in Milton Keynes and Alex Alderton, Roberto Amato, Jeffrey Barrett, Sonia Goncalves, Ewan Harrison, David K. Jackson, Ian Johnston, Dominic Kwiatkowski, Cordelia Langford, John Sillitoe on behalf of the Wellcome Sanger Institute COVID-19 Surveillance Team |
| EPI_ISL_3440687 | Lighthouse Laboratory Plymouth | Wellcome Sanger Institute for the COVID-19 Genomics UK (COG-UK) Consortium | Lighthouse Laboratory Plymouth and Alex Alderton, Roberto Amato, Jeffrey Barrett, Sonia Goncalves, Ewan Harrison, David K. Jackson, Ian Johnston, Dominic Kwiatkowski, Cordelia Langford, John Sillitoe on behalf of the Wellcome Sanger Institute COVID-19 Surveillance Team |
| EPI_ISL_3440688 | Lighthouse Lab in Glasgow | Wellcome Sanger Institute for the COVID-19 Genomics UK (COG-UK) Consortium | Harper VanSteenhouse, Yumi Kasai, David Gray, Carol Clugston, Anna Dominiczak and Alex Alderton, Roberto Amato, Jeffrey Barrett, Sonia Goncalves, Ewan Harrison, David K. Jackson, Ian Johnston, Dominic Kwiatkowski, Cordelia Langford, John Sillitoe on behalf of the Wellcome Sanger Institute COVID-19 Surveillance Team |
| EPI_ISL_3440689 | Lighthouse Laboratory Plymouth | Wellcome Sanger Institute for the COVID-19 Genomics UK (COG-UK) Consortium | Lighthouse Laboratory Plymouth and Alex Alderton, Roberto Amato, Jeffrey Barrett, Sonia Goncalves, Ewan Harrison, David K. Jackson, Ian Johnston, Dominic Kwiatkowski, Cordelia Langford, John Sillitoe on behalf of the Wellcome Sanger Institute COVID-19 Surveillance Team |
| EPI_ISL_3440690, EPI_ISL_3440691, EPI_ISL_3440692, | Lighthouse Lab in Milton Keynes | Wellcome Sanger Institute for the COVID-19 Genomics UK (COG-UK) Consortium | The Lighthouse Lab in Milton Keynes and Alex Alderton, Roberto Amato, Jeffrey Barrett, Sonia Goncalves, Ewan Harrison, David K. Jackson, Ian Johnston, Dominic Kwiatkowski, Cordelia Langford, John Sillitoe on behalf of the Wellcome Sanger Institute COVID-19 Surveillance Team |

| | | | |
|---|---|---|---|
| EPI_ISL_3440693, EPI_ISL_3440694, EPI_ISL_3440695 | | | |
| EPI_ISL_3440696 | Lighthouse Lab in Glasgow | Wellcome Sanger Institute for the COVID-19 Genomics UK (COG-UK) Consortium | Harper VanSteenhouse, Yumi Kasai, David Gray, Carol Clugston, Anna Dominiczak and Alex Alderton, Roberto Amato, Jeffrey Barrett, Sonia Goncalves, Ewan Harrison, David K. Jackson, Ian Johnston, Dominic Kwiatkowski, Cordelia Langford, John Sillitoe on behalf of the Wellcome Sanger Institute COVID-19 Surveillance Team |
| EPI_ISL_3440697 | Lighthouse Lab in Milton Keynes | Wellcome Sanger Institute for the COVID-19 Genomics UK (COG-UK) Consortium | The Lighthouse Lab in Milton Keynes and Alex Alderton, Roberto Amato, Jeffrey Barrett, Sonia Goncalves, Ewan Harrison, David K. Jackson, Ian Johnston, Dominic Kwiatkowski, Cordelia Langford, John Sillitoe on behalf of the Wellcome Sanger Institute COVID-19 Surveillance Team |
| EPI_ISL_3440698 | Lighthouse Lab in Glasgow | Wellcome Sanger Institute for the COVID-19 Genomics UK (COG-UK) Consortium | Harper VanSteenhouse, Yumi Kasai, David Gray, Carol Clugston, Anna Dominiczak and Alex Alderton, Roberto Amato, Jeffrey Barrett, Sonia Goncalves, Ewan Harrison, David K. Jackson, Ian Johnston, Dominic Kwiatkowski, Cordelia Langford, John Sillitoe on behalf of the Wellcome Sanger Institute COVID-19 Surveillance Team |
| EPI_ISL_3440699 | Lighthouse Laboratory Plymouth | Wellcome Sanger Institute for the COVID-19 Genomics UK (COG-UK) Consortium | Lighthouse Laboratory Plymouth and Alex Alderton, Roberto Amato, Jeffrey Barrett, Sonia Goncalves, Ewan Harrison, David K. Jackson, Ian Johnston, Dominic Kwiatkowski, Cordelia Langford, John Sillitoe on behalf of the Wellcome Sanger Institute COVID-19 Surveillance Team |
| EPI_ISL_3440700, EPI_ISL_3440701, EPI_ISL_3440702 | Lighthouse Lab in Milton Keynes | Wellcome Sanger Institute for the COVID-19 Genomics UK (COG-UK) Consortium | The Lighthouse Lab in Milton Keynes and Alex Alderton, Roberto Amato, Jeffrey Barrett, Sonia Goncalves, Ewan Harrison, David K. Jackson, Ian Johnston, Dominic Kwiatkowski, Cordelia Langford, John Sillitoe on behalf of the Wellcome Sanger Institute COVID-19 Surveillance Team |
| EPI_ISL_3440703 | Lighthouse Lab in Glasgow | Wellcome Sanger Institute for the COVID-19 Genomics UK (COG-UK) Consortium | Harper VanSteenhouse, Yumi Kasai, David Gray, Carol Clugston, Anna Dominiczak and Alex Alderton, Roberto Amato, Jeffrey Barrett, Sonia Goncalves, Ewan Harrison, David K. Jackson, Ian Johnston, Dominic Kwiatkowski, Cordelia Langford, John Sillitoe on behalf of the Wellcome Sanger Institute COVID-19 Surveillance Team |
| EPI_ISL_3440704, EPI_ISL_3440705 | Lighthouse Lab in Milton Keynes | Wellcome Sanger Institute for the COVID-19 Genomics UK (COG-UK) Consortium | The Lighthouse Lab in Milton Keynes and Alex Alderton, Roberto Amato, Jeffrey Barrett, Sonia Goncalves, Ewan Harrison, David K. Jackson, Ian Johnston, Dominic Kwiatkowski, Cordelia Langford, John Sillitoe on behalf of the Wellcome Sanger Institute COVID-19 Surveillance Team |
| EPI_ISL_3440706, EPI_ISL_3440707 | Lighthouse Laboratory Plymouth | Wellcome Sanger Institute for the COVID-19 Genomics UK (COG-UK) Consortium | Lighthouse Laboratory Plymouth and Alex Alderton, Roberto Amato, Jeffrey Barrett, Sonia Goncalves, Ewan Harrison, David K. Jackson, Ian Johnston, Dominic Kwiatkowski, Cordelia Langford, John Sillitoe on behalf of the Wellcome Sanger Institute COVID-19 Surveillance Team |
| EPI_ISL_3440708 | Lighthouse Lab in Milton Keynes | Wellcome Sanger Institute for the COVID-19 Genomics UK (COG-UK) Consortium | The Lighthouse Lab in Milton Keynes and Alex Alderton, Roberto Amato, Jeffrey Barrett, Sonia Goncalves, Ewan Harrison, David K. Jackson, Ian Johnston, Dominic Kwiatkowski, Cordelia Langford, John Sillitoe on behalf of the Wellcome Sanger Institute COVID-19 Surveillance Team |
| EPI_ISL_3440709 | Lighthouse Laboratory Plymouth | Wellcome Sanger Institute for the COVID-19 Genomics UK (COG-UK) Consortium | Lighthouse Laboratory Plymouth and Alex Alderton, Roberto Amato, Jeffrey Barrett, Sonia Goncalves, Ewan Harrison, David K. Jackson, Ian Johnston, Dominic Kwiatkowski, Cordelia Langford, John Sillitoe on behalf of the Wellcome Sanger Institute COVID-19 Surveillance Team |
| EPI_ISL_3440710, EPI_ISL_3440711, EPI_ISL_3440712, EPI_ISL_3440713, EPI_ISL_3440714 | Lighthouse Lab in Milton Keynes | Wellcome Sanger Institute for the COVID-19 Genomics UK (COG-UK) Consortium | The Lighthouse Lab in Milton Keynes and Alex Alderton, Roberto Amato, Jeffrey Barrett, Sonia Goncalves, Ewan Harrison, David K. Jackson, Ian Johnston, Dominic Kwiatkowski, Cordelia Langford, John Sillitoe on behalf of the Wellcome Sanger Institute COVID-19 Surveillance Team |
| EPI_ISL_3440715, EPI_ISL_3440716 | Lighthouse Lab in Glasgow | Wellcome Sanger Institute for the COVID-19 Genomics UK (COG-UK) Consortium | Harper VanSteenhouse, Yumi Kasai, David Gray, Carol Clugston, Anna Dominiczak and Alex Alderton, Roberto Amato, Jeffrey Barrett, Sonia Goncalves, Ewan Harrison, David K. Jackson, Ian Johnston, Dominic Kwiatkowski, Cordelia Langford, John Sillitoe on behalf of the Wellcome Sanger Institute COVID-19 Surveillance Team |
| EPI_ISL_3440717 | Lighthouse Lab in Milton Keynes | Wellcome Sanger Institute for the COVID-19 Genomics UK (COG-UK) Consortium | The Lighthouse Lab in Milton Keynes and Alex Alderton, Roberto Amato, Jeffrey Barrett, Sonia Goncalves, Ewan Harrison, David K. Jackson, Ian Johnston, Dominic Kwiatkowski, Cordelia Langford, John Sillitoe on behalf of the Wellcome Sanger Institute COVID-19 Surveillance Team |
| EPI_ISL_3440718 | Lighthouse Lab in Glasgow | Wellcome Sanger Institute for the COVID-19 Genomics UK (COG-UK) Consortium | Harper VanSteenhouse, Yumi Kasai, David Gray, Carol Clugston, Anna Dominiczak and Alex Alderton, Roberto Amato, Jeffrey Barrett, Sonia Goncalves, Ewan Harrison, David K. Jackson, Ian Johnston, Dominic Kwiatkowski, Cordelia Langford, John Sillitoe on behalf of the Wellcome Sanger Institute COVID-19 Surveillance Team |
| EPI_ISL_3440719, EPI_ISL_3440720 | Lighthouse Lab in Milton Keynes | Wellcome Sanger Institute for the COVID-19 Genomics UK (COG-UK) Consortium | The Lighthouse Lab in Milton Keynes and Alex Alderton, Roberto Amato, Jeffrey Barrett, Sonia Goncalves, Ewan Harrison, David K. Jackson, Ian Johnston, Dominic Kwiatkowski, Cordelia Langford, John Sillitoe on behalf of the Wellcome Sanger Institute COVID-19 Surveillance Team |
| EPI_ISL_3440721, EPI_ISL_3440722, EPI_ISL_3440723 | Lighthouse Laboratory Plymouth | Wellcome Sanger Institute for the COVID-19 Genomics UK (COG-UK) Consortium | Lighthouse Laboratory Plymouth and Alex Alderton, Roberto Amato, Jeffrey Barrett, Sonia Goncalves, Ewan Harrison, David K. Jackson, Ian Johnston, Dominic Kwiatkowski, Cordelia Langford, John Sillitoe on behalf of the Wellcome Sanger Institute COVID-19 Surveillance Team |
| EPI_ISL_3440724 | Lighthouse Lab in Milton Keynes | Wellcome Sanger Institute for the COVID-19 Genomics UK (COG-UK) Consortium | The Lighthouse Lab in Milton Keynes and Alex Alderton, Roberto Amato, Jeffrey Barrett, Sonia Goncalves, Ewan Harrison, David K. Jackson, Ian Johnston, Dominic Kwiatkowski, Cordelia Langford, John Sillitoe on behalf of the Wellcome Sanger Institute COVID-19 Surveillance Team |
| EPI_ISL_3440725 | Lighthouse Laboratory Plymouth | Wellcome Sanger Institute for the COVID-19 Genomics UK (COG-UK) Consortium | Lighthouse Laboratory Plymouth and Alex Alderton, Roberto Amato, Jeffrey Barrett, Sonia Goncalves, Ewan Harrison, David K. Jackson, Ian Johnston, Dominic Kwiatkowski, Cordelia Langford, John Sillitoe on behalf of the Wellcome Sanger Institute COVID-19 Surveillance Team |
| EPI_ISL_3440726, EPI_ISL_3440727 | Lighthouse Lab in Glasgow | Wellcome Sanger Institute for the COVID-19 Genomics UK (COG-UK) Consortium | Harper VanSteenhouse, Yumi Kasai, David Gray, Carol Clugston, Anna Dominiczak and Alex Alderton, Roberto Amato, Jeffrey Barrett, Sonia Goncalves, Ewan Harrison, David K. Jackson, Ian Johnston, Dominic Kwiatkowski, Cordelia Langford, John Sillitoe on behalf of the Wellcome Sanger Institute COVID-19 Surveillance Team |
| EPI_ISL_3440728, EPI_ISL_3440729, EPI_ISL_3440730, EPI_ISL_3440731, EPI_ISL_3440732, EPI_ISL_3440733, EPI_ISL_3440734, EPI_ISL_3440735 | Lighthouse Lab in Milton Keynes | Wellcome Sanger Institute for the COVID-19 Genomics UK (COG-UK) Consortium | The Lighthouse Lab in Milton Keynes and Alex Alderton, Roberto Amato, Jeffrey Barrett, Sonia Goncalves, Ewan Harrison, David K. Jackson, Ian Johnston, Dominic Kwiatkowski, Cordelia Langford, John Sillitoe on behalf of the Wellcome Sanger Institute COVID-19 Surveillance Team |
| EPI_ISL_3440736, EPI_ISL_3440737 | Lighthouse Laboratory Plymouth | Wellcome Sanger Institute for the COVID-19 Genomics UK (COG-UK) Consortium | Lighthouse Laboratory Plymouth and Alex Alderton, Roberto Amato, Jeffrey Barrett, Sonia Goncalves, Ewan Harrison, David K. Jackson, Ian Johnston, Dominic Kwiatkowski, Cordelia Langford, John Sillitoe on behalf of the Wellcome Sanger Institute COVID-19 Surveillance Team |
| EPI_ISL_3440738, EPI_ISL_3440739, EPI_ISL_3440740 | Lighthouse Lab in Milton Keynes | Wellcome Sanger Institute for the COVID-19 Genomics UK (COG-UK) Consortium | The Lighthouse Lab in Milton Keynes and Alex Alderton, Roberto Amato, Jeffrey Barrett, Sonia Goncalves, Ewan Harrison, David K. Jackson, Ian Johnston, Dominic Kwiatkowski, Cordelia Langford, John Sillitoe on behalf of the Wellcome Sanger Institute COVID-19 Surveillance Team |
| EPI_ISL_3440741, EPI_ISL_3440742 | Lighthouse Lab in Glasgow | Wellcome Sanger Institute for the COVID-19 Genomics UK (COG-UK) Consortium | Harper VanSteenhouse, Yumi Kasai, David Gray, Carol Clugston, Anna Dominiczak and Alex Alderton, Roberto Amato, Jeffrey Barrett, Sonia Goncalves, Ewan Harrison, David K. Jackson, Ian Johnston, Dominic Kwiatkowski, Cordelia Langford, John Sillitoe on behalf of the Wellcome Sanger Institute COVID-19 Surveillance Team |
| EPI_ISL_3440743, EPI_ISL_3440744, EPI_ISL_3440745 | Lighthouse Lab in Milton Keynes | Wellcome Sanger Institute for the COVID-19 Genomics UK (COG-UK) Consortium | The Lighthouse Lab in Milton Keynes and Alex Alderton, Roberto Amato, Jeffrey Barrett, Sonia Goncalves, Ewan Harrison, David K. Jackson, Ian Johnston, Dominic Kwiatkowski, Cordelia Langford, John Sillitoe on behalf of the Wellcome Sanger Institute COVID-19 Surveillance Team |
| EPI_ISL_3440746 | Lighthouse Laboratory Plymouth | Wellcome Sanger Institute for the COVID-19 Genomics UK (COG-UK) Consortium | Lighthouse Laboratory Plymouth and Alex Alderton, Roberto Amato, Jeffrey Barrett, Sonia Goncalves, Ewan Harrison, David K. Jackson, Ian Johnston, Dominic Kwiatkowski, Cordelia Langford, John Sillitoe on behalf of the Wellcome Sanger Institute COVID-19 Surveillance Team |
| EPI_ISL_3440747, EPI_ISL_3440748 | Lighthouse Lab in Milton Keynes | Wellcome Sanger Institute for the COVID-19 Genomics UK (COG-UK) Consortium | The Lighthouse Lab in Milton Keynes and Alex Alderton, Roberto Amato, Jeffrey Barrett, Sonia Goncalves, Ewan Harrison, David K. Jackson, Ian Johnston, Dominic Kwiatkowski, Cordelia Langford, John Sillitoe on behalf of the Wellcome Sanger Institute COVID-19 Surveillance Team |
| EPI_ISL_3440749 | Lighthouse Laboratory Plymouth | Wellcome Sanger Institute for the COVID-19 Genomics UK (COG-UK) Consortium | Lighthouse Laboratory Plymouth and Alex Alderton, Roberto Amato, Jeffrey Barrett, Sonia Goncalves, Ewan Harrison, David K. Jackson, Ian Johnston, Dominic Kwiatkowski, Cordelia Langford, John Sillitoe on behalf of the Wellcome Sanger Institute COVID-19 Surveillance Team |
| EPI_ISL_3440750, EPI_ISL_3440751, EPI_ISL_3440752, EPI_ISL_3440753 | Lighthouse Lab in Milton Keynes | Wellcome Sanger Institute for the COVID-19 Genomics UK (COG-UK) Consortium | The Lighthouse Lab in Milton Keynes and Alex Alderton, Roberto Amato, Jeffrey Barrett, Sonia Goncalves, Ewan Harrison, David K. Jackson, Ian Johnston, Dominic Kwiatkowski, Cordelia Langford, John Sillitoe on behalf of the Wellcome Sanger Institute COVID-19 Surveillance Team |
| EPI_ISL_3440754 | Lighthouse Laboratory Plymouth | Wellcome Sanger Institute for the COVID-19 Genomics UK (COG-UK) Consortium | Lighthouse Laboratory Plymouth and Alex Alderton, Roberto Amato, Jeffrey Barrett, Sonia Goncalves, Ewan Harrison, David K. Jackson, Ian Johnston, Dominic Kwiatkowski, Cordelia Langford, John Sillitoe on behalf of the Wellcome Sanger Institute COVID-19 Surveillance Team |
| EPI_ISL_3440755, EPI_ISL_3440756 | Lighthouse Lab in Glasgow | Wellcome Sanger Institute for the COVID-19 Genomics UK (COG-UK) Consortium | Harper VanSteenhouse, Yumi Kasai, David Gray, Carol Clugston, Anna Dominiczak and Alex Alderton, Roberto Amato, Jeffrey Barrett, Sonia Goncalves, Ewan Harrison, David K. Jackson, Ian Johnston, Dominic Kwiatkowski, Cordelia Langford, John Sillitoe on behalf of the Wellcome Sanger Institute COVID-19 Surveillance Team |
| EPI_ISL_3440757 | Lighthouse Lab in Milton Keynes | Wellcome Sanger Institute for the COVID-19 Genomics UK (COG-UK) Consortium | The Lighthouse Lab in Milton Keynes and Alex Alderton, Roberto Amato, Jeffrey Barrett, Sonia Goncalves, Ewan Harrison, David K. Jackson, Ian Johnston, Dominic Kwiatkowski, Cordelia Langford, John Sillitoe on behalf of the Wellcome Sanger Institute COVID-19 Surveillance Team |
| EPI_ISL_3440758 | Lighthouse Lab in Glasgow | Wellcome Sanger Institute for the COVID-19 Genomics UK (COG-UK) Consortium | Harper VanSteenhouse, Yumi Kasai, David Gray, Carol Clugston, Anna Dominiczak and Alex Alderton, Roberto Amato, Jeffrey Barrett, Sonia Goncalves, Ewan Harrison, David K. Jackson, Ian Johnston, Dominic Kwiatkowski, Cordelia Langford, John Sillitoe on behalf of the Wellcome Sanger Institute COVID-19 Surveillance Team |
| EPI_ISL_3440759, EPI_ISL_3440760 | Lighthouse Laboratory Plymouth | Wellcome Sanger Institute for the COVID-19 Genomics UK (COG-UK) Consortium | Lighthouse Laboratory Plymouth and Alex Alderton, Roberto Amato, Jeffrey Barrett, Sonia Goncalves, Ewan Harrison, David K. Jackson, Ian Johnston, Dominic Kwiatkowski, Cordelia Langford, John Sillitoe on behalf of the Wellcome Sanger Institute COVID-19 Surveillance Team |
| EPI_ISL_3440761, EPI_ISL_3440762, EPI_ISL_3440763 | Lighthouse Lab in Milton Keynes | Wellcome Sanger Institute for the COVID-19 Genomics UK (COG-UK) Consortium | The Lighthouse Lab in Milton Keynes and Alex Alderton, Roberto Amato, Jeffrey Barrett, Sonia Goncalves, Ewan Harrison, David K. Jackson, Ian Johnston, Dominic Kwiatkowski, Cordelia Langford, John Sillitoe on behalf of the Wellcome Sanger Institute COVID-19 Surveillance Team |
| EPI_ISL_3440764 | Lighthouse Lab in Glasgow | Wellcome Sanger Institute for the COVID-19 Genomics UK (COG-UK) Consortium | Harper VanSteenhouse, Yumi Kasai, David Gray, Carol Clugston, Anna Dominiczak and Alex Alderton, Roberto Amato, Jeffrey Barrett, Sonia Goncalves, Ewan Harrison, David K. Jackson, Ian Johnston, Dominic Kwiatkowski, Cordelia Langford, John Sillitoe on behalf of the Wellcome Sanger Institute COVID-19 Surveillance Team |
| EPI_ISL_3440765 | Lighthouse Lab in Milton Keynes | Wellcome Sanger Institute for the COVID-19 Genomics UK (COG-UK) Consortium | The Lighthouse Lab in Milton Keynes and Alex Alderton, Roberto Amato, Jeffrey Barrett, Sonia Goncalves, Ewan Harrison, David K. Jackson, Ian Johnston, Dominic Kwiatkowski, Cordelia Langford, John Sillitoe on behalf of the Wellcome Sanger Institute COVID-19 Surveillance Team |
| EPI_ISL_3440766 | Lighthouse Lab in Glasgow | Wellcome Sanger Institute for the COVID-19 Genomics UK (COG-UK) Consortium | Harper VanSteenhouse, Yumi Kasai, David Gray, Carol Clugston, Anna Dominiczak and Alex Alderton, Roberto Amato, Jeffrey Barrett, Sonia Goncalves, Ewan Harrison, David K. Jackson, Ian Johnston, Dominic Kwiatkowski, Cordelia Langford, John Sillitoe on behalf of the Wellcome Sanger Institute COVID-19 Surveillance Team |

| | | | |
|---|---|---|---|
| EPI_ISL_3440767 | Lighthouse Laboratory Plymouth | Wellcome Sanger Institute for the COVID-19 Genomics UK (COG-UK) Consortium | Lighthouse Laboratory Plymouth and Alex Alderton, Roberto Amato, Jeffrey Barrett, Sonia Goncalves, Ewan Harrison, David K. Jackson, Ian Johnston, Dominic Kwiatkowski, Cordelia Langford, John Sillitoe on behalf of the Wellcome Sanger Institute COVID-19 Surveillance Team |
| EPI_ISL_3440768 | Lighthouse Lab in Milton Keynes | Wellcome Sanger Institute for the COVID-19 Genomics UK (COG-UK) Consortium | The Lighthouse Lab in Milton Keynes and Alex Alderton, Roberto Amato, Jeffrey Barrett, Sonia Goncalves, Ewan Harrison, David K. Jackson, Ian Johnston, Dominic Kwiatkowski, Cordelia Langford, John Sillitoe on behalf of the Wellcome Sanger Institute COVID-19 Surveillance Team |
| EPI_ISL_3440769 | Lighthouse Lab in Glasgow | Wellcome Sanger Institute for the COVID-19 Genomics UK (COG-UK) Consortium | Harper VanSteenhouse, Yumi Kasai, David Gray, Carol Clugston, Anna Dominiczak and Alex Alderton, Roberto Amato, Jeffrey Barrett, Sonia Goncalves, Ewan Harrison, David K. Jackson, Ian Johnston, Dominic Kwiatkowski, Cordelia Langford, John Sillitoe on behalf of the Wellcome Sanger Institute COVID-19 Surveillance Team |
| EPI_ISL_3440770, EPI_ISL_3440771 | Lighthouse Lab in Milton Keynes | Wellcome Sanger Institute for the COVID-19 Genomics UK (COG-UK) Consortium | The Lighthouse Lab in Milton Keynes and Alex Alderton, Roberto Amato, Jeffrey Barrett, Sonia Goncalves, Ewan Harrison, David K. Jackson, Ian Johnston, Dominic Kwiatkowski, Cordelia Langford, John Sillitoe on behalf of the Wellcome Sanger Institute COVID-19 Surveillance Team |
| EPI_ISL_3440772, EPI_ISL_3440773 | Lighthouse Laboratory Plymouth | Wellcome Sanger Institute for the COVID-19 Genomics UK (COG-UK) Consortium | Lighthouse Laboratory Plymouth and Alex Alderton, Roberto Amato, Jeffrey Barrett, Sonia Goncalves, Ewan Harrison, David K. Jackson, Ian Johnston, Dominic Kwiatkowski, Cordelia Langford, John Sillitoe on behalf of the Wellcome Sanger Institute COVID-19 Surveillance Team |
| EPI_ISL_3440774, EPI_ISL_3440775 | Lighthouse Lab in Glasgow | Wellcome Sanger Institute for the COVID-19 Genomics UK (COG-UK) Consortium | Harper VanSteenhouse, Yumi Kasai, David Gray, Carol Clugston, Anna Dominiczak and Alex Alderton, Roberto Amato, Jeffrey Barrett, Sonia Goncalves, Ewan Harrison, David K. Jackson, Ian Johnston, Dominic Kwiatkowski, Cordelia Langford, John Sillitoe on behalf of the Wellcome Sanger Institute COVID-19 Surveillance Team |
| EPI_ISL_3440776, EPI_ISL_3440777, EPI_ISL_3440778 | Lighthouse Lab in Milton Keynes | Wellcome Sanger Institute for the COVID-19 Genomics UK (COG-UK) Consortium | The Lighthouse Lab in Milton Keynes and Alex Alderton, Roberto Amato, Jeffrey Barrett, Sonia Goncalves, Ewan Harrison, David K. Jackson, Ian Johnston, Dominic Kwiatkowski, Cordelia Langford, John Sillitoe on behalf of the Wellcome Sanger Institute COVID-19 Surveillance Team |
| EPI_ISL_3440779 | Lighthouse Laboratory Plymouth | Wellcome Sanger Institute for the COVID-19 Genomics UK (COG-UK) Consortium | Lighthouse Laboratory Plymouth and Alex Alderton, Roberto Amato, Jeffrey Barrett, Sonia Goncalves, Ewan Harrison, David K. Jackson, Ian Johnston, Dominic Kwiatkowski, Cordelia Langford, John Sillitoe on behalf of the Wellcome Sanger Institute COVID-19 Surveillance Team |
| EPI_ISL_3440780, EPI_ISL_3440781, EPI_ISL_3440782 | Lighthouse Lab in Milton Keynes | Wellcome Sanger Institute for the COVID-19 Genomics UK (COG-UK) Consortium | The Lighthouse Lab in Milton Keynes and Alex Alderton, Roberto Amato, Jeffrey Barrett, Sonia Goncalves, Ewan Harrison, David K. Jackson, Ian Johnston, Dominic Kwiatkowski, Cordelia Langford, John Sillitoe on behalf of the Wellcome Sanger Institute COVID-19 Surveillance Team |
| EPI_ISL_3440783 | Lighthouse Laboratory Plymouth | Wellcome Sanger Institute for the COVID-19 Genomics UK (COG-UK) Consortium | Lighthouse Laboratory Plymouth and Alex Alderton, Roberto Amato, Jeffrey Barrett, Sonia Goncalves, Ewan Harrison, David K. Jackson, Ian Johnston, Dominic Kwiatkowski, Cordelia Langford, John Sillitoe on behalf of the Wellcome Sanger Institute COVID-19 Surveillance Team |
| EPI_ISL_3440784 | Lighthouse Lab in Glasgow | Wellcome Sanger Institute for the COVID-19 Genomics UK (COG-UK) Consortium | Harper VanSteenhouse, Yumi Kasai, David Gray, Carol Clugston, Anna Dominiczak and Alex Alderton, Roberto Amato, Jeffrey Barrett, Sonia Goncalves, Ewan Harrison, David K. Jackson, Ian Johnston, Dominic Kwiatkowski, Cordelia Langford, John Sillitoe on behalf of the Wellcome Sanger Institute COVID-19 Surveillance Team |
| EPI_ISL_3440785, EPI_ISL_3440786 | Lighthouse Lab in Milton Keynes | Wellcome Sanger Institute for the COVID-19 Genomics UK (COG-UK) Consortium | The Lighthouse Lab in Milton Keynes and Alex Alderton, Roberto Amato, Jeffrey Barrett, Sonia Goncalves, Ewan Harrison, David K. Jackson, Ian Johnston, Dominic Kwiatkowski, Cordelia Langford, John Sillitoe on behalf of the Wellcome Sanger Institute COVID-19 Surveillance Team |
| EPI_ISL_3440787, EPI_ISL_3440788 | Lighthouse Lab in Glasgow | Wellcome Sanger Institute for the COVID-19 Genomics UK (COG-UK) Consortium | Harper VanSteenhouse, Yumi Kasai, David Gray, Carol Clugston, Anna Dominiczak and Alex Alderton, Roberto Amato, Jeffrey Barrett, Sonia Goncalves, Ewan Harrison, David K. Jackson, Ian Johnston, Dominic Kwiatkowski, Cordelia Langford, John Sillitoe on behalf of the Wellcome Sanger Institute COVID-19 Surveillance Team |
| EPI_ISL_3440789, EPI_ISL_3440790, EPI_ISL_3440791 | Lighthouse Lab in Milton Keynes | Wellcome Sanger Institute for the COVID-19 Genomics UK (COG-UK) Consortium | The Lighthouse Lab in Milton Keynes and Alex Alderton, Roberto Amato, Jeffrey Barrett, Sonia Goncalves, Ewan Harrison, David K. Jackson, Ian Johnston, Dominic Kwiatkowski, Cordelia Langford, John Sillitoe on behalf of the Wellcome Sanger Institute COVID-19 Surveillance Team |
| EPI_ISL_3440792 | Lighthouse Lab in Glasgow | Wellcome Sanger Institute for the COVID-19 Genomics UK (COG-UK) Consortium | Harper VanSteenhouse, Yumi Kasai, David Gray, Carol Clugston, Anna Dominiczak and Alex Alderton, Roberto Amato, Jeffrey Barrett, Sonia Goncalves, Ewan Harrison, David K. Jackson, Ian Johnston, Dominic Kwiatkowski, Cordelia Langford, John Sillitoe on behalf of the Wellcome Sanger Institute COVID-19 Surveillance Team |
| EPI_ISL_3440793 | Lighthouse Laboratory Plymouth | Wellcome Sanger Institute for the COVID-19 Genomics UK (COG-UK) Consortium | Lighthouse Laboratory Plymouth and Alex Alderton, Roberto Amato, Jeffrey Barrett, Sonia Goncalves, Ewan Harrison, David K. Jackson, Ian Johnston, Dominic Kwiatkowski, Cordelia Langford, John Sillitoe on behalf of the Wellcome Sanger Institute COVID-19 Surveillance Team |
| EPI_ISL_3440794, EPI_ISL_3440795 | Lighthouse Lab in Milton Keynes | Wellcome Sanger Institute for the COVID-19 Genomics UK (COG-UK) Consortium | The Lighthouse Lab in Milton Keynes and Alex Alderton, Roberto Amato, Jeffrey Barrett, Sonia Goncalves, Ewan Harrison, David K. Jackson, Ian Johnston, Dominic Kwiatkowski, Cordelia Langford, John Sillitoe on behalf of the Wellcome Sanger Institute COVID-19 Surveillance Team |
| EPI_ISL_3440796 | Lighthouse Laboratory Plymouth | Wellcome Sanger Institute for the COVID-19 Genomics UK (COG-UK) Consortium | Lighthouse Laboratory Plymouth and Alex Alderton, Roberto Amato, Jeffrey Barrett, Sonia Goncalves, Ewan Harrison, David K. Jackson, Ian Johnston, Dominic Kwiatkowski, Cordelia Langford, John Sillitoe on behalf of the Wellcome Sanger Institute COVID-19 Surveillance Team |
| EPI_ISL_3440797 | Lighthouse Lab in Milton Keynes | Wellcome Sanger Institute for the COVID-19 Genomics UK (COG-UK) Consortium | The Lighthouse Lab in Milton Keynes and Alex Alderton, Roberto Amato, Jeffrey Barrett, Sonia Goncalves, Ewan Harrison, David K. Jackson, Ian Johnston, Dominic Kwiatkowski, Cordelia Langford, John Sillitoe on behalf of the Wellcome Sanger Institute COVID-19 Surveillance Team |
| EPI_ISL_3440798 | Lighthouse Laboratory Plymouth | Wellcome Sanger Institute for the COVID-19 Genomics UK (COG-UK) Consortium | Lighthouse Laboratory Plymouth and Alex Alderton, Roberto Amato, Jeffrey Barrett, Sonia Goncalves, Ewan Harrison, David K. Jackson, Ian Johnston, Dominic Kwiatkowski, Cordelia Langford, John Sillitoe on behalf of the Wellcome Sanger Institute COVID-19 Surveillance Team |
| EPI_ISL_3440799, EPI_ISL_3440800, EPI_ISL_3440801, EPI_ISL_3440802, EPI_ISL_3440803 | Lighthouse Lab in Milton Keynes | Wellcome Sanger Institute for the COVID-19 Genomics UK (COG-UK) Consortium | The Lighthouse Lab in Milton Keynes and Alex Alderton, Roberto Amato, Jeffrey Barrett, Sonia Goncalves, Ewan Harrison, David K. Jackson, Ian Johnston, Dominic Kwiatkowski, Cordelia Langford, John Sillitoe on behalf of the Wellcome Sanger Institute COVID-19 Surveillance Team |
| EPI_ISL_3440804 | Lighthouse Lab in Glasgow | Wellcome Sanger Institute for the COVID-19 Genomics UK (COG-UK) Consortium | Harper VanSteenhouse, Yumi Kasai, David Gray, Carol Clugston, Anna Dominiczak and Alex Alderton, Roberto Amato, Jeffrey Barrett, Sonia Goncalves, Ewan Harrison, David K. Jackson, Ian Johnston, Dominic Kwiatkowski, Cordelia Langford, John Sillitoe on behalf of the Wellcome Sanger Institute COVID-19 Surveillance Team |
| EPI_ISL_3440805 | Lighthouse Laboratory Plymouth | Wellcome Sanger Institute for the COVID-19 Genomics UK (COG-UK) Consortium | Lighthouse Laboratory Plymouth and Alex Alderton, Roberto Amato, Jeffrey Barrett, Sonia Goncalves, Ewan Harrison, David K. Jackson, Ian Johnston, Dominic Kwiatkowski, Cordelia Langford, John Sillitoe on behalf of the Wellcome Sanger Institute COVID-19 Surveillance Team |
| EPI_ISL_3440806 | Lighthouse Lab in Glasgow | Wellcome Sanger Institute for the COVID-19 Genomics UK (COG-UK) Consortium | Harper VanSteenhouse, Yumi Kasai, David Gray, Carol Clugston, Anna Dominiczak and Alex Alderton, Roberto Amato, Jeffrey Barrett, Sonia Goncalves, Ewan Harrison, David K. Jackson, Ian Johnston, Dominic Kwiatkowski, Cordelia Langford, John Sillitoe on behalf of the Wellcome Sanger Institute COVID-19 Surveillance Team |
| EPI_ISL_3440807 | Lighthouse Laboratory Plymouth | Wellcome Sanger Institute for the COVID-19 Genomics UK (COG-UK) Consortium | Lighthouse Laboratory Plymouth and Alex Alderton, Roberto Amato, Jeffrey Barrett, Sonia Goncalves, Ewan Harrison, David K. Jackson, Ian Johnston, Dominic Kwiatkowski, Cordelia Langford, John Sillitoe on behalf of the Wellcome Sanger Institute COVID-19 Surveillance Team |
| EPI_ISL_3440808 | Lighthouse Lab in Milton Keynes | Wellcome Sanger Institute for the COVID-19 Genomics UK (COG-UK) Consortium | The Lighthouse Lab in Milton Keynes and Alex Alderton, Roberto Amato, Jeffrey Barrett, Sonia Goncalves, Ewan Harrison, David K. Jackson, Ian Johnston, Dominic Kwiatkowski, Cordelia Langford, John Sillitoe on behalf of the Wellcome Sanger Institute COVID-19 Surveillance Team |
| EPI_ISL_3440809, EPI_ISL_3440810 | Lighthouse Lab in Glasgow | Wellcome Sanger Institute for the COVID-19 Genomics UK (COG-UK) Consortium | Harper VanSteenhouse, Yumi Kasai, David Gray, Carol Clugston, Anna Dominiczak and Alex Alderton, Roberto Amato, Jeffrey Barrett, Sonia Goncalves, Ewan Harrison, David K. Jackson, Ian Johnston, Dominic Kwiatkowski, Cordelia Langford, John Sillitoe on behalf of the Wellcome Sanger Institute COVID-19 Surveillance Team |
| EPI_ISL_3440811 | Lighthouse Laboratory Plymouth | Wellcome Sanger Institute for the COVID-19 Genomics UK (COG-UK) Consortium | Lighthouse Laboratory Plymouth and Alex Alderton, Roberto Amato, Jeffrey Barrett, Sonia Goncalves, Ewan Harrison, David K. Jackson, Ian Johnston, Dominic Kwiatkowski, Cordelia Langford, John Sillitoe on behalf of the Wellcome Sanger Institute COVID-19 Surveillance Team |
| EPI_ISL_3440812 | Lighthouse Lab in Glasgow | Wellcome Sanger Institute for the COVID-19 Genomics UK (COG-UK) Consortium | Harper VanSteenhouse, Yumi Kasai, David Gray, Carol Clugston, Anna Dominiczak and Alex Alderton, Roberto Amato, Jeffrey Barrett, Sonia Goncalves, Ewan Harrison, David K. Jackson, Ian Johnston, Dominic Kwiatkowski, Cordelia Langford, John Sillitoe on behalf of the Wellcome Sanger Institute COVID-19 Surveillance Team |
| EPI_ISL_3440813, EPI_ISL_3440814, EPI_ISL_3440815 | Lighthouse Lab in Milton Keynes | Wellcome Sanger Institute for the COVID-19 Genomics UK (COG-UK) Consortium | The Lighthouse Lab in Milton Keynes and Alex Alderton, Roberto Amato, Jeffrey Barrett, Sonia Goncalves, Ewan Harrison, David K. Jackson, Ian Johnston, Dominic Kwiatkowski, Cordelia Langford, John Sillitoe on behalf of the Wellcome Sanger Institute COVID-19 Surveillance Team |
| EPI_ISL_3440816 | Lighthouse Lab in Glasgow | Wellcome Sanger Institute for the COVID-19 Genomics UK (COG-UK) Consortium | Harper VanSteenhouse, Yumi Kasai, David Gray, Carol Clugston, Anna Dominiczak and Alex Alderton, Roberto Amato, Jeffrey Barrett, Sonia Goncalves, Ewan Harrison, David K. Jackson, Ian Johnston, Dominic Kwiatkowski, Cordelia Langford, John Sillitoe on behalf of the Wellcome Sanger Institute COVID-19 Surveillance Team |
| EPI_ISL_3440817 | Lighthouse Lab in Milton Keynes | Wellcome Sanger Institute for the COVID-19 Genomics UK (COG-UK) Consortium | The Lighthouse Lab in Milton Keynes and Alex Alderton, Roberto Amato, Jeffrey Barrett, Sonia Goncalves, Ewan Harrison, David K. Jackson, Ian Johnston, Dominic Kwiatkowski, Cordelia Langford, John Sillitoe on behalf of the Wellcome Sanger Institute COVID-19 Surveillance Team |
| EPI_ISL_3440818 | Lighthouse Laboratory Plymouth | Wellcome Sanger Institute for the COVID-19 Genomics UK (COG-UK) Consortium | Lighthouse Laboratory Plymouth and Alex Alderton, Roberto Amato, Jeffrey Barrett, Sonia Goncalves, Ewan Harrison, David K. Jackson, Ian Johnston, Dominic Kwiatkowski, Cordelia Langford, John Sillitoe on behalf of the Wellcome Sanger Institute COVID-19 Surveillance Team |
| EPI_ISL_3440819, EPI_ISL_3440820 | Lighthouse Lab in Milton Keynes | Wellcome Sanger Institute for the COVID-19 Genomics UK (COG-UK) Consortium | The Lighthouse Lab in Milton Keynes and Alex Alderton, Roberto Amato, Jeffrey Barrett, Sonia Goncalves, Ewan Harrison, David K. Jackson, Ian Johnston, Dominic Kwiatkowski, Cordelia Langford, John Sillitoe on behalf of the Wellcome Sanger Institute COVID-19 Surveillance Team |
| EPI_ISL_3440821 | Lighthouse Laboratory Plymouth | Wellcome Sanger Institute for the COVID-19 Genomics UK (COG-UK) Consortium | Lighthouse Laboratory Plymouth and Alex Alderton, Roberto Amato, Jeffrey Barrett, Sonia Goncalves, Ewan Harrison, David K. Jackson, Ian Johnston, Dominic Kwiatkowski, Cordelia Langford, John Sillitoe on behalf of the Wellcome Sanger Institute COVID-19 Surveillance Team |
| EPI_ISL_3440822, EPI_ISL_3440823, EPI_ISL_3440824 | Lighthouse Lab in Milton Keynes | Wellcome Sanger Institute for the COVID-19 Genomics UK (COG-UK) Consortium | The Lighthouse Lab in Milton Keynes and Alex Alderton, Roberto Amato, Jeffrey Barrett, Sonia Goncalves, Ewan Harrison, David K. Jackson, Ian Johnston, Dominic Kwiatkowski, Cordelia Langford, John Sillitoe on behalf of the Wellcome Sanger Institute COVID-19 Surveillance Team |
| EPI_ISL_3440825, EPI_ISL_3440826 | Lighthouse Lab in Glasgow | Wellcome Sanger Institute for the COVID-19 Genomics UK (COG-UK) Consortium | Harper VanSteenhouse, Yumi Kasai, David Gray, Carol Clugston, Anna Dominiczak and Alex Alderton, Roberto Amato, Jeffrey Barrett, Sonia Goncalves, Ewan Harrison, David K. Jackson, Ian Johnston, Dominic Kwiatkowski, Cordelia Langford, John Sillitoe on behalf of the Wellcome Sanger Institute COVID-19 Surveillance Team |
| EPI_ISL_3440827 | Lighthouse Lab in Milton Keynes | Wellcome Sanger Institute for the COVID-19 Genomics UK (COG-UK) Consortium | The Lighthouse Lab in Milton Keynes and Alex Alderton, Roberto Amato, Jeffrey Barrett, Sonia Goncalves, Ewan Harrison, David K. Jackson, Ian Johnston, Dominic Kwiatkowski, Cordelia Langford, John Sillitoe on behalf of the Wellcome Sanger Institute COVID-19 Surveillance Team |
| EPI_ISL_3440828 | Lighthouse Lab in Glasgow | Wellcome Sanger Institute for the COVID-19 Genomics UK (COG-UK) Consortium | Harper VanSteenhouse, Yumi Kasai, David Gray, Carol Clugston, Anna Dominiczak and Alex Alderton, Roberto Amato, Jeffrey Barrett, Sonia Goncalves, Ewan Harrison, David K. Jackson, Ian Johnston, Dominic Kwiatkowski, Cordelia Langford, John Sillitoe on behalf of the Wellcome Sanger Institute COVID-19 Surveillance Team |
| EPI_ISL_3440829, EPI_ISL_3440830, EPI_ISL_3440831, EPI_ISL_3440832, EPI_ISL_3440833 | Lighthouse Lab in Milton Keynes | Wellcome Sanger Institute for the COVID-19 Genomics UK (COG-UK) Consortium | The Lighthouse Lab in Milton Keynes and Alex Alderton, Roberto Amato, Jeffrey Barrett, Sonia Goncalves, Ewan Harrison, David K. Jackson, Ian Johnston, Dominic Kwiatkowski, Cordelia Langford, John Sillitoe on behalf of the Wellcome Sanger Institute COVID-19 Surveillance Team |
| EPI_ISL_3440834 | Lighthouse Laboratory Plymouth | Wellcome Sanger Institute for the COVID-19 Genomics UK (COG-UK) | Lighthouse Laboratory Plymouth and Alex Alderton, Roberto Amato, Jeffrey Barrett, Sonia Goncalves, Ewan Harrison, David K. Jackson, Ian Johnston, Dominic Kwiatkowski, Cordelia Langford, John Sillitoe on behalf of the Wellcome Sanger |

| | | | |
|---|---|---|---|
| | | Consortium | Institute COVID-19 Surveillance Team |
| EPI_ISL_3440835 | Lighthouse Lab in Milton Keynes | Wellcome Sanger Institute for the COVID-19 Genomics UK (COG-UK) Consortium | The Lighthouse Lab in Milton Keynes and Alex Alderton, Roberto Amato, Jeffrey Barrett, Sonia Goncalves, Ewan Harrison, David K. Jackson, Ian Johnston, Dominic Kwiatkowski, Cordelia Langford, John Sillitoe on behalf of the Wellcome Sanger Institute COVID-19 Surveillance Team |
| EPI_ISL_3440836 | Lighthouse Laboratory Plymouth | Wellcome Sanger Institute for the COVID-19 Genomics UK (COG-UK) Consortium | Lighthouse Laboratory Plymouth and Alex Alderton, Roberto Amato, Jeffrey Barrett, Sonia Goncalves, Ewan Harrison, David K. Jackson, Ian Johnston, Dominic Kwiatkowski, Cordelia Langford, John Sillitoe on behalf of the Wellcome Sanger Institute COVID-19 Surveillance Team |
| EPI_ISL_3440837 | Lighthouse Lab in Glasgow | Wellcome Sanger Institute for the COVID-19 Genomics UK (COG-UK) Consortium | Harper VanSteenhouse, Yumi Kasai, David Gray, Carol Clugston, Anna Dominiczak and Alex Alderton, Roberto Amato, Jeffrey Barrett, Sonia Goncalves, Ewan Harrison, David K. Jackson, Ian Johnston, Dominic Kwiatkowski, Cordelia Langford, John Sillitoe on behalf of the Wellcome Sanger Institute COVID-19 Surveillance Team |
| EPI_ISL_3440838, EPI_ISL_3440839, EPI_ISL_3440840 | Lighthouse Lab in Milton Keynes | Wellcome Sanger Institute for the COVID-19 Genomics UK (COG-UK) Consortium | The Lighthouse Lab in Milton Keynes and Alex Alderton, Roberto Amato, Jeffrey Barrett, Sonia Goncalves, Ewan Harrison, David K. Jackson, Ian Johnston, Dominic Kwiatkowski, Cordelia Langford, John Sillitoe on behalf of the Wellcome Sanger Institute COVID-19 Surveillance Team |
| EPI_ISL_3440841 | Lighthouse Lab in Glasgow | Wellcome Sanger Institute for the COVID-19 Genomics UK (COG-UK) Consortium | Harper VanSteenhouse, Yumi Kasai, David Gray, Carol Clugston, Anna Dominiczak and Alex Alderton, Roberto Amato, Jeffrey Barrett, Sonia Goncalves, Ewan Harrison, David K. Jackson, Ian Johnston, Dominic Kwiatkowski, Cordelia Langford, John Sillitoe on behalf of the Wellcome Sanger Institute COVID-19 Surveillance Team |
| EPI_ISL_3440842, EPI_ISL_3440843 | Lighthouse Lab in Milton Keynes | Wellcome Sanger Institute for the COVID-19 Genomics UK (COG-UK) Consortium | The Lighthouse Lab in Milton Keynes and Alex Alderton, Roberto Amato, Jeffrey Barrett, Sonia Goncalves, Ewan Harrison, David K. Jackson, Ian Johnston, Dominic Kwiatkowski, Cordelia Langford, John Sillitoe on behalf of the Wellcome Sanger Institute COVID-19 Surveillance Team |
| EPI_ISL_3440844 | Lighthouse Lab in Glasgow | Wellcome Sanger Institute for the COVID-19 Genomics UK (COG-UK) Consortium | Harper VanSteenhouse, Yumi Kasai, David Gray, Carol Clugston, Anna Dominiczak and Alex Alderton, Roberto Amato, Jeffrey Barrett, Sonia Goncalves, Ewan Harrison, David K. Jackson, Ian Johnston, Dominic Kwiatkowski, Cordelia Langford, John Sillitoe on behalf of the Wellcome Sanger Institute COVID-19 Surveillance Team |
| EPI_ISL_3440845, EPI_ISL_3440846, EPI_ISL_3440847, EPI_ISL_3440848, EPI_ISL_3440849, EPI_ISL_3440850 | Lighthouse Lab in Milton Keynes | Wellcome Sanger Institute for the COVID-19 Genomics UK (COG-UK) Consortium | The Lighthouse Lab in Milton Keynes and Alex Alderton, Roberto Amato, Jeffrey Barrett, Sonia Goncalves, Ewan Harrison, David K. Jackson, Ian Johnston, Dominic Kwiatkowski, Cordelia Langford, John Sillitoe on behalf of the Wellcome Sanger Institute COVID-19 Surveillance Team |
| EPI_ISL_3440851 | Lighthouse Laboratory Plymouth | Wellcome Sanger Institute for the COVID-19 Genomics UK (COG-UK) Consortium | Lighthouse Laboratory Plymouth and Alex Alderton, Roberto Amato, Jeffrey Barrett, Sonia Goncalves, Ewan Harrison, David K. Jackson, Ian Johnston, Dominic Kwiatkowski, Cordelia Langford, John Sillitoe on behalf of the Wellcome Sanger Institute COVID-19 Surveillance Team |
| EPI_ISL_3440852, EPI_ISL_3440853 | Lighthouse Lab in Milton Keynes | Wellcome Sanger Institute for the COVID-19 Genomics UK (COG-UK) Consortium | The Lighthouse Lab in Milton Keynes and Alex Alderton, Roberto Amato, Jeffrey Barrett, Sonia Goncalves, Ewan Harrison, David K. Jackson, Ian Johnston, Dominic Kwiatkowski, Cordelia Langford, John Sillitoe on behalf of the Wellcome Sanger Institute COVID-19 Surveillance Team |
| EPI_ISL_3440854 | Lighthouse Laboratory Plymouth | Wellcome Sanger Institute for the COVID-19 Genomics UK (COG-UK) Consortium | Lighthouse Laboratory Plymouth and Alex Alderton, Roberto Amato, Jeffrey Barrett, Sonia Goncalves, Ewan Harrison, David K. Jackson, Ian Johnston, Dominic Kwiatkowski, Cordelia Langford, John Sillitoe on behalf of the Wellcome Sanger Institute COVID-19 Surveillance Team |
| EPI_ISL_3440855 | Lighthouse Lab in Milton Keynes | Wellcome Sanger Institute for the COVID-19 Genomics UK (COG-UK) Consortium | The Lighthouse Lab in Milton Keynes and Alex Alderton, Roberto Amato, Jeffrey Barrett, Sonia Goncalves, Ewan Harrison, David K. Jackson, Ian Johnston, Dominic Kwiatkowski, Cordelia Langford, John Sillitoe on behalf of the Wellcome Sanger Institute COVID-19 Surveillance Team |
| EPI_ISL_3440856 | Lighthouse Laboratory Plymouth | Wellcome Sanger Institute for the COVID-19 Genomics UK (COG-UK) Consortium | Lighthouse Laboratory Plymouth and Alex Alderton, Roberto Amato, Jeffrey Barrett, Sonia Goncalves, Ewan Harrison, David K. Jackson, Ian Johnston, Dominic Kwiatkowski, Cordelia Langford, John Sillitoe on behalf of the Wellcome Sanger Institute COVID-19 Surveillance Team |
| EPI_ISL_3440857 | Lighthouse Lab in Milton Keynes | Wellcome Sanger Institute for the COVID-19 Genomics UK (COG-UK) Consortium | The Lighthouse Lab in Milton Keynes and Alex Alderton, Roberto Amato, Jeffrey Barrett, Sonia Goncalves, Ewan Harrison, David K. Jackson, Ian Johnston, Dominic Kwiatkowski, Cordelia Langford, John Sillitoe on behalf of the Wellcome Sanger Institute COVID-19 Surveillance Team |
| EPI_ISL_3440858 | Lighthouse Laboratory Plymouth | Wellcome Sanger Institute for the COVID-19 Genomics UK (COG-UK) Consortium | Lighthouse Laboratory Plymouth and Alex Alderton, Roberto Amato, Jeffrey Barrett, Sonia Goncalves, Ewan Harrison, David K. Jackson, Ian Johnston, Dominic Kwiatkowski, Cordelia Langford, John Sillitoe on behalf of the Wellcome Sanger Institute COVID-19 Surveillance Team |
| EPI_ISL_3440859 | Lighthouse Lab in Milton Keynes | Wellcome Sanger Institute for the COVID-19 Genomics UK (COG-UK) Consortium | The Lighthouse Lab in Milton Keynes and Alex Alderton, Roberto Amato, Jeffrey Barrett, Sonia Goncalves, Ewan Harrison, David K. Jackson, Ian Johnston, Dominic Kwiatkowski, Cordelia Langford, John Sillitoe on behalf of the Wellcome Sanger Institute COVID-19 Surveillance Team |
| EPI_ISL_3440860 | Lighthouse Laboratory Plymouth | Wellcome Sanger Institute for the COVID-19 Genomics UK (COG-UK) Consortium | Lighthouse Laboratory Plymouth and Alex Alderton, Roberto Amato, Jeffrey Barrett, Sonia Goncalves, Ewan Harrison, David K. Jackson, Ian Johnston, Dominic Kwiatkowski, Cordelia Langford, John Sillitoe on behalf of the Wellcome Sanger Institute COVID-19 Surveillance Team |
| EPI_ISL_3440861, EPI_ISL_3440862, EPI_ISL_3440863, EPI_ISL_3440864, EPI_ISL_3440865 | Lighthouse Lab in Milton Keynes | Wellcome Sanger Institute for the COVID-19 Genomics UK (COG-UK) Consortium | The Lighthouse Lab in Milton Keynes and Alex Alderton, Roberto Amato, Jeffrey Barrett, Sonia Goncalves, Ewan Harrison, David K. Jackson, Ian Johnston, Dominic Kwiatkowski, Cordelia Langford, John Sillitoe on behalf of the Wellcome Sanger Institute COVID-19 Surveillance Team |
| EPI_ISL_3440866 | Lighthouse Laboratory Plymouth | Wellcome Sanger Institute for the COVID-19 Genomics UK (COG-UK) Consortium | Lighthouse Laboratory Plymouth and Alex Alderton, Roberto Amato, Jeffrey Barrett, Sonia Goncalves, Ewan Harrison, David K. Jackson, Ian Johnston, Dominic Kwiatkowski, Cordelia Langford, John Sillitoe on behalf of the Wellcome Sanger Institute COVID-19 Surveillance Team |
| EPI_ISL_3440867 | Lighthouse Lab in Milton Keynes | Wellcome Sanger Institute for the COVID-19 Genomics UK (COG-UK) Consortium | The Lighthouse Lab in Milton Keynes and Alex Alderton, Roberto Amato, Jeffrey Barrett, Sonia Goncalves, Ewan Harrison, David K. Jackson, Ian Johnston, Dominic Kwiatkowski, Cordelia Langford, John Sillitoe on behalf of the Wellcome Sanger Institute COVID-19 Surveillance Team |
| EPI_ISL_3440868, EPI_ISL_3440869, EPI_ISL_3440870 | Lighthouse Laboratory Plymouth | Wellcome Sanger Institute for the COVID-19 Genomics UK (COG-UK) Consortium | Lighthouse Laboratory Plymouth and Alex Alderton, Roberto Amato, Jeffrey Barrett, Sonia Goncalves, Ewan Harrison, David K. Jackson, Ian Johnston, Dominic Kwiatkowski, Cordelia Langford, John Sillitoe on behalf of the Wellcome Sanger Institute COVID-19 Surveillance Team |
| EPI_ISL_3440871, EPI_ISL_3440872 | Lighthouse Lab in Milton Keynes | Wellcome Sanger Institute for the COVID-19 Genomics UK (COG-UK) Consortium | The Lighthouse Lab in Milton Keynes and Alex Alderton, Roberto Amato, Jeffrey Barrett, Sonia Goncalves, Ewan Harrison, David K. Jackson, Ian Johnston, Dominic Kwiatkowski, Cordelia Langford, John Sillitoe on behalf of the Wellcome Sanger Institute COVID-19 Surveillance Team |
| EPI_ISL_3440873 | Lighthouse Lab in Glasgow | Wellcome Sanger Institute for the COVID-19 Genomics UK (COG-UK) Consortium | Harper VanSteenhouse, Yumi Kasai, David Gray, Carol Clugston, Anna Dominiczak and Alex Alderton, Roberto Amato, Jeffrey Barrett, Sonia Goncalves, Ewan Harrison, David K. Jackson, Ian Johnston, Dominic Kwiatkowski, Cordelia Langford, John Sillitoe on behalf of the Wellcome Sanger Institute COVID-19 Surveillance Team |
| EPI_ISL_3440874, EPI_ISL_3440875, EPI_ISL_3440876 | Lighthouse Laboratory Plymouth | Wellcome Sanger Institute for the COVID-19 Genomics UK (COG-UK) Consortium | Lighthouse Laboratory Plymouth and Alex Alderton, Roberto Amato, Jeffrey Barrett, Sonia Goncalves, Ewan Harrison, David K. Jackson, Ian Johnston, Dominic Kwiatkowski, Cordelia Langford, John Sillitoe on behalf of the Wellcome Sanger Institute COVID-19 Surveillance Team |
| EPI_ISL_3440877 | Lighthouse Lab in Glasgow | Wellcome Sanger Institute for the COVID-19 Genomics UK (COG-UK) Consortium | Harper VanSteenhouse, Yumi Kasai, David Gray, Carol Clugston, Anna Dominiczak and Alex Alderton, Roberto Amato, Jeffrey Barrett, Sonia Goncalves, Ewan Harrison, David K. Jackson, Ian Johnston, Dominic Kwiatkowski, Cordelia Langford, John Sillitoe on behalf of the Wellcome Sanger Institute COVID-19 Surveillance Team |
| EPI_ISL_3440878, EPI_ISL_3440879 | Lighthouse Laboratory Plymouth | Wellcome Sanger Institute for the COVID-19 Genomics UK (COG-UK) Consortium | Lighthouse Laboratory Plymouth and Alex Alderton, Roberto Amato, Jeffrey Barrett, Sonia Goncalves, Ewan Harrison, David K. Jackson, Ian Johnston, Dominic Kwiatkowski, Cordelia Langford, John Sillitoe on behalf of the Wellcome Sanger Institute COVID-19 Surveillance Team |
| EPI_ISL_3440880, EPI_ISL_3440881 | Lighthouse Lab in Milton Keynes | Wellcome Sanger Institute for the COVID-19 Genomics UK (COG-UK) Consortium | The Lighthouse Lab in Milton Keynes and Alex Alderton, Roberto Amato, Jeffrey Barrett, Sonia Goncalves, Ewan Harrison, David K. Jackson, Ian Johnston, Dominic Kwiatkowski, Cordelia Langford, John Sillitoe on behalf of the Wellcome Sanger Institute COVID-19 Surveillance Team |
| EPI_ISL_3440882 | Lighthouse Lab in Glasgow | Wellcome Sanger Institute for the COVID-19 Genomics UK (COG-UK) Consortium | Harper VanSteenhouse, Yumi Kasai, David Gray, Carol Clugston, Anna Dominiczak and Alex Alderton, Roberto Amato, Jeffrey Barrett, Sonia Goncalves, Ewan Harrison, David K. Jackson, Ian Johnston, Dominic Kwiatkowski, Cordelia Langford, John Sillitoe on behalf of the Wellcome Sanger Institute COVID-19 Surveillance Team |
| EPI_ISL_3440883, EPI_ISL_3440884 | Lighthouse Lab in Milton Keynes | Wellcome Sanger Institute for the COVID-19 Genomics UK (COG-UK) Consortium | The Lighthouse Lab in Milton Keynes and Alex Alderton, Roberto Amato, Jeffrey Barrett, Sonia Goncalves, Ewan Harrison, David K. Jackson, Ian Johnston, Dominic Kwiatkowski, Cordelia Langford, John Sillitoe on behalf of the Wellcome Sanger Institute COVID-19 Surveillance Team |
| EPI_ISL_3440885, EPI_ISL_3440886 | Lighthouse Lab in Glasgow | Wellcome Sanger Institute for the COVID-19 Genomics UK (COG-UK) Consortium | Harper VanSteenhouse, Yumi Kasai, David Gray, Carol Clugston, Anna Dominiczak and Alex Alderton, Roberto Amato, Jeffrey Barrett, Sonia Goncalves, Ewan Harrison, David K. Jackson, Ian Johnston, Dominic Kwiatkowski, Cordelia Langford, John Sillitoe on behalf of the Wellcome Sanger Institute COVID-19 Surveillance Team |
| EPI_ISL_3440887 | Lighthouse Lab in Milton Keynes | Wellcome Sanger Institute for the COVID-19 Genomics UK (COG-UK) Consortium | The Lighthouse Lab in Milton Keynes and Alex Alderton, Roberto Amato, Jeffrey Barrett, Sonia Goncalves, Ewan Harrison, David K. Jackson, Ian Johnston, Dominic Kwiatkowski, Cordelia Langford, John Sillitoe on behalf of the Wellcome Sanger Institute COVID-19 Surveillance Team |
| EPI_ISL_3440888 | Lighthouse Lab in Glasgow | Wellcome Sanger Institute for the COVID-19 Genomics UK (COG-UK) Consortium | Harper VanSteenhouse, Yumi Kasai, David Gray, Carol Clugston, Anna Dominiczak and Alex Alderton, Roberto Amato, Jeffrey Barrett, Sonia Goncalves, Ewan Harrison, David K. Jackson, Ian Johnston, Dominic Kwiatkowski, Cordelia Langford, John Sillitoe on behalf of the Wellcome Sanger Institute COVID-19 Surveillance Team |
| EPI_ISL_3440889 | Lighthouse Lab in Milton Keynes | Wellcome Sanger Institute for the COVID-19 Genomics UK (COG-UK) Consortium | The Lighthouse Lab in Milton Keynes and Alex Alderton, Roberto Amato, Jeffrey Barrett, Sonia Goncalves, Ewan Harrison, David K. Jackson, Ian Johnston, Dominic Kwiatkowski, Cordelia Langford, John Sillitoe on behalf of the Wellcome Sanger Institute COVID-19 Surveillance Team |
| EPI_ISL_3440890, EPI_ISL_3440891, EPI_ISL_3440892, EPI_ISL_3440893, EPI_ISL_3440894 | Lighthouse Laboratory Plymouth | Wellcome Sanger Institute for the COVID-19 Genomics UK (COG-UK) Consortium | Lighthouse Laboratory Plymouth and Alex Alderton, Roberto Amato, Jeffrey Barrett, Sonia Goncalves, Ewan Harrison, David K. Jackson, Ian Johnston, Dominic Kwiatkowski, Cordelia Langford, John Sillitoe on behalf of the Wellcome Sanger Institute COVID-19 Surveillance Team |
| EPI_ISL_3440895, EPI_ISL_3440896, EPI_ISL_3440897, EPI_ISL_3440898, EPI_ISL_3440899, EPI_ISL_3440900, EPI_ISL_3440901 | Lighthouse Lab in Milton Keynes | Wellcome Sanger Institute for the COVID-19 Genomics UK (COG-UK) Consortium | The Lighthouse Lab in Milton Keynes and Alex Alderton, Roberto Amato, Jeffrey Barrett, Sonia Goncalves, Ewan Harrison, David K. Jackson, Ian Johnston, Dominic Kwiatkowski, Cordelia Langford, John Sillitoe on behalf of the Wellcome Sanger Institute COVID-19 Surveillance Team |
| EPI_ISL_3440902 | Lighthouse Laboratory Plymouth | Wellcome Sanger Institute for the COVID-19 Genomics UK (COG-UK) Consortium | Lighthouse Laboratory Plymouth and Alex Alderton, Roberto Amato, Jeffrey Barrett, Sonia Goncalves, Ewan Harrison, David K. Jackson, Ian Johnston, Dominic Kwiatkowski, Cordelia Langford, John Sillitoe on behalf of the Wellcome Sanger Institute COVID-19 Surveillance Team |
| EPI_ISL_3440903, EPI_ISL_3440904, EPI_ISL_3440905 | Lighthouse Lab in Milton Keynes | Wellcome Sanger Institute for the COVID-19 Genomics UK (COG-UK) Consortium | The Lighthouse Lab in Milton Keynes and Alex Alderton, Roberto Amato, Jeffrey Barrett, Sonia Goncalves, Ewan Harrison, David K. Jackson, Ian Johnston, Dominic Kwiatkowski, Cordelia Langford, John Sillitoe on behalf of the Wellcome Sanger Institute COVID-19 Surveillance Team |

| | | | |
|---|---|---|---|
| EPI_ISL_3440906 | Lighthouse Laboratory Plymouth | Wellcome Sanger Institute for the COVID-19 Genomics UK (COG-UK) Consortium | Lighthouse Laboratory Plymouth and Alex Alderton, Roberto Amato, Jeffrey Barrett, Sonia Goncalves, Ewan Harrison, David K. Jackson, Ian Johnston, Dominic Kwiatkowski, Cordelia Langford, John Sillitoe on behalf of the Wellcome Sanger Institute COVID-19 Surveillance Team |
| EPI_ISL_3440907, EPI_ISL_3440908 | Lighthouse Lab in Milton Keynes | Wellcome Sanger Institute for the COVID-19 Genomics UK (COG-UK) Consortium | The Lighthouse Lab in Milton Keynes and Alex Alderton, Roberto Amato, Jeffrey Barrett, Sonia Goncalves, Ewan Harrison, David K. Jackson, Ian Johnston, Dominic Kwiatkowski, Cordelia Langford, John Sillitoe on behalf of the Wellcome Sanger Institute COVID-19 Surveillance Team |
| EPI_ISL_3440909 | Lighthouse Lab in Glasgow | Wellcome Sanger Institute for the COVID-19 Genomics UK (COG-UK) Consortium | Harper VanSteenhouse, Yumi Kasai, David Gray, Carol Clugston, Anna Dominiczak and Alex Alderton, Roberto Amato, Jeffrey Barrett, Sonia Goncalves, Ewan Harrison, David K. Jackson, Ian Johnston, Dominic Kwiatkowski, Cordelia Langford, John Sillitoe on behalf of the Wellcome Sanger Institute COVID-19 Surveillance Team |
| EPI_ISL_3440910 | Lighthouse Lab in Milton Keynes | Wellcome Sanger Institute for the COVID-19 Genomics UK (COG-UK) Consortium | The Lighthouse Lab in Milton Keynes and Alex Alderton, Roberto Amato, Jeffrey Barrett, Sonia Goncalves, Ewan Harrison, David K. Jackson, Ian Johnston, Dominic Kwiatkowski, Cordelia Langford, John Sillitoe on behalf of the Wellcome Sanger Institute COVID-19 Surveillance Team |
| EPI_ISL_3440911 | Lighthouse Laboratory Plymouth | Wellcome Sanger Institute for the COVID-19 Genomics UK (COG-UK) Consortium | Lighthouse Laboratory Plymouth and Alex Alderton, Roberto Amato, Jeffrey Barrett, Sonia Goncalves, Ewan Harrison, David K. Jackson, Ian Johnston, Dominic Kwiatkowski, Cordelia Langford, John Sillitoe on behalf of the Wellcome Sanger Institute COVID-19 Surveillance Team |
| EPI_ISL_3440912 | Lighthouse Lab in Milton Keynes | Wellcome Sanger Institute for the COVID-19 Genomics UK (COG-UK) Consortium | The Lighthouse Lab in Milton Keynes and Alex Alderton, Roberto Amato, Jeffrey Barrett, Sonia Goncalves, Ewan Harrison, David K. Jackson, Ian Johnston, Dominic Kwiatkowski, Cordelia Langford, John Sillitoe on behalf of the Wellcome Sanger Institute COVID-19 Surveillance Team |
| EPI_ISL_3440913 | Lighthouse Laboratory Plymouth | Wellcome Sanger Institute for the COVID-19 Genomics UK (COG-UK) Consortium | Lighthouse Laboratory Plymouth and Alex Alderton, Roberto Amato, Jeffrey Barrett, Sonia Goncalves, Ewan Harrison, David K. Jackson, Ian Johnston, Dominic Kwiatkowski, Cordelia Langford, John Sillitoe on behalf of the Wellcome Sanger Institute COVID-19 Surveillance Team |
| EPI_ISL_3440914 | Lighthouse Lab in Glasgow | Wellcome Sanger Institute for the COVID-19 Genomics UK (COG-UK) Consortium | Harper VanSteenhouse, Yumi Kasai, David Gray, Carol Clugston, Anna Dominiczak and Alex Alderton, Roberto Amato, Jeffrey Barrett, Sonia Goncalves, Ewan Harrison, David K. Jackson, Ian Johnston, Dominic Kwiatkowski, Cordelia Langford, John Sillitoe on behalf of the Wellcome Sanger Institute COVID-19 Surveillance Team |
| EPI_ISL_3440915 | Lighthouse Lab in Milton Keynes | Wellcome Sanger Institute for the COVID-19 Genomics UK (COG-UK) Consortium | The Lighthouse Lab in Milton Keynes and Alex Alderton, Roberto Amato, Jeffrey Barrett, Sonia Goncalves, Ewan Harrison, David K. Jackson, Ian Johnston, Dominic Kwiatkowski, Cordelia Langford, John Sillitoe on behalf of the Wellcome Sanger Institute COVID-19 Surveillance Team |
| EPI_ISL_3440916 | Lighthouse Laboratory Plymouth | Wellcome Sanger Institute for the COVID-19 Genomics UK (COG-UK) Consortium | Lighthouse Laboratory Plymouth and Alex Alderton, Roberto Amato, Jeffrey Barrett, Sonia Goncalves, Ewan Harrison, David K. Jackson, Ian Johnston, Dominic Kwiatkowski, Cordelia Langford, John Sillitoe on behalf of the Wellcome Sanger Institute COVID-19 Surveillance Team |
| EPI_ISL_3440917, EPI_ISL_3440918 | Lighthouse Lab in Glasgow | Wellcome Sanger Institute for the COVID-19 Genomics UK (COG-UK) Consortium | Harper VanSteenhouse, Yumi Kasai, David Gray, Carol Clugston, Anna Dominiczak and Alex Alderton, Roberto Amato, Jeffrey Barrett, Sonia Goncalves, Ewan Harrison, David K. Jackson, Ian Johnston, Dominic Kwiatkowski, Cordelia Langford, John Sillitoe on behalf of the Wellcome Sanger Institute COVID-19 Surveillance Team |
| EPI_ISL_3440919, EPI_ISL_3440920, EPI_ISL_3440921 | Lighthouse Laboratory Plymouth | Wellcome Sanger Institute for the COVID-19 Genomics UK (COG-UK) Consortium | Lighthouse Laboratory Plymouth and Alex Alderton, Roberto Amato, Jeffrey Barrett, Sonia Goncalves, Ewan Harrison, David K. Jackson, Ian Johnston, Dominic Kwiatkowski, Cordelia Langford, John Sillitoe on behalf of the Wellcome Sanger Institute COVID-19 Surveillance Team |
| EPI_ISL_3440922 | Lighthouse Lab in Glasgow | Wellcome Sanger Institute for the COVID-19 Genomics UK (COG-UK) Consortium | Harper VanSteenhouse, Yumi Kasai, David Gray, Carol Clugston, Anna Dominiczak and Alex Alderton, Roberto Amato, Jeffrey Barrett, Sonia Goncalves, Ewan Harrison, David K. Jackson, Ian Johnston, Dominic Kwiatkowski, Cordelia Langford, John Sillitoe on behalf of the Wellcome Sanger Institute COVID-19 Surveillance Team |
| EPI_ISL_3440923 | Lighthouse Laboratory Plymouth | Wellcome Sanger Institute for the COVID-19 Genomics UK (COG-UK) Consortium | Lighthouse Laboratory Plymouth and Alex Alderton, Roberto Amato, Jeffrey Barrett, Sonia Goncalves, Ewan Harrison, David K. Jackson, Ian Johnston, Dominic Kwiatkowski, Cordelia Langford, John Sillitoe on behalf of the Wellcome Sanger Institute COVID-19 Surveillance Team |
| EPI_ISL_3440924 | Lighthouse Lab in Glasgow | Wellcome Sanger Institute for the COVID-19 Genomics UK (COG-UK) Consortium | Harper VanSteenhouse, Yumi Kasai, David Gray, Carol Clugston, Anna Dominiczak and Alex Alderton, Roberto Amato, Jeffrey Barrett, Sonia Goncalves, Ewan Harrison, David K. Jackson, Ian Johnston, Dominic Kwiatkowski, Cordelia Langford, John Sillitoe on behalf of the Wellcome Sanger Institute COVID-19 Surveillance Team |
| EPI_ISL_3440925 | Lighthouse Lab in Milton Keynes | Wellcome Sanger Institute for the COVID-19 Genomics UK (COG-UK) Consortium | The Lighthouse Lab in Milton Keynes and Alex Alderton, Roberto Amato, Jeffrey Barrett, Sonia Goncalves, Ewan Harrison, David K. Jackson, Ian Johnston, Dominic Kwiatkowski, Cordelia Langford, John Sillitoe on behalf of the Wellcome Sanger Institute COVID-19 Surveillance Team |
| EPI_ISL_3440926 | Lighthouse Laboratory Plymouth | Wellcome Sanger Institute for the COVID-19 Genomics UK (COG-UK) Consortium | Lighthouse Laboratory Plymouth and Alex Alderton, Roberto Amato, Jeffrey Barrett, Sonia Goncalves, Ewan Harrison, David K. Jackson, Ian Johnston, Dominic Kwiatkowski, Cordelia Langford, John Sillitoe on behalf of the Wellcome Sanger Institute COVID-19 Surveillance Team |
| EPI_ISL_3440927 | Lighthouse Lab in Milton Keynes | Wellcome Sanger Institute for the COVID-19 Genomics UK (COG-UK) Consortium | The Lighthouse Lab in Milton Keynes and Alex Alderton, Roberto Amato, Jeffrey Barrett, Sonia Goncalves, Ewan Harrison, David K. Jackson, Ian Johnston, Dominic Kwiatkowski, Cordelia Langford, John Sillitoe on behalf of the Wellcome Sanger Institute COVID-19 Surveillance Team |
| EPI_ISL_3440928, EPI_ISL_3440929 | Lighthouse Lab in Glasgow | Wellcome Sanger Institute for the COVID-19 Genomics UK (COG-UK) Consortium | Harper VanSteenhouse, Yumi Kasai, David Gray, Carol Clugston, Anna Dominiczak and Alex Alderton, Roberto Amato, Jeffrey Barrett, Sonia Goncalves, Ewan Harrison, David K. Jackson, Ian Johnston, Dominic Kwiatkowski, Cordelia Langford, John Sillitoe on behalf of the Wellcome Sanger Institute COVID-19 Surveillance Team |
| EPI_ISL_3440930, EPI_ISL_3440931 | Lighthouse Laboratory Plymouth | Wellcome Sanger Institute for the COVID-19 Genomics UK (COG-UK) Consortium | Lighthouse Laboratory Plymouth and Alex Alderton, Roberto Amato, Jeffrey Barrett, Sonia Goncalves, Ewan Harrison, David K. Jackson, Ian Johnston, Dominic Kwiatkowski, Cordelia Langford, John Sillitoe on behalf of the Wellcome Sanger Institute COVID-19 Surveillance Team |
| EPI_ISL_3440932 | Lighthouse Lab in Glasgow | Wellcome Sanger Institute for the COVID-19 Genomics UK (COG-UK) Consortium | Harper VanSteenhouse, Yumi Kasai, David Gray, Carol Clugston, Anna Dominiczak and Alex Alderton, Roberto Amato, Jeffrey Barrett, Sonia Goncalves, Ewan Harrison, David K. Jackson, Ian Johnston, Dominic Kwiatkowski, Cordelia Langford, John Sillitoe on behalf of the Wellcome Sanger Institute COVID-19 Surveillance Team |
| EPI_ISL_3440933, EPI_ISL_3440934 | Lighthouse Lab in Milton Keynes | Wellcome Sanger Institute for the COVID-19 Genomics UK (COG-UK) Consortium | The Lighthouse Lab in Milton Keynes and Alex Alderton, Roberto Amato, Jeffrey Barrett, Sonia Goncalves, Ewan Harrison, David K. Jackson, Ian Johnston, Dominic Kwiatkowski, Cordelia Langford, John Sillitoe on behalf of the Wellcome Sanger Institute COVID-19 Surveillance Team |
| EPI_ISL_3440935 | Lighthouse Lab in Glasgow | Wellcome Sanger Institute for the COVID-19 Genomics UK (COG-UK) Consortium | Harper VanSteenhouse, Yumi Kasai, David Gray, Carol Clugston, Anna Dominiczak and Alex Alderton, Roberto Amato, Jeffrey Barrett, Sonia Goncalves, Ewan Harrison, David K. Jackson, Ian Johnston, Dominic Kwiatkowski, Cordelia Langford, John Sillitoe on behalf of the Wellcome Sanger Institute COVID-19 Surveillance Team |
| EPI_ISL_3440936 | Lighthouse Lab in Milton Keynes | Wellcome Sanger Institute for the COVID-19 Genomics UK (COG-UK) Consortium | The Lighthouse Lab in Milton Keynes and Alex Alderton, Roberto Amato, Jeffrey Barrett, Sonia Goncalves, Ewan Harrison, David K. Jackson, Ian Johnston, Dominic Kwiatkowski, Cordelia Langford, John Sillitoe on behalf of the Wellcome Sanger Institute COVID-19 Surveillance Team |
| EPI_ISL_3440937 | Health Services Laboratories | Wellcome Sanger Institute for the COVID-19 Genomics UK (COG-UK) Consortium | Health Services Laboratories and Alex Alderton, Roberto Amato, Jeffrey Barrett, Sonia Goncalves, Ewan Harrison, David K. Jackson, Ian Johnston, Dominic Kwiatkowski, Cordelia Langford, John Sillitoe on behalf of the Wellcome Sanger Institute COVID-19 Surveillance Team |
| EPI_ISL_3440938 | Lighthouse Lab in Milton Keynes | Wellcome Sanger Institute for the COVID-19 Genomics UK (COG-UK) Consortium | The Lighthouse Lab in Milton Keynes and Alex Alderton, Roberto Amato, Jeffrey Barrett, Sonia Goncalves, Ewan Harrison, David K. Jackson, Ian Johnston, Dominic Kwiatkowski, Cordelia Langford, John Sillitoe on behalf of the Wellcome Sanger Institute COVID-19 Surveillance Team |
| EPI_ISL_3440939 | Health Services Laboratories | Wellcome Sanger Institute for the COVID-19 Genomics UK (COG-UK) Consortium | Health Services Laboratories and Alex Alderton, Roberto Amato, Jeffrey Barrett, Sonia Goncalves, Ewan Harrison, David K. Jackson, Ian Johnston, Dominic Kwiatkowski, Cordelia Langford, John Sillitoe on behalf of the Wellcome Sanger Institute COVID-19 Surveillance Team |
| EPI_ISL_3440940 | Lighthouse Lab in Glasgow | Wellcome Sanger Institute for the COVID-19 Genomics UK (COG-UK) Consortium | Harper VanSteenhouse, Yumi Kasai, David Gray, Carol Clugston, Anna Dominiczak and Alex Alderton, Roberto Amato, Jeffrey Barrett, Sonia Goncalves, Ewan Harrison, David K. Jackson, Ian Johnston, Dominic Kwiatkowski, Cordelia Langford, John Sillitoe on behalf of the Wellcome Sanger Institute COVID-19 Surveillance Team |
| EPI_ISL_3440941, EPI_ISL_3440942 | Health Services Laboratories | Wellcome Sanger Institute for the COVID-19 Genomics UK (COG-UK) Consortium | Health Services Laboratories and Alex Alderton, Roberto Amato, Jeffrey Barrett, Sonia Goncalves, Ewan Harrison, David K. Jackson, Ian Johnston, Dominic Kwiatkowski, Cordelia Langford, John Sillitoe on behalf of the Wellcome Sanger Institute COVID-19 Surveillance Team |
| EPI_ISL_3440943, EPI_ISL_3440944 | Lighthouse Laboratory Plymouth | Wellcome Sanger Institute for the COVID-19 Genomics UK (COG-UK) Consortium | Lighthouse Laboratory Plymouth and Alex Alderton, Roberto Amato, Jeffrey Barrett, Sonia Goncalves, Ewan Harrison, David K. Jackson, Ian Johnston, Dominic Kwiatkowski, Cordelia Langford, John Sillitoe on behalf of the Wellcome Sanger Institute COVID-19 Surveillance Team |
| EPI_ISL_3440945 | Lighthouse Lab in Glasgow | Wellcome Sanger Institute for the COVID-19 Genomics UK (COG-UK) Consortium | Harper VanSteenhouse, Yumi Kasai, David Gray, Carol Clugston, Anna Dominiczak and Alex Alderton, Roberto Amato, Jeffrey Barrett, Sonia Goncalves, Ewan Harrison, David K. Jackson, Ian Johnston, Dominic Kwiatkowski, Cordelia Langford, John Sillitoe on behalf of the Wellcome Sanger Institute COVID-19 Surveillance Team |
| EPI_ISL_3440946, EPI_ISL_3440947 | Lighthouse Lab in Milton Keynes | Wellcome Sanger Institute for the COVID-19 Genomics UK (COG-UK) Consortium | The Lighthouse Lab in Milton Keynes and Alex Alderton, Roberto Amato, Jeffrey Barrett, Sonia Goncalves, Ewan Harrison, David K. Jackson, Ian Johnston, Dominic Kwiatkowski, Cordelia Langford, John Sillitoe on behalf of the Wellcome Sanger Institute COVID-19 Surveillance Team |
| EPI_ISL_3440948 | Lighthouse Lab in Glasgow | Wellcome Sanger Institute for the COVID-19 Genomics UK (COG-UK) Consortium | Harper VanSteenhouse, Yumi Kasai, David Gray, Carol Clugston, Anna Dominiczak and Alex Alderton, Roberto Amato, Jeffrey Barrett, Sonia Goncalves, Ewan Harrison, David K. Jackson, Ian Johnston, Dominic Kwiatkowski, Cordelia Langford, John Sillitoe on behalf of the Wellcome Sanger Institute COVID-19 Surveillance Team |
| EPI_ISL_3440949, EPI_ISL_3440950 | Lighthouse Lab in Milton Keynes | Wellcome Sanger Institute for the COVID-19 Genomics UK (COG-UK) Consortium | The Lighthouse Lab in Milton Keynes and Alex Alderton, Roberto Amato, Jeffrey Barrett, Sonia Goncalves, Ewan Harrison, David K. Jackson, Ian Johnston, Dominic Kwiatkowski, Cordelia Langford, John Sillitoe on behalf of the Wellcome Sanger Institute COVID-19 Surveillance Team |
| EPI_ISL_3440951 | Lighthouse Lab in Glasgow | Wellcome Sanger Institute for the COVID-19 Genomics UK (COG-UK) Consortium | Harper VanSteenhouse, Yumi Kasai, David Gray, Carol Clugston, Anna Dominiczak and Alex Alderton, Roberto Amato, Jeffrey Barrett, Sonia Goncalves, Ewan Harrison, David K. Jackson, Ian Johnston, Dominic Kwiatkowski, Cordelia Langford, John Sillitoe on behalf of the Wellcome Sanger Institute COVID-19 Surveillance Team |
| EPI_ISL_3440952, EPI_ISL_3440953 | Berkshire and Surrey Pathology Services Lighthouse Laboratory | Wellcome Sanger Institute for the COVID-19 Genomics UK (COG-UK) Consortium | Berkshire and Surrey Pathology Services Lighthouse Laboratory and Alex Alderton, Roberto Amato, Jeffrey Barrett, Sonia Goncalves, Ewan Harrison, David K. Jackson, Ian Johnston, Dominic Kwiatkowski, Cordelia Langford, John Sillitoe on behalf of the Wellcome Sanger Institute COVID-19 Surveillance Team |
| EPI_ISL_3440954 | Lighthouse Lab in Glasgow | Wellcome Sanger Institute for the COVID-19 Genomics UK (COG-UK) Consortium | Harper VanSteenhouse, Yumi Kasai, David Gray, Carol Clugston, Anna Dominiczak and Alex Alderton, Roberto Amato, Jeffrey Barrett, Sonia Goncalves, Ewan Harrison, David K. Jackson, Ian Johnston, Dominic Kwiatkowski, Cordelia Langford, John Sillitoe on behalf of the Wellcome Sanger Institute COVID-19 Surveillance Team |
| EPI_ISL_3440955 | Lighthouse Laboratory Plymouth | Wellcome Sanger Institute for the COVID-19 Genomics UK (COG-UK) Consortium | Lighthouse Laboratory Plymouth and Alex Alderton, Roberto Amato, Jeffrey Barrett, Sonia Goncalves, Ewan Harrison, David K. Jackson, Ian Johnston, Dominic Kwiatkowski, Cordelia Langford, John Sillitoe on behalf of the Wellcome Sanger Institute COVID-19 Surveillance Team |
| EPI_ISL_3440956 | Lighthouse Lab in Milton Keynes | Wellcome Sanger Institute for the COVID-19 Genomics UK (COG-UK) Consortium | The Lighthouse Lab in Milton Keynes and Alex Alderton, Roberto Amato, Jeffrey Barrett, Sonia Goncalves, Ewan Harrison, David K. Jackson, Ian Johnston, Dominic Kwiatkowski, Cordelia Langford, John Sillitoe on behalf of the Wellcome Sanger Institute COVID-19 Surveillance Team |
| EPI_ISL_3440957 | Lighthouse Lab in Glasgow | Wellcome Sanger Institute for the COVID-19 Genomics UK (COG-UK) Consortium | Harper VanSteenhouse, Yumi Kasai, David Gray, Carol Clugston, Anna Dominiczak and Alex Alderton, Roberto Amato, Jeffrey Barrett, Sonia Goncalves, Ewan Harrison, David K. Jackson, Ian Johnston, Dominic Kwiatkowski, Cordelia Langford, John Sillitoe on behalf of the Wellcome Sanger Institute COVID-19 Surveillance Team |
| EPI_ISL_3440958, EPI_ISL_3440959 | Lighthouse Lab in Milton Keynes | Wellcome Sanger Institute for the COVID-19 Genomics UK (COG-UK) Consortium | The Lighthouse Lab in Milton Keynes and Alex Alderton, Roberto Amato, Jeffrey Barrett, Sonia Goncalves, Ewan Harrison, David K. Jackson, Ian Johnston, Dominic Kwiatkowski, Cordelia Langford, John Sillitoe on behalf of the Wellcome Sanger Institute COVID-19 Surveillance Team |
| EPI_ISL_3440960 | Health Services Laboratories | Wellcome Sanger Institute for the COVID-19 Genomics UK (COG-UK) Consortium | Health Services Laboratories and Alex Alderton, Roberto Amato, Jeffrey Barrett, Sonia Goncalves, Ewan Harrison, David K. Jackson, Ian Johnston, Dominic Kwiatkowski, Cordelia Langford, John Sillitoe on behalf of the Wellcome Sanger Institute COVID-19 Surveillance Team |
| EPI_ISL_3440961 | Lighthouse Lab in Milton Keynes | Wellcome Sanger Institute for the COVID-19 Genomics UK (COG-UK) Consortium | The Lighthouse Lab in Milton Keynes and Alex Alderton, Roberto Amato, Jeffrey Barrett, Sonia Goncalves, Ewan Harrison, David K. Jackson, Ian Johnston, Dominic Kwiatkowski, Cordelia Langford, John Sillitoe on behalf of the Wellcome Sanger Institute COVID-19 Surveillance Team |
| EPI_ISL_3440962 | Lighthouse Lab in Glasgow | Wellcome Sanger Institute for the COVID-19 Genomics UK (COG-UK) Consortium | Harper VanSteenhouse, Yumi Kasai, David Gray, Carol Clugston, Anna Dominiczak and Alex Alderton, Roberto Amato, Jeffrey Barrett, Sonia Goncalves, Ewan Harrison, David K. Jackson, Ian Johnston, Dominic Kwiatkowski, Cordelia Langford, John Sillitoe on behalf of the Wellcome Sanger Institute COVID-19 Surveillance Team |
| EPI_ISL_3440963 | Lighthouse Lab in Milton Keynes | Wellcome Sanger Institute for the COVID-19 Genomics UK (COG-UK) | The Lighthouse Lab in Milton Keynes and Alex Alderton, Roberto Amato, Jeffrey Barrett, Sonia Goncalves, Ewan Harrison, David K. Jackson, Ian Johnston, Dominic Kwiatkowski, Cordelia Langford, John Sillitoe on behalf of the Wellcome Sanger |

| | | | |
|---|---|---|---|
| | | Consortium | Institute COVID-19 Surveillance Team |
| EPI_ISL_3440964 | Berkshire and Surrey Pathology Services Lighthouse Laboratory | Wellcome Sanger Institute for the COVID-19 Genomics UK (COG-UK) Consortium | Berkshire and Surrey Pathology Services Lighthouse Laboratory and Alex Alderton, Roberto Amato, Jeffrey Barrett, Sonia Goncalves, Ewan Harrison, David K. Jackson, Ian Johnston, Dominic Kwiatkowski, Cordelia Langford, John Sillitoe on behalf of the Wellcome Sanger Institute COVID-19 Surveillance Team |
| EPI_ISL_3440965 | Lighthouse Lab in Milton Keynes | Wellcome Sanger Institute for the COVID-19 Genomics UK (COG-UK) Consortium | The Lighthouse Lab in Milton Keynes and Alex Alderton, Roberto Amato, Jeffrey Barrett, Sonia Goncalves, Ewan Harrison, David K. Jackson, Ian Johnston, Dominic Kwiatkowski, Cordelia Langford, John Sillitoe on behalf of the Wellcome Sanger Institute COVID-19 Surveillance Team |
| EPI_ISL_3440966, EPI_ISL_3440967 | Lighthouse Laboratory Plymouth | Wellcome Sanger Institute for the COVID-19 Genomics UK (COG-UK) Consortium | Lighthouse Laboratory Plymouth and Alex Alderton, Roberto Amato, Jeffrey Barrett, Sonia Goncalves, Ewan Harrison, David K. Jackson, Ian Johnston, Dominic Kwiatkowski, Cordelia Langford, John Sillitoe on behalf of the Wellcome Sanger Institute COVID-19 Surveillance Team |
| EPI_ISL_3440968 | Lighthouse Lab in Milton Keynes | Wellcome Sanger Institute for the COVID-19 Genomics UK (COG-UK) Consortium | The Lighthouse Lab in Milton Keynes and Alex Alderton, Roberto Amato, Jeffrey Barrett, Sonia Goncalves, Ewan Harrison, David K. Jackson, Ian Johnston, Dominic Kwiatkowski, Cordelia Langford, John Sillitoe on behalf of the Wellcome Sanger Institute COVID-19 Surveillance Team |
| EPI_ISL_3440969, EPI_ISL_3440970, EPI_ISL_3440971 | Lighthouse Lab in Glasgow | Wellcome Sanger Institute for the COVID-19 Genomics UK (COG-UK) Consortium | Harper VanSteenhouse, Yumi Kasai, David Gray, Carol Clugston, Anna Dominiczak and Alex Alderton, Roberto Amato, Jeffrey Barrett, Sonia Goncalves, Ewan Harrison, David K. Jackson, Ian Johnston, Dominic Kwiatkowski, Cordelia Langford, John Sillitoe on behalf of the Wellcome Sanger Institute COVID-19 Surveillance Team |
| EPI_ISL_3440972, EPI_ISL_3440973, EPI_ISL_3440974 | Lighthouse Laboratory Plymouth | Wellcome Sanger Institute for the COVID-19 Genomics UK (COG-UK) Consortium | Lighthouse Laboratory Plymouth and Alex Alderton, Roberto Amato, Jeffrey Barrett, Sonia Goncalves, Ewan Harrison, David K. Jackson, Ian Johnston, Dominic Kwiatkowski, Cordelia Langford, John Sillitoe on behalf of the Wellcome Sanger Institute COVID-19 Surveillance Team |
| EPI_ISL_3440975, EPI_ISL_3440976 | Lighthouse Lab in Milton Keynes | Wellcome Sanger Institute for the COVID-19 Genomics UK (COG-UK) Consortium | The Lighthouse Lab in Milton Keynes and Alex Alderton, Roberto Amato, Jeffrey Barrett, Sonia Goncalves, Ewan Harrison, David K. Jackson, Ian Johnston, Dominic Kwiatkowski, Cordelia Langford, John Sillitoe on behalf of the Wellcome Sanger Institute COVID-19 Surveillance Team |
| EPI_ISL_3440977 | Health Services Laboratories | Wellcome Sanger Institute for the COVID-19 Genomics UK (COG-UK) Consortium | Health Services Laboratories and Alex Alderton, Roberto Amato, Jeffrey Barrett, Sonia Goncalves, Ewan Harrison, David K. Jackson, Ian Johnston, Dominic Kwiatkowski, Cordelia Langford, John Sillitoe on behalf of the Wellcome Sanger Institute COVID-19 Surveillance Team |
| EPI_ISL_3440978 | Lighthouse Lab in Milton Keynes | Wellcome Sanger Institute for the COVID-19 Genomics UK (COG-UK) Consortium | The Lighthouse Lab in Milton Keynes and Alex Alderton, Roberto Amato, Jeffrey Barrett, Sonia Goncalves, Ewan Harrison, David K. Jackson, Ian Johnston, Dominic Kwiatkowski, Cordelia Langford, John Sillitoe on behalf of the Wellcome Sanger Institute COVID-19 Surveillance Team |
| EPI_ISL_3440979 | Lighthouse Laboratory Plymouth | Wellcome Sanger Institute for the COVID-19 Genomics UK (COG-UK) Consortium | Lighthouse Laboratory Plymouth and Alex Alderton, Roberto Amato, Jeffrey Barrett, Sonia Goncalves, Ewan Harrison, David K. Jackson, Ian Johnston, Dominic Kwiatkowski, Cordelia Langford, John Sillitoe on behalf of the Wellcome Sanger Institute COVID-19 Surveillance Team |
| EPI_ISL_3440980 | Lighthouse Lab in Glasgow | Wellcome Sanger Institute for the COVID-19 Genomics UK (COG-UK) Consortium | Harper VanSteenhouse, Yumi Kasai, David Gray, Carol Clugston, Anna Dominiczak and Alex Alderton, Roberto Amato, Jeffrey Barrett, Sonia Goncalves, Ewan Harrison, David K. Jackson, Ian Johnston, Dominic Kwiatkowski, Cordelia Langford, John Sillitoe on behalf of the Wellcome Sanger Institute COVID-19 Surveillance Team |
| EPI_ISL_3440981, EPI_ISL_3440982 | Lighthouse Lab in Milton Keynes | Wellcome Sanger Institute for the COVID-19 Genomics UK (COG-UK) Consortium | The Lighthouse Lab in Milton Keynes and Alex Alderton, Roberto Amato, Jeffrey Barrett, Sonia Goncalves, Ewan Harrison, David K. Jackson, Ian Johnston, Dominic Kwiatkowski, Cordelia Langford, John Sillitoe on behalf of the Wellcome Sanger Institute COVID-19 Surveillance Team |
| EPI_ISL_3440983 | Lighthouse Laboratory Plymouth | Wellcome Sanger Institute for the COVID-19 Genomics UK (COG-UK) Consortium | Lighthouse Laboratory Plymouth and Alex Alderton, Roberto Amato, Jeffrey Barrett, Sonia Goncalves, Ewan Harrison, David K. Jackson, Ian Johnston, Dominic Kwiatkowski, Cordelia Langford, John Sillitoe on behalf of the Wellcome Sanger Institute COVID-19 Surveillance Team |
| EPI_ISL_3440984, EPI_ISL_3440985, EPI_ISL_3440986 | Lighthouse Lab in Milton Keynes | Wellcome Sanger Institute for the COVID-19 Genomics UK (COG-UK) Consortium | The Lighthouse Lab in Milton Keynes and Alex Alderton, Roberto Amato, Jeffrey Barrett, Sonia Goncalves, Ewan Harrison, David K. Jackson, Ian Johnston, Dominic Kwiatkowski, Cordelia Langford, John Sillitoe on behalf of the Wellcome Sanger Institute COVID-19 Surveillance Team |
| EPI_ISL_3440987 | Health Services Laboratories | Wellcome Sanger Institute for the COVID-19 Genomics UK (COG-UK) Consortium | Health Services Laboratories and Alex Alderton, Roberto Amato, Jeffrey Barrett, Sonia Goncalves, Ewan Harrison, David K. Jackson, Ian Johnston, Dominic Kwiatkowski, Cordelia Langford, John Sillitoe on behalf of the Wellcome Sanger Institute COVID-19 Surveillance Team |
| EPI_ISL_3440988, EPI_ISL_3440989 | Lighthouse Laboratory Plymouth | Wellcome Sanger Institute for the COVID-19 Genomics UK (COG-UK) Consortium | Lighthouse Laboratory Plymouth and Alex Alderton, Roberto Amato, Jeffrey Barrett, Sonia Goncalves, Ewan Harrison, David K. Jackson, Ian Johnston, Dominic Kwiatkowski, Cordelia Langford, John Sillitoe on behalf of the Wellcome Sanger Institute COVID-19 Surveillance Team |
| EPI_ISL_3440990 | Health Services Laboratories | Wellcome Sanger Institute for the COVID-19 Genomics UK (COG-UK) Consortium | Health Services Laboratories and Alex Alderton, Roberto Amato, Jeffrey Barrett, Sonia Goncalves, Ewan Harrison, David K. Jackson, Ian Johnston, Dominic Kwiatkowski, Cordelia Langford, John Sillitoe on behalf of the Wellcome Sanger Institute COVID-19 Surveillance Team |
| EPI_ISL_3440991 | Lighthouse Lab in Glasgow | Wellcome Sanger Institute for the COVID-19 Genomics UK (COG-UK) Consortium | Harper VanSteenhouse, Yumi Kasai, David Gray, Carol Clugston, Anna Dominiczak and Alex Alderton, Roberto Amato, Jeffrey Barrett, Sonia Goncalves, Ewan Harrison, David K. Jackson, Ian Johnston, Dominic Kwiatkowski, Cordelia Langford, John Sillitoe on behalf of the Wellcome Sanger Institute COVID-19 Surveillance Team |
| EPI_ISL_3440992 | Lighthouse Lab in Milton Keynes | Wellcome Sanger Institute for the COVID-19 Genomics UK (COG-UK) Consortium | The Lighthouse Lab in Milton Keynes and Alex Alderton, Roberto Amato, Jeffrey Barrett, Sonia Goncalves, Ewan Harrison, David K. Jackson, Ian Johnston, Dominic Kwiatkowski, Cordelia Langford, John Sillitoe on behalf of the Wellcome Sanger Institute COVID-19 Surveillance Team |
| EPI_ISL_3440993 | Lighthouse Lab in Glasgow | Wellcome Sanger Institute for the COVID-19 Genomics UK (COG-UK) Consortium | Harper VanSteenhouse, Yumi Kasai, David Gray, Carol Clugston, Anna Dominiczak and Alex Alderton, Roberto Amato, Jeffrey Barrett, Sonia Goncalves, Ewan Harrison, David K. Jackson, Ian Johnston, Dominic Kwiatkowski, Cordelia Langford, John Sillitoe on behalf of the Wellcome Sanger Institute COVID-19 Surveillance Team |
| EPI_ISL_3440994 | Lighthouse Lab in Milton Keynes | Wellcome Sanger Institute for the COVID-19 Genomics UK (COG-UK) Consortium | The Lighthouse Lab in Milton Keynes and Alex Alderton, Roberto Amato, Jeffrey Barrett, Sonia Goncalves, Ewan Harrison, David K. Jackson, Ian Johnston, Dominic Kwiatkowski, Cordelia Langford, John Sillitoe on behalf of the Wellcome Sanger Institute COVID-19 Surveillance Team |
| EPI_ISL_3440995 | Lighthouse Laboratory Plymouth | Wellcome Sanger Institute for the COVID-19 Genomics UK (COG-UK) Consortium | Lighthouse Laboratory Plymouth and Alex Alderton, Roberto Amato, Jeffrey Barrett, Sonia Goncalves, Ewan Harrison, David K. Jackson, Ian Johnston, Dominic Kwiatkowski, Cordelia Langford, John Sillitoe on behalf of the Wellcome Sanger Institute COVID-19 Surveillance Team |
| EPI_ISL_3440996 | Lighthouse Lab in Glasgow | Wellcome Sanger Institute for the COVID-19 Genomics UK (COG-UK) Consortium | Harper VanSteenhouse, Yumi Kasai, David Gray, Carol Clugston, Anna Dominiczak and Alex Alderton, Roberto Amato, Jeffrey Barrett, Sonia Goncalves, Ewan Harrison, David K. Jackson, Ian Johnston, Dominic Kwiatkowski, Cordelia Langford, John Sillitoe on behalf of the Wellcome Sanger Institute COVID-19 Surveillance Team |
| EPI_ISL_3440997 | Lighthouse Lab in Milton Keynes | Wellcome Sanger Institute for the COVID-19 Genomics UK (COG-UK) Consortium | The Lighthouse Lab in Milton Keynes and Alex Alderton, Roberto Amato, Jeffrey Barrett, Sonia Goncalves, Ewan Harrison, David K. Jackson, Ian Johnston, Dominic Kwiatkowski, Cordelia Langford, John Sillitoe on behalf of the Wellcome Sanger Institute COVID-19 Surveillance Team |
| EPI_ISL_3440998, EPI_ISL_3440999 | Lighthouse Lab in Glasgow | Wellcome Sanger Institute for the COVID-19 Genomics UK (COG-UK) Consortium | Harper VanSteenhouse, Yumi Kasai, David Gray, Carol Clugston, Anna Dominiczak and Alex Alderton, Roberto Amato, Jeffrey Barrett, Sonia Goncalves, Ewan Harrison, David K. Jackson, Ian Johnston, Dominic Kwiatkowski, Cordelia Langford, John Sillitoe on behalf of the Wellcome Sanger Institute COVID-19 Surveillance Team |
| EPI_ISL_3441000 | Lighthouse Laboratory Plymouth | Wellcome Sanger Institute for the COVID-19 Genomics UK (COG-UK) Consortium | Lighthouse Laboratory Plymouth and Alex Alderton, Roberto Amato, Jeffrey Barrett, Sonia Goncalves, Ewan Harrison, David K. Jackson, Ian Johnston, Dominic Kwiatkowski, Cordelia Langford, John Sillitoe on behalf of the Wellcome Sanger Institute COVID-19 Surveillance Team |
| EPI_ISL_3441001, EPI_ISL_3441002, EPI_ISL_3441003 | Lighthouse Lab in Glasgow | Wellcome Sanger Institute for the COVID-19 Genomics UK (COG-UK) Consortium | Harper VanSteenhouse, Yumi Kasai, David Gray, Carol Clugston, Anna Dominiczak and Alex Alderton, Roberto Amato, Jeffrey Barrett, Sonia Goncalves, Ewan Harrison, David K. Jackson, Ian Johnston, Dominic Kwiatkowski, Cordelia Langford, John Sillitoe on behalf of the Wellcome Sanger Institute COVID-19 Surveillance Team |
| EPI_ISL_3441004 | Berkshire and Surrey Pathology Services Lighthouse Laboratory | Wellcome Sanger Institute for the COVID-19 Genomics UK (COG-UK) Consortium | Berkshire and Surrey Pathology Services Lighthouse Laboratory and Alex Alderton, Roberto Amato, Jeffrey Barrett, Sonia Goncalves, Ewan Harrison, David K. Jackson, Ian Johnston, Dominic Kwiatkowski, Cordelia Langford, John Sillitoe on behalf of the Wellcome Sanger Institute COVID-19 Surveillance Team |
| EPI_ISL_3441005 | Lighthouse Lab in Glasgow | Wellcome Sanger Institute for the COVID-19 Genomics UK (COG-UK) Consortium | Harper VanSteenhouse, Yumi Kasai, David Gray, Carol Clugston, Anna Dominiczak and Alex Alderton, Roberto Amato, Jeffrey Barrett, Sonia Goncalves, Ewan Harrison, David K. Jackson, Ian Johnston, Dominic Kwiatkowski, Cordelia Langford, John Sillitoe on behalf of the Wellcome Sanger Institute COVID-19 Surveillance Team |
| EPI_ISL_3441006, EPI_ISL_3441007, EPI_ISL_3441008 | Berkshire and Surrey Pathology Services Lighthouse Laboratory | Wellcome Sanger Institute for the COVID-19 Genomics UK (COG-UK) Consortium | Berkshire and Surrey Pathology Services Lighthouse Laboratory and Alex Alderton, Roberto Amato, Jeffrey Barrett, Sonia Goncalves, Ewan Harrison, David K. Jackson, Ian Johnston, Dominic Kwiatkowski, Cordelia Langford, John Sillitoe on behalf of the Wellcome Sanger Institute COVID-19 Surveillance Team |
| EPI_ISL_3441009 | Lighthouse Laboratory Plymouth | Wellcome Sanger Institute for the COVID-19 Genomics UK (COG-UK) Consortium | Lighthouse Laboratory Plymouth and Alex Alderton, Roberto Amato, Jeffrey Barrett, Sonia Goncalves, Ewan Harrison, David K. Jackson, Ian Johnston, Dominic Kwiatkowski, Cordelia Langford, John Sillitoe on behalf of the Wellcome Sanger Institute COVID-19 Surveillance Team |
| EPI_ISL_3441010 | Lighthouse Lab in Milton Keynes | Wellcome Sanger Institute for the COVID-19 Genomics UK (COG-UK) Consortium | The Lighthouse Lab in Milton Keynes and Alex Alderton, Roberto Amato, Jeffrey Barrett, Sonia Goncalves, Ewan Harrison, David K. Jackson, Ian Johnston, Dominic Kwiatkowski, Cordelia Langford, John Sillitoe on behalf of the Wellcome Sanger Institute COVID-19 Surveillance Team |
| EPI_ISL_3441011 | Lighthouse Laboratory Plymouth | Wellcome Sanger Institute for the COVID-19 Genomics UK (COG-UK) Consortium | Lighthouse Laboratory Plymouth and Alex Alderton, Roberto Amato, Jeffrey Barrett, Sonia Goncalves, Ewan Harrison, David K. Jackson, Ian Johnston, Dominic Kwiatkowski, Cordelia Langford, John Sillitoe on behalf of the Wellcome Sanger Institute COVID-19 Surveillance Team |
| EPI_ISL_3441012 | Health Services Laboratories | Wellcome Sanger Institute for the COVID-19 Genomics UK (COG-UK) Consortium | Health Services Laboratories and Alex Alderton, Roberto Amato, Jeffrey Barrett, Sonia Goncalves, Ewan Harrison, David K. Jackson, Ian Johnston, Dominic Kwiatkowski, Cordelia Langford, John Sillitoe on behalf of the Wellcome Sanger Institute COVID-19 Surveillance Team |
| EPI_ISL_3441013, EPI_ISL_3441014 | Lighthouse Lab in Milton Keynes | Wellcome Sanger Institute for the COVID-19 Genomics UK (COG-UK) Consortium | The Lighthouse Lab in Milton Keynes and Alex Alderton, Roberto Amato, Jeffrey Barrett, Sonia Goncalves, Ewan Harrison, David K. Jackson, Ian Johnston, Dominic Kwiatkowski, Cordelia Langford, John Sillitoe on behalf of the Wellcome Sanger Institute COVID-19 Surveillance Team |
| EPI_ISL_3441015 | Berkshire and Surrey Pathology Services Lighthouse Laboratory | Wellcome Sanger Institute for the COVID-19 Genomics UK (COG-UK) Consortium | Berkshire and Surrey Pathology Services Lighthouse Laboratory and Alex Alderton, Roberto Amato, Jeffrey Barrett, Sonia Goncalves, Ewan Harrison, David K. Jackson, Ian Johnston, Dominic Kwiatkowski, Cordelia Langford, John Sillitoe on behalf of the Wellcome Sanger Institute COVID-19 Surveillance Team |
| EPI_ISL_3441016 | Lighthouse Laboratory Plymouth | Wellcome Sanger Institute for the COVID-19 Genomics UK (COG-UK) Consortium | Lighthouse Laboratory Plymouth and Alex Alderton, Roberto Amato, Jeffrey Barrett, Sonia Goncalves, Ewan Harrison, David K. Jackson, Ian Johnston, Dominic Kwiatkowski, Cordelia Langford, John Sillitoe on behalf of the Wellcome Sanger Institute COVID-19 Surveillance Team |
| EPI_ISL_3441017 | Health Services Laboratories | Wellcome Sanger Institute for the COVID-19 Genomics UK (COG-UK) Consortium | Health Services Laboratories and Alex Alderton, Roberto Amato, Jeffrey Barrett, Sonia Goncalves, Ewan Harrison, David K. Jackson, Ian Johnston, Dominic Kwiatkowski, Cordelia Langford, John Sillitoe on behalf of the Wellcome Sanger Institute COVID-19 Surveillance Team |
| EPI_ISL_3441018 | Lighthouse Lab in Milton Keynes | Wellcome Sanger Institute for the COVID-19 Genomics UK (COG-UK) Consortium | The Lighthouse Lab in Milton Keynes and Alex Alderton, Roberto Amato, Jeffrey Barrett, Sonia Goncalves, Ewan Harrison, David K. Jackson, Ian Johnston, Dominic Kwiatkowski, Cordelia Langford, John Sillitoe on behalf of the Wellcome Sanger Institute COVID-19 Surveillance Team |
| EPI_ISL_3441019 | Lighthouse Lab in Glasgow | Wellcome Sanger Institute for the COVID-19 Genomics UK (COG-UK) Consortium | Harper VanSteenhouse, Yumi Kasai, David Gray, Carol Clugston, Anna Dominiczak and Alex Alderton, Roberto Amato, Jeffrey Barrett, Sonia Goncalves, Ewan Harrison, David K. Jackson, Ian Johnston, Dominic Kwiatkowski, Cordelia Langford, John Sillitoe on behalf of the Wellcome Sanger Institute COVID-19 Surveillance Team |
| EPI_ISL_3441020, EPI_ISL_3441021 | Lighthouse Lab in Milton Keynes | Wellcome Sanger Institute for the COVID-19 Genomics UK (COG-UK) Consortium | The Lighthouse Lab in Milton Keynes and Alex Alderton, Roberto Amato, Jeffrey Barrett, Sonia Goncalves, Ewan Harrison, David K. Jackson, Ian Johnston, Dominic Kwiatkowski, Cordelia Langford, John Sillitoe on behalf of the Wellcome Sanger Institute COVID-19 Surveillance Team |
| EPI_ISL_3441022 | Lighthouse Lab in Glasgow | Wellcome Sanger Institute for the COVID-19 Genomics UK (COG-UK) Consortium | Harper VanSteenhouse, Yumi Kasai, David Gray, Carol Clugston, Anna Dominiczak and Alex Alderton, Roberto Amato, Jeffrey Barrett, Sonia Goncalves, Ewan Harrison, David K. Jackson, Ian Johnston, Dominic Kwiatkowski, Cordelia Langford, John Sillitoe on behalf of the Wellcome Sanger Institute COVID-19 Surveillance Team |
| EPI_ISL_3441023 | Lighthouse Lab in Milton Keynes | Wellcome Sanger Institute for the COVID-19 Genomics UK (COG-UK) | The Lighthouse Lab in Milton Keynes and Alex Alderton, Roberto Amato, Jeffrey Barrett, Sonia Goncalves, Ewan Harrison, David K. Jackson, Ian Johnston, Dominic Kwiatkowski, Cordelia Langford, John Sillitoe on behalf of the Wellcome Sanger |

| | | | |
|---|---|---|---|
| | | Consortium | Institute COVID-19 Surveillance Team |
| EPI_ISL_3441024 | Health Services Laboratories | Wellcome Sanger Institute for the COVID-19 Genomics UK (COG-UK) Consortium | Health Services Laboratories and Alex Alderton, Roberto Amato, Jeffrey Barrett, Sonia Goncalves, Ewan Harrison, David K. Jackson, Ian Johnston, Dominic Kwiatkowski, Cordelia Langford, John Sillitoe on behalf of the Wellcome Sanger Institute COVID-19 Surveillance Team |
| EPI_ISL_3441025 | Lighthouse Laboratory Plymouth | Wellcome Sanger Institute for the COVID-19 Genomics UK (COG-UK) Consortium | Lighthouse Laboratory Plymouth and Alex Alderton, Roberto Amato, Jeffrey Barrett, Sonia Goncalves, Ewan Harrison, David K. Jackson, Ian Johnston, Dominic Kwiatkowski, Cordelia Langford, John Sillitoe on behalf of the Wellcome Sanger Institute COVID-19 Surveillance Team |
| EPI_ISL_3441026 | Lighthouse Lab in Milton Keynes | Wellcome Sanger Institute for the COVID-19 Genomics UK (COG-UK) Consortium | The Lighthouse Lab in Milton Keynes and Alex Alderton, Roberto Amato, Jeffrey Barrett, Sonia Goncalves, Ewan Harrison, David K. Jackson, Ian Johnston, Dominic Kwiatkowski, Cordelia Langford, John Sillitoe on behalf of the Wellcome Sanger Institute COVID-19 Surveillance Team |
| EPI_ISL_3441027 | Lighthouse Laboratory Plymouth | Wellcome Sanger Institute for the COVID-19 Genomics UK (COG-UK) Consortium | Lighthouse Laboratory Plymouth and Alex Alderton, Roberto Amato, Jeffrey Barrett, Sonia Goncalves, Ewan Harrison, David K. Jackson, Ian Johnston, Dominic Kwiatkowski, Cordelia Langford, John Sillitoe on behalf of the Wellcome Sanger Institute COVID-19 Surveillance Team |
| EPI_ISL_3441028 | Health Services Laboratories | Wellcome Sanger Institute for the COVID-19 Genomics UK (COG-UK) Consortium | Health Services Laboratories and Alex Alderton, Roberto Amato, Jeffrey Barrett, Sonia Goncalves, Ewan Harrison, David K. Jackson, Ian Johnston, Dominic Kwiatkowski, Cordelia Langford, John Sillitoe on behalf of the Wellcome Sanger Institute COVID-19 Surveillance Team |
| EPI_ISL_3441029, EPI_ISL_3441030 | Lighthouse Lab in Milton Keynes | Wellcome Sanger Institute for the COVID-19 Genomics UK (COG-UK) Consortium | The Lighthouse Lab in Milton Keynes and Alex Alderton, Roberto Amato, Jeffrey Barrett, Sonia Goncalves, Ewan Harrison, David K. Jackson, Ian Johnston, Dominic Kwiatkowski, Cordelia Langford, John Sillitoe on behalf of the Wellcome Sanger Institute COVID-19 Surveillance Team |
| EPI_ISL_3441031 | Lighthouse Laboratory Plymouth | Wellcome Sanger Institute for the COVID-19 Genomics UK (COG-UK) Consortium | Lighthouse Laboratory Plymouth and Alex Alderton, Roberto Amato, Jeffrey Barrett, Sonia Goncalves, Ewan Harrison, David K. Jackson, Ian Johnston, Dominic Kwiatkowski, Cordelia Langford, John Sillitoe on behalf of the Wellcome Sanger Institute COVID-19 Surveillance Team |
| EPI_ISL_3441032 | Lighthouse Lab in Milton Keynes | Wellcome Sanger Institute for the COVID-19 Genomics UK (COG-UK) Consortium | The Lighthouse Lab in Milton Keynes and Alex Alderton, Roberto Amato, Jeffrey Barrett, Sonia Goncalves, Ewan Harrison, David K. Jackson, Ian Johnston, Dominic Kwiatkowski, Cordelia Langford, John Sillitoe on behalf of the Wellcome Sanger Institute COVID-19 Surveillance Team |
| EPI_ISL_3441033 | Lighthouse Laboratory Plymouth | Wellcome Sanger Institute for the COVID-19 Genomics UK (COG-UK) Consortium | Lighthouse Laboratory Plymouth and Alex Alderton, Roberto Amato, Jeffrey Barrett, Sonia Goncalves, Ewan Harrison, David K. Jackson, Ian Johnston, Dominic Kwiatkowski, Cordelia Langford, John Sillitoe on behalf of the Wellcome Sanger Institute COVID-19 Surveillance Team |
| EPI_ISL_3441034 | Lighthouse Lab in Milton Keynes | Wellcome Sanger Institute for the COVID-19 Genomics UK (COG-UK) Consortium | The Lighthouse Lab in Milton Keynes and Alex Alderton, Roberto Amato, Jeffrey Barrett, Sonia Goncalves, Ewan Harrison, David K. Jackson, Ian Johnston, Dominic Kwiatkowski, Cordelia Langford, John Sillitoe on behalf of the Wellcome Sanger Institute COVID-19 Surveillance Team |
| EPI_ISL_3441035 | Health Services Laboratories | Wellcome Sanger Institute for the COVID-19 Genomics UK (COG-UK) Consortium | Health Services Laboratories and Alex Alderton, Roberto Amato, Jeffrey Barrett, Sonia Goncalves, Ewan Harrison, David K. Jackson, Ian Johnston, Dominic Kwiatkowski, Cordelia Langford, John Sillitoe on behalf of the Wellcome Sanger Institute COVID-19 Surveillance Team |
| EPI_ISL_3441036 | Lighthouse Laboratory Plymouth | Wellcome Sanger Institute for the COVID-19 Genomics UK (COG-UK) Consortium | Lighthouse Laboratory Plymouth and Alex Alderton, Roberto Amato, Jeffrey Barrett, Sonia Goncalves, Ewan Harrison, David K. Jackson, Ian Johnston, Dominic Kwiatkowski, Cordelia Langford, John Sillitoe on behalf of the Wellcome Sanger Institute COVID-19 Surveillance Team |
| EPI_ISL_3441037 | Lighthouse Lab in Glasgow | Wellcome Sanger Institute for the COVID-19 Genomics UK (COG-UK) Consortium | Harper VanSteenhouse, Yumi Kasai, David Gray, Carol Clugston, Anna Dominiczak and Alex Alderton, Roberto Amato, Jeffrey Barrett, Sonia Goncalves, Ewan Harrison, David K. Jackson, Ian Johnston, Dominic Kwiatkowski, Cordelia Langford, John Sillitoe on behalf of the Wellcome Sanger Institute COVID-19 Surveillance Team |
| EPI_ISL_3441038 | Lighthouse Laboratory Plymouth | Wellcome Sanger Institute for the COVID-19 Genomics UK (COG-UK) Consortium | Lighthouse Laboratory Plymouth and Alex Alderton, Roberto Amato, Jeffrey Barrett, Sonia Goncalves, Ewan Harrison, David K. Jackson, Ian Johnston, Dominic Kwiatkowski, Cordelia Langford, John Sillitoe on behalf of the Wellcome Sanger Institute COVID-19 Surveillance Team |
| EPI_ISL_3441039 | Lighthouse Lab in Glasgow | Wellcome Sanger Institute for the COVID-19 Genomics UK (COG-UK) Consortium | Harper VanSteenhouse, Yumi Kasai, David Gray, Carol Clugston, Anna Dominiczak and Alex Alderton, Roberto Amato, Jeffrey Barrett, Sonia Goncalves, Ewan Harrison, David K. Jackson, Ian Johnston, Dominic Kwiatkowski, Cordelia Langford, John Sillitoe on behalf of the Wellcome Sanger Institute COVID-19 Surveillance Team |
| EPI_ISL_3441040 | Lighthouse Lab in Milton Keynes | Wellcome Sanger Institute for the COVID-19 Genomics UK (COG-UK) Consortium | The Lighthouse Lab in Milton Keynes and Alex Alderton, Roberto Amato, Jeffrey Barrett, Sonia Goncalves, Ewan Harrison, David K. Jackson, Ian Johnston, Dominic Kwiatkowski, Cordelia Langford, John Sillitoe on behalf of the Wellcome Sanger Institute COVID-19 Surveillance Team |
| EPI_ISL_3441041, EPI_ISL_3441042 | Lighthouse Lab in Glasgow | Wellcome Sanger Institute for the COVID-19 Genomics UK (COG-UK) Consortium | Harper VanSteenhouse, Yumi Kasai, David Gray, Carol Clugston, Anna Dominiczak and Alex Alderton, Roberto Amato, Jeffrey Barrett, Sonia Goncalves, Ewan Harrison, David K. Jackson, Ian Johnston, Dominic Kwiatkowski, Cordelia Langford, John Sillitoe on behalf of the Wellcome Sanger Institute COVID-19 Surveillance Team |
| EPI_ISL_3441043 | Berkshire and Surrey Pathology Services Lighthouse Laboratory | Wellcome Sanger Institute for the COVID-19 Genomics UK (COG-UK) Consortium | Berkshire and Surrey Pathology Services Lighthouse Laboratory and Alex Alderton, Roberto Amato, Jeffrey Barrett, Sonia Goncalves, Ewan Harrison, David K. Jackson, Ian Johnston, Dominic Kwiatkowski, Cordelia Langford, John Sillitoe on behalf of the Wellcome Sanger Institute COVID-19 Surveillance Team |
| EPI_ISL_3441044 | Lighthouse Lab in Milton Keynes | Wellcome Sanger Institute for the COVID-19 Genomics UK (COG-UK) Consortium | The Lighthouse Lab in Milton Keynes and Alex Alderton, Roberto Amato, Jeffrey Barrett, Sonia Goncalves, Ewan Harrison, David K. Jackson, Ian Johnston, Dominic Kwiatkowski, Cordelia Langford, John Sillitoe on behalf of the Wellcome Sanger Institute COVID-19 Surveillance Team |
| EPI_ISL_3441045 | Health Services Laboratories | Wellcome Sanger Institute for the COVID-19 Genomics UK (COG-UK) Consortium | Health Services Laboratories and Alex Alderton, Roberto Amato, Jeffrey Barrett, Sonia Goncalves, Ewan Harrison, David K. Jackson, Ian Johnston, Dominic Kwiatkowski, Cordelia Langford, John Sillitoe on behalf of the Wellcome Sanger Institute COVID-19 Surveillance Team |
| EPI_ISL_3441046, EPI_ISL_3441047 | Lighthouse Laboratory Plymouth | Wellcome Sanger Institute for the COVID-19 Genomics UK (COG-UK) Consortium | Lighthouse Laboratory Plymouth and Alex Alderton, Roberto Amato, Jeffrey Barrett, Sonia Goncalves, Ewan Harrison, David K. Jackson, Ian Johnston, Dominic Kwiatkowski, Cordelia Langford, John Sillitoe on behalf of the Wellcome Sanger Institute COVID-19 Surveillance Team |
| EPI_ISL_3441048 | Lighthouse Lab in Glasgow | Wellcome Sanger Institute for the COVID-19 Genomics UK (COG-UK) Consortium | Harper VanSteenhouse, Yumi Kasai, David Gray, Carol Clugston, Anna Dominiczak and Alex Alderton, Roberto Amato, Jeffrey Barrett, Sonia Goncalves, Ewan Harrison, David K. Jackson, Ian Johnston, Dominic Kwiatkowski, Cordelia Langford, John Sillitoe on behalf of the Wellcome Sanger Institute COVID-19 Surveillance Team |
| EPI_ISL_3441049, EPI_ISL_3441050 | Lighthouse Laboratory Plymouth | Wellcome Sanger Institute for the COVID-19 Genomics UK (COG-UK) Consortium | Lighthouse Laboratory Plymouth and Alex Alderton, Roberto Amato, Jeffrey Barrett, Sonia Goncalves, Ewan Harrison, David K. Jackson, Ian Johnston, Dominic Kwiatkowski, Cordelia Langford, John Sillitoe on behalf of the Wellcome Sanger Institute COVID-19 Surveillance Team |
| EPI_ISL_3441051 | Lighthouse Lab in Milton Keynes | Wellcome Sanger Institute for the COVID-19 Genomics UK (COG-UK) Consortium | The Lighthouse Lab in Milton Keynes and Alex Alderton, Roberto Amato, Jeffrey Barrett, Sonia Goncalves, Ewan Harrison, David K. Jackson, Ian Johnston, Dominic Kwiatkowski, Cordelia Langford, John Sillitoe on behalf of the Wellcome Sanger Institute COVID-19 Surveillance Team |
| EPI_ISL_3441052 | Lighthouse Laboratory Plymouth | Wellcome Sanger Institute for the COVID-19 Genomics UK (COG-UK) Consortium | Lighthouse Laboratory Plymouth and Alex Alderton, Roberto Amato, Jeffrey Barrett, Sonia Goncalves, Ewan Harrison, David K. Jackson, Ian Johnston, Dominic Kwiatkowski, Cordelia Langford, John Sillitoe on behalf of the Wellcome Sanger Institute COVID-19 Surveillance Team |
| EPI_ISL_3441053, EPI_ISL_3441054 | Lighthouse Lab in Glasgow | Wellcome Sanger Institute for the COVID-19 Genomics UK (COG-UK) Consortium | Harper VanSteenhouse, Yumi Kasai, David Gray, Carol Clugston, Anna Dominiczak and Alex Alderton, Roberto Amato, Jeffrey Barrett, Sonia Goncalves, Ewan Harrison, David K. Jackson, Ian Johnston, Dominic Kwiatkowski, Cordelia Langford, John Sillitoe on behalf of the Wellcome Sanger Institute COVID-19 Surveillance Team |
| EPI_ISL_3441055 | Lighthouse Laboratory Plymouth | Wellcome Sanger Institute for the COVID-19 Genomics UK (COG-UK) Consortium | Lighthouse Laboratory Plymouth and Alex Alderton, Roberto Amato, Jeffrey Barrett, Sonia Goncalves, Ewan Harrison, David K. Jackson, Ian Johnston, Dominic Kwiatkowski, Cordelia Langford, John Sillitoe on behalf of the Wellcome Sanger Institute COVID-19 Surveillance Team |
| EPI_ISL_3441056 | Lighthouse Lab in Milton Keynes | Wellcome Sanger Institute for the COVID-19 Genomics UK (COG-UK) Consortium | The Lighthouse Lab in Milton Keynes and Alex Alderton, Roberto Amato, Jeffrey Barrett, Sonia Goncalves, Ewan Harrison, David K. Jackson, Ian Johnston, Dominic Kwiatkowski, Cordelia Langford, John Sillitoe on behalf of the Wellcome Sanger Institute COVID-19 Surveillance Team |
| EPI_ISL_3441057 | Lighthouse Lab in Glasgow | Wellcome Sanger Institute for the COVID-19 Genomics UK (COG-UK) Consortium | Harper VanSteenhouse, Yumi Kasai, David Gray, Carol Clugston, Anna Dominiczak and Alex Alderton, Roberto Amato, Jeffrey Barrett, Sonia Goncalves, Ewan Harrison, David K. Jackson, Ian Johnston, Dominic Kwiatkowski, Cordelia Langford, John Sillitoe on behalf of the Wellcome Sanger Institute COVID-19 Surveillance Team |
| EPI_ISL_3441058 | Lighthouse Laboratory Plymouth | Wellcome Sanger Institute for the COVID-19 Genomics UK (COG-UK) Consortium | Lighthouse Laboratory Plymouth and Alex Alderton, Roberto Amato, Jeffrey Barrett, Sonia Goncalves, Ewan Harrison, David K. Jackson, Ian Johnston, Dominic Kwiatkowski, Cordelia Langford, John Sillitoe on behalf of the Wellcome Sanger Institute COVID-19 Surveillance Team |
| EPI_ISL_3441059 | Lighthouse Lab in Glasgow | Wellcome Sanger Institute for the COVID-19 Genomics UK (COG-UK) Consortium | Harper VanSteenhouse, Yumi Kasai, David Gray, Carol Clugston, Anna Dominiczak and Alex Alderton, Roberto Amato, Jeffrey Barrett, Sonia Goncalves, Ewan Harrison, David K. Jackson, Ian Johnston, Dominic Kwiatkowski, Cordelia Langford, John Sillitoe on behalf of the Wellcome Sanger Institute COVID-19 Surveillance Team |
| EPI_ISL_3441060 | Lighthouse Lab in Milton Keynes | Wellcome Sanger Institute for the COVID-19 Genomics UK (COG-UK) Consortium | The Lighthouse Lab in Milton Keynes and Alex Alderton, Roberto Amato, Jeffrey Barrett, Sonia Goncalves, Ewan Harrison, David K. Jackson, Ian Johnston, Dominic Kwiatkowski, Cordelia Langford, John Sillitoe on behalf of the Wellcome Sanger Institute COVID-19 Surveillance Team |
| EPI_ISL_3441061, EPI_ISL_3441062, EPI_ISL_3441063 | Lighthouse Laboratory Plymouth | Wellcome Sanger Institute for the COVID-19 Genomics UK (COG-UK) Consortium | Lighthouse Laboratory Plymouth and Alex Alderton, Roberto Amato, Jeffrey Barrett, Sonia Goncalves, Ewan Harrison, David K. Jackson, Ian Johnston, Dominic Kwiatkowski, Cordelia Langford, John Sillitoe on behalf of the Wellcome Sanger Institute COVID-19 Surveillance Team |
| EPI_ISL_3441064 | Lighthouse Lab in Milton Keynes | Wellcome Sanger Institute for the COVID-19 Genomics UK (COG-UK) Consortium | The Lighthouse Lab in Milton Keynes and Alex Alderton, Roberto Amato, Jeffrey Barrett, Sonia Goncalves, Ewan Harrison, David K. Jackson, Ian Johnston, Dominic Kwiatkowski, Cordelia Langford, John Sillitoe on behalf of the Wellcome Sanger Institute COVID-19 Surveillance Team |
| EPI_ISL_3441065 | Berkshire and Surrey Pathology Services Lighthouse Laboratory | Wellcome Sanger Institute for the COVID-19 Genomics UK (COG-UK) Consortium | Berkshire and Surrey Pathology Services Lighthouse Laboratory and Alex Alderton, Roberto Amato, Jeffrey Barrett, Sonia Goncalves, Ewan Harrison, David K. Jackson, Ian Johnston, Dominic Kwiatkowski, Cordelia Langford, John Sillitoe on behalf of the Wellcome Sanger Institute COVID-19 Surveillance Team |
| EPI_ISL_3441066 | Lighthouse Lab in Milton Keynes | Wellcome Sanger Institute for the COVID-19 Genomics UK (COG-UK) Consortium | The Lighthouse Lab in Milton Keynes and Alex Alderton, Roberto Amato, Jeffrey Barrett, Sonia Goncalves, Ewan Harrison, David K. Jackson, Ian Johnston, Dominic Kwiatkowski, Cordelia Langford, John Sillitoe on behalf of the Wellcome Sanger Institute COVID-19 Surveillance Team |
| EPI_ISL_3441067 | Health Services Laboratories | Wellcome Sanger Institute for the COVID-19 Genomics UK (COG-UK) Consortium | Health Services Laboratories and Alex Alderton, Roberto Amato, Jeffrey Barrett, Sonia Goncalves, Ewan Harrison, David K. Jackson, Ian Johnston, Dominic Kwiatkowski, Cordelia Langford, John Sillitoe on behalf of the Wellcome Sanger Institute COVID-19 Surveillance Team |
| EPI_ISL_3441068, EPI_ISL_3441069, EPI_ISL_3441070 | Lighthouse Lab in Glasgow | Wellcome Sanger Institute for the COVID-19 Genomics UK (COG-UK) Consortium | Harper VanSteenhouse, Yumi Kasai, David Gray, Carol Clugston, Anna Dominiczak and Alex Alderton, Roberto Amato, Jeffrey Barrett, Sonia Goncalves, Ewan Harrison, David K. Jackson, Ian Johnston, Dominic Kwiatkowski, Cordelia Langford, John Sillitoe on behalf of the Wellcome Sanger Institute COVID-19 Surveillance Team |
| EPI_ISL_3441071 | Berkshire and Surrey Pathology Services Lighthouse Laboratory | Wellcome Sanger Institute for the COVID-19 Genomics UK (COG-UK) Consortium | Berkshire and Surrey Pathology Services Lighthouse Laboratory and Alex Alderton, Roberto Amato, Jeffrey Barrett, Sonia Goncalves, Ewan Harrison, David K. Jackson, Ian Johnston, Dominic Kwiatkowski, Cordelia Langford, John Sillitoe on behalf of the Wellcome Sanger Institute COVID-19 Surveillance Team |
| EPI_ISL_3441072 | Lighthouse Lab in Milton Keynes | Wellcome Sanger Institute for the COVID-19 Genomics UK (COG-UK) Consortium | The Lighthouse Lab in Milton Keynes and Alex Alderton, Roberto Amato, Jeffrey Barrett, Sonia Goncalves, Ewan Harrison, David K. Jackson, Ian Johnston, Dominic Kwiatkowski, Cordelia Langford, John Sillitoe on behalf of the Wellcome Sanger Institute COVID-19 Surveillance Team |
| EPI_ISL_3441073, EPI_ISL_3441074, EPI_ISL_3441075 | Lighthouse Laboratory Plymouth | Wellcome Sanger Institute for the COVID-19 Genomics UK (COG-UK) Consortium | Lighthouse Laboratory Plymouth and Alex Alderton, Roberto Amato, Jeffrey Barrett, Sonia Goncalves, Ewan Harrison, David K. Jackson, Ian Johnston, Dominic Kwiatkowski, Cordelia Langford, John Sillitoe on behalf of the Wellcome Sanger Institute COVID-19 Surveillance Team |
| EPI_ISL_3441076 | Lighthouse Lab in Glasgow | Wellcome Sanger Institute for the COVID-19 Genomics UK (COG-UK) Consortium | Harper VanSteenhouse, Yumi Kasai, David Gray, Carol Clugston, Anna Dominiczak and Alex Alderton, Roberto Amato, Jeffrey Barrett, Sonia Goncalves, Ewan Harrison, David K. Jackson, Ian Johnston, Dominic Kwiatkowski, Cordelia Langford, John Sillitoe on behalf of the Wellcome Sanger Institute COVID-19 Surveillance Team |
| EPI_ISL_3441077 | Berkshire and Surrey Pathology Services Lighthouse Laboratory | Wellcome Sanger Institute for the COVID-19 Genomics UK (COG-UK) Consortium | Berkshire and Surrey Pathology Services Lighthouse Laboratory and Alex Alderton, Roberto Amato, Jeffrey Barrett, Sonia Goncalves, Ewan Harrison, David K. Jackson, Ian Johnston, Dominic Kwiatkowski, Cordelia Langford, John Sillitoe on behalf of the Wellcome Sanger Institute COVID-19 Surveillance Team |
| EPI_ISL_3441078 | Lighthouse Lab in Glasgow | Wellcome Sanger Institute for the COVID-19 Genomics UK (COG-UK) | Harper VanSteenhouse, Yumi Kasai, David Gray, Carol Clugston, Anna Dominiczak and Alex Alderton, Roberto Amato, Jeffrey Barrett, Sonia Goncalves, Ewan Harrison, David K. Jackson, Ian Johnston, Dominic Kwiatkowski, Cordelia Langford, |

| | | | |
|---|---|---|---|
| | | Consortium | John Sillitoe on behalf of the Wellcome Sanger Institute COVID-19 Surveillance Team |
| EPI_ISL_3441079, EPI_ISL_3441080 | Lighthouse Laboratory Plymouth | Wellcome Sanger Institute for the COVID-19 Genomics UK (COG-UK) Consortium | Lighthouse Laboratory Plymouth and Alex Alderton, Roberto Amato, Jeffrey Barrett, Sonia Goncalves, Ewan Harrison, David K. Jackson, Ian Johnston, Dominic Kwiatkowski, Cordelia Langford, John Sillitoe on behalf of the Wellcome Sanger Institute COVID-19 Surveillance Team |
| EPI_ISL_3441081, EPI_ISL_3441082, EPI_ISL_3441083 | Lighthouse Lab in Glasgow | Wellcome Sanger Institute for the COVID-19 Genomics UK (COG-UK) Consortium | Harper VanSteenhouse, Yumi Kasai, David Gray, Carol Clugston, Anna Dominiczak and Alex Alderton, Roberto Amato, Jeffrey Barrett, Sonia Goncalves, Ewan Harrison, David K. Jackson, Ian Johnston, Dominic Kwiatkowski, Cordelia Langford, John Sillitoe on behalf of the Wellcome Sanger Institute COVID-19 Surveillance Team |
| EPI_ISL_3441084 | Health Services Laboratories | Wellcome Sanger Institute for the COVID-19 Genomics UK (COG-UK) Consortium | Health Services Laboratories and Alex Alderton, Roberto Amato, Jeffrey Barrett, Sonia Goncalves, Ewan Harrison, David K. Jackson, Ian Johnston, Dominic Kwiatkowski, Cordelia Langford, John Sillitoe on behalf of the Wellcome Sanger Institute COVID-19 Surveillance Team |
| EPI_ISL_3441085, EPI_ISL_3441086 | Lighthouse Lab in Milton Keynes | Wellcome Sanger Institute for the COVID-19 Genomics UK (COG-UK) Consortium | The Lighthouse Lab in Milton Keynes and Alex Alderton, Roberto Amato, Jeffrey Barrett, Sonia Goncalves, Ewan Harrison, David K. Jackson, Ian Johnston, Dominic Kwiatkowski, Cordelia Langford, John Sillitoe on behalf of the Wellcome Sanger Institute COVID-19 Surveillance Team |
| EPI_ISL_3441087, EPI_ISL_3441088 | Health Services Laboratories | Wellcome Sanger Institute for the COVID-19 Genomics UK (COG-UK) Consortium | Health Services Laboratories and Alex Alderton, Roberto Amato, Jeffrey Barrett, Sonia Goncalves, Ewan Harrison, David K. Jackson, Ian Johnston, Dominic Kwiatkowski, Cordelia Langford, John Sillitoe on behalf of the Wellcome Sanger Institute COVID-19 Surveillance Team |
| EPI_ISL_3441089 | Lighthouse Lab in Milton Keynes | Wellcome Sanger Institute for the COVID-19 Genomics UK (COG-UK) Consortium | The Lighthouse Lab in Milton Keynes and Alex Alderton, Roberto Amato, Jeffrey Barrett, Sonia Goncalves, Ewan Harrison, David K. Jackson, Ian Johnston, Dominic Kwiatkowski, Cordelia Langford, John Sillitoe on behalf of the Wellcome Sanger Institute COVID-19 Surveillance Team |
| EPI_ISL_3441090 | Lighthouse Laboratory Plymouth | Wellcome Sanger Institute for the COVID-19 Genomics UK (COG-UK) Consortium | Lighthouse Laboratory Plymouth and Alex Alderton, Roberto Amato, Jeffrey Barrett, Sonia Goncalves, Ewan Harrison, David K. Jackson, Ian Johnston, Dominic Kwiatkowski, Cordelia Langford, John Sillitoe on behalf of the Wellcome Sanger Institute COVID-19 Surveillance Team |
| EPI_ISL_3441091, EPI_ISL_3441092 | Lighthouse Lab in Glasgow | Wellcome Sanger Institute for the COVID-19 Genomics UK (COG-UK) Consortium | Harper VanSteenhouse, Yumi Kasai, David Gray, Carol Clugston, Anna Dominiczak and Alex Alderton, Roberto Amato, Jeffrey Barrett, Sonia Goncalves, Ewan Harrison, David K. Jackson, Ian Johnston, Dominic Kwiatkowski, Cordelia Langford, John Sillitoe on behalf of the Wellcome Sanger Institute COVID-19 Surveillance Team |
| EPI_ISL_3441093 | Lighthouse Lab in Milton Keynes | Wellcome Sanger Institute for the COVID-19 Genomics UK (COG-UK) Consortium | The Lighthouse Lab in Milton Keynes and Alex Alderton, Roberto Amato, Jeffrey Barrett, Sonia Goncalves, Ewan Harrison, David K. Jackson, Ian Johnston, Dominic Kwiatkowski, Cordelia Langford, John Sillitoe on behalf of the Wellcome Sanger Institute COVID-19 Surveillance Team |
| EPI_ISL_3441094, EPI_ISL_3441095 | Lighthouse Lab in Glasgow | Wellcome Sanger Institute for the COVID-19 Genomics UK (COG-UK) Consortium | Harper VanSteenhouse, Yumi Kasai, David Gray, Carol Clugston, Anna Dominiczak and Alex Alderton, Roberto Amato, Jeffrey Barrett, Sonia Goncalves, Ewan Harrison, David K. Jackson, Ian Johnston, Dominic Kwiatkowski, Cordelia Langford, John Sillitoe on behalf of the Wellcome Sanger Institute COVID-19 Surveillance Team |
| EPI_ISL_3441096 | Lighthouse Laboratory Plymouth | Wellcome Sanger Institute for the COVID-19 Genomics UK (COG-UK) Consortium | Lighthouse Laboratory Plymouth and Alex Alderton, Roberto Amato, Jeffrey Barrett, Sonia Goncalves, Ewan Harrison, David K. Jackson, Ian Johnston, Dominic Kwiatkowski, Cordelia Langford, John Sillitoe on behalf of the Wellcome Sanger Institute COVID-19 Surveillance Team |
| EPI_ISL_3441097 | Berkshire and Surrey Pathology Services Lighthouse Laboratory | Wellcome Sanger Institute for the COVID-19 Genomics UK (COG-UK) Consortium | Berkshire and Surrey Pathology Services Lighthouse Laboratory and Alex Alderton, Roberto Amato, Jeffrey Barrett, Sonia Goncalves, Ewan Harrison, David K. Jackson, Ian Johnston, Dominic Kwiatkowski, Cordelia Langford, John Sillitoe on behalf of the Wellcome Sanger Institute COVID-19 Surveillance Team |
| EPI_ISL_3441098, EPI_ISL_3441099 | Lighthouse Lab in Milton Keynes | Wellcome Sanger Institute for the COVID-19 Genomics UK (COG-UK) Consortium | The Lighthouse Lab in Milton Keynes and Alex Alderton, Roberto Amato, Jeffrey Barrett, Sonia Goncalves, Ewan Harrison, David K. Jackson, Ian Johnston, Dominic Kwiatkowski, Cordelia Langford, John Sillitoe on behalf of the Wellcome Sanger Institute COVID-19 Surveillance Team |
| EPI_ISL_3441100 | Health Services Laboratories | Wellcome Sanger Institute for the COVID-19 Genomics UK (COG-UK) Consortium | Health Services Laboratories and Alex Alderton, Roberto Amato, Jeffrey Barrett, Sonia Goncalves, Ewan Harrison, David K. Jackson, Ian Johnston, Dominic Kwiatkowski, Cordelia Langford, John Sillitoe on behalf of the Wellcome Sanger Institute COVID-19 Surveillance Team |
| EPI_ISL_3441101 | Berkshire and Surrey Pathology Services Lighthouse Laboratory | Wellcome Sanger Institute for the COVID-19 Genomics UK (COG-UK) Consortium | Berkshire and Surrey Pathology Services Lighthouse Laboratory and Alex Alderton, Roberto Amato, Jeffrey Barrett, Sonia Goncalves, Ewan Harrison, David K. Jackson, Ian Johnston, Dominic Kwiatkowski, Cordelia Langford, John Sillitoe on behalf of the Wellcome Sanger Institute COVID-19 Surveillance Team |
| EPI_ISL_3441102 | Lighthouse Lab in Glasgow | Wellcome Sanger Institute for the COVID-19 Genomics UK (COG-UK) Consortium | Harper VanSteenhouse, Yumi Kasai, David Gray, Carol Clugston, Anna Dominiczak and Alex Alderton, Roberto Amato, Jeffrey Barrett, Sonia Goncalves, Ewan Harrison, David K. Jackson, Ian Johnston, Dominic Kwiatkowski, Cordelia Langford, John Sillitoe on behalf of the Wellcome Sanger Institute COVID-19 Surveillance Team |
| EPI_ISL_3441103 | Lighthouse Lab in Milton Keynes | Wellcome Sanger Institute for the COVID-19 Genomics UK (COG-UK) Consortium | The Lighthouse Lab in Milton Keynes and Alex Alderton, Roberto Amato, Jeffrey Barrett, Sonia Goncalves, Ewan Harrison, David K. Jackson, Ian Johnston, Dominic Kwiatkowski, Cordelia Langford, John Sillitoe on behalf of the Wellcome Sanger Institute COVID-19 Surveillance Team |
| EPI_ISL_3441104 | Lighthouse Laboratory Plymouth | Wellcome Sanger Institute for the COVID-19 Genomics UK (COG-UK) Consortium | Lighthouse Laboratory Plymouth and Alex Alderton, Roberto Amato, Jeffrey Barrett, Sonia Goncalves, Ewan Harrison, David K. Jackson, Ian Johnston, Dominic Kwiatkowski, Cordelia Langford, John Sillitoe on behalf of the Wellcome Sanger Institute COVID-19 Surveillance Team |
| EPI_ISL_3441105, EPI_ISL_3441106 | Lighthouse Lab in Milton Keynes | Wellcome Sanger Institute for the COVID-19 Genomics UK (COG-UK) Consortium | The Lighthouse Lab in Milton Keynes and Alex Alderton, Roberto Amato, Jeffrey Barrett, Sonia Goncalves, Ewan Harrison, David K. Jackson, Ian Johnston, Dominic Kwiatkowski, Cordelia Langford, John Sillitoe on behalf of the Wellcome Sanger Institute COVID-19 Surveillance Team |
| EPI_ISL_3441107 | Lighthouse Laboratory Plymouth | Wellcome Sanger Institute for the COVID-19 Genomics UK (COG-UK) Consortium | Lighthouse Laboratory Plymouth and Alex Alderton, Roberto Amato, Jeffrey Barrett, Sonia Goncalves, Ewan Harrison, David K. Jackson, Ian Johnston, Dominic Kwiatkowski, Cordelia Langford, John Sillitoe on behalf of the Wellcome Sanger Institute COVID-19 Surveillance Team |
| EPI_ISL_3441108, EPI_ISL_3441109 | Health Services Laboratories | Wellcome Sanger Institute for the COVID-19 Genomics UK (COG-UK) Consortium | Health Services Laboratories and Alex Alderton, Roberto Amato, Jeffrey Barrett, Sonia Goncalves, Ewan Harrison, David K. Jackson, Ian Johnston, Dominic Kwiatkowski, Cordelia Langford, John Sillitoe on behalf of the Wellcome Sanger Institute COVID-19 Surveillance Team |
| EPI_ISL_3441110 | Lighthouse Laboratory Plymouth | Wellcome Sanger Institute for the COVID-19 Genomics UK (COG-UK) Consortium | Lighthouse Laboratory Plymouth and Alex Alderton, Roberto Amato, Jeffrey Barrett, Sonia Goncalves, Ewan Harrison, David K. Jackson, Ian Johnston, Dominic Kwiatkowski, Cordelia Langford, John Sillitoe on behalf of the Wellcome Sanger Institute COVID-19 Surveillance Team |
| EPI_ISL_3441111, EPI_ISL_3441112 | Lighthouse Lab in Glasgow | Wellcome Sanger Institute for the COVID-19 Genomics UK (COG-UK) Consortium | Harper VanSteenhouse, Yumi Kasai, David Gray, Carol Clugston, Anna Dominiczak and Alex Alderton, Roberto Amato, Jeffrey Barrett, Sonia Goncalves, Ewan Harrison, David K. Jackson, Ian Johnston, Dominic Kwiatkowski, Cordelia Langford, John Sillitoe on behalf of the Wellcome Sanger Institute COVID-19 Surveillance Team |
| EPI_ISL_3441113, EPI_ISL_3441114 | Lighthouse Laboratory Plymouth | Wellcome Sanger Institute for the COVID-19 Genomics UK (COG-UK) Consortium | Lighthouse Laboratory Plymouth and Alex Alderton, Roberto Amato, Jeffrey Barrett, Sonia Goncalves, Ewan Harrison, David K. Jackson, Ian Johnston, Dominic Kwiatkowski, Cordelia Langford, John Sillitoe on behalf of the Wellcome Sanger Institute COVID-19 Surveillance Team |
| EPI_ISL_3441115, EPI_ISL_3441116, EPI_ISL_3441117 | Lighthouse Lab in Milton Keynes | Wellcome Sanger Institute for the COVID-19 Genomics UK (COG-UK) Consortium | The Lighthouse Lab in Milton Keynes and Alex Alderton, Roberto Amato, Jeffrey Barrett, Sonia Goncalves, Ewan Harrison, David K. Jackson, Ian Johnston, Dominic Kwiatkowski, Cordelia Langford, John Sillitoe on behalf of the Wellcome Sanger Institute COVID-19 Surveillance Team |
| EPI_ISL_3441118 | Lighthouse Lab in Glasgow | Wellcome Sanger Institute for the COVID-19 Genomics UK (COG-UK) Consortium | Harper VanSteenhouse, Yumi Kasai, David Gray, Carol Clugston, Anna Dominiczak and Alex Alderton, Roberto Amato, Jeffrey Barrett, Sonia Goncalves, Ewan Harrison, David K. Jackson, Ian Johnston, Dominic Kwiatkowski, Cordelia Langford, John Sillitoe on behalf of the Wellcome Sanger Institute COVID-19 Surveillance Team |
| EPI_ISL_3441119 | Berkshire and Surrey Pathology Services Lighthouse Laboratory | Wellcome Sanger Institute for the COVID-19 Genomics UK (COG-UK) Consortium | Berkshire and Surrey Pathology Services Lighthouse Laboratory and Alex Alderton, Roberto Amato, Jeffrey Barrett, Sonia Goncalves, Ewan Harrison, David K. Jackson, Ian Johnston, Dominic Kwiatkowski, Cordelia Langford, John Sillitoe on behalf of the Wellcome Sanger Institute COVID-19 Surveillance Team |
| EPI_ISL_3441120 | Lighthouse Laboratory Plymouth | Wellcome Sanger Institute for the COVID-19 Genomics UK (COG-UK) Consortium | Lighthouse Laboratory Plymouth and Alex Alderton, Roberto Amato, Jeffrey Barrett, Sonia Goncalves, Ewan Harrison, David K. Jackson, Ian Johnston, Dominic Kwiatkowski, Cordelia Langford, John Sillitoe on behalf of the Wellcome Sanger Institute COVID-19 Surveillance Team |
| EPI_ISL_3441121, EPI_ISL_3441122, EPI_ISL_3441123 | Lighthouse Lab in Glasgow | Wellcome Sanger Institute for the COVID-19 Genomics UK (COG-UK) Consortium | Harper VanSteenhouse, Yumi Kasai, David Gray, Carol Clugston, Anna Dominiczak and Alex Alderton, Roberto Amato, Jeffrey Barrett, Sonia Goncalves, Ewan Harrison, David K. Jackson, Ian Johnston, Dominic Kwiatkowski, Cordelia Langford, John Sillitoe on behalf of the Wellcome Sanger Institute COVID-19 Surveillance Team |
| EPI_ISL_3441124, EPI_ISL_3441125 | Lighthouse Lab in Milton Keynes | Wellcome Sanger Institute for the COVID-19 Genomics UK (COG-UK) Consortium | The Lighthouse Lab in Milton Keynes and Alex Alderton, Roberto Amato, Jeffrey Barrett, Sonia Goncalves, Ewan Harrison, David K. Jackson, Ian Johnston, Dominic Kwiatkowski, Cordelia Langford, John Sillitoe on behalf of the Wellcome Sanger Institute COVID-19 Surveillance Team |
| EPI_ISL_3441126 | Lighthouse Lab in Glasgow | Wellcome Sanger Institute for the COVID-19 Genomics UK (COG-UK) Consortium | Harper VanSteenhouse, Yumi Kasai, David Gray, Carol Clugston, Anna Dominiczak and Alex Alderton, Roberto Amato, Jeffrey Barrett, Sonia Goncalves, Ewan Harrison, David K. Jackson, Ian Johnston, Dominic Kwiatkowski, Cordelia Langford, John Sillitoe on behalf of the Wellcome Sanger Institute COVID-19 Surveillance Team |
| EPI_ISL_3441127, EPI_ISL_3441128 | Lighthouse Laboratory Plymouth | Wellcome Sanger Institute for the COVID-19 Genomics UK (COG-UK) Consortium | Lighthouse Laboratory Plymouth and Alex Alderton, Roberto Amato, Jeffrey Barrett, Sonia Goncalves, Ewan Harrison, David K. Jackson, Ian Johnston, Dominic Kwiatkowski, Cordelia Langford, John Sillitoe on behalf of the Wellcome Sanger Institute COVID-19 Surveillance Team |
| EPI_ISL_3441129 | Lighthouse Lab in Milton Keynes | Wellcome Sanger Institute for the COVID-19 Genomics UK (COG-UK) Consortium | The Lighthouse Lab in Milton Keynes and Alex Alderton, Roberto Amato, Jeffrey Barrett, Sonia Goncalves, Ewan Harrison, David K. Jackson, Ian Johnston, Dominic Kwiatkowski, Cordelia Langford, John Sillitoe on behalf of the Wellcome Sanger Institute COVID-19 Surveillance Team |
| EPI_ISL_3441130 | Lighthouse Lab in Glasgow | Wellcome Sanger Institute for the COVID-19 Genomics UK (COG-UK) Consortium | Harper VanSteenhouse, Yumi Kasai, David Gray, Carol Clugston, Anna Dominiczak and Alex Alderton, Roberto Amato, Jeffrey Barrett, Sonia Goncalves, Ewan Harrison, David K. Jackson, Ian Johnston, Dominic Kwiatkowski, Cordelia Langford, John Sillitoe on behalf of the Wellcome Sanger Institute COVID-19 Surveillance Team |
| EPI_ISL_3441131 | Health Services Laboratories | Wellcome Sanger Institute for the COVID-19 Genomics UK (COG-UK) Consortium | Health Services Laboratories and Alex Alderton, Roberto Amato, Jeffrey Barrett, Sonia Goncalves, Ewan Harrison, David K. Jackson, Ian Johnston, Dominic Kwiatkowski, Cordelia Langford, John Sillitoe on behalf of the Wellcome Sanger Institute COVID-19 Surveillance Team |
| EPI_ISL_3441132 | Lighthouse Lab in Glasgow | Wellcome Sanger Institute for the COVID-19 Genomics UK (COG-UK) Consortium | Harper VanSteenhouse, Yumi Kasai, David Gray, Carol Clugston, Anna Dominiczak and Alex Alderton, Roberto Amato, Jeffrey Barrett, Sonia Goncalves, Ewan Harrison, David K. Jackson, Ian Johnston, Dominic Kwiatkowski, Cordelia Langford, John Sillitoe on behalf of the Wellcome Sanger Institute COVID-19 Surveillance Team |
| EPI_ISL_3441133 | Lighthouse Lab in Milton Keynes | Wellcome Sanger Institute for the COVID-19 Genomics UK (COG-UK) Consortium | The Lighthouse Lab in Milton Keynes and Alex Alderton, Roberto Amato, Jeffrey Barrett, Sonia Goncalves, Ewan Harrison, David K. Jackson, Ian Johnston, Dominic Kwiatkowski, Cordelia Langford, John Sillitoe on behalf of the Wellcome Sanger Institute COVID-19 Surveillance Team |
| EPI_ISL_3441134, EPI_ISL_3441135 | Lighthouse Laboratory Plymouth | Wellcome Sanger Institute for the COVID-19 Genomics UK (COG-UK) Consortium | Lighthouse Laboratory Plymouth and Alex Alderton, Roberto Amato, Jeffrey Barrett, Sonia Goncalves, Ewan Harrison, David K. Jackson, Ian Johnston, Dominic Kwiatkowski, Cordelia Langford, John Sillitoe on behalf of the Wellcome Sanger Institute COVID-19 Surveillance Team |
| EPI_ISL_3441136 | Lighthouse Lab in Milton Keynes | Wellcome Sanger Institute for the COVID-19 Genomics UK (COG-UK) Consortium | The Lighthouse Lab in Milton Keynes and Alex Alderton, Roberto Amato, Jeffrey Barrett, Sonia Goncalves, Ewan Harrison, David K. Jackson, Ian Johnston, Dominic Kwiatkowski, Cordelia Langford, John Sillitoe on behalf of the Wellcome Sanger Institute COVID-19 Surveillance Team |
| EPI_ISL_3441137, EPI_ISL_3441138 | Health Services Laboratories | Wellcome Sanger Institute for the COVID-19 Genomics UK (COG-UK) Consortium | Health Services Laboratories and Alex Alderton, Roberto Amato, Jeffrey Barrett, Sonia Goncalves, Ewan Harrison, David K. Jackson, Ian Johnston, Dominic Kwiatkowski, Cordelia Langford, John Sillitoe on behalf of the Wellcome Sanger Institute COVID-19 Surveillance Team |
| EPI_ISL_3441139, EPI_ISL_3441140, EPI_ISL_3441141 | Lighthouse Lab in Milton Keynes | Wellcome Sanger Institute for the COVID-19 Genomics UK (COG-UK) Consortium | The Lighthouse Lab in Milton Keynes and Alex Alderton, Roberto Amato, Jeffrey Barrett, Sonia Goncalves, Ewan Harrison, David K. Jackson, Ian Johnston, Dominic Kwiatkowski, Cordelia Langford, John Sillitoe on behalf of the Wellcome Sanger Institute COVID-19 Surveillance Team |
| EPI_ISL_3441142 | Lighthouse Laboratory Plymouth | Wellcome Sanger Institute for the COVID-19 Genomics UK (COG-UK) Consortium | Lighthouse Laboratory Plymouth and Alex Alderton, Roberto Amato, Jeffrey Barrett, Sonia Goncalves, Ewan Harrison, David K. Jackson, Ian Johnston, Dominic Kwiatkowski, Cordelia Langford, John Sillitoe on behalf of the Wellcome Sanger Institute COVID-19 Surveillance Team |
| EPI_ISL_3441143 | Lighthouse Lab in Glasgow | Wellcome Sanger Institute for the COVID-19 Genomics UK (COG-UK) | Harper VanSteenhouse, Yumi Kasai, David Gray, Carol Clugston, Anna Dominiczak and Alex Alderton, Roberto Amato, Jeffrey Barrett, Sonia Goncalves, Ewan Harrison, David K. Jackson, Ian Johnston, Dominic Kwiatkowski, Cordelia Langford, |

| | | | |
|---|---|---|---|
| | | Consortium | John Sillitoe on behalf of the Wellcome Sanger Institute COVID-19 Surveillance Team |
| EPI_ISL_3441144 | Lighthouse Lab in Milton Keynes | Wellcome Sanger Institute for the COVID-19 Genomics UK (COG-UK) Consortium | The Lighthouse Lab in Milton Keynes and Alex Alderton, Roberto Amato, Jeffrey Barrett, Sonia Goncalves, Ewan Harrison, David K. Jackson, Ian Johnston, Dominic Kwiatkowski, Cordelia Langford, John Sillitoe on behalf of the Wellcome Sanger Institute COVID-19 Surveillance Team |
| EPI_ISL_3441145 | Health Services Laboratories | Wellcome Sanger Institute for the COVID-19 Genomics UK (COG-UK) Consortium | Health Services Laboratories and Alex Alderton, Roberto Amato, Jeffrey Barrett, Sonia Goncalves, Ewan Harrison, David K. Jackson, Ian Johnston, Dominic Kwiatkowski, Cordelia Langford, John Sillitoe on behalf of the Wellcome Sanger Institute COVID-19 Surveillance Team |
| EPI_ISL_3441146 | Lighthouse Laboratory Plymouth | Wellcome Sanger Institute for the COVID-19 Genomics UK (COG-UK) Consortium | Lighthouse Laboratory Plymouth and Alex Alderton, Roberto Amato, Jeffrey Barrett, Sonia Goncalves, Ewan Harrison, David K. Jackson, Ian Johnston, Dominic Kwiatkowski, Cordelia Langford, John Sillitoe on behalf of the Wellcome Sanger Institute COVID-19 Surveillance Team |
| EPI_ISL_3441147 | Lighthouse Lab in Milton Keynes | Wellcome Sanger Institute for the COVID-19 Genomics UK (COG-UK) Consortium | The Lighthouse Lab in Milton Keynes and Alex Alderton, Roberto Amato, Jeffrey Barrett, Sonia Goncalves, Ewan Harrison, David K. Jackson, Ian Johnston, Dominic Kwiatkowski, Cordelia Langford, John Sillitoe on behalf of the Wellcome Sanger Institute COVID-19 Surveillance Team |
| EPI_ISL_3441148 | Lighthouse Lab in Glasgow | Wellcome Sanger Institute for the COVID-19 Genomics UK (COG-UK) Consortium | Harper VanSteenhouse, Yumi Kasai, David Gray, Carol Clugston, Anna Dominiczak and Alex Alderton, Roberto Amato, Jeffrey Barrett, Sonia Goncalves, Ewan Harrison, David K. Jackson, Ian Johnston, Dominic Kwiatkowski, Cordelia Langford, John Sillitoe on behalf of the Wellcome Sanger Institute COVID-19 Surveillance Team |
| EPI_ISL_3441149 | Berkshire and Surrey Pathology Services Lighthouse Laboratory | Wellcome Sanger Institute for the COVID-19 Genomics UK (COG-UK) Consortium | Berkshire and Surrey Pathology Services Lighthouse Laboratory and Alex Alderton, Roberto Amato, Jeffrey Barrett, Sonia Goncalves, Ewan Harrison, David K. Jackson, Ian Johnston, Dominic Kwiatkowski, Cordelia Langford, John Sillitoe on behalf of the Wellcome Sanger Institute COVID-19 Surveillance Team |
| EPI_ISL_3441150 | Health Services Laboratories | Wellcome Sanger Institute for the COVID-19 Genomics UK (COG-UK) Consortium | Health Services Laboratories and Alex Alderton, Roberto Amato, Jeffrey Barrett, Sonia Goncalves, Ewan Harrison, David K. Jackson, Ian Johnston, Dominic Kwiatkowski, Cordelia Langford, John Sillitoe on behalf of the Wellcome Sanger Institute COVID-19 Surveillance Team |
| EPI_ISL_3441151, EPI_ISL_3441152 | Lighthouse Laboratory Plymouth | Wellcome Sanger Institute for the COVID-19 Genomics UK (COG-UK) Consortium | Lighthouse Laboratory Plymouth and Alex Alderton, Roberto Amato, Jeffrey Barrett, Sonia Goncalves, Ewan Harrison, David K. Jackson, Ian Johnston, Dominic Kwiatkowski, Cordelia Langford, John Sillitoe on behalf of the Wellcome Sanger Institute COVID-19 Surveillance Team |
| EPI_ISL_3441153 | Lighthouse Lab in Milton Keynes | Wellcome Sanger Institute for the COVID-19 Genomics UK (COG-UK) Consortium | The Lighthouse Lab in Milton Keynes and Alex Alderton, Roberto Amato, Jeffrey Barrett, Sonia Goncalves, Ewan Harrison, David K. Jackson, Ian Johnston, Dominic Kwiatkowski, Cordelia Langford, John Sillitoe on behalf of the Wellcome Sanger Institute COVID-19 Surveillance Team |
| EPI_ISL_3441154 | Health Services Laboratories | Wellcome Sanger Institute for the COVID-19 Genomics UK (COG-UK) Consortium | Health Services Laboratories and Alex Alderton, Roberto Amato, Jeffrey Barrett, Sonia Goncalves, Ewan Harrison, David K. Jackson, Ian Johnston, Dominic Kwiatkowski, Cordelia Langford, John Sillitoe on behalf of the Wellcome Sanger Institute COVID-19 Surveillance Team |
| EPI_ISL_3441155 | Berkshire and Surrey Pathology Services Lighthouse Laboratory | Wellcome Sanger Institute for the COVID-19 Genomics UK (COG-UK) Consortium | Berkshire and Surrey Pathology Services Lighthouse Laboratory and Alex Alderton, Roberto Amato, Jeffrey Barrett, Sonia Goncalves, Ewan Harrison, David K. Jackson, Ian Johnston, Dominic Kwiatkowski, Cordelia Langford, John Sillitoe on behalf of the Wellcome Sanger Institute COVID-19 Surveillance Team |
| EPI_ISL_3441156, EPI_ISL_3441157 | Lighthouse Lab in Glasgow | Wellcome Sanger Institute for the COVID-19 Genomics UK (COG-UK) Consortium | Harper VanSteenhouse, Yumi Kasai, David Gray, Carol Clugston, Anna Dominiczak and Alex Alderton, Roberto Amato, Jeffrey Barrett, Sonia Goncalves, Ewan Harrison, David K. Jackson, Ian Johnston, Dominic Kwiatkowski, Cordelia Langford, John Sillitoe on behalf of the Wellcome Sanger Institute COVID-19 Surveillance Team |
| EPI_ISL_3441158, EPI_ISL_3441159 | Lighthouse Lab in Milton Keynes | Wellcome Sanger Institute for the COVID-19 Genomics UK (COG-UK) Consortium | The Lighthouse Lab in Milton Keynes and Alex Alderton, Roberto Amato, Jeffrey Barrett, Sonia Goncalves, Ewan Harrison, David K. Jackson, Ian Johnston, Dominic Kwiatkowski, Cordelia Langford, John Sillitoe on behalf of the Wellcome Sanger Institute COVID-19 Surveillance Team |
| EPI_ISL_3441160, EPI_ISL_3441161 | Lighthouse Laboratory Plymouth | Wellcome Sanger Institute for the COVID-19 Genomics UK (COG-UK) Consortium | Lighthouse Laboratory Plymouth and Alex Alderton, Roberto Amato, Jeffrey Barrett, Sonia Goncalves, Ewan Harrison, David K. Jackson, Ian Johnston, Dominic Kwiatkowski, Cordelia Langford, John Sillitoe on behalf of the Wellcome Sanger Institute COVID-19 Surveillance Team |
| EPI_ISL_3441162 | Lighthouse Lab in Milton Keynes | Wellcome Sanger Institute for the COVID-19 Genomics UK (COG-UK) Consortium | The Lighthouse Lab in Milton Keynes and Alex Alderton, Roberto Amato, Jeffrey Barrett, Sonia Goncalves, Ewan Harrison, David K. Jackson, Ian Johnston, Dominic Kwiatkowski, Cordelia Langford, John Sillitoe on behalf of the Wellcome Sanger Institute COVID-19 Surveillance Team |
| EPI_ISL_3441163 | Lighthouse Laboratory Plymouth | Wellcome Sanger Institute for the COVID-19 Genomics UK (COG-UK) Consortium | Lighthouse Laboratory Plymouth and Alex Alderton, Roberto Amato, Jeffrey Barrett, Sonia Goncalves, Ewan Harrison, David K. Jackson, Ian Johnston, Dominic Kwiatkowski, Cordelia Langford, John Sillitoe on behalf of the Wellcome Sanger Institute COVID-19 Surveillance Team |
| EPI_ISL_3441164 | Lighthouse Lab in Milton Keynes | Wellcome Sanger Institute for the COVID-19 Genomics UK (COG-UK) Consortium | The Lighthouse Lab in Milton Keynes and Alex Alderton, Roberto Amato, Jeffrey Barrett, Sonia Goncalves, Ewan Harrison, David K. Jackson, Ian Johnston, Dominic Kwiatkowski, Cordelia Langford, John Sillitoe on behalf of the Wellcome Sanger Institute COVID-19 Surveillance Team |
| EPI_ISL_3441165, EPI_ISL_3441166 | Lighthouse Laboratory Plymouth | Wellcome Sanger Institute for the COVID-19 Genomics UK (COG-UK) Consortium | Lighthouse Laboratory Plymouth and Alex Alderton, Roberto Amato, Jeffrey Barrett, Sonia Goncalves, Ewan Harrison, David K. Jackson, Ian Johnston, Dominic Kwiatkowski, Cordelia Langford, John Sillitoe on behalf of the Wellcome Sanger Institute COVID-19 Surveillance Team |
| EPI_ISL_3441167, EPI_ISL_3441168 | Lighthouse Lab in Glasgow | Wellcome Sanger Institute for the COVID-19 Genomics UK (COG-UK) Consortium | Harper VanSteenhouse, Yumi Kasai, David Gray, Carol Clugston, Anna Dominiczak and Alex Alderton, Roberto Amato, Jeffrey Barrett, Sonia Goncalves, Ewan Harrison, David K. Jackson, Ian Johnston, Dominic Kwiatkowski, Cordelia Langford, John Sillitoe on behalf of the Wellcome Sanger Institute COVID-19 Surveillance Team |
| EPI_ISL_3441169 | Lighthouse Lab in Milton Keynes | Wellcome Sanger Institute for the COVID-19 Genomics UK (COG-UK) Consortium | The Lighthouse Lab in Milton Keynes and Alex Alderton, Roberto Amato, Jeffrey Barrett, Sonia Goncalves, Ewan Harrison, David K. Jackson, Ian Johnston, Dominic Kwiatkowski, Cordelia Langford, John Sillitoe on behalf of the Wellcome Sanger Institute COVID-19 Surveillance Team |
| EPI_ISL_3441170 | Lighthouse Laboratory Plymouth | Wellcome Sanger Institute for the COVID-19 Genomics UK (COG-UK) Consortium | Lighthouse Laboratory Plymouth and Alex Alderton, Roberto Amato, Jeffrey Barrett, Sonia Goncalves, Ewan Harrison, David K. Jackson, Ian Johnston, Dominic Kwiatkowski, Cordelia Langford, John Sillitoe on behalf of the Wellcome Sanger Institute COVID-19 Surveillance Team |
| EPI_ISL_3441171 | Health Services Laboratories | Wellcome Sanger Institute for the COVID-19 Genomics UK (COG-UK) Consortium | Health Services Laboratories and Alex Alderton, Roberto Amato, Jeffrey Barrett, Sonia Goncalves, Ewan Harrison, David K. Jackson, Ian Johnston, Dominic Kwiatkowski, Cordelia Langford, John Sillitoe on behalf of the Wellcome Sanger Institute COVID-19 Surveillance Team |
| EPI_ISL_3441172, EPI_ISL_3441173 | Lighthouse Lab in Milton Keynes | Wellcome Sanger Institute for the COVID-19 Genomics UK (COG-UK) Consortium | The Lighthouse Lab in Milton Keynes and Alex Alderton, Roberto Amato, Jeffrey Barrett, Sonia Goncalves, Ewan Harrison, David K. Jackson, Ian Johnston, Dominic Kwiatkowski, Cordelia Langford, John Sillitoe on behalf of the Wellcome Sanger Institute COVID-19 Surveillance Team |
| EPI_ISL_3441174 | Lighthouse Lab in Glasgow | Wellcome Sanger Institute for the COVID-19 Genomics UK (COG-UK) Consortium | Harper VanSteenhouse, Yumi Kasai, David Gray, Carol Clugston, Anna Dominiczak and Alex Alderton, Roberto Amato, Jeffrey Barrett, Sonia Goncalves, Ewan Harrison, David K. Jackson, Ian Johnston, Dominic Kwiatkowski, Cordelia Langford, John Sillitoe on behalf of the Wellcome Sanger Institute COVID-19 Surveillance Team |
| EPI_ISL_3441175, EPI_ISL_3441176, EPI_ISL_3441177 | Lighthouse Laboratory Plymouth | Wellcome Sanger Institute for the COVID-19 Genomics UK (COG-UK) Consortium | Lighthouse Laboratory Plymouth and Alex Alderton, Roberto Amato, Jeffrey Barrett, Sonia Goncalves, Ewan Harrison, David K. Jackson, Ian Johnston, Dominic Kwiatkowski, Cordelia Langford, John Sillitoe on behalf of the Wellcome Sanger Institute COVID-19 Surveillance Team |
| EPI_ISL_3441178 | Lighthouse Lab in Milton Keynes | Wellcome Sanger Institute for the COVID-19 Genomics UK (COG-UK) Consortium | The Lighthouse Lab in Milton Keynes and Alex Alderton, Roberto Amato, Jeffrey Barrett, Sonia Goncalves, Ewan Harrison, David K. Jackson, Ian Johnston, Dominic Kwiatkowski, Cordelia Langford, John Sillitoe on behalf of the Wellcome Sanger Institute COVID-19 Surveillance Team |
| EPI_ISL_3441179 | Lighthouse Laboratory Plymouth | Wellcome Sanger Institute for the COVID-19 Genomics UK (COG-UK) Consortium | Lighthouse Laboratory Plymouth and Alex Alderton, Roberto Amato, Jeffrey Barrett, Sonia Goncalves, Ewan Harrison, David K. Jackson, Ian Johnston, Dominic Kwiatkowski, Cordelia Langford, John Sillitoe on behalf of the Wellcome Sanger Institute COVID-19 Surveillance Team |
| EPI_ISL_3441180, EPI_ISL_3441181, EPI_ISL_3441182 | Lighthouse Lab in Glasgow | Wellcome Sanger Institute for the COVID-19 Genomics UK (COG-UK) Consortium | Harper VanSteenhouse, Yumi Kasai, David Gray, Carol Clugston, Anna Dominiczak and Alex Alderton, Roberto Amato, Jeffrey Barrett, Sonia Goncalves, Ewan Harrison, David K. Jackson, Ian Johnston, Dominic Kwiatkowski, Cordelia Langford, John Sillitoe on behalf of the Wellcome Sanger Institute COVID-19 Surveillance Team |
| EPI_ISL_3441183 | Lighthouse Laboratory Plymouth | Wellcome Sanger Institute for the COVID-19 Genomics UK (COG-UK) Consortium | Lighthouse Laboratory Plymouth and Alex Alderton, Roberto Amato, Jeffrey Barrett, Sonia Goncalves, Ewan Harrison, David K. Jackson, Ian Johnston, Dominic Kwiatkowski, Cordelia Langford, John Sillitoe on behalf of the Wellcome Sanger Institute COVID-19 Surveillance Team |
| EPI_ISL_3441184 | Lighthouse Lab in Milton Keynes | Wellcome Sanger Institute for the COVID-19 Genomics UK (COG-UK) Consortium | The Lighthouse Lab in Milton Keynes and Alex Alderton, Roberto Amato, Jeffrey Barrett, Sonia Goncalves, Ewan Harrison, David K. Jackson, Ian Johnston, Dominic Kwiatkowski, Cordelia Langford, John Sillitoe on behalf of the Wellcome Sanger Institute COVID-19 Surveillance Team |
| EPI_ISL_3441185 | Lighthouse Laboratory Plymouth | Wellcome Sanger Institute for the COVID-19 Genomics UK (COG-UK) Consortium | Lighthouse Laboratory Plymouth and Alex Alderton, Roberto Amato, Jeffrey Barrett, Sonia Goncalves, Ewan Harrison, David K. Jackson, Ian Johnston, Dominic Kwiatkowski, Cordelia Langford, John Sillitoe on behalf of the Wellcome Sanger Institute COVID-19 Surveillance Team |
| EPI_ISL_3441186 | Health Services Laboratories | Wellcome Sanger Institute for the COVID-19 Genomics UK (COG-UK) Consortium | Health Services Laboratories and Alex Alderton, Roberto Amato, Jeffrey Barrett, Sonia Goncalves, Ewan Harrison, David K. Jackson, Ian Johnston, Dominic Kwiatkowski, Cordelia Langford, John Sillitoe on behalf of the Wellcome Sanger Institute COVID-19 Surveillance Team |
| EPI_ISL_3441187, EPI_ISL_3441188 | Lighthouse Laboratory Plymouth | Wellcome Sanger Institute for the COVID-19 Genomics UK (COG-UK) Consortium | Lighthouse Laboratory Plymouth and Alex Alderton, Roberto Amato, Jeffrey Barrett, Sonia Goncalves, Ewan Harrison, David K. Jackson, Ian Johnston, Dominic Kwiatkowski, Cordelia Langford, John Sillitoe on behalf of the Wellcome Sanger Institute COVID-19 Surveillance Team |
| EPI_ISL_3441189 | Lighthouse Lab in Glasgow | Wellcome Sanger Institute for the COVID-19 Genomics UK (COG-UK) Consortium | Harper VanSteenhouse, Yumi Kasai, David Gray, Carol Clugston, Anna Dominiczak and Alex Alderton, Roberto Amato, Jeffrey Barrett, Sonia Goncalves, Ewan Harrison, David K. Jackson, Ian Johnston, Dominic Kwiatkowski, Cordelia Langford, John Sillitoe on behalf of the Wellcome Sanger Institute COVID-19 Surveillance Team |
| EPI_ISL_3441190 | Berkshire and Surrey Pathology Services Lighthouse Laboratory | Wellcome Sanger Institute for the COVID-19 Genomics UK (COG-UK) Consortium | Berkshire and Surrey Pathology Services Lighthouse Laboratory and Alex Alderton, Roberto Amato, Jeffrey Barrett, Sonia Goncalves, Ewan Harrison, David K. Jackson, Ian Johnston, Dominic Kwiatkowski, Cordelia Langford, John Sillitoe on behalf of the Wellcome Sanger Institute COVID-19 Surveillance Team |
| EPI_ISL_3441191 | Lighthouse Lab in Glasgow | Wellcome Sanger Institute for the COVID-19 Genomics UK (COG-UK) Consortium | Harper VanSteenhouse, Yumi Kasai, David Gray, Carol Clugston, Anna Dominiczak and Alex Alderton, Roberto Amato, Jeffrey Barrett, Sonia Goncalves, Ewan Harrison, David K. Jackson, Ian Johnston, Dominic Kwiatkowski, Cordelia Langford, John Sillitoe on behalf of the Wellcome Sanger Institute COVID-19 Surveillance Team |
| EPI_ISL_3441192 | Health Services Laboratories | Wellcome Sanger Institute for the COVID-19 Genomics UK (COG-UK) Consortium | Health Services Laboratories and Alex Alderton, Roberto Amato, Jeffrey Barrett, Sonia Goncalves, Ewan Harrison, David K. Jackson, Ian Johnston, Dominic Kwiatkowski, Cordelia Langford, John Sillitoe on behalf of the Wellcome Sanger Institute COVID-19 Surveillance Team |
| EPI_ISL_3441193, EPI_ISL_3441194 | Lighthouse Lab in Glasgow | Wellcome Sanger Institute for the COVID-19 Genomics UK (COG-UK) Consortium | Harper VanSteenhouse, Yumi Kasai, David Gray, Carol Clugston, Anna Dominiczak and Alex Alderton, Roberto Amato, Jeffrey Barrett, Sonia Goncalves, Ewan Harrison, David K. Jackson, Ian Johnston, Dominic Kwiatkowski, Cordelia Langford, John Sillitoe on behalf of the Wellcome Sanger Institute COVID-19 Surveillance Team |
| EPI_ISL_3441195, EPI_ISL_3441196, EPI_ISL_3441197 | Lighthouse Lab in Milton Keynes | Wellcome Sanger Institute for the COVID-19 Genomics UK (COG-UK) Consortium | The Lighthouse Lab in Milton Keynes and Alex Alderton, Roberto Amato, Jeffrey Barrett, Sonia Goncalves, Ewan Harrison, David K. Jackson, Ian Johnston, Dominic Kwiatkowski, Cordelia Langford, John Sillitoe on behalf of the Wellcome Sanger Institute COVID-19 Surveillance Team |
| EPI_ISL_3441198 | Health Services Laboratories | Wellcome Sanger Institute for the COVID-19 Genomics UK (COG-UK) Consortium | Health Services Laboratories and Alex Alderton, Roberto Amato, Jeffrey Barrett, Sonia Goncalves, Ewan Harrison, David K. Jackson, Ian Johnston, Dominic Kwiatkowski, Cordelia Langford, John Sillitoe on behalf of the Wellcome Sanger Institute COVID-19 Surveillance Team |
| EPI_ISL_3441199, EPI_ISL_3441200 | Lighthouse Lab in Glasgow | Wellcome Sanger Institute for the COVID-19 Genomics UK (COG-UK) Consortium | Harper VanSteenhouse, Yumi Kasai, David Gray, Carol Clugston, Anna Dominiczak and Alex Alderton, Roberto Amato, Jeffrey Barrett, Sonia Goncalves, Ewan Harrison, David K. Jackson, Ian Johnston, Dominic Kwiatkowski, Cordelia Langford, John Sillitoe on behalf of the Wellcome Sanger Institute COVID-19 Surveillance Team |
| EPI_ISL_3441201, EPI_ISL_3441202 | Lighthouse Laboratory Plymouth | Wellcome Sanger Institute for the COVID-19 Genomics UK (COG-UK) Consortium | Lighthouse Laboratory Plymouth and Alex Alderton, Roberto Amato, Jeffrey Barrett, Sonia Goncalves, Ewan Harrison, David K. Jackson, Ian Johnston, Dominic Kwiatkowski, Cordelia Langford, John Sillitoe on behalf of the Wellcome Sanger Institute COVID-19 Surveillance Team |
| EPI_ISL_3441203 | Health Services Laboratories | Wellcome Sanger Institute for the COVID-19 Genomics UK (COG-UK) Consortium | Health Services Laboratories and Alex Alderton, Roberto Amato, Jeffrey Barrett, Sonia Goncalves, Ewan Harrison, David K. Jackson, Ian Johnston, Dominic Kwiatkowski, Cordelia Langford, John Sillitoe on behalf of the Wellcome Sanger Institute COVID-19 Surveillance Team |
| EPI_ISL_3441204 | Lighthouse Lab in Glasgow | Wellcome Sanger Institute for the COVID-19 Genomics UK (COG-UK) | Harper VanSteenhouse, Yumi Kasai, David Gray, Carol Clugston, Anna Dominiczak and Alex Alderton, Roberto Amato, Jeffrey Barrett, Sonia Goncalves, Ewan Harrison, David K. Jackson, Ian Johnston, Dominic Kwiatkowski, Cordelia Langford, |

| | | | |
|---|---|---|---|
| | | Consortium | John Sillitoe on behalf of the Wellcome Sanger Institute COVID-19 Surveillance Team |
| EPI_ISL_3441205, EPI_ISL_3441206 | Lighthouse Laboratory Plymouth | Wellcome Sanger Institute for the COVID-19 Genomics UK (COG-UK) Consortium | Lighthouse Laboratory Plymouth and Alex Alderton, Roberto Amato, Jeffrey Barrett, Sonia Goncalves, Ewan Harrison, David K. Jackson, Ian Johnston, Dominic Kwiatkowski, Cordelia Langford, John Sillitoe on behalf of the Wellcome Sanger Institute COVID-19 Surveillance Team |
| EPI_ISL_3441207 | Berkshire and Surrey Pathology Services Lighthouse Laboratory | Wellcome Sanger Institute for the COVID-19 Genomics UK (COG-UK) Consortium | Berkshire and Surrey Pathology Services Lighthouse Laboratory and Alex Alderton, Roberto Amato, Jeffrey Barrett, Sonia Goncalves, Ewan Harrison, David K. Jackson, Ian Johnston, Dominic Kwiatkowski, Cordelia Langford, John Sillitoe on behalf of the Wellcome Sanger Institute COVID-19 Surveillance Team |
| EPI_ISL_3441208 | Lighthouse Lab in Milton Keynes | Wellcome Sanger Institute for the COVID-19 Genomics UK (COG-UK) Consortium | The Lighthouse Lab in Milton Keynes and Alex Alderton, Roberto Amato, Jeffrey Barrett, Sonia Goncalves, Ewan Harrison, David K. Jackson, Ian Johnston, Dominic Kwiatkowski, Cordelia Langford, John Sillitoe on behalf of the Wellcome Sanger Institute COVID-19 Surveillance Team |
| EPI_ISL_3441209 | Berkshire and Surrey Pathology Services Lighthouse Laboratory | Wellcome Sanger Institute for the COVID-19 Genomics UK (COG-UK) Consortium | Berkshire and Surrey Pathology Services Lighthouse Laboratory and Alex Alderton, Roberto Amato, Jeffrey Barrett, Sonia Goncalves, Ewan Harrison, David K. Jackson, Ian Johnston, Dominic Kwiatkowski, Cordelia Langford, John Sillitoe on behalf of the Wellcome Sanger Institute COVID-19 Surveillance Team |
| EPI_ISL_3441210 | Lighthouse Lab in Glasgow | Wellcome Sanger Institute for the COVID-19 Genomics UK (COG-UK) Consortium | Harper VanSteenhouse, Yumi Kasai, David Gray, Carol Clugston, Anna Dominiczak and Alex Alderton, Roberto Amato, Jeffrey Barrett, Sonia Goncalves, Ewan Harrison, David K. Jackson, Ian Johnston, Dominic Kwiatkowski, Cordelia Langford, John Sillitoe on behalf of the Wellcome Sanger Institute COVID-19 Surveillance Team |
| EPI_ISL_3441211, EPI_ISL_3441212 | Lighthouse Lab in Milton Keynes | Wellcome Sanger Institute for the COVID-19 Genomics UK (COG-UK) Consortium | The Lighthouse Lab in Milton Keynes and Alex Alderton, Roberto Amato, Jeffrey Barrett, Sonia Goncalves, Ewan Harrison, David K. Jackson, Ian Johnston, Dominic Kwiatkowski, Cordelia Langford, John Sillitoe on behalf of the Wellcome Sanger Institute COVID-19 Surveillance Team |
| EPI_ISL_3441213 | Berkshire and Surrey Pathology Services Lighthouse Laboratory | Wellcome Sanger Institute for the COVID-19 Genomics UK (COG-UK) Consortium | Berkshire and Surrey Pathology Services Lighthouse Laboratory and Alex Alderton, Roberto Amato, Jeffrey Barrett, Sonia Goncalves, Ewan Harrison, David K. Jackson, Ian Johnston, Dominic Kwiatkowski, Cordelia Langford, John Sillitoe on behalf of the Wellcome Sanger Institute COVID-19 Surveillance Team |
| EPI_ISL_3441214, EPI_ISL_3441215 | Lighthouse Lab in Glasgow | Wellcome Sanger Institute for the COVID-19 Genomics UK (COG-UK) Consortium | Harper VanSteenhouse, Yumi Kasai, David Gray, Carol Clugston, Anna Dominiczak and Alex Alderton, Roberto Amato, Jeffrey Barrett, Sonia Goncalves, Ewan Harrison, David K. Jackson, Ian Johnston, Dominic Kwiatkowski, Cordelia Langford, John Sillitoe on behalf of the Wellcome Sanger Institute COVID-19 Surveillance Team |
| EPI_ISL_3441216 | Lighthouse Laboratory Plymouth | Wellcome Sanger Institute for the COVID-19 Genomics UK (COG-UK) Consortium | Lighthouse Laboratory Plymouth and Alex Alderton, Roberto Amato, Jeffrey Barrett, Sonia Goncalves, Ewan Harrison, David K. Jackson, Ian Johnston, Dominic Kwiatkowski, Cordelia Langford, John Sillitoe on behalf of the Wellcome Sanger Institute COVID-19 Surveillance Team |
| EPI_ISL_3441217 | Lighthouse Lab in Milton Keynes | Wellcome Sanger Institute for the COVID-19 Genomics UK (COG-UK) Consortium | The Lighthouse Lab in Milton Keynes and Alex Alderton, Roberto Amato, Jeffrey Barrett, Sonia Goncalves, Ewan Harrison, David K. Jackson, Ian Johnston, Dominic Kwiatkowski, Cordelia Langford, John Sillitoe on behalf of the Wellcome Sanger Institute COVID-19 Surveillance Team |
| EPI_ISL_3441218 | Lighthouse Laboratory Plymouth | Wellcome Sanger Institute for the COVID-19 Genomics UK (COG-UK) Consortium | Lighthouse Laboratory Plymouth and Alex Alderton, Roberto Amato, Jeffrey Barrett, Sonia Goncalves, Ewan Harrison, David K. Jackson, Ian Johnston, Dominic Kwiatkowski, Cordelia Langford, John Sillitoe on behalf of the Wellcome Sanger Institute COVID-19 Surveillance Team |
| EPI_ISL_3441219 | Lighthouse Lab in Milton Keynes | Wellcome Sanger Institute for the COVID-19 Genomics UK (COG-UK) Consortium | The Lighthouse Lab in Milton Keynes and Alex Alderton, Roberto Amato, Jeffrey Barrett, Sonia Goncalves, Ewan Harrison, David K. Jackson, Ian Johnston, Dominic Kwiatkowski, Cordelia Langford, John Sillitoe on behalf of the Wellcome Sanger Institute COVID-19 Surveillance Team |
| EPI_ISL_3441220, EPI_ISL_3441221 | Lighthouse Lab in Glasgow | Wellcome Sanger Institute for the COVID-19 Genomics UK (COG-UK) Consortium | Harper VanSteenhouse, Yumi Kasai, David Gray, Carol Clugston, Anna Dominiczak and Alex Alderton, Roberto Amato, Jeffrey Barrett, Sonia Goncalves, Ewan Harrison, David K. Jackson, Ian Johnston, Dominic Kwiatkowski, Cordelia Langford, John Sillitoe on behalf of the Wellcome Sanger Institute COVID-19 Surveillance Team |
| EPI_ISL_3441222 | Berkshire and Surrey Pathology Services Lighthouse Laboratory | Wellcome Sanger Institute for the COVID-19 Genomics UK (COG-UK) Consortium | Berkshire and Surrey Pathology Services Lighthouse Laboratory and Alex Alderton, Roberto Amato, Jeffrey Barrett, Sonia Goncalves, Ewan Harrison, David K. Jackson, Ian Johnston, Dominic Kwiatkowski, Cordelia Langford, John Sillitoe on behalf of the Wellcome Sanger Institute COVID-19 Surveillance Team |
| EPI_ISL_3441223 | Lighthouse Laboratory Plymouth | Wellcome Sanger Institute for the COVID-19 Genomics UK (COG-UK) Consortium | Lighthouse Laboratory Plymouth and Alex Alderton, Roberto Amato, Jeffrey Barrett, Sonia Goncalves, Ewan Harrison, David K. Jackson, Ian Johnston, Dominic Kwiatkowski, Cordelia Langford, John Sillitoe on behalf of the Wellcome Sanger Institute COVID-19 Surveillance Team |
| EPI_ISL_3441224, EPI_ISL_3441225 | Lighthouse Lab in Glasgow | Wellcome Sanger Institute for the COVID-19 Genomics UK (COG-UK) Consortium | Harper VanSteenhouse, Yumi Kasai, David Gray, Carol Clugston, Anna Dominiczak and Alex Alderton, Roberto Amato, Jeffrey Barrett, Sonia Goncalves, Ewan Harrison, David K. Jackson, Ian Johnston, Dominic Kwiatkowski, Cordelia Langford, John Sillitoe on behalf of the Wellcome Sanger Institute COVID-19 Surveillance Team |
| EPI_ISL_3441226 | Berkshire and Surrey Pathology Services Lighthouse Laboratory | Wellcome Sanger Institute for the COVID-19 Genomics UK (COG-UK) Consortium | Berkshire and Surrey Pathology Services Lighthouse Laboratory and Alex Alderton, Roberto Amato, Jeffrey Barrett, Sonia Goncalves, Ewan Harrison, David K. Jackson, Ian Johnston, Dominic Kwiatkowski, Cordelia Langford, John Sillitoe on behalf of the Wellcome Sanger Institute COVID-19 Surveillance Team |
| EPI_ISL_3441227 | Lighthouse Laboratory Plymouth | Wellcome Sanger Institute for the COVID-19 Genomics UK (COG-UK) Consortium | Lighthouse Laboratory Plymouth and Alex Alderton, Roberto Amato, Jeffrey Barrett, Sonia Goncalves, Ewan Harrison, David K. Jackson, Ian Johnston, Dominic Kwiatkowski, Cordelia Langford, John Sillitoe on behalf of the Wellcome Sanger Institute COVID-19 Surveillance Team |
| EPI_ISL_3441228, EPI_ISL_3441229, EPI_ISL_3441230 | Lighthouse Lab in Glasgow | Wellcome Sanger Institute for the COVID-19 Genomics UK (COG-UK) Consortium | Harper VanSteenhouse, Yumi Kasai, David Gray, Carol Clugston, Anna Dominiczak and Alex Alderton, Roberto Amato, Jeffrey Barrett, Sonia Goncalves, Ewan Harrison, David K. Jackson, Ian Johnston, Dominic Kwiatkowski, Cordelia Langford, John Sillitoe on behalf of the Wellcome Sanger Institute COVID-19 Surveillance Team |
| EPI_ISL_3441231 | Lighthouse Lab in Milton Keynes | Wellcome Sanger Institute for the COVID-19 Genomics UK (COG-UK) Consortium | The Lighthouse Lab in Milton Keynes and Alex Alderton, Roberto Amato, Jeffrey Barrett, Sonia Goncalves, Ewan Harrison, David K. Jackson, Ian Johnston, Dominic Kwiatkowski, Cordelia Langford, John Sillitoe on behalf of the Wellcome Sanger Institute COVID-19 Surveillance Team |
| EPI_ISL_3441232 | Lighthouse Laboratory Plymouth | Wellcome Sanger Institute for the COVID-19 Genomics UK (COG-UK) Consortium | Lighthouse Laboratory Plymouth and Alex Alderton, Roberto Amato, Jeffrey Barrett, Sonia Goncalves, Ewan Harrison, David K. Jackson, Ian Johnston, Dominic Kwiatkowski, Cordelia Langford, John Sillitoe on behalf of the Wellcome Sanger Institute COVID-19 Surveillance Team |
| EPI_ISL_3441233 | Berkshire and Surrey Pathology Services Lighthouse Laboratory | Wellcome Sanger Institute for the COVID-19 Genomics UK (COG-UK) Consortium | Berkshire and Surrey Pathology Services Lighthouse Laboratory and Alex Alderton, Roberto Amato, Jeffrey Barrett, Sonia Goncalves, Ewan Harrison, David K. Jackson, Ian Johnston, Dominic Kwiatkowski, Cordelia Langford, John Sillitoe on behalf of the Wellcome Sanger Institute COVID-19 Surveillance Team |
| EPI_ISL_3441234 | Lighthouse Laboratory Plymouth | Wellcome Sanger Institute for the COVID-19 Genomics UK (COG-UK) Consortium | Lighthouse Laboratory Plymouth and Alex Alderton, Roberto Amato, Jeffrey Barrett, Sonia Goncalves, Ewan Harrison, David K. Jackson, Ian Johnston, Dominic Kwiatkowski, Cordelia Langford, John Sillitoe on behalf of the Wellcome Sanger Institute COVID-19 Surveillance Team |
| EPI_ISL_3441235, EPI_ISL_3441236 | Lighthouse Lab in Glasgow | Wellcome Sanger Institute for the COVID-19 Genomics UK (COG-UK) Consortium | Harper VanSteenhouse, Yumi Kasai, David Gray, Carol Clugston, Anna Dominiczak and Alex Alderton, Roberto Amato, Jeffrey Barrett, Sonia Goncalves, Ewan Harrison, David K. Jackson, Ian Johnston, Dominic Kwiatkowski, Cordelia Langford, John Sillitoe on behalf of the Wellcome Sanger Institute COVID-19 Surveillance Team |
| EPI_ISL_3441237 | Lighthouse Laboratory Plymouth | Wellcome Sanger Institute for the COVID-19 Genomics UK (COG-UK) Consortium | Lighthouse Laboratory Plymouth and Alex Alderton, Roberto Amato, Jeffrey Barrett, Sonia Goncalves, Ewan Harrison, David K. Jackson, Ian Johnston, Dominic Kwiatkowski, Cordelia Langford, John Sillitoe on behalf of the Wellcome Sanger Institute COVID-19 Surveillance Team |
| EPI_ISL_3441238, EPI_ISL_3441239, EPI_ISL_3441240, EPI_ISL_3441241, EPI_ISL_3441242 | Berkshire and Surrey Pathology Services Lighthouse Laboratory | Wellcome Sanger Institute for the COVID-19 Genomics UK (COG-UK) Consortium | Berkshire and Surrey Pathology Services Lighthouse Laboratory and Alex Alderton, Roberto Amato, Jeffrey Barrett, Sonia Goncalves, Ewan Harrison, David K. Jackson, Ian Johnston, Dominic Kwiatkowski, Cordelia Langford, John Sillitoe on behalf of the Wellcome Sanger Institute COVID-19 Surveillance Team |
| EPI_ISL_3441243 | Lighthouse Lab in Glasgow | Wellcome Sanger Institute for the COVID-19 Genomics UK (COG-UK) Consortium | Harper VanSteenhouse, Yumi Kasai, David Gray, Carol Clugston, Anna Dominiczak and Alex Alderton, Roberto Amato, Jeffrey Barrett, Sonia Goncalves, Ewan Harrison, David K. Jackson, Ian Johnston, Dominic Kwiatkowski, Cordelia Langford, John Sillitoe on behalf of the Wellcome Sanger Institute COVID-19 Surveillance Team |
| EPI_ISL_3441244 | Lighthouse Lab in Milton Keynes | Wellcome Sanger Institute for the COVID-19 Genomics UK (COG-UK) Consortium | The Lighthouse Lab in Milton Keynes and Alex Alderton, Roberto Amato, Jeffrey Barrett, Sonia Goncalves, Ewan Harrison, David K. Jackson, Ian Johnston, Dominic Kwiatkowski, Cordelia Langford, John Sillitoe on behalf of the Wellcome Sanger Institute COVID-19 Surveillance Team |
| EPI_ISL_3441245 | Lighthouse Lab in Glasgow | Wellcome Sanger Institute for the COVID-19 Genomics UK (COG-UK) Consortium | Harper VanSteenhouse, Yumi Kasai, David Gray, Carol Clugston, Anna Dominiczak and Alex Alderton, Roberto Amato, Jeffrey Barrett, Sonia Goncalves, Ewan Harrison, David K. Jackson, Ian Johnston, Dominic Kwiatkowski, Cordelia Langford, John Sillitoe on behalf of the Wellcome Sanger Institute COVID-19 Surveillance Team |
| EPI_ISL_3441246 | Lighthouse Laboratory Plymouth | Wellcome Sanger Institute for the COVID-19 Genomics UK (COG-UK) Consortium | Lighthouse Laboratory Plymouth and Alex Alderton, Roberto Amato, Jeffrey Barrett, Sonia Goncalves, Ewan Harrison, David K. Jackson, Ian Johnston, Dominic Kwiatkowski, Cordelia Langford, John Sillitoe on behalf of the Wellcome Sanger Institute COVID-19 Surveillance Team |
| EPI_ISL_3441247 | Lighthouse Lab in Glasgow | Wellcome Sanger Institute for the COVID-19 Genomics UK (COG-UK) Consortium | Harper VanSteenhouse, Yumi Kasai, David Gray, Carol Clugston, Anna Dominiczak and Alex Alderton, Roberto Amato, Jeffrey Barrett, Sonia Goncalves, Ewan Harrison, David K. Jackson, Ian Johnston, Dominic Kwiatkowski, Cordelia Langford, John Sillitoe on behalf of the Wellcome Sanger Institute COVID-19 Surveillance Team |
| EPI_ISL_3441248 | Lighthouse Laboratory Plymouth | Wellcome Sanger Institute for the COVID-19 Genomics UK (COG-UK) Consortium | Lighthouse Laboratory Plymouth and Alex Alderton, Roberto Amato, Jeffrey Barrett, Sonia Goncalves, Ewan Harrison, David K. Jackson, Ian Johnston, Dominic Kwiatkowski, Cordelia Langford, John Sillitoe on behalf of the Wellcome Sanger Institute COVID-19 Surveillance Team |
| EPI_ISL_3441249 | Lighthouse Lab in Milton Keynes | Wellcome Sanger Institute for the COVID-19 Genomics UK (COG-UK) Consortium | The Lighthouse Lab in Milton Keynes and Alex Alderton, Roberto Amato, Jeffrey Barrett, Sonia Goncalves, Ewan Harrison, David K. Jackson, Ian Johnston, Dominic Kwiatkowski, Cordelia Langford, John Sillitoe on behalf of the Wellcome Sanger Institute COVID-19 Surveillance Team |
| EPI_ISL_3441250 | Lighthouse Laboratory Plymouth | Wellcome Sanger Institute for the COVID-19 Genomics UK (COG-UK) Consortium | Lighthouse Laboratory Plymouth and Alex Alderton, Roberto Amato, Jeffrey Barrett, Sonia Goncalves, Ewan Harrison, David K. Jackson, Ian Johnston, Dominic Kwiatkowski, Cordelia Langford, John Sillitoe on behalf of the Wellcome Sanger Institute COVID-19 Surveillance Team |
| EPI_ISL_3441251, EPI_ISL_3441252 | Lighthouse Lab in Glasgow | Wellcome Sanger Institute for the COVID-19 Genomics UK (COG-UK) Consortium | Harper VanSteenhouse, Yumi Kasai, David Gray, Carol Clugston, Anna Dominiczak and Alex Alderton, Roberto Amato, Jeffrey Barrett, Sonia Goncalves, Ewan Harrison, David K. Jackson, Ian Johnston, Dominic Kwiatkowski, Cordelia Langford, John Sillitoe on behalf of the Wellcome Sanger Institute COVID-19 Surveillance Team |
| EPI_ISL_3441253 | Berkshire and Surrey Pathology Services Lighthouse Laboratory | Wellcome Sanger Institute for the COVID-19 Genomics UK (COG-UK) Consortium | Berkshire and Surrey Pathology Services Lighthouse Laboratory and Alex Alderton, Roberto Amato, Jeffrey Barrett, Sonia Goncalves, Ewan Harrison, David K. Jackson, Ian Johnston, Dominic Kwiatkowski, Cordelia Langford, John Sillitoe on behalf of the Wellcome Sanger Institute COVID-19 Surveillance Team |
| EPI_ISL_3441254 | Lighthouse Laboratory Plymouth | Wellcome Sanger Institute for the COVID-19 Genomics UK (COG-UK) Consortium | Lighthouse Laboratory Plymouth and Alex Alderton, Roberto Amato, Jeffrey Barrett, Sonia Goncalves, Ewan Harrison, David K. Jackson, Ian Johnston, Dominic Kwiatkowski, Cordelia Langford, John Sillitoe on behalf of the Wellcome Sanger Institute COVID-19 Surveillance Team |
| EPI_ISL_3441255, EPI_ISL_3441256, EPI_ISL_3441257, EPI_ISL_3441258 | Lighthouse Lab in Milton Keynes | Wellcome Sanger Institute for the COVID-19 Genomics UK (COG-UK) Consortium | The Lighthouse Lab in Milton Keynes and Alex Alderton, Roberto Amato, Jeffrey Barrett, Sonia Goncalves, Ewan Harrison, David K. Jackson, Ian Johnston, Dominic Kwiatkowski, Cordelia Langford, John Sillitoe on behalf of the Wellcome Sanger Institute COVID-19 Surveillance Team |
| EPI_ISL_3441259 | Lighthouse Lab in Glasgow | Wellcome Sanger Institute for the COVID-19 Genomics UK (COG-UK) Consortium | Harper VanSteenhouse, Yumi Kasai, David Gray, Carol Clugston, Anna Dominiczak and Alex Alderton, Roberto Amato, Jeffrey Barrett, Sonia Goncalves, Ewan Harrison, David K. Jackson, Ian Johnston, Dominic Kwiatkowski, Cordelia Langford, John Sillitoe on behalf of the Wellcome Sanger Institute COVID-19 Surveillance Team |
| EPI_ISL_3441260 | Berkshire and Surrey Pathology Services Lighthouse Laboratory | Wellcome Sanger Institute for the COVID-19 Genomics UK (COG-UK) Consortium | Berkshire and Surrey Pathology Services Lighthouse Laboratory and Alex Alderton, Roberto Amato, Jeffrey Barrett, Sonia Goncalves, Ewan Harrison, David K. Jackson, Ian Johnston, Dominic Kwiatkowski, Cordelia Langford, John Sillitoe on behalf of the Wellcome Sanger Institute COVID-19 Surveillance Team |
| EPI_ISL_3441261, EPI_ISL_3441262 | Lighthouse Lab in Glasgow | Wellcome Sanger Institute for the COVID-19 Genomics UK (COG-UK) Consortium | Harper VanSteenhouse, Yumi Kasai, David Gray, Carol Clugston, Anna Dominiczak and Alex Alderton, Roberto Amato, Jeffrey Barrett, Sonia Goncalves, Ewan Harrison, David K. Jackson, Ian Johnston, Dominic Kwiatkowski, Cordelia Langford, John Sillitoe on behalf of the Wellcome Sanger Institute COVID-19 Surveillance Team |
| EPI_ISL_3441263 | Lighthouse Laboratory Plymouth | Wellcome Sanger Institute for the COVID-19 Genomics UK (COG-UK) Consortium | Lighthouse Laboratory Plymouth and Alex Alderton, Roberto Amato, Jeffrey Barrett, Sonia Goncalves, Ewan Harrison, David K. Jackson, Ian Johnston, Dominic Kwiatkowski, Cordelia Langford, John Sillitoe on behalf of the Wellcome Sanger Institute COVID-19 Surveillance Team |

| | | | |
|---|---|---|---|
| EPI_ISL_3441264, EPI_ISL_3441265 | Lighthouse Lab in Milton Keynes | Wellcome Sanger Institute for the COVID-19 Genomics UK (COG-UK) Consortium | The Lighthouse Lab in Milton Keynes and Alex Alderton, Roberto Amato, Jeffrey Barrett, Sonia Goncalves, Ewan Harrison, David K. Jackson, Ian Johnston, Dominic Kwiatkowski, Cordelia Langford, John Sillitoe on behalf of the Wellcome Sanger Institute COVID-19 Surveillance Team |
| EPI_ISL_3441266 | Lighthouse Lab in Glasgow | Wellcome Sanger Institute for the COVID-19 Genomics UK (COG-UK) Consortium | Harper VanSteenhouse, Yumi Kasai, David Gray, Carol Clugston, Anna Dominiczak and Alex Alderton, Roberto Amato, Jeffrey Barrett, Sonia Goncalves, Ewan Harrison, David K. Jackson, Ian Johnston, Dominic Kwiatkowski, Cordelia Langford, John Sillitoe on behalf of the Wellcome Sanger Institute COVID-19 Surveillance Team |
| EPI_ISL_3441267, EPI_ISL_3441268 | Lighthouse Lab in Milton Keynes | Wellcome Sanger Institute for the COVID-19 Genomics UK (COG-UK) Consortium | The Lighthouse Lab in Milton Keynes and Alex Alderton, Roberto Amato, Jeffrey Barrett, Sonia Goncalves, Ewan Harrison, David K. Jackson, Ian Johnston, Dominic Kwiatkowski, Cordelia Langford, John Sillitoe on behalf of the Wellcome Sanger Institute COVID-19 Surveillance Team |
| EPI_ISL_3441269, EPI_ISL_3441270 | Lighthouse Lab in Glasgow | Wellcome Sanger Institute for the COVID-19 Genomics UK (COG-UK) Consortium | Harper VanSteenhouse, Yumi Kasai, David Gray, Carol Clugston, Anna Dominiczak and Alex Alderton, Roberto Amato, Jeffrey Barrett, Sonia Goncalves, Ewan Harrison, David K. Jackson, Ian Johnston, Dominic Kwiatkowski, Cordelia Langford, John Sillitoe on behalf of the Wellcome Sanger Institute COVID-19 Surveillance Team |
| EPI_ISL_3441271, EPI_ISL_3441272, EPI_ISL_3441273 | Lighthouse Laboratory Plymouth | Wellcome Sanger Institute for the COVID-19 Genomics UK (COG-UK) Consortium | Lighthouse Laboratory Plymouth and Alex Alderton, Roberto Amato, Jeffrey Barrett, Sonia Goncalves, Ewan Harrison, David K. Jackson, Ian Johnston, Dominic Kwiatkowski, Cordelia Langford, John Sillitoe on behalf of the Wellcome Sanger Institute COVID-19 Surveillance Team |
| EPI_ISL_3441274 | Lighthouse Lab in Milton Keynes | Wellcome Sanger Institute for the COVID-19 Genomics UK (COG-UK) Consortium | The Lighthouse Lab in Milton Keynes and Alex Alderton, Roberto Amato, Jeffrey Barrett, Sonia Goncalves, Ewan Harrison, David K. Jackson, Ian Johnston, Dominic Kwiatkowski, Cordelia Langford, John Sillitoe on behalf of the Wellcome Sanger Institute COVID-19 Surveillance Team |
| EPI_ISL_3441275, EPI_ISL_3441276 | Berkshire and Surrey Pathology Services Lighthouse Laboratory | Wellcome Sanger Institute for the COVID-19 Genomics UK (COG-UK) Consortium | Berkshire and Surrey Pathology Services Lighthouse Laboratory and Alex Alderton, Roberto Amato, Jeffrey Barrett, Sonia Goncalves, Ewan Harrison, David K. Jackson, Ian Johnston, Dominic Kwiatkowski, Cordelia Langford, John Sillitoe on behalf of the Wellcome Sanger Institute COVID-19 Surveillance Team |
| EPI_ISL_3441277 | Lighthouse Laboratory Plymouth | Wellcome Sanger Institute for the COVID-19 Genomics UK (COG-UK) Consortium | Lighthouse Laboratory Plymouth and Alex Alderton, Roberto Amato, Jeffrey Barrett, Sonia Goncalves, Ewan Harrison, David K. Jackson, Ian Johnston, Dominic Kwiatkowski, Cordelia Langford, John Sillitoe on behalf of the Wellcome Sanger Institute COVID-19 Surveillance Team |
| EPI_ISL_3441278 | Lighthouse Lab in Glasgow | Wellcome Sanger Institute for the COVID-19 Genomics UK (COG-UK) Consortium | Harper VanSteenhouse, Yumi Kasai, David Gray, Carol Clugston, Anna Dominiczak and Alex Alderton, Roberto Amato, Jeffrey Barrett, Sonia Goncalves, Ewan Harrison, David K. Jackson, Ian Johnston, Dominic Kwiatkowski, Cordelia Langford, John Sillitoe on behalf of the Wellcome Sanger Institute COVID-19 Surveillance Team |
| EPI_ISL_3441279 | Berkshire and Surrey Pathology Services Lighthouse Laboratory | Wellcome Sanger Institute for the COVID-19 Genomics UK (COG-UK) Consortium | Berkshire and Surrey Pathology Services Lighthouse Laboratory and Alex Alderton, Roberto Amato, Jeffrey Barrett, Sonia Goncalves, Ewan Harrison, David K. Jackson, Ian Johnston, Dominic Kwiatkowski, Cordelia Langford, John Sillitoe on behalf of the Wellcome Sanger Institute COVID-19 Surveillance Team |
| EPI_ISL_3441280, EPI_ISL_3441281, EPI_ISL_3441282 | Lighthouse Laboratory Plymouth | Wellcome Sanger Institute for the COVID-19 Genomics UK (COG-UK) Consortium | Lighthouse Laboratory Plymouth and Alex Alderton, Roberto Amato, Jeffrey Barrett, Sonia Goncalves, Ewan Harrison, David K. Jackson, Ian Johnston, Dominic Kwiatkowski, Cordelia Langford, John Sillitoe on behalf of the Wellcome Sanger Institute COVID-19 Surveillance Team |
| EPI_ISL_3441283 | Lighthouse Lab in Glasgow | Wellcome Sanger Institute for the COVID-19 Genomics UK (COG-UK) Consortium | Harper VanSteenhouse, Yumi Kasai, David Gray, Carol Clugston, Anna Dominiczak and Alex Alderton, Roberto Amato, Jeffrey Barrett, Sonia Goncalves, Ewan Harrison, David K. Jackson, Ian Johnston, Dominic Kwiatkowski, Cordelia Langford, John Sillitoe on behalf of the Wellcome Sanger Institute COVID-19 Surveillance Team |
| EPI_ISL_3441284 | Lighthouse Laboratory Plymouth | Wellcome Sanger Institute for the COVID-19 Genomics UK (COG-UK) Consortium | Lighthouse Laboratory Plymouth and Alex Alderton, Roberto Amato, Jeffrey Barrett, Sonia Goncalves, Ewan Harrison, David K. Jackson, Ian Johnston, Dominic Kwiatkowski, Cordelia Langford, John Sillitoe on behalf of the Wellcome Sanger Institute COVID-19 Surveillance Team |
| EPI_ISL_3441285 | Lighthouse Lab in Milton Keynes | Wellcome Sanger Institute for the COVID-19 Genomics UK (COG-UK) Consortium | The Lighthouse Lab in Milton Keynes and Alex Alderton, Roberto Amato, Jeffrey Barrett, Sonia Goncalves, Ewan Harrison, David K. Jackson, Ian Johnston, Dominic Kwiatkowski, Cordelia Langford, John Sillitoe on behalf of the Wellcome Sanger Institute COVID-19 Surveillance Team |
| EPI_ISL_3441286, EPI_ISL_3441287 | Lighthouse Lab in Glasgow | Wellcome Sanger Institute for the COVID-19 Genomics UK (COG-UK) Consortium | Harper VanSteenhouse, Yumi Kasai, David Gray, Carol Clugston, Anna Dominiczak and Alex Alderton, Roberto Amato, Jeffrey Barrett, Sonia Goncalves, Ewan Harrison, David K. Jackson, Ian Johnston, Dominic Kwiatkowski, Cordelia Langford, John Sillitoe on behalf of the Wellcome Sanger Institute COVID-19 Surveillance Team |
| EPI_ISL_3441288 | Lighthouse Laboratory Plymouth | Wellcome Sanger Institute for the COVID-19 Genomics UK (COG-UK) Consortium | Lighthouse Laboratory Plymouth and Alex Alderton, Roberto Amato, Jeffrey Barrett, Sonia Goncalves, Ewan Harrison, David K. Jackson, Ian Johnston, Dominic Kwiatkowski, Cordelia Langford, John Sillitoe on behalf of the Wellcome Sanger Institute COVID-19 Surveillance Team |
| EPI_ISL_3441289 | Berkshire and Surrey Pathology Services Lighthouse Laboratory | Wellcome Sanger Institute for the COVID-19 Genomics UK (COG-UK) Consortium | Berkshire and Surrey Pathology Services Lighthouse Laboratory and Alex Alderton, Roberto Amato, Jeffrey Barrett, Sonia Goncalves, Ewan Harrison, David K. Jackson, Ian Johnston, Dominic Kwiatkowski, Cordelia Langford, John Sillitoe on behalf of the Wellcome Sanger Institute COVID-19 Surveillance Team |
| EPI_ISL_3441290 | Lighthouse Laboratory Plymouth | Wellcome Sanger Institute for the COVID-19 Genomics UK (COG-UK) Consortium | Lighthouse Laboratory Plymouth and Alex Alderton, Roberto Amato, Jeffrey Barrett, Sonia Goncalves, Ewan Harrison, David K. Jackson, Ian Johnston, Dominic Kwiatkowski, Cordelia Langford, John Sillitoe on behalf of the Wellcome Sanger Institute COVID-19 Surveillance Team |
| EPI_ISL_3441291, EPI_ISL_3441292 | Berkshire and Surrey Pathology Services Lighthouse Laboratory | Wellcome Sanger Institute for the COVID-19 Genomics UK (COG-UK) Consortium | Berkshire and Surrey Pathology Services Lighthouse Laboratory and Alex Alderton, Roberto Amato, Jeffrey Barrett, Sonia Goncalves, Ewan Harrison, David K. Jackson, Ian Johnston, Dominic Kwiatkowski, Cordelia Langford, John Sillitoe on behalf of the Wellcome Sanger Institute COVID-19 Surveillance Team |
| EPI_ISL_3441293, EPI_ISL_3441294 | Lighthouse Lab in Milton Keynes | Wellcome Sanger Institute for the COVID-19 Genomics UK (COG-UK) Consortium | The Lighthouse Lab in Milton Keynes and Alex Alderton, Roberto Amato, Jeffrey Barrett, Sonia Goncalves, Ewan Harrison, David K. Jackson, Ian Johnston, Dominic Kwiatkowski, Cordelia Langford, John Sillitoe on behalf of the Wellcome Sanger Institute COVID-19 Surveillance Team |
| EPI_ISL_3441295 | Lighthouse Laboratory Plymouth | Wellcome Sanger Institute for the COVID-19 Genomics UK (COG-UK) Consortium | Lighthouse Laboratory Plymouth and Alex Alderton, Roberto Amato, Jeffrey Barrett, Sonia Goncalves, Ewan Harrison, David K. Jackson, Ian Johnston, Dominic Kwiatkowski, Cordelia Langford, John Sillitoe on behalf of the Wellcome Sanger Institute COVID-19 Surveillance Team |
| EPI_ISL_3441296 | Lighthouse Lab in Milton Keynes | Wellcome Sanger Institute for the COVID-19 Genomics UK (COG-UK) Consortium | The Lighthouse Lab in Milton Keynes and Alex Alderton, Roberto Amato, Jeffrey Barrett, Sonia Goncalves, Ewan Harrison, David K. Jackson, Ian Johnston, Dominic Kwiatkowski, Cordelia Langford, John Sillitoe on behalf of the Wellcome Sanger Institute COVID-19 Surveillance Team |
| EPI_ISL_3441297, EPI_ISL_3441298 | Lighthouse Laboratory Plymouth | Wellcome Sanger Institute for the COVID-19 Genomics UK (COG-UK) Consortium | Lighthouse Laboratory Plymouth and Alex Alderton, Roberto Amato, Jeffrey Barrett, Sonia Goncalves, Ewan Harrison, David K. Jackson, Ian Johnston, Dominic Kwiatkowski, Cordelia Langford, John Sillitoe on behalf of the Wellcome Sanger Institute COVID-19 Surveillance Team |
| EPI_ISL_3441299 | Lighthouse Lab in Milton Keynes | Wellcome Sanger Institute for the COVID-19 Genomics UK (COG-UK) Consortium | The Lighthouse Lab in Milton Keynes and Alex Alderton, Roberto Amato, Jeffrey Barrett, Sonia Goncalves, Ewan Harrison, David K. Jackson, Ian Johnston, Dominic Kwiatkowski, Cordelia Langford, John Sillitoe on behalf of the Wellcome Sanger Institute COVID-19 Surveillance Team |
| EPI_ISL_3441300 | Lighthouse Lab in Glasgow | Wellcome Sanger Institute for the COVID-19 Genomics UK (COG-UK) Consortium | Harper VanSteenhouse, Yumi Kasai, David Gray, Carol Clugston, Anna Dominiczak and Alex Alderton, Roberto Amato, Jeffrey Barrett, Sonia Goncalves, Ewan Harrison, David K. Jackson, Ian Johnston, Dominic Kwiatkowski, Cordelia Langford, John Sillitoe on behalf of the Wellcome Sanger Institute COVID-19 Surveillance Team |
| EPI_ISL_3441301 | Berkshire and Surrey Pathology Services Lighthouse Laboratory | Wellcome Sanger Institute for the COVID-19 Genomics UK (COG-UK) Consortium | Berkshire and Surrey Pathology Services Lighthouse Laboratory and Alex Alderton, Roberto Amato, Jeffrey Barrett, Sonia Goncalves, Ewan Harrison, David K. Jackson, Ian Johnston, Dominic Kwiatkowski, Cordelia Langford, John Sillitoe on behalf of the Wellcome Sanger Institute COVID-19 Surveillance Team |
| EPI_ISL_3441302 | Lighthouse Lab in Glasgow | Wellcome Sanger Institute for the COVID-19 Genomics UK (COG-UK) Consortium | Harper VanSteenhouse, Yumi Kasai, David Gray, Carol Clugston, Anna Dominiczak and Alex Alderton, Roberto Amato, Jeffrey Barrett, Sonia Goncalves, Ewan Harrison, David K. Jackson, Ian Johnston, Dominic Kwiatkowski, Cordelia Langford, John Sillitoe on behalf of the Wellcome Sanger Institute COVID-19 Surveillance Team |
| EPI_ISL_3441303 | Berkshire and Surrey Pathology Services Lighthouse Laboratory | Wellcome Sanger Institute for the COVID-19 Genomics UK (COG-UK) Consortium | Berkshire and Surrey Pathology Services Lighthouse Laboratory and Alex Alderton, Roberto Amato, Jeffrey Barrett, Sonia Goncalves, Ewan Harrison, David K. Jackson, Ian Johnston, Dominic Kwiatkowski, Cordelia Langford, John Sillitoe on behalf of the Wellcome Sanger Institute COVID-19 Surveillance Team |
| EPI_ISL_3441304 | Lighthouse Laboratory Plymouth | Wellcome Sanger Institute for the COVID-19 Genomics UK (COG-UK) Consortium | Lighthouse Laboratory Plymouth and Alex Alderton, Roberto Amato, Jeffrey Barrett, Sonia Goncalves, Ewan Harrison, David K. Jackson, Ian Johnston, Dominic Kwiatkowski, Cordelia Langford, John Sillitoe on behalf of the Wellcome Sanger Institute COVID-19 Surveillance Team |
| EPI_ISL_3441305 | Lighthouse Lab in Milton Keynes | Wellcome Sanger Institute for the COVID-19 Genomics UK (COG-UK) Consortium | The Lighthouse Lab in Milton Keynes and Alex Alderton, Roberto Amato, Jeffrey Barrett, Sonia Goncalves, Ewan Harrison, David K. Jackson, Ian Johnston, Dominic Kwiatkowski, Cordelia Langford, John Sillitoe on behalf of the Wellcome Sanger Institute COVID-19 Surveillance Team |
| EPI_ISL_3441306 | Lighthouse Lab in Glasgow | Wellcome Sanger Institute for the COVID-19 Genomics UK (COG-UK) Consortium | Harper VanSteenhouse, Yumi Kasai, David Gray, Carol Clugston, Anna Dominiczak and Alex Alderton, Roberto Amato, Jeffrey Barrett, Sonia Goncalves, Ewan Harrison, David K. Jackson, Ian Johnston, Dominic Kwiatkowski, Cordelia Langford, John Sillitoe on behalf of the Wellcome Sanger Institute COVID-19 Surveillance Team |
| EPI_ISL_3441307 | Lighthouse Laboratory Plymouth | Wellcome Sanger Institute for the COVID-19 Genomics UK (COG-UK) Consortium | Lighthouse Laboratory Plymouth and Alex Alderton, Roberto Amato, Jeffrey Barrett, Sonia Goncalves, Ewan Harrison, David K. Jackson, Ian Johnston, Dominic Kwiatkowski, Cordelia Langford, John Sillitoe on behalf of the Wellcome Sanger Institute COVID-19 Surveillance Team |
| EPI_ISL_3441308 | Lighthouse Lab in Milton Keynes | Wellcome Sanger Institute for the COVID-19 Genomics UK (COG-UK) Consortium | The Lighthouse Lab in Milton Keynes and Alex Alderton, Roberto Amato, Jeffrey Barrett, Sonia Goncalves, Ewan Harrison, David K. Jackson, Ian Johnston, Dominic Kwiatkowski, Cordelia Langford, John Sillitoe on behalf of the Wellcome Sanger Institute COVID-19 Surveillance Team |
| EPI_ISL_3441309, EPI_ISL_3441310 | Lighthouse Laboratory Plymouth | Wellcome Sanger Institute for the COVID-19 Genomics UK (COG-UK) Consortium | Lighthouse Laboratory Plymouth and Alex Alderton, Roberto Amato, Jeffrey Barrett, Sonia Goncalves, Ewan Harrison, David K. Jackson, Ian Johnston, Dominic Kwiatkowski, Cordelia Langford, John Sillitoe on behalf of the Wellcome Sanger Institute COVID-19 Surveillance Team |
| EPI_ISL_3441311 | Lighthouse Lab in Milton Keynes | Wellcome Sanger Institute for the COVID-19 Genomics UK (COG-UK) Consortium | The Lighthouse Lab in Milton Keynes and Alex Alderton, Roberto Amato, Jeffrey Barrett, Sonia Goncalves, Ewan Harrison, David K. Jackson, Ian Johnston, Dominic Kwiatkowski, Cordelia Langford, John Sillitoe on behalf of the Wellcome Sanger Institute COVID-19 Surveillance Team |
| EPI_ISL_3441312 | Lighthouse Lab in Glasgow | Wellcome Sanger Institute for the COVID-19 Genomics UK (COG-UK) Consortium | Harper VanSteenhouse, Yumi Kasai, David Gray, Carol Clugston, Anna Dominiczak and Alex Alderton, Roberto Amato, Jeffrey Barrett, Sonia Goncalves, Ewan Harrison, David K. Jackson, Ian Johnston, Dominic Kwiatkowski, Cordelia Langford, John Sillitoe on behalf of the Wellcome Sanger Institute COVID-19 Surveillance Team |
| EPI_ISL_3441313 | Lighthouse Lab in Milton Keynes | Wellcome Sanger Institute for the COVID-19 Genomics UK (COG-UK) Consortium | The Lighthouse Lab in Milton Keynes and Alex Alderton, Roberto Amato, Jeffrey Barrett, Sonia Goncalves, Ewan Harrison, David K. Jackson, Ian Johnston, Dominic Kwiatkowski, Cordelia Langford, John Sillitoe on behalf of the Wellcome Sanger Institute COVID-19 Surveillance Team |
| EPI_ISL_3441314 | Berkshire and Surrey Pathology Services Lighthouse Laboratory | Wellcome Sanger Institute for the COVID-19 Genomics UK (COG-UK) Consortium | Berkshire and Surrey Pathology Services Lighthouse Laboratory and Alex Alderton, Roberto Amato, Jeffrey Barrett, Sonia Goncalves, Ewan Harrison, David K. Jackson, Ian Johnston, Dominic Kwiatkowski, Cordelia Langford, John Sillitoe on behalf of the Wellcome Sanger Institute COVID-19 Surveillance Team |
| EPI_ISL_3441315 | Lighthouse Lab in Glasgow | Wellcome Sanger Institute for the COVID-19 Genomics UK (COG-UK) Consortium | Harper VanSteenhouse, Yumi Kasai, David Gray, Carol Clugston, Anna Dominiczak and Alex Alderton, Roberto Amato, Jeffrey Barrett, Sonia Goncalves, Ewan Harrison, David K. Jackson, Ian Johnston, Dominic Kwiatkowski, Cordelia Langford, John Sillitoe on behalf of the Wellcome Sanger Institute COVID-19 Surveillance Team |
| EPI_ISL_3441316 | Berkshire and Surrey Pathology Services Lighthouse Laboratory | Wellcome Sanger Institute for the COVID-19 Genomics UK (COG-UK) Consortium | Berkshire and Surrey Pathology Services Lighthouse Laboratory and Alex Alderton, Roberto Amato, Jeffrey Barrett, Sonia Goncalves, Ewan Harrison, David K. Jackson, Ian Johnston, Dominic Kwiatkowski, Cordelia Langford, John Sillitoe on behalf of the Wellcome Sanger Institute COVID-19 Surveillance Team |
| EPI_ISL_3441317 | Lighthouse Lab in Milton Keynes | Wellcome Sanger Institute for the COVID-19 Genomics UK (COG-UK) Consortium | The Lighthouse Lab in Milton Keynes and Alex Alderton, Roberto Amato, Jeffrey Barrett, Sonia Goncalves, Ewan Harrison, David K. Jackson, Ian Johnston, Dominic Kwiatkowski, Cordelia Langford, John Sillitoe on behalf of the Wellcome Sanger Institute COVID-19 Surveillance Team |
| EPI_ISL_3441318 | Berkshire and Surrey Pathology Services Lighthouse Laboratory | Wellcome Sanger Institute for the COVID-19 Genomics UK (COG-UK) Consortium | Berkshire and Surrey Pathology Services Lighthouse Laboratory and Alex Alderton, Roberto Amato, Jeffrey Barrett, Sonia Goncalves, Ewan Harrison, David K. Jackson, Ian Johnston, Dominic Kwiatkowski, Cordelia Langford, John Sillitoe on behalf of the Wellcome Sanger Institute COVID-19 Surveillance Team |
| EPI_ISL_3441319 | Lighthouse Lab in Glasgow | Wellcome Sanger Institute for the COVID-19 Genomics UK (COG-UK) Consortium | Harper VanSteenhouse, Yumi Kasai, David Gray, Carol Clugston, Anna Dominiczak and Alex Alderton, Roberto Amato, Jeffrey Barrett, Sonia Goncalves, Ewan Harrison, David K. Jackson, Ian Johnston, Dominic Kwiatkowski, Cordelia Langford, John Sillitoe on behalf of the Wellcome Sanger Institute COVID-19 Surveillance Team |
| EPI_ISL_3441320, EPI_ISL_3441321 | Berkshire and Surrey Pathology Services Lighthouse Laboratory | Wellcome Sanger Institute for the COVID-19 Genomics UK (COG-UK) Consortium | Berkshire and Surrey Pathology Services Lighthouse Laboratory and Alex Alderton, Roberto Amato, Jeffrey Barrett, Sonia Goncalves, Ewan Harrison, David K. Jackson, Ian Johnston, Dominic Kwiatkowski, Cordelia Langford, John Sillitoe on behalf of the Wellcome Sanger Institute COVID-19 Surveillance Team |

| | | | |
|---|---|---|---|
| EPI_ISL_3441322 | Lighthouse Lab in Glasgow | Wellcome Sanger Institute for the COVID-19 Genomics UK (COG-UK) Consortium | Harper VanSteenhouse, Yumi Kasai, David Gray, Carol Clugston, Anna Dominiczak and Alex Alderton, Roberto Amato, Jeffrey Barrett, Sonia Goncalves, Ewan Harrison, David K. Jackson, Ian Johnston, Dominic Kwiatkowski, Cordelia Langford, John Sillitoe on behalf of the Wellcome Sanger Institute COVID-19 Surveillance Team |
| EPI_ISL_3441323 | Lighthouse Laboratory Plymouth | Wellcome Sanger Institute for the COVID-19 Genomics UK (COG-UK) Consortium | Lighthouse Laboratory Plymouth and Alex Alderton, Roberto Amato, Jeffrey Barrett, Sonia Goncalves, Ewan Harrison, David K. Jackson, Ian Johnston, Dominic Kwiatkowski, Cordelia Langford, John Sillitoe on behalf of the Wellcome Sanger Institute COVID-19 Surveillance Team |
| EPI_ISL_3441324, EPI_ISL_3441325 | Lighthouse Lab in Glasgow | Wellcome Sanger Institute for the COVID-19 Genomics UK (COG-UK) Consortium | Harper VanSteenhouse, Yumi Kasai, David Gray, Carol Clugston, Anna Dominiczak and Alex Alderton, Roberto Amato, Jeffrey Barrett, Sonia Goncalves, Ewan Harrison, David K. Jackson, Ian Johnston, Dominic Kwiatkowski, Cordelia Langford, John Sillitoe on behalf of the Wellcome Sanger Institute COVID-19 Surveillance Team |
| EPI_ISL_3441326, EPI_ISL_3441327 | Lighthouse Laboratory Plymouth | Wellcome Sanger Institute for the COVID-19 Genomics UK (COG-UK) Consortium | Lighthouse Laboratory Plymouth and Alex Alderton, Roberto Amato, Jeffrey Barrett, Sonia Goncalves, Ewan Harrison, David K. Jackson, Ian Johnston, Dominic Kwiatkowski, Cordelia Langford, John Sillitoe on behalf of the Wellcome Sanger Institute COVID-19 Surveillance Team |
| EPI_ISL_3441328, EPI_ISL_3441329 | Berkshire and Surrey Pathology Services Lighthouse Laboratory | Wellcome Sanger Institute for the COVID-19 Genomics UK (COG-UK) Consortium | Berkshire and Surrey Pathology Services Lighthouse Laboratory and Alex Alderton, Roberto Amato, Jeffrey Barrett, Sonia Goncalves, Ewan Harrison, David K. Jackson, Ian Johnston, Dominic Kwiatkowski, Cordelia Langford, John Sillitoe on behalf of the Wellcome Sanger Institute COVID-19 Surveillance Team |
| EPI_ISL_3441330 | Lighthouse Laboratory Plymouth | Wellcome Sanger Institute for the COVID-19 Genomics UK (COG-UK) Consortium | Lighthouse Laboratory Plymouth and Alex Alderton, Roberto Amato, Jeffrey Barrett, Sonia Goncalves, Ewan Harrison, David K. Jackson, Ian Johnston, Dominic Kwiatkowski, Cordelia Langford, John Sillitoe on behalf of the Wellcome Sanger Institute COVID-19 Surveillance Team |
| EPI_ISL_3441331 | Lighthouse Lab in Glasgow | Wellcome Sanger Institute for the COVID-19 Genomics UK (COG-UK) Consortium | Harper VanSteenhouse, Yumi Kasai, David Gray, Carol Clugston, Anna Dominiczak and Alex Alderton, Roberto Amato, Jeffrey Barrett, Sonia Goncalves, Ewan Harrison, David K. Jackson, Ian Johnston, Dominic Kwiatkowski, Cordelia Langford, John Sillitoe on behalf of the Wellcome Sanger Institute COVID-19 Surveillance Team |
| EPI_ISL_3441332 | Lighthouse Laboratory Plymouth | Wellcome Sanger Institute for the COVID-19 Genomics UK (COG-UK) Consortium | Lighthouse Laboratory Plymouth and Alex Alderton, Roberto Amato, Jeffrey Barrett, Sonia Goncalves, Ewan Harrison, David K. Jackson, Ian Johnston, Dominic Kwiatkowski, Cordelia Langford, John Sillitoe on behalf of the Wellcome Sanger Institute COVID-19 Surveillance Team |
| EPI_ISL_3441333 | Berkshire and Surrey Pathology Services Lighthouse Laboratory | Wellcome Sanger Institute for the COVID-19 Genomics UK (COG-UK) Consortium | Berkshire and Surrey Pathology Services Lighthouse Laboratory and Alex Alderton, Roberto Amato, Jeffrey Barrett, Sonia Goncalves, Ewan Harrison, David K. Jackson, Ian Johnston, Dominic Kwiatkowski, Cordelia Langford, John Sillitoe on behalf of the Wellcome Sanger Institute COVID-19 Surveillance Team |
| EPI_ISL_3441334, EPI_ISL_3441335 | Lighthouse Laboratory Plymouth | Wellcome Sanger Institute for the COVID-19 Genomics UK (COG-UK) Consortium | Lighthouse Laboratory Plymouth and Alex Alderton, Roberto Amato, Jeffrey Barrett, Sonia Goncalves, Ewan Harrison, David K. Jackson, Ian Johnston, Dominic Kwiatkowski, Cordelia Langford, John Sillitoe on behalf of the Wellcome Sanger Institute COVID-19 Surveillance Team |
| EPI_ISL_3441336 | Berkshire and Surrey Pathology Services Lighthouse Laboratory | Wellcome Sanger Institute for the COVID-19 Genomics UK (COG-UK) Consortium | Berkshire and Surrey Pathology Services Lighthouse Laboratory and Alex Alderton, Roberto Amato, Jeffrey Barrett, Sonia Goncalves, Ewan Harrison, David K. Jackson, Ian Johnston, Dominic Kwiatkowski, Cordelia Langford, John Sillitoe on behalf of the Wellcome Sanger Institute COVID-19 Surveillance Team |
| EPI_ISL_3441337, EPI_ISL_3441338 | Lighthouse Lab in Glasgow | Wellcome Sanger Institute for the COVID-19 Genomics UK (COG-UK) Consortium | Harper VanSteenhouse, Yumi Kasai, David Gray, Carol Clugston, Anna Dominiczak and Alex Alderton, Roberto Amato, Jeffrey Barrett, Sonia Goncalves, Ewan Harrison, David K. Jackson, Ian Johnston, Dominic Kwiatkowski, Cordelia Langford, John Sillitoe on behalf of the Wellcome Sanger Institute COVID-19 Surveillance Team |
| EPI_ISL_3441339 | Berkshire and Surrey Pathology Services Lighthouse Laboratory | Wellcome Sanger Institute for the COVID-19 Genomics UK (COG-UK) Consortium | Berkshire and Surrey Pathology Services Lighthouse Laboratory and Alex Alderton, Roberto Amato, Jeffrey Barrett, Sonia Goncalves, Ewan Harrison, David K. Jackson, Ian Johnston, Dominic Kwiatkowski, Cordelia Langford, John Sillitoe on behalf of the Wellcome Sanger Institute COVID-19 Surveillance Team |
| EPI_ISL_3441340 | Lighthouse Lab in Glasgow | Wellcome Sanger Institute for the COVID-19 Genomics UK (COG-UK) Consortium | Harper VanSteenhouse, Yumi Kasai, David Gray, Carol Clugston, Anna Dominiczak and Alex Alderton, Roberto Amato, Jeffrey Barrett, Sonia Goncalves, Ewan Harrison, David K. Jackson, Ian Johnston, Dominic Kwiatkowski, Cordelia Langford, John Sillitoe on behalf of the Wellcome Sanger Institute COVID-19 Surveillance Team |
| EPI_ISL_3441341, EPI_ISL_3441342 | Berkshire and Surrey Pathology Services Lighthouse Laboratory | Wellcome Sanger Institute for the COVID-19 Genomics UK (COG-UK) Consortium | Berkshire and Surrey Pathology Services Lighthouse Laboratory and Alex Alderton, Roberto Amato, Jeffrey Barrett, Sonia Goncalves, Ewan Harrison, David K. Jackson, Ian Johnston, Dominic Kwiatkowski, Cordelia Langford, John Sillitoe on behalf of the Wellcome Sanger Institute COVID-19 Surveillance Team |
| EPI_ISL_3441343 | Lighthouse Lab in Milton Keynes | Wellcome Sanger Institute for the COVID-19 Genomics UK (COG-UK) Consortium | The Lighthouse Lab in Milton Keynes and Alex Alderton, Roberto Amato, Jeffrey Barrett, Sonia Goncalves, Ewan Harrison, David K. Jackson, Ian Johnston, Dominic Kwiatkowski, Cordelia Langford, John Sillitoe on behalf of the Wellcome Sanger Institute COVID-19 Surveillance Team |
| EPI_ISL_3441344 | Berkshire and Surrey Pathology Services Lighthouse Laboratory | Wellcome Sanger Institute for the COVID-19 Genomics UK (COG-UK) Consortium | Berkshire and Surrey Pathology Services Lighthouse Laboratory and Alex Alderton, Roberto Amato, Jeffrey Barrett, Sonia Goncalves, Ewan Harrison, David K. Jackson, Ian Johnston, Dominic Kwiatkowski, Cordelia Langford, John Sillitoe on behalf of the Wellcome Sanger Institute COVID-19 Surveillance Team |
| EPI_ISL_3441345 | Lighthouse Lab in Milton Keynes | Wellcome Sanger Institute for the COVID-19 Genomics UK (COG-UK) Consortium | The Lighthouse Lab in Milton Keynes and Alex Alderton, Roberto Amato, Jeffrey Barrett, Sonia Goncalves, Ewan Harrison, David K. Jackson, Ian Johnston, Dominic Kwiatkowski, Cordelia Langford, John Sillitoe on behalf of the Wellcome Sanger Institute COVID-19 Surveillance Team |
| EPI_ISL_3441346 | Lighthouse Lab in Glasgow | Wellcome Sanger Institute for the COVID-19 Genomics UK (COG-UK) Consortium | Harper VanSteenhouse, Yumi Kasai, David Gray, Carol Clugston, Anna Dominiczak and Alex Alderton, Roberto Amato, Jeffrey Barrett, Sonia Goncalves, Ewan Harrison, David K. Jackson, Ian Johnston, Dominic Kwiatkowski, Cordelia Langford, John Sillitoe on behalf of the Wellcome Sanger Institute COVID-19 Surveillance Team |
| EPI_ISL_3441347 | Berkshire and Surrey Pathology Services Lighthouse Laboratory | Wellcome Sanger Institute for the COVID-19 Genomics UK (COG-UK) Consortium | Berkshire and Surrey Pathology Services Lighthouse Laboratory and Alex Alderton, Roberto Amato, Jeffrey Barrett, Sonia Goncalves, Ewan Harrison, David K. Jackson, Ian Johnston, Dominic Kwiatkowski, Cordelia Langford, John Sillitoe on behalf of the Wellcome Sanger Institute COVID-19 Surveillance Team |
| EPI_ISL_3441348, EPI_ISL_3441349 | Lighthouse Lab in Milton Keynes | Wellcome Sanger Institute for the COVID-19 Genomics UK (COG-UK) Consortium | The Lighthouse Lab in Milton Keynes and Alex Alderton, Roberto Amato, Jeffrey Barrett, Sonia Goncalves, Ewan Harrison, David K. Jackson, Ian Johnston, Dominic Kwiatkowski, Cordelia Langford, John Sillitoe on behalf of the Wellcome Sanger Institute COVID-19 Surveillance Team |
| EPI_ISL_3441350 | Berkshire and Surrey Pathology Services Lighthouse Laboratory | Wellcome Sanger Institute for the COVID-19 Genomics UK (COG-UK) Consortium | Berkshire and Surrey Pathology Services Lighthouse Laboratory and Alex Alderton, Roberto Amato, Jeffrey Barrett, Sonia Goncalves, Ewan Harrison, David K. Jackson, Ian Johnston, Dominic Kwiatkowski, Cordelia Langford, John Sillitoe on behalf of the Wellcome Sanger Institute COVID-19 Surveillance Team |
| EPI_ISL_3441351 | Lighthouse Lab in Glasgow | Wellcome Sanger Institute for the COVID-19 Genomics UK (COG-UK) Consortium | Harper VanSteenhouse, Yumi Kasai, David Gray, Carol Clugston, Anna Dominiczak and Alex Alderton, Roberto Amato, Jeffrey Barrett, Sonia Goncalves, Ewan Harrison, David K. Jackson, Ian Johnston, Dominic Kwiatkowski, Cordelia Langford, John Sillitoe on behalf of the Wellcome Sanger Institute COVID-19 Surveillance Team |
| EPI_ISL_3441352 | Lighthouse Lab in Milton Keynes | Wellcome Sanger Institute for the COVID-19 Genomics UK (COG-UK) Consortium | The Lighthouse Lab in Milton Keynes and Alex Alderton, Roberto Amato, Jeffrey Barrett, Sonia Goncalves, Ewan Harrison, David K. Jackson, Ian Johnston, Dominic Kwiatkowski, Cordelia Langford, John Sillitoe on behalf of the Wellcome Sanger Institute COVID-19 Surveillance Team |
| EPI_ISL_3441353, EPI_ISL_3441354 | Berkshire and Surrey Pathology Services Lighthouse Laboratory | Wellcome Sanger Institute for the COVID-19 Genomics UK (COG-UK) Consortium | Berkshire and Surrey Pathology Services Lighthouse Laboratory and Alex Alderton, Roberto Amato, Jeffrey Barrett, Sonia Goncalves, Ewan Harrison, David K. Jackson, Ian Johnston, Dominic Kwiatkowski, Cordelia Langford, John Sillitoe on behalf of the Wellcome Sanger Institute COVID-19 Surveillance Team |
| EPI_ISL_3441355 | Lighthouse Lab in Glasgow | Wellcome Sanger Institute for the COVID-19 Genomics UK (COG-UK) Consortium | Harper VanSteenhouse, Yumi Kasai, David Gray, Carol Clugston, Anna Dominiczak and Alex Alderton, Roberto Amato, Jeffrey Barrett, Sonia Goncalves, Ewan Harrison, David K. Jackson, Ian Johnston, Dominic Kwiatkowski, Cordelia Langford, John Sillitoe on behalf of the Wellcome Sanger Institute COVID-19 Surveillance Team |
| EPI_ISL_3441356 | Berkshire and Surrey Pathology Services Lighthouse Laboratory | Wellcome Sanger Institute for the COVID-19 Genomics UK (COG-UK) Consortium | Berkshire and Surrey Pathology Services Lighthouse Laboratory and Alex Alderton, Roberto Amato, Jeffrey Barrett, Sonia Goncalves, Ewan Harrison, David K. Jackson, Ian Johnston, Dominic Kwiatkowski, Cordelia Langford, John Sillitoe on behalf of the Wellcome Sanger Institute COVID-19 Surveillance Team |
| EPI_ISL_3441357 | Lighthouse Lab in Glasgow | Wellcome Sanger Institute for the COVID-19 Genomics UK (COG-UK) Consortium | Harper VanSteenhouse, Yumi Kasai, David Gray, Carol Clugston, Anna Dominiczak and Alex Alderton, Roberto Amato, Jeffrey Barrett, Sonia Goncalves, Ewan Harrison, David K. Jackson, Ian Johnston, Dominic Kwiatkowski, Cordelia Langford, John Sillitoe on behalf of the Wellcome Sanger Institute COVID-19 Surveillance Team |
| EPI_ISL_3441358, EPI_ISL_3441359 | Lighthouse Laboratory Plymouth | Wellcome Sanger Institute for the COVID-19 Genomics UK (COG-UK) Consortium | Lighthouse Laboratory Plymouth and Alex Alderton, Roberto Amato, Jeffrey Barrett, Sonia Goncalves, Ewan Harrison, David K. Jackson, Ian Johnston, Dominic Kwiatkowski, Cordelia Langford, John Sillitoe on behalf of the Wellcome Sanger Institute COVID-19 Surveillance Team |
| EPI_ISL_3441360, EPI_ISL_3441361 | Lighthouse Lab in Milton Keynes | Wellcome Sanger Institute for the COVID-19 Genomics UK (COG-UK) Consortium | The Lighthouse Lab in Milton Keynes and Alex Alderton, Roberto Amato, Jeffrey Barrett, Sonia Goncalves, Ewan Harrison, David K. Jackson, Ian Johnston, Dominic Kwiatkowski, Cordelia Langford, John Sillitoe on behalf of the Wellcome Sanger Institute COVID-19 Surveillance Team |
| EPI_ISL_3441362 | Lighthouse Lab in Glasgow | Wellcome Sanger Institute for the COVID-19 Genomics UK (COG-UK) Consortium | Harper VanSteenhouse, Yumi Kasai, David Gray, Carol Clugston, Anna Dominiczak and Alex Alderton, Roberto Amato, Jeffrey Barrett, Sonia Goncalves, Ewan Harrison, David K. Jackson, Ian Johnston, Dominic Kwiatkowski, Cordelia Langford, John Sillitoe on behalf of the Wellcome Sanger Institute COVID-19 Surveillance Team |
| EPI_ISL_3441363, EPI_ISL_3441364, EPI_ISL_3441365 | Lighthouse Laboratory Plymouth | Wellcome Sanger Institute for the COVID-19 Genomics UK (COG-UK) Consortium | Lighthouse Laboratory Plymouth and Alex Alderton, Roberto Amato, Jeffrey Barrett, Sonia Goncalves, Ewan Harrison, David K. Jackson, Ian Johnston, Dominic Kwiatkowski, Cordelia Langford, John Sillitoe on behalf of the Wellcome Sanger Institute COVID-19 Surveillance Team |
| EPI_ISL_3441366 | Lighthouse Lab in Milton Keynes | Wellcome Sanger Institute for the COVID-19 Genomics UK (COG-UK) Consortium | The Lighthouse Lab in Milton Keynes and Alex Alderton, Roberto Amato, Jeffrey Barrett, Sonia Goncalves, Ewan Harrison, David K. Jackson, Ian Johnston, Dominic Kwiatkowski, Cordelia Langford, John Sillitoe on behalf of the Wellcome Sanger Institute COVID-19 Surveillance Team |
| EPI_ISL_3441367 | Lighthouse Laboratory Plymouth | Wellcome Sanger Institute for the COVID-19 Genomics UK (COG-UK) Consortium | Lighthouse Laboratory Plymouth and Alex Alderton, Roberto Amato, Jeffrey Barrett, Sonia Goncalves, Ewan Harrison, David K. Jackson, Ian Johnston, Dominic Kwiatkowski, Cordelia Langford, John Sillitoe on behalf of the Wellcome Sanger Institute COVID-19 Surveillance Team |
| EPI_ISL_3441368 | Berkshire and Surrey Pathology Services Lighthouse Laboratory | Wellcome Sanger Institute for the COVID-19 Genomics UK (COG-UK) Consortium | Berkshire and Surrey Pathology Services Lighthouse Laboratory and Alex Alderton, Roberto Amato, Jeffrey Barrett, Sonia Goncalves, Ewan Harrison, David K. Jackson, Ian Johnston, Dominic Kwiatkowski, Cordelia Langford, John Sillitoe on behalf of the Wellcome Sanger Institute COVID-19 Surveillance Team |
| EPI_ISL_3441369, EPI_ISL_3441370 | Lighthouse Laboratory Plymouth | Wellcome Sanger Institute for the COVID-19 Genomics UK (COG-UK) Consortium | Lighthouse Laboratory Plymouth and Alex Alderton, Roberto Amato, Jeffrey Barrett, Sonia Goncalves, Ewan Harrison, David K. Jackson, Ian Johnston, Dominic Kwiatkowski, Cordelia Langford, John Sillitoe on behalf of the Wellcome Sanger Institute COVID-19 Surveillance Team |
| EPI_ISL_3441371 | Lighthouse Lab in Milton Keynes | Wellcome Sanger Institute for the COVID-19 Genomics UK (COG-UK) Consortium | The Lighthouse Lab in Milton Keynes and Alex Alderton, Roberto Amato, Jeffrey Barrett, Sonia Goncalves, Ewan Harrison, David K. Jackson, Ian Johnston, Dominic Kwiatkowski, Cordelia Langford, John Sillitoe on behalf of the Wellcome Sanger Institute COVID-19 Surveillance Team |
| EPI_ISL_3441372 | Lighthouse Laboratory Plymouth | Wellcome Sanger Institute for the COVID-19 Genomics UK (COG-UK) Consortium | Lighthouse Laboratory Plymouth and Alex Alderton, Roberto Amato, Jeffrey Barrett, Sonia Goncalves, Ewan Harrison, David K. Jackson, Ian Johnston, Dominic Kwiatkowski, Cordelia Langford, John Sillitoe on behalf of the Wellcome Sanger Institute COVID-19 Surveillance Team |
| EPI_ISL_3441373 | Berkshire and Surrey Pathology Services Lighthouse Laboratory | Wellcome Sanger Institute for the COVID-19 Genomics UK (COG-UK) Consortium | Berkshire and Surrey Pathology Services Lighthouse Laboratory and Alex Alderton, Roberto Amato, Jeffrey Barrett, Sonia Goncalves, Ewan Harrison, David K. Jackson, Ian Johnston, Dominic Kwiatkowski, Cordelia Langford, John Sillitoe on behalf of the Wellcome Sanger Institute COVID-19 Surveillance Team |
| EPI_ISL_3441374 | Lighthouse Lab in Milton Keynes | Wellcome Sanger Institute for the COVID-19 Genomics UK (COG-UK) Consortium | The Lighthouse Lab in Milton Keynes and Alex Alderton, Roberto Amato, Jeffrey Barrett, Sonia Goncalves, Ewan Harrison, David K. Jackson, Ian Johnston, Dominic Kwiatkowski, Cordelia Langford, John Sillitoe on behalf of the Wellcome Sanger Institute COVID-19 Surveillance Team |
| EPI_ISL_3441375 | Lighthouse Laboratory Plymouth | Wellcome Sanger Institute for the COVID-19 Genomics UK (COG-UK) Consortium | Lighthouse Laboratory Plymouth and Alex Alderton, Roberto Amato, Jeffrey Barrett, Sonia Goncalves, Ewan Harrison, David K. Jackson, Ian Johnston, Dominic Kwiatkowski, Cordelia Langford, John Sillitoe on behalf of the Wellcome Sanger Institute COVID-19 Surveillance Team |
| EPI_ISL_3441376 | Lighthouse Lab in Milton Keynes | Wellcome Sanger Institute for the COVID-19 Genomics UK (COG-UK) Consortium | The Lighthouse Lab in Milton Keynes and Alex Alderton, Roberto Amato, Jeffrey Barrett, Sonia Goncalves, Ewan Harrison, David K. Jackson, Ian Johnston, Dominic Kwiatkowski, Cordelia Langford, John Sillitoe on behalf of the Wellcome Sanger Institute COVID-19 Surveillance Team |
| EPI_ISL_3441377, EPI_ISL_3441378, EPI_ISL_3441379 | Lighthouse Laboratory Plymouth | Wellcome Sanger Institute for the COVID-19 Genomics UK (COG-UK) Consortium | Lighthouse Laboratory Plymouth and Alex Alderton, Roberto Amato, Jeffrey Barrett, Sonia Goncalves, Ewan Harrison, David K. Jackson, Ian Johnston, Dominic Kwiatkowski, Cordelia Langford, John Sillitoe on behalf of the Wellcome Sanger Institute COVID-19 Surveillance Team |
| EPI_ISL_3441380 | Lighthouse Lab in Milton Keynes | Wellcome Sanger Institute for the COVID-19 Genomics UK (COG-UK) Consortium | The Lighthouse Lab in Milton Keynes and Alex Alderton, Roberto Amato, Jeffrey Barrett, Sonia Goncalves, Ewan Harrison, David K. Jackson, Ian Johnston, Dominic Kwiatkowski, Cordelia Langford, John Sillitoe on behalf of the Wellcome Sanger Institute COVID-19 Surveillance Team |

| | | | |
|---|---|---|---|
| EPI_ISL_3441381 | Lighthouse Laboratory Plymouth | Wellcome Sanger Institute for the COVID-19 Genomics UK (COG-UK) Consortium | Lighthouse Laboratory Plymouth and Alex Alderton, Roberto Amato, Jeffrey Barrett, Sonia Goncalves, Ewan Harrison, David K. Jackson, Ian Johnston, Dominic Kwiatkowski, Cordelia Langford, John Sillitoe on behalf of the Wellcome Sanger Institute COVID-19 Surveillance Team |
| EPI_ISL_3441382, EPI_ISL_3441383, EPI_ISL_3441384 | Berkshire and Surrey Pathology Services Lighthouse Laboratory | Wellcome Sanger Institute for the COVID-19 Genomics UK (COG-UK) Consortium | Berkshire and Surrey Pathology Services Lighthouse Laboratory and Alex Alderton, Roberto Amato, Jeffrey Barrett, Sonia Goncalves, Ewan Harrison, David K. Jackson, Ian Johnston, Dominic Kwiatkowski, Cordelia Langford, John Sillitoe on behalf of the Wellcome Sanger Institute COVID-19 Surveillance Team |
| EPI_ISL_3441385 | Lighthouse Lab in Milton Keynes | Wellcome Sanger Institute for the COVID-19 Genomics UK (COG-UK) Consortium | The Lighthouse Lab in Milton Keynes and Alex Alderton, Roberto Amato, Jeffrey Barrett, Sonia Goncalves, Ewan Harrison, David K. Jackson, Ian Johnston, Dominic Kwiatkowski, Cordelia Langford, John Sillitoe on behalf of the Wellcome Sanger Institute COVID-19 Surveillance Team |
| EPI_ISL_3441386 | Lighthouse Laboratory Plymouth | Wellcome Sanger Institute for the COVID-19 Genomics UK (COG-UK) Consortium | Lighthouse Laboratory Plymouth and Alex Alderton, Roberto Amato, Jeffrey Barrett, Sonia Goncalves, Ewan Harrison, David K. Jackson, Ian Johnston, Dominic Kwiatkowski, Cordelia Langford, John Sillitoe on behalf of the Wellcome Sanger Institute COVID-19 Surveillance Team |
| EPI_ISL_3441387 | Berkshire and Surrey Pathology Services Lighthouse Laboratory | Wellcome Sanger Institute for the COVID-19 Genomics UK (COG-UK) Consortium | Berkshire and Surrey Pathology Services Lighthouse Laboratory and Alex Alderton, Roberto Amato, Jeffrey Barrett, Sonia Goncalves, Ewan Harrison, David K. Jackson, Ian Johnston, Dominic Kwiatkowski, Cordelia Langford, John Sillitoe on behalf of the Wellcome Sanger Institute COVID-19 Surveillance Team |
| EPI_ISL_3441388 | Lighthouse Lab in Glasgow | Wellcome Sanger Institute for the COVID-19 Genomics UK (COG-UK) Consortium | Harper VanSteenhouse, Yumi Kasai, David Gray, Carol Clugston, Anna Dominiczak and Alex Alderton, Roberto Amato, Jeffrey Barrett, Sonia Goncalves, Ewan Harrison, David K. Jackson, Ian Johnston, Dominic Kwiatkowski, Cordelia Langford, John Sillitoe on behalf of the Wellcome Sanger Institute COVID-19 Surveillance Team |
| EPI_ISL_3441389 | Lighthouse Lab in Milton Keynes | Wellcome Sanger Institute for the COVID-19 Genomics UK (COG-UK) Consortium | The Lighthouse Lab in Milton Keynes and Alex Alderton, Roberto Amato, Jeffrey Barrett, Sonia Goncalves, Ewan Harrison, David K. Jackson, Ian Johnston, Dominic Kwiatkowski, Cordelia Langford, John Sillitoe on behalf of the Wellcome Sanger Institute COVID-19 Surveillance Team |
| EPI_ISL_3441390 | Lighthouse Lab in Glasgow | Wellcome Sanger Institute for the COVID-19 Genomics UK (COG-UK) Consortium | Harper VanSteenhouse, Yumi Kasai, David Gray, Carol Clugston, Anna Dominiczak and Alex Alderton, Roberto Amato, Jeffrey Barrett, Sonia Goncalves, Ewan Harrison, David K. Jackson, Ian Johnston, Dominic Kwiatkowski, Cordelia Langford, John Sillitoe on behalf of the Wellcome Sanger Institute COVID-19 Surveillance Team |
| EPI_ISL_3441391, EPI_ISL_3441392, EPI_ISL_3441393 | Lighthouse Laboratory Plymouth | Wellcome Sanger Institute for the COVID-19 Genomics UK (COG-UK) Consortium | Lighthouse Laboratory Plymouth and Alex Alderton, Roberto Amato, Jeffrey Barrett, Sonia Goncalves, Ewan Harrison, David K. Jackson, Ian Johnston, Dominic Kwiatkowski, Cordelia Langford, John Sillitoe on behalf of the Wellcome Sanger Institute COVID-19 Surveillance Team |
| EPI_ISL_3441394 | Lighthouse Lab in Glasgow | Wellcome Sanger Institute for the COVID-19 Genomics UK (COG-UK) Consortium | Harper VanSteenhouse, Yumi Kasai, David Gray, Carol Clugston, Anna Dominiczak and Alex Alderton, Roberto Amato, Jeffrey Barrett, Sonia Goncalves, Ewan Harrison, David K. Jackson, Ian Johnston, Dominic Kwiatkowski, Cordelia Langford, John Sillitoe on behalf of the Wellcome Sanger Institute COVID-19 Surveillance Team |
| EPI_ISL_3441395 | Berkshire and Surrey Pathology Services Lighthouse Laboratory | Wellcome Sanger Institute for the COVID-19 Genomics UK (COG-UK) Consortium | Berkshire and Surrey Pathology Services Lighthouse Laboratory and Alex Alderton, Roberto Amato, Jeffrey Barrett, Sonia Goncalves, Ewan Harrison, David K. Jackson, Ian Johnston, Dominic Kwiatkowski, Cordelia Langford, John Sillitoe on behalf of the Wellcome Sanger Institute COVID-19 Surveillance Team |
| EPI_ISL_3441396, EPI_ISL_3441397 | Lighthouse Lab in Milton Keynes | Wellcome Sanger Institute for the COVID-19 Genomics UK (COG-UK) Consortium | The Lighthouse Lab in Milton Keynes and Alex Alderton, Roberto Amato, Jeffrey Barrett, Sonia Goncalves, Ewan Harrison, David K. Jackson, Ian Johnston, Dominic Kwiatkowski, Cordelia Langford, John Sillitoe on behalf of the Wellcome Sanger Institute COVID-19 Surveillance Team |
| EPI_ISL_3441398, EPI_ISL_3441399 | Berkshire and Surrey Pathology Services Lighthouse Laboratory | Wellcome Sanger Institute for the COVID-19 Genomics UK (COG-UK) Consortium | Berkshire and Surrey Pathology Services Lighthouse Laboratory and Alex Alderton, Roberto Amato, Jeffrey Barrett, Sonia Goncalves, Ewan Harrison, David K. Jackson, Ian Johnston, Dominic Kwiatkowski, Cordelia Langford, John Sillitoe on behalf of the Wellcome Sanger Institute COVID-19 Surveillance Team |
| EPI_ISL_3441400 | Lighthouse Lab in Glasgow | Wellcome Sanger Institute for the COVID-19 Genomics UK (COG-UK) Consortium | Harper VanSteenhouse, Yumi Kasai, David Gray, Carol Clugston, Anna Dominiczak and Alex Alderton, Roberto Amato, Jeffrey Barrett, Sonia Goncalves, Ewan Harrison, David K. Jackson, Ian Johnston, Dominic Kwiatkowski, Cordelia Langford, John Sillitoe on behalf of the Wellcome Sanger Institute COVID-19 Surveillance Team |
| EPI_ISL_3441401 | Lighthouse Lab in Milton Keynes | Wellcome Sanger Institute for the COVID-19 Genomics UK (COG-UK) Consortium | The Lighthouse Lab in Milton Keynes and Alex Alderton, Roberto Amato, Jeffrey Barrett, Sonia Goncalves, Ewan Harrison, David K. Jackson, Ian Johnston, Dominic Kwiatkowski, Cordelia Langford, John Sillitoe on behalf of the Wellcome Sanger Institute COVID-19 Surveillance Team |
| EPI_ISL_3441402 | Lighthouse Laboratory Plymouth | Wellcome Sanger Institute for the COVID-19 Genomics UK (COG-UK) Consortium | Lighthouse Laboratory Plymouth and Alex Alderton, Roberto Amato, Jeffrey Barrett, Sonia Goncalves, Ewan Harrison, David K. Jackson, Ian Johnston, Dominic Kwiatkowski, Cordelia Langford, John Sillitoe on behalf of the Wellcome Sanger Institute COVID-19 Surveillance Team |
| EPI_ISL_3441403 | Berkshire and Surrey Pathology Services Lighthouse Laboratory | Wellcome Sanger Institute for the COVID-19 Genomics UK (COG-UK) Consortium | Berkshire and Surrey Pathology Services Lighthouse Laboratory and Alex Alderton, Roberto Amato, Jeffrey Barrett, Sonia Goncalves, Ewan Harrison, David K. Jackson, Ian Johnston, Dominic Kwiatkowski, Cordelia Langford, John Sillitoe on behalf of the Wellcome Sanger Institute COVID-19 Surveillance Team |
| EPI_ISL_3441404 | Lighthouse Laboratory Plymouth | Wellcome Sanger Institute for the COVID-19 Genomics UK (COG-UK) Consortium | Lighthouse Laboratory Plymouth and Alex Alderton, Roberto Amato, Jeffrey Barrett, Sonia Goncalves, Ewan Harrison, David K. Jackson, Ian Johnston, Dominic Kwiatkowski, Cordelia Langford, John Sillitoe on behalf of the Wellcome Sanger Institute COVID-19 Surveillance Team |
| EPI_ISL_3441405 | Berkshire and Surrey Pathology Services Lighthouse Laboratory | Wellcome Sanger Institute for the COVID-19 Genomics UK (COG-UK) Consortium | Berkshire and Surrey Pathology Services Lighthouse Laboratory and Alex Alderton, Roberto Amato, Jeffrey Barrett, Sonia Goncalves, Ewan Harrison, David K. Jackson, Ian Johnston, Dominic Kwiatkowski, Cordelia Langford, John Sillitoe on behalf of the Wellcome Sanger Institute COVID-19 Surveillance Team |
| EPI_ISL_3441406 | Lighthouse Lab in Milton Keynes | Wellcome Sanger Institute for the COVID-19 Genomics UK (COG-UK) Consortium | The Lighthouse Lab in Milton Keynes and Alex Alderton, Roberto Amato, Jeffrey Barrett, Sonia Goncalves, Ewan Harrison, David K. Jackson, Ian Johnston, Dominic Kwiatkowski, Cordelia Langford, John Sillitoe on behalf of the Wellcome Sanger Institute COVID-19 Surveillance Team |
| EPI_ISL_3441407 | Berkshire and Surrey Pathology Services Lighthouse Laboratory | Wellcome Sanger Institute for the COVID-19 Genomics UK (COG-UK) Consortium | Berkshire and Surrey Pathology Services Lighthouse Laboratory and Alex Alderton, Roberto Amato, Jeffrey Barrett, Sonia Goncalves, Ewan Harrison, David K. Jackson, Ian Johnston, Dominic Kwiatkowski, Cordelia Langford, John Sillitoe on behalf of the Wellcome Sanger Institute COVID-19 Surveillance Team |
| EPI_ISL_3441408 | Lighthouse Lab in Glasgow | Wellcome Sanger Institute for the COVID-19 Genomics UK (COG-UK) Consortium | Harper VanSteenhouse, Yumi Kasai, David Gray, Carol Clugston, Anna Dominiczak and Alex Alderton, Roberto Amato, Jeffrey Barrett, Sonia Goncalves, Ewan Harrison, David K. Jackson, Ian Johnston, Dominic Kwiatkowski, Cordelia Langford, John Sillitoe on behalf of the Wellcome Sanger Institute COVID-19 Surveillance Team |
| EPI_ISL_3441409, EPI_ISL_3441410 | Lighthouse Lab in Milton Keynes | Wellcome Sanger Institute for the COVID-19 Genomics UK (COG-UK) Consortium | The Lighthouse Lab in Milton Keynes and Alex Alderton, Roberto Amato, Jeffrey Barrett, Sonia Goncalves, Ewan Harrison, David K. Jackson, Ian Johnston, Dominic Kwiatkowski, Cordelia Langford, John Sillitoe on behalf of the Wellcome Sanger Institute COVID-19 Surveillance Team |
| EPI_ISL_3441411, EPI_ISL_3441412 | Lighthouse Laboratory Plymouth | Wellcome Sanger Institute for the COVID-19 Genomics UK (COG-UK) Consortium | Lighthouse Laboratory Plymouth and Alex Alderton, Roberto Amato, Jeffrey Barrett, Sonia Goncalves, Ewan Harrison, David K. Jackson, Ian Johnston, Dominic Kwiatkowski, Cordelia Langford, John Sillitoe on behalf of the Wellcome Sanger Institute COVID-19 Surveillance Team |
| EPI_ISL_3441413, EPI_ISL_3441414 | Lighthouse Lab in Milton Keynes | Wellcome Sanger Institute for the COVID-19 Genomics UK (COG-UK) Consortium | The Lighthouse Lab in Milton Keynes and Alex Alderton, Roberto Amato, Jeffrey Barrett, Sonia Goncalves, Ewan Harrison, David K. Jackson, Ian Johnston, Dominic Kwiatkowski, Cordelia Langford, John Sillitoe on behalf of the Wellcome Sanger Institute COVID-19 Surveillance Team |
| EPI_ISL_3441415, EPI_ISL_3441416, EPI_ISL_3441417 | Lighthouse Lab in Glasgow | Wellcome Sanger Institute for the COVID-19 Genomics UK (COG-UK) Consortium | Harper VanSteenhouse, Yumi Kasai, David Gray, Carol Clugston, Anna Dominiczak and Alex Alderton, Roberto Amato, Jeffrey Barrett, Sonia Goncalves, Ewan Harrison, David K. Jackson, Ian Johnston, Dominic Kwiatkowski, Cordelia Langford, John Sillitoe on behalf of the Wellcome Sanger Institute COVID-19 Surveillance Team |
| EPI_ISL_3441418, EPI_ISL_3441419 | Berkshire and Surrey Pathology Services Lighthouse Laboratory | Wellcome Sanger Institute for the COVID-19 Genomics UK (COG-UK) Consortium | Berkshire and Surrey Pathology Services Lighthouse Laboratory and Alex Alderton, Roberto Amato, Jeffrey Barrett, Sonia Goncalves, Ewan Harrison, David K. Jackson, Ian Johnston, Dominic Kwiatkowski, Cordelia Langford, John Sillitoe on behalf of the Wellcome Sanger Institute COVID-19 Surveillance Team |
| EPI_ISL_3441420 | Lighthouse Lab in Glasgow | Wellcome Sanger Institute for the COVID-19 Genomics UK (COG-UK) Consortium | Harper VanSteenhouse, Yumi Kasai, David Gray, Carol Clugston, Anna Dominiczak and Alex Alderton, Roberto Amato, Jeffrey Barrett, Sonia Goncalves, Ewan Harrison, David K. Jackson, Ian Johnston, Dominic Kwiatkowski, Cordelia Langford, John Sillitoe on behalf of the Wellcome Sanger Institute COVID-19 Surveillance Team |
| EPI_ISL_3441421, EPI_ISL_3441422 | Berkshire and Surrey Pathology Services Lighthouse Laboratory | Wellcome Sanger Institute for the COVID-19 Genomics UK (COG-UK) Consortium | Berkshire and Surrey Pathology Services Lighthouse Laboratory and Alex Alderton, Roberto Amato, Jeffrey Barrett, Sonia Goncalves, Ewan Harrison, David K. Jackson, Ian Johnston, Dominic Kwiatkowski, Cordelia Langford, John Sillitoe on behalf of the Wellcome Sanger Institute COVID-19 Surveillance Team |
| EPI_ISL_3441423 | Lighthouse Lab in Glasgow | Wellcome Sanger Institute for the COVID-19 Genomics UK (COG-UK) Consortium | Harper VanSteenhouse, Yumi Kasai, David Gray, Carol Clugston, Anna Dominiczak and Alex Alderton, Roberto Amato, Jeffrey Barrett, Sonia Goncalves, Ewan Harrison, David K. Jackson, Ian Johnston, Dominic Kwiatkowski, Cordelia Langford, John Sillitoe on behalf of the Wellcome Sanger Institute COVID-19 Surveillance Team |
| EPI_ISL_3441424 | Lighthouse Lab in Milton Keynes | Wellcome Sanger Institute for the COVID-19 Genomics UK (COG-UK) Consortium | The Lighthouse Lab in Milton Keynes and Alex Alderton, Roberto Amato, Jeffrey Barrett, Sonia Goncalves, Ewan Harrison, David K. Jackson, Ian Johnston, Dominic Kwiatkowski, Cordelia Langford, John Sillitoe on behalf of the Wellcome Sanger Institute COVID-19 Surveillance Team |
| EPI_ISL_3441425 | Lighthouse Laboratory Plymouth | Wellcome Sanger Institute for the COVID-19 Genomics UK (COG-UK) Consortium | Lighthouse Laboratory Plymouth and Alex Alderton, Roberto Amato, Jeffrey Barrett, Sonia Goncalves, Ewan Harrison, David K. Jackson, Ian Johnston, Dominic Kwiatkowski, Cordelia Langford, John Sillitoe on behalf of the Wellcome Sanger Institute COVID-19 Surveillance Team |
| EPI_ISL_3441426 | Lighthouse Lab in Glasgow | Wellcome Sanger Institute for the COVID-19 Genomics UK (COG-UK) Consortium | Harper VanSteenhouse, Yumi Kasai, David Gray, Carol Clugston, Anna Dominiczak and Alex Alderton, Roberto Amato, Jeffrey Barrett, Sonia Goncalves, Ewan Harrison, David K. Jackson, Ian Johnston, Dominic Kwiatkowski, Cordelia Langford, John Sillitoe on behalf of the Wellcome Sanger Institute COVID-19 Surveillance Team |
| EPI_ISL_3441427 | Lighthouse Lab in Milton Keynes | Wellcome Sanger Institute for the COVID-19 Genomics UK (COG-UK) Consortium | The Lighthouse Lab in Milton Keynes and Alex Alderton, Roberto Amato, Jeffrey Barrett, Sonia Goncalves, Ewan Harrison, David K. Jackson, Ian Johnston, Dominic Kwiatkowski, Cordelia Langford, John Sillitoe on behalf of the Wellcome Sanger Institute COVID-19 Surveillance Team |
| EPI_ISL_3441428 | Lighthouse Lab in Glasgow | Wellcome Sanger Institute for the COVID-19 Genomics UK (COG-UK) Consortium | Harper VanSteenhouse, Yumi Kasai, David Gray, Carol Clugston, Anna Dominiczak and Alex Alderton, Roberto Amato, Jeffrey Barrett, Sonia Goncalves, Ewan Harrison, David K. Jackson, Ian Johnston, Dominic Kwiatkowski, Cordelia Langford, John Sillitoe on behalf of the Wellcome Sanger Institute COVID-19 Surveillance Team |
| EPI_ISL_3441429 | Berkshire and Surrey Pathology Services Lighthouse Laboratory | Wellcome Sanger Institute for the COVID-19 Genomics UK (COG-UK) Consortium | Berkshire and Surrey Pathology Services Lighthouse Laboratory and Alex Alderton, Roberto Amato, Jeffrey Barrett, Sonia Goncalves, Ewan Harrison, David K. Jackson, Ian Johnston, Dominic Kwiatkowski, Cordelia Langford, John Sillitoe on behalf of the Wellcome Sanger Institute COVID-19 Surveillance Team |
| EPI_ISL_3441430 | Lighthouse Lab in Glasgow | Wellcome Sanger Institute for the COVID-19 Genomics UK (COG-UK) Consortium | Harper VanSteenhouse, Yumi Kasai, David Gray, Carol Clugston, Anna Dominiczak and Alex Alderton, Roberto Amato, Jeffrey Barrett, Sonia Goncalves, Ewan Harrison, David K. Jackson, Ian Johnston, Dominic Kwiatkowski, Cordelia Langford, John Sillitoe on behalf of the Wellcome Sanger Institute COVID-19 Surveillance Team |
| EPI_ISL_3441431, EPI_ISL_3441432 | Berkshire and Surrey Pathology Services Lighthouse Laboratory | Wellcome Sanger Institute for the COVID-19 Genomics UK (COG-UK) Consortium | Berkshire and Surrey Pathology Services Lighthouse Laboratory and Alex Alderton, Roberto Amato, Jeffrey Barrett, Sonia Goncalves, Ewan Harrison, David K. Jackson, Ian Johnston, Dominic Kwiatkowski, Cordelia Langford, John Sillitoe on behalf of the Wellcome Sanger Institute COVID-19 Surveillance Team |
| EPI_ISL_3441433 | Lighthouse Lab in Glasgow | Wellcome Sanger Institute for the COVID-19 Genomics UK (COG-UK) Consortium | Harper VanSteenhouse, Yumi Kasai, David Gray, Carol Clugston, Anna Dominiczak and Alex Alderton, Roberto Amato, Jeffrey Barrett, Sonia Goncalves, Ewan Harrison, David K. Jackson, Ian Johnston, Dominic Kwiatkowski, Cordelia Langford, John Sillitoe on behalf of the Wellcome Sanger Institute COVID-19 Surveillance Team |
| EPI_ISL_3441434, EPI_ISL_3441435 | Lighthouse Lab in Milton Keynes | Wellcome Sanger Institute for the COVID-19 Genomics UK (COG-UK) Consortium | The Lighthouse Lab in Milton Keynes and Alex Alderton, Roberto Amato, Jeffrey Barrett, Sonia Goncalves, Ewan Harrison, David K. Jackson, Ian Johnston, Dominic Kwiatkowski, Cordelia Langford, John Sillitoe on behalf of the Wellcome Sanger Institute COVID-19 Surveillance Team |
| EPI_ISL_3441436 | Lighthouse Lab in Glasgow | Wellcome Sanger Institute for the COVID-19 Genomics UK (COG-UK) Consortium | Harper VanSteenhouse, Yumi Kasai, David Gray, Carol Clugston, Anna Dominiczak and Alex Alderton, Roberto Amato, Jeffrey Barrett, Sonia Goncalves, Ewan Harrison, David K. Jackson, Ian Johnston, Dominic Kwiatkowski, Cordelia Langford, John Sillitoe on behalf of the Wellcome Sanger Institute COVID-19 Surveillance Team |
| EPI_ISL_3441437, EPI_ISL_3441438 | Lighthouse Laboratory Plymouth | Wellcome Sanger Institute for the COVID-19 Genomics UK (COG-UK) Consortium | Lighthouse Laboratory Plymouth and Alex Alderton, Roberto Amato, Jeffrey Barrett, Sonia Goncalves, Ewan Harrison, David K. Jackson, Ian Johnston, Dominic Kwiatkowski, Cordelia Langford, John Sillitoe on behalf of the Wellcome Sanger Institute COVID-19 Surveillance Team |
| EPI_ISL_3441439, EPI_ISL_3441440 | Lighthouse Lab in Glasgow | Wellcome Sanger Institute for the COVID-19 Genomics UK (COG-UK) Consortium | Harper VanSteenhouse, Yumi Kasai, David Gray, Carol Clugston, Anna Dominiczak and Alex Alderton, Roberto Amato, Jeffrey Barrett, Sonia Goncalves, Ewan Harrison, David K. Jackson, Ian Johnston, Dominic Kwiatkowski, Cordelia Langford, John Sillitoe on behalf of the Wellcome Sanger Institute COVID-19 Surveillance Team |
| EPI_ISL_3441441 | Berkshire and Surrey Pathology Services Lighthouse Laboratory | Wellcome Sanger Institute for the COVID-19 Genomics UK (COG-UK) Consortium | Berkshire and Surrey Pathology Services Lighthouse Laboratory and Alex Alderton, Roberto Amato, Jeffrey Barrett, Sonia Goncalves, Ewan Harrison, David K. Jackson, Ian Johnston, Dominic Kwiatkowski, Cordelia Langford, John Sillitoe on behalf of the Wellcome Sanger Institute COVID-19 Surveillance Team |

| | | | |
|---|---|---|---|
| EPI_ISL_3441442 | Lighthouse Lab in Milton Keynes | Wellcome Sanger Institute for the COVID-19 Genomics UK (COG-UK) Consortium | The Lighthouse Lab in Milton Keynes and Alex Alderton, Roberto Amato, Jeffrey Barrett, Sonia Goncalves, Ewan Harrison, David K. Jackson, Ian Johnston, Dominic Kwiatkowski, Cordelia Langford, John Sillitoe on behalf of the Wellcome Sanger Institute COVID-19 Surveillance Team |
| EPI_ISL_3441443 | Lighthouse Lab in Glasgow | Wellcome Sanger Institute for the COVID-19 Genomics UK (COG-UK) Consortium | Harper VanSteenhouse, Yumi Kasai, David Gray, Carol Clugston, Anna Dominiczak and Alex Alderton, Roberto Amato, Jeffrey Barrett, Sonia Goncalves, Ewan Harrison, David K. Jackson, Ian Johnston, Dominic Kwiatkowski, Cordelia Langford, John Sillitoe on behalf of the Wellcome Sanger Institute COVID-19 Surveillance Team |
| EPI_ISL_3441444, EPI_ISL_3441445 | Lighthouse Laboratory Plymouth | Wellcome Sanger Institute for the COVID-19 Genomics UK (COG-UK) Consortium | Lighthouse Laboratory Plymouth and Alex Alderton, Roberto Amato, Jeffrey Barrett, Sonia Goncalves, Ewan Harrison, David K. Jackson, Ian Johnston, Dominic Kwiatkowski, Cordelia Langford, John Sillitoe on behalf of the Wellcome Sanger Institute COVID-19 Surveillance Team |
| EPI_ISL_3441446 | Lighthouse Lab in Glasgow | Wellcome Sanger Institute for the COVID-19 Genomics UK (COG-UK) Consortium | Harper VanSteenhouse, Yumi Kasai, David Gray, Carol Clugston, Anna Dominiczak and Alex Alderton, Roberto Amato, Jeffrey Barrett, Sonia Goncalves, Ewan Harrison, David K. Jackson, Ian Johnston, Dominic Kwiatkowski, Cordelia Langford, John Sillitoe on behalf of the Wellcome Sanger Institute COVID-19 Surveillance Team |
| EPI_ISL_3441447 | Lighthouse Laboratory Plymouth | Wellcome Sanger Institute for the COVID-19 Genomics UK (COG-UK) Consortium | Lighthouse Laboratory Plymouth and Alex Alderton, Roberto Amato, Jeffrey Barrett, Sonia Goncalves, Ewan Harrison, David K. Jackson, Ian Johnston, Dominic Kwiatkowski, Cordelia Langford, John Sillitoe on behalf of the Wellcome Sanger Institute COVID-19 Surveillance Team |
| EPI_ISL_3441448 | Lighthouse Lab in Milton Keynes | Wellcome Sanger Institute for the COVID-19 Genomics UK (COG-UK) Consortium | The Lighthouse Lab in Milton Keynes and Alex Alderton, Roberto Amato, Jeffrey Barrett, Sonia Goncalves, Ewan Harrison, David K. Jackson, Ian Johnston, Dominic Kwiatkowski, Cordelia Langford, John Sillitoe on behalf of the Wellcome Sanger Institute COVID-19 Surveillance Team |
| EPI_ISL_3441449 | Berkshire and Surrey Pathology Services Lighthouse Laboratory | Wellcome Sanger Institute for the COVID-19 Genomics UK (COG-UK) Consortium | Berkshire and Surrey Pathology Services Lighthouse Laboratory and Alex Alderton, Roberto Amato, Jeffrey Barrett, Sonia Goncalves, Ewan Harrison, David K. Jackson, Ian Johnston, Dominic Kwiatkowski, Cordelia Langford, John Sillitoe on behalf of the Wellcome Sanger Institute COVID-19 Surveillance Team |
| EPI_ISL_3441450 | Lighthouse Laboratory Plymouth | Wellcome Sanger Institute for the COVID-19 Genomics UK (COG-UK) Consortium | Lighthouse Laboratory Plymouth and Alex Alderton, Roberto Amato, Jeffrey Barrett, Sonia Goncalves, Ewan Harrison, David K. Jackson, Ian Johnston, Dominic Kwiatkowski, Cordelia Langford, John Sillitoe on behalf of the Wellcome Sanger Institute COVID-19 Surveillance Team |
| EPI_ISL_3441451 | Lighthouse Lab in Glasgow | Wellcome Sanger Institute for the COVID-19 Genomics UK (COG-UK) Consortium | Harper VanSteenhouse, Yumi Kasai, David Gray, Carol Clugston, Anna Dominiczak and Alex Alderton, Roberto Amato, Jeffrey Barrett, Sonia Goncalves, Ewan Harrison, David K. Jackson, Ian Johnston, Dominic Kwiatkowski, Cordelia Langford, John Sillitoe on behalf of the Wellcome Sanger Institute COVID-19 Surveillance Team |
| EPI_ISL_3441452 | Berkshire and Surrey Pathology Services Lighthouse Laboratory | Wellcome Sanger Institute for the COVID-19 Genomics UK (COG-UK) Consortium | Berkshire and Surrey Pathology Services Lighthouse Laboratory and Alex Alderton, Roberto Amato, Jeffrey Barrett, Sonia Goncalves, Ewan Harrison, David K. Jackson, Ian Johnston, Dominic Kwiatkowski, Cordelia Langford, John Sillitoe on behalf of the Wellcome Sanger Institute COVID-19 Surveillance Team |
| EPI_ISL_3441453, EPI_ISL_3441454 | Lighthouse Laboratory Plymouth | Wellcome Sanger Institute for the COVID-19 Genomics UK (COG-UK) Consortium | Lighthouse Laboratory Plymouth and Alex Alderton, Roberto Amato, Jeffrey Barrett, Sonia Goncalves, Ewan Harrison, David K. Jackson, Ian Johnston, Dominic Kwiatkowski, Cordelia Langford, John Sillitoe on behalf of the Wellcome Sanger Institute COVID-19 Surveillance Team |
| EPI_ISL_3441455 | Lighthouse Lab in Milton Keynes | Wellcome Sanger Institute for the COVID-19 Genomics UK (COG-UK) Consortium | The Lighthouse Lab in Milton Keynes and Alex Alderton, Roberto Amato, Jeffrey Barrett, Sonia Goncalves, Ewan Harrison, David K. Jackson, Ian Johnston, Dominic Kwiatkowski, Cordelia Langford, John Sillitoe on behalf of the Wellcome Sanger Institute COVID-19 Surveillance Team |
| EPI_ISL_3441456 | Lighthouse Laboratory Plymouth | Wellcome Sanger Institute for the COVID-19 Genomics UK (COG-UK) Consortium | Lighthouse Laboratory Plymouth and Alex Alderton, Roberto Amato, Jeffrey Barrett, Sonia Goncalves, Ewan Harrison, David K. Jackson, Ian Johnston, Dominic Kwiatkowski, Cordelia Langford, John Sillitoe on behalf of the Wellcome Sanger Institute COVID-19 Surveillance Team |
| EPI_ISL_3441457, EPI_ISL_3441458, EPI_ISL_3441459 | Lighthouse Lab in Milton Keynes | Wellcome Sanger Institute for the COVID-19 Genomics UK (COG-UK) Consortium | The Lighthouse Lab in Milton Keynes and Alex Alderton, Roberto Amato, Jeffrey Barrett, Sonia Goncalves, Ewan Harrison, David K. Jackson, Ian Johnston, Dominic Kwiatkowski, Cordelia Langford, John Sillitoe on behalf of the Wellcome Sanger Institute COVID-19 Surveillance Team |
| EPI_ISL_3441460 | Berkshire and Surrey Pathology Services Lighthouse Laboratory | Wellcome Sanger Institute for the COVID-19 Genomics UK (COG-UK) Consortium | Berkshire and Surrey Pathology Services Lighthouse Laboratory and Alex Alderton, Roberto Amato, Jeffrey Barrett, Sonia Goncalves, Ewan Harrison, David K. Jackson, Ian Johnston, Dominic Kwiatkowski, Cordelia Langford, John Sillitoe on behalf of the Wellcome Sanger Institute COVID-19 Surveillance Team |
| EPI_ISL_3441461 | Lighthouse Laboratory Plymouth | Wellcome Sanger Institute for the COVID-19 Genomics UK (COG-UK) Consortium | Lighthouse Laboratory Plymouth and Alex Alderton, Roberto Amato, Jeffrey Barrett, Sonia Goncalves, Ewan Harrison, David K. Jackson, Ian Johnston, Dominic Kwiatkowski, Cordelia Langford, John Sillitoe on behalf of the Wellcome Sanger Institute COVID-19 Surveillance Team |
| EPI_ISL_3441462 | Lighthouse Lab in Milton Keynes | Wellcome Sanger Institute for the COVID-19 Genomics UK (COG-UK) Consortium | The Lighthouse Lab in Milton Keynes and Alex Alderton, Roberto Amato, Jeffrey Barrett, Sonia Goncalves, Ewan Harrison, David K. Jackson, Ian Johnston, Dominic Kwiatkowski, Cordelia Langford, John Sillitoe on behalf of the Wellcome Sanger Institute COVID-19 Surveillance Team |
| EPI_ISL_3441463, EPI_ISL_3441464, EPI_ISL_3441465 | Lighthouse Lab in Glasgow | Wellcome Sanger Institute for the COVID-19 Genomics UK (COG-UK) Consortium | Harper VanSteenhouse, Yumi Kasai, David Gray, Carol Clugston, Anna Dominiczak and Alex Alderton, Roberto Amato, Jeffrey Barrett, Sonia Goncalves, Ewan Harrison, David K. Jackson, Ian Johnston, Dominic Kwiatkowski, Cordelia Langford, John Sillitoe on behalf of the Wellcome Sanger Institute COVID-19 Surveillance Team |
| EPI_ISL_3441466, EPI_ISL_3441467, EPI_ISL_3441468 | Berkshire and Surrey Pathology Services Lighthouse Laboratory | Wellcome Sanger Institute for the COVID-19 Genomics UK (COG-UK) Consortium | Berkshire and Surrey Pathology Services Lighthouse Laboratory and Alex Alderton, Roberto Amato, Jeffrey Barrett, Sonia Goncalves, Ewan Harrison, David K. Jackson, Ian Johnston, Dominic Kwiatkowski, Cordelia Langford, John Sillitoe on behalf of the Wellcome Sanger Institute COVID-19 Surveillance Team |
| EPI_ISL_3441469 | Lighthouse Lab in Glasgow | Wellcome Sanger Institute for the COVID-19 Genomics UK (COG-UK) Consortium | Harper VanSteenhouse, Yumi Kasai, David Gray, Carol Clugston, Anna Dominiczak and Alex Alderton, Roberto Amato, Jeffrey Barrett, Sonia Goncalves, Ewan Harrison, David K. Jackson, Ian Johnston, Dominic Kwiatkowski, Cordelia Langford, John Sillitoe on behalf of the Wellcome Sanger Institute COVID-19 Surveillance Team |
| EPI_ISL_3441470 | Lighthouse Laboratory Plymouth | Wellcome Sanger Institute for the COVID-19 Genomics UK (COG-UK) Consortium | Lighthouse Laboratory Plymouth and Alex Alderton, Roberto Amato, Jeffrey Barrett, Sonia Goncalves, Ewan Harrison, David K. Jackson, Ian Johnston, Dominic Kwiatkowski, Cordelia Langford, John Sillitoe on behalf of the Wellcome Sanger Institute COVID-19 Surveillance Team |
| EPI_ISL_3441471 | Lighthouse Lab in Milton Keynes | Wellcome Sanger Institute for the COVID-19 Genomics UK (COG-UK) Consortium | The Lighthouse Lab in Milton Keynes and Alex Alderton, Roberto Amato, Jeffrey Barrett, Sonia Goncalves, Ewan Harrison, David K. Jackson, Ian Johnston, Dominic Kwiatkowski, Cordelia Langford, John Sillitoe on behalf of the Wellcome Sanger Institute COVID-19 Surveillance Team |
| EPI_ISL_3441472 | Berkshire and Surrey Pathology Services Lighthouse Laboratory | Wellcome Sanger Institute for the COVID-19 Genomics UK (COG-UK) Consortium | Berkshire and Surrey Pathology Services Lighthouse Laboratory and Alex Alderton, Roberto Amato, Jeffrey Barrett, Sonia Goncalves, Ewan Harrison, David K. Jackson, Ian Johnston, Dominic Kwiatkowski, Cordelia Langford, John Sillitoe on behalf of the Wellcome Sanger Institute COVID-19 Surveillance Team |
| EPI_ISL_3441473 | Lighthouse Laboratory Plymouth | Wellcome Sanger Institute for the COVID-19 Genomics UK (COG-UK) Consortium | Lighthouse Laboratory Plymouth and Alex Alderton, Roberto Amato, Jeffrey Barrett, Sonia Goncalves, Ewan Harrison, David K. Jackson, Ian Johnston, Dominic Kwiatkowski, Cordelia Langford, John Sillitoe on behalf of the Wellcome Sanger Institute COVID-19 Surveillance Team |
| EPI_ISL_3441474 | Lighthouse Lab in Glasgow | Wellcome Sanger Institute for the COVID-19 Genomics UK (COG-UK) Consortium | Harper VanSteenhouse, Yumi Kasai, David Gray, Carol Clugston, Anna Dominiczak and Alex Alderton, Roberto Amato, Jeffrey Barrett, Sonia Goncalves, Ewan Harrison, David K. Jackson, Ian Johnston, Dominic Kwiatkowski, Cordelia Langford, John Sillitoe on behalf of the Wellcome Sanger Institute COVID-19 Surveillance Team |
| EPI_ISL_3441475 | Berkshire and Surrey Pathology Services Lighthouse Laboratory | Wellcome Sanger Institute for the COVID-19 Genomics UK (COG-UK) Consortium | Berkshire and Surrey Pathology Services Lighthouse Laboratory and Alex Alderton, Roberto Amato, Jeffrey Barrett, Sonia Goncalves, Ewan Harrison, David K. Jackson, Ian Johnston, Dominic Kwiatkowski, Cordelia Langford, John Sillitoe on behalf of the Wellcome Sanger Institute COVID-19 Surveillance Team |
| EPI_ISL_3441476 | Lighthouse Lab in Milton Keynes | Wellcome Sanger Institute for the COVID-19 Genomics UK (COG-UK) Consortium | The Lighthouse Lab in Milton Keynes and Alex Alderton, Roberto Amato, Jeffrey Barrett, Sonia Goncalves, Ewan Harrison, David K. Jackson, Ian Johnston, Dominic Kwiatkowski, Cordelia Langford, John Sillitoe on behalf of the Wellcome Sanger Institute COVID-19 Surveillance Team |
| EPI_ISL_3441477 | Lighthouse Laboratory Plymouth | Wellcome Sanger Institute for the COVID-19 Genomics UK (COG-UK) Consortium | Lighthouse Laboratory Plymouth and Alex Alderton, Roberto Amato, Jeffrey Barrett, Sonia Goncalves, Ewan Harrison, David K. Jackson, Ian Johnston, Dominic Kwiatkowski, Cordelia Langford, John Sillitoe on behalf of the Wellcome Sanger Institute COVID-19 Surveillance Team |
| EPI_ISL_3441478 | Lighthouse Lab in Milton Keynes | Wellcome Sanger Institute for the COVID-19 Genomics UK (COG-UK) Consortium | The Lighthouse Lab in Milton Keynes and Alex Alderton, Roberto Amato, Jeffrey Barrett, Sonia Goncalves, Ewan Harrison, David K. Jackson, Ian Johnston, Dominic Kwiatkowski, Cordelia Langford, John Sillitoe on behalf of the Wellcome Sanger Institute COVID-19 Surveillance Team |
| EPI_ISL_3441479 | Lighthouse Lab in Glasgow | Wellcome Sanger Institute for the COVID-19 Genomics UK (COG-UK) Consortium | Harper VanSteenhouse, Yumi Kasai, David Gray, Carol Clugston, Anna Dominiczak and Alex Alderton, Roberto Amato, Jeffrey Barrett, Sonia Goncalves, Ewan Harrison, David K. Jackson, Ian Johnston, Dominic Kwiatkowski, Cordelia Langford, John Sillitoe on behalf of the Wellcome Sanger Institute COVID-19 Surveillance Team |
| EPI_ISL_3441480 | Lighthouse Lab in Milton Keynes | Wellcome Sanger Institute for the COVID-19 Genomics UK (COG-UK) Consortium | The Lighthouse Lab in Milton Keynes and Alex Alderton, Roberto Amato, Jeffrey Barrett, Sonia Goncalves, Ewan Harrison, David K. Jackson, Ian Johnston, Dominic Kwiatkowski, Cordelia Langford, John Sillitoe on behalf of the Wellcome Sanger Institute COVID-19 Surveillance Team |
| EPI_ISL_3441481 | Lighthouse Lab in Glasgow | Wellcome Sanger Institute for the COVID-19 Genomics UK (COG-UK) Consortium | Harper VanSteenhouse, Yumi Kasai, David Gray, Carol Clugston, Anna Dominiczak and Alex Alderton, Roberto Amato, Jeffrey Barrett, Sonia Goncalves, Ewan Harrison, David K. Jackson, Ian Johnston, Dominic Kwiatkowski, Cordelia Langford, John Sillitoe on behalf of the Wellcome Sanger Institute COVID-19 Surveillance Team |
| EPI_ISL_3441482 | Berkshire and Surrey Pathology Services Lighthouse Laboratory | Wellcome Sanger Institute for the COVID-19 Genomics UK (COG-UK) Consortium | Berkshire and Surrey Pathology Services Lighthouse Laboratory and Alex Alderton, Roberto Amato, Jeffrey Barrett, Sonia Goncalves, Ewan Harrison, David K. Jackson, Ian Johnston, Dominic Kwiatkowski, Cordelia Langford, John Sillitoe on behalf of the Wellcome Sanger Institute COVID-19 Surveillance Team |
| EPI_ISL_3441483 | Lighthouse Lab in Milton Keynes | Wellcome Sanger Institute for the COVID-19 Genomics UK (COG-UK) Consortium | The Lighthouse Lab in Milton Keynes and Alex Alderton, Roberto Amato, Jeffrey Barrett, Sonia Goncalves, Ewan Harrison, David K. Jackson, Ian Johnston, Dominic Kwiatkowski, Cordelia Langford, John Sillitoe on behalf of the Wellcome Sanger Institute COVID-19 Surveillance Team |
| EPI_ISL_3441484, EPI_ISL_3441485 | Lighthouse Lab in Glasgow | Wellcome Sanger Institute for the COVID-19 Genomics UK (COG-UK) Consortium | Harper VanSteenhouse, Yumi Kasai, David Gray, Carol Clugston, Anna Dominiczak and Alex Alderton, Roberto Amato, Jeffrey Barrett, Sonia Goncalves, Ewan Harrison, David K. Jackson, Ian Johnston, Dominic Kwiatkowski, Cordelia Langford, John Sillitoe on behalf of the Wellcome Sanger Institute COVID-19 Surveillance Team |
| EPI_ISL_3441486 | Berkshire and Surrey Pathology Services Lighthouse Laboratory | Wellcome Sanger Institute for the COVID-19 Genomics UK (COG-UK) Consortium | Berkshire and Surrey Pathology Services Lighthouse Laboratory and Alex Alderton, Roberto Amato, Jeffrey Barrett, Sonia Goncalves, Ewan Harrison, David K. Jackson, Ian Johnston, Dominic Kwiatkowski, Cordelia Langford, John Sillitoe on behalf of the Wellcome Sanger Institute COVID-19 Surveillance Team |
| EPI_ISL_3441487 | Lighthouse Lab in Glasgow | Wellcome Sanger Institute for the COVID-19 Genomics UK (COG-UK) Consortium | Harper VanSteenhouse, Yumi Kasai, David Gray, Carol Clugston, Anna Dominiczak and Alex Alderton, Roberto Amato, Jeffrey Barrett, Sonia Goncalves, Ewan Harrison, David K. Jackson, Ian Johnston, Dominic Kwiatkowski, Cordelia Langford, John Sillitoe on behalf of the Wellcome Sanger Institute COVID-19 Surveillance Team |
| EPI_ISL_3441488 | Berkshire and Surrey Pathology Services Lighthouse Laboratory | Wellcome Sanger Institute for the COVID-19 Genomics UK (COG-UK) Consortium | Berkshire and Surrey Pathology Services Lighthouse Laboratory and Alex Alderton, Roberto Amato, Jeffrey Barrett, Sonia Goncalves, Ewan Harrison, David K. Jackson, Ian Johnston, Dominic Kwiatkowski, Cordelia Langford, John Sillitoe on behalf of the Wellcome Sanger Institute COVID-19 Surveillance Team |
| EPI_ISL_3441489 | Lighthouse Lab in Glasgow | Wellcome Sanger Institute for the COVID-19 Genomics UK (COG-UK) Consortium | Harper VanSteenhouse, Yumi Kasai, David Gray, Carol Clugston, Anna Dominiczak and Alex Alderton, Roberto Amato, Jeffrey Barrett, Sonia Goncalves, Ewan Harrison, David K. Jackson, Ian Johnston, Dominic Kwiatkowski, Cordelia Langford, John Sillitoe on behalf of the Wellcome Sanger Institute COVID-19 Surveillance Team |
| EPI_ISL_3441490 | Berkshire and Surrey Pathology Services Lighthouse Laboratory | Wellcome Sanger Institute for the COVID-19 Genomics UK (COG-UK) Consortium | Berkshire and Surrey Pathology Services Lighthouse Laboratory and Alex Alderton, Roberto Amato, Jeffrey Barrett, Sonia Goncalves, Ewan Harrison, David K. Jackson, Ian Johnston, Dominic Kwiatkowski, Cordelia Langford, John Sillitoe on behalf of the Wellcome Sanger Institute COVID-19 Surveillance Team |
| EPI_ISL_3441491 | Lighthouse Lab in Glasgow | Wellcome Sanger Institute for the COVID-19 Genomics UK (COG-UK) Consortium | Harper VanSteenhouse, Yumi Kasai, David Gray, Carol Clugston, Anna Dominiczak and Alex Alderton, Roberto Amato, Jeffrey Barrett, Sonia Goncalves, Ewan Harrison, David K. Jackson, Ian Johnston, Dominic Kwiatkowski, Cordelia Langford, John Sillitoe on behalf of the Wellcome Sanger Institute COVID-19 Surveillance Team |
| EPI_ISL_3441492 | Lighthouse Lab in Milton Keynes | Wellcome Sanger Institute for the COVID-19 Genomics UK (COG-UK) Consortium | The Lighthouse Lab in Milton Keynes and Alex Alderton, Roberto Amato, Jeffrey Barrett, Sonia Goncalves, Ewan Harrison, David K. Jackson, Ian Johnston, Dominic Kwiatkowski, Cordelia Langford, John Sillitoe on behalf of the Wellcome Sanger Institute COVID-19 Surveillance Team |
| EPI_ISL_3441493 | Lighthouse Lab in Glasgow | Wellcome Sanger Institute for the COVID-19 Genomics UK (COG-UK) Consortium | Harper VanSteenhouse, Yumi Kasai, David Gray, Carol Clugston, Anna Dominiczak and Alex Alderton, Roberto Amato, Jeffrey Barrett, Sonia Goncalves, Ewan Harrison, David K. Jackson, Ian Johnston, Dominic Kwiatkowski, Cordelia Langford, John Sillitoe on behalf of the Wellcome Sanger Institute COVID-19 Surveillance Team |
| EPI_ISL_3441494 | Lighthouse Laboratory Plymouth | Wellcome Sanger Institute for the COVID-19 Genomics UK (COG-UK) Consortium | Lighthouse Laboratory Plymouth and Alex Alderton, Roberto Amato, Jeffrey Barrett, Sonia Goncalves, Ewan Harrison, David K. Jackson, Ian Johnston, Dominic Kwiatkowski, Cordelia Langford, John Sillitoe on behalf of the Wellcome Sanger Institute COVID-19 Surveillance Team |

| | | | |
|---|---|---|---|
| EPI_ISL_3441495, EPI_ISL_3441496 | Lighthouse Lab in Milton Keynes | Wellcome Sanger Institute for the COVID-19 Genomics UK (COG-UK) Consortium | The Lighthouse Lab in Milton Keynes and Alex Alderton, Roberto Amato, Jeffrey Barrett, Sonia Goncalves, Ewan Harrison, David K. Jackson, Ian Johnston, Dominic Kwiatkowski, Cordelia Langford, John Sillitoe on behalf of the Wellcome Sanger Institute COVID-19 Surveillance Team |
| EPI_ISL_3441497 | Berkshire and Surrey Pathology Services Lighthouse Laboratory | Wellcome Sanger Institute for the COVID-19 Genomics UK (COG-UK) Consortium | Berkshire and Surrey Pathology Services Lighthouse Laboratory and Alex Alderton, Roberto Amato, Jeffrey Barrett, Sonia Goncalves, Ewan Harrison, David K. Jackson, Ian Johnston, Dominic Kwiatkowski, Cordelia Langford, John Sillitoe on behalf of the Wellcome Sanger Institute COVID-19 Surveillance Team |
| EPI_ISL_3441498, EPI_ISL_3441499 | Lighthouse Lab in Glasgow | Wellcome Sanger Institute for the COVID-19 Genomics UK (COG-UK) Consortium | Harper VanSteenhouse, Yumi Kasai, David Gray, Carol Clugston, Anna Dominiczak and Alex Alderton, Roberto Amato, Jeffrey Barrett, Sonia Goncalves, Ewan Harrison, David K. Jackson, Ian Johnston, Dominic Kwiatkowski, Cordelia Langford, John Sillitoe on behalf of the Wellcome Sanger Institute COVID-19 Surveillance Team |
| EPI_ISL_3441500, EPI_ISL_3441501, EPI_ISL_3441502 | Lighthouse Laboratory Plymouth | Wellcome Sanger Institute for the COVID-19 Genomics UK (COG-UK) Consortium | Lighthouse Laboratory Plymouth and Alex Alderton, Roberto Amato, Jeffrey Barrett, Sonia Goncalves, Ewan Harrison, David K. Jackson, Ian Johnston, Dominic Kwiatkowski, Cordelia Langford, John Sillitoe on behalf of the Wellcome Sanger Institute COVID-19 Surveillance Team |
| EPI_ISL_3441503 | Berkshire and Surrey Pathology Services Lighthouse Laboratory | Wellcome Sanger Institute for the COVID-19 Genomics UK (COG-UK) Consortium | Berkshire and Surrey Pathology Services Lighthouse Laboratory and Alex Alderton, Roberto Amato, Jeffrey Barrett, Sonia Goncalves, Ewan Harrison, David K. Jackson, Ian Johnston, Dominic Kwiatkowski, Cordelia Langford, John Sillitoe on behalf of the Wellcome Sanger Institute COVID-19 Surveillance Team |
| EPI_ISL_3441504 | Lighthouse Lab in Glasgow | Wellcome Sanger Institute for the COVID-19 Genomics UK (COG-UK) Consortium | Harper VanSteenhouse, Yumi Kasai, David Gray, Carol Clugston, Anna Dominiczak and Alex Alderton, Roberto Amato, Jeffrey Barrett, Sonia Goncalves, Ewan Harrison, David K. Jackson, Ian Johnston, Dominic Kwiatkowski, Cordelia Langford, John Sillitoe on behalf of the Wellcome Sanger Institute COVID-19 Surveillance Team |
| EPI_ISL_3441505 | Lighthouse Lab in Milton Keynes | Wellcome Sanger Institute for the COVID-19 Genomics UK (COG-UK) Consortium | The Lighthouse Lab in Milton Keynes and Alex Alderton, Roberto Amato, Jeffrey Barrett, Sonia Goncalves, Ewan Harrison, David K. Jackson, Ian Johnston, Dominic Kwiatkowski, Cordelia Langford, John Sillitoe on behalf of the Wellcome Sanger Institute COVID-19 Surveillance Team |
| EPI_ISL_3441506 | Berkshire and Surrey Pathology Services Lighthouse Laboratory | Wellcome Sanger Institute for the COVID-19 Genomics UK (COG-UK) Consortium | Berkshire and Surrey Pathology Services Lighthouse Laboratory and Alex Alderton, Roberto Amato, Jeffrey Barrett, Sonia Goncalves, Ewan Harrison, David K. Jackson, Ian Johnston, Dominic Kwiatkowski, Cordelia Langford, John Sillitoe on behalf of the Wellcome Sanger Institute COVID-19 Surveillance Team |
| EPI_ISL_3441507, EPI_ISL_3441508 | Lighthouse Lab in Milton Keynes | Wellcome Sanger Institute for the COVID-19 Genomics UK (COG-UK) Consortium | The Lighthouse Lab in Milton Keynes and Alex Alderton, Roberto Amato, Jeffrey Barrett, Sonia Goncalves, Ewan Harrison, David K. Jackson, Ian Johnston, Dominic Kwiatkowski, Cordelia Langford, John Sillitoe on behalf of the Wellcome Sanger Institute COVID-19 Surveillance Team |
| EPI_ISL_3441509 | Berkshire and Surrey Pathology Services Lighthouse Laboratory | Wellcome Sanger Institute for the COVID-19 Genomics UK (COG-UK) Consortium | Berkshire and Surrey Pathology Services Lighthouse Laboratory and Alex Alderton, Roberto Amato, Jeffrey Barrett, Sonia Goncalves, Ewan Harrison, David K. Jackson, Ian Johnston, Dominic Kwiatkowski, Cordelia Langford, John Sillitoe on behalf of the Wellcome Sanger Institute COVID-19 Surveillance Team |
| EPI_ISL_3441510 | Lighthouse Laboratory Plymouth | Wellcome Sanger Institute for the COVID-19 Genomics UK (COG-UK) Consortium | Lighthouse Laboratory Plymouth and Alex Alderton, Roberto Amato, Jeffrey Barrett, Sonia Goncalves, Ewan Harrison, David K. Jackson, Ian Johnston, Dominic Kwiatkowski, Cordelia Langford, John Sillitoe on behalf of the Wellcome Sanger Institute COVID-19 Surveillance Team |
| EPI_ISL_3441511, EPI_ISL_3441512 | Lighthouse Lab in Glasgow | Wellcome Sanger Institute for the COVID-19 Genomics UK (COG-UK) Consortium | Harper VanSteenhouse, Yumi Kasai, David Gray, Carol Clugston, Anna Dominiczak and Alex Alderton, Roberto Amato, Jeffrey Barrett, Sonia Goncalves, Ewan Harrison, David K. Jackson, Ian Johnston, Dominic Kwiatkowski, Cordelia Langford, John Sillitoe on behalf of the Wellcome Sanger Institute COVID-19 Surveillance Team |
| EPI_ISL_3441513, EPI_ISL_3441514 | Lighthouse Lab in Milton Keynes | Wellcome Sanger Institute for the COVID-19 Genomics UK (COG-UK) Consortium | The Lighthouse Lab in Milton Keynes and Alex Alderton, Roberto Amato, Jeffrey Barrett, Sonia Goncalves, Ewan Harrison, David K. Jackson, Ian Johnston, Dominic Kwiatkowski, Cordelia Langford, John Sillitoe on behalf of the Wellcome Sanger Institute COVID-19 Surveillance Team |
| EPI_ISL_3441515 | Berkshire and Surrey Pathology Services Lighthouse Laboratory | Wellcome Sanger Institute for the COVID-19 Genomics UK (COG-UK) Consortium | Berkshire and Surrey Pathology Services Lighthouse Laboratory and Alex Alderton, Roberto Amato, Jeffrey Barrett, Sonia Goncalves, Ewan Harrison, David K. Jackson, Ian Johnston, Dominic Kwiatkowski, Cordelia Langford, John Sillitoe on behalf of the Wellcome Sanger Institute COVID-19 Surveillance Team |
| EPI_ISL_3441516 | Lighthouse Lab in Glasgow | Wellcome Sanger Institute for the COVID-19 Genomics UK (COG-UK) Consortium | Harper VanSteenhouse, Yumi Kasai, David Gray, Carol Clugston, Anna Dominiczak and Alex Alderton, Roberto Amato, Jeffrey Barrett, Sonia Goncalves, Ewan Harrison, David K. Jackson, Ian Johnston, Dominic Kwiatkowski, Cordelia Langford, John Sillitoe on behalf of the Wellcome Sanger Institute COVID-19 Surveillance Team |
| EPI_ISL_3441517 | Lighthouse Lab in Milton Keynes | Wellcome Sanger Institute for the COVID-19 Genomics UK (COG-UK) Consortium | The Lighthouse Lab in Milton Keynes and Alex Alderton, Roberto Amato, Jeffrey Barrett, Sonia Goncalves, Ewan Harrison, David K. Jackson, Ian Johnston, Dominic Kwiatkowski, Cordelia Langford, John Sillitoe on behalf of the Wellcome Sanger Institute COVID-19 Surveillance Team |
| EPI_ISL_3441518 | Berkshire and Surrey Pathology Services Lighthouse Laboratory | Wellcome Sanger Institute for the COVID-19 Genomics UK (COG-UK) Consortium | Berkshire and Surrey Pathology Services Lighthouse Laboratory and Alex Alderton, Roberto Amato, Jeffrey Barrett, Sonia Goncalves, Ewan Harrison, David K. Jackson, Ian Johnston, Dominic Kwiatkowski, Cordelia Langford, John Sillitoe on behalf of the Wellcome Sanger Institute COVID-19 Surveillance Team |
| EPI_ISL_3441519 | Lighthouse Lab in Glasgow | Wellcome Sanger Institute for the COVID-19 Genomics UK (COG-UK) Consortium | Harper VanSteenhouse, Yumi Kasai, David Gray, Carol Clugston, Anna Dominiczak and Alex Alderton, Roberto Amato, Jeffrey Barrett, Sonia Goncalves, Ewan Harrison, David K. Jackson, Ian Johnston, Dominic Kwiatkowski, Cordelia Langford, John Sillitoe on behalf of the Wellcome Sanger Institute COVID-19 Surveillance Team |
| EPI_ISL_3441520, EPI_ISL_3441521 | Lighthouse Lab in Milton Keynes | Wellcome Sanger Institute for the COVID-19 Genomics UK (COG-UK) Consortium | The Lighthouse Lab in Milton Keynes and Alex Alderton, Roberto Amato, Jeffrey Barrett, Sonia Goncalves, Ewan Harrison, David K. Jackson, Ian Johnston, Dominic Kwiatkowski, Cordelia Langford, John Sillitoe on behalf of the Wellcome Sanger Institute COVID-19 Surveillance Team |
| EPI_ISL_3441522 | Lighthouse Laboratory Plymouth | Wellcome Sanger Institute for the COVID-19 Genomics UK (COG-UK) Consortium | Lighthouse Laboratory Plymouth and Alex Alderton, Roberto Amato, Jeffrey Barrett, Sonia Goncalves, Ewan Harrison, David K. Jackson, Ian Johnston, Dominic Kwiatkowski, Cordelia Langford, John Sillitoe on behalf of the Wellcome Sanger Institute COVID-19 Surveillance Team |
| EPI_ISL_3441523 | Berkshire and Surrey Pathology Services Lighthouse Laboratory | Wellcome Sanger Institute for the COVID-19 Genomics UK (COG-UK) Consortium | Berkshire and Surrey Pathology Services Lighthouse Laboratory and Alex Alderton, Roberto Amato, Jeffrey Barrett, Sonia Goncalves, Ewan Harrison, David K. Jackson, Ian Johnston, Dominic Kwiatkowski, Cordelia Langford, John Sillitoe on behalf of the Wellcome Sanger Institute COVID-19 Surveillance Team |
| EPI_ISL_3441524 | Lighthouse Lab in Milton Keynes | Wellcome Sanger Institute for the COVID-19 Genomics UK (COG-UK) Consortium | The Lighthouse Lab in Milton Keynes and Alex Alderton, Roberto Amato, Jeffrey Barrett, Sonia Goncalves, Ewan Harrison, David K. Jackson, Ian Johnston, Dominic Kwiatkowski, Cordelia Langford, John Sillitoe on behalf of the Wellcome Sanger Institute COVID-19 Surveillance Team |
| EPI_ISL_3441525, EPI_ISL_3441526, EPI_ISL_3441527 | Lighthouse Lab in Glasgow | Wellcome Sanger Institute for the COVID-19 Genomics UK (COG-UK) Consortium | Harper VanSteenhouse, Yumi Kasai, David Gray, Carol Clugston, Anna Dominiczak and Alex Alderton, Roberto Amato, Jeffrey Barrett, Sonia Goncalves, Ewan Harrison, David K. Jackson, Ian Johnston, Dominic Kwiatkowski, Cordelia Langford, John Sillitoe on behalf of the Wellcome Sanger Institute COVID-19 Surveillance Team |
| EPI_ISL_3441528, EPI_ISL_3441529 | Berkshire and Surrey Pathology Services Lighthouse Laboratory | Wellcome Sanger Institute for the COVID-19 Genomics UK (COG-UK) Consortium | Berkshire and Surrey Pathology Services Lighthouse Laboratory and Alex Alderton, Roberto Amato, Jeffrey Barrett, Sonia Goncalves, Ewan Harrison, David K. Jackson, Ian Johnston, Dominic Kwiatkowski, Cordelia Langford, John Sillitoe on behalf of the Wellcome Sanger Institute COVID-19 Surveillance Team |
| EPI_ISL_3441530, EPI_ISL_3441531 | Lighthouse Lab in Glasgow | Wellcome Sanger Institute for the COVID-19 Genomics UK (COG-UK) Consortium | Harper VanSteenhouse, Yumi Kasai, David Gray, Carol Clugston, Anna Dominiczak and Alex Alderton, Roberto Amato, Jeffrey Barrett, Sonia Goncalves, Ewan Harrison, David K. Jackson, Ian Johnston, Dominic Kwiatkowski, Cordelia Langford, John Sillitoe on behalf of the Wellcome Sanger Institute COVID-19 Surveillance Team |
| EPI_ISL_3441532 | Lighthouse Lab in Milton Keynes | Wellcome Sanger Institute for the COVID-19 Genomics UK (COG-UK) Consortium | The Lighthouse Lab in Milton Keynes and Alex Alderton, Roberto Amato, Jeffrey Barrett, Sonia Goncalves, Ewan Harrison, David K. Jackson, Ian Johnston, Dominic Kwiatkowski, Cordelia Langford, John Sillitoe on behalf of the Wellcome Sanger Institute COVID-19 Surveillance Team |
| EPI_ISL_3441533 | Lighthouse Laboratory Plymouth | Wellcome Sanger Institute for the COVID-19 Genomics UK (COG-UK) Consortium | Lighthouse Laboratory Plymouth and Alex Alderton, Roberto Amato, Jeffrey Barrett, Sonia Goncalves, Ewan Harrison, David K. Jackson, Ian Johnston, Dominic Kwiatkowski, Cordelia Langford, John Sillitoe on behalf of the Wellcome Sanger Institute COVID-19 Surveillance Team |
| EPI_ISL_3441534, EPI_ISL_3441535 | Berkshire and Surrey Pathology Services Lighthouse Laboratory | Wellcome Sanger Institute for the COVID-19 Genomics UK (COG-UK) Consortium | Berkshire and Surrey Pathology Services Lighthouse Laboratory and Alex Alderton, Roberto Amato, Jeffrey Barrett, Sonia Goncalves, Ewan Harrison, David K. Jackson, Ian Johnston, Dominic Kwiatkowski, Cordelia Langford, John Sillitoe on behalf of the Wellcome Sanger Institute COVID-19 Surveillance Team |
| EPI_ISL_3441536, EPI_ISL_3441537 | Lighthouse Lab in Milton Keynes | Wellcome Sanger Institute for the COVID-19 Genomics UK (COG-UK) Consortium | The Lighthouse Lab in Milton Keynes and Alex Alderton, Roberto Amato, Jeffrey Barrett, Sonia Goncalves, Ewan Harrison, David K. Jackson, Ian Johnston, Dominic Kwiatkowski, Cordelia Langford, John Sillitoe on behalf of the Wellcome Sanger Institute COVID-19 Surveillance Team |
| EPI_ISL_3441538 | Lighthouse Laboratory Plymouth | Wellcome Sanger Institute for the COVID-19 Genomics UK (COG-UK) Consortium | Lighthouse Laboratory Plymouth and Alex Alderton, Roberto Amato, Jeffrey Barrett, Sonia Goncalves, Ewan Harrison, David K. Jackson, Ian Johnston, Dominic Kwiatkowski, Cordelia Langford, John Sillitoe on behalf of the Wellcome Sanger Institute COVID-19 Surveillance Team |
| EPI_ISL_3441539, EPI_ISL_3441540 | Berkshire and Surrey Pathology Services Lighthouse Laboratory | Wellcome Sanger Institute for the COVID-19 Genomics UK (COG-UK) Consortium | Berkshire and Surrey Pathology Services Lighthouse Laboratory and Alex Alderton, Roberto Amato, Jeffrey Barrett, Sonia Goncalves, Ewan Harrison, David K. Jackson, Ian Johnston, Dominic Kwiatkowski, Cordelia Langford, John Sillitoe on behalf of the Wellcome Sanger Institute COVID-19 Surveillance Team |
| EPI_ISL_3441541, EPI_ISL_3441542, EPI_ISL_3441543, EPI_ISL_3441544, EPI_ISL_3441545 | Lighthouse Lab in Glasgow | Wellcome Sanger Institute for the COVID-19 Genomics UK (COG-UK) Consortium | Harper VanSteenhouse, Yumi Kasai, David Gray, Carol Clugston, Anna Dominiczak and Alex Alderton, Roberto Amato, Jeffrey Barrett, Sonia Goncalves, Ewan Harrison, David K. Jackson, Ian Johnston, Dominic Kwiatkowski, Cordelia Langford, John Sillitoe on behalf of the Wellcome Sanger Institute COVID-19 Surveillance Team |
| EPI_ISL_3441546 | Berkshire and Surrey Pathology Services Lighthouse Laboratory | Wellcome Sanger Institute for the COVID-19 Genomics UK (COG-UK) Consortium | Berkshire and Surrey Pathology Services Lighthouse Laboratory and Alex Alderton, Roberto Amato, Jeffrey Barrett, Sonia Goncalves, Ewan Harrison, David K. Jackson, Ian Johnston, Dominic Kwiatkowski, Cordelia Langford, John Sillitoe on behalf of the Wellcome Sanger Institute COVID-19 Surveillance Team |
| EPI_ISL_3441547 | Lighthouse Lab in Glasgow | Wellcome Sanger Institute for the COVID-19 Genomics UK (COG-UK) Consortium | Harper VanSteenhouse, Yumi Kasai, David Gray, Carol Clugston, Anna Dominiczak and Alex Alderton, Roberto Amato, Jeffrey Barrett, Sonia Goncalves, Ewan Harrison, David K. Jackson, Ian Johnston, Dominic Kwiatkowski, Cordelia Langford, John Sillitoe on behalf of the Wellcome Sanger Institute COVID-19 Surveillance Team |
| EPI_ISL_3441548 | Berkshire and Surrey Pathology Services Lighthouse Laboratory | Wellcome Sanger Institute for the COVID-19 Genomics UK (COG-UK) Consortium | Berkshire and Surrey Pathology Services Lighthouse Laboratory and Alex Alderton, Roberto Amato, Jeffrey Barrett, Sonia Goncalves, Ewan Harrison, David K. Jackson, Ian Johnston, Dominic Kwiatkowski, Cordelia Langford, John Sillitoe on behalf of the Wellcome Sanger Institute COVID-19 Surveillance Team |
| EPI_ISL_3441549 | Lighthouse Lab in Glasgow | Wellcome Sanger Institute for the COVID-19 Genomics UK (COG-UK) Consortium | Harper VanSteenhouse, Yumi Kasai, David Gray, Carol Clugston, Anna Dominiczak and Alex Alderton, Roberto Amato, Jeffrey Barrett, Sonia Goncalves, Ewan Harrison, David K. Jackson, Ian Johnston, Dominic Kwiatkowski, Cordelia Langford, John Sillitoe on behalf of the Wellcome Sanger Institute COVID-19 Surveillance Team |
| EPI_ISL_3441550, EPI_ISL_3441551 | Berkshire and Surrey Pathology Services Lighthouse Laboratory | Wellcome Sanger Institute for the COVID-19 Genomics UK (COG-UK) Consortium | Berkshire and Surrey Pathology Services Lighthouse Laboratory and Alex Alderton, Roberto Amato, Jeffrey Barrett, Sonia Goncalves, Ewan Harrison, David K. Jackson, Ian Johnston, Dominic Kwiatkowski, Cordelia Langford, John Sillitoe on behalf of the Wellcome Sanger Institute COVID-19 Surveillance Team |
| EPI_ISL_3441552 | Lighthouse Lab in Glasgow | Wellcome Sanger Institute for the COVID-19 Genomics UK (COG-UK) Consortium | Harper VanSteenhouse, Yumi Kasai, David Gray, Carol Clugston, Anna Dominiczak and Alex Alderton, Roberto Amato, Jeffrey Barrett, Sonia Goncalves, Ewan Harrison, David K. Jackson, Ian Johnston, Dominic Kwiatkowski, Cordelia Langford, John Sillitoe on behalf of the Wellcome Sanger Institute COVID-19 Surveillance Team |
| EPI_ISL_3441553, EPI_ISL_3441554, EPI_ISL_3441555, EPI_ISL_3441556 | Berkshire and Surrey Pathology Services Lighthouse Laboratory | Wellcome Sanger Institute for the COVID-19 Genomics UK (COG-UK) Consortium | Berkshire and Surrey Pathology Services Lighthouse Laboratory and Alex Alderton, Roberto Amato, Jeffrey Barrett, Sonia Goncalves, Ewan Harrison, David K. Jackson, Ian Johnston, Dominic Kwiatkowski, Cordelia Langford, John Sillitoe on behalf of the Wellcome Sanger Institute COVID-19 Surveillance Team |
| EPI_ISL_3441557 | Lighthouse Lab in Glasgow | Wellcome Sanger Institute for the COVID-19 Genomics UK (COG-UK) Consortium | Harper VanSteenhouse, Yumi Kasai, David Gray, Carol Clugston, Anna Dominiczak and Alex Alderton, Roberto Amato, Jeffrey Barrett, Sonia Goncalves, Ewan Harrison, David K. Jackson, Ian Johnston, Dominic Kwiatkowski, Cordelia Langford, John Sillitoe on behalf of the Wellcome Sanger Institute COVID-19 Surveillance Team |
| EPI_ISL_3441558, EPI_ISL_3441559, EPI_ISL_3441560 | Berkshire and Surrey Pathology Services Lighthouse Laboratory | Wellcome Sanger Institute for the COVID-19 Genomics UK (COG-UK) Consortium | Berkshire and Surrey Pathology Services Lighthouse Laboratory and Alex Alderton, Roberto Amato, Jeffrey Barrett, Sonia Goncalves, Ewan Harrison, David K. Jackson, Ian Johnston, Dominic Kwiatkowski, Cordelia Langford, John Sillitoe on behalf of the Wellcome Sanger Institute COVID-19 Surveillance Team |
| EPI_ISL_3441561 | Lighthouse Lab in Glasgow | Wellcome Sanger Institute for the COVID-19 Genomics UK (COG-UK) Consortium | Harper VanSteenhouse, Yumi Kasai, David Gray, Carol Clugston, Anna Dominiczak and Alex Alderton, Roberto Amato, Jeffrey Barrett, Sonia Goncalves, Ewan Harrison, David K. Jackson, Ian Johnston, Dominic Kwiatkowski, Cordelia Langford, John Sillitoe on behalf of the Wellcome Sanger Institute COVID-19 Surveillance Team |

| | | | |
|---|---|---|---|
| EPI_ISL_3441562 | Berkshire and Surrey Pathology Services Lighthouse Laboratory | Wellcome Sanger Institute for the COVID-19 Genomics UK (COG-UK) Consortium | Berkshire and Surrey Pathology Services Lighthouse Laboratory and Alex Alderton, Roberto Amato, Jeffrey Barrett, Sonia Goncalves, Ewan Harrison, David K. Jackson, Ian Johnston, Dominic Kwiatkowski, Cordelia Langford, John Sillitoe on behalf of the Wellcome Sanger Institute COVID-19 Surveillance Team |
| EPI_ISL_3441563 | Lighthouse Lab in Glasgow | Wellcome Sanger Institute for the COVID-19 Genomics UK (COG-UK) Consortium | Harper VanSteenhouse, Yumi Kasai, David Gray, Carol Clugston, Anna Dominiczak and Alex Alderton, Roberto Amato, Jeffrey Barrett, Sonia Goncalves, Ewan Harrison, David K. Jackson, Ian Johnston, Dominic Kwiatkowski, Cordelia Langford, John Sillitoe on behalf of the Wellcome Sanger Institute COVID-19 Surveillance Team |
| EPI_ISL_3441564, EPI_ISL_3441565 | Berkshire and Surrey Pathology Services Lighthouse Laboratory | Wellcome Sanger Institute for the COVID-19 Genomics UK (COG-UK) Consortium | Berkshire and Surrey Pathology Services Lighthouse Laboratory and Alex Alderton, Roberto Amato, Jeffrey Barrett, Sonia Goncalves, Ewan Harrison, David K. Jackson, Ian Johnston, Dominic Kwiatkowski, Cordelia Langford, John Sillitoe on behalf of the Wellcome Sanger Institute COVID-19 Surveillance Team |
| EPI_ISL_3441566, EPI_ISL_3441567 | Lighthouse Lab in Glasgow | Wellcome Sanger Institute for the COVID-19 Genomics UK (COG-UK) Consortium | Harper VanSteenhouse, Yumi Kasai, David Gray, Carol Clugston, Anna Dominiczak and Alex Alderton, Roberto Amato, Jeffrey Barrett, Sonia Goncalves, Ewan Harrison, David K. Jackson, Ian Johnston, Dominic Kwiatkowski, Cordelia Langford, John Sillitoe on behalf of the Wellcome Sanger Institute COVID-19 Surveillance Team |
| EPI_ISL_3441568 | Berkshire and Surrey Pathology Services Lighthouse Laboratory | Wellcome Sanger Institute for the COVID-19 Genomics UK (COG-UK) Consortium | Berkshire and Surrey Pathology Services Lighthouse Laboratory and Alex Alderton, Roberto Amato, Jeffrey Barrett, Sonia Goncalves, Ewan Harrison, David K. Jackson, Ian Johnston, Dominic Kwiatkowski, Cordelia Langford, John Sillitoe on behalf of the Wellcome Sanger Institute COVID-19 Surveillance Team |
| EPI_ISL_3441569 | Lighthouse Lab in Glasgow | Wellcome Sanger Institute for the COVID-19 Genomics UK (COG-UK) Consortium | Harper VanSteenhouse, Yumi Kasai, David Gray, Carol Clugston, Anna Dominiczak and Alex Alderton, Roberto Amato, Jeffrey Barrett, Sonia Goncalves, Ewan Harrison, David K. Jackson, Ian Johnston, Dominic Kwiatkowski, Cordelia Langford, John Sillitoe on behalf of the Wellcome Sanger Institute COVID-19 Surveillance Team |
| EPI_ISL_3441570, EPI_ISL_3441571, EPI_ISL_3441572 | Berkshire and Surrey Pathology Services Lighthouse Laboratory | Wellcome Sanger Institute for the COVID-19 Genomics UK (COG-UK) Consortium | Berkshire and Surrey Pathology Services Lighthouse Laboratory and Alex Alderton, Roberto Amato, Jeffrey Barrett, Sonia Goncalves, Ewan Harrison, David K. Jackson, Ian Johnston, Dominic Kwiatkowski, Cordelia Langford, John Sillitoe on behalf of the Wellcome Sanger Institute COVID-19 Surveillance Team |
| EPI_ISL_3441573, EPI_ISL_3441574 | Lighthouse Lab in Glasgow | Wellcome Sanger Institute for the COVID-19 Genomics UK (COG-UK) Consortium | Harper VanSteenhouse, Yumi Kasai, David Gray, Carol Clugston, Anna Dominiczak and Alex Alderton, Roberto Amato, Jeffrey Barrett, Sonia Goncalves, Ewan Harrison, David K. Jackson, Ian Johnston, Dominic Kwiatkowski, Cordelia Langford, John Sillitoe on behalf of the Wellcome Sanger Institute COVID-19 Surveillance Team |
| EPI_ISL_3441575 | Berkshire and Surrey Pathology Services Lighthouse Laboratory | Wellcome Sanger Institute for the COVID-19 Genomics UK (COG-UK) Consortium | Berkshire and Surrey Pathology Services Lighthouse Laboratory and Alex Alderton, Roberto Amato, Jeffrey Barrett, Sonia Goncalves, Ewan Harrison, David K. Jackson, Ian Johnston, Dominic Kwiatkowski, Cordelia Langford, John Sillitoe on behalf of the Wellcome Sanger Institute COVID-19 Surveillance Team |
| EPI_ISL_3441576, EPI_ISL_3441577, EPI_ISL_3441578 | Lighthouse Lab in Glasgow | Wellcome Sanger Institute for the COVID-19 Genomics UK (COG-UK) Consortium | Harper VanSteenhouse, Yumi Kasai, David Gray, Carol Clugston, Anna Dominiczak and Alex Alderton, Roberto Amato, Jeffrey Barrett, Sonia Goncalves, Ewan Harrison, David K. Jackson, Ian Johnston, Dominic Kwiatkowski, Cordelia Langford, John Sillitoe on behalf of the Wellcome Sanger Institute COVID-19 Surveillance Team |
| EPI_ISL_3441579 | Berkshire and Surrey Pathology Services Lighthouse Laboratory | Wellcome Sanger Institute for the COVID-19 Genomics UK (COG-UK) Consortium | Berkshire and Surrey Pathology Services Lighthouse Laboratory and Alex Alderton, Roberto Amato, Jeffrey Barrett, Sonia Goncalves, Ewan Harrison, David K. Jackson, Ian Johnston, Dominic Kwiatkowski, Cordelia Langford, John Sillitoe on behalf of the Wellcome Sanger Institute COVID-19 Surveillance Team |
| EPI_ISL_3441580 | Lighthouse Lab in Glasgow | Wellcome Sanger Institute for the COVID-19 Genomics UK (COG-UK) Consortium | Harper VanSteenhouse, Yumi Kasai, David Gray, Carol Clugston, Anna Dominiczak and Alex Alderton, Roberto Amato, Jeffrey Barrett, Sonia Goncalves, Ewan Harrison, David K. Jackson, Ian Johnston, Dominic Kwiatkowski, Cordelia Langford, John Sillitoe on behalf of the Wellcome Sanger Institute COVID-19 Surveillance Team |
| EPI_ISL_3441581 | Berkshire and Surrey Pathology Services Lighthouse Laboratory | Wellcome Sanger Institute for the COVID-19 Genomics UK (COG-UK) Consortium | Berkshire and Surrey Pathology Services Lighthouse Laboratory and Alex Alderton, Roberto Amato, Jeffrey Barrett, Sonia Goncalves, Ewan Harrison, David K. Jackson, Ian Johnston, Dominic Kwiatkowski, Cordelia Langford, John Sillitoe on behalf of the Wellcome Sanger Institute COVID-19 Surveillance Team |
| EPI_ISL_3441582, EPI_ISL_3441583, EPI_ISL_3441584 | Lighthouse Lab in Glasgow | Wellcome Sanger Institute for the COVID-19 Genomics UK (COG-UK) Consortium | Harper VanSteenhouse, Yumi Kasai, David Gray, Carol Clugston, Anna Dominiczak and Alex Alderton, Roberto Amato, Jeffrey Barrett, Sonia Goncalves, Ewan Harrison, David K. Jackson, Ian Johnston, Dominic Kwiatkowski, Cordelia Langford, John Sillitoe on behalf of the Wellcome Sanger Institute COVID-19 Surveillance Team |
| EPI_ISL_3441585, EPI_ISL_3441586 | Berkshire and Surrey Pathology Services Lighthouse Laboratory | Wellcome Sanger Institute for the COVID-19 Genomics UK (COG-UK) Consortium | Berkshire and Surrey Pathology Services Lighthouse Laboratory and Alex Alderton, Roberto Amato, Jeffrey Barrett, Sonia Goncalves, Ewan Harrison, David K. Jackson, Ian Johnston, Dominic Kwiatkowski, Cordelia Langford, John Sillitoe on behalf of the Wellcome Sanger Institute COVID-19 Surveillance Team |
| EPI_ISL_3441587, EPI_ISL_3441588, EPI_ISL_3441589 | Lighthouse Lab in Glasgow | Wellcome Sanger Institute for the COVID-19 Genomics UK (COG-UK) Consortium | Harper VanSteenhouse, Yumi Kasai, David Gray, Carol Clugston, Anna Dominiczak and Alex Alderton, Roberto Amato, Jeffrey Barrett, Sonia Goncalves, Ewan Harrison, David K. Jackson, Ian Johnston, Dominic Kwiatkowski, Cordelia Langford, John Sillitoe on behalf of the Wellcome Sanger Institute COVID-19 Surveillance Team |
| EPI_ISL_3441590, EPI_ISL_3441591, EPI_ISL_3441592 | Berkshire and Surrey Pathology Services Lighthouse Laboratory | Wellcome Sanger Institute for the COVID-19 Genomics UK (COG-UK) Consortium | Berkshire and Surrey Pathology Services Lighthouse Laboratory and Alex Alderton, Roberto Amato, Jeffrey Barrett, Sonia Goncalves, Ewan Harrison, David K. Jackson, Ian Johnston, Dominic Kwiatkowski, Cordelia Langford, John Sillitoe on behalf of the Wellcome Sanger Institute COVID-19 Surveillance Team |
| EPI_ISL_3441593, EPI_ISL_3441594, EPI_ISL_3441595, EPI_ISL_3441596, EPI_ISL_3441597, EPI_ISL_3441598 | Lighthouse Lab in Glasgow | Wellcome Sanger Institute for the COVID-19 Genomics UK (COG-UK) Consortium | Harper VanSteenhouse, Yumi Kasai, David Gray, Carol Clugston, Anna Dominiczak and Alex Alderton, Roberto Amato, Jeffrey Barrett, Sonia Goncalves, Ewan Harrison, David K. Jackson, Ian Johnston, Dominic Kwiatkowski, Cordelia Langford, John Sillitoe on behalf of the Wellcome Sanger Institute COVID-19 Surveillance Team |
| EPI_ISL_3441599, EPI_ISL_3441600, EPI_ISL_3441601, EPI_ISL_3441602 | Berkshire and Surrey Pathology Services Lighthouse Laboratory | Wellcome Sanger Institute for the COVID-19 Genomics UK (COG-UK) Consortium | Berkshire and Surrey Pathology Services Lighthouse Laboratory and Alex Alderton, Roberto Amato, Jeffrey Barrett, Sonia Goncalves, Ewan Harrison, David K. Jackson, Ian Johnston, Dominic Kwiatkowski, Cordelia Langford, John Sillitoe on behalf of the Wellcome Sanger Institute COVID-19 Surveillance Team |
| EPI_ISL_3441603, EPI_ISL_3441604, EPI_ISL_3441605 | Lighthouse Lab in Glasgow | Wellcome Sanger Institute for the COVID-19 Genomics UK (COG-UK) Consortium | Harper VanSteenhouse, Yumi Kasai, David Gray, Carol Clugston, Anna Dominiczak and Alex Alderton, Roberto Amato, Jeffrey Barrett, Sonia Goncalves, Ewan Harrison, David K. Jackson, Ian Johnston, Dominic Kwiatkowski, Cordelia Langford, John Sillitoe on behalf of the Wellcome Sanger Institute COVID-19 Surveillance Team |
| EPI_ISL_3441606 | Berkshire and Surrey Pathology Services Lighthouse Laboratory | Wellcome Sanger Institute for the COVID-19 Genomics UK (COG-UK) Consortium | Berkshire and Surrey Pathology Services Lighthouse Laboratory and Alex Alderton, Roberto Amato, Jeffrey Barrett, Sonia Goncalves, Ewan Harrison, David K. Jackson, Ian Johnston, Dominic Kwiatkowski, Cordelia Langford, John Sillitoe on behalf of the Wellcome Sanger Institute COVID-19 Surveillance Team |
| EPI_ISL_3441607, EPI_ISL_3441608, EPI_ISL_3441609, EPI_ISL_3441610 | Lighthouse Lab in Glasgow | Wellcome Sanger Institute for the COVID-19 Genomics UK (COG-UK) Consortium | Harper VanSteenhouse, Yumi Kasai, David Gray, Carol Clugston, Anna Dominiczak and Alex Alderton, Roberto Amato, Jeffrey Barrett, Sonia Goncalves, Ewan Harrison, David K. Jackson, Ian Johnston, Dominic Kwiatkowski, Cordelia Langford, John Sillitoe on behalf of the Wellcome Sanger Institute COVID-19 Surveillance Team |
| EPI_ISL_3441611, EPI_ISL_3441612, EPI_ISL_3441613, EPI_ISL_3441614, EPI_ISL_3441615 | Berkshire and Surrey Pathology Services Lighthouse Laboratory | Wellcome Sanger Institute for the COVID-19 Genomics UK (COG-UK) Consortium | Berkshire and Surrey Pathology Services Lighthouse Laboratory and Alex Alderton, Roberto Amato, Jeffrey Barrett, Sonia Goncalves, Ewan Harrison, David K. Jackson, Ian Johnston, Dominic Kwiatkowski, Cordelia Langford, John Sillitoe on behalf of the Wellcome Sanger Institute COVID-19 Surveillance Team |
| EPI_ISL_3441616 | Lighthouse Lab in Glasgow | Wellcome Sanger Institute for the COVID-19 Genomics UK (COG-UK) Consortium | Harper VanSteenhouse, Yumi Kasai, David Gray, Carol Clugston, Anna Dominiczak and Alex Alderton, Roberto Amato, Jeffrey Barrett, Sonia Goncalves, Ewan Harrison, David K. Jackson, Ian Johnston, Dominic Kwiatkowski, Cordelia Langford, John Sillitoe on behalf of the Wellcome Sanger Institute COVID-19 Surveillance Team |
| EPI_ISL_3441617 | Berkshire and Surrey Pathology Services Lighthouse Laboratory | Wellcome Sanger Institute for the COVID-19 Genomics UK (COG-UK) Consortium | Berkshire and Surrey Pathology Services Lighthouse Laboratory and Alex Alderton, Roberto Amato, Jeffrey Barrett, Sonia Goncalves, Ewan Harrison, David K. Jackson, Ian Johnston, Dominic Kwiatkowski, Cordelia Langford, John Sillitoe on behalf of the Wellcome Sanger Institute COVID-19 Surveillance Team |
| EPI_ISL_3441618, EPI_ISL_3441619 | Lighthouse Lab in Glasgow | Wellcome Sanger Institute for the COVID-19 Genomics UK (COG-UK) Consortium | Harper VanSteenhouse, Yumi Kasai, David Gray, Carol Clugston, Anna Dominiczak and Alex Alderton, Roberto Amato, Jeffrey Barrett, Sonia Goncalves, Ewan Harrison, David K. Jackson, Ian Johnston, Dominic Kwiatkowski, Cordelia Langford, John Sillitoe on behalf of the Wellcome Sanger Institute COVID-19 Surveillance Team |
| EPI_ISL_3441620 | Berkshire and Surrey Pathology Services Lighthouse Laboratory | Wellcome Sanger Institute for the COVID-19 Genomics UK (COG-UK) Consortium | Berkshire and Surrey Pathology Services Lighthouse Laboratory and Alex Alderton, Roberto Amato, Jeffrey Barrett, Sonia Goncalves, Ewan Harrison, David K. Jackson, Ian Johnston, Dominic Kwiatkowski, Cordelia Langford, John Sillitoe on behalf of the Wellcome Sanger Institute COVID-19 Surveillance Team |
| EPI_ISL_3441621, EPI_ISL_3441622, EPI_ISL_3441623 | Lighthouse Lab in Glasgow | Wellcome Sanger Institute for the COVID-19 Genomics UK (COG-UK) Consortium | Harper VanSteenhouse, Yumi Kasai, David Gray, Carol Clugston, Anna Dominiczak and Alex Alderton, Roberto Amato, Jeffrey Barrett, Sonia Goncalves, Ewan Harrison, David K. Jackson, Ian Johnston, Dominic Kwiatkowski, Cordelia Langford, John Sillitoe on behalf of the Wellcome Sanger Institute COVID-19 Surveillance Team |
| EPI_ISL_3441624, EPI_ISL_3441625, EPI_ISL_3441626, EPI_ISL_3441627, EPI_ISL_3441628, EPI_ISL_3441629 | Berkshire and Surrey Pathology Services Lighthouse Laboratory | Wellcome Sanger Institute for the COVID-19 Genomics UK (COG-UK) Consortium | Berkshire and Surrey Pathology Services Lighthouse Laboratory and Alex Alderton, Roberto Amato, Jeffrey Barrett, Sonia Goncalves, Ewan Harrison, David K. Jackson, Ian Johnston, Dominic Kwiatkowski, Cordelia Langford, John Sillitoe on behalf of the Wellcome Sanger Institute COVID-19 Surveillance Team |
| EPI_ISL_3441630 | Lighthouse Lab in Glasgow | Wellcome Sanger Institute for the COVID-19 Genomics UK (COG-UK) Consortium | Harper VanSteenhouse, Yumi Kasai, David Gray, Carol Clugston, Anna Dominiczak and Alex Alderton, Roberto Amato, Jeffrey Barrett, Sonia Goncalves, Ewan Harrison, David K. Jackson, Ian Johnston, Dominic Kwiatkowski, Cordelia Langford, John Sillitoe on behalf of the Wellcome Sanger Institute COVID-19 Surveillance Team |
| EPI_ISL_3441631 | Berkshire and Surrey Pathology Services Lighthouse Laboratory | Wellcome Sanger Institute for the COVID-19 Genomics UK (COG-UK) Consortium | Berkshire and Surrey Pathology Services Lighthouse Laboratory and Alex Alderton, Roberto Amato, Jeffrey Barrett, Sonia Goncalves, Ewan Harrison, David K. Jackson, Ian Johnston, Dominic Kwiatkowski, Cordelia Langford, John Sillitoe on behalf of the Wellcome Sanger Institute COVID-19 Surveillance Team |
| EPI_ISL_3441632 | Lighthouse Lab in Glasgow | Wellcome Sanger Institute for the COVID-19 Genomics UK (COG-UK) Consortium | Harper VanSteenhouse, Yumi Kasai, David Gray, Carol Clugston, Anna Dominiczak and Alex Alderton, Roberto Amato, Jeffrey Barrett, Sonia Goncalves, Ewan Harrison, David K. Jackson, Ian Johnston, Dominic Kwiatkowski, Cordelia Langford, John Sillitoe on behalf of the Wellcome Sanger Institute COVID-19 Surveillance Team |
| EPI_ISL_3441633 | Berkshire and Surrey Pathology Services Lighthouse Laboratory | Wellcome Sanger Institute for the COVID-19 Genomics UK (COG-UK) Consortium | Berkshire and Surrey Pathology Services Lighthouse Laboratory and Alex Alderton, Roberto Amato, Jeffrey Barrett, Sonia Goncalves, Ewan Harrison, David K. Jackson, Ian Johnston, Dominic Kwiatkowski, Cordelia Langford, John Sillitoe on behalf of the Wellcome Sanger Institute COVID-19 Surveillance Team |
| EPI_ISL_3441634 | Lighthouse Lab in Glasgow | Wellcome Sanger Institute for the COVID-19 Genomics UK (COG-UK) Consortium | Harper VanSteenhouse, Yumi Kasai, David Gray, Carol Clugston, Anna Dominiczak and Alex Alderton, Roberto Amato, Jeffrey Barrett, Sonia Goncalves, Ewan Harrison, David K. Jackson, Ian Johnston, Dominic Kwiatkowski, Cordelia Langford, John Sillitoe on behalf of the Wellcome Sanger Institute COVID-19 Surveillance Team |
| EPI_ISL_3441635, EPI_ISL_3441636 | Berkshire and Surrey Pathology Services Lighthouse Laboratory | Wellcome Sanger Institute for the COVID-19 Genomics UK (COG-UK) Consortium | Berkshire and Surrey Pathology Services Lighthouse Laboratory and Alex Alderton, Roberto Amato, Jeffrey Barrett, Sonia Goncalves, Ewan Harrison, David K. Jackson, Ian Johnston, Dominic Kwiatkowski, Cordelia Langford, John Sillitoe on behalf of the Wellcome Sanger Institute COVID-19 Surveillance Team |
| EPI_ISL_3441637, EPI_ISL_3441638 | Lighthouse Lab in Glasgow | Wellcome Sanger Institute for the COVID-19 Genomics UK (COG-UK) Consortium | Harper VanSteenhouse, Yumi Kasai, David Gray, Carol Clugston, Anna Dominiczak and Alex Alderton, Roberto Amato, Jeffrey Barrett, Sonia Goncalves, Ewan Harrison, David K. Jackson, Ian Johnston, Dominic Kwiatkowski, Cordelia Langford, John Sillitoe on behalf of the Wellcome Sanger Institute COVID-19 Surveillance Team |

| | | | |
|---|---|---|---|
| EPI_ISL_3441639, EPI_ISL_3441640 | Berkshire and Surrey Pathology Services Lighthouse Laboratory | Wellcome Sanger Institute for the COVID-19 Genomics UK (COG-UK) Consortium | Berkshire and Surrey Pathology Services Lighthouse Laboratory and Alex Alderton, Roberto Amato, Jeffrey Barrett, Sonia Goncalves, Ewan Harrison, David K. Jackson, Ian Johnston, Dominic Kwiatkowski, Cordelia Langford, John Sillitoe on behalf of the Wellcome Sanger Institute COVID-19 Surveillance Team |
| EPI_ISL_3441641 | Lighthouse Lab in Glasgow | Wellcome Sanger Institute for the COVID-19 Genomics UK (COG-UK) Consortium | Harper VanSteenhouse, Yumi Kasai, David Gray, Carol Clugston, Anna Dominiczak and Alex Alderton, Roberto Amato, Jeffrey Barrett, Sonia Goncalves, Ewan Harrison, David K. Jackson, Ian Johnston, Dominic Kwiatkowski, Cordelia Langford, John Sillitoe on behalf of the Wellcome Sanger Institute COVID-19 Surveillance Team |
| EPI_ISL_3441642, EPI_ISL_3441643 | Berkshire and Surrey Pathology Services Lighthouse Laboratory | Wellcome Sanger Institute for the COVID-19 Genomics UK (COG-UK) Consortium | Berkshire and Surrey Pathology Services Lighthouse Laboratory and Alex Alderton, Roberto Amato, Jeffrey Barrett, Sonia Goncalves, Ewan Harrison, David K. Jackson, Ian Johnston, Dominic Kwiatkowski, Cordelia Langford, John Sillitoe on behalf of the Wellcome Sanger Institute COVID-19 Surveillance Team |
| EPI_ISL_3441644, EPI_ISL_3441645 | Lighthouse Lab in Glasgow | Wellcome Sanger Institute for the COVID-19 Genomics UK (COG-UK) Consortium | Harper VanSteenhouse, Yumi Kasai, David Gray, Carol Clugston, Anna Dominiczak and Alex Alderton, Roberto Amato, Jeffrey Barrett, Sonia Goncalves, Ewan Harrison, David K. Jackson, Ian Johnston, Dominic Kwiatkowski, Cordelia Langford, John Sillitoe on behalf of the Wellcome Sanger Institute COVID-19 Surveillance Team |
| EPI_ISL_3441646, EPI_ISL_3441647, EPI_ISL_3441648 | Berkshire and Surrey Pathology Services Lighthouse Laboratory | Wellcome Sanger Institute for the COVID-19 Genomics UK (COG-UK) Consortium | Berkshire and Surrey Pathology Services Lighthouse Laboratory and Alex Alderton, Roberto Amato, Jeffrey Barrett, Sonia Goncalves, Ewan Harrison, David K. Jackson, Ian Johnston, Dominic Kwiatkowski, Cordelia Langford, John Sillitoe on behalf of the Wellcome Sanger Institute COVID-19 Surveillance Team |
| EPI_ISL_3441649, EPI_ISL_3441650 | Lighthouse Lab in Glasgow | Wellcome Sanger Institute for the COVID-19 Genomics UK (COG-UK) Consortium | Harper VanSteenhouse, Yumi Kasai, David Gray, Carol Clugston, Anna Dominiczak and Alex Alderton, Roberto Amato, Jeffrey Barrett, Sonia Goncalves, Ewan Harrison, David K. Jackson, Ian Johnston, Dominic Kwiatkowski, Cordelia Langford, John Sillitoe on behalf of the Wellcome Sanger Institute COVID-19 Surveillance Team |
| EPI_ISL_3441651, EPI_ISL_3441652 | Berkshire and Surrey Pathology Services Lighthouse Laboratory | Wellcome Sanger Institute for the COVID-19 Genomics UK (COG-UK) Consortium | Berkshire and Surrey Pathology Services Lighthouse Laboratory and Alex Alderton, Roberto Amato, Jeffrey Barrett, Sonia Goncalves, Ewan Harrison, David K. Jackson, Ian Johnston, Dominic Kwiatkowski, Cordelia Langford, John Sillitoe on behalf of the Wellcome Sanger Institute COVID-19 Surveillance Team |
| EPI_ISL_3441653, EPI_ISL_3441654, EPI_ISL_3441655 | Lighthouse Lab in Glasgow | Wellcome Sanger Institute for the COVID-19 Genomics UK (COG-UK) Consortium | Harper VanSteenhouse, Yumi Kasai, David Gray, Carol Clugston, Anna Dominiczak and Alex Alderton, Roberto Amato, Jeffrey Barrett, Sonia Goncalves, Ewan Harrison, David K. Jackson, Ian Johnston, Dominic Kwiatkowski, Cordelia Langford, John Sillitoe on behalf of the Wellcome Sanger Institute COVID-19 Surveillance Team |
| EPI_ISL_3441656 | Berkshire and Surrey Pathology Services Lighthouse Laboratory | Wellcome Sanger Institute for the COVID-19 Genomics UK (COG-UK) Consortium | Berkshire and Surrey Pathology Services Lighthouse Laboratory and Alex Alderton, Roberto Amato, Jeffrey Barrett, Sonia Goncalves, Ewan Harrison, David K. Jackson, Ian Johnston, Dominic Kwiatkowski, Cordelia Langford, John Sillitoe on behalf of the Wellcome Sanger Institute COVID-19 Surveillance Team |
| EPI_ISL_3441657, EPI_ISL_3441658 | Lighthouse Lab in Glasgow | Wellcome Sanger Institute for the COVID-19 Genomics UK (COG-UK) Consortium | Harper VanSteenhouse, Yumi Kasai, David Gray, Carol Clugston, Anna Dominiczak and Alex Alderton, Roberto Amato, Jeffrey Barrett, Sonia Goncalves, Ewan Harrison, David K. Jackson, Ian Johnston, Dominic Kwiatkowski, Cordelia Langford, John Sillitoe on behalf of the Wellcome Sanger Institute COVID-19 Surveillance Team |
| EPI_ISL_3441659 | Berkshire and Surrey Pathology Services Lighthouse Laboratory | Wellcome Sanger Institute for the COVID-19 Genomics UK (COG-UK) Consortium | Berkshire and Surrey Pathology Services Lighthouse Laboratory and Alex Alderton, Roberto Amato, Jeffrey Barrett, Sonia Goncalves, Ewan Harrison, David K. Jackson, Ian Johnston, Dominic Kwiatkowski, Cordelia Langford, John Sillitoe on behalf of the Wellcome Sanger Institute COVID-19 Surveillance Team |
| EPI_ISL_3441660, EPI_ISL_3441661, EPI_ISL_3441662 | Lighthouse Lab in Glasgow | Wellcome Sanger Institute for the COVID-19 Genomics UK (COG-UK) Consortium | Harper VanSteenhouse, Yumi Kasai, David Gray, Carol Clugston, Anna Dominiczak and Alex Alderton, Roberto Amato, Jeffrey Barrett, Sonia Goncalves, Ewan Harrison, David K. Jackson, Ian Johnston, Dominic Kwiatkowski, Cordelia Langford, John Sillitoe on behalf of the Wellcome Sanger Institute COVID-19 Surveillance Team |
| EPI_ISL_3441663, EPI_ISL_3441664 | Berkshire and Surrey Pathology Services Lighthouse Laboratory | Wellcome Sanger Institute for the COVID-19 Genomics UK (COG-UK) Consortium | Berkshire and Surrey Pathology Services Lighthouse Laboratory and Alex Alderton, Roberto Amato, Jeffrey Barrett, Sonia Goncalves, Ewan Harrison, David K. Jackson, Ian Johnston, Dominic Kwiatkowski, Cordelia Langford, John Sillitoe on behalf of the Wellcome Sanger Institute COVID-19 Surveillance Team |
| EPI_ISL_3441665, EPI_ISL_3441666 | Lighthouse Lab in Glasgow | Wellcome Sanger Institute for the COVID-19 Genomics UK (COG-UK) Consortium | Harper VanSteenhouse, Yumi Kasai, David Gray, Carol Clugston, Anna Dominiczak and Alex Alderton, Roberto Amato, Jeffrey Barrett, Sonia Goncalves, Ewan Harrison, David K. Jackson, Ian Johnston, Dominic Kwiatkowski, Cordelia Langford, John Sillitoe on behalf of the Wellcome Sanger Institute COVID-19 Surveillance Team |
| EPI_ISL_3441667 | Berkshire and Surrey Pathology Services Lighthouse Laboratory | Wellcome Sanger Institute for the COVID-19 Genomics UK (COG-UK) Consortium | Berkshire and Surrey Pathology Services Lighthouse Laboratory and Alex Alderton, Roberto Amato, Jeffrey Barrett, Sonia Goncalves, Ewan Harrison, David K. Jackson, Ian Johnston, Dominic Kwiatkowski, Cordelia Langford, John Sillitoe on behalf of the Wellcome Sanger Institute COVID-19 Surveillance Team |
| EPI_ISL_3441668, EPI_ISL_3441669 | Lighthouse Lab in Glasgow | Wellcome Sanger Institute for the COVID-19 Genomics UK (COG-UK) Consortium | Harper VanSteenhouse, Yumi Kasai, David Gray, Carol Clugston, Anna Dominiczak and Alex Alderton, Roberto Amato, Jeffrey Barrett, Sonia Goncalves, Ewan Harrison, David K. Jackson, Ian Johnston, Dominic Kwiatkowski, Cordelia Langford, John Sillitoe on behalf of the Wellcome Sanger Institute COVID-19 Surveillance Team |
| EPI_ISL_3441670, EPI_ISL_3441671, EPI_ISL_3441672 | Berkshire and Surrey Pathology Services Lighthouse Laboratory | Wellcome Sanger Institute for the COVID-19 Genomics UK (COG-UK) Consortium | Berkshire and Surrey Pathology Services Lighthouse Laboratory and Alex Alderton, Roberto Amato, Jeffrey Barrett, Sonia Goncalves, Ewan Harrison, David K. Jackson, Ian Johnston, Dominic Kwiatkowski, Cordelia Langford, John Sillitoe on behalf of the Wellcome Sanger Institute COVID-19 Surveillance Team |
| EPI_ISL_3441673, EPI_ISL_3441674, EPI_ISL_3441675, EPI_ISL_3441676, EPI_ISL_3441677 | Lighthouse Lab in Glasgow | Wellcome Sanger Institute for the COVID-19 Genomics UK (COG-UK) Consortium | Harper VanSteenhouse, Yumi Kasai, David Gray, Carol Clugston, Anna Dominiczak and Alex Alderton, Roberto Amato, Jeffrey Barrett, Sonia Goncalves, Ewan Harrison, David K. Jackson, Ian Johnston, Dominic Kwiatkowski, Cordelia Langford, John Sillitoe on behalf of the Wellcome Sanger Institute COVID-19 Surveillance Team |
| EPI_ISL_3441678 | Berkshire and Surrey Pathology Services Lighthouse Laboratory | Wellcome Sanger Institute for the COVID-19 Genomics UK (COG-UK) Consortium | Berkshire and Surrey Pathology Services Lighthouse Laboratory and Alex Alderton, Roberto Amato, Jeffrey Barrett, Sonia Goncalves, Ewan Harrison, David K. Jackson, Ian Johnston, Dominic Kwiatkowski, Cordelia Langford, John Sillitoe on behalf of the Wellcome Sanger Institute COVID-19 Surveillance Team |
| EPI_ISL_3441679 | Lighthouse Lab in Glasgow | Wellcome Sanger Institute for the COVID-19 Genomics UK (COG-UK) Consortium | Harper VanSteenhouse, Yumi Kasai, David Gray, Carol Clugston, Anna Dominiczak and Alex Alderton, Roberto Amato, Jeffrey Barrett, Sonia Goncalves, Ewan Harrison, David K. Jackson, Ian Johnston, Dominic Kwiatkowski, Cordelia Langford, John Sillitoe on behalf of the Wellcome Sanger Institute COVID-19 Surveillance Team |
| EPI_ISL_3441680 | Berkshire and Surrey Pathology Services Lighthouse Laboratory | Wellcome Sanger Institute for the COVID-19 Genomics UK (COG-UK) Consortium | Berkshire and Surrey Pathology Services Lighthouse Laboratory and Alex Alderton, Roberto Amato, Jeffrey Barrett, Sonia Goncalves, Ewan Harrison, David K. Jackson, Ian Johnston, Dominic Kwiatkowski, Cordelia Langford, John Sillitoe on behalf of the Wellcome Sanger Institute COVID-19 Surveillance Team |
| EPI_ISL_3441681, EPI_ISL_3441682, EPI_ISL_3441683, EPI_ISL_3441684, EPI_ISL_3441685, EPI_ISL_3441686, EPI_ISL_3441687 | Lighthouse Lab in Glasgow | Wellcome Sanger Institute for the COVID-19 Genomics UK (COG-UK) Consortium | Harper VanSteenhouse, Yumi Kasai, David Gray, Carol Clugston, Anna Dominiczak and Alex Alderton, Roberto Amato, Jeffrey Barrett, Sonia Goncalves, Ewan Harrison, David K. Jackson, Ian Johnston, Dominic Kwiatkowski, Cordelia Langford, John Sillitoe on behalf of the Wellcome Sanger Institute COVID-19 Surveillance Team |
| EPI_ISL_3441688, EPI_ISL_3441689 | Berkshire and Surrey Pathology Services Lighthouse Laboratory | Wellcome Sanger Institute for the COVID-19 Genomics UK (COG-UK) Consortium | Berkshire and Surrey Pathology Services Lighthouse Laboratory and Alex Alderton, Roberto Amato, Jeffrey Barrett, Sonia Goncalves, Ewan Harrison, David K. Jackson, Ian Johnston, Dominic Kwiatkowski, Cordelia Langford, John Sillitoe on behalf of the Wellcome Sanger Institute COVID-19 Surveillance Team |
| EPI_ISL_3441690 | Lighthouse Lab in Glasgow | Wellcome Sanger Institute for the COVID-19 Genomics UK (COG-UK) Consortium | Harper VanSteenhouse, Yumi Kasai, David Gray, Carol Clugston, Anna Dominiczak and Alex Alderton, Roberto Amato, Jeffrey Barrett, Sonia Goncalves, Ewan Harrison, David K. Jackson, Ian Johnston, Dominic Kwiatkowski, Cordelia Langford, John Sillitoe on behalf of the Wellcome Sanger Institute COVID-19 Surveillance Team |
| EPI_ISL_3441691, EPI_ISL_3441692 | Berkshire and Surrey Pathology Services Lighthouse Laboratory | Wellcome Sanger Institute for the COVID-19 Genomics UK (COG-UK) Consortium | Berkshire and Surrey Pathology Services Lighthouse Laboratory and Alex Alderton, Roberto Amato, Jeffrey Barrett, Sonia Goncalves, Ewan Harrison, David K. Jackson, Ian Johnston, Dominic Kwiatkowski, Cordelia Langford, John Sillitoe on behalf of the Wellcome Sanger Institute COVID-19 Surveillance Team |
| EPI_ISL_3441693, EPI_ISL_3441694 | Lighthouse Lab in Glasgow | Wellcome Sanger Institute for the COVID-19 Genomics UK (COG-UK) Consortium | Harper VanSteenhouse, Yumi Kasai, David Gray, Carol Clugston, Anna Dominiczak and Alex Alderton, Roberto Amato, Jeffrey Barrett, Sonia Goncalves, Ewan Harrison, David K. Jackson, Ian Johnston, Dominic Kwiatkowski, Cordelia Langford, John Sillitoe on behalf of the Wellcome Sanger Institute COVID-19 Surveillance Team |
| EPI_ISL_3441695, EPI_ISL_3441696 | Berkshire and Surrey Pathology Services Lighthouse Laboratory | Wellcome Sanger Institute for the COVID-19 Genomics UK (COG-UK) Consortium | Berkshire and Surrey Pathology Services Lighthouse Laboratory and Alex Alderton, Roberto Amato, Jeffrey Barrett, Sonia Goncalves, Ewan Harrison, David K. Jackson, Ian Johnston, Dominic Kwiatkowski, Cordelia Langford, John Sillitoe on behalf of the Wellcome Sanger Institute COVID-19 Surveillance Team |
| EPI_ISL_3441697 | Lighthouse Lab in Glasgow | Wellcome Sanger Institute for the COVID-19 Genomics UK (COG-UK) Consortium | Harper VanSteenhouse, Yumi Kasai, David Gray, Carol Clugston, Anna Dominiczak and Alex Alderton, Roberto Amato, Jeffrey Barrett, Sonia Goncalves, Ewan Harrison, David K. Jackson, Ian Johnston, Dominic Kwiatkowski, Cordelia Langford, John Sillitoe on behalf of the Wellcome Sanger Institute COVID-19 Surveillance Team |
| EPI_ISL_3441698, EPI_ISL_3441699, EPI_ISL_3441700 | Berkshire and Surrey Pathology Services Lighthouse Laboratory | Wellcome Sanger Institute for the COVID-19 Genomics UK (COG-UK) Consortium | Berkshire and Surrey Pathology Services Lighthouse Laboratory and Alex Alderton, Roberto Amato, Jeffrey Barrett, Sonia Goncalves, Ewan Harrison, David K. Jackson, Ian Johnston, Dominic Kwiatkowski, Cordelia Langford, John Sillitoe on behalf of the Wellcome Sanger Institute COVID-19 Surveillance Team |
| EPI_ISL_3441701, EPI_ISL_3441702, EPI_ISL_3441703 | Lighthouse Lab in Glasgow | Wellcome Sanger Institute for the COVID-19 Genomics UK (COG-UK) Consortium | Harper VanSteenhouse, Yumi Kasai, David Gray, Carol Clugston, Anna Dominiczak and Alex Alderton, Roberto Amato, Jeffrey Barrett, Sonia Goncalves, Ewan Harrison, David K. Jackson, Ian Johnston, Dominic Kwiatkowski, Cordelia Langford, John Sillitoe on behalf of the Wellcome Sanger Institute COVID-19 Surveillance Team |
| EPI_ISL_3441704, EPI_ISL_3441705 | Berkshire and Surrey Pathology Services Lighthouse Laboratory | Wellcome Sanger Institute for the COVID-19 Genomics UK (COG-UK) Consortium | Berkshire and Surrey Pathology Services Lighthouse Laboratory and Alex Alderton, Roberto Amato, Jeffrey Barrett, Sonia Goncalves, Ewan Harrison, David K. Jackson, Ian Johnston, Dominic Kwiatkowski, Cordelia Langford, John Sillitoe on behalf of the Wellcome Sanger Institute COVID-19 Surveillance Team |
| EPI_ISL_3441706 | Lighthouse Lab in Glasgow | Wellcome Sanger Institute for the COVID-19 Genomics UK (COG-UK) Consortium | Harper VanSteenhouse, Yumi Kasai, David Gray, Carol Clugston, Anna Dominiczak and Alex Alderton, Roberto Amato, Jeffrey Barrett, Sonia Goncalves, Ewan Harrison, David K. Jackson, Ian Johnston, Dominic Kwiatkowski, Cordelia Langford, John Sillitoe on behalf of the Wellcome Sanger Institute COVID-19 Surveillance Team |
| EPI_ISL_3441707 | Berkshire and Surrey Pathology Services Lighthouse Laboratory | Wellcome Sanger Institute for the COVID-19 Genomics UK (COG-UK) Consortium | Berkshire and Surrey Pathology Services Lighthouse Laboratory and Alex Alderton, Roberto Amato, Jeffrey Barrett, Sonia Goncalves, Ewan Harrison, David K. Jackson, Ian Johnston, Dominic Kwiatkowski, Cordelia Langford, John Sillitoe on behalf of the Wellcome Sanger Institute COVID-19 Surveillance Team |
| EPI_ISL_3441708 | Lighthouse Lab in Glasgow | Wellcome Sanger Institute for the COVID-19 Genomics UK (COG-UK) Consortium | Harper VanSteenhouse, Yumi Kasai, David Gray, Carol Clugston, Anna Dominiczak and Alex Alderton, Roberto Amato, Jeffrey Barrett, Sonia Goncalves, Ewan Harrison, David K. Jackson, Ian Johnston, Dominic Kwiatkowski, Cordelia Langford, John Sillitoe on behalf of the Wellcome Sanger Institute COVID-19 Surveillance Team |
| EPI_ISL_3441709 | Berkshire and Surrey Pathology Services Lighthouse Laboratory | Wellcome Sanger Institute for the COVID-19 Genomics UK (COG-UK) Consortium | Berkshire and Surrey Pathology Services Lighthouse Laboratory and Alex Alderton, Roberto Amato, Jeffrey Barrett, Sonia Goncalves, Ewan Harrison, David K. Jackson, Ian Johnston, Dominic Kwiatkowski, Cordelia Langford, John Sillitoe on behalf of the Wellcome Sanger Institute COVID-19 Surveillance Team |
| EPI_ISL_3441710 | Lighthouse Lab in Glasgow | Wellcome Sanger Institute for the COVID-19 Genomics UK (COG-UK) Consortium | Harper VanSteenhouse, Yumi Kasai, David Gray, Carol Clugston, Anna Dominiczak and Alex Alderton, Roberto Amato, Jeffrey Barrett, Sonia Goncalves, Ewan Harrison, David K. Jackson, Ian Johnston, Dominic Kwiatkowski, Cordelia Langford, John Sillitoe on behalf of the Wellcome Sanger Institute COVID-19 Surveillance Team |
| EPI_ISL_3441711 | Berkshire and Surrey Pathology Services Lighthouse Laboratory | Wellcome Sanger Institute for the COVID-19 Genomics UK (COG-UK) Consortium | Berkshire and Surrey Pathology Services Lighthouse Laboratory and Alex Alderton, Roberto Amato, Jeffrey Barrett, Sonia Goncalves, Ewan Harrison, David K. Jackson, Ian Johnston, Dominic Kwiatkowski, Cordelia Langford, John Sillitoe on behalf of the Wellcome Sanger Institute COVID-19 Surveillance Team |
| EPI_ISL_3441712 | Lighthouse Lab in Glasgow | Wellcome Sanger Institute for the COVID-19 Genomics UK (COG-UK) Consortium | Harper VanSteenhouse, Yumi Kasai, David Gray, Carol Clugston, Anna Dominiczak and Alex Alderton, Roberto Amato, Jeffrey Barrett, Sonia Goncalves, Ewan Harrison, David K. Jackson, Ian Johnston, Dominic Kwiatkowski, Cordelia Langford, John Sillitoe on behalf of the Wellcome Sanger Institute COVID-19 Surveillance Team |
| EPI_ISL_3441713, EPI_ISL_3441714, EPI_ISL_3441715 | Berkshire and Surrey Pathology Services Lighthouse Laboratory | Wellcome Sanger Institute for the COVID-19 Genomics UK (COG-UK) Consortium | Berkshire and Surrey Pathology Services Lighthouse Laboratory and Alex Alderton, Roberto Amato, Jeffrey Barrett, Sonia Goncalves, Ewan Harrison, David K. Jackson, Ian Johnston, Dominic Kwiatkowski, Cordelia Langford, John Sillitoe on behalf of the Wellcome Sanger Institute COVID-19 Surveillance Team |

| | | | |
|---|---|---|---|
| EPI_ISL_3441716 | Lighthouse Lab in Glasgow | Wellcome Sanger Institute for the COVID-19 Genomics UK (COG-UK) Consortium | Harper VanSteenhouse, Yumi Kasai, David Gray, Carol Clugston, Anna Dominiczak and Alex Alderton, Roberto Amato, Jeffrey Barrett, Sonia Goncalves, Ewan Harrison, David K. Jackson, Ian Johnston, Dominic Kwiatkowski, Cordelia Langford, John Sillitoe on behalf of the Wellcome Sanger Institute COVID-19 Surveillance Team |
| EPI_ISL_3441717 | Berkshire and Surrey Pathology Services Lighthouse Laboratory | Wellcome Sanger Institute for the COVID-19 Genomics UK (COG-UK) Consortium | Berkshire and Surrey Pathology Services Lighthouse Laboratory and Alex Alderton, Roberto Amato, Jeffrey Barrett, Sonia Goncalves, Ewan Harrison, David K. Jackson, Ian Johnston, Dominic Kwiatkowski, Cordelia Langford, John Sillitoe on behalf of the Wellcome Sanger Institute COVID-19 Surveillance Team |
| EPI_ISL_3441718 | Lighthouse Lab in Glasgow | Wellcome Sanger Institute for the COVID-19 Genomics UK (COG-UK) Consortium | Harper VanSteenhouse, Yumi Kasai, David Gray, Carol Clugston, Anna Dominiczak and Alex Alderton, Roberto Amato, Jeffrey Barrett, Sonia Goncalves, Ewan Harrison, David K. Jackson, Ian Johnston, Dominic Kwiatkowski, Cordelia Langford, John Sillitoe on behalf of the Wellcome Sanger Institute COVID-19 Surveillance Team |
| EPI_ISL_3441719, EPI_ISL_3441720 | Berkshire and Surrey Pathology Services Lighthouse Laboratory | Wellcome Sanger Institute for the COVID-19 Genomics UK (COG-UK) Consortium | Berkshire and Surrey Pathology Services Lighthouse Laboratory and Alex Alderton, Roberto Amato, Jeffrey Barrett, Sonia Goncalves, Ewan Harrison, David K. Jackson, Ian Johnston, Dominic Kwiatkowski, Cordelia Langford, John Sillitoe on behalf of the Wellcome Sanger Institute COVID-19 Surveillance Team |
| EPI_ISL_3441721, EPI_ISL_3441722, EPI_ISL_3441723 | Lighthouse Lab in Glasgow | Wellcome Sanger Institute for the COVID-19 Genomics UK (COG-UK) Consortium | Harper VanSteenhouse, Yumi Kasai, David Gray, Carol Clugston, Anna Dominiczak and Alex Alderton, Roberto Amato, Jeffrey Barrett, Sonia Goncalves, Ewan Harrison, David K. Jackson, Ian Johnston, Dominic Kwiatkowski, Cordelia Langford, John Sillitoe on behalf of the Wellcome Sanger Institute COVID-19 Surveillance Team |
| EPI_ISL_3441724, EPI_ISL_3441725, EPI_ISL_3441726 | Berkshire and Surrey Pathology Services Lighthouse Laboratory | Wellcome Sanger Institute for the COVID-19 Genomics UK (COG-UK) Consortium | Berkshire and Surrey Pathology Services Lighthouse Laboratory and Alex Alderton, Roberto Amato, Jeffrey Barrett, Sonia Goncalves, Ewan Harrison, David K. Jackson, Ian Johnston, Dominic Kwiatkowski, Cordelia Langford, John Sillitoe on behalf of the Wellcome Sanger Institute COVID-19 Surveillance Team |
| EPI_ISL_3441727 | Lighthouse Lab in Glasgow | Wellcome Sanger Institute for the COVID-19 Genomics UK (COG-UK) Consortium | Harper VanSteenhouse, Yumi Kasai, David Gray, Carol Clugston, Anna Dominiczak and Alex Alderton, Roberto Amato, Jeffrey Barrett, Sonia Goncalves, Ewan Harrison, David K. Jackson, Ian Johnston, Dominic Kwiatkowski, Cordelia Langford, John Sillitoe on behalf of the Wellcome Sanger Institute COVID-19 Surveillance Team |
| EPI_ISL_3441728, EPI_ISL_3441729, EPI_ISL_3441730 | Berkshire and Surrey Pathology Services Lighthouse Laboratory | Wellcome Sanger Institute for the COVID-19 Genomics UK (COG-UK) Consortium | Berkshire and Surrey Pathology Services Lighthouse Laboratory and Alex Alderton, Roberto Amato, Jeffrey Barrett, Sonia Goncalves, Ewan Harrison, David K. Jackson, Ian Johnston, Dominic Kwiatkowski, Cordelia Langford, John Sillitoe on behalf of the Wellcome Sanger Institute COVID-19 Surveillance Team |
| EPI_ISL_3441731 | Lighthouse Lab in Glasgow | Wellcome Sanger Institute for the COVID-19 Genomics UK (COG-UK) Consortium | Harper VanSteenhouse, Yumi Kasai, David Gray, Carol Clugston, Anna Dominiczak and Alex Alderton, Roberto Amato, Jeffrey Barrett, Sonia Goncalves, Ewan Harrison, David K. Jackson, Ian Johnston, Dominic Kwiatkowski, Cordelia Langford, John Sillitoe on behalf of the Wellcome Sanger Institute COVID-19 Surveillance Team |
| EPI_ISL_3441732, EPI_ISL_3441733, EPI_ISL_3441734 | Berkshire and Surrey Pathology Services Lighthouse Laboratory | Wellcome Sanger Institute for the COVID-19 Genomics UK (COG-UK) Consortium | Berkshire and Surrey Pathology Services Lighthouse Laboratory and Alex Alderton, Roberto Amato, Jeffrey Barrett, Sonia Goncalves, Ewan Harrison, David K. Jackson, Ian Johnston, Dominic Kwiatkowski, Cordelia Langford, John Sillitoe on behalf of the Wellcome Sanger Institute COVID-19 Surveillance Team |
| EPI_ISL_3441735, EPI_ISL_3441736 | Lighthouse Lab in Glasgow | Wellcome Sanger Institute for the COVID-19 Genomics UK (COG-UK) Consortium | Harper VanSteenhouse, Yumi Kasai, David Gray, Carol Clugston, Anna Dominiczak and Alex Alderton, Roberto Amato, Jeffrey Barrett, Sonia Goncalves, Ewan Harrison, David K. Jackson, Ian Johnston, Dominic Kwiatkowski, Cordelia Langford, John Sillitoe on behalf of the Wellcome Sanger Institute COVID-19 Surveillance Team |
| EPI_ISL_3441737, EPI_ISL_3441738, EPI_ISL_3441739, EPI_ISL_3441740, EPI_ISL_3441741 | Berkshire and Surrey Pathology Services Lighthouse Laboratory | Wellcome Sanger Institute for the COVID-19 Genomics UK (COG-UK) Consortium | Berkshire and Surrey Pathology Services Lighthouse Laboratory and Alex Alderton, Roberto Amato, Jeffrey Barrett, Sonia Goncalves, Ewan Harrison, David K. Jackson, Ian Johnston, Dominic Kwiatkowski, Cordelia Langford, John Sillitoe on behalf of the Wellcome Sanger Institute COVID-19 Surveillance Team |
| EPI_ISL_3441742 | Lighthouse Lab in Glasgow | Wellcome Sanger Institute for the COVID-19 Genomics UK (COG-UK) Consortium | Harper VanSteenhouse, Yumi Kasai, David Gray, Carol Clugston, Anna Dominiczak and Alex Alderton, Roberto Amato, Jeffrey Barrett, Sonia Goncalves, Ewan Harrison, David K. Jackson, Ian Johnston, Dominic Kwiatkowski, Cordelia Langford, John Sillitoe on behalf of the Wellcome Sanger Institute COVID-19 Surveillance Team |
| EPI_ISL_3441743, EPI_ISL_3441744, EPI_ISL_3441745, EPI_ISL_3441746 | Berkshire and Surrey Pathology Services Lighthouse Laboratory | Wellcome Sanger Institute for the COVID-19 Genomics UK (COG-UK) Consortium | Berkshire and Surrey Pathology Services Lighthouse Laboratory and Alex Alderton, Roberto Amato, Jeffrey Barrett, Sonia Goncalves, Ewan Harrison, David K. Jackson, Ian Johnston, Dominic Kwiatkowski, Cordelia Langford, John Sillitoe on behalf of the Wellcome Sanger Institute COVID-19 Surveillance Team |
| EPI_ISL_3441747 | Lighthouse Lab in Glasgow | Wellcome Sanger Institute for the COVID-19 Genomics UK (COG-UK) Consortium | Harper VanSteenhouse, Yumi Kasai, David Gray, Carol Clugston, Anna Dominiczak and Alex Alderton, Roberto Amato, Jeffrey Barrett, Sonia Goncalves, Ewan Harrison, David K. Jackson, Ian Johnston, Dominic Kwiatkowski, Cordelia Langford, John Sillitoe on behalf of the Wellcome Sanger Institute COVID-19 Surveillance Team |
| EPI_ISL_3441748 | Berkshire and Surrey Pathology Services Lighthouse Laboratory | Wellcome Sanger Institute for the COVID-19 Genomics UK (COG-UK) Consortium | Berkshire and Surrey Pathology Services Lighthouse Laboratory and Alex Alderton, Roberto Amato, Jeffrey Barrett, Sonia Goncalves, Ewan Harrison, David K. Jackson, Ian Johnston, Dominic Kwiatkowski, Cordelia Langford, John Sillitoe on behalf of the Wellcome Sanger Institute COVID-19 Surveillance Team |
| EPI_ISL_3441749 | Lighthouse Lab in Glasgow | Wellcome Sanger Institute for the COVID-19 Genomics UK (COG-UK) Consortium | Harper VanSteenhouse, Yumi Kasai, David Gray, Carol Clugston, Anna Dominiczak and Alex Alderton, Roberto Amato, Jeffrey Barrett, Sonia Goncalves, Ewan Harrison, David K. Jackson, Ian Johnston, Dominic Kwiatkowski, Cordelia Langford, John Sillitoe on behalf of the Wellcome Sanger Institute COVID-19 Surveillance Team |
| EPI_ISL_3441750, EPI_ISL_3441751, EPI_ISL_3441752, EPI_ISL_3441753 | Berkshire and Surrey Pathology Services Lighthouse Laboratory | Wellcome Sanger Institute for the COVID-19 Genomics UK (COG-UK) Consortium | Berkshire and Surrey Pathology Services Lighthouse Laboratory and Alex Alderton, Roberto Amato, Jeffrey Barrett, Sonia Goncalves, Ewan Harrison, David K. Jackson, Ian Johnston, Dominic Kwiatkowski, Cordelia Langford, John Sillitoe on behalf of the Wellcome Sanger Institute COVID-19 Surveillance Team |
| EPI_ISL_3441754 | Lighthouse Lab in Glasgow | Wellcome Sanger Institute for the COVID-19 Genomics UK (COG-UK) Consortium | Harper VanSteenhouse, Yumi Kasai, David Gray, Carol Clugston, Anna Dominiczak and Alex Alderton, Roberto Amato, Jeffrey Barrett, Sonia Goncalves, Ewan Harrison, David K. Jackson, Ian Johnston, Dominic Kwiatkowski, Cordelia Langford, John Sillitoe on behalf of the Wellcome Sanger Institute COVID-19 Surveillance Team |
| EPI_ISL_3441755, EPI_ISL_3441756, EPI_ISL_3441757, EPI_ISL_3441758, EPI_ISL_3441759, EPI_ISL_3441760, EPI_ISL_3441761, EPI_ISL_3441762, EPI_ISL_3441763, EPI_ISL_3441764 | Berkshire and Surrey Pathology Services Lighthouse Laboratory | Wellcome Sanger Institute for the COVID-19 Genomics UK (COG-UK) Consortium | Berkshire and Surrey Pathology Services Lighthouse Laboratory and Alex Alderton, Roberto Amato, Jeffrey Barrett, Sonia Goncalves, Ewan Harrison, David K. Jackson, Ian Johnston, Dominic Kwiatkowski, Cordelia Langford, John Sillitoe on behalf of the Wellcome Sanger Institute COVID-19 Surveillance Team |
| EPI_ISL_3441765 | Lighthouse Lab in Glasgow | Wellcome Sanger Institute for the COVID-19 Genomics UK (COG-UK) Consortium | Harper VanSteenhouse, Yumi Kasai, David Gray, Carol Clugston, Anna Dominiczak and Alex Alderton, Roberto Amato, Jeffrey Barrett, Sonia Goncalves, Ewan Harrison, David K. Jackson, Ian Johnston, Dominic Kwiatkowski, Cordelia Langford, John Sillitoe on behalf of the Wellcome Sanger Institute COVID-19 Surveillance Team |
| EPI_ISL_3441766, EPI_ISL_3441767, EPI_ISL_3441768 | Berkshire and Surrey Pathology Services Lighthouse Laboratory | Wellcome Sanger Institute for the COVID-19 Genomics UK (COG-UK) Consortium | Berkshire and Surrey Pathology Services Lighthouse Laboratory and Alex Alderton, Roberto Amato, Jeffrey Barrett, Sonia Goncalves, Ewan Harrison, David K. Jackson, Ian Johnston, Dominic Kwiatkowski, Cordelia Langford, John Sillitoe on behalf of the Wellcome Sanger Institute COVID-19 Surveillance Team |
| EPI_ISL_3441769 | Lighthouse Lab in Glasgow | Wellcome Sanger Institute for the COVID-19 Genomics UK (COG-UK) Consortium | Harper VanSteenhouse, Yumi Kasai, David Gray, Carol Clugston, Anna Dominiczak and Alex Alderton, Roberto Amato, Jeffrey Barrett, Sonia Goncalves, Ewan Harrison, David K. Jackson, Ian Johnston, Dominic Kwiatkowski, Cordelia Langford, John Sillitoe on behalf of the Wellcome Sanger Institute COVID-19 Surveillance Team |
| EPI_ISL_3441770 | Berkshire and Surrey Pathology Services Lighthouse Laboratory | Wellcome Sanger Institute for the COVID-19 Genomics UK (COG-UK) Consortium | Berkshire and Surrey Pathology Services Lighthouse Laboratory and Alex Alderton, Roberto Amato, Jeffrey Barrett, Sonia Goncalves, Ewan Harrison, David K. Jackson, Ian Johnston, Dominic Kwiatkowski, Cordelia Langford, John Sillitoe on behalf of the Wellcome Sanger Institute COVID-19 Surveillance Team |
| EPI_ISL_3441771, EPI_ISL_3441772 | Lighthouse Lab in Glasgow | Wellcome Sanger Institute for the COVID-19 Genomics UK (COG-UK) Consortium | Harper VanSteenhouse, Yumi Kasai, David Gray, Carol Clugston, Anna Dominiczak and Alex Alderton, Roberto Amato, Jeffrey Barrett, Sonia Goncalves, Ewan Harrison, David K. Jackson, Ian Johnston, Dominic Kwiatkowski, Cordelia Langford, John Sillitoe on behalf of the Wellcome Sanger Institute COVID-19 Surveillance Team |
| EPI_ISL_3441773, EPI_ISL_3441774 | Berkshire and Surrey Pathology Services Lighthouse Laboratory | Wellcome Sanger Institute for the COVID-19 Genomics UK (COG-UK) Consortium | Berkshire and Surrey Pathology Services Lighthouse Laboratory and Alex Alderton, Roberto Amato, Jeffrey Barrett, Sonia Goncalves, Ewan Harrison, David K. Jackson, Ian Johnston, Dominic Kwiatkowski, Cordelia Langford, John Sillitoe on behalf of the Wellcome Sanger Institute COVID-19 Surveillance Team |
| EPI_ISL_3441775 | Lighthouse Lab in Glasgow | Wellcome Sanger Institute for the COVID-19 Genomics UK (COG-UK) Consortium | Harper VanSteenhouse, Yumi Kasai, David Gray, Carol Clugston, Anna Dominiczak and Alex Alderton, Roberto Amato, Jeffrey Barrett, Sonia Goncalves, Ewan Harrison, David K. Jackson, Ian Johnston, Dominic Kwiatkowski, Cordelia Langford, John Sillitoe on behalf of the Wellcome Sanger Institute COVID-19 Surveillance Team |
| EPI_ISL_3441776 | Berkshire and Surrey Pathology Services Lighthouse Laboratory | Wellcome Sanger Institute for the COVID-19 Genomics UK (COG-UK) Consortium | Berkshire and Surrey Pathology Services Lighthouse Laboratory and Alex Alderton, Roberto Amato, Jeffrey Barrett, Sonia Goncalves, Ewan Harrison, David K. Jackson, Ian Johnston, Dominic Kwiatkowski, Cordelia Langford, John Sillitoe on behalf of the Wellcome Sanger Institute COVID-19 Surveillance Team |
| EPI_ISL_3441777 | Lighthouse Lab in Glasgow | Wellcome Sanger Institute for the COVID-19 Genomics UK (COG-UK) Consortium | Harper VanSteenhouse, Yumi Kasai, David Gray, Carol Clugston, Anna Dominiczak and Alex Alderton, Roberto Amato, Jeffrey Barrett, Sonia Goncalves, Ewan Harrison, David K. Jackson, Ian Johnston, Dominic Kwiatkowski, Cordelia Langford, John Sillitoe on behalf of the Wellcome Sanger Institute COVID-19 Surveillance Team |
| EPI_ISL_3441778 | Berkshire and Surrey Pathology Services Lighthouse Laboratory | Wellcome Sanger Institute for the COVID-19 Genomics UK (COG-UK) Consortium | Berkshire and Surrey Pathology Services Lighthouse Laboratory and Alex Alderton, Roberto Amato, Jeffrey Barrett, Sonia Goncalves, Ewan Harrison, David K. Jackson, Ian Johnston, Dominic Kwiatkowski, Cordelia Langford, John Sillitoe on behalf of the Wellcome Sanger Institute COVID-19 Surveillance Team |
| EPI_ISL_3441779 | Lighthouse Lab in Glasgow | Wellcome Sanger Institute for the COVID-19 Genomics UK (COG-UK) Consortium | Harper VanSteenhouse, Yumi Kasai, David Gray, Carol Clugston, Anna Dominiczak and Alex Alderton, Roberto Amato, Jeffrey Barrett, Sonia Goncalves, Ewan Harrison, David K. Jackson, Ian Johnston, Dominic Kwiatkowski, Cordelia Langford, John Sillitoe on behalf of the Wellcome Sanger Institute COVID-19 Surveillance Team |
| EPI_ISL_3441780 | Berkshire and Surrey Pathology Services Lighthouse Laboratory | Wellcome Sanger Institute for the COVID-19 Genomics UK (COG-UK) Consortium | Berkshire and Surrey Pathology Services Lighthouse Laboratory and Alex Alderton, Roberto Amato, Jeffrey Barrett, Sonia Goncalves, Ewan Harrison, David K. Jackson, Ian Johnston, Dominic Kwiatkowski, Cordelia Langford, John Sillitoe on behalf of the Wellcome Sanger Institute COVID-19 Surveillance Team |
| EPI_ISL_3441781, EPI_ISL_3441782, EPI_ISL_3441783 | Lighthouse Lab in Glasgow | Wellcome Sanger Institute for the COVID-19 Genomics UK (COG-UK) Consortium | Harper VanSteenhouse, Yumi Kasai, David Gray, Carol Clugston, Anna Dominiczak and Alex Alderton, Roberto Amato, Jeffrey Barrett, Sonia Goncalves, Ewan Harrison, David K. Jackson, Ian Johnston, Dominic Kwiatkowski, Cordelia Langford, John Sillitoe on behalf of the Wellcome Sanger Institute COVID-19 Surveillance Team |
| EPI_ISL_3441784, EPI_ISL_3441785, EPI_ISL_3441786, EPI_ISL_3441787, EPI_ISL_3441788, EPI_ISL_3441789, EPI_ISL_3441790 | Berkshire and Surrey Pathology Services Lighthouse Laboratory | Wellcome Sanger Institute for the COVID-19 Genomics UK (COG-UK) Consortium | Berkshire and Surrey Pathology Services Lighthouse Laboratory and Alex Alderton, Roberto Amato, Jeffrey Barrett, Sonia Goncalves, Ewan Harrison, David K. Jackson, Ian Johnston, Dominic Kwiatkowski, Cordelia Langford, John Sillitoe on behalf of the Wellcome Sanger Institute COVID-19 Surveillance Team |
| EPI_ISL_3441791 | Lighthouse Lab in Glasgow | Wellcome Sanger Institute for the COVID-19 Genomics UK (COG-UK) | Harper VanSteenhouse, Yumi Kasai, David Gray, Carol Clugston, Anna Dominiczak and Alex Alderton, Roberto Amato, Jeffrey Barrett, Sonia Goncalves, Ewan Harrison, David K. Jackson, Ian Johnston, Dominic Kwiatkowski, Cordelia Langford, |

| | | | |
|---|---|---|---|
| | | Consortium | John Sillitoe on behalf of the Wellcome Sanger Institute COVID-19 Surveillance Team |
| EPI_ISL_3441792, EPI_ISL_3441793 | Berkshire and Surrey Pathology Services Lighthouse Laboratory | Wellcome Sanger Institute for the COVID-19 Genomics UK (COG-UK) Consortium | Berkshire and Surrey Pathology Services Lighthouse Laboratory and Alex Alderton, Roberto Amato, Jeffrey Barrett, Sonia Goncalves, Ewan Harrison, David K. Jackson, Ian Johnston, Dominic Kwiatkowski, Cordelia Langford, John Sillitoe on behalf of the Wellcome Sanger Institute COVID-19 Surveillance Team |
| EPI_ISL_3441794 | Lighthouse Lab in Glasgow | Wellcome Sanger Institute for the COVID-19 Genomics UK (COG-UK) Consortium | Harper VanSteenhouse, Yumi Kasai, David Gray, Carol Clugston, Anna Dominiczak and Alex Alderton, Roberto Amato, Jeffrey Barrett, Sonia Goncalves, Ewan Harrison, David K. Jackson, Ian Johnston, Dominic Kwiatkowski, Cordelia Langford, John Sillitoe on behalf of the Wellcome Sanger Institute COVID-19 Surveillance Team |
| EPI_ISL_3441795, EPI_ISL_3441796 | Berkshire and Surrey Pathology Services Lighthouse Laboratory | Wellcome Sanger Institute for the COVID-19 Genomics UK (COG-UK) Consortium | Berkshire and Surrey Pathology Services Lighthouse Laboratory and Alex Alderton, Roberto Amato, Jeffrey Barrett, Sonia Goncalves, Ewan Harrison, David K. Jackson, Ian Johnston, Dominic Kwiatkowski, Cordelia Langford, John Sillitoe on behalf of the Wellcome Sanger Institute COVID-19 Surveillance Team |
| EPI_ISL_3441797 | Lighthouse Lab in Glasgow | Wellcome Sanger Institute for the COVID-19 Genomics UK (COG-UK) Consortium | Harper VanSteenhouse, Yumi Kasai, David Gray, Carol Clugston, Anna Dominiczak and Alex Alderton, Roberto Amato, Jeffrey Barrett, Sonia Goncalves, Ewan Harrison, David K. Jackson, Ian Johnston, Dominic Kwiatkowski, Cordelia Langford, John Sillitoe on behalf of the Wellcome Sanger Institute COVID-19 Surveillance Team |
| EPI_ISL_3441798, EPI_ISL_3441799 | Berkshire and Surrey Pathology Services Lighthouse Laboratory | Wellcome Sanger Institute for the COVID-19 Genomics UK (COG-UK) Consortium | Berkshire and Surrey Pathology Services Lighthouse Laboratory and Alex Alderton, Roberto Amato, Jeffrey Barrett, Sonia Goncalves, Ewan Harrison, David K. Jackson, Ian Johnston, Dominic Kwiatkowski, Cordelia Langford, John Sillitoe on behalf of the Wellcome Sanger Institute COVID-19 Surveillance Team |
| EPI_ISL_3441800 | Lighthouse Lab in Glasgow | Wellcome Sanger Institute for the COVID-19 Genomics UK (COG-UK) Consortium | Harper VanSteenhouse, Yumi Kasai, David Gray, Carol Clugston, Anna Dominiczak and Alex Alderton, Roberto Amato, Jeffrey Barrett, Sonia Goncalves, Ewan Harrison, David K. Jackson, Ian Johnston, Dominic Kwiatkowski, Cordelia Langford, John Sillitoe on behalf of the Wellcome Sanger Institute COVID-19 Surveillance Team |
| EPI_ISL_3441801 | Berkshire and Surrey Pathology Services Lighthouse Laboratory | Wellcome Sanger Institute for the COVID-19 Genomics UK (COG-UK) Consortium | Berkshire and Surrey Pathology Services Lighthouse Laboratory and Alex Alderton, Roberto Amato, Jeffrey Barrett, Sonia Goncalves, Ewan Harrison, David K. Jackson, Ian Johnston, Dominic Kwiatkowski, Cordelia Langford, John Sillitoe on behalf of the Wellcome Sanger Institute COVID-19 Surveillance Team |
| EPI_ISL_3441802, EPI_ISL_3441803 | Lighthouse Lab in Glasgow | Wellcome Sanger Institute for the COVID-19 Genomics UK (COG-UK) Consortium | Harper VanSteenhouse, Yumi Kasai, David Gray, Carol Clugston, Anna Dominiczak and Alex Alderton, Roberto Amato, Jeffrey Barrett, Sonia Goncalves, Ewan Harrison, David K. Jackson, Ian Johnston, Dominic Kwiatkowski, Cordelia Langford, John Sillitoe on behalf of the Wellcome Sanger Institute COVID-19 Surveillance Team |
| EPI_ISL_3441804 | Berkshire and Surrey Pathology Services Lighthouse Laboratory | Wellcome Sanger Institute for the COVID-19 Genomics UK (COG-UK) Consortium | Berkshire and Surrey Pathology Services Lighthouse Laboratory and Alex Alderton, Roberto Amato, Jeffrey Barrett, Sonia Goncalves, Ewan Harrison, David K. Jackson, Ian Johnston, Dominic Kwiatkowski, Cordelia Langford, John Sillitoe on behalf of the Wellcome Sanger Institute COVID-19 Surveillance Team |
| EPI_ISL_3441805 | Lighthouse Lab in Glasgow | Wellcome Sanger Institute for the COVID-19 Genomics UK (COG-UK) Consortium | Harper VanSteenhouse, Yumi Kasai, David Gray, Carol Clugston, Anna Dominiczak and Alex Alderton, Roberto Amato, Jeffrey Barrett, Sonia Goncalves, Ewan Harrison, David K. Jackson, Ian Johnston, Dominic Kwiatkowski, Cordelia Langford, John Sillitoe on behalf of the Wellcome Sanger Institute COVID-19 Surveillance Team |
| EPI_ISL_3441806 | Berkshire and Surrey Pathology Services Lighthouse Laboratory | Wellcome Sanger Institute for the COVID-19 Genomics UK (COG-UK) Consortium | Berkshire and Surrey Pathology Services Lighthouse Laboratory and Alex Alderton, Roberto Amato, Jeffrey Barrett, Sonia Goncalves, Ewan Harrison, David K. Jackson, Ian Johnston, Dominic Kwiatkowski, Cordelia Langford, John Sillitoe on behalf of the Wellcome Sanger Institute COVID-19 Surveillance Team |
| EPI_ISL_3441807 | Lighthouse Lab in Glasgow | Wellcome Sanger Institute for the COVID-19 Genomics UK (COG-UK) Consortium | Harper VanSteenhouse, Yumi Kasai, David Gray, Carol Clugston, Anna Dominiczak and Alex Alderton, Roberto Amato, Jeffrey Barrett, Sonia Goncalves, Ewan Harrison, David K. Jackson, Ian Johnston, Dominic Kwiatkowski, Cordelia Langford, John Sillitoe on behalf of the Wellcome Sanger Institute COVID-19 Surveillance Team |
| EPI_ISL_3441808, EPI_ISL_3441809, EPI_ISL_3441810 | Berkshire and Surrey Pathology Services Lighthouse Laboratory | Wellcome Sanger Institute for the COVID-19 Genomics UK (COG-UK) Consortium | Berkshire and Surrey Pathology Services Lighthouse Laboratory and Alex Alderton, Roberto Amato, Jeffrey Barrett, Sonia Goncalves, Ewan Harrison, David K. Jackson, Ian Johnston, Dominic Kwiatkowski, Cordelia Langford, John Sillitoe on behalf of the Wellcome Sanger Institute COVID-19 Surveillance Team |
| EPI_ISL_3441811 | Lighthouse Lab in Glasgow | Wellcome Sanger Institute for the COVID-19 Genomics UK (COG-UK) Consortium | Harper VanSteenhouse, Yumi Kasai, David Gray, Carol Clugston, Anna Dominiczak and Alex Alderton, Roberto Amato, Jeffrey Barrett, Sonia Goncalves, Ewan Harrison, David K. Jackson, Ian Johnston, Dominic Kwiatkowski, Cordelia Langford, John Sillitoe on behalf of the Wellcome Sanger Institute COVID-19 Surveillance Team |
| EPI_ISL_3441812 | Berkshire and Surrey Pathology Services Lighthouse Laboratory | Wellcome Sanger Institute for the COVID-19 Genomics UK (COG-UK) Consortium | Berkshire and Surrey Pathology Services Lighthouse Laboratory and Alex Alderton, Roberto Amato, Jeffrey Barrett, Sonia Goncalves, Ewan Harrison, David K. Jackson, Ian Johnston, Dominic Kwiatkowski, Cordelia Langford, John Sillitoe on behalf of the Wellcome Sanger Institute COVID-19 Surveillance Team |
| EPI_ISL_3441813 | Lighthouse Lab in Glasgow | Wellcome Sanger Institute for the COVID-19 Genomics UK (COG-UK) Consortium | Harper VanSteenhouse, Yumi Kasai, David Gray, Carol Clugston, Anna Dominiczak and Alex Alderton, Roberto Amato, Jeffrey Barrett, Sonia Goncalves, Ewan Harrison, David K. Jackson, Ian Johnston, Dominic Kwiatkowski, Cordelia Langford, John Sillitoe on behalf of the Wellcome Sanger Institute COVID-19 Surveillance Team |
| EPI_ISL_3441814 | Berkshire and Surrey Pathology Services Lighthouse Laboratory | Wellcome Sanger Institute for the COVID-19 Genomics UK (COG-UK) Consortium | Berkshire and Surrey Pathology Services Lighthouse Laboratory and Alex Alderton, Roberto Amato, Jeffrey Barrett, Sonia Goncalves, Ewan Harrison, David K. Jackson, Ian Johnston, Dominic Kwiatkowski, Cordelia Langford, John Sillitoe on behalf of the Wellcome Sanger Institute COVID-19 Surveillance Team |
| EPI_ISL_3441815, EPI_ISL_3441816 | Lighthouse Lab in Glasgow | Wellcome Sanger Institute for the COVID-19 Genomics UK (COG-UK) Consortium | Harper VanSteenhouse, Yumi Kasai, David Gray, Carol Clugston, Anna Dominiczak and Alex Alderton, Roberto Amato, Jeffrey Barrett, Sonia Goncalves, Ewan Harrison, David K. Jackson, Ian Johnston, Dominic Kwiatkowski, Cordelia Langford, John Sillitoe on behalf of the Wellcome Sanger Institute COVID-19 Surveillance Team |
| EPI_ISL_3441817 | Berkshire and Surrey Pathology Services Lighthouse Laboratory | Wellcome Sanger Institute for the COVID-19 Genomics UK (COG-UK) Consortium | Berkshire and Surrey Pathology Services Lighthouse Laboratory and Alex Alderton, Roberto Amato, Jeffrey Barrett, Sonia Goncalves, Ewan Harrison, David K. Jackson, Ian Johnston, Dominic Kwiatkowski, Cordelia Langford, John Sillitoe on behalf of the Wellcome Sanger Institute COVID-19 Surveillance Team |
| EPI_ISL_3441818 | Lighthouse Lab in Glasgow | Wellcome Sanger Institute for the COVID-19 Genomics UK (COG-UK) Consortium | Harper VanSteenhouse, Yumi Kasai, David Gray, Carol Clugston, Anna Dominiczak and Alex Alderton, Roberto Amato, Jeffrey Barrett, Sonia Goncalves, Ewan Harrison, David K. Jackson, Ian Johnston, Dominic Kwiatkowski, Cordelia Langford, John Sillitoe on behalf of the Wellcome Sanger Institute COVID-19 Surveillance Team |
| EPI_ISL_3441819, EPI_ISL_3441820, EPI_ISL_3441821, EPI_ISL_3441822 | Berkshire and Surrey Pathology Services Lighthouse Laboratory | Wellcome Sanger Institute for the COVID-19 Genomics UK (COG-UK) Consortium | Berkshire and Surrey Pathology Services Lighthouse Laboratory and Alex Alderton, Roberto Amato, Jeffrey Barrett, Sonia Goncalves, Ewan Harrison, David K. Jackson, Ian Johnston, Dominic Kwiatkowski, Cordelia Langford, John Sillitoe on behalf of the Wellcome Sanger Institute COVID-19 Surveillance Team |
| EPI_ISL_3441823 | Lighthouse Lab in Glasgow | Wellcome Sanger Institute for the COVID-19 Genomics UK (COG-UK) Consortium | Harper VanSteenhouse, Yumi Kasai, David Gray, Carol Clugston, Anna Dominiczak and Alex Alderton, Roberto Amato, Jeffrey Barrett, Sonia Goncalves, Ewan Harrison, David K. Jackson, Ian Johnston, Dominic Kwiatkowski, Cordelia Langford, John Sillitoe on behalf of the Wellcome Sanger Institute COVID-19 Surveillance Team |
| EPI_ISL_3441824, EPI_ISL_3441825, EPI_ISL_3441826, EPI_ISL_3441827, EPI_ISL_3441828, EPI_ISL_3441829, EPI_ISL_3441830 | Berkshire and Surrey Pathology Services Lighthouse Laboratory | Wellcome Sanger Institute for the COVID-19 Genomics UK (COG-UK) Consortium | Berkshire and Surrey Pathology Services Lighthouse Laboratory and Alex Alderton, Roberto Amato, Jeffrey Barrett, Sonia Goncalves, Ewan Harrison, David K. Jackson, Ian Johnston, Dominic Kwiatkowski, Cordelia Langford, John Sillitoe on behalf of the Wellcome Sanger Institute COVID-19 Surveillance Team |
| EPI_ISL_3441831, EPI_ISL_3441832, EPI_ISL_3441833 | Lighthouse Lab in Glasgow | Wellcome Sanger Institute for the COVID-19 Genomics UK (COG-UK) Consortium | Harper VanSteenhouse, Yumi Kasai, David Gray, Carol Clugston, Anna Dominiczak and Alex Alderton, Roberto Amato, Jeffrey Barrett, Sonia Goncalves, Ewan Harrison, David K. Jackson, Ian Johnston, Dominic Kwiatkowski, Cordelia Langford, John Sillitoe on behalf of the Wellcome Sanger Institute COVID-19 Surveillance Team |
| EPI_ISL_3441834, EPI_ISL_3441835, EPI_ISL_3441836, EPI_ISL_3441837, EPI_ISL_3441838 | Berkshire and Surrey Pathology Services Lighthouse Laboratory | Wellcome Sanger Institute for the COVID-19 Genomics UK (COG-UK) Consortium | Berkshire and Surrey Pathology Services Lighthouse Laboratory and Alex Alderton, Roberto Amato, Jeffrey Barrett, Sonia Goncalves, Ewan Harrison, David K. Jackson, Ian Johnston, Dominic Kwiatkowski, Cordelia Langford, John Sillitoe on behalf of the Wellcome Sanger Institute COVID-19 Surveillance Team |
| EPI_ISL_3441839, EPI_ISL_3441840 | Lighthouse Lab in Glasgow | Wellcome Sanger Institute for the COVID-19 Genomics UK (COG-UK) Consortium | Harper VanSteenhouse, Yumi Kasai, David Gray, Carol Clugston, Anna Dominiczak and Alex Alderton, Roberto Amato, Jeffrey Barrett, Sonia Goncalves, Ewan Harrison, David K. Jackson, Ian Johnston, Dominic Kwiatkowski, Cordelia Langford, John Sillitoe on behalf of the Wellcome Sanger Institute COVID-19 Surveillance Team |
| EPI_ISL_3441841, EPI_ISL_3441842 | Berkshire and Surrey Pathology Services Lighthouse Laboratory | Wellcome Sanger Institute for the COVID-19 Genomics UK (COG-UK) Consortium | Berkshire and Surrey Pathology Services Lighthouse Laboratory and Alex Alderton, Roberto Amato, Jeffrey Barrett, Sonia Goncalves, Ewan Harrison, David K. Jackson, Ian Johnston, Dominic Kwiatkowski, Cordelia Langford, John Sillitoe on behalf of the Wellcome Sanger Institute COVID-19 Surveillance Team |
| EPI_ISL_3441843 | Lighthouse Lab in Glasgow | Wellcome Sanger Institute for the COVID-19 Genomics UK (COG-UK) Consortium | Harper VanSteenhouse, Yumi Kasai, David Gray, Carol Clugston, Anna Dominiczak and Alex Alderton, Roberto Amato, Jeffrey Barrett, Sonia Goncalves, Ewan Harrison, David K. Jackson, Ian Johnston, Dominic Kwiatkowski, Cordelia Langford, John Sillitoe on behalf of the Wellcome Sanger Institute COVID-19 Surveillance Team |
| EPI_ISL_3441844, EPI_ISL_3441845 | Berkshire and Surrey Pathology Services Lighthouse Laboratory | Wellcome Sanger Institute for the COVID-19 Genomics UK (COG-UK) Consortium | Berkshire and Surrey Pathology Services Lighthouse Laboratory and Alex Alderton, Roberto Amato, Jeffrey Barrett, Sonia Goncalves, Ewan Harrison, David K. Jackson, Ian Johnston, Dominic Kwiatkowski, Cordelia Langford, John Sillitoe on behalf of the Wellcome Sanger Institute COVID-19 Surveillance Team |
| EPI_ISL_3441846 | Lighthouse Lab in Glasgow | Wellcome Sanger Institute for the COVID-19 Genomics UK (COG-UK) Consortium | Harper VanSteenhouse, Yumi Kasai, David Gray, Carol Clugston, Anna Dominiczak and Alex Alderton, Roberto Amato, Jeffrey Barrett, Sonia Goncalves, Ewan Harrison, David K. Jackson, Ian Johnston, Dominic Kwiatkowski, Cordelia Langford, John Sillitoe on behalf of the Wellcome Sanger Institute COVID-19 Surveillance Team |
| EPI_ISL_3441847 | Berkshire and Surrey Pathology Services Lighthouse Laboratory | Wellcome Sanger Institute for the COVID-19 Genomics UK (COG-UK) Consortium | Berkshire and Surrey Pathology Services Lighthouse Laboratory and Alex Alderton, Roberto Amato, Jeffrey Barrett, Sonia Goncalves, Ewan Harrison, David K. Jackson, Ian Johnston, Dominic Kwiatkowski, Cordelia Langford, John Sillitoe on behalf of the Wellcome Sanger Institute COVID-19 Surveillance Team |
| EPI_ISL_3441848 | Lighthouse Lab in Glasgow | Wellcome Sanger Institute for the COVID-19 Genomics UK (COG-UK) Consortium | Harper VanSteenhouse, Yumi Kasai, David Gray, Carol Clugston, Anna Dominiczak and Alex Alderton, Roberto Amato, Jeffrey Barrett, Sonia Goncalves, Ewan Harrison, David K. Jackson, Ian Johnston, Dominic Kwiatkowski, Cordelia Langford, John Sillitoe on behalf of the Wellcome Sanger Institute COVID-19 Surveillance Team |
| EPI_ISL_3441849 | Berkshire and Surrey Pathology Services Lighthouse Laboratory | Wellcome Sanger Institute for the COVID-19 Genomics UK (COG-UK) Consortium | Berkshire and Surrey Pathology Services Lighthouse Laboratory and Alex Alderton, Roberto Amato, Jeffrey Barrett, Sonia Goncalves, Ewan Harrison, David K. Jackson, Ian Johnston, Dominic Kwiatkowski, Cordelia Langford, John Sillitoe on behalf of the Wellcome Sanger Institute COVID-19 Surveillance Team |
| EPI_ISL_3441850 | Lighthouse Lab in Glasgow | Wellcome Sanger Institute for the COVID-19 Genomics UK (COG-UK) Consortium | Harper VanSteenhouse, Yumi Kasai, David Gray, Carol Clugston, Anna Dominiczak and Alex Alderton, Roberto Amato, Jeffrey Barrett, Sonia Goncalves, Ewan Harrison, David K. Jackson, Ian Johnston, Dominic Kwiatkowski, Cordelia Langford, John Sillitoe on behalf of the Wellcome Sanger Institute COVID-19 Surveillance Team |
| EPI_ISL_3441851, EPI_ISL_3441852, EPI_ISL_3441853, EPI_ISL_3441854, EPI_ISL_3441855 | Berkshire and Surrey Pathology Services Lighthouse Laboratory | Wellcome Sanger Institute for the COVID-19 Genomics UK (COG-UK) Consortium | Berkshire and Surrey Pathology Services Lighthouse Laboratory and Alex Alderton, Roberto Amato, Jeffrey Barrett, Sonia Goncalves, Ewan Harrison, David K. Jackson, Ian Johnston, Dominic Kwiatkowski, Cordelia Langford, John Sillitoe on behalf of the Wellcome Sanger Institute COVID-19 Surveillance Team |
| EPI_ISL_3441856 | Lighthouse Lab in Glasgow | Wellcome Sanger Institute for the COVID-19 Genomics UK (COG-UK) Consortium | Harper VanSteenhouse, Yumi Kasai, David Gray, Carol Clugston, Anna Dominiczak and Alex Alderton, Roberto Amato, Jeffrey Barrett, Sonia Goncalves, Ewan Harrison, David K. Jackson, Ian Johnston, Dominic Kwiatkowski, Cordelia Langford, John Sillitoe on behalf of the Wellcome Sanger Institute COVID-19 Surveillance Team |
| EPI_ISL_3441857, EPI_ISL_3441858 | Berkshire and Surrey Pathology Services Lighthouse Laboratory | Wellcome Sanger Institute for the COVID-19 Genomics UK (COG-UK) Consortium | Berkshire and Surrey Pathology Services Lighthouse Laboratory and Alex Alderton, Roberto Amato, Jeffrey Barrett, Sonia Goncalves, Ewan Harrison, David K. Jackson, Ian Johnston, Dominic Kwiatkowski, Cordelia Langford, John Sillitoe on behalf of the Wellcome Sanger Institute COVID-19 Surveillance Team |
| EPI_ISL_3441859 | Lighthouse Lab in Glasgow | Wellcome Sanger Institute for the COVID-19 Genomics UK (COG-UK) Consortium | Harper VanSteenhouse, Yumi Kasai, David Gray, Carol Clugston, Anna Dominiczak and Alex Alderton, Roberto Amato, Jeffrey Barrett, Sonia Goncalves, Ewan Harrison, David K. Jackson, Ian Johnston, Dominic Kwiatkowski, Cordelia Langford, John Sillitoe on behalf of the Wellcome Sanger Institute COVID-19 Surveillance Team |
| EPI_ISL_3441860 | Berkshire and Surrey Pathology Services Lighthouse Laboratory | Wellcome Sanger Institute for the COVID-19 Genomics UK (COG-UK) | Berkshire and Surrey Pathology Services Lighthouse Laboratory and Alex Alderton, Roberto Amato, Jeffrey Barrett, Sonia Goncalves, Ewan Harrison, David K. Jackson, Ian Johnston, Dominic Kwiatkowski, Cordelia Langford, John Sillitoe on behalf |

| | | | |
|---|---|---|---|
| | | Consortium | of the Wellcome Sanger Institute COVID-19 Surveillance Team |
| EPI_ISL_3441861 | Lighthouse Lab in Glasgow | Wellcome Sanger Institute for the COVID-19 Genomics UK (COG-UK) Consortium | Harper VanSteenhouse, Yumi Kasai, David Gray, Carol Clugston, Anna Dominiczak and Alex Alderton, Roberto Amato, Jeffrey Barrett, Sonia Goncalves, Ewan Harrison, David K. Jackson, Ian Johnston, Dominic Kwiatkowski, Cordelia Langford, John Sillitoe on behalf of the Wellcome Sanger Institute COVID-19 Surveillance Team |
| EPI_ISL_3441862, EPI_ISL_3441863, EPI_ISL_3441864, EPI_ISL_3441865 | Berkshire and Surrey Pathology Services Lighthouse Laboratory | Wellcome Sanger Institute for the COVID-19 Genomics UK (COG-UK) Consortium | Berkshire and Surrey Pathology Services Lighthouse Laboratory and Alex Alderton, Roberto Amato, Jeffrey Barrett, Sonia Goncalves, Ewan Harrison, David K. Jackson, Ian Johnston, Dominic Kwiatkowski, Cordelia Langford, John Sillitoe on behalf of the Wellcome Sanger Institute COVID-19 Surveillance Team |
| EPI_ISL_3441866, EPI_ISL_3441867 | Lighthouse Lab in Glasgow | Wellcome Sanger Institute for the COVID-19 Genomics UK (COG-UK) Consortium | Harper VanSteenhouse, Yumi Kasai, David Gray, Carol Clugston, Anna Dominiczak and Alex Alderton, Roberto Amato, Jeffrey Barrett, Sonia Goncalves, Ewan Harrison, David K. Jackson, Ian Johnston, Dominic Kwiatkowski, Cordelia Langford, John Sillitoe on behalf of the Wellcome Sanger Institute COVID-19 Surveillance Team |
| EPI_ISL_3441868 | Berkshire and Surrey Pathology Services Lighthouse Laboratory | Wellcome Sanger Institute for the COVID-19 Genomics UK (COG-UK) Consortium | Berkshire and Surrey Pathology Services Lighthouse Laboratory and Alex Alderton, Roberto Amato, Jeffrey Barrett, Sonia Goncalves, Ewan Harrison, David K. Jackson, Ian Johnston, Dominic Kwiatkowski, Cordelia Langford, John Sillitoe on behalf of the Wellcome Sanger Institute COVID-19 Surveillance Team |
| EPI_ISL_3441869, EPI_ISL_3441870 | Lighthouse Lab in Glasgow | Wellcome Sanger Institute for the COVID-19 Genomics UK (COG-UK) Consortium | Harper VanSteenhouse, Yumi Kasai, David Gray, Carol Clugston, Anna Dominiczak and Alex Alderton, Roberto Amato, Jeffrey Barrett, Sonia Goncalves, Ewan Harrison, David K. Jackson, Ian Johnston, Dominic Kwiatkowski, Cordelia Langford, John Sillitoe on behalf of the Wellcome Sanger Institute COVID-19 Surveillance Team |
| EPI_ISL_3441871 | Berkshire and Surrey Pathology Services Lighthouse Laboratory | Wellcome Sanger Institute for the COVID-19 Genomics UK (COG-UK) Consortium | Berkshire and Surrey Pathology Services Lighthouse Laboratory and Alex Alderton, Roberto Amato, Jeffrey Barrett, Sonia Goncalves, Ewan Harrison, David K. Jackson, Ian Johnston, Dominic Kwiatkowski, Cordelia Langford, John Sillitoe on behalf of the Wellcome Sanger Institute COVID-19 Surveillance Team |
| EPI_ISL_3441872 | Lighthouse Lab in Glasgow | Wellcome Sanger Institute for the COVID-19 Genomics UK (COG-UK) Consortium | Harper VanSteenhouse, Yumi Kasai, David Gray, Carol Clugston, Anna Dominiczak and Alex Alderton, Roberto Amato, Jeffrey Barrett, Sonia Goncalves, Ewan Harrison, David K. Jackson, Ian Johnston, Dominic Kwiatkowski, Cordelia Langford, John Sillitoe on behalf of the Wellcome Sanger Institute COVID-19 Surveillance Team |
| EPI_ISL_3441873, EPI_ISL_3441874, EPI_ISL_3441875, EPI_ISL_3441876, EPI_ISL_3441877 | Berkshire and Surrey Pathology Services Lighthouse Laboratory | Wellcome Sanger Institute for the COVID-19 Genomics UK (COG-UK) Consortium | Berkshire and Surrey Pathology Services Lighthouse Laboratory and Alex Alderton, Roberto Amato, Jeffrey Barrett, Sonia Goncalves, Ewan Harrison, David K. Jackson, Ian Johnston, Dominic Kwiatkowski, Cordelia Langford, John Sillitoe on behalf of the Wellcome Sanger Institute COVID-19 Surveillance Team |
| EPI_ISL_3441878 | Lighthouse Lab in Glasgow | Wellcome Sanger Institute for the COVID-19 Genomics UK (COG-UK) Consortium | Harper VanSteenhouse, Yumi Kasai, David Gray, Carol Clugston, Anna Dominiczak and Alex Alderton, Roberto Amato, Jeffrey Barrett, Sonia Goncalves, Ewan Harrison, David K. Jackson, Ian Johnston, Dominic Kwiatkowski, Cordelia Langford, John Sillitoe on behalf of the Wellcome Sanger Institute COVID-19 Surveillance Team |
| EPI_ISL_3441879 | Berkshire and Surrey Pathology Services Lighthouse Laboratory | Wellcome Sanger Institute for the COVID-19 Genomics UK (COG-UK) Consortium | Berkshire and Surrey Pathology Services Lighthouse Laboratory and Alex Alderton, Roberto Amato, Jeffrey Barrett, Sonia Goncalves, Ewan Harrison, David K. Jackson, Ian Johnston, Dominic Kwiatkowski, Cordelia Langford, John Sillitoe on behalf of the Wellcome Sanger Institute COVID-19 Surveillance Team |
| EPI_ISL_3441880, EPI_ISL_3441881, EPI_ISL_3441882, EPI_ISL_3441883, EPI_ISL_3441884 | Lighthouse Lab in Glasgow | Wellcome Sanger Institute for the COVID-19 Genomics UK (COG-UK) Consortium | Harper VanSteenhouse, Yumi Kasai, David Gray, Carol Clugston, Anna Dominiczak and Alex Alderton, Roberto Amato, Jeffrey Barrett, Sonia Goncalves, Ewan Harrison, David K. Jackson, Ian Johnston, Dominic Kwiatkowski, Cordelia Langford, John Sillitoe on behalf of the Wellcome Sanger Institute COVID-19 Surveillance Team |
| EPI_ISL_3441885 | Berkshire and Surrey Pathology Services Lighthouse Laboratory | Wellcome Sanger Institute for the COVID-19 Genomics UK (COG-UK) Consortium | Berkshire and Surrey Pathology Services Lighthouse Laboratory and Alex Alderton, Roberto Amato, Jeffrey Barrett, Sonia Goncalves, Ewan Harrison, David K. Jackson, Ian Johnston, Dominic Kwiatkowski, Cordelia Langford, John Sillitoe on behalf of the Wellcome Sanger Institute COVID-19 Surveillance Team |
| EPI_ISL_3441886 | Lighthouse Lab in Glasgow | Wellcome Sanger Institute for the COVID-19 Genomics UK (COG-UK) Consortium | Harper VanSteenhouse, Yumi Kasai, David Gray, Carol Clugston, Anna Dominiczak and Alex Alderton, Roberto Amato, Jeffrey Barrett, Sonia Goncalves, Ewan Harrison, David K. Jackson, Ian Johnston, Dominic Kwiatkowski, Cordelia Langford, John Sillitoe on behalf of the Wellcome Sanger Institute COVID-19 Surveillance Team |
| EPI_ISL_3441887, EPI_ISL_3441888 | Berkshire and Surrey Pathology Services Lighthouse Laboratory | Wellcome Sanger Institute for the COVID-19 Genomics UK (COG-UK) Consortium | Berkshire and Surrey Pathology Services Lighthouse Laboratory and Alex Alderton, Roberto Amato, Jeffrey Barrett, Sonia Goncalves, Ewan Harrison, David K. Jackson, Ian Johnston, Dominic Kwiatkowski, Cordelia Langford, John Sillitoe on behalf of the Wellcome Sanger Institute COVID-19 Surveillance Team |
| EPI_ISL_3441889, EPI_ISL_3441890, EPI_ISL_3441891 | Lighthouse Lab in Glasgow | Wellcome Sanger Institute for the COVID-19 Genomics UK (COG-UK) Consortium | Harper VanSteenhouse, Yumi Kasai, David Gray, Carol Clugston, Anna Dominiczak and Alex Alderton, Roberto Amato, Jeffrey Barrett, Sonia Goncalves, Ewan Harrison, David K. Jackson, Ian Johnston, Dominic Kwiatkowski, Cordelia Langford, John Sillitoe on behalf of the Wellcome Sanger Institute COVID-19 Surveillance Team |
| EPI_ISL_3441892 | Berkshire and Surrey Pathology Services Lighthouse Laboratory | Wellcome Sanger Institute for the COVID-19 Genomics UK (COG-UK) Consortium | Berkshire and Surrey Pathology Services Lighthouse Laboratory and Alex Alderton, Roberto Amato, Jeffrey Barrett, Sonia Goncalves, Ewan Harrison, David K. Jackson, Ian Johnston, Dominic Kwiatkowski, Cordelia Langford, John Sillitoe on behalf of the Wellcome Sanger Institute COVID-19 Surveillance Team |
| EPI_ISL_3441893 | Lighthouse Lab in Glasgow | Wellcome Sanger Institute for the COVID-19 Genomics UK (COG-UK) Consortium | Harper VanSteenhouse, Yumi Kasai, David Gray, Carol Clugston, Anna Dominiczak and Alex Alderton, Roberto Amato, Jeffrey Barrett, Sonia Goncalves, Ewan Harrison, David K. Jackson, Ian Johnston, Dominic Kwiatkowski, Cordelia Langford, John Sillitoe on behalf of the Wellcome Sanger Institute COVID-19 Surveillance Team |
| EPI_ISL_3441894, EPI_ISL_3441895 | Berkshire and Surrey Pathology Services Lighthouse Laboratory | Wellcome Sanger Institute for the COVID-19 Genomics UK (COG-UK) Consortium | Berkshire and Surrey Pathology Services Lighthouse Laboratory and Alex Alderton, Roberto Amato, Jeffrey Barrett, Sonia Goncalves, Ewan Harrison, David K. Jackson, Ian Johnston, Dominic Kwiatkowski, Cordelia Langford, John Sillitoe on behalf of the Wellcome Sanger Institute COVID-19 Surveillance Team |
| EPI_ISL_3441896 | Lighthouse Lab in Glasgow | Wellcome Sanger Institute for the COVID-19 Genomics UK (COG-UK) Consortium | Harper VanSteenhouse, Yumi Kasai, David Gray, Carol Clugston, Anna Dominiczak and Alex Alderton, Roberto Amato, Jeffrey Barrett, Sonia Goncalves, Ewan Harrison, David K. Jackson, Ian Johnston, Dominic Kwiatkowski, Cordelia Langford, John Sillitoe on behalf of the Wellcome Sanger Institute COVID-19 Surveillance Team |
| EPI_ISL_3441897 | Berkshire and Surrey Pathology Services Lighthouse Laboratory | Wellcome Sanger Institute for the COVID-19 Genomics UK (COG-UK) Consortium | Berkshire and Surrey Pathology Services Lighthouse Laboratory and Alex Alderton, Roberto Amato, Jeffrey Barrett, Sonia Goncalves, Ewan Harrison, David K. Jackson, Ian Johnston, Dominic Kwiatkowski, Cordelia Langford, John Sillitoe on behalf of the Wellcome Sanger Institute COVID-19 Surveillance Team |
| EPI_ISL_3441898 | Lighthouse Lab in Glasgow | Wellcome Sanger Institute for the COVID-19 Genomics UK (COG-UK) Consortium | Harper VanSteenhouse, Yumi Kasai, David Gray, Carol Clugston, Anna Dominiczak and Alex Alderton, Roberto Amato, Jeffrey Barrett, Sonia Goncalves, Ewan Harrison, David K. Jackson, Ian Johnston, Dominic Kwiatkowski, Cordelia Langford, John Sillitoe on behalf of the Wellcome Sanger Institute COVID-19 Surveillance Team |
| EPI_ISL_3441899, EPI_ISL_3441900 | Berkshire and Surrey Pathology Services Lighthouse Laboratory | Wellcome Sanger Institute for the COVID-19 Genomics UK (COG-UK) Consortium | Berkshire and Surrey Pathology Services Lighthouse Laboratory and Alex Alderton, Roberto Amato, Jeffrey Barrett, Sonia Goncalves, Ewan Harrison, David K. Jackson, Ian Johnston, Dominic Kwiatkowski, Cordelia Langford, John Sillitoe on behalf of the Wellcome Sanger Institute COVID-19 Surveillance Team |
| EPI_ISL_3441901 | Lighthouse Lab in Glasgow | Wellcome Sanger Institute for the COVID-19 Genomics UK (COG-UK) Consortium | Harper VanSteenhouse, Yumi Kasai, David Gray, Carol Clugston, Anna Dominiczak and Alex Alderton, Roberto Amato, Jeffrey Barrett, Sonia Goncalves, Ewan Harrison, David K. Jackson, Ian Johnston, Dominic Kwiatkowski, Cordelia Langford, John Sillitoe on behalf of the Wellcome Sanger Institute COVID-19 Surveillance Team |
| EPI_ISL_3441902, EPI_ISL_3441903, EPI_ISL_3441904 | Berkshire and Surrey Pathology Services Lighthouse Laboratory | Wellcome Sanger Institute for the COVID-19 Genomics UK (COG-UK) Consortium | Berkshire and Surrey Pathology Services Lighthouse Laboratory and Alex Alderton, Roberto Amato, Jeffrey Barrett, Sonia Goncalves, Ewan Harrison, David K. Jackson, Ian Johnston, Dominic Kwiatkowski, Cordelia Langford, John Sillitoe on behalf of the Wellcome Sanger Institute COVID-19 Surveillance Team |
| EPI_ISL_3441905 | Lighthouse Lab in Glasgow | Wellcome Sanger Institute for the COVID-19 Genomics UK (COG-UK) Consortium | Harper VanSteenhouse, Yumi Kasai, David Gray, Carol Clugston, Anna Dominiczak and Alex Alderton, Roberto Amato, Jeffrey Barrett, Sonia Goncalves, Ewan Harrison, David K. Jackson, Ian Johnston, Dominic Kwiatkowski, Cordelia Langford, John Sillitoe on behalf of the Wellcome Sanger Institute COVID-19 Surveillance Team |
| EPI_ISL_3441906, EPI_ISL_3441907, EPI_ISL_3441908, EPI_ISL_3441909, EPI_ISL_3441910, EPI_ISL_3441911, EPI_ISL_3441912, EPI_ISL_3441913, EPI_ISL_3441914 | Berkshire and Surrey Pathology Services Lighthouse Laboratory | Wellcome Sanger Institute for the COVID-19 Genomics UK (COG-UK) Consortium | Berkshire and Surrey Pathology Services Lighthouse Laboratory and Alex Alderton, Roberto Amato, Jeffrey Barrett, Sonia Goncalves, Ewan Harrison, David K. Jackson, Ian Johnston, Dominic Kwiatkowski, Cordelia Langford, John Sillitoe on behalf of the Wellcome Sanger Institute COVID-19 Surveillance Team |
| EPI_ISL_3441915 | Lighthouse Lab in Glasgow | Wellcome Sanger Institute for the COVID-19 Genomics UK (COG-UK) Consortium | Harper VanSteenhouse, Yumi Kasai, David Gray, Carol Clugston, Anna Dominiczak and Alex Alderton, Roberto Amato, Jeffrey Barrett, Sonia Goncalves, Ewan Harrison, David K. Jackson, Ian Johnston, Dominic Kwiatkowski, Cordelia Langford, John Sillitoe on behalf of the Wellcome Sanger Institute COVID-19 Surveillance Team |
| EPI_ISL_3441916, EPI_ISL_3441917 | Berkshire and Surrey Pathology Services Lighthouse Laboratory | Wellcome Sanger Institute for the COVID-19 Genomics UK (COG-UK) Consortium | Berkshire and Surrey Pathology Services Lighthouse Laboratory and Alex Alderton, Roberto Amato, Jeffrey Barrett, Sonia Goncalves, Ewan Harrison, David K. Jackson, Ian Johnston, Dominic Kwiatkowski, Cordelia Langford, John Sillitoe on behalf of the Wellcome Sanger Institute COVID-19 Surveillance Team |
| EPI_ISL_3441918 | Lighthouse Lab in Glasgow | Wellcome Sanger Institute for the COVID-19 Genomics UK (COG-UK) Consortium | Harper VanSteenhouse, Yumi Kasai, David Gray, Carol Clugston, Anna Dominiczak and Alex Alderton, Roberto Amato, Jeffrey Barrett, Sonia Goncalves, Ewan Harrison, David K. Jackson, Ian Johnston, Dominic Kwiatkowski, Cordelia Langford, John Sillitoe on behalf of the Wellcome Sanger Institute COVID-19 Surveillance Team |
| EPI_ISL_3441919 | Berkshire and Surrey Pathology Services Lighthouse Laboratory | Wellcome Sanger Institute for the COVID-19 Genomics UK (COG-UK) Consortium | Berkshire and Surrey Pathology Services Lighthouse Laboratory and Alex Alderton, Roberto Amato, Jeffrey Barrett, Sonia Goncalves, Ewan Harrison, David K. Jackson, Ian Johnston, Dominic Kwiatkowski, Cordelia Langford, John Sillitoe on behalf of the Wellcome Sanger Institute COVID-19 Surveillance Team |
| EPI_ISL_3441920, EPI_ISL_3441921, EPI_ISL_3441922 | Lighthouse Lab in Glasgow | Wellcome Sanger Institute for the COVID-19 Genomics UK (COG-UK) Consortium | Harper VanSteenhouse, Yumi Kasai, David Gray, Carol Clugston, Anna Dominiczak and Alex Alderton, Roberto Amato, Jeffrey Barrett, Sonia Goncalves, Ewan Harrison, David K. Jackson, Ian Johnston, Dominic Kwiatkowski, Cordelia Langford, John Sillitoe on behalf of the Wellcome Sanger Institute COVID-19 Surveillance Team |
| EPI_ISL_3441923, EPI_ISL_3441924 | Berkshire and Surrey Pathology Services Lighthouse Laboratory | Wellcome Sanger Institute for the COVID-19 Genomics UK (COG-UK) Consortium | Berkshire and Surrey Pathology Services Lighthouse Laboratory and Alex Alderton, Roberto Amato, Jeffrey Barrett, Sonia Goncalves, Ewan Harrison, David K. Jackson, Ian Johnston, Dominic Kwiatkowski, Cordelia Langford, John Sillitoe on behalf of the Wellcome Sanger Institute COVID-19 Surveillance Team |
| EPI_ISL_3441925, EPI_ISL_3441926 | Lighthouse Lab in Glasgow | Wellcome Sanger Institute for the COVID-19 Genomics UK (COG-UK) Consortium | Harper VanSteenhouse, Yumi Kasai, David Gray, Carol Clugston, Anna Dominiczak and Alex Alderton, Roberto Amato, Jeffrey Barrett, Sonia Goncalves, Ewan Harrison, David K. Jackson, Ian Johnston, Dominic Kwiatkowski, Cordelia Langford, John Sillitoe on behalf of the Wellcome Sanger Institute COVID-19 Surveillance Team |
| EPI_ISL_3441927, EPI_ISL_3441928, EPI_ISL_3441929, EPI_ISL_3441930, EPI_ISL_3441931 | Berkshire and Surrey Pathology Services Lighthouse Laboratory | Wellcome Sanger Institute for the COVID-19 Genomics UK (COG-UK) Consortium | Berkshire and Surrey Pathology Services Lighthouse Laboratory and Alex Alderton, Roberto Amato, Jeffrey Barrett, Sonia Goncalves, Ewan Harrison, David K. Jackson, Ian Johnston, Dominic Kwiatkowski, Cordelia Langford, John Sillitoe on behalf of the Wellcome Sanger Institute COVID-19 Surveillance Team |
| EPI_ISL_3441932 | Lighthouse Lab in Glasgow | Wellcome Sanger Institute for the COVID-19 Genomics UK (COG-UK) Consortium | Harper VanSteenhouse, Yumi Kasai, David Gray, Carol Clugston, Anna Dominiczak and Alex Alderton, Roberto Amato, Jeffrey Barrett, Sonia Goncalves, Ewan Harrison, David K. Jackson, Ian Johnston, Dominic Kwiatkowski, Cordelia Langford, John Sillitoe on behalf of the Wellcome Sanger Institute COVID-19 Surveillance Team |
| EPI_ISL_3441933, EPI_ISL_3441934, | Berkshire and Surrey Pathology Services Lighthouse Laboratory | Wellcome Sanger Institute for the COVID-19 Genomics UK (COG-UK) Consortium | Berkshire and Surrey Pathology Services Lighthouse Laboratory and Alex Alderton, Roberto Amato, Jeffrey Barrett, Sonia Goncalves, Ewan Harrison, David K. Jackson, Ian Johnston, Dominic Kwiatkowski, Cordelia Langford, John Sillitoe on behalf of the Wellcome Sanger Institute COVID-19 Surveillance Team |

| | | | |
|---|---|---|---|
| EPI_ISL_3441935 | | | |
| EPI_ISL_3441936 | Lighthouse Lab in Glasgow | Wellcome Sanger Institute for the COVID-19 Genomics UK (COG-UK) Consortium | Harper VanSteenhouse, Yumi Kasai, David Gray, Carol Clugston, Anna Dominiczak and Alex Alderton, Roberto Amato, Jeffrey Barrett, Sonia Goncalves, Ewan Harrison, David K. Jackson, Ian Johnston, Dominic Kwiatkowski, Cordelia Langford, John Sillitoe on behalf of the Wellcome Sanger Institute COVID-19 Surveillance Team |
| EPI_ISL_3441937, EPI_ISL_3441938 | Berkshire and Surrey Pathology Services Lighthouse Laboratory | Wellcome Sanger Institute for the COVID-19 Genomics UK (COG-UK) Consortium | Berkshire and Surrey Pathology Services Lighthouse Laboratory and Alex Alderton, Roberto Amato, Jeffrey Barrett, Sonia Goncalves, Ewan Harrison, David K. Jackson, Ian Johnston, Dominic Kwiatkowski, Cordelia Langford, John Sillitoe on behalf of the Wellcome Sanger Institute COVID-19 Surveillance Team |
| EPI_ISL_3441939 | Lighthouse Lab in Glasgow | Wellcome Sanger Institute for the COVID-19 Genomics UK (COG-UK) Consortium | Harper VanSteenhouse, Yumi Kasai, David Gray, Carol Clugston, Anna Dominiczak and Alex Alderton, Roberto Amato, Jeffrey Barrett, Sonia Goncalves, Ewan Harrison, David K. Jackson, Ian Johnston, Dominic Kwiatkowski, Cordelia Langford, John Sillitoe on behalf of the Wellcome Sanger Institute COVID-19 Surveillance Team |
| EPI_ISL_3441940, EPI_ISL_3441941 | Berkshire and Surrey Pathology Services Lighthouse Laboratory | Wellcome Sanger Institute for the COVID-19 Genomics UK (COG-UK) Consortium | Berkshire and Surrey Pathology Services Lighthouse Laboratory and Alex Alderton, Roberto Amato, Jeffrey Barrett, Sonia Goncalves, Ewan Harrison, David K. Jackson, Ian Johnston, Dominic Kwiatkowski, Cordelia Langford, John Sillitoe on behalf of the Wellcome Sanger Institute COVID-19 Surveillance Team |
| EPI_ISL_3441942 | Lighthouse Lab in Glasgow | Wellcome Sanger Institute for the COVID-19 Genomics UK (COG-UK) Consortium | Harper VanSteenhouse, Yumi Kasai, David Gray, Carol Clugston, Anna Dominiczak and Alex Alderton, Roberto Amato, Jeffrey Barrett, Sonia Goncalves, Ewan Harrison, David K. Jackson, Ian Johnston, Dominic Kwiatkowski, Cordelia Langford, John Sillitoe on behalf of the Wellcome Sanger Institute COVID-19 Surveillance Team |
| EPI_ISL_3441943, EPI_ISL_3441944 | Berkshire and Surrey Pathology Services Lighthouse Laboratory | Wellcome Sanger Institute for the COVID-19 Genomics UK (COG-UK) Consortium | Berkshire and Surrey Pathology Services Lighthouse Laboratory and Alex Alderton, Roberto Amato, Jeffrey Barrett, Sonia Goncalves, Ewan Harrison, David K. Jackson, Ian Johnston, Dominic Kwiatkowski, Cordelia Langford, John Sillitoe on behalf of the Wellcome Sanger Institute COVID-19 Surveillance Team |
| EPI_ISL_3441945 | Lighthouse Lab in Glasgow | Wellcome Sanger Institute for the COVID-19 Genomics UK (COG-UK) Consortium | Harper VanSteenhouse, Yumi Kasai, David Gray, Carol Clugston, Anna Dominiczak and Alex Alderton, Roberto Amato, Jeffrey Barrett, Sonia Goncalves, Ewan Harrison, David K. Jackson, Ian Johnston, Dominic Kwiatkowski, Cordelia Langford, John Sillitoe on behalf of the Wellcome Sanger Institute COVID-19 Surveillance Team |
| EPI_ISL_3441946, EPI_ISL_3441947, EPI_ISL_3441948, EPI_ISL_3441949, EPI_ISL_3441950 | Berkshire and Surrey Pathology Services Lighthouse Laboratory | Wellcome Sanger Institute for the COVID-19 Genomics UK (COG-UK) Consortium | Berkshire and Surrey Pathology Services Lighthouse Laboratory and Alex Alderton, Roberto Amato, Jeffrey Barrett, Sonia Goncalves, Ewan Harrison, David K. Jackson, Ian Johnston, Dominic Kwiatkowski, Cordelia Langford, John Sillitoe on behalf of the Wellcome Sanger Institute COVID-19 Surveillance Team |
| EPI_ISL_3441951, EPI_ISL_3441952 | Lighthouse Lab in Glasgow | Wellcome Sanger Institute for the COVID-19 Genomics UK (COG-UK) Consortium | Harper VanSteenhouse, Yumi Kasai, David Gray, Carol Clugston, Anna Dominiczak and Alex Alderton, Roberto Amato, Jeffrey Barrett, Sonia Goncalves, Ewan Harrison, David K. Jackson, Ian Johnston, Dominic Kwiatkowski, Cordelia Langford, John Sillitoe on behalf of the Wellcome Sanger Institute COVID-19 Surveillance Team |
| EPI_ISL_3441953, EPI_ISL_3441954, EPI_ISL_3441955, EPI_ISL_3441956 | Lighthouse Laboratory Plymouth | Wellcome Sanger Institute for the COVID-19 Genomics UK (COG-UK) Consortium | Lighthouse Laboratory Plymouth and Alex Alderton, Roberto Amato, Jeffrey Barrett, Sonia Goncalves, Ewan Harrison, David K. Jackson, Ian Johnston, Dominic Kwiatkowski, Cordelia Langford, John Sillitoe on behalf of the Wellcome Sanger Institute COVID-19 Surveillance Team |
| EPI_ISL_3441957, EPI_ISL_3441958, EPI_ISL_3441959 | Lighthouse Lab in Glasgow | Wellcome Sanger Institute for the COVID-19 Genomics UK (COG-UK) Consortium | Harper VanSteenhouse, Yumi Kasai, David Gray, Carol Clugston, Anna Dominiczak and Alex Alderton, Roberto Amato, Jeffrey Barrett, Sonia Goncalves, Ewan Harrison, David K. Jackson, Ian Johnston, Dominic Kwiatkowski, Cordelia Langford, John Sillitoe on behalf of the Wellcome Sanger Institute COVID-19 Surveillance Team |
| EPI_ISL_3441960, EPI_ISL_3441961 | Lighthouse Laboratory Plymouth | Wellcome Sanger Institute for the COVID-19 Genomics UK (COG-UK) Consortium | Lighthouse Laboratory Plymouth and Alex Alderton, Roberto Amato, Jeffrey Barrett, Sonia Goncalves, Ewan Harrison, David K. Jackson, Ian Johnston, Dominic Kwiatkowski, Cordelia Langford, John Sillitoe on behalf of the Wellcome Sanger Institute COVID-19 Surveillance Team |
| EPI_ISL_3441962 | Lighthouse Lab in Glasgow | Wellcome Sanger Institute for the COVID-19 Genomics UK (COG-UK) Consortium | Harper VanSteenhouse, Yumi Kasai, David Gray, Carol Clugston, Anna Dominiczak and Alex Alderton, Roberto Amato, Jeffrey Barrett, Sonia Goncalves, Ewan Harrison, David K. Jackson, Ian Johnston, Dominic Kwiatkowski, Cordelia Langford, John Sillitoe on behalf of the Wellcome Sanger Institute COVID-19 Surveillance Team |
| EPI_ISL_3441963, EPI_ISL_3441964 | Lighthouse Laboratory Plymouth | Wellcome Sanger Institute for the COVID-19 Genomics UK (COG-UK) Consortium | Lighthouse Laboratory Plymouth and Alex Alderton, Roberto Amato, Jeffrey Barrett, Sonia Goncalves, Ewan Harrison, David K. Jackson, Ian Johnston, Dominic Kwiatkowski, Cordelia Langford, John Sillitoe on behalf of the Wellcome Sanger Institute COVID-19 Surveillance Team |
| EPI_ISL_3441965 | Lighthouse Lab in Glasgow | Wellcome Sanger Institute for the COVID-19 Genomics UK (COG-UK) Consortium | Harper VanSteenhouse, Yumi Kasai, David Gray, Carol Clugston, Anna Dominiczak and Alex Alderton, Roberto Amato, Jeffrey Barrett, Sonia Goncalves, Ewan Harrison, David K. Jackson, Ian Johnston, Dominic Kwiatkowski, Cordelia Langford, John Sillitoe on behalf of the Wellcome Sanger Institute COVID-19 Surveillance Team |
| EPI_ISL_3441966, EPI_ISL_3441967 | Lighthouse Laboratory Plymouth | Wellcome Sanger Institute for the COVID-19 Genomics UK (COG-UK) Consortium | Lighthouse Laboratory Plymouth and Alex Alderton, Roberto Amato, Jeffrey Barrett, Sonia Goncalves, Ewan Harrison, David K. Jackson, Ian Johnston, Dominic Kwiatkowski, Cordelia Langford, John Sillitoe on behalf of the Wellcome Sanger Institute COVID-19 Surveillance Team |
| EPI_ISL_3441968 | Lighthouse Lab in Glasgow | Wellcome Sanger Institute for the COVID-19 Genomics UK (COG-UK) Consortium | Harper VanSteenhouse, Yumi Kasai, David Gray, Carol Clugston, Anna Dominiczak and Alex Alderton, Roberto Amato, Jeffrey Barrett, Sonia Goncalves, Ewan Harrison, David K. Jackson, Ian Johnston, Dominic Kwiatkowski, Cordelia Langford, John Sillitoe on behalf of the Wellcome Sanger Institute COVID-19 Surveillance Team |
| EPI_ISL_3441969 | Lighthouse Laboratory Plymouth | Wellcome Sanger Institute for the COVID-19 Genomics UK (COG-UK) Consortium | Lighthouse Laboratory Plymouth and Alex Alderton, Roberto Amato, Jeffrey Barrett, Sonia Goncalves, Ewan Harrison, David K. Jackson, Ian Johnston, Dominic Kwiatkowski, Cordelia Langford, John Sillitoe on behalf of the Wellcome Sanger Institute COVID-19 Surveillance Team |
| EPI_ISL_3441970, EPI_ISL_3441971 | Lighthouse Lab in Glasgow | Wellcome Sanger Institute for the COVID-19 Genomics UK (COG-UK) Consortium | Harper VanSteenhouse, Yumi Kasai, David Gray, Carol Clugston, Anna Dominiczak and Alex Alderton, Roberto Amato, Jeffrey Barrett, Sonia Goncalves, Ewan Harrison, David K. Jackson, Ian Johnston, Dominic Kwiatkowski, Cordelia Langford, John Sillitoe on behalf of the Wellcome Sanger Institute COVID-19 Surveillance Team |
| EPI_ISL_3441972, EPI_ISL_3441973 | Lighthouse Laboratory Plymouth | Wellcome Sanger Institute for the COVID-19 Genomics UK (COG-UK) Consortium | Lighthouse Laboratory Plymouth and Alex Alderton, Roberto Amato, Jeffrey Barrett, Sonia Goncalves, Ewan Harrison, David K. Jackson, Ian Johnston, Dominic Kwiatkowski, Cordelia Langford, John Sillitoe on behalf of the Wellcome Sanger Institute COVID-19 Surveillance Team |
| EPI_ISL_3441974 | Lighthouse Lab in Glasgow | Wellcome Sanger Institute for the COVID-19 Genomics UK (COG-UK) Consortium | Harper VanSteenhouse, Yumi Kasai, David Gray, Carol Clugston, Anna Dominiczak and Alex Alderton, Roberto Amato, Jeffrey Barrett, Sonia Goncalves, Ewan Harrison, David K. Jackson, Ian Johnston, Dominic Kwiatkowski, Cordelia Langford, John Sillitoe on behalf of the Wellcome Sanger Institute COVID-19 Surveillance Team |
| EPI_ISL_3441975, EPI_ISL_3441976, EPI_ISL_3441977 | Lighthouse Laboratory Plymouth | Wellcome Sanger Institute for the COVID-19 Genomics UK (COG-UK) Consortium | Lighthouse Laboratory Plymouth and Alex Alderton, Roberto Amato, Jeffrey Barrett, Sonia Goncalves, Ewan Harrison, David K. Jackson, Ian Johnston, Dominic Kwiatkowski, Cordelia Langford, John Sillitoe on behalf of the Wellcome Sanger Institute COVID-19 Surveillance Team |
| EPI_ISL_3441978, EPI_ISL_3441979, EPI_ISL_3441980 | Lighthouse Lab in Glasgow | Wellcome Sanger Institute for the COVID-19 Genomics UK (COG-UK) Consortium | Harper VanSteenhouse, Yumi Kasai, David Gray, Carol Clugston, Anna Dominiczak and Alex Alderton, Roberto Amato, Jeffrey Barrett, Sonia Goncalves, Ewan Harrison, David K. Jackson, Ian Johnston, Dominic Kwiatkowski, Cordelia Langford, John Sillitoe on behalf of the Wellcome Sanger Institute COVID-19 Surveillance Team |
| EPI_ISL_3441981, EPI_ISL_3441982, EPI_ISL_3441983, EPI_ISL_3441984 | Lighthouse Laboratory Plymouth | Wellcome Sanger Institute for the COVID-19 Genomics UK (COG-UK) Consortium | Lighthouse Laboratory Plymouth and Alex Alderton, Roberto Amato, Jeffrey Barrett, Sonia Goncalves, Ewan Harrison, David K. Jackson, Ian Johnston, Dominic Kwiatkowski, Cordelia Langford, John Sillitoe on behalf of the Wellcome Sanger Institute COVID-19 Surveillance Team |
| EPI_ISL_3441985, EPI_ISL_3441986 | Lighthouse Lab in Glasgow | Wellcome Sanger Institute for the COVID-19 Genomics UK (COG-UK) Consortium | Harper VanSteenhouse, Yumi Kasai, David Gray, Carol Clugston, Anna Dominiczak and Alex Alderton, Roberto Amato, Jeffrey Barrett, Sonia Goncalves, Ewan Harrison, David K. Jackson, Ian Johnston, Dominic Kwiatkowski, Cordelia Langford, John Sillitoe on behalf of the Wellcome Sanger Institute COVID-19 Surveillance Team |
| EPI_ISL_3441987 | Lighthouse Laboratory Plymouth | Wellcome Sanger Institute for the COVID-19 Genomics UK (COG-UK) Consortium | Lighthouse Laboratory Plymouth and Alex Alderton, Roberto Amato, Jeffrey Barrett, Sonia Goncalves, Ewan Harrison, David K. Jackson, Ian Johnston, Dominic Kwiatkowski, Cordelia Langford, John Sillitoe on behalf of the Wellcome Sanger Institute COVID-19 Surveillance Team |
| EPI_ISL_3441988 | Lighthouse Lab in Glasgow | Wellcome Sanger Institute for the COVID-19 Genomics UK (COG-UK) Consortium | Harper VanSteenhouse, Yumi Kasai, David Gray, Carol Clugston, Anna Dominiczak and Alex Alderton, Roberto Amato, Jeffrey Barrett, Sonia Goncalves, Ewan Harrison, David K. Jackson, Ian Johnston, Dominic Kwiatkowski, Cordelia Langford, John Sillitoe on behalf of the Wellcome Sanger Institute COVID-19 Surveillance Team |
| EPI_ISL_3441989, EPI_ISL_3441990, EPI_ISL_3441991 | Lighthouse Laboratory Plymouth | Wellcome Sanger Institute for the COVID-19 Genomics UK (COG-UK) Consortium | Lighthouse Laboratory Plymouth and Alex Alderton, Roberto Amato, Jeffrey Barrett, Sonia Goncalves, Ewan Harrison, David K. Jackson, Ian Johnston, Dominic Kwiatkowski, Cordelia Langford, John Sillitoe on behalf of the Wellcome Sanger Institute COVID-19 Surveillance Team |
| EPI_ISL_3441992, EPI_ISL_3441993, EPI_ISL_3441994, EPI_ISL_3441995 | Lighthouse Lab in Glasgow | Wellcome Sanger Institute for the COVID-19 Genomics UK (COG-UK) Consortium | Harper VanSteenhouse, Yumi Kasai, David Gray, Carol Clugston, Anna Dominiczak and Alex Alderton, Roberto Amato, Jeffrey Barrett, Sonia Goncalves, Ewan Harrison, David K. Jackson, Ian Johnston, Dominic Kwiatkowski, Cordelia Langford, John Sillitoe on behalf of the Wellcome Sanger Institute COVID-19 Surveillance Team |
| EPI_ISL_3441996 | Lighthouse Laboratory Plymouth | Wellcome Sanger Institute for the COVID-19 Genomics UK (COG-UK) Consortium | Lighthouse Laboratory Plymouth and Alex Alderton, Roberto Amato, Jeffrey Barrett, Sonia Goncalves, Ewan Harrison, David K. Jackson, Ian Johnston, Dominic Kwiatkowski, Cordelia Langford, John Sillitoe on behalf of the Wellcome Sanger Institute COVID-19 Surveillance Team |
| EPI_ISL_3441997 | Lighthouse Lab in Glasgow | Wellcome Sanger Institute for the COVID-19 Genomics UK (COG-UK) Consortium | Harper VanSteenhouse, Yumi Kasai, David Gray, Carol Clugston, Anna Dominiczak and Alex Alderton, Roberto Amato, Jeffrey Barrett, Sonia Goncalves, Ewan Harrison, David K. Jackson, Ian Johnston, Dominic Kwiatkowski, Cordelia Langford, John Sillitoe on behalf of the Wellcome Sanger Institute COVID-19 Surveillance Team |
| EPI_ISL_3441998 | Lighthouse Laboratory Plymouth | Wellcome Sanger Institute for the COVID-19 Genomics UK (COG-UK) Consortium | Lighthouse Laboratory Plymouth and Alex Alderton, Roberto Amato, Jeffrey Barrett, Sonia Goncalves, Ewan Harrison, David K. Jackson, Ian Johnston, Dominic Kwiatkowski, Cordelia Langford, John Sillitoe on behalf of the Wellcome Sanger Institute COVID-19 Surveillance Team |
| EPI_ISL_3441999 | Lighthouse Lab in Glasgow | Wellcome Sanger Institute for the COVID-19 Genomics UK (COG-UK) Consortium | Harper VanSteenhouse, Yumi Kasai, David Gray, Carol Clugston, Anna Dominiczak and Alex Alderton, Roberto Amato, Jeffrey Barrett, Sonia Goncalves, Ewan Harrison, David K. Jackson, Ian Johnston, Dominic Kwiatkowski, Cordelia Langford, John Sillitoe on behalf of the Wellcome Sanger Institute COVID-19 Surveillance Team |
| EPI_ISL_3442000 | Lighthouse Laboratory Plymouth | Wellcome Sanger Institute for the COVID-19 Genomics UK (COG-UK) Consortium | Lighthouse Laboratory Plymouth and Alex Alderton, Roberto Amato, Jeffrey Barrett, Sonia Goncalves, Ewan Harrison, David K. Jackson, Ian Johnston, Dominic Kwiatkowski, Cordelia Langford, John Sillitoe on behalf of the Wellcome Sanger Institute COVID-19 Surveillance Team |
| EPI_ISL_3442001 | Lighthouse Lab in Glasgow | Wellcome Sanger Institute for the COVID-19 Genomics UK (COG-UK) Consortium | Harper VanSteenhouse, Yumi Kasai, David Gray, Carol Clugston, Anna Dominiczak and Alex Alderton, Roberto Amato, Jeffrey Barrett, Sonia Goncalves, Ewan Harrison, David K. Jackson, Ian Johnston, Dominic Kwiatkowski, Cordelia Langford, John Sillitoe on behalf of the Wellcome Sanger Institute COVID-19 Surveillance Team |
| EPI_ISL_3442002, EPI_ISL_3442003, EPI_ISL_3442004 | Lighthouse Laboratory Plymouth | Wellcome Sanger Institute for the COVID-19 Genomics UK (COG-UK) Consortium | Lighthouse Laboratory Plymouth and Alex Alderton, Roberto Amato, Jeffrey Barrett, Sonia Goncalves, Ewan Harrison, David K. Jackson, Ian Johnston, Dominic Kwiatkowski, Cordelia Langford, John Sillitoe on behalf of the Wellcome Sanger Institute COVID-19 Surveillance Team |
| EPI_ISL_3442005 | Lighthouse Lab in Glasgow | Wellcome Sanger Institute for the COVID-19 Genomics UK (COG-UK) Consortium | Harper VanSteenhouse, Yumi Kasai, David Gray, Carol Clugston, Anna Dominiczak and Alex Alderton, Roberto Amato, Jeffrey Barrett, Sonia Goncalves, Ewan Harrison, David K. Jackson, Ian Johnston, Dominic Kwiatkowski, Cordelia Langford, John Sillitoe on behalf of the Wellcome Sanger Institute COVID-19 Surveillance Team |
| EPI_ISL_3442006, EPI_ISL_3442007 | Lighthouse Laboratory Plymouth | Wellcome Sanger Institute for the COVID-19 Genomics UK (COG-UK) Consortium | Lighthouse Laboratory Plymouth and Alex Alderton, Roberto Amato, Jeffrey Barrett, Sonia Goncalves, Ewan Harrison, David K. Jackson, Ian Johnston, Dominic Kwiatkowski, Cordelia Langford, John Sillitoe on behalf of the Wellcome Sanger Institute COVID-19 Surveillance Team |
| EPI_ISL_3442008, EPI_ISL_3442009 | Lighthouse Lab in Glasgow | Wellcome Sanger Institute for the COVID-19 Genomics UK (COG-UK) Consortium | Harper VanSteenhouse, Yumi Kasai, David Gray, Carol Clugston, Anna Dominiczak and Alex Alderton, Roberto Amato, Jeffrey Barrett, Sonia Goncalves, Ewan Harrison, David K. Jackson, Ian Johnston, Dominic Kwiatkowski, Cordelia Langford, John Sillitoe on behalf of the Wellcome Sanger Institute COVID-19 Surveillance Team |

| | | | |
|---|---|---|---|
| EPI_ISL_3442010, EPI_ISL_3442011 | Lighthouse Laboratory Plymouth | Wellcome Sanger Institute for the COVID-19 Genomics UK (COG-UK) Consortium | Lighthouse Laboratory Plymouth and Alex Alderton, Roberto Amato, Jeffrey Barrett, Sonia Goncalves, Ewan Harrison, David K. Jackson, Ian Johnston, Dominic Kwiatkowski, Cordelia Langford, John Sillitoe on behalf of the Wellcome Sanger Institute COVID-19 Surveillance Team |
| EPI_ISL_3442012 | Lighthouse Lab in Glasgow | Wellcome Sanger Institute for the COVID-19 Genomics UK (COG-UK) Consortium | Harper VanSteenhouse, Yumi Kasai, David Gray, Carol Clugston, Anna Dominiczak and Alex Alderton, Roberto Amato, Jeffrey Barrett, Sonia Goncalves, Ewan Harrison, David K. Jackson, Ian Johnston, Dominic Kwiatkowski, Cordelia Langford, John Sillitoe on behalf of the Wellcome Sanger Institute COVID-19 Surveillance Team |
| EPI_ISL_3442013, EPI_ISL_3442014, EPI_ISL_3442015 | Lighthouse Laboratory Plymouth | Wellcome Sanger Institute for the COVID-19 Genomics UK (COG-UK) Consortium | Lighthouse Laboratory Plymouth and Alex Alderton, Roberto Amato, Jeffrey Barrett, Sonia Goncalves, Ewan Harrison, David K. Jackson, Ian Johnston, Dominic Kwiatkowski, Cordelia Langford, John Sillitoe on behalf of the Wellcome Sanger Institute COVID-19 Surveillance Team |
| EPI_ISL_3442016 | Lighthouse Lab in Glasgow | Wellcome Sanger Institute for the COVID-19 Genomics UK (COG-UK) Consortium | Harper VanSteenhouse, Yumi Kasai, David Gray, Carol Clugston, Anna Dominiczak and Alex Alderton, Roberto Amato, Jeffrey Barrett, Sonia Goncalves, Ewan Harrison, David K. Jackson, Ian Johnston, Dominic Kwiatkowski, Cordelia Langford, John Sillitoe on behalf of the Wellcome Sanger Institute COVID-19 Surveillance Team |
| EPI_ISL_3442017 | Lighthouse Laboratory Plymouth | Wellcome Sanger Institute for the COVID-19 Genomics UK (COG-UK) Consortium | Lighthouse Laboratory Plymouth and Alex Alderton, Roberto Amato, Jeffrey Barrett, Sonia Goncalves, Ewan Harrison, David K. Jackson, Ian Johnston, Dominic Kwiatkowski, Cordelia Langford, John Sillitoe on behalf of the Wellcome Sanger Institute COVID-19 Surveillance Team |
| EPI_ISL_3442018, EPI_ISL_3442019, EPI_ISL_3442020 | Lighthouse Lab in Glasgow | Wellcome Sanger Institute for the COVID-19 Genomics UK (COG-UK) Consortium | Harper VanSteenhouse, Yumi Kasai, David Gray, Carol Clugston, Anna Dominiczak and Alex Alderton, Roberto Amato, Jeffrey Barrett, Sonia Goncalves, Ewan Harrison, David K. Jackson, Ian Johnston, Dominic Kwiatkowski, Cordelia Langford, John Sillitoe on behalf of the Wellcome Sanger Institute COVID-19 Surveillance Team |
| EPI_ISL_3442021, EPI_ISL_3442022 | Lighthouse Laboratory Plymouth | Wellcome Sanger Institute for the COVID-19 Genomics UK (COG-UK) Consortium | Lighthouse Laboratory Plymouth and Alex Alderton, Roberto Amato, Jeffrey Barrett, Sonia Goncalves, Ewan Harrison, David K. Jackson, Ian Johnston, Dominic Kwiatkowski, Cordelia Langford, John Sillitoe on behalf of the Wellcome Sanger Institute COVID-19 Surveillance Team |
| EPI_ISL_3442023 | Lighthouse Lab in Glasgow | Wellcome Sanger Institute for the COVID-19 Genomics UK (COG-UK) Consortium | Harper VanSteenhouse, Yumi Kasai, David Gray, Carol Clugston, Anna Dominiczak and Alex Alderton, Roberto Amato, Jeffrey Barrett, Sonia Goncalves, Ewan Harrison, David K. Jackson, Ian Johnston, Dominic Kwiatkowski, Cordelia Langford, John Sillitoe on behalf of the Wellcome Sanger Institute COVID-19 Surveillance Team |
| EPI_ISL_3442024, EPI_ISL_3442025 | Lighthouse Laboratory Plymouth | Wellcome Sanger Institute for the COVID-19 Genomics UK (COG-UK) Consortium | Lighthouse Laboratory Plymouth and Alex Alderton, Roberto Amato, Jeffrey Barrett, Sonia Goncalves, Ewan Harrison, David K. Jackson, Ian Johnston, Dominic Kwiatkowski, Cordelia Langford, John Sillitoe on behalf of the Wellcome Sanger Institute COVID-19 Surveillance Team |
| EPI_ISL_3442026 | Lighthouse Lab in Glasgow | Wellcome Sanger Institute for the COVID-19 Genomics UK (COG-UK) Consortium | Harper VanSteenhouse, Yumi Kasai, David Gray, Carol Clugston, Anna Dominiczak and Alex Alderton, Roberto Amato, Jeffrey Barrett, Sonia Goncalves, Ewan Harrison, David K. Jackson, Ian Johnston, Dominic Kwiatkowski, Cordelia Langford, John Sillitoe on behalf of the Wellcome Sanger Institute COVID-19 Surveillance Team |
| EPI_ISL_3442027 | Lighthouse Laboratory Plymouth | Wellcome Sanger Institute for the COVID-19 Genomics UK (COG-UK) Consortium | Lighthouse Laboratory Plymouth and Alex Alderton, Roberto Amato, Jeffrey Barrett, Sonia Goncalves, Ewan Harrison, David K. Jackson, Ian Johnston, Dominic Kwiatkowski, Cordelia Langford, John Sillitoe on behalf of the Wellcome Sanger Institute COVID-19 Surveillance Team |
| EPI_ISL_3442028, EPI_ISL_3442029 | Lighthouse Lab in Glasgow | Wellcome Sanger Institute for the COVID-19 Genomics UK (COG-UK) Consortium | Harper VanSteenhouse, Yumi Kasai, David Gray, Carol Clugston, Anna Dominiczak and Alex Alderton, Roberto Amato, Jeffrey Barrett, Sonia Goncalves, Ewan Harrison, David K. Jackson, Ian Johnston, Dominic Kwiatkowski, Cordelia Langford, John Sillitoe on behalf of the Wellcome Sanger Institute COVID-19 Surveillance Team |
| EPI_ISL_3442030, EPI_ISL_3442031 | Lighthouse Laboratory Plymouth | Wellcome Sanger Institute for the COVID-19 Genomics UK (COG-UK) Consortium | Lighthouse Laboratory Plymouth and Alex Alderton, Roberto Amato, Jeffrey Barrett, Sonia Goncalves, Ewan Harrison, David K. Jackson, Ian Johnston, Dominic Kwiatkowski, Cordelia Langford, John Sillitoe on behalf of the Wellcome Sanger Institute COVID-19 Surveillance Team |
| EPI_ISL_3442032 | Lighthouse Lab in Glasgow | Wellcome Sanger Institute for the COVID-19 Genomics UK (COG-UK) Consortium | Harper VanSteenhouse, Yumi Kasai, David Gray, Carol Clugston, Anna Dominiczak and Alex Alderton, Roberto Amato, Jeffrey Barrett, Sonia Goncalves, Ewan Harrison, David K. Jackson, Ian Johnston, Dominic Kwiatkowski, Cordelia Langford, John Sillitoe on behalf of the Wellcome Sanger Institute COVID-19 Surveillance Team |
| EPI_ISL_3442033, EPI_ISL_3442034, EPI_ISL_3442035, EPI_ISL_3442036, EPI_ISL_3442037 | Lighthouse Laboratory Plymouth | Wellcome Sanger Institute for the COVID-19 Genomics UK (COG-UK) Consortium | Lighthouse Laboratory Plymouth and Alex Alderton, Roberto Amato, Jeffrey Barrett, Sonia Goncalves, Ewan Harrison, David K. Jackson, Ian Johnston, Dominic Kwiatkowski, Cordelia Langford, John Sillitoe on behalf of the Wellcome Sanger Institute COVID-19 Surveillance Team |
| EPI_ISL_3442038 | Lighthouse Lab in Glasgow | Wellcome Sanger Institute for the COVID-19 Genomics UK (COG-UK) Consortium | Harper VanSteenhouse, Yumi Kasai, David Gray, Carol Clugston, Anna Dominiczak and Alex Alderton, Roberto Amato, Jeffrey Barrett, Sonia Goncalves, Ewan Harrison, David K. Jackson, Ian Johnston, Dominic Kwiatkowski, Cordelia Langford, John Sillitoe on behalf of the Wellcome Sanger Institute COVID-19 Surveillance Team |
| EPI_ISL_3442039 | Lighthouse Laboratory Plymouth | Wellcome Sanger Institute for the COVID-19 Genomics UK (COG-UK) Consortium | Lighthouse Laboratory Plymouth and Alex Alderton, Roberto Amato, Jeffrey Barrett, Sonia Goncalves, Ewan Harrison, David K. Jackson, Ian Johnston, Dominic Kwiatkowski, Cordelia Langford, John Sillitoe on behalf of the Wellcome Sanger Institute COVID-19 Surveillance Team |
| EPI_ISL_3442040, EPI_ISL_3442041, EPI_ISL_3442042, EPI_ISL_3442043, EPI_ISL_3442044, EPI_ISL_3442045 | Lighthouse Lab in Glasgow | Wellcome Sanger Institute for the COVID-19 Genomics UK (COG-UK) Consortium | Harper VanSteenhouse, Yumi Kasai, David Gray, Carol Clugston, Anna Dominiczak and Alex Alderton, Roberto Amato, Jeffrey Barrett, Sonia Goncalves, Ewan Harrison, David K. Jackson, Ian Johnston, Dominic Kwiatkowski, Cordelia Langford, John Sillitoe on behalf of the Wellcome Sanger Institute COVID-19 Surveillance Team |
| EPI_ISL_3442046, EPI_ISL_3442047 | Lighthouse Laboratory Plymouth | Wellcome Sanger Institute for the COVID-19 Genomics UK (COG-UK) Consortium | Lighthouse Laboratory Plymouth and Alex Alderton, Roberto Amato, Jeffrey Barrett, Sonia Goncalves, Ewan Harrison, David K. Jackson, Ian Johnston, Dominic Kwiatkowski, Cordelia Langford, John Sillitoe on behalf of the Wellcome Sanger Institute COVID-19 Surveillance Team |
| EPI_ISL_3442048, EPI_ISL_3442049, EPI_ISL_3442050 | Lighthouse Lab in Glasgow | Wellcome Sanger Institute for the COVID-19 Genomics UK (COG-UK) Consortium | Harper VanSteenhouse, Yumi Kasai, David Gray, Carol Clugston, Anna Dominiczak and Alex Alderton, Roberto Amato, Jeffrey Barrett, Sonia Goncalves, Ewan Harrison, David K. Jackson, Ian Johnston, Dominic Kwiatkowski, Cordelia Langford, John Sillitoe on behalf of the Wellcome Sanger Institute COVID-19 Surveillance Team |
| EPI_ISL_3442051, EPI_ISL_3442052 | Lighthouse Laboratory Plymouth | Wellcome Sanger Institute for the COVID-19 Genomics UK (COG-UK) Consortium | Lighthouse Laboratory Plymouth and Alex Alderton, Roberto Amato, Jeffrey Barrett, Sonia Goncalves, Ewan Harrison, David K. Jackson, Ian Johnston, Dominic Kwiatkowski, Cordelia Langford, John Sillitoe on behalf of the Wellcome Sanger Institute COVID-19 Surveillance Team |
| EPI_ISL_3442053 | Lighthouse Lab in Glasgow | Wellcome Sanger Institute for the COVID-19 Genomics UK (COG-UK) Consortium | Harper VanSteenhouse, Yumi Kasai, David Gray, Carol Clugston, Anna Dominiczak and Alex Alderton, Roberto Amato, Jeffrey Barrett, Sonia Goncalves, Ewan Harrison, David K. Jackson, Ian Johnston, Dominic Kwiatkowski, Cordelia Langford, John Sillitoe on behalf of the Wellcome Sanger Institute COVID-19 Surveillance Team |
| EPI_ISL_3442054, EPI_ISL_3442055 | Lighthouse Laboratory Plymouth | Wellcome Sanger Institute for the COVID-19 Genomics UK (COG-UK) Consortium | Lighthouse Laboratory Plymouth and Alex Alderton, Roberto Amato, Jeffrey Barrett, Sonia Goncalves, Ewan Harrison, David K. Jackson, Ian Johnston, Dominic Kwiatkowski, Cordelia Langford, John Sillitoe on behalf of the Wellcome Sanger Institute COVID-19 Surveillance Team |
| EPI_ISL_3442056 | Lighthouse Lab in Glasgow | Wellcome Sanger Institute for the COVID-19 Genomics UK (COG-UK) Consortium | Harper VanSteenhouse, Yumi Kasai, David Gray, Carol Clugston, Anna Dominiczak and Alex Alderton, Roberto Amato, Jeffrey Barrett, Sonia Goncalves, Ewan Harrison, David K. Jackson, Ian Johnston, Dominic Kwiatkowski, Cordelia Langford, John Sillitoe on behalf of the Wellcome Sanger Institute COVID-19 Surveillance Team |
| EPI_ISL_3442057 | Lighthouse Laboratory Plymouth | Wellcome Sanger Institute for the COVID-19 Genomics UK (COG-UK) Consortium | Lighthouse Laboratory Plymouth and Alex Alderton, Roberto Amato, Jeffrey Barrett, Sonia Goncalves, Ewan Harrison, David K. Jackson, Ian Johnston, Dominic Kwiatkowski, Cordelia Langford, John Sillitoe on behalf of the Wellcome Sanger Institute COVID-19 Surveillance Team |
| EPI_ISL_3442058 | Lighthouse Lab in Glasgow | Wellcome Sanger Institute for the COVID-19 Genomics UK (COG-UK) Consortium | Harper VanSteenhouse, Yumi Kasai, David Gray, Carol Clugston, Anna Dominiczak and Alex Alderton, Roberto Amato, Jeffrey Barrett, Sonia Goncalves, Ewan Harrison, David K. Jackson, Ian Johnston, Dominic Kwiatkowski, Cordelia Langford, John Sillitoe on behalf of the Wellcome Sanger Institute COVID-19 Surveillance Team |
| EPI_ISL_3442059, EPI_ISL_3442060 | Lighthouse Laboratory Plymouth | Wellcome Sanger Institute for the COVID-19 Genomics UK (COG-UK) Consortium | Lighthouse Laboratory Plymouth and Alex Alderton, Roberto Amato, Jeffrey Barrett, Sonia Goncalves, Ewan Harrison, David K. Jackson, Ian Johnston, Dominic Kwiatkowski, Cordelia Langford, John Sillitoe on behalf of the Wellcome Sanger Institute COVID-19 Surveillance Team |
| EPI_ISL_3442061 | Lighthouse Lab in Glasgow | Wellcome Sanger Institute for the COVID-19 Genomics UK (COG-UK) Consortium | Harper VanSteenhouse, Yumi Kasai, David Gray, Carol Clugston, Anna Dominiczak and Alex Alderton, Roberto Amato, Jeffrey Barrett, Sonia Goncalves, Ewan Harrison, David K. Jackson, Ian Johnston, Dominic Kwiatkowski, Cordelia Langford, John Sillitoe on behalf of the Wellcome Sanger Institute COVID-19 Surveillance Team |
| EPI_ISL_3442062 | Lighthouse Laboratory Plymouth | Wellcome Sanger Institute for the COVID-19 Genomics UK (COG-UK) Consortium | Lighthouse Laboratory Plymouth and Alex Alderton, Roberto Amato, Jeffrey Barrett, Sonia Goncalves, Ewan Harrison, David K. Jackson, Ian Johnston, Dominic Kwiatkowski, Cordelia Langford, John Sillitoe on behalf of the Wellcome Sanger Institute COVID-19 Surveillance Team |
| EPI_ISL_3442063, EPI_ISL_3442064, EPI_ISL_3442065, EPI_ISL_3442066 | Lighthouse Lab in Glasgow | Wellcome Sanger Institute for the COVID-19 Genomics UK (COG-UK) Consortium | Harper VanSteenhouse, Yumi Kasai, David Gray, Carol Clugston, Anna Dominiczak and Alex Alderton, Roberto Amato, Jeffrey Barrett, Sonia Goncalves, Ewan Harrison, David K. Jackson, Ian Johnston, Dominic Kwiatkowski, Cordelia Langford, John Sillitoe on behalf of the Wellcome Sanger Institute COVID-19 Surveillance Team |
| EPI_ISL_3442067, EPI_ISL_3442068 | Lighthouse Laboratory Plymouth | Wellcome Sanger Institute for the COVID-19 Genomics UK (COG-UK) Consortium | Lighthouse Laboratory Plymouth and Alex Alderton, Roberto Amato, Jeffrey Barrett, Sonia Goncalves, Ewan Harrison, David K. Jackson, Ian Johnston, Dominic Kwiatkowski, Cordelia Langford, John Sillitoe on behalf of the Wellcome Sanger Institute COVID-19 Surveillance Team |
| EPI_ISL_3442069 | Lighthouse Lab in Glasgow | Wellcome Sanger Institute for the COVID-19 Genomics UK (COG-UK) Consortium | Harper VanSteenhouse, Yumi Kasai, David Gray, Carol Clugston, Anna Dominiczak and Alex Alderton, Roberto Amato, Jeffrey Barrett, Sonia Goncalves, Ewan Harrison, David K. Jackson, Ian Johnston, Dominic Kwiatkowski, Cordelia Langford, John Sillitoe on behalf of the Wellcome Sanger Institute COVID-19 Surveillance Team |
| EPI_ISL_3442070, EPI_ISL_3442071, EPI_ISL_3442072 | Lighthouse Laboratory Plymouth | Wellcome Sanger Institute for the COVID-19 Genomics UK (COG-UK) Consortium | Lighthouse Laboratory Plymouth and Alex Alderton, Roberto Amato, Jeffrey Barrett, Sonia Goncalves, Ewan Harrison, David K. Jackson, Ian Johnston, Dominic Kwiatkowski, Cordelia Langford, John Sillitoe on behalf of the Wellcome Sanger Institute COVID-19 Surveillance Team |
| EPI_ISL_3442073 | Lighthouse Lab in Glasgow | Wellcome Sanger Institute for the COVID-19 Genomics UK (COG-UK) Consortium | Harper VanSteenhouse, Yumi Kasai, David Gray, Carol Clugston, Anna Dominiczak and Alex Alderton, Roberto Amato, Jeffrey Barrett, Sonia Goncalves, Ewan Harrison, David K. Jackson, Ian Johnston, Dominic Kwiatkowski, Cordelia Langford, John Sillitoe on behalf of the Wellcome Sanger Institute COVID-19 Surveillance Team |
| EPI_ISL_3442074, EPI_ISL_3442075, EPI_ISL_3442076, EPI_ISL_3442077 | Lighthouse Laboratory Plymouth | Wellcome Sanger Institute for the COVID-19 Genomics UK (COG-UK) Consortium | Lighthouse Laboratory Plymouth and Alex Alderton, Roberto Amato, Jeffrey Barrett, Sonia Goncalves, Ewan Harrison, David K. Jackson, Ian Johnston, Dominic Kwiatkowski, Cordelia Langford, John Sillitoe on behalf of the Wellcome Sanger Institute COVID-19 Surveillance Team |
| EPI_ISL_3442078 | Lighthouse Lab in Glasgow | Wellcome Sanger Institute for the COVID-19 Genomics UK (COG-UK) Consortium | Harper VanSteenhouse, Yumi Kasai, David Gray, Carol Clugston, Anna Dominiczak and Alex Alderton, Roberto Amato, Jeffrey Barrett, Sonia Goncalves, Ewan Harrison, David K. Jackson, Ian Johnston, Dominic Kwiatkowski, Cordelia Langford, John Sillitoe on behalf of the Wellcome Sanger Institute COVID-19 Surveillance Team |
| EPI_ISL_3442079, EPI_ISL_3442080 | Lighthouse Laboratory Plymouth | Wellcome Sanger Institute for the COVID-19 Genomics UK (COG-UK) Consortium | Lighthouse Laboratory Plymouth and Alex Alderton, Roberto Amato, Jeffrey Barrett, Sonia Goncalves, Ewan Harrison, David K. Jackson, Ian Johnston, Dominic Kwiatkowski, Cordelia Langford, John Sillitoe on behalf of the Wellcome Sanger Institute COVID-19 Surveillance Team |
| EPI_ISL_3442081, EPI_ISL_3442082 | Lighthouse Lab in Glasgow | Wellcome Sanger Institute for the COVID-19 Genomics UK (COG-UK) Consortium | Harper VanSteenhouse, Yumi Kasai, David Gray, Carol Clugston, Anna Dominiczak and Alex Alderton, Roberto Amato, Jeffrey Barrett, Sonia Goncalves, Ewan Harrison, David K. Jackson, Ian Johnston, Dominic Kwiatkowski, Cordelia Langford, John Sillitoe on behalf of the Wellcome Sanger Institute COVID-19 Surveillance Team |
| EPI_ISL_3442083, EPI_ISL_3442084 | Lighthouse Laboratory Plymouth | Wellcome Sanger Institute for the COVID-19 Genomics UK (COG-UK) Consortium | Lighthouse Laboratory Plymouth and Alex Alderton, Roberto Amato, Jeffrey Barrett, Sonia Goncalves, Ewan Harrison, David K. Jackson, Ian Johnston, Dominic Kwiatkowski, Cordelia Langford, John Sillitoe on behalf of the Wellcome Sanger Institute COVID-19 Surveillance Team |

| | | | |
|---|---|---|---|
| EPI_ISL_3442085 | Lighthouse Lab in Glasgow | Wellcome Sanger Institute for the COVID-19 Genomics UK (COG-UK) Consortium | Harper VanSteenhouse, Yumi Kasai, David Gray, Carol Clugston, Anna Dominiczak and Alex Alderton, Roberto Amato, Jeffrey Barrett, Sonia Goncalves, Ewan Harrison, David K. Jackson, Ian Johnston, Dominic Kwiatkowski, Cordelia Langford, John Sillitoe on behalf of the Wellcome Sanger Institute COVID-19 Surveillance Team |
| EPI_ISL_3442086 | Lighthouse Laboratory Plymouth | Wellcome Sanger Institute for the COVID-19 Genomics UK (COG-UK) Consortium | Lighthouse Laboratory Plymouth and Alex Alderton, Roberto Amato, Jeffrey Barrett, Sonia Goncalves, Ewan Harrison, David K. Jackson, Ian Johnston, Dominic Kwiatkowski, Cordelia Langford, John Sillitoe on behalf of the Wellcome Sanger Institute COVID-19 Surveillance Team |
| EPI_ISL_3442087 | Lighthouse Lab in Glasgow | Wellcome Sanger Institute for the COVID-19 Genomics UK (COG-UK) Consortium | Harper VanSteenhouse, Yumi Kasai, David Gray, Carol Clugston, Anna Dominiczak and Alex Alderton, Roberto Amato, Jeffrey Barrett, Sonia Goncalves, Ewan Harrison, David K. Jackson, Ian Johnston, Dominic Kwiatkowski, Cordelia Langford, John Sillitoe on behalf of the Wellcome Sanger Institute COVID-19 Surveillance Team |
| EPI_ISL_3442088, EPI_ISL_3442089, EPI_ISL_3442090, EPI_ISL_3442091, EPI_ISL_3442092 | Lighthouse Laboratory Plymouth | Wellcome Sanger Institute for the COVID-19 Genomics UK (COG-UK) Consortium | Lighthouse Laboratory Plymouth and Alex Alderton, Roberto Amato, Jeffrey Barrett, Sonia Goncalves, Ewan Harrison, David K. Jackson, Ian Johnston, Dominic Kwiatkowski, Cordelia Langford, John Sillitoe on behalf of the Wellcome Sanger Institute COVID-19 Surveillance Team |
| EPI_ISL_3442093, EPI_ISL_3442094, EPI_ISL_3442095, EPI_ISL_3442096, EPI_ISL_3442097, EPI_ISL_3442098, EPI_ISL_3442099, EPI_ISL_3442100 | Lighthouse Lab in Glasgow | Wellcome Sanger Institute for the COVID-19 Genomics UK (COG-UK) Consortium | Harper VanSteenhouse, Yumi Kasai, David Gray, Carol Clugston, Anna Dominiczak and Alex Alderton, Roberto Amato, Jeffrey Barrett, Sonia Goncalves, Ewan Harrison, David K. Jackson, Ian Johnston, Dominic Kwiatkowski, Cordelia Langford, John Sillitoe on behalf of the Wellcome Sanger Institute COVID-19 Surveillance Team |
| EPI_ISL_3442101 | Lighthouse Laboratory Plymouth | Wellcome Sanger Institute for the COVID-19 Genomics UK (COG-UK) Consortium | Lighthouse Laboratory Plymouth and Alex Alderton, Roberto Amato, Jeffrey Barrett, Sonia Goncalves, Ewan Harrison, David K. Jackson, Ian Johnston, Dominic Kwiatkowski, Cordelia Langford, John Sillitoe on behalf of the Wellcome Sanger Institute COVID-19 Surveillance Team |
| EPI_ISL_3442102, EPI_ISL_3442103 | Lighthouse Lab in Glasgow | Wellcome Sanger Institute for the COVID-19 Genomics UK (COG-UK) Consortium | Harper VanSteenhouse, Yumi Kasai, David Gray, Carol Clugston, Anna Dominiczak and Alex Alderton, Roberto Amato, Jeffrey Barrett, Sonia Goncalves, Ewan Harrison, David K. Jackson, Ian Johnston, Dominic Kwiatkowski, Cordelia Langford, John Sillitoe on behalf of the Wellcome Sanger Institute COVID-19 Surveillance Team |
| EPI_ISL_3442104 | Lighthouse Laboratory Plymouth | Wellcome Sanger Institute for the COVID-19 Genomics UK (COG-UK) Consortium | Lighthouse Laboratory Plymouth and Alex Alderton, Roberto Amato, Jeffrey Barrett, Sonia Goncalves, Ewan Harrison, David K. Jackson, Ian Johnston, Dominic Kwiatkowski, Cordelia Langford, John Sillitoe on behalf of the Wellcome Sanger Institute COVID-19 Surveillance Team |
| EPI_ISL_3442105 | Lighthouse Lab in Glasgow | Wellcome Sanger Institute for the COVID-19 Genomics UK (COG-UK) Consortium | Harper VanSteenhouse, Yumi Kasai, David Gray, Carol Clugston, Anna Dominiczak and Alex Alderton, Roberto Amato, Jeffrey Barrett, Sonia Goncalves, Ewan Harrison, David K. Jackson, Ian Johnston, Dominic Kwiatkowski, Cordelia Langford, John Sillitoe on behalf of the Wellcome Sanger Institute COVID-19 Surveillance Team |
| EPI_ISL_3442106, EPI_ISL_3442107, EPI_ISL_3442108, EPI_ISL_3442109, EPI_ISL_3442110, EPI_ISL_3442111, EPI_ISL_3442112, EPI_ISL_3442113, EPI_ISL_3442114, EPI_ISL_3442115, EPI_ISL_3442116 | | | |
| see above | Lighthouse Laboratory Plymouth | Wellcome Sanger Institute for the COVID-19 Genomics UK (COG-UK) Consortium | Lighthouse Laboratory Plymouth and Alex Alderton, Roberto Amato, Jeffrey Barrett, Sonia Goncalves, Ewan Harrison, David K. Jackson, Ian Johnston, Dominic Kwiatkowski, Cordelia Langford, John Sillitoe on behalf of the Wellcome Sanger Institute COVID-19 Surveillance Team |
| EPI_ISL_3442117, EPI_ISL_3442118, EPI_ISL_3442119 | Lighthouse Lab in Glasgow | Wellcome Sanger Institute for the COVID-19 Genomics UK (COG-UK) Consortium | Harper VanSteenhouse, Yumi Kasai, David Gray, Carol Clugston, Anna Dominiczak and Alex Alderton, Roberto Amato, Jeffrey Barrett, Sonia Goncalves, Ewan Harrison, David K. Jackson, Ian Johnston, Dominic Kwiatkowski, Cordelia Langford, John Sillitoe on behalf of the Wellcome Sanger Institute COVID-19 Surveillance Team |
| EPI_ISL_3442120, EPI_ISL_3442121 | Lighthouse Laboratory Plymouth | Wellcome Sanger Institute for the COVID-19 Genomics UK (COG-UK) Consortium | Lighthouse Laboratory Plymouth and Alex Alderton, Roberto Amato, Jeffrey Barrett, Sonia Goncalves, Ewan Harrison, David K. Jackson, Ian Johnston, Dominic Kwiatkowski, Cordelia Langford, John Sillitoe on behalf of the Wellcome Sanger Institute COVID-19 Surveillance Team |
| EPI_ISL_3442122 | Lighthouse Lab in Glasgow | Wellcome Sanger Institute for the COVID-19 Genomics UK (COG-UK) Consortium | Harper VanSteenhouse, Yumi Kasai, David Gray, Carol Clugston, Anna Dominiczak and Alex Alderton, Roberto Amato, Jeffrey Barrett, Sonia Goncalves, Ewan Harrison, David K. Jackson, Ian Johnston, Dominic Kwiatkowski, Cordelia Langford, John Sillitoe on behalf of the Wellcome Sanger Institute COVID-19 Surveillance Team |
| EPI_ISL_3442123, EPI_ISL_3442124 | Lighthouse Laboratory Plymouth | Wellcome Sanger Institute for the COVID-19 Genomics UK (COG-UK) Consortium | Lighthouse Laboratory Plymouth and Alex Alderton, Roberto Amato, Jeffrey Barrett, Sonia Goncalves, Ewan Harrison, David K. Jackson, Ian Johnston, Dominic Kwiatkowski, Cordelia Langford, John Sillitoe on behalf of the Wellcome Sanger Institute COVID-19 Surveillance Team |
| EPI_ISL_3442125, EPI_ISL_3442126 | Lighthouse Lab in Glasgow | Wellcome Sanger Institute for the COVID-19 Genomics UK (COG-UK) Consortium | Harper VanSteenhouse, Yumi Kasai, David Gray, Carol Clugston, Anna Dominiczak and Alex Alderton, Roberto Amato, Jeffrey Barrett, Sonia Goncalves, Ewan Harrison, David K. Jackson, Ian Johnston, Dominic Kwiatkowski, Cordelia Langford, John Sillitoe on behalf of the Wellcome Sanger Institute COVID-19 Surveillance Team |
| EPI_ISL_3442127 | Lighthouse Laboratory Plymouth | Wellcome Sanger Institute for the COVID-19 Genomics UK (COG-UK) Consortium | Lighthouse Laboratory Plymouth and Alex Alderton, Roberto Amato, Jeffrey Barrett, Sonia Goncalves, Ewan Harrison, David K. Jackson, Ian Johnston, Dominic Kwiatkowski, Cordelia Langford, John Sillitoe on behalf of the Wellcome Sanger Institute COVID-19 Surveillance Team |
| EPI_ISL_3442128, EPI_ISL_3442129, EPI_ISL_3442130 | Lighthouse Lab in Glasgow | Wellcome Sanger Institute for the COVID-19 Genomics UK (COG-UK) Consortium | Harper VanSteenhouse, Yumi Kasai, David Gray, Carol Clugston, Anna Dominiczak and Alex Alderton, Roberto Amato, Jeffrey Barrett, Sonia Goncalves, Ewan Harrison, David K. Jackson, Ian Johnston, Dominic Kwiatkowski, Cordelia Langford, John Sillitoe on behalf of the Wellcome Sanger Institute COVID-19 Surveillance Team |
| EPI_ISL_3442131 | Lighthouse Laboratory Plymouth | Wellcome Sanger Institute for the COVID-19 Genomics UK (COG-UK) Consortium | Lighthouse Laboratory Plymouth and Alex Alderton, Roberto Amato, Jeffrey Barrett, Sonia Goncalves, Ewan Harrison, David K. Jackson, Ian Johnston, Dominic Kwiatkowski, Cordelia Langford, John Sillitoe on behalf of the Wellcome Sanger Institute COVID-19 Surveillance Team |
| EPI_ISL_3442132, EPI_ISL_3442133, EPI_ISL_3442134 | Lighthouse Lab in Glasgow | Wellcome Sanger Institute for the COVID-19 Genomics UK (COG-UK) Consortium | Harper VanSteenhouse, Yumi Kasai, David Gray, Carol Clugston, Anna Dominiczak and Alex Alderton, Roberto Amato, Jeffrey Barrett, Sonia Goncalves, Ewan Harrison, David K. Jackson, Ian Johnston, Dominic Kwiatkowski, Cordelia Langford, John Sillitoe on behalf of the Wellcome Sanger Institute COVID-19 Surveillance Team |
| EPI_ISL_3442135 | Lighthouse Laboratory Plymouth | Wellcome Sanger Institute for the COVID-19 Genomics UK (COG-UK) Consortium | Lighthouse Laboratory Plymouth and Alex Alderton, Roberto Amato, Jeffrey Barrett, Sonia Goncalves, Ewan Harrison, David K. Jackson, Ian Johnston, Dominic Kwiatkowski, Cordelia Langford, John Sillitoe on behalf of the Wellcome Sanger Institute COVID-19 Surveillance Team |
| EPI_ISL_3442136, EPI_ISL_3442137 | Lighthouse Lab in Glasgow | Wellcome Sanger Institute for the COVID-19 Genomics UK (COG-UK) Consortium | Harper VanSteenhouse, Yumi Kasai, David Gray, Carol Clugston, Anna Dominiczak and Alex Alderton, Roberto Amato, Jeffrey Barrett, Sonia Goncalves, Ewan Harrison, David K. Jackson, Ian Johnston, Dominic Kwiatkowski, Cordelia Langford, John Sillitoe on behalf of the Wellcome Sanger Institute COVID-19 Surveillance Team |
| EPI_ISL_3442138, EPI_ISL_3442139 | Lighthouse Laboratory Plymouth | Wellcome Sanger Institute for the COVID-19 Genomics UK (COG-UK) Consortium | Lighthouse Laboratory Plymouth and Alex Alderton, Roberto Amato, Jeffrey Barrett, Sonia Goncalves, Ewan Harrison, David K. Jackson, Ian Johnston, Dominic Kwiatkowski, Cordelia Langford, John Sillitoe on behalf of the Wellcome Sanger Institute COVID-19 Surveillance Team |
| EPI_ISL_3442140 | Lighthouse Lab in Glasgow | Wellcome Sanger Institute for the COVID-19 Genomics UK (COG-UK) Consortium | Harper VanSteenhouse, Yumi Kasai, David Gray, Carol Clugston, Anna Dominiczak and Alex Alderton, Roberto Amato, Jeffrey Barrett, Sonia Goncalves, Ewan Harrison, David K. Jackson, Ian Johnston, Dominic Kwiatkowski, Cordelia Langford, John Sillitoe on behalf of the Wellcome Sanger Institute COVID-19 Surveillance Team |
| EPI_ISL_3442141, EPI_ISL_3442142, EPI_ISL_3442143, EPI_ISL_3442144, EPI_ISL_3442145 | Lighthouse Laboratory Plymouth | Wellcome Sanger Institute for the COVID-19 Genomics UK (COG-UK) Consortium | Lighthouse Laboratory Plymouth and Alex Alderton, Roberto Amato, Jeffrey Barrett, Sonia Goncalves, Ewan Harrison, David K. Jackson, Ian Johnston, Dominic Kwiatkowski, Cordelia Langford, John Sillitoe on behalf of the Wellcome Sanger Institute COVID-19 Surveillance Team |
| EPI_ISL_3442146 | Lighthouse Lab in Glasgow | Wellcome Sanger Institute for the COVID-19 Genomics UK (COG-UK) Consortium | Harper VanSteenhouse, Yumi Kasai, David Gray, Carol Clugston, Anna Dominiczak and Alex Alderton, Roberto Amato, Jeffrey Barrett, Sonia Goncalves, Ewan Harrison, David K. Jackson, Ian Johnston, Dominic Kwiatkowski, Cordelia Langford, John Sillitoe on behalf of the Wellcome Sanger Institute COVID-19 Surveillance Team |
| EPI_ISL_3442147 | Lighthouse Laboratory Plymouth | Wellcome Sanger Institute for the COVID-19 Genomics UK (COG-UK) Consortium | Lighthouse Laboratory Plymouth and Alex Alderton, Roberto Amato, Jeffrey Barrett, Sonia Goncalves, Ewan Harrison, David K. Jackson, Ian Johnston, Dominic Kwiatkowski, Cordelia Langford, John Sillitoe on behalf of the Wellcome Sanger Institute COVID-19 Surveillance Team |
| EPI_ISL_3442148 | Lighthouse Lab in Glasgow | Wellcome Sanger Institute for the COVID-19 Genomics UK (COG-UK) Consortium | Harper VanSteenhouse, Yumi Kasai, David Gray, Carol Clugston, Anna Dominiczak and Alex Alderton, Roberto Amato, Jeffrey Barrett, Sonia Goncalves, Ewan Harrison, David K. Jackson, Ian Johnston, Dominic Kwiatkowski, Cordelia Langford, John Sillitoe on behalf of the Wellcome Sanger Institute COVID-19 Surveillance Team |
| EPI_ISL_3442149, EPI_ISL_3442150, EPI_ISL_3442151, EPI_ISL_3442152 | Lighthouse Laboratory Plymouth | Wellcome Sanger Institute for the COVID-19 Genomics UK (COG-UK) Consortium | Lighthouse Laboratory Plymouth and Alex Alderton, Roberto Amato, Jeffrey Barrett, Sonia Goncalves, Ewan Harrison, David K. Jackson, Ian Johnston, Dominic Kwiatkowski, Cordelia Langford, John Sillitoe on behalf of the Wellcome Sanger Institute COVID-19 Surveillance Team |
| EPI_ISL_3442153, EPI_ISL_3442154 | Lighthouse Lab in Glasgow | Wellcome Sanger Institute for the COVID-19 Genomics UK (COG-UK) Consortium | Harper VanSteenhouse, Yumi Kasai, David Gray, Carol Clugston, Anna Dominiczak and Alex Alderton, Roberto Amato, Jeffrey Barrett, Sonia Goncalves, Ewan Harrison, David K. Jackson, Ian Johnston, Dominic Kwiatkowski, Cordelia Langford, John Sillitoe on behalf of the Wellcome Sanger Institute COVID-19 Surveillance Team |
| EPI_ISL_3442155 | Lighthouse Laboratory Plymouth | Wellcome Sanger Institute for the COVID-19 Genomics UK (COG-UK) Consortium | Lighthouse Laboratory Plymouth and Alex Alderton, Roberto Amato, Jeffrey Barrett, Sonia Goncalves, Ewan Harrison, David K. Jackson, Ian Johnston, Dominic Kwiatkowski, Cordelia Langford, John Sillitoe on behalf of the Wellcome Sanger Institute COVID-19 Surveillance Team |
| EPI_ISL_3442156, EPI_ISL_3442157 | Lighthouse Lab in Glasgow | Wellcome Sanger Institute for the COVID-19 Genomics UK (COG-UK) Consortium | Harper VanSteenhouse, Yumi Kasai, David Gray, Carol Clugston, Anna Dominiczak and Alex Alderton, Roberto Amato, Jeffrey Barrett, Sonia Goncalves, Ewan Harrison, David K. Jackson, Ian Johnston, Dominic Kwiatkowski, Cordelia Langford, John Sillitoe on behalf of the Wellcome Sanger Institute COVID-19 Surveillance Team |
| EPI_ISL_3442158 | Lighthouse Laboratory Plymouth | Wellcome Sanger Institute for the COVID-19 Genomics UK (COG-UK) Consortium | Lighthouse Laboratory Plymouth and Alex Alderton, Roberto Amato, Jeffrey Barrett, Sonia Goncalves, Ewan Harrison, David K. Jackson, Ian Johnston, Dominic Kwiatkowski, Cordelia Langford, John Sillitoe on behalf of the Wellcome Sanger Institute COVID-19 Surveillance Team |
| EPI_ISL_3442159, EPI_ISL_3442160, EPI_ISL_3442161, EPI_ISL_3442162 | Lighthouse Lab in Glasgow | Wellcome Sanger Institute for the COVID-19 Genomics UK (COG-UK) Consortium | Harper VanSteenhouse, Yumi Kasai, David Gray, Carol Clugston, Anna Dominiczak and Alex Alderton, Roberto Amato, Jeffrey Barrett, Sonia Goncalves, Ewan Harrison, David K. Jackson, Ian Johnston, Dominic Kwiatkowski, Cordelia Langford, John Sillitoe on behalf of the Wellcome Sanger Institute COVID-19 Surveillance Team |
| EPI_ISL_3442163, EPI_ISL_3442164 | Lighthouse Laboratory Plymouth | Wellcome Sanger Institute for the COVID-19 Genomics UK (COG-UK) Consortium | Lighthouse Laboratory Plymouth and Alex Alderton, Roberto Amato, Jeffrey Barrett, Sonia Goncalves, Ewan Harrison, David K. Jackson, Ian Johnston, Dominic Kwiatkowski, Cordelia Langford, John Sillitoe on behalf of the Wellcome Sanger Institute COVID-19 Surveillance Team |
| EPI_ISL_3442165, EPI_ISL_3442166, EPI_ISL_3442167 | Lighthouse Lab in Glasgow | Wellcome Sanger Institute for the COVID-19 Genomics UK (COG-UK) Consortium | Harper VanSteenhouse, Yumi Kasai, David Gray, Carol Clugston, Anna Dominiczak and Alex Alderton, Roberto Amato, Jeffrey Barrett, Sonia Goncalves, Ewan Harrison, David K. Jackson, Ian Johnston, Dominic Kwiatkowski, Cordelia Langford, John Sillitoe on behalf of the Wellcome Sanger Institute COVID-19 Surveillance Team |
| EPI_ISL_3442168, EPI_ISL_3442169, EPI_ISL_3442170, EPI_ISL_3442171, EPI_ISL_3442172, EPI_ISL_3442173, EPI_ISL_3442174, EPI_ISL_3442175, EPI_ISL_3442176, EPI_ISL_3442177, EPI_ISL_3442178, EPI_ISL_3442179 | | | |
| see above | Lighthouse Laboratory Plymouth | Wellcome Sanger Institute for the COVID-19 Genomics UK (COG-UK) | Lighthouse Laboratory Plymouth and Alex Alderton, Roberto Amato, Jeffrey Barrett, Sonia Goncalves, Ewan Harrison, David K. Jackson, Ian Johnston, Dominic Kwiatkowski, Cordelia Langford, John Sillitoe on behalf of the Wellcome Sanger |

| | | | |
|---|---|---|---|
| | | Consortium | Institute COVID-19 Surveillance Team |
| EPI_ISL_3442180 | Lighthouse Lab in Glasgow | Wellcome Sanger Institute for the COVID-19 Genomics UK (COG-UK) Consortium | Harper VanSteenhouse, Yumi Kasai, David Gray, Carol Clugston, Anna Dominiczak and Alex Alderton, Roberto Amato, Jeffrey Barrett, Sonia Goncalves, Ewan Harrison, David K. Jackson, Ian Johnston, Dominic Kwiatkowski, Cordelia Langford, John Sillitoe on behalf of the Wellcome Sanger Institute COVID-19 Surveillance Team |
| EPI_ISL_3442181, EPI_ISL_3442182 | Lighthouse Laboratory Plymouth | Wellcome Sanger Institute for the COVID-19 Genomics UK (COG-UK) Consortium | Lighthouse Laboratory Plymouth and Alex Alderton, Roberto Amato, Jeffrey Barrett, Sonia Goncalves, Ewan Harrison, David K. Jackson, Ian Johnston, Dominic Kwiatkowski, Cordelia Langford, John Sillitoe on behalf of the Wellcome Sanger Institute COVID-19 Surveillance Team |
| EPI_ISL_3442183 | Lighthouse Lab in Glasgow | Wellcome Sanger Institute for the COVID-19 Genomics UK (COG-UK) Consortium | Harper VanSteenhouse, Yumi Kasai, David Gray, Carol Clugston, Anna Dominiczak and Alex Alderton, Roberto Amato, Jeffrey Barrett, Sonia Goncalves, Ewan Harrison, David K. Jackson, Ian Johnston, Dominic Kwiatkowski, Cordelia Langford, John Sillitoe on behalf of the Wellcome Sanger Institute COVID-19 Surveillance Team |
| EPI_ISL_3442184, EPI_ISL_3442185 | Lighthouse Laboratory Plymouth | Wellcome Sanger Institute for the COVID-19 Genomics UK (COG-UK) Consortium | Lighthouse Laboratory Plymouth and Alex Alderton, Roberto Amato, Jeffrey Barrett, Sonia Goncalves, Ewan Harrison, David K. Jackson, Ian Johnston, Dominic Kwiatkowski, Cordelia Langford, John Sillitoe on behalf of the Wellcome Sanger Institute COVID-19 Surveillance Team |
| EPI_ISL_3442186, EPI_ISL_3442187 | Lighthouse Lab in Glasgow | Wellcome Sanger Institute for the COVID-19 Genomics UK (COG-UK) Consortium | Harper VanSteenhouse, Yumi Kasai, David Gray, Carol Clugston, Anna Dominiczak and Alex Alderton, Roberto Amato, Jeffrey Barrett, Sonia Goncalves, Ewan Harrison, David K. Jackson, Ian Johnston, Dominic Kwiatkowski, Cordelia Langford, John Sillitoe on behalf of the Wellcome Sanger Institute COVID-19 Surveillance Team |
| EPI_ISL_3442188, EPI_ISL_3442189 | Lighthouse Laboratory Plymouth | Wellcome Sanger Institute for the COVID-19 Genomics UK (COG-UK) Consortium | Lighthouse Laboratory Plymouth and Alex Alderton, Roberto Amato, Jeffrey Barrett, Sonia Goncalves, Ewan Harrison, David K. Jackson, Ian Johnston, Dominic Kwiatkowski, Cordelia Langford, John Sillitoe on behalf of the Wellcome Sanger Institute COVID-19 Surveillance Team |
| EPI_ISL_3442190, EPI_ISL_3442191, EPI_ISL_3442192 | Lighthouse Lab in Glasgow | Wellcome Sanger Institute for the COVID-19 Genomics UK (COG-UK) Consortium | Harper VanSteenhouse, Yumi Kasai, David Gray, Carol Clugston, Anna Dominiczak and Alex Alderton, Roberto Amato, Jeffrey Barrett, Sonia Goncalves, Ewan Harrison, David K. Jackson, Ian Johnston, Dominic Kwiatkowski, Cordelia Langford, John Sillitoe on behalf of the Wellcome Sanger Institute COVID-19 Surveillance Team |
| EPI_ISL_3442193, EPI_ISL_3442194, EPI_ISL_3442195, EPI_ISL_3442196 | Lighthouse Laboratory Plymouth | Wellcome Sanger Institute for the COVID-19 Genomics UK (COG-UK) Consortium | Lighthouse Laboratory Plymouth and Alex Alderton, Roberto Amato, Jeffrey Barrett, Sonia Goncalves, Ewan Harrison, David K. Jackson, Ian Johnston, Dominic Kwiatkowski, Cordelia Langford, John Sillitoe on behalf of the Wellcome Sanger Institute COVID-19 Surveillance Team |
| EPI_ISL_3442197 | Lighthouse Lab in Glasgow | Wellcome Sanger Institute for the COVID-19 Genomics UK (COG-UK) Consortium | Harper VanSteenhouse, Yumi Kasai, David Gray, Carol Clugston, Anna Dominiczak and Alex Alderton, Roberto Amato, Jeffrey Barrett, Sonia Goncalves, Ewan Harrison, David K. Jackson, Ian Johnston, Dominic Kwiatkowski, Cordelia Langford, John Sillitoe on behalf of the Wellcome Sanger Institute COVID-19 Surveillance Team |
| EPI_ISL_3442198 | Lighthouse Laboratory Plymouth | Wellcome Sanger Institute for the COVID-19 Genomics UK (COG-UK) Consortium | Lighthouse Laboratory Plymouth and Alex Alderton, Roberto Amato, Jeffrey Barrett, Sonia Goncalves, Ewan Harrison, David K. Jackson, Ian Johnston, Dominic Kwiatkowski, Cordelia Langford, John Sillitoe on behalf of the Wellcome Sanger Institute COVID-19 Surveillance Team |
| EPI_ISL_3442199, EPI_ISL_3442200, EPI_ISL_3442201 | Lighthouse Lab in Glasgow | Wellcome Sanger Institute for the COVID-19 Genomics UK (COG-UK) Consortium | Harper VanSteenhouse, Yumi Kasai, David Gray, Carol Clugston, Anna Dominiczak and Alex Alderton, Roberto Amato, Jeffrey Barrett, Sonia Goncalves, Ewan Harrison, David K. Jackson, Ian Johnston, Dominic Kwiatkowski, Cordelia Langford, John Sillitoe on behalf of the Wellcome Sanger Institute COVID-19 Surveillance Team |
| EPI_ISL_3442202, EPI_ISL_3442203 | Lighthouse Laboratory Plymouth | Wellcome Sanger Institute for the COVID-19 Genomics UK (COG-UK) Consortium | Lighthouse Laboratory Plymouth and Alex Alderton, Roberto Amato, Jeffrey Barrett, Sonia Goncalves, Ewan Harrison, David K. Jackson, Ian Johnston, Dominic Kwiatkowski, Cordelia Langford, John Sillitoe on behalf of the Wellcome Sanger Institute COVID-19 Surveillance Team |
| EPI_ISL_3442204, EPI_ISL_3442205, EPI_ISL_3442206, EPI_ISL_3442207 | Lighthouse Lab in Glasgow | Wellcome Sanger Institute for the COVID-19 Genomics UK (COG-UK) Consortium | Harper VanSteenhouse, Yumi Kasai, David Gray, Carol Clugston, Anna Dominiczak and Alex Alderton, Roberto Amato, Jeffrey Barrett, Sonia Goncalves, Ewan Harrison, David K. Jackson, Ian Johnston, Dominic Kwiatkowski, Cordelia Langford, John Sillitoe on behalf of the Wellcome Sanger Institute COVID-19 Surveillance Team |
| EPI_ISL_3442208 | Lighthouse Laboratory Plymouth | Wellcome Sanger Institute for the COVID-19 Genomics UK (COG-UK) Consortium | Lighthouse Laboratory Plymouth and Alex Alderton, Roberto Amato, Jeffrey Barrett, Sonia Goncalves, Ewan Harrison, David K. Jackson, Ian Johnston, Dominic Kwiatkowski, Cordelia Langford, John Sillitoe on behalf of the Wellcome Sanger Institute COVID-19 Surveillance Team |
| EPI_ISL_3442209, EPI_ISL_3442210 | Lighthouse Lab in Glasgow | Wellcome Sanger Institute for the COVID-19 Genomics UK (COG-UK) Consortium | Harper VanSteenhouse, Yumi Kasai, David Gray, Carol Clugston, Anna Dominiczak and Alex Alderton, Roberto Amato, Jeffrey Barrett, Sonia Goncalves, Ewan Harrison, David K. Jackson, Ian Johnston, Dominic Kwiatkowski, Cordelia Langford, John Sillitoe on behalf of the Wellcome Sanger Institute COVID-19 Surveillance Team |
| EPI_ISL_3442211 | Lighthouse Laboratory Plymouth | Wellcome Sanger Institute for the COVID-19 Genomics UK (COG-UK) Consortium | Lighthouse Laboratory Plymouth and Alex Alderton, Roberto Amato, Jeffrey Barrett, Sonia Goncalves, Ewan Harrison, David K. Jackson, Ian Johnston, Dominic Kwiatkowski, Cordelia Langford, John Sillitoe on behalf of the Wellcome Sanger Institute COVID-19 Surveillance Team |
| EPI_ISL_3442212 | Lighthouse Lab in Glasgow | Wellcome Sanger Institute for the COVID-19 Genomics UK (COG-UK) Consortium | Harper VanSteenhouse, Yumi Kasai, David Gray, Carol Clugston, Anna Dominiczak and Alex Alderton, Roberto Amato, Jeffrey Barrett, Sonia Goncalves, Ewan Harrison, David K. Jackson, Ian Johnston, Dominic Kwiatkowski, Cordelia Langford, John Sillitoe on behalf of the Wellcome Sanger Institute COVID-19 Surveillance Team |
| EPI_ISL_3442213 | Lighthouse Laboratory Plymouth | Wellcome Sanger Institute for the COVID-19 Genomics UK (COG-UK) Consortium | Lighthouse Laboratory Plymouth and Alex Alderton, Roberto Amato, Jeffrey Barrett, Sonia Goncalves, Ewan Harrison, David K. Jackson, Ian Johnston, Dominic Kwiatkowski, Cordelia Langford, John Sillitoe on behalf of the Wellcome Sanger Institute COVID-19 Surveillance Team |
| EPI_ISL_3442214 | Lighthouse Lab in Glasgow | Wellcome Sanger Institute for the COVID-19 Genomics UK (COG-UK) Consortium | Harper VanSteenhouse, Yumi Kasai, David Gray, Carol Clugston, Anna Dominiczak and Alex Alderton, Roberto Amato, Jeffrey Barrett, Sonia Goncalves, Ewan Harrison, David K. Jackson, Ian Johnston, Dominic Kwiatkowski, Cordelia Langford, John Sillitoe on behalf of the Wellcome Sanger Institute COVID-19 Surveillance Team |
| EPI_ISL_3442215, EPI_ISL_3442216, EPI_ISL_3442217 | Lighthouse Laboratory Plymouth | Wellcome Sanger Institute for the COVID-19 Genomics UK (COG-UK) Consortium | Lighthouse Laboratory Plymouth and Alex Alderton, Roberto Amato, Jeffrey Barrett, Sonia Goncalves, Ewan Harrison, David K. Jackson, Ian Johnston, Dominic Kwiatkowski, Cordelia Langford, John Sillitoe on behalf of the Wellcome Sanger Institute COVID-19 Surveillance Team |
| EPI_ISL_3442218 | Lighthouse Lab in Glasgow | Wellcome Sanger Institute for the COVID-19 Genomics UK (COG-UK) Consortium | Harper VanSteenhouse, Yumi Kasai, David Gray, Carol Clugston, Anna Dominiczak and Alex Alderton, Roberto Amato, Jeffrey Barrett, Sonia Goncalves, Ewan Harrison, David K. Jackson, Ian Johnston, Dominic Kwiatkowski, Cordelia Langford, John Sillitoe on behalf of the Wellcome Sanger Institute COVID-19 Surveillance Team |
| EPI_ISL_3442219, EPI_ISL_3442220 | Lighthouse Laboratory Plymouth | Wellcome Sanger Institute for the COVID-19 Genomics UK (COG-UK) Consortium | Lighthouse Laboratory Plymouth and Alex Alderton, Roberto Amato, Jeffrey Barrett, Sonia Goncalves, Ewan Harrison, David K. Jackson, Ian Johnston, Dominic Kwiatkowski, Cordelia Langford, John Sillitoe on behalf of the Wellcome Sanger Institute COVID-19 Surveillance Team |
| EPI_ISL_3442221 | Lighthouse Lab in Glasgow | Wellcome Sanger Institute for the COVID-19 Genomics UK (COG-UK) Consortium | Harper VanSteenhouse, Yumi Kasai, David Gray, Carol Clugston, Anna Dominiczak and Alex Alderton, Roberto Amato, Jeffrey Barrett, Sonia Goncalves, Ewan Harrison, David K. Jackson, Ian Johnston, Dominic Kwiatkowski, Cordelia Langford, John Sillitoe on behalf of the Wellcome Sanger Institute COVID-19 Surveillance Team |
| EPI_ISL_3442222, EPI_ISL_3442223, EPI_ISL_3442224, EPI_ISL_3442225 | Lighthouse Laboratory Plymouth | Wellcome Sanger Institute for the COVID-19 Genomics UK (COG-UK) Consortium | Lighthouse Laboratory Plymouth and Alex Alderton, Roberto Amato, Jeffrey Barrett, Sonia Goncalves, Ewan Harrison, David K. Jackson, Ian Johnston, Dominic Kwiatkowski, Cordelia Langford, John Sillitoe on behalf of the Wellcome Sanger Institute COVID-19 Surveillance Team |
| EPI_ISL_3442226 | Lighthouse Lab in Glasgow | Wellcome Sanger Institute for the COVID-19 Genomics UK (COG-UK) Consortium | Harper VanSteenhouse, Yumi Kasai, David Gray, Carol Clugston, Anna Dominiczak and Alex Alderton, Roberto Amato, Jeffrey Barrett, Sonia Goncalves, Ewan Harrison, David K. Jackson, Ian Johnston, Dominic Kwiatkowski, Cordelia Langford, John Sillitoe on behalf of the Wellcome Sanger Institute COVID-19 Surveillance Team |
| EPI_ISL_3442227, EPI_ISL_3442228 | Lighthouse Laboratory Plymouth | Wellcome Sanger Institute for the COVID-19 Genomics UK (COG-UK) Consortium | Lighthouse Laboratory Plymouth and Alex Alderton, Roberto Amato, Jeffrey Barrett, Sonia Goncalves, Ewan Harrison, David K. Jackson, Ian Johnston, Dominic Kwiatkowski, Cordelia Langford, John Sillitoe on behalf of the Wellcome Sanger Institute COVID-19 Surveillance Team |
| EPI_ISL_3442229 | Lighthouse Lab in Glasgow | Wellcome Sanger Institute for the COVID-19 Genomics UK (COG-UK) Consortium | Harper VanSteenhouse, Yumi Kasai, David Gray, Carol Clugston, Anna Dominiczak and Alex Alderton, Roberto Amato, Jeffrey Barrett, Sonia Goncalves, Ewan Harrison, David K. Jackson, Ian Johnston, Dominic Kwiatkowski, Cordelia Langford, John Sillitoe on behalf of the Wellcome Sanger Institute COVID-19 Surveillance Team |
| EPI_ISL_3442230, EPI_ISL_3442231, EPI_ISL_3442232, EPI_ISL_3442233 | Lighthouse Laboratory Plymouth | Wellcome Sanger Institute for the COVID-19 Genomics UK (COG-UK) Consortium | Lighthouse Laboratory Plymouth and Alex Alderton, Roberto Amato, Jeffrey Barrett, Sonia Goncalves, Ewan Harrison, David K. Jackson, Ian Johnston, Dominic Kwiatkowski, Cordelia Langford, John Sillitoe on behalf of the Wellcome Sanger Institute COVID-19 Surveillance Team |
| EPI_ISL_3442234, EPI_ISL_3442235, EPI_ISL_3442236, EPI_ISL_3442237 | Lighthouse Lab in Glasgow | Wellcome Sanger Institute for the COVID-19 Genomics UK (COG-UK) Consortium | Harper VanSteenhouse, Yumi Kasai, David Gray, Carol Clugston, Anna Dominiczak and Alex Alderton, Roberto Amato, Jeffrey Barrett, Sonia Goncalves, Ewan Harrison, David K. Jackson, Ian Johnston, Dominic Kwiatkowski, Cordelia Langford, John Sillitoe on behalf of the Wellcome Sanger Institute COVID-19 Surveillance Team |
| EPI_ISL_3442238, EPI_ISL_3442239, EPI_ISL_3442240 | Lighthouse Laboratory Plymouth | Wellcome Sanger Institute for the COVID-19 Genomics UK (COG-UK) Consortium | Lighthouse Laboratory Plymouth and Alex Alderton, Roberto Amato, Jeffrey Barrett, Sonia Goncalves, Ewan Harrison, David K. Jackson, Ian Johnston, Dominic Kwiatkowski, Cordelia Langford, John Sillitoe on behalf of the Wellcome Sanger Institute COVID-19 Surveillance Team |
| EPI_ISL_3442241, EPI_ISL_3442242, EPI_ISL_3442243, EPI_ISL_3442244, EPI_ISL_3442245 | Lighthouse Lab in Glasgow | Wellcome Sanger Institute for the COVID-19 Genomics UK (COG-UK) Consortium | Harper VanSteenhouse, Yumi Kasai, David Gray, Carol Clugston, Anna Dominiczak and Alex Alderton, Roberto Amato, Jeffrey Barrett, Sonia Goncalves, Ewan Harrison, David K. Jackson, Ian Johnston, Dominic Kwiatkowski, Cordelia Langford, John Sillitoe on behalf of the Wellcome Sanger Institute COVID-19 Surveillance Team |
| EPI_ISL_3442246 | Lighthouse Laboratory Plymouth | Wellcome Sanger Institute for the COVID-19 Genomics UK (COG-UK) Consortium | Lighthouse Laboratory Plymouth and Alex Alderton, Roberto Amato, Jeffrey Barrett, Sonia Goncalves, Ewan Harrison, David K. Jackson, Ian Johnston, Dominic Kwiatkowski, Cordelia Langford, John Sillitoe on behalf of the Wellcome Sanger Institute COVID-19 Surveillance Team |
| EPI_ISL_3442247, EPI_ISL_3442248, EPI_ISL_3442249, EPI_ISL_3442250 | Lighthouse Lab in Glasgow | Wellcome Sanger Institute for the COVID-19 Genomics UK (COG-UK) Consortium | Harper VanSteenhouse, Yumi Kasai, David Gray, Carol Clugston, Anna Dominiczak and Alex Alderton, Roberto Amato, Jeffrey Barrett, Sonia Goncalves, Ewan Harrison, David K. Jackson, Ian Johnston, Dominic Kwiatkowski, Cordelia Langford, John Sillitoe on behalf of the Wellcome Sanger Institute COVID-19 Surveillance Team |
| EPI_ISL_3442251, EPI_ISL_3442252 | Lighthouse Laboratory Plymouth | Wellcome Sanger Institute for the COVID-19 Genomics UK (COG-UK) Consortium | Lighthouse Laboratory Plymouth and Alex Alderton, Roberto Amato, Jeffrey Barrett, Sonia Goncalves, Ewan Harrison, David K. Jackson, Ian Johnston, Dominic Kwiatkowski, Cordelia Langford, John Sillitoe on behalf of the Wellcome Sanger Institute COVID-19 Surveillance Team |
| EPI_ISL_3442253, EPI_ISL_3442254, EPI_ISL_3442255 | Lighthouse Lab in Glasgow | Wellcome Sanger Institute for the COVID-19 Genomics UK (COG-UK) Consortium | Harper VanSteenhouse, Yumi Kasai, David Gray, Carol Clugston, Anna Dominiczak and Alex Alderton, Roberto Amato, Jeffrey Barrett, Sonia Goncalves, Ewan Harrison, David K. Jackson, Ian Johnston, Dominic Kwiatkowski, Cordelia Langford, John Sillitoe on behalf of the Wellcome Sanger Institute COVID-19 Surveillance Team |
| EPI_ISL_3442256, EPI_ISL_3442257 | Lighthouse Laboratory Plymouth | Wellcome Sanger Institute for the COVID-19 Genomics UK (COG-UK) Consortium | Lighthouse Laboratory Plymouth and Alex Alderton, Roberto Amato, Jeffrey Barrett, Sonia Goncalves, Ewan Harrison, David K. Jackson, Ian Johnston, Dominic Kwiatkowski, Cordelia Langford, John Sillitoe on behalf of the Wellcome Sanger Institute COVID-19 Surveillance Team |

| | | | |
|---|---|---|---|
| EPI_ISL_3442258 | Lighthouse Lab in Glasgow | Wellcome Sanger Institute for the COVID-19 Genomics UK (COG-UK) Consortium | Harper VanSteenhouse, Yumi Kasai, David Gray, Carol Clugston, Anna Dominiczak and Alex Alderton, Roberto Amato, Jeffrey Barrett, Sonia Goncalves, Ewan Harrison, David K. Jackson, Ian Johnston, Dominic Kwiatkowski, Cordelia Langford, John Sillitoe on behalf of the Wellcome Sanger Institute COVID-19 Surveillance Team |
| EPI_ISL_3442259, EPI_ISL_3442260 | Lighthouse Laboratory Plymouth | Wellcome Sanger Institute for the COVID-19 Genomics UK (COG-UK) Consortium | Lighthouse Laboratory Plymouth and Alex Alderton, Roberto Amato, Jeffrey Barrett, Sonia Goncalves, Ewan Harrison, David K. Jackson, Ian Johnston, Dominic Kwiatkowski, Cordelia Langford, John Sillitoe on behalf of the Wellcome Sanger Institute COVID-19 Surveillance Team |
| EPI_ISL_3442261 | Lighthouse Lab in Glasgow | Wellcome Sanger Institute for the COVID-19 Genomics UK (COG-UK) Consortium | Harper VanSteenhouse, Yumi Kasai, David Gray, Carol Clugston, Anna Dominiczak and Alex Alderton, Roberto Amato, Jeffrey Barrett, Sonia Goncalves, Ewan Harrison, David K. Jackson, Ian Johnston, Dominic Kwiatkowski, Cordelia Langford, John Sillitoe on behalf of the Wellcome Sanger Institute COVID-19 Surveillance Team |
| EPI_ISL_3442262 | Lighthouse Laboratory Plymouth | Wellcome Sanger Institute for the COVID-19 Genomics UK (COG-UK) Consortium | Lighthouse Laboratory Plymouth and Alex Alderton, Roberto Amato, Jeffrey Barrett, Sonia Goncalves, Ewan Harrison, David K. Jackson, Ian Johnston, Dominic Kwiatkowski, Cordelia Langford, John Sillitoe on behalf of the Wellcome Sanger Institute COVID-19 Surveillance Team |
| EPI_ISL_3442263, EPI_ISL_3442264, EPI_ISL_3442265, EPI_ISL_3442266 | Lighthouse Lab in Glasgow | Wellcome Sanger Institute for the COVID-19 Genomics UK (COG-UK) Consortium | Harper VanSteenhouse, Yumi Kasai, David Gray, Carol Clugston, Anna Dominiczak and Alex Alderton, Roberto Amato, Jeffrey Barrett, Sonia Goncalves, Ewan Harrison, David K. Jackson, Ian Johnston, Dominic Kwiatkowski, Cordelia Langford, John Sillitoe on behalf of the Wellcome Sanger Institute COVID-19 Surveillance Team |
| EPI_ISL_3442267 | Lighthouse Laboratory Plymouth | Wellcome Sanger Institute for the COVID-19 Genomics UK (COG-UK) Consortium | Lighthouse Laboratory Plymouth and Alex Alderton, Roberto Amato, Jeffrey Barrett, Sonia Goncalves, Ewan Harrison, David K. Jackson, Ian Johnston, Dominic Kwiatkowski, Cordelia Langford, John Sillitoe on behalf of the Wellcome Sanger Institute COVID-19 Surveillance Team |
| EPI_ISL_3442268, EPI_ISL_3442269, EPI_ISL_3442270, EPI_ISL_3442271 | Lighthouse Lab in Glasgow | Wellcome Sanger Institute for the COVID-19 Genomics UK (COG-UK) Consortium | Harper VanSteenhouse, Yumi Kasai, David Gray, Carol Clugston, Anna Dominiczak and Alex Alderton, Roberto Amato, Jeffrey Barrett, Sonia Goncalves, Ewan Harrison, David K. Jackson, Ian Johnston, Dominic Kwiatkowski, Cordelia Langford, John Sillitoe on behalf of the Wellcome Sanger Institute COVID-19 Surveillance Team |
| EPI_ISL_3442272, EPI_ISL_3442273 | Lighthouse Laboratory Plymouth | Wellcome Sanger Institute for the COVID-19 Genomics UK (COG-UK) Consortium | Lighthouse Laboratory Plymouth and Alex Alderton, Roberto Amato, Jeffrey Barrett, Sonia Goncalves, Ewan Harrison, David K. Jackson, Ian Johnston, Dominic Kwiatkowski, Cordelia Langford, John Sillitoe on behalf of the Wellcome Sanger Institute COVID-19 Surveillance Team |
| EPI_ISL_3442274, EPI_ISL_3442275 | Lighthouse Lab in Glasgow | Wellcome Sanger Institute for the COVID-19 Genomics UK (COG-UK) Consortium | Harper VanSteenhouse, Yumi Kasai, David Gray, Carol Clugston, Anna Dominiczak and Alex Alderton, Roberto Amato, Jeffrey Barrett, Sonia Goncalves, Ewan Harrison, David K. Jackson, Ian Johnston, Dominic Kwiatkowski, Cordelia Langford, John Sillitoe on behalf of the Wellcome Sanger Institute COVID-19 Surveillance Team |
| EPI_ISL_3442276 | Lighthouse Laboratory Plymouth | Wellcome Sanger Institute for the COVID-19 Genomics UK (COG-UK) Consortium | Lighthouse Laboratory Plymouth and Alex Alderton, Roberto Amato, Jeffrey Barrett, Sonia Goncalves, Ewan Harrison, David K. Jackson, Ian Johnston, Dominic Kwiatkowski, Cordelia Langford, John Sillitoe on behalf of the Wellcome Sanger Institute COVID-19 Surveillance Team |
| EPI_ISL_3442277 | Lighthouse Lab in Glasgow | Wellcome Sanger Institute for the COVID-19 Genomics UK (COG-UK) Consortium | Harper VanSteenhouse, Yumi Kasai, David Gray, Carol Clugston, Anna Dominiczak and Alex Alderton, Roberto Amato, Jeffrey Barrett, Sonia Goncalves, Ewan Harrison, David K. Jackson, Ian Johnston, Dominic Kwiatkowski, Cordelia Langford, John Sillitoe on behalf of the Wellcome Sanger Institute COVID-19 Surveillance Team |
| EPI_ISL_3442278, EPI_ISL_3442279 | Lighthouse Laboratory Plymouth | Wellcome Sanger Institute for the COVID-19 Genomics UK (COG-UK) Consortium | Lighthouse Laboratory Plymouth and Alex Alderton, Roberto Amato, Jeffrey Barrett, Sonia Goncalves, Ewan Harrison, David K. Jackson, Ian Johnston, Dominic Kwiatkowski, Cordelia Langford, John Sillitoe on behalf of the Wellcome Sanger Institute COVID-19 Surveillance Team |
| EPI_ISL_3442280 | Lighthouse Lab in Glasgow | Wellcome Sanger Institute for the COVID-19 Genomics UK (COG-UK) Consortium | Harper VanSteenhouse, Yumi Kasai, David Gray, Carol Clugston, Anna Dominiczak and Alex Alderton, Roberto Amato, Jeffrey Barrett, Sonia Goncalves, Ewan Harrison, David K. Jackson, Ian Johnston, Dominic Kwiatkowski, Cordelia Langford, John Sillitoe on behalf of the Wellcome Sanger Institute COVID-19 Surveillance Team |
| EPI_ISL_3442281, EPI_ISL_3442282 | Lighthouse Laboratory Plymouth | Wellcome Sanger Institute for the COVID-19 Genomics UK (COG-UK) Consortium | Lighthouse Laboratory Plymouth and Alex Alderton, Roberto Amato, Jeffrey Barrett, Sonia Goncalves, Ewan Harrison, David K. Jackson, Ian Johnston, Dominic Kwiatkowski, Cordelia Langford, John Sillitoe on behalf of the Wellcome Sanger Institute COVID-19 Surveillance Team |
| EPI_ISL_3442283 | Lighthouse Lab in Glasgow | Wellcome Sanger Institute for the COVID-19 Genomics UK (COG-UK) Consortium | Harper VanSteenhouse, Yumi Kasai, David Gray, Carol Clugston, Anna Dominiczak and Alex Alderton, Roberto Amato, Jeffrey Barrett, Sonia Goncalves, Ewan Harrison, David K. Jackson, Ian Johnston, Dominic Kwiatkowski, Cordelia Langford, John Sillitoe on behalf of the Wellcome Sanger Institute COVID-19 Surveillance Team |
| EPI_ISL_3442284 | Lighthouse Laboratory Plymouth | Wellcome Sanger Institute for the COVID-19 Genomics UK (COG-UK) Consortium | Lighthouse Laboratory Plymouth and Alex Alderton, Roberto Amato, Jeffrey Barrett, Sonia Goncalves, Ewan Harrison, David K. Jackson, Ian Johnston, Dominic Kwiatkowski, Cordelia Langford, John Sillitoe on behalf of the Wellcome Sanger Institute COVID-19 Surveillance Team |
| EPI_ISL_3442285 | Lighthouse Lab in Glasgow | Wellcome Sanger Institute for the COVID-19 Genomics UK (COG-UK) Consortium | Harper VanSteenhouse, Yumi Kasai, David Gray, Carol Clugston, Anna Dominiczak and Alex Alderton, Roberto Amato, Jeffrey Barrett, Sonia Goncalves, Ewan Harrison, David K. Jackson, Ian Johnston, Dominic Kwiatkowski, Cordelia Langford, John Sillitoe on behalf of the Wellcome Sanger Institute COVID-19 Surveillance Team |
| EPI_ISL_3442286, EPI_ISL_3442287, EPI_ISL_3442288, EPI_ISL_3442289, EPI_ISL_3442290 | Lighthouse Laboratory Plymouth | Wellcome Sanger Institute for the COVID-19 Genomics UK (COG-UK) Consortium | Lighthouse Laboratory Plymouth and Alex Alderton, Roberto Amato, Jeffrey Barrett, Sonia Goncalves, Ewan Harrison, David K. Jackson, Ian Johnston, Dominic Kwiatkowski, Cordelia Langford, John Sillitoe on behalf of the Wellcome Sanger Institute COVID-19 Surveillance Team |
| EPI_ISL_3442291, EPI_ISL_3442292 | Lighthouse Lab in Glasgow | Wellcome Sanger Institute for the COVID-19 Genomics UK (COG-UK) Consortium | Harper VanSteenhouse, Yumi Kasai, David Gray, Carol Clugston, Anna Dominiczak and Alex Alderton, Roberto Amato, Jeffrey Barrett, Sonia Goncalves, Ewan Harrison, David K. Jackson, Ian Johnston, Dominic Kwiatkowski, Cordelia Langford, John Sillitoe on behalf of the Wellcome Sanger Institute COVID-19 Surveillance Team |
| EPI_ISL_3442293, EPI_ISL_3442294 | Lighthouse Laboratory Plymouth | Wellcome Sanger Institute for the COVID-19 Genomics UK (COG-UK) Consortium | Lighthouse Laboratory Plymouth and Alex Alderton, Roberto Amato, Jeffrey Barrett, Sonia Goncalves, Ewan Harrison, David K. Jackson, Ian Johnston, Dominic Kwiatkowski, Cordelia Langford, John Sillitoe on behalf of the Wellcome Sanger Institute COVID-19 Surveillance Team |
| EPI_ISL_3442295 | Lighthouse Lab in Glasgow | Wellcome Sanger Institute for the COVID-19 Genomics UK (COG-UK) Consortium | Harper VanSteenhouse, Yumi Kasai, David Gray, Carol Clugston, Anna Dominiczak and Alex Alderton, Roberto Amato, Jeffrey Barrett, Sonia Goncalves, Ewan Harrison, David K. Jackson, Ian Johnston, Dominic Kwiatkowski, Cordelia Langford, John Sillitoe on behalf of the Wellcome Sanger Institute COVID-19 Surveillance Team |
| EPI_ISL_3442296, EPI_ISL_3442297, EPI_ISL_3442298 | Lighthouse Laboratory Plymouth | Wellcome Sanger Institute for the COVID-19 Genomics UK (COG-UK) Consortium | Lighthouse Laboratory Plymouth and Alex Alderton, Roberto Amato, Jeffrey Barrett, Sonia Goncalves, Ewan Harrison, David K. Jackson, Ian Johnston, Dominic Kwiatkowski, Cordelia Langford, John Sillitoe on behalf of the Wellcome Sanger Institute COVID-19 Surveillance Team |
| EPI_ISL_3442299 | Lighthouse Lab in Glasgow | Wellcome Sanger Institute for the COVID-19 Genomics UK (COG-UK) Consortium | Harper VanSteenhouse, Yumi Kasai, David Gray, Carol Clugston, Anna Dominiczak and Alex Alderton, Roberto Amato, Jeffrey Barrett, Sonia Goncalves, Ewan Harrison, David K. Jackson, Ian Johnston, Dominic Kwiatkowski, Cordelia Langford, John Sillitoe on behalf of the Wellcome Sanger Institute COVID-19 Surveillance Team |
| EPI_ISL_3442300 | Lighthouse Laboratory Plymouth | Wellcome Sanger Institute for the COVID-19 Genomics UK (COG-UK) Consortium | Lighthouse Laboratory Plymouth and Alex Alderton, Roberto Amato, Jeffrey Barrett, Sonia Goncalves, Ewan Harrison, David K. Jackson, Ian Johnston, Dominic Kwiatkowski, Cordelia Langford, John Sillitoe on behalf of the Wellcome Sanger Institute COVID-19 Surveillance Team |
| EPI_ISL_3442301, EPI_ISL_3442302, EPI_ISL_3442303, EPI_ISL_3442304, EPI_ISL_3442305 | Lighthouse Lab in Glasgow | Wellcome Sanger Institute for the COVID-19 Genomics UK (COG-UK) Consortium | Harper VanSteenhouse, Yumi Kasai, David Gray, Carol Clugston, Anna Dominiczak and Alex Alderton, Roberto Amato, Jeffrey Barrett, Sonia Goncalves, Ewan Harrison, David K. Jackson, Ian Johnston, Dominic Kwiatkowski, Cordelia Langford, John Sillitoe on behalf of the Wellcome Sanger Institute COVID-19 Surveillance Team |
| EPI_ISL_3442306 | Lighthouse Laboratory Plymouth | Wellcome Sanger Institute for the COVID-19 Genomics UK (COG-UK) Consortium | Lighthouse Laboratory Plymouth and Alex Alderton, Roberto Amato, Jeffrey Barrett, Sonia Goncalves, Ewan Harrison, David K. Jackson, Ian Johnston, Dominic Kwiatkowski, Cordelia Langford, John Sillitoe on behalf of the Wellcome Sanger Institute COVID-19 Surveillance Team |
| EPI_ISL_3442307 | Lighthouse Lab in Glasgow | Wellcome Sanger Institute for the COVID-19 Genomics UK (COG-UK) Consortium | Harper VanSteenhouse, Yumi Kasai, David Gray, Carol Clugston, Anna Dominiczak and Alex Alderton, Roberto Amato, Jeffrey Barrett, Sonia Goncalves, Ewan Harrison, David K. Jackson, Ian Johnston, Dominic Kwiatkowski, Cordelia Langford, John Sillitoe on behalf of the Wellcome Sanger Institute COVID-19 Surveillance Team |
| EPI_ISL_3442308 | Lighthouse Laboratory Plymouth | Wellcome Sanger Institute for the COVID-19 Genomics UK (COG-UK) Consortium | Lighthouse Laboratory Plymouth and Alex Alderton, Roberto Amato, Jeffrey Barrett, Sonia Goncalves, Ewan Harrison, David K. Jackson, Ian Johnston, Dominic Kwiatkowski, Cordelia Langford, John Sillitoe on behalf of the Wellcome Sanger Institute COVID-19 Surveillance Team |
| EPI_ISL_3442309, EPI_ISL_3442310 | Lighthouse Lab in Glasgow | Wellcome Sanger Institute for the COVID-19 Genomics UK (COG-UK) Consortium | Harper VanSteenhouse, Yumi Kasai, David Gray, Carol Clugston, Anna Dominiczak and Alex Alderton, Roberto Amato, Jeffrey Barrett, Sonia Goncalves, Ewan Harrison, David K. Jackson, Ian Johnston, Dominic Kwiatkowski, Cordelia Langford, John Sillitoe on behalf of the Wellcome Sanger Institute COVID-19 Surveillance Team |
| EPI_ISL_3442311 | Lighthouse Laboratory Plymouth | Wellcome Sanger Institute for the COVID-19 Genomics UK (COG-UK) Consortium | Lighthouse Laboratory Plymouth and Alex Alderton, Roberto Amato, Jeffrey Barrett, Sonia Goncalves, Ewan Harrison, David K. Jackson, Ian Johnston, Dominic Kwiatkowski, Cordelia Langford, John Sillitoe on behalf of the Wellcome Sanger Institute COVID-19 Surveillance Team |
| EPI_ISL_3442312, EPI_ISL_3442313, EPI_ISL_3442314 | Lighthouse Lab in Glasgow | Wellcome Sanger Institute for the COVID-19 Genomics UK (COG-UK) Consortium | Harper VanSteenhouse, Yumi Kasai, David Gray, Carol Clugston, Anna Dominiczak and Alex Alderton, Roberto Amato, Jeffrey Barrett, Sonia Goncalves, Ewan Harrison, David K. Jackson, Ian Johnston, Dominic Kwiatkowski, Cordelia Langford, John Sillitoe on behalf of the Wellcome Sanger Institute COVID-19 Surveillance Team |
| EPI_ISL_3442315, EPI_ISL_3442316, EPI_ISL_3442317, EPI_ISL_3442318 | Lighthouse Laboratory Plymouth | Wellcome Sanger Institute for the COVID-19 Genomics UK (COG-UK) Consortium | Lighthouse Laboratory Plymouth and Alex Alderton, Roberto Amato, Jeffrey Barrett, Sonia Goncalves, Ewan Harrison, David K. Jackson, Ian Johnston, Dominic Kwiatkowski, Cordelia Langford, John Sillitoe on behalf of the Wellcome Sanger Institute COVID-19 Surveillance Team |
| EPI_ISL_3442319 | Lighthouse Lab in Glasgow | Wellcome Sanger Institute for the COVID-19 Genomics UK (COG-UK) Consortium | Harper VanSteenhouse, Yumi Kasai, David Gray, Carol Clugston, Anna Dominiczak and Alex Alderton, Roberto Amato, Jeffrey Barrett, Sonia Goncalves, Ewan Harrison, David K. Jackson, Ian Johnston, Dominic Kwiatkowski, Cordelia Langford, John Sillitoe on behalf of the Wellcome Sanger Institute COVID-19 Surveillance Team |
| EPI_ISL_3442320, EPI_ISL_3442321, EPI_ISL_3442322 | Lighthouse Laboratory Plymouth | Wellcome Sanger Institute for the COVID-19 Genomics UK (COG-UK) Consortium | Lighthouse Laboratory Plymouth and Alex Alderton, Roberto Amato, Jeffrey Barrett, Sonia Goncalves, Ewan Harrison, David K. Jackson, Ian Johnston, Dominic Kwiatkowski, Cordelia Langford, John Sillitoe on behalf of the Wellcome Sanger Institute COVID-19 Surveillance Team |
| EPI_ISL_3442323 | Lighthouse Lab in Glasgow | Wellcome Sanger Institute for the COVID-19 Genomics UK (COG-UK) Consortium | Harper VanSteenhouse, Yumi Kasai, David Gray, Carol Clugston, Anna Dominiczak and Alex Alderton, Roberto Amato, Jeffrey Barrett, Sonia Goncalves, Ewan Harrison, David K. Jackson, Ian Johnston, Dominic Kwiatkowski, Cordelia Langford, John Sillitoe on behalf of the Wellcome Sanger Institute COVID-19 Surveillance Team |
| EPI_ISL_3442324, EPI_ISL_3442325, EPI_ISL_3442326, EPI_ISL_3442327 | Lighthouse Laboratory Plymouth | Wellcome Sanger Institute for the COVID-19 Genomics UK (COG-UK) Consortium | Lighthouse Laboratory Plymouth and Alex Alderton, Roberto Amato, Jeffrey Barrett, Sonia Goncalves, Ewan Harrison, David K. Jackson, Ian Johnston, Dominic Kwiatkowski, Cordelia Langford, John Sillitoe on behalf of the Wellcome Sanger Institute COVID-19 Surveillance Team |
| EPI_ISL_3442328 | Lighthouse Lab in Glasgow | Wellcome Sanger Institute for the COVID-19 Genomics UK (COG-UK) Consortium | Harper VanSteenhouse, Yumi Kasai, David Gray, Carol Clugston, Anna Dominiczak and Alex Alderton, Roberto Amato, Jeffrey Barrett, Sonia Goncalves, Ewan Harrison, David K. Jackson, Ian Johnston, Dominic Kwiatkowski, Cordelia Langford, John Sillitoe on behalf of the Wellcome Sanger Institute COVID-19 Surveillance Team |
| EPI_ISL_3442329, EPI_ISL_3442330, EPI_ISL_3442331 | Lighthouse Laboratory Plymouth | Wellcome Sanger Institute for the COVID-19 Genomics UK (COG-UK) Consortium | Lighthouse Laboratory Plymouth and Alex Alderton, Roberto Amato, Jeffrey Barrett, Sonia Goncalves, Ewan Harrison, David K. Jackson, Ian Johnston, Dominic Kwiatkowski, Cordelia Langford, John Sillitoe on behalf of the Wellcome Sanger Institute COVID-19 Surveillance Team |

| | | | |
|---|---|---|---|
| EPI_ISL_3442332 | Lighthouse Lab in Glasgow | Wellcome Sanger Institute for the COVID-19 Genomics UK (COG-UK) Consortium | Harper VanSteenhouse, Yumi Kasai, David Gray, Carol Clugston, Anna Dominiczak and Alex Alderton, Roberto Amato, Jeffrey Barrett, Sonia Goncalves, Ewan Harrison, David K. Jackson, Ian Johnston, Dominic Kwiatkowski, Cordelia Langford, John Sillitoe on behalf of the Wellcome Sanger Institute COVID-19 Surveillance Team |
| EPI_ISL_3442333, EPI_ISL_3442334, EPI_ISL_3442335, EPI_ISL_3442336 | Lighthouse Laboratory Plymouth | Wellcome Sanger Institute for the COVID-19 Genomics UK (COG-UK) Consortium | Lighthouse Laboratory Plymouth and Alex Alderton, Roberto Amato, Jeffrey Barrett, Sonia Goncalves, Ewan Harrison, David K. Jackson, Ian Johnston, Dominic Kwiatkowski, Cordelia Langford, John Sillitoe on behalf of the Wellcome Sanger Institute COVID-19 Surveillance Team |
| EPI_ISL_3442337 | Lighthouse Lab in Glasgow | Wellcome Sanger Institute for the COVID-19 Genomics UK (COG-UK) Consortium | Harper VanSteenhouse, Yumi Kasai, David Gray, Carol Clugston, Anna Dominiczak and Alex Alderton, Roberto Amato, Jeffrey Barrett, Sonia Goncalves, Ewan Harrison, David K. Jackson, Ian Johnston, Dominic Kwiatkowski, Cordelia Langford, John Sillitoe on behalf of the Wellcome Sanger Institute COVID-19 Surveillance Team |
| EPI_ISL_3442338, EPI_ISL_3442339, EPI_ISL_3442340 | Lighthouse Laboratory Plymouth | Wellcome Sanger Institute for the COVID-19 Genomics UK (COG-UK) Consortium | Lighthouse Laboratory Plymouth and Alex Alderton, Roberto Amato, Jeffrey Barrett, Sonia Goncalves, Ewan Harrison, David K. Jackson, Ian Johnston, Dominic Kwiatkowski, Cordelia Langford, John Sillitoe on behalf of the Wellcome Sanger Institute COVID-19 Surveillance Team |
| EPI_ISL_3442341, EPI_ISL_3442342 | Lighthouse Lab in Glasgow | Wellcome Sanger Institute for the COVID-19 Genomics UK (COG-UK) Consortium | Harper VanSteenhouse, Yumi Kasai, David Gray, Carol Clugston, Anna Dominiczak and Alex Alderton, Roberto Amato, Jeffrey Barrett, Sonia Goncalves, Ewan Harrison, David K. Jackson, Ian Johnston, Dominic Kwiatkowski, Cordelia Langford, John Sillitoe on behalf of the Wellcome Sanger Institute COVID-19 Surveillance Team |
| EPI_ISL_3442343, EPI_ISL_3442344, EPI_ISL_3442345 | Lighthouse Laboratory Plymouth | Wellcome Sanger Institute for the COVID-19 Genomics UK (COG-UK) Consortium | Lighthouse Laboratory Plymouth and Alex Alderton, Roberto Amato, Jeffrey Barrett, Sonia Goncalves, Ewan Harrison, David K. Jackson, Ian Johnston, Dominic Kwiatkowski, Cordelia Langford, John Sillitoe on behalf of the Wellcome Sanger Institute COVID-19 Surveillance Team |
| EPI_ISL_3442346, EPI_ISL_3442347, EPI_ISL_3442348 | Lighthouse Lab in Glasgow | Wellcome Sanger Institute for the COVID-19 Genomics UK (COG-UK) Consortium | Harper VanSteenhouse, Yumi Kasai, David Gray, Carol Clugston, Anna Dominiczak and Alex Alderton, Roberto Amato, Jeffrey Barrett, Sonia Goncalves, Ewan Harrison, David K. Jackson, Ian Johnston, Dominic Kwiatkowski, Cordelia Langford, John Sillitoe on behalf of the Wellcome Sanger Institute COVID-19 Surveillance Team |
| EPI_ISL_3442349, EPI_ISL_3442350, EPI_ISL_3442351 | Lighthouse Laboratory Plymouth | Wellcome Sanger Institute for the COVID-19 Genomics UK (COG-UK) Consortium | Lighthouse Laboratory Plymouth and Alex Alderton, Roberto Amato, Jeffrey Barrett, Sonia Goncalves, Ewan Harrison, David K. Jackson, Ian Johnston, Dominic Kwiatkowski, Cordelia Langford, John Sillitoe on behalf of the Wellcome Sanger Institute COVID-19 Surveillance Team |
| EPI_ISL_3442352 | Lighthouse Lab in Glasgow | Wellcome Sanger Institute for the COVID-19 Genomics UK (COG-UK) Consortium | Harper VanSteenhouse, Yumi Kasai, David Gray, Carol Clugston, Anna Dominiczak and Alex Alderton, Roberto Amato, Jeffrey Barrett, Sonia Goncalves, Ewan Harrison, David K. Jackson, Ian Johnston, Dominic Kwiatkowski, Cordelia Langford, John Sillitoe on behalf of the Wellcome Sanger Institute COVID-19 Surveillance Team |
| EPI_ISL_3442353, EPI_ISL_3442354 | Lighthouse Laboratory Plymouth | Wellcome Sanger Institute for the COVID-19 Genomics UK (COG-UK) Consortium | Lighthouse Laboratory Plymouth and Alex Alderton, Roberto Amato, Jeffrey Barrett, Sonia Goncalves, Ewan Harrison, David K. Jackson, Ian Johnston, Dominic Kwiatkowski, Cordelia Langford, John Sillitoe on behalf of the Wellcome Sanger Institute COVID-19 Surveillance Team |
| EPI_ISL_3442355 | Lighthouse Lab in Glasgow | Wellcome Sanger Institute for the COVID-19 Genomics UK (COG-UK) Consortium | Harper VanSteenhouse, Yumi Kasai, David Gray, Carol Clugston, Anna Dominiczak and Alex Alderton, Roberto Amato, Jeffrey Barrett, Sonia Goncalves, Ewan Harrison, David K. Jackson, Ian Johnston, Dominic Kwiatkowski, Cordelia Langford, John Sillitoe on behalf of the Wellcome Sanger Institute COVID-19 Surveillance Team |
| EPI_ISL_3442356, EPI_ISL_3442357, EPI_ISL_3442358 | Lighthouse Laboratory Plymouth | Wellcome Sanger Institute for the COVID-19 Genomics UK (COG-UK) Consortium | Lighthouse Laboratory Plymouth and Alex Alderton, Roberto Amato, Jeffrey Barrett, Sonia Goncalves, Ewan Harrison, David K. Jackson, Ian Johnston, Dominic Kwiatkowski, Cordelia Langford, John Sillitoe on behalf of the Wellcome Sanger Institute COVID-19 Surveillance Team |
| EPI_ISL_3442359, EPI_ISL_3442360, EPI_ISL_3442361 | Lighthouse Lab in Glasgow | Wellcome Sanger Institute for the COVID-19 Genomics UK (COG-UK) Consortium | Harper VanSteenhouse, Yumi Kasai, David Gray, Carol Clugston, Anna Dominiczak and Alex Alderton, Roberto Amato, Jeffrey Barrett, Sonia Goncalves, Ewan Harrison, David K. Jackson, Ian Johnston, Dominic Kwiatkowski, Cordelia Langford, John Sillitoe on behalf of the Wellcome Sanger Institute COVID-19 Surveillance Team |
| EPI_ISL_3442362, EPI_ISL_3442363, EPI_ISL_3442364 | Lighthouse Laboratory Plymouth | Wellcome Sanger Institute for the COVID-19 Genomics UK (COG-UK) Consortium | Lighthouse Laboratory Plymouth and Alex Alderton, Roberto Amato, Jeffrey Barrett, Sonia Goncalves, Ewan Harrison, David K. Jackson, Ian Johnston, Dominic Kwiatkowski, Cordelia Langford, John Sillitoe on behalf of the Wellcome Sanger Institute COVID-19 Surveillance Team |
| EPI_ISL_3442365, EPI_ISL_3442366, EPI_ISL_3442367, EPI_ISL_3442368 | Lighthouse Lab in Glasgow | Wellcome Sanger Institute for the COVID-19 Genomics UK (COG-UK) Consortium | Harper VanSteenhouse, Yumi Kasai, David Gray, Carol Clugston, Anna Dominiczak and Alex Alderton, Roberto Amato, Jeffrey Barrett, Sonia Goncalves, Ewan Harrison, David K. Jackson, Ian Johnston, Dominic Kwiatkowski, Cordelia Langford, John Sillitoe on behalf of the Wellcome Sanger Institute COVID-19 Surveillance Team |
| EPI_ISL_3442369, EPI_ISL_3442370, EPI_ISL_3442371, EPI_ISL_3442372, EPI_ISL_3442373, EPI_ISL_3442374, EPI_ISL_3442375, EPI_ISL_3442376, EPI_ISL_3442377, EPI_ISL_3442378, EPI_ISL_3442379 | | | |
| see above | Lighthouse Laboratory Plymouth | Wellcome Sanger Institute for the COVID-19 Genomics UK (COG-UK) Consortium | Lighthouse Laboratory Plymouth and Alex Alderton, Roberto Amato, Jeffrey Barrett, Sonia Goncalves, Ewan Harrison, David K. Jackson, Ian Johnston, Dominic Kwiatkowski, Cordelia Langford, John Sillitoe on behalf of the Wellcome Sanger Institute COVID-19 Surveillance Team |
| EPI_ISL_3442380, EPI_ISL_3442381 | Lighthouse Lab in Glasgow | Wellcome Sanger Institute for the COVID-19 Genomics UK (COG-UK) Consortium | Harper VanSteenhouse, Yumi Kasai, David Gray, Carol Clugston, Anna Dominiczak and Alex Alderton, Roberto Amato, Jeffrey Barrett, Sonia Goncalves, Ewan Harrison, David K. Jackson, Ian Johnston, Dominic Kwiatkowski, Cordelia Langford, John Sillitoe on behalf of the Wellcome Sanger Institute COVID-19 Surveillance Team |
| EPI_ISL_3442382, EPI_ISL_3442383 | Lighthouse Laboratory Plymouth | Wellcome Sanger Institute for the COVID-19 Genomics UK (COG-UK) Consortium | Lighthouse Laboratory Plymouth and Alex Alderton, Roberto Amato, Jeffrey Barrett, Sonia Goncalves, Ewan Harrison, David K. Jackson, Ian Johnston, Dominic Kwiatkowski, Cordelia Langford, John Sillitoe on behalf of the Wellcome Sanger Institute COVID-19 Surveillance Team |
| EPI_ISL_3442384 | Lighthouse Lab in Glasgow | Wellcome Sanger Institute for the COVID-19 Genomics UK (COG-UK) Consortium | Harper VanSteenhouse, Yumi Kasai, David Gray, Carol Clugston, Anna Dominiczak and Alex Alderton, Roberto Amato, Jeffrey Barrett, Sonia Goncalves, Ewan Harrison, David K. Jackson, Ian Johnston, Dominic Kwiatkowski, Cordelia Langford, John Sillitoe on behalf of the Wellcome Sanger Institute COVID-19 Surveillance Team |
| EPI_ISL_3442385, EPI_ISL_3442386 | Lighthouse Laboratory Plymouth | Wellcome Sanger Institute for the COVID-19 Genomics UK (COG-UK) Consortium | Lighthouse Laboratory Plymouth and Alex Alderton, Roberto Amato, Jeffrey Barrett, Sonia Goncalves, Ewan Harrison, David K. Jackson, Ian Johnston, Dominic Kwiatkowski, Cordelia Langford, John Sillitoe on behalf of the Wellcome Sanger Institute COVID-19 Surveillance Team |
| EPI_ISL_3442387 | Lighthouse Lab in Glasgow | Wellcome Sanger Institute for the COVID-19 Genomics UK (COG-UK) Consortium | Harper VanSteenhouse, Yumi Kasai, David Gray, Carol Clugston, Anna Dominiczak and Alex Alderton, Roberto Amato, Jeffrey Barrett, Sonia Goncalves, Ewan Harrison, David K. Jackson, Ian Johnston, Dominic Kwiatkowski, Cordelia Langford, John Sillitoe on behalf of the Wellcome Sanger Institute COVID-19 Surveillance Team |
| EPI_ISL_3442388 | Lighthouse Laboratory Plymouth | Wellcome Sanger Institute for the COVID-19 Genomics UK (COG-UK) Consortium | Lighthouse Laboratory Plymouth and Alex Alderton, Roberto Amato, Jeffrey Barrett, Sonia Goncalves, Ewan Harrison, David K. Jackson, Ian Johnston, Dominic Kwiatkowski, Cordelia Langford, John Sillitoe on behalf of the Wellcome Sanger Institute COVID-19 Surveillance Team |
| EPI_ISL_3442389 | Lighthouse Lab in Glasgow | Wellcome Sanger Institute for the COVID-19 Genomics UK (COG-UK) Consortium | Harper VanSteenhouse, Yumi Kasai, David Gray, Carol Clugston, Anna Dominiczak and Alex Alderton, Roberto Amato, Jeffrey Barrett, Sonia Goncalves, Ewan Harrison, David K. Jackson, Ian Johnston, Dominic Kwiatkowski, Cordelia Langford, John Sillitoe on behalf of the Wellcome Sanger Institute COVID-19 Surveillance Team |
| EPI_ISL_3442390, EPI_ISL_3442391 | Lighthouse Laboratory Plymouth | Wellcome Sanger Institute for the COVID-19 Genomics UK (COG-UK) Consortium | Lighthouse Laboratory Plymouth and Alex Alderton, Roberto Amato, Jeffrey Barrett, Sonia Goncalves, Ewan Harrison, David K. Jackson, Ian Johnston, Dominic Kwiatkowski, Cordelia Langford, John Sillitoe on behalf of the Wellcome Sanger Institute COVID-19 Surveillance Team |
| EPI_ISL_3442392, EPI_ISL_3442393 | Lighthouse Lab in Glasgow | Wellcome Sanger Institute for the COVID-19 Genomics UK (COG-UK) Consortium | Harper VanSteenhouse, Yumi Kasai, David Gray, Carol Clugston, Anna Dominiczak and Alex Alderton, Roberto Amato, Jeffrey Barrett, Sonia Goncalves, Ewan Harrison, David K. Jackson, Ian Johnston, Dominic Kwiatkowski, Cordelia Langford, John Sillitoe on behalf of the Wellcome Sanger Institute COVID-19 Surveillance Team |
| EPI_ISL_3442394, EPI_ISL_3442395, EPI_ISL_3442396 | Lighthouse Laboratory Plymouth | Wellcome Sanger Institute for the COVID-19 Genomics UK (COG-UK) Consortium | Lighthouse Laboratory Plymouth and Alex Alderton, Roberto Amato, Jeffrey Barrett, Sonia Goncalves, Ewan Harrison, David K. Jackson, Ian Johnston, Dominic Kwiatkowski, Cordelia Langford, John Sillitoe on behalf of the Wellcome Sanger Institute COVID-19 Surveillance Team |
| EPI_ISL_3442397, EPI_ISL_3442398, EPI_ISL_3442399 | Lighthouse Lab in Glasgow | Wellcome Sanger Institute for the COVID-19 Genomics UK (COG-UK) Consortium | Harper VanSteenhouse, Yumi Kasai, David Gray, Carol Clugston, Anna Dominiczak and Alex Alderton, Roberto Amato, Jeffrey Barrett, Sonia Goncalves, Ewan Harrison, David K. Jackson, Ian Johnston, Dominic Kwiatkowski, Cordelia Langford, John Sillitoe on behalf of the Wellcome Sanger Institute COVID-19 Surveillance Team |
| EPI_ISL_3442400 | Lighthouse Laboratory Plymouth | Wellcome Sanger Institute for the COVID-19 Genomics UK (COG-UK) Consortium | Lighthouse Laboratory Plymouth and Alex Alderton, Roberto Amato, Jeffrey Barrett, Sonia Goncalves, Ewan Harrison, David K. Jackson, Ian Johnston, Dominic Kwiatkowski, Cordelia Langford, John Sillitoe on behalf of the Wellcome Sanger Institute COVID-19 Surveillance Team |
| EPI_ISL_3442401, EPI_ISL_3442402, EPI_ISL_3442403 | Lighthouse Lab in Glasgow | Wellcome Sanger Institute for the COVID-19 Genomics UK (COG-UK) Consortium | Harper VanSteenhouse, Yumi Kasai, David Gray, Carol Clugston, Anna Dominiczak and Alex Alderton, Roberto Amato, Jeffrey Barrett, Sonia Goncalves, Ewan Harrison, David K. Jackson, Ian Johnston, Dominic Kwiatkowski, Cordelia Langford, John Sillitoe on behalf of the Wellcome Sanger Institute COVID-19 Surveillance Team |
| EPI_ISL_3442404, EPI_ISL_3442405 | Lighthouse Laboratory Plymouth | Wellcome Sanger Institute for the COVID-19 Genomics UK (COG-UK) Consortium | Lighthouse Laboratory Plymouth and Alex Alderton, Roberto Amato, Jeffrey Barrett, Sonia Goncalves, Ewan Harrison, David K. Jackson, Ian Johnston, Dominic Kwiatkowski, Cordelia Langford, John Sillitoe on behalf of the Wellcome Sanger Institute COVID-19 Surveillance Team |
| EPI_ISL_3442406 | Lighthouse Lab in Glasgow | Wellcome Sanger Institute for the COVID-19 Genomics UK (COG-UK) Consortium | Harper VanSteenhouse, Yumi Kasai, David Gray, Carol Clugston, Anna Dominiczak and Alex Alderton, Roberto Amato, Jeffrey Barrett, Sonia Goncalves, Ewan Harrison, David K. Jackson, Ian Johnston, Dominic Kwiatkowski, Cordelia Langford, John Sillitoe on behalf of the Wellcome Sanger Institute COVID-19 Surveillance Team |
| EPI_ISL_3442407 | Lighthouse Laboratory Plymouth | Wellcome Sanger Institute for the COVID-19 Genomics UK (COG-UK) Consortium | Lighthouse Laboratory Plymouth and Alex Alderton, Roberto Amato, Jeffrey Barrett, Sonia Goncalves, Ewan Harrison, David K. Jackson, Ian Johnston, Dominic Kwiatkowski, Cordelia Langford, John Sillitoe on behalf of the Wellcome Sanger Institute COVID-19 Surveillance Team |
| EPI_ISL_3442408, EPI_ISL_3442409, EPI_ISL_3442410 | Lighthouse Lab in Glasgow | Wellcome Sanger Institute for the COVID-19 Genomics UK (COG-UK) Consortium | Harper VanSteenhouse, Yumi Kasai, David Gray, Carol Clugston, Anna Dominiczak and Alex Alderton, Roberto Amato, Jeffrey Barrett, Sonia Goncalves, Ewan Harrison, David K. Jackson, Ian Johnston, Dominic Kwiatkowski, Cordelia Langford, John Sillitoe on behalf of the Wellcome Sanger Institute COVID-19 Surveillance Team |
| EPI_ISL_3442411 | Lighthouse Laboratory Plymouth | Wellcome Sanger Institute for the COVID-19 Genomics UK (COG-UK) Consortium | Lighthouse Laboratory Plymouth and Alex Alderton, Roberto Amato, Jeffrey Barrett, Sonia Goncalves, Ewan Harrison, David K. Jackson, Ian Johnston, Dominic Kwiatkowski, Cordelia Langford, John Sillitoe on behalf of the Wellcome Sanger Institute COVID-19 Surveillance Team |
| EPI_ISL_3442412 | Lighthouse Lab in Glasgow | Wellcome Sanger Institute for the COVID-19 Genomics UK (COG-UK) Consortium | Harper VanSteenhouse, Yumi Kasai, David Gray, Carol Clugston, Anna Dominiczak and Alex Alderton, Roberto Amato, Jeffrey Barrett, Sonia Goncalves, Ewan Harrison, David K. Jackson, Ian Johnston, Dominic Kwiatkowski, Cordelia Langford, John Sillitoe on behalf of the Wellcome Sanger Institute COVID-19 Surveillance Team |
| EPI_ISL_3442413 | Berkshire and Surrey Pathology Services Lighthouse Laboratory | Wellcome Sanger Institute for the COVID-19 Genomics UK (COG-UK) Consortium | Berkshire and Surrey Pathology Services Lighthouse Laboratory and Alex Alderton, Roberto Amato, Jeffrey Barrett, Sonia Goncalves, Ewan Harrison, David K. Jackson, Ian Johnston, Dominic Kwiatkowski, Cordelia Langford, John Sillitoe on behalf of the Wellcome Sanger Institute COVID-19 Surveillance Team |
| EPI_ISL_3442414, EPI_ISL_3442415, EPI_ISL_3442416 | Lighthouse Lab in Glasgow | Wellcome Sanger Institute for the COVID-19 Genomics UK (COG-UK) Consortium | Harper VanSteenhouse, Yumi Kasai, David Gray, Carol Clugston, Anna Dominiczak and Alex Alderton, Roberto Amato, Jeffrey Barrett, Sonia Goncalves, Ewan Harrison, David K. Jackson, Ian Johnston, Dominic Kwiatkowski, Cordelia Langford, John Sillitoe on behalf of the Wellcome Sanger Institute COVID-19 Surveillance Team |
| EPI_ISL_3442417 | Berkshire and Surrey Pathology Services Lighthouse Laboratory | Wellcome Sanger Institute for the COVID-19 Genomics UK (COG-UK) Consortium | Berkshire and Surrey Pathology Services Lighthouse Laboratory and Alex Alderton, Roberto Amato, Jeffrey Barrett, Sonia Goncalves, Ewan Harrison, David K. Jackson, Ian Johnston, Dominic Kwiatkowski, Cordelia Langford, John Sillitoe on behalf of the Wellcome Sanger Institute COVID-19 Surveillance Team |

| | | | |
|---|---|---|---|
| EPI_ISL_3442418 | Lighthouse Lab in Glasgow | Wellcome Sanger Institute for the COVID-19 Genomics UK (COG-UK) Consortium | Harper VanSteenhouse, Yumi Kasai, David Gray, Carol Clugston, Anna Dominiczak and Alex Alderton, Roberto Amato, Jeffrey Barrett, Sonia Goncalves, Ewan Harrison, David K. Jackson, Ian Johnston, Dominic Kwiatkowski, Cordelia Langford, John Sillitoe on behalf of the Wellcome Sanger Institute COVID-19 Surveillance Team |
| EPI_ISL_3442419 | Berkshire and Surrey Pathology Services Lighthouse Laboratory | Wellcome Sanger Institute for the COVID-19 Genomics UK (COG-UK) Consortium | Berkshire and Surrey Pathology Services Lighthouse Laboratory and Alex Alderton, Roberto Amato, Jeffrey Barrett, Sonia Goncalves, Ewan Harrison, David K. Jackson, Ian Johnston, Dominic Kwiatkowski, Cordelia Langford, John Sillitoe on behalf of the Wellcome Sanger Institute COVID-19 Surveillance Team |
| EPI_ISL_3442420 | Lighthouse Lab in Glasgow | Wellcome Sanger Institute for the COVID-19 Genomics UK (COG-UK) Consortium | Harper VanSteenhouse, Yumi Kasai, David Gray, Carol Clugston, Anna Dominiczak and Alex Alderton, Roberto Amato, Jeffrey Barrett, Sonia Goncalves, Ewan Harrison, David K. Jackson, Ian Johnston, Dominic Kwiatkowski, Cordelia Langford, John Sillitoe on behalf of the Wellcome Sanger Institute COVID-19 Surveillance Team |
| EPI_ISL_3442421 | Berkshire and Surrey Pathology Services Lighthouse Laboratory | Wellcome Sanger Institute for the COVID-19 Genomics UK (COG-UK) Consortium | Berkshire and Surrey Pathology Services Lighthouse Laboratory and Alex Alderton, Roberto Amato, Jeffrey Barrett, Sonia Goncalves, Ewan Harrison, David K. Jackson, Ian Johnston, Dominic Kwiatkowski, Cordelia Langford, John Sillitoe on behalf of the Wellcome Sanger Institute COVID-19 Surveillance Team |
| EPI_ISL_3442422, EPI_ISL_3442423, EPI_ISL_3442424, EPI_ISL_3442425 | Lighthouse Lab in Glasgow | Wellcome Sanger Institute for the COVID-19 Genomics UK (COG-UK) Consortium | Harper VanSteenhouse, Yumi Kasai, David Gray, Carol Clugston, Anna Dominiczak and Alex Alderton, Roberto Amato, Jeffrey Barrett, Sonia Goncalves, Ewan Harrison, David K. Jackson, Ian Johnston, Dominic Kwiatkowski, Cordelia Langford, John Sillitoe on behalf of the Wellcome Sanger Institute COVID-19 Surveillance Team |
| EPI_ISL_3442426, EPI_ISL_3442427 | Lighthouse Laboratory Plymouth | Wellcome Sanger Institute for the COVID-19 Genomics UK (COG-UK) Consortium | Lighthouse Laboratory Plymouth and Alex Alderton, Roberto Amato, Jeffrey Barrett, Sonia Goncalves, Ewan Harrison, David K. Jackson, Ian Johnston, Dominic Kwiatkowski, Cordelia Langford, John Sillitoe on behalf of the Wellcome Sanger Institute COVID-19 Surveillance Team |
| EPI_ISL_3442428 | Berkshire and Surrey Pathology Services Lighthouse Laboratory | Wellcome Sanger Institute for the COVID-19 Genomics UK (COG-UK) Consortium | Berkshire and Surrey Pathology Services Lighthouse Laboratory and Alex Alderton, Roberto Amato, Jeffrey Barrett, Sonia Goncalves, Ewan Harrison, David K. Jackson, Ian Johnston, Dominic Kwiatkowski, Cordelia Langford, John Sillitoe on behalf of the Wellcome Sanger Institute COVID-19 Surveillance Team |
| EPI_ISL_3442429, EPI_ISL_3442430 | Lighthouse Laboratory Plymouth | Wellcome Sanger Institute for the COVID-19 Genomics UK (COG-UK) Consortium | Lighthouse Laboratory Plymouth and Alex Alderton, Roberto Amato, Jeffrey Barrett, Sonia Goncalves, Ewan Harrison, David K. Jackson, Ian Johnston, Dominic Kwiatkowski, Cordelia Langford, John Sillitoe on behalf of the Wellcome Sanger Institute COVID-19 Surveillance Team |
| EPI_ISL_3442431, EPI_ISL_3442432, EPI_ISL_3442433 | Lighthouse Lab in Glasgow | Wellcome Sanger Institute for the COVID-19 Genomics UK (COG-UK) Consortium | Harper VanSteenhouse, Yumi Kasai, David Gray, Carol Clugston, Anna Dominiczak and Alex Alderton, Roberto Amato, Jeffrey Barrett, Sonia Goncalves, Ewan Harrison, David K. Jackson, Ian Johnston, Dominic Kwiatkowski, Cordelia Langford, John Sillitoe on behalf of the Wellcome Sanger Institute COVID-19 Surveillance Team |
| EPI_ISL_3442434, EPI_ISL_3442435 | Berkshire and Surrey Pathology Services Lighthouse Laboratory | Wellcome Sanger Institute for the COVID-19 Genomics UK (COG-UK) Consortium | Berkshire and Surrey Pathology Services Lighthouse Laboratory and Alex Alderton, Roberto Amato, Jeffrey Barrett, Sonia Goncalves, Ewan Harrison, David K. Jackson, Ian Johnston, Dominic Kwiatkowski, Cordelia Langford, John Sillitoe on behalf of the Wellcome Sanger Institute COVID-19 Surveillance Team |
| EPI_ISL_3442436 | Lighthouse Lab in Glasgow | Wellcome Sanger Institute for the COVID-19 Genomics UK (COG-UK) Consortium | Harper VanSteenhouse, Yumi Kasai, David Gray, Carol Clugston, Anna Dominiczak and Alex Alderton, Roberto Amato, Jeffrey Barrett, Sonia Goncalves, Ewan Harrison, David K. Jackson, Ian Johnston, Dominic Kwiatkowski, Cordelia Langford, John Sillitoe on behalf of the Wellcome Sanger Institute COVID-19 Surveillance Team |
| EPI_ISL_3442437 | Lighthouse Laboratory Plymouth | Wellcome Sanger Institute for the COVID-19 Genomics UK (COG-UK) Consortium | Lighthouse Laboratory Plymouth and Alex Alderton, Roberto Amato, Jeffrey Barrett, Sonia Goncalves, Ewan Harrison, David K. Jackson, Ian Johnston, Dominic Kwiatkowski, Cordelia Langford, John Sillitoe on behalf of the Wellcome Sanger Institute COVID-19 Surveillance Team |
| EPI_ISL_3442438, EPI_ISL_3442439, EPI_ISL_3442440, EPI_ISL_3442441 | Lighthouse Lab in Glasgow | Wellcome Sanger Institute for the COVID-19 Genomics UK (COG-UK) Consortium | Harper VanSteenhouse, Yumi Kasai, David Gray, Carol Clugston, Anna Dominiczak and Alex Alderton, Roberto Amato, Jeffrey Barrett, Sonia Goncalves, Ewan Harrison, David K. Jackson, Ian Johnston, Dominic Kwiatkowski, Cordelia Langford, John Sillitoe on behalf of the Wellcome Sanger Institute COVID-19 Surveillance Team |
| EPI_ISL_3442442, EPI_ISL_3442443 | Berkshire and Surrey Pathology Services Lighthouse Laboratory | Wellcome Sanger Institute for the COVID-19 Genomics UK (COG-UK) Consortium | Berkshire and Surrey Pathology Services Lighthouse Laboratory and Alex Alderton, Roberto Amato, Jeffrey Barrett, Sonia Goncalves, Ewan Harrison, David K. Jackson, Ian Johnston, Dominic Kwiatkowski, Cordelia Langford, John Sillitoe on behalf of the Wellcome Sanger Institute COVID-19 Surveillance Team |
| EPI_ISL_3442444 | Lighthouse Laboratory Plymouth | Wellcome Sanger Institute for the COVID-19 Genomics UK (COG-UK) Consortium | Lighthouse Laboratory Plymouth and Alex Alderton, Roberto Amato, Jeffrey Barrett, Sonia Goncalves, Ewan Harrison, David K. Jackson, Ian Johnston, Dominic Kwiatkowski, Cordelia Langford, John Sillitoe on behalf of the Wellcome Sanger Institute COVID-19 Surveillance Team |
| EPI_ISL_3442445 | Lighthouse Lab in Glasgow | Wellcome Sanger Institute for the COVID-19 Genomics UK (COG-UK) Consortium | Harper VanSteenhouse, Yumi Kasai, David Gray, Carol Clugston, Anna Dominiczak and Alex Alderton, Roberto Amato, Jeffrey Barrett, Sonia Goncalves, Ewan Harrison, David K. Jackson, Ian Johnston, Dominic Kwiatkowski, Cordelia Langford, John Sillitoe on behalf of the Wellcome Sanger Institute COVID-19 Surveillance Team |
| EPI_ISL_3442446 | Lighthouse Laboratory Plymouth | Wellcome Sanger Institute for the COVID-19 Genomics UK (COG-UK) Consortium | Lighthouse Laboratory Plymouth and Alex Alderton, Roberto Amato, Jeffrey Barrett, Sonia Goncalves, Ewan Harrison, David K. Jackson, Ian Johnston, Dominic Kwiatkowski, Cordelia Langford, John Sillitoe on behalf of the Wellcome Sanger Institute COVID-19 Surveillance Team |
| EPI_ISL_3442447, EPI_ISL_3442448, EPI_ISL_3442449, EPI_ISL_3442450 | Lighthouse Lab in Glasgow | Wellcome Sanger Institute for the COVID-19 Genomics UK (COG-UK) Consortium | Harper VanSteenhouse, Yumi Kasai, David Gray, Carol Clugston, Anna Dominiczak and Alex Alderton, Roberto Amato, Jeffrey Barrett, Sonia Goncalves, Ewan Harrison, David K. Jackson, Ian Johnston, Dominic Kwiatkowski, Cordelia Langford, John Sillitoe on behalf of the Wellcome Sanger Institute COVID-19 Surveillance Team |
| EPI_ISL_3442451 | Lighthouse Laboratory Plymouth | Wellcome Sanger Institute for the COVID-19 Genomics UK (COG-UK) Consortium | Lighthouse Laboratory Plymouth and Alex Alderton, Roberto Amato, Jeffrey Barrett, Sonia Goncalves, Ewan Harrison, David K. Jackson, Ian Johnston, Dominic Kwiatkowski, Cordelia Langford, John Sillitoe on behalf of the Wellcome Sanger Institute COVID-19 Surveillance Team |
| EPI_ISL_3442452, EPI_ISL_3442453 | Lighthouse Lab in Glasgow | Wellcome Sanger Institute for the COVID-19 Genomics UK (COG-UK) Consortium | Harper VanSteenhouse, Yumi Kasai, David Gray, Carol Clugston, Anna Dominiczak and Alex Alderton, Roberto Amato, Jeffrey Barrett, Sonia Goncalves, Ewan Harrison, David K. Jackson, Ian Johnston, Dominic Kwiatkowski, Cordelia Langford, John Sillitoe on behalf of the Wellcome Sanger Institute COVID-19 Surveillance Team |
| EPI_ISL_3442454 | Berkshire and Surrey Pathology Services Lighthouse Laboratory | Wellcome Sanger Institute for the COVID-19 Genomics UK (COG-UK) Consortium | Berkshire and Surrey Pathology Services Lighthouse Laboratory and Alex Alderton, Roberto Amato, Jeffrey Barrett, Sonia Goncalves, Ewan Harrison, David K. Jackson, Ian Johnston, Dominic Kwiatkowski, Cordelia Langford, John Sillitoe on behalf of the Wellcome Sanger Institute COVID-19 Surveillance Team |
| EPI_ISL_3442455, EPI_ISL_3442456, EPI_ISL_3442457, EPI_ISL_3442458, EPI_ISL_3442459, EPI_ISL_3442460, EPI_ISL_3442461 | Lighthouse Lab in Glasgow | Wellcome Sanger Institute for the COVID-19 Genomics UK (COG-UK) Consortium | Harper VanSteenhouse, Yumi Kasai, David Gray, Carol Clugston, Anna Dominiczak and Alex Alderton, Roberto Amato, Jeffrey Barrett, Sonia Goncalves, Ewan Harrison, David K. Jackson, Ian Johnston, Dominic Kwiatkowski, Cordelia Langford, John Sillitoe on behalf of the Wellcome Sanger Institute COVID-19 Surveillance Team |
| EPI_ISL_3442462 | Lighthouse Laboratory Plymouth | Wellcome Sanger Institute for the COVID-19 Genomics UK (COG-UK) Consortium | Lighthouse Laboratory Plymouth and Alex Alderton, Roberto Amato, Jeffrey Barrett, Sonia Goncalves, Ewan Harrison, David K. Jackson, Ian Johnston, Dominic Kwiatkowski, Cordelia Langford, John Sillitoe on behalf of the Wellcome Sanger Institute COVID-19 Surveillance Team |
| EPI_ISL_3442463, EPI_ISL_3442464 | Lighthouse Lab in Glasgow | Wellcome Sanger Institute for the COVID-19 Genomics UK (COG-UK) Consortium | Harper VanSteenhouse, Yumi Kasai, David Gray, Carol Clugston, Anna Dominiczak and Alex Alderton, Roberto Amato, Jeffrey Barrett, Sonia Goncalves, Ewan Harrison, David K. Jackson, Ian Johnston, Dominic Kwiatkowski, Cordelia Langford, John Sillitoe on behalf of the Wellcome Sanger Institute COVID-19 Surveillance Team |
| EPI_ISL_3442465 | Berkshire and Surrey Pathology Services Lighthouse Laboratory | Wellcome Sanger Institute for the COVID-19 Genomics UK (COG-UK) Consortium | Berkshire and Surrey Pathology Services Lighthouse Laboratory and Alex Alderton, Roberto Amato, Jeffrey Barrett, Sonia Goncalves, Ewan Harrison, David K. Jackson, Ian Johnston, Dominic Kwiatkowski, Cordelia Langford, John Sillitoe on behalf of the Wellcome Sanger Institute COVID-19 Surveillance Team |
| EPI_ISL_3442466 | Lighthouse Lab in Glasgow | Wellcome Sanger Institute for the COVID-19 Genomics UK (COG-UK) Consortium | Harper VanSteenhouse, Yumi Kasai, David Gray, Carol Clugston, Anna Dominiczak and Alex Alderton, Roberto Amato, Jeffrey Barrett, Sonia Goncalves, Ewan Harrison, David K. Jackson, Ian Johnston, Dominic Kwiatkowski, Cordelia Langford, John Sillitoe on behalf of the Wellcome Sanger Institute COVID-19 Surveillance Team |
| EPI_ISL_3442467 | Lighthouse Laboratory Plymouth | Wellcome Sanger Institute for the COVID-19 Genomics UK (COG-UK) Consortium | Lighthouse Laboratory Plymouth and Alex Alderton, Roberto Amato, Jeffrey Barrett, Sonia Goncalves, Ewan Harrison, David K. Jackson, Ian Johnston, Dominic Kwiatkowski, Cordelia Langford, John Sillitoe on behalf of the Wellcome Sanger Institute COVID-19 Surveillance Team |
| EPI_ISL_3442468, EPI_ISL_3442469 | Berkshire and Surrey Pathology Services Lighthouse Laboratory | Wellcome Sanger Institute for the COVID-19 Genomics UK (COG-UK) Consortium | Berkshire and Surrey Pathology Services Lighthouse Laboratory and Alex Alderton, Roberto Amato, Jeffrey Barrett, Sonia Goncalves, Ewan Harrison, David K. Jackson, Ian Johnston, Dominic Kwiatkowski, Cordelia Langford, John Sillitoe on behalf of the Wellcome Sanger Institute COVID-19 Surveillance Team |
| EPI_ISL_3442470 | Lighthouse Lab in Glasgow | Wellcome Sanger Institute for the COVID-19 Genomics UK (COG-UK) Consortium | Harper VanSteenhouse, Yumi Kasai, David Gray, Carol Clugston, Anna Dominiczak and Alex Alderton, Roberto Amato, Jeffrey Barrett, Sonia Goncalves, Ewan Harrison, David K. Jackson, Ian Johnston, Dominic Kwiatkowski, Cordelia Langford, John Sillitoe on behalf of the Wellcome Sanger Institute COVID-19 Surveillance Team |
| EPI_ISL_3442471 | Lighthouse Laboratory Plymouth | Wellcome Sanger Institute for the COVID-19 Genomics UK (COG-UK) Consortium | Lighthouse Laboratory Plymouth and Alex Alderton, Roberto Amato, Jeffrey Barrett, Sonia Goncalves, Ewan Harrison, David K. Jackson, Ian Johnston, Dominic Kwiatkowski, Cordelia Langford, John Sillitoe on behalf of the Wellcome Sanger Institute COVID-19 Surveillance Team |
| EPI_ISL_3442472, EPI_ISL_3442473, EPI_ISL_3442474 | Lighthouse Lab in Glasgow | Wellcome Sanger Institute for the COVID-19 Genomics UK (COG-UK) Consortium | Harper VanSteenhouse, Yumi Kasai, David Gray, Carol Clugston, Anna Dominiczak and Alex Alderton, Roberto Amato, Jeffrey Barrett, Sonia Goncalves, Ewan Harrison, David K. Jackson, Ian Johnston, Dominic Kwiatkowski, Cordelia Langford, John Sillitoe on behalf of the Wellcome Sanger Institute COVID-19 Surveillance Team |
| EPI_ISL_3442475 | Lighthouse Laboratory Plymouth | Wellcome Sanger Institute for the COVID-19 Genomics UK (COG-UK) Consortium | Lighthouse Laboratory Plymouth and Alex Alderton, Roberto Amato, Jeffrey Barrett, Sonia Goncalves, Ewan Harrison, David K. Jackson, Ian Johnston, Dominic Kwiatkowski, Cordelia Langford, John Sillitoe on behalf of the Wellcome Sanger Institute COVID-19 Surveillance Team |
| EPI_ISL_3442476 | Lighthouse Lab in Glasgow | Wellcome Sanger Institute for the COVID-19 Genomics UK (COG-UK) Consortium | Harper VanSteenhouse, Yumi Kasai, David Gray, Carol Clugston, Anna Dominiczak and Alex Alderton, Roberto Amato, Jeffrey Barrett, Sonia Goncalves, Ewan Harrison, David K. Jackson, Ian Johnston, Dominic Kwiatkowski, Cordelia Langford, John Sillitoe on behalf of the Wellcome Sanger Institute COVID-19 Surveillance Team |
| EPI_ISL_3442477 | Berkshire and Surrey Pathology Services Lighthouse Laboratory | Wellcome Sanger Institute for the COVID-19 Genomics UK (COG-UK) Consortium | Berkshire and Surrey Pathology Services Lighthouse Laboratory and Alex Alderton, Roberto Amato, Jeffrey Barrett, Sonia Goncalves, Ewan Harrison, David K. Jackson, Ian Johnston, Dominic Kwiatkowski, Cordelia Langford, John Sillitoe on behalf of the Wellcome Sanger Institute COVID-19 Surveillance Team |
| EPI_ISL_3442478, EPI_ISL_3442479, EPI_ISL_3442480 | Lighthouse Lab in Glasgow | Wellcome Sanger Institute for the COVID-19 Genomics UK (COG-UK) Consortium | Harper VanSteenhouse, Yumi Kasai, David Gray, Carol Clugston, Anna Dominiczak and Alex Alderton, Roberto Amato, Jeffrey Barrett, Sonia Goncalves, Ewan Harrison, David K. Jackson, Ian Johnston, Dominic Kwiatkowski, Cordelia Langford, John Sillitoe on behalf of the Wellcome Sanger Institute COVID-19 Surveillance Team |
| EPI_ISL_3442481 | Lighthouse Laboratory Plymouth | Wellcome Sanger Institute for the COVID-19 Genomics UK (COG-UK) Consortium | Lighthouse Laboratory Plymouth and Alex Alderton, Roberto Amato, Jeffrey Barrett, Sonia Goncalves, Ewan Harrison, David K. Jackson, Ian Johnston, Dominic Kwiatkowski, Cordelia Langford, John Sillitoe on behalf of the Wellcome Sanger Institute COVID-19 Surveillance Team |
| EPI_ISL_3442482 | Berkshire and Surrey Pathology Services Lighthouse Laboratory | Wellcome Sanger Institute for the COVID-19 Genomics UK (COG-UK) Consortium | Berkshire and Surrey Pathology Services Lighthouse Laboratory and Alex Alderton, Roberto Amato, Jeffrey Barrett, Sonia Goncalves, Ewan Harrison, David K. Jackson, Ian Johnston, Dominic Kwiatkowski, Cordelia Langford, John Sillitoe on behalf of the Wellcome Sanger Institute COVID-19 Surveillance Team |
| EPI_ISL_3442483 | Lighthouse Lab in Glasgow | Wellcome Sanger Institute for the COVID-19 Genomics UK (COG-UK) Consortium | Harper VanSteenhouse, Yumi Kasai, David Gray, Carol Clugston, Anna Dominiczak and Alex Alderton, Roberto Amato, Jeffrey Barrett, Sonia Goncalves, Ewan Harrison, David K. Jackson, Ian Johnston, Dominic Kwiatkowski, Cordelia Langford, John Sillitoe on behalf of the Wellcome Sanger Institute COVID-19 Surveillance Team |
| EPI_ISL_3442484 | Lighthouse Laboratory Plymouth | Wellcome Sanger Institute for the COVID-19 Genomics UK (COG-UK) Consortium | Lighthouse Laboratory Plymouth and Alex Alderton, Roberto Amato, Jeffrey Barrett, Sonia Goncalves, Ewan Harrison, David K. Jackson, Ian Johnston, Dominic Kwiatkowski, Cordelia Langford, John Sillitoe on behalf of the Wellcome Sanger Institute COVID-19 Surveillance Team |
| EPI_ISL_3442485 | Lighthouse Lab in Glasgow | Wellcome Sanger Institute for the COVID-19 Genomics UK (COG-UK) | Harper VanSteenhouse, Yumi Kasai, David Gray, Carol Clugston, Anna Dominiczak and Alex Alderton, Roberto Amato, Jeffrey Barrett, Sonia Goncalves, Ewan Harrison, David K. Jackson, Ian Johnston, Dominic Kwiatkowski, Cordelia Langford, |

| | | | |
|---|---|---|---|
| | | Consortium | John Sillitoe on behalf of the Wellcome Sanger Institute COVID-19 Surveillance Team |
| EPI_ISL_3442486 | Lighthouse Laboratory Plymouth | Wellcome Sanger Institute for the COVID-19 Genomics UK (COG-UK) Consortium | Lighthouse Laboratory Plymouth and Alex Alderton, Roberto Amato, Jeffrey Barrett, Sonia Goncalves, Ewan Harrison, David K. Jackson, Ian Johnston, Dominic Kwiatkowski, Cordelia Langford, John Sillitoe on behalf of the Wellcome Sanger Institute COVID-19 Surveillance Team |
| EPI_ISL_3442487 | Lighthouse Lab in Glasgow | Wellcome Sanger Institute for the COVID-19 Genomics UK (COG-UK) Consortium | Harper VanSteenhouse, Yumi Kasai, David Gray, Carol Clugston, Anna Dominiczak and Alex Alderton, Roberto Amato, Jeffrey Barrett, Sonia Goncalves, Ewan Harrison, David K. Jackson, Ian Johnston, Dominic Kwiatkowski, Cordelia Langford, John Sillitoe on behalf of the Wellcome Sanger Institute COVID-19 Surveillance Team |
| EPI_ISL_3442488 | Berkshire and Surrey Pathology Services Lighthouse Laboratory | Wellcome Sanger Institute for the COVID-19 Genomics UK (COG-UK) Consortium | Berkshire and Surrey Pathology Services Lighthouse Laboratory and Alex Alderton, Roberto Amato, Jeffrey Barrett, Sonia Goncalves, Ewan Harrison, David K. Jackson, Ian Johnston, Dominic Kwiatkowski, Cordelia Langford, John Sillitoe on behalf of the Wellcome Sanger Institute COVID-19 Surveillance Team |
| EPI_ISL_3442489, EPI_ISL_3442490 | Lighthouse Lab in Glasgow | Wellcome Sanger Institute for the COVID-19 Genomics UK (COG-UK) Consortium | Harper VanSteenhouse, Yumi Kasai, David Gray, Carol Clugston, Anna Dominiczak and Alex Alderton, Roberto Amato, Jeffrey Barrett, Sonia Goncalves, Ewan Harrison, David K. Jackson, Ian Johnston, Dominic Kwiatkowski, Cordelia Langford, John Sillitoe on behalf of the Wellcome Sanger Institute COVID-19 Surveillance Team |
| EPI_ISL_3442491 | Lighthouse Laboratory Plymouth | Wellcome Sanger Institute for the COVID-19 Genomics UK (COG-UK) Consortium | Lighthouse Laboratory Plymouth and Alex Alderton, Roberto Amato, Jeffrey Barrett, Sonia Goncalves, Ewan Harrison, David K. Jackson, Ian Johnston, Dominic Kwiatkowski, Cordelia Langford, John Sillitoe on behalf of the Wellcome Sanger Institute COVID-19 Surveillance Team |
| EPI_ISL_3442492 | Lighthouse Lab in Glasgow | Wellcome Sanger Institute for the COVID-19 Genomics UK (COG-UK) Consortium | Harper VanSteenhouse, Yumi Kasai, David Gray, Carol Clugston, Anna Dominiczak and Alex Alderton, Roberto Amato, Jeffrey Barrett, Sonia Goncalves, Ewan Harrison, David K. Jackson, Ian Johnston, Dominic Kwiatkowski, Cordelia Langford, John Sillitoe on behalf of the Wellcome Sanger Institute COVID-19 Surveillance Team |
| EPI_ISL_3442493 | Lighthouse Laboratory Plymouth | Wellcome Sanger Institute for the COVID-19 Genomics UK (COG-UK) Consortium | Lighthouse Laboratory Plymouth and Alex Alderton, Roberto Amato, Jeffrey Barrett, Sonia Goncalves, Ewan Harrison, David K. Jackson, Ian Johnston, Dominic Kwiatkowski, Cordelia Langford, John Sillitoe on behalf of the Wellcome Sanger Institute COVID-19 Surveillance Team |
| EPI_ISL_3442494 | Lighthouse Lab in Glasgow | Wellcome Sanger Institute for the COVID-19 Genomics UK (COG-UK) Consortium | Harper VanSteenhouse, Yumi Kasai, David Gray, Carol Clugston, Anna Dominiczak and Alex Alderton, Roberto Amato, Jeffrey Barrett, Sonia Goncalves, Ewan Harrison, David K. Jackson, Ian Johnston, Dominic Kwiatkowski, Cordelia Langford, John Sillitoe on behalf of the Wellcome Sanger Institute COVID-19 Surveillance Team |
| EPI_ISL_3442495 | Lighthouse Laboratory Plymouth | Wellcome Sanger Institute for the COVID-19 Genomics UK (COG-UK) Consortium | Lighthouse Laboratory Plymouth and Alex Alderton, Roberto Amato, Jeffrey Barrett, Sonia Goncalves, Ewan Harrison, David K. Jackson, Ian Johnston, Dominic Kwiatkowski, Cordelia Langford, John Sillitoe on behalf of the Wellcome Sanger Institute COVID-19 Surveillance Team |
| EPI_ISL_3442496 | Berkshire and Surrey Pathology Services Lighthouse Laboratory | Wellcome Sanger Institute for the COVID-19 Genomics UK (COG-UK) Consortium | Berkshire and Surrey Pathology Services Lighthouse Laboratory and Alex Alderton, Roberto Amato, Jeffrey Barrett, Sonia Goncalves, Ewan Harrison, David K. Jackson, Ian Johnston, Dominic Kwiatkowski, Cordelia Langford, John Sillitoe on behalf of the Wellcome Sanger Institute COVID-19 Surveillance Team |
| EPI_ISL_3442497, EPI_ISL_3442498 | Lighthouse Lab in Glasgow | Wellcome Sanger Institute for the COVID-19 Genomics UK (COG-UK) Consortium | Harper VanSteenhouse, Yumi Kasai, David Gray, Carol Clugston, Anna Dominiczak and Alex Alderton, Roberto Amato, Jeffrey Barrett, Sonia Goncalves, Ewan Harrison, David K. Jackson, Ian Johnston, Dominic Kwiatkowski, Cordelia Langford, John Sillitoe on behalf of the Wellcome Sanger Institute COVID-19 Surveillance Team |
| EPI_ISL_3442499, EPI_ISL_3442500 | Lighthouse Laboratory Plymouth | Wellcome Sanger Institute for the COVID-19 Genomics UK (COG-UK) Consortium | Lighthouse Laboratory Plymouth and Alex Alderton, Roberto Amato, Jeffrey Barrett, Sonia Goncalves, Ewan Harrison, David K. Jackson, Ian Johnston, Dominic Kwiatkowski, Cordelia Langford, John Sillitoe on behalf of the Wellcome Sanger Institute COVID-19 Surveillance Team |
| EPI_ISL_3442501, EPI_ISL_3442502 | Lighthouse Lab in Glasgow | Wellcome Sanger Institute for the COVID-19 Genomics UK (COG-UK) Consortium | Harper VanSteenhouse, Yumi Kasai, David Gray, Carol Clugston, Anna Dominiczak and Alex Alderton, Roberto Amato, Jeffrey Barrett, Sonia Goncalves, Ewan Harrison, David K. Jackson, Ian Johnston, Dominic Kwiatkowski, Cordelia Langford, John Sillitoe on behalf of the Wellcome Sanger Institute COVID-19 Surveillance Team |
| EPI_ISL_3442503 | Lighthouse Laboratory Plymouth | Wellcome Sanger Institute for the COVID-19 Genomics UK (COG-UK) Consortium | Lighthouse Laboratory Plymouth and Alex Alderton, Roberto Amato, Jeffrey Barrett, Sonia Goncalves, Ewan Harrison, David K. Jackson, Ian Johnston, Dominic Kwiatkowski, Cordelia Langford, John Sillitoe on behalf of the Wellcome Sanger Institute COVID-19 Surveillance Team |
| EPI_ISL_3442504 | Lighthouse Lab in Glasgow | Wellcome Sanger Institute for the COVID-19 Genomics UK (COG-UK) Consortium | Harper VanSteenhouse, Yumi Kasai, David Gray, Carol Clugston, Anna Dominiczak and Alex Alderton, Roberto Amato, Jeffrey Barrett, Sonia Goncalves, Ewan Harrison, David K. Jackson, Ian Johnston, Dominic Kwiatkowski, Cordelia Langford, John Sillitoe on behalf of the Wellcome Sanger Institute COVID-19 Surveillance Team |
| EPI_ISL_3442505, EPI_ISL_3442506 | Lighthouse Laboratory Plymouth | Wellcome Sanger Institute for the COVID-19 Genomics UK (COG-UK) Consortium | Lighthouse Laboratory Plymouth and Alex Alderton, Roberto Amato, Jeffrey Barrett, Sonia Goncalves, Ewan Harrison, David K. Jackson, Ian Johnston, Dominic Kwiatkowski, Cordelia Langford, John Sillitoe on behalf of the Wellcome Sanger Institute COVID-19 Surveillance Team |
| EPI_ISL_3442507 | Lighthouse Lab in Glasgow | Wellcome Sanger Institute for the COVID-19 Genomics UK (COG-UK) Consortium | Harper VanSteenhouse, Yumi Kasai, David Gray, Carol Clugston, Anna Dominiczak and Alex Alderton, Roberto Amato, Jeffrey Barrett, Sonia Goncalves, Ewan Harrison, David K. Jackson, Ian Johnston, Dominic Kwiatkowski, Cordelia Langford, John Sillitoe on behalf of the Wellcome Sanger Institute COVID-19 Surveillance Team |
| EPI_ISL_3442508 | Berkshire and Surrey Pathology Services Lighthouse Laboratory | Wellcome Sanger Institute for the COVID-19 Genomics UK (COG-UK) Consortium | Berkshire and Surrey Pathology Services Lighthouse Laboratory and Alex Alderton, Roberto Amato, Jeffrey Barrett, Sonia Goncalves, Ewan Harrison, David K. Jackson, Ian Johnston, Dominic Kwiatkowski, Cordelia Langford, John Sillitoe on behalf of the Wellcome Sanger Institute COVID-19 Surveillance Team |
| EPI_ISL_3442509, EPI_ISL_3442510 | Lighthouse Lab in Glasgow | Wellcome Sanger Institute for the COVID-19 Genomics UK (COG-UK) Consortium | Harper VanSteenhouse, Yumi Kasai, David Gray, Carol Clugston, Anna Dominiczak and Alex Alderton, Roberto Amato, Jeffrey Barrett, Sonia Goncalves, Ewan Harrison, David K. Jackson, Ian Johnston, Dominic Kwiatkowski, Cordelia Langford, John Sillitoe on behalf of the Wellcome Sanger Institute COVID-19 Surveillance Team |
| EPI_ISL_3442511 | Lighthouse Laboratory Plymouth | Wellcome Sanger Institute for the COVID-19 Genomics UK (COG-UK) Consortium | Lighthouse Laboratory Plymouth and Alex Alderton, Roberto Amato, Jeffrey Barrett, Sonia Goncalves, Ewan Harrison, David K. Jackson, Ian Johnston, Dominic Kwiatkowski, Cordelia Langford, John Sillitoe on behalf of the Wellcome Sanger Institute COVID-19 Surveillance Team |
| EPI_ISL_3442512, EPI_ISL_3442513 | Lighthouse Lab in Glasgow | Wellcome Sanger Institute for the COVID-19 Genomics UK (COG-UK) Consortium | Harper VanSteenhouse, Yumi Kasai, David Gray, Carol Clugston, Anna Dominiczak and Alex Alderton, Roberto Amato, Jeffrey Barrett, Sonia Goncalves, Ewan Harrison, David K. Jackson, Ian Johnston, Dominic Kwiatkowski, Cordelia Langford, John Sillitoe on behalf of the Wellcome Sanger Institute COVID-19 Surveillance Team |
| EPI_ISL_3442514, EPI_ISL_3442515 | Lighthouse Laboratory Plymouth | Wellcome Sanger Institute for the COVID-19 Genomics UK (COG-UK) Consortium | Lighthouse Laboratory Plymouth and Alex Alderton, Roberto Amato, Jeffrey Barrett, Sonia Goncalves, Ewan Harrison, David K. Jackson, Ian Johnston, Dominic Kwiatkowski, Cordelia Langford, John Sillitoe on behalf of the Wellcome Sanger Institute COVID-19 Surveillance Team |
| EPI_ISL_3442516, EPI_ISL_3442517 | Lighthouse Lab in Glasgow | Wellcome Sanger Institute for the COVID-19 Genomics UK (COG-UK) Consortium | Harper VanSteenhouse, Yumi Kasai, David Gray, Carol Clugston, Anna Dominiczak and Alex Alderton, Roberto Amato, Jeffrey Barrett, Sonia Goncalves, Ewan Harrison, David K. Jackson, Ian Johnston, Dominic Kwiatkowski, Cordelia Langford, John Sillitoe on behalf of the Wellcome Sanger Institute COVID-19 Surveillance Team |
| EPI_ISL_3442518, EPI_ISL_3442519 | Lighthouse Laboratory Plymouth | Wellcome Sanger Institute for the COVID-19 Genomics UK (COG-UK) Consortium | Lighthouse Laboratory Plymouth and Alex Alderton, Roberto Amato, Jeffrey Barrett, Sonia Goncalves, Ewan Harrison, David K. Jackson, Ian Johnston, Dominic Kwiatkowski, Cordelia Langford, John Sillitoe on behalf of the Wellcome Sanger Institute COVID-19 Surveillance Team |
| EPI_ISL_3442520 | Lighthouse Lab in Glasgow | Wellcome Sanger Institute for the COVID-19 Genomics UK (COG-UK) Consortium | Harper VanSteenhouse, Yumi Kasai, David Gray, Carol Clugston, Anna Dominiczak and Alex Alderton, Roberto Amato, Jeffrey Barrett, Sonia Goncalves, Ewan Harrison, David K. Jackson, Ian Johnston, Dominic Kwiatkowski, Cordelia Langford, John Sillitoe on behalf of the Wellcome Sanger Institute COVID-19 Surveillance Team |
| EPI_ISL_3442521 | Lighthouse Laboratory Plymouth | Wellcome Sanger Institute for the COVID-19 Genomics UK (COG-UK) Consortium | Lighthouse Laboratory Plymouth and Alex Alderton, Roberto Amato, Jeffrey Barrett, Sonia Goncalves, Ewan Harrison, David K. Jackson, Ian Johnston, Dominic Kwiatkowski, Cordelia Langford, John Sillitoe on behalf of the Wellcome Sanger Institute COVID-19 Surveillance Team |
| EPI_ISL_3442522, EPI_ISL_3442523 | Lighthouse Lab in Glasgow | Wellcome Sanger Institute for the COVID-19 Genomics UK (COG-UK) Consortium | Harper VanSteenhouse, Yumi Kasai, David Gray, Carol Clugston, Anna Dominiczak and Alex Alderton, Roberto Amato, Jeffrey Barrett, Sonia Goncalves, Ewan Harrison, David K. Jackson, Ian Johnston, Dominic Kwiatkowski, Cordelia Langford, John Sillitoe on behalf of the Wellcome Sanger Institute COVID-19 Surveillance Team |
| EPI_ISL_3442524 | Berkshire and Surrey Pathology Services Lighthouse Laboratory | Wellcome Sanger Institute for the COVID-19 Genomics UK (COG-UK) Consortium | Berkshire and Surrey Pathology Services Lighthouse Laboratory and Alex Alderton, Roberto Amato, Jeffrey Barrett, Sonia Goncalves, Ewan Harrison, David K. Jackson, Ian Johnston, Dominic Kwiatkowski, Cordelia Langford, John Sillitoe on behalf of the Wellcome Sanger Institute COVID-19 Surveillance Team |
| EPI_ISL_3442525 | Lighthouse Laboratory Plymouth | Wellcome Sanger Institute for the COVID-19 Genomics UK (COG-UK) Consortium | Lighthouse Laboratory Plymouth and Alex Alderton, Roberto Amato, Jeffrey Barrett, Sonia Goncalves, Ewan Harrison, David K. Jackson, Ian Johnston, Dominic Kwiatkowski, Cordelia Langford, John Sillitoe on behalf of the Wellcome Sanger Institute COVID-19 Surveillance Team |
| EPI_ISL_3442526 | Lighthouse Lab in Glasgow | Wellcome Sanger Institute for the COVID-19 Genomics UK (COG-UK) Consortium | Harper VanSteenhouse, Yumi Kasai, David Gray, Carol Clugston, Anna Dominiczak and Alex Alderton, Roberto Amato, Jeffrey Barrett, Sonia Goncalves, Ewan Harrison, David K. Jackson, Ian Johnston, Dominic Kwiatkowski, Cordelia Langford, John Sillitoe on behalf of the Wellcome Sanger Institute COVID-19 Surveillance Team |
| EPI_ISL_3442527 | Berkshire and Surrey Pathology Services Lighthouse Laboratory | Wellcome Sanger Institute for the COVID-19 Genomics UK (COG-UK) Consortium | Berkshire and Surrey Pathology Services Lighthouse Laboratory and Alex Alderton, Roberto Amato, Jeffrey Barrett, Sonia Goncalves, Ewan Harrison, David K. Jackson, Ian Johnston, Dominic Kwiatkowski, Cordelia Langford, John Sillitoe on behalf of the Wellcome Sanger Institute COVID-19 Surveillance Team |
| EPI_ISL_3442528 | Lighthouse Lab in Glasgow | Wellcome Sanger Institute for the COVID-19 Genomics UK (COG-UK) Consortium | Harper VanSteenhouse, Yumi Kasai, David Gray, Carol Clugston, Anna Dominiczak and Alex Alderton, Roberto Amato, Jeffrey Barrett, Sonia Goncalves, Ewan Harrison, David K. Jackson, Ian Johnston, Dominic Kwiatkowski, Cordelia Langford, John Sillitoe on behalf of the Wellcome Sanger Institute COVID-19 Surveillance Team |
| EPI_ISL_3442529 | Berkshire and Surrey Pathology Services Lighthouse Laboratory | Wellcome Sanger Institute for the COVID-19 Genomics UK (COG-UK) Consortium | Berkshire and Surrey Pathology Services Lighthouse Laboratory and Alex Alderton, Roberto Amato, Jeffrey Barrett, Sonia Goncalves, Ewan Harrison, David K. Jackson, Ian Johnston, Dominic Kwiatkowski, Cordelia Langford, John Sillitoe on behalf of the Wellcome Sanger Institute COVID-19 Surveillance Team |
| EPI_ISL_3442530 | Lighthouse Laboratory Plymouth | Wellcome Sanger Institute for the COVID-19 Genomics UK (COG-UK) Consortium | Lighthouse Laboratory Plymouth and Alex Alderton, Roberto Amato, Jeffrey Barrett, Sonia Goncalves, Ewan Harrison, David K. Jackson, Ian Johnston, Dominic Kwiatkowski, Cordelia Langford, John Sillitoe on behalf of the Wellcome Sanger Institute COVID-19 Surveillance Team |
| EPI_ISL_3442531, EPI_ISL_3442532, EPI_ISL_3442533 | Lighthouse Lab in Glasgow | Wellcome Sanger Institute for the COVID-19 Genomics UK (COG-UK) Consortium | Harper VanSteenhouse, Yumi Kasai, David Gray, Carol Clugston, Anna Dominiczak and Alex Alderton, Roberto Amato, Jeffrey Barrett, Sonia Goncalves, Ewan Harrison, David K. Jackson, Ian Johnston, Dominic Kwiatkowski, Cordelia Langford, John Sillitoe on behalf of the Wellcome Sanger Institute COVID-19 Surveillance Team |
| EPI_ISL_3442534 | Berkshire and Surrey Pathology Services Lighthouse Laboratory | Wellcome Sanger Institute for the COVID-19 Genomics UK (COG-UK) Consortium | Berkshire and Surrey Pathology Services Lighthouse Laboratory and Alex Alderton, Roberto Amato, Jeffrey Barrett, Sonia Goncalves, Ewan Harrison, David K. Jackson, Ian Johnston, Dominic Kwiatkowski, Cordelia Langford, John Sillitoe on behalf of the Wellcome Sanger Institute COVID-19 Surveillance Team |
| EPI_ISL_3442535 | Lighthouse Lab in Glasgow | Wellcome Sanger Institute for the COVID-19 Genomics UK (COG-UK) Consortium | Harper VanSteenhouse, Yumi Kasai, David Gray, Carol Clugston, Anna Dominiczak and Alex Alderton, Roberto Amato, Jeffrey Barrett, Sonia Goncalves, Ewan Harrison, David K. Jackson, Ian Johnston, Dominic Kwiatkowski, Cordelia Langford, John Sillitoe on behalf of the Wellcome Sanger Institute COVID-19 Surveillance Team |
| EPI_ISL_3442536 | Lighthouse Laboratory Plymouth | Wellcome Sanger Institute for the COVID-19 Genomics UK (COG-UK) Consortium | Lighthouse Laboratory Plymouth and Alex Alderton, Roberto Amato, Jeffrey Barrett, Sonia Goncalves, Ewan Harrison, David K. Jackson, Ian Johnston, Dominic Kwiatkowski, Cordelia Langford, John Sillitoe on behalf of the Wellcome Sanger Institute COVID-19 Surveillance Team |
| EPI_ISL_3442537 | Berkshire and Surrey Pathology Services Lighthouse Laboratory | Wellcome Sanger Institute for the COVID-19 Genomics UK (COG-UK) Consortium | Berkshire and Surrey Pathology Services Lighthouse Laboratory and Alex Alderton, Roberto Amato, Jeffrey Barrett, Sonia Goncalves, Ewan Harrison, David K. Jackson, Ian Johnston, Dominic Kwiatkowski, Cordelia Langford, John Sillitoe on behalf of the Wellcome Sanger Institute COVID-19 Surveillance Team |
| EPI_ISL_3442538 | Lighthouse Laboratory Plymouth | Wellcome Sanger Institute for the COVID-19 Genomics UK (COG-UK) Consortium | Lighthouse Laboratory Plymouth and Alex Alderton, Roberto Amato, Jeffrey Barrett, Sonia Goncalves, Ewan Harrison, David K. Jackson, Ian Johnston, Dominic Kwiatkowski, Cordelia Langford, John Sillitoe on behalf of the Wellcome Sanger Institute COVID-19 Surveillance Team |
| EPI_ISL_3442539 | Lighthouse Lab in Glasgow | Wellcome Sanger Institute for the COVID-19 Genomics UK (COG-UK) Consortium | Harper VanSteenhouse, Yumi Kasai, David Gray, Carol Clugston, Anna Dominiczak and Alex Alderton, Roberto Amato, Jeffrey Barrett, Sonia Goncalves, Ewan Harrison, David K. Jackson, Ian Johnston, Dominic Kwiatkowski, Cordelia Langford, John Sillitoe on behalf of the Wellcome Sanger Institute COVID-19 Surveillance Team |
| EPI_ISL_3442540 | Lighthouse Laboratory Plymouth | Wellcome Sanger Institute for the COVID-19 Genomics UK (COG-UK) Consortium | Lighthouse Laboratory Plymouth and Alex Alderton, Roberto Amato, Jeffrey Barrett, Sonia Goncalves, Ewan Harrison, David K. Jackson, Ian Johnston, Dominic Kwiatkowski, Cordelia Langford, John Sillitoe on behalf of the Wellcome Sanger Institute COVID-19 Surveillance Team |
| EPI_ISL_3442541 | Lighthouse Lab in Glasgow | Wellcome Sanger Institute for the COVID-19 Genomics UK (COG-UK) Consortium | Harper VanSteenhouse, Yumi Kasai, David Gray, Carol Clugston, Anna Dominiczak and Alex Alderton, Roberto Amato, Jeffrey Barrett, Sonia Goncalves, Ewan Harrison, David K. Jackson, Ian Johnston, Dominic Kwiatkowski, Cordelia Langford, John Sillitoe on behalf of the Wellcome Sanger Institute COVID-19 Surveillance Team |
| EPI_ISL_3442542 | Berkshire and Surrey Pathology Services Lighthouse Laboratory | Wellcome Sanger Institute for the COVID-19 Genomics UK (COG-UK) Consortium | Berkshire and Surrey Pathology Services Lighthouse Laboratory and Alex Alderton, Roberto Amato, Jeffrey Barrett, Sonia Goncalves, Ewan Harrison, David K. Jackson, Ian Johnston, Dominic Kwiatkowski, Cordelia Langford, John Sillitoe on behalf of the Wellcome Sanger Institute COVID-19 Surveillance Team |

| | | | |
|---|---|---|---|
| EPI_ISL_3442543 | Lighthouse Lab in Glasgow | Wellcome Sanger Institute for the COVID-19 Genomics UK (COG-UK) Consortium | Harper VanSteenhouse, Yumi Kasai, David Gray, Carol Clugston, Anna Dominiczak and Alex Alderton, Roberto Amato, Jeffrey Barrett, Sonia Goncalves, Ewan Harrison, David K. Jackson, Ian Johnston, Dominic Kwiatkowski, Cordelia Langford, John Sillitoe on behalf of the Wellcome Sanger Institute COVID-19 Surveillance Team |
| EPI_ISL_3442544 | Lighthouse Laboratory Plymouth | Wellcome Sanger Institute for the COVID-19 Genomics UK (COG-UK) Consortium | Lighthouse Laboratory Plymouth and Alex Alderton, Roberto Amato, Jeffrey Barrett, Sonia Goncalves, Ewan Harrison, David K. Jackson, Ian Johnston, Dominic Kwiatkowski, Cordelia Langford, John Sillitoe on behalf of the Wellcome Sanger Institute COVID-19 Surveillance Team |
| EPI_ISL_3442545, EPI_ISL_3442546, EPI_ISL_3442547, EPI_ISL_3442548 | Lighthouse Lab in Glasgow | Wellcome Sanger Institute for the COVID-19 Genomics UK (COG-UK) Consortium | Harper VanSteenhouse, Yumi Kasai, David Gray, Carol Clugston, Anna Dominiczak and Alex Alderton, Roberto Amato, Jeffrey Barrett, Sonia Goncalves, Ewan Harrison, David K. Jackson, Ian Johnston, Dominic Kwiatkowski, Cordelia Langford, John Sillitoe on behalf of the Wellcome Sanger Institute COVID-19 Surveillance Team |
| EPI_ISL_3442549 | Lighthouse Laboratory Plymouth | Wellcome Sanger Institute for the COVID-19 Genomics UK (COG-UK) Consortium | Lighthouse Laboratory Plymouth and Alex Alderton, Roberto Amato, Jeffrey Barrett, Sonia Goncalves, Ewan Harrison, David K. Jackson, Ian Johnston, Dominic Kwiatkowski, Cordelia Langford, John Sillitoe on behalf of the Wellcome Sanger Institute COVID-19 Surveillance Team |
| EPI_ISL_3442550, EPI_ISL_3442551 | Berkshire and Surrey Pathology Services Lighthouse Laboratory | Wellcome Sanger Institute for the COVID-19 Genomics UK (COG-UK) Consortium | Berkshire and Surrey Pathology Services Lighthouse Laboratory and Alex Alderton, Roberto Amato, Jeffrey Barrett, Sonia Goncalves, Ewan Harrison, David K. Jackson, Ian Johnston, Dominic Kwiatkowski, Cordelia Langford, John Sillitoe on behalf of the Wellcome Sanger Institute COVID-19 Surveillance Team |
| EPI_ISL_3442552 | Lighthouse Lab in Glasgow | Wellcome Sanger Institute for the COVID-19 Genomics UK (COG-UK) Consortium | Harper VanSteenhouse, Yumi Kasai, David Gray, Carol Clugston, Anna Dominiczak and Alex Alderton, Roberto Amato, Jeffrey Barrett, Sonia Goncalves, Ewan Harrison, David K. Jackson, Ian Johnston, Dominic Kwiatkowski, Cordelia Langford, John Sillitoe on behalf of the Wellcome Sanger Institute COVID-19 Surveillance Team |
| EPI_ISL_3442553 | Lighthouse Laboratory Plymouth | Wellcome Sanger Institute for the COVID-19 Genomics UK (COG-UK) Consortium | Lighthouse Laboratory Plymouth and Alex Alderton, Roberto Amato, Jeffrey Barrett, Sonia Goncalves, Ewan Harrison, David K. Jackson, Ian Johnston, Dominic Kwiatkowski, Cordelia Langford, John Sillitoe on behalf of the Wellcome Sanger Institute COVID-19 Surveillance Team |
| EPI_ISL_3442554, EPI_ISL_3442555, EPI_ISL_3442556, EPI_ISL_3442557 | Lighthouse Lab in Glasgow | Wellcome Sanger Institute for the COVID-19 Genomics UK (COG-UK) Consortium | Harper VanSteenhouse, Yumi Kasai, David Gray, Carol Clugston, Anna Dominiczak and Alex Alderton, Roberto Amato, Jeffrey Barrett, Sonia Goncalves, Ewan Harrison, David K. Jackson, Ian Johnston, Dominic Kwiatkowski, Cordelia Langford, John Sillitoe on behalf of the Wellcome Sanger Institute COVID-19 Surveillance Team |
| EPI_ISL_3442558, EPI_ISL_3442559 | Lighthouse Laboratory Plymouth | Wellcome Sanger Institute for the COVID-19 Genomics UK (COG-UK) Consortium | Lighthouse Laboratory Plymouth and Alex Alderton, Roberto Amato, Jeffrey Barrett, Sonia Goncalves, Ewan Harrison, David K. Jackson, Ian Johnston, Dominic Kwiatkowski, Cordelia Langford, John Sillitoe on behalf of the Wellcome Sanger Institute COVID-19 Surveillance Team |
| EPI_ISL_3442560, EPI_ISL_3442561, EPI_ISL_3442562 | Lighthouse Lab in Glasgow | Wellcome Sanger Institute for the COVID-19 Genomics UK (COG-UK) Consortium | Harper VanSteenhouse, Yumi Kasai, David Gray, Carol Clugston, Anna Dominiczak and Alex Alderton, Roberto Amato, Jeffrey Barrett, Sonia Goncalves, Ewan Harrison, David K. Jackson, Ian Johnston, Dominic Kwiatkowski, Cordelia Langford, John Sillitoe on behalf of the Wellcome Sanger Institute COVID-19 Surveillance Team |
| EPI_ISL_3442563, EPI_ISL_3442564 | Lighthouse Laboratory Plymouth | Wellcome Sanger Institute for the COVID-19 Genomics UK (COG-UK) Consortium | Lighthouse Laboratory Plymouth and Alex Alderton, Roberto Amato, Jeffrey Barrett, Sonia Goncalves, Ewan Harrison, David K. Jackson, Ian Johnston, Dominic Kwiatkowski, Cordelia Langford, John Sillitoe on behalf of the Wellcome Sanger Institute COVID-19 Surveillance Team |
| EPI_ISL_3442565 | Lighthouse Lab in Glasgow | Wellcome Sanger Institute for the COVID-19 Genomics UK (COG-UK) Consortium | Harper VanSteenhouse, Yumi Kasai, David Gray, Carol Clugston, Anna Dominiczak and Alex Alderton, Roberto Amato, Jeffrey Barrett, Sonia Goncalves, Ewan Harrison, David K. Jackson, Ian Johnston, Dominic Kwiatkowski, Cordelia Langford, John Sillitoe on behalf of the Wellcome Sanger Institute COVID-19 Surveillance Team |
| EPI_ISL_3442566 | Lighthouse Laboratory Plymouth | Wellcome Sanger Institute for the COVID-19 Genomics UK (COG-UK) Consortium | Lighthouse Laboratory Plymouth and Alex Alderton, Roberto Amato, Jeffrey Barrett, Sonia Goncalves, Ewan Harrison, David K. Jackson, Ian Johnston, Dominic Kwiatkowski, Cordelia Langford, John Sillitoe on behalf of the Wellcome Sanger Institute COVID-19 Surveillance Team |
| EPI_ISL_3442567, EPI_ISL_3442568, EPI_ISL_3442569, EPI_ISL_3442570, EPI_ISL_3442571 | Lighthouse Lab in Glasgow | Wellcome Sanger Institute for the COVID-19 Genomics UK (COG-UK) Consortium | Harper VanSteenhouse, Yumi Kasai, David Gray, Carol Clugston, Anna Dominiczak and Alex Alderton, Roberto Amato, Jeffrey Barrett, Sonia Goncalves, Ewan Harrison, David K. Jackson, Ian Johnston, Dominic Kwiatkowski, Cordelia Langford, John Sillitoe on behalf of the Wellcome Sanger Institute COVID-19 Surveillance Team |
| EPI_ISL_3442572 | Lighthouse Laboratory Plymouth | Wellcome Sanger Institute for the COVID-19 Genomics UK (COG-UK) Consortium | Lighthouse Laboratory Plymouth and Alex Alderton, Roberto Amato, Jeffrey Barrett, Sonia Goncalves, Ewan Harrison, David K. Jackson, Ian Johnston, Dominic Kwiatkowski, Cordelia Langford, John Sillitoe on behalf of the Wellcome Sanger Institute COVID-19 Surveillance Team |
| EPI_ISL_3442573 | Lighthouse Lab in Glasgow | Wellcome Sanger Institute for the COVID-19 Genomics UK (COG-UK) Consortium | Harper VanSteenhouse, Yumi Kasai, David Gray, Carol Clugston, Anna Dominiczak and Alex Alderton, Roberto Amato, Jeffrey Barrett, Sonia Goncalves, Ewan Harrison, David K. Jackson, Ian Johnston, Dominic Kwiatkowski, Cordelia Langford, John Sillitoe on behalf of the Wellcome Sanger Institute COVID-19 Surveillance Team |
| EPI_ISL_3442574, EPI_ISL_3442575 | Lighthouse Laboratory Plymouth | Wellcome Sanger Institute for the COVID-19 Genomics UK (COG-UK) Consortium | Lighthouse Laboratory Plymouth and Alex Alderton, Roberto Amato, Jeffrey Barrett, Sonia Goncalves, Ewan Harrison, David K. Jackson, Ian Johnston, Dominic Kwiatkowski, Cordelia Langford, John Sillitoe on behalf of the Wellcome Sanger Institute COVID-19 Surveillance Team |
| EPI_ISL_3442576, EPI_ISL_3442577, EPI_ISL_3442578, EPI_ISL_3442579, EPI_ISL_3442580, EPI_ISL_3442581 | Lighthouse Lab in Glasgow | Wellcome Sanger Institute for the COVID-19 Genomics UK (COG-UK) Consortium | Harper VanSteenhouse, Yumi Kasai, David Gray, Carol Clugston, Anna Dominiczak and Alex Alderton, Roberto Amato, Jeffrey Barrett, Sonia Goncalves, Ewan Harrison, David K. Jackson, Ian Johnston, Dominic Kwiatkowski, Cordelia Langford, John Sillitoe on behalf of the Wellcome Sanger Institute COVID-19 Surveillance Team |
| EPI_ISL_3442582 | Lighthouse Laboratory Plymouth | Wellcome Sanger Institute for the COVID-19 Genomics UK (COG-UK) Consortium | Lighthouse Laboratory Plymouth and Alex Alderton, Roberto Amato, Jeffrey Barrett, Sonia Goncalves, Ewan Harrison, David K. Jackson, Ian Johnston, Dominic Kwiatkowski, Cordelia Langford, John Sillitoe on behalf of the Wellcome Sanger Institute COVID-19 Surveillance Team |
| EPI_ISL_3442583, EPI_ISL_3442584, EPI_ISL_3442585 | Lighthouse Lab in Glasgow | Wellcome Sanger Institute for the COVID-19 Genomics UK (COG-UK) Consortium | Harper VanSteenhouse, Yumi Kasai, David Gray, Carol Clugston, Anna Dominiczak and Alex Alderton, Roberto Amato, Jeffrey Barrett, Sonia Goncalves, Ewan Harrison, David K. Jackson, Ian Johnston, Dominic Kwiatkowski, Cordelia Langford, John Sillitoe on behalf of the Wellcome Sanger Institute COVID-19 Surveillance Team |
| EPI_ISL_3442586 | Lighthouse Laboratory Plymouth | Wellcome Sanger Institute for the COVID-19 Genomics UK (COG-UK) Consortium | Lighthouse Laboratory Plymouth and Alex Alderton, Roberto Amato, Jeffrey Barrett, Sonia Goncalves, Ewan Harrison, David K. Jackson, Ian Johnston, Dominic Kwiatkowski, Cordelia Langford, John Sillitoe on behalf of the Wellcome Sanger Institute COVID-19 Surveillance Team |
| EPI_ISL_3442587, EPI_ISL_3442588, EPI_ISL_3442589 | Lighthouse Lab in Glasgow | Wellcome Sanger Institute for the COVID-19 Genomics UK (COG-UK) Consortium | Harper VanSteenhouse, Yumi Kasai, David Gray, Carol Clugston, Anna Dominiczak and Alex Alderton, Roberto Amato, Jeffrey Barrett, Sonia Goncalves, Ewan Harrison, David K. Jackson, Ian Johnston, Dominic Kwiatkowski, Cordelia Langford, John Sillitoe on behalf of the Wellcome Sanger Institute COVID-19 Surveillance Team |
| EPI_ISL_3442590, EPI_ISL_3442591 | Lighthouse Laboratory Plymouth | Wellcome Sanger Institute for the COVID-19 Genomics UK (COG-UK) Consortium | Lighthouse Laboratory Plymouth and Alex Alderton, Roberto Amato, Jeffrey Barrett, Sonia Goncalves, Ewan Harrison, David K. Jackson, Ian Johnston, Dominic Kwiatkowski, Cordelia Langford, John Sillitoe on behalf of the Wellcome Sanger Institute COVID-19 Surveillance Team |
| EPI_ISL_3442592, EPI_ISL_3442593, EPI_ISL_3442594, EPI_ISL_3442595 | Lighthouse Lab in Glasgow | Wellcome Sanger Institute for the COVID-19 Genomics UK (COG-UK) Consortium | Harper VanSteenhouse, Yumi Kasai, David Gray, Carol Clugston, Anna Dominiczak and Alex Alderton, Roberto Amato, Jeffrey Barrett, Sonia Goncalves, Ewan Harrison, David K. Jackson, Ian Johnston, Dominic Kwiatkowski, Cordelia Langford, John Sillitoe on behalf of the Wellcome Sanger Institute COVID-19 Surveillance Team |
| EPI_ISL_3442596 | Lighthouse Laboratory Plymouth | Wellcome Sanger Institute for the COVID-19 Genomics UK (COG-UK) Consortium | Lighthouse Laboratory Plymouth and Alex Alderton, Roberto Amato, Jeffrey Barrett, Sonia Goncalves, Ewan Harrison, David K. Jackson, Ian Johnston, Dominic Kwiatkowski, Cordelia Langford, John Sillitoe on behalf of the Wellcome Sanger Institute COVID-19 Surveillance Team |
| EPI_ISL_3442597, EPI_ISL_3442598, EPI_ISL_3442599, EPI_ISL_3442600, EPI_ISL_3442601, EPI_ISL_3442602, EPI_ISL_3442603, EPI_ISL_3442604 | Lighthouse Lab in Glasgow | Wellcome Sanger Institute for the COVID-19 Genomics UK (COG-UK) Consortium | Harper VanSteenhouse, Yumi Kasai, David Gray, Carol Clugston, Anna Dominiczak and Alex Alderton, Roberto Amato, Jeffrey Barrett, Sonia Goncalves, Ewan Harrison, David K. Jackson, Ian Johnston, Dominic Kwiatkowski, Cordelia Langford, John Sillitoe on behalf of the Wellcome Sanger Institute COVID-19 Surveillance Team |
| EPI_ISL_3442605 | Lighthouse Laboratory Plymouth | Wellcome Sanger Institute for the COVID-19 Genomics UK (COG-UK) Consortium | Lighthouse Laboratory Plymouth and Alex Alderton, Roberto Amato, Jeffrey Barrett, Sonia Goncalves, Ewan Harrison, David K. Jackson, Ian Johnston, Dominic Kwiatkowski, Cordelia Langford, John Sillitoe on behalf of the Wellcome Sanger Institute COVID-19 Surveillance Team |
| EPI_ISL_3442606 | Lighthouse Lab in Glasgow | Wellcome Sanger Institute for the COVID-19 Genomics UK (COG-UK) Consortium | Harper VanSteenhouse, Yumi Kasai, David Gray, Carol Clugston, Anna Dominiczak and Alex Alderton, Roberto Amato, Jeffrey Barrett, Sonia Goncalves, Ewan Harrison, David K. Jackson, Ian Johnston, Dominic Kwiatkowski, Cordelia Langford, John Sillitoe on behalf of the Wellcome Sanger Institute COVID-19 Surveillance Team |
| EPI_ISL_3442607 | Lighthouse Laboratory Plymouth | Wellcome Sanger Institute for the COVID-19 Genomics UK (COG-UK) Consortium | Lighthouse Laboratory Plymouth and Alex Alderton, Roberto Amato, Jeffrey Barrett, Sonia Goncalves, Ewan Harrison, David K. Jackson, Ian Johnston, Dominic Kwiatkowski, Cordelia Langford, John Sillitoe on behalf of the Wellcome Sanger Institute COVID-19 Surveillance Team |
| EPI_ISL_3442608, EPI_ISL_3442609 | Lighthouse Lab in Glasgow | Wellcome Sanger Institute for the COVID-19 Genomics UK (COG-UK) Consortium | Harper VanSteenhouse, Yumi Kasai, David Gray, Carol Clugston, Anna Dominiczak and Alex Alderton, Roberto Amato, Jeffrey Barrett, Sonia Goncalves, Ewan Harrison, David K. Jackson, Ian Johnston, Dominic Kwiatkowski, Cordelia Langford, John Sillitoe on behalf of the Wellcome Sanger Institute COVID-19 Surveillance Team |
| EPI_ISL_3442610, EPI_ISL_3442611 | Lighthouse Laboratory Plymouth | Wellcome Sanger Institute for the COVID-19 Genomics UK (COG-UK) Consortium | Lighthouse Laboratory Plymouth and Alex Alderton, Roberto Amato, Jeffrey Barrett, Sonia Goncalves, Ewan Harrison, David K. Jackson, Ian Johnston, Dominic Kwiatkowski, Cordelia Langford, John Sillitoe on behalf of the Wellcome Sanger Institute COVID-19 Surveillance Team |
| EPI_ISL_3442612 | Lighthouse Lab in Glasgow | Wellcome Sanger Institute for the COVID-19 Genomics UK (COG-UK) Consortium | Harper VanSteenhouse, Yumi Kasai, David Gray, Carol Clugston, Anna Dominiczak and Alex Alderton, Roberto Amato, Jeffrey Barrett, Sonia Goncalves, Ewan Harrison, David K. Jackson, Ian Johnston, Dominic Kwiatkowski, Cordelia Langford, John Sillitoe on behalf of the Wellcome Sanger Institute COVID-19 Surveillance Team |
| EPI_ISL_3442613, EPI_ISL_3442614 | Lighthouse Laboratory Plymouth | Wellcome Sanger Institute for the COVID-19 Genomics UK (COG-UK) Consortium | Lighthouse Laboratory Plymouth and Alex Alderton, Roberto Amato, Jeffrey Barrett, Sonia Goncalves, Ewan Harrison, David K. Jackson, Ian Johnston, Dominic Kwiatkowski, Cordelia Langford, John Sillitoe on behalf of the Wellcome Sanger Institute COVID-19 Surveillance Team |
| EPI_ISL_3442615, EPI_ISL_3442616, EPI_ISL_3442617, EPI_ISL_3442618, EPI_ISL_3442619, EPI_ISL_3442620, EPI_ISL_3442621, | Lighthouse Lab in Glasgow | Wellcome Sanger Institute for the COVID-19 Genomics UK (COG-UK) Consortium | Harper VanSteenhouse, Yumi Kasai, David Gray, Carol Clugston, Anna Dominiczak and Alex Alderton, Roberto Amato, Jeffrey Barrett, Sonia Goncalves, Ewan Harrison, David K. Jackson, Ian Johnston, Dominic Kwiatkowski, Cordelia Langford, John Sillitoe on behalf of the Wellcome Sanger Institute COVID-19 Surveillance Team |

| | | | |
|---|---|---|---|
| EPI_ISL_3442622, EPI_ISL_3442623 | | | |
| EPI_ISL_3442624 | Lighthouse Laboratory Plymouth | Wellcome Sanger Institute for the COVID-19 Genomics UK (COG-UK) Consortium | Lighthouse Laboratory Plymouth and Alex Alderton, Roberto Amato, Jeffrey Barrett, Sonia Goncalves, Ewan Harrison, David K. Jackson, Ian Johnston, Dominic Kwiatkowski, Cordelia Langford, John Sillitoe on behalf of the Wellcome Sanger Institute COVID-19 Surveillance Team |
| EPI_ISL_3442625, EPI_ISL_3442626, EPI_ISL_3442627, EPI_ISL_3442628, EPI_ISL_3442629 | Lighthouse Lab in Glasgow | Wellcome Sanger Institute for the COVID-19 Genomics UK (COG-UK) Consortium | Harper VanSteenhouse, Yumi Kasai, David Gray, Carol Clugston, Anna Dominiczak and Alex Alderton, Roberto Amato, Jeffrey Barrett, Sonia Goncalves, Ewan Harrison, David K. Jackson, Ian Johnston, Dominic Kwiatkowski, Cordelia Langford, John Sillitoe on behalf of the Wellcome Sanger Institute COVID-19 Surveillance Team |
| EPI_ISL_3442630 | Lighthouse Laboratory Plymouth | Wellcome Sanger Institute for the COVID-19 Genomics UK (COG-UK) Consortium | Lighthouse Laboratory Plymouth and Alex Alderton, Roberto Amato, Jeffrey Barrett, Sonia Goncalves, Ewan Harrison, David K. Jackson, Ian Johnston, Dominic Kwiatkowski, Cordelia Langford, John Sillitoe on behalf of the Wellcome Sanger Institute COVID-19 Surveillance Team |
| EPI_ISL_3442631, EPI_ISL_3442632, EPI_ISL_3442633 | Lighthouse Lab in Glasgow | Wellcome Sanger Institute for the COVID-19 Genomics UK (COG-UK) Consortium | Harper VanSteenhouse, Yumi Kasai, David Gray, Carol Clugston, Anna Dominiczak and Alex Alderton, Roberto Amato, Jeffrey Barrett, Sonia Goncalves, Ewan Harrison, David K. Jackson, Ian Johnston, Dominic Kwiatkowski, Cordelia Langford, John Sillitoe on behalf of the Wellcome Sanger Institute COVID-19 Surveillance Team |
| EPI_ISL_3442634 | Lighthouse Laboratory Plymouth | Wellcome Sanger Institute for the COVID-19 Genomics UK (COG-UK) Consortium | Lighthouse Laboratory Plymouth and Alex Alderton, Roberto Amato, Jeffrey Barrett, Sonia Goncalves, Ewan Harrison, David K. Jackson, Ian Johnston, Dominic Kwiatkowski, Cordelia Langford, John Sillitoe on behalf of the Wellcome Sanger Institute COVID-19 Surveillance Team |
| EPI_ISL_3442635, EPI_ISL_3442636 | Lighthouse Lab in Glasgow | Wellcome Sanger Institute for the COVID-19 Genomics UK (COG-UK) Consortium | Harper VanSteenhouse, Yumi Kasai, David Gray, Carol Clugston, Anna Dominiczak and Alex Alderton, Roberto Amato, Jeffrey Barrett, Sonia Goncalves, Ewan Harrison, David K. Jackson, Ian Johnston, Dominic Kwiatkowski, Cordelia Langford, John Sillitoe on behalf of the Wellcome Sanger Institute COVID-19 Surveillance Team |
| EPI_ISL_3442637 | Lighthouse Laboratory Plymouth | Wellcome Sanger Institute for the COVID-19 Genomics UK (COG-UK) Consortium | Lighthouse Laboratory Plymouth and Alex Alderton, Roberto Amato, Jeffrey Barrett, Sonia Goncalves, Ewan Harrison, David K. Jackson, Ian Johnston, Dominic Kwiatkowski, Cordelia Langford, John Sillitoe on behalf of the Wellcome Sanger Institute COVID-19 Surveillance Team |
| EPI_ISL_3442638 | Lighthouse Lab in Glasgow | Wellcome Sanger Institute for the COVID-19 Genomics UK (COG-UK) Consortium | Harper VanSteenhouse, Yumi Kasai, David Gray, Carol Clugston, Anna Dominiczak and Alex Alderton, Roberto Amato, Jeffrey Barrett, Sonia Goncalves, Ewan Harrison, David K. Jackson, Ian Johnston, Dominic Kwiatkowski, Cordelia Langford, John Sillitoe on behalf of the Wellcome Sanger Institute COVID-19 Surveillance Team |
| EPI_ISL_3442639, EPI_ISL_3442640 | Lighthouse Laboratory Plymouth | Wellcome Sanger Institute for the COVID-19 Genomics UK (COG-UK) Consortium | Lighthouse Laboratory Plymouth and Alex Alderton, Roberto Amato, Jeffrey Barrett, Sonia Goncalves, Ewan Harrison, David K. Jackson, Ian Johnston, Dominic Kwiatkowski, Cordelia Langford, John Sillitoe on behalf of the Wellcome Sanger Institute COVID-19 Surveillance Team |
| EPI_ISL_3442641, EPI_ISL_3442642, EPI_ISL_3442643 | Lighthouse Lab in Glasgow | Wellcome Sanger Institute for the COVID-19 Genomics UK (COG-UK) Consortium | Harper VanSteenhouse, Yumi Kasai, David Gray, Carol Clugston, Anna Dominiczak and Alex Alderton, Roberto Amato, Jeffrey Barrett, Sonia Goncalves, Ewan Harrison, David K. Jackson, Ian Johnston, Dominic Kwiatkowski, Cordelia Langford, John Sillitoe on behalf of the Wellcome Sanger Institute COVID-19 Surveillance Team |
| EPI_ISL_3442644 | Lighthouse Laboratory Plymouth | Wellcome Sanger Institute for the COVID-19 Genomics UK (COG-UK) Consortium | Lighthouse Laboratory Plymouth and Alex Alderton, Roberto Amato, Jeffrey Barrett, Sonia Goncalves, Ewan Harrison, David K. Jackson, Ian Johnston, Dominic Kwiatkowski, Cordelia Langford, John Sillitoe on behalf of the Wellcome Sanger Institute COVID-19 Surveillance Team |
| EPI_ISL_3442645, EPI_ISL_3442646 | Lighthouse Lab in Glasgow | Wellcome Sanger Institute for the COVID-19 Genomics UK (COG-UK) Consortium | Harper VanSteenhouse, Yumi Kasai, David Gray, Carol Clugston, Anna Dominiczak and Alex Alderton, Roberto Amato, Jeffrey Barrett, Sonia Goncalves, Ewan Harrison, David K. Jackson, Ian Johnston, Dominic Kwiatkowski, Cordelia Langford, John Sillitoe on behalf of the Wellcome Sanger Institute COVID-19 Surveillance Team |
| EPI_ISL_3442647 | Lighthouse Laboratory Plymouth | Wellcome Sanger Institute for the COVID-19 Genomics UK (COG-UK) Consortium | Lighthouse Laboratory Plymouth and Alex Alderton, Roberto Amato, Jeffrey Barrett, Sonia Goncalves, Ewan Harrison, David K. Jackson, Ian Johnston, Dominic Kwiatkowski, Cordelia Langford, John Sillitoe on behalf of the Wellcome Sanger Institute COVID-19 Surveillance Team |
| EPI_ISL_3442648 | Lighthouse Lab in Glasgow | Wellcome Sanger Institute for the COVID-19 Genomics UK (COG-UK) Consortium | Harper VanSteenhouse, Yumi Kasai, David Gray, Carol Clugston, Anna Dominiczak and Alex Alderton, Roberto Amato, Jeffrey Barrett, Sonia Goncalves, Ewan Harrison, David K. Jackson, Ian Johnston, Dominic Kwiatkowski, Cordelia Langford, John Sillitoe on behalf of the Wellcome Sanger Institute COVID-19 Surveillance Team |
| EPI_ISL_3442649 | Lighthouse Laboratory Plymouth | Wellcome Sanger Institute for the COVID-19 Genomics UK (COG-UK) Consortium | Lighthouse Laboratory Plymouth and Alex Alderton, Roberto Amato, Jeffrey Barrett, Sonia Goncalves, Ewan Harrison, David K. Jackson, Ian Johnston, Dominic Kwiatkowski, Cordelia Langford, John Sillitoe on behalf of the Wellcome Sanger Institute COVID-19 Surveillance Team |
| EPI_ISL_3442650 | Lighthouse Lab in Glasgow | Wellcome Sanger Institute for the COVID-19 Genomics UK (COG-UK) Consortium | Harper VanSteenhouse, Yumi Kasai, David Gray, Carol Clugston, Anna Dominiczak and Alex Alderton, Roberto Amato, Jeffrey Barrett, Sonia Goncalves, Ewan Harrison, David K. Jackson, Ian Johnston, Dominic Kwiatkowski, Cordelia Langford, John Sillitoe on behalf of the Wellcome Sanger Institute COVID-19 Surveillance Team |
| EPI_ISL_3442651, EPI_ISL_3442652 | Lighthouse Laboratory Plymouth | Wellcome Sanger Institute for the COVID-19 Genomics UK (COG-UK) Consortium | Lighthouse Laboratory Plymouth and Alex Alderton, Roberto Amato, Jeffrey Barrett, Sonia Goncalves, Ewan Harrison, David K. Jackson, Ian Johnston, Dominic Kwiatkowski, Cordelia Langford, John Sillitoe on behalf of the Wellcome Sanger Institute COVID-19 Surveillance Team |
| EPI_ISL_3442653, EPI_ISL_3442654 | Lighthouse Lab in Glasgow | Wellcome Sanger Institute for the COVID-19 Genomics UK (COG-UK) Consortium | Harper VanSteenhouse, Yumi Kasai, David Gray, Carol Clugston, Anna Dominiczak and Alex Alderton, Roberto Amato, Jeffrey Barrett, Sonia Goncalves, Ewan Harrison, David K. Jackson, Ian Johnston, Dominic Kwiatkowski, Cordelia Langford, John Sillitoe on behalf of the Wellcome Sanger Institute COVID-19 Surveillance Team |
| EPI_ISL_3442655, EPI_ISL_3442656, EPI_ISL_3442657, EPI_ISL_3442658, EPI_ISL_3442659, EPI_ISL_3442660 | Lighthouse Laboratory Plymouth | Wellcome Sanger Institute for the COVID-19 Genomics UK (COG-UK) Consortium | Lighthouse Laboratory Plymouth and Alex Alderton, Roberto Amato, Jeffrey Barrett, Sonia Goncalves, Ewan Harrison, David K. Jackson, Ian Johnston, Dominic Kwiatkowski, Cordelia Langford, John Sillitoe on behalf of the Wellcome Sanger Institute COVID-19 Surveillance Team |
| EPI_ISL_3442661, EPI_ISL_3442662 | Lighthouse Lab in Glasgow | Wellcome Sanger Institute for the COVID-19 Genomics UK (COG-UK) Consortium | Harper VanSteenhouse, Yumi Kasai, David Gray, Carol Clugston, Anna Dominiczak and Alex Alderton, Roberto Amato, Jeffrey Barrett, Sonia Goncalves, Ewan Harrison, David K. Jackson, Ian Johnston, Dominic Kwiatkowski, Cordelia Langford, John Sillitoe on behalf of the Wellcome Sanger Institute COVID-19 Surveillance Team |
| EPI_ISL_3442663 | Lighthouse Laboratory Plymouth | Wellcome Sanger Institute for the COVID-19 Genomics UK (COG-UK) Consortium | Lighthouse Laboratory Plymouth and Alex Alderton, Roberto Amato, Jeffrey Barrett, Sonia Goncalves, Ewan Harrison, David K. Jackson, Ian Johnston, Dominic Kwiatkowski, Cordelia Langford, John Sillitoe on behalf of the Wellcome Sanger Institute COVID-19 Surveillance Team |
| EPI_ISL_3442664, EPI_ISL_3442665, EPI_ISL_3442666, EPI_ISL_3442667, EPI_ISL_3442668 | Lighthouse Lab in Glasgow | Wellcome Sanger Institute for the COVID-19 Genomics UK (COG-UK) Consortium | Harper VanSteenhouse, Yumi Kasai, David Gray, Carol Clugston, Anna Dominiczak and Alex Alderton, Roberto Amato, Jeffrey Barrett, Sonia Goncalves, Ewan Harrison, David K. Jackson, Ian Johnston, Dominic Kwiatkowski, Cordelia Langford, John Sillitoe on behalf of the Wellcome Sanger Institute COVID-19 Surveillance Team |
| EPI_ISL_3442669 | Lighthouse Laboratory Plymouth | Wellcome Sanger Institute for the COVID-19 Genomics UK (COG-UK) Consortium | Lighthouse Laboratory Plymouth and Alex Alderton, Roberto Amato, Jeffrey Barrett, Sonia Goncalves, Ewan Harrison, David K. Jackson, Ian Johnston, Dominic Kwiatkowski, Cordelia Langford, John Sillitoe on behalf of the Wellcome Sanger Institute COVID-19 Surveillance Team |
| EPI_ISL_3442670, EPI_ISL_3442671, EPI_ISL_3442672, EPI_ISL_3442673, EPI_ISL_3442674, EPI_ISL_3442675 | Lighthouse Lab in Glasgow | Wellcome Sanger Institute for the COVID-19 Genomics UK (COG-UK) Consortium | Harper VanSteenhouse, Yumi Kasai, David Gray, Carol Clugston, Anna Dominiczak and Alex Alderton, Roberto Amato, Jeffrey Barrett, Sonia Goncalves, Ewan Harrison, David K. Jackson, Ian Johnston, Dominic Kwiatkowski, Cordelia Langford, John Sillitoe on behalf of the Wellcome Sanger Institute COVID-19 Surveillance Team |
| EPI_ISL_3442676, EPI_ISL_3442677 | Lighthouse Laboratory Plymouth | Wellcome Sanger Institute for the COVID-19 Genomics UK (COG-UK) Consortium | Lighthouse Laboratory Plymouth and Alex Alderton, Roberto Amato, Jeffrey Barrett, Sonia Goncalves, Ewan Harrison, David K. Jackson, Ian Johnston, Dominic Kwiatkowski, Cordelia Langford, John Sillitoe on behalf of the Wellcome Sanger Institute COVID-19 Surveillance Team |
| EPI_ISL_3442678, EPI_ISL_3442679 | Lighthouse Lab in Glasgow | Wellcome Sanger Institute for the COVID-19 Genomics UK (COG-UK) Consortium | Harper VanSteenhouse, Yumi Kasai, David Gray, Carol Clugston, Anna Dominiczak and Alex Alderton, Roberto Amato, Jeffrey Barrett, Sonia Goncalves, Ewan Harrison, David K. Jackson, Ian Johnston, Dominic Kwiatkowski, Cordelia Langford, John Sillitoe on behalf of the Wellcome Sanger Institute COVID-19 Surveillance Team |
| EPI_ISL_3442680, EPI_ISL_3442681 | Lighthouse Laboratory Plymouth | Wellcome Sanger Institute for the COVID-19 Genomics UK (COG-UK) Consortium | Lighthouse Laboratory Plymouth and Alex Alderton, Roberto Amato, Jeffrey Barrett, Sonia Goncalves, Ewan Harrison, David K. Jackson, Ian Johnston, Dominic Kwiatkowski, Cordelia Langford, John Sillitoe on behalf of the Wellcome Sanger Institute COVID-19 Surveillance Team |
| EPI_ISL_3442682, EPI_ISL_3442683, EPI_ISL_3442684 | Lighthouse Lab in Glasgow | Wellcome Sanger Institute for the COVID-19 Genomics UK (COG-UK) Consortium | Harper VanSteenhouse, Yumi Kasai, David Gray, Carol Clugston, Anna Dominiczak and Alex Alderton, Roberto Amato, Jeffrey Barrett, Sonia Goncalves, Ewan Harrison, David K. Jackson, Ian Johnston, Dominic Kwiatkowski, Cordelia Langford, John Sillitoe on behalf of the Wellcome Sanger Institute COVID-19 Surveillance Team |
| EPI_ISL_3442685 | Lighthouse Laboratory Plymouth | Wellcome Sanger Institute for the COVID-19 Genomics UK (COG-UK) Consortium | Lighthouse Laboratory Plymouth and Alex Alderton, Roberto Amato, Jeffrey Barrett, Sonia Goncalves, Ewan Harrison, David K. Jackson, Ian Johnston, Dominic Kwiatkowski, Cordelia Langford, John Sillitoe on behalf of the Wellcome Sanger Institute COVID-19 Surveillance Team |
| EPI_ISL_3442686 | Lighthouse Lab in Glasgow | Wellcome Sanger Institute for the COVID-19 Genomics UK (COG-UK) Consortium | Harper VanSteenhouse, Yumi Kasai, David Gray, Carol Clugston, Anna Dominiczak and Alex Alderton, Roberto Amato, Jeffrey Barrett, Sonia Goncalves, Ewan Harrison, David K. Jackson, Ian Johnston, Dominic Kwiatkowski, Cordelia Langford, John Sillitoe on behalf of the Wellcome Sanger Institute COVID-19 Surveillance Team |
| EPI_ISL_3442687 | Lighthouse Laboratory Plymouth | Wellcome Sanger Institute for the COVID-19 Genomics UK (COG-UK) Consortium | Lighthouse Laboratory Plymouth and Alex Alderton, Roberto Amato, Jeffrey Barrett, Sonia Goncalves, Ewan Harrison, David K. Jackson, Ian Johnston, Dominic Kwiatkowski, Cordelia Langford, John Sillitoe on behalf of the Wellcome Sanger Institute COVID-19 Surveillance Team |
| EPI_ISL_3442688 | Lighthouse Lab in Glasgow | Wellcome Sanger Institute for the COVID-19 Genomics UK (COG-UK) Consortium | Harper VanSteenhouse, Yumi Kasai, David Gray, Carol Clugston, Anna Dominiczak and Alex Alderton, Roberto Amato, Jeffrey Barrett, Sonia Goncalves, Ewan Harrison, David K. Jackson, Ian Johnston, Dominic Kwiatkowski, Cordelia Langford, John Sillitoe on behalf of the Wellcome Sanger Institute COVID-19 Surveillance Team |
| EPI_ISL_3442689 | Lighthouse Laboratory Plymouth | Wellcome Sanger Institute for the COVID-19 Genomics UK (COG-UK) Consortium | Lighthouse Laboratory Plymouth and Alex Alderton, Roberto Amato, Jeffrey Barrett, Sonia Goncalves, Ewan Harrison, David K. Jackson, Ian Johnston, Dominic Kwiatkowski, Cordelia Langford, John Sillitoe on behalf of the Wellcome Sanger Institute COVID-19 Surveillance Team |
| EPI_ISL_3442690, EPI_ISL_3442691, EPI_ISL_3442692, EPI_ISL_3442693, EPI_ISL_3442694 | Lighthouse Lab in Glasgow | Wellcome Sanger Institute for the COVID-19 Genomics UK (COG-UK) Consortium | Harper VanSteenhouse, Yumi Kasai, David Gray, Carol Clugston, Anna Dominiczak and Alex Alderton, Roberto Amato, Jeffrey Barrett, Sonia Goncalves, Ewan Harrison, David K. Jackson, Ian Johnston, Dominic Kwiatkowski, Cordelia Langford, John Sillitoe on behalf of the Wellcome Sanger Institute COVID-19 Surveillance Team |
| EPI_ISL_3442695, EPI_ISL_3442696 | Lighthouse Laboratory Plymouth | Wellcome Sanger Institute for the COVID-19 Genomics UK (COG-UK) Consortium | Lighthouse Laboratory Plymouth and Alex Alderton, Roberto Amato, Jeffrey Barrett, Sonia Goncalves, Ewan Harrison, David K. Jackson, Ian Johnston, Dominic Kwiatkowski, Cordelia Langford, John Sillitoe on behalf of the Wellcome Sanger Institute COVID-19 Surveillance Team |
| EPI_ISL_3442697, EPI_ISL_3442698, | Lighthouse Lab in Glasgow | Wellcome Sanger Institute for the COVID-19 Genomics UK (COG-UK) Consortium | Harper VanSteenhouse, Yumi Kasai, David Gray, Carol Clugston, Anna Dominiczak and Alex Alderton, Roberto Amato, Jeffrey Barrett, Sonia Goncalves, Ewan Harrison, David K. Jackson, Ian Johnston, Dominic Kwiatkowski, Cordelia Langford, John Sillitoe on behalf of the Wellcome Sanger Institute COVID-19 Surveillance Team |

| | | | |
|---|---|---|---|
| EPI_ISL_3442699, EPI_ISL_3442700, EPI_ISL_3442701 | | | |
| EPI_ISL_3442702 | Lighthouse Laboratory Plymouth | Wellcome Sanger Institute for the COVID-19 Genomics UK (COG-UK) Consortium | Lighthouse Laboratory Plymouth and Alex Alderton, Roberto Amato, Jeffrey Barrett, Sonia Goncalves, Ewan Harrison, David K. Jackson, Ian Johnston, Dominic Kwiatkowski, Cordelia Langford, John Sillitoe on behalf of the Wellcome Sanger Institute COVID-19 Surveillance Team |
| EPI_ISL_3442703 | Lighthouse Lab in Glasgow | Wellcome Sanger Institute for the COVID-19 Genomics UK (COG-UK) Consortium | Harper VanSteenhouse, Yumi Kasai, David Gray, Carol Clugston, Anna Dominiczak and Alex Alderton, Roberto Amato, Jeffrey Barrett, Sonia Goncalves, Ewan Harrison, David K. Jackson, Ian Johnston, Dominic Kwiatkowski, Cordelia Langford, John Sillitoe on behalf of the Wellcome Sanger Institute COVID-19 Surveillance Team |
| EPI_ISL_3442704 | Lighthouse Laboratory Plymouth | Wellcome Sanger Institute for the COVID-19 Genomics UK (COG-UK) Consortium | Lighthouse Laboratory Plymouth and Alex Alderton, Roberto Amato, Jeffrey Barrett, Sonia Goncalves, Ewan Harrison, David K. Jackson, Ian Johnston, Dominic Kwiatkowski, Cordelia Langford, John Sillitoe on behalf of the Wellcome Sanger Institute COVID-19 Surveillance Team |
| EPI_ISL_3442705 | Lighthouse Lab in Glasgow | Wellcome Sanger Institute for the COVID-19 Genomics UK (COG-UK) Consortium | Harper VanSteenhouse, Yumi Kasai, David Gray, Carol Clugston, Anna Dominiczak and Alex Alderton, Roberto Amato, Jeffrey Barrett, Sonia Goncalves, Ewan Harrison, David K. Jackson, Ian Johnston, Dominic Kwiatkowski, Cordelia Langford, John Sillitoe on behalf of the Wellcome Sanger Institute COVID-19 Surveillance Team |
| EPI_ISL_3442706, EPI_ISL_3442707 | Lighthouse Laboratory Plymouth | Wellcome Sanger Institute for the COVID-19 Genomics UK (COG-UK) Consortium | Lighthouse Laboratory Plymouth and Alex Alderton, Roberto Amato, Jeffrey Barrett, Sonia Goncalves, Ewan Harrison, David K. Jackson, Ian Johnston, Dominic Kwiatkowski, Cordelia Langford, John Sillitoe on behalf of the Wellcome Sanger Institute COVID-19 Surveillance Team |
| EPI_ISL_3442708, EPI_ISL_3442709, EPI_ISL_3442710 | Lighthouse Lab in Glasgow | Wellcome Sanger Institute for the COVID-19 Genomics UK (COG-UK) Consortium | Harper VanSteenhouse, Yumi Kasai, David Gray, Carol Clugston, Anna Dominiczak and Alex Alderton, Roberto Amato, Jeffrey Barrett, Sonia Goncalves, Ewan Harrison, David K. Jackson, Ian Johnston, Dominic Kwiatkowski, Cordelia Langford, John Sillitoe on behalf of the Wellcome Sanger Institute COVID-19 Surveillance Team |
| EPI_ISL_3442711, EPI_ISL_3442712 | Lighthouse Laboratory Plymouth | Wellcome Sanger Institute for the COVID-19 Genomics UK (COG-UK) Consortium | Lighthouse Laboratory Plymouth and Alex Alderton, Roberto Amato, Jeffrey Barrett, Sonia Goncalves, Ewan Harrison, David K. Jackson, Ian Johnston, Dominic Kwiatkowski, Cordelia Langford, John Sillitoe on behalf of the Wellcome Sanger Institute COVID-19 Surveillance Team |
| EPI_ISL_3442713, EPI_ISL_3442714, EPI_ISL_3442715, EPI_ISL_3442716, EPI_ISL_3442717, EPI_ISL_3442718 | Lighthouse Lab in Glasgow | Wellcome Sanger Institute for the COVID-19 Genomics UK (COG-UK) Consortium | Harper VanSteenhouse, Yumi Kasai, David Gray, Carol Clugston, Anna Dominiczak and Alex Alderton, Roberto Amato, Jeffrey Barrett, Sonia Goncalves, Ewan Harrison, David K. Jackson, Ian Johnston, Dominic Kwiatkowski, Cordelia Langford, John Sillitoe on behalf of the Wellcome Sanger Institute COVID-19 Surveillance Team |
| EPI_ISL_3442719, EPI_ISL_3442720 | Lighthouse Laboratory Plymouth | Wellcome Sanger Institute for the COVID-19 Genomics UK (COG-UK) Consortium | Lighthouse Laboratory Plymouth and Alex Alderton, Roberto Amato, Jeffrey Barrett, Sonia Goncalves, Ewan Harrison, David K. Jackson, Ian Johnston, Dominic Kwiatkowski, Cordelia Langford, John Sillitoe on behalf of the Wellcome Sanger Institute COVID-19 Surveillance Team |
| EPI_ISL_3442721, EPI_ISL_3442722, EPI_ISL_3442723, EPI_ISL_3442724 | Lighthouse Lab in Glasgow | Wellcome Sanger Institute for the COVID-19 Genomics UK (COG-UK) Consortium | Harper VanSteenhouse, Yumi Kasai, David Gray, Carol Clugston, Anna Dominiczak and Alex Alderton, Roberto Amato, Jeffrey Barrett, Sonia Goncalves, Ewan Harrison, David K. Jackson, Ian Johnston, Dominic Kwiatkowski, Cordelia Langford, John Sillitoe on behalf of the Wellcome Sanger Institute COVID-19 Surveillance Team |
| EPI_ISL_3442725, EPI_ISL_3442726, EPI_ISL_3442727 | Lighthouse Laboratory Plymouth | Wellcome Sanger Institute for the COVID-19 Genomics UK (COG-UK) Consortium | Lighthouse Laboratory Plymouth and Alex Alderton, Roberto Amato, Jeffrey Barrett, Sonia Goncalves, Ewan Harrison, David K. Jackson, Ian Johnston, Dominic Kwiatkowski, Cordelia Langford, John Sillitoe on behalf of the Wellcome Sanger Institute COVID-19 Surveillance Team |
| EPI_ISL_3442728, EPI_ISL_3442729, EPI_ISL_3442730, EPI_ISL_3442731, EPI_ISL_3442732, EPI_ISL_3442733, EPI_ISL_3442734 | Lighthouse Lab in Glasgow | Wellcome Sanger Institute for the COVID-19 Genomics UK (COG-UK) Consortium | Harper VanSteenhouse, Yumi Kasai, David Gray, Carol Clugston, Anna Dominiczak and Alex Alderton, Roberto Amato, Jeffrey Barrett, Sonia Goncalves, Ewan Harrison, David K. Jackson, Ian Johnston, Dominic Kwiatkowski, Cordelia Langford, John Sillitoe on behalf of the Wellcome Sanger Institute COVID-19 Surveillance Team |
| EPI_ISL_3442735 | Lighthouse Laboratory Plymouth | Wellcome Sanger Institute for the COVID-19 Genomics UK (COG-UK) Consortium | Lighthouse Laboratory Plymouth and Alex Alderton, Roberto Amato, Jeffrey Barrett, Sonia Goncalves, Ewan Harrison, David K. Jackson, Ian Johnston, Dominic Kwiatkowski, Cordelia Langford, John Sillitoe on behalf of the Wellcome Sanger Institute COVID-19 Surveillance Team |
| EPI_ISL_3442736, EPI_ISL_3442737, EPI_ISL_3442738, EPI_ISL_3442739, EPI_ISL_3442740 | Lighthouse Lab in Glasgow | Wellcome Sanger Institute for the COVID-19 Genomics UK (COG-UK) Consortium | Harper VanSteenhouse, Yumi Kasai, David Gray, Carol Clugston, Anna Dominiczak and Alex Alderton, Roberto Amato, Jeffrey Barrett, Sonia Goncalves, Ewan Harrison, David K. Jackson, Ian Johnston, Dominic Kwiatkowski, Cordelia Langford, John Sillitoe on behalf of the Wellcome Sanger Institute COVID-19 Surveillance Team |
| EPI_ISL_3442741 | Lighthouse Laboratory Plymouth | Wellcome Sanger Institute for the COVID-19 Genomics UK (COG-UK) Consortium | Lighthouse Laboratory Plymouth and Alex Alderton, Roberto Amato, Jeffrey Barrett, Sonia Goncalves, Ewan Harrison, David K. Jackson, Ian Johnston, Dominic Kwiatkowski, Cordelia Langford, John Sillitoe on behalf of the Wellcome Sanger Institute COVID-19 Surveillance Team |
| EPI_ISL_3442742 | Lighthouse Lab in Glasgow | Wellcome Sanger Institute for the COVID-19 Genomics UK (COG-UK) Consortium | Harper VanSteenhouse, Yumi Kasai, David Gray, Carol Clugston, Anna Dominiczak and Alex Alderton, Roberto Amato, Jeffrey Barrett, Sonia Goncalves, Ewan Harrison, David K. Jackson, Ian Johnston, Dominic Kwiatkowski, Cordelia Langford, John Sillitoe on behalf of the Wellcome Sanger Institute COVID-19 Surveillance Team |
| EPI_ISL_3442743 | Lighthouse Laboratory Plymouth | Wellcome Sanger Institute for the COVID-19 Genomics UK (COG-UK) Consortium | Lighthouse Laboratory Plymouth and Alex Alderton, Roberto Amato, Jeffrey Barrett, Sonia Goncalves, Ewan Harrison, David K. Jackson, Ian Johnston, Dominic Kwiatkowski, Cordelia Langford, John Sillitoe on behalf of the Wellcome Sanger Institute COVID-19 Surveillance Team |
| EPI_ISL_3442744 | Lighthouse Lab in Glasgow | Wellcome Sanger Institute for the COVID-19 Genomics UK (COG-UK) Consortium | Harper VanSteenhouse, Yumi Kasai, David Gray, Carol Clugston, Anna Dominiczak and Alex Alderton, Roberto Amato, Jeffrey Barrett, Sonia Goncalves, Ewan Harrison, David K. Jackson, Ian Johnston, Dominic Kwiatkowski, Cordelia Langford, John Sillitoe on behalf of the Wellcome Sanger Institute COVID-19 Surveillance Team |
| EPI_ISL_3442745 | Lighthouse Laboratory Plymouth | Wellcome Sanger Institute for the COVID-19 Genomics UK (COG-UK) Consortium | Lighthouse Laboratory Plymouth and Alex Alderton, Roberto Amato, Jeffrey Barrett, Sonia Goncalves, Ewan Harrison, David K. Jackson, Ian Johnston, Dominic Kwiatkowski, Cordelia Langford, John Sillitoe on behalf of the Wellcome Sanger Institute COVID-19 Surveillance Team |
| EPI_ISL_3442746, EPI_ISL_3442747 | Lighthouse Lab in Glasgow | Wellcome Sanger Institute for the COVID-19 Genomics UK (COG-UK) Consortium | Harper VanSteenhouse, Yumi Kasai, David Gray, Carol Clugston, Anna Dominiczak and Alex Alderton, Roberto Amato, Jeffrey Barrett, Sonia Goncalves, Ewan Harrison, David K. Jackson, Ian Johnston, Dominic Kwiatkowski, Cordelia Langford, John Sillitoe on behalf of the Wellcome Sanger Institute COVID-19 Surveillance Team |
| EPI_ISL_3442748 | Lighthouse Laboratory Plymouth | Wellcome Sanger Institute for the COVID-19 Genomics UK (COG-UK) Consortium | Lighthouse Laboratory Plymouth and Alex Alderton, Roberto Amato, Jeffrey Barrett, Sonia Goncalves, Ewan Harrison, David K. Jackson, Ian Johnston, Dominic Kwiatkowski, Cordelia Langford, John Sillitoe on behalf of the Wellcome Sanger Institute COVID-19 Surveillance Team |
| EPI_ISL_3442749 | Lighthouse Lab in Glasgow | Wellcome Sanger Institute for the COVID-19 Genomics UK (COG-UK) Consortium | Harper VanSteenhouse, Yumi Kasai, David Gray, Carol Clugston, Anna Dominiczak and Alex Alderton, Roberto Amato, Jeffrey Barrett, Sonia Goncalves, Ewan Harrison, David K. Jackson, Ian Johnston, Dominic Kwiatkowski, Cordelia Langford, John Sillitoe on behalf of the Wellcome Sanger Institute COVID-19 Surveillance Team |
| EPI_ISL_3442750, EPI_ISL_3442751, EPI_ISL_3442752 | Lighthouse Laboratory Plymouth | Wellcome Sanger Institute for the COVID-19 Genomics UK (COG-UK) Consortium | Lighthouse Laboratory Plymouth and Alex Alderton, Roberto Amato, Jeffrey Barrett, Sonia Goncalves, Ewan Harrison, David K. Jackson, Ian Johnston, Dominic Kwiatkowski, Cordelia Langford, John Sillitoe on behalf of the Wellcome Sanger Institute COVID-19 Surveillance Team |
| EPI_ISL_3442753 | Lighthouse Lab in Glasgow | Wellcome Sanger Institute for the COVID-19 Genomics UK (COG-UK) Consortium | Harper VanSteenhouse, Yumi Kasai, David Gray, Carol Clugston, Anna Dominiczak and Alex Alderton, Roberto Amato, Jeffrey Barrett, Sonia Goncalves, Ewan Harrison, David K. Jackson, Ian Johnston, Dominic Kwiatkowski, Cordelia Langford, John Sillitoe on behalf of the Wellcome Sanger Institute COVID-19 Surveillance Team |
| EPI_ISL_3442754 | Lighthouse Laboratory Plymouth | Wellcome Sanger Institute for the COVID-19 Genomics UK (COG-UK) Consortium | Lighthouse Laboratory Plymouth and Alex Alderton, Roberto Amato, Jeffrey Barrett, Sonia Goncalves, Ewan Harrison, David K. Jackson, Ian Johnston, Dominic Kwiatkowski, Cordelia Langford, John Sillitoe on behalf of the Wellcome Sanger Institute COVID-19 Surveillance Team |
| EPI_ISL_3442755 | Lighthouse Lab in Glasgow | Wellcome Sanger Institute for the COVID-19 Genomics UK (COG-UK) Consortium | Harper VanSteenhouse, Yumi Kasai, David Gray, Carol Clugston, Anna Dominiczak and Alex Alderton, Roberto Amato, Jeffrey Barrett, Sonia Goncalves, Ewan Harrison, David K. Jackson, Ian Johnston, Dominic Kwiatkowski, Cordelia Langford, John Sillitoe on behalf of the Wellcome Sanger Institute COVID-19 Surveillance Team |
| EPI_ISL_3442756 | Lighthouse Laboratory Plymouth | Wellcome Sanger Institute for the COVID-19 Genomics UK (COG-UK) Consortium | Lighthouse Laboratory Plymouth and Alex Alderton, Roberto Amato, Jeffrey Barrett, Sonia Goncalves, Ewan Harrison, David K. Jackson, Ian Johnston, Dominic Kwiatkowski, Cordelia Langford, John Sillitoe on behalf of the Wellcome Sanger Institute COVID-19 Surveillance Team |
| EPI_ISL_3442757 | Lighthouse Lab in Glasgow | Wellcome Sanger Institute for the COVID-19 Genomics UK (COG-UK) Consortium | Harper VanSteenhouse, Yumi Kasai, David Gray, Carol Clugston, Anna Dominiczak and Alex Alderton, Roberto Amato, Jeffrey Barrett, Sonia Goncalves, Ewan Harrison, David K. Jackson, Ian Johnston, Dominic Kwiatkowski, Cordelia Langford, John Sillitoe on behalf of the Wellcome Sanger Institute COVID-19 Surveillance Team |
| EPI_ISL_3442758, EPI_ISL_3442759 | Lighthouse Laboratory Plymouth | Wellcome Sanger Institute for the COVID-19 Genomics UK (COG-UK) Consortium | Lighthouse Laboratory Plymouth and Alex Alderton, Roberto Amato, Jeffrey Barrett, Sonia Goncalves, Ewan Harrison, David K. Jackson, Ian Johnston, Dominic Kwiatkowski, Cordelia Langford, John Sillitoe on behalf of the Wellcome Sanger Institute COVID-19 Surveillance Team |
| EPI_ISL_3442760, EPI_ISL_3442761 | Lighthouse Lab in Glasgow | Wellcome Sanger Institute for the COVID-19 Genomics UK (COG-UK) Consortium | Harper VanSteenhouse, Yumi Kasai, David Gray, Carol Clugston, Anna Dominiczak and Alex Alderton, Roberto Amato, Jeffrey Barrett, Sonia Goncalves, Ewan Harrison, David K. Jackson, Ian Johnston, Dominic Kwiatkowski, Cordelia Langford, John Sillitoe on behalf of the Wellcome Sanger Institute COVID-19 Surveillance Team |
| EPI_ISL_3442762 | Lighthouse Laboratory Plymouth | Wellcome Sanger Institute for the COVID-19 Genomics UK (COG-UK) Consortium | Lighthouse Laboratory Plymouth and Alex Alderton, Roberto Amato, Jeffrey Barrett, Sonia Goncalves, Ewan Harrison, David K. Jackson, Ian Johnston, Dominic Kwiatkowski, Cordelia Langford, John Sillitoe on behalf of the Wellcome Sanger Institute COVID-19 Surveillance Team |
| EPI_ISL_3442763, EPI_ISL_3442764, EPI_ISL_3442765 | Lighthouse Lab in Glasgow | Wellcome Sanger Institute for the COVID-19 Genomics UK (COG-UK) Consortium | Harper VanSteenhouse, Yumi Kasai, David Gray, Carol Clugston, Anna Dominiczak and Alex Alderton, Roberto Amato, Jeffrey Barrett, Sonia Goncalves, Ewan Harrison, David K. Jackson, Ian Johnston, Dominic Kwiatkowski, Cordelia Langford, John Sillitoe on behalf of the Wellcome Sanger Institute COVID-19 Surveillance Team |
| EPI_ISL_3442766 | Lighthouse Laboratory Plymouth | Wellcome Sanger Institute for the COVID-19 Genomics UK (COG-UK) Consortium | Lighthouse Laboratory Plymouth and Alex Alderton, Roberto Amato, Jeffrey Barrett, Sonia Goncalves, Ewan Harrison, David K. Jackson, Ian Johnston, Dominic Kwiatkowski, Cordelia Langford, John Sillitoe on behalf of the Wellcome Sanger Institute COVID-19 Surveillance Team |
| EPI_ISL_3442767, EPI_ISL_3442768, EPI_ISL_3442769 | Lighthouse Lab in Glasgow | Wellcome Sanger Institute for the COVID-19 Genomics UK (COG-UK) Consortium | Harper VanSteenhouse, Yumi Kasai, David Gray, Carol Clugston, Anna Dominiczak and Alex Alderton, Roberto Amato, Jeffrey Barrett, Sonia Goncalves, Ewan Harrison, David K. Jackson, Ian Johnston, Dominic Kwiatkowski, Cordelia Langford, John Sillitoe on behalf of the Wellcome Sanger Institute COVID-19 Surveillance Team |
| EPI_ISL_3442770 | Lighthouse Laboratory Plymouth | Wellcome Sanger Institute for the COVID-19 Genomics UK (COG-UK) Consortium | Lighthouse Laboratory Plymouth and Alex Alderton, Roberto Amato, Jeffrey Barrett, Sonia Goncalves, Ewan Harrison, David K. Jackson, Ian Johnston, Dominic Kwiatkowski, Cordelia Langford, John Sillitoe on behalf of the Wellcome Sanger Institute COVID-19 Surveillance Team |

EPI_ISL_3442771, EPI_ISL_3442772, EPI_ISL_3442773, EPI_ISL_3442774, EPI_ISL_3442775, EPI_ISL_3442776, EPI_ISL_3442777, EPI_ISL_3442778, EPI_ISL_3442779, EPI_ISL_3442780, EPI_ISL_3442781, EPI_ISL_3442782

| | | | |
|---|---|---|---|
| see above | Lighthouse Lab in Glasgow | Wellcome Sanger Institute for the COVID-19 Genomics UK (COG-UK) Consortium | Harper VanSteenhouse, Yumi Kasai, David Gray, Carol Clugston, Anna Dominiczak and Alex Alderton, Roberto Amato, Jeffrey Barrett, Sonia Goncalves, Ewan Harrison, David K. Jackson, Ian Johnston, Dominic Kwiatkowski, Cordelia Langford, John Sillitoe on behalf of the Wellcome Sanger Institute COVID-19 Surveillance Team |
| EPI_ISL_3442783 | Lighthouse Laboratory Plymouth | Wellcome Sanger Institute for the COVID-19 Genomics UK (COG-UK) Consortium | Lighthouse Laboratory Plymouth and Alex Alderton, Roberto Amato, Jeffrey Barrett, Sonia Goncalves, Ewan Harrison, David K. Jackson, Ian Johnston, Dominic Kwiatkowski, Cordelia Langford, John Sillitoe on behalf of the Wellcome Sanger Institute COVID-19 Surveillance Team |
| EPI_ISL_3442784, EPI_ISL_3442785, EPI_ISL_3442786, EPI_ISL_3442787, EPI_ISL_3442788 | Lighthouse Lab in Glasgow | Wellcome Sanger Institute for the COVID-19 Genomics UK (COG-UK) Consortium | Harper VanSteenhouse, Yumi Kasai, David Gray, Carol Clugston, Anna Dominiczak and Alex Alderton, Roberto Amato, Jeffrey Barrett, Sonia Goncalves, Ewan Harrison, David K. Jackson, Ian Johnston, Dominic Kwiatkowski, Cordelia Langford, John Sillitoe on behalf of the Wellcome Sanger Institute COVID-19 Surveillance Team |
| EPI_ISL_3442789 | Lighthouse Laboratory Plymouth | Wellcome Sanger Institute for the COVID-19 Genomics UK (COG-UK) Consortium | Lighthouse Laboratory Plymouth and Alex Alderton, Roberto Amato, Jeffrey Barrett, Sonia Goncalves, Ewan Harrison, David K. Jackson, Ian Johnston, Dominic Kwiatkowski, Cordelia Langford, John Sillitoe on behalf of the Wellcome Sanger Institute COVID-19 Surveillance Team |
| EPI_ISL_3442790, EPI_ISL_3442791 | Lighthouse Lab in Glasgow | Wellcome Sanger Institute for the COVID-19 Genomics UK (COG-UK) Consortium | Harper VanSteenhouse, Yumi Kasai, David Gray, Carol Clugston, Anna Dominiczak and Alex Alderton, Roberto Amato, Jeffrey Barrett, Sonia Goncalves, Ewan Harrison, David K. Jackson, Ian Johnston, Dominic Kwiatkowski, Cordelia Langford, John Sillitoe on behalf of the Wellcome Sanger Institute COVID-19 Surveillance Team |
| EPI_ISL_3442792 | Lighthouse Laboratory Plymouth | Wellcome Sanger Institute for the COVID-19 Genomics UK (COG-UK) Consortium | Lighthouse Laboratory Plymouth and Alex Alderton, Roberto Amato, Jeffrey Barrett, Sonia Goncalves, Ewan Harrison, David K. Jackson, Ian Johnston, Dominic Kwiatkowski, Cordelia Langford, John Sillitoe on behalf of the Wellcome Sanger Institute COVID-19 Surveillance Team |
| EPI_ISL_3442793 | Berkshire and Surrey Pathology Services Lighthouse Laboratory | Wellcome Sanger Institute for the COVID-19 Genomics UK (COG-UK) Consortium | Berkshire and Surrey Pathology Services Lighthouse Laboratory and Alex Alderton, Roberto Amato, Jeffrey Barrett, Sonia Goncalves, Ewan Harrison, David K. Jackson, Ian Johnston, Dominic Kwiatkowski, Cordelia Langford, John Sillitoe on behalf of the Wellcome Sanger Institute COVID-19 Surveillance Team |
| EPI_ISL_3442794, EPI_ISL_3442795, EPI_ISL_3442796 | Health Services Laboratories | Wellcome Sanger Institute for the COVID-19 Genomics UK (COG-UK) Consortium | Health Services Laboratories and Alex Alderton, Roberto Amato, Jeffrey Barrett, Sonia Goncalves, Ewan Harrison, David K. Jackson, Ian Johnston, Dominic Kwiatkowski, Cordelia Langford, John Sillitoe on behalf of the Wellcome Sanger Institute COVID-19 Surveillance Team |
| EPI_ISL_3442797 | Berkshire and Surrey Pathology Services Lighthouse Laboratory | Wellcome Sanger Institute for the COVID-19 Genomics UK (COG-UK) Consortium | Berkshire and Surrey Pathology Services Lighthouse Laboratory and Alex Alderton, Roberto Amato, Jeffrey Barrett, Sonia Goncalves, Ewan Harrison, David K. Jackson, Ian Johnston, Dominic Kwiatkowski, Cordelia Langford, John Sillitoe on behalf of the Wellcome Sanger Institute COVID-19 Surveillance Team |
| EPI_ISL_3442798, EPI_ISL_3442799 | Health Services Laboratories | Wellcome Sanger Institute for the COVID-19 Genomics UK (COG-UK) Consortium | Health Services Laboratories and Alex Alderton, Roberto Amato, Jeffrey Barrett, Sonia Goncalves, Ewan Harrison, David K. Jackson, Ian Johnston, Dominic Kwiatkowski, Cordelia Langford, John Sillitoe on behalf of the Wellcome Sanger Institute COVID-19 Surveillance Team |
| EPI_ISL_3442800 | Berkshire and Surrey Pathology Services Lighthouse Laboratory | Wellcome Sanger Institute for the COVID-19 Genomics UK (COG-UK) Consortium | Berkshire and Surrey Pathology Services Lighthouse Laboratory and Alex Alderton, Roberto Amato, Jeffrey Barrett, Sonia Goncalves, Ewan Harrison, David K. Jackson, Ian Johnston, Dominic Kwiatkowski, Cordelia Langford, John Sillitoe on behalf of the Wellcome Sanger Institute COVID-19 Surveillance Team |
| EPI_ISL_3442801, EPI_ISL_3442802 | Health Services Laboratories | Wellcome Sanger Institute for the COVID-19 Genomics UK (COG-UK) Consortium | Health Services Laboratories and Alex Alderton, Roberto Amato, Jeffrey Barrett, Sonia Goncalves, Ewan Harrison, David K. Jackson, Ian Johnston, Dominic Kwiatkowski, Cordelia Langford, John Sillitoe on behalf of the Wellcome Sanger Institute COVID-19 Surveillance Team |
| EPI_ISL_3442803, EPI_ISL_3442804 | Berkshire and Surrey Pathology Services Lighthouse Laboratory | Wellcome Sanger Institute for the COVID-19 Genomics UK (COG-UK) Consortium | Berkshire and Surrey Pathology Services Lighthouse Laboratory and Alex Alderton, Roberto Amato, Jeffrey Barrett, Sonia Goncalves, Ewan Harrison, David K. Jackson, Ian Johnston, Dominic Kwiatkowski, Cordelia Langford, John Sillitoe on behalf of the Wellcome Sanger Institute COVID-19 Surveillance Team |
| EPI_ISL_3442805, EPI_ISL_3442806 | Health Services Laboratories | Wellcome Sanger Institute for the COVID-19 Genomics UK (COG-UK) Consortium | Health Services Laboratories and Alex Alderton, Roberto Amato, Jeffrey Barrett, Sonia Goncalves, Ewan Harrison, David K. Jackson, Ian Johnston, Dominic Kwiatkowski, Cordelia Langford, John Sillitoe on behalf of the Wellcome Sanger Institute COVID-19 Surveillance Team |
| EPI_ISL_3442807, EPI_ISL_3442808, EPI_ISL_3442809, EPI_ISL_3442810, EPI_ISL_3442811, EPI_ISL_3442812, EPI_ISL_3442813 | Berkshire and Surrey Pathology Services Lighthouse Laboratory | Wellcome Sanger Institute for the COVID-19 Genomics UK (COG-UK) Consortium | Berkshire and Surrey Pathology Services Lighthouse Laboratory and Alex Alderton, Roberto Amato, Jeffrey Barrett, Sonia Goncalves, Ewan Harrison, David K. Jackson, Ian Johnston, Dominic Kwiatkowski, Cordelia Langford, John Sillitoe on behalf of the Wellcome Sanger Institute COVID-19 Surveillance Team |
| EPI_ISL_3442814 | Health Services Laboratories | Wellcome Sanger Institute for the COVID-19 Genomics UK (COG-UK) Consortium | Health Services Laboratories and Alex Alderton, Roberto Amato, Jeffrey Barrett, Sonia Goncalves, Ewan Harrison, David K. Jackson, Ian Johnston, Dominic Kwiatkowski, Cordelia Langford, John Sillitoe on behalf of the Wellcome Sanger Institute COVID-19 Surveillance Team |
| EPI_ISL_3442815, EPI_ISL_3442816, EPI_ISL_3442817 | Berkshire and Surrey Pathology Services Lighthouse Laboratory | Wellcome Sanger Institute for the COVID-19 Genomics UK (COG-UK) Consortium | Berkshire and Surrey Pathology Services Lighthouse Laboratory and Alex Alderton, Roberto Amato, Jeffrey Barrett, Sonia Goncalves, Ewan Harrison, David K. Jackson, Ian Johnston, Dominic Kwiatkowski, Cordelia Langford, John Sillitoe on behalf of the Wellcome Sanger Institute COVID-19 Surveillance Team |
| EPI_ISL_3442818 | Health Services Laboratories | Wellcome Sanger Institute for the COVID-19 Genomics UK (COG-UK) Consortium | Health Services Laboratories and Alex Alderton, Roberto Amato, Jeffrey Barrett, Sonia Goncalves, Ewan Harrison, David K. Jackson, Ian Johnston, Dominic Kwiatkowski, Cordelia Langford, John Sillitoe on behalf of the Wellcome Sanger Institute COVID-19 Surveillance Team |
| EPI_ISL_3442819, EPI_ISL_3442820 | Berkshire and Surrey Pathology Services Lighthouse Laboratory | Wellcome Sanger Institute for the COVID-19 Genomics UK (COG-UK) Consortium | Berkshire and Surrey Pathology Services Lighthouse Laboratory and Alex Alderton, Roberto Amato, Jeffrey Barrett, Sonia Goncalves, Ewan Harrison, David K. Jackson, Ian Johnston, Dominic Kwiatkowski, Cordelia Langford, John Sillitoe on behalf of the Wellcome Sanger Institute COVID-19 Surveillance Team |
| EPI_ISL_3442821 | Health Services Laboratories | Wellcome Sanger Institute for the COVID-19 Genomics UK (COG-UK) Consortium | Health Services Laboratories and Alex Alderton, Roberto Amato, Jeffrey Barrett, Sonia Goncalves, Ewan Harrison, David K. Jackson, Ian Johnston, Dominic Kwiatkowski, Cordelia Langford, John Sillitoe on behalf of the Wellcome Sanger Institute COVID-19 Surveillance Team |
| EPI_ISL_3442822, EPI_ISL_3442823, EPI_ISL_3442824, EPI_ISL_3442825, EPI_ISL_3442826, EPI_ISL_3442827, EPI_ISL_3442828, EPI_ISL_3442829, EPI_ISL_3442830 | Berkshire and Surrey Pathology Services Lighthouse Laboratory | Wellcome Sanger Institute for the COVID-19 Genomics UK (COG-UK) Consortium | Berkshire and Surrey Pathology Services Lighthouse Laboratory and Alex Alderton, Roberto Amato, Jeffrey Barrett, Sonia Goncalves, Ewan Harrison, David K. Jackson, Ian Johnston, Dominic Kwiatkowski, Cordelia Langford, John Sillitoe on behalf of the Wellcome Sanger Institute COVID-19 Surveillance Team |
| EPI_ISL_3442831 | Health Services Laboratories | Wellcome Sanger Institute for the COVID-19 Genomics UK (COG-UK) Consortium | Health Services Laboratories and Alex Alderton, Roberto Amato, Jeffrey Barrett, Sonia Goncalves, Ewan Harrison, David K. Jackson, Ian Johnston, Dominic Kwiatkowski, Cordelia Langford, John Sillitoe on behalf of the Wellcome Sanger Institute COVID-19 Surveillance Team |
| EPI_ISL_3442832, EPI_ISL_3442833, EPI_ISL_3442834, EPI_ISL_3442835, EPI_ISL_3442836, EPI_ISL_3442837, EPI_ISL_3442838, EPI_ISL_3442839 | Berkshire and Surrey Pathology Services Lighthouse Laboratory | Wellcome Sanger Institute for the COVID-19 Genomics UK (COG-UK) Consortium | Berkshire and Surrey Pathology Services Lighthouse Laboratory and Alex Alderton, Roberto Amato, Jeffrey Barrett, Sonia Goncalves, Ewan Harrison, David K. Jackson, Ian Johnston, Dominic Kwiatkowski, Cordelia Langford, John Sillitoe on behalf of the Wellcome Sanger Institute COVID-19 Surveillance Team |
| EPI_ISL_3442840, EPI_ISL_3442841 | Health Services Laboratories | Wellcome Sanger Institute for the COVID-19 Genomics UK (COG-UK) Consortium | Health Services Laboratories and Alex Alderton, Roberto Amato, Jeffrey Barrett, Sonia Goncalves, Ewan Harrison, David K. Jackson, Ian Johnston, Dominic Kwiatkowski, Cordelia Langford, John Sillitoe on behalf of the Wellcome Sanger Institute COVID-19 Surveillance Team |
| EPI_ISL_3442842, EPI_ISL_3442843 | Berkshire and Surrey Pathology Services Lighthouse Laboratory | Wellcome Sanger Institute for the COVID-19 Genomics UK (COG-UK) Consortium | Berkshire and Surrey Pathology Services Lighthouse Laboratory and Alex Alderton, Roberto Amato, Jeffrey Barrett, Sonia Goncalves, Ewan Harrison, David K. Jackson, Ian Johnston, Dominic Kwiatkowski, Cordelia Langford, John Sillitoe on behalf of the Wellcome Sanger Institute COVID-19 Surveillance Team |
| EPI_ISL_3442844 | Health Services Laboratories | Wellcome Sanger Institute for the COVID-19 Genomics UK (COG-UK) Consortium | Health Services Laboratories and Alex Alderton, Roberto Amato, Jeffrey Barrett, Sonia Goncalves, Ewan Harrison, David K. Jackson, Ian Johnston, Dominic Kwiatkowski, Cordelia Langford, John Sillitoe on behalf of the Wellcome Sanger Institute COVID-19 Surveillance Team |
| EPI_ISL_3442845, EPI_ISL_3442846, EPI_ISL_3442847, EPI_ISL_3442848, EPI_ISL_3442849 | Berkshire and Surrey Pathology Services Lighthouse Laboratory | Wellcome Sanger Institute for the COVID-19 Genomics UK (COG-UK) Consortium | Berkshire and Surrey Pathology Services Lighthouse Laboratory and Alex Alderton, Roberto Amato, Jeffrey Barrett, Sonia Goncalves, Ewan Harrison, David K. Jackson, Ian Johnston, Dominic Kwiatkowski, Cordelia Langford, John Sillitoe on behalf of the Wellcome Sanger Institute COVID-19 Surveillance Team |
| EPI_ISL_3442850 | Health Services Laboratories | Wellcome Sanger Institute for the COVID-19 Genomics UK (COG-UK) Consortium | Health Services Laboratories and Alex Alderton, Roberto Amato, Jeffrey Barrett, Sonia Goncalves, Ewan Harrison, David K. Jackson, Ian Johnston, Dominic Kwiatkowski, Cordelia Langford, John Sillitoe on behalf of the Wellcome Sanger Institute COVID-19 Surveillance Team |
| EPI_ISL_3442851, EPI_ISL_3442852, EPI_ISL_3442853 | Berkshire and Surrey Pathology Services Lighthouse Laboratory | Wellcome Sanger Institute for the COVID-19 Genomics UK (COG-UK) Consortium | Berkshire and Surrey Pathology Services Lighthouse Laboratory and Alex Alderton, Roberto Amato, Jeffrey Barrett, Sonia Goncalves, Ewan Harrison, David K. Jackson, Ian Johnston, Dominic Kwiatkowski, Cordelia Langford, John Sillitoe on behalf of the Wellcome Sanger Institute COVID-19 Surveillance Team |
| EPI_ISL_3442854, EPI_ISL_3442855 | Health Services Laboratories | Wellcome Sanger Institute for the COVID-19 Genomics UK (COG-UK) Consortium | Health Services Laboratories and Alex Alderton, Roberto Amato, Jeffrey Barrett, Sonia Goncalves, Ewan Harrison, David K. Jackson, Ian Johnston, Dominic Kwiatkowski, Cordelia Langford, John Sillitoe on behalf of the Wellcome Sanger Institute COVID-19 Surveillance Team |
| EPI_ISL_3442856, EPI_ISL_3442857, EPI_ISL_3442858, EPI_ISL_3442859, EPI_ISL_3442860 | Berkshire and Surrey Pathology Services Lighthouse Laboratory | Wellcome Sanger Institute for the COVID-19 Genomics UK (COG-UK) Consortium | Berkshire and Surrey Pathology Services Lighthouse Laboratory and Alex Alderton, Roberto Amato, Jeffrey Barrett, Sonia Goncalves, Ewan Harrison, David K. Jackson, Ian Johnston, Dominic Kwiatkowski, Cordelia Langford, John Sillitoe on behalf of the Wellcome Sanger Institute COVID-19 Surveillance Team |
| EPI_ISL_3442861 | Health Services Laboratories | Wellcome Sanger Institute for the COVID-19 Genomics UK (COG-UK) Consortium | Health Services Laboratories and Alex Alderton, Roberto Amato, Jeffrey Barrett, Sonia Goncalves, Ewan Harrison, David K. Jackson, Ian Johnston, Dominic Kwiatkowski, Cordelia Langford, John Sillitoe on behalf of the Wellcome Sanger Institute COVID-19 Surveillance Team |
| EPI_ISL_3442862 | Berkshire and Surrey Pathology Services Lighthouse Laboratory | Wellcome Sanger Institute for the COVID-19 Genomics UK (COG-UK) | Berkshire and Surrey Pathology Services Lighthouse Laboratory and Alex Alderton, Roberto Amato, Jeffrey Barrett, Sonia Goncalves, Ewan Harrison, David K. Jackson, Ian Johnston, Dominic Kwiatkowski, Cordelia Langford, John Sillitoe on behalf |

| | | | |
|---|---|---|---|
| | | Consortium | of the Wellcome Sanger Institute COVID-19 Surveillance Team |
| EPI_ISL_3442863 | Health Services Laboratories | Wellcome Sanger Institute for the COVID-19 Genomics UK (COG-UK) Consortium | Health Services Laboratories and Alex Alderton, Roberto Amato, Jeffrey Barrett, Sonia Goncalves, Ewan Harrison, David K. Jackson, Ian Johnston, Dominic Kwiatkowski, Cordelia Langford, John Sillitoe on behalf of the Wellcome Sanger Institute COVID-19 Surveillance Team |
| EPI_ISL_3442864, EPI_ISL_3442865, EPI_ISL_3442866 | Berkshire and Surrey Pathology Services Lighthouse Laboratory | Wellcome Sanger Institute for the COVID-19 Genomics UK (COG-UK) Consortium | Berkshire and Surrey Pathology Services Lighthouse Laboratory and Alex Alderton, Roberto Amato, Jeffrey Barrett, Sonia Goncalves, Ewan Harrison, David K. Jackson, Ian Johnston, Dominic Kwiatkowski, Cordelia Langford, John Sillitoe on behalf of the Wellcome Sanger Institute COVID-19 Surveillance Team |
| EPI_ISL_3442867 | Health Services Laboratories | Wellcome Sanger Institute for the COVID-19 Genomics UK (COG-UK) Consortium | Health Services Laboratories and Alex Alderton, Roberto Amato, Jeffrey Barrett, Sonia Goncalves, Ewan Harrison, David K. Jackson, Ian Johnston, Dominic Kwiatkowski, Cordelia Langford, John Sillitoe on behalf of the Wellcome Sanger Institute COVID-19 Surveillance Team |
| EPI_ISL_3442868, EPI_ISL_3442869, EPI_ISL_3442870 | Berkshire and Surrey Pathology Services Lighthouse Laboratory | Wellcome Sanger Institute for the COVID-19 Genomics UK (COG-UK) Consortium | Berkshire and Surrey Pathology Services Lighthouse Laboratory and Alex Alderton, Roberto Amato, Jeffrey Barrett, Sonia Goncalves, Ewan Harrison, David K. Jackson, Ian Johnston, Dominic Kwiatkowski, Cordelia Langford, John Sillitoe on behalf of the Wellcome Sanger Institute COVID-19 Surveillance Team |
| EPI_ISL_3442871, EPI_ISL_3442872, EPI_ISL_3442873, EPI_ISL_3442874 | Health Services Laboratories | Wellcome Sanger Institute for the COVID-19 Genomics UK (COG-UK) Consortium | Health Services Laboratories and Alex Alderton, Roberto Amato, Jeffrey Barrett, Sonia Goncalves, Ewan Harrison, David K. Jackson, Ian Johnston, Dominic Kwiatkowski, Cordelia Langford, John Sillitoe on behalf of the Wellcome Sanger Institute COVID-19 Surveillance Team |
| EPI_ISL_3442875, EPI_ISL_3442876, EPI_ISL_3442877 | Berkshire and Surrey Pathology Services Lighthouse Laboratory | Wellcome Sanger Institute for the COVID-19 Genomics UK (COG-UK) Consortium | Berkshire and Surrey Pathology Services Lighthouse Laboratory and Alex Alderton, Roberto Amato, Jeffrey Barrett, Sonia Goncalves, Ewan Harrison, David K. Jackson, Ian Johnston, Dominic Kwiatkowski, Cordelia Langford, John Sillitoe on behalf of the Wellcome Sanger Institute COVID-19 Surveillance Team |
| EPI_ISL_3442878 | Health Services Laboratories | Wellcome Sanger Institute for the COVID-19 Genomics UK (COG-UK) Consortium | Health Services Laboratories and Alex Alderton, Roberto Amato, Jeffrey Barrett, Sonia Goncalves, Ewan Harrison, David K. Jackson, Ian Johnston, Dominic Kwiatkowski, Cordelia Langford, John Sillitoe on behalf of the Wellcome Sanger Institute COVID-19 Surveillance Team |
| EPI_ISL_3442879 | Berkshire and Surrey Pathology Services Lighthouse Laboratory | Wellcome Sanger Institute for the COVID-19 Genomics UK (COG-UK) Consortium | Berkshire and Surrey Pathology Services Lighthouse Laboratory and Alex Alderton, Roberto Amato, Jeffrey Barrett, Sonia Goncalves, Ewan Harrison, David K. Jackson, Ian Johnston, Dominic Kwiatkowski, Cordelia Langford, John Sillitoe on behalf of the Wellcome Sanger Institute COVID-19 Surveillance Team |
| EPI_ISL_3442880 | Health Services Laboratories | Wellcome Sanger Institute for the COVID-19 Genomics UK (COG-UK) Consortium | Health Services Laboratories and Alex Alderton, Roberto Amato, Jeffrey Barrett, Sonia Goncalves, Ewan Harrison, David K. Jackson, Ian Johnston, Dominic Kwiatkowski, Cordelia Langford, John Sillitoe on behalf of the Wellcome Sanger Institute COVID-19 Surveillance Team |
| EPI_ISL_3442881, EPI_ISL_3442882, EPI_ISL_3442883 | Berkshire and Surrey Pathology Services Lighthouse Laboratory | Wellcome Sanger Institute for the COVID-19 Genomics UK (COG-UK) Consortium | Berkshire and Surrey Pathology Services Lighthouse Laboratory and Alex Alderton, Roberto Amato, Jeffrey Barrett, Sonia Goncalves, Ewan Harrison, David K. Jackson, Ian Johnston, Dominic Kwiatkowski, Cordelia Langford, John Sillitoe on behalf of the Wellcome Sanger Institute COVID-19 Surveillance Team |
| EPI_ISL_3442884, EPI_ISL_3442885 | Health Services Laboratories | Wellcome Sanger Institute for the COVID-19 Genomics UK (COG-UK) Consortium | Health Services Laboratories and Alex Alderton, Roberto Amato, Jeffrey Barrett, Sonia Goncalves, Ewan Harrison, David K. Jackson, Ian Johnston, Dominic Kwiatkowski, Cordelia Langford, John Sillitoe on behalf of the Wellcome Sanger Institute COVID-19 Surveillance Team |
| EPI_ISL_3442886 | Berkshire and Surrey Pathology Services Lighthouse Laboratory | Wellcome Sanger Institute for the COVID-19 Genomics UK (COG-UK) Consortium | Berkshire and Surrey Pathology Services Lighthouse Laboratory and Alex Alderton, Roberto Amato, Jeffrey Barrett, Sonia Goncalves, Ewan Harrison, David K. Jackson, Ian Johnston, Dominic Kwiatkowski, Cordelia Langford, John Sillitoe on behalf of the Wellcome Sanger Institute COVID-19 Surveillance Team |
| EPI_ISL_3442887 | Health Services Laboratories | Wellcome Sanger Institute for the COVID-19 Genomics UK (COG-UK) Consortium | Health Services Laboratories and Alex Alderton, Roberto Amato, Jeffrey Barrett, Sonia Goncalves, Ewan Harrison, David K. Jackson, Ian Johnston, Dominic Kwiatkowski, Cordelia Langford, John Sillitoe on behalf of the Wellcome Sanger Institute COVID-19 Surveillance Team |
| EPI_ISL_3442888 | Berkshire and Surrey Pathology Services Lighthouse Laboratory | Wellcome Sanger Institute for the COVID-19 Genomics UK (COG-UK) Consortium | Berkshire and Surrey Pathology Services Lighthouse Laboratory and Alex Alderton, Roberto Amato, Jeffrey Barrett, Sonia Goncalves, Ewan Harrison, David K. Jackson, Ian Johnston, Dominic Kwiatkowski, Cordelia Langford, John Sillitoe on behalf of the Wellcome Sanger Institute COVID-19 Surveillance Team |
| EPI_ISL_3442889, EPI_ISL_3442890 | Health Services Laboratories | Wellcome Sanger Institute for the COVID-19 Genomics UK (COG-UK) Consortium | Health Services Laboratories and Alex Alderton, Roberto Amato, Jeffrey Barrett, Sonia Goncalves, Ewan Harrison, David K. Jackson, Ian Johnston, Dominic Kwiatkowski, Cordelia Langford, John Sillitoe on behalf of the Wellcome Sanger Institute COVID-19 Surveillance Team |
| EPI_ISL_3442891 | Berkshire and Surrey Pathology Services Lighthouse Laboratory | Wellcome Sanger Institute for the COVID-19 Genomics UK (COG-UK) Consortium | Berkshire and Surrey Pathology Services Lighthouse Laboratory and Alex Alderton, Roberto Amato, Jeffrey Barrett, Sonia Goncalves, Ewan Harrison, David K. Jackson, Ian Johnston, Dominic Kwiatkowski, Cordelia Langford, John Sillitoe on behalf of the Wellcome Sanger Institute COVID-19 Surveillance Team |
| EPI_ISL_3442892 | Health Services Laboratories | Wellcome Sanger Institute for the COVID-19 Genomics UK (COG-UK) Consortium | Health Services Laboratories and Alex Alderton, Roberto Amato, Jeffrey Barrett, Sonia Goncalves, Ewan Harrison, David K. Jackson, Ian Johnston, Dominic Kwiatkowski, Cordelia Langford, John Sillitoe on behalf of the Wellcome Sanger Institute COVID-19 Surveillance Team |
| EPI_ISL_3442893, EPI_ISL_3442894, EPI_ISL_3442895 | Berkshire and Surrey Pathology Services Lighthouse Laboratory | Wellcome Sanger Institute for the COVID-19 Genomics UK (COG-UK) Consortium | Berkshire and Surrey Pathology Services Lighthouse Laboratory and Alex Alderton, Roberto Amato, Jeffrey Barrett, Sonia Goncalves, Ewan Harrison, David K. Jackson, Ian Johnston, Dominic Kwiatkowski, Cordelia Langford, John Sillitoe on behalf of the Wellcome Sanger Institute COVID-19 Surveillance Team |
| EPI_ISL_3442896, EPI_ISL_3442897 | Health Services Laboratories | Wellcome Sanger Institute for the COVID-19 Genomics UK (COG-UK) Consortium | Health Services Laboratories and Alex Alderton, Roberto Amato, Jeffrey Barrett, Sonia Goncalves, Ewan Harrison, David K. Jackson, Ian Johnston, Dominic Kwiatkowski, Cordelia Langford, John Sillitoe on behalf of the Wellcome Sanger Institute COVID-19 Surveillance Team |
| EPI_ISL_3442898, EPI_ISL_3442899, EPI_ISL_3442900, EPI_ISL_3442901, EPI_ISL_3442902, EPI_ISL_3442903, EPI_ISL_3442904 | Berkshire and Surrey Pathology Services Lighthouse Laboratory | Wellcome Sanger Institute for the COVID-19 Genomics UK (COG-UK) Consortium | Berkshire and Surrey Pathology Services Lighthouse Laboratory and Alex Alderton, Roberto Amato, Jeffrey Barrett, Sonia Goncalves, Ewan Harrison, David K. Jackson, Ian Johnston, Dominic Kwiatkowski, Cordelia Langford, John Sillitoe on behalf of the Wellcome Sanger Institute COVID-19 Surveillance Team |
| EPI_ISL_3442905 | Health Services Laboratories | Wellcome Sanger Institute for the COVID-19 Genomics UK (COG-UK) Consortium | Health Services Laboratories and Alex Alderton, Roberto Amato, Jeffrey Barrett, Sonia Goncalves, Ewan Harrison, David K. Jackson, Ian Johnston, Dominic Kwiatkowski, Cordelia Langford, John Sillitoe on behalf of the Wellcome Sanger Institute COVID-19 Surveillance Team |
| EPI_ISL_3442906, EPI_ISL_3442907, EPI_ISL_3442908, EPI_ISL_3442909 | Berkshire and Surrey Pathology Services Lighthouse Laboratory | Wellcome Sanger Institute for the COVID-19 Genomics UK (COG-UK) Consortium | Berkshire and Surrey Pathology Services Lighthouse Laboratory and Alex Alderton, Roberto Amato, Jeffrey Barrett, Sonia Goncalves, Ewan Harrison, David K. Jackson, Ian Johnston, Dominic Kwiatkowski, Cordelia Langford, John Sillitoe on behalf of the Wellcome Sanger Institute COVID-19 Surveillance Team |
| EPI_ISL_3442910, EPI_ISL_3442911 | Health Services Laboratories | Wellcome Sanger Institute for the COVID-19 Genomics UK (COG-UK) Consortium | Health Services Laboratories and Alex Alderton, Roberto Amato, Jeffrey Barrett, Sonia Goncalves, Ewan Harrison, David K. Jackson, Ian Johnston, Dominic Kwiatkowski, Cordelia Langford, John Sillitoe on behalf of the Wellcome Sanger Institute COVID-19 Surveillance Team |
| EPI_ISL_3442912 | Berkshire and Surrey Pathology Services Lighthouse Laboratory | Wellcome Sanger Institute for the COVID-19 Genomics UK (COG-UK) Consortium | Berkshire and Surrey Pathology Services Lighthouse Laboratory and Alex Alderton, Roberto Amato, Jeffrey Barrett, Sonia Goncalves, Ewan Harrison, David K. Jackson, Ian Johnston, Dominic Kwiatkowski, Cordelia Langford, John Sillitoe on behalf of the Wellcome Sanger Institute COVID-19 Surveillance Team |
| EPI_ISL_3442913 | Health Services Laboratories | Wellcome Sanger Institute for the COVID-19 Genomics UK (COG-UK) Consortium | Health Services Laboratories and Alex Alderton, Roberto Amato, Jeffrey Barrett, Sonia Goncalves, Ewan Harrison, David K. Jackson, Ian Johnston, Dominic Kwiatkowski, Cordelia Langford, John Sillitoe on behalf of the Wellcome Sanger Institute COVID-19 Surveillance Team |
| EPI_ISL_3442914, EPI_ISL_3442915, EPI_ISL_3442916 | Berkshire and Surrey Pathology Services Lighthouse Laboratory | Wellcome Sanger Institute for the COVID-19 Genomics UK (COG-UK) Consortium | Berkshire and Surrey Pathology Services Lighthouse Laboratory and Alex Alderton, Roberto Amato, Jeffrey Barrett, Sonia Goncalves, Ewan Harrison, David K. Jackson, Ian Johnston, Dominic Kwiatkowski, Cordelia Langford, John Sillitoe on behalf of the Wellcome Sanger Institute COVID-19 Surveillance Team |
| EPI_ISL_3442917, EPI_ISL_3442918 | Health Services Laboratories | Wellcome Sanger Institute for the COVID-19 Genomics UK (COG-UK) Consortium | Health Services Laboratories and Alex Alderton, Roberto Amato, Jeffrey Barrett, Sonia Goncalves, Ewan Harrison, David K. Jackson, Ian Johnston, Dominic Kwiatkowski, Cordelia Langford, John Sillitoe on behalf of the Wellcome Sanger Institute COVID-19 Surveillance Team |
| EPI_ISL_3442919, EPI_ISL_3442920, EPI_ISL_3442921, EPI_ISL_3442922 | Berkshire and Surrey Pathology Services Lighthouse Laboratory | Wellcome Sanger Institute for the COVID-19 Genomics UK (COG-UK) Consortium | Berkshire and Surrey Pathology Services Lighthouse Laboratory and Alex Alderton, Roberto Amato, Jeffrey Barrett, Sonia Goncalves, Ewan Harrison, David K. Jackson, Ian Johnston, Dominic Kwiatkowski, Cordelia Langford, John Sillitoe on behalf of the Wellcome Sanger Institute COVID-19 Surveillance Team |
| EPI_ISL_3442923 | Health Services Laboratories | Wellcome Sanger Institute for the COVID-19 Genomics UK (COG-UK) Consortium | Health Services Laboratories and Alex Alderton, Roberto Amato, Jeffrey Barrett, Sonia Goncalves, Ewan Harrison, David K. Jackson, Ian Johnston, Dominic Kwiatkowski, Cordelia Langford, John Sillitoe on behalf of the Wellcome Sanger Institute COVID-19 Surveillance Team |
| EPI_ISL_3442924, EPI_ISL_3442925, EPI_ISL_3442926, EPI_ISL_3442927 | Berkshire and Surrey Pathology Services Lighthouse Laboratory | Wellcome Sanger Institute for the COVID-19 Genomics UK (COG-UK) Consortium | Berkshire and Surrey Pathology Services Lighthouse Laboratory and Alex Alderton, Roberto Amato, Jeffrey Barrett, Sonia Goncalves, Ewan Harrison, David K. Jackson, Ian Johnston, Dominic Kwiatkowski, Cordelia Langford, John Sillitoe on behalf of the Wellcome Sanger Institute COVID-19 Surveillance Team |
| EPI_ISL_3442928, EPI_ISL_3442929 | Health Services Laboratories | Wellcome Sanger Institute for the COVID-19 Genomics UK (COG-UK) Consortium | Health Services Laboratories and Alex Alderton, Roberto Amato, Jeffrey Barrett, Sonia Goncalves, Ewan Harrison, David K. Jackson, Ian Johnston, Dominic Kwiatkowski, Cordelia Langford, John Sillitoe on behalf of the Wellcome Sanger Institute COVID-19 Surveillance Team |
| EPI_ISL_3442930, EPI_ISL_3442931 | Berkshire and Surrey Pathology Services Lighthouse Laboratory | Wellcome Sanger Institute for the COVID-19 Genomics UK (COG-UK) Consortium | Berkshire and Surrey Pathology Services Lighthouse Laboratory and Alex Alderton, Roberto Amato, Jeffrey Barrett, Sonia Goncalves, Ewan Harrison, David K. Jackson, Ian Johnston, Dominic Kwiatkowski, Cordelia Langford, John Sillitoe on behalf of the Wellcome Sanger Institute COVID-19 Surveillance Team |
| EPI_ISL_3442932 | Health Services Laboratories | Wellcome Sanger Institute for the COVID-19 Genomics UK (COG-UK) Consortium | Health Services Laboratories and Alex Alderton, Roberto Amato, Jeffrey Barrett, Sonia Goncalves, Ewan Harrison, David K. Jackson, Ian Johnston, Dominic Kwiatkowski, Cordelia Langford, John Sillitoe on behalf of the Wellcome Sanger Institute COVID-19 Surveillance Team |
| EPI_ISL_3442933, EPI_ISL_3442934, EPI_ISL_3442935, EPI_ISL_3442936 | Berkshire and Surrey Pathology Services Lighthouse Laboratory | Wellcome Sanger Institute for the COVID-19 Genomics UK (COG-UK) Consortium | Berkshire and Surrey Pathology Services Lighthouse Laboratory and Alex Alderton, Roberto Amato, Jeffrey Barrett, Sonia Goncalves, Ewan Harrison, David K. Jackson, Ian Johnston, Dominic Kwiatkowski, Cordelia Langford, John Sillitoe on behalf of the Wellcome Sanger Institute COVID-19 Surveillance Team |
| EPI_ISL_3442937, EPI_ISL_3442938, EPI_ISL_3442939 | Health Services Laboratories | Wellcome Sanger Institute for the COVID-19 Genomics UK (COG-UK) Consortium | Health Services Laboratories and Alex Alderton, Roberto Amato, Jeffrey Barrett, Sonia Goncalves, Ewan Harrison, David K. Jackson, Ian Johnston, Dominic Kwiatkowski, Cordelia Langford, John Sillitoe on behalf of the Wellcome Sanger Institute COVID-19 Surveillance Team |
| EPI_ISL_3442940, | Berkshire and Surrey Pathology Services Lighthouse Laboratory | Wellcome Sanger Institute for the COVID-19 Genomics UK (COG-UK) | Berkshire and Surrey Pathology Services Lighthouse Laboratory and Alex Alderton, Roberto Amato, Jeffrey Barrett, Sonia Goncalves, Ewan Harrison, David K. Jackson, Ian Johnston, Dominic Kwiatkowski, Cordelia Langford, John Sillitoe on behalf |

| | | | |
|---|---|---|---|
| EPI_ISL_3442941, EPI_ISL_3442942 | | Consortium | of the Wellcome Sanger Institute COVID-19 Surveillance Team |
| EPI_ISL_3442943 | Health Services Laboratories | Wellcome Sanger Institute for the COVID-19 Genomics UK (COG-UK) Consortium | Health Services Laboratories and Alex Alderton, Roberto Amato, Jeffrey Barrett, Sonia Goncalves, Ewan Harrison, David K. Jackson, Ian Johnston, Dominic Kwiatkowski, Cordelia Langford, John Sillitoe on behalf of the Wellcome Sanger Institute COVID-19 Surveillance Team |
| EPI_ISL_3442944 | Berkshire and Surrey Pathology Services Lighthouse Laboratory | Wellcome Sanger Institute for the COVID-19 Genomics UK (COG-UK) Consortium | Berkshire and Surrey Pathology Services Lighthouse Laboratory and Alex Alderton, Roberto Amato, Jeffrey Barrett, Sonia Goncalves, Ewan Harrison, David K. Jackson, Ian Johnston, Dominic Kwiatkowski, Cordelia Langford, John Sillitoe on behalf of the Wellcome Sanger Institute COVID-19 Surveillance Team |
| EPI_ISL_3442945 | Health Services Laboratories | Wellcome Sanger Institute for the COVID-19 Genomics UK (COG-UK) Consortium | Health Services Laboratories and Alex Alderton, Roberto Amato, Jeffrey Barrett, Sonia Goncalves, Ewan Harrison, David K. Jackson, Ian Johnston, Dominic Kwiatkowski, Cordelia Langford, John Sillitoe on behalf of the Wellcome Sanger Institute COVID-19 Surveillance Team |
| EPI_ISL_3442946, EPI_ISL_3442947, EPI_ISL_3442948, EPI_ISL_3442949 | Berkshire and Surrey Pathology Services Lighthouse Laboratory | Wellcome Sanger Institute for the COVID-19 Genomics UK (COG-UK) Consortium | Berkshire and Surrey Pathology Services Lighthouse Laboratory and Alex Alderton, Roberto Amato, Jeffrey Barrett, Sonia Goncalves, Ewan Harrison, David K. Jackson, Ian Johnston, Dominic Kwiatkowski, Cordelia Langford, John Sillitoe on behalf of the Wellcome Sanger Institute COVID-19 Surveillance Team |
| EPI_ISL_3442950 | Health Services Laboratories | Wellcome Sanger Institute for the COVID-19 Genomics UK (COG-UK) Consortium | Health Services Laboratories and Alex Alderton, Roberto Amato, Jeffrey Barrett, Sonia Goncalves, Ewan Harrison, David K. Jackson, Ian Johnston, Dominic Kwiatkowski, Cordelia Langford, John Sillitoe on behalf of the Wellcome Sanger Institute COVID-19 Surveillance Team |
| EPI_ISL_3442951, EPI_ISL_3442952 | Berkshire and Surrey Pathology Services Lighthouse Laboratory | Wellcome Sanger Institute for the COVID-19 Genomics UK (COG-UK) Consortium | Berkshire and Surrey Pathology Services Lighthouse Laboratory and Alex Alderton, Roberto Amato, Jeffrey Barrett, Sonia Goncalves, Ewan Harrison, David K. Jackson, Ian Johnston, Dominic Kwiatkowski, Cordelia Langford, John Sillitoe on behalf of the Wellcome Sanger Institute COVID-19 Surveillance Team |
| EPI_ISL_3442953 | Health Services Laboratories | Wellcome Sanger Institute for the COVID-19 Genomics UK (COG-UK) Consortium | Health Services Laboratories and Alex Alderton, Roberto Amato, Jeffrey Barrett, Sonia Goncalves, Ewan Harrison, David K. Jackson, Ian Johnston, Dominic Kwiatkowski, Cordelia Langford, John Sillitoe on behalf of the Wellcome Sanger Institute COVID-19 Surveillance Team |
| EPI_ISL_3442954, EPI_ISL_3442955, EPI_ISL_3442956, EPI_ISL_3442957 | Berkshire and Surrey Pathology Services Lighthouse Laboratory | Wellcome Sanger Institute for the COVID-19 Genomics UK (COG-UK) Consortium | Berkshire and Surrey Pathology Services Lighthouse Laboratory and Alex Alderton, Roberto Amato, Jeffrey Barrett, Sonia Goncalves, Ewan Harrison, David K. Jackson, Ian Johnston, Dominic Kwiatkowski, Cordelia Langford, John Sillitoe on behalf of the Wellcome Sanger Institute COVID-19 Surveillance Team |
| EPI_ISL_3442958 | Health Services Laboratories | Wellcome Sanger Institute for the COVID-19 Genomics UK (COG-UK) Consortium | Health Services Laboratories and Alex Alderton, Roberto Amato, Jeffrey Barrett, Sonia Goncalves, Ewan Harrison, David K. Jackson, Ian Johnston, Dominic Kwiatkowski, Cordelia Langford, John Sillitoe on behalf of the Wellcome Sanger Institute COVID-19 Surveillance Team |
| EPI_ISL_3442959, EPI_ISL_3442960, EPI_ISL_3442961, EPI_ISL_3442962, EPI_ISL_3442963 | Berkshire and Surrey Pathology Services Lighthouse Laboratory | Wellcome Sanger Institute for the COVID-19 Genomics UK (COG-UK) Consortium | Berkshire and Surrey Pathology Services Lighthouse Laboratory and Alex Alderton, Roberto Amato, Jeffrey Barrett, Sonia Goncalves, Ewan Harrison, David K. Jackson, Ian Johnston, Dominic Kwiatkowski, Cordelia Langford, John Sillitoe on behalf of the Wellcome Sanger Institute COVID-19 Surveillance Team |
| EPI_ISL_3442964 | Health Services Laboratories | Wellcome Sanger Institute for the COVID-19 Genomics UK (COG-UK) Consortium | Health Services Laboratories and Alex Alderton, Roberto Amato, Jeffrey Barrett, Sonia Goncalves, Ewan Harrison, David K. Jackson, Ian Johnston, Dominic Kwiatkowski, Cordelia Langford, John Sillitoe on behalf of the Wellcome Sanger Institute COVID-19 Surveillance Team |
| EPI_ISL_3442965, EPI_ISL_3442966, EPI_ISL_3442967, EPI_ISL_3442968, EPI_ISL_3442969, EPI_ISL_3442970, EPI_ISL_3442971, EPI_ISL_3442972 | Berkshire and Surrey Pathology Services Lighthouse Laboratory | Wellcome Sanger Institute for the COVID-19 Genomics UK (COG-UK) Consortium | Berkshire and Surrey Pathology Services Lighthouse Laboratory and Alex Alderton, Roberto Amato, Jeffrey Barrett, Sonia Goncalves, Ewan Harrison, David K. Jackson, Ian Johnston, Dominic Kwiatkowski, Cordelia Langford, John Sillitoe on behalf of the Wellcome Sanger Institute COVID-19 Surveillance Team |
| EPI_ISL_3442973, EPI_ISL_3442974, EPI_ISL_3442975 | Health Services Laboratories | Wellcome Sanger Institute for the COVID-19 Genomics UK (COG-UK) Consortium | Health Services Laboratories and Alex Alderton, Roberto Amato, Jeffrey Barrett, Sonia Goncalves, Ewan Harrison, David K. Jackson, Ian Johnston, Dominic Kwiatkowski, Cordelia Langford, John Sillitoe on behalf of the Wellcome Sanger Institute COVID-19 Surveillance Team |
| EPI_ISL_3442976, EPI_ISL_3442977 | Berkshire and Surrey Pathology Services Lighthouse Laboratory | Wellcome Sanger Institute for the COVID-19 Genomics UK (COG-UK) Consortium | Berkshire and Surrey Pathology Services Lighthouse Laboratory and Alex Alderton, Roberto Amato, Jeffrey Barrett, Sonia Goncalves, Ewan Harrison, David K. Jackson, Ian Johnston, Dominic Kwiatkowski, Cordelia Langford, John Sillitoe on behalf of the Wellcome Sanger Institute COVID-19 Surveillance Team |
| EPI_ISL_3442978, EPI_ISL_3442979 | Health Services Laboratories | Wellcome Sanger Institute for the COVID-19 Genomics UK (COG-UK) Consortium | Health Services Laboratories and Alex Alderton, Roberto Amato, Jeffrey Barrett, Sonia Goncalves, Ewan Harrison, David K. Jackson, Ian Johnston, Dominic Kwiatkowski, Cordelia Langford, John Sillitoe on behalf of the Wellcome Sanger Institute COVID-19 Surveillance Team |
| EPI_ISL_3442980 | Berkshire and Surrey Pathology Services Lighthouse Laboratory | Wellcome Sanger Institute for the COVID-19 Genomics UK (COG-UK) Consortium | Berkshire and Surrey Pathology Services Lighthouse Laboratory and Alex Alderton, Roberto Amato, Jeffrey Barrett, Sonia Goncalves, Ewan Harrison, David K. Jackson, Ian Johnston, Dominic Kwiatkowski, Cordelia Langford, John Sillitoe on behalf of the Wellcome Sanger Institute COVID-19 Surveillance Team |
| EPI_ISL_3442981 | Health Services Laboratories | Wellcome Sanger Institute for the COVID-19 Genomics UK (COG-UK) Consortium | Health Services Laboratories and Alex Alderton, Roberto Amato, Jeffrey Barrett, Sonia Goncalves, Ewan Harrison, David K. Jackson, Ian Johnston, Dominic Kwiatkowski, Cordelia Langford, John Sillitoe on behalf of the Wellcome Sanger Institute COVID-19 Surveillance Team |
| EPI_ISL_3442982, EPI_ISL_3442983, EPI_ISL_3442984, EPI_ISL_3442985, EPI_ISL_3442986 | Berkshire and Surrey Pathology Services Lighthouse Laboratory | Wellcome Sanger Institute for the COVID-19 Genomics UK (COG-UK) Consortium | Berkshire and Surrey Pathology Services Lighthouse Laboratory and Alex Alderton, Roberto Amato, Jeffrey Barrett, Sonia Goncalves, Ewan Harrison, David K. Jackson, Ian Johnston, Dominic Kwiatkowski, Cordelia Langford, John Sillitoe on behalf of the Wellcome Sanger Institute COVID-19 Surveillance Team |
| EPI_ISL_3442987 | Health Services Laboratories | Wellcome Sanger Institute for the COVID-19 Genomics UK (COG-UK) Consortium | Health Services Laboratories and Alex Alderton, Roberto Amato, Jeffrey Barrett, Sonia Goncalves, Ewan Harrison, David K. Jackson, Ian Johnston, Dominic Kwiatkowski, Cordelia Langford, John Sillitoe on behalf of the Wellcome Sanger Institute COVID-19 Surveillance Team |
| EPI_ISL_3442988, EPI_ISL_3442989 | Berkshire and Surrey Pathology Services Lighthouse Laboratory | Wellcome Sanger Institute for the COVID-19 Genomics UK (COG-UK) Consortium | Berkshire and Surrey Pathology Services Lighthouse Laboratory and Alex Alderton, Roberto Amato, Jeffrey Barrett, Sonia Goncalves, Ewan Harrison, David K. Jackson, Ian Johnston, Dominic Kwiatkowski, Cordelia Langford, John Sillitoe on behalf of the Wellcome Sanger Institute COVID-19 Surveillance Team |
| EPI_ISL_3442990 | Health Services Laboratories | Wellcome Sanger Institute for the COVID-19 Genomics UK (COG-UK) Consortium | Health Services Laboratories and Alex Alderton, Roberto Amato, Jeffrey Barrett, Sonia Goncalves, Ewan Harrison, David K. Jackson, Ian Johnston, Dominic Kwiatkowski, Cordelia Langford, John Sillitoe on behalf of the Wellcome Sanger Institute COVID-19 Surveillance Team |
| EPI_ISL_3442991, EPI_ISL_3442992, EPI_ISL_3442993 | Berkshire and Surrey Pathology Services Lighthouse Laboratory | Wellcome Sanger Institute for the COVID-19 Genomics UK (COG-UK) Consortium | Berkshire and Surrey Pathology Services Lighthouse Laboratory and Alex Alderton, Roberto Amato, Jeffrey Barrett, Sonia Goncalves, Ewan Harrison, David K. Jackson, Ian Johnston, Dominic Kwiatkowski, Cordelia Langford, John Sillitoe on behalf of the Wellcome Sanger Institute COVID-19 Surveillance Team |
| EPI_ISL_3442994, EPI_ISL_3442995 | Health Services Laboratories | Wellcome Sanger Institute for the COVID-19 Genomics UK (COG-UK) Consortium | Health Services Laboratories and Alex Alderton, Roberto Amato, Jeffrey Barrett, Sonia Goncalves, Ewan Harrison, David K. Jackson, Ian Johnston, Dominic Kwiatkowski, Cordelia Langford, John Sillitoe on behalf of the Wellcome Sanger Institute COVID-19 Surveillance Team |
| EPI_ISL_3442996, EPI_ISL_3442997, EPI_ISL_3442998, EPI_ISL_3442999 | Berkshire and Surrey Pathology Services Lighthouse Laboratory | Wellcome Sanger Institute for the COVID-19 Genomics UK (COG-UK) Consortium | Berkshire and Surrey Pathology Services Lighthouse Laboratory and Alex Alderton, Roberto Amato, Jeffrey Barrett, Sonia Goncalves, Ewan Harrison, David K. Jackson, Ian Johnston, Dominic Kwiatkowski, Cordelia Langford, John Sillitoe on behalf of the Wellcome Sanger Institute COVID-19 Surveillance Team |
| EPI_ISL_3443000, EPI_ISL_3443001 | Health Services Laboratories | Wellcome Sanger Institute for the COVID-19 Genomics UK (COG-UK) Consortium | Health Services Laboratories and Alex Alderton, Roberto Amato, Jeffrey Barrett, Sonia Goncalves, Ewan Harrison, David K. Jackson, Ian Johnston, Dominic Kwiatkowski, Cordelia Langford, John Sillitoe on behalf of the Wellcome Sanger Institute COVID-19 Surveillance Team |
| EPI_ISL_3443002, EPI_ISL_3443003 | Berkshire and Surrey Pathology Services Lighthouse Laboratory | Wellcome Sanger Institute for the COVID-19 Genomics UK (COG-UK) Consortium | Berkshire and Surrey Pathology Services Lighthouse Laboratory and Alex Alderton, Roberto Amato, Jeffrey Barrett, Sonia Goncalves, Ewan Harrison, David K. Jackson, Ian Johnston, Dominic Kwiatkowski, Cordelia Langford, John Sillitoe on behalf of the Wellcome Sanger Institute COVID-19 Surveillance Team |
| EPI_ISL_3443004 | Health Services Laboratories | Wellcome Sanger Institute for the COVID-19 Genomics UK (COG-UK) Consortium | Health Services Laboratories and Alex Alderton, Roberto Amato, Jeffrey Barrett, Sonia Goncalves, Ewan Harrison, David K. Jackson, Ian Johnston, Dominic Kwiatkowski, Cordelia Langford, John Sillitoe on behalf of the Wellcome Sanger Institute COVID-19 Surveillance Team |
| EPI_ISL_3443005, EPI_ISL_3443006, EPI_ISL_3443007, EPI_ISL_3443008, EPI_ISL_3443009, EPI_ISL_3443010, EPI_ISL_3443011, EPI_ISL_3443012, EPI_ISL_3443013, EPI_ISL_3443014, EPI_ISL_3443015, EPI_ISL_3443016, EPI_ISL_3443017, EPI_ISL_3443018, EPI_ISL_3443019, EPI_ISL_3443020, EPI_ISL_3443021, EPI_ISL_3443022, EPI_ISL_3443023, EPI_ISL_3443024, EPI_ISL_3443025, EPI_ISL_3443026, EPI_ISL_3443027, EPI_ISL_3443028, EPI_ISL_3443029, EPI_ISL_3443030, EPI_ISL_3443031, EPI_ISL_3443032, EPI_ISL_3443033, EPI_ISL_3443034, EPI_ISL_3443035, EPI_ISL_3443036, EPI_ISL_3443037, EPI_ISL_3443038, EPI_ISL_3443039, EPI_ISL_3443040, EPI_ISL_3443041, EPI_ISL_3443042, EPI_ISL_3443043, EPI_ISL_3443044, EPI_ISL_3443045, EPI_ISL_3443046, EPI_ISL_3443047, EPI_ISL_3443048, EPI_ISL_3443049, EPI_ISL_3443050, EPI_ISL_3443051, EPI_ISL_3443052, EPI_ISL_3443053, EPI_ISL_3443054, EPI_ISL_3443055, EPI_ISL_3443056, EPI_ISL_3443057, EPI_ISL_3443058, EPI_ISL_3443059, EPI_ISL_3443060, EPI_ISL_3443061, EPI_ISL_3443062, EPI_ISL_3443063, EPI_ISL_3443064, EPI_ISL_3443065, EPI_ISL_3443066, EPI_ISL_3443067, EPI_ISL_3443068, EPI_ISL_3443069, EPI_ISL_3443070, EPI_ISL_3443071, EPI_ISL_3443072, EPI_ISL_3443073, EPI_ISL_3443074, EPI_ISL_3443075, EPI_ISL_3443076, EPI_ISL_3443078, EPI_ISL_3443079, EPI_ISL_3443080, EPI_ISL_3443081 | | | |
| see above | Berkshire and Surrey Pathology Services Lighthouse Laboratory | Wellcome Sanger Institute for the COVID-19 Genomics UK (COG-UK) Consortium | Berkshire and Surrey Pathology Services Lighthouse Laboratory and Alex Alderton, Roberto Amato, Jeffrey Barrett, Sonia Goncalves, Ewan Harrison, David K. Jackson, Ian Johnston, Dominic Kwiatkowski, Cordelia Langford, John Sillitoe on behalf of the Wellcome Sanger Institute COVID-19 Surveillance Team |
| EPI_ISL_3443082 | Lighthouse Lab in Glasgow | Wellcome Sanger Institute for the COVID-19 Genomics UK (COG-UK) Consortium | Harper VanSteenhouse, Yumi Kasai, David Gray, Carol Clugston, Anna Dominiczak and Alex Alderton, Roberto Amato, Jeffrey Barrett, Sonia Goncalves, Ewan Harrison, David K. Jackson, Ian Johnston, Dominic Kwiatkowski, Cordelia Langford, John Sillitoe on behalf of the Wellcome Sanger Institute COVID-19 Surveillance Team |
| EPI_ISL_3443083 | Lighthouse Lab in Alderley Park | Wellcome Sanger Institute for the COVID-19 Genomics UK (COG-UK) Consortium | Jacquelyn Wynn, Mairead Hyland, The Lighthouse Lab in Alderley Park and Alex Alderton, Roberto Amato, Jeffrey Barrett, Sonia Goncalves, Ewan Harrison, David K. Jackson, Ian Johnston, Dominic Kwiatkowski, Cordelia Langford, John Sillitoe on behalf of the Wellcome Sanger Institute COVID-19 Surveillance Team |
| EPI_ISL_3443084, EPI_ISL_3443085 | Lighthouse Lab in Glasgow | Wellcome Sanger Institute for the COVID-19 Genomics UK (COG-UK) Consortium | Harper VanSteenhouse, Yumi Kasai, David Gray, Carol Clugston, Anna Dominiczak and Alex Alderton, Roberto Amato, Jeffrey Barrett, Sonia Goncalves, Ewan Harrison, David K. Jackson, Ian Johnston, Dominic Kwiatkowski, Cordelia Langford, John Sillitoe on behalf of the Wellcome Sanger Institute COVID-19 Surveillance Team |
| EPI_ISL_3443086 | Lighthouse Lab in Alderley Park | Wellcome Sanger Institute for the COVID-19 Genomics UK (COG-UK) Consortium | Jacquelyn Wynn, Mairead Hyland, The Lighthouse Lab in Alderley Park and Alex Alderton, Roberto Amato, Jeffrey Barrett, Sonia Goncalves, Ewan Harrison, David K. Jackson, Ian Johnston, Dominic Kwiatkowski, Cordelia Langford, John Sillitoe on behalf of the Wellcome Sanger Institute COVID-19 Surveillance Team |
| EPI_ISL_3443087, EPI_ISL_3443088, EPI_ISL_3443089 | Lighthouse Lab in Glasgow | Wellcome Sanger Institute for the COVID-19 Genomics UK (COG-UK) Consortium | Harper VanSteenhouse, Yumi Kasai, David Gray, Carol Clugston, Anna Dominiczak and Alex Alderton, Roberto Amato, Jeffrey Barrett, Sonia Goncalves, Ewan Harrison, David K. Jackson, Ian Johnston, Dominic Kwiatkowski, Cordelia Langford, John Sillitoe on behalf of the Wellcome Sanger Institute COVID-19 Surveillance Team |
| EPI_ISL_3443090 | Lighthouse Lab in Alderley Park | Wellcome Sanger Institute for the COVID-19 Genomics UK (COG-UK) | Jacquelyn Wynn, Mairead Hyland, The Lighthouse Lab in Alderley Park and Alex Alderton, Roberto Amato, Jeffrey Barrett, Sonia Goncalves, Ewan Harrison, David K. Jackson, Ian Johnston, Dominic Kwiatkowski, Cordelia Langford, John Sillitoe on |

| | | | |
|---|---|---|---|
| | | Consortium | behalf of the Wellcome Sanger Institute COVID-19 Surveillance Team |
| EPI_ISL_3443091, EPI_ISL_3443092, EPI_ISL_3443093, EPI_ISL_3443094, EPI_ISL_3443095, EPI_ISL_3443096, EPI_ISL_3443097, EPI_ISL_3443098 | Lighthouse Lab in Glasgow | Wellcome Sanger Institute for the COVID-19 Genomics UK (COG-UK) Consortium | Harper VanSteenhouse, Yumi Kasai, David Gray, Carol Clugston, Anna Dominiczak and Alex Alderton, Roberto Amato, Jeffrey Barrett, Sonia Goncalves, Ewan Harrison, David K. Jackson, Ian Johnston, Dominic Kwiatkowski, Cordelia Langford, John Sillitoe on behalf of the Wellcome Sanger Institute COVID-19 Surveillance Team |
| EPI_ISL_3443099 | Lighthouse Lab in Alderley Park | Wellcome Sanger Institute for the COVID-19 Genomics UK (COG-UK) Consortium | Jacquelyn Wynn, Mairead Hyland, The Lighthouse Lab in Alderley Park and Alex Alderton, Roberto Amato, Jeffrey Barrett, Sonia Goncalves, Ewan Harrison, David K. Jackson, Ian Johnston, Dominic Kwiatkowski, Cordelia Langford, John Sillitoe on behalf of the Wellcome Sanger Institute COVID-19 Surveillance Team |
| EPI_ISL_3443100, EPI_ISL_3443101 | Lighthouse Lab in Glasgow | Wellcome Sanger Institute for the COVID-19 Genomics UK (COG-UK) Consortium | Harper VanSteenhouse, Yumi Kasai, David Gray, Carol Clugston, Anna Dominiczak and Alex Alderton, Roberto Amato, Jeffrey Barrett, Sonia Goncalves, Ewan Harrison, David K. Jackson, Ian Johnston, Dominic Kwiatkowski, Cordelia Langford, John Sillitoe on behalf of the Wellcome Sanger Institute COVID-19 Surveillance Team |
| EPI_ISL_3443102 | Lighthouse Lab in Alderley Park | Wellcome Sanger Institute for the COVID-19 Genomics UK (COG-UK) Consortium | Jacquelyn Wynn, Mairead Hyland, The Lighthouse Lab in Alderley Park and Alex Alderton, Roberto Amato, Jeffrey Barrett, Sonia Goncalves, Ewan Harrison, David K. Jackson, Ian Johnston, Dominic Kwiatkowski, Cordelia Langford, John Sillitoe on behalf of the Wellcome Sanger Institute COVID-19 Surveillance Team |
| EPI_ISL_3443103, EPI_ISL_3443104 | Lighthouse Lab in Glasgow | Wellcome Sanger Institute for the COVID-19 Genomics UK (COG-UK) Consortium | Harper VanSteenhouse, Yumi Kasai, David Gray, Carol Clugston, Anna Dominiczak and Alex Alderton, Roberto Amato, Jeffrey Barrett, Sonia Goncalves, Ewan Harrison, David K. Jackson, Ian Johnston, Dominic Kwiatkowski, Cordelia Langford, John Sillitoe on behalf of the Wellcome Sanger Institute COVID-19 Surveillance Team |
| EPI_ISL_3443105 | Lighthouse Lab in Alderley Park | Wellcome Sanger Institute for the COVID-19 Genomics UK (COG-UK) Consortium | Jacquelyn Wynn, Mairead Hyland, The Lighthouse Lab in Alderley Park and Alex Alderton, Roberto Amato, Jeffrey Barrett, Sonia Goncalves, Ewan Harrison, David K. Jackson, Ian Johnston, Dominic Kwiatkowski, Cordelia Langford, John Sillitoe on behalf of the Wellcome Sanger Institute COVID-19 Surveillance Team |
| EPI_ISL_3443106 | Lighthouse Lab in Glasgow | Wellcome Sanger Institute for the COVID-19 Genomics UK (COG-UK) Consortium | Harper VanSteenhouse, Yumi Kasai, David Gray, Carol Clugston, Anna Dominiczak and Alex Alderton, Roberto Amato, Jeffrey Barrett, Sonia Goncalves, Ewan Harrison, David K. Jackson, Ian Johnston, Dominic Kwiatkowski, Cordelia Langford, John Sillitoe on behalf of the Wellcome Sanger Institute COVID-19 Surveillance Team |
| EPI_ISL_3443107 | Lighthouse Lab in Alderley Park | Wellcome Sanger Institute for the COVID-19 Genomics UK (COG-UK) Consortium | Jacquelyn Wynn, Mairead Hyland, The Lighthouse Lab in Alderley Park and Alex Alderton, Roberto Amato, Jeffrey Barrett, Sonia Goncalves, Ewan Harrison, David K. Jackson, Ian Johnston, Dominic Kwiatkowski, Cordelia Langford, John Sillitoe on behalf of the Wellcome Sanger Institute COVID-19 Surveillance Team |
| EPI_ISL_3443108, EPI_ISL_3443109, EPI_ISL_3443110 | Lighthouse Lab in Glasgow | Wellcome Sanger Institute for the COVID-19 Genomics UK (COG-UK) Consortium | Harper VanSteenhouse, Yumi Kasai, David Gray, Carol Clugston, Anna Dominiczak and Alex Alderton, Roberto Amato, Jeffrey Barrett, Sonia Goncalves, Ewan Harrison, David K. Jackson, Ian Johnston, Dominic Kwiatkowski, Cordelia Langford, John Sillitoe on behalf of the Wellcome Sanger Institute COVID-19 Surveillance Team |
| EPI_ISL_3443111 | Lighthouse Lab in Alderley Park | Wellcome Sanger Institute for the COVID-19 Genomics UK (COG-UK) Consortium | Jacquelyn Wynn, Mairead Hyland, The Lighthouse Lab in Alderley Park and Alex Alderton, Roberto Amato, Jeffrey Barrett, Sonia Goncalves, Ewan Harrison, David K. Jackson, Ian Johnston, Dominic Kwiatkowski, Cordelia Langford, John Sillitoe on behalf of the Wellcome Sanger Institute COVID-19 Surveillance Team |
| EPI_ISL_3443112, EPI_ISL_3443113 | Lighthouse Lab in Glasgow | Wellcome Sanger Institute for the COVID-19 Genomics UK (COG-UK) Consortium | Harper VanSteenhouse, Yumi Kasai, David Gray, Carol Clugston, Anna Dominiczak and Alex Alderton, Roberto Amato, Jeffrey Barrett, Sonia Goncalves, Ewan Harrison, David K. Jackson, Ian Johnston, Dominic Kwiatkowski, Cordelia Langford, John Sillitoe on behalf of the Wellcome Sanger Institute COVID-19 Surveillance Team |
| EPI_ISL_3443114, EPI_ISL_3443115, EPI_ISL_3443116, EPI_ISL_3443117 | Lighthouse Lab in Alderley Park | Wellcome Sanger Institute for the COVID-19 Genomics UK (COG-UK) Consortium | Jacquelyn Wynn, Mairead Hyland, The Lighthouse Lab in Alderley Park and Alex Alderton, Roberto Amato, Jeffrey Barrett, Sonia Goncalves, Ewan Harrison, David K. Jackson, Ian Johnston, Dominic Kwiatkowski, Cordelia Langford, John Sillitoe on behalf of the Wellcome Sanger Institute COVID-19 Surveillance Team |
| EPI_ISL_3443118 | Lighthouse Lab in Glasgow | Wellcome Sanger Institute for the COVID-19 Genomics UK (COG-UK) Consortium | Harper VanSteenhouse, Yumi Kasai, David Gray, Carol Clugston, Anna Dominiczak and Alex Alderton, Roberto Amato, Jeffrey Barrett, Sonia Goncalves, Ewan Harrison, David K. Jackson, Ian Johnston, Dominic Kwiatkowski, Cordelia Langford, John Sillitoe on behalf of the Wellcome Sanger Institute COVID-19 Surveillance Team |
| EPI_ISL_3443119 | Lighthouse Lab in Alderley Park | Wellcome Sanger Institute for the COVID-19 Genomics UK (COG-UK) Consortium | Jacquelyn Wynn, Mairead Hyland, The Lighthouse Lab in Alderley Park and Alex Alderton, Roberto Amato, Jeffrey Barrett, Sonia Goncalves, Ewan Harrison, David K. Jackson, Ian Johnston, Dominic Kwiatkowski, Cordelia Langford, John Sillitoe on behalf of the Wellcome Sanger Institute COVID-19 Surveillance Team |
| EPI_ISL_3443120, EPI_ISL_3443121, EPI_ISL_3443122 | Lighthouse Lab in Glasgow | Wellcome Sanger Institute for the COVID-19 Genomics UK (COG-UK) Consortium | Harper VanSteenhouse, Yumi Kasai, David Gray, Carol Clugston, Anna Dominiczak and Alex Alderton, Roberto Amato, Jeffrey Barrett, Sonia Goncalves, Ewan Harrison, David K. Jackson, Ian Johnston, Dominic Kwiatkowski, Cordelia Langford, John Sillitoe on behalf of the Wellcome Sanger Institute COVID-19 Surveillance Team |
| EPI_ISL_3443123 | Lighthouse Lab in Alderley Park | Wellcome Sanger Institute for the COVID-19 Genomics UK (COG-UK) Consortium | Jacquelyn Wynn, Mairead Hyland, The Lighthouse Lab in Alderley Park and Alex Alderton, Roberto Amato, Jeffrey Barrett, Sonia Goncalves, Ewan Harrison, David K. Jackson, Ian Johnston, Dominic Kwiatkowski, Cordelia Langford, John Sillitoe on behalf of the Wellcome Sanger Institute COVID-19 Surveillance Team |
| EPI_ISL_3443124, EPI_ISL_3443125 | Lighthouse Lab in Glasgow | Wellcome Sanger Institute for the COVID-19 Genomics UK (COG-UK) Consortium | Harper VanSteenhouse, Yumi Kasai, David Gray, Carol Clugston, Anna Dominiczak and Alex Alderton, Roberto Amato, Jeffrey Barrett, Sonia Goncalves, Ewan Harrison, David K. Jackson, Ian Johnston, Dominic Kwiatkowski, Cordelia Langford, John Sillitoe on behalf of the Wellcome Sanger Institute COVID-19 Surveillance Team |
| EPI_ISL_3443126, EPI_ISL_3443127, EPI_ISL_3443128 | Lighthouse Lab in Alderley Park | Wellcome Sanger Institute for the COVID-19 Genomics UK (COG-UK) Consortium | Jacquelyn Wynn, Mairead Hyland, The Lighthouse Lab in Alderley Park and Alex Alderton, Roberto Amato, Jeffrey Barrett, Sonia Goncalves, Ewan Harrison, David K. Jackson, Ian Johnston, Dominic Kwiatkowski, Cordelia Langford, John Sillitoe on behalf of the Wellcome Sanger Institute COVID-19 Surveillance Team |
| EPI_ISL_3443129 | Lighthouse Lab in Glasgow | Wellcome Sanger Institute for the COVID-19 Genomics UK (COG-UK) Consortium | Harper VanSteenhouse, Yumi Kasai, David Gray, Carol Clugston, Anna Dominiczak and Alex Alderton, Roberto Amato, Jeffrey Barrett, Sonia Goncalves, Ewan Harrison, David K. Jackson, Ian Johnston, Dominic Kwiatkowski, Cordelia Langford, John Sillitoe on behalf of the Wellcome Sanger Institute COVID-19 Surveillance Team |
| EPI_ISL_3443130 | Lighthouse Lab in Alderley Park | Wellcome Sanger Institute for the COVID-19 Genomics UK (COG-UK) Consortium | Jacquelyn Wynn, Mairead Hyland, The Lighthouse Lab in Alderley Park and Alex Alderton, Roberto Amato, Jeffrey Barrett, Sonia Goncalves, Ewan Harrison, David K. Jackson, Ian Johnston, Dominic Kwiatkowski, Cordelia Langford, John Sillitoe on behalf of the Wellcome Sanger Institute COVID-19 Surveillance Team |
| EPI_ISL_3443131, EPI_ISL_3443132 | Lighthouse Lab in Glasgow | Wellcome Sanger Institute for the COVID-19 Genomics UK (COG-UK) Consortium | Harper VanSteenhouse, Yumi Kasai, David Gray, Carol Clugston, Anna Dominiczak and Alex Alderton, Roberto Amato, Jeffrey Barrett, Sonia Goncalves, Ewan Harrison, David K. Jackson, Ian Johnston, Dominic Kwiatkowski, Cordelia Langford, John Sillitoe on behalf of the Wellcome Sanger Institute COVID-19 Surveillance Team |
| EPI_ISL_3443133, EPI_ISL_3443134 | Lighthouse Lab in Alderley Park | Wellcome Sanger Institute for the COVID-19 Genomics UK (COG-UK) Consortium | Jacquelyn Wynn, Mairead Hyland, The Lighthouse Lab in Alderley Park and Alex Alderton, Roberto Amato, Jeffrey Barrett, Sonia Goncalves, Ewan Harrison, David K. Jackson, Ian Johnston, Dominic Kwiatkowski, Cordelia Langford, John Sillitoe on behalf of the Wellcome Sanger Institute COVID-19 Surveillance Team |
| EPI_ISL_3443135 | Lighthouse Lab in Glasgow | Wellcome Sanger Institute for the COVID-19 Genomics UK (COG-UK) Consortium | Harper VanSteenhouse, Yumi Kasai, David Gray, Carol Clugston, Anna Dominiczak and Alex Alderton, Roberto Amato, Jeffrey Barrett, Sonia Goncalves, Ewan Harrison, David K. Jackson, Ian Johnston, Dominic Kwiatkowski, Cordelia Langford, John Sillitoe on behalf of the Wellcome Sanger Institute COVID-19 Surveillance Team |
| EPI_ISL_3443136 | Lighthouse Lab in Alderley Park | Wellcome Sanger Institute for the COVID-19 Genomics UK (COG-UK) Consortium | Jacquelyn Wynn, Mairead Hyland, The Lighthouse Lab in Alderley Park and Alex Alderton, Roberto Amato, Jeffrey Barrett, Sonia Goncalves, Ewan Harrison, David K. Jackson, Ian Johnston, Dominic Kwiatkowski, Cordelia Langford, John Sillitoe on behalf of the Wellcome Sanger Institute COVID-19 Surveillance Team |
| EPI_ISL_3443137, EPI_ISL_3443138, EPI_ISL_3443139 | Lighthouse Lab in Glasgow | Wellcome Sanger Institute for the COVID-19 Genomics UK (COG-UK) Consortium | Harper VanSteenhouse, Yumi Kasai, David Gray, Carol Clugston, Anna Dominiczak and Alex Alderton, Roberto Amato, Jeffrey Barrett, Sonia Goncalves, Ewan Harrison, David K. Jackson, Ian Johnston, Dominic Kwiatkowski, Cordelia Langford, John Sillitoe on behalf of the Wellcome Sanger Institute COVID-19 Surveillance Team |
| EPI_ISL_3443140, EPI_ISL_3443141 | Lighthouse Lab in Alderley Park | Wellcome Sanger Institute for the COVID-19 Genomics UK (COG-UK) Consortium | Jacquelyn Wynn, Mairead Hyland, The Lighthouse Lab in Alderley Park and Alex Alderton, Roberto Amato, Jeffrey Barrett, Sonia Goncalves, Ewan Harrison, David K. Jackson, Ian Johnston, Dominic Kwiatkowski, Cordelia Langford, John Sillitoe on behalf of the Wellcome Sanger Institute COVID-19 Surveillance Team |
| EPI_ISL_3443142 | Lighthouse Lab in Glasgow | Wellcome Sanger Institute for the COVID-19 Genomics UK (COG-UK) Consortium | Harper VanSteenhouse, Yumi Kasai, David Gray, Carol Clugston, Anna Dominiczak and Alex Alderton, Roberto Amato, Jeffrey Barrett, Sonia Goncalves, Ewan Harrison, David K. Jackson, Ian Johnston, Dominic Kwiatkowski, Cordelia Langford, John Sillitoe on behalf of the Wellcome Sanger Institute COVID-19 Surveillance Team |
| EPI_ISL_3443143 | Lighthouse Lab in Alderley Park | Wellcome Sanger Institute for the COVID-19 Genomics UK (COG-UK) Consortium | Jacquelyn Wynn, Mairead Hyland, The Lighthouse Lab in Alderley Park and Alex Alderton, Roberto Amato, Jeffrey Barrett, Sonia Goncalves, Ewan Harrison, David K. Jackson, Ian Johnston, Dominic Kwiatkowski, Cordelia Langford, John Sillitoe on behalf of the Wellcome Sanger Institute COVID-19 Surveillance Team |
| EPI_ISL_3443144, EPI_ISL_3443145, EPI_ISL_3443146, EPI_ISL_3443147, EPI_ISL_3443148, EPI_ISL_3443149, EPI_ISL_3443150 | Lighthouse Lab in Glasgow | Wellcome Sanger Institute for the COVID-19 Genomics UK (COG-UK) Consortium | Harper VanSteenhouse, Yumi Kasai, David Gray, Carol Clugston, Anna Dominiczak and Alex Alderton, Roberto Amato, Jeffrey Barrett, Sonia Goncalves, Ewan Harrison, David K. Jackson, Ian Johnston, Dominic Kwiatkowski, Cordelia Langford, John Sillitoe on behalf of the Wellcome Sanger Institute COVID-19 Surveillance Team |
| EPI_ISL_3443151 | Lighthouse Lab in Alderley Park | Wellcome Sanger Institute for the COVID-19 Genomics UK (COG-UK) Consortium | Jacquelyn Wynn, Mairead Hyland, The Lighthouse Lab in Alderley Park and Alex Alderton, Roberto Amato, Jeffrey Barrett, Sonia Goncalves, Ewan Harrison, David K. Jackson, Ian Johnston, Dominic Kwiatkowski, Cordelia Langford, John Sillitoe on behalf of the Wellcome Sanger Institute COVID-19 Surveillance Team |
| EPI_ISL_3443152, EPI_ISL_3443153, EPI_ISL_3443154, EPI_ISL_3443155, EPI_ISL_3443156, EPI_ISL_3443157 | Lighthouse Lab in Glasgow | Wellcome Sanger Institute for the COVID-19 Genomics UK (COG-UK) Consortium | Harper VanSteenhouse, Yumi Kasai, David Gray, Carol Clugston, Anna Dominiczak and Alex Alderton, Roberto Amato, Jeffrey Barrett, Sonia Goncalves, Ewan Harrison, David K. Jackson, Ian Johnston, Dominic Kwiatkowski, Cordelia Langford, John Sillitoe on behalf of the Wellcome Sanger Institute COVID-19 Surveillance Team |
| EPI_ISL_3443158 | Lighthouse Lab in Alderley Park | Wellcome Sanger Institute for the COVID-19 Genomics UK (COG-UK) Consortium | Jacquelyn Wynn, Mairead Hyland, The Lighthouse Lab in Alderley Park and Alex Alderton, Roberto Amato, Jeffrey Barrett, Sonia Goncalves, Ewan Harrison, David K. Jackson, Ian Johnston, Dominic Kwiatkowski, Cordelia Langford, John Sillitoe on behalf of the Wellcome Sanger Institute COVID-19 Surveillance Team |
| EPI_ISL_3443159 | Lighthouse Lab in Glasgow | Wellcome Sanger Institute for the COVID-19 Genomics UK (COG-UK) Consortium | Harper VanSteenhouse, Yumi Kasai, David Gray, Carol Clugston, Anna Dominiczak and Alex Alderton, Roberto Amato, Jeffrey Barrett, Sonia Goncalves, Ewan Harrison, David K. Jackson, Ian Johnston, Dominic Kwiatkowski, Cordelia Langford, John Sillitoe on behalf of the Wellcome Sanger Institute COVID-19 Surveillance Team |
| EPI_ISL_3443160 | Lighthouse Lab in Alderley Park | Wellcome Sanger Institute for the COVID-19 Genomics UK (COG-UK) Consortium | Jacquelyn Wynn, Mairead Hyland, The Lighthouse Lab in Alderley Park and Alex Alderton, Roberto Amato, Jeffrey Barrett, Sonia Goncalves, Ewan Harrison, David K. Jackson, Ian Johnston, Dominic Kwiatkowski, Cordelia Langford, John Sillitoe on behalf of the Wellcome Sanger Institute COVID-19 Surveillance Team |
| EPI_ISL_3443161, EPI_ISL_3443162, EPI_ISL_3443163, EPI_ISL_3443164, EPI_ISL_3443165, | Lighthouse Lab in Glasgow | Wellcome Sanger Institute for the COVID-19 Genomics UK (COG-UK) Consortium | Harper VanSteenhouse, Yumi Kasai, David Gray, Carol Clugston, Anna Dominiczak and Alex Alderton, Roberto Amato, Jeffrey Barrett, Sonia Goncalves, Ewan Harrison, David K. Jackson, Ian Johnston, Dominic Kwiatkowski, Cordelia Langford, John Sillitoe on behalf of the Wellcome Sanger Institute COVID-19 Surveillance Team |

| | | | |
|---|---|---|---|
| EPI_ISL_3443166, EPI_ISL_3443167, EPI_ISL_3443168, EPI_ISL_3443169 | | | |
| EPI_ISL_3443170, EPI_ISL_3443171 | Lighthouse Lab in Alderley Park | Wellcome Sanger Institute for the COVID-19 Genomics UK (COG-UK) Consortium | Jacquelyn Wynn, Mairead Hyland, The Lighthouse Lab in Alderley Park and Alex Alderton, Roberto Amato, Jeffrey Barrett, Sonia Goncalves, Ewan Harrison, David K. Jackson, Ian Johnston, Dominic Kwiatkowski, Cordelia Langford, John Sillitoe on behalf of the Wellcome Sanger Institute COVID-19 Surveillance Team |
| EPI_ISL_3443172, EPI_ISL_3443173, EPI_ISL_3443174, EPI_ISL_3443175, EPI_ISL_3443176 | Lighthouse Lab in Glasgow | Wellcome Sanger Institute for the COVID-19 Genomics UK (COG-UK) Consortium | Harper VanSteenhouse, Yumi Kasai, David Gray, Carol Clugston, Anna Dominiczak and Alex Alderton, Roberto Amato, Jeffrey Barrett, Sonia Goncalves, Ewan Harrison, David K. Jackson, Ian Johnston, Dominic Kwiatkowski, Cordelia Langford, John Sillitoe on behalf of the Wellcome Sanger Institute COVID-19 Surveillance Team |
| EPI_ISL_3443177, EPI_ISL_3443178 | Lighthouse Lab in Alderley Park | Wellcome Sanger Institute for the COVID-19 Genomics UK (COG-UK) Consortium | Jacquelyn Wynn, Mairead Hyland, The Lighthouse Lab in Alderley Park and Alex Alderton, Roberto Amato, Jeffrey Barrett, Sonia Goncalves, Ewan Harrison, David K. Jackson, Ian Johnston, Dominic Kwiatkowski, Cordelia Langford, John Sillitoe on behalf of the Wellcome Sanger Institute COVID-19 Surveillance Team |
| EPI_ISL_3443179, EPI_ISL_3443180, EPI_ISL_3443181, EPI_ISL_3443182, EPI_ISL_3443183, EPI_ISL_3443184, EPI_ISL_3443185, EPI_ISL_3443186, EPI_ISL_3443187, EPI_ISL_3443188 | Lighthouse Lab in Glasgow | Wellcome Sanger Institute for the COVID-19 Genomics UK (COG-UK) Consortium | Harper VanSteenhouse, Yumi Kasai, David Gray, Carol Clugston, Anna Dominiczak and Alex Alderton, Roberto Amato, Jeffrey Barrett, Sonia Goncalves, Ewan Harrison, David K. Jackson, Ian Johnston, Dominic Kwiatkowski, Cordelia Langford, John Sillitoe on behalf of the Wellcome Sanger Institute COVID-19 Surveillance Team |
| EPI_ISL_3443189, EPI_ISL_3443190 | Lighthouse Lab in Alderley Park | Wellcome Sanger Institute for the COVID-19 Genomics UK (COG-UK) Consortium | Jacquelyn Wynn, Mairead Hyland, The Lighthouse Lab in Alderley Park and Alex Alderton, Roberto Amato, Jeffrey Barrett, Sonia Goncalves, Ewan Harrison, David K. Jackson, Ian Johnston, Dominic Kwiatkowski, Cordelia Langford, John Sillitoe on behalf of the Wellcome Sanger Institute COVID-19 Surveillance Team |
| EPI_ISL_3443191, EPI_ISL_3443192, EPI_ISL_3443193, EPI_ISL_3443194, EPI_ISL_3443195, EPI_ISL_3443196, EPI_ISL_3443197, EPI_ISL_3443198, EPI_ISL_3443199, EPI_ISL_3443200, EPI_ISL_3443201, EPI_ISL_3443202, EPI_ISL_3443203, EPI_ISL_3443204, EPI_ISL_3443205, EPI_ISL_3443206 | | | |
| see above | Lighthouse Lab in Glasgow | Wellcome Sanger Institute for the COVID-19 Genomics UK (COG-UK) Consortium | Harper VanSteenhouse, Yumi Kasai, David Gray, Carol Clugston, Anna Dominiczak and Alex Alderton, Roberto Amato, Jeffrey Barrett, Sonia Goncalves, Ewan Harrison, David K. Jackson, Ian Johnston, Dominic Kwiatkowski, Cordelia Langford, John Sillitoe on behalf of the Wellcome Sanger Institute COVID-19 Surveillance Team |
| EPI_ISL_3443207 | Lighthouse Lab in Alderley Park | Wellcome Sanger Institute for the COVID-19 Genomics UK (COG-UK) Consortium | Jacquelyn Wynn, Mairead Hyland, The Lighthouse Lab in Alderley Park and Alex Alderton, Roberto Amato, Jeffrey Barrett, Sonia Goncalves, Ewan Harrison, David K. Jackson, Ian Johnston, Dominic Kwiatkowski, Cordelia Langford, John Sillitoe on behalf of the Wellcome Sanger Institute COVID-19 Surveillance Team |
| EPI_ISL_3443208, EPI_ISL_3443209, EPI_ISL_3443210, EPI_ISL_3443211, EPI_ISL_3443212 | Lighthouse Lab in Glasgow | Wellcome Sanger Institute for the COVID-19 Genomics UK (COG-UK) Consortium | Harper VanSteenhouse, Yumi Kasai, David Gray, Carol Clugston, Anna Dominiczak and Alex Alderton, Roberto Amato, Jeffrey Barrett, Sonia Goncalves, Ewan Harrison, David K. Jackson, Ian Johnston, Dominic Kwiatkowski, Cordelia Langford, John Sillitoe on behalf of the Wellcome Sanger Institute COVID-19 Surveillance Team |
| EPI_ISL_3443213 | Lighthouse Lab in Alderley Park | Wellcome Sanger Institute for the COVID-19 Genomics UK (COG-UK) Consortium | Jacquelyn Wynn, Mairead Hyland, The Lighthouse Lab in Alderley Park and Alex Alderton, Roberto Amato, Jeffrey Barrett, Sonia Goncalves, Ewan Harrison, David K. Jackson, Ian Johnston, Dominic Kwiatkowski, Cordelia Langford, John Sillitoe on behalf of the Wellcome Sanger Institute COVID-19 Surveillance Team |
| EPI_ISL_3443214, EPI_ISL_3443215, EPI_ISL_3443216, EPI_ISL_3443217, EPI_ISL_3443218, EPI_ISL_3443219 | Lighthouse Lab in Glasgow | Wellcome Sanger Institute for the COVID-19 Genomics UK (COG-UK) Consortium | Harper VanSteenhouse, Yumi Kasai, David Gray, Carol Clugston, Anna Dominiczak and Alex Alderton, Roberto Amato, Jeffrey Barrett, Sonia Goncalves, Ewan Harrison, David K. Jackson, Ian Johnston, Dominic Kwiatkowski, Cordelia Langford, John Sillitoe on behalf of the Wellcome Sanger Institute COVID-19 Surveillance Team |
| EPI_ISL_3443220 | Lighthouse Lab in Alderley Park | Wellcome Sanger Institute for the COVID-19 Genomics UK (COG-UK) Consortium | Jacquelyn Wynn, Mairead Hyland, The Lighthouse Lab in Alderley Park and Alex Alderton, Roberto Amato, Jeffrey Barrett, Sonia Goncalves, Ewan Harrison, David K. Jackson, Ian Johnston, Dominic Kwiatkowski, Cordelia Langford, John Sillitoe on behalf of the Wellcome Sanger Institute COVID-19 Surveillance Team |
| EPI_ISL_3443221, EPI_ISL_3443222, EPI_ISL_3443223, EPI_ISL_3443224, EPI_ISL_3443225, EPI_ISL_3443226, EPI_ISL_3443227, EPI_ISL_3443228, EPI_ISL_3443229, EPI_ISL_3443230 | Lighthouse Lab in Glasgow | Wellcome Sanger Institute for the COVID-19 Genomics UK (COG-UK) Consortium | Harper VanSteenhouse, Yumi Kasai, David Gray, Carol Clugston, Anna Dominiczak and Alex Alderton, Roberto Amato, Jeffrey Barrett, Sonia Goncalves, Ewan Harrison, David K. Jackson, Ian Johnston, Dominic Kwiatkowski, Cordelia Langford, John Sillitoe on behalf of the Wellcome Sanger Institute COVID-19 Surveillance Team |
| EPI_ISL_3443231, EPI_ISL_3443232 | Lighthouse Lab in Alderley Park | Wellcome Sanger Institute for the COVID-19 Genomics UK (COG-UK) Consortium | Jacquelyn Wynn, Mairead Hyland, The Lighthouse Lab in Alderley Park and Alex Alderton, Roberto Amato, Jeffrey Barrett, Sonia Goncalves, Ewan Harrison, David K. Jackson, Ian Johnston, Dominic Kwiatkowski, Cordelia Langford, John Sillitoe on behalf of the Wellcome Sanger Institute COVID-19 Surveillance Team |
| EPI_ISL_3443233, EPI_ISL_3443234, EPI_ISL_3443235, EPI_ISL_3443236, EPI_ISL_3443237 | Lighthouse Lab in Glasgow | Wellcome Sanger Institute for the COVID-19 Genomics UK (COG-UK) Consortium | Harper VanSteenhouse, Yumi Kasai, David Gray, Carol Clugston, Anna Dominiczak and Alex Alderton, Roberto Amato, Jeffrey Barrett, Sonia Goncalves, Ewan Harrison, David K. Jackson, Ian Johnston, Dominic Kwiatkowski, Cordelia Langford, John Sillitoe on behalf of the Wellcome Sanger Institute COVID-19 Surveillance Team |
| EPI_ISL_3443238 | Lighthouse Lab in Alderley Park | Wellcome Sanger Institute for the COVID-19 Genomics UK (COG-UK) Consortium | Jacquelyn Wynn, Mairead Hyland, The Lighthouse Lab in Alderley Park and Alex Alderton, Roberto Amato, Jeffrey Barrett, Sonia Goncalves, Ewan Harrison, David K. Jackson, Ian Johnston, Dominic Kwiatkowski, Cordelia Langford, John Sillitoe on behalf of the Wellcome Sanger Institute COVID-19 Surveillance Team |
| EPI_ISL_3443239, EPI_ISL_3443240, EPI_ISL_3443241, EPI_ISL_3443242, EPI_ISL_3443243 | Lighthouse Lab in Glasgow | Wellcome Sanger Institute for the COVID-19 Genomics UK (COG-UK) Consortium | Harper VanSteenhouse, Yumi Kasai, David Gray, Carol Clugston, Anna Dominiczak and Alex Alderton, Roberto Amato, Jeffrey Barrett, Sonia Goncalves, Ewan Harrison, David K. Jackson, Ian Johnston, Dominic Kwiatkowski, Cordelia Langford, John Sillitoe on behalf of the Wellcome Sanger Institute COVID-19 Surveillance Team |
| EPI_ISL_3443244 | Lighthouse Lab in Alderley Park | Wellcome Sanger Institute for the COVID-19 Genomics UK (COG-UK) Consortium | Jacquelyn Wynn, Mairead Hyland, The Lighthouse Lab in Alderley Park and Alex Alderton, Roberto Amato, Jeffrey Barrett, Sonia Goncalves, Ewan Harrison, David K. Jackson, Ian Johnston, Dominic Kwiatkowski, Cordelia Langford, John Sillitoe on behalf of the Wellcome Sanger Institute COVID-19 Surveillance Team |
| EPI_ISL_3443245, EPI_ISL_3443246 | Lighthouse Lab in Glasgow | Wellcome Sanger Institute for the COVID-19 Genomics UK (COG-UK) Consortium | Harper VanSteenhouse, Yumi Kasai, David Gray, Carol Clugston, Anna Dominiczak and Alex Alderton, Roberto Amato, Jeffrey Barrett, Sonia Goncalves, Ewan Harrison, David K. Jackson, Ian Johnston, Dominic Kwiatkowski, Cordelia Langford, John Sillitoe on behalf of the Wellcome Sanger Institute COVID-19 Surveillance Team |
| EPI_ISL_3443247 | Lighthouse Lab in Alderley Park | Wellcome Sanger Institute for the COVID-19 Genomics UK (COG-UK) Consortium | Jacquelyn Wynn, Mairead Hyland, The Lighthouse Lab in Alderley Park and Alex Alderton, Roberto Amato, Jeffrey Barrett, Sonia Goncalves, Ewan Harrison, David K. Jackson, Ian Johnston, Dominic Kwiatkowski, Cordelia Langford, John Sillitoe on behalf of the Wellcome Sanger Institute COVID-19 Surveillance Team |
| EPI_ISL_3443248, EPI_ISL_3443249, EPI_ISL_3443250, EPI_ISL_3443251, EPI_ISL_3443252, EPI_ISL_3443253, EPI_ISL_3443254, EPI_ISL_3443255, EPI_ISL_3443256 | Lighthouse Lab in Glasgow | Wellcome Sanger Institute for the COVID-19 Genomics UK (COG-UK) Consortium | Harper VanSteenhouse, Yumi Kasai, David Gray, Carol Clugston, Anna Dominiczak and Alex Alderton, Roberto Amato, Jeffrey Barrett, Sonia Goncalves, Ewan Harrison, David K. Jackson, Ian Johnston, Dominic Kwiatkowski, Cordelia Langford, John Sillitoe on behalf of the Wellcome Sanger Institute COVID-19 Surveillance Team |
| EPI_ISL_3443257 | Lighthouse Lab in Alderley Park | Wellcome Sanger Institute for the COVID-19 Genomics UK (COG-UK) Consortium | Jacquelyn Wynn, Mairead Hyland, The Lighthouse Lab in Alderley Park and Alex Alderton, Roberto Amato, Jeffrey Barrett, Sonia Goncalves, Ewan Harrison, David K. Jackson, Ian Johnston, Dominic Kwiatkowski, Cordelia Langford, John Sillitoe on behalf of the Wellcome Sanger Institute COVID-19 Surveillance Team |
| EPI_ISL_3443258, EPI_ISL_3443259, EPI_ISL_3443260 | Lighthouse Lab in Glasgow | Wellcome Sanger Institute for the COVID-19 Genomics UK (COG-UK) Consortium | Harper VanSteenhouse, Yumi Kasai, David Gray, Carol Clugston, Anna Dominiczak and Alex Alderton, Roberto Amato, Jeffrey Barrett, Sonia Goncalves, Ewan Harrison, David K. Jackson, Ian Johnston, Dominic Kwiatkowski, Cordelia Langford, John Sillitoe on behalf of the Wellcome Sanger Institute COVID-19 Surveillance Team |
| EPI_ISL_3443261 | Lighthouse Lab in Alderley Park | Wellcome Sanger Institute for the COVID-19 Genomics UK (COG-UK) Consortium | Jacquelyn Wynn, Mairead Hyland, The Lighthouse Lab in Alderley Park and Alex Alderton, Roberto Amato, Jeffrey Barrett, Sonia Goncalves, Ewan Harrison, David K. Jackson, Ian Johnston, Dominic Kwiatkowski, Cordelia Langford, John Sillitoe on behalf of the Wellcome Sanger Institute COVID-19 Surveillance Team |
| EPI_ISL_3443262, EPI_ISL_3443263 | Lighthouse Lab in Glasgow | Wellcome Sanger Institute for the COVID-19 Genomics UK (COG-UK) Consortium | Harper VanSteenhouse, Yumi Kasai, David Gray, Carol Clugston, Anna Dominiczak and Alex Alderton, Roberto Amato, Jeffrey Barrett, Sonia Goncalves, Ewan Harrison, David K. Jackson, Ian Johnston, Dominic Kwiatkowski, Cordelia Langford, John Sillitoe on behalf of the Wellcome Sanger Institute COVID-19 Surveillance Team |
| EPI_ISL_3443264 | Lighthouse Lab in Alderley Park | Wellcome Sanger Institute for the COVID-19 Genomics UK (COG-UK) Consortium | Jacquelyn Wynn, Mairead Hyland, The Lighthouse Lab in Alderley Park and Alex Alderton, Roberto Amato, Jeffrey Barrett, Sonia Goncalves, Ewan Harrison, David K. Jackson, Ian Johnston, Dominic Kwiatkowski, Cordelia Langford, John Sillitoe on behalf of the Wellcome Sanger Institute COVID-19 Surveillance Team |
| EPI_ISL_3443265, EPI_ISL_3443266 | Lighthouse Lab in Glasgow | Wellcome Sanger Institute for the COVID-19 Genomics UK (COG-UK) Consortium | Harper VanSteenhouse, Yumi Kasai, David Gray, Carol Clugston, Anna Dominiczak and Alex Alderton, Roberto Amato, Jeffrey Barrett, Sonia Goncalves, Ewan Harrison, David K. Jackson, Ian Johnston, Dominic Kwiatkowski, Cordelia Langford, John Sillitoe on behalf of the Wellcome Sanger Institute COVID-19 Surveillance Team |
| EPI_ISL_3443267 | Lighthouse Lab in Alderley Park | Wellcome Sanger Institute for the COVID-19 Genomics UK (COG-UK) | Jacquelyn Wynn, Mairead Hyland, The Lighthouse Lab in Alderley Park and Alex Alderton, Roberto Amato, Jeffrey Barrett, Sonia Goncalves, Ewan Harrison, David K. Jackson, Ian Johnston, Dominic Kwiatkowski, Cordelia Langford, John Sillitoe on |

| | | | |
|---|---|---|---|
| | | Consortium | behalf of the Wellcome Sanger Institute COVID-19 Surveillance Team |
| EPI_ISL_3443268, EPI_ISL_3443269, EPI_ISL_3443270 | Lighthouse Lab in Glasgow | Wellcome Sanger Institute for the COVID-19 Genomics UK (COG-UK) Consortium | Harper VanSteenhouse, Yumi Kasai, David Gray, Carol Clugston, Anna Dominiczak and Alex Alderton, Roberto Amato, Jeffrey Barrett, Sonia Goncalves, Ewan Harrison, David K. Jackson, Ian Johnston, Dominic Kwiatkowski, Cordelia Langford, John Sillitoe on behalf of the Wellcome Sanger Institute COVID-19 Surveillance Team |
| EPI_ISL_3443271 | Lighthouse Lab in Alderley Park | Wellcome Sanger Institute for the COVID-19 Genomics UK (COG-UK) Consortium | Jacquelyn Wynn, Mairead Hyland, The Lighthouse Lab in Alderley Park and Alex Alderton, Roberto Amato, Jeffrey Barrett, Sonia Goncalves, Ewan Harrison, David K. Jackson, Ian Johnston, Dominic Kwiatkowski, Cordelia Langford, John Sillitoe on behalf of the Wellcome Sanger Institute COVID-19 Surveillance Team |
| EPI_ISL_3443272, EPI_ISL_3443273 | Lighthouse Lab in Glasgow | Wellcome Sanger Institute for the COVID-19 Genomics UK (COG-UK) Consortium | Harper VanSteenhouse, Yumi Kasai, David Gray, Carol Clugston, Anna Dominiczak and Alex Alderton, Roberto Amato, Jeffrey Barrett, Sonia Goncalves, Ewan Harrison, David K. Jackson, Ian Johnston, Dominic Kwiatkowski, Cordelia Langford, John Sillitoe on behalf of the Wellcome Sanger Institute COVID-19 Surveillance Team |
| EPI_ISL_3443274 | Lighthouse Lab in Alderley Park | Wellcome Sanger Institute for the COVID-19 Genomics UK (COG-UK) Consortium | Jacquelyn Wynn, Mairead Hyland, The Lighthouse Lab in Alderley Park and Alex Alderton, Roberto Amato, Jeffrey Barrett, Sonia Goncalves, Ewan Harrison, David K. Jackson, Ian Johnston, Dominic Kwiatkowski, Cordelia Langford, John Sillitoe on behalf of the Wellcome Sanger Institute COVID-19 Surveillance Team |
| EPI_ISL_3443275 | Lighthouse Lab in Glasgow | Wellcome Sanger Institute for the COVID-19 Genomics UK (COG-UK) Consortium | Harper VanSteenhouse, Yumi Kasai, David Gray, Carol Clugston, Anna Dominiczak and Alex Alderton, Roberto Amato, Jeffrey Barrett, Sonia Goncalves, Ewan Harrison, David K. Jackson, Ian Johnston, Dominic Kwiatkowski, Cordelia Langford, John Sillitoe on behalf of the Wellcome Sanger Institute COVID-19 Surveillance Team |
| EPI_ISL_3443276 | Lighthouse Lab in Alderley Park | Wellcome Sanger Institute for the COVID-19 Genomics UK (COG-UK) Consortium | Jacquelyn Wynn, Mairead Hyland, The Lighthouse Lab in Alderley Park and Alex Alderton, Roberto Amato, Jeffrey Barrett, Sonia Goncalves, Ewan Harrison, David K. Jackson, Ian Johnston, Dominic Kwiatkowski, Cordelia Langford, John Sillitoe on behalf of the Wellcome Sanger Institute COVID-19 Surveillance Team |
| EPI_ISL_3443277, EPI_ISL_3443278, EPI_ISL_3443279, EPI_ISL_3443280 | Lighthouse Lab in Glasgow | Wellcome Sanger Institute for the COVID-19 Genomics UK (COG-UK) Consortium | Harper VanSteenhouse, Yumi Kasai, David Gray, Carol Clugston, Anna Dominiczak and Alex Alderton, Roberto Amato, Jeffrey Barrett, Sonia Goncalves, Ewan Harrison, David K. Jackson, Ian Johnston, Dominic Kwiatkowski, Cordelia Langford, John Sillitoe on behalf of the Wellcome Sanger Institute COVID-19 Surveillance Team |
| EPI_ISL_3443281, EPI_ISL_3443282, EPI_ISL_3443283, EPI_ISL_3443284 | Lighthouse Lab in Alderley Park | Wellcome Sanger Institute for the COVID-19 Genomics UK (COG-UK) Consortium | Jacquelyn Wynn, Mairead Hyland, The Lighthouse Lab in Alderley Park and Alex Alderton, Roberto Amato, Jeffrey Barrett, Sonia Goncalves, Ewan Harrison, David K. Jackson, Ian Johnston, Dominic Kwiatkowski, Cordelia Langford, John Sillitoe on behalf of the Wellcome Sanger Institute COVID-19 Surveillance Team |
| EPI_ISL_3443285, EPI_ISL_3443286, EPI_ISL_3443287, EPI_ISL_3443288, EPI_ISL_3443289 | Lighthouse Lab in Glasgow | Wellcome Sanger Institute for the COVID-19 Genomics UK (COG-UK) Consortium | Harper VanSteenhouse, Yumi Kasai, David Gray, Carol Clugston, Anna Dominiczak and Alex Alderton, Roberto Amato, Jeffrey Barrett, Sonia Goncalves, Ewan Harrison, David K. Jackson, Ian Johnston, Dominic Kwiatkowski, Cordelia Langford, John Sillitoe on behalf of the Wellcome Sanger Institute COVID-19 Surveillance Team |
| EPI_ISL_3443290, EPI_ISL_3443291 | Lighthouse Lab in Alderley Park | Wellcome Sanger Institute for the COVID-19 Genomics UK (COG-UK) Consortium | Jacquelyn Wynn, Mairead Hyland, The Lighthouse Lab in Alderley Park and Alex Alderton, Roberto Amato, Jeffrey Barrett, Sonia Goncalves, Ewan Harrison, David K. Jackson, Ian Johnston, Dominic Kwiatkowski, Cordelia Langford, John Sillitoe on behalf of the Wellcome Sanger Institute COVID-19 Surveillance Team |
| EPI_ISL_3443292, EPI_ISL_3443293, EPI_ISL_3443294, EPI_ISL_3443295 | Lighthouse Lab in Glasgow | Wellcome Sanger Institute for the COVID-19 Genomics UK (COG-UK) Consortium | Harper VanSteenhouse, Yumi Kasai, David Gray, Carol Clugston, Anna Dominiczak and Alex Alderton, Roberto Amato, Jeffrey Barrett, Sonia Goncalves, Ewan Harrison, David K. Jackson, Ian Johnston, Dominic Kwiatkowski, Cordelia Langford, John Sillitoe on behalf of the Wellcome Sanger Institute COVID-19 Surveillance Team |
| EPI_ISL_3443296 | Lighthouse Lab in Alderley Park | Wellcome Sanger Institute for the COVID-19 Genomics UK (COG-UK) Consortium | Jacquelyn Wynn, Mairead Hyland, The Lighthouse Lab in Alderley Park and Alex Alderton, Roberto Amato, Jeffrey Barrett, Sonia Goncalves, Ewan Harrison, David K. Jackson, Ian Johnston, Dominic Kwiatkowski, Cordelia Langford, John Sillitoe on behalf of the Wellcome Sanger Institute COVID-19 Surveillance Team |
| EPI_ISL_3443297 | Lighthouse Lab in Glasgow | Wellcome Sanger Institute for the COVID-19 Genomics UK (COG-UK) Consortium | Harper VanSteenhouse, Yumi Kasai, David Gray, Carol Clugston, Anna Dominiczak and Alex Alderton, Roberto Amato, Jeffrey Barrett, Sonia Goncalves, Ewan Harrison, David K. Jackson, Ian Johnston, Dominic Kwiatkowski, Cordelia Langford, John Sillitoe on behalf of the Wellcome Sanger Institute COVID-19 Surveillance Team |
| EPI_ISL_3443298 | Lighthouse Lab in Alderley Park | Wellcome Sanger Institute for the COVID-19 Genomics UK (COG-UK) Consortium | Jacquelyn Wynn, Mairead Hyland, The Lighthouse Lab in Alderley Park and Alex Alderton, Roberto Amato, Jeffrey Barrett, Sonia Goncalves, Ewan Harrison, David K. Jackson, Ian Johnston, Dominic Kwiatkowski, Cordelia Langford, John Sillitoe on behalf of the Wellcome Sanger Institute COVID-19 Surveillance Team |
| EPI_ISL_3443299, EPI_ISL_3443300 | Lighthouse Lab in Glasgow | Wellcome Sanger Institute for the COVID-19 Genomics UK (COG-UK) Consortium | Harper VanSteenhouse, Yumi Kasai, David Gray, Carol Clugston, Anna Dominiczak and Alex Alderton, Roberto Amato, Jeffrey Barrett, Sonia Goncalves, Ewan Harrison, David K. Jackson, Ian Johnston, Dominic Kwiatkowski, Cordelia Langford, John Sillitoe on behalf of the Wellcome Sanger Institute COVID-19 Surveillance Team |
| EPI_ISL_3443301 | Lighthouse Lab in Alderley Park | Wellcome Sanger Institute for the COVID-19 Genomics UK (COG-UK) Consortium | Jacquelyn Wynn, Mairead Hyland, The Lighthouse Lab in Alderley Park and Alex Alderton, Roberto Amato, Jeffrey Barrett, Sonia Goncalves, Ewan Harrison, David K. Jackson, Ian Johnston, Dominic Kwiatkowski, Cordelia Langford, John Sillitoe on behalf of the Wellcome Sanger Institute COVID-19 Surveillance Team |
| EPI_ISL_3443302, EPI_ISL_3443303, EPI_ISL_3443304 | Lighthouse Lab in Glasgow | Wellcome Sanger Institute for the COVID-19 Genomics UK (COG-UK) Consortium | Harper VanSteenhouse, Yumi Kasai, David Gray, Carol Clugston, Anna Dominiczak and Alex Alderton, Roberto Amato, Jeffrey Barrett, Sonia Goncalves, Ewan Harrison, David K. Jackson, Ian Johnston, Dominic Kwiatkowski, Cordelia Langford, John Sillitoe on behalf of the Wellcome Sanger Institute COVID-19 Surveillance Team |
| EPI_ISL_3443305 | Lighthouse Lab in Alderley Park | Wellcome Sanger Institute for the COVID-19 Genomics UK (COG-UK) Consortium | Jacquelyn Wynn, Mairead Hyland, The Lighthouse Lab in Alderley Park and Alex Alderton, Roberto Amato, Jeffrey Barrett, Sonia Goncalves, Ewan Harrison, David K. Jackson, Ian Johnston, Dominic Kwiatkowski, Cordelia Langford, John Sillitoe on behalf of the Wellcome Sanger Institute COVID-19 Surveillance Team |
| EPI_ISL_3443306, EPI_ISL_3443307, EPI_ISL_3443308, EPI_ISL_3443309, EPI_ISL_3443310 | Lighthouse Lab in Glasgow | Wellcome Sanger Institute for the COVID-19 Genomics UK (COG-UK) Consortium | Harper VanSteenhouse, Yumi Kasai, David Gray, Carol Clugston, Anna Dominiczak and Alex Alderton, Roberto Amato, Jeffrey Barrett, Sonia Goncalves, Ewan Harrison, David K. Jackson, Ian Johnston, Dominic Kwiatkowski, Cordelia Langford, John Sillitoe on behalf of the Wellcome Sanger Institute COVID-19 Surveillance Team |
| EPI_ISL_3443311, EPI_ISL_3443312, EPI_ISL_3443313 | Lighthouse Lab in Alderley Park | Wellcome Sanger Institute for the COVID-19 Genomics UK (COG-UK) Consortium | Jacquelyn Wynn, Mairead Hyland, The Lighthouse Lab in Alderley Park and Alex Alderton, Roberto Amato, Jeffrey Barrett, Sonia Goncalves, Ewan Harrison, David K. Jackson, Ian Johnston, Dominic Kwiatkowski, Cordelia Langford, John Sillitoe on behalf of the Wellcome Sanger Institute COVID-19 Surveillance Team |
| EPI_ISL_3443314, EPI_ISL_3443315, EPI_ISL_3443316 | Lighthouse Lab in Glasgow | Wellcome Sanger Institute for the COVID-19 Genomics UK (COG-UK) Consortium | Harper VanSteenhouse, Yumi Kasai, David Gray, Carol Clugston, Anna Dominiczak and Alex Alderton, Roberto Amato, Jeffrey Barrett, Sonia Goncalves, Ewan Harrison, David K. Jackson, Ian Johnston, Dominic Kwiatkowski, Cordelia Langford, John Sillitoe on behalf of the Wellcome Sanger Institute COVID-19 Surveillance Team |
| EPI_ISL_3443317 | Lighthouse Lab in Alderley Park | Wellcome Sanger Institute for the COVID-19 Genomics UK (COG-UK) Consortium | Jacquelyn Wynn, Mairead Hyland, The Lighthouse Lab in Alderley Park and Alex Alderton, Roberto Amato, Jeffrey Barrett, Sonia Goncalves, Ewan Harrison, David K. Jackson, Ian Johnston, Dominic Kwiatkowski, Cordelia Langford, John Sillitoe on behalf of the Wellcome Sanger Institute COVID-19 Surveillance Team |
| EPI_ISL_3443318, EPI_ISL_3443319, EPI_ISL_3443320 | Lighthouse Lab in Glasgow | Wellcome Sanger Institute for the COVID-19 Genomics UK (COG-UK) Consortium | Harper VanSteenhouse, Yumi Kasai, David Gray, Carol Clugston, Anna Dominiczak and Alex Alderton, Roberto Amato, Jeffrey Barrett, Sonia Goncalves, Ewan Harrison, David K. Jackson, Ian Johnston, Dominic Kwiatkowski, Cordelia Langford, John Sillitoe on behalf of the Wellcome Sanger Institute COVID-19 Surveillance Team |
| EPI_ISL_3443321 | Lighthouse Lab in Alderley Park | Wellcome Sanger Institute for the COVID-19 Genomics UK (COG-UK) Consortium | Jacquelyn Wynn, Mairead Hyland, The Lighthouse Lab in Alderley Park and Alex Alderton, Roberto Amato, Jeffrey Barrett, Sonia Goncalves, Ewan Harrison, David K. Jackson, Ian Johnston, Dominic Kwiatkowski, Cordelia Langford, John Sillitoe on behalf of the Wellcome Sanger Institute COVID-19 Surveillance Team |
| EPI_ISL_3443322, EPI_ISL_3443323 | Lighthouse Lab in Glasgow | Wellcome Sanger Institute for the COVID-19 Genomics UK (COG-UK) Consortium | Harper VanSteenhouse, Yumi Kasai, David Gray, Carol Clugston, Anna Dominiczak and Alex Alderton, Roberto Amato, Jeffrey Barrett, Sonia Goncalves, Ewan Harrison, David K. Jackson, Ian Johnston, Dominic Kwiatkowski, Cordelia Langford, John Sillitoe on behalf of the Wellcome Sanger Institute COVID-19 Surveillance Team |
| EPI_ISL_3443324 | Lighthouse Lab in Alderley Park | Wellcome Sanger Institute for the COVID-19 Genomics UK (COG-UK) Consortium | Jacquelyn Wynn, Mairead Hyland, The Lighthouse Lab in Alderley Park and Alex Alderton, Roberto Amato, Jeffrey Barrett, Sonia Goncalves, Ewan Harrison, David K. Jackson, Ian Johnston, Dominic Kwiatkowski, Cordelia Langford, John Sillitoe on behalf of the Wellcome Sanger Institute COVID-19 Surveillance Team |
| EPI_ISL_3443325 | Lighthouse Lab in Glasgow | Wellcome Sanger Institute for the COVID-19 Genomics UK (COG-UK) Consortium | Harper VanSteenhouse, Yumi Kasai, David Gray, Carol Clugston, Anna Dominiczak and Alex Alderton, Roberto Amato, Jeffrey Barrett, Sonia Goncalves, Ewan Harrison, David K. Jackson, Ian Johnston, Dominic Kwiatkowski, Cordelia Langford, John Sillitoe on behalf of the Wellcome Sanger Institute COVID-19 Surveillance Team |
| EPI_ISL_3443326, EPI_ISL_3443327, EPI_ISL_3443328 | Lighthouse Lab in Alderley Park | Wellcome Sanger Institute for the COVID-19 Genomics UK (COG-UK) Consortium | Jacquelyn Wynn, Mairead Hyland, The Lighthouse Lab in Alderley Park and Alex Alderton, Roberto Amato, Jeffrey Barrett, Sonia Goncalves, Ewan Harrison, David K. Jackson, Ian Johnston, Dominic Kwiatkowski, Cordelia Langford, John Sillitoe on behalf of the Wellcome Sanger Institute COVID-19 Surveillance Team |
| EPI_ISL_3443329, EPI_ISL_3443330, EPI_ISL_3443331, EPI_ISL_3443332, EPI_ISL_3443333, EPI_ISL_3443334 | Lighthouse Lab in Glasgow | Wellcome Sanger Institute for the COVID-19 Genomics UK (COG-UK) Consortium | Harper VanSteenhouse, Yumi Kasai, David Gray, Carol Clugston, Anna Dominiczak and Alex Alderton, Roberto Amato, Jeffrey Barrett, Sonia Goncalves, Ewan Harrison, David K. Jackson, Ian Johnston, Dominic Kwiatkowski, Cordelia Langford, John Sillitoe on behalf of the Wellcome Sanger Institute COVID-19 Surveillance Team |
| EPI_ISL_3443335, EPI_ISL_3443336 | Lighthouse Lab in Alderley Park | Wellcome Sanger Institute for the COVID-19 Genomics UK (COG-UK) Consortium | Jacquelyn Wynn, Mairead Hyland, The Lighthouse Lab in Alderley Park and Alex Alderton, Roberto Amato, Jeffrey Barrett, Sonia Goncalves, Ewan Harrison, David K. Jackson, Ian Johnston, Dominic Kwiatkowski, Cordelia Langford, John Sillitoe on behalf of the Wellcome Sanger Institute COVID-19 Surveillance Team |
| EPI_ISL_3443337, EPI_ISL_3443338 | Lighthouse Lab in Glasgow | Wellcome Sanger Institute for the COVID-19 Genomics UK (COG-UK) Consortium | Harper VanSteenhouse, Yumi Kasai, David Gray, Carol Clugston, Anna Dominiczak and Alex Alderton, Roberto Amato, Jeffrey Barrett, Sonia Goncalves, Ewan Harrison, David K. Jackson, Ian Johnston, Dominic Kwiatkowski, Cordelia Langford, John Sillitoe on behalf of the Wellcome Sanger Institute COVID-19 Surveillance Team |
| EPI_ISL_3443339 | Lighthouse Lab in Alderley Park | Wellcome Sanger Institute for the COVID-19 Genomics UK (COG-UK) Consortium | Jacquelyn Wynn, Mairead Hyland, The Lighthouse Lab in Alderley Park and Alex Alderton, Roberto Amato, Jeffrey Barrett, Sonia Goncalves, Ewan Harrison, David K. Jackson, Ian Johnston, Dominic Kwiatkowski, Cordelia Langford, John Sillitoe on behalf of the Wellcome Sanger Institute COVID-19 Surveillance Team |
| EPI_ISL_3443340, EPI_ISL_3443341, EPI_ISL_3443342 | Lighthouse Lab in Glasgow | Wellcome Sanger Institute for the COVID-19 Genomics UK (COG-UK) Consortium | Harper VanSteenhouse, Yumi Kasai, David Gray, Carol Clugston, Anna Dominiczak and Alex Alderton, Roberto Amato, Jeffrey Barrett, Sonia Goncalves, Ewan Harrison, David K. Jackson, Ian Johnston, Dominic Kwiatkowski, Cordelia Langford, John Sillitoe on behalf of the Wellcome Sanger Institute COVID-19 Surveillance Team |
| EPI_ISL_3443343 | Lighthouse Lab in Alderley Park | Wellcome Sanger Institute for the COVID-19 Genomics UK (COG-UK) Consortium | Jacquelyn Wynn, Mairead Hyland, The Lighthouse Lab in Alderley Park and Alex Alderton, Roberto Amato, Jeffrey Barrett, Sonia Goncalves, Ewan Harrison, David K. Jackson, Ian Johnston, Dominic Kwiatkowski, Cordelia Langford, John Sillitoe on behalf of the Wellcome Sanger Institute COVID-19 Surveillance Team |
| EPI_ISL_3443344, EPI_ISL_3443345, EPI_ISL_3443346, EPI_ISL_3443347, EPI_ISL_3443348, EPI_ISL_3443349, EPI_ISL_3443350, EPI_ISL_3443351, EPI_ISL_3443352, EPI_ISL_3443353, EPI_ISL_3443354 | | | |
| see above | Lighthouse Lab in Glasgow | Wellcome Sanger Institute for the COVID-19 Genomics UK (COG-UK) | Harper VanSteenhouse, Yumi Kasai, David Gray, Carol Clugston, Anna Dominiczak and Alex Alderton, Roberto Amato, Jeffrey Barrett, Sonia Goncalves, Ewan Harrison, David K. Jackson, Ian Johnston, Dominic Kwiatkowski, Cordelia Langford, |

| | | | |
|---|---|---|---|
| | | Consortium | John Sillitoe on behalf of the Wellcome Sanger Institute COVID-19 Surveillance Team |
| EPI_ISL_3443355 | Lighthouse Lab in Alderley Park | Wellcome Sanger Institute for the COVID-19 Genomics UK (COG-UK) Consortium | Jacquelyn Wynn, Mairead Hyland, The Lighthouse Lab in Alderley Park and Alex Alderton, Roberto Amato, Jeffrey Barrett, Sonia Goncalves, Ewan Harrison, David K. Jackson, Ian Johnston, Dominic Kwiatkowski, Cordelia Langford, John Sillitoe on behalf of the Wellcome Sanger Institute COVID-19 Surveillance Team |
| EPI_ISL_3443356, EPI_ISL_3443357 | Lighthouse Lab in Glasgow | Wellcome Sanger Institute for the COVID-19 Genomics UK (COG-UK) Consortium | Harper VanSteenhouse, Yumi Kasai, David Gray, Carol Clugston, Anna Dominiczak and Alex Alderton, Roberto Amato, Jeffrey Barrett, Sonia Goncalves, Ewan Harrison, David K. Jackson, Ian Johnston, Dominic Kwiatkowski, Cordelia Langford, John Sillitoe on behalf of the Wellcome Sanger Institute COVID-19 Surveillance Team |
| EPI_ISL_3443358 | Lighthouse Lab in Alderley Park | Wellcome Sanger Institute for the COVID-19 Genomics UK (COG-UK) Consortium | Jacquelyn Wynn, Mairead Hyland, The Lighthouse Lab in Alderley Park and Alex Alderton, Roberto Amato, Jeffrey Barrett, Sonia Goncalves, Ewan Harrison, David K. Jackson, Ian Johnston, Dominic Kwiatkowski, Cordelia Langford, John Sillitoe on behalf of the Wellcome Sanger Institute COVID-19 Surveillance Team |
| EPI_ISL_3443359, EPI_ISL_3443360, EPI_ISL_3443361 | Lighthouse Lab in Glasgow | Wellcome Sanger Institute for the COVID-19 Genomics UK (COG-UK) Consortium | Harper VanSteenhouse, Yumi Kasai, David Gray, Carol Clugston, Anna Dominiczak and Alex Alderton, Roberto Amato, Jeffrey Barrett, Sonia Goncalves, Ewan Harrison, David K. Jackson, Ian Johnston, Dominic Kwiatkowski, Cordelia Langford, John Sillitoe on behalf of the Wellcome Sanger Institute COVID-19 Surveillance Team |
| EPI_ISL_3443362, EPI_ISL_3443363 | Lighthouse Lab in Alderley Park | Wellcome Sanger Institute for the COVID-19 Genomics UK (COG-UK) Consortium | Jacquelyn Wynn, Mairead Hyland, The Lighthouse Lab in Alderley Park and Alex Alderton, Roberto Amato, Jeffrey Barrett, Sonia Goncalves, Ewan Harrison, David K. Jackson, Ian Johnston, Dominic Kwiatkowski, Cordelia Langford, John Sillitoe on behalf of the Wellcome Sanger Institute COVID-19 Surveillance Team |
| EPI_ISL_3443364, EPI_ISL_3443365, EPI_ISL_3443366, EPI_ISL_3443367, EPI_ISL_3443368, EPI_ISL_3443369, EPI_ISL_3443370 | Lighthouse Lab in Glasgow | Wellcome Sanger Institute for the COVID-19 Genomics UK (COG-UK) Consortium | Harper VanSteenhouse, Yumi Kasai, David Gray, Carol Clugston, Anna Dominiczak and Alex Alderton, Roberto Amato, Jeffrey Barrett, Sonia Goncalves, Ewan Harrison, David K. Jackson, Ian Johnston, Dominic Kwiatkowski, Cordelia Langford, John Sillitoe on behalf of the Wellcome Sanger Institute COVID-19 Surveillance Team |
| EPI_ISL_3443371 | Lighthouse Lab in Alderley Park | Wellcome Sanger Institute for the COVID-19 Genomics UK (COG-UK) Consortium | Jacquelyn Wynn, Mairead Hyland, The Lighthouse Lab in Alderley Park and Alex Alderton, Roberto Amato, Jeffrey Barrett, Sonia Goncalves, Ewan Harrison, David K. Jackson, Ian Johnston, Dominic Kwiatkowski, Cordelia Langford, John Sillitoe on behalf of the Wellcome Sanger Institute COVID-19 Surveillance Team |
| EPI_ISL_3443372, EPI_ISL_3443373, EPI_ISL_3443374, EPI_ISL_3443375, EPI_ISL_3443376, EPI_ISL_3443377, EPI_ISL_3443378, EPI_ISL_3443379, EPI_ISL_3443380, EPI_ISL_3443381, EPI_ISL_3443382 | | | |
| see above | Lighthouse Lab in Glasgow | Wellcome Sanger Institute for the COVID-19 Genomics UK (COG-UK) Consortium | Harper VanSteenhouse, Yumi Kasai, David Gray, Carol Clugston, Anna Dominiczak and Alex Alderton, Roberto Amato, Jeffrey Barrett, Sonia Goncalves, Ewan Harrison, David K. Jackson, Ian Johnston, Dominic Kwiatkowski, Cordelia Langford, John Sillitoe on behalf of the Wellcome Sanger Institute COVID-19 Surveillance Team |
| EPI_ISL_3443383, EPI_ISL_3443384, EPI_ISL_3443385 | Lighthouse Lab in Alderley Park | Wellcome Sanger Institute for the COVID-19 Genomics UK (COG-UK) Consortium | Jacquelyn Wynn, Mairead Hyland, The Lighthouse Lab in Alderley Park and Alex Alderton, Roberto Amato, Jeffrey Barrett, Sonia Goncalves, Ewan Harrison, David K. Jackson, Ian Johnston, Dominic Kwiatkowski, Cordelia Langford, John Sillitoe on behalf of the Wellcome Sanger Institute COVID-19 Surveillance Team |
| EPI_ISL_3443386, EPI_ISL_3443387 | Lighthouse Lab in Glasgow | Wellcome Sanger Institute for the COVID-19 Genomics UK (COG-UK) Consortium | Harper VanSteenhouse, Yumi Kasai, David Gray, Carol Clugston, Anna Dominiczak and Alex Alderton, Roberto Amato, Jeffrey Barrett, Sonia Goncalves, Ewan Harrison, David K. Jackson, Ian Johnston, Dominic Kwiatkowski, Cordelia Langford, John Sillitoe on behalf of the Wellcome Sanger Institute COVID-19 Surveillance Team |
| EPI_ISL_3443388 | Lighthouse Lab in Alderley Park | Wellcome Sanger Institute for the COVID-19 Genomics UK (COG-UK) Consortium | Jacquelyn Wynn, Mairead Hyland, The Lighthouse Lab in Alderley Park and Alex Alderton, Roberto Amato, Jeffrey Barrett, Sonia Goncalves, Ewan Harrison, David K. Jackson, Ian Johnston, Dominic Kwiatkowski, Cordelia Langford, John Sillitoe on behalf of the Wellcome Sanger Institute COVID-19 Surveillance Team |
| EPI_ISL_3443389 | Lighthouse Lab in Glasgow | Wellcome Sanger Institute for the COVID-19 Genomics UK (COG-UK) Consortium | Harper VanSteenhouse, Yumi Kasai, David Gray, Carol Clugston, Anna Dominiczak and Alex Alderton, Roberto Amato, Jeffrey Barrett, Sonia Goncalves, Ewan Harrison, David K. Jackson, Ian Johnston, Dominic Kwiatkowski, Cordelia Langford, John Sillitoe on behalf of the Wellcome Sanger Institute COVID-19 Surveillance Team |
| EPI_ISL_3443390 | Lighthouse Lab in Alderley Park | Wellcome Sanger Institute for the COVID-19 Genomics UK (COG-UK) Consortium | Jacquelyn Wynn, Mairead Hyland, The Lighthouse Lab in Alderley Park and Alex Alderton, Roberto Amato, Jeffrey Barrett, Sonia Goncalves, Ewan Harrison, David K. Jackson, Ian Johnston, Dominic Kwiatkowski, Cordelia Langford, John Sillitoe on behalf of the Wellcome Sanger Institute COVID-19 Surveillance Team |
| EPI_ISL_3443391, EPI_ISL_3443392, EPI_ISL_3443393, EPI_ISL_3443394, EPI_ISL_3443395, EPI_ISL_3443396, EPI_ISL_3443397, EPI_ISL_3443398, EPI_ISL_3443399, EPI_ISL_3443400, EPI_ISL_3443401, EPI_ISL_3443402, EPI_ISL_3443403, EPI_ISL_3443404, EPI_ISL_3443405, EPI_ISL_3443406, EPI_ISL_3443407, EPI_ISL_3443408, EPI_ISL_3443409, EPI_ISL_3443410, EPI_ISL_3443411, EPI_ISL_3443412, EPI_ISL_3443413, EPI_ISL_3443414, EPI_ISL_3443415, EPI_ISL_3443416, EPI_ISL_3443417, EPI_ISL_3443418, EPI_ISL_3443419, EPI_ISL_3443420, EPI_ISL_3443421, EPI_ISL_3443422, EPI_ISL_3443423, EPI_ISL_3443424, EPI_ISL_3443425, EPI_ISL_3443426, EPI_ISL_3443427, EPI_ISL_3443428, EPI_ISL_3443429, EPI_ISL_3443430, EPI_ISL_3443431, EPI_ISL_3443432, EPI_ISL_3443433, EPI_ISL_3443434, EPI_ISL_3443435, EPI_ISL_3443436, EPI_ISL_3443437, EPI_ISL_3443438, EPI_ISL_3443439, EPI_ISL_3443440, EPI_ISL_3443441, EPI_ISL_3443442, EPI_ISL_3443443, EPI_ISL_3443444, EPI_ISL_3443445, EPI_ISL_3443446, EPI_ISL_3443447, EPI_ISL_3443448, EPI_ISL_3443449, EPI_ISL_3443450, EPI_ISL_3443451, EPI_ISL_3443452, EPI_ISL_3443453, EPI_ISL_3443454, EPI_ISL_3443455, EPI_ISL_3443456, EPI_ISL_3443457, EPI_ISL_3443458, EPI_ISL_3443459, EPI_ISL_3443460, EPI_ISL_3443461, EPI_ISL_3443462, EPI_ISL_3443463, EPI_ISL_3443464, EPI_ISL_3443465, EPI_ISL_3443466, EPI_ISL_3443467, EPI_ISL_3443468, EPI_ISL_3443469, EPI_ISL_3443470, EPI_ISL_3443471, EPI_ISL_3443472, EPI_ISL_3443473, EPI_ISL_3443474, EPI_ISL_3443475, EPI_ISL_3443476, EPI_ISL_3443477, EPI_ISL_3443478, EPI_ISL_3443479, EPI_ISL_3443480, EPI_ISL_3443481, EPI_ISL_3443482, EPI_ISL_3443483, EPI_ISL_3443484, EPI_ISL_3443485, EPI_ISL_3443486, EPI_ISL_3443487, EPI_ISL_3443488, EPI_ISL_3443489, EPI_ISL_3443490, EPI_ISL_3443491, EPI_ISL_3443492, EPI_ISL_3443493, EPI_ISL_3443494, EPI_ISL_3443495, EPI_ISL_3443496, EPI_ISL_3443497, EPI_ISL_3443498, EPI_ISL_3443499, EPI_ISL_3443500, EPI_ISL_3443501, EPI_ISL_3443502, EPI_ISL_3443503, EPI_ISL_3443504, EPI_ISL_3443505, EPI_ISL_3443506, EPI_ISL_3443507, EPI_ISL_3443508, EPI_ISL_3443509, EPI_ISL_3443510, EPI_ISL_3443511, EPI_ISL_3443512, EPI_ISL_3443513, EPI_ISL_3443514, EPI_ISL_3443515, EPI_ISL_3443516, EPI_ISL_3443517, EPI_ISL_3443518, EPI_ISL_3443519, EPI_ISL_3443520, EPI_ISL_3443521, EPI_ISL_3443522, EPI_ISL_3443523, EPI_ISL_3443524, EPI_ISL_3443525, EPI_ISL_3443526, EPI_ISL_3443527, EPI_ISL_3443528, EPI_ISL_3443529, EPI_ISL_3443530, EPI_ISL_3443531, EPI_ISL_3443532, EPI_ISL_3443533, EPI_ISL_3443534, EPI_ISL_3443535, EPI_ISL_3443536, EPI_ISL_3443537, EPI_ISL_3443538, EPI_ISL_3443539, EPI_ISL_3443540, EPI_ISL_3443541, EPI_ISL_3443542, EPI_ISL_3443543, EPI_ISL_3443544, EPI_ISL_3443545, EPI_ISL_3443546, EPI_ISL_3443547, EPI_ISL_3443548, EPI_ISL_3443549, EPI_ISL_3443550, EPI_ISL_3443551, EPI_ISL_3443552, EPI_ISL_3443553, EPI_ISL_3443554, EPI_ISL_3443555, EPI_ISL_3443556, EPI_ISL_3443557, EPI_ISL_3443558, EPI_ISL_3443559, EPI_ISL_3443560, EPI_ISL_3443561, EPI_ISL_3443562, EPI_ISL_3443563, EPI_ISL_3443564, EPI_ISL_3443565, EPI_ISL_3443566, EPI_ISL_3443567, EPI_ISL_3443568, EPI_ISL_3443569, EPI_ISL_3443570, EPI_ISL_3443571, EPI_ISL_3443572, EPI_ISL_3443573, EPI_ISL_3443574, EPI_ISL_3443575, EPI_ISL_3443576, EPI_ISL_3443577, EPI_ISL_3443578, EPI_ISL_3443579, EPI_ISL_3443580, EPI_ISL_3443581, EPI_ISL_3443582, EPI_ISL_3443583, EPI_ISL_3443584, EPI_ISL_3443585, EPI_ISL_3443586, EPI_ISL_3443587, EPI_ISL_3443588, EPI_ISL_3443589, EPI_ISL_3443590, EPI_ISL_3443591, EPI_ISL_3443592, EPI_ISL_3443593, EPI_ISL_3443594, EPI_ISL_3443595, EPI_ISL_3443596, EPI_ISL_3443597, EPI_ISL_3443598, EPI_ISL_3443599, EPI_ISL_3443600, EPI_ISL_3443601, EPI_ISL_3443602, EPI_ISL_3443603, EPI_ISL_3443604, EPI_ISL_3443605, EPI_ISL_3443606, EPI_ISL_3443607, EPI_ISL_3443608, EPI_ISL_3443609, EPI_ISL_3443610, EPI_ISL_3443611, EPI_ISL_3443612, EPI_ISL_3443613, EPI_ISL_3443614, EPI_ISL_3443615, EPI_ISL_3443616, EPI_ISL_3443617, EPI_ISL_3443618, EPI_ISL_3443619, EPI_ISL_3443620, EPI_ISL_3443621, EPI_ISL_3443622, EPI_ISL_3443623, EPI_ISL_3443624, EPI_ISL_3443625, EPI_ISL_3443626, EPI_ISL_3443627, EPI_ISL_3443628, EPI_ISL_3443629, EPI_ISL_3443630, EPI_ISL_3443631, EPI_ISL_3443632, EPI_ISL_3443633, EPI_ISL_3443634, EPI_ISL_3443635, EPI_ISL_3443636, EPI_ISL_3443637, EPI_ISL_3443638, EPI_ISL_3443639, EPI_ISL_3443640, EPI_ISL_3443641, EPI_ISL_3443642, EPI_ISL_3443643, EPI_ISL_3443644, EPI_ISL_3443645, EPI_ISL_3443646, EPI_ISL_3443647, EPI_ISL_3443648, EPI_ISL_3443649, EPI_ISL_3443650, EPI_ISL_3443651, EPI_ISL_3443652, EPI_ISL_3443653, EPI_ISL_3443654, EPI_ISL_3443655, EPI_ISL_3443656, EPI_ISL_3443657, EPI_ISL_3443658, EPI_ISL_3443659, EPI_ISL_3443660, EPI_ISL_3443661, EPI_ISL_3443662, EPI_ISL_3443663, EPI_ISL_3443664, EPI_ISL_3443665, EPI_ISL_3443666, EPI_ISL_3443667, EPI_ISL_3443668, EPI_ISL_3443669, EPI_ISL_3443670, EPI_ISL_3443671, EPI_ISL_3443672, EPI_ISL_3443673, EPI_ISL_3443674, EPI_ISL_3443675, EPI_ISL_3443676, EPI_ISL_3443677, EPI_ISL_3443678, EPI_ISL_3443679, EPI_ISL_3443680, EPI_ISL_3443681, EPI_ISL_3443682, EPI_ISL_3443683, EPI_ISL_3443684, EPI_ISL_3443685, EPI_ISL_3443686, EPI_ISL_3443687, EPI_ISL_3443688, EPI_ISL_3443689 | | | |
| see above | Lighthouse Lab in Glasgow | Wellcome Sanger Institute for the COVID-19 Genomics UK (COG-UK) Consortium | Harper VanSteenhouse, Yumi Kasai, David Gray, Carol Clugston, Anna Dominiczak and Alex Alderton, Roberto Amato, Jeffrey Barrett, Sonia Goncalves, Ewan Harrison, David K. Jackson, Ian Johnston, Dominic Kwiatkowski, Cordelia Langford, John Sillitoe on behalf of the Wellcome Sanger Institute COVID-19 Surveillance Team |
| EPI_ISL_3443690, EPI_ISL_3443691, EPI_ISL_3443692, EPI_ISL_3443693, EPI_ISL_3443694 | Lighthouse Lab in Milton Keynes | Wellcome Sanger Institute for the COVID-19 Genomics UK (COG-UK) Consortium | The Lighthouse Lab in Milton Keynes and Alex Alderton, Roberto Amato, Jeffrey Barrett, Sonia Goncalves, Ewan Harrison, David K. Jackson, Ian Johnston, Dominic Kwiatkowski, Cordelia Langford, John Sillitoe on behalf of the Wellcome Sanger Institute COVID-19 Surveillance Team |
| EPI_ISL_3443695, EPI_ISL_3443696, EPI_ISL_3443697 | Lighthouse Lab in Glasgow | Wellcome Sanger Institute for the COVID-19 Genomics UK (COG-UK) Consortium | Harper VanSteenhouse, Yumi Kasai, David Gray, Carol Clugston, Anna Dominiczak and Alex Alderton, Roberto Amato, Jeffrey Barrett, Sonia Goncalves, Ewan Harrison, David K. Jackson, Ian Johnston, Dominic Kwiatkowski, Cordelia Langford, John Sillitoe on behalf of the Wellcome Sanger Institute COVID-19 Surveillance Team |
| EPI_ISL_3443698 | Lighthouse Lab in Milton Keynes | Wellcome Sanger Institute for the COVID-19 Genomics UK (COG-UK) Consortium | The Lighthouse Lab in Milton Keynes and Alex Alderton, Roberto Amato, Jeffrey Barrett, Sonia Goncalves, Ewan Harrison, David K. Jackson, Ian Johnston, Dominic Kwiatkowski, Cordelia Langford, John Sillitoe on behalf of the Wellcome Sanger Institute COVID-19 Surveillance Team |
| EPI_ISL_3443699 | Lighthouse Lab in Glasgow | Wellcome Sanger Institute for the COVID-19 Genomics UK (COG-UK) Consortium | Harper VanSteenhouse, Yumi Kasai, David Gray, Carol Clugston, Anna Dominiczak and Alex Alderton, Roberto Amato, Jeffrey Barrett, Sonia Goncalves, Ewan Harrison, David K. Jackson, Ian Johnston, Dominic Kwiatkowski, Cordelia Langford, John Sillitoe on behalf of the Wellcome Sanger Institute COVID-19 Surveillance Team |
| EPI_ISL_3443700 | Lighthouse Lab in Milton Keynes | Wellcome Sanger Institute for the COVID-19 Genomics UK (COG-UK) Consortium | The Lighthouse Lab in Milton Keynes and Alex Alderton, Roberto Amato, Jeffrey Barrett, Sonia Goncalves, Ewan Harrison, David K. Jackson, Ian Johnston, Dominic Kwiatkowski, Cordelia Langford, John Sillitoe on behalf of the Wellcome Sanger Institute COVID-19 Surveillance Team |
| EPI_ISL_3443701 | Lighthouse Lab in Glasgow | Wellcome Sanger Institute for the COVID-19 Genomics UK (COG-UK) Consortium | Harper VanSteenhouse, Yumi Kasai, David Gray, Carol Clugston, Anna Dominiczak and Alex Alderton, Roberto Amato, Jeffrey Barrett, Sonia Goncalves, Ewan Harrison, David K. Jackson, Ian Johnston, Dominic Kwiatkowski, Cordelia Langford, John Sillitoe on behalf of the Wellcome Sanger Institute COVID-19 Surveillance Team |
| EPI_ISL_3443702 | Lighthouse Lab in Milton Keynes | Wellcome Sanger Institute for the COVID-19 Genomics UK (COG-UK) Consortium | The Lighthouse Lab in Milton Keynes and Alex Alderton, Roberto Amato, Jeffrey Barrett, Sonia Goncalves, Ewan Harrison, David K. Jackson, Ian Johnston, Dominic Kwiatkowski, Cordelia Langford, John Sillitoe on behalf of the Wellcome Sanger Institute COVID-19 Surveillance Team |
| EPI_ISL_3443703, EPI_ISL_3443704 | Lighthouse Lab in Glasgow | Wellcome Sanger Institute for the COVID-19 Genomics UK (COG-UK) Consortium | Harper VanSteenhouse, Yumi Kasai, David Gray, Carol Clugston, Anna Dominiczak and Alex Alderton, Roberto Amato, Jeffrey Barrett, Sonia Goncalves, Ewan Harrison, David K. Jackson, Ian Johnston, Dominic Kwiatkowski, Cordelia Langford, John Sillitoe on behalf of the Wellcome Sanger Institute COVID-19 Surveillance Team |
| EPI_ISL_3443705 | Lighthouse Lab in Milton Keynes | Wellcome Sanger Institute for the COVID-19 Genomics UK (COG-UK) Consortium | The Lighthouse Lab in Milton Keynes and Alex Alderton, Roberto Amato, Jeffrey Barrett, Sonia Goncalves, Ewan Harrison, David K. Jackson, Ian Johnston, Dominic Kwiatkowski, Cordelia Langford, John Sillitoe on behalf of the Wellcome Sanger Institute COVID-19 Surveillance Team |
| EPI_ISL_3443706, EPI_ISL_3443707 | Lighthouse Lab in Glasgow | Wellcome Sanger Institute for the COVID-19 Genomics UK (COG-UK) Consortium | Harper VanSteenhouse, Yumi Kasai, David Gray, Carol Clugston, Anna Dominiczak and Alex Alderton, Roberto Amato, Jeffrey Barrett, Sonia Goncalves, Ewan Harrison, David K. Jackson, Ian Johnston, Dominic Kwiatkowski, Cordelia Langford, John Sillitoe on behalf of the Wellcome Sanger Institute COVID-19 Surveillance Team |
| EPI_ISL_3443708, EPI_ISL_3443709, EPI_ISL_3443710 | Lighthouse Lab in Milton Keynes | Wellcome Sanger Institute for the COVID-19 Genomics UK (COG-UK) Consortium | The Lighthouse Lab in Milton Keynes and Alex Alderton, Roberto Amato, Jeffrey Barrett, Sonia Goncalves, Ewan Harrison, David K. Jackson, Ian Johnston, Dominic Kwiatkowski, Cordelia Langford, John Sillitoe on behalf of the Wellcome Sanger Institute COVID-19 Surveillance Team |
| EPI_ISL_3443711 | Lighthouse Lab in Glasgow | Wellcome Sanger Institute for the COVID-19 Genomics UK (COG-UK) Consortium | Harper VanSteenhouse, Yumi Kasai, David Gray, Carol Clugston, Anna Dominiczak and Alex Alderton, Roberto Amato, Jeffrey Barrett, Sonia Goncalves, Ewan Harrison, David K. Jackson, Ian Johnston, Dominic Kwiatkowski, Cordelia Langford, John Sillitoe on behalf of the Wellcome Sanger Institute COVID-19 Surveillance Team |
| EPI_ISL_3443712, EPI_ISL_3443713, EPI_ISL_3443714 | Lighthouse Lab in Milton Keynes | Wellcome Sanger Institute for the COVID-19 Genomics UK (COG-UK) Consortium | The Lighthouse Lab in Milton Keynes and Alex Alderton, Roberto Amato, Jeffrey Barrett, Sonia Goncalves, Ewan Harrison, David K. Jackson, Ian Johnston, Dominic Kwiatkowski, Cordelia Langford, John Sillitoe on behalf of the Wellcome Sanger Institute COVID-19 Surveillance Team |
| EPI_ISL_3443715, EPI_ISL_3443716, EPI_ISL_3443717 | Lighthouse Lab in Glasgow | Wellcome Sanger Institute for the COVID-19 Genomics UK (COG-UK) Consortium | Harper VanSteenhouse, Yumi Kasai, David Gray, Carol Clugston, Anna Dominiczak and Alex Alderton, Roberto Amato, Jeffrey Barrett, Sonia Goncalves, Ewan Harrison, David K. Jackson, Ian Johnston, Dominic Kwiatkowski, Cordelia Langford, John Sillitoe on behalf of the Wellcome Sanger Institute COVID-19 Surveillance Team |
| EPI_ISL_3443718, EPI_ISL_3443719, EPI_ISL_3443720, EPI_ISL_3443721 | Lighthouse Lab in Milton Keynes | Wellcome Sanger Institute for the COVID-19 Genomics UK (COG-UK) Consortium | The Lighthouse Lab in Milton Keynes and Alex Alderton, Roberto Amato, Jeffrey Barrett, Sonia Goncalves, Ewan Harrison, David K. Jackson, Ian Johnston, Dominic Kwiatkowski, Cordelia Langford, John Sillitoe on behalf of the Wellcome Sanger Institute COVID-19 Surveillance Team |
| EPI_ISL_3443722, EPI_ISL_3443723, EPI_ISL_3443724, EPI_ISL_3443725, EPI_ISL_3443726 | Lighthouse Lab in Glasgow | Wellcome Sanger Institute for the COVID-19 Genomics UK (COG-UK) Consortium | Harper VanSteenhouse, Yumi Kasai, David Gray, Carol Clugston, Anna Dominiczak and Alex Alderton, Roberto Amato, Jeffrey Barrett, Sonia Goncalves, Ewan Harrison, David K. Jackson, Ian Johnston, Dominic Kwiatkowski, Cordelia Langford, John Sillitoe on behalf of the Wellcome Sanger Institute COVID-19 Surveillance Team |
| EPI_ISL_3443727 | Lighthouse Lab in Milton Keynes | Wellcome Sanger Institute for the COVID-19 Genomics UK (COG-UK) | The Lighthouse Lab in Milton Keynes and Alex Alderton, Roberto Amato, Jeffrey Barrett, Sonia Goncalves, Ewan Harrison, David K. Jackson, Ian Johnston, Dominic Kwiatkowski, Cordelia Langford, John Sillitoe on behalf of the Wellcome Sanger |

| | | | |
|---|---|---|---|
| | | Consortium | Institute COVID-19 Surveillance Team |
| EPI_ISL_3443728, EPI_ISL_3443729, EPI_ISL_3443730 | Lighthouse Lab in Glasgow | Wellcome Sanger Institute for the COVID-19 Genomics UK (COG-UK) Consortium | Harper VanSteenhouse, Yumi Kasai, David Gray, Carol Clugston, Anna Dominiczak and Alex Alderton, Roberto Amato, Jeffrey Barrett, Sonia Goncalves, Ewan Harrison, David K. Jackson, Ian Johnston, Dominic Kwiatkowski, Cordelia Langford, John Sillitoe on behalf of the Wellcome Sanger Institute COVID-19 Surveillance Team |
| EPI_ISL_3443731, EPI_ISL_3443732, EPI_ISL_3443733 | Lighthouse Lab in Milton Keynes | Wellcome Sanger Institute for the COVID-19 Genomics UK (COG-UK) Consortium | The Lighthouse Lab in Milton Keynes and Alex Alderton, Roberto Amato, Jeffrey Barrett, Sonia Goncalves, Ewan Harrison, David K. Jackson, Ian Johnston, Dominic Kwiatkowski, Cordelia Langford, John Sillitoe on behalf of the Wellcome Sanger Institute COVID-19 Surveillance Team |
| EPI_ISL_3443734 | Lighthouse Lab in Glasgow | Wellcome Sanger Institute for the COVID-19 Genomics UK (COG-UK) Consortium | Harper VanSteenhouse, Yumi Kasai, David Gray, Carol Clugston, Anna Dominiczak and Alex Alderton, Roberto Amato, Jeffrey Barrett, Sonia Goncalves, Ewan Harrison, David K. Jackson, Ian Johnston, Dominic Kwiatkowski, Cordelia Langford, John Sillitoe on behalf of the Wellcome Sanger Institute COVID-19 Surveillance Team |
| EPI_ISL_3443735, EPI_ISL_3443736, EPI_ISL_3443737, EPI_ISL_3443738, EPI_ISL_3443739 | Lighthouse Lab in Milton Keynes | Wellcome Sanger Institute for the COVID-19 Genomics UK (COG-UK) Consortium | The Lighthouse Lab in Milton Keynes and Alex Alderton, Roberto Amato, Jeffrey Barrett, Sonia Goncalves, Ewan Harrison, David K. Jackson, Ian Johnston, Dominic Kwiatkowski, Cordelia Langford, John Sillitoe on behalf of the Wellcome Sanger Institute COVID-19 Surveillance Team |
| EPI_ISL_3443740, EPI_ISL_3443741 | Lighthouse Lab in Glasgow | Wellcome Sanger Institute for the COVID-19 Genomics UK (COG-UK) Consortium | Harper VanSteenhouse, Yumi Kasai, David Gray, Carol Clugston, Anna Dominiczak and Alex Alderton, Roberto Amato, Jeffrey Barrett, Sonia Goncalves, Ewan Harrison, David K. Jackson, Ian Johnston, Dominic Kwiatkowski, Cordelia Langford, John Sillitoe on behalf of the Wellcome Sanger Institute COVID-19 Surveillance Team |
| EPI_ISL_3443742 | Lighthouse Lab in Milton Keynes | Wellcome Sanger Institute for the COVID-19 Genomics UK (COG-UK) Consortium | The Lighthouse Lab in Milton Keynes and Alex Alderton, Roberto Amato, Jeffrey Barrett, Sonia Goncalves, Ewan Harrison, David K. Jackson, Ian Johnston, Dominic Kwiatkowski, Cordelia Langford, John Sillitoe on behalf of the Wellcome Sanger Institute COVID-19 Surveillance Team |
| EPI_ISL_3443743 | Lighthouse Lab in Glasgow | Wellcome Sanger Institute for the COVID-19 Genomics UK (COG-UK) Consortium | Harper VanSteenhouse, Yumi Kasai, David Gray, Carol Clugston, Anna Dominiczak and Alex Alderton, Roberto Amato, Jeffrey Barrett, Sonia Goncalves, Ewan Harrison, David K. Jackson, Ian Johnston, Dominic Kwiatkowski, Cordelia Langford, John Sillitoe on behalf of the Wellcome Sanger Institute COVID-19 Surveillance Team |
| EPI_ISL_3443744, EPI_ISL_3443745 | Lighthouse Lab in Milton Keynes | Wellcome Sanger Institute for the COVID-19 Genomics UK (COG-UK) Consortium | The Lighthouse Lab in Milton Keynes and Alex Alderton, Roberto Amato, Jeffrey Barrett, Sonia Goncalves, Ewan Harrison, David K. Jackson, Ian Johnston, Dominic Kwiatkowski, Cordelia Langford, John Sillitoe on behalf of the Wellcome Sanger Institute COVID-19 Surveillance Team |
| EPI_ISL_3443746, EPI_ISL_3443747, EPI_ISL_3443748 | Lighthouse Lab in Glasgow | Wellcome Sanger Institute for the COVID-19 Genomics UK (COG-UK) Consortium | Harper VanSteenhouse, Yumi Kasai, David Gray, Carol Clugston, Anna Dominiczak and Alex Alderton, Roberto Amato, Jeffrey Barrett, Sonia Goncalves, Ewan Harrison, David K. Jackson, Ian Johnston, Dominic Kwiatkowski, Cordelia Langford, John Sillitoe on behalf of the Wellcome Sanger Institute COVID-19 Surveillance Team |
| EPI_ISL_3443749 | Lighthouse Lab in Milton Keynes | Wellcome Sanger Institute for the COVID-19 Genomics UK (COG-UK) Consortium | The Lighthouse Lab in Milton Keynes and Alex Alderton, Roberto Amato, Jeffrey Barrett, Sonia Goncalves, Ewan Harrison, David K. Jackson, Ian Johnston, Dominic Kwiatkowski, Cordelia Langford, John Sillitoe on behalf of the Wellcome Sanger Institute COVID-19 Surveillance Team |
| EPI_ISL_3443750, EPI_ISL_3443751 | Lighthouse Lab in Glasgow | Wellcome Sanger Institute for the COVID-19 Genomics UK (COG-UK) Consortium | Harper VanSteenhouse, Yumi Kasai, David Gray, Carol Clugston, Anna Dominiczak and Alex Alderton, Roberto Amato, Jeffrey Barrett, Sonia Goncalves, Ewan Harrison, David K. Jackson, Ian Johnston, Dominic Kwiatkowski, Cordelia Langford, John Sillitoe on behalf of the Wellcome Sanger Institute COVID-19 Surveillance Team |
| EPI_ISL_3443752 | Lighthouse Lab in Milton Keynes | Wellcome Sanger Institute for the COVID-19 Genomics UK (COG-UK) Consortium | The Lighthouse Lab in Milton Keynes and Alex Alderton, Roberto Amato, Jeffrey Barrett, Sonia Goncalves, Ewan Harrison, David K. Jackson, Ian Johnston, Dominic Kwiatkowski, Cordelia Langford, John Sillitoe on behalf of the Wellcome Sanger Institute COVID-19 Surveillance Team |
| EPI_ISL_3443753 | Lighthouse Lab in Glasgow | Wellcome Sanger Institute for the COVID-19 Genomics UK (COG-UK) Consortium | Harper VanSteenhouse, Yumi Kasai, David Gray, Carol Clugston, Anna Dominiczak and Alex Alderton, Roberto Amato, Jeffrey Barrett, Sonia Goncalves, Ewan Harrison, David K. Jackson, Ian Johnston, Dominic Kwiatkowski, Cordelia Langford, John Sillitoe on behalf of the Wellcome Sanger Institute COVID-19 Surveillance Team |
| EPI_ISL_3443754, EPI_ISL_3443755, EPI_ISL_3443756 | Lighthouse Lab in Milton Keynes | Wellcome Sanger Institute for the COVID-19 Genomics UK (COG-UK) Consortium | The Lighthouse Lab in Milton Keynes and Alex Alderton, Roberto Amato, Jeffrey Barrett, Sonia Goncalves, Ewan Harrison, David K. Jackson, Ian Johnston, Dominic Kwiatkowski, Cordelia Langford, John Sillitoe on behalf of the Wellcome Sanger Institute COVID-19 Surveillance Team |
| EPI_ISL_3443757 | Lighthouse Lab in Glasgow | Wellcome Sanger Institute for the COVID-19 Genomics UK (COG-UK) Consortium | Harper VanSteenhouse, Yumi Kasai, David Gray, Carol Clugston, Anna Dominiczak and Alex Alderton, Roberto Amato, Jeffrey Barrett, Sonia Goncalves, Ewan Harrison, David K. Jackson, Ian Johnston, Dominic Kwiatkowski, Cordelia Langford, John Sillitoe on behalf of the Wellcome Sanger Institute COVID-19 Surveillance Team |
| EPI_ISL_3443758 | Lighthouse Lab in Milton Keynes | Wellcome Sanger Institute for the COVID-19 Genomics UK (COG-UK) Consortium | The Lighthouse Lab in Milton Keynes and Alex Alderton, Roberto Amato, Jeffrey Barrett, Sonia Goncalves, Ewan Harrison, David K. Jackson, Ian Johnston, Dominic Kwiatkowski, Cordelia Langford, John Sillitoe on behalf of the Wellcome Sanger Institute COVID-19 Surveillance Team |
| EPI_ISL_3443759, EPI_ISL_3443760 | Lighthouse Lab in Glasgow | Wellcome Sanger Institute for the COVID-19 Genomics UK (COG-UK) Consortium | Harper VanSteenhouse, Yumi Kasai, David Gray, Carol Clugston, Anna Dominiczak and Alex Alderton, Roberto Amato, Jeffrey Barrett, Sonia Goncalves, Ewan Harrison, David K. Jackson, Ian Johnston, Dominic Kwiatkowski, Cordelia Langford, John Sillitoe on behalf of the Wellcome Sanger Institute COVID-19 Surveillance Team |
| EPI_ISL_3443761, EPI_ISL_3443762, EPI_ISL_3443763 | Lighthouse Lab in Milton Keynes | Wellcome Sanger Institute for the COVID-19 Genomics UK (COG-UK) Consortium | The Lighthouse Lab in Milton Keynes and Alex Alderton, Roberto Amato, Jeffrey Barrett, Sonia Goncalves, Ewan Harrison, David K. Jackson, Ian Johnston, Dominic Kwiatkowski, Cordelia Langford, John Sillitoe on behalf of the Wellcome Sanger Institute COVID-19 Surveillance Team |
| EPI_ISL_3443764 | Lighthouse Lab in Glasgow | Wellcome Sanger Institute for the COVID-19 Genomics UK (COG-UK) Consortium | Harper VanSteenhouse, Yumi Kasai, David Gray, Carol Clugston, Anna Dominiczak and Alex Alderton, Roberto Amato, Jeffrey Barrett, Sonia Goncalves, Ewan Harrison, David K. Jackson, Ian Johnston, Dominic Kwiatkowski, Cordelia Langford, John Sillitoe on behalf of the Wellcome Sanger Institute COVID-19 Surveillance Team |
| EPI_ISL_3443765 | Lighthouse Lab in Milton Keynes | Wellcome Sanger Institute for the COVID-19 Genomics UK (COG-UK) Consortium | The Lighthouse Lab in Milton Keynes and Alex Alderton, Roberto Amato, Jeffrey Barrett, Sonia Goncalves, Ewan Harrison, David K. Jackson, Ian Johnston, Dominic Kwiatkowski, Cordelia Langford, John Sillitoe on behalf of the Wellcome Sanger Institute COVID-19 Surveillance Team |
| EPI_ISL_3443766 | Lighthouse Lab in Glasgow | Wellcome Sanger Institute for the COVID-19 Genomics UK (COG-UK) Consortium | Harper VanSteenhouse, Yumi Kasai, David Gray, Carol Clugston, Anna Dominiczak and Alex Alderton, Roberto Amato, Jeffrey Barrett, Sonia Goncalves, Ewan Harrison, David K. Jackson, Ian Johnston, Dominic Kwiatkowski, Cordelia Langford, John Sillitoe on behalf of the Wellcome Sanger Institute COVID-19 Surveillance Team |
| EPI_ISL_3443767 | Lighthouse Lab in Milton Keynes | Wellcome Sanger Institute for the COVID-19 Genomics UK (COG-UK) Consortium | The Lighthouse Lab in Milton Keynes and Alex Alderton, Roberto Amato, Jeffrey Barrett, Sonia Goncalves, Ewan Harrison, David K. Jackson, Ian Johnston, Dominic Kwiatkowski, Cordelia Langford, John Sillitoe on behalf of the Wellcome Sanger Institute COVID-19 Surveillance Team |
| EPI_ISL_3443768 | Lighthouse Lab in Glasgow | Wellcome Sanger Institute for the COVID-19 Genomics UK (COG-UK) Consortium | Harper VanSteenhouse, Yumi Kasai, David Gray, Carol Clugston, Anna Dominiczak and Alex Alderton, Roberto Amato, Jeffrey Barrett, Sonia Goncalves, Ewan Harrison, David K. Jackson, Ian Johnston, Dominic Kwiatkowski, Cordelia Langford, John Sillitoe on behalf of the Wellcome Sanger Institute COVID-19 Surveillance Team |
| EPI_ISL_3443769, EPI_ISL_3443770 | Lighthouse Lab in Milton Keynes | Wellcome Sanger Institute for the COVID-19 Genomics UK (COG-UK) Consortium | The Lighthouse Lab in Milton Keynes and Alex Alderton, Roberto Amato, Jeffrey Barrett, Sonia Goncalves, Ewan Harrison, David K. Jackson, Ian Johnston, Dominic Kwiatkowski, Cordelia Langford, John Sillitoe on behalf of the Wellcome Sanger Institute COVID-19 Surveillance Team |
| EPI_ISL_3443771, EPI_ISL_3443772 | Lighthouse Lab in Glasgow | Wellcome Sanger Institute for the COVID-19 Genomics UK (COG-UK) Consortium | Harper VanSteenhouse, Yumi Kasai, David Gray, Carol Clugston, Anna Dominiczak and Alex Alderton, Roberto Amato, Jeffrey Barrett, Sonia Goncalves, Ewan Harrison, David K. Jackson, Ian Johnston, Dominic Kwiatkowski, Cordelia Langford, John Sillitoe on behalf of the Wellcome Sanger Institute COVID-19 Surveillance Team |
| EPI_ISL_3443773 | Lighthouse Lab in Milton Keynes | Wellcome Sanger Institute for the COVID-19 Genomics UK (COG-UK) Consortium | The Lighthouse Lab in Milton Keynes and Alex Alderton, Roberto Amato, Jeffrey Barrett, Sonia Goncalves, Ewan Harrison, David K. Jackson, Ian Johnston, Dominic Kwiatkowski, Cordelia Langford, John Sillitoe on behalf of the Wellcome Sanger Institute COVID-19 Surveillance Team |
| EPI_ISL_3443774 | Lighthouse Lab in Glasgow | Wellcome Sanger Institute for the COVID-19 Genomics UK (COG-UK) Consortium | Harper VanSteenhouse, Yumi Kasai, David Gray, Carol Clugston, Anna Dominiczak and Alex Alderton, Roberto Amato, Jeffrey Barrett, Sonia Goncalves, Ewan Harrison, David K. Jackson, Ian Johnston, Dominic Kwiatkowski, Cordelia Langford, John Sillitoe on behalf of the Wellcome Sanger Institute COVID-19 Surveillance Team |
| EPI_ISL_3443775 | Lighthouse Lab in Milton Keynes | Wellcome Sanger Institute for the COVID-19 Genomics UK (COG-UK) Consortium | The Lighthouse Lab in Milton Keynes and Alex Alderton, Roberto Amato, Jeffrey Barrett, Sonia Goncalves, Ewan Harrison, David K. Jackson, Ian Johnston, Dominic Kwiatkowski, Cordelia Langford, John Sillitoe on behalf of the Wellcome Sanger Institute COVID-19 Surveillance Team |
| EPI_ISL_3443776, EPI_ISL_3443777 | Lighthouse Lab in Glasgow | Wellcome Sanger Institute for the COVID-19 Genomics UK (COG-UK) Consortium | Harper VanSteenhouse, Yumi Kasai, David Gray, Carol Clugston, Anna Dominiczak and Alex Alderton, Roberto Amato, Jeffrey Barrett, Sonia Goncalves, Ewan Harrison, David K. Jackson, Ian Johnston, Dominic Kwiatkowski, Cordelia Langford, John Sillitoe on behalf of the Wellcome Sanger Institute COVID-19 Surveillance Team |
| EPI_ISL_3443778, EPI_ISL_3443779, EPI_ISL_3443780 | Lighthouse Lab in Milton Keynes | Wellcome Sanger Institute for the COVID-19 Genomics UK (COG-UK) Consortium | The Lighthouse Lab in Milton Keynes and Alex Alderton, Roberto Amato, Jeffrey Barrett, Sonia Goncalves, Ewan Harrison, David K. Jackson, Ian Johnston, Dominic Kwiatkowski, Cordelia Langford, John Sillitoe on behalf of the Wellcome Sanger Institute COVID-19 Surveillance Team |
| EPI_ISL_3443781 | Lighthouse Lab in Glasgow | Wellcome Sanger Institute for the COVID-19 Genomics UK (COG-UK) Consortium | Harper VanSteenhouse, Yumi Kasai, David Gray, Carol Clugston, Anna Dominiczak and Alex Alderton, Roberto Amato, Jeffrey Barrett, Sonia Goncalves, Ewan Harrison, David K. Jackson, Ian Johnston, Dominic Kwiatkowski, Cordelia Langford, John Sillitoe on behalf of the Wellcome Sanger Institute COVID-19 Surveillance Team |
| EPI_ISL_3443782 | Lighthouse Lab in Milton Keynes | Wellcome Sanger Institute for the COVID-19 Genomics UK (COG-UK) Consortium | The Lighthouse Lab in Milton Keynes and Alex Alderton, Roberto Amato, Jeffrey Barrett, Sonia Goncalves, Ewan Harrison, David K. Jackson, Ian Johnston, Dominic Kwiatkowski, Cordelia Langford, John Sillitoe on behalf of the Wellcome Sanger Institute COVID-19 Surveillance Team |
| EPI_ISL_3443783, EPI_ISL_3443784 | Lighthouse Lab in Glasgow | Wellcome Sanger Institute for the COVID-19 Genomics UK (COG-UK) Consortium | Harper VanSteenhouse, Yumi Kasai, David Gray, Carol Clugston, Anna Dominiczak and Alex Alderton, Roberto Amato, Jeffrey Barrett, Sonia Goncalves, Ewan Harrison, David K. Jackson, Ian Johnston, Dominic Kwiatkowski, Cordelia Langford, John Sillitoe on behalf of the Wellcome Sanger Institute COVID-19 Surveillance Team |
| EPI_ISL_3443785, EPI_ISL_3443786, EPI_ISL_3443787, EPI_ISL_3443788 | Lighthouse Lab in Milton Keynes | Wellcome Sanger Institute for the COVID-19 Genomics UK (COG-UK) Consortium | The Lighthouse Lab in Milton Keynes and Alex Alderton, Roberto Amato, Jeffrey Barrett, Sonia Goncalves, Ewan Harrison, David K. Jackson, Ian Johnston, Dominic Kwiatkowski, Cordelia Langford, John Sillitoe on behalf of the Wellcome Sanger Institute COVID-19 Surveillance Team |
| EPI_ISL_3443789 | Lighthouse Lab in Glasgow | Wellcome Sanger Institute for the COVID-19 Genomics UK (COG-UK) Consortium | Harper VanSteenhouse, Yumi Kasai, David Gray, Carol Clugston, Anna Dominiczak and Alex Alderton, Roberto Amato, Jeffrey Barrett, Sonia Goncalves, Ewan Harrison, David K. Jackson, Ian Johnston, Dominic Kwiatkowski, Cordelia Langford, John Sillitoe on behalf of the Wellcome Sanger Institute COVID-19 Surveillance Team |
| EPI_ISL_3443790, EPI_ISL_3443791 | Lighthouse Lab in Milton Keynes | Wellcome Sanger Institute for the COVID-19 Genomics UK (COG-UK) Consortium | The Lighthouse Lab in Milton Keynes and Alex Alderton, Roberto Amato, Jeffrey Barrett, Sonia Goncalves, Ewan Harrison, David K. Jackson, Ian Johnston, Dominic Kwiatkowski, Cordelia Langford, John Sillitoe on behalf of the Wellcome Sanger Institute COVID-19 Surveillance Team |
| EPI_ISL_3443792, EPI_ISL_3443793 | Lighthouse Lab in Glasgow | Wellcome Sanger Institute for the COVID-19 Genomics UK (COG-UK) Consortium | Harper VanSteenhouse, Yumi Kasai, David Gray, Carol Clugston, Anna Dominiczak and Alex Alderton, Roberto Amato, Jeffrey Barrett, Sonia Goncalves, Ewan Harrison, David K. Jackson, Ian Johnston, Dominic Kwiatkowski, Cordelia Langford, John Sillitoe on behalf of the Wellcome Sanger Institute COVID-19 Surveillance Team |
| EPI_ISL_3443794 | Lighthouse Lab in Milton Keynes | Wellcome Sanger Institute for the COVID-19 Genomics UK (COG-UK) Consortium | The Lighthouse Lab in Milton Keynes and Alex Alderton, Roberto Amato, Jeffrey Barrett, Sonia Goncalves, Ewan Harrison, David K. Jackson, Ian Johnston, Dominic Kwiatkowski, Cordelia Langford, John Sillitoe on behalf of the Wellcome Sanger Institute COVID-19 Surveillance Team |
| EPI_ISL_3443795 | Lighthouse Lab in Glasgow | Wellcome Sanger Institute for the COVID-19 Genomics UK (COG-UK) Consortium | Harper VanSteenhouse, Yumi Kasai, David Gray, Carol Clugston, Anna Dominiczak and Alex Alderton, Roberto Amato, Jeffrey Barrett, Sonia Goncalves, Ewan Harrison, David K. Jackson, Ian Johnston, Dominic Kwiatkowski, Cordelia Langford, John Sillitoe on behalf of the Wellcome Sanger Institute COVID-19 Surveillance Team |
| EPI_ISL_3443796 | Lighthouse Lab in Milton Keynes | Wellcome Sanger Institute for the COVID-19 Genomics UK (COG-UK) Consortium | The Lighthouse Lab in Milton Keynes and Alex Alderton, Roberto Amato, Jeffrey Barrett, Sonia Goncalves, Ewan Harrison, David K. Jackson, Ian Johnston, Dominic Kwiatkowski, Cordelia Langford, John Sillitoe on behalf of the Wellcome Sanger Institute COVID-19 Surveillance Team |

| | | | |
|---|---|---|---|
| EPI_ISL_3443797, EPI_ISL_3443798 | Lighthouse Lab in Glasgow | Wellcome Sanger Institute for the COVID-19 Genomics UK (COG-UK) Consortium | Harper VanSteenhouse, Yumi Kasai, David Gray, Carol Clugston, Anna Dominiczak and Alex Alderton, Roberto Amato, Jeffrey Barrett, Sonia Goncalves, Ewan Harrison, David K. Jackson, Ian Johnston, Dominic Kwiatkowski, Cordelia Langford, John Sillitoe on behalf of the Wellcome Sanger Institute COVID-19 Surveillance Team |
| EPI_ISL_3443799, EPI_ISL_3443800, EPI_ISL_3443801 | Lighthouse Lab in Milton Keynes | Wellcome Sanger Institute for the COVID-19 Genomics UK (COG-UK) Consortium | The Lighthouse Lab in Milton Keynes and Alex Alderton, Roberto Amato, Jeffrey Barrett, Sonia Goncalves, Ewan Harrison, David K. Jackson, Ian Johnston, Dominic Kwiatkowski, Cordelia Langford, John Sillitoe on behalf of the Wellcome Sanger Institute COVID-19 Surveillance Team |
| EPI_ISL_3443802, EPI_ISL_3443803, EPI_ISL_3443804, EPI_ISL_3443805 | Lighthouse Lab in Glasgow | Wellcome Sanger Institute for the COVID-19 Genomics UK (COG-UK) Consortium | Harper VanSteenhouse, Yumi Kasai, David Gray, Carol Clugston, Anna Dominiczak and Alex Alderton, Roberto Amato, Jeffrey Barrett, Sonia Goncalves, Ewan Harrison, David K. Jackson, Ian Johnston, Dominic Kwiatkowski, Cordelia Langford, John Sillitoe on behalf of the Wellcome Sanger Institute COVID-19 Surveillance Team |
| EPI_ISL_3443806, EPI_ISL_3443807, EPI_ISL_3443808, EPI_ISL_3443809, EPI_ISL_3443810, EPI_ISL_3443811, EPI_ISL_3443812, EPI_ISL_3443813, EPI_ISL_3443814, EPI_ISL_3443815, EPI_ISL_3443816, EPI_ISL_3443817 | | | |
| see above | Lighthouse Lab in Milton Keynes | Wellcome Sanger Institute for the COVID-19 Genomics UK (COG-UK) Consortium | The Lighthouse Lab in Milton Keynes and Alex Alderton, Roberto Amato, Jeffrey Barrett, Sonia Goncalves, Ewan Harrison, David K. Jackson, Ian Johnston, Dominic Kwiatkowski, Cordelia Langford, John Sillitoe on behalf of the Wellcome Sanger Institute COVID-19 Surveillance Team |
| EPI_ISL_3443818 | Lighthouse Lab in Glasgow | Wellcome Sanger Institute for the COVID-19 Genomics UK (COG-UK) Consortium | Harper VanSteenhouse, Yumi Kasai, David Gray, Carol Clugston, Anna Dominiczak and Alex Alderton, Roberto Amato, Jeffrey Barrett, Sonia Goncalves, Ewan Harrison, David K. Jackson, Ian Johnston, Dominic Kwiatkowski, Cordelia Langford, John Sillitoe on behalf of the Wellcome Sanger Institute COVID-19 Surveillance Team |
| EPI_ISL_3443819, EPI_ISL_3443820 | Lighthouse Lab in Milton Keynes | Wellcome Sanger Institute for the COVID-19 Genomics UK (COG-UK) Consortium | The Lighthouse Lab in Milton Keynes and Alex Alderton, Roberto Amato, Jeffrey Barrett, Sonia Goncalves, Ewan Harrison, David K. Jackson, Ian Johnston, Dominic Kwiatkowski, Cordelia Langford, John Sillitoe on behalf of the Wellcome Sanger Institute COVID-19 Surveillance Team |
| EPI_ISL_3443821, EPI_ISL_3443822, EPI_ISL_3443823 | Lighthouse Lab in Glasgow | Wellcome Sanger Institute for the COVID-19 Genomics UK (COG-UK) Consortium | Harper VanSteenhouse, Yumi Kasai, David Gray, Carol Clugston, Anna Dominiczak and Alex Alderton, Roberto Amato, Jeffrey Barrett, Sonia Goncalves, Ewan Harrison, David K. Jackson, Ian Johnston, Dominic Kwiatkowski, Cordelia Langford, John Sillitoe on behalf of the Wellcome Sanger Institute COVID-19 Surveillance Team |
| EPI_ISL_3443824, EPI_ISL_3443825, EPI_ISL_3443826, EPI_ISL_3443827 | Lighthouse Lab in Milton Keynes | Wellcome Sanger Institute for the COVID-19 Genomics UK (COG-UK) Consortium | The Lighthouse Lab in Milton Keynes and Alex Alderton, Roberto Amato, Jeffrey Barrett, Sonia Goncalves, Ewan Harrison, David K. Jackson, Ian Johnston, Dominic Kwiatkowski, Cordelia Langford, John Sillitoe on behalf of the Wellcome Sanger Institute COVID-19 Surveillance Team |
| EPI_ISL_3443828 | Lighthouse Lab in Glasgow | Wellcome Sanger Institute for the COVID-19 Genomics UK (COG-UK) Consortium | Harper VanSteenhouse, Yumi Kasai, David Gray, Carol Clugston, Anna Dominiczak and Alex Alderton, Roberto Amato, Jeffrey Barrett, Sonia Goncalves, Ewan Harrison, David K. Jackson, Ian Johnston, Dominic Kwiatkowski, Cordelia Langford, John Sillitoe on behalf of the Wellcome Sanger Institute COVID-19 Surveillance Team |
| EPI_ISL_3443829, EPI_ISL_3443830 | Lighthouse Lab in Milton Keynes | Wellcome Sanger Institute for the COVID-19 Genomics UK (COG-UK) Consortium | The Lighthouse Lab in Milton Keynes and Alex Alderton, Roberto Amato, Jeffrey Barrett, Sonia Goncalves, Ewan Harrison, David K. Jackson, Ian Johnston, Dominic Kwiatkowski, Cordelia Langford, John Sillitoe on behalf of the Wellcome Sanger Institute COVID-19 Surveillance Team |
| EPI_ISL_3443831 | Lighthouse Lab in Glasgow | Wellcome Sanger Institute for the COVID-19 Genomics UK (COG-UK) Consortium | Harper VanSteenhouse, Yumi Kasai, David Gray, Carol Clugston, Anna Dominiczak and Alex Alderton, Roberto Amato, Jeffrey Barrett, Sonia Goncalves, Ewan Harrison, David K. Jackson, Ian Johnston, Dominic Kwiatkowski, Cordelia Langford, John Sillitoe on behalf of the Wellcome Sanger Institute COVID-19 Surveillance Team |
| EPI_ISL_3443832, EPI_ISL_3443833 | Lighthouse Lab in Milton Keynes | Wellcome Sanger Institute for the COVID-19 Genomics UK (COG-UK) Consortium | The Lighthouse Lab in Milton Keynes and Alex Alderton, Roberto Amato, Jeffrey Barrett, Sonia Goncalves, Ewan Harrison, David K. Jackson, Ian Johnston, Dominic Kwiatkowski, Cordelia Langford, John Sillitoe on behalf of the Wellcome Sanger Institute COVID-19 Surveillance Team |
| EPI_ISL_3443834 | Lighthouse Lab in Glasgow | Wellcome Sanger Institute for the COVID-19 Genomics UK (COG-UK) Consortium | Harper VanSteenhouse, Yumi Kasai, David Gray, Carol Clugston, Anna Dominiczak and Alex Alderton, Roberto Amato, Jeffrey Barrett, Sonia Goncalves, Ewan Harrison, David K. Jackson, Ian Johnston, Dominic Kwiatkowski, Cordelia Langford, John Sillitoe on behalf of the Wellcome Sanger Institute COVID-19 Surveillance Team |
| EPI_ISL_3443835 | Lighthouse Lab in Milton Keynes | Wellcome Sanger Institute for the COVID-19 Genomics UK (COG-UK) Consortium | The Lighthouse Lab in Milton Keynes and Alex Alderton, Roberto Amato, Jeffrey Barrett, Sonia Goncalves, Ewan Harrison, David K. Jackson, Ian Johnston, Dominic Kwiatkowski, Cordelia Langford, John Sillitoe on behalf of the Wellcome Sanger Institute COVID-19 Surveillance Team |
| EPI_ISL_3443836, EPI_ISL_3443837, EPI_ISL_3443838 | Lighthouse Lab in Glasgow | Wellcome Sanger Institute for the COVID-19 Genomics UK (COG-UK) Consortium | Harper VanSteenhouse, Yumi Kasai, David Gray, Carol Clugston, Anna Dominiczak and Alex Alderton, Roberto Amato, Jeffrey Barrett, Sonia Goncalves, Ewan Harrison, David K. Jackson, Ian Johnston, Dominic Kwiatkowski, Cordelia Langford, John Sillitoe on behalf of the Wellcome Sanger Institute COVID-19 Surveillance Team |
| EPI_ISL_3443839 | Lighthouse Lab in Milton Keynes | Wellcome Sanger Institute for the COVID-19 Genomics UK (COG-UK) Consortium | The Lighthouse Lab in Milton Keynes and Alex Alderton, Roberto Amato, Jeffrey Barrett, Sonia Goncalves, Ewan Harrison, David K. Jackson, Ian Johnston, Dominic Kwiatkowski, Cordelia Langford, John Sillitoe on behalf of the Wellcome Sanger Institute COVID-19 Surveillance Team |
| EPI_ISL_3443840, EPI_ISL_3443841, EPI_ISL_3443842 | Lighthouse Lab in Glasgow | Wellcome Sanger Institute for the COVID-19 Genomics UK (COG-UK) Consortium | Harper VanSteenhouse, Yumi Kasai, David Gray, Carol Clugston, Anna Dominiczak and Alex Alderton, Roberto Amato, Jeffrey Barrett, Sonia Goncalves, Ewan Harrison, David K. Jackson, Ian Johnston, Dominic Kwiatkowski, Cordelia Langford, John Sillitoe on behalf of the Wellcome Sanger Institute COVID-19 Surveillance Team |
| EPI_ISL_3443843, EPI_ISL_3443844, EPI_ISL_3443845, EPI_ISL_3443846, EPI_ISL_3443847 | Lighthouse Lab in Milton Keynes | Wellcome Sanger Institute for the COVID-19 Genomics UK (COG-UK) Consortium | The Lighthouse Lab in Milton Keynes and Alex Alderton, Roberto Amato, Jeffrey Barrett, Sonia Goncalves, Ewan Harrison, David K. Jackson, Ian Johnston, Dominic Kwiatkowski, Cordelia Langford, John Sillitoe on behalf of the Wellcome Sanger Institute COVID-19 Surveillance Team |
| EPI_ISL_3443848 | Lighthouse Lab in Glasgow | Wellcome Sanger Institute for the COVID-19 Genomics UK (COG-UK) Consortium | Harper VanSteenhouse, Yumi Kasai, David Gray, Carol Clugston, Anna Dominiczak and Alex Alderton, Roberto Amato, Jeffrey Barrett, Sonia Goncalves, Ewan Harrison, David K. Jackson, Ian Johnston, Dominic Kwiatkowski, Cordelia Langford, John Sillitoe on behalf of the Wellcome Sanger Institute COVID-19 Surveillance Team |
| EPI_ISL_3443849 | Lighthouse Lab in Milton Keynes | Wellcome Sanger Institute for the COVID-19 Genomics UK (COG-UK) Consortium | The Lighthouse Lab in Milton Keynes and Alex Alderton, Roberto Amato, Jeffrey Barrett, Sonia Goncalves, Ewan Harrison, David K. Jackson, Ian Johnston, Dominic Kwiatkowski, Cordelia Langford, John Sillitoe on behalf of the Wellcome Sanger Institute COVID-19 Surveillance Team |
| EPI_ISL_3443850 | Lighthouse Lab in Glasgow | Wellcome Sanger Institute for the COVID-19 Genomics UK (COG-UK) Consortium | Harper VanSteenhouse, Yumi Kasai, David Gray, Carol Clugston, Anna Dominiczak and Alex Alderton, Roberto Amato, Jeffrey Barrett, Sonia Goncalves, Ewan Harrison, David K. Jackson, Ian Johnston, Dominic Kwiatkowski, Cordelia Langford, John Sillitoe on behalf of the Wellcome Sanger Institute COVID-19 Surveillance Team |
| EPI_ISL_3443851 | Lighthouse Lab in Milton Keynes | Wellcome Sanger Institute for the COVID-19 Genomics UK (COG-UK) Consortium | The Lighthouse Lab in Milton Keynes and Alex Alderton, Roberto Amato, Jeffrey Barrett, Sonia Goncalves, Ewan Harrison, David K. Jackson, Ian Johnston, Dominic Kwiatkowski, Cordelia Langford, John Sillitoe on behalf of the Wellcome Sanger Institute COVID-19 Surveillance Team |
| EPI_ISL_3443852, EPI_ISL_3443853 | Lighthouse Lab in Glasgow | Wellcome Sanger Institute for the COVID-19 Genomics UK (COG-UK) Consortium | Harper VanSteenhouse, Yumi Kasai, David Gray, Carol Clugston, Anna Dominiczak and Alex Alderton, Roberto Amato, Jeffrey Barrett, Sonia Goncalves, Ewan Harrison, David K. Jackson, Ian Johnston, Dominic Kwiatkowski, Cordelia Langford, John Sillitoe on behalf of the Wellcome Sanger Institute COVID-19 Surveillance Team |
| EPI_ISL_3443854 | Lighthouse Lab in Milton Keynes | Wellcome Sanger Institute for the COVID-19 Genomics UK (COG-UK) Consortium | The Lighthouse Lab in Milton Keynes and Alex Alderton, Roberto Amato, Jeffrey Barrett, Sonia Goncalves, Ewan Harrison, David K. Jackson, Ian Johnston, Dominic Kwiatkowski, Cordelia Langford, John Sillitoe on behalf of the Wellcome Sanger Institute COVID-19 Surveillance Team |
| EPI_ISL_3443855, EPI_ISL_3443856 | Lighthouse Lab in Glasgow | Wellcome Sanger Institute for the COVID-19 Genomics UK (COG-UK) Consortium | Harper VanSteenhouse, Yumi Kasai, David Gray, Carol Clugston, Anna Dominiczak and Alex Alderton, Roberto Amato, Jeffrey Barrett, Sonia Goncalves, Ewan Harrison, David K. Jackson, Ian Johnston, Dominic Kwiatkowski, Cordelia Langford, John Sillitoe on behalf of the Wellcome Sanger Institute COVID-19 Surveillance Team |
| EPI_ISL_3443857 | Lighthouse Lab in Milton Keynes | Wellcome Sanger Institute for the COVID-19 Genomics UK (COG-UK) Consortium | The Lighthouse Lab in Milton Keynes and Alex Alderton, Roberto Amato, Jeffrey Barrett, Sonia Goncalves, Ewan Harrison, David K. Jackson, Ian Johnston, Dominic Kwiatkowski, Cordelia Langford, John Sillitoe on behalf of the Wellcome Sanger Institute COVID-19 Surveillance Team |
| EPI_ISL_3443858, EPI_ISL_3443859, EPI_ISL_3443860 | Lighthouse Lab in Glasgow | Wellcome Sanger Institute for the COVID-19 Genomics UK (COG-UK) Consortium | Harper VanSteenhouse, Yumi Kasai, David Gray, Carol Clugston, Anna Dominiczak and Alex Alderton, Roberto Amato, Jeffrey Barrett, Sonia Goncalves, Ewan Harrison, David K. Jackson, Ian Johnston, Dominic Kwiatkowski, Cordelia Langford, John Sillitoe on behalf of the Wellcome Sanger Institute COVID-19 Surveillance Team |
| EPI_ISL_3443861 | Lighthouse Lab in Milton Keynes | Wellcome Sanger Institute for the COVID-19 Genomics UK (COG-UK) Consortium | The Lighthouse Lab in Milton Keynes and Alex Alderton, Roberto Amato, Jeffrey Barrett, Sonia Goncalves, Ewan Harrison, David K. Jackson, Ian Johnston, Dominic Kwiatkowski, Cordelia Langford, John Sillitoe on behalf of the Wellcome Sanger Institute COVID-19 Surveillance Team |
| EPI_ISL_3443862, EPI_ISL_3443863, EPI_ISL_3443864 | Lighthouse Lab in Glasgow | Wellcome Sanger Institute for the COVID-19 Genomics UK (COG-UK) Consortium | Harper VanSteenhouse, Yumi Kasai, David Gray, Carol Clugston, Anna Dominiczak and Alex Alderton, Roberto Amato, Jeffrey Barrett, Sonia Goncalves, Ewan Harrison, David K. Jackson, Ian Johnston, Dominic Kwiatkowski, Cordelia Langford, John Sillitoe on behalf of the Wellcome Sanger Institute COVID-19 Surveillance Team |
| EPI_ISL_3443865 | Lighthouse Lab in Milton Keynes | Wellcome Sanger Institute for the COVID-19 Genomics UK (COG-UK) Consortium | The Lighthouse Lab in Milton Keynes and Alex Alderton, Roberto Amato, Jeffrey Barrett, Sonia Goncalves, Ewan Harrison, David K. Jackson, Ian Johnston, Dominic Kwiatkowski, Cordelia Langford, John Sillitoe on behalf of the Wellcome Sanger Institute COVID-19 Surveillance Team |
| EPI_ISL_3443866, EPI_ISL_3443867 | Lighthouse Lab in Glasgow | Wellcome Sanger Institute for the COVID-19 Genomics UK (COG-UK) Consortium | Harper VanSteenhouse, Yumi Kasai, David Gray, Carol Clugston, Anna Dominiczak and Alex Alderton, Roberto Amato, Jeffrey Barrett, Sonia Goncalves, Ewan Harrison, David K. Jackson, Ian Johnston, Dominic Kwiatkowski, Cordelia Langford, John Sillitoe on behalf of the Wellcome Sanger Institute COVID-19 Surveillance Team |
| EPI_ISL_3443868, EPI_ISL_3443869, EPI_ISL_3443870, EPI_ISL_3443871, EPI_ISL_3443872, EPI_ISL_3443873, EPI_ISL_3443874 | Lighthouse Lab in Milton Keynes | Wellcome Sanger Institute for the COVID-19 Genomics UK (COG-UK) Consortium | The Lighthouse Lab in Milton Keynes and Alex Alderton, Roberto Amato, Jeffrey Barrett, Sonia Goncalves, Ewan Harrison, David K. Jackson, Ian Johnston, Dominic Kwiatkowski, Cordelia Langford, John Sillitoe on behalf of the Wellcome Sanger Institute COVID-19 Surveillance Team |
| EPI_ISL_3443875 | Lighthouse Lab in Glasgow | Wellcome Sanger Institute for the COVID-19 Genomics UK (COG-UK) Consortium | Harper VanSteenhouse, Yumi Kasai, David Gray, Carol Clugston, Anna Dominiczak and Alex Alderton, Roberto Amato, Jeffrey Barrett, Sonia Goncalves, Ewan Harrison, David K. Jackson, Ian Johnston, Dominic Kwiatkowski, Cordelia Langford, John Sillitoe on behalf of the Wellcome Sanger Institute COVID-19 Surveillance Team |
| EPI_ISL_3443876, EPI_ISL_3443877 | Lighthouse Lab in Milton Keynes | Wellcome Sanger Institute for the COVID-19 Genomics UK (COG-UK) Consortium | The Lighthouse Lab in Milton Keynes and Alex Alderton, Roberto Amato, Jeffrey Barrett, Sonia Goncalves, Ewan Harrison, David K. Jackson, Ian Johnston, Dominic Kwiatkowski, Cordelia Langford, John Sillitoe on behalf of the Wellcome Sanger Institute COVID-19 Surveillance Team |
| EPI_ISL_3443878 | Lighthouse Lab in Glasgow | Wellcome Sanger Institute for the COVID-19 Genomics UK (COG-UK) Consortium | Harper VanSteenhouse, Yumi Kasai, David Gray, Carol Clugston, Anna Dominiczak and Alex Alderton, Roberto Amato, Jeffrey Barrett, Sonia Goncalves, Ewan Harrison, David K. Jackson, Ian Johnston, Dominic Kwiatkowski, Cordelia Langford, John Sillitoe on behalf of the Wellcome Sanger Institute COVID-19 Surveillance Team |
| EPI_ISL_3443879, EPI_ISL_3443880, EPI_ISL_3443881, EPI_ISL_3443882 | Lighthouse Lab in Milton Keynes | Wellcome Sanger Institute for the COVID-19 Genomics UK (COG-UK) Consortium | The Lighthouse Lab in Milton Keynes and Alex Alderton, Roberto Amato, Jeffrey Barrett, Sonia Goncalves, Ewan Harrison, David K. Jackson, Ian Johnston, Dominic Kwiatkowski, Cordelia Langford, John Sillitoe on behalf of the Wellcome Sanger Institute COVID-19 Surveillance Team |

| | | | |
|---|---|---|---|
| EPI_ISL_3443883, EPI_ISL_3443884 | Lighthouse Lab in Glasgow | Wellcome Sanger Institute for the COVID-19 Genomics UK (COG-UK) Consortium | Harper VanSteenhouse, Yumi Kasai, David Gray, Carol Clugston, Anna Dominiczak and Alex Alderton, Roberto Amato, Jeffrey Barrett, Sonia Goncalves, Ewan Harrison, David K. Jackson, Ian Johnston, Dominic Kwiatkowski, Cordelia Langford, John Sillitoe on behalf of the Wellcome Sanger Institute COVID-19 Surveillance Team |
| EPI_ISL_3443885, EPI_ISL_3443886, EPI_ISL_3443887 | Lighthouse Lab in Milton Keynes | Wellcome Sanger Institute for the COVID-19 Genomics UK (COG-UK) Consortium | The Lighthouse Lab in Milton Keynes and Alex Alderton, Roberto Amato, Jeffrey Barrett, Sonia Goncalves, Ewan Harrison, David K. Jackson, Ian Johnston, Dominic Kwiatkowski, Cordelia Langford, John Sillitoe on behalf of the Wellcome Sanger Institute COVID-19 Surveillance Team |
| EPI_ISL_3443888, EPI_ISL_3443889, EPI_ISL_3443890 | Lighthouse Lab in Glasgow | Wellcome Sanger Institute for the COVID-19 Genomics UK (COG-UK) Consortium | Harper VanSteenhouse, Yumi Kasai, David Gray, Carol Clugston, Anna Dominiczak and Alex Alderton, Roberto Amato, Jeffrey Barrett, Sonia Goncalves, Ewan Harrison, David K. Jackson, Ian Johnston, Dominic Kwiatkowski, Cordelia Langford, John Sillitoe on behalf of the Wellcome Sanger Institute COVID-19 Surveillance Team |
| EPI_ISL_3443891 | Lighthouse Lab in Milton Keynes | Wellcome Sanger Institute for the COVID-19 Genomics UK (COG-UK) Consortium | The Lighthouse Lab in Milton Keynes and Alex Alderton, Roberto Amato, Jeffrey Barrett, Sonia Goncalves, Ewan Harrison, David K. Jackson, Ian Johnston, Dominic Kwiatkowski, Cordelia Langford, John Sillitoe on behalf of the Wellcome Sanger Institute COVID-19 Surveillance Team |
| EPI_ISL_3443892, EPI_ISL_3443893 | Lighthouse Lab in Glasgow | Wellcome Sanger Institute for the COVID-19 Genomics UK (COG-UK) Consortium | Harper VanSteenhouse, Yumi Kasai, David Gray, Carol Clugston, Anna Dominiczak and Alex Alderton, Roberto Amato, Jeffrey Barrett, Sonia Goncalves, Ewan Harrison, David K. Jackson, Ian Johnston, Dominic Kwiatkowski, Cordelia Langford, John Sillitoe on behalf of the Wellcome Sanger Institute COVID-19 Surveillance Team |
| EPI_ISL_3443894, EPI_ISL_3443895 | Lighthouse Lab in Milton Keynes | Wellcome Sanger Institute for the COVID-19 Genomics UK (COG-UK) Consortium | The Lighthouse Lab in Milton Keynes and Alex Alderton, Roberto Amato, Jeffrey Barrett, Sonia Goncalves, Ewan Harrison, David K. Jackson, Ian Johnston, Dominic Kwiatkowski, Cordelia Langford, John Sillitoe on behalf of the Wellcome Sanger Institute COVID-19 Surveillance Team |
| EPI_ISL_3443896 | Lighthouse Lab in Glasgow | Wellcome Sanger Institute for the COVID-19 Genomics UK (COG-UK) Consortium | Harper VanSteenhouse, Yumi Kasai, David Gray, Carol Clugston, Anna Dominiczak and Alex Alderton, Roberto Amato, Jeffrey Barrett, Sonia Goncalves, Ewan Harrison, David K. Jackson, Ian Johnston, Dominic Kwiatkowski, Cordelia Langford, John Sillitoe on behalf of the Wellcome Sanger Institute COVID-19 Surveillance Team |
| EPI_ISL_3443897, EPI_ISL_3443898 | Lighthouse Lab in Milton Keynes | Wellcome Sanger Institute for the COVID-19 Genomics UK (COG-UK) Consortium | The Lighthouse Lab in Milton Keynes and Alex Alderton, Roberto Amato, Jeffrey Barrett, Sonia Goncalves, Ewan Harrison, David K. Jackson, Ian Johnston, Dominic Kwiatkowski, Cordelia Langford, John Sillitoe on behalf of the Wellcome Sanger Institute COVID-19 Surveillance Team |
| EPI_ISL_3443899, EPI_ISL_3443900, EPI_ISL_3443901, EPI_ISL_3443902, EPI_ISL_3443903 | Lighthouse Lab in Glasgow | Wellcome Sanger Institute for the COVID-19 Genomics UK (COG-UK) Consortium | Harper VanSteenhouse, Yumi Kasai, David Gray, Carol Clugston, Anna Dominiczak and Alex Alderton, Roberto Amato, Jeffrey Barrett, Sonia Goncalves, Ewan Harrison, David K. Jackson, Ian Johnston, Dominic Kwiatkowski, Cordelia Langford, John Sillitoe on behalf of the Wellcome Sanger Institute COVID-19 Surveillance Team |
| EPI_ISL_3443904, EPI_ISL_3443905, EPI_ISL_3443906, EPI_ISL_3443907, EPI_ISL_3443908, EPI_ISL_3443909, EPI_ISL_3443910, EPI_ISL_3443911 | Lighthouse Lab in Milton Keynes | Wellcome Sanger Institute for the COVID-19 Genomics UK (COG-UK) Consortium | The Lighthouse Lab in Milton Keynes and Alex Alderton, Roberto Amato, Jeffrey Barrett, Sonia Goncalves, Ewan Harrison, David K. Jackson, Ian Johnston, Dominic Kwiatkowski, Cordelia Langford, John Sillitoe on behalf of the Wellcome Sanger Institute COVID-19 Surveillance Team |
| EPI_ISL_3443912 | Lighthouse Lab in Glasgow | Wellcome Sanger Institute for the COVID-19 Genomics UK (COG-UK) Consortium | Harper VanSteenhouse, Yumi Kasai, David Gray, Carol Clugston, Anna Dominiczak and Alex Alderton, Roberto Amato, Jeffrey Barrett, Sonia Goncalves, Ewan Harrison, David K. Jackson, Ian Johnston, Dominic Kwiatkowski, Cordelia Langford, John Sillitoe on behalf of the Wellcome Sanger Institute COVID-19 Surveillance Team |
| EPI_ISL_3443913, EPI_ISL_3443914 | Lighthouse Lab in Milton Keynes | Wellcome Sanger Institute for the COVID-19 Genomics UK (COG-UK) Consortium | The Lighthouse Lab in Milton Keynes and Alex Alderton, Roberto Amato, Jeffrey Barrett, Sonia Goncalves, Ewan Harrison, David K. Jackson, Ian Johnston, Dominic Kwiatkowski, Cordelia Langford, John Sillitoe on behalf of the Wellcome Sanger Institute COVID-19 Surveillance Team |
| EPI_ISL_3443915, EPI_ISL_3443916, EPI_ISL_3443917, EPI_ISL_3443918, EPI_ISL_3443919 | Lighthouse Lab in Glasgow | Wellcome Sanger Institute for the COVID-19 Genomics UK (COG-UK) Consortium | Harper VanSteenhouse, Yumi Kasai, David Gray, Carol Clugston, Anna Dominiczak and Alex Alderton, Roberto Amato, Jeffrey Barrett, Sonia Goncalves, Ewan Harrison, David K. Jackson, Ian Johnston, Dominic Kwiatkowski, Cordelia Langford, John Sillitoe on behalf of the Wellcome Sanger Institute COVID-19 Surveillance Team |
| EPI_ISL_3443920, EPI_ISL_3443921 | Lighthouse Lab in Milton Keynes | Wellcome Sanger Institute for the COVID-19 Genomics UK (COG-UK) Consortium | The Lighthouse Lab in Milton Keynes and Alex Alderton, Roberto Amato, Jeffrey Barrett, Sonia Goncalves, Ewan Harrison, David K. Jackson, Ian Johnston, Dominic Kwiatkowski, Cordelia Langford, John Sillitoe on behalf of the Wellcome Sanger Institute COVID-19 Surveillance Team |
| EPI_ISL_3443922 | Lighthouse Lab in Glasgow | Wellcome Sanger Institute for the COVID-19 Genomics UK (COG-UK) Consortium | Harper VanSteenhouse, Yumi Kasai, David Gray, Carol Clugston, Anna Dominiczak and Alex Alderton, Roberto Amato, Jeffrey Barrett, Sonia Goncalves, Ewan Harrison, David K. Jackson, Ian Johnston, Dominic Kwiatkowski, Cordelia Langford, John Sillitoe on behalf of the Wellcome Sanger Institute COVID-19 Surveillance Team |
| EPI_ISL_3443923 | Lighthouse Lab in Milton Keynes | Wellcome Sanger Institute for the COVID-19 Genomics UK (COG-UK) Consortium | The Lighthouse Lab in Milton Keynes and Alex Alderton, Roberto Amato, Jeffrey Barrett, Sonia Goncalves, Ewan Harrison, David K. Jackson, Ian Johnston, Dominic Kwiatkowski, Cordelia Langford, John Sillitoe on behalf of the Wellcome Sanger Institute COVID-19 Surveillance Team |
| EPI_ISL_3443924, EPI_ISL_3443925, EPI_ISL_3443926 | Lighthouse Lab in Glasgow | Wellcome Sanger Institute for the COVID-19 Genomics UK (COG-UK) Consortium | Harper VanSteenhouse, Yumi Kasai, David Gray, Carol Clugston, Anna Dominiczak and Alex Alderton, Roberto Amato, Jeffrey Barrett, Sonia Goncalves, Ewan Harrison, David K. Jackson, Ian Johnston, Dominic Kwiatkowski, Cordelia Langford, John Sillitoe on behalf of the Wellcome Sanger Institute COVID-19 Surveillance Team |
| EPI_ISL_3443927, EPI_ISL_3443928 | Lighthouse Lab in Milton Keynes | Wellcome Sanger Institute for the COVID-19 Genomics UK (COG-UK) Consortium | The Lighthouse Lab in Milton Keynes and Alex Alderton, Roberto Amato, Jeffrey Barrett, Sonia Goncalves, Ewan Harrison, David K. Jackson, Ian Johnston, Dominic Kwiatkowski, Cordelia Langford, John Sillitoe on behalf of the Wellcome Sanger Institute COVID-19 Surveillance Team |
| EPI_ISL_3443929 | Lighthouse Lab in Glasgow | Wellcome Sanger Institute for the COVID-19 Genomics UK (COG-UK) Consortium | Harper VanSteenhouse, Yumi Kasai, David Gray, Carol Clugston, Anna Dominiczak and Alex Alderton, Roberto Amato, Jeffrey Barrett, Sonia Goncalves, Ewan Harrison, David K. Jackson, Ian Johnston, Dominic Kwiatkowski, Cordelia Langford, John Sillitoe on behalf of the Wellcome Sanger Institute COVID-19 Surveillance Team |
| EPI_ISL_3443930, EPI_ISL_3443931 | Lighthouse Lab in Milton Keynes | Wellcome Sanger Institute for the COVID-19 Genomics UK (COG-UK) Consortium | The Lighthouse Lab in Milton Keynes and Alex Alderton, Roberto Amato, Jeffrey Barrett, Sonia Goncalves, Ewan Harrison, David K. Jackson, Ian Johnston, Dominic Kwiatkowski, Cordelia Langford, John Sillitoe on behalf of the Wellcome Sanger Institute COVID-19 Surveillance Team |
| EPI_ISL_3443932 | Lighthouse Lab in Glasgow | Wellcome Sanger Institute for the COVID-19 Genomics UK (COG-UK) Consortium | Harper VanSteenhouse, Yumi Kasai, David Gray, Carol Clugston, Anna Dominiczak and Alex Alderton, Roberto Amato, Jeffrey Barrett, Sonia Goncalves, Ewan Harrison, David K. Jackson, Ian Johnston, Dominic Kwiatkowski, Cordelia Langford, John Sillitoe on behalf of the Wellcome Sanger Institute COVID-19 Surveillance Team |
| EPI_ISL_3443933, EPI_ISL_3443934 | Lighthouse Lab in Milton Keynes | Wellcome Sanger Institute for the COVID-19 Genomics UK (COG-UK) Consortium | The Lighthouse Lab in Milton Keynes and Alex Alderton, Roberto Amato, Jeffrey Barrett, Sonia Goncalves, Ewan Harrison, David K. Jackson, Ian Johnston, Dominic Kwiatkowski, Cordelia Langford, John Sillitoe on behalf of the Wellcome Sanger Institute COVID-19 Surveillance Team |
| EPI_ISL_3443935, EPI_ISL_3443936 | Lighthouse Lab in Glasgow | Wellcome Sanger Institute for the COVID-19 Genomics UK (COG-UK) Consortium | Harper VanSteenhouse, Yumi Kasai, David Gray, Carol Clugston, Anna Dominiczak and Alex Alderton, Roberto Amato, Jeffrey Barrett, Sonia Goncalves, Ewan Harrison, David K. Jackson, Ian Johnston, Dominic Kwiatkowski, Cordelia Langford, John Sillitoe on behalf of the Wellcome Sanger Institute COVID-19 Surveillance Team |
| EPI_ISL_3443937, EPI_ISL_3443938, EPI_ISL_3443939 | Lighthouse Lab in Milton Keynes | Wellcome Sanger Institute for the COVID-19 Genomics UK (COG-UK) Consortium | The Lighthouse Lab in Milton Keynes and Alex Alderton, Roberto Amato, Jeffrey Barrett, Sonia Goncalves, Ewan Harrison, David K. Jackson, Ian Johnston, Dominic Kwiatkowski, Cordelia Langford, John Sillitoe on behalf of the Wellcome Sanger Institute COVID-19 Surveillance Team |
| EPI_ISL_3443940 | Lighthouse Lab in Glasgow | Wellcome Sanger Institute for the COVID-19 Genomics UK (COG-UK) Consortium | Harper VanSteenhouse, Yumi Kasai, David Gray, Carol Clugston, Anna Dominiczak and Alex Alderton, Roberto Amato, Jeffrey Barrett, Sonia Goncalves, Ewan Harrison, David K. Jackson, Ian Johnston, Dominic Kwiatkowski, Cordelia Langford, John Sillitoe on behalf of the Wellcome Sanger Institute COVID-19 Surveillance Team |
| EPI_ISL_3443941, EPI_ISL_3443942 | Lighthouse Lab in Milton Keynes | Wellcome Sanger Institute for the COVID-19 Genomics UK (COG-UK) Consortium | The Lighthouse Lab in Milton Keynes and Alex Alderton, Roberto Amato, Jeffrey Barrett, Sonia Goncalves, Ewan Harrison, David K. Jackson, Ian Johnston, Dominic Kwiatkowski, Cordelia Langford, John Sillitoe on behalf of the Wellcome Sanger Institute COVID-19 Surveillance Team |
| EPI_ISL_3443943, EPI_ISL_3443944 | Lighthouse Lab in Glasgow | Wellcome Sanger Institute for the COVID-19 Genomics UK (COG-UK) Consortium | Harper VanSteenhouse, Yumi Kasai, David Gray, Carol Clugston, Anna Dominiczak and Alex Alderton, Roberto Amato, Jeffrey Barrett, Sonia Goncalves, Ewan Harrison, David K. Jackson, Ian Johnston, Dominic Kwiatkowski, Cordelia Langford, John Sillitoe on behalf of the Wellcome Sanger Institute COVID-19 Surveillance Team |
| EPI_ISL_3443945 | Lighthouse Lab in Milton Keynes | Wellcome Sanger Institute for the COVID-19 Genomics UK (COG-UK) Consortium | The Lighthouse Lab in Milton Keynes and Alex Alderton, Roberto Amato, Jeffrey Barrett, Sonia Goncalves, Ewan Harrison, David K. Jackson, Ian Johnston, Dominic Kwiatkowski, Cordelia Langford, John Sillitoe on behalf of the Wellcome Sanger Institute COVID-19 Surveillance Team |
| EPI_ISL_3443946, EPI_ISL_3443947, EPI_ISL_3443948 | Lighthouse Lab in Glasgow | Wellcome Sanger Institute for the COVID-19 Genomics UK (COG-UK) Consortium | Harper VanSteenhouse, Yumi Kasai, David Gray, Carol Clugston, Anna Dominiczak and Alex Alderton, Roberto Amato, Jeffrey Barrett, Sonia Goncalves, Ewan Harrison, David K. Jackson, Ian Johnston, Dominic Kwiatkowski, Cordelia Langford, John Sillitoe on behalf of the Wellcome Sanger Institute COVID-19 Surveillance Team |
| EPI_ISL_3443949, EPI_ISL_3443950 | Lighthouse Lab in Milton Keynes | Wellcome Sanger Institute for the COVID-19 Genomics UK (COG-UK) Consortium | The Lighthouse Lab in Milton Keynes and Alex Alderton, Roberto Amato, Jeffrey Barrett, Sonia Goncalves, Ewan Harrison, David K. Jackson, Ian Johnston, Dominic Kwiatkowski, Cordelia Langford, John Sillitoe on behalf of the Wellcome Sanger Institute COVID-19 Surveillance Team |
| EPI_ISL_3443951, EPI_ISL_3443952 | Lighthouse Lab in Glasgow | Wellcome Sanger Institute for the COVID-19 Genomics UK (COG-UK) Consortium | Harper VanSteenhouse, Yumi Kasai, David Gray, Carol Clugston, Anna Dominiczak and Alex Alderton, Roberto Amato, Jeffrey Barrett, Sonia Goncalves, Ewan Harrison, David K. Jackson, Ian Johnston, Dominic Kwiatkowski, Cordelia Langford, John Sillitoe on behalf of the Wellcome Sanger Institute COVID-19 Surveillance Team |
| EPI_ISL_3443953 | Lighthouse Lab in Milton Keynes | Wellcome Sanger Institute for the COVID-19 Genomics UK (COG-UK) Consortium | The Lighthouse Lab in Milton Keynes and Alex Alderton, Roberto Amato, Jeffrey Barrett, Sonia Goncalves, Ewan Harrison, David K. Jackson, Ian Johnston, Dominic Kwiatkowski, Cordelia Langford, John Sillitoe on behalf of the Wellcome Sanger Institute COVID-19 Surveillance Team |
| EPI_ISL_3443954, EPI_ISL_3443955, EPI_ISL_3443956 | Lighthouse Lab in Glasgow | Wellcome Sanger Institute for the COVID-19 Genomics UK (COG-UK) Consortium | Harper VanSteenhouse, Yumi Kasai, David Gray, Carol Clugston, Anna Dominiczak and Alex Alderton, Roberto Amato, Jeffrey Barrett, Sonia Goncalves, Ewan Harrison, David K. Jackson, Ian Johnston, Dominic Kwiatkowski, Cordelia Langford, John Sillitoe on behalf of the Wellcome Sanger Institute COVID-19 Surveillance Team |
| EPI_ISL_3443957 | Lighthouse Lab in Milton Keynes | Wellcome Sanger Institute for the COVID-19 Genomics UK (COG-UK) Consortium | The Lighthouse Lab in Milton Keynes and Alex Alderton, Roberto Amato, Jeffrey Barrett, Sonia Goncalves, Ewan Harrison, David K. Jackson, Ian Johnston, Dominic Kwiatkowski, Cordelia Langford, John Sillitoe on behalf of the Wellcome Sanger Institute COVID-19 Surveillance Team |
| EPI_ISL_3443958 | Lighthouse Lab in Glasgow | Wellcome Sanger Institute for the COVID-19 Genomics UK (COG-UK) Consortium | Harper VanSteenhouse, Yumi Kasai, David Gray, Carol Clugston, Anna Dominiczak and Alex Alderton, Roberto Amato, Jeffrey Barrett, Sonia Goncalves, Ewan Harrison, David K. Jackson, Ian Johnston, Dominic Kwiatkowski, Cordelia Langford, John Sillitoe on behalf of the Wellcome Sanger Institute COVID-19 Surveillance Team |
| EPI_ISL_3443959, EPI_ISL_3443960, EPI_ISL_3443961 | Lighthouse Lab in Milton Keynes | Wellcome Sanger Institute for the COVID-19 Genomics UK (COG-UK) Consortium | The Lighthouse Lab in Milton Keynes and Alex Alderton, Roberto Amato, Jeffrey Barrett, Sonia Goncalves, Ewan Harrison, David K. Jackson, Ian Johnston, Dominic Kwiatkowski, Cordelia Langford, John Sillitoe on behalf of the Wellcome Sanger Institute COVID-19 Surveillance Team |
| EPI_ISL_3443962 | Lighthouse Lab in Glasgow | Wellcome Sanger Institute for the COVID-19 Genomics UK (COG-UK) Consortium | Harper VanSteenhouse, Yumi Kasai, David Gray, Carol Clugston, Anna Dominiczak and Alex Alderton, Roberto Amato, Jeffrey Barrett, Sonia Goncalves, Ewan Harrison, David K. Jackson, Ian Johnston, Dominic Kwiatkowski, Cordelia Langford, John Sillitoe on behalf of the Wellcome Sanger Institute COVID-19 Surveillance Team |
| EPI_ISL_3443963 | Lighthouse Lab in Milton Keynes | Wellcome Sanger Institute for the COVID-19 Genomics UK (COG-UK) | The Lighthouse Lab in Milton Keynes and Alex Alderton, Roberto Amato, Jeffrey Barrett, Sonia Goncalves, Ewan Harrison, David K. Jackson, Ian Johnston, Dominic Kwiatkowski, Cordelia Langford, John Sillitoe on behalf of the Wellcome Sanger |

| | | | |
|---|---|---|---|
| | | Consortium | Institute COVID-19 Surveillance Team |
| EPI_ISL_3443964 | Lighthouse Lab in Glasgow | Wellcome Sanger Institute for the COVID-19 Genomics UK (COG-UK) Consortium | Harper VanSteenhouse, Yumi Kasai, David Gray, Carol Clugston, Anna Dominiczak and Alex Alderton, Roberto Amato, Jeffrey Barrett, Sonia Goncalves, Ewan Harrison, David K. Jackson, Ian Johnston, Dominic Kwiatkowski, Cordelia Langford, John Sillitoe on behalf of the Wellcome Sanger Institute COVID-19 Surveillance Team |
| EPI_ISL_3443965 | Lighthouse Lab in Milton Keynes | Wellcome Sanger Institute for the COVID-19 Genomics UK (COG-UK) Consortium | The Lighthouse Lab in Milton Keynes and Alex Alderton, Roberto Amato, Jeffrey Barrett, Sonia Goncalves, Ewan Harrison, David K. Jackson, Ian Johnston, Dominic Kwiatkowski, Cordelia Langford, John Sillitoe on behalf of the Wellcome Sanger Institute COVID-19 Surveillance Team |
| EPI_ISL_3443966 | Lighthouse Lab in Glasgow | Wellcome Sanger Institute for the COVID-19 Genomics UK (COG-UK) Consortium | Harper VanSteenhouse, Yumi Kasai, David Gray, Carol Clugston, Anna Dominiczak and Alex Alderton, Roberto Amato, Jeffrey Barrett, Sonia Goncalves, Ewan Harrison, David K. Jackson, Ian Johnston, Dominic Kwiatkowski, Cordelia Langford, John Sillitoe on behalf of the Wellcome Sanger Institute COVID-19 Surveillance Team |
| EPI_ISL_3443967, EPI_ISL_3443968, EPI_ISL_3443969 | Lighthouse Lab in Milton Keynes | Wellcome Sanger Institute for the COVID-19 Genomics UK (COG-UK) Consortium | The Lighthouse Lab in Milton Keynes and Alex Alderton, Roberto Amato, Jeffrey Barrett, Sonia Goncalves, Ewan Harrison, David K. Jackson, Ian Johnston, Dominic Kwiatkowski, Cordelia Langford, John Sillitoe on behalf of the Wellcome Sanger Institute COVID-19 Surveillance Team |
| EPI_ISL_3443970, EPI_ISL_3443971, EPI_ISL_3443972 | Lighthouse Lab in Glasgow | Wellcome Sanger Institute for the COVID-19 Genomics UK (COG-UK) Consortium | Harper VanSteenhouse, Yumi Kasai, David Gray, Carol Clugston, Anna Dominiczak and Alex Alderton, Roberto Amato, Jeffrey Barrett, Sonia Goncalves, Ewan Harrison, David K. Jackson, Ian Johnston, Dominic Kwiatkowski, Cordelia Langford, John Sillitoe on behalf of the Wellcome Sanger Institute COVID-19 Surveillance Team |
| EPI_ISL_3443973 | Lighthouse Lab in Milton Keynes | Wellcome Sanger Institute for the COVID-19 Genomics UK (COG-UK) Consortium | The Lighthouse Lab in Milton Keynes and Alex Alderton, Roberto Amato, Jeffrey Barrett, Sonia Goncalves, Ewan Harrison, David K. Jackson, Ian Johnston, Dominic Kwiatkowski, Cordelia Langford, John Sillitoe on behalf of the Wellcome Sanger Institute COVID-19 Surveillance Team |
| EPI_ISL_3443974 | Lighthouse Lab in Glasgow | Wellcome Sanger Institute for the COVID-19 Genomics UK (COG-UK) Consortium | Harper VanSteenhouse, Yumi Kasai, David Gray, Carol Clugston, Anna Dominiczak and Alex Alderton, Roberto Amato, Jeffrey Barrett, Sonia Goncalves, Ewan Harrison, David K. Jackson, Ian Johnston, Dominic Kwiatkowski, Cordelia Langford, John Sillitoe on behalf of the Wellcome Sanger Institute COVID-19 Surveillance Team |
| EPI_ISL_3443975 | Lighthouse Lab in Milton Keynes | Wellcome Sanger Institute for the COVID-19 Genomics UK (COG-UK) Consortium | The Lighthouse Lab in Milton Keynes and Alex Alderton, Roberto Amato, Jeffrey Barrett, Sonia Goncalves, Ewan Harrison, David K. Jackson, Ian Johnston, Dominic Kwiatkowski, Cordelia Langford, John Sillitoe on behalf of the Wellcome Sanger Institute COVID-19 Surveillance Team |
| EPI_ISL_3443976 | Lighthouse Lab in Glasgow | Wellcome Sanger Institute for the COVID-19 Genomics UK (COG-UK) Consortium | Harper VanSteenhouse, Yumi Kasai, David Gray, Carol Clugston, Anna Dominiczak and Alex Alderton, Roberto Amato, Jeffrey Barrett, Sonia Goncalves, Ewan Harrison, David K. Jackson, Ian Johnston, Dominic Kwiatkowski, Cordelia Langford, John Sillitoe on behalf of the Wellcome Sanger Institute COVID-19 Surveillance Team |
| EPI_ISL_3443977, EPI_ISL_3443978, EPI_ISL_3443979 | Lighthouse Lab in Milton Keynes | Wellcome Sanger Institute for the COVID-19 Genomics UK (COG-UK) Consortium | The Lighthouse Lab in Milton Keynes and Alex Alderton, Roberto Amato, Jeffrey Barrett, Sonia Goncalves, Ewan Harrison, David K. Jackson, Ian Johnston, Dominic Kwiatkowski, Cordelia Langford, John Sillitoe on behalf of the Wellcome Sanger Institute COVID-19 Surveillance Team |
| EPI_ISL_3443980 | Lighthouse Lab in Glasgow | Wellcome Sanger Institute for the COVID-19 Genomics UK (COG-UK) Consortium | Harper VanSteenhouse, Yumi Kasai, David Gray, Carol Clugston, Anna Dominiczak and Alex Alderton, Roberto Amato, Jeffrey Barrett, Sonia Goncalves, Ewan Harrison, David K. Jackson, Ian Johnston, Dominic Kwiatkowski, Cordelia Langford, John Sillitoe on behalf of the Wellcome Sanger Institute COVID-19 Surveillance Team |
| EPI_ISL_3443981, EPI_ISL_3443982, EPI_ISL_3443983, EPI_ISL_3443984 | Lighthouse Lab in Milton Keynes | Wellcome Sanger Institute for the COVID-19 Genomics UK (COG-UK) Consortium | The Lighthouse Lab in Milton Keynes and Alex Alderton, Roberto Amato, Jeffrey Barrett, Sonia Goncalves, Ewan Harrison, David K. Jackson, Ian Johnston, Dominic Kwiatkowski, Cordelia Langford, John Sillitoe on behalf of the Wellcome Sanger Institute COVID-19 Surveillance Team |
| EPI_ISL_3443985, EPI_ISL_3443986 | Lighthouse Lab in Glasgow | Wellcome Sanger Institute for the COVID-19 Genomics UK (COG-UK) Consortium | Harper VanSteenhouse, Yumi Kasai, David Gray, Carol Clugston, Anna Dominiczak and Alex Alderton, Roberto Amato, Jeffrey Barrett, Sonia Goncalves, Ewan Harrison, David K. Jackson, Ian Johnston, Dominic Kwiatkowski, Cordelia Langford, John Sillitoe on behalf of the Wellcome Sanger Institute COVID-19 Surveillance Team |
| EPI_ISL_3443987 | Lighthouse Lab in Milton Keynes | Wellcome Sanger Institute for the COVID-19 Genomics UK (COG-UK) Consortium | The Lighthouse Lab in Milton Keynes and Alex Alderton, Roberto Amato, Jeffrey Barrett, Sonia Goncalves, Ewan Harrison, David K. Jackson, Ian Johnston, Dominic Kwiatkowski, Cordelia Langford, John Sillitoe on behalf of the Wellcome Sanger Institute COVID-19 Surveillance Team |
| EPI_ISL_3443988 | Lighthouse Lab in Glasgow | Wellcome Sanger Institute for the COVID-19 Genomics UK (COG-UK) Consortium | Harper VanSteenhouse, Yumi Kasai, David Gray, Carol Clugston, Anna Dominiczak and Alex Alderton, Roberto Amato, Jeffrey Barrett, Sonia Goncalves, Ewan Harrison, David K. Jackson, Ian Johnston, Dominic Kwiatkowski, Cordelia Langford, John Sillitoe on behalf of the Wellcome Sanger Institute COVID-19 Surveillance Team |
| EPI_ISL_3443989 | Lighthouse Lab in Milton Keynes | Wellcome Sanger Institute for the COVID-19 Genomics UK (COG-UK) Consortium | The Lighthouse Lab in Milton Keynes and Alex Alderton, Roberto Amato, Jeffrey Barrett, Sonia Goncalves, Ewan Harrison, David K. Jackson, Ian Johnston, Dominic Kwiatkowski, Cordelia Langford, John Sillitoe on behalf of the Wellcome Sanger Institute COVID-19 Surveillance Team |
| EPI_ISL_3443990 | Lighthouse Lab in Glasgow | Wellcome Sanger Institute for the COVID-19 Genomics UK (COG-UK) Consortium | Harper VanSteenhouse, Yumi Kasai, David Gray, Carol Clugston, Anna Dominiczak and Alex Alderton, Roberto Amato, Jeffrey Barrett, Sonia Goncalves, Ewan Harrison, David K. Jackson, Ian Johnston, Dominic Kwiatkowski, Cordelia Langford, John Sillitoe on behalf of the Wellcome Sanger Institute COVID-19 Surveillance Team |
| EPI_ISL_3443991 | Lighthouse Lab in Milton Keynes | Wellcome Sanger Institute for the COVID-19 Genomics UK (COG-UK) Consortium | The Lighthouse Lab in Milton Keynes and Alex Alderton, Roberto Amato, Jeffrey Barrett, Sonia Goncalves, Ewan Harrison, David K. Jackson, Ian Johnston, Dominic Kwiatkowski, Cordelia Langford, John Sillitoe on behalf of the Wellcome Sanger Institute COVID-19 Surveillance Team |
| EPI_ISL_3443992 | Lighthouse Lab in Glasgow | Wellcome Sanger Institute for the COVID-19 Genomics UK (COG-UK) Consortium | Harper VanSteenhouse, Yumi Kasai, David Gray, Carol Clugston, Anna Dominiczak and Alex Alderton, Roberto Amato, Jeffrey Barrett, Sonia Goncalves, Ewan Harrison, David K. Jackson, Ian Johnston, Dominic Kwiatkowski, Cordelia Langford, John Sillitoe on behalf of the Wellcome Sanger Institute COVID-19 Surveillance Team |
| EPI_ISL_3443993 | Lighthouse Lab in Milton Keynes | Wellcome Sanger Institute for the COVID-19 Genomics UK (COG-UK) Consortium | The Lighthouse Lab in Milton Keynes and Alex Alderton, Roberto Amato, Jeffrey Barrett, Sonia Goncalves, Ewan Harrison, David K. Jackson, Ian Johnston, Dominic Kwiatkowski, Cordelia Langford, John Sillitoe on behalf of the Wellcome Sanger Institute COVID-19 Surveillance Team |
| EPI_ISL_3443994 | Lighthouse Lab in Glasgow | Wellcome Sanger Institute for the COVID-19 Genomics UK (COG-UK) Consortium | Harper VanSteenhouse, Yumi Kasai, David Gray, Carol Clugston, Anna Dominiczak and Alex Alderton, Roberto Amato, Jeffrey Barrett, Sonia Goncalves, Ewan Harrison, David K. Jackson, Ian Johnston, Dominic Kwiatkowski, Cordelia Langford, John Sillitoe on behalf of the Wellcome Sanger Institute COVID-19 Surveillance Team |
| EPI_ISL_3443995 | Lighthouse Lab in Milton Keynes | Wellcome Sanger Institute for the COVID-19 Genomics UK (COG-UK) Consortium | The Lighthouse Lab in Milton Keynes and Alex Alderton, Roberto Amato, Jeffrey Barrett, Sonia Goncalves, Ewan Harrison, David K. Jackson, Ian Johnston, Dominic Kwiatkowski, Cordelia Langford, John Sillitoe on behalf of the Wellcome Sanger Institute COVID-19 Surveillance Team |
| EPI_ISL_3443996 | Lighthouse Lab in Glasgow | Wellcome Sanger Institute for the COVID-19 Genomics UK (COG-UK) Consortium | Harper VanSteenhouse, Yumi Kasai, David Gray, Carol Clugston, Anna Dominiczak and Alex Alderton, Roberto Amato, Jeffrey Barrett, Sonia Goncalves, Ewan Harrison, David K. Jackson, Ian Johnston, Dominic Kwiatkowski, Cordelia Langford, John Sillitoe on behalf of the Wellcome Sanger Institute COVID-19 Surveillance Team |
| EPI_ISL_3443997 | Lighthouse Lab in Milton Keynes | Wellcome Sanger Institute for the COVID-19 Genomics UK (COG-UK) Consortium | The Lighthouse Lab in Milton Keynes and Alex Alderton, Roberto Amato, Jeffrey Barrett, Sonia Goncalves, Ewan Harrison, David K. Jackson, Ian Johnston, Dominic Kwiatkowski, Cordelia Langford, John Sillitoe on behalf of the Wellcome Sanger Institute COVID-19 Surveillance Team |
| EPI_ISL_3443998, EPI_ISL_3443999, EPI_ISL_3444000, EPI_ISL_3444001 | Lighthouse Lab in Glasgow | Wellcome Sanger Institute for the COVID-19 Genomics UK (COG-UK) Consortium | Harper VanSteenhouse, Yumi Kasai, David Gray, Carol Clugston, Anna Dominiczak and Alex Alderton, Roberto Amato, Jeffrey Barrett, Sonia Goncalves, Ewan Harrison, David K. Jackson, Ian Johnston, Dominic Kwiatkowski, Cordelia Langford, John Sillitoe on behalf of the Wellcome Sanger Institute COVID-19 Surveillance Team |
| EPI_ISL_3444002, EPI_ISL_3444003 | Lighthouse Lab in Milton Keynes | Wellcome Sanger Institute for the COVID-19 Genomics UK (COG-UK) Consortium | The Lighthouse Lab in Milton Keynes and Alex Alderton, Roberto Amato, Jeffrey Barrett, Sonia Goncalves, Ewan Harrison, David K. Jackson, Ian Johnston, Dominic Kwiatkowski, Cordelia Langford, John Sillitoe on behalf of the Wellcome Sanger Institute COVID-19 Surveillance Team |
| EPI_ISL_3444004, EPI_ISL_3444005 | Lighthouse Lab in Glasgow | Wellcome Sanger Institute for the COVID-19 Genomics UK (COG-UK) Consortium | Harper VanSteenhouse, Yumi Kasai, David Gray, Carol Clugston, Anna Dominiczak and Alex Alderton, Roberto Amato, Jeffrey Barrett, Sonia Goncalves, Ewan Harrison, David K. Jackson, Ian Johnston, Dominic Kwiatkowski, Cordelia Langford, John Sillitoe on behalf of the Wellcome Sanger Institute COVID-19 Surveillance Team |
| EPI_ISL_3444006 | Lighthouse Lab in Milton Keynes | Wellcome Sanger Institute for the COVID-19 Genomics UK (COG-UK) Consortium | The Lighthouse Lab in Milton Keynes and Alex Alderton, Roberto Amato, Jeffrey Barrett, Sonia Goncalves, Ewan Harrison, David K. Jackson, Ian Johnston, Dominic Kwiatkowski, Cordelia Langford, John Sillitoe on behalf of the Wellcome Sanger Institute COVID-19 Surveillance Team |
| EPI_ISL_3444007, EPI_ISL_3444008, EPI_ISL_3444009, EPI_ISL_3444010, EPI_ISL_3444011, EPI_ISL_3444012, EPI_ISL_3444013, EPI_ISL_3444014, EPI_ISL_3444015, EPI_ISL_3444016, EPI_ISL_3444017 | | | |
| see above | Lighthouse Lab in Glasgow | Wellcome Sanger Institute for the COVID-19 Genomics UK (COG-UK) Consortium | Harper VanSteenhouse, Yumi Kasai, David Gray, Carol Clugston, Anna Dominiczak and Alex Alderton, Roberto Amato, Jeffrey Barrett, Sonia Goncalves, Ewan Harrison, David K. Jackson, Ian Johnston, Dominic Kwiatkowski, Cordelia Langford, John Sillitoe on behalf of the Wellcome Sanger Institute COVID-19 Surveillance Team |
| EPI_ISL_3444018 | Lighthouse Lab in Milton Keynes | Wellcome Sanger Institute for the COVID-19 Genomics UK (COG-UK) Consortium | The Lighthouse Lab in Milton Keynes and Alex Alderton, Roberto Amato, Jeffrey Barrett, Sonia Goncalves, Ewan Harrison, David K. Jackson, Ian Johnston, Dominic Kwiatkowski, Cordelia Langford, John Sillitoe on behalf of the Wellcome Sanger Institute COVID-19 Surveillance Team |
| EPI_ISL_3444019, EPI_ISL_3444020, EPI_ISL_3444021, EPI_ISL_3444022, EPI_ISL_3444023, EPI_ISL_3444024, EPI_ISL_3444025, EPI_ISL_3444026, EPI_ISL_3444027, EPI_ISL_3444028, EPI_ISL_3444029, EPI_ISL_3444030, EPI_ISL_3444031, EPI_ISL_3444032 | | | |
| see above | Lighthouse Lab in Glasgow | Wellcome Sanger Institute for the COVID-19 Genomics UK (COG-UK) Consortium | Harper VanSteenhouse, Yumi Kasai, David Gray, Carol Clugston, Anna Dominiczak and Alex Alderton, Roberto Amato, Jeffrey Barrett, Sonia Goncalves, Ewan Harrison, David K. Jackson, Ian Johnston, Dominic Kwiatkowski, Cordelia Langford, John Sillitoe on behalf of the Wellcome Sanger Institute COVID-19 Surveillance Team |
| EPI_ISL_3444033 | Lighthouse Lab in Milton Keynes | Wellcome Sanger Institute for the COVID-19 Genomics UK (COG-UK) Consortium | The Lighthouse Lab in Milton Keynes and Alex Alderton, Roberto Amato, Jeffrey Barrett, Sonia Goncalves, Ewan Harrison, David K. Jackson, Ian Johnston, Dominic Kwiatkowski, Cordelia Langford, John Sillitoe on behalf of the Wellcome Sanger Institute COVID-19 Surveillance Team |
| EPI_ISL_3444034, EPI_ISL_3444035, EPI_ISL_3444036, EPI_ISL_3444037, EPI_ISL_3444038, EPI_ISL_3444039, EPI_ISL_3444040, EPI_ISL_3444041, EPI_ISL_3444042, EPI_ISL_3444043, EPI_ISL_3444044, EPI_ISL_3444045, EPI_ISL_3444046, EPI_ISL_3444047, EPI_ISL_3444048, EPI_ISL_3444049, EPI_ISL_3444050, EPI_ISL_3444051, EPI_ISL_3444052, EPI_ISL_3444053, EPI_ISL_3444054, EPI_ISL_3444055, EPI_ISL_3444056, EPI_ISL_3444057, EPI_ISL_3444058, EPI_ISL_3444059, EPI_ISL_3444060, EPI_ISL_3444061, EPI_ISL_3444062, EPI_ISL_3444063 | | | |
| see above | Lighthouse Lab in Glasgow | Wellcome Sanger Institute for the COVID-19 Genomics UK (COG-UK) Consortium | Harper VanSteenhouse, Yumi Kasai, David Gray, Carol Clugston, Anna Dominiczak and Alex Alderton, Roberto Amato, Jeffrey Barrett, Sonia Goncalves, Ewan Harrison, David K. Jackson, Ian Johnston, Dominic Kwiatkowski, Cordelia Langford, John Sillitoe on behalf of the Wellcome Sanger Institute COVID-19 Surveillance Team |
| EPI_ISL_3444064 | Lighthouse Lab in Milton Keynes | Wellcome Sanger Institute for the COVID-19 Genomics UK (COG-UK) Consortium | The Lighthouse Lab in Milton Keynes and Alex Alderton, Roberto Amato, Jeffrey Barrett, Sonia Goncalves, Ewan Harrison, David K. Jackson, Ian Johnston, Dominic Kwiatkowski, Cordelia Langford, John Sillitoe on behalf of the Wellcome Sanger Institute COVID-19 Surveillance Team |
| EPI_ISL_3444065, EPI_ISL_3444066, EPI_ISL_3444067, EPI_ISL_3444068, EPI_ISL_3444069 | Lighthouse Lab in Glasgow | Wellcome Sanger Institute for the COVID-19 Genomics UK (COG-UK) Consortium | Harper VanSteenhouse, Yumi Kasai, David Gray, Carol Clugston, Anna Dominiczak and Alex Alderton, Roberto Amato, Jeffrey Barrett, Sonia Goncalves, Ewan Harrison, David K. Jackson, Ian Johnston, Dominic Kwiatkowski, Cordelia Langford, John Sillitoe on behalf of the Wellcome Sanger Institute COVID-19 Surveillance Team |
| EPI_ISL_3444070 | Lighthouse Lab in Milton Keynes | Wellcome Sanger Institute for the COVID-19 Genomics UK (COG-UK) Consortium | The Lighthouse Lab in Milton Keynes and Alex Alderton, Roberto Amato, Jeffrey Barrett, Sonia Goncalves, Ewan Harrison, David K. Jackson, Ian Johnston, Dominic Kwiatkowski, Cordelia Langford, John Sillitoe on behalf of the Wellcome Sanger Institute COVID-19 Surveillance Team |
| EPI_ISL_3444071, EPI_ISL_3444072, EPI_ISL_3444073, EPI_ISL_3444074, EPI_ISL_3444075, EPI_ISL_3444076, EPI_ISL_3444077, EPI_ISL_3444078, EPI_ISL_3444079, EPI_ISL_3444080, EPI_ISL_3444081, EPI_ISL_3444082, EPI_ISL_3444083, EPI_ISL_3444084, EPI_ISL_3444085, EPI_ISL_3444086, EPI_ISL_3444087, EPI_ISL_3444088, EPI_ISL_3444089, EPI_ISL_3444090, EPI_ISL_3444091, EPI_ISL_3444092, EPI_ISL_3444093, EPI_ISL_3444094, EPI_ISL_3444095, EPI_ISL_3444096, EPI_ISL_3444097, EPI_ISL_3444098, EPI_ISL_3444099, EPI_ISL_3444100, EPI_ISL_3444101, EPI_ISL_3444102, EPI_ISL_3444103, EPI_ISL_3444104, EPI_ISL_3444105, EPI_ISL_3444106, EPI_ISL_3444107, EPI_ISL_3444108, EPI_ISL_3444109, EPI_ISL_3444110, EPI_ISL_3444111, EPI_ISL_3444112, EPI_ISL_3444113, EPI_ISL_3444114, EPI_ISL_3444115, EPI_ISL_3444116, EPI_ISL_3444117, EPI_ISL_3444118, EPI_ISL_3444119, EPI_ISL_3444120, EPI_ISL_3444121, EPI_ISL_3444122, EPI_ISL_3444123, EPI_ISL_3444124, EPI_ISL_3444125, EPI_ISL_3444126, EPI_ISL_3444127, EPI_ISL_3444128, EPI_ISL_3444129, EPI_ISL_3444130, EPI_ISL_3444131, EPI_ISL_3444132, EPI_ISL_3444133, EPI_ISL_3444134, EPI_ISL_3444135, EPI_ISL_3444136, EPI_ISL_3444137, EPI_ISL_3444138, EPI_ISL_3444139, EPI_ISL_3444140, EPI_ISL_3444141, EPI_ISL_3444142, EPI_ISL_3444143, EPI_ISL_3444144, EPI_ISL_3444145, EPI_ISL_3444146, EPI_ISL_3444147, EPI_ISL_3444148, EPI_ISL_3444149, EPI_ISL_3444150, EPI_ISL_3444151, EPI_ISL_3444152, EPI_ISL_3444153, EPI_ISL_3444154, EPI_ISL_3444155, EPI_ISL_3444156, EPI_ISL_3444157, EPI_ISL_3444158, EPI_ISL_3444159, EPI_ISL_3444160 | | | |

| | | | |
|---|---|---|---|
| see above | Lighthouse Lab in Glasgow | Wellcome Sanger Institute for the COVID-19 Genomics UK (COG-UK) Consortium | Harper VanSteenhouse, Yumi Kasai, David Gray, Carol Clugston, Anna Dominiczak and Alex Alderton, Roberto Amato, Jeffrey Barrett, Sonia Goncalves, Ewan Harrison, David K. Jackson, Ian Johnston, Dominic Kwiatkowski, Cordelia Langford, John Sillitoe on behalf of the Wellcome Sanger Institute COVID-19 Surveillance Team |
| EPI_ISL_3444161 | Lighthouse Lab in Milton Keynes | Wellcome Sanger Institute for the COVID-19 Genomics UK (COG-UK) Consortium | The Lighthouse Lab in Milton Keynes and Alex Alderton, Roberto Amato, Jeffrey Barrett, Sonia Goncalves, Ewan Harrison, David K. Jackson, Ian Johnston, Dominic Kwiatkowski, Cordelia Langford, John Sillitoe on behalf of the Wellcome Sanger Institute COVID-19 Surveillance Team |
| EPI_ISL_3444162, EPI_ISL_3444163, EPI_ISL_3444164, EPI_ISL_3444165, EPI_ISL_3444166, EPI_ISL_3444167, EPI_ISL_3444168, EPI_ISL_3444169, EPI_ISL_3444170, EPI_ISL_3444171, EPI_ISL_3444172, EPI_ISL_3444173, EPI_ISL_3444174, EPI_ISL_3444175, EPI_ISL_3444176, EPI_ISL_3444177, EPI_ISL_3444178, EPI_ISL_3444179, EPI_ISL_3444180, EPI_ISL_3444181, EPI_ISL_3444182, EPI_ISL_3444183, EPI_ISL_3444184, EPI_ISL_3444185, EPI_ISL_3444186, EPI_ISL_3444187, EPI_ISL_3444188, EPI_ISL_3444189, EPI_ISL_3444190, EPI_ISL_3444191, EPI_ISL_3444192, EPI_ISL_3444193, EPI_ISL_3444194, EPI_ISL_3444195, EPI_ISL_3444196, EPI_ISL_3444197, EPI_ISL_3444198, EPI_ISL_3444199, EPI_ISL_3444200, EPI_ISL_3444201, EPI_ISL_3444202, EPI_ISL_3444203, EPI_ISL_3444204, EPI_ISL_3444205, EPI_ISL_3444206, EPI_ISL_3444207, EPI_ISL_3444208, EPI_ISL_3444209, EPI_ISL_3444210, EPI_ISL_3444211, EPI_ISL_3444212, EPI_ISL_3444213, EPI_ISL_3444214, EPI_ISL_3444215, EPI_ISL_3444216, EPI_ISL_3444217, EPI_ISL_3444218, EPI_ISL_3444219, EPI_ISL_3444220, EPI_ISL_3444221, EPI_ISL_3444222, EPI_ISL_3444223, EPI_ISL_3444224, EPI_ISL_3444225, EPI_ISL_3444226, EPI_ISL_3444227, EPI_ISL_3444228, EPI_ISL_3444229, EPI_ISL_3444230, EPI_ISL_3444231, EPI_ISL_3444232, EPI_ISL_3444233, EPI_ISL_3444234, EPI_ISL_3444235, EPI_ISL_3444236, EPI_ISL_3444237, EPI_ISL_3444238, EPI_ISL_3444239, EPI_ISL_3444240, EPI_ISL_3444241, EPI_ISL_3444242, EPI_ISL_3444243, EPI_ISL_3444244, EPI_ISL_3444245, EPI_ISL_3444246, EPI_ISL_3444247, EPI_ISL_3444248, EPI_ISL_3444249, EPI_ISL_3444250, EPI_ISL_3444251, EPI_ISL_3444252, EPI_ISL_3444253, EPI_ISL_3444254 | | | |
| see above | Lighthouse Lab in Glasgow | Wellcome Sanger Institute for the COVID-19 Genomics UK (COG-UK) Consortium | Harper VanSteenhouse, Yumi Kasai, David Gray, Carol Clugston, Anna Dominiczak and Alex Alderton, Roberto Amato, Jeffrey Barrett, Sonia Goncalves, Ewan Harrison, David K. Jackson, Ian Johnston, Dominic Kwiatkowski, Cordelia Langford, John Sillitoe on behalf of the Wellcome Sanger Institute COVID-19 Surveillance Team |
| EPI_ISL_3444255, EPI_ISL_3444256, EPI_ISL_3444257, EPI_ISL_3444258 | Lighthouse Lab in Milton Keynes | Wellcome Sanger Institute for the COVID-19 Genomics UK (COG-UK) Consortium | The Lighthouse Lab in Milton Keynes and Alex Alderton, Roberto Amato, Jeffrey Barrett, Sonia Goncalves, Ewan Harrison, David K. Jackson, Ian Johnston, Dominic Kwiatkowski, Cordelia Langford, John Sillitoe on behalf of the Wellcome Sanger Institute COVID-19 Surveillance Team |
| EPI_ISL_3444259 | Lighthouse Lab in Glasgow | Wellcome Sanger Institute for the COVID-19 Genomics UK (COG-UK) Consortium | Harper VanSteenhouse, Yumi Kasai, David Gray, Carol Clugston, Anna Dominiczak and Alex Alderton, Roberto Amato, Jeffrey Barrett, Sonia Goncalves, Ewan Harrison, David K. Jackson, Ian Johnston, Dominic Kwiatkowski, Cordelia Langford, John Sillitoe on behalf of the Wellcome Sanger Institute COVID-19 Surveillance Team |
| EPI_ISL_3444260 | Lighthouse Lab in Milton Keynes | Wellcome Sanger Institute for the COVID-19 Genomics UK (COG-UK) Consortium | The Lighthouse Lab in Milton Keynes and Alex Alderton, Roberto Amato, Jeffrey Barrett, Sonia Goncalves, Ewan Harrison, David K. Jackson, Ian Johnston, Dominic Kwiatkowski, Cordelia Langford, John Sillitoe on behalf of the Wellcome Sanger Institute COVID-19 Surveillance Team |
| EPI_ISL_3444261, EPI_ISL_3444262 | Lighthouse Lab in Glasgow | Wellcome Sanger Institute for the COVID-19 Genomics UK (COG-UK) Consortium | Harper VanSteenhouse, Yumi Kasai, David Gray, Carol Clugston, Anna Dominiczak and Alex Alderton, Roberto Amato, Jeffrey Barrett, Sonia Goncalves, Ewan Harrison, David K. Jackson, Ian Johnston, Dominic Kwiatkowski, Cordelia Langford, John Sillitoe on behalf of the Wellcome Sanger Institute COVID-19 Surveillance Team |
| EPI_ISL_3444263 | Lighthouse Lab in Milton Keynes | Wellcome Sanger Institute for the COVID-19 Genomics UK (COG-UK) Consortium | The Lighthouse Lab in Milton Keynes and Alex Alderton, Roberto Amato, Jeffrey Barrett, Sonia Goncalves, Ewan Harrison, David K. Jackson, Ian Johnston, Dominic Kwiatkowski, Cordelia Langford, John Sillitoe on behalf of the Wellcome Sanger Institute COVID-19 Surveillance Team |
| EPI_ISL_3444264, EPI_ISL_3444265, EPI_ISL_3444266 | Lighthouse Lab in Glasgow | Wellcome Sanger Institute for the COVID-19 Genomics UK (COG-UK) Consortium | Harper VanSteenhouse, Yumi Kasai, David Gray, Carol Clugston, Anna Dominiczak and Alex Alderton, Roberto Amato, Jeffrey Barrett, Sonia Goncalves, Ewan Harrison, David K. Jackson, Ian Johnston, Dominic Kwiatkowski, Cordelia Langford, John Sillitoe on behalf of the Wellcome Sanger Institute COVID-19 Surveillance Team |
| EPI_ISL_3444267, EPI_ISL_3444268 | Lighthouse Lab in Milton Keynes | Wellcome Sanger Institute for the COVID-19 Genomics UK (COG-UK) Consortium | The Lighthouse Lab in Milton Keynes and Alex Alderton, Roberto Amato, Jeffrey Barrett, Sonia Goncalves, Ewan Harrison, David K. Jackson, Ian Johnston, Dominic Kwiatkowski, Cordelia Langford, John Sillitoe on behalf of the Wellcome Sanger Institute COVID-19 Surveillance Team |
| EPI_ISL_3444269, EPI_ISL_3444270 | Lighthouse Lab in Glasgow | Wellcome Sanger Institute for the COVID-19 Genomics UK (COG-UK) Consortium | Harper VanSteenhouse, Yumi Kasai, David Gray, Carol Clugston, Anna Dominiczak and Alex Alderton, Roberto Amato, Jeffrey Barrett, Sonia Goncalves, Ewan Harrison, David K. Jackson, Ian Johnston, Dominic Kwiatkowski, Cordelia Langford, John Sillitoe on behalf of the Wellcome Sanger Institute COVID-19 Surveillance Team |
| EPI_ISL_3444271 | Lighthouse Lab in Milton Keynes | Wellcome Sanger Institute for the COVID-19 Genomics UK (COG-UK) Consortium | The Lighthouse Lab in Milton Keynes and Alex Alderton, Roberto Amato, Jeffrey Barrett, Sonia Goncalves, Ewan Harrison, David K. Jackson, Ian Johnston, Dominic Kwiatkowski, Cordelia Langford, John Sillitoe on behalf of the Wellcome Sanger Institute COVID-19 Surveillance Team |
| EPI_ISL_3444272 | Lighthouse Lab in Glasgow | Wellcome Sanger Institute for the COVID-19 Genomics UK (COG-UK) Consortium | Harper VanSteenhouse, Yumi Kasai, David Gray, Carol Clugston, Anna Dominiczak and Alex Alderton, Roberto Amato, Jeffrey Barrett, Sonia Goncalves, Ewan Harrison, David K. Jackson, Ian Johnston, Dominic Kwiatkowski, Cordelia Langford, John Sillitoe on behalf of the Wellcome Sanger Institute COVID-19 Surveillance Team |
| EPI_ISL_3444273, EPI_ISL_3444274 | Lighthouse Lab in Milton Keynes | Wellcome Sanger Institute for the COVID-19 Genomics UK (COG-UK) Consortium | The Lighthouse Lab in Milton Keynes and Alex Alderton, Roberto Amato, Jeffrey Barrett, Sonia Goncalves, Ewan Harrison, David K. Jackson, Ian Johnston, Dominic Kwiatkowski, Cordelia Langford, John Sillitoe on behalf of the Wellcome Sanger Institute COVID-19 Surveillance Team |
| EPI_ISL_3444275, EPI_ISL_3444276 | Lighthouse Lab in Glasgow | Wellcome Sanger Institute for the COVID-19 Genomics UK (COG-UK) Consortium | Harper VanSteenhouse, Yumi Kasai, David Gray, Carol Clugston, Anna Dominiczak and Alex Alderton, Roberto Amato, Jeffrey Barrett, Sonia Goncalves, Ewan Harrison, David K. Jackson, Ian Johnston, Dominic Kwiatkowski, Cordelia Langford, John Sillitoe on behalf of the Wellcome Sanger Institute COVID-19 Surveillance Team |
| EPI_ISL_3444277, EPI_ISL_3444278 | Lighthouse Lab in Milton Keynes | Wellcome Sanger Institute for the COVID-19 Genomics UK (COG-UK) Consortium | The Lighthouse Lab in Milton Keynes and Alex Alderton, Roberto Amato, Jeffrey Barrett, Sonia Goncalves, Ewan Harrison, David K. Jackson, Ian Johnston, Dominic Kwiatkowski, Cordelia Langford, John Sillitoe on behalf of the Wellcome Sanger Institute COVID-19 Surveillance Team |
| EPI_ISL_3444279 | Lighthouse Lab in Glasgow | Wellcome Sanger Institute for the COVID-19 Genomics UK (COG-UK) Consortium | Harper VanSteenhouse, Yumi Kasai, David Gray, Carol Clugston, Anna Dominiczak and Alex Alderton, Roberto Amato, Jeffrey Barrett, Sonia Goncalves, Ewan Harrison, David K. Jackson, Ian Johnston, Dominic Kwiatkowski, Cordelia Langford, John Sillitoe on behalf of the Wellcome Sanger Institute COVID-19 Surveillance Team |
| EPI_ISL_3444280 | Lighthouse Lab in Milton Keynes | Wellcome Sanger Institute for the COVID-19 Genomics UK (COG-UK) Consortium | The Lighthouse Lab in Milton Keynes and Alex Alderton, Roberto Amato, Jeffrey Barrett, Sonia Goncalves, Ewan Harrison, David K. Jackson, Ian Johnston, Dominic Kwiatkowski, Cordelia Langford, John Sillitoe on behalf of the Wellcome Sanger Institute COVID-19 Surveillance Team |
| EPI_ISL_3444281, EPI_ISL_3444282, EPI_ISL_3444283, EPI_ISL_3444284 | Lighthouse Lab in Glasgow | Wellcome Sanger Institute for the COVID-19 Genomics UK (COG-UK) Consortium | Harper VanSteenhouse, Yumi Kasai, David Gray, Carol Clugston, Anna Dominiczak and Alex Alderton, Roberto Amato, Jeffrey Barrett, Sonia Goncalves, Ewan Harrison, David K. Jackson, Ian Johnston, Dominic Kwiatkowski, Cordelia Langford, John Sillitoe on behalf of the Wellcome Sanger Institute COVID-19 Surveillance Team |
| EPI_ISL_3444285, EPI_ISL_3444286 | Lighthouse Lab in Milton Keynes | Wellcome Sanger Institute for the COVID-19 Genomics UK (COG-UK) Consortium | The Lighthouse Lab in Milton Keynes and Alex Alderton, Roberto Amato, Jeffrey Barrett, Sonia Goncalves, Ewan Harrison, David K. Jackson, Ian Johnston, Dominic Kwiatkowski, Cordelia Langford, John Sillitoe on behalf of the Wellcome Sanger Institute COVID-19 Surveillance Team |
| EPI_ISL_3444287 | Lighthouse Lab in Glasgow | Wellcome Sanger Institute for the COVID-19 Genomics UK (COG-UK) Consortium | Harper VanSteenhouse, Yumi Kasai, David Gray, Carol Clugston, Anna Dominiczak and Alex Alderton, Roberto Amato, Jeffrey Barrett, Sonia Goncalves, Ewan Harrison, David K. Jackson, Ian Johnston, Dominic Kwiatkowski, Cordelia Langford, John Sillitoe on behalf of the Wellcome Sanger Institute COVID-19 Surveillance Team |
| EPI_ISL_3444288 | Lighthouse Lab in Milton Keynes | Wellcome Sanger Institute for the COVID-19 Genomics UK (COG-UK) Consortium | The Lighthouse Lab in Milton Keynes and Alex Alderton, Roberto Amato, Jeffrey Barrett, Sonia Goncalves, Ewan Harrison, David K. Jackson, Ian Johnston, Dominic Kwiatkowski, Cordelia Langford, John Sillitoe on behalf of the Wellcome Sanger Institute COVID-19 Surveillance Team |
| EPI_ISL_3444289, EPI_ISL_3444290 | Lighthouse Lab in Glasgow | Wellcome Sanger Institute for the COVID-19 Genomics UK (COG-UK) Consortium | Harper VanSteenhouse, Yumi Kasai, David Gray, Carol Clugston, Anna Dominiczak and Alex Alderton, Roberto Amato, Jeffrey Barrett, Sonia Goncalves, Ewan Harrison, David K. Jackson, Ian Johnston, Dominic Kwiatkowski, Cordelia Langford, John Sillitoe on behalf of the Wellcome Sanger Institute COVID-19 Surveillance Team |
| EPI_ISL_3444291 | Lighthouse Lab in Milton Keynes | Wellcome Sanger Institute for the COVID-19 Genomics UK (COG-UK) Consortium | The Lighthouse Lab in Milton Keynes and Alex Alderton, Roberto Amato, Jeffrey Barrett, Sonia Goncalves, Ewan Harrison, David K. Jackson, Ian Johnston, Dominic Kwiatkowski, Cordelia Langford, John Sillitoe on behalf of the Wellcome Sanger Institute COVID-19 Surveillance Team |
| EPI_ISL_3444292, EPI_ISL_3444293, EPI_ISL_3444294, EPI_ISL_3444295, EPI_ISL_3444296 | Lighthouse Lab in Glasgow | Wellcome Sanger Institute for the COVID-19 Genomics UK (COG-UK) Consortium | Harper VanSteenhouse, Yumi Kasai, David Gray, Carol Clugston, Anna Dominiczak and Alex Alderton, Roberto Amato, Jeffrey Barrett, Sonia Goncalves, Ewan Harrison, David K. Jackson, Ian Johnston, Dominic Kwiatkowski, Cordelia Langford, John Sillitoe on behalf of the Wellcome Sanger Institute COVID-19 Surveillance Team |
| EPI_ISL_3444297, EPI_ISL_3444298, EPI_ISL_3444299, EPI_ISL_3444300, EPI_ISL_3444301, EPI_ISL_3444302, EPI_ISL_3444303, EPI_ISL_3444304 | Lighthouse Lab in Milton Keynes | Wellcome Sanger Institute for the COVID-19 Genomics UK (COG-UK) Consortium | The Lighthouse Lab in Milton Keynes and Alex Alderton, Roberto Amato, Jeffrey Barrett, Sonia Goncalves, Ewan Harrison, David K. Jackson, Ian Johnston, Dominic Kwiatkowski, Cordelia Langford, John Sillitoe on behalf of the Wellcome Sanger Institute COVID-19 Surveillance Team |
| EPI_ISL_3444305, EPI_ISL_3444306 | Lighthouse Lab in Glasgow | Wellcome Sanger Institute for the COVID-19 Genomics UK (COG-UK) Consortium | Harper VanSteenhouse, Yumi Kasai, David Gray, Carol Clugston, Anna Dominiczak and Alex Alderton, Roberto Amato, Jeffrey Barrett, Sonia Goncalves, Ewan Harrison, David K. Jackson, Ian Johnston, Dominic Kwiatkowski, Cordelia Langford, John Sillitoe on behalf of the Wellcome Sanger Institute COVID-19 Surveillance Team |
| EPI_ISL_3444307, EPI_ISL_3444308, EPI_ISL_3444309 | Lighthouse Lab in Milton Keynes | Wellcome Sanger Institute for the COVID-19 Genomics UK (COG-UK) Consortium | The Lighthouse Lab in Milton Keynes and Alex Alderton, Roberto Amato, Jeffrey Barrett, Sonia Goncalves, Ewan Harrison, David K. Jackson, Ian Johnston, Dominic Kwiatkowski, Cordelia Langford, John Sillitoe on behalf of the Wellcome Sanger Institute COVID-19 Surveillance Team |
| EPI_ISL_3444310, EPI_ISL_3444311 | Lighthouse Lab in Glasgow | Wellcome Sanger Institute for the COVID-19 Genomics UK (COG-UK) Consortium | Harper VanSteenhouse, Yumi Kasai, David Gray, Carol Clugston, Anna Dominiczak and Alex Alderton, Roberto Amato, Jeffrey Barrett, Sonia Goncalves, Ewan Harrison, David K. Jackson, Ian Johnston, Dominic Kwiatkowski, Cordelia Langford, John Sillitoe on behalf of the Wellcome Sanger Institute COVID-19 Surveillance Team |
| EPI_ISL_3444312, EPI_ISL_3444313, EPI_ISL_3444314, EPI_ISL_3444315, EPI_ISL_3444316, EPI_ISL_3444317 | Lighthouse Lab in Milton Keynes | Wellcome Sanger Institute for the COVID-19 Genomics UK (COG-UK) Consortium | The Lighthouse Lab in Milton Keynes and Alex Alderton, Roberto Amato, Jeffrey Barrett, Sonia Goncalves, Ewan Harrison, David K. Jackson, Ian Johnston, Dominic Kwiatkowski, Cordelia Langford, John Sillitoe on behalf of the Wellcome Sanger Institute COVID-19 Surveillance Team |
| EPI_ISL_3444318, EPI_ISL_3444319 | Lighthouse Lab in Glasgow | Wellcome Sanger Institute for the COVID-19 Genomics UK (COG-UK) Consortium | Harper VanSteenhouse, Yumi Kasai, David Gray, Carol Clugston, Anna Dominiczak and Alex Alderton, Roberto Amato, Jeffrey Barrett, Sonia Goncalves, Ewan Harrison, David K. Jackson, Ian Johnston, Dominic Kwiatkowski, Cordelia Langford, John Sillitoe on behalf of the Wellcome Sanger Institute COVID-19 Surveillance Team |
| EPI_ISL_3444320 | Lighthouse Lab in Milton Keynes | Wellcome Sanger Institute for the COVID-19 Genomics UK (COG-UK) Consortium | The Lighthouse Lab in Milton Keynes and Alex Alderton, Roberto Amato, Jeffrey Barrett, Sonia Goncalves, Ewan Harrison, David K. Jackson, Ian Johnston, Dominic Kwiatkowski, Cordelia Langford, John Sillitoe on behalf of the Wellcome Sanger Institute COVID-19 Surveillance Team |
| EPI_ISL_3444321 | Lighthouse Lab in Glasgow | Wellcome Sanger Institute for the COVID-19 Genomics UK (COG-UK) Consortium | Harper VanSteenhouse, Yumi Kasai, David Gray, Carol Clugston, Anna Dominiczak and Alex Alderton, Roberto Amato, Jeffrey Barrett, Sonia Goncalves, Ewan Harrison, David K. Jackson, Ian Johnston, Dominic Kwiatkowski, Cordelia Langford, John Sillitoe on behalf of the Wellcome Sanger Institute COVID-19 Surveillance Team |
| EPI_ISL_3444322, EPI_ISL_3444323, EPI_ISL_3444324, EPI_ISL_3444325, EPI_ISL_3444326 | Lighthouse Lab in Milton Keynes | Wellcome Sanger Institute for the COVID-19 Genomics UK (COG-UK) Consortium | The Lighthouse Lab in Milton Keynes and Alex Alderton, Roberto Amato, Jeffrey Barrett, Sonia Goncalves, Ewan Harrison, David K. Jackson, Ian Johnston, Dominic Kwiatkowski, Cordelia Langford, John Sillitoe on behalf of the Wellcome Sanger Institute COVID-19 Surveillance Team |

| | | | |
|---|---|---|---|
| EPI_ISL_3444327, EPI_ISL_3444328, EPI_ISL_3444329 | Lighthouse Lab in Glasgow | Wellcome Sanger Institute for the COVID-19 Genomics UK (COG-UK) Consortium | Harper VanSteenhouse, Yumi Kasai, David Gray, Carol Clugston, Anna Dominiczak and Alex Alderton, Roberto Amato, Jeffrey Barrett, Sonia Goncalves, Ewan Harrison, David K. Jackson, Ian Johnston, Dominic Kwiatkowski, Cordelia Langford, John Sillitoe on behalf of the Wellcome Sanger Institute COVID-19 Surveillance Team |
| EPI_ISL_3444330, EPI_ISL_3444331 | Lighthouse Lab in Milton Keynes | Wellcome Sanger Institute for the COVID-19 Genomics UK (COG-UK) Consortium | The Lighthouse Lab in Milton Keynes and Alex Alderton, Roberto Amato, Jeffrey Barrett, Sonia Goncalves, Ewan Harrison, David K. Jackson, Ian Johnston, Dominic Kwiatkowski, Cordelia Langford, John Sillitoe on behalf of the Wellcome Sanger Institute COVID-19 Surveillance Team |
| EPI_ISL_3444332 | Lighthouse Lab in Glasgow | Wellcome Sanger Institute for the COVID-19 Genomics UK (COG-UK) Consortium | Harper VanSteenhouse, Yumi Kasai, David Gray, Carol Clugston, Anna Dominiczak and Alex Alderton, Roberto Amato, Jeffrey Barrett, Sonia Goncalves, Ewan Harrison, David K. Jackson, Ian Johnston, Dominic Kwiatkowski, Cordelia Langford, John Sillitoe on behalf of the Wellcome Sanger Institute COVID-19 Surveillance Team |
| EPI_ISL_3444333, EPI_ISL_3444334, EPI_ISL_3444335 | Lighthouse Lab in Milton Keynes | Wellcome Sanger Institute for the COVID-19 Genomics UK (COG-UK) Consortium | The Lighthouse Lab in Milton Keynes and Alex Alderton, Roberto Amato, Jeffrey Barrett, Sonia Goncalves, Ewan Harrison, David K. Jackson, Ian Johnston, Dominic Kwiatkowski, Cordelia Langford, John Sillitoe on behalf of the Wellcome Sanger Institute COVID-19 Surveillance Team |
| EPI_ISL_3444336, EPI_ISL_3444337 | Lighthouse Lab in Glasgow | Wellcome Sanger Institute for the COVID-19 Genomics UK (COG-UK) Consortium | Harper VanSteenhouse, Yumi Kasai, David Gray, Carol Clugston, Anna Dominiczak and Alex Alderton, Roberto Amato, Jeffrey Barrett, Sonia Goncalves, Ewan Harrison, David K. Jackson, Ian Johnston, Dominic Kwiatkowski, Cordelia Langford, John Sillitoe on behalf of the Wellcome Sanger Institute COVID-19 Surveillance Team |
| EPI_ISL_3444338, EPI_ISL_3444339 | Lighthouse Lab in Milton Keynes | Wellcome Sanger Institute for the COVID-19 Genomics UK (COG-UK) Consortium | The Lighthouse Lab in Milton Keynes and Alex Alderton, Roberto Amato, Jeffrey Barrett, Sonia Goncalves, Ewan Harrison, David K. Jackson, Ian Johnston, Dominic Kwiatkowski, Cordelia Langford, John Sillitoe on behalf of the Wellcome Sanger Institute COVID-19 Surveillance Team |
| EPI_ISL_3444340 | Lighthouse Lab in Glasgow | Wellcome Sanger Institute for the COVID-19 Genomics UK (COG-UK) Consortium | Harper VanSteenhouse, Yumi Kasai, David Gray, Carol Clugston, Anna Dominiczak and Alex Alderton, Roberto Amato, Jeffrey Barrett, Sonia Goncalves, Ewan Harrison, David K. Jackson, Ian Johnston, Dominic Kwiatkowski, Cordelia Langford, John Sillitoe on behalf of the Wellcome Sanger Institute COVID-19 Surveillance Team |
| EPI_ISL_3444341 | Lighthouse Lab in Milton Keynes | Wellcome Sanger Institute for the COVID-19 Genomics UK (COG-UK) Consortium | The Lighthouse Lab in Milton Keynes and Alex Alderton, Roberto Amato, Jeffrey Barrett, Sonia Goncalves, Ewan Harrison, David K. Jackson, Ian Johnston, Dominic Kwiatkowski, Cordelia Langford, John Sillitoe on behalf of the Wellcome Sanger Institute COVID-19 Surveillance Team |
| EPI_ISL_3444342, EPI_ISL_3444343 | Lighthouse Lab in Glasgow | Wellcome Sanger Institute for the COVID-19 Genomics UK (COG-UK) Consortium | Harper VanSteenhouse, Yumi Kasai, David Gray, Carol Clugston, Anna Dominiczak and Alex Alderton, Roberto Amato, Jeffrey Barrett, Sonia Goncalves, Ewan Harrison, David K. Jackson, Ian Johnston, Dominic Kwiatkowski, Cordelia Langford, John Sillitoe on behalf of the Wellcome Sanger Institute COVID-19 Surveillance Team |
| EPI_ISL_3444344 | Lighthouse Lab in Milton Keynes | Wellcome Sanger Institute for the COVID-19 Genomics UK (COG-UK) Consortium | The Lighthouse Lab in Milton Keynes and Alex Alderton, Roberto Amato, Jeffrey Barrett, Sonia Goncalves, Ewan Harrison, David K. Jackson, Ian Johnston, Dominic Kwiatkowski, Cordelia Langford, John Sillitoe on behalf of the Wellcome Sanger Institute COVID-19 Surveillance Team |
| EPI_ISL_3444345, EPI_ISL_3444346 | Lighthouse Lab in Glasgow | Wellcome Sanger Institute for the COVID-19 Genomics UK (COG-UK) Consortium | Harper VanSteenhouse, Yumi Kasai, David Gray, Carol Clugston, Anna Dominiczak and Alex Alderton, Roberto Amato, Jeffrey Barrett, Sonia Goncalves, Ewan Harrison, David K. Jackson, Ian Johnston, Dominic Kwiatkowski, Cordelia Langford, John Sillitoe on behalf of the Wellcome Sanger Institute COVID-19 Surveillance Team |
| EPI_ISL_3444347, EPI_ISL_3444348 | Lighthouse Lab in Milton Keynes | Wellcome Sanger Institute for the COVID-19 Genomics UK (COG-UK) Consortium | The Lighthouse Lab in Milton Keynes and Alex Alderton, Roberto Amato, Jeffrey Barrett, Sonia Goncalves, Ewan Harrison, David K. Jackson, Ian Johnston, Dominic Kwiatkowski, Cordelia Langford, John Sillitoe on behalf of the Wellcome Sanger Institute COVID-19 Surveillance Team |
| EPI_ISL_3444349 | Lighthouse Lab in Glasgow | Wellcome Sanger Institute for the COVID-19 Genomics UK (COG-UK) Consortium | Harper VanSteenhouse, Yumi Kasai, David Gray, Carol Clugston, Anna Dominiczak and Alex Alderton, Roberto Amato, Jeffrey Barrett, Sonia Goncalves, Ewan Harrison, David K. Jackson, Ian Johnston, Dominic Kwiatkowski, Cordelia Langford, John Sillitoe on behalf of the Wellcome Sanger Institute COVID-19 Surveillance Team |
| EPI_ISL_3444350 | Lighthouse Lab in Milton Keynes | Wellcome Sanger Institute for the COVID-19 Genomics UK (COG-UK) Consortium | The Lighthouse Lab in Milton Keynes and Alex Alderton, Roberto Amato, Jeffrey Barrett, Sonia Goncalves, Ewan Harrison, David K. Jackson, Ian Johnston, Dominic Kwiatkowski, Cordelia Langford, John Sillitoe on behalf of the Wellcome Sanger Institute COVID-19 Surveillance Team |
| EPI_ISL_3444351 | Lighthouse Lab in Glasgow | Wellcome Sanger Institute for the COVID-19 Genomics UK (COG-UK) Consortium | Harper VanSteenhouse, Yumi Kasai, David Gray, Carol Clugston, Anna Dominiczak and Alex Alderton, Roberto Amato, Jeffrey Barrett, Sonia Goncalves, Ewan Harrison, David K. Jackson, Ian Johnston, Dominic Kwiatkowski, Cordelia Langford, John Sillitoe on behalf of the Wellcome Sanger Institute COVID-19 Surveillance Team |
| EPI_ISL_3444352, EPI_ISL_3444353 | Lighthouse Lab in Milton Keynes | Wellcome Sanger Institute for the COVID-19 Genomics UK (COG-UK) Consortium | The Lighthouse Lab in Milton Keynes and Alex Alderton, Roberto Amato, Jeffrey Barrett, Sonia Goncalves, Ewan Harrison, David K. Jackson, Ian Johnston, Dominic Kwiatkowski, Cordelia Langford, John Sillitoe on behalf of the Wellcome Sanger Institute COVID-19 Surveillance Team |
| EPI_ISL_3444354, EPI_ISL_3444355 | Lighthouse Lab in Glasgow | Wellcome Sanger Institute for the COVID-19 Genomics UK (COG-UK) Consortium | Harper VanSteenhouse, Yumi Kasai, David Gray, Carol Clugston, Anna Dominiczak and Alex Alderton, Roberto Amato, Jeffrey Barrett, Sonia Goncalves, Ewan Harrison, David K. Jackson, Ian Johnston, Dominic Kwiatkowski, Cordelia Langford, John Sillitoe on behalf of the Wellcome Sanger Institute COVID-19 Surveillance Team |
| EPI_ISL_3444356, EPI_ISL_3444357, EPI_ISL_3444358, EPI_ISL_3444359, EPI_ISL_3444360, EPI_ISL_3444361 | Lighthouse Lab in Milton Keynes | Wellcome Sanger Institute for the COVID-19 Genomics UK (COG-UK) Consortium | The Lighthouse Lab in Milton Keynes and Alex Alderton, Roberto Amato, Jeffrey Barrett, Sonia Goncalves, Ewan Harrison, David K. Jackson, Ian Johnston, Dominic Kwiatkowski, Cordelia Langford, John Sillitoe on behalf of the Wellcome Sanger Institute COVID-19 Surveillance Team |
| EPI_ISL_3444362, EPI_ISL_3444363 | Lighthouse Lab in Glasgow | Wellcome Sanger Institute for the COVID-19 Genomics UK (COG-UK) Consortium | Harper VanSteenhouse, Yumi Kasai, David Gray, Carol Clugston, Anna Dominiczak and Alex Alderton, Roberto Amato, Jeffrey Barrett, Sonia Goncalves, Ewan Harrison, David K. Jackson, Ian Johnston, Dominic Kwiatkowski, Cordelia Langford, John Sillitoe on behalf of the Wellcome Sanger Institute COVID-19 Surveillance Team |
| EPI_ISL_3444364 | Lighthouse Lab in Milton Keynes | Wellcome Sanger Institute for the COVID-19 Genomics UK (COG-UK) Consortium | The Lighthouse Lab in Milton Keynes and Alex Alderton, Roberto Amato, Jeffrey Barrett, Sonia Goncalves, Ewan Harrison, David K. Jackson, Ian Johnston, Dominic Kwiatkowski, Cordelia Langford, John Sillitoe on behalf of the Wellcome Sanger Institute COVID-19 Surveillance Team |
| EPI_ISL_3444365, EPI_ISL_3444366, EPI_ISL_3444367, EPI_ISL_3444368 | Lighthouse Lab in Glasgow | Wellcome Sanger Institute for the COVID-19 Genomics UK (COG-UK) Consortium | Harper VanSteenhouse, Yumi Kasai, David Gray, Carol Clugston, Anna Dominiczak and Alex Alderton, Roberto Amato, Jeffrey Barrett, Sonia Goncalves, Ewan Harrison, David K. Jackson, Ian Johnston, Dominic Kwiatkowski, Cordelia Langford, John Sillitoe on behalf of the Wellcome Sanger Institute COVID-19 Surveillance Team |
| EPI_ISL_3444369, EPI_ISL_3444370 | Lighthouse Lab in Milton Keynes | Wellcome Sanger Institute for the COVID-19 Genomics UK (COG-UK) Consortium | The Lighthouse Lab in Milton Keynes and Alex Alderton, Roberto Amato, Jeffrey Barrett, Sonia Goncalves, Ewan Harrison, David K. Jackson, Ian Johnston, Dominic Kwiatkowski, Cordelia Langford, John Sillitoe on behalf of the Wellcome Sanger Institute COVID-19 Surveillance Team |
| EPI_ISL_3444371 | Lighthouse Lab in Glasgow | Wellcome Sanger Institute for the COVID-19 Genomics UK (COG-UK) Consortium | Harper VanSteenhouse, Yumi Kasai, David Gray, Carol Clugston, Anna Dominiczak and Alex Alderton, Roberto Amato, Jeffrey Barrett, Sonia Goncalves, Ewan Harrison, David K. Jackson, Ian Johnston, Dominic Kwiatkowski, Cordelia Langford, John Sillitoe on behalf of the Wellcome Sanger Institute COVID-19 Surveillance Team |
| EPI_ISL_3444372, EPI_ISL_3444373, EPI_ISL_3444374 | Lighthouse Lab in Milton Keynes | Wellcome Sanger Institute for the COVID-19 Genomics UK (COG-UK) Consortium | The Lighthouse Lab in Milton Keynes and Alex Alderton, Roberto Amato, Jeffrey Barrett, Sonia Goncalves, Ewan Harrison, David K. Jackson, Ian Johnston, Dominic Kwiatkowski, Cordelia Langford, John Sillitoe on behalf of the Wellcome Sanger Institute COVID-19 Surveillance Team |
| EPI_ISL_3444375 | Lighthouse Lab in Glasgow | Wellcome Sanger Institute for the COVID-19 Genomics UK (COG-UK) Consortium | Harper VanSteenhouse, Yumi Kasai, David Gray, Carol Clugston, Anna Dominiczak and Alex Alderton, Roberto Amato, Jeffrey Barrett, Sonia Goncalves, Ewan Harrison, David K. Jackson, Ian Johnston, Dominic Kwiatkowski, Cordelia Langford, John Sillitoe on behalf of the Wellcome Sanger Institute COVID-19 Surveillance Team |
| EPI_ISL_3444376, EPI_ISL_3444377, EPI_ISL_3444378 | Lighthouse Lab in Milton Keynes | Wellcome Sanger Institute for the COVID-19 Genomics UK (COG-UK) Consortium | The Lighthouse Lab in Milton Keynes and Alex Alderton, Roberto Amato, Jeffrey Barrett, Sonia Goncalves, Ewan Harrison, David K. Jackson, Ian Johnston, Dominic Kwiatkowski, Cordelia Langford, John Sillitoe on behalf of the Wellcome Sanger Institute COVID-19 Surveillance Team |
| EPI_ISL_3444379 | Lighthouse Lab in Glasgow | Wellcome Sanger Institute for the COVID-19 Genomics UK (COG-UK) Consortium | Harper VanSteenhouse, Yumi Kasai, David Gray, Carol Clugston, Anna Dominiczak and Alex Alderton, Roberto Amato, Jeffrey Barrett, Sonia Goncalves, Ewan Harrison, David K. Jackson, Ian Johnston, Dominic Kwiatkowski, Cordelia Langford, John Sillitoe on behalf of the Wellcome Sanger Institute COVID-19 Surveillance Team |
| EPI_ISL_3444380, EPI_ISL_3444381 | Lighthouse Lab in Milton Keynes | Wellcome Sanger Institute for the COVID-19 Genomics UK (COG-UK) Consortium | The Lighthouse Lab in Milton Keynes and Alex Alderton, Roberto Amato, Jeffrey Barrett, Sonia Goncalves, Ewan Harrison, David K. Jackson, Ian Johnston, Dominic Kwiatkowski, Cordelia Langford, John Sillitoe on behalf of the Wellcome Sanger Institute COVID-19 Surveillance Team |
| EPI_ISL_3444382 | Lighthouse Lab in Glasgow | Wellcome Sanger Institute for the COVID-19 Genomics UK (COG-UK) Consortium | Harper VanSteenhouse, Yumi Kasai, David Gray, Carol Clugston, Anna Dominiczak and Alex Alderton, Roberto Amato, Jeffrey Barrett, Sonia Goncalves, Ewan Harrison, David K. Jackson, Ian Johnston, Dominic Kwiatkowski, Cordelia Langford, John Sillitoe on behalf of the Wellcome Sanger Institute COVID-19 Surveillance Team |
| EPI_ISL_3444383, EPI_ISL_3444384, EPI_ISL_3444385, EPI_ISL_3444386, EPI_ISL_3444387, EPI_ISL_3444388 | Lighthouse Lab in Milton Keynes | Wellcome Sanger Institute for the COVID-19 Genomics UK (COG-UK) Consortium | The Lighthouse Lab in Milton Keynes and Alex Alderton, Roberto Amato, Jeffrey Barrett, Sonia Goncalves, Ewan Harrison, David K. Jackson, Ian Johnston, Dominic Kwiatkowski, Cordelia Langford, John Sillitoe on behalf of the Wellcome Sanger Institute COVID-19 Surveillance Team |
| EPI_ISL_3444389 | Lighthouse Lab in Glasgow | Wellcome Sanger Institute for the COVID-19 Genomics UK (COG-UK) Consortium | Harper VanSteenhouse, Yumi Kasai, David Gray, Carol Clugston, Anna Dominiczak and Alex Alderton, Roberto Amato, Jeffrey Barrett, Sonia Goncalves, Ewan Harrison, David K. Jackson, Ian Johnston, Dominic Kwiatkowski, Cordelia Langford, John Sillitoe on behalf of the Wellcome Sanger Institute COVID-19 Surveillance Team |
| EPI_ISL_3444390, EPI_ISL_3444391 | Lighthouse Lab in Milton Keynes | Wellcome Sanger Institute for the COVID-19 Genomics UK (COG-UK) Consortium | The Lighthouse Lab in Milton Keynes and Alex Alderton, Roberto Amato, Jeffrey Barrett, Sonia Goncalves, Ewan Harrison, David K. Jackson, Ian Johnston, Dominic Kwiatkowski, Cordelia Langford, John Sillitoe on behalf of the Wellcome Sanger Institute COVID-19 Surveillance Team |
| EPI_ISL_3444392 | Lighthouse Lab in Glasgow | Wellcome Sanger Institute for the COVID-19 Genomics UK (COG-UK) Consortium | Harper VanSteenhouse, Yumi Kasai, David Gray, Carol Clugston, Anna Dominiczak and Alex Alderton, Roberto Amato, Jeffrey Barrett, Sonia Goncalves, Ewan Harrison, David K. Jackson, Ian Johnston, Dominic Kwiatkowski, Cordelia Langford, John Sillitoe on behalf of the Wellcome Sanger Institute COVID-19 Surveillance Team |
| EPI_ISL_3444393 | Lighthouse Lab in Milton Keynes | Wellcome Sanger Institute for the COVID-19 Genomics UK (COG-UK) Consortium | The Lighthouse Lab in Milton Keynes and Alex Alderton, Roberto Amato, Jeffrey Barrett, Sonia Goncalves, Ewan Harrison, David K. Jackson, Ian Johnston, Dominic Kwiatkowski, Cordelia Langford, John Sillitoe on behalf of the Wellcome Sanger Institute COVID-19 Surveillance Team |
| EPI_ISL_3444394, EPI_ISL_3444395, EPI_ISL_3444396, EPI_ISL_3444397 | Lighthouse Lab in Glasgow | Wellcome Sanger Institute for the COVID-19 Genomics UK (COG-UK) Consortium | Harper VanSteenhouse, Yumi Kasai, David Gray, Carol Clugston, Anna Dominiczak and Alex Alderton, Roberto Amato, Jeffrey Barrett, Sonia Goncalves, Ewan Harrison, David K. Jackson, Ian Johnston, Dominic Kwiatkowski, Cordelia Langford, John Sillitoe on behalf of the Wellcome Sanger Institute COVID-19 Surveillance Team |
| EPI_ISL_3444398 | Lighthouse Lab in Milton Keynes | Wellcome Sanger Institute for the COVID-19 Genomics UK (COG-UK) Consortium | The Lighthouse Lab in Milton Keynes and Alex Alderton, Roberto Amato, Jeffrey Barrett, Sonia Goncalves, Ewan Harrison, David K. Jackson, Ian Johnston, Dominic Kwiatkowski, Cordelia Langford, John Sillitoe on behalf of the Wellcome Sanger Institute COVID-19 Surveillance Team |
| EPI_ISL_3444399, EPI_ISL_3444400 | Lighthouse Lab in Glasgow | Wellcome Sanger Institute for the COVID-19 Genomics UK (COG-UK) Consortium | Harper VanSteenhouse, Yumi Kasai, David Gray, Carol Clugston, Anna Dominiczak and Alex Alderton, Roberto Amato, Jeffrey Barrett, Sonia Goncalves, Ewan Harrison, David K. Jackson, Ian Johnston, Dominic Kwiatkowski, Cordelia Langford, John Sillitoe on behalf of the Wellcome Sanger Institute COVID-19 Surveillance Team |
| EPI_ISL_3444401, EPI_ISL_3444402, EPI_ISL_3444403, | Lighthouse Lab in Milton Keynes | Wellcome Sanger Institute for the COVID-19 Genomics UK (COG-UK) Consortium | The Lighthouse Lab in Milton Keynes and Alex Alderton, Roberto Amato, Jeffrey Barrett, Sonia Goncalves, Ewan Harrison, David K. Jackson, Ian Johnston, Dominic Kwiatkowski, Cordelia Langford, John Sillitoe on behalf of the Wellcome Sanger Institute COVID-19 Surveillance Team |

| | | | |
|---|---|---|---|
| EPI_ISL_3444404 | | | |
| EPI_ISL_3444405 | Lighthouse Lab in Glasgow | Wellcome Sanger Institute for the COVID-19 Genomics UK (COG-UK) Consortium | Harper VanSteenhouse, Yumi Kasai, David Gray, Carol Clugston, Anna Dominiczak and Alex Alderton, Roberto Amato, Jeffrey Barrett, Sonia Goncalves, Ewan Harrison, David K. Jackson, Ian Johnston, Dominic Kwiatkowski, Cordelia Langford, John Sillitoe on behalf of the Wellcome Sanger Institute COVID-19 Surveillance Team |
| EPI_ISL_3444406 | Lighthouse Lab in Milton Keynes | Wellcome Sanger Institute for the COVID-19 Genomics UK (COG-UK) Consortium | The Lighthouse Lab in Milton Keynes and Alex Alderton, Roberto Amato, Jeffrey Barrett, Sonia Goncalves, Ewan Harrison, David K. Jackson, Ian Johnston, Dominic Kwiatkowski, Cordelia Langford, John Sillitoe on behalf of the Wellcome Sanger Institute COVID-19 Surveillance Team |
| EPI_ISL_3444407, EPI_ISL_3444408 | Lighthouse Lab in Glasgow | Wellcome Sanger Institute for the COVID-19 Genomics UK (COG-UK) Consortium | Harper VanSteenhouse, Yumi Kasai, David Gray, Carol Clugston, Anna Dominiczak and Alex Alderton, Roberto Amato, Jeffrey Barrett, Sonia Goncalves, Ewan Harrison, David K. Jackson, Ian Johnston, Dominic Kwiatkowski, Cordelia Langford, John Sillitoe on behalf of the Wellcome Sanger Institute COVID-19 Surveillance Team |
| EPI_ISL_3444409 | Lighthouse Lab in Milton Keynes | Wellcome Sanger Institute for the COVID-19 Genomics UK (COG-UK) Consortium | The Lighthouse Lab in Milton Keynes and Alex Alderton, Roberto Amato, Jeffrey Barrett, Sonia Goncalves, Ewan Harrison, David K. Jackson, Ian Johnston, Dominic Kwiatkowski, Cordelia Langford, John Sillitoe on behalf of the Wellcome Sanger Institute COVID-19 Surveillance Team |
| EPI_ISL_3444410, EPI_ISL_3444411, EPI_ISL_3444412 | Lighthouse Lab in Glasgow | Wellcome Sanger Institute for the COVID-19 Genomics UK (COG-UK) Consortium | Harper VanSteenhouse, Yumi Kasai, David Gray, Carol Clugston, Anna Dominiczak and Alex Alderton, Roberto Amato, Jeffrey Barrett, Sonia Goncalves, Ewan Harrison, David K. Jackson, Ian Johnston, Dominic Kwiatkowski, Cordelia Langford, John Sillitoe on behalf of the Wellcome Sanger Institute COVID-19 Surveillance Team |
| EPI_ISL_3444413 | Lighthouse Lab in Milton Keynes | Wellcome Sanger Institute for the COVID-19 Genomics UK (COG-UK) Consortium | The Lighthouse Lab in Milton Keynes and Alex Alderton, Roberto Amato, Jeffrey Barrett, Sonia Goncalves, Ewan Harrison, David K. Jackson, Ian Johnston, Dominic Kwiatkowski, Cordelia Langford, John Sillitoe on behalf of the Wellcome Sanger Institute COVID-19 Surveillance Team |
| EPI_ISL_3444414 | Lighthouse Lab in Glasgow | Wellcome Sanger Institute for the COVID-19 Genomics UK (COG-UK) Consortium | Harper VanSteenhouse, Yumi Kasai, David Gray, Carol Clugston, Anna Dominiczak and Alex Alderton, Roberto Amato, Jeffrey Barrett, Sonia Goncalves, Ewan Harrison, David K. Jackson, Ian Johnston, Dominic Kwiatkowski, Cordelia Langford, John Sillitoe on behalf of the Wellcome Sanger Institute COVID-19 Surveillance Team |
| EPI_ISL_3444415 | Lighthouse Lab in Milton Keynes | Wellcome Sanger Institute for the COVID-19 Genomics UK (COG-UK) Consortium | The Lighthouse Lab in Milton Keynes and Alex Alderton, Roberto Amato, Jeffrey Barrett, Sonia Goncalves, Ewan Harrison, David K. Jackson, Ian Johnston, Dominic Kwiatkowski, Cordelia Langford, John Sillitoe on behalf of the Wellcome Sanger Institute COVID-19 Surveillance Team |
| EPI_ISL_3444416 | Lighthouse Lab in Glasgow | Wellcome Sanger Institute for the COVID-19 Genomics UK (COG-UK) Consortium | Harper VanSteenhouse, Yumi Kasai, David Gray, Carol Clugston, Anna Dominiczak and Alex Alderton, Roberto Amato, Jeffrey Barrett, Sonia Goncalves, Ewan Harrison, David K. Jackson, Ian Johnston, Dominic Kwiatkowski, Cordelia Langford, John Sillitoe on behalf of the Wellcome Sanger Institute COVID-19 Surveillance Team |
| EPI_ISL_3444417 | Lighthouse Lab in Milton Keynes | Wellcome Sanger Institute for the COVID-19 Genomics UK (COG-UK) Consortium | The Lighthouse Lab in Milton Keynes and Alex Alderton, Roberto Amato, Jeffrey Barrett, Sonia Goncalves, Ewan Harrison, David K. Jackson, Ian Johnston, Dominic Kwiatkowski, Cordelia Langford, John Sillitoe on behalf of the Wellcome Sanger Institute COVID-19 Surveillance Team |
| EPI_ISL_3444418 | Lighthouse Lab in Glasgow | Wellcome Sanger Institute for the COVID-19 Genomics UK (COG-UK) Consortium | Harper VanSteenhouse, Yumi Kasai, David Gray, Carol Clugston, Anna Dominiczak and Alex Alderton, Roberto Amato, Jeffrey Barrett, Sonia Goncalves, Ewan Harrison, David K. Jackson, Ian Johnston, Dominic Kwiatkowski, Cordelia Langford, John Sillitoe on behalf of the Wellcome Sanger Institute COVID-19 Surveillance Team |
| EPI_ISL_3444419 | Lighthouse Lab in Milton Keynes | Wellcome Sanger Institute for the COVID-19 Genomics UK (COG-UK) Consortium | The Lighthouse Lab in Milton Keynes and Alex Alderton, Roberto Amato, Jeffrey Barrett, Sonia Goncalves, Ewan Harrison, David K. Jackson, Ian Johnston, Dominic Kwiatkowski, Cordelia Langford, John Sillitoe on behalf of the Wellcome Sanger Institute COVID-19 Surveillance Team |
| EPI_ISL_3444420, EPI_ISL_3444421, EPI_ISL_3444422, EPI_ISL_3444423 | Lighthouse Lab in Glasgow | Wellcome Sanger Institute for the COVID-19 Genomics UK (COG-UK) Consortium | Harper VanSteenhouse, Yumi Kasai, David Gray, Carol Clugston, Anna Dominiczak and Alex Alderton, Roberto Amato, Jeffrey Barrett, Sonia Goncalves, Ewan Harrison, David K. Jackson, Ian Johnston, Dominic Kwiatkowski, Cordelia Langford, John Sillitoe on behalf of the Wellcome Sanger Institute COVID-19 Surveillance Team |
| EPI_ISL_3444424, EPI_ISL_3444425 | Lighthouse Lab in Milton Keynes | Wellcome Sanger Institute for the COVID-19 Genomics UK (COG-UK) Consortium | The Lighthouse Lab in Milton Keynes and Alex Alderton, Roberto Amato, Jeffrey Barrett, Sonia Goncalves, Ewan Harrison, David K. Jackson, Ian Johnston, Dominic Kwiatkowski, Cordelia Langford, John Sillitoe on behalf of the Wellcome Sanger Institute COVID-19 Surveillance Team |
| EPI_ISL_3444426, EPI_ISL_3444427, EPI_ISL_3444428 | Lighthouse Lab in Glasgow | Wellcome Sanger Institute for the COVID-19 Genomics UK (COG-UK) Consortium | Harper VanSteenhouse, Yumi Kasai, David Gray, Carol Clugston, Anna Dominiczak and Alex Alderton, Roberto Amato, Jeffrey Barrett, Sonia Goncalves, Ewan Harrison, David K. Jackson, Ian Johnston, Dominic Kwiatkowski, Cordelia Langford, John Sillitoe on behalf of the Wellcome Sanger Institute COVID-19 Surveillance Team |
| EPI_ISL_3444429, EPI_ISL_3444430, EPI_ISL_3444431, EPI_ISL_3444432, EPI_ISL_3444433, EPI_ISL_3444434 | Lighthouse Lab in Milton Keynes | Wellcome Sanger Institute for the COVID-19 Genomics UK (COG-UK) Consortium | The Lighthouse Lab in Milton Keynes and Alex Alderton, Roberto Amato, Jeffrey Barrett, Sonia Goncalves, Ewan Harrison, David K. Jackson, Ian Johnston, Dominic Kwiatkowski, Cordelia Langford, John Sillitoe on behalf of the Wellcome Sanger Institute COVID-19 Surveillance Team |
| EPI_ISL_3444435, EPI_ISL_3444436 | Lighthouse Lab in Glasgow | Wellcome Sanger Institute for the COVID-19 Genomics UK (COG-UK) Consortium | Harper VanSteenhouse, Yumi Kasai, David Gray, Carol Clugston, Anna Dominiczak and Alex Alderton, Roberto Amato, Jeffrey Barrett, Sonia Goncalves, Ewan Harrison, David K. Jackson, Ian Johnston, Dominic Kwiatkowski, Cordelia Langford, John Sillitoe on behalf of the Wellcome Sanger Institute COVID-19 Surveillance Team |
| EPI_ISL_3444437 | Lighthouse Lab in Milton Keynes | Wellcome Sanger Institute for the COVID-19 Genomics UK (COG-UK) Consortium | The Lighthouse Lab in Milton Keynes and Alex Alderton, Roberto Amato, Jeffrey Barrett, Sonia Goncalves, Ewan Harrison, David K. Jackson, Ian Johnston, Dominic Kwiatkowski, Cordelia Langford, John Sillitoe on behalf of the Wellcome Sanger Institute COVID-19 Surveillance Team |
| EPI_ISL_3444438 | Lighthouse Lab in Glasgow | Wellcome Sanger Institute for the COVID-19 Genomics UK (COG-UK) Consortium | Harper VanSteenhouse, Yumi Kasai, David Gray, Carol Clugston, Anna Dominiczak and Alex Alderton, Roberto Amato, Jeffrey Barrett, Sonia Goncalves, Ewan Harrison, David K. Jackson, Ian Johnston, Dominic Kwiatkowski, Cordelia Langford, John Sillitoe on behalf of the Wellcome Sanger Institute COVID-19 Surveillance Team |
| EPI_ISL_3444439, EPI_ISL_3444440 | Lighthouse Lab in Milton Keynes | Wellcome Sanger Institute for the COVID-19 Genomics UK (COG-UK) Consortium | The Lighthouse Lab in Milton Keynes and Alex Alderton, Roberto Amato, Jeffrey Barrett, Sonia Goncalves, Ewan Harrison, David K. Jackson, Ian Johnston, Dominic Kwiatkowski, Cordelia Langford, John Sillitoe on behalf of the Wellcome Sanger Institute COVID-19 Surveillance Team |
| EPI_ISL_3444441 | Lighthouse Lab in Glasgow | Wellcome Sanger Institute for the COVID-19 Genomics UK (COG-UK) Consortium | Harper VanSteenhouse, Yumi Kasai, David Gray, Carol Clugston, Anna Dominiczak and Alex Alderton, Roberto Amato, Jeffrey Barrett, Sonia Goncalves, Ewan Harrison, David K. Jackson, Ian Johnston, Dominic Kwiatkowski, Cordelia Langford, John Sillitoe on behalf of the Wellcome Sanger Institute COVID-19 Surveillance Team |
| EPI_ISL_3444442, EPI_ISL_3444443, EPI_ISL_3444444, EPI_ISL_3444445 | Lighthouse Lab in Milton Keynes | Wellcome Sanger Institute for the COVID-19 Genomics UK (COG-UK) Consortium | The Lighthouse Lab in Milton Keynes and Alex Alderton, Roberto Amato, Jeffrey Barrett, Sonia Goncalves, Ewan Harrison, David K. Jackson, Ian Johnston, Dominic Kwiatkowski, Cordelia Langford, John Sillitoe on behalf of the Wellcome Sanger Institute COVID-19 Surveillance Team |
| EPI_ISL_3444446 | Lighthouse Lab in Glasgow | Wellcome Sanger Institute for the COVID-19 Genomics UK (COG-UK) Consortium | Harper VanSteenhouse, Yumi Kasai, David Gray, Carol Clugston, Anna Dominiczak and Alex Alderton, Roberto Amato, Jeffrey Barrett, Sonia Goncalves, Ewan Harrison, David K. Jackson, Ian Johnston, Dominic Kwiatkowski, Cordelia Langford, John Sillitoe on behalf of the Wellcome Sanger Institute COVID-19 Surveillance Team |
| EPI_ISL_3444447 | Lighthouse Lab in Milton Keynes | Wellcome Sanger Institute for the COVID-19 Genomics UK (COG-UK) Consortium | The Lighthouse Lab in Milton Keynes and Alex Alderton, Roberto Amato, Jeffrey Barrett, Sonia Goncalves, Ewan Harrison, David K. Jackson, Ian Johnston, Dominic Kwiatkowski, Cordelia Langford, John Sillitoe on behalf of the Wellcome Sanger Institute COVID-19 Surveillance Team |
| EPI_ISL_3444448, EPI_ISL_3444449 | Lighthouse Lab in Glasgow | Wellcome Sanger Institute for the COVID-19 Genomics UK (COG-UK) Consortium | Harper VanSteenhouse, Yumi Kasai, David Gray, Carol Clugston, Anna Dominiczak and Alex Alderton, Roberto Amato, Jeffrey Barrett, Sonia Goncalves, Ewan Harrison, David K. Jackson, Ian Johnston, Dominic Kwiatkowski, Cordelia Langford, John Sillitoe on behalf of the Wellcome Sanger Institute COVID-19 Surveillance Team |
| EPI_ISL_3444450, EPI_ISL_3444451, EPI_ISL_3444452 | Lighthouse Lab in Milton Keynes | Wellcome Sanger Institute for the COVID-19 Genomics UK (COG-UK) Consortium | The Lighthouse Lab in Milton Keynes and Alex Alderton, Roberto Amato, Jeffrey Barrett, Sonia Goncalves, Ewan Harrison, David K. Jackson, Ian Johnston, Dominic Kwiatkowski, Cordelia Langford, John Sillitoe on behalf of the Wellcome Sanger Institute COVID-19 Surveillance Team |
| EPI_ISL_3444453, EPI_ISL_3444454 | Lighthouse Lab in Glasgow | Wellcome Sanger Institute for the COVID-19 Genomics UK (COG-UK) Consortium | Harper VanSteenhouse, Yumi Kasai, David Gray, Carol Clugston, Anna Dominiczak and Alex Alderton, Roberto Amato, Jeffrey Barrett, Sonia Goncalves, Ewan Harrison, David K. Jackson, Ian Johnston, Dominic Kwiatkowski, Cordelia Langford, John Sillitoe on behalf of the Wellcome Sanger Institute COVID-19 Surveillance Team |
| EPI_ISL_3444455 | Lighthouse Lab in Milton Keynes | Wellcome Sanger Institute for the COVID-19 Genomics UK (COG-UK) Consortium | The Lighthouse Lab in Milton Keynes and Alex Alderton, Roberto Amato, Jeffrey Barrett, Sonia Goncalves, Ewan Harrison, David K. Jackson, Ian Johnston, Dominic Kwiatkowski, Cordelia Langford, John Sillitoe on behalf of the Wellcome Sanger Institute COVID-19 Surveillance Team |
| EPI_ISL_3444456, EPI_ISL_3444457 | Lighthouse Lab in Glasgow | Wellcome Sanger Institute for the COVID-19 Genomics UK (COG-UK) Consortium | Harper VanSteenhouse, Yumi Kasai, David Gray, Carol Clugston, Anna Dominiczak and Alex Alderton, Roberto Amato, Jeffrey Barrett, Sonia Goncalves, Ewan Harrison, David K. Jackson, Ian Johnston, Dominic Kwiatkowski, Cordelia Langford, John Sillitoe on behalf of the Wellcome Sanger Institute COVID-19 Surveillance Team |
| EPI_ISL_3444458 | Lighthouse Lab in Milton Keynes | Wellcome Sanger Institute for the COVID-19 Genomics UK (COG-UK) Consortium | The Lighthouse Lab in Milton Keynes and Alex Alderton, Roberto Amato, Jeffrey Barrett, Sonia Goncalves, Ewan Harrison, David K. Jackson, Ian Johnston, Dominic Kwiatkowski, Cordelia Langford, John Sillitoe on behalf of the Wellcome Sanger Institute COVID-19 Surveillance Team |
| EPI_ISL_3444459, EPI_ISL_3444460, EPI_ISL_3444461 | Lighthouse Lab in Glasgow | Wellcome Sanger Institute for the COVID-19 Genomics UK (COG-UK) Consortium | Harper VanSteenhouse, Yumi Kasai, David Gray, Carol Clugston, Anna Dominiczak and Alex Alderton, Roberto Amato, Jeffrey Barrett, Sonia Goncalves, Ewan Harrison, David K. Jackson, Ian Johnston, Dominic Kwiatkowski, Cordelia Langford, John Sillitoe on behalf of the Wellcome Sanger Institute COVID-19 Surveillance Team |
| EPI_ISL_3444462, EPI_ISL_3444463 | Lighthouse Lab in Milton Keynes | Wellcome Sanger Institute for the COVID-19 Genomics UK (COG-UK) Consortium | The Lighthouse Lab in Milton Keynes and Alex Alderton, Roberto Amato, Jeffrey Barrett, Sonia Goncalves, Ewan Harrison, David K. Jackson, Ian Johnston, Dominic Kwiatkowski, Cordelia Langford, John Sillitoe on behalf of the Wellcome Sanger Institute COVID-19 Surveillance Team |
| EPI_ISL_3444464, EPI_ISL_3444465, EPI_ISL_3444466, EPI_ISL_3444467, EPI_ISL_3444468 | Lighthouse Lab in Glasgow | Wellcome Sanger Institute for the COVID-19 Genomics UK (COG-UK) Consortium | Harper VanSteenhouse, Yumi Kasai, David Gray, Carol Clugston, Anna Dominiczak and Alex Alderton, Roberto Amato, Jeffrey Barrett, Sonia Goncalves, Ewan Harrison, David K. Jackson, Ian Johnston, Dominic Kwiatkowski, Cordelia Langford, John Sillitoe on behalf of the Wellcome Sanger Institute COVID-19 Surveillance Team |
| EPI_ISL_3444469 | Lighthouse Lab in Milton Keynes | Wellcome Sanger Institute for the COVID-19 Genomics UK (COG-UK) Consortium | The Lighthouse Lab in Milton Keynes and Alex Alderton, Roberto Amato, Jeffrey Barrett, Sonia Goncalves, Ewan Harrison, David K. Jackson, Ian Johnston, Dominic Kwiatkowski, Cordelia Langford, John Sillitoe on behalf of the Wellcome Sanger Institute COVID-19 Surveillance Team |
| EPI_ISL_3444470, EPI_ISL_3444471 | Lighthouse Lab in Glasgow | Wellcome Sanger Institute for the COVID-19 Genomics UK (COG-UK) Consortium | Harper VanSteenhouse, Yumi Kasai, David Gray, Carol Clugston, Anna Dominiczak and Alex Alderton, Roberto Amato, Jeffrey Barrett, Sonia Goncalves, Ewan Harrison, David K. Jackson, Ian Johnston, Dominic Kwiatkowski, Cordelia Langford, John Sillitoe on behalf of the Wellcome Sanger Institute COVID-19 Surveillance Team |
| EPI_ISL_3444472, EPI_ISL_3444473, EPI_ISL_3444474, EPI_ISL_3444475, EPI_ISL_3444476 | Lighthouse Lab in Milton Keynes | Wellcome Sanger Institute for the COVID-19 Genomics UK (COG-UK) Consortium | The Lighthouse Lab in Milton Keynes and Alex Alderton, Roberto Amato, Jeffrey Barrett, Sonia Goncalves, Ewan Harrison, David K. Jackson, Ian Johnston, Dominic Kwiatkowski, Cordelia Langford, John Sillitoe on behalf of the Wellcome Sanger Institute COVID-19 Surveillance Team |
| EPI_ISL_3444477 | Lighthouse Lab in Glasgow | Wellcome Sanger Institute for the COVID-19 Genomics UK (COG-UK) Consortium | Harper VanSteenhouse, Yumi Kasai, David Gray, Carol Clugston, Anna Dominiczak and Alex Alderton, Roberto Amato, Jeffrey Barrett, Sonia Goncalves, Ewan Harrison, David K. Jackson, Ian Johnston, Dominic Kwiatkowski, Cordelia Langford, John Sillitoe on behalf of the Wellcome Sanger Institute COVID-19 Surveillance Team |

| | | | |
|---|---|---|---|
| EPI_ISL_3444478, EPI_ISL_3444479, EPI_ISL_3444480 | Lighthouse Lab in Milton Keynes | Wellcome Sanger Institute for the COVID-19 Genomics UK (COG-UK) Consortium | The Lighthouse Lab in Milton Keynes and Alex Alderton, Roberto Amato, Jeffrey Barrett, Sonia Goncalves, Ewan Harrison, David K. Jackson, Ian Johnston, Dominic Kwiatkowski, Cordelia Langford, John Sillitoe on behalf of the Wellcome Sanger Institute COVID-19 Surveillance Team |
| EPI_ISL_3444481, EPI_ISL_3444482 | Lighthouse Lab in Glasgow | Wellcome Sanger Institute for the COVID-19 Genomics UK (COG-UK) Consortium | Harper VanSteenhouse, Yumi Kasai, David Gray, Carol Clugston, Anna Dominiczak and Alex Alderton, Roberto Amato, Jeffrey Barrett, Sonia Goncalves, Ewan Harrison, David K. Jackson, Ian Johnston, Dominic Kwiatkowski, Cordelia Langford, John Sillitoe on behalf of the Wellcome Sanger Institute COVID-19 Surveillance Team |
| EPI_ISL_3444483 | Lighthouse Lab in Milton Keynes | Wellcome Sanger Institute for the COVID-19 Genomics UK (COG-UK) Consortium | The Lighthouse Lab in Milton Keynes and Alex Alderton, Roberto Amato, Jeffrey Barrett, Sonia Goncalves, Ewan Harrison, David K. Jackson, Ian Johnston, Dominic Kwiatkowski, Cordelia Langford, John Sillitoe on behalf of the Wellcome Sanger Institute COVID-19 Surveillance Team |
| EPI_ISL_3444484, EPI_ISL_3444485 | Lighthouse Lab in Glasgow | Wellcome Sanger Institute for the COVID-19 Genomics UK (COG-UK) Consortium | Harper VanSteenhouse, Yumi Kasai, David Gray, Carol Clugston, Anna Dominiczak and Alex Alderton, Roberto Amato, Jeffrey Barrett, Sonia Goncalves, Ewan Harrison, David K. Jackson, Ian Johnston, Dominic Kwiatkowski, Cordelia Langford, John Sillitoe on behalf of the Wellcome Sanger Institute COVID-19 Surveillance Team |
| EPI_ISL_3444486 | Lighthouse Lab in Milton Keynes | Wellcome Sanger Institute for the COVID-19 Genomics UK (COG-UK) Consortium | The Lighthouse Lab in Milton Keynes and Alex Alderton, Roberto Amato, Jeffrey Barrett, Sonia Goncalves, Ewan Harrison, David K. Jackson, Ian Johnston, Dominic Kwiatkowski, Cordelia Langford, John Sillitoe on behalf of the Wellcome Sanger Institute COVID-19 Surveillance Team |
| EPI_ISL_3444487, EPI_ISL_3444488 | Lighthouse Lab in Glasgow | Wellcome Sanger Institute for the COVID-19 Genomics UK (COG-UK) Consortium | Harper VanSteenhouse, Yumi Kasai, David Gray, Carol Clugston, Anna Dominiczak and Alex Alderton, Roberto Amato, Jeffrey Barrett, Sonia Goncalves, Ewan Harrison, David K. Jackson, Ian Johnston, Dominic Kwiatkowski, Cordelia Langford, John Sillitoe on behalf of the Wellcome Sanger Institute COVID-19 Surveillance Team |
| EPI_ISL_3444489, EPI_ISL_3444490, EPI_ISL_3444491 | Lighthouse Lab in Milton Keynes | Wellcome Sanger Institute for the COVID-19 Genomics UK (COG-UK) Consortium | The Lighthouse Lab in Milton Keynes and Alex Alderton, Roberto Amato, Jeffrey Barrett, Sonia Goncalves, Ewan Harrison, David K. Jackson, Ian Johnston, Dominic Kwiatkowski, Cordelia Langford, John Sillitoe on behalf of the Wellcome Sanger Institute COVID-19 Surveillance Team |
| EPI_ISL_3444492, EPI_ISL_3444493, EPI_ISL_3444494, EPI_ISL_3444495, EPI_ISL_3444496, EPI_ISL_3444497, EPI_ISL_3444498, EPI_ISL_3444499 | Lighthouse Lab in Glasgow | Wellcome Sanger Institute for the COVID-19 Genomics UK (COG-UK) Consortium | Harper VanSteenhouse, Yumi Kasai, David Gray, Carol Clugston, Anna Dominiczak and Alex Alderton, Roberto Amato, Jeffrey Barrett, Sonia Goncalves, Ewan Harrison, David K. Jackson, Ian Johnston, Dominic Kwiatkowski, Cordelia Langford, John Sillitoe on behalf of the Wellcome Sanger Institute COVID-19 Surveillance Team |
| EPI_ISL_3444500, EPI_ISL_3444501, EPI_ISL_3444502 | Lighthouse Lab in Milton Keynes | Wellcome Sanger Institute for the COVID-19 Genomics UK (COG-UK) Consortium | The Lighthouse Lab in Milton Keynes and Alex Alderton, Roberto Amato, Jeffrey Barrett, Sonia Goncalves, Ewan Harrison, David K. Jackson, Ian Johnston, Dominic Kwiatkowski, Cordelia Langford, John Sillitoe on behalf of the Wellcome Sanger Institute COVID-19 Surveillance Team |
| EPI_ISL_3444503, EPI_ISL_3444504 | Lighthouse Lab in Glasgow | Wellcome Sanger Institute for the COVID-19 Genomics UK (COG-UK) Consortium | Harper VanSteenhouse, Yumi Kasai, David Gray, Carol Clugston, Anna Dominiczak and Alex Alderton, Roberto Amato, Jeffrey Barrett, Sonia Goncalves, Ewan Harrison, David K. Jackson, Ian Johnston, Dominic Kwiatkowski, Cordelia Langford, John Sillitoe on behalf of the Wellcome Sanger Institute COVID-19 Surveillance Team |
| EPI_ISL_3444505, EPI_ISL_3444506, EPI_ISL_3444507, EPI_ISL_3444508, EPI_ISL_3444509 | Lighthouse Lab in Milton Keynes | Wellcome Sanger Institute for the COVID-19 Genomics UK (COG-UK) Consortium | The Lighthouse Lab in Milton Keynes and Alex Alderton, Roberto Amato, Jeffrey Barrett, Sonia Goncalves, Ewan Harrison, David K. Jackson, Ian Johnston, Dominic Kwiatkowski, Cordelia Langford, John Sillitoe on behalf of the Wellcome Sanger Institute COVID-19 Surveillance Team |
| EPI_ISL_3444510, EPI_ISL_3444511 | Lighthouse Lab in Glasgow | Wellcome Sanger Institute for the COVID-19 Genomics UK (COG-UK) Consortium | Harper VanSteenhouse, Yumi Kasai, David Gray, Carol Clugston, Anna Dominiczak and Alex Alderton, Roberto Amato, Jeffrey Barrett, Sonia Goncalves, Ewan Harrison, David K. Jackson, Ian Johnston, Dominic Kwiatkowski, Cordelia Langford, John Sillitoe on behalf of the Wellcome Sanger Institute COVID-19 Surveillance Team |
| EPI_ISL_3444512 | Lighthouse Lab in Milton Keynes | Wellcome Sanger Institute for the COVID-19 Genomics UK (COG-UK) Consortium | The Lighthouse Lab in Milton Keynes and Alex Alderton, Roberto Amato, Jeffrey Barrett, Sonia Goncalves, Ewan Harrison, David K. Jackson, Ian Johnston, Dominic Kwiatkowski, Cordelia Langford, John Sillitoe on behalf of the Wellcome Sanger Institute COVID-19 Surveillance Team |
| EPI_ISL_3444513, EPI_ISL_3444514, EPI_ISL_3444515 | Lighthouse Lab in Glasgow | Wellcome Sanger Institute for the COVID-19 Genomics UK (COG-UK) Consortium | Harper VanSteenhouse, Yumi Kasai, David Gray, Carol Clugston, Anna Dominiczak and Alex Alderton, Roberto Amato, Jeffrey Barrett, Sonia Goncalves, Ewan Harrison, David K. Jackson, Ian Johnston, Dominic Kwiatkowski, Cordelia Langford, John Sillitoe on behalf of the Wellcome Sanger Institute COVID-19 Surveillance Team |
| EPI_ISL_3444516, EPI_ISL_3444517 | Lighthouse Lab in Milton Keynes | Wellcome Sanger Institute for the COVID-19 Genomics UK (COG-UK) Consortium | The Lighthouse Lab in Milton Keynes and Alex Alderton, Roberto Amato, Jeffrey Barrett, Sonia Goncalves, Ewan Harrison, David K. Jackson, Ian Johnston, Dominic Kwiatkowski, Cordelia Langford, John Sillitoe on behalf of the Wellcome Sanger Institute COVID-19 Surveillance Team |
| EPI_ISL_3444518 | Lighthouse Lab in Glasgow | Wellcome Sanger Institute for the COVID-19 Genomics UK (COG-UK) Consortium | Harper VanSteenhouse, Yumi Kasai, David Gray, Carol Clugston, Anna Dominiczak and Alex Alderton, Roberto Amato, Jeffrey Barrett, Sonia Goncalves, Ewan Harrison, David K. Jackson, Ian Johnston, Dominic Kwiatkowski, Cordelia Langford, John Sillitoe on behalf of the Wellcome Sanger Institute COVID-19 Surveillance Team |
| EPI_ISL_3444519, EPI_ISL_3444520 | Lighthouse Lab in Milton Keynes | Wellcome Sanger Institute for the COVID-19 Genomics UK (COG-UK) Consortium | The Lighthouse Lab in Milton Keynes and Alex Alderton, Roberto Amato, Jeffrey Barrett, Sonia Goncalves, Ewan Harrison, David K. Jackson, Ian Johnston, Dominic Kwiatkowski, Cordelia Langford, John Sillitoe on behalf of the Wellcome Sanger Institute COVID-19 Surveillance Team |
| EPI_ISL_3444521, EPI_ISL_3444522, EPI_ISL_3444523, EPI_ISL_3444524, EPI_ISL_3444525 | Lighthouse Lab in Glasgow | Wellcome Sanger Institute for the COVID-19 Genomics UK (COG-UK) Consortium | Harper VanSteenhouse, Yumi Kasai, David Gray, Carol Clugston, Anna Dominiczak and Alex Alderton, Roberto Amato, Jeffrey Barrett, Sonia Goncalves, Ewan Harrison, David K. Jackson, Ian Johnston, Dominic Kwiatkowski, Cordelia Langford, John Sillitoe on behalf of the Wellcome Sanger Institute COVID-19 Surveillance Team |
| EPI_ISL_3444526 | Lighthouse Lab in Milton Keynes | Wellcome Sanger Institute for the COVID-19 Genomics UK (COG-UK) Consortium | The Lighthouse Lab in Milton Keynes and Alex Alderton, Roberto Amato, Jeffrey Barrett, Sonia Goncalves, Ewan Harrison, David K. Jackson, Ian Johnston, Dominic Kwiatkowski, Cordelia Langford, John Sillitoe on behalf of the Wellcome Sanger Institute COVID-19 Surveillance Team |
| EPI_ISL_3444527, EPI_ISL_3444528, EPI_ISL_3444529, EPI_ISL_3444530 | Lighthouse Lab in Glasgow | Wellcome Sanger Institute for the COVID-19 Genomics UK (COG-UK) Consortium | Harper VanSteenhouse, Yumi Kasai, David Gray, Carol Clugston, Anna Dominiczak and Alex Alderton, Roberto Amato, Jeffrey Barrett, Sonia Goncalves, Ewan Harrison, David K. Jackson, Ian Johnston, Dominic Kwiatkowski, Cordelia Langford, John Sillitoe on behalf of the Wellcome Sanger Institute COVID-19 Surveillance Team |
| EPI_ISL_3444531 | Lighthouse Lab in Milton Keynes | Wellcome Sanger Institute for the COVID-19 Genomics UK (COG-UK) Consortium | The Lighthouse Lab in Milton Keynes and Alex Alderton, Roberto Amato, Jeffrey Barrett, Sonia Goncalves, Ewan Harrison, David K. Jackson, Ian Johnston, Dominic Kwiatkowski, Cordelia Langford, John Sillitoe on behalf of the Wellcome Sanger Institute COVID-19 Surveillance Team |
| EPI_ISL_3444532 | Lighthouse Lab in Glasgow | Wellcome Sanger Institute for the COVID-19 Genomics UK (COG-UK) Consortium | Harper VanSteenhouse, Yumi Kasai, David Gray, Carol Clugston, Anna Dominiczak and Alex Alderton, Roberto Amato, Jeffrey Barrett, Sonia Goncalves, Ewan Harrison, David K. Jackson, Ian Johnston, Dominic Kwiatkowski, Cordelia Langford, John Sillitoe on behalf of the Wellcome Sanger Institute COVID-19 Surveillance Team |
| EPI_ISL_3444533 | Lighthouse Lab in Milton Keynes | Wellcome Sanger Institute for the COVID-19 Genomics UK (COG-UK) Consortium | The Lighthouse Lab in Milton Keynes and Alex Alderton, Roberto Amato, Jeffrey Barrett, Sonia Goncalves, Ewan Harrison, David K. Jackson, Ian Johnston, Dominic Kwiatkowski, Cordelia Langford, John Sillitoe on behalf of the Wellcome Sanger Institute COVID-19 Surveillance Team |
| EPI_ISL_3444534, EPI_ISL_3444535, EPI_ISL_3444536 | Lighthouse Lab in Glasgow | Wellcome Sanger Institute for the COVID-19 Genomics UK (COG-UK) Consortium | Harper VanSteenhouse, Yumi Kasai, David Gray, Carol Clugston, Anna Dominiczak and Alex Alderton, Roberto Amato, Jeffrey Barrett, Sonia Goncalves, Ewan Harrison, David K. Jackson, Ian Johnston, Dominic Kwiatkowski, Cordelia Langford, John Sillitoe on behalf of the Wellcome Sanger Institute COVID-19 Surveillance Team |
| EPI_ISL_3444537, EPI_ISL_3444538 | Lighthouse Lab in Milton Keynes | Wellcome Sanger Institute for the COVID-19 Genomics UK (COG-UK) Consortium | The Lighthouse Lab in Milton Keynes and Alex Alderton, Roberto Amato, Jeffrey Barrett, Sonia Goncalves, Ewan Harrison, David K. Jackson, Ian Johnston, Dominic Kwiatkowski, Cordelia Langford, John Sillitoe on behalf of the Wellcome Sanger Institute COVID-19 Surveillance Team |
| EPI_ISL_3444539 | Lighthouse Lab in Glasgow | Wellcome Sanger Institute for the COVID-19 Genomics UK (COG-UK) Consortium | Harper VanSteenhouse, Yumi Kasai, David Gray, Carol Clugston, Anna Dominiczak and Alex Alderton, Roberto Amato, Jeffrey Barrett, Sonia Goncalves, Ewan Harrison, David K. Jackson, Ian Johnston, Dominic Kwiatkowski, Cordelia Langford, John Sillitoe on behalf of the Wellcome Sanger Institute COVID-19 Surveillance Team |
| EPI_ISL_3444540 | Lighthouse Lab in Milton Keynes | Wellcome Sanger Institute for the COVID-19 Genomics UK (COG-UK) Consortium | The Lighthouse Lab in Milton Keynes and Alex Alderton, Roberto Amato, Jeffrey Barrett, Sonia Goncalves, Ewan Harrison, David K. Jackson, Ian Johnston, Dominic Kwiatkowski, Cordelia Langford, John Sillitoe on behalf of the Wellcome Sanger Institute COVID-19 Surveillance Team |
| EPI_ISL_3444541, EPI_ISL_3444542, EPI_ISL_3444543, EPI_ISL_3444544, EPI_ISL_3444545, EPI_ISL_3444546 | Lighthouse Lab in Glasgow | Wellcome Sanger Institute for the COVID-19 Genomics UK (COG-UK) Consortium | Harper VanSteenhouse, Yumi Kasai, David Gray, Carol Clugston, Anna Dominiczak and Alex Alderton, Roberto Amato, Jeffrey Barrett, Sonia Goncalves, Ewan Harrison, David K. Jackson, Ian Johnston, Dominic Kwiatkowski, Cordelia Langford, John Sillitoe on behalf of the Wellcome Sanger Institute COVID-19 Surveillance Team |
| EPI_ISL_3444547, EPI_ISL_3444548, EPI_ISL_3444549, EPI_ISL_3444550 | Lighthouse Lab in Milton Keynes | Wellcome Sanger Institute for the COVID-19 Genomics UK (COG-UK) Consortium | The Lighthouse Lab in Milton Keynes and Alex Alderton, Roberto Amato, Jeffrey Barrett, Sonia Goncalves, Ewan Harrison, David K. Jackson, Ian Johnston, Dominic Kwiatkowski, Cordelia Langford, John Sillitoe on behalf of the Wellcome Sanger Institute COVID-19 Surveillance Team |
| EPI_ISL_3444551 | Lighthouse Lab in Glasgow | Wellcome Sanger Institute for the COVID-19 Genomics UK (COG-UK) Consortium | Harper VanSteenhouse, Yumi Kasai, David Gray, Carol Clugston, Anna Dominiczak and Alex Alderton, Roberto Amato, Jeffrey Barrett, Sonia Goncalves, Ewan Harrison, David K. Jackson, Ian Johnston, Dominic Kwiatkowski, Cordelia Langford, John Sillitoe on behalf of the Wellcome Sanger Institute COVID-19 Surveillance Team |
| EPI_ISL_3444552, EPI_ISL_3444553, EPI_ISL_3444554, EPI_ISL_3444555, EPI_ISL_3444556, EPI_ISL_3444557 | Lighthouse Lab in Alderley Park | Wellcome Sanger Institute for the COVID-19 Genomics UK (COG-UK) Consortium | Jacquelyn Wynn, Mairead Hyland, The Lighthouse Lab in Alderley Park and Alex Alderton, Roberto Amato, Jeffrey Barrett, Sonia Goncalves, Ewan Harrison, David K. Jackson, Ian Johnston, Dominic Kwiatkowski, Cordelia Langford, John Sillitoe on behalf of the Wellcome Sanger Institute COVID-19 Surveillance Team |
| EPI_ISL_3444558, EPI_ISL_3444559 | Lighthouse Lab in Glasgow | Wellcome Sanger Institute for the COVID-19 Genomics UK (COG-UK) Consortium | Harper VanSteenhouse, Yumi Kasai, David Gray, Carol Clugston, Anna Dominiczak and Alex Alderton, Roberto Amato, Jeffrey Barrett, Sonia Goncalves, Ewan Harrison, David K. Jackson, Ian Johnston, Dominic Kwiatkowski, Cordelia Langford, John Sillitoe on behalf of the Wellcome Sanger Institute COVID-19 Surveillance Team |
| EPI_ISL_3444560, EPI_ISL_3444561 | Lighthouse Lab in Alderley Park | Wellcome Sanger Institute for the COVID-19 Genomics UK (COG-UK) Consortium | Jacquelyn Wynn, Mairead Hyland, The Lighthouse Lab in Alderley Park and Alex Alderton, Roberto Amato, Jeffrey Barrett, Sonia Goncalves, Ewan Harrison, David K. Jackson, Ian Johnston, Dominic Kwiatkowski, Cordelia Langford, John Sillitoe on behalf of the Wellcome Sanger Institute COVID-19 Surveillance Team |

| | | | |
|---|---|---|---|
| EPI_ISL_3444562, EPI_ISL_3444563 | Lighthouse Lab in Glasgow | Wellcome Sanger Institute for the COVID-19 Genomics UK (COG-UK) Consortium | Harper VanSteenhouse, Yumi Kasai, David Gray, Carol Clugston, Anna Dominiczak and Alex Alderton, Roberto Amato, Jeffrey Barrett, Sonia Goncalves, Ewan Harrison, David K. Jackson, Ian Johnston, Dominic Kwiatkowski, Cordelia Langford, John Sillitoe on behalf of the Wellcome Sanger Institute COVID-19 Surveillance Team |
| EPI_ISL_3444564 | Lighthouse Lab in Alderley Park | Wellcome Sanger Institute for the COVID-19 Genomics UK (COG-UK) Consortium | Jacquelyn Wynn, Mairead Hyland, The Lighthouse Lab in Alderley Park and Alex Alderton, Roberto Amato, Jeffrey Barrett, Sonia Goncalves, Ewan Harrison, David K. Jackson, Ian Johnston, Dominic Kwiatkowski, Cordelia Langford, John Sillitoe on behalf of the Wellcome Sanger Institute COVID-19 Surveillance Team |
| EPI_ISL_3444565 | Lighthouse Lab in Glasgow | Wellcome Sanger Institute for the COVID-19 Genomics UK (COG-UK) Consortium | Harper VanSteenhouse, Yumi Kasai, David Gray, Carol Clugston, Anna Dominiczak and Alex Alderton, Roberto Amato, Jeffrey Barrett, Sonia Goncalves, Ewan Harrison, David K. Jackson, Ian Johnston, Dominic Kwiatkowski, Cordelia Langford, John Sillitoe on behalf of the Wellcome Sanger Institute COVID-19 Surveillance Team |
| EPI_ISL_3444566, EPI_ISL_3444567, EPI_ISL_3444568 | Lighthouse Lab in Alderley Park | Wellcome Sanger Institute for the COVID-19 Genomics UK (COG-UK) Consortium | Jacquelyn Wynn, Mairead Hyland, The Lighthouse Lab in Alderley Park and Alex Alderton, Roberto Amato, Jeffrey Barrett, Sonia Goncalves, Ewan Harrison, David K. Jackson, Ian Johnston, Dominic Kwiatkowski, Cordelia Langford, John Sillitoe on behalf of the Wellcome Sanger Institute COVID-19 Surveillance Team |
| EPI_ISL_3444569 | Lighthouse Lab in Glasgow | Wellcome Sanger Institute for the COVID-19 Genomics UK (COG-UK) Consortium | Harper VanSteenhouse, Yumi Kasai, David Gray, Carol Clugston, Anna Dominiczak and Alex Alderton, Roberto Amato, Jeffrey Barrett, Sonia Goncalves, Ewan Harrison, David K. Jackson, Ian Johnston, Dominic Kwiatkowski, Cordelia Langford, John Sillitoe on behalf of the Wellcome Sanger Institute COVID-19 Surveillance Team |
| EPI_ISL_3444570, EPI_ISL_3444571, EPI_ISL_3444572 | Lighthouse Lab in Alderley Park | Wellcome Sanger Institute for the COVID-19 Genomics UK (COG-UK) Consortium | Jacquelyn Wynn, Mairead Hyland, The Lighthouse Lab in Alderley Park and Alex Alderton, Roberto Amato, Jeffrey Barrett, Sonia Goncalves, Ewan Harrison, David K. Jackson, Ian Johnston, Dominic Kwiatkowski, Cordelia Langford, John Sillitoe on behalf of the Wellcome Sanger Institute COVID-19 Surveillance Team |
| EPI_ISL_3444573, EPI_ISL_3444574 | Lighthouse Lab in Glasgow | Wellcome Sanger Institute for the COVID-19 Genomics UK (COG-UK) Consortium | Harper VanSteenhouse, Yumi Kasai, David Gray, Carol Clugston, Anna Dominiczak and Alex Alderton, Roberto Amato, Jeffrey Barrett, Sonia Goncalves, Ewan Harrison, David K. Jackson, Ian Johnston, Dominic Kwiatkowski, Cordelia Langford, John Sillitoe on behalf of the Wellcome Sanger Institute COVID-19 Surveillance Team |
| EPI_ISL_3444575, EPI_ISL_3444576 | Lighthouse Lab in Alderley Park | Wellcome Sanger Institute for the COVID-19 Genomics UK (COG-UK) Consortium | Jacquelyn Wynn, Mairead Hyland, The Lighthouse Lab in Alderley Park and Alex Alderton, Roberto Amato, Jeffrey Barrett, Sonia Goncalves, Ewan Harrison, David K. Jackson, Ian Johnston, Dominic Kwiatkowski, Cordelia Langford, John Sillitoe on behalf of the Wellcome Sanger Institute COVID-19 Surveillance Team |
| EPI_ISL_3444577, EPI_ISL_3444578 | Lighthouse Lab in Glasgow | Wellcome Sanger Institute for the COVID-19 Genomics UK (COG-UK) Consortium | Harper VanSteenhouse, Yumi Kasai, David Gray, Carol Clugston, Anna Dominiczak and Alex Alderton, Roberto Amato, Jeffrey Barrett, Sonia Goncalves, Ewan Harrison, David K. Jackson, Ian Johnston, Dominic Kwiatkowski, Cordelia Langford, John Sillitoe on behalf of the Wellcome Sanger Institute COVID-19 Surveillance Team |
| EPI_ISL_3444579, EPI_ISL_3444580 | Lighthouse Lab in Alderley Park | Wellcome Sanger Institute for the COVID-19 Genomics UK (COG-UK) Consortium | Jacquelyn Wynn, Mairead Hyland, The Lighthouse Lab in Alderley Park and Alex Alderton, Roberto Amato, Jeffrey Barrett, Sonia Goncalves, Ewan Harrison, David K. Jackson, Ian Johnston, Dominic Kwiatkowski, Cordelia Langford, John Sillitoe on behalf of the Wellcome Sanger Institute COVID-19 Surveillance Team |
| EPI_ISL_3444581 | Lighthouse Lab in Glasgow | Wellcome Sanger Institute for the COVID-19 Genomics UK (COG-UK) Consortium | Harper VanSteenhouse, Yumi Kasai, David Gray, Carol Clugston, Anna Dominiczak and Alex Alderton, Roberto Amato, Jeffrey Barrett, Sonia Goncalves, Ewan Harrison, David K. Jackson, Ian Johnston, Dominic Kwiatkowski, Cordelia Langford, John Sillitoe on behalf of the Wellcome Sanger Institute COVID-19 Surveillance Team |
| EPI_ISL_3444582 | Lighthouse Lab in Alderley Park | Wellcome Sanger Institute for the COVID-19 Genomics UK (COG-UK) Consortium | Jacquelyn Wynn, Mairead Hyland, The Lighthouse Lab in Alderley Park and Alex Alderton, Roberto Amato, Jeffrey Barrett, Sonia Goncalves, Ewan Harrison, David K. Jackson, Ian Johnston, Dominic Kwiatkowski, Cordelia Langford, John Sillitoe on behalf of the Wellcome Sanger Institute COVID-19 Surveillance Team |
| EPI_ISL_3444583 | Lighthouse Lab in Glasgow | Wellcome Sanger Institute for the COVID-19 Genomics UK (COG-UK) Consortium | Harper VanSteenhouse, Yumi Kasai, David Gray, Carol Clugston, Anna Dominiczak and Alex Alderton, Roberto Amato, Jeffrey Barrett, Sonia Goncalves, Ewan Harrison, David K. Jackson, Ian Johnston, Dominic Kwiatkowski, Cordelia Langford, John Sillitoe on behalf of the Wellcome Sanger Institute COVID-19 Surveillance Team |
| EPI_ISL_3444584, EPI_ISL_3444585, EPI_ISL_3444586, EPI_ISL_3444587, EPI_ISL_3444588, EPI_ISL_3444589 | Lighthouse Lab in Alderley Park | Wellcome Sanger Institute for the COVID-19 Genomics UK (COG-UK) Consortium | Jacquelyn Wynn, Mairead Hyland, The Lighthouse Lab in Alderley Park and Alex Alderton, Roberto Amato, Jeffrey Barrett, Sonia Goncalves, Ewan Harrison, David K. Jackson, Ian Johnston, Dominic Kwiatkowski, Cordelia Langford, John Sillitoe on behalf of the Wellcome Sanger Institute COVID-19 Surveillance Team |
| EPI_ISL_3444590 | Lighthouse Lab in Glasgow | Wellcome Sanger Institute for the COVID-19 Genomics UK (COG-UK) Consortium | Harper VanSteenhouse, Yumi Kasai, David Gray, Carol Clugston, Anna Dominiczak and Alex Alderton, Roberto Amato, Jeffrey Barrett, Sonia Goncalves, Ewan Harrison, David K. Jackson, Ian Johnston, Dominic Kwiatkowski, Cordelia Langford, John Sillitoe on behalf of the Wellcome Sanger Institute COVID-19 Surveillance Team |
| EPI_ISL_3444591, EPI_ISL_3444592 | Lighthouse Lab in Alderley Park | Wellcome Sanger Institute for the COVID-19 Genomics UK (COG-UK) Consortium | Jacquelyn Wynn, Mairead Hyland, The Lighthouse Lab in Alderley Park and Alex Alderton, Roberto Amato, Jeffrey Barrett, Sonia Goncalves, Ewan Harrison, David K. Jackson, Ian Johnston, Dominic Kwiatkowski, Cordelia Langford, John Sillitoe on behalf of the Wellcome Sanger Institute COVID-19 Surveillance Team |
| EPI_ISL_3444593 | Lighthouse Lab in Glasgow | Wellcome Sanger Institute for the COVID-19 Genomics UK (COG-UK) Consortium | Harper VanSteenhouse, Yumi Kasai, David Gray, Carol Clugston, Anna Dominiczak and Alex Alderton, Roberto Amato, Jeffrey Barrett, Sonia Goncalves, Ewan Harrison, David K. Jackson, Ian Johnston, Dominic Kwiatkowski, Cordelia Langford, John Sillitoe on behalf of the Wellcome Sanger Institute COVID-19 Surveillance Team |
| EPI_ISL_3444594, EPI_ISL_3444595 | Lighthouse Lab in Alderley Park | Wellcome Sanger Institute for the COVID-19 Genomics UK (COG-UK) Consortium | Jacquelyn Wynn, Mairead Hyland, The Lighthouse Lab in Alderley Park and Alex Alderton, Roberto Amato, Jeffrey Barrett, Sonia Goncalves, Ewan Harrison, David K. Jackson, Ian Johnston, Dominic Kwiatkowski, Cordelia Langford, John Sillitoe on behalf of the Wellcome Sanger Institute COVID-19 Surveillance Team |
| EPI_ISL_3444596, EPI_ISL_3444597 | Lighthouse Lab in Glasgow | Wellcome Sanger Institute for the COVID-19 Genomics UK (COG-UK) Consortium | Harper VanSteenhouse, Yumi Kasai, David Gray, Carol Clugston, Anna Dominiczak and Alex Alderton, Roberto Amato, Jeffrey Barrett, Sonia Goncalves, Ewan Harrison, David K. Jackson, Ian Johnston, Dominic Kwiatkowski, Cordelia Langford, John Sillitoe on behalf of the Wellcome Sanger Institute COVID-19 Surveillance Team |
| EPI_ISL_3444598, EPI_ISL_3444599, EPI_ISL_3444600, EPI_ISL_3444601 | Lighthouse Lab in Alderley Park | Wellcome Sanger Institute for the COVID-19 Genomics UK (COG-UK) Consortium | Jacquelyn Wynn, Mairead Hyland, The Lighthouse Lab in Alderley Park and Alex Alderton, Roberto Amato, Jeffrey Barrett, Sonia Goncalves, Ewan Harrison, David K. Jackson, Ian Johnston, Dominic Kwiatkowski, Cordelia Langford, John Sillitoe on behalf of the Wellcome Sanger Institute COVID-19 Surveillance Team |
| EPI_ISL_3444602 | Lighthouse Lab in Glasgow | Wellcome Sanger Institute for the COVID-19 Genomics UK (COG-UK) Consortium | Harper VanSteenhouse, Yumi Kasai, David Gray, Carol Clugston, Anna Dominiczak and Alex Alderton, Roberto Amato, Jeffrey Barrett, Sonia Goncalves, Ewan Harrison, David K. Jackson, Ian Johnston, Dominic Kwiatkowski, Cordelia Langford, John Sillitoe on behalf of the Wellcome Sanger Institute COVID-19 Surveillance Team |
| EPI_ISL_3444603 | Lighthouse Lab in Alderley Park | Wellcome Sanger Institute for the COVID-19 Genomics UK (COG-UK) Consortium | Jacquelyn Wynn, Mairead Hyland, The Lighthouse Lab in Alderley Park and Alex Alderton, Roberto Amato, Jeffrey Barrett, Sonia Goncalves, Ewan Harrison, David K. Jackson, Ian Johnston, Dominic Kwiatkowski, Cordelia Langford, John Sillitoe on behalf of the Wellcome Sanger Institute COVID-19 Surveillance Team |
| EPI_ISL_3444604 | Lighthouse Lab in Glasgow | Wellcome Sanger Institute for the COVID-19 Genomics UK (COG-UK) Consortium | Harper VanSteenhouse, Yumi Kasai, David Gray, Carol Clugston, Anna Dominiczak and Alex Alderton, Roberto Amato, Jeffrey Barrett, Sonia Goncalves, Ewan Harrison, David K. Jackson, Ian Johnston, Dominic Kwiatkowski, Cordelia Langford, John Sillitoe on behalf of the Wellcome Sanger Institute COVID-19 Surveillance Team |
| EPI_ISL_3444605 | Lighthouse Lab in Alderley Park | Wellcome Sanger Institute for the COVID-19 Genomics UK (COG-UK) Consortium | Jacquelyn Wynn, Mairead Hyland, The Lighthouse Lab in Alderley Park and Alex Alderton, Roberto Amato, Jeffrey Barrett, Sonia Goncalves, Ewan Harrison, David K. Jackson, Ian Johnston, Dominic Kwiatkowski, Cordelia Langford, John Sillitoe on behalf of the Wellcome Sanger Institute COVID-19 Surveillance Team |
| EPI_ISL_3444606 | Lighthouse Lab in Glasgow | Wellcome Sanger Institute for the COVID-19 Genomics UK (COG-UK) Consortium | Harper VanSteenhouse, Yumi Kasai, David Gray, Carol Clugston, Anna Dominiczak and Alex Alderton, Roberto Amato, Jeffrey Barrett, Sonia Goncalves, Ewan Harrison, David K. Jackson, Ian Johnston, Dominic Kwiatkowski, Cordelia Langford, John Sillitoe on behalf of the Wellcome Sanger Institute COVID-19 Surveillance Team |
| EPI_ISL_3444607, EPI_ISL_3444608 | Lighthouse Lab in Alderley Park | Wellcome Sanger Institute for the COVID-19 Genomics UK (COG-UK) Consortium | Jacquelyn Wynn, Mairead Hyland, The Lighthouse Lab in Alderley Park and Alex Alderton, Roberto Amato, Jeffrey Barrett, Sonia Goncalves, Ewan Harrison, David K. Jackson, Ian Johnston, Dominic Kwiatkowski, Cordelia Langford, John Sillitoe on behalf of the Wellcome Sanger Institute COVID-19 Surveillance Team |
| EPI_ISL_3444609, EPI_ISL_3444610, EPI_ISL_3444611 | Lighthouse Lab in Glasgow | Wellcome Sanger Institute for the COVID-19 Genomics UK (COG-UK) Consortium | Harper VanSteenhouse, Yumi Kasai, David Gray, Carol Clugston, Anna Dominiczak and Alex Alderton, Roberto Amato, Jeffrey Barrett, Sonia Goncalves, Ewan Harrison, David K. Jackson, Ian Johnston, Dominic Kwiatkowski, Cordelia Langford, John Sillitoe on behalf of the Wellcome Sanger Institute COVID-19 Surveillance Team |
| EPI_ISL_3444612, EPI_ISL_3444613, EPI_ISL_3444614, EPI_ISL_3444615, EPI_ISL_3444616, EPI_ISL_3444617, EPI_ISL_3444618 | Lighthouse Lab in Alderley Park | Wellcome Sanger Institute for the COVID-19 Genomics UK (COG-UK) Consortium | Jacquelyn Wynn, Mairead Hyland, The Lighthouse Lab in Alderley Park and Alex Alderton, Roberto Amato, Jeffrey Barrett, Sonia Goncalves, Ewan Harrison, David K. Jackson, Ian Johnston, Dominic Kwiatkowski, Cordelia Langford, John Sillitoe on behalf of the Wellcome Sanger Institute COVID-19 Surveillance Team |
| EPI_ISL_3444619 | Lighthouse Lab in Glasgow | Wellcome Sanger Institute for the COVID-19 Genomics UK (COG-UK) Consortium | Harper VanSteenhouse, Yumi Kasai, David Gray, Carol Clugston, Anna Dominiczak and Alex Alderton, Roberto Amato, Jeffrey Barrett, Sonia Goncalves, Ewan Harrison, David K. Jackson, Ian Johnston, Dominic Kwiatkowski, Cordelia Langford, John Sillitoe on behalf of the Wellcome Sanger Institute COVID-19 Surveillance Team |
| EPI_ISL_3444620 | Lighthouse Lab in Alderley Park | Wellcome Sanger Institute for the COVID-19 Genomics UK (COG-UK) Consortium | Jacquelyn Wynn, Mairead Hyland, The Lighthouse Lab in Alderley Park and Alex Alderton, Roberto Amato, Jeffrey Barrett, Sonia Goncalves, Ewan Harrison, David K. Jackson, Ian Johnston, Dominic Kwiatkowski, Cordelia Langford, John Sillitoe on behalf of the Wellcome Sanger Institute COVID-19 Surveillance Team |
| EPI_ISL_3444621, EPI_ISL_3444622 | Lighthouse Lab in Glasgow | Wellcome Sanger Institute for the COVID-19 Genomics UK (COG-UK) Consortium | Harper VanSteenhouse, Yumi Kasai, David Gray, Carol Clugston, Anna Dominiczak and Alex Alderton, Roberto Amato, Jeffrey Barrett, Sonia Goncalves, Ewan Harrison, David K. Jackson, Ian Johnston, Dominic Kwiatkowski, Cordelia Langford, John Sillitoe on behalf of the Wellcome Sanger Institute COVID-19 Surveillance Team |
| EPI_ISL_3444623, EPI_ISL_3444624, EPI_ISL_3444625 | Lighthouse Lab in Alderley Park | Wellcome Sanger Institute for the COVID-19 Genomics UK (COG-UK) Consortium | Jacquelyn Wynn, Mairead Hyland, The Lighthouse Lab in Alderley Park and Alex Alderton, Roberto Amato, Jeffrey Barrett, Sonia Goncalves, Ewan Harrison, David K. Jackson, Ian Johnston, Dominic Kwiatkowski, Cordelia Langford, John Sillitoe on behalf of the Wellcome Sanger Institute COVID-19 Surveillance Team |
| EPI_ISL_3444626, EPI_ISL_3444627 | Lighthouse Lab in Glasgow | Wellcome Sanger Institute for the COVID-19 Genomics UK (COG-UK) Consortium | Harper VanSteenhouse, Yumi Kasai, David Gray, Carol Clugston, Anna Dominiczak and Alex Alderton, Roberto Amato, Jeffrey Barrett, Sonia Goncalves, Ewan Harrison, David K. Jackson, Ian Johnston, Dominic Kwiatkowski, Cordelia Langford, John Sillitoe on behalf of the Wellcome Sanger Institute COVID-19 Surveillance Team |
| EPI_ISL_3444628 | Lighthouse Lab in Alderley Park | Wellcome Sanger Institute for the COVID-19 Genomics UK (COG-UK) Consortium | Jacquelyn Wynn, Mairead Hyland, The Lighthouse Lab in Alderley Park and Alex Alderton, Roberto Amato, Jeffrey Barrett, Sonia Goncalves, Ewan Harrison, David K. Jackson, Ian Johnston, Dominic Kwiatkowski, Cordelia Langford, John Sillitoe on behalf of the Wellcome Sanger Institute COVID-19 Surveillance Team |
| EPI_ISL_3444629 | Lighthouse Lab in Glasgow | Wellcome Sanger Institute for the COVID-19 Genomics UK (COG-UK) Consortium | Harper VanSteenhouse, Yumi Kasai, David Gray, Carol Clugston, Anna Dominiczak and Alex Alderton, Roberto Amato, Jeffrey Barrett, Sonia Goncalves, Ewan Harrison, David K. Jackson, Ian Johnston, Dominic Kwiatkowski, Cordelia Langford, John Sillitoe on behalf of the Wellcome Sanger Institute COVID-19 Surveillance Team |
| EPI_ISL_3444630, EPI_ISL_3444631, EPI_ISL_3444632, EPI_ISL_3444633, EPI_ISL_3444634, EPI_ISL_3444635, EPI_ISL_3444636, EPI_ISL_3444637, EPI_ISL_3444638, EPI_ISL_3444639 | Lighthouse Lab in Alderley Park | Wellcome Sanger Institute for the COVID-19 Genomics UK (COG-UK) Consortium | Jacquelyn Wynn, Mairead Hyland, The Lighthouse Lab in Alderley Park and Alex Alderton, Roberto Amato, Jeffrey Barrett, Sonia Goncalves, Ewan Harrison, David K. Jackson, Ian Johnston, Dominic Kwiatkowski, Cordelia Langford, John Sillitoe on behalf of the Wellcome Sanger Institute COVID-19 Surveillance Team |

| | | | |
|---|---|---|---|
| EPI_ISL_3444640 | Lighthouse Lab in Glasgow | Wellcome Sanger Institute for the COVID-19 Genomics UK (COG-UK) Consortium | Harper VanSteenhouse, Yumi Kasai, David Gray, Carol Clugston, Anna Dominiczak and Alex Alderton, Roberto Amato, Jeffrey Barrett, Sonia Goncalves, Ewan Harrison, David K. Jackson, Ian Johnston, Dominic Kwiatkowski, Cordelia Langford, John Sillitoe on behalf of the Wellcome Sanger Institute COVID-19 Surveillance Team |
| EPI_ISL_3444641, EPI_ISL_3444642, EPI_ISL_3444643 | Lighthouse Lab in Alderley Park | Wellcome Sanger Institute for the COVID-19 Genomics UK (COG-UK) Consortium | Jacquelyn Wynn, Mairead Hyland, The Lighthouse Lab in Alderley Park and Alex Alderton, Roberto Amato, Jeffrey Barrett, Sonia Goncalves, Ewan Harrison, David K. Jackson, Ian Johnston, Dominic Kwiatkowski, Cordelia Langford, John Sillitoe on behalf of the Wellcome Sanger Institute COVID-19 Surveillance Team |
| EPI_ISL_3444644 | Lighthouse Lab in Glasgow | Wellcome Sanger Institute for the COVID-19 Genomics UK (COG-UK) Consortium | Harper VanSteenhouse, Yumi Kasai, David Gray, Carol Clugston, Anna Dominiczak and Alex Alderton, Roberto Amato, Jeffrey Barrett, Sonia Goncalves, Ewan Harrison, David K. Jackson, Ian Johnston, Dominic Kwiatkowski, Cordelia Langford, John Sillitoe on behalf of the Wellcome Sanger Institute COVID-19 Surveillance Team |
| EPI_ISL_3444645, EPI_ISL_3444646 | Lighthouse Lab in Alderley Park | Wellcome Sanger Institute for the COVID-19 Genomics UK (COG-UK) Consortium | Jacquelyn Wynn, Mairead Hyland, The Lighthouse Lab in Alderley Park and Alex Alderton, Roberto Amato, Jeffrey Barrett, Sonia Goncalves, Ewan Harrison, David K. Jackson, Ian Johnston, Dominic Kwiatkowski, Cordelia Langford, John Sillitoe on behalf of the Wellcome Sanger Institute COVID-19 Surveillance Team |
| EPI_ISL_3444647 | Lighthouse Lab in Glasgow | Wellcome Sanger Institute for the COVID-19 Genomics UK (COG-UK) Consortium | Harper VanSteenhouse, Yumi Kasai, David Gray, Carol Clugston, Anna Dominiczak and Alex Alderton, Roberto Amato, Jeffrey Barrett, Sonia Goncalves, Ewan Harrison, David K. Jackson, Ian Johnston, Dominic Kwiatkowski, Cordelia Langford, John Sillitoe on behalf of the Wellcome Sanger Institute COVID-19 Surveillance Team |
| EPI_ISL_3444648, EPI_ISL_3444649, EPI_ISL_3444650, EPI_ISL_3444651 | Lighthouse Lab in Alderley Park | Wellcome Sanger Institute for the COVID-19 Genomics UK (COG-UK) Consortium | Jacquelyn Wynn, Mairead Hyland, The Lighthouse Lab in Alderley Park and Alex Alderton, Roberto Amato, Jeffrey Barrett, Sonia Goncalves, Ewan Harrison, David K. Jackson, Ian Johnston, Dominic Kwiatkowski, Cordelia Langford, John Sillitoe on behalf of the Wellcome Sanger Institute COVID-19 Surveillance Team |
| EPI_ISL_3444652 | Lighthouse Lab in Glasgow | Wellcome Sanger Institute for the COVID-19 Genomics UK (COG-UK) Consortium | Harper VanSteenhouse, Yumi Kasai, David Gray, Carol Clugston, Anna Dominiczak and Alex Alderton, Roberto Amato, Jeffrey Barrett, Sonia Goncalves, Ewan Harrison, David K. Jackson, Ian Johnston, Dominic Kwiatkowski, Cordelia Langford, John Sillitoe on behalf of the Wellcome Sanger Institute COVID-19 Surveillance Team |
| EPI_ISL_3444653, EPI_ISL_3444654, EPI_ISL_3444655 | Lighthouse Lab in Alderley Park | Wellcome Sanger Institute for the COVID-19 Genomics UK (COG-UK) Consortium | Jacquelyn Wynn, Mairead Hyland, The Lighthouse Lab in Alderley Park and Alex Alderton, Roberto Amato, Jeffrey Barrett, Sonia Goncalves, Ewan Harrison, David K. Jackson, Ian Johnston, Dominic Kwiatkowski, Cordelia Langford, John Sillitoe on behalf of the Wellcome Sanger Institute COVID-19 Surveillance Team |
| EPI_ISL_3444656 | Lighthouse Lab in Glasgow | Wellcome Sanger Institute for the COVID-19 Genomics UK (COG-UK) Consortium | Harper VanSteenhouse, Yumi Kasai, David Gray, Carol Clugston, Anna Dominiczak and Alex Alderton, Roberto Amato, Jeffrey Barrett, Sonia Goncalves, Ewan Harrison, David K. Jackson, Ian Johnston, Dominic Kwiatkowski, Cordelia Langford, John Sillitoe on behalf of the Wellcome Sanger Institute COVID-19 Surveillance Team |
| EPI_ISL_3444657, EPI_ISL_3444658, EPI_ISL_3444659 | Lighthouse Lab in Alderley Park | Wellcome Sanger Institute for the COVID-19 Genomics UK (COG-UK) Consortium | Jacquelyn Wynn, Mairead Hyland, The Lighthouse Lab in Alderley Park and Alex Alderton, Roberto Amato, Jeffrey Barrett, Sonia Goncalves, Ewan Harrison, David K. Jackson, Ian Johnston, Dominic Kwiatkowski, Cordelia Langford, John Sillitoe on behalf of the Wellcome Sanger Institute COVID-19 Surveillance Team |
| EPI_ISL_3444660 | Lighthouse Lab in Glasgow | Wellcome Sanger Institute for the COVID-19 Genomics UK (COG-UK) Consortium | Harper VanSteenhouse, Yumi Kasai, David Gray, Carol Clugston, Anna Dominiczak and Alex Alderton, Roberto Amato, Jeffrey Barrett, Sonia Goncalves, Ewan Harrison, David K. Jackson, Ian Johnston, Dominic Kwiatkowski, Cordelia Langford, John Sillitoe on behalf of the Wellcome Sanger Institute COVID-19 Surveillance Team |
| EPI_ISL_3444661 | Lighthouse Lab in Alderley Park | Wellcome Sanger Institute for the COVID-19 Genomics UK (COG-UK) Consortium | Jacquelyn Wynn, Mairead Hyland, The Lighthouse Lab in Alderley Park and Alex Alderton, Roberto Amato, Jeffrey Barrett, Sonia Goncalves, Ewan Harrison, David K. Jackson, Ian Johnston, Dominic Kwiatkowski, Cordelia Langford, John Sillitoe on behalf of the Wellcome Sanger Institute COVID-19 Surveillance Team |
| EPI_ISL_3444662, EPI_ISL_3444663 | Lighthouse Lab in Glasgow | Wellcome Sanger Institute for the COVID-19 Genomics UK (COG-UK) Consortium | Harper VanSteenhouse, Yumi Kasai, David Gray, Carol Clugston, Anna Dominiczak and Alex Alderton, Roberto Amato, Jeffrey Barrett, Sonia Goncalves, Ewan Harrison, David K. Jackson, Ian Johnston, Dominic Kwiatkowski, Cordelia Langford, John Sillitoe on behalf of the Wellcome Sanger Institute COVID-19 Surveillance Team |
| EPI_ISL_3444664, EPI_ISL_3444665 | Lighthouse Lab in Alderley Park | Wellcome Sanger Institute for the COVID-19 Genomics UK (COG-UK) Consortium | Jacquelyn Wynn, Mairead Hyland, The Lighthouse Lab in Alderley Park and Alex Alderton, Roberto Amato, Jeffrey Barrett, Sonia Goncalves, Ewan Harrison, David K. Jackson, Ian Johnston, Dominic Kwiatkowski, Cordelia Langford, John Sillitoe on behalf of the Wellcome Sanger Institute COVID-19 Surveillance Team |
| EPI_ISL_3444666, EPI_ISL_3444667 | Lighthouse Lab in Glasgow | Wellcome Sanger Institute for the COVID-19 Genomics UK (COG-UK) Consortium | Harper VanSteenhouse, Yumi Kasai, David Gray, Carol Clugston, Anna Dominiczak and Alex Alderton, Roberto Amato, Jeffrey Barrett, Sonia Goncalves, Ewan Harrison, David K. Jackson, Ian Johnston, Dominic Kwiatkowski, Cordelia Langford, John Sillitoe on behalf of the Wellcome Sanger Institute COVID-19 Surveillance Team |
| EPI_ISL_3444668, EPI_ISL_3444669 | Lighthouse Lab in Alderley Park | Wellcome Sanger Institute for the COVID-19 Genomics UK (COG-UK) Consortium | Jacquelyn Wynn, Mairead Hyland, The Lighthouse Lab in Alderley Park and Alex Alderton, Roberto Amato, Jeffrey Barrett, Sonia Goncalves, Ewan Harrison, David K. Jackson, Ian Johnston, Dominic Kwiatkowski, Cordelia Langford, John Sillitoe on behalf of the Wellcome Sanger Institute COVID-19 Surveillance Team |
| EPI_ISL_3444670 | Lighthouse Lab in Glasgow | Wellcome Sanger Institute for the COVID-19 Genomics UK (COG-UK) Consortium | Harper VanSteenhouse, Yumi Kasai, David Gray, Carol Clugston, Anna Dominiczak and Alex Alderton, Roberto Amato, Jeffrey Barrett, Sonia Goncalves, Ewan Harrison, David K. Jackson, Ian Johnston, Dominic Kwiatkowski, Cordelia Langford, John Sillitoe on behalf of the Wellcome Sanger Institute COVID-19 Surveillance Team |
| EPI_ISL_3444671, EPI_ISL_3444672, EPI_ISL_3444673, EPI_ISL_3444674, EPI_ISL_3444675 | Lighthouse Lab in Alderley Park | Wellcome Sanger Institute for the COVID-19 Genomics UK (COG-UK) Consortium | Jacquelyn Wynn, Mairead Hyland, The Lighthouse Lab in Alderley Park and Alex Alderton, Roberto Amato, Jeffrey Barrett, Sonia Goncalves, Ewan Harrison, David K. Jackson, Ian Johnston, Dominic Kwiatkowski, Cordelia Langford, John Sillitoe on behalf of the Wellcome Sanger Institute COVID-19 Surveillance Team |
| EPI_ISL_3444676, EPI_ISL_3444677 | Lighthouse Lab in Glasgow | Wellcome Sanger Institute for the COVID-19 Genomics UK (COG-UK) Consortium | Harper VanSteenhouse, Yumi Kasai, David Gray, Carol Clugston, Anna Dominiczak and Alex Alderton, Roberto Amato, Jeffrey Barrett, Sonia Goncalves, Ewan Harrison, David K. Jackson, Ian Johnston, Dominic Kwiatkowski, Cordelia Langford, John Sillitoe on behalf of the Wellcome Sanger Institute COVID-19 Surveillance Team |
| EPI_ISL_3444678, EPI_ISL_3444679, EPI_ISL_3444680, EPI_ISL_3444681, EPI_ISL_3444682, EPI_ISL_3444683 | Lighthouse Lab in Alderley Park | Wellcome Sanger Institute for the COVID-19 Genomics UK (COG-UK) Consortium | Jacquelyn Wynn, Mairead Hyland, The Lighthouse Lab in Alderley Park and Alex Alderton, Roberto Amato, Jeffrey Barrett, Sonia Goncalves, Ewan Harrison, David K. Jackson, Ian Johnston, Dominic Kwiatkowski, Cordelia Langford, John Sillitoe on behalf of the Wellcome Sanger Institute COVID-19 Surveillance Team |
| EPI_ISL_3444684 | Lighthouse Lab in Glasgow | Wellcome Sanger Institute for the COVID-19 Genomics UK (COG-UK) Consortium | Harper VanSteenhouse, Yumi Kasai, David Gray, Carol Clugston, Anna Dominiczak and Alex Alderton, Roberto Amato, Jeffrey Barrett, Sonia Goncalves, Ewan Harrison, David K. Jackson, Ian Johnston, Dominic Kwiatkowski, Cordelia Langford, John Sillitoe on behalf of the Wellcome Sanger Institute COVID-19 Surveillance Team |
| EPI_ISL_3444685, EPI_ISL_3444686, EPI_ISL_3444687, EPI_ISL_3444688, EPI_ISL_3444689, EPI_ISL_3444690, EPI_ISL_3444691 | Lighthouse Lab in Alderley Park | Wellcome Sanger Institute for the COVID-19 Genomics UK (COG-UK) Consortium | Jacquelyn Wynn, Mairead Hyland, The Lighthouse Lab in Alderley Park and Alex Alderton, Roberto Amato, Jeffrey Barrett, Sonia Goncalves, Ewan Harrison, David K. Jackson, Ian Johnston, Dominic Kwiatkowski, Cordelia Langford, John Sillitoe on behalf of the Wellcome Sanger Institute COVID-19 Surveillance Team |
| EPI_ISL_3444692, EPI_ISL_3444693 | Lighthouse Lab in Glasgow | Wellcome Sanger Institute for the COVID-19 Genomics UK (COG-UK) Consortium | Harper VanSteenhouse, Yumi Kasai, David Gray, Carol Clugston, Anna Dominiczak and Alex Alderton, Roberto Amato, Jeffrey Barrett, Sonia Goncalves, Ewan Harrison, David K. Jackson, Ian Johnston, Dominic Kwiatkowski, Cordelia Langford, John Sillitoe on behalf of the Wellcome Sanger Institute COVID-19 Surveillance Team |
| EPI_ISL_3444694, EPI_ISL_3444695, EPI_ISL_3444696 | Lighthouse Lab in Alderley Park | Wellcome Sanger Institute for the COVID-19 Genomics UK (COG-UK) Consortium | Jacquelyn Wynn, Mairead Hyland, The Lighthouse Lab in Alderley Park and Alex Alderton, Roberto Amato, Jeffrey Barrett, Sonia Goncalves, Ewan Harrison, David K. Jackson, Ian Johnston, Dominic Kwiatkowski, Cordelia Langford, John Sillitoe on behalf of the Wellcome Sanger Institute COVID-19 Surveillance Team |
| EPI_ISL_3444697 | Lighthouse Lab in Glasgow | Wellcome Sanger Institute for the COVID-19 Genomics UK (COG-UK) Consortium | Harper VanSteenhouse, Yumi Kasai, David Gray, Carol Clugston, Anna Dominiczak and Alex Alderton, Roberto Amato, Jeffrey Barrett, Sonia Goncalves, Ewan Harrison, David K. Jackson, Ian Johnston, Dominic Kwiatkowski, Cordelia Langford, John Sillitoe on behalf of the Wellcome Sanger Institute COVID-19 Surveillance Team |
| EPI_ISL_3444698, EPI_ISL_3444699 | Lighthouse Lab in Alderley Park | Wellcome Sanger Institute for the COVID-19 Genomics UK (COG-UK) Consortium | Jacquelyn Wynn, Mairead Hyland, The Lighthouse Lab in Alderley Park and Alex Alderton, Roberto Amato, Jeffrey Barrett, Sonia Goncalves, Ewan Harrison, David K. Jackson, Ian Johnston, Dominic Kwiatkowski, Cordelia Langford, John Sillitoe on behalf of the Wellcome Sanger Institute COVID-19 Surveillance Team |
| EPI_ISL_3444700, EPI_ISL_3444701 | Lighthouse Lab in Glasgow | Wellcome Sanger Institute for the COVID-19 Genomics UK (COG-UK) Consortium | Harper VanSteenhouse, Yumi Kasai, David Gray, Carol Clugston, Anna Dominiczak and Alex Alderton, Roberto Amato, Jeffrey Barrett, Sonia Goncalves, Ewan Harrison, David K. Jackson, Ian Johnston, Dominic Kwiatkowski, Cordelia Langford, John Sillitoe on behalf of the Wellcome Sanger Institute COVID-19 Surveillance Team |
| EPI_ISL_3444702, EPI_ISL_3444703, EPI_ISL_3444704, EPI_ISL_3444705 | Lighthouse Lab in Alderley Park | Wellcome Sanger Institute for the COVID-19 Genomics UK (COG-UK) Consortium | Jacquelyn Wynn, Mairead Hyland, The Lighthouse Lab in Alderley Park and Alex Alderton, Roberto Amato, Jeffrey Barrett, Sonia Goncalves, Ewan Harrison, David K. Jackson, Ian Johnston, Dominic Kwiatkowski, Cordelia Langford, John Sillitoe on behalf of the Wellcome Sanger Institute COVID-19 Surveillance Team |
| EPI_ISL_3444706 | Lighthouse Lab in Glasgow | Wellcome Sanger Institute for the COVID-19 Genomics UK (COG-UK) Consortium | Harper VanSteenhouse, Yumi Kasai, David Gray, Carol Clugston, Anna Dominiczak and Alex Alderton, Roberto Amato, Jeffrey Barrett, Sonia Goncalves, Ewan Harrison, David K. Jackson, Ian Johnston, Dominic Kwiatkowski, Cordelia Langford, John Sillitoe on behalf of the Wellcome Sanger Institute COVID-19 Surveillance Team |
| EPI_ISL_3444707, EPI_ISL_3444708, EPI_ISL_3444709, EPI_ISL_3444710, EPI_ISL_3444711, EPI_ISL_3444712 | Lighthouse Lab in Alderley Park | Wellcome Sanger Institute for the COVID-19 Genomics UK (COG-UK) Consortium | Jacquelyn Wynn, Mairead Hyland, The Lighthouse Lab in Alderley Park and Alex Alderton, Roberto Amato, Jeffrey Barrett, Sonia Goncalves, Ewan Harrison, David K. Jackson, Ian Johnston, Dominic Kwiatkowski, Cordelia Langford, John Sillitoe on behalf of the Wellcome Sanger Institute COVID-19 Surveillance Team |
| EPI_ISL_3444713 | Lighthouse Lab in Glasgow | Wellcome Sanger Institute for the COVID-19 Genomics UK (COG-UK) Consortium | Harper VanSteenhouse, Yumi Kasai, David Gray, Carol Clugston, Anna Dominiczak and Alex Alderton, Roberto Amato, Jeffrey Barrett, Sonia Goncalves, Ewan Harrison, David K. Jackson, Ian Johnston, Dominic Kwiatkowski, Cordelia Langford, John Sillitoe on behalf of the Wellcome Sanger Institute COVID-19 Surveillance Team |
| EPI_ISL_3444714, EPI_ISL_3444715 | Lighthouse Lab in Alderley Park | Wellcome Sanger Institute for the COVID-19 Genomics UK (COG-UK) Consortium | Jacquelyn Wynn, Mairead Hyland, The Lighthouse Lab in Alderley Park and Alex Alderton, Roberto Amato, Jeffrey Barrett, Sonia Goncalves, Ewan Harrison, David K. Jackson, Ian Johnston, Dominic Kwiatkowski, Cordelia Langford, John Sillitoe on behalf of the Wellcome Sanger Institute COVID-19 Surveillance Team |
| EPI_ISL_3444716, EPI_ISL_3444717, EPI_ISL_3444718 | Lighthouse Lab in Glasgow | Wellcome Sanger Institute for the COVID-19 Genomics UK (COG-UK) Consortium | Harper VanSteenhouse, Yumi Kasai, David Gray, Carol Clugston, Anna Dominiczak and Alex Alderton, Roberto Amato, Jeffrey Barrett, Sonia Goncalves, Ewan Harrison, David K. Jackson, Ian Johnston, Dominic Kwiatkowski, Cordelia Langford, John Sillitoe on behalf of the Wellcome Sanger Institute COVID-19 Surveillance Team |
| EPI_ISL_3444719, EPI_ISL_3444720, EPI_ISL_3444721, EPI_ISL_3444722, | Lighthouse Lab in Alderley Park | Wellcome Sanger Institute for the COVID-19 Genomics UK (COG-UK) Consortium | Jacquelyn Wynn, Mairead Hyland, The Lighthouse Lab in Alderley Park and Alex Alderton, Roberto Amato, Jeffrey Barrett, Sonia Goncalves, Ewan Harrison, David K. Jackson, Ian Johnston, Dominic Kwiatkowski, Cordelia Langford, John Sillitoe on behalf of the Wellcome Sanger Institute COVID-19 Surveillance Team |

| | | | |
|---|---|---|---|
| EPI_ISL_3444723 | | | |
| EPI_ISL_3444724, EPI_ISL_3444725, EPI_ISL_3444726 | Lighthouse Lab in Glasgow | Wellcome Sanger Institute for the COVID-19 Genomics UK (COG-UK) Consortium | Harper VanSteenhouse, Yumi Kasai, David Gray, Carol Clugston, Anna Dominiczak and Alex Alderton, Roberto Amato, Jeffrey Barrett, Sonia Goncalves, Ewan Harrison, David K. Jackson, Ian Johnston, Dominic Kwiatkowski, Cordelia Langford, John Sillitoe on behalf of the Wellcome Sanger Institute COVID-19 Surveillance Team |
| EPI_ISL_3444727, EPI_ISL_3444728, EPI_ISL_3444729 | Lighthouse Lab in Alderley Park | Wellcome Sanger Institute for the COVID-19 Genomics UK (COG-UK) Consortium | Jacquelyn Wynn, Mairead Hyland, The Lighthouse Lab in Alderley Park and Alex Alderton, Roberto Amato, Jeffrey Barrett, Sonia Goncalves, Ewan Harrison, David K. Jackson, Ian Johnston, Dominic Kwiatkowski, Cordelia Langford, John Sillitoe on behalf of the Wellcome Sanger Institute COVID-19 Surveillance Team |
| EPI_ISL_3444730 | Lighthouse Lab in Glasgow | Wellcome Sanger Institute for the COVID-19 Genomics UK (COG-UK) Consortium | Harper VanSteenhouse, Yumi Kasai, David Gray, Carol Clugston, Anna Dominiczak and Alex Alderton, Roberto Amato, Jeffrey Barrett, Sonia Goncalves, Ewan Harrison, David K. Jackson, Ian Johnston, Dominic Kwiatkowski, Cordelia Langford, John Sillitoe on behalf of the Wellcome Sanger Institute COVID-19 Surveillance Team |
| EPI_ISL_3444731, EPI_ISL_3444732, EPI_ISL_3444733, EPI_ISL_3444734, EPI_ISL_3444735 | Lighthouse Lab in Alderley Park | Wellcome Sanger Institute for the COVID-19 Genomics UK (COG-UK) Consortium | Jacquelyn Wynn, Mairead Hyland, The Lighthouse Lab in Alderley Park and Alex Alderton, Roberto Amato, Jeffrey Barrett, Sonia Goncalves, Ewan Harrison, David K. Jackson, Ian Johnston, Dominic Kwiatkowski, Cordelia Langford, John Sillitoe on behalf of the Wellcome Sanger Institute COVID-19 Surveillance Team |
| EPI_ISL_3444736 | Lighthouse Lab in Glasgow | Wellcome Sanger Institute for the COVID-19 Genomics UK (COG-UK) Consortium | Harper VanSteenhouse, Yumi Kasai, David Gray, Carol Clugston, Anna Dominiczak and Alex Alderton, Roberto Amato, Jeffrey Barrett, Sonia Goncalves, Ewan Harrison, David K. Jackson, Ian Johnston, Dominic Kwiatkowski, Cordelia Langford, John Sillitoe on behalf of the Wellcome Sanger Institute COVID-19 Surveillance Team |
| EPI_ISL_3444737, EPI_ISL_3444738, EPI_ISL_3444739, EPI_ISL_3444740, EPI_ISL_3444741 | Lighthouse Lab in Alderley Park | Wellcome Sanger Institute for the COVID-19 Genomics UK (COG-UK) Consortium | Jacquelyn Wynn, Mairead Hyland, The Lighthouse Lab in Alderley Park and Alex Alderton, Roberto Amato, Jeffrey Barrett, Sonia Goncalves, Ewan Harrison, David K. Jackson, Ian Johnston, Dominic Kwiatkowski, Cordelia Langford, John Sillitoe on behalf of the Wellcome Sanger Institute COVID-19 Surveillance Team |
| EPI_ISL_3444742 | Lighthouse Lab in Glasgow | Wellcome Sanger Institute for the COVID-19 Genomics UK (COG-UK) Consortium | Harper VanSteenhouse, Yumi Kasai, David Gray, Carol Clugston, Anna Dominiczak and Alex Alderton, Roberto Amato, Jeffrey Barrett, Sonia Goncalves, Ewan Harrison, David K. Jackson, Ian Johnston, Dominic Kwiatkowski, Cordelia Langford, John Sillitoe on behalf of the Wellcome Sanger Institute COVID-19 Surveillance Team |
| EPI_ISL_3444743, EPI_ISL_3444744, EPI_ISL_3444745, EPI_ISL_3444746 | Lighthouse Lab in Alderley Park | Wellcome Sanger Institute for the COVID-19 Genomics UK (COG-UK) Consortium | Jacquelyn Wynn, Mairead Hyland, The Lighthouse Lab in Alderley Park and Alex Alderton, Roberto Amato, Jeffrey Barrett, Sonia Goncalves, Ewan Harrison, David K. Jackson, Ian Johnston, Dominic Kwiatkowski, Cordelia Langford, John Sillitoe on behalf of the Wellcome Sanger Institute COVID-19 Surveillance Team |
| EPI_ISL_3444747, EPI_ISL_3444748 | Lighthouse Lab in Glasgow | Wellcome Sanger Institute for the COVID-19 Genomics UK (COG-UK) Consortium | Harper VanSteenhouse, Yumi Kasai, David Gray, Carol Clugston, Anna Dominiczak and Alex Alderton, Roberto Amato, Jeffrey Barrett, Sonia Goncalves, Ewan Harrison, David K. Jackson, Ian Johnston, Dominic Kwiatkowski, Cordelia Langford, John Sillitoe on behalf of the Wellcome Sanger Institute COVID-19 Surveillance Team |
| EPI_ISL_3444749 | Lighthouse Lab in Alderley Park | Wellcome Sanger Institute for the COVID-19 Genomics UK (COG-UK) Consortium | Jacquelyn Wynn, Mairead Hyland, The Lighthouse Lab in Alderley Park and Alex Alderton, Roberto Amato, Jeffrey Barrett, Sonia Goncalves, Ewan Harrison, David K. Jackson, Ian Johnston, Dominic Kwiatkowski, Cordelia Langford, John Sillitoe on behalf of the Wellcome Sanger Institute COVID-19 Surveillance Team |
| EPI_ISL_3444750, EPI_ISL_3444751 | Lighthouse Lab in Glasgow | Wellcome Sanger Institute for the COVID-19 Genomics UK (COG-UK) Consortium | Harper VanSteenhouse, Yumi Kasai, David Gray, Carol Clugston, Anna Dominiczak and Alex Alderton, Roberto Amato, Jeffrey Barrett, Sonia Goncalves, Ewan Harrison, David K. Jackson, Ian Johnston, Dominic Kwiatkowski, Cordelia Langford, John Sillitoe on behalf of the Wellcome Sanger Institute COVID-19 Surveillance Team |
| EPI_ISL_3444752, EPI_ISL_3444753 | Lighthouse Lab in Alderley Park | Wellcome Sanger Institute for the COVID-19 Genomics UK (COG-UK) Consortium | Jacquelyn Wynn, Mairead Hyland, The Lighthouse Lab in Alderley Park and Alex Alderton, Roberto Amato, Jeffrey Barrett, Sonia Goncalves, Ewan Harrison, David K. Jackson, Ian Johnston, Dominic Kwiatkowski, Cordelia Langford, John Sillitoe on behalf of the Wellcome Sanger Institute COVID-19 Surveillance Team |
| EPI_ISL_3444754 | Lighthouse Lab in Glasgow | Wellcome Sanger Institute for the COVID-19 Genomics UK (COG-UK) Consortium | Harper VanSteenhouse, Yumi Kasai, David Gray, Carol Clugston, Anna Dominiczak and Alex Alderton, Roberto Amato, Jeffrey Barrett, Sonia Goncalves, Ewan Harrison, David K. Jackson, Ian Johnston, Dominic Kwiatkowski, Cordelia Langford, John Sillitoe on behalf of the Wellcome Sanger Institute COVID-19 Surveillance Team |
| EPI_ISL_3444755, EPI_ISL_3444756, EPI_ISL_3444757 | Lighthouse Lab in Alderley Park | Wellcome Sanger Institute for the COVID-19 Genomics UK (COG-UK) Consortium | Jacquelyn Wynn, Mairead Hyland, The Lighthouse Lab in Alderley Park and Alex Alderton, Roberto Amato, Jeffrey Barrett, Sonia Goncalves, Ewan Harrison, David K. Jackson, Ian Johnston, Dominic Kwiatkowski, Cordelia Langford, John Sillitoe on behalf of the Wellcome Sanger Institute COVID-19 Surveillance Team |
| EPI_ISL_3444758 | Lighthouse Lab in Glasgow | Wellcome Sanger Institute for the COVID-19 Genomics UK (COG-UK) Consortium | Harper VanSteenhouse, Yumi Kasai, David Gray, Carol Clugston, Anna Dominiczak and Alex Alderton, Roberto Amato, Jeffrey Barrett, Sonia Goncalves, Ewan Harrison, David K. Jackson, Ian Johnston, Dominic Kwiatkowski, Cordelia Langford, John Sillitoe on behalf of the Wellcome Sanger Institute COVID-19 Surveillance Team |
| EPI_ISL_3444759, EPI_ISL_3444760 | Lighthouse Lab in Alderley Park | Wellcome Sanger Institute for the COVID-19 Genomics UK (COG-UK) Consortium | Jacquelyn Wynn, Mairead Hyland, The Lighthouse Lab in Alderley Park and Alex Alderton, Roberto Amato, Jeffrey Barrett, Sonia Goncalves, Ewan Harrison, David K. Jackson, Ian Johnston, Dominic Kwiatkowski, Cordelia Langford, John Sillitoe on behalf of the Wellcome Sanger Institute COVID-19 Surveillance Team |
| EPI_ISL_3444761 | Lighthouse Lab in Glasgow | Wellcome Sanger Institute for the COVID-19 Genomics UK (COG-UK) Consortium | Harper VanSteenhouse, Yumi Kasai, David Gray, Carol Clugston, Anna Dominiczak and Alex Alderton, Roberto Amato, Jeffrey Barrett, Sonia Goncalves, Ewan Harrison, David K. Jackson, Ian Johnston, Dominic Kwiatkowski, Cordelia Langford, John Sillitoe on behalf of the Wellcome Sanger Institute COVID-19 Surveillance Team |
| EPI_ISL_3444762 | Lighthouse Lab in Alderley Park | Wellcome Sanger Institute for the COVID-19 Genomics UK (COG-UK) Consortium | Jacquelyn Wynn, Mairead Hyland, The Lighthouse Lab in Alderley Park and Alex Alderton, Roberto Amato, Jeffrey Barrett, Sonia Goncalves, Ewan Harrison, David K. Jackson, Ian Johnston, Dominic Kwiatkowski, Cordelia Langford, John Sillitoe on behalf of the Wellcome Sanger Institute COVID-19 Surveillance Team |
| EPI_ISL_3444763 | Lighthouse Lab in Glasgow | Wellcome Sanger Institute for the COVID-19 Genomics UK (COG-UK) Consortium | Harper VanSteenhouse, Yumi Kasai, David Gray, Carol Clugston, Anna Dominiczak and Alex Alderton, Roberto Amato, Jeffrey Barrett, Sonia Goncalves, Ewan Harrison, David K. Jackson, Ian Johnston, Dominic Kwiatkowski, Cordelia Langford, John Sillitoe on behalf of the Wellcome Sanger Institute COVID-19 Surveillance Team |
| EPI_ISL_3444764 | Lighthouse Lab in Alderley Park | Wellcome Sanger Institute for the COVID-19 Genomics UK (COG-UK) Consortium | Jacquelyn Wynn, Mairead Hyland, The Lighthouse Lab in Alderley Park and Alex Alderton, Roberto Amato, Jeffrey Barrett, Sonia Goncalves, Ewan Harrison, David K. Jackson, Ian Johnston, Dominic Kwiatkowski, Cordelia Langford, John Sillitoe on behalf of the Wellcome Sanger Institute COVID-19 Surveillance Team |
| EPI_ISL_3444765 | Lighthouse Lab in Glasgow | Wellcome Sanger Institute for the COVID-19 Genomics UK (COG-UK) Consortium | Harper VanSteenhouse, Yumi Kasai, David Gray, Carol Clugston, Anna Dominiczak and Alex Alderton, Roberto Amato, Jeffrey Barrett, Sonia Goncalves, Ewan Harrison, David K. Jackson, Ian Johnston, Dominic Kwiatkowski, Cordelia Langford, John Sillitoe on behalf of the Wellcome Sanger Institute COVID-19 Surveillance Team |
| EPI_ISL_3444766 | Lighthouse Lab in Alderley Park | Wellcome Sanger Institute for the COVID-19 Genomics UK (COG-UK) Consortium | Jacquelyn Wynn, Mairead Hyland, The Lighthouse Lab in Alderley Park and Alex Alderton, Roberto Amato, Jeffrey Barrett, Sonia Goncalves, Ewan Harrison, David K. Jackson, Ian Johnston, Dominic Kwiatkowski, Cordelia Langford, John Sillitoe on behalf of the Wellcome Sanger Institute COVID-19 Surveillance Team |
| EPI_ISL_3444767 | Lighthouse Lab in Glasgow | Wellcome Sanger Institute for the COVID-19 Genomics UK (COG-UK) Consortium | Harper VanSteenhouse, Yumi Kasai, David Gray, Carol Clugston, Anna Dominiczak and Alex Alderton, Roberto Amato, Jeffrey Barrett, Sonia Goncalves, Ewan Harrison, David K. Jackson, Ian Johnston, Dominic Kwiatkowski, Cordelia Langford, John Sillitoe on behalf of the Wellcome Sanger Institute COVID-19 Surveillance Team |
| EPI_ISL_3444768, EPI_ISL_3444769 | Lighthouse Lab in Alderley Park | Wellcome Sanger Institute for the COVID-19 Genomics UK (COG-UK) Consortium | Jacquelyn Wynn, Mairead Hyland, The Lighthouse Lab in Alderley Park and Alex Alderton, Roberto Amato, Jeffrey Barrett, Sonia Goncalves, Ewan Harrison, David K. Jackson, Ian Johnston, Dominic Kwiatkowski, Cordelia Langford, John Sillitoe on behalf of the Wellcome Sanger Institute COVID-19 Surveillance Team |
| EPI_ISL_3444770, EPI_ISL_3444771 | Lighthouse Lab in Glasgow | Wellcome Sanger Institute for the COVID-19 Genomics UK (COG-UK) Consortium | Harper VanSteenhouse, Yumi Kasai, David Gray, Carol Clugston, Anna Dominiczak and Alex Alderton, Roberto Amato, Jeffrey Barrett, Sonia Goncalves, Ewan Harrison, David K. Jackson, Ian Johnston, Dominic Kwiatkowski, Cordelia Langford, John Sillitoe on behalf of the Wellcome Sanger Institute COVID-19 Surveillance Team |
| EPI_ISL_3444772, EPI_ISL_3444773, EPI_ISL_3444774, EPI_ISL_3444775, EPI_ISL_3444776, EPI_ISL_3444777, EPI_ISL_3444778 | Lighthouse Lab in Alderley Park | Wellcome Sanger Institute for the COVID-19 Genomics UK (COG-UK) Consortium | Jacquelyn Wynn, Mairead Hyland, The Lighthouse Lab in Alderley Park and Alex Alderton, Roberto Amato, Jeffrey Barrett, Sonia Goncalves, Ewan Harrison, David K. Jackson, Ian Johnston, Dominic Kwiatkowski, Cordelia Langford, John Sillitoe on behalf of the Wellcome Sanger Institute COVID-19 Surveillance Team |
| EPI_ISL_3444779, EPI_ISL_3444780, EPI_ISL_3444781 | Lighthouse Lab in Glasgow | Wellcome Sanger Institute for the COVID-19 Genomics UK (COG-UK) Consortium | Harper VanSteenhouse, Yumi Kasai, David Gray, Carol Clugston, Anna Dominiczak and Alex Alderton, Roberto Amato, Jeffrey Barrett, Sonia Goncalves, Ewan Harrison, David K. Jackson, Ian Johnston, Dominic Kwiatkowski, Cordelia Langford, John Sillitoe on behalf of the Wellcome Sanger Institute COVID-19 Surveillance Team |
| EPI_ISL_3444782 | Lighthouse Lab in Alderley Park | Wellcome Sanger Institute for the COVID-19 Genomics UK (COG-UK) Consortium | Jacquelyn Wynn, Mairead Hyland, The Lighthouse Lab in Alderley Park and Alex Alderton, Roberto Amato, Jeffrey Barrett, Sonia Goncalves, Ewan Harrison, David K. Jackson, Ian Johnston, Dominic Kwiatkowski, Cordelia Langford, John Sillitoe on behalf of the Wellcome Sanger Institute COVID-19 Surveillance Team |
| EPI_ISL_3444783 | Lighthouse Lab in Glasgow | Wellcome Sanger Institute for the COVID-19 Genomics UK (COG-UK) Consortium | Harper VanSteenhouse, Yumi Kasai, David Gray, Carol Clugston, Anna Dominiczak and Alex Alderton, Roberto Amato, Jeffrey Barrett, Sonia Goncalves, Ewan Harrison, David K. Jackson, Ian Johnston, Dominic Kwiatkowski, Cordelia Langford, John Sillitoe on behalf of the Wellcome Sanger Institute COVID-19 Surveillance Team |
| EPI_ISL_3444784 | Lighthouse Lab in Alderley Park | Wellcome Sanger Institute for the COVID-19 Genomics UK (COG-UK) Consortium | Jacquelyn Wynn, Mairead Hyland, The Lighthouse Lab in Alderley Park and Alex Alderton, Roberto Amato, Jeffrey Barrett, Sonia Goncalves, Ewan Harrison, David K. Jackson, Ian Johnston, Dominic Kwiatkowski, Cordelia Langford, John Sillitoe on behalf of the Wellcome Sanger Institute COVID-19 Surveillance Team |
| EPI_ISL_3444785 | Lighthouse Lab in Glasgow | Wellcome Sanger Institute for the COVID-19 Genomics UK (COG-UK) Consortium | Harper VanSteenhouse, Yumi Kasai, David Gray, Carol Clugston, Anna Dominiczak and Alex Alderton, Roberto Amato, Jeffrey Barrett, Sonia Goncalves, Ewan Harrison, David K. Jackson, Ian Johnston, Dominic Kwiatkowski, Cordelia Langford, John Sillitoe on behalf of the Wellcome Sanger Institute COVID-19 Surveillance Team |
| EPI_ISL_3444786, EPI_ISL_3444787, EPI_ISL_3444788 | Lighthouse Lab in Alderley Park | Wellcome Sanger Institute for the COVID-19 Genomics UK (COG-UK) Consortium | Jacquelyn Wynn, Mairead Hyland, The Lighthouse Lab in Alderley Park and Alex Alderton, Roberto Amato, Jeffrey Barrett, Sonia Goncalves, Ewan Harrison, David K. Jackson, Ian Johnston, Dominic Kwiatkowski, Cordelia Langford, John Sillitoe on behalf of the Wellcome Sanger Institute COVID-19 Surveillance Team |
| EPI_ISL_3444789, EPI_ISL_3444790, EPI_ISL_3444791 | Lighthouse Lab in Glasgow | Wellcome Sanger Institute for the COVID-19 Genomics UK (COG-UK) Consortium | Harper VanSteenhouse, Yumi Kasai, David Gray, Carol Clugston, Anna Dominiczak and Alex Alderton, Roberto Amato, Jeffrey Barrett, Sonia Goncalves, Ewan Harrison, David K. Jackson, Ian Johnston, Dominic Kwiatkowski, Cordelia Langford, John Sillitoe on behalf of the Wellcome Sanger Institute COVID-19 Surveillance Team |
| EPI_ISL_3444792, EPI_ISL_3444793, EPI_ISL_3444794, EPI_ISL_3444795, EPI_ISL_3444796, EPI_ISL_3444797, EPI_ISL_3444798, EPI_ISL_3444799 | Lighthouse Lab in Alderley Park | Wellcome Sanger Institute for the COVID-19 Genomics UK (COG-UK) Consortium | Jacquelyn Wynn, Mairead Hyland, The Lighthouse Lab in Alderley Park and Alex Alderton, Roberto Amato, Jeffrey Barrett, Sonia Goncalves, Ewan Harrison, David K. Jackson, Ian Johnston, Dominic Kwiatkowski, Cordelia Langford, John Sillitoe on behalf of the Wellcome Sanger Institute COVID-19 Surveillance Team |
| EPI_ISL_3444800 | Lighthouse Lab in Glasgow | Wellcome Sanger Institute for the COVID-19 Genomics UK (COG-UK) | Harper VanSteenhouse, Yumi Kasai, David Gray, Carol Clugston, Anna Dominiczak and Alex Alderton, Roberto Amato, Jeffrey Barrett, Sonia Goncalves, Ewan Harrison, David K. Jackson, Ian Johnston, Dominic Kwiatkowski, Cordelia Langford, |

| | | | |
|---|---|---|---|
| | | Consortium | John Sillitoe on behalf of the Wellcome Sanger Institute COVID-19 Surveillance Team |
| EPI_ISL_3444801 | Lighthouse Lab in Alderley Park | Wellcome Sanger Institute for the COVID-19 Genomics UK (COG-UK) Consortium | Jacquelyn Wynn, Mairead Hyland, The Lighthouse Lab in Alderley Park and Alex Alderton, Roberto Amato, Jeffrey Barrett, Sonia Goncalves, Ewan Harrison, David K. Jackson, Ian Johnston, Dominic Kwiatkowski, Cordelia Langford, John Sillitoe on behalf of the Wellcome Sanger Institute COVID-19 Surveillance Team |
| EPI_ISL_3444802, EPI_ISL_3444803, EPI_ISL_3444804, EPI_ISL_3444805, EPI_ISL_3444806, EPI_ISL_3444807, EPI_ISL_3444808, EPI_ISL_3444809, EPI_ISL_3444810, EPI_ISL_3444811, EPI_ISL_3444812, EPI_ISL_3444813, EPI_ISL_3444814, EPI_ISL_3444815, EPI_ISL_3444816, EPI_ISL_3444817, EPI_ISL_3444818, EPI_ISL_3444819, EPI_ISL_3444820, EPI_ISL_3444821, EPI_ISL_3444822, EPI_ISL_3444823, EPI_ISL_3444824, EPI_ISL_3444825, EPI_ISL_3444826, EPI_ISL_3444827, EPI_ISL_3444828, EPI_ISL_3444829, EPI_ISL_3444830, EPI_ISL_3444831, EPI_ISL_3444832, EPI_ISL_3444833, EPI_ISL_3444834, EPI_ISL_3444835, EPI_ISL_3444836, EPI_ISL_3444837, EPI_ISL_3444838, EPI_ISL_3444839, EPI_ISL_3444840, EPI_ISL_3444841, EPI_ISL_3444842, EPI_ISL_3444843, EPI_ISL_3444844, EPI_ISL_3444845, EPI_ISL_3444846, EPI_ISL_3444847, EPI_ISL_3444848, EPI_ISL_3444849, EPI_ISL_3444850, EPI_ISL_3444851, EPI_ISL_3444852, EPI_ISL_3444853, EPI_ISL_3444854, EPI_ISL_3444855, EPI_ISL_3444856, EPI_ISL_3444857, EPI_ISL_3444858, EPI_ISL_3444859, EPI_ISL_3444860, EPI_ISL_3444861, EPI_ISL_3444862, EPI_ISL_3444863, EPI_ISL_3444864, EPI_ISL_3444865, EPI_ISL_3444866, EPI_ISL_3444867, EPI_ISL_3444868, EPI_ISL_3444869, EPI_ISL_3444870, EPI_ISL_3444871, EPI_ISL_3444872, EPI_ISL_3444873, EPI_ISL_3444874, EPI_ISL_3444875, EPI_ISL_3444876, EPI_ISL_3444877, EPI_ISL_3444878, EPI_ISL_3444879, EPI_ISL_3444880, EPI_ISL_3444881, EPI_ISL_3444882, EPI_ISL_3444883, EPI_ISL_3444884, EPI_ISL_3444885 | | | |
| see above | Lighthouse Lab in Glasgow | Wellcome Sanger Institute for the COVID-19 Genomics UK (COG-UK) Consortium | Harper VanSteenhouse, Yumi Kasai, David Gray, Carol Clugston, Anna Dominiczak and Alex Alderton, Roberto Amato, Jeffrey Barrett, Sonia Goncalves, Ewan Harrison, David K. Jackson, Ian Johnston, Dominic Kwiatkowski, Cordelia Langford, John Sillitoe on behalf of the Wellcome Sanger Institute COVID-19 Surveillance Team |
| EPI_ISL_3444886 | Lighthouse Lab in Milton Keynes | Wellcome Sanger Institute for the COVID-19 Genomics UK (COG-UK) Consortium | The Lighthouse Lab in Milton Keynes and Alex Alderton, Roberto Amato, Jeffrey Barrett, Sonia Goncalves, Ewan Harrison, David K. Jackson, Ian Johnston, Dominic Kwiatkowski, Cordelia Langford, John Sillitoe on behalf of the Wellcome Sanger Institute COVID-19 Surveillance Team |
| EPI_ISL_3444887, EPI_ISL_3444888, EPI_ISL_3444889 | Lighthouse Lab in Glasgow | Wellcome Sanger Institute for the COVID-19 Genomics UK (COG-UK) Consortium | Harper VanSteenhouse, Yumi Kasai, David Gray, Carol Clugston, Anna Dominiczak and Alex Alderton, Roberto Amato, Jeffrey Barrett, Sonia Goncalves, Ewan Harrison, David K. Jackson, Ian Johnston, Dominic Kwiatkowski, Cordelia Langford, John Sillitoe on behalf of the Wellcome Sanger Institute COVID-19 Surveillance Team |
| EPI_ISL_3444890 | Lighthouse Lab in Milton Keynes | Wellcome Sanger Institute for the COVID-19 Genomics UK (COG-UK) Consortium | The Lighthouse Lab in Milton Keynes and Alex Alderton, Roberto Amato, Jeffrey Barrett, Sonia Goncalves, Ewan Harrison, David K. Jackson, Ian Johnston, Dominic Kwiatkowski, Cordelia Langford, John Sillitoe on behalf of the Wellcome Sanger Institute COVID-19 Surveillance Team |
| EPI_ISL_3444891, EPI_ISL_3444892, EPI_ISL_3444893 | Lighthouse Lab in Glasgow | Wellcome Sanger Institute for the COVID-19 Genomics UK (COG-UK) Consortium | Harper VanSteenhouse, Yumi Kasai, David Gray, Carol Clugston, Anna Dominiczak and Alex Alderton, Roberto Amato, Jeffrey Barrett, Sonia Goncalves, Ewan Harrison, David K. Jackson, Ian Johnston, Dominic Kwiatkowski, Cordelia Langford, John Sillitoe on behalf of the Wellcome Sanger Institute COVID-19 Surveillance Team |
| EPI_ISL_3444894 | Lighthouse Lab in Milton Keynes | Wellcome Sanger Institute for the COVID-19 Genomics UK (COG-UK) Consortium | The Lighthouse Lab in Milton Keynes and Alex Alderton, Roberto Amato, Jeffrey Barrett, Sonia Goncalves, Ewan Harrison, David K. Jackson, Ian Johnston, Dominic Kwiatkowski, Cordelia Langford, John Sillitoe on behalf of the Wellcome Sanger Institute COVID-19 Surveillance Team |
| EPI_ISL_3444895, EPI_ISL_3444896, EPI_ISL_3444897 | Lighthouse Lab in Glasgow | Wellcome Sanger Institute for the COVID-19 Genomics UK (COG-UK) Consortium | Harper VanSteenhouse, Yumi Kasai, David Gray, Carol Clugston, Anna Dominiczak and Alex Alderton, Roberto Amato, Jeffrey Barrett, Sonia Goncalves, Ewan Harrison, David K. Jackson, Ian Johnston, Dominic Kwiatkowski, Cordelia Langford, John Sillitoe on behalf of the Wellcome Sanger Institute COVID-19 Surveillance Team |
| EPI_ISL_3444898, EPI_ISL_3444899 | Lighthouse Lab in Milton Keynes | Wellcome Sanger Institute for the COVID-19 Genomics UK (COG-UK) Consortium | The Lighthouse Lab in Milton Keynes and Alex Alderton, Roberto Amato, Jeffrey Barrett, Sonia Goncalves, Ewan Harrison, David K. Jackson, Ian Johnston, Dominic Kwiatkowski, Cordelia Langford, John Sillitoe on behalf of the Wellcome Sanger Institute COVID-19 Surveillance Team |
| EPI_ISL_3444900 | Lighthouse Lab in Glasgow | Wellcome Sanger Institute for the COVID-19 Genomics UK (COG-UK) Consortium | Harper VanSteenhouse, Yumi Kasai, David Gray, Carol Clugston, Anna Dominiczak and Alex Alderton, Roberto Amato, Jeffrey Barrett, Sonia Goncalves, Ewan Harrison, David K. Jackson, Ian Johnston, Dominic Kwiatkowski, Cordelia Langford, John Sillitoe on behalf of the Wellcome Sanger Institute COVID-19 Surveillance Team |
| EPI_ISL_3444901 | Lighthouse Lab in Milton Keynes | Wellcome Sanger Institute for the COVID-19 Genomics UK (COG-UK) Consortium | The Lighthouse Lab in Milton Keynes and Alex Alderton, Roberto Amato, Jeffrey Barrett, Sonia Goncalves, Ewan Harrison, David K. Jackson, Ian Johnston, Dominic Kwiatkowski, Cordelia Langford, John Sillitoe on behalf of the Wellcome Sanger Institute COVID-19 Surveillance Team |
| EPI_ISL_3444902 | Lighthouse Lab in Glasgow | Wellcome Sanger Institute for the COVID-19 Genomics UK (COG-UK) Consortium | Harper VanSteenhouse, Yumi Kasai, David Gray, Carol Clugston, Anna Dominiczak and Alex Alderton, Roberto Amato, Jeffrey Barrett, Sonia Goncalves, Ewan Harrison, David K. Jackson, Ian Johnston, Dominic Kwiatkowski, Cordelia Langford, John Sillitoe on behalf of the Wellcome Sanger Institute COVID-19 Surveillance Team |
| EPI_ISL_3444903, EPI_ISL_3444904, EPI_ISL_3444905, EPI_ISL_3444906 | Lighthouse Lab in Milton Keynes | Wellcome Sanger Institute for the COVID-19 Genomics UK (COG-UK) Consortium | The Lighthouse Lab in Milton Keynes and Alex Alderton, Roberto Amato, Jeffrey Barrett, Sonia Goncalves, Ewan Harrison, David K. Jackson, Ian Johnston, Dominic Kwiatkowski, Cordelia Langford, John Sillitoe on behalf of the Wellcome Sanger Institute COVID-19 Surveillance Team |
| EPI_ISL_3444907 | Lighthouse Lab in Glasgow | Wellcome Sanger Institute for the COVID-19 Genomics UK (COG-UK) Consortium | Harper VanSteenhouse, Yumi Kasai, David Gray, Carol Clugston, Anna Dominiczak and Alex Alderton, Roberto Amato, Jeffrey Barrett, Sonia Goncalves, Ewan Harrison, David K. Jackson, Ian Johnston, Dominic Kwiatkowski, Cordelia Langford, John Sillitoe on behalf of the Wellcome Sanger Institute COVID-19 Surveillance Team |
| EPI_ISL_3444908 | Lighthouse Lab in Milton Keynes | Wellcome Sanger Institute for the COVID-19 Genomics UK (COG-UK) Consortium | The Lighthouse Lab in Milton Keynes and Alex Alderton, Roberto Amato, Jeffrey Barrett, Sonia Goncalves, Ewan Harrison, David K. Jackson, Ian Johnston, Dominic Kwiatkowski, Cordelia Langford, John Sillitoe on behalf of the Wellcome Sanger Institute COVID-19 Surveillance Team |
| EPI_ISL_3444909 | Lighthouse Lab in Glasgow | Wellcome Sanger Institute for the COVID-19 Genomics UK (COG-UK) Consortium | Harper VanSteenhouse, Yumi Kasai, David Gray, Carol Clugston, Anna Dominiczak and Alex Alderton, Roberto Amato, Jeffrey Barrett, Sonia Goncalves, Ewan Harrison, David K. Jackson, Ian Johnston, Dominic Kwiatkowski, Cordelia Langford, John Sillitoe on behalf of the Wellcome Sanger Institute COVID-19 Surveillance Team |
| EPI_ISL_3444910 | Lighthouse Lab in Milton Keynes | Wellcome Sanger Institute for the COVID-19 Genomics UK (COG-UK) Consortium | The Lighthouse Lab in Milton Keynes and Alex Alderton, Roberto Amato, Jeffrey Barrett, Sonia Goncalves, Ewan Harrison, David K. Jackson, Ian Johnston, Dominic Kwiatkowski, Cordelia Langford, John Sillitoe on behalf of the Wellcome Sanger Institute COVID-19 Surveillance Team |
| EPI_ISL_3444911, EPI_ISL_3444912, EPI_ISL_3444913, EPI_ISL_3444914, EPI_ISL_3444915, EPI_ISL_3444916 | Lighthouse Lab in Glasgow | Wellcome Sanger Institute for the COVID-19 Genomics UK (COG-UK) Consortium | Harper VanSteenhouse, Yumi Kasai, David Gray, Carol Clugston, Anna Dominiczak and Alex Alderton, Roberto Amato, Jeffrey Barrett, Sonia Goncalves, Ewan Harrison, David K. Jackson, Ian Johnston, Dominic Kwiatkowski, Cordelia Langford, John Sillitoe on behalf of the Wellcome Sanger Institute COVID-19 Surveillance Team |
| EPI_ISL_3444917, EPI_ISL_3444918 | Lighthouse Lab in Milton Keynes | Wellcome Sanger Institute for the COVID-19 Genomics UK (COG-UK) Consortium | The Lighthouse Lab in Milton Keynes and Alex Alderton, Roberto Amato, Jeffrey Barrett, Sonia Goncalves, Ewan Harrison, David K. Jackson, Ian Johnston, Dominic Kwiatkowski, Cordelia Langford, John Sillitoe on behalf of the Wellcome Sanger Institute COVID-19 Surveillance Team |
| EPI_ISL_3444919 | Lighthouse Lab in Glasgow | Wellcome Sanger Institute for the COVID-19 Genomics UK (COG-UK) Consortium | Harper VanSteenhouse, Yumi Kasai, David Gray, Carol Clugston, Anna Dominiczak and Alex Alderton, Roberto Amato, Jeffrey Barrett, Sonia Goncalves, Ewan Harrison, David K. Jackson, Ian Johnston, Dominic Kwiatkowski, Cordelia Langford, John Sillitoe on behalf of the Wellcome Sanger Institute COVID-19 Surveillance Team |
| EPI_ISL_3444920 | Lighthouse Lab in Milton Keynes | Wellcome Sanger Institute for the COVID-19 Genomics UK (COG-UK) Consortium | The Lighthouse Lab in Milton Keynes and Alex Alderton, Roberto Amato, Jeffrey Barrett, Sonia Goncalves, Ewan Harrison, David K. Jackson, Ian Johnston, Dominic Kwiatkowski, Cordelia Langford, John Sillitoe on behalf of the Wellcome Sanger Institute COVID-19 Surveillance Team |
| EPI_ISL_3444921, EPI_ISL_3444922 | Lighthouse Lab in Glasgow | Wellcome Sanger Institute for the COVID-19 Genomics UK (COG-UK) Consortium | Harper VanSteenhouse, Yumi Kasai, David Gray, Carol Clugston, Anna Dominiczak and Alex Alderton, Roberto Amato, Jeffrey Barrett, Sonia Goncalves, Ewan Harrison, David K. Jackson, Ian Johnston, Dominic Kwiatkowski, Cordelia Langford, John Sillitoe on behalf of the Wellcome Sanger Institute COVID-19 Surveillance Team |
| EPI_ISL_3444923 | Lighthouse Lab in Milton Keynes | Wellcome Sanger Institute for the COVID-19 Genomics UK (COG-UK) Consortium | The Lighthouse Lab in Milton Keynes and Alex Alderton, Roberto Amato, Jeffrey Barrett, Sonia Goncalves, Ewan Harrison, David K. Jackson, Ian Johnston, Dominic Kwiatkowski, Cordelia Langford, John Sillitoe on behalf of the Wellcome Sanger Institute COVID-19 Surveillance Team |
| EPI_ISL_3444924, EPI_ISL_3444925, EPI_ISL_3444926, EPI_ISL_3444927, EPI_ISL_3444928 | Lighthouse Lab in Glasgow | Wellcome Sanger Institute for the COVID-19 Genomics UK (COG-UK) Consortium | Harper VanSteenhouse, Yumi Kasai, David Gray, Carol Clugston, Anna Dominiczak and Alex Alderton, Roberto Amato, Jeffrey Barrett, Sonia Goncalves, Ewan Harrison, David K. Jackson, Ian Johnston, Dominic Kwiatkowski, Cordelia Langford, John Sillitoe on behalf of the Wellcome Sanger Institute COVID-19 Surveillance Team |
| EPI_ISL_3444929, EPI_ISL_3444930 | Lighthouse Lab in Milton Keynes | Wellcome Sanger Institute for the COVID-19 Genomics UK (COG-UK) Consortium | The Lighthouse Lab in Milton Keynes and Alex Alderton, Roberto Amato, Jeffrey Barrett, Sonia Goncalves, Ewan Harrison, David K. Jackson, Ian Johnston, Dominic Kwiatkowski, Cordelia Langford, John Sillitoe on behalf of the Wellcome Sanger Institute COVID-19 Surveillance Team |
| EPI_ISL_3444931 | Lighthouse Lab in Glasgow | Wellcome Sanger Institute for the COVID-19 Genomics UK (COG-UK) Consortium | Harper VanSteenhouse, Yumi Kasai, David Gray, Carol Clugston, Anna Dominiczak and Alex Alderton, Roberto Amato, Jeffrey Barrett, Sonia Goncalves, Ewan Harrison, David K. Jackson, Ian Johnston, Dominic Kwiatkowski, Cordelia Langford, John Sillitoe on behalf of the Wellcome Sanger Institute COVID-19 Surveillance Team |
| EPI_ISL_3444932 | Lighthouse Lab in Milton Keynes | Wellcome Sanger Institute for the COVID-19 Genomics UK (COG-UK) Consortium | The Lighthouse Lab in Milton Keynes and Alex Alderton, Roberto Amato, Jeffrey Barrett, Sonia Goncalves, Ewan Harrison, David K. Jackson, Ian Johnston, Dominic Kwiatkowski, Cordelia Langford, John Sillitoe on behalf of the Wellcome Sanger Institute COVID-19 Surveillance Team |
| EPI_ISL_3444933, EPI_ISL_3444934, EPI_ISL_3444935 | Lighthouse Lab in Glasgow | Wellcome Sanger Institute for the COVID-19 Genomics UK (COG-UK) Consortium | Harper VanSteenhouse, Yumi Kasai, David Gray, Carol Clugston, Anna Dominiczak and Alex Alderton, Roberto Amato, Jeffrey Barrett, Sonia Goncalves, Ewan Harrison, David K. Jackson, Ian Johnston, Dominic Kwiatkowski, Cordelia Langford, John Sillitoe on behalf of the Wellcome Sanger Institute COVID-19 Surveillance Team |
| EPI_ISL_3444936 | Lighthouse Lab in Milton Keynes | Wellcome Sanger Institute for the COVID-19 Genomics UK (COG-UK) Consortium | The Lighthouse Lab in Milton Keynes and Alex Alderton, Roberto Amato, Jeffrey Barrett, Sonia Goncalves, Ewan Harrison, David K. Jackson, Ian Johnston, Dominic Kwiatkowski, Cordelia Langford, John Sillitoe on behalf of the Wellcome Sanger Institute COVID-19 Surveillance Team |
| EPI_ISL_3444937, EPI_ISL_3444938, EPI_ISL_3444939, EPI_ISL_3444940 | Lighthouse Lab in Glasgow | Wellcome Sanger Institute for the COVID-19 Genomics UK (COG-UK) Consortium | Harper VanSteenhouse, Yumi Kasai, David Gray, Carol Clugston, Anna Dominiczak and Alex Alderton, Roberto Amato, Jeffrey Barrett, Sonia Goncalves, Ewan Harrison, David K. Jackson, Ian Johnston, Dominic Kwiatkowski, Cordelia Langford, John Sillitoe on behalf of the Wellcome Sanger Institute COVID-19 Surveillance Team |
| EPI_ISL_3444941, EPI_ISL_3444942 | Lighthouse Lab in Milton Keynes | Wellcome Sanger Institute for the COVID-19 Genomics UK (COG-UK) Consortium | The Lighthouse Lab in Milton Keynes and Alex Alderton, Roberto Amato, Jeffrey Barrett, Sonia Goncalves, Ewan Harrison, David K. Jackson, Ian Johnston, Dominic Kwiatkowski, Cordelia Langford, John Sillitoe on behalf of the Wellcome Sanger Institute COVID-19 Surveillance Team |
| EPI_ISL_3444943 | Lighthouse Lab in Glasgow | Wellcome Sanger Institute for the COVID-19 Genomics UK (COG-UK) Consortium | Harper VanSteenhouse, Yumi Kasai, David Gray, Carol Clugston, Anna Dominiczak and Alex Alderton, Roberto Amato, Jeffrey Barrett, Sonia Goncalves, Ewan Harrison, David K. Jackson, Ian Johnston, Dominic Kwiatkowski, Cordelia Langford, John Sillitoe on behalf of the Wellcome Sanger Institute COVID-19 Surveillance Team |
| EPI_ISL_3444944 | Lighthouse Lab in Milton Keynes | Wellcome Sanger Institute for the COVID-19 Genomics UK (COG-UK) Consortium | The Lighthouse Lab in Milton Keynes and Alex Alderton, Roberto Amato, Jeffrey Barrett, Sonia Goncalves, Ewan Harrison, David K. Jackson, Ian Johnston, Dominic Kwiatkowski, Cordelia Langford, John Sillitoe on behalf of the Wellcome Sanger Institute COVID-19 Surveillance Team |
| EPI_ISL_3444945 | Lighthouse Lab in Glasgow | Wellcome Sanger Institute for the COVID-19 Genomics UK (COG-UK) Consortium | Harper VanSteenhouse, Yumi Kasai, David Gray, Carol Clugston, Anna Dominiczak and Alex Alderton, Roberto Amato, Jeffrey Barrett, Sonia Goncalves, Ewan Harrison, David K. Jackson, Ian Johnston, Dominic Kwiatkowski, Cordelia Langford, John Sillitoe on behalf of the Wellcome Sanger Institute COVID-19 Surveillance Team |
| EPI_ISL_3444946, EPI_ISL_3444947 | Lighthouse Lab in Milton Keynes | Wellcome Sanger Institute for the COVID-19 Genomics UK (COG-UK) Consortium | The Lighthouse Lab in Milton Keynes and Alex Alderton, Roberto Amato, Jeffrey Barrett, Sonia Goncalves, Ewan Harrison, David K. Jackson, Ian Johnston, Dominic Kwiatkowski, Cordelia Langford, John Sillitoe on behalf of the Wellcome Sanger Institute COVID-19 Surveillance Team |
| EPI_ISL_3444948, EPI_ISL_3444949 | Lighthouse Lab in Glasgow | Wellcome Sanger Institute for the COVID-19 Genomics UK (COG-UK) Consortium | Harper VanSteenhouse, Yumi Kasai, David Gray, Carol Clugston, Anna Dominiczak and Alex Alderton, Roberto Amato, Jeffrey Barrett, Sonia Goncalves, Ewan Harrison, David K. Jackson, Ian Johnston, Dominic Kwiatkowski, Cordelia Langford, John Sillitoe on behalf of the Wellcome Sanger Institute COVID-19 Surveillance Team |
| EPI_ISL_3444950 | Lighthouse Lab in Milton Keynes | Wellcome Sanger Institute for the COVID-19 Genomics UK (COG-UK) | The Lighthouse Lab in Milton Keynes and Alex Alderton, Roberto Amato, Jeffrey Barrett, Sonia Goncalves, Ewan Harrison, David K. Jackson, Ian Johnston, Dominic Kwiatkowski, Cordelia Langford, John Sillitoe on behalf of the Wellcome Sanger |

| | | | |
|---|---|---|---|
| | | Consortium | Institute COVID-19 Surveillance Team |
| EPI_ISL_3444951 | Lighthouse Lab in Glasgow | Wellcome Sanger Institute for the COVID-19 Genomics UK (COG-UK) Consortium | Harper VanSteenhouse, Yumi Kasai, David Gray, Carol Clugston, Anna Dominiczak and Alex Alderton, Roberto Amato, Jeffrey Barrett, Sonia Goncalves, Ewan Harrison, David K. Jackson, Ian Johnston, Dominic Kwiatkowski, Cordelia Langford, John Sillitoe on behalf of the Wellcome Sanger Institute COVID-19 Surveillance Team |
| EPI_ISL_3444952 | Lighthouse Lab in Milton Keynes | Wellcome Sanger Institute for the COVID-19 Genomics UK (COG-UK) Consortium | The Lighthouse Lab in Milton Keynes and Alex Alderton, Roberto Amato, Jeffrey Barrett, Sonia Goncalves, Ewan Harrison, David K. Jackson, Ian Johnston, Dominic Kwiatkowski, Cordelia Langford, John Sillitoe on behalf of the Wellcome Sanger Institute COVID-19 Surveillance Team |
| EPI_ISL_3444953, EPI_ISL_3444954 | Lighthouse Lab in Glasgow | Wellcome Sanger Institute for the COVID-19 Genomics UK (COG-UK) Consortium | Harper VanSteenhouse, Yumi Kasai, David Gray, Carol Clugston, Anna Dominiczak and Alex Alderton, Roberto Amato, Jeffrey Barrett, Sonia Goncalves, Ewan Harrison, David K. Jackson, Ian Johnston, Dominic Kwiatkowski, Cordelia Langford, John Sillitoe on behalf of the Wellcome Sanger Institute COVID-19 Surveillance Team |
| EPI_ISL_3444955 | Lighthouse Lab in Milton Keynes | Wellcome Sanger Institute for the COVID-19 Genomics UK (COG-UK) Consortium | The Lighthouse Lab in Milton Keynes and Alex Alderton, Roberto Amato, Jeffrey Barrett, Sonia Goncalves, Ewan Harrison, David K. Jackson, Ian Johnston, Dominic Kwiatkowski, Cordelia Langford, John Sillitoe on behalf of the Wellcome Sanger Institute COVID-19 Surveillance Team |
| EPI_ISL_3444956, EPI_ISL_3444957 | Lighthouse Lab in Glasgow | Wellcome Sanger Institute for the COVID-19 Genomics UK (COG-UK) Consortium | Harper VanSteenhouse, Yumi Kasai, David Gray, Carol Clugston, Anna Dominiczak and Alex Alderton, Roberto Amato, Jeffrey Barrett, Sonia Goncalves, Ewan Harrison, David K. Jackson, Ian Johnston, Dominic Kwiatkowski, Cordelia Langford, John Sillitoe on behalf of the Wellcome Sanger Institute COVID-19 Surveillance Team |
| EPI_ISL_3444958 | Lighthouse Lab in Milton Keynes | Wellcome Sanger Institute for the COVID-19 Genomics UK (COG-UK) Consortium | The Lighthouse Lab in Milton Keynes and Alex Alderton, Roberto Amato, Jeffrey Barrett, Sonia Goncalves, Ewan Harrison, David K. Jackson, Ian Johnston, Dominic Kwiatkowski, Cordelia Langford, John Sillitoe on behalf of the Wellcome Sanger Institute COVID-19 Surveillance Team |
| EPI_ISL_3444959, EPI_ISL_3444960 | Lighthouse Lab in Glasgow | Wellcome Sanger Institute for the COVID-19 Genomics UK (COG-UK) Consortium | Harper VanSteenhouse, Yumi Kasai, David Gray, Carol Clugston, Anna Dominiczak and Alex Alderton, Roberto Amato, Jeffrey Barrett, Sonia Goncalves, Ewan Harrison, David K. Jackson, Ian Johnston, Dominic Kwiatkowski, Cordelia Langford, John Sillitoe on behalf of the Wellcome Sanger Institute COVID-19 Surveillance Team |
| EPI_ISL_3444961 | Lighthouse Lab in Milton Keynes | Wellcome Sanger Institute for the COVID-19 Genomics UK (COG-UK) Consortium | The Lighthouse Lab in Milton Keynes and Alex Alderton, Roberto Amato, Jeffrey Barrett, Sonia Goncalves, Ewan Harrison, David K. Jackson, Ian Johnston, Dominic Kwiatkowski, Cordelia Langford, John Sillitoe on behalf of the Wellcome Sanger Institute COVID-19 Surveillance Team |
| EPI_ISL_3444962, EPI_ISL_3444963, EPI_ISL_3444964, EPI_ISL_3444965, EPI_ISL_3444966, EPI_ISL_3444967, EPI_ISL_3444968, EPI_ISL_3444969, EPI_ISL_3444970, EPI_ISL_3444971, EPI_ISL_3444972, EPI_ISL_3444973, EPI_ISL_3444974, EPI_ISL_3444975 | | | |
| see above | Lighthouse Lab in Glasgow | Wellcome Sanger Institute for the COVID-19 Genomics UK (COG-UK) Consortium | Harper VanSteenhouse, Yumi Kasai, David Gray, Carol Clugston, Anna Dominiczak and Alex Alderton, Roberto Amato, Jeffrey Barrett, Sonia Goncalves, Ewan Harrison, David K. Jackson, Ian Johnston, Dominic Kwiatkowski, Cordelia Langford, John Sillitoe on behalf of the Wellcome Sanger Institute COVID-19 Surveillance Team |
| EPI_ISL_3444976 | Lighthouse Lab in Milton Keynes | Wellcome Sanger Institute for the COVID-19 Genomics UK (COG-UK) Consortium | The Lighthouse Lab in Milton Keynes and Alex Alderton, Roberto Amato, Jeffrey Barrett, Sonia Goncalves, Ewan Harrison, David K. Jackson, Ian Johnston, Dominic Kwiatkowski, Cordelia Langford, John Sillitoe on behalf of the Wellcome Sanger Institute COVID-19 Surveillance Team |
| EPI_ISL_3444977 | Lighthouse Lab in Glasgow | Wellcome Sanger Institute for the COVID-19 Genomics UK (COG-UK) Consortium | Harper VanSteenhouse, Yumi Kasai, David Gray, Carol Clugston, Anna Dominiczak and Alex Alderton, Roberto Amato, Jeffrey Barrett, Sonia Goncalves, Ewan Harrison, David K. Jackson, Ian Johnston, Dominic Kwiatkowski, Cordelia Langford, John Sillitoe on behalf of the Wellcome Sanger Institute COVID-19 Surveillance Team |
| EPI_ISL_3444978, EPI_ISL_3444979 | Lighthouse Lab in Milton Keynes | Wellcome Sanger Institute for the COVID-19 Genomics UK (COG-UK) Consortium | The Lighthouse Lab in Milton Keynes and Alex Alderton, Roberto Amato, Jeffrey Barrett, Sonia Goncalves, Ewan Harrison, David K. Jackson, Ian Johnston, Dominic Kwiatkowski, Cordelia Langford, John Sillitoe on behalf of the Wellcome Sanger Institute COVID-19 Surveillance Team |
| EPI_ISL_3444980, EPI_ISL_3444981, EPI_ISL_3444982, EPI_ISL_3444983, EPI_ISL_3444984 | Lighthouse Lab in Glasgow | Wellcome Sanger Institute for the COVID-19 Genomics UK (COG-UK) Consortium | Harper VanSteenhouse, Yumi Kasai, David Gray, Carol Clugston, Anna Dominiczak and Alex Alderton, Roberto Amato, Jeffrey Barrett, Sonia Goncalves, Ewan Harrison, David K. Jackson, Ian Johnston, Dominic Kwiatkowski, Cordelia Langford, John Sillitoe on behalf of the Wellcome Sanger Institute COVID-19 Surveillance Team |
| EPI_ISL_3444985 | Lighthouse Lab in Milton Keynes | Wellcome Sanger Institute for the COVID-19 Genomics UK (COG-UK) Consortium | The Lighthouse Lab in Milton Keynes and Alex Alderton, Roberto Amato, Jeffrey Barrett, Sonia Goncalves, Ewan Harrison, David K. Jackson, Ian Johnston, Dominic Kwiatkowski, Cordelia Langford, John Sillitoe on behalf of the Wellcome Sanger Institute COVID-19 Surveillance Team |
| EPI_ISL_3444986 | Lighthouse Lab in Glasgow | Wellcome Sanger Institute for the COVID-19 Genomics UK (COG-UK) Consortium | Harper VanSteenhouse, Yumi Kasai, David Gray, Carol Clugston, Anna Dominiczak and Alex Alderton, Roberto Amato, Jeffrey Barrett, Sonia Goncalves, Ewan Harrison, David K. Jackson, Ian Johnston, Dominic Kwiatkowski, Cordelia Langford, John Sillitoe on behalf of the Wellcome Sanger Institute COVID-19 Surveillance Team |
| EPI_ISL_3444987, EPI_ISL_3444988, EPI_ISL_3444989 | Lighthouse Lab in Milton Keynes | Wellcome Sanger Institute for the COVID-19 Genomics UK (COG-UK) Consortium | The Lighthouse Lab in Milton Keynes and Alex Alderton, Roberto Amato, Jeffrey Barrett, Sonia Goncalves, Ewan Harrison, David K. Jackson, Ian Johnston, Dominic Kwiatkowski, Cordelia Langford, John Sillitoe on behalf of the Wellcome Sanger Institute COVID-19 Surveillance Team |
| EPI_ISL_3444990, EPI_ISL_3444991, EPI_ISL_3444992, EPI_ISL_3444993, EPI_ISL_3444994 | Lighthouse Lab in Glasgow | Wellcome Sanger Institute for the COVID-19 Genomics UK (COG-UK) Consortium | Harper VanSteenhouse, Yumi Kasai, David Gray, Carol Clugston, Anna Dominiczak and Alex Alderton, Roberto Amato, Jeffrey Barrett, Sonia Goncalves, Ewan Harrison, David K. Jackson, Ian Johnston, Dominic Kwiatkowski, Cordelia Langford, John Sillitoe on behalf of the Wellcome Sanger Institute COVID-19 Surveillance Team |
| EPI_ISL_3444995, EPI_ISL_3444996 | Lighthouse Lab in Milton Keynes | Wellcome Sanger Institute for the COVID-19 Genomics UK (COG-UK) Consortium | The Lighthouse Lab in Milton Keynes and Alex Alderton, Roberto Amato, Jeffrey Barrett, Sonia Goncalves, Ewan Harrison, David K. Jackson, Ian Johnston, Dominic Kwiatkowski, Cordelia Langford, John Sillitoe on behalf of the Wellcome Sanger Institute COVID-19 Surveillance Team |
| EPI_ISL_3444997, EPI_ISL_3444998 | Lighthouse Lab in Glasgow | Wellcome Sanger Institute for the COVID-19 Genomics UK (COG-UK) Consortium | Harper VanSteenhouse, Yumi Kasai, David Gray, Carol Clugston, Anna Dominiczak and Alex Alderton, Roberto Amato, Jeffrey Barrett, Sonia Goncalves, Ewan Harrison, David K. Jackson, Ian Johnston, Dominic Kwiatkowski, Cordelia Langford, John Sillitoe on behalf of the Wellcome Sanger Institute COVID-19 Surveillance Team |
| EPI_ISL_3444999 | Lighthouse Lab in Milton Keynes | Wellcome Sanger Institute for the COVID-19 Genomics UK (COG-UK) Consortium | The Lighthouse Lab in Milton Keynes and Alex Alderton, Roberto Amato, Jeffrey Barrett, Sonia Goncalves, Ewan Harrison, David K. Jackson, Ian Johnston, Dominic Kwiatkowski, Cordelia Langford, John Sillitoe on behalf of the Wellcome Sanger Institute COVID-19 Surveillance Team |
| EPI_ISL_3445000, EPI_ISL_3445001, EPI_ISL_3445002, EPI_ISL_3445003 | Lighthouse Lab in Glasgow | Wellcome Sanger Institute for the COVID-19 Genomics UK (COG-UK) Consortium | Harper VanSteenhouse, Yumi Kasai, David Gray, Carol Clugston, Anna Dominiczak and Alex Alderton, Roberto Amato, Jeffrey Barrett, Sonia Goncalves, Ewan Harrison, David K. Jackson, Ian Johnston, Dominic Kwiatkowski, Cordelia Langford, John Sillitoe on behalf of the Wellcome Sanger Institute COVID-19 Surveillance Team |
| EPI_ISL_3445004 | Lighthouse Lab in Milton Keynes | Wellcome Sanger Institute for the COVID-19 Genomics UK (COG-UK) Consortium | The Lighthouse Lab in Milton Keynes and Alex Alderton, Roberto Amato, Jeffrey Barrett, Sonia Goncalves, Ewan Harrison, David K. Jackson, Ian Johnston, Dominic Kwiatkowski, Cordelia Langford, John Sillitoe on behalf of the Wellcome Sanger Institute COVID-19 Surveillance Team |
| EPI_ISL_3445005, EPI_ISL_3445006, EPI_ISL_3445007, EPI_ISL_3445008, EPI_ISL_3445009, EPI_ISL_3445010, EPI_ISL_3445011, EPI_ISL_3445012, EPI_ISL_3445013 | Lighthouse Lab in Glasgow | Wellcome Sanger Institute for the COVID-19 Genomics UK (COG-UK) Consortium | Harper VanSteenhouse, Yumi Kasai, David Gray, Carol Clugston, Anna Dominiczak and Alex Alderton, Roberto Amato, Jeffrey Barrett, Sonia Goncalves, Ewan Harrison, David K. Jackson, Ian Johnston, Dominic Kwiatkowski, Cordelia Langford, John Sillitoe on behalf of the Wellcome Sanger Institute COVID-19 Surveillance Team |
| EPI_ISL_3445014 | Lighthouse Lab in Milton Keynes | Wellcome Sanger Institute for the COVID-19 Genomics UK (COG-UK) Consortium | The Lighthouse Lab in Milton Keynes and Alex Alderton, Roberto Amato, Jeffrey Barrett, Sonia Goncalves, Ewan Harrison, David K. Jackson, Ian Johnston, Dominic Kwiatkowski, Cordelia Langford, John Sillitoe on behalf of the Wellcome Sanger Institute COVID-19 Surveillance Team |
| EPI_ISL_3445015, EPI_ISL_3445016, EPI_ISL_3445017, EPI_ISL_3445018 | Lighthouse Lab in Glasgow | Wellcome Sanger Institute for the COVID-19 Genomics UK (COG-UK) Consortium | Harper VanSteenhouse, Yumi Kasai, David Gray, Carol Clugston, Anna Dominiczak and Alex Alderton, Roberto Amato, Jeffrey Barrett, Sonia Goncalves, Ewan Harrison, David K. Jackson, Ian Johnston, Dominic Kwiatkowski, Cordelia Langford, John Sillitoe on behalf of the Wellcome Sanger Institute COVID-19 Surveillance Team |
| EPI_ISL_3445019 | Lighthouse Lab in Milton Keynes | Wellcome Sanger Institute for the COVID-19 Genomics UK (COG-UK) Consortium | The Lighthouse Lab in Milton Keynes and Alex Alderton, Roberto Amato, Jeffrey Barrett, Sonia Goncalves, Ewan Harrison, David K. Jackson, Ian Johnston, Dominic Kwiatkowski, Cordelia Langford, John Sillitoe on behalf of the Wellcome Sanger Institute COVID-19 Surveillance Team |
| EPI_ISL_3445020, EPI_ISL_3445021, EPI_ISL_3445022, EPI_ISL_3445023, EPI_ISL_3445024, EPI_ISL_3445025, EPI_ISL_3445026, EPI_ISL_3445027 | Lighthouse Lab in Glasgow | Wellcome Sanger Institute for the COVID-19 Genomics UK (COG-UK) Consortium | Harper VanSteenhouse, Yumi Kasai, David Gray, Carol Clugston, Anna Dominiczak and Alex Alderton, Roberto Amato, Jeffrey Barrett, Sonia Goncalves, Ewan Harrison, David K. Jackson, Ian Johnston, Dominic Kwiatkowski, Cordelia Langford, John Sillitoe on behalf of the Wellcome Sanger Institute COVID-19 Surveillance Team |
| EPI_ISL_3445028, EPI_ISL_3445029 | Lighthouse Lab in Milton Keynes | Wellcome Sanger Institute for the COVID-19 Genomics UK (COG-UK) Consortium | The Lighthouse Lab in Milton Keynes and Alex Alderton, Roberto Amato, Jeffrey Barrett, Sonia Goncalves, Ewan Harrison, David K. Jackson, Ian Johnston, Dominic Kwiatkowski, Cordelia Langford, John Sillitoe on behalf of the Wellcome Sanger Institute COVID-19 Surveillance Team |
| EPI_ISL_3445030, EPI_ISL_3445031, EPI_ISL_3445032 | Lighthouse Lab in Glasgow | Wellcome Sanger Institute for the COVID-19 Genomics UK (COG-UK) Consortium | Harper VanSteenhouse, Yumi Kasai, David Gray, Carol Clugston, Anna Dominiczak and Alex Alderton, Roberto Amato, Jeffrey Barrett, Sonia Goncalves, Ewan Harrison, David K. Jackson, Ian Johnston, Dominic Kwiatkowski, Cordelia Langford, John Sillitoe on behalf of the Wellcome Sanger Institute COVID-19 Surveillance Team |
| EPI_ISL_3445033 | Lighthouse Lab in Milton Keynes | Wellcome Sanger Institute for the COVID-19 Genomics UK (COG-UK) Consortium | The Lighthouse Lab in Milton Keynes and Alex Alderton, Roberto Amato, Jeffrey Barrett, Sonia Goncalves, Ewan Harrison, David K. Jackson, Ian Johnston, Dominic Kwiatkowski, Cordelia Langford, John Sillitoe on behalf of the Wellcome Sanger Institute COVID-19 Surveillance Team |
| EPI_ISL_3445034, EPI_ISL_3445035 | Lighthouse Lab in Glasgow | Wellcome Sanger Institute for the COVID-19 Genomics UK (COG-UK) Consortium | Harper VanSteenhouse, Yumi Kasai, David Gray, Carol Clugston, Anna Dominiczak and Alex Alderton, Roberto Amato, Jeffrey Barrett, Sonia Goncalves, Ewan Harrison, David K. Jackson, Ian Johnston, Dominic Kwiatkowski, Cordelia Langford, John Sillitoe on behalf of the Wellcome Sanger Institute COVID-19 Surveillance Team |
| EPI_ISL_3445036 | Lighthouse Lab in Milton Keynes | Wellcome Sanger Institute for the COVID-19 Genomics UK (COG-UK) Consortium | The Lighthouse Lab in Milton Keynes and Alex Alderton, Roberto Amato, Jeffrey Barrett, Sonia Goncalves, Ewan Harrison, David K. Jackson, Ian Johnston, Dominic Kwiatkowski, Cordelia Langford, John Sillitoe on behalf of the Wellcome Sanger Institute COVID-19 Surveillance Team |
| EPI_ISL_3445037, EPI_ISL_3445038, EPI_ISL_3445039, EPI_ISL_3445040, | Lighthouse Lab in Glasgow | Wellcome Sanger Institute for the COVID-19 Genomics UK (COG-UK) Consortium | Harper VanSteenhouse, Yumi Kasai, David Gray, Carol Clugston, Anna Dominiczak and Alex Alderton, Roberto Amato, Jeffrey Barrett, Sonia Goncalves, Ewan Harrison, David K. Jackson, Ian Johnston, Dominic Kwiatkowski, Cordelia Langford, John Sillitoe on behalf of the Wellcome Sanger Institute COVID-19 Surveillance Team |

| | | | |
|---|---|---|---|
| EPI_ISL_3445041, EPI_ISL_3445042, EPI_ISL_3445043, EPI_ISL_3445044, EPI_ISL_3445045 | | | |
| EPI_ISL_3445046 | Lighthouse Lab in Milton Keynes | Wellcome Sanger Institute for the COVID-19 Genomics UK (COG-UK) Consortium | The Lighthouse Lab in Milton Keynes and Alex Alderton, Roberto Amato, Jeffrey Barrett, Sonia Goncalves, Ewan Harrison, David K. Jackson, Ian Johnston, Dominic Kwiatkowski, Cordelia Langford, John Sillitoe on behalf of the Wellcome Sanger Institute COVID-19 Surveillance Team |
| EPI_ISL_3445047, EPI_ISL_3445048, EPI_ISL_3445049, EPI_ISL_3445050 | Lighthouse Lab in Glasgow | Wellcome Sanger Institute for the COVID-19 Genomics UK (COG-UK) Consortium | Harper VanSteenhouse, Yumi Kasai, David Gray, Carol Clugston, Anna Dominiczak and Alex Alderton, Roberto Amato, Jeffrey Barrett, Sonia Goncalves, Ewan Harrison, David K. Jackson, Ian Johnston, Dominic Kwiatkowski, Cordelia Langford, John Sillitoe on behalf of the Wellcome Sanger Institute COVID-19 Surveillance Team |
| EPI_ISL_3445051 | Lighthouse Lab in Milton Keynes | Wellcome Sanger Institute for the COVID-19 Genomics UK (COG-UK) Consortium | The Lighthouse Lab in Milton Keynes and Alex Alderton, Roberto Amato, Jeffrey Barrett, Sonia Goncalves, Ewan Harrison, David K. Jackson, Ian Johnston, Dominic Kwiatkowski, Cordelia Langford, John Sillitoe on behalf of the Wellcome Sanger Institute COVID-19 Surveillance Team |
| EPI_ISL_3445052, EPI_ISL_3445053, EPI_ISL_3445054, EPI_ISL_3445055, EPI_ISL_3445056, EPI_ISL_3445057, EPI_ISL_3445058 | Lighthouse Lab in Glasgow | Wellcome Sanger Institute for the COVID-19 Genomics UK (COG-UK) Consortium | Harper VanSteenhouse, Yumi Kasai, David Gray, Carol Clugston, Anna Dominiczak and Alex Alderton, Roberto Amato, Jeffrey Barrett, Sonia Goncalves, Ewan Harrison, David K. Jackson, Ian Johnston, Dominic Kwiatkowski, Cordelia Langford, John Sillitoe on behalf of the Wellcome Sanger Institute COVID-19 Surveillance Team |
| EPI_ISL_3445059 | Lighthouse Lab in Milton Keynes | Wellcome Sanger Institute for the COVID-19 Genomics UK (COG-UK) Consortium | The Lighthouse Lab in Milton Keynes and Alex Alderton, Roberto Amato, Jeffrey Barrett, Sonia Goncalves, Ewan Harrison, David K. Jackson, Ian Johnston, Dominic Kwiatkowski, Cordelia Langford, John Sillitoe on behalf of the Wellcome Sanger Institute COVID-19 Surveillance Team |
| EPI_ISL_3445060, EPI_ISL_3445061, EPI_ISL_3445062, EPI_ISL_3445063, EPI_ISL_3445064, EPI_ISL_3445065 | Lighthouse Lab in Glasgow | Wellcome Sanger Institute for the COVID-19 Genomics UK (COG-UK) Consortium | Harper VanSteenhouse, Yumi Kasai, David Gray, Carol Clugston, Anna Dominiczak and Alex Alderton, Roberto Amato, Jeffrey Barrett, Sonia Goncalves, Ewan Harrison, David K. Jackson, Ian Johnston, Dominic Kwiatkowski, Cordelia Langford, John Sillitoe on behalf of the Wellcome Sanger Institute COVID-19 Surveillance Team |
| EPI_ISL_3445066 | Lighthouse Lab in Milton Keynes | Wellcome Sanger Institute for the COVID-19 Genomics UK (COG-UK) Consortium | The Lighthouse Lab in Milton Keynes and Alex Alderton, Roberto Amato, Jeffrey Barrett, Sonia Goncalves, Ewan Harrison, David K. Jackson, Ian Johnston, Dominic Kwiatkowski, Cordelia Langford, John Sillitoe on behalf of the Wellcome Sanger Institute COVID-19 Surveillance Team |
| EPI_ISL_3445067 | Lighthouse Lab in Glasgow | Wellcome Sanger Institute for the COVID-19 Genomics UK (COG-UK) Consortium | Harper VanSteenhouse, Yumi Kasai, David Gray, Carol Clugston, Anna Dominiczak and Alex Alderton, Roberto Amato, Jeffrey Barrett, Sonia Goncalves, Ewan Harrison, David K. Jackson, Ian Johnston, Dominic Kwiatkowski, Cordelia Langford, John Sillitoe on behalf of the Wellcome Sanger Institute COVID-19 Surveillance Team |
| EPI_ISL_3445068 | Lighthouse Lab in Milton Keynes | Wellcome Sanger Institute for the COVID-19 Genomics UK (COG-UK) Consortium | The Lighthouse Lab in Milton Keynes and Alex Alderton, Roberto Amato, Jeffrey Barrett, Sonia Goncalves, Ewan Harrison, David K. Jackson, Ian Johnston, Dominic Kwiatkowski, Cordelia Langford, John Sillitoe on behalf of the Wellcome Sanger Institute COVID-19 Surveillance Team |
| EPI_ISL_3445069 | Lighthouse Lab in Glasgow | Wellcome Sanger Institute for the COVID-19 Genomics UK (COG-UK) Consortium | Harper VanSteenhouse, Yumi Kasai, David Gray, Carol Clugston, Anna Dominiczak and Alex Alderton, Roberto Amato, Jeffrey Barrett, Sonia Goncalves, Ewan Harrison, David K. Jackson, Ian Johnston, Dominic Kwiatkowski, Cordelia Langford, John Sillitoe on behalf of the Wellcome Sanger Institute COVID-19 Surveillance Team |
| EPI_ISL_3445070, EPI_ISL_3445071, EPI_ISL_3445072 | Lighthouse Lab in Milton Keynes | Wellcome Sanger Institute for the COVID-19 Genomics UK (COG-UK) Consortium | The Lighthouse Lab in Milton Keynes and Alex Alderton, Roberto Amato, Jeffrey Barrett, Sonia Goncalves, Ewan Harrison, David K. Jackson, Ian Johnston, Dominic Kwiatkowski, Cordelia Langford, John Sillitoe on behalf of the Wellcome Sanger Institute COVID-19 Surveillance Team |
| EPI_ISL_3445073, EPI_ISL_3445074, EPI_ISL_3445075, EPI_ISL_3445076, EPI_ISL_3445077, EPI_ISL_3445078, EPI_ISL_3445079, EPI_ISL_3445080 | Lighthouse Lab in Glasgow | Wellcome Sanger Institute for the COVID-19 Genomics UK (COG-UK) Consortium | Harper VanSteenhouse, Yumi Kasai, David Gray, Carol Clugston, Anna Dominiczak and Alex Alderton, Roberto Amato, Jeffrey Barrett, Sonia Goncalves, Ewan Harrison, David K. Jackson, Ian Johnston, Dominic Kwiatkowski, Cordelia Langford, John Sillitoe on behalf of the Wellcome Sanger Institute COVID-19 Surveillance Team |
| EPI_ISL_3445081 | Lighthouse Lab in Milton Keynes | Wellcome Sanger Institute for the COVID-19 Genomics UK (COG-UK) Consortium | The Lighthouse Lab in Milton Keynes and Alex Alderton, Roberto Amato, Jeffrey Barrett, Sonia Goncalves, Ewan Harrison, David K. Jackson, Ian Johnston, Dominic Kwiatkowski, Cordelia Langford, John Sillitoe on behalf of the Wellcome Sanger Institute COVID-19 Surveillance Team |
| EPI_ISL_3445082 | Lighthouse Lab in Glasgow | Wellcome Sanger Institute for the COVID-19 Genomics UK (COG-UK) Consortium | Harper VanSteenhouse, Yumi Kasai, David Gray, Carol Clugston, Anna Dominiczak and Alex Alderton, Roberto Amato, Jeffrey Barrett, Sonia Goncalves, Ewan Harrison, David K. Jackson, Ian Johnston, Dominic Kwiatkowski, Cordelia Langford, John Sillitoe on behalf of the Wellcome Sanger Institute COVID-19 Surveillance Team |
| EPI_ISL_3445083, EPI_ISL_3445084, EPI_ISL_3445085 | Lighthouse Lab in Milton Keynes | Wellcome Sanger Institute for the COVID-19 Genomics UK (COG-UK) Consortium | The Lighthouse Lab in Milton Keynes and Alex Alderton, Roberto Amato, Jeffrey Barrett, Sonia Goncalves, Ewan Harrison, David K. Jackson, Ian Johnston, Dominic Kwiatkowski, Cordelia Langford, John Sillitoe on behalf of the Wellcome Sanger Institute COVID-19 Surveillance Team |
| EPI_ISL_3445086, EPI_ISL_3445087, EPI_ISL_3445088, EPI_ISL_3445089, EPI_ISL_3445090, EPI_ISL_3445091 | Lighthouse Lab in Glasgow | Wellcome Sanger Institute for the COVID-19 Genomics UK (COG-UK) Consortium | Harper VanSteenhouse, Yumi Kasai, David Gray, Carol Clugston, Anna Dominiczak and Alex Alderton, Roberto Amato, Jeffrey Barrett, Sonia Goncalves, Ewan Harrison, David K. Jackson, Ian Johnston, Dominic Kwiatkowski, Cordelia Langford, John Sillitoe on behalf of the Wellcome Sanger Institute COVID-19 Surveillance Team |
| EPI_ISL_3445092 | Lighthouse Lab in Milton Keynes | Wellcome Sanger Institute for the COVID-19 Genomics UK (COG-UK) Consortium | The Lighthouse Lab in Milton Keynes and Alex Alderton, Roberto Amato, Jeffrey Barrett, Sonia Goncalves, Ewan Harrison, David K. Jackson, Ian Johnston, Dominic Kwiatkowski, Cordelia Langford, John Sillitoe on behalf of the Wellcome Sanger Institute COVID-19 Surveillance Team |
| EPI_ISL_3445093, EPI_ISL_3445094, EPI_ISL_3445095 | Lighthouse Lab in Glasgow | Wellcome Sanger Institute for the COVID-19 Genomics UK (COG-UK) Consortium | Harper VanSteenhouse, Yumi Kasai, David Gray, Carol Clugston, Anna Dominiczak and Alex Alderton, Roberto Amato, Jeffrey Barrett, Sonia Goncalves, Ewan Harrison, David K. Jackson, Ian Johnston, Dominic Kwiatkowski, Cordelia Langford, John Sillitoe on behalf of the Wellcome Sanger Institute COVID-19 Surveillance Team |
| EPI_ISL_3445096 | Lighthouse Lab in Milton Keynes | Wellcome Sanger Institute for the COVID-19 Genomics UK (COG-UK) Consortium | The Lighthouse Lab in Milton Keynes and Alex Alderton, Roberto Amato, Jeffrey Barrett, Sonia Goncalves, Ewan Harrison, David K. Jackson, Ian Johnston, Dominic Kwiatkowski, Cordelia Langford, John Sillitoe on behalf of the Wellcome Sanger Institute COVID-19 Surveillance Team |
| EPI_ISL_3445097 | Lighthouse Lab in Glasgow | Wellcome Sanger Institute for the COVID-19 Genomics UK (COG-UK) Consortium | Harper VanSteenhouse, Yumi Kasai, David Gray, Carol Clugston, Anna Dominiczak and Alex Alderton, Roberto Amato, Jeffrey Barrett, Sonia Goncalves, Ewan Harrison, David K. Jackson, Ian Johnston, Dominic Kwiatkowski, Cordelia Langford, John Sillitoe on behalf of the Wellcome Sanger Institute COVID-19 Surveillance Team |
| EPI_ISL_3445098 | Lighthouse Lab in Milton Keynes | Wellcome Sanger Institute for the COVID-19 Genomics UK (COG-UK) Consortium | The Lighthouse Lab in Milton Keynes and Alex Alderton, Roberto Amato, Jeffrey Barrett, Sonia Goncalves, Ewan Harrison, David K. Jackson, Ian Johnston, Dominic Kwiatkowski, Cordelia Langford, John Sillitoe on behalf of the Wellcome Sanger Institute COVID-19 Surveillance Team |
| EPI_ISL_3445099 | Lighthouse Lab in Glasgow | Wellcome Sanger Institute for the COVID-19 Genomics UK (COG-UK) Consortium | Harper VanSteenhouse, Yumi Kasai, David Gray, Carol Clugston, Anna Dominiczak and Alex Alderton, Roberto Amato, Jeffrey Barrett, Sonia Goncalves, Ewan Harrison, David K. Jackson, Ian Johnston, Dominic Kwiatkowski, Cordelia Langford, John Sillitoe on behalf of the Wellcome Sanger Institute COVID-19 Surveillance Team |
| EPI_ISL_3445100 | Lighthouse Lab in Milton Keynes | Wellcome Sanger Institute for the COVID-19 Genomics UK (COG-UK) Consortium | The Lighthouse Lab in Milton Keynes and Alex Alderton, Roberto Amato, Jeffrey Barrett, Sonia Goncalves, Ewan Harrison, David K. Jackson, Ian Johnston, Dominic Kwiatkowski, Cordelia Langford, John Sillitoe on behalf of the Wellcome Sanger Institute COVID-19 Surveillance Team |
| EPI_ISL_3445101, EPI_ISL_3445102 | Lighthouse Lab in Glasgow | Wellcome Sanger Institute for the COVID-19 Genomics UK (COG-UK) Consortium | Harper VanSteenhouse, Yumi Kasai, David Gray, Carol Clugston, Anna Dominiczak and Alex Alderton, Roberto Amato, Jeffrey Barrett, Sonia Goncalves, Ewan Harrison, David K. Jackson, Ian Johnston, Dominic Kwiatkowski, Cordelia Langford, John Sillitoe on behalf of the Wellcome Sanger Institute COVID-19 Surveillance Team |
| EPI_ISL_3445103, EPI_ISL_3445104, EPI_ISL_3445105 | Lighthouse Lab in Milton Keynes | Wellcome Sanger Institute for the COVID-19 Genomics UK (COG-UK) Consortium | The Lighthouse Lab in Milton Keynes and Alex Alderton, Roberto Amato, Jeffrey Barrett, Sonia Goncalves, Ewan Harrison, David K. Jackson, Ian Johnston, Dominic Kwiatkowski, Cordelia Langford, John Sillitoe on behalf of the Wellcome Sanger Institute COVID-19 Surveillance Team |
| EPI_ISL_3445106, EPI_ISL_3445107, EPI_ISL_3445108, EPI_ISL_3445109, EPI_ISL_3445110, EPI_ISL_3445111 | Lighthouse Lab in Glasgow | Wellcome Sanger Institute for the COVID-19 Genomics UK (COG-UK) Consortium | Harper VanSteenhouse, Yumi Kasai, David Gray, Carol Clugston, Anna Dominiczak and Alex Alderton, Roberto Amato, Jeffrey Barrett, Sonia Goncalves, Ewan Harrison, David K. Jackson, Ian Johnston, Dominic Kwiatkowski, Cordelia Langford, John Sillitoe on behalf of the Wellcome Sanger Institute COVID-19 Surveillance Team |
| EPI_ISL_3445112 | Lighthouse Lab in Milton Keynes | Wellcome Sanger Institute for the COVID-19 Genomics UK (COG-UK) Consortium | The Lighthouse Lab in Milton Keynes and Alex Alderton, Roberto Amato, Jeffrey Barrett, Sonia Goncalves, Ewan Harrison, David K. Jackson, Ian Johnston, Dominic Kwiatkowski, Cordelia Langford, John Sillitoe on behalf of the Wellcome Sanger Institute COVID-19 Surveillance Team |
| EPI_ISL_3445113, EPI_ISL_3445114, EPI_ISL_3445115, EPI_ISL_3445116 | Lighthouse Lab in Glasgow | Wellcome Sanger Institute for the COVID-19 Genomics UK (COG-UK) Consortium | Harper VanSteenhouse, Yumi Kasai, David Gray, Carol Clugston, Anna Dominiczak and Alex Alderton, Roberto Amato, Jeffrey Barrett, Sonia Goncalves, Ewan Harrison, David K. Jackson, Ian Johnston, Dominic Kwiatkowski, Cordelia Langford, John Sillitoe on behalf of the Wellcome Sanger Institute COVID-19 Surveillance Team |
| EPI_ISL_3445117, EPI_ISL_3445118, EPI_ISL_3445119, EPI_ISL_3445120 | Lighthouse Lab in Milton Keynes | Wellcome Sanger Institute for the COVID-19 Genomics UK (COG-UK) Consortium | The Lighthouse Lab in Milton Keynes and Alex Alderton, Roberto Amato, Jeffrey Barrett, Sonia Goncalves, Ewan Harrison, David K. Jackson, Ian Johnston, Dominic Kwiatkowski, Cordelia Langford, John Sillitoe on behalf of the Wellcome Sanger Institute COVID-19 Surveillance Team |
| EPI_ISL_3445121, | Lighthouse Lab in Glasgow | Wellcome Sanger Institute for the COVID-19 Genomics UK (COG-UK) | Harper VanSteenhouse, Yumi Kasai, David Gray, Carol Clugston, Anna Dominiczak and Alex Alderton, Roberto Amato, Jeffrey Barrett, Sonia Goncalves, Ewan Harrison, David K. Jackson, Ian Johnston, Dominic Kwiatkowski, Cordelia Langford, |

| | | | |
|---|---|---|---|
| EPI_ISL_3445122, EPI_ISL_3445123, EPI_ISL_3445124, EPI_ISL_3445125, EPI_ISL_3445126, EPI_ISL_3445127, EPI_ISL_3445128 | | Consortium | John Sillitoe on behalf of the Wellcome Sanger Institute COVID-19 Surveillance Team |
| EPI_ISL_3445129 | Lighthouse Lab in Milton Keynes | Wellcome Sanger Institute for the COVID-19 Genomics UK (COG-UK) Consortium | The Lighthouse Lab in Milton Keynes and Alex Alderton, Roberto Amato, Jeffrey Barrett, Sonia Goncalves, Ewan Harrison, David K. Jackson, Ian Johnston, Dominic Kwiatkowski, Cordelia Langford, John Sillitoe on behalf of the Wellcome Sanger Institute COVID-19 Surveillance Team |
| EPI_ISL_3445130, EPI_ISL_3445131, EPI_ISL_3445132 | Lighthouse Lab in Glasgow | Wellcome Sanger Institute for the COVID-19 Genomics UK (COG-UK) Consortium | Harper VanSteenhouse, Yumi Kasai, David Gray, Carol Clugston, Anna Dominiczak and Alex Alderton, Roberto Amato, Jeffrey Barrett, Sonia Goncalves, Ewan Harrison, David K. Jackson, Ian Johnston, Dominic Kwiatkowski, Cordelia Langford, John Sillitoe on behalf of the Wellcome Sanger Institute COVID-19 Surveillance Team |
| EPI_ISL_3445133 | Lighthouse Lab in Milton Keynes | Wellcome Sanger Institute for the COVID-19 Genomics UK (COG-UK) Consortium | The Lighthouse Lab in Milton Keynes and Alex Alderton, Roberto Amato, Jeffrey Barrett, Sonia Goncalves, Ewan Harrison, David K. Jackson, Ian Johnston, Dominic Kwiatkowski, Cordelia Langford, John Sillitoe on behalf of the Wellcome Sanger Institute COVID-19 Surveillance Team |
| EPI_ISL_3445134 | Lighthouse Lab in Glasgow | Wellcome Sanger Institute for the COVID-19 Genomics UK (COG-UK) Consortium | Harper VanSteenhouse, Yumi Kasai, David Gray, Carol Clugston, Anna Dominiczak and Alex Alderton, Roberto Amato, Jeffrey Barrett, Sonia Goncalves, Ewan Harrison, David K. Jackson, Ian Johnston, Dominic Kwiatkowski, Cordelia Langford, John Sillitoe on behalf of the Wellcome Sanger Institute COVID-19 Surveillance Team |
| EPI_ISL_3445135 | Berkshire and Surrey Pathology Services Lighthouse Laboratory | Wellcome Sanger Institute for the COVID-19 Genomics UK (COG-UK) Consortium | Berkshire and Surrey Pathology Services Lighthouse Laboratory and Alex Alderton, Roberto Amato, Jeffrey Barrett, Sonia Goncalves, Ewan Harrison, David K. Jackson, Ian Johnston, Dominic Kwiatkowski, Cordelia Langford, John Sillitoe on behalf of the Wellcome Sanger Institute COVID-19 Surveillance Team |
| EPI_ISL_3445136 | Lighthouse Laboratory Plymouth | Wellcome Sanger Institute for the COVID-19 Genomics UK (COG-UK) Consortium | Lighthouse Laboratory Plymouth and Alex Alderton, Roberto Amato, Jeffrey Barrett, Sonia Goncalves, Ewan Harrison, David K. Jackson, Ian Johnston, Dominic Kwiatkowski, Cordelia Langford, John Sillitoe on behalf of the Wellcome Sanger Institute COVID-19 Surveillance Team |
| EPI_ISL_3445137, EPI_ISL_3445138 | Berkshire and Surrey Pathology Services Lighthouse Laboratory | Wellcome Sanger Institute for the COVID-19 Genomics UK (COG-UK) Consortium | Berkshire and Surrey Pathology Services Lighthouse Laboratory and Alex Alderton, Roberto Amato, Jeffrey Barrett, Sonia Goncalves, Ewan Harrison, David K. Jackson, Ian Johnston, Dominic Kwiatkowski, Cordelia Langford, John Sillitoe on behalf of the Wellcome Sanger Institute COVID-19 Surveillance Team |
| EPI_ISL_3445139 | Lighthouse Laboratory Plymouth | Wellcome Sanger Institute for the COVID-19 Genomics UK (COG-UK) Consortium | Lighthouse Laboratory Plymouth and Alex Alderton, Roberto Amato, Jeffrey Barrett, Sonia Goncalves, Ewan Harrison, David K. Jackson, Ian Johnston, Dominic Kwiatkowski, Cordelia Langford, John Sillitoe on behalf of the Wellcome Sanger Institute COVID-19 Surveillance Team |
| EPI_ISL_3445140, EPI_ISL_3445141, EPI_ISL_3445142, EPI_ISL_3445143, EPI_ISL_3445144 | Berkshire and Surrey Pathology Services Lighthouse Laboratory | Wellcome Sanger Institute for the COVID-19 Genomics UK (COG-UK) Consortium | Berkshire and Surrey Pathology Services Lighthouse Laboratory and Alex Alderton, Roberto Amato, Jeffrey Barrett, Sonia Goncalves, Ewan Harrison, David K. Jackson, Ian Johnston, Dominic Kwiatkowski, Cordelia Langford, John Sillitoe on behalf of the Wellcome Sanger Institute COVID-19 Surveillance Team |
| EPI_ISL_3445145, EPI_ISL_3445146, EPI_ISL_3445147 | Lighthouse Laboratory Plymouth | Wellcome Sanger Institute for the COVID-19 Genomics UK (COG-UK) Consortium | Lighthouse Laboratory Plymouth and Alex Alderton, Roberto Amato, Jeffrey Barrett, Sonia Goncalves, Ewan Harrison, David K. Jackson, Ian Johnston, Dominic Kwiatkowski, Cordelia Langford, John Sillitoe on behalf of the Wellcome Sanger Institute COVID-19 Surveillance Team |
| EPI_ISL_3445148, EPI_ISL_3445149, EPI_ISL_3445150, EPI_ISL_3445151, EPI_ISL_3445152, EPI_ISL_3445153 | Berkshire and Surrey Pathology Services Lighthouse Laboratory | Wellcome Sanger Institute for the COVID-19 Genomics UK (COG-UK) Consortium | Berkshire and Surrey Pathology Services Lighthouse Laboratory and Alex Alderton, Roberto Amato, Jeffrey Barrett, Sonia Goncalves, Ewan Harrison, David K. Jackson, Ian Johnston, Dominic Kwiatkowski, Cordelia Langford, John Sillitoe on behalf of the Wellcome Sanger Institute COVID-19 Surveillance Team |
| EPI_ISL_3445154, EPI_ISL_3445155 | Lighthouse Laboratory Plymouth | Wellcome Sanger Institute for the COVID-19 Genomics UK (COG-UK) Consortium | Lighthouse Laboratory Plymouth and Alex Alderton, Roberto Amato, Jeffrey Barrett, Sonia Goncalves, Ewan Harrison, David K. Jackson, Ian Johnston, Dominic Kwiatkowski, Cordelia Langford, John Sillitoe on behalf of the Wellcome Sanger Institute COVID-19 Surveillance Team |
| EPI_ISL_3445156 | Berkshire and Surrey Pathology Services Lighthouse Laboratory | Wellcome Sanger Institute for the COVID-19 Genomics UK (COG-UK) Consortium | Berkshire and Surrey Pathology Services Lighthouse Laboratory and Alex Alderton, Roberto Amato, Jeffrey Barrett, Sonia Goncalves, Ewan Harrison, David K. Jackson, Ian Johnston, Dominic Kwiatkowski, Cordelia Langford, John Sillitoe on behalf of the Wellcome Sanger Institute COVID-19 Surveillance Team |
| EPI_ISL_3445157, EPI_ISL_3445158 | Lighthouse Laboratory Plymouth | Wellcome Sanger Institute for the COVID-19 Genomics UK (COG-UK) Consortium | Lighthouse Laboratory Plymouth and Alex Alderton, Roberto Amato, Jeffrey Barrett, Sonia Goncalves, Ewan Harrison, David K. Jackson, Ian Johnston, Dominic Kwiatkowski, Cordelia Langford, John Sillitoe on behalf of the Wellcome Sanger Institute COVID-19 Surveillance Team |
| EPI_ISL_3445159 | Berkshire and Surrey Pathology Services Lighthouse Laboratory | Wellcome Sanger Institute for the COVID-19 Genomics UK (COG-UK) Consortium | Berkshire and Surrey Pathology Services Lighthouse Laboratory and Alex Alderton, Roberto Amato, Jeffrey Barrett, Sonia Goncalves, Ewan Harrison, David K. Jackson, Ian Johnston, Dominic Kwiatkowski, Cordelia Langford, John Sillitoe on behalf of the Wellcome Sanger Institute COVID-19 Surveillance Team |
| EPI_ISL_3445160, EPI_ISL_3445161 | Lighthouse Laboratory Plymouth | Wellcome Sanger Institute for the COVID-19 Genomics UK (COG-UK) Consortium | Lighthouse Laboratory Plymouth and Alex Alderton, Roberto Amato, Jeffrey Barrett, Sonia Goncalves, Ewan Harrison, David K. Jackson, Ian Johnston, Dominic Kwiatkowski, Cordelia Langford, John Sillitoe on behalf of the Wellcome Sanger Institute COVID-19 Surveillance Team |
| EPI_ISL_3445162 | Berkshire and Surrey Pathology Services Lighthouse Laboratory | Wellcome Sanger Institute for the COVID-19 Genomics UK (COG-UK) Consortium | Berkshire and Surrey Pathology Services Lighthouse Laboratory and Alex Alderton, Roberto Amato, Jeffrey Barrett, Sonia Goncalves, Ewan Harrison, David K. Jackson, Ian Johnston, Dominic Kwiatkowski, Cordelia Langford, John Sillitoe on behalf of the Wellcome Sanger Institute COVID-19 Surveillance Team |
| EPI_ISL_3445163, EPI_ISL_3445164, EPI_ISL_3445165, EPI_ISL_3445166, EPI_ISL_3445167, EPI_ISL_3445168, EPI_ISL_3445169 | Lighthouse Laboratory Plymouth | Wellcome Sanger Institute for the COVID-19 Genomics UK (COG-UK) Consortium | Lighthouse Laboratory Plymouth and Alex Alderton, Roberto Amato, Jeffrey Barrett, Sonia Goncalves, Ewan Harrison, David K. Jackson, Ian Johnston, Dominic Kwiatkowski, Cordelia Langford, John Sillitoe on behalf of the Wellcome Sanger Institute COVID-19 Surveillance Team |
| EPI_ISL_3445170 | Berkshire and Surrey Pathology Services Lighthouse Laboratory | Wellcome Sanger Institute for the COVID-19 Genomics UK (COG-UK) Consortium | Berkshire and Surrey Pathology Services Lighthouse Laboratory and Alex Alderton, Roberto Amato, Jeffrey Barrett, Sonia Goncalves, Ewan Harrison, David K. Jackson, Ian Johnston, Dominic Kwiatkowski, Cordelia Langford, John Sillitoe on behalf of the Wellcome Sanger Institute COVID-19 Surveillance Team |
| EPI_ISL_3445171, EPI_ISL_3445172, EPI_ISL_3445173 | Lighthouse Laboratory Plymouth | Wellcome Sanger Institute for the COVID-19 Genomics UK (COG-UK) Consortium | Lighthouse Laboratory Plymouth and Alex Alderton, Roberto Amato, Jeffrey Barrett, Sonia Goncalves, Ewan Harrison, David K. Jackson, Ian Johnston, Dominic Kwiatkowski, Cordelia Langford, John Sillitoe on behalf of the Wellcome Sanger Institute COVID-19 Surveillance Team |
| EPI_ISL_3445174, EPI_ISL_3445175 | Berkshire and Surrey Pathology Services Lighthouse Laboratory | Wellcome Sanger Institute for the COVID-19 Genomics UK (COG-UK) Consortium | Berkshire and Surrey Pathology Services Lighthouse Laboratory and Alex Alderton, Roberto Amato, Jeffrey Barrett, Sonia Goncalves, Ewan Harrison, David K. Jackson, Ian Johnston, Dominic Kwiatkowski, Cordelia Langford, John Sillitoe on behalf of the Wellcome Sanger Institute COVID-19 Surveillance Team |
| EPI_ISL_3445176 | Lighthouse Laboratory Plymouth | Wellcome Sanger Institute for the COVID-19 Genomics UK (COG-UK) Consortium | Lighthouse Laboratory Plymouth and Alex Alderton, Roberto Amato, Jeffrey Barrett, Sonia Goncalves, Ewan Harrison, David K. Jackson, Ian Johnston, Dominic Kwiatkowski, Cordelia Langford, John Sillitoe on behalf of the Wellcome Sanger Institute COVID-19 Surveillance Team |
| EPI_ISL_3445177 | Berkshire and Surrey Pathology Services Lighthouse Laboratory | Wellcome Sanger Institute for the COVID-19 Genomics UK (COG-UK) Consortium | Berkshire and Surrey Pathology Services Lighthouse Laboratory and Alex Alderton, Roberto Amato, Jeffrey Barrett, Sonia Goncalves, Ewan Harrison, David K. Jackson, Ian Johnston, Dominic Kwiatkowski, Cordelia Langford, John Sillitoe on behalf of the Wellcome Sanger Institute COVID-19 Surveillance Team |
| EPI_ISL_3445178, EPI_ISL_3445179 | Lighthouse Laboratory Plymouth | Wellcome Sanger Institute for the COVID-19 Genomics UK (COG-UK) Consortium | Lighthouse Laboratory Plymouth and Alex Alderton, Roberto Amato, Jeffrey Barrett, Sonia Goncalves, Ewan Harrison, David K. Jackson, Ian Johnston, Dominic Kwiatkowski, Cordelia Langford, John Sillitoe on behalf of the Wellcome Sanger Institute COVID-19 Surveillance Team |
| EPI_ISL_3445180, EPI_ISL_3445181, EPI_ISL_3445182, EPI_ISL_3445183 | Berkshire and Surrey Pathology Services Lighthouse Laboratory | Wellcome Sanger Institute for the COVID-19 Genomics UK (COG-UK) Consortium | Berkshire and Surrey Pathology Services Lighthouse Laboratory and Alex Alderton, Roberto Amato, Jeffrey Barrett, Sonia Goncalves, Ewan Harrison, David K. Jackson, Ian Johnston, Dominic Kwiatkowski, Cordelia Langford, John Sillitoe on behalf of the Wellcome Sanger Institute COVID-19 Surveillance Team |
| EPI_ISL_3445184 | Lighthouse Laboratory Plymouth | Wellcome Sanger Institute for the COVID-19 Genomics UK (COG-UK) Consortium | Lighthouse Laboratory Plymouth and Alex Alderton, Roberto Amato, Jeffrey Barrett, Sonia Goncalves, Ewan Harrison, David K. Jackson, Ian Johnston, Dominic Kwiatkowski, Cordelia Langford, John Sillitoe on behalf of the Wellcome Sanger Institute COVID-19 Surveillance Team |
| EPI_ISL_3445185, EPI_ISL_3445186 | Berkshire and Surrey Pathology Services Lighthouse Laboratory | Wellcome Sanger Institute for the COVID-19 Genomics UK (COG-UK) Consortium | Berkshire and Surrey Pathology Services Lighthouse Laboratory and Alex Alderton, Roberto Amato, Jeffrey Barrett, Sonia Goncalves, Ewan Harrison, David K. Jackson, Ian Johnston, Dominic Kwiatkowski, Cordelia Langford, John Sillitoe on behalf of the Wellcome Sanger Institute COVID-19 Surveillance Team |
| EPI_ISL_3445187 | Lighthouse Laboratory Plymouth | Wellcome Sanger Institute for the COVID-19 Genomics UK (COG-UK) Consortium | Lighthouse Laboratory Plymouth and Alex Alderton, Roberto Amato, Jeffrey Barrett, Sonia Goncalves, Ewan Harrison, David K. Jackson, Ian Johnston, Dominic Kwiatkowski, Cordelia Langford, John Sillitoe on behalf of the Wellcome Sanger Institute COVID-19 Surveillance Team |
| EPI_ISL_3445188 | Berkshire and Surrey Pathology Services Lighthouse Laboratory | Wellcome Sanger Institute for the COVID-19 Genomics UK (COG-UK) Consortium | Berkshire and Surrey Pathology Services Lighthouse Laboratory and Alex Alderton, Roberto Amato, Jeffrey Barrett, Sonia Goncalves, Ewan Harrison, David K. Jackson, Ian Johnston, Dominic Kwiatkowski, Cordelia Langford, John Sillitoe on behalf of the Wellcome Sanger Institute COVID-19 Surveillance Team |
| EPI_ISL_3445189, EPI_ISL_3445190 | Lighthouse Laboratory Plymouth | Wellcome Sanger Institute for the COVID-19 Genomics UK (COG-UK) Consortium | Lighthouse Laboratory Plymouth and Alex Alderton, Roberto Amato, Jeffrey Barrett, Sonia Goncalves, Ewan Harrison, David K. Jackson, Ian Johnston, Dominic Kwiatkowski, Cordelia Langford, John Sillitoe on behalf of the Wellcome Sanger Institute COVID-19 Surveillance Team |
| EPI_ISL_3445191, EPI_ISL_3445192 | Berkshire and Surrey Pathology Services Lighthouse Laboratory | Wellcome Sanger Institute for the COVID-19 Genomics UK (COG-UK) Consortium | Berkshire and Surrey Pathology Services Lighthouse Laboratory and Alex Alderton, Roberto Amato, Jeffrey Barrett, Sonia Goncalves, Ewan Harrison, David K. Jackson, Ian Johnston, Dominic Kwiatkowski, Cordelia Langford, John Sillitoe on behalf of the Wellcome Sanger Institute COVID-19 Surveillance Team |
| EPI_ISL_3445193, EPI_ISL_3445194, EPI_ISL_3445195 | Lighthouse Laboratory Plymouth | Wellcome Sanger Institute for the COVID-19 Genomics UK (COG-UK) Consortium | Lighthouse Laboratory Plymouth and Alex Alderton, Roberto Amato, Jeffrey Barrett, Sonia Goncalves, Ewan Harrison, David K. Jackson, Ian Johnston, Dominic Kwiatkowski, Cordelia Langford, John Sillitoe on behalf of the Wellcome Sanger Institute COVID-19 Surveillance Team |
| EPI_ISL_3445196 | Berkshire and Surrey Pathology Services Lighthouse Laboratory | Wellcome Sanger Institute for the COVID-19 Genomics UK (COG-UK) Consortium | Berkshire and Surrey Pathology Services Lighthouse Laboratory and Alex Alderton, Roberto Amato, Jeffrey Barrett, Sonia Goncalves, Ewan Harrison, David K. Jackson, Ian Johnston, Dominic Kwiatkowski, Cordelia Langford, John Sillitoe on behalf of the Wellcome Sanger Institute COVID-19 Surveillance Team |
| EPI_ISL_3445197, EPI_ISL_3445198 | Lighthouse Laboratory Plymouth | Wellcome Sanger Institute for the COVID-19 Genomics UK (COG-UK) Consortium | Lighthouse Laboratory Plymouth and Alex Alderton, Roberto Amato, Jeffrey Barrett, Sonia Goncalves, Ewan Harrison, David K. Jackson, Ian Johnston, Dominic Kwiatkowski, Cordelia Langford, John Sillitoe on behalf of the Wellcome Sanger Institute COVID-19 Surveillance Team |
| EPI_ISL_3445199 | Berkshire and Surrey Pathology Services Lighthouse Laboratory | Wellcome Sanger Institute for the COVID-19 Genomics UK (COG-UK) | Berkshire and Surrey Pathology Services Lighthouse Laboratory and Alex Alderton, Roberto Amato, Jeffrey Barrett, Sonia Goncalves, Ewan Harrison, David K. Jackson, Ian Johnston, Dominic Kwiatkowski, Cordelia Langford, John Sillitoe on behalf |

| | | | |
|---|---|---|---|
| | | Consortium | of the Wellcome Sanger Institute COVID-19 Surveillance Team |
| EPI_ISL_3445200, EPI_ISL_3445201 | Lighthouse Laboratory Plymouth | Wellcome Sanger Institute for the COVID-19 Genomics UK (COG-UK) Consortium | Lighthouse Laboratory Plymouth and Alex Alderton, Roberto Amato, Jeffrey Barrett, Sonia Goncalves, Ewan Harrison, David K. Jackson, Ian Johnston, Dominic Kwiatkowski, Cordelia Langford, John Sillitoe on behalf of the Wellcome Sanger Institute COVID-19 Surveillance Team |
| EPI_ISL_3445202, EPI_ISL_3445203 | Berkshire and Surrey Pathology Services Lighthouse Laboratory | Wellcome Sanger Institute for the COVID-19 Genomics UK (COG-UK) Consortium | Berkshire and Surrey Pathology Services Lighthouse Laboratory and Alex Alderton, Roberto Amato, Jeffrey Barrett, Sonia Goncalves, Ewan Harrison, David K. Jackson, Ian Johnston, Dominic Kwiatkowski, Cordelia Langford, John Sillitoe on behalf of the Wellcome Sanger Institute COVID-19 Surveillance Team |
| EPI_ISL_3445204 | Lighthouse Laboratory Plymouth | Wellcome Sanger Institute for the COVID-19 Genomics UK (COG-UK) Consortium | Lighthouse Laboratory Plymouth and Alex Alderton, Roberto Amato, Jeffrey Barrett, Sonia Goncalves, Ewan Harrison, David K. Jackson, Ian Johnston, Dominic Kwiatkowski, Cordelia Langford, John Sillitoe on behalf of the Wellcome Sanger Institute COVID-19 Surveillance Team |
| EPI_ISL_3445205 | Berkshire and Surrey Pathology Services Lighthouse Laboratory | Wellcome Sanger Institute for the COVID-19 Genomics UK (COG-UK) Consortium | Berkshire and Surrey Pathology Services Lighthouse Laboratory and Alex Alderton, Roberto Amato, Jeffrey Barrett, Sonia Goncalves, Ewan Harrison, David K. Jackson, Ian Johnston, Dominic Kwiatkowski, Cordelia Langford, John Sillitoe on behalf of the Wellcome Sanger Institute COVID-19 Surveillance Team |
| EPI_ISL_3445206 | Lighthouse Laboratory Plymouth | Wellcome Sanger Institute for the COVID-19 Genomics UK (COG-UK) Consortium | Lighthouse Laboratory Plymouth and Alex Alderton, Roberto Amato, Jeffrey Barrett, Sonia Goncalves, Ewan Harrison, David K. Jackson, Ian Johnston, Dominic Kwiatkowski, Cordelia Langford, John Sillitoe on behalf of the Wellcome Sanger Institute COVID-19 Surveillance Team |
| EPI_ISL_3445207 | Berkshire and Surrey Pathology Services Lighthouse Laboratory | Wellcome Sanger Institute for the COVID-19 Genomics UK (COG-UK) Consortium | Berkshire and Surrey Pathology Services Lighthouse Laboratory and Alex Alderton, Roberto Amato, Jeffrey Barrett, Sonia Goncalves, Ewan Harrison, David K. Jackson, Ian Johnston, Dominic Kwiatkowski, Cordelia Langford, John Sillitoe on behalf of the Wellcome Sanger Institute COVID-19 Surveillance Team |
| EPI_ISL_3445208, EPI_ISL_3445209, EPI_ISL_3445210, EPI_ISL_3445211, EPI_ISL_3445212, EPI_ISL_3445213 | Lighthouse Laboratory Plymouth | Wellcome Sanger Institute for the COVID-19 Genomics UK (COG-UK) Consortium | Lighthouse Laboratory Plymouth and Alex Alderton, Roberto Amato, Jeffrey Barrett, Sonia Goncalves, Ewan Harrison, David K. Jackson, Ian Johnston, Dominic Kwiatkowski, Cordelia Langford, John Sillitoe on behalf of the Wellcome Sanger Institute COVID-19 Surveillance Team |
| EPI_ISL_3445214 | Berkshire and Surrey Pathology Services Lighthouse Laboratory | Wellcome Sanger Institute for the COVID-19 Genomics UK (COG-UK) Consortium | Berkshire and Surrey Pathology Services Lighthouse Laboratory and Alex Alderton, Roberto Amato, Jeffrey Barrett, Sonia Goncalves, Ewan Harrison, David K. Jackson, Ian Johnston, Dominic Kwiatkowski, Cordelia Langford, John Sillitoe on behalf of the Wellcome Sanger Institute COVID-19 Surveillance Team |
| EPI_ISL_3445215 | Lighthouse Laboratory Plymouth | Wellcome Sanger Institute for the COVID-19 Genomics UK (COG-UK) Consortium | Lighthouse Laboratory Plymouth and Alex Alderton, Roberto Amato, Jeffrey Barrett, Sonia Goncalves, Ewan Harrison, David K. Jackson, Ian Johnston, Dominic Kwiatkowski, Cordelia Langford, John Sillitoe on behalf of the Wellcome Sanger Institute COVID-19 Surveillance Team |
| EPI_ISL_3445216, EPI_ISL_3445217 | Berkshire and Surrey Pathology Services Lighthouse Laboratory | Wellcome Sanger Institute for the COVID-19 Genomics UK (COG-UK) Consortium | Berkshire and Surrey Pathology Services Lighthouse Laboratory and Alex Alderton, Roberto Amato, Jeffrey Barrett, Sonia Goncalves, Ewan Harrison, David K. Jackson, Ian Johnston, Dominic Kwiatkowski, Cordelia Langford, John Sillitoe on behalf of the Wellcome Sanger Institute COVID-19 Surveillance Team |
| EPI_ISL_3445218 | Lighthouse Laboratory Plymouth | Wellcome Sanger Institute for the COVID-19 Genomics UK (COG-UK) Consortium | Lighthouse Laboratory Plymouth and Alex Alderton, Roberto Amato, Jeffrey Barrett, Sonia Goncalves, Ewan Harrison, David K. Jackson, Ian Johnston, Dominic Kwiatkowski, Cordelia Langford, John Sillitoe on behalf of the Wellcome Sanger Institute COVID-19 Surveillance Team |
| EPI_ISL_3445219 | Berkshire and Surrey Pathology Services Lighthouse Laboratory | Wellcome Sanger Institute for the COVID-19 Genomics UK (COG-UK) Consortium | Berkshire and Surrey Pathology Services Lighthouse Laboratory and Alex Alderton, Roberto Amato, Jeffrey Barrett, Sonia Goncalves, Ewan Harrison, David K. Jackson, Ian Johnston, Dominic Kwiatkowski, Cordelia Langford, John Sillitoe on behalf of the Wellcome Sanger Institute COVID-19 Surveillance Team |
| EPI_ISL_3445220 | Lighthouse Laboratory Plymouth | Wellcome Sanger Institute for the COVID-19 Genomics UK (COG-UK) Consortium | Lighthouse Laboratory Plymouth and Alex Alderton, Roberto Amato, Jeffrey Barrett, Sonia Goncalves, Ewan Harrison, David K. Jackson, Ian Johnston, Dominic Kwiatkowski, Cordelia Langford, John Sillitoe on behalf of the Wellcome Sanger Institute COVID-19 Surveillance Team |
| EPI_ISL_3445221 | Berkshire and Surrey Pathology Services Lighthouse Laboratory | Wellcome Sanger Institute for the COVID-19 Genomics UK (COG-UK) Consortium | Berkshire and Surrey Pathology Services Lighthouse Laboratory and Alex Alderton, Roberto Amato, Jeffrey Barrett, Sonia Goncalves, Ewan Harrison, David K. Jackson, Ian Johnston, Dominic Kwiatkowski, Cordelia Langford, John Sillitoe on behalf of the Wellcome Sanger Institute COVID-19 Surveillance Team |
| EPI_ISL_3445222 | Lighthouse Laboratory Plymouth | Wellcome Sanger Institute for the COVID-19 Genomics UK (COG-UK) Consortium | Lighthouse Laboratory Plymouth and Alex Alderton, Roberto Amato, Jeffrey Barrett, Sonia Goncalves, Ewan Harrison, David K. Jackson, Ian Johnston, Dominic Kwiatkowski, Cordelia Langford, John Sillitoe on behalf of the Wellcome Sanger Institute COVID-19 Surveillance Team |
| EPI_ISL_3445223 | Berkshire and Surrey Pathology Services Lighthouse Laboratory | Wellcome Sanger Institute for the COVID-19 Genomics UK (COG-UK) Consortium | Berkshire and Surrey Pathology Services Lighthouse Laboratory and Alex Alderton, Roberto Amato, Jeffrey Barrett, Sonia Goncalves, Ewan Harrison, David K. Jackson, Ian Johnston, Dominic Kwiatkowski, Cordelia Langford, John Sillitoe on behalf of the Wellcome Sanger Institute COVID-19 Surveillance Team |
| EPI_ISL_3445224, EPI_ISL_3445225, EPI_ISL_3445226 | Lighthouse Laboratory Plymouth | Wellcome Sanger Institute for the COVID-19 Genomics UK (COG-UK) Consortium | Lighthouse Laboratory Plymouth and Alex Alderton, Roberto Amato, Jeffrey Barrett, Sonia Goncalves, Ewan Harrison, David K. Jackson, Ian Johnston, Dominic Kwiatkowski, Cordelia Langford, John Sillitoe on behalf of the Wellcome Sanger Institute COVID-19 Surveillance Team |
| EPI_ISL_3445227 | Berkshire and Surrey Pathology Services Lighthouse Laboratory | Wellcome Sanger Institute for the COVID-19 Genomics UK (COG-UK) Consortium | Berkshire and Surrey Pathology Services Lighthouse Laboratory and Alex Alderton, Roberto Amato, Jeffrey Barrett, Sonia Goncalves, Ewan Harrison, David K. Jackson, Ian Johnston, Dominic Kwiatkowski, Cordelia Langford, John Sillitoe on behalf of the Wellcome Sanger Institute COVID-19 Surveillance Team |
| EPI_ISL_3445228 | Lighthouse Laboratory Plymouth | Wellcome Sanger Institute for the COVID-19 Genomics UK (COG-UK) Consortium | Lighthouse Laboratory Plymouth and Alex Alderton, Roberto Amato, Jeffrey Barrett, Sonia Goncalves, Ewan Harrison, David K. Jackson, Ian Johnston, Dominic Kwiatkowski, Cordelia Langford, John Sillitoe on behalf of the Wellcome Sanger Institute COVID-19 Surveillance Team |
| EPI_ISL_3445229, EPI_ISL_3445230 | Berkshire and Surrey Pathology Services Lighthouse Laboratory | Wellcome Sanger Institute for the COVID-19 Genomics UK (COG-UK) Consortium | Berkshire and Surrey Pathology Services Lighthouse Laboratory and Alex Alderton, Roberto Amato, Jeffrey Barrett, Sonia Goncalves, Ewan Harrison, David K. Jackson, Ian Johnston, Dominic Kwiatkowski, Cordelia Langford, John Sillitoe on behalf of the Wellcome Sanger Institute COVID-19 Surveillance Team |
| EPI_ISL_3445231, EPI_ISL_3445232 | Lighthouse Laboratory Plymouth | Wellcome Sanger Institute for the COVID-19 Genomics UK (COG-UK) Consortium | Lighthouse Laboratory Plymouth and Alex Alderton, Roberto Amato, Jeffrey Barrett, Sonia Goncalves, Ewan Harrison, David K. Jackson, Ian Johnston, Dominic Kwiatkowski, Cordelia Langford, John Sillitoe on behalf of the Wellcome Sanger Institute COVID-19 Surveillance Team |
| EPI_ISL_3445233 | Berkshire and Surrey Pathology Services Lighthouse Laboratory | Wellcome Sanger Institute for the COVID-19 Genomics UK (COG-UK) Consortium | Berkshire and Surrey Pathology Services Lighthouse Laboratory and Alex Alderton, Roberto Amato, Jeffrey Barrett, Sonia Goncalves, Ewan Harrison, David K. Jackson, Ian Johnston, Dominic Kwiatkowski, Cordelia Langford, John Sillitoe on behalf of the Wellcome Sanger Institute COVID-19 Surveillance Team |
| EPI_ISL_3445234, EPI_ISL_3445235, EPI_ISL_3445236, EPI_ISL_3445237, EPI_ISL_3445238, EPI_ISL_3445239 | Lighthouse Laboratory Plymouth | Wellcome Sanger Institute for the COVID-19 Genomics UK (COG-UK) Consortium | Lighthouse Laboratory Plymouth and Alex Alderton, Roberto Amato, Jeffrey Barrett, Sonia Goncalves, Ewan Harrison, David K. Jackson, Ian Johnston, Dominic Kwiatkowski, Cordelia Langford, John Sillitoe on behalf of the Wellcome Sanger Institute COVID-19 Surveillance Team |
| EPI_ISL_3445240, EPI_ISL_3445241, EPI_ISL_3445242, EPI_ISL_3445243, EPI_ISL_3445244 | Berkshire and Surrey Pathology Services Lighthouse Laboratory | Wellcome Sanger Institute for the COVID-19 Genomics UK (COG-UK) Consortium | Berkshire and Surrey Pathology Services Lighthouse Laboratory and Alex Alderton, Roberto Amato, Jeffrey Barrett, Sonia Goncalves, Ewan Harrison, David K. Jackson, Ian Johnston, Dominic Kwiatkowski, Cordelia Langford, John Sillitoe on behalf of the Wellcome Sanger Institute COVID-19 Surveillance Team |
| EPI_ISL_3445245 | Lighthouse Laboratory Plymouth | Wellcome Sanger Institute for the COVID-19 Genomics UK (COG-UK) Consortium | Lighthouse Laboratory Plymouth and Alex Alderton, Roberto Amato, Jeffrey Barrett, Sonia Goncalves, Ewan Harrison, David K. Jackson, Ian Johnston, Dominic Kwiatkowski, Cordelia Langford, John Sillitoe on behalf of the Wellcome Sanger Institute COVID-19 Surveillance Team |
| EPI_ISL_3445246 | Berkshire and Surrey Pathology Services Lighthouse Laboratory | Wellcome Sanger Institute for the COVID-19 Genomics UK (COG-UK) Consortium | Berkshire and Surrey Pathology Services Lighthouse Laboratory and Alex Alderton, Roberto Amato, Jeffrey Barrett, Sonia Goncalves, Ewan Harrison, David K. Jackson, Ian Johnston, Dominic Kwiatkowski, Cordelia Langford, John Sillitoe on behalf of the Wellcome Sanger Institute COVID-19 Surveillance Team |
| EPI_ISL_3445247, EPI_ISL_3445248 | Lighthouse Laboratory Plymouth | Wellcome Sanger Institute for the COVID-19 Genomics UK (COG-UK) Consortium | Lighthouse Laboratory Plymouth and Alex Alderton, Roberto Amato, Jeffrey Barrett, Sonia Goncalves, Ewan Harrison, David K. Jackson, Ian Johnston, Dominic Kwiatkowski, Cordelia Langford, John Sillitoe on behalf of the Wellcome Sanger Institute COVID-19 Surveillance Team |
| EPI_ISL_3445249, EPI_ISL_3445250, EPI_ISL_3445251, EPI_ISL_3445252 | Berkshire and Surrey Pathology Services Lighthouse Laboratory | Wellcome Sanger Institute for the COVID-19 Genomics UK (COG-UK) Consortium | Berkshire and Surrey Pathology Services Lighthouse Laboratory and Alex Alderton, Roberto Amato, Jeffrey Barrett, Sonia Goncalves, Ewan Harrison, David K. Jackson, Ian Johnston, Dominic Kwiatkowski, Cordelia Langford, John Sillitoe on behalf of the Wellcome Sanger Institute COVID-19 Surveillance Team |
| EPI_ISL_3445253, EPI_ISL_3445254 | Lighthouse Laboratory Plymouth | Wellcome Sanger Institute for the COVID-19 Genomics UK (COG-UK) Consortium | Lighthouse Laboratory Plymouth and Alex Alderton, Roberto Amato, Jeffrey Barrett, Sonia Goncalves, Ewan Harrison, David K. Jackson, Ian Johnston, Dominic Kwiatkowski, Cordelia Langford, John Sillitoe on behalf of the Wellcome Sanger Institute COVID-19 Surveillance Team |
| EPI_ISL_3445255 | Berkshire and Surrey Pathology Services Lighthouse Laboratory | Wellcome Sanger Institute for the COVID-19 Genomics UK (COG-UK) Consortium | Berkshire and Surrey Pathology Services Lighthouse Laboratory and Alex Alderton, Roberto Amato, Jeffrey Barrett, Sonia Goncalves, Ewan Harrison, David K. Jackson, Ian Johnston, Dominic Kwiatkowski, Cordelia Langford, John Sillitoe on behalf of the Wellcome Sanger Institute COVID-19 Surveillance Team |
| EPI_ISL_3445256, EPI_ISL_3445257, EPI_ISL_3445258 | Lighthouse Laboratory Plymouth | Wellcome Sanger Institute for the COVID-19 Genomics UK (COG-UK) Consortium | Lighthouse Laboratory Plymouth and Alex Alderton, Roberto Amato, Jeffrey Barrett, Sonia Goncalves, Ewan Harrison, David K. Jackson, Ian Johnston, Dominic Kwiatkowski, Cordelia Langford, John Sillitoe on behalf of the Wellcome Sanger Institute COVID-19 Surveillance Team |
| EPI_ISL_3445259, EPI_ISL_3445260 | Berkshire and Surrey Pathology Services Lighthouse Laboratory | Wellcome Sanger Institute for the COVID-19 Genomics UK (COG-UK) Consortium | Berkshire and Surrey Pathology Services Lighthouse Laboratory and Alex Alderton, Roberto Amato, Jeffrey Barrett, Sonia Goncalves, Ewan Harrison, David K. Jackson, Ian Johnston, Dominic Kwiatkowski, Cordelia Langford, John Sillitoe on behalf of the Wellcome Sanger Institute COVID-19 Surveillance Team |
| EPI_ISL_3445261, EPI_ISL_3445262 | Lighthouse Laboratory Plymouth | Wellcome Sanger Institute for the COVID-19 Genomics UK (COG-UK) Consortium | Lighthouse Laboratory Plymouth and Alex Alderton, Roberto Amato, Jeffrey Barrett, Sonia Goncalves, Ewan Harrison, David K. Jackson, Ian Johnston, Dominic Kwiatkowski, Cordelia Langford, John Sillitoe on behalf of the Wellcome Sanger Institute COVID-19 Surveillance Team |
| EPI_ISL_3445263, EPI_ISL_3445264, EPI_ISL_3445265, EPI_ISL_3445266, EPI_ISL_3445267, EPI_ISL_3445268 | Berkshire and Surrey Pathology Services Lighthouse Laboratory | Wellcome Sanger Institute for the COVID-19 Genomics UK (COG-UK) Consortium | Berkshire and Surrey Pathology Services Lighthouse Laboratory and Alex Alderton, Roberto Amato, Jeffrey Barrett, Sonia Goncalves, Ewan Harrison, David K. Jackson, Ian Johnston, Dominic Kwiatkowski, Cordelia Langford, John Sillitoe on behalf of the Wellcome Sanger Institute COVID-19 Surveillance Team |
| EPI_ISL_3445269, EPI_ISL_3445270, EPI_ISL_3445271, EPI_ISL_3445272, EPI_ISL_3445273, EPI_ISL_3445274 | Lighthouse Laboratory Plymouth | Wellcome Sanger Institute for the COVID-19 Genomics UK (COG-UK) Consortium | Lighthouse Laboratory Plymouth and Alex Alderton, Roberto Amato, Jeffrey Barrett, Sonia Goncalves, Ewan Harrison, David K. Jackson, Ian Johnston, Dominic Kwiatkowski, Cordelia Langford, John Sillitoe on behalf of the Wellcome Sanger Institute COVID-19 Surveillance Team |

| | | | |
|---|---|---|---|
| EPI_ISL_3445275 | Berkshire and Surrey Pathology Services Lighthouse Laboratory | Wellcome Sanger Institute for the COVID-19 Genomics UK (COG-UK) Consortium | Berkshire and Surrey Pathology Services Lighthouse Laboratory and Alex Alderton, Roberto Amato, Jeffrey Barrett, Sonia Goncalves, Ewan Harrison, David K. Jackson, Ian Johnston, Dominic Kwiatkowski, Cordelia Langford, John Sillitoe on behalf of the Wellcome Sanger Institute COVID-19 Surveillance Team |
| EPI_ISL_3445276 | Lighthouse Laboratory Plymouth | Wellcome Sanger Institute for the COVID-19 Genomics UK (COG-UK) Consortium | Lighthouse Laboratory Plymouth and Alex Alderton, Roberto Amato, Jeffrey Barrett, Sonia Goncalves, Ewan Harrison, David K. Jackson, Ian Johnston, Dominic Kwiatkowski, Cordelia Langford, John Sillitoe on behalf of the Wellcome Sanger Institute COVID-19 Surveillance Team |
| EPI_ISL_3445277, EPI_ISL_3445278 | Berkshire and Surrey Pathology Services Lighthouse Laboratory | Wellcome Sanger Institute for the COVID-19 Genomics UK (COG-UK) Consortium | Berkshire and Surrey Pathology Services Lighthouse Laboratory and Alex Alderton, Roberto Amato, Jeffrey Barrett, Sonia Goncalves, Ewan Harrison, David K. Jackson, Ian Johnston, Dominic Kwiatkowski, Cordelia Langford, John Sillitoe on behalf of the Wellcome Sanger Institute COVID-19 Surveillance Team |
| EPI_ISL_3445279 | Lighthouse Laboratory Plymouth | Wellcome Sanger Institute for the COVID-19 Genomics UK (COG-UK) Consortium | Lighthouse Laboratory Plymouth and Alex Alderton, Roberto Amato, Jeffrey Barrett, Sonia Goncalves, Ewan Harrison, David K. Jackson, Ian Johnston, Dominic Kwiatkowski, Cordelia Langford, John Sillitoe on behalf of the Wellcome Sanger Institute COVID-19 Surveillance Team |
| EPI_ISL_3445280 | Berkshire and Surrey Pathology Services Lighthouse Laboratory | Wellcome Sanger Institute for the COVID-19 Genomics UK (COG-UK) Consortium | Berkshire and Surrey Pathology Services Lighthouse Laboratory and Alex Alderton, Roberto Amato, Jeffrey Barrett, Sonia Goncalves, Ewan Harrison, David K. Jackson, Ian Johnston, Dominic Kwiatkowski, Cordelia Langford, John Sillitoe on behalf of the Wellcome Sanger Institute COVID-19 Surveillance Team |
| EPI_ISL_3445281, EPI_ISL_3445282 | Lighthouse Laboratory Plymouth | Wellcome Sanger Institute for the COVID-19 Genomics UK (COG-UK) Consortium | Lighthouse Laboratory Plymouth and Alex Alderton, Roberto Amato, Jeffrey Barrett, Sonia Goncalves, Ewan Harrison, David K. Jackson, Ian Johnston, Dominic Kwiatkowski, Cordelia Langford, John Sillitoe on behalf of the Wellcome Sanger Institute COVID-19 Surveillance Team |
| EPI_ISL_3445283 | Berkshire and Surrey Pathology Services Lighthouse Laboratory | Wellcome Sanger Institute for the COVID-19 Genomics UK (COG-UK) Consortium | Berkshire and Surrey Pathology Services Lighthouse Laboratory and Alex Alderton, Roberto Amato, Jeffrey Barrett, Sonia Goncalves, Ewan Harrison, David K. Jackson, Ian Johnston, Dominic Kwiatkowski, Cordelia Langford, John Sillitoe on behalf of the Wellcome Sanger Institute COVID-19 Surveillance Team |
| EPI_ISL_3445284, EPI_ISL_3445285 | Lighthouse Laboratory Plymouth | Wellcome Sanger Institute for the COVID-19 Genomics UK (COG-UK) Consortium | Lighthouse Laboratory Plymouth and Alex Alderton, Roberto Amato, Jeffrey Barrett, Sonia Goncalves, Ewan Harrison, David K. Jackson, Ian Johnston, Dominic Kwiatkowski, Cordelia Langford, John Sillitoe on behalf of the Wellcome Sanger Institute COVID-19 Surveillance Team |
| EPI_ISL_3445286 | Berkshire and Surrey Pathology Services Lighthouse Laboratory | Wellcome Sanger Institute for the COVID-19 Genomics UK (COG-UK) Consortium | Berkshire and Surrey Pathology Services Lighthouse Laboratory and Alex Alderton, Roberto Amato, Jeffrey Barrett, Sonia Goncalves, Ewan Harrison, David K. Jackson, Ian Johnston, Dominic Kwiatkowski, Cordelia Langford, John Sillitoe on behalf of the Wellcome Sanger Institute COVID-19 Surveillance Team |
| EPI_ISL_3445287, EPI_ISL_3445288 | Lighthouse Laboratory Plymouth | Wellcome Sanger Institute for the COVID-19 Genomics UK (COG-UK) Consortium | Lighthouse Laboratory Plymouth and Alex Alderton, Roberto Amato, Jeffrey Barrett, Sonia Goncalves, Ewan Harrison, David K. Jackson, Ian Johnston, Dominic Kwiatkowski, Cordelia Langford, John Sillitoe on behalf of the Wellcome Sanger Institute COVID-19 Surveillance Team |
| EPI_ISL_3445289 | Berkshire and Surrey Pathology Services Lighthouse Laboratory | Wellcome Sanger Institute for the COVID-19 Genomics UK (COG-UK) Consortium | Berkshire and Surrey Pathology Services Lighthouse Laboratory and Alex Alderton, Roberto Amato, Jeffrey Barrett, Sonia Goncalves, Ewan Harrison, David K. Jackson, Ian Johnston, Dominic Kwiatkowski, Cordelia Langford, John Sillitoe on behalf of the Wellcome Sanger Institute COVID-19 Surveillance Team |
| EPI_ISL_3445290, EPI_ISL_3445291, EPI_ISL_3445292 | Lighthouse Laboratory Plymouth | Wellcome Sanger Institute for the COVID-19 Genomics UK (COG-UK) Consortium | Lighthouse Laboratory Plymouth and Alex Alderton, Roberto Amato, Jeffrey Barrett, Sonia Goncalves, Ewan Harrison, David K. Jackson, Ian Johnston, Dominic Kwiatkowski, Cordelia Langford, John Sillitoe on behalf of the Wellcome Sanger Institute COVID-19 Surveillance Team |
| EPI_ISL_3445293 | Berkshire and Surrey Pathology Services Lighthouse Laboratory | Wellcome Sanger Institute for the COVID-19 Genomics UK (COG-UK) Consortium | Berkshire and Surrey Pathology Services Lighthouse Laboratory and Alex Alderton, Roberto Amato, Jeffrey Barrett, Sonia Goncalves, Ewan Harrison, David K. Jackson, Ian Johnston, Dominic Kwiatkowski, Cordelia Langford, John Sillitoe on behalf of the Wellcome Sanger Institute COVID-19 Surveillance Team |
| EPI_ISL_3445294 | Lighthouse Laboratory Plymouth | Wellcome Sanger Institute for the COVID-19 Genomics UK (COG-UK) Consortium | Lighthouse Laboratory Plymouth and Alex Alderton, Roberto Amato, Jeffrey Barrett, Sonia Goncalves, Ewan Harrison, David K. Jackson, Ian Johnston, Dominic Kwiatkowski, Cordelia Langford, John Sillitoe on behalf of the Wellcome Sanger Institute COVID-19 Surveillance Team |
| EPI_ISL_3445295, EPI_ISL_3445296 | Berkshire and Surrey Pathology Services Lighthouse Laboratory | Wellcome Sanger Institute for the COVID-19 Genomics UK (COG-UK) Consortium | Berkshire and Surrey Pathology Services Lighthouse Laboratory and Alex Alderton, Roberto Amato, Jeffrey Barrett, Sonia Goncalves, Ewan Harrison, David K. Jackson, Ian Johnston, Dominic Kwiatkowski, Cordelia Langford, John Sillitoe on behalf of the Wellcome Sanger Institute COVID-19 Surveillance Team |
| EPI_ISL_3445297 | Lighthouse Laboratory Plymouth | Wellcome Sanger Institute for the COVID-19 Genomics UK (COG-UK) Consortium | Lighthouse Laboratory Plymouth and Alex Alderton, Roberto Amato, Jeffrey Barrett, Sonia Goncalves, Ewan Harrison, David K. Jackson, Ian Johnston, Dominic Kwiatkowski, Cordelia Langford, John Sillitoe on behalf of the Wellcome Sanger Institute COVID-19 Surveillance Team |
| EPI_ISL_3445298 | Berkshire and Surrey Pathology Services Lighthouse Laboratory | Wellcome Sanger Institute for the COVID-19 Genomics UK (COG-UK) Consortium | Berkshire and Surrey Pathology Services Lighthouse Laboratory and Alex Alderton, Roberto Amato, Jeffrey Barrett, Sonia Goncalves, Ewan Harrison, David K. Jackson, Ian Johnston, Dominic Kwiatkowski, Cordelia Langford, John Sillitoe on behalf of the Wellcome Sanger Institute COVID-19 Surveillance Team |
| EPI_ISL_3445299 | Lighthouse Laboratory Plymouth | Wellcome Sanger Institute for the COVID-19 Genomics UK (COG-UK) Consortium | Lighthouse Laboratory Plymouth and Alex Alderton, Roberto Amato, Jeffrey Barrett, Sonia Goncalves, Ewan Harrison, David K. Jackson, Ian Johnston, Dominic Kwiatkowski, Cordelia Langford, John Sillitoe on behalf of the Wellcome Sanger Institute COVID-19 Surveillance Team |
| EPI_ISL_3445300 | Berkshire and Surrey Pathology Services Lighthouse Laboratory | Wellcome Sanger Institute for the COVID-19 Genomics UK (COG-UK) Consortium | Berkshire and Surrey Pathology Services Lighthouse Laboratory and Alex Alderton, Roberto Amato, Jeffrey Barrett, Sonia Goncalves, Ewan Harrison, David K. Jackson, Ian Johnston, Dominic Kwiatkowski, Cordelia Langford, John Sillitoe on behalf of the Wellcome Sanger Institute COVID-19 Surveillance Team |
| EPI_ISL_3445301 | Lighthouse Laboratory Plymouth | Wellcome Sanger Institute for the COVID-19 Genomics UK (COG-UK) Consortium | Lighthouse Laboratory Plymouth and Alex Alderton, Roberto Amato, Jeffrey Barrett, Sonia Goncalves, Ewan Harrison, David K. Jackson, Ian Johnston, Dominic Kwiatkowski, Cordelia Langford, John Sillitoe on behalf of the Wellcome Sanger Institute COVID-19 Surveillance Team |
| EPI_ISL_3445302 | Berkshire and Surrey Pathology Services Lighthouse Laboratory | Wellcome Sanger Institute for the COVID-19 Genomics UK (COG-UK) Consortium | Berkshire and Surrey Pathology Services Lighthouse Laboratory and Alex Alderton, Roberto Amato, Jeffrey Barrett, Sonia Goncalves, Ewan Harrison, David K. Jackson, Ian Johnston, Dominic Kwiatkowski, Cordelia Langford, John Sillitoe on behalf of the Wellcome Sanger Institute COVID-19 Surveillance Team |
| EPI_ISL_3445303 | Lighthouse Laboratory Plymouth | Wellcome Sanger Institute for the COVID-19 Genomics UK (COG-UK) Consortium | Lighthouse Laboratory Plymouth and Alex Alderton, Roberto Amato, Jeffrey Barrett, Sonia Goncalves, Ewan Harrison, David K. Jackson, Ian Johnston, Dominic Kwiatkowski, Cordelia Langford, John Sillitoe on behalf of the Wellcome Sanger Institute COVID-19 Surveillance Team |
| EPI_ISL_3445304, EPI_ISL_3445305 | Berkshire and Surrey Pathology Services Lighthouse Laboratory | Wellcome Sanger Institute for the COVID-19 Genomics UK (COG-UK) Consortium | Berkshire and Surrey Pathology Services Lighthouse Laboratory and Alex Alderton, Roberto Amato, Jeffrey Barrett, Sonia Goncalves, Ewan Harrison, David K. Jackson, Ian Johnston, Dominic Kwiatkowski, Cordelia Langford, John Sillitoe on behalf of the Wellcome Sanger Institute COVID-19 Surveillance Team |
| EPI_ISL_3445306, EPI_ISL_3445307 | Lighthouse Laboratory Plymouth | Wellcome Sanger Institute for the COVID-19 Genomics UK (COG-UK) Consortium | Lighthouse Laboratory Plymouth and Alex Alderton, Roberto Amato, Jeffrey Barrett, Sonia Goncalves, Ewan Harrison, David K. Jackson, Ian Johnston, Dominic Kwiatkowski, Cordelia Langford, John Sillitoe on behalf of the Wellcome Sanger Institute COVID-19 Surveillance Team |
| EPI_ISL_3445308, EPI_ISL_3445309 | Berkshire and Surrey Pathology Services Lighthouse Laboratory | Wellcome Sanger Institute for the COVID-19 Genomics UK (COG-UK) Consortium | Berkshire and Surrey Pathology Services Lighthouse Laboratory and Alex Alderton, Roberto Amato, Jeffrey Barrett, Sonia Goncalves, Ewan Harrison, David K. Jackson, Ian Johnston, Dominic Kwiatkowski, Cordelia Langford, John Sillitoe on behalf of the Wellcome Sanger Institute COVID-19 Surveillance Team |
| EPI_ISL_3445310, EPI_ISL_3445311, EPI_ISL_3445312 | Lighthouse Laboratory Plymouth | Wellcome Sanger Institute for the COVID-19 Genomics UK (COG-UK) Consortium | Lighthouse Laboratory Plymouth and Alex Alderton, Roberto Amato, Jeffrey Barrett, Sonia Goncalves, Ewan Harrison, David K. Jackson, Ian Johnston, Dominic Kwiatkowski, Cordelia Langford, John Sillitoe on behalf of the Wellcome Sanger Institute COVID-19 Surveillance Team |
| EPI_ISL_3445313, EPI_ISL_3445314, EPI_ISL_3445315 | Berkshire and Surrey Pathology Services Lighthouse Laboratory | Wellcome Sanger Institute for the COVID-19 Genomics UK (COG-UK) Consortium | Berkshire and Surrey Pathology Services Lighthouse Laboratory and Alex Alderton, Roberto Amato, Jeffrey Barrett, Sonia Goncalves, Ewan Harrison, David K. Jackson, Ian Johnston, Dominic Kwiatkowski, Cordelia Langford, John Sillitoe on behalf of the Wellcome Sanger Institute COVID-19 Surveillance Team |
| EPI_ISL_3445316 | Lighthouse Laboratory Plymouth | Wellcome Sanger Institute for the COVID-19 Genomics UK (COG-UK) Consortium | Lighthouse Laboratory Plymouth and Alex Alderton, Roberto Amato, Jeffrey Barrett, Sonia Goncalves, Ewan Harrison, David K. Jackson, Ian Johnston, Dominic Kwiatkowski, Cordelia Langford, John Sillitoe on behalf of the Wellcome Sanger Institute COVID-19 Surveillance Team |
| EPI_ISL_3445317 | Berkshire and Surrey Pathology Services Lighthouse Laboratory | Wellcome Sanger Institute for the COVID-19 Genomics UK (COG-UK) Consortium | Berkshire and Surrey Pathology Services Lighthouse Laboratory and Alex Alderton, Roberto Amato, Jeffrey Barrett, Sonia Goncalves, Ewan Harrison, David K. Jackson, Ian Johnston, Dominic Kwiatkowski, Cordelia Langford, John Sillitoe on behalf of the Wellcome Sanger Institute COVID-19 Surveillance Team |
| EPI_ISL_3445318, EPI_ISL_3445319, EPI_ISL_3445320 | Lighthouse Laboratory Plymouth | Wellcome Sanger Institute for the COVID-19 Genomics UK (COG-UK) Consortium | Lighthouse Laboratory Plymouth and Alex Alderton, Roberto Amato, Jeffrey Barrett, Sonia Goncalves, Ewan Harrison, David K. Jackson, Ian Johnston, Dominic Kwiatkowski, Cordelia Langford, John Sillitoe on behalf of the Wellcome Sanger Institute COVID-19 Surveillance Team |
| EPI_ISL_3445321, EPI_ISL_3445322 | Berkshire and Surrey Pathology Services Lighthouse Laboratory | Wellcome Sanger Institute for the COVID-19 Genomics UK (COG-UK) Consortium | Berkshire and Surrey Pathology Services Lighthouse Laboratory and Alex Alderton, Roberto Amato, Jeffrey Barrett, Sonia Goncalves, Ewan Harrison, David K. Jackson, Ian Johnston, Dominic Kwiatkowski, Cordelia Langford, John Sillitoe on behalf of the Wellcome Sanger Institute COVID-19 Surveillance Team |
| EPI_ISL_3445323, EPI_ISL_3445324 | Lighthouse Laboratory Plymouth | Wellcome Sanger Institute for the COVID-19 Genomics UK (COG-UK) Consortium | Lighthouse Laboratory Plymouth and Alex Alderton, Roberto Amato, Jeffrey Barrett, Sonia Goncalves, Ewan Harrison, David K. Jackson, Ian Johnston, Dominic Kwiatkowski, Cordelia Langford, John Sillitoe on behalf of the Wellcome Sanger Institute COVID-19 Surveillance Team |
| EPI_ISL_3445325, EPI_ISL_3445326 | Berkshire and Surrey Pathology Services Lighthouse Laboratory | Wellcome Sanger Institute for the COVID-19 Genomics UK (COG-UK) Consortium | Berkshire and Surrey Pathology Services Lighthouse Laboratory and Alex Alderton, Roberto Amato, Jeffrey Barrett, Sonia Goncalves, Ewan Harrison, David K. Jackson, Ian Johnston, Dominic Kwiatkowski, Cordelia Langford, John Sillitoe on behalf of the Wellcome Sanger Institute COVID-19 Surveillance Team |
| EPI_ISL_3445327 | Lighthouse Laboratory Plymouth | Wellcome Sanger Institute for the COVID-19 Genomics UK (COG-UK) Consortium | Lighthouse Laboratory Plymouth and Alex Alderton, Roberto Amato, Jeffrey Barrett, Sonia Goncalves, Ewan Harrison, David K. Jackson, Ian Johnston, Dominic Kwiatkowski, Cordelia Langford, John Sillitoe on behalf of the Wellcome Sanger Institute COVID-19 Surveillance Team |
| EPI_ISL_3445328 | Berkshire and Surrey Pathology Services Lighthouse Laboratory | Wellcome Sanger Institute for the COVID-19 Genomics UK (COG-UK) Consortium | Berkshire and Surrey Pathology Services Lighthouse Laboratory and Alex Alderton, Roberto Amato, Jeffrey Barrett, Sonia Goncalves, Ewan Harrison, David K. Jackson, Ian Johnston, Dominic Kwiatkowski, Cordelia Langford, John Sillitoe on behalf of the Wellcome Sanger Institute COVID-19 Surveillance Team |
| EPI_ISL_3445329, EPI_ISL_3445330, EPI_ISL_3445331, EPI_ISL_3445332 | Lighthouse Laboratory Plymouth | Wellcome Sanger Institute for the COVID-19 Genomics UK (COG-UK) Consortium | Lighthouse Laboratory Plymouth and Alex Alderton, Roberto Amato, Jeffrey Barrett, Sonia Goncalves, Ewan Harrison, David K. Jackson, Ian Johnston, Dominic Kwiatkowski, Cordelia Langford, John Sillitoe on behalf of the Wellcome Sanger Institute COVID-19 Surveillance Team |
| EPI_ISL_3445333, EPI_ISL_3445334 | Berkshire and Surrey Pathology Services Lighthouse Laboratory | Wellcome Sanger Institute for the COVID-19 Genomics UK (COG-UK) Consortium | Berkshire and Surrey Pathology Services Lighthouse Laboratory and Alex Alderton, Roberto Amato, Jeffrey Barrett, Sonia Goncalves, Ewan Harrison, David K. Jackson, Ian Johnston, Dominic Kwiatkowski, Cordelia Langford, John Sillitoe on behalf of the Wellcome Sanger Institute COVID-19 Surveillance Team |
| EPI_ISL_3445335 | Lighthouse Laboratory Plymouth | Wellcome Sanger Institute for the COVID-19 Genomics UK (COG-UK) Consortium | Lighthouse Laboratory Plymouth and Alex Alderton, Roberto Amato, Jeffrey Barrett, Sonia Goncalves, Ewan Harrison, David K. Jackson, Ian Johnston, Dominic Kwiatkowski, Cordelia Langford, John Sillitoe on behalf of the Wellcome Sanger Institute COVID-19 Surveillance Team |
| EPI_ISL_3445336 | Berkshire and Surrey Pathology Services Lighthouse Laboratory | Wellcome Sanger Institute for the COVID-19 Genomics UK (COG-UK) Consortium | Berkshire and Surrey Pathology Services Lighthouse Laboratory and Alex Alderton, Roberto Amato, Jeffrey Barrett, Sonia Goncalves, Ewan Harrison, David K. Jackson, Ian Johnston, Dominic Kwiatkowski, Cordelia Langford, John Sillitoe on behalf of the Wellcome Sanger Institute COVID-19 Surveillance Team |
| EPI_ISL_3445337 | Lighthouse Laboratory Plymouth | Wellcome Sanger Institute for the COVID-19 Genomics UK (COG-UK) Consortium | Lighthouse Laboratory Plymouth and Alex Alderton, Roberto Amato, Jeffrey Barrett, Sonia Goncalves, Ewan Harrison, David K. Jackson, Ian Johnston, Dominic Kwiatkowski, Cordelia Langford, John Sillitoe on behalf of the Wellcome Sanger Institute COVID-19 Surveillance Team |
| EPI_ISL_3445338 | Berkshire and Surrey Pathology Services Lighthouse Laboratory | Wellcome Sanger Institute for the COVID-19 Genomics UK (COG-UK) Consortium | Berkshire and Surrey Pathology Services Lighthouse Laboratory and Alex Alderton, Roberto Amato, Jeffrey Barrett, Sonia Goncalves, Ewan Harrison, David K. Jackson, Ian Johnston, Dominic Kwiatkowski, Cordelia Langford, John Sillitoe on behalf of the Wellcome Sanger Institute COVID-19 Surveillance Team |
| EPI_ISL_3445339, EPI_ISL_3445340, EPI_ISL_3445341, | Lighthouse Laboratory Plymouth | Wellcome Sanger Institute for the COVID-19 Genomics UK (COG-UK) Consortium | Lighthouse Laboratory Plymouth and Alex Alderton, Roberto Amato, Jeffrey Barrett, Sonia Goncalves, Ewan Harrison, David K. Jackson, Ian Johnston, Dominic Kwiatkowski, Cordelia Langford, John Sillitoe on behalf of the Wellcome Sanger Institute COVID-19 Surveillance Team |

| | | | |
|---|---|---|---|
| EPI_ISL_3445342 | | | |
| EPI_ISL_3445343, EPI_ISL_3445344, EPI_ISL_3445345, EPI_ISL_3445346 | Berkshire and Surrey Pathology Services Lighthouse Laboratory | Wellcome Sanger Institute for the COVID-19 Genomics UK (COG-UK) Consortium | Berkshire and Surrey Pathology Services Lighthouse Laboratory and Alex Alderton, Roberto Amato, Jeffrey Barrett, Sonia Goncalves, Ewan Harrison, David K. Jackson, Ian Johnston, Dominic Kwiatkowski, Cordelia Langford, John Sillitoe on behalf of the Wellcome Sanger Institute COVID-19 Surveillance Team |
| EPI_ISL_3445347 | Lighthouse Laboratory Plymouth | Wellcome Sanger Institute for the COVID-19 Genomics UK (COG-UK) Consortium | Lighthouse Laboratory Plymouth and Alex Alderton, Roberto Amato, Jeffrey Barrett, Sonia Goncalves, Ewan Harrison, David K. Jackson, Ian Johnston, Dominic Kwiatkowski, Cordelia Langford, John Sillitoe on behalf of the Wellcome Sanger Institute COVID-19 Surveillance Team |
| EPI_ISL_3445348, EPI_ISL_3445349 | Berkshire and Surrey Pathology Services Lighthouse Laboratory | Wellcome Sanger Institute for the COVID-19 Genomics UK (COG-UK) Consortium | Berkshire and Surrey Pathology Services Lighthouse Laboratory and Alex Alderton, Roberto Amato, Jeffrey Barrett, Sonia Goncalves, Ewan Harrison, David K. Jackson, Ian Johnston, Dominic Kwiatkowski, Cordelia Langford, John Sillitoe on behalf of the Wellcome Sanger Institute COVID-19 Surveillance Team |
| EPI_ISL_3445350 | Lighthouse Laboratory Plymouth | Wellcome Sanger Institute for the COVID-19 Genomics UK (COG-UK) Consortium | Lighthouse Laboratory Plymouth and Alex Alderton, Roberto Amato, Jeffrey Barrett, Sonia Goncalves, Ewan Harrison, David K. Jackson, Ian Johnston, Dominic Kwiatkowski, Cordelia Langford, John Sillitoe on behalf of the Wellcome Sanger Institute COVID-19 Surveillance Team |
| EPI_ISL_3445351 | Berkshire and Surrey Pathology Services Lighthouse Laboratory | Wellcome Sanger Institute for the COVID-19 Genomics UK (COG-UK) Consortium | Berkshire and Surrey Pathology Services Lighthouse Laboratory and Alex Alderton, Roberto Amato, Jeffrey Barrett, Sonia Goncalves, Ewan Harrison, David K. Jackson, Ian Johnston, Dominic Kwiatkowski, Cordelia Langford, John Sillitoe on behalf of the Wellcome Sanger Institute COVID-19 Surveillance Team |
| EPI_ISL_3445352, EPI_ISL_3445353, EPI_ISL_3445354, EPI_ISL_3445355 | Lighthouse Laboratory Plymouth | Wellcome Sanger Institute for the COVID-19 Genomics UK (COG-UK) Consortium | Lighthouse Laboratory Plymouth and Alex Alderton, Roberto Amato, Jeffrey Barrett, Sonia Goncalves, Ewan Harrison, David K. Jackson, Ian Johnston, Dominic Kwiatkowski, Cordelia Langford, John Sillitoe on behalf of the Wellcome Sanger Institute COVID-19 Surveillance Team |
| EPI_ISL_3445356, EPI_ISL_3445357 | Berkshire and Surrey Pathology Services Lighthouse Laboratory | Wellcome Sanger Institute for the COVID-19 Genomics UK (COG-UK) Consortium | Berkshire and Surrey Pathology Services Lighthouse Laboratory and Alex Alderton, Roberto Amato, Jeffrey Barrett, Sonia Goncalves, Ewan Harrison, David K. Jackson, Ian Johnston, Dominic Kwiatkowski, Cordelia Langford, John Sillitoe on behalf of the Wellcome Sanger Institute COVID-19 Surveillance Team |
| EPI_ISL_3445358 | Lighthouse Laboratory Plymouth | Wellcome Sanger Institute for the COVID-19 Genomics UK (COG-UK) Consortium | Lighthouse Laboratory Plymouth and Alex Alderton, Roberto Amato, Jeffrey Barrett, Sonia Goncalves, Ewan Harrison, David K. Jackson, Ian Johnston, Dominic Kwiatkowski, Cordelia Langford, John Sillitoe on behalf of the Wellcome Sanger Institute COVID-19 Surveillance Team |
| EPI_ISL_3445359 | Berkshire and Surrey Pathology Services Lighthouse Laboratory | Wellcome Sanger Institute for the COVID-19 Genomics UK (COG-UK) Consortium | Berkshire and Surrey Pathology Services Lighthouse Laboratory and Alex Alderton, Roberto Amato, Jeffrey Barrett, Sonia Goncalves, Ewan Harrison, David K. Jackson, Ian Johnston, Dominic Kwiatkowski, Cordelia Langford, John Sillitoe on behalf of the Wellcome Sanger Institute COVID-19 Surveillance Team |
| EPI_ISL_3445360, EPI_ISL_3445361 | Lighthouse Laboratory Plymouth | Wellcome Sanger Institute for the COVID-19 Genomics UK (COG-UK) Consortium | Lighthouse Laboratory Plymouth and Alex Alderton, Roberto Amato, Jeffrey Barrett, Sonia Goncalves, Ewan Harrison, David K. Jackson, Ian Johnston, Dominic Kwiatkowski, Cordelia Langford, John Sillitoe on behalf of the Wellcome Sanger Institute COVID-19 Surveillance Team |
| EPI_ISL_3445362 | Berkshire and Surrey Pathology Services Lighthouse Laboratory | Wellcome Sanger Institute for the COVID-19 Genomics UK (COG-UK) Consortium | Berkshire and Surrey Pathology Services Lighthouse Laboratory and Alex Alderton, Roberto Amato, Jeffrey Barrett, Sonia Goncalves, Ewan Harrison, David K. Jackson, Ian Johnston, Dominic Kwiatkowski, Cordelia Langford, John Sillitoe on behalf of the Wellcome Sanger Institute COVID-19 Surveillance Team |
| EPI_ISL_3445363, EPI_ISL_3445364, EPI_ISL_3445365 | Lighthouse Laboratory Plymouth | Wellcome Sanger Institute for the COVID-19 Genomics UK (COG-UK) Consortium | Lighthouse Laboratory Plymouth and Alex Alderton, Roberto Amato, Jeffrey Barrett, Sonia Goncalves, Ewan Harrison, David K. Jackson, Ian Johnston, Dominic Kwiatkowski, Cordelia Langford, John Sillitoe on behalf of the Wellcome Sanger Institute COVID-19 Surveillance Team |
| EPI_ISL_3445366, EPI_ISL_3445367 | Berkshire and Surrey Pathology Services Lighthouse Laboratory | Wellcome Sanger Institute for the COVID-19 Genomics UK (COG-UK) Consortium | Berkshire and Surrey Pathology Services Lighthouse Laboratory and Alex Alderton, Roberto Amato, Jeffrey Barrett, Sonia Goncalves, Ewan Harrison, David K. Jackson, Ian Johnston, Dominic Kwiatkowski, Cordelia Langford, John Sillitoe on behalf of the Wellcome Sanger Institute COVID-19 Surveillance Team |
| EPI_ISL_3445368 | Lighthouse Laboratory Plymouth | Wellcome Sanger Institute for the COVID-19 Genomics UK (COG-UK) Consortium | Lighthouse Laboratory Plymouth and Alex Alderton, Roberto Amato, Jeffrey Barrett, Sonia Goncalves, Ewan Harrison, David K. Jackson, Ian Johnston, Dominic Kwiatkowski, Cordelia Langford, John Sillitoe on behalf of the Wellcome Sanger Institute COVID-19 Surveillance Team |
| EPI_ISL_3445369 | Berkshire and Surrey Pathology Services Lighthouse Laboratory | Wellcome Sanger Institute for the COVID-19 Genomics UK (COG-UK) Consortium | Berkshire and Surrey Pathology Services Lighthouse Laboratory and Alex Alderton, Roberto Amato, Jeffrey Barrett, Sonia Goncalves, Ewan Harrison, David K. Jackson, Ian Johnston, Dominic Kwiatkowski, Cordelia Langford, John Sillitoe on behalf of the Wellcome Sanger Institute COVID-19 Surveillance Team |
| EPI_ISL_3445370, EPI_ISL_3445371, EPI_ISL_3445372 | Lighthouse Laboratory Plymouth | Wellcome Sanger Institute for the COVID-19 Genomics UK (COG-UK) Consortium | Lighthouse Laboratory Plymouth and Alex Alderton, Roberto Amato, Jeffrey Barrett, Sonia Goncalves, Ewan Harrison, David K. Jackson, Ian Johnston, Dominic Kwiatkowski, Cordelia Langford, John Sillitoe on behalf of the Wellcome Sanger Institute COVID-19 Surveillance Team |
| EPI_ISL_3445373, EPI_ISL_3445374 | Berkshire and Surrey Pathology Services Lighthouse Laboratory | Wellcome Sanger Institute for the COVID-19 Genomics UK (COG-UK) Consortium | Berkshire and Surrey Pathology Services Lighthouse Laboratory and Alex Alderton, Roberto Amato, Jeffrey Barrett, Sonia Goncalves, Ewan Harrison, David K. Jackson, Ian Johnston, Dominic Kwiatkowski, Cordelia Langford, John Sillitoe on behalf of the Wellcome Sanger Institute COVID-19 Surveillance Team |
| EPI_ISL_3445375 | Lighthouse Laboratory Plymouth | Wellcome Sanger Institute for the COVID-19 Genomics UK (COG-UK) Consortium | Lighthouse Laboratory Plymouth and Alex Alderton, Roberto Amato, Jeffrey Barrett, Sonia Goncalves, Ewan Harrison, David K. Jackson, Ian Johnston, Dominic Kwiatkowski, Cordelia Langford, John Sillitoe on behalf of the Wellcome Sanger Institute COVID-19 Surveillance Team |
| EPI_ISL_3445376 | Berkshire and Surrey Pathology Services Lighthouse Laboratory | Wellcome Sanger Institute for the COVID-19 Genomics UK (COG-UK) Consortium | Berkshire and Surrey Pathology Services Lighthouse Laboratory and Alex Alderton, Roberto Amato, Jeffrey Barrett, Sonia Goncalves, Ewan Harrison, David K. Jackson, Ian Johnston, Dominic Kwiatkowski, Cordelia Langford, John Sillitoe on behalf of the Wellcome Sanger Institute COVID-19 Surveillance Team |
| EPI_ISL_3445377, EPI_ISL_3445378, EPI_ISL_3445379, EPI_ISL_3445380, EPI_ISL_3445381 | Lighthouse Laboratory Plymouth | Wellcome Sanger Institute for the COVID-19 Genomics UK (COG-UK) Consortium | Lighthouse Laboratory Plymouth and Alex Alderton, Roberto Amato, Jeffrey Barrett, Sonia Goncalves, Ewan Harrison, David K. Jackson, Ian Johnston, Dominic Kwiatkowski, Cordelia Langford, John Sillitoe on behalf of the Wellcome Sanger Institute COVID-19 Surveillance Team |
| EPI_ISL_3445382 | Berkshire and Surrey Pathology Services Lighthouse Laboratory | Wellcome Sanger Institute for the COVID-19 Genomics UK (COG-UK) Consortium | Berkshire and Surrey Pathology Services Lighthouse Laboratory and Alex Alderton, Roberto Amato, Jeffrey Barrett, Sonia Goncalves, Ewan Harrison, David K. Jackson, Ian Johnston, Dominic Kwiatkowski, Cordelia Langford, John Sillitoe on behalf of the Wellcome Sanger Institute COVID-19 Surveillance Team |
| EPI_ISL_3445383 | Lighthouse Laboratory Plymouth | Wellcome Sanger Institute for the COVID-19 Genomics UK (COG-UK) Consortium | Lighthouse Laboratory Plymouth and Alex Alderton, Roberto Amato, Jeffrey Barrett, Sonia Goncalves, Ewan Harrison, David K. Jackson, Ian Johnston, Dominic Kwiatkowski, Cordelia Langford, John Sillitoe on behalf of the Wellcome Sanger Institute COVID-19 Surveillance Team |
| EPI_ISL_3445384 | Berkshire and Surrey Pathology Services Lighthouse Laboratory | Wellcome Sanger Institute for the COVID-19 Genomics UK (COG-UK) Consortium | Berkshire and Surrey Pathology Services Lighthouse Laboratory and Alex Alderton, Roberto Amato, Jeffrey Barrett, Sonia Goncalves, Ewan Harrison, David K. Jackson, Ian Johnston, Dominic Kwiatkowski, Cordelia Langford, John Sillitoe on behalf of the Wellcome Sanger Institute COVID-19 Surveillance Team |
| EPI_ISL_3445385, EPI_ISL_3445386, EPI_ISL_3445387 | Lighthouse Laboratory Plymouth | Wellcome Sanger Institute for the COVID-19 Genomics UK (COG-UK) Consortium | Lighthouse Laboratory Plymouth and Alex Alderton, Roberto Amato, Jeffrey Barrett, Sonia Goncalves, Ewan Harrison, David K. Jackson, Ian Johnston, Dominic Kwiatkowski, Cordelia Langford, John Sillitoe on behalf of the Wellcome Sanger Institute COVID-19 Surveillance Team |
| EPI_ISL_3445388 | Berkshire and Surrey Pathology Services Lighthouse Laboratory | Wellcome Sanger Institute for the COVID-19 Genomics UK (COG-UK) Consortium | Berkshire and Surrey Pathology Services Lighthouse Laboratory and Alex Alderton, Roberto Amato, Jeffrey Barrett, Sonia Goncalves, Ewan Harrison, David K. Jackson, Ian Johnston, Dominic Kwiatkowski, Cordelia Langford, John Sillitoe on behalf of the Wellcome Sanger Institute COVID-19 Surveillance Team |
| EPI_ISL_3445389, EPI_ISL_3445390 | Lighthouse Laboratory Plymouth | Wellcome Sanger Institute for the COVID-19 Genomics UK (COG-UK) Consortium | Lighthouse Laboratory Plymouth and Alex Alderton, Roberto Amato, Jeffrey Barrett, Sonia Goncalves, Ewan Harrison, David K. Jackson, Ian Johnston, Dominic Kwiatkowski, Cordelia Langford, John Sillitoe on behalf of the Wellcome Sanger Institute COVID-19 Surveillance Team |
| EPI_ISL_3445391, EPI_ISL_3445392 | Berkshire and Surrey Pathology Services Lighthouse Laboratory | Wellcome Sanger Institute for the COVID-19 Genomics UK (COG-UK) Consortium | Berkshire and Surrey Pathology Services Lighthouse Laboratory and Alex Alderton, Roberto Amato, Jeffrey Barrett, Sonia Goncalves, Ewan Harrison, David K. Jackson, Ian Johnston, Dominic Kwiatkowski, Cordelia Langford, John Sillitoe on behalf of the Wellcome Sanger Institute COVID-19 Surveillance Team |
| EPI_ISL_3445393, EPI_ISL_3445394 | Lighthouse Laboratory Plymouth | Wellcome Sanger Institute for the COVID-19 Genomics UK (COG-UK) Consortium | Lighthouse Laboratory Plymouth and Alex Alderton, Roberto Amato, Jeffrey Barrett, Sonia Goncalves, Ewan Harrison, David K. Jackson, Ian Johnston, Dominic Kwiatkowski, Cordelia Langford, John Sillitoe on behalf of the Wellcome Sanger Institute COVID-19 Surveillance Team |
| EPI_ISL_3445395 | Berkshire and Surrey Pathology Services Lighthouse Laboratory | Wellcome Sanger Institute for the COVID-19 Genomics UK (COG-UK) Consortium | Berkshire and Surrey Pathology Services Lighthouse Laboratory and Alex Alderton, Roberto Amato, Jeffrey Barrett, Sonia Goncalves, Ewan Harrison, David K. Jackson, Ian Johnston, Dominic Kwiatkowski, Cordelia Langford, John Sillitoe on behalf of the Wellcome Sanger Institute COVID-19 Surveillance Team |
| EPI_ISL_3445396, EPI_ISL_3445397, EPI_ISL_3445398 | Lighthouse Laboratory Plymouth | Wellcome Sanger Institute for the COVID-19 Genomics UK (COG-UK) Consortium | Lighthouse Laboratory Plymouth and Alex Alderton, Roberto Amato, Jeffrey Barrett, Sonia Goncalves, Ewan Harrison, David K. Jackson, Ian Johnston, Dominic Kwiatkowski, Cordelia Langford, John Sillitoe on behalf of the Wellcome Sanger Institute COVID-19 Surveillance Team |
| EPI_ISL_3445399 | Berkshire and Surrey Pathology Services Lighthouse Laboratory | Wellcome Sanger Institute for the COVID-19 Genomics UK (COG-UK) Consortium | Berkshire and Surrey Pathology Services Lighthouse Laboratory and Alex Alderton, Roberto Amato, Jeffrey Barrett, Sonia Goncalves, Ewan Harrison, David K. Jackson, Ian Johnston, Dominic Kwiatkowski, Cordelia Langford, John Sillitoe on behalf of the Wellcome Sanger Institute COVID-19 Surveillance Team |
| EPI_ISL_3445400 | Lighthouse Laboratory Plymouth | Wellcome Sanger Institute for the COVID-19 Genomics UK (COG-UK) Consortium | Lighthouse Laboratory Plymouth and Alex Alderton, Roberto Amato, Jeffrey Barrett, Sonia Goncalves, Ewan Harrison, David K. Jackson, Ian Johnston, Dominic Kwiatkowski, Cordelia Langford, John Sillitoe on behalf of the Wellcome Sanger Institute COVID-19 Surveillance Team |
| EPI_ISL_3445401, EPI_ISL_3445402, EPI_ISL_3445403 | Berkshire and Surrey Pathology Services Lighthouse Laboratory | Wellcome Sanger Institute for the COVID-19 Genomics UK (COG-UK) Consortium | Berkshire and Surrey Pathology Services Lighthouse Laboratory and Alex Alderton, Roberto Amato, Jeffrey Barrett, Sonia Goncalves, Ewan Harrison, David K. Jackson, Ian Johnston, Dominic Kwiatkowski, Cordelia Langford, John Sillitoe on behalf of the Wellcome Sanger Institute COVID-19 Surveillance Team |
| EPI_ISL_3445404 | Lighthouse Laboratory Plymouth | Wellcome Sanger Institute for the COVID-19 Genomics UK (COG-UK) Consortium | Lighthouse Laboratory Plymouth and Alex Alderton, Roberto Amato, Jeffrey Barrett, Sonia Goncalves, Ewan Harrison, David K. Jackson, Ian Johnston, Dominic Kwiatkowski, Cordelia Langford, John Sillitoe on behalf of the Wellcome Sanger Institute COVID-19 Surveillance Team |
| EPI_ISL_3445405 | Berkshire and Surrey Pathology Services Lighthouse Laboratory | Wellcome Sanger Institute for the COVID-19 Genomics UK (COG-UK) Consortium | Berkshire and Surrey Pathology Services Lighthouse Laboratory and Alex Alderton, Roberto Amato, Jeffrey Barrett, Sonia Goncalves, Ewan Harrison, David K. Jackson, Ian Johnston, Dominic Kwiatkowski, Cordelia Langford, John Sillitoe on behalf of the Wellcome Sanger Institute COVID-19 Surveillance Team |
| EPI_ISL_3445406, EPI_ISL_3445407 | Lighthouse Laboratory Plymouth | Wellcome Sanger Institute for the COVID-19 Genomics UK (COG-UK) Consortium | Lighthouse Laboratory Plymouth and Alex Alderton, Roberto Amato, Jeffrey Barrett, Sonia Goncalves, Ewan Harrison, David K. Jackson, Ian Johnston, Dominic Kwiatkowski, Cordelia Langford, John Sillitoe on behalf of the Wellcome Sanger Institute COVID-19 Surveillance Team |
| EPI_ISL_3445408, EPI_ISL_3445409, EPI_ISL_3445410, EPI_ISL_3445411 | Berkshire and Surrey Pathology Services Lighthouse Laboratory | Wellcome Sanger Institute for the COVID-19 Genomics UK (COG-UK) Consortium | Berkshire and Surrey Pathology Services Lighthouse Laboratory and Alex Alderton, Roberto Amato, Jeffrey Barrett, Sonia Goncalves, Ewan Harrison, David K. Jackson, Ian Johnston, Dominic Kwiatkowski, Cordelia Langford, John Sillitoe on behalf of the Wellcome Sanger Institute COVID-19 Surveillance Team |
| EPI_ISL_3445412, EPI_ISL_3445413 | Lighthouse Laboratory Plymouth | Wellcome Sanger Institute for the COVID-19 Genomics UK (COG-UK) Consortium | Lighthouse Laboratory Plymouth and Alex Alderton, Roberto Amato, Jeffrey Barrett, Sonia Goncalves, Ewan Harrison, David K. Jackson, Ian Johnston, Dominic Kwiatkowski, Cordelia Langford, John Sillitoe on behalf of the Wellcome Sanger Institute COVID-19 Surveillance Team |
| EPI_ISL_3445414, EPI_ISL_3445415, | Berkshire and Surrey Pathology Services Lighthouse Laboratory | Wellcome Sanger Institute for the COVID-19 Genomics UK (COG-UK) Consortium | Berkshire and Surrey Pathology Services Lighthouse Laboratory and Alex Alderton, Roberto Amato, Jeffrey Barrett, Sonia Goncalves, Ewan Harrison, David K. Jackson, Ian Johnston, Dominic Kwiatkowski, Cordelia Langford, John Sillitoe on behalf of the Wellcome Sanger Institute COVID-19 Surveillance Team |

| | | | |
|---|---|---|---|
| EPI_ISL_3445416, EPI_ISL_3445417, EPI_ISL_3445418, EPI_ISL_3445419, EPI_ISL_3445420 | | | |
| EPI_ISL_3445421 | Lighthouse Laboratory Plymouth | Wellcome Sanger Institute for the COVID-19 Genomics UK (COG-UK) Consortium | Lighthouse Laboratory Plymouth and Alex Alderton, Roberto Amato, Jeffrey Barrett, Sonia Goncalves, Ewan Harrison, David K. Jackson, Ian Johnston, Dominic Kwiatkowski, Cordelia Langford, John Sillitoe on behalf of the Wellcome Sanger Institute COVID-19 Surveillance Team |
| EPI_ISL_3445422 | Berkshire and Surrey Pathology Services Lighthouse Laboratory | Wellcome Sanger Institute for the COVID-19 Genomics UK (COG-UK) Consortium | Berkshire and Surrey Pathology Services Lighthouse Laboratory and Alex Alderton, Roberto Amato, Jeffrey Barrett, Sonia Goncalves, Ewan Harrison, David K. Jackson, Ian Johnston, Dominic Kwiatkowski, Cordelia Langford, John Sillitoe on behalf of the Wellcome Sanger Institute COVID-19 Surveillance Team |
| EPI_ISL_3445423, EPI_ISL_3445424 | Lighthouse Laboratory Plymouth | Wellcome Sanger Institute for the COVID-19 Genomics UK (COG-UK) Consortium | Lighthouse Laboratory Plymouth and Alex Alderton, Roberto Amato, Jeffrey Barrett, Sonia Goncalves, Ewan Harrison, David K. Jackson, Ian Johnston, Dominic Kwiatkowski, Cordelia Langford, John Sillitoe on behalf of the Wellcome Sanger Institute COVID-19 Surveillance Team |
| EPI_ISL_3445425, EPI_ISL_3445426, EPI_ISL_3445427, EPI_ISL_3445428, EPI_ISL_3445429, EPI_ISL_3445430 | Berkshire and Surrey Pathology Services Lighthouse Laboratory | Wellcome Sanger Institute for the COVID-19 Genomics UK (COG-UK) Consortium | Berkshire and Surrey Pathology Services Lighthouse Laboratory and Alex Alderton, Roberto Amato, Jeffrey Barrett, Sonia Goncalves, Ewan Harrison, David K. Jackson, Ian Johnston, Dominic Kwiatkowski, Cordelia Langford, John Sillitoe on behalf of the Wellcome Sanger Institute COVID-19 Surveillance Team |
| EPI_ISL_3445431, EPI_ISL_3445432 | Lighthouse Laboratory Plymouth | Wellcome Sanger Institute for the COVID-19 Genomics UK (COG-UK) Consortium | Lighthouse Laboratory Plymouth and Alex Alderton, Roberto Amato, Jeffrey Barrett, Sonia Goncalves, Ewan Harrison, David K. Jackson, Ian Johnston, Dominic Kwiatkowski, Cordelia Langford, John Sillitoe on behalf of the Wellcome Sanger Institute COVID-19 Surveillance Team |
| EPI_ISL_3445433, EPI_ISL_3445434 | Lighthouse Lab in Glasgow | Wellcome Sanger Institute for the COVID-19 Genomics UK (COG-UK) Consortium | Harper VanSteenhouse, Yumi Kasai, David Gray, Carol Clugston, Anna Dominiczak and Alex Alderton, Roberto Amato, Jeffrey Barrett, Sonia Goncalves, Ewan Harrison, David K. Jackson, Ian Johnston, Dominic Kwiatkowski, Cordelia Langford, John Sillitoe on behalf of the Wellcome Sanger Institute COVID-19 Surveillance Team |
| EPI_ISL_3445435 | Lighthouse Lab in Alderley Park | Wellcome Sanger Institute for the COVID-19 Genomics UK (COG-UK) Consortium | Jacquelyn Wynn, Mairead Hyland, The Lighthouse Lab in Alderley Park and Alex Alderton, Roberto Amato, Jeffrey Barrett, Sonia Goncalves, Ewan Harrison, David K. Jackson, Ian Johnston, Dominic Kwiatkowski, Cordelia Langford, John Sillitoe on behalf of the Wellcome Sanger Institute COVID-19 Surveillance Team |
| EPI_ISL_3445436, EPI_ISL_3445437, EPI_ISL_3445438, EPI_ISL_3445439, EPI_ISL_3445440, EPI_ISL_3445441, EPI_ISL_3445442 | Lighthouse Lab in Glasgow | Wellcome Sanger Institute for the COVID-19 Genomics UK (COG-UK) Consortium | Harper VanSteenhouse, Yumi Kasai, David Gray, Carol Clugston, Anna Dominiczak and Alex Alderton, Roberto Amato, Jeffrey Barrett, Sonia Goncalves, Ewan Harrison, David K. Jackson, Ian Johnston, Dominic Kwiatkowski, Cordelia Langford, John Sillitoe on behalf of the Wellcome Sanger Institute COVID-19 Surveillance Team |
| EPI_ISL_3445443 | Berkshire and Surrey Pathology Services Lighthouse Laboratory | Wellcome Sanger Institute for the COVID-19 Genomics UK (COG-UK) Consortium | Berkshire and Surrey Pathology Services Lighthouse Laboratory and Alex Alderton, Roberto Amato, Jeffrey Barrett, Sonia Goncalves, Ewan Harrison, David K. Jackson, Ian Johnston, Dominic Kwiatkowski, Cordelia Langford, John Sillitoe on behalf of the Wellcome Sanger Institute COVID-19 Surveillance Team |
| EPI_ISL_3445444, EPI_ISL_3445445, EPI_ISL_3445446 | Lighthouse Lab in Glasgow | Wellcome Sanger Institute for the COVID-19 Genomics UK (COG-UK) Consortium | Harper VanSteenhouse, Yumi Kasai, David Gray, Carol Clugston, Anna Dominiczak and Alex Alderton, Roberto Amato, Jeffrey Barrett, Sonia Goncalves, Ewan Harrison, David K. Jackson, Ian Johnston, Dominic Kwiatkowski, Cordelia Langford, John Sillitoe on behalf of the Wellcome Sanger Institute COVID-19 Surveillance Team |
| EPI_ISL_3445447 | Lighthouse Lab in Alderley Park | Wellcome Sanger Institute for the COVID-19 Genomics UK (COG-UK) Consortium | Jacquelyn Wynn, Mairead Hyland, The Lighthouse Lab in Alderley Park and Alex Alderton, Roberto Amato, Jeffrey Barrett, Sonia Goncalves, Ewan Harrison, David K. Jackson, Ian Johnston, Dominic Kwiatkowski, Cordelia Langford, John Sillitoe on behalf of the Wellcome Sanger Institute COVID-19 Surveillance Team |
| EPI_ISL_3445448, EPI_ISL_3445449, EPI_ISL_3445450, EPI_ISL_3445451, EPI_ISL_3445452, EPI_ISL_3445453, EPI_ISL_3445454, EPI_ISL_3445455, EPI_ISL_3445456, EPI_ISL_3445457, EPI_ISL_3445458, EPI_ISL_3445459, EPI_ISL_3445460, EPI_ISL_3445461, EPI_ISL_3445462 | | | |
| see above | Lighthouse Lab in Glasgow | Wellcome Sanger Institute for the COVID-19 Genomics UK (COG-UK) Consortium | Harper VanSteenhouse, Yumi Kasai, David Gray, Carol Clugston, Anna Dominiczak and Alex Alderton, Roberto Amato, Jeffrey Barrett, Sonia Goncalves, Ewan Harrison, David K. Jackson, Ian Johnston, Dominic Kwiatkowski, Cordelia Langford, John Sillitoe on behalf of the Wellcome Sanger Institute COVID-19 Surveillance Team |
| EPI_ISL_3445463 | Lighthouse Lab in Alderley Park | Wellcome Sanger Institute for the COVID-19 Genomics UK (COG-UK) Consortium | Jacquelyn Wynn, Mairead Hyland, The Lighthouse Lab in Alderley Park and Alex Alderton, Roberto Amato, Jeffrey Barrett, Sonia Goncalves, Ewan Harrison, David K. Jackson, Ian Johnston, Dominic Kwiatkowski, Cordelia Langford, John Sillitoe on behalf of the Wellcome Sanger Institute COVID-19 Surveillance Team |
| EPI_ISL_3445464, EPI_ISL_3445465, EPI_ISL_3445466, EPI_ISL_3445467, EPI_ISL_3445468, EPI_ISL_3445469, EPI_ISL_3445470 | Lighthouse Lab in Glasgow | Wellcome Sanger Institute for the COVID-19 Genomics UK (COG-UK) Consortium | Harper VanSteenhouse, Yumi Kasai, David Gray, Carol Clugston, Anna Dominiczak and Alex Alderton, Roberto Amato, Jeffrey Barrett, Sonia Goncalves, Ewan Harrison, David K. Jackson, Ian Johnston, Dominic Kwiatkowski, Cordelia Langford, John Sillitoe on behalf of the Wellcome Sanger Institute COVID-19 Surveillance Team |
| EPI_ISL_3445471, EPI_ISL_3445472 | Berkshire and Surrey Pathology Services Lighthouse Laboratory | Wellcome Sanger Institute for the COVID-19 Genomics UK (COG-UK) Consortium | Berkshire and Surrey Pathology Services Lighthouse Laboratory and Alex Alderton, Roberto Amato, Jeffrey Barrett, Sonia Goncalves, Ewan Harrison, David K. Jackson, Ian Johnston, Dominic Kwiatkowski, Cordelia Langford, John Sillitoe on behalf of the Wellcome Sanger Institute COVID-19 Surveillance Team |
| EPI_ISL_3445473 | Lighthouse Lab in Milton Keynes | Wellcome Sanger Institute for the COVID-19 Genomics UK (COG-UK) Consortium | The Lighthouse Lab in Milton Keynes and Alex Alderton, Roberto Amato, Jeffrey Barrett, Sonia Goncalves, Ewan Harrison, David K. Jackson, Ian Johnston, Dominic Kwiatkowski, Cordelia Langford, John Sillitoe on behalf of the Wellcome Sanger Institute COVID-19 Surveillance Team |
| EPI_ISL_3445474, EPI_ISL_3445475 | Berkshire and Surrey Pathology Services Lighthouse Laboratory | Wellcome Sanger Institute for the COVID-19 Genomics UK (COG-UK) Consortium | Berkshire and Surrey Pathology Services Lighthouse Laboratory and Alex Alderton, Roberto Amato, Jeffrey Barrett, Sonia Goncalves, Ewan Harrison, David K. Jackson, Ian Johnston, Dominic Kwiatkowski, Cordelia Langford, John Sillitoe on behalf of the Wellcome Sanger Institute COVID-19 Surveillance Team |
| EPI_ISL_3445476 | Lighthouse Lab in Milton Keynes | Wellcome Sanger Institute for the COVID-19 Genomics UK (COG-UK) Consortium | The Lighthouse Lab in Milton Keynes and Alex Alderton, Roberto Amato, Jeffrey Barrett, Sonia Goncalves, Ewan Harrison, David K. Jackson, Ian Johnston, Dominic Kwiatkowski, Cordelia Langford, John Sillitoe on behalf of the Wellcome Sanger Institute COVID-19 Surveillance Team |
| EPI_ISL_3445477 | Berkshire and Surrey Pathology Services Lighthouse Laboratory | Wellcome Sanger Institute for the COVID-19 Genomics UK (COG-UK) Consortium | Berkshire and Surrey Pathology Services Lighthouse Laboratory and Alex Alderton, Roberto Amato, Jeffrey Barrett, Sonia Goncalves, Ewan Harrison, David K. Jackson, Ian Johnston, Dominic Kwiatkowski, Cordelia Langford, John Sillitoe on behalf of the Wellcome Sanger Institute COVID-19 Surveillance Team |
| EPI_ISL_3445478 | Lighthouse Lab in Glasgow | Wellcome Sanger Institute for the COVID-19 Genomics UK (COG-UK) Consortium | Harper VanSteenhouse, Yumi Kasai, David Gray, Carol Clugston, Anna Dominiczak and Alex Alderton, Roberto Amato, Jeffrey Barrett, Sonia Goncalves, Ewan Harrison, David K. Jackson, Ian Johnston, Dominic Kwiatkowski, Cordelia Langford, John Sillitoe on behalf of the Wellcome Sanger Institute COVID-19 Surveillance Team |
| EPI_ISL_3445479 | Lighthouse Lab in Milton Keynes | Wellcome Sanger Institute for the COVID-19 Genomics UK (COG-UK) Consortium | The Lighthouse Lab in Milton Keynes and Alex Alderton, Roberto Amato, Jeffrey Barrett, Sonia Goncalves, Ewan Harrison, David K. Jackson, Ian Johnston, Dominic Kwiatkowski, Cordelia Langford, John Sillitoe on behalf of the Wellcome Sanger Institute COVID-19 Surveillance Team |
| EPI_ISL_3445480 | Lighthouse Lab in Glasgow | Wellcome Sanger Institute for the COVID-19 Genomics UK (COG-UK) Consortium | Harper VanSteenhouse, Yumi Kasai, David Gray, Carol Clugston, Anna Dominiczak and Alex Alderton, Roberto Amato, Jeffrey Barrett, Sonia Goncalves, Ewan Harrison, David K. Jackson, Ian Johnston, Dominic Kwiatkowski, Cordelia Langford, John Sillitoe on behalf of the Wellcome Sanger Institute COVID-19 Surveillance Team |
| EPI_ISL_3445481, EPI_ISL_3445482, EPI_ISL_3445483 | Lighthouse Lab in Milton Keynes | Wellcome Sanger Institute for the COVID-19 Genomics UK (COG-UK) Consortium | The Lighthouse Lab in Milton Keynes and Alex Alderton, Roberto Amato, Jeffrey Barrett, Sonia Goncalves, Ewan Harrison, David K. Jackson, Ian Johnston, Dominic Kwiatkowski, Cordelia Langford, John Sillitoe on behalf of the Wellcome Sanger Institute COVID-19 Surveillance Team |
| EPI_ISL_3445484, EPI_ISL_3445485, EPI_ISL_3445486, EPI_ISL_3445487 | Lighthouse Lab in Glasgow | Wellcome Sanger Institute for the COVID-19 Genomics UK (COG-UK) Consortium | Harper VanSteenhouse, Yumi Kasai, David Gray, Carol Clugston, Anna Dominiczak and Alex Alderton, Roberto Amato, Jeffrey Barrett, Sonia Goncalves, Ewan Harrison, David K. Jackson, Ian Johnston, Dominic Kwiatkowski, Cordelia Langford, John Sillitoe on behalf of the Wellcome Sanger Institute COVID-19 Surveillance Team |
| EPI_ISL_3445488 | Lighthouse Lab in Milton Keynes | Wellcome Sanger Institute for the COVID-19 Genomics UK (COG-UK) Consortium | The Lighthouse Lab in Milton Keynes and Alex Alderton, Roberto Amato, Jeffrey Barrett, Sonia Goncalves, Ewan Harrison, David K. Jackson, Ian Johnston, Dominic Kwiatkowski, Cordelia Langford, John Sillitoe on behalf of the Wellcome Sanger Institute COVID-19 Surveillance Team |
| EPI_ISL_3445489, EPI_ISL_3445490, EPI_ISL_3445491, EPI_ISL_3445492, EPI_ISL_3445493, EPI_ISL_3445494 | Lighthouse Lab in Glasgow | Wellcome Sanger Institute for the COVID-19 Genomics UK (COG-UK) Consortium | Harper VanSteenhouse, Yumi Kasai, David Gray, Carol Clugston, Anna Dominiczak and Alex Alderton, Roberto Amato, Jeffrey Barrett, Sonia Goncalves, Ewan Harrison, David K. Jackson, Ian Johnston, Dominic Kwiatkowski, Cordelia Langford, John Sillitoe on behalf of the Wellcome Sanger Institute COVID-19 Surveillance Team |
| EPI_ISL_3445495 | Lighthouse Lab in Milton Keynes | Wellcome Sanger Institute for the COVID-19 Genomics UK (COG-UK) Consortium | The Lighthouse Lab in Milton Keynes and Alex Alderton, Roberto Amato, Jeffrey Barrett, Sonia Goncalves, Ewan Harrison, David K. Jackson, Ian Johnston, Dominic Kwiatkowski, Cordelia Langford, John Sillitoe on behalf of the Wellcome Sanger Institute COVID-19 Surveillance Team |
| EPI_ISL_3445496, EPI_ISL_3445497, EPI_ISL_3445498, EPI_ISL_3445499, EPI_ISL_3445500 | Lighthouse Lab in Glasgow | Wellcome Sanger Institute for the COVID-19 Genomics UK (COG-UK) Consortium | Harper VanSteenhouse, Yumi Kasai, David Gray, Carol Clugston, Anna Dominiczak and Alex Alderton, Roberto Amato, Jeffrey Barrett, Sonia Goncalves, Ewan Harrison, David K. Jackson, Ian Johnston, Dominic Kwiatkowski, Cordelia Langford, John Sillitoe on behalf of the Wellcome Sanger Institute COVID-19 Surveillance Team |
| EPI_ISL_3445501, EPI_ISL_3445502 | Lighthouse Lab in Milton Keynes | Wellcome Sanger Institute for the COVID-19 Genomics UK (COG-UK) Consortium | The Lighthouse Lab in Milton Keynes and Alex Alderton, Roberto Amato, Jeffrey Barrett, Sonia Goncalves, Ewan Harrison, David K. Jackson, Ian Johnston, Dominic Kwiatkowski, Cordelia Langford, John Sillitoe on behalf of the Wellcome Sanger Institute COVID-19 Surveillance Team |
| EPI_ISL_3445503, EPI_ISL_3445504, EPI_ISL_3445505, EPI_ISL_3445506, EPI_ISL_3445507, EPI_ISL_3445508, EPI_ISL_3445509 | Lighthouse Lab in Glasgow | Wellcome Sanger Institute for the COVID-19 Genomics UK (COG-UK) Consortium | Harper VanSteenhouse, Yumi Kasai, David Gray, Carol Clugston, Anna Dominiczak and Alex Alderton, Roberto Amato, Jeffrey Barrett, Sonia Goncalves, Ewan Harrison, David K. Jackson, Ian Johnston, Dominic Kwiatkowski, Cordelia Langford, John Sillitoe on behalf of the Wellcome Sanger Institute COVID-19 Surveillance Team |

| | | | |
|---|---|---|---|
| EPI_ISL_3445510 | Lighthouse Lab in Milton Keynes | Wellcome Sanger Institute for the COVID-19 Genomics UK (COG-UK) Consortium | The Lighthouse Lab in Milton Keynes and Alex Alderton, Roberto Amato, Jeffrey Barrett, Sonia Goncalves, Ewan Harrison, David K. Jackson, Ian Johnston, Dominic Kwiatkowski, Cordelia Langford, John Sillitoe on behalf of the Wellcome Sanger Institute COVID-19 Surveillance Team |
| EPI_ISL_3445511, EPI_ISL_3445512, EPI_ISL_3445513, EPI_ISL_3445514, EPI_ISL_3445515, EPI_ISL_3445516, EPI_ISL_3445517 | Lighthouse Lab in Glasgow | Wellcome Sanger Institute for the COVID-19 Genomics UK (COG-UK) Consortium | Harper VanSteenhouse, Yumi Kasai, David Gray, Carol Clugston, Anna Dominiczak and Alex Alderton, Roberto Amato, Jeffrey Barrett, Sonia Goncalves, Ewan Harrison, David K. Jackson, Ian Johnston, Dominic Kwiatkowski, Cordelia Langford, John Sillitoe on behalf of the Wellcome Sanger Institute COVID-19 Surveillance Team |
| EPI_ISL_3445518 | Lighthouse Lab in Milton Keynes | Wellcome Sanger Institute for the COVID-19 Genomics UK (COG-UK) Consortium | The Lighthouse Lab in Milton Keynes and Alex Alderton, Roberto Amato, Jeffrey Barrett, Sonia Goncalves, Ewan Harrison, David K. Jackson, Ian Johnston, Dominic Kwiatkowski, Cordelia Langford, John Sillitoe on behalf of the Wellcome Sanger Institute COVID-19 Surveillance Team |
| EPI_ISL_3445519 | Lighthouse Lab in Glasgow | Wellcome Sanger Institute for the COVID-19 Genomics UK (COG-UK) Consortium | Harper VanSteenhouse, Yumi Kasai, David Gray, Carol Clugston, Anna Dominiczak and Alex Alderton, Roberto Amato, Jeffrey Barrett, Sonia Goncalves, Ewan Harrison, David K. Jackson, Ian Johnston, Dominic Kwiatkowski, Cordelia Langford, John Sillitoe on behalf of the Wellcome Sanger Institute COVID-19 Surveillance Team |
| EPI_ISL_3445520 | Lighthouse Lab in Milton Keynes | Wellcome Sanger Institute for the COVID-19 Genomics UK (COG-UK) Consortium | The Lighthouse Lab in Milton Keynes and Alex Alderton, Roberto Amato, Jeffrey Barrett, Sonia Goncalves, Ewan Harrison, David K. Jackson, Ian Johnston, Dominic Kwiatkowski, Cordelia Langford, John Sillitoe on behalf of the Wellcome Sanger Institute COVID-19 Surveillance Team |
| EPI_ISL_3445521, EPI_ISL_3445522 | Lighthouse Lab in Glasgow | Wellcome Sanger Institute for the COVID-19 Genomics UK (COG-UK) Consortium | Harper VanSteenhouse, Yumi Kasai, David Gray, Carol Clugston, Anna Dominiczak and Alex Alderton, Roberto Amato, Jeffrey Barrett, Sonia Goncalves, Ewan Harrison, David K. Jackson, Ian Johnston, Dominic Kwiatkowski, Cordelia Langford, John Sillitoe on behalf of the Wellcome Sanger Institute COVID-19 Surveillance Team |
| EPI_ISL_3445523, EPI_ISL_3445524 | Lighthouse Lab in Milton Keynes | Wellcome Sanger Institute for the COVID-19 Genomics UK (COG-UK) Consortium | The Lighthouse Lab in Milton Keynes and Alex Alderton, Roberto Amato, Jeffrey Barrett, Sonia Goncalves, Ewan Harrison, David K. Jackson, Ian Johnston, Dominic Kwiatkowski, Cordelia Langford, John Sillitoe on behalf of the Wellcome Sanger Institute COVID-19 Surveillance Team |
| EPI_ISL_3445525, EPI_ISL_3445526 | Lighthouse Lab in Glasgow | Wellcome Sanger Institute for the COVID-19 Genomics UK (COG-UK) Consortium | Harper VanSteenhouse, Yumi Kasai, David Gray, Carol Clugston, Anna Dominiczak and Alex Alderton, Roberto Amato, Jeffrey Barrett, Sonia Goncalves, Ewan Harrison, David K. Jackson, Ian Johnston, Dominic Kwiatkowski, Cordelia Langford, John Sillitoe on behalf of the Wellcome Sanger Institute COVID-19 Surveillance Team |
| EPI_ISL_3445527, EPI_ISL_3445528 | Lighthouse Lab in Milton Keynes | Wellcome Sanger Institute for the COVID-19 Genomics UK (COG-UK) Consortium | The Lighthouse Lab in Milton Keynes and Alex Alderton, Roberto Amato, Jeffrey Barrett, Sonia Goncalves, Ewan Harrison, David K. Jackson, Ian Johnston, Dominic Kwiatkowski, Cordelia Langford, John Sillitoe on behalf of the Wellcome Sanger Institute COVID-19 Surveillance Team |
| EPI_ISL_3445529, EPI_ISL_3445530, EPI_ISL_3445531, EPI_ISL_3445532, EPI_ISL_3445533, EPI_ISL_3445534 | Lighthouse Lab in Glasgow | Wellcome Sanger Institute for the COVID-19 Genomics UK (COG-UK) Consortium | Harper VanSteenhouse, Yumi Kasai, David Gray, Carol Clugston, Anna Dominiczak and Alex Alderton, Roberto Amato, Jeffrey Barrett, Sonia Goncalves, Ewan Harrison, David K. Jackson, Ian Johnston, Dominic Kwiatkowski, Cordelia Langford, John Sillitoe on behalf of the Wellcome Sanger Institute COVID-19 Surveillance Team |
| EPI_ISL_3445535 | Lighthouse Lab in Milton Keynes | Wellcome Sanger Institute for the COVID-19 Genomics UK (COG-UK) Consortium | The Lighthouse Lab in Milton Keynes and Alex Alderton, Roberto Amato, Jeffrey Barrett, Sonia Goncalves, Ewan Harrison, David K. Jackson, Ian Johnston, Dominic Kwiatkowski, Cordelia Langford, John Sillitoe on behalf of the Wellcome Sanger Institute COVID-19 Surveillance Team |
| EPI_ISL_3445536 | Lighthouse Lab in Glasgow | Wellcome Sanger Institute for the COVID-19 Genomics UK (COG-UK) Consortium | Harper VanSteenhouse, Yumi Kasai, David Gray, Carol Clugston, Anna Dominiczak and Alex Alderton, Roberto Amato, Jeffrey Barrett, Sonia Goncalves, Ewan Harrison, David K. Jackson, Ian Johnston, Dominic Kwiatkowski, Cordelia Langford, John Sillitoe on behalf of the Wellcome Sanger Institute COVID-19 Surveillance Team |
| EPI_ISL_3445537 | Lighthouse Lab in Milton Keynes | Wellcome Sanger Institute for the COVID-19 Genomics UK (COG-UK) Consortium | The Lighthouse Lab in Milton Keynes and Alex Alderton, Roberto Amato, Jeffrey Barrett, Sonia Goncalves, Ewan Harrison, David K. Jackson, Ian Johnston, Dominic Kwiatkowski, Cordelia Langford, John Sillitoe on behalf of the Wellcome Sanger Institute COVID-19 Surveillance Team |
| EPI_ISL_3445538 | Lighthouse Lab in Glasgow | Wellcome Sanger Institute for the COVID-19 Genomics UK (COG-UK) Consortium | Harper VanSteenhouse, Yumi Kasai, David Gray, Carol Clugston, Anna Dominiczak and Alex Alderton, Roberto Amato, Jeffrey Barrett, Sonia Goncalves, Ewan Harrison, David K. Jackson, Ian Johnston, Dominic Kwiatkowski, Cordelia Langford, John Sillitoe on behalf of the Wellcome Sanger Institute COVID-19 Surveillance Team |
| EPI_ISL_3445539 | Lighthouse Lab in Milton Keynes | Wellcome Sanger Institute for the COVID-19 Genomics UK (COG-UK) Consortium | The Lighthouse Lab in Milton Keynes and Alex Alderton, Roberto Amato, Jeffrey Barrett, Sonia Goncalves, Ewan Harrison, David K. Jackson, Ian Johnston, Dominic Kwiatkowski, Cordelia Langford, John Sillitoe on behalf of the Wellcome Sanger Institute COVID-19 Surveillance Team |
| EPI_ISL_3445540 | Lighthouse Lab in Glasgow | Wellcome Sanger Institute for the COVID-19 Genomics UK (COG-UK) Consortium | Harper VanSteenhouse, Yumi Kasai, David Gray, Carol Clugston, Anna Dominiczak and Alex Alderton, Roberto Amato, Jeffrey Barrett, Sonia Goncalves, Ewan Harrison, David K. Jackson, Ian Johnston, Dominic Kwiatkowski, Cordelia Langford, John Sillitoe on behalf of the Wellcome Sanger Institute COVID-19 Surveillance Team |
| EPI_ISL_3445541 | Lighthouse Lab in Milton Keynes | Wellcome Sanger Institute for the COVID-19 Genomics UK (COG-UK) Consortium | The Lighthouse Lab in Milton Keynes and Alex Alderton, Roberto Amato, Jeffrey Barrett, Sonia Goncalves, Ewan Harrison, David K. Jackson, Ian Johnston, Dominic Kwiatkowski, Cordelia Langford, John Sillitoe on behalf of the Wellcome Sanger Institute COVID-19 Surveillance Team |
| EPI_ISL_3445542, EPI_ISL_3445543, EPI_ISL_3445544 | Lighthouse Lab in Glasgow | Wellcome Sanger Institute for the COVID-19 Genomics UK (COG-UK) Consortium | Harper VanSteenhouse, Yumi Kasai, David Gray, Carol Clugston, Anna Dominiczak and Alex Alderton, Roberto Amato, Jeffrey Barrett, Sonia Goncalves, Ewan Harrison, David K. Jackson, Ian Johnston, Dominic Kwiatkowski, Cordelia Langford, John Sillitoe on behalf of the Wellcome Sanger Institute COVID-19 Surveillance Team |
| EPI_ISL_3445545, EPI_ISL_3445546, EPI_ISL_3445547, EPI_ISL_3445548 | Lighthouse Lab in Milton Keynes | Wellcome Sanger Institute for the COVID-19 Genomics UK (COG-UK) Consortium | The Lighthouse Lab in Milton Keynes and Alex Alderton, Roberto Amato, Jeffrey Barrett, Sonia Goncalves, Ewan Harrison, David K. Jackson, Ian Johnston, Dominic Kwiatkowski, Cordelia Langford, John Sillitoe on behalf of the Wellcome Sanger Institute COVID-19 Surveillance Team |
| EPI_ISL_3445549 | Lighthouse Lab in Glasgow | Wellcome Sanger Institute for the COVID-19 Genomics UK (COG-UK) Consortium | Harper VanSteenhouse, Yumi Kasai, David Gray, Carol Clugston, Anna Dominiczak and Alex Alderton, Roberto Amato, Jeffrey Barrett, Sonia Goncalves, Ewan Harrison, David K. Jackson, Ian Johnston, Dominic Kwiatkowski, Cordelia Langford, John Sillitoe on behalf of the Wellcome Sanger Institute COVID-19 Surveillance Team |
| EPI_ISL_3445550 | Lighthouse Lab in Milton Keynes | Wellcome Sanger Institute for the COVID-19 Genomics UK (COG-UK) Consortium | The Lighthouse Lab in Milton Keynes and Alex Alderton, Roberto Amato, Jeffrey Barrett, Sonia Goncalves, Ewan Harrison, David K. Jackson, Ian Johnston, Dominic Kwiatkowski, Cordelia Langford, John Sillitoe on behalf of the Wellcome Sanger Institute COVID-19 Surveillance Team |
| EPI_ISL_3445551 | Lighthouse Lab in Glasgow | Wellcome Sanger Institute for the COVID-19 Genomics UK (COG-UK) Consortium | Harper VanSteenhouse, Yumi Kasai, David Gray, Carol Clugston, Anna Dominiczak and Alex Alderton, Roberto Amato, Jeffrey Barrett, Sonia Goncalves, Ewan Harrison, David K. Jackson, Ian Johnston, Dominic Kwiatkowski, Cordelia Langford, John Sillitoe on behalf of the Wellcome Sanger Institute COVID-19 Surveillance Team |
| EPI_ISL_3445552, EPI_ISL_3445553 | Lighthouse Lab in Milton Keynes | Wellcome Sanger Institute for the COVID-19 Genomics UK (COG-UK) Consortium | The Lighthouse Lab in Milton Keynes and Alex Alderton, Roberto Amato, Jeffrey Barrett, Sonia Goncalves, Ewan Harrison, David K. Jackson, Ian Johnston, Dominic Kwiatkowski, Cordelia Langford, John Sillitoe on behalf of the Wellcome Sanger Institute COVID-19 Surveillance Team |
| EPI_ISL_3445554, EPI_ISL_3445555, EPI_ISL_3445556 | Lighthouse Lab in Glasgow | Wellcome Sanger Institute for the COVID-19 Genomics UK (COG-UK) Consortium | Harper VanSteenhouse, Yumi Kasai, David Gray, Carol Clugston, Anna Dominiczak and Alex Alderton, Roberto Amato, Jeffrey Barrett, Sonia Goncalves, Ewan Harrison, David K. Jackson, Ian Johnston, Dominic Kwiatkowski, Cordelia Langford, John Sillitoe on behalf of the Wellcome Sanger Institute COVID-19 Surveillance Team |
| EPI_ISL_3445557 | Lighthouse Lab in Milton Keynes | Wellcome Sanger Institute for the COVID-19 Genomics UK (COG-UK) Consortium | The Lighthouse Lab in Milton Keynes and Alex Alderton, Roberto Amato, Jeffrey Barrett, Sonia Goncalves, Ewan Harrison, David K. Jackson, Ian Johnston, Dominic Kwiatkowski, Cordelia Langford, John Sillitoe on behalf of the Wellcome Sanger Institute COVID-19 Surveillance Team |
| EPI_ISL_3445558, EPI_ISL_3445559 | Lighthouse Lab in Glasgow | Wellcome Sanger Institute for the COVID-19 Genomics UK (COG-UK) Consortium | Harper VanSteenhouse, Yumi Kasai, David Gray, Carol Clugston, Anna Dominiczak and Alex Alderton, Roberto Amato, Jeffrey Barrett, Sonia Goncalves, Ewan Harrison, David K. Jackson, Ian Johnston, Dominic Kwiatkowski, Cordelia Langford, John Sillitoe on behalf of the Wellcome Sanger Institute COVID-19 Surveillance Team |
| EPI_ISL_3445560 | Lighthouse Lab in Milton Keynes | Wellcome Sanger Institute for the COVID-19 Genomics UK (COG-UK) Consortium | The Lighthouse Lab in Milton Keynes and Alex Alderton, Roberto Amato, Jeffrey Barrett, Sonia Goncalves, Ewan Harrison, David K. Jackson, Ian Johnston, Dominic Kwiatkowski, Cordelia Langford, John Sillitoe on behalf of the Wellcome Sanger Institute COVID-19 Surveillance Team |
| EPI_ISL_3445561, EPI_ISL_3445562, EPI_ISL_3445563 | Lighthouse Lab in Glasgow | Wellcome Sanger Institute for the COVID-19 Genomics UK (COG-UK) Consortium | Harper VanSteenhouse, Yumi Kasai, David Gray, Carol Clugston, Anna Dominiczak and Alex Alderton, Roberto Amato, Jeffrey Barrett, Sonia Goncalves, Ewan Harrison, David K. Jackson, Ian Johnston, Dominic Kwiatkowski, Cordelia Langford, John Sillitoe on behalf of the Wellcome Sanger Institute COVID-19 Surveillance Team |
| EPI_ISL_3445564, EPI_ISL_3445565 | Lighthouse Lab in Milton Keynes | Wellcome Sanger Institute for the COVID-19 Genomics UK (COG-UK) Consortium | The Lighthouse Lab in Milton Keynes and Alex Alderton, Roberto Amato, Jeffrey Barrett, Sonia Goncalves, Ewan Harrison, David K. Jackson, Ian Johnston, Dominic Kwiatkowski, Cordelia Langford, John Sillitoe on behalf of the Wellcome Sanger Institute COVID-19 Surveillance Team |
| EPI_ISL_3445566, EPI_ISL_3445567 | Lighthouse Lab in Glasgow | Wellcome Sanger Institute for the COVID-19 Genomics UK (COG-UK) Consortium | Harper VanSteenhouse, Yumi Kasai, David Gray, Carol Clugston, Anna Dominiczak and Alex Alderton, Roberto Amato, Jeffrey Barrett, Sonia Goncalves, Ewan Harrison, David K. Jackson, Ian Johnston, Dominic Kwiatkowski, Cordelia Langford, John Sillitoe on behalf of the Wellcome Sanger Institute COVID-19 Surveillance Team |
| EPI_ISL_3445568 | Lighthouse Lab in Milton Keynes | Wellcome Sanger Institute for the COVID-19 Genomics UK (COG-UK) Consortium | The Lighthouse Lab in Milton Keynes and Alex Alderton, Roberto Amato, Jeffrey Barrett, Sonia Goncalves, Ewan Harrison, David K. Jackson, Ian Johnston, Dominic Kwiatkowski, Cordelia Langford, John Sillitoe on behalf of the Wellcome Sanger Institute COVID-19 Surveillance Team |
| EPI_ISL_3445569 | Lighthouse Lab in Glasgow | Wellcome Sanger Institute for the COVID-19 Genomics UK (COG-UK) Consortium | Harper VanSteenhouse, Yumi Kasai, David Gray, Carol Clugston, Anna Dominiczak and Alex Alderton, Roberto Amato, Jeffrey Barrett, Sonia Goncalves, Ewan Harrison, David K. Jackson, Ian Johnston, Dominic Kwiatkowski, Cordelia Langford, John Sillitoe on behalf of the Wellcome Sanger Institute COVID-19 Surveillance Team |
| EPI_ISL_3445570 | Lighthouse Lab in Milton Keynes | Wellcome Sanger Institute for the COVID-19 Genomics UK (COG-UK) Consortium | The Lighthouse Lab in Milton Keynes and Alex Alderton, Roberto Amato, Jeffrey Barrett, Sonia Goncalves, Ewan Harrison, David K. Jackson, Ian Johnston, Dominic Kwiatkowski, Cordelia Langford, John Sillitoe on behalf of the Wellcome Sanger Institute COVID-19 Surveillance Team |
| EPI_ISL_3445571, EPI_ISL_3445572, EPI_ISL_3445573 | Lighthouse Lab in Glasgow | Wellcome Sanger Institute for the COVID-19 Genomics UK (COG-UK) Consortium | Harper VanSteenhouse, Yumi Kasai, David Gray, Carol Clugston, Anna Dominiczak and Alex Alderton, Roberto Amato, Jeffrey Barrett, Sonia Goncalves, Ewan Harrison, David K. Jackson, Ian Johnston, Dominic Kwiatkowski, Cordelia Langford, John Sillitoe on behalf of the Wellcome Sanger Institute COVID-19 Surveillance Team |
| EPI_ISL_3445574 | Lighthouse Lab in Milton Keynes | Wellcome Sanger Institute for the COVID-19 Genomics UK (COG-UK) Consortium | The Lighthouse Lab in Milton Keynes and Alex Alderton, Roberto Amato, Jeffrey Barrett, Sonia Goncalves, Ewan Harrison, David K. Jackson, Ian Johnston, Dominic Kwiatkowski, Cordelia Langford, John Sillitoe on behalf of the Wellcome Sanger Institute COVID-19 Surveillance Team |
| EPI_ISL_3445575, EPI_ISL_3445576, EPI_ISL_3445577, EPI_ISL_3445578, EPI_ISL_3445579, EPI_ISL_3445580 | Lighthouse Lab in Glasgow | Wellcome Sanger Institute for the COVID-19 Genomics UK (COG-UK) Consortium | Harper VanSteenhouse, Yumi Kasai, David Gray, Carol Clugston, Anna Dominiczak and Alex Alderton, Roberto Amato, Jeffrey Barrett, Sonia Goncalves, Ewan Harrison, David K. Jackson, Ian Johnston, Dominic Kwiatkowski, Cordelia Langford, John Sillitoe on behalf of the Wellcome Sanger Institute COVID-19 Surveillance Team |
| EPI_ISL_3445581 | Lighthouse Lab in Milton Keynes | Wellcome Sanger Institute for the COVID-19 Genomics UK (COG-UK) Consortium | The Lighthouse Lab in Milton Keynes and Alex Alderton, Roberto Amato, Jeffrey Barrett, Sonia Goncalves, Ewan Harrison, David K. Jackson, Ian Johnston, Dominic Kwiatkowski, Cordelia Langford, John Sillitoe on behalf of the Wellcome Sanger Institute COVID-19 Surveillance Team |
| EPI_ISL_3445582, | Lighthouse Lab in Glasgow | Wellcome Sanger Institute for the COVID-19 Genomics UK (COG-UK) | Harper VanSteenhouse, Yumi Kasai, David Gray, Carol Clugston, Anna Dominiczak and Alex Alderton, Roberto Amato, Jeffrey Barrett, Sonia Goncalves, Ewan Harrison, David K. Jackson, Ian Johnston, Dominic Kwiatkowski, Cordelia Langford, |

| | | | |
|---|---|---|---|
| EPI_ISL_3445583, EPI_ISL_3445584 | | Consortium | John Sillitoe on behalf of the Wellcome Sanger Institute COVID-19 Surveillance Team |
| EPI_ISL_3445585 | Lighthouse Lab in Milton Keynes | Wellcome Sanger Institute for the COVID-19 Genomics UK (COG-UK) Consortium | The Lighthouse Lab in Milton Keynes and Alex Alderton, Roberto Amato, Jeffrey Barrett, Sonia Goncalves, Ewan Harrison, David K. Jackson, Ian Johnston, Dominic Kwiatkowski, Cordelia Langford, John Sillitoe on behalf of the Wellcome Sanger Institute COVID-19 Surveillance Team |
| EPI_ISL_3445586, EPI_ISL_3445587, EPI_ISL_3445588, EPI_ISL_3445589, EPI_ISL_3445590, EPI_ISL_3445591, EPI_ISL_3445592 | Lighthouse Lab in Glasgow | Wellcome Sanger Institute for the COVID-19 Genomics UK (COG-UK) Consortium | Harper VanSteenhouse, Yumi Kasai, David Gray, Carol Clugston, Anna Dominiczak and Alex Alderton, Roberto Amato, Jeffrey Barrett, Sonia Goncalves, Ewan Harrison, David K. Jackson, Ian Johnston, Dominic Kwiatkowski, Cordelia Langford, John Sillitoe on behalf of the Wellcome Sanger Institute COVID-19 Surveillance Team |
| EPI_ISL_3445593 | Lighthouse Lab in Milton Keynes | Wellcome Sanger Institute for the COVID-19 Genomics UK (COG-UK) Consortium | The Lighthouse Lab in Milton Keynes and Alex Alderton, Roberto Amato, Jeffrey Barrett, Sonia Goncalves, Ewan Harrison, David K. Jackson, Ian Johnston, Dominic Kwiatkowski, Cordelia Langford, John Sillitoe on behalf of the Wellcome Sanger Institute COVID-19 Surveillance Team |
| EPI_ISL_3445594, EPI_ISL_3445595, EPI_ISL_3445596, EPI_ISL_3445597 | Lighthouse Lab in Glasgow | Wellcome Sanger Institute for the COVID-19 Genomics UK (COG-UK) Consortium | Harper VanSteenhouse, Yumi Kasai, David Gray, Carol Clugston, Anna Dominiczak and Alex Alderton, Roberto Amato, Jeffrey Barrett, Sonia Goncalves, Ewan Harrison, David K. Jackson, Ian Johnston, Dominic Kwiatkowski, Cordelia Langford, John Sillitoe on behalf of the Wellcome Sanger Institute COVID-19 Surveillance Team |
| EPI_ISL_3445598 | Lighthouse Lab in Milton Keynes | Wellcome Sanger Institute for the COVID-19 Genomics UK (COG-UK) Consortium | The Lighthouse Lab in Milton Keynes and Alex Alderton, Roberto Amato, Jeffrey Barrett, Sonia Goncalves, Ewan Harrison, David K. Jackson, Ian Johnston, Dominic Kwiatkowski, Cordelia Langford, John Sillitoe on behalf of the Wellcome Sanger Institute COVID-19 Surveillance Team |
| EPI_ISL_3445599, EPI_ISL_3445600, EPI_ISL_3445601, EPI_ISL_3445602, EPI_ISL_3445603, EPI_ISL_3445604, EPI_ISL_3445605, EPI_ISL_3445606, EPI_ISL_3445607 | Lighthouse Lab in Glasgow | Wellcome Sanger Institute for the COVID-19 Genomics UK (COG-UK) Consortium | Harper VanSteenhouse, Yumi Kasai, David Gray, Carol Clugston, Anna Dominiczak and Alex Alderton, Roberto Amato, Jeffrey Barrett, Sonia Goncalves, Ewan Harrison, David K. Jackson, Ian Johnston, Dominic Kwiatkowski, Cordelia Langford, John Sillitoe on behalf of the Wellcome Sanger Institute COVID-19 Surveillance Team |
| EPI_ISL_3445608 | Lighthouse Lab in Milton Keynes | Wellcome Sanger Institute for the COVID-19 Genomics UK (COG-UK) Consortium | The Lighthouse Lab in Milton Keynes and Alex Alderton, Roberto Amato, Jeffrey Barrett, Sonia Goncalves, Ewan Harrison, David K. Jackson, Ian Johnston, Dominic Kwiatkowski, Cordelia Langford, John Sillitoe on behalf of the Wellcome Sanger Institute COVID-19 Surveillance Team |
| EPI_ISL_3445609 | Lighthouse Lab in Glasgow | Wellcome Sanger Institute for the COVID-19 Genomics UK (COG-UK) Consortium | Harper VanSteenhouse, Yumi Kasai, David Gray, Carol Clugston, Anna Dominiczak and Alex Alderton, Roberto Amato, Jeffrey Barrett, Sonia Goncalves, Ewan Harrison, David K. Jackson, Ian Johnston, Dominic Kwiatkowski, Cordelia Langford, John Sillitoe on behalf of the Wellcome Sanger Institute COVID-19 Surveillance Team |
| EPI_ISL_3445610, EPI_ISL_3445611 | Lighthouse Lab in Milton Keynes | Wellcome Sanger Institute for the COVID-19 Genomics UK (COG-UK) Consortium | The Lighthouse Lab in Milton Keynes and Alex Alderton, Roberto Amato, Jeffrey Barrett, Sonia Goncalves, Ewan Harrison, David K. Jackson, Ian Johnston, Dominic Kwiatkowski, Cordelia Langford, John Sillitoe on behalf of the Wellcome Sanger Institute COVID-19 Surveillance Team |
| EPI_ISL_3445612, EPI_ISL_3445613, EPI_ISL_3445614, EPI_ISL_3445615, EPI_ISL_3445616 | Lighthouse Lab in Glasgow | Wellcome Sanger Institute for the COVID-19 Genomics UK (COG-UK) Consortium | Harper VanSteenhouse, Yumi Kasai, David Gray, Carol Clugston, Anna Dominiczak and Alex Alderton, Roberto Amato, Jeffrey Barrett, Sonia Goncalves, Ewan Harrison, David K. Jackson, Ian Johnston, Dominic Kwiatkowski, Cordelia Langford, John Sillitoe on behalf of the Wellcome Sanger Institute COVID-19 Surveillance Team |
| EPI_ISL_3445617 | Lighthouse Lab in Milton Keynes | Wellcome Sanger Institute for the COVID-19 Genomics UK (COG-UK) Consortium | The Lighthouse Lab in Milton Keynes and Alex Alderton, Roberto Amato, Jeffrey Barrett, Sonia Goncalves, Ewan Harrison, David K. Jackson, Ian Johnston, Dominic Kwiatkowski, Cordelia Langford, John Sillitoe on behalf of the Wellcome Sanger Institute COVID-19 Surveillance Team |
| EPI_ISL_3445618 | Lighthouse Lab in Glasgow | Wellcome Sanger Institute for the COVID-19 Genomics UK (COG-UK) Consortium | Harper VanSteenhouse, Yumi Kasai, David Gray, Carol Clugston, Anna Dominiczak and Alex Alderton, Roberto Amato, Jeffrey Barrett, Sonia Goncalves, Ewan Harrison, David K. Jackson, Ian Johnston, Dominic Kwiatkowski, Cordelia Langford, John Sillitoe on behalf of the Wellcome Sanger Institute COVID-19 Surveillance Team |
| EPI_ISL_3445619 | Lighthouse Lab in Milton Keynes | Wellcome Sanger Institute for the COVID-19 Genomics UK (COG-UK) Consortium | The Lighthouse Lab in Milton Keynes and Alex Alderton, Roberto Amato, Jeffrey Barrett, Sonia Goncalves, Ewan Harrison, David K. Jackson, Ian Johnston, Dominic Kwiatkowski, Cordelia Langford, John Sillitoe on behalf of the Wellcome Sanger Institute COVID-19 Surveillance Team |
| EPI_ISL_3445620, EPI_ISL_3445621, EPI_ISL_3445622, EPI_ISL_3445623, EPI_ISL_3445624 | Lighthouse Lab in Glasgow | Wellcome Sanger Institute for the COVID-19 Genomics UK (COG-UK) Consortium | Harper VanSteenhouse, Yumi Kasai, David Gray, Carol Clugston, Anna Dominiczak and Alex Alderton, Roberto Amato, Jeffrey Barrett, Sonia Goncalves, Ewan Harrison, David K. Jackson, Ian Johnston, Dominic Kwiatkowski, Cordelia Langford, John Sillitoe on behalf of the Wellcome Sanger Institute COVID-19 Surveillance Team |
| EPI_ISL_3445625, EPI_ISL_3445626, EPI_ISL_3445627 | Lighthouse Lab in Milton Keynes | Wellcome Sanger Institute for the COVID-19 Genomics UK (COG-UK) Consortium | The Lighthouse Lab in Milton Keynes and Alex Alderton, Roberto Amato, Jeffrey Barrett, Sonia Goncalves, Ewan Harrison, David K. Jackson, Ian Johnston, Dominic Kwiatkowski, Cordelia Langford, John Sillitoe on behalf of the Wellcome Sanger Institute COVID-19 Surveillance Team |
| EPI_ISL_3445628 | Lighthouse Lab in Glasgow | Wellcome Sanger Institute for the COVID-19 Genomics UK (COG-UK) Consortium | Harper VanSteenhouse, Yumi Kasai, David Gray, Carol Clugston, Anna Dominiczak and Alex Alderton, Roberto Amato, Jeffrey Barrett, Sonia Goncalves, Ewan Harrison, David K. Jackson, Ian Johnston, Dominic Kwiatkowski, Cordelia Langford, John Sillitoe on behalf of the Wellcome Sanger Institute COVID-19 Surveillance Team |
| EPI_ISL_3445629 | Lighthouse Lab in Milton Keynes | Wellcome Sanger Institute for the COVID-19 Genomics UK (COG-UK) Consortium | The Lighthouse Lab in Milton Keynes and Alex Alderton, Roberto Amato, Jeffrey Barrett, Sonia Goncalves, Ewan Harrison, David K. Jackson, Ian Johnston, Dominic Kwiatkowski, Cordelia Langford, John Sillitoe on behalf of the Wellcome Sanger Institute COVID-19 Surveillance Team |
| EPI_ISL_3445630, EPI_ISL_3445631, EPI_ISL_3445632, EPI_ISL_3445633 | Lighthouse Lab in Glasgow | Wellcome Sanger Institute for the COVID-19 Genomics UK (COG-UK) Consortium | Harper VanSteenhouse, Yumi Kasai, David Gray, Carol Clugston, Anna Dominiczak and Alex Alderton, Roberto Amato, Jeffrey Barrett, Sonia Goncalves, Ewan Harrison, David K. Jackson, Ian Johnston, Dominic Kwiatkowski, Cordelia Langford, John Sillitoe on behalf of the Wellcome Sanger Institute COVID-19 Surveillance Team |
| EPI_ISL_3445634, EPI_ISL_3445635 | Lighthouse Lab in Milton Keynes | Wellcome Sanger Institute for the COVID-19 Genomics UK (COG-UK) Consortium | The Lighthouse Lab in Milton Keynes and Alex Alderton, Roberto Amato, Jeffrey Barrett, Sonia Goncalves, Ewan Harrison, David K. Jackson, Ian Johnston, Dominic Kwiatkowski, Cordelia Langford, John Sillitoe on behalf of the Wellcome Sanger Institute COVID-19 Surveillance Team |
| EPI_ISL_3445636 | Lighthouse Lab in Glasgow | Wellcome Sanger Institute for the COVID-19 Genomics UK (COG-UK) Consortium | Harper VanSteenhouse, Yumi Kasai, David Gray, Carol Clugston, Anna Dominiczak and Alex Alderton, Roberto Amato, Jeffrey Barrett, Sonia Goncalves, Ewan Harrison, David K. Jackson, Ian Johnston, Dominic Kwiatkowski, Cordelia Langford, John Sillitoe on behalf of the Wellcome Sanger Institute COVID-19 Surveillance Team |
| EPI_ISL_3445637 | Lighthouse Lab in Milton Keynes | Wellcome Sanger Institute for the COVID-19 Genomics UK (COG-UK) Consortium | The Lighthouse Lab in Milton Keynes and Alex Alderton, Roberto Amato, Jeffrey Barrett, Sonia Goncalves, Ewan Harrison, David K. Jackson, Ian Johnston, Dominic Kwiatkowski, Cordelia Langford, John Sillitoe on behalf of the Wellcome Sanger Institute COVID-19 Surveillance Team |
| EPI_ISL_3445638 | Lighthouse Lab in Glasgow | Wellcome Sanger Institute for the COVID-19 Genomics UK (COG-UK) Consortium | Harper VanSteenhouse, Yumi Kasai, David Gray, Carol Clugston, Anna Dominiczak and Alex Alderton, Roberto Amato, Jeffrey Barrett, Sonia Goncalves, Ewan Harrison, David K. Jackson, Ian Johnston, Dominic Kwiatkowski, Cordelia Langford, John Sillitoe on behalf of the Wellcome Sanger Institute COVID-19 Surveillance Team |
| EPI_ISL_3445639 | Lighthouse Lab in Milton Keynes | Wellcome Sanger Institute for the COVID-19 Genomics UK (COG-UK) Consortium | The Lighthouse Lab in Milton Keynes and Alex Alderton, Roberto Amato, Jeffrey Barrett, Sonia Goncalves, Ewan Harrison, David K. Jackson, Ian Johnston, Dominic Kwiatkowski, Cordelia Langford, John Sillitoe on behalf of the Wellcome Sanger Institute COVID-19 Surveillance Team |
| EPI_ISL_3445640, EPI_ISL_3445641 | Lighthouse Lab in Glasgow | Wellcome Sanger Institute for the COVID-19 Genomics UK (COG-UK) Consortium | Harper VanSteenhouse, Yumi Kasai, David Gray, Carol Clugston, Anna Dominiczak and Alex Alderton, Roberto Amato, Jeffrey Barrett, Sonia Goncalves, Ewan Harrison, David K. Jackson, Ian Johnston, Dominic Kwiatkowski, Cordelia Langford, John Sillitoe on behalf of the Wellcome Sanger Institute COVID-19 Surveillance Team |
| EPI_ISL_3445642 | Lighthouse Lab in Milton Keynes | Wellcome Sanger Institute for the COVID-19 Genomics UK (COG-UK) Consortium | The Lighthouse Lab in Milton Keynes and Alex Alderton, Roberto Amato, Jeffrey Barrett, Sonia Goncalves, Ewan Harrison, David K. Jackson, Ian Johnston, Dominic Kwiatkowski, Cordelia Langford, John Sillitoe on behalf of the Wellcome Sanger Institute COVID-19 Surveillance Team |
| EPI_ISL_3445643, EPI_ISL_3445644, EPI_ISL_3445645, EPI_ISL_3445646, EPI_ISL_3445647, EPI_ISL_3445648, EPI_ISL_3445649, EPI_ISL_3445650 | Lighthouse Lab in Glasgow | Wellcome Sanger Institute for the COVID-19 Genomics UK (COG-UK) Consortium | Harper VanSteenhouse, Yumi Kasai, David Gray, Carol Clugston, Anna Dominiczak and Alex Alderton, Roberto Amato, Jeffrey Barrett, Sonia Goncalves, Ewan Harrison, David K. Jackson, Ian Johnston, Dominic Kwiatkowski, Cordelia Langford, John Sillitoe on behalf of the Wellcome Sanger Institute COVID-19 Surveillance Team |
| EPI_ISL_3445651 | Lighthouse Lab in Milton Keynes | Wellcome Sanger Institute for the COVID-19 Genomics UK (COG-UK) Consortium | The Lighthouse Lab in Milton Keynes and Alex Alderton, Roberto Amato, Jeffrey Barrett, Sonia Goncalves, Ewan Harrison, David K. Jackson, Ian Johnston, Dominic Kwiatkowski, Cordelia Langford, John Sillitoe on behalf of the Wellcome Sanger Institute COVID-19 Surveillance Team |
| EPI_ISL_3445652, EPI_ISL_3445653, EPI_ISL_3445654, EPI_ISL_3445655 | Lighthouse Lab in Glasgow | Wellcome Sanger Institute for the COVID-19 Genomics UK (COG-UK) Consortium | Harper VanSteenhouse, Yumi Kasai, David Gray, Carol Clugston, Anna Dominiczak and Alex Alderton, Roberto Amato, Jeffrey Barrett, Sonia Goncalves, Ewan Harrison, David K. Jackson, Ian Johnston, Dominic Kwiatkowski, Cordelia Langford, John Sillitoe on behalf of the Wellcome Sanger Institute COVID-19 Surveillance Team |
| EPI_ISL_3445656 | Lighthouse Lab in Milton Keynes | Wellcome Sanger Institute for the COVID-19 Genomics UK (COG-UK) Consortium | The Lighthouse Lab in Milton Keynes and Alex Alderton, Roberto Amato, Jeffrey Barrett, Sonia Goncalves, Ewan Harrison, David K. Jackson, Ian Johnston, Dominic Kwiatkowski, Cordelia Langford, John Sillitoe on behalf of the Wellcome Sanger Institute COVID-19 Surveillance Team |
| EPI_ISL_3445657 | Lighthouse Lab in Glasgow | Wellcome Sanger Institute for the COVID-19 Genomics UK (COG-UK) Consortium | Harper VanSteenhouse, Yumi Kasai, David Gray, Carol Clugston, Anna Dominiczak and Alex Alderton, Roberto Amato, Jeffrey Barrett, Sonia Goncalves, Ewan Harrison, David K. Jackson, Ian Johnston, Dominic Kwiatkowski, Cordelia Langford, John Sillitoe on behalf of the Wellcome Sanger Institute COVID-19 Surveillance Team |
| EPI_ISL_3445658 | Lighthouse Lab in Milton Keynes | Wellcome Sanger Institute for the COVID-19 Genomics UK (COG-UK) Consortium | The Lighthouse Lab in Milton Keynes and Alex Alderton, Roberto Amato, Jeffrey Barrett, Sonia Goncalves, Ewan Harrison, David K. Jackson, Ian Johnston, Dominic Kwiatkowski, Cordelia Langford, John Sillitoe on behalf of the Wellcome Sanger Institute COVID-19 Surveillance Team |
| EPI_ISL_3445659, | Lighthouse Lab in Glasgow | Wellcome Sanger Institute for the COVID-19 Genomics UK (COG-UK) | Harper VanSteenhouse, Yumi Kasai, David Gray, Carol Clugston, Anna Dominiczak and Alex Alderton, Roberto Amato, Jeffrey Barrett, Sonia Goncalves, Ewan Harrison, David K. Jackson, Ian Johnston, Dominic Kwiatkowski, Cordelia Langford, |

| | | | |
|---|---|---|---|
| EPI_ISL_3445660, EPI_ISL_3445661, EPI_ISL_3445662, EPI_ISL_3445663, EPI_ISL_3445664, EPI_ISL_3445665, EPI_ISL_3445666, EPI_ISL_3445667 | | Consortium | John Sillitoe on behalf of the Wellcome Sanger Institute COVID-19 Surveillance Team |
| EPI_ISL_3445668 | Lighthouse Lab in Milton Keynes | Wellcome Sanger Institute for the COVID-19 Genomics UK (COG-UK) Consortium | The Lighthouse Lab in Milton Keynes and Alex Alderton, Roberto Amato, Jeffrey Barrett, Sonia Goncalves, Ewan Harrison, David K. Jackson, Ian Johnston, Dominic Kwiatkowski, Cordelia Langford, John Sillitoe on behalf of the Wellcome Sanger Institute COVID-19 Surveillance Team |
| EPI_ISL_3445669, EPI_ISL_3445670 | Lighthouse Lab in Glasgow | Wellcome Sanger Institute for the COVID-19 Genomics UK (COG-UK) Consortium | Harper VanSteenhouse, Yumi Kasai, David Gray, Carol Clugston, Anna Dominiczak and Alex Alderton, Roberto Amato, Jeffrey Barrett, Sonia Goncalves, Ewan Harrison, David K. Jackson, Ian Johnston, Dominic Kwiatkowski, Cordelia Langford, John Sillitoe on behalf of the Wellcome Sanger Institute COVID-19 Surveillance Team |
| EPI_ISL_3445671 | Lighthouse Lab in Milton Keynes | Wellcome Sanger Institute for the COVID-19 Genomics UK (COG-UK) Consortium | The Lighthouse Lab in Milton Keynes and Alex Alderton, Roberto Amato, Jeffrey Barrett, Sonia Goncalves, Ewan Harrison, David K. Jackson, Ian Johnston, Dominic Kwiatkowski, Cordelia Langford, John Sillitoe on behalf of the Wellcome Sanger Institute COVID-19 Surveillance Team |
| EPI_ISL_3445672, EPI_ISL_3445673 | Lighthouse Lab in Glasgow | Wellcome Sanger Institute for the COVID-19 Genomics UK (COG-UK) Consortium | Harper VanSteenhouse, Yumi Kasai, David Gray, Carol Clugston, Anna Dominiczak and Alex Alderton, Roberto Amato, Jeffrey Barrett, Sonia Goncalves, Ewan Harrison, David K. Jackson, Ian Johnston, Dominic Kwiatkowski, Cordelia Langford, John Sillitoe on behalf of the Wellcome Sanger Institute COVID-19 Surveillance Team |
| EPI_ISL_3445674, EPI_ISL_3445675, EPI_ISL_3445676, EPI_ISL_3445677, EPI_ISL_3445678 | Lighthouse Lab in Milton Keynes | Wellcome Sanger Institute for the COVID-19 Genomics UK (COG-UK) Consortium | The Lighthouse Lab in Milton Keynes and Alex Alderton, Roberto Amato, Jeffrey Barrett, Sonia Goncalves, Ewan Harrison, David K. Jackson, Ian Johnston, Dominic Kwiatkowski, Cordelia Langford, John Sillitoe on behalf of the Wellcome Sanger Institute COVID-19 Surveillance Team |
| EPI_ISL_3445679, EPI_ISL_3445680, EPI_ISL_3445681, EPI_ISL_3445682, EPI_ISL_3445683, EPI_ISL_3445684, EPI_ISL_3445685, EPI_ISL_3445686, EPI_ISL_3445687 | Lighthouse Lab in Glasgow | Wellcome Sanger Institute for the COVID-19 Genomics UK (COG-UK) Consortium | Harper VanSteenhouse, Yumi Kasai, David Gray, Carol Clugston, Anna Dominiczak and Alex Alderton, Roberto Amato, Jeffrey Barrett, Sonia Goncalves, Ewan Harrison, David K. Jackson, Ian Johnston, Dominic Kwiatkowski, Cordelia Langford, John Sillitoe on behalf of the Wellcome Sanger Institute COVID-19 Surveillance Team |
| EPI_ISL_3445688, EPI_ISL_3445689 | Lighthouse Lab in Milton Keynes | Wellcome Sanger Institute for the COVID-19 Genomics UK (COG-UK) Consortium | The Lighthouse Lab in Milton Keynes and Alex Alderton, Roberto Amato, Jeffrey Barrett, Sonia Goncalves, Ewan Harrison, David K. Jackson, Ian Johnston, Dominic Kwiatkowski, Cordelia Langford, John Sillitoe on behalf of the Wellcome Sanger Institute COVID-19 Surveillance Team |
| EPI_ISL_3445690, EPI_ISL_3445691, EPI_ISL_3445692 | Lighthouse Lab in Glasgow | Wellcome Sanger Institute for the COVID-19 Genomics UK (COG-UK) Consortium | Harper VanSteenhouse, Yumi Kasai, David Gray, Carol Clugston, Anna Dominiczak and Alex Alderton, Roberto Amato, Jeffrey Barrett, Sonia Goncalves, Ewan Harrison, David K. Jackson, Ian Johnston, Dominic Kwiatkowski, Cordelia Langford, John Sillitoe on behalf of the Wellcome Sanger Institute COVID-19 Surveillance Team |
| EPI_ISL_3445693 | Lighthouse Lab in Milton Keynes | Wellcome Sanger Institute for the COVID-19 Genomics UK (COG-UK) Consortium | The Lighthouse Lab in Milton Keynes and Alex Alderton, Roberto Amato, Jeffrey Barrett, Sonia Goncalves, Ewan Harrison, David K. Jackson, Ian Johnston, Dominic Kwiatkowski, Cordelia Langford, John Sillitoe on behalf of the Wellcome Sanger Institute COVID-19 Surveillance Team |
| EPI_ISL_3445694 | Lighthouse Lab in Glasgow | Wellcome Sanger Institute for the COVID-19 Genomics UK (COG-UK) Consortium | Harper VanSteenhouse, Yumi Kasai, David Gray, Carol Clugston, Anna Dominiczak and Alex Alderton, Roberto Amato, Jeffrey Barrett, Sonia Goncalves, Ewan Harrison, David K. Jackson, Ian Johnston, Dominic Kwiatkowski, Cordelia Langford, John Sillitoe on behalf of the Wellcome Sanger Institute COVID-19 Surveillance Team |
| EPI_ISL_3445695 | Lighthouse Lab in Milton Keynes | Wellcome Sanger Institute for the COVID-19 Genomics UK (COG-UK) Consortium | The Lighthouse Lab in Milton Keynes and Alex Alderton, Roberto Amato, Jeffrey Barrett, Sonia Goncalves, Ewan Harrison, David K. Jackson, Ian Johnston, Dominic Kwiatkowski, Cordelia Langford, John Sillitoe on behalf of the Wellcome Sanger Institute COVID-19 Surveillance Team |
| EPI_ISL_3445696 | Lighthouse Lab in Glasgow | Wellcome Sanger Institute for the COVID-19 Genomics UK (COG-UK) Consortium | Harper VanSteenhouse, Yumi Kasai, David Gray, Carol Clugston, Anna Dominiczak and Alex Alderton, Roberto Amato, Jeffrey Barrett, Sonia Goncalves, Ewan Harrison, David K. Jackson, Ian Johnston, Dominic Kwiatkowski, Cordelia Langford, John Sillitoe on behalf of the Wellcome Sanger Institute COVID-19 Surveillance Team |
| EPI_ISL_3445697, EPI_ISL_3445698 | Lighthouse Lab in Milton Keynes | Wellcome Sanger Institute for the COVID-19 Genomics UK (COG-UK) Consortium | The Lighthouse Lab in Milton Keynes and Alex Alderton, Roberto Amato, Jeffrey Barrett, Sonia Goncalves, Ewan Harrison, David K. Jackson, Ian Johnston, Dominic Kwiatkowski, Cordelia Langford, John Sillitoe on behalf of the Wellcome Sanger Institute COVID-19 Surveillance Team |
| EPI_ISL_3445699 | Lighthouse Lab in Glasgow | Wellcome Sanger Institute for the COVID-19 Genomics UK (COG-UK) Consortium | Harper VanSteenhouse, Yumi Kasai, David Gray, Carol Clugston, Anna Dominiczak and Alex Alderton, Roberto Amato, Jeffrey Barrett, Sonia Goncalves, Ewan Harrison, David K. Jackson, Ian Johnston, Dominic Kwiatkowski, Cordelia Langford, John Sillitoe on behalf of the Wellcome Sanger Institute COVID-19 Surveillance Team |
| EPI_ISL_3445700, EPI_ISL_3445701 | Lighthouse Lab in Milton Keynes | Wellcome Sanger Institute for the COVID-19 Genomics UK (COG-UK) Consortium | The Lighthouse Lab in Milton Keynes and Alex Alderton, Roberto Amato, Jeffrey Barrett, Sonia Goncalves, Ewan Harrison, David K. Jackson, Ian Johnston, Dominic Kwiatkowski, Cordelia Langford, John Sillitoe on behalf of the Wellcome Sanger Institute COVID-19 Surveillance Team |
| EPI_ISL_3445702 | Lighthouse Lab in Glasgow | Wellcome Sanger Institute for the COVID-19 Genomics UK (COG-UK) Consortium | Harper VanSteenhouse, Yumi Kasai, David Gray, Carol Clugston, Anna Dominiczak and Alex Alderton, Roberto Amato, Jeffrey Barrett, Sonia Goncalves, Ewan Harrison, David K. Jackson, Ian Johnston, Dominic Kwiatkowski, Cordelia Langford, John Sillitoe on behalf of the Wellcome Sanger Institute COVID-19 Surveillance Team |
| EPI_ISL_3445703 | Lighthouse Lab in Milton Keynes | Wellcome Sanger Institute for the COVID-19 Genomics UK (COG-UK) Consortium | The Lighthouse Lab in Milton Keynes and Alex Alderton, Roberto Amato, Jeffrey Barrett, Sonia Goncalves, Ewan Harrison, David K. Jackson, Ian Johnston, Dominic Kwiatkowski, Cordelia Langford, John Sillitoe on behalf of the Wellcome Sanger Institute COVID-19 Surveillance Team |
| EPI_ISL_3445704, EPI_ISL_3445705, EPI_ISL_3445706 | Lighthouse Lab in Glasgow | Wellcome Sanger Institute for the COVID-19 Genomics UK (COG-UK) Consortium | Harper VanSteenhouse, Yumi Kasai, David Gray, Carol Clugston, Anna Dominiczak and Alex Alderton, Roberto Amato, Jeffrey Barrett, Sonia Goncalves, Ewan Harrison, David K. Jackson, Ian Johnston, Dominic Kwiatkowski, Cordelia Langford, John Sillitoe on behalf of the Wellcome Sanger Institute COVID-19 Surveillance Team |
| EPI_ISL_3445707 | Lighthouse Lab in Milton Keynes | Wellcome Sanger Institute for the COVID-19 Genomics UK (COG-UK) Consortium | The Lighthouse Lab in Milton Keynes and Alex Alderton, Roberto Amato, Jeffrey Barrett, Sonia Goncalves, Ewan Harrison, David K. Jackson, Ian Johnston, Dominic Kwiatkowski, Cordelia Langford, John Sillitoe on behalf of the Wellcome Sanger Institute COVID-19 Surveillance Team |
| EPI_ISL_3445708 | Lighthouse Lab in Glasgow | Wellcome Sanger Institute for the COVID-19 Genomics UK (COG-UK) Consortium | Harper VanSteenhouse, Yumi Kasai, David Gray, Carol Clugston, Anna Dominiczak and Alex Alderton, Roberto Amato, Jeffrey Barrett, Sonia Goncalves, Ewan Harrison, David K. Jackson, Ian Johnston, Dominic Kwiatkowski, Cordelia Langford, John Sillitoe on behalf of the Wellcome Sanger Institute COVID-19 Surveillance Team |
| EPI_ISL_3445709 | Lighthouse Lab in Milton Keynes | Wellcome Sanger Institute for the COVID-19 Genomics UK (COG-UK) Consortium | The Lighthouse Lab in Milton Keynes and Alex Alderton, Roberto Amato, Jeffrey Barrett, Sonia Goncalves, Ewan Harrison, David K. Jackson, Ian Johnston, Dominic Kwiatkowski, Cordelia Langford, John Sillitoe on behalf of the Wellcome Sanger Institute COVID-19 Surveillance Team |
| EPI_ISL_3445710, EPI_ISL_3445711 | Lighthouse Lab in Glasgow | Wellcome Sanger Institute for the COVID-19 Genomics UK (COG-UK) Consortium | Harper VanSteenhouse, Yumi Kasai, David Gray, Carol Clugston, Anna Dominiczak and Alex Alderton, Roberto Amato, Jeffrey Barrett, Sonia Goncalves, Ewan Harrison, David K. Jackson, Ian Johnston, Dominic Kwiatkowski, Cordelia Langford, John Sillitoe on behalf of the Wellcome Sanger Institute COVID-19 Surveillance Team |
| EPI_ISL_3445712 | Lighthouse Lab in Milton Keynes | Wellcome Sanger Institute for the COVID-19 Genomics UK (COG-UK) Consortium | The Lighthouse Lab in Milton Keynes and Alex Alderton, Roberto Amato, Jeffrey Barrett, Sonia Goncalves, Ewan Harrison, David K. Jackson, Ian Johnston, Dominic Kwiatkowski, Cordelia Langford, John Sillitoe on behalf of the Wellcome Sanger Institute COVID-19 Surveillance Team |
| EPI_ISL_3445713 | Lighthouse Lab in Glasgow | Wellcome Sanger Institute for the COVID-19 Genomics UK (COG-UK) Consortium | Harper VanSteenhouse, Yumi Kasai, David Gray, Carol Clugston, Anna Dominiczak and Alex Alderton, Roberto Amato, Jeffrey Barrett, Sonia Goncalves, Ewan Harrison, David K. Jackson, Ian Johnston, Dominic Kwiatkowski, Cordelia Langford, John Sillitoe on behalf of the Wellcome Sanger Institute COVID-19 Surveillance Team |
| EPI_ISL_3445714, EPI_ISL_3445715 | Lighthouse Lab in Milton Keynes | Wellcome Sanger Institute for the COVID-19 Genomics UK (COG-UK) Consortium | The Lighthouse Lab in Milton Keynes and Alex Alderton, Roberto Amato, Jeffrey Barrett, Sonia Goncalves, Ewan Harrison, David K. Jackson, Ian Johnston, Dominic Kwiatkowski, Cordelia Langford, John Sillitoe on behalf of the Wellcome Sanger Institute COVID-19 Surveillance Team |
| EPI_ISL_3445716, EPI_ISL_3445717 | Lighthouse Lab in Glasgow | Wellcome Sanger Institute for the COVID-19 Genomics UK (COG-UK) Consortium | Harper VanSteenhouse, Yumi Kasai, David Gray, Carol Clugston, Anna Dominiczak and Alex Alderton, Roberto Amato, Jeffrey Barrett, Sonia Goncalves, Ewan Harrison, David K. Jackson, Ian Johnston, Dominic Kwiatkowski, Cordelia Langford, John Sillitoe on behalf of the Wellcome Sanger Institute COVID-19 Surveillance Team |
| EPI_ISL_3445718 | Lighthouse Lab in Milton Keynes | Wellcome Sanger Institute for the COVID-19 Genomics UK (COG-UK) Consortium | The Lighthouse Lab in Milton Keynes and Alex Alderton, Roberto Amato, Jeffrey Barrett, Sonia Goncalves, Ewan Harrison, David K. Jackson, Ian Johnston, Dominic Kwiatkowski, Cordelia Langford, John Sillitoe on behalf of the Wellcome Sanger Institute COVID-19 Surveillance Team |
| EPI_ISL_3445719, EPI_ISL_3445720, EPI_ISL_3445721, EPI_ISL_3445722, EPI_ISL_3445723, EPI_ISL_3445724, EPI_ISL_3445725 | Lighthouse Lab in Glasgow | Wellcome Sanger Institute for the COVID-19 Genomics UK (COG-UK) Consortium | Harper VanSteenhouse, Yumi Kasai, David Gray, Carol Clugston, Anna Dominiczak and Alex Alderton, Roberto Amato, Jeffrey Barrett, Sonia Goncalves, Ewan Harrison, David K. Jackson, Ian Johnston, Dominic Kwiatkowski, Cordelia Langford, John Sillitoe on behalf of the Wellcome Sanger Institute COVID-19 Surveillance Team |
| EPI_ISL_3445726 | Lighthouse Lab in Milton Keynes | Wellcome Sanger Institute for the COVID-19 Genomics UK (COG-UK) Consortium | The Lighthouse Lab in Milton Keynes and Alex Alderton, Roberto Amato, Jeffrey Barrett, Sonia Goncalves, Ewan Harrison, David K. Jackson, Ian Johnston, Dominic Kwiatkowski, Cordelia Langford, John Sillitoe on behalf of the Wellcome Sanger Institute COVID-19 Surveillance Team |
| EPI_ISL_3445727, EPI_ISL_3445728, EPI_ISL_3445729, EPI_ISL_3445730, EPI_ISL_3445731 | Lighthouse Lab in Glasgow | Wellcome Sanger Institute for the COVID-19 Genomics UK (COG-UK) Consortium | Harper VanSteenhouse, Yumi Kasai, David Gray, Carol Clugston, Anna Dominiczak and Alex Alderton, Roberto Amato, Jeffrey Barrett, Sonia Goncalves, Ewan Harrison, David K. Jackson, Ian Johnston, Dominic Kwiatkowski, Cordelia Langford, John Sillitoe on behalf of the Wellcome Sanger Institute COVID-19 Surveillance Team |
| EPI_ISL_3445732 | Lighthouse Lab in Milton Keynes | Wellcome Sanger Institute for the COVID-19 Genomics UK (COG-UK) Consortium | The Lighthouse Lab in Milton Keynes and Alex Alderton, Roberto Amato, Jeffrey Barrett, Sonia Goncalves, Ewan Harrison, David K. Jackson, Ian Johnston, Dominic Kwiatkowski, Cordelia Langford, John Sillitoe on behalf of the Wellcome Sanger Institute COVID-19 Surveillance Team |
| EPI_ISL_3445733 | Lighthouse Lab in Glasgow | Wellcome Sanger Institute for the COVID-19 Genomics UK (COG-UK) Consortium | Harper VanSteenhouse, Yumi Kasai, David Gray, Carol Clugston, Anna Dominiczak and Alex Alderton, Roberto Amato, Jeffrey Barrett, Sonia Goncalves, Ewan Harrison, David K. Jackson, Ian Johnston, Dominic Kwiatkowski, Cordelia Langford, John Sillitoe on behalf of the Wellcome Sanger Institute COVID-19 Surveillance Team |
| EPI_ISL_3445734 | Lighthouse Laboratory Plymouth | Wellcome Sanger Institute for the COVID-19 Genomics UK (COG-UK) Consortium | Lighthouse Laboratory Plymouth and Alex Alderton, Roberto Amato, Jeffrey Barrett, Sonia Goncalves, Ewan Harrison, David K. Jackson, Ian Johnston, Dominic Kwiatkowski, Cordelia Langford, John Sillitoe on behalf of the Wellcome Sanger Institute COVID-19 Surveillance Team |
| EPI_ISL_3445735, | Lighthouse Lab in Glasgow | Wellcome Sanger Institute for the COVID-19 Genomics UK (COG-UK) | Harper VanSteenhouse, Yumi Kasai, David Gray, Carol Clugston, Anna Dominiczak and Alex Alderton, Roberto Amato, Jeffrey Barrett, Sonia Goncalves, Ewan Harrison, David K. Jackson, Ian Johnston, Dominic Kwiatkowski, Cordelia Langford, |

| | | | |
|---|---|---|---|
| EPI_ISL_3445736 | | Consortium | John Sillitoe on behalf of the Wellcome Sanger Institute COVID-19 Surveillance Team |
| EPI_ISL_3445737 | Lighthouse Lab in Milton Keynes | Wellcome Sanger Institute for the COVID-19 Genomics UK (COG-UK) Consortium | The Lighthouse Lab in Milton Keynes and Alex Alderton, Roberto Amato, Jeffrey Barrett, Sonia Goncalves, Ewan Harrison, David K. Jackson, Ian Johnston, Dominic Kwiatkowski, Cordelia Langford, John Sillitoe on behalf of the Wellcome Sanger Institute COVID-19 Surveillance Team |
| EPI_ISL_3445738 | Lighthouse Laboratory Plymouth | Wellcome Sanger Institute for the COVID-19 Genomics UK (COG-UK) Consortium | Lighthouse Laboratory Plymouth and Alex Alderton, Roberto Amato, Jeffrey Barrett, Sonia Goncalves, Ewan Harrison, David K. Jackson, Ian Johnston, Dominic Kwiatkowski, Cordelia Langford, John Sillitoe on behalf of the Wellcome Sanger Institute COVID-19 Surveillance Team |
| EPI_ISL_3445739 | Lighthouse Lab in Glasgow | Wellcome Sanger Institute for the COVID-19 Genomics UK (COG-UK) Consortium | Harper VanSteenhouse, Yumi Kasai, David Gray, Carol Clugston, Anna Dominiczak and Alex Alderton, Roberto Amato, Jeffrey Barrett, Sonia Goncalves, Ewan Harrison, David K. Jackson, Ian Johnston, Dominic Kwiatkowski, Cordelia Langford, John Sillitoe on behalf of the Wellcome Sanger Institute COVID-19 Surveillance Team |
| EPI_ISL_3445740 | Lighthouse Lab in Milton Keynes | Wellcome Sanger Institute for the COVID-19 Genomics UK (COG-UK) Consortium | The Lighthouse Lab in Milton Keynes and Alex Alderton, Roberto Amato, Jeffrey Barrett, Sonia Goncalves, Ewan Harrison, David K. Jackson, Ian Johnston, Dominic Kwiatkowski, Cordelia Langford, John Sillitoe on behalf of the Wellcome Sanger Institute COVID-19 Surveillance Team |
| EPI_ISL_3445741, EPI_ISL_3445742, EPI_ISL_3445743 | Lighthouse Lab in Glasgow | Wellcome Sanger Institute for the COVID-19 Genomics UK (COG-UK) Consortium | Harper VanSteenhouse, Yumi Kasai, David Gray, Carol Clugston, Anna Dominiczak and Alex Alderton, Roberto Amato, Jeffrey Barrett, Sonia Goncalves, Ewan Harrison, David K. Jackson, Ian Johnston, Dominic Kwiatkowski, Cordelia Langford, John Sillitoe on behalf of the Wellcome Sanger Institute COVID-19 Surveillance Team |
| EPI_ISL_3445744 | Lighthouse Lab in Milton Keynes | Wellcome Sanger Institute for the COVID-19 Genomics UK (COG-UK) Consortium | The Lighthouse Lab in Milton Keynes and Alex Alderton, Roberto Amato, Jeffrey Barrett, Sonia Goncalves, Ewan Harrison, David K. Jackson, Ian Johnston, Dominic Kwiatkowski, Cordelia Langford, John Sillitoe on behalf of the Wellcome Sanger Institute COVID-19 Surveillance Team |
| EPI_ISL_3445745, EPI_ISL_3445746 | Lighthouse Lab in Glasgow | Wellcome Sanger Institute for the COVID-19 Genomics UK (COG-UK) Consortium | Harper VanSteenhouse, Yumi Kasai, David Gray, Carol Clugston, Anna Dominiczak and Alex Alderton, Roberto Amato, Jeffrey Barrett, Sonia Goncalves, Ewan Harrison, David K. Jackson, Ian Johnston, Dominic Kwiatkowski, Cordelia Langford, John Sillitoe on behalf of the Wellcome Sanger Institute COVID-19 Surveillance Team |
| EPI_ISL_3445747 | Lighthouse Laboratory Plymouth | Wellcome Sanger Institute for the COVID-19 Genomics UK (COG-UK) Consortium | Lighthouse Laboratory Plymouth and Alex Alderton, Roberto Amato, Jeffrey Barrett, Sonia Goncalves, Ewan Harrison, David K. Jackson, Ian Johnston, Dominic Kwiatkowski, Cordelia Langford, John Sillitoe on behalf of the Wellcome Sanger Institute COVID-19 Surveillance Team |
| EPI_ISL_3445748 | Lighthouse Lab in Milton Keynes | Wellcome Sanger Institute for the COVID-19 Genomics UK (COG-UK) Consortium | The Lighthouse Lab in Milton Keynes and Alex Alderton, Roberto Amato, Jeffrey Barrett, Sonia Goncalves, Ewan Harrison, David K. Jackson, Ian Johnston, Dominic Kwiatkowski, Cordelia Langford, John Sillitoe on behalf of the Wellcome Sanger Institute COVID-19 Surveillance Team |
| EPI_ISL_3445749, EPI_ISL_3445750 | Lighthouse Lab in Glasgow | Wellcome Sanger Institute for the COVID-19 Genomics UK (COG-UK) Consortium | Harper VanSteenhouse, Yumi Kasai, David Gray, Carol Clugston, Anna Dominiczak and Alex Alderton, Roberto Amato, Jeffrey Barrett, Sonia Goncalves, Ewan Harrison, David K. Jackson, Ian Johnston, Dominic Kwiatkowski, Cordelia Langford, John Sillitoe on behalf of the Wellcome Sanger Institute COVID-19 Surveillance Team |
| EPI_ISL_3445751 | Lighthouse Lab in Milton Keynes | Wellcome Sanger Institute for the COVID-19 Genomics UK (COG-UK) Consortium | The Lighthouse Lab in Milton Keynes and Alex Alderton, Roberto Amato, Jeffrey Barrett, Sonia Goncalves, Ewan Harrison, David K. Jackson, Ian Johnston, Dominic Kwiatkowski, Cordelia Langford, John Sillitoe on behalf of the Wellcome Sanger Institute COVID-19 Surveillance Team |
| EPI_ISL_3445752, EPI_ISL_3445753 | Lighthouse Lab in Glasgow | Wellcome Sanger Institute for the COVID-19 Genomics UK (COG-UK) Consortium | Harper VanSteenhouse, Yumi Kasai, David Gray, Carol Clugston, Anna Dominiczak and Alex Alderton, Roberto Amato, Jeffrey Barrett, Sonia Goncalves, Ewan Harrison, David K. Jackson, Ian Johnston, Dominic Kwiatkowski, Cordelia Langford, John Sillitoe on behalf of the Wellcome Sanger Institute COVID-19 Surveillance Team |
| EPI_ISL_3445754 | Lighthouse Lab in Milton Keynes | Wellcome Sanger Institute for the COVID-19 Genomics UK (COG-UK) Consortium | The Lighthouse Lab in Milton Keynes and Alex Alderton, Roberto Amato, Jeffrey Barrett, Sonia Goncalves, Ewan Harrison, David K. Jackson, Ian Johnston, Dominic Kwiatkowski, Cordelia Langford, John Sillitoe on behalf of the Wellcome Sanger Institute COVID-19 Surveillance Team |
| EPI_ISL_3445755 | Lighthouse Lab in Glasgow | Wellcome Sanger Institute for the COVID-19 Genomics UK (COG-UK) Consortium | Harper VanSteenhouse, Yumi Kasai, David Gray, Carol Clugston, Anna Dominiczak and Alex Alderton, Roberto Amato, Jeffrey Barrett, Sonia Goncalves, Ewan Harrison, David K. Jackson, Ian Johnston, Dominic Kwiatkowski, Cordelia Langford, John Sillitoe on behalf of the Wellcome Sanger Institute COVID-19 Surveillance Team |
| EPI_ISL_3445756 | Lighthouse Lab in Milton Keynes | Wellcome Sanger Institute for the COVID-19 Genomics UK (COG-UK) Consortium | The Lighthouse Lab in Milton Keynes and Alex Alderton, Roberto Amato, Jeffrey Barrett, Sonia Goncalves, Ewan Harrison, David K. Jackson, Ian Johnston, Dominic Kwiatkowski, Cordelia Langford, John Sillitoe on behalf of the Wellcome Sanger Institute COVID-19 Surveillance Team |
| EPI_ISL_3445757, EPI_ISL_3445758 | Lighthouse Lab in Glasgow | Wellcome Sanger Institute for the COVID-19 Genomics UK (COG-UK) Consortium | Harper VanSteenhouse, Yumi Kasai, David Gray, Carol Clugston, Anna Dominiczak and Alex Alderton, Roberto Amato, Jeffrey Barrett, Sonia Goncalves, Ewan Harrison, David K. Jackson, Ian Johnston, Dominic Kwiatkowski, Cordelia Langford, John Sillitoe on behalf of the Wellcome Sanger Institute COVID-19 Surveillance Team |
| EPI_ISL_3445759 | Lighthouse Lab in Milton Keynes | Wellcome Sanger Institute for the COVID-19 Genomics UK (COG-UK) Consortium | The Lighthouse Lab in Milton Keynes and Alex Alderton, Roberto Amato, Jeffrey Barrett, Sonia Goncalves, Ewan Harrison, David K. Jackson, Ian Johnston, Dominic Kwiatkowski, Cordelia Langford, John Sillitoe on behalf of the Wellcome Sanger Institute COVID-19 Surveillance Team |
| EPI_ISL_3445760 | Lighthouse Lab in Glasgow | Wellcome Sanger Institute for the COVID-19 Genomics UK (COG-UK) Consortium | Harper VanSteenhouse, Yumi Kasai, David Gray, Carol Clugston, Anna Dominiczak and Alex Alderton, Roberto Amato, Jeffrey Barrett, Sonia Goncalves, Ewan Harrison, David K. Jackson, Ian Johnston, Dominic Kwiatkowski, Cordelia Langford, John Sillitoe on behalf of the Wellcome Sanger Institute COVID-19 Surveillance Team |
| EPI_ISL_3445761 | Lighthouse Laboratory Plymouth | Wellcome Sanger Institute for the COVID-19 Genomics UK (COG-UK) Consortium | Lighthouse Laboratory Plymouth and Alex Alderton, Roberto Amato, Jeffrey Barrett, Sonia Goncalves, Ewan Harrison, David K. Jackson, Ian Johnston, Dominic Kwiatkowski, Cordelia Langford, John Sillitoe on behalf of the Wellcome Sanger Institute COVID-19 Surveillance Team |
| EPI_ISL_3445762 | Lighthouse Lab in Milton Keynes | Wellcome Sanger Institute for the COVID-19 Genomics UK (COG-UK) Consortium | The Lighthouse Lab in Milton Keynes and Alex Alderton, Roberto Amato, Jeffrey Barrett, Sonia Goncalves, Ewan Harrison, David K. Jackson, Ian Johnston, Dominic Kwiatkowski, Cordelia Langford, John Sillitoe on behalf of the Wellcome Sanger Institute COVID-19 Surveillance Team |
| EPI_ISL_3445763 | Lighthouse Lab in Glasgow | Wellcome Sanger Institute for the COVID-19 Genomics UK (COG-UK) Consortium | Harper VanSteenhouse, Yumi Kasai, David Gray, Carol Clugston, Anna Dominiczak and Alex Alderton, Roberto Amato, Jeffrey Barrett, Sonia Goncalves, Ewan Harrison, David K. Jackson, Ian Johnston, Dominic Kwiatkowski, Cordelia Langford, John Sillitoe on behalf of the Wellcome Sanger Institute COVID-19 Surveillance Team |
| EPI_ISL_3445764 | Lighthouse Lab in Milton Keynes | Wellcome Sanger Institute for the COVID-19 Genomics UK (COG-UK) Consortium | The Lighthouse Lab in Milton Keynes and Alex Alderton, Roberto Amato, Jeffrey Barrett, Sonia Goncalves, Ewan Harrison, David K. Jackson, Ian Johnston, Dominic Kwiatkowski, Cordelia Langford, John Sillitoe on behalf of the Wellcome Sanger Institute COVID-19 Surveillance Team |
| EPI_ISL_3445765 | Lighthouse Laboratory Plymouth | Wellcome Sanger Institute for the COVID-19 Genomics UK (COG-UK) Consortium | Lighthouse Laboratory Plymouth and Alex Alderton, Roberto Amato, Jeffrey Barrett, Sonia Goncalves, Ewan Harrison, David K. Jackson, Ian Johnston, Dominic Kwiatkowski, Cordelia Langford, John Sillitoe on behalf of the Wellcome Sanger Institute COVID-19 Surveillance Team |
| EPI_ISL_3445766 | Lighthouse Lab in Milton Keynes | Wellcome Sanger Institute for the COVID-19 Genomics UK (COG-UK) Consortium | The Lighthouse Lab in Milton Keynes and Alex Alderton, Roberto Amato, Jeffrey Barrett, Sonia Goncalves, Ewan Harrison, David K. Jackson, Ian Johnston, Dominic Kwiatkowski, Cordelia Langford, John Sillitoe on behalf of the Wellcome Sanger Institute COVID-19 Surveillance Team |
| EPI_ISL_3445767 | Lighthouse Laboratory Plymouth | Wellcome Sanger Institute for the COVID-19 Genomics UK (COG-UK) Consortium | Lighthouse Laboratory Plymouth and Alex Alderton, Roberto Amato, Jeffrey Barrett, Sonia Goncalves, Ewan Harrison, David K. Jackson, Ian Johnston, Dominic Kwiatkowski, Cordelia Langford, John Sillitoe on behalf of the Wellcome Sanger Institute COVID-19 Surveillance Team |
| EPI_ISL_3445768, EPI_ISL_3445769 | Lighthouse Lab in Glasgow | Wellcome Sanger Institute for the COVID-19 Genomics UK (COG-UK) Consortium | Harper VanSteenhouse, Yumi Kasai, David Gray, Carol Clugston, Anna Dominiczak and Alex Alderton, Roberto Amato, Jeffrey Barrett, Sonia Goncalves, Ewan Harrison, David K. Jackson, Ian Johnston, Dominic Kwiatkowski, Cordelia Langford, John Sillitoe on behalf of the Wellcome Sanger Institute COVID-19 Surveillance Team |
| EPI_ISL_3445770 | Lighthouse Lab in Milton Keynes | Wellcome Sanger Institute for the COVID-19 Genomics UK (COG-UK) Consortium | The Lighthouse Lab in Milton Keynes and Alex Alderton, Roberto Amato, Jeffrey Barrett, Sonia Goncalves, Ewan Harrison, David K. Jackson, Ian Johnston, Dominic Kwiatkowski, Cordelia Langford, John Sillitoe on behalf of the Wellcome Sanger Institute COVID-19 Surveillance Team |
| EPI_ISL_3445771, EPI_ISL_3445772, EPI_ISL_3445773 | Lighthouse Lab in Glasgow | Wellcome Sanger Institute for the COVID-19 Genomics UK (COG-UK) Consortium | Harper VanSteenhouse, Yumi Kasai, David Gray, Carol Clugston, Anna Dominiczak and Alex Alderton, Roberto Amato, Jeffrey Barrett, Sonia Goncalves, Ewan Harrison, David K. Jackson, Ian Johnston, Dominic Kwiatkowski, Cordelia Langford, John Sillitoe on behalf of the Wellcome Sanger Institute COVID-19 Surveillance Team |
| EPI_ISL_3445774 | Lighthouse Laboratory Plymouth | Wellcome Sanger Institute for the COVID-19 Genomics UK (COG-UK) Consortium | Lighthouse Laboratory Plymouth and Alex Alderton, Roberto Amato, Jeffrey Barrett, Sonia Goncalves, Ewan Harrison, David K. Jackson, Ian Johnston, Dominic Kwiatkowski, Cordelia Langford, John Sillitoe on behalf of the Wellcome Sanger Institute COVID-19 Surveillance Team |
| EPI_ISL_3445775, EPI_ISL_3445776 | Lighthouse Lab in Glasgow | Wellcome Sanger Institute for the COVID-19 Genomics UK (COG-UK) Consortium | Harper VanSteenhouse, Yumi Kasai, David Gray, Carol Clugston, Anna Dominiczak and Alex Alderton, Roberto Amato, Jeffrey Barrett, Sonia Goncalves, Ewan Harrison, David K. Jackson, Ian Johnston, Dominic Kwiatkowski, Cordelia Langford, John Sillitoe on behalf of the Wellcome Sanger Institute COVID-19 Surveillance Team |
| EPI_ISL_3445777 | Lighthouse Laboratory Plymouth | Wellcome Sanger Institute for the COVID-19 Genomics UK (COG-UK) Consortium | Lighthouse Laboratory Plymouth and Alex Alderton, Roberto Amato, Jeffrey Barrett, Sonia Goncalves, Ewan Harrison, David K. Jackson, Ian Johnston, Dominic Kwiatkowski, Cordelia Langford, John Sillitoe on behalf of the Wellcome Sanger Institute COVID-19 Surveillance Team |
| EPI_ISL_3445778 | Lighthouse Lab in Glasgow | Wellcome Sanger Institute for the COVID-19 Genomics UK (COG-UK) Consortium | Harper VanSteenhouse, Yumi Kasai, David Gray, Carol Clugston, Anna Dominiczak and Alex Alderton, Roberto Amato, Jeffrey Barrett, Sonia Goncalves, Ewan Harrison, David K. Jackson, Ian Johnston, Dominic Kwiatkowski, Cordelia Langford, John Sillitoe on behalf of the Wellcome Sanger Institute COVID-19 Surveillance Team |
| EPI_ISL_3445779 | Lighthouse Lab in Milton Keynes | Wellcome Sanger Institute for the COVID-19 Genomics UK (COG-UK) Consortium | The Lighthouse Lab in Milton Keynes and Alex Alderton, Roberto Amato, Jeffrey Barrett, Sonia Goncalves, Ewan Harrison, David K. Jackson, Ian Johnston, Dominic Kwiatkowski, Cordelia Langford, John Sillitoe on behalf of the Wellcome Sanger Institute COVID-19 Surveillance Team |
| EPI_ISL_3445780 | Lighthouse Lab in Glasgow | Wellcome Sanger Institute for the COVID-19 Genomics UK (COG-UK) Consortium | Harper VanSteenhouse, Yumi Kasai, David Gray, Carol Clugston, Anna Dominiczak and Alex Alderton, Roberto Amato, Jeffrey Barrett, Sonia Goncalves, Ewan Harrison, David K. Jackson, Ian Johnston, Dominic Kwiatkowski, Cordelia Langford, John Sillitoe on behalf of the Wellcome Sanger Institute COVID-19 Surveillance Team |
| EPI_ISL_3445781 | Lighthouse Lab in Milton Keynes | Wellcome Sanger Institute for the COVID-19 Genomics UK (COG-UK) Consortium | The Lighthouse Lab in Milton Keynes and Alex Alderton, Roberto Amato, Jeffrey Barrett, Sonia Goncalves, Ewan Harrison, David K. Jackson, Ian Johnston, Dominic Kwiatkowski, Cordelia Langford, John Sillitoe on behalf of the Wellcome Sanger Institute COVID-19 Surveillance Team |
| EPI_ISL_3445782, EPI_ISL_3445783 | Lighthouse Lab in Glasgow | Wellcome Sanger Institute for the COVID-19 Genomics UK (COG-UK) Consortium | Harper VanSteenhouse, Yumi Kasai, David Gray, Carol Clugston, Anna Dominiczak and Alex Alderton, Roberto Amato, Jeffrey Barrett, Sonia Goncalves, Ewan Harrison, David K. Jackson, Ian Johnston, Dominic Kwiatkowski, Cordelia Langford, John Sillitoe on behalf of the Wellcome Sanger Institute COVID-19 Surveillance Team |
| EPI_ISL_3445784 | Lighthouse Lab in Milton Keynes | Wellcome Sanger Institute for the COVID-19 Genomics UK (COG-UK) Consortium | The Lighthouse Lab in Milton Keynes and Alex Alderton, Roberto Amato, Jeffrey Barrett, Sonia Goncalves, Ewan Harrison, David K. Jackson, Ian Johnston, Dominic Kwiatkowski, Cordelia Langford, John Sillitoe on behalf of the Wellcome Sanger Institute COVID-19 Surveillance Team |
| EPI_ISL_3445785 | Lighthouse Laboratory Plymouth | Wellcome Sanger Institute for the COVID-19 Genomics UK (COG-UK) Consortium | Lighthouse Laboratory Plymouth and Alex Alderton, Roberto Amato, Jeffrey Barrett, Sonia Goncalves, Ewan Harrison, David K. Jackson, Ian Johnston, Dominic Kwiatkowski, Cordelia Langford, John Sillitoe on behalf of the Wellcome Sanger Institute COVID-19 Surveillance Team |
| EPI_ISL_3445786 | Lighthouse Lab in Milton Keynes | Wellcome Sanger Institute for the COVID-19 Genomics UK (COG-UK) Consortium | The Lighthouse Lab in Milton Keynes and Alex Alderton, Roberto Amato, Jeffrey Barrett, Sonia Goncalves, Ewan Harrison, David K. Jackson, Ian Johnston, Dominic Kwiatkowski, Cordelia Langford, John Sillitoe on behalf of the Wellcome Sanger Institute COVID-19 Surveillance Team |
| EPI_ISL_3445787 | Lighthouse Laboratory Plymouth | Wellcome Sanger Institute for the COVID-19 Genomics UK (COG-UK) Consortium | Lighthouse Laboratory Plymouth and Alex Alderton, Roberto Amato, Jeffrey Barrett, Sonia Goncalves, Ewan Harrison, David K. Jackson, Ian Johnston, Dominic Kwiatkowski, Cordelia Langford, John Sillitoe on behalf of the Wellcome Sanger Institute COVID-19 Surveillance Team |
| EPI_ISL_3445788, EPI_ISL_3445789, EPI_ISL_3445790, EPI_ISL_3445791 | Lighthouse Lab in Glasgow | Wellcome Sanger Institute for the COVID-19 Genomics UK (COG-UK) Consortium | Harper VanSteenhouse, Yumi Kasai, David Gray, Carol Clugston, Anna Dominiczak and Alex Alderton, Roberto Amato, Jeffrey Barrett, Sonia Goncalves, Ewan Harrison, David K. Jackson, Ian Johnston, Dominic Kwiatkowski, Cordelia Langford, John Sillitoe on behalf of the Wellcome Sanger Institute COVID-19 Surveillance Team |
| EPI_ISL_3445792 | Lighthouse Laboratory Plymouth | Wellcome Sanger Institute for the COVID-19 Genomics UK (COG-UK) Consortium | Lighthouse Laboratory Plymouth and Alex Alderton, Roberto Amato, Jeffrey Barrett, Sonia Goncalves, Ewan Harrison, David K. Jackson, Ian Johnston, Dominic Kwiatkowski, Cordelia Langford, John Sillitoe on behalf of the Wellcome Sanger Institute COVID-19 Surveillance Team |
| EPI_ISL_3445793, EPI_ISL_3445794 | Lighthouse Lab in Glasgow | Wellcome Sanger Institute for the COVID-19 Genomics UK (COG-UK) Consortium | Harper VanSteenhouse, Yumi Kasai, David Gray, Carol Clugston, Anna Dominiczak and Alex Alderton, Roberto Amato, Jeffrey Barrett, Sonia Goncalves, Ewan Harrison, David K. Jackson, Ian Johnston, Dominic Kwiatkowski, Cordelia Langford, John Sillitoe on behalf of the Wellcome Sanger Institute COVID-19 Surveillance Team |

| | | | |
|---|---|---|---|
| EPI_ISL_3445795 | Lighthouse Lab in Milton Keynes | Wellcome Sanger Institute for the COVID-19 Genomics UK (COG-UK) Consortium | The Lighthouse Lab in Milton Keynes and Alex Alderton, Roberto Amato, Jeffrey Barrett, Sonia Goncalves, Ewan Harrison, David K. Jackson, Ian Johnston, Dominic Kwiatkowski, Cordelia Langford, John Sillitoe on behalf of the Wellcome Sanger Institute COVID-19 Surveillance Team |
| EPI_ISL_3445796, EPI_ISL_3445797, EPI_ISL_3445798, EPI_ISL_3445799 | Lighthouse Lab in Glasgow | Wellcome Sanger Institute for the COVID-19 Genomics UK (COG-UK) Consortium | Harper VanSteenhouse, Yumi Kasai, David Gray, Carol Clugston, Anna Dominiczak and Alex Alderton, Roberto Amato, Jeffrey Barrett, Sonia Goncalves, Ewan Harrison, David K. Jackson, Ian Johnston, Dominic Kwiatkowski, Cordelia Langford, John Sillitoe on behalf of the Wellcome Sanger Institute COVID-19 Surveillance Team |
| EPI_ISL_3445800 | Lighthouse Lab in Milton Keynes | Wellcome Sanger Institute for the COVID-19 Genomics UK (COG-UK) Consortium | The Lighthouse Lab in Milton Keynes and Alex Alderton, Roberto Amato, Jeffrey Barrett, Sonia Goncalves, Ewan Harrison, David K. Jackson, Ian Johnston, Dominic Kwiatkowski, Cordelia Langford, John Sillitoe on behalf of the Wellcome Sanger Institute COVID-19 Surveillance Team |
| EPI_ISL_3445801, EPI_ISL_3445802, EPI_ISL_3445803 | Lighthouse Lab in Glasgow | Wellcome Sanger Institute for the COVID-19 Genomics UK (COG-UK) Consortium | Harper VanSteenhouse, Yumi Kasai, David Gray, Carol Clugston, Anna Dominiczak and Alex Alderton, Roberto Amato, Jeffrey Barrett, Sonia Goncalves, Ewan Harrison, David K. Jackson, Ian Johnston, Dominic Kwiatkowski, Cordelia Langford, John Sillitoe on behalf of the Wellcome Sanger Institute COVID-19 Surveillance Team |
| EPI_ISL_3445804, EPI_ISL_3445805 | Lighthouse Lab in Milton Keynes | Wellcome Sanger Institute for the COVID-19 Genomics UK (COG-UK) Consortium | The Lighthouse Lab in Milton Keynes and Alex Alderton, Roberto Amato, Jeffrey Barrett, Sonia Goncalves, Ewan Harrison, David K. Jackson, Ian Johnston, Dominic Kwiatkowski, Cordelia Langford, John Sillitoe on behalf of the Wellcome Sanger Institute COVID-19 Surveillance Team |
| EPI_ISL_3445806, EPI_ISL_3445807 | Lighthouse Lab in Glasgow | Wellcome Sanger Institute for the COVID-19 Genomics UK (COG-UK) Consortium | Harper VanSteenhouse, Yumi Kasai, David Gray, Carol Clugston, Anna Dominiczak and Alex Alderton, Roberto Amato, Jeffrey Barrett, Sonia Goncalves, Ewan Harrison, David K. Jackson, Ian Johnston, Dominic Kwiatkowski, Cordelia Langford, John Sillitoe on behalf of the Wellcome Sanger Institute COVID-19 Surveillance Team |
| EPI_ISL_3445808 | Lighthouse Lab in Milton Keynes | Wellcome Sanger Institute for the COVID-19 Genomics UK (COG-UK) Consortium | The Lighthouse Lab in Milton Keynes and Alex Alderton, Roberto Amato, Jeffrey Barrett, Sonia Goncalves, Ewan Harrison, David K. Jackson, Ian Johnston, Dominic Kwiatkowski, Cordelia Langford, John Sillitoe on behalf of the Wellcome Sanger Institute COVID-19 Surveillance Team |
| EPI_ISL_3445809 | Lighthouse Lab in Glasgow | Wellcome Sanger Institute for the COVID-19 Genomics UK (COG-UK) Consortium | Harper VanSteenhouse, Yumi Kasai, David Gray, Carol Clugston, Anna Dominiczak and Alex Alderton, Roberto Amato, Jeffrey Barrett, Sonia Goncalves, Ewan Harrison, David K. Jackson, Ian Johnston, Dominic Kwiatkowski, Cordelia Langford, John Sillitoe on behalf of the Wellcome Sanger Institute COVID-19 Surveillance Team |
| EPI_ISL_3445810 | Lighthouse Lab in Milton Keynes | Wellcome Sanger Institute for the COVID-19 Genomics UK (COG-UK) Consortium | The Lighthouse Lab in Milton Keynes and Alex Alderton, Roberto Amato, Jeffrey Barrett, Sonia Goncalves, Ewan Harrison, David K. Jackson, Ian Johnston, Dominic Kwiatkowski, Cordelia Langford, John Sillitoe on behalf of the Wellcome Sanger Institute COVID-19 Surveillance Team |
| EPI_ISL_3445811 | Lighthouse Lab in Glasgow | Wellcome Sanger Institute for the COVID-19 Genomics UK (COG-UK) Consortium | Harper VanSteenhouse, Yumi Kasai, David Gray, Carol Clugston, Anna Dominiczak and Alex Alderton, Roberto Amato, Jeffrey Barrett, Sonia Goncalves, Ewan Harrison, David K. Jackson, Ian Johnston, Dominic Kwiatkowski, Cordelia Langford, John Sillitoe on behalf of the Wellcome Sanger Institute COVID-19 Surveillance Team |
| EPI_ISL_3445812 | Lighthouse Lab in Milton Keynes | Wellcome Sanger Institute for the COVID-19 Genomics UK (COG-UK) Consortium | The Lighthouse Lab in Milton Keynes and Alex Alderton, Roberto Amato, Jeffrey Barrett, Sonia Goncalves, Ewan Harrison, David K. Jackson, Ian Johnston, Dominic Kwiatkowski, Cordelia Langford, John Sillitoe on behalf of the Wellcome Sanger Institute COVID-19 Surveillance Team |
| EPI_ISL_3445813 | Lighthouse Lab in Glasgow | Wellcome Sanger Institute for the COVID-19 Genomics UK (COG-UK) Consortium | Harper VanSteenhouse, Yumi Kasai, David Gray, Carol Clugston, Anna Dominiczak and Alex Alderton, Roberto Amato, Jeffrey Barrett, Sonia Goncalves, Ewan Harrison, David K. Jackson, Ian Johnston, Dominic Kwiatkowski, Cordelia Langford, John Sillitoe on behalf of the Wellcome Sanger Institute COVID-19 Surveillance Team |
| EPI_ISL_3445814 | Lighthouse Lab in Milton Keynes | Wellcome Sanger Institute for the COVID-19 Genomics UK (COG-UK) Consortium | The Lighthouse Lab in Milton Keynes and Alex Alderton, Roberto Amato, Jeffrey Barrett, Sonia Goncalves, Ewan Harrison, David K. Jackson, Ian Johnston, Dominic Kwiatkowski, Cordelia Langford, John Sillitoe on behalf of the Wellcome Sanger Institute COVID-19 Surveillance Team |
| EPI_ISL_3445815 | Lighthouse Laboratory Plymouth | Wellcome Sanger Institute for the COVID-19 Genomics UK (COG-UK) Consortium | Lighthouse Laboratory Plymouth and Alex Alderton, Roberto Amato, Jeffrey Barrett, Sonia Goncalves, Ewan Harrison, David K. Jackson, Ian Johnston, Dominic Kwiatkowski, Cordelia Langford, John Sillitoe on behalf of the Wellcome Sanger Institute COVID-19 Surveillance Team |
| EPI_ISL_3445816, EPI_ISL_3445817 | Lighthouse Lab in Glasgow | Wellcome Sanger Institute for the COVID-19 Genomics UK (COG-UK) Consortium | Harper VanSteenhouse, Yumi Kasai, David Gray, Carol Clugston, Anna Dominiczak and Alex Alderton, Roberto Amato, Jeffrey Barrett, Sonia Goncalves, Ewan Harrison, David K. Jackson, Ian Johnston, Dominic Kwiatkowski, Cordelia Langford, John Sillitoe on behalf of the Wellcome Sanger Institute COVID-19 Surveillance Team |
| EPI_ISL_3445818 | Lighthouse Laboratory Plymouth | Wellcome Sanger Institute for the COVID-19 Genomics UK (COG-UK) Consortium | Lighthouse Laboratory Plymouth and Alex Alderton, Roberto Amato, Jeffrey Barrett, Sonia Goncalves, Ewan Harrison, David K. Jackson, Ian Johnston, Dominic Kwiatkowski, Cordelia Langford, John Sillitoe on behalf of the Wellcome Sanger Institute COVID-19 Surveillance Team |
| EPI_ISL_3445819 | Lighthouse Lab in Milton Keynes | Wellcome Sanger Institute for the COVID-19 Genomics UK (COG-UK) Consortium | The Lighthouse Lab in Milton Keynes and Alex Alderton, Roberto Amato, Jeffrey Barrett, Sonia Goncalves, Ewan Harrison, David K. Jackson, Ian Johnston, Dominic Kwiatkowski, Cordelia Langford, John Sillitoe on behalf of the Wellcome Sanger Institute COVID-19 Surveillance Team |
| EPI_ISL_3445820, EPI_ISL_3445821, EPI_ISL_3445822, EPI_ISL_3445823, EPI_ISL_3445824, EPI_ISL_3445825, EPI_ISL_3445826, EPI_ISL_3445827, EPI_ISL_3445828, EPI_ISL_3445829, EPI_ISL_3445830, EPI_ISL_3445831, EPI_ISL_3445832 | | | |
| see above | Lighthouse Lab in Glasgow | Wellcome Sanger Institute for the COVID-19 Genomics UK (COG-UK) Consortium | Harper VanSteenhouse, Yumi Kasai, David Gray, Carol Clugston, Anna Dominiczak and Alex Alderton, Roberto Amato, Jeffrey Barrett, Sonia Goncalves, Ewan Harrison, David K. Jackson, Ian Johnston, Dominic Kwiatkowski, Cordelia Langford, John Sillitoe on behalf of the Wellcome Sanger Institute COVID-19 Surveillance Team |
| EPI_ISL_3445833 | Lighthouse Laboratory Plymouth | Wellcome Sanger Institute for the COVID-19 Genomics UK (COG-UK) Consortium | Lighthouse Laboratory Plymouth and Alex Alderton, Roberto Amato, Jeffrey Barrett, Sonia Goncalves, Ewan Harrison, David K. Jackson, Ian Johnston, Dominic Kwiatkowski, Cordelia Langford, John Sillitoe on behalf of the Wellcome Sanger Institute COVID-19 Surveillance Team |
| EPI_ISL_3445834 | Lighthouse Lab in Glasgow | Wellcome Sanger Institute for the COVID-19 Genomics UK (COG-UK) Consortium | Harper VanSteenhouse, Yumi Kasai, David Gray, Carol Clugston, Anna Dominiczak and Alex Alderton, Roberto Amato, Jeffrey Barrett, Sonia Goncalves, Ewan Harrison, David K. Jackson, Ian Johnston, Dominic Kwiatkowski, Cordelia Langford, John Sillitoe on behalf of the Wellcome Sanger Institute COVID-19 Surveillance Team |
| EPI_ISL_3445835 | Lighthouse Lab in Milton Keynes | Wellcome Sanger Institute for the COVID-19 Genomics UK (COG-UK) Consortium | The Lighthouse Lab in Milton Keynes and Alex Alderton, Roberto Amato, Jeffrey Barrett, Sonia Goncalves, Ewan Harrison, David K. Jackson, Ian Johnston, Dominic Kwiatkowski, Cordelia Langford, John Sillitoe on behalf of the Wellcome Sanger Institute COVID-19 Surveillance Team |
| EPI_ISL_3445836, EPI_ISL_3445837 | Lighthouse Lab in Glasgow | Wellcome Sanger Institute for the COVID-19 Genomics UK (COG-UK) Consortium | Harper VanSteenhouse, Yumi Kasai, David Gray, Carol Clugston, Anna Dominiczak and Alex Alderton, Roberto Amato, Jeffrey Barrett, Sonia Goncalves, Ewan Harrison, David K. Jackson, Ian Johnston, Dominic Kwiatkowski, Cordelia Langford, John Sillitoe on behalf of the Wellcome Sanger Institute COVID-19 Surveillance Team |
| EPI_ISL_3445838, EPI_ISL_3445839 | Lighthouse Lab in Milton Keynes | Wellcome Sanger Institute for the COVID-19 Genomics UK (COG-UK) Consortium | The Lighthouse Lab in Milton Keynes and Alex Alderton, Roberto Amato, Jeffrey Barrett, Sonia Goncalves, Ewan Harrison, David K. Jackson, Ian Johnston, Dominic Kwiatkowski, Cordelia Langford, John Sillitoe on behalf of the Wellcome Sanger Institute COVID-19 Surveillance Team |
| EPI_ISL_3445840, EPI_ISL_3445841, EPI_ISL_3445842 | Lighthouse Lab in Glasgow | Wellcome Sanger Institute for the COVID-19 Genomics UK (COG-UK) Consortium | Harper VanSteenhouse, Yumi Kasai, David Gray, Carol Clugston, Anna Dominiczak and Alex Alderton, Roberto Amato, Jeffrey Barrett, Sonia Goncalves, Ewan Harrison, David K. Jackson, Ian Johnston, Dominic Kwiatkowski, Cordelia Langford, John Sillitoe on behalf of the Wellcome Sanger Institute COVID-19 Surveillance Team |
| EPI_ISL_3445843 | Lighthouse Laboratory Plymouth | Wellcome Sanger Institute for the COVID-19 Genomics UK (COG-UK) Consortium | Lighthouse Laboratory Plymouth and Alex Alderton, Roberto Amato, Jeffrey Barrett, Sonia Goncalves, Ewan Harrison, David K. Jackson, Ian Johnston, Dominic Kwiatkowski, Cordelia Langford, John Sillitoe on behalf of the Wellcome Sanger Institute COVID-19 Surveillance Team |
| EPI_ISL_3445844, EPI_ISL_3445845 | Lighthouse Lab in Glasgow | Wellcome Sanger Institute for the COVID-19 Genomics UK (COG-UK) Consortium | Harper VanSteenhouse, Yumi Kasai, David Gray, Carol Clugston, Anna Dominiczak and Alex Alderton, Roberto Amato, Jeffrey Barrett, Sonia Goncalves, Ewan Harrison, David K. Jackson, Ian Johnston, Dominic Kwiatkowski, Cordelia Langford, John Sillitoe on behalf of the Wellcome Sanger Institute COVID-19 Surveillance Team |
| EPI_ISL_3445846, EPI_ISL_3445847 | Lighthouse Laboratory Plymouth | Wellcome Sanger Institute for the COVID-19 Genomics UK (COG-UK) Consortium | Lighthouse Laboratory Plymouth and Alex Alderton, Roberto Amato, Jeffrey Barrett, Sonia Goncalves, Ewan Harrison, David K. Jackson, Ian Johnston, Dominic Kwiatkowski, Cordelia Langford, John Sillitoe on behalf of the Wellcome Sanger Institute COVID-19 Surveillance Team |
| EPI_ISL_3445848, EPI_ISL_3445849 | Lighthouse Lab in Milton Keynes | Wellcome Sanger Institute for the COVID-19 Genomics UK (COG-UK) Consortium | The Lighthouse Lab in Milton Keynes and Alex Alderton, Roberto Amato, Jeffrey Barrett, Sonia Goncalves, Ewan Harrison, David K. Jackson, Ian Johnston, Dominic Kwiatkowski, Cordelia Langford, John Sillitoe on behalf of the Wellcome Sanger Institute COVID-19 Surveillance Team |
| EPI_ISL_3445850, EPI_ISL_3445851, EPI_ISL_3445852, EPI_ISL_3445853, EPI_ISL_3445854, EPI_ISL_3445855, EPI_ISL_3445856 | Lighthouse Lab in Glasgow | Wellcome Sanger Institute for the COVID-19 Genomics UK (COG-UK) Consortium | Harper VanSteenhouse, Yumi Kasai, David Gray, Carol Clugston, Anna Dominiczak and Alex Alderton, Roberto Amato, Jeffrey Barrett, Sonia Goncalves, Ewan Harrison, David K. Jackson, Ian Johnston, Dominic Kwiatkowski, Cordelia Langford, John Sillitoe on behalf of the Wellcome Sanger Institute COVID-19 Surveillance Team |
| EPI_ISL_3445857 | Lighthouse Lab in Milton Keynes | Wellcome Sanger Institute for the COVID-19 Genomics UK (COG-UK) Consortium | The Lighthouse Lab in Milton Keynes and Alex Alderton, Roberto Amato, Jeffrey Barrett, Sonia Goncalves, Ewan Harrison, David K. Jackson, Ian Johnston, Dominic Kwiatkowski, Cordelia Langford, John Sillitoe on behalf of the Wellcome Sanger Institute COVID-19 Surveillance Team |
| EPI_ISL_3445858, EPI_ISL_3445859 | Lighthouse Lab in Glasgow | Wellcome Sanger Institute for the COVID-19 Genomics UK (COG-UK) Consortium | Harper VanSteenhouse, Yumi Kasai, David Gray, Carol Clugston, Anna Dominiczak and Alex Alderton, Roberto Amato, Jeffrey Barrett, Sonia Goncalves, Ewan Harrison, David K. Jackson, Ian Johnston, Dominic Kwiatkowski, Cordelia Langford, John Sillitoe on behalf of the Wellcome Sanger Institute COVID-19 Surveillance Team |
| EPI_ISL_3445860 | Lighthouse Laboratory Plymouth | Wellcome Sanger Institute for the COVID-19 Genomics UK (COG-UK) Consortium | Lighthouse Laboratory Plymouth and Alex Alderton, Roberto Amato, Jeffrey Barrett, Sonia Goncalves, Ewan Harrison, David K. Jackson, Ian Johnston, Dominic Kwiatkowski, Cordelia Langford, John Sillitoe on behalf of the Wellcome Sanger Institute COVID-19 Surveillance Team |
| EPI_ISL_3445861, EPI_ISL_3445862, EPI_ISL_3445863, EPI_ISL_3445864 | Lighthouse Lab in Glasgow | Wellcome Sanger Institute for the COVID-19 Genomics UK (COG-UK) Consortium | Harper VanSteenhouse, Yumi Kasai, David Gray, Carol Clugston, Anna Dominiczak and Alex Alderton, Roberto Amato, Jeffrey Barrett, Sonia Goncalves, Ewan Harrison, David K. Jackson, Ian Johnston, Dominic Kwiatkowski, Cordelia Langford, John Sillitoe on behalf of the Wellcome Sanger Institute COVID-19 Surveillance Team |
| EPI_ISL_3445865 | Lighthouse Lab in Milton Keynes | Wellcome Sanger Institute for the COVID-19 Genomics UK (COG-UK) Consortium | The Lighthouse Lab in Milton Keynes and Alex Alderton, Roberto Amato, Jeffrey Barrett, Sonia Goncalves, Ewan Harrison, David K. Jackson, Ian Johnston, Dominic Kwiatkowski, Cordelia Langford, John Sillitoe on behalf of the Wellcome Sanger Institute COVID-19 Surveillance Team |
| EPI_ISL_3445866 | Lighthouse Lab in Glasgow | Wellcome Sanger Institute for the COVID-19 Genomics UK (COG-UK) Consortium | Harper VanSteenhouse, Yumi Kasai, David Gray, Carol Clugston, Anna Dominiczak and Alex Alderton, Roberto Amato, Jeffrey Barrett, Sonia Goncalves, Ewan Harrison, David K. Jackson, Ian Johnston, Dominic Kwiatkowski, Cordelia Langford, John Sillitoe on behalf of the Wellcome Sanger Institute COVID-19 Surveillance Team |
| EPI_ISL_3445867 | Lighthouse Lab in Milton Keynes | Wellcome Sanger Institute for the COVID-19 Genomics UK (COG-UK) Consortium | The Lighthouse Lab in Milton Keynes and Alex Alderton, Roberto Amato, Jeffrey Barrett, Sonia Goncalves, Ewan Harrison, David K. Jackson, Ian Johnston, Dominic Kwiatkowski, Cordelia Langford, John Sillitoe on behalf of the Wellcome Sanger Institute COVID-19 Surveillance Team |
| EPI_ISL_3445868 | Lighthouse Lab in Glasgow | Wellcome Sanger Institute for the COVID-19 Genomics UK (COG-UK) Consortium | Harper VanSteenhouse, Yumi Kasai, David Gray, Carol Clugston, Anna Dominiczak and Alex Alderton, Roberto Amato, Jeffrey Barrett, Sonia Goncalves, Ewan Harrison, David K. Jackson, Ian Johnston, Dominic Kwiatkowski, Cordelia Langford, John Sillitoe on behalf of the Wellcome Sanger Institute COVID-19 Surveillance Team |
| EPI_ISL_3445869, EPI_ISL_3445870 | Lighthouse Laboratory Plymouth | Wellcome Sanger Institute for the COVID-19 Genomics UK (COG-UK) Consortium | Lighthouse Laboratory Plymouth and Alex Alderton, Roberto Amato, Jeffrey Barrett, Sonia Goncalves, Ewan Harrison, David K. Jackson, Ian Johnston, Dominic Kwiatkowski, Cordelia Langford, John Sillitoe on behalf of the Wellcome Sanger Institute COVID-19 Surveillance Team |
| EPI_ISL_3445871 | Lighthouse Lab in Glasgow | Wellcome Sanger Institute for the COVID-19 Genomics UK (COG-UK) Consortium | Harper VanSteenhouse, Yumi Kasai, David Gray, Carol Clugston, Anna Dominiczak and Alex Alderton, Roberto Amato, Jeffrey Barrett, Sonia Goncalves, Ewan Harrison, David K. Jackson, Ian Johnston, Dominic Kwiatkowski, Cordelia Langford, John Sillitoe on behalf of the Wellcome Sanger Institute COVID-19 Surveillance Team |
| EPI_ISL_3445872 | Lighthouse Lab in Milton Keynes | Wellcome Sanger Institute for the COVID-19 Genomics UK (COG-UK) Consortium | The Lighthouse Lab in Milton Keynes and Alex Alderton, Roberto Amato, Jeffrey Barrett, Sonia Goncalves, Ewan Harrison, David K. Jackson, Ian Johnston, Dominic Kwiatkowski, Cordelia Langford, John Sillitoe on behalf of the Wellcome Sanger Institute COVID-19 Surveillance Team |

| | | | |
|---|---|---|---|
| EPI_ISL_3445873 | Lighthouse Lab in Glasgow | Wellcome Sanger Institute for the COVID-19 Genomics UK (COG-UK) Consortium | Harper VanSteenhouse, Yumi Kasai, David Gray, Carol Clugston, Anna Dominiczak and Alex Alderton, Roberto Amato, Jeffrey Barrett, Sonia Goncalves, Ewan Harrison, David K. Jackson, Ian Johnston, Dominic Kwiatkowski, Cordelia Langford, John Sillitoe on behalf of the Wellcome Sanger Institute COVID-19 Surveillance Team |
| EPI_ISL_3445874 | Lighthouse Laboratory Plymouth | Wellcome Sanger Institute for the COVID-19 Genomics UK (COG-UK) Consortium | Lighthouse Laboratory Plymouth and Alex Alderton, Roberto Amato, Jeffrey Barrett, Sonia Goncalves, Ewan Harrison, David K. Jackson, Ian Johnston, Dominic Kwiatkowski, Cordelia Langford, John Sillitoe on behalf of the Wellcome Sanger Institute COVID-19 Surveillance Team |
| EPI_ISL_3445875 | Lighthouse Lab in Glasgow | Wellcome Sanger Institute for the COVID-19 Genomics UK (COG-UK) Consortium | Harper VanSteenhouse, Yumi Kasai, David Gray, Carol Clugston, Anna Dominiczak and Alex Alderton, Roberto Amato, Jeffrey Barrett, Sonia Goncalves, Ewan Harrison, David K. Jackson, Ian Johnston, Dominic Kwiatkowski, Cordelia Langford, John Sillitoe on behalf of the Wellcome Sanger Institute COVID-19 Surveillance Team |
| EPI_ISL_3445876 | Lighthouse Laboratory Plymouth | Wellcome Sanger Institute for the COVID-19 Genomics UK (COG-UK) Consortium | Lighthouse Laboratory Plymouth and Alex Alderton, Roberto Amato, Jeffrey Barrett, Sonia Goncalves, Ewan Harrison, David K. Jackson, Ian Johnston, Dominic Kwiatkowski, Cordelia Langford, John Sillitoe on behalf of the Wellcome Sanger Institute COVID-19 Surveillance Team |
| EPI_ISL_3445877 | Lighthouse Lab in Glasgow | Wellcome Sanger Institute for the COVID-19 Genomics UK (COG-UK) Consortium | Harper VanSteenhouse, Yumi Kasai, David Gray, Carol Clugston, Anna Dominiczak and Alex Alderton, Roberto Amato, Jeffrey Barrett, Sonia Goncalves, Ewan Harrison, David K. Jackson, Ian Johnston, Dominic Kwiatkowski, Cordelia Langford, John Sillitoe on behalf of the Wellcome Sanger Institute COVID-19 Surveillance Team |
| EPI_ISL_3445878 | Lighthouse Lab in Milton Keynes | Wellcome Sanger Institute for the COVID-19 Genomics UK (COG-UK) Consortium | The Lighthouse Lab in Milton Keynes and Alex Alderton, Roberto Amato, Jeffrey Barrett, Sonia Goncalves, Ewan Harrison, David K. Jackson, Ian Johnston, Dominic Kwiatkowski, Cordelia Langford, John Sillitoe on behalf of the Wellcome Sanger Institute COVID-19 Surveillance Team |
| EPI_ISL_3445879 | Lighthouse Lab in Glasgow | Wellcome Sanger Institute for the COVID-19 Genomics UK (COG-UK) Consortium | Harper VanSteenhouse, Yumi Kasai, David Gray, Carol Clugston, Anna Dominiczak and Alex Alderton, Roberto Amato, Jeffrey Barrett, Sonia Goncalves, Ewan Harrison, David K. Jackson, Ian Johnston, Dominic Kwiatkowski, Cordelia Langford, John Sillitoe on behalf of the Wellcome Sanger Institute COVID-19 Surveillance Team |
| EPI_ISL_3445880, EPI_ISL_3445881, EPI_ISL_3445882 | Lighthouse Laboratory Plymouth | Wellcome Sanger Institute for the COVID-19 Genomics UK (COG-UK) Consortium | Lighthouse Laboratory Plymouth and Alex Alderton, Roberto Amato, Jeffrey Barrett, Sonia Goncalves, Ewan Harrison, David K. Jackson, Ian Johnston, Dominic Kwiatkowski, Cordelia Langford, John Sillitoe on behalf of the Wellcome Sanger Institute COVID-19 Surveillance Team |
| EPI_ISL_3445883 | Lighthouse Lab in Glasgow | Wellcome Sanger Institute for the COVID-19 Genomics UK (COG-UK) Consortium | Harper VanSteenhouse, Yumi Kasai, David Gray, Carol Clugston, Anna Dominiczak and Alex Alderton, Roberto Amato, Jeffrey Barrett, Sonia Goncalves, Ewan Harrison, David K. Jackson, Ian Johnston, Dominic Kwiatkowski, Cordelia Langford, John Sillitoe on behalf of the Wellcome Sanger Institute COVID-19 Surveillance Team |
| EPI_ISL_3445884 | Lighthouse Lab in Milton Keynes | Wellcome Sanger Institute for the COVID-19 Genomics UK (COG-UK) Consortium | The Lighthouse Lab in Milton Keynes and Alex Alderton, Roberto Amato, Jeffrey Barrett, Sonia Goncalves, Ewan Harrison, David K. Jackson, Ian Johnston, Dominic Kwiatkowski, Cordelia Langford, John Sillitoe on behalf of the Wellcome Sanger Institute COVID-19 Surveillance Team |
| EPI_ISL_3445885 | Lighthouse Laboratory Plymouth | Wellcome Sanger Institute for the COVID-19 Genomics UK (COG-UK) Consortium | Lighthouse Laboratory Plymouth and Alex Alderton, Roberto Amato, Jeffrey Barrett, Sonia Goncalves, Ewan Harrison, David K. Jackson, Ian Johnston, Dominic Kwiatkowski, Cordelia Langford, John Sillitoe on behalf of the Wellcome Sanger Institute COVID-19 Surveillance Team |
| EPI_ISL_3445886 | Lighthouse Lab in Milton Keynes | Wellcome Sanger Institute for the COVID-19 Genomics UK (COG-UK) Consortium | The Lighthouse Lab in Milton Keynes and Alex Alderton, Roberto Amato, Jeffrey Barrett, Sonia Goncalves, Ewan Harrison, David K. Jackson, Ian Johnston, Dominic Kwiatkowski, Cordelia Langford, John Sillitoe on behalf of the Wellcome Sanger Institute COVID-19 Surveillance Team |
| EPI_ISL_3445887, EPI_ISL_3445888, EPI_ISL_3445889, EPI_ISL_3445890, EPI_ISL_3445891, EPI_ISL_3445892, EPI_ISL_3445893, EPI_ISL_3445894 | Lighthouse Lab in Glasgow | Wellcome Sanger Institute for the COVID-19 Genomics UK (COG-UK) Consortium | Harper VanSteenhouse, Yumi Kasai, David Gray, Carol Clugston, Anna Dominiczak and Alex Alderton, Roberto Amato, Jeffrey Barrett, Sonia Goncalves, Ewan Harrison, David K. Jackson, Ian Johnston, Dominic Kwiatkowski, Cordelia Langford, John Sillitoe on behalf of the Wellcome Sanger Institute COVID-19 Surveillance Team |
| EPI_ISL_3445895 | Lighthouse Laboratory Plymouth | Wellcome Sanger Institute for the COVID-19 Genomics UK (COG-UK) Consortium | Lighthouse Laboratory Plymouth and Alex Alderton, Roberto Amato, Jeffrey Barrett, Sonia Goncalves, Ewan Harrison, David K. Jackson, Ian Johnston, Dominic Kwiatkowski, Cordelia Langford, John Sillitoe on behalf of the Wellcome Sanger Institute COVID-19 Surveillance Team |
| EPI_ISL_3445896 | Lighthouse Lab in Milton Keynes | Wellcome Sanger Institute for the COVID-19 Genomics UK (COG-UK) Consortium | The Lighthouse Lab in Milton Keynes and Alex Alderton, Roberto Amato, Jeffrey Barrett, Sonia Goncalves, Ewan Harrison, David K. Jackson, Ian Johnston, Dominic Kwiatkowski, Cordelia Langford, John Sillitoe on behalf of the Wellcome Sanger Institute COVID-19 Surveillance Team |
| EPI_ISL_3445897, EPI_ISL_3445898, EPI_ISL_3445899 | Lighthouse Lab in Glasgow | Wellcome Sanger Institute for the COVID-19 Genomics UK (COG-UK) Consortium | Harper VanSteenhouse, Yumi Kasai, David Gray, Carol Clugston, Anna Dominiczak and Alex Alderton, Roberto Amato, Jeffrey Barrett, Sonia Goncalves, Ewan Harrison, David K. Jackson, Ian Johnston, Dominic Kwiatkowski, Cordelia Langford, John Sillitoe on behalf of the Wellcome Sanger Institute COVID-19 Surveillance Team |
| EPI_ISL_3445900 | Lighthouse Laboratory Plymouth | Wellcome Sanger Institute for the COVID-19 Genomics UK (COG-UK) Consortium | Lighthouse Laboratory Plymouth and Alex Alderton, Roberto Amato, Jeffrey Barrett, Sonia Goncalves, Ewan Harrison, David K. Jackson, Ian Johnston, Dominic Kwiatkowski, Cordelia Langford, John Sillitoe on behalf of the Wellcome Sanger Institute COVID-19 Surveillance Team |
| EPI_ISL_3445901 | Lighthouse Lab in Glasgow | Wellcome Sanger Institute for the COVID-19 Genomics UK (COG-UK) Consortium | Harper VanSteenhouse, Yumi Kasai, David Gray, Carol Clugston, Anna Dominiczak and Alex Alderton, Roberto Amato, Jeffrey Barrett, Sonia Goncalves, Ewan Harrison, David K. Jackson, Ian Johnston, Dominic Kwiatkowski, Cordelia Langford, John Sillitoe on behalf of the Wellcome Sanger Institute COVID-19 Surveillance Team |
| EPI_ISL_3445902 | Lighthouse Lab in Milton Keynes | Wellcome Sanger Institute for the COVID-19 Genomics UK (COG-UK) Consortium | The Lighthouse Lab in Milton Keynes and Alex Alderton, Roberto Amato, Jeffrey Barrett, Sonia Goncalves, Ewan Harrison, David K. Jackson, Ian Johnston, Dominic Kwiatkowski, Cordelia Langford, John Sillitoe on behalf of the Wellcome Sanger Institute COVID-19 Surveillance Team |
| EPI_ISL_3445903, EPI_ISL_3445904, EPI_ISL_3445905, EPI_ISL_3445906 | Lighthouse Lab in Glasgow | Wellcome Sanger Institute for the COVID-19 Genomics UK (COG-UK) Consortium | Harper VanSteenhouse, Yumi Kasai, David Gray, Carol Clugston, Anna Dominiczak and Alex Alderton, Roberto Amato, Jeffrey Barrett, Sonia Goncalves, Ewan Harrison, David K. Jackson, Ian Johnston, Dominic Kwiatkowski, Cordelia Langford, John Sillitoe on behalf of the Wellcome Sanger Institute COVID-19 Surveillance Team |
| EPI_ISL_3445907 | Lighthouse Lab in Milton Keynes | Wellcome Sanger Institute for the COVID-19 Genomics UK (COG-UK) Consortium | The Lighthouse Lab in Milton Keynes and Alex Alderton, Roberto Amato, Jeffrey Barrett, Sonia Goncalves, Ewan Harrison, David K. Jackson, Ian Johnston, Dominic Kwiatkowski, Cordelia Langford, John Sillitoe on behalf of the Wellcome Sanger Institute COVID-19 Surveillance Team |
| EPI_ISL_3445908 | Lighthouse Laboratory Plymouth | Wellcome Sanger Institute for the COVID-19 Genomics UK (COG-UK) Consortium | Lighthouse Laboratory Plymouth and Alex Alderton, Roberto Amato, Jeffrey Barrett, Sonia Goncalves, Ewan Harrison, David K. Jackson, Ian Johnston, Dominic Kwiatkowski, Cordelia Langford, John Sillitoe on behalf of the Wellcome Sanger Institute COVID-19 Surveillance Team |
| EPI_ISL_3445909 | Lighthouse Lab in Glasgow | Wellcome Sanger Institute for the COVID-19 Genomics UK (COG-UK) Consortium | Harper VanSteenhouse, Yumi Kasai, David Gray, Carol Clugston, Anna Dominiczak and Alex Alderton, Roberto Amato, Jeffrey Barrett, Sonia Goncalves, Ewan Harrison, David K. Jackson, Ian Johnston, Dominic Kwiatkowski, Cordelia Langford, John Sillitoe on behalf of the Wellcome Sanger Institute COVID-19 Surveillance Team |
| EPI_ISL_3445910 | Lighthouse Lab in Milton Keynes | Wellcome Sanger Institute for the COVID-19 Genomics UK (COG-UK) Consortium | The Lighthouse Lab in Milton Keynes and Alex Alderton, Roberto Amato, Jeffrey Barrett, Sonia Goncalves, Ewan Harrison, David K. Jackson, Ian Johnston, Dominic Kwiatkowski, Cordelia Langford, John Sillitoe on behalf of the Wellcome Sanger Institute COVID-19 Surveillance Team |
| EPI_ISL_3445911, EPI_ISL_3445912 | Lighthouse Lab in Glasgow | Wellcome Sanger Institute for the COVID-19 Genomics UK (COG-UK) Consortium | Harper VanSteenhouse, Yumi Kasai, David Gray, Carol Clugston, Anna Dominiczak and Alex Alderton, Roberto Amato, Jeffrey Barrett, Sonia Goncalves, Ewan Harrison, David K. Jackson, Ian Johnston, Dominic Kwiatkowski, Cordelia Langford, John Sillitoe on behalf of the Wellcome Sanger Institute COVID-19 Surveillance Team |
| EPI_ISL_3445913 | Lighthouse Lab in Milton Keynes | Wellcome Sanger Institute for the COVID-19 Genomics UK (COG-UK) Consortium | The Lighthouse Lab in Milton Keynes and Alex Alderton, Roberto Amato, Jeffrey Barrett, Sonia Goncalves, Ewan Harrison, David K. Jackson, Ian Johnston, Dominic Kwiatkowski, Cordelia Langford, John Sillitoe on behalf of the Wellcome Sanger Institute COVID-19 Surveillance Team |
| EPI_ISL_3445914 | Lighthouse Laboratory Plymouth | Wellcome Sanger Institute for the COVID-19 Genomics UK (COG-UK) Consortium | Lighthouse Laboratory Plymouth and Alex Alderton, Roberto Amato, Jeffrey Barrett, Sonia Goncalves, Ewan Harrison, David K. Jackson, Ian Johnston, Dominic Kwiatkowski, Cordelia Langford, John Sillitoe on behalf of the Wellcome Sanger Institute COVID-19 Surveillance Team |
| EPI_ISL_3445915, EPI_ISL_3445916 | Lighthouse Lab in Glasgow | Wellcome Sanger Institute for the COVID-19 Genomics UK (COG-UK) Consortium | Harper VanSteenhouse, Yumi Kasai, David Gray, Carol Clugston, Anna Dominiczak and Alex Alderton, Roberto Amato, Jeffrey Barrett, Sonia Goncalves, Ewan Harrison, David K. Jackson, Ian Johnston, Dominic Kwiatkowski, Cordelia Langford, John Sillitoe on behalf of the Wellcome Sanger Institute COVID-19 Surveillance Team |
| EPI_ISL_3445917 | Lighthouse Lab in Milton Keynes | Wellcome Sanger Institute for the COVID-19 Genomics UK (COG-UK) Consortium | The Lighthouse Lab in Milton Keynes and Alex Alderton, Roberto Amato, Jeffrey Barrett, Sonia Goncalves, Ewan Harrison, David K. Jackson, Ian Johnston, Dominic Kwiatkowski, Cordelia Langford, John Sillitoe on behalf of the Wellcome Sanger Institute COVID-19 Surveillance Team |
| EPI_ISL_3445918, EPI_ISL_3445919 | Lighthouse Lab in Glasgow | Wellcome Sanger Institute for the COVID-19 Genomics UK (COG-UK) Consortium | Harper VanSteenhouse, Yumi Kasai, David Gray, Carol Clugston, Anna Dominiczak and Alex Alderton, Roberto Amato, Jeffrey Barrett, Sonia Goncalves, Ewan Harrison, David K. Jackson, Ian Johnston, Dominic Kwiatkowski, Cordelia Langford, John Sillitoe on behalf of the Wellcome Sanger Institute COVID-19 Surveillance Team |
| EPI_ISL_3445920 | Lighthouse Laboratory Plymouth | Wellcome Sanger Institute for the COVID-19 Genomics UK (COG-UK) Consortium | Lighthouse Laboratory Plymouth and Alex Alderton, Roberto Amato, Jeffrey Barrett, Sonia Goncalves, Ewan Harrison, David K. Jackson, Ian Johnston, Dominic Kwiatkowski, Cordelia Langford, John Sillitoe on behalf of the Wellcome Sanger Institute COVID-19 Surveillance Team |
| EPI_ISL_3445921 | Lighthouse Lab in Milton Keynes | Wellcome Sanger Institute for the COVID-19 Genomics UK (COG-UK) Consortium | The Lighthouse Lab in Milton Keynes and Alex Alderton, Roberto Amato, Jeffrey Barrett, Sonia Goncalves, Ewan Harrison, David K. Jackson, Ian Johnston, Dominic Kwiatkowski, Cordelia Langford, John Sillitoe on behalf of the Wellcome Sanger Institute COVID-19 Surveillance Team |
| EPI_ISL_3445922, EPI_ISL_3445923 | Lighthouse Lab in Glasgow | Wellcome Sanger Institute for the COVID-19 Genomics UK (COG-UK) Consortium | Harper VanSteenhouse, Yumi Kasai, David Gray, Carol Clugston, Anna Dominiczak and Alex Alderton, Roberto Amato, Jeffrey Barrett, Sonia Goncalves, Ewan Harrison, David K. Jackson, Ian Johnston, Dominic Kwiatkowski, Cordelia Langford, John Sillitoe on behalf of the Wellcome Sanger Institute COVID-19 Surveillance Team |
| EPI_ISL_3445924 | Lighthouse Lab in Milton Keynes | Wellcome Sanger Institute for the COVID-19 Genomics UK (COG-UK) Consortium | The Lighthouse Lab in Milton Keynes and Alex Alderton, Roberto Amato, Jeffrey Barrett, Sonia Goncalves, Ewan Harrison, David K. Jackson, Ian Johnston, Dominic Kwiatkowski, Cordelia Langford, John Sillitoe on behalf of the Wellcome Sanger Institute COVID-19 Surveillance Team |
| EPI_ISL_3445925, EPI_ISL_3445926 | Lighthouse Lab in Glasgow | Wellcome Sanger Institute for the COVID-19 Genomics UK (COG-UK) Consortium | Harper VanSteenhouse, Yumi Kasai, David Gray, Carol Clugston, Anna Dominiczak and Alex Alderton, Roberto Amato, Jeffrey Barrett, Sonia Goncalves, Ewan Harrison, David K. Jackson, Ian Johnston, Dominic Kwiatkowski, Cordelia Langford, John Sillitoe on behalf of the Wellcome Sanger Institute COVID-19 Surveillance Team |
| EPI_ISL_3445927 | Lighthouse Laboratory Plymouth | Wellcome Sanger Institute for the COVID-19 Genomics UK (COG-UK) Consortium | Lighthouse Laboratory Plymouth and Alex Alderton, Roberto Amato, Jeffrey Barrett, Sonia Goncalves, Ewan Harrison, David K. Jackson, Ian Johnston, Dominic Kwiatkowski, Cordelia Langford, John Sillitoe on behalf of the Wellcome Sanger Institute COVID-19 Surveillance Team |
| EPI_ISL_3445928 | Lighthouse Lab in Milton Keynes | Wellcome Sanger Institute for the COVID-19 Genomics UK (COG-UK) Consortium | The Lighthouse Lab in Milton Keynes and Alex Alderton, Roberto Amato, Jeffrey Barrett, Sonia Goncalves, Ewan Harrison, David K. Jackson, Ian Johnston, Dominic Kwiatkowski, Cordelia Langford, John Sillitoe on behalf of the Wellcome Sanger Institute COVID-19 Surveillance Team |
| EPI_ISL_3445929 | Lighthouse Laboratory Plymouth | Wellcome Sanger Institute for the COVID-19 Genomics UK (COG-UK) Consortium | Lighthouse Laboratory Plymouth and Alex Alderton, Roberto Amato, Jeffrey Barrett, Sonia Goncalves, Ewan Harrison, David K. Jackson, Ian Johnston, Dominic Kwiatkowski, Cordelia Langford, John Sillitoe on behalf of the Wellcome Sanger Institute COVID-19 Surveillance Team |
| EPI_ISL_3445930, EPI_ISL_3445931 | Lighthouse Lab in Glasgow | Wellcome Sanger Institute for the COVID-19 Genomics UK (COG-UK) Consortium | Harper VanSteenhouse, Yumi Kasai, David Gray, Carol Clugston, Anna Dominiczak and Alex Alderton, Roberto Amato, Jeffrey Barrett, Sonia Goncalves, Ewan Harrison, David K. Jackson, Ian Johnston, Dominic Kwiatkowski, Cordelia Langford, John Sillitoe on behalf of the Wellcome Sanger Institute COVID-19 Surveillance Team |
| EPI_ISL_3445932 | Lighthouse Laboratory Plymouth | Wellcome Sanger Institute for the COVID-19 Genomics UK (COG-UK) Consortium | Lighthouse Laboratory Plymouth and Alex Alderton, Roberto Amato, Jeffrey Barrett, Sonia Goncalves, Ewan Harrison, David K. Jackson, Ian Johnston, Dominic Kwiatkowski, Cordelia Langford, John Sillitoe on behalf of the Wellcome Sanger Institute COVID-19 Surveillance Team |
| EPI_ISL_3445933, EPI_ISL_3445934, EPI_ISL_3445935, | Lighthouse Lab in Milton Keynes | Wellcome Sanger Institute for the COVID-19 Genomics UK (COG-UK) Consortium | The Lighthouse Lab in Milton Keynes and Alex Alderton, Roberto Amato, Jeffrey Barrett, Sonia Goncalves, Ewan Harrison, David K. Jackson, Ian Johnston, Dominic Kwiatkowski, Cordelia Langford, John Sillitoe on behalf of the Wellcome Sanger Institute COVID-19 Surveillance Team |

| | | | |
|---|---|---|---|
| EPI_ISL_3445936 | | | |
| EPI_ISL_3445937 | Lighthouse Lab in Glasgow | Wellcome Sanger Institute for the COVID-19 Genomics UK (COG-UK) Consortium | Harper VanSteenhouse, Yumi Kasai, David Gray, Carol Clugston, Anna Dominiczak and Alex Alderton, Roberto Amato, Jeffrey Barrett, Sonia Goncalves, Ewan Harrison, David K. Jackson, Ian Johnston, Dominic Kwiatkowski, Cordelia Langford, John Sillitoe on behalf of the Wellcome Sanger Institute COVID-19 Surveillance Team |
| EPI_ISL_3445938 | Lighthouse Lab in Milton Keynes | Wellcome Sanger Institute for the COVID-19 Genomics UK (COG-UK) Consortium | The Lighthouse Lab in Milton Keynes and Alex Alderton, Roberto Amato, Jeffrey Barrett, Sonia Goncalves, Ewan Harrison, David K. Jackson, Ian Johnston, Dominic Kwiatkowski, Cordelia Langford, John Sillitoe on behalf of the Wellcome Sanger Institute COVID-19 Surveillance Team |
| EPI_ISL_3445939, EPI_ISL_3445940 | Lighthouse Laboratory Plymouth | Wellcome Sanger Institute for the COVID-19 Genomics UK (COG-UK) Consortium | Lighthouse Laboratory Plymouth and Alex Alderton, Roberto Amato, Jeffrey Barrett, Sonia Goncalves, Ewan Harrison, David K. Jackson, Ian Johnston, Dominic Kwiatkowski, Cordelia Langford, John Sillitoe on behalf of the Wellcome Sanger Institute COVID-19 Surveillance Team |
| EPI_ISL_3445941, EPI_ISL_3445942, EPI_ISL_3445943, EPI_ISL_3445944 | Lighthouse Lab in Milton Keynes | Wellcome Sanger Institute for the COVID-19 Genomics UK (COG-UK) Consortium | The Lighthouse Lab in Milton Keynes and Alex Alderton, Roberto Amato, Jeffrey Barrett, Sonia Goncalves, Ewan Harrison, David K. Jackson, Ian Johnston, Dominic Kwiatkowski, Cordelia Langford, John Sillitoe on behalf of the Wellcome Sanger Institute COVID-19 Surveillance Team |
| EPI_ISL_3445945 | Lighthouse Lab in Glasgow | Wellcome Sanger Institute for the COVID-19 Genomics UK (COG-UK) Consortium | Harper VanSteenhouse, Yumi Kasai, David Gray, Carol Clugston, Anna Dominiczak and Alex Alderton, Roberto Amato, Jeffrey Barrett, Sonia Goncalves, Ewan Harrison, David K. Jackson, Ian Johnston, Dominic Kwiatkowski, Cordelia Langford, John Sillitoe on behalf of the Wellcome Sanger Institute COVID-19 Surveillance Team |
| EPI_ISL_3445946, EPI_ISL_3445947 | Lighthouse Lab in Milton Keynes | Wellcome Sanger Institute for the COVID-19 Genomics UK (COG-UK) Consortium | The Lighthouse Lab in Milton Keynes and Alex Alderton, Roberto Amato, Jeffrey Barrett, Sonia Goncalves, Ewan Harrison, David K. Jackson, Ian Johnston, Dominic Kwiatkowski, Cordelia Langford, John Sillitoe on behalf of the Wellcome Sanger Institute COVID-19 Surveillance Team |
| EPI_ISL_3445948 | Lighthouse Lab in Glasgow | Wellcome Sanger Institute for the COVID-19 Genomics UK (COG-UK) Consortium | Harper VanSteenhouse, Yumi Kasai, David Gray, Carol Clugston, Anna Dominiczak and Alex Alderton, Roberto Amato, Jeffrey Barrett, Sonia Goncalves, Ewan Harrison, David K. Jackson, Ian Johnston, Dominic Kwiatkowski, Cordelia Langford, John Sillitoe on behalf of the Wellcome Sanger Institute COVID-19 Surveillance Team |
| EPI_ISL_3445949, EPI_ISL_3445950 | Lighthouse Lab in Milton Keynes | Wellcome Sanger Institute for the COVID-19 Genomics UK (COG-UK) Consortium | The Lighthouse Lab in Milton Keynes and Alex Alderton, Roberto Amato, Jeffrey Barrett, Sonia Goncalves, Ewan Harrison, David K. Jackson, Ian Johnston, Dominic Kwiatkowski, Cordelia Langford, John Sillitoe on behalf of the Wellcome Sanger Institute COVID-19 Surveillance Team |
| EPI_ISL_3445951, EPI_ISL_3445952 | Lighthouse Lab in Glasgow | Wellcome Sanger Institute for the COVID-19 Genomics UK (COG-UK) Consortium | Harper VanSteenhouse, Yumi Kasai, David Gray, Carol Clugston, Anna Dominiczak and Alex Alderton, Roberto Amato, Jeffrey Barrett, Sonia Goncalves, Ewan Harrison, David K. Jackson, Ian Johnston, Dominic Kwiatkowski, Cordelia Langford, John Sillitoe on behalf of the Wellcome Sanger Institute COVID-19 Surveillance Team |
| EPI_ISL_3445953 | Lighthouse Laboratory Plymouth | Wellcome Sanger Institute for the COVID-19 Genomics UK (COG-UK) Consortium | Lighthouse Laboratory Plymouth and Alex Alderton, Roberto Amato, Jeffrey Barrett, Sonia Goncalves, Ewan Harrison, David K. Jackson, Ian Johnston, Dominic Kwiatkowski, Cordelia Langford, John Sillitoe on behalf of the Wellcome Sanger Institute COVID-19 Surveillance Team |
| EPI_ISL_3445954, EPI_ISL_3445955 | Lighthouse Lab in Milton Keynes | Wellcome Sanger Institute for the COVID-19 Genomics UK (COG-UK) Consortium | The Lighthouse Lab in Milton Keynes and Alex Alderton, Roberto Amato, Jeffrey Barrett, Sonia Goncalves, Ewan Harrison, David K. Jackson, Ian Johnston, Dominic Kwiatkowski, Cordelia Langford, John Sillitoe on behalf of the Wellcome Sanger Institute COVID-19 Surveillance Team |
| EPI_ISL_3445956, EPI_ISL_3445957 | Lighthouse Lab in Glasgow | Wellcome Sanger Institute for the COVID-19 Genomics UK (COG-UK) Consortium | Harper VanSteenhouse, Yumi Kasai, David Gray, Carol Clugston, Anna Dominiczak and Alex Alderton, Roberto Amato, Jeffrey Barrett, Sonia Goncalves, Ewan Harrison, David K. Jackson, Ian Johnston, Dominic Kwiatkowski, Cordelia Langford, John Sillitoe on behalf of the Wellcome Sanger Institute COVID-19 Surveillance Team |
| EPI_ISL_3445958, EPI_ISL_3445959 | Lighthouse Lab in Milton Keynes | Wellcome Sanger Institute for the COVID-19 Genomics UK (COG-UK) Consortium | The Lighthouse Lab in Milton Keynes and Alex Alderton, Roberto Amato, Jeffrey Barrett, Sonia Goncalves, Ewan Harrison, David K. Jackson, Ian Johnston, Dominic Kwiatkowski, Cordelia Langford, John Sillitoe on behalf of the Wellcome Sanger Institute COVID-19 Surveillance Team |
| EPI_ISL_3445960 | Lighthouse Laboratory Plymouth | Wellcome Sanger Institute for the COVID-19 Genomics UK (COG-UK) Consortium | Lighthouse Laboratory Plymouth and Alex Alderton, Roberto Amato, Jeffrey Barrett, Sonia Goncalves, Ewan Harrison, David K. Jackson, Ian Johnston, Dominic Kwiatkowski, Cordelia Langford, John Sillitoe on behalf of the Wellcome Sanger Institute COVID-19 Surveillance Team |
| EPI_ISL_3445961, EPI_ISL_3445962 | Lighthouse Lab in Glasgow | Wellcome Sanger Institute for the COVID-19 Genomics UK (COG-UK) Consortium | Harper VanSteenhouse, Yumi Kasai, David Gray, Carol Clugston, Anna Dominiczak and Alex Alderton, Roberto Amato, Jeffrey Barrett, Sonia Goncalves, Ewan Harrison, David K. Jackson, Ian Johnston, Dominic Kwiatkowski, Cordelia Langford, John Sillitoe on behalf of the Wellcome Sanger Institute COVID-19 Surveillance Team |
| EPI_ISL_3445963, EPI_ISL_3445964 | Lighthouse Lab in Milton Keynes | Wellcome Sanger Institute for the COVID-19 Genomics UK (COG-UK) Consortium | The Lighthouse Lab in Milton Keynes and Alex Alderton, Roberto Amato, Jeffrey Barrett, Sonia Goncalves, Ewan Harrison, David K. Jackson, Ian Johnston, Dominic Kwiatkowski, Cordelia Langford, John Sillitoe on behalf of the Wellcome Sanger Institute COVID-19 Surveillance Team |
| EPI_ISL_3445965 | Lighthouse Lab in Glasgow | Wellcome Sanger Institute for the COVID-19 Genomics UK (COG-UK) Consortium | Harper VanSteenhouse, Yumi Kasai, David Gray, Carol Clugston, Anna Dominiczak and Alex Alderton, Roberto Amato, Jeffrey Barrett, Sonia Goncalves, Ewan Harrison, David K. Jackson, Ian Johnston, Dominic Kwiatkowski, Cordelia Langford, John Sillitoe on behalf of the Wellcome Sanger Institute COVID-19 Surveillance Team |
| EPI_ISL_3445966 | Lighthouse Lab in Milton Keynes | Wellcome Sanger Institute for the COVID-19 Genomics UK (COG-UK) Consortium | The Lighthouse Lab in Milton Keynes and Alex Alderton, Roberto Amato, Jeffrey Barrett, Sonia Goncalves, Ewan Harrison, David K. Jackson, Ian Johnston, Dominic Kwiatkowski, Cordelia Langford, John Sillitoe on behalf of the Wellcome Sanger Institute COVID-19 Surveillance Team |
| EPI_ISL_3445967, EPI_ISL_3445968, EPI_ISL_3445969, EPI_ISL_3445970 | Lighthouse Lab in Glasgow | Wellcome Sanger Institute for the COVID-19 Genomics UK (COG-UK) Consortium | Harper VanSteenhouse, Yumi Kasai, David Gray, Carol Clugston, Anna Dominiczak and Alex Alderton, Roberto Amato, Jeffrey Barrett, Sonia Goncalves, Ewan Harrison, David K. Jackson, Ian Johnston, Dominic Kwiatkowski, Cordelia Langford, John Sillitoe on behalf of the Wellcome Sanger Institute COVID-19 Surveillance Team |
| EPI_ISL_3445971 | Lighthouse Laboratory Plymouth | Wellcome Sanger Institute for the COVID-19 Genomics UK (COG-UK) Consortium | Lighthouse Laboratory Plymouth and Alex Alderton, Roberto Amato, Jeffrey Barrett, Sonia Goncalves, Ewan Harrison, David K. Jackson, Ian Johnston, Dominic Kwiatkowski, Cordelia Langford, John Sillitoe on behalf of the Wellcome Sanger Institute COVID-19 Surveillance Team |
| EPI_ISL_3445972 | Lighthouse Lab in Milton Keynes | Wellcome Sanger Institute for the COVID-19 Genomics UK (COG-UK) Consortium | The Lighthouse Lab in Milton Keynes and Alex Alderton, Roberto Amato, Jeffrey Barrett, Sonia Goncalves, Ewan Harrison, David K. Jackson, Ian Johnston, Dominic Kwiatkowski, Cordelia Langford, John Sillitoe on behalf of the Wellcome Sanger Institute COVID-19 Surveillance Team |
| EPI_ISL_3445973, EPI_ISL_3445974, EPI_ISL_3445975 | Lighthouse Lab in Glasgow | Wellcome Sanger Institute for the COVID-19 Genomics UK (COG-UK) Consortium | Harper VanSteenhouse, Yumi Kasai, David Gray, Carol Clugston, Anna Dominiczak and Alex Alderton, Roberto Amato, Jeffrey Barrett, Sonia Goncalves, Ewan Harrison, David K. Jackson, Ian Johnston, Dominic Kwiatkowski, Cordelia Langford, John Sillitoe on behalf of the Wellcome Sanger Institute COVID-19 Surveillance Team |
| EPI_ISL_3445976 | Lighthouse Laboratory Plymouth | Wellcome Sanger Institute for the COVID-19 Genomics UK (COG-UK) Consortium | Lighthouse Laboratory Plymouth and Alex Alderton, Roberto Amato, Jeffrey Barrett, Sonia Goncalves, Ewan Harrison, David K. Jackson, Ian Johnston, Dominic Kwiatkowski, Cordelia Langford, John Sillitoe on behalf of the Wellcome Sanger Institute COVID-19 Surveillance Team |
| EPI_ISL_3445977 | Lighthouse Lab in Glasgow | Wellcome Sanger Institute for the COVID-19 Genomics UK (COG-UK) Consortium | Harper VanSteenhouse, Yumi Kasai, David Gray, Carol Clugston, Anna Dominiczak and Alex Alderton, Roberto Amato, Jeffrey Barrett, Sonia Goncalves, Ewan Harrison, David K. Jackson, Ian Johnston, Dominic Kwiatkowski, Cordelia Langford, John Sillitoe on behalf of the Wellcome Sanger Institute COVID-19 Surveillance Team |
| EPI_ISL_3445978, EPI_ISL_3445979 | Lighthouse Lab in Milton Keynes | Wellcome Sanger Institute for the COVID-19 Genomics UK (COG-UK) Consortium | The Lighthouse Lab in Milton Keynes and Alex Alderton, Roberto Amato, Jeffrey Barrett, Sonia Goncalves, Ewan Harrison, David K. Jackson, Ian Johnston, Dominic Kwiatkowski, Cordelia Langford, John Sillitoe on behalf of the Wellcome Sanger Institute COVID-19 Surveillance Team |
| EPI_ISL_3445980 | Lighthouse Lab in Glasgow | Wellcome Sanger Institute for the COVID-19 Genomics UK (COG-UK) Consortium | Harper VanSteenhouse, Yumi Kasai, David Gray, Carol Clugston, Anna Dominiczak and Alex Alderton, Roberto Amato, Jeffrey Barrett, Sonia Goncalves, Ewan Harrison, David K. Jackson, Ian Johnston, Dominic Kwiatkowski, Cordelia Langford, John Sillitoe on behalf of the Wellcome Sanger Institute COVID-19 Surveillance Team |
| EPI_ISL_3445981, EPI_ISL_3445982 | Lighthouse Lab in Milton Keynes | Wellcome Sanger Institute for the COVID-19 Genomics UK (COG-UK) Consortium | The Lighthouse Lab in Milton Keynes and Alex Alderton, Roberto Amato, Jeffrey Barrett, Sonia Goncalves, Ewan Harrison, David K. Jackson, Ian Johnston, Dominic Kwiatkowski, Cordelia Langford, John Sillitoe on behalf of the Wellcome Sanger Institute COVID-19 Surveillance Team |
| EPI_ISL_3445983, EPI_ISL_3445984, EPI_ISL_3445985, EPI_ISL_3445986, EPI_ISL_3445987 | Lighthouse Lab in Glasgow | Wellcome Sanger Institute for the COVID-19 Genomics UK (COG-UK) Consortium | Harper VanSteenhouse, Yumi Kasai, David Gray, Carol Clugston, Anna Dominiczak and Alex Alderton, Roberto Amato, Jeffrey Barrett, Sonia Goncalves, Ewan Harrison, David K. Jackson, Ian Johnston, Dominic Kwiatkowski, Cordelia Langford, John Sillitoe on behalf of the Wellcome Sanger Institute COVID-19 Surveillance Team |
| EPI_ISL_3445988 | Lighthouse Lab in Milton Keynes | Wellcome Sanger Institute for the COVID-19 Genomics UK (COG-UK) Consortium | The Lighthouse Lab in Milton Keynes and Alex Alderton, Roberto Amato, Jeffrey Barrett, Sonia Goncalves, Ewan Harrison, David K. Jackson, Ian Johnston, Dominic Kwiatkowski, Cordelia Langford, John Sillitoe on behalf of the Wellcome Sanger Institute COVID-19 Surveillance Team |
| EPI_ISL_3445989, EPI_ISL_3445990, EPI_ISL_3445991, EPI_ISL_3445992 | Lighthouse Lab in Glasgow | Wellcome Sanger Institute for the COVID-19 Genomics UK (COG-UK) Consortium | Harper VanSteenhouse, Yumi Kasai, David Gray, Carol Clugston, Anna Dominiczak and Alex Alderton, Roberto Amato, Jeffrey Barrett, Sonia Goncalves, Ewan Harrison, David K. Jackson, Ian Johnston, Dominic Kwiatkowski, Cordelia Langford, John Sillitoe on behalf of the Wellcome Sanger Institute COVID-19 Surveillance Team |
| EPI_ISL_3445993 | Lighthouse Laboratory Plymouth | Wellcome Sanger Institute for the COVID-19 Genomics UK (COG-UK) Consortium | Lighthouse Laboratory Plymouth and Alex Alderton, Roberto Amato, Jeffrey Barrett, Sonia Goncalves, Ewan Harrison, David K. Jackson, Ian Johnston, Dominic Kwiatkowski, Cordelia Langford, John Sillitoe on behalf of the Wellcome Sanger Institute COVID-19 Surveillance Team |
| EPI_ISL_3445994, EPI_ISL_3445995 | Lighthouse Lab in Milton Keynes | Wellcome Sanger Institute for the COVID-19 Genomics UK (COG-UK) Consortium | The Lighthouse Lab in Milton Keynes and Alex Alderton, Roberto Amato, Jeffrey Barrett, Sonia Goncalves, Ewan Harrison, David K. Jackson, Ian Johnston, Dominic Kwiatkowski, Cordelia Langford, John Sillitoe on behalf of the Wellcome Sanger Institute COVID-19 Surveillance Team |
| EPI_ISL_3445996, EPI_ISL_3445997 | Lighthouse Lab in Glasgow | Wellcome Sanger Institute for the COVID-19 Genomics UK (COG-UK) Consortium | Harper VanSteenhouse, Yumi Kasai, David Gray, Carol Clugston, Anna Dominiczak and Alex Alderton, Roberto Amato, Jeffrey Barrett, Sonia Goncalves, Ewan Harrison, David K. Jackson, Ian Johnston, Dominic Kwiatkowski, Cordelia Langford, John Sillitoe on behalf of the Wellcome Sanger Institute COVID-19 Surveillance Team |
| EPI_ISL_3445998, EPI_ISL_3445999, EPI_ISL_3446000 | Lighthouse Lab in Milton Keynes | Wellcome Sanger Institute for the COVID-19 Genomics UK (COG-UK) Consortium | The Lighthouse Lab in Milton Keynes and Alex Alderton, Roberto Amato, Jeffrey Barrett, Sonia Goncalves, Ewan Harrison, David K. Jackson, Ian Johnston, Dominic Kwiatkowski, Cordelia Langford, John Sillitoe on behalf of the Wellcome Sanger Institute COVID-19 Surveillance Team |
| EPI_ISL_3446001, EPI_ISL_3446002, EPI_ISL_3446003, EPI_ISL_3446004, EPI_ISL_3446005, EPI_ISL_3446006, EPI_ISL_3446007, EPI_ISL_3446008, EPI_ISL_3446009, EPI_ISL_3446010, EPI_ISL_3446011, EPI_ISL_3446012, EPI_ISL_3446013, EPI_ISL_3446014, EPI_ISL_3446015, EPI_ISL_3446016, EPI_ISL_3446017, EPI_ISL_3446018, EPI_ISL_3446019, EPI_ISL_3446020, EPI_ISL_3446021, EPI_ISL_3446022, EPI_ISL_3446023, EPI_ISL_3446024, EPI_ISL_3446025, EPI_ISL_3446026, EPI_ISL_3446027, EPI_ISL_3446028, EPI_ISL_3446029, EPI_ISL_3446030, EPI_ISL_3446031, EPI_ISL_3446032, EPI_ISL_3446033, EPI_ISL_3446034, EPI_ISL_3446035, EPI_ISL_3446036, EPI_ISL_3446037, EPI_ISL_3446038, EPI_ISL_3446039, EPI_ISL_3446040, EPI_ISL_3446041, EPI_ISL_3446042, EPI_ISL_3446043, EPI_ISL_3446044, EPI_ISL_3446045, EPI_ISL_3446046, EPI_ISL_3446047, EPI_ISL_3446048, EPI_ISL_3446049, EPI_ISL_3446050, EPI_ISL_3446051, EPI_ISL_3446052, EPI_ISL_3446053, EPI_ISL_3446054, EPI_ISL_3446055, EPI_ISL_3446056, EPI_ISL_3446057, EPI_ISL_3446058, EPI_ISL_3446059, EPI_ISL_3446060, EPI_ISL_3446061, EPI_ISL_3446062, EPI_ISL_3446063, EPI_ISL_3446064, EPI_ISL_3446065, EPI_ISL_3446066, EPI_ISL_3446067, EPI_ISL_3446068, EPI_ISL_3446069, EPI_ISL_3446070, EPI_ISL_3446071, EPI_ISL_3446072, EPI_ISL_3446073, EPI_ISL_3446074 | | | |
| see above | Berkshire and Surrey Pathology Services Lighthouse Laboratory | Wellcome Sanger Institute for the COVID-19 Genomics UK (COG-UK) Consortium | Berkshire and Surrey Pathology Services Lighthouse Laboratory and Alex Alderton, Roberto Amato, Jeffrey Barrett, Sonia Goncalves, Ewan Harrison, David K. Jackson, Ian Johnston, Dominic Kwiatkowski, Cordelia Langford, John Sillitoe on behalf of the Wellcome Sanger Institute COVID-19 Surveillance Team |
| EPI_ISL_3446075, EPI_ISL_3446076 | Lighthouse Lab in Alderley Park | Wellcome Sanger Institute for the COVID-19 Genomics UK (COG-UK) Consortium | Jacquelyn Wynn, Mairead Hyland, The Lighthouse Lab in Alderley Park and Alex Alderton, Roberto Amato, Jeffrey Barrett, Sonia Goncalves, Ewan Harrison, David K. Jackson, Ian Johnston, Dominic Kwiatkowski, Cordelia Langford, John Sillitoe on behalf of the Wellcome Sanger Institute COVID-19 Surveillance Team |
| EPI_ISL_3446077 | Lighthouse Lab in Milton Keynes | Wellcome Sanger Institute for the COVID-19 Genomics UK (COG-UK) Consortium | The Lighthouse Lab in Milton Keynes and Alex Alderton, Roberto Amato, Jeffrey Barrett, Sonia Goncalves, Ewan Harrison, David K. Jackson, Ian Johnston, Dominic Kwiatkowski, Cordelia Langford, John Sillitoe on behalf of the Wellcome Sanger Institute COVID-19 Surveillance Team |
| EPI_ISL_3446078 | Lighthouse Lab in Glasgow | Wellcome Sanger Institute for the COVID-19 Genomics UK (COG-UK) Consortium | Harper VanSteenhouse, Yumi Kasai, David Gray, Carol Clugston, Anna Dominiczak and Alex Alderton, Roberto Amato, Jeffrey Barrett, Sonia Goncalves, Ewan Harrison, David K. Jackson, Ian Johnston, Dominic Kwiatkowski, Cordelia Langford, John Sillitoe on behalf of the Wellcome Sanger Institute COVID-19 Surveillance Team |

| | | | |
|---|---|---|---|
| EPI_ISL_3446079, EPI_ISL_3446080, EPI_ISL_3446081 | Lighthouse Lab in Alderley Park | Wellcome Sanger Institute for the COVID-19 Genomics UK (COG-UK) Consortium | Jacquelyn Wynn, Mairead Hyland, The Lighthouse Lab in Alderley Park and Alex Alderton, Roberto Amato, Jeffrey Barrett, Sonia Goncalves, Ewan Harrison, David K. Jackson, Ian Johnston, Dominic Kwiatkowski, Cordelia Langford, John Sillitoe on behalf of the Wellcome Sanger Institute COVID-19 Surveillance Team |
| EPI_ISL_3446082 | Lighthouse Lab in Milton Keynes | Wellcome Sanger Institute for the COVID-19 Genomics UK (COG-UK) Consortium | The Lighthouse Lab in Milton Keynes and Alex Alderton, Roberto Amato, Jeffrey Barrett, Sonia Goncalves, Ewan Harrison, David K. Jackson, Ian Johnston, Dominic Kwiatkowski, Cordelia Langford, John Sillitoe on behalf of the Wellcome Sanger Institute COVID-19 Surveillance Team |
| EPI_ISL_3446083, EPI_ISL_3446084, EPI_ISL_3446085 | Lighthouse Lab in Glasgow | Wellcome Sanger Institute for the COVID-19 Genomics UK (COG-UK) Consortium | Harper VanSteenhouse, Yumi Kasai, David Gray, Carol Clugston, Anna Dominiczak and Alex Alderton, Roberto Amato, Jeffrey Barrett, Sonia Goncalves, Ewan Harrison, David K. Jackson, Ian Johnston, Dominic Kwiatkowski, Cordelia Langford, John Sillitoe on behalf of the Wellcome Sanger Institute COVID-19 Surveillance Team |
| EPI_ISL_3446086, EPI_ISL_3446087 | Lighthouse Lab in Milton Keynes | Wellcome Sanger Institute for the COVID-19 Genomics UK (COG-UK) Consortium | The Lighthouse Lab in Milton Keynes and Alex Alderton, Roberto Amato, Jeffrey Barrett, Sonia Goncalves, Ewan Harrison, David K. Jackson, Ian Johnston, Dominic Kwiatkowski, Cordelia Langford, John Sillitoe on behalf of the Wellcome Sanger Institute COVID-19 Surveillance Team |
| EPI_ISL_3446088 | Lighthouse Lab in Glasgow | Wellcome Sanger Institute for the COVID-19 Genomics UK (COG-UK) Consortium | Harper VanSteenhouse, Yumi Kasai, David Gray, Carol Clugston, Anna Dominiczak and Alex Alderton, Roberto Amato, Jeffrey Barrett, Sonia Goncalves, Ewan Harrison, David K. Jackson, Ian Johnston, Dominic Kwiatkowski, Cordelia Langford, John Sillitoe on behalf of the Wellcome Sanger Institute COVID-19 Surveillance Team |
| EPI_ISL_3446089 | Lighthouse Lab in Alderley Park | Wellcome Sanger Institute for the COVID-19 Genomics UK (COG-UK) Consortium | Jacquelyn Wynn, Mairead Hyland, The Lighthouse Lab in Alderley Park and Alex Alderton, Roberto Amato, Jeffrey Barrett, Sonia Goncalves, Ewan Harrison, David K. Jackson, Ian Johnston, Dominic Kwiatkowski, Cordelia Langford, John Sillitoe on behalf of the Wellcome Sanger Institute COVID-19 Surveillance Team |
| EPI_ISL_3446090, EPI_ISL_3446091 | Lighthouse Lab in Glasgow | Wellcome Sanger Institute for the COVID-19 Genomics UK (COG-UK) Consortium | Harper VanSteenhouse, Yumi Kasai, David Gray, Carol Clugston, Anna Dominiczak and Alex Alderton, Roberto Amato, Jeffrey Barrett, Sonia Goncalves, Ewan Harrison, David K. Jackson, Ian Johnston, Dominic Kwiatkowski, Cordelia Langford, John Sillitoe on behalf of the Wellcome Sanger Institute COVID-19 Surveillance Team |
| EPI_ISL_3446092 | Lighthouse Lab in Milton Keynes | Wellcome Sanger Institute for the COVID-19 Genomics UK (COG-UK) Consortium | The Lighthouse Lab in Milton Keynes and Alex Alderton, Roberto Amato, Jeffrey Barrett, Sonia Goncalves, Ewan Harrison, David K. Jackson, Ian Johnston, Dominic Kwiatkowski, Cordelia Langford, John Sillitoe on behalf of the Wellcome Sanger Institute COVID-19 Surveillance Team |
| EPI_ISL_3446093 | Lighthouse Lab in Glasgow | Wellcome Sanger Institute for the COVID-19 Genomics UK (COG-UK) Consortium | Harper VanSteenhouse, Yumi Kasai, David Gray, Carol Clugston, Anna Dominiczak and Alex Alderton, Roberto Amato, Jeffrey Barrett, Sonia Goncalves, Ewan Harrison, David K. Jackson, Ian Johnston, Dominic Kwiatkowski, Cordelia Langford, John Sillitoe on behalf of the Wellcome Sanger Institute COVID-19 Surveillance Team |
| EPI_ISL_3446094, EPI_ISL_3446095 | Lighthouse Lab in Alderley Park | Wellcome Sanger Institute for the COVID-19 Genomics UK (COG-UK) Consortium | Jacquelyn Wynn, Mairead Hyland, The Lighthouse Lab in Alderley Park and Alex Alderton, Roberto Amato, Jeffrey Barrett, Sonia Goncalves, Ewan Harrison, David K. Jackson, Ian Johnston, Dominic Kwiatkowski, Cordelia Langford, John Sillitoe on behalf of the Wellcome Sanger Institute COVID-19 Surveillance Team |
| EPI_ISL_3446096, EPI_ISL_3446097, EPI_ISL_3446098, EPI_ISL_3446099, EPI_ISL_3446100 | Lighthouse Lab in Glasgow | Wellcome Sanger Institute for the COVID-19 Genomics UK (COG-UK) Consortium | Harper VanSteenhouse, Yumi Kasai, David Gray, Carol Clugston, Anna Dominiczak and Alex Alderton, Roberto Amato, Jeffrey Barrett, Sonia Goncalves, Ewan Harrison, David K. Jackson, Ian Johnston, Dominic Kwiatkowski, Cordelia Langford, John Sillitoe on behalf of the Wellcome Sanger Institute COVID-19 Surveillance Team |
| EPI_ISL_3446101 | Lighthouse Lab in Alderley Park | Wellcome Sanger Institute for the COVID-19 Genomics UK (COG-UK) Consortium | Jacquelyn Wynn, Mairead Hyland, The Lighthouse Lab in Alderley Park and Alex Alderton, Roberto Amato, Jeffrey Barrett, Sonia Goncalves, Ewan Harrison, David K. Jackson, Ian Johnston, Dominic Kwiatkowski, Cordelia Langford, John Sillitoe on behalf of the Wellcome Sanger Institute COVID-19 Surveillance Team |
| EPI_ISL_3446102, EPI_ISL_3446103 | Lighthouse Lab in Glasgow | Wellcome Sanger Institute for the COVID-19 Genomics UK (COG-UK) Consortium | Harper VanSteenhouse, Yumi Kasai, David Gray, Carol Clugston, Anna Dominiczak and Alex Alderton, Roberto Amato, Jeffrey Barrett, Sonia Goncalves, Ewan Harrison, David K. Jackson, Ian Johnston, Dominic Kwiatkowski, Cordelia Langford, John Sillitoe on behalf of the Wellcome Sanger Institute COVID-19 Surveillance Team |
| EPI_ISL_3446104 | Lighthouse Lab in Milton Keynes | Wellcome Sanger Institute for the COVID-19 Genomics UK (COG-UK) Consortium | The Lighthouse Lab in Milton Keynes and Alex Alderton, Roberto Amato, Jeffrey Barrett, Sonia Goncalves, Ewan Harrison, David K. Jackson, Ian Johnston, Dominic Kwiatkowski, Cordelia Langford, John Sillitoe on behalf of the Wellcome Sanger Institute COVID-19 Surveillance Team |
| EPI_ISL_3446105, EPI_ISL_3446106 | Lighthouse Lab in Glasgow | Wellcome Sanger Institute for the COVID-19 Genomics UK (COG-UK) Consortium | Harper VanSteenhouse, Yumi Kasai, David Gray, Carol Clugston, Anna Dominiczak and Alex Alderton, Roberto Amato, Jeffrey Barrett, Sonia Goncalves, Ewan Harrison, David K. Jackson, Ian Johnston, Dominic Kwiatkowski, Cordelia Langford, John Sillitoe on behalf of the Wellcome Sanger Institute COVID-19 Surveillance Team |
| EPI_ISL_3446107 | Lighthouse Lab in Alderley Park | Wellcome Sanger Institute for the COVID-19 Genomics UK (COG-UK) Consortium | Jacquelyn Wynn, Mairead Hyland, The Lighthouse Lab in Alderley Park and Alex Alderton, Roberto Amato, Jeffrey Barrett, Sonia Goncalves, Ewan Harrison, David K. Jackson, Ian Johnston, Dominic Kwiatkowski, Cordelia Langford, John Sillitoe on behalf of the Wellcome Sanger Institute COVID-19 Surveillance Team |
| EPI_ISL_3446108, EPI_ISL_3446109, EPI_ISL_3446110, EPI_ISL_3446111 | Lighthouse Lab in Glasgow | Wellcome Sanger Institute for the COVID-19 Genomics UK (COG-UK) Consortium | Harper VanSteenhouse, Yumi Kasai, David Gray, Carol Clugston, Anna Dominiczak and Alex Alderton, Roberto Amato, Jeffrey Barrett, Sonia Goncalves, Ewan Harrison, David K. Jackson, Ian Johnston, Dominic Kwiatkowski, Cordelia Langford, John Sillitoe on behalf of the Wellcome Sanger Institute COVID-19 Surveillance Team |
| EPI_ISL_3446112, EPI_ISL_3446113 | Lighthouse Lab in Milton Keynes | Wellcome Sanger Institute for the COVID-19 Genomics UK (COG-UK) Consortium | The Lighthouse Lab in Milton Keynes and Alex Alderton, Roberto Amato, Jeffrey Barrett, Sonia Goncalves, Ewan Harrison, David K. Jackson, Ian Johnston, Dominic Kwiatkowski, Cordelia Langford, John Sillitoe on behalf of the Wellcome Sanger Institute COVID-19 Surveillance Team |
| EPI_ISL_3446114 | Lighthouse Lab in Alderley Park | Wellcome Sanger Institute for the COVID-19 Genomics UK (COG-UK) Consortium | Jacquelyn Wynn, Mairead Hyland, The Lighthouse Lab in Alderley Park and Alex Alderton, Roberto Amato, Jeffrey Barrett, Sonia Goncalves, Ewan Harrison, David K. Jackson, Ian Johnston, Dominic Kwiatkowski, Cordelia Langford, John Sillitoe on behalf of the Wellcome Sanger Institute COVID-19 Surveillance Team |
| EPI_ISL_3446115 | Lighthouse Lab in Glasgow | Wellcome Sanger Institute for the COVID-19 Genomics UK (COG-UK) Consortium | Harper VanSteenhouse, Yumi Kasai, David Gray, Carol Clugston, Anna Dominiczak and Alex Alderton, Roberto Amato, Jeffrey Barrett, Sonia Goncalves, Ewan Harrison, David K. Jackson, Ian Johnston, Dominic Kwiatkowski, Cordelia Langford, John Sillitoe on behalf of the Wellcome Sanger Institute COVID-19 Surveillance Team |
| EPI_ISL_3446116 | Lighthouse Lab in Alderley Park | Wellcome Sanger Institute for the COVID-19 Genomics UK (COG-UK) Consortium | Jacquelyn Wynn, Mairead Hyland, The Lighthouse Lab in Alderley Park and Alex Alderton, Roberto Amato, Jeffrey Barrett, Sonia Goncalves, Ewan Harrison, David K. Jackson, Ian Johnston, Dominic Kwiatkowski, Cordelia Langford, John Sillitoe on behalf of the Wellcome Sanger Institute COVID-19 Surveillance Team |
| EPI_ISL_3446117 | Lighthouse Lab in Glasgow | Wellcome Sanger Institute for the COVID-19 Genomics UK (COG-UK) Consortium | Harper VanSteenhouse, Yumi Kasai, David Gray, Carol Clugston, Anna Dominiczak and Alex Alderton, Roberto Amato, Jeffrey Barrett, Sonia Goncalves, Ewan Harrison, David K. Jackson, Ian Johnston, Dominic Kwiatkowski, Cordelia Langford, John Sillitoe on behalf of the Wellcome Sanger Institute COVID-19 Surveillance Team |
| EPI_ISL_3446118, EPI_ISL_3446119 | Lighthouse Lab in Alderley Park | Wellcome Sanger Institute for the COVID-19 Genomics UK (COG-UK) Consortium | Jacquelyn Wynn, Mairead Hyland, The Lighthouse Lab in Alderley Park and Alex Alderton, Roberto Amato, Jeffrey Barrett, Sonia Goncalves, Ewan Harrison, David K. Jackson, Ian Johnston, Dominic Kwiatkowski, Cordelia Langford, John Sillitoe on behalf of the Wellcome Sanger Institute COVID-19 Surveillance Team |
| EPI_ISL_3446120 | Lighthouse Lab in Glasgow | Wellcome Sanger Institute for the COVID-19 Genomics UK (COG-UK) Consortium | Harper VanSteenhouse, Yumi Kasai, David Gray, Carol Clugston, Anna Dominiczak and Alex Alderton, Roberto Amato, Jeffrey Barrett, Sonia Goncalves, Ewan Harrison, David K. Jackson, Ian Johnston, Dominic Kwiatkowski, Cordelia Langford, John Sillitoe on behalf of the Wellcome Sanger Institute COVID-19 Surveillance Team |
| EPI_ISL_3446121, EPI_ISL_3446122, EPI_ISL_3446123 | Lighthouse Lab in Milton Keynes | Wellcome Sanger Institute for the COVID-19 Genomics UK (COG-UK) Consortium | The Lighthouse Lab in Milton Keynes and Alex Alderton, Roberto Amato, Jeffrey Barrett, Sonia Goncalves, Ewan Harrison, David K. Jackson, Ian Johnston, Dominic Kwiatkowski, Cordelia Langford, John Sillitoe on behalf of the Wellcome Sanger Institute COVID-19 Surveillance Team |
| EPI_ISL_3446124 | Lighthouse Lab in Glasgow | Wellcome Sanger Institute for the COVID-19 Genomics UK (COG-UK) Consortium | Harper VanSteenhouse, Yumi Kasai, David Gray, Carol Clugston, Anna Dominiczak and Alex Alderton, Roberto Amato, Jeffrey Barrett, Sonia Goncalves, Ewan Harrison, David K. Jackson, Ian Johnston, Dominic Kwiatkowski, Cordelia Langford, John Sillitoe on behalf of the Wellcome Sanger Institute COVID-19 Surveillance Team |
| EPI_ISL_3446125 | Lighthouse Lab in Milton Keynes | Wellcome Sanger Institute for the COVID-19 Genomics UK (COG-UK) Consortium | The Lighthouse Lab in Milton Keynes and Alex Alderton, Roberto Amato, Jeffrey Barrett, Sonia Goncalves, Ewan Harrison, David K. Jackson, Ian Johnston, Dominic Kwiatkowski, Cordelia Langford, John Sillitoe on behalf of the Wellcome Sanger Institute COVID-19 Surveillance Team |
| EPI_ISL_3446126 | Lighthouse Lab in Glasgow | Wellcome Sanger Institute for the COVID-19 Genomics UK (COG-UK) Consortium | Harper VanSteenhouse, Yumi Kasai, David Gray, Carol Clugston, Anna Dominiczak and Alex Alderton, Roberto Amato, Jeffrey Barrett, Sonia Goncalves, Ewan Harrison, David K. Jackson, Ian Johnston, Dominic Kwiatkowski, Cordelia Langford, John Sillitoe on behalf of the Wellcome Sanger Institute COVID-19 Surveillance Team |
| EPI_ISL_3446127 | Lighthouse Lab in Alderley Park | Wellcome Sanger Institute for the COVID-19 Genomics UK (COG-UK) Consortium | Jacquelyn Wynn, Mairead Hyland, The Lighthouse Lab in Alderley Park and Alex Alderton, Roberto Amato, Jeffrey Barrett, Sonia Goncalves, Ewan Harrison, David K. Jackson, Ian Johnston, Dominic Kwiatkowski, Cordelia Langford, John Sillitoe on behalf of the Wellcome Sanger Institute COVID-19 Surveillance Team |
| EPI_ISL_3446128, EPI_ISL_3446129 | Lighthouse Lab in Glasgow | Wellcome Sanger Institute for the COVID-19 Genomics UK (COG-UK) Consortium | Harper VanSteenhouse, Yumi Kasai, David Gray, Carol Clugston, Anna Dominiczak and Alex Alderton, Roberto Amato, Jeffrey Barrett, Sonia Goncalves, Ewan Harrison, David K. Jackson, Ian Johnston, Dominic Kwiatkowski, Cordelia Langford, John Sillitoe on behalf of the Wellcome Sanger Institute COVID-19 Surveillance Team |
| EPI_ISL_3446130 | Lighthouse Lab in Alderley Park | Wellcome Sanger Institute for the COVID-19 Genomics UK (COG-UK) Consortium | Jacquelyn Wynn, Mairead Hyland, The Lighthouse Lab in Alderley Park and Alex Alderton, Roberto Amato, Jeffrey Barrett, Sonia Goncalves, Ewan Harrison, David K. Jackson, Ian Johnston, Dominic Kwiatkowski, Cordelia Langford, John Sillitoe on behalf of the Wellcome Sanger Institute COVID-19 Surveillance Team |
| EPI_ISL_3446131, EPI_ISL_3446132, EPI_ISL_3446133 | Lighthouse Lab in Glasgow | Wellcome Sanger Institute for the COVID-19 Genomics UK (COG-UK) Consortium | Harper VanSteenhouse, Yumi Kasai, David Gray, Carol Clugston, Anna Dominiczak and Alex Alderton, Roberto Amato, Jeffrey Barrett, Sonia Goncalves, Ewan Harrison, David K. Jackson, Ian Johnston, Dominic Kwiatkowski, Cordelia Langford, John Sillitoe on behalf of the Wellcome Sanger Institute COVID-19 Surveillance Team |
| EPI_ISL_3446134 | Lighthouse Lab in Milton Keynes | Wellcome Sanger Institute for the COVID-19 Genomics UK (COG-UK) Consortium | The Lighthouse Lab in Milton Keynes and Alex Alderton, Roberto Amato, Jeffrey Barrett, Sonia Goncalves, Ewan Harrison, David K. Jackson, Ian Johnston, Dominic Kwiatkowski, Cordelia Langford, John Sillitoe on behalf of the Wellcome Sanger Institute COVID-19 Surveillance Team |
| EPI_ISL_3446135, EPI_ISL_3446136 | Lighthouse Lab in Glasgow | Wellcome Sanger Institute for the COVID-19 Genomics UK (COG-UK) Consortium | Harper VanSteenhouse, Yumi Kasai, David Gray, Carol Clugston, Anna Dominiczak and Alex Alderton, Roberto Amato, Jeffrey Barrett, Sonia Goncalves, Ewan Harrison, David K. Jackson, Ian Johnston, Dominic Kwiatkowski, Cordelia Langford, John Sillitoe on behalf of the Wellcome Sanger Institute COVID-19 Surveillance Team |
| EPI_ISL_3446137, EPI_ISL_3446138, EPI_ISL_3446139 | Lighthouse Lab in Alderley Park | Wellcome Sanger Institute for the COVID-19 Genomics UK (COG-UK) Consortium | Jacquelyn Wynn, Mairead Hyland, The Lighthouse Lab in Alderley Park and Alex Alderton, Roberto Amato, Jeffrey Barrett, Sonia Goncalves, Ewan Harrison, David K. Jackson, Ian Johnston, Dominic Kwiatkowski, Cordelia Langford, John Sillitoe on behalf of the Wellcome Sanger Institute COVID-19 Surveillance Team |
| EPI_ISL_3446140 | Lighthouse Lab in Milton Keynes | Wellcome Sanger Institute for the COVID-19 Genomics UK (COG-UK) Consortium | The Lighthouse Lab in Milton Keynes and Alex Alderton, Roberto Amato, Jeffrey Barrett, Sonia Goncalves, Ewan Harrison, David K. Jackson, Ian Johnston, Dominic Kwiatkowski, Cordelia Langford, John Sillitoe on behalf of the Wellcome Sanger Institute COVID-19 Surveillance Team |
| EPI_ISL_3446141 | Lighthouse Lab in Glasgow | Wellcome Sanger Institute for the COVID-19 Genomics UK (COG-UK) Consortium | Harper VanSteenhouse, Yumi Kasai, David Gray, Carol Clugston, Anna Dominiczak and Alex Alderton, Roberto Amato, Jeffrey Barrett, Sonia Goncalves, Ewan Harrison, David K. Jackson, Ian Johnston, Dominic Kwiatkowski, Cordelia Langford, John Sillitoe on behalf of the Wellcome Sanger Institute COVID-19 Surveillance Team |
| EPI_ISL_3446142 | Lighthouse Lab in Alderley Park | Wellcome Sanger Institute for the COVID-19 Genomics UK (COG-UK) Consortium | Jacquelyn Wynn, Mairead Hyland, The Lighthouse Lab in Alderley Park and Alex Alderton, Roberto Amato, Jeffrey Barrett, Sonia Goncalves, Ewan Harrison, David K. Jackson, Ian Johnston, Dominic Kwiatkowski, Cordelia Langford, John Sillitoe on behalf of the Wellcome Sanger Institute COVID-19 Surveillance Team |
| EPI_ISL_3446143 | Lighthouse Lab in Glasgow | Wellcome Sanger Institute for the COVID-19 Genomics UK (COG-UK) Consortium | Harper VanSteenhouse, Yumi Kasai, David Gray, Carol Clugston, Anna Dominiczak and Alex Alderton, Roberto Amato, Jeffrey Barrett, Sonia Goncalves, Ewan Harrison, David K. Jackson, Ian Johnston, Dominic Kwiatkowski, Cordelia Langford, John Sillitoe on behalf of the Wellcome Sanger Institute COVID-19 Surveillance Team |
| EPI_ISL_3446144 | Lighthouse Lab in Alderley Park | Wellcome Sanger Institute for the COVID-19 Genomics UK (COG-UK) Consortium | Jacquelyn Wynn, Mairead Hyland, The Lighthouse Lab in Alderley Park and Alex Alderton, Roberto Amato, Jeffrey Barrett, Sonia Goncalves, Ewan Harrison, David K. Jackson, Ian Johnston, Dominic Kwiatkowski, Cordelia Langford, John Sillitoe on behalf of the Wellcome Sanger Institute COVID-19 Surveillance Team |
| EPI_ISL_3446145, EPI_ISL_3446146, | Lighthouse Lab in Glasgow | Wellcome Sanger Institute for the COVID-19 Genomics UK (COG-UK) Consortium | Harper VanSteenhouse, Yumi Kasai, David Gray, Carol Clugston, Anna Dominiczak and Alex Alderton, Roberto Amato, Jeffrey Barrett, Sonia Goncalves, Ewan Harrison, David K. Jackson, Ian Johnston, Dominic Kwiatkowski, Cordelia Langford, John Sillitoe on behalf of the Wellcome Sanger Institute COVID-19 Surveillance Team |

| | | | |
|---|---|---|---|
| EPI_ISL_3446147, EPI_ISL_3446148 | | | |
| EPI_ISL_3446149 | Lighthouse Lab in Milton Keynes | Wellcome Sanger Institute for the COVID-19 Genomics UK (COG-UK) Consortium | The Lighthouse Lab in Milton Keynes and Alex Alderton, Roberto Amato, Jeffrey Barrett, Sonia Goncalves, Ewan Harrison, David K. Jackson, Ian Johnston, Dominic Kwiatkowski, Cordelia Langford, John Sillitoe on behalf of the Wellcome Sanger Institute COVID-19 Surveillance Team |
| EPI_ISL_3446150 | Lighthouse Lab in Glasgow | Wellcome Sanger Institute for the COVID-19 Genomics UK (COG-UK) Consortium | Harper VanSteenhouse, Yumi Kasai, David Gray, Carol Clugston, Anna Dominiczak and Alex Alderton, Roberto Amato, Jeffrey Barrett, Sonia Goncalves, Ewan Harrison, David K. Jackson, Ian Johnston, Dominic Kwiatkowski, Cordelia Langford, John Sillitoe on behalf of the Wellcome Sanger Institute COVID-19 Surveillance Team |
| EPI_ISL_3446151, EPI_ISL_3446152 | Lighthouse Lab in Milton Keynes | Wellcome Sanger Institute for the COVID-19 Genomics UK (COG-UK) Consortium | The Lighthouse Lab in Milton Keynes and Alex Alderton, Roberto Amato, Jeffrey Barrett, Sonia Goncalves, Ewan Harrison, David K. Jackson, Ian Johnston, Dominic Kwiatkowski, Cordelia Langford, John Sillitoe on behalf of the Wellcome Sanger Institute COVID-19 Surveillance Team |
| EPI_ISL_3446153 | Lighthouse Lab in Glasgow | Wellcome Sanger Institute for the COVID-19 Genomics UK (COG-UK) Consortium | Harper VanSteenhouse, Yumi Kasai, David Gray, Carol Clugston, Anna Dominiczak and Alex Alderton, Roberto Amato, Jeffrey Barrett, Sonia Goncalves, Ewan Harrison, David K. Jackson, Ian Johnston, Dominic Kwiatkowski, Cordelia Langford, John Sillitoe on behalf of the Wellcome Sanger Institute COVID-19 Surveillance Team |
| EPI_ISL_3446154 | Lighthouse Lab in Milton Keynes | Wellcome Sanger Institute for the COVID-19 Genomics UK (COG-UK) Consortium | The Lighthouse Lab in Milton Keynes and Alex Alderton, Roberto Amato, Jeffrey Barrett, Sonia Goncalves, Ewan Harrison, David K. Jackson, Ian Johnston, Dominic Kwiatkowski, Cordelia Langford, John Sillitoe on behalf of the Wellcome Sanger Institute COVID-19 Surveillance Team |
| EPI_ISL_3446155 | Lighthouse Lab in Alderley Park | Wellcome Sanger Institute for the COVID-19 Genomics UK (COG-UK) Consortium | Jacquelyn Wynn, Mairead Hyland, The Lighthouse Lab in Alderley Park and Alex Alderton, Roberto Amato, Jeffrey Barrett, Sonia Goncalves, Ewan Harrison, David K. Jackson, Ian Johnston, Dominic Kwiatkowski, Cordelia Langford, John Sillitoe on behalf of the Wellcome Sanger Institute COVID-19 Surveillance Team |
| EPI_ISL_3446156, EPI_ISL_3446157 | Lighthouse Lab in Glasgow | Wellcome Sanger Institute for the COVID-19 Genomics UK (COG-UK) Consortium | Harper VanSteenhouse, Yumi Kasai, David Gray, Carol Clugston, Anna Dominiczak and Alex Alderton, Roberto Amato, Jeffrey Barrett, Sonia Goncalves, Ewan Harrison, David K. Jackson, Ian Johnston, Dominic Kwiatkowski, Cordelia Langford, John Sillitoe on behalf of the Wellcome Sanger Institute COVID-19 Surveillance Team |
| EPI_ISL_3446158, EPI_ISL_3446159 | Lighthouse Lab in Alderley Park | Wellcome Sanger Institute for the COVID-19 Genomics UK (COG-UK) Consortium | Jacquelyn Wynn, Mairead Hyland, The Lighthouse Lab in Alderley Park and Alex Alderton, Roberto Amato, Jeffrey Barrett, Sonia Goncalves, Ewan Harrison, David K. Jackson, Ian Johnston, Dominic Kwiatkowski, Cordelia Langford, John Sillitoe on behalf of the Wellcome Sanger Institute COVID-19 Surveillance Team |
| EPI_ISL_3446160 | Lighthouse Lab in Milton Keynes | Wellcome Sanger Institute for the COVID-19 Genomics UK (COG-UK) Consortium | The Lighthouse Lab in Milton Keynes and Alex Alderton, Roberto Amato, Jeffrey Barrett, Sonia Goncalves, Ewan Harrison, David K. Jackson, Ian Johnston, Dominic Kwiatkowski, Cordelia Langford, John Sillitoe on behalf of the Wellcome Sanger Institute COVID-19 Surveillance Team |
| EPI_ISL_3446161, EPI_ISL_3446162 | Lighthouse Lab in Glasgow | Wellcome Sanger Institute for the COVID-19 Genomics UK (COG-UK) Consortium | Harper VanSteenhouse, Yumi Kasai, David Gray, Carol Clugston, Anna Dominiczak and Alex Alderton, Roberto Amato, Jeffrey Barrett, Sonia Goncalves, Ewan Harrison, David K. Jackson, Ian Johnston, Dominic Kwiatkowski, Cordelia Langford, John Sillitoe on behalf of the Wellcome Sanger Institute COVID-19 Surveillance Team |
| EPI_ISL_3446163 | Lighthouse Lab in Milton Keynes | Wellcome Sanger Institute for the COVID-19 Genomics UK (COG-UK) Consortium | The Lighthouse Lab in Milton Keynes and Alex Alderton, Roberto Amato, Jeffrey Barrett, Sonia Goncalves, Ewan Harrison, David K. Jackson, Ian Johnston, Dominic Kwiatkowski, Cordelia Langford, John Sillitoe on behalf of the Wellcome Sanger Institute COVID-19 Surveillance Team |
| EPI_ISL_3446164 | Lighthouse Lab in Alderley Park | Wellcome Sanger Institute for the COVID-19 Genomics UK (COG-UK) Consortium | Jacquelyn Wynn, Mairead Hyland, The Lighthouse Lab in Alderley Park and Alex Alderton, Roberto Amato, Jeffrey Barrett, Sonia Goncalves, Ewan Harrison, David K. Jackson, Ian Johnston, Dominic Kwiatkowski, Cordelia Langford, John Sillitoe on behalf of the Wellcome Sanger Institute COVID-19 Surveillance Team |
| EPI_ISL_3446165, EPI_ISL_3446166 | Lighthouse Lab in Glasgow | Wellcome Sanger Institute for the COVID-19 Genomics UK (COG-UK) Consortium | Harper VanSteenhouse, Yumi Kasai, David Gray, Carol Clugston, Anna Dominiczak and Alex Alderton, Roberto Amato, Jeffrey Barrett, Sonia Goncalves, Ewan Harrison, David K. Jackson, Ian Johnston, Dominic Kwiatkowski, Cordelia Langford, John Sillitoe on behalf of the Wellcome Sanger Institute COVID-19 Surveillance Team |
| EPI_ISL_3446167, EPI_ISL_3446168 | Lighthouse Lab in Alderley Park | Wellcome Sanger Institute for the COVID-19 Genomics UK (COG-UK) Consortium | Jacquelyn Wynn, Mairead Hyland, The Lighthouse Lab in Alderley Park and Alex Alderton, Roberto Amato, Jeffrey Barrett, Sonia Goncalves, Ewan Harrison, David K. Jackson, Ian Johnston, Dominic Kwiatkowski, Cordelia Langford, John Sillitoe on behalf of the Wellcome Sanger Institute COVID-19 Surveillance Team |
| EPI_ISL_3446169 | Lighthouse Lab in Milton Keynes | Wellcome Sanger Institute for the COVID-19 Genomics UK (COG-UK) Consortium | The Lighthouse Lab in Milton Keynes and Alex Alderton, Roberto Amato, Jeffrey Barrett, Sonia Goncalves, Ewan Harrison, David K. Jackson, Ian Johnston, Dominic Kwiatkowski, Cordelia Langford, John Sillitoe on behalf of the Wellcome Sanger Institute COVID-19 Surveillance Team |
| EPI_ISL_3446170, EPI_ISL_3446171 | Lighthouse Lab in Alderley Park | Wellcome Sanger Institute for the COVID-19 Genomics UK (COG-UK) Consortium | Jacquelyn Wynn, Mairead Hyland, The Lighthouse Lab in Alderley Park and Alex Alderton, Roberto Amato, Jeffrey Barrett, Sonia Goncalves, Ewan Harrison, David K. Jackson, Ian Johnston, Dominic Kwiatkowski, Cordelia Langford, John Sillitoe on behalf of the Wellcome Sanger Institute COVID-19 Surveillance Team |
| EPI_ISL_3446172 | Lighthouse Lab in Glasgow | Wellcome Sanger Institute for the COVID-19 Genomics UK (COG-UK) Consortium | Harper VanSteenhouse, Yumi Kasai, David Gray, Carol Clugston, Anna Dominiczak and Alex Alderton, Roberto Amato, Jeffrey Barrett, Sonia Goncalves, Ewan Harrison, David K. Jackson, Ian Johnston, Dominic Kwiatkowski, Cordelia Langford, John Sillitoe on behalf of the Wellcome Sanger Institute COVID-19 Surveillance Team |
| EPI_ISL_3446173 | Lighthouse Lab in Milton Keynes | Wellcome Sanger Institute for the COVID-19 Genomics UK (COG-UK) Consortium | The Lighthouse Lab in Milton Keynes and Alex Alderton, Roberto Amato, Jeffrey Barrett, Sonia Goncalves, Ewan Harrison, David K. Jackson, Ian Johnston, Dominic Kwiatkowski, Cordelia Langford, John Sillitoe on behalf of the Wellcome Sanger Institute COVID-19 Surveillance Team |
| EPI_ISL_3446174 | Lighthouse Lab in Glasgow | Wellcome Sanger Institute for the COVID-19 Genomics UK (COG-UK) Consortium | Harper VanSteenhouse, Yumi Kasai, David Gray, Carol Clugston, Anna Dominiczak and Alex Alderton, Roberto Amato, Jeffrey Barrett, Sonia Goncalves, Ewan Harrison, David K. Jackson, Ian Johnston, Dominic Kwiatkowski, Cordelia Langford, John Sillitoe on behalf of the Wellcome Sanger Institute COVID-19 Surveillance Team |
| EPI_ISL_3446175 | Lighthouse Lab in Alderley Park | Wellcome Sanger Institute for the COVID-19 Genomics UK (COG-UK) Consortium | Jacquelyn Wynn, Mairead Hyland, The Lighthouse Lab in Alderley Park and Alex Alderton, Roberto Amato, Jeffrey Barrett, Sonia Goncalves, Ewan Harrison, David K. Jackson, Ian Johnston, Dominic Kwiatkowski, Cordelia Langford, John Sillitoe on behalf of the Wellcome Sanger Institute COVID-19 Surveillance Team |
| EPI_ISL_3446176 | Lighthouse Lab in Milton Keynes | Wellcome Sanger Institute for the COVID-19 Genomics UK (COG-UK) Consortium | The Lighthouse Lab in Milton Keynes and Alex Alderton, Roberto Amato, Jeffrey Barrett, Sonia Goncalves, Ewan Harrison, David K. Jackson, Ian Johnston, Dominic Kwiatkowski, Cordelia Langford, John Sillitoe on behalf of the Wellcome Sanger Institute COVID-19 Surveillance Team |
| EPI_ISL_3446177 | Lighthouse Lab in Glasgow | Wellcome Sanger Institute for the COVID-19 Genomics UK (COG-UK) Consortium | Harper VanSteenhouse, Yumi Kasai, David Gray, Carol Clugston, Anna Dominiczak and Alex Alderton, Roberto Amato, Jeffrey Barrett, Sonia Goncalves, Ewan Harrison, David K. Jackson, Ian Johnston, Dominic Kwiatkowski, Cordelia Langford, John Sillitoe on behalf of the Wellcome Sanger Institute COVID-19 Surveillance Team |
| EPI_ISL_3446178 | Lighthouse Lab in Milton Keynes | Wellcome Sanger Institute for the COVID-19 Genomics UK (COG-UK) Consortium | The Lighthouse Lab in Milton Keynes and Alex Alderton, Roberto Amato, Jeffrey Barrett, Sonia Goncalves, Ewan Harrison, David K. Jackson, Ian Johnston, Dominic Kwiatkowski, Cordelia Langford, John Sillitoe on behalf of the Wellcome Sanger Institute COVID-19 Surveillance Team |
| EPI_ISL_3446179 | Lighthouse Lab in Alderley Park | Wellcome Sanger Institute for the COVID-19 Genomics UK (COG-UK) Consortium | Jacquelyn Wynn, Mairead Hyland, The Lighthouse Lab in Alderley Park and Alex Alderton, Roberto Amato, Jeffrey Barrett, Sonia Goncalves, Ewan Harrison, David K. Jackson, Ian Johnston, Dominic Kwiatkowski, Cordelia Langford, John Sillitoe on behalf of the Wellcome Sanger Institute COVID-19 Surveillance Team |
| EPI_ISL_3446180, EPI_ISL_3446181 | Lighthouse Lab in Milton Keynes | Wellcome Sanger Institute for the COVID-19 Genomics UK (COG-UK) Consortium | The Lighthouse Lab in Milton Keynes and Alex Alderton, Roberto Amato, Jeffrey Barrett, Sonia Goncalves, Ewan Harrison, David K. Jackson, Ian Johnston, Dominic Kwiatkowski, Cordelia Langford, John Sillitoe on behalf of the Wellcome Sanger Institute COVID-19 Surveillance Team |
| EPI_ISL_3446182, EPI_ISL_3446183 | Lighthouse Lab in Alderley Park | Wellcome Sanger Institute for the COVID-19 Genomics UK (COG-UK) Consortium | Jacquelyn Wynn, Mairead Hyland, The Lighthouse Lab in Alderley Park and Alex Alderton, Roberto Amato, Jeffrey Barrett, Sonia Goncalves, Ewan Harrison, David K. Jackson, Ian Johnston, Dominic Kwiatkowski, Cordelia Langford, John Sillitoe on behalf of the Wellcome Sanger Institute COVID-19 Surveillance Team |
| EPI_ISL_3446184, EPI_ISL_3446185 | Lighthouse Lab in Glasgow | Wellcome Sanger Institute for the COVID-19 Genomics UK (COG-UK) Consortium | Harper VanSteenhouse, Yumi Kasai, David Gray, Carol Clugston, Anna Dominiczak and Alex Alderton, Roberto Amato, Jeffrey Barrett, Sonia Goncalves, Ewan Harrison, David K. Jackson, Ian Johnston, Dominic Kwiatkowski, Cordelia Langford, John Sillitoe on behalf of the Wellcome Sanger Institute COVID-19 Surveillance Team |
| EPI_ISL_3446186, EPI_ISL_3446187 | Lighthouse Lab in Alderley Park | Wellcome Sanger Institute for the COVID-19 Genomics UK (COG-UK) Consortium | Jacquelyn Wynn, Mairead Hyland, The Lighthouse Lab in Alderley Park and Alex Alderton, Roberto Amato, Jeffrey Barrett, Sonia Goncalves, Ewan Harrison, David K. Jackson, Ian Johnston, Dominic Kwiatkowski, Cordelia Langford, John Sillitoe on behalf of the Wellcome Sanger Institute COVID-19 Surveillance Team |
| EPI_ISL_3446188, EPI_ISL_3446189 | Lighthouse Lab in Glasgow | Wellcome Sanger Institute for the COVID-19 Genomics UK (COG-UK) Consortium | Harper VanSteenhouse, Yumi Kasai, David Gray, Carol Clugston, Anna Dominiczak and Alex Alderton, Roberto Amato, Jeffrey Barrett, Sonia Goncalves, Ewan Harrison, David K. Jackson, Ian Johnston, Dominic Kwiatkowski, Cordelia Langford, John Sillitoe on behalf of the Wellcome Sanger Institute COVID-19 Surveillance Team |
| EPI_ISL_3446190 | Lighthouse Lab in Alderley Park | Wellcome Sanger Institute for the COVID-19 Genomics UK (COG-UK) Consortium | Jacquelyn Wynn, Mairead Hyland, The Lighthouse Lab in Alderley Park and Alex Alderton, Roberto Amato, Jeffrey Barrett, Sonia Goncalves, Ewan Harrison, David K. Jackson, Ian Johnston, Dominic Kwiatkowski, Cordelia Langford, John Sillitoe on behalf of the Wellcome Sanger Institute COVID-19 Surveillance Team |
| EPI_ISL_3446191 | Lighthouse Lab in Glasgow | Wellcome Sanger Institute for the COVID-19 Genomics UK (COG-UK) Consortium | Harper VanSteenhouse, Yumi Kasai, David Gray, Carol Clugston, Anna Dominiczak and Alex Alderton, Roberto Amato, Jeffrey Barrett, Sonia Goncalves, Ewan Harrison, David K. Jackson, Ian Johnston, Dominic Kwiatkowski, Cordelia Langford, John Sillitoe on behalf of the Wellcome Sanger Institute COVID-19 Surveillance Team |
| EPI_ISL_3446192, EPI_ISL_3446193, EPI_ISL_3446194 | Lighthouse Lab in Milton Keynes | Wellcome Sanger Institute for the COVID-19 Genomics UK (COG-UK) Consortium | The Lighthouse Lab in Milton Keynes and Alex Alderton, Roberto Amato, Jeffrey Barrett, Sonia Goncalves, Ewan Harrison, David K. Jackson, Ian Johnston, Dominic Kwiatkowski, Cordelia Langford, John Sillitoe on behalf of the Wellcome Sanger Institute COVID-19 Surveillance Team |
| EPI_ISL_3446195 | Lighthouse Lab in Glasgow | Wellcome Sanger Institute for the COVID-19 Genomics UK (COG-UK) Consortium | Harper VanSteenhouse, Yumi Kasai, David Gray, Carol Clugston, Anna Dominiczak and Alex Alderton, Roberto Amato, Jeffrey Barrett, Sonia Goncalves, Ewan Harrison, David K. Jackson, Ian Johnston, Dominic Kwiatkowski, Cordelia Langford, John Sillitoe on behalf of the Wellcome Sanger Institute COVID-19 Surveillance Team |
| EPI_ISL_3446196 | Lighthouse Lab in Alderley Park | Wellcome Sanger Institute for the COVID-19 Genomics UK (COG-UK) Consortium | Jacquelyn Wynn, Mairead Hyland, The Lighthouse Lab in Alderley Park and Alex Alderton, Roberto Amato, Jeffrey Barrett, Sonia Goncalves, Ewan Harrison, David K. Jackson, Ian Johnston, Dominic Kwiatkowski, Cordelia Langford, John Sillitoe on behalf of the Wellcome Sanger Institute COVID-19 Surveillance Team |
| EPI_ISL_3446197 | Lighthouse Lab in Glasgow | Wellcome Sanger Institute for the COVID-19 Genomics UK (COG-UK) Consortium | Harper VanSteenhouse, Yumi Kasai, David Gray, Carol Clugston, Anna Dominiczak and Alex Alderton, Roberto Amato, Jeffrey Barrett, Sonia Goncalves, Ewan Harrison, David K. Jackson, Ian Johnston, Dominic Kwiatkowski, Cordelia Langford, John Sillitoe on behalf of the Wellcome Sanger Institute COVID-19 Surveillance Team |
| EPI_ISL_3446198 | Lighthouse Lab in Alderley Park | Wellcome Sanger Institute for the COVID-19 Genomics UK (COG-UK) Consortium | Jacquelyn Wynn, Mairead Hyland, The Lighthouse Lab in Alderley Park and Alex Alderton, Roberto Amato, Jeffrey Barrett, Sonia Goncalves, Ewan Harrison, David K. Jackson, Ian Johnston, Dominic Kwiatkowski, Cordelia Langford, John Sillitoe on behalf of the Wellcome Sanger Institute COVID-19 Surveillance Team |
| EPI_ISL_3446199, EPI_ISL_3446200, EPI_ISL_3446201 | Lighthouse Lab in Glasgow | Wellcome Sanger Institute for the COVID-19 Genomics UK (COG-UK) Consortium | Harper VanSteenhouse, Yumi Kasai, David Gray, Carol Clugston, Anna Dominiczak and Alex Alderton, Roberto Amato, Jeffrey Barrett, Sonia Goncalves, Ewan Harrison, David K. Jackson, Ian Johnston, Dominic Kwiatkowski, Cordelia Langford, John Sillitoe on behalf of the Wellcome Sanger Institute COVID-19 Surveillance Team |
| EPI_ISL_3446202, EPI_ISL_3446203 | Lighthouse Lab in Alderley Park | Wellcome Sanger Institute for the COVID-19 Genomics UK (COG-UK) Consortium | Jacquelyn Wynn, Mairead Hyland, The Lighthouse Lab in Alderley Park and Alex Alderton, Roberto Amato, Jeffrey Barrett, Sonia Goncalves, Ewan Harrison, David K. Jackson, Ian Johnston, Dominic Kwiatkowski, Cordelia Langford, John Sillitoe on behalf of the Wellcome Sanger Institute COVID-19 Surveillance Team |
| EPI_ISL_3446204, EPI_ISL_3446205 | Lighthouse Lab in Milton Keynes | Wellcome Sanger Institute for the COVID-19 Genomics UK (COG-UK) Consortium | The Lighthouse Lab in Milton Keynes and Alex Alderton, Roberto Amato, Jeffrey Barrett, Sonia Goncalves, Ewan Harrison, David K. Jackson, Ian Johnston, Dominic Kwiatkowski, Cordelia Langford, John Sillitoe on behalf of the Wellcome Sanger Institute COVID-19 Surveillance Team |
| EPI_ISL_3446206, EPI_ISL_3446207, EPI_ISL_3446208 | Lighthouse Lab in Glasgow | Wellcome Sanger Institute for the COVID-19 Genomics UK (COG-UK) Consortium | Harper VanSteenhouse, Yumi Kasai, David Gray, Carol Clugston, Anna Dominiczak and Alex Alderton, Roberto Amato, Jeffrey Barrett, Sonia Goncalves, Ewan Harrison, David K. Jackson, Ian Johnston, Dominic Kwiatkowski, Cordelia Langford, John Sillitoe on behalf of the Wellcome Sanger Institute COVID-19 Surveillance Team |
| EPI_ISL_3446209 | Lighthouse Lab in Milton Keynes | Wellcome Sanger Institute for the COVID-19 Genomics UK (COG-UK) Consortium | The Lighthouse Lab in Milton Keynes and Alex Alderton, Roberto Amato, Jeffrey Barrett, Sonia Goncalves, Ewan Harrison, David K. Jackson, Ian Johnston, Dominic Kwiatkowski, Cordelia Langford, John Sillitoe on behalf of the Wellcome Sanger Institute COVID-19 Surveillance Team |
| EPI_ISL_3446210 | Lighthouse Lab in Glasgow | Wellcome Sanger Institute for the COVID-19 Genomics UK (COG-UK) Consortium | Harper VanSteenhouse, Yumi Kasai, David Gray, Carol Clugston, Anna Dominiczak and Alex Alderton, Roberto Amato, Jeffrey Barrett, Sonia Goncalves, Ewan Harrison, David K. Jackson, Ian Johnston, Dominic Kwiatkowski, Cordelia Langford, John Sillitoe on behalf of the Wellcome Sanger Institute COVID-19 Surveillance Team |
| EPI_ISL_3446211 | Lighthouse Lab in Milton Keynes | Wellcome Sanger Institute for the COVID-19 Genomics UK (COG-UK) | The Lighthouse Lab in Milton Keynes and Alex Alderton, Roberto Amato, Jeffrey Barrett, Sonia Goncalves, Ewan Harrison, David K. Jackson, Ian Johnston, Dominic Kwiatkowski, Cordelia Langford, John Sillitoe on behalf of the Wellcome Sanger |

| | | | |
|---|---|---|---|
| | | Consortium | Institute COVID-19 Surveillance Team |
| EPI_ISL_3446212, EPI_ISL_3446213 | Lighthouse Lab in Glasgow | Wellcome Sanger Institute for the COVID-19 Genomics UK (COG-UK) Consortium | Harper VanSteenhouse, Yumi Kasai, David Gray, Carol Clugston, Anna Dominiczak and Alex Alderton, Roberto Amato, Jeffrey Barrett, Sonia Goncalves, Ewan Harrison, David K. Jackson, Ian Johnston, Dominic Kwiatkowski, Cordelia Langford, John Sillitoe on behalf of the Wellcome Sanger Institute COVID-19 Surveillance Team |
| EPI_ISL_3446214, EPI_ISL_3446215, EPI_ISL_3446216 | Lighthouse Lab in Milton Keynes | Wellcome Sanger Institute for the COVID-19 Genomics UK (COG-UK) Consortium | The Lighthouse Lab in Milton Keynes and Alex Alderton, Roberto Amato, Jeffrey Barrett, Sonia Goncalves, Ewan Harrison, David K. Jackson, Ian Johnston, Dominic Kwiatkowski, Cordelia Langford, John Sillitoe on behalf of the Wellcome Sanger Institute COVID-19 Surveillance Team |
| EPI_ISL_3446217, EPI_ISL_3446218, EPI_ISL_3446219 | Lighthouse Lab in Alderley Park | Wellcome Sanger Institute for the COVID-19 Genomics UK (COG-UK) Consortium | Jacquelyn Wynn, Mairead Hyland, The Lighthouse Lab in Alderley Park and Alex Alderton, Roberto Amato, Jeffrey Barrett, Sonia Goncalves, Ewan Harrison, David K. Jackson, Ian Johnston, Dominic Kwiatkowski, Cordelia Langford, John Sillitoe on behalf of the Wellcome Sanger Institute COVID-19 Surveillance Team |
| EPI_ISL_3446220, EPI_ISL_3446221 | Lighthouse Lab in Glasgow | Wellcome Sanger Institute for the COVID-19 Genomics UK (COG-UK) Consortium | Harper VanSteenhouse, Yumi Kasai, David Gray, Carol Clugston, Anna Dominiczak and Alex Alderton, Roberto Amato, Jeffrey Barrett, Sonia Goncalves, Ewan Harrison, David K. Jackson, Ian Johnston, Dominic Kwiatkowski, Cordelia Langford, John Sillitoe on behalf of the Wellcome Sanger Institute COVID-19 Surveillance Team |
| EPI_ISL_3446222 | Lighthouse Lab in Alderley Park | Wellcome Sanger Institute for the COVID-19 Genomics UK (COG-UK) Consortium | Jacquelyn Wynn, Mairead Hyland, The Lighthouse Lab in Alderley Park and Alex Alderton, Roberto Amato, Jeffrey Barrett, Sonia Goncalves, Ewan Harrison, David K. Jackson, Ian Johnston, Dominic Kwiatkowski, Cordelia Langford, John Sillitoe on behalf of the Wellcome Sanger Institute COVID-19 Surveillance Team |
| EPI_ISL_3446223 | Lighthouse Lab in Milton Keynes | Wellcome Sanger Institute for the COVID-19 Genomics UK (COG-UK) Consortium | The Lighthouse Lab in Milton Keynes and Alex Alderton, Roberto Amato, Jeffrey Barrett, Sonia Goncalves, Ewan Harrison, David K. Jackson, Ian Johnston, Dominic Kwiatkowski, Cordelia Langford, John Sillitoe on behalf of the Wellcome Sanger Institute COVID-19 Surveillance Team |
| EPI_ISL_3446224, EPI_ISL_3446225, EPI_ISL_3446226 | Lighthouse Lab in Glasgow | Wellcome Sanger Institute for the COVID-19 Genomics UK (COG-UK) Consortium | Harper VanSteenhouse, Yumi Kasai, David Gray, Carol Clugston, Anna Dominiczak and Alex Alderton, Roberto Amato, Jeffrey Barrett, Sonia Goncalves, Ewan Harrison, David K. Jackson, Ian Johnston, Dominic Kwiatkowski, Cordelia Langford, John Sillitoe on behalf of the Wellcome Sanger Institute COVID-19 Surveillance Team |
| EPI_ISL_3446227 | Lighthouse Lab in Milton Keynes | Wellcome Sanger Institute for the COVID-19 Genomics UK (COG-UK) Consortium | The Lighthouse Lab in Milton Keynes and Alex Alderton, Roberto Amato, Jeffrey Barrett, Sonia Goncalves, Ewan Harrison, David K. Jackson, Ian Johnston, Dominic Kwiatkowski, Cordelia Langford, John Sillitoe on behalf of the Wellcome Sanger Institute COVID-19 Surveillance Team |
| EPI_ISL_3446228, EPI_ISL_3446229, EPI_ISL_3446230, EPI_ISL_3446231 | Lighthouse Lab in Glasgow | Wellcome Sanger Institute for the COVID-19 Genomics UK (COG-UK) Consortium | Harper VanSteenhouse, Yumi Kasai, David Gray, Carol Clugston, Anna Dominiczak and Alex Alderton, Roberto Amato, Jeffrey Barrett, Sonia Goncalves, Ewan Harrison, David K. Jackson, Ian Johnston, Dominic Kwiatkowski, Cordelia Langford, John Sillitoe on behalf of the Wellcome Sanger Institute COVID-19 Surveillance Team |
| EPI_ISL_3446232, EPI_ISL_3446233 | Lighthouse Lab in Alderley Park | Wellcome Sanger Institute for the COVID-19 Genomics UK (COG-UK) Consortium | Jacquelyn Wynn, Mairead Hyland, The Lighthouse Lab in Alderley Park and Alex Alderton, Roberto Amato, Jeffrey Barrett, Sonia Goncalves, Ewan Harrison, David K. Jackson, Ian Johnston, Dominic Kwiatkowski, Cordelia Langford, John Sillitoe on behalf of the Wellcome Sanger Institute COVID-19 Surveillance Team |
| EPI_ISL_3446234, EPI_ISL_3446235 | Lighthouse Lab in Glasgow | Wellcome Sanger Institute for the COVID-19 Genomics UK (COG-UK) Consortium | Harper VanSteenhouse, Yumi Kasai, David Gray, Carol Clugston, Anna Dominiczak and Alex Alderton, Roberto Amato, Jeffrey Barrett, Sonia Goncalves, Ewan Harrison, David K. Jackson, Ian Johnston, Dominic Kwiatkowski, Cordelia Langford, John Sillitoe on behalf of the Wellcome Sanger Institute COVID-19 Surveillance Team |
| EPI_ISL_3446236 | Lighthouse Lab in Milton Keynes | Wellcome Sanger Institute for the COVID-19 Genomics UK (COG-UK) Consortium | The Lighthouse Lab in Milton Keynes and Alex Alderton, Roberto Amato, Jeffrey Barrett, Sonia Goncalves, Ewan Harrison, David K. Jackson, Ian Johnston, Dominic Kwiatkowski, Cordelia Langford, John Sillitoe on behalf of the Wellcome Sanger Institute COVID-19 Surveillance Team |
| EPI_ISL_3446237 | Lighthouse Lab in Alderley Park | Wellcome Sanger Institute for the COVID-19 Genomics UK (COG-UK) Consortium | Jacquelyn Wynn, Mairead Hyland, The Lighthouse Lab in Alderley Park and Alex Alderton, Roberto Amato, Jeffrey Barrett, Sonia Goncalves, Ewan Harrison, David K. Jackson, Ian Johnston, Dominic Kwiatkowski, Cordelia Langford, John Sillitoe on behalf of the Wellcome Sanger Institute COVID-19 Surveillance Team |
| EPI_ISL_3446238 | Lighthouse Lab in Glasgow | Wellcome Sanger Institute for the COVID-19 Genomics UK (COG-UK) Consortium | Harper VanSteenhouse, Yumi Kasai, David Gray, Carol Clugston, Anna Dominiczak and Alex Alderton, Roberto Amato, Jeffrey Barrett, Sonia Goncalves, Ewan Harrison, David K. Jackson, Ian Johnston, Dominic Kwiatkowski, Cordelia Langford, John Sillitoe on behalf of the Wellcome Sanger Institute COVID-19 Surveillance Team |
| EPI_ISL_3446239 | Lighthouse Lab in Milton Keynes | Wellcome Sanger Institute for the COVID-19 Genomics UK (COG-UK) Consortium | The Lighthouse Lab in Milton Keynes and Alex Alderton, Roberto Amato, Jeffrey Barrett, Sonia Goncalves, Ewan Harrison, David K. Jackson, Ian Johnston, Dominic Kwiatkowski, Cordelia Langford, John Sillitoe on behalf of the Wellcome Sanger Institute COVID-19 Surveillance Team |
| EPI_ISL_3446240, EPI_ISL_3446241 | Lighthouse Lab in Alderley Park | Wellcome Sanger Institute for the COVID-19 Genomics UK (COG-UK) Consortium | Jacquelyn Wynn, Mairead Hyland, The Lighthouse Lab in Alderley Park and Alex Alderton, Roberto Amato, Jeffrey Barrett, Sonia Goncalves, Ewan Harrison, David K. Jackson, Ian Johnston, Dominic Kwiatkowski, Cordelia Langford, John Sillitoe on behalf of the Wellcome Sanger Institute COVID-19 Surveillance Team |
| EPI_ISL_3446242 | Lighthouse Lab in Milton Keynes | Wellcome Sanger Institute for the COVID-19 Genomics UK (COG-UK) Consortium | The Lighthouse Lab in Milton Keynes and Alex Alderton, Roberto Amato, Jeffrey Barrett, Sonia Goncalves, Ewan Harrison, David K. Jackson, Ian Johnston, Dominic Kwiatkowski, Cordelia Langford, John Sillitoe on behalf of the Wellcome Sanger Institute COVID-19 Surveillance Team |
| EPI_ISL_3446243 | Lighthouse Lab in Alderley Park | Wellcome Sanger Institute for the COVID-19 Genomics UK (COG-UK) Consortium | Jacquelyn Wynn, Mairead Hyland, The Lighthouse Lab in Alderley Park and Alex Alderton, Roberto Amato, Jeffrey Barrett, Sonia Goncalves, Ewan Harrison, David K. Jackson, Ian Johnston, Dominic Kwiatkowski, Cordelia Langford, John Sillitoe on behalf of the Wellcome Sanger Institute COVID-19 Surveillance Team |
| EPI_ISL_3446244, EPI_ISL_3446245, EPI_ISL_3446246, EPI_ISL_3446247, EPI_ISL_3446248, EPI_ISL_3446249 | Lighthouse Lab in Glasgow | Wellcome Sanger Institute for the COVID-19 Genomics UK (COG-UK) Consortium | Harper VanSteenhouse, Yumi Kasai, David Gray, Carol Clugston, Anna Dominiczak and Alex Alderton, Roberto Amato, Jeffrey Barrett, Sonia Goncalves, Ewan Harrison, David K. Jackson, Ian Johnston, Dominic Kwiatkowski, Cordelia Langford, John Sillitoe on behalf of the Wellcome Sanger Institute COVID-19 Surveillance Team |
| EPI_ISL_3446250 | Lighthouse Lab in Milton Keynes | Wellcome Sanger Institute for the COVID-19 Genomics UK (COG-UK) Consortium | The Lighthouse Lab in Milton Keynes and Alex Alderton, Roberto Amato, Jeffrey Barrett, Sonia Goncalves, Ewan Harrison, David K. Jackson, Ian Johnston, Dominic Kwiatkowski, Cordelia Langford, John Sillitoe on behalf of the Wellcome Sanger Institute COVID-19 Surveillance Team |
| EPI_ISL_3446251 | Lighthouse Lab in Glasgow | Wellcome Sanger Institute for the COVID-19 Genomics UK (COG-UK) Consortium | Harper VanSteenhouse, Yumi Kasai, David Gray, Carol Clugston, Anna Dominiczak and Alex Alderton, Roberto Amato, Jeffrey Barrett, Sonia Goncalves, Ewan Harrison, David K. Jackson, Ian Johnston, Dominic Kwiatkowski, Cordelia Langford, John Sillitoe on behalf of the Wellcome Sanger Institute COVID-19 Surveillance Team |
| EPI_ISL_3446252, EPI_ISL_3446253 | Lighthouse Lab in Alderley Park | Wellcome Sanger Institute for the COVID-19 Genomics UK (COG-UK) Consortium | Jacquelyn Wynn, Mairead Hyland, The Lighthouse Lab in Alderley Park and Alex Alderton, Roberto Amato, Jeffrey Barrett, Sonia Goncalves, Ewan Harrison, David K. Jackson, Ian Johnston, Dominic Kwiatkowski, Cordelia Langford, John Sillitoe on behalf of the Wellcome Sanger Institute COVID-19 Surveillance Team |
| EPI_ISL_3446254 | Lighthouse Lab in Milton Keynes | Wellcome Sanger Institute for the COVID-19 Genomics UK (COG-UK) Consortium | The Lighthouse Lab in Milton Keynes and Alex Alderton, Roberto Amato, Jeffrey Barrett, Sonia Goncalves, Ewan Harrison, David K. Jackson, Ian Johnston, Dominic Kwiatkowski, Cordelia Langford, John Sillitoe on behalf of the Wellcome Sanger Institute COVID-19 Surveillance Team |
| EPI_ISL_3446255 | Lighthouse Lab in Glasgow | Wellcome Sanger Institute for the COVID-19 Genomics UK (COG-UK) Consortium | Harper VanSteenhouse, Yumi Kasai, David Gray, Carol Clugston, Anna Dominiczak and Alex Alderton, Roberto Amato, Jeffrey Barrett, Sonia Goncalves, Ewan Harrison, David K. Jackson, Ian Johnston, Dominic Kwiatkowski, Cordelia Langford, John Sillitoe on behalf of the Wellcome Sanger Institute COVID-19 Surveillance Team |
| EPI_ISL_3446256 | Lighthouse Lab in Alderley Park | Wellcome Sanger Institute for the COVID-19 Genomics UK (COG-UK) Consortium | Jacquelyn Wynn, Mairead Hyland, The Lighthouse Lab in Alderley Park and Alex Alderton, Roberto Amato, Jeffrey Barrett, Sonia Goncalves, Ewan Harrison, David K. Jackson, Ian Johnston, Dominic Kwiatkowski, Cordelia Langford, John Sillitoe on behalf of the Wellcome Sanger Institute COVID-19 Surveillance Team |
| EPI_ISL_3446257, EPI_ISL_3446258 | Lighthouse Lab in Glasgow | Wellcome Sanger Institute for the COVID-19 Genomics UK (COG-UK) Consortium | Harper VanSteenhouse, Yumi Kasai, David Gray, Carol Clugston, Anna Dominiczak and Alex Alderton, Roberto Amato, Jeffrey Barrett, Sonia Goncalves, Ewan Harrison, David K. Jackson, Ian Johnston, Dominic Kwiatkowski, Cordelia Langford, John Sillitoe on behalf of the Wellcome Sanger Institute COVID-19 Surveillance Team |
| EPI_ISL_3446259 | Lighthouse Lab in Milton Keynes | Wellcome Sanger Institute for the COVID-19 Genomics UK (COG-UK) Consortium | The Lighthouse Lab in Milton Keynes and Alex Alderton, Roberto Amato, Jeffrey Barrett, Sonia Goncalves, Ewan Harrison, David K. Jackson, Ian Johnston, Dominic Kwiatkowski, Cordelia Langford, John Sillitoe on behalf of the Wellcome Sanger Institute COVID-19 Surveillance Team |
| EPI_ISL_3446260, EPI_ISL_3446261 | Lighthouse Lab in Alderley Park | Wellcome Sanger Institute for the COVID-19 Genomics UK (COG-UK) Consortium | Jacquelyn Wynn, Mairead Hyland, The Lighthouse Lab in Alderley Park and Alex Alderton, Roberto Amato, Jeffrey Barrett, Sonia Goncalves, Ewan Harrison, David K. Jackson, Ian Johnston, Dominic Kwiatkowski, Cordelia Langford, John Sillitoe on behalf of the Wellcome Sanger Institute COVID-19 Surveillance Team |
| EPI_ISL_3446262, EPI_ISL_3446263, EPI_ISL_3446264 | Lighthouse Lab in Glasgow | Wellcome Sanger Institute for the COVID-19 Genomics UK (COG-UK) Consortium | Harper VanSteenhouse, Yumi Kasai, David Gray, Carol Clugston, Anna Dominiczak and Alex Alderton, Roberto Amato, Jeffrey Barrett, Sonia Goncalves, Ewan Harrison, David K. Jackson, Ian Johnston, Dominic Kwiatkowski, Cordelia Langford, John Sillitoe on behalf of the Wellcome Sanger Institute COVID-19 Surveillance Team |
| EPI_ISL_3446265 | Lighthouse Lab in Milton Keynes | Wellcome Sanger Institute for the COVID-19 Genomics UK (COG-UK) Consortium | The Lighthouse Lab in Milton Keynes and Alex Alderton, Roberto Amato, Jeffrey Barrett, Sonia Goncalves, Ewan Harrison, David K. Jackson, Ian Johnston, Dominic Kwiatkowski, Cordelia Langford, John Sillitoe on behalf of the Wellcome Sanger Institute COVID-19 Surveillance Team |
| EPI_ISL_3446266, EPI_ISL_3446267, EPI_ISL_3446268 | Lighthouse Lab in Glasgow | Wellcome Sanger Institute for the COVID-19 Genomics UK (COG-UK) Consortium | Harper VanSteenhouse, Yumi Kasai, David Gray, Carol Clugston, Anna Dominiczak and Alex Alderton, Roberto Amato, Jeffrey Barrett, Sonia Goncalves, Ewan Harrison, David K. Jackson, Ian Johnston, Dominic Kwiatkowski, Cordelia Langford, John Sillitoe on behalf of the Wellcome Sanger Institute COVID-19 Surveillance Team |
| EPI_ISL_3446269 | Lighthouse Lab in Milton Keynes | Wellcome Sanger Institute for the COVID-19 Genomics UK (COG-UK) Consortium | The Lighthouse Lab in Milton Keynes and Alex Alderton, Roberto Amato, Jeffrey Barrett, Sonia Goncalves, Ewan Harrison, David K. Jackson, Ian Johnston, Dominic Kwiatkowski, Cordelia Langford, John Sillitoe on behalf of the Wellcome Sanger Institute COVID-19 Surveillance Team |
| EPI_ISL_3446270, EPI_ISL_3446271 | Lighthouse Lab in Glasgow | Wellcome Sanger Institute for the COVID-19 Genomics UK (COG-UK) Consortium | Harper VanSteenhouse, Yumi Kasai, David Gray, Carol Clugston, Anna Dominiczak and Alex Alderton, Roberto Amato, Jeffrey Barrett, Sonia Goncalves, Ewan Harrison, David K. Jackson, Ian Johnston, Dominic Kwiatkowski, Cordelia Langford, John Sillitoe on behalf of the Wellcome Sanger Institute COVID-19 Surveillance Team |
| EPI_ISL_3446272 | Lighthouse Lab in Alderley Park | Wellcome Sanger Institute for the COVID-19 Genomics UK (COG-UK) Consortium | Jacquelyn Wynn, Mairead Hyland, The Lighthouse Lab in Alderley Park and Alex Alderton, Roberto Amato, Jeffrey Barrett, Sonia Goncalves, Ewan Harrison, David K. Jackson, Ian Johnston, Dominic Kwiatkowski, Cordelia Langford, John Sillitoe on behalf of the Wellcome Sanger Institute COVID-19 Surveillance Team |
| EPI_ISL_3446273 | Lighthouse Lab in Glasgow | Wellcome Sanger Institute for the COVID-19 Genomics UK (COG-UK) Consortium | Harper VanSteenhouse, Yumi Kasai, David Gray, Carol Clugston, Anna Dominiczak and Alex Alderton, Roberto Amato, Jeffrey Barrett, Sonia Goncalves, Ewan Harrison, David K. Jackson, Ian Johnston, Dominic Kwiatkowski, Cordelia Langford, John Sillitoe on behalf of the Wellcome Sanger Institute COVID-19 Surveillance Team |
| EPI_ISL_3446274 | Lighthouse Lab in Alderley Park | Wellcome Sanger Institute for the COVID-19 Genomics UK (COG-UK) Consortium | Jacquelyn Wynn, Mairead Hyland, The Lighthouse Lab in Alderley Park and Alex Alderton, Roberto Amato, Jeffrey Barrett, Sonia Goncalves, Ewan Harrison, David K. Jackson, Ian Johnston, Dominic Kwiatkowski, Cordelia Langford, John Sillitoe on behalf of the Wellcome Sanger Institute COVID-19 Surveillance Team |
| EPI_ISL_3446275, EPI_ISL_3446276, EPI_ISL_3446277, EPI_ISL_3446278, EPI_ISL_3446279 | Lighthouse Lab in Glasgow | Wellcome Sanger Institute for the COVID-19 Genomics UK (COG-UK) Consortium | Harper VanSteenhouse, Yumi Kasai, David Gray, Carol Clugston, Anna Dominiczak and Alex Alderton, Roberto Amato, Jeffrey Barrett, Sonia Goncalves, Ewan Harrison, David K. Jackson, Ian Johnston, Dominic Kwiatkowski, Cordelia Langford, John Sillitoe on behalf of the Wellcome Sanger Institute COVID-19 Surveillance Team |
| EPI_ISL_3446280 | Lighthouse Lab in Alderley Park | Wellcome Sanger Institute for the COVID-19 Genomics UK (COG-UK) Consortium | Jacquelyn Wynn, Mairead Hyland, The Lighthouse Lab in Alderley Park and Alex Alderton, Roberto Amato, Jeffrey Barrett, Sonia Goncalves, Ewan Harrison, David K. Jackson, Ian Johnston, Dominic Kwiatkowski, Cordelia Langford, John Sillitoe on behalf of the Wellcome Sanger Institute COVID-19 Surveillance Team |
| EPI_ISL_3446281 | Lighthouse Lab in Milton Keynes | Wellcome Sanger Institute for the COVID-19 Genomics UK (COG-UK) | The Lighthouse Lab in Milton Keynes and Alex Alderton, Roberto Amato, Jeffrey Barrett, Sonia Goncalves, Ewan Harrison, David K. Jackson, Ian Johnston, Dominic Kwiatkowski, Cordelia Langford, John Sillitoe on behalf of the Wellcome Sanger |

| | | | |
|---|---|---|---|
| | | Consortium | Institute COVID-19 Surveillance Team |
| EPI_ISL_3446282 | Lighthouse Lab in Glasgow | Wellcome Sanger Institute for the COVID-19 Genomics UK (COG-UK) Consortium | Harper VanSteenhouse, Yumi Kasai, David Gray, Carol Clugston, Anna Dominiczak and Alex Alderton, Roberto Amato, Jeffrey Barrett, Sonia Goncalves, Ewan Harrison, David K. Jackson, Ian Johnston, Dominic Kwiatkowski, Cordelia Langford, John Sillitoe on behalf of the Wellcome Sanger Institute COVID-19 Surveillance Team |
| EPI_ISL_3446283, EPI_ISL_3446284 | Lighthouse Lab in Alderley Park | Wellcome Sanger Institute for the COVID-19 Genomics UK (COG-UK) Consortium | Jacquelyn Wynn, Mairead Hyland, The Lighthouse Lab in Alderley Park and Alex Alderton, Roberto Amato, Jeffrey Barrett, Sonia Goncalves, Ewan Harrison, David K. Jackson, Ian Johnston, Dominic Kwiatkowski, Cordelia Langford, John Sillitoe on behalf of the Wellcome Sanger Institute COVID-19 Surveillance Team |
| EPI_ISL_3446285 | Lighthouse Lab in Glasgow | Wellcome Sanger Institute for the COVID-19 Genomics UK (COG-UK) Consortium | Harper VanSteenhouse, Yumi Kasai, David Gray, Carol Clugston, Anna Dominiczak and Alex Alderton, Roberto Amato, Jeffrey Barrett, Sonia Goncalves, Ewan Harrison, David K. Jackson, Ian Johnston, Dominic Kwiatkowski, Cordelia Langford, John Sillitoe on behalf of the Wellcome Sanger Institute COVID-19 Surveillance Team |
| EPI_ISL_3446286 | Lighthouse Lab in Milton Keynes | Wellcome Sanger Institute for the COVID-19 Genomics UK (COG-UK) Consortium | The Lighthouse Lab in Milton Keynes and Alex Alderton, Roberto Amato, Jeffrey Barrett, Sonia Goncalves, Ewan Harrison, David K. Jackson, Ian Johnston, Dominic Kwiatkowski, Cordelia Langford, John Sillitoe on behalf of the Wellcome Sanger Institute COVID-19 Surveillance Team |
| EPI_ISL_3446287 | Lighthouse Lab in Glasgow | Wellcome Sanger Institute for the COVID-19 Genomics UK (COG-UK) Consortium | Harper VanSteenhouse, Yumi Kasai, David Gray, Carol Clugston, Anna Dominiczak and Alex Alderton, Roberto Amato, Jeffrey Barrett, Sonia Goncalves, Ewan Harrison, David K. Jackson, Ian Johnston, Dominic Kwiatkowski, Cordelia Langford, John Sillitoe on behalf of the Wellcome Sanger Institute COVID-19 Surveillance Team |
| EPI_ISL_3446288, EPI_ISL_3446289 | Lighthouse Lab in Milton Keynes | Wellcome Sanger Institute for the COVID-19 Genomics UK (COG-UK) Consortium | The Lighthouse Lab in Milton Keynes and Alex Alderton, Roberto Amato, Jeffrey Barrett, Sonia Goncalves, Ewan Harrison, David K. Jackson, Ian Johnston, Dominic Kwiatkowski, Cordelia Langford, John Sillitoe on behalf of the Wellcome Sanger Institute COVID-19 Surveillance Team |
| EPI_ISL_3446290 | Lighthouse Lab in Glasgow | Wellcome Sanger Institute for the COVID-19 Genomics UK (COG-UK) Consortium | Harper VanSteenhouse, Yumi Kasai, David Gray, Carol Clugston, Anna Dominiczak and Alex Alderton, Roberto Amato, Jeffrey Barrett, Sonia Goncalves, Ewan Harrison, David K. Jackson, Ian Johnston, Dominic Kwiatkowski, Cordelia Langford, John Sillitoe on behalf of the Wellcome Sanger Institute COVID-19 Surveillance Team |
| EPI_ISL_3446291 | Lighthouse Lab in Alderley Park | Wellcome Sanger Institute for the COVID-19 Genomics UK (COG-UK) Consortium | Jacquelyn Wynn, Mairead Hyland, The Lighthouse Lab in Alderley Park and Alex Alderton, Roberto Amato, Jeffrey Barrett, Sonia Goncalves, Ewan Harrison, David K. Jackson, Ian Johnston, Dominic Kwiatkowski, Cordelia Langford, John Sillitoe on behalf of the Wellcome Sanger Institute COVID-19 Surveillance Team |
| EPI_ISL_3446292 | Lighthouse Lab in Glasgow | Wellcome Sanger Institute for the COVID-19 Genomics UK (COG-UK) Consortium | Harper VanSteenhouse, Yumi Kasai, David Gray, Carol Clugston, Anna Dominiczak and Alex Alderton, Roberto Amato, Jeffrey Barrett, Sonia Goncalves, Ewan Harrison, David K. Jackson, Ian Johnston, Dominic Kwiatkowski, Cordelia Langford, John Sillitoe on behalf of the Wellcome Sanger Institute COVID-19 Surveillance Team |
| EPI_ISL_3446293 | Lighthouse Lab in Alderley Park | Wellcome Sanger Institute for the COVID-19 Genomics UK (COG-UK) Consortium | Jacquelyn Wynn, Mairead Hyland, The Lighthouse Lab in Alderley Park and Alex Alderton, Roberto Amato, Jeffrey Barrett, Sonia Goncalves, Ewan Harrison, David K. Jackson, Ian Johnston, Dominic Kwiatkowski, Cordelia Langford, John Sillitoe on behalf of the Wellcome Sanger Institute COVID-19 Surveillance Team |
| EPI_ISL_3446294 | Lighthouse Lab in Glasgow | Wellcome Sanger Institute for the COVID-19 Genomics UK (COG-UK) Consortium | Harper VanSteenhouse, Yumi Kasai, David Gray, Carol Clugston, Anna Dominiczak and Alex Alderton, Roberto Amato, Jeffrey Barrett, Sonia Goncalves, Ewan Harrison, David K. Jackson, Ian Johnston, Dominic Kwiatkowski, Cordelia Langford, John Sillitoe on behalf of the Wellcome Sanger Institute COVID-19 Surveillance Team |
| EPI_ISL_3446295, EPI_ISL_3446296, EPI_ISL_3446297, EPI_ISL_3446298 | Lighthouse Lab in Milton Keynes | Wellcome Sanger Institute for the COVID-19 Genomics UK (COG-UK) Consortium | The Lighthouse Lab in Milton Keynes and Alex Alderton, Roberto Amato, Jeffrey Barrett, Sonia Goncalves, Ewan Harrison, David K. Jackson, Ian Johnston, Dominic Kwiatkowski, Cordelia Langford, John Sillitoe on behalf of the Wellcome Sanger Institute COVID-19 Surveillance Team |
| EPI_ISL_3446299 | Lighthouse Lab in Glasgow | Wellcome Sanger Institute for the COVID-19 Genomics UK (COG-UK) Consortium | Harper VanSteenhouse, Yumi Kasai, David Gray, Carol Clugston, Anna Dominiczak and Alex Alderton, Roberto Amato, Jeffrey Barrett, Sonia Goncalves, Ewan Harrison, David K. Jackson, Ian Johnston, Dominic Kwiatkowski, Cordelia Langford, John Sillitoe on behalf of the Wellcome Sanger Institute COVID-19 Surveillance Team |
| EPI_ISL_3446300 | Lighthouse Lab in Alderley Park | Wellcome Sanger Institute for the COVID-19 Genomics UK (COG-UK) Consortium | Jacquelyn Wynn, Mairead Hyland, The Lighthouse Lab in Alderley Park and Alex Alderton, Roberto Amato, Jeffrey Barrett, Sonia Goncalves, Ewan Harrison, David K. Jackson, Ian Johnston, Dominic Kwiatkowski, Cordelia Langford, John Sillitoe on behalf of the Wellcome Sanger Institute COVID-19 Surveillance Team |
| EPI_ISL_3446301 | Lighthouse Lab in Milton Keynes | Wellcome Sanger Institute for the COVID-19 Genomics UK (COG-UK) Consortium | The Lighthouse Lab in Milton Keynes and Alex Alderton, Roberto Amato, Jeffrey Barrett, Sonia Goncalves, Ewan Harrison, David K. Jackson, Ian Johnston, Dominic Kwiatkowski, Cordelia Langford, John Sillitoe on behalf of the Wellcome Sanger Institute COVID-19 Surveillance Team |
| EPI_ISL_3446302, EPI_ISL_3446303, EPI_ISL_3446304 | Lighthouse Lab in Glasgow | Wellcome Sanger Institute for the COVID-19 Genomics UK (COG-UK) Consortium | Harper VanSteenhouse, Yumi Kasai, David Gray, Carol Clugston, Anna Dominiczak and Alex Alderton, Roberto Amato, Jeffrey Barrett, Sonia Goncalves, Ewan Harrison, David K. Jackson, Ian Johnston, Dominic Kwiatkowski, Cordelia Langford, John Sillitoe on behalf of the Wellcome Sanger Institute COVID-19 Surveillance Team |
| EPI_ISL_3446305 | Lighthouse Lab in Milton Keynes | Wellcome Sanger Institute for the COVID-19 Genomics UK (COG-UK) Consortium | The Lighthouse Lab in Milton Keynes and Alex Alderton, Roberto Amato, Jeffrey Barrett, Sonia Goncalves, Ewan Harrison, David K. Jackson, Ian Johnston, Dominic Kwiatkowski, Cordelia Langford, John Sillitoe on behalf of the Wellcome Sanger Institute COVID-19 Surveillance Team |
| EPI_ISL_3446306 | Lighthouse Lab in Alderley Park | Wellcome Sanger Institute for the COVID-19 Genomics UK (COG-UK) Consortium | Jacquelyn Wynn, Mairead Hyland, The Lighthouse Lab in Alderley Park and Alex Alderton, Roberto Amato, Jeffrey Barrett, Sonia Goncalves, Ewan Harrison, David K. Jackson, Ian Johnston, Dominic Kwiatkowski, Cordelia Langford, John Sillitoe on behalf of the Wellcome Sanger Institute COVID-19 Surveillance Team |
| EPI_ISL_3446307, EPI_ISL_3446308, EPI_ISL_3446309, EPI_ISL_3446310, EPI_ISL_3446311 | Lighthouse Lab in Glasgow | Wellcome Sanger Institute for the COVID-19 Genomics UK (COG-UK) Consortium | Harper VanSteenhouse, Yumi Kasai, David Gray, Carol Clugston, Anna Dominiczak and Alex Alderton, Roberto Amato, Jeffrey Barrett, Sonia Goncalves, Ewan Harrison, David K. Jackson, Ian Johnston, Dominic Kwiatkowski, Cordelia Langford, John Sillitoe on behalf of the Wellcome Sanger Institute COVID-19 Surveillance Team |
| EPI_ISL_3446312 | Lighthouse Lab in Alderley Park | Wellcome Sanger Institute for the COVID-19 Genomics UK (COG-UK) Consortium | Jacquelyn Wynn, Mairead Hyland, The Lighthouse Lab in Alderley Park and Alex Alderton, Roberto Amato, Jeffrey Barrett, Sonia Goncalves, Ewan Harrison, David K. Jackson, Ian Johnston, Dominic Kwiatkowski, Cordelia Langford, John Sillitoe on behalf of the Wellcome Sanger Institute COVID-19 Surveillance Team |
| EPI_ISL_3446313 | Lighthouse Lab in Milton Keynes | Wellcome Sanger Institute for the COVID-19 Genomics UK (COG-UK) Consortium | The Lighthouse Lab in Milton Keynes and Alex Alderton, Roberto Amato, Jeffrey Barrett, Sonia Goncalves, Ewan Harrison, David K. Jackson, Ian Johnston, Dominic Kwiatkowski, Cordelia Langford, John Sillitoe on behalf of the Wellcome Sanger Institute COVID-19 Surveillance Team |
| EPI_ISL_3446314 | Lighthouse Lab in Alderley Park | Wellcome Sanger Institute for the COVID-19 Genomics UK (COG-UK) Consortium | Jacquelyn Wynn, Mairead Hyland, The Lighthouse Lab in Alderley Park and Alex Alderton, Roberto Amato, Jeffrey Barrett, Sonia Goncalves, Ewan Harrison, David K. Jackson, Ian Johnston, Dominic Kwiatkowski, Cordelia Langford, John Sillitoe on behalf of the Wellcome Sanger Institute COVID-19 Surveillance Team |
| EPI_ISL_3446315, EPI_ISL_3446316, EPI_ISL_3446317 | Lighthouse Lab in Glasgow | Wellcome Sanger Institute for the COVID-19 Genomics UK (COG-UK) Consortium | Harper VanSteenhouse, Yumi Kasai, David Gray, Carol Clugston, Anna Dominiczak and Alex Alderton, Roberto Amato, Jeffrey Barrett, Sonia Goncalves, Ewan Harrison, David K. Jackson, Ian Johnston, Dominic Kwiatkowski, Cordelia Langford, John Sillitoe on behalf of the Wellcome Sanger Institute COVID-19 Surveillance Team |
| EPI_ISL_3446318 | Lighthouse Lab in Milton Keynes | Wellcome Sanger Institute for the COVID-19 Genomics UK (COG-UK) Consortium | The Lighthouse Lab in Milton Keynes and Alex Alderton, Roberto Amato, Jeffrey Barrett, Sonia Goncalves, Ewan Harrison, David K. Jackson, Ian Johnston, Dominic Kwiatkowski, Cordelia Langford, John Sillitoe on behalf of the Wellcome Sanger Institute COVID-19 Surveillance Team |
| EPI_ISL_3446319 | Lighthouse Lab in Glasgow | Wellcome Sanger Institute for the COVID-19 Genomics UK (COG-UK) Consortium | Harper VanSteenhouse, Yumi Kasai, David Gray, Carol Clugston, Anna Dominiczak and Alex Alderton, Roberto Amato, Jeffrey Barrett, Sonia Goncalves, Ewan Harrison, David K. Jackson, Ian Johnston, Dominic Kwiatkowski, Cordelia Langford, John Sillitoe on behalf of the Wellcome Sanger Institute COVID-19 Surveillance Team |
| EPI_ISL_3446320 | Lighthouse Lab in Alderley Park | Wellcome Sanger Institute for the COVID-19 Genomics UK (COG-UK) Consortium | Jacquelyn Wynn, Mairead Hyland, The Lighthouse Lab in Alderley Park and Alex Alderton, Roberto Amato, Jeffrey Barrett, Sonia Goncalves, Ewan Harrison, David K. Jackson, Ian Johnston, Dominic Kwiatkowski, Cordelia Langford, John Sillitoe on behalf of the Wellcome Sanger Institute COVID-19 Surveillance Team |
| EPI_ISL_3446321 | Lighthouse Lab in Glasgow | Wellcome Sanger Institute for the COVID-19 Genomics UK (COG-UK) Consortium | Harper VanSteenhouse, Yumi Kasai, David Gray, Carol Clugston, Anna Dominiczak and Alex Alderton, Roberto Amato, Jeffrey Barrett, Sonia Goncalves, Ewan Harrison, David K. Jackson, Ian Johnston, Dominic Kwiatkowski, Cordelia Langford, John Sillitoe on behalf of the Wellcome Sanger Institute COVID-19 Surveillance Team |
| EPI_ISL_3446322 | Lighthouse Lab in Alderley Park | Wellcome Sanger Institute for the COVID-19 Genomics UK (COG-UK) Consortium | Jacquelyn Wynn, Mairead Hyland, The Lighthouse Lab in Alderley Park and Alex Alderton, Roberto Amato, Jeffrey Barrett, Sonia Goncalves, Ewan Harrison, David K. Jackson, Ian Johnston, Dominic Kwiatkowski, Cordelia Langford, John Sillitoe on behalf of the Wellcome Sanger Institute COVID-19 Surveillance Team |
| EPI_ISL_3446323 | Lighthouse Lab in Glasgow | Wellcome Sanger Institute for the COVID-19 Genomics UK (COG-UK) Consortium | Harper VanSteenhouse, Yumi Kasai, David Gray, Carol Clugston, Anna Dominiczak and Alex Alderton, Roberto Amato, Jeffrey Barrett, Sonia Goncalves, Ewan Harrison, David K. Jackson, Ian Johnston, Dominic Kwiatkowski, Cordelia Langford, John Sillitoe on behalf of the Wellcome Sanger Institute COVID-19 Surveillance Team |
| EPI_ISL_3446324 | Lighthouse Lab in Alderley Park | Wellcome Sanger Institute for the COVID-19 Genomics UK (COG-UK) Consortium | Jacquelyn Wynn, Mairead Hyland, The Lighthouse Lab in Alderley Park and Alex Alderton, Roberto Amato, Jeffrey Barrett, Sonia Goncalves, Ewan Harrison, David K. Jackson, Ian Johnston, Dominic Kwiatkowski, Cordelia Langford, John Sillitoe on behalf of the Wellcome Sanger Institute COVID-19 Surveillance Team |
| EPI_ISL_3446325, EPI_ISL_3446326, EPI_ISL_3446327 | Lighthouse Lab in Glasgow | Wellcome Sanger Institute for the COVID-19 Genomics UK (COG-UK) Consortium | Harper VanSteenhouse, Yumi Kasai, David Gray, Carol Clugston, Anna Dominiczak and Alex Alderton, Roberto Amato, Jeffrey Barrett, Sonia Goncalves, Ewan Harrison, David K. Jackson, Ian Johnston, Dominic Kwiatkowski, Cordelia Langford, John Sillitoe on behalf of the Wellcome Sanger Institute COVID-19 Surveillance Team |
| EPI_ISL_3446328 | Lighthouse Lab in Milton Keynes | Wellcome Sanger Institute for the COVID-19 Genomics UK (COG-UK) Consortium | The Lighthouse Lab in Milton Keynes and Alex Alderton, Roberto Amato, Jeffrey Barrett, Sonia Goncalves, Ewan Harrison, David K. Jackson, Ian Johnston, Dominic Kwiatkowski, Cordelia Langford, John Sillitoe on behalf of the Wellcome Sanger Institute COVID-19 Surveillance Team |
| EPI_ISL_3446329 | Lighthouse Lab in Glasgow | Wellcome Sanger Institute for the COVID-19 Genomics UK (COG-UK) Consortium | Harper VanSteenhouse, Yumi Kasai, David Gray, Carol Clugston, Anna Dominiczak and Alex Alderton, Roberto Amato, Jeffrey Barrett, Sonia Goncalves, Ewan Harrison, David K. Jackson, Ian Johnston, Dominic Kwiatkowski, Cordelia Langford, John Sillitoe on behalf of the Wellcome Sanger Institute COVID-19 Surveillance Team |
| EPI_ISL_3446330 | Lighthouse Lab in Milton Keynes | Wellcome Sanger Institute for the COVID-19 Genomics UK (COG-UK) Consortium | The Lighthouse Lab in Milton Keynes and Alex Alderton, Roberto Amato, Jeffrey Barrett, Sonia Goncalves, Ewan Harrison, David K. Jackson, Ian Johnston, Dominic Kwiatkowski, Cordelia Langford, John Sillitoe on behalf of the Wellcome Sanger Institute COVID-19 Surveillance Team |
| EPI_ISL_3446331 | Lighthouse Lab in Alderley Park | Wellcome Sanger Institute for the COVID-19 Genomics UK (COG-UK) Consortium | Jacquelyn Wynn, Mairead Hyland, The Lighthouse Lab in Alderley Park and Alex Alderton, Roberto Amato, Jeffrey Barrett, Sonia Goncalves, Ewan Harrison, David K. Jackson, Ian Johnston, Dominic Kwiatkowski, Cordelia Langford, John Sillitoe on behalf of the Wellcome Sanger Institute COVID-19 Surveillance Team |
| EPI_ISL_3446332 | Lighthouse Lab in Glasgow | Wellcome Sanger Institute for the COVID-19 Genomics UK (COG-UK) Consortium | Harper VanSteenhouse, Yumi Kasai, David Gray, Carol Clugston, Anna Dominiczak and Alex Alderton, Roberto Amato, Jeffrey Barrett, Sonia Goncalves, Ewan Harrison, David K. Jackson, Ian Johnston, Dominic Kwiatkowski, Cordelia Langford, John Sillitoe on behalf of the Wellcome Sanger Institute COVID-19 Surveillance Team |
| EPI_ISL_3446333 | Lighthouse Lab in Alderley Park | Wellcome Sanger Institute for the COVID-19 Genomics UK (COG-UK) Consortium | Jacquelyn Wynn, Mairead Hyland, The Lighthouse Lab in Alderley Park and Alex Alderton, Roberto Amato, Jeffrey Barrett, Sonia Goncalves, Ewan Harrison, David K. Jackson, Ian Johnston, Dominic Kwiatkowski, Cordelia Langford, John Sillitoe on behalf of the Wellcome Sanger Institute COVID-19 Surveillance Team |
| EPI_ISL_3446334 | Lighthouse Lab in Milton Keynes | Wellcome Sanger Institute for the COVID-19 Genomics UK (COG-UK) Consortium | The Lighthouse Lab in Milton Keynes and Alex Alderton, Roberto Amato, Jeffrey Barrett, Sonia Goncalves, Ewan Harrison, David K. Jackson, Ian Johnston, Dominic Kwiatkowski, Cordelia Langford, John Sillitoe on behalf of the Wellcome Sanger Institute COVID-19 Surveillance Team |
| EPI_ISL_3446335, EPI_ISL_3446336 | Lighthouse Lab in Glasgow | Wellcome Sanger Institute for the COVID-19 Genomics UK (COG-UK) Consortium | Harper VanSteenhouse, Yumi Kasai, David Gray, Carol Clugston, Anna Dominiczak and Alex Alderton, Roberto Amato, Jeffrey Barrett, Sonia Goncalves, Ewan Harrison, David K. Jackson, Ian Johnston, Dominic Kwiatkowski, Cordelia Langford, John Sillitoe on behalf of the Wellcome Sanger Institute COVID-19 Surveillance Team |
| EPI_ISL_3446337 | Lighthouse Lab in Milton Keynes | Wellcome Sanger Institute for the COVID-19 Genomics UK (COG-UK) Consortium | The Lighthouse Lab in Milton Keynes and Alex Alderton, Roberto Amato, Jeffrey Barrett, Sonia Goncalves, Ewan Harrison, David K. Jackson, Ian Johnston, Dominic Kwiatkowski, Cordelia Langford, John Sillitoe on behalf of the Wellcome Sanger Institute COVID-19 Surveillance Team |
| EPI_ISL_3446338 | Lighthouse Lab in Alderley Park | Wellcome Sanger Institute for the COVID-19 Genomics UK (COG-UK) Consortium | Jacquelyn Wynn, Mairead Hyland, The Lighthouse Lab in Alderley Park and Alex Alderton, Roberto Amato, Jeffrey Barrett, Sonia Goncalves, Ewan Harrison, David K. Jackson, Ian Johnston, Dominic Kwiatkowski, Cordelia Langford, John Sillitoe on behalf of the Wellcome Sanger Institute COVID-19 Surveillance Team |

| | | | |
|---|---|---|---|
| EPI_ISL_3446339, EPI_ISL_3446340, EPI_ISL_3446341, EPI_ISL_3446342 | Lighthouse Lab in Glasgow | Wellcome Sanger Institute for the COVID-19 Genomics UK (COG-UK) Consortium | Harper VanSteenhouse, Yumi Kasai, David Gray, Carol Clugston, Anna Dominiczak and Alex Alderton, Roberto Amato, Jeffrey Barrett, Sonia Goncalves, Ewan Harrison, David K. Jackson, Ian Johnston, Dominic Kwiatkowski, Cordelia Langford, John Sillitoe on behalf of the Wellcome Sanger Institute COVID-19 Surveillance Team |
| EPI_ISL_3446343 | Lighthouse Lab in Alderley Park | Wellcome Sanger Institute for the COVID-19 Genomics UK (COG-UK) Consortium | Jacquelyn Wynn, Mairead Hyland, The Lighthouse Lab in Alderley Park and Alex Alderton, Roberto Amato, Jeffrey Barrett, Sonia Goncalves, Ewan Harrison, David K. Jackson, Ian Johnston, Dominic Kwiatkowski, Cordelia Langford, John Sillitoe on behalf of the Wellcome Sanger Institute COVID-19 Surveillance Team |
| EPI_ISL_3446344 | Lighthouse Lab in Milton Keynes | Wellcome Sanger Institute for the COVID-19 Genomics UK (COG-UK) Consortium | The Lighthouse Lab in Milton Keynes and Alex Alderton, Roberto Amato, Jeffrey Barrett, Sonia Goncalves, Ewan Harrison, David K. Jackson, Ian Johnston, Dominic Kwiatkowski, Cordelia Langford, John Sillitoe on behalf of the Wellcome Sanger Institute COVID-19 Surveillance Team |
| EPI_ISL_3446345, EPI_ISL_3446346 | Lighthouse Lab in Glasgow | Wellcome Sanger Institute for the COVID-19 Genomics UK (COG-UK) Consortium | Harper VanSteenhouse, Yumi Kasai, David Gray, Carol Clugston, Anna Dominiczak and Alex Alderton, Roberto Amato, Jeffrey Barrett, Sonia Goncalves, Ewan Harrison, David K. Jackson, Ian Johnston, Dominic Kwiatkowski, Cordelia Langford, John Sillitoe on behalf of the Wellcome Sanger Institute COVID-19 Surveillance Team |
| EPI_ISL_3446347 | Lighthouse Lab in Milton Keynes | Wellcome Sanger Institute for the COVID-19 Genomics UK (COG-UK) Consortium | The Lighthouse Lab in Milton Keynes and Alex Alderton, Roberto Amato, Jeffrey Barrett, Sonia Goncalves, Ewan Harrison, David K. Jackson, Ian Johnston, Dominic Kwiatkowski, Cordelia Langford, John Sillitoe on behalf of the Wellcome Sanger Institute COVID-19 Surveillance Team |
| EPI_ISL_3446348 | Lighthouse Lab in Alderley Park | Wellcome Sanger Institute for the COVID-19 Genomics UK (COG-UK) Consortium | Jacquelyn Wynn, Mairead Hyland, The Lighthouse Lab in Alderley Park and Alex Alderton, Roberto Amato, Jeffrey Barrett, Sonia Goncalves, Ewan Harrison, David K. Jackson, Ian Johnston, Dominic Kwiatkowski, Cordelia Langford, John Sillitoe on behalf of the Wellcome Sanger Institute COVID-19 Surveillance Team |
| EPI_ISL_3446349 | Lighthouse Lab in Glasgow | Wellcome Sanger Institute for the COVID-19 Genomics UK (COG-UK) Consortium | Harper VanSteenhouse, Yumi Kasai, David Gray, Carol Clugston, Anna Dominiczak and Alex Alderton, Roberto Amato, Jeffrey Barrett, Sonia Goncalves, Ewan Harrison, David K. Jackson, Ian Johnston, Dominic Kwiatkowski, Cordelia Langford, John Sillitoe on behalf of the Wellcome Sanger Institute COVID-19 Surveillance Team |
| EPI_ISL_3446350, EPI_ISL_3446351 | Lighthouse Lab in Milton Keynes | Wellcome Sanger Institute for the COVID-19 Genomics UK (COG-UK) Consortium | The Lighthouse Lab in Milton Keynes and Alex Alderton, Roberto Amato, Jeffrey Barrett, Sonia Goncalves, Ewan Harrison, David K. Jackson, Ian Johnston, Dominic Kwiatkowski, Cordelia Langford, John Sillitoe on behalf of the Wellcome Sanger Institute COVID-19 Surveillance Team |
| EPI_ISL_3446352 | Lighthouse Lab in Alderley Park | Wellcome Sanger Institute for the COVID-19 Genomics UK (COG-UK) Consortium | Jacquelyn Wynn, Mairead Hyland, The Lighthouse Lab in Alderley Park and Alex Alderton, Roberto Amato, Jeffrey Barrett, Sonia Goncalves, Ewan Harrison, David K. Jackson, Ian Johnston, Dominic Kwiatkowski, Cordelia Langford, John Sillitoe on behalf of the Wellcome Sanger Institute COVID-19 Surveillance Team |
| EPI_ISL_3446353, EPI_ISL_3446354 | Lighthouse Lab in Glasgow | Wellcome Sanger Institute for the COVID-19 Genomics UK (COG-UK) Consortium | Harper VanSteenhouse, Yumi Kasai, David Gray, Carol Clugston, Anna Dominiczak and Alex Alderton, Roberto Amato, Jeffrey Barrett, Sonia Goncalves, Ewan Harrison, David K. Jackson, Ian Johnston, Dominic Kwiatkowski, Cordelia Langford, John Sillitoe on behalf of the Wellcome Sanger Institute COVID-19 Surveillance Team |
| EPI_ISL_3446355 | Lighthouse Lab in Alderley Park | Wellcome Sanger Institute for the COVID-19 Genomics UK (COG-UK) Consortium | Jacquelyn Wynn, Mairead Hyland, The Lighthouse Lab in Alderley Park and Alex Alderton, Roberto Amato, Jeffrey Barrett, Sonia Goncalves, Ewan Harrison, David K. Jackson, Ian Johnston, Dominic Kwiatkowski, Cordelia Langford, John Sillitoe on behalf of the Wellcome Sanger Institute COVID-19 Surveillance Team |
| EPI_ISL_3446356, EPI_ISL_3446357, EPI_ISL_3446358, EPI_ISL_3446359 | Lighthouse Lab in Glasgow | Wellcome Sanger Institute for the COVID-19 Genomics UK (COG-UK) Consortium | Harper VanSteenhouse, Yumi Kasai, David Gray, Carol Clugston, Anna Dominiczak and Alex Alderton, Roberto Amato, Jeffrey Barrett, Sonia Goncalves, Ewan Harrison, David K. Jackson, Ian Johnston, Dominic Kwiatkowski, Cordelia Langford, John Sillitoe on behalf of the Wellcome Sanger Institute COVID-19 Surveillance Team |
| EPI_ISL_3446360 | Lighthouse Lab in Alderley Park | Wellcome Sanger Institute for the COVID-19 Genomics UK (COG-UK) Consortium | Jacquelyn Wynn, Mairead Hyland, The Lighthouse Lab in Alderley Park and Alex Alderton, Roberto Amato, Jeffrey Barrett, Sonia Goncalves, Ewan Harrison, David K. Jackson, Ian Johnston, Dominic Kwiatkowski, Cordelia Langford, John Sillitoe on behalf of the Wellcome Sanger Institute COVID-19 Surveillance Team |
| EPI_ISL_3446361, EPI_ISL_3446362 | Lighthouse Lab in Glasgow | Wellcome Sanger Institute for the COVID-19 Genomics UK (COG-UK) Consortium | Harper VanSteenhouse, Yumi Kasai, David Gray, Carol Clugston, Anna Dominiczak and Alex Alderton, Roberto Amato, Jeffrey Barrett, Sonia Goncalves, Ewan Harrison, David K. Jackson, Ian Johnston, Dominic Kwiatkowski, Cordelia Langford, John Sillitoe on behalf of the Wellcome Sanger Institute COVID-19 Surveillance Team |
| EPI_ISL_3446363 | Lighthouse Lab in Alderley Park | Wellcome Sanger Institute for the COVID-19 Genomics UK (COG-UK) Consortium | Jacquelyn Wynn, Mairead Hyland, The Lighthouse Lab in Alderley Park and Alex Alderton, Roberto Amato, Jeffrey Barrett, Sonia Goncalves, Ewan Harrison, David K. Jackson, Ian Johnston, Dominic Kwiatkowski, Cordelia Langford, John Sillitoe on behalf of the Wellcome Sanger Institute COVID-19 Surveillance Team |
| EPI_ISL_3446364 | Lighthouse Lab in Milton Keynes | Wellcome Sanger Institute for the COVID-19 Genomics UK (COG-UK) Consortium | The Lighthouse Lab in Milton Keynes and Alex Alderton, Roberto Amato, Jeffrey Barrett, Sonia Goncalves, Ewan Harrison, David K. Jackson, Ian Johnston, Dominic Kwiatkowski, Cordelia Langford, John Sillitoe on behalf of the Wellcome Sanger Institute COVID-19 Surveillance Team |
| EPI_ISL_3446365 | Lighthouse Lab in Glasgow | Wellcome Sanger Institute for the COVID-19 Genomics UK (COG-UK) Consortium | Harper VanSteenhouse, Yumi Kasai, David Gray, Carol Clugston, Anna Dominiczak and Alex Alderton, Roberto Amato, Jeffrey Barrett, Sonia Goncalves, Ewan Harrison, David K. Jackson, Ian Johnston, Dominic Kwiatkowski, Cordelia Langford, John Sillitoe on behalf of the Wellcome Sanger Institute COVID-19 Surveillance Team |
| EPI_ISL_3446366 | Lighthouse Lab in Milton Keynes | Wellcome Sanger Institute for the COVID-19 Genomics UK (COG-UK) Consortium | The Lighthouse Lab in Milton Keynes and Alex Alderton, Roberto Amato, Jeffrey Barrett, Sonia Goncalves, Ewan Harrison, David K. Jackson, Ian Johnston, Dominic Kwiatkowski, Cordelia Langford, John Sillitoe on behalf of the Wellcome Sanger Institute COVID-19 Surveillance Team |
| EPI_ISL_3446367, EPI_ISL_3446368 | Lighthouse Lab in Alderley Park | Wellcome Sanger Institute for the COVID-19 Genomics UK (COG-UK) Consortium | Jacquelyn Wynn, Mairead Hyland, The Lighthouse Lab in Alderley Park and Alex Alderton, Roberto Amato, Jeffrey Barrett, Sonia Goncalves, Ewan Harrison, David K. Jackson, Ian Johnston, Dominic Kwiatkowski, Cordelia Langford, John Sillitoe on behalf of the Wellcome Sanger Institute COVID-19 Surveillance Team |
| EPI_ISL_3446369 | Lighthouse Lab in Milton Keynes | Wellcome Sanger Institute for the COVID-19 Genomics UK (COG-UK) Consortium | The Lighthouse Lab in Milton Keynes and Alex Alderton, Roberto Amato, Jeffrey Barrett, Sonia Goncalves, Ewan Harrison, David K. Jackson, Ian Johnston, Dominic Kwiatkowski, Cordelia Langford, John Sillitoe on behalf of the Wellcome Sanger Institute COVID-19 Surveillance Team |
| EPI_ISL_3446370, EPI_ISL_3446371, EPI_ISL_3446372, EPI_ISL_3446373 | Lighthouse Lab in Alderley Park | Wellcome Sanger Institute for the COVID-19 Genomics UK (COG-UK) Consortium | Jacquelyn Wynn, Mairead Hyland, The Lighthouse Lab in Alderley Park and Alex Alderton, Roberto Amato, Jeffrey Barrett, Sonia Goncalves, Ewan Harrison, David K. Jackson, Ian Johnston, Dominic Kwiatkowski, Cordelia Langford, John Sillitoe on behalf of the Wellcome Sanger Institute COVID-19 Surveillance Team |
| EPI_ISL_3446374 | Lighthouse Lab in Glasgow | Wellcome Sanger Institute for the COVID-19 Genomics UK (COG-UK) Consortium | Harper VanSteenhouse, Yumi Kasai, David Gray, Carol Clugston, Anna Dominiczak and Alex Alderton, Roberto Amato, Jeffrey Barrett, Sonia Goncalves, Ewan Harrison, David K. Jackson, Ian Johnston, Dominic Kwiatkowski, Cordelia Langford, John Sillitoe on behalf of the Wellcome Sanger Institute COVID-19 Surveillance Team |
| EPI_ISL_3446375, EPI_ISL_3446376, EPI_ISL_3446377 | Lighthouse Lab in Milton Keynes | Wellcome Sanger Institute for the COVID-19 Genomics UK (COG-UK) Consortium | The Lighthouse Lab in Milton Keynes and Alex Alderton, Roberto Amato, Jeffrey Barrett, Sonia Goncalves, Ewan Harrison, David K. Jackson, Ian Johnston, Dominic Kwiatkowski, Cordelia Langford, John Sillitoe on behalf of the Wellcome Sanger Institute COVID-19 Surveillance Team |
| EPI_ISL_3446378 | Lighthouse Lab in Glasgow | Wellcome Sanger Institute for the COVID-19 Genomics UK (COG-UK) Consortium | Harper VanSteenhouse, Yumi Kasai, David Gray, Carol Clugston, Anna Dominiczak and Alex Alderton, Roberto Amato, Jeffrey Barrett, Sonia Goncalves, Ewan Harrison, David K. Jackson, Ian Johnston, Dominic Kwiatkowski, Cordelia Langford, John Sillitoe on behalf of the Wellcome Sanger Institute COVID-19 Surveillance Team |
| EPI_ISL_3446379, EPI_ISL_3446380 | Lighthouse Lab in Alderley Park | Wellcome Sanger Institute for the COVID-19 Genomics UK (COG-UK) Consortium | Jacquelyn Wynn, Mairead Hyland, The Lighthouse Lab in Alderley Park and Alex Alderton, Roberto Amato, Jeffrey Barrett, Sonia Goncalves, Ewan Harrison, David K. Jackson, Ian Johnston, Dominic Kwiatkowski, Cordelia Langford, John Sillitoe on behalf of the Wellcome Sanger Institute COVID-19 Surveillance Team |
| EPI_ISL_3446381 | Lighthouse Lab in Glasgow | Wellcome Sanger Institute for the COVID-19 Genomics UK (COG-UK) Consortium | Harper VanSteenhouse, Yumi Kasai, David Gray, Carol Clugston, Anna Dominiczak and Alex Alderton, Roberto Amato, Jeffrey Barrett, Sonia Goncalves, Ewan Harrison, David K. Jackson, Ian Johnston, Dominic Kwiatkowski, Cordelia Langford, John Sillitoe on behalf of the Wellcome Sanger Institute COVID-19 Surveillance Team |
| EPI_ISL_3446382 | Lighthouse Lab in Milton Keynes | Wellcome Sanger Institute for the COVID-19 Genomics UK (COG-UK) Consortium | The Lighthouse Lab in Milton Keynes and Alex Alderton, Roberto Amato, Jeffrey Barrett, Sonia Goncalves, Ewan Harrison, David K. Jackson, Ian Johnston, Dominic Kwiatkowski, Cordelia Langford, John Sillitoe on behalf of the Wellcome Sanger Institute COVID-19 Surveillance Team |
| EPI_ISL_3446383, EPI_ISL_3446384, EPI_ISL_3446385 | Lighthouse Lab in Alderley Park | Wellcome Sanger Institute for the COVID-19 Genomics UK (COG-UK) Consortium | Jacquelyn Wynn, Mairead Hyland, The Lighthouse Lab in Alderley Park and Alex Alderton, Roberto Amato, Jeffrey Barrett, Sonia Goncalves, Ewan Harrison, David K. Jackson, Ian Johnston, Dominic Kwiatkowski, Cordelia Langford, John Sillitoe on behalf of the Wellcome Sanger Institute COVID-19 Surveillance Team |
| EPI_ISL_3446386, EPI_ISL_3446387 | Lighthouse Lab in Glasgow | Wellcome Sanger Institute for the COVID-19 Genomics UK (COG-UK) Consortium | Harper VanSteenhouse, Yumi Kasai, David Gray, Carol Clugston, Anna Dominiczak and Alex Alderton, Roberto Amato, Jeffrey Barrett, Sonia Goncalves, Ewan Harrison, David K. Jackson, Ian Johnston, Dominic Kwiatkowski, Cordelia Langford, John Sillitoe on behalf of the Wellcome Sanger Institute COVID-19 Surveillance Team |
| EPI_ISL_3446388 | Lighthouse Lab in Alderley Park | Wellcome Sanger Institute for the COVID-19 Genomics UK (COG-UK) Consortium | Jacquelyn Wynn, Mairead Hyland, The Lighthouse Lab in Alderley Park and Alex Alderton, Roberto Amato, Jeffrey Barrett, Sonia Goncalves, Ewan Harrison, David K. Jackson, Ian Johnston, Dominic Kwiatkowski, Cordelia Langford, John Sillitoe on behalf of the Wellcome Sanger Institute COVID-19 Surveillance Team |
| EPI_ISL_3446389 | Lighthouse Lab in Milton Keynes | Wellcome Sanger Institute for the COVID-19 Genomics UK (COG-UK) Consortium | The Lighthouse Lab in Milton Keynes and Alex Alderton, Roberto Amato, Jeffrey Barrett, Sonia Goncalves, Ewan Harrison, David K. Jackson, Ian Johnston, Dominic Kwiatkowski, Cordelia Langford, John Sillitoe on behalf of the Wellcome Sanger Institute COVID-19 Surveillance Team |
| EPI_ISL_3446390 | Lighthouse Lab in Glasgow | Wellcome Sanger Institute for the COVID-19 Genomics UK (COG-UK) Consortium | Harper VanSteenhouse, Yumi Kasai, David Gray, Carol Clugston, Anna Dominiczak and Alex Alderton, Roberto Amato, Jeffrey Barrett, Sonia Goncalves, Ewan Harrison, David K. Jackson, Ian Johnston, Dominic Kwiatkowski, Cordelia Langford, John Sillitoe on behalf of the Wellcome Sanger Institute COVID-19 Surveillance Team |
| EPI_ISL_3446391 | Lighthouse Lab in Alderley Park | Wellcome Sanger Institute for the COVID-19 Genomics UK (COG-UK) Consortium | Jacquelyn Wynn, Mairead Hyland, The Lighthouse Lab in Alderley Park and Alex Alderton, Roberto Amato, Jeffrey Barrett, Sonia Goncalves, Ewan Harrison, David K. Jackson, Ian Johnston, Dominic Kwiatkowski, Cordelia Langford, John Sillitoe on behalf of the Wellcome Sanger Institute COVID-19 Surveillance Team |
| EPI_ISL_3446392, EPI_ISL_3446393 | Lighthouse Lab in Milton Keynes | Wellcome Sanger Institute for the COVID-19 Genomics UK (COG-UK) Consortium | The Lighthouse Lab in Milton Keynes and Alex Alderton, Roberto Amato, Jeffrey Barrett, Sonia Goncalves, Ewan Harrison, David K. Jackson, Ian Johnston, Dominic Kwiatkowski, Cordelia Langford, John Sillitoe on behalf of the Wellcome Sanger Institute COVID-19 Surveillance Team |
| EPI_ISL_3446394, EPI_ISL_3446395, EPI_ISL_3446396, EPI_ISL_3446397 | Lighthouse Lab in Glasgow | Wellcome Sanger Institute for the COVID-19 Genomics UK (COG-UK) Consortium | Harper VanSteenhouse, Yumi Kasai, David Gray, Carol Clugston, Anna Dominiczak and Alex Alderton, Roberto Amato, Jeffrey Barrett, Sonia Goncalves, Ewan Harrison, David K. Jackson, Ian Johnston, Dominic Kwiatkowski, Cordelia Langford, John Sillitoe on behalf of the Wellcome Sanger Institute COVID-19 Surveillance Team |
| EPI_ISL_3446398 | Lighthouse Lab in Milton Keynes | Wellcome Sanger Institute for the COVID-19 Genomics UK (COG-UK) Consortium | The Lighthouse Lab in Milton Keynes and Alex Alderton, Roberto Amato, Jeffrey Barrett, Sonia Goncalves, Ewan Harrison, David K. Jackson, Ian Johnston, Dominic Kwiatkowski, Cordelia Langford, John Sillitoe on behalf of the Wellcome Sanger Institute COVID-19 Surveillance Team |
| EPI_ISL_3446399, EPI_ISL_3446400, EPI_ISL_3446401, EPI_ISL_3446402, EPI_ISL_3446403 | Lighthouse Lab in Glasgow | Wellcome Sanger Institute for the COVID-19 Genomics UK (COG-UK) Consortium | Harper VanSteenhouse, Yumi Kasai, David Gray, Carol Clugston, Anna Dominiczak and Alex Alderton, Roberto Amato, Jeffrey Barrett, Sonia Goncalves, Ewan Harrison, David K. Jackson, Ian Johnston, Dominic Kwiatkowski, Cordelia Langford, John Sillitoe on behalf of the Wellcome Sanger Institute COVID-19 Surveillance Team |
| EPI_ISL_3446404, EPI_ISL_3446405, EPI_ISL_3446406 | Lighthouse Lab in Alderley Park | Wellcome Sanger Institute for the COVID-19 Genomics UK (COG-UK) Consortium | Jacquelyn Wynn, Mairead Hyland, The Lighthouse Lab in Alderley Park and Alex Alderton, Roberto Amato, Jeffrey Barrett, Sonia Goncalves, Ewan Harrison, David K. Jackson, Ian Johnston, Dominic Kwiatkowski, Cordelia Langford, John Sillitoe on behalf of the Wellcome Sanger Institute COVID-19 Surveillance Team |
| EPI_ISL_3446407 | Lighthouse Lab in Glasgow | Wellcome Sanger Institute for the COVID-19 Genomics UK (COG-UK) Consortium | Harper VanSteenhouse, Yumi Kasai, David Gray, Carol Clugston, Anna Dominiczak and Alex Alderton, Roberto Amato, Jeffrey Barrett, Sonia Goncalves, Ewan Harrison, David K. Jackson, Ian Johnston, Dominic Kwiatkowski, Cordelia Langford, John Sillitoe on behalf of the Wellcome Sanger Institute COVID-19 Surveillance Team |

| | | | |
|---|---|---|---|
| EPI_ISL_3446408 | Lighthouse Lab in Alderley Park | Wellcome Sanger Institute for the COVID-19 Genomics UK (COG-UK) Consortium | Jacquelyn Wynn, Mairead Hyland, The Lighthouse Lab in Alderley Park and Alex Alderton, Roberto Amato, Jeffrey Barrett, Sonia Goncalves, Ewan Harrison, David K. Jackson, Ian Johnston, Dominic Kwiatkowski, Cordelia Langford, John Sillitoe on behalf of the Wellcome Sanger Institute COVID-19 Surveillance Team |
| EPI_ISL_3446409 | Lighthouse Lab in Glasgow | Wellcome Sanger Institute for the COVID-19 Genomics UK (COG-UK) Consortium | Harper VanSteenhouse, Yumi Kasai, David Gray, Carol Clugston, Anna Dominiczak and Alex Alderton, Roberto Amato, Jeffrey Barrett, Sonia Goncalves, Ewan Harrison, David K. Jackson, Ian Johnston, Dominic Kwiatkowski, Cordelia Langford, John Sillitoe on behalf of the Wellcome Sanger Institute COVID-19 Surveillance Team |
| EPI_ISL_3446410 | Lighthouse Lab in Alderley Park | Wellcome Sanger Institute for the COVID-19 Genomics UK (COG-UK) Consortium | Jacquelyn Wynn, Mairead Hyland, The Lighthouse Lab in Alderley Park and Alex Alderton, Roberto Amato, Jeffrey Barrett, Sonia Goncalves, Ewan Harrison, David K. Jackson, Ian Johnston, Dominic Kwiatkowski, Cordelia Langford, John Sillitoe on behalf of the Wellcome Sanger Institute COVID-19 Surveillance Team |
| EPI_ISL_3446411 | Lighthouse Lab in Glasgow | Wellcome Sanger Institute for the COVID-19 Genomics UK (COG-UK) Consortium | Harper VanSteenhouse, Yumi Kasai, David Gray, Carol Clugston, Anna Dominiczak and Alex Alderton, Roberto Amato, Jeffrey Barrett, Sonia Goncalves, Ewan Harrison, David K. Jackson, Ian Johnston, Dominic Kwiatkowski, Cordelia Langford, John Sillitoe on behalf of the Wellcome Sanger Institute COVID-19 Surveillance Team |
| EPI_ISL_3446412 | Lighthouse Lab in Alderley Park | Wellcome Sanger Institute for the COVID-19 Genomics UK (COG-UK) Consortium | Jacquelyn Wynn, Mairead Hyland, The Lighthouse Lab in Alderley Park and Alex Alderton, Roberto Amato, Jeffrey Barrett, Sonia Goncalves, Ewan Harrison, David K. Jackson, Ian Johnston, Dominic Kwiatkowski, Cordelia Langford, John Sillitoe on behalf of the Wellcome Sanger Institute COVID-19 Surveillance Team |
| EPI_ISL_3446413 | Lighthouse Lab in Glasgow | Wellcome Sanger Institute for the COVID-19 Genomics UK (COG-UK) Consortium | Harper VanSteenhouse, Yumi Kasai, David Gray, Carol Clugston, Anna Dominiczak and Alex Alderton, Roberto Amato, Jeffrey Barrett, Sonia Goncalves, Ewan Harrison, David K. Jackson, Ian Johnston, Dominic Kwiatkowski, Cordelia Langford, John Sillitoe on behalf of the Wellcome Sanger Institute COVID-19 Surveillance Team |
| EPI_ISL_3446414 | Lighthouse Lab in Alderley Park | Wellcome Sanger Institute for the COVID-19 Genomics UK (COG-UK) Consortium | Jacquelyn Wynn, Mairead Hyland, The Lighthouse Lab in Alderley Park and Alex Alderton, Roberto Amato, Jeffrey Barrett, Sonia Goncalves, Ewan Harrison, David K. Jackson, Ian Johnston, Dominic Kwiatkowski, Cordelia Langford, John Sillitoe on behalf of the Wellcome Sanger Institute COVID-19 Surveillance Team |
| EPI_ISL_3446415, EPI_ISL_3446416 | Lighthouse Lab in Glasgow | Wellcome Sanger Institute for the COVID-19 Genomics UK (COG-UK) Consortium | Harper VanSteenhouse, Yumi Kasai, David Gray, Carol Clugston, Anna Dominiczak and Alex Alderton, Roberto Amato, Jeffrey Barrett, Sonia Goncalves, Ewan Harrison, David K. Jackson, Ian Johnston, Dominic Kwiatkowski, Cordelia Langford, John Sillitoe on behalf of the Wellcome Sanger Institute COVID-19 Surveillance Team |
| EPI_ISL_3446417, EPI_ISL_3446418, EPI_ISL_3446419, EPI_ISL_3446420 | Lighthouse Lab in Alderley Park | Wellcome Sanger Institute for the COVID-19 Genomics UK (COG-UK) Consortium | Jacquelyn Wynn, Mairead Hyland, The Lighthouse Lab in Alderley Park and Alex Alderton, Roberto Amato, Jeffrey Barrett, Sonia Goncalves, Ewan Harrison, David K. Jackson, Ian Johnston, Dominic Kwiatkowski, Cordelia Langford, John Sillitoe on behalf of the Wellcome Sanger Institute COVID-19 Surveillance Team |
| EPI_ISL_3446421 | Lighthouse Lab in Milton Keynes | Wellcome Sanger Institute for the COVID-19 Genomics UK (COG-UK) Consortium | The Lighthouse Lab in Milton Keynes and Alex Alderton, Roberto Amato, Jeffrey Barrett, Sonia Goncalves, Ewan Harrison, David K. Jackson, Ian Johnston, Dominic Kwiatkowski, Cordelia Langford, John Sillitoe on behalf of the Wellcome Sanger Institute COVID-19 Surveillance Team |
| EPI_ISL_3446422 | Lighthouse Lab in Alderley Park | Wellcome Sanger Institute for the COVID-19 Genomics UK (COG-UK) Consortium | Jacquelyn Wynn, Mairead Hyland, The Lighthouse Lab in Alderley Park and Alex Alderton, Roberto Amato, Jeffrey Barrett, Sonia Goncalves, Ewan Harrison, David K. Jackson, Ian Johnston, Dominic Kwiatkowski, Cordelia Langford, John Sillitoe on behalf of the Wellcome Sanger Institute COVID-19 Surveillance Team |
| EPI_ISL_3446423 | Lighthouse Lab in Milton Keynes | Wellcome Sanger Institute for the COVID-19 Genomics UK (COG-UK) Consortium | The Lighthouse Lab in Milton Keynes and Alex Alderton, Roberto Amato, Jeffrey Barrett, Sonia Goncalves, Ewan Harrison, David K. Jackson, Ian Johnston, Dominic Kwiatkowski, Cordelia Langford, John Sillitoe on behalf of the Wellcome Sanger Institute COVID-19 Surveillance Team |
| EPI_ISL_3446424, EPI_ISL_3446425 | Lighthouse Lab in Alderley Park | Wellcome Sanger Institute for the COVID-19 Genomics UK (COG-UK) Consortium | Jacquelyn Wynn, Mairead Hyland, The Lighthouse Lab in Alderley Park and Alex Alderton, Roberto Amato, Jeffrey Barrett, Sonia Goncalves, Ewan Harrison, David K. Jackson, Ian Johnston, Dominic Kwiatkowski, Cordelia Langford, John Sillitoe on behalf of the Wellcome Sanger Institute COVID-19 Surveillance Team |
| EPI_ISL_3446426, EPI_ISL_3446427 | Lighthouse Lab in Milton Keynes | Wellcome Sanger Institute for the COVID-19 Genomics UK (COG-UK) Consortium | The Lighthouse Lab in Milton Keynes and Alex Alderton, Roberto Amato, Jeffrey Barrett, Sonia Goncalves, Ewan Harrison, David K. Jackson, Ian Johnston, Dominic Kwiatkowski, Cordelia Langford, John Sillitoe on behalf of the Wellcome Sanger Institute COVID-19 Surveillance Team |
| EPI_ISL_3446428 | Lighthouse Lab in Alderley Park | Wellcome Sanger Institute for the COVID-19 Genomics UK (COG-UK) Consortium | Jacquelyn Wynn, Mairead Hyland, The Lighthouse Lab in Alderley Park and Alex Alderton, Roberto Amato, Jeffrey Barrett, Sonia Goncalves, Ewan Harrison, David K. Jackson, Ian Johnston, Dominic Kwiatkowski, Cordelia Langford, John Sillitoe on behalf of the Wellcome Sanger Institute COVID-19 Surveillance Team |
| EPI_ISL_3446429, EPI_ISL_3446430, EPI_ISL_3446431 | Lighthouse Lab in Milton Keynes | Wellcome Sanger Institute for the COVID-19 Genomics UK (COG-UK) Consortium | The Lighthouse Lab in Milton Keynes and Alex Alderton, Roberto Amato, Jeffrey Barrett, Sonia Goncalves, Ewan Harrison, David K. Jackson, Ian Johnston, Dominic Kwiatkowski, Cordelia Langford, John Sillitoe on behalf of the Wellcome Sanger Institute COVID-19 Surveillance Team |
| EPI_ISL_3446432 | Lighthouse Lab in Alderley Park | Wellcome Sanger Institute for the COVID-19 Genomics UK (COG-UK) Consortium | Jacquelyn Wynn, Mairead Hyland, The Lighthouse Lab in Alderley Park and Alex Alderton, Roberto Amato, Jeffrey Barrett, Sonia Goncalves, Ewan Harrison, David K. Jackson, Ian Johnston, Dominic Kwiatkowski, Cordelia Langford, John Sillitoe on behalf of the Wellcome Sanger Institute COVID-19 Surveillance Team |
| EPI_ISL_3446433, EPI_ISL_3446434 | Lighthouse Lab in Milton Keynes | Wellcome Sanger Institute for the COVID-19 Genomics UK (COG-UK) Consortium | The Lighthouse Lab in Milton Keynes and Alex Alderton, Roberto Amato, Jeffrey Barrett, Sonia Goncalves, Ewan Harrison, David K. Jackson, Ian Johnston, Dominic Kwiatkowski, Cordelia Langford, John Sillitoe on behalf of the Wellcome Sanger Institute COVID-19 Surveillance Team |
| EPI_ISL_3446435, EPI_ISL_3446436 | Lighthouse Lab in Alderley Park | Wellcome Sanger Institute for the COVID-19 Genomics UK (COG-UK) Consortium | Jacquelyn Wynn, Mairead Hyland, The Lighthouse Lab in Alderley Park and Alex Alderton, Roberto Amato, Jeffrey Barrett, Sonia Goncalves, Ewan Harrison, David K. Jackson, Ian Johnston, Dominic Kwiatkowski, Cordelia Langford, John Sillitoe on behalf of the Wellcome Sanger Institute COVID-19 Surveillance Team |
| EPI_ISL_3446437 | Lighthouse Lab in Milton Keynes | Wellcome Sanger Institute for the COVID-19 Genomics UK (COG-UK) Consortium | The Lighthouse Lab in Milton Keynes and Alex Alderton, Roberto Amato, Jeffrey Barrett, Sonia Goncalves, Ewan Harrison, David K. Jackson, Ian Johnston, Dominic Kwiatkowski, Cordelia Langford, John Sillitoe on behalf of the Wellcome Sanger Institute COVID-19 Surveillance Team |
| EPI_ISL_3446438 | Berkshire and Surrey Pathology Services Lighthouse Laboratory | Wellcome Sanger Institute for the COVID-19 Genomics UK (COG-UK) Consortium | Berkshire and Surrey Pathology Services Lighthouse Laboratory and Alex Alderton, Roberto Amato, Jeffrey Barrett, Sonia Goncalves, Ewan Harrison, David K. Jackson, Ian Johnston, Dominic Kwiatkowski, Cordelia Langford, John Sillitoe on behalf of the Wellcome Sanger Institute COVID-19 Surveillance Team |
| EPI_ISL_3446439 | Health Services Laboratories | Wellcome Sanger Institute for the COVID-19 Genomics UK (COG-UK) Consortium | Health Services Laboratories and Alex Alderton, Roberto Amato, Jeffrey Barrett, Sonia Goncalves, Ewan Harrison, David K. Jackson, Ian Johnston, Dominic Kwiatkowski, Cordelia Langford, John Sillitoe on behalf of the Wellcome Sanger Institute COVID-19 Surveillance Team |
| EPI_ISL_3446440, EPI_ISL_3446441 | Berkshire and Surrey Pathology Services Lighthouse Laboratory | Wellcome Sanger Institute for the COVID-19 Genomics UK (COG-UK) Consortium | Berkshire and Surrey Pathology Services Lighthouse Laboratory and Alex Alderton, Roberto Amato, Jeffrey Barrett, Sonia Goncalves, Ewan Harrison, David K. Jackson, Ian Johnston, Dominic Kwiatkowski, Cordelia Langford, John Sillitoe on behalf of the Wellcome Sanger Institute COVID-19 Surveillance Team |
| EPI_ISL_3446442 | Health Services Laboratories | Wellcome Sanger Institute for the COVID-19 Genomics UK (COG-UK) Consortium | Health Services Laboratories and Alex Alderton, Roberto Amato, Jeffrey Barrett, Sonia Goncalves, Ewan Harrison, David K. Jackson, Ian Johnston, Dominic Kwiatkowski, Cordelia Langford, John Sillitoe on behalf of the Wellcome Sanger Institute COVID-19 Surveillance Team |
| EPI_ISL_3446443 | Berkshire and Surrey Pathology Services Lighthouse Laboratory | Wellcome Sanger Institute for the COVID-19 Genomics UK (COG-UK) Consortium | Berkshire and Surrey Pathology Services Lighthouse Laboratory and Alex Alderton, Roberto Amato, Jeffrey Barrett, Sonia Goncalves, Ewan Harrison, David K. Jackson, Ian Johnston, Dominic Kwiatkowski, Cordelia Langford, John Sillitoe on behalf of the Wellcome Sanger Institute COVID-19 Surveillance Team |
| EPI_ISL_3446444 | Health Services Laboratories | Wellcome Sanger Institute for the COVID-19 Genomics UK (COG-UK) Consortium | Health Services Laboratories and Alex Alderton, Roberto Amato, Jeffrey Barrett, Sonia Goncalves, Ewan Harrison, David K. Jackson, Ian Johnston, Dominic Kwiatkowski, Cordelia Langford, John Sillitoe on behalf of the Wellcome Sanger Institute COVID-19 Surveillance Team |
| EPI_ISL_3446445 | Berkshire and Surrey Pathology Services Lighthouse Laboratory | Wellcome Sanger Institute for the COVID-19 Genomics UK (COG-UK) Consortium | Berkshire and Surrey Pathology Services Lighthouse Laboratory and Alex Alderton, Roberto Amato, Jeffrey Barrett, Sonia Goncalves, Ewan Harrison, David K. Jackson, Ian Johnston, Dominic Kwiatkowski, Cordelia Langford, John Sillitoe on behalf of the Wellcome Sanger Institute COVID-19 Surveillance Team |
| EPI_ISL_3446446, EPI_ISL_3446447 | Health Services Laboratories | Wellcome Sanger Institute for the COVID-19 Genomics UK (COG-UK) Consortium | Health Services Laboratories and Alex Alderton, Roberto Amato, Jeffrey Barrett, Sonia Goncalves, Ewan Harrison, David K. Jackson, Ian Johnston, Dominic Kwiatkowski, Cordelia Langford, John Sillitoe on behalf of the Wellcome Sanger Institute COVID-19 Surveillance Team |
| EPI_ISL_3446448, EPI_ISL_3446449 | Berkshire and Surrey Pathology Services Lighthouse Laboratory | Wellcome Sanger Institute for the COVID-19 Genomics UK (COG-UK) Consortium | Berkshire and Surrey Pathology Services Lighthouse Laboratory and Alex Alderton, Roberto Amato, Jeffrey Barrett, Sonia Goncalves, Ewan Harrison, David K. Jackson, Ian Johnston, Dominic Kwiatkowski, Cordelia Langford, John Sillitoe on behalf of the Wellcome Sanger Institute COVID-19 Surveillance Team |
| EPI_ISL_3446450, EPI_ISL_3446451, EPI_ISL_3446452 | Health Services Laboratories | Wellcome Sanger Institute for the COVID-19 Genomics UK (COG-UK) Consortium | Health Services Laboratories and Alex Alderton, Roberto Amato, Jeffrey Barrett, Sonia Goncalves, Ewan Harrison, David K. Jackson, Ian Johnston, Dominic Kwiatkowski, Cordelia Langford, John Sillitoe on behalf of the Wellcome Sanger Institute COVID-19 Surveillance Team |
| EPI_ISL_3446453 | Berkshire and Surrey Pathology Services Lighthouse Laboratory | Wellcome Sanger Institute for the COVID-19 Genomics UK (COG-UK) Consortium | Berkshire and Surrey Pathology Services Lighthouse Laboratory and Alex Alderton, Roberto Amato, Jeffrey Barrett, Sonia Goncalves, Ewan Harrison, David K. Jackson, Ian Johnston, Dominic Kwiatkowski, Cordelia Langford, John Sillitoe on behalf of the Wellcome Sanger Institute COVID-19 Surveillance Team |
| EPI_ISL_3446454, EPI_ISL_3446455, EPI_ISL_3446456, EPI_ISL_3446457 | Health Services Laboratories | Wellcome Sanger Institute for the COVID-19 Genomics UK (COG-UK) Consortium | Health Services Laboratories and Alex Alderton, Roberto Amato, Jeffrey Barrett, Sonia Goncalves, Ewan Harrison, David K. Jackson, Ian Johnston, Dominic Kwiatkowski, Cordelia Langford, John Sillitoe on behalf of the Wellcome Sanger Institute COVID-19 Surveillance Team |
| EPI_ISL_3446458, EPI_ISL_3446459, EPI_ISL_3446460, EPI_ISL_3446461 | Berkshire and Surrey Pathology Services Lighthouse Laboratory | Wellcome Sanger Institute for the COVID-19 Genomics UK (COG-UK) Consortium | Berkshire and Surrey Pathology Services Lighthouse Laboratory and Alex Alderton, Roberto Amato, Jeffrey Barrett, Sonia Goncalves, Ewan Harrison, David K. Jackson, Ian Johnston, Dominic Kwiatkowski, Cordelia Langford, John Sillitoe on behalf of the Wellcome Sanger Institute COVID-19 Surveillance Team |
| EPI_ISL_3446462, EPI_ISL_3446463 | Health Services Laboratories | Wellcome Sanger Institute for the COVID-19 Genomics UK (COG-UK) Consortium | Health Services Laboratories and Alex Alderton, Roberto Amato, Jeffrey Barrett, Sonia Goncalves, Ewan Harrison, David K. Jackson, Ian Johnston, Dominic Kwiatkowski, Cordelia Langford, John Sillitoe on behalf of the Wellcome Sanger Institute COVID-19 Surveillance Team |
| EPI_ISL_3446464, EPI_ISL_3446465 | Berkshire and Surrey Pathology Services Lighthouse Laboratory | Wellcome Sanger Institute for the COVID-19 Genomics UK (COG-UK) Consortium | Berkshire and Surrey Pathology Services Lighthouse Laboratory and Alex Alderton, Roberto Amato, Jeffrey Barrett, Sonia Goncalves, Ewan Harrison, David K. Jackson, Ian Johnston, Dominic Kwiatkowski, Cordelia Langford, John Sillitoe on behalf of the Wellcome Sanger Institute COVID-19 Surveillance Team |
| EPI_ISL_3446466, EPI_ISL_3446467, EPI_ISL_3446468 | Health Services Laboratories | Wellcome Sanger Institute for the COVID-19 Genomics UK (COG-UK) Consortium | Health Services Laboratories and Alex Alderton, Roberto Amato, Jeffrey Barrett, Sonia Goncalves, Ewan Harrison, David K. Jackson, Ian Johnston, Dominic Kwiatkowski, Cordelia Langford, John Sillitoe on behalf of the Wellcome Sanger Institute COVID-19 Surveillance Team |
| EPI_ISL_3446469 | Berkshire and Surrey Pathology Services Lighthouse Laboratory | Wellcome Sanger Institute for the COVID-19 Genomics UK (COG-UK) Consortium | Berkshire and Surrey Pathology Services Lighthouse Laboratory and Alex Alderton, Roberto Amato, Jeffrey Barrett, Sonia Goncalves, Ewan Harrison, David K. Jackson, Ian Johnston, Dominic Kwiatkowski, Cordelia Langford, John Sillitoe on behalf of the Wellcome Sanger Institute COVID-19 Surveillance Team |
| EPI_ISL_3446470, EPI_ISL_3446471, EPI_ISL_3446472, EPI_ISL_3446473, EPI_ISL_3446474, EPI_ISL_3446475, EPI_ISL_3446476, EPI_ISL_3446477, EPI_ISL_3446478, EPI_ISL_3446479, EPI_ISL_3446480 | | | |
| see above | Health Services Laboratories | Wellcome Sanger Institute for the COVID-19 Genomics UK (COG-UK) Consortium | Health Services Laboratories and Alex Alderton, Roberto Amato, Jeffrey Barrett, Sonia Goncalves, Ewan Harrison, David K. Jackson, Ian Johnston, Dominic Kwiatkowski, Cordelia Langford, John Sillitoe on behalf of the Wellcome Sanger Institute COVID-19 Surveillance Team |
| EPI_ISL_3446481 | Berkshire and Surrey Pathology Services Lighthouse Laboratory | Wellcome Sanger Institute for the COVID-19 Genomics UK (COG-UK) Consortium | Berkshire and Surrey Pathology Services Lighthouse Laboratory and Alex Alderton, Roberto Amato, Jeffrey Barrett, Sonia Goncalves, Ewan Harrison, David K. Jackson, Ian Johnston, Dominic Kwiatkowski, Cordelia Langford, John Sillitoe on behalf of the Wellcome Sanger Institute COVID-19 Surveillance Team |
| EPI_ISL_3446482, EPI_ISL_3446483, EPI_ISL_3446484 | Lighthouse Lab in Alderley Park | Wellcome Sanger Institute for the COVID-19 Genomics UK (COG-UK) Consortium | Jacquelyn Wynn, Mairead Hyland, The Lighthouse Lab in Alderley Park and Alex Alderton, Roberto Amato, Jeffrey Barrett, Sonia Goncalves, Ewan Harrison, David K. Jackson, Ian Johnston, Dominic Kwiatkowski, Cordelia Langford, John Sillitoe on behalf of the Wellcome Sanger Institute COVID-19 Surveillance Team |
| EPI_ISL_3446485 | Lighthouse Lab in Glasgow | Wellcome Sanger Institute for the COVID-19 Genomics UK (COG-UK) | Harper VanSteenhouse, Yumi Kasai, David Gray, Carol Clugston, Anna Dominiczak and Alex Alderton, Roberto Amato, Jeffrey Barrett, Sonia Goncalves, Ewan Harrison, David K. Jackson, Ian Johnston, Dominic Kwiatkowski, Cordelia Langford, |

| | | | |
|---|---|---|---|
| | | Consortium | John Sillitoe on behalf of the Wellcome Sanger Institute COVID-19 Surveillance Team |
| EPI_ISL_3446486 | Lighthouse Lab in Alderley Park | Wellcome Sanger Institute for the COVID-19 Genomics UK (COG-UK) Consortium | Jacquelyn Wynn, Mairead Hyland, The Lighthouse Lab in Alderley Park and Alex Alderton, Roberto Amato, Jeffrey Barrett, Sonia Goncalves, Ewan Harrison, David K. Jackson, Ian Johnston, Dominic Kwiatkowski, Cordelia Langford, John Sillitoe on behalf of the Wellcome Sanger Institute COVID-19 Surveillance Team |
| EPI_ISL_3446487, EPI_ISL_3446488 | Lighthouse Lab in Milton Keynes | Wellcome Sanger Institute for the COVID-19 Genomics UK (COG-UK) Consortium | The Lighthouse Lab in Milton Keynes and Alex Alderton, Roberto Amato, Jeffrey Barrett, Sonia Goncalves, Ewan Harrison, David K. Jackson, Ian Johnston, Dominic Kwiatkowski, Cordelia Langford, John Sillitoe on behalf of the Wellcome Sanger Institute COVID-19 Surveillance Team |
| EPI_ISL_3446489, EPI_ISL_3446490, EPI_ISL_3446491 | Lighthouse Lab in Alderley Park | Wellcome Sanger Institute for the COVID-19 Genomics UK (COG-UK) Consortium | Jacquelyn Wynn, Mairead Hyland, The Lighthouse Lab in Alderley Park and Alex Alderton, Roberto Amato, Jeffrey Barrett, Sonia Goncalves, Ewan Harrison, David K. Jackson, Ian Johnston, Dominic Kwiatkowski, Cordelia Langford, John Sillitoe on behalf of the Wellcome Sanger Institute COVID-19 Surveillance Team |
| EPI_ISL_3446492, EPI_ISL_3446493 | Berkshire and Surrey Pathology Services Lighthouse Laboratory | Wellcome Sanger Institute for the COVID-19 Genomics UK (COG-UK) Consortium | Berkshire and Surrey Pathology Services Lighthouse Laboratory and Alex Alderton, Roberto Amato, Jeffrey Barrett, Sonia Goncalves, Ewan Harrison, David K. Jackson, Ian Johnston, Dominic Kwiatkowski, Cordelia Langford, John Sillitoe on behalf of the Wellcome Sanger Institute COVID-19 Surveillance Team |
| EPI_ISL_3446494, EPI_ISL_3446495 | Lighthouse Lab in Milton Keynes | Wellcome Sanger Institute for the COVID-19 Genomics UK (COG-UK) Consortium | The Lighthouse Lab in Milton Keynes and Alex Alderton, Roberto Amato, Jeffrey Barrett, Sonia Goncalves, Ewan Harrison, David K. Jackson, Ian Johnston, Dominic Kwiatkowski, Cordelia Langford, John Sillitoe on behalf of the Wellcome Sanger Institute COVID-19 Surveillance Team |
| EPI_ISL_3446496 | Lighthouse Lab in Alderley Park | Wellcome Sanger Institute for the COVID-19 Genomics UK (COG-UK) Consortium | Jacquelyn Wynn, Mairead Hyland, The Lighthouse Lab in Alderley Park and Alex Alderton, Roberto Amato, Jeffrey Barrett, Sonia Goncalves, Ewan Harrison, David K. Jackson, Ian Johnston, Dominic Kwiatkowski, Cordelia Langford, John Sillitoe on behalf of the Wellcome Sanger Institute COVID-19 Surveillance Team |
| EPI_ISL_3446497, EPI_ISL_3446498, EPI_ISL_3446499 | Lighthouse Lab in Milton Keynes | Wellcome Sanger Institute for the COVID-19 Genomics UK (COG-UK) Consortium | The Lighthouse Lab in Milton Keynes and Alex Alderton, Roberto Amato, Jeffrey Barrett, Sonia Goncalves, Ewan Harrison, David K. Jackson, Ian Johnston, Dominic Kwiatkowski, Cordelia Langford, John Sillitoe on behalf of the Wellcome Sanger Institute COVID-19 Surveillance Team |
| EPI_ISL_3446500 | Berkshire and Surrey Pathology Services Lighthouse Laboratory | Wellcome Sanger Institute for the COVID-19 Genomics UK (COG-UK) Consortium | Berkshire and Surrey Pathology Services Lighthouse Laboratory and Alex Alderton, Roberto Amato, Jeffrey Barrett, Sonia Goncalves, Ewan Harrison, David K. Jackson, Ian Johnston, Dominic Kwiatkowski, Cordelia Langford, John Sillitoe on behalf of the Wellcome Sanger Institute COVID-19 Surveillance Team |
| EPI_ISL_3446501 | Lighthouse Lab in Milton Keynes | Wellcome Sanger Institute for the COVID-19 Genomics UK (COG-UK) Consortium | The Lighthouse Lab in Milton Keynes and Alex Alderton, Roberto Amato, Jeffrey Barrett, Sonia Goncalves, Ewan Harrison, David K. Jackson, Ian Johnston, Dominic Kwiatkowski, Cordelia Langford, John Sillitoe on behalf of the Wellcome Sanger Institute COVID-19 Surveillance Team |
| EPI_ISL_3446502 | Lighthouse Lab in Alderley Park | Wellcome Sanger Institute for the COVID-19 Genomics UK (COG-UK) Consortium | Jacquelyn Wynn, Mairead Hyland, The Lighthouse Lab in Alderley Park and Alex Alderton, Roberto Amato, Jeffrey Barrett, Sonia Goncalves, Ewan Harrison, David K. Jackson, Ian Johnston, Dominic Kwiatkowski, Cordelia Langford, John Sillitoe on behalf of the Wellcome Sanger Institute COVID-19 Surveillance Team |
| EPI_ISL_3446503, EPI_ISL_3446504, EPI_ISL_3446505 | Lighthouse Lab in Milton Keynes | Wellcome Sanger Institute for the COVID-19 Genomics UK (COG-UK) Consortium | The Lighthouse Lab in Milton Keynes and Alex Alderton, Roberto Amato, Jeffrey Barrett, Sonia Goncalves, Ewan Harrison, David K. Jackson, Ian Johnston, Dominic Kwiatkowski, Cordelia Langford, John Sillitoe on behalf of the Wellcome Sanger Institute COVID-19 Surveillance Team |
| EPI_ISL_3446506, EPI_ISL_3446507, EPI_ISL_3446508, EPI_ISL_3446509, EPI_ISL_3446510 | Lighthouse Lab in Alderley Park | Wellcome Sanger Institute for the COVID-19 Genomics UK (COG-UK) Consortium | Jacquelyn Wynn, Mairead Hyland, The Lighthouse Lab in Alderley Park and Alex Alderton, Roberto Amato, Jeffrey Barrett, Sonia Goncalves, Ewan Harrison, David K. Jackson, Ian Johnston, Dominic Kwiatkowski, Cordelia Langford, John Sillitoe on behalf of the Wellcome Sanger Institute COVID-19 Surveillance Team |
| EPI_ISL_3446511, EPI_ISL_3446512, EPI_ISL_3446513 | Lighthouse Lab in Glasgow | Wellcome Sanger Institute for the COVID-19 Genomics UK (COG-UK) Consortium | Harper VanSteenhouse, Yumi Kasai, David Gray, Carol Clugston, Anna Dominiczak and Alex Alderton, Roberto Amato, Jeffrey Barrett, Sonia Goncalves, Ewan Harrison, David K. Jackson, Ian Johnston, Dominic Kwiatkowski, Cordelia Langford, John Sillitoe on behalf of the Wellcome Sanger Institute COVID-19 Surveillance Team |
| EPI_ISL_3446514 | Lighthouse Lab in Alderley Park | Wellcome Sanger Institute for the COVID-19 Genomics UK (COG-UK) Consortium | Jacquelyn Wynn, Mairead Hyland, The Lighthouse Lab in Alderley Park and Alex Alderton, Roberto Amato, Jeffrey Barrett, Sonia Goncalves, Ewan Harrison, David K. Jackson, Ian Johnston, Dominic Kwiatkowski, Cordelia Langford, John Sillitoe on behalf of the Wellcome Sanger Institute COVID-19 Surveillance Team |
| EPI_ISL_3446515, EPI_ISL_3446516, EPI_ISL_3446517, EPI_ISL_3446518 | Lighthouse Lab in Glasgow | Wellcome Sanger Institute for the COVID-19 Genomics UK (COG-UK) Consortium | Harper VanSteenhouse, Yumi Kasai, David Gray, Carol Clugston, Anna Dominiczak and Alex Alderton, Roberto Amato, Jeffrey Barrett, Sonia Goncalves, Ewan Harrison, David K. Jackson, Ian Johnston, Dominic Kwiatkowski, Cordelia Langford, John Sillitoe on behalf of the Wellcome Sanger Institute COVID-19 Surveillance Team |
| EPI_ISL_3446519 | Randox Laboratories | Wellcome Sanger Institute for the COVID-19 Genomics UK (COG-UK) Consortium | Randox Laboratories and Alex Alderton, Roberto Amato, Jeffrey Barrett, Sonia Goncalves, Ewan Harrison, David K. Jackson, Ian Johnston, Dominic Kwiatkowski, Cordelia Langford, John Sillitoe on behalf of the Wellcome Sanger Institute COVID-19 Surveillance Team |
| EPI_ISL_3446520 | Lighthouse Laboratory Plymouth | Wellcome Sanger Institute for the COVID-19 Genomics UK (COG-UK) Consortium | Lighthouse Laboratory Plymouth and Alex Alderton, Roberto Amato, Jeffrey Barrett, Sonia Goncalves, Ewan Harrison, David K. Jackson, Ian Johnston, Dominic Kwiatkowski, Cordelia Langford, John Sillitoe on behalf of the Wellcome Sanger Institute COVID-19 Surveillance Team |
| EPI_ISL_3446521 | Berkshire and Surrey Pathology Services Lighthouse Laboratory | Wellcome Sanger Institute for the COVID-19 Genomics UK (COG-UK) Consortium | Berkshire and Surrey Pathology Services Lighthouse Laboratory and Alex Alderton, Roberto Amato, Jeffrey Barrett, Sonia Goncalves, Ewan Harrison, David K. Jackson, Ian Johnston, Dominic Kwiatkowski, Cordelia Langford, John Sillitoe on behalf of the Wellcome Sanger Institute COVID-19 Surveillance Team |
| EPI_ISL_3446522 | Lighthouse Lab in Alderley Park | Wellcome Sanger Institute for the COVID-19 Genomics UK (COG-UK) Consortium | Jacquelyn Wynn, Mairead Hyland, The Lighthouse Lab in Alderley Park and Alex Alderton, Roberto Amato, Jeffrey Barrett, Sonia Goncalves, Ewan Harrison, David K. Jackson, Ian Johnston, Dominic Kwiatkowski, Cordelia Langford, John Sillitoe on behalf of the Wellcome Sanger Institute COVID-19 Surveillance Team |
| EPI_ISL_3446523, EPI_ISL_3446524, EPI_ISL_3446525, EPI_ISL_3446526, EPI_ISL_3446527, EPI_ISL_3446528, EPI_ISL_3446529 | Lighthouse Lab in Glasgow | Wellcome Sanger Institute for the COVID-19 Genomics UK (COG-UK) Consortium | Harper VanSteenhouse, Yumi Kasai, David Gray, Carol Clugston, Anna Dominiczak and Alex Alderton, Roberto Amato, Jeffrey Barrett, Sonia Goncalves, Ewan Harrison, David K. Jackson, Ian Johnston, Dominic Kwiatkowski, Cordelia Langford, John Sillitoe on behalf of the Wellcome Sanger Institute COVID-19 Surveillance Team |
| EPI_ISL_3446546, EPI_ISL_3446547, EPI_ISL_3446548, EPI_ISL_3446549, EPI_ISL_3446550, EPI_ISL_3446551, EPI_ISL_3446552, EPI_ISL_3446553, EPI_ISL_3446554, EPI_ISL_3446555, EPI_ISL_3446556, EPI_ISL_3446557, EPI_ISL_3446558, EPI_ISL_3446559, EPI_ISL_3446560, EPI_ISL_3446561, EPI_ISL_3446562, EPI_ISL_3446563, EPI_ISL_3446564, EPI_ISL_3446565, EPI_ISL_3446566, EPI_ISL_3446567, EPI_ISL_3446568, EPI_ISL_3446569, EPI_ISL_3446570, EPI_ISL_3446571, EPI_ISL_3446572, EPI_ISL_3446573, EPI_ISL_3446574, EPI_ISL_3446575, EPI_ISL_3446576, EPI_ISL_3446577, EPI_ISL_3446579, EPI_ISL_3446580, EPI_ISL_3446581, EPI_ISL_3446582, EPI_ISL_3446583, EPI_ISL_3446584, EPI_ISL_3446586, EPI_ISL_3446587, EPI_ISL_3446588, EPI_ISL_3446589, EPI_ISL_3446590, EPI_ISL_3446591 | | | |
| see above | Michigan Department of Health and Human Services, Bureau of Laboratories | Michigan Department of Health and Human Services, Bureau of Laboratories | Blankenship HM, Riner D, Soehnlen MK |
| EPI_ISL_3446592, EPI_ISL_3446593 | Hospital Universitari Joan XXIII | Hospital Universitari Joan XXIII | Gemma Recio, Cristina Gutiérrez, Clara Benavent, Carla Martín, Margarida Terrón, Ester Picó, Natalia Bastón |
| EPI_ISL_3446597, EPI_ISL_3446598, EPI_ISL_3446599 | St. Anne´s University Hospital Brno | University Hospital Brno, CMBG | Bezdicek Matej, Svaton Jan, Dufkova Kristyna, Lengerova Martina, Volfova Pavlina |
| EPI_ISL_3446600, EPI_ISL_3446601 | SYNLAB | University Hospital Brno, CMBG | Bezdicek Matej, Svaton Jan, Dufkova Kristyna, Lengerova Martina, Volfova Pavlina |
| EPI_ISL_3446602, EPI_ISL_3446603, EPI_ISL_3446604 | St. Anne´s University Hospital Brno | University Hospital Brno, CMBG | Bezdicek Matej, Svaton Jan, Dufkova Kristyna, Lengerova Martina, Volfova Pavlina |
| EPI_ISL_3446605 | Nemocnice Breclav | University Hospital Brno, CMBG | Bezdicek Matej, Svaton Jan, Dufkova Kristyna, Lengerova Martina, Volfova Pavlina |
| EPI_ISL_3446606 | IFCOR Brno | University Hospital Brno, CMBG | Bezdicek Matej, Svaton Jan, Dufkova Kristyna, Lengerova Martina, Volfova Pavlina |
| EPI_ISL_3446607, EPI_ISL_3446608, EPI_ISL_3446609, EPI_ISL_3446610, EPI_ISL_3446611, EPI_ISL_3446612, EPI_ISL_3446613 | St. Anne´s University Hospital Brno | University Hospital Brno, CMBG | Bezdicek Matej, Svaton Jan, Dufkova Kristyna, Lengerova Martina, Volfova Pavlina |
| EPI_ISL_3446614 | Nemocnice Vyskov | University Hospital Brno, CMBG | Bezdicek Matej, Svaton Jan, Dufkova Kristyna, Lengerova Martina, Volfova Pavlina |
| EPI_ISL_3446615 | Nemocnice TGM Hodonin | University Hospital Brno, CMBG | Bezdicek Matej, Svaton Jan, Dufkova Kristyna, Lengerova Martina, Volfova Pavlina |
| EPI_ISL_3446616 | SYNLAB | University Hospital Brno, CMBG | Bezdicek Matej, Svaton Jan, Dufkova Kristyna, Lengerova Martina, Volfova Pavlina |
| EPI_ISL_3446617 | IFCOR Brno | University Hospital Brno, CMBG | Bezdicek Matej, Svaton Jan, Dufkova Kristyna, Lengerova Martina, Volfova Pavlina |
| EPI_ISL_3446618, EPI_ISL_3446619 | St. Anne´s University Hospital Brno | University Hospital Brno, CMBG | Bezdicek Matej, Svaton Jan, Dufkova Kristyna, Lengerova Martina, Volfova Pavlina |
| EPI_ISL_3446620 | Nemocnice Vyskov | University Hospital Brno, CMBG | Bezdicek Matej, Svaton Jan, Dufkova Kristyna, Lengerova Martina, Volfova Pavlina |
| EPI_ISL_3446621, EPI_ISL_3446622 | SYNLAB | University Hospital Brno, CMBG | Bezdicek Matej, Svaton Jan, Dufkova Kristyna, Lengerova Martina, Volfova Pavlina |
| EPI_ISL_3446623 | Nemocnice Havlickuv Brod | University Hospital Brno, CMBG | Bezdicek Matej, Svaton Jan, Dufkova Kristyna, Lengerova Martina, Volfova Pavlina |
| EPI_ISL_3446624, EPI_ISL_3446625, EPI_ISL_3446626 | Nemocnice Jihlava | University Hospital Brno, CMBG | Bezdicek Matej, Svaton Jan, Dufkova Kristyna, Lengerova Martina, Volfova Pavlina |
| EPI_ISL_3446627, | Nemocnice Havlickuv Brod | University Hospital Brno, CMBG | Bezdicek Matej, Svaton Jan, Dufkova Kristyna, Lengerova Martina, Volfova Pavlina |

| | | | |
|---|---|---|---|
| EPI_ISL_3446628 | | | |
| EPI_ISL_3446629, EPI_ISL_3446630, EPI_ISL_3446631, EPI_ISL_3446632, EPI_ISL_3446633, EPI_ISL_3446634 | Nemocnice Jihlava | University Hospital Brno, CMBG | Bezdicek Matej, Svaton Jan, Dufkova Kristyna, Lengerova Martina, Volfova Pavlina |
| EPI_ISL_3446635 | Nemocnice Trebic | University Hospital Brno, CMBG | Bezdicek Matej, Svaton Jan, Dufkova Kristyna, Lengerova Martina, Volfova Pavlina |
| EPI_ISL_3446636 | Cytogeneticka laborator Brno | University Hospital Brno, CMBG | Bezdicek Matej, Svaton Jan, Dufkova Kristyna, Lengerova Martina, Volfova Pavlina |
| EPI_ISL_3446637, EPI_ISL_3446638, EPI_ISL_3446639, EPI_ISL_3446640, EPI_ISL_3446641 | Nemocnice Jihlava | University Hospital Brno, CMBG | Bezdicek Matej, Svaton Jan, Dufkova Kristyna, Lengerova Martina, Volfova Pavlina |
| EPI_ISL_3446642 | Nemocnice Havlickuv Brod | University Hospital Brno, CMBG | Bezdicek Matej, Svaton Jan, Dufkova Kristyna, Lengerova Martina, Volfova Pavlina |
| EPI_ISL_3446643 | Cytogeneticka laborator Brno | University Hospital Brno, CMBG | Bezdicek Matej, Svaton Jan, Dufkova Kristyna, Lengerova Martina, Volfova Pavlina |
| EPI_ISL_3446644 | Nemocnice Jihlava | University Hospital Brno, CMBG | Bezdicek Matej, Svaton Jan, Dufkova Kristyna, Lengerova Martina, Volfova Pavlina |
| EPI_ISL_3446645 | IFCOR Jihlava | University Hospital Brno, CMBG | Bezdicek Matej, Svaton Jan, Dufkova Kristyna, Lengerova Martina, Volfova Pavlina |
| EPI_ISL_3446646 | Nemocnice Havlickuv Brod | University Hospital Brno, CMBG | Bezdicek Matej, Svaton Jan, Dufkova Kristyna, Lengerova Martina, Volfova Pavlina |
| EPI_ISL_3446647, EPI_ISL_3446648, EPI_ISL_3446649, EPI_ISL_3446650, EPI_ISL_3446651 | Nemocnice Trebic | University Hospital Brno, CMBG | Bezdicek Matej, Svaton Jan, Dufkova Kristyna, Lengerova Martina, Volfova Pavlina |
| EPI_ISL_3446652 | IFCOR Jihlava | University Hospital Brno, CMBG | Bezdicek Matej, Svaton Jan, Dufkova Kristyna, Lengerova Martina, Volfova Pavlina |
| EPI_ISL_3446653 | Nemocnice Jihlava | University Hospital Brno, CMBG | Bezdicek Matej, Svaton Jan, Dufkova Kristyna, Lengerova Martina, Volfova Pavlina |
| EPI_ISL_3446654 | Hola Biolabs | University Hospital Brno, CMBG | Bezdicek Matej, Svaton Jan, Dufkova Kristyna, Lengerova Martina, Volfova Pavlina |
| EPI_ISL_3446655 | Dr. Yoshitaka Tamura Department of Clinical Laboratory, Osaka Habikino Medical Center | Department of Developmental Medicine, Research Institute, Osaka Women's and Children's Hospital | Itaru Yanagihara, Yukiko Nakura |
| EPI_ISL_3446656 | Port Klang Health Clinic | Institute for Medical Research, Infectious Disease Research Centre, National Institutes of Health, Ministry of Health Malaysia | Suppiah J, Kamel K, Mohd Zawawi Z, Azizan MA, Ramly N, Robert F, Anasir MI, Thayan R |
| EPI_ISL_3446657 | Kuala Kubu Bharu Hospital | Institute for Medical Research, Infectious Disease Research Centre, National Institutes of Health, Ministry of Health Malaysia | Suppiah J, Kamel K, Mohd Zawawi Z, Azizan MA, Ramly N, Robert F, Anasir MI, Thayan R |
| EPI_ISL_3446658 | Kuala kubu Bharu Hospital | Institute for Medical Research, Infectious Disease Research Centre, National Institutes of Health, Ministry of Health Malaysia | Suppiah J, Kamel K, Mohd Zawawi Z, Azizan MA, Ramly N, Robert F, Anasir MI, Thayan R |
| EPI_ISL_3446659, EPI_ISL_3446660, EPI_ISL_3446661 | Tengku Ampuan Rahimah Hospital | Institute for Medical Research, Infectious Disease Research Centre, National Institutes of Health, Ministry of Health Malaysia | Suppiah J, Kamel K, Mohd Zawawi Z, Azizan MA, Ramly N, Robert F, Anasir MI, Thayan R |
| EPI_ISL_3446663, EPI_ISL_3446664 | Kampung Bandar Health Clinic | Institute for Medical Research, Infectious Disease Research Centre, National Institutes of Health, Ministry of Health Malaysia | Suppiah J, Kamel K, Mohd Zawawi Z, Azizan MA, Ramly N, Robert F, Anasir MI, Thayan R |
| EPI_ISL_3446665, EPI_ISL_3446666, EPI_ISL_3446668 | Batu Health Clinic | Institute for Medical Research, Infectious Disease Research Centre, National Institutes of Health, Ministry of Health Malaysia | Suppiah J, Kamel K, Mohd Zawawi Z, Azizan MA, Ramly N, Robert F, Anasir MI, Thayan R |
| EPI_ISL_3446671, EPI_ISL_3446673 | Bukit Changgang Health Clinic | Institute for Medical Research, Infectious Disease Research Centre, National Institutes of Health, Ministry of Health Malaysia | Suppiah J, Kamel K, Mohd Zawawi Z, Azizan MA, Ramly N, Robert F, Anasir MI, Thayan R |
| EPI_ISL_3446674 | Bukit Kuda Heath Clinic | Institute for Medical Research, Infectious Disease Research Centre, National Institutes of Health, Ministry of Health Malaysia | Suppiah J, Kamel K, Mohd Zawawi Z, Azizan MA, Ramly N, Robert F, Anasir MI, Thayan R |
| EPI_ISL_3446676 | Bukit Kuda Health Clinic | Institute for Medical Research, Infectious Disease Research Centre, National Institutes of Health, Ministry of Health Malaysia | Suppiah J, Kamel K, Mohd Zawawi Z, Azizan MA, Ramly N, Robert F, Anasir MI, Thayan R |
| EPI_ISL_3446678 | Dengkil Health Clinic | Institute for Medical Research, Infectious Disease Research Centre, National Institutes of Health, Ministry of Health Malaysia | Suppiah J, Kamel K, Mohd Zawawi Z, Azizan MA, Ramly N, Robert F, Anasir MI, Thayan R |
| EPI_ISL_3446679 | Jenjarom Health Clinic | Institute for Medical Research, Infectious Disease Research Centre, National Institutes of Health, Ministry of Health Malaysia | Suppiah J, Kamel K, Mohd Zawawi Z, Azizan MA, Ramly N, Robert F, Anasir MI, Thayan R |
| EPI_ISL_3446681, EPI_ISL_3446683 | Jeram Health Clinic | Institute for Medical Research, Infectious Disease Research Centre, National Institutes of Health, Ministry of Health Malaysia | Suppiah J, Kamel K, Mohd Zawawi Z, Azizan MA, Ramly N, Robert F, Anasir MI, Thayan R |
| EPI_ISL_3446686, EPI_ISL_3446688 | Kebun Baru Health Clinic | Institute for Medical Research, Infectious Disease Research Centre, National Institutes of Health, Ministry of Health Malaysia | Suppiah J, Kamel K, Mohd Zawawi Z, Azizan MA, Ramly N, Robert F, Anasir MI, Thayan R |
| EPI_ISL_3446692, EPI_ISL_3446695 | Kuala Selangor Health Clinic | Institute for Medical Research, Infectious Disease Research Centre, National Institutes of Health, Ministry of Health Malaysia | Suppiah J, Kamel K, Mohd Zawawi Z, Azizan MA, Ramly N, Robert F, Anasir MI, Thayan R |
| EPI_ISL_3446696, EPI_ISL_3446697 | Pandamaran Health Clinic | Institute for Medical Research, Infectious Disease Research Centre, National Institutes of Health, Ministry of Health Malaysia | Suppiah J, Kamel K, Mohd Zawawi Z, Azizan MA, Ramly N, Robert F, Anasir MI, Thayan R |
| EPI_ISL_3446702 | Pulau Ketam Health Clinic | Institute for Medical Research, Infectious Disease Research Centre, National Institutes of Health, Ministry of Health Malaysia | Suppiah J, Kamel K, Mohd Zawawi Z, Azizan MA, Ramly N, Robert F, Anasir MI, Thayan R |
| EPI_ISL_3446703 | Sijangkang Health Clinic | Institute for Medical Research, Infectious Disease Research Centre, National Institutes of Health, Ministry of Health Malaysia | Suppiah J, Kamel K, Mohd Zawawi Z, Azizan MA, Ramly N, Robert F, Anasir MI, Thayan R |
| EPI_ISL_3446704, EPI_ISL_3446706 | Tanjung Sepat Health Clinic | Institute for Medical Research, Infectious Disease Research Centre, National Institutes of Health, Ministry of Health Malaysia | Suppiah J, Kamel K, Mohd Zawawi Z, Azizan MA, Ramly N, Robert F, Anasir MI, Thayan R |
| EPI_ISL_3446709, EPI_ISL_3446712 | Teluk Datok Health Clinic | Institute for Medical Research, Infectious Disease Research Centre, National Institutes of Health, Ministry of Health Malaysia | Suppiah J, Kamel K, Mohd Zawawi Z, Azizan MA, Ramly N, Robert F, Anasir MI, Thayan R |
| EPI_ISL_3446714 | Teluk Panglima Garang Health Clinic | Institute for Medical Research, Infectious Disease Research Centre, National Institutes of Health, Ministry of Health Malaysia | Suppiah J, Kamel K, Mohd Zawawi Z, Azizan MA, Ramly N, Robert F, Anasir MI, Thayan R |
| EPI_ISL_3446716 | Telok Panglima Garang Health Clinic | Institute for Medical Research, Infectious Disease Research Centre, National Institutes of Health, Ministry of Health Malaysia | Suppiah J, Kamel K, Mohd Zawawi Z, Azizan MA, Ramly N, Robert F, Anasir MI, Thayan R |
| EPI_ISL_3446717, EPI_ISL_3446719, EPI_ISL_3446726 | Kuala Langat District Health Office | Institute for Medical Research, Infectious Disease Research Centre, National Institutes of Health, Ministry of Health Malaysia | Suppiah J, Kamel K, Mohd Zawawi Z, Azizan MA, Ramly N, Robert F, Anasir MI, Thayan R |
| EPI_ISL_3446727 | Laboratorio Nacional de Salud | Asociación de Salud Integral/Clínica Familiar "Luis Ángel García" | Eduardo Arathoon, Ana S. Gonzalez-Reiche, Claudia Rangel, Luis Rivas, Oscar Bonilla, Narda Medina, Luis Aguirre, Danicela Mercado, Hilda Ruiz, Osmar Gamboa. |
| EPI_ISL_3446728 | Kuala Langat District Health Office | Institute for Medical Research, Infectious Disease Research Centre, National Institutes of Health, Ministry of Health Malaysia | Suppiah J, Kamel K, Mohd Zawawi Z, Azizan MA, Ramly N, Robert F, Anasir MI, Thayan R |
| EPI_ISL_3446730, EPI_ISL_3446731, EPI_ISL_3446732, EPI_ISL_3446733, EPI_ISL_3446734, EPI_ISL_3446735, EPI_ISL_3446736, EPI_ISL_3446737, EPI_ISL_3446738, EPI_ISL_3446739, EPI_ISL_3446740, EPI_ISL_3446741, EPI_ISL_3446742, EPI_ISL_3446743, EPI_ISL_3446744, EPI_ISL_3446745, EPI_ISL_3446746, EPI_ISL_3446747, EPI_ISL_3446748, EPI_ISL_3446749, EPI_ISL_3446750, EPI_ISL_3446751, EPI_ISL_3446752, EPI_ISL_3446753, EPI_ISL_3446754, EPI_ISL_3446755, EPI_ISL_3446756, EPI_ISL_3446757, EPI_ISL_3446758, EPI_ISL_3446759, EPI_ISL_3446760, EPI_ISL_3446761, EPI_ISL_3446762, EPI_ISL_3446763, EPI_ISL_3446764, EPI_ISL_3446765, EPI_ISL_3446766 | | | |
| see above | Fondazione IRCCS Ca' Granda Ospedale Maggiore Policlinico | Fondazione IRCCS Ca' Granda Ospedale Maggiore Policlinico | Ferruccio Ceriotti, Sara Uceda Renteria |
| EPI_ISL_3446768, EPI_ISL_3446769 | Biological Hazard and Health Research Laboratory, Centre for Advanced Research in Science (CARS), University of Dhaka | Genomic Research Lab, BCSIR | Md. Murshed Hasan Sarkar, Md. Ahasan Habib, Shahina Akter, Tanjina Akhter Banu, Barna Goswami, Iffat Jahan, Mohammad Mohi Uddin,Abu Sayeed Mohammad Mahmud, Mohammad Samir Uzzaman, Eshrar Osman, Md. Kamrul Islam, Md. Shaminur Rahman, Latiful Bari, Md Abdul Malek, Md Mizanur Rahaman, Md. Salim Khan |
| EPI_ISL_3446770, EPI_ISL_3446771 | IstitutoAuxologicoItaliano | Istituto Auxologico Italiano | Erminio Torresani |
| EPI_ISL_3446772 | Istituto Auxologico Italiano | Istituto Auxologico Italiano | Erminio Torresani |
| EPI_ISL_3446773, EPI_ISL_3446774, EPI_ISL_3446775, EPI_ISL_3446776, EPI_ISL_3446777, EPI_ISL_3446778, EPI_ISL_3446779, EPI_ISL_3446780, EPI_ISL_3446781, EPI_ISL_3446782, EPI_ISL_3446783 | | | |
| see above | IstitutoAuxologicoItaliano | Istituto Auxologico Italiano | Erminio Torresani |
| EPI_ISL_3446784, | Biological Hazard and Health Research Laboratory, Centre for | Genomic Research Lab, BCSIR | Md. Murshed Hasan Sarkar, Md. Ahasan Habib, Shahina Akter, Tanjina Akhter Banu, Barna Goswami, Iffat Jahan, Mohammad Mohi Uddin,Abu Sayeed Mohammad Mahmud, Mohammad Samir Uzzaman, Eshrar Osman, Md. Kamrul Islam, Md. |

| | | | |
|---|---|---|---|
| EPI_ISL_3446794 | Advanced Research in Science (CARS), University of Dhaka | | Shaminur Rahman, Latiful Bari, Md Abdul Malek, Md Mizanur Rahaman, Md. Salim Khan |
| EPI_ISL_3446795 | AULSS 9 Scaligera - Villafranca | UOSD Genetica e Citogenetica - Azienda ULSS 3 Serenissima; Istituto Zooprofilattico Sperimentale delle Venezie | Mosé Favarato, Elisa Squarcina, Laura Squarzon, Claudia Perini, Laura Bevilacqua, Luca Sorino, Adelaide Milani, Alessia Schivo, Annalisa Salviato, Elisa Palumbo, Erika Giorgia Quaranta, Luca Tassoni, Ambra Pastori, Edoardo Giussani, Alice Fusaro, Isabella Monne, Calogero Terregino, Antonia Ricci |
| EPI_ISL_3446796, EPI_ISL_3446797, EPI_ISL_3446798, EPI_ISL_3446800, EPI_ISL_3446801, EPI_ISL_3446802, EPI_ISL_3446803, EPI_ISL_3446804, EPI_ISL_3446805, EPI_ISL_3446806, EPI_ISL_3446807, EPI_ISL_3446808, EPI_ISL_3446809, EPI_ISL_3446810, EPI_ISL_3446811, EPI_ISL_3446812, EPI_ISL_3446813, EPI_ISL_3446814 | | | |
| see above | AULSS 3 Venezia | UOSD Genetica e Citogenetica - Azienda ULSS 3 Serenissima; Istituto Zooprofilattico Sperimentale delle Venezie | Mosé Favarato, Elisa Squarcina, Laura Squarzon, Claudia Perini, Laura Bevilacqua, Luca Sorino, Adelaide Milani, Alessia Schivo, Annalisa Salviato, Elisa Palumbo, Erika Giorgia Quaranta, Luca Tassoni, Ambra Pastori, Edoardo Giussani, Alice Fusaro, Isabella Monne, Calogero Terregino, Antonia Ricci |
| EPI_ISL_3446815, EPI_ISL_3446816, EPI_ISL_3446817, EPI_ISL_3446818, EPI_ISL_3446819, EPI_ISL_3446820, EPI_ISL_3446821, EPI_ISL_3446822, EPI_ISL_3446823, EPI_ISL_3446824, EPI_ISL_3446825 | | | |
| see above | Area of Virology, Serology and Virology Division (SAViD), New South Wales Health Pathology Randwick | Virology Research Laboratory; Area of Virology, Serology and Virology Division (SAViD), New South Wales Health Pathology Randwick | Foster, C.; Au, J.; Ruiz Silva, M.; Deveson, I.; Bull, R.; Van Hal, S.; Rawlinson, W. |
| EPI_ISL_3446900, EPI_ISL_3446901 | RM3/IRCCS INMI Lazzaro Spallanzani | National Institute for Infectious Diseases (INMI) L. Spallanzani I.R.C.C.S | E Giombini, B Bartolini, M Rueca, G Bonfiglio, O Butera, CEM Gruber, F Santini, F Messina, A Di Caro, MR Capobianchi |
| EPI_ISL_3446902, EPI_ISL_3446903 | RI/Osp. S. Camillo De Lellis | National Institute for Infectious Diseases (INMI) L. Spallanzani I.R.C.C.S | E Giombini, B Bartolini, M Rueca, G Bonfiglio, O Butera, CEM Gruber, F Santini, F Messina, A Di Caro, MR Capobianchi |
| EPI_ISL_3446904 | FR/Osp. F.Spaziani | National Institute for Infectious Diseases (INMI) L. Spallanzani I.R.C.C.S | E Giombini, B Bartolini, M Rueca, G Bonfiglio, O Butera, CEM Gruber, F Santini, F Messina, A Di Caro, MR Capobianchi |
| EPI_ISL_3446905 | RM1/Osp.C. S.Pietro Fatebenefratelli | National Institute for Infectious Diseases (INMI) L. Spallanzani I.R.C.C.S | E Giombini, B Bartolini, M Rueca, G Bonfiglio, O Butera, CEM Gruber, F Santini, F Messina, A Di Caro, MR Capobianchi |
| EPI_ISL_3446906 | RM3/IFO | National Institute for Infectious Diseases (INMI) L. Spallanzani I.R.C.C.S | E Giombini, B Bartolini, M Rueca, G Bonfiglio, O Butera, CEM Gruber, F Santini, F Messina, A Di Caro, MR Capobianchi |
| EPI_ISL_3446907 | RM3/IRCCS INMI Lazzaro Spallanzani | National Institute for Infectious Diseases (INMI) L. Spallanzani I.R.C.C.S | E Giombini, B Bartolini, M Rueca, G Bonfiglio, O Butera, CEM Gruber, F Santini, F Messina, A Di Caro, MR Capobianchi |
| EPI_ISL_3446908 | RM3/A.O. S.Camillo | National Institute for Infectious Diseases (INMI) L. Spallanzani I.R.C.C.S | E Giombini, B Bartolini, M Rueca, G Bonfiglio, O Butera, CEM Gruber, F Santini, F Messina, A Di Caro, MR Capobianchi |
| EPI_ISL_3446909 | RM1/Osp.C. S.Pietro Fatebenefratelli | National Institute for Infectious Diseases (INMI) L. Spallanzani I.R.C.C.S | E Giombini, B Bartolini, M Rueca, G Bonfiglio, O Butera, CEM Gruber, F Santini, F Messina, A Di Caro, MR Capobianchi |
| EPI_ISL_3446910 | RM3/A.O. S.Camillo | National Institute for Infectious Diseases (INMI) L. Spallanzani I.R.C.C.S | E Giombini, B Bartolini, M Rueca, G Bonfiglio, O Butera, CEM Gruber, F Santini, F Messina, A Di Caro, MR Capobianchi |
| EPI_ISL_3446911, EPI_ISL_3446912 | FR/Osp. F.Spaziani | National Institute for Infectious Diseases (INMI) L. Spallanzani I.R.C.C.S | E Giombini, B Bartolini, M Rueca, G Bonfiglio, O Butera, CEM Gruber, F Santini, F Messina, A Di Caro, MR Capobianchi |
| EPI_ISL_3446913 | RM3/IRCCS INMI Lazzaro Spallanzani | National Institute for Infectious Diseases (INMI) L. Spallanzani I.R.C.C.S | E Giombini, B Bartolini, M Rueca, G Bonfiglio, O Butera, CEM Gruber, F Santini, F Messina, A Di Caro, MR Capobianchi |
| EPI_ISL_3446914 | RM1/Osp.C. S.Pietro Fatebenefratelli | National Institute for Infectious Diseases (INMI) L. Spallanzani I.R.C.C.S | E Giombini, B Bartolini, M Rueca, G Bonfiglio, O Butera, CEM Gruber, F Santini, F Messina, A Di Caro, MR Capobianchi |
| EPI_ISL_3446915 | RM2/IRCCS S.Lucia | National Institute for Infectious Diseases (INMI) L. Spallanzani I.R.C.C.S | E Giombini, B Bartolini, M Rueca, G Bonfiglio, O Butera, CEM Gruber, F Santini, F Messina, A Di Caro, MR Capobianchi |
| EPI_ISL_3446916 | RM3/A.O. S.Camillo | National Institute for Infectious Diseases (INMI) L. Spallanzani I.R.C.C.S | E Giombini, B Bartolini, M Rueca, G Bonfiglio, O Butera, CEM Gruber, F Santini, F Messina, A Di Caro, MR Capobianchi |
| EPI_ISL_3446917 | RM1/A.U. S.Andrea | National Institute for Infectious Diseases (INMI) L. Spallanzani I.R.C.C.S | E Giombini, B Bartolini, M Rueca, G Bonfiglio, O Butera, CEM Gruber, F Santini, F Messina, A Di Caro, MR Capobianchi |
| EPI_ISL_3446919 | RM3/A.O. S.Camillo | National Institute for Infectious Diseases (INMI) L. Spallanzani I.R.C.C.S | E Giombini, B Bartolini, M Rueca, G Bonfiglio, O Butera, CEM Gruber, F Santini, F Messina, A Di Caro, MR Capobianchi |
| EPI_ISL_3446920, EPI_ISL_3446923 | RM1/Osp. S.Filippo Neri | National Institute for Infectious Diseases (INMI) L. Spallanzani I.R.C.C.S | E Giombini, B Bartolini, M Rueca, G Bonfiglio, O Butera, CEM Gruber, F Santini, F Messina, A Di Caro, MR Capobianchi |
| EPI_ISL_3446925 | RI/Osp. S. Camillo De Lellis | National Institute for Infectious Diseases (INMI) L. Spallanzani I.R.C.C.S | E Giombini, B Bartolini, M Rueca, G Bonfiglio, O Butera, CEM Gruber, F Santini, F Messina, A Di Caro, MR Capobianchi |
| EPI_ISL_3446926 | FR/Osp. F.Spaziani | National Institute for Infectious Diseases (INMI) L. Spallanzani I.R.C.C.S | E Giombini, B Bartolini, M Rueca, G Bonfiglio, O Butera, CEM Gruber, F Santini, F Messina, A Di Caro, MR Capobianchi |
| EPI_ISL_3446927 | RM1/Osp. S.Filippo Neri | National Institute for Infectious Diseases (INMI) L. Spallanzani I.R.C.C.S | E Giombini, B Bartolini, M Rueca, G Bonfiglio, O Butera, CEM Gruber, F Santini, F Messina, A Di Caro, MR Capobianchi |
| EPI_ISL_3446928 | RI/Osp. S. Camillo De Lellis | National Institute for Infectious Diseases (INMI) L. Spallanzani I.R.C.C.S | E Giombini, B Bartolini, M Rueca, G Bonfiglio, O Butera, CEM Gruber, F Santini, F Messina, A Di Caro, MR Capobianchi |
| EPI_ISL_3446929 | RM3/IRCCS INMI Lazzaro Spallanzani | National Institute for Infectious Diseases (INMI) L. Spallanzani I.R.C.C.S | E Giombini, B Bartolini, M Rueca, G Bonfiglio, O Butera, CEM Gruber, F Santini, F Messina, A Di Caro, MR Capobianchi |
| EPI_ISL_3446930 | RM1/Osp. S.Filippo Neri | National Institute for Infectious Diseases (INMI) L. Spallanzani I.R.C.C.S | E Giombini, B Bartolini, M Rueca, G Bonfiglio, O Butera, CEM Gruber, F Santini, F Messina, A Di Caro, MR Capobianchi |
| EPI_ISL_3446931 | RM3/IRCCS INMI Lazzaro Spallanzani | National Institute for Infectious Diseases (INMI) L. Spallanzani I.R.C.C.S | E Giombini, B Bartolini, M Rueca, G Bonfiglio, O Butera, CEM Gruber, F Santini, F Messina, A Di Caro, MR Capobianchi |
| EPI_ISL_3446932 | RM3/A.O. S.Camillo | National Institute for Infectious Diseases (INMI) L. Spallanzani I.R.C.C.S | E Giombini, B Bartolini, M Rueca, G Bonfiglio, O Butera, CEM Gruber, F Santini, F Messina, A Di Caro, MR Capobianchi |
| EPI_ISL_3446933, EPI_ISL_3446934 | RM1/Osp. S.Filippo Neri | National Institute for Infectious Diseases (INMI) L. Spallanzani I.R.C.C.S | E Giombini, B Bartolini, M Rueca, G Bonfiglio, O Butera, CEM Gruber, F Santini, F Messina, A Di Caro, MR Capobianchi |
| EPI_ISL_3446935 | RM3/IFO | National Institute for Infectious Diseases (INMI) L. Spallanzani I.R.C.C.S | E Giombini, B Bartolini, M Rueca, G Bonfiglio, O Butera, CEM Gruber, F Santini, F Messina, A Di Caro, MR Capobianchi |
| EPI_ISL_3446936, EPI_ISL_3446937, EPI_ISL_3446938 | RM5/Osp. S.Giovanni Evangelista | National Institute for Infectious Diseases (INMI) L. Spallanzani I.R.C.C.S | E Giombini, B Bartolini, M Rueca, G Bonfiglio, O Butera, CEM Gruber, F Santini, F Messina, A Di Caro, MR Capobianchi |
| EPI_ISL_3446939 | RM3/IFO | National Institute for Infectious Diseases (INMI) L. Spallanzani I.R.C.C.S | E Giombini, B Bartolini, M Rueca, G Bonfiglio, O Butera, CEM Gruber, F Santini, F Messina, A Di Caro, MR Capobianchi |
| EPI_ISL_3446940, EPI_ISL_3446941 | RM5/Osp. S.Giovanni Evangelista | National Institute for Infectious Diseases (INMI) L. Spallanzani I.R.C.C.S | E Giombini, B Bartolini, M Rueca, G Bonfiglio, O Butera, CEM Gruber, F Santini, F Messina, A Di Caro, MR Capobianchi |
| EPI_ISL_3446942, EPI_ISL_3446943, EPI_ISL_3446944, EPI_ISL_3446945 | RM3/IFO | National Institute for Infectious Diseases (INMI) L. Spallanzani I.R.C.C.S | E Giombini, B Bartolini, M Rueca, G Bonfiglio, O Butera, CEM Gruber, F Santini, F Messina, A Di Caro, MR Capobianchi |
| EPI_ISL_3446946 | RM1/Osp. S.Filippo Neri | National Institute for Infectious Diseases (INMI) L. Spallanzani I.R.C.C.S | E Giombini, B Bartolini, M Rueca, G Bonfiglio, O Butera, CEM Gruber, F Santini, F Messina, A Di Caro, MR Capobianchi |
| EPI_ISL_3446947, EPI_ISL_3446948 | RM3/IFO | National Institute for Infectious Diseases (INMI) L. Spallanzani I.R.C.C.S | E Giombini, B Bartolini, M Rueca, G Bonfiglio, O Butera, CEM Gruber, F Santini, F Messina, A Di Caro, MR Capobianchi |
| EPI_ISL_3446949, EPI_ISL_3446950 | RM5/Osp. S.Giovanni Evangelista | National Institute for Infectious Diseases (INMI) L. Spallanzani I.R.C.C.S | E Giombini, B Bartolini, M Rueca, G Bonfiglio, O Butera, CEM Gruber, F Santini, F Messina, A Di Caro, MR Capobianchi |
| EPI_ISL_3446952, EPI_ISL_3446953, EPI_ISL_3446954, EPI_ISL_3446955, EPI_ISL_3446956, EPI_ISL_3446957, EPI_ISL_3446958 | RM3/IFO | National Institute for Infectious Diseases (INMI) L. Spallanzani I.R.C.C.S | E Giombini, B Bartolini, M Rueca, G Bonfiglio, O Butera, CEM Gruber, F Santini, F Messina, A Di Caro, MR Capobianchi |
| EPI_ISL_3446959 | IFO | National Institute for Infectious Diseases (INMI) L. Spallanzani I.R.C.C.S | E Giombini, B Bartolini, M Rueca, G Bonfiglio, O Butera, CEM Gruber, F Santini, F Messina, A Di Caro, MR Capobianchi |
| EPI_ISL_3446960, | RM3/IFO | National Institute for Infectious Diseases (INMI) L. Spallanzani | E Giombini, B Bartolini, M Rueca, G Bonfiglio, O Butera, CEM Gruber, F Santini, F Messina, A Di Caro, MR Capobianchi |

| | | | |
|---|---|---|---|
| EPI_ISL_3446961, EPI_ISL_3446962, EPI_ISL_3446963, EPI_ISL_3446964, EPI_ISL_3446965, EPI_ISL_3446966, EPI_ISL_3446967, EPI_ISL_3446968 | | I.R.C.C.S | |
| EPI_ISL_3446969, EPI_ISL_3446970, EPI_ISL_3446971 | RM2/Osp. S.Eugenio | National Institute for Infectious Diseases (INMI) L. Spallanzani I.R.C.C.S | E Giombini, B Bartolini, M Rueca, G Bonfiglio, O Butera, CEM Gruber, F Santini, F Messina, A Di Caro, MR Capobianchi |
| EPI_ISL_3446972 | RM1/Osp. S.Filippo Neri | National Institute for Infectious Diseases (INMI) L. Spallanzani I.R.C.C.S | E Giombini, B Bartolini, M Rueca, G Bonfiglio, O Butera, CEM Gruber, F Santini, F Messina, A Di Caro, MR Capobianchi |
| EPI_ISL_3446973 | RM2/Osp. S.Eugenio | National Institute for Infectious Diseases (INMI) L. Spallanzani I.R.C.C.S | E Giombini, B Bartolini, M Rueca, G Bonfiglio, O Butera, CEM Gruber, F Santini, F Messina, A Di Caro, MR Capobianchi |
| EPI_ISL_3446974 | RM1/Osp. S.Filippo Neri | National Institute for Infectious Diseases (INMI) L. Spallanzani I.R.C.C.S | E Giombini, B Bartolini, M Rueca, G Bonfiglio, O Butera, CEM Gruber, F Santini, F Messina, A Di Caro, MR Capobianchi |
| EPI_ISL_3446975 | RM2/Osp. S.Eugenio | National Institute for Infectious Diseases (INMI) L. Spallanzani I.R.C.C.S | E Giombini, B Bartolini, M Rueca, G Bonfiglio, O Butera, CEM Gruber, F Santini, F Messina, A Di Caro, MR Capobianchi |
| EPI_ISL_3446976 | RM1/Osp. S.Filippo Neri | National Institute for Infectious Diseases (INMI) L. Spallanzani I.R.C.C.S | E Giombini, B Bartolini, M Rueca, G Bonfiglio, O Butera, CEM Gruber, F Santini, F Messina, A Di Caro, MR Capobianchi |
| EPI_ISL_3446977 | RM2/Osp. S.Eugenio | National Institute for Infectious Diseases (INMI) L. Spallanzani I.R.C.C.S | E Giombini, B Bartolini, M Rueca, G Bonfiglio, O Butera, CEM Gruber, F Santini, F Messina, A Di Caro, MR Capobianchi |
| EPI_ISL_3446978, EPI_ISL_3446979 | RM1/Osp. S.Filippo Neri | National Institute for Infectious Diseases (INMI) L. Spallanzani I.R.C.C.S | E Giombini, B Bartolini, M Rueca, G Bonfiglio, O Butera, CEM Gruber, F Santini, F Messina, A Di Caro, MR Capobianchi |
| EPI_ISL_3446980, EPI_ISL_3446981 | RM2/Osp. S.Eugenio | National Institute for Infectious Diseases (INMI) L. Spallanzani I.R.C.C.S | E Giombini, B Bartolini, M Rueca, G Bonfiglio, O Butera, CEM Gruber, F Santini, F Messina, A Di Caro, MR Capobianchi |
| EPI_ISL_3446982, EPI_ISL_3446983, EPI_ISL_3446984, EPI_ISL_3446985, EPI_ISL_3446986, EPI_ISL_3446987, EPI_ISL_3446988, EPI_ISL_3446989, EPI_ISL_3446990, EPI_ISL_3446991, EPI_ISL_3446992, EPI_ISL_3446993, EPI_ISL_3446994, EPI_ISL_3446995, EPI_ISL_3446996, EPI_ISL_3446997, EPI_ISL_3446998, EPI_ISL_3446999, EPI_ISL_3447000, EPI_ISL_3447001, EPI_ISL_3447002, EPI_ISL_3447003, EPI_ISL_3447004, EPI_ISL_3447005, EPI_ISL_3447006, EPI_ISL_3447007, EPI_ISL_3447008, EPI_ISL_3447009, EPI_ISL_3447010, EPI_ISL_3447011, EPI_ISL_3447012, EPI_ISL_3447013, EPI_ISL_3447014, EPI_ISL_3447015 | | | |
| see above | RM1/Osp. S.Filippo Neri | National Institute for Infectious Diseases (INMI) L. Spallanzani I.R.C.C.S | E Giombini, B Bartolini, M Rueca, G Bonfiglio, O Butera, CEM Gruber, F Santini, F Messina, A Di Caro, MR Capobianchi |
| EPI_ISL_3447016, EPI_ISL_3447017, EPI_ISL_3447018, EPI_ISL_3447019, EPI_ISL_3447020, EPI_ISL_3447021, EPI_ISL_3447022, EPI_ISL_3447024, EPI_ISL_3447025, EPI_ISL_3447026, EPI_ISL_3447027, EPI_ISL_3447028, EPI_ISL_3447029, EPI_ISL_3447030, EPI_ISL_3447031, EPI_ISL_3447033, EPI_ISL_3447034, EPI_ISL_3447035, EPI_ISL_3447036, EPI_ISL_3447037, EPI_ISL_3447038, EPI_ISL_3447039, EPI_ISL_3447040, EPI_ISL_3447041, EPI_ISL_3447042, EPI_ISL_3447043, EPI_ISL_3447044, EPI_ISL_3447045, EPI_ISL_3447047, EPI_ISL_3447048, EPI_ISL_3447049, EPI_ISL_3447050, EPI_ISL_3447051, EPI_ISL_3447052, EPI_ISL_3447053, EPI_ISL_3447054, EPI_ISL_3447055, EPI_ISL_3447056, EPI_ISL_3447057, EPI_ISL_3447058, EPI_ISL_3447059, EPI_ISL_3447060, EPI_ISL_3447061, EPI_ISL_3447062, EPI_ISL_3447063, EPI_ISL_3447064, EPI_ISL_3447065, EPI_ISL_3447066 | | | |
| see above | RM3/IFO | National Institute for Infectious Diseases (INMI) L. Spallanzani I.R.C.C.S | E Giombini, B Bartolini, M Rueca, G Bonfiglio, O Butera, CEM Gruber, F Santini, F Messina, A Di Caro, MR Capobianchi |
| EPI_ISL_3447068 | Biological Hazard and Health Research Laboratory, Centre for Advanced Research in Science (CARS), University of Dhaka | Genomic Research Lab, BCSIR | Md. Murshed Hasan Sarkar, Md. Ahasan Habib, Shahina Akter, Tanjina Akhter Banu, Barna Goswami, Iffat Jahan, Mohammad Mohi Uddin,Abu Sayeed Mohammad Mahmud, Mohammad Samir Uzzaman, Eshrar Osman, Md. Kamrul Islam, Md. Shaminur Rahman, Latiful Bari, Md Abdul Malek, Md Mizanur Rahaman, Md. Salim Khan |
| EPI_ISL_3447070, EPI_ISL_3447071, EPI_ISL_3447072, EPI_ISL_3447073, EPI_ISL_3447074, EPI_ISL_3447075, EPI_ISL_3447076, EPI_ISL_3447077, EPI_ISL_3447078, EPI_ISL_3447079, EPI_ISL_3447080, EPI_ISL_3447081, EPI_ISL_3447082, EPI_ISL_3447083 | | | |
| see above | IstitutoAuxologicoItaliano | Istituto Auxologico Italiano | Erminio Torresani |
| EPI_ISL_3447085, EPI_ISL_3447086, EPI_ISL_3447087, EPI_ISL_3447088, EPI_ISL_3447089, EPI_ISL_3447090, EPI_ISL_3447091, EPI_ISL_3447092, EPI_ISL_3447093, EPI_ISL_3447094, EPI_ISL_3447095, EPI_ISL_3447096, EPI_ISL_3447097, EPI_ISL_3447098, EPI_ISL_3447099, EPI_ISL_3447100, EPI_ISL_3447101, EPI_ISL_3447102, EPI_ISL_3447103, EPI_ISL_3447104, EPI_ISL_3447105, EPI_ISL_3447106, EPI_ISL_3447107, EPI_ISL_3447108, EPI_ISL_3447109, EPI_ISL_3447110, EPI_ISL_3447111, EPI_ISL_3447112, EPI_ISL_3447113, EPI_ISL_3447114, EPI_ISL_3447115, EPI_ISL_3447116, EPI_ISL_3447117, EPI_ISL_3447118, EPI_ISL_3447119, EPI_ISL_3447120, EPI_ISL_3447121, EPI_ISL_3447122, EPI_ISL_3447123, EPI_ISL_3447124, EPI_ISL_3447125, EPI_ISL_3447126, EPI_ISL_3447127, EPI_ISL_3447128, EPI_ISL_3447129, EPI_ISL_3447130, EPI_ISL_3447131, EPI_ISL_3447132, EPI_ISL_3447133, EPI_ISL_3447134, EPI_ISL_3447135, EPI_ISL_3447136, EPI_ISL_3447137, EPI_ISL_3447138, EPI_ISL_3447139, EPI_ISL_3447140, EPI_ISL_3447141, EPI_ISL_3447142, EPI_ISL_3447143, EPI_ISL_3447144, EPI_ISL_3447146, EPI_ISL_3447147, EPI_ISL_3447148, EPI_ISL_3447149, EPI_ISL_3447150, EPI_ISL_3447151, EPI_ISL_3447152, EPI_ISL_3447153, EPI_ISL_3447154, EPI_ISL_3447155, EPI_ISL_3447156, EPI_ISL_3447157, EPI_ISL_3447158, EPI_ISL_3447159, EPI_ISL_3447160, EPI_ISL_3447161, EPI_ISL_3447162, EPI_ISL_3447163, EPI_ISL_3447164, EPI_ISL_3447165, EPI_ISL_3447166, EPI_ISL_3447167, EPI_ISL_3447168, EPI_ISL_3447169, EPI_ISL_3447170, EPI_ISL_3447171, EPI_ISL_3447172, EPI_ISL_3447173, EPI_ISL_3447174, EPI_ISL_3447175, EPI_ISL_3447176, EPI_ISL_3447177, EPI_ISL_3447178, EPI_ISL_3447179, EPI_ISL_3447180, EPI_ISL_3447181, EPI_ISL_3447182, EPI_ISL_3447183, EPI_ISL_3447184, EPI_ISL_3447185, EPI_ISL_3447186, EPI_ISL_3447187, EPI_ISL_3447188, EPI_ISL_3447189, EPI_ISL_3447190, EPI_ISL_3447191, EPI_ISL_3447192, EPI_ISL_3447193, EPI_ISL_3447194, EPI_ISL_3447195, EPI_ISL_3447196, EPI_ISL_3447197, EPI_ISL_3447198, EPI_ISL_3447199, EPI_ISL_3447200, EPI_ISL_3447201, EPI_ISL_3447202, EPI_ISL_3447203, EPI_ISL_3447204, EPI_ISL_3447206, EPI_ISL_3447207, EPI_ISL_3447208, EPI_ISL_3447209, EPI_ISL_3447210, EPI_ISL_3447211, EPI_ISL_3447212, EPI_ISL_3447213, EPI_ISL_3447214, EPI_ISL_3447215, EPI_ISL_3447216, EPI_ISL_3447217, EPI_ISL_3447218, EPI_ISL_3447219, EPI_ISL_3447220, EPI_ISL_3447221, EPI_ISL_3447222, EPI_ISL_3447223, EPI_ISL_3447224, EPI_ISL_3447225, EPI_ISL_3447226 | | | |
| see above | Labor Berlin Charité Vivantes GmbH / Institut für Virologie | Charité Universitätsmedizin Berlin, Institut für Virologie/Labor Berlin | Peter Menzel, Christine Stephan, Rolf Schwarzer, Victor M Corman, Barbara Mühlemann, Terry Jones, Christian Drosten |
| EPI_ISL_3447227 | Biological Hazard and Health Research Laboratory, Centre for Advanced Research in Science (CARS), University of Dhaka | Genomic Research Lab, BCSIR | Md. Murshed Hasan Sarkar, Md. Ahasan Habib, Shahina Akter, Tanjina Akhter Banu, Barna Goswami, Iffat Jahan, Mohammad Mohi Uddin,Abu Sayeed Mohammad Mahmud, Mohammad Samir Uzzaman, Eshrar Osman, Md. Kamrul Islam, Md. Shaminur Rahman, Latiful Bari, Md Abdul Malek, Md Mizanur Rahaman, Md. Salim Khan |
| EPI_ISL_3447228 | Biological Hazard and Health Research Laboratory, Centre for Advanced Research in Science (CARS), University of Dhaka | Genomic Research Lab, BCSIR | Dr. Qudrat-I-Khuda Road, Dhaka-1205, Bangladesh |
| EPI_ISL_3447230, EPI_ISL_3447231, EPI_ISL_3447232, EPI_ISL_3447233, EPI_ISL_3447234, EPI_ISL_3447235, EPI_ISL_3447236, EPI_ISL_3447237, EPI_ISL_3447238, EPI_ISL_3447239, EPI_ISL_3447240, EPI_ISL_3447241, EPI_ISL_3447242, EPI_ISL_3447243, EPI_ISL_3447244, EPI_ISL_3447245, EPI_ISL_3447246, EPI_ISL_3447247, EPI_ISL_3447248, EPI_ISL_3447249, EPI_ISL_3447250 | | | |
| see above | IstitutoAuxologicoItaliano | Istituto Auxologico Italiano | Erminio Torresani |
| EPI_ISL_3447251 | Istituto Auxologico Italiano | Istituto Auxologico Italiano | Erminio Torresani |
| EPI_ISL_3447252, EPI_ISL_3447253, EPI_ISL_3447254, EPI_ISL_3447255, EPI_ISL_3447256, EPI_ISL_3447257, EPI_ISL_3447258, EPI_ISL_3447259 | IstitutoAuxologicoItaliano | Istituto Auxologico Italiano | Erminio Torresani |
| EPI_ISL_3447261, EPI_ISL_3447262, EPI_ISL_3447263, EPI_ISL_3447264, EPI_ISL_3447265, EPI_ISL_3447266, EPI_ISL_3447267, EPI_ISL_3447268, EPI_ISL_3447269, EPI_ISL_3447270, EPI_ISL_3447271, EPI_ISL_3447272, EPI_ISL_3447273, EPI_ISL_3447274, EPI_ISL_3447275, EPI_ISL_3447276, EPI_ISL_3447277, EPI_ISL_3447278, EPI_ISL_3447279, EPI_ISL_3447280, EPI_ISL_3447281, EPI_ISL_3447282, EPI_ISL_3447283, EPI_ISL_3447284, EPI_ISL_3447285, EPI_ISL_3447286, EPI_ISL_3447287, EPI_ISL_3447288, EPI_ISL_3447289, EPI_ISL_3447290, EPI_ISL_3447291, EPI_ISL_3447292, EPI_ISL_3447293, EPI_ISL_3447294, EPI_ISL_3447295, EPI_ISL_3447296, EPI_ISL_3447297, EPI_ISL_3447298, EPI_ISL_3447299, EPI_ISL_3447300, EPI_ISL_3447301, EPI_ISL_3447302, EPI_ISL_3447303, EPI_ISL_3447304, EPI_ISL_3447305, EPI_ISL_3447306, EPI_ISL_3447307, EPI_ISL_3447308, EPI_ISL_3447309, EPI_ISL_3447310, EPI_ISL_3447311, EPI_ISL_3447312, EPI_ISL_3447313, EPI_ISL_3447314, EPI_ISL_3447315, EPI_ISL_3447316, EPI_ISL_3447317, EPI_ISL_3447318, EPI_ISL_3447319, EPI_ISL_3447320, EPI_ISL_3447321, EPI_ISL_3447322, EPI_ISL_3447324, EPI_ISL_3447325, EPI_ISL_3447326, EPI_ISL_3447328, EPI_ISL_3447329, EPI_ISL_3447330, EPI_ISL_3447331, EPI_ISL_3447332, EPI_ISL_3447333, EPI_ISL_3447334, EPI_ISL_3447335, EPI_ISL_3447336, EPI_ISL_3447337, EPI_ISL_3447338, EPI_ISL_3447339, EPI_ISL_3447340, EPI_ISL_3447341 | | | |
| see above | Illinois Department of Public Health - Springfield Lab | Illinois Department of Public Health - Springfield Lab | Bryan Sim, Marshall Smock |
| EPI_ISL_3447342 | Biological Hazard and Health Research Laboratory, Centre for Advanced Research in Science (CARS), University of Dhaka | Genomic Research Lab, BCSIR | Md. Murshed Hasan Sarkar, Md. Ahasan Habib, Shahina Akter, Tanjina Akhter Banu, Barna Goswami, Iffat Jahan, Mohammad Mohi Uddin,Abu Sayeed Mohammad Mahmud, Mohammad Samir Uzzaman, Eshrar Osman, Md. Kamrul Islam, Md. Shaminur Rahman, Latiful Bari, Md Abdul Malek, Md Mizanur Rahaman, Md. Salim Khan |
| EPI_ISL_3447343, EPI_ISL_3447344, EPI_ISL_3447345, EPI_ISL_3447346, EPI_ISL_3447347, EPI_ISL_3447348, EPI_ISL_3447349, EPI_ISL_3447350, EPI_ISL_3447351, EPI_ISL_3447352, EPI_ISL_3447353, EPI_ISL_3447354, EPI_ISL_3447355, EPI_ISL_3447356, EPI_ISL_3447357, EPI_ISL_3447358, EPI_ISL_3447359, EPI_ISL_3447360, EPI_ISL_3447361, EPI_ISL_3447362, EPI_ISL_3447363, EPI_ISL_3447364, EPI_ISL_3447365, EPI_ISL_3447366, EPI_ISL_3447367, EPI_ISL_3447368, EPI_ISL_3447369, EPI_ISL_3447370 | | | |
| see above | Nebraska Public Health Laboratory | NPHL COVID-19 Response Team | NPHL COVID-19 Response Team |
| EPI_ISL_3447371, EPI_ISL_3447372 | Biological Hazard and Health Research Laboratory, Centre for Advanced Research in Science (CARS), University of Dhaka | Genomic Research Lab, BCSIR | Md. Murshed Hasan Sarkar, Md. Ahasan Habib, Shahina Akter, Tanjina Akhter Banu, Barna Goswami, Iffat Jahan, Mohammad Mohi Uddin,Abu Sayeed Mohammad Mahmud, Mohammad Samir Uzzaman, Eshrar Osman, Md. Kamrul Islam, Md. Shaminur Rahman, Latiful Bari, Md Abdul Malek, Md Mizanur Rahaman, Md. Salim Khan |
| EPI_ISL_3447373, EPI_ISL_3447374, EPI_ISL_3447375, EPI_ISL_3447376, EPI_ISL_3447377, EPI_ISL_3447378, EPI_ISL_3447379, EPI_ISL_3447380, EPI_ISL_3447381, EPI_ISL_3447382, EPI_ISL_3447383, EPI_ISL_3447384, EPI_ISL_3447385, EPI_ISL_3447386, EPI_ISL_3447387, EPI_ISL_3447388, EPI_ISL_3447389, EPI_ISL_3447390, EPI_ISL_3447391, EPI_ISL_3447392, EPI_ISL_3447393, EPI_ISL_3447394, EPI_ISL_3447395, EPI_ISL_3447396, EPI_ISL_3447397, EPI_ISL_3447398, EPI_ISL_3447399, EPI_ISL_3447400, EPI_ISL_3447401, EPI_ISL_3447402, EPI_ISL_3447403, EPI_ISL_3447404, EPI_ISL_3447405, EPI_ISL_3447406, EPI_ISL_3447407, EPI_ISL_3447408, EPI_ISL_3447409 | | | |
| see above | Virginia Division of Consolidated Laboratory Services | Virginia Division of Consolidated Laboratory Services | Virginia Division of Consolidated Laboratory Services |
| EPI_ISL_3447410 | IstitutoAuxologicoItaliano | Istituto Auxologico Italiano | Erminio Torresani |
| EPI_ISL_3447445, EPI_ISL_3447452 | LACEN/PE | WallauLab on behalf of Fiocruz COVID-19 Genomic Surveillance Network | Marcelo Henrique dos Santos Paiva, Cassia Docena, Matheus Filgueira Bezerra, Filipe Zimmer Dezordi, Lais Ceschini Machado, Lilian Caroliny Amorim Silva, Gustavo Barbosa de Lima, Alexandre Freitas da Silva, Sinval Pinto Brandão Filho, Constância Flávia Junqueira Ayres, Gabriel Luz Wallau |
| EPI_ISL_3447601, EPI_ISL_3447602, EPI_ISL_3447603, EPI_ISL_3447604, EPI_ISL_3447605, EPI_ISL_3447606, EPI_ISL_3447607, EPI_ISL_3447608, EPI_ISL_3447609, EPI_ISL_3447610, EPI_ISL_3447611, EPI_ISL_3447612, EPI_ISL_3447613, EPI_ISL_3447614, EPI_ISL_3447615, EPI_ISL_3447616, EPI_ISL_3447617, EPI_ISL_3447618, EPI_ISL_3447619, EPI_ISL_3447620, EPI_ISL_3447621, EPI_ISL_3447622, EPI_ISL_3447623, EPI_ISL_3447624, EPI_ISL_3447625, EPI_ISL_3447626, EPI_ISL_3447627, EPI_ISL_3447628, EPI_ISL_3447629, EPI_ISL_3447630, EPI_ISL_3447631, EPI_ISL_3447632, EPI_ISL_3447633, EPI_ISL_3447634, EPI_ISL_3447635, EPI_ISL_3447636, EPI_ISL_3447637, EPI_ISL_3447638, EPI_ISL_3447639, EPI_ISL_3447640, EPI_ISL_3447641, EPI_ISL_3447642, EPI_ISL_3447643, EPI_ISL_3447644, EPI_ISL_3447645, EPI_ISL_3447646, EPI_ISL_3447647, EPI_ISL_3447648, EPI_ISL_3447649, EPI_ISL_3447650, EPI_ISL_3447651, EPI_ISL_3447652, EPI_ISL_3447653, EPI_ISL_3447654, EPI_ISL_3447655, EPI_ISL_3447656, EPI_ISL_3447657, EPI_ISL_3447658, EPI_ISL_3447659, EPI_ISL_3447660, EPI_ISL_3447661, EPI_ISL_3447662, EPI_ISL_3447663, EPI_ISL_3447664, EPI_ISL_3447665, EPI_ISL_3447666, EPI_ISL_3447667, EPI_ISL_3447668, EPI_ISL_3447669, EPI_ISL_3447670, EPI_ISL_3447671, EPI_ISL_3447672, EPI_ISL_3447673 | | | |
| see above | Minnesota Department of Health, Public Health Laboratory | Minnesota Department of Health, Public Health Laboratory | Alexandra Lorentz, Jacob Garfin, Matt Plumb, and Xiong Wang |
| EPI_ISL_3447674, EPI_ISL_3447675, EPI_ISL_3447676, EPI_ISL_3447677, EPI_ISL_3447678, EPI_ISL_3447679 | IstitutoAuxologicoItaliano | Istituto Auxologico Italiano | Erminio Torresani |
| EPI_ISL_3447680 | Istituto Auxologico Italiano | Istituto Auxologico Italiano | Erminio Torresani |

| | | | |
|---|---|---|---|
| EPI_ISL_3447681, EPI_ISL_3447682, EPI_ISL_3447683, EPI_ISL_3447684, EPI_ISL_3447685, EPI_ISL_3447686, EPI_ISL_3447687, EPI_ISL_3447688, EPI_ISL_3447689, EPI_ISL_3447690, EPI_ISL_3447691, EPI_ISL_3447692, EPI_ISL_3447693, EPI_ISL_3447694, EPI_ISL_3447695, EPI_ISL_3447696, EPI_ISL_3447697, EPI_ISL_3447698, EPI_ISL_3447699, EPI_ISL_3447700, EPI_ISL_3447701, EPI_ISL_3447702, EPI_ISL_3447703, EPI_ISL_3447704, EPI_ISL_3447705 | | | |
| see above | Tokyo Metropolitan Institute of Public Health | Tokyo Metropolitan Institute of Public Health | Masaki Hayashi, Takako Yamazaki, Akane Negishi, Hirofumi Miyake, Kenshirou Kuroki, Maya Isogai, Takushi Fujiwara, Mami Nagashima, Jun Suzuki, Kenji Sadamasu |
| EPI_ISL_3447706, EPI_ISL_3447707, EPI_ISL_3447708 | NHLS_JHB | KRISP, KZN Research Innovation and Sequencing Platform | Florette Treurnicht, Bulelani Manene, Kathleen Subramoney,Giandhari Jennifer, Pillay Sureshnee, Yajna Ramphal, Naidoo Yeshnee, Tshiabuila Derek,Tegally Houriiyah, San James, Wilkinson Eduan, de Oliveira Tulio |
| EPI_ISL_3447709, EPI_ISL_3447710, EPI_ISL_3447711, EPI_ISL_3447716 | NHLS_IALCH | KRISP, KZN Research Innovation and Sequencing Platform | Giandhari Jennifer, Pillay Sureshnee, Yajna Ramphal, Naidoo Yeshnee, Tshiabuila Derek,Tegally Houriiyah, San James, Wilkinson Eduan, de Oliveira Tulio |
| EPI_ISL_3447718 | NHLS_JHB | KRISP, KZN Research Innovation and Sequencing Platform | Florette Treurnicht, Bulelani Manene, Kathleen Subramoney,Giandhari Jennifer, Pillay Sureshnee, Yajna Ramphal, Naidoo Yeshnee, Tshiabuila Derek,Tegally Houriiyah, San James, Wilkinson Eduan, de Oliveira Tulio |
| EPI_ISL_3447719, EPI_ISL_3447720, EPI_ISL_3447721, EPI_ISL_3447722, EPI_ISL_3447723, EPI_ISL_3447724, EPI_ISL_3447725, EPI_ISL_3447726, EPI_ISL_3447727 | NHLS_IALCH | KRISP, KZN Research Innovation and Sequencing Platform | Giandhari Jennifer, Pillay Sureshnee, Yajna Ramphal, Naidoo Yeshnee, Tshiabuila Derek,Tegally Houriiyah, San James, Wilkinson Eduan, de Oliveira Tulio |
| EPI_ISL_3447728, EPI_ISL_3447729 | NHLS_JHB | KRISP, KZN Research Innovation and Sequencing Platform | Florette Treurnicht, Bulelani Manene, Kathleen Subramoney,Giandhari Jennifer, Pillay Sureshnee, Yajna Ramphal, Naidoo Yeshnee, Tshiabuila Derek,Tegally Houriiyah, San James, Wilkinson Eduan, de Oliveira Tulio |
| EPI_ISL_3447730, EPI_ISL_3447731, EPI_ISL_3447732, EPI_ISL_3447733 | NHLS_IALCH | KRISP, KZN Research Innovation and Sequencing Platform | Giandhari Jennifer, Pillay Sureshnee, Yajna Ramphal, Naidoo Yeshnee, Tshiabuila Derek,Tegally Houriiyah, San James, Wilkinson Eduan, de Oliveira Tulio |
| EPI_ISL_3447734 | NHLS_JHB | KRISP, KZN Research Innovation and Sequencing Platform | Florette Treurnicht, Bulelani Manene, Kathleen Subramoney,Giandhari Jennifer, Pillay Sureshnee, Yajna Ramphal, Naidoo Yeshnee, Tshiabuila Derek,Tegally Houriiyah, San James, Wilkinson Eduan, de Oliveira Tulio |
| EPI_ISL_3447735, EPI_ISL_3447736 | NHLS_IALCH | KRISP, KZN Research Innovation and Sequencing Platform | Giandhari Jennifer, Pillay Sureshnee, Yajna Ramphal, Naidoo Yeshnee, Tshiabuila Derek,Tegally Houriiyah, San James, Wilkinson Eduan, de Oliveira Tulio |
| EPI_ISL_3447737, EPI_ISL_3447738, EPI_ISL_3447739 | NHLS_JHB | KRISP, KZN Research Innovation and Sequencing Platform | Florette Treurnicht, Bulelani Manene, Kathleen Subramoney,Giandhari Jennifer, Pillay Sureshnee, Yajna Ramphal, Naidoo Yeshnee, Tshiabuila Derek,Tegally Houriiyah, San James, Wilkinson Eduan, de Oliveira Tulio |
| EPI_ISL_3447740, EPI_ISL_3447741, EPI_ISL_3447742 | NHLS_IALCH | KRISP, KZN Research Innovation and Sequencing Platform | Giandhari Jennifer, Pillay Sureshnee, Yajna Ramphal, Naidoo Yeshnee, Tshiabuila Derek,Tegally Houriiyah, San James, Wilkinson Eduan, de Oliveira Tulio |
| EPI_ISL_3447743, EPI_ISL_3447744 | NHLS_JHB | KRISP, KZN Research Innovation and Sequencing Platform | Florette Treurnicht, Bulelani Manene, Kathleen Subramoney,Giandhari Jennifer, Pillay Sureshnee, Yajna Ramphal, Naidoo Yeshnee, Tshiabuila Derek,Tegally Houriiyah, San James, Wilkinson Eduan, de Oliveira Tulio |
| EPI_ISL_3447745 | NHLS_IALCH | KRISP, KZN Research Innovation and Sequencing Platform | Giandhari Jennifer, Pillay Sureshnee, Yajna Ramphal, Naidoo Yeshnee, Tshiabuila Derek,Tegally Houriiyah, San James, Wilkinson Eduan, de Oliveira Tulio |
| EPI_ISL_3447746 | NHLS_JHB | KRISP, KZN Research Innovation and Sequencing Platform | Florette Treurnicht, Bulelani Manene, Kathleen Subramoney,Giandhari Jennifer, Pillay Sureshnee, Yajna Ramphal, Naidoo Yeshnee, Tshiabuila Derek,Tegally Houriiyah, San James, Wilkinson Eduan, de Oliveira Tulio |
| EPI_ISL_3447747 | NHLS_IALCH | KRISP, KZN Research Innovation and Sequencing Platform | Giandhari Jennifer, Pillay Sureshnee, Yajna Ramphal, Naidoo Yeshnee, Tshiabuila Derek,Tegally Houriiyah, San James, Wilkinson Eduan, de Oliveira Tulio |
| EPI_ISL_3447748 | NHLS_JHB | KRISP, KZN Research Innovation and Sequencing Platform | Florette Treurnicht, Bulelani Manene, Kathleen Subramoney,Giandhari Jennifer, Pillay Sureshnee, Yajna Ramphal, Naidoo Yeshnee, Tshiabuila Derek,Tegally Houriiyah, San James, Wilkinson Eduan, de Oliveira Tulio |
| EPI_ISL_3447749, EPI_ISL_3447750, EPI_ISL_3447751 | NHLS_IALCH | KRISP, KZN Research Innovation and Sequencing Platform | Giandhari Jennifer, Pillay Sureshnee, Yajna Ramphal, Naidoo Yeshnee, Tshiabuila Derek,Tegally Houriiyah, San James, Wilkinson Eduan, de Oliveira Tulio |
| EPI_ISL_3447752 | AHRI- Alex Sigal Lab | KRISP, KZN Research Innovation and Sequencing Platform | Alex Sigal, Sandile Cele, Mallory Bernstein, Giandhari Jennifer, Pillay Sureshnee, Yajna Ramphal, Naidoo Yeshnee, Tshabuila Derek,Tegally Houriiyah, San E. James, Wilkinson Eduan, de Oliveira Tulio |
| EPI_ISL_3447753 | NHLS_IALCH | KRISP, KZN Research Innovation and Sequencing Platform | Giandhari Jennifer, Pillay Sureshnee, Yajna Ramphal, Naidoo Yeshnee, Tshiabuila Derek,Tegally Houriiyah, San James, Wilkinson Eduan, de Oliveira Tulio |
| EPI_ISL_3447754 | NHLS_JHB | KRISP, KZN Research Innovation and Sequencing Platform | Florette Treurnicht, Bulelani Manene, Kathleen Subramoney,Giandhari Jennifer, Pillay Sureshnee, Yajna Ramphal, Naidoo Yeshnee, Tshiabuila Derek,Tegally Houriiyah, San James, Wilkinson Eduan, de Oliveira Tulio |
| EPI_ISL_3447755, EPI_ISL_3447756, EPI_ISL_3447757, EPI_ISL_3447759 | NHLS_IALCH | KRISP, KZN Research Innovation and Sequencing Platform | Giandhari Jennifer, Pillay Sureshnee, Yajna Ramphal, Naidoo Yeshnee, Tshiabuila Derek,Tegally Houriiyah, San James, Wilkinson Eduan, de Oliveira Tulio |
| EPI_ISL_3447760, EPI_ISL_3447761, EPI_ISL_3447762, EPI_ISL_3447763, EPI_ISL_3447764, EPI_ISL_3447765, EPI_ISL_3447766, EPI_ISL_3447767, EPI_ISL_3447768, EPI_ISL_3447769, EPI_ISL_3447770, EPI_ISL_3447771, EPI_ISL_3447772, EPI_ISL_3447774, EPI_ISL_3447775, EPI_ISL_3447776, EPI_ISL_3447777, EPI_ISL_3447778 | | | |
| see above | NHLS_JHB | KRISP, KZN Research Innovation and Sequencing Platform | Florette Treurnicht, Bulelani Manene, Kathleen Subramoney,Giandhari Jennifer, Pillay Sureshnee, Yajna Ramphal, Naidoo Yeshnee, Tshiabuila Derek,Tegally Houriiyah, San James, Wilkinson Eduan, de Oliveira Tulio |
| EPI_ISL_3447779 | AHRI- Alex Sigal Lab | KRISP, KZN Research Innovation and Sequencing Platform | Alex Sigal, Sandile Cele, Mallory Bernstein, Giandhari Jennifer, Pillay Sureshnee, Yajna Ramphal, Naidoo Yeshnee, Tshabuila Derek,Tegally Houriiyah, San E. James, Wilkinson Eduan, de Oliveira Tulio |
| EPI_ISL_3447780, EPI_ISL_3447781 | NHLS_JHB | KRISP, KZN Research Innovation and Sequencing Platform | Florette Treurnicht, Bulelani Manene, Kathleen Subramoney,Giandhari Jennifer, Pillay Sureshnee, Yajna Ramphal, Naidoo Yeshnee, Tshiabuila Derek,Tegally Houriiyah, San James, Wilkinson Eduan, de Oliveira Tulio |
| EPI_ISL_3447782 | NHLS_IALCH | KRISP, KZN Research Innovation and Sequencing Platform | Giandhari Jennifer, Pillay Sureshnee, Yajna Ramphal, Naidoo Yeshnee, Tshiabuila Derek,Tegally Houriiyah, San James, Wilkinson Eduan, de Oliveira Tulio |
| EPI_ISL_3447783, EPI_ISL_3447784, EPI_ISL_3447785, EPI_ISL_3447786, EPI_ISL_3447787, EPI_ISL_3447788, EPI_ISL_3447789, EPI_ISL_3447790, EPI_ISL_3447791, EPI_ISL_3447792, EPI_ISL_3447793, EPI_ISL_3447794, EPI_ISL_3447795, EPI_ISL_3447796 | | | |
| see above | NHLS_JHB | KRISP, KZN Research Innovation and Sequencing Platform | Florette Treurnicht, Bulelani Manene, Kathleen Subramoney,Giandhari Jennifer, Pillay Sureshnee, Yajna Ramphal, Naidoo Yeshnee, Tshiabuila Derek,Tegally Houriiyah, San James, Wilkinson Eduan, de Oliveira Tulio |
| EPI_ISL_3447797 | NHLS_IALCH | KRISP, KZN Research Innovation and Sequencing Platform | Giandhari Jennifer, Pillay Sureshnee, Yajna Ramphal, Naidoo Yeshnee, Tshiabuila Derek,Tegally Houriiyah, San James, Wilkinson Eduan, de Oliveira Tulio |
| EPI_ISL_3447798, EPI_ISL_3447799, EPI_ISL_3447800 | NHLS_JHB | KRISP, KZN Research Innovation and Sequencing Platform | Florette Treurnicht, Bulelani Manene, Kathleen Subramoney,Giandhari Jennifer, Pillay Sureshnee, Yajna Ramphal, Naidoo Yeshnee, Tshiabuila Derek,Tegally Houriiyah, San James, Wilkinson Eduan, de Oliveira Tulio |
| EPI_ISL_3447801 | NHLS_IALCH | KRISP, KZN Research Innovation and Sequencing Platform | Giandhari Jennifer, Pillay Sureshnee, Yajna Ramphal, Naidoo Yeshnee, Tshiabuila Derek,Tegally Houriiyah, San James, Wilkinson Eduan, de Oliveira Tulio |
| EPI_ISL_3447802 | AHRI- Alex Sigal Lab | KRISP, KZN Research Innovation and Sequencing Platform | Alex Sigal, Sandile Cele, Mallory Bernstein, Giandhari Jennifer, Pillay Sureshnee, Yajna Ramphal, Naidoo Yeshnee, Tshabuila Derek,Tegally Houriiyah, San E. James, Wilkinson Eduan, de Oliveira Tulio |
| EPI_ISL_3447803, EPI_ISL_3447804 | NHLS_JHB | KRISP, KZN Research Innovation and Sequencing Platform | Florette Treurnicht, Bulelani Manene, Kathleen Subramoney,Giandhari Jennifer, Pillay Sureshnee, Yajna Ramphal, Naidoo Yeshnee, Tshiabuila Derek,Tegally Houriiyah, San James, Wilkinson Eduan, de Oliveira Tulio |
| EPI_ISL_3447805 | NHLS_IALCH | KRISP, KZN Research Innovation and Sequencing Platform | Giandhari Jennifer, Pillay Sureshnee, Yajna Ramphal, Naidoo Yeshnee, Tshiabuila Derek,Tegally Houriiyah, San James, Wilkinson Eduan, de Oliveira Tulio |
| EPI_ISL_3447806, EPI_ISL_3447807, EPI_ISL_3447808, EPI_ISL_3447809 | NHLS_JHB | KRISP, KZN Research Innovation and Sequencing Platform | Florette Treurnicht, Bulelani Manene, Kathleen Subramoney,Giandhari Jennifer, Pillay Sureshnee, Yajna Ramphal, Naidoo Yeshnee, Tshiabuila Derek,Tegally Houriiyah, San James, Wilkinson Eduan, de Oliveira Tulio |
| EPI_ISL_3447810, EPI_ISL_3447811 | NHLS_IALCH | KRISP, KZN Research Innovation and Sequencing Platform | Giandhari Jennifer, Pillay Sureshnee, Yajna Ramphal, Naidoo Yeshnee, Tshabuila Derek,Tegally Houriiyah, San James, Wilkinson Eduan, de Oliveira Tulio |
| EPI_ISL_3447812 | Vaccines and Infectious Diseases Analytics Research Unit (VIDA) | KRISP, KZN Research Innovation and Sequencing Platform | Baillie Vicky, du Plessis Jeanine, Giandhari Jennifer, Pillay Sureshnee, Yajna Ramphal, Naidoo Yeshnee, Tshabuila Derek,Tegally Houriiyah, San James, Wilkinson Eduan, de Oliveira Tulio, Madhi Shabir |
| EPI_ISL_3447813 | NHLS_IALCH | KRISP, KZN Research Innovation and Sequencing Platform | Giandhari Jennifer, Pillay Sureshnee, Yajna Ramphal, Naidoo Yeshnee, Tshabuila Derek,Tegally Houriiyah, San James, Wilkinson Eduan, de Oliveira Tulio |
| EPI_ISL_3447814, EPI_ISL_3447815 | Vaccines and Infectious Diseases Analytics Research Unit (VIDA) | KRISP, KZN Research Innovation and Sequencing Platform | Baillie Vicky, du Plessis Jeanine, Giandhari Jennifer, Pillay Sureshnee, Yajna Ramphal, Naidoo Yeshnee, Tshabuila Derek,Tegally Houriiyah, San James, Wilkinson Eduan, de Oliveira Tulio, Madhi Shabir |
| EPI_ISL_3447817 | NHLS_IALCH | KRISP, KZN Research Innovation and Sequencing Platform | Giandhari Jennifer, Pillay Sureshnee, Yajna Ramphal, Naidoo Yeshnee, Tshabuila Derek,Tegally Houriiyah, San James, Wilkinson Eduan, de Oliveira Tulio |
| EPI_ISL_3447818 | NHLS Port Elizabeth Laboratory | KRISP, KZN Research Innovation and Sequencing Platform | Giandhari Jennifer, Pillay Sureshnee, Yajna Ramphal, Naidoo Yeshnee, Tshabuila Derek,Tegally Houriiyah, San James, Wilkinson Eduan, de Oliveira Tulio |
| EPI_ISL_3447819 | NHLS_IALCH | KRISP, KZN Research Innovation and Sequencing Platform | Giandhari Jennifer, Pillay Sureshnee, Yajna Ramphal, Naidoo Yeshnee, Tshabuila Derek,Tegally Houriiyah, San James, Wilkinson Eduan, de Oliveira Tulio |
| EPI_ISL_3447820, EPI_ISL_3447821 | NHLS Port Elizabeth Laboratory | KRISP, KZN Research Innovation and Sequencing Platform | Giandhari Jennifer, Pillay Sureshnee, Yajna Ramphal, Naidoo Yeshnee, Tshabuila Derek,Tegally Houriiyah, San James, Wilkinson Eduan, de Oliveira Tulio |
| EPI_ISL_3447822, EPI_ISL_3447824 | NHLS_IALCH | KRISP, KZN Research Innovation and Sequencing Platform | Giandhari Jennifer, Pillay Sureshnee, Yajna Ramphal, Naidoo Yeshnee, Tshabuila Derek,Tegally Houriiyah, San James, Wilkinson Eduan, de Oliveira Tulio |
| EPI_ISL_3447825 | Vaccines and Infectious Diseases Analytics Research Unit (VIDA) | KRISP, KZN Research Innovation and Sequencing Platform | Baillie Vicky, du Plessis Jeanine, Giandhari Jennifer, Pillay Sureshnee, Yajna Ramphal, Naidoo Yeshnee, Tshabuila Derek,Tegally Houriiyah, San James, Wilkinson Eduan, de Oliveira Tulio, Madhi Shabir |
| EPI_ISL_3447826, EPI_ISL_3447827, EPI_ISL_3447828, EPI_ISL_3447829, EPI_ISL_3447830, EPI_ISL_3447831, EPI_ISL_3447832, EPI_ISL_3447833, EPI_ISL_3447834, EPI_ISL_3447835, EPI_ISL_3447836 | | | |
| see above | NHLS_IALCH | KRISP, KZN Research Innovation and Sequencing Platform | Giandhari Jennifer, Pillay Sureshnee, Yajna Ramphal, Naidoo Yeshnee, Tshabuila Derek,Tegally Houriiyah, San James, Wilkinson Eduan, de Oliveira Tulio |
| EPI_ISL_3447837 | NHLS Port Elizabeth Laboratory | KRISP, KZN Research Innovation and Sequencing Platform | Giandhari Jennifer, Pillay Sureshnee, Yajna Ramphal, Naidoo Yeshnee, Tshabuila Derek,Tegally Houriiyah, San James, Wilkinson Eduan, de Oliveira Tulio |

| | | | |
|---|---|---|---|
| EPI_ISL_3447838, EPI_ISL_3447840, EPI_ISL_3447841, EPI_ISL_3447842, EPI_ISL_3447855 | NHLS_IALCH | KRISP, KZN Research Innovation and Sequencing Platform | Giandhari Jennifer, Pillay Sureshnee, Yajna Ramphal, Naidoo Yeshnee, Tshabuila Derek,Tegally Houriiyah, San James, Wilkinson Eduan, de Oliveira Tulio |
| EPI_ISL_3447856, EPI_ISL_3447857, EPI_ISL_3447858, EPI_ISL_3447859, EPI_ISL_3447860, EPI_ISL_3447861, EPI_ISL_3447862 | NHLS Port Elizabeth Laboratory | KRISP, KZN Research Innovation and Sequencing Platform | Giandhari Jennifer, Pillay Sureshnee, Yajna Ramphal, Naidoo Yeshnee, Tshabuila Derek,Tegally Houriiyah, San James, Wilkinson Eduan, de Oliveira Tulio |
| EPI_ISL_3447863, EPI_ISL_3447864, EPI_ISL_3447865, EPI_ISL_3447866, EPI_ISL_3447867, EPI_ISL_3447868 | Vaccines and Infectious Diseases Analytics Research Unit (VIDA) | KRISP, KZN Research Innovation and Sequencing Platform | Baillie Vicky, du Plessis Jeanine, Giandhari Jennifer, Pillay Sureshnee, Yajna Ramphal, Naidoo Yeshnee, Tshabuila Derek,Tegally Houriiyah, San James, Wilkinson Eduan, de Oliveira Tulio, Madhi Shabir |
| EPI_ISL_3447869, EPI_ISL_3447870, EPI_ISL_3447871, EPI_ISL_3447872, EPI_ISL_3447873, EPI_ISL_3447874, EPI_ISL_3447875, EPI_ISL_3447876, EPI_ISL_3447877, EPI_ISL_3447878, EPI_ISL_3447879, EPI_ISL_3447880 | | | |
| see above | NHLS_IALCH | KRISP, KZN Research Innovation and Sequencing Platform | Giandhari Jennifer, Pillay Sureshnee, Yajna Ramphal, Naidoo Yeshnee, Tshabuila Derek,Tegally Houriiyah, San James, Wilkinson Eduan, de Oliveira Tulio |
| EPI_ISL_3447882, EPI_ISL_3447883, EPI_ISL_3447884, EPI_ISL_3447885, EPI_ISL_3447886, EPI_ISL_3447887, EPI_ISL_3447888, EPI_ISL_3447889, EPI_ISL_3447890, EPI_ISL_3447891, EPI_ISL_3447892, EPI_ISL_3447893, EPI_ISL_3447894, EPI_ISL_3447895, EPI_ISL_3447896, EPI_ISL_3447897, EPI_ISL_3447898, EPI_ISL_3447899, EPI_ISL_3447900, EPI_ISL_3447901, EPI_ISL_3447902, EPI_ISL_3447903, EPI_ISL_3447904, EPI_ISL_3447905, EPI_ISL_3447906, EPI_ISL_3447907, EPI_ISL_3447908, EPI_ISL_3447909, EPI_ISL_3447910, EPI_ISL_3447911, EPI_ISL_3447912, EPI_ISL_3447913, EPI_ISL_3447914, EPI_ISL_3447915, EPI_ISL_3447916, EPI_ISL_3447917, EPI_ISL_3447918, EPI_ISL_3447919, EPI_ISL_3447920, EPI_ISL_3447921, EPI_ISL_3447922, EPI_ISL_3447923, EPI_ISL_3447924, EPI_ISL_3447925, EPI_ISL_3447926, EPI_ISL_3447927, EPI_ISL_3447928, EPI_ISL_3447929, EPI_ISL_3447930, EPI_ISL_3447931, EPI_ISL_3447932, EPI_ISL_3447933, EPI_ISL_3447934, EPI_ISL_3447935, EPI_ISL_3447936, EPI_ISL_3447937, EPI_ISL_3447938, EPI_ISL_3447939, EPI_ISL_3447940, EPI_ISL_3447941, EPI_ISL_3447942, EPI_ISL_3447943, EPI_ISL_3447944, EPI_ISL_3447945, EPI_ISL_3447946, EPI_ISL_3447947, EPI_ISL_3447948, EPI_ISL_3447949, EPI_ISL_3447950, EPI_ISL_3447951, EPI_ISL_3447952, EPI_ISL_3447953, EPI_ISL_3447954, EPI_ISL_3447955, EPI_ISL_3447956, EPI_ISL_3447957, EPI_ISL_3447958, EPI_ISL_3447959, EPI_ISL_3447960, EPI_ISL_3447961, EPI_ISL_3447962, EPI_ISL_3447963, EPI_ISL_3447964, EPI_ISL_3447965, EPI_ISL_3447967, EPI_ISL_3447968, EPI_ISL_3447969, EPI_ISL_3447970, EPI_ISL_3447971, EPI_ISL_3447972, EPI_ISL_3447973 | | | |
| see above | OHSU Lab Services Molecular Microbiology Lab | Oregon SARS-CoV-2 Genome Sequencing Center | Brendan L. O'Connell, Sonia Acharya, Cierra LaBlanc, Ruth V. Nichols, Alec J. Hirsch, Donna Hansel, Guang Fan, Xuan Qin, Daniel N. Streblow, William B. Messer, Andrew C. Adey, Benjamin N. Bimber, Brian J. O'Roak |
| EPI_ISL_3447999, EPI_ISL_3448000, EPI_ISL_3448001, EPI_ISL_3448002, EPI_ISL_3448003, EPI_ISL_3448004, EPI_ISL_3448005, EPI_ISL_3448006, EPI_ISL_3448007, EPI_ISL_3448008, EPI_ISL_3448009 | | | |
| see above | Minnesota Department of Health, Public Health Laboratory | Minnesota Department of Health, Public Health Laboratory | Alexandra Lorentz, Jacob Garfin, Matt Plumb, and Xiong Wang |
| EPI_ISL_3448010, EPI_ISL_3448011, EPI_ISL_3448012, EPI_ISL_3448013, EPI_ISL_3448014, EPI_ISL_3448015 | Centracare Laboratory Services | Minnesota Department of Health, Public Health Laboratory | Alexandra Lorentz, Jacob Garfin, Matt Plumb, and Xiong Wang |
| EPI_ISL_3448016, EPI_ISL_3448017, EPI_ISL_3448018, EPI_ISL_3448019, EPI_ISL_3448020, EPI_ISL_3448021, EPI_ISL_3448022, EPI_ISL_3448023, EPI_ISL_3448024, EPI_ISL_3448025, EPI_ISL_3448028, EPI_ISL_3448029 | | | |
| see above | Minnesota Department of Health, Public Health Laboratory | Minnesota Department of Health, Public Health Laboratory | Alexandra Lorentz, Jacob Garfin, Matt Plumb, and Xiong Wang |
| EPI_ISL_3448030 | Centracare Laboratory Services | Minnesota Department of Health, Public Health Laboratory | Alexandra Lorentz, Jacob Garfin, Matt Plumb, and Xiong Wang |
| EPI_ISL_3448031, EPI_ISL_3448032, EPI_ISL_3448033, EPI_ISL_3448034 | Minnesota Department of Health, Public Health Laboratory | Minnesota Department of Health, Public Health Laboratory | Alexandra Lorentz, Jacob Garfin, Matt Plumb, and Xiong Wang |
| EPI_ISL_3448035 | Centracare Laboratory Services | Minnesota Department of Health, Public Health Laboratory | Alexandra Lorentz, Jacob Garfin, Matt Plumb, and Xiong Wang |
| EPI_ISL_3448036, EPI_ISL_3448037, EPI_ISL_3448038, EPI_ISL_3448039, EPI_ISL_3448040, EPI_ISL_3448041, EPI_ISL_3448042, EPI_ISL_3448043, EPI_ISL_3448044, EPI_ISL_3448045, EPI_ISL_3448046, EPI_ISL_3448047, EPI_ISL_3448048 | | | |
| see above | Minnesota Department of Health, Public Health Laboratory | Minnesota Department of Health, Public Health Laboratory | Alexandra Lorentz, Jacob Garfin, Matt Plumb, and Xiong Wang |
| EPI_ISL_3448049, EPI_ISL_3448050, EPI_ISL_3448051 | Centracare Laboratory Services | Minnesota Department of Health, Public Health Laboratory | Alexandra Lorentz, Jacob Garfin, Matt Plumb, and Xiong Wang |
| EPI_ISL_3448052, EPI_ISL_3448053, EPI_ISL_3448054, EPI_ISL_3448055, EPI_ISL_3448056, EPI_ISL_3448057, EPI_ISL_3448058, EPI_ISL_3448059, EPI_ISL_3448060, EPI_ISL_3448061, EPI_ISL_3448062, EPI_ISL_3448063, EPI_ISL_3448064, EPI_ISL_3448065 | | | |
| see above | Minnesota Department of Health, Public Health Laboratory | Minnesota Department of Health, Public Health Laboratory | Alexandra Lorentz, Jacob Garfin, Matt Plumb, and Xiong Wang |
| EPI_ISL_3448066 | Centracare Laboratory Services | Minnesota Department of Health, Public Health Laboratory | Alexandra Lorentz, Jacob Garfin, Matt Plumb, and Xiong Wang |
| EPI_ISL_3448067, EPI_ISL_3448068 | Minnesota Department of Health, Public Health Laboratory | Minnesota Department of Health, Public Health Laboratory | Alexandra Lorentz, Jacob Garfin, Matt Plumb, and Xiong Wang |
| EPI_ISL_3448069, EPI_ISL_3448070 | Centracare Laboratory Services | Minnesota Department of Health, Public Health Laboratory | Alexandra Lorentz, Jacob Garfin, Matt Plumb, and Xiong Wang |
| EPI_ISL_3448071, EPI_ISL_3448072, EPI_ISL_3448073, EPI_ISL_3448074, EPI_ISL_3448075, EPI_ISL_3448077, EPI_ISL_3448078, EPI_ISL_3448079, EPI_ISL_3448080, EPI_ISL_3448081, EPI_ISL_3448082, EPI_ISL_3448083, EPI_ISL_3448084, EPI_ISL_3448085, EPI_ISL_3448086, EPI_ISL_3448087, EPI_ISL_3448088, EPI_ISL_3448089, EPI_ISL_3448090, EPI_ISL_3448091, EPI_ISL_3448092, EPI_ISL_3448093, EPI_ISL_3448094, EPI_ISL_3448095, EPI_ISL_3448096, EPI_ISL_3448097, EPI_ISL_3448098, EPI_ISL_3448099, EPI_ISL_3448100, EPI_ISL_3448101, EPI_ISL_3448102, EPI_ISL_3448103, EPI_ISL_3448104, EPI_ISL_3448106, EPI_ISL_3448107, EPI_ISL_3448108, EPI_ISL_3448109, EPI_ISL_3448110, EPI_ISL_3448111, EPI_ISL_3448112, EPI_ISL_3448113, EPI_ISL_3448114, EPI_ISL_3448115, EPI_ISL_3448116, EPI_ISL_3448117, EPI_ISL_3448118, EPI_ISL_3448119, EPI_ISL_3448120, EPI_ISL_3448121, EPI_ISL_3448122, EPI_ISL_3448123, EPI_ISL_3448124, EPI_ISL_3448125, EPI_ISL_3448126, EPI_ISL_3448127, EPI_ISL_3448128, EPI_ISL_3448129, EPI_ISL_3448130, EPI_ISL_3448131, EPI_ISL_3448132, EPI_ISL_3448133, EPI_ISL_3448134, EPI_ISL_3448135, EPI_ISL_3448136, EPI_ISL_3448137, EPI_ISL_3448138, EPI_ISL_3448139, EPI_ISL_3448140, EPI_ISL_3448141, EPI_ISL_3448142, EPI_ISL_3448143, EPI_ISL_3448144, EPI_ISL_3448145, EPI_ISL_3448146, EPI_ISL_3448147, EPI_ISL_3448148, EPI_ISL_3448149, EPI_ISL_3448150, EPI_ISL_3448151, EPI_ISL_3448152, EPI_ISL_3448153, EPI_ISL_3448154, EPI_ISL_3448155, EPI_ISL_3448156, EPI_ISL_3448158, EPI_ISL_3448159, EPI_ISL_3448160, EPI_ISL_3448161, EPI_ISL_3448162, EPI_ISL_3448163, EPI_ISL_3448164, EPI_ISL_3448165, EPI_ISL_3448166, EPI_ISL_3448167, EPI_ISL_3448168, EPI_ISL_3448169, EPI_ISL_3448170, EPI_ISL_3448171, EPI_ISL_3448172, EPI_ISL_3448173, EPI_ISL_3448174, EPI_ISL_3448175, EPI_ISL_3448177, EPI_ISL_3448178, EPI_ISL_3448179, EPI_ISL_3448180, EPI_ISL_3448181, EPI_ISL_3448182, EPI_ISL_3448183, EPI_ISL_3448184, EPI_ISL_3448187, EPI_ISL_3448188, EPI_ISL_3448189, EPI_ISL_3448191, EPI_ISL_3448192, EPI_ISL_3448193, EPI_ISL_3448194, EPI_ISL_3448195, EPI_ISL_3448196, EPI_ISL_3448197, EPI_ISL_3448198, EPI_ISL_3448199, EPI_ISL_3448200, EPI_ISL_3448201, EPI_ISL_3448203, EPI_ISL_3448204, EPI_ISL_3448205, EPI_ISL_3448206, EPI_ISL_3448207, EPI_ISL_3448208, EPI_ISL_3448209, EPI_ISL_3448210, EPI_ISL_3448211, EPI_ISL_3448212, EPI_ISL_3448213, EPI_ISL_3448215, EPI_ISL_3448216, EPI_ISL_3448217, EPI_ISL_3448218, EPI_ISL_3448219, EPI_ISL_3448220, EPI_ISL_3448221, EPI_ISL_3448222, EPI_ISL_3448223, EPI_ISL_3448224, EPI_ISL_3448225, EPI_ISL_3448226, EPI_ISL_3448227, EPI_ISL_3448228, EPI_ISL_3448229, EPI_ISL_3448230, EPI_ISL_3448231, EPI_ISL_3448232, EPI_ISL_3448233, EPI_ISL_3448234, EPI_ISL_3448235, EPI_ISL_3448236, EPI_ISL_3448237, EPI_ISL_3448238, EPI_ISL_3448239, EPI_ISL_3448240, EPI_ISL_3448241, EPI_ISL_3448242, EPI_ISL_3448243, EPI_ISL_3448244, EPI_ISL_3448245, EPI_ISL_3448246, EPI_ISL_3448247, EPI_ISL_3448248, EPI_ISL_3448249, EPI_ISL_3448250, EPI_ISL_3448251, EPI_ISL_3448252, EPI_ISL_3448253, EPI_ISL_3448254, EPI_ISL_3448255, EPI_ISL_3448256, EPI_ISL_3448257, EPI_ISL_3448258, EPI_ISL_3448259, EPI_ISL_3448260, EPI_ISL_3448261, EPI_ISL_3448262, EPI_ISL_3448263, EPI_ISL_3448264, EPI_ISL_3448265, EPI_ISL_3448266, EPI_ISL_3448267, EPI_ISL_3448268, EPI_ISL_3448269, EPI_ISL_3448270, EPI_ISL_3448271, EPI_ISL_3448272, EPI_ISL_3448273, EPI_ISL_3448274, EPI_ISL_3448275, EPI_ISL_3448276, EPI_ISL_3448277, EPI_ISL_3448278, EPI_ISL_3448279, EPI_ISL_3448280, EPI_ISL_3448281, EPI_ISL_3448282, EPI_ISL_3448283, EPI_ISL_3448284, EPI_ISL_3448285, EPI_ISL_3448286, EPI_ISL_3448287, EPI_ISL_3448288, EPI_ISL_3448289, EPI_ISL_3448290, EPI_ISL_3448291, EPI_ISL_3448292, EPI_ISL_3448293, EPI_ISL_3448294, EPI_ISL_3448295, EPI_ISL_3448296, EPI_ISL_3448297, EPI_ISL_3448298, EPI_ISL_3448299, EPI_ISL_3448300, EPI_ISL_3448301, EPI_ISL_3448302, EPI_ISL_3448303, EPI_ISL_3448304, EPI_ISL_3448305, EPI_ISL_3448306, EPI_ISL_3448307, EPI_ISL_3448308, EPI_ISL_3448309, EPI_ISL_3448310, EPI_ISL_3448311, EPI_ISL_3448312, EPI_ISL_3448313, EPI_ISL_3448314, EPI_ISL_3448315, EPI_ISL_3448316, EPI_ISL_3448317, EPI_ISL_3448318, EPI_ISL_3448319, EPI_ISL_3448320, EPI_ISL_3448321, EPI_ISL_3448322, EPI_ISL_3448323, EPI_ISL_3448324, EPI_ISL_3448325, EPI_ISL_3448326, EPI_ISL_3448327, EPI_ISL_3448328, EPI_ISL_3448329, EPI_ISL_3448330, EPI_ISL_3448331, EPI_ISL_3448332, EPI_ISL_3448333, EPI_ISL_3448334, EPI_ISL_3448335, EPI_ISL_3448336, EPI_ISL_3448337, EPI_ISL_3448338, EPI_ISL_3448339, EPI_ISL_3448340, EPI_ISL_3448341, EPI_ISL_3448342, EPI_ISL_3448343, EPI_ISL_3448344, EPI_ISL_3448345, EPI_ISL_3448346, EPI_ISL_3448347, EPI_ISL_3448348, EPI_ISL_3448349, EPI_ISL_3448350, EPI_ISL_3448351, EPI_ISL_3448353, EPI_ISL_3448354, EPI_ISL_3448355, EPI_ISL_3448356, EPI_ISL_3448357, EPI_ISL_3448358, EPI_ISL_3448360, EPI_ISL_3448361, EPI_ISL_3448362, EPI_ISL_3448363, EPI_ISL_3448364, EPI_ISL_3448365, EPI_ISL_3448366, EPI_ISL_3448367, EPI_ISL_3448368, EPI_ISL_3448369, EPI_ISL_3448370, EPI_ISL_3448371, EPI_ISL_3448372, EPI_ISL_3448373, EPI_ISL_3448374, EPI_ISL_3448375, EPI_ISL_3448376, EPI_ISL_3448377, EPI_ISL_3448378, EPI_ISL_3448379, EPI_ISL_3448380, EPI_ISL_3448381, EPI_ISL_3448382, EPI_ISL_3448383, EPI_ISL_3448384, EPI_ISL_3448385, EPI_ISL_3448386, EPI_ISL_3448388, EPI_ISL_3448389, EPI_ISL_3448390, EPI_ISL_3448391, EPI_ISL_3448392, EPI_ISL_3448393, EPI_ISL_3448394, EPI_ISL_3448395, EPI_ISL_3448396, EPI_ISL_3448397, EPI_ISL_3448398, EPI_ISL_3448399, EPI_ISL_3448400, EPI_ISL_3448401, EPI_ISL_3448402, EPI_ISL_3448403, EPI_ISL_3448404, EPI_ISL_3448405, EPI_ISL_3448406, EPI_ISL_3448407, EPI_ISL_3448408, EPI_ISL_3448412, EPI_ISL_3448413, EPI_ISL_3448425, EPI_ISL_3448426, EPI_ISL_3448427, EPI_ISL_3448428, EPI_ISL_3448429, EPI_ISL_3448430, EPI_ISL_3448431, EPI_ISL_3448432, EPI_ISL_3448433, EPI_ISL_3448434, EPI_ISL_3448435, EPI_ISL_3448436, EPI_ISL_3448437, EPI_ISL_3448438, EPI_ISL_3448439, EPI_ISL_3448440, EPI_ISL_3448442, EPI_ISL_3448443, EPI_ISL_3448444, EPI_ISL_3448445, EPI_ISL_3448446, EPI_ISL_3448447, EPI_ISL_3448448, EPI_ISL_3448449, EPI_ISL_3448450, EPI_ISL_3448451, EPI_ISL_3448452, EPI_ISL_3448453, EPI_ISL_3448454, EPI_ISL_3448455, EPI_ISL_3448456, EPI_ISL_3448457, EPI_ISL_3448458, EPI_ISL_3448459, EPI_ISL_3448460, EPI_ISL_3448461, EPI_ISL_3448462, EPI_ISL_3448463, EPI_ISL_3448464, EPI_ISL_3448465, EPI_ISL_3448466, EPI_ISL_3448467, EPI_ISL_3448468, EPI_ISL_3448469, EPI_ISL_3448470, EPI_ISL_3448471, EPI_ISL_3448472, EPI_ISL_3448473, EPI_ISL_3448474, EPI_ISL_3448475, EPI_ISL_3448476, EPI_ISL_3448477, EPI_ISL_3448478, EPI_ISL_3448479, EPI_ISL_3448480, EPI_ISL_3448481, EPI_ISL_3448482, EPI_ISL_3448483, EPI_ISL_3448484, EPI_ISL_3448486, EPI_ISL_3448487, EPI_ISL_3448488, EPI_ISL_3448489, EPI_ISL_3448490, EPI_ISL_3448491, EPI_ISL_3448492, EPI_ISL_3448493, EPI_ISL_3448494, EPI_ISL_3448495, EPI_ISL_3448496, EPI_ISL_3448497, EPI_ISL_3448498, EPI_ISL_3448499, EPI_ISL_3448500, EPI_ISL_3448502, EPI_ISL_3448503, EPI_ISL_3448504, EPI_ISL_3448505, EPI_ISL_3448506, EPI_ISL_3448509, EPI_ISL_3448510, EPI_ISL_3448511, EPI_ISL_3448512, EPI_ISL_3448523, EPI_ISL_3448525, EPI_ISL_3448526, EPI_ISL_3448527, EPI_ISL_3448528, EPI_ISL_3448529, EPI_ISL_3448530, EPI_ISL_3448531, EPI_ISL_3448532, EPI_ISL_3448533, EPI_ISL_3448534, EPI_ISL_3448535, EPI_ISL_3448536, EPI_ISL_3448537, EPI_ISL_3448538, EPI_ISL_3448539, EPI_ISL_3448540, EPI_ISL_3448541, EPI_ISL_3448542, EPI_ISL_3448543, EPI_ISL_3448546, EPI_ISL_3448547, EPI_ISL_3448548, EPI_ISL_3448549, EPI_ISL_3448550, EPI_ISL_3448551, EPI_ISL_3448552, EPI_ISL_3448573, EPI_ISL_3448574, EPI_ISL_3448575, EPI_ISL_3448576, EPI_ISL_3448577, EPI_ISL_3448578, EPI_ISL_3448579, EPI_ISL_3448580, EPI_ISL_3448581, EPI_ISL_3448582, EPI_ISL_3448583, EPI_ISL_3448584, EPI_ISL_3448585, EPI_ISL_3448586, EPI_ISL_3448587, EPI_ISL_3448588, EPI_ISL_3448589, EPI_ISL_3448590, EPI_ISL_3448591, EPI_ISL_3448592, EPI_ISL_3448593, EPI_ISL_3448594, EPI_ISL_3448595, EPI_ISL_3448596, EPI_ISL_3448597, EPI_ISL_3448598, EPI_ISL_3448599, EPI_ISL_3448600, EPI_ISL_3448601, EPI_ISL_3448602, EPI_ISL_3448603, EPI_ISL_3448604, EPI_ISL_3448605, EPI_ISL_3448606, EPI_ISL_3448607, EPI_ISL_3448608, EPI_ISL_3448609, EPI_ISL_3448610, EPI_ISL_3448611, EPI_ISL_3448612, EPI_ISL_3448613, EPI_ISL_3448614, EPI_ISL_3448615, EPI_ISL_3448616, EPI_ISL_3448617, EPI_ISL_3448618, EPI_ISL_3448619, EPI_ISL_3448620, EPI_ISL_3448621, EPI_ISL_3448622, EPI_ISL_3448623, EPI_ISL_3448624, EPI_ISL_3448625, EPI_ISL_3448626, EPI_ISL_3448627, EPI_ISL_3448628, EPI_ISL_3448629, EPI_ISL_3448630, EPI_ISL_3448631, EPI_ISL_3448632, EPI_ISL_3448633, EPI_ISL_3448634, EPI_ISL_3448635, EPI_ISL_3448636, EPI_ISL_3448637, EPI_ISL_3448638, EPI_ISL_3448639, EPI_ISL_3448640, EPI_ISL_3448641, EPI_ISL_3448642, EPI_ISL_3448643, EPI_ISL_3448644, EPI_ISL_3448645, EPI_ISL_3448646, EPI_ISL_3448647, EPI_ISL_3448648, EPI_ISL_3448649, EPI_ISL_3448650, EPI_ISL_3448651, EPI_ISL_3448652, EPI_ISL_3448653, EPI_ISL_3448654, EPI_ISL_3448655, EPI_ISL_3448656, EPI_ISL_3448657, EPI_ISL_3448658, EPI_ISL_3448659, EPI_ISL_3448660, EPI_ISL_3448661, EPI_ISL_3448662, EPI_ISL_3448663, EPI_ISL_3448664, EPI_ISL_3448665, EPI_ISL_3448666, EPI_ISL_3448667, EPI_ISL_3448668, EPI_ISL_3448669, EPI_ISL_3448670, EPI_ISL_3448671, EPI_ISL_3448672, EPI_ISL_3448673, EPI_ISL_3448674, EPI_ISL_3448675, EPI_ISL_3448676, EPI_ISL_3448677, EPI_ISL_3448678, EPI_ISL_3448679, EPI_ISL_3448680, EPI_ISL_3448681, EPI_ISL_3448682, EPI_ISL_3448683, EPI_ISL_3448684, EPI_ISL_3448685, EPI_ISL_3448686, EPI_ISL_3448687, EPI_ISL_3448688, EPI_ISL_3448689, EPI_ISL_3448690, EPI_ISL_3448691, EPI_ISL_3448692, EPI_ISL_3448693, EPI_ISL_3448694, EPI_ISL_3448695, EPI_ISL_3448696, EPI_ISL_3448697, EPI_ISL_3448698, EPI_ISL_3448699, EPI_ISL_3448700, EPI_ISL_3448701, EPI_ISL_3448702, EPI_ISL_3448703, EPI_ISL_3448704, EPI_ISL_3448705, EPI_ISL_3448706, EPI_ISL_3448707, EPI_ISL_3448708, EPI_ISL_3448709, EPI_ISL_3448710, EPI_ISL_3448711, EPI_ISL_3448712, EPI_ISL_3448713, EPI_ISL_3448714, EPI_ISL_3448715, EPI_ISL_3448716, EPI_ISL_3448717, EPI_ISL_3448718, EPI_ISL_3448719, EPI_ISL_3448720, EPI_ISL_3448721, EPI_ISL_3448722, EPI_ISL_3448723, EPI_ISL_3448724, EPI_ISL_3448725, EPI_ISL_3448726, EPI_ISL_3448727, EPI_ISL_3448728, EPI_ISL_3448729, EPI_ISL_3448730, EPI_ISL_3448731, EPI_ISL_3448732, EPI_ISL_3448733, EPI_ISL_3448734, EPI_ISL_3448735, EPI_ISL_3448736, EPI_ISL_3448737, EPI_ISL_3448738, EPI_ISL_3448739, EPI_ISL_3448740, EPI_ISL_3448741, EPI_ISL_3448742, EPI_ISL_3448743, EPI_ISL_3448744, EPI_ISL_3448745, EPI_ISL_3448746, EPI_ISL_3448747, EPI_ISL_3448748, EPI_ISL_3448749, EPI_ISL_3448750, EPI_ISL_3448751, EPI_ISL_3448752, EPI_ISL_3448753, EPI_ISL_3448754, EPI_ISL_3448755, EPI_ISL_3448756, EPI_ISL_3448757, EPI_ISL_3448758, EPI_ISL_3448759, EPI_ISL_3448760, EPI_ISL_3448761, EPI_ISL_3448762, EPI_ISL_3448763, EPI_ISL_3448764, EPI_ISL_3448765, EPI_ISL_3448766, EPI_ISL_3448767, EPI_ISL_3448768, EPI_ISL_3448769, EPI_ISL_3448770, EPI_ISL_3448771, EPI_ISL_3448772, EPI_ISL_3448773, EPI_ISL_3448774, EPI_ISL_3448775, EPI_ISL_3448776, EPI_ISL_3448777, EPI_ISL_3448778, EPI_ISL_3448779, EPI_ISL_3448780, EPI_ISL_3448781, EPI_ISL_3448782, EPI_ISL_3448783, EPI_ISL_3448784, EPI_ISL_3448785, EPI_ISL_3448786, EPI_ISL_3448787, EPI_ISL_3448788, EPI_ISL_3448789, EPI_ISL_3448790, EPI_ISL_3448791, EPI_ISL_3448792, EPI_ISL_3448793, EPI_ISL_3448794, EPI_ISL_3448795, EPI_ISL_3448796, EPI_ISL_3448797, EPI_ISL_3448798, EPI_ISL_3448799, EPI_ISL_3448800, EPI_ISL_3448801, EPI_ISL_3448802, EPI_ISL_3448803, EPI_ISL_3448804, EPI_ISL_3448805, EPI_ISL_3448806, EPI_ISL_3448807, EPI_ISL_3448808, EPI_ISL_3448809, EPI_ISL_3448810, EPI_ISL_3448811, EPI_ISL_3448812, EPI_ISL_3448813, EPI_ISL_3448814, EPI_ISL_3448815, EPI_ISL_3448816, EPI_ISL_3448817, EPI_ISL_3448818, EPI_ISL_3448819, EPI_ISL_3448820, EPI_ISL_3448821, EPI_ISL_3448822, EPI_ISL_3448823, EPI_ISL_3448824, EPI_ISL_3448825, EPI_ISL_3448826, EPI_ISL_3448827, EPI_ISL_3448828, EPI_ISL_3448829, EPI_ISL_3448830, EPI_ISL_3448831, EPI_ISL_3448832, EPI_ISL_3448833, EPI_ISL_3448834, EPI_ISL_3448835, EPI_ISL_3448836, EPI_ISL_3448837, EPI_ISL_3448838, EPI_ISL_3448839, EPI_ISL_3448840, EPI_ISL_3448841, EPI_ISL_3448842, EPI_ISL_3448843, EPI_ISL_3448844, EPI_ISL_3448845, EPI_ISL_3448846, EPI_ISL_3448847, EPI_ISL_3448848 | | | |
| see above | Public Health Ontario Laboratory | Public Health Ontario Laboratory | Vanessa G Allen, Philip Banh, Yao Chen, Richard de Borja, Alireza Eshaghi, Nahuel Fittipaldi, Christine Frantz, Jonathan B Gubbay, Jennifer L Guthrie, Lawrence Heisler, Esha Joshi, Michael Laszloffy, Aimin Li, Michael CY Li, Dean Maxwell, Sandeep Nagra, Samir N Patel, Jared Simpson, Karthikeyan Sivaraman, Ashleigh Sullivan, Yogi Sundaravadanam, Sarah Teatero, Andre Villegas, Matthew Watson, Sandra Zittermann |
| EPI_ISL_3448953, EPI_ISL_3448954, EPI_ISL_3448955 | Division of Emerging Infectious Diseases, Bureau of Infectious Diseases Diagnosis Control, Korea Disease Control and Prevention Agency | Division of Emerging Infectious Diseases, Bureau of Infectious Diseases Diagnosis Control, Korea Disease Control and Prevention Agency | Ae Kyung Park, Il-Hwan Kim, Heui Man Kim, Jeong-Ah Kim, Chae Young Lee, Eun-Jin Kim |
| EPI_ISL_3448956, EPI_ISL_3448957, EPI_ISL_3448958, EPI_ISL_3448959, EPI_ISL_3448960, EPI_ISL_3448961, EPI_ISL_3448962, EPI_ISL_3448963, EPI_ISL_3448964, EPI_ISL_3448965, EPI_ISL_3448966, EPI_ISL_3448967, EPI_ISL_3448968, EPI_ISL_3448971, EPI_ISL_3448972, EPI_ISL_3448973, EPI_ISL_3448974, EPI_ISL_3448975, EPI_ISL_3448976, EPI_ISL_3448977, EPI_ISL_3448978, EPI_ISL_3448979, EPI_ISL_3448980, EPI_ISL_3448981, EPI_ISL_3448982, EPI_ISL_3448983, EPI_ISL_3448984, EPI_ISL_3448985, EPI_ISL_3448986, EPI_ISL_3448987, EPI_ISL_3448988, EPI_ISL_3448989, EPI_ISL_3448990, EPI_ISL_3448991, EPI_ISL_3448992, EPI_ISL_3448993, EPI_ISL_3448994, EPI_ISL_3448995, EPI_ISL_3448996, EPI_ISL_3448997, EPI_ISL_3448998, EPI_ISL_3448999, EPI_ISL_3449000, EPI_ISL_3449001, EPI_ISL_3449002, EPI_ISL_3449003, EPI_ISL_3449004, EPI_ISL_3449005, EPI_ISL_3449006, EPI_ISL_3449007, EPI_ISL_3449008, EPI_ISL_3449009, EPI_ISL_3449010, EPI_ISL_3449011, EPI_ISL_3449012, EPI_ISL_3449013, EPI_ISL_3449014, EPI_ISL_3449015, EPI_ISL_3449016, EPI_ISL_3449019, EPI_ISL_3449020, EPI_ISL_3449021, EPI_ISL_3449022, EPI_ISL_3449023, EPI_ISL_3449024, EPI_ISL_3449025, EPI_ISL_3449026, EPI_ISL_3449027, EPI_ISL_3449028, EPI_ISL_3449029, EPI_ISL_3449030, EPI_ISL_3449031, EPI_ISL_3449032, EPI_ISL_3449033, EPI_ISL_3449034, EPI_ISL_3449035, EPI_ISL_3449036, EPI_ISL_3449037, EPI_ISL_3449038, EPI_ISL_3449039, EPI_ISL_3449040, EPI_ISL_3449041, EPI_ISL_3449042, EPI_ISL_3449043, EPI_ISL_3449044, EPI_ISL_3449045, EPI_ISL_3449046, EPI_ISL_3449047, EPI_ISL_3449048, EPI_ISL_3449049, EPI_ISL_3449050, EPI_ISL_3449051, EPI_ISL_3449052, EPI_ISL_3449053, EPI_ISL_3449054, EPI_ISL_3449055, EPI_ISL_3449056, EPI_ISL_3449057, EPI_ISL_3449058, EPI_ISL_3449059, EPI_ISL_3449060, EPI_ISL_3449061, EPI_ISL_3449062, EPI_ISL_3449063, EPI_ISL_3449064, EPI_ISL_3449065, EPI_ISL_3449066, EPI_ISL_3449067, EPI_ISL_3449068, EPI_ISL_3449069, EPI_ISL_3449070, EPI_ISL_3449071, EPI_ISL_3449072, EPI_ISL_3449073, EPI_ISL_3449074, EPI_ISL_3449075, EPI_ISL_3449076, EPI_ISL_3449077, EPI_ISL_3449078, EPI_ISL_3449079, EPI_ISL_3449080, EPI_ISL_3449081, EPI_ISL_3449082, EPI_ISL_3449083, EPI_ISL_3449084, EPI_ISL_3449085, EPI_ISL_3449086, EPI_ISL_3449087, EPI_ISL_3449088, EPI_ISL_3449089, EPI_ISL_3449090, EPI_ISL_3449091, | | | |

| | | | |
|---|---|---|---|
| EPI_ISL_3449092, EPI_ISL_3449093, EPI_ISL_3449094, EPI_ISL_3449095, EPI_ISL_3449096, EPI_ISL_3449097, EPI_ISL_3449098, EPI_ISL_3449099, EPI_ISL_3449100, EPI_ISL_3449101, EPI_ISL_3449102, EPI_ISL_3449103, EPI_ISL_3449104, EPI_ISL_3449105, EPI_ISL_3449106, EPI_ISL_3449107, EPI_ISL_3449108, EPI_ISL_3449109, EPI_ISL_3449110, EPI_ISL_3449111, EPI_ISL_3449112, EPI_ISL_3449113, EPI_ISL_3449114, EPI_ISL_3449115, EPI_ISL_3449116, EPI_ISL_3449117, EPI_ISL_3449118, EPI_ISL_3449119, EPI_ISL_3449120, EPI_ISL_3449121, EPI_ISL_3449122, EPI_ISL_3449123, EPI_ISL_3449124, EPI_ISL_3449125, EPI_ISL_3449126, EPI_ISL_3449127, EPI_ISL_3449128, EPI_ISL_3449129, EPI_ISL_3449130, EPI_ISL_3449131, EPI_ISL_3449132, EPI_ISL_3449133, EPI_ISL_3449134, EPI_ISL_3449135, EPI_ISL_3449136, EPI_ISL_3449137, EPI_ISL_3449138, EPI_ISL_3449139, EPI_ISL_3449140, EPI_ISL_3449141, EPI_ISL_3449142, EPI_ISL_3449143, EPI_ISL_3449144, EPI_ISL_3449145, EPI_ISL_3449146, EPI_ISL_3449147, EPI_ISL_3449148, EPI_ISL_3449149, EPI_ISL_3449150, EPI_ISL_3449151, EPI_ISL_3449152, EPI_ISL_3449153, EPI_ISL_3449154, EPI_ISL_3449155, EPI_ISL_3449156, EPI_ISL_3449157, EPI_ISL_3449158, EPI_ISL_3449159, EPI_ISL_3449160, EPI_ISL_3449161, EPI_ISL_3449162, EPI_ISL_3449163, EPI_ISL_3449164, EPI_ISL_3449165, EPI_ISL_3449166, EPI_ISL_3449167, EPI_ISL_3449168, EPI_ISL_3449169, EPI_ISL_3449170, EPI_ISL_3449171, EPI_ISL_3449172, EPI_ISL_3449173, EPI_ISL_3449174, EPI_ISL_3449175, EPI_ISL_3449176, EPI_ISL_3449177, EPI_ISL_3449178, EPI_ISL_3449179, EPI_ISL_3449180, EPI_ISL_3449181, EPI_ISL_3449182, EPI_ISL_3449183, EPI_ISL_3449184, EPI_ISL_3449185, EPI_ISL_3449186, EPI_ISL_3449187, EPI_ISL_3449188, EPI_ISL_3449189, EPI_ISL_3449190, EPI_ISL_3449191, EPI_ISL_3449192, EPI_ISL_3449193, EPI_ISL_3449194, EPI_ISL_3449195, EPI_ISL_3449196, EPI_ISL_3449197, EPI_ISL_3449198, EPI_ISL_3449199, EPI_ISL_3449200, EPI_ISL_3449201, EPI_ISL_3449202, EPI_ISL_3449203, EPI_ISL_3449204, EPI_ISL_3449205, EPI_ISL_3449206, EPI_ISL_3449207, EPI_ISL_3449208, EPI_ISL_3449209, EPI_ISL_3449210, EPI_ISL_3449211, EPI_ISL_3449212, EPI_ISL_3449213, EPI_ISL_3449214, EPI_ISL_3449215, EPI_ISL_3449216, EPI_ISL_3449217, EPI_ISL_3449218, EPI_ISL_3449219, EPI_ISL_3449220, EPI_ISL_3449221, EPI_ISL_3449222, EPI_ISL_3449223, EPI_ISL_3449224, EPI_ISL_3449225, EPI_ISL_3449226, EPI_ISL_3449227, EPI_ISL_3449228, EPI_ISL_3449229, EPI_ISL_3449230, EPI_ISL_3449231, EPI_ISL_3449232 | | | |
| see above | UW Virology Lab | UW Virology Lab | Pavitra Roychoudhury, Shah Mohamed Bakhash, Ricardo Perez, Maria Lukes, Hong Xie, Tien V. Nguyen, Savanna S. Carmack, Sean Ellis, Nathan Breit, Robert J. Livingston, Meei-Li Huang, Keith R Jerome, Patrick Mathias, Alexander Greninger |
| EPI_ISL_3449234, EPI_ISL_3449235, EPI_ISL_3449236, EPI_ISL_3449237, EPI_ISL_3449238, EPI_ISL_3449239, EPI_ISL_3449240, EPI_ISL_3449241, EPI_ISL_3449242, EPI_ISL_3449243, EPI_ISL_3449244, EPI_ISL_3449245, EPI_ISL_3449246, EPI_ISL_3449247, EPI_ISL_3449248, EPI_ISL_3449249, EPI_ISL_3449250, EPI_ISL_3449251, EPI_ISL_3449252, EPI_ISL_3449253, EPI_ISL_3449254, EPI_ISL_3449255, EPI_ISL_3449256, EPI_ISL_3449257, EPI_ISL_3449258, EPI_ISL_3449259, EPI_ISL_3449260, EPI_ISL_3449261, EPI_ISL_3449262, EPI_ISL_3449263, EPI_ISL_3449264, EPI_ISL_3449265, EPI_ISL_3449266, EPI_ISL_3449267, EPI_ISL_3449268, EPI_ISL_3449269 | | | |
| see above | Oregon State Public Health Laboratory | Oregon State Public Health Laboratory | Rafia Razzaque, Eugene Yeboah, Vanda Makris, Laura Tsaknaridis, John Fontana and Shane Sevey |
| EPI_ISL_3449270, EPI_ISL_3449271, EPI_ISL_3449272, EPI_ISL_3449273, EPI_ISL_3449274, EPI_ISL_3449275, EPI_ISL_3449276, EPI_ISL_3449277, EPI_ISL_3449278, EPI_ISL_3449279, EPI_ISL_3449280, EPI_ISL_3449281, EPI_ISL_3449282, EPI_ISL_3449283, EPI_ISL_3449284, EPI_ISL_3449285, EPI_ISL_3449286, EPI_ISL_3449287, EPI_ISL_3449288, EPI_ISL_3449289, EPI_ISL_3449290, EPI_ISL_3449291, EPI_ISL_3449292, EPI_ISL_3449293, EPI_ISL_3449294, EPI_ISL_3449295, EPI_ISL_3449296, EPI_ISL_3449297, EPI_ISL_3449298, EPI_ISL_3449299, EPI_ISL_3449300, EPI_ISL_3449301, EPI_ISL_3449302, EPI_ISL_3449303 | | | |
| see above | Plateforme de testing Namuroise | Plateforme de testing Namuroise | Lesly Nyinkeu Kemamen; Nicolas Gilliard; Demars Aurore ; Maschietto Céline ; Nobis Chloé ; Otto Gaetan ; Denis Olivier ; Degosserie Jonathan ; Mullier François |
| EPI_ISL_3449306, EPI_ISL_3449307, EPI_ISL_3449308, EPI_ISL_3449309, EPI_ISL_3449310, EPI_ISL_3449311, EPI_ISL_3449312, EPI_ISL_3449313, EPI_ISL_3449315, EPI_ISL_3449316, EPI_ISL_3449317, EPI_ISL_3449318, EPI_ISL_3449319 | | | |
| see above | Department of Microbiology, Yamanashi Institute for Public Health and Environment | Genome Analysis Center, Yamanashi Central Hospital | Yosuke Hirotsu |
| EPI_ISL_3449320, EPI_ISL_3449321, EPI_ISL_3449322, EPI_ISL_3449323, EPI_ISL_3449324, EPI_ISL_3449325, EPI_ISL_3449326, EPI_ISL_3449327, EPI_ISL_3449328, EPI_ISL_3449329, EPI_ISL_3449330, EPI_ISL_3449331, EPI_ISL_3449332, EPI_ISL_3449333, EPI_ISL_3449334, EPI_ISL_3449335, EPI_ISL_3449336, EPI_ISL_3449337, EPI_ISL_3449338, EPI_ISL_3449339, EPI_ISL_3449340, EPI_ISL_3449341, EPI_ISL_3449342, EPI_ISL_3449343, EPI_ISL_3449344, EPI_ISL_3449345, EPI_ISL_3449346, EPI_ISL_3449347, EPI_ISL_3449348, EPI_ISL_3449349, EPI_ISL_3449350, EPI_ISL_3449351, EPI_ISL_3449352, EPI_ISL_3449353 | | | |
| see above | Labo Analyses Med | CERBA HealthCare | Bénédicte Roquebert, Laura Verdurme, Sabine Trombert, Stéphanie Haim-Boukobza |
| EPI_ISL_3449354, EPI_ISL_3449355, EPI_ISL_3449356, EPI_ISL_3449357, EPI_ISL_3449358, EPI_ISL_3449359, EPI_ISL_3449360, EPI_ISL_3449361, EPI_ISL_3449362, EPI_ISL_3449363, EPI_ISL_3449364, EPI_ISL_3449365, EPI_ISL_3449366, EPI_ISL_3449367, EPI_ISL_3449368, EPI_ISL_3449369, EPI_ISL_3449370, EPI_ISL_3449371, EPI_ISL_3449372, EPI_ISL_3449373, EPI_ISL_3449374, EPI_ISL_3449375, EPI_ISL_3449376, EPI_ISL_3449377, EPI_ISL_3449378, EPI_ISL_3449379, EPI_ISL_3449380, EPI_ISL_3449381, EPI_ISL_3449382, EPI_ISL_3449383, EPI_ISL_3449384, EPI_ISL_3449385, EPI_ISL_3449386, EPI_ISL_3449387, EPI_ISL_3449388, EPI_ISL_3449389, EPI_ISL_3449390, EPI_ISL_3449391, EPI_ISL_3449392, EPI_ISL_3449393, EPI_ISL_3449394, EPI_ISL_3449395, EPI_ISL_3449396, EPI_ISL_3449397, EPI_ISL_3449398, EPI_ISL_3449399, EPI_ISL_3449400, EPI_ISL_3449401, EPI_ISL_3449402, EPI_ISL_3449403, EPI_ISL_3449404, EPI_ISL_3449405, EPI_ISL_3449406, EPI_ISL_3449407, EPI_ISL_3449408, EPI_ISL_3449409, EPI_ISL_3449410, EPI_ISL_3449411, EPI_ISL_3449412, EPI_ISL_3449413, EPI_ISL_3449414, EPI_ISL_3449415, EPI_ISL_3449416, EPI_ISL_3449417 | | | |
| see above | Department of Bacteria, Parasites and Fungi, Statens Serum Institut, Copenhagen, Denmark | Statens Serum Institut Bioinformatics and Microbial Genomics | Danish Covid-19 Genome Consortium |
| EPI_ISL_3449418 | Tulare County Public Health Laboratory | Tulare County Public Health Laboratory | Denise L Lopez, Joshua Kootstra |
| EPI_ISL_3449419, EPI_ISL_3449420 | Department of Bacteria, Parasites and Fungi, Statens Serum Institut, Copenhagen, Denmark | Statens Serum Institut Bioinformatics and Microbial Genomics | Danish Covid-19 Genome Consortium |
| EPI_ISL_3449421 | Tulare County Public Health Laboratory | Tulare County Public Health Laboratory | Denise L Lopez, Joshua Kootstra |
| EPI_ISL_3449422 | Department of Bacteria, Parasites and Fungi, Statens Serum Institut, Copenhagen, Denmark | Statens Serum Institut Bioinformatics and Microbial Genomics | Danish Covid-19 Genome Consortium |
| EPI_ISL_3449423 | Tulare County Public Health Laboratory | Tulare County Public Health Laboratory | Denise L Lopez, Joshua Kootstra |
| EPI_ISL_3449424, EPI_ISL_3449426 | Department of Bacteria, Parasites and Fungi, Statens Serum Institut, Copenhagen, Denmark | Statens Serum Institut Bioinformatics and Microbial Genomics | Danish Covid-19 Genome Consortium |
| EPI_ISL_3449427 | Tulare County Public Health Laboratory | Tulare County Public Health Laboratory | Denise L Lopez, Joshua Kootstra |
| EPI_ISL_3449428 | Department of Bacteria, Parasites and Fungi, Statens Serum Institut, Copenhagen, Denmark | Statens Serum Institut Bioinformatics and Microbial Genomics | Danish Covid-19 Genome Consortium |
| EPI_ISL_3449429 | Tulare County Public Health Laboratory | Tulare County Public Health Laboratory | Denise L Lopez, Joshua Kootstra |
| EPI_ISL_3449430, EPI_ISL_3449431 | Department of Bacteria, Parasites and Fungi, Statens Serum Institut, Copenhagen, Denmark | Statens Serum Institut Bioinformatics and Microbial Genomics | Danish Covid-19 Genome Consortium |
| EPI_ISL_3449432 | Tulare County Public Health Laboratory | Tulare County Public Health Laboratory | Denise L Lopez, Joshua Kootstra |
| EPI_ISL_3449433 | Department of Bacteria, Parasites and Fungi, Statens Serum Institut, Copenhagen, Denmark | Statens Serum Institut Bioinformatics and Microbial Genomics | Danish Covid-19 Genome Consortium |
| EPI_ISL_3449434 | Tulare County Public Health Laboratory | Tulare County Public Health Laboratory | Denise L Lopez, Joshua Kootstra |
| EPI_ISL_3449435 | Department of Bacteria, Parasites and Fungi, Statens Serum Institut, Copenhagen, Denmark | Statens Serum Institut Bioinformatics and Microbial Genomics | Danish Covid-19 Genome Consortium |
| EPI_ISL_3449436 | Tulare County Public Health Laboratory | Tulare County Public Health Laboratory | Denise L Lopez, Joshua Kootstra |
| EPI_ISL_3449437 | Department of Bacteria, Parasites and Fungi, Statens Serum Institut, Copenhagen, Denmark | Statens Serum Institut Bioinformatics and Microbial Genomics | Danish Covid-19 Genome Consortium |
| EPI_ISL_3449438 | Tulare County Public Health Laboratory | Tulare County Public Health Laboratory | Denise L Lopez, Joshua Kootstra |
| EPI_ISL_3449439 | Department of Bacteria, Parasites and Fungi, Statens Serum Institut, Copenhagen, Denmark | Statens Serum Institut Bioinformatics and Microbial Genomics | Danish Covid-19 Genome Consortium |
| EPI_ISL_3449440 | Tulare County Public Health Laboratory | Tulare County Public Health Laboratory | Denise L Lopez, Joshua Kootstra |
| EPI_ISL_3449441, EPI_ISL_3449442 | Department of Bacteria, Parasites and Fungi, Statens Serum Institut, Copenhagen, Denmark | Statens Serum Institut Bioinformatics and Microbial Genomics | Danish Covid-19 Genome Consortium |
| EPI_ISL_3449443 | Tulare County Public Health Laboratory | Tulare County Public Health Laboratory | Denise L Lopez, Joshua Kootstra |
| EPI_ISL_3449444 | Department of Bacteria, Parasites and Fungi, Statens Serum Institut, Copenhagen, Denmark | Statens Serum Institut Bioinformatics and Microbial Genomics | Danish Covid-19 Genome Consortium |
| EPI_ISL_3449445 | Tulare County Public Health Laboratory | Tulare County Public Health Laboratory | Denise L Lopez, Joshua Kootstra |
| EPI_ISL_3449446 | Department of Bacteria, Parasites and Fungi, Statens Serum Institut, Copenhagen, Denmark | Statens Serum Institut Bioinformatics and Microbial Genomics | Danish Covid-19 Genome Consortium |
| EPI_ISL_3449447 | Tulare County Public Health Laboratory | Tulare County Public Health Laboratory | Denise L Lopez, Joshua Kootstra |
| EPI_ISL_3449448 | Department of Bacteria, Parasites and Fungi, Statens Serum Institut, Copenhagen, Denmark | Statens Serum Institut Bioinformatics and Microbial Genomics | Danish Covid-19 Genome Consortium |
| EPI_ISL_3449449 | Tulare County Public Health Laboratory | Tulare County Public Health Laboratory | Denise L Lopez, Joshua Kootstra |
| EPI_ISL_3449450 | Department of Bacteria, Parasites and Fungi, Statens Serum Institut, Copenhagen, Denmark | Statens Serum Institut Bioinformatics and Microbial Genomics | Danish Covid-19 Genome Consortium |
| EPI_ISL_3449451 | Tulare County Public Health Laboratory | Tulare County Public Health Laboratory | Denise L Lopez, Joshua Kootstra |
| EPI_ISL_3449452 | Department of Bacteria, Parasites and Fungi, Statens Serum Institut, Copenhagen, Denmark | Statens Serum Institut Bioinformatics and Microbial Genomics | Danish Covid-19 Genome Consortium |
| EPI_ISL_3449453 | Tulare County Public Health Laboratory | Tulare County Public Health Laboratory | Denise L Lopez, Joshua Kootstra |
| EPI_ISL_3449454 | Department of Bacteria, Parasites and Fungi, Statens Serum Institut, Copenhagen, Denmark | Statens Serum Institut Bioinformatics and Microbial Genomics | Danish Covid-19 Genome Consortium |
| EPI_ISL_3449455 | Tulare County Public Health Laboratory | Tulare County Public Health Laboratory | Denise L Lopez, Joshua Kootstra |
| EPI_ISL_3449456 | Department of Bacteria, Parasites and Fungi, Statens Serum Institut, Copenhagen, Denmark | Statens Serum Institut Bioinformatics and Microbial Genomics | Danish Covid-19 Genome Consortium |
| EPI_ISL_3449457 | Tulare County Public Health Laboratory | Tulare County Public Health Laboratory | Denise L Lopez, Joshua Kootstra |
| EPI_ISL_3449458 | Department of Bacteria, Parasites and Fungi, Statens Serum Institut, Copenhagen, Denmark | Statens Serum Institut Bioinformatics and Microbial Genomics | Danish Covid-19 Genome Consortium |
| EPI_ISL_3449459 | Tulare County Public Health Laboratory | Tulare County Public Health Laboratory | Denise L Lopez, Joshua Kootstra |
| EPI_ISL_3449460 | Department of Bacteria, Parasites and Fungi, Statens Serum Institut, Copenhagen, Denmark | Statens Serum Institut Bioinformatics and Microbial Genomics | Danish Covid-19 Genome Consortium |
| EPI_ISL_3449461 | Tulare County Public Health Laboratory | Tulare County Public Health Laboratory | Denise L Lopez, Joshua Kootstra |
| EPI_ISL_3449462 | Department of Bacteria, Parasites and Fungi, Statens Serum Institut, Copenhagen, Denmark | Statens Serum Institut Bioinformatics and Microbial Genomics | Danish Covid-19 Genome Consortium |
| EPI_ISL_3449463 | Tulare County Public Health Laboratory | Tulare County Public Health Laboratory | Denise L Lopez, Joshua Kootstra |
| EPI_ISL_3449464 | Department of Bacteria, Parasites and Fungi, Statens Serum | Statens Serum Institut Bioinformatics and Microbial Genomics | Danish Covid-19 Genome Consortium |

| | | | |
|---|---|---|---|
| | Institut, Copenhagen, Denmark | | |
| EPI_ISL_3449465 | Tulare County Public Health Laboratory | Tulare County Public Health Laboratory | Denise L Lopez, Joshua Kootstra |
| EPI_ISL_3449466 | Department of Bacteria, Parasites and Fungi, Statens Serum Institut, Copenhagen, Denmark | Statens Serum Institut Bioinformatics and Microbial Genomics | Danish Covid-19 Genome Consortium |
| EPI_ISL_3449467 | Tulare County Public Health Laboratory | Tulare County Public Health Laboratory | Denise L Lopez, Joshua Kootstra |
| EPI_ISL_3449468 | Department of Bacteria, Parasites and Fungi, Statens Serum Institut, Copenhagen, Denmark | Statens Serum Institut Bioinformatics and Microbial Genomics | Danish Covid-19 Genome Consortium |
| EPI_ISL_3449469 | Tulare County Public Health Laboratory | Tulare County Public Health Laboratory | Denise L Lopez, Joshua Kootstra |
| EPI_ISL_3449470 | Department of Bacteria, Parasites and Fungi, Statens Serum Institut, Copenhagen, Denmark | Statens Serum Institut Bioinformatics and Microbial Genomics | Danish Covid-19 Genome Consortium |
| EPI_ISL_3449471 | Tulare County Public Health Laboratory | Tulare County Public Health Laboratory | Denise L Lopez, Joshua Kootstra |
| EPI_ISL_3449472, EPI_ISL_3449473 | Department of Bacteria, Parasites and Fungi, Statens Serum Institut, Copenhagen, Denmark | Statens Serum Institut Bioinformatics and Microbial Genomics | Danish Covid-19 Genome Consortium |
| EPI_ISL_3449474 | Tulare County Public Health Laboratory | Tulare County Public Health Laboratory | Denise L Lopez, Joshua Kootstra |
| EPI_ISL_3449475 | Department of Bacteria, Parasites and Fungi, Statens Serum Institut, Copenhagen, Denmark | Statens Serum Institut Bioinformatics and Microbial Genomics | Danish Covid-19 Genome Consortium |
| EPI_ISL_3449476 | Tulare County Public Health Laboratory | Tulare County Public Health Laboratory | Denise L Lopez, Joshua Kootstra |
| EPI_ISL_3449477 | Department of Bacteria, Parasites and Fungi, Statens Serum Institut, Copenhagen, Denmark | Statens Serum Institut Bioinformatics and Microbial Genomics | Danish Covid-19 Genome Consortium |
| EPI_ISL_3449478 | Tulare County Public Health Laboratory | Tulare County Public Health Laboratory | Denise L Lopez, Joshua Kootstra |
| EPI_ISL_3449479, EPI_ISL_3449480, EPI_ISL_3449481, EPI_ISL_3449482, EPI_ISL_3449483, EPI_ISL_3449484, EPI_ISL_3449485, EPI_ISL_3449486, EPI_ISL_3449487, EPI_ISL_3449488, EPI_ISL_3449489, EPI_ISL_3449490, EPI_ISL_3449491, EPI_ISL_3449492, EPI_ISL_3449493, EPI_ISL_3449494, EPI_ISL_3449495, EPI_ISL_3449496, EPI_ISL_3449497, EPI_ISL_3449498, EPI_ISL_3449499, EPI_ISL_3449500, EPI_ISL_3449501, EPI_ISL_3449502, EPI_ISL_3449503, EPI_ISL_3449504, EPI_ISL_3449505, EPI_ISL_3449506, EPI_ISL_3449507, EPI_ISL_3449508, EPI_ISL_3449509, EPI_ISL_3449510, EPI_ISL_3449511, EPI_ISL_3449512, EPI_ISL_3449513, EPI_ISL_3449514, EPI_ISL_3449515, EPI_ISL_3449516, EPI_ISL_3449517, EPI_ISL_3449518, EPI_ISL_3449519, EPI_ISL_3449520, EPI_ISL_3449521, EPI_ISL_3449522, EPI_ISL_3449523, EPI_ISL_3449524, EPI_ISL_3449525, EPI_ISL_3449526, EPI_ISL_3449527, EPI_ISL_3449528, EPI_ISL_3449530, EPI_ISL_3449532, EPI_ISL_3449534, EPI_ISL_3449536, EPI_ISL_3449538, EPI_ISL_3449540, EPI_ISL_3449541, EPI_ISL_3449543, EPI_ISL_3449545, EPI_ISL_3449547, EPI_ISL_3449549, EPI_ISL_3449551, EPI_ISL_3449553, EPI_ISL_3449555, EPI_ISL_3449557, EPI_ISL_3449558, EPI_ISL_3449560, EPI_ISL_3449562, EPI_ISL_3449564, EPI_ISL_3449566, EPI_ISL_3449568, EPI_ISL_3449570, EPI_ISL_3449572, EPI_ISL_3449573, EPI_ISL_3449575, EPI_ISL_3449577, EPI_ISL_3449579, EPI_ISL_3449581, EPI_ISL_3449583, EPI_ISL_3449585, EPI_ISL_3449587, EPI_ISL_3449590, EPI_ISL_3449592, EPI_ISL_3449594, EPI_ISL_3449596, EPI_ISL_3449598, EPI_ISL_3449600, EPI_ISL_3449602, EPI_ISL_3449604, EPI_ISL_3449605, EPI_ISL_3449607, EPI_ISL_3449609, EPI_ISL_3449611, EPI_ISL_3449613, EPI_ISL_3449615, EPI_ISL_3449617, EPI_ISL_3449619, EPI_ISL_3449621, EPI_ISL_3449623, EPI_ISL_3449625, EPI_ISL_3449627, EPI_ISL_3449629, EPI_ISL_3449630, EPI_ISL_3449632, EPI_ISL_3449634 | | | |
| see above | Department of Bacteria, Parasites and Fungi, Statens Serum Institut, Copenhagen, Denmark | Statens Serum Institut Bioinformatics and Microbial Genomics | Danish Covid-19 Genome Consortium |
| EPI_ISL_3449635 | Virology Laboratory, Scientific Department, Army Medical Center | Virology Laboratory, Scientific Department, Army Medical Center | Silvia Fillo, Riccardo De Sanctis, Antonella Fortunato, Anella Monte, Rossella Brandi, Giulia Campoli, Marzia Cavalli, Lucia Nicosia, Anna Anselmo, Vanessa Vera Fain, Francesco Giordani, Giandomenico Cerreto, Filippo Molinari, Giancarlo Petralito, Florigio Lista |
| EPI_ISL_3449636 | Department of Bacteria, Parasites and Fungi, Statens Serum Institut, Copenhagen, Denmark | Statens Serum Institut Bioinformatics and Microbial Genomics | Danish Covid-19 Genome Consortium |
| EPI_ISL_3449637 | Virology Laboratory, Scientific Department, Army Medical Center | Virology Laboratory, Scientific Department, Army Medical Center | Silvia Fillo, Riccardo De Sanctis, Antonella Fortunato, Anella Monte, Rossella Brandi, Giulia Campoli, Marzia Cavalli, Lucia Nicosia, Anna Anselmo, Vanessa Vera Fain, Francesco Giordani, Giandomenico Cerreto, Filippo Molinari, Giancarlo Petralito, Florigio Lista |
| EPI_ISL_3449638 | Department of Bacteria, Parasites and Fungi, Statens Serum Institut, Copenhagen, Denmark | Statens Serum Institut Bioinformatics and Microbial Genomics | Danish Covid-19 Genome Consortium |
| EPI_ISL_3449639 | Virology Laboratory, Scientific Department, Army Medical Center | Virology Laboratory, Scientific Department, Army Medical Center | Silvia Fillo, Riccardo De Sanctis, Antonella Fortunato, Anella Monte, Rossella Brandi, Giulia Campoli, Marzia Cavalli, Lucia Nicosia, Anna Anselmo, Vanessa Vera Fain, Francesco Giordani, Giandomenico Cerreto, Filippo Molinari, Giancarlo Petralito, Florigio Lista |
| EPI_ISL_3449640 | Department of Bacteria, Parasites and Fungi, Statens Serum Institut, Copenhagen, Denmark | Statens Serum Institut Bioinformatics and Microbial Genomics | Danish Covid-19 Genome Consortium |
| EPI_ISL_3449641 | Virology Laboratory, Scientific Department, Army Medical Center | Virology Laboratory, Scientific Department, Army Medical Center | Silvia Fillo, Riccardo De Sanctis, Antonella Fortunato, Anella Monte, Rossella Brandi, Giulia Campoli, Marzia Cavalli, Lucia Nicosia, Anna Anselmo, Vanessa Vera Fain, Francesco Giordani, Giandomenico Cerreto, Filippo Molinari, Giancarlo Petralito, Florigio Lista |
| EPI_ISL_3449642 | Department of Bacteria, Parasites and Fungi, Statens Serum Institut, Copenhagen, Denmark | Statens Serum Institut Bioinformatics and Microbial Genomics | Danish Covid-19 Genome Consortium |
| EPI_ISL_3449643 | Virology Laboratory, Scientific Department, Army Medical Center | Virology Laboratory, Scientific Department, Army Medical Center | Silvia Fillo, Riccardo De Sanctis, Antonella Fortunato, Anella Monte, Rossella Brandi, Giulia Campoli, Marzia Cavalli, Lucia Nicosia, Anna Anselmo, Vanessa Vera Fain, Francesco Giordani, Giandomenico Cerreto, Filippo Molinari, Giancarlo Petralito, Florigio Lista |
| EPI_ISL_3449644 | Department of Bacteria, Parasites and Fungi, Statens Serum Institut, Copenhagen, Denmark | Statens Serum Institut Bioinformatics and Microbial Genomics | Danish Covid-19 Genome Consortium |
| EPI_ISL_3449645 | Virology Laboratory, Scientific Department, Army Medical Center | Virology Laboratory, Scientific Department, Army Medical Center | Silvia Fillo, Riccardo De Sanctis, Antonella Fortunato, Anella Monte, Rossella Brandi, Giulia Campoli, Marzia Cavalli, Lucia Nicosia, Anna Anselmo, Vanessa Vera Fain, Francesco Giordani, Giandomenico Cerreto, Filippo Molinari, Giancarlo Petralito, Florigio Lista |
| EPI_ISL_3449646 | Department of Bacteria, Parasites and Fungi, Statens Serum Institut, Copenhagen, Denmark | Statens Serum Institut Bioinformatics and Microbial Genomics | Danish Covid-19 Genome Consortium |
| EPI_ISL_3449647 | Virology Laboratory, Scientific Department, Army Medical Center | Virology Laboratory, Scientific Department, Army Medical Center | Silvia Fillo, Riccardo De Sanctis, Antonella Fortunato, Anella Monte, Rossella Brandi, Giulia Campoli, Marzia Cavalli, Lucia Nicosia, Anna Anselmo, Vanessa Vera Fain, Francesco Giordani, Giandomenico Cerreto, Filippo Molinari, Giancarlo Petralito, Florigio Lista |
| EPI_ISL_3449648, EPI_ISL_3449649 | Department of Bacteria, Parasites and Fungi, Statens Serum Institut, Copenhagen, Denmark | Statens Serum Institut Bioinformatics and Microbial Genomics | Danish Covid-19 Genome Consortium |
| EPI_ISL_3449650 | Virology Laboratory, Scientific Department, Army Medical Center | Virology Laboratory, Scientific Department, Army Medical Center | Silvia Fillo, Riccardo De Sanctis, Antonella Fortunato, Anella Monte, Rossella Brandi, Giulia Campoli, Marzia Cavalli, Lucia Nicosia, Anna Anselmo, Vanessa Vera Fain, Francesco Giordani, Giandomenico Cerreto, Filippo Molinari, Giancarlo Petralito, Florigio Lista |
| EPI_ISL_3449651 | Department of Bacteria, Parasites and Fungi, Statens Serum Institut, Copenhagen, Denmark | Statens Serum Institut Bioinformatics and Microbial Genomics | Danish Covid-19 Genome Consortium |
| EPI_ISL_3449652 | Virology Laboratory, Scientific Department, Army Medical Center | Virology Laboratory, Scientific Department, Army Medical Center | Silvia Fillo, Riccardo De Sanctis, Antonella Fortunato, Anella Monte, Rossella Brandi, Giulia Campoli, Marzia Cavalli, Lucia Nicosia, Anna Anselmo, Vanessa Vera Fain, Francesco Giordani, Giandomenico Cerreto, Filippo Molinari, Giancarlo Petralito, Florigio Lista |
| EPI_ISL_3449653, EPI_ISL_3449655, EPI_ISL_3449657, EPI_ISL_3449659, EPI_ISL_3449661, EPI_ISL_3449663, EPI_ISL_3449665, EPI_ISL_3449667, EPI_ISL_3449668, EPI_ISL_3449670, EPI_ISL_3449672, EPI_ISL_3449674, EPI_ISL_3449676, EPI_ISL_3449678, EPI_ISL_3449680, EPI_ISL_3449682, EPI_ISL_3449684, EPI_ISL_3449686, EPI_ISL_3449688, EPI_ISL_3449689, EPI_ISL_3449691, EPI_ISL_3449693, EPI_ISL_3449695, EPI_ISL_3449697, EPI_ISL_3449699, EPI_ISL_3449701, EPI_ISL_3449704, EPI_ISL_3449705, EPI_ISL_3449707, EPI_ISL_3449710, EPI_ISL_3449712, EPI_ISL_3449713, EPI_ISL_3449716, EPI_ISL_3449718, EPI_ISL_3449720, EPI_ISL_3449722, EPI_ISL_3449724 | | | |
| see above | Department of Bacteria, Parasites and Fungi, Statens Serum Institut, Copenhagen, Denmark | Statens Serum Institut Bioinformatics and Microbial Genomics | Danish Covid-19 Genome Consortium |
| EPI_ISL_3449725 | Virology Laboratory, Scientific Department, Army Medical Center | Virology Laboratory, Scientific Department, Army Medical Center | Silvia Fillo, Riccardo De Sanctis, Antonella Fortunato, Anella Monte, Rossella Brandi, Giulia Campoli, Marzia Cavalli, Lucia Nicosia, Anna Anselmo, Vanessa Vera Fain, Francesco Giordani, Giandomenico Cerreto, Filippo Molinari, Giancarlo Petralito, Florigio Lista |
| EPI_ISL_3449726 | Department of Bacteria, Parasites and Fungi, Statens Serum Institut, Copenhagen, Denmark | Statens Serum Institut Bioinformatics and Microbial Genomics | Danish Covid-19 Genome Consortium |
| EPI_ISL_3449727 | Virology Laboratory, Scientific Department, Army Medical Center | Virology Laboratory, Scientific Department, Army Medical Center | Silvia Fillo, Riccardo De Sanctis, Antonella Fortunato, Anella Monte, Rossella Brandi, Giulia Campoli, Marzia Cavalli, Lucia Nicosia, Anna Anselmo, Vanessa Vera Fain, Francesco Giordani, Giandomenico Cerreto, Filippo Molinari, Giancarlo Petralito, Florigio Lista |
| EPI_ISL_3449728, EPI_ISL_3449730 | Department of Bacteria, Parasites and Fungi, Statens Serum Institut, Copenhagen, Denmark | Statens Serum Institut Bioinformatics and Microbial Genomics | Danish Covid-19 Genome Consortium |
| EPI_ISL_3449731 | Laboratorio di Diagnostica Molecolare | Virology Laboratory, Scientific Department, Army Medical Center | Silvia Fillo, Riccardo De Sanctis, Antonella Fortunato, Anella Monte, Rossella Brandi, Giulia Campoli, Marzia Cavalli, Lucia Nicosia, Anna Anselmo, Vanessa Vera Fain, Francesco Giordani, Giandomenico Cerreto, Filippo Molinari, Giancarlo Petralito, Florigio Lista, Manuel Francesca, Federica Nuccio, Francesco Colaci, Mario Tucciarone. |
| EPI_ISL_3449732 | Department of Bacteria, Parasites and Fungi, Statens Serum Institut, Copenhagen, Denmark | Statens Serum Institut Bioinformatics and Microbial Genomics | Danish Covid-19 Genome Consortium |
| EPI_ISL_3449733 | Virology Laboratory, Scientific Department, Army Medical Center | Virology Laboratory, Scientific Department, Army Medical Center | Silvia Fillo, Riccardo De Sanctis, Antonella Fortunato, Anella Monte, Rossella Brandi, Giulia Campoli, Marzia Cavalli, Lucia Nicosia, Anna Anselmo, Vanessa Vera Fain, Francesco Giordani, Giandomenico Cerreto, Filippo Molinari, Giancarlo Petralito, Florigio Lista |
| EPI_ISL_3449734, EPI_ISL_3449735 | Department of Bacteria, Parasites and Fungi, Statens Serum Institut, Copenhagen, Denmark | Statens Serum Institut Bioinformatics and Microbial Genomics | Danish Covid-19 Genome Consortium |
| EPI_ISL_3449736 | Virology Laboratory, Scientific Department, Army Medical Center | Virology Laboratory, Scientific Department, Army Medical Center | Silvia Fillo, Riccardo De Sanctis, Antonella Fortunato, Anella Monte, Rossella Brandi, Giulia Campoli, Marzia Cavalli, Lucia Nicosia, Anna Anselmo, Vanessa Vera Fain, Francesco Giordani, Giandomenico Cerreto, Filippo Molinari, Giancarlo Petralito, Florigio Lista |
| EPI_ISL_3449737 | Department of Bacteria, Parasites and Fungi, Statens Serum Institut, Copenhagen, Denmark | Statens Serum Institut Bioinformatics and Microbial Genomics | Danish Covid-19 Genome Consortium |
| EPI_ISL_3449738 | Virology Laboratory, Scientific Department, Army Medical Center | Virology Laboratory, Scientific Department, Army Medical Center | Silvia Fillo, Riccardo De Sanctis, Antonella Fortunato, Anella Monte, Rossella Brandi, Giulia Campoli, Marzia Cavalli, Lucia Nicosia, Anna Anselmo, Vanessa Vera Fain, Francesco Giordani, Giandomenico Cerreto, Filippo Molinari, Giancarlo Petralito, Florigio Lista |
| EPI_ISL_3449739 | Department of Bacteria, Parasites and Fungi, Statens Serum Institut, Copenhagen, Denmark | Statens Serum Institut Bioinformatics and Microbial Genomics | Danish Covid-19 Genome Consortium |

| | | | |
|---|---|---|---|
| EPI_ISL_3449740 | Virology Laboratory, Scientific Department, Army Medical Center | Virology Laboratory, Scientific Department, Army Medical Center | Silvia Fillo, Riccardo De Sanctis, Antonella Fortunato, Anella Monte, Rossella Brandi, Giulia Campoli, Marzia Cavalli, Lucia Nicosia, Anna Anselmo, Vanessa Vera Fain, Francesco Giordani, Giandomenico Cerreto, Filippo Molinari, Giancarlo Petralito, Florigio Lista |
| EPI_ISL_3449741, EPI_ISL_3449742, EPI_ISL_3449743, EPI_ISL_3449744, EPI_ISL_3449745, EPI_ISL_3449746, EPI_ISL_3449747, EPI_ISL_3449748, EPI_ISL_3449749, EPI_ISL_3449750, EPI_ISL_3449751, EPI_ISL_3449752, EPI_ISL_3449753, EPI_ISL_3449754, EPI_ISL_3449755, EPI_ISL_3449756, EPI_ISL_3449757, EPI_ISL_3449758, EPI_ISL_3449759, EPI_ISL_3449760, EPI_ISL_3449761, EPI_ISL_3449762, EPI_ISL_3449763, EPI_ISL_3449764, EPI_ISL_3449765, EPI_ISL_3449766, EPI_ISL_3449767, EPI_ISL_3449768, EPI_ISL_3449769, EPI_ISL_3449770, EPI_ISL_3449771, EPI_ISL_3449773, EPI_ISL_3449774, EPI_ISL_3449775, EPI_ISL_3449776, EPI_ISL_3449777, EPI_ISL_3449778, EPI_ISL_3449780, EPI_ISL_3449781, EPI_ISL_3449782, EPI_ISL_3449783, EPI_ISL_3449784, EPI_ISL_3449785, EPI_ISL_3449786, EPI_ISL_3449787, EPI_ISL_3449788, EPI_ISL_3449789, EPI_ISL_3449790, EPI_ISL_3449791, EPI_ISL_3449792, EPI_ISL_3449793, EPI_ISL_3449794, EPI_ISL_3449795, EPI_ISL_3449796, EPI_ISL_3449797, EPI_ISL_3449798, EPI_ISL_3449799, EPI_ISL_3449801, EPI_ISL_3449802, EPI_ISL_3449803, EPI_ISL_3449804, EPI_ISL_3449805, EPI_ISL_3449806, EPI_ISL_3449807, EPI_ISL_3449808, EPI_ISL_3449809, EPI_ISL_3449810, EPI_ISL_3449811, EPI_ISL_3449812, EPI_ISL_3449813, EPI_ISL_3449814, EPI_ISL_3449815, EPI_ISL_3449816, EPI_ISL_3449817, EPI_ISL_3449818, EPI_ISL_3449819, EPI_ISL_3449820, EPI_ISL_3449821, EPI_ISL_3449822, EPI_ISL_3449823, EPI_ISL_3449824, EPI_ISL_3449825, EPI_ISL_3449826, EPI_ISL_3449827, EPI_ISL_3449828, EPI_ISL_3449829, EPI_ISL_3449830, EPI_ISL_3449831, EPI_ISL_3449832, EPI_ISL_3449833, EPI_ISL_3449834, EPI_ISL_3449835, EPI_ISL_3449836, EPI_ISL_3449837, EPI_ISL_3449839, EPI_ISL_3449840, EPI_ISL_3449841, EPI_ISL_3449842, EPI_ISL_3449843, EPI_ISL_3449844, EPI_ISL_3449845, EPI_ISL_3449846, EPI_ISL_3449847, EPI_ISL_3449848, EPI_ISL_3449849 | | | |
| see above | Department of Bacteria, Parasites and Fungi, Statens Serum Institut, Copenhagen, Denmark | Statens Serum Institut Bioinformatics and Microbial Genomics | Danish Covid-19 Genome Consortium |
| EPI_ISL_3449850 | Utah Public Health Laboratory | Utah Public Health Laboratory | Erin L. Young, Kelly F. Oakeson, Pooja Gupta, Olinto Linares-Perdomo |
| EPI_ISL_3449851, EPI_ISL_3449852 | Department of Bacteria, Parasites and Fungi, Statens Serum Institut, Copenhagen, Denmark | Statens Serum Institut Bioinformatics and Microbial Genomics | Danish Covid-19 Genome Consortium |
| EPI_ISL_3449853 | Utah Public Health Laboratory | Utah Public Health Laboratory | Erin L. Young, Kelly F. Oakeson, Pooja Gupta, Olinto Linares-Perdomo |
| EPI_ISL_3449854 | Department of Bacteria, Parasites and Fungi, Statens Serum Institut, Copenhagen, Denmark | Statens Serum Institut Bioinformatics and Microbial Genomics | Danish Covid-19 Genome Consortium |
| EPI_ISL_3449855 | Utah Public Health Laboratory | Utah Public Health Laboratory | Erin L. Young, Kelly F. Oakeson, Pooja Gupta, Olinto Linares-Perdomo |
| EPI_ISL_3449856 | Department of Bacteria, Parasites and Fungi, Statens Serum Institut, Copenhagen, Denmark | Statens Serum Institut Bioinformatics and Microbial Genomics | Danish Covid-19 Genome Consortium |
| EPI_ISL_3449857, EPI_ISL_3449858 | Utah Public Health Laboratory | Utah Public Health Laboratory | Erin L. Young, Kelly F. Oakeson, Pooja Gupta, Olinto Linares-Perdomo |
| EPI_ISL_3449859 | Department of Bacteria, Parasites and Fungi, Statens Serum Institut, Copenhagen, Denmark | Statens Serum Institut Bioinformatics and Microbial Genomics | Danish Covid-19 Genome Consortium |
| EPI_ISL_3449860 | Utah Public Health Laboratory | Utah Public Health Laboratory | Erin L. Young, Kelly F. Oakeson, Pooja Gupta, Olinto Linares-Perdomo |
| EPI_ISL_3449861 | Department of Bacteria, Parasites and Fungi, Statens Serum Institut, Copenhagen, Denmark | Statens Serum Institut Bioinformatics and Microbial Genomics | Danish Covid-19 Genome Consortium |
| EPI_ISL_3449862 | Utah Public Health Laboratory | Utah Public Health Laboratory | Erin L. Young, Kelly F. Oakeson, Pooja Gupta, Olinto Linares-Perdomo |
| EPI_ISL_3449863 | Department of Bacteria, Parasites and Fungi, Statens Serum Institut, Copenhagen, Denmark | Statens Serum Institut Bioinformatics and Microbial Genomics | Danish Covid-19 Genome Consortium |
| EPI_ISL_3449864 | Utah Public Health Laboratory | Utah Public Health Laboratory | Erin L. Young, Kelly F. Oakeson, Pooja Gupta, Olinto Linares-Perdomo |
| EPI_ISL_3449865 | Department of Bacteria, Parasites and Fungi, Statens Serum Institut, Copenhagen, Denmark | Statens Serum Institut Bioinformatics and Microbial Genomics | Danish Covid-19 Genome Consortium |
| EPI_ISL_3449866 | Utah Public Health Laboratory | Utah Public Health Laboratory | Erin L. Young, Kelly F. Oakeson, Pooja Gupta, Olinto Linares-Perdomo |
| EPI_ISL_3449867 | Department of Bacteria, Parasites and Fungi, Statens Serum Institut, Copenhagen, Denmark | Statens Serum Institut Bioinformatics and Microbial Genomics | Danish Covid-19 Genome Consortium |
| EPI_ISL_3449868 | Utah Public Health Laboratory | Utah Public Health Laboratory | Erin L. Young, Kelly F. Oakeson, Pooja Gupta, Olinto Linares-Perdomo |
| EPI_ISL_3449869 | Department of Bacteria, Parasites and Fungi, Statens Serum Institut, Copenhagen, Denmark | Statens Serum Institut Bioinformatics and Microbial Genomics | Danish Covid-19 Genome Consortium |
| EPI_ISL_3449870 | Utah Public Health Laboratory | Utah Public Health Laboratory | Erin L. Young, Kelly F. Oakeson, Pooja Gupta, Olinto Linares-Perdomo |
| EPI_ISL_3449871, EPI_ISL_3449872 | Department of Bacteria, Parasites and Fungi, Statens Serum Institut, Copenhagen, Denmark | Statens Serum Institut Bioinformatics and Microbial Genomics | Danish Covid-19 Genome Consortium |
| EPI_ISL_3449873, EPI_ISL_3449874 | Utah Public Health Laboratory | Utah Public Health Laboratory | Erin L. Young, Kelly F. Oakeson, Pooja Gupta, Olinto Linares-Perdomo |
| EPI_ISL_3449875, EPI_ISL_3449876 | Department of Bacteria, Parasites and Fungi, Statens Serum Institut, Copenhagen, Denmark | Statens Serum Institut Bioinformatics and Microbial Genomics | Danish Covid-19 Genome Consortium |
| EPI_ISL_3449877 | Utah Public Health Laboratory | Utah Public Health Laboratory | Erin L. Young, Kelly F. Oakeson, Pooja Gupta, Olinto Linares-Perdomo |
| EPI_ISL_3449878 | Department of Bacteria, Parasites and Fungi, Statens Serum Institut, Copenhagen, Denmark | Statens Serum Institut Bioinformatics and Microbial Genomics | Danish Covid-19 Genome Consortium |
| EPI_ISL_3449879 | Utah Public Health Laboratory | Utah Public Health Laboratory | Erin L. Young, Kelly F. Oakeson, Pooja Gupta, Olinto Linares-Perdomo |
| EPI_ISL_3449880 | Department of Bacteria, Parasites and Fungi, Statens Serum Institut, Copenhagen, Denmark | Statens Serum Institut Bioinformatics and Microbial Genomics | Danish Covid-19 Genome Consortium |
| EPI_ISL_3449881 | Utah Public Health Laboratory | Utah Public Health Laboratory | Erin L. Young, Kelly F. Oakeson, Pooja Gupta, Olinto Linares-Perdomo |
| EPI_ISL_3449882 | Department of Bacteria, Parasites and Fungi, Statens Serum Institut, Copenhagen, Denmark | Statens Serum Institut Bioinformatics and Microbial Genomics | Danish Covid-19 Genome Consortium |
| EPI_ISL_3449883 | Utah Public Health Laboratory | Utah Public Health Laboratory | Erin L. Young, Kelly F. Oakeson, Pooja Gupta, Olinto Linares-Perdomo |
| EPI_ISL_3449884 | Department of Bacteria, Parasites and Fungi, Statens Serum Institut, Copenhagen, Denmark | Statens Serum Institut Bioinformatics and Microbial Genomics | Danish Covid-19 Genome Consortium |
| EPI_ISL_3449885 | Utah Public Health Laboratory | Utah Public Health Laboratory | Erin L. Young, Kelly F. Oakeson, Pooja Gupta, Olinto Linares-Perdomo |
| EPI_ISL_3449886, EPI_ISL_3449887 | Department of Bacteria, Parasites and Fungi, Statens Serum Institut, Copenhagen, Denmark | Statens Serum Institut Bioinformatics and Microbial Genomics | Danish Covid-19 Genome Consortium |
| EPI_ISL_3449888 | Utah Public Health Laboratory | Utah Public Health Laboratory | Erin L. Young, Kelly F. Oakeson, Pooja Gupta, Olinto Linares-Perdomo |
| EPI_ISL_3449889 | Department of Bacteria, Parasites and Fungi, Statens Serum Institut, Copenhagen, Denmark | Statens Serum Institut Bioinformatics and Microbial Genomics | Danish Covid-19 Genome Consortium |
| EPI_ISL_3449890 | Utah Public Health Laboratory | Utah Public Health Laboratory | Erin L. Young, Kelly F. Oakeson, Pooja Gupta, Olinto Linares-Perdomo |
| EPI_ISL_3449891 | Department of Bacteria, Parasites and Fungi, Statens Serum Institut, Copenhagen, Denmark | Statens Serum Institut Bioinformatics and Microbial Genomics | Danish Covid-19 Genome Consortium |
| EPI_ISL_3449892 | Utah Public Health Laboratory | Utah Public Health Laboratory | Erin L. Young, Kelly F. Oakeson, Pooja Gupta, Olinto Linares-Perdomo |
| EPI_ISL_3449893 | Department of Bacteria, Parasites and Fungi, Statens Serum Institut, Copenhagen, Denmark | Statens Serum Institut Bioinformatics and Microbial Genomics | Danish Covid-19 Genome Consortium |
| EPI_ISL_3449894 | Utah Public Health Laboratory | Utah Public Health Laboratory | Erin L. Young, Kelly F. Oakeson, Pooja Gupta, Olinto Linares-Perdomo |
| EPI_ISL_3449895 | Department of Bacteria, Parasites and Fungi, Statens Serum Institut, Copenhagen, Denmark | Statens Serum Institut Bioinformatics and Microbial Genomics | Danish Covid-19 Genome Consortium |
| EPI_ISL_3449896 | Utah Public Health Laboratory | Utah Public Health Laboratory | Erin L. Young, Kelly F. Oakeson, Pooja Gupta, Olinto Linares-Perdomo |
| EPI_ISL_3449897 | Department of Bacteria, Parasites and Fungi, Statens Serum Institut, Copenhagen, Denmark | Statens Serum Institut Bioinformatics and Microbial Genomics | Danish Covid-19 Genome Consortium |
| EPI_ISL_3449898 | Utah Public Health Laboratory | Utah Public Health Laboratory | Erin L. Young, Kelly F. Oakeson, Pooja Gupta, Olinto Linares-Perdomo |
| EPI_ISL_3449899 | Department of Bacteria, Parasites and Fungi, Statens Serum Institut, Copenhagen, Denmark | Statens Serum Institut Bioinformatics and Microbial Genomics | Danish Covid-19 Genome Consortium |
| EPI_ISL_3449900 | Utah Public Health Laboratory | Utah Public Health Laboratory | Erin L. Young, Kelly F. Oakeson, Pooja Gupta, Olinto Linares-Perdomo |
| EPI_ISL_3449901, EPI_ISL_3449902 | Department of Bacteria, Parasites and Fungi, Statens Serum Institut, Copenhagen, Denmark | Statens Serum Institut Bioinformatics and Microbial Genomics | Danish Covid-19 Genome Consortium |
| EPI_ISL_3449903 | Utah Public Health Laboratory | Utah Public Health Laboratory | Erin L. Young, Kelly F. Oakeson, Pooja Gupta, Olinto Linares-Perdomo |
| EPI_ISL_3449904 | Department of Bacteria, Parasites and Fungi, Statens Serum Institut, Copenhagen, Denmark | Statens Serum Institut Bioinformatics and Microbial Genomics | Danish Covid-19 Genome Consortium |
| EPI_ISL_3449905 | Utah Public Health Laboratory | Utah Public Health Laboratory | Erin L. Young, Kelly F. Oakeson, Pooja Gupta, Olinto Linares-Perdomo |
| EPI_ISL_3449906 | Department of Bacteria, Parasites and Fungi, Statens Serum Institut, Copenhagen, Denmark | Statens Serum Institut Bioinformatics and Microbial Genomics | Danish Covid-19 Genome Consortium |
| EPI_ISL_3449907 | Utah Public Health Laboratory | Utah Public Health Laboratory | Erin L. Young, Kelly F. Oakeson, Pooja Gupta, Olinto Linares-Perdomo |

| | | | |
|---|---|---|---|
| EPI_ISL_3449908 | Department of Bacteria, Parasites and Fungi, Statens Serum Institut, Copenhagen, Denmark | Statens Serum Institut Bioinformatics and Microbial Genomics | Danish Covid-19 Genome Consortium |
| EPI_ISL_3449909 | Utah Public Health Laboratory | Utah Public Health Laboratory | Erin L. Young, Kelly F. Oakeson, Pooja Gupta, Olinto Linares-Perdomo |
| EPI_ISL_3449910 | Department of Bacteria, Parasites and Fungi, Statens Serum Institut, Copenhagen, Denmark | Statens Serum Institut Bioinformatics and Microbial Genomics | Danish Covid-19 Genome Consortium |
| EPI_ISL_3449911 | Utah Public Health Laboratory | Utah Public Health Laboratory | Erin L. Young, Kelly F. Oakeson, Pooja Gupta, Olinto Linares-Perdomo |
| EPI_ISL_3449912 | Department of Bacteria, Parasites and Fungi, Statens Serum Institut, Copenhagen, Denmark | Statens Serum Institut Bioinformatics and Microbial Genomics | Danish Covid-19 Genome Consortium |
| EPI_ISL_3449913, EPI_ISL_3449915 | Utah Public Health Laboratory | Utah Public Health Laboratory | Erin L. Young, Kelly F. Oakeson, Pooja Gupta, Olinto Linares-Perdomo |
| EPI_ISL_3449916 | Department of Bacteria, Parasites and Fungi, Statens Serum Institut, Copenhagen, Denmark | Statens Serum Institut Bioinformatics and Microbial Genomics | Danish Covid-19 Genome Consortium |
| EPI_ISL_3449917, EPI_ISL_3449919 | Utah Public Health Laboratory | Utah Public Health Laboratory | Erin L. Young, Kelly F. Oakeson, Pooja Gupta, Olinto Linares-Perdomo |
| EPI_ISL_3449920, EPI_ISL_3449922, EPI_ISL_3449924 | Department of Bacteria, Parasites and Fungi, Statens Serum Institut, Copenhagen, Denmark | Statens Serum Institut Bioinformatics and Microbial Genomics | Danish Covid-19 Genome Consortium |
| EPI_ISL_3449925 | Utah Public Health Laboratory | Utah Public Health Laboratory | Erin L. Young, Kelly F. Oakeson, Pooja Gupta, Olinto Linares-Perdomo |
| EPI_ISL_3449926 | Department of Bacteria, Parasites and Fungi, Statens Serum Institut, Copenhagen, Denmark | Statens Serum Institut Bioinformatics and Microbial Genomics | Danish Covid-19 Genome Consortium |
| EPI_ISL_3449927 | Utah Public Health Laboratory | Utah Public Health Laboratory | Erin L. Young, Kelly F. Oakeson, Pooja Gupta, Olinto Linares-Perdomo |
| EPI_ISL_3449928, EPI_ISL_3449929 | Department of Bacteria, Parasites and Fungi, Statens Serum Institut, Copenhagen, Denmark | Statens Serum Institut Bioinformatics and Microbial Genomics | Danish Covid-19 Genome Consortium |
| EPI_ISL_3449930, EPI_ISL_3449931 | Utah Public Health Laboratory | Utah Public Health Laboratory | Erin L. Young, Kelly F. Oakeson, Pooja Gupta, Olinto Linares-Perdomo |
| EPI_ISL_3449932 | Department of Bacteria, Parasites and Fungi, Statens Serum Institut, Copenhagen, Denmark | Statens Serum Institut Bioinformatics and Microbial Genomics | Danish Covid-19 Genome Consortium |
| EPI_ISL_3449933 | Utah Public Health Laboratory | Utah Public Health Laboratory | Erin L. Young, Kelly F. Oakeson, Pooja Gupta, Olinto Linares-Perdomo |
| EPI_ISL_3449934 | Department of Bacteria, Parasites and Fungi, Statens Serum Institut, Copenhagen, Denmark | Statens Serum Institut Bioinformatics and Microbial Genomics | Danish Covid-19 Genome Consortium |
| EPI_ISL_3449935 | Utah Public Health Laboratory | Utah Public Health Laboratory | Erin L. Young, Kelly F. Oakeson, Pooja Gupta, Olinto Linares-Perdomo |
| EPI_ISL_3449936 | Department of Bacteria, Parasites and Fungi, Statens Serum Institut, Copenhagen, Denmark | Statens Serum Institut Bioinformatics and Microbial Genomics | Danish Covid-19 Genome Consortium |
| EPI_ISL_3449937 | Utah Public Health Laboratory | Utah Public Health Laboratory | Erin L. Young, Kelly F. Oakeson, Pooja Gupta, Olinto Linares-Perdomo |
| EPI_ISL_3449939 | Department of Bacteria, Parasites and Fungi, Statens Serum Institut, Copenhagen, Denmark | Statens Serum Institut Bioinformatics and Microbial Genomics | Danish Covid-19 Genome Consortium |
| EPI_ISL_3449940 | Utah Public Health Laboratory | Utah Public Health Laboratory | Erin L. Young, Kelly F. Oakeson, Pooja Gupta, Olinto Linares-Perdomo |
| EPI_ISL_3449941 | Department of Bacteria, Parasites and Fungi, Statens Serum Institut, Copenhagen, Denmark | Statens Serum Institut Bioinformatics and Microbial Genomics | Danish Covid-19 Genome Consortium |
| EPI_ISL_3449942 | Utah Public Health Laboratory | Utah Public Health Laboratory | Erin L. Young, Kelly F. Oakeson, Pooja Gupta, Olinto Linares-Perdomo |
| EPI_ISL_3449943 | Department of Bacteria, Parasites and Fungi, Statens Serum Institut, Copenhagen, Denmark | Statens Serum Institut Bioinformatics and Microbial Genomics | Danish Covid-19 Genome Consortium |
| EPI_ISL_3449944 | Utah Public Health Laboratory | Utah Public Health Laboratory | Erin L. Young, Kelly F. Oakeson, Pooja Gupta, Olinto Linares-Perdomo |
| EPI_ISL_3449945 | Department of Bacteria, Parasites and Fungi, Statens Serum Institut, Copenhagen, Denmark | Statens Serum Institut Bioinformatics and Microbial Genomics | Danish Covid-19 Genome Consortium |
| EPI_ISL_3449946, EPI_ISL_3449947 | Utah Public Health Laboratory | Utah Public Health Laboratory | Erin L. Young, Kelly F. Oakeson, Pooja Gupta, Olinto Linares-Perdomo |
| EPI_ISL_3449948, EPI_ISL_3449949 | Department of Bacteria, Parasites and Fungi, Statens Serum Institut, Copenhagen, Denmark | Statens Serum Institut Bioinformatics and Microbial Genomics | Danish Covid-19 Genome Consortium |
| EPI_ISL_3449950 | Utah Public Health Laboratory | Utah Public Health Laboratory | Erin L. Young, Kelly F. Oakeson, Pooja Gupta, Olinto Linares-Perdomo |
| EPI_ISL_3449951 | Department of Bacteria, Parasites and Fungi, Statens Serum Institut, Copenhagen, Denmark | Statens Serum Institut Bioinformatics and Microbial Genomics | Danish Covid-19 Genome Consortium |
| EPI_ISL_3449952 | Utah Public Health Laboratory | Utah Public Health Laboratory | Erin L. Young, Kelly F. Oakeson, Pooja Gupta, Olinto Linares-Perdomo |
| EPI_ISL_3449953 | Department of Bacteria, Parasites and Fungi, Statens Serum Institut, Copenhagen, Denmark | Statens Serum Institut Bioinformatics and Microbial Genomics | Danish Covid-19 Genome Consortium |
| EPI_ISL_3449954 | Utah Public Health Laboratory | Utah Public Health Laboratory | Erin L. Young, Kelly F. Oakeson, Pooja Gupta, Olinto Linares-Perdomo |
| EPI_ISL_3449955, EPI_ISL_3449957 | Department of Bacteria, Parasites and Fungi, Statens Serum Institut, Copenhagen, Denmark | Statens Serum Institut Bioinformatics and Microbial Genomics | Danish Covid-19 Genome Consortium |
| EPI_ISL_3449958 | Utah Public Health Laboratory | Utah Public Health Laboratory | Erin L. Young, Kelly F. Oakeson, Pooja Gupta, Olinto Linares-Perdomo |
| EPI_ISL_3449959 | Department of Bacteria, Parasites and Fungi, Statens Serum Institut, Copenhagen, Denmark | Statens Serum Institut Bioinformatics and Microbial Genomics | Danish Covid-19 Genome Consortium |
| EPI_ISL_3449960 | Utah Public Health Laboratory | Utah Public Health Laboratory | Erin L. Young, Kelly F. Oakeson, Pooja Gupta, Olinto Linares-Perdomo |
| EPI_ISL_3449961 | Department of Bacteria, Parasites and Fungi, Statens Serum Institut, Copenhagen, Denmark | Statens Serum Institut Bioinformatics and Microbial Genomics | Danish Covid-19 Genome Consortium |
| EPI_ISL_3449962 | Utah Public Health Laboratory | Utah Public Health Laboratory | Erin L. Young, Kelly F. Oakeson, Pooja Gupta, Olinto Linares-Perdomo |
| EPI_ISL_3449963 | Department of Bacteria, Parasites and Fungi, Statens Serum Institut, Copenhagen, Denmark | Statens Serum Institut Bioinformatics and Microbial Genomics | Danish Covid-19 Genome Consortium |
| EPI_ISL_3449964 | Utah Public Health Laboratory | Utah Public Health Laboratory | Erin L. Young, Kelly F. Oakeson, Pooja Gupta, Olinto Linares-Perdomo |
| EPI_ISL_3449965 | Department of Bacteria, Parasites and Fungi, Statens Serum Institut, Copenhagen, Denmark | Statens Serum Institut Bioinformatics and Microbial Genomics | Danish Covid-19 Genome Consortium |
| EPI_ISL_3449966 | Utah Public Health Laboratory | Utah Public Health Laboratory | Erin L. Young, Kelly F. Oakeson, Pooja Gupta, Olinto Linares-Perdomo |
| EPI_ISL_3449967 | Department of Bacteria, Parasites and Fungi, Statens Serum Institut, Copenhagen, Denmark | Statens Serum Institut Bioinformatics and Microbial Genomics | Danish Covid-19 Genome Consortium |
| EPI_ISL_3449968 | Utah Public Health Laboratory | Utah Public Health Laboratory | Erin L. Young, Kelly F. Oakeson, Pooja Gupta, Olinto Linares-Perdomo |
| EPI_ISL_3449969, EPI_ISL_3449970, EPI_ISL_3449972 | Department of Bacteria, Parasites and Fungi, Statens Serum Institut, Copenhagen, Denmark | Statens Serum Institut Bioinformatics and Microbial Genomics | Danish Covid-19 Genome Consortium |
| EPI_ISL_3449973 | Utah Public Health Laboratory | Utah Public Health Laboratory | Erin L. Young, Kelly F. Oakeson, Pooja Gupta, Olinto Linares-Perdomo |
| EPI_ISL_3449974 | Department of Bacteria, Parasites and Fungi, Statens Serum Institut, Copenhagen, Denmark | Statens Serum Institut Bioinformatics and Microbial Genomics | Danish Covid-19 Genome Consortium |
| EPI_ISL_3449975 | Utah Public Health Laboratory | Utah Public Health Laboratory | Erin L. Young, Kelly F. Oakeson, Pooja Gupta, Olinto Linares-Perdomo |
| EPI_ISL_3449976 | Department of Bacteria, Parasites and Fungi, Statens Serum Institut, Copenhagen, Denmark | Statens Serum Institut Bioinformatics and Microbial Genomics | Danish Covid-19 Genome Consortium |
| EPI_ISL_3449977, EPI_ISL_3449979 | Utah Public Health Laboratory | Utah Public Health Laboratory | Erin L. Young, Kelly F. Oakeson, Pooja Gupta, Olinto Linares-Perdomo |
| EPI_ISL_3449980, | Department of Bacteria, Parasites and Fungi, Statens Serum | Statens Serum Institut Bioinformatics and Microbial Genomics | Danish Covid-19 Genome Consortium |

| | | | |
|---|---|---|---|
| EPI_ISL_3449981 | Institut, Copenhagen, Denmark | | |
| EPI_ISL_3449982 | Utah Public Health Laboratory | Utah Public Health Laboratory | Erin L. Young, Kelly F. Oakeson, Pooja Gupta, Olinto Linares-Perdomo |
| EPI_ISL_3449983 | Department of Bacteria, Parasites and Fungi, Statens Serum Institut, Copenhagen, Denmark | Statens Serum Institut Bioinformatics and Microbial Genomics | Danish Covid-19 Genome Consortium |
| EPI_ISL_3449984 | Utah Public Health Laboratory | Utah Public Health Laboratory | Erin L. Young, Kelly F. Oakeson, Pooja Gupta, Olinto Linares-Perdomo |
| EPI_ISL_3449985 | Department of Bacteria, Parasites and Fungi, Statens Serum Institut, Copenhagen, Denmark | Statens Serum Institut Bioinformatics and Microbial Genomics | Danish Covid-19 Genome Consortium |
| EPI_ISL_3449986 | Utah Public Health Laboratory | Utah Public Health Laboratory | Erin L. Young, Kelly F. Oakeson, Pooja Gupta, Olinto Linares-Perdomo |
| EPI_ISL_3449987 | Department of Bacteria, Parasites and Fungi, Statens Serum Institut, Copenhagen, Denmark | Statens Serum Institut Bioinformatics and Microbial Genomics | Danish Covid-19 Genome Consortium |
| EPI_ISL_3449988 | Utah Public Health Laboratory | Utah Public Health Laboratory | Erin L. Young, Kelly F. Oakeson, Pooja Gupta, Olinto Linares-Perdomo |
| EPI_ISL_3449989 | Department of Bacteria, Parasites and Fungi, Statens Serum Institut, Copenhagen, Denmark | Statens Serum Institut Bioinformatics and Microbial Genomics | Danish Covid-19 Genome Consortium |
| EPI_ISL_3449990 | Utah Public Health Laboratory | Utah Public Health Laboratory | Erin L. Young, Kelly F. Oakeson, Pooja Gupta, Olinto Linares-Perdomo |
| EPI_ISL_3449991 | Department of Bacteria, Parasites and Fungi, Statens Serum Institut, Copenhagen, Denmark | Statens Serum Institut Bioinformatics and Microbial Genomics | Danish Covid-19 Genome Consortium |
| EPI_ISL_3449992 | Utah Public Health Laboratory | Utah Public Health Laboratory | Erin L. Young, Kelly F. Oakeson, Pooja Gupta, Olinto Linares-Perdomo |
| EPI_ISL_3449993 | Department of Bacteria, Parasites and Fungi, Statens Serum Institut, Copenhagen, Denmark | Statens Serum Institut Bioinformatics and Microbial Genomics | Danish Covid-19 Genome Consortium |
| EPI_ISL_3449994 | Utah Public Health Laboratory | Utah Public Health Laboratory | Erin L. Young, Kelly F. Oakeson, Pooja Gupta, Olinto Linares-Perdomo |
| EPI_ISL_3449995, EPI_ISL_3449996 | Department of Bacteria, Parasites and Fungi, Statens Serum Institut, Copenhagen, Denmark | Statens Serum Institut Bioinformatics and Microbial Genomics | Danish Covid-19 Genome Consortium |
| EPI_ISL_3449997 | Utah Public Health Laboratory | Utah Public Health Laboratory | Erin L. Young, Kelly F. Oakeson, Pooja Gupta, Olinto Linares-Perdomo |
| EPI_ISL_3449998 | Department of Bacteria, Parasites and Fungi, Statens Serum Institut, Copenhagen, Denmark | Statens Serum Institut Bioinformatics and Microbial Genomics | Danish Covid-19 Genome Consortium |
| EPI_ISL_3449999 | Utah Public Health Laboratory | Utah Public Health Laboratory | Erin L. Young, Kelly F. Oakeson, Pooja Gupta, Olinto Linares-Perdomo |
| EPI_ISL_3450000 | Department of Bacteria, Parasites and Fungi, Statens Serum Institut, Copenhagen, Denmark | Statens Serum Institut Bioinformatics and Microbial Genomics | Danish Covid-19 Genome Consortium |
| EPI_ISL_3450001 | Utah Public Health Laboratory | Utah Public Health Laboratory | Erin L. Young, Kelly F. Oakeson, Pooja Gupta, Olinto Linares-Perdomo |
| EPI_ISL_3450002 | Department of Bacteria, Parasites and Fungi, Statens Serum Institut, Copenhagen, Denmark | Statens Serum Institut Bioinformatics and Microbial Genomics | Danish Covid-19 Genome Consortium |
| EPI_ISL_3450003 | Utah Public Health Laboratory | Utah Public Health Laboratory | Erin L. Young, Kelly F. Oakeson, Pooja Gupta, Olinto Linares-Perdomo |
| EPI_ISL_3450004, EPI_ISL_3450005 | Department of Bacteria, Parasites and Fungi, Statens Serum Institut, Copenhagen, Denmark | Statens Serum Institut Bioinformatics and Microbial Genomics | Danish Covid-19 Genome Consortium |
| EPI_ISL_3450006 | Utah Public Health Laboratory | Utah Public Health Laboratory | Erin L. Young, Kelly F. Oakeson, Pooja Gupta, Olinto Linares-Perdomo |
| EPI_ISL_3450007 | Department of Bacteria, Parasites and Fungi, Statens Serum Institut, Copenhagen, Denmark | Statens Serum Institut Bioinformatics and Microbial Genomics | Danish Covid-19 Genome Consortium |
| EPI_ISL_3450008 | Utah Public Health Laboratory | Utah Public Health Laboratory | Erin L. Young, Kelly F. Oakeson, Pooja Gupta, Olinto Linares-Perdomo |
| EPI_ISL_3450009 | Department of Bacteria, Parasites and Fungi, Statens Serum Institut, Copenhagen, Denmark | Statens Serum Institut Bioinformatics and Microbial Genomics | Danish Covid-19 Genome Consortium |
| EPI_ISL_3450010 | Utah Public Health Laboratory | Utah Public Health Laboratory | Erin L. Young, Kelly F. Oakeson, Pooja Gupta, Olinto Linares-Perdomo |
| EPI_ISL_3450011 | Department of Bacteria, Parasites and Fungi, Statens Serum Institut, Copenhagen, Denmark | Statens Serum Institut Bioinformatics and Microbial Genomics | Danish Covid-19 Genome Consortium |
| EPI_ISL_3450012 | Utah Public Health Laboratory | Utah Public Health Laboratory | Erin L. Young, Kelly F. Oakeson, Pooja Gupta, Olinto Linares-Perdomo |
| EPI_ISL_3450013 | Department of Bacteria, Parasites and Fungi, Statens Serum Institut, Copenhagen, Denmark | Statens Serum Institut Bioinformatics and Microbial Genomics | Danish Covid-19 Genome Consortium |
| EPI_ISL_3450014 | Utah Public Health Laboratory | Utah Public Health Laboratory | Erin L. Young, Kelly F. Oakeson, Pooja Gupta, Olinto Linares-Perdomo |
| EPI_ISL_3450015 | Department of Bacteria, Parasites and Fungi, Statens Serum Institut, Copenhagen, Denmark | Statens Serum Institut Bioinformatics and Microbial Genomics | Danish Covid-19 Genome Consortium |
| EPI_ISL_3450016 | Utah Public Health Laboratory | Utah Public Health Laboratory | Erin L. Young, Kelly F. Oakeson, Pooja Gupta, Olinto Linares-Perdomo |
| EPI_ISL_3450017 | Department of Bacteria, Parasites and Fungi, Statens Serum Institut, Copenhagen, Denmark | Statens Serum Institut Bioinformatics and Microbial Genomics | Danish Covid-19 Genome Consortium |
| EPI_ISL_3450018 | Utah Public Health Laboratory | Utah Public Health Laboratory | Erin L. Young, Kelly F. Oakeson, Pooja Gupta, Olinto Linares-Perdomo |
| EPI_ISL_3450019 | Department of Bacteria, Parasites and Fungi, Statens Serum Institut, Copenhagen, Denmark | Statens Serum Institut Bioinformatics and Microbial Genomics | Danish Covid-19 Genome Consortium |
| EPI_ISL_3450020 | Utah Public Health Laboratory | Utah Public Health Laboratory | Erin L. Young, Kelly F. Oakeson, Pooja Gupta, Olinto Linares-Perdomo |
| EPI_ISL_3450021 | Department of Bacteria, Parasites and Fungi, Statens Serum Institut, Copenhagen, Denmark | Statens Serum Institut Bioinformatics and Microbial Genomics | Danish Covid-19 Genome Consortium |
| EPI_ISL_3450022 | Utah Public Health Laboratory | Utah Public Health Laboratory | Erin L. Young, Kelly F. Oakeson, Pooja Gupta, Olinto Linares-Perdomo |
| EPI_ISL_3450023 | Department of Bacteria, Parasites and Fungi, Statens Serum Institut, Copenhagen, Denmark | Statens Serum Institut Bioinformatics and Microbial Genomics | Danish Covid-19 Genome Consortium |
| EPI_ISL_3450024 | Utah Public Health Laboratory | Utah Public Health Laboratory | Erin L. Young, Kelly F. Oakeson, Pooja Gupta, Olinto Linares-Perdomo |
| EPI_ISL_3450025, EPI_ISL_3450026, EPI_ISL_3450027 | Department of Bacteria, Parasites and Fungi, Statens Serum Institut, Copenhagen, Denmark | Statens Serum Institut Bioinformatics and Microbial Genomics | Danish Covid-19 Genome Consortium |
| EPI_ISL_3450028 | Utah Public Health Laboratory | Utah Public Health Laboratory | Erin L. Young, Kelly F. Oakeson, Pooja Gupta, Olinto Linares-Perdomo |
| EPI_ISL_3450029, EPI_ISL_3450030 | Department of Bacteria, Parasites and Fungi, Statens Serum Institut, Copenhagen, Denmark | Statens Serum Institut Bioinformatics and Microbial Genomics | Danish Covid-19 Genome Consortium |
| EPI_ISL_3450031 | Utah Public Health Laboratory | Utah Public Health Laboratory | Erin L. Young, Kelly F. Oakeson, Pooja Gupta, Olinto Linares-Perdomo |
| EPI_ISL_3450032 | Department of Bacteria, Parasites and Fungi, Statens Serum Institut, Copenhagen, Denmark | Statens Serum Institut Bioinformatics and Microbial Genomics | Danish Covid-19 Genome Consortium |
| EPI_ISL_3450033 | Utah Public Health Laboratory | Utah Public Health Laboratory | Erin L. Young, Kelly F. Oakeson, Pooja Gupta, Olinto Linares-Perdomo |
| EPI_ISL_3450034 | Department of Bacteria, Parasites and Fungi, Statens Serum Institut, Copenhagen, Denmark | Statens Serum Institut Bioinformatics and Microbial Genomics | Danish Covid-19 Genome Consortium |
| EPI_ISL_3450035 | Utah Public Health Laboratory | Utah Public Health Laboratory | Erin L. Young, Kelly F. Oakeson, Pooja Gupta, Olinto Linares-Perdomo |
| EPI_ISL_3450036 | Department of Bacteria, Parasites and Fungi, Statens Serum Institut, Copenhagen, Denmark | Statens Serum Institut Bioinformatics and Microbial Genomics | Danish Covid-19 Genome Consortium |
| EPI_ISL_3450037 | Utah Public Health Laboratory | Utah Public Health Laboratory | Erin L. Young, Kelly F. Oakeson, Pooja Gupta, Olinto Linares-Perdomo |
| EPI_ISL_3450038 | Department of Bacteria, Parasites and Fungi, Statens Serum Institut, Copenhagen, Denmark | Statens Serum Institut Bioinformatics and Microbial Genomics | Danish Covid-19 Genome Consortium |
| EPI_ISL_3450039 | Utah Public Health Laboratory | Utah Public Health Laboratory | Erin L. Young, Kelly F. Oakeson, Pooja Gupta, Olinto Linares-Perdomo |
| EPI_ISL_3450040, EPI_ISL_3450041 | Department of Bacteria, Parasites and Fungi, Statens Serum Institut, Copenhagen, Denmark | Statens Serum Institut Bioinformatics and Microbial Genomics | Danish Covid-19 Genome Consortium |
| EPI_ISL_3450042 | Utah Public Health Laboratory | Utah Public Health Laboratory | Erin L. Young, Kelly F. Oakeson, Pooja Gupta, Olinto Linares-Perdomo |
| EPI_ISL_3450043 | Department of Bacteria, Parasites and Fungi, Statens Serum | Statens Serum Institut Bioinformatics and Microbial Genomics | Danish Covid-19 Genome Consortium |

| | | | |
|---|---|---|---|
| | Institut, Copenhagen, Denmark | | |
| EPI_ISL_3450044 | Utah Public Health Laboratory | Utah Public Health Laboratory | Erin L. Young, Kelly F. Oakeson, Pooja Gupta, Olinto Linares-Perdomo |
| EPI_ISL_3450045 | Department of Bacteria, Parasites and Fungi, Statens Serum Institut, Copenhagen, Denmark | Statens Serum Institut Bioinformatics and Microbial Genomics | Danish Covid-19 Genome Consortium |
| EPI_ISL_3450046 | Utah Public Health Laboratory | Utah Public Health Laboratory | Erin L. Young, Kelly F. Oakeson, Pooja Gupta, Olinto Linares-Perdomo |
| EPI_ISL_3450047 | Department of Bacteria, Parasites and Fungi, Statens Serum Institut, Copenhagen, Denmark | Statens Serum Institut Bioinformatics and Microbial Genomics | Danish Covid-19 Genome Consortium |
| EPI_ISL_3450048 | Utah Public Health Laboratory | Utah Public Health Laboratory | Erin L. Young, Kelly F. Oakeson, Pooja Gupta, Olinto Linares-Perdomo |
| EPI_ISL_3450049 | Department of Bacteria, Parasites and Fungi, Statens Serum Institut, Copenhagen, Denmark | Statens Serum Institut Bioinformatics and Microbial Genomics | Danish Covid-19 Genome Consortium |
| EPI_ISL_3450050 | Utah Public Health Laboratory | Utah Public Health Laboratory | Erin L. Young, Kelly F. Oakeson, Pooja Gupta, Olinto Linares-Perdomo |
| EPI_ISL_3450051 | Department of Bacteria, Parasites and Fungi, Statens Serum Institut, Copenhagen, Denmark | Statens Serum Institut Bioinformatics and Microbial Genomics | Danish Covid-19 Genome Consortium |
| EPI_ISL_3450052 | Utah Public Health Laboratory | Utah Public Health Laboratory | Erin L. Young, Kelly F. Oakeson, Pooja Gupta, Olinto Linares-Perdomo |
| EPI_ISL_3450053, EPI_ISL_3450054 | Department of Bacteria, Parasites and Fungi, Statens Serum Institut, Copenhagen, Denmark | Statens Serum Institut Bioinformatics and Microbial Genomics | Danish Covid-19 Genome Consortium |
| EPI_ISL_3450055, EPI_ISL_3450057, EPI_ISL_3450058 | Utah Public Health Laboratory | Utah Public Health Laboratory | Erin L. Young, Kelly F. Oakeson, Pooja Gupta, Olinto Linares-Perdomo |
| EPI_ISL_3450059, EPI_ISL_3450060 | Department of Bacteria, Parasites and Fungi, Statens Serum Institut, Copenhagen, Denmark | Statens Serum Institut Bioinformatics and Microbial Genomics | Danish Covid-19 Genome Consortium |
| EPI_ISL_3450061, EPI_ISL_3450062 | Utah Public Health Laboratory | Utah Public Health Laboratory | Erin L. Young, Kelly F. Oakeson, Pooja Gupta, Olinto Linares-Perdomo |
| EPI_ISL_3450063 | Department of Bacteria, Parasites and Fungi, Statens Serum Institut, Copenhagen, Denmark | Statens Serum Institut Bioinformatics and Microbial Genomics | Danish Covid-19 Genome Consortium |
| EPI_ISL_3450064 | Utah Public Health Laboratory | Utah Public Health Laboratory | Erin L. Young, Kelly F. Oakeson, Pooja Gupta, Olinto Linares-Perdomo |
| EPI_ISL_3450065 | Department of Bacteria, Parasites and Fungi, Statens Serum Institut, Copenhagen, Denmark | Statens Serum Institut Bioinformatics and Microbial Genomics | Danish Covid-19 Genome Consortium |
| EPI_ISL_3450066 | Utah Public Health Laboratory | Utah Public Health Laboratory | Erin L. Young, Kelly F. Oakeson, Pooja Gupta, Olinto Linares-Perdomo |
| EPI_ISL_3450067 | Department of Bacteria, Parasites and Fungi, Statens Serum Institut, Copenhagen, Denmark | Statens Serum Institut Bioinformatics and Microbial Genomics | Danish Covid-19 Genome Consortium |
| EPI_ISL_3450068 | Utah Public Health Laboratory | Utah Public Health Laboratory | Erin L. Young, Kelly F. Oakeson, Pooja Gupta, Olinto Linares-Perdomo |
| EPI_ISL_3450069 | Department of Bacteria, Parasites and Fungi, Statens Serum Institut, Copenhagen, Denmark | Statens Serum Institut Bioinformatics and Microbial Genomics | Danish Covid-19 Genome Consortium |
| EPI_ISL_3450071 | Utah Public Health Laboratory | Utah Public Health Laboratory | Erin L. Young, Kelly F. Oakeson, Pooja Gupta, Olinto Linares-Perdomo |
| EPI_ISL_3450072 | Department of Bacteria, Parasites and Fungi, Statens Serum Institut, Copenhagen, Denmark | Statens Serum Institut Bioinformatics and Microbial Genomics | Danish Covid-19 Genome Consortium |
| EPI_ISL_3450073 | Utah Public Health Laboratory | Utah Public Health Laboratory | Erin L. Young, Kelly F. Oakeson, Pooja Gupta, Olinto Linares-Perdomo |
| EPI_ISL_3450074 | Department of Bacteria, Parasites and Fungi, Statens Serum Institut, Copenhagen, Denmark | Statens Serum Institut Bioinformatics and Microbial Genomics | Danish Covid-19 Genome Consortium |
| EPI_ISL_3450075 | Utah Public Health Laboratory | Utah Public Health Laboratory | Erin L. Young, Kelly F. Oakeson, Pooja Gupta, Olinto Linares-Perdomo |
| EPI_ISL_3450076 | Department of Bacteria, Parasites and Fungi, Statens Serum Institut, Copenhagen, Denmark | Statens Serum Institut Bioinformatics and Microbial Genomics | Danish Covid-19 Genome Consortium |
| EPI_ISL_3450077 | Utah Public Health Laboratory | Utah Public Health Laboratory | Erin L. Young, Kelly F. Oakeson, Pooja Gupta, Olinto Linares-Perdomo |
| EPI_ISL_3450078, EPI_ISL_3450079 | Department of Bacteria, Parasites and Fungi, Statens Serum Institut, Copenhagen, Denmark | Statens Serum Institut Bioinformatics and Microbial Genomics | Danish Covid-19 Genome Consortium |
| EPI_ISL_3450080 | Utah Public Health Laboratory | Utah Public Health Laboratory | Erin L. Young, Kelly F. Oakeson, Pooja Gupta, Olinto Linares-Perdomo |
| EPI_ISL_3450081 | Department of Bacteria, Parasites and Fungi, Statens Serum Institut, Copenhagen, Denmark | Statens Serum Institut Bioinformatics and Microbial Genomics | Danish Covid-19 Genome Consortium |
| EPI_ISL_3450082 | Utah Public Health Laboratory | Utah Public Health Laboratory | Erin L. Young, Kelly F. Oakeson, Pooja Gupta, Olinto Linares-Perdomo |
| EPI_ISL_3450083 | Department of Bacteria, Parasites and Fungi, Statens Serum Institut, Copenhagen, Denmark | Statens Serum Institut Bioinformatics and Microbial Genomics | Danish Covid-19 Genome Consortium |
| EPI_ISL_3450084 | Utah Public Health Laboratory | Utah Public Health Laboratory | Erin L. Young, Kelly F. Oakeson, Pooja Gupta, Olinto Linares-Perdomo |
| EPI_ISL_3450085, EPI_ISL_3450086 | Department of Bacteria, Parasites and Fungi, Statens Serum Institut, Copenhagen, Denmark | Statens Serum Institut Bioinformatics and Microbial Genomics | Danish Covid-19 Genome Consortium |
| EPI_ISL_3450087 | Utah Public Health Laboratory | Utah Public Health Laboratory | Erin L. Young, Kelly F. Oakeson, Pooja Gupta, Olinto Linares-Perdomo |
| EPI_ISL_3450088 | Department of Bacteria, Parasites and Fungi, Statens Serum Institut, Copenhagen, Denmark | Statens Serum Institut Bioinformatics and Microbial Genomics | Danish Covid-19 Genome Consortium |
| EPI_ISL_3450089 | Utah Public Health Laboratory | Utah Public Health Laboratory | Erin L. Young, Kelly F. Oakeson, Pooja Gupta, Olinto Linares-Perdomo |
| EPI_ISL_3450090 | Department of Bacteria, Parasites and Fungi, Statens Serum Institut, Copenhagen, Denmark | Statens Serum Institut Bioinformatics and Microbial Genomics | Danish Covid-19 Genome Consortium |
| EPI_ISL_3450091 | Utah Public Health Laboratory | Utah Public Health Laboratory | Erin L. Young, Kelly F. Oakeson, Pooja Gupta, Olinto Linares-Perdomo |
| EPI_ISL_3450092 | Department of Bacteria, Parasites and Fungi, Statens Serum Institut, Copenhagen, Denmark | Statens Serum Institut Bioinformatics and Microbial Genomics | Danish Covid-19 Genome Consortium |
| EPI_ISL_3450093 | Utah Public Health Laboratory | Utah Public Health Laboratory | Erin L. Young, Kelly F. Oakeson, Pooja Gupta, Olinto Linares-Perdomo |
| EPI_ISL_3450094 | Department of Bacteria, Parasites and Fungi, Statens Serum Institut, Copenhagen, Denmark | Statens Serum Institut Bioinformatics and Microbial Genomics | Danish Covid-19 Genome Consortium |
| EPI_ISL_3450095 | Utah Public Health Laboratory | Utah Public Health Laboratory | Erin L. Young, Kelly F. Oakeson, Pooja Gupta, Olinto Linares-Perdomo |
| EPI_ISL_3450096 | Department of Bacteria, Parasites and Fungi, Statens Serum Institut, Copenhagen, Denmark | Statens Serum Institut Bioinformatics and Microbial Genomics | Danish Covid-19 Genome Consortium |
| EPI_ISL_3450097 | Utah Public Health Laboratory | Utah Public Health Laboratory | Erin L. Young, Kelly F. Oakeson, Pooja Gupta, Olinto Linares-Perdomo |
| EPI_ISL_3450098 | Department of Bacteria, Parasites and Fungi, Statens Serum Institut, Copenhagen, Denmark | Statens Serum Institut Bioinformatics and Microbial Genomics | Danish Covid-19 Genome Consortium |
| EPI_ISL_3450099 | Utah Public Health Laboratory | Utah Public Health Laboratory | Erin L. Young, Kelly F. Oakeson, Pooja Gupta, Olinto Linares-Perdomo |
| EPI_ISL_3450100 | Department of Bacteria, Parasites and Fungi, Statens Serum Institut, Copenhagen, Denmark | Statens Serum Institut Bioinformatics and Microbial Genomics | Danish Covid-19 Genome Consortium |
| EPI_ISL_3450101 | Utah Public Health Laboratory | Utah Public Health Laboratory | Erin L. Young, Kelly F. Oakeson, Pooja Gupta, Olinto Linares-Perdomo |
| EPI_ISL_3450102, EPI_ISL_3450103 | Department of Bacteria, Parasites and Fungi, Statens Serum Institut, Copenhagen, Denmark | Statens Serum Institut Bioinformatics and Microbial Genomics | Danish Covid-19 Genome Consortium |
| EPI_ISL_3450104, EPI_ISL_3450105, EPI_ISL_3450106, EPI_ISL_3450107 | Utah Public Health Laboratory | Utah Public Health Laboratory | Erin L. Young, Kelly F. Oakeson, Pooja Gupta, Olinto Linares-Perdomo |
| EPI_ISL_3450108 | Department of Bacteria, Parasites and Fungi, Statens Serum Institut, Copenhagen, Denmark | Statens Serum Institut Bioinformatics and Microbial Genomics | Danish Covid-19 Genome Consortium |
| EPI_ISL_3450109 | Utah Public Health Laboratory | Utah Public Health Laboratory | Erin L. Young, Kelly F. Oakeson, Pooja Gupta, Olinto Linares-Perdomo |

| | | | |
|---|---|---|---|
| EPI_ISL_3450110, EPI_ISL_3450111 | Department of Bacteria, Parasites and Fungi, Statens Serum Institut, Copenhagen, Denmark | Statens Serum Institut Bioinformatics and Microbial Genomics | Danish Covid-19 Genome Consortium |
| EPI_ISL_3450112 | Utah Public Health Laboratory | Utah Public Health Laboratory | Erin L. Young, Kelly F. Oakeson, Pooja Gupta, Olinto Linares-Perdomo |
| EPI_ISL_3450113 | Department of Bacteria, Parasites and Fungi, Statens Serum Institut, Copenhagen, Denmark | Statens Serum Institut Bioinformatics and Microbial Genomics | Danish Covid-19 Genome Consortium |
| EPI_ISL_3450114 | Utah Public Health Laboratory | Utah Public Health Laboratory | Erin L. Young, Kelly F. Oakeson, Pooja Gupta, Olinto Linares-Perdomo |
| EPI_ISL_3450115 | Department of Bacteria, Parasites and Fungi, Statens Serum Institut, Copenhagen, Denmark | Statens Serum Institut Bioinformatics and Microbial Genomics | Danish Covid-19 Genome Consortium |
| EPI_ISL_3450116 | Utah Public Health Laboratory | Utah Public Health Laboratory | Erin L. Young, Kelly F. Oakeson, Pooja Gupta, Olinto Linares-Perdomo |
| EPI_ISL_3450117 | Department of Bacteria, Parasites and Fungi, Statens Serum Institut, Copenhagen, Denmark | Statens Serum Institut Bioinformatics and Microbial Genomics | Danish Covid-19 Genome Consortium |
| EPI_ISL_3450118 | Utah Public Health Laboratory | Utah Public Health Laboratory | Erin L. Young, Kelly F. Oakeson, Pooja Gupta, Olinto Linares-Perdomo |
| EPI_ISL_3450119 | Department of Bacteria, Parasites and Fungi, Statens Serum Institut, Copenhagen, Denmark | Statens Serum Institut Bioinformatics and Microbial Genomics | Danish Covid-19 Genome Consortium |
| EPI_ISL_3450120 | Utah Public Health Laboratory | Utah Public Health Laboratory | Erin L. Young, Kelly F. Oakeson, Pooja Gupta, Olinto Linares-Perdomo |
| EPI_ISL_3450121 | Department of Bacteria, Parasites and Fungi, Statens Serum Institut, Copenhagen, Denmark | Statens Serum Institut Bioinformatics and Microbial Genomics | Danish Covid-19 Genome Consortium |
| EPI_ISL_3450122 | Utah Public Health Laboratory | Utah Public Health Laboratory | Erin L. Young, Kelly F. Oakeson, Pooja Gupta, Olinto Linares-Perdomo |
| EPI_ISL_3450123 | Department of Bacteria, Parasites and Fungi, Statens Serum Institut, Copenhagen, Denmark | Statens Serum Institut Bioinformatics and Microbial Genomics | Danish Covid-19 Genome Consortium |
| EPI_ISL_3450124 | Utah Public Health Laboratory | Utah Public Health Laboratory | Erin L. Young, Kelly F. Oakeson, Pooja Gupta, Olinto Linares-Perdomo |
| EPI_ISL_3450125 | Department of Bacteria, Parasites and Fungi, Statens Serum Institut, Copenhagen, Denmark | Statens Serum Institut Bioinformatics and Microbial Genomics | Danish Covid-19 Genome Consortium |
| EPI_ISL_3450126 | Utah Public Health Laboratory | Utah Public Health Laboratory | Erin L. Young, Kelly F. Oakeson, Pooja Gupta, Olinto Linares-Perdomo |
| EPI_ISL_3450127 | Department of Bacteria, Parasites and Fungi, Statens Serum Institut, Copenhagen, Denmark | Statens Serum Institut Bioinformatics and Microbial Genomics | Danish Covid-19 Genome Consortium |
| EPI_ISL_3450128 | Utah Public Health Laboratory | Utah Public Health Laboratory | Erin L. Young, Kelly F. Oakeson, Pooja Gupta, Olinto Linares-Perdomo |
| EPI_ISL_3450129 | Department of Bacteria, Parasites and Fungi, Statens Serum Institut, Copenhagen, Denmark | Statens Serum Institut Bioinformatics and Microbial Genomics | Danish Covid-19 Genome Consortium |
| EPI_ISL_3450130 | Utah Public Health Laboratory | Utah Public Health Laboratory | Erin L. Young, Kelly F. Oakeson, Pooja Gupta, Olinto Linares-Perdomo |
| EPI_ISL_3450131, EPI_ISL_3450132 | Department of Bacteria, Parasites and Fungi, Statens Serum Institut, Copenhagen, Denmark | Statens Serum Institut Bioinformatics and Microbial Genomics | Danish Covid-19 Genome Consortium |
| EPI_ISL_3450133 | Utah Public Health Laboratory | Utah Public Health Laboratory | Erin L. Young, Kelly F. Oakeson, Pooja Gupta, Olinto Linares-Perdomo |
| EPI_ISL_3450134 | Department of Bacteria, Parasites and Fungi, Statens Serum Institut, Copenhagen, Denmark | Statens Serum Institut Bioinformatics and Microbial Genomics | Danish Covid-19 Genome Consortium |
| EPI_ISL_3450135 | Utah Public Health Laboratory | Utah Public Health Laboratory | Erin L. Young, Kelly F. Oakeson, Pooja Gupta, Olinto Linares-Perdomo |
| EPI_ISL_3450136 | Department of Bacteria, Parasites and Fungi, Statens Serum Institut, Copenhagen, Denmark | Statens Serum Institut Bioinformatics and Microbial Genomics | Danish Covid-19 Genome Consortium |
| EPI_ISL_3450137 | Utah Public Health Laboratory | Utah Public Health Laboratory | Erin L. Young, Kelly F. Oakeson, Pooja Gupta, Olinto Linares-Perdomo |
| EPI_ISL_3450138 | Department of Bacteria, Parasites and Fungi, Statens Serum Institut, Copenhagen, Denmark | Statens Serum Institut Bioinformatics and Microbial Genomics | Danish Covid-19 Genome Consortium |
| EPI_ISL_3450139 | Utah Public Health Laboratory | Utah Public Health Laboratory | Erin L. Young, Kelly F. Oakeson, Pooja Gupta, Olinto Linares-Perdomo |
| EPI_ISL_3450140 | Department of Bacteria, Parasites and Fungi, Statens Serum Institut, Copenhagen, Denmark | Statens Serum Institut Bioinformatics and Microbial Genomics | Danish Covid-19 Genome Consortium |
| EPI_ISL_3450141 | Utah Public Health Laboratory | Utah Public Health Laboratory | Erin L. Young, Kelly F. Oakeson, Pooja Gupta, Olinto Linares-Perdomo |
| EPI_ISL_3450142 | Department of Bacteria, Parasites and Fungi, Statens Serum Institut, Copenhagen, Denmark | Statens Serum Institut Bioinformatics and Microbial Genomics | Danish Covid-19 Genome Consortium |
| EPI_ISL_3450143 | Utah Public Health Laboratory | Utah Public Health Laboratory | Erin L. Young, Kelly F. Oakeson, Pooja Gupta, Olinto Linares-Perdomo |
| EPI_ISL_3450144 | Department of Bacteria, Parasites and Fungi, Statens Serum Institut, Copenhagen, Denmark | Statens Serum Institut Bioinformatics and Microbial Genomics | Danish Covid-19 Genome Consortium |
| EPI_ISL_3450145 | Utah Public Health Laboratory | Utah Public Health Laboratory | Erin L. Young, Kelly F. Oakeson, Pooja Gupta, Olinto Linares-Perdomo |
| EPI_ISL_3450146 | Department of Bacteria, Parasites and Fungi, Statens Serum Institut, Copenhagen, Denmark | Statens Serum Institut Bioinformatics and Microbial Genomics | Danish Covid-19 Genome Consortium |
| EPI_ISL_3450147 | Utah Public Health Laboratory | Utah Public Health Laboratory | Erin L. Young, Kelly F. Oakeson, Pooja Gupta, Olinto Linares-Perdomo |
| EPI_ISL_3450148 | Department of Bacteria, Parasites and Fungi, Statens Serum Institut, Copenhagen, Denmark | Statens Serum Institut Bioinformatics and Microbial Genomics | Danish Covid-19 Genome Consortium |
| EPI_ISL_3450149 | Utah Public Health Laboratory | Utah Public Health Laboratory | Erin L. Young, Kelly F. Oakeson, Pooja Gupta, Olinto Linares-Perdomo |
| EPI_ISL_3450150 | Department of Bacteria, Parasites and Fungi, Statens Serum Institut, Copenhagen, Denmark | Statens Serum Institut Bioinformatics and Microbial Genomics | Danish Covid-19 Genome Consortium |
| EPI_ISL_3450151 | Utah Public Health Laboratory | Utah Public Health Laboratory | Erin L. Young, Kelly F. Oakeson, Pooja Gupta, Olinto Linares-Perdomo |
| EPI_ISL_3450152 | Department of Bacteria, Parasites and Fungi, Statens Serum Institut, Copenhagen, Denmark | Statens Serum Institut Bioinformatics and Microbial Genomics | Danish Covid-19 Genome Consortium |
| EPI_ISL_3450153 | Utah Public Health Laboratory | Utah Public Health Laboratory | Erin L. Young, Kelly F. Oakeson, Pooja Gupta, Olinto Linares-Perdomo |
| EPI_ISL_3450154 | Department of Bacteria, Parasites and Fungi, Statens Serum Institut, Copenhagen, Denmark | Statens Serum Institut Bioinformatics and Microbial Genomics | Danish Covid-19 Genome Consortium |
| EPI_ISL_3450155 | Utah Public Health Laboratory | Utah Public Health Laboratory | Erin L. Young, Kelly F. Oakeson, Pooja Gupta, Olinto Linares-Perdomo |
| EPI_ISL_3450156 | Department of Bacteria, Parasites and Fungi, Statens Serum Institut, Copenhagen, Denmark | Statens Serum Institut Bioinformatics and Microbial Genomics | Danish Covid-19 Genome Consortium |
| EPI_ISL_3450157 | Utah Public Health Laboratory | Utah Public Health Laboratory | Erin L. Young, Kelly F. Oakeson, Pooja Gupta, Olinto Linares-Perdomo |
| EPI_ISL_3450158 | Department of Bacteria, Parasites and Fungi, Statens Serum Institut, Copenhagen, Denmark | Statens Serum Institut Bioinformatics and Microbial Genomics | Danish Covid-19 Genome Consortium |
| EPI_ISL_3450159 | Utah Public Health Laboratory | Utah Public Health Laboratory | Erin L. Young, Kelly F. Oakeson, Pooja Gupta, Olinto Linares-Perdomo |
| EPI_ISL_3450160, EPI_ISL_3450161 | Department of Bacteria, Parasites and Fungi, Statens Serum Institut, Copenhagen, Denmark | Statens Serum Institut Bioinformatics and Microbial Genomics | Danish Covid-19 Genome Consortium |
| EPI_ISL_3450162 | Utah Public Health Laboratory | Utah Public Health Laboratory | Erin L. Young, Kelly F. Oakeson, Pooja Gupta, Olinto Linares-Perdomo |
| EPI_ISL_3450163 | Department of Bacteria, Parasites and Fungi, Statens Serum Institut, Copenhagen, Denmark | Statens Serum Institut Bioinformatics and Microbial Genomics | Danish Covid-19 Genome Consortium |
| EPI_ISL_3450164 | Utah Public Health Laboratory | Utah Public Health Laboratory | Erin L. Young, Kelly F. Oakeson, Pooja Gupta, Olinto Linares-Perdomo |
| EPI_ISL_3450165 | Department of Bacteria, Parasites and Fungi, Statens Serum Institut, Copenhagen, Denmark | Statens Serum Institut Bioinformatics and Microbial Genomics | Danish Covid-19 Genome Consortium |
| EPI_ISL_3450166, EPI_ISL_3450167 | Utah Public Health Laboratory | Utah Public Health Laboratory | Erin L. Young, Kelly F. Oakeson, Pooja Gupta, Olinto Linares-Perdomo |
| EPI_ISL_3450168 | Department of Bacteria, Parasites and Fungi, Statens Serum Institut, Copenhagen, Denmark | Statens Serum Institut Bioinformatics and Microbial Genomics | Danish Covid-19 Genome Consortium |
| EPI_ISL_3450169 | Utah Public Health Laboratory | Utah Public Health Laboratory | Erin L. Young, Kelly F. Oakeson, Pooja Gupta, Olinto Linares-Perdomo |

| | | | |
|---|---|---|---|
| EPI_ISL_3450170 | Department of Bacteria, Parasites and Fungi, Statens Serum Institut, Copenhagen, Denmark | Statens Serum Institut Bioinformatics and Microbial Genomics | Danish Covid-19 Genome Consortium |
| EPI_ISL_3450171 | Utah Public Health Laboratory | Utah Public Health Laboratory | Erin L. Young, Kelly F. Oakeson, Pooja Gupta, Olinto Linares-Perdomo |
| EPI_ISL_3450172, EPI_ISL_3450173 | Department of Bacteria, Parasites and Fungi, Statens Serum Institut, Copenhagen, Denmark | Statens Serum Institut Bioinformatics and Microbial Genomics | Danish Covid-19 Genome Consortium |
| EPI_ISL_3450174 | Utah Public Health Laboratory | Utah Public Health Laboratory | Erin L. Young, Kelly F. Oakeson, Pooja Gupta, Olinto Linares-Perdomo |
| EPI_ISL_3450175 | Department of Bacteria, Parasites and Fungi, Statens Serum Institut, Copenhagen, Denmark | Statens Serum Institut Bioinformatics and Microbial Genomics | Danish Covid-19 Genome Consortium |
| EPI_ISL_3450176 | Utah Public Health Laboratory | Utah Public Health Laboratory | Erin L. Young, Kelly F. Oakeson, Pooja Gupta, Olinto Linares-Perdomo |
| EPI_ISL_3450177, EPI_ISL_3450178 | Department of Bacteria, Parasites and Fungi, Statens Serum Institut, Copenhagen, Denmark | Statens Serum Institut Bioinformatics and Microbial Genomics | Danish Covid-19 Genome Consortium |
| EPI_ISL_3450179 | Utah Public Health Laboratory | Utah Public Health Laboratory | Erin L. Young, Kelly F. Oakeson, Pooja Gupta, Olinto Linares-Perdomo |
| EPI_ISL_3450180, EPI_ISL_3450181 | Department of Bacteria, Parasites and Fungi, Statens Serum Institut, Copenhagen, Denmark | Statens Serum Institut Bioinformatics and Microbial Genomics | Danish Covid-19 Genome Consortium |
| EPI_ISL_3450182 | Utah Public Health Laboratory | Utah Public Health Laboratory | Erin L. Young, Kelly F. Oakeson, Pooja Gupta, Olinto Linares-Perdomo |
| EPI_ISL_3450183 | Department of Bacteria, Parasites and Fungi, Statens Serum Institut, Copenhagen, Denmark | Statens Serum Institut Bioinformatics and Microbial Genomics | Danish Covid-19 Genome Consortium |
| EPI_ISL_3450184 | Utah Public Health Laboratory | Utah Public Health Laboratory | Erin L. Young, Kelly F. Oakeson, Pooja Gupta, Olinto Linares-Perdomo |
| EPI_ISL_3450185 | Department of Bacteria, Parasites and Fungi, Statens Serum Institut, Copenhagen, Denmark | Statens Serum Institut Bioinformatics and Microbial Genomics | Danish Covid-19 Genome Consortium |
| EPI_ISL_3450186 | Utah Public Health Laboratory | Utah Public Health Laboratory | Erin L. Young, Kelly F. Oakeson, Pooja Gupta, Olinto Linares-Perdomo |
| EPI_ISL_3450187 | Department of Bacteria, Parasites and Fungi, Statens Serum Institut, Copenhagen, Denmark | Statens Serum Institut Bioinformatics and Microbial Genomics | Danish Covid-19 Genome Consortium |
| EPI_ISL_3450188 | Utah Public Health Laboratory | Utah Public Health Laboratory | Erin L. Young, Kelly F. Oakeson, Pooja Gupta, Olinto Linares-Perdomo |
| EPI_ISL_3450189 | Department of Bacteria, Parasites and Fungi, Statens Serum Institut, Copenhagen, Denmark | Statens Serum Institut Bioinformatics and Microbial Genomics | Danish Covid-19 Genome Consortium |
| EPI_ISL_3450190 | Utah Public Health Laboratory | Utah Public Health Laboratory | Erin L. Young, Kelly F. Oakeson, Pooja Gupta, Olinto Linares-Perdomo |
| EPI_ISL_3450191 | Department of Bacteria, Parasites and Fungi, Statens Serum Institut, Copenhagen, Denmark | Statens Serum Institut Bioinformatics and Microbial Genomics | Danish Covid-19 Genome Consortium |
| EPI_ISL_3450192 | Utah Public Health Laboratory | Utah Public Health Laboratory | Erin L. Young, Kelly F. Oakeson, Pooja Gupta, Olinto Linares-Perdomo |
| EPI_ISL_3450193 | Department of Bacteria, Parasites and Fungi, Statens Serum Institut, Copenhagen, Denmark | Statens Serum Institut Bioinformatics and Microbial Genomics | Danish Covid-19 Genome Consortium |
| EPI_ISL_3450194 | Utah Public Health Laboratory | Utah Public Health Laboratory | Erin L. Young, Kelly F. Oakeson, Pooja Gupta, Olinto Linares-Perdomo |
| EPI_ISL_3450195, EPI_ISL_3450196 | Department of Bacteria, Parasites and Fungi, Statens Serum Institut, Copenhagen, Denmark | Statens Serum Institut Bioinformatics and Microbial Genomics | Danish Covid-19 Genome Consortium |
| EPI_ISL_3450197 | Utah Public Health Laboratory | Utah Public Health Laboratory | Erin L. Young, Kelly F. Oakeson, Pooja Gupta, Olinto Linares-Perdomo |
| EPI_ISL_3450198 | Department of Bacteria, Parasites and Fungi, Statens Serum Institut, Copenhagen, Denmark | Statens Serum Institut Bioinformatics and Microbial Genomics | Danish Covid-19 Genome Consortium |
| EPI_ISL_3450199 | Utah Public Health Laboratory | Utah Public Health Laboratory | Erin L. Young, Kelly F. Oakeson, Pooja Gupta, Olinto Linares-Perdomo |
| EPI_ISL_3450200 | Department of Bacteria, Parasites and Fungi, Statens Serum Institut, Copenhagen, Denmark | Statens Serum Institut Bioinformatics and Microbial Genomics | Danish Covid-19 Genome Consortium |
| EPI_ISL_3450201 | Utah Public Health Laboratory | Utah Public Health Laboratory | Erin L. Young, Kelly F. Oakeson, Pooja Gupta, Olinto Linares-Perdomo |
| EPI_ISL_3450202 | Department of Bacteria, Parasites and Fungi, Statens Serum Institut, Copenhagen, Denmark | Statens Serum Institut Bioinformatics and Microbial Genomics | Danish Covid-19 Genome Consortium |
| EPI_ISL_3450203 | Utah Public Health Laboratory | Utah Public Health Laboratory | Erin L. Young, Kelly F. Oakeson, Pooja Gupta, Olinto Linares-Perdomo |
| EPI_ISL_3450204 | Department of Bacteria, Parasites and Fungi, Statens Serum Institut, Copenhagen, Denmark | Statens Serum Institut Bioinformatics and Microbial Genomics | Danish Covid-19 Genome Consortium |
| EPI_ISL_3450205 | Utah Public Health Laboratory | Utah Public Health Laboratory | Erin L. Young, Kelly F. Oakeson, Pooja Gupta, Olinto Linares-Perdomo |
| EPI_ISL_3450206 | Department of Bacteria, Parasites and Fungi, Statens Serum Institut, Copenhagen, Denmark | Statens Serum Institut Bioinformatics and Microbial Genomics | Danish Covid-19 Genome Consortium |
| EPI_ISL_3450207 | Utah Public Health Laboratory | Utah Public Health Laboratory | Erin L. Young, Kelly F. Oakeson, Pooja Gupta, Olinto Linares-Perdomo |
| EPI_ISL_3450208, EPI_ISL_3450209 | Department of Bacteria, Parasites and Fungi, Statens Serum Institut, Copenhagen, Denmark | Statens Serum Institut Bioinformatics and Microbial Genomics | Danish Covid-19 Genome Consortium |
| EPI_ISL_3450210 | Utah Public Health Laboratory | Utah Public Health Laboratory | Erin L. Young, Kelly F. Oakeson, Pooja Gupta, Olinto Linares-Perdomo |
| EPI_ISL_3450211 | Department of Bacteria, Parasites and Fungi, Statens Serum Institut, Copenhagen, Denmark | Statens Serum Institut Bioinformatics and Microbial Genomics | Danish Covid-19 Genome Consortium |
| EPI_ISL_3450212 | Utah Public Health Laboratory | Utah Public Health Laboratory | Erin L. Young, Kelly F. Oakeson, Pooja Gupta, Olinto Linares-Perdomo |
| EPI_ISL_3450213 | Department of Bacteria, Parasites and Fungi, Statens Serum Institut, Copenhagen, Denmark | Statens Serum Institut Bioinformatics and Microbial Genomics | Danish Covid-19 Genome Consortium |
| EPI_ISL_3450214 | Utah Public Health Laboratory | Utah Public Health Laboratory | Erin L. Young, Kelly F. Oakeson, Pooja Gupta, Olinto Linares-Perdomo |
| EPI_ISL_3450215 | Department of Bacteria, Parasites and Fungi, Statens Serum Institut, Copenhagen, Denmark | Statens Serum Institut Bioinformatics and Microbial Genomics | Danish Covid-19 Genome Consortium |
| EPI_ISL_3450216 | Utah Public Health Laboratory | Utah Public Health Laboratory | Erin L. Young, Kelly F. Oakeson, Pooja Gupta, Olinto Linares-Perdomo |
| EPI_ISL_3450217 | Department of Bacteria, Parasites and Fungi, Statens Serum Institut, Copenhagen, Denmark | Statens Serum Institut Bioinformatics and Microbial Genomics | Danish Covid-19 Genome Consortium |
| EPI_ISL_3450218 | Utah Public Health Laboratory | Utah Public Health Laboratory | Erin L. Young, Kelly F. Oakeson, Pooja Gupta, Olinto Linares-Perdomo |
| EPI_ISL_3450219 | Department of Bacteria, Parasites and Fungi, Statens Serum Institut, Copenhagen, Denmark | Statens Serum Institut Bioinformatics and Microbial Genomics | Danish Covid-19 Genome Consortium |
| EPI_ISL_3450220 | Utah Public Health Laboratory | Utah Public Health Laboratory | Erin L. Young, Kelly F. Oakeson, Pooja Gupta, Olinto Linares-Perdomo |
| EPI_ISL_3450221 | Department of Bacteria, Parasites and Fungi, Statens Serum Institut, Copenhagen, Denmark | Statens Serum Institut Bioinformatics and Microbial Genomics | Danish Covid-19 Genome Consortium |
| EPI_ISL_3450222 | Utah Public Health Laboratory | Utah Public Health Laboratory | Erin L. Young, Kelly F. Oakeson, Pooja Gupta, Olinto Linares-Perdomo |
| EPI_ISL_3450223 | Department of Bacteria, Parasites and Fungi, Statens Serum Institut, Copenhagen, Denmark | Statens Serum Institut Bioinformatics and Microbial Genomics | Danish Covid-19 Genome Consortium |
| EPI_ISL_3450224 | Utah Public Health Laboratory | Utah Public Health Laboratory | Erin L. Young, Kelly F. Oakeson, Pooja Gupta, Olinto Linares-Perdomo |
| EPI_ISL_3450225 | Department of Bacteria, Parasites and Fungi, Statens Serum Institut, Copenhagen, Denmark | Statens Serum Institut Bioinformatics and Microbial Genomics | Danish Covid-19 Genome Consortium |
| EPI_ISL_3450226 | Utah Public Health Laboratory | Utah Public Health Laboratory | Erin L. Young, Kelly F. Oakeson, Pooja Gupta, Olinto Linares-Perdomo |
| EPI_ISL_3450227 | Department of Bacteria, Parasites and Fungi, Statens Serum Institut, Copenhagen, Denmark | Statens Serum Institut Bioinformatics and Microbial Genomics | Danish Covid-19 Genome Consortium |
| EPI_ISL_3450228 | Utah Public Health Laboratory | Utah Public Health Laboratory | Erin L. Young, Kelly F. Oakeson, Pooja Gupta, Olinto Linares-Perdomo |
| EPI_ISL_3450229 | Department of Bacteria, Parasites and Fungi, Statens Serum Institut, Copenhagen, Denmark | Statens Serum Institut Bioinformatics and Microbial Genomics | Danish Covid-19 Genome Consortium |
| EPI_ISL_3450230 | Utah Public Health Laboratory | Utah Public Health Laboratory | Erin L. Young, Kelly F. Oakeson, Pooja Gupta, Olinto Linares-Perdomo |
| EPI_ISL_3450231 | Department of Bacteria, Parasites and Fungi, Statens Serum | Statens Serum Institut Bioinformatics and Microbial Genomics | Danish Covid-19 Genome Consortium |

| | | | |
|---|---|---|---|
| | Institut, Copenhagen, Denmark | | |
| EPI_ISL_3450232 | Utah Public Health Laboratory | Utah Public Health Laboratory | Erin L. Young, Kelly F. Oakeson, Pooja Gupta, Olinto Linares-Perdomo |
| EPI_ISL_3450233, EPI_ISL_3450234 | Department of Bacteria, Parasites and Fungi, Statens Serum Institut, Copenhagen, Denmark | Statens Serum Institut Bioinformatics and Microbial Genomics | Danish Covid-19 Genome Consortium |
| EPI_ISL_3450235 | Utah Public Health Laboratory | Utah Public Health Laboratory | Erin L. Young, Kelly F. Oakeson, Pooja Gupta, Olinto Linares-Perdomo |
| EPI_ISL_3450236 | Department of Bacteria, Parasites and Fungi, Statens Serum Institut, Copenhagen, Denmark | Statens Serum Institut Bioinformatics and Microbial Genomics | Danish Covid-19 Genome Consortium |
| EPI_ISL_3450237 | Utah Public Health Laboratory | Utah Public Health Laboratory | Erin L. Young, Kelly F. Oakeson, Pooja Gupta, Olinto Linares-Perdomo |
| EPI_ISL_3450238 | Department of Bacteria, Parasites and Fungi, Statens Serum Institut, Copenhagen, Denmark | Statens Serum Institut Bioinformatics and Microbial Genomics | Danish Covid-19 Genome Consortium |
| EPI_ISL_3450239 | Utah Public Health Laboratory | Utah Public Health Laboratory | Erin L. Young, Kelly F. Oakeson, Pooja Gupta, Olinto Linares-Perdomo |
| EPI_ISL_3450240 | Department of Bacteria, Parasites and Fungi, Statens Serum Institut, Copenhagen, Denmark | Statens Serum Institut Bioinformatics and Microbial Genomics | Danish Covid-19 Genome Consortium |
| EPI_ISL_3450241 | Utah Public Health Laboratory | Utah Public Health Laboratory | Erin L. Young, Kelly F. Oakeson, Pooja Gupta, Olinto Linares-Perdomo |
| EPI_ISL_3450242 | Department of Bacteria, Parasites and Fungi, Statens Serum Institut, Copenhagen, Denmark | Statens Serum Institut Bioinformatics and Microbial Genomics | Danish Covid-19 Genome Consortium |
| EPI_ISL_3450243 | Utah Public Health Laboratory | Utah Public Health Laboratory | Erin L. Young, Kelly F. Oakeson, Pooja Gupta, Olinto Linares-Perdomo |
| EPI_ISL_3450244, EPI_ISL_3450245 | Department of Bacteria, Parasites and Fungi, Statens Serum Institut, Copenhagen, Denmark | Statens Serum Institut Bioinformatics and Microbial Genomics | Danish Covid-19 Genome Consortium |
| EPI_ISL_3450246 | Utah Public Health Laboratory | Utah Public Health Laboratory | Erin L. Young, Kelly F. Oakeson, Pooja Gupta, Olinto Linares-Perdomo |
| EPI_ISL_3450247 | Department of Bacteria, Parasites and Fungi, Statens Serum Institut, Copenhagen, Denmark | Statens Serum Institut Bioinformatics and Microbial Genomics | Danish Covid-19 Genome Consortium |
| EPI_ISL_3450248 | Utah Public Health Laboratory | Utah Public Health Laboratory | Erin L. Young, Kelly F. Oakeson, Pooja Gupta, Olinto Linares-Perdomo |
| EPI_ISL_3450249 | Department of Bacteria, Parasites and Fungi, Statens Serum Institut, Copenhagen, Denmark | Statens Serum Institut Bioinformatics and Microbial Genomics | Danish Covid-19 Genome Consortium |
| EPI_ISL_3450250 | Utah Public Health Laboratory | Utah Public Health Laboratory | Erin L. Young, Kelly F. Oakeson, Pooja Gupta, Olinto Linares-Perdomo |
| EPI_ISL_3450251 | Department of Bacteria, Parasites and Fungi, Statens Serum Institut, Copenhagen, Denmark | Statens Serum Institut Bioinformatics and Microbial Genomics | Danish Covid-19 Genome Consortium |
| EPI_ISL_3450252 | Utah Public Health Laboratory | Utah Public Health Laboratory | Erin L. Young, Kelly F. Oakeson, Pooja Gupta, Olinto Linares-Perdomo |
| EPI_ISL_3450253 | Department of Bacteria, Parasites and Fungi, Statens Serum Institut, Copenhagen, Denmark | Statens Serum Institut Bioinformatics and Microbial Genomics | Danish Covid-19 Genome Consortium |
| EPI_ISL_3450254 | Utah Public Health Laboratory | Utah Public Health Laboratory | Erin L. Young, Kelly F. Oakeson, Pooja Gupta, Olinto Linares-Perdomo |
| EPI_ISL_3450255 | Department of Bacteria, Parasites and Fungi, Statens Serum Institut, Copenhagen, Denmark | Statens Serum Institut Bioinformatics and Microbial Genomics | Danish Covid-19 Genome Consortium |
| EPI_ISL_3450256 | Utah Public Health Laboratory | Utah Public Health Laboratory | Erin L. Young, Kelly F. Oakeson, Pooja Gupta, Olinto Linares-Perdomo |
| EPI_ISL_3450257 | Department of Bacteria, Parasites and Fungi, Statens Serum Institut, Copenhagen, Denmark | Statens Serum Institut Bioinformatics and Microbial Genomics | Danish Covid-19 Genome Consortium |
| EPI_ISL_3450258 | Utah Public Health Laboratory | Utah Public Health Laboratory | Erin L. Young, Kelly F. Oakeson, Pooja Gupta, Olinto Linares-Perdomo |
| EPI_ISL_3450259 | Department of Bacteria, Parasites and Fungi, Statens Serum Institut, Copenhagen, Denmark | Statens Serum Institut Bioinformatics and Microbial Genomics | Danish Covid-19 Genome Consortium |
| EPI_ISL_3450260 | Utah Public Health Laboratory | Utah Public Health Laboratory | Erin L. Young, Kelly F. Oakeson, Pooja Gupta, Olinto Linares-Perdomo |
| EPI_ISL_3450261 | Department of Bacteria, Parasites and Fungi, Statens Serum Institut, Copenhagen, Denmark | Statens Serum Institut Bioinformatics and Microbial Genomics | Danish Covid-19 Genome Consortium |
| EPI_ISL_3450262 | Utah Public Health Laboratory | Utah Public Health Laboratory | Erin L. Young, Kelly F. Oakeson, Pooja Gupta, Olinto Linares-Perdomo |
| EPI_ISL_3450263 | Department of Bacteria, Parasites and Fungi, Statens Serum Institut, Copenhagen, Denmark | Statens Serum Institut Bioinformatics and Microbial Genomics | Danish Covid-19 Genome Consortium |
| EPI_ISL_3450264 | Utah Public Health Laboratory | Utah Public Health Laboratory | Erin L. Young, Kelly F. Oakeson, Pooja Gupta, Olinto Linares-Perdomo |
| EPI_ISL_3450265 | Department of Bacteria, Parasites and Fungi, Statens Serum Institut, Copenhagen, Denmark | Statens Serum Institut Bioinformatics and Microbial Genomics | Danish Covid-19 Genome Consortium |
| EPI_ISL_3450266 | Utah Public Health Laboratory | Utah Public Health Laboratory | Erin L. Young, Kelly F. Oakeson, Pooja Gupta, Olinto Linares-Perdomo |
| EPI_ISL_3450267 | Department of Bacteria, Parasites and Fungi, Statens Serum Institut, Copenhagen, Denmark | Statens Serum Institut Bioinformatics and Microbial Genomics | Danish Covid-19 Genome Consortium |
| EPI_ISL_3450268 | Utah Public Health Laboratory | Utah Public Health Laboratory | Erin L. Young, Kelly F. Oakeson, Pooja Gupta, Olinto Linares-Perdomo |
| EPI_ISL_3450269 | Department of Bacteria, Parasites and Fungi, Statens Serum Institut, Copenhagen, Denmark | Statens Serum Institut Bioinformatics and Microbial Genomics | Danish Covid-19 Genome Consortium |
| EPI_ISL_3450270 | Utah Public Health Laboratory | Utah Public Health Laboratory | Erin L. Young, Kelly F. Oakeson, Pooja Gupta, Olinto Linares-Perdomo |
| EPI_ISL_3450271, EPI_ISL_3450272 | Department of Bacteria, Parasites and Fungi, Statens Serum Institut, Copenhagen, Denmark | Statens Serum Institut Bioinformatics and Microbial Genomics | Danish Covid-19 Genome Consortium |
| EPI_ISL_3450273 | Utah Public Health Laboratory | Utah Public Health Laboratory | Erin L. Young, Kelly F. Oakeson, Pooja Gupta, Olinto Linares-Perdomo |
| EPI_ISL_3450274 | Department of Bacteria, Parasites and Fungi, Statens Serum Institut, Copenhagen, Denmark | Statens Serum Institut Bioinformatics and Microbial Genomics | Danish Covid-19 Genome Consortium |
| EPI_ISL_3450275 | Utah Public Health Laboratory | Utah Public Health Laboratory | Erin L. Young, Kelly F. Oakeson, Pooja Gupta, Olinto Linares-Perdomo |
| EPI_ISL_3450276 | Department of Bacteria, Parasites and Fungi, Statens Serum Institut, Copenhagen, Denmark | Statens Serum Institut Bioinformatics and Microbial Genomics | Danish Covid-19 Genome Consortium |
| EPI_ISL_3450277 | Utah Public Health Laboratory | Utah Public Health Laboratory | Erin L. Young, Kelly F. Oakeson, Pooja Gupta, Olinto Linares-Perdomo |
| EPI_ISL_3450278 | Department of Bacteria, Parasites and Fungi, Statens Serum Institut, Copenhagen, Denmark | Statens Serum Institut Bioinformatics and Microbial Genomics | Danish Covid-19 Genome Consortium |
| EPI_ISL_3450279 | Utah Public Health Laboratory | Utah Public Health Laboratory | Erin L. Young, Kelly F. Oakeson, Pooja Gupta, Olinto Linares-Perdomo |
| EPI_ISL_3450280 | Department of Bacteria, Parasites and Fungi, Statens Serum Institut, Copenhagen, Denmark | Statens Serum Institut Bioinformatics and Microbial Genomics | Danish Covid-19 Genome Consortium |
| EPI_ISL_3450281 | Utah Public Health Laboratory | Utah Public Health Laboratory | Erin L. Young, Kelly F. Oakeson, Pooja Gupta, Olinto Linares-Perdomo |
| EPI_ISL_3450282 | Department of Bacteria, Parasites and Fungi, Statens Serum Institut, Copenhagen, Denmark | Statens Serum Institut Bioinformatics and Microbial Genomics | Danish Covid-19 Genome Consortium |
| EPI_ISL_3450283 | Utah Public Health Laboratory | Utah Public Health Laboratory | Erin L. Young, Kelly F. Oakeson, Pooja Gupta, Olinto Linares-Perdomo |
| EPI_ISL_3450284 | Department of Bacteria, Parasites and Fungi, Statens Serum Institut, Copenhagen, Denmark | Statens Serum Institut Bioinformatics and Microbial Genomics | Danish Covid-19 Genome Consortium |
| EPI_ISL_3450285 | Utah Public Health Laboratory | Utah Public Health Laboratory | Erin L. Young, Kelly F. Oakeson, Pooja Gupta, Olinto Linares-Perdomo |
| EPI_ISL_3450286, EPI_ISL_3450287 | Department of Bacteria, Parasites and Fungi, Statens Serum Institut, Copenhagen, Denmark | Statens Serum Institut Bioinformatics and Microbial Genomics | Danish Covid-19 Genome Consortium |
| EPI_ISL_3450288 | Utah Public Health Laboratory | Utah Public Health Laboratory | Erin L. Young, Kelly F. Oakeson, Pooja Gupta, Olinto Linares-Perdomo |
| EPI_ISL_3450289 | Department of Bacteria, Parasites and Fungi, Statens Serum Institut, Copenhagen, Denmark | Statens Serum Institut Bioinformatics and Microbial Genomics | Danish Covid-19 Genome Consortium |
| EPI_ISL_3450290 | Utah Public Health Laboratory | Utah Public Health Laboratory | Erin L. Young, Kelly F. Oakeson, Pooja Gupta, Olinto Linares-Perdomo |
| EPI_ISL_3450291 | Department of Bacteria, Parasites and Fungi, Statens Serum Institut, Copenhagen, Denmark | Statens Serum Institut Bioinformatics and Microbial Genomics | Danish Covid-19 Genome Consortium |

| | | | |
|---|---|---|---|
| EPI_ISL_3450292 | Utah Public Health Laboratory | Utah Public Health Laboratory | Erin L. Young, Kelly F. Oakeson, Pooja Gupta, Olinto Linares-Perdomo |
| EPI_ISL_3450293 | Department of Bacteria, Parasites and Fungi, Statens Serum Institut, Copenhagen, Denmark | Statens Serum Institut Bioinformatics and Microbial Genomics | Danish Covid-19 Genome Consortium |
| EPI_ISL_3450294, EPI_ISL_3450295 | Utah Public Health Laboratory | Utah Public Health Laboratory | Erin L. Young, Kelly F. Oakeson, Pooja Gupta, Olinto Linares-Perdomo |
| EPI_ISL_3450296, EPI_ISL_3450297 | Department of Bacteria, Parasites and Fungi, Statens Serum Institut, Copenhagen, Denmark | Statens Serum Institut Bioinformatics and Microbial Genomics | Danish Covid-19 Genome Consortium |
| EPI_ISL_3450298 | Utah Public Health Laboratory | Utah Public Health Laboratory | Erin L. Young, Kelly F. Oakeson, Pooja Gupta, Olinto Linares-Perdomo |
| EPI_ISL_3450299 | Department of Bacteria, Parasites and Fungi, Statens Serum Institut, Copenhagen, Denmark | Statens Serum Institut Bioinformatics and Microbial Genomics | Danish Covid-19 Genome Consortium |
| EPI_ISL_3450300 | Utah Public Health Laboratory | Utah Public Health Laboratory | Erin L. Young, Kelly F. Oakeson, Pooja Gupta, Olinto Linares-Perdomo |
| EPI_ISL_3450301 | Department of Bacteria, Parasites and Fungi, Statens Serum Institut, Copenhagen, Denmark | Statens Serum Institut Bioinformatics and Microbial Genomics | Danish Covid-19 Genome Consortium |
| EPI_ISL_3450302 | Utah Public Health Laboratory | Utah Public Health Laboratory | Erin L. Young, Kelly F. Oakeson, Pooja Gupta, Olinto Linares-Perdomo |
| EPI_ISL_3450303 | Department of Bacteria, Parasites and Fungi, Statens Serum Institut, Copenhagen, Denmark | Statens Serum Institut Bioinformatics and Microbial Genomics | Danish Covid-19 Genome Consortium |
| EPI_ISL_3450304 | Utah Public Health Laboratory | Utah Public Health Laboratory | Erin L. Young, Kelly F. Oakeson, Pooja Gupta, Olinto Linares-Perdomo |
| EPI_ISL_3450305 | Department of Bacteria, Parasites and Fungi, Statens Serum Institut, Copenhagen, Denmark | Statens Serum Institut Bioinformatics and Microbial Genomics | Danish Covid-19 Genome Consortium |
| EPI_ISL_3450306 | Utah Public Health Laboratory | Utah Public Health Laboratory | Erin L. Young, Kelly F. Oakeson, Pooja Gupta, Olinto Linares-Perdomo |
| EPI_ISL_3450307 | Department of Bacteria, Parasites and Fungi, Statens Serum Institut, Copenhagen, Denmark | Statens Serum Institut Bioinformatics and Microbial Genomics | Danish Covid-19 Genome Consortium |
| EPI_ISL_3450308 | Utah Public Health Laboratory | Utah Public Health Laboratory | Erin L. Young, Kelly F. Oakeson, Pooja Gupta, Olinto Linares-Perdomo |
| EPI_ISL_3450309 | Department of Bacteria, Parasites and Fungi, Statens Serum Institut, Copenhagen, Denmark | Statens Serum Institut Bioinformatics and Microbial Genomics | Danish Covid-19 Genome Consortium |
| EPI_ISL_3450310 | Utah Public Health Laboratory | Utah Public Health Laboratory | Erin L. Young, Kelly F. Oakeson, Pooja Gupta, Olinto Linares-Perdomo |
| EPI_ISL_3450311 | Department of Bacteria, Parasites and Fungi, Statens Serum Institut, Copenhagen, Denmark | Statens Serum Institut Bioinformatics and Microbial Genomics | Danish Covid-19 Genome Consortium |
| EPI_ISL_3450312 | Utah Public Health Laboratory | Utah Public Health Laboratory | Erin L. Young, Kelly F. Oakeson, Pooja Gupta, Olinto Linares-Perdomo |
| EPI_ISL_3450313 | Department of Bacteria, Parasites and Fungi, Statens Serum Institut, Copenhagen, Denmark | Statens Serum Institut Bioinformatics and Microbial Genomics | Danish Covid-19 Genome Consortium |
| EPI_ISL_3450314 | Utah Public Health Laboratory | Utah Public Health Laboratory | Erin L. Young, Kelly F. Oakeson, Pooja Gupta, Olinto Linares-Perdomo |
| EPI_ISL_3450315 | Department of Bacteria, Parasites and Fungi, Statens Serum Institut, Copenhagen, Denmark | Statens Serum Institut Bioinformatics and Microbial Genomics | Danish Covid-19 Genome Consortium |
| EPI_ISL_3450316 | Utah Public Health Laboratory | Utah Public Health Laboratory | Erin L. Young, Kelly F. Oakeson, Pooja Gupta, Olinto Linares-Perdomo |
| EPI_ISL_3450317, EPI_ISL_3450318 | Department of Bacteria, Parasites and Fungi, Statens Serum Institut, Copenhagen, Denmark | Statens Serum Institut Bioinformatics and Microbial Genomics | Danish Covid-19 Genome Consortium |
| EPI_ISL_3450319, EPI_ISL_3450320 | Utah Public Health Laboratory | Utah Public Health Laboratory | Erin L. Young, Kelly F. Oakeson, Pooja Gupta, Olinto Linares-Perdomo |
| EPI_ISL_3450321, EPI_ISL_3450322 | Department of Bacteria, Parasites and Fungi, Statens Serum Institut, Copenhagen, Denmark | Statens Serum Institut Bioinformatics and Microbial Genomics | Danish Covid-19 Genome Consortium |
| EPI_ISL_3450323 | Utah Public Health Laboratory | Utah Public Health Laboratory | Erin L. Young, Kelly F. Oakeson, Pooja Gupta, Olinto Linares-Perdomo |
| EPI_ISL_3450324 | Department of Bacteria, Parasites and Fungi, Statens Serum Institut, Copenhagen, Denmark | Statens Serum Institut Bioinformatics and Microbial Genomics | Danish Covid-19 Genome Consortium |
| EPI_ISL_3450325 | Utah Public Health Laboratory | Utah Public Health Laboratory | Erin L. Young, Kelly F. Oakeson, Pooja Gupta, Olinto Linares-Perdomo |
| EPI_ISL_3450326 | Department of Bacteria, Parasites and Fungi, Statens Serum Institut, Copenhagen, Denmark | Statens Serum Institut Bioinformatics and Microbial Genomics | Danish Covid-19 Genome Consortium |
| EPI_ISL_3450327 | Utah Public Health Laboratory | Utah Public Health Laboratory | Erin L. Young, Kelly F. Oakeson, Pooja Gupta, Olinto Linares-Perdomo |
| EPI_ISL_3450328 | Department of Bacteria, Parasites and Fungi, Statens Serum Institut, Copenhagen, Denmark | Statens Serum Institut Bioinformatics and Microbial Genomics | Danish Covid-19 Genome Consortium |
| EPI_ISL_3450329 | Utah Public Health Laboratory | Utah Public Health Laboratory | Erin L. Young, Kelly F. Oakeson, Pooja Gupta, Olinto Linares-Perdomo |
| EPI_ISL_3450330 | Department of Bacteria, Parasites and Fungi, Statens Serum Institut, Copenhagen, Denmark | Statens Serum Institut Bioinformatics and Microbial Genomics | Danish Covid-19 Genome Consortium |
| EPI_ISL_3450331 | Utah Public Health Laboratory | Utah Public Health Laboratory | Erin L. Young, Kelly F. Oakeson, Pooja Gupta, Olinto Linares-Perdomo |
| EPI_ISL_3450332 | Department of Bacteria, Parasites and Fungi, Statens Serum Institut, Copenhagen, Denmark | Statens Serum Institut Bioinformatics and Microbial Genomics | Danish Covid-19 Genome Consortium |
| EPI_ISL_3450333 | Utah Public Health Laboratory | Utah Public Health Laboratory | Erin L. Young, Kelly F. Oakeson, Pooja Gupta, Olinto Linares-Perdomo |
| EPI_ISL_3450334 | Department of Bacteria, Parasites and Fungi, Statens Serum Institut, Copenhagen, Denmark | Statens Serum Institut Bioinformatics and Microbial Genomics | Danish Covid-19 Genome Consortium |
| EPI_ISL_3450335 | Utah Public Health Laboratory | Utah Public Health Laboratory | Erin L. Young, Kelly F. Oakeson, Pooja Gupta, Olinto Linares-Perdomo |
| EPI_ISL_3450336 | Department of Bacteria, Parasites and Fungi, Statens Serum Institut, Copenhagen, Denmark | Statens Serum Institut Bioinformatics and Microbial Genomics | Danish Covid-19 Genome Consortium |
| EPI_ISL_3450337 | Utah Public Health Laboratory | Utah Public Health Laboratory | Erin L. Young, Kelly F. Oakeson, Pooja Gupta, Olinto Linares-Perdomo |
| EPI_ISL_3450338, EPI_ISL_3450339 | Department of Bacteria, Parasites and Fungi, Statens Serum Institut, Copenhagen, Denmark | Statens Serum Institut Bioinformatics and Microbial Genomics | Danish Covid-19 Genome Consortium |
| EPI_ISL_3450340 | Utah Public Health Laboratory | Utah Public Health Laboratory | Erin L. Young, Kelly F. Oakeson, Pooja Gupta, Olinto Linares-Perdomo |
| EPI_ISL_3450341 | Department of Bacteria, Parasites and Fungi, Statens Serum Institut, Copenhagen, Denmark | Statens Serum Institut Bioinformatics and Microbial Genomics | Danish Covid-19 Genome Consortium |
| EPI_ISL_3450342 | Utah Public Health Laboratory | Utah Public Health Laboratory | Erin L. Young, Kelly F. Oakeson, Pooja Gupta, Olinto Linares-Perdomo |
| EPI_ISL_3450343 | Department of Bacteria, Parasites and Fungi, Statens Serum Institut, Copenhagen, Denmark | Statens Serum Institut Bioinformatics and Microbial Genomics | Danish Covid-19 Genome Consortium |
| EPI_ISL_3450344 | Utah Public Health Laboratory | Utah Public Health Laboratory | Erin L. Young, Kelly F. Oakeson, Pooja Gupta, Olinto Linares-Perdomo |
| EPI_ISL_3450345 | Department of Bacteria, Parasites and Fungi, Statens Serum Institut, Copenhagen, Denmark | Statens Serum Institut Bioinformatics and Microbial Genomics | Danish Covid-19 Genome Consortium |
| EPI_ISL_3450346, EPI_ISL_3450347 | Utah Public Health Laboratory | Utah Public Health Laboratory | Erin L. Young, Kelly F. Oakeson, Pooja Gupta, Olinto Linares-Perdomo |
| EPI_ISL_3450349 | Department of Bacteria, Parasites and Fungi, Statens Serum Institut, Copenhagen, Denmark | Statens Serum Institut Bioinformatics and Microbial Genomics | Danish Covid-19 Genome Consortium |
| EPI_ISL_3450350 | Utah Public Health Laboratory | Utah Public Health Laboratory | Erin L. Young, Kelly F. Oakeson, Pooja Gupta, Olinto Linares-Perdomo |
| EPI_ISL_3450351 | Department of Bacteria, Parasites and Fungi, Statens Serum Institut, Copenhagen, Denmark | Statens Serum Institut Bioinformatics and Microbial Genomics | Danish Covid-19 Genome Consortium |
| EPI_ISL_3450352 | Utah Public Health Laboratory | Utah Public Health Laboratory | Erin L. Young, Kelly F. Oakeson, Pooja Gupta, Olinto Linares-Perdomo |
| EPI_ISL_3450353 | Department of Bacteria, Parasites and Fungi, Statens Serum Institut, Copenhagen, Denmark | Statens Serum Institut Bioinformatics and Microbial Genomics | Danish Covid-19 Genome Consortium |
| EPI_ISL_3450354 | Utah Public Health Laboratory | Utah Public Health Laboratory | Erin L. Young, Kelly F. Oakeson, Pooja Gupta, Olinto Linares-Perdomo |
| EPI_ISL_3450355, | Department of Bacteria, Parasites and Fungi, Statens Serum | Statens Serum Institut Bioinformatics and Microbial Genomics | Danish Covid-19 Genome Consortium |

| | | | |
|---|---|---|---|
| EPI_ISL_3450356 | Institut, Copenhagen, Denmark | | |
| EPI_ISL_3450357 | Utah Public Health Laboratory | Utah Public Health Laboratory | Erin L. Young, Kelly F. Oakeson, Pooja Gupta, Olinto Linares-Perdomo |
| EPI_ISL_3450358 | Department of Bacteria, Parasites and Fungi, Statens Serum Institut, Copenhagen, Denmark | Statens Serum Institut Bioinformatics and Microbial Genomics | Danish Covid-19 Genome Consortium |
| EPI_ISL_3450359, EPI_ISL_3450360 | Utah Public Health Laboratory | Utah Public Health Laboratory | Erin L. Young, Kelly F. Oakeson, Pooja Gupta, Olinto Linares-Perdomo |
| EPI_ISL_3450361 | Department of Bacteria, Parasites and Fungi, Statens Serum Institut, Copenhagen, Denmark | Statens Serum Institut Bioinformatics and Microbial Genomics | Danish Covid-19 Genome Consortium |
| EPI_ISL_3450362 | Utah Public Health Laboratory | Utah Public Health Laboratory | Erin L. Young, Kelly F. Oakeson, Pooja Gupta, Olinto Linares-Perdomo |
| EPI_ISL_3450363 | Department of Bacteria, Parasites and Fungi, Statens Serum Institut, Copenhagen, Denmark | Statens Serum Institut Bioinformatics and Microbial Genomics | Danish Covid-19 Genome Consortium |
| EPI_ISL_3450364 | Utah Public Health Laboratory | Utah Public Health Laboratory | Erin L. Young, Kelly F. Oakeson, Pooja Gupta, Olinto Linares-Perdomo |
| EPI_ISL_3450365 | Department of Bacteria, Parasites and Fungi, Statens Serum Institut, Copenhagen, Denmark | Statens Serum Institut Bioinformatics and Microbial Genomics | Danish Covid-19 Genome Consortium |
| EPI_ISL_3450366 | Utah Public Health Laboratory | Utah Public Health Laboratory | Erin L. Young, Kelly F. Oakeson, Pooja Gupta, Olinto Linares-Perdomo |
| EPI_ISL_3450367 | Department of Bacteria, Parasites and Fungi, Statens Serum Institut, Copenhagen, Denmark | Statens Serum Institut Bioinformatics and Microbial Genomics | Danish Covid-19 Genome Consortium |
| EPI_ISL_3450368 | Utah Public Health Laboratory | Utah Public Health Laboratory | Erin L. Young, Kelly F. Oakeson, Pooja Gupta, Olinto Linares-Perdomo |
| EPI_ISL_3450369 | Department of Bacteria, Parasites and Fungi, Statens Serum Institut, Copenhagen, Denmark | Statens Serum Institut Bioinformatics and Microbial Genomics | Danish Covid-19 Genome Consortium |
| EPI_ISL_3450370 | Utah Public Health Laboratory | Utah Public Health Laboratory | Erin L. Young, Kelly F. Oakeson, Pooja Gupta, Olinto Linares-Perdomo |
| EPI_ISL_3450371 | Department of Bacteria, Parasites and Fungi, Statens Serum Institut, Copenhagen, Denmark | Statens Serum Institut Bioinformatics and Microbial Genomics | Danish Covid-19 Genome Consortium |
| EPI_ISL_3450372 | Utah Public Health Laboratory | Utah Public Health Laboratory | Erin L. Young, Kelly F. Oakeson, Pooja Gupta, Olinto Linares-Perdomo |
| EPI_ISL_3450373 | Department of Bacteria, Parasites and Fungi, Statens Serum Institut, Copenhagen, Denmark | Statens Serum Institut Bioinformatics and Microbial Genomics | Danish Covid-19 Genome Consortium |
| EPI_ISL_3450374 | Utah Public Health Laboratory | Utah Public Health Laboratory | Erin L. Young, Kelly F. Oakeson, Pooja Gupta, Olinto Linares-Perdomo |
| EPI_ISL_3450375 | Department of Bacteria, Parasites and Fungi, Statens Serum Institut, Copenhagen, Denmark | Statens Serum Institut Bioinformatics and Microbial Genomics | Danish Covid-19 Genome Consortium |
| EPI_ISL_3450376 | Utah Public Health Laboratory | Utah Public Health Laboratory | Erin L. Young, Kelly F. Oakeson, Pooja Gupta, Olinto Linares-Perdomo |
| EPI_ISL_3450377 | Department of Bacteria, Parasites and Fungi, Statens Serum Institut, Copenhagen, Denmark | Statens Serum Institut Bioinformatics and Microbial Genomics | Danish Covid-19 Genome Consortium |
| EPI_ISL_3450378 | Utah Public Health Laboratory | Utah Public Health Laboratory | Erin L. Young, Kelly F. Oakeson, Pooja Gupta, Olinto Linares-Perdomo |
| EPI_ISL_3450379 | Department of Bacteria, Parasites and Fungi, Statens Serum Institut, Copenhagen, Denmark | Statens Serum Institut Bioinformatics and Microbial Genomics | Danish Covid-19 Genome Consortium |
| EPI_ISL_3450380 | Utah Public Health Laboratory | Utah Public Health Laboratory | Erin L. Young, Kelly F. Oakeson, Pooja Gupta, Olinto Linares-Perdomo |
| EPI_ISL_3450381 | Department of Bacteria, Parasites and Fungi, Statens Serum Institut, Copenhagen, Denmark | Statens Serum Institut Bioinformatics and Microbial Genomics | Danish Covid-19 Genome Consortium |
| EPI_ISL_3450382, EPI_ISL_3450384 | Utah Public Health Laboratory | Utah Public Health Laboratory | Erin L. Young, Kelly F. Oakeson, Pooja Gupta, Olinto Linares-Perdomo |
| EPI_ISL_3450385 | Department of Bacteria, Parasites and Fungi, Statens Serum Institut, Copenhagen, Denmark | Statens Serum Institut Bioinformatics and Microbial Genomics | Danish Covid-19 Genome Consortium |
| EPI_ISL_3450386 | Utah Public Health Laboratory | Utah Public Health Laboratory | Erin L. Young, Kelly F. Oakeson, Pooja Gupta, Olinto Linares-Perdomo |
| EPI_ISL_3450387 | Department of Bacteria, Parasites and Fungi, Statens Serum Institut, Copenhagen, Denmark | Statens Serum Institut Bioinformatics and Microbial Genomics | Danish Covid-19 Genome Consortium |
| EPI_ISL_3450388 | Utah Public Health Laboratory | Utah Public Health Laboratory | Erin L. Young, Kelly F. Oakeson, Pooja Gupta, Olinto Linares-Perdomo |
| EPI_ISL_3450389, EPI_ISL_3450390 | Department of Bacteria, Parasites and Fungi, Statens Serum Institut, Copenhagen, Denmark | Statens Serum Institut Bioinformatics and Microbial Genomics | Danish Covid-19 Genome Consortium |
| EPI_ISL_3450391 | Utah Public Health Laboratory | Utah Public Health Laboratory | Erin L. Young, Kelly F. Oakeson, Pooja Gupta, Olinto Linares-Perdomo |
| EPI_ISL_3450392 | Department of Bacteria, Parasites and Fungi, Statens Serum Institut, Copenhagen, Denmark | Statens Serum Institut Bioinformatics and Microbial Genomics | Danish Covid-19 Genome Consortium |
| EPI_ISL_3450393 | Utah Public Health Laboratory | Utah Public Health Laboratory | Erin L. Young, Kelly F. Oakeson, Pooja Gupta, Olinto Linares-Perdomo |
| EPI_ISL_3450394 | Department of Bacteria, Parasites and Fungi, Statens Serum Institut, Copenhagen, Denmark | Statens Serum Institut Bioinformatics and Microbial Genomics | Danish Covid-19 Genome Consortium |
| EPI_ISL_3450395 | Utah Public Health Laboratory | Utah Public Health Laboratory | Erin L. Young, Kelly F. Oakeson, Pooja Gupta, Olinto Linares-Perdomo |
| EPI_ISL_3450396 | Department of Bacteria, Parasites and Fungi, Statens Serum Institut, Copenhagen, Denmark | Statens Serum Institut Bioinformatics and Microbial Genomics | Danish Covid-19 Genome Consortium |
| EPI_ISL_3450397 | Utah Public Health Laboratory | Utah Public Health Laboratory | Erin L. Young, Kelly F. Oakeson, Pooja Gupta, Olinto Linares-Perdomo |
| EPI_ISL_3450398 | Department of Bacteria, Parasites and Fungi, Statens Serum Institut, Copenhagen, Denmark | Statens Serum Institut Bioinformatics and Microbial Genomics | Danish Covid-19 Genome Consortium |
| EPI_ISL_3450399 | Utah Public Health Laboratory | Utah Public Health Laboratory | Erin L. Young, Kelly F. Oakeson, Pooja Gupta, Olinto Linares-Perdomo |
| EPI_ISL_3450400 | Department of Bacteria, Parasites and Fungi, Statens Serum Institut, Copenhagen, Denmark | Statens Serum Institut Bioinformatics and Microbial Genomics | Danish Covid-19 Genome Consortium |
| EPI_ISL_3450401 | Utah Public Health Laboratory | Utah Public Health Laboratory | Erin L. Young, Kelly F. Oakeson, Pooja Gupta, Olinto Linares-Perdomo |
| EPI_ISL_3450402, EPI_ISL_3450403 | Department of Bacteria, Parasites and Fungi, Statens Serum Institut, Copenhagen, Denmark | Statens Serum Institut Bioinformatics and Microbial Genomics | Danish Covid-19 Genome Consortium |
| EPI_ISL_3450404 | Utah Public Health Laboratory | Utah Public Health Laboratory | Erin L. Young, Kelly F. Oakeson, Pooja Gupta, Olinto Linares-Perdomo |
| EPI_ISL_3450405 | Department of Bacteria, Parasites and Fungi, Statens Serum Institut, Copenhagen, Denmark | Statens Serum Institut Bioinformatics and Microbial Genomics | Danish Covid-19 Genome Consortium |
| EPI_ISL_3450406 | Utah Public Health Laboratory | Utah Public Health Laboratory | Erin L. Young, Kelly F. Oakeson, Pooja Gupta, Olinto Linares-Perdomo |
| EPI_ISL_3450407 | Department of Bacteria, Parasites and Fungi, Statens Serum Institut, Copenhagen, Denmark | Statens Serum Institut Bioinformatics and Microbial Genomics | Danish Covid-19 Genome Consortium |
| EPI_ISL_3450408 | Utah Public Health Laboratory | Utah Public Health Laboratory | Erin L. Young, Kelly F. Oakeson, Pooja Gupta, Olinto Linares-Perdomo |
| EPI_ISL_3450409 | Department of Bacteria, Parasites and Fungi, Statens Serum Institut, Copenhagen, Denmark | Statens Serum Institut Bioinformatics and Microbial Genomics | Danish Covid-19 Genome Consortium |
| EPI_ISL_3450410 | Utah Public Health Laboratory | Utah Public Health Laboratory | Erin L. Young, Kelly F. Oakeson, Pooja Gupta, Olinto Linares-Perdomo |
| EPI_ISL_3450411 | Department of Bacteria, Parasites and Fungi, Statens Serum Institut, Copenhagen, Denmark | Statens Serum Institut Bioinformatics and Microbial Genomics | Danish Covid-19 Genome Consortium |
| EPI_ISL_3450412 | Utah Public Health Laboratory | Utah Public Health Laboratory | Erin L. Young, Kelly F. Oakeson, Pooja Gupta, Olinto Linares-Perdomo |
| EPI_ISL_3450413 | Department of Bacteria, Parasites and Fungi, Statens Serum Institut, Copenhagen, Denmark | Statens Serum Institut Bioinformatics and Microbial Genomics | Danish Covid-19 Genome Consortium |
| EPI_ISL_3450414 | Utah Public Health Laboratory | Utah Public Health Laboratory | Erin L. Young, Kelly F. Oakeson, Pooja Gupta, Olinto Linares-Perdomo |
| EPI_ISL_3450415 | Department of Bacteria, Parasites and Fungi, Statens Serum Institut, Copenhagen, Denmark | Statens Serum Institut Bioinformatics and Microbial Genomics | Danish Covid-19 Genome Consortium |
| EPI_ISL_3450416 | Utah Public Health Laboratory | Utah Public Health Laboratory | Erin L. Young, Kelly F. Oakeson, Pooja Gupta, Olinto Linares-Perdomo |

| | | | |
|---|---|---|---|
| EPI_ISL_3450417 | Department of Bacteria, Parasites and Fungi, Statens Serum Institut, Copenhagen, Denmark | Statens Serum Institut Bioinformatics and Microbial Genomics | Danish Covid-19 Genome Consortium |
| EPI_ISL_3450418 | Utah Public Health Laboratory | Utah Public Health Laboratory | Erin L. Young, Kelly F. Oakeson, Pooja Gupta, Olinto Linares-Perdomo |
| EPI_ISL_3450419 | Department of Bacteria, Parasites and Fungi, Statens Serum Institut, Copenhagen, Denmark | Statens Serum Institut Bioinformatics and Microbial Genomics | Danish Covid-19 Genome Consortium |
| EPI_ISL_3450420 | Utah Public Health Laboratory | Utah Public Health Laboratory | Erin L. Young, Kelly F. Oakeson, Pooja Gupta, Olinto Linares-Perdomo |
| EPI_ISL_3450421, EPI_ISL_3450422 | Department of Bacteria, Parasites and Fungi, Statens Serum Institut, Copenhagen, Denmark | Statens Serum Institut Bioinformatics and Microbial Genomics | Danish Covid-19 Genome Consortium |
| EPI_ISL_3450423 | Utah Public Health Laboratory | Utah Public Health Laboratory | Erin L. Young, Kelly F. Oakeson, Pooja Gupta, Olinto Linares-Perdomo |
| EPI_ISL_3450424 | Department of Bacteria, Parasites and Fungi, Statens Serum Institut, Copenhagen, Denmark | Statens Serum Institut Bioinformatics and Microbial Genomics | Danish Covid-19 Genome Consortium |
| EPI_ISL_3450425 | Utah Public Health Laboratory | Utah Public Health Laboratory | Erin L. Young, Kelly F. Oakeson, Pooja Gupta, Olinto Linares-Perdomo |
| EPI_ISL_3450426 | Department of Bacteria, Parasites and Fungi, Statens Serum Institut, Copenhagen, Denmark | Statens Serum Institut Bioinformatics and Microbial Genomics | Danish Covid-19 Genome Consortium |
| EPI_ISL_3450427 | Utah Public Health Laboratory | Utah Public Health Laboratory | Erin L. Young, Kelly F. Oakeson, Pooja Gupta, Olinto Linares-Perdomo |
| EPI_ISL_3450428 | Department of Bacteria, Parasites and Fungi, Statens Serum Institut, Copenhagen, Denmark | Statens Serum Institut Bioinformatics and Microbial Genomics | Danish Covid-19 Genome Consortium |
| EPI_ISL_3450429 | Utah Public Health Laboratory | Utah Public Health Laboratory | Erin L. Young, Kelly F. Oakeson, Pooja Gupta, Olinto Linares-Perdomo |
| EPI_ISL_3450430, EPI_ISL_3450431 | Department of Bacteria, Parasites and Fungi, Statens Serum Institut, Copenhagen, Denmark | Statens Serum Institut Bioinformatics and Microbial Genomics | Danish Covid-19 Genome Consortium |
| EPI_ISL_3450432 | Utah Public Health Laboratory | Utah Public Health Laboratory | Erin L. Young, Kelly F. Oakeson, Pooja Gupta, Olinto Linares-Perdomo |
| EPI_ISL_3450433 | Department of Bacteria, Parasites and Fungi, Statens Serum Institut, Copenhagen, Denmark | Statens Serum Institut Bioinformatics and Microbial Genomics | Danish Covid-19 Genome Consortium |
| EPI_ISL_3450434 | Utah Public Health Laboratory | Utah Public Health Laboratory | Erin L. Young, Kelly F. Oakeson, Pooja Gupta, Olinto Linares-Perdomo |
| EPI_ISL_3450435 | Department of Bacteria, Parasites and Fungi, Statens Serum Institut, Copenhagen, Denmark | Statens Serum Institut Bioinformatics and Microbial Genomics | Danish Covid-19 Genome Consortium |
| EPI_ISL_3450436 | Utah Public Health Laboratory | Utah Public Health Laboratory | Erin L. Young, Kelly F. Oakeson, Pooja Gupta, Olinto Linares-Perdomo |
| EPI_ISL_3450437 | Department of Bacteria, Parasites and Fungi, Statens Serum Institut, Copenhagen, Denmark | Statens Serum Institut Bioinformatics and Microbial Genomics | Danish Covid-19 Genome Consortium |
| EPI_ISL_3450438 | Utah Public Health Laboratory | Utah Public Health Laboratory | Erin L. Young, Kelly F. Oakeson, Pooja Gupta, Olinto Linares-Perdomo |
| EPI_ISL_3450439 | Department of Bacteria, Parasites and Fungi, Statens Serum Institut, Copenhagen, Denmark | Statens Serum Institut Bioinformatics and Microbial Genomics | Danish Covid-19 Genome Consortium |
| EPI_ISL_3450440 | Utah Public Health Laboratory | Utah Public Health Laboratory | Erin L. Young, Kelly F. Oakeson, Pooja Gupta, Olinto Linares-Perdomo |
| EPI_ISL_3450441 | Department of Bacteria, Parasites and Fungi, Statens Serum Institut, Copenhagen, Denmark | Statens Serum Institut Bioinformatics and Microbial Genomics | Danish Covid-19 Genome Consortium |
| EPI_ISL_3450442 | Utah Public Health Laboratory | Utah Public Health Laboratory | Erin L. Young, Kelly F. Oakeson, Pooja Gupta, Olinto Linares-Perdomo |
| EPI_ISL_3450443, EPI_ISL_3450444 | Department of Bacteria, Parasites and Fungi, Statens Serum Institut, Copenhagen, Denmark | Statens Serum Institut Bioinformatics and Microbial Genomics | Danish Covid-19 Genome Consortium |
| EPI_ISL_3450445 | Utah Public Health Laboratory | Utah Public Health Laboratory | Erin L. Young, Kelly F. Oakeson, Pooja Gupta, Olinto Linares-Perdomo |
| EPI_ISL_3450446 | Department of Bacteria, Parasites and Fungi, Statens Serum Institut, Copenhagen, Denmark | Statens Serum Institut Bioinformatics and Microbial Genomics | Danish Covid-19 Genome Consortium |
| EPI_ISL_3450447 | Utah Public Health Laboratory | Utah Public Health Laboratory | Erin L. Young, Kelly F. Oakeson, Pooja Gupta, Olinto Linares-Perdomo |
| EPI_ISL_3450448 | Department of Bacteria, Parasites and Fungi, Statens Serum Institut, Copenhagen, Denmark | Statens Serum Institut Bioinformatics and Microbial Genomics | Danish Covid-19 Genome Consortium |
| EPI_ISL_3450449 | Utah Public Health Laboratory | Utah Public Health Laboratory | Erin L. Young, Kelly F. Oakeson, Pooja Gupta, Olinto Linares-Perdomo |
| EPI_ISL_3450450, EPI_ISL_3450451 | Department of Bacteria, Parasites and Fungi, Statens Serum Institut, Copenhagen, Denmark | Statens Serum Institut Bioinformatics and Microbial Genomics | Danish Covid-19 Genome Consortium |
| EPI_ISL_3450452 | Utah Public Health Laboratory | Utah Public Health Laboratory | Erin L. Young, Kelly F. Oakeson, Pooja Gupta, Olinto Linares-Perdomo |
| EPI_ISL_3450453 | Department of Bacteria, Parasites and Fungi, Statens Serum Institut, Copenhagen, Denmark | Statens Serum Institut Bioinformatics and Microbial Genomics | Danish Covid-19 Genome Consortium |
| EPI_ISL_3450454 | Utah Public Health Laboratory | Utah Public Health Laboratory | Erin L. Young, Kelly F. Oakeson, Pooja Gupta, Olinto Linares-Perdomo |
| EPI_ISL_3450455 | Department of Bacteria, Parasites and Fungi, Statens Serum Institut, Copenhagen, Denmark | Statens Serum Institut Bioinformatics and Microbial Genomics | Danish Covid-19 Genome Consortium |
| EPI_ISL_3450456 | Utah Public Health Laboratory | Utah Public Health Laboratory | Erin L. Young, Kelly F. Oakeson, Pooja Gupta, Olinto Linares-Perdomo |
| EPI_ISL_3450457 | Department of Bacteria, Parasites and Fungi, Statens Serum Institut, Copenhagen, Denmark | Statens Serum Institut Bioinformatics and Microbial Genomics | Danish Covid-19 Genome Consortium |
| EPI_ISL_3450458 | Utah Public Health Laboratory | Utah Public Health Laboratory | Erin L. Young, Kelly F. Oakeson, Pooja Gupta, Olinto Linares-Perdomo |
| EPI_ISL_3450459 | Department of Bacteria, Parasites and Fungi, Statens Serum Institut, Copenhagen, Denmark | Statens Serum Institut Bioinformatics and Microbial Genomics | Danish Covid-19 Genome Consortium |
| EPI_ISL_3450460 | Utah Public Health Laboratory | Utah Public Health Laboratory | Erin L. Young, Kelly F. Oakeson, Pooja Gupta, Olinto Linares-Perdomo |
| EPI_ISL_3450461 | Department of Bacteria, Parasites and Fungi, Statens Serum Institut, Copenhagen, Denmark | Statens Serum Institut Bioinformatics and Microbial Genomics | Danish Covid-19 Genome Consortium |
| EPI_ISL_3450462 | Utah Public Health Laboratory | Utah Public Health Laboratory | Erin L. Young, Kelly F. Oakeson, Pooja Gupta, Olinto Linares-Perdomo |
| EPI_ISL_3450463 | Department of Bacteria, Parasites and Fungi, Statens Serum Institut, Copenhagen, Denmark | Statens Serum Institut Bioinformatics and Microbial Genomics | Danish Covid-19 Genome Consortium |
| EPI_ISL_3450464 | Utah Public Health Laboratory | Utah Public Health Laboratory | Erin L. Young, Kelly F. Oakeson, Pooja Gupta, Olinto Linares-Perdomo |
| EPI_ISL_3450465 | Department of Bacteria, Parasites and Fungi, Statens Serum Institut, Copenhagen, Denmark | Statens Serum Institut Bioinformatics and Microbial Genomics | Danish Covid-19 Genome Consortium |
| EPI_ISL_3450466 | Utah Public Health Laboratory | Utah Public Health Laboratory | Erin L. Young, Kelly F. Oakeson, Pooja Gupta, Olinto Linares-Perdomo |
| EPI_ISL_3450467, EPI_ISL_3450468 | Department of Bacteria, Parasites and Fungi, Statens Serum Institut, Copenhagen, Denmark | Statens Serum Institut Bioinformatics and Microbial Genomics | Danish Covid-19 Genome Consortium |
| EPI_ISL_3450469 | Utah Public Health Laboratory | Utah Public Health Laboratory | Erin L. Young, Kelly F. Oakeson, Pooja Gupta, Olinto Linares-Perdomo |
| EPI_ISL_3450470 | Department of Bacteria, Parasites and Fungi, Statens Serum Institut, Copenhagen, Denmark | Statens Serum Institut Bioinformatics and Microbial Genomics | Danish Covid-19 Genome Consortium |
| EPI_ISL_3450471 | Utah Public Health Laboratory | Utah Public Health Laboratory | Erin L. Young, Kelly F. Oakeson, Pooja Gupta, Olinto Linares-Perdomo |
| EPI_ISL_3450472 | Department of Bacteria, Parasites and Fungi, Statens Serum Institut, Copenhagen, Denmark | Statens Serum Institut Bioinformatics and Microbial Genomics | Danish Covid-19 Genome Consortium |
| EPI_ISL_3450473 | Utah Public Health Laboratory | Utah Public Health Laboratory | Erin L. Young, Kelly F. Oakeson, Pooja Gupta, Olinto Linares-Perdomo |
| EPI_ISL_3450474 | Department of Bacteria, Parasites and Fungi, Statens Serum Institut, Copenhagen, Denmark | Statens Serum Institut Bioinformatics and Microbial Genomics | Danish Covid-19 Genome Consortium |
| EPI_ISL_3450475 | Utah Public Health Laboratory | Utah Public Health Laboratory | Erin L. Young, Kelly F. Oakeson, Pooja Gupta, Olinto Linares-Perdomo |
| EPI_ISL_3450476 | Department of Bacteria, Parasites and Fungi, Statens Serum Institut, Copenhagen, Denmark | Statens Serum Institut Bioinformatics and Microbial Genomics | Danish Covid-19 Genome Consortium |
| EPI_ISL_3450477 | Utah Public Health Laboratory | Utah Public Health Laboratory | Erin L. Young, Kelly F. Oakeson, Pooja Gupta, Olinto Linares-Perdomo |
| EPI_ISL_3450478, | Department of Bacteria, Parasites and Fungi, Statens Serum | Statens Serum Institut Bioinformatics and Microbial Genomics | Danish Covid-19 Genome Consortium |

| | | | |
|---|---|---|---|
| EPI_ISL_3450479 | Institut, Copenhagen, Denmark | | |
| EPI_ISL_3450480 | Utah Public Health Laboratory | Utah Public Health Laboratory | Erin L. Young, Kelly F. Oakeson, Pooja Gupta, Olinto Linares-Perdomo |
| EPI_ISL_3450481 | Department of Bacteria, Parasites and Fungi, Statens Serum Institut, Copenhagen, Denmark | Statens Serum Institut Bioinformatics and Microbial Genomics | Danish Covid-19 Genome Consortium |
| EPI_ISL_3450482 | Utah Public Health Laboratory | Utah Public Health Laboratory | Erin L. Young, Kelly F. Oakeson, Pooja Gupta, Olinto Linares-Perdomo |
| EPI_ISL_3450483 | Department of Bacteria, Parasites and Fungi, Statens Serum Institut, Copenhagen, Denmark | Statens Serum Institut Bioinformatics and Microbial Genomics | Danish Covid-19 Genome Consortium |
| EPI_ISL_3450484 | Utah Public Health Laboratory | Utah Public Health Laboratory | Erin L. Young, Kelly F. Oakeson, Pooja Gupta, Olinto Linares-Perdomo |
| EPI_ISL_3450485 | Department of Bacteria, Parasites and Fungi, Statens Serum Institut, Copenhagen, Denmark | Statens Serum Institut Bioinformatics and Microbial Genomics | Danish Covid-19 Genome Consortium |
| EPI_ISL_3450486 | Utah Public Health Laboratory | Utah Public Health Laboratory | Erin L. Young, Kelly F. Oakeson, Pooja Gupta, Olinto Linares-Perdomo |
| EPI_ISL_3450487 | Department of Bacteria, Parasites and Fungi, Statens Serum Institut, Copenhagen, Denmark | Statens Serum Institut Bioinformatics and Microbial Genomics | Danish Covid-19 Genome Consortium |
| EPI_ISL_3450488 | Utah Public Health Laboratory | Utah Public Health Laboratory | Erin L. Young, Kelly F. Oakeson, Pooja Gupta, Olinto Linares-Perdomo |
| EPI_ISL_3450489, EPI_ISL_3450490 | Department of Bacteria, Parasites and Fungi, Statens Serum Institut, Copenhagen, Denmark | Statens Serum Institut Bioinformatics and Microbial Genomics | Danish Covid-19 Genome Consortium |
| EPI_ISL_3450491 | Utah Public Health Laboratory | Utah Public Health Laboratory | Erin L. Young, Kelly F. Oakeson, Pooja Gupta, Olinto Linares-Perdomo |
| EPI_ISL_3450492 | Department of Bacteria, Parasites and Fungi, Statens Serum Institut, Copenhagen, Denmark | Statens Serum Institut Bioinformatics and Microbial Genomics | Danish Covid-19 Genome Consortium |
| EPI_ISL_3450493 | Utah Public Health Laboratory | Utah Public Health Laboratory | Erin L. Young, Kelly F. Oakeson, Pooja Gupta, Olinto Linares-Perdomo |
| EPI_ISL_3450494 | Department of Bacteria, Parasites and Fungi, Statens Serum Institut, Copenhagen, Denmark | Statens Serum Institut Bioinformatics and Microbial Genomics | Danish Covid-19 Genome Consortium |
| EPI_ISL_3450495 | Utah Public Health Laboratory | Utah Public Health Laboratory | Erin L. Young, Kelly F. Oakeson, Pooja Gupta, Olinto Linares-Perdomo |
| EPI_ISL_3450496 | Department of Bacteria, Parasites and Fungi, Statens Serum Institut, Copenhagen, Denmark | Statens Serum Institut Bioinformatics and Microbial Genomics | Danish Covid-19 Genome Consortium |
| EPI_ISL_3450497 | Utah Public Health Laboratory | Utah Public Health Laboratory | Erin L. Young, Kelly F. Oakeson, Pooja Gupta, Olinto Linares-Perdomo |
| EPI_ISL_3450498, EPI_ISL_3450499 | Department of Bacteria, Parasites and Fungi, Statens Serum Institut, Copenhagen, Denmark | Statens Serum Institut Bioinformatics and Microbial Genomics | Danish Covid-19 Genome Consortium |
| EPI_ISL_3450500 | Utah Public Health Laboratory | Utah Public Health Laboratory | Erin L. Young, Kelly F. Oakeson, Pooja Gupta, Olinto Linares-Perdomo |
| EPI_ISL_3450501 | Department of Bacteria, Parasites and Fungi, Statens Serum Institut, Copenhagen, Denmark | Statens Serum Institut Bioinformatics and Microbial Genomics | Danish Covid-19 Genome Consortium |
| EPI_ISL_3450502 | Utah Public Health Laboratory | Utah Public Health Laboratory | Erin L. Young, Kelly F. Oakeson, Pooja Gupta, Olinto Linares-Perdomo |
| EPI_ISL_3450503 | Department of Bacteria, Parasites and Fungi, Statens Serum Institut, Copenhagen, Denmark | Statens Serum Institut Bioinformatics and Microbial Genomics | Danish Covid-19 Genome Consortium |
| EPI_ISL_3450504 | Utah Public Health Laboratory | Utah Public Health Laboratory | Erin L. Young, Kelly F. Oakeson, Pooja Gupta, Olinto Linares-Perdomo |
| EPI_ISL_3450505, EPI_ISL_3450506 | Department of Bacteria, Parasites and Fungi, Statens Serum Institut, Copenhagen, Denmark | Statens Serum Institut Bioinformatics and Microbial Genomics | Danish Covid-19 Genome Consortium |
| EPI_ISL_3450507 | Utah Public Health Laboratory | Utah Public Health Laboratory | Erin L. Young, Kelly F. Oakeson, Pooja Gupta, Olinto Linares-Perdomo |
| EPI_ISL_3450508 | Department of Bacteria, Parasites and Fungi, Statens Serum Institut, Copenhagen, Denmark | Statens Serum Institut Bioinformatics and Microbial Genomics | Danish Covid-19 Genome Consortium |
| EPI_ISL_3450509 | Utah Public Health Laboratory | Utah Public Health Laboratory | Erin L. Young, Kelly F. Oakeson, Pooja Gupta, Olinto Linares-Perdomo |
| EPI_ISL_3450510 | Department of Bacteria, Parasites and Fungi, Statens Serum Institut, Copenhagen, Denmark | Statens Serum Institut Bioinformatics and Microbial Genomics | Danish Covid-19 Genome Consortium |
| EPI_ISL_3450511 | Utah Public Health Laboratory | Utah Public Health Laboratory | Erin L. Young, Kelly F. Oakeson, Pooja Gupta, Olinto Linares-Perdomo |
| EPI_ISL_3450512 | Department of Bacteria, Parasites and Fungi, Statens Serum Institut, Copenhagen, Denmark | Statens Serum Institut Bioinformatics and Microbial Genomics | Danish Covid-19 Genome Consortium |
| EPI_ISL_3450513 | Utah Public Health Laboratory | Utah Public Health Laboratory | Erin L. Young, Kelly F. Oakeson, Pooja Gupta, Olinto Linares-Perdomo |
| EPI_ISL_3450514 | Department of Bacteria, Parasites and Fungi, Statens Serum Institut, Copenhagen, Denmark | Statens Serum Institut Bioinformatics and Microbial Genomics | Danish Covid-19 Genome Consortium |
| EPI_ISL_3450515 | Utah Public Health Laboratory | Utah Public Health Laboratory | Erin L. Young, Kelly F. Oakeson, Pooja Gupta, Olinto Linares-Perdomo |
| EPI_ISL_3450516, EPI_ISL_3450517 | Department of Bacteria, Parasites and Fungi, Statens Serum Institut, Copenhagen, Denmark | Statens Serum Institut Bioinformatics and Microbial Genomics | Danish Covid-19 Genome Consortium |
| EPI_ISL_3450518 | Utah Public Health Laboratory | Utah Public Health Laboratory | Erin L. Young, Kelly F. Oakeson, Pooja Gupta, Olinto Linares-Perdomo |
| EPI_ISL_3450519 | Department of Bacteria, Parasites and Fungi, Statens Serum Institut, Copenhagen, Denmark | Statens Serum Institut Bioinformatics and Microbial Genomics | Danish Covid-19 Genome Consortium |
| EPI_ISL_3450520 | Utah Public Health Laboratory | Utah Public Health Laboratory | Erin L. Young, Kelly F. Oakeson, Pooja Gupta, Olinto Linares-Perdomo |
| EPI_ISL_3450521 | Department of Bacteria, Parasites and Fungi, Statens Serum Institut, Copenhagen, Denmark | Statens Serum Institut Bioinformatics and Microbial Genomics | Danish Covid-19 Genome Consortium |
| EPI_ISL_3450522 | Utah Public Health Laboratory | Utah Public Health Laboratory | Erin L. Young, Kelly F. Oakeson, Pooja Gupta, Olinto Linares-Perdomo |
| EPI_ISL_3450523 | Department of Bacteria, Parasites and Fungi, Statens Serum Institut, Copenhagen, Denmark | Statens Serum Institut Bioinformatics and Microbial Genomics | Danish Covid-19 Genome Consortium |
| EPI_ISL_3450524 | Utah Public Health Laboratory | Utah Public Health Laboratory | Erin L. Young, Kelly F. Oakeson, Pooja Gupta, Olinto Linares-Perdomo |
| EPI_ISL_3450525 | Department of Bacteria, Parasites and Fungi, Statens Serum Institut, Copenhagen, Denmark | Statens Serum Institut Bioinformatics and Microbial Genomics | Danish Covid-19 Genome Consortium |
| EPI_ISL_3450526 | Utah Public Health Laboratory | Utah Public Health Laboratory | Erin L. Young, Kelly F. Oakeson, Pooja Gupta, Olinto Linares-Perdomo |
| EPI_ISL_3450527 | Department of Bacteria, Parasites and Fungi, Statens Serum Institut, Copenhagen, Denmark | Statens Serum Institut Bioinformatics and Microbial Genomics | Danish Covid-19 Genome Consortium |
| EPI_ISL_3450528 | Utah Public Health Laboratory | Utah Public Health Laboratory | Erin L. Young, Kelly F. Oakeson, Pooja Gupta, Olinto Linares-Perdomo |
| EPI_ISL_3450529 | Department of Bacteria, Parasites and Fungi, Statens Serum Institut, Copenhagen, Denmark | Statens Serum Institut Bioinformatics and Microbial Genomics | Danish Covid-19 Genome Consortium |
| EPI_ISL_3450530 | Utah Public Health Laboratory | Utah Public Health Laboratory | Erin L. Young, Kelly F. Oakeson, Pooja Gupta, Olinto Linares-Perdomo |
| EPI_ISL_3450531 | Department of Bacteria, Parasites and Fungi, Statens Serum Institut, Copenhagen, Denmark | Statens Serum Institut Bioinformatics and Microbial Genomics | Danish Covid-19 Genome Consortium |
| EPI_ISL_3450532 | Utah Public Health Laboratory | Utah Public Health Laboratory | Erin L. Young, Kelly F. Oakeson, Pooja Gupta, Olinto Linares-Perdomo |
| EPI_ISL_3450533 | Department of Bacteria, Parasites and Fungi, Statens Serum Institut, Copenhagen, Denmark | Statens Serum Institut Bioinformatics and Microbial Genomics | Danish Covid-19 Genome Consortium |
| EPI_ISL_3450534 | Utah Public Health Laboratory | Utah Public Health Laboratory | Erin L. Young, Kelly F. Oakeson, Pooja Gupta, Olinto Linares-Perdomo |
| EPI_ISL_3450535 | Department of Bacteria, Parasites and Fungi, Statens Serum Institut, Copenhagen, Denmark | Statens Serum Institut Bioinformatics and Microbial Genomics | Danish Covid-19 Genome Consortium |
| EPI_ISL_3450536 | Utah Public Health Laboratory | Utah Public Health Laboratory | Erin L. Young, Kelly F. Oakeson, Pooja Gupta, Olinto Linares-Perdomo |
| EPI_ISL_3450537 | Department of Bacteria, Parasites and Fungi, Statens Serum Institut, Copenhagen, Denmark | Statens Serum Institut Bioinformatics and Microbial Genomics | Danish Covid-19 Genome Consortium |
| EPI_ISL_3450538 | Utah Public Health Laboratory | Utah Public Health Laboratory | Erin L. Young, Kelly F. Oakeson, Pooja Gupta, Olinto Linares-Perdomo |
| EPI_ISL_3450539 | Department of Bacteria, Parasites and Fungi, Statens Serum Institut, Copenhagen, Denmark | Statens Serum Institut Bioinformatics and Microbial Genomics | Danish Covid-19 Genome Consortium |

| | | | |
|---|---|---|---|
| EPI_ISL_3450540 | Utah Public Health Laboratory | Utah Public Health Laboratory | Erin L. Young, Kelly F. Oakeson, Pooja Gupta, Olinto Linares-Perdomo |
| EPI_ISL_3450541, EPI_ISL_3450542 | Department of Bacteria, Parasites and Fungi, Statens Serum Institut, Copenhagen, Denmark | Statens Serum Institut Bioinformatics and Microbial Genomics | Danish Covid-19 Genome Consortium |
| EPI_ISL_3450543 | Utah Public Health Laboratory | Utah Public Health Laboratory | Erin L. Young, Kelly F. Oakeson, Pooja Gupta, Olinto Linares-Perdomo |
| EPI_ISL_3450544 | Department of Bacteria, Parasites and Fungi, Statens Serum Institut, Copenhagen, Denmark | Statens Serum Institut Bioinformatics and Microbial Genomics | Danish Covid-19 Genome Consortium |
| EPI_ISL_3450545 | Utah Public Health Laboratory | Utah Public Health Laboratory | Erin L. Young, Kelly F. Oakeson, Pooja Gupta, Olinto Linares-Perdomo |
| EPI_ISL_3450546 | Department of Bacteria, Parasites and Fungi, Statens Serum Institut, Copenhagen, Denmark | Statens Serum Institut Bioinformatics and Microbial Genomics | Danish Covid-19 Genome Consortium |
| EPI_ISL_3450547 | Utah Public Health Laboratory | Utah Public Health Laboratory | Erin L. Young, Kelly F. Oakeson, Pooja Gupta, Olinto Linares-Perdomo |
| EPI_ISL_3450548 | Department of Bacteria, Parasites and Fungi, Statens Serum Institut, Copenhagen, Denmark | Statens Serum Institut Bioinformatics and Microbial Genomics | Danish Covid-19 Genome Consortium |
| EPI_ISL_3450549 | Utah Public Health Laboratory | Utah Public Health Laboratory | Erin L. Young, Kelly F. Oakeson, Pooja Gupta, Olinto Linares-Perdomo |
| EPI_ISL_3450550 | Department of Bacteria, Parasites and Fungi, Statens Serum Institut, Copenhagen, Denmark | Statens Serum Institut Bioinformatics and Microbial Genomics | Danish Covid-19 Genome Consortium |
| EPI_ISL_3450551 | Utah Public Health Laboratory | Utah Public Health Laboratory | Erin L. Young, Kelly F. Oakeson, Pooja Gupta, Olinto Linares-Perdomo |
| EPI_ISL_3450552 | Department of Bacteria, Parasites and Fungi, Statens Serum Institut, Copenhagen, Denmark | Statens Serum Institut Bioinformatics and Microbial Genomics | Danish Covid-19 Genome Consortium |
| EPI_ISL_3450553 | Utah Public Health Laboratory | Utah Public Health Laboratory | Erin L. Young, Kelly F. Oakeson, Pooja Gupta, Olinto Linares-Perdomo |
| EPI_ISL_3450554 | Department of Bacteria, Parasites and Fungi, Statens Serum Institut, Copenhagen, Denmark | Statens Serum Institut Bioinformatics and Microbial Genomics | Danish Covid-19 Genome Consortium |
| EPI_ISL_3450555 | Utah Public Health Laboratory | Utah Public Health Laboratory | Erin L. Young, Kelly F. Oakeson, Pooja Gupta, Olinto Linares-Perdomo |
| EPI_ISL_3450556, EPI_ISL_3450557 | Department of Bacteria, Parasites and Fungi, Statens Serum Institut, Copenhagen, Denmark | Statens Serum Institut Bioinformatics and Microbial Genomics | Danish Covid-19 Genome Consortium |
| EPI_ISL_3450558, EPI_ISL_3450559 | Utah Public Health Laboratory | Utah Public Health Laboratory | Erin L. Young, Kelly F. Oakeson, Pooja Gupta, Olinto Linares-Perdomo |
| EPI_ISL_3450560 | Department of Bacteria, Parasites and Fungi, Statens Serum Institut, Copenhagen, Denmark | Statens Serum Institut Bioinformatics and Microbial Genomics | Danish Covid-19 Genome Consortium |
| EPI_ISL_3450561 | Utah Public Health Laboratory | Utah Public Health Laboratory | Erin L. Young, Kelly F. Oakeson, Pooja Gupta, Olinto Linares-Perdomo |
| EPI_ISL_3450562, EPI_ISL_3450563 | Department of Bacteria, Parasites and Fungi, Statens Serum Institut, Copenhagen, Denmark | Statens Serum Institut Bioinformatics and Microbial Genomics | Danish Covid-19 Genome Consortium |
| EPI_ISL_3450564 | Utah Public Health Laboratory | Utah Public Health Laboratory | Erin L. Young, Kelly F. Oakeson, Pooja Gupta, Olinto Linares-Perdomo |
| EPI_ISL_3450565 | Department of Bacteria, Parasites and Fungi, Statens Serum Institut, Copenhagen, Denmark | Statens Serum Institut Bioinformatics and Microbial Genomics | Danish Covid-19 Genome Consortium |
| EPI_ISL_3450566 | Utah Public Health Laboratory | Utah Public Health Laboratory | Erin L. Young, Kelly F. Oakeson, Pooja Gupta, Olinto Linares-Perdomo |
| EPI_ISL_3450567 | Department of Bacteria, Parasites and Fungi, Statens Serum Institut, Copenhagen, Denmark | Statens Serum Institut Bioinformatics and Microbial Genomics | Danish Covid-19 Genome Consortium |
| EPI_ISL_3450568 | Utah Public Health Laboratory | Utah Public Health Laboratory | Erin L. Young, Kelly F. Oakeson, Pooja Gupta, Olinto Linares-Perdomo |
| EPI_ISL_3450569 | Department of Bacteria, Parasites and Fungi, Statens Serum Institut, Copenhagen, Denmark | Statens Serum Institut Bioinformatics and Microbial Genomics | Danish Covid-19 Genome Consortium |
| EPI_ISL_3450570 | Utah Public Health Laboratory | Utah Public Health Laboratory | Erin L. Young, Kelly F. Oakeson, Pooja Gupta, Olinto Linares-Perdomo |
| EPI_ISL_3450571 | Department of Bacteria, Parasites and Fungi, Statens Serum Institut, Copenhagen, Denmark | Statens Serum Institut Bioinformatics and Microbial Genomics | Danish Covid-19 Genome Consortium |
| EPI_ISL_3450572 | Utah Public Health Laboratory | Utah Public Health Laboratory | Erin L. Young, Kelly F. Oakeson, Pooja Gupta, Olinto Linares-Perdomo |
| EPI_ISL_3450573, EPI_ISL_3450574 | Department of Bacteria, Parasites and Fungi, Statens Serum Institut, Copenhagen, Denmark | Statens Serum Institut Bioinformatics and Microbial Genomics | Danish Covid-19 Genome Consortium |
| EPI_ISL_3450575 | Utah Public Health Laboratory | Utah Public Health Laboratory | Erin L. Young, Kelly F. Oakeson, Pooja Gupta, Olinto Linares-Perdomo |
| EPI_ISL_3450576 | Department of Bacteria, Parasites and Fungi, Statens Serum Institut, Copenhagen, Denmark | Statens Serum Institut Bioinformatics and Microbial Genomics | Danish Covid-19 Genome Consortium |
| EPI_ISL_3450577 | Utah Public Health Laboratory | Utah Public Health Laboratory | Erin L. Young, Kelly F. Oakeson, Pooja Gupta, Olinto Linares-Perdomo |
| EPI_ISL_3450578 | Department of Bacteria, Parasites and Fungi, Statens Serum Institut, Copenhagen, Denmark | Statens Serum Institut Bioinformatics and Microbial Genomics | Danish Covid-19 Genome Consortium |
| EPI_ISL_3450579 | Utah Public Health Laboratory | Utah Public Health Laboratory | Erin L. Young, Kelly F. Oakeson, Pooja Gupta, Olinto Linares-Perdomo |
| EPI_ISL_3450580 | Department of Bacteria, Parasites and Fungi, Statens Serum Institut, Copenhagen, Denmark | Statens Serum Institut Bioinformatics and Microbial Genomics | Danish Covid-19 Genome Consortium |
| EPI_ISL_3450581 | Utah Public Health Laboratory | Utah Public Health Laboratory | Erin L. Young, Kelly F. Oakeson, Pooja Gupta, Olinto Linares-Perdomo |
| EPI_ISL_3450582 | Department of Bacteria, Parasites and Fungi, Statens Serum Institut, Copenhagen, Denmark | Statens Serum Institut Bioinformatics and Microbial Genomics | Danish Covid-19 Genome Consortium |
| EPI_ISL_3450583 | Utah Public Health Laboratory | Utah Public Health Laboratory | Erin L. Young, Kelly F. Oakeson, Pooja Gupta, Olinto Linares-Perdomo |
| EPI_ISL_3450584 | Department of Bacteria, Parasites and Fungi, Statens Serum Institut, Copenhagen, Denmark | Statens Serum Institut Bioinformatics and Microbial Genomics | Danish Covid-19 Genome Consortium |
| EPI_ISL_3450585 | Utah Public Health Laboratory | Utah Public Health Laboratory | Erin L. Young, Kelly F. Oakeson, Pooja Gupta, Olinto Linares-Perdomo |
| EPI_ISL_3450586 | Department of Bacteria, Parasites and Fungi, Statens Serum Institut, Copenhagen, Denmark | Statens Serum Institut Bioinformatics and Microbial Genomics | Danish Covid-19 Genome Consortium |
| EPI_ISL_3450587 | Utah Public Health Laboratory | Utah Public Health Laboratory | Erin L. Young, Kelly F. Oakeson, Pooja Gupta, Olinto Linares-Perdomo |
| EPI_ISL_3450588 | Department of Bacteria, Parasites and Fungi, Statens Serum Institut, Copenhagen, Denmark | Statens Serum Institut Bioinformatics and Microbial Genomics | Danish Covid-19 Genome Consortium |
| EPI_ISL_3450589 | Utah Public Health Laboratory | Utah Public Health Laboratory | Erin L. Young, Kelly F. Oakeson, Pooja Gupta, Olinto Linares-Perdomo |
| EPI_ISL_3450590, EPI_ISL_3450591 | Department of Bacteria, Parasites and Fungi, Statens Serum Institut, Copenhagen, Denmark | Statens Serum Institut Bioinformatics and Microbial Genomics | Danish Covid-19 Genome Consortium |
| EPI_ISL_3450592 | Utah Public Health Laboratory | Utah Public Health Laboratory | Erin L. Young, Kelly F. Oakeson, Pooja Gupta, Olinto Linares-Perdomo |
| EPI_ISL_3450593 | Department of Bacteria, Parasites and Fungi, Statens Serum Institut, Copenhagen, Denmark | Statens Serum Institut Bioinformatics and Microbial Genomics | Danish Covid-19 Genome Consortium |
| EPI_ISL_3450594 | Utah Public Health Laboratory | Utah Public Health Laboratory | Erin L. Young, Kelly F. Oakeson, Pooja Gupta, Olinto Linares-Perdomo |
| EPI_ISL_3450595 | Department of Bacteria, Parasites and Fungi, Statens Serum Institut, Copenhagen, Denmark | Statens Serum Institut Bioinformatics and Microbial Genomics | Danish Covid-19 Genome Consortium |
| EPI_ISL_3450596, EPI_ISL_3450597 | Utah Public Health Laboratory | Utah Public Health Laboratory | Erin L. Young, Kelly F. Oakeson, Pooja Gupta, Olinto Linares-Perdomo |
| EPI_ISL_3450598, EPI_ISL_3450599 | Department of Bacteria, Parasites and Fungi, Statens Serum Institut, Copenhagen, Denmark | Statens Serum Institut Bioinformatics and Microbial Genomics | Danish Covid-19 Genome Consortium |
| EPI_ISL_3450600 | Utah Public Health Laboratory | Utah Public Health Laboratory | Erin L. Young, Kelly F. Oakeson, Pooja Gupta, Olinto Linares-Perdomo |
| EPI_ISL_3450601 | Department of Bacteria, Parasites and Fungi, Statens Serum Institut, Copenhagen, Denmark | Statens Serum Institut Bioinformatics and Microbial Genomics | Danish Covid-19 Genome Consortium |
| EPI_ISL_3450602, EPI_ISL_3450603 | Utah Public Health Laboratory | Utah Public Health Laboratory | Erin L. Young, Kelly F. Oakeson, Pooja Gupta, Olinto Linares-Perdomo |
| EPI_ISL_3450604, | Department of Bacteria, Parasites and Fungi, Statens Serum | Statens Serum Institut Bioinformatics and Microbial Genomics | Danish Covid-19 Genome Consortium |

| | | | |
|---|---|---|---|
| EPI_ISL_3450605 | Institut, Copenhagen, Denmark | | |
| EPI_ISL_3450606 | Utah Public Health Laboratory | Utah Public Health Laboratory | Erin L. Young, Kelly F. Oakeson, Pooja Gupta, Olinto Linares-Perdomo |
| EPI_ISL_3450607 | Department of Bacteria, Parasites and Fungi, Statens Serum Institut, Copenhagen, Denmark | Statens Serum Institut Bioinformatics and Microbial Genomics | Danish Covid-19 Genome Consortium |
| EPI_ISL_3450608 | Utah Public Health Laboratory | Utah Public Health Laboratory | Erin L. Young, Kelly F. Oakeson, Pooja Gupta, Olinto Linares-Perdomo |
| EPI_ISL_3450609, EPI_ISL_3450610 | Department of Bacteria, Parasites and Fungi, Statens Serum Institut, Copenhagen, Denmark | Statens Serum Institut Bioinformatics and Microbial Genomics | Danish Covid-19 Genome Consortium |
| EPI_ISL_3450611 | Utah Public Health Laboratory | Utah Public Health Laboratory | Erin L. Young, Kelly F. Oakeson, Pooja Gupta, Olinto Linares-Perdomo |
| EPI_ISL_3450612 | Department of Bacteria, Parasites and Fungi, Statens Serum Institut, Copenhagen, Denmark | Statens Serum Institut Bioinformatics and Microbial Genomics | Danish Covid-19 Genome Consortium |
| EPI_ISL_3450613 | Utah Public Health Laboratory | Utah Public Health Laboratory | Erin L. Young, Kelly F. Oakeson, Pooja Gupta, Olinto Linares-Perdomo |
| EPI_ISL_3450614 | Department of Bacteria, Parasites and Fungi, Statens Serum Institut, Copenhagen, Denmark | Statens Serum Institut Bioinformatics and Microbial Genomics | Danish Covid-19 Genome Consortium |
| EPI_ISL_3450615 | Utah Public Health Laboratory | Utah Public Health Laboratory | Erin L. Young, Kelly F. Oakeson, Pooja Gupta, Olinto Linares-Perdomo |
| EPI_ISL_3450616, EPI_ISL_3450617 | Department of Bacteria, Parasites and Fungi, Statens Serum Institut, Copenhagen, Denmark | Statens Serum Institut Bioinformatics and Microbial Genomics | Danish Covid-19 Genome Consortium |
| EPI_ISL_3450618 | Utah Public Health Laboratory | Utah Public Health Laboratory | Erin L. Young, Kelly F. Oakeson, Pooja Gupta, Olinto Linares-Perdomo |
| EPI_ISL_3450619, EPI_ISL_3450620 | Department of Bacteria, Parasites and Fungi, Statens Serum Institut, Copenhagen, Denmark | Statens Serum Institut Bioinformatics and Microbial Genomics | Danish Covid-19 Genome Consortium |
| EPI_ISL_3450621 | Utah Public Health Laboratory | Utah Public Health Laboratory | Erin L. Young, Kelly F. Oakeson, Pooja Gupta, Olinto Linares-Perdomo |
| EPI_ISL_3450622, EPI_ISL_3450623, EPI_ISL_3450624 | Department of Bacteria, Parasites and Fungi, Statens Serum Institut, Copenhagen, Denmark | Statens Serum Institut Bioinformatics and Microbial Genomics | Danish Covid-19 Genome Consortium |
| EPI_ISL_3450625 | Utah Public Health Laboratory | Utah Public Health Laboratory | Erin L. Young, Kelly F. Oakeson, Pooja Gupta, Olinto Linares-Perdomo |
| EPI_ISL_3450626 | Department of Bacteria, Parasites and Fungi, Statens Serum Institut, Copenhagen, Denmark | Statens Serum Institut Bioinformatics and Microbial Genomics | Danish Covid-19 Genome Consortium |
| EPI_ISL_3450627 | Utah Public Health Laboratory | Utah Public Health Laboratory | Erin L. Young, Kelly F. Oakeson, Pooja Gupta, Olinto Linares-Perdomo |
| EPI_ISL_3450628 | Department of Bacteria, Parasites and Fungi, Statens Serum Institut, Copenhagen, Denmark | Statens Serum Institut Bioinformatics and Microbial Genomics | Danish Covid-19 Genome Consortium |
| EPI_ISL_3450629 | Utah Public Health Laboratory | Utah Public Health Laboratory | Erin L. Young, Kelly F. Oakeson, Pooja Gupta, Olinto Linares-Perdomo |
| EPI_ISL_3450630 | Department of Bacteria, Parasites and Fungi, Statens Serum Institut, Copenhagen, Denmark | Statens Serum Institut Bioinformatics and Microbial Genomics | Danish Covid-19 Genome Consortium |
| EPI_ISL_3450631 | Utah Public Health Laboratory | Utah Public Health Laboratory | Erin L. Young, Kelly F. Oakeson, Pooja Gupta, Olinto Linares-Perdomo |
| EPI_ISL_3450632 | Department of Bacteria, Parasites and Fungi, Statens Serum Institut, Copenhagen, Denmark | Statens Serum Institut Bioinformatics and Microbial Genomics | Danish Covid-19 Genome Consortium |
| EPI_ISL_3450633 | Utah Public Health Laboratory | Utah Public Health Laboratory | Erin L. Young, Kelly F. Oakeson, Pooja Gupta, Olinto Linares-Perdomo |
| EPI_ISL_3450634 | Department of Bacteria, Parasites and Fungi, Statens Serum Institut, Copenhagen, Denmark | Statens Serum Institut Bioinformatics and Microbial Genomics | Danish Covid-19 Genome Consortium |
| EPI_ISL_3450635 | Utah Public Health Laboratory | Utah Public Health Laboratory | Erin L. Young, Kelly F. Oakeson, Pooja Gupta, Olinto Linares-Perdomo |
| EPI_ISL_3450636 | Department of Bacteria, Parasites and Fungi, Statens Serum Institut, Copenhagen, Denmark | Statens Serum Institut Bioinformatics and Microbial Genomics | Danish Covid-19 Genome Consortium |
| EPI_ISL_3450637 | Utah Public Health Laboratory | Utah Public Health Laboratory | Erin L. Young, Kelly F. Oakeson, Pooja Gupta, Olinto Linares-Perdomo |
| EPI_ISL_3450638 | Department of Bacteria, Parasites and Fungi, Statens Serum Institut, Copenhagen, Denmark | Statens Serum Institut Bioinformatics and Microbial Genomics | Danish Covid-19 Genome Consortium |
| EPI_ISL_3450639 | Utah Public Health Laboratory | Utah Public Health Laboratory | Erin L. Young, Kelly F. Oakeson, Pooja Gupta, Olinto Linares-Perdomo |
| EPI_ISL_3450640 | Department of Bacteria, Parasites and Fungi, Statens Serum Institut, Copenhagen, Denmark | Statens Serum Institut Bioinformatics and Microbial Genomics | Danish Covid-19 Genome Consortium |
| EPI_ISL_3450641 | Utah Public Health Laboratory | Utah Public Health Laboratory | Erin L. Young, Kelly F. Oakeson, Pooja Gupta, Olinto Linares-Perdomo |
| EPI_ISL_3450642 | Department of Bacteria, Parasites and Fungi, Statens Serum Institut, Copenhagen, Denmark | Statens Serum Institut Bioinformatics and Microbial Genomics | Danish Covid-19 Genome Consortium |
| EPI_ISL_3450643 | Utah Public Health Laboratory | Utah Public Health Laboratory | Erin L. Young, Kelly F. Oakeson, Pooja Gupta, Olinto Linares-Perdomo |
| EPI_ISL_3450644 | Department of Bacteria, Parasites and Fungi, Statens Serum Institut, Copenhagen, Denmark | Statens Serum Institut Bioinformatics and Microbial Genomics | Danish Covid-19 Genome Consortium |
| EPI_ISL_3450645 | Utah Public Health Laboratory | Utah Public Health Laboratory | Erin L. Young, Kelly F. Oakeson, Pooja Gupta, Olinto Linares-Perdomo |
| EPI_ISL_3450646, EPI_ISL_3450647 | Department of Bacteria, Parasites and Fungi, Statens Serum Institut, Copenhagen, Denmark | Statens Serum Institut Bioinformatics and Microbial Genomics | Danish Covid-19 Genome Consortium |
| EPI_ISL_3450648 | Utah Public Health Laboratory | Utah Public Health Laboratory | Erin L. Young, Kelly F. Oakeson, Pooja Gupta, Olinto Linares-Perdomo |
| EPI_ISL_3450649 | Department of Bacteria, Parasites and Fungi, Statens Serum Institut, Copenhagen, Denmark | Statens Serum Institut Bioinformatics and Microbial Genomics | Danish Covid-19 Genome Consortium |
| EPI_ISL_3450650 | Utah Public Health Laboratory | Utah Public Health Laboratory | Erin L. Young, Kelly F. Oakeson, Pooja Gupta, Olinto Linares-Perdomo |
| EPI_ISL_3450651 | Department of Bacteria, Parasites and Fungi, Statens Serum Institut, Copenhagen, Denmark | Statens Serum Institut Bioinformatics and Microbial Genomics | Danish Covid-19 Genome Consortium |
| EPI_ISL_3450652 | Utah Public Health Laboratory | Utah Public Health Laboratory | Erin L. Young, Kelly F. Oakeson, Pooja Gupta, Olinto Linares-Perdomo |
| EPI_ISL_3450653, EPI_ISL_3450654, EPI_ISL_3450655, EPI_ISL_3450656, EPI_ISL_3450657, EPI_ISL_3450658, EPI_ISL_3450659, EPI_ISL_3450660, EPI_ISL_3450661, EPI_ISL_3450662, EPI_ISL_3450663, EPI_ISL_3450664, EPI_ISL_3450665, EPI_ISL_3450666, EPI_ISL_3450667, EPI_ISL_3450668, EPI_ISL_3450669, EPI_ISL_3450670, EPI_ISL_3450671, EPI_ISL_3450672, EPI_ISL_3450673, EPI_ISL_3450674, EPI_ISL_3450675, EPI_ISL_3450677, EPI_ISL_3450678, EPI_ISL_3450679, EPI_ISL_3450680, EPI_ISL_3450681, EPI_ISL_3450682, EPI_ISL_3450683, EPI_ISL_3450684, EPI_ISL_3450685, EPI_ISL_3450686, EPI_ISL_3450687, EPI_ISL_3450688, EPI_ISL_3450689, EPI_ISL_3450690, EPI_ISL_3450691, EPI_ISL_3450692, EPI_ISL_3450693, EPI_ISL_3450694, EPI_ISL_3450695, EPI_ISL_3450696, EPI_ISL_3450697, EPI_ISL_3450698, EPI_ISL_3450699, EPI_ISL_3450700, EPI_ISL_3450701, EPI_ISL_3450702, EPI_ISL_3450703, EPI_ISL_3450704, EPI_ISL_3450705, EPI_ISL_3450706, EPI_ISL_3450707, EPI_ISL_3450708, EPI_ISL_3450709, EPI_ISL_3450710, EPI_ISL_3450711, EPI_ISL_3450712, EPI_ISL_3450713, EPI_ISL_3450714, EPI_ISL_3450715, EPI_ISL_3450716, EPI_ISL_3450717, EPI_ISL_3450718, EPI_ISL_3450719, EPI_ISL_3450720, EPI_ISL_3450721, EPI_ISL_3450722, EPI_ISL_3450723, EPI_ISL_3450724, EPI_ISL_3450725, EPI_ISL_3450726, EPI_ISL_3450727, EPI_ISL_3450728, EPI_ISL_3450729, EPI_ISL_3450730, EPI_ISL_3450731, EPI_ISL_3450732, EPI_ISL_3450733, EPI_ISL_3450734, EPI_ISL_3450735, EPI_ISL_3450736, EPI_ISL_3450737, EPI_ISL_3450738, EPI_ISL_3450739, EPI_ISL_3450740, EPI_ISL_3450741, EPI_ISL_3450742, EPI_ISL_3450743, EPI_ISL_3450744, EPI_ISL_3450745, EPI_ISL_3450746, EPI_ISL_3450747, EPI_ISL_3450748, EPI_ISL_3450750, EPI_ISL_3450751, EPI_ISL_3450752, EPI_ISL_3450753, EPI_ISL_3450754, EPI_ISL_3450755, EPI_ISL_3450756, EPI_ISL_3450757, EPI_ISL_3450758, EPI_ISL_3450759, EPI_ISL_3450760, EPI_ISL_3450761, EPI_ISL_3450762, EPI_ISL_3450763, EPI_ISL_3450764, EPI_ISL_3450765, EPI_ISL_3450766, EPI_ISL_3450767, EPI_ISL_3450768, EPI_ISL_3450769, EPI_ISL_3450770, EPI_ISL_3450771, EPI_ISL_3450772, EPI_ISL_3450773, EPI_ISL_3450774, EPI_ISL_3450775, EPI_ISL_3450776, EPI_ISL_3450777, EPI_ISL_3450778, EPI_ISL_3450779, EPI_ISL_3450780, EPI_ISL_3450781, EPI_ISL_3450782, EPI_ISL_3450783, EPI_ISL_3450784, EPI_ISL_3450785, EPI_ISL_3450786, EPI_ISL_3450787, EPI_ISL_3450788, EPI_ISL_3450789, EPI_ISL_3450790, EPI_ISL_3450791, EPI_ISL_3450792, EPI_ISL_3450793, EPI_ISL_3450794, EPI_ISL_3450795, EPI_ISL_3450796, EPI_ISL_3450797, EPI_ISL_3450798, EPI_ISL_3450799, EPI_ISL_3450800, EPI_ISL_3450801, EPI_ISL_3450802 | | | |
| see above | Department of Bacteria, Parasites and Fungi, Statens Serum Institut, Copenhagen, Denmark | Statens Serum Institut Bioinformatics and Microbial Genomics | Danish Covid-19 Genome Consortium |
| EPI_ISL_3450804, EPI_ISL_3450805, EPI_ISL_3450806, EPI_ISL_3450807, EPI_ISL_3450808, EPI_ISL_3450809, EPI_ISL_3450810, EPI_ISL_3450811, EPI_ISL_3450812, EPI_ISL_3450813, EPI_ISL_3450814, EPI_ISL_3450815, EPI_ISL_3450816, EPI_ISL_3450817, EPI_ISL_3450818, EPI_ISL_3450819, EPI_ISL_3450820, EPI_ISL_3450821, EPI_ISL_3450822, EPI_ISL_3450823, EPI_ISL_3450824, EPI_ISL_3450825, EPI_ISL_3450826, EPI_ISL_3450827, EPI_ISL_3450828, EPI_ISL_3450829, EPI_ISL_3450830, EPI_ISL_3450831, EPI_ISL_3450832, EPI_ISL_3450833, EPI_ISL_3450834, EPI_ISL_3450835, EPI_ISL_3450836, EPI_ISL_3450837, EPI_ISL_3450838, EPI_ISL_3450839, EPI_ISL_3450840, EPI_ISL_3450841, EPI_ISL_3450842, EPI_ISL_3450843, EPI_ISL_3450844, EPI_ISL_3450845, EPI_ISL_3450846, EPI_ISL_3450847, EPI_ISL_3450848, EPI_ISL_3450849, EPI_ISL_3450850, EPI_ISL_3450851, EPI_ISL_3450852, EPI_ISL_3450853, EPI_ISL_3450854, EPI_ISL_3450855, EPI_ISL_3450856, EPI_ISL_3450857, EPI_ISL_3450858, EPI_ISL_3450859, EPI_ISL_3450860, EPI_ISL_3450861, EPI_ISL_3450862, EPI_ISL_3450863, EPI_ISL_3450864, EPI_ISL_3450865, EPI_ISL_3450866, EPI_ISL_3450867, EPI_ISL_3450868, EPI_ISL_3450869, EPI_ISL_3450870, EPI_ISL_3450871, EPI_ISL_3450872, EPI_ISL_3450873, EPI_ISL_3450874, EPI_ISL_3450875, EPI_ISL_3450876, EPI_ISL_3450877, EPI_ISL_3450878, EPI_ISL_3450879, EPI_ISL_3450880, EPI_ISL_3450881, EPI_ISL_3450882, EPI_ISL_3450883, EPI_ISL_3450884, EPI_ISL_3450885, EPI_ISL_3450886, EPI_ISL_3450887, EPI_ISL_3450888, EPI_ISL_3450889, EPI_ISL_3450890, EPI_ISL_3450891, EPI_ISL_3450892, EPI_ISL_3450893, EPI_ISL_3450894, EPI_ISL_3450895, EPI_ISL_3450896, EPI_ISL_3450897, EPI_ISL_3450898, EPI_ISL_3450899, EPI_ISL_3450900, EPI_ISL_3450901, EPI_ISL_3450902, EPI_ISL_3450903, EPI_ISL_3450904, EPI_ISL_3450905, EPI_ISL_3450906, EPI_ISL_3450907, EPI_ISL_3450908, EPI_ISL_3450909, EPI_ISL_3450910, EPI_ISL_3450911, EPI_ISL_3450912, EPI_ISL_3450913, EPI_ISL_3450914, EPI_ISL_3450915, EPI_ISL_3450916, EPI_ISL_3450917, EPI_ISL_3450918, EPI_ISL_3450919, EPI_ISL_3450920, EPI_ISL_3450921, EPI_ISL_3450922, EPI_ISL_3450923, EPI_ISL_3450924, EPI_ISL_3450925, EPI_ISL_3450926, EPI_ISL_3450927, EPI_ISL_3450928, EPI_ISL_3450929, EPI_ISL_3450930, EPI_ISL_3450931, EPI_ISL_3450932, EPI_ISL_3450933, EPI_ISL_3450934, EPI_ISL_3450935, EPI_ISL_3450936, EPI_ISL_3450937, EPI_ISL_3450938, EPI_ISL_3450939, EPI_ISL_3450940, EPI_ISL_3450941, EPI_ISL_3450942, EPI_ISL_3450943, EPI_ISL_3450944, EPI_ISL_3450945, EPI_ISL_3450946, EPI_ISL_3450947, EPI_ISL_3450948, EPI_ISL_3450949, EPI_ISL_3450950, EPI_ISL_3450951, EPI_ISL_3450952, EPI_ISL_3450953, EPI_ISL_3450954, EPI_ISL_3450955, EPI_ISL_3450956, EPI_ISL_3450957, EPI_ISL_3450958, EPI_ISL_3450959, EPI_ISL_3450960, EPI_ISL_3450961, EPI_ISL_3450962, EPI_ISL_3450963, EPI_ISL_3450964, EPI_ISL_3450965, EPI_ISL_3450966, EPI_ISL_3450967, EPI_ISL_3450968, EPI_ISL_3450969, EPI_ISL_3450970, EPI_ISL_3450971, EPI_ISL_3450972, EPI_ISL_3450973, EPI_ISL_3450974, EPI_ISL_3450975, EPI_ISL_3450976, EPI_ISL_3450977, EPI_ISL_3450978, EPI_ISL_3450979, EPI_ISL_3450980, EPI_ISL_3450981, EPI_ISL_3450982, EPI_ISL_3450983, EPI_ISL_3450984, EPI_ISL_3450985, EPI_ISL_3450986, EPI_ISL_3450987, EPI_ISL_3450988, EPI_ISL_3450989, EPI_ISL_3450990, EPI_ISL_3450991, EPI_ISL_3450992, EPI_ISL_3450993, EPI_ISL_3450994, EPI_ISL_3450995, EPI_ISL_3450996, EPI_ISL_3450997, EPI_ISL_3450998, EPI_ISL_3450999, EPI_ISL_3451000, EPI_ISL_3451001, EPI_ISL_3451002, EPI_ISL_3451003, EPI_ISL_3451004, EPI_ISL_3451005, EPI_ISL_3451006, EPI_ISL_3451007, EPI_ISL_3451008, EPI_ISL_3451011, EPI_ISL_3451012, EPI_ISL_3451013, EPI_ISL_3451014, EPI_ISL_3451015, EPI_ISL_3451016, EPI_ISL_3451018, EPI_ISL_3451019, EPI_ISL_3451020, EPI_ISL_3451021, EPI_ISL_3451022, EPI_ISL_3451023, EPI_ISL_3451024, EPI_ISL_3451025, EPI_ISL_3451026, EPI_ISL_3451028, EPI_ISL_3451029, EPI_ISL_3451030, EPI_ISL_3451034, EPI_ISL_3451035, EPI_ISL_3451036, EPI_ISL_3451037, EPI_ISL_3451038, EPI_ISL_3451039, EPI_ISL_3451040, EPI_ISL_3451041, EPI_ISL_3451042, EPI_ISL_3451043, EPI_ISL_3451044, EPI_ISL_3451045, EPI_ISL_3451046, EPI_ISL_3451047, EPI_ISL_3451048, EPI_ISL_3451049, EPI_ISL_3451050, EPI_ISL_3451051, EPI_ISL_3451052, EPI_ISL_3451053, EPI_ISL_3451054, EPI_ISL_3451055, EPI_ISL_3451056, EPI_ISL_3451057, EPI_ISL_3451058, EPI_ISL_3451059, EPI_ISL_3451060, EPI_ISL_3451061, EPI_ISL_3451062, EPI_ISL_3451063, EPI_ISL_3451064, EPI_ISL_3451065, EPI_ISL_3451067, EPI_ISL_3451068, EPI_ISL_3451069, EPI_ISL_3451070, EPI_ISL_3451071, EPI_ISL_3451072, EPI_ISL_3451073 | | | |
| see above | Washington State Department of Health Public Health Laboratories | Washington State Department of Health Public Health Laboratories | Drew MacKellar, Philip Dykema, Denny Russell, Joenice Gonzalez, Hannah Gray, Geoff Melly, Vanessa De Los Santos, Darren Lucas, JohnAric Peterson, Avi Singh, Rebecca Cao |
| EPI_ISL_3451074 | MANKWENG LABORATORY | National Institute for Communicable Diseases of the National Health Laboratory Service | Amoako DG, Everatt J, Scheepers C, Mohale T, Ntuli N, Mahlangu B, Mnguni A, Ismail A, Bhiman JN |
| EPI_ISL_3451076, EPI_ISL_3451077, EPI_ISL_3451079, EPI_ISL_3451080, EPI_ISL_3451081, EPI_ISL_3451082, EPI_ISL_3451083, EPI_ISL_3451084, EPI_ISL_3451085, EPI_ISL_3451086, EPI_ISL_3451087, EPI_ISL_3451088 | | | |

| | | | |
|---|---|---|---|
| see above | PORT ELIZABETH LABORATORY | National Institute for Communicable Diseases of the National Health Laboratory Service | Amoako DG, Everatt J, Scheepers C, Mohale T, Ntuli N, Mahlangu B, Mnguni A, Ismail A, Bhiman JN |
| EPI_ISL_3451095, EPI_ISL_3451096, EPI_ISL_3451097, EPI_ISL_3451098, EPI_ISL_3451099, EPI_ISL_3451100 | RUSTENBURG LABORATORY | National Institute for Communicable Diseases of the National Health Laboratory Service | Amoako DG, Everatt J, Scheepers C, Mohale T, Ntuli N, Mahlangu B, Mnguni A, Ismail A, Bhiman JN |
| EPI_ISL_3451101, EPI_ISL_3451102, EPI_ISL_3451103, EPI_ISL_3451104, EPI_ISL_3451105, EPI_ISL_3451106, EPI_ISL_3451107, EPI_ISL_3451108, EPI_ISL_3451109, EPI_ISL_3451110, EPI_ISL_3451111, EPI_ISL_3451112, EPI_ISL_3451113, EPI_ISL_3451114, EPI_ISL_3451115, EPI_ISL_3451116, EPI_ISL_3451117, EPI_ISL_3451118, EPI_ISL_3451119, EPI_ISL_3451120, EPI_ISL_3451121, EPI_ISL_3451122, EPI_ISL_3451123, EPI_ISL_3451124, EPI_ISL_3451125, EPI_ISL_3451126, EPI_ISL_3451127, EPI_ISL_3451128, EPI_ISL_3451129, EPI_ISL_3451130, EPI_ISL_3451131 | | | |
| see above | PORT ELIZABETH LABORATORY | National Institute for Communicable Diseases of the National Health Laboratory Service | Amoako DG, Everatt J, Scheepers C, Mohale T, Ntuli N, Mahlangu B, Mnguni A, Ismail A, Bhiman JN |
| EPI_ISL_3451132, EPI_ISL_3451133, EPI_ISL_3451134, EPI_ISL_3451135, EPI_ISL_3451136, EPI_ISL_3451137, EPI_ISL_3451138, EPI_ISL_3451139, EPI_ISL_3451140, EPI_ISL_3451141, EPI_ISL_3451142, EPI_ISL_3451143, EPI_ISL_3451145, EPI_ISL_3451146, EPI_ISL_3451147, EPI_ISL_3451148, EPI_ISL_3451149, EPI_ISL_3451151, EPI_ISL_3451152, EPI_ISL_3451153, EPI_ISL_3451154 | | | |
| see above | MANKWENG LABORATORY | National Institute for Communicable Diseases of the National Health Laboratory Service | Amoako DG, Everatt J, Scheepers C, Mohale T, Ntuli N, Mahlangu B, Mnguni A, Ismail A, Bhiman JN |
| EPI_ISL_3451155, EPI_ISL_3451156, EPI_ISL_3451157, EPI_ISL_3451158, EPI_ISL_3451159, EPI_ISL_3451160, EPI_ISL_3451161, EPI_ISL_3451162, EPI_ISL_3451163, EPI_ISL_3451164, EPI_ISL_3451165, EPI_ISL_3451166, EPI_ISL_3451167, EPI_ISL_3451168, EPI_ISL_3451169, EPI_ISL_3451170, EPI_ISL_3451171, EPI_ISL_3451172, EPI_ISL_3451173, EPI_ISL_3451174, EPI_ISL_3451175, EPI_ISL_3451176, EPI_ISL_3451177, EPI_ISL_3451178, EPI_ISL_3451179, EPI_ISL_3451180, EPI_ISL_3451181, EPI_ISL_3451182, EPI_ISL_3451183 | | | |
| see above | PORT ELIZABETH LABORATORY | National Institute for Communicable Diseases of the National Health Laboratory Service | Amoako DG, Everatt J, Scheepers C, Mohale T, Ntuli N, Mahlangu B, Mnguni A, Ismail A, Bhiman JN |
| EPI_ISL_3451184, EPI_ISL_3451185, EPI_ISL_3451186, EPI_ISL_3451187, EPI_ISL_3451188, EPI_ISL_3451189, EPI_ISL_3451190, EPI_ISL_3451191, EPI_ISL_3451192, EPI_ISL_3451193, EPI_ISL_3451194, EPI_ISL_3451196, EPI_ISL_3451197, EPI_ISL_3451198, EPI_ISL_3451199, EPI_ISL_3451200, EPI_ISL_3451201, EPI_ISL_3451202, EPI_ISL_3451203, EPI_ISL_3451204, EPI_ISL_3451205, EPI_ISL_3451206, EPI_ISL_3451207, EPI_ISL_3451208, EPI_ISL_3451209, EPI_ISL_3451210, EPI_ISL_3451211, EPI_ISL_3451212, EPI_ISL_3451213, EPI_ISL_3451215, EPI_ISL_3451216 | | | |
| see above | TAMBO MEMORIAL LABORATORY | National Institute for Communicable Diseases of the National Health Laboratory Service | Amoako DG, Everatt J, Scheepers C, Mohale T, Ntuli N, Mahlangu B, Mnguni A, Ismail A, Bhiman JN |
| EPI_ISL_3451217, EPI_ISL_3451218, EPI_ISL_3451219, EPI_ISL_3451220, EPI_ISL_3451221, EPI_ISL_3451223, EPI_ISL_3451224, EPI_ISL_3451225, EPI_ISL_3451226, EPI_ISL_3451227, EPI_ISL_3451228, EPI_ISL_3451229, EPI_ISL_3451230, EPI_ISL_3451231, EPI_ISL_3451232, EPI_ISL_3451233, EPI_ISL_3451234, EPI_ISL_3451235, EPI_ISL_3451236, EPI_ISL_3451237, EPI_ISL_3451238, EPI_ISL_3451239, EPI_ISL_3451240, EPI_ISL_3451241, EPI_ISL_3451242, EPI_ISL_3451243, EPI_ISL_3451244, EPI_ISL_3451245, EPI_ISL_3451246, EPI_ISL_3451247, EPI_ISL_3451250, EPI_ISL_3451251, EPI_ISL_3451252, EPI_ISL_3451253, EPI_ISL_3451254 | | | |
| see above | POLOKWANE LABORATORY | National Institute for Communicable Diseases of the National Health Laboratory Service | Amoako DG, Everatt J, Scheepers C, Mohale T, Ntuli N, Mahlangu B, Mnguni A, Ismail A, Bhiman JN |
| EPI_ISL_3451255, EPI_ISL_3451256 | KOPANONG LABORATORY | National Institute for Communicable Diseases of the National Health Laboratory Service | Amoako DG, Everatt J, Scheepers C, Mohale T, Ntuli N, Mahlangu B, Mnguni A, Ismail A, Bhiman JN |
| EPI_ISL_3451262, EPI_ISL_3451263, EPI_ISL_3451264, EPI_ISL_3451266, EPI_ISL_3451269, EPI_ISL_3451271, EPI_ISL_3451272, EPI_ISL_3451275, EPI_ISL_3451277, EPI_ISL_3451278, EPI_ISL_3451280, EPI_ISL_3451281, EPI_ISL_3451282, EPI_ISL_3451284, EPI_ISL_3451285, EPI_ISL_3451286, EPI_ISL_3451287, EPI_ISL_3451288, EPI_ISL_3451289, EPI_ISL_3451290, EPI_ISL_3451291, EPI_ISL_3451292, EPI_ISL_3451293, EPI_ISL_3451294, EPI_ISL_3451296, EPI_ISL_3451297, EPI_ISL_3451298, EPI_ISL_3451299, EPI_ISL_3451302, EPI_ISL_3451303 | | | |
| see above | KIMBERLEY LABORATORY | National Institute for Communicable Diseases of the National Health Laboratory Service | Amoako DG, Everatt J, Scheepers C, Mohale T, Ntuli N, Mahlangu B, Mnguni A, Ismail A, Bhiman JN |
| EPI_ISL_3451304, EPI_ISL_3451305, EPI_ISL_3451306, EPI_ISL_3451307, EPI_ISL_3451308, EPI_ISL_3451309, EPI_ISL_3451310, EPI_ISL_3451311, EPI_ISL_3451312, EPI_ISL_3451313, EPI_ISL_3451314, EPI_ISL_3451317, EPI_ISL_3451318, EPI_ISL_3451319, EPI_ISL_3451320, EPI_ISL_3451321, EPI_ISL_3451322, EPI_ISL_3451323, EPI_ISL_3451324, EPI_ISL_3451325, EPI_ISL_3451326, EPI_ISL_3451327, EPI_ISL_3451328, EPI_ISL_3451329 | | | |
| see above | CHRIS HANI BARAGWANATH LABORATORY | National Institute for Communicable Diseases of the National Health Laboratory Service | Amoako DG, Everatt J, Scheepers C, Mohale T, Ntuli N, Mahlangu B, Mnguni A, Ismail A, Bhiman JN |
| EPI_ISL_3451330, EPI_ISL_3451331, EPI_ISL_3451332, EPI_ISL_3451333, EPI_ISL_3451334, EPI_ISL_3451336, EPI_ISL_3451337, EPI_ISL_3451338, EPI_ISL_3451339, EPI_ISL_3451340 | PORT ELIZABETH LABORATORY | National Institute for Communicable Diseases of the National Health Laboratory Service | Amoako DG, Everatt J, Scheepers C, Mohale T, Ntuli N, Mahlangu B, Mnguni A, Ismail A, Bhiman JN |
| EPI_ISL_3451341, EPI_ISL_3451342, EPI_ISL_3451343, EPI_ISL_3451344, EPI_ISL_3451345, EPI_ISL_3451346, EPI_ISL_3451347, EPI_ISL_3451348, EPI_ISL_3451349, EPI_ISL_3451350, EPI_ISL_3451351, EPI_ISL_3451352, EPI_ISL_3451353, EPI_ISL_3451354, EPI_ISL_3451355, EPI_ISL_3451356, EPI_ISL_3451357, EPI_ISL_3451359, EPI_ISL_3451360, EPI_ISL_3451361, EPI_ISL_3451363, EPI_ISL_3451364, EPI_ISL_3451365, EPI_ISL_3451367, EPI_ISL_3451368, EPI_ISL_3451370, EPI_ISL_3451371, EPI_ISL_3451372, EPI_ISL_3451374, EPI_ISL_3451375, EPI_ISL_3451376, EPI_ISL_3451377, EPI_ISL_3451379, EPI_ISL_3451380, EPI_ISL_3451382, EPI_ISL_3451383, EPI_ISL_3451384, EPI_ISL_3451385, EPI_ISL_3451386, EPI_ISL_3451387, EPI_ISL_3451388, EPI_ISL_3451389, EPI_ISL_3451390, EPI_ISL_3451392, EPI_ISL_3451393, EPI_ISL_3451394, EPI_ISL_3451395, EPI_ISL_3451397, EPI_ISL_3451398, EPI_ISL_3451399, EPI_ISL_3451400, EPI_ISL_3451401 | | | |
| see above | LEBOWAKGOMO LABORATORY | National Institute for Communicable Diseases of the National Health Laboratory Service | Amoako DG, Everatt J, Scheepers C, Mohale T, Ntuli N, Mahlangu B, Mnguni A, Ismail A, Bhiman JN |
| EPI_ISL_3451402, EPI_ISL_3451403, EPI_ISL_3451404, EPI_ISL_3451405, EPI_ISL_3451406, EPI_ISL_3451407, EPI_ISL_3451408, EPI_ISL_3451409, EPI_ISL_3451410, EPI_ISL_3451411, EPI_ISL_3451412, EPI_ISL_3451413, EPI_ISL_3451414, EPI_ISL_3451415, EPI_ISL_3451416, EPI_ISL_3451417, EPI_ISL_3451418, EPI_ISL_3451419, EPI_ISL_3451420, EPI_ISL_3451421, EPI_ISL_3451422, EPI_ISL_3451423, EPI_ISL_3451424, EPI_ISL_3451425, EPI_ISL_3451426, EPI_ISL_3451427, EPI_ISL_3451428, EPI_ISL_3451429, EPI_ISL_3451430, EPI_ISL_3451431, EPI_ISL_3451432, EPI_ISL_3451433, EPI_ISL_3451434, EPI_ISL_3451435, EPI_ISL_3451436, EPI_ISL_3451437, EPI_ISL_3451438 | | | |
| see above | PORT ELIZABETH LABORATORY | National Institute for Communicable Diseases of the National Health Laboratory Service | Amoako DG, Everatt J, Scheepers C, Mohale T, Ntuli N, Mahlangu B, Mnguni A, Ismail A, Bhiman JN |
| EPI_ISL_3451455, EPI_ISL_3451478 | ROB FERREIRA LABORATORY | National Institute for Communicable Diseases of the National Health Laboratory Service | Amoako DG, Everatt J, Scheepers C, Mohale T, Ntuli N, Mahlangu B, Mnguni A, Ismail A, Bhiman JN |
| EPI_ISL_3451498, EPI_ISL_3451500, EPI_ISL_3451502 | KOPANONG LABORATORY | National Institute for Communicable Diseases of the National Health Laboratory Service | Amoako DG, Everatt J, Scheepers C, Mohale T, Ntuli N, Mahlangu B, Mnguni A, Ismail A, Bhiman JN |
| EPI_ISL_3451505, EPI_ISL_3451506, EPI_ISL_3451507, EPI_ISL_3451508, EPI_ISL_3451509, EPI_ISL_3451511, EPI_ISL_3451512, EPI_ISL_3451515, EPI_ISL_3451517, EPI_ISL_3451523, EPI_ISL_3451526, EPI_ISL_3451528, EPI_ISL_3451529 | | | |
| see above | TAMBO MEMORIAL LABORATORY | National Institute for Communicable Diseases of the National Health Laboratory Service | Amoako DG, Everatt J, Scheepers C, Mohale T, Ntuli N, Mahlangu B, Mnguni A, Ismail A, Bhiman JN |
| EPI_ISL_3451530, EPI_ISL_3451531 | HELEN JOSEPH LABORATORY | National Institute for Communicable Diseases of the National Health Laboratory Service | Amoako DG, Everatt J, Scheepers C, Mohale T, Ntuli N, Mahlangu B, Mnguni A, Ismail A, Bhiman JN |
| EPI_ISL_3451532, EPI_ISL_3451533 | MANKWENG LABORATORY | National Institute for Communicable Diseases of the National Health Laboratory Service | Amoako DG, Everatt J, Scheepers C, Mohale T, Ntuli N, Mahlangu B, Mnguni A, Ismail A, Bhiman JN |
| EPI_ISL_3451534, EPI_ISL_3451535, EPI_ISL_3451536, EPI_ISL_3451537, EPI_ISL_3451538, EPI_ISL_3451539, EPI_ISL_3451540, EPI_ISL_3451541, EPI_ISL_3451542, EPI_ISL_3451543, EPI_ISL_3451545, EPI_ISL_3451546, EPI_ISL_3451547, EPI_ISL_3451548, EPI_ISL_3451549, EPI_ISL_3451550, EPI_ISL_3451551, EPI_ISL_3451552, EPI_ISL_3451553, EPI_ISL_3451554, EPI_ISL_3451555, EPI_ISL_3451556, EPI_ISL_3451557, EPI_ISL_3451558, EPI_ISL_3451559, EPI_ISL_3451560, EPI_ISL_3451561, EPI_ISL_3451562, EPI_ISL_3451563, EPI_ISL_3451564, EPI_ISL_3451565, EPI_ISL_3451566, EPI_ISL_3451567, EPI_ISL_3451568, EPI_ISL_3451570, EPI_ISL_3451571, EPI_ISL_3451572, EPI_ISL_3451574, EPI_ISL_3451575, EPI_ISL_3451576, EPI_ISL_3451577, EPI_ISL_3451578, EPI_ISL_3451579, EPI_ISL_3451580, EPI_ISL_3451581, EPI_ISL_3451582, EPI_ISL_3451583, EPI_ISL_3451584, EPI_ISL_3451585, EPI_ISL_3451586, EPI_ISL_3451587, EPI_ISL_3451588, EPI_ISL_3451589, EPI_ISL_3451590, EPI_ISL_3451591, EPI_ISL_3451592, EPI_ISL_3451593, EPI_ISL_3451594, EPI_ISL_3451595, EPI_ISL_3451596, EPI_ISL_3451597, EPI_ISL_3451598, EPI_ISL_3451599, EPI_ISL_3451600, EPI_ISL_3451601, EPI_ISL_3451602, EPI_ISL_3451603, EPI_ISL_3451604, EPI_ISL_3451605, EPI_ISL_3451606, EPI_ISL_3451607, EPI_ISL_3451608, EPI_ISL_3451609, EPI_ISL_3451610, EPI_ISL_3451611, EPI_ISL_3451612, EPI_ISL_3451613, EPI_ISL_3451614, EPI_ISL_3451615, EPI_ISL_3451616, EPI_ISL_3451617, EPI_ISL_3451618, EPI_ISL_3451620, EPI_ISL_3451621, EPI_ISL_3451622, EPI_ISL_3451623, EPI_ISL_3451624, EPI_ISL_3451625, EPI_ISL_3451626, EPI_ISL_3451628, EPI_ISL_3451629, EPI_ISL_3451630, EPI_ISL_3451631, EPI_ISL_3451632, EPI_ISL_3451633, EPI_ISL_3451635, EPI_ISL_3451636, EPI_ISL_3451637 | | | |
| see above | National Institute for Communicable Diseases of the National Health Laboratory Service | National Institute for Communicable Diseases of the National Health Laboratory Service | Amoako DG, Everatt J, Scheepers C, Mohale T, Ntuli N, Mahlangu B, Mnguni A, Ismail A, Bhiman JN |
| EPI_ISL_3451638, EPI_ISL_3451639, EPI_ISL_3451640, EPI_ISL_3451641, EPI_ISL_3451642, EPI_ISL_3451643, EPI_ISL_3451644, EPI_ISL_3451645, EPI_ISL_3451646, EPI_ISL_3451647, EPI_ISL_3451648, EPI_ISL_3451649, EPI_ISL_3451650, EPI_ISL_3451651, EPI_ISL_3451652, EPI_ISL_3451653, EPI_ISL_3451654, EPI_ISL_3451655, EPI_ISL_3451656, EPI_ISL_3451657, EPI_ISL_3451658, EPI_ISL_3451659, EPI_ISL_3451660, EPI_ISL_3451661, EPI_ISL_3451662, EPI_ISL_3451663, EPI_ISL_3451664, EPI_ISL_3451665, EPI_ISL_3451666, EPI_ISL_3451667, EPI_ISL_3451668, EPI_ISL_3451669 | | | |
| see above | Klinisch Laboratorium ZNA | Klinisch Laboratorium ZNA | Verstrepen et al. |
| EPI_ISL_3451670, EPI_ISL_3451671, EPI_ISL_3451673, EPI_ISL_3451674, EPI_ISL_3451675, EPI_ISL_3451676, EPI_ISL_3451677, EPI_ISL_3451678, EPI_ISL_3451679, EPI_ISL_3451680, EPI_ISL_3451681, EPI_ISL_3451683, EPI_ISL_3451684, EPI_ISL_3451685 | | | |
| see above | Clinical Virology | Clinical Bacteriology | Tim Roloff, Fanny Wegner, Helena MB Seth-Smith, Alfredo Mari, Karoline Leuzinger, Julia Bielicki, Manuel Battegay, Hans Hirsch, Adrian Egli |
| EPI_ISL_3451686, EPI_ISL_3451687, EPI_ISL_3451688, EPI_ISL_3451689 | Medics Labor AG | Clinical Bacteriology | Tim Roloff, Fanny Wegner, Helena MB Seth-Smith, Alfredo Mari, Karoline Leuzinger, Julia Bielicki, Manuel Battegay, Hans Hirsch, Adrian Egli |
| EPI_ISL_3451690 | Synlab Suisse SA | Clinical Bacteriology | Tim Roloff, Fanny Wegner, Helena MB Seth-Smith, Alfredo Mari, Karoline Leuzinger, Julia Bielicki, Manuel Battegay, Hans Hirsch, Adrian Egli |
| EPI_ISL_3451691, EPI_ISL_3451692, EPI_ISL_3451693, EPI_ISL_3451694, EPI_ISL_3451695, EPI_ISL_3451696, EPI_ISL_3451697, EPI_ISL_3451698, EPI_ISL_3451699, EPI_ISL_3451700, EPI_ISL_3451701, EPI_ISL_3451702, EPI_ISL_3451703, EPI_ISL_3451704, EPI_ISL_3451705, EPI_ISL_3451706, EPI_ISL_3451707, EPI_ISL_3451708, EPI_ISL_3451709, EPI_ISL_3451710, EPI_ISL_3451711, EPI_ISL_3451712, EPI_ISL_3451713, EPI_ISL_3451714 | | | |
| see above | Medics Labor AG | Clinical Bacteriology | Tim Roloff, Fanny Wegner, Helena MB Seth-Smith, Alfredo Mari, Karoline Leuzinger, Julia Bielicki, Manuel Battegay, Hans Hirsch, Adrian Egli |
| EPI_ISL_3451715, EPI_ISL_3451716, EPI_ISL_3451717, EPI_ISL_3451718 | Synlab Suisse SA | Clinical Bacteriology | Tim Roloff, Fanny Wegner, Helena MB Seth-Smith, Alfredo Mari, Karoline Leuzinger, Julia Bielicki, Manuel Battegay, Hans Hirsch, Adrian Egli |
| EPI_ISL_3451719, EPI_ISL_3451720, EPI_ISL_3451721, EPI_ISL_3451722, EPI_ISL_3451723, EPI_ISL_3451724, EPI_ISL_3451725, EPI_ISL_3451726, EPI_ISL_3451727, EPI_ISL_3451728, EPI_ISL_3451729, EPI_ISL_3451730 | | | |

| | | | |
|---|---|---|---|
| see above | Medics Labor AG | Clinical Bacteriology | Tim Roloff, Fanny Wegner, Helena MB Seth-Smith, Alfredo Mari, Karoline Leuzinger, Julia Bielicki, Manuel Battegay, Hans Hirsch, Adrian Egli |
| EPI_ISL_3451732, EPI_ISL_3451733 | Rothen Medizinische Laboratorien AG | Clinical Bacteriology | Tim Roloff, Fanny Wegner, Helena MB Seth-Smith, Alfredo Mari, Karoline Leuzinger, Julia Bielicki, Ingrid Steffen, Manuel Battegay, Hans Hirsch, Adrian Egli |
| EPI_ISL_3451734, EPI_ISL_3451735, EPI_ISL_3451736 | Synlab Suisse SA | Clinical Bacteriology | Tim Roloff, Fanny Wegner, Helena MB Seth-Smith, Alfredo Mari, Karoline Leuzinger, Julia Bielicki, Manuel Battegay, Hans Hirsch, Adrian Egli |
| EPI_ISL_3451737 | Clinical Virology | Clinical Bacteriology | Tim Roloff, Fanny Wegner, Helena MB Seth-Smith, Alfredo Mari, Karoline Leuzinger, Julia Bielicki, Manuel Battegay, Hans Hirsch, Adrian Egli |
| EPI_ISL_3451738 | Unilabs | Institute of Medical Virology | Daniel Ehrsam, Isabel Stürmer, Catharine Aquino, Joel Wirz, Weihong Qi, Hubert Rehrauer, Verena Kufner, Gabriela Ziltener, Maryam Zaheri, Stefan Schmutz, Annette Audigé, Maria Grünberg, Kevin Steiner, Jon Huder, Cyril Shah, Riccarda Capaul, Guido Bloemberg, Jürg Böni, Michael Huber, Alexandra Trkola |
| EPI_ISL_3451739 | UniversitätsSpital Zürich | Institute of Medical Virology | Daniel Ehrsam, Isabel Stürmer, Catharine Aquino, Joel Wirz, Weihong Qi, Hubert Rehrauer, Verena Kufner, Gabriela Ziltener, Maryam Zaheri, Stefan Schmutz, Annette Audigé, Maria Grünberg, Kevin Steiner, Jon Huder, Cyril Shah, Riccarda Capaul, Guido Bloemberg, Jürg Böni, Michael Huber, Alexandra Trkola |
| EPI_ISL_3451740, EPI_ISL_3451741 | Stadtspital Triemli | Institute of Medical Virology | Daniel Ehrsam, Isabel Stürmer, Catharine Aquino, Joel Wirz, Weihong Qi, Hubert Rehrauer, Verena Kufner, Gabriela Ziltener, Maryam Zaheri, Stefan Schmutz, Annette Audigé, Maria Grünberg, Kevin Steiner, Jon Huder, Cyril Shah, Riccarda Capaul, Guido Bloemberg, Jürg Böni, Michael Huber, Alexandra Trkola |
| EPI_ISL_3451742, EPI_ISL_3451743, EPI_ISL_3451744, EPI_ISL_3451745 | UniversitätsSpital Zürich | Institute of Medical Virology | Daniel Ehrsam, Isabel Stürmer, Catharine Aquino, Joel Wirz, Weihong Qi, Hubert Rehrauer, Verena Kufner, Gabriela Ziltener, Maryam Zaheri, Stefan Schmutz, Annette Audigé, Maria Grünberg, Kevin Steiner, Jon Huder, Cyril Shah, Riccarda Capaul, Guido Bloemberg, Jürg Böni, Michael Huber, Alexandra Trkola |
| EPI_ISL_3451746 | Universitäts-Kinderspital Zürich | Institute of Medical Virology | Daniel Ehrsam, Isabel Stürmer, Catharine Aquino, Joel Wirz, Weihong Qi, Hubert Rehrauer, Verena Kufner, Gabriela Ziltener, Maryam Zaheri, Stefan Schmutz, Annette Audigé, Maria Grünberg, Kevin Steiner, Jon Huder, Cyril Shah, Riccarda Capaul, Guido Bloemberg, Jürg Böni, Michael Huber, Alexandra Trkola |
| EPI_ISL_3451747, EPI_ISL_3451748, EPI_ISL_3451749, EPI_ISL_3451750 | Universität Zürich | Institute of Medical Virology | Daniel Ehrsam, Isabel Stürmer, Catharine Aquino, Joel Wirz, Weihong Qi, Hubert Rehrauer, Verena Kufner, Gabriela Ziltener, Maryam Zaheri, Stefan Schmutz, Annette Audigé, Maria Grünberg, Kevin Steiner, Jon Huder, Cyril Shah, Riccarda Capaul, Guido Bloemberg, Jürg Böni, Michael Huber, Alexandra Trkola |
| EPI_ISL_3451752 | UniversitätsSpital Zürich | Institute of Medical Virology | Daniel Ehrsam, Isabel Stürmer, Catharine Aquino, Joel Wirz, Weihong Qi, Hubert Rehrauer, Verena Kufner, Gabriela Ziltener, Maryam Zaheri, Stefan Schmutz, Annette Audigé, Maria Grünberg, Kevin Steiner, Jon Huder, Cyril Shah, Riccarda Capaul, Guido Bloemberg, Jürg Böni, Michael Huber, Alexandra Trkola |
| EPI_ISL_3451753, EPI_ISL_3451754 | Zentrallabor Zürich | Institute of Medical Virology | Daniel Ehrsam, Isabel Stürmer, Catharine Aquino, Joel Wirz, Weihong Qi, Hubert Rehrauer, Verena Kufner, Gabriela Ziltener, Maryam Zaheri, Stefan Schmutz, Annette Audigé, Maria Grünberg, Kevin Steiner, Jon Huder, Cyril Shah, Riccarda Capaul, Guido Bloemberg, Jürg Böni, Michael Huber, Alexandra Trkola |
| EPI_ISL_3451755, EPI_ISL_3451756, EPI_ISL_3451757, EPI_ISL_3451758, EPI_ISL_3451759 | Stadtspital Triemli | Institute of Medical Virology | Daniel Ehrsam, Isabel Stürmer, Catharine Aquino, Joel Wirz, Weihong Qi, Hubert Rehrauer, Verena Kufner, Gabriela Ziltener, Maryam Zaheri, Stefan Schmutz, Annette Audigé, Maria Grünberg, Kevin Steiner, Jon Huder, Cyril Shah, Riccarda Capaul, Guido Bloemberg, Jürg Böni, Michael Huber, Alexandra Trkola |
| EPI_ISL_3451760 | Universität Zürich | Institute of Medical Virology | Daniel Ehrsam, Isabel Stürmer, Catharine Aquino, Joel Wirz, Weihong Qi, Hubert Rehrauer, Verena Kufner, Gabriela Ziltener, Maryam Zaheri, Stefan Schmutz, Annette Audigé, Maria Grünberg, Kevin Steiner, Jon Huder, Cyril Shah, Riccarda Capaul, Guido Bloemberg, Jürg Böni, Michael Huber, Alexandra Trkola |
| EPI_ISL_3451761, EPI_ISL_3451762 | Universitäts-Kinderspital Zürich | Institute of Medical Virology | Daniel Ehrsam, Isabel Stürmer, Catharine Aquino, Joel Wirz, Weihong Qi, Hubert Rehrauer, Verena Kufner, Gabriela Ziltener, Maryam Zaheri, Stefan Schmutz, Annette Audigé, Maria Grünberg, Kevin Steiner, Jon Huder, Cyril Shah, Riccarda Capaul, Guido Bloemberg, Jürg Böni, Michael Huber, Alexandra Trkola |
| EPI_ISL_3451763, EPI_ISL_3451764, EPI_ISL_3451765 | Universität Zürich | Institute of Medical Virology | Daniel Ehrsam, Isabel Stürmer, Catharine Aquino, Joel Wirz, Weihong Qi, Hubert Rehrauer, Verena Kufner, Gabriela Ziltener, Maryam Zaheri, Stefan Schmutz, Annette Audigé, Maria Grünberg, Kevin Steiner, Jon Huder, Cyril Shah, Riccarda Capaul, Guido Bloemberg, Jürg Böni, Michael Huber, Alexandra Trkola |
| EPI_ISL_3451766, EPI_ISL_3451767 | UniversitätsSpital Zürich | Institute of Medical Virology | Daniel Ehrsam, Isabel Stürmer, Catharine Aquino, Joel Wirz, Weihong Qi, Hubert Rehrauer, Verena Kufner, Gabriela Ziltener, Maryam Zaheri, Stefan Schmutz, Annette Audigé, Maria Grünberg, Kevin Steiner, Jon Huder, Cyril Shah, Riccarda Capaul, Guido Bloemberg, Jürg Böni, Michael Huber, Alexandra Trkola |
| EPI_ISL_3451768 | Universitäts-Kinderspital Zürich | Institute of Medical Virology | Daniel Ehrsam, Isabel Stürmer, Catharine Aquino, Joel Wirz, Weihong Qi, Hubert Rehrauer, Verena Kufner, Gabriela Ziltener, Maryam Zaheri, Stefan Schmutz, Annette Audigé, Maria Grünberg, Kevin Steiner, Jon Huder, Cyril Shah, Riccarda Capaul, Guido Bloemberg, Jürg Böni, Michael Huber, Alexandra Trkola |
| EPI_ISL_3451769, EPI_ISL_3451770, EPI_ISL_3451771, EPI_ISL_3451772, EPI_ISL_3451773, EPI_ISL_3451774, EPI_ISL_3451775, EPI_ISL_3451776, EPI_ISL_3451777, EPI_ISL_3451778, EPI_ISL_3451779, EPI_ISL_3451780, EPI_ISL_3451781, EPI_ISL_3451782, EPI_ISL_3451783, EPI_ISL_3451784, EPI_ISL_3451785, EPI_ISL_3451786, EPI_ISL_3451787, EPI_ISL_3451788, EPI_ISL_3451789, EPI_ISL_3451791, EPI_ISL_3451792, EPI_ISL_3451793, EPI_ISL_3451794, EPI_ISL_3451795, EPI_ISL_3451796, EPI_ISL_3451797, EPI_ISL_3451798, EPI_ISL_3451799, EPI_ISL_3451801, EPI_ISL_3451802 | | | |
| see above | Kantonsspital Winterthur | Institute of Medical Virology | Daniel Ehrsam, Isabel Stürmer, Catharine Aquino, Joel Wirz, Weihong Qi, Hubert Rehrauer, Verena Kufner, Gabriela Ziltener, Maryam Zaheri, Stefan Schmutz, Annette Audigé, Maria Grünberg, Kevin Steiner, Jon Huder, Cyril Shah, Riccarda Capaul, Guido Bloemberg, Jürg Böni, Michael Huber, Alexandra Trkola |
| EPI_ISL_3451803, EPI_ISL_3451804, EPI_ISL_3451805, EPI_ISL_3451806, EPI_ISL_3451807 | Stadtspital Triemli | Institute of Medical Virology | Daniel Ehrsam, Isabel Stürmer, Catharine Aquino, Joel Wirz, Weihong Qi, Hubert Rehrauer, Verena Kufner, Gabriela Ziltener, Maryam Zaheri, Stefan Schmutz, Annette Audigé, Maria Grünberg, Kevin Steiner, Jon Huder, Cyril Shah, Riccarda Capaul, Guido Bloemberg, Jürg Böni, Michael Huber, Alexandra Trkola |
| EPI_ISL_3451808, EPI_ISL_3451809, EPI_ISL_3451810 | UniversitätsSpital Zürich | Institute of Medical Virology | Daniel Ehrsam, Isabel Stürmer, Catharine Aquino, Joel Wirz, Weihong Qi, Hubert Rehrauer, Verena Kufner, Gabriela Ziltener, Maryam Zaheri, Stefan Schmutz, Annette Audigé, Maria Grünberg, Kevin Steiner, Jon Huder, Cyril Shah, Riccarda Capaul, Guido Bloemberg, Jürg Böni, Michael Huber, Alexandra Trkola |
| EPI_ISL_3451811, EPI_ISL_3451812, EPI_ISL_3451813, EPI_ISL_3451814, EPI_ISL_3451815 | Universität Zürich | Institute of Medical Virology | Daniel Ehrsam, Isabel Stürmer, Catharine Aquino, Joel Wirz, Weihong Qi, Hubert Rehrauer, Verena Kufner, Gabriela Ziltener, Maryam Zaheri, Stefan Schmutz, Annette Audigé, Maria Grünberg, Kevin Steiner, Jon Huder, Cyril Shah, Riccarda Capaul, Guido Bloemberg, Jürg Böni, Michael Huber, Alexandra Trkola |
| EPI_ISL_3451816 | Zentrallabor Zürich | Institute of Medical Virology | Daniel Ehrsam, Isabel Stürmer, Catharine Aquino, Joel Wirz, Weihong Qi, Hubert Rehrauer, Verena Kufner, Gabriela Ziltener, Maryam Zaheri, Stefan Schmutz, Annette Audigé, Maria Grünberg, Kevin Steiner, Jon Huder, Cyril Shah, Riccarda Capaul, Guido Bloemberg, Jürg Böni, Michael Huber, Alexandra Trkola |
| EPI_ISL_3451817, EPI_ISL_3451818, EPI_ISL_3451819 | Universitäts-Kinderspital Zürich | Institute of Medical Virology | Daniel Ehrsam, Isabel Stürmer, Catharine Aquino, Joel Wirz, Weihong Qi, Hubert Rehrauer, Verena Kufner, Gabriela Ziltener, Maryam Zaheri, Stefan Schmutz, Annette Audigé, Maria Grünberg, Kevin Steiner, Jon Huder, Cyril Shah, Riccarda Capaul, Guido Bloemberg, Jürg Böni, Michael Huber, Alexandra Trkola |
| EPI_ISL_3451828, EPI_ISL_3451829, EPI_ISL_3451830, EPI_ISL_3451832, EPI_ISL_3451833, EPI_ISL_3451834, EPI_ISL_3451839, EPI_ISL_3451840, EPI_ISL_3451841 | Pennsylvania Department of Health Bureau of Laboratories | Pennsylvania Department of Health Bureau of Laboratories | Dongxiang Xia |
| EPI_ISL_3451848, EPI_ISL_3451849, EPI_ISL_3451850, EPI_ISL_3451851, EPI_ISL_3451852, EPI_ISL_3451853 | Stadtspital Triemli | Institute of Medical Virology | Daniel Ehrsam, Isabel Stürmer, Catharine Aquino, Joel Wirz, Weihong Qi, Hubert Rehrauer, Verena Kufner, Gabriela Ziltener, Maryam Zaheri, Stefan Schmutz, Annette Audigé, Maria Grünberg, Kevin Steiner, Jon Huder, Cyril Shah, Riccarda Capaul, Guido Bloemberg, Jürg Böni, Michael Huber, Alexandra Trkola |
| EPI_ISL_3451854 | Universitäts-Kinderspital Zürich | Institute of Medical Virology | Daniel Ehrsam, Isabel Stürmer, Catharine Aquino, Joel Wirz, Weihong Qi, Hubert Rehrauer, Verena Kufner, Gabriela Ziltener, Maryam Zaheri, Stefan Schmutz, Annette Audigé, Maria Grünberg, Kevin Steiner, Jon Huder, Cyril Shah, Riccarda Capaul, Guido Bloemberg, Jürg Böni, Michael Huber, Alexandra Trkola |
| EPI_ISL_3451855 | Universität Zürich | Institute of Medical Virology | Daniel Ehrsam, Isabel Stürmer, Catharine Aquino, Joel Wirz, Weihong Qi, Hubert Rehrauer, Verena Kufner, Gabriela Ziltener, Maryam Zaheri, Stefan Schmutz, Annette Audigé, Maria Grünberg, Kevin Steiner, Jon Huder, Cyril Shah, Riccarda Capaul, Guido Bloemberg, Jürg Böni, Michael Huber, Alexandra Trkola |
| EPI_ISL_3451856, EPI_ISL_3451857 | UniversitätsSpital Zürich | Institute of Medical Virology | Daniel Ehrsam, Isabel Stürmer, Catharine Aquino, Joel Wirz, Weihong Qi, Hubert Rehrauer, Verena Kufner, Gabriela Ziltener, Maryam Zaheri, Stefan Schmutz, Annette Audigé, Maria Grünberg, Kevin Steiner, Jon Huder, Cyril Shah, Riccarda Capaul, Guido Bloemberg, Jürg Böni, Michael Huber, Alexandra Trkola |
| EPI_ISL_3451858, EPI_ISL_3451859, EPI_ISL_3451860, EPI_ISL_3451861, EPI_ISL_3451862, EPI_ISL_3451863, EPI_ISL_3451864, EPI_ISL_3451865 | Stadtspital Triemli | Institute of Medical Virology | Daniel Ehrsam, Isabel Stürmer, Catharine Aquino, Joel Wirz, Weihong Qi, Hubert Rehrauer, Verena Kufner, Gabriela Ziltener, Maryam Zaheri, Stefan Schmutz, Annette Audigé, Maria Grünberg, Kevin Steiner, Jon Huder, Cyril Shah, Riccarda Capaul, Guido Bloemberg, Jürg Böni, Michael Huber, Alexandra Trkola |
| EPI_ISL_3451866 | Universität Zürich | Institute of Medical Virology | Daniel Ehrsam, Isabel Stürmer, Catharine Aquino, Joel Wirz, Weihong Qi, Hubert Rehrauer, Verena Kufner, Gabriela Ziltener, Maryam Zaheri, Stefan Schmutz, Annette Audigé, Maria Grünberg, Kevin Steiner, Jon Huder, Cyril Shah, Riccarda Capaul, Guido Bloemberg, Jürg Böni, Michael Huber, Alexandra Trkola |

| | | | |
|---|---|---|---|
| EPI_ISL_3451867, EPI_ISL_3451868, EPI_ISL_3451869, EPI_ISL_3451870, EPI_ISL_3451871 | UniversitätsSpital Zürich | Institute of Medical Virology | Daniel Ehrsam, Isabel Stürmer, Catharine Aquino, Joel Wirz, Weihong Qi, Hubert Rehrauer, Verena Kufner, Gabriela Ziltener, Maryam Zaheri, Stefan Schmutz, Annette Audigé, Maria Grünberg, Kevin Steiner, Jon Huder, Cyril Shah, Riccarda Capaul, Guido Bloemberg, Jürg Böni, Michael Huber, Alexandra Trkola |
| EPI_ISL_3451872, EPI_ISL_3451873, EPI_ISL_3451874 | Universität Zürich | Institute of Medical Virology | Daniel Ehrsam, Isabel Stürmer, Catharine Aquino, Joel Wirz, Weihong Qi, Hubert Rehrauer, Verena Kufner, Gabriela Ziltener, Maryam Zaheri, Stefan Schmutz, Annette Audigé, Maria Grünberg, Kevin Steiner, Jon Huder, Cyril Shah, Riccarda Capaul, Guido Bloemberg, Jürg Böni, Michael Huber, Alexandra Trkola |
| EPI_ISL_3451875, EPI_ISL_3451876, EPI_ISL_3451877, EPI_ISL_3451880, EPI_ISL_3451881, EPI_ISL_3451882, EPI_ISL_3451883, EPI_ISL_3451884, EPI_ISL_3451885, EPI_ISL_3451886, EPI_ISL_3451887, EPI_ISL_3451888, EPI_ISL_3451890, EPI_ISL_3451891, EPI_ISL_3451892, EPI_ISL_3451893, EPI_ISL_3451894, EPI_ISL_3451895, EPI_ISL_3451896, EPI_ISL_3451897, EPI_ISL_3451898, EPI_ISL_3451900, EPI_ISL_3451901, EPI_ISL_3451903, EPI_ISL_3451904, EPI_ISL_3451907, EPI_ISL_3451908, EPI_ISL_3451909, EPI_ISL_3451911, EPI_ISL_3451912, EPI_ISL_3451913, EPI_ISL_3451914, EPI_ISL_3451915, EPI_ISL_3451917, EPI_ISL_3451919, EPI_ISL_3451920, EPI_ISL_3451921, EPI_ISL_3451924, EPI_ISL_3451925, EPI_ISL_3451926, EPI_ISL_3451927, EPI_ISL_3451928, EPI_ISL_3451951, EPI_ISL_3451952, EPI_ISL_3451954, EPI_ISL_3451959, EPI_ISL_3451960, EPI_ISL_3451961, EPI_ISL_3451962, EPI_ISL_3451967, EPI_ISL_3451968, EPI_ISL_3451969, EPI_ISL_3451970, EPI_ISL_3451971, EPI_ISL_3451972, EPI_ISL_3451973, EPI_ISL_3451974, EPI_ISL_3451975, EPI_ISL_3451976, EPI_ISL_3451977, EPI_ISL_3451979, EPI_ISL_3451980, EPI_ISL_3451981, EPI_ISL_3451983, EPI_ISL_3451984, EPI_ISL_3451985, EPI_ISL_3451989, EPI_ISL_3451990, EPI_ISL_3451991, EPI_ISL_3451992, EPI_ISL_3451993, EPI_ISL_3451997, EPI_ISL_3451999, EPI_ISL_3452002, EPI_ISL_3452003, EPI_ISL_3452004, EPI_ISL_3452005, EPI_ISL_3452006, EPI_ISL_3452007, EPI_ISL_3452008, EPI_ISL_3452010, EPI_ISL_3452011, EPI_ISL_3452013, EPI_ISL_3452014, EPI_ISL_3452015, EPI_ISL_3452016, EPI_ISL_3452018, EPI_ISL_3452019, EPI_ISL_3452021, EPI_ISL_3452022, EPI_ISL_3452023, EPI_ISL_3452025, EPI_ISL_3452026, EPI_ISL_3452027, EPI_ISL_3452028, EPI_ISL_3452029, EPI_ISL_3452030, EPI_ISL_3452031, EPI_ISL_3452032, EPI_ISL_3452033, EPI_ISL_3452034, EPI_ISL_3452035, EPI_ISL_3452036, EPI_ISL_3452037, EPI_ISL_3452038, EPI_ISL_3452039, EPI_ISL_3452040, EPI_ISL_3452041, EPI_ISL_3452042, EPI_ISL_3452043, EPI_ISL_3452044, EPI_ISL_3452045, EPI_ISL_3452046, EPI_ISL_3452047, EPI_ISL_3452048, EPI_ISL_3452049, EPI_ISL_3452050, EPI_ISL_3452051, EPI_ISL_3452052, EPI_ISL_3452053, EPI_ISL_3452054, EPI_ISL_3452055, EPI_ISL_3452057, EPI_ISL_3452059, EPI_ISL_3452060, EPI_ISL_3452061, EPI_ISL_3452062, EPI_ISL_3452064, EPI_ISL_3452065, EPI_ISL_3452066, EPI_ISL_3452068, EPI_ISL_3452069, EPI_ISL_3452070, EPI_ISL_3452071, EPI_ISL_3452072, EPI_ISL_3452073, EPI_ISL_3452074, EPI_ISL_3452076, EPI_ISL_3452077, EPI_ISL_3452078, EPI_ISL_3452079, EPI_ISL_3452080, EPI_ISL_3452081, EPI_ISL_3452083, EPI_ISL_3452084, EPI_ISL_3452086, EPI_ISL_3452087, EPI_ISL_3452088, EPI_ISL_3452089, EPI_ISL_3452090, EPI_ISL_3452092, EPI_ISL_3452095, EPI_ISL_3452100, EPI_ISL_3452102, EPI_ISL_3452103, EPI_ISL_3452105, EPI_ISL_3452116, EPI_ISL_3452119, EPI_ISL_3452122, EPI_ISL_3452125, EPI_ISL_3452126, EPI_ISL_3452127, EPI_ISL_3452131, EPI_ISL_3452139, EPI_ISL_3452140, EPI_ISL_3452143, EPI_ISL_3452145, EPI_ISL_3452146, EPI_ISL_3452148, EPI_ISL_3452149, EPI_ISL_3452152, EPI_ISL_3452154, EPI_ISL_3452155, EPI_ISL_3452158, EPI_ISL_3452160, EPI_ISL_3452162, EPI_ISL_3452163, EPI_ISL_3452165, EPI_ISL_3452169, EPI_ISL_3452171, EPI_ISL_3452172, EPI_ISL_3452174, EPI_ISL_3452176, EPI_ISL_3452177, EPI_ISL_3452179, EPI_ISL_3452180, EPI_ISL_3452182, EPI_ISL_3452183, EPI_ISL_3452185, EPI_ISL_3452187, EPI_ISL_3452188, EPI_ISL_3452189, EPI_ISL_3452191, EPI_ISL_3452193, EPI_ISL_3452194, EPI_ISL_3452197, EPI_ISL_3452199, EPI_ISL_3452200, EPI_ISL_3452203, EPI_ISL_3452205, EPI_ISL_3452206, EPI_ISL_3452208, EPI_ISL_3452210, EPI_ISL_3452211, EPI_ISL_3452213, EPI_ISL_3452214, EPI_ISL_3452216, EPI_ISL_3452218, EPI_ISL_3452219, EPI_ISL_3452221, EPI_ISL_3452224, EPI_ISL_3452227, EPI_ISL_3452228, EPI_ISL_3452230, EPI_ISL_3452231, EPI_ISL_3452234, EPI_ISL_3452236, EPI_ISL_3452237, EPI_ISL_3452239, EPI_ISL_3452242, EPI_ISL_3452244, EPI_ISL_3452245, EPI_ISL_3452247, EPI_ISL_3452248, EPI_ISL_3452250, EPI_ISL_3452251, EPI_ISL_3452256, EPI_ISL_3452257, EPI_ISL_3452259, EPI_ISL_3452263, EPI_ISL_3452265, EPI_ISL_3452268, EPI_ISL_3452270, EPI_ISL_3452271, EPI_ISL_3452273, EPI_ISL_3452275, EPI_ISL_3452276, EPI_ISL_3452278, EPI_ISL_3452279, EPI_ISL_3452281, EPI_ISL_3452283, EPI_ISL_3452284, EPI_ISL_3452286, EPI_ISL_3452287, EPI_ISL_3452289, EPI_ISL_3452290, EPI_ISL_3452291, EPI_ISL_3452293, EPI_ISL_3452295, EPI_ISL_3452296, EPI_ISL_3452298, EPI_ISL_3452299, EPI_ISL_3452301, EPI_ISL_3452304, EPI_ISL_3452306, EPI_ISL_3452307, EPI_ISL_3452309, EPI_ISL_3452311, EPI_ISL_3452312, EPI_ISL_3452314, EPI_ISL_3452316, EPI_ISL_3452318, EPI_ISL_3452319, EPI_ISL_3452321, EPI_ISL_3452324, EPI_ISL_3452325, EPI_ISL_3452327, EPI_ISL_3452329, EPI_ISL_3452331, EPI_ISL_3452334, EPI_ISL_3452337, EPI_ISL_3452339, EPI_ISL_3452340, EPI_ISL_3452342, EPI_ISL_3452345, EPI_ISL_3452348, EPI_ISL_3452349, EPI_ISL_3452351, EPI_ISL_3452353, EPI_ISL_3452354, EPI_ISL_3452356, EPI_ISL_3452359, EPI_ISL_3452362, EPI_ISL_3452364, EPI_ISL_3452365, EPI_ISL_3452367, EPI_ISL_3452373, EPI_ISL_3452376, EPI_ISL_3452378, EPI_ISL_3452379, EPI_ISL_3452381, EPI_ISL_3452384, EPI_ISL_3452385, EPI_ISL_3452387, EPI_ISL_3452389, EPI_ISL_3452390, EPI_ISL_3452392, EPI_ISL_3452393, EPI_ISL_3452396, EPI_ISL_3452401, EPI_ISL_3452402, EPI_ISL_3452404, EPI_ISL_3452405, EPI_ISL_3452407, EPI_ISL_3452411, EPI_ISL_3452413, EPI_ISL_3452416, EPI_ISL_3452419, EPI_ISL_3452421, EPI_ISL_3452422, EPI_ISL_3452425, EPI_ISL_3452427, EPI_ISL_3452429, EPI_ISL_3452430, EPI_ISL_3452432, EPI_ISL_3452433, EPI_ISL_3452435, EPI_ISL_3452436, EPI_ISL_3452438, EPI_ISL_3452439, EPI_ISL_3452440, EPI_ISL_3452441, EPI_ISL_3452442, EPI_ISL_3452443, EPI_ISL_3452444, EPI_ISL_3452445, EPI_ISL_3452446, EPI_ISL_3452447, EPI_ISL_3452448, EPI_ISL_3452449, EPI_ISL_3452450, EPI_ISL_3452451, EPI_ISL_3452452, EPI_ISL_3452453, EPI_ISL_3452454, EPI_ISL_3452455, EPI_ISL_3452456, EPI_ISL_3452457, EPI_ISL_3452458, EPI_ISL_3452461, EPI_ISL_3452462, EPI_ISL_3452463, EPI_ISL_3452464, EPI_ISL_3452465, EPI_ISL_3452468, EPI_ISL_3452469, EPI_ISL_3452470, EPI_ISL_3452471, EPI_ISL_3452472, EPI_ISL_3452473, EPI_ISL_3452474, EPI_ISL_3452475, EPI_ISL_3452476, EPI_ISL_3452477, EPI_ISL_3452478, EPI_ISL_3452479, EPI_ISL_3452480, EPI_ISL_3452481, EPI_ISL_3452482, EPI_ISL_3452483, EPI_ISL_3452484, EPI_ISL_3452485, EPI_ISL_3452486, EPI_ISL_3452489, EPI_ISL_3452490, EPI_ISL_3452491, EPI_ISL_3452492, EPI_ISL_3452494, EPI_ISL_3452495, EPI_ISL_3452496 | | | |
| see above | Division of Emerging Infectious Diseases, Bureau of Infectious Diseases Diagnosis Control, Korea Disease Control and Prevention Agency | Division of Emerging Infectious Diseases, Bureau of Infectious Diseases Diagnosis Control, Korea Disease Control and Prevention Agency | Ae Kyung Park, Il-Hwan Kim, Heui Man Kim, Jeong-Ah Kim, Chae Young Lee, Eun-Jin Kim |
| EPI_ISL_3452497 | Laboratori de Referencia de Catalunya | Laboratori de Referencia de Catalunya | Ramirez A., Bellosillo B., Padilla E., Vilas A., Hernandez JJ., Canal M. |
| EPI_ISL_3452499, EPI_ISL_3452500, EPI_ISL_3452501, EPI_ISL_3452502, EPI_ISL_3452503, EPI_ISL_3452504, EPI_ISL_3452505, EPI_ISL_3452506, EPI_ISL_3452507, EPI_ISL_3452508, EPI_ISL_3452509, EPI_ISL_3452510, EPI_ISL_3452511, EPI_ISL_3452512, EPI_ISL_3452513, EPI_ISL_3452514, EPI_ISL_3452515, EPI_ISL_3452516, EPI_ISL_3452517, EPI_ISL_3452518, EPI_ISL_3452519, EPI_ISL_3452520, EPI_ISL_3452521, EPI_ISL_3452522, EPI_ISL_3452523, EPI_ISL_3452524, EPI_ISL_3452525, EPI_ISL_3452526, EPI_ISL_3452527, EPI_ISL_3452528, EPI_ISL_3452529, EPI_ISL_3452530, EPI_ISL_3452531, EPI_ISL_3452533, EPI_ISL_3452534, EPI_ISL_3452535, EPI_ISL_3452536, EPI_ISL_3452537, EPI_ISL_3452538, EPI_ISL_3452539, EPI_ISL_3452540, EPI_ISL_3452541, EPI_ISL_3452542, EPI_ISL_3452543, EPI_ISL_3452544, EPI_ISL_3452545, EPI_ISL_3452546 | | | |
| see above | MD PHL | Maryland Department of Health Laboratories Administration | Eric N. Keller, Jillian Loomis, Kwang Low, Terence L. Moore, Ami Patel, and Robert Myers |
| EPI_ISL_3452547, EPI_ISL_3452548, EPI_ISL_3452549, EPI_ISL_3452550, EPI_ISL_3452551, EPI_ISL_3452552, EPI_ISL_3452553, EPI_ISL_3452554, EPI_ISL_3452555, EPI_ISL_3452556, EPI_ISL_3452557, EPI_ISL_3452558, EPI_ISL_3452559, EPI_ISL_3452560, EPI_ISL_3452561, EPI_ISL_3452562, EPI_ISL_3452563, EPI_ISL_3452564, EPI_ISL_3452565, EPI_ISL_3452566, EPI_ISL_3452567, EPI_ISL_3452568, EPI_ISL_3452569, EPI_ISL_3452570, EPI_ISL_3452571, EPI_ISL_3452572, EPI_ISL_3452573, EPI_ISL_3452574, EPI_ISL_3452575, EPI_ISL_3452576, EPI_ISL_3452577, EPI_ISL_3452578, EPI_ISL_3452579, EPI_ISL_3452580, EPI_ISL_3452581, EPI_ISL_3452582, EPI_ISL_3452583, EPI_ISL_3452584, EPI_ISL_3452585, EPI_ISL_3452586, EPI_ISL_3452587, EPI_ISL_3452588, EPI_ISL_3452589, EPI_ISL_3452590, EPI_ISL_3452591, EPI_ISL_3452592, EPI_ISL_3452593, EPI_ISL_3452594, EPI_ISL_3452595, EPI_ISL_3452596, EPI_ISL_3452597, EPI_ISL_3452598, EPI_ISL_3452599, EPI_ISL_3452600, EPI_ISL_3452601, EPI_ISL_3452602, EPI_ISL_3452603, EPI_ISL_3452604, EPI_ISL_3452605, EPI_ISL_3452606, EPI_ISL_3452607, EPI_ISL_3452608, EPI_ISL_3452609, EPI_ISL_3452610, EPI_ISL_3452611, EPI_ISL_3452612, EPI_ISL_3452613, EPI_ISL_3452614, EPI_ISL_3452615, EPI_ISL_3452616, EPI_ISL_3452617, EPI_ISL_3452618, EPI_ISL_3452619, EPI_ISL_3452620, EPI_ISL_3452621, EPI_ISL_3452622, EPI_ISL_3452623, EPI_ISL_3452624, EPI_ISL_3452625, EPI_ISL_3452626, EPI_ISL_3452627, EPI_ISL_3452628, EPI_ISL_3452629, EPI_ISL_3452631, EPI_ISL_3452632, EPI_ISL_3452633, EPI_ISL_3452634, EPI_ISL_3452635, EPI_ISL_3452636, EPI_ISL_3452637, EPI_ISL_3452638, EPI_ISL_3452639, EPI_ISL_3452640, EPI_ISL_3452641, EPI_ISL_3452642, EPI_ISL_3452643, EPI_ISL_3452644, EPI_ISL_3452645, EPI_ISL_3452646, EPI_ISL_3452647, EPI_ISL_3452648, EPI_ISL_3452649, EPI_ISL_3452650, EPI_ISL_3452651, EPI_ISL_3452652, EPI_ISL_3452653, EPI_ISL_3452654, EPI_ISL_3452655, EPI_ISL_3452656, EPI_ISL_3452657, EPI_ISL_3452658, EPI_ISL_3452659, EPI_ISL_3452660, EPI_ISL_3452661, EPI_ISL_3452662, EPI_ISL_3452663, EPI_ISL_3452664, EPI_ISL_3452665, EPI_ISL_3452666, EPI_ISL_3452667, EPI_ISL_3452668, EPI_ISL_3452669, EPI_ISL_3452670, EPI_ISL_3452671, EPI_ISL_3452672, EPI_ISL_3452673, EPI_ISL_3452674, EPI_ISL_3452675, EPI_ISL_3452676, EPI_ISL_3452677, EPI_ISL_3452678, EPI_ISL_3452679, EPI_ISL_3452680, EPI_ISL_3452681, EPI_ISL_3452682, EPI_ISL_3452683, EPI_ISL_3452684, EPI_ISL_3452685, EPI_ISL_3452686, EPI_ISL_3452687, EPI_ISL_3452688, EPI_ISL_3452689, EPI_ISL_3452690, EPI_ISL_3452691, EPI_ISL_3452692, EPI_ISL_3452693, EPI_ISL_3452694, EPI_ISL_3452695, EPI_ISL_3452696, EPI_ISL_3452697, EPI_ISL_3452698, EPI_ISL_3452699, EPI_ISL_3452700, EPI_ISL_3452701, EPI_ISL_3452702, EPI_ISL_3452703, EPI_ISL_3452704, EPI_ISL_3452705, EPI_ISL_3452706, EPI_ISL_3452707, EPI_ISL_3452708, EPI_ISL_3452709, EPI_ISL_3452710, EPI_ISL_3452711, EPI_ISL_3452712 | | | |
| see above | Public Health Authority of the Slovak Republic | Laboratory of Genomics and Bioinformatics, Comenius University Science Park | Tatiana Sedláčková, Diana Rusňáková, Miroslav Böhmer, Anna Gičová, Jaroslav Budiš, Tomáš Szemes, Jakub Styk |
| EPI_ISL_3452713, EPI_ISL_3452714, EPI_ISL_3452715, EPI_ISL_3452716, EPI_ISL_3452717, EPI_ISL_3452718 | Laboratori de Referencia de Catalunya | Laboratori de Referencia de Catalunya | Ramirez A., Bellosillo B., Padilla E., Vilas A., Hernandez JJ., Canal M. |
| EPI_ISL_3452735, EPI_ISL_3452736, EPI_ISL_3452737, EPI_ISL_3452738, EPI_ISL_3452739, EPI_ISL_3452740, EPI_ISL_3452741, EPI_ISL_3452742, EPI_ISL_3452743, EPI_ISL_3452744, EPI_ISL_3452745, EPI_ISL_3452746, EPI_ISL_3452747, EPI_ISL_3452748, EPI_ISL_3452749, EPI_ISL_3452750, EPI_ISL_3452751, EPI_ISL_3452752, EPI_ISL_3452753, EPI_ISL_3452754, EPI_ISL_3452755, EPI_ISL_3452756, EPI_ISL_3452757, EPI_ISL_3452758, EPI_ISL_3452759 | | | |
| see above | CHU Purpan - Laboratoire de Virologie - Institut Fédératif de Biologie | CHU Purpan - Laboratoire de Virologie - Institut Fédératif de Biologie | Latour J., Jeanne N., Ranger N., Carcenac R., Harter A., Boyer P., Ferrer V., Tremeaux R., Izopet J. |
| EPI_ISL_3452760, EPI_ISL_3452761, EPI_ISL_3452762, EPI_ISL_3452763, EPI_ISL_3452764, EPI_ISL_3452765, EPI_ISL_3452766, EPI_ISL_3452767, EPI_ISL_3452768, EPI_ISL_3452769, EPI_ISL_3452770, EPI_ISL_3452771, EPI_ISL_3452772, EPI_ISL_3452773, EPI_ISL_3452774, EPI_ISL_3452775, EPI_ISL_3452776, EPI_ISL_3452777, EPI_ISL_3452778, EPI_ISL_3452779, EPI_ISL_3452780, EPI_ISL_3452781, EPI_ISL_3452782, EPI_ISL_3452783, EPI_ISL_3452784, EPI_ISL_3452785, EPI_ISL_3452786, EPI_ISL_3452787, EPI_ISL_3452788, EPI_ISL_3452789, EPI_ISL_3452790, EPI_ISL_3452791, EPI_ISL_3452792, EPI_ISL_3452793, EPI_ISL_3452794, EPI_ISL_3452795, EPI_ISL_3452796, EPI_ISL_3452797, EPI_ISL_3452798, EPI_ISL_3452799, EPI_ISL_3452800, EPI_ISL_3452801, EPI_ISL_3452802, EPI_ISL_3452803, EPI_ISL_3452804, EPI_ISL_3452805, EPI_ISL_3452806 | | | |
| see above | LABO BIO DOC | CHU Purpan - Laboratoire de Virologie - Institut Fédératif de Biologie | Latour J., Jeanne N., Ranger N., Carcenac R., Harter A., Boyer P., Ferrer V., Tremeaux R., Izopet J. |
| EPI_ISL_3452807, EPI_ISL_3452808 | CHU Purpan - Laboratoire de Virologie - Institut Fédératif de Biologie | CHU Purpan - Laboratoire de Virologie - Institut Fédératif de Biologie | Latour J., Jeanne N., Ranger N., Carcenac R., Harter A., Boyer P., Ferrer V., Tremeaux R., Izopet J. |
| EPI_ISL_3452810, EPI_ISL_3452811 | HOPITAL NOSTRA SENYORA MERITXELL | CHU Purpan - Laboratoire de Virologie - Institut Fédératif de Biologie | Latour J., Jeanne N., Ranger N., Carcenac R., Harter A., Boyer P., Ferrer V., Tremeaux R., Izopet J. |
| EPI_ISL_3452812, EPI_ISL_3452813, EPI_ISL_3452814, EPI_ISL_3452816, EPI_ISL_3452817, EPI_ISL_3452818, EPI_ISL_3452819, EPI_ISL_3452820, EPI_ISL_3452821, EPI_ISL_3452822, EPI_ISL_3452823, EPI_ISL_3452824, EPI_ISL_3452825, EPI_ISL_3452826, EPI_ISL_3452827, EPI_ISL_3452828, EPI_ISL_3452829, EPI_ISL_3452830, EPI_ISL_3452831, EPI_ISL_3452832, EPI_ISL_3452833, EPI_ISL_3452834, EPI_ISL_3452835, EPI_ISL_3452836, EPI_ISL_3452837 | | | |
| see above | CHU Purpan - Laboratoire de Virologie - Institut Fédératif de Biologie | CHU Purpan - Laboratoire de Virologie - Institut Fédératif de Biologie | Latour J., Jeanne N., Ranger N., Carcenac R., Harter A., Boyer P., Ferrer V., Tremeaux R., Izopet J. |
| EPI_ISL_3452838 | Biogroup Bio Lam-LCD Saint-Denis | Department of Virology, Henri Mondor University Hospital, Assistance Publique Hôpitaux de Paris, Université Paris-Est Créteil, INSERM U955 | Christophe Rodriguez, Slim Fourati, Vanessa Demontant, Guillaume Gricourt, Melissa N'Debi, Alexandre Soulier, Elisabeth Trawinski, Jean-Michel Pawlotsky |
| EPI_ISL_3452839 | SELARL BIO | Department of Virology, Henri Mondor University Hospital, Assistance Publique Hôpitaux de Paris, Université Paris-Est Créteil, INSERM U955 | Christophe Rodriguez, Slim Fourati, Vanessa Demontant, Guillaume Gricourt, Melissa N'Debi, Alexandre Soulier, Elisabeth Trawinski, Jean-Michel Pawlotsky |
| EPI_ISL_3452840 | SELAS BC-LAB | Department of Virology, Henri Mondor University Hospital, Assistance Publique Hôpitaux de Paris, Université Paris-Est Créteil, INSERM U955 | Christophe Rodriguez, Slim Fourati, Vanessa Demontant, Guillaume Gricourt, Melissa N'Debi, Alexandre Soulier, Elisabeth Trawinski, Jean-Michel Pawlotsky |
| EPI_ISL_3452841, EPI_ISL_3452842 | BROUSSAIS | Department of Virology, Henri Mondor University Hospital, Assistance Publique Hôpitaux de Paris, Université Paris-Est Créteil, INSERM U955 | Christophe Rodriguez, Slim Fourati, Vanessa Demontant, Guillaume Gricourt, Melissa N'Debi, Alexandre Soulier, Elisabeth Trawinski, Jean-Michel Pawlotsky |
| EPI_ISL_3452843 | SELAS BC-LAB | Department of Virology, Henri Mondor University Hospital, Assistance Publique Hôpitaux de Paris, Université Paris-Est Créteil, INSERM U955 | Christophe Rodriguez, Slim Fourati, Vanessa Demontant, Guillaume Gricourt, Melissa N'Debi, Alexandre Soulier, Elisabeth Trawinski, Jean-Michel Pawlotsky |
| EPI_ISL_3452844 | CERBALLIANCE Valmy | Department of Virology, Henri Mondor University Hospital, Assistance Publique Hôpitaux de Paris, Université Paris-Est Créteil, INSERM U955 | Christophe Rodriguez, Slim Fourati, Vanessa Demontant, Guillaume Gricourt, Melissa N'Debi, Alexandre Soulier, Elisabeth Trawinski, Jean-Michel Pawlotsky |
| EPI_ISL_3452845 | BROUSSAIS | Department of Virology, Henri Mondor University Hospital, Assistance Publique Hôpitaux de Paris, Université Paris-Est Créteil, INSERM U955 | Christophe Rodriguez, Slim Fourati, Vanessa Demontant, Guillaume Gricourt, Melissa N'Debi, Alexandre Soulier, Elisabeth Trawinski, Jean-Michel Pawlotsky |
| EPI_ISL_3452846 | SELAS BC-LAB | Department of Virology, Henri Mondor University Hospital, Assistance Publique Hôpitaux de Paris, Université Paris-Est Créteil, INSERM U955 | Christophe Rodriguez, Slim Fourati, Vanessa Demontant, Guillaume Gricourt, Melissa N'Debi, Alexandre Soulier, Elisabeth Trawinski, Jean-Michel Pawlotsky |
| EPI_ISL_3452847 | BROUSSAIS | Department of Virology, Henri Mondor University Hospital, Assistance Publique Hôpitaux de Paris, Université Paris-Est Créteil, INSERM U955 | Christophe Rodriguez, Slim Fourati, Vanessa Demontant, Guillaume Gricourt, Melissa N'Debi, Alexandre Soulier, Elisabeth Trawinski, Jean-Michel Pawlotsky |
| EPI_ISL_3452848, EPI_ISL_3452849 | SELAS BC-LAB | Department of Virology, Henri Mondor University Hospital, Assistance Publique Hôpitaux de Paris, Université Paris-Est Créteil, INSERM U955 | Christophe Rodriguez, Slim Fourati, Vanessa Demontant, Guillaume Gricourt, Melissa N'Debi, Alexandre Soulier, Elisabeth Trawinski, Jean-Michel Pawlotsky |
| EPI_ISL_3452850, EPI_ISL_3452851 | BROUSSAIS | Department of Virology, Henri Mondor University Hospital, Assistance Publique Hôpitaux de Paris, Université Paris-Est Créteil, | Christophe Rodriguez, Slim Fourati, Vanessa Demontant, Guillaume Gricourt, Melissa N'Debi, Alexandre Soulier, Elisabeth Trawinski, Jean-Michel Pawlotsky |

| | | | |
|---|---|---|---|
| | | INSERM U955 | |
| EPI_ISL_3452852 | BIOLLAN Brognard | Department of Virology, Henri Mondor University Hospital, Assistance Publique Hôpitaux de Paris, Université Paris-Est Créteil, INSERM U955 | Christophe Rodriguez, Slim Fourati, Vanessa Demontant, Guillaume Gricourt, Melissa N'Debi, Alexandre Soulier, Elisabeth Trawinski, Jean-Michel Pawlotsky |
| EPI_ISL_3452853, EPI_ISL_3452854, EPI_ISL_3452855 | Biogroup Bio Lam-LCD Saint-Denis | Department of Virology, Henri Mondor University Hospital, Assistance Publique Hôpitaux de Paris, Université Paris-Est Créteil, INSERM U955 | Christophe Rodriguez, Slim Fourati, Vanessa Demontant, Guillaume Gricourt, Melissa N'Debi, Alexandre Soulier, Elisabeth Trawinski, Jean-Michel Pawlotsky |
| EPI_ISL_3452856 | BROUSSAIS | Department of Virology, Henri Mondor University Hospital, Assistance Publique Hôpitaux de Paris, Université Paris-Est Créteil, INSERM U955 | Christophe Rodriguez, Slim Fourati, Vanessa Demontant, Guillaume Gricourt, Melissa N'Debi, Alexandre Soulier, Elisabeth Trawinski, Jean-Michel Pawlotsky |
| EPI_ISL_3452857 | CH. ANDRE GREGOIRE | Department of Virology, Henri Mondor University Hospital, Assistance Publique Hôpitaux de Paris, Université Paris-Est Créteil, INSERM U955 | Christophe Rodriguez, Slim Fourati, Vanessa Demontant, Guillaume Gricourt, Melissa N'Debi, Alexandre Soulier, Elisabeth Trawinski, Jean-Michel Pawlotsky |
| EPI_ISL_3452859 | Biogroup Bio Lam-LCD Saint-Denis | Department of Virology, Henri Mondor University Hospital, Assistance Publique Hôpitaux de Paris, Université Paris-Est Créteil, INSERM U955 | Christophe Rodriguez, Slim Fourati, Vanessa Demontant, Guillaume Gricourt, Melissa N'Debi, Alexandre Soulier, Elisabeth Trawinski, Jean-Michel Pawlotsky |
| EPI_ISL_3452860 | Laboratoire BIOTOV | Department of Virology, Henri Mondor University Hospital, Assistance Publique Hôpitaux de Paris, Université Paris-Est Créteil, INSERM U955 | Christophe Rodriguez, Slim Fourati, Vanessa Demontant, Guillaume Gricourt, Melissa N'Debi, Alexandre Soulier, Elisabeth Trawinski, Jean-Michel Pawlotsky |
| EPI_ISL_3452861 | Hôpital Paul Brousse | Department of Virology, Henri Mondor University Hospital, Assistance Publique Hôpitaux de Paris, Université Paris-Est Créteil, INSERM U955 | Christophe Rodriguez, Slim Fourati, Vanessa Demontant, Guillaume Gricourt, Melissa N'Debi, Alexandre Soulier, Elisabeth Trawinski, Jean-Michel Pawlotsky |
| EPI_ISL_3452862 | SELAS BC-LAB | Department of Virology, Henri Mondor University Hospital, Assistance Publique Hôpitaux de Paris, Université Paris-Est Créteil, INSERM U955 | Christophe Rodriguez, Slim Fourati, Vanessa Demontant, Guillaume Gricourt, Melissa N'Debi, Alexandre Soulier, Elisabeth Trawinski, Jean-Michel Pawlotsky |
| EPI_ISL_3452863 | Laboratoire CBM 25 TERRE ROUGE | Department of Virology, Henri Mondor University Hospital, Assistance Publique Hôpitaux de Paris, Université Paris-Est Créteil, INSERM U955 | Christophe Rodriguez, Slim Fourati, Vanessa Demontant, Guillaume Gricourt, Melissa N'Debi, Alexandre Soulier, Elisabeth Trawinski, Jean-Michel Pawlotsky |
| EPI_ISL_3452865 | CH. ANDRE GREGOIRE | Department of Virology, Henri Mondor University Hospital, Assistance Publique Hôpitaux de Paris, Université Paris-Est Créteil, INSERM U955 | Christophe Rodriguez, Slim Fourati, Vanessa Demontant, Guillaume Gricourt, Melissa N'Debi, Alexandre Soulier, Elisabeth Trawinski, Jean-Michel Pawlotsky |
| EPI_ISL_3452866 | BIOLLAN Brognard | Department of Virology, Henri Mondor University Hospital, Assistance Publique Hôpitaux de Paris, Université Paris-Est Créteil, INSERM U955 | Christophe Rodriguez, Slim Fourati, Vanessa Demontant, Guillaume Gricourt, Melissa N'Debi, Alexandre Soulier, Elisabeth Trawinski, Jean-Michel Pawlotsky |
| EPI_ISL_3452867, EPI_ISL_3452868 | Hôpital Avicenne | Department of Virology, Henri Mondor University Hospital, Assistance Publique Hôpitaux de Paris, Université Paris-Est Créteil, INSERM U955 | Christophe Rodriguez, Slim Fourati, Vanessa Demontant, Guillaume Gricourt, Melissa N'Debi, Alexandre Soulier, Elisabeth Trawinski, Jean-Michel Pawlotsky |
| EPI_ISL_3452869 | Hôpital Ambroise-Paré | Department of Virology, Henri Mondor University Hospital, Assistance Publique Hôpitaux de Paris, Université Paris-Est Créteil, INSERM U955 | Christophe Rodriguez, Slim Fourati, Vanessa Demontant, Guillaume Gricourt, Melissa N'Debi, Alexandre Soulier, Elisabeth Trawinski, Jean-Michel Pawlotsky |
| EPI_ISL_3452870 | SELAS BC-LAB | Department of Virology, Henri Mondor University Hospital, Assistance Publique Hôpitaux de Paris, Université Paris-Est Créteil, INSERM U955 | Christophe Rodriguez, Slim Fourati, Vanessa Demontant, Guillaume Gricourt, Melissa N'Debi, Alexandre Soulier, Elisabeth Trawinski, Jean-Michel Pawlotsky |
| EPI_ISL_3452871 | SELARL BIO | Department of Virology, Henri Mondor University Hospital, Assistance Publique Hôpitaux de Paris, Université Paris-Est Créteil, INSERM U955 | Christophe Rodriguez, Slim Fourati, Vanessa Demontant, Guillaume Gricourt, Melissa N'Debi, Alexandre Soulier, Elisabeth Trawinski, Jean-Michel Pawlotsky |
| EPI_ISL_3452872, EPI_ISL_3452873 | Biogroup Bio Lam-LCD Saint-Denis | Department of Virology, Henri Mondor University Hospital, Assistance Publique Hôpitaux de Paris, Université Paris-Est Créteil, INSERM U955 | Christophe Rodriguez, Slim Fourati, Vanessa Demontant, Guillaume Gricourt, Melissa N'Debi, Alexandre Soulier, Elisabeth Trawinski, Jean-Michel Pawlotsky |
| EPI_ISL_3452874 | SELAFA SANTE LABO | Department of Virology, Henri Mondor University Hospital, Assistance Publique Hôpitaux de Paris, Université Paris-Est Créteil, INSERM U955 | Christophe Rodriguez, Slim Fourati, Vanessa Demontant, Guillaume Gricourt, Melissa N'Debi, Alexandre Soulier, Elisabeth Trawinski, Jean-Michel Pawlotsky |
| EPI_ISL_3452875 | BIOLLAN Brognard | Department of Virology, Henri Mondor University Hospital, Assistance Publique Hôpitaux de Paris, Université Paris-Est Créteil, INSERM U955 | Christophe Rodriguez, Slim Fourati, Vanessa Demontant, Guillaume Gricourt, Melissa N'Debi, Alexandre Soulier, Elisabeth Trawinski, Jean-Michel Pawlotsky |
| EPI_ISL_3452876 | SELAS BC-LAB | Department of Virology, Henri Mondor University Hospital, Assistance Publique Hôpitaux de Paris, Université Paris-Est Créteil, INSERM U955 | Christophe Rodriguez, Slim Fourati, Vanessa Demontant, Guillaume Gricourt, Melissa N'Debi, Alexandre Soulier, Elisabeth Trawinski, Jean-Michel Pawlotsky |
| EPI_ISL_3452877 | GH A.CHENEVIER-H.MONDOR | Department of Virology, Henri Mondor University Hospital, Assistance Publique Hôpitaux de Paris, Université Paris-Est Créteil, INSERM U955 | Christophe Rodriguez, Slim Fourati, Vanessa Demontant, Guillaume Gricourt, Melissa N'Debi, Alexandre Soulier, Elisabeth Trawinski, Jean-Michel Pawlotsky |
| EPI_ISL_3452878 | Hôpital Avicenne | Department of Virology, Henri Mondor University Hospital, Assistance Publique Hôpitaux de Paris, Université Paris-Est Créteil, INSERM U955 | Christophe Rodriguez, Slim Fourati, Vanessa Demontant, Guillaume Gricourt, Melissa N'Debi, Alexandre Soulier, Elisabeth Trawinski, Jean-Michel Pawlotsky |
| EPI_ISL_3452879 | Biogroup Bio Lam-LCD Saint-Denis | Department of Virology, Henri Mondor University Hospital, Assistance Publique Hôpitaux de Paris, Université Paris-Est Créteil, INSERM U955 | Christophe Rodriguez, Slim Fourati, Vanessa Demontant, Guillaume Gricourt, Melissa N'Debi, Alexandre Soulier, Elisabeth Trawinski, Jean-Michel Pawlotsky |
| EPI_ISL_3452880 | SELAFA SANTE LABO | Department of Virology, Henri Mondor University Hospital, Assistance Publique Hôpitaux de Paris, Université Paris-Est Créteil, INSERM U955 | Christophe Rodriguez, Slim Fourati, Vanessa Demontant, Guillaume Gricourt, Melissa N'Debi, Alexandre Soulier, Elisabeth Trawinski, Jean-Michel Pawlotsky |
| EPI_ISL_3452881 | GH A.CHENEVIER-H.MONDOR | Department of Virology, Henri Mondor University Hospital, Assistance Publique Hôpitaux de Paris, Université Paris-Est Créteil, INSERM U955 | Christophe Rodriguez, Slim Fourati, Vanessa Demontant, Guillaume Gricourt, Melissa N'Debi, Alexandre Soulier, Elisabeth Trawinski, Jean-Michel Pawlotsky |
| EPI_ISL_3452882 | LABM DEFRANCE | Department of Virology, Henri Mondor University Hospital, Assistance Publique Hôpitaux de Paris, Université Paris-Est Créteil, INSERM U955 | Christophe Rodriguez, Slim Fourati, Vanessa Demontant, Guillaume Gricourt, Melissa N'Debi, Alexandre Soulier, Elisabeth Trawinski, Jean-Michel Pawlotsky |
| EPI_ISL_3452883 | GH A.CHENEVIER-H.MONDOR | Department of Virology, Henri Mondor University Hospital, Assistance Publique Hôpitaux de Paris, Université Paris-Est Créteil, INSERM U955 | Christophe Rodriguez, Slim Fourati, Vanessa Demontant, Guillaume Gricourt, Melissa N'Debi, Alexandre Soulier, Elisabeth Trawinski, Jean-Michel Pawlotsky |
| EPI_ISL_3452884 | SELAS BC-LAB | Department of Virology, Henri Mondor University Hospital, Assistance Publique Hôpitaux de Paris, Université Paris-Est Créteil, INSERM U955 | Christophe Rodriguez, Slim Fourati, Vanessa Demontant, Guillaume Gricourt, Melissa N'Debi, Alexandre Soulier, Elisabeth Trawinski, Jean-Michel Pawlotsky |
| EPI_ISL_3452886 | Biogroup Bio Lam-LCD Saint-Denis | Department of Virology, Henri Mondor University Hospital, Assistance Publique Hôpitaux de Paris, Université Paris-Est Créteil, INSERM U955 | Christophe Rodriguez, Slim Fourati, Vanessa Demontant, Guillaume Gricourt, Melissa N'Debi, Alexandre Soulier, Elisabeth Trawinski, Jean-Michel Pawlotsky |
| EPI_ISL_3452887 | SELAS BC-LAB | Department of Virology, Henri Mondor University Hospital, Assistance Publique Hôpitaux de Paris, Université Paris-Est Créteil, INSERM U955 | Christophe Rodriguez, Slim Fourati, Vanessa Demontant, Guillaume Gricourt, Melissa N'Debi, Alexandre Soulier, Elisabeth Trawinski, Jean-Michel Pawlotsky |
| EPI_ISL_3452888, EPI_ISL_3452889 | SELARL BIO | Department of Virology, Henri Mondor University Hospital, Assistance Publique Hôpitaux de Paris, Université Paris-Est Créteil, INSERM U955 | Christophe Rodriguez, Slim Fourati, Vanessa Demontant, Guillaume Gricourt, Melissa N'Debi, Alexandre Soulier, Elisabeth Trawinski, Jean-Michel Pawlotsky |
| EPI_ISL_3452890 | SELAS BC-LAB | Department of Virology, Henri Mondor University Hospital, Assistance Publique Hôpitaux de Paris, Université Paris-Est Créteil, INSERM U955 | Christophe Rodriguez, Slim Fourati, Vanessa Demontant, Guillaume Gricourt, Melissa N'Debi, Alexandre Soulier, Elisabeth Trawinski, Jean-Michel Pawlotsky |
| EPI_ISL_3452891 | CH. ANDRE GREGOIRE | Department of Virology, Henri Mondor University Hospital, Assistance Publique Hôpitaux de Paris, Université Paris-Est Créteil, | Christophe Rodriguez, Slim Fourati, Vanessa Demontant, Guillaume Gricourt, Melissa N'Debi, Alexandre Soulier, Elisabeth Trawinski, Jean-Michel Pawlotsky |

| | | | |
|---|---|---|---|
| | | INSERM U955 | |
| EPI_ISL_3452892 | SELAS BC-LAB | Department of Virology, Henri Mondor University Hospital, Assistance Publique Hôpitaux de Paris, Université Paris-Est Créteil, INSERM U955 | Christophe Rodriguez, Slim Fourati, Vanessa Demontant, Guillaume Gricourt, Melissa N'Debi, Alexandre Soulier, Elisabeth Trawinski, Jean-Michel Pawlotsky |
| EPI_ISL_3452893, EPI_ISL_3452894, EPI_ISL_3452895 | Biogroup Bio Lam-LCD Saint-Denis | Department of Virology, Henri Mondor University Hospital, Assistance Publique Hôpitaux de Paris, Université Paris-Est Créteil, INSERM U955 | Christophe Rodriguez, Slim Fourati, Vanessa Demontant, Guillaume Gricourt, Melissa N'Debi, Alexandre Soulier, Elisabeth Trawinski, Jean-Michel Pawlotsky |
| EPI_ISL_3452896 | SELARL BIO | Department of Virology, Henri Mondor University Hospital, Assistance Publique Hôpitaux de Paris, Université Paris-Est Créteil, INSERM U955 | Christophe Rodriguez, Slim Fourati, Vanessa Demontant, Guillaume Gricourt, Melissa N'Debi, Alexandre Soulier, Elisabeth Trawinski, Jean-Michel Pawlotsky |
| EPI_ISL_3452897 | SELAS BC-LAB | Department of Virology, Henri Mondor University Hospital, Assistance Publique Hôpitaux de Paris, Université Paris-Est Créteil, INSERM U955 | Christophe Rodriguez, Slim Fourati, Vanessa Demontant, Guillaume Gricourt, Melissa N'Debi, Alexandre Soulier, Elisabeth Trawinski, Jean-Michel Pawlotsky |
| EPI_ISL_3452898 | GH A.CHENEVIER-H.MONDOR | Department of Virology, Henri Mondor University Hospital, Assistance Publique Hôpitaux de Paris, Université Paris-Est Créteil, INSERM U955 | Christophe Rodriguez, Slim Fourati, Vanessa Demontant, Guillaume Gricourt, Melissa N'Debi, Alexandre Soulier, Elisabeth Trawinski, Jean-Michel Pawlotsky |
| EPI_ISL_3452899 | ESPACEBIO | Department of Virology, Henri Mondor University Hospital, Assistance Publique Hôpitaux de Paris, Université Paris-Est Créteil, INSERM U955 | Christophe Rodriguez, Slim Fourati, Vanessa Demontant, Guillaume Gricourt, Melissa N'Debi, Alexandre Soulier, Elisabeth Trawinski, Jean-Michel Pawlotsky |
| EPI_ISL_3452900 | Biogroup Bio Lam-LCD Saint-Denis | Department of Virology, Henri Mondor University Hospital, Assistance Publique Hôpitaux de Paris, Université Paris-Est Créteil, INSERM U955 | Christophe Rodriguez, Slim Fourati, Vanessa Demontant, Guillaume Gricourt, Melissa N'Debi, Alexandre Soulier, Elisabeth Trawinski, Jean-Michel Pawlotsky |
| EPI_ISL_3452901 | ESPACEBIO | Department of Virology, Henri Mondor University Hospital, Assistance Publique Hôpitaux de Paris, Université Paris-Est Créteil, INSERM U955 | Christophe Rodriguez, Slim Fourati, Vanessa Demontant, Guillaume Gricourt, Melissa N'Debi, Alexandre Soulier, Elisabeth Trawinski, Jean-Michel Pawlotsky |
| EPI_ISL_3452902 | SELARL BIO | Department of Virology, Henri Mondor University Hospital, Assistance Publique Hôpitaux de Paris, Université Paris-Est Créteil, INSERM U955 | Christophe Rodriguez, Slim Fourati, Vanessa Demontant, Guillaume Gricourt, Melissa N'Debi, Alexandre Soulier, Elisabeth Trawinski, Jean-Michel Pawlotsky |
| EPI_ISL_3452903 | BIOLLAN Brognard | Department of Virology, Henri Mondor University Hospital, Assistance Publique Hôpitaux de Paris, Université Paris-Est Créteil, INSERM U955 | Christophe Rodriguez, Slim Fourati, Vanessa Demontant, Guillaume Gricourt, Melissa N'Debi, Alexandre Soulier, Elisabeth Trawinski, Jean-Michel Pawlotsky |
| EPI_ISL_3452904 | Biogroup Bio Lam-LCD Saint-Denis | Department of Virology, Henri Mondor University Hospital, Assistance Publique Hôpitaux de Paris, Université Paris-Est Créteil, INSERM U955 | Christophe Rodriguez, Slim Fourati, Vanessa Demontant, Guillaume Gricourt, Melissa N'Debi, Alexandre Soulier, Elisabeth Trawinski, Jean-Michel Pawlotsky |
| EPI_ISL_3452905 | ESPACEBIO | Department of Virology, Henri Mondor University Hospital, Assistance Publique Hôpitaux de Paris, Université Paris-Est Créteil, INSERM U955 | Christophe Rodriguez, Slim Fourati, Vanessa Demontant, Guillaume Gricourt, Melissa N'Debi, Alexandre Soulier, Elisabeth Trawinski, Jean-Michel Pawlotsky |
| EPI_ISL_3452906 | SELAFA SANTE LABO | Department of Virology, Henri Mondor University Hospital, Assistance Publique Hôpitaux de Paris, Université Paris-Est Créteil, INSERM U955 | Christophe Rodriguez, Slim Fourati, Vanessa Demontant, Guillaume Gricourt, Melissa N'Debi, Alexandre Soulier, Elisabeth Trawinski, Jean-Michel Pawlotsky |
| EPI_ISL_3452907 | Biogroup Bio Lam-LCD Saint-Denis | Department of Virology, Henri Mondor University Hospital, Assistance Publique Hôpitaux de Paris, Université Paris-Est Créteil, INSERM U955 | Christophe Rodriguez, Slim Fourati, Vanessa Demontant, Guillaume Gricourt, Melissa N'Debi, Alexandre Soulier, Elisabeth Trawinski, Jean-Michel Pawlotsky |
| EPI_ISL_3452908 | Hôpital Ambroise-Paré | Department of Virology, Henri Mondor University Hospital, Assistance Publique Hôpitaux de Paris, Université Paris-Est Créteil, INSERM U955 | Christophe Rodriguez, Slim Fourati, Vanessa Demontant, Guillaume Gricourt, Melissa N'Debi, Alexandre Soulier, Elisabeth Trawinski, Jean-Michel Pawlotsky |
| EPI_ISL_3452909 | Biogroup Bio Lam-LCD Saint-Denis | Department of Virology, Henri Mondor University Hospital, Assistance Publique Hôpitaux de Paris, Université Paris-Est Créteil, INSERM U955 | Christophe Rodriguez, Slim Fourati, Vanessa Demontant, Guillaume Gricourt, Melissa N'Debi, Alexandre Soulier, Elisabeth Trawinski, Jean-Michel Pawlotsky |
| EPI_ISL_3452910 | SELARL BIO | Department of Virology, Henri Mondor University Hospital, Assistance Publique Hôpitaux de Paris, Université Paris-Est Créteil, INSERM U955 | Christophe Rodriguez, Slim Fourati, Vanessa Demontant, Guillaume Gricourt, Melissa N'Debi, Alexandre Soulier, Elisabeth Trawinski, Jean-Michel Pawlotsky |
| EPI_ISL_3452911 | Laboratoire CBM 25 TERRE ROUGE | Department of Virology, Henri Mondor University Hospital, Assistance Publique Hôpitaux de Paris, Université Paris-Est Créteil, INSERM U955 | Christophe Rodriguez, Slim Fourati, Vanessa Demontant, Guillaume Gricourt, Melissa N'Debi, Alexandre Soulier, Elisabeth Trawinski, Jean-Michel Pawlotsky |
| EPI_ISL_3452912 | Biogroup Bio Lam-LCD Saint-Denis | Department of Virology, Henri Mondor University Hospital, Assistance Publique Hôpitaux de Paris, Université Paris-Est Créteil, INSERM U955 | Christophe Rodriguez, Slim Fourati, Vanessa Demontant, Guillaume Gricourt, Melissa N'Debi, Alexandre Soulier, Elisabeth Trawinski, Jean-Michel Pawlotsky |
| EPI_ISL_3452913 | Laboratoire CBM 25 TERRE ROUGE | Department of Virology, Henri Mondor University Hospital, Assistance Publique Hôpitaux de Paris, Université Paris-Est Créteil, INSERM U955 | Christophe Rodriguez, Slim Fourati, Vanessa Demontant, Guillaume Gricourt, Melissa N'Debi, Alexandre Soulier, Elisabeth Trawinski, Jean-Michel Pawlotsky |
| EPI_ISL_3452914 | CH. CHU DE REIMS | Department of Virology, Henri Mondor University Hospital, Assistance Publique Hôpitaux de Paris, Université Paris-Est Créteil, INSERM U955 | Christophe Rodriguez, Slim Fourati, Vanessa Demontant, Guillaume Gricourt, Melissa N'Debi, Alexandre Soulier, Elisabeth Trawinski, Jean-Michel Pawlotsky |
| EPI_ISL_3452915 | Biogroup Bio Lam-LCD Saint-Denis | Department of Virology, Henri Mondor University Hospital, Assistance Publique Hôpitaux de Paris, Université Paris-Est Créteil, INSERM U955 | Christophe Rodriguez, Slim Fourati, Vanessa Demontant, Guillaume Gricourt, Melissa N'Debi, Alexandre Soulier, Elisabeth Trawinski, Jean-Michel Pawlotsky |
| EPI_ISL_3452916 | SELAS BC-LAB | Department of Virology, Henri Mondor University Hospital, Assistance Publique Hôpitaux de Paris, Université Paris-Est Créteil, INSERM U955 | Christophe Rodriguez, Slim Fourati, Vanessa Demontant, Guillaume Gricourt, Melissa N'Debi, Alexandre Soulier, Elisabeth Trawinski, Jean-Michel Pawlotsky |
| EPI_ISL_3452917 | SELARL BIO | Department of Virology, Henri Mondor University Hospital, Assistance Publique Hôpitaux de Paris, Université Paris-Est Créteil, INSERM U955 | Christophe Rodriguez, Slim Fourati, Vanessa Demontant, Guillaume Gricourt, Melissa N'Debi, Alexandre Soulier, Elisabeth Trawinski, Jean-Michel Pawlotsky |
| EPI_ISL_3452918 | GH A.CHENEVIER-H.MONDOR | Department of Virology, Henri Mondor University Hospital, Assistance Publique Hôpitaux de Paris, Université Paris-Est Créteil, INSERM U955 | Christophe Rodriguez, Slim Fourati, Vanessa Demontant, Guillaume Gricourt, Melissa N'Debi, Alexandre Soulier, Elisabeth Trawinski, Jean-Michel Pawlotsky |
| EPI_ISL_3452920 | Biogroup Bio Lam-LCD Saint-Denis | Department of Virology, Henri Mondor University Hospital, Assistance Publique Hôpitaux de Paris, Université Paris-Est Créteil, INSERM U955 | Christophe Rodriguez, Slim Fourati, Vanessa Demontant, Guillaume Gricourt, Melissa N'Debi, Alexandre Soulier, Elisabeth Trawinski, Jean-Michel Pawlotsky |
| EPI_ISL_3452921, EPI_ISL_3452922 | BIOLLAN Brognard | Department of Virology, Henri Mondor University Hospital, Assistance Publique Hôpitaux de Paris, Université Paris-Est Créteil, INSERM U955 | Christophe Rodriguez, Slim Fourati, Vanessa Demontant, Guillaume Gricourt, Melissa N'Debi, Alexandre Soulier, Elisabeth Trawinski, Jean-Michel Pawlotsky |
| EPI_ISL_3452923 | ESPACEBIO | Department of Virology, Henri Mondor University Hospital, Assistance Publique Hôpitaux de Paris, Université Paris-Est Créteil, INSERM U955 | Christophe Rodriguez, Slim Fourati, Vanessa Demontant, Guillaume Gricourt, Melissa N'Debi, Alexandre Soulier, Elisabeth Trawinski, Jean-Michel Pawlotsky |
| EPI_ISL_3452924 | Biogroup Bio Lam-LCD Saint-Denis | Department of Virology, Henri Mondor University Hospital, Assistance Publique Hôpitaux de Paris, Université Paris-Est Créteil, INSERM U955 | Christophe Rodriguez, Slim Fourati, Vanessa Demontant, Guillaume Gricourt, Melissa N'Debi, Alexandre Soulier, Elisabeth Trawinski, Jean-Michel Pawlotsky |
| EPI_ISL_3452925 | Laboratoire MAYMAT | Department of Virology, Henri Mondor University Hospital, Assistance Publique Hôpitaux de Paris, Université Paris-Est Créteil, INSERM U955 | Christophe Rodriguez, Slim Fourati, Vanessa Demontant, Guillaume Gricourt, Melissa N'Debi, Alexandre Soulier, Elisabeth Trawinski, Jean-Michel Pawlotsky |
| EPI_ISL_3452926 | Hôpital Armand-Trousseau | Department of Virology, Henri Mondor University Hospital, Assistance Publique Hôpitaux de Paris, Université Paris-Est Créteil, INSERM U955 | Christophe Rodriguez, Slim Fourati, Vanessa Demontant, Guillaume Gricourt, Melissa N'Debi, Alexandre Soulier, Elisabeth Trawinski, Jean-Michel Pawlotsky |
| EPI_ISL_3452927 | BIOLLAN Brognard | Department of Virology, Henri Mondor University Hospital, Assistance Publique Hôpitaux de Paris, Université Paris-Est Créteil, | Christophe Rodriguez, Slim Fourati, Vanessa Demontant, Guillaume Gricourt, Melissa N'Debi, Alexandre Soulier, Elisabeth Trawinski, Jean-Michel Pawlotsky |

| | | | |
|---|---|---|---|
| | | INSERM U955 | |
| EPI_ISL_3452928 | SELAFA SANTE LABO | Department of Virology, Henri Mondor University Hospital, Assistance Publique Hôpitaux de Paris, Université Paris-Est Créteil, INSERM U955 | Christophe Rodriguez, Slim Fourati, Vanessa Demontant, Guillaume Gricourt, Melissa N'Debi, Alexandre Soulier, Elisabeth Trawinski, Jean-Michel Pawlotsky |
| EPI_ISL_3452929 | Biogroup Bio Lam-LCD Saint-Denis | Department of Virology, Henri Mondor University Hospital, Assistance Publique Hôpitaux de Paris, Université Paris-Est Créteil, INSERM U955 | Christophe Rodriguez, Slim Fourati, Vanessa Demontant, Guillaume Gricourt, Melissa N'Debi, Alexandre Soulier, Elisabeth Trawinski, Jean-Michel Pawlotsky |
| EPI_ISL_3452930 | Laboratoire MAYMAT | Department of Virology, Henri Mondor University Hospital, Assistance Publique Hôpitaux de Paris, Université Paris-Est Créteil, INSERM U955 | Christophe Rodriguez, Slim Fourati, Vanessa Demontant, Guillaume Gricourt, Melissa N'Debi, Alexandre Soulier, Elisabeth Trawinski, Jean-Michel Pawlotsky |
| EPI_ISL_3452931 | Biogroup Bio Lam-LCD Saint-Denis | Department of Virology, Henri Mondor University Hospital, Assistance Publique Hôpitaux de Paris, Université Paris-Est Créteil, INSERM U955 | Christophe Rodriguez, Slim Fourati, Vanessa Demontant, Guillaume Gricourt, Melissa N'Debi, Alexandre Soulier, Elisabeth Trawinski, Jean-Michel Pawlotsky |
| EPI_ISL_3452932 | CH. CHU DE REIMS | Department of Virology, Henri Mondor University Hospital, Assistance Publique Hôpitaux de Paris, Université Paris-Est Créteil, INSERM U955 | Christophe Rodriguez, Slim Fourati, Vanessa Demontant, Guillaume Gricourt, Melissa N'Debi, Alexandre Soulier, Elisabeth Trawinski, Jean-Michel Pawlotsky |
| EPI_ISL_3452933, EPI_ISL_3452934 | Biogroup Bio Lam-LCD Saint-Denis | Department of Virology, Henri Mondor University Hospital, Assistance Publique Hôpitaux de Paris, Université Paris-Est Créteil, INSERM U955 | Christophe Rodriguez, Slim Fourati, Vanessa Demontant, Guillaume Gricourt, Melissa N'Debi, Alexandre Soulier, Elisabeth Trawinski, Jean-Michel Pawlotsky |
| EPI_ISL_3452935 | ESPACEBIO | Department of Virology, Henri Mondor University Hospital, Assistance Publique Hôpitaux de Paris, Université Paris-Est Créteil, INSERM U955 | Christophe Rodriguez, Slim Fourati, Vanessa Demontant, Guillaume Gricourt, Melissa N'Debi, Alexandre Soulier, Elisabeth Trawinski, Jean-Michel Pawlotsky |
| EPI_ISL_3452936 | SELAS ANALYSIS | Department of Virology, Henri Mondor University Hospital, Assistance Publique Hôpitaux de Paris, Université Paris-Est Créteil, INSERM U955 | Christophe Rodriguez, Slim Fourati, Vanessa Demontant, Guillaume Gricourt, Melissa N'Debi, Alexandre Soulier, Elisabeth Trawinski, Jean-Michel Pawlotsky |
| EPI_ISL_3452937 | ESPACEBIO | Department of Virology, Henri Mondor University Hospital, Assistance Publique Hôpitaux de Paris, Université Paris-Est Créteil, INSERM U955 | Christophe Rodriguez, Slim Fourati, Vanessa Demontant, Guillaume Gricourt, Melissa N'Debi, Alexandre Soulier, Elisabeth Trawinski, Jean-Michel Pawlotsky |
| EPI_ISL_3452938 | Biogroup Bio Lam-LCD Saint-Denis | Department of Virology, Henri Mondor University Hospital, Assistance Publique Hôpitaux de Paris, Université Paris-Est Créteil, INSERM U955 | Christophe Rodriguez, Slim Fourati, Vanessa Demontant, Guillaume Gricourt, Melissa N'Debi, Alexandre Soulier, Elisabeth Trawinski, Jean-Michel Pawlotsky |
| EPI_ISL_3452952 | BROUSSAIS | Department of Virology, Henri Mondor University Hospital, Assistance Publique Hôpitaux de Paris, Université Paris-Est Créteil, INSERM U955 | Christophe Rodriguez, Slim Fourati, Vanessa Demontant, Guillaume Gricourt, Melissa N'Debi, Alexandre Soulier, Elisabeth Trawinski, Jean-Michel Pawlotsky |
| EPI_ISL_3452954, EPI_ISL_3452955 | SELAS BC-LAB | Department of Virology, Henri Mondor University Hospital, Assistance Publique Hôpitaux de Paris, Université Paris-Est Créteil, INSERM U955 | Christophe Rodriguez, Slim Fourati, Vanessa Demontant, Guillaume Gricourt, Melissa N'Debi, Alexandre Soulier, Elisabeth Trawinski, Jean-Michel Pawlotsky |
| EPI_ISL_3452956 | CH Eure-Seine EVREUX | Department of Virology, Henri Mondor University Hospital, Assistance Publique Hôpitaux de Paris, Université Paris-Est Créteil, INSERM U955 | Christophe Rodriguez, Slim Fourati, Vanessa Demontant, Guillaume Gricourt, Melissa N'Debi, Alexandre Soulier, Elisabeth Trawinski, Jean-Michel Pawlotsky |
| EPI_ISL_3452957 | BROUSSAIS | Department of Virology, Henri Mondor University Hospital, Assistance Publique Hôpitaux de Paris, Université Paris-Est Créteil, INSERM U955 | Christophe Rodriguez, Slim Fourati, Vanessa Demontant, Guillaume Gricourt, Melissa N'Debi, Alexandre Soulier, Elisabeth Trawinski, Jean-Michel Pawlotsky |
| EPI_ISL_3452958 | CH. ANDRE GREGOIRE | Department of Virology, Henri Mondor University Hospital, Assistance Publique Hôpitaux de Paris, Université Paris-Est Créteil, INSERM U955 | Christophe Rodriguez, Slim Fourati, Vanessa Demontant, Guillaume Gricourt, Melissa N'Debi, Alexandre Soulier, Elisabeth Trawinski, Jean-Michel Pawlotsky |
| EPI_ISL_3452959 | GH A.CHENEVIER-H.MONDOR | Department of Virology, Henri Mondor University Hospital, Assistance Publique Hôpitaux de Paris, Université Paris-Est Créteil, INSERM U955 | Christophe Rodriguez, Slim Fourati, Vanessa Demontant, Guillaume Gricourt, Melissa N'Debi, Alexandre Soulier, Elisabeth Trawinski, Jean-Michel Pawlotsky |
| EPI_ISL_3452960 | Laboratoire CBM 25 TERRE ROUGE | Department of Virology, Henri Mondor University Hospital, Assistance Publique Hôpitaux de Paris, Université Paris-Est Créteil, INSERM U955 | Christophe Rodriguez, Slim Fourati, Vanessa Demontant, Guillaume Gricourt, Melissa N'Debi, Alexandre Soulier, Elisabeth Trawinski, Jean-Michel Pawlotsky |
| EPI_ISL_3452961, EPI_ISL_3452962, EPI_ISL_3452963, EPI_ISL_3452964, EPI_ISL_3452965, EPI_ISL_3452966, EPI_ISL_3452967 | BROUSSAIS | Department of Virology, Henri Mondor University Hospital, Assistance Publique Hôpitaux de Paris, Université Paris-Est Créteil, INSERM U955 | Christophe Rodriguez, Slim Fourati, Vanessa Demontant, Guillaume Gricourt, Melissa N'Debi, Alexandre Soulier, Elisabeth Trawinski, Jean-Michel Pawlotsky |
| EPI_ISL_3452968 | SELAS BC-LAB | Department of Virology, Henri Mondor University Hospital, Assistance Publique Hôpitaux de Paris, Université Paris-Est Créteil, INSERM U955 | Christophe Rodriguez, Slim Fourati, Vanessa Demontant, Guillaume Gricourt, Melissa N'Debi, Alexandre Soulier, Elisabeth Trawinski, Jean-Michel Pawlotsky |
| EPI_ISL_3452969 | BROUSSAIS | Department of Virology, Henri Mondor University Hospital, Assistance Publique Hôpitaux de Paris, Université Paris-Est Créteil, INSERM U955 | Christophe Rodriguez, Slim Fourati, Vanessa Demontant, Guillaume Gricourt, Melissa N'Debi, Alexandre Soulier, Elisabeth Trawinski, Jean-Michel Pawlotsky |
| EPI_ISL_3452970 | SELAS BC-LAB | Department of Virology, Henri Mondor University Hospital, Assistance Publique Hôpitaux de Paris, Université Paris-Est Créteil, INSERM U955 | Christophe Rodriguez, Slim Fourati, Vanessa Demontant, Guillaume Gricourt, Melissa N'Debi, Alexandre Soulier, Elisabeth Trawinski, Jean-Michel Pawlotsky |
| EPI_ISL_3452971, EPI_ISL_3452972, EPI_ISL_3452973, EPI_ISL_3452974, EPI_ISL_3452975 | BROUSSAIS | Department of Virology, Henri Mondor University Hospital, Assistance Publique Hôpitaux de Paris, Université Paris-Est Créteil, INSERM U955 | Christophe Rodriguez, Slim Fourati, Vanessa Demontant, Guillaume Gricourt, Melissa N'Debi, Alexandre Soulier, Elisabeth Trawinski, Jean-Michel Pawlotsky |
| EPI_ISL_3452976 | SELAS BC-LAB | Department of Virology, Henri Mondor University Hospital, Assistance Publique Hôpitaux de Paris, Université Paris-Est Créteil, INSERM U955 | Christophe Rodriguez, Slim Fourati, Vanessa Demontant, Guillaume Gricourt, Melissa N'Debi, Alexandre Soulier, Elisabeth Trawinski, Jean-Michel Pawlotsky |
| EPI_ISL_3452977 | BIOLLAN Brognard | Department of Virology, Henri Mondor University Hospital, Assistance Publique Hôpitaux de Paris, Université Paris-Est Créteil, INSERM U955 | Christophe Rodriguez, Slim Fourati, Vanessa Demontant, Guillaume Gricourt, Melissa N'Debi, Alexandre Soulier, Elisabeth Trawinski, Jean-Michel Pawlotsky |
| EPI_ISL_3452978 | BROUSSAIS | Department of Virology, Henri Mondor University Hospital, Assistance Publique Hôpitaux de Paris, Université Paris-Est Créteil, INSERM U955 | Christophe Rodriguez, Slim Fourati, Vanessa Demontant, Guillaume Gricourt, Melissa N'Debi, Alexandre Soulier, Elisabeth Trawinski, Jean-Michel Pawlotsky |
| EPI_ISL_3452983 | GH A.CHENEVIER-H.MONDOR | Department of Virology, Henri Mondor University Hospital, Assistance Publique Hôpitaux de Paris, Université Paris-Est Créteil, INSERM U955 | Christophe Rodriguez, Slim Fourati, Vanessa Demontant, Guillaume Gricourt, Melissa N'Debi, Alexandre Soulier, Elisabeth Trawinski, Jean-Michel Pawlotsky |
| EPI_ISL_3452984 | SELAS BC-LAB | Department of Virology, Henri Mondor University Hospital, Assistance Publique Hôpitaux de Paris, Université Paris-Est Créteil, INSERM U955 | Christophe Rodriguez, Slim Fourati, Vanessa Demontant, Guillaume Gricourt, Melissa N'Debi, Alexandre Soulier, Elisabeth Trawinski, Jean-Michel Pawlotsky |
| EPI_ISL_3452987 | Laboratoire CBM 25 TERRE ROUGE | Department of Virology, Henri Mondor University Hospital, Assistance Publique Hôpitaux de Paris, Université Paris-Est Créteil, INSERM U955 | Christophe Rodriguez, Slim Fourati, Vanessa Demontant, Guillaume Gricourt, Melissa N'Debi, Alexandre Soulier, Elisabeth Trawinski, Jean-Michel Pawlotsky |
| EPI_ISL_3452988 | SELAS BC-LAB | Department of Virology, Henri Mondor University Hospital, Assistance Publique Hôpitaux de Paris, Université Paris-Est Créteil, INSERM U955 | Christophe Rodriguez, Slim Fourati, Vanessa Demontant, Guillaume Gricourt, Melissa N'Debi, Alexandre Soulier, Elisabeth Trawinski, Jean-Michel Pawlotsky |
| EPI_ISL_3452990 | BROUSSAIS | Department of Virology, Henri Mondor University Hospital, Assistance Publique Hôpitaux de Paris, Université Paris-Est Créteil, | Christophe Rodriguez, Slim Fourati, Vanessa Demontant, Guillaume Gricourt, Melissa N'Debi, Alexandre Soulier, Elisabeth Trawinski, Jean-Michel Pawlotsky |

| | | | |
|---|---|---|---|
| | | INSERM U955 | |
| EPI_ISL_3452991 | Laboratoire CBM 25 TERRE ROUGE | Department of Virology, Henri Mondor University Hospital, Assistance Publique Hôpitaux de Paris, Université Paris-Est Créteil, INSERM U955 | Christophe Rodriguez, Slim Fourati, Vanessa Demontant, Guillaume Gricourt, Melissa N'Debi, Alexandre Soulier, Elisabeth Trawinski, Jean-Michel Pawlotsky |
| EPI_ISL_3452992, EPI_ISL_3452993, EPI_ISL_3452994, EPI_ISL_3452995 | SELAS BC-LAB | Department of Virology, Henri Mondor University Hospital, Assistance Publique Hôpitaux de Paris, Université Paris-Est Créteil, INSERM U955 | Christophe Rodriguez, Slim Fourati, Vanessa Demontant, Guillaume Gricourt, Melissa N'Debi, Alexandre Soulier, Elisabeth Trawinski, Jean-Michel Pawlotsky |
| EPI_ISL_3452996, EPI_ISL_3452997 | BROUSSAIS | Department of Virology, Henri Mondor University Hospital, Assistance Publique Hôpitaux de Paris, Université Paris-Est Créteil, INSERM U955 | Christophe Rodriguez, Slim Fourati, Vanessa Demontant, Guillaume Gricourt, Melissa N'Debi, Alexandre Soulier, Elisabeth Trawinski, Jean-Michel Pawlotsky |
| EPI_ISL_3452998, EPI_ISL_3452999, EPI_ISL_3453000 | SELAS BC-LAB | Department of Virology, Henri Mondor University Hospital, Assistance Publique Hôpitaux de Paris, Université Paris-Est Créteil, INSERM U955 | Christophe Rodriguez, Slim Fourati, Vanessa Demontant, Guillaume Gricourt, Melissa N'Debi, Alexandre Soulier, Elisabeth Trawinski, Jean-Michel Pawlotsky |
| EPI_ISL_3453001 | Biogroup Bio Lam-LCD Saint-Denis | Department of Virology, Henri Mondor University Hospital, Assistance Publique Hôpitaux de Paris, Université Paris-Est Créteil, INSERM U955 | Christophe Rodriguez, Slim Fourati, Vanessa Demontant, Guillaume Gricourt, Melissa N'Debi, Alexandre Soulier, Elisabeth Trawinski, Jean-Michel Pawlotsky |
| EPI_ISL_3453002 | BROUSSAIS | Department of Virology, Henri Mondor University Hospital, Assistance Publique Hôpitaux de Paris, Université Paris-Est Créteil, INSERM U955 | Christophe Rodriguez, Slim Fourati, Vanessa Demontant, Guillaume Gricourt, Melissa N'Debi, Alexandre Soulier, Elisabeth Trawinski, Jean-Michel Pawlotsky |
| EPI_ISL_3453004 | BIOLLAN Brognard | Department of Virology, Henri Mondor University Hospital, Assistance Publique Hôpitaux de Paris, Université Paris-Est Créteil, INSERM U955 | Christophe Rodriguez, Slim Fourati, Vanessa Demontant, Guillaume Gricourt, Melissa N'Debi, Alexandre Soulier, Elisabeth Trawinski, Jean-Michel Pawlotsky |
| EPI_ISL_3453005 | CH. ANDRE GREGOIRE | Department of Virology, Henri Mondor University Hospital, Assistance Publique Hôpitaux de Paris, Université Paris-Est Créteil, INSERM U955 | Christophe Rodriguez, Slim Fourati, Vanessa Demontant, Guillaume Gricourt, Melissa N'Debi, Alexandre Soulier, Elisabeth Trawinski, Jean-Michel Pawlotsky |
| EPI_ISL_3453006 | Biogroup Bio Lam-LCD Saint-Denis | Department of Virology, Henri Mondor University Hospital, Assistance Publique Hôpitaux de Paris, Université Paris-Est Créteil, INSERM U955 | Christophe Rodriguez, Slim Fourati, Vanessa Demontant, Guillaume Gricourt, Melissa N'Debi, Alexandre Soulier, Elisabeth Trawinski, Jean-Michel Pawlotsky |
| EPI_ISL_3453007 | CH. ANDRE GREGOIRE | Department of Virology, Henri Mondor University Hospital, Assistance Publique Hôpitaux de Paris, Université Paris-Est Créteil, INSERM U955 | Christophe Rodriguez, Slim Fourati, Vanessa Demontant, Guillaume Gricourt, Melissa N'Debi, Alexandre Soulier, Elisabeth Trawinski, Jean-Michel Pawlotsky |
| EPI_ISL_3453008, EPI_ISL_3453009, EPI_ISL_3453010 | Biogroup Bio Lam-LCD Saint-Denis | Department of Virology, Henri Mondor University Hospital, Assistance Publique Hôpitaux de Paris, Université Paris-Est Créteil, INSERM U955 | Christophe Rodriguez, Slim Fourati, Vanessa Demontant, Guillaume Gricourt, Melissa N'Debi, Alexandre Soulier, Elisabeth Trawinski, Jean-Michel Pawlotsky |
| EPI_ISL_3453011 | Laboratoire MAYMAT | Department of Virology, Henri Mondor University Hospital, Assistance Publique Hôpitaux de Paris, Université Paris-Est Créteil, INSERM U955 | Christophe Rodriguez, Slim Fourati, Vanessa Demontant, Guillaume Gricourt, Melissa N'Debi, Alexandre Soulier, Elisabeth Trawinski, Jean-Michel Pawlotsky |
| EPI_ISL_3453012 | BIO ARD'AISNE | Department of Virology, Henri Mondor University Hospital, Assistance Publique Hôpitaux de Paris, Université Paris-Est Créteil, INSERM U955 | Christophe Rodriguez, Slim Fourati, Vanessa Demontant, Guillaume Gricourt, Melissa N'Debi, Alexandre Soulier, Elisabeth Trawinski, Jean-Michel Pawlotsky |
| EPI_ISL_3453013 | BIOLLAN Brognard | Department of Virology, Henri Mondor University Hospital, Assistance Publique Hôpitaux de Paris, Université Paris-Est Créteil, INSERM U955 | Christophe Rodriguez, Slim Fourati, Vanessa Demontant, Guillaume Gricourt, Melissa N'Debi, Alexandre Soulier, Elisabeth Trawinski, Jean-Michel Pawlotsky |
| EPI_ISL_3453014 | Laboratoire Bioconcorde | Department of Virology, Henri Mondor University Hospital, Assistance Publique Hôpitaux de Paris, Université Paris-Est Créteil, INSERM U955 | Christophe Rodriguez, Slim Fourati, Vanessa Demontant, Guillaume Gricourt, Melissa N'Debi, Alexandre Soulier, Elisabeth Trawinski, Jean-Michel Pawlotsky |
| EPI_ISL_3453017 | SELARL BIO | Department of Virology, Henri Mondor University Hospital, Assistance Publique Hôpitaux de Paris, Université Paris-Est Créteil, INSERM U955 | Christophe Rodriguez, Slim Fourati, Vanessa Demontant, Guillaume Gricourt, Melissa N'Debi, Alexandre Soulier, Elisabeth Trawinski, Jean-Michel Pawlotsky |
| EPI_ISL_3453018 | SELAFA SANTE LABO | Department of Virology, Henri Mondor University Hospital, Assistance Publique Hôpitaux de Paris, Université Paris-Est Créteil, INSERM U955 | Christophe Rodriguez, Slim Fourati, Vanessa Demontant, Guillaume Gricourt, Melissa N'Debi, Alexandre Soulier, Elisabeth Trawinski, Jean-Michel Pawlotsky |
| EPI_ISL_3453019 | CERBALLIANCE Valmy | Department of Virology, Henri Mondor University Hospital, Assistance Publique Hôpitaux de Paris, Université Paris-Est Créteil, INSERM U955 | Christophe Rodriguez, Slim Fourati, Vanessa Demontant, Guillaume Gricourt, Melissa N'Debi, Alexandre Soulier, Elisabeth Trawinski, Jean-Michel Pawlotsky |
| EPI_ISL_3453020 | Laboratoire CBM 25 TERRE ROUGE | Department of Virology, Henri Mondor University Hospital, Assistance Publique Hôpitaux de Paris, Université Paris-Est Créteil, INSERM U955 | Christophe Rodriguez, Slim Fourati, Vanessa Demontant, Guillaume Gricourt, Melissa N'Debi, Alexandre Soulier, Elisabeth Trawinski, Jean-Michel Pawlotsky |
| EPI_ISL_3453021 | Laboratoire MAYMAT | Department of Virology, Henri Mondor University Hospital, Assistance Publique Hôpitaux de Paris, Université Paris-Est Créteil, INSERM U955 | Christophe Rodriguez, Slim Fourati, Vanessa Demontant, Guillaume Gricourt, Melissa N'Debi, Alexandre Soulier, Elisabeth Trawinski, Jean-Michel Pawlotsky |
| EPI_ISL_3453022 | CERBALLIANCE Valmy | Department of Virology, Henri Mondor University Hospital, Assistance Publique Hôpitaux de Paris, Université Paris-Est Créteil, INSERM U955 | Christophe Rodriguez, Slim Fourati, Vanessa Demontant, Guillaume Gricourt, Melissa N'Debi, Alexandre Soulier, Elisabeth Trawinski, Jean-Michel Pawlotsky |
| EPI_ISL_3453023 | Laboratoire CBM 25 TERRE ROUGE | Department of Virology, Henri Mondor University Hospital, Assistance Publique Hôpitaux de Paris, Université Paris-Est Créteil, INSERM U955 | Christophe Rodriguez, Slim Fourati, Vanessa Demontant, Guillaume Gricourt, Melissa N'Debi, Alexandre Soulier, Elisabeth Trawinski, Jean-Michel Pawlotsky |
| EPI_ISL_3453024 | BIOLAB UNILABS | Department of Virology, Henri Mondor University Hospital, Assistance Publique Hôpitaux de Paris, Université Paris-Est Créteil, INSERM U955 | Christophe Rodriguez, Slim Fourati, Vanessa Demontant, Guillaume Gricourt, Melissa N'Debi, Alexandre Soulier, Elisabeth Trawinski, Jean-Michel Pawlotsky |
| EPI_ISL_3453025 | SELAFA SANTE LABO | Department of Virology, Henri Mondor University Hospital, Assistance Publique Hôpitaux de Paris, Université Paris-Est Créteil, INSERM U955 | Christophe Rodriguez, Slim Fourati, Vanessa Demontant, Guillaume Gricourt, Melissa N'Debi, Alexandre Soulier, Elisabeth Trawinski, Jean-Michel Pawlotsky |
| EPI_ISL_3453026 | Laboratoire MAYMAT | Department of Virology, Henri Mondor University Hospital, Assistance Publique Hôpitaux de Paris, Université Paris-Est Créteil, INSERM U955 | Christophe Rodriguez, Slim Fourati, Vanessa Demontant, Guillaume Gricourt, Melissa N'Debi, Alexandre Soulier, Elisabeth Trawinski, Jean-Michel Pawlotsky |
| EPI_ISL_3453027 | Laboratoire CBM 25 TERRE ROUGE | Department of Virology, Henri Mondor University Hospital, Assistance Publique Hôpitaux de Paris, Université Paris-Est Créteil, INSERM U955 | Christophe Rodriguez, Slim Fourati, Vanessa Demontant, Guillaume Gricourt, Melissa N'Debi, Alexandre Soulier, Elisabeth Trawinski, Jean-Michel Pawlotsky |
| EPI_ISL_3453028 | SELAS BC-LAB | Department of Virology, Henri Mondor University Hospital, Assistance Publique Hôpitaux de Paris, Université Paris-Est Créteil, INSERM U955 | Christophe Rodriguez, Slim Fourati, Vanessa Demontant, Guillaume Gricourt, Melissa N'Debi, Alexandre Soulier, Elisabeth Trawinski, Jean-Michel Pawlotsky |
| EPI_ISL_3453029 | Laboratoire MAYMAT | Department of Virology, Henri Mondor University Hospital, Assistance Publique Hôpitaux de Paris, Université Paris-Est Créteil, INSERM U955 | Christophe Rodriguez, Slim Fourati, Vanessa Demontant, Guillaume Gricourt, Melissa N'Debi, Alexandre Soulier, Elisabeth Trawinski, Jean-Michel Pawlotsky |
| EPI_ISL_3453030 | BIOLLAN Brognard | Department of Virology, Henri Mondor University Hospital, Assistance Publique Hôpitaux de Paris, Université Paris-Est Créteil, INSERM U955 | Christophe Rodriguez, Slim Fourati, Vanessa Demontant, Guillaume Gricourt, Melissa N'Debi, Alexandre Soulier, Elisabeth Trawinski, Jean-Michel Pawlotsky |
| EPI_ISL_3453031 | CERBALLIANCE Valmy | Department of Virology, Henri Mondor University Hospital, Assistance Publique Hôpitaux de Paris, Université Paris-Est Créteil, INSERM U955 | Christophe Rodriguez, Slim Fourati, Vanessa Demontant, Guillaume Gricourt, Melissa N'Debi, Alexandre Soulier, Elisabeth Trawinski, Jean-Michel Pawlotsky |
| EPI_ISL_3453032 | Laboratoire MAYMAT | Department of Virology, Henri Mondor University Hospital, Assistance Publique Hôpitaux de Paris, Université Paris-Est Créteil, INSERM U955 | Christophe Rodriguez, Slim Fourati, Vanessa Demontant, Guillaume Gricourt, Melissa N'Debi, Alexandre Soulier, Elisabeth Trawinski, Jean-Michel Pawlotsky |
| EPI_ISL_3453033, | Laboratoire CBM 25 TERRE ROUGE | Department of Virology, Henri Mondor University Hospital, | Christophe Rodriguez, Slim Fourati, Vanessa Demontant, Guillaume Gricourt, Melissa N'Debi, Alexandre Soulier, Elisabeth Trawinski, Jean-Michel Pawlotsky |

| | | | |
|---|---|---|---|
| EPI_ISL_3453034 | | Assistance Publique Hôpitaux de Paris, Université Paris-Est Créteil, INSERM U955 | |
| EPI_ISL_3453035 | CERBALLIANCE Valmy | Department of Virology, Henri Mondor University Hospital, Assistance Publique Hôpitaux de Paris, Université Paris-Est Créteil, INSERM U955 | Christophe Rodriguez, Slim Fourati, Vanessa Demontant, Guillaume Gricourt, Melissa N'Debi, Alexandre Soulier, Elisabeth Trawinski, Jean-Michel Pawlotsky |
| EPI_ISL_3453036 | SELAFA SANTE LABO | Department of Virology, Henri Mondor University Hospital, Assistance Publique Hôpitaux de Paris, Université Paris-Est Créteil, INSERM U955 | Christophe Rodriguez, Slim Fourati, Vanessa Demontant, Guillaume Gricourt, Melissa N'Debi, Alexandre Soulier, Elisabeth Trawinski, Jean-Michel Pawlotsky |
| EPI_ISL_3453037 | CERBALLIANCE Valmy | Department of Virology, Henri Mondor University Hospital, Assistance Publique Hôpitaux de Paris, Université Paris-Est Créteil, INSERM U955 | Christophe Rodriguez, Slim Fourati, Vanessa Demontant, Guillaume Gricourt, Melissa N'Debi, Alexandre Soulier, Elisabeth Trawinski, Jean-Michel Pawlotsky |
| EPI_ISL_3453038, EPI_ISL_3453039, EPI_ISL_3453040 | BIOLLAN Brognard | Department of Virology, Henri Mondor University Hospital, Assistance Publique Hôpitaux de Paris, Université Paris-Est Créteil, INSERM U955 | Christophe Rodriguez, Slim Fourati, Vanessa Demontant, Guillaume Gricourt, Melissa N'Debi, Alexandre Soulier, Elisabeth Trawinski, Jean-Michel Pawlotsky |
| EPI_ISL_3453041, EPI_ISL_3453042 | SELAFA SANTE LABO | Department of Virology, Henri Mondor University Hospital, Assistance Publique Hôpitaux de Paris, Université Paris-Est Créteil, INSERM U955 | Christophe Rodriguez, Slim Fourati, Vanessa Demontant, Guillaume Gricourt, Melissa N'Debi, Alexandre Soulier, Elisabeth Trawinski, Jean-Michel Pawlotsky |
| EPI_ISL_3453043, EPI_ISL_3453044 | Laboratoire CBM 25 TERRE ROUGE | Department of Virology, Henri Mondor University Hospital, Assistance Publique Hôpitaux de Paris, Université Paris-Est Créteil, INSERM U955 | Christophe Rodriguez, Slim Fourati, Vanessa Demontant, Guillaume Gricourt, Melissa N'Debi, Alexandre Soulier, Elisabeth Trawinski, Jean-Michel Pawlotsky |
| EPI_ISL_3453045 | SELAFA SANTE LABO | Department of Virology, Henri Mondor University Hospital, Assistance Publique Hôpitaux de Paris, Université Paris-Est Créteil, INSERM U955 | Christophe Rodriguez, Slim Fourati, Vanessa Demontant, Guillaume Gricourt, Melissa N'Debi, Alexandre Soulier, Elisabeth Trawinski, Jean-Michel Pawlotsky |
| EPI_ISL_3453046 | Laboratoire CBM 25 TERRE ROUGE | Department of Virology, Henri Mondor University Hospital, Assistance Publique Hôpitaux de Paris, Université Paris-Est Créteil, INSERM U955 | Christophe Rodriguez, Slim Fourati, Vanessa Demontant, Guillaume Gricourt, Melissa N'Debi, Alexandre Soulier, Elisabeth Trawinski, Jean-Michel Pawlotsky |
| EPI_ISL_3453047 | CERBALLIANCE Valmy | Department of Virology, Henri Mondor University Hospital, Assistance Publique Hôpitaux de Paris, Université Paris-Est Créteil, INSERM U955 | Christophe Rodriguez, Slim Fourati, Vanessa Demontant, Guillaume Gricourt, Melissa N'Debi, Alexandre Soulier, Elisabeth Trawinski, Jean-Michel Pawlotsky |
| EPI_ISL_3453048 | Laboratoire MAYMAT | Department of Virology, Henri Mondor University Hospital, Assistance Publique Hôpitaux de Paris, Université Paris-Est Créteil, INSERM U955 | Christophe Rodriguez, Slim Fourati, Vanessa Demontant, Guillaume Gricourt, Melissa N'Debi, Alexandre Soulier, Elisabeth Trawinski, Jean-Michel Pawlotsky |
| EPI_ISL_3453049 | CERBALLIANCE Valmy | Department of Virology, Henri Mondor University Hospital, Assistance Publique Hôpitaux de Paris, Université Paris-Est Créteil, INSERM U955 | Christophe Rodriguez, Slim Fourati, Vanessa Demontant, Guillaume Gricourt, Melissa N'Debi, Alexandre Soulier, Elisabeth Trawinski, Jean-Michel Pawlotsky |
| EPI_ISL_3453050 | Laboratoire CBM 25 TERRE ROUGE | Department of Virology, Henri Mondor University Hospital, Assistance Publique Hôpitaux de Paris, Université Paris-Est Créteil, INSERM U955 | Christophe Rodriguez, Slim Fourati, Vanessa Demontant, Guillaume Gricourt, Melissa N'Debi, Alexandre Soulier, Elisabeth Trawinski, Jean-Michel Pawlotsky |
| EPI_ISL_3453051 | Laboratoire MAYMAT | Department of Virology, Henri Mondor University Hospital, Assistance Publique Hôpitaux de Paris, Université Paris-Est Créteil, INSERM U955 | Christophe Rodriguez, Slim Fourati, Vanessa Demontant, Guillaume Gricourt, Melissa N'Debi, Alexandre Soulier, Elisabeth Trawinski, Jean-Michel Pawlotsky |
| EPI_ISL_3453052 | SELAS BC-LAB | Department of Virology, Henri Mondor University Hospital, Assistance Publique Hôpitaux de Paris, Université Paris-Est Créteil, INSERM U955 | Christophe Rodriguez, Slim Fourati, Vanessa Demontant, Guillaume Gricourt, Melissa N'Debi, Alexandre Soulier, Elisabeth Trawinski, Jean-Michel Pawlotsky |
| EPI_ISL_3453053 | Laboratoire MAYMAT | Department of Virology, Henri Mondor University Hospital, Assistance Publique Hôpitaux de Paris, Université Paris-Est Créteil, INSERM U955 | Christophe Rodriguez, Slim Fourati, Vanessa Demontant, Guillaume Gricourt, Melissa N'Debi, Alexandre Soulier, Elisabeth Trawinski, Jean-Michel Pawlotsky |
| EPI_ISL_3453054 | BIOLLAN Brognard | Department of Virology, Henri Mondor University Hospital, Assistance Publique Hôpitaux de Paris, Université Paris-Est Créteil, INSERM U955 | Christophe Rodriguez, Slim Fourati, Vanessa Demontant, Guillaume Gricourt, Melissa N'Debi, Alexandre Soulier, Elisabeth Trawinski, Jean-Michel Pawlotsky |
| EPI_ISL_3453055, EPI_ISL_3453056 | SELAFA SANTE LABO | Department of Virology, Henri Mondor University Hospital, Assistance Publique Hôpitaux de Paris, Université Paris-Est Créteil, INSERM U955 | Christophe Rodriguez, Slim Fourati, Vanessa Demontant, Guillaume Gricourt, Melissa N'Debi, Alexandre Soulier, Elisabeth Trawinski, Jean-Michel Pawlotsky |
| EPI_ISL_3453057 | BIOLLAN Brognard | Department of Virology, Henri Mondor University Hospital, Assistance Publique Hôpitaux de Paris, Université Paris-Est Créteil, INSERM U955 | Christophe Rodriguez, Slim Fourati, Vanessa Demontant, Guillaume Gricourt, Melissa N'Debi, Alexandre Soulier, Elisabeth Trawinski, Jean-Michel Pawlotsky |
| EPI_ISL_3453058 | CERBALLIANCE Valmy | Department of Virology, Henri Mondor University Hospital, Assistance Publique Hôpitaux de Paris, Université Paris-Est Créteil, INSERM U955 | Christophe Rodriguez, Slim Fourati, Vanessa Demontant, Guillaume Gricourt, Melissa N'Debi, Alexandre Soulier, Elisabeth Trawinski, Jean-Michel Pawlotsky |
| EPI_ISL_3453059 | Laboratoire MAYMAT | Department of Virology, Henri Mondor University Hospital, Assistance Publique Hôpitaux de Paris, Université Paris-Est Créteil, INSERM U955 | Christophe Rodriguez, Slim Fourati, Vanessa Demontant, Guillaume Gricourt, Melissa N'Debi, Alexandre Soulier, Elisabeth Trawinski, Jean-Michel Pawlotsky |
| EPI_ISL_3453060 | BIOLLAN Brognard | Department of Virology, Henri Mondor University Hospital, Assistance Publique Hôpitaux de Paris, Université Paris-Est Créteil, INSERM U955 | Christophe Rodriguez, Slim Fourati, Vanessa Demontant, Guillaume Gricourt, Melissa N'Debi, Alexandre Soulier, Elisabeth Trawinski, Jean-Michel Pawlotsky |
| EPI_ISL_3453061 | CERBALLIANCE Valmy | Department of Virology, Henri Mondor University Hospital, Assistance Publique Hôpitaux de Paris, Université Paris-Est Créteil, INSERM U955 | Christophe Rodriguez, Slim Fourati, Vanessa Demontant, Guillaume Gricourt, Melissa N'Debi, Alexandre Soulier, Elisabeth Trawinski, Jean-Michel Pawlotsky |
| EPI_ISL_3453062 | BIOLLAN Brognard | Department of Virology, Henri Mondor University Hospital, Assistance Publique Hôpitaux de Paris, Université Paris-Est Créteil, INSERM U955 | Christophe Rodriguez, Slim Fourati, Vanessa Demontant, Guillaume Gricourt, Melissa N'Debi, Alexandre Soulier, Elisabeth Trawinski, Jean-Michel Pawlotsky |
| EPI_ISL_3453063 | Laboratoire CBM 25 TERRE ROUGE | Department of Virology, Henri Mondor University Hospital, Assistance Publique Hôpitaux de Paris, Université Paris-Est Créteil, INSERM U955 | Christophe Rodriguez, Slim Fourati, Vanessa Demontant, Guillaume Gricourt, Melissa N'Debi, Alexandre Soulier, Elisabeth Trawinski, Jean-Michel Pawlotsky |
| EPI_ISL_3453064 | SELAS BC-LAB | Department of Virology, Henri Mondor University Hospital, Assistance Publique Hôpitaux de Paris, Université Paris-Est Créteil, INSERM U955 | Christophe Rodriguez, Slim Fourati, Vanessa Demontant, Guillaume Gricourt, Melissa N'Debi, Alexandre Soulier, Elisabeth Trawinski, Jean-Michel Pawlotsky |
| EPI_ISL_3453065 | BIOLAB UNILABS | Department of Virology, Henri Mondor University Hospital, Assistance Publique Hôpitaux de Paris, Université Paris-Est Créteil, INSERM U955 | Christophe Rodriguez, Slim Fourati, Vanessa Demontant, Guillaume Gricourt, Melissa N'Debi, Alexandre Soulier, Elisabeth Trawinski, Jean-Michel Pawlotsky |
| EPI_ISL_3453066 | ESPACEBIO | Department of Virology, Henri Mondor University Hospital, Assistance Publique Hôpitaux de Paris, Université Paris-Est Créteil, INSERM U955 | Christophe Rodriguez, Slim Fourati, Vanessa Demontant, Guillaume Gricourt, Melissa N'Debi, Alexandre Soulier, Elisabeth Trawinski, Jean-Michel Pawlotsky |
| EPI_ISL_3453067 | SELAS ANALYSIS | Department of Virology, Henri Mondor University Hospital, Assistance Publique Hôpitaux de Paris, Université Paris-Est Créteil, INSERM U955 | Christophe Rodriguez, Slim Fourati, Vanessa Demontant, Guillaume Gricourt, Melissa N'Debi, Alexandre Soulier, Elisabeth Trawinski, Jean-Michel Pawlotsky |
| EPI_ISL_3453068 | ESPACEBIO | Department of Virology, Henri Mondor University Hospital, Assistance Publique Hôpitaux de Paris, Université Paris-Est Créteil, INSERM U955 | Christophe Rodriguez, Slim Fourati, Vanessa Demontant, Guillaume Gricourt, Melissa N'Debi, Alexandre Soulier, Elisabeth Trawinski, Jean-Michel Pawlotsky |
| EPI_ISL_3453069, EPI_ISL_3453070 | SELAS ANALYSIS | Department of Virology, Henri Mondor University Hospital, Assistance Publique Hôpitaux de Paris, Université Paris-Est Créteil, INSERM U955 | Christophe Rodriguez, Slim Fourati, Vanessa Demontant, Guillaume Gricourt, Melissa N'Debi, Alexandre Soulier, Elisabeth Trawinski, Jean-Michel Pawlotsky |
| EPI_ISL_3453071 | ESPACEBIO | Department of Virology, Henri Mondor University Hospital, Assistance Publique Hôpitaux de Paris, Université Paris-Est Créteil, INSERM U955 | Christophe Rodriguez, Slim Fourati, Vanessa Demontant, Guillaume Gricourt, Melissa N'Debi, Alexandre Soulier, Elisabeth Trawinski, Jean-Michel Pawlotsky |
| EPI_ISL_3453072, | SELAS ANALYSIS | Department of Virology, Henri Mondor University Hospital, | Christophe Rodriguez, Slim Fourati, Vanessa Demontant, Guillaume Gricourt, Melissa N'Debi, Alexandre Soulier, Elisabeth Trawinski, Jean-Michel Pawlotsky |

| | | | |
|---|---|---|---|
| EPI_ISL_3453073 | | Assistance Publique Hôpitaux de Paris, Université Paris-Est Créteil, INSERM U955 | |
| EPI_ISL_3453074, EPI_ISL_3453075 | ESPACEBIO | Department of Virology, Henri Mondor University Hospital, Assistance Publique Hôpitaux de Paris, Université Paris-Est Créteil, INSERM U955 | Christophe Rodriguez, Slim Fourati, Vanessa Demontant, Guillaume Gricourt, Melissa N'Debi, Alexandre Soulier, Elisabeth Trawinski, Jean-Michel Pawlotsky |
| EPI_ISL_3453076 | SELAS ANALYSIS | Department of Virology, Henri Mondor University Hospital, Assistance Publique Hôpitaux de Paris, Université Paris-Est Créteil, INSERM U955 | Christophe Rodriguez, Slim Fourati, Vanessa Demontant, Guillaume Gricourt, Melissa N'Debi, Alexandre Soulier, Elisabeth Trawinski, Jean-Michel Pawlotsky |
| EPI_ISL_3453077 | ESPACEBIO | Department of Virology, Henri Mondor University Hospital, Assistance Publique Hôpitaux de Paris, Université Paris-Est Créteil, INSERM U955 | Christophe Rodriguez, Slim Fourati, Vanessa Demontant, Guillaume Gricourt, Melissa N'Debi, Alexandre Soulier, Elisabeth Trawinski, Jean-Michel Pawlotsky |
| EPI_ISL_3453078 | SELAS ANALYSIS | Department of Virology, Henri Mondor University Hospital, Assistance Publique Hôpitaux de Paris, Université Paris-Est Créteil, INSERM U955 | Christophe Rodriguez, Slim Fourati, Vanessa Demontant, Guillaume Gricourt, Melissa N'Debi, Alexandre Soulier, Elisabeth Trawinski, Jean-Michel Pawlotsky |
| EPI_ISL_3453079, EPI_ISL_3453080, EPI_ISL_3453081, EPI_ISL_3453082 | ESPACEBIO | Department of Virology, Henri Mondor University Hospital, Assistance Publique Hôpitaux de Paris, Université Paris-Est Créteil, INSERM U955 | Christophe Rodriguez, Slim Fourati, Vanessa Demontant, Guillaume Gricourt, Melissa N'Debi, Alexandre Soulier, Elisabeth Trawinski, Jean-Michel Pawlotsky |
| EPI_ISL_3453083 | SELAS ANALYSIS | Department of Virology, Henri Mondor University Hospital, Assistance Publique Hôpitaux de Paris, Université Paris-Est Créteil, INSERM U955 | Christophe Rodriguez, Slim Fourati, Vanessa Demontant, Guillaume Gricourt, Melissa N'Debi, Alexandre Soulier, Elisabeth Trawinski, Jean-Michel Pawlotsky |
| EPI_ISL_3453084, EPI_ISL_3453085 | ESPACEBIO | Department of Virology, Henri Mondor University Hospital, Assistance Publique Hôpitaux de Paris, Université Paris-Est Créteil, INSERM U955 | Christophe Rodriguez, Slim Fourati, Vanessa Demontant, Guillaume Gricourt, Melissa N'Debi, Alexandre Soulier, Elisabeth Trawinski, Jean-Michel Pawlotsky |
| EPI_ISL_3453086 | CH. CHU DE REIMS | Department of Virology, Henri Mondor University Hospital, Assistance Publique Hôpitaux de Paris, Université Paris-Est Créteil, INSERM U955 | Christophe Rodriguez, Slim Fourati, Vanessa Demontant, Guillaume Gricourt, Melissa N'Debi, Alexandre Soulier, Elisabeth Trawinski, Jean-Michel Pawlotsky |
| EPI_ISL_3453087, EPI_ISL_3453088, EPI_ISL_3453089, EPI_ISL_3453090, EPI_ISL_3453091, EPI_ISL_3453092 | Biogroup Bio Lam-LCD Saint-Denis | Department of Virology, Henri Mondor University Hospital, Assistance Publique Hôpitaux de Paris, Université Paris-Est Créteil, INSERM U955 | Christophe Rodriguez, Slim Fourati, Vanessa Demontant, Guillaume Gricourt, Melissa N'Debi, Alexandre Soulier, Elisabeth Trawinski, Jean-Michel Pawlotsky |
| EPI_ISL_3453093 | CH. ANDRE GREGOIRE | Department of Virology, Henri Mondor University Hospital, Assistance Publique Hôpitaux de Paris, Université Paris-Est Créteil, INSERM U955 | Christophe Rodriguez, Slim Fourati, Vanessa Demontant, Guillaume Gricourt, Melissa N'Debi, Alexandre Soulier, Elisabeth Trawinski, Jean-Michel Pawlotsky |
| EPI_ISL_3453094, EPI_ISL_3453095, EPI_ISL_3453096, EPI_ISL_3453097, EPI_ISL_3453098, EPI_ISL_3453099, EPI_ISL_3453100, EPI_ISL_3453101, EPI_ISL_3453102, EPI_ISL_3453103, EPI_ISL_3453104, EPI_ISL_3453105, EPI_ISL_3453106, EPI_ISL_3453107, EPI_ISL_3453108, EPI_ISL_3453109, EPI_ISL_3453110, EPI_ISL_3453111, EPI_ISL_3453112, EPI_ISL_3453113 | | | |
| see above | Biogroup Bio Lam-LCD Saint-Denis | Department of Virology, Henri Mondor University Hospital, Assistance Publique Hôpitaux de Paris, Université Paris-Est Créteil, INSERM U955 | Christophe Rodriguez, Slim Fourati, Vanessa Demontant, Guillaume Gricourt, Melissa N'Debi, Alexandre Soulier, Elisabeth Trawinski, Jean-Michel Pawlotsky |
| EPI_ISL_3453114, EPI_ISL_3453115 | CH. ANDRE GREGOIRE | Department of Virology, Henri Mondor University Hospital, Assistance Publique Hôpitaux de Paris, Université Paris-Est Créteil, INSERM U955 | Christophe Rodriguez, Slim Fourati, Vanessa Demontant, Guillaume Gricourt, Melissa N'Debi, Alexandre Soulier, Elisabeth Trawinski, Jean-Michel Pawlotsky |
| EPI_ISL_3453116, EPI_ISL_3453117 | Biogroup Bio Lam-LCD Saint-Denis | Department of Virology, Henri Mondor University Hospital, Assistance Publique Hôpitaux de Paris, Université Paris-Est Créteil, INSERM U955 | Christophe Rodriguez, Slim Fourati, Vanessa Demontant, Guillaume Gricourt, Melissa N'Debi, Alexandre Soulier, Elisabeth Trawinski, Jean-Michel Pawlotsky |
| EPI_ISL_3453118 | CH. ANDRE GREGOIRE | Department of Virology, Henri Mondor University Hospital, Assistance Publique Hôpitaux de Paris, Université Paris-Est Créteil, INSERM U955 | Christophe Rodriguez, Slim Fourati, Vanessa Demontant, Guillaume Gricourt, Melissa N'Debi, Alexandre Soulier, Elisabeth Trawinski, Jean-Michel Pawlotsky |
| EPI_ISL_3453119, EPI_ISL_3453120, EPI_ISL_3453121, EPI_ISL_3453122, EPI_ISL_3453123, EPI_ISL_3453124, EPI_ISL_3453125, EPI_ISL_3453126 | Biogroup Bio Lam-LCD Saint-Denis | Department of Virology, Henri Mondor University Hospital, Assistance Publique Hôpitaux de Paris, Université Paris-Est Créteil, INSERM U955 | Christophe Rodriguez, Slim Fourati, Vanessa Demontant, Guillaume Gricourt, Melissa N'Debi, Alexandre Soulier, Elisabeth Trawinski, Jean-Michel Pawlotsky |
| EPI_ISL_3453127 | GH de l'Est Francilien | Department of Virology, Henri Mondor University Hospital, Assistance Publique Hôpitaux de Paris, Université Paris-Est Créteil, INSERM U955 | Christophe Rodriguez, Slim Fourati, Vanessa Demontant, Guillaume Gricourt, Melissa N'Debi, Alexandre Soulier, Elisabeth Trawinski, Jean-Michel Pawlotsky |
| EPI_ISL_3453128, EPI_ISL_3453129, EPI_ISL_3453130, EPI_ISL_3453131, EPI_ISL_3453132, EPI_ISL_3453133, EPI_ISL_3453134, EPI_ISL_3453135, EPI_ISL_3453136, EPI_ISL_3453137, EPI_ISL_3453138, EPI_ISL_3453139, EPI_ISL_3453140, EPI_ISL_3453141, EPI_ISL_3453142, EPI_ISL_3453143, EPI_ISL_3453144, EPI_ISL_3453145, EPI_ISL_3453146 | | | |
| see above | Biogroup Bio Lam-LCD Saint-Denis | Department of Virology, Henri Mondor University Hospital, Assistance Publique Hôpitaux de Paris, Université Paris-Est Créteil, INSERM U955 | Christophe Rodriguez, Slim Fourati, Vanessa Demontant, Guillaume Gricourt, Melissa N'Debi, Alexandre Soulier, Elisabeth Trawinski, Jean-Michel Pawlotsky |
| EPI_ISL_3453147 | CH. BEGIN | Department of Virology, Henri Mondor University Hospital, Assistance Publique Hôpitaux de Paris, Université Paris-Est Créteil, INSERM U955 | Christophe Rodriguez, Slim Fourati, Vanessa Demontant, Guillaume Gricourt, Melissa N'Debi, Alexandre Soulier, Elisabeth Trawinski, Jean-Michel Pawlotsky |
| EPI_ISL_3453148, EPI_ISL_3453149, EPI_ISL_3453150, EPI_ISL_3453151, EPI_ISL_3453152, EPI_ISL_3453153, EPI_ISL_3453154, EPI_ISL_3453155, EPI_ISL_3453156 | Biogroup Bio Lam-LCD Saint-Denis | Department of Virology, Henri Mondor University Hospital, Assistance Publique Hôpitaux de Paris, Université Paris-Est Créteil, INSERM U955 | Christophe Rodriguez, Slim Fourati, Vanessa Demontant, Guillaume Gricourt, Melissa N'Debi, Alexandre Soulier, Elisabeth Trawinski, Jean-Michel Pawlotsky |
| EPI_ISL_3453157 | LABM DEFRANCE | Department of Virology, Henri Mondor University Hospital, Assistance Publique Hôpitaux de Paris, Université Paris-Est Créteil, INSERM U955 | Christophe Rodriguez, Slim Fourati, Vanessa Demontant, Guillaume Gricourt, Melissa N'Debi, Alexandre Soulier, Elisabeth Trawinski, Jean-Michel Pawlotsky |
| EPI_ISL_3453158 | SELARL BIO | Department of Virology, Henri Mondor University Hospital, Assistance Publique Hôpitaux de Paris, Université Paris-Est Créteil, INSERM U955 | Christophe Rodriguez, Slim Fourati, Vanessa Demontant, Guillaume Gricourt, Melissa N'Debi, Alexandre Soulier, Elisabeth Trawinski, Jean-Michel Pawlotsky |
| EPI_ISL_3453159 | LISIEUX Cerballiance | Department of Virology, Henri Mondor University Hospital, Assistance Publique Hôpitaux de Paris, Université Paris-Est Créteil, INSERM U955 | Christophe Rodriguez, Slim Fourati, Vanessa Demontant, Guillaume Gricourt, Melissa N'Debi, Alexandre Soulier, Elisabeth Trawinski, Jean-Michel Pawlotsky |
| EPI_ISL_3453160 | LABM DEFRANCE | Department of Virology, Henri Mondor University Hospital, Assistance Publique Hôpitaux de Paris, Université Paris-Est Créteil, INSERM U955 | Christophe Rodriguez, Slim Fourati, Vanessa Demontant, Guillaume Gricourt, Melissa N'Debi, Alexandre Soulier, Elisabeth Trawinski, Jean-Michel Pawlotsky |
| EPI_ISL_3453161 | LISIEUX Cerballiance | Department of Virology, Henri Mondor University Hospital, Assistance Publique Hôpitaux de Paris, Université Paris-Est Créteil, INSERM U955 | Christophe Rodriguez, Slim Fourati, Vanessa Demontant, Guillaume Gricourt, Melissa N'Debi, Alexandre Soulier, Elisabeth Trawinski, Jean-Michel Pawlotsky |
| EPI_ISL_3453162 | LABM DEFRANCE | Department of Virology, Henri Mondor University Hospital, Assistance Publique Hôpitaux de Paris, Université Paris-Est Créteil, INSERM U955 | Christophe Rodriguez, Slim Fourati, Vanessa Demontant, Guillaume Gricourt, Melissa N'Debi, Alexandre Soulier, Elisabeth Trawinski, Jean-Michel Pawlotsky |
| EPI_ISL_3453163 | SELARL BIO | Department of Virology, Henri Mondor University Hospital, Assistance Publique Hôpitaux de Paris, Université Paris-Est Créteil, INSERM U955 | Christophe Rodriguez, Slim Fourati, Vanessa Demontant, Guillaume Gricourt, Melissa N'Debi, Alexandre Soulier, Elisabeth Trawinski, Jean-Michel Pawlotsky |

| | | | |
|---|---|---|---|
| EPI_ISL_3453164, EPI_ISL_3453165, EPI_ISL_3453167, EPI_ISL_3453168, EPI_ISL_3453169, EPI_ISL_3453170, EPI_ISL_3453171, EPI_ISL_3453172, EPI_ISL_3453173, EPI_ISL_3453174, EPI_ISL_3453175, EPI_ISL_3453176, EPI_ISL_3453177, EPI_ISL_3453179 | | | |
| see above | Laboratory of Microbiology and Virology, Ospedale Amedeo di Savoia, ASL "Città di Torino" | Laboratory of Microbiology and Virology, Ospedale Amedeo di Savoia, ASL "Città di Torino" | Tiziano Allice, Francesco Cerutti, Maria Grazia Milia, Gabriella Gregori, Elisa Burdino, Paola Tremante, Antonella Bottoni, Valeria Ghisetti |
| EPI_ISL_3453180, EPI_ISL_3453181, EPI_ISL_3453182, EPI_ISL_3453183, EPI_ISL_3453184, EPI_ISL_3453185, EPI_ISL_3453186, EPI_ISL_3453187, EPI_ISL_3453188, EPI_ISL_3453189, EPI_ISL_3453190, EPI_ISL_3453191, EPI_ISL_3453192, EPI_ISL_3453193, EPI_ISL_3453194, EPI_ISL_3453196, EPI_ISL_3453197, EPI_ISL_3453198, EPI_ISL_3453200, EPI_ISL_3453201, EPI_ISL_3453202, EPI_ISL_3453203, EPI_ISL_3453204, EPI_ISL_3453205, EPI_ISL_3453206, EPI_ISL_3453207, EPI_ISL_3453208, EPI_ISL_3453209, EPI_ISL_3453210, EPI_ISL_3453211, EPI_ISL_3453212, EPI_ISL_3453213, EPI_ISL_3453214, EPI_ISL_3453215, EPI_ISL_3453216, EPI_ISL_3453217 | | | |
| see above | National Platform bis UMONS/Jolimont | National Platform bis UMONS/Jolimont | François Dufrasne, Guillaume Bayon-Vicente, Florian Juszczak, Laetitia Gheysen, Eric Tarantino, Gautier Detry, Ruddy Wattiez |
| EPI_ISL_3453218, EPI_ISL_3453220, EPI_ISL_3453221, EPI_ISL_3453223, EPI_ISL_3453224, EPI_ISL_3453225, EPI_ISL_3453226, EPI_ISL_3453227, EPI_ISL_3453228, EPI_ISL_3453229, EPI_ISL_3453230, EPI_ISL_3453231, EPI_ISL_3453232, EPI_ISL_3453233, EPI_ISL_3453234, EPI_ISL_3453236, EPI_ISL_3453237, EPI_ISL_3453238, EPI_ISL_3453239, EPI_ISL_3453240, EPI_ISL_3453241, EPI_ISL_3453242, EPI_ISL_3453243, EPI_ISL_3453244, EPI_ISL_3453247, EPI_ISL_3453248, EPI_ISL_3453249, EPI_ISL_3453251, EPI_ISL_3453252, EPI_ISL_3453253, EPI_ISL_3453254, EPI_ISL_3453255, EPI_ISL_3453257 | | | |
| see above | Fondazione IRCCS Ca' Granda Ospedale Maggiore Policlinico | Fondazione IRCCS Ca' Granda Ospedale Maggiore Policlinico | Ferruccio Ceriotti, Sara Uceda Renteria |
| EPI_ISL_3453270, EPI_ISL_3453272, EPI_ISL_3453276, EPI_ISL_3453278, EPI_ISL_3453280, EPI_ISL_3453281, EPI_ISL_3453284, EPI_ISL_3453285, EPI_ISL_3453286, EPI_ISL_3453287, EPI_ISL_3453288, EPI_ISL_3453290, EPI_ISL_3453291, EPI_ISL_3453292, EPI_ISL_3453293, EPI_ISL_3453294, EPI_ISL_3453295, EPI_ISL_3453297, EPI_ISL_3453299, EPI_ISL_3453300, EPI_ISL_3453301, EPI_ISL_3453302, EPI_ISL_3453303, EPI_ISL_3453304, EPI_ISL_3453305, EPI_ISL_3453306, EPI_ISL_3453307, EPI_ISL_3453308, EPI_ISL_3453309, EPI_ISL_3453311, EPI_ISL_3453312, EPI_ISL_3453315, EPI_ISL_3453316, EPI_ISL_3453317, EPI_ISL_3453318, EPI_ISL_3453319, EPI_ISL_3453320, EPI_ISL_3453321, EPI_ISL_3453322, EPI_ISL_3453323, EPI_ISL_3453324, EPI_ISL_3453325, EPI_ISL_3453327, EPI_ISL_3453328, EPI_ISL_3453329, EPI_ISL_3453331, EPI_ISL_3453332, EPI_ISL_3453333, EPI_ISL_3453334, EPI_ISL_3453336, EPI_ISL_3453337, EPI_ISL_3453338, EPI_ISL_3453339, EPI_ISL_3453340, EPI_ISL_3453341, EPI_ISL_3453342, EPI_ISL_3453343, EPI_ISL_3453344, EPI_ISL_3453345, EPI_ISL_3453346, EPI_ISL_3453347, EPI_ISL_3453348, EPI_ISL_3453349, EPI_ISL_3453350, EPI_ISL_3453351, EPI_ISL_3453352, EPI_ISL_3453354, EPI_ISL_3453355, EPI_ISL_3453356, EPI_ISL_3453358, EPI_ISL_3453359, EPI_ISL_3453360, EPI_ISL_3453362 | | | |
| see above | Institute of Microbiology and Immunology, Faculty of Medicine, University of Ljubljana | Institute of Microbiology and Immunology, Faculty of Medicine, University of Ljubljana | Alen Suljič, Samo Zakotnik, Tomaž Mark Zorec, Matic Brvar, Doroteja Vlaj, Andraž Celar, Dominika Šturm, Patricija Pozvek, Špela Pleh, Miša Korva, Mario Poljak, Tatjana Avšič – Županc |
| EPI_ISL_3453363 | AP SSO | CSIR-Centre for Cellular and Molecular Biology-INSACOG | Payel Mukherjee, Lamuk Zaveri, Onkar Kulkarni,Tulasi Nagabandi, Sofia Banu, Ara Sreenivas,Shreekant Verma, Amareshwar Vodapalli ,B Himasri,Valli Nagalakshmi Undamatla, Sumedha Avadhanula, Vidhyadhari Methuku, Priya Nurkurthy, Archana Bharadwaj Siva, Karthik Bharadwaj Tallapaka,Krishna Khairnar,Rakesh K Mishra,Divya Tej Sowpati |
| EPI_ISL_3453364 | AP SSO | CSIR-Centre for Cellular and Molecular Biology-INSACOG | Amareshwar Vodapalli,Ara Sreenivas,Shreekant Verma, B Himasri,Valli Nagalakshmi Undamatla,Payel Mukherjee, Lamuk Zaveri, Onkar Kulkarni,Tulasi Nagabandi, Sofia Banu, Jandhyala Sai Krishna,Priya Nurkurthy, Vidhyadhari Methuku, Sumedha Avadhanula, Archana Bharadwaj Siva, Karthik Bharadwaj Tallapaka,Rakesh K Mishra,Divya Tej Sowpati |
| EPI_ISL_3453366 | Mapmygenome | CSIR-Centre for Cellular and Molecular Biology-INSACOG | Tulasi Nagabandi,Lamuk Zaveri,Payel Mukherjee,Onkar Kulkarni,Jandhyala Sai Krishna, Sofia Banu, Ara Sreenivas,Shreekant Verma, Amareshwar Vodapalli ,B Himasri,Valli Nagalakshmi Undamatla, Priya Nurkurthy,Sumedha Avadhanula, Vidhyadhari Methuku, Archana Bharadwaj Siva, Karthik Bharadwaj Tallapaka,Rakesh K Mishra,Divya Tej Sowpati |
| EPI_ISL_3453370 | AP SSO | CSIR-Centre for Cellular and Molecular Biology-INSACOG | Sumedha Avadhanula, Lamuk Zaveri, Tulasi Nagabandi, Ara Sreenivas,Shreekant Verma, Amareshwar Vodapalli ,B Himasri,Valli Nagalakshmi Undamatla, Vidhyadhari Methuku, Priya Nurkurthy, Payel Mukherjee,Jandhyala Sai Krishna,Onkar Kulkarni, Sofia Banu, Archana Bharadwaj Siva, Karthik Bharadwaj Tallapaka,Rakesh K Mishra,Divya Tej Sowpati |
| EPI_ISL_3453375 | AP SSO | CSIR-Centre for Cellular and Molecular Biology-INSACOG | B Himasri,Amareshwar Vodapalli,Ara Sreenivas,Shreekant Verma,Valli Nagalakshmi Undamatla,Payel Mukherjee, Lamuk Zaveri, Onkar Kulkarni,Tulasi Nagabandi, Sofia Banu, Vidhyadhari Methuku,Priya Nurkurthy, Jandhyala Sai Krishna,Sumedha Avadhanula, Archana Bharadwaj Siva, Karthik Bharadwaj Tallapaka,Rakesh K Mishra,Divya Tej Sowpati |
| EPI_ISL_3453376 | RUHS College of Medical Sciences | CSIR-Centre for Cellular and Molecular Biology - INSACOG | Dr. Gaurav Dalela,Dr. Vaseem Naheed Baig,Dr. Sonali Sharma,Dr. Ramesh Sharma,Dr. Nilofer Khayyam,Dr. Jitendra Panda,Dr. Sudhanshu Kacker,Dr. Raja Babu Panwar,Dr. Rajeev Gupta, Lamuk Zaveri,Tulasi Nagabandi,Payel Mukherjee,Onkar Kulkarni,Jandhyala Sai Krishna, Sofia Banu, Ara Sreenivas,Shreekant Verma, Amareshwar Vodapalli ,B Himasri,Valli Nagalakshmi Undamatla, Sumedha Avadhanula, Priya Nurkurthy, Vidhyadhari Methuku, Archana Bharadwaj Siva, Karthik Bharadwaj Tallapaka,Rakesh K Mishra,Divya Tej Sowpati |
| EPI_ISL_3453379 | AP SSO | CSIR-Centre for Cellular and Molecular Biology-INSACOG | Tulasi Nagabandi,Lamuk Zaveri,Payel Mukherjee,Onkar Kulkarni,Jandhyala Sai Krishna, Sofia Banu, Ara Sreenivas,Shreekant Verma, Amareshwar Vodapalli ,B Himasri,Valli Nagalakshmi Undamatla, Priya Nurkurthy,Sumedha Avadhanula, Vidhyadhari Methuku, Archana Bharadwaj Siva, Karthik Bharadwaj Tallapaka,Rakesh K Mishra,Divya Tej Sowpati |
| EPI_ISL_3453381 | Mapmygenome | CSIR-Centre for Cellular and Molecular Biology-INSACOG | Priya Nurkurthy,Ara Sreenivas,Shreekant Verma, Amareshwar Vodapalli ,Lamuk Zaveri,Tulasi Nagabandi,B Himasri,Valli Nagalakshmi Undamatla, Sumedha Avadhanula, Vidhyadhari Methuku,Payel Mukherjee,Onkar Kulkarni, Jandhyala Sai Krishna,Sofia Banu, Archana Bharadwaj Siva, Karthik Bharadwaj Tallapaka,Rakesh K Mishra,Divya Tej Sowpati |
| EPI_ISL_3453383 | Mapmygenome | CSIR-Centre for Cellular and Molecular Biology-INSACOG | Shreekant Verma, Amareshwar Vodapalli ,Ara Sreenivas,B Himasri,Valli Nagalakshmi Undamatla,Payel Mukherjee, Lamuk Zaveri, Onkar Kulkarni,Tulasi Nagabandi, Sofia Banu, Jandhyala Sai Krishna,Sumedha Avadhanula, Priya Nurkurthy, Vidhyadhari Methuku, Archana Bharadwaj Siva, Karthik Bharadwaj Tallapaka,Rakesh K Mishra,Divya Tej Sowpati |
| EPI_ISL_3453385 | AP SSO | CSIR-Centre for Cellular and Molecular Biology-INSACOG | B Himasri,Amareshwar Vodapalli,Ara Sreenivas,Shreekant Verma,Valli Nagalakshmi Undamatla,Payel Mukherjee, Lamuk Zaveri, Onkar Kulkarni,Tulasi Nagabandi, Sofia Banu, Vidhyadhari Methuku,Priya Nurkurthy, Jandhyala Sai Krishna,Sumedha Avadhanula, Archana Bharadwaj Siva, Karthik Bharadwaj Tallapaka,Rakesh K Mishra,Divya Tej Sowpati |
| EPI_ISL_3453390 | AP SSO | CSIR-Centre for Cellular and Molecular Biology-INSACOG | Sumedha Avadhanula, Lamuk Zaveri, Tulasi Nagabandi, Ara Sreenivas,Shreekant Verma, Amareshwar Vodapalli ,B Himasri,Valli Nagalakshmi Undamatla, Vidhyadhari Methuku, Priya Nurkurthy, Payel Mukherjee,Jandhyala Sai Krishna,Onkar Kulkarni, Sofia Banu, Archana Bharadwaj Siva, Karthik Bharadwaj Tallapaka,Rakesh K Mishra,Divya Tej Sowpati |
| EPI_ISL_3453391 | Mapmygenome | CSIR-Centre for Cellular and Molecular Biology-INSACOG | Shreekant Verma, Amareshwar Vodapalli ,Ara Sreenivas,B Himasri,Valli Nagalakshmi Undamatla,Payel Mukherjee, Lamuk Zaveri, Onkar Kulkarni,Tulasi Nagabandi, Sofia Banu, Jandhyala Sai Krishna,Sumedha Avadhanula, Priya Nurkurthy, Vidhyadhari Methuku, Archana Bharadwaj Siva, Karthik Bharadwaj Tallapaka,Rakesh K Mishra,Divya Tej Sowpati |
| EPI_ISL_3453393, EPI_ISL_3453396 | AP SSO | CSIR-Centre for Cellular and Molecular Biology-INSACOG | B Himasri,Amareshwar Vodapalli,Ara Sreenivas,Shreekant Verma,Valli Nagalakshmi Undamatla,Payel Mukherjee, Lamuk Zaveri, Onkar Kulkarni,Tulasi Nagabandi, Sofia Banu, Vidhyadhari Methuku,Priya Nurkurthy, Jandhyala Sai Krishna,Sumedha Avadhanula, Archana Bharadwaj Siva, Karthik Bharadwaj Tallapaka,Rakesh K Mishra,Divya Tej Sowpati |
| EPI_ISL_3453403 | AP SSO | CSIR-Centre for Cellular and Molecular Biology-INSACOG | Sumedha Avadhanula, Lamuk Zaveri, Tulasi Nagabandi, Ara Sreenivas,Shreekant Verma, Amareshwar Vodapalli ,B Himasri,Valli Nagalakshmi Undamatla, Vidhyadhari Methuku, Priya Nurkurthy, Payel Mukherjee,Jandhyala Sai Krishna,Onkar Kulkarni, Sofia Banu, Archana Bharadwaj Siva, Karthik Bharadwaj Tallapaka,Rakesh K Mishra,Divya Tej Sowpati |
| EPI_ISL_3453404 | AP SSO | CSIR-Centre for Cellular and Molecular Biology-INSACOG | Amareshwar Vodapalli,Ara Sreenivas,Shreekant Verma, B Himasri,Valli Nagalakshmi Undamatla,Payel Mukherjee, Lamuk Zaveri, Onkar Kulkarni,Tulasi Nagabandi, Sofia Banu, Jandhyala Sai Krishna,Priya Nurkurthy, Vidhyadhari Methuku, Sumedha Avadhanula, Archana Bharadwaj Siva, Karthik Bharadwaj Tallapaka,Rakesh K Mishra,Divya Tej Sowpati |
| EPI_ISL_3453406 | AP SSO | CSIR-Centre for Cellular and Molecular Biology-INSACOG | Priya Nurkurthy,Ara Sreenivas,Shreekant Verma, Amareshwar Vodapalli ,Lamuk Zaveri,Tulasi Nagabandi,B Himasri,Valli Nagalakshmi Undamatla, Sumedha Avadhanula, Vidhyadhari Methuku,Payel Mukherjee,Onkar Kulkarni, Jandhyala Sai Krishna,Sofia Banu, Archana Bharadwaj Siva, Karthik Bharadwaj Tallapaka,Rakesh K Mishra,Divya Tej Sowpati |
| EPI_ISL_3453409 | Mapmygenome | CSIR-Centre for Cellular and Molecular Biology-INSACOG | Tulasi Nagabandi,Lamuk Zaveri,Payel Mukherjee,Onkar Kulkarni,Jandhyala Sai Krishna, Sofia Banu, Ara Sreenivas,Shreekant Verma, Amareshwar Vodapalli ,B Himasri,Valli Nagalakshmi Undamatla, Priya Nurkurthy,Sumedha Avadhanula, Vidhyadhari Methuku, Archana Bharadwaj Siva, Karthik Bharadwaj Tallapaka,Rakesh K Mishra,Divya Tej Sowpati |
| EPI_ISL_3453410 | Mapmygenome | CSIR-Centre for Cellular and Molecular Biology-INSACOG | Vidhyadhari Methuku,Tulasi Nagabandi,Lamuk Zaveri, Ara Sreenivas,Shreekant Verma, Amareshwar Vodapalli ,B Himasri,Valli Nagalakshmi Undamatla, Priya Nurkurthy,Sumedha Avadhanula, Archana Bharadwaj Siva,Payel Mukherjee,Onkar Kulkarni, Jandhyala Sai Krishna,Sofia Banu, Karthik Bharadwaj Tallapaka,Rakesh K Mishra,Divya Tej Sowpati |
| EPI_ISL_3453411 | AP SSO | CSIR-Centre for Cellular and Molecular Biology-INSACOG | Amareshwar Vodapalli,Ara Sreenivas,Shreekant Verma, B Himasri,Valli Nagalakshmi Undamatla,Payel Mukherjee, Lamuk Zaveri, Onkar Kulkarni,Tulasi Nagabandi, Sofia Banu, Jandhyala Sai Krishna,Priya Nurkurthy, Vidhyadhari Methuku, Sumedha Avadhanula, Archana Bharadwaj Siva, Karthik Bharadwaj Tallapaka,Rakesh K Mishra,Divya Tej Sowpati |
| EPI_ISL_3453412 | AP SSO | CSIR-Centre for Cellular and Molecular Biology-INSACOG | Lamuk Zaveri,Tulasi Nagabandi,Payel Mukherjee,Onkar Kulkarni, Sofia Banu, Jandhyala Sai Krishna,Ara Sreenivas,Shreekant Verma, Amareshwar Vodapalli ,B Himasri,Valli Nagalakshmi Undamatla, Sumedha Avadhanula, Priya Nurkurthy, Vidhyadhari Methuku, Archana Bharadwaj Siva, Karthik Bharadwaj Tallapaka,Rakesh K Mishra,Divya Tej Sowpati |
| EPI_ISL_3453413 | AP SSO | CSIR-Centre for Cellular and Molecular Biology-INSACOG | Shreekant Verma, Amareshwar Vodapalli ,Ara Sreenivas,B Himasri,Valli Nagalakshmi Undamatla,Payel Mukherjee, Lamuk Zaveri, Onkar Kulkarni,Tulasi Nagabandi, Sofia Banu, Jandhyala Sai Krishna,Sumedha Avadhanula, Priya Nurkurthy, Vidhyadhari Methuku, Archana Bharadwaj Siva, Karthik Bharadwaj Tallapaka,Rakesh K Mishra,Divya Tej Sowpati |
| EPI_ISL_3453414 | Mapmygenome | CSIR-Centre for Cellular and Molecular Biology-INSACOG | Priya Nurkurthy,Ara Sreenivas,Shreekant Verma, Amareshwar Vodapalli ,Lamuk Zaveri,Tulasi Nagabandi,B Himasri,Valli Nagalakshmi Undamatla, Sumedha Avadhanula, Vidhyadhari Methuku,Payel Mukherjee,Onkar Kulkarni, Jandhyala Sai Krishna,Sofia Banu, Archana Bharadwaj Siva, Karthik Bharadwaj Tallapaka,Rakesh K Mishra,Divya Tej Sowpati |
| EPI_ISL_3453416 | AP SSO | CSIR-Centre for Cellular and Molecular Biology-INSACOG | Payel Mukherjee, Lamuk Zaveri, Onkar Kulkarni,Jandhyala Sai Krishna,Tulasi Nagabandi, Sofia Banu, Ara Sreenivas,Shreekant Verma, Amareshwar Vodapalli ,B Himasri,Valli Nagalakshmi Undamatla, Sumedha Avadhanula, Vidhyadhari Methuku, Priya Nurkurthy, Archana Bharadwaj Siva, Karthik Bharadwaj Tallapaka,Rakesh K Mishra,Divya Tej Sowpati |
| EPI_ISL_3453417 | AP SSO | CSIR-Centre for Cellular and Molecular Biology-INSACOG | Lamuk Zaveri,Tulasi Nagabandi,Payel Mukherjee,Onkar Kulkarni, Sofia Banu, Jandhyala Sai Krishna,Ara Sreenivas,Shreekant Verma, Amareshwar Vodapalli ,B Himasri,Valli Nagalakshmi Undamatla, Sumedha Avadhanula, Priya Nurkurthy, Vidhyadhari Methuku, Archana Bharadwaj Siva, Karthik Bharadwaj Tallapaka,Rakesh K Mishra,Divya Tej Sowpati |
| EPI_ISL_3453419 | AP SSO | CSIR-Centre for Cellular and Molecular Biology-INSACOG | Sumedha Avadhanula, Lamuk Zaveri, Tulasi Nagabandi, Ara Sreenivas,Shreekant Verma, Amareshwar Vodapalli ,B Himasri,Valli Nagalakshmi Undamatla, Vidhyadhari Methuku, Priya Nurkurthy, Payel Mukherjee,Jandhyala Sai Krishna,Onkar Kulkarni, Sofia Banu, Archana Bharadwaj Siva, Karthik Bharadwaj Tallapaka,Rakesh K Mishra,Divya Tej Sowpati |
| EPI_ISL_3453420 | AP SSO | CSIR-Centre for Cellular and Molecular Biology-INSACOG | Amareshwar Vodapalli,Ara Sreenivas,Shreekant Verma, B Himasri,Valli Nagalakshmi Undamatla,Payel Mukherjee, Lamuk Zaveri, Onkar Kulkarni,Tulasi Nagabandi, Sofia Banu, Jandhyala Sai Krishna,Priya Nurkurthy, Vidhyadhari Methuku, Sumedha Avadhanula, Archana Bharadwaj Siva, Karthik Bharadwaj Tallapaka,Rakesh K Mishra,Divya Tej Sowpati |
| EPI_ISL_3453421 | AP SSO | CSIR-Centre for Cellular and Molecular Biology-INSACOG | Tulasi Nagabandi,Lamuk Zaveri,Payel Mukherjee,Onkar Kulkarni,Jandhyala Sai Krishna, Sofia Banu, Ara Sreenivas,Shreekant Verma, Amareshwar Vodapalli ,B Himasri,Valli Nagalakshmi Undamatla, Priya Nurkurthy,Sumedha Avadhanula, Vidhyadhari Methuku, Archana Bharadwaj Siva, Karthik Bharadwaj Tallapaka,Rakesh K Mishra,Divya Tej Sowpati |
| EPI_ISL_3453422 | AP SSO | CSIR-Centre for Cellular and Molecular Biology-INSACOG | Sumedha Avadhanula, Lamuk Zaveri, Tulasi Nagabandi, Ara Sreenivas,Shreekant Verma, Amareshwar Vodapalli ,B Himasri,Valli Nagalakshmi Undamatla, Vidhyadhari Methuku, Priya Nurkurthy, Payel Mukherjee,Jandhyala Sai Krishna,Onkar Kulkarni, Sofia Banu, Archana Bharadwaj Siva, Karthik Bharadwaj Tallapaka,Rakesh K Mishra,Divya Tej Sowpati |
| EPI_ISL_3453423 | AP SSO | CSIR-Centre for Cellular and Molecular Biology-INSACOG | Tulasi Nagabandi,Lamuk Zaveri,Payel Mukherjee,Onkar Kulkarni,Jandhyala Sai Krishna, Sofia Banu, Ara Sreenivas,Shreekant Verma, Amareshwar Vodapalli ,B Himasri,Valli Nagalakshmi Undamatla, Priya Nurkurthy,Sumedha Avadhanula, Vidhyadhari Methuku, Archana Bharadwaj Siva, Karthik Bharadwaj Tallapaka,Rakesh K Mishra,Divya Tej Sowpati |
| EPI_ISL_3453425 | AP SSO | CSIR-Centre for Cellular and Molecular Biology-INSACOG | Lamuk Zaveri,Tulasi Nagabandi,Payel Mukherjee,Onkar Kulkarni, Sofia Banu, Jandhyala Sai Krishna,Ara Sreenivas,Shreekant Verma, Amareshwar Vodapalli ,B Himasri,Valli Nagalakshmi Undamatla, Sumedha Avadhanula, Priya Nurkurthy, Vidhyadhari Methuku, Archana Bharadwaj Siva, Karthik Bharadwaj Tallapaka,Rakesh K Mishra,Divya Tej Sowpati |
| EPI_ISL_3453427 | AP SSO | CSIR-Centre for Cellular and Molecular Biology-INSACOG | Sumedha Avadhanula, Lamuk Zaveri, Tulasi Nagabandi, Ara Sreenivas,Shreekant Verma, Amareshwar Vodapalli ,B Himasri,Valli Nagalakshmi Undamatla, Vidhyadhari Methuku, Priya Nurkurthy, Payel Mukherjee,Jandhyala Sai Krishna,Onkar Kulkarni, Sofia Banu, Archana Bharadwaj Siva, Karthik Bharadwaj Tallapaka,Rakesh K Mishra,Divya Tej Sowpati |
| EPI_ISL_3453428 | AP SSO | CSIR-Centre for Cellular and Molecular Biology-INSACOG | Priya Nurkurthy,Ara Sreenivas,Shreekant Verma, Amareshwar Vodapalli ,Lamuk Zaveri,Tulasi Nagabandi,B Himasri,Valli Nagalakshmi Undamatla, Sumedha Avadhanula, Vidhyadhari Methuku,Payel Mukherjee,Onkar Kulkarni, Jandhyala Sai Krishna,Sofia Banu, Archana Bharadwaj Siva, Karthik Bharadwaj Tallapaka,Rakesh K Mishra,Divya Tej Sowpati |
| EPI_ISL_3453431 | AP SSO | CSIR-Centre for Cellular and Molecular Biology-INSACOG | B Himasri,Amareshwar Vodapalli,Ara Sreenivas,Shreekant Verma,Valli Nagalakshmi Undamatla,Payel Mukherjee, Lamuk Zaveri, Onkar Kulkarni,Tulasi Nagabandi, Sofia Banu, Vidhyadhari Methuku,Priya Nurkurthy, Jandhyala Sai Krishna,Sumedha Avadhanula, Archana Bharadwaj Siva, Karthik Bharadwaj Tallapaka,Rakesh K Mishra,Divya Tej Sowpati |
| EPI_ISL_3453433 | AP SSO | CSIR-Centre for Cellular and Molecular Biology-INSACOG | Sumedha Avadhanula, Lamuk Zaveri, Tulasi Nagabandi, Ara Sreenivas,Shreekant Verma, Amareshwar Vodapalli ,B Himasri,Valli Nagalakshmi Undamatla, Vidhyadhari Methuku, Priya Nurkurthy, Payel Mukherjee,Jandhyala Sai Krishna,Onkar Kulkarni, Sofia Banu, Archana Bharadwaj Siva, Karthik Bharadwaj Tallapaka,Rakesh K Mishra,Divya Tej Sowpati |
| EPI_ISL_3453434 | AP SSO | CSIR-Centre for Cellular and Molecular Biology-INSACOG | Shreekant Verma, Amareshwar Vodapalli ,Ara Sreenivas,B Himasri,Valli Nagalakshmi Undamatla,Payel Mukherjee, Lamuk Zaveri, Onkar Kulkarni,Tulasi Nagabandi, Sofia Banu, Jandhyala Sai Krishna,Sumedha Avadhanula, Priya Nurkurthy, Vidhyadhari Methuku, Archana Bharadwaj Siva, Karthik Bharadwaj Tallapaka,Rakesh K Mishra,Divya Tej Sowpati |
| EPI_ISL_3453435 | AP SSO | CSIR-Centre for Cellular and Molecular Biology-INSACOG | Lamuk Zaveri,Tulasi Nagabandi,Payel Mukherjee,Onkar Kulkarni, Sofia Banu, Jandhyala Sai Krishna,Ara Sreenivas,Shreekant Verma, Amareshwar Vodapalli ,B Himasri,Valli Nagalakshmi Undamatla, Sumedha Avadhanula, Priya Nurkurthy, Vidhyadhari Methuku, Archana Bharadwaj Siva, Karthik Bharadwaj Tallapaka,Rakesh K Mishra,Divya Tej Sowpati |
| EPI_ISL_3453436 | AP SSO | CSIR-Centre for Cellular and Molecular Biology-INSACOG | Priya Nurkurthy,Ara Sreenivas,Shreekant Verma, Amareshwar Vodapalli ,Lamuk Zaveri,Tulasi Nagabandi,B Himasri,Valli Nagalakshmi Undamatla, Sumedha Avadhanula, Vidhyadhari Methuku,Payel Mukherjee,Onkar Kulkarni, Jandhyala Sai Krishna,Sofia Banu, Archana Bharadwaj Siva, Karthik Bharadwaj Tallapaka,Rakesh K Mishra,Divya Tej Sowpati |

| | | | |
|---|---|---|---|
| EPI_ISL_3453437 | AP SSO | CSIR-Centre for Cellular and Molecular Biology-INSACOG | Payel Mukherjee, Lamuk Zaveri, Onkar Kulkarni,Jandhyala Sai Krishna,Tulasi Nagabandi, Sofia Banu, Ara Sreenivas,Shreekant Verma, Amareshwar Vodapalli ,B Himasri,Valli Nagalakshmi Undamatla, Sumedha Avadhanula, Vidhyadhari Methuku, Priya Nurkurthy, Archana Bharadwaj Siva, Karthik Bharadwaj Tallapaka,Rakesh K Mishra,Divya Tej Sowpati |
| EPI_ISL_3453438 | AP SSO | CSIR-Centre for Cellular and Molecular Biology-INSACOG | Sumedha Avadhanula, Lamuk Zaveri, Tulasi Nagabandi, Ara Sreenivas,Shreekant Verma, Amareshwar Vodapalli ,B Himasri,Valli Nagalakshmi Undamatla, Vidhyadhari Methuku, Priya Nurkurthy, Payel Mukherjee,Jandhyala Sai Krishna,Onkar Kulkarni, Sofia Banu, Archana Bharadwaj Siva, Karthik Bharadwaj Tallapaka,Rakesh K Mishra,Divya Tej Sowpati |
| EPI_ISL_3453439 | Mapmygenome | CSIR-Centre for Cellular and Molecular Biology-INSACOG | Sumedha Avadhanula, Lamuk Zaveri, Tulasi Nagabandi, Ara Sreenivas,Shreekant Verma, Amareshwar Vodapalli ,B Himasri,Valli Nagalakshmi Undamatla, Vidhyadhari Methuku, Priya Nurkurthy, Payel Mukherjee,Jandhyala Sai Krishna,Onkar Kulkarni, Sofia Banu, Archana Bharadwaj Siva, Karthik Bharadwaj Tallapaka,Rakesh K Mishra,Divya Tej Sowpati |
| EPI_ISL_3453440 | RUHS College of Medical Sciences | CSIR-Centre for Cellular and Molecular Biology - INSACOG | Dr. Gaurav Dalela,Dr. Vaseem Naheed Baig,Dr. Sonali Sharma,Dr. Ramesh Sharma,Dr. Nilofer Khayyam,Dr. Jitendra Panda,Dr. Sudhanshu Kacker,Dr. Raja Babu Panwar,Dr. Rajeev Gupta, Jandhyala Sai Krishna,Onkar Kulkarni, Lamuk Zaveri, Tulasi Nagabandi,Sofia Banu,Payel Mukherjee, Ara Sreenivas,Shreekant Verma, Amareshwar Vodapalli ,B Himasri,Valli Nagalakshmi Undamatla, Sumedha Avadhanula, Vidhyadhari Methuku, Priya Nurkurthy, Archana Bharadwaj Siva, Karthik Bharadwaj Tallapaka,Rakesh K Mishra,Divya Tej Sowpati |
| EPI_ISL_3453441 | Mapmygenome | CSIR-Centre for Cellular and Molecular Biology-INSACOG | Shreekant Verma, Amareshwar Vodapalli ,Ara Sreenivas,B Himasri,Valli Nagalakshmi Undamatla,Payel Mukherjee, Lamuk Zaveri, Onkar Kulkarni,Tulasi Nagabandi, Sofia Banu, Jandhyala Sai Krishna,Sumedha Avadhanula, Priya Nurkurthy, Vidhyadhari Methuku, Archana Bharadwaj Siva, Karthik Bharadwaj Tallapaka,Rakesh K Mishra,Divya Tej Sowpati |
| EPI_ISL_3453442 | AP SSO | CSIR-Centre for Cellular and Molecular Biology-INSACOG | Vidhyadhari Methuku,Tulasi Nagabandi,Lamuk Zaveri, Ara Sreenivas,Shreekant Verma, Amareshwar Vodapalli ,B Himasri,Valli Nagalakshmi Undamatla, Priya Nurkurthy,Sumedha Avadhanula, Archana Bharadwaj Siva,Payel Mukherjee,Onkar Kulkarni, Jandhyala Sai Krishna,Sofia Banu, Karthik Bharadwaj Tallapaka,Rakesh K Mishra,Divya Tej Sowpati |
| EPI_ISL_3453443 | AP SSO | CSIR-Centre for Cellular and Molecular Biology-INSACOG | Lamuk Zaveri,Tulasi Nagabandi,Payel Mukherjee,Onkar Kulkarni, Sofia Banu, Jandhyala Sai Krishna,Ara Sreenivas,Shreekant Verma, Amareshwar Vodapalli ,B Himasri,Valli Nagalakshmi Undamatla, Sumedha Avadhanula, Priya Nurkurthy, Vidhyadhari Methuku, Archana Bharadwaj Siva, Karthik Bharadwaj Tallapaka,Rakesh K Mishra,Divya Tej Sowpati |
| EPI_ISL_3453444 | AP SSO | CSIR-Centre for Cellular and Molecular Biology-INSACOG | Sumedha Avadhanula, Lamuk Zaveri, Tulasi Nagabandi, Ara Sreenivas,Shreekant Verma, Amareshwar Vodapalli ,B Himasri,Valli Nagalakshmi Undamatla, Vidhyadhari Methuku, Priya Nurkurthy, Payel Mukherjee,Jandhyala Sai Krishna,Onkar Kulkarni, Sofia Banu, Archana Bharadwaj Siva, Karthik Bharadwaj Tallapaka,Rakesh K Mishra,Divya Tej Sowpati |
| EPI_ISL_3453445 | AP SSO | CSIR-Centre for Cellular and Molecular Biology-INSACOG | Lamuk Zaveri,Tulasi Nagabandi,Payel Mukherjee,Onkar Kulkarni, Sofia Banu, Jandhyala Sai Krishna,Ara Sreenivas,Shreekant Verma, Amareshwar Vodapalli ,B Himasri,Valli Nagalakshmi Undamatla, Sumedha Avadhanula, Priya Nurkurthy, Vidhyadhari Methuku, Archana Bharadwaj Siva, Karthik Bharadwaj Tallapaka,Rakesh K Mishra,Divya Tej Sowpati |
| EPI_ISL_3453446 | AP SSO | CSIR-Centre for Cellular and Molecular Biology-INSACOG | Sofia Banu,Payel Mukherjee, Lamuk Zaveri, Onkar Kulkarni,Tulasi Nagabandi, Jandhyala Sai Krishna, Ara Sreenivas,Shreekant Verma, Amareshwar Vodapalli ,B Himasri,Valli Nagalakshmi Undamatla, Sumedha Avadhanula, Vidhyadhari Methuku, Priya Nurkurthy, Archana Bharadwaj Siva, Karthik Bharadwaj Tallapaka,Rakesh K Mishra,Divya Tej Sowpati |
| EPI_ISL_3453447 | AP SSO | CSIR-Centre for Cellular and Molecular Biology-INSACOG | Tulasi Nagabandi,Lamuk Zaveri,Payel Mukherjee,Onkar Kulkarni,Jandhyala Sai Krishna, Sofia Banu, Ara Sreenivas,Shreekant Verma, Amareshwar Vodapalli ,B Himasri,Valli Nagalakshmi Undamatla, Priya Nurkurthy,Sumedha Avadhanula, Vidhyadhari Methuku, Archana Bharadwaj Siva, Karthik Bharadwaj Tallapaka,Rakesh K Mishra,Divya Tej Sowpati |
| EPI_ISL_3453448, EPI_ISL_3453449 | AP SSO | CSIR-Centre for Cellular and Molecular Biology-INSACOG | Sumedha Avadhanula, Lamuk Zaveri, Tulasi Nagabandi, Ara Sreenivas,Shreekant Verma, Amareshwar Vodapalli ,B Himasri,Valli Nagalakshmi Undamatla, Vidhyadhari Methuku, Priya Nurkurthy, Payel Mukherjee,Jandhyala Sai Krishna,Onkar Kulkarni, Sofia Banu, Archana Bharadwaj Siva, Karthik Bharadwaj Tallapaka,Rakesh K Mishra,Divya Tej Sowpati |
| EPI_ISL_3453451 | Mapmygenome | CSIR-Centre for Cellular and Molecular Biology-INSACOG | Vidhyadhari Methuku,Tulasi Nagabandi,Lamuk Zaveri, Ara Sreenivas,Shreekant Verma, Amareshwar Vodapalli ,B Himasri,Valli Nagalakshmi Undamatla, Priya Nurkurthy,Sumedha Avadhanula, Archana Bharadwaj Siva,Payel Mukherjee,Onkar Kulkarni, Jandhyala Sai Krishna,Sofia Banu, Karthik Bharadwaj Tallapaka,Rakesh K Mishra,Divya Tej Sowpati |
| EPI_ISL_3453452 | Mapmygenome | CSIR-Centre for Cellular and Molecular Biology-INSACOG | Amareshwar Vodapalli,Ara Sreenivas,Shreekant Verma, B Himasri,Valli Nagalakshmi Undamatla,Payel Mukherjee, Lamuk Zaveri, Onkar Kulkarni,Tulasi Nagabandi, Sofia Banu, Jandhyala Sai Krishna,Priya Nurkurthy, Vidhyadhari Methuku, Sumedha Avadhanula, Archana Bharadwaj Siva, Karthik Bharadwaj Tallapaka,Rakesh K Mishra,Divya Tej Sowpati |
| EPI_ISL_3453453 | AP SSO | CSIR-Centre for Cellular and Molecular Biology-INSACOG | Sofia Banu,Payel Mukherjee, Lamuk Zaveri, Onkar Kulkarni,Tulasi Nagabandi, Jandhyala Sai Krishna, Ara Sreenivas,Shreekant Verma, Amareshwar Vodapalli ,B Himasri,Valli Nagalakshmi Undamatla, Sumedha Avadhanula, Vidhyadhari Methuku, Priya Nurkurthy, Archana Bharadwaj Siva, Karthik Bharadwaj Tallapaka,Rakesh K Mishra,Divya Tej Sowpati |
| EPI_ISL_3453454 | AP SSO | CSIR-Centre for Cellular and Molecular Biology-INSACOG | Ara Sreenivas,Shreekant Verma, Amareshwar Vodapalli ,B Himasri,Valli Nagalakshmi Undamatla,Payel Mukherjee, Lamuk Zaveri, Onkar Kulkarni,Jandhyala Sai Krishna,Tulasi Nagabandi, Sofia Banu, ,Priya Nurkurthy, Vidhyadhari Methuku, Sumedha Avadhanula, Archana Bharadwaj Siva, Karthik Bharadwaj Tallapaka,Rakesh K Mishra,Divya Tej Sowpati |
| EPI_ISL_3453455 | AP SSO | CSIR-Centre for Cellular and Molecular Biology-INSACOG | Shreekant Verma, Amareshwar Vodapalli ,Ara Sreenivas,B Himasri,Valli Nagalakshmi Undamatla,Payel Mukherjee, Lamuk Zaveri, Onkar Kulkarni,Tulasi Nagabandi, Sofia Banu, Jandhyala Sai Krishna,Sumedha Avadhanula, Priya Nurkurthy, Vidhyadhari Methuku, Archana Bharadwaj Siva, Karthik Bharadwaj Tallapaka,Rakesh K Mishra,Divya Tej Sowpati |
| EPI_ISL_3453456 | Mapmygenome | CSIR-Centre for Cellular and Molecular Biology-INSACOG | Vidhyadhari Methuku,Tulasi Nagabandi,Lamuk Zaveri, Ara Sreenivas,Shreekant Verma, Amareshwar Vodapalli ,B Himasri,Valli Nagalakshmi Undamatla, Priya Nurkurthy,Sumedha Avadhanula, Archana Bharadwaj Siva,Payel Mukherjee,Onkar Kulkarni, Jandhyala Sai Krishna,Sofia Banu, Karthik Bharadwaj Tallapaka,Rakesh K Mishra,Divya Tej Sowpati |
| EPI_ISL_3453457 | AP SSO | CSIR-Centre for Cellular and Molecular Biology-INSACOG | Amareshwar Vodapalli,Ara Sreenivas,Shreekant Verma, B Himasri,Valli Nagalakshmi Undamatla,Payel Mukherjee, Lamuk Zaveri, Onkar Kulkarni,Tulasi Nagabandi, Sofia Banu, Jandhyala Sai Krishna,Priya Nurkurthy, Vidhyadhari Methuku, Sumedha Avadhanula, Archana Bharadwaj Siva, Karthik Bharadwaj Tallapaka,Rakesh K Mishra,Divya Tej Sowpati |
| EPI_ISL_3453458 | AP SSO | CSIR-Centre for Cellular and Molecular Biology-INSACOG | Shreekant Verma, Amareshwar Vodapalli ,Ara Sreenivas,B Himasri,Valli Nagalakshmi Undamatla,Payel Mukherjee, Lamuk Zaveri, Onkar Kulkarni,Tulasi Nagabandi, Sofia Banu, Jandhyala Sai Krishna,Sumedha Avadhanula, Priya Nurkurthy, Vidhyadhari Methuku, Archana Bharadwaj Siva, Karthik Bharadwaj Tallapaka,Rakesh K Mishra,Divya Tej Sowpati |
| EPI_ISL_3453459 | AP SSO | CSIR-Centre for Cellular and Molecular Biology-INSACOG | Payel Mukherjee, Lamuk Zaveri, Onkar Kulkarni,Tulasi Nagabandi, Sofia Banu, Ara Sreenivas,Shreekant Verma, Amareshwar Vodapalli ,B Himasri,Valli Nagalakshmi Undamatla, Sumedha Avadhanula, Vidhyadhari Methuku, Priya Nurkurthy, Archana Bharadwaj Siva, Karthik Bharadwaj Tallapaka,Krishna Khairnar,Rakesh K Mishra,Divya Tej Sowpati |
| EPI_ISL_3453460 | AP SSO | CSIR-Centre for Cellular and Molecular Biology-INSACOG | Lamuk Zaveri,Tulasi Nagabandi,Payel Mukherjee,Onkar Kulkarni, Sofia Banu, Jandhyala Sai Krishna,Ara Sreenivas,Shreekant Verma, Amareshwar Vodapalli ,B Himasri,Valli Nagalakshmi Undamatla, Sumedha Avadhanula, Priya Nurkurthy, Vidhyadhari Methuku, Archana Bharadwaj Siva, Karthik Bharadwaj Tallapaka,Rakesh K Mishra,Divya Tej Sowpati |
| EPI_ISL_3453461 | AP SSO | CSIR-Centre for Cellular and Molecular Biology-INSACOG | Vidhyadhari Methuku,Tulasi Nagabandi,Lamuk Zaveri, Ara Sreenivas,Shreekant Verma, Amareshwar Vodapalli ,B Himasri,Valli Nagalakshmi Undamatla, Priya Nurkurthy,Sumedha Avadhanula, Archana Bharadwaj Siva,Payel Mukherjee,Onkar Kulkarni, Jandhyala Sai Krishna,Sofia Banu, Karthik Bharadwaj Tallapaka,Rakesh K Mishra,Divya Tej Sowpati |
| EPI_ISL_3453462 | AP SSO | CSIR-Centre for Cellular and Molecular Biology-INSACOG | Payel Mukherjee, Lamuk Zaveri, Onkar Kulkarni,Jandhyala Sai Krishna,Tulasi Nagabandi, Sofia Banu, Ara Sreenivas,Shreekant Verma, Amareshwar Vodapalli ,B Himasri,Valli Nagalakshmi Undamatla, Sumedha Avadhanula, Vidhyadhari Methuku, Priya Nurkurthy, Archana Bharadwaj Siva, Karthik Bharadwaj Tallapaka,Rakesh K Mishra,Divya Tej Sowpati |
| EPI_ISL_3453463 | AP SSO | CSIR-Centre for Cellular and Molecular Biology-INSACOG | Sumedha Avadhanula, Lamuk Zaveri, Tulasi Nagabandi, Ara Sreenivas,Shreekant Verma, Amareshwar Vodapalli ,B Himasri,Valli Nagalakshmi Undamatla, Vidhyadhari Methuku, Priya Nurkurthy, Payel Mukherjee,Jandhyala Sai Krishna,Onkar Kulkarni, Sofia Banu, Archana Bharadwaj Siva, Karthik Bharadwaj Tallapaka,Rakesh K Mishra,Divya Tej Sowpati |
| EPI_ISL_3453465 | AP SSO | CSIR-Centre for Cellular and Molecular Biology-INSACOG | B Himasri,Amareshwar Vodapalli,Ara Sreenivas,Shreekant Verma,Valli Nagalakshmi Undamatla,Payel Mukherjee, Lamuk Zaveri, Onkar Kulkarni,Tulasi Nagabandi, Sofia Banu, Vidhyadhari Methuku,Priya Nurkurthy, Jandhyala Sai Krishna,Sumedha Avadhanula, Archana Bharadwaj Siva, Karthik Bharadwaj Tallapaka,Rakesh K Mishra,Divya Tej Sowpati |
| EPI_ISL_3453469 | AP SSO | CSIR-Centre for Cellular and Molecular Biology-INSACOG | Sumedha Avadhanula, Lamuk Zaveri, Tulasi Nagabandi, Ara Sreenivas,Shreekant Verma, Amareshwar Vodapalli ,B Himasri,Valli Nagalakshmi Undamatla, Vidhyadhari Methuku, Priya Nurkurthy, Payel Mukherjee,Jandhyala Sai Krishna,Onkar Kulkarni, Sofia Banu, Archana Bharadwaj Siva, Karthik Bharadwaj Tallapaka,Rakesh K Mishra,Divya Tej Sowpati |
| EPI_ISL_3453471 | Mapmygenome | CSIR-Centre for Cellular and Molecular Biology-INSACOG | Tulasi Nagabandi,Lamuk Zaveri,Payel Mukherjee,Onkar Kulkarni,Jandhyala Sai Krishna, Sofia Banu, Ara Sreenivas,Shreekant Verma, Amareshwar Vodapalli ,B Himasri,Valli Nagalakshmi Undamatla, Priya Nurkurthy,Sumedha Avadhanula, Vidhyadhari Methuku, Archana Bharadwaj Siva, Karthik Bharadwaj Tallapaka,Rakesh K Mishra,Divya Tej Sowpati |
| EPI_ISL_3453472 | AP SSO | CSIR-Centre for Cellular and Molecular Biology-INSACOG | Payel Mukherjee, Lamuk Zaveri, Onkar Kulkarni,Jandhyala Sai Krishna,Tulasi Nagabandi, Sofia Banu, Ara Sreenivas,Shreekant Verma, Amareshwar Vodapalli ,B Himasri,Valli Nagalakshmi Undamatla, Sumedha Avadhanula, Vidhyadhari Methuku, Priya Nurkurthy, Archana Bharadwaj Siva, Karthik Bharadwaj Tallapaka,Rakesh K Mishra,Divya Tej Sowpati |
| EPI_ISL_3453473 | AP SSO | CSIR-Centre for Cellular and Molecular Biology-INSACOG | Sumedha Avadhanula, Lamuk Zaveri, Tulasi Nagabandi, Ara Sreenivas,Shreekant Verma, Amareshwar Vodapalli ,B Himasri,Valli Nagalakshmi Undamatla, Vidhyadhari Methuku, Priya Nurkurthy, Payel Mukherjee,Jandhyala Sai Krishna,Onkar Kulkarni, Sofia Banu, Archana Bharadwaj Siva, Karthik Bharadwaj Tallapaka,Rakesh K Mishra,Divya Tej Sowpati |
| EPI_ISL_3453474 | AP SSO | CSIR-Centre for Cellular and Molecular Biology-INSACOG | Tulasi Nagabandi,Lamuk Zaveri,Payel Mukherjee,Onkar Kulkarni,Jandhyala Sai Krishna, Sofia Banu, Ara Sreenivas,Shreekant Verma, Amareshwar Vodapalli ,B Himasri,Valli Nagalakshmi Undamatla, Priya Nurkurthy,Sumedha Avadhanula, Vidhyadhari Methuku, Archana Bharadwaj Siva, Karthik Bharadwaj Tallapaka,Rakesh K Mishra,Divya Tej Sowpati |
| EPI_ISL_3453475 | AP SSO | CSIR-Centre for Cellular and Molecular Biology-INSACOG | Sumedha Avadhanula, Lamuk Zaveri, Tulasi Nagabandi, Ara Sreenivas,Shreekant Verma, Amareshwar Vodapalli ,B Himasri,Valli Nagalakshmi Undamatla, Vidhyadhari Methuku, Priya Nurkurthy, Payel Mukherjee,Jandhyala Sai Krishna,Onkar Kulkarni, Sofia Banu, Archana Bharadwaj Siva, Karthik Bharadwaj Tallapaka,Rakesh K Mishra,Divya Tej Sowpati |
| EPI_ISL_3453477 | AP SSO | CSIR-Centre for Cellular and Molecular Biology-INSACOG | Tulasi Nagabandi,Lamuk Zaveri,Payel Mukherjee,Onkar Kulkarni,Jandhyala Sai Krishna, Sofia Banu, Ara Sreenivas,Shreekant Verma, Amareshwar Vodapalli ,B Himasri,Valli Nagalakshmi Undamatla, Priya Nurkurthy,Sumedha Avadhanula, Vidhyadhari Methuku, Archana Bharadwaj Siva, Karthik Bharadwaj Tallapaka,Rakesh K Mishra,Divya Tej Sowpati |
| EPI_ISL_3453479 | AP SSO | CSIR-Centre for Cellular and Molecular Biology-INSACOG | Amareshwar Vodapalli,Ara Sreenivas,Shreekant Verma, B Himasri,Valli Nagalakshmi Undamatla,Payel Mukherjee, Lamuk Zaveri, Onkar Kulkarni,Tulasi Nagabandi, Sofia Banu, Jandhyala Sai Krishna,Priya Nurkurthy, Vidhyadhari Methuku, Sumedha Avadhanula, Archana Bharadwaj Siva, Karthik Bharadwaj Tallapaka,Rakesh K Mishra,Divya Tej Sowpati |
| EPI_ISL_3453480 | AP SSO | CSIR-Centre for Cellular and Molecular Biology-INSACOG | Vidhyadhari Methuku,Tulasi Nagabandi,Lamuk Zaveri, Ara Sreenivas,Shreekant Verma, Amareshwar Vodapalli ,B Himasri,Valli Nagalakshmi Undamatla, Priya Nurkurthy,Sumedha Avadhanula, Archana Bharadwaj Siva,Payel Mukherjee,Onkar Kulkarni, Jandhyala Sai Krishna,Sofia Banu, Karthik Bharadwaj Tallapaka,Rakesh K Mishra,Divya Tej Sowpati |
| EPI_ISL_3453483 | Mapmygenome | CSIR-Centre for Cellular and Molecular Biology-INSACOG | Vidhyadhari Methuku,Tulasi Nagabandi,Lamuk Zaveri, Ara Sreenivas,Shreekant Verma, Amareshwar Vodapalli ,B Himasri,Valli Nagalakshmi Undamatla, Priya Nurkurthy,Sumedha Avadhanula, Archana Bharadwaj Siva,Payel Mukherjee,Onkar Kulkarni, Jandhyala Sai Krishna,Sofia Banu, Karthik Bharadwaj Tallapaka,Rakesh K Mishra,Divya Tej Sowpati |
| EPI_ISL_3453486 | AP SSO | CSIR-Centre for Cellular and Molecular Biology-INSACOG | Sumedha Avadhanula, Lamuk Zaveri, Tulasi Nagabandi, Ara Sreenivas,Shreekant Verma, Amareshwar Vodapalli ,B Himasri,Valli Nagalakshmi Undamatla, Vidhyadhari Methuku, Priya Nurkurthy, Payel Mukherjee,Jandhyala Sai Krishna,Onkar Kulkarni, Sofia Banu, Archana Bharadwaj Siva, Karthik Bharadwaj Tallapaka,Rakesh K Mishra,Divya Tej Sowpati |
| EPI_ISL_3453488 | AP SSO | CSIR-Centre for Cellular and Molecular Biology-INSACOG | Ara Sreenivas,Shreekant Verma, Amareshwar Vodapalli ,B Himasri,Valli Nagalakshmi Undamatla,Payel Mukherjee, Lamuk Zaveri, Onkar Kulkarni,Jandhyala Sai Krishna,Tulasi Nagabandi, Sofia Banu, ,Priya Nurkurthy, Vidhyadhari Methuku, Sumedha Avadhanula, Archana Bharadwaj Siva, Karthik Bharadwaj Tallapaka,Rakesh K Mishra,Divya Tej Sowpati |
| EPI_ISL_3453491 | AP SSO | CSIR-Centre for Cellular and Molecular Biology-INSACOG | Lamuk Zaveri,Tulasi Nagabandi,Payel Mukherjee,Onkar Kulkarni, Sofia Banu, Jandhyala Sai Krishna,Ara Sreenivas,Shreekant Verma, Amareshwar Vodapalli ,B Himasri,Valli Nagalakshmi Undamatla, Sumedha Avadhanula, Priya Nurkurthy, Vidhyadhari Methuku, Archana Bharadwaj Siva, Karthik Bharadwaj Tallapaka,Rakesh K Mishra,Divya Tej Sowpati |
| EPI_ISL_3453492 | Mapmygenome | CSIR-Centre for Cellular and Molecular Biology-INSACOG | Payel Mukherjee, Lamuk Zaveri, Onkar Kulkarni,Jandhyala Sai Krishna,Tulasi Nagabandi, Sofia Banu, Ara Sreenivas,Shreekant Verma, Amareshwar Vodapalli ,B Himasri,Valli Nagalakshmi Undamatla, Sumedha Avadhanula, Vidhyadhari Methuku, Priya Nurkurthy, Archana Bharadwaj Siva, Karthik Bharadwaj Tallapaka,Rakesh K Mishra,Divya Tej Sowpati |
| EPI_ISL_3453493 | AP SSO | CSIR-Centre for Cellular and Molecular Biology-INSACOG | B Himasri,Amareshwar Vodapalli,Ara Sreenivas,Shreekant Verma,Valli Nagalakshmi Undamatla,Payel Mukherjee, Lamuk Zaveri, Onkar Kulkarni,Tulasi Nagabandi, Sofia Banu, Vidhyadhari Methuku,Priya Nurkurthy, Jandhyala Sai Krishna,Sumedha Avadhanula, Archana Bharadwaj Siva, Karthik Bharadwaj Tallapaka,Rakesh K Mishra,Divya Tej Sowpati |
| EPI_ISL_3453494 | AP SSO | CSIR-Centre for Cellular and Molecular Biology-INSACOG | Shreekant Verma, Amareshwar Vodapalli ,Ara Sreenivas,B Himasri,Valli Nagalakshmi Undamatla,Payel Mukherjee, Lamuk Zaveri, Onkar Kulkarni,Tulasi Nagabandi, Sofia Banu, Jandhyala Sai Krishna,Sumedha Avadhanula, Priya Nurkurthy, Vidhyadhari Methuku, Archana Bharadwaj Siva, Karthik Bharadwaj Tallapaka,Rakesh K Mishra,Divya Tej Sowpati |
| EPI_ISL_3453495 | AP SSO | CSIR-Centre for Cellular and Molecular Biology-INSACOG | Tulasi Nagabandi,Lamuk Zaveri,Payel Mukherjee,Onkar Kulkarni,Jandhyala Sai Krishna, Sofia Banu, Ara Sreenivas,Shreekant Verma, Amareshwar Vodapalli ,B Himasri,Valli Nagalakshmi Undamatla, Priya Nurkurthy,Sumedha Avadhanula, Vidhyadhari Methuku, Archana Bharadwaj Siva, Karthik Bharadwaj Tallapaka,Rakesh K Mishra,Divya Tej Sowpati |
| EPI_ISL_3453496 | AP SSO | CSIR-Centre for Cellular and Molecular Biology-INSACOG | Lamuk Zaveri,Tulasi Nagabandi,Payel Mukherjee,Onkar Kulkarni, Sofia Banu, Jandhyala Sai Krishna,Ara Sreenivas,Shreekant Verma, Amareshwar Vodapalli ,B Himasri,Valli Nagalakshmi Undamatla, Sumedha Avadhanula, Priya Nurkurthy, Vidhyadhari Methuku, Archana Bharadwaj Siva, Karthik Bharadwaj Tallapaka,Rakesh K Mishra,Divya Tej Sowpati |
| EPI_ISL_3453497 | AP SSO | CSIR-Centre for Cellular and Molecular Biology-INSACOG | B Himasri,Amareshwar Vodapalli,Ara Sreenivas,Shreekant Verma,Valli Nagalakshmi Undamatla,Payel Mukherjee, Lamuk Zaveri, Onkar Kulkarni,Tulasi Nagabandi, Sofia Banu, Vidhyadhari Methuku,Priya Nurkurthy, Jandhyala Sai |

| | | | |
|---|---|---|---|
| | | | Krishna,Sumedha Avadhanula, Archana Bharadwaj Siva, Karthik Bharadwaj Tallapaka,Rakesh K Mishra,Divya Tej Sowpati |
| EPI_ISL_3453498 | AP SSO | CSIR-Centre for Cellular and Molecular Biology-INSACOG | Payel Mukherjee, Lamuk Zaveri, Onkar Kulkarni,Jandhyala Sai Krishna,Tulasi Nagabandi, Sofia Banu, Ara Sreenivas,Shreekant Verma, Amareshwar Vodapalli ,B Himasri,Valli Nagalakshmi Undamatla, Sumedha Avadhanula, Vidhyadhari Methuku, Priya Nurkurthy, Archana Bharadwaj Siva, Karthik Bharadwaj Tallapaka,Rakesh K Mishra,Divya Tej Sowpati |
| EPI_ISL_3453499, EPI_ISL_3453500 | AP SSO | CSIR-Centre for Cellular and Molecular Biology-INSACOG | Tulasi Nagabandi,Lamuk Zaveri,Payel Mukherjee,Onkar Kulkarni,Jandhyala Sai Krishna, Sofia Banu, Ara Sreenivas,Shreekant Verma, Amareshwar Vodapalli ,B Himasri,Valli Nagalakshmi Undamatla, Priya Nurkurthy,Sumedha Avadhanula, Vidhyadhari Methuku, Archana Bharadwaj Siva, Karthik Bharadwaj Tallapaka,Rakesh K Mishra,Divya Tej Sowpati |
| EPI_ISL_3453501 | AP SSO | CSIR-Centre for Cellular and Molecular Biology-INSACOG | Payel Mukherjee, Lamuk Zaveri, Onkar Kulkarni,Jandhyala Sai Krishna,Tulasi Nagabandi, Sofia Banu, Ara Sreenivas,Shreekant Verma, Amareshwar Vodapalli ,B Himasri,Valli Nagalakshmi Undamatla, Sumedha Avadhanula, Vidhyadhari Methuku, Priya Nurkurthy, Archana Bharadwaj Siva, Karthik Bharadwaj Tallapaka,Rakesh K Mishra,Divya Tej Sowpati |
| EPI_ISL_3453502 | AP SSO | CSIR-Centre for Cellular and Molecular Biology-INSACOG | Lamuk Zaveri,Tulasi Nagabandi,Payel Mukherjee,Onkar Kulkarni, Sofia Banu, Jandhyala Sai Krishna,Ara Sreenivas,Shreekant Verma, Amareshwar Vodapalli ,B Himasri,Valli Nagalakshmi Undamatla, Sumedha Avadhanula, Priya Nurkurthy, Vidhyadhari Methuku, Archana Bharadwaj Siva, Karthik Bharadwaj Tallapaka,Rakesh K Mishra,Divya Tej Sowpati |
| EPI_ISL_3453503 | AP SSO | CSIR-Centre for Cellular and Molecular Biology-INSACOG | Shreekant Verma, Amareshwar Vodapalli ,Ara Sreenivas,B Himasri,Valli Nagalakshmi Undamatla,Payel Mukherjee, Lamuk Zaveri, Onkar Kulkarni,Tulasi Nagabandi, Sofia Banu, Jandhyala Sai Krishna,Sumedha Avadhanula, Priya Nurkurthy, Vidhyadhari Methuku, Archana Bharadwaj Siva, Karthik Bharadwaj Tallapaka,Rakesh K Mishra,Divya Tej Sowpati |
| EPI_ISL_3453504 | AP SSO | CSIR-Centre for Cellular and Molecular Biology-INSACOG | Vidhyadhari Methuku,Tulasi Nagabandi,Lamuk Zaveri, Ara Sreenivas,Shreekant Verma, Amareshwar Vodapalli ,B Himasri,Valli Nagalakshmi Undamatla, Priya Nurkurthy,Sumedha Avadhanula, Archana Bharadwaj Siva,Payel Mukherjee,Onkar Kulkarni, Jandhyala Sai Krishna,Sofia Banu, Karthik Bharadwaj Tallapaka,Rakesh K Mishra,Divya Tej Sowpati |
| EPI_ISL_3453506, EPI_ISL_3453519 | AP SSO | CSIR-Centre for Cellular and Molecular Biology-INSACOG | Tulasi Nagabandi,Lamuk Zaveri,Payel Mukherjee,Onkar Kulkarni,Jandhyala Sai Krishna, Sofia Banu, Ara Sreenivas,Shreekant Verma, Amareshwar Vodapalli ,B Himasri,Valli Nagalakshmi Undamatla, Priya Nurkurthy,Sumedha Avadhanula, Vidhyadhari Methuku, Archana Bharadwaj Siva, Karthik Bharadwaj Tallapaka,Rakesh K Mishra,Divya Tej Sowpati |
| EPI_ISL_3453520 | AP SSO | CSIR-Centre for Cellular and Molecular Biology-INSACOG | Sofia Banu,Payel Mukherjee, Lamuk Zaveri, Onkar Kulkarni,Tulasi Nagabandi, Jandhyala Sai Krishna, Ara Sreenivas,Shreekant Verma, Amareshwar Vodapalli ,B Himasri,Valli Nagalakshmi Undamatla, Sumedha Avadhanula, Vidhyadhari Methuku, Priya Nurkurthy, Archana Bharadwaj Siva, Karthik Bharadwaj Tallapaka,Rakesh K Mishra,Divya Tej Sowpati |
| EPI_ISL_3453521 | AP SSO | CSIR-Centre for Cellular and Molecular Biology-INSACOG | Priya Nurkurthy,Ara Sreenivas,Shreekant Verma, Amareshwar Vodapalli ,Lamuk Zaveri,Tulasi Nagabandi,B Himasri,Valli Nagalakshmi Undamatla, Sumedha Avadhanula, Vidhyadhari Methuku,Payel Mukherjee,Onkar Kulkarni, Jandhyala Sai Krishna,Sofia Banu, Archana Bharadwaj Siva, Karthik Bharadwaj Tallapaka,Rakesh K Mishra,Divya Tej Sowpati |
| EPI_ISL_3453522 | AP SSO | CSIR-Centre for Cellular and Molecular Biology-INSACOG | Sumedha Avadhanula, Lamuk Zaveri, Tulasi Nagabandi, Ara Sreenivas,Shreekant Verma, Amareshwar Vodapalli ,B Himasri,Valli Nagalakshmi Undamatla, Vidhyadhari Methuku, Priya Nurkurthy, Payel Mukherjee,Jandhyala Sai Krishna,Onkar Kulkarni, Sofia Banu, Archana Bharadwaj Siva, Karthik Bharadwaj Tallapaka,Rakesh K Mishra,Divya Tej Sowpati |
| EPI_ISL_3453523 | AP SSO | CSIR-Centre for Cellular and Molecular Biology-INSACOG | Amareshwar Vodapalli,Ara Sreenivas,Shreekant Verma, B Himasri,Valli Nagalakshmi Undamatla,Payel Mukherjee, Lamuk Zaveri, Onkar Kulkarni,Tulasi Nagabandi, Sofia Banu, Jandhyala Sai Krishna,Priya Nurkurthy, Vidhyadhari Methuku, Sumedha Avadhanula, Archana Bharadwaj Siva, Karthik Bharadwaj Tallapaka,Rakesh K Mishra,Divya Tej Sowpati |
| EPI_ISL_3453526 | AP SSO | CSIR-Centre for Cellular and Molecular Biology-INSACOG | Shreekant Verma, Amareshwar Vodapalli ,Ara Sreenivas,B Himasri,Valli Nagalakshmi Undamatla,Payel Mukherjee, Lamuk Zaveri, Onkar Kulkarni,Tulasi Nagabandi, Sofia Banu, Jandhyala Sai Krishna,Sumedha Avadhanula, Priya Nurkurthy, Vidhyadhari Methuku, Archana Bharadwaj Siva, Karthik Bharadwaj Tallapaka,Rakesh K Mishra,Divya Tej Sowpati |
| EPI_ISL_3453530 | RUHS College of Medical Sciences | CSIR-Centre for Cellular and Molecular Biology - INSACOG | Dr. Gaurav Dalela,Dr. Vaseem Naheed Baig,Dr. Sonali Sharma,Dr. Ramesh Sharma,Dr. Nilofer Khayyam,Dr. Jitendra Panda,Dr. Sudhanshu Kacker,Dr. Raja Babu Panwar,Dr. Rajeev Gupta, Ara Sreenivas,Shreekant Verma, Amareshwar Vodapalli ,B Himasri,Jandhyala Sai Krishna,Valli Nagalakshmi Undamatla,Payel Mukherjee, Lamuk Zaveri, Onkar Kulkarni,Tulasi Nagabandi, Sofia Banu, Priya Nurkurthy, Vidhyadhari Methuku, Sumedha Avadhanula, Archana Bharadwaj Siva, Karthik Bharadwaj Tallapaka,Rakesh K Mishra,Divya Tej Sowpati |
| EPI_ISL_3453540 | AP SSO | CSIR-Centre for Cellular and Molecular Biology-INSACOG | Sumedha Avadhanula, Lamuk Zaveri, Tulasi Nagabandi, Ara Sreenivas,Shreekant Verma, Amareshwar Vodapalli ,B Himasri,Valli Nagalakshmi Undamatla, Vidhyadhari Methuku, Priya Nurkurthy, Payel Mukherjee,Jandhyala Sai Krishna,Onkar Kulkarni, Sofia Banu, Archana Bharadwaj Siva, Karthik Bharadwaj Tallapaka,Rakesh K Mishra,Divya Tej Sowpati |
| EPI_ISL_3453544 | RUHS College of Medical Sciences | CSIR-Centre for Cellular and Molecular Biology - INSACOG | Dr. Gaurav Dalela,Dr. Vaseem Naheed Baig,Dr. Sonali Sharma,Dr. Ramesh Sharma,Dr. Nilofer Khayyam,Dr. Jitendra Panda,Dr. Sudhanshu Kacker,Dr. Raja Babu Panwar,Dr. Rajeev Gupta, Payel Mukherjee, Lamuk Zaveri, Jandhyala Sai Krishna,Onkar Kulkarni,Tulasi Nagabandi, Sofia Banu, Ara Sreenivas,Shreekant Verma, Amareshwar Vodapalli ,B Himasri,Valli Nagalakshmi Undamatla, Sumedha Avadhanula, Vidhyadhari Methuku, Priya Nurkurthy, Archana Bharadwaj Siva, Karthik Bharadwaj Tallapaka,Rakesh K Mishra,Divya Tej Sowpati |
| EPI_ISL_3453547 | AP SSO | CSIR-Centre for Cellular and Molecular Biology-INSACOG | Shreekant Verma, Amareshwar Vodapalli ,Ara Sreenivas,B Himasri,Valli Nagalakshmi Undamatla,Payel Mukherjee, Lamuk Zaveri, Onkar Kulkarni,Tulasi Nagabandi, Sofia Banu, Jandhyala Sai Krishna,Sumedha Avadhanula, Priya Nurkurthy, Vidhyadhari Methuku, Archana Bharadwaj Siva, Karthik Bharadwaj Tallapaka,Rakesh K Mishra,Divya Tej Sowpati |
| EPI_ISL_3453549 | RUHS College of Medical Sciences | CSIR-Centre for Cellular and Molecular Biology - INSACOG | Dr. Gaurav Dalela,Dr. Vaseem Naheed Baig,Dr. Sonali Sharma,Dr. Ramesh Sharma,Dr. Nilofer Khayyam,Dr. Jitendra Panda,Dr. Sudhanshu Kacker,Dr. Raja Babu Panwar,Dr. Rajeev Gupta, Amareshwar Vodapalli,Ara Sreenivas,Shreekant Verma, B Himasri,Valli Nagalakshmi Undamatla,Payel Mukherjee, Lamuk Zaveri, Onkar Kulkarni,Tulasi Nagabandi, Sofia Banu,Jandhyala Sai Krishna ,Priya Nurkurthy, Vidhyadhari Methuku, Sumedha Avadhanula, Archana Bharadwaj Siva, Karthik Bharadwaj Tallapaka,Rakesh K Mishra,Divya Tej Sowpati |
| EPI_ISL_3453559 | RUHS College of Medical Sciences | CSIR-Centre for Cellular and Molecular Biology - INSACOG | Dr. Gaurav Dalela,Dr. Vaseem Naheed Baig,Dr. Sonali Sharma,Dr. Ramesh Sharma,Dr. Nilofer Khayyam,Dr. Jitendra Panda,Dr. Sudhanshu Kacker,Dr. Raja Babu Panwar,Dr. Rajeev Gupta, Shreekant Verma, Amareshwar Vodapalli ,Ara Sreenivas,B Himasri,Valli Nagalakshmi Undamatla,Payel Mukherjee, Lamuk Zaveri, Onkar Kulkarni,Tulasi Nagabandi,Jandhyala Sai Krishna, Sofia Banu, Sumedha Avadhanula, Priya Nurkurthy, Vidhyadhari Methuku, Archana Bharadwaj Siva, Karthik Bharadwaj Tallapaka,Rakesh K Mishra,Divya Tej Sowpati |
| EPI_ISL_3453562 | RUHS College of Medical Sciences | CSIR-Centre for Cellular and Molecular Biology - INSACOG | Dr. Gaurav Dalela,Dr. Vaseem Naheed Baig,Dr. Sonali Sharma,Dr. Ramesh Sharma,Dr. Nilofer Khayyam,Dr. Jitendra Panda,Dr. Sudhanshu Kacker,Dr. Raja Babu Panwar,Dr. Rajeev Gupta, Lamuk Zaveri,Tulasi Nagabandi,Payel Mukherjee,Onkar Kulkarni,Jandhyala Sai Krishna, Sofia Banu, Ara Sreenivas,Shreekant Verma, Amareshwar Vodapalli ,B Himasri,Valli Nagalakshmi Undamatla, Sumedha Avadhanula, Priya Nurkurthy, Vidhyadhari Methuku, Archana Bharadwaj Siva, Karthik Bharadwaj Tallapaka,Rakesh K Mishra,Divya Tej Sowpati |
| EPI_ISL_3453563 | AP SSO | CSIR-Centre for Cellular and Molecular Biology-INSACOG | Lamuk Zaveri,Tulasi Nagabandi,Payel Mukherjee,Onkar Kulkarni, Sofia Banu, Jandhyala Sai Krishna,Ara Sreenivas,Shreekant Verma, Amareshwar Vodapalli ,B Himasri,Valli Nagalakshmi Undamatla, Sumedha Avadhanula, Priya Nurkurthy, Vidhyadhari Methuku, Archana Bharadwaj Siva, Karthik Bharadwaj Tallapaka,Rakesh K Mishra,Divya Tej Sowpati |
| EPI_ISL_3453566 | AP SSO | CSIR-Centre for Cellular and Molecular Biology-INSACOG | Sumedha Avadhanula, Lamuk Zaveri, Tulasi Nagabandi, Ara Sreenivas,Shreekant Verma, Amareshwar Vodapalli ,B Himasri,Valli Nagalakshmi Undamatla, Vidhyadhari Methuku, Priya Nurkurthy, Payel Mukherjee,Jandhyala Sai Krishna,Onkar Kulkarni, Sofia Banu, Archana Bharadwaj Siva, Karthik Bharadwaj Tallapaka,Rakesh K Mishra,Divya Tej Sowpati |
| EPI_ISL_3453568 | RUHS College of Medical Sciences | CSIR-Centre for Cellular and Molecular Biology - INSACOG | Dr. Gaurav Dalela,Dr. Vaseem Naheed Baig,Dr. Sonali Sharma,Dr. Ramesh Sharma,Dr. Nilofer Khayyam,Dr. Jitendra Panda,Dr. Sudhanshu Kacker,Dr. Raja Babu Panwar,Dr. Rajeev Gupta, Ara Sreenivas,Shreekant Verma, Amareshwar Vodapalli ,B Himasri,Jandhyala Sai Krishna,Valli Nagalakshmi Undamatla,Payel Mukherjee, Lamuk Zaveri, Onkar Kulkarni,Tulasi Nagabandi, Sofia Banu, Priya Nurkurthy, Vidhyadhari Methuku, Sumedha Avadhanula, Archana Bharadwaj Siva, Karthik Bharadwaj Tallapaka,Rakesh K Mishra,Divya Tej Sowpati |
| EPI_ISL_3453572 | AP SSO | CSIR-Centre for Cellular and Molecular Biology-INSACOG | B Himasri,Amareshwar Vodapalli,Ara Sreenivas,Shreekant Verma,Valli Nagalakshmi Undamatla,Payel Mukherjee, Lamuk Zaveri, Onkar Kulkarni,Tulasi Nagabandi, Sofia Banu, Vidhyadhari Methuku,Priya Nurkurthy, Jandhyala Sai Krishna,Sumedha Avadhanula, Archana Bharadwaj Siva, Karthik Bharadwaj Tallapaka,Rakesh K Mishra,Divya Tej Sowpati |
| EPI_ISL_3453575 | RUHS College of Medical Sciences | CSIR-Centre for Cellular and Molecular Biology - INSACOG | Dr. Gaurav Dalela,Dr. Vaseem Naheed Baig,Dr. Sonali Sharma,Dr. Ramesh Sharma,Dr. Nilofer Khayyam,Dr. Jitendra Panda,Dr. Sudhanshu Kacker,Dr. Raja Babu Panwar,Dr. Rajeev Gupta, Shreekant Verma, Amareshwar Vodapalli ,Ara Sreenivas,B Himasri,Valli Nagalakshmi Undamatla,Payel Mukherjee, Lamuk Zaveri, Onkar Kulkarni,Tulasi Nagabandi,Jandhyala Sai Krishna, Sofia Banu, Sumedha Avadhanula, Priya Nurkurthy, Vidhyadhari Methuku, Archana Bharadwaj Siva, Karthik Bharadwaj Tallapaka,Rakesh K Mishra,Divya Tej Sowpati |
| EPI_ISL_3453576 | Mapmygenome | CSIR-Centre for Cellular and Molecular Biology-INSACOG | Tulasi Nagabandi,Lamuk Zaveri,Payel Mukherjee,Onkar Kulkarni,Jandhyala Sai Krishna, Sofia Banu, Ara Sreenivas,Shreekant Verma, Amareshwar Vodapalli ,B Himasri,Valli Nagalakshmi Undamatla, Priya Nurkurthy,Sumedha Avadhanula, Vidhyadhari Methuku, Archana Bharadwaj Siva, Karthik Bharadwaj Tallapaka,Rakesh K Mishra,Divya Tej Sowpati |
| EPI_ISL_3453577 | AP SSO | CSIR-Centre for Cellular and Molecular Biology-INSACOG | Amareshwar Vodapalli,Ara Sreenivas,Shreekant Verma, B Himasri,Valli Nagalakshmi Undamatla,Payel Mukherjee, Lamuk Zaveri, Onkar Kulkarni,Tulasi Nagabandi, Sofia Banu, Jandhyala Sai Krishna,Priya Nurkurthy, Vidhyadhari Methuku, Sumedha Avadhanula, Archana Bharadwaj Siva, Karthik Bharadwaj Tallapaka,Rakesh K Mishra,Divya Tej Sowpati |
| EPI_ISL_3453578 | CSIR-Centre for Cellular and Molecular Biology | CSIR-Centre for Cellular and Molecular Biology-INSACOG | Shreekant Verma, Amareshwar Vodapalli ,Ara Sreenivas,B Himasri,Valli Nagalakshmi Undamatla,Payel Mukherjee, Lamuk Zaveri, Onkar Kulkarni,Tulasi Nagabandi, Sofia Banu, Jandhyala Sai Krishna,Sumedha Avadhanula, Priya Nurkurthy, Vidhyadhari Methuku, Archana Bharadwaj Siva, Karthik Bharadwaj Tallapaka,Rakesh K Mishra,Divya Tej Sowpati |
| EPI_ISL_3453581 | RUHS College of Medical Sciences | CSIR-Centre for Cellular and Molecular Biology - INSACOG | Dr. Gaurav Dalela,Dr. Vaseem Naheed Baig,Dr. Sonali Sharma,Dr. Ramesh Sharma,Dr. Nilofer Khayyam,Dr. Jitendra Panda,Dr. Sudhanshu Kacker,Dr. Raja Babu Panwar,Dr. Rajeev Gupta, Payel Mukherjee, Lamuk Zaveri, Jandhyala Sai Krishna,Onkar Kulkarni,Tulasi Nagabandi, Sofia Banu, Ara Sreenivas,Shreekant Verma, Amareshwar Vodapalli ,B Himasri,Valli Nagalakshmi Undamatla, Sumedha Avadhanula, Vidhyadhari Methuku, Priya Nurkurthy, Archana Bharadwaj Siva, Karthik Bharadwaj Tallapaka,Rakesh K Mishra,Divya Tej Sowpati |
| EPI_ISL_3453586 | AP SSO | CSIR-Centre for Cellular and Molecular Biology-INSACOG | Shreekant Verma, Amareshwar Vodapalli ,Ara Sreenivas,B Himasri,Valli Nagalakshmi Undamatla,Payel Mukherjee, Lamuk Zaveri, Onkar Kulkarni,Tulasi Nagabandi, Sofia Banu, Jandhyala Sai Krishna,Sumedha Avadhanula, Priya Nurkurthy, Vidhyadhari Methuku, Archana Bharadwaj Siva, Karthik Bharadwaj Tallapaka,Rakesh K Mishra,Divya Tej Sowpati |
| EPI_ISL_3453588 | RUHS College of Medical Sciences | CSIR-Centre for Cellular and Molecular Biology - INSACOG | Dr. Gaurav Dalela,Dr. Vaseem Naheed Baig,Dr. Sonali Sharma,Dr. Ramesh Sharma,Dr. Nilofer Khayyam,Dr. Jitendra Panda,Dr. Sudhanshu Kacker,Dr. Raja Babu Panwar,Dr. Rajeev Gupta, Amareshwar Vodapalli,Ara Sreenivas,Shreekant Verma, B Himasri,Valli Nagalakshmi Undamatla,Payel Mukherjee, Lamuk Zaveri, Onkar Kulkarni,Tulasi Nagabandi, Sofia Banu,Jandhyala Sai Krishna ,Priya Nurkurthy, Vidhyadhari Methuku, Sumedha Avadhanula, Archana Bharadwaj Siva, Karthik Bharadwaj Tallapaka,Rakesh K Mishra,Divya Tej Sowpati |
| EPI_ISL_3453589 | AP SSO | CSIR-Centre for Cellular and Molecular Biology-INSACOG | Amareshwar Vodapalli,Ara Sreenivas,Shreekant Verma, B Himasri,Valli Nagalakshmi Undamatla,Payel Mukherjee, Lamuk Zaveri, Onkar Kulkarni,Tulasi Nagabandi, Sofia Banu, Jandhyala Sai Krishna,Priya Nurkurthy, Vidhyadhari Methuku, Sumedha Avadhanula, Archana Bharadwaj Siva, Karthik Bharadwaj Tallapaka,Rakesh K Mishra,Divya Tej Sowpati |
| EPI_ISL_3453590 | RUHS College of Medical Sciences | CSIR-Centre for Cellular and Molecular Biology - INSACOG | Dr. Gaurav Dalela,Dr. Vaseem Naheed Baig,Dr. Sonali Sharma,Dr. Ramesh Sharma,Dr. Nilofer Khayyam,Dr. Jitendra Panda,Dr. Sudhanshu Kacker,Dr. Raja Babu Panwar,Dr. Rajeev Gupta, Tulasi Nagabandi,Lamuk Zaveri,Jandhyala Sai Krishna,Payel Mukherjee,Onkar Kulkarni, Sofia Banu, Ara Sreenivas,Shreekant Verma, Amareshwar Vodapalli ,B Himasri,Valli Nagalakshmi Undamatla, Priya Nurkurthy,Sumedha Avadhanula, Vidhyadhari Methuku, Archana Bharadwaj Siva, Karthik Bharadwaj Tallapaka,Rakesh K Mishra,Divya Tej Sowpati |
| EPI_ISL_3453592 | AP SSO | CSIR-Centre for Cellular and Molecular Biology-INSACOG | Ara Sreenivas,Shreekant Verma, Amareshwar Vodapalli ,B Himasri,Valli Nagalakshmi Undamatla,Payel Mukherjee, Lamuk Zaveri, Onkar Kulkarni,Jandhyala Sai Krishna,Tulasi Nagabandi, Sofia Banu, ,Priya Nurkurthy, Vidhyadhari Methuku, Sumedha Avadhanula, Archana Bharadwaj Siva, Karthik Bharadwaj Tallapaka,Rakesh K Mishra,Divya Tej Sowpati |
| EPI_ISL_3453593 | Mapmygenome | CSIR-Centre for Cellular and Molecular Biology-INSACOG | Shreekant Verma, Amareshwar Vodapalli ,Ara Sreenivas,B Himasri,Valli Nagalakshmi Undamatla,Payel Mukherjee, Lamuk Zaveri, Onkar Kulkarni,Tulasi Nagabandi, Sofia Banu, Jandhyala Sai Krishna,Sumedha Avadhanula, Priya Nurkurthy, Vidhyadhari Methuku, Archana Bharadwaj Siva, Karthik Bharadwaj Tallapaka,Rakesh K Mishra,Divya Tej Sowpati |
| EPI_ISL_3453594 | AP SSO | CSIR-Centre for Cellular and Molecular Biology-INSACOG | Payel Mukherjee, Lamuk Zaveri, Onkar Kulkarni,Tulasi Nagabandi, Sofia Banu, Ara Sreenivas,Shreekant Verma, Amareshwar Vodapalli ,B Himasri,Valli Nagalakshmi Undamatla, Sumedha Avadhanula, Vidhyadhari Methuku, Priya Nurkurthy, Archana Bharadwaj Siva, Karthik Bharadwaj Tallapaka,Krishna Khairnar,Rakesh K Mishra,Divya Tej Sowpati |
| EPI_ISL_3453595 | Mapmygenome | CSIR-Centre for Cellular and Molecular Biology-INSACOG | Amareshwar Vodapalli,Ara Sreenivas,Shreekant Verma, B Himasri,Valli Nagalakshmi Undamatla,Payel Mukherjee, Lamuk Zaveri, Onkar Kulkarni,Tulasi Nagabandi, Sofia Banu, Jandhyala Sai Krishna,Priya Nurkurthy, Vidhyadhari Methuku, Sumedha Avadhanula, Archana Bharadwaj Siva, Karthik Bharadwaj Tallapaka,Rakesh K Mishra,Divya Tej Sowpati |
| EPI_ISL_3453598 | RUHS College of Medical Sciences | CSIR-Centre for Cellular and Molecular Biology - INSACOG | Dr. Gaurav Dalela,Dr. Vaseem Naheed Baig,Dr. Sonali Sharma,Dr. Ramesh Sharma,Dr. Nilofer Khayyam,Dr. Jitendra Panda,Dr. Sudhanshu Kacker,Dr. Raja Babu Panwar,Dr. Rajeev Gupta, Shreekant Verma, Amareshwar Vodapalli ,Ara Sreenivas,B Himasri,Valli Nagalakshmi Undamatla,Payel Mukherjee, Lamuk Zaveri, Onkar Kulkarni,Tulasi Nagabandi,Jandhyala Sai Krishna, Sofia Banu, Sumedha Avadhanula, Priya Nurkurthy, Vidhyadhari Methuku, Archana Bharadwaj Siva, Karthik Bharadwaj Tallapaka,Rakesh K Mishra,Divya Tej Sowpati |
| EPI_ISL_3453600 | AP SSO | CSIR-Centre for Cellular and Molecular Biology-INSACOG | Shreekant Verma, Amareshwar Vodapalli ,Ara Sreenivas,B Himasri,Valli Nagalakshmi Undamatla,Payel Mukherjee, Lamuk Zaveri, Onkar Kulkarni,Tulasi Nagabandi, Sofia Banu, Jandhyala Sai Krishna,Sumedha Avadhanula, Priya Nurkurthy, Vidhyadhari Methuku, Archana Bharadwaj Siva, Karthik Bharadwaj Tallapaka,Rakesh K Mishra,Divya Tej Sowpati |
| EPI_ISL_3453603 | Mapmygenome | CSIR-Centre for Cellular and Molecular Biology-INSACOG | Vidhyadhari Methuku,Tulasi Nagabandi,Lamuk Zaveri, Ara Sreenivas,Shreekant Verma, Amareshwar Vodapalli ,B Himasri,Valli Nagalakshmi Undamatla, Priya Nurkurthy,Sumedha Avadhanula, Archana Bharadwaj Siva,Payel Mukherjee,Onkar Kulkarni, Jandhyala Sai Krishna,Sofia Banu, Karthik Bharadwaj Tallapaka,Rakesh K Mishra,Divya Tej Sowpati |
| EPI_ISL_3453608 | AP SSO | CSIR-Centre for Cellular and Molecular Biology-INSACOG | Sumedha Avadhanula, Lamuk Zaveri, Tulasi Nagabandi, Ara Sreenivas,Shreekant Verma, Amareshwar Vodapalli ,B Himasri,Valli Nagalakshmi Undamatla, Vidhyadhari Methuku, Priya Nurkurthy, Payel Mukherjee,Jandhyala Sai Krishna,Onkar Kulkarni, Sofia Banu, Archana Bharadwaj Siva, Karthik Bharadwaj Tallapaka,Rakesh K Mishra,Divya Tej Sowpati |

| | | | |
|---|---|---|---|
| EPI_ISL_3453613 | AP SSO | CSIR-Centre for Cellular and Molecular Biology-INSACOG | Lamuk Zaveri,Tulasi Nagabandi,Payel Mukherjee,Onkar Kulkarni, Sofia Banu, Jandhyala Sai Krishna,Ara Sreenivas,Shreekant Verma, Amareshwar Vodapalli ,B Himasri,Valli Nagalakshmi Undamatla, Sumedha Avadhanula, Priya Nurkurthy, Vidhyadhari Methuku, Archana Bharadwaj Siva, Karthik Bharadwaj Tallapaka,Rakesh K Mishra,Divya Tej Sowpati |
| EPI_ISL_3453615 | RUHS College of Medical Sciences | CSIR-Centre for Cellular and Molecular Biology - INSACOG | Dr. Gaurav Dalela,Dr. Vaseem Naheed Baig,Dr. Sonali Sharma,Dr. Ramesh Sharma,Dr. Nilofer Khayyam,Dr. Jitendra Panda,Dr. Sudhanshu Kacker,Dr. Raja Babu Panwar,Dr. Rajeev Gupta, Ara Sreenivas,Shreekant Verma, Amareshwar Vodapalli ,B Himasri,Jandhyala Sai Krishna,Valli Nagalakshmi Undamatla,Payel Mukherjee, Lamuk Zaveri, Onkar Kulkarni,Tulasi Nagabandi, Sofia Banu, Priya Nurkurthy, Vidhyadhari Methuku, Sumedha Avadhanula, Archana Bharadwaj Siva, Karthik Bharadwaj Tallapaka,Rakesh K Mishra,Divya Tej Sowpati |
| EPI_ISL_3453621 | Mapmygenome | CSIR-Centre for Cellular and Molecular Biology-INSACOG | Shreekant Verma, Amareshwar Vodapalli ,Ara Sreenivas,B Himasri,Valli Nagalakshmi Undamatla,Payel Mukherjee, Lamuk Zaveri, Onkar Kulkarni,Tulasi Nagabandi, Sofia Banu, Jandhyala Sai Krishna,Sumedha Avadhanula, Priya Nurkurthy, Vidhyadhari Methuku, Archana Bharadwaj Siva, Karthik Bharadwaj Tallapaka,Rakesh K Mishra,Divya Tej Sowpati |
| EPI_ISL_3453623 | AP SSO | CSIR-Centre for Cellular and Molecular Biology-INSACOG | Lamuk Zaveri,Tulasi Nagabandi,Payel Mukherjee,Onkar Kulkarni, Sofia Banu, Jandhyala Sai Krishna,Ara Sreenivas,Shreekant Verma, Amareshwar Vodapalli ,B Himasri,Valli Nagalakshmi Undamatla, Sumedha Avadhanula, Priya Nurkurthy, Vidhyadhari Methuku, Archana Bharadwaj Siva, Karthik Bharadwaj Tallapaka,Rakesh K Mishra,Divya Tej Sowpati |
| EPI_ISL_3453624 | RUHS College of Medical Sciences | CSIR-Centre for Cellular and Molecular Biology - INSACOG | Dr. Gaurav Dalela,Dr. Vaseem Naheed Baig,Dr. Sonali Sharma,Dr. Ramesh Sharma,Dr. Nilofer Khayyam,Dr. Jitendra Panda,Dr. Sudhanshu Kacker,Dr. Raja Babu Panwar,Dr. Rajeev Gupta, Ara Sreenivas,Shreekant Verma, Amareshwar Vodapalli ,B Himasri,Jandhyala Sai Krishna,Valli Nagalakshmi Undamatla,Payel Mukherjee, Lamuk Zaveri, Onkar Kulkarni,Tulasi Nagabandi, Sofia Banu, Priya Nurkurthy, Vidhyadhari Methuku, Sumedha Avadhanula, Archana Bharadwaj Siva, Karthik Bharadwaj Tallapaka,Rakesh K Mishra,Divya Tej Sowpati |
| EPI_ISL_3453628 | RUHS College of Medical Sciences | CSIR-Centre for Cellular and Molecular Biology - INSACOG | Dr. Gaurav Dalela,Dr. Vaseem Naheed Baig,Dr. Sonali Sharma,Dr. Ramesh Sharma,Dr. Nilofer Khayyam,Dr. Jitendra Panda,Dr. Sudhanshu Kacker,Dr. Raja Babu Panwar,Dr. Rajeev Gupta, Lamuk Zaveri,Tulasi Nagabandi,Payel Mukherjee,Onkar Kulkarni,Jandhyala Sai Krishna, Sofia Banu, Ara Sreenivas,Shreekant Verma, Amareshwar Vodapalli ,B Himasri,Valli Nagalakshmi Undamatla, Sumedha Avadhanula, Priya Nurkurthy, Vidhyadhari Methuku, Archana Bharadwaj Siva, Karthik Bharadwaj Tallapaka,Rakesh K Mishra,Divya Tej Sowpati |
| EPI_ISL_3453629 | AP SSO | CSIR-Centre for Cellular and Molecular Biology-INSACOG | Tulasi Nagabandi,Lamuk Zaveri,Payel Mukherjee,Onkar Kulkarni,Jandhyala Sai Krishna, Sofia Banu, Ara Sreenivas,Shreekant Verma, Amareshwar Vodapalli ,B Himasri,Valli Nagalakshmi Undamatla, Priya Nurkurthy,Sumedha Avadhanula, Vidhyadhari Methuku, Archana Bharadwaj Siva, Karthik Bharadwaj Tallapaka,Rakesh K Mishra,Divya Tej Sowpati |
| EPI_ISL_3453631 | AP SSO | CSIR-Centre for Cellular and Molecular Biology-INSACOG | Sumedha Avadhanula, Lamuk Zaveri, Tulasi Nagabandi, Ara Sreenivas,Shreekant Verma, Amareshwar Vodapalli ,B Himasri,Valli Nagalakshmi Undamatla, Vidhyadhari Methuku, Priya Nurkurthy, Payel Mukherjee,Jandhyala Sai Krishna,Onkar Kulkarni, Sofia Banu, Archana Bharadwaj Siva, Karthik Bharadwaj Tallapaka,Rakesh K Mishra,Divya Tej Sowpati |
| EPI_ISL_3453633 | AP SSO | CSIR-Centre for Cellular and Molecular Biology-INSACOG | Amareshwar Vodapalli,Ara Sreenivas,Shreekant Verma, B Himasri,Valli Nagalakshmi Undamatla,Payel Mukherjee, Lamuk Zaveri, Onkar Kulkarni,Tulasi Nagabandi, Sofia Banu, Jandhyala Sai Krishna,Priya Nurkurthy, Vidhyadhari Methuku, Sumedha Avadhanula, Archana Bharadwaj Siva, Karthik Bharadwaj Tallapaka,Rakesh K Mishra,Divya Tej Sowpati |
| EPI_ISL_3453659 | AP SSO | CSIR-Centre for Cellular and Molecular Biology-INSACOG | Lamuk Zaveri,Tulasi Nagabandi,Payel Mukherjee,Onkar Kulkarni, Sofia Banu, Jandhyala Sai Krishna,Ara Sreenivas,Shreekant Verma, Amareshwar Vodapalli ,B Himasri,Valli Nagalakshmi Undamatla, Sumedha Avadhanula, Priya Nurkurthy, Vidhyadhari Methuku, Archana Bharadwaj Siva, Karthik Bharadwaj Tallapaka,Rakesh K Mishra,Divya Tej Sowpati |
| EPI_ISL_3453668 | AP SSO | CSIR-Centre for Cellular and Molecular Biology-INSACOG | Sumedha Avadhanula, Lamuk Zaveri, Tulasi Nagabandi, Ara Sreenivas,Shreekant Verma, Amareshwar Vodapalli ,B Himasri,Valli Nagalakshmi Undamatla, Vidhyadhari Methuku, Priya Nurkurthy, Payel Mukherjee,Jandhyala Sai Krishna,Onkar Kulkarni, Sofia Banu, Archana Bharadwaj Siva, Karthik Bharadwaj Tallapaka,Rakesh K Mishra,Divya Tej Sowpati |
| EPI_ISL_3453677 | AP SSO | CSIR-Centre for Cellular and Molecular Biology-INSACOG | Amareshwar Vodapalli,Ara Sreenivas,Shreekant Verma, B Himasri,Valli Nagalakshmi Undamatla,Payel Mukherjee, Lamuk Zaveri, Onkar Kulkarni,Tulasi Nagabandi, Sofia Banu, Jandhyala Sai Krishna,Priya Nurkurthy, Vidhyadhari Methuku, Sumedha Avadhanula, Archana Bharadwaj Siva, Karthik Bharadwaj Tallapaka,Rakesh K Mishra,Divya Tej Sowpati |
| EPI_ISL_3453684 | AP SSO | CSIR-Centre for Cellular and Molecular Biology-INSACOG | Sumedha Avadhanula, Lamuk Zaveri, Tulasi Nagabandi, Ara Sreenivas,Shreekant Verma, Amareshwar Vodapalli ,B Himasri,Valli Nagalakshmi Undamatla, Vidhyadhari Methuku, Priya Nurkurthy, Payel Mukherjee,Jandhyala Sai Krishna,Onkar Kulkarni, Sofia Banu, Archana Bharadwaj Siva, Karthik Bharadwaj Tallapaka,Rakesh K Mishra,Divya Tej Sowpati |
| EPI_ISL_3453688, EPI_ISL_3453689, EPI_ISL_3453690, EPI_ISL_3453691 | AP SSO | CSIR-Centre for Cellular and Molecular Biology-INSACOG | Payel Mukherjee, Lamuk Zaveri, Onkar Kulkarni,Jandhyala Sai Krishna,Tulasi Nagabandi, Sofia Banu, Ara Sreenivas,Shreekant Verma, Amareshwar Vodapalli ,B Himasri,Valli Nagalakshmi Undamatla, Sumedha Avadhanula, Vidhyadhari Methuku, Priya Nurkurthy, Archana Bharadwaj Siva, Karthik Bharadwaj Tallapaka,Rakesh K Mishra,Divya Tej Sowpati |
| EPI_ISL_3453692, EPI_ISL_3453693, EPI_ISL_3453694, EPI_ISL_3453695, EPI_ISL_3453696 | AP SSO | CSIR-Centre for Cellular and Molecular Biology-INSACOG | Lamuk Zaveri,Tulasi Nagabandi,Payel Mukherjee,Onkar Kulkarni, Sofia Banu, Jandhyala Sai Krishna,Ara Sreenivas,Shreekant Verma, Amareshwar Vodapalli ,B Himasri,Valli Nagalakshmi Undamatla, Sumedha Avadhanula, Priya Nurkurthy, Vidhyadhari Methuku, Archana Bharadwaj Siva, Karthik Bharadwaj Tallapaka,Rakesh K Mishra,Divya Tej Sowpati |
| EPI_ISL_3453697 | AP SSO | CSIR-Centre for Cellular and Molecular Biology-INSACOG | Priya Nurkurthy,Ara Sreenivas,Shreekant Verma, Amareshwar Vodapalli ,Lamuk Zaveri,Tulasi Nagabandi,B Himasri,Valli Nagalakshmi Undamatla, Sumedha Avadhanula, Vidhyadhari Methuku,Payel Mukherjee,Onkar Kulkarni, Jandhyala Sai Krishna,Sofia Banu, Archana Bharadwaj Siva, Karthik Bharadwaj Tallapaka,Rakesh K Mishra,Divya Tej Sowpati |
| EPI_ISL_3453698, EPI_ISL_3453699 | AP SSO | CSIR-Centre for Cellular and Molecular Biology-INSACOG | Vidhyadhari Methuku,Tulasi Nagabandi,Lamuk Zaveri, Ara Sreenivas,Shreekant Verma, Amareshwar Vodapalli ,B Himasri,Valli Nagalakshmi Undamatla, Priya Nurkurthy,Sumedha Avadhanula, Archana Bharadwaj Siva,Payel Mukherjee,Onkar Kulkarni, Jandhyala Sai Krishna,Sofia Banu, Karthik Bharadwaj Tallapaka,Rakesh K Mishra,Divya Tej Sowpati |
| EPI_ISL_3453700, EPI_ISL_3453701 | AP SSO | CSIR-Centre for Cellular and Molecular Biology-INSACOG | Payel Mukherjee, Lamuk Zaveri, Onkar Kulkarni,Jandhyala Sai Krishna,Tulasi Nagabandi, Sofia Banu, Ara Sreenivas,Shreekant Verma, Amareshwar Vodapalli ,B Himasri,Valli Nagalakshmi Undamatla, Sumedha Avadhanula, Vidhyadhari Methuku, Priya Nurkurthy, Archana Bharadwaj Siva, Karthik Bharadwaj Tallapaka,Rakesh K Mishra,Divya Tej Sowpati |
| EPI_ISL_3453702 | AP SSO | CSIR-Centre for Cellular and Molecular Biology-INSACOG | Amareshwar Vodapalli,Ara Sreenivas,Shreekant Verma, B Himasri,Valli Nagalakshmi Undamatla,Payel Mukherjee, Lamuk Zaveri, Onkar Kulkarni,Tulasi Nagabandi, Sofia Banu, Jandhyala Sai Krishna,Priya Nurkurthy, Vidhyadhari Methuku, Sumedha Avadhanula, Archana Bharadwaj Siva, Karthik Bharadwaj Tallapaka,Rakesh K Mishra,Divya Tej Sowpati |
| EPI_ISL_3453703, EPI_ISL_3453704, EPI_ISL_3453705, EPI_ISL_3453706, EPI_ISL_3453707, EPI_ISL_3453708, EPI_ISL_3453709 | AP SSO | CSIR-Centre for Cellular and Molecular Biology-INSACOG | B Himasri,Amareshwar Vodapalli,Ara Sreenivas,Shreekant Verma,Valli Nagalakshmi Undamatla,Payel Mukherjee, Lamuk Zaveri, Onkar Kulkarni,Tulasi Nagabandi, Sofia Banu, Vidhyadhari Methuku,Priya Nurkurthy, Jandhyala Sai Krishna,Sumedha Avadhanula, Archana Bharadwaj Siva, Karthik Bharadwaj Tallapaka,Rakesh K Mishra,Divya Tej Sowpati |
| EPI_ISL_3453710, EPI_ISL_3453711, EPI_ISL_3453712 | AP SSO | CSIR-Centre for Cellular and Molecular Biology-INSACOG | Sumedha Avadhanula, Lamuk Zaveri, Tulasi Nagabandi, Ara Sreenivas,Shreekant Verma, Amareshwar Vodapalli ,B Himasri,Valli Nagalakshmi Undamatla, Vidhyadhari Methuku, Priya Nurkurthy, Payel Mukherjee,Jandhyala Sai Krishna,Onkar Kulkarni, Sofia Banu, Archana Bharadwaj Siva, Karthik Bharadwaj Tallapaka,Rakesh K Mishra,Divya Tej Sowpati |
| EPI_ISL_3453713, EPI_ISL_3453714, EPI_ISL_3453715, EPI_ISL_3453716 | AP SSO | CSIR-Centre for Cellular and Molecular Biology-INSACOG | Shreekant Verma, Amareshwar Vodapalli ,Ara Sreenivas,B Himasri,Valli Nagalakshmi Undamatla,Payel Mukherjee, Lamuk Zaveri, Onkar Kulkarni,Tulasi Nagabandi, Sofia Banu, Jandhyala Sai Krishna,Sumedha Avadhanula, Priya Nurkurthy, Vidhyadhari Methuku, Archana Bharadwaj Siva, Karthik Bharadwaj Tallapaka,Rakesh K Mishra,Divya Tej Sowpati |
| EPI_ISL_3453717, EPI_ISL_3453718, EPI_ISL_3453719, EPI_ISL_3453720, EPI_ISL_3453721, EPI_ISL_3453722, EPI_ISL_3453723 | AP SSO | CSIR-Centre for Cellular and Molecular Biology-INSACOG | Amareshwar Vodapalli,Ara Sreenivas,Shreekant Verma, B Himasri,Valli Nagalakshmi Undamatla,Payel Mukherjee, Lamuk Zaveri, Onkar Kulkarni,Tulasi Nagabandi, Sofia Banu, Jandhyala Sai Krishna,Priya Nurkurthy, Vidhyadhari Methuku, Sumedha Avadhanula, Archana Bharadwaj Siva, Karthik Bharadwaj Tallapaka,Rakesh K Mishra,Divya Tej Sowpati |
| EPI_ISL_3453724, EPI_ISL_3453725 | AP SSO | CSIR-Centre for Cellular and Molecular Biology-INSACOG | B Himasri,Amareshwar Vodapalli,Ara Sreenivas,Shreekant Verma,Valli Nagalakshmi Undamatla,Payel Mukherjee, Lamuk Zaveri, Onkar Kulkarni,Tulasi Nagabandi, Sofia Banu, Vidhyadhari Methuku,Priya Nurkurthy, Jandhyala Sai Krishna,Sumedha Avadhanula, Archana Bharadwaj Siva, Karthik Bharadwaj Tallapaka,Rakesh K Mishra,Divya Tej Sowpati |
| EPI_ISL_3453726, EPI_ISL_3453727, EPI_ISL_3453728 | AP SSO | CSIR-Centre for Cellular and Molecular Biology-INSACOG | Sofia Banu,Payel Mukherjee, Lamuk Zaveri, Onkar Kulkarni,Tulasi Nagabandi, Jandhyala Sai Krishna, Ara Sreenivas,Shreekant Verma, Amareshwar Vodapalli ,B Himasri,Valli Nagalakshmi Undamatla, Sumedha Avadhanula, Vidhyadhari Methuku, Priya Nurkurthy, Archana Bharadwaj Siva, Karthik Bharadwaj Tallapaka,Rakesh K Mishra,Divya Tej Sowpati |
| EPI_ISL_3453729, EPI_ISL_3453730, EPI_ISL_3453731, EPI_ISL_3453732 | AP SSO | CSIR-Centre for Cellular and Molecular Biology-INSACOG | Sumedha Avadhanula, Lamuk Zaveri, Tulasi Nagabandi, Ara Sreenivas,Shreekant Verma, Amareshwar Vodapalli ,B Himasri,Valli Nagalakshmi Undamatla, Vidhyadhari Methuku, Priya Nurkurthy, Payel Mukherjee,Jandhyala Sai Krishna,Onkar Kulkarni, Sofia Banu, Archana Bharadwaj Siva, Karthik Bharadwaj Tallapaka,Rakesh K Mishra,Divya Tej Sowpati |
| EPI_ISL_3453733, EPI_ISL_3453734, EPI_ISL_3453735, EPI_ISL_3453736, EPI_ISL_3453737 | AP SSO | CSIR-Centre for Cellular and Molecular Biology-INSACOG | Priya Nurkurthy,Ara Sreenivas,Shreekant Verma, Amareshwar Vodapalli ,Lamuk Zaveri,Tulasi Nagabandi,B Himasri,Valli Nagalakshmi Undamatla, Sumedha Avadhanula, Vidhyadhari Methuku,Payel Mukherjee,Onkar Kulkarni, Jandhyala Sai Krishna,Sofia Banu, Archana Bharadwaj Siva, Karthik Bharadwaj Tallapaka,Rakesh K Mishra,Divya Tej Sowpati |
| EPI_ISL_3453738, EPI_ISL_3453739, EPI_ISL_3453740, EPI_ISL_3453741, EPI_ISL_3453742 | AP SSO | CSIR-Centre for Cellular and Molecular Biology-INSACOG | Lamuk Zaveri,Tulasi Nagabandi,Payel Mukherjee,Onkar Kulkarni, Sofia Banu, Jandhyala Sai Krishna,Ara Sreenivas,Shreekant Verma, Amareshwar Vodapalli ,B Himasri,Valli Nagalakshmi Undamatla, Sumedha Avadhanula, Priya Nurkurthy, Vidhyadhari Methuku, Archana Bharadwaj Siva, Karthik Bharadwaj Tallapaka,Rakesh K Mishra,Divya Tej Sowpati |
| EPI_ISL_3453743, EPI_ISL_3453744, EPI_ISL_3453745, EPI_ISL_3453746, EPI_ISL_3453747, EPI_ISL_3453748 | AP SSO | CSIR-Centre for Cellular and Molecular Biology-INSACOG | Tulasi Nagabandi,Lamuk Zaveri,Payel Mukherjee,Onkar Kulkarni,Jandhyala Sai Krishna, Sofia Banu, Ara Sreenivas,Shreekant Verma, Amareshwar Vodapalli ,B Himasri,Valli Nagalakshmi Undamatla, Priya Nurkurthy,Sumedha Avadhanula, Vidhyadhari Methuku, Archana Bharadwaj Siva, Karthik Bharadwaj Tallapaka,Rakesh K Mishra,Divya Tej Sowpati |
| EPI_ISL_3453749, EPI_ISL_3453750 | AP SSO | CSIR-Centre for Cellular and Molecular Biology-INSACOG | Sumedha Avadhanula, Lamuk Zaveri, Tulasi Nagabandi, Ara Sreenivas,Shreekant Verma, Amareshwar Vodapalli ,B Himasri,Valli Nagalakshmi Undamatla, Vidhyadhari Methuku, Priya Nurkurthy, Payel Mukherjee,Jandhyala Sai Krishna,Onkar Kulkarni, Sofia Banu, Archana Bharadwaj Siva, Karthik Bharadwaj Tallapaka,Rakesh K Mishra,Divya Tej Sowpati |
| EPI_ISL_3453751 | AP SSO | CSIR-Centre for Cellular and Molecular Biology-INSACOG | Shreekant Verma, Amareshwar Vodapalli ,Ara Sreenivas,B Himasri,Valli Nagalakshmi Undamatla,Payel Mukherjee, Lamuk Zaveri, Onkar Kulkarni,Tulasi Nagabandi, Sofia Banu, Jandhyala Sai Krishna,Sumedha Avadhanula, Priya Nurkurthy, Vidhyadhari Methuku, Archana Bharadwaj Siva, Karthik Bharadwaj Tallapaka,Rakesh K Mishra,Divya Tej Sowpati |
| EPI_ISL_3453752, | AP SSO | CSIR-Centre for Cellular and Molecular Biology-INSACOG | Tulasi Nagabandi,Lamuk Zaveri,Payel Mukherjee,Onkar Kulkarni,Jandhyala Sai Krishna, Sofia Banu, Ara Sreenivas,Shreekant Verma, Amareshwar Vodapalli ,B Himasri,Valli Nagalakshmi Undamatla, Priya Nurkurthy,Sumedha Avadhanula, |

| | | | |
|---|---|---|---|
| EPI_ISL_3453753 | | | Vidhyadhari Methuku, Archana Bharadwaj Siva, Karthik Bharadwaj Tallapaka,Rakesh K Mishra,Divya Tej Sowpati |
| EPI_ISL_3453754, EPI_ISL_3453755, EPI_ISL_3453756 | AP SSO | CSIR-Centre for Cellular and Molecular Biology-INSACOG | Shreekant Verma, Amareshwar Vodapalli ,Ara Sreenivas,B Himasri,Valli Nagalakshmi Undamatla,Payel Mukherjee, Lamuk Zaveri, Onkar Kulkarni,Tulasi Nagabandi, Sofia Banu, Jandhyala Sai Krishna,Sumedha Avadhanula, Priya Nurkurthy, Vidhyadhari Methuku, Archana Bharadwaj Siva, Karthik Bharadwaj Tallapaka,Rakesh K Mishra,Divya Tej Sowpati |
| EPI_ISL_3453757 | AP SSO | CSIR-Centre for Cellular and Molecular Biology-INSACOG | Amareshwar Vodapalli,Ara Sreenivas,Shreekant Verma, B Himasri,Valli Nagalakshmi Undamatla,Payel Mukherjee, Lamuk Zaveri, Onkar Kulkarni,Tulasi Nagabandi, Sofia Banu, Jandhyala Sai Krishna,Priya Nurkurthy, Vidhyadhari Methuku, Sumedha Avadhanula, Archana Bharadwaj Siva, Karthik Bharadwaj Tallapaka,Rakesh K Mishra,Divya Tej Sowpati |
| EPI_ISL_3453758, EPI_ISL_3453759 | AP SSO | CSIR-Centre for Cellular and Molecular Biology-INSACOG | B Himasri,Amareshwar Vodapalli,Ara Sreenivas,Shreekant Verma,Valli Nagalakshmi Undamatla,Payel Mukherjee, Lamuk Zaveri, Onkar Kulkarni,Tulasi Nagabandi, Sofia Banu, Vidhyadhari Methuku,Priya Nurkurthy, Jandhyala Sai Krishna,Sumedha Avadhanula, Archana Bharadwaj Siva, Karthik Bharadwaj Tallapaka,Rakesh K Mishra,Divya Tej Sowpati |
| EPI_ISL_3453760, EPI_ISL_3453761 | AP SSO | CSIR-Centre for Cellular and Molecular Biology-INSACOG | Sumedha Avadhanula, Lamuk Zaveri, Tulasi Nagabandi, Ara Sreenivas,Shreekant Verma, Amareshwar Vodapalli ,B Himasri,Valli Nagalakshmi Undamatla, Vidhyadhari Methuku, Priya Nurkurthy, Payel Mukherjee,Jandhyala Sai Krishna,Onkar Kulkarni, Sofia Banu, Archana Bharadwaj Siva, Karthik Bharadwaj Tallapaka,Rakesh K Mishra,Divya Tej Sowpati |
| EPI_ISL_3453762 | AP SSO | CSIR-Centre for Cellular and Molecular Biology-INSACOG | Payel Mukherjee, Lamuk Zaveri, Onkar Kulkarni,Tulasi Nagabandi, Sofia Banu, Ara Sreenivas,Shreekant Verma, Amareshwar Vodapalli ,B Himasri,Valli Nagalakshmi Undamatla, Sumedha Avadhanula, Vidhyadhari Methuku, Priya Nurkurthy, Archana Bharadwaj Siva, Karthik Bharadwaj Tallapaka,Krishna Khairnar,Rakesh K Mishra,Divya Tej Sowpati |
| EPI_ISL_3453763 | AP SSO | CSIR-Centre for Cellular and Molecular Biology-INSACOG | Tulasi Nagabandi,Lamuk Zaveri,Payel Mukherjee,Onkar Kulkarni,Jandhyala Sai Krishna, Sofia Banu, Ara Sreenivas,Shreekant Verma, Amareshwar Vodapalli ,B Himasri,Valli Nagalakshmi Undamatla, Priya Nurkurthy,Sumedha Avadhanula, Vidhyadhari Methuku, Archana Bharadwaj Siva, Karthik Bharadwaj Tallapaka,Rakesh K Mishra,Divya Tej Sowpati |
| EPI_ISL_3453764, EPI_ISL_3453765 | AP SSO | CSIR-Centre for Cellular and Molecular Biology-INSACOG | Lamuk Zaveri,Tulasi Nagabandi,Payel Mukherjee,Onkar Kulkarni, Sofia Banu, Jandhyala Sai Krishna,Ara Sreenivas,Shreekant Verma, Amareshwar Vodapalli ,B Himasri,Valli Nagalakshmi Undamatla, Sumedha Avadhanula, Priya Nurkurthy, Vidhyadhari Methuku, Archana Bharadwaj Siva, Karthik Bharadwaj Tallapaka,Rakesh K Mishra,Divya Tej Sowpati |
| EPI_ISL_3453766, EPI_ISL_3453767, EPI_ISL_3453768, EPI_ISL_3453769, EPI_ISL_3453770 | AP SSO | CSIR-Centre for Cellular and Molecular Biology-INSACOG | Shreekant Verma, Amareshwar Vodapalli ,Ara Sreenivas,B Himasri,Valli Nagalakshmi Undamatla,Payel Mukherjee, Lamuk Zaveri, Onkar Kulkarni,Tulasi Nagabandi, Sofia Banu, Jandhyala Sai Krishna,Sumedha Avadhanula, Priya Nurkurthy, Vidhyadhari Methuku, Archana Bharadwaj Siva, Karthik Bharadwaj Tallapaka,Rakesh K Mishra,Divya Tej Sowpati |
| EPI_ISL_3453771 | AP SSO | CSIR-Centre for Cellular and Molecular Biology-INSACOG | Amareshwar Vodapalli,Ara Sreenivas,Shreekant Verma, B Himasri,Valli Nagalakshmi Undamatla,Payel Mukherjee, Lamuk Zaveri, Onkar Kulkarni,Tulasi Nagabandi, Sofia Banu, Jandhyala Sai Krishna,Priya Nurkurthy, Vidhyadhari Methuku, Sumedha Avadhanula, Archana Bharadwaj Siva, Karthik Bharadwaj Tallapaka,Rakesh K Mishra,Divya Tej Sowpati |
| EPI_ISL_3453772, EPI_ISL_3453773 | AP SSO | CSIR-Centre for Cellular and Molecular Biology-INSACOG | B Himasri,Amareshwar Vodapalli,Ara Sreenivas,Shreekant Verma,Valli Nagalakshmi Undamatla,Payel Mukherjee, Lamuk Zaveri, Onkar Kulkarni,Tulasi Nagabandi, Sofia Banu, Vidhyadhari Methuku,Priya Nurkurthy, Jandhyala Sai Krishna,Sumedha Avadhanula, Archana Bharadwaj Siva, Karthik Bharadwaj Tallapaka,Rakesh K Mishra,Divya Tej Sowpati |
| EPI_ISL_3453774, EPI_ISL_3453775 | AP SSO | CSIR-Centre for Cellular and Molecular Biology-INSACOG | Sofia Banu,Payel Mukherjee, Lamuk Zaveri, Onkar Kulkarni,Tulasi Nagabandi, Jandhyala Sai Krishna, Ara Sreenivas,Shreekant Verma, Amareshwar Vodapalli ,B Himasri,Valli Nagalakshmi Undamatla, Sumedha Avadhanula, Vidhyadhari Methuku, Priya Nurkurthy, Archana Bharadwaj Siva, Karthik Bharadwaj Tallapaka,Rakesh K Mishra,Divya Tej Sowpati |
| EPI_ISL_3453776, EPI_ISL_3453777, EPI_ISL_3453778, EPI_ISL_3453779 | AP SSO | CSIR-Centre for Cellular and Molecular Biology-INSACOG | Sumedha Avadhanula, Lamuk Zaveri, Tulasi Nagabandi, Ara Sreenivas,Shreekant Verma, Amareshwar Vodapalli ,B Himasri,Valli Nagalakshmi Undamatla, Vidhyadhari Methuku, Priya Nurkurthy, Payel Mukherjee,Jandhyala Sai Krishna,Onkar Kulkarni, Sofia Banu, Archana Bharadwaj Siva, Karthik Bharadwaj Tallapaka,Rakesh K Mishra,Divya Tej Sowpati |
| EPI_ISL_3453780, EPI_ISL_3453781 | AP SSO | CSIR-Centre for Cellular and Molecular Biology-INSACOG | Priya Nurkurthy,Ara Sreenivas,Shreekant Verma, Amareshwar Vodapalli ,Lamuk Zaveri,Tulasi Nagabandi,B Himasri,Valli Nagalakshmi Undamatla, Sumedha Avadhanula, Vidhyadhari Methuku,Payel Mukherjee,Onkar Kulkarni, Jandhyala Sai Krishna,Sofia Banu, Archana Bharadwaj Siva, Karthik Bharadwaj Tallapaka,Rakesh K Mishra,Divya Tej Sowpati |
| EPI_ISL_3453782, EPI_ISL_3453783 | AP SSO | CSIR-Centre for Cellular and Molecular Biology-INSACOG | Tulasi Nagabandi,Lamuk Zaveri,Payel Mukherjee,Onkar Kulkarni,Jandhyala Sai Krishna, Sofia Banu, Ara Sreenivas,Shreekant Verma, Amareshwar Vodapalli ,B Himasri,Valli Nagalakshmi Undamatla, Priya Nurkurthy,Sumedha Avadhanula, Vidhyadhari Methuku, Archana Bharadwaj Siva, Karthik Bharadwaj Tallapaka,Rakesh K Mishra,Divya Tej Sowpati |
| EPI_ISL_3453784 | AP SSO | CSIR-Centre for Cellular and Molecular Biology-INSACOG | Sumedha Avadhanula, Lamuk Zaveri, Tulasi Nagabandi, Ara Sreenivas,Shreekant Verma, Amareshwar Vodapalli ,B Himasri,Valli Nagalakshmi Undamatla, Vidhyadhari Methuku, Priya Nurkurthy, Payel Mukherjee,Jandhyala Sai Krishna,Onkar Kulkarni, Sofia Banu, Archana Bharadwaj Siva, Karthik Bharadwaj Tallapaka,Rakesh K Mishra,Divya Tej Sowpati |
| EPI_ISL_3453785, EPI_ISL_3453786, EPI_ISL_3453787 | AP SSO | CSIR-Centre for Cellular and Molecular Biology-INSACOG | Payel Mukherjee, Lamuk Zaveri, Onkar Kulkarni,Jandhyala Sai Krishna,Tulasi Nagabandi, Sofia Banu, Ara Sreenivas,Shreekant Verma, Amareshwar Vodapalli ,B Himasri,Valli Nagalakshmi Undamatla, Sumedha Avadhanula, Vidhyadhari Methuku, Priya Nurkurthy, Archana Bharadwaj Siva, Karthik Bharadwaj Tallapaka,Rakesh K Mishra,Divya Tej Sowpati |
| EPI_ISL_3453788, EPI_ISL_3453789, EPI_ISL_3453790, EPI_ISL_3453791, EPI_ISL_3453792, EPI_ISL_3453793, EPI_ISL_3453794, EPI_ISL_3453795, EPI_ISL_3453796, EPI_ISL_3453797 | AP SSO | CSIR-Centre for Cellular and Molecular Biology-INSACOG | Lamuk Zaveri,Tulasi Nagabandi,Payel Mukherjee,Onkar Kulkarni, Sofia Banu, Jandhyala Sai Krishna,Ara Sreenivas,Shreekant Verma, Amareshwar Vodapalli ,B Himasri,Valli Nagalakshmi Undamatla, Sumedha Avadhanula, Priya Nurkurthy, Vidhyadhari Methuku, Archana Bharadwaj Siva, Karthik Bharadwaj Tallapaka,Rakesh K Mishra,Divya Tej Sowpati |
| EPI_ISL_3453798 | AP SSO | CSIR-Centre for Cellular and Molecular Biology-INSACOG | Tulasi Nagabandi,Lamuk Zaveri,Payel Mukherjee,Onkar Kulkarni,Jandhyala Sai Krishna, Sofia Banu, Ara Sreenivas,Shreekant Verma, Amareshwar Vodapalli ,B Himasri,Valli Nagalakshmi Undamatla, Priya Nurkurthy,Sumedha Avadhanula, Vidhyadhari Methuku, Archana Bharadwaj Siva, Karthik Bharadwaj Tallapaka,Rakesh K Mishra,Divya Tej Sowpati |
| EPI_ISL_3453836 | RUHS College of Medical Sciences | CSIR-Centre for Cellular and Molecular Biology - INSACOG | Dr. Gaurav Dalela,Dr. Vaseem Naheed Baig,Dr. Sonali Sharma,Dr. Ramesh Sharma,Dr. Nilofer Khayyam,Dr. Jitendra Panda,Dr. Sudhanshu Kacker,Dr. Raja Babu Panwar,Dr. Rajeev Gupta, Sofia Banu,Payel Mukherjee, Lamuk Zaveri, Jandhyala Sai Krishna,Onkar Kulkarni,Tulasi Nagabandi, Ara Sreenivas,Shreekant Verma, Amareshwar Vodapalli ,B Himasri,Valli Nagalakshmi Undamatla, Sumedha Avadhanula, Vidhyadhari Methuku, Priya Nurkurthy, Archana Bharadwaj Siva, Karthik Bharadwaj Tallapaka,Rakesh K Mishra,Divya Tej Sowpati |
| EPI_ISL_3453837 | RUHS College of Medical Sciences | CSIR-Centre for Cellular and Molecular Biology - INSACOG | Dr. Gaurav Dalela,Dr. Vaseem Naheed Baig,Dr. Sonali Sharma,Dr. Ramesh Sharma,Dr. Nilofer Khayyam,Dr. Jitendra Panda,Dr. Sudhanshu Kacker,Dr. Raja Babu Panwar,Dr. Rajeev Gupta, Onkar Kulkarni, Lamuk Zaveri, Jandhyala Sai Krishna,Tulasi Nagabandi,Sofia Banu,Payel Mukherjee, Ara Sreenivas,Shreekant Verma, Amareshwar Vodapalli ,B Himasri,Valli Nagalakshmi Undamatla, Sumedha Avadhanula, Vidhyadhari Methuku, Priya Nurkurthy, Archana Bharadwaj Siva, Karthik Bharadwaj Tallapaka,Rakesh K Mishra,Divya Tej Sowpati |
| EPI_ISL_3453840, EPI_ISL_3453841 | Mapmygenome | CSIR-Centre for Cellular and Molecular Biology-INSACOG | Vidhyadhari Methuku,Tulasi Nagabandi,Lamuk Zaveri, Ara Sreenivas,Shreekant Verma, Amareshwar Vodapalli ,B Himasri,Valli Nagalakshmi Undamatla, Priya Nurkurthy,Sumedha Avadhanula, Archana Bharadwaj Siva,Payel Mukherjee,Onkar Kulkarni, Jandhyala Sai Krishna,Sofia Banu, Karthik Bharadwaj Tallapaka,Rakesh K Mishra,Divya Tej Sowpati |
| EPI_ISL_3453842, EPI_ISL_3453843, EPI_ISL_3453844, EPI_ISL_3453845 | Mapmygenome | CSIR-Centre for Cellular and Molecular Biology-INSACOG | Payel Mukherjee, Lamuk Zaveri, Onkar Kulkarni,Jandhyala Sai Krishna,Tulasi Nagabandi, Sofia Banu, Ara Sreenivas,Shreekant Verma, Amareshwar Vodapalli ,B Himasri,Valli Nagalakshmi Undamatla, Sumedha Avadhanula, Vidhyadhari Methuku, Priya Nurkurthy, Archana Bharadwaj Siva, Karthik Bharadwaj Tallapaka,Rakesh K Mishra,Divya Tej Sowpati |
| EPI_ISL_3453846 | Mapmygenome | CSIR-Centre for Cellular and Molecular Biology-INSACOG | Shreekant Verma, Amareshwar Vodapalli ,Ara Sreenivas,B Himasri,Valli Nagalakshmi Undamatla,Payel Mukherjee, Lamuk Zaveri, Onkar Kulkarni,Tulasi Nagabandi, Sofia Banu, Jandhyala Sai Krishna,Sumedha Avadhanula, Priya Nurkurthy, Vidhyadhari Methuku, Archana Bharadwaj Siva, Karthik Bharadwaj Tallapaka,Rakesh K Mishra,Divya Tej Sowpati |
| EPI_ISL_3453847, EPI_ISL_3453848 | Mapmygenome | CSIR-Centre for Cellular and Molecular Biology-INSACOG | Tulasi Nagabandi,Lamuk Zaveri,Payel Mukherjee,Onkar Kulkarni,Jandhyala Sai Krishna, Sofia Banu, Ara Sreenivas,Shreekant Verma, Amareshwar Vodapalli ,B Himasri,Valli Nagalakshmi Undamatla, Priya Nurkurthy,Sumedha Avadhanula, Vidhyadhari Methuku, Archana Bharadwaj Siva, Karthik Bharadwaj Tallapaka,Rakesh K Mishra,Divya Tej Sowpati |
| EPI_ISL_3453849, EPI_ISL_3453850 | Mapmygenome | CSIR-Centre for Cellular and Molecular Biology-INSACOG | Ara Sreenivas,Shreekant Verma, Amareshwar Vodapalli ,B Himasri,Valli Nagalakshmi Undamatla,Payel Mukherjee, Lamuk Zaveri, Onkar Kulkarni,Jandhyala Sai Krishna,Tulasi Nagabandi, Sofia Banu, ,Priya Nurkurthy, Vidhyadhari Methuku, Sumedha Avadhanula, Archana Bharadwaj Siva, Karthik Bharadwaj Tallapaka,Rakesh K Mishra,Divya Tej Sowpati |
| EPI_ISL_3453851, EPI_ISL_3453852 | Mapmygenome | CSIR-Centre for Cellular and Molecular Biology-INSACOG | Shreekant Verma, Amareshwar Vodapalli ,Ara Sreenivas,B Himasri,Valli Nagalakshmi Undamatla,Payel Mukherjee, Lamuk Zaveri, Onkar Kulkarni,Tulasi Nagabandi, Sofia Banu, Jandhyala Sai Krishna,Sumedha Avadhanula, Priya Nurkurthy, Vidhyadhari Methuku, Archana Bharadwaj Siva, Karthik Bharadwaj Tallapaka,Rakesh K Mishra,Divya Tej Sowpati |
| EPI_ISL_3453853 | Mapmygenome | CSIR-Centre for Cellular and Molecular Biology-INSACOG | Amareshwar Vodapalli,Ara Sreenivas,Shreekant Verma, B Himasri,Valli Nagalakshmi Undamatla,Payel Mukherjee, Lamuk Zaveri, Onkar Kulkarni,Tulasi Nagabandi, Sofia Banu, Jandhyala Sai Krishna,Priya Nurkurthy, Vidhyadhari Methuku, Sumedha Avadhanula, Archana Bharadwaj Siva, Karthik Bharadwaj Tallapaka,Rakesh K Mishra,Divya Tej Sowpati |
| EPI_ISL_3453855, EPI_ISL_3453856, EPI_ISL_3453857, EPI_ISL_3453858, EPI_ISL_3453859, EPI_ISL_3453860, EPI_ISL_3453862, EPI_ISL_3453863, EPI_ISL_3453865, EPI_ISL_3453866, EPI_ISL_3453867, EPI_ISL_3453868, EPI_ISL_3453869, EPI_ISL_3453870, EPI_ISL_3453871, EPI_ISL_3453873, EPI_ISL_3453874, EPI_ISL_3453875, EPI_ISL_3453876, EPI_ISL_3453879, EPI_ISL_3453880 | | | |
| see above | Palapye Primary Hospital Laboratory | Botswana Harvard HIV Reference Laboratory | Sikhulile Moyo, Wonderful T. Choga, Dorcas Maruapula, Thongbotho Mphoyakgosi, Boitumelo J.L Zuze, Botshelo Radibe, Legodile Kooepile, Ontlametse T. Bareng, Letsibogo Gaoraelwe, Thela Tefelo, Keoratile Ntshambiwa, Modisa Motswaledi, Madisa Mine, Joseph Makhema, Roger Shapiro, Shahin Lockman, Mosepele Mosepele, Simani Gaseitsiwe |
| EPI_ISL_3453882, EPI_ISL_3453883, EPI_ISL_3453884, EPI_ISL_3453885, EPI_ISL_3453886, EPI_ISL_3453887, EPI_ISL_3453888, EPI_ISL_3453889, EPI_ISL_3453890, EPI_ISL_3453891, EPI_ISL_3453892, EPI_ISL_3453893, EPI_ISL_3453894, EPI_ISL_3453895, EPI_ISL_3453896, EPI_ISL_3453897, EPI_ISL_3453898, EPI_ISL_3453899, EPI_ISL_3453900, EPI_ISL_3453901, EPI_ISL_3453902, EPI_ISL_3453903, EPI_ISL_3453904, EPI_ISL_3453905, EPI_ISL_3453906, EPI_ISL_3453907, EPI_ISL_3453908, EPI_ISL_3453909 | | | |
| see above | Maun Reference Hospital laboratory | Botswana Harvard HIV Reference Laboratory | Sikhulile Moyo, Wonderful T. Choga, Dorcas Maruapula, Thongbotho Mphoyakgosi, Boitumelo J.L Zuze, Botshelo Radibe, Legodile Kooepile, Ontlametse T. Bareng, Sandra Maripe, Timothy Odirile Matsuokwane, Keoratile Ntshambiwa, Modisa Motswaledi, Madisa Mine, Joseph Makhema, Roger Shapiro, Shahin Lockman, Mosepele Mosepele, Simani Gaseitsiwe |
| EPI_ISL_3453911, EPI_ISL_3453912, EPI_ISL_3453913, EPI_ISL_3453914, EPI_ISL_3453915, EPI_ISL_3453916, EPI_ISL_3453917 | Botswana Harvard HIV Reference Laboratory | Botswana Harvard HIV Reference Laboratory | Sikhulile Moyo, Wonderful T. Choga, Dorcas Maruapula, Thongbotho Mphoyakgosi, Boitumelo J.L Zuze, Botshelo Radibe, Legodile Kooepile, Ontlametse T. Bareng, Keoratile Ntshambiwa, Modisa Motswaledi, Pamela Lawrence-Smith, Joseph Makhema, Roger Shapiro, Shahin Lockman, Mosepele Mosepele, Simani Gaseitsiwe |
| EPI_ISL_3453918 | Lighthouse Laboratory Plymouth | Wellcome Sanger Institute for the COVID-19 Genomics UK (COG-UK) Consortium | Lighthouse Laboratory Plymouth and Alex Alderton, Roberto Amato, Jeffrey Barrett, Sonia Goncalves, Ewan Harrison, David K. Jackson, Ian Johnston, Dominic Kwiatkowski, Cordelia Langford, John Sillitoe on behalf of the Wellcome Sanger Institute COVID-19 Surveillance Team |
| EPI_ISL_3453919 | Lighthouse Lab in Milton Keynes | Wellcome Sanger Institute for the COVID-19 Genomics UK (COG-UK) Consortium | The Lighthouse Lab in Milton Keynes and Alex Alderton, Roberto Amato, Jeffrey Barrett, Sonia Goncalves, Ewan Harrison, David K. Jackson, Ian Johnston, Dominic Kwiatkowski, Cordelia Langford, John Sillitoe on behalf of the Wellcome Sanger Institute COVID-19 Surveillance Team |
| EPI_ISL_3453920, EPI_ISL_3453921 | Lighthouse Lab in Alderley Park | Wellcome Sanger Institute for the COVID-19 Genomics UK (COG-UK) Consortium | Jacquelyn Wynn, Mairead Hyland, The Lighthouse Lab in Alderley Park and Alex Alderton, Roberto Amato, Jeffrey Barrett, Sonia Goncalves, Ewan Harrison, David K. Jackson, Ian Johnston, Dominic Kwiatkowski, Cordelia Langford, John Sillitoe on behalf of the Wellcome Sanger Institute COVID-19 Surveillance Team |

| | | | |
|---|---|---|---|
| EPI_ISL_3453922 | Lighthouse Laboratory Plymouth | Wellcome Sanger Institute for the COVID-19 Genomics UK (COG-UK) Consortium | Lighthouse Laboratory Plymouth and Alex Alderton, Roberto Amato, Jeffrey Barrett, Sonia Goncalves, Ewan Harrison, David K. Jackson, Ian Johnston, Dominic Kwiatkowski, Cordelia Langford, John Sillitoe on behalf of the Wellcome Sanger Institute COVID-19 Surveillance Team |
| EPI_ISL_3453923 | Lighthouse Lab in Alderley Park | Wellcome Sanger Institute for the COVID-19 Genomics UK (COG-UK) Consortium | Jacquelyn Wynn, Mairead Hyland, The Lighthouse Lab in Alderley Park and Alex Alderton, Roberto Amato, Jeffrey Barrett, Sonia Goncalves, Ewan Harrison, David K. Jackson, Ian Johnston, Dominic Kwiatkowski, Cordelia Langford, John Sillitoe on behalf of the Wellcome Sanger Institute COVID-19 Surveillance Team |
| EPI_ISL_3453924 | Lighthouse Lab in Milton Keynes | Wellcome Sanger Institute for the COVID-19 Genomics UK (COG-UK) Consortium | The Lighthouse Lab in Milton Keynes and Alex Alderton, Roberto Amato, Jeffrey Barrett, Sonia Goncalves, Ewan Harrison, David K. Jackson, Ian Johnston, Dominic Kwiatkowski, Cordelia Langford, John Sillitoe on behalf of the Wellcome Sanger Institute COVID-19 Surveillance Team |
| EPI_ISL_3453925 | Lighthouse Lab in Alderley Park | Wellcome Sanger Institute for the COVID-19 Genomics UK (COG-UK) Consortium | Jacquelyn Wynn, Mairead Hyland, The Lighthouse Lab in Alderley Park and Alex Alderton, Roberto Amato, Jeffrey Barrett, Sonia Goncalves, Ewan Harrison, David K. Jackson, Ian Johnston, Dominic Kwiatkowski, Cordelia Langford, John Sillitoe on behalf of the Wellcome Sanger Institute COVID-19 Surveillance Team |
| EPI_ISL_3453926 | Lighthouse Laboratory Plymouth | Wellcome Sanger Institute for the COVID-19 Genomics UK (COG-UK) Consortium | Lighthouse Laboratory Plymouth and Alex Alderton, Roberto Amato, Jeffrey Barrett, Sonia Goncalves, Ewan Harrison, David K. Jackson, Ian Johnston, Dominic Kwiatkowski, Cordelia Langford, John Sillitoe on behalf of the Wellcome Sanger Institute COVID-19 Surveillance Team |
| EPI_ISL_3453927 | Lighthouse Lab in Alderley Park | Wellcome Sanger Institute for the COVID-19 Genomics UK (COG-UK) Consortium | Jacquelyn Wynn, Mairead Hyland, The Lighthouse Lab in Alderley Park and Alex Alderton, Roberto Amato, Jeffrey Barrett, Sonia Goncalves, Ewan Harrison, David K. Jackson, Ian Johnston, Dominic Kwiatkowski, Cordelia Langford, John Sillitoe on behalf of the Wellcome Sanger Institute COVID-19 Surveillance Team |
| EPI_ISL_3453928 | Lighthouse Laboratory Plymouth | Wellcome Sanger Institute for the COVID-19 Genomics UK (COG-UK) Consortium | Lighthouse Laboratory Plymouth and Alex Alderton, Roberto Amato, Jeffrey Barrett, Sonia Goncalves, Ewan Harrison, David K. Jackson, Ian Johnston, Dominic Kwiatkowski, Cordelia Langford, John Sillitoe on behalf of the Wellcome Sanger Institute COVID-19 Surveillance Team |
| EPI_ISL_3453929, EPI_ISL_3453930, EPI_ISL_3453931, EPI_ISL_3453932, EPI_ISL_3453933 | Lighthouse Lab in Alderley Park | Wellcome Sanger Institute for the COVID-19 Genomics UK (COG-UK) Consortium | Jacquelyn Wynn, Mairead Hyland, The Lighthouse Lab in Alderley Park and Alex Alderton, Roberto Amato, Jeffrey Barrett, Sonia Goncalves, Ewan Harrison, David K. Jackson, Ian Johnston, Dominic Kwiatkowski, Cordelia Langford, John Sillitoe on behalf of the Wellcome Sanger Institute COVID-19 Surveillance Team |
| EPI_ISL_3453934, EPI_ISL_3453935 | Lighthouse Lab in Milton Keynes | Wellcome Sanger Institute for the COVID-19 Genomics UK (COG-UK) Consortium | The Lighthouse Lab in Milton Keynes and Alex Alderton, Roberto Amato, Jeffrey Barrett, Sonia Goncalves, Ewan Harrison, David K. Jackson, Ian Johnston, Dominic Kwiatkowski, Cordelia Langford, John Sillitoe on behalf of the Wellcome Sanger Institute COVID-19 Surveillance Team |
| EPI_ISL_3453936, EPI_ISL_3453937 | Lighthouse Lab in Alderley Park | Wellcome Sanger Institute for the COVID-19 Genomics UK (COG-UK) Consortium | Jacquelyn Wynn, Mairead Hyland, The Lighthouse Lab in Alderley Park and Alex Alderton, Roberto Amato, Jeffrey Barrett, Sonia Goncalves, Ewan Harrison, David K. Jackson, Ian Johnston, Dominic Kwiatkowski, Cordelia Langford, John Sillitoe on behalf of the Wellcome Sanger Institute COVID-19 Surveillance Team |
| EPI_ISL_3453938 | Lighthouse Laboratory Plymouth | Wellcome Sanger Institute for the COVID-19 Genomics UK (COG-UK) Consortium | Lighthouse Laboratory Plymouth and Alex Alderton, Roberto Amato, Jeffrey Barrett, Sonia Goncalves, Ewan Harrison, David K. Jackson, Ian Johnston, Dominic Kwiatkowski, Cordelia Langford, John Sillitoe on behalf of the Wellcome Sanger Institute COVID-19 Surveillance Team |
| EPI_ISL_3453939 | Lighthouse Lab in Milton Keynes | Wellcome Sanger Institute for the COVID-19 Genomics UK (COG-UK) Consortium | The Lighthouse Lab in Milton Keynes and Alex Alderton, Roberto Amato, Jeffrey Barrett, Sonia Goncalves, Ewan Harrison, David K. Jackson, Ian Johnston, Dominic Kwiatkowski, Cordelia Langford, John Sillitoe on behalf of the Wellcome Sanger Institute COVID-19 Surveillance Team |
| EPI_ISL_3453940, EPI_ISL_3453941, EPI_ISL_3453942 | Lighthouse Lab in Alderley Park | Wellcome Sanger Institute for the COVID-19 Genomics UK (COG-UK) Consortium | Jacquelyn Wynn, Mairead Hyland, The Lighthouse Lab in Alderley Park and Alex Alderton, Roberto Amato, Jeffrey Barrett, Sonia Goncalves, Ewan Harrison, David K. Jackson, Ian Johnston, Dominic Kwiatkowski, Cordelia Langford, John Sillitoe on behalf of the Wellcome Sanger Institute COVID-19 Surveillance Team |
| EPI_ISL_3453943 | Lighthouse Laboratory Plymouth | Wellcome Sanger Institute for the COVID-19 Genomics UK (COG-UK) Consortium | Lighthouse Laboratory Plymouth and Alex Alderton, Roberto Amato, Jeffrey Barrett, Sonia Goncalves, Ewan Harrison, David K. Jackson, Ian Johnston, Dominic Kwiatkowski, Cordelia Langford, John Sillitoe on behalf of the Wellcome Sanger Institute COVID-19 Surveillance Team |
| EPI_ISL_3453944, EPI_ISL_3453945 | Lighthouse Lab in Milton Keynes | Wellcome Sanger Institute for the COVID-19 Genomics UK (COG-UK) Consortium | The Lighthouse Lab in Milton Keynes and Alex Alderton, Roberto Amato, Jeffrey Barrett, Sonia Goncalves, Ewan Harrison, David K. Jackson, Ian Johnston, Dominic Kwiatkowski, Cordelia Langford, John Sillitoe on behalf of the Wellcome Sanger Institute COVID-19 Surveillance Team |
| EPI_ISL_3453946, EPI_ISL_3453947, EPI_ISL_3453948, EPI_ISL_3453949 | Lighthouse Laboratory Plymouth | Wellcome Sanger Institute for the COVID-19 Genomics UK (COG-UK) Consortium | Lighthouse Laboratory Plymouth and Alex Alderton, Roberto Amato, Jeffrey Barrett, Sonia Goncalves, Ewan Harrison, David K. Jackson, Ian Johnston, Dominic Kwiatkowski, Cordelia Langford, John Sillitoe on behalf of the Wellcome Sanger Institute COVID-19 Surveillance Team |
| EPI_ISL_3453950 | Lighthouse Lab in Alderley Park | Wellcome Sanger Institute for the COVID-19 Genomics UK (COG-UK) Consortium | Jacquelyn Wynn, Mairead Hyland, The Lighthouse Lab in Alderley Park and Alex Alderton, Roberto Amato, Jeffrey Barrett, Sonia Goncalves, Ewan Harrison, David K. Jackson, Ian Johnston, Dominic Kwiatkowski, Cordelia Langford, John Sillitoe on behalf of the Wellcome Sanger Institute COVID-19 Surveillance Team |
| EPI_ISL_3453951 | Lighthouse Lab in Milton Keynes | Wellcome Sanger Institute for the COVID-19 Genomics UK (COG-UK) Consortium | The Lighthouse Lab in Milton Keynes and Alex Alderton, Roberto Amato, Jeffrey Barrett, Sonia Goncalves, Ewan Harrison, David K. Jackson, Ian Johnston, Dominic Kwiatkowski, Cordelia Langford, John Sillitoe on behalf of the Wellcome Sanger Institute COVID-19 Surveillance Team |
| EPI_ISL_3453952, EPI_ISL_3453953 | Lighthouse Laboratory Plymouth | Wellcome Sanger Institute for the COVID-19 Genomics UK (COG-UK) Consortium | Lighthouse Laboratory Plymouth and Alex Alderton, Roberto Amato, Jeffrey Barrett, Sonia Goncalves, Ewan Harrison, David K. Jackson, Ian Johnston, Dominic Kwiatkowski, Cordelia Langford, John Sillitoe on behalf of the Wellcome Sanger Institute COVID-19 Surveillance Team |
| EPI_ISL_3453954, EPI_ISL_3453955, EPI_ISL_3453956 | Lighthouse Lab in Alderley Park | Wellcome Sanger Institute for the COVID-19 Genomics UK (COG-UK) Consortium | Jacquelyn Wynn, Mairead Hyland, The Lighthouse Lab in Alderley Park and Alex Alderton, Roberto Amato, Jeffrey Barrett, Sonia Goncalves, Ewan Harrison, David K. Jackson, Ian Johnston, Dominic Kwiatkowski, Cordelia Langford, John Sillitoe on behalf of the Wellcome Sanger Institute COVID-19 Surveillance Team |
| EPI_ISL_3453957 | Lighthouse Lab in Milton Keynes | Wellcome Sanger Institute for the COVID-19 Genomics UK (COG-UK) Consortium | The Lighthouse Lab in Milton Keynes and Alex Alderton, Roberto Amato, Jeffrey Barrett, Sonia Goncalves, Ewan Harrison, David K. Jackson, Ian Johnston, Dominic Kwiatkowski, Cordelia Langford, John Sillitoe on behalf of the Wellcome Sanger Institute COVID-19 Surveillance Team |
| EPI_ISL_3453958 | Lighthouse Laboratory Plymouth | Wellcome Sanger Institute for the COVID-19 Genomics UK (COG-UK) Consortium | Lighthouse Laboratory Plymouth and Alex Alderton, Roberto Amato, Jeffrey Barrett, Sonia Goncalves, Ewan Harrison, David K. Jackson, Ian Johnston, Dominic Kwiatkowski, Cordelia Langford, John Sillitoe on behalf of the Wellcome Sanger Institute COVID-19 Surveillance Team |
| EPI_ISL_3453959 | Lighthouse Lab in Milton Keynes | Wellcome Sanger Institute for the COVID-19 Genomics UK (COG-UK) Consortium | The Lighthouse Lab in Milton Keynes and Alex Alderton, Roberto Amato, Jeffrey Barrett, Sonia Goncalves, Ewan Harrison, David K. Jackson, Ian Johnston, Dominic Kwiatkowski, Cordelia Langford, John Sillitoe on behalf of the Wellcome Sanger Institute COVID-19 Surveillance Team |
| EPI_ISL_3453960 | Lighthouse Lab in Alderley Park | Wellcome Sanger Institute for the COVID-19 Genomics UK (COG-UK) Consortium | Jacquelyn Wynn, Mairead Hyland, The Lighthouse Lab in Alderley Park and Alex Alderton, Roberto Amato, Jeffrey Barrett, Sonia Goncalves, Ewan Harrison, David K. Jackson, Ian Johnston, Dominic Kwiatkowski, Cordelia Langford, John Sillitoe on behalf of the Wellcome Sanger Institute COVID-19 Surveillance Team |
| EPI_ISL_3453961 | Lighthouse Laboratory Plymouth | Wellcome Sanger Institute for the COVID-19 Genomics UK (COG-UK) Consortium | Lighthouse Laboratory Plymouth and Alex Alderton, Roberto Amato, Jeffrey Barrett, Sonia Goncalves, Ewan Harrison, David K. Jackson, Ian Johnston, Dominic Kwiatkowski, Cordelia Langford, John Sillitoe on behalf of the Wellcome Sanger Institute COVID-19 Surveillance Team |
| EPI_ISL_3453962 | Lighthouse Lab in Alderley Park | Wellcome Sanger Institute for the COVID-19 Genomics UK (COG-UK) Consortium | Jacquelyn Wynn, Mairead Hyland, The Lighthouse Lab in Alderley Park and Alex Alderton, Roberto Amato, Jeffrey Barrett, Sonia Goncalves, Ewan Harrison, David K. Jackson, Ian Johnston, Dominic Kwiatkowski, Cordelia Langford, John Sillitoe on behalf of the Wellcome Sanger Institute COVID-19 Surveillance Team |
| EPI_ISL_3453963 | Lighthouse Lab in Milton Keynes | Wellcome Sanger Institute for the COVID-19 Genomics UK (COG-UK) Consortium | The Lighthouse Lab in Milton Keynes and Alex Alderton, Roberto Amato, Jeffrey Barrett, Sonia Goncalves, Ewan Harrison, David K. Jackson, Ian Johnston, Dominic Kwiatkowski, Cordelia Langford, John Sillitoe on behalf of the Wellcome Sanger Institute COVID-19 Surveillance Team |
| EPI_ISL_3453964 | Lighthouse Laboratory Plymouth | Wellcome Sanger Institute for the COVID-19 Genomics UK (COG-UK) Consortium | Lighthouse Laboratory Plymouth and Alex Alderton, Roberto Amato, Jeffrey Barrett, Sonia Goncalves, Ewan Harrison, David K. Jackson, Ian Johnston, Dominic Kwiatkowski, Cordelia Langford, John Sillitoe on behalf of the Wellcome Sanger Institute COVID-19 Surveillance Team |
| EPI_ISL_3453965 | Lighthouse Lab in Milton Keynes | Wellcome Sanger Institute for the COVID-19 Genomics UK (COG-UK) Consortium | The Lighthouse Lab in Milton Keynes and Alex Alderton, Roberto Amato, Jeffrey Barrett, Sonia Goncalves, Ewan Harrison, David K. Jackson, Ian Johnston, Dominic Kwiatkowski, Cordelia Langford, John Sillitoe on behalf of the Wellcome Sanger Institute COVID-19 Surveillance Team |
| EPI_ISL_3453966, EPI_ISL_3453967, EPI_ISL_3453968 | Lighthouse Lab in Alderley Park | Wellcome Sanger Institute for the COVID-19 Genomics UK (COG-UK) Consortium | Jacquelyn Wynn, Mairead Hyland, The Lighthouse Lab in Alderley Park and Alex Alderton, Roberto Amato, Jeffrey Barrett, Sonia Goncalves, Ewan Harrison, David K. Jackson, Ian Johnston, Dominic Kwiatkowski, Cordelia Langford, John Sillitoe on behalf of the Wellcome Sanger Institute COVID-19 Surveillance Team |
| EPI_ISL_3453969 | Lighthouse Laboratory Plymouth | Wellcome Sanger Institute for the COVID-19 Genomics UK (COG-UK) Consortium | Lighthouse Laboratory Plymouth and Alex Alderton, Roberto Amato, Jeffrey Barrett, Sonia Goncalves, Ewan Harrison, David K. Jackson, Ian Johnston, Dominic Kwiatkowski, Cordelia Langford, John Sillitoe on behalf of the Wellcome Sanger Institute COVID-19 Surveillance Team |
| EPI_ISL_3453970 | Lighthouse Lab in Alderley Park | Wellcome Sanger Institute for the COVID-19 Genomics UK (COG-UK) Consortium | Jacquelyn Wynn, Mairead Hyland, The Lighthouse Lab in Alderley Park and Alex Alderton, Roberto Amato, Jeffrey Barrett, Sonia Goncalves, Ewan Harrison, David K. Jackson, Ian Johnston, Dominic Kwiatkowski, Cordelia Langford, John Sillitoe on behalf of the Wellcome Sanger Institute COVID-19 Surveillance Team |
| EPI_ISL_3453971, EPI_ISL_3453972 | Lighthouse Laboratory Plymouth | Wellcome Sanger Institute for the COVID-19 Genomics UK (COG-UK) Consortium | Lighthouse Laboratory Plymouth and Alex Alderton, Roberto Amato, Jeffrey Barrett, Sonia Goncalves, Ewan Harrison, David K. Jackson, Ian Johnston, Dominic Kwiatkowski, Cordelia Langford, John Sillitoe on behalf of the Wellcome Sanger Institute COVID-19 Surveillance Team |
| EPI_ISL_3453973 | Lighthouse Lab in Milton Keynes | Wellcome Sanger Institute for the COVID-19 Genomics UK (COG-UK) Consortium | The Lighthouse Lab in Milton Keynes and Alex Alderton, Roberto Amato, Jeffrey Barrett, Sonia Goncalves, Ewan Harrison, David K. Jackson, Ian Johnston, Dominic Kwiatkowski, Cordelia Langford, John Sillitoe on behalf of the Wellcome Sanger Institute COVID-19 Surveillance Team |
| EPI_ISL_3453974, EPI_ISL_3453975 | Lighthouse Laboratory Plymouth | Wellcome Sanger Institute for the COVID-19 Genomics UK (COG-UK) Consortium | Lighthouse Laboratory Plymouth and Alex Alderton, Roberto Amato, Jeffrey Barrett, Sonia Goncalves, Ewan Harrison, David K. Jackson, Ian Johnston, Dominic Kwiatkowski, Cordelia Langford, John Sillitoe on behalf of the Wellcome Sanger Institute COVID-19 Surveillance Team |
| EPI_ISL_3453976, EPI_ISL_3453977, EPI_ISL_3453978 | Lighthouse Lab in Alderley Park | Wellcome Sanger Institute for the COVID-19 Genomics UK (COG-UK) Consortium | Jacquelyn Wynn, Mairead Hyland, The Lighthouse Lab in Alderley Park and Alex Alderton, Roberto Amato, Jeffrey Barrett, Sonia Goncalves, Ewan Harrison, David K. Jackson, Ian Johnston, Dominic Kwiatkowski, Cordelia Langford, John Sillitoe on behalf of the Wellcome Sanger Institute COVID-19 Surveillance Team |
| EPI_ISL_3453979 | Lighthouse Laboratory Plymouth | Wellcome Sanger Institute for the COVID-19 Genomics UK (COG-UK) Consortium | Lighthouse Laboratory Plymouth and Alex Alderton, Roberto Amato, Jeffrey Barrett, Sonia Goncalves, Ewan Harrison, David K. Jackson, Ian Johnston, Dominic Kwiatkowski, Cordelia Langford, John Sillitoe on behalf of the Wellcome Sanger Institute COVID-19 Surveillance Team |
| EPI_ISL_3453980 | Lighthouse Lab in Alderley Park | Wellcome Sanger Institute for the COVID-19 Genomics UK (COG-UK) Consortium | Jacquelyn Wynn, Mairead Hyland, The Lighthouse Lab in Alderley Park and Alex Alderton, Roberto Amato, Jeffrey Barrett, Sonia Goncalves, Ewan Harrison, David K. Jackson, Ian Johnston, Dominic Kwiatkowski, Cordelia Langford, John Sillitoe on behalf of the Wellcome Sanger Institute COVID-19 Surveillance Team |
| EPI_ISL_3453981 | Lighthouse Lab in Milton Keynes | Wellcome Sanger Institute for the COVID-19 Genomics UK (COG-UK) Consortium | The Lighthouse Lab in Milton Keynes and Alex Alderton, Roberto Amato, Jeffrey Barrett, Sonia Goncalves, Ewan Harrison, David K. Jackson, Ian Johnston, Dominic Kwiatkowski, Cordelia Langford, John Sillitoe on behalf of the Wellcome Sanger Institute COVID-19 Surveillance Team |
| EPI_ISL_3453982 | Lighthouse Laboratory Plymouth | Wellcome Sanger Institute for the COVID-19 Genomics UK (COG-UK) Consortium | Lighthouse Laboratory Plymouth and Alex Alderton, Roberto Amato, Jeffrey Barrett, Sonia Goncalves, Ewan Harrison, David K. Jackson, Ian Johnston, Dominic Kwiatkowski, Cordelia Langford, John Sillitoe on behalf of the Wellcome Sanger Institute COVID-19 Surveillance Team |
| EPI_ISL_3453983, EPI_ISL_3453984 | Lighthouse Lab in Milton Keynes | Wellcome Sanger Institute for the COVID-19 Genomics UK (COG-UK) Consortium | The Lighthouse Lab in Milton Keynes and Alex Alderton, Roberto Amato, Jeffrey Barrett, Sonia Goncalves, Ewan Harrison, David K. Jackson, Ian Johnston, Dominic Kwiatkowski, Cordelia Langford, John Sillitoe on behalf of the Wellcome Sanger Institute COVID-19 Surveillance Team |

| | | | |
|---|---|---|---|
| EPI_ISL_3453985 | Lighthouse Lab in Alderley Park | Wellcome Sanger Institute for the COVID-19 Genomics UK (COG-UK) Consortium | Jacquelyn Wynn, Mairead Hyland, The Lighthouse Lab in Alderley Park and Alex Alderton, Roberto Amato, Jeffrey Barrett, Sonia Goncalves, Ewan Harrison, David K. Jackson, Ian Johnston, Dominic Kwiatkowski, Cordelia Langford, John Sillitoe on behalf of the Wellcome Sanger Institute COVID-19 Surveillance Team |
| EPI_ISL_3453986 | Lighthouse Lab in Milton Keynes | Wellcome Sanger Institute for the COVID-19 Genomics UK (COG-UK) Consortium | The Lighthouse Lab in Milton Keynes and Alex Alderton, Roberto Amato, Jeffrey Barrett, Sonia Goncalves, Ewan Harrison, David K. Jackson, Ian Johnston, Dominic Kwiatkowski, Cordelia Langford, John Sillitoe on behalf of the Wellcome Sanger Institute COVID-19 Surveillance Team |
| EPI_ISL_3453987 | Lighthouse Laboratory Plymouth | Wellcome Sanger Institute for the COVID-19 Genomics UK (COG-UK) Consortium | Lighthouse Laboratory Plymouth and Alex Alderton, Roberto Amato, Jeffrey Barrett, Sonia Goncalves, Ewan Harrison, David K. Jackson, Ian Johnston, Dominic Kwiatkowski, Cordelia Langford, John Sillitoe on behalf of the Wellcome Sanger Institute COVID-19 Surveillance Team |
| EPI_ISL_3453988 | Lighthouse Lab in Alderley Park | Wellcome Sanger Institute for the COVID-19 Genomics UK (COG-UK) Consortium | Jacquelyn Wynn, Mairead Hyland, The Lighthouse Lab in Alderley Park and Alex Alderton, Roberto Amato, Jeffrey Barrett, Sonia Goncalves, Ewan Harrison, David K. Jackson, Ian Johnston, Dominic Kwiatkowski, Cordelia Langford, John Sillitoe on behalf of the Wellcome Sanger Institute COVID-19 Surveillance Team |
| EPI_ISL_3453989, EPI_ISL_3453990 | Lighthouse Laboratory Plymouth | Wellcome Sanger Institute for the COVID-19 Genomics UK (COG-UK) Consortium | Lighthouse Laboratory Plymouth and Alex Alderton, Roberto Amato, Jeffrey Barrett, Sonia Goncalves, Ewan Harrison, David K. Jackson, Ian Johnston, Dominic Kwiatkowski, Cordelia Langford, John Sillitoe on behalf of the Wellcome Sanger Institute COVID-19 Surveillance Team |
| EPI_ISL_3453991 | Lighthouse Lab in Milton Keynes | Wellcome Sanger Institute for the COVID-19 Genomics UK (COG-UK) Consortium | The Lighthouse Lab in Milton Keynes and Alex Alderton, Roberto Amato, Jeffrey Barrett, Sonia Goncalves, Ewan Harrison, David K. Jackson, Ian Johnston, Dominic Kwiatkowski, Cordelia Langford, John Sillitoe on behalf of the Wellcome Sanger Institute COVID-19 Surveillance Team |
| EPI_ISL_3453992, EPI_ISL_3453993 | Lighthouse Lab in Alderley Park | Wellcome Sanger Institute for the COVID-19 Genomics UK (COG-UK) Consortium | Jacquelyn Wynn, Mairead Hyland, The Lighthouse Lab in Alderley Park and Alex Alderton, Roberto Amato, Jeffrey Barrett, Sonia Goncalves, Ewan Harrison, David K. Jackson, Ian Johnston, Dominic Kwiatkowski, Cordelia Langford, John Sillitoe on behalf of the Wellcome Sanger Institute COVID-19 Surveillance Team |
| EPI_ISL_3453994, EPI_ISL_3453995 | Lighthouse Lab in Milton Keynes | Wellcome Sanger Institute for the COVID-19 Genomics UK (COG-UK) Consortium | The Lighthouse Lab in Milton Keynes and Alex Alderton, Roberto Amato, Jeffrey Barrett, Sonia Goncalves, Ewan Harrison, David K. Jackson, Ian Johnston, Dominic Kwiatkowski, Cordelia Langford, John Sillitoe on behalf of the Wellcome Sanger Institute COVID-19 Surveillance Team |
| EPI_ISL_3453996 | Lighthouse Lab in Alderley Park | Wellcome Sanger Institute for the COVID-19 Genomics UK (COG-UK) Consortium | Jacquelyn Wynn, Mairead Hyland, The Lighthouse Lab in Alderley Park and Alex Alderton, Roberto Amato, Jeffrey Barrett, Sonia Goncalves, Ewan Harrison, David K. Jackson, Ian Johnston, Dominic Kwiatkowski, Cordelia Langford, John Sillitoe on behalf of the Wellcome Sanger Institute COVID-19 Surveillance Team |
| EPI_ISL_3453997 | Lighthouse Lab in Milton Keynes | Wellcome Sanger Institute for the COVID-19 Genomics UK (COG-UK) Consortium | The Lighthouse Lab in Milton Keynes and Alex Alderton, Roberto Amato, Jeffrey Barrett, Sonia Goncalves, Ewan Harrison, David K. Jackson, Ian Johnston, Dominic Kwiatkowski, Cordelia Langford, John Sillitoe on behalf of the Wellcome Sanger Institute COVID-19 Surveillance Team |
| EPI_ISL_3453998 | Kuzbass Clinical Dermatovenerologic Dispensary | WHO National Influenza Centre Russian Federation | Andrey Komissarov, Artem Fadeev, Kseniya Komissarova, Oula Mansour, Kirill Varchenko, Mikhail Bakaev, Tamila Musaeva, Veronika Eder, Maria Pisareva, Nikita Yolshin, Daria Danilenko, Ksenia Safina, Elena Nabieva, Georgii Bazykin, Dmitry Lioznov |
| EPI_ISL_3453999, EPI_ISL_3454000 | Lighthouse Laboratory Plymouth | Wellcome Sanger Institute for the COVID-19 Genomics UK (COG-UK) Consortium | Lighthouse Laboratory Plymouth and Alex Alderton, Roberto Amato, Jeffrey Barrett, Sonia Goncalves, Ewan Harrison, David K. Jackson, Ian Johnston, Dominic Kwiatkowski, Cordelia Langford, John Sillitoe on behalf of the Wellcome Sanger Institute COVID-19 Surveillance Team |
| EPI_ISL_3454001 | Kuzbass Clinical Dermatovenerologic Dispensary | WHO National Influenza Centre Russian Federation | Andrey Komissarov, Artem Fadeev, Kseniya Komissarova, Oula Mansour, Kirill Varchenko, Mikhail Bakaev, Tamila Musaeva, Veronika Eder, Maria Pisareva, Nikita Yolshin, Daria Danilenko, Ksenia Safina, Elena Nabieva, Georgii Bazykin, Dmitry Lioznov |
| EPI_ISL_3454002 | Lighthouse Lab in Alderley Park | Wellcome Sanger Institute for the COVID-19 Genomics UK (COG-UK) Consortium | Jacquelyn Wynn, Mairead Hyland, The Lighthouse Lab in Alderley Park and Alex Alderton, Roberto Amato, Jeffrey Barrett, Sonia Goncalves, Ewan Harrison, David K. Jackson, Ian Johnston, Dominic Kwiatkowski, Cordelia Langford, John Sillitoe on behalf of the Wellcome Sanger Institute COVID-19 Surveillance Team |
| EPI_ISL_3454003 | Lighthouse Lab in Milton Keynes | Wellcome Sanger Institute for the COVID-19 Genomics UK (COG-UK) Consortium | The Lighthouse Lab in Milton Keynes and Alex Alderton, Roberto Amato, Jeffrey Barrett, Sonia Goncalves, Ewan Harrison, David K. Jackson, Ian Johnston, Dominic Kwiatkowski, Cordelia Langford, John Sillitoe on behalf of the Wellcome Sanger Institute COVID-19 Surveillance Team |
| EPI_ISL_3454004 | Lighthouse Laboratory Plymouth | Wellcome Sanger Institute for the COVID-19 Genomics UK (COG-UK) Consortium | Lighthouse Laboratory Plymouth and Alex Alderton, Roberto Amato, Jeffrey Barrett, Sonia Goncalves, Ewan Harrison, David K. Jackson, Ian Johnston, Dominic Kwiatkowski, Cordelia Langford, John Sillitoe on behalf of the Wellcome Sanger Institute COVID-19 Surveillance Team |
| EPI_ISL_3454005 | Kuzbass Clinical Dermatovenerologic Dispensary | WHO National Influenza Centre Russian Federation | Andrey Komissarov, Artem Fadeev, Kseniya Komissarova, Oula Mansour, Kirill Varchenko, Mikhail Bakaev, Tamila Musaeva, Veronika Eder, Maria Pisareva, Nikita Yolshin, Daria Danilenko, Ksenia Safina, Elena Nabieva, Georgii Bazykin, Dmitry Lioznov |
| EPI_ISL_3454006, EPI_ISL_3454007 | Lighthouse Lab in Alderley Park | Wellcome Sanger Institute for the COVID-19 Genomics UK (COG-UK) Consortium | Jacquelyn Wynn, Mairead Hyland, The Lighthouse Lab in Alderley Park and Alex Alderton, Roberto Amato, Jeffrey Barrett, Sonia Goncalves, Ewan Harrison, David K. Jackson, Ian Johnston, Dominic Kwiatkowski, Cordelia Langford, John Sillitoe on behalf of the Wellcome Sanger Institute COVID-19 Surveillance Team |
| EPI_ISL_3454009 | Lighthouse Laboratory Plymouth | Wellcome Sanger Institute for the COVID-19 Genomics UK (COG-UK) Consortium | Lighthouse Laboratory Plymouth and Alex Alderton, Roberto Amato, Jeffrey Barrett, Sonia Goncalves, Ewan Harrison, David K. Jackson, Ian Johnston, Dominic Kwiatkowski, Cordelia Langford, John Sillitoe on behalf of the Wellcome Sanger Institute COVID-19 Surveillance Team |
| EPI_ISL_3454010 | Lighthouse Lab in Alderley Park | Wellcome Sanger Institute for the COVID-19 Genomics UK (COG-UK) Consortium | Jacquelyn Wynn, Mairead Hyland, The Lighthouse Lab in Alderley Park and Alex Alderton, Roberto Amato, Jeffrey Barrett, Sonia Goncalves, Ewan Harrison, David K. Jackson, Ian Johnston, Dominic Kwiatkowski, Cordelia Langford, John Sillitoe on behalf of the Wellcome Sanger Institute COVID-19 Surveillance Team |
| EPI_ISL_3454011 | Lighthouse Lab in Milton Keynes | Wellcome Sanger Institute for the COVID-19 Genomics UK (COG-UK) Consortium | The Lighthouse Lab in Milton Keynes and Alex Alderton, Roberto Amato, Jeffrey Barrett, Sonia Goncalves, Ewan Harrison, David K. Jackson, Ian Johnston, Dominic Kwiatkowski, Cordelia Langford, John Sillitoe on behalf of the Wellcome Sanger Institute COVID-19 Surveillance Team |
| EPI_ISL_3454012 | Kuzbass Clinical Dermatovenerologic Dispensary | WHO National Influenza Centre Russian Federation | Andrey Komissarov, Artem Fadeev, Kseniya Komissarova, Oula Mansour, Kirill Varchenko, Mikhail Bakaev, Tamila Musaeva, Veronika Eder, Maria Pisareva, Nikita Yolshin, Daria Danilenko, Ksenia Safina, Elena Nabieva, Georgii Bazykin, Dmitry Lioznov |
| EPI_ISL_3454013, EPI_ISL_3454014 | Lighthouse Lab in Alderley Park | Wellcome Sanger Institute for the COVID-19 Genomics UK (COG-UK) Consortium | Jacquelyn Wynn, Mairead Hyland, The Lighthouse Lab in Alderley Park and Alex Alderton, Roberto Amato, Jeffrey Barrett, Sonia Goncalves, Ewan Harrison, David K. Jackson, Ian Johnston, Dominic Kwiatkowski, Cordelia Langford, John Sillitoe on behalf of the Wellcome Sanger Institute COVID-19 Surveillance Team |
| EPI_ISL_3454015 | Kuzbass Clinical Dermatovenerologic Dispensary | WHO National Influenza Centre Russian Federation | Andrey Komissarov, Artem Fadeev, Kseniya Komissarova, Oula Mansour, Kirill Varchenko, Mikhail Bakaev, Tamila Musaeva, Veronika Eder, Maria Pisareva, Nikita Yolshin, Daria Danilenko, Ksenia Safina, Elena Nabieva, Georgii Bazykin, Dmitry Lioznov |
| EPI_ISL_3454016 | Lighthouse Lab in Milton Keynes | Wellcome Sanger Institute for the COVID-19 Genomics UK (COG-UK) Consortium | The Lighthouse Lab in Milton Keynes and Alex Alderton, Roberto Amato, Jeffrey Barrett, Sonia Goncalves, Ewan Harrison, David K. Jackson, Ian Johnston, Dominic Kwiatkowski, Cordelia Langford, John Sillitoe on behalf of the Wellcome Sanger Institute COVID-19 Surveillance Team |
| EPI_ISL_3454017 | Lighthouse Lab in Alderley Park | Wellcome Sanger Institute for the COVID-19 Genomics UK (COG-UK) Consortium | Jacquelyn Wynn, Mairead Hyland, The Lighthouse Lab in Alderley Park and Alex Alderton, Roberto Amato, Jeffrey Barrett, Sonia Goncalves, Ewan Harrison, David K. Jackson, Ian Johnston, Dominic Kwiatkowski, Cordelia Langford, John Sillitoe on behalf of the Wellcome Sanger Institute COVID-19 Surveillance Team |
| EPI_ISL_3454018 | Kuzbass Clinical Dermatovenerologic Dispensary | WHO National Influenza Centre Russian Federation | Andrey Komissarov, Artem Fadeev, Kseniya Komissarova, Oula Mansour, Kirill Varchenko, Mikhail Bakaev, Tamila Musaeva, Veronika Eder, Maria Pisareva, Nikita Yolshin, Daria Danilenko, Ksenia Safina, Elena Nabieva, Georgii Bazykin, Dmitry Lioznov |
| EPI_ISL_3454019, EPI_ISL_3454020 | Lighthouse Laboratory Plymouth | Wellcome Sanger Institute for the COVID-19 Genomics UK (COG-UK) Consortium | Lighthouse Laboratory Plymouth and Alex Alderton, Roberto Amato, Jeffrey Barrett, Sonia Goncalves, Ewan Harrison, David K. Jackson, Ian Johnston, Dominic Kwiatkowski, Cordelia Langford, John Sillitoe on behalf of the Wellcome Sanger Institute COVID-19 Surveillance Team |
| EPI_ISL_3454021 | Lighthouse Lab in Alderley Park | Wellcome Sanger Institute for the COVID-19 Genomics UK (COG-UK) Consortium | Jacquelyn Wynn, Mairead Hyland, The Lighthouse Lab in Alderley Park and Alex Alderton, Roberto Amato, Jeffrey Barrett, Sonia Goncalves, Ewan Harrison, David K. Jackson, Ian Johnston, Dominic Kwiatkowski, Cordelia Langford, John Sillitoe on behalf of the Wellcome Sanger Institute COVID-19 Surveillance Team |
| EPI_ISL_3454022 | Kuzbass Clinical Dermatovenerologic Dispensary | WHO National Influenza Centre Russian Federation | Andrey Komissarov, Artem Fadeev, Kseniya Komissarova, Oula Mansour, Kirill Varchenko, Mikhail Bakaev, Tamila Musaeva, Veronika Eder, Maria Pisareva, Nikita Yolshin, Daria Danilenko, Ksenia Safina, Elena Nabieva, Georgii Bazykin, Dmitry Lioznov |
| EPI_ISL_3454023, EPI_ISL_3454024 | Lighthouse Lab in Alderley Park | Wellcome Sanger Institute for the COVID-19 Genomics UK (COG-UK) Consortium | Jacquelyn Wynn, Mairead Hyland, The Lighthouse Lab in Alderley Park and Alex Alderton, Roberto Amato, Jeffrey Barrett, Sonia Goncalves, Ewan Harrison, David K. Jackson, Ian Johnston, Dominic Kwiatkowski, Cordelia Langford, John Sillitoe on behalf of the Wellcome Sanger Institute COVID-19 Surveillance Team |
| EPI_ISL_3454025 | Kuzbass Clinical Dermatovenerologic Dispensary | WHO National Influenza Centre Russian Federation | Andrey Komissarov, Artem Fadeev, Kseniya Komissarova, Oula Mansour, Kirill Varchenko, Mikhail Bakaev, Tamila Musaeva, Veronika Eder, Maria Pisareva, Nikita Yolshin, Daria Danilenko, Ksenia Safina, Elena Nabieva, Georgii Bazykin, Dmitry Lioznov |
| EPI_ISL_3454026, EPI_ISL_3454027 | Lighthouse Laboratory Plymouth | Wellcome Sanger Institute for the COVID-19 Genomics UK (COG-UK) Consortium | Lighthouse Laboratory Plymouth and Alex Alderton, Roberto Amato, Jeffrey Barrett, Sonia Goncalves, Ewan Harrison, David K. Jackson, Ian Johnston, Dominic Kwiatkowski, Cordelia Langford, John Sillitoe on behalf of the Wellcome Sanger Institute COVID-19 Surveillance Team |
| EPI_ISL_3454028 | Lighthouse Lab in Milton Keynes | Wellcome Sanger Institute for the COVID-19 Genomics UK (COG-UK) Consortium | The Lighthouse Lab in Milton Keynes and Alex Alderton, Roberto Amato, Jeffrey Barrett, Sonia Goncalves, Ewan Harrison, David K. Jackson, Ian Johnston, Dominic Kwiatkowski, Cordelia Langford, John Sillitoe on behalf of the Wellcome Sanger Institute COVID-19 Surveillance Team |
| EPI_ISL_3454029 | Perm Krai Blood Transfusion Station | WHO National Influenza Centre Russian Federation | Andrey Komissarov, Artem Fadeev, Kseniya Komissarova, Oula Mansour, Kirill Varchenko, Mikhail Bakaev, Tamila Musaeva, Veronika Eder, Maria Pisareva, Nikita Yolshin, Daria Danilenko, Ksenia Safina, Elena Nabieva, Georgii Bazykin, Dmitry Lioznov |
| EPI_ISL_3454030 | Lighthouse Laboratory Plymouth | Wellcome Sanger Institute for the COVID-19 Genomics UK (COG-UK) Consortium | Lighthouse Laboratory Plymouth and Alex Alderton, Roberto Amato, Jeffrey Barrett, Sonia Goncalves, Ewan Harrison, David K. Jackson, Ian Johnston, Dominic Kwiatkowski, Cordelia Langford, John Sillitoe on behalf of the Wellcome Sanger Institute COVID-19 Surveillance Team |
| EPI_ISL_3454031 | Lighthouse Lab in Milton Keynes | Wellcome Sanger Institute for the COVID-19 Genomics UK (COG-UK) Consortium | The Lighthouse Lab in Milton Keynes and Alex Alderton, Roberto Amato, Jeffrey Barrett, Sonia Goncalves, Ewan Harrison, David K. Jackson, Ian Johnston, Dominic Kwiatkowski, Cordelia Langford, John Sillitoe on behalf of the Wellcome Sanger Institute COVID-19 Surveillance Team |
| EPI_ISL_3454032 | Lighthouse Lab in Alderley Park | Wellcome Sanger Institute for the COVID-19 Genomics UK (COG-UK) Consortium | Jacquelyn Wynn, Mairead Hyland, The Lighthouse Lab in Alderley Park and Alex Alderton, Roberto Amato, Jeffrey Barrett, Sonia Goncalves, Ewan Harrison, David K. Jackson, Ian Johnston, Dominic Kwiatkowski, Cordelia Langford, John Sillitoe on behalf of the Wellcome Sanger Institute COVID-19 Surveillance Team |
| EPI_ISL_3454033 | Perm Krai Blood Transfusion Station | WHO National Influenza Centre Russian Federation | Andrey Komissarov, Artem Fadeev, Kseniya Komissarova, Oula Mansour, Kirill Varchenko, Mikhail Bakaev, Tamila Musaeva, Veronika Eder, Maria Pisareva, Nikita Yolshin, Daria Danilenko, Ksenia Safina, Elena Nabieva, Georgii Bazykin, Dmitry Lioznov |
| EPI_ISL_3454034 | Lighthouse Laboratory Plymouth | Wellcome Sanger Institute for the COVID-19 Genomics UK (COG-UK) Consortium | Lighthouse Laboratory Plymouth and Alex Alderton, Roberto Amato, Jeffrey Barrett, Sonia Goncalves, Ewan Harrison, David K. Jackson, Ian Johnston, Dominic Kwiatkowski, Cordelia Langford, John Sillitoe on behalf of the Wellcome Sanger Institute COVID-19 Surveillance Team |
| EPI_ISL_3454035 | Lighthouse Lab in Milton Keynes | Wellcome Sanger Institute for the COVID-19 Genomics UK (COG-UK) Consortium | The Lighthouse Lab in Milton Keynes and Alex Alderton, Roberto Amato, Jeffrey Barrett, Sonia Goncalves, Ewan Harrison, David K. Jackson, Ian Johnston, Dominic Kwiatkowski, Cordelia Langford, John Sillitoe on behalf of the Wellcome Sanger Institute COVID-19 Surveillance Team |
| EPI_ISL_3454036 | Perm Krai Blood Transfusion Station | WHO National Influenza Centre Russian Federation | Andrey Komissarov, Artem Fadeev, Kseniya Komissarova, Oula Mansour, Kirill Varchenko, Mikhail Bakaev, Tamila Musaeva, Veronika Eder, Maria Pisareva, Nikita Yolshin, Daria Danilenko, Ksenia Safina, Elena Nabieva, Georgii Bazykin, Dmitry Lioznov |
| EPI_ISL_3454037 | Lighthouse Lab in Alderley Park | Wellcome Sanger Institute for the COVID-19 Genomics UK (COG-UK) Consortium | Jacquelyn Wynn, Mairead Hyland, The Lighthouse Lab in Alderley Park and Alex Alderton, Roberto Amato, Jeffrey Barrett, Sonia Goncalves, Ewan Harrison, David K. Jackson, Ian Johnston, Dominic Kwiatkowski, Cordelia Langford, John Sillitoe on behalf of the Wellcome Sanger Institute COVID-19 Surveillance Team |
| EPI_ISL_3454038 | Lighthouse Laboratory Plymouth | Wellcome Sanger Institute for the COVID-19 Genomics UK (COG-UK) Consortium | Lighthouse Laboratory Plymouth and Alex Alderton, Roberto Amato, Jeffrey Barrett, Sonia Goncalves, Ewan Harrison, David K. Jackson, Ian Johnston, Dominic Kwiatkowski, Cordelia Langford, John Sillitoe on behalf of the Wellcome Sanger Institute COVID-19 Surveillance Team |
| EPI_ISL_3454039 | Lighthouse Lab in Alderley Park | Wellcome Sanger Institute for the COVID-19 Genomics UK (COG-UK) Consortium | Jacquelyn Wynn, Mairead Hyland, The Lighthouse Lab in Alderley Park and Alex Alderton, Roberto Amato, Jeffrey Barrett, Sonia Goncalves, Ewan Harrison, David K. Jackson, Ian Johnston, Dominic Kwiatkowski, Cordelia Langford, John Sillitoe on behalf of the Wellcome Sanger Institute COVID-19 Surveillance Team |

| | | | |
|---|---|---|---|
| EPI_ISL_3454040 | Perm Krai Blood Transfusion Station | WHO National Influenza Centre Russian Federation | Andrey Komissarov, Artem Fadeev, Kseniya Komissarova, Oula Mansour, Kirill Varchenko, Mikhail Bakaev, Tamila Musaeva, Veronika Eder, Maria Pisareva, Nikita Yolshin, Daria Danilenko, Ksenia Safina, Elena Nabieva, Georgii Bazykin, Dmitry Lioznov |
| EPI_ISL_3454041 | Lighthouse Laboratory Plymouth | Wellcome Sanger Institute for the COVID-19 Genomics UK (COG-UK) Consortium | Lighthouse Laboratory Plymouth and Alex Alderton, Roberto Amato, Jeffrey Barrett, Sonia Goncalves, Ewan Harrison, David K. Jackson, Ian Johnston, Dominic Kwiatkowski, Cordelia Langford, John Sillitoe on behalf of the Wellcome Sanger Institute COVID-19 Surveillance Team |
| EPI_ISL_3454042 | Lighthouse Lab in Alderley Park | Wellcome Sanger Institute for the COVID-19 Genomics UK (COG-UK) Consortium | Jacquelyn Wynn, Mairead Hyland, The Lighthouse Lab in Alderley Park and Alex Alderton, Roberto Amato, Jeffrey Barrett, Sonia Goncalves, Ewan Harrison, David K. Jackson, Ian Johnston, Dominic Kwiatkowski, Cordelia Langford, John Sillitoe on behalf of the Wellcome Sanger Institute COVID-19 Surveillance Team |
| EPI_ISL_3454043 | Perm Krai Blood Transfusion Station | WHO National Influenza Centre Russian Federation | Andrey Komissarov, Artem Fadeev, Kseniya Komissarova, Oula Mansour, Kirill Varchenko, Mikhail Bakaev, Tamila Musaeva, Veronika Eder, Maria Pisareva, Nikita Yolshin, Daria Danilenko, Ksenia Safina, Elena Nabieva, Georgii Bazykin, Dmitry Lioznov |
| EPI_ISL_3454044, EPI_ISL_3454045 | Lighthouse Lab in Milton Keynes | Wellcome Sanger Institute for the COVID-19 Genomics UK (COG-UK) Consortium | The Lighthouse Lab in Milton Keynes and Alex Alderton, Roberto Amato, Jeffrey Barrett, Sonia Goncalves, Ewan Harrison, David K. Jackson, Ian Johnston, Dominic Kwiatkowski, Cordelia Langford, John Sillitoe on behalf of the Wellcome Sanger Institute COVID-19 Surveillance Team |
| EPI_ISL_3454046 | Perm Krai Blood Transfusion Station | WHO National Influenza Centre Russian Federation | Andrey Komissarov, Artem Fadeev, Kseniya Komissarova, Oula Mansour, Kirill Varchenko, Mikhail Bakaev, Tamila Musaeva, Veronika Eder, Maria Pisareva, Nikita Yolshin, Daria Danilenko, Ksenia Safina, Elena Nabieva, Georgii Bazykin, Dmitry Lioznov |
| EPI_ISL_3454047 | Lighthouse Lab in Alderley Park | Wellcome Sanger Institute for the COVID-19 Genomics UK (COG-UK) Consortium | Jacquelyn Wynn, Mairead Hyland, The Lighthouse Lab in Alderley Park and Alex Alderton, Roberto Amato, Jeffrey Barrett, Sonia Goncalves, Ewan Harrison, David K. Jackson, Ian Johnston, Dominic Kwiatkowski, Cordelia Langford, John Sillitoe on behalf of the Wellcome Sanger Institute COVID-19 Surveillance Team |
| EPI_ISL_3454048 | Lighthouse Lab in Milton Keynes | Wellcome Sanger Institute for the COVID-19 Genomics UK (COG-UK) Consortium | The Lighthouse Lab in Milton Keynes and Alex Alderton, Roberto Amato, Jeffrey Barrett, Sonia Goncalves, Ewan Harrison, David K. Jackson, Ian Johnston, Dominic Kwiatkowski, Cordelia Langford, John Sillitoe on behalf of the Wellcome Sanger Institute COVID-19 Surveillance Team |
| EPI_ISL_3454049 | Lighthouse Lab in Alderley Park | Wellcome Sanger Institute for the COVID-19 Genomics UK (COG-UK) Consortium | Jacquelyn Wynn, Mairead Hyland, The Lighthouse Lab in Alderley Park and Alex Alderton, Roberto Amato, Jeffrey Barrett, Sonia Goncalves, Ewan Harrison, David K. Jackson, Ian Johnston, Dominic Kwiatkowski, Cordelia Langford, John Sillitoe on behalf of the Wellcome Sanger Institute COVID-19 Surveillance Team |
| EPI_ISL_3454050 | Perm Krai Blood Transfusion Station | WHO National Influenza Centre Russian Federation | Andrey Komissarov, Artem Fadeev, Kseniya Komissarova, Oula Mansour, Kirill Varchenko, Mikhail Bakaev, Tamila Musaeva, Veronika Eder, Maria Pisareva, Nikita Yolshin, Daria Danilenko, Ksenia Safina, Elena Nabieva, Georgii Bazykin, Dmitry Lioznov |
| EPI_ISL_3454051 | Lighthouse Lab in Alderley Park | Wellcome Sanger Institute for the COVID-19 Genomics UK (COG-UK) Consortium | Jacquelyn Wynn, Mairead Hyland, The Lighthouse Lab in Alderley Park and Alex Alderton, Roberto Amato, Jeffrey Barrett, Sonia Goncalves, Ewan Harrison, David K. Jackson, Ian Johnston, Dominic Kwiatkowski, Cordelia Langford, John Sillitoe on behalf of the Wellcome Sanger Institute COVID-19 Surveillance Team |
| EPI_ISL_3454052 | Lighthouse Laboratory Plymouth | Wellcome Sanger Institute for the COVID-19 Genomics UK (COG-UK) Consortium | Lighthouse Laboratory Plymouth and Alex Alderton, Roberto Amato, Jeffrey Barrett, Sonia Goncalves, Ewan Harrison, David K. Jackson, Ian Johnston, Dominic Kwiatkowski, Cordelia Langford, John Sillitoe on behalf of the Wellcome Sanger Institute COVID-19 Surveillance Team |
| EPI_ISL_3454053 | Perm Krai Blood Transfusion Station | WHO National Influenza Centre Russian Federation | Andrey Komissarov, Artem Fadeev, Kseniya Komissarova, Oula Mansour, Kirill Varchenko, Mikhail Bakaev, Tamila Musaeva, Veronika Eder, Maria Pisareva, Nikita Yolshin, Daria Danilenko, Ksenia Safina, Elena Nabieva, Georgii Bazykin, Dmitry Lioznov |
| EPI_ISL_3454054 | Lighthouse Lab in Milton Keynes | Wellcome Sanger Institute for the COVID-19 Genomics UK (COG-UK) Consortium | The Lighthouse Lab in Milton Keynes and Alex Alderton, Roberto Amato, Jeffrey Barrett, Sonia Goncalves, Ewan Harrison, David K. Jackson, Ian Johnston, Dominic Kwiatkowski, Cordelia Langford, John Sillitoe on behalf of the Wellcome Sanger Institute COVID-19 Surveillance Team |
| EPI_ISL_3454055, EPI_ISL_3454056 | Lighthouse Laboratory Plymouth | Wellcome Sanger Institute for the COVID-19 Genomics UK (COG-UK) Consortium | Lighthouse Laboratory Plymouth and Alex Alderton, Roberto Amato, Jeffrey Barrett, Sonia Goncalves, Ewan Harrison, David K. Jackson, Ian Johnston, Dominic Kwiatkowski, Cordelia Langford, John Sillitoe on behalf of the Wellcome Sanger Institute COVID-19 Surveillance Team |
| EPI_ISL_3454057 | Perm Krai Blood Transfusion Station | WHO National Influenza Centre Russian Federation | Andrey Komissarov, Artem Fadeev, Kseniya Komissarova, Oula Mansour, Kirill Varchenko, Mikhail Bakaev, Tamila Musaeva, Veronika Eder, Maria Pisareva, Nikita Yolshin, Daria Danilenko, Ksenia Safina, Elena Nabieva, Georgii Bazykin, Dmitry Lioznov |
| EPI_ISL_3454058 | Lighthouse Lab in Milton Keynes | Wellcome Sanger Institute for the COVID-19 Genomics UK (COG-UK) Consortium | The Lighthouse Lab in Milton Keynes and Alex Alderton, Roberto Amato, Jeffrey Barrett, Sonia Goncalves, Ewan Harrison, David K. Jackson, Ian Johnston, Dominic Kwiatkowski, Cordelia Langford, John Sillitoe on behalf of the Wellcome Sanger Institute COVID-19 Surveillance Team |
| EPI_ISL_3454059 | Lighthouse Lab in Alderley Park | Wellcome Sanger Institute for the COVID-19 Genomics UK (COG-UK) Consortium | Jacquelyn Wynn, Mairead Hyland, The Lighthouse Lab in Alderley Park and Alex Alderton, Roberto Amato, Jeffrey Barrett, Sonia Goncalves, Ewan Harrison, David K. Jackson, Ian Johnston, Dominic Kwiatkowski, Cordelia Langford, John Sillitoe on behalf of the Wellcome Sanger Institute COVID-19 Surveillance Team |
| EPI_ISL_3454060 | N. N. Burdenko Penza Regional Clinical Hospital | WHO National Influenza Centre Russian Federation | Andrey Komissarov, Artem Fadeev, Kseniya Komissarova, Oula Mansour, Kirill Varchenko, Mikhail Bakaev, Tamila Musaeva, Veronika Eder, Maria Pisareva, Nikita Yolshin, Daria Danilenko, Ksenia Safina, Elena Nabieva, Georgii Bazykin, Dmitry Lioznov |
| EPI_ISL_3454061, EPI_ISL_3454062, EPI_ISL_3454063 | Lighthouse Lab in Milton Keynes | Wellcome Sanger Institute for the COVID-19 Genomics UK (COG-UK) Consortium | The Lighthouse Lab in Milton Keynes and Alex Alderton, Roberto Amato, Jeffrey Barrett, Sonia Goncalves, Ewan Harrison, David K. Jackson, Ian Johnston, Dominic Kwiatkowski, Cordelia Langford, John Sillitoe on behalf of the Wellcome Sanger Institute COVID-19 Surveillance Team |
| EPI_ISL_3454064 | N. N. Burdenko Penza Regional Clinical Hospital | WHO National Influenza Centre Russian Federation | Andrey Komissarov, Artem Fadeev, Kseniya Komissarova, Oula Mansour, Kirill Varchenko, Mikhail Bakaev, Tamila Musaeva, Veronika Eder, Maria Pisareva, Nikita Yolshin, Daria Danilenko, Ksenia Safina, Elena Nabieva, Georgii Bazykin, Dmitry Lioznov |
| EPI_ISL_3454065 | Lighthouse Lab in Alderley Park | Wellcome Sanger Institute for the COVID-19 Genomics UK (COG-UK) Consortium | Jacquelyn Wynn, Mairead Hyland, The Lighthouse Lab in Alderley Park and Alex Alderton, Roberto Amato, Jeffrey Barrett, Sonia Goncalves, Ewan Harrison, David K. Jackson, Ian Johnston, Dominic Kwiatkowski, Cordelia Langford, John Sillitoe on behalf of the Wellcome Sanger Institute COVID-19 Surveillance Team |
| EPI_ISL_3454066 | Lighthouse Laboratory Plymouth | Wellcome Sanger Institute for the COVID-19 Genomics UK (COG-UK) Consortium | Lighthouse Laboratory Plymouth and Alex Alderton, Roberto Amato, Jeffrey Barrett, Sonia Goncalves, Ewan Harrison, David K. Jackson, Ian Johnston, Dominic Kwiatkowski, Cordelia Langford, John Sillitoe on behalf of the Wellcome Sanger Institute COVID-19 Surveillance Team |
| EPI_ISL_3454067 | Lighthouse Lab in Milton Keynes | Wellcome Sanger Institute for the COVID-19 Genomics UK (COG-UK) Consortium | The Lighthouse Lab in Milton Keynes and Alex Alderton, Roberto Amato, Jeffrey Barrett, Sonia Goncalves, Ewan Harrison, David K. Jackson, Ian Johnston, Dominic Kwiatkowski, Cordelia Langford, John Sillitoe on behalf of the Wellcome Sanger Institute COVID-19 Surveillance Team |
| EPI_ISL_3454068 | N. N. Burdenko Penza Regional Clinical Hospital | WHO National Influenza Centre Russian Federation | Andrey Komissarov, Artem Fadeev, Kseniya Komissarova, Oula Mansour, Kirill Varchenko, Mikhail Bakaev, Tamila Musaeva, Veronika Eder, Maria Pisareva, Nikita Yolshin, Daria Danilenko, Ksenia Safina, Elena Nabieva, Georgii Bazykin, Dmitry Lioznov |
| EPI_ISL_3454069 | Lighthouse Lab in Alderley Park | Wellcome Sanger Institute for the COVID-19 Genomics UK (COG-UK) Consortium | Jacquelyn Wynn, Mairead Hyland, The Lighthouse Lab in Alderley Park and Alex Alderton, Roberto Amato, Jeffrey Barrett, Sonia Goncalves, Ewan Harrison, David K. Jackson, Ian Johnston, Dominic Kwiatkowski, Cordelia Langford, John Sillitoe on behalf of the Wellcome Sanger Institute COVID-19 Surveillance Team |
| EPI_ISL_3454070 | Lighthouse Laboratory Plymouth | Wellcome Sanger Institute for the COVID-19 Genomics UK (COG-UK) Consortium | Lighthouse Laboratory Plymouth and Alex Alderton, Roberto Amato, Jeffrey Barrett, Sonia Goncalves, Ewan Harrison, David K. Jackson, Ian Johnston, Dominic Kwiatkowski, Cordelia Langford, John Sillitoe on behalf of the Wellcome Sanger Institute COVID-19 Surveillance Team |
| EPI_ISL_3454071 | N. N. Burdenko Penza Regional Clinical Hospital | WHO National Influenza Centre Russian Federation | Andrey Komissarov, Artem Fadeev, Kseniya Komissarova, Oula Mansour, Kirill Varchenko, Mikhail Bakaev, Tamila Musaeva, Veronika Eder, Maria Pisareva, Nikita Yolshin, Daria Danilenko, Ksenia Safina, Elena Nabieva, Georgii Bazykin, Dmitry Lioznov |
| EPI_ISL_3454072, EPI_ISL_3454073 | Lighthouse Lab in Alderley Park | Wellcome Sanger Institute for the COVID-19 Genomics UK (COG-UK) Consortium | Jacquelyn Wynn, Mairead Hyland, The Lighthouse Lab in Alderley Park and Alex Alderton, Roberto Amato, Jeffrey Barrett, Sonia Goncalves, Ewan Harrison, David K. Jackson, Ian Johnston, Dominic Kwiatkowski, Cordelia Langford, John Sillitoe on behalf of the Wellcome Sanger Institute COVID-19 Surveillance Team |
| EPI_ISL_3454074 | Lighthouse Laboratory Plymouth | Wellcome Sanger Institute for the COVID-19 Genomics UK (COG-UK) Consortium | Lighthouse Laboratory Plymouth and Alex Alderton, Roberto Amato, Jeffrey Barrett, Sonia Goncalves, Ewan Harrison, David K. Jackson, Ian Johnston, Dominic Kwiatkowski, Cordelia Langford, John Sillitoe on behalf of the Wellcome Sanger Institute COVID-19 Surveillance Team |
| EPI_ISL_3454075 | N. N. Burdenko Penza Regional Clinical Hospital | WHO National Influenza Centre Russian Federation | Andrey Komissarov, Artem Fadeev, Kseniya Komissarova, Oula Mansour, Kirill Varchenko, Mikhail Bakaev, Tamila Musaeva, Veronika Eder, Maria Pisareva, Nikita Yolshin, Daria Danilenko, Ksenia Safina, Elena Nabieva, Georgii Bazykin, Dmitry Lioznov |
| EPI_ISL_3454076 | Lighthouse Lab in Alderley Park | Wellcome Sanger Institute for the COVID-19 Genomics UK (COG-UK) Consortium | Jacquelyn Wynn, Mairead Hyland, The Lighthouse Lab in Alderley Park and Alex Alderton, Roberto Amato, Jeffrey Barrett, Sonia Goncalves, Ewan Harrison, David K. Jackson, Ian Johnston, Dominic Kwiatkowski, Cordelia Langford, John Sillitoe on behalf of the Wellcome Sanger Institute COVID-19 Surveillance Team |
| EPI_ISL_3454077 | Lighthouse Laboratory Plymouth | Wellcome Sanger Institute for the COVID-19 Genomics UK (COG-UK) Consortium | Lighthouse Laboratory Plymouth and Alex Alderton, Roberto Amato, Jeffrey Barrett, Sonia Goncalves, Ewan Harrison, David K. Jackson, Ian Johnston, Dominic Kwiatkowski, Cordelia Langford, John Sillitoe on behalf of the Wellcome Sanger Institute COVID-19 Surveillance Team |
| EPI_ISL_3454078 | N. N. Burdenko Penza Regional Clinical Hospital | WHO National Influenza Centre Russian Federation | Andrey Komissarov, Artem Fadeev, Kseniya Komissarova, Oula Mansour, Kirill Varchenko, Mikhail Bakaev, Tamila Musaeva, Veronika Eder, Maria Pisareva, Nikita Yolshin, Daria Danilenko, Ksenia Safina, Elena Nabieva, Georgii Bazykin, Dmitry Lioznov |
| EPI_ISL_3454079 | Lighthouse Lab in Alderley Park | Wellcome Sanger Institute for the COVID-19 Genomics UK (COG-UK) Consortium | Jacquelyn Wynn, Mairead Hyland, The Lighthouse Lab in Alderley Park and Alex Alderton, Roberto Amato, Jeffrey Barrett, Sonia Goncalves, Ewan Harrison, David K. Jackson, Ian Johnston, Dominic Kwiatkowski, Cordelia Langford, John Sillitoe on behalf of the Wellcome Sanger Institute COVID-19 Surveillance Team |
| EPI_ISL_3454080 | Lighthouse Laboratory Plymouth | Wellcome Sanger Institute for the COVID-19 Genomics UK (COG-UK) Consortium | Lighthouse Laboratory Plymouth and Alex Alderton, Roberto Amato, Jeffrey Barrett, Sonia Goncalves, Ewan Harrison, David K. Jackson, Ian Johnston, Dominic Kwiatkowski, Cordelia Langford, John Sillitoe on behalf of the Wellcome Sanger Institute COVID-19 Surveillance Team |
| EPI_ISL_3454081 | Lighthouse Lab in Alderley Park | Wellcome Sanger Institute for the COVID-19 Genomics UK (COG-UK) Consortium | Jacquelyn Wynn, Mairead Hyland, The Lighthouse Lab in Alderley Park and Alex Alderton, Roberto Amato, Jeffrey Barrett, Sonia Goncalves, Ewan Harrison, David K. Jackson, Ian Johnston, Dominic Kwiatkowski, Cordelia Langford, John Sillitoe on behalf of the Wellcome Sanger Institute COVID-19 Surveillance Team |
| EPI_ISL_3454082 | N. N. Burdenko Penza Regional Clinical Hospital | WHO National Influenza Centre Russian Federation | Andrey Komissarov, Artem Fadeev, Kseniya Komissarova, Oula Mansour, Kirill Varchenko, Mikhail Bakaev, Tamila Musaeva, Veronika Eder, Maria Pisareva, Nikita Yolshin, Daria Danilenko, Ksenia Safina, Elena Nabieva, Georgii Bazykin, Dmitry Lioznov |
| EPI_ISL_3454083 | Lighthouse Laboratory Plymouth | Wellcome Sanger Institute for the COVID-19 Genomics UK (COG-UK) Consortium | Lighthouse Laboratory Plymouth and Alex Alderton, Roberto Amato, Jeffrey Barrett, Sonia Goncalves, Ewan Harrison, David K. Jackson, Ian Johnston, Dominic Kwiatkowski, Cordelia Langford, John Sillitoe on behalf of the Wellcome Sanger Institute COVID-19 Surveillance Team |
| EPI_ISL_3454084 | Lighthouse Lab in Milton Keynes | Wellcome Sanger Institute for the COVID-19 Genomics UK (COG-UK) Consortium | The Lighthouse Lab in Milton Keynes and Alex Alderton, Roberto Amato, Jeffrey Barrett, Sonia Goncalves, Ewan Harrison, David K. Jackson, Ian Johnston, Dominic Kwiatkowski, Cordelia Langford, John Sillitoe on behalf of the Wellcome Sanger Institute COVID-19 Surveillance Team |
| EPI_ISL_3454085 | Lighthouse Lab in Alderley Park | Wellcome Sanger Institute for the COVID-19 Genomics UK (COG-UK) Consortium | Jacquelyn Wynn, Mairead Hyland, The Lighthouse Lab in Alderley Park and Alex Alderton, Roberto Amato, Jeffrey Barrett, Sonia Goncalves, Ewan Harrison, David K. Jackson, Ian Johnston, Dominic Kwiatkowski, Cordelia Langford, John Sillitoe on behalf of the Wellcome Sanger Institute COVID-19 Surveillance Team |
| EPI_ISL_3454086 | Lighthouse Lab in Milton Keynes | Wellcome Sanger Institute for the COVID-19 Genomics UK (COG-UK) Consortium | The Lighthouse Lab in Milton Keynes and Alex Alderton, Roberto Amato, Jeffrey Barrett, Sonia Goncalves, Ewan Harrison, David K. Jackson, Ian Johnston, Dominic Kwiatkowski, Cordelia Langford, John Sillitoe on behalf of the Wellcome Sanger Institute COVID-19 Surveillance Team |
| EPI_ISL_3454087, EPI_ISL_3454088 | Lighthouse Lab in Alderley Park | Wellcome Sanger Institute for the COVID-19 Genomics UK (COG-UK) Consortium | Jacquelyn Wynn, Mairead Hyland, The Lighthouse Lab in Alderley Park and Alex Alderton, Roberto Amato, Jeffrey Barrett, Sonia Goncalves, Ewan Harrison, David K. Jackson, Ian Johnston, Dominic Kwiatkowski, Cordelia Langford, John Sillitoe on behalf of the Wellcome Sanger Institute COVID-19 Surveillance Team |
| EPI_ISL_3454089, EPI_ISL_3454090, EPI_ISL_3454091 | Lighthouse Laboratory Plymouth | Wellcome Sanger Institute for the COVID-19 Genomics UK (COG-UK) Consortium | Lighthouse Laboratory Plymouth and Alex Alderton, Roberto Amato, Jeffrey Barrett, Sonia Goncalves, Ewan Harrison, David K. Jackson, Ian Johnston, Dominic Kwiatkowski, Cordelia Langford, John Sillitoe on behalf of the Wellcome Sanger Institute COVID-19 Surveillance Team |

| | | | |
|---|---|---|---|
| EPI_ISL_3454092 | Lighthouse Lab in Alderley Park | Wellcome Sanger Institute for the COVID-19 Genomics UK (COG-UK) Consortium | Jacquelyn Wynn, Mairead Hyland, The Lighthouse Lab in Alderley Park and Alex Alderton, Roberto Amato, Jeffrey Barrett, Sonia Goncalves, Ewan Harrison, David K. Jackson, Ian Johnston, Dominic Kwiatkowski, Cordelia Langford, John Sillitoe on behalf of the Wellcome Sanger Institute COVID-19 Surveillance Team |
| EPI_ISL_3454093, EPI_ISL_3454094 | Lighthouse Laboratory Plymouth | Wellcome Sanger Institute for the COVID-19 Genomics UK (COG-UK) Consortium | Lighthouse Laboratory Plymouth and Alex Alderton, Roberto Amato, Jeffrey Barrett, Sonia Goncalves, Ewan Harrison, David K. Jackson, Ian Johnston, Dominic Kwiatkowski, Cordelia Langford, John Sillitoe on behalf of the Wellcome Sanger Institute COVID-19 Surveillance Team |
| EPI_ISL_3454095 | Lighthouse Lab in Alderley Park | Wellcome Sanger Institute for the COVID-19 Genomics UK (COG-UK) Consortium | Jacquelyn Wynn, Mairead Hyland, The Lighthouse Lab in Alderley Park and Alex Alderton, Roberto Amato, Jeffrey Barrett, Sonia Goncalves, Ewan Harrison, David K. Jackson, Ian Johnston, Dominic Kwiatkowski, Cordelia Langford, John Sillitoe on behalf of the Wellcome Sanger Institute COVID-19 Surveillance Team |
| EPI_ISL_3454096, EPI_ISL_3454097 | Lighthouse Laboratory Plymouth | Wellcome Sanger Institute for the COVID-19 Genomics UK (COG-UK) Consortium | Lighthouse Laboratory Plymouth and Alex Alderton, Roberto Amato, Jeffrey Barrett, Sonia Goncalves, Ewan Harrison, David K. Jackson, Ian Johnston, Dominic Kwiatkowski, Cordelia Langford, John Sillitoe on behalf of the Wellcome Sanger Institute COVID-19 Surveillance Team |
| EPI_ISL_3454098 | Lighthouse Lab in Alderley Park | Wellcome Sanger Institute for the COVID-19 Genomics UK (COG-UK) Consortium | Jacquelyn Wynn, Mairead Hyland, The Lighthouse Lab in Alderley Park and Alex Alderton, Roberto Amato, Jeffrey Barrett, Sonia Goncalves, Ewan Harrison, David K. Jackson, Ian Johnston, Dominic Kwiatkowski, Cordelia Langford, John Sillitoe on behalf of the Wellcome Sanger Institute COVID-19 Surveillance Team |
| EPI_ISL_3454099 | Lighthouse Laboratory Plymouth | Wellcome Sanger Institute for the COVID-19 Genomics UK (COG-UK) Consortium | Lighthouse Laboratory Plymouth and Alex Alderton, Roberto Amato, Jeffrey Barrett, Sonia Goncalves, Ewan Harrison, David K. Jackson, Ian Johnston, Dominic Kwiatkowski, Cordelia Langford, John Sillitoe on behalf of the Wellcome Sanger Institute COVID-19 Surveillance Team |
| EPI_ISL_3454100 | Lighthouse Lab in Milton Keynes | Wellcome Sanger Institute for the COVID-19 Genomics UK (COG-UK) Consortium | The Lighthouse Lab in Milton Keynes and Alex Alderton, Roberto Amato, Jeffrey Barrett, Sonia Goncalves, Ewan Harrison, David K. Jackson, Ian Johnston, Dominic Kwiatkowski, Cordelia Langford, John Sillitoe on behalf of the Wellcome Sanger Institute COVID-19 Surveillance Team |
| EPI_ISL_3454101 | Lighthouse Laboratory Plymouth | Wellcome Sanger Institute for the COVID-19 Genomics UK (COG-UK) Consortium | Lighthouse Laboratory Plymouth and Alex Alderton, Roberto Amato, Jeffrey Barrett, Sonia Goncalves, Ewan Harrison, David K. Jackson, Ian Johnston, Dominic Kwiatkowski, Cordelia Langford, John Sillitoe on behalf of the Wellcome Sanger Institute COVID-19 Surveillance Team |
| EPI_ISL_3454102, EPI_ISL_3454103, EPI_ISL_3454104, EPI_ISL_3454105 | Lighthouse Lab in Alderley Park | Wellcome Sanger Institute for the COVID-19 Genomics UK (COG-UK) Consortium | Jacquelyn Wynn, Mairead Hyland, The Lighthouse Lab in Alderley Park and Alex Alderton, Roberto Amato, Jeffrey Barrett, Sonia Goncalves, Ewan Harrison, David K. Jackson, Ian Johnston, Dominic Kwiatkowski, Cordelia Langford, John Sillitoe on behalf of the Wellcome Sanger Institute COVID-19 Surveillance Team |
| EPI_ISL_3454106, EPI_ISL_3454107, EPI_ISL_3454108 | Lighthouse Lab in Milton Keynes | Wellcome Sanger Institute for the COVID-19 Genomics UK (COG-UK) Consortium | The Lighthouse Lab in Milton Keynes and Alex Alderton, Roberto Amato, Jeffrey Barrett, Sonia Goncalves, Ewan Harrison, David K. Jackson, Ian Johnston, Dominic Kwiatkowski, Cordelia Langford, John Sillitoe on behalf of the Wellcome Sanger Institute COVID-19 Surveillance Team |
| EPI_ISL_3454109 | Lighthouse Lab in Alderley Park | Wellcome Sanger Institute for the COVID-19 Genomics UK (COG-UK) Consortium | Jacquelyn Wynn, Mairead Hyland, The Lighthouse Lab in Alderley Park and Alex Alderton, Roberto Amato, Jeffrey Barrett, Sonia Goncalves, Ewan Harrison, David K. Jackson, Ian Johnston, Dominic Kwiatkowski, Cordelia Langford, John Sillitoe on behalf of the Wellcome Sanger Institute COVID-19 Surveillance Team |
| EPI_ISL_3454110 | Lighthouse Lab in Milton Keynes | Wellcome Sanger Institute for the COVID-19 Genomics UK (COG-UK) Consortium | The Lighthouse Lab in Milton Keynes and Alex Alderton, Roberto Amato, Jeffrey Barrett, Sonia Goncalves, Ewan Harrison, David K. Jackson, Ian Johnston, Dominic Kwiatkowski, Cordelia Langford, John Sillitoe on behalf of the Wellcome Sanger Institute COVID-19 Surveillance Team |
| EPI_ISL_3454111 | Lighthouse Laboratory Plymouth | Wellcome Sanger Institute for the COVID-19 Genomics UK (COG-UK) Consortium | Lighthouse Laboratory Plymouth and Alex Alderton, Roberto Amato, Jeffrey Barrett, Sonia Goncalves, Ewan Harrison, David K. Jackson, Ian Johnston, Dominic Kwiatkowski, Cordelia Langford, John Sillitoe on behalf of the Wellcome Sanger Institute COVID-19 Surveillance Team |
| EPI_ISL_3454112 | Lighthouse Lab in Alderley Park | Wellcome Sanger Institute for the COVID-19 Genomics UK (COG-UK) Consortium | Jacquelyn Wynn, Mairead Hyland, The Lighthouse Lab in Alderley Park and Alex Alderton, Roberto Amato, Jeffrey Barrett, Sonia Goncalves, Ewan Harrison, David K. Jackson, Ian Johnston, Dominic Kwiatkowski, Cordelia Langford, John Sillitoe on behalf of the Wellcome Sanger Institute COVID-19 Surveillance Team |
| EPI_ISL_3454113 | Lighthouse Laboratory Plymouth | Wellcome Sanger Institute for the COVID-19 Genomics UK (COG-UK) Consortium | Lighthouse Laboratory Plymouth and Alex Alderton, Roberto Amato, Jeffrey Barrett, Sonia Goncalves, Ewan Harrison, David K. Jackson, Ian Johnston, Dominic Kwiatkowski, Cordelia Langford, John Sillitoe on behalf of the Wellcome Sanger Institute COVID-19 Surveillance Team |
| EPI_ISL_3454114, EPI_ISL_3454115 | Lighthouse Lab in Milton Keynes | Wellcome Sanger Institute for the COVID-19 Genomics UK (COG-UK) Consortium | The Lighthouse Lab in Milton Keynes and Alex Alderton, Roberto Amato, Jeffrey Barrett, Sonia Goncalves, Ewan Harrison, David K. Jackson, Ian Johnston, Dominic Kwiatkowski, Cordelia Langford, John Sillitoe on behalf of the Wellcome Sanger Institute COVID-19 Surveillance Team |
| EPI_ISL_3454116 | Lighthouse Laboratory Plymouth | Wellcome Sanger Institute for the COVID-19 Genomics UK (COG-UK) Consortium | Lighthouse Laboratory Plymouth and Alex Alderton, Roberto Amato, Jeffrey Barrett, Sonia Goncalves, Ewan Harrison, David K. Jackson, Ian Johnston, Dominic Kwiatkowski, Cordelia Langford, John Sillitoe on behalf of the Wellcome Sanger Institute COVID-19 Surveillance Team |
| EPI_ISL_3454117 | Lighthouse Lab in Milton Keynes | Wellcome Sanger Institute for the COVID-19 Genomics UK (COG-UK) Consortium | The Lighthouse Lab in Milton Keynes and Alex Alderton, Roberto Amato, Jeffrey Barrett, Sonia Goncalves, Ewan Harrison, David K. Jackson, Ian Johnston, Dominic Kwiatkowski, Cordelia Langford, John Sillitoe on behalf of the Wellcome Sanger Institute COVID-19 Surveillance Team |
| EPI_ISL_3454118 | Lighthouse Laboratory Plymouth | Wellcome Sanger Institute for the COVID-19 Genomics UK (COG-UK) Consortium | Lighthouse Laboratory Plymouth and Alex Alderton, Roberto Amato, Jeffrey Barrett, Sonia Goncalves, Ewan Harrison, David K. Jackson, Ian Johnston, Dominic Kwiatkowski, Cordelia Langford, John Sillitoe on behalf of the Wellcome Sanger Institute COVID-19 Surveillance Team |
| EPI_ISL_3454119 | Lighthouse Lab in Alderley Park | Wellcome Sanger Institute for the COVID-19 Genomics UK (COG-UK) Consortium | Jacquelyn Wynn, Mairead Hyland, The Lighthouse Lab in Alderley Park and Alex Alderton, Roberto Amato, Jeffrey Barrett, Sonia Goncalves, Ewan Harrison, David K. Jackson, Ian Johnston, Dominic Kwiatkowski, Cordelia Langford, John Sillitoe on behalf of the Wellcome Sanger Institute COVID-19 Surveillance Team |
| EPI_ISL_3454120 | Lighthouse Lab in Milton Keynes | Wellcome Sanger Institute for the COVID-19 Genomics UK (COG-UK) Consortium | The Lighthouse Lab in Milton Keynes and Alex Alderton, Roberto Amato, Jeffrey Barrett, Sonia Goncalves, Ewan Harrison, David K. Jackson, Ian Johnston, Dominic Kwiatkowski, Cordelia Langford, John Sillitoe on behalf of the Wellcome Sanger Institute COVID-19 Surveillance Team |
| EPI_ISL_3454121 | Lighthouse Lab in Alderley Park | Wellcome Sanger Institute for the COVID-19 Genomics UK (COG-UK) Consortium | Jacquelyn Wynn, Mairead Hyland, The Lighthouse Lab in Alderley Park and Alex Alderton, Roberto Amato, Jeffrey Barrett, Sonia Goncalves, Ewan Harrison, David K. Jackson, Ian Johnston, Dominic Kwiatkowski, Cordelia Langford, John Sillitoe on behalf of the Wellcome Sanger Institute COVID-19 Surveillance Team |
| EPI_ISL_3454122 | Lighthouse Lab in Milton Keynes | Wellcome Sanger Institute for the COVID-19 Genomics UK (COG-UK) Consortium | The Lighthouse Lab in Milton Keynes and Alex Alderton, Roberto Amato, Jeffrey Barrett, Sonia Goncalves, Ewan Harrison, David K. Jackson, Ian Johnston, Dominic Kwiatkowski, Cordelia Langford, John Sillitoe on behalf of the Wellcome Sanger Institute COVID-19 Surveillance Team |
| EPI_ISL_3454123, EPI_ISL_3454124 | Lighthouse Lab in Alderley Park | Wellcome Sanger Institute for the COVID-19 Genomics UK (COG-UK) Consortium | Jacquelyn Wynn, Mairead Hyland, The Lighthouse Lab in Alderley Park and Alex Alderton, Roberto Amato, Jeffrey Barrett, Sonia Goncalves, Ewan Harrison, David K. Jackson, Ian Johnston, Dominic Kwiatkowski, Cordelia Langford, John Sillitoe on behalf of the Wellcome Sanger Institute COVID-19 Surveillance Team |
| EPI_ISL_3454125 | Lighthouse Laboratory Plymouth | Wellcome Sanger Institute for the COVID-19 Genomics UK (COG-UK) Consortium | Lighthouse Laboratory Plymouth and Alex Alderton, Roberto Amato, Jeffrey Barrett, Sonia Goncalves, Ewan Harrison, David K. Jackson, Ian Johnston, Dominic Kwiatkowski, Cordelia Langford, John Sillitoe on behalf of the Wellcome Sanger Institute COVID-19 Surveillance Team |
| EPI_ISL_3454126, EPI_ISL_3454127 | Lighthouse Lab in Alderley Park | Wellcome Sanger Institute for the COVID-19 Genomics UK (COG-UK) Consortium | Jacquelyn Wynn, Mairead Hyland, The Lighthouse Lab in Alderley Park and Alex Alderton, Roberto Amato, Jeffrey Barrett, Sonia Goncalves, Ewan Harrison, David K. Jackson, Ian Johnston, Dominic Kwiatkowski, Cordelia Langford, John Sillitoe on behalf of the Wellcome Sanger Institute COVID-19 Surveillance Team |
| EPI_ISL_3454128, EPI_ISL_3454129 | Lighthouse Lab in Milton Keynes | Wellcome Sanger Institute for the COVID-19 Genomics UK (COG-UK) Consortium | The Lighthouse Lab in Milton Keynes and Alex Alderton, Roberto Amato, Jeffrey Barrett, Sonia Goncalves, Ewan Harrison, David K. Jackson, Ian Johnston, Dominic Kwiatkowski, Cordelia Langford, John Sillitoe on behalf of the Wellcome Sanger Institute COVID-19 Surveillance Team |
| EPI_ISL_3454130, EPI_ISL_3454131, EPI_ISL_3454132 | Lighthouse Lab in Alderley Park | Wellcome Sanger Institute for the COVID-19 Genomics UK (COG-UK) Consortium | Jacquelyn Wynn, Mairead Hyland, The Lighthouse Lab in Alderley Park and Alex Alderton, Roberto Amato, Jeffrey Barrett, Sonia Goncalves, Ewan Harrison, David K. Jackson, Ian Johnston, Dominic Kwiatkowski, Cordelia Langford, John Sillitoe on behalf of the Wellcome Sanger Institute COVID-19 Surveillance Team |
| EPI_ISL_3454133, EPI_ISL_3454134 | Lighthouse Lab in Milton Keynes | Wellcome Sanger Institute for the COVID-19 Genomics UK (COG-UK) Consortium | The Lighthouse Lab in Milton Keynes and Alex Alderton, Roberto Amato, Jeffrey Barrett, Sonia Goncalves, Ewan Harrison, David K. Jackson, Ian Johnston, Dominic Kwiatkowski, Cordelia Langford, John Sillitoe on behalf of the Wellcome Sanger Institute COVID-19 Surveillance Team |
| EPI_ISL_3454135 | Lighthouse Lab in Alderley Park | Wellcome Sanger Institute for the COVID-19 Genomics UK (COG-UK) Consortium | Jacquelyn Wynn, Mairead Hyland, The Lighthouse Lab in Alderley Park and Alex Alderton, Roberto Amato, Jeffrey Barrett, Sonia Goncalves, Ewan Harrison, David K. Jackson, Ian Johnston, Dominic Kwiatkowski, Cordelia Langford, John Sillitoe on behalf of the Wellcome Sanger Institute COVID-19 Surveillance Team |
| EPI_ISL_3454136, EPI_ISL_3454137 | Lighthouse Lab in Milton Keynes | Wellcome Sanger Institute for the COVID-19 Genomics UK (COG-UK) Consortium | The Lighthouse Lab in Milton Keynes and Alex Alderton, Roberto Amato, Jeffrey Barrett, Sonia Goncalves, Ewan Harrison, David K. Jackson, Ian Johnston, Dominic Kwiatkowski, Cordelia Langford, John Sillitoe on behalf of the Wellcome Sanger Institute COVID-19 Surveillance Team |
| EPI_ISL_3454138 | Lighthouse Laboratory Plymouth | Wellcome Sanger Institute for the COVID-19 Genomics UK (COG-UK) Consortium | Lighthouse Laboratory Plymouth and Alex Alderton, Roberto Amato, Jeffrey Barrett, Sonia Goncalves, Ewan Harrison, David K. Jackson, Ian Johnston, Dominic Kwiatkowski, Cordelia Langford, John Sillitoe on behalf of the Wellcome Sanger Institute COVID-19 Surveillance Team |
| EPI_ISL_3454139, EPI_ISL_3454140 | Lighthouse Lab in Alderley Park | Wellcome Sanger Institute for the COVID-19 Genomics UK (COG-UK) Consortium | Jacquelyn Wynn, Mairead Hyland, The Lighthouse Lab in Alderley Park and Alex Alderton, Roberto Amato, Jeffrey Barrett, Sonia Goncalves, Ewan Harrison, David K. Jackson, Ian Johnston, Dominic Kwiatkowski, Cordelia Langford, John Sillitoe on behalf of the Wellcome Sanger Institute COVID-19 Surveillance Team |
| EPI_ISL_3454141 | Lighthouse Lab in Milton Keynes | Wellcome Sanger Institute for the COVID-19 Genomics UK (COG-UK) Consortium | The Lighthouse Lab in Milton Keynes and Alex Alderton, Roberto Amato, Jeffrey Barrett, Sonia Goncalves, Ewan Harrison, David K. Jackson, Ian Johnston, Dominic Kwiatkowski, Cordelia Langford, John Sillitoe on behalf of the Wellcome Sanger Institute COVID-19 Surveillance Team |
| EPI_ISL_3454142 | Lighthouse Lab in Alderley Park | Wellcome Sanger Institute for the COVID-19 Genomics UK (COG-UK) Consortium | Jacquelyn Wynn, Mairead Hyland, The Lighthouse Lab in Alderley Park and Alex Alderton, Roberto Amato, Jeffrey Barrett, Sonia Goncalves, Ewan Harrison, David K. Jackson, Ian Johnston, Dominic Kwiatkowski, Cordelia Langford, John Sillitoe on behalf of the Wellcome Sanger Institute COVID-19 Surveillance Team |
| EPI_ISL_3454143 | Lighthouse Laboratory Plymouth | Wellcome Sanger Institute for the COVID-19 Genomics UK (COG-UK) Consortium | Lighthouse Laboratory Plymouth and Alex Alderton, Roberto Amato, Jeffrey Barrett, Sonia Goncalves, Ewan Harrison, David K. Jackson, Ian Johnston, Dominic Kwiatkowski, Cordelia Langford, John Sillitoe on behalf of the Wellcome Sanger Institute COVID-19 Surveillance Team |
| EPI_ISL_3454144 | Lighthouse Lab in Alderley Park | Wellcome Sanger Institute for the COVID-19 Genomics UK (COG-UK) Consortium | Jacquelyn Wynn, Mairead Hyland, The Lighthouse Lab in Alderley Park and Alex Alderton, Roberto Amato, Jeffrey Barrett, Sonia Goncalves, Ewan Harrison, David K. Jackson, Ian Johnston, Dominic Kwiatkowski, Cordelia Langford, John Sillitoe on behalf of the Wellcome Sanger Institute COVID-19 Surveillance Team |
| EPI_ISL_3454145 | Lighthouse Lab in Milton Keynes | Wellcome Sanger Institute for the COVID-19 Genomics UK (COG-UK) Consortium | The Lighthouse Lab in Milton Keynes and Alex Alderton, Roberto Amato, Jeffrey Barrett, Sonia Goncalves, Ewan Harrison, David K. Jackson, Ian Johnston, Dominic Kwiatkowski, Cordelia Langford, John Sillitoe on behalf of the Wellcome Sanger Institute COVID-19 Surveillance Team |
| EPI_ISL_3454146 | Lighthouse Lab in Alderley Park | Wellcome Sanger Institute for the COVID-19 Genomics UK (COG-UK) Consortium | Jacquelyn Wynn, Mairead Hyland, The Lighthouse Lab in Alderley Park and Alex Alderton, Roberto Amato, Jeffrey Barrett, Sonia Goncalves, Ewan Harrison, David K. Jackson, Ian Johnston, Dominic Kwiatkowski, Cordelia Langford, John Sillitoe on behalf of the Wellcome Sanger Institute COVID-19 Surveillance Team |
| EPI_ISL_3454147 | Lighthouse Lab in Milton Keynes | Wellcome Sanger Institute for the COVID-19 Genomics UK (COG-UK) Consortium | The Lighthouse Lab in Milton Keynes and Alex Alderton, Roberto Amato, Jeffrey Barrett, Sonia Goncalves, Ewan Harrison, David K. Jackson, Ian Johnston, Dominic Kwiatkowski, Cordelia Langford, John Sillitoe on behalf of the Wellcome Sanger Institute COVID-19 Surveillance Team |
| EPI_ISL_3454148, EPI_ISL_3454149, EPI_ISL_3454150, EPI_ISL_3454151 | Lighthouse Laboratory Plymouth | Wellcome Sanger Institute for the COVID-19 Genomics UK (COG-UK) Consortium | Lighthouse Laboratory Plymouth and Alex Alderton, Roberto Amato, Jeffrey Barrett, Sonia Goncalves, Ewan Harrison, David K. Jackson, Ian Johnston, Dominic Kwiatkowski, Cordelia Langford, John Sillitoe on behalf of the Wellcome Sanger Institute COVID-19 Surveillance Team |
| EPI_ISL_3454152, EPI_ISL_3454153 | Lighthouse Lab in Milton Keynes | Wellcome Sanger Institute for the COVID-19 Genomics UK (COG-UK) Consortium | The Lighthouse Lab in Milton Keynes and Alex Alderton, Roberto Amato, Jeffrey Barrett, Sonia Goncalves, Ewan Harrison, David K. Jackson, Ian Johnston, Dominic Kwiatkowski, Cordelia Langford, John Sillitoe on behalf of the Wellcome Sanger Institute COVID-19 Surveillance Team |
| EPI_ISL_3454154, | Lighthouse Lab in Alderley Park | Wellcome Sanger Institute for the COVID-19 Genomics UK (COG-UK) | Jacquelyn Wynn, Mairead Hyland, The Lighthouse Lab in Alderley Park and Alex Alderton, Roberto Amato, Jeffrey Barrett, Sonia Goncalves, Ewan Harrison, David K. Jackson, Ian Johnston, Dominic Kwiatkowski, Cordelia Langford, John Sillitoe on |

| | | | |
|---|---|---|---|
| EPI_ISL_3454155 | | Consortium | behalf of the Wellcome Sanger Institute COVID-19 Surveillance Team |
| EPI_ISL_3454156 | Lighthouse Lab in Milton Keynes | Wellcome Sanger Institute for the COVID-19 Genomics UK (COG-UK) Consortium | The Lighthouse Lab in Milton Keynes and Alex Alderton, Roberto Amato, Jeffrey Barrett, Sonia Goncalves, Ewan Harrison, David K. Jackson, Ian Johnston, Dominic Kwiatkowski, Cordelia Langford, John Sillitoe on behalf of the Wellcome Sanger Institute COVID-19 Surveillance Team |
| EPI_ISL_3454157 | Lighthouse Lab in Alderley Park | Wellcome Sanger Institute for the COVID-19 Genomics UK (COG-UK) Consortium | Jacquelyn Wynn, Mairead Hyland, The Lighthouse Lab in Alderley Park and Alex Alderton, Roberto Amato, Jeffrey Barrett, Sonia Goncalves, Ewan Harrison, David K. Jackson, Ian Johnston, Dominic Kwiatkowski, Cordelia Langford, John Sillitoe on behalf of the Wellcome Sanger Institute COVID-19 Surveillance Team |
| EPI_ISL_3454158 | Lighthouse Lab in Milton Keynes | Wellcome Sanger Institute for the COVID-19 Genomics UK (COG-UK) Consortium | The Lighthouse Lab in Milton Keynes and Alex Alderton, Roberto Amato, Jeffrey Barrett, Sonia Goncalves, Ewan Harrison, David K. Jackson, Ian Johnston, Dominic Kwiatkowski, Cordelia Langford, John Sillitoe on behalf of the Wellcome Sanger Institute COVID-19 Surveillance Team |
| EPI_ISL_3454159 | Lighthouse Laboratory Plymouth | Wellcome Sanger Institute for the COVID-19 Genomics UK (COG-UK) Consortium | Lighthouse Laboratory Plymouth and Alex Alderton, Roberto Amato, Jeffrey Barrett, Sonia Goncalves, Ewan Harrison, David K. Jackson, Ian Johnston, Dominic Kwiatkowski, Cordelia Langford, John Sillitoe on behalf of the Wellcome Sanger Institute COVID-19 Surveillance Team |
| EPI_ISL_3454160 | Lighthouse Lab in Milton Keynes | Wellcome Sanger Institute for the COVID-19 Genomics UK (COG-UK) Consortium | The Lighthouse Lab in Milton Keynes and Alex Alderton, Roberto Amato, Jeffrey Barrett, Sonia Goncalves, Ewan Harrison, David K. Jackson, Ian Johnston, Dominic Kwiatkowski, Cordelia Langford, John Sillitoe on behalf of the Wellcome Sanger Institute COVID-19 Surveillance Team |
| EPI_ISL_3454161 | Lighthouse Lab in Alderley Park | Wellcome Sanger Institute for the COVID-19 Genomics UK (COG-UK) Consortium | Jacquelyn Wynn, Mairead Hyland, The Lighthouse Lab in Alderley Park and Alex Alderton, Roberto Amato, Jeffrey Barrett, Sonia Goncalves, Ewan Harrison, David K. Jackson, Ian Johnston, Dominic Kwiatkowski, Cordelia Langford, John Sillitoe on behalf of the Wellcome Sanger Institute COVID-19 Surveillance Team |
| EPI_ISL_3454162 | Lighthouse Lab in Milton Keynes | Wellcome Sanger Institute for the COVID-19 Genomics UK (COG-UK) Consortium | The Lighthouse Lab in Milton Keynes and Alex Alderton, Roberto Amato, Jeffrey Barrett, Sonia Goncalves, Ewan Harrison, David K. Jackson, Ian Johnston, Dominic Kwiatkowski, Cordelia Langford, John Sillitoe on behalf of the Wellcome Sanger Institute COVID-19 Surveillance Team |
| EPI_ISL_3454163, EPI_ISL_3454164, EPI_ISL_3454165 | Lighthouse Lab in Alderley Park | Wellcome Sanger Institute for the COVID-19 Genomics UK (COG-UK) Consortium | Jacquelyn Wynn, Mairead Hyland, The Lighthouse Lab in Alderley Park and Alex Alderton, Roberto Amato, Jeffrey Barrett, Sonia Goncalves, Ewan Harrison, David K. Jackson, Ian Johnston, Dominic Kwiatkowski, Cordelia Langford, John Sillitoe on behalf of the Wellcome Sanger Institute COVID-19 Surveillance Team |
| EPI_ISL_3454166 | Lighthouse Laboratory Plymouth | Wellcome Sanger Institute for the COVID-19 Genomics UK (COG-UK) Consortium | Lighthouse Laboratory Plymouth and Alex Alderton, Roberto Amato, Jeffrey Barrett, Sonia Goncalves, Ewan Harrison, David K. Jackson, Ian Johnston, Dominic Kwiatkowski, Cordelia Langford, John Sillitoe on behalf of the Wellcome Sanger Institute COVID-19 Surveillance Team |
| EPI_ISL_3454167, EPI_ISL_3454168, EPI_ISL_3454169 | Lighthouse Lab in Alderley Park | Wellcome Sanger Institute for the COVID-19 Genomics UK (COG-UK) Consortium | Jacquelyn Wynn, Mairead Hyland, The Lighthouse Lab in Alderley Park and Alex Alderton, Roberto Amato, Jeffrey Barrett, Sonia Goncalves, Ewan Harrison, David K. Jackson, Ian Johnston, Dominic Kwiatkowski, Cordelia Langford, John Sillitoe on behalf of the Wellcome Sanger Institute COVID-19 Surveillance Team |
| EPI_ISL_3454170, EPI_ISL_3454171, EPI_ISL_3454172 | Lighthouse Laboratory Plymouth | Wellcome Sanger Institute for the COVID-19 Genomics UK (COG-UK) Consortium | Lighthouse Laboratory Plymouth and Alex Alderton, Roberto Amato, Jeffrey Barrett, Sonia Goncalves, Ewan Harrison, David K. Jackson, Ian Johnston, Dominic Kwiatkowski, Cordelia Langford, John Sillitoe on behalf of the Wellcome Sanger Institute COVID-19 Surveillance Team |
| EPI_ISL_3454173 | Lighthouse Lab in Milton Keynes | Wellcome Sanger Institute for the COVID-19 Genomics UK (COG-UK) Consortium | The Lighthouse Lab in Milton Keynes and Alex Alderton, Roberto Amato, Jeffrey Barrett, Sonia Goncalves, Ewan Harrison, David K. Jackson, Ian Johnston, Dominic Kwiatkowski, Cordelia Langford, John Sillitoe on behalf of the Wellcome Sanger Institute COVID-19 Surveillance Team |
| EPI_ISL_3454174 | Lighthouse Laboratory Plymouth | Wellcome Sanger Institute for the COVID-19 Genomics UK (COG-UK) Consortium | Lighthouse Laboratory Plymouth and Alex Alderton, Roberto Amato, Jeffrey Barrett, Sonia Goncalves, Ewan Harrison, David K. Jackson, Ian Johnston, Dominic Kwiatkowski, Cordelia Langford, John Sillitoe on behalf of the Wellcome Sanger Institute COVID-19 Surveillance Team |
| EPI_ISL_3454175 | Lighthouse Lab in Alderley Park | Wellcome Sanger Institute for the COVID-19 Genomics UK (COG-UK) Consortium | Jacquelyn Wynn, Mairead Hyland, The Lighthouse Lab in Alderley Park and Alex Alderton, Roberto Amato, Jeffrey Barrett, Sonia Goncalves, Ewan Harrison, David K. Jackson, Ian Johnston, Dominic Kwiatkowski, Cordelia Langford, John Sillitoe on behalf of the Wellcome Sanger Institute COVID-19 Surveillance Team |
| EPI_ISL_3454176 | Lighthouse Lab in Milton Keynes | Wellcome Sanger Institute for the COVID-19 Genomics UK (COG-UK) Consortium | The Lighthouse Lab in Milton Keynes and Alex Alderton, Roberto Amato, Jeffrey Barrett, Sonia Goncalves, Ewan Harrison, David K. Jackson, Ian Johnston, Dominic Kwiatkowski, Cordelia Langford, John Sillitoe on behalf of the Wellcome Sanger Institute COVID-19 Surveillance Team |
| EPI_ISL_3454177, EPI_ISL_3454178 | Lighthouse Lab in Alderley Park | Wellcome Sanger Institute for the COVID-19 Genomics UK (COG-UK) Consortium | Jacquelyn Wynn, Mairead Hyland, The Lighthouse Lab in Alderley Park and Alex Alderton, Roberto Amato, Jeffrey Barrett, Sonia Goncalves, Ewan Harrison, David K. Jackson, Ian Johnston, Dominic Kwiatkowski, Cordelia Langford, John Sillitoe on behalf of the Wellcome Sanger Institute COVID-19 Surveillance Team |
| EPI_ISL_3454179 | Lighthouse Lab in Milton Keynes | Wellcome Sanger Institute for the COVID-19 Genomics UK (COG-UK) Consortium | The Lighthouse Lab in Milton Keynes and Alex Alderton, Roberto Amato, Jeffrey Barrett, Sonia Goncalves, Ewan Harrison, David K. Jackson, Ian Johnston, Dominic Kwiatkowski, Cordelia Langford, John Sillitoe on behalf of the Wellcome Sanger Institute COVID-19 Surveillance Team |
| EPI_ISL_3454180 | Lighthouse Lab in Alderley Park | Wellcome Sanger Institute for the COVID-19 Genomics UK (COG-UK) Consortium | Jacquelyn Wynn, Mairead Hyland, The Lighthouse Lab in Alderley Park and Alex Alderton, Roberto Amato, Jeffrey Barrett, Sonia Goncalves, Ewan Harrison, David K. Jackson, Ian Johnston, Dominic Kwiatkowski, Cordelia Langford, John Sillitoe on behalf of the Wellcome Sanger Institute COVID-19 Surveillance Team |
| EPI_ISL_3454181 | Lighthouse Lab in Milton Keynes | Wellcome Sanger Institute for the COVID-19 Genomics UK (COG-UK) Consortium | The Lighthouse Lab in Milton Keynes and Alex Alderton, Roberto Amato, Jeffrey Barrett, Sonia Goncalves, Ewan Harrison, David K. Jackson, Ian Johnston, Dominic Kwiatkowski, Cordelia Langford, John Sillitoe on behalf of the Wellcome Sanger Institute COVID-19 Surveillance Team |
| EPI_ISL_3454182 | Lighthouse Lab in Alderley Park | Wellcome Sanger Institute for the COVID-19 Genomics UK (COG-UK) Consortium | Jacquelyn Wynn, Mairead Hyland, The Lighthouse Lab in Alderley Park and Alex Alderton, Roberto Amato, Jeffrey Barrett, Sonia Goncalves, Ewan Harrison, David K. Jackson, Ian Johnston, Dominic Kwiatkowski, Cordelia Langford, John Sillitoe on behalf of the Wellcome Sanger Institute COVID-19 Surveillance Team |
| EPI_ISL_3454183 | Lighthouse Lab in Milton Keynes | Wellcome Sanger Institute for the COVID-19 Genomics UK (COG-UK) Consortium | The Lighthouse Lab in Milton Keynes and Alex Alderton, Roberto Amato, Jeffrey Barrett, Sonia Goncalves, Ewan Harrison, David K. Jackson, Ian Johnston, Dominic Kwiatkowski, Cordelia Langford, John Sillitoe on behalf of the Wellcome Sanger Institute COVID-19 Surveillance Team |
| EPI_ISL_3454184 | Lighthouse Lab in Alderley Park | Wellcome Sanger Institute for the COVID-19 Genomics UK (COG-UK) Consortium | Jacquelyn Wynn, Mairead Hyland, The Lighthouse Lab in Alderley Park and Alex Alderton, Roberto Amato, Jeffrey Barrett, Sonia Goncalves, Ewan Harrison, David K. Jackson, Ian Johnston, Dominic Kwiatkowski, Cordelia Langford, John Sillitoe on behalf of the Wellcome Sanger Institute COVID-19 Surveillance Team |
| EPI_ISL_3454185 | Lighthouse Lab in Milton Keynes | Wellcome Sanger Institute for the COVID-19 Genomics UK (COG-UK) Consortium | The Lighthouse Lab in Milton Keynes and Alex Alderton, Roberto Amato, Jeffrey Barrett, Sonia Goncalves, Ewan Harrison, David K. Jackson, Ian Johnston, Dominic Kwiatkowski, Cordelia Langford, John Sillitoe on behalf of the Wellcome Sanger Institute COVID-19 Surveillance Team |
| EPI_ISL_3454186 | Lighthouse Laboratory Plymouth | Wellcome Sanger Institute for the COVID-19 Genomics UK (COG-UK) Consortium | Lighthouse Laboratory Plymouth and Alex Alderton, Roberto Amato, Jeffrey Barrett, Sonia Goncalves, Ewan Harrison, David K. Jackson, Ian Johnston, Dominic Kwiatkowski, Cordelia Langford, John Sillitoe on behalf of the Wellcome Sanger Institute COVID-19 Surveillance Team |
| EPI_ISL_3454187, EPI_ISL_3454188 | Lighthouse Lab in Milton Keynes | Wellcome Sanger Institute for the COVID-19 Genomics UK (COG-UK) Consortium | The Lighthouse Lab in Milton Keynes and Alex Alderton, Roberto Amato, Jeffrey Barrett, Sonia Goncalves, Ewan Harrison, David K. Jackson, Ian Johnston, Dominic Kwiatkowski, Cordelia Langford, John Sillitoe on behalf of the Wellcome Sanger Institute COVID-19 Surveillance Team |
| EPI_ISL_3454189, EPI_ISL_3454190, EPI_ISL_3454191 | Lighthouse Lab in Alderley Park | Wellcome Sanger Institute for the COVID-19 Genomics UK (COG-UK) Consortium | Jacquelyn Wynn, Mairead Hyland, The Lighthouse Lab in Alderley Park and Alex Alderton, Roberto Amato, Jeffrey Barrett, Sonia Goncalves, Ewan Harrison, David K. Jackson, Ian Johnston, Dominic Kwiatkowski, Cordelia Langford, John Sillitoe on behalf of the Wellcome Sanger Institute COVID-19 Surveillance Team |
| EPI_ISL_3454192, EPI_ISL_3454193 | Lighthouse Laboratory Plymouth | Wellcome Sanger Institute for the COVID-19 Genomics UK (COG-UK) Consortium | Lighthouse Laboratory Plymouth and Alex Alderton, Roberto Amato, Jeffrey Barrett, Sonia Goncalves, Ewan Harrison, David K. Jackson, Ian Johnston, Dominic Kwiatkowski, Cordelia Langford, John Sillitoe on behalf of the Wellcome Sanger Institute COVID-19 Surveillance Team |
| EPI_ISL_3454194 | Lighthouse Lab in Milton Keynes | Wellcome Sanger Institute for the COVID-19 Genomics UK (COG-UK) Consortium | The Lighthouse Lab in Milton Keynes and Alex Alderton, Roberto Amato, Jeffrey Barrett, Sonia Goncalves, Ewan Harrison, David K. Jackson, Ian Johnston, Dominic Kwiatkowski, Cordelia Langford, John Sillitoe on behalf of the Wellcome Sanger Institute COVID-19 Surveillance Team |
| EPI_ISL_3454195 | Lighthouse Laboratory Plymouth | Wellcome Sanger Institute for the COVID-19 Genomics UK (COG-UK) Consortium | Lighthouse Laboratory Plymouth and Alex Alderton, Roberto Amato, Jeffrey Barrett, Sonia Goncalves, Ewan Harrison, David K. Jackson, Ian Johnston, Dominic Kwiatkowski, Cordelia Langford, John Sillitoe on behalf of the Wellcome Sanger Institute COVID-19 Surveillance Team |
| EPI_ISL_3454196 | Lighthouse Lab in Alderley Park | Wellcome Sanger Institute for the COVID-19 Genomics UK (COG-UK) Consortium | Jacquelyn Wynn, Mairead Hyland, The Lighthouse Lab in Alderley Park and Alex Alderton, Roberto Amato, Jeffrey Barrett, Sonia Goncalves, Ewan Harrison, David K. Jackson, Ian Johnston, Dominic Kwiatkowski, Cordelia Langford, John Sillitoe on behalf of the Wellcome Sanger Institute COVID-19 Surveillance Team |
| EPI_ISL_3454197, EPI_ISL_3454198 | Lighthouse Lab in Milton Keynes | Wellcome Sanger Institute for the COVID-19 Genomics UK (COG-UK) Consortium | The Lighthouse Lab in Milton Keynes and Alex Alderton, Roberto Amato, Jeffrey Barrett, Sonia Goncalves, Ewan Harrison, David K. Jackson, Ian Johnston, Dominic Kwiatkowski, Cordelia Langford, John Sillitoe on behalf of the Wellcome Sanger Institute COVID-19 Surveillance Team |
| EPI_ISL_3454199 | Lighthouse Lab in Alderley Park | Wellcome Sanger Institute for the COVID-19 Genomics UK (COG-UK) Consortium | Jacquelyn Wynn, Mairead Hyland, The Lighthouse Lab in Alderley Park and Alex Alderton, Roberto Amato, Jeffrey Barrett, Sonia Goncalves, Ewan Harrison, David K. Jackson, Ian Johnston, Dominic Kwiatkowski, Cordelia Langford, John Sillitoe on behalf of the Wellcome Sanger Institute COVID-19 Surveillance Team |
| EPI_ISL_3454200 | Lighthouse Lab in Milton Keynes | Wellcome Sanger Institute for the COVID-19 Genomics UK (COG-UK) Consortium | The Lighthouse Lab in Milton Keynes and Alex Alderton, Roberto Amato, Jeffrey Barrett, Sonia Goncalves, Ewan Harrison, David K. Jackson, Ian Johnston, Dominic Kwiatkowski, Cordelia Langford, John Sillitoe on behalf of the Wellcome Sanger Institute COVID-19 Surveillance Team |
| EPI_ISL_3454201, EPI_ISL_3454202 | Lighthouse Lab in Alderley Park | Wellcome Sanger Institute for the COVID-19 Genomics UK (COG-UK) Consortium | Jacquelyn Wynn, Mairead Hyland, The Lighthouse Lab in Alderley Park and Alex Alderton, Roberto Amato, Jeffrey Barrett, Sonia Goncalves, Ewan Harrison, David K. Jackson, Ian Johnston, Dominic Kwiatkowski, Cordelia Langford, John Sillitoe on behalf of the Wellcome Sanger Institute COVID-19 Surveillance Team |
| EPI_ISL_3454203 | Lighthouse Lab in Milton Keynes | Wellcome Sanger Institute for the COVID-19 Genomics UK (COG-UK) Consortium | The Lighthouse Lab in Milton Keynes and Alex Alderton, Roberto Amato, Jeffrey Barrett, Sonia Goncalves, Ewan Harrison, David K. Jackson, Ian Johnston, Dominic Kwiatkowski, Cordelia Langford, John Sillitoe on behalf of the Wellcome Sanger Institute COVID-19 Surveillance Team |
| EPI_ISL_3454204 | Lighthouse Lab in Alderley Park | Wellcome Sanger Institute for the COVID-19 Genomics UK (COG-UK) Consortium | Jacquelyn Wynn, Mairead Hyland, The Lighthouse Lab in Alderley Park and Alex Alderton, Roberto Amato, Jeffrey Barrett, Sonia Goncalves, Ewan Harrison, David K. Jackson, Ian Johnston, Dominic Kwiatkowski, Cordelia Langford, John Sillitoe on behalf of the Wellcome Sanger Institute COVID-19 Surveillance Team |
| EPI_ISL_3454205, EPI_ISL_3454206 | Lighthouse Laboratory Plymouth | Wellcome Sanger Institute for the COVID-19 Genomics UK (COG-UK) Consortium | Lighthouse Laboratory Plymouth and Alex Alderton, Roberto Amato, Jeffrey Barrett, Sonia Goncalves, Ewan Harrison, David K. Jackson, Ian Johnston, Dominic Kwiatkowski, Cordelia Langford, John Sillitoe on behalf of the Wellcome Sanger Institute COVID-19 Surveillance Team |
| EPI_ISL_3454207, EPI_ISL_3454208, EPI_ISL_3454209 | Lighthouse Lab in Alderley Park | Wellcome Sanger Institute for the COVID-19 Genomics UK (COG-UK) Consortium | Jacquelyn Wynn, Mairead Hyland, The Lighthouse Lab in Alderley Park and Alex Alderton, Roberto Amato, Jeffrey Barrett, Sonia Goncalves, Ewan Harrison, David K. Jackson, Ian Johnston, Dominic Kwiatkowski, Cordelia Langford, John Sillitoe on behalf of the Wellcome Sanger Institute COVID-19 Surveillance Team |
| EPI_ISL_3454210 | Lighthouse Lab in Milton Keynes | Wellcome Sanger Institute for the COVID-19 Genomics UK (COG-UK) Consortium | The Lighthouse Lab in Milton Keynes and Alex Alderton, Roberto Amato, Jeffrey Barrett, Sonia Goncalves, Ewan Harrison, David K. Jackson, Ian Johnston, Dominic Kwiatkowski, Cordelia Langford, John Sillitoe on behalf of the Wellcome Sanger Institute COVID-19 Surveillance Team |
| EPI_ISL_3454211 | Lighthouse Laboratory Plymouth | Wellcome Sanger Institute for the COVID-19 Genomics UK (COG-UK) Consortium | Lighthouse Laboratory Plymouth and Alex Alderton, Roberto Amato, Jeffrey Barrett, Sonia Goncalves, Ewan Harrison, David K. Jackson, Ian Johnston, Dominic Kwiatkowski, Cordelia Langford, John Sillitoe on behalf of the Wellcome Sanger Institute COVID-19 Surveillance Team |
| EPI_ISL_3454212 | Lighthouse Lab in Milton Keynes | Wellcome Sanger Institute for the COVID-19 Genomics UK (COG-UK) Consortium | The Lighthouse Lab in Milton Keynes and Alex Alderton, Roberto Amato, Jeffrey Barrett, Sonia Goncalves, Ewan Harrison, David K. Jackson, Ian Johnston, Dominic Kwiatkowski, Cordelia Langford, John Sillitoe on behalf of the Wellcome Sanger Institute COVID-19 Surveillance Team |
| EPI_ISL_3454213 | Lighthouse Lab in Alderley Park | Wellcome Sanger Institute for the COVID-19 Genomics UK (COG-UK) Consortium | Jacquelyn Wynn, Mairead Hyland, The Lighthouse Lab in Alderley Park and Alex Alderton, Roberto Amato, Jeffrey Barrett, Sonia Goncalves, Ewan Harrison, David K. Jackson, Ian Johnston, Dominic Kwiatkowski, Cordelia Langford, John Sillitoe on behalf of the Wellcome Sanger Institute COVID-19 Surveillance Team |
| EPI_ISL_3454214 | Lighthouse Lab in Milton Keynes | Wellcome Sanger Institute for the COVID-19 Genomics UK (COG-UK) | The Lighthouse Lab in Milton Keynes and Alex Alderton, Roberto Amato, Jeffrey Barrett, Sonia Goncalves, Ewan Harrison, David K. Jackson, Ian Johnston, Dominic Kwiatkowski, Cordelia Langford, John Sillitoe on behalf of the Wellcome Sanger |

| | | | |
|---|---|---|---|
| | | Consortium | Institute COVID-19 Surveillance Team |
| EPI_ISL_3454215, EPI_ISL_3454216 | Lighthouse Laboratory Plymouth | Wellcome Sanger Institute for the COVID-19 Genomics UK (COG-UK) Consortium | Lighthouse Laboratory Plymouth and Alex Alderton, Roberto Amato, Jeffrey Barrett, Sonia Goncalves, Ewan Harrison, David K. Jackson, Ian Johnston, Dominic Kwiatkowski, Cordelia Langford, John Sillitoe on behalf of the Wellcome Sanger Institute COVID-19 Surveillance Team |
| EPI_ISL_3454217, EPI_ISL_3454218, EPI_ISL_3454219 | Lighthouse Lab in Alderley Park | Wellcome Sanger Institute for the COVID-19 Genomics UK (COG-UK) Consortium | Jacquelyn Wynn, Mairead Hyland, The Lighthouse Lab in Alderley Park and Alex Alderton, Roberto Amato, Jeffrey Barrett, Sonia Goncalves, Ewan Harrison, David K. Jackson, Ian Johnston, Dominic Kwiatkowski, Cordelia Langford, John Sillitoe on behalf of the Wellcome Sanger Institute COVID-19 Surveillance Team |
| EPI_ISL_3454220 | Lighthouse Laboratory Plymouth | Wellcome Sanger Institute for the COVID-19 Genomics UK (COG-UK) Consortium | Lighthouse Laboratory Plymouth and Alex Alderton, Roberto Amato, Jeffrey Barrett, Sonia Goncalves, Ewan Harrison, David K. Jackson, Ian Johnston, Dominic Kwiatkowski, Cordelia Langford, John Sillitoe on behalf of the Wellcome Sanger Institute COVID-19 Surveillance Team |
| EPI_ISL_3454221, EPI_ISL_3454222 | Lighthouse Lab in Milton Keynes | Wellcome Sanger Institute for the COVID-19 Genomics UK (COG-UK) Consortium | The Lighthouse Lab in Milton Keynes and Alex Alderton, Roberto Amato, Jeffrey Barrett, Sonia Goncalves, Ewan Harrison, David K. Jackson, Ian Johnston, Dominic Kwiatkowski, Cordelia Langford, John Sillitoe on behalf of the Wellcome Sanger Institute COVID-19 Surveillance Team |
| EPI_ISL_3454223, EPI_ISL_3454224 | Lighthouse Laboratory Plymouth | Wellcome Sanger Institute for the COVID-19 Genomics UK (COG-UK) Consortium | Lighthouse Laboratory Plymouth and Alex Alderton, Roberto Amato, Jeffrey Barrett, Sonia Goncalves, Ewan Harrison, David K. Jackson, Ian Johnston, Dominic Kwiatkowski, Cordelia Langford, John Sillitoe on behalf of the Wellcome Sanger Institute COVID-19 Surveillance Team |
| EPI_ISL_3454225, EPI_ISL_3454226, EPI_ISL_3454227, EPI_ISL_3454228, EPI_ISL_3454229, EPI_ISL_3454230, EPI_ISL_3454231, EPI_ISL_3454232, EPI_ISL_3454233, EPI_ISL_3454234, EPI_ISL_3454235, EPI_ISL_3454236, EPI_ISL_3454237, EPI_ISL_3454238, EPI_ISL_3454239, EPI_ISL_3454240, EPI_ISL_3454241, EPI_ISL_3454242, EPI_ISL_3454243, EPI_ISL_3454244, EPI_ISL_3454245, EPI_ISL_3454246, EPI_ISL_3454247, EPI_ISL_3454248, EPI_ISL_3454249, EPI_ISL_3454250, EPI_ISL_3454251, EPI_ISL_3454252, EPI_ISL_3454253, EPI_ISL_3454254, EPI_ISL_3454255, EPI_ISL_3454256, EPI_ISL_3454257, EPI_ISL_3454258, EPI_ISL_3454259, EPI_ISL_3454260, EPI_ISL_3454261, EPI_ISL_3454262, EPI_ISL_3454263, EPI_ISL_3454264, EPI_ISL_3454265, EPI_ISL_3454266, EPI_ISL_3454267, EPI_ISL_3454268, EPI_ISL_3454269, EPI_ISL_3454270, EPI_ISL_3454271, EPI_ISL_3454272, EPI_ISL_3454273, EPI_ISL_3454274, EPI_ISL_3454275, EPI_ISL_3454276, EPI_ISL_3454277, EPI_ISL_3454278, EPI_ISL_3454279, EPI_ISL_3454280, EPI_ISL_3454281, EPI_ISL_3454282, EPI_ISL_3454283, EPI_ISL_3454284, EPI_ISL_3454285, EPI_ISL_3454286, EPI_ISL_3454287, EPI_ISL_3454288, EPI_ISL_3454289, EPI_ISL_3454290, EPI_ISL_3454291, EPI_ISL_3454292, EPI_ISL_3454293, EPI_ISL_3454294, EPI_ISL_3454295, EPI_ISL_3454296, EPI_ISL_3454297, EPI_ISL_3454298, EPI_ISL_3454299, EPI_ISL_3454300, EPI_ISL_3454301, EPI_ISL_3454302, EPI_ISL_3454303, EPI_ISL_3454304, EPI_ISL_3454305, EPI_ISL_3454306, EPI_ISL_3454307, EPI_ISL_3454308, EPI_ISL_3454309, EPI_ISL_3454310, EPI_ISL_3454311, EPI_ISL_3454312, EPI_ISL_3454313, EPI_ISL_3454314, EPI_ISL_3454315, EPI_ISL_3454316, EPI_ISL_3454317, EPI_ISL_3454318, EPI_ISL_3454319, EPI_ISL_3454320, EPI_ISL_3454321, EPI_ISL_3454322, EPI_ISL_3454323, EPI_ISL_3454324, EPI_ISL_3454325, EPI_ISL_3454326, EPI_ISL_3454327, EPI_ISL_3454328, EPI_ISL_3454329, EPI_ISL_3454330, EPI_ISL_3454331, EPI_ISL_3454332, EPI_ISL_3454333, EPI_ISL_3454334, EPI_ISL_3454335, EPI_ISL_3454336, EPI_ISL_3454337, EPI_ISL_3454338, EPI_ISL_3454339, EPI_ISL_3454340, EPI_ISL_3454341, EPI_ISL_3454342, EPI_ISL_3454343, EPI_ISL_3454344, EPI_ISL_3454345, EPI_ISL_3454346, EPI_ISL_3454347, EPI_ISL_3454348, EPI_ISL_3454349, EPI_ISL_3454350, EPI_ISL_3454351, EPI_ISL_3454352, EPI_ISL_3454353, EPI_ISL_3454354, EPI_ISL_3454355, EPI_ISL_3454356, EPI_ISL_3454357, EPI_ISL_3454358, EPI_ISL_3454359, EPI_ISL_3454360, EPI_ISL_3454361, EPI_ISL_3454362, EPI_ISL_3454363, EPI_ISL_3454364, EPI_ISL_3454365, EPI_ISL_3454366, EPI_ISL_3454367, EPI_ISL_3454368, EPI_ISL_3454369, EPI_ISL_3454370, EPI_ISL_3454371, EPI_ISL_3454372, EPI_ISL_3454373, EPI_ISL_3454374, EPI_ISL_3454375, EPI_ISL_3454376, EPI_ISL_3454377, EPI_ISL_3454378, EPI_ISL_3454379, EPI_ISL_3454380, EPI_ISL_3454381, EPI_ISL_3454382, EPI_ISL_3454383, EPI_ISL_3454384, EPI_ISL_3454385, EPI_ISL_3454386, EPI_ISL_3454387, EPI_ISL_3454388, EPI_ISL_3454389, EPI_ISL_3454390, EPI_ISL_3454391, EPI_ISL_3454392, EPI_ISL_3454393, EPI_ISL_3454394, EPI_ISL_3454395, EPI_ISL_3454396, EPI_ISL_3454397, EPI_ISL_3454398, EPI_ISL_3454399, EPI_ISL_3454400, EPI_ISL_3454401, EPI_ISL_3454402, EPI_ISL_3454403, EPI_ISL_3454404, EPI_ISL_3454405, EPI_ISL_3454406, EPI_ISL_3454407, EPI_ISL_3454408, EPI_ISL_3454409, EPI_ISL_3454410, EPI_ISL_3454411, EPI_ISL_3454412, EPI_ISL_3454413, EPI_ISL_3454414, EPI_ISL_3454415, EPI_ISL_3454416, EPI_ISL_3454417, EPI_ISL_3454418, EPI_ISL_3454419, EPI_ISL_3454420, EPI_ISL_3454421, EPI_ISL_3454422, EPI_ISL_3454423, EPI_ISL_3454424, EPI_ISL_3454425, EPI_ISL_3454426, EPI_ISL_3454427, EPI_ISL_3454428, EPI_ISL_3454429, EPI_ISL_3454430, EPI_ISL_3454431, EPI_ISL_3454432, EPI_ISL_3454433, EPI_ISL_3454434, EPI_ISL_3454435, EPI_ISL_3454436, EPI_ISL_3454437, EPI_ISL_3454438, EPI_ISL_3454439, EPI_ISL_3454440, EPI_ISL_3454441, EPI_ISL_3454442, EPI_ISL_3454443, EPI_ISL_3454444, EPI_ISL_3454445, EPI_ISL_3454446, EPI_ISL_3454447, EPI_ISL_3454448, EPI_ISL_3454449, EPI_ISL_3454450, EPI_ISL_3454451, EPI_ISL_3454452, EPI_ISL_3454453, EPI_ISL_3454454, EPI_ISL_3454455, EPI_ISL_3454456, EPI_ISL_3454457, EPI_ISL_3454458, EPI_ISL_3454459, EPI_ISL_3454460, EPI_ISL_3454461, EPI_ISL_3454462, EPI_ISL_3454463, EPI_ISL_3454464, EPI_ISL_3454465, EPI_ISL_3454466, EPI_ISL_3454467, EPI_ISL_3454468, EPI_ISL_3454469, EPI_ISL_3454470, EPI_ISL_3454471, EPI_ISL_3454472, EPI_ISL_3454473, EPI_ISL_3454474, EPI_ISL_3454475, EPI_ISL_3454476, EPI_ISL_3454477, EPI_ISL_3454478, EPI_ISL_3454479, EPI_ISL_3454480, EPI_ISL_3454481, EPI_ISL_3454482, EPI_ISL_3454483, EPI_ISL_3454484, EPI_ISL_3454485, EPI_ISL_3454486, EPI_ISL_3454487, EPI_ISL_3454488, EPI_ISL_3454489, EPI_ISL_3454490, EPI_ISL_3454491, EPI_ISL_3454492, EPI_ISL_3454493, EPI_ISL_3454494, EPI_ISL_3454495, EPI_ISL_3454496, EPI_ISL_3454497, EPI_ISL_3454498, EPI_ISL_3454499, EPI_ISL_3454500, EPI_ISL_3454501, EPI_ISL_3454502, EPI_ISL_3454503, EPI_ISL_3454504, EPI_ISL_3454505, EPI_ISL_3454506, EPI_ISL_3454507, EPI_ISL_3454508, EPI_ISL_3454509, EPI_ISL_3454510, EPI_ISL_3454511, EPI_ISL_3454512, EPI_ISL_3454513, EPI_ISL_3454514, EPI_ISL_3454515, EPI_ISL_3454516, EPI_ISL_3454517, EPI_ISL_3454518, EPI_ISL_3454519, EPI_ISL_3454520, EPI_ISL_3454521, EPI_ISL_3454522, EPI_ISL_3454523, EPI_ISL_3454524, EPI_ISL_3454525, EPI_ISL_3454526, EPI_ISL_3454527, EPI_ISL_3454528, EPI_ISL_3454529, EPI_ISL_3454530, EPI_ISL_3454531, EPI_ISL_3454532, EPI_ISL_3454533, EPI_ISL_3454534, EPI_ISL_3454535, EPI_ISL_3454536, EPI_ISL_3454537, EPI_ISL_3454538, EPI_ISL_3454539, EPI_ISL_3454540 | | | |
| see above | Lighthouse Lab in Milton Keynes | Wellcome Sanger Institute for the COVID-19 Genomics UK (COG-UK) Consortium | The Lighthouse Lab in Milton Keynes and Alex Alderton, Roberto Amato, Jeffrey Barrett, Sonia Goncalves, Ewan Harrison, David K. Jackson, Ian Johnston, Dominic Kwiatkowski, Cordelia Langford, John Sillitoe on behalf of the Wellcome Sanger Institute COVID-19 Surveillance Team |
| EPI_ISL_3454541 | Lighthouse Lab in Alderley Park | Wellcome Sanger Institute for the COVID-19 Genomics UK (COG-UK) Consortium | Jacquelyn Wynn, Mairead Hyland, The Lighthouse Lab in Alderley Park and Alex Alderton, Roberto Amato, Jeffrey Barrett, Sonia Goncalves, Ewan Harrison, David K. Jackson, Ian Johnston, Dominic Kwiatkowski, Cordelia Langford, John Sillitoe on behalf of the Wellcome Sanger Institute COVID-19 Surveillance Team |
| EPI_ISL_3454542 | Lighthouse Lab in Glasgow | Wellcome Sanger Institute for the COVID-19 Genomics UK (COG-UK) Consortium | Harper VanSteenhouse, Yumi Kasai, David Gray, Carol Clugston, Anna Dominiczak and Alex Alderton, Roberto Amato, Jeffrey Barrett, Sonia Goncalves, Ewan Harrison, David K. Jackson, Ian Johnston, Dominic Kwiatkowski, Cordelia Langford, John Sillitoe on behalf of the Wellcome Sanger Institute COVID-19 Surveillance Team |
| EPI_ISL_3454543 | Lighthouse Lab in Alderley Park | Wellcome Sanger Institute for the COVID-19 Genomics UK (COG-UK) Consortium | Jacquelyn Wynn, Mairead Hyland, The Lighthouse Lab in Alderley Park and Alex Alderton, Roberto Amato, Jeffrey Barrett, Sonia Goncalves, Ewan Harrison, David K. Jackson, Ian Johnston, Dominic Kwiatkowski, Cordelia Langford, John Sillitoe on behalf of the Wellcome Sanger Institute COVID-19 Surveillance Team |
| EPI_ISL_3454544 | Lighthouse Lab in Glasgow | Wellcome Sanger Institute for the COVID-19 Genomics UK (COG-UK) Consortium | Harper VanSteenhouse, Yumi Kasai, David Gray, Carol Clugston, Anna Dominiczak and Alex Alderton, Roberto Amato, Jeffrey Barrett, Sonia Goncalves, Ewan Harrison, David K. Jackson, Ian Johnston, Dominic Kwiatkowski, Cordelia Langford, John Sillitoe on behalf of the Wellcome Sanger Institute COVID-19 Surveillance Team |
| EPI_ISL_3454545, EPI_ISL_3454546, EPI_ISL_3454547 | Lighthouse Lab in Alderley Park | Wellcome Sanger Institute for the COVID-19 Genomics UK (COG-UK) Consortium | Jacquelyn Wynn, Mairead Hyland, The Lighthouse Lab in Alderley Park and Alex Alderton, Roberto Amato, Jeffrey Barrett, Sonia Goncalves, Ewan Harrison, David K. Jackson, Ian Johnston, Dominic Kwiatkowski, Cordelia Langford, John Sillitoe on behalf of the Wellcome Sanger Institute COVID-19 Surveillance Team |
| EPI_ISL_3454548, EPI_ISL_3454549 | Lighthouse Lab in Glasgow | Wellcome Sanger Institute for the COVID-19 Genomics UK (COG-UK) Consortium | Harper VanSteenhouse, Yumi Kasai, David Gray, Carol Clugston, Anna Dominiczak and Alex Alderton, Roberto Amato, Jeffrey Barrett, Sonia Goncalves, Ewan Harrison, David K. Jackson, Ian Johnston, Dominic Kwiatkowski, Cordelia Langford, John Sillitoe on behalf of the Wellcome Sanger Institute COVID-19 Surveillance Team |
| EPI_ISL_3454550, EPI_ISL_3454551 | Lighthouse Lab in Alderley Park | Wellcome Sanger Institute for the COVID-19 Genomics UK (COG-UK) Consortium | Jacquelyn Wynn, Mairead Hyland, The Lighthouse Lab in Alderley Park and Alex Alderton, Roberto Amato, Jeffrey Barrett, Sonia Goncalves, Ewan Harrison, David K. Jackson, Ian Johnston, Dominic Kwiatkowski, Cordelia Langford, John Sillitoe on behalf of the Wellcome Sanger Institute COVID-19 Surveillance Team |
| EPI_ISL_3454552 | Lighthouse Lab in Glasgow | Wellcome Sanger Institute for the COVID-19 Genomics UK (COG-UK) Consortium | Harper VanSteenhouse, Yumi Kasai, David Gray, Carol Clugston, Anna Dominiczak and Alex Alderton, Roberto Amato, Jeffrey Barrett, Sonia Goncalves, Ewan Harrison, David K. Jackson, Ian Johnston, Dominic Kwiatkowski, Cordelia Langford, John Sillitoe on behalf of the Wellcome Sanger Institute COVID-19 Surveillance Team |
| EPI_ISL_3454553 | Lighthouse Lab in Alderley Park | Wellcome Sanger Institute for the COVID-19 Genomics UK (COG-UK) Consortium | Jacquelyn Wynn, Mairead Hyland, The Lighthouse Lab in Alderley Park and Alex Alderton, Roberto Amato, Jeffrey Barrett, Sonia Goncalves, Ewan Harrison, David K. Jackson, Ian Johnston, Dominic Kwiatkowski, Cordelia Langford, John Sillitoe on behalf of the Wellcome Sanger Institute COVID-19 Surveillance Team |
| EPI_ISL_3454554 | Lighthouse Lab in Glasgow | Wellcome Sanger Institute for the COVID-19 Genomics UK (COG-UK) Consortium | Harper VanSteenhouse, Yumi Kasai, David Gray, Carol Clugston, Anna Dominiczak and Alex Alderton, Roberto Amato, Jeffrey Barrett, Sonia Goncalves, Ewan Harrison, David K. Jackson, Ian Johnston, Dominic Kwiatkowski, Cordelia Langford, John Sillitoe on behalf of the Wellcome Sanger Institute COVID-19 Surveillance Team |
| EPI_ISL_3454555, EPI_ISL_3454556, EPI_ISL_3454557, EPI_ISL_3454558, EPI_ISL_3454559, EPI_ISL_3454560, EPI_ISL_3454561 | Lighthouse Lab in Alderley Park | Wellcome Sanger Institute for the COVID-19 Genomics UK (COG-UK) Consortium | Jacquelyn Wynn, Mairead Hyland, The Lighthouse Lab in Alderley Park and Alex Alderton, Roberto Amato, Jeffrey Barrett, Sonia Goncalves, Ewan Harrison, David K. Jackson, Ian Johnston, Dominic Kwiatkowski, Cordelia Langford, John Sillitoe on behalf of the Wellcome Sanger Institute COVID-19 Surveillance Team |
| EPI_ISL_3454562 | Lighthouse Lab in Glasgow | Wellcome Sanger Institute for the COVID-19 Genomics UK (COG-UK) Consortium | Harper VanSteenhouse, Yumi Kasai, David Gray, Carol Clugston, Anna Dominiczak and Alex Alderton, Roberto Amato, Jeffrey Barrett, Sonia Goncalves, Ewan Harrison, David K. Jackson, Ian Johnston, Dominic Kwiatkowski, Cordelia Langford, John Sillitoe on behalf of the Wellcome Sanger Institute COVID-19 Surveillance Team |
| EPI_ISL_3454563 | Lighthouse Lab in Alderley Park | Wellcome Sanger Institute for the COVID-19 Genomics UK (COG-UK) Consortium | Jacquelyn Wynn, Mairead Hyland, The Lighthouse Lab in Alderley Park and Alex Alderton, Roberto Amato, Jeffrey Barrett, Sonia Goncalves, Ewan Harrison, David K. Jackson, Ian Johnston, Dominic Kwiatkowski, Cordelia Langford, John Sillitoe on behalf of the Wellcome Sanger Institute COVID-19 Surveillance Team |
| EPI_ISL_3454564 | Lighthouse Lab in Glasgow | Wellcome Sanger Institute for the COVID-19 Genomics UK (COG-UK) Consortium | Harper VanSteenhouse, Yumi Kasai, David Gray, Carol Clugston, Anna Dominiczak and Alex Alderton, Roberto Amato, Jeffrey Barrett, Sonia Goncalves, Ewan Harrison, David K. Jackson, Ian Johnston, Dominic Kwiatkowski, Cordelia Langford, John Sillitoe on behalf of the Wellcome Sanger Institute COVID-19 Surveillance Team |
| EPI_ISL_3454565, EPI_ISL_3454566 | Lighthouse Lab in Alderley Park | Wellcome Sanger Institute for the COVID-19 Genomics UK (COG-UK) Consortium | Jacquelyn Wynn, Mairead Hyland, The Lighthouse Lab in Alderley Park and Alex Alderton, Roberto Amato, Jeffrey Barrett, Sonia Goncalves, Ewan Harrison, David K. Jackson, Ian Johnston, Dominic Kwiatkowski, Cordelia Langford, John Sillitoe on behalf of the Wellcome Sanger Institute COVID-19 Surveillance Team |
| EPI_ISL_3454567 | Lighthouse Lab in Glasgow | Wellcome Sanger Institute for the COVID-19 Genomics UK (COG-UK) Consortium | Harper VanSteenhouse, Yumi Kasai, David Gray, Carol Clugston, Anna Dominiczak and Alex Alderton, Roberto Amato, Jeffrey Barrett, Sonia Goncalves, Ewan Harrison, David K. Jackson, Ian Johnston, Dominic Kwiatkowski, Cordelia Langford, John Sillitoe on behalf of the Wellcome Sanger Institute COVID-19 Surveillance Team |
| EPI_ISL_3454568, EPI_ISL_3454569, EPI_ISL_3454570, EPI_ISL_3454571 | Lighthouse Lab in Alderley Park | Wellcome Sanger Institute for the COVID-19 Genomics UK (COG-UK) Consortium | Jacquelyn Wynn, Mairead Hyland, The Lighthouse Lab in Alderley Park and Alex Alderton, Roberto Amato, Jeffrey Barrett, Sonia Goncalves, Ewan Harrison, David K. Jackson, Ian Johnston, Dominic Kwiatkowski, Cordelia Langford, John Sillitoe on behalf of the Wellcome Sanger Institute COVID-19 Surveillance Team |
| EPI_ISL_3454572, EPI_ISL_3454573, EPI_ISL_3454574, EPI_ISL_3454575 | Lighthouse Lab in Milton Keynes | Wellcome Sanger Institute for the COVID-19 Genomics UK (COG-UK) Consortium | The Lighthouse Lab in Milton Keynes and Alex Alderton, Roberto Amato, Jeffrey Barrett, Sonia Goncalves, Ewan Harrison, David K. Jackson, Ian Johnston, Dominic Kwiatkowski, Cordelia Langford, John Sillitoe on behalf of the Wellcome Sanger Institute COVID-19 Surveillance Team |
| EPI_ISL_3454576 | Berkshire and Surrey Pathology Services Lighthouse Laboratory | Wellcome Sanger Institute for the COVID-19 Genomics UK (COG-UK) Consortium | Berkshire and Surrey Pathology Services Lighthouse Laboratory and Alex Alderton, Roberto Amato, Jeffrey Barrett, Sonia Goncalves, Ewan Harrison, David K. Jackson, Ian Johnston, Dominic Kwiatkowski, Cordelia Langford, John Sillitoe on behalf of the Wellcome Sanger Institute COVID-19 Surveillance Team |
| EPI_ISL_3454577 | Lighthouse Lab in Milton Keynes | Wellcome Sanger Institute for the COVID-19 Genomics UK (COG-UK) Consortium | The Lighthouse Lab in Milton Keynes and Alex Alderton, Roberto Amato, Jeffrey Barrett, Sonia Goncalves, Ewan Harrison, David K. Jackson, Ian Johnston, Dominic Kwiatkowski, Cordelia Langford, John Sillitoe on behalf of the Wellcome Sanger Institute COVID-19 Surveillance Team |
| EPI_ISL_3454578 | Berkshire and Surrey Pathology Services Lighthouse Laboratory | Wellcome Sanger Institute for the COVID-19 Genomics UK (COG-UK) Consortium | Berkshire and Surrey Pathology Services Lighthouse Laboratory and Alex Alderton, Roberto Amato, Jeffrey Barrett, Sonia Goncalves, Ewan Harrison, David K. Jackson, Ian Johnston, Dominic Kwiatkowski, Cordelia Langford, John Sillitoe on behalf of the Wellcome Sanger Institute COVID-19 Surveillance Team |
| EPI_ISL_3454579 | Lighthouse Lab in Milton Keynes | Wellcome Sanger Institute for the COVID-19 Genomics UK (COG-UK) Consortium | The Lighthouse Lab in Milton Keynes and Alex Alderton, Roberto Amato, Jeffrey Barrett, Sonia Goncalves, Ewan Harrison, David K. Jackson, Ian Johnston, Dominic Kwiatkowski, Cordelia Langford, John Sillitoe on behalf of the Wellcome Sanger Institute COVID-19 Surveillance Team |
| EPI_ISL_3454580 | Berkshire and Surrey Pathology Services Lighthouse Laboratory | Wellcome Sanger Institute for the COVID-19 Genomics UK (COG-UK) Consortium | Berkshire and Surrey Pathology Services Lighthouse Laboratory and Alex Alderton, Roberto Amato, Jeffrey Barrett, Sonia Goncalves, Ewan Harrison, David K. Jackson, Ian Johnston, Dominic Kwiatkowski, Cordelia Langford, John Sillitoe on behalf of the Wellcome Sanger Institute COVID-19 Surveillance Team |
| EPI_ISL_3454581, EPI_ISL_3454582 | Lighthouse Lab in Milton Keynes | Wellcome Sanger Institute for the COVID-19 Genomics UK (COG-UK) Consortium | The Lighthouse Lab in Milton Keynes and Alex Alderton, Roberto Amato, Jeffrey Barrett, Sonia Goncalves, Ewan Harrison, David K. Jackson, Ian Johnston, Dominic Kwiatkowski, Cordelia Langford, John Sillitoe on behalf of the Wellcome Sanger Institute COVID-19 Surveillance Team |
| EPI_ISL_3454583 | Berkshire and Surrey Pathology Services Lighthouse Laboratory | Wellcome Sanger Institute for the COVID-19 Genomics UK (COG-UK) Consortium | Berkshire and Surrey Pathology Services Lighthouse Laboratory and Alex Alderton, Roberto Amato, Jeffrey Barrett, Sonia Goncalves, Ewan Harrison, David K. Jackson, Ian Johnston, Dominic Kwiatkowski, Cordelia Langford, John Sillitoe on behalf of the Wellcome Sanger Institute COVID-19 Surveillance Team |
| EPI_ISL_3454584, EPI_ISL_3454585, EPI_ISL_3454586, EPI_ISL_3454587 | Lighthouse Lab in Milton Keynes | Wellcome Sanger Institute for the COVID-19 Genomics UK (COG-UK) Consortium | The Lighthouse Lab in Milton Keynes and Alex Alderton, Roberto Amato, Jeffrey Barrett, Sonia Goncalves, Ewan Harrison, David K. Jackson, Ian Johnston, Dominic Kwiatkowski, Cordelia Langford, John Sillitoe on behalf of the Wellcome Sanger Institute COVID-19 Surveillance Team |
| EPI_ISL_3454588 | Lighthouse Lab in Alderley Park | Wellcome Sanger Institute for the COVID-19 Genomics UK (COG-UK) Consortium | Jacquelyn Wynn, Mairead Hyland, The Lighthouse Lab in Alderley Park and Alex Alderton, Roberto Amato, Jeffrey Barrett, Sonia Goncalves, Ewan Harrison, David K. Jackson, Ian Johnston, Dominic Kwiatkowski, Cordelia Langford, John Sillitoe on behalf of the Wellcome Sanger Institute COVID-19 Surveillance Team |

| | | | |
|---|---|---|---|
| EPI_ISL_3454589 | Lighthouse Lab in Milton Keynes | Wellcome Sanger Institute for the COVID-19 Genomics UK (COG-UK) Consortium | The Lighthouse Lab in Milton Keynes and Alex Alderton, Roberto Amato, Jeffrey Barrett, Sonia Goncalves, Ewan Harrison, David K. Jackson, Ian Johnston, Dominic Kwiatkowski, Cordelia Langford, John Sillitoe on behalf of the Wellcome Sanger Institute COVID-19 Surveillance Team |
| EPI_ISL_3454590 | Berkshire and Surrey Pathology Services Lighthouse Laboratory | Wellcome Sanger Institute for the COVID-19 Genomics UK (COG-UK) Consortium | Berkshire and Surrey Pathology Services Lighthouse Laboratory and Alex Alderton, Roberto Amato, Jeffrey Barrett, Sonia Goncalves, Ewan Harrison, David K. Jackson, Ian Johnston, Dominic Kwiatkowski, Cordelia Langford, John Sillitoe on behalf of the Wellcome Sanger Institute COVID-19 Surveillance Team |
| EPI_ISL_3454591, EPI_ISL_3454592, EPI_ISL_3454593 | Lighthouse Lab in Milton Keynes | Wellcome Sanger Institute for the COVID-19 Genomics UK (COG-UK) Consortium | The Lighthouse Lab in Milton Keynes and Alex Alderton, Roberto Amato, Jeffrey Barrett, Sonia Goncalves, Ewan Harrison, David K. Jackson, Ian Johnston, Dominic Kwiatkowski, Cordelia Langford, John Sillitoe on behalf of the Wellcome Sanger Institute COVID-19 Surveillance Team |
| EPI_ISL_3454594 | Lighthouse Lab in Alderley Park | Wellcome Sanger Institute for the COVID-19 Genomics UK (COG-UK) Consortium | Jacquelyn Wynn, Mairead Hyland, The Lighthouse Lab in Alderley Park and Alex Alderton, Roberto Amato, Jeffrey Barrett, Sonia Goncalves, Ewan Harrison, David K. Jackson, Ian Johnston, Dominic Kwiatkowski, Cordelia Langford, John Sillitoe on behalf of the Wellcome Sanger Institute COVID-19 Surveillance Team |
| EPI_ISL_3454595, EPI_ISL_3454596, EPI_ISL_3454597 | Lighthouse Lab in Milton Keynes | Wellcome Sanger Institute for the COVID-19 Genomics UK (COG-UK) Consortium | The Lighthouse Lab in Milton Keynes and Alex Alderton, Roberto Amato, Jeffrey Barrett, Sonia Goncalves, Ewan Harrison, David K. Jackson, Ian Johnston, Dominic Kwiatkowski, Cordelia Langford, John Sillitoe on behalf of the Wellcome Sanger Institute COVID-19 Surveillance Team |
| EPI_ISL_3454598 | Lighthouse Lab in Alderley Park | Wellcome Sanger Institute for the COVID-19 Genomics UK (COG-UK) Consortium | Jacquelyn Wynn, Mairead Hyland, The Lighthouse Lab in Alderley Park and Alex Alderton, Roberto Amato, Jeffrey Barrett, Sonia Goncalves, Ewan Harrison, David K. Jackson, Ian Johnston, Dominic Kwiatkowski, Cordelia Langford, John Sillitoe on behalf of the Wellcome Sanger Institute COVID-19 Surveillance Team |
| EPI_ISL_3454599 | Lighthouse Lab in Milton Keynes | Wellcome Sanger Institute for the COVID-19 Genomics UK (COG-UK) Consortium | The Lighthouse Lab in Milton Keynes and Alex Alderton, Roberto Amato, Jeffrey Barrett, Sonia Goncalves, Ewan Harrison, David K. Jackson, Ian Johnston, Dominic Kwiatkowski, Cordelia Langford, John Sillitoe on behalf of the Wellcome Sanger Institute COVID-19 Surveillance Team |
| EPI_ISL_3454600, EPI_ISL_3454601, EPI_ISL_3454602, EPI_ISL_3454603 | Lighthouse Lab in Alderley Park | Wellcome Sanger Institute for the COVID-19 Genomics UK (COG-UK) Consortium | Jacquelyn Wynn, Mairead Hyland, The Lighthouse Lab in Alderley Park and Alex Alderton, Roberto Amato, Jeffrey Barrett, Sonia Goncalves, Ewan Harrison, David K. Jackson, Ian Johnston, Dominic Kwiatkowski, Cordelia Langford, John Sillitoe on behalf of the Wellcome Sanger Institute COVID-19 Surveillance Team |
| EPI_ISL_3454604, EPI_ISL_3454605, EPI_ISL_3454606 | Lighthouse Lab in Milton Keynes | Wellcome Sanger Institute for the COVID-19 Genomics UK (COG-UK) Consortium | The Lighthouse Lab in Milton Keynes and Alex Alderton, Roberto Amato, Jeffrey Barrett, Sonia Goncalves, Ewan Harrison, David K. Jackson, Ian Johnston, Dominic Kwiatkowski, Cordelia Langford, John Sillitoe on behalf of the Wellcome Sanger Institute COVID-19 Surveillance Team |
| EPI_ISL_3454607, EPI_ISL_3454608 | Lighthouse Lab in Alderley Park | Wellcome Sanger Institute for the COVID-19 Genomics UK (COG-UK) Consortium | Jacquelyn Wynn, Mairead Hyland, The Lighthouse Lab in Alderley Park and Alex Alderton, Roberto Amato, Jeffrey Barrett, Sonia Goncalves, Ewan Harrison, David K. Jackson, Ian Johnston, Dominic Kwiatkowski, Cordelia Langford, John Sillitoe on behalf of the Wellcome Sanger Institute COVID-19 Surveillance Team |
| EPI_ISL_3454609, EPI_ISL_3454610 | Lighthouse Lab in Milton Keynes | Wellcome Sanger Institute for the COVID-19 Genomics UK (COG-UK) Consortium | The Lighthouse Lab in Milton Keynes and Alex Alderton, Roberto Amato, Jeffrey Barrett, Sonia Goncalves, Ewan Harrison, David K. Jackson, Ian Johnston, Dominic Kwiatkowski, Cordelia Langford, John Sillitoe on behalf of the Wellcome Sanger Institute COVID-19 Surveillance Team |
| EPI_ISL_3454611, EPI_ISL_3454612 | Lighthouse Lab in Alderley Park | Wellcome Sanger Institute for the COVID-19 Genomics UK (COG-UK) Consortium | Jacquelyn Wynn, Mairead Hyland, The Lighthouse Lab in Alderley Park and Alex Alderton, Roberto Amato, Jeffrey Barrett, Sonia Goncalves, Ewan Harrison, David K. Jackson, Ian Johnston, Dominic Kwiatkowski, Cordelia Langford, John Sillitoe on behalf of the Wellcome Sanger Institute COVID-19 Surveillance Team |
| EPI_ISL_3454613, EPI_ISL_3454614, EPI_ISL_3454615, EPI_ISL_3454616 | Lighthouse Lab in Milton Keynes | Wellcome Sanger Institute for the COVID-19 Genomics UK (COG-UK) Consortium | The Lighthouse Lab in Milton Keynes and Alex Alderton, Roberto Amato, Jeffrey Barrett, Sonia Goncalves, Ewan Harrison, David K. Jackson, Ian Johnston, Dominic Kwiatkowski, Cordelia Langford, John Sillitoe on behalf of the Wellcome Sanger Institute COVID-19 Surveillance Team |
| EPI_ISL_3454617, EPI_ISL_3454618 | Lighthouse Lab in Alderley Park | Wellcome Sanger Institute for the COVID-19 Genomics UK (COG-UK) Consortium | Jacquelyn Wynn, Mairead Hyland, The Lighthouse Lab in Alderley Park and Alex Alderton, Roberto Amato, Jeffrey Barrett, Sonia Goncalves, Ewan Harrison, David K. Jackson, Ian Johnston, Dominic Kwiatkowski, Cordelia Langford, John Sillitoe on behalf of the Wellcome Sanger Institute COVID-19 Surveillance Team |
| EPI_ISL_3454619 | Berkshire and Surrey Pathology Services Lighthouse Laboratory | Wellcome Sanger Institute for the COVID-19 Genomics UK (COG-UK) Consortium | Berkshire and Surrey Pathology Services Lighthouse Laboratory and Alex Alderton, Roberto Amato, Jeffrey Barrett, Sonia Goncalves, Ewan Harrison, David K. Jackson, Ian Johnston, Dominic Kwiatkowski, Cordelia Langford, John Sillitoe on behalf of the Wellcome Sanger Institute COVID-19 Surveillance Team |
| EPI_ISL_3454620 | Lighthouse Lab in Milton Keynes | Wellcome Sanger Institute for the COVID-19 Genomics UK (COG-UK) Consortium | The Lighthouse Lab in Milton Keynes and Alex Alderton, Roberto Amato, Jeffrey Barrett, Sonia Goncalves, Ewan Harrison, David K. Jackson, Ian Johnston, Dominic Kwiatkowski, Cordelia Langford, John Sillitoe on behalf of the Wellcome Sanger Institute COVID-19 Surveillance Team |
| EPI_ISL_3454621 | Berkshire and Surrey Pathology Services Lighthouse Laboratory | Wellcome Sanger Institute for the COVID-19 Genomics UK (COG-UK) Consortium | Berkshire and Surrey Pathology Services Lighthouse Laboratory and Alex Alderton, Roberto Amato, Jeffrey Barrett, Sonia Goncalves, Ewan Harrison, David K. Jackson, Ian Johnston, Dominic Kwiatkowski, Cordelia Langford, John Sillitoe on behalf of the Wellcome Sanger Institute COVID-19 Surveillance Team |
| EPI_ISL_3454622, EPI_ISL_3454623 | Lighthouse Lab in Milton Keynes | Wellcome Sanger Institute for the COVID-19 Genomics UK (COG-UK) Consortium | The Lighthouse Lab in Milton Keynes and Alex Alderton, Roberto Amato, Jeffrey Barrett, Sonia Goncalves, Ewan Harrison, David K. Jackson, Ian Johnston, Dominic Kwiatkowski, Cordelia Langford, John Sillitoe on behalf of the Wellcome Sanger Institute COVID-19 Surveillance Team |
| EPI_ISL_3454624 | Berkshire and Surrey Pathology Services Lighthouse Laboratory | Wellcome Sanger Institute for the COVID-19 Genomics UK (COG-UK) Consortium | Berkshire and Surrey Pathology Services Lighthouse Laboratory and Alex Alderton, Roberto Amato, Jeffrey Barrett, Sonia Goncalves, Ewan Harrison, David K. Jackson, Ian Johnston, Dominic Kwiatkowski, Cordelia Langford, John Sillitoe on behalf of the Wellcome Sanger Institute COVID-19 Surveillance Team |
| EPI_ISL_3454625 | Lighthouse Lab in Alderley Park | Wellcome Sanger Institute for the COVID-19 Genomics UK (COG-UK) Consortium | Jacquelyn Wynn, Mairead Hyland, The Lighthouse Lab in Alderley Park and Alex Alderton, Roberto Amato, Jeffrey Barrett, Sonia Goncalves, Ewan Harrison, David K. Jackson, Ian Johnston, Dominic Kwiatkowski, Cordelia Langford, John Sillitoe on behalf of the Wellcome Sanger Institute COVID-19 Surveillance Team |
| EPI_ISL_3454626 | Lighthouse Lab in Milton Keynes | Wellcome Sanger Institute for the COVID-19 Genomics UK (COG-UK) Consortium | The Lighthouse Lab in Milton Keynes and Alex Alderton, Roberto Amato, Jeffrey Barrett, Sonia Goncalves, Ewan Harrison, David K. Jackson, Ian Johnston, Dominic Kwiatkowski, Cordelia Langford, John Sillitoe on behalf of the Wellcome Sanger Institute COVID-19 Surveillance Team |
| EPI_ISL_3454627 | Berkshire and Surrey Pathology Services Lighthouse Laboratory | Wellcome Sanger Institute for the COVID-19 Genomics UK (COG-UK) Consortium | Berkshire and Surrey Pathology Services Lighthouse Laboratory and Alex Alderton, Roberto Amato, Jeffrey Barrett, Sonia Goncalves, Ewan Harrison, David K. Jackson, Ian Johnston, Dominic Kwiatkowski, Cordelia Langford, John Sillitoe on behalf of the Wellcome Sanger Institute COVID-19 Surveillance Team |
| EPI_ISL_3454628, EPI_ISL_3454629 | Lighthouse Lab in Milton Keynes | Wellcome Sanger Institute for the COVID-19 Genomics UK (COG-UK) Consortium | The Lighthouse Lab in Milton Keynes and Alex Alderton, Roberto Amato, Jeffrey Barrett, Sonia Goncalves, Ewan Harrison, David K. Jackson, Ian Johnston, Dominic Kwiatkowski, Cordelia Langford, John Sillitoe on behalf of the Wellcome Sanger Institute COVID-19 Surveillance Team |
| EPI_ISL_3454630, EPI_ISL_3454631 | Lighthouse Lab in Alderley Park | Wellcome Sanger Institute for the COVID-19 Genomics UK (COG-UK) Consortium | Jacquelyn Wynn, Mairead Hyland, The Lighthouse Lab in Alderley Park and Alex Alderton, Roberto Amato, Jeffrey Barrett, Sonia Goncalves, Ewan Harrison, David K. Jackson, Ian Johnston, Dominic Kwiatkowski, Cordelia Langford, John Sillitoe on behalf of the Wellcome Sanger Institute COVID-19 Surveillance Team |
| EPI_ISL_3454632 | Lighthouse Lab in Milton Keynes | Wellcome Sanger Institute for the COVID-19 Genomics UK (COG-UK) Consortium | The Lighthouse Lab in Milton Keynes and Alex Alderton, Roberto Amato, Jeffrey Barrett, Sonia Goncalves, Ewan Harrison, David K. Jackson, Ian Johnston, Dominic Kwiatkowski, Cordelia Langford, John Sillitoe on behalf of the Wellcome Sanger Institute COVID-19 Surveillance Team |
| EPI_ISL_3454633 | Lighthouse Lab in Alderley Park | Wellcome Sanger Institute for the COVID-19 Genomics UK (COG-UK) Consortium | Jacquelyn Wynn, Mairead Hyland, The Lighthouse Lab in Alderley Park and Alex Alderton, Roberto Amato, Jeffrey Barrett, Sonia Goncalves, Ewan Harrison, David K. Jackson, Ian Johnston, Dominic Kwiatkowski, Cordelia Langford, John Sillitoe on behalf of the Wellcome Sanger Institute COVID-19 Surveillance Team |
| EPI_ISL_3454634, EPI_ISL_3454635 | Lighthouse Lab in Milton Keynes | Wellcome Sanger Institute for the COVID-19 Genomics UK (COG-UK) Consortium | The Lighthouse Lab in Milton Keynes and Alex Alderton, Roberto Amato, Jeffrey Barrett, Sonia Goncalves, Ewan Harrison, David K. Jackson, Ian Johnston, Dominic Kwiatkowski, Cordelia Langford, John Sillitoe on behalf of the Wellcome Sanger Institute COVID-19 Surveillance Team |
| EPI_ISL_3454636 | Lighthouse Lab in Alderley Park | Wellcome Sanger Institute for the COVID-19 Genomics UK (COG-UK) Consortium | Jacquelyn Wynn, Mairead Hyland, The Lighthouse Lab in Alderley Park and Alex Alderton, Roberto Amato, Jeffrey Barrett, Sonia Goncalves, Ewan Harrison, David K. Jackson, Ian Johnston, Dominic Kwiatkowski, Cordelia Langford, John Sillitoe on behalf of the Wellcome Sanger Institute COVID-19 Surveillance Team |
| EPI_ISL_3454637, EPI_ISL_3454638 | Berkshire and Surrey Pathology Services Lighthouse Laboratory | Wellcome Sanger Institute for the COVID-19 Genomics UK (COG-UK) Consortium | Berkshire and Surrey Pathology Services Lighthouse Laboratory and Alex Alderton, Roberto Amato, Jeffrey Barrett, Sonia Goncalves, Ewan Harrison, David K. Jackson, Ian Johnston, Dominic Kwiatkowski, Cordelia Langford, John Sillitoe on behalf of the Wellcome Sanger Institute COVID-19 Surveillance Team |
| EPI_ISL_3454639, EPI_ISL_3454640 | Lighthouse Lab in Milton Keynes | Wellcome Sanger Institute for the COVID-19 Genomics UK (COG-UK) Consortium | The Lighthouse Lab in Milton Keynes and Alex Alderton, Roberto Amato, Jeffrey Barrett, Sonia Goncalves, Ewan Harrison, David K. Jackson, Ian Johnston, Dominic Kwiatkowski, Cordelia Langford, John Sillitoe on behalf of the Wellcome Sanger Institute COVID-19 Surveillance Team |
| EPI_ISL_3454641 | Lighthouse Lab in Alderley Park | Wellcome Sanger Institute for the COVID-19 Genomics UK (COG-UK) Consortium | Jacquelyn Wynn, Mairead Hyland, The Lighthouse Lab in Alderley Park and Alex Alderton, Roberto Amato, Jeffrey Barrett, Sonia Goncalves, Ewan Harrison, David K. Jackson, Ian Johnston, Dominic Kwiatkowski, Cordelia Langford, John Sillitoe on behalf of the Wellcome Sanger Institute COVID-19 Surveillance Team |
| EPI_ISL_3454642 | Berkshire and Surrey Pathology Services Lighthouse Laboratory | Wellcome Sanger Institute for the COVID-19 Genomics UK (COG-UK) Consortium | Berkshire and Surrey Pathology Services Lighthouse Laboratory and Alex Alderton, Roberto Amato, Jeffrey Barrett, Sonia Goncalves, Ewan Harrison, David K. Jackson, Ian Johnston, Dominic Kwiatkowski, Cordelia Langford, John Sillitoe on behalf of the Wellcome Sanger Institute COVID-19 Surveillance Team |
| EPI_ISL_3454643, EPI_ISL_3454644 | Lighthouse Lab in Milton Keynes | Wellcome Sanger Institute for the COVID-19 Genomics UK (COG-UK) Consortium | The Lighthouse Lab in Milton Keynes and Alex Alderton, Roberto Amato, Jeffrey Barrett, Sonia Goncalves, Ewan Harrison, David K. Jackson, Ian Johnston, Dominic Kwiatkowski, Cordelia Langford, John Sillitoe on behalf of the Wellcome Sanger Institute COVID-19 Surveillance Team |
| EPI_ISL_3454645 | Lighthouse Lab in Alderley Park | Wellcome Sanger Institute for the COVID-19 Genomics UK (COG-UK) Consortium | Jacquelyn Wynn, Mairead Hyland, The Lighthouse Lab in Alderley Park and Alex Alderton, Roberto Amato, Jeffrey Barrett, Sonia Goncalves, Ewan Harrison, David K. Jackson, Ian Johnston, Dominic Kwiatkowski, Cordelia Langford, John Sillitoe on behalf of the Wellcome Sanger Institute COVID-19 Surveillance Team |
| EPI_ISL_3454646 | Lighthouse Lab in Milton Keynes | Wellcome Sanger Institute for the COVID-19 Genomics UK (COG-UK) Consortium | The Lighthouse Lab in Milton Keynes and Alex Alderton, Roberto Amato, Jeffrey Barrett, Sonia Goncalves, Ewan Harrison, David K. Jackson, Ian Johnston, Dominic Kwiatkowski, Cordelia Langford, John Sillitoe on behalf of the Wellcome Sanger Institute COVID-19 Surveillance Team |
| EPI_ISL_3454647 | Lighthouse Lab in Alderley Park | Wellcome Sanger Institute for the COVID-19 Genomics UK (COG-UK) Consortium | Jacquelyn Wynn, Mairead Hyland, The Lighthouse Lab in Alderley Park and Alex Alderton, Roberto Amato, Jeffrey Barrett, Sonia Goncalves, Ewan Harrison, David K. Jackson, Ian Johnston, Dominic Kwiatkowski, Cordelia Langford, John Sillitoe on behalf of the Wellcome Sanger Institute COVID-19 Surveillance Team |
| EPI_ISL_3454648 | Berkshire and Surrey Pathology Services Lighthouse Laboratory | Wellcome Sanger Institute for the COVID-19 Genomics UK (COG-UK) Consortium | Berkshire and Surrey Pathology Services Lighthouse Laboratory and Alex Alderton, Roberto Amato, Jeffrey Barrett, Sonia Goncalves, Ewan Harrison, David K. Jackson, Ian Johnston, Dominic Kwiatkowski, Cordelia Langford, John Sillitoe on behalf of the Wellcome Sanger Institute COVID-19 Surveillance Team |
| EPI_ISL_3454649 | Lighthouse Lab in Alderley Park | Wellcome Sanger Institute for the COVID-19 Genomics UK (COG-UK) Consortium | Jacquelyn Wynn, Mairead Hyland, The Lighthouse Lab in Alderley Park and Alex Alderton, Roberto Amato, Jeffrey Barrett, Sonia Goncalves, Ewan Harrison, David K. Jackson, Ian Johnston, Dominic Kwiatkowski, Cordelia Langford, John Sillitoe on behalf of the Wellcome Sanger Institute COVID-19 Surveillance Team |
| EPI_ISL_3454650, EPI_ISL_3454651, EPI_ISL_3454652 | Lighthouse Lab in Milton Keynes | Wellcome Sanger Institute for the COVID-19 Genomics UK (COG-UK) Consortium | The Lighthouse Lab in Milton Keynes and Alex Alderton, Roberto Amato, Jeffrey Barrett, Sonia Goncalves, Ewan Harrison, David K. Jackson, Ian Johnston, Dominic Kwiatkowski, Cordelia Langford, John Sillitoe on behalf of the Wellcome Sanger Institute COVID-19 Surveillance Team |
| EPI_ISL_3454653, EPI_ISL_3454654 | Lighthouse Lab in Alderley Park | Wellcome Sanger Institute for the COVID-19 Genomics UK (COG-UK) Consortium | Jacquelyn Wynn, Mairead Hyland, The Lighthouse Lab in Alderley Park and Alex Alderton, Roberto Amato, Jeffrey Barrett, Sonia Goncalves, Ewan Harrison, David K. Jackson, Ian Johnston, Dominic Kwiatkowski, Cordelia Langford, John Sillitoe on behalf of the Wellcome Sanger Institute COVID-19 Surveillance Team |
| EPI_ISL_3454655, EPI_ISL_3454656 | Lighthouse Lab in Milton Keynes | Wellcome Sanger Institute for the COVID-19 Genomics UK (COG-UK) Consortium | The Lighthouse Lab in Milton Keynes and Alex Alderton, Roberto Amato, Jeffrey Barrett, Sonia Goncalves, Ewan Harrison, David K. Jackson, Ian Johnston, Dominic Kwiatkowski, Cordelia Langford, John Sillitoe on behalf of the Wellcome Sanger Institute COVID-19 Surveillance Team |
| EPI_ISL_3454657 | Berkshire and Surrey Pathology Services Lighthouse Laboratory | Wellcome Sanger Institute for the COVID-19 Genomics UK (COG-UK) | Berkshire and Surrey Pathology Services Lighthouse Laboratory and Alex Alderton, Roberto Amato, Jeffrey Barrett, Sonia Goncalves, Ewan Harrison, David K. Jackson, Ian Johnston, Dominic Kwiatkowski, Cordelia Langford, John Sillitoe on behalf |

| | | | |
|---|---|---|---|
| | | Consortium | of the Wellcome Sanger Institute COVID-19 Surveillance Team |
| EPI_ISL_3454658, EPI_ISL_3454659 | Lighthouse Lab in Milton Keynes | Wellcome Sanger Institute for the COVID-19 Genomics UK (COG-UK) Consortium | The Lighthouse Lab in Milton Keynes and Alex Alderton, Roberto Amato, Jeffrey Barrett, Sonia Goncalves, Ewan Harrison, David K. Jackson, Ian Johnston, Dominic Kwiatkowski, Cordelia Langford, John Sillitoe on behalf of the Wellcome Sanger Institute COVID-19 Surveillance Team |
| EPI_ISL_3454660 | Lighthouse Lab in Alderley Park | Wellcome Sanger Institute for the COVID-19 Genomics UK (COG-UK) Consortium | Jacquelyn Wynn, Mairead Hyland, The Lighthouse Lab in Alderley Park and Alex Alderton, Roberto Amato, Jeffrey Barrett, Sonia Goncalves, Ewan Harrison, David K. Jackson, Ian Johnston, Dominic Kwiatkowski, Cordelia Langford, John Sillitoe on behalf of the Wellcome Sanger Institute COVID-19 Surveillance Team |
| EPI_ISL_3454661 | Berkshire and Surrey Pathology Services Lighthouse Laboratory | Wellcome Sanger Institute for the COVID-19 Genomics UK (COG-UK) Consortium | Berkshire and Surrey Pathology Services Lighthouse Laboratory and Alex Alderton, Roberto Amato, Jeffrey Barrett, Sonia Goncalves, Ewan Harrison, David K. Jackson, Ian Johnston, Dominic Kwiatkowski, Cordelia Langford, John Sillitoe on behalf of the Wellcome Sanger Institute COVID-19 Surveillance Team |
| EPI_ISL_3454662 | Lighthouse Lab in Milton Keynes | Wellcome Sanger Institute for the COVID-19 Genomics UK (COG-UK) Consortium | The Lighthouse Lab in Milton Keynes and Alex Alderton, Roberto Amato, Jeffrey Barrett, Sonia Goncalves, Ewan Harrison, David K. Jackson, Ian Johnston, Dominic Kwiatkowski, Cordelia Langford, John Sillitoe on behalf of the Wellcome Sanger Institute COVID-19 Surveillance Team |
| EPI_ISL_3454663 | Lighthouse Lab in Alderley Park | Wellcome Sanger Institute for the COVID-19 Genomics UK (COG-UK) Consortium | Jacquelyn Wynn, Mairead Hyland, The Lighthouse Lab in Alderley Park and Alex Alderton, Roberto Amato, Jeffrey Barrett, Sonia Goncalves, Ewan Harrison, David K. Jackson, Ian Johnston, Dominic Kwiatkowski, Cordelia Langford, John Sillitoe on behalf of the Wellcome Sanger Institute COVID-19 Surveillance Team |
| EPI_ISL_3454664, EPI_ISL_3454665 | Lighthouse Lab in Milton Keynes | Wellcome Sanger Institute for the COVID-19 Genomics UK (COG-UK) Consortium | The Lighthouse Lab in Milton Keynes and Alex Alderton, Roberto Amato, Jeffrey Barrett, Sonia Goncalves, Ewan Harrison, David K. Jackson, Ian Johnston, Dominic Kwiatkowski, Cordelia Langford, John Sillitoe on behalf of the Wellcome Sanger Institute COVID-19 Surveillance Team |
| EPI_ISL_3454666 | Berkshire and Surrey Pathology Services Lighthouse Laboratory | Wellcome Sanger Institute for the COVID-19 Genomics UK (COG-UK) Consortium | Berkshire and Surrey Pathology Services Lighthouse Laboratory and Alex Alderton, Roberto Amato, Jeffrey Barrett, Sonia Goncalves, Ewan Harrison, David K. Jackson, Ian Johnston, Dominic Kwiatkowski, Cordelia Langford, John Sillitoe on behalf of the Wellcome Sanger Institute COVID-19 Surveillance Team |
| EPI_ISL_3454667, EPI_ISL_3454668 | Lighthouse Lab in Alderley Park | Wellcome Sanger Institute for the COVID-19 Genomics UK (COG-UK) Consortium | Jacquelyn Wynn, Mairead Hyland, The Lighthouse Lab in Alderley Park and Alex Alderton, Roberto Amato, Jeffrey Barrett, Sonia Goncalves, Ewan Harrison, David K. Jackson, Ian Johnston, Dominic Kwiatkowski, Cordelia Langford, John Sillitoe on behalf of the Wellcome Sanger Institute COVID-19 Surveillance Team |
| EPI_ISL_3454669 | Lighthouse Lab in Milton Keynes | Wellcome Sanger Institute for the COVID-19 Genomics UK (COG-UK) Consortium | The Lighthouse Lab in Milton Keynes and Alex Alderton, Roberto Amato, Jeffrey Barrett, Sonia Goncalves, Ewan Harrison, David K. Jackson, Ian Johnston, Dominic Kwiatkowski, Cordelia Langford, John Sillitoe on behalf of the Wellcome Sanger Institute COVID-19 Surveillance Team |
| EPI_ISL_3454670 | Berkshire and Surrey Pathology Services Lighthouse Laboratory | Wellcome Sanger Institute for the COVID-19 Genomics UK (COG-UK) Consortium | Berkshire and Surrey Pathology Services Lighthouse Laboratory and Alex Alderton, Roberto Amato, Jeffrey Barrett, Sonia Goncalves, Ewan Harrison, David K. Jackson, Ian Johnston, Dominic Kwiatkowski, Cordelia Langford, John Sillitoe on behalf of the Wellcome Sanger Institute COVID-19 Surveillance Team |
| EPI_ISL_3454671 | Lighthouse Lab in Alderley Park | Wellcome Sanger Institute for the COVID-19 Genomics UK (COG-UK) Consortium | Jacquelyn Wynn, Mairead Hyland, The Lighthouse Lab in Alderley Park and Alex Alderton, Roberto Amato, Jeffrey Barrett, Sonia Goncalves, Ewan Harrison, David K. Jackson, Ian Johnston, Dominic Kwiatkowski, Cordelia Langford, John Sillitoe on behalf of the Wellcome Sanger Institute COVID-19 Surveillance Team |
| EPI_ISL_3454672, EPI_ISL_3454673 | Lighthouse Lab in Milton Keynes | Wellcome Sanger Institute for the COVID-19 Genomics UK (COG-UK) Consortium | The Lighthouse Lab in Milton Keynes and Alex Alderton, Roberto Amato, Jeffrey Barrett, Sonia Goncalves, Ewan Harrison, David K. Jackson, Ian Johnston, Dominic Kwiatkowski, Cordelia Langford, John Sillitoe on behalf of the Wellcome Sanger Institute COVID-19 Surveillance Team |
| EPI_ISL_3454674 | Lighthouse Lab in Alderley Park | Wellcome Sanger Institute for the COVID-19 Genomics UK (COG-UK) Consortium | Jacquelyn Wynn, Mairead Hyland, The Lighthouse Lab in Alderley Park and Alex Alderton, Roberto Amato, Jeffrey Barrett, Sonia Goncalves, Ewan Harrison, David K. Jackson, Ian Johnston, Dominic Kwiatkowski, Cordelia Langford, John Sillitoe on behalf of the Wellcome Sanger Institute COVID-19 Surveillance Team |
| EPI_ISL_3454675, EPI_ISL_3454676 | Lighthouse Lab in Milton Keynes | Wellcome Sanger Institute for the COVID-19 Genomics UK (COG-UK) Consortium | The Lighthouse Lab in Milton Keynes and Alex Alderton, Roberto Amato, Jeffrey Barrett, Sonia Goncalves, Ewan Harrison, David K. Jackson, Ian Johnston, Dominic Kwiatkowski, Cordelia Langford, John Sillitoe on behalf of the Wellcome Sanger Institute COVID-19 Surveillance Team |
| EPI_ISL_3454677 | Lighthouse Lab in Alderley Park | Wellcome Sanger Institute for the COVID-19 Genomics UK (COG-UK) Consortium | Jacquelyn Wynn, Mairead Hyland, The Lighthouse Lab in Alderley Park and Alex Alderton, Roberto Amato, Jeffrey Barrett, Sonia Goncalves, Ewan Harrison, David K. Jackson, Ian Johnston, Dominic Kwiatkowski, Cordelia Langford, John Sillitoe on behalf of the Wellcome Sanger Institute COVID-19 Surveillance Team |
| EPI_ISL_3454678 | Lighthouse Lab in Milton Keynes | Wellcome Sanger Institute for the COVID-19 Genomics UK (COG-UK) Consortium | The Lighthouse Lab in Milton Keynes and Alex Alderton, Roberto Amato, Jeffrey Barrett, Sonia Goncalves, Ewan Harrison, David K. Jackson, Ian Johnston, Dominic Kwiatkowski, Cordelia Langford, John Sillitoe on behalf of the Wellcome Sanger Institute COVID-19 Surveillance Team |
| EPI_ISL_3454679 | Lighthouse Lab in Alderley Park | Wellcome Sanger Institute for the COVID-19 Genomics UK (COG-UK) Consortium | Jacquelyn Wynn, Mairead Hyland, The Lighthouse Lab in Alderley Park and Alex Alderton, Roberto Amato, Jeffrey Barrett, Sonia Goncalves, Ewan Harrison, David K. Jackson, Ian Johnston, Dominic Kwiatkowski, Cordelia Langford, John Sillitoe on behalf of the Wellcome Sanger Institute COVID-19 Surveillance Team |
| EPI_ISL_3454680 | Lighthouse Lab in Milton Keynes | Wellcome Sanger Institute for the COVID-19 Genomics UK (COG-UK) Consortium | The Lighthouse Lab in Milton Keynes and Alex Alderton, Roberto Amato, Jeffrey Barrett, Sonia Goncalves, Ewan Harrison, David K. Jackson, Ian Johnston, Dominic Kwiatkowski, Cordelia Langford, John Sillitoe on behalf of the Wellcome Sanger Institute COVID-19 Surveillance Team |
| EPI_ISL_3454681 | Berkshire and Surrey Pathology Services Lighthouse Laboratory | Wellcome Sanger Institute for the COVID-19 Genomics UK (COG-UK) Consortium | Berkshire and Surrey Pathology Services Lighthouse Laboratory and Alex Alderton, Roberto Amato, Jeffrey Barrett, Sonia Goncalves, Ewan Harrison, David K. Jackson, Ian Johnston, Dominic Kwiatkowski, Cordelia Langford, John Sillitoe on behalf of the Wellcome Sanger Institute COVID-19 Surveillance Team |
| EPI_ISL_3454682 | Lighthouse Lab in Milton Keynes | Wellcome Sanger Institute for the COVID-19 Genomics UK (COG-UK) Consortium | The Lighthouse Lab in Milton Keynes and Alex Alderton, Roberto Amato, Jeffrey Barrett, Sonia Goncalves, Ewan Harrison, David K. Jackson, Ian Johnston, Dominic Kwiatkowski, Cordelia Langford, John Sillitoe on behalf of the Wellcome Sanger Institute COVID-19 Surveillance Team |
| EPI_ISL_3454683 | Lighthouse Lab in Alderley Park | Wellcome Sanger Institute for the COVID-19 Genomics UK (COG-UK) Consortium | Jacquelyn Wynn, Mairead Hyland, The Lighthouse Lab in Alderley Park and Alex Alderton, Roberto Amato, Jeffrey Barrett, Sonia Goncalves, Ewan Harrison, David K. Jackson, Ian Johnston, Dominic Kwiatkowski, Cordelia Langford, John Sillitoe on behalf of the Wellcome Sanger Institute COVID-19 Surveillance Team |
| EPI_ISL_3454684, EPI_ISL_3454685, EPI_ISL_3454686, EPI_ISL_3454687, EPI_ISL_3454688 | Lighthouse Lab in Milton Keynes | Wellcome Sanger Institute for the COVID-19 Genomics UK (COG-UK) Consortium | The Lighthouse Lab in Milton Keynes and Alex Alderton, Roberto Amato, Jeffrey Barrett, Sonia Goncalves, Ewan Harrison, David K. Jackson, Ian Johnston, Dominic Kwiatkowski, Cordelia Langford, John Sillitoe on behalf of the Wellcome Sanger Institute COVID-19 Surveillance Team |
| EPI_ISL_3454689 | Lighthouse Lab in Alderley Park | Wellcome Sanger Institute for the COVID-19 Genomics UK (COG-UK) Consortium | Jacquelyn Wynn, Mairead Hyland, The Lighthouse Lab in Alderley Park and Alex Alderton, Roberto Amato, Jeffrey Barrett, Sonia Goncalves, Ewan Harrison, David K. Jackson, Ian Johnston, Dominic Kwiatkowski, Cordelia Langford, John Sillitoe on behalf of the Wellcome Sanger Institute COVID-19 Surveillance Team |
| EPI_ISL_3454690 | Lighthouse Lab in Milton Keynes | Wellcome Sanger Institute for the COVID-19 Genomics UK (COG-UK) Consortium | The Lighthouse Lab in Milton Keynes and Alex Alderton, Roberto Amato, Jeffrey Barrett, Sonia Goncalves, Ewan Harrison, David K. Jackson, Ian Johnston, Dominic Kwiatkowski, Cordelia Langford, John Sillitoe on behalf of the Wellcome Sanger Institute COVID-19 Surveillance Team |
| EPI_ISL_3454691 | Lighthouse Lab in Alderley Park | Wellcome Sanger Institute for the COVID-19 Genomics UK (COG-UK) Consortium | Jacquelyn Wynn, Mairead Hyland, The Lighthouse Lab in Alderley Park and Alex Alderton, Roberto Amato, Jeffrey Barrett, Sonia Goncalves, Ewan Harrison, David K. Jackson, Ian Johnston, Dominic Kwiatkowski, Cordelia Langford, John Sillitoe on behalf of the Wellcome Sanger Institute COVID-19 Surveillance Team |
| EPI_ISL_3454692, EPI_ISL_3454693 | Lighthouse Lab in Milton Keynes | Wellcome Sanger Institute for the COVID-19 Genomics UK (COG-UK) Consortium | The Lighthouse Lab in Milton Keynes and Alex Alderton, Roberto Amato, Jeffrey Barrett, Sonia Goncalves, Ewan Harrison, David K. Jackson, Ian Johnston, Dominic Kwiatkowski, Cordelia Langford, John Sillitoe on behalf of the Wellcome Sanger Institute COVID-19 Surveillance Team |
| EPI_ISL_3454694 | Berkshire and Surrey Pathology Services Lighthouse Laboratory | Wellcome Sanger Institute for the COVID-19 Genomics UK (COG-UK) Consortium | Berkshire and Surrey Pathology Services Lighthouse Laboratory and Alex Alderton, Roberto Amato, Jeffrey Barrett, Sonia Goncalves, Ewan Harrison, David K. Jackson, Ian Johnston, Dominic Kwiatkowski, Cordelia Langford, John Sillitoe on behalf of the Wellcome Sanger Institute COVID-19 Surveillance Team |
| EPI_ISL_3454695 | Lighthouse Lab in Milton Keynes | Wellcome Sanger Institute for the COVID-19 Genomics UK (COG-UK) Consortium | The Lighthouse Lab in Milton Keynes and Alex Alderton, Roberto Amato, Jeffrey Barrett, Sonia Goncalves, Ewan Harrison, David K. Jackson, Ian Johnston, Dominic Kwiatkowski, Cordelia Langford, John Sillitoe on behalf of the Wellcome Sanger Institute COVID-19 Surveillance Team |
| EPI_ISL_3454696 | Berkshire and Surrey Pathology Services Lighthouse Laboratory | Wellcome Sanger Institute for the COVID-19 Genomics UK (COG-UK) Consortium | Berkshire and Surrey Pathology Services Lighthouse Laboratory and Alex Alderton, Roberto Amato, Jeffrey Barrett, Sonia Goncalves, Ewan Harrison, David K. Jackson, Ian Johnston, Dominic Kwiatkowski, Cordelia Langford, John Sillitoe on behalf of the Wellcome Sanger Institute COVID-19 Surveillance Team |
| EPI_ISL_3454697 | Lighthouse Lab in Alderley Park | Wellcome Sanger Institute for the COVID-19 Genomics UK (COG-UK) Consortium | Jacquelyn Wynn, Mairead Hyland, The Lighthouse Lab in Alderley Park and Alex Alderton, Roberto Amato, Jeffrey Barrett, Sonia Goncalves, Ewan Harrison, David K. Jackson, Ian Johnston, Dominic Kwiatkowski, Cordelia Langford, John Sillitoe on behalf of the Wellcome Sanger Institute COVID-19 Surveillance Team |
| EPI_ISL_3454698 | Lighthouse Lab in Milton Keynes | Wellcome Sanger Institute for the COVID-19 Genomics UK (COG-UK) Consortium | The Lighthouse Lab in Milton Keynes and Alex Alderton, Roberto Amato, Jeffrey Barrett, Sonia Goncalves, Ewan Harrison, David K. Jackson, Ian Johnston, Dominic Kwiatkowski, Cordelia Langford, John Sillitoe on behalf of the Wellcome Sanger Institute COVID-19 Surveillance Team |
| EPI_ISL_3454699 | Lighthouse Lab in Alderley Park | Wellcome Sanger Institute for the COVID-19 Genomics UK (COG-UK) Consortium | Jacquelyn Wynn, Mairead Hyland, The Lighthouse Lab in Alderley Park and Alex Alderton, Roberto Amato, Jeffrey Barrett, Sonia Goncalves, Ewan Harrison, David K. Jackson, Ian Johnston, Dominic Kwiatkowski, Cordelia Langford, John Sillitoe on behalf of the Wellcome Sanger Institute COVID-19 Surveillance Team |
| EPI_ISL_3454700 | Lighthouse Lab in Milton Keynes | Wellcome Sanger Institute for the COVID-19 Genomics UK (COG-UK) Consortium | The Lighthouse Lab in Milton Keynes and Alex Alderton, Roberto Amato, Jeffrey Barrett, Sonia Goncalves, Ewan Harrison, David K. Jackson, Ian Johnston, Dominic Kwiatkowski, Cordelia Langford, John Sillitoe on behalf of the Wellcome Sanger Institute COVID-19 Surveillance Team |
| EPI_ISL_3454701 | Berkshire and Surrey Pathology Services Lighthouse Laboratory | Wellcome Sanger Institute for the COVID-19 Genomics UK (COG-UK) Consortium | Berkshire and Surrey Pathology Services Lighthouse Laboratory and Alex Alderton, Roberto Amato, Jeffrey Barrett, Sonia Goncalves, Ewan Harrison, David K. Jackson, Ian Johnston, Dominic Kwiatkowski, Cordelia Langford, John Sillitoe on behalf of the Wellcome Sanger Institute COVID-19 Surveillance Team |
| EPI_ISL_3454702 | Lighthouse Lab in Alderley Park | Wellcome Sanger Institute for the COVID-19 Genomics UK (COG-UK) Consortium | Jacquelyn Wynn, Mairead Hyland, The Lighthouse Lab in Alderley Park and Alex Alderton, Roberto Amato, Jeffrey Barrett, Sonia Goncalves, Ewan Harrison, David K. Jackson, Ian Johnston, Dominic Kwiatkowski, Cordelia Langford, John Sillitoe on behalf of the Wellcome Sanger Institute COVID-19 Surveillance Team |
| EPI_ISL_3454703 | Lighthouse Lab in Milton Keynes | Wellcome Sanger Institute for the COVID-19 Genomics UK (COG-UK) Consortium | The Lighthouse Lab in Milton Keynes and Alex Alderton, Roberto Amato, Jeffrey Barrett, Sonia Goncalves, Ewan Harrison, David K. Jackson, Ian Johnston, Dominic Kwiatkowski, Cordelia Langford, John Sillitoe on behalf of the Wellcome Sanger Institute COVID-19 Surveillance Team |
| EPI_ISL_3454704 | Lighthouse Lab in Alderley Park | Wellcome Sanger Institute for the COVID-19 Genomics UK (COG-UK) Consortium | Jacquelyn Wynn, Mairead Hyland, The Lighthouse Lab in Alderley Park and Alex Alderton, Roberto Amato, Jeffrey Barrett, Sonia Goncalves, Ewan Harrison, David K. Jackson, Ian Johnston, Dominic Kwiatkowski, Cordelia Langford, John Sillitoe on behalf of the Wellcome Sanger Institute COVID-19 Surveillance Team |
| EPI_ISL_3454705, EPI_ISL_3454706, EPI_ISL_3454707 | Lighthouse Lab in Milton Keynes | Wellcome Sanger Institute for the COVID-19 Genomics UK (COG-UK) Consortium | The Lighthouse Lab in Milton Keynes and Alex Alderton, Roberto Amato, Jeffrey Barrett, Sonia Goncalves, Ewan Harrison, David K. Jackson, Ian Johnston, Dominic Kwiatkowski, Cordelia Langford, John Sillitoe on behalf of the Wellcome Sanger Institute COVID-19 Surveillance Team |
| EPI_ISL_3454708 | Berkshire and Surrey Pathology Services Lighthouse Laboratory | Wellcome Sanger Institute for the COVID-19 Genomics UK (COG-UK) Consortium | Berkshire and Surrey Pathology Services Lighthouse Laboratory and Alex Alderton, Roberto Amato, Jeffrey Barrett, Sonia Goncalves, Ewan Harrison, David K. Jackson, Ian Johnston, Dominic Kwiatkowski, Cordelia Langford, John Sillitoe on behalf of the Wellcome Sanger Institute COVID-19 Surveillance Team |
| EPI_ISL_3454709 | Lighthouse Lab in Alderley Park | Wellcome Sanger Institute for the COVID-19 Genomics UK (COG-UK) Consortium | Jacquelyn Wynn, Mairead Hyland, The Lighthouse Lab in Alderley Park and Alex Alderton, Roberto Amato, Jeffrey Barrett, Sonia Goncalves, Ewan Harrison, David K. Jackson, Ian Johnston, Dominic Kwiatkowski, Cordelia Langford, John Sillitoe on behalf of the Wellcome Sanger Institute COVID-19 Surveillance Team |
| EPI_ISL_3454710, EPI_ISL_3454711, EPI_ISL_3454712, EPI_ISL_3454713, EPI_ISL_3454714, EPI_ISL_3454715, | Lighthouse Lab in Milton Keynes | Wellcome Sanger Institute for the COVID-19 Genomics UK (COG-UK) Consortium | The Lighthouse Lab in Milton Keynes and Alex Alderton, Roberto Amato, Jeffrey Barrett, Sonia Goncalves, Ewan Harrison, David K. Jackson, Ian Johnston, Dominic Kwiatkowski, Cordelia Langford, John Sillitoe on behalf of the Wellcome Sanger Institute COVID-19 Surveillance Team |

| | | | |
|---|---|---|---|
| EPI_ISL_3454716 | | | |
| EPI_ISL_3454717 | Berkshire and Surrey Pathology Services Lighthouse Laboratory | Wellcome Sanger Institute for the COVID-19 Genomics UK (COG-UK) Consortium | Berkshire and Surrey Pathology Services Lighthouse Laboratory and Alex Alderton, Roberto Amato, Jeffrey Barrett, Sonia Goncalves, Ewan Harrison, David K. Jackson, Ian Johnston, Dominic Kwiatkowski, Cordelia Langford, John Sillitoe on behalf of the Wellcome Sanger Institute COVID-19 Surveillance Team |
| EPI_ISL_3454718 | Lighthouse Lab in Milton Keynes | Wellcome Sanger Institute for the COVID-19 Genomics UK (COG-UK) Consortium | The Lighthouse Lab in Milton Keynes and Alex Alderton, Roberto Amato, Jeffrey Barrett, Sonia Goncalves, Ewan Harrison, David K. Jackson, Ian Johnston, Dominic Kwiatkowski, Cordelia Langford, John Sillitoe on behalf of the Wellcome Sanger Institute COVID-19 Surveillance Team |
| EPI_ISL_3454719, EPI_ISL_3454720, EPI_ISL_3454721, EPI_ISL_3454722 | Lighthouse Lab in Alderley Park | Wellcome Sanger Institute for the COVID-19 Genomics UK (COG-UK) Consortium | Jacquelyn Wynn, Mairead Hyland, The Lighthouse Lab in Alderley Park and Alex Alderton, Roberto Amato, Jeffrey Barrett, Sonia Goncalves, Ewan Harrison, David K. Jackson, Ian Johnston, Dominic Kwiatkowski, Cordelia Langford, John Sillitoe on behalf of the Wellcome Sanger Institute COVID-19 Surveillance Team |
| EPI_ISL_3454723 | Berkshire and Surrey Pathology Services Lighthouse Laboratory | Wellcome Sanger Institute for the COVID-19 Genomics UK (COG-UK) Consortium | Berkshire and Surrey Pathology Services Lighthouse Laboratory and Alex Alderton, Roberto Amato, Jeffrey Barrett, Sonia Goncalves, Ewan Harrison, David K. Jackson, Ian Johnston, Dominic Kwiatkowski, Cordelia Langford, John Sillitoe on behalf of the Wellcome Sanger Institute COVID-19 Surveillance Team |
| EPI_ISL_3454724, EPI_ISL_3454725 | Lighthouse Lab in Milton Keynes | Wellcome Sanger Institute for the COVID-19 Genomics UK (COG-UK) Consortium | The Lighthouse Lab in Milton Keynes and Alex Alderton, Roberto Amato, Jeffrey Barrett, Sonia Goncalves, Ewan Harrison, David K. Jackson, Ian Johnston, Dominic Kwiatkowski, Cordelia Langford, John Sillitoe on behalf of the Wellcome Sanger Institute COVID-19 Surveillance Team |
| EPI_ISL_3454726 | Lighthouse Lab in Alderley Park | Wellcome Sanger Institute for the COVID-19 Genomics UK (COG-UK) Consortium | Jacquelyn Wynn, Mairead Hyland, The Lighthouse Lab in Alderley Park and Alex Alderton, Roberto Amato, Jeffrey Barrett, Sonia Goncalves, Ewan Harrison, David K. Jackson, Ian Johnston, Dominic Kwiatkowski, Cordelia Langford, John Sillitoe on behalf of the Wellcome Sanger Institute COVID-19 Surveillance Team |
| EPI_ISL_3454727, EPI_ISL_3454728 | Lighthouse Lab in Milton Keynes | Wellcome Sanger Institute for the COVID-19 Genomics UK (COG-UK) Consortium | The Lighthouse Lab in Milton Keynes and Alex Alderton, Roberto Amato, Jeffrey Barrett, Sonia Goncalves, Ewan Harrison, David K. Jackson, Ian Johnston, Dominic Kwiatkowski, Cordelia Langford, John Sillitoe on behalf of the Wellcome Sanger Institute COVID-19 Surveillance Team |
| EPI_ISL_3454729 | Berkshire and Surrey Pathology Services Lighthouse Laboratory | Wellcome Sanger Institute for the COVID-19 Genomics UK (COG-UK) Consortium | Berkshire and Surrey Pathology Services Lighthouse Laboratory and Alex Alderton, Roberto Amato, Jeffrey Barrett, Sonia Goncalves, Ewan Harrison, David K. Jackson, Ian Johnston, Dominic Kwiatkowski, Cordelia Langford, John Sillitoe on behalf of the Wellcome Sanger Institute COVID-19 Surveillance Team |
| EPI_ISL_3454730, EPI_ISL_3454731, EPI_ISL_3454732 | Lighthouse Lab in Milton Keynes | Wellcome Sanger Institute for the COVID-19 Genomics UK (COG-UK) Consortium | The Lighthouse Lab in Milton Keynes and Alex Alderton, Roberto Amato, Jeffrey Barrett, Sonia Goncalves, Ewan Harrison, David K. Jackson, Ian Johnston, Dominic Kwiatkowski, Cordelia Langford, John Sillitoe on behalf of the Wellcome Sanger Institute COVID-19 Surveillance Team |
| EPI_ISL_3454733 | Lighthouse Lab in Alderley Park | Wellcome Sanger Institute for the COVID-19 Genomics UK (COG-UK) Consortium | Jacquelyn Wynn, Mairead Hyland, The Lighthouse Lab in Alderley Park and Alex Alderton, Roberto Amato, Jeffrey Barrett, Sonia Goncalves, Ewan Harrison, David K. Jackson, Ian Johnston, Dominic Kwiatkowski, Cordelia Langford, John Sillitoe on behalf of the Wellcome Sanger Institute COVID-19 Surveillance Team |
| EPI_ISL_3454734, EPI_ISL_3454735 | Lighthouse Lab in Milton Keynes | Wellcome Sanger Institute for the COVID-19 Genomics UK (COG-UK) Consortium | The Lighthouse Lab in Milton Keynes and Alex Alderton, Roberto Amato, Jeffrey Barrett, Sonia Goncalves, Ewan Harrison, David K. Jackson, Ian Johnston, Dominic Kwiatkowski, Cordelia Langford, John Sillitoe on behalf of the Wellcome Sanger Institute COVID-19 Surveillance Team |
| EPI_ISL_3454736 | Berkshire and Surrey Pathology Services Lighthouse Laboratory | Wellcome Sanger Institute for the COVID-19 Genomics UK (COG-UK) Consortium | Berkshire and Surrey Pathology Services Lighthouse Laboratory and Alex Alderton, Roberto Amato, Jeffrey Barrett, Sonia Goncalves, Ewan Harrison, David K. Jackson, Ian Johnston, Dominic Kwiatkowski, Cordelia Langford, John Sillitoe on behalf of the Wellcome Sanger Institute COVID-19 Surveillance Team |
| EPI_ISL_3454737 | Lighthouse Lab in Alderley Park | Wellcome Sanger Institute for the COVID-19 Genomics UK (COG-UK) Consortium | Jacquelyn Wynn, Mairead Hyland, The Lighthouse Lab in Alderley Park and Alex Alderton, Roberto Amato, Jeffrey Barrett, Sonia Goncalves, Ewan Harrison, David K. Jackson, Ian Johnston, Dominic Kwiatkowski, Cordelia Langford, John Sillitoe on behalf of the Wellcome Sanger Institute COVID-19 Surveillance Team |
| EPI_ISL_3454738, EPI_ISL_3454739, EPI_ISL_3454740 | Lighthouse Lab in Milton Keynes | Wellcome Sanger Institute for the COVID-19 Genomics UK (COG-UK) Consortium | The Lighthouse Lab in Milton Keynes and Alex Alderton, Roberto Amato, Jeffrey Barrett, Sonia Goncalves, Ewan Harrison, David K. Jackson, Ian Johnston, Dominic Kwiatkowski, Cordelia Langford, John Sillitoe on behalf of the Wellcome Sanger Institute COVID-19 Surveillance Team |
| EPI_ISL_3454741 | Lighthouse Lab in Alderley Park | Wellcome Sanger Institute for the COVID-19 Genomics UK (COG-UK) Consortium | Jacquelyn Wynn, Mairead Hyland, The Lighthouse Lab in Alderley Park and Alex Alderton, Roberto Amato, Jeffrey Barrett, Sonia Goncalves, Ewan Harrison, David K. Jackson, Ian Johnston, Dominic Kwiatkowski, Cordelia Langford, John Sillitoe on behalf of the Wellcome Sanger Institute COVID-19 Surveillance Team |
| EPI_ISL_3454742, EPI_ISL_3454743, EPI_ISL_3454744, EPI_ISL_3454745 | Lighthouse Lab in Milton Keynes | Wellcome Sanger Institute for the COVID-19 Genomics UK (COG-UK) Consortium | The Lighthouse Lab in Milton Keynes and Alex Alderton, Roberto Amato, Jeffrey Barrett, Sonia Goncalves, Ewan Harrison, David K. Jackson, Ian Johnston, Dominic Kwiatkowski, Cordelia Langford, John Sillitoe on behalf of the Wellcome Sanger Institute COVID-19 Surveillance Team |
| EPI_ISL_3454746 | Lighthouse Lab in Alderley Park | Wellcome Sanger Institute for the COVID-19 Genomics UK (COG-UK) Consortium | Jacquelyn Wynn, Mairead Hyland, The Lighthouse Lab in Alderley Park and Alex Alderton, Roberto Amato, Jeffrey Barrett, Sonia Goncalves, Ewan Harrison, David K. Jackson, Ian Johnston, Dominic Kwiatkowski, Cordelia Langford, John Sillitoe on behalf of the Wellcome Sanger Institute COVID-19 Surveillance Team |
| EPI_ISL_3454747 | Lighthouse Lab in Milton Keynes | Wellcome Sanger Institute for the COVID-19 Genomics UK (COG-UK) Consortium | The Lighthouse Lab in Milton Keynes and Alex Alderton, Roberto Amato, Jeffrey Barrett, Sonia Goncalves, Ewan Harrison, David K. Jackson, Ian Johnston, Dominic Kwiatkowski, Cordelia Langford, John Sillitoe on behalf of the Wellcome Sanger Institute COVID-19 Surveillance Team |
| EPI_ISL_3454748, EPI_ISL_3454749 | Lighthouse Lab in Alderley Park | Wellcome Sanger Institute for the COVID-19 Genomics UK (COG-UK) Consortium | Jacquelyn Wynn, Mairead Hyland, The Lighthouse Lab in Alderley Park and Alex Alderton, Roberto Amato, Jeffrey Barrett, Sonia Goncalves, Ewan Harrison, David K. Jackson, Ian Johnston, Dominic Kwiatkowski, Cordelia Langford, John Sillitoe on behalf of the Wellcome Sanger Institute COVID-19 Surveillance Team |
| EPI_ISL_3454750, EPI_ISL_3454751 | Lighthouse Lab in Milton Keynes | Wellcome Sanger Institute for the COVID-19 Genomics UK (COG-UK) Consortium | The Lighthouse Lab in Milton Keynes and Alex Alderton, Roberto Amato, Jeffrey Barrett, Sonia Goncalves, Ewan Harrison, David K. Jackson, Ian Johnston, Dominic Kwiatkowski, Cordelia Langford, John Sillitoe on behalf of the Wellcome Sanger Institute COVID-19 Surveillance Team |
| EPI_ISL_3454752 | Lighthouse Lab in Alderley Park | Wellcome Sanger Institute for the COVID-19 Genomics UK (COG-UK) Consortium | Jacquelyn Wynn, Mairead Hyland, The Lighthouse Lab in Alderley Park and Alex Alderton, Roberto Amato, Jeffrey Barrett, Sonia Goncalves, Ewan Harrison, David K. Jackson, Ian Johnston, Dominic Kwiatkowski, Cordelia Langford, John Sillitoe on behalf of the Wellcome Sanger Institute COVID-19 Surveillance Team |
| EPI_ISL_3454753 | Lighthouse Lab in Milton Keynes | Wellcome Sanger Institute for the COVID-19 Genomics UK (COG-UK) Consortium | The Lighthouse Lab in Milton Keynes and Alex Alderton, Roberto Amato, Jeffrey Barrett, Sonia Goncalves, Ewan Harrison, David K. Jackson, Ian Johnston, Dominic Kwiatkowski, Cordelia Langford, John Sillitoe on behalf of the Wellcome Sanger Institute COVID-19 Surveillance Team |
| EPI_ISL_3454754 | Lighthouse Lab in Alderley Park | Wellcome Sanger Institute for the COVID-19 Genomics UK (COG-UK) Consortium | Jacquelyn Wynn, Mairead Hyland, The Lighthouse Lab in Alderley Park and Alex Alderton, Roberto Amato, Jeffrey Barrett, Sonia Goncalves, Ewan Harrison, David K. Jackson, Ian Johnston, Dominic Kwiatkowski, Cordelia Langford, John Sillitoe on behalf of the Wellcome Sanger Institute COVID-19 Surveillance Team |
| EPI_ISL_3454755, EPI_ISL_3454756 | Lighthouse Lab in Milton Keynes | Wellcome Sanger Institute for the COVID-19 Genomics UK (COG-UK) Consortium | The Lighthouse Lab in Milton Keynes and Alex Alderton, Roberto Amato, Jeffrey Barrett, Sonia Goncalves, Ewan Harrison, David K. Jackson, Ian Johnston, Dominic Kwiatkowski, Cordelia Langford, John Sillitoe on behalf of the Wellcome Sanger Institute COVID-19 Surveillance Team |
| EPI_ISL_3454757 | Berkshire and Surrey Pathology Services Lighthouse Laboratory | Wellcome Sanger Institute for the COVID-19 Genomics UK (COG-UK) Consortium | Berkshire and Surrey Pathology Services Lighthouse Laboratory and Alex Alderton, Roberto Amato, Jeffrey Barrett, Sonia Goncalves, Ewan Harrison, David K. Jackson, Ian Johnston, Dominic Kwiatkowski, Cordelia Langford, John Sillitoe on behalf of the Wellcome Sanger Institute COVID-19 Surveillance Team |
| EPI_ISL_3454758 | Lighthouse Lab in Alderley Park | Wellcome Sanger Institute for the COVID-19 Genomics UK (COG-UK) Consortium | Jacquelyn Wynn, Mairead Hyland, The Lighthouse Lab in Alderley Park and Alex Alderton, Roberto Amato, Jeffrey Barrett, Sonia Goncalves, Ewan Harrison, David K. Jackson, Ian Johnston, Dominic Kwiatkowski, Cordelia Langford, John Sillitoe on behalf of the Wellcome Sanger Institute COVID-19 Surveillance Team |
| EPI_ISL_3454759, EPI_ISL_3454760, EPI_ISL_3454761, EPI_ISL_3454762, EPI_ISL_3454763 | Lighthouse Lab in Milton Keynes | Wellcome Sanger Institute for the COVID-19 Genomics UK (COG-UK) Consortium | The Lighthouse Lab in Milton Keynes and Alex Alderton, Roberto Amato, Jeffrey Barrett, Sonia Goncalves, Ewan Harrison, David K. Jackson, Ian Johnston, Dominic Kwiatkowski, Cordelia Langford, John Sillitoe on behalf of the Wellcome Sanger Institute COVID-19 Surveillance Team |
| EPI_ISL_3454764 | Lighthouse Lab in Alderley Park | Wellcome Sanger Institute for the COVID-19 Genomics UK (COG-UK) Consortium | Jacquelyn Wynn, Mairead Hyland, The Lighthouse Lab in Alderley Park and Alex Alderton, Roberto Amato, Jeffrey Barrett, Sonia Goncalves, Ewan Harrison, David K. Jackson, Ian Johnston, Dominic Kwiatkowski, Cordelia Langford, John Sillitoe on behalf of the Wellcome Sanger Institute COVID-19 Surveillance Team |
| EPI_ISL_3454765 | Lighthouse Lab in Milton Keynes | Wellcome Sanger Institute for the COVID-19 Genomics UK (COG-UK) Consortium | The Lighthouse Lab in Milton Keynes and Alex Alderton, Roberto Amato, Jeffrey Barrett, Sonia Goncalves, Ewan Harrison, David K. Jackson, Ian Johnston, Dominic Kwiatkowski, Cordelia Langford, John Sillitoe on behalf of the Wellcome Sanger Institute COVID-19 Surveillance Team |
| EPI_ISL_3454766 | Berkshire and Surrey Pathology Services Lighthouse Laboratory | Wellcome Sanger Institute for the COVID-19 Genomics UK (COG-UK) Consortium | Berkshire and Surrey Pathology Services Lighthouse Laboratory and Alex Alderton, Roberto Amato, Jeffrey Barrett, Sonia Goncalves, Ewan Harrison, David K. Jackson, Ian Johnston, Dominic Kwiatkowski, Cordelia Langford, John Sillitoe on behalf of the Wellcome Sanger Institute COVID-19 Surveillance Team |
| EPI_ISL_3454767 | Lighthouse Lab in Alderley Park | Wellcome Sanger Institute for the COVID-19 Genomics UK (COG-UK) Consortium | Jacquelyn Wynn, Mairead Hyland, The Lighthouse Lab in Alderley Park and Alex Alderton, Roberto Amato, Jeffrey Barrett, Sonia Goncalves, Ewan Harrison, David K. Jackson, Ian Johnston, Dominic Kwiatkowski, Cordelia Langford, John Sillitoe on behalf of the Wellcome Sanger Institute COVID-19 Surveillance Team |
| EPI_ISL_3454768, EPI_ISL_3454769, EPI_ISL_3454770, EPI_ISL_3454771 | Lighthouse Lab in Milton Keynes | Wellcome Sanger Institute for the COVID-19 Genomics UK (COG-UK) Consortium | The Lighthouse Lab in Milton Keynes and Alex Alderton, Roberto Amato, Jeffrey Barrett, Sonia Goncalves, Ewan Harrison, David K. Jackson, Ian Johnston, Dominic Kwiatkowski, Cordelia Langford, John Sillitoe on behalf of the Wellcome Sanger Institute COVID-19 Surveillance Team |
| EPI_ISL_3454772 | Berkshire and Surrey Pathology Services Lighthouse Laboratory | Wellcome Sanger Institute for the COVID-19 Genomics UK (COG-UK) Consortium | Berkshire and Surrey Pathology Services Lighthouse Laboratory and Alex Alderton, Roberto Amato, Jeffrey Barrett, Sonia Goncalves, Ewan Harrison, David K. Jackson, Ian Johnston, Dominic Kwiatkowski, Cordelia Langford, John Sillitoe on behalf of the Wellcome Sanger Institute COVID-19 Surveillance Team |
| EPI_ISL_3454773, EPI_ISL_3454774, EPI_ISL_3454775 | Lighthouse Lab in Milton Keynes | Wellcome Sanger Institute for the COVID-19 Genomics UK (COG-UK) Consortium | The Lighthouse Lab in Milton Keynes and Alex Alderton, Roberto Amato, Jeffrey Barrett, Sonia Goncalves, Ewan Harrison, David K. Jackson, Ian Johnston, Dominic Kwiatkowski, Cordelia Langford, John Sillitoe on behalf of the Wellcome Sanger Institute COVID-19 Surveillance Team |
| EPI_ISL_3454777 | Lighthouse Lab in Alderley Park | Wellcome Sanger Institute for the COVID-19 Genomics UK (COG-UK) Consortium | Jacquelyn Wynn, Mairead Hyland, The Lighthouse Lab in Alderley Park and Alex Alderton, Roberto Amato, Jeffrey Barrett, Sonia Goncalves, Ewan Harrison, David K. Jackson, Ian Johnston, Dominic Kwiatkowski, Cordelia Langford, John Sillitoe on behalf of the Wellcome Sanger Institute COVID-19 Surveillance Team |
| EPI_ISL_3454778 | Berkshire and Surrey Pathology Services Lighthouse Laboratory | Wellcome Sanger Institute for the COVID-19 Genomics UK (COG-UK) Consortium | Berkshire and Surrey Pathology Services Lighthouse Laboratory and Alex Alderton, Roberto Amato, Jeffrey Barrett, Sonia Goncalves, Ewan Harrison, David K. Jackson, Ian Johnston, Dominic Kwiatkowski, Cordelia Langford, John Sillitoe on behalf of the Wellcome Sanger Institute COVID-19 Surveillance Team |
| EPI_ISL_3454780, EPI_ISL_3454781, EPI_ISL_3454783, EPI_ISL_3454784 | Lighthouse Lab in Milton Keynes | Wellcome Sanger Institute for the COVID-19 Genomics UK (COG-UK) Consortium | The Lighthouse Lab in Milton Keynes and Alex Alderton, Roberto Amato, Jeffrey Barrett, Sonia Goncalves, Ewan Harrison, David K. Jackson, Ian Johnston, Dominic Kwiatkowski, Cordelia Langford, John Sillitoe on behalf of the Wellcome Sanger Institute COVID-19 Surveillance Team |
| EPI_ISL_3454785 | Lighthouse Lab in Alderley Park | Wellcome Sanger Institute for the COVID-19 Genomics UK (COG-UK) Consortium | Jacquelyn Wynn, Mairead Hyland, The Lighthouse Lab in Alderley Park and Alex Alderton, Roberto Amato, Jeffrey Barrett, Sonia Goncalves, Ewan Harrison, David K. Jackson, Ian Johnston, Dominic Kwiatkowski, Cordelia Langford, John Sillitoe on behalf of the Wellcome Sanger Institute COVID-19 Surveillance Team |
| EPI_ISL_3454786 | WHO National Influenza Centre Russian Federation | WHO National Influenza Centre Russian Federation | Andrey Komissarov, Artem Fadeev, Kseniya Komissarova, Oula Mansour, Kirill Varchenko, Mikhail Bakaev, Tamila Musaeva, Veronika Eder, Maria Pisareva, Nikita Yolshin, Daria Danilenko, Oksana Stanevich, Ksenia Safina, Elena Nabieva, Georgii Bazykin, Dmitry Lioznov |

| | | | |
|---|---|---|---|
| EPI_ISL_3454787 | Lighthouse Lab in Milton Keynes | Wellcome Sanger Institute for the COVID-19 Genomics UK (COG-UK) Consortium | The Lighthouse Lab in Milton Keynes and Alex Alderton, Roberto Amato, Jeffrey Barrett, Sonia Goncalves, Ewan Harrison, David K. Jackson, Ian Johnston, Dominic Kwiatkowski, Cordelia Langford, John Sillitoe on behalf of the Wellcome Sanger Institute COVID-19 Surveillance Team |
| EPI_ISL_3454788 | Berkshire and Surrey Pathology Services Lighthouse Laboratory | Wellcome Sanger Institute for the COVID-19 Genomics UK (COG-UK) Consortium | Berkshire and Surrey Pathology Services Lighthouse Laboratory and Alex Alderton, Roberto Amato, Jeffrey Barrett, Sonia Goncalves, Ewan Harrison, David K. Jackson, Ian Johnston, Dominic Kwiatkowski, Cordelia Langford, John Sillitoe on behalf of the Wellcome Sanger Institute COVID-19 Surveillance Team |
| EPI_ISL_3454789 | Lighthouse Lab in Milton Keynes | Wellcome Sanger Institute for the COVID-19 Genomics UK (COG-UK) Consortium | The Lighthouse Lab in Milton Keynes and Alex Alderton, Roberto Amato, Jeffrey Barrett, Sonia Goncalves, Ewan Harrison, David K. Jackson, Ian Johnston, Dominic Kwiatkowski, Cordelia Langford, John Sillitoe on behalf of the Wellcome Sanger Institute COVID-19 Surveillance Team |
| EPI_ISL_3454791 | Lighthouse Lab in Alderley Park | Wellcome Sanger Institute for the COVID-19 Genomics UK (COG-UK) Consortium | Jacquelyn Wynn, Mairead Hyland, The Lighthouse Lab in Alderley Park and Alex Alderton, Roberto Amato, Jeffrey Barrett, Sonia Goncalves, Ewan Harrison, David K. Jackson, Ian Johnston, Dominic Kwiatkowski, Cordelia Langford, John Sillitoe on behalf of the Wellcome Sanger Institute COVID-19 Surveillance Team |
| EPI_ISL_3454792 | Lighthouse Lab in Milton Keynes | Wellcome Sanger Institute for the COVID-19 Genomics UK (COG-UK) Consortium | The Lighthouse Lab in Milton Keynes and Alex Alderton, Roberto Amato, Jeffrey Barrett, Sonia Goncalves, Ewan Harrison, David K. Jackson, Ian Johnston, Dominic Kwiatkowski, Cordelia Langford, John Sillitoe on behalf of the Wellcome Sanger Institute COVID-19 Surveillance Team |
| EPI_ISL_3454793 | WHO National Influenza Centre Russian Federation | WHO National Influenza Centre Russian Federation | Andrey Komissarov, Artem Fadeev, Kseniya Komissarova, Oula Mansour, Kirill Varchenko, Mikhail Bakaev, Veronika Eder, Maria Pisareva, Nikita Yolshin, Daria Danilenko, Oksana Stanevich, Ksenia Safina, Elena Nabieva, Georgii Bazykin, Dmitry Lioznov |
| EPI_ISL_3454794 | Lighthouse Lab in Alderley Park | Wellcome Sanger Institute for the COVID-19 Genomics UK (COG-UK) Consortium | Jacquelyn Wynn, Mairead Hyland, The Lighthouse Lab in Alderley Park and Alex Alderton, Roberto Amato, Jeffrey Barrett, Sonia Goncalves, Ewan Harrison, David K. Jackson, Ian Johnston, Dominic Kwiatkowski, Cordelia Langford, John Sillitoe on behalf of the Wellcome Sanger Institute COVID-19 Surveillance Team |
| EPI_ISL_3454795 | Berkshire and Surrey Pathology Services Lighthouse Laboratory | Wellcome Sanger Institute for the COVID-19 Genomics UK (COG-UK) Consortium | Berkshire and Surrey Pathology Services Lighthouse Laboratory and Alex Alderton, Roberto Amato, Jeffrey Barrett, Sonia Goncalves, Ewan Harrison, David K. Jackson, Ian Johnston, Dominic Kwiatkowski, Cordelia Langford, John Sillitoe on behalf of the Wellcome Sanger Institute COVID-19 Surveillance Team |
| EPI_ISL_3454796, EPI_ISL_3454798, EPI_ISL_3454799, EPI_ISL_3454800 | Lighthouse Lab in Milton Keynes | Wellcome Sanger Institute for the COVID-19 Genomics UK (COG-UK) Consortium | The Lighthouse Lab in Milton Keynes and Alex Alderton, Roberto Amato, Jeffrey Barrett, Sonia Goncalves, Ewan Harrison, David K. Jackson, Ian Johnston, Dominic Kwiatkowski, Cordelia Langford, John Sillitoe on behalf of the Wellcome Sanger Institute COVID-19 Surveillance Team |
| EPI_ISL_3454802 | Lighthouse Lab in Alderley Park | Wellcome Sanger Institute for the COVID-19 Genomics UK (COG-UK) Consortium | Jacquelyn Wynn, Mairead Hyland, The Lighthouse Lab in Alderley Park and Alex Alderton, Roberto Amato, Jeffrey Barrett, Sonia Goncalves, Ewan Harrison, David K. Jackson, Ian Johnston, Dominic Kwiatkowski, Cordelia Langford, John Sillitoe on behalf of the Wellcome Sanger Institute COVID-19 Surveillance Team |
| EPI_ISL_3454803 | Lighthouse Lab in Milton Keynes | Wellcome Sanger Institute for the COVID-19 Genomics UK (COG-UK) Consortium | The Lighthouse Lab in Milton Keynes and Alex Alderton, Roberto Amato, Jeffrey Barrett, Sonia Goncalves, Ewan Harrison, David K. Jackson, Ian Johnston, Dominic Kwiatkowski, Cordelia Langford, John Sillitoe on behalf of the Wellcome Sanger Institute COVID-19 Surveillance Team |
| EPI_ISL_3454804 | WHO National Influenza Centre Russian Federation | WHO National Influenza Centre Russian Federation | Andrey Komissarov, Artem Fadeev, Kseniya Komissarova, Oula Mansour, Kirill Varchenko, Mikhail Bakaev, Tamila Musaeva, Veronika Eder, Maria Pisareva, Nikita Yolshin, Daria Danilenko, Ksenia Safina, Elena Nabieva, Georgii Bazykin, Dmitry Lioznov |
| EPI_ISL_3454805 | Lighthouse Lab in Alderley Park | Wellcome Sanger Institute for the COVID-19 Genomics UK (COG-UK) Consortium | Jacquelyn Wynn, Mairead Hyland, The Lighthouse Lab in Alderley Park and Alex Alderton, Roberto Amato, Jeffrey Barrett, Sonia Goncalves, Ewan Harrison, David K. Jackson, Ian Johnston, Dominic Kwiatkowski, Cordelia Langford, John Sillitoe on behalf of the Wellcome Sanger Institute COVID-19 Surveillance Team |
| EPI_ISL_3454806 | Lighthouse Lab in Milton Keynes | Wellcome Sanger Institute for the COVID-19 Genomics UK (COG-UK) Consortium | The Lighthouse Lab in Milton Keynes and Alex Alderton, Roberto Amato, Jeffrey Barrett, Sonia Goncalves, Ewan Harrison, David K. Jackson, Ian Johnston, Dominic Kwiatkowski, Cordelia Langford, John Sillitoe on behalf of the Wellcome Sanger Institute COVID-19 Surveillance Team |
| EPI_ISL_3454807, EPI_ISL_3454809 | Lighthouse Lab in Alderley Park | Wellcome Sanger Institute for the COVID-19 Genomics UK (COG-UK) Consortium | Jacquelyn Wynn, Mairead Hyland, The Lighthouse Lab in Alderley Park and Alex Alderton, Roberto Amato, Jeffrey Barrett, Sonia Goncalves, Ewan Harrison, David K. Jackson, Ian Johnston, Dominic Kwiatkowski, Cordelia Langford, John Sillitoe on behalf of the Wellcome Sanger Institute COVID-19 Surveillance Team |
| EPI_ISL_3454810, EPI_ISL_3454811, EPI_ISL_3454813, EPI_ISL_3454814, EPI_ISL_3454816, EPI_ISL_3454817, EPI_ISL_3454819, EPI_ISL_3454820 | Lighthouse Lab in Milton Keynes | Wellcome Sanger Institute for the COVID-19 Genomics UK (COG-UK) Consortium | The Lighthouse Lab in Milton Keynes and Alex Alderton, Roberto Amato, Jeffrey Barrett, Sonia Goncalves, Ewan Harrison, David K. Jackson, Ian Johnston, Dominic Kwiatkowski, Cordelia Langford, John Sillitoe on behalf of the Wellcome Sanger Institute COVID-19 Surveillance Team |
| EPI_ISL_3454821, EPI_ISL_3454823 | Lighthouse Lab in Alderley Park | Wellcome Sanger Institute for the COVID-19 Genomics UK (COG-UK) Consortium | Jacquelyn Wynn, Mairead Hyland, The Lighthouse Lab in Alderley Park and Alex Alderton, Roberto Amato, Jeffrey Barrett, Sonia Goncalves, Ewan Harrison, David K. Jackson, Ian Johnston, Dominic Kwiatkowski, Cordelia Langford, John Sillitoe on behalf of the Wellcome Sanger Institute COVID-19 Surveillance Team |
| EPI_ISL_3454824, EPI_ISL_3454826 | Lighthouse Lab in Milton Keynes | Wellcome Sanger Institute for the COVID-19 Genomics UK (COG-UK) Consortium | The Lighthouse Lab in Milton Keynes and Alex Alderton, Roberto Amato, Jeffrey Barrett, Sonia Goncalves, Ewan Harrison, David K. Jackson, Ian Johnston, Dominic Kwiatkowski, Cordelia Langford, John Sillitoe on behalf of the Wellcome Sanger Institute COVID-19 Surveillance Team |
| EPI_ISL_3454827, EPI_ISL_3454828 | Lighthouse Lab in Alderley Park | Wellcome Sanger Institute for the COVID-19 Genomics UK (COG-UK) Consortium | Jacquelyn Wynn, Mairead Hyland, The Lighthouse Lab in Alderley Park and Alex Alderton, Roberto Amato, Jeffrey Barrett, Sonia Goncalves, Ewan Harrison, David K. Jackson, Ian Johnston, Dominic Kwiatkowski, Cordelia Langford, John Sillitoe on behalf of the Wellcome Sanger Institute COVID-19 Surveillance Team |
| EPI_ISL_3454830, EPI_ISL_3454831, EPI_ISL_3454832 | Lighthouse Lab in Milton Keynes | Wellcome Sanger Institute for the COVID-19 Genomics UK (COG-UK) Consortium | The Lighthouse Lab in Milton Keynes and Alex Alderton, Roberto Amato, Jeffrey Barrett, Sonia Goncalves, Ewan Harrison, David K. Jackson, Ian Johnston, Dominic Kwiatkowski, Cordelia Langford, John Sillitoe on behalf of the Wellcome Sanger Institute COVID-19 Surveillance Team |
| EPI_ISL_3454834 | Lighthouse Lab in Alderley Park | Wellcome Sanger Institute for the COVID-19 Genomics UK (COG-UK) Consortium | Jacquelyn Wynn, Mairead Hyland, The Lighthouse Lab in Alderley Park and Alex Alderton, Roberto Amato, Jeffrey Barrett, Sonia Goncalves, Ewan Harrison, David K. Jackson, Ian Johnston, Dominic Kwiatkowski, Cordelia Langford, John Sillitoe on behalf of the Wellcome Sanger Institute COVID-19 Surveillance Team |
| EPI_ISL_3454835 | Berkshire and Surrey Pathology Services Lighthouse Laboratory | Wellcome Sanger Institute for the COVID-19 Genomics UK (COG-UK) Consortium | Berkshire and Surrey Pathology Services Lighthouse Laboratory and Alex Alderton, Roberto Amato, Jeffrey Barrett, Sonia Goncalves, Ewan Harrison, David K. Jackson, Ian Johnston, Dominic Kwiatkowski, Cordelia Langford, John Sillitoe on behalf of the Wellcome Sanger Institute COVID-19 Surveillance Team |
| EPI_ISL_3454836, EPI_ISL_3454838, EPI_ISL_3454839, EPI_ISL_3454840, EPI_ISL_3454842, EPI_ISL_3454843 | Lighthouse Lab in Milton Keynes | Wellcome Sanger Institute for the COVID-19 Genomics UK (COG-UK) Consortium | The Lighthouse Lab in Milton Keynes and Alex Alderton, Roberto Amato, Jeffrey Barrett, Sonia Goncalves, Ewan Harrison, David K. Jackson, Ian Johnston, Dominic Kwiatkowski, Cordelia Langford, John Sillitoe on behalf of the Wellcome Sanger Institute COVID-19 Surveillance Team |
| EPI_ISL_3454844 | Lighthouse Lab in Alderley Park | Wellcome Sanger Institute for the COVID-19 Genomics UK (COG-UK) Consortium | Jacquelyn Wynn, Mairead Hyland, The Lighthouse Lab in Alderley Park and Alex Alderton, Roberto Amato, Jeffrey Barrett, Sonia Goncalves, Ewan Harrison, David K. Jackson, Ian Johnston, Dominic Kwiatkowski, Cordelia Langford, John Sillitoe on behalf of the Wellcome Sanger Institute COVID-19 Surveillance Team |
| EPI_ISL_3454845 | Lighthouse Lab in Milton Keynes | Wellcome Sanger Institute for the COVID-19 Genomics UK (COG-UK) Consortium | The Lighthouse Lab in Milton Keynes and Alex Alderton, Roberto Amato, Jeffrey Barrett, Sonia Goncalves, Ewan Harrison, David K. Jackson, Ian Johnston, Dominic Kwiatkowski, Cordelia Langford, John Sillitoe on behalf of the Wellcome Sanger Institute COVID-19 Surveillance Team |
| EPI_ISL_3454846 | Berkshire and Surrey Pathology Services Lighthouse Laboratory | Wellcome Sanger Institute for the COVID-19 Genomics UK (COG-UK) Consortium | Berkshire and Surrey Pathology Services Lighthouse Laboratory and Alex Alderton, Roberto Amato, Jeffrey Barrett, Sonia Goncalves, Ewan Harrison, David K. Jackson, Ian Johnston, Dominic Kwiatkowski, Cordelia Langford, John Sillitoe on behalf of the Wellcome Sanger Institute COVID-19 Surveillance Team |
| EPI_ISL_3454847, EPI_ISL_3454848 | Lighthouse Lab in Milton Keynes | Wellcome Sanger Institute for the COVID-19 Genomics UK (COG-UK) Consortium | The Lighthouse Lab in Milton Keynes and Alex Alderton, Roberto Amato, Jeffrey Barrett, Sonia Goncalves, Ewan Harrison, David K. Jackson, Ian Johnston, Dominic Kwiatkowski, Cordelia Langford, John Sillitoe on behalf of the Wellcome Sanger Institute COVID-19 Surveillance Team |
| EPI_ISL_3454849 | Berkshire and Surrey Pathology Services Lighthouse Laboratory | Wellcome Sanger Institute for the COVID-19 Genomics UK (COG-UK) Consortium | Berkshire and Surrey Pathology Services Lighthouse Laboratory and Alex Alderton, Roberto Amato, Jeffrey Barrett, Sonia Goncalves, Ewan Harrison, David K. Jackson, Ian Johnston, Dominic Kwiatkowski, Cordelia Langford, John Sillitoe on behalf of the Wellcome Sanger Institute COVID-19 Surveillance Team |
| EPI_ISL_3454850 | Lighthouse Lab in Alderley Park | Wellcome Sanger Institute for the COVID-19 Genomics UK (COG-UK) Consortium | Jacquelyn Wynn, Mairead Hyland, The Lighthouse Lab in Alderley Park and Alex Alderton, Roberto Amato, Jeffrey Barrett, Sonia Goncalves, Ewan Harrison, David K. Jackson, Ian Johnston, Dominic Kwiatkowski, Cordelia Langford, John Sillitoe on behalf of the Wellcome Sanger Institute COVID-19 Surveillance Team |
| EPI_ISL_3454851 | Lighthouse Lab in Milton Keynes | Wellcome Sanger Institute for the COVID-19 Genomics UK (COG-UK) Consortium | The Lighthouse Lab in Milton Keynes and Alex Alderton, Roberto Amato, Jeffrey Barrett, Sonia Goncalves, Ewan Harrison, David K. Jackson, Ian Johnston, Dominic Kwiatkowski, Cordelia Langford, John Sillitoe on behalf of the Wellcome Sanger Institute COVID-19 Surveillance Team |
| EPI_ISL_3454852, EPI_ISL_3454853 | Lighthouse Lab in Alderley Park | Wellcome Sanger Institute for the COVID-19 Genomics UK (COG-UK) Consortium | Jacquelyn Wynn, Mairead Hyland, The Lighthouse Lab in Alderley Park and Alex Alderton, Roberto Amato, Jeffrey Barrett, Sonia Goncalves, Ewan Harrison, David K. Jackson, Ian Johnston, Dominic Kwiatkowski, Cordelia Langford, John Sillitoe on behalf of the Wellcome Sanger Institute COVID-19 Surveillance Team |
| EPI_ISL_3454854 | Lighthouse Lab in Milton Keynes | Wellcome Sanger Institute for the COVID-19 Genomics UK (COG-UK) Consortium | The Lighthouse Lab in Milton Keynes and Alex Alderton, Roberto Amato, Jeffrey Barrett, Sonia Goncalves, Ewan Harrison, David K. Jackson, Ian Johnston, Dominic Kwiatkowski, Cordelia Langford, John Sillitoe on behalf of the Wellcome Sanger Institute COVID-19 Surveillance Team |
| EPI_ISL_3454855 | Lighthouse Lab in Alderley Park | Wellcome Sanger Institute for the COVID-19 Genomics UK (COG-UK) Consortium | Jacquelyn Wynn, Mairead Hyland, The Lighthouse Lab in Alderley Park and Alex Alderton, Roberto Amato, Jeffrey Barrett, Sonia Goncalves, Ewan Harrison, David K. Jackson, Ian Johnston, Dominic Kwiatkowski, Cordelia Langford, John Sillitoe on behalf of the Wellcome Sanger Institute COVID-19 Surveillance Team |
| EPI_ISL_3454856, EPI_ISL_3454857 | Lighthouse Lab in Milton Keynes | Wellcome Sanger Institute for the COVID-19 Genomics UK (COG-UK) Consortium | The Lighthouse Lab in Milton Keynes and Alex Alderton, Roberto Amato, Jeffrey Barrett, Sonia Goncalves, Ewan Harrison, David K. Jackson, Ian Johnston, Dominic Kwiatkowski, Cordelia Langford, John Sillitoe on behalf of the Wellcome Sanger Institute COVID-19 Surveillance Team |
| EPI_ISL_3454858, EPI_ISL_3454859, EPI_ISL_3454860 | Lighthouse Lab in Alderley Park | Wellcome Sanger Institute for the COVID-19 Genomics UK (COG-UK) Consortium | Jacquelyn Wynn, Mairead Hyland, The Lighthouse Lab in Alderley Park and Alex Alderton, Roberto Amato, Jeffrey Barrett, Sonia Goncalves, Ewan Harrison, David K. Jackson, Ian Johnston, Dominic Kwiatkowski, Cordelia Langford, John Sillitoe on behalf of the Wellcome Sanger Institute COVID-19 Surveillance Team |
| EPI_ISL_3454861 | Lighthouse Lab in Milton Keynes | Wellcome Sanger Institute for the COVID-19 Genomics UK (COG-UK) Consortium | The Lighthouse Lab in Milton Keynes and Alex Alderton, Roberto Amato, Jeffrey Barrett, Sonia Goncalves, Ewan Harrison, David K. Jackson, Ian Johnston, Dominic Kwiatkowski, Cordelia Langford, John Sillitoe on behalf of the Wellcome Sanger Institute COVID-19 Surveillance Team |
| EPI_ISL_3454862 | Berkshire and Surrey Pathology Services Lighthouse Laboratory | Wellcome Sanger Institute for the COVID-19 Genomics UK (COG-UK) Consortium | Berkshire and Surrey Pathology Services Lighthouse Laboratory and Alex Alderton, Roberto Amato, Jeffrey Barrett, Sonia Goncalves, Ewan Harrison, David K. Jackson, Ian Johnston, Dominic Kwiatkowski, Cordelia Langford, John Sillitoe on behalf of the Wellcome Sanger Institute COVID-19 Surveillance Team |
| EPI_ISL_3454863 | Lighthouse Lab in Milton Keynes | Wellcome Sanger Institute for the COVID-19 Genomics UK (COG-UK) Consortium | The Lighthouse Lab in Milton Keynes and Alex Alderton, Roberto Amato, Jeffrey Barrett, Sonia Goncalves, Ewan Harrison, David K. Jackson, Ian Johnston, Dominic Kwiatkowski, Cordelia Langford, John Sillitoe on behalf of the Wellcome Sanger Institute COVID-19 Surveillance Team |
| EPI_ISL_3454864 | Lighthouse Lab in Alderley Park | Wellcome Sanger Institute for the COVID-19 Genomics UK (COG-UK) Consortium | Jacquelyn Wynn, Mairead Hyland, The Lighthouse Lab in Alderley Park and Alex Alderton, Roberto Amato, Jeffrey Barrett, Sonia Goncalves, Ewan Harrison, David K. Jackson, Ian Johnston, Dominic Kwiatkowski, Cordelia Langford, John Sillitoe on behalf of the Wellcome Sanger Institute COVID-19 Surveillance Team |
| EPI_ISL_3454865, | Lighthouse Lab in Milton Keynes | Wellcome Sanger Institute for the COVID-19 Genomics UK (COG-UK) | The Lighthouse Lab in Milton Keynes and Alex Alderton, Roberto Amato, Jeffrey Barrett, Sonia Goncalves, Ewan Harrison, David K. Jackson, Ian Johnston, Dominic Kwiatkowski, Cordelia Langford, John Sillitoe on behalf of the Wellcome Sanger |

| | | | |
|---|---|---|---|
| EPI_ISL_3454866 | | Consortium | Institute COVID-19 Surveillance Team |
| EPI_ISL_3454867 | Lighthouse Lab in Alderley Park | Wellcome Sanger Institute for the COVID-19 Genomics UK (COG-UK) Consortium | Jacquelyn Wynn, Mairead Hyland, The Lighthouse Lab in Alderley Park and Alex Alderton, Roberto Amato, Jeffrey Barrett, Sonia Goncalves, Ewan Harrison, David K. Jackson, Ian Johnston, Dominic Kwiatkowski, Cordelia Langford, John Sillitoe on behalf of the Wellcome Sanger Institute COVID-19 Surveillance Team |
| EPI_ISL_3454868, EPI_ISL_3454869, EPI_ISL_3454870, EPI_ISL_3454871, EPI_ISL_3454872, EPI_ISL_3454873 | Lighthouse Lab in Milton Keynes | Wellcome Sanger Institute for the COVID-19 Genomics UK (COG-UK) Consortium | The Lighthouse Lab in Milton Keynes and Alex Alderton, Roberto Amato, Jeffrey Barrett, Sonia Goncalves, Ewan Harrison, David K. Jackson, Ian Johnston, Dominic Kwiatkowski, Cordelia Langford, John Sillitoe on behalf of the Wellcome Sanger Institute COVID-19 Surveillance Team |
| EPI_ISL_3454874, EPI_ISL_3454875 | Lighthouse Lab in Alderley Park | Wellcome Sanger Institute for the COVID-19 Genomics UK (COG-UK) Consortium | Jacquelyn Wynn, Mairead Hyland, The Lighthouse Lab in Alderley Park and Alex Alderton, Roberto Amato, Jeffrey Barrett, Sonia Goncalves, Ewan Harrison, David K. Jackson, Ian Johnston, Dominic Kwiatkowski, Cordelia Langford, John Sillitoe on behalf of the Wellcome Sanger Institute COVID-19 Surveillance Team |
| EPI_ISL_3454876, EPI_ISL_3454877, EPI_ISL_3454878, EPI_ISL_3454879, EPI_ISL_3454880 | Lighthouse Lab in Milton Keynes | Wellcome Sanger Institute for the COVID-19 Genomics UK (COG-UK) Consortium | The Lighthouse Lab in Milton Keynes and Alex Alderton, Roberto Amato, Jeffrey Barrett, Sonia Goncalves, Ewan Harrison, David K. Jackson, Ian Johnston, Dominic Kwiatkowski, Cordelia Langford, John Sillitoe on behalf of the Wellcome Sanger Institute COVID-19 Surveillance Team |
| EPI_ISL_3454881 | Lighthouse Lab in Alderley Park | Wellcome Sanger Institute for the COVID-19 Genomics UK (COG-UK) Consortium | Jacquelyn Wynn, Mairead Hyland, The Lighthouse Lab in Alderley Park and Alex Alderton, Roberto Amato, Jeffrey Barrett, Sonia Goncalves, Ewan Harrison, David K. Jackson, Ian Johnston, Dominic Kwiatkowski, Cordelia Langford, John Sillitoe on behalf of the Wellcome Sanger Institute COVID-19 Surveillance Team |
| EPI_ISL_3454882 | Lighthouse Lab in Milton Keynes | Wellcome Sanger Institute for the COVID-19 Genomics UK (COG-UK) Consortium | The Lighthouse Lab in Milton Keynes and Alex Alderton, Roberto Amato, Jeffrey Barrett, Sonia Goncalves, Ewan Harrison, David K. Jackson, Ian Johnston, Dominic Kwiatkowski, Cordelia Langford, John Sillitoe on behalf of the Wellcome Sanger Institute COVID-19 Surveillance Team |
| EPI_ISL_3454883 | Lighthouse Lab in Alderley Park | Wellcome Sanger Institute for the COVID-19 Genomics UK (COG-UK) Consortium | Jacquelyn Wynn, Mairead Hyland, The Lighthouse Lab in Alderley Park and Alex Alderton, Roberto Amato, Jeffrey Barrett, Sonia Goncalves, Ewan Harrison, David K. Jackson, Ian Johnston, Dominic Kwiatkowski, Cordelia Langford, John Sillitoe on behalf of the Wellcome Sanger Institute COVID-19 Surveillance Team |
| EPI_ISL_3454884, EPI_ISL_3454885, EPI_ISL_3454886, EPI_ISL_3454887, EPI_ISL_3454888, EPI_ISL_3454889, EPI_ISL_3454890, EPI_ISL_3454891, EPI_ISL_3454892 | Lighthouse Lab in Milton Keynes | Wellcome Sanger Institute for the COVID-19 Genomics UK (COG-UK) Consortium | The Lighthouse Lab in Milton Keynes and Alex Alderton, Roberto Amato, Jeffrey Barrett, Sonia Goncalves, Ewan Harrison, David K. Jackson, Ian Johnston, Dominic Kwiatkowski, Cordelia Langford, John Sillitoe on behalf of the Wellcome Sanger Institute COVID-19 Surveillance Team |
| EPI_ISL_3454893 | Lighthouse Lab in Alderley Park | Wellcome Sanger Institute for the COVID-19 Genomics UK (COG-UK) Consortium | Jacquelyn Wynn, Mairead Hyland, The Lighthouse Lab in Alderley Park and Alex Alderton, Roberto Amato, Jeffrey Barrett, Sonia Goncalves, Ewan Harrison, David K. Jackson, Ian Johnston, Dominic Kwiatkowski, Cordelia Langford, John Sillitoe on behalf of the Wellcome Sanger Institute COVID-19 Surveillance Team |
| EPI_ISL_3454894, EPI_ISL_3454895, EPI_ISL_3454896, EPI_ISL_3454897, EPI_ISL_3454898, EPI_ISL_3454899, EPI_ISL_3454900, EPI_ISL_3454901, EPI_ISL_3454902, EPI_ISL_3454903, EPI_ISL_3454904, EPI_ISL_3454905, EPI_ISL_3454906, EPI_ISL_3454907, EPI_ISL_3454908, EPI_ISL_3454909, EPI_ISL_3454910, EPI_ISL_3454911, EPI_ISL_3454912, EPI_ISL_3454913, EPI_ISL_3454914, EPI_ISL_3454915, EPI_ISL_3454916, EPI_ISL_3454917, EPI_ISL_3454918, EPI_ISL_3454919, EPI_ISL_3454920, EPI_ISL_3454921, EPI_ISL_3454922, EPI_ISL_3454923, EPI_ISL_3454924, EPI_ISL_3454925, EPI_ISL_3454926, EPI_ISL_3454927, EPI_ISL_3454928, EPI_ISL_3454929, EPI_ISL_3454930, EPI_ISL_3454931, EPI_ISL_3454932, EPI_ISL_3454933, EPI_ISL_3454934, EPI_ISL_3454935, EPI_ISL_3454936, EPI_ISL_3454937, EPI_ISL_3454938, EPI_ISL_3454939, EPI_ISL_3454940, EPI_ISL_3454941, EPI_ISL_3454942, EPI_ISL_3454943, EPI_ISL_3454944, EPI_ISL_3454945, EPI_ISL_3454946, EPI_ISL_3454947, EPI_ISL_3454948, EPI_ISL_3454949, EPI_ISL_3454950, EPI_ISL_3454951, EPI_ISL_3454952, EPI_ISL_3454953, EPI_ISL_3454954, EPI_ISL_3454955, EPI_ISL_3454956, EPI_ISL_3454957, EPI_ISL_3454958, EPI_ISL_3454959, EPI_ISL_3454960, EPI_ISL_3454961, EPI_ISL_3454962, EPI_ISL_3454963, EPI_ISL_3454964, EPI_ISL_3454965, EPI_ISL_3454966, EPI_ISL_3454967, EPI_ISL_3454968, EPI_ISL_3454969, EPI_ISL_3454970, EPI_ISL_3454971, EPI_ISL_3454972, EPI_ISL_3454973, EPI_ISL_3454974, EPI_ISL_3454975, EPI_ISL_3454976, EPI_ISL_3454977, EPI_ISL_3454978, EPI_ISL_3454979, EPI_ISL_3454980, EPI_ISL_3454981, EPI_ISL_3454982, EPI_ISL_3454983, EPI_ISL_3454984, EPI_ISL_3454985, EPI_ISL_3454986, EPI_ISL_3454987, EPI_ISL_3454988, EPI_ISL_3454989, EPI_ISL_3454990, EPI_ISL_3454991, EPI_ISL_3454992, EPI_ISL_3454993, EPI_ISL_3454994, EPI_ISL_3454995, EPI_ISL_3454996, EPI_ISL_3454997, EPI_ISL_3454998, EPI_ISL_3454999, EPI_ISL_3455000, EPI_ISL_3455001, EPI_ISL_3455002, EPI_ISL_3455003, EPI_ISL_3455004, EPI_ISL_3455005, EPI_ISL_3455006, EPI_ISL_3455007, EPI_ISL_3455008, EPI_ISL_3455009, EPI_ISL_3455010, EPI_ISL_3455011, EPI_ISL_3455012, EPI_ISL_3455013, EPI_ISL_3455014, EPI_ISL_3455015, EPI_ISL_3455016, EPI_ISL_3455017, EPI_ISL_3455018, EPI_ISL_3455019, EPI_ISL_3455020, EPI_ISL_3455021, EPI_ISL_3455022, EPI_ISL_3455023, EPI_ISL_3455024, EPI_ISL_3455025, EPI_ISL_3455026, EPI_ISL_3455027, EPI_ISL_3455028, EPI_ISL_3455029, EPI_ISL_3455030, EPI_ISL_3455031, EPI_ISL_3455032, EPI_ISL_3455033, EPI_ISL_3455034, EPI_ISL_3455035, EPI_ISL_3455036, EPI_ISL_3455037, EPI_ISL_3455038, EPI_ISL_3455039, EPI_ISL_3455040, EPI_ISL_3455041, EPI_ISL_3455042, EPI_ISL_3455043, EPI_ISL_3455044, EPI_ISL_3455045, EPI_ISL_3455046, EPI_ISL_3455047, EPI_ISL_3455048, EPI_ISL_3455049, EPI_ISL_3455050, EPI_ISL_3455051, EPI_ISL_3455052, EPI_ISL_3455053, EPI_ISL_3455054, EPI_ISL_3455055, EPI_ISL_3455056, EPI_ISL_3455057, EPI_ISL_3455058, EPI_ISL_3455059, EPI_ISL_3455060, EPI_ISL_3455061, EPI_ISL_3455062, EPI_ISL_3455063, EPI_ISL_3455064, EPI_ISL_3455065, EPI_ISL_3455066, EPI_ISL_3455067, EPI_ISL_3455068, EPI_ISL_3455069, EPI_ISL_3455070, EPI_ISL_3455071, EPI_ISL_3455072, EPI_ISL_3455073, EPI_ISL_3455074, EPI_ISL_3455075, EPI_ISL_3455076, EPI_ISL_3455077, EPI_ISL_3455078, EPI_ISL_3455079, EPI_ISL_3455080, EPI_ISL_3455081, EPI_ISL_3455082, EPI_ISL_3455083, EPI_ISL_3455084, EPI_ISL_3455085, EPI_ISL_3455086, EPI_ISL_3455087, EPI_ISL_3455088, EPI_ISL_3455089, EPI_ISL_3455090, EPI_ISL_3455091, EPI_ISL_3455092, EPI_ISL_3455093, EPI_ISL_3455094, EPI_ISL_3455095, EPI_ISL_3455096, EPI_ISL_3455097, EPI_ISL_3455098, EPI_ISL_3455099, EPI_ISL_3455100, EPI_ISL_3455101, EPI_ISL_3455102, EPI_ISL_3455103, EPI_ISL_3455104, EPI_ISL_3455105, EPI_ISL_3455106, EPI_ISL_3455107, EPI_ISL_3455108, EPI_ISL_3455109, EPI_ISL_3455110, EPI_ISL_3455111, EPI_ISL_3455112, EPI_ISL_3455113, EPI_ISL_3455114, EPI_ISL_3455115, EPI_ISL_3455116, EPI_ISL_3455117, EPI_ISL_3455118, EPI_ISL_3455119, EPI_ISL_3455120, EPI_ISL_3455121, EPI_ISL_3455122, EPI_ISL_3455123, EPI_ISL_3455124, EPI_ISL_3455125, EPI_ISL_3455126, EPI_ISL_3455127, EPI_ISL_3455128, EPI_ISL_3455129, EPI_ISL_3455130, EPI_ISL_3455131, EPI_ISL_3455132, EPI_ISL_3455133, EPI_ISL_3455134, EPI_ISL_3455135, EPI_ISL_3455136, EPI_ISL_3455137, EPI_ISL_3455138, EPI_ISL_3455139, EPI_ISL_3455140, EPI_ISL_3455141, EPI_ISL_3455142, EPI_ISL_3455143, EPI_ISL_3455144, EPI_ISL_3455145, EPI_ISL_3455146, EPI_ISL_3455147, EPI_ISL_3455148, EPI_ISL_3455149, EPI_ISL_3455150, EPI_ISL_3455151, EPI_ISL_3455152, EPI_ISL_3455153, EPI_ISL_3455154, EPI_ISL_3455155, EPI_ISL_3455156, EPI_ISL_3455157, EPI_ISL_3455158, EPI_ISL_3455159 | | | |
| see above | Lighthouse Lab in Milton Keynes | Wellcome Sanger Institute for the COVID-19 Genomics UK (COG-UK) Consortium | The Lighthouse Lab in Milton Keynes and Alex Alderton, Roberto Amato, Jeffrey Barrett, Sonia Goncalves, Ewan Harrison, David K. Jackson, Ian Johnston, Dominic Kwiatkowski, Cordelia Langford, John Sillitoe on behalf of the Wellcome Sanger Institute COVID-19 Surveillance Team |
| EPI_ISL_3455160 | Lighthouse Lab in Glasgow | Wellcome Sanger Institute for the COVID-19 Genomics UK (COG-UK) Consortium | Harper VanSteenhouse, Yumi Kasai, David Gray, Carol Clugston, Anna Dominiczak and Alex Alderton, Roberto Amato, Jeffrey Barrett, Sonia Goncalves, Ewan Harrison, David K. Jackson, Ian Johnston, Dominic Kwiatkowski, Cordelia Langford, John Sillitoe on behalf of the Wellcome Sanger Institute COVID-19 Surveillance Team |
| EPI_ISL_3455161, EPI_ISL_3455162, EPI_ISL_3455163, EPI_ISL_3455164, EPI_ISL_3455165, EPI_ISL_3455166, EPI_ISL_3455167 | Lighthouse Lab in Milton Keynes | Wellcome Sanger Institute for the COVID-19 Genomics UK (COG-UK) Consortium | The Lighthouse Lab in Milton Keynes and Alex Alderton, Roberto Amato, Jeffrey Barrett, Sonia Goncalves, Ewan Harrison, David K. Jackson, Ian Johnston, Dominic Kwiatkowski, Cordelia Langford, John Sillitoe on behalf of the Wellcome Sanger Institute COVID-19 Surveillance Team |
| EPI_ISL_3455168 | Lighthouse Lab in Alderley Park | Wellcome Sanger Institute for the COVID-19 Genomics UK (COG-UK) Consortium | Jacquelyn Wynn, Mairead Hyland, The Lighthouse Lab in Alderley Park and Alex Alderton, Roberto Amato, Jeffrey Barrett, Sonia Goncalves, Ewan Harrison, David K. Jackson, Ian Johnston, Dominic Kwiatkowski, Cordelia Langford, John Sillitoe on behalf of the Wellcome Sanger Institute COVID-19 Surveillance Team |
| EPI_ISL_3455169, EPI_ISL_3455170, EPI_ISL_3455171 | Lighthouse Lab in Milton Keynes | Wellcome Sanger Institute for the COVID-19 Genomics UK (COG-UK) Consortium | The Lighthouse Lab in Milton Keynes and Alex Alderton, Roberto Amato, Jeffrey Barrett, Sonia Goncalves, Ewan Harrison, David K. Jackson, Ian Johnston, Dominic Kwiatkowski, Cordelia Langford, John Sillitoe on behalf of the Wellcome Sanger Institute COVID-19 Surveillance Team |
| EPI_ISL_3455172 | Lighthouse Lab in Glasgow | Wellcome Sanger Institute for the COVID-19 Genomics UK (COG-UK) Consortium | Harper VanSteenhouse, Yumi Kasai, David Gray, Carol Clugston, Anna Dominiczak and Alex Alderton, Roberto Amato, Jeffrey Barrett, Sonia Goncalves, Ewan Harrison, David K. Jackson, Ian Johnston, Dominic Kwiatkowski, Cordelia Langford, John Sillitoe on behalf of the Wellcome Sanger Institute COVID-19 Surveillance Team |
| EPI_ISL_3455173, EPI_ISL_3455174, EPI_ISL_3455175 | Lighthouse Lab in Milton Keynes | Wellcome Sanger Institute for the COVID-19 Genomics UK (COG-UK) Consortium | The Lighthouse Lab in Milton Keynes and Alex Alderton, Roberto Amato, Jeffrey Barrett, Sonia Goncalves, Ewan Harrison, David K. Jackson, Ian Johnston, Dominic Kwiatkowski, Cordelia Langford, John Sillitoe on behalf of the Wellcome Sanger Institute COVID-19 Surveillance Team |
| EPI_ISL_3455176 | Lighthouse Lab in Glasgow | Wellcome Sanger Institute for the COVID-19 Genomics UK (COG-UK) Consortium | Harper VanSteenhouse, Yumi Kasai, David Gray, Carol Clugston, Anna Dominiczak and Alex Alderton, Roberto Amato, Jeffrey Barrett, Sonia Goncalves, Ewan Harrison, David K. Jackson, Ian Johnston, Dominic Kwiatkowski, Cordelia Langford, John Sillitoe on behalf of the Wellcome Sanger Institute COVID-19 Surveillance Team |
| EPI_ISL_3455177, EPI_ISL_3455178, EPI_ISL_3455179, EPI_ISL_3455180 | Lighthouse Lab in Milton Keynes | Wellcome Sanger Institute for the COVID-19 Genomics UK (COG-UK) Consortium | The Lighthouse Lab in Milton Keynes and Alex Alderton, Roberto Amato, Jeffrey Barrett, Sonia Goncalves, Ewan Harrison, David K. Jackson, Ian Johnston, Dominic Kwiatkowski, Cordelia Langford, John Sillitoe on behalf of the Wellcome Sanger Institute COVID-19 Surveillance Team |
| EPI_ISL_3455181 | Lighthouse Lab in Glasgow | Wellcome Sanger Institute for the COVID-19 Genomics UK (COG-UK) Consortium | Harper VanSteenhouse, Yumi Kasai, David Gray, Carol Clugston, Anna Dominiczak and Alex Alderton, Roberto Amato, Jeffrey Barrett, Sonia Goncalves, Ewan Harrison, David K. Jackson, Ian Johnston, Dominic Kwiatkowski, Cordelia Langford, John Sillitoe on behalf of the Wellcome Sanger Institute COVID-19 Surveillance Team |
| EPI_ISL_3455182, EPI_ISL_3455183 | Lighthouse Lab in Milton Keynes | Wellcome Sanger Institute for the COVID-19 Genomics UK (COG-UK) Consortium | The Lighthouse Lab in Milton Keynes and Alex Alderton, Roberto Amato, Jeffrey Barrett, Sonia Goncalves, Ewan Harrison, David K. Jackson, Ian Johnston, Dominic Kwiatkowski, Cordelia Langford, John Sillitoe on behalf of the Wellcome Sanger Institute COVID-19 Surveillance Team |
| EPI_ISL_3455184 | Lighthouse Lab in Glasgow | Wellcome Sanger Institute for the COVID-19 Genomics UK (COG-UK) Consortium | Harper VanSteenhouse, Yumi Kasai, David Gray, Carol Clugston, Anna Dominiczak and Alex Alderton, Roberto Amato, Jeffrey Barrett, Sonia Goncalves, Ewan Harrison, David K. Jackson, Ian Johnston, Dominic Kwiatkowski, Cordelia Langford, John Sillitoe on behalf of the Wellcome Sanger Institute COVID-19 Surveillance Team |
| EPI_ISL_3455185, EPI_ISL_3455186, EPI_ISL_3455187 | Lighthouse Lab in Alderley Park | Wellcome Sanger Institute for the COVID-19 Genomics UK (COG-UK) Consortium | Jacquelyn Wynn, Mairead Hyland, The Lighthouse Lab in Alderley Park and Alex Alderton, Roberto Amato, Jeffrey Barrett, Sonia Goncalves, Ewan Harrison, David K. Jackson, Ian Johnston, Dominic Kwiatkowski, Cordelia Langford, John Sillitoe on behalf of the Wellcome Sanger Institute COVID-19 Surveillance Team |
| EPI_ISL_3455188 | Lighthouse Lab in Milton Keynes | Wellcome Sanger Institute for the COVID-19 Genomics UK (COG-UK) Consortium | The Lighthouse Lab in Milton Keynes and Alex Alderton, Roberto Amato, Jeffrey Barrett, Sonia Goncalves, Ewan Harrison, David K. Jackson, Ian Johnston, Dominic Kwiatkowski, Cordelia Langford, John Sillitoe on behalf of the Wellcome Sanger Institute COVID-19 Surveillance Team |
| EPI_ISL_3455189 | Lighthouse Lab in Alderley Park | Wellcome Sanger Institute for the COVID-19 Genomics UK (COG-UK) Consortium | Jacquelyn Wynn, Mairead Hyland, The Lighthouse Lab in Alderley Park and Alex Alderton, Roberto Amato, Jeffrey Barrett, Sonia Goncalves, Ewan Harrison, David K. Jackson, Ian Johnston, Dominic Kwiatkowski, Cordelia Langford, John Sillitoe on behalf of the Wellcome Sanger Institute COVID-19 Surveillance Team |
| EPI_ISL_3455190 | Lighthouse Lab in Milton Keynes | Wellcome Sanger Institute for the COVID-19 Genomics UK (COG-UK) Consortium | The Lighthouse Lab in Milton Keynes and Alex Alderton, Roberto Amato, Jeffrey Barrett, Sonia Goncalves, Ewan Harrison, David K. Jackson, Ian Johnston, Dominic Kwiatkowski, Cordelia Langford, John Sillitoe on behalf of the Wellcome Sanger Institute COVID-19 Surveillance Team |
| EPI_ISL_3455191, EPI_ISL_3455192, EPI_ISL_3455193 | Berkshire and Surrey Pathology Services Lighthouse Laboratory | Wellcome Sanger Institute for the COVID-19 Genomics UK (COG-UK) Consortium | Berkshire and Surrey Pathology Services Lighthouse Laboratory and Alex Alderton, Roberto Amato, Jeffrey Barrett, Sonia Goncalves, Ewan Harrison, David K. Jackson, Ian Johnston, Dominic Kwiatkowski, Cordelia Langford, John Sillitoe on behalf of the Wellcome Sanger Institute COVID-19 Surveillance Team |
| EPI_ISL_3455194 | Lighthouse Lab in Alderley Park | Wellcome Sanger Institute for the COVID-19 Genomics UK (COG-UK) Consortium | Jacquelyn Wynn, Mairead Hyland, The Lighthouse Lab in Alderley Park and Alex Alderton, Roberto Amato, Jeffrey Barrett, Sonia Goncalves, Ewan Harrison, David K. Jackson, Ian Johnston, Dominic Kwiatkowski, Cordelia Langford, John Sillitoe on behalf of the Wellcome Sanger Institute COVID-19 Surveillance Team |
| EPI_ISL_3455195 | Berkshire and Surrey Pathology Services Lighthouse Laboratory | Wellcome Sanger Institute for the COVID-19 Genomics UK (COG-UK) Consortium | Berkshire and Surrey Pathology Services Lighthouse Laboratory and Alex Alderton, Roberto Amato, Jeffrey Barrett, Sonia Goncalves, Ewan Harrison, David K. Jackson, Ian Johnston, Dominic Kwiatkowski, Cordelia Langford, John Sillitoe on behalf of the Wellcome Sanger Institute COVID-19 Surveillance Team |
| EPI_ISL_3455196, | Lighthouse Lab in Alderley Park | Wellcome Sanger Institute for the COVID-19 Genomics UK (COG-UK) | Jacquelyn Wynn, Mairead Hyland, The Lighthouse Lab in Alderley Park and Alex Alderton, Roberto Amato, Jeffrey Barrett, Sonia Goncalves, Ewan Harrison, David K. Jackson, Ian Johnston, Dominic Kwiatkowski, Cordelia Langford, John Sillitoe on |

| | | | |
|---|---|---|---|
| EPI_ISL_3455197, EPI_ISL_3455198 | | Consortium | behalf of the Wellcome Sanger Institute COVID-19 Surveillance Team |
| EPI_ISL_3455199, EPI_ISL_3455200, EPI_ISL_3455201, EPI_ISL_3455202 | Lighthouse Lab in Glasgow | Wellcome Sanger Institute for the COVID-19 Genomics UK (COG-UK) Consortium | Harper VanSteenhouse, Yumi Kasai, David Gray, Carol Clugston, Anna Dominiczak and Alex Alderton, Roberto Amato, Jeffrey Barrett, Sonia Goncalves, Ewan Harrison, David K. Jackson, Ian Johnston, Dominic Kwiatkowski, Cordelia Langford, John Sillitoe on behalf of the Wellcome Sanger Institute COVID-19 Surveillance Team |
| EPI_ISL_3455203 | Lighthouse Lab in Alderley Park | Wellcome Sanger Institute for the COVID-19 Genomics UK (COG-UK) Consortium | Jacquelyn Wynn, Mairead Hyland, The Lighthouse Lab in Alderley Park and Alex Alderton, Roberto Amato, Jeffrey Barrett, Sonia Goncalves, Ewan Harrison, David K. Jackson, Ian Johnston, Dominic Kwiatkowski, Cordelia Langford, John Sillitoe on behalf of the Wellcome Sanger Institute COVID-19 Surveillance Team |
| EPI_ISL_3455204, EPI_ISL_3455205, EPI_ISL_3455206 | Lighthouse Lab in Glasgow | Wellcome Sanger Institute for the COVID-19 Genomics UK (COG-UK) Consortium | Harper VanSteenhouse, Yumi Kasai, David Gray, Carol Clugston, Anna Dominiczak and Alex Alderton, Roberto Amato, Jeffrey Barrett, Sonia Goncalves, Ewan Harrison, David K. Jackson, Ian Johnston, Dominic Kwiatkowski, Cordelia Langford, John Sillitoe on behalf of the Wellcome Sanger Institute COVID-19 Surveillance Team |
| EPI_ISL_3455207 | Lighthouse Lab in Alderley Park | Wellcome Sanger Institute for the COVID-19 Genomics UK (COG-UK) Consortium | Jacquelyn Wynn, Mairead Hyland, The Lighthouse Lab in Alderley Park and Alex Alderton, Roberto Amato, Jeffrey Barrett, Sonia Goncalves, Ewan Harrison, David K. Jackson, Ian Johnston, Dominic Kwiatkowski, Cordelia Langford, John Sillitoe on behalf of the Wellcome Sanger Institute COVID-19 Surveillance Team |
| EPI_ISL_3455208, EPI_ISL_3455209 | Lighthouse Lab in Glasgow | Wellcome Sanger Institute for the COVID-19 Genomics UK (COG-UK) Consortium | Harper VanSteenhouse, Yumi Kasai, David Gray, Carol Clugston, Anna Dominiczak and Alex Alderton, Roberto Amato, Jeffrey Barrett, Sonia Goncalves, Ewan Harrison, David K. Jackson, Ian Johnston, Dominic Kwiatkowski, Cordelia Langford, John Sillitoe on behalf of the Wellcome Sanger Institute COVID-19 Surveillance Team |
| EPI_ISL_3455210 | Lighthouse Lab in Alderley Park | Wellcome Sanger Institute for the COVID-19 Genomics UK (COG-UK) Consortium | Jacquelyn Wynn, Mairead Hyland, The Lighthouse Lab in Alderley Park and Alex Alderton, Roberto Amato, Jeffrey Barrett, Sonia Goncalves, Ewan Harrison, David K. Jackson, Ian Johnston, Dominic Kwiatkowski, Cordelia Langford, John Sillitoe on behalf of the Wellcome Sanger Institute COVID-19 Surveillance Team |
| EPI_ISL_3455211, EPI_ISL_3455212, EPI_ISL_3455213, EPI_ISL_3455214, EPI_ISL_3455215, EPI_ISL_3455216, EPI_ISL_3455217, EPI_ISL_3455218, EPI_ISL_3455219 | Lighthouse Lab in Glasgow | Wellcome Sanger Institute for the COVID-19 Genomics UK (COG-UK) Consortium | Harper VanSteenhouse, Yumi Kasai, David Gray, Carol Clugston, Anna Dominiczak and Alex Alderton, Roberto Amato, Jeffrey Barrett, Sonia Goncalves, Ewan Harrison, David K. Jackson, Ian Johnston, Dominic Kwiatkowski, Cordelia Langford, John Sillitoe on behalf of the Wellcome Sanger Institute COVID-19 Surveillance Team |
| EPI_ISL_3455220 | Lighthouse Lab in Alderley Park | Wellcome Sanger Institute for the COVID-19 Genomics UK (COG-UK) Consortium | Jacquelyn Wynn, Mairead Hyland, The Lighthouse Lab in Alderley Park and Alex Alderton, Roberto Amato, Jeffrey Barrett, Sonia Goncalves, Ewan Harrison, David K. Jackson, Ian Johnston, Dominic Kwiatkowski, Cordelia Langford, John Sillitoe on behalf of the Wellcome Sanger Institute COVID-19 Surveillance Team |
| EPI_ISL_3455221 | Berkshire and Surrey Pathology Services Lighthouse Laboratory | Wellcome Sanger Institute for the COVID-19 Genomics UK (COG-UK) Consortium | Berkshire and Surrey Pathology Services Lighthouse Laboratory and Alex Alderton, Roberto Amato, Jeffrey Barrett, Sonia Goncalves, Ewan Harrison, David K. Jackson, Ian Johnston, Dominic Kwiatkowski, Cordelia Langford, John Sillitoe on behalf of the Wellcome Sanger Institute COVID-19 Surveillance Team |
| EPI_ISL_3455222 | Health Services Laboratories | Wellcome Sanger Institute for the COVID-19 Genomics UK (COG-UK) Consortium | Health Services Laboratories and Alex Alderton, Roberto Amato, Jeffrey Barrett, Sonia Goncalves, Ewan Harrison, David K. Jackson, Ian Johnston, Dominic Kwiatkowski, Cordelia Langford, John Sillitoe on behalf of the Wellcome Sanger Institute COVID-19 Surveillance Team |
| EPI_ISL_3455223 | Lighthouse Lab in Alderley Park | Wellcome Sanger Institute for the COVID-19 Genomics UK (COG-UK) Consortium | Jacquelyn Wynn, Mairead Hyland, The Lighthouse Lab in Alderley Park and Alex Alderton, Roberto Amato, Jeffrey Barrett, Sonia Goncalves, Ewan Harrison, David K. Jackson, Ian Johnston, Dominic Kwiatkowski, Cordelia Langford, John Sillitoe on behalf of the Wellcome Sanger Institute COVID-19 Surveillance Team |
| EPI_ISL_3455224 | Berkshire and Surrey Pathology Services Lighthouse Laboratory | Wellcome Sanger Institute for the COVID-19 Genomics UK (COG-UK) Consortium | Berkshire and Surrey Pathology Services Lighthouse Laboratory and Alex Alderton, Roberto Amato, Jeffrey Barrett, Sonia Goncalves, Ewan Harrison, David K. Jackson, Ian Johnston, Dominic Kwiatkowski, Cordelia Langford, John Sillitoe on behalf of the Wellcome Sanger Institute COVID-19 Surveillance Team |
| EPI_ISL_3455225 | Health Services Laboratories | Wellcome Sanger Institute for the COVID-19 Genomics UK (COG-UK) Consortium | Health Services Laboratories and Alex Alderton, Roberto Amato, Jeffrey Barrett, Sonia Goncalves, Ewan Harrison, David K. Jackson, Ian Johnston, Dominic Kwiatkowski, Cordelia Langford, John Sillitoe on behalf of the Wellcome Sanger Institute COVID-19 Surveillance Team |
| EPI_ISL_3455226, EPI_ISL_3455227 | Lighthouse Lab in Alderley Park | Wellcome Sanger Institute for the COVID-19 Genomics UK (COG-UK) Consortium | Jacquelyn Wynn, Mairead Hyland, The Lighthouse Lab in Alderley Park and Alex Alderton, Roberto Amato, Jeffrey Barrett, Sonia Goncalves, Ewan Harrison, David K. Jackson, Ian Johnston, Dominic Kwiatkowski, Cordelia Langford, John Sillitoe on behalf of the Wellcome Sanger Institute COVID-19 Surveillance Team |
| EPI_ISL_3455228 | Health Services Laboratories | Wellcome Sanger Institute for the COVID-19 Genomics UK (COG-UK) Consortium | Health Services Laboratories and Alex Alderton, Roberto Amato, Jeffrey Barrett, Sonia Goncalves, Ewan Harrison, David K. Jackson, Ian Johnston, Dominic Kwiatkowski, Cordelia Langford, John Sillitoe on behalf of the Wellcome Sanger Institute COVID-19 Surveillance Team |
| EPI_ISL_3455229 | Berkshire and Surrey Pathology Services Lighthouse Laboratory | Wellcome Sanger Institute for the COVID-19 Genomics UK (COG-UK) Consortium | Berkshire and Surrey Pathology Services Lighthouse Laboratory and Alex Alderton, Roberto Amato, Jeffrey Barrett, Sonia Goncalves, Ewan Harrison, David K. Jackson, Ian Johnston, Dominic Kwiatkowski, Cordelia Langford, John Sillitoe on behalf of the Wellcome Sanger Institute COVID-19 Surveillance Team |
| EPI_ISL_3455230, EPI_ISL_3455231 | Health Services Laboratories | Wellcome Sanger Institute for the COVID-19 Genomics UK (COG-UK) Consortium | Health Services Laboratories and Alex Alderton, Roberto Amato, Jeffrey Barrett, Sonia Goncalves, Ewan Harrison, David K. Jackson, Ian Johnston, Dominic Kwiatkowski, Cordelia Langford, John Sillitoe on behalf of the Wellcome Sanger Institute COVID-19 Surveillance Team |
| EPI_ISL_3455232 | Berkshire and Surrey Pathology Services Lighthouse Laboratory | Wellcome Sanger Institute for the COVID-19 Genomics UK (COG-UK) Consortium | Berkshire and Surrey Pathology Services Lighthouse Laboratory and Alex Alderton, Roberto Amato, Jeffrey Barrett, Sonia Goncalves, Ewan Harrison, David K. Jackson, Ian Johnston, Dominic Kwiatkowski, Cordelia Langford, John Sillitoe on behalf of the Wellcome Sanger Institute COVID-19 Surveillance Team |
| EPI_ISL_3455233 | Health Services Laboratories | Wellcome Sanger Institute for the COVID-19 Genomics UK (COG-UK) Consortium | Health Services Laboratories and Alex Alderton, Roberto Amato, Jeffrey Barrett, Sonia Goncalves, Ewan Harrison, David K. Jackson, Ian Johnston, Dominic Kwiatkowski, Cordelia Langford, John Sillitoe on behalf of the Wellcome Sanger Institute COVID-19 Surveillance Team |
| EPI_ISL_3455234 | Lighthouse Lab in Alderley Park | Wellcome Sanger Institute for the COVID-19 Genomics UK (COG-UK) Consortium | Jacquelyn Wynn, Mairead Hyland, The Lighthouse Lab in Alderley Park and Alex Alderton, Roberto Amato, Jeffrey Barrett, Sonia Goncalves, Ewan Harrison, David K. Jackson, Ian Johnston, Dominic Kwiatkowski, Cordelia Langford, John Sillitoe on behalf of the Wellcome Sanger Institute COVID-19 Surveillance Team |
| EPI_ISL_3455235, EPI_ISL_3455236, EPI_ISL_3455237, EPI_ISL_3455238, EPI_ISL_3455239, EPI_ISL_3455240, EPI_ISL_3455241 | Health Services Laboratories | Wellcome Sanger Institute for the COVID-19 Genomics UK (COG-UK) Consortium | Health Services Laboratories and Alex Alderton, Roberto Amato, Jeffrey Barrett, Sonia Goncalves, Ewan Harrison, David K. Jackson, Ian Johnston, Dominic Kwiatkowski, Cordelia Langford, John Sillitoe on behalf of the Wellcome Sanger Institute COVID-19 Surveillance Team |
| EPI_ISL_3455242 | Lighthouse Laboratory Plymouth | Wellcome Sanger Institute for the COVID-19 Genomics UK (COG-UK) Consortium | Lighthouse Laboratory Plymouth and Alex Alderton, Roberto Amato, Jeffrey Barrett, Sonia Goncalves, Ewan Harrison, David K. Jackson, Ian Johnston, Dominic Kwiatkowski, Cordelia Langford, John Sillitoe on behalf of the Wellcome Sanger Institute COVID-19 Surveillance Team |
| EPI_ISL_3455243, EPI_ISL_3455244, EPI_ISL_3455245, EPI_ISL_3455246, EPI_ISL_3455247, EPI_ISL_3455248, EPI_ISL_3455249 | Health Services Laboratories | Wellcome Sanger Institute for the COVID-19 Genomics UK (COG-UK) Consortium | Health Services Laboratories and Alex Alderton, Roberto Amato, Jeffrey Barrett, Sonia Goncalves, Ewan Harrison, David K. Jackson, Ian Johnston, Dominic Kwiatkowski, Cordelia Langford, John Sillitoe on behalf of the Wellcome Sanger Institute COVID-19 Surveillance Team |
| EPI_ISL_3455250 | Berkshire and Surrey Pathology Services Lighthouse Laboratory | Wellcome Sanger Institute for the COVID-19 Genomics UK (COG-UK) Consortium | Berkshire and Surrey Pathology Services Lighthouse Laboratory and Alex Alderton, Roberto Amato, Jeffrey Barrett, Sonia Goncalves, Ewan Harrison, David K. Jackson, Ian Johnston, Dominic Kwiatkowski, Cordelia Langford, John Sillitoe on behalf of the Wellcome Sanger Institute COVID-19 Surveillance Team |
| EPI_ISL_3455251, EPI_ISL_3455252 | Health Services Laboratories | Wellcome Sanger Institute for the COVID-19 Genomics UK (COG-UK) Consortium | Health Services Laboratories and Alex Alderton, Roberto Amato, Jeffrey Barrett, Sonia Goncalves, Ewan Harrison, David K. Jackson, Ian Johnston, Dominic Kwiatkowski, Cordelia Langford, John Sillitoe on behalf of the Wellcome Sanger Institute COVID-19 Surveillance Team |
| EPI_ISL_3455253, EPI_ISL_3455254 | Lighthouse Lab in Alderley Park | Wellcome Sanger Institute for the COVID-19 Genomics UK (COG-UK) Consortium | Jacquelyn Wynn, Mairead Hyland, The Lighthouse Lab in Alderley Park and Alex Alderton, Roberto Amato, Jeffrey Barrett, Sonia Goncalves, Ewan Harrison, David K. Jackson, Ian Johnston, Dominic Kwiatkowski, Cordelia Langford, John Sillitoe on behalf of the Wellcome Sanger Institute COVID-19 Surveillance Team |
| EPI_ISL_3455255, EPI_ISL_3455256, EPI_ISL_3455257 | Berkshire and Surrey Pathology Services Lighthouse Laboratory | Wellcome Sanger Institute for the COVID-19 Genomics UK (COG-UK) Consortium | Berkshire and Surrey Pathology Services Lighthouse Laboratory and Alex Alderton, Roberto Amato, Jeffrey Barrett, Sonia Goncalves, Ewan Harrison, David K. Jackson, Ian Johnston, Dominic Kwiatkowski, Cordelia Langford, John Sillitoe on behalf of the Wellcome Sanger Institute COVID-19 Surveillance Team |
| EPI_ISL_3455258, EPI_ISL_3455259 | Lighthouse Lab in Milton Keynes | Wellcome Sanger Institute for the COVID-19 Genomics UK (COG-UK) Consortium | The Lighthouse Lab in Milton Keynes and Alex Alderton, Roberto Amato, Jeffrey Barrett, Sonia Goncalves, Ewan Harrison, David K. Jackson, Ian Johnston, Dominic Kwiatkowski, Cordelia Langford, John Sillitoe on behalf of the Wellcome Sanger Institute COVID-19 Surveillance Team |
| EPI_ISL_3455260 | Lighthouse Lab in Alderley Park | Wellcome Sanger Institute for the COVID-19 Genomics UK (COG-UK) Consortium | Jacquelyn Wynn, Mairead Hyland, The Lighthouse Lab in Alderley Park and Alex Alderton, Roberto Amato, Jeffrey Barrett, Sonia Goncalves, Ewan Harrison, David K. Jackson, Ian Johnston, Dominic Kwiatkowski, Cordelia Langford, John Sillitoe on behalf of the Wellcome Sanger Institute COVID-19 Surveillance Team |
| EPI_ISL_3455261 | Lighthouse Lab in Milton Keynes | Wellcome Sanger Institute for the COVID-19 Genomics UK (COG-UK) Consortium | The Lighthouse Lab in Milton Keynes and Alex Alderton, Roberto Amato, Jeffrey Barrett, Sonia Goncalves, Ewan Harrison, David K. Jackson, Ian Johnston, Dominic Kwiatkowski, Cordelia Langford, John Sillitoe on behalf of the Wellcome Sanger Institute COVID-19 Surveillance Team |
| EPI_ISL_3455262 | Lighthouse Lab in Alderley Park | Wellcome Sanger Institute for the COVID-19 Genomics UK (COG-UK) Consortium | Jacquelyn Wynn, Mairead Hyland, The Lighthouse Lab in Alderley Park and Alex Alderton, Roberto Amato, Jeffrey Barrett, Sonia Goncalves, Ewan Harrison, David K. Jackson, Ian Johnston, Dominic Kwiatkowski, Cordelia Langford, John Sillitoe on behalf of the Wellcome Sanger Institute COVID-19 Surveillance Team |
| EPI_ISL_3455263 | Lighthouse Lab in Milton Keynes | Wellcome Sanger Institute for the COVID-19 Genomics UK (COG-UK) Consortium | The Lighthouse Lab in Milton Keynes and Alex Alderton, Roberto Amato, Jeffrey Barrett, Sonia Goncalves, Ewan Harrison, David K. Jackson, Ian Johnston, Dominic Kwiatkowski, Cordelia Langford, John Sillitoe on behalf of the Wellcome Sanger Institute COVID-19 Surveillance Team |
| EPI_ISL_3455264 | Lighthouse Lab in Alderley Park | Wellcome Sanger Institute for the COVID-19 Genomics UK (COG-UK) Consortium | Jacquelyn Wynn, Mairead Hyland, The Lighthouse Lab in Alderley Park and Alex Alderton, Roberto Amato, Jeffrey Barrett, Sonia Goncalves, Ewan Harrison, David K. Jackson, Ian Johnston, Dominic Kwiatkowski, Cordelia Langford, John Sillitoe on behalf of the Wellcome Sanger Institute COVID-19 Surveillance Team |
| EPI_ISL_3455265, EPI_ISL_3455266, EPI_ISL_3455267, EPI_ISL_3455268 | Lighthouse Lab in Milton Keynes | Wellcome Sanger Institute for the COVID-19 Genomics UK (COG-UK) Consortium | The Lighthouse Lab in Milton Keynes and Alex Alderton, Roberto Amato, Jeffrey Barrett, Sonia Goncalves, Ewan Harrison, David K. Jackson, Ian Johnston, Dominic Kwiatkowski, Cordelia Langford, John Sillitoe on behalf of the Wellcome Sanger Institute COVID-19 Surveillance Team |
| EPI_ISL_3455269, | Lighthouse Lab in Alderley Park | Wellcome Sanger Institute for the COVID-19 Genomics UK (COG-UK) | Jacquelyn Wynn, Mairead Hyland, The Lighthouse Lab in Alderley Park and Alex Alderton, Roberto Amato, Jeffrey Barrett, Sonia Goncalves, Ewan Harrison, David K. Jackson, Ian Johnston, Dominic Kwiatkowski, Cordelia Langford, John Sillitoe on |

| | | | |
|---|---|---|---|
| EPI_ISL_3455270 | | Consortium | behalf of the Wellcome Sanger Institute COVID-19 Surveillance Team |
| EPI_ISL_3455271, EPI_ISL_3455272, EPI_ISL_3455273, EPI_ISL_3455274, EPI_ISL_3455275, EPI_ISL_3455276, EPI_ISL_3455277, EPI_ISL_3455278, EPI_ISL_3455279, EPI_ISL_3455280, EPI_ISL_3455281 | | | |
| see above | Lighthouse Lab in Milton Keynes | Wellcome Sanger Institute for the COVID-19 Genomics UK (COG-UK) Consortium | The Lighthouse Lab in Milton Keynes and Alex Alderton, Roberto Amato, Jeffrey Barrett, Sonia Goncalves, Ewan Harrison, David K. Jackson, Ian Johnston, Dominic Kwiatkowski, Cordelia Langford, John Sillitoe on behalf of the Wellcome Sanger Institute COVID-19 Surveillance Team |
| EPI_ISL_3455282 | Lighthouse Lab in Glasgow | Wellcome Sanger Institute for the COVID-19 Genomics UK (COG-UK) Consortium | Harper VanSteenhouse, Yumi Kasai, David Gray, Carol Clugston, Anna Dominiczak and Alex Alderton, Roberto Amato, Jeffrey Barrett, Sonia Goncalves, Ewan Harrison, David K. Jackson, Ian Johnston, Dominic Kwiatkowski, Cordelia Langford, John Sillitoe on behalf of the Wellcome Sanger Institute COVID-19 Surveillance Team |
| EPI_ISL_3455283, EPI_ISL_3455284, EPI_ISL_3455285 | Lighthouse Lab in Milton Keynes | Wellcome Sanger Institute for the COVID-19 Genomics UK (COG-UK) Consortium | The Lighthouse Lab in Milton Keynes and Alex Alderton, Roberto Amato, Jeffrey Barrett, Sonia Goncalves, Ewan Harrison, David K. Jackson, Ian Johnston, Dominic Kwiatkowski, Cordelia Langford, John Sillitoe on behalf of the Wellcome Sanger Institute COVID-19 Surveillance Team |
| EPI_ISL_3455286, EPI_ISL_3455287, EPI_ISL_3455288, EPI_ISL_3455289, EPI_ISL_3455290, EPI_ISL_3455291 | Lighthouse Lab in Glasgow | Wellcome Sanger Institute for the COVID-19 Genomics UK (COG-UK) Consortium | Harper VanSteenhouse, Yumi Kasai, David Gray, Carol Clugston, Anna Dominiczak and Alex Alderton, Roberto Amato, Jeffrey Barrett, Sonia Goncalves, Ewan Harrison, David K. Jackson, Ian Johnston, Dominic Kwiatkowski, Cordelia Langford, John Sillitoe on behalf of the Wellcome Sanger Institute COVID-19 Surveillance Team |
| EPI_ISL_3455292, EPI_ISL_3455293 | Lighthouse Lab in Alderley Park | Wellcome Sanger Institute for the COVID-19 Genomics UK (COG-UK) Consortium | Jacquelyn Wynn, Mairead Hyland, The Lighthouse Lab in Alderley Park and Alex Alderton, Roberto Amato, Jeffrey Barrett, Sonia Goncalves, Ewan Harrison, David K. Jackson, Ian Johnston, Dominic Kwiatkowski, Cordelia Langford, John Sillitoe on behalf of the Wellcome Sanger Institute COVID-19 Surveillance Team |
| EPI_ISL_3455294 | Lighthouse Lab in Glasgow | Wellcome Sanger Institute for the COVID-19 Genomics UK (COG-UK) Consortium | Harper VanSteenhouse, Yumi Kasai, David Gray, Carol Clugston, Anna Dominiczak and Alex Alderton, Roberto Amato, Jeffrey Barrett, Sonia Goncalves, Ewan Harrison, David K. Jackson, Ian Johnston, Dominic Kwiatkowski, Cordelia Langford, John Sillitoe on behalf of the Wellcome Sanger Institute COVID-19 Surveillance Team |
| EPI_ISL_3455295 | Lighthouse Lab in Alderley Park | Wellcome Sanger Institute for the COVID-19 Genomics UK (COG-UK) Consortium | Jacquelyn Wynn, Mairead Hyland, The Lighthouse Lab in Alderley Park and Alex Alderton, Roberto Amato, Jeffrey Barrett, Sonia Goncalves, Ewan Harrison, David K. Jackson, Ian Johnston, Dominic Kwiatkowski, Cordelia Langford, John Sillitoe on behalf of the Wellcome Sanger Institute COVID-19 Surveillance Team |
| EPI_ISL_3455296, EPI_ISL_3455297, EPI_ISL_3455298, EPI_ISL_3455299, EPI_ISL_3455300 | Lighthouse Lab in Glasgow | Wellcome Sanger Institute for the COVID-19 Genomics UK (COG-UK) Consortium | Harper VanSteenhouse, Yumi Kasai, David Gray, Carol Clugston, Anna Dominiczak and Alex Alderton, Roberto Amato, Jeffrey Barrett, Sonia Goncalves, Ewan Harrison, David K. Jackson, Ian Johnston, Dominic Kwiatkowski, Cordelia Langford, John Sillitoe on behalf of the Wellcome Sanger Institute COVID-19 Surveillance Team |
| EPI_ISL_3455301 | Lighthouse Lab in Alderley Park | Wellcome Sanger Institute for the COVID-19 Genomics UK (COG-UK) Consortium | Jacquelyn Wynn, Mairead Hyland, The Lighthouse Lab in Alderley Park and Alex Alderton, Roberto Amato, Jeffrey Barrett, Sonia Goncalves, Ewan Harrison, David K. Jackson, Ian Johnston, Dominic Kwiatkowski, Cordelia Langford, John Sillitoe on behalf of the Wellcome Sanger Institute COVID-19 Surveillance Team |
| EPI_ISL_3455302, EPI_ISL_3455303 | Lighthouse Lab in Glasgow | Wellcome Sanger Institute for the COVID-19 Genomics UK (COG-UK) Consortium | Harper VanSteenhouse, Yumi Kasai, David Gray, Carol Clugston, Anna Dominiczak and Alex Alderton, Roberto Amato, Jeffrey Barrett, Sonia Goncalves, Ewan Harrison, David K. Jackson, Ian Johnston, Dominic Kwiatkowski, Cordelia Langford, John Sillitoe on behalf of the Wellcome Sanger Institute COVID-19 Surveillance Team |
| EPI_ISL_3455304, EPI_ISL_3455305, EPI_ISL_3455306, EPI_ISL_3455307 | Health Services Laboratories | Wellcome Sanger Institute for the COVID-19 Genomics UK (COG-UK) Consortium | Health Services Laboratories and Alex Alderton, Roberto Amato, Jeffrey Barrett, Sonia Goncalves, Ewan Harrison, David K. Jackson, Ian Johnston, Dominic Kwiatkowski, Cordelia Langford, John Sillitoe on behalf of the Wellcome Sanger Institute COVID-19 Surveillance Team |
| EPI_ISL_3455308 | Berkshire and Surrey Pathology Services Lighthouse Laboratory | Wellcome Sanger Institute for the COVID-19 Genomics UK (COG-UK) Consortium | Berkshire and Surrey Pathology Services Lighthouse Laboratory and Alex Alderton, Roberto Amato, Jeffrey Barrett, Sonia Goncalves, Ewan Harrison, David K. Jackson, Ian Johnston, Dominic Kwiatkowski, Cordelia Langford, John Sillitoe on behalf of the Wellcome Sanger Institute COVID-19 Surveillance Team |
| EPI_ISL_3455309 | Health Services Laboratories | Wellcome Sanger Institute for the COVID-19 Genomics UK (COG-UK) Consortium | Health Services Laboratories and Alex Alderton, Roberto Amato, Jeffrey Barrett, Sonia Goncalves, Ewan Harrison, David K. Jackson, Ian Johnston, Dominic Kwiatkowski, Cordelia Langford, John Sillitoe on behalf of the Wellcome Sanger Institute COVID-19 Surveillance Team |
| EPI_ISL_3455310 | Lighthouse Lab in Alderley Park | Wellcome Sanger Institute for the COVID-19 Genomics UK (COG-UK) Consortium | Jacquelyn Wynn, Mairead Hyland, The Lighthouse Lab in Alderley Park and Alex Alderton, Roberto Amato, Jeffrey Barrett, Sonia Goncalves, Ewan Harrison, David K. Jackson, Ian Johnston, Dominic Kwiatkowski, Cordelia Langford, John Sillitoe on behalf of the Wellcome Sanger Institute COVID-19 Surveillance Team |
| EPI_ISL_3455311, EPI_ISL_3455312, EPI_ISL_3455313 | Health Services Laboratories | Wellcome Sanger Institute for the COVID-19 Genomics UK (COG-UK) Consortium | Health Services Laboratories and Alex Alderton, Roberto Amato, Jeffrey Barrett, Sonia Goncalves, Ewan Harrison, David K. Jackson, Ian Johnston, Dominic Kwiatkowski, Cordelia Langford, John Sillitoe on behalf of the Wellcome Sanger Institute COVID-19 Surveillance Team |
| EPI_ISL_3455314 | Lighthouse Lab in Alderley Park | Wellcome Sanger Institute for the COVID-19 Genomics UK (COG-UK) Consortium | Jacquelyn Wynn, Mairead Hyland, The Lighthouse Lab in Alderley Park and Alex Alderton, Roberto Amato, Jeffrey Barrett, Sonia Goncalves, Ewan Harrison, David K. Jackson, Ian Johnston, Dominic Kwiatkowski, Cordelia Langford, John Sillitoe on behalf of the Wellcome Sanger Institute COVID-19 Surveillance Team |
| EPI_ISL_3455315 | Health Services Laboratories | Wellcome Sanger Institute for the COVID-19 Genomics UK (COG-UK) Consortium | Health Services Laboratories and Alex Alderton, Roberto Amato, Jeffrey Barrett, Sonia Goncalves, Ewan Harrison, David K. Jackson, Ian Johnston, Dominic Kwiatkowski, Cordelia Langford, John Sillitoe on behalf of the Wellcome Sanger Institute COVID-19 Surveillance Team |
| EPI_ISL_3455316 | Lighthouse Lab in Alderley Park | Wellcome Sanger Institute for the COVID-19 Genomics UK (COG-UK) Consortium | Jacquelyn Wynn, Mairead Hyland, The Lighthouse Lab in Alderley Park and Alex Alderton, Roberto Amato, Jeffrey Barrett, Sonia Goncalves, Ewan Harrison, David K. Jackson, Ian Johnston, Dominic Kwiatkowski, Cordelia Langford, John Sillitoe on behalf of the Wellcome Sanger Institute COVID-19 Surveillance Team |
| EPI_ISL_3455317, EPI_ISL_3455318 | Health Services Laboratories | Wellcome Sanger Institute for the COVID-19 Genomics UK (COG-UK) Consortium | Health Services Laboratories and Alex Alderton, Roberto Amato, Jeffrey Barrett, Sonia Goncalves, Ewan Harrison, David K. Jackson, Ian Johnston, Dominic Kwiatkowski, Cordelia Langford, John Sillitoe on behalf of the Wellcome Sanger Institute COVID-19 Surveillance Team |
| EPI_ISL_3455319 | Lighthouse Lab in Alderley Park | Wellcome Sanger Institute for the COVID-19 Genomics UK (COG-UK) Consortium | Jacquelyn Wynn, Mairead Hyland, The Lighthouse Lab in Alderley Park and Alex Alderton, Roberto Amato, Jeffrey Barrett, Sonia Goncalves, Ewan Harrison, David K. Jackson, Ian Johnston, Dominic Kwiatkowski, Cordelia Langford, John Sillitoe on behalf of the Wellcome Sanger Institute COVID-19 Surveillance Team |
| EPI_ISL_3455320 | Health Services Laboratories | Wellcome Sanger Institute for the COVID-19 Genomics UK (COG-UK) Consortium | Health Services Laboratories and Alex Alderton, Roberto Amato, Jeffrey Barrett, Sonia Goncalves, Ewan Harrison, David K. Jackson, Ian Johnston, Dominic Kwiatkowski, Cordelia Langford, John Sillitoe on behalf of the Wellcome Sanger Institute COVID-19 Surveillance Team |
| EPI_ISL_3455321 | Lighthouse Lab in Alderley Park | Wellcome Sanger Institute for the COVID-19 Genomics UK (COG-UK) Consortium | Jacquelyn Wynn, Mairead Hyland, The Lighthouse Lab in Alderley Park and Alex Alderton, Roberto Amato, Jeffrey Barrett, Sonia Goncalves, Ewan Harrison, David K. Jackson, Ian Johnston, Dominic Kwiatkowski, Cordelia Langford, John Sillitoe on behalf of the Wellcome Sanger Institute COVID-19 Surveillance Team |
| EPI_ISL_3455322 | Health Services Laboratories | Wellcome Sanger Institute for the COVID-19 Genomics UK (COG-UK) Consortium | Health Services Laboratories and Alex Alderton, Roberto Amato, Jeffrey Barrett, Sonia Goncalves, Ewan Harrison, David K. Jackson, Ian Johnston, Dominic Kwiatkowski, Cordelia Langford, John Sillitoe on behalf of the Wellcome Sanger Institute COVID-19 Surveillance Team |
| EPI_ISL_3455323 | Lighthouse Lab in Alderley Park | Wellcome Sanger Institute for the COVID-19 Genomics UK (COG-UK) Consortium | Jacquelyn Wynn, Mairead Hyland, The Lighthouse Lab in Alderley Park and Alex Alderton, Roberto Amato, Jeffrey Barrett, Sonia Goncalves, Ewan Harrison, David K. Jackson, Ian Johnston, Dominic Kwiatkowski, Cordelia Langford, John Sillitoe on behalf of the Wellcome Sanger Institute COVID-19 Surveillance Team |
| EPI_ISL_3455324 | Berkshire and Surrey Pathology Services Lighthouse Laboratory | Wellcome Sanger Institute for the COVID-19 Genomics UK (COG-UK) Consortium | Berkshire and Surrey Pathology Services Lighthouse Laboratory and Alex Alderton, Roberto Amato, Jeffrey Barrett, Sonia Goncalves, Ewan Harrison, David K. Jackson, Ian Johnston, Dominic Kwiatkowski, Cordelia Langford, John Sillitoe on behalf of the Wellcome Sanger Institute COVID-19 Surveillance Team |
| EPI_ISL_3455325, EPI_ISL_3455326, EPI_ISL_3455327 | Lighthouse Lab in Alderley Park | Wellcome Sanger Institute for the COVID-19 Genomics UK (COG-UK) Consortium | Jacquelyn Wynn, Mairead Hyland, The Lighthouse Lab in Alderley Park and Alex Alderton, Roberto Amato, Jeffrey Barrett, Sonia Goncalves, Ewan Harrison, David K. Jackson, Ian Johnston, Dominic Kwiatkowski, Cordelia Langford, John Sillitoe on behalf of the Wellcome Sanger Institute COVID-19 Surveillance Team |
| EPI_ISL_3455328, EPI_ISL_3455329 | Lighthouse Laboratory Plymouth | Wellcome Sanger Institute for the COVID-19 Genomics UK (COG-UK) Consortium | Lighthouse Laboratory Plymouth and Alex Alderton, Roberto Amato, Jeffrey Barrett, Sonia Goncalves, Ewan Harrison, David K. Jackson, Ian Johnston, Dominic Kwiatkowski, Cordelia Langford, John Sillitoe on behalf of the Wellcome Sanger Institute COVID-19 Surveillance Team |
| EPI_ISL_3455330, EPI_ISL_3455331 | Berkshire and Surrey Pathology Services Lighthouse Laboratory | Wellcome Sanger Institute for the COVID-19 Genomics UK (COG-UK) Consortium | Berkshire and Surrey Pathology Services Lighthouse Laboratory and Alex Alderton, Roberto Amato, Jeffrey Barrett, Sonia Goncalves, Ewan Harrison, David K. Jackson, Ian Johnston, Dominic Kwiatkowski, Cordelia Langford, John Sillitoe on behalf of the Wellcome Sanger Institute COVID-19 Surveillance Team |
| EPI_ISL_3455332, EPI_ISL_3455333, EPI_ISL_3455334, EPI_ISL_3455335, EPI_ISL_3455336, EPI_ISL_3455337 | Health Services Laboratories | Wellcome Sanger Institute for the COVID-19 Genomics UK (COG-UK) Consortium | Health Services Laboratories and Alex Alderton, Roberto Amato, Jeffrey Barrett, Sonia Goncalves, Ewan Harrison, David K. Jackson, Ian Johnston, Dominic Kwiatkowski, Cordelia Langford, John Sillitoe on behalf of the Wellcome Sanger Institute COVID-19 Surveillance Team |
| EPI_ISL_3455338 | Berkshire and Surrey Pathology Services Lighthouse Laboratory | Wellcome Sanger Institute for the COVID-19 Genomics UK (COG-UK) Consortium | Berkshire and Surrey Pathology Services Lighthouse Laboratory and Alex Alderton, Roberto Amato, Jeffrey Barrett, Sonia Goncalves, Ewan Harrison, David K. Jackson, Ian Johnston, Dominic Kwiatkowski, Cordelia Langford, John Sillitoe on behalf of the Wellcome Sanger Institute COVID-19 Surveillance Team |
| EPI_ISL_3455339 | Health Services Laboratories | Wellcome Sanger Institute for the COVID-19 Genomics UK (COG-UK) Consortium | Health Services Laboratories and Alex Alderton, Roberto Amato, Jeffrey Barrett, Sonia Goncalves, Ewan Harrison, David K. Jackson, Ian Johnston, Dominic Kwiatkowski, Cordelia Langford, John Sillitoe on behalf of the Wellcome Sanger Institute COVID-19 Surveillance Team |
| EPI_ISL_3455340 | Berkshire and Surrey Pathology Services Lighthouse Laboratory | Wellcome Sanger Institute for the COVID-19 Genomics UK (COG-UK) Consortium | Berkshire and Surrey Pathology Services Lighthouse Laboratory and Alex Alderton, Roberto Amato, Jeffrey Barrett, Sonia Goncalves, Ewan Harrison, David K. Jackson, Ian Johnston, Dominic Kwiatkowski, Cordelia Langford, John Sillitoe on behalf of the Wellcome Sanger Institute COVID-19 Surveillance Team |
| EPI_ISL_3455341, EPI_ISL_3455342, EPI_ISL_3455343, EPI_ISL_3455344, EPI_ISL_3455345 | Health Services Laboratories | Wellcome Sanger Institute for the COVID-19 Genomics UK (COG-UK) Consortium | Health Services Laboratories and Alex Alderton, Roberto Amato, Jeffrey Barrett, Sonia Goncalves, Ewan Harrison, David K. Jackson, Ian Johnston, Dominic Kwiatkowski, Cordelia Langford, John Sillitoe on behalf of the Wellcome Sanger Institute COVID-19 Surveillance Team |
| EPI_ISL_3455346, EPI_ISL_3455347 | Berkshire and Surrey Pathology Services Lighthouse Laboratory | Wellcome Sanger Institute for the COVID-19 Genomics UK (COG-UK) Consortium | Berkshire and Surrey Pathology Services Lighthouse Laboratory and Alex Alderton, Roberto Amato, Jeffrey Barrett, Sonia Goncalves, Ewan Harrison, David K. Jackson, Ian Johnston, Dominic Kwiatkowski, Cordelia Langford, John Sillitoe on behalf of the Wellcome Sanger Institute COVID-19 Surveillance Team |
| EPI_ISL_3455348 | Health Services Laboratories | Wellcome Sanger Institute for the COVID-19 Genomics UK (COG-UK) Consortium | Health Services Laboratories and Alex Alderton, Roberto Amato, Jeffrey Barrett, Sonia Goncalves, Ewan Harrison, David K. Jackson, Ian Johnston, Dominic Kwiatkowski, Cordelia Langford, John Sillitoe on behalf of the Wellcome Sanger Institute COVID-19 Surveillance Team |
| EPI_ISL_3455349 | Berkshire and Surrey Pathology Services Lighthouse Laboratory | Wellcome Sanger Institute for the COVID-19 Genomics UK (COG-UK) Consortium | Berkshire and Surrey Pathology Services Lighthouse Laboratory and Alex Alderton, Roberto Amato, Jeffrey Barrett, Sonia Goncalves, Ewan Harrison, David K. Jackson, Ian Johnston, Dominic Kwiatkowski, Cordelia Langford, John Sillitoe on behalf of the Wellcome Sanger Institute COVID-19 Surveillance Team |

| | | | |
|---|---|---|---|
| EPI_ISL_3455350, EPI_ISL_3455351 | Health Services Laboratories | Wellcome Sanger Institute for the COVID-19 Genomics UK (COG-UK) Consortium | Health Services Laboratories and Alex Alderton, Roberto Amato, Jeffrey Barrett, Sonia Goncalves, Ewan Harrison, David K. Jackson, Ian Johnston, Dominic Kwiatkowski, Cordelia Langford, John Sillitoe on behalf of the Wellcome Sanger Institute COVID-19 Surveillance Team |
| EPI_ISL_3455352, EPI_ISL_3455353, EPI_ISL_3455354 | Berkshire and Surrey Pathology Services Lighthouse Laboratory | Wellcome Sanger Institute for the COVID-19 Genomics UK (COG-UK) Consortium | Berkshire and Surrey Pathology Services Lighthouse Laboratory and Alex Alderton, Roberto Amato, Jeffrey Barrett, Sonia Goncalves, Ewan Harrison, David K. Jackson, Ian Johnston, Dominic Kwiatkowski, Cordelia Langford, John Sillitoe on behalf of the Wellcome Sanger Institute COVID-19 Surveillance Team |
| EPI_ISL_3455355, EPI_ISL_3455356 | Lighthouse Lab in Glasgow | Wellcome Sanger Institute for the COVID-19 Genomics UK (COG-UK) Consortium | Harper VanSteenhouse, Yumi Kasai, David Gray, Carol Clugston, Anna Dominiczak and Alex Alderton, Roberto Amato, Jeffrey Barrett, Sonia Goncalves, Ewan Harrison, David K. Jackson, Ian Johnston, Dominic Kwiatkowski, Cordelia Langford, John Sillitoe on behalf of the Wellcome Sanger Institute COVID-19 Surveillance Team |
| EPI_ISL_3455357, EPI_ISL_3455358 | Lighthouse Lab in Alderley Park | Wellcome Sanger Institute for the COVID-19 Genomics UK (COG-UK) Consortium | Jacquelyn Wynn, Mairead Hyland, The Lighthouse Lab in Alderley Park and Alex Alderton, Roberto Amato, Jeffrey Barrett, Sonia Goncalves, Ewan Harrison, David K. Jackson, Ian Johnston, Dominic Kwiatkowski, Cordelia Langford, John Sillitoe on behalf of the Wellcome Sanger Institute COVID-19 Surveillance Team |
| EPI_ISL_3455359, EPI_ISL_3455360, EPI_ISL_3455361, EPI_ISL_3455362, EPI_ISL_3455363, EPI_ISL_3455364, EPI_ISL_3455365, EPI_ISL_3455366, EPI_ISL_3455367, EPI_ISL_3455368, EPI_ISL_3455369 | | | |
| see above | Lighthouse Lab in Glasgow | Wellcome Sanger Institute for the COVID-19 Genomics UK (COG-UK) Consortium | Harper VanSteenhouse, Yumi Kasai, David Gray, Carol Clugston, Anna Dominiczak and Alex Alderton, Roberto Amato, Jeffrey Barrett, Sonia Goncalves, Ewan Harrison, David K. Jackson, Ian Johnston, Dominic Kwiatkowski, Cordelia Langford, John Sillitoe on behalf of the Wellcome Sanger Institute COVID-19 Surveillance Team |
| EPI_ISL_3455370 | Lighthouse Laboratory Plymouth | Wellcome Sanger Institute for the COVID-19 Genomics UK (COG-UK) Consortium | Lighthouse Laboratory Plymouth and Alex Alderton, Roberto Amato, Jeffrey Barrett, Sonia Goncalves, Ewan Harrison, David K. Jackson, Ian Johnston, Dominic Kwiatkowski, Cordelia Langford, John Sillitoe on behalf of the Wellcome Sanger Institute COVID-19 Surveillance Team |
| EPI_ISL_3455371, EPI_ISL_3455372, EPI_ISL_3455373, EPI_ISL_3455374, EPI_ISL_3455375, EPI_ISL_3455376, EPI_ISL_3455377, EPI_ISL_3455378, EPI_ISL_3455379, EPI_ISL_3455380, EPI_ISL_3455381, EPI_ISL_3455382, EPI_ISL_3455383, EPI_ISL_3455384, EPI_ISL_3455385, EPI_ISL_3455386, EPI_ISL_3455387, EPI_ISL_3455388, EPI_ISL_3455389, EPI_ISL_3455390, EPI_ISL_3455391, EPI_ISL_3455392, EPI_ISL_3455393, EPI_ISL_3455394, EPI_ISL_3455395, EPI_ISL_3455396, EPI_ISL_3455397 | | | |
| see above | Lighthouse Lab in Glasgow | Wellcome Sanger Institute for the COVID-19 Genomics UK (COG-UK) Consortium | Harper VanSteenhouse, Yumi Kasai, David Gray, Carol Clugston, Anna Dominiczak and Alex Alderton, Roberto Amato, Jeffrey Barrett, Sonia Goncalves, Ewan Harrison, David K. Jackson, Ian Johnston, Dominic Kwiatkowski, Cordelia Langford, John Sillitoe on behalf of the Wellcome Sanger Institute COVID-19 Surveillance Team |
| EPI_ISL_3455398, EPI_ISL_3455399, EPI_ISL_3455400, EPI_ISL_3455401 | Lighthouse Lab in Alderley Park | Wellcome Sanger Institute for the COVID-19 Genomics UK (COG-UK) Consortium | Jacquelyn Wynn, Mairead Hyland, The Lighthouse Lab in Alderley Park and Alex Alderton, Roberto Amato, Jeffrey Barrett, Sonia Goncalves, Ewan Harrison, David K. Jackson, Ian Johnston, Dominic Kwiatkowski, Cordelia Langford, John Sillitoe on behalf of the Wellcome Sanger Institute COVID-19 Surveillance Team |
| EPI_ISL_3455402 | Lighthouse Lab in Glasgow | Wellcome Sanger Institute for the COVID-19 Genomics UK (COG-UK) Consortium | Harper VanSteenhouse, Yumi Kasai, David Gray, Carol Clugston, Anna Dominiczak and Alex Alderton, Roberto Amato, Jeffrey Barrett, Sonia Goncalves, Ewan Harrison, David K. Jackson, Ian Johnston, Dominic Kwiatkowski, Cordelia Langford, John Sillitoe on behalf of the Wellcome Sanger Institute COVID-19 Surveillance Team |
| EPI_ISL_3455403, EPI_ISL_3455404 | ZOL | Jessa | Berden et al. on behalf of the Jessa_cmdLab |
| EPI_ISL_3455405 | LKO | Jessa | Berden et al. on behalf of the Jessa_cmdLab |
| EPI_ISL_3455406 | Jessa | Jessa | Berden et al. on behalf of the Jessa_cmdLab |
| EPI_ISL_3455407, EPI_ISL_3455408, EPI_ISL_3455409, EPI_ISL_3455410, EPI_ISL_3455411 | ZOL | Jessa | Berden et al. on behalf of the Jessa_cmdLab |
| EPI_ISL_3455413 | Jessa | Jessa | Berden et al. on behalf of the Jessa_cmdLab |
| EPI_ISL_3455414 | ZOL | Jessa | Berden et al. on behalf of the Jessa_cmdLab |
| EPI_ISL_3455415 | Jessa | Jessa | Berden et al. on behalf of the Jessa_cmdLab |
| EPI_ISL_3455416, EPI_ISL_3455417 | ZOL | Jessa | Berden et al. on behalf of the Jessa_cmdLab |
| EPI_ISL_3455418, EPI_ISL_3455419 | Jessa | Jessa | Berden et al. on behalf of the Jessa_cmdLab |
| EPI_ISL_3455420, EPI_ISL_3455421, EPI_ISL_3455422 | ZOL | Jessa | Berden et al. on behalf of the Jessa_cmdLab |
| EPI_ISL_3455423 | Jessa | Jessa | Berden et al. on behalf of the Jessa_cmdLab |
| EPI_ISL_3455424, EPI_ISL_3455425 | ZOL | Jessa | Berden et al. on behalf of the Jessa_cmdLab |
| EPI_ISL_3455426, EPI_ISL_3455427 | Jessa | Jessa | Berden et al. on behalf of the Jessa_cmdLab |
| EPI_ISL_3455428 | ZOL | Jessa | Berden et al. on behalf of the Jessa_cmdLab |
| EPI_ISL_3455429 | Jessa | Jessa | Berden et al. on behalf of the Jessa_cmdLab |
| EPI_ISL_3455430 | ZOL | Jessa | Berden et al. on behalf of the Jessa_cmdLab |
| EPI_ISL_3455431 | LKO | Jessa | Berden et al. on behalf of the Jessa_cmdLab |
| EPI_ISL_3455432, EPI_ISL_3455433, EPI_ISL_3455434, EPI_ISL_3455435, EPI_ISL_3455436 | ZOL | Jessa | Berden et al. on behalf of the Jessa_cmdLab |
| EPI_ISL_3455437, EPI_ISL_3455438, EPI_ISL_3455439, EPI_ISL_3455440, EPI_ISL_3455441, EPI_ISL_3455442, EPI_ISL_3455443, EPI_ISL_3455444, EPI_ISL_3455445 | Jessa | Jessa | Berden et al. on behalf of the Jessa_cmdLab |
| EPI_ISL_3455446, EPI_ISL_3455447, EPI_ISL_3455448, EPI_ISL_3455449, EPI_ISL_3455450, EPI_ISL_3455451, EPI_ISL_3455452, EPI_ISL_3455453, EPI_ISL_3455454, EPI_ISL_3455455, EPI_ISL_3455456, EPI_ISL_3455457, EPI_ISL_3455458, EPI_ISL_3455459, EPI_ISL_3455460, EPI_ISL_3455461, EPI_ISL_3455462, EPI_ISL_3455463, EPI_ISL_3455464, EPI_ISL_3455465, EPI_ISL_3455466, EPI_ISL_3455467, EPI_ISL_3455468 | | | |
| see above | The Jackson Laboratory | The Jackson Laboratory | Long J, Renzette N, Adams M, Omerza G, Kelly K, Li L |
| EPI_ISL_3455469, EPI_ISL_3455470, EPI_ISL_3455471, EPI_ISL_3455472, EPI_ISL_3455473, EPI_ISL_3455474, EPI_ISL_3455476, EPI_ISL_3455477, EPI_ISL_3455478, EPI_ISL_3455479, EPI_ISL_3455480, EPI_ISL_3455481, EPI_ISL_3455482, EPI_ISL_3455483, EPI_ISL_3455484, EPI_ISL_3455486, EPI_ISL_3455487, EPI_ISL_3455488, EPI_ISL_3455489, EPI_ISL_3455490, EPI_ISL_3455491, EPI_ISL_3455492, EPI_ISL_3455493, EPI_ISL_3455494, EPI_ISL_3455495, EPI_ISL_3455496, EPI_ISL_3455497, EPI_ISL_3455498, EPI_ISL_3455499, EPI_ISL_3455500, EPI_ISL_3455501, EPI_ISL_3455502, EPI_ISL_3455503, EPI_ISL_3455504, EPI_ISL_3455505, EPI_ISL_3455506, EPI_ISL_3455507, EPI_ISL_3455508, EPI_ISL_3455509, EPI_ISL_3455510, EPI_ISL_3455511, EPI_ISL_3455512, EPI_ISL_3455513, EPI_ISL_3455514, EPI_ISL_3455515, EPI_ISL_3455516, EPI_ISL_3455517, EPI_ISL_3455518, EPI_ISL_3455519, EPI_ISL_3455520, EPI_ISL_3455521, EPI_ISL_3455522, EPI_ISL_3455523, EPI_ISL_3455524, EPI_ISL_3455525, EPI_ISL_3455526, EPI_ISL_3455527, EPI_ISL_3455528, EPI_ISL_3455529, EPI_ISL_3455530, EPI_ISL_3455531, EPI_ISL_3455532, EPI_ISL_3455533, EPI_ISL_3455534, EPI_ISL_3455535, EPI_ISL_3455536, EPI_ISL_3455537, EPI_ISL_3455538, EPI_ISL_3455539, EPI_ISL_3455540, EPI_ISL_3455541, EPI_ISL_3455542, EPI_ISL_3455543, EPI_ISL_3455544, EPI_ISL_3455545, EPI_ISL_3455546, EPI_ISL_3455547, EPI_ISL_3455548, EPI_ISL_3455549, EPI_ISL_3455550, EPI_ISL_3455551, EPI_ISL_3455552, EPI_ISL_3455553, EPI_ISL_3455554, EPI_ISL_3455555, EPI_ISL_3455556, EPI_ISL_3455557, EPI_ISL_3455558, EPI_ISL_3455559, EPI_ISL_3455560, EPI_ISL_3455561, EPI_ISL_3455563, EPI_ISL_3455565, EPI_ISL_3455566, EPI_ISL_3455570, EPI_ISL_3455572, EPI_ISL_3455573, EPI_ISL_3455575 | | | |
| see above | Azienda Sanitaria dell'Alto Adige - Laboratorio Aziendale di Microbiologia e Virologia | Azienda Sanitaria dell'Alto Adige | Irene Bianconi |
| EPI_ISL_3455577, EPI_ISL_3455578, EPI_ISL_3455579, EPI_ISL_3455580, EPI_ISL_3455581, EPI_ISL_3455582, EPI_ISL_3455583, EPI_ISL_3455584, EPI_ISL_3455585, EPI_ISL_3455586, EPI_ISL_3455587, EPI_ISL_3455588, EPI_ISL_3455589, EPI_ISL_3455590, EPI_ISL_3455591, EPI_ISL_3455592, EPI_ISL_3455593, EPI_ISL_3455594, EPI_ISL_3455595, EPI_ISL_3455596, EPI_ISL_3455597, EPI_ISL_3455598, EPI_ISL_3455599, EPI_ISL_3455600, EPI_ISL_3455601, EPI_ISL_3455602, EPI_ISL_3455603, EPI_ISL_3455604, EPI_ISL_3455605, EPI_ISL_3455606, EPI_ISL_3455607, EPI_ISL_3455608, EPI_ISL_3455610, EPI_ISL_3455612, EPI_ISL_3455614, EPI_ISL_3455615, EPI_ISL_3455616, EPI_ISL_3455617, EPI_ISL_3455618, EPI_ISL_3455619, EPI_ISL_3455620, EPI_ISL_3455621, EPI_ISL_3455622, EPI_ISL_3455623, EPI_ISL_3455624, EPI_ISL_3455625, EPI_ISL_3455626, EPI_ISL_3455627, EPI_ISL_3455628, EPI_ISL_3455629, EPI_ISL_3455630, EPI_ISL_3455631, EPI_ISL_3455632, EPI_ISL_3455633, EPI_ISL_3455634, EPI_ISL_3455635, EPI_ISL_3455636, EPI_ISL_3455637, EPI_ISL_3455638, EPI_ISL_3455639, EPI_ISL_3455640, EPI_ISL_3455641, EPI_ISL_3455642, EPI_ISL_3455643, EPI_ISL_3455644, EPI_ISL_3455645, EPI_ISL_3455646, EPI_ISL_3455647, EPI_ISL_3455648, EPI_ISL_3455649, EPI_ISL_3455650, EPI_ISL_3455651, EPI_ISL_3455652, EPI_ISL_3455653, EPI_ISL_3455654, EPI_ISL_3455655, EPI_ISL_3455656, EPI_ISL_3455657, EPI_ISL_3455658, EPI_ISL_3455659, EPI_ISL_3455660, EPI_ISL_3455661, EPI_ISL_3455662, EPI_ISL_3455663, EPI_ISL_3455664 | | | |
| see above | TXDSHS | TXDSHS | Rashmi Tuladhar, Bonnie Oh, Jenny Zhang, Maliha Rahman, Mayela Pedrueza, Anita Pokharel, Karen Bobier, Lorraine Rodriguez, Myong Koag, Chun Wang, Rachel Lee, Grace Kubin |
| EPI_ISL_3455665, EPI_ISL_3455666, EPI_ISL_3455667, EPI_ISL_3455668, EPI_ISL_3455669, EPI_ISL_3455670, EPI_ISL_3455671 | Indian Council of Medical Research-National Institute of Virology, Microbial Containment Complex | Indian Council of Medical Research-National Institute of Virology, Microbial Containment Complex | Pragya D Yadav |
| EPI_ISL_3455672, EPI_ISL_3455673, EPI_ISL_3455674, EPI_ISL_3455675, EPI_ISL_3455676, EPI_ISL_3455677, EPI_ISL_3455678, EPI_ISL_3455679, EPI_ISL_3455680, EPI_ISL_3455681, EPI_ISL_3455682, EPI_ISL_3455683, EPI_ISL_3455684, EPI_ISL_3455686, EPI_ISL_3455687, EPI_ISL_3455688, EPI_ISL_3455689, EPI_ISL_3455691, EPI_ISL_3455692, EPI_ISL_3455693, EPI_ISL_3455694, EPI_ISL_3455695, EPI_ISL_3455696, EPI_ISL_3455697, EPI_ISL_3455698, EPI_ISL_3455700, EPI_ISL_3455701, EPI_ISL_3455702, EPI_ISL_3455703, EPI_ISL_3455704, EPI_ISL_3455705, EPI_ISL_3455706, EPI_ISL_3455707, EPI_ISL_3455708, EPI_ISL_3455709, EPI_ISL_3455710, EPI_ISL_3455711, EPI_ISL_3455712, EPI_ISL_3455713, EPI_ISL_3455714, EPI_ISL_3455715, EPI_ISL_3455716, EPI_ISL_3455717, EPI_ISL_3455718, EPI_ISL_3455719, EPI_ISL_3455720, EPI_ISL_3455721, EPI_ISL_3455722, EPI_ISL_3455723, EPI_ISL_3455724, EPI_ISL_3455725, EPI_ISL_3455726 | | | |
| see above | TXDSHS | TXDSHS | Rashmi Tuladhar, Bonnie Oh, Jenny Zhang, Maliha Rahman, Mayela Pedrueza, Anita Pokharel, Karen Bobier, Lorraine Rodriguez, Myong Koag, Chun Wang, Rachel Lee, Grace Kubin |
| EPI_ISL_3455727, | Laboratori de Referencia de Catalunya | Laboratori de Referencia de Catalunya | Ramirez A., Bellosillo B., Padilla E., Vilas A., Hernandez JJ., Canal M. |

| | | | |
|---|---|---|---|
| EPI_ISL_3455728, EPI_ISL_3455729, EPI_ISL_3455730, EPI_ISL_3455731 | | | |
| EPI_ISL_3455732, EPI_ISL_3455733, EPI_ISL_3455734, EPI_ISL_3455735, EPI_ISL_3455736, EPI_ISL_3455737, EPI_ISL_3455738, EPI_ISL_3455739 | Microbiology Department, Laboratori Clínic Metropolitana Nord. Hospital Universitari Germans Trias i Pujol | Can Ruti SARS-CoV-2 Sequencing Hub (HUGTiP/IrsiCaixa/IGTP) | Marc Noguera-Julian, Pilar Armengol, Ignacio Blanco, Antoni E Bordoy, Francesc Catala-Moll, Pere-Joan Cardona, Maria Casadellà, Cristina Casañ, Gemma Clara, Bonaventura Clotet, Montserrat Giménez, Mercedes Guerrero, Anna Not, David Panisello, Roger Paredes, Mariona Parera, Alexia París, Verónica Saludes, Alba Sánchez, Laia Soler, Lauro Sumoy, and Elisa Martró on behalf of the Can Ruti SARS-CoV-2 Sequencing Hub |
| EPI_ISL_3455740 | Microbiologia CATLAB | Can Ruti SARS-CoV-2 Sequencing Hub (HUGTiP/IrsiCaixa/IGTP) | Marc Noguera-Julian, Pilar Armengol, Ignacio Blanco, Antoni E Bordoy, Francesc Catala-Moll, Pere-Joan Cardona, Maria Casadellà, Cristina Casañ, Gemma Clara, Bonaventura Clotet, Montserrat Giménez, Mercedes Guerrero, Anna Not, David Panisello, Roger Paredes, Mariona Parera, Alexia París, Verónica Saludes, Alba Sánchez, Laia Soler, Lauro Sumoy, and Elisa Martró on behalf of the Can Ruti SARS-CoV-2 Sequencing Hub |
| EPI_ISL_3455741, EPI_ISL_3455742, EPI_ISL_3455743, EPI_ISL_3455744 | Microbiology Department, Laboratori Clínic Metropolitana Nord. Hospital Universitari Germans Trias i Pujol | Can Ruti SARS-CoV-2 Sequencing Hub (HUGTiP/IrsiCaixa/IGTP) | Marc Noguera-Julian, Pilar Armengol, Ignacio Blanco, Antoni E Bordoy, Francesc Catala-Moll, Pere-Joan Cardona, Maria Casadellà, Cristina Casañ, Gemma Clara, Bonaventura Clotet, Montserrat Giménez, Mercedes Guerrero, Anna Not, David Panisello, Roger Paredes, Mariona Parera, Alexia París, Verónica Saludes, Alba Sánchez, Laia Soler, Lauro Sumoy, and Elisa Martró on behalf of the Can Ruti SARS-CoV-2 Sequencing Hub |
| EPI_ISL_3455745 | Microbiologia CATLAB | Can Ruti SARS-CoV-2 Sequencing Hub (HUGTiP/IrsiCaixa/IGTP) | Marc Noguera-Julian, Pilar Armengol, Ignacio Blanco, Antoni E Bordoy, Francesc Catala-Moll, Pere-Joan Cardona, Maria Casadellà, Cristina Casañ, Gemma Clara, Bonaventura Clotet, Montserrat Giménez, Mercedes Guerrero, Anna Not, David Panisello, Roger Paredes, Mariona Parera, Alexia París, Verónica Saludes, Alba Sánchez, Laia Soler, Lauro Sumoy, and Elisa Martró on behalf of the Can Ruti SARS-CoV-2 Sequencing Hub |
| EPI_ISL_3455746, EPI_ISL_3455747, EPI_ISL_3455748, EPI_ISL_3455749 | Microbiology Department, Laboratori Clínic Metropolitana Nord. Hospital Universitari Germans Trias i Pujol | Can Ruti SARS-CoV-2 Sequencing Hub (HUGTiP/IrsiCaixa/IGTP) | Marc Noguera-Julian, Pilar Armengol, Ignacio Blanco, Antoni E Bordoy, Francesc Catala-Moll, Pere-Joan Cardona, Maria Casadellà, Cristina Casañ, Gemma Clara, Bonaventura Clotet, Montserrat Giménez, Mercedes Guerrero, Anna Not, David Panisello, Roger Paredes, Mariona Parera, Alexia París, Verónica Saludes, Alba Sánchez, Laia Soler, Lauro Sumoy, and Elisa Martró on behalf of the Can Ruti SARS-CoV-2 Sequencing Hub |
| EPI_ISL_3455750 | Microbiologia CATLAB | Can Ruti SARS-CoV-2 Sequencing Hub (HUGTiP/IrsiCaixa/IGTP) | Marc Noguera-Julian, Pilar Armengol, Ignacio Blanco, Antoni E Bordoy, Francesc Catala-Moll, Pere-Joan Cardona, Maria Casadellà, Cristina Casañ, Gemma Clara, Bonaventura Clotet, Montserrat Giménez, Mercedes Guerrero, Anna Not, David Panisello, Roger Paredes, Mariona Parera, Alexia París, Verónica Saludes, Alba Sánchez, Laia Soler, Lauro Sumoy, and Elisa Martró on behalf of the Can Ruti SARS-CoV-2 Sequencing Hub |
| EPI_ISL_3455751, EPI_ISL_3455753, EPI_ISL_3455754, EPI_ISL_3455755, EPI_ISL_3455756, EPI_ISL_3455758, EPI_ISL_3455759, EPI_ISL_3455760, EPI_ISL_3455761, EPI_ISL_3455762, EPI_ISL_3455763, EPI_ISL_3455765, EPI_ISL_3455766, EPI_ISL_3455767, EPI_ISL_3455768 | | | |
| see above | Microbiology Department, Laboratori Clínic Metropolitana Nord. Hospital Universitari Germans Trias i Pujol | Can Ruti SARS-CoV-2 Sequencing Hub (HUGTiP/IrsiCaixa/IGTP) | Marc Noguera-Julian, Pilar Armengol, Ignacio Blanco, Antoni E Bordoy, Francesc Catala-Moll, Pere-Joan Cardona, Maria Casadellà, Cristina Casañ, Gemma Clara, Bonaventura Clotet, Montserrat Giménez, Mercedes Guerrero, Anna Not, David Panisello, Roger Paredes, Mariona Parera, Alexia París, Verónica Saludes, Alba Sánchez, Laia Soler, Lauro Sumoy, and Elisa Martró on behalf of the Can Ruti SARS-CoV-2 Sequencing Hub |
| EPI_ISL_3455769, EPI_ISL_3455770 | Microbiologia CATLAB | Can Ruti SARS-CoV-2 Sequencing Hub (HUGTiP/IrsiCaixa/IGTP) | Marc Noguera-Julian, Pilar Armengol, Ignacio Blanco, Antoni E Bordoy, Francesc Catala-Moll, Pere-Joan Cardona, Maria Casadellà, Cristina Casañ, Gemma Clara, Bonaventura Clotet, Montserrat Giménez, Mercedes Guerrero, Anna Not, David Panisello, Roger Paredes, Mariona Parera, Alexia París, Verónica Saludes, Alba Sánchez, Laia Soler, Lauro Sumoy, and Elisa Martró on behalf of the Can Ruti SARS-CoV-2 Sequencing Hub |
| EPI_ISL_3455771, EPI_ISL_3455772, EPI_ISL_3455774, EPI_ISL_3455775, EPI_ISL_3455776, EPI_ISL_3455777, EPI_ISL_3455778, EPI_ISL_3455779, EPI_ISL_3455780, EPI_ISL_3455781, EPI_ISL_3455782, EPI_ISL_3455783, EPI_ISL_3455784, EPI_ISL_3455785, EPI_ISL_3455786, EPI_ISL_3455787, EPI_ISL_3455788, EPI_ISL_3455789, EPI_ISL_3455790, EPI_ISL_3455791, EPI_ISL_3455792, EPI_ISL_3455793, EPI_ISL_3455794, EPI_ISL_3455795, EPI_ISL_3455796, EPI_ISL_3455797, EPI_ISL_3455798, EPI_ISL_3455800, EPI_ISL_3455801, EPI_ISL_3455802, EPI_ISL_3455804, EPI_ISL_3455805, EPI_ISL_3455806, EPI_ISL_3455807, EPI_ISL_3455808, EPI_ISL_3455809, EPI_ISL_3455810, EPI_ISL_3455811, EPI_ISL_3455812, EPI_ISL_3455813, EPI_ISL_3455814, EPI_ISL_3455815, EPI_ISL_3455816, EPI_ISL_3455817, EPI_ISL_3455818, EPI_ISL_3455819, EPI_ISL_3455820, EPI_ISL_3455821, EPI_ISL_3455822, EPI_ISL_3455823, EPI_ISL_3455824, EPI_ISL_3455825, EPI_ISL_3455826, EPI_ISL_3455827, EPI_ISL_3455828, EPI_ISL_3455829, EPI_ISL_3455830, EPI_ISL_3455831, EPI_ISL_3455832, EPI_ISL_3455833, EPI_ISL_3455834, EPI_ISL_3455835, EPI_ISL_3455836, EPI_ISL_3455837, EPI_ISL_3455838, EPI_ISL_3455839, EPI_ISL_3455840, EPI_ISL_3455841, EPI_ISL_3455842, EPI_ISL_3455843, EPI_ISL_3455844, EPI_ISL_3455845, EPI_ISL_3455846, EPI_ISL_3455847, EPI_ISL_3455848, EPI_ISL_3455849, EPI_ISL_3455850, EPI_ISL_3455851, EPI_ISL_3455852 | | | |
| see above | Microbiology Department, Laboratori Clínic Metropolitana Nord. Hospital Universitari Germans Trias i Pujol | Can Ruti SARS-CoV-2 Sequencing Hub (HUGTiP/IrsiCaixa/IGTP) | Marc Noguera-Julian, Pilar Armengol, Ignacio Blanco, Antoni E Bordoy, Francesc Catala-Moll, Pere-Joan Cardona, Maria Casadellà, Cristina Casañ, Gemma Clara, Bonaventura Clotet, Montserrat Giménez, Mercedes Guerrero, Anna Not, David Panisello, Roger Paredes, Mariona Parera, Alexia París, Verónica Saludes, Alba Sánchez, Laia Soler, Lauro Sumoy, and Elisa Martró on behalf of the Can Ruti SARS-CoV-2 Sequencing Hub |
| EPI_ISL_3455853 | Microbiologia CATLAB | Can Ruti SARS-CoV-2 Sequencing Hub (HUGTiP/IrsiCaixa/IGTP) | Marc Noguera-Julian, Pilar Armengol, Ignacio Blanco, Antoni E Bordoy, Francesc Catala-Moll, Pere-Joan Cardona, Maria Casadellà, Cristina Casañ, Gemma Clara, Bonaventura Clotet, Montserrat Giménez, Mercedes Guerrero, Anna Not, David Panisello, Roger Paredes, Mariona Parera, Alexia París, Verónica Saludes, Alba Sánchez, Laia Soler, Lauro Sumoy, and Elisa Martró on behalf of the Can Ruti SARS-CoV-2 Sequencing Hub |
| EPI_ISL_3455854, EPI_ISL_3455855, EPI_ISL_3455856, EPI_ISL_3455857, EPI_ISL_3455859, EPI_ISL_3455860, EPI_ISL_3455861 | Microbiology Department, Laboratori Clínic Metropolitana Nord. Hospital Universitari Germans Trias i Pujol | Can Ruti SARS-CoV-2 Sequencing Hub (HUGTiP/IrsiCaixa/IGTP) | Marc Noguera-Julian, Pilar Armengol, Ignacio Blanco, Antoni E Bordoy, Francesc Catala-Moll, Pere-Joan Cardona, Maria Casadellà, Cristina Casañ, Gemma Clara, Bonaventura Clotet, Montserrat Giménez, Mercedes Guerrero, Anna Not, David Panisello, Roger Paredes, Mariona Parera, Alexia París, Verónica Saludes, Alba Sánchez, Laia Soler, Lauro Sumoy, and Elisa Martró on behalf of the Can Ruti SARS-CoV-2 Sequencing Hub |
| EPI_ISL_3456029 | HOSPITAL UNIVERSITARIO HERNANDO MONCALEANO PERDOMO | Instituto Nacional de Salud | Katherine Laiton-Donato, Diego A. Álvarez-Díaz, Carlos Franco-Muñoz, Hector Alejandro Ruiz-Moreno, Paola Rojas, Maria T. Herrera-Sepúlveda, Diego Andrés Prada, Jhonnatan Reales-González, Sheryll Corchuelo, Julian Naizaque, Jorge Rivera, Gerardo Santamaría, Sergio Gomez, Lisseth Pardo, Juan Camilo Martinez, Marta Lopez Blanco, Ángela Alarcon Cruz, Diana Malo, Carmen Osorio, Magdalena Wiesner, Martha Lucia Ospina Martinez, Marcela Mercado-Reyes |
| EPI_ISL_3456042, EPI_ISL_3456044, EPI_ISL_3456045, EPI_ISL_3456046, EPI_ISL_3456047, EPI_ISL_3456048, EPI_ISL_3456049, EPI_ISL_3456050, EPI_ISL_3456051, EPI_ISL_3456052, EPI_ISL_3456053, EPI_ISL_3456054, EPI_ISL_3456056, EPI_ISL_3456057, EPI_ISL_3456058, EPI_ISL_3456059 | | | |
| see above | CHU Pontchaillou | CHU Pontchaillou | GROLHIER Claire, DENOUAL Florent, ETCHEVERRY Amandine, SASSI Mohamed, JAGLINE Steven, FEBREAU Christine, PRONIER Charlotte, GALIBERT Marie Dominique, DE TAYRAC Marie, THIBAULT Vincent |
| EPI_ISL_3456060 | Clinical Microbiology Laboratory, Faculty of Medicine, Universitas Indonesia | Clinical Microbiology Laboratory, Faculty of Medicine, Universitas Indonesia | Ibrahim F, Natasha A, Yasmon A, Fitrhiyah, Saharman YR, Karuniawati A, Sudarmono P. |
| EPI_ISL_3456061 | Laboratório de Microbiologia Molecular - Universidade FEEVALE | Molecular Microbiology Laboratory | Alana Witt Hansen, Fágner Henrique Heldt, Fernando Rosado Spilki, Juliana Schons Gularte, Juliana Deise Fleck, Mariana Soares da Silva, Meriane Demoliner, Matheus Nunes Weber, Paula Rodrigues de Almeida, Micheli Filippi. |
| EPI_ISL_3456062 | Laboratório de Microbiologia Molecular - Universidade FEEVALE | Molecular Microbiology Laboratory | Alana Witt Hansen, Fágner Henrique Heldt, Fernando Rosado Spilki, Juliana Schons Gulart, Juliane Deise Fleck, Mariana Soares da Silva, Meriane Demoliner, Matheus Nunes Weber, Paula Rodrigues de Almeida, Michele Filippi. |
| EPI_ISL_3456071, EPI_ISL_3456072, EPI_ISL_3456073, EPI_ISL_3456074, EPI_ISL_3456075, EPI_ISL_3456076 | National Platform bis UMONS/Jolimont | National Platform bis UMONS/Jolimont | François Dufrasne, Guillaume Bayon-Vicente, Florian Juszczak, Laetitia Gheysen, Eric Tarantino, Gautier Detry, Ruddy Wattiez |
| EPI_ISL_3456077, EPI_ISL_3456078, EPI_ISL_3456079, EPI_ISL_3456080, EPI_ISL_3456081, EPI_ISL_3456082, EPI_ISL_3456083, EPI_ISL_3456084, EPI_ISL_3456085, EPI_ISL_3456086, EPI_ISL_3456087, EPI_ISL_3456088, EPI_ISL_3456089, EPI_ISL_3456090, EPI_ISL_3456091, EPI_ISL_3456092, EPI_ISL_3456093, EPI_ISL_3456094, EPI_ISL_3456095, EPI_ISL_3456096, EPI_ISL_3456097, EPI_ISL_3456099, EPI_ISL_3456100, EPI_ISL_3456101, EPI_ISL_3456102, EPI_ISL_3456103, EPI_ISL_3456104, EPI_ISL_3456105, EPI_ISL_3456106, EPI_ISL_3456107, EPI_ISL_3456108, EPI_ISL_3456109, EPI_ISL_3456110, EPI_ISL_3456111, EPI_ISL_3456112, EPI_ISL_3456113, EPI_ISL_3456114, EPI_ISL_3456115, EPI_ISL_3456116, EPI_ISL_3456117, EPI_ISL_3456118, EPI_ISL_3456119, EPI_ISL_3456120, EPI_ISL_3456121, EPI_ISL_3456122, EPI_ISL_3456123, EPI_ISL_3456124, EPI_ISL_3456125, EPI_ISL_3456126, EPI_ISL_3456127, EPI_ISL_3456128, EPI_ISL_3456129, EPI_ISL_3456130, EPI_ISL_3456131, EPI_ISL_3456132, EPI_ISL_3456133, EPI_ISL_3456134, EPI_ISL_3456135, EPI_ISL_3456136, EPI_ISL_3456137, EPI_ISL_3456138, EPI_ISL_3456139, EPI_ISL_3456140, EPI_ISL_3456141, EPI_ISL_3456142, EPI_ISL_3456143, EPI_ISL_3456144, EPI_ISL_3456145, EPI_ISL_3456146, EPI_ISL_3456147, EPI_ISL_3456148, EPI_ISL_3456149, EPI_ISL_3456150, EPI_ISL_3456151, EPI_ISL_3456152, EPI_ISL_3456153, EPI_ISL_3456154, EPI_ISL_3456155, EPI_ISL_3456156, EPI_ISL_3456157, EPI_ISL_3456158, EPI_ISL_3456159, EPI_ISL_3456160, EPI_ISL_3456161, EPI_ISL_3456162, EPI_ISL_3456163, EPI_ISL_3456164, EPI_ISL_3456165, EPI_ISL_3456166, EPI_ISL_3456167, EPI_ISL_3456168, EPI_ISL_3456169, EPI_ISL_3456170, EPI_ISL_3456171, EPI_ISL_3456172, EPI_ISL_3456173, EPI_ISL_3456174, EPI_ISL_3456175, EPI_ISL_3456176, EPI_ISL_3456177, EPI_ISL_3456178, EPI_ISL_3456179, EPI_ISL_3456180, EPI_ISL_3456181, EPI_ISL_3456182, EPI_ISL_3456183, EPI_ISL_3456184, EPI_ISL_3456186, EPI_ISL_3456187, EPI_ISL_3456188, EPI_ISL_3456189, EPI_ISL_3456190, EPI_ISL_3456191, EPI_ISL_3456192, EPI_ISL_3456193, EPI_ISL_3456194, EPI_ISL_3456195, EPI_ISL_3456196, EPI_ISL_3456197, EPI_ISL_3456198, EPI_ISL_3456199, EPI_ISL_3456200, EPI_ISL_3456202, EPI_ISL_3456203, EPI_ISL_3456204, EPI_ISL_3456205, EPI_ISL_3456206, EPI_ISL_3456207, EPI_ISL_3456208, EPI_ISL_3456209, EPI_ISL_3456210, EPI_ISL_3456211, EPI_ISL_3456212, EPI_ISL_3456213, EPI_ISL_3456214, EPI_ISL_3456215, EPI_ISL_3456216, EPI_ISL_3456217, EPI_ISL_3456218, EPI_ISL_3456219, EPI_ISL_3456220, EPI_ISL_3456221, EPI_ISL_3456222, EPI_ISL_3456223, EPI_ISL_3456224, EPI_ISL_3456225, EPI_ISL_3456226, EPI_ISL_3456227, EPI_ISL_3456228, EPI_ISL_3456229, EPI_ISL_3456230, EPI_ISL_3456231, EPI_ISL_3456232, EPI_ISL_3456233, EPI_ISL_3456234, EPI_ISL_3456235, EPI_ISL_3456236, EPI_ISL_3456237, EPI_ISL_3456238, EPI_ISL_3456239, EPI_ISL_3456240, EPI_ISL_3456241, EPI_ISL_3456242, EPI_ISL_3456243, EPI_ISL_3456244, EPI_ISL_3456245, EPI_ISL_3456246, EPI_ISL_3456247, EPI_ISL_3456248, EPI_ISL_3456249, EPI_ISL_3456250, EPI_ISL_3456251, EPI_ISL_3456252, EPI_ISL_3456253, EPI_ISL_3456254, EPI_ISL_3456255, EPI_ISL_3456256, EPI_ISL_3456257, EPI_ISL_3456258, EPI_ISL_3456259, EPI_ISL_3456260, EPI_ISL_3456262, EPI_ISL_3456263, EPI_ISL_3456264, EPI_ISL_3456265, EPI_ISL_3456266, EPI_ISL_3456267, EPI_ISL_3456268, EPI_ISL_3456269, EPI_ISL_3456270, EPI_ISL_3456271, EPI_ISL_3456272, EPI_ISL_3456273, EPI_ISL_3456274, EPI_ISL_3456276, EPI_ISL_3456277, EPI_ISL_3456278, EPI_ISL_3456279, EPI_ISL_3456280, EPI_ISL_3456281, EPI_ISL_3456282, EPI_ISL_3456283, EPI_ISL_3456284, EPI_ISL_3456285, EPI_ISL_3456286, EPI_ISL_3456287, EPI_ISL_3456288, EPI_ISL_3456289, EPI_ISL_3456290, EPI_ISL_3456291, EPI_ISL_3456292, EPI_ISL_3456293, EPI_ISL_3456294, EPI_ISL_3456295, EPI_ISL_3456296, EPI_ISL_3456297, EPI_ISL_3456298, EPI_ISL_3456299, EPI_ISL_3456300, EPI_ISL_3456301, EPI_ISL_3456302, EPI_ISL_3456303, EPI_ISL_3456304, EPI_ISL_3456305, EPI_ISL_3456306, EPI_ISL_3456307, EPI_ISL_3456308, EPI_ISL_3456309, EPI_ISL_3456310, EPI_ISL_3456311, EPI_ISL_3456312, EPI_ISL_3456313, EPI_ISL_3456314, EPI_ISL_3456315, EPI_ISL_3456316, EPI_ISL_3456317, EPI_ISL_3456318, EPI_ISL_3456319, EPI_ISL_3456320, EPI_ISL_3456321, EPI_ISL_3456322, EPI_ISL_3456323, EPI_ISL_3456324, EPI_ISL_3456325, EPI_ISL_3456326, EPI_ISL_3456327, EPI_ISL_3456328, EPI_ISL_3456329, EPI_ISL_3456330, EPI_ISL_3456331, EPI_ISL_3456332, EPI_ISL_3456333, EPI_ISL_3456334, EPI_ISL_3456335, EPI_ISL_3456336, EPI_ISL_3456337, EPI_ISL_3456338, EPI_ISL_3456339, EPI_ISL_3456340, EPI_ISL_3456341, EPI_ISL_3456342, EPI_ISL_3456343, EPI_ISL_3456344, EPI_ISL_3456345, EPI_ISL_3456346, EPI_ISL_3456347, EPI_ISL_3456348, EPI_ISL_3456349, EPI_ISL_3456350, EPI_ISL_3456351, EPI_ISL_3456352, EPI_ISL_3456353, EPI_ISL_3456354, EPI_ISL_3456355, EPI_ISL_3456356, EPI_ISL_3456357, EPI_ISL_3456358, EPI_ISL_3456359, EPI_ISL_3456360, EPI_ISL_3456361, EPI_ISL_3456362, EPI_ISL_3456363, EPI_ISL_3456364, EPI_ISL_3456365, EPI_ISL_3456366, EPI_ISL_3456367, EPI_ISL_3456368, EPI_ISL_3456369, EPI_ISL_3456370, EPI_ISL_3456371, EPI_ISL_3456372, EPI_ISL_3456373, EPI_ISL_3456374, EPI_ISL_3456375, EPI_ISL_3456376, EPI_ISL_3456377, EPI_ISL_3456378, EPI_ISL_3456379, EPI_ISL_3456380, EPI_ISL_3456381, EPI_ISL_3456382, EPI_ISL_3456383, EPI_ISL_3456384, EPI_ISL_3456385, EPI_ISL_3456386, EPI_ISL_3456387, EPI_ISL_3456388, EPI_ISL_3456389, EPI_ISL_3456390, EPI_ISL_3456391, EPI_ISL_3456392, EPI_ISL_3456393, EPI_ISL_3456394, EPI_ISL_3456395, EPI_ISL_3456396, EPI_ISL_3456397, EPI_ISL_3456398, EPI_ISL_3456399, EPI_ISL_3456400, EPI_ISL_3456401, EPI_ISL_3456402, EPI_ISL_3456403, EPI_ISL_3456404, EPI_ISL_3456405, EPI_ISL_3456406, EPI_ISL_3456407, EPI_ISL_3456408, EPI_ISL_3456409, EPI_ISL_3456410, EPI_ISL_3456411, EPI_ISL_3456412, EPI_ISL_3456413, EPI_ISL_3456414, EPI_ISL_3456415, EPI_ISL_3456416, EPI_ISL_3456417, EPI_ISL_3456418, EPI_ISL_3456419, EPI_ISL_3456420, EPI_ISL_3456421, EPI_ISL_3456422, EPI_ISL_3456423, EPI_ISL_3456424, EPI_ISL_3456425, EPI_ISL_3456426, EPI_ISL_3456427, EPI_ISL_3456428, EPI_ISL_3456429, EPI_ISL_3456430, EPI_ISL_3456431, EPI_ISL_3456432, EPI_ISL_3456433, EPI_ISL_3456434, EPI_ISL_3456436, EPI_ISL_3456437, EPI_ISL_3456438, EPI_ISL_3456439, EPI_ISL_3456440, EPI_ISL_3456441, EPI_ISL_3456442, EPI_ISL_3456443, EPI_ISL_3456444, EPI_ISL_3456445, EPI_ISL_3456447, EPI_ISL_3456448, EPI_ISL_3456449, EPI_ISL_3456450, EPI_ISL_3456451, EPI_ISL_3456452, EPI_ISL_3456453, EPI_ISL_3456454, EPI_ISL_3456455, EPI_ISL_3456456, EPI_ISL_3456457, EPI_ISL_3456458, EPI_ISL_3456460, EPI_ISL_3456461, EPI_ISL_3456462, EPI_ISL_3456463, EPI_ISL_3456464, EPI_ISL_3456465, EPI_ISL_3456466, EPI_ISL_3456467, EPI_ISL_3456468, EPI_ISL_3456469, EPI_ISL_3456470, EPI_ISL_3456471, EPI_ISL_3456472, EPI_ISL_3456473, EPI_ISL_3456474, EPI_ISL_3456475, EPI_ISL_3456476, EPI_ISL_3456477, EPI_ISL_3456478, EPI_ISL_3456479, EPI_ISL_3456480, EPI_ISL_3456481, EPI_ISL_3456482, EPI_ISL_3456483, EPI_ISL_3456484, EPI_ISL_3456485, EPI_ISL_3456486, EPI_ISL_3456487, EPI_ISL_3456488, EPI_ISL_3456489, EPI_ISL_3456490, EPI_ISL_3456491, EPI_ISL_3456492, EPI_ISL_3456493, EPI_ISL_3456494, EPI_ISL_3456495, EPI_ISL_3456496, EPI_ISL_3456497, EPI_ISL_3456498, EPI_ISL_3456499, EPI_ISL_3456500, EPI_ISL_3456501, EPI_ISL_3456502, EPI_ISL_3456503, EPI_ISL_3456504, EPI_ISL_3456506, EPI_ISL_3456507, EPI_ISL_3456508, EPI_ISL_3456509, EPI_ISL_3456510, EPI_ISL_3456511, EPI_ISL_3456513, EPI_ISL_3456514, EPI_ISL_3456515, EPI_ISL_3456516, EPI_ISL_3456517, EPI_ISL_3456518, EPI_ISL_3456519, EPI_ISL_3456520, EPI_ISL_3456521, EPI_ISL_3456522, EPI_ISL_3456523, EPI_ISL_3456524, EPI_ISL_3456525, EPI_ISL_3456526, EPI_ISL_3456527, EPI_ISL_3456528, EPI_ISL_3456529, EPI_ISL_3456530, EPI_ISL_3456531, EPI_ISL_3456532, EPI_ISL_3456533, EPI_ISL_3456534, EPI_ISL_3456535, EPI_ISL_3456536, EPI_ISL_3456537, EPI_ISL_3456538, EPI_ISL_3456539, EPI_ISL_3456540, EPI_ISL_3456541, EPI_ISL_3456542, EPI_ISL_3456543, EPI_ISL_3456544, EPI_ISL_3456545, EPI_ISL_3456546, EPI_ISL_3456548, EPI_ISL_3456549, EPI_ISL_3456550, EPI_ISL_3456551, EPI_ISL_3456552, EPI_ISL_3456553, EPI_ISL_3456554, EPI_ISL_3456555, EPI_ISL_3456556, EPI_ISL_3456557, EPI_ISL_3456558, EPI_ISL_3456559, EPI_ISL_3456561, EPI_ISL_3456562, EPI_ISL_3456563, EPI_ISL_3456564, EPI_ISL_3456565, EPI_ISL_3456566, EPI_ISL_3456567, EPI_ISL_3456568, EPI_ISL_3456569, EPI_ISL_3456570, EPI_ISL_3456571, EPI_ISL_3456572, EPI_ISL_3456573, EPI_ISL_3456574, EPI_ISL_3456575, EPI_ISL_3456576, EPI_ISL_3456577, EPI_ISL_3456578, EPI_ISL_3456579, EPI_ISL_3456580, EPI_ISL_3456581, EPI_ISL_3456582, EPI_ISL_3456583, EPI_ISL_3456584, EPI_ISL_3456585, EPI_ISL_3456586, EPI_ISL_3456587, EPI_ISL_3456588, EPI_ISL_3456589, EPI_ISL_3456590, EPI_ISL_3456591, EPI_ISL_3456592, EPI_ISL_3456593, EPI_ISL_3456594, EPI_ISL_3456595, EPI_ISL_3456596, EPI_ISL_3456597, EPI_ISL_3456598, EPI_ISL_3456599, EPI_ISL_3456600, EPI_ISL_3456601, EPI_ISL_3456602 | | | |
| see above | Quest Diagnostics Incorporated | Centers for Disease Control and Prevention Division of Viral Diseases, Pathogen Discovery | Dakota Howard, Dhwani Batra, Peter W. Cook, Kara Moser, Adrian Paskey, Jason Caravas, Benjamin Rambo-Martin, Shatavia Morrison, Christopher Gulvick, Scott Sammons, Yvette Unoarumhi, Darlene Wagner, Matthew Schmerer, S. H. Rosenthal, A. Gerasimova, R. M. Kagan, B. Anderson, M. Hua, Y. Liu, L.E. Bernstein, K.E. Livingston, A. Perez, I. A. Shlyakhter, R. V. Rolando, R. Owen, P. Tanpaiboon, F. Lacbawan, Clinton R. Paden, Duncan MacCannell |
| EPI_ISL_3456603, EPI_ISL_3456604, EPI_ISL_3456605, EPI_ISL_3456606, EPI_ISL_3456607, EPI_ISL_3456608, EPI_ISL_3456609, EPI_ISL_3456610, EPI_ISL_3456611, EPI_ISL_3456612, EPI_ISL_3456613, EPI_ISL_3456614, EPI_ISL_3456615, EPI_ISL_3456616, EPI_ISL_3456617, EPI_ISL_3456618, EPI_ISL_3456619, EPI_ISL_3456620, EPI_ISL_3456621, EPI_ISL_3456622, EPI_ISL_3456623, EPI_ISL_3456624, EPI_ISL_3456625, EPI_ISL_3456626, EPI_ISL_3456627, EPI_ISL_3456628, EPI_ISL_3456630, EPI_ISL_3456631, EPI_ISL_3456633, EPI_ISL_3456634, EPI_ISL_3456635, EPI_ISL_3456636, EPI_ISL_3456637, EPI_ISL_3456638, EPI_ISL_3456639, EPI_ISL_3456640, EPI_ISL_3456641, EPI_ISL_3456642, EPI_ISL_3456643, EPI_ISL_3456644, EPI_ISL_3456645, EPI_ISL_3456646, EPI_ISL_3456647, EPI_ISL_3456648, EPI_ISL_3456649, EPI_ISL_3456650, EPI_ISL_3456651, EPI_ISL_3456652, EPI_ISL_3456653, EPI_ISL_3456654, EPI_ISL_3456655, EPI_ISL_3456656, EPI_ISL_3456657, EPI_ISL_3456658, EPI_ISL_3456659, EPI_ISL_3456660, EPI_ISL_3456661, EPI_ISL_3456662, EPI_ISL_3456663, EPI_ISL_3456664, EPI_ISL_3456665, EPI_ISL_3456666, EPI_ISL_3456667, EPI_ISL_3456668, EPI_ISL_3456669, EPI_ISL_3456670, EPI_ISL_3456671, EPI_ISL_3456672, EPI_ISL_3456673, EPI_ISL_3456674, EPI_ISL_3456675, EPI_ISL_3456676, EPI_ISL_3456677, EPI_ISL_3456678, EPI_ISL_3456679, EPI_ISL_3456681, EPI_ISL_3456682, EPI_ISL_3456683, EPI_ISL_3456684, EPI_ISL_3456685, EPI_ISL_3456686, EPI_ISL_3456687, EPI_ISL_3456688, EPI_ISL_3456689, EPI_ISL_3456690, EPI_ISL_3456691, EPI_ISL_3456692, EPI_ISL_3456693, EPI_ISL_3456694, EPI_ISL_3456695, EPI_ISL_3456696, EPI_ISL_3456697, EPI_ISL_3456698, EPI_ISL_3456699, EPI_ISL_3456700, EPI_ISL_3456701, EPI_ISL_3456702, EPI_ISL_3456703, EPI_ISL_3456704, EPI_ISL_3456705, EPI_ISL_3456706, EPI_ISL_3456707, EPI_ISL_3456708, EPI_ISL_3456710, EPI_ISL_3456711, EPI_ISL_3456712, EPI_ISL_3456713, EPI_ISL_3456714, EPI_ISL_3456715, EPI_ISL_3456716, EPI_ISL_3456717, EPI_ISL_3456718, EPI_ISL_3456719, EPI_ISL_3456720, EPI_ISL_3456721, EPI_ISL_3456722, EPI_ISL_3456723, EPI_ISL_3456724, EPI_ISL_3456725, EPI_ISL_3456726, EPI_ISL_3456727, EPI_ISL_3456728, EPI_ISL_3456729, EPI_ISL_3456730, EPI_ISL_3456731, EPI_ISL_3456732, EPI_ISL_3456733, EPI_ISL_3456734, EPI_ISL_3456735, EPI_ISL_3456736, EPI_ISL_3456737, EPI_ISL_3456738, EPI_ISL_3456739, EPI_ISL_3456740, EPI_ISL_3456743, EPI_ISL_3456744, EPI_ISL_3456745, EPI_ISL_3456746, EPI_ISL_3456747, EPI_ISL_3456748, EPI_ISL_3456749, EPI_ISL_3456750, EPI_ISL_3456751, EPI_ISL_3456752, EPI_ISL_3456753, EPI_ISL_3456754, EPI_ISL_3456755, EPI_ISL_3456756, EPI_ISL_3456757, EPI_ISL_3456758, EPI_ISL_3456759, EPI_ISL_3456760, EPI_ISL_3456761, EPI_ISL_3456762, EPI_ISL_3456763, EPI_ISL_3456764, EPI_ISL_3456765, EPI_ISL_3456766, EPI_ISL_3456767, EPI_ISL_3456768, EPI_ISL_3456769, EPI_ISL_3456770, EPI_ISL_3456771, EPI_ISL_3456772, EPI_ISL_3456773, EPI_ISL_3456774, EPI_ISL_3456775, EPI_ISL_3456776, EPI_ISL_3456777, EPI_ISL_3456778, EPI_ISL_3456779, EPI_ISL_3456780, EPI_ISL_3456781, EPI_ISL_3456782, EPI_ISL_3456783, EPI_ISL_3456784, EPI_ISL_3456785, EPI_ISL_3456786, EPI_ISL_3456787, EPI_ISL_3456788, EPI_ISL_3456789, EPI_ISL_3456790, EPI_ISL_3456791, EPI_ISL_3456792, EPI_ISL_3456793, EPI_ISL_3456794, EPI_ISL_3456795, EPI_ISL_3456796, EPI_ISL_3456797, EPI_ISL_3456798, EPI_ISL_3456799, EPI_ISL_3456800, EPI_ISL_3456801, EPI_ISL_3456802, EPI_ISL_3456803, EPI_ISL_3456804, EPI_ISL_3456805, EPI_ISL_3456806, EPI_ISL_3456807, EPI_ISL_3456808, EPI_ISL_3456809, EPI_ISL_3456810, EPI_ISL_3456811, EPI_ISL_3456812, EPI_ISL_3456813, EPI_ISL_3456815, EPI_ISL_3456816, EPI_ISL_3456817, EPI_ISL_3456818, EPI_ISL_3456819, EPI_ISL_3456820, EPI_ISL_3456821, EPI_ISL_3456822, EPI_ISL_3456823, EPI_ISL_3456824, EPI_ISL_3456825, EPI_ISL_3456826, EPI_ISL_3456827, EPI_ISL_3456828, EPI_ISL_3456829, EPI_ISL_3456830, EPI_ISL_3456831, EPI_ISL_3456832, EPI_ISL_3456833, EPI_ISL_3456834, EPI_ISL_3456835, EPI_ISL_3456836, EPI_ISL_3456837, EPI_ISL_3456838, EPI_ISL_3456839, EPI_ISL_3456840, EPI_ISL_3456841, EPI_ISL_3456842, EPI_ISL_3456843, EPI_ISL_3456844, EPI_ISL_3456845, EPI_ISL_3456846, EPI_ISL_3456847, EPI_ISL_3456848, EPI_ISL_3456849, EPI_ISL_3456850, EPI_ISL_3456851, EPI_ISL_3456852, EPI_ISL_3456853, EPI_ISL_3456854, EPI_ISL_3456855, EPI_ISL_3456856, EPI_ISL_3456857, EPI_ISL_3456858, EPI_ISL_3456859, EPI_ISL_3456860, EPI_ISL_3456861, EPI_ISL_3456862, EPI_ISL_3456863, EPI_ISL_3456864, EPI_ISL_3456865, EPI_ISL_3456866, EPI_ISL_3456867, EPI_ISL_3456868, EPI_ISL_3456869, EPI_ISL_3456870, EPI_ISL_3456871, EPI_ISL_3456873, EPI_ISL_3456874, EPI_ISL_3456875, EPI_ISL_3456876, EPI_ISL_3456877, EPI_ISL_3456878, EPI_ISL_3456879, EPI_ISL_3456880, EPI_ISL_3456881, EPI_ISL_3456882, EPI_ISL_3456883, EPI_ISL_3456884, EPI_ISL_3456885, EPI_ISL_3456886, EPI_ISL_3456887, EPI_ISL_3456888, EPI_ISL_3456889, EPI_ISL_3456890, EPI_ISL_3456891, EPI_ISL_3456892, EPI_ISL_3456893, EPI_ISL_3456894, EPI_ISL_3456895, EPI_ISL_3456896, EPI_ISL_3456897, EPI_ISL_3456898, EPI_ISL_3456899, EPI_ISL_3456900, EPI_ISL_3456901, EPI_ISL_3456902, EPI_ISL_3456903, EPI_ISL_3456904, EPI_ISL_3456905, EPI_ISL_3456906, EPI_ISL_3456907, EPI_ISL_3456908, EPI_ISL_3456909, EPI_ISL_3456910, EPI_ISL_3456911, EPI_ISL_3456912, EPI_ISL_3456913, EPI_ISL_3456914, EPI_ISL_3456915, EPI_ISL_3456916, EPI_ISL_3456917, EPI_ISL_3456918, | | | |

| | | | |
|---|---|---|---|
| EPI_ISL_3456919, EPI_ISL_3456920, EPI_ISL_3456921, EPI_ISL_3456922, EPI_ISL_3456923, EPI_ISL_3456924, EPI_ISL_3456925, EPI_ISL_3456926, EPI_ISL_3456927, EPI_ISL_3456928, EPI_ISL_3456929, EPI_ISL_3456930, EPI_ISL_3456931, EPI_ISL_3456932 | | | |
| see above | Fulgent Genetics | Centers for Disease Control and Prevention Division of Viral Diseases, Pathogen Discovery | Dakota Howard, Dhwani Batra, Peter W. Cook, Kara Moser, Adrian Paskey, Jason Caravas, Benjamin Rambo-Martin, Shatavia Morrison, Christopher Gulvick, Scott Sammons, Yvette Unoarumhi, Darlene Wagner, Matthew Schmerer, Harry Gao, Mickey Li, John Gao, Joseph Fierro, Benafsh Sapra, Becky Tsai, Yan Meng, Doreen Ng, James Xie, Clinton R. Paden, Duncan MacCannell |
| EPI_ISL_3456933 | Azienda Sanitaria dell'Alto Adige - Laboratorio Aziendale di Microbiologia e Virologia | Azienda Sanitaria dell'Alto Adige | Irene Bianconi |
| EPI_ISL_3456934 | Fulgent Genetics | Centers for Disease Control and Prevention Division of Viral Diseases, Pathogen Discovery | Dakota Howard, Dhwani Batra, Peter W. Cook, Kara Moser, Adrian Paskey, Jason Caravas, Benjamin Rambo-Martin, Shatavia Morrison, Christopher Gulvick, Scott Sammons, Yvette Unoarumhi, Darlene Wagner, Matthew Schmerer, Harry Gao, Mickey Li, John Gao, Joseph Fierro, Benafsh Sapra, Becky Tsai, Yan Meng, Doreen Ng, James Xie, Clinton R. Paden, Duncan MacCannell |
| EPI_ISL_3456935 | Azienda Sanitaria dell'Alto Adige - Laboratorio Aziendale di Microbiologia e Virologia | Azienda Sanitaria dell'Alto Adige | Irene Bianconi |
| EPI_ISL_3456936 | Fulgent Genetics | Centers for Disease Control and Prevention Division of Viral Diseases, Pathogen Discovery | Dakota Howard, Dhwani Batra, Peter W. Cook, Kara Moser, Adrian Paskey, Jason Caravas, Benjamin Rambo-Martin, Shatavia Morrison, Christopher Gulvick, Scott Sammons, Yvette Unoarumhi, Darlene Wagner, Matthew Schmerer, Harry Gao, Mickey Li, John Gao, Joseph Fierro, Benafsh Sapra, Becky Tsai, Yan Meng, Doreen Ng, James Xie, Clinton R. Paden, Duncan MacCannell |
| EPI_ISL_3456937, EPI_ISL_3456938 | Azienda Sanitaria dell'Alto Adige - Laboratorio Aziendale di Microbiologia e Virologia | Azienda Sanitaria dell'Alto Adige | Irene Bianconi |
| EPI_ISL_3456939 | Fulgent Genetics | Centers for Disease Control and Prevention Division of Viral Diseases, Pathogen Discovery | Dakota Howard, Dhwani Batra, Peter W. Cook, Kara Moser, Adrian Paskey, Jason Caravas, Benjamin Rambo-Martin, Shatavia Morrison, Christopher Gulvick, Scott Sammons, Yvette Unoarumhi, Darlene Wagner, Matthew Schmerer, Harry Gao, Mickey Li, John Gao, Joseph Fierro, Benafsh Sapra, Becky Tsai, Yan Meng, Doreen Ng, James Xie, Clinton R. Paden, Duncan MacCannell |
| EPI_ISL_3456940 | Azienda Sanitaria dell'Alto Adige - Laboratorio Aziendale di Microbiologia e Virologia | Azienda Sanitaria dell'Alto Adige | Irene Bianconi |
| EPI_ISL_3456941, EPI_ISL_3456942, EPI_ISL_3456943, EPI_ISL_3456944, EPI_ISL_3456945, EPI_ISL_3456946, EPI_ISL_3456947, EPI_ISL_3456948, EPI_ISL_3456949, EPI_ISL_3456950, EPI_ISL_3456951, EPI_ISL_3456952, EPI_ISL_3456953, EPI_ISL_3456954, EPI_ISL_3456955, EPI_ISL_3456956, EPI_ISL_3456957, EPI_ISL_3456958, EPI_ISL_3456959, EPI_ISL_3456960, EPI_ISL_3456961, EPI_ISL_3456962, EPI_ISL_3456963, EPI_ISL_3456964, EPI_ISL_3456965, EPI_ISL_3456966, EPI_ISL_3456967, EPI_ISL_3456968, EPI_ISL_3456969, EPI_ISL_3456970, EPI_ISL_3456971, EPI_ISL_3456972, EPI_ISL_3456974, EPI_ISL_3456975, EPI_ISL_3456976, EPI_ISL_3456977, EPI_ISL_3456978, EPI_ISL_3456979, EPI_ISL_3456980, EPI_ISL_3456981, EPI_ISL_3456982, EPI_ISL_3456983, EPI_ISL_3456984, EPI_ISL_3456985, EPI_ISL_3456986, EPI_ISL_3456987, EPI_ISL_3456988, EPI_ISL_3456989, EPI_ISL_3456990, EPI_ISL_3456991, EPI_ISL_3456992, EPI_ISL_3456993, EPI_ISL_3456994, EPI_ISL_3456995, EPI_ISL_3456996, EPI_ISL_3456997, EPI_ISL_3456998, EPI_ISL_3456999, EPI_ISL_3457000, EPI_ISL_3457001, EPI_ISL_3457002, EPI_ISL_3457003, EPI_ISL_3457004, EPI_ISL_3457006, EPI_ISL_3457007, EPI_ISL_3457008, EPI_ISL_3457009, EPI_ISL_3457010, EPI_ISL_3457011, EPI_ISL_3457012, EPI_ISL_3457013, EPI_ISL_3457014, EPI_ISL_3457015, EPI_ISL_3457017, EPI_ISL_3457018, EPI_ISL_3457019, EPI_ISL_3457020, EPI_ISL_3457021, EPI_ISL_3457022, EPI_ISL_3457023, EPI_ISL_3457024, EPI_ISL_3457025, EPI_ISL_3457026, EPI_ISL_3457027, EPI_ISL_3457028, EPI_ISL_3457029, EPI_ISL_3457030, EPI_ISL_3457031, EPI_ISL_3457032, EPI_ISL_3457033, EPI_ISL_3457034, EPI_ISL_3457035, EPI_ISL_3457036, EPI_ISL_3457037, EPI_ISL_3457038, EPI_ISL_3457039, EPI_ISL_3457040, EPI_ISL_3457041, EPI_ISL_3457042, EPI_ISL_3457043, EPI_ISL_3457045, EPI_ISL_3457046, EPI_ISL_3457047, EPI_ISL_3457048, EPI_ISL_3457049, EPI_ISL_3457050, EPI_ISL_3457051, EPI_ISL_3457052, EPI_ISL_3457053, EPI_ISL_3457055, EPI_ISL_3457058, EPI_ISL_3457061, EPI_ISL_3457063, EPI_ISL_3457064, EPI_ISL_3457065, EPI_ISL_3457066, EPI_ISL_3457067, EPI_ISL_3457068, EPI_ISL_3457069, EPI_ISL_3457070, EPI_ISL_3457071, EPI_ISL_3457072, EPI_ISL_3457073, EPI_ISL_3457074, EPI_ISL_3457075, EPI_ISL_3457076, EPI_ISL_3457077, EPI_ISL_3457078, EPI_ISL_3457079, EPI_ISL_3457080, EPI_ISL_3457081, EPI_ISL_3457082, EPI_ISL_3457083, EPI_ISL_3457084, EPI_ISL_3457085, EPI_ISL_3457086, EPI_ISL_3457087, EPI_ISL_3457088, EPI_ISL_3457089, EPI_ISL_3457090, EPI_ISL_3457091, EPI_ISL_3457092, EPI_ISL_3457093, EPI_ISL_3457094, EPI_ISL_3457095, EPI_ISL_3457096, EPI_ISL_3457097, EPI_ISL_3457098, EPI_ISL_3457099, EPI_ISL_3457100, EPI_ISL_3457101, EPI_ISL_3457102, EPI_ISL_3457103, EPI_ISL_3457104, EPI_ISL_3457105, EPI_ISL_3457106, EPI_ISL_3457107, EPI_ISL_3457108, EPI_ISL_3457109, EPI_ISL_3457110, EPI_ISL_3457111, EPI_ISL_3457112, EPI_ISL_3457113, EPI_ISL_3457114, EPI_ISL_3457115, EPI_ISL_3457116, EPI_ISL_3457117, EPI_ISL_3457118, EPI_ISL_3457119, EPI_ISL_3457120, EPI_ISL_3457121, EPI_ISL_3457122, EPI_ISL_3457123, EPI_ISL_3457124, EPI_ISL_3457125, EPI_ISL_3457126, EPI_ISL_3457127, EPI_ISL_3457128, EPI_ISL_3457129, EPI_ISL_3457130, EPI_ISL_3457131, EPI_ISL_3457132, EPI_ISL_3457133, EPI_ISL_3457134, EPI_ISL_3457135, EPI_ISL_3457136, EPI_ISL_3457137, EPI_ISL_3457138, EPI_ISL_3457139, EPI_ISL_3457140, EPI_ISL_3457141, EPI_ISL_3457142, EPI_ISL_3457143, EPI_ISL_3457144, EPI_ISL_3457145, EPI_ISL_3457146, EPI_ISL_3457147, EPI_ISL_3457148, EPI_ISL_3457149, EPI_ISL_3457150, EPI_ISL_3457151, EPI_ISL_3457152, EPI_ISL_3457153, EPI_ISL_3457154, EPI_ISL_3457155, EPI_ISL_3457156, EPI_ISL_3457157, EPI_ISL_3457158, EPI_ISL_3457159, EPI_ISL_3457160, EPI_ISL_3457161, EPI_ISL_3457162, EPI_ISL_3457163, EPI_ISL_3457164, EPI_ISL_3457165, EPI_ISL_3457166, EPI_ISL_3457167, EPI_ISL_3457168, EPI_ISL_3457169, EPI_ISL_3457170, EPI_ISL_3457171, EPI_ISL_3457172, EPI_ISL_3457173, EPI_ISL_3457174, EPI_ISL_3457175, EPI_ISL_3457176, EPI_ISL_3457177, EPI_ISL_3457178, EPI_ISL_3457179, EPI_ISL_3457180, EPI_ISL_3457181, EPI_ISL_3457182, EPI_ISL_3457183, EPI_ISL_3457184, EPI_ISL_3457185, EPI_ISL_3457186, EPI_ISL_3457187, EPI_ISL_3457188, EPI_ISL_3457189, EPI_ISL_3457191, EPI_ISL_3457192, EPI_ISL_3457193, EPI_ISL_3457194, EPI_ISL_3457195, EPI_ISL_3457196, EPI_ISL_3457197, EPI_ISL_3457198, EPI_ISL_3457199, EPI_ISL_3457200, EPI_ISL_3457201, EPI_ISL_3457202, EPI_ISL_3457203, EPI_ISL_3457204, EPI_ISL_3457205, EPI_ISL_3457206, EPI_ISL_3457207, EPI_ISL_3457208, EPI_ISL_3457209, EPI_ISL_3457210, EPI_ISL_3457211, EPI_ISL_3457212, EPI_ISL_3457213, EPI_ISL_3457214, EPI_ISL_3457215, EPI_ISL_3457216, EPI_ISL_3457217, EPI_ISL_3457218, EPI_ISL_3457219, EPI_ISL_3457220, EPI_ISL_3457221, EPI_ISL_3457222, EPI_ISL_3457223, EPI_ISL_3457224, EPI_ISL_3457225, EPI_ISL_3457226, EPI_ISL_3457227, EPI_ISL_3457228, EPI_ISL_3457229, EPI_ISL_3457230, EPI_ISL_3457231, EPI_ISL_3457232, EPI_ISL_3457233, EPI_ISL_3457234, EPI_ISL_3457235, EPI_ISL_3457236, EPI_ISL_3457237, EPI_ISL_3457238, EPI_ISL_3457239, EPI_ISL_3457240, EPI_ISL_3457241, EPI_ISL_3457242, EPI_ISL_3457243, EPI_ISL_3457244, EPI_ISL_3457245, EPI_ISL_3457246, EPI_ISL_3457247, EPI_ISL_3457248, EPI_ISL_3457249, EPI_ISL_3457250, EPI_ISL_3457251, EPI_ISL_3457252, EPI_ISL_3457253, EPI_ISL_3457254, EPI_ISL_3457255, EPI_ISL_3457256, EPI_ISL_3457257, EPI_ISL_3457258, EPI_ISL_3457259, EPI_ISL_3457260, EPI_ISL_3457261, EPI_ISL_3457262, EPI_ISL_3457263, EPI_ISL_3457264, EPI_ISL_3457265, EPI_ISL_3457266, EPI_ISL_3457267, EPI_ISL_3457268, EPI_ISL_3457269, EPI_ISL_3457270, EPI_ISL_3457271, EPI_ISL_3457272, EPI_ISL_3457273, EPI_ISL_3457274, EPI_ISL_3457275, EPI_ISL_3457276, EPI_ISL_3457277, EPI_ISL_3457278, EPI_ISL_3457279, EPI_ISL_3457280, EPI_ISL_3457281, EPI_ISL_3457282, EPI_ISL_3457283, EPI_ISL_3457284, EPI_ISL_3457285, EPI_ISL_3457286, EPI_ISL_3457287, EPI_ISL_3457288, EPI_ISL_3457289, EPI_ISL_3457290, EPI_ISL_3457291, EPI_ISL_3457292, EPI_ISL_3457293, EPI_ISL_3457294, EPI_ISL_3457295, EPI_ISL_3457296, EPI_ISL_3457297, EPI_ISL_3457298, EPI_ISL_3457299, EPI_ISL_3457300, EPI_ISL_3457301, EPI_ISL_3457302, EPI_ISL_3457303, EPI_ISL_3457304, EPI_ISL_3457305, EPI_ISL_3457306, EPI_ISL_3457307, EPI_ISL_3457308, EPI_ISL_3457309, EPI_ISL_3457310, EPI_ISL_3457311, EPI_ISL_3457312, EPI_ISL_3457313, EPI_ISL_3457314, EPI_ISL_3457315, EPI_ISL_3457316, EPI_ISL_3457317, EPI_ISL_3457318, EPI_ISL_3457319, EPI_ISL_3457320, EPI_ISL_3457321, EPI_ISL_3457322, EPI_ISL_3457323, EPI_ISL_3457324, EPI_ISL_3457325, EPI_ISL_3457326, EPI_ISL_3457327, EPI_ISL_3457329, EPI_ISL_3457330, EPI_ISL_3457331, EPI_ISL_3457332, EPI_ISL_3457333, EPI_ISL_3457334, EPI_ISL_3457335, EPI_ISL_3457336, EPI_ISL_3457337, EPI_ISL_3457338, EPI_ISL_3457339, EPI_ISL_3457340, EPI_ISL_3457341, EPI_ISL_3457342, EPI_ISL_3457343, EPI_ISL_3457344, EPI_ISL_3457345, EPI_ISL_3457346, EPI_ISL_3457347, EPI_ISL_3457348, EPI_ISL_3457349, EPI_ISL_3457350, EPI_ISL_3457351, EPI_ISL_3457352, EPI_ISL_3457353, EPI_ISL_3457354, EPI_ISL_3457355, EPI_ISL_3457356, EPI_ISL_3457357, EPI_ISL_3457358, EPI_ISL_3457359, EPI_ISL_3457360, EPI_ISL_3457361, EPI_ISL_3457362, EPI_ISL_3457363, EPI_ISL_3457364, EPI_ISL_3457365, EPI_ISL_3457366, EPI_ISL_3457367, EPI_ISL_3457368, EPI_ISL_3457369, EPI_ISL_3457370, EPI_ISL_3457371, EPI_ISL_3457372, EPI_ISL_3457373, EPI_ISL_3457376, EPI_ISL_3457377, EPI_ISL_3457378, EPI_ISL_3457379, EPI_ISL_3457380, EPI_ISL_3457381, EPI_ISL_3457382, EPI_ISL_3457383, EPI_ISL_3457384, EPI_ISL_3457385, EPI_ISL_3457386, EPI_ISL_3457387, EPI_ISL_3457388, EPI_ISL_3457389, EPI_ISL_3457390, EPI_ISL_3457391, EPI_ISL_3457392, EPI_ISL_3457393, EPI_ISL_3457394, EPI_ISL_3457395, EPI_ISL_3457396, EPI_ISL_3457397, EPI_ISL_3457398, EPI_ISL_3457399, EPI_ISL_3457400, EPI_ISL_3457401, EPI_ISL_3457402, EPI_ISL_3457403, EPI_ISL_3457404, EPI_ISL_3457405, EPI_ISL_3457406, EPI_ISL_3457407, EPI_ISL_3457408, EPI_ISL_3457409, EPI_ISL_3457410, EPI_ISL_3457411, EPI_ISL_3457412, EPI_ISL_3457413, EPI_ISL_3457414, EPI_ISL_3457415, EPI_ISL_3457416, EPI_ISL_3457417, EPI_ISL_3457418, EPI_ISL_3457419, EPI_ISL_3457420, EPI_ISL_3457421, EPI_ISL_3457422, EPI_ISL_3457424, EPI_ISL_3457425, EPI_ISL_3457426, EPI_ISL_3457427, EPI_ISL_3457428, EPI_ISL_3457429 | | | |
| see above | Fulgent Genetics | Centers for Disease Control and Prevention Division of Viral Diseases, Pathogen Discovery | Dakota Howard, Dhwani Batra, Peter W. Cook, Kara Moser, Adrian Paskey, Jason Caravas, Benjamin Rambo-Martin, Shatavia Morrison, Christopher Gulvick, Scott Sammons, Yvette Unoarumhi, Darlene Wagner, Matthew Schmerer, Harry Gao, Mickey Li, John Gao, Joseph Fierro, Benafsh Sapra, Becky Tsai, Yan Meng, Doreen Ng, James Xie, Clinton R. Paden, Duncan MacCannell |
| EPI_ISL_3457430 | Plateforme de testing Namuroise | Plateforme de testing Namuroise | Lesly Nyinkeu Kemamen: Nicolas Gilliard; Demars Aurore ; Maschietto Céline ; Nobis Chloé ; Otto Gaetan ; Denis Olivier ; Degosserie Jonathan ; Mullier François |
| EPI_ISL_3457431, EPI_ISL_3457432, EPI_ISL_3457433, EPI_ISL_3457434, EPI_ISL_3457435, EPI_ISL_3457437, EPI_ISL_3457438, EPI_ISL_3457439, EPI_ISL_3457440, EPI_ISL_3457441, EPI_ISL_3457442, EPI_ISL_3457443, EPI_ISL_3457444, EPI_ISL_3457445, EPI_ISL_3457446, EPI_ISL_3457447, EPI_ISL_3457448, EPI_ISL_3457449, EPI_ISL_3457450, EPI_ISL_3457451, EPI_ISL_3457452, EPI_ISL_3457453, EPI_ISL_3457454, EPI_ISL_3457455, EPI_ISL_3457456, EPI_ISL_3457457, EPI_ISL_3457458, EPI_ISL_3457459, EPI_ISL_3457460, EPI_ISL_3457461, EPI_ISL_3457462, EPI_ISL_3457463, EPI_ISL_3457464, EPI_ISL_3457465, EPI_ISL_3457466, EPI_ISL_3457467, EPI_ISL_3457468, EPI_ISL_3457469, EPI_ISL_3457470, EPI_ISL_3457471, EPI_ISL_3457472, EPI_ISL_3457473, EPI_ISL_3457474, EPI_ISL_3457475, EPI_ISL_3457476, EPI_ISL_3457477, EPI_ISL_3457478, EPI_ISL_3457479, EPI_ISL_3457480, EPI_ISL_3457481, EPI_ISL_3457482, EPI_ISL_3457483, EPI_ISL_3457485, EPI_ISL_3457486, EPI_ISL_3457487, EPI_ISL_3457488, EPI_ISL_3457489, EPI_ISL_3457490, EPI_ISL_3457491, EPI_ISL_3457492, EPI_ISL_3457493, EPI_ISL_3457494, EPI_ISL_3457495, EPI_ISL_3457496, EPI_ISL_3457497, EPI_ISL_3457498, EPI_ISL_3457499, EPI_ISL_3457500, EPI_ISL_3457501, EPI_ISL_3457502, EPI_ISL_3457503, EPI_ISL_3457504, EPI_ISL_3457505, EPI_ISL_3457506, EPI_ISL_3457507, EPI_ISL_3457508, EPI_ISL_3457509, EPI_ISL_3457510, EPI_ISL_3457511, EPI_ISL_3457512, EPI_ISL_3457513, EPI_ISL_3457514, EPI_ISL_3457515, EPI_ISL_3457516, EPI_ISL_3457517, EPI_ISL_3457518, EPI_ISL_3457519, EPI_ISL_3457520, EPI_ISL_3457521, EPI_ISL_3457522, EPI_ISL_3457523, EPI_ISL_3457524, EPI_ISL_3457525, EPI_ISL_3457526, EPI_ISL_3457527, EPI_ISL_3457528, EPI_ISL_3457529, EPI_ISL_3457530, EPI_ISL_3457531, EPI_ISL_3457532, EPI_ISL_3457533, EPI_ISL_3457534, EPI_ISL_3457535, EPI_ISL_3457536, EPI_ISL_3457537, EPI_ISL_3457538, EPI_ISL_3457539, EPI_ISL_3457540, EPI_ISL_3457541, EPI_ISL_3457542, EPI_ISL_3457543, EPI_ISL_3457544, EPI_ISL_3457545, EPI_ISL_3457546, EPI_ISL_3457547, EPI_ISL_3457548, EPI_ISL_3457549, EPI_ISL_3457550, EPI_ISL_3457551, EPI_ISL_3457552, EPI_ISL_3457553, EPI_ISL_3457554, EPI_ISL_3457555, EPI_ISL_3457556, EPI_ISL_3457557, EPI_ISL_3457558, EPI_ISL_3457559, EPI_ISL_3457560, EPI_ISL_3457561, EPI_ISL_3457562, EPI_ISL_3457563, EPI_ISL_3457564, EPI_ISL_3457565, EPI_ISL_3457566, EPI_ISL_3457567, EPI_ISL_3457568, EPI_ISL_3457569, EPI_ISL_3457570, EPI_ISL_3457571, EPI_ISL_3457572, EPI_ISL_3457573, EPI_ISL_3457574, EPI_ISL_3457575, EPI_ISL_3457576, EPI_ISL_3457577, EPI_ISL_3457578, EPI_ISL_3457579, EPI_ISL_3457580, EPI_ISL_3457581, EPI_ISL_3457582, EPI_ISL_3457583, EPI_ISL_3457584, EPI_ISL_3457585, EPI_ISL_3457586, EPI_ISL_3457587, EPI_ISL_3457588, EPI_ISL_3457589, EPI_ISL_3457590, EPI_ISL_3457591, EPI_ISL_3457592, EPI_ISL_3457593, EPI_ISL_3457594, EPI_ISL_3457595, EPI_ISL_3457596, EPI_ISL_3457597, EPI_ISL_3457598, EPI_ISL_3457599, EPI_ISL_3457600, EPI_ISL_3457601, EPI_ISL_3457602, EPI_ISL_3457603, EPI_ISL_3457604, EPI_ISL_3457605, EPI_ISL_3457606, EPI_ISL_3457607, EPI_ISL_3457608, EPI_ISL_3457609, EPI_ISL_3457610, EPI_ISL_3457611, EPI_ISL_3457612, EPI_ISL_3457613, EPI_ISL_3457614, EPI_ISL_3457615, EPI_ISL_3457616, EPI_ISL_3457617, EPI_ISL_3457618, EPI_ISL_3457619, EPI_ISL_3457620, EPI_ISL_3457621, EPI_ISL_3457622, EPI_ISL_3457623, EPI_ISL_3457624, EPI_ISL_3457625, EPI_ISL_3457626, EPI_ISL_3457627, EPI_ISL_3457628, EPI_ISL_3457629, EPI_ISL_3457630, EPI_ISL_3457631, EPI_ISL_3457632, EPI_ISL_3457633, EPI_ISL_3457634, EPI_ISL_3457635, EPI_ISL_3457636, EPI_ISL_3457637, EPI_ISL_3457638, EPI_ISL_3457639, EPI_ISL_3457640, EPI_ISL_3457641, EPI_ISL_3457642, EPI_ISL_3457643, EPI_ISL_3457644, EPI_ISL_3457645, EPI_ISL_3457646, EPI_ISL_3457647, EPI_ISL_3457648, EPI_ISL_3457649, EPI_ISL_3457650, EPI_ISL_3457651, EPI_ISL_3457652, EPI_ISL_3457653, EPI_ISL_3457654, EPI_ISL_3457655, EPI_ISL_3457656, EPI_ISL_3457657, EPI_ISL_3457658, EPI_ISL_3457659, EPI_ISL_3457660, EPI_ISL_3457661, EPI_ISL_3457662, EPI_ISL_3457663, EPI_ISL_3457664, EPI_ISL_3457665, EPI_ISL_3457666, EPI_ISL_3457667, EPI_ISL_3457668, EPI_ISL_3457669, EPI_ISL_3457670, EPI_ISL_3457671, EPI_ISL_3457672, EPI_ISL_3457673, EPI_ISL_3457674, EPI_ISL_3457675, EPI_ISL_3457676, EPI_ISL_3457677, EPI_ISL_3457678, EPI_ISL_3457679, EPI_ISL_3457680, EPI_ISL_3457681, EPI_ISL_3457682, EPI_ISL_3457683, EPI_ISL_3457684, EPI_ISL_3457685, EPI_ISL_3457686, EPI_ISL_3457687, EPI_ISL_3457688, EPI_ISL_3457689, EPI_ISL_3457690, EPI_ISL_3457691, EPI_ISL_3457692, EPI_ISL_3457693, EPI_ISL_3457694, EPI_ISL_3457695, EPI_ISL_3457696, EPI_ISL_3457697, EPI_ISL_3457698, EPI_ISL_3457699, EPI_ISL_3457700, EPI_ISL_3457701, EPI_ISL_3457702, EPI_ISL_3457703, EPI_ISL_3457704, EPI_ISL_3457705, EPI_ISL_3457706, EPI_ISL_3457707, EPI_ISL_3457708, EPI_ISL_3457709, EPI_ISL_3457710, EPI_ISL_3457711, EPI_ISL_3457712, EPI_ISL_3457713, EPI_ISL_3457714, EPI_ISL_3457715, EPI_ISL_3457716, EPI_ISL_3457717, EPI_ISL_3457718, EPI_ISL_3457719, EPI_ISL_3457720, EPI_ISL_3457721, EPI_ISL_3457722, EPI_ISL_3457723, EPI_ISL_3457724, EPI_ISL_3457725, EPI_ISL_3457726, EPI_ISL_3457727, EPI_ISL_3457728, EPI_ISL_3457729, EPI_ISL_3457730, EPI_ISL_3457731, EPI_ISL_3457732, EPI_ISL_3457733, EPI_ISL_3457734, EPI_ISL_3457735, EPI_ISL_3457736, EPI_ISL_3457737, EPI_ISL_3457738, EPI_ISL_3457739, EPI_ISL_3457740, EPI_ISL_3457741, EPI_ISL_3457742, EPI_ISL_3457743, EPI_ISL_3457744, EPI_ISL_3457745, EPI_ISL_3457746, EPI_ISL_3457747, EPI_ISL_3457748, EPI_ISL_3457749, EPI_ISL_3457750, EPI_ISL_3457751, EPI_ISL_3457752, EPI_ISL_3457753, EPI_ISL_3457754, EPI_ISL_3457755, EPI_ISL_3457756, EPI_ISL_3457757, EPI_ISL_3457758, EPI_ISL_3457759, EPI_ISL_3457760, EPI_ISL_3457761, EPI_ISL_3457762, EPI_ISL_3457763, EPI_ISL_3457764, EPI_ISL_3457765, EPI_ISL_3457766, EPI_ISL_3457767, EPI_ISL_3457768, EPI_ISL_3457769, EPI_ISL_3457770, EPI_ISL_3457771, EPI_ISL_3457772, EPI_ISL_3457773, EPI_ISL_3457774, EPI_ISL_3457775, EPI_ISL_3457776, EPI_ISL_3457777, EPI_ISL_3457778, EPI_ISL_3457779, EPI_ISL_3457780, EPI_ISL_3457781, EPI_ISL_3457782, EPI_ISL_3457783, EPI_ISL_3457784, EPI_ISL_3457785, EPI_ISL_3457786, EPI_ISL_3457787, EPI_ISL_3457789, EPI_ISL_3457790, EPI_ISL_3457791, EPI_ISL_3457792, EPI_ISL_3457793, EPI_ISL_3457794, EPI_ISL_3457795, EPI_ISL_3457796, EPI_ISL_3457797, EPI_ISL_3457798, EPI_ISL_3457799, EPI_ISL_3457800, EPI_ISL_3457801, EPI_ISL_3457803, EPI_ISL_3457804, EPI_ISL_3457806, EPI_ISL_3457809, EPI_ISL_3457811, EPI_ISL_3457814, EPI_ISL_3457817, EPI_ISL_3457820, EPI_ISL_3457822, EPI_ISL_3457825, EPI_ISL_3457828, EPI_ISL_3457830, EPI_ISL_3457833, EPI_ISL_3457835, EPI_ISL_3457838, EPI_ISL_3457840, EPI_ISL_3457846, EPI_ISL_3457848, EPI_ISL_3457851, EPI_ISL_3457853, EPI_ISL_3457855, EPI_ISL_3457858, EPI_ISL_3457860, EPI_ISL_3457863, EPI_ISL_3457865, EPI_ISL_3457868, EPI_ISL_3457871, EPI_ISL_3457873, EPI_ISL_3457876, EPI_ISL_3457878, EPI_ISL_3457881, EPI_ISL_3457884, EPI_ISL_3457886, EPI_ISL_3457889, EPI_ISL_3457892, EPI_ISL_3457894, EPI_ISL_3457897, EPI_ISL_3457899, EPI_ISL_3457902, EPI_ISL_3457905, EPI_ISL_3457907, EPI_ISL_3457910, EPI_ISL_3457912, EPI_ISL_3457914, EPI_ISL_3457916, EPI_ISL_3457919, EPI_ISL_3457922, EPI_ISL_3457924, EPI_ISL_3457927, EPI_ISL_3457930, EPI_ISL_3457933, EPI_ISL_3457935, EPI_ISL_3457938, EPI_ISL_3457940, EPI_ISL_3457943, EPI_ISL_3457945, EPI_ISL_3457948, EPI_ISL_3457950, EPI_ISL_3457953, EPI_ISL_3457956, EPI_ISL_3457958, EPI_ISL_3457961, EPI_ISL_3457963, EPI_ISL_3457966, EPI_ISL_3457968, EPI_ISL_3457971, EPI_ISL_3457973, EPI_ISL_3457976, EPI_ISL_3457978, EPI_ISL_3457981, EPI_ISL_3457984, EPI_ISL_3457986, EPI_ISL_3457988, EPI_ISL_3457991, EPI_ISL_3457993, EPI_ISL_3457996, EPI_ISL_3457998, EPI_ISL_3458001, EPI_ISL_3458003, EPI_ISL_3458006, EPI_ISL_3458009, EPI_ISL_3458012, EPI_ISL_3458015, EPI_ISL_3458017, EPI_ISL_3458020, EPI_ISL_3458022, EPI_ISL_3458025, EPI_ISL_3458028, EPI_ISL_3458031, EPI_ISL_3458033, EPI_ISL_3458035, EPI_ISL_3458038, EPI_ISL_3458041, EPI_ISL_3458043, EPI_ISL_3458046, EPI_ISL_3458049, EPI_ISL_3458051, EPI_ISL_3458054, EPI_ISL_3458056, EPI_ISL_3458058, EPI_ISL_3458061, EPI_ISL_3458063, EPI_ISL_3458065, EPI_ISL_3458068, EPI_ISL_3458070, EPI_ISL_3458073, EPI_ISL_3458075, EPI_ISL_3458078, EPI_ISL_3458080, EPI_ISL_3458083, EPI_ISL_3458085, EPI_ISL_3458087, EPI_ISL_3458090, EPI_ISL_3458092 | | | |
| see above | Fulgent Genetics | Centers for Disease Control and Prevention Division of Viral Diseases, Pathogen Discovery | Dakota Howard, Dhwani Batra, Peter W. Cook, Kara Moser, Adrian Paskey, Jason Caravas, Benjamin Rambo-Martin, Shatavia Morrison, Christopher Gulvick, Scott Sammons, Yvette Unoarumhi, Darlene Wagner, Matthew Schmerer, Harry Gao, Mickey Li, John Gao, Joseph Fierro, Benafsh Sapra, Becky Tsai, Yan Meng, Doreen Ng, James Xie, Clinton R. Paden, Duncan MacCannell |
| EPI_ISL_3458577, EPI_ISL_3458631, EPI_ISL_3458676, EPI_ISL_3458745, EPI_ISL_3458746, EPI_ISL_3458754, EPI_ISL_3458758, EPI_ISL_3458759, EPI_ISL_3458761, EPI_ISL_3458764, EPI_ISL_3458765, EPI_ISL_3458766, EPI_ISL_3458770, EPI_ISL_3458771, EPI_ISL_3458773, EPI_ISL_3458774, EPI_ISL_3458775, EPI_ISL_3458776, EPI_ISL_3458777, EPI_ISL_3458790, EPI_ISL_3458792, EPI_ISL_3458795, EPI_ISL_3458802, EPI_ISL_3458807, EPI_ISL_3458818, EPI_ISL_3458829, EPI_ISL_3458830, EPI_ISL_3458831, EPI_ISL_3458832, EPI_ISL_3458833, EPI_ISL_3458834, EPI_ISL_3458835, EPI_ISL_3458836, EPI_ISL_3458837, EPI_ISL_3458838, EPI_ISL_3458839, EPI_ISL_3458843, EPI_ISL_3458846, EPI_ISL_3458852, EPI_ISL_3458863, EPI_ISL_3458868, EPI_ISL_3458869, EPI_ISL_3458872, EPI_ISL_3458874, EPI_ISL_3458882, EPI_ISL_3458883, EPI_ISL_3458890, EPI_ISL_3458902, EPI_ISL_3458903, EPI_ISL_3458908, EPI_ISL_3458911, EPI_ISL_3458913, EPI_ISL_3458915, EPI_ISL_3458917, EPI_ISL_3458918, EPI_ISL_3458919, EPI_ISL_3458920, EPI_ISL_3458922, EPI_ISL_3458923, EPI_ISL_3458926, EPI_ISL_3458934, EPI_ISL_3458936, EPI_ISL_3458937, EPI_ISL_3458938, EPI_ISL_3458939, EPI_ISL_3458940, EPI_ISL_3458941, EPI_ISL_3458942, EPI_ISL_3458943, EPI_ISL_3458944, EPI_ISL_3458946, EPI_ISL_3458947, EPI_ISL_3458951, EPI_ISL_3458953, EPI_ISL_3458955, EPI_ISL_3458956, EPI_ISL_3458957, EPI_ISL_3458958, EPI_ISL_3458959, EPI_ISL_3458960, EPI_ISL_3458961, EPI_ISL_3458962, EPI_ISL_3458963, EPI_ISL_3458964, EPI_ISL_3458965, EPI_ISL_3458972, EPI_ISL_3458977, EPI_ISL_3458979, EPI_ISL_3458984, EPI_ISL_3458991, EPI_ISL_3458996, EPI_ISL_3459000, EPI_ISL_3459001, EPI_ISL_3459003, EPI_ISL_3459004, EPI_ISL_3459005, EPI_ISL_3459010, EPI_ISL_3459012, EPI_ISL_3459016, EPI_ISL_3459018, EPI_ISL_3459019, EPI_ISL_3459020, EPI_ISL_3459022, EPI_ISL_3459023, EPI_ISL_3459024, EPI_ISL_3459026, EPI_ISL_3459028, EPI_ISL_3459029, EPI_ISL_3459030, EPI_ISL_3459038, EPI_ISL_3459039, EPI_ISL_3459040, EPI_ISL_3459043, EPI_ISL_3459044, EPI_ISL_3459045, EPI_ISL_3459047, EPI_ISL_3459052, EPI_ISL_3459054, EPI_ISL_3459055, EPI_ISL_3459059, EPI_ISL_3459060, EPI_ISL_3459064, EPI_ISL_3459065, EPI_ISL_3459066, EPI_ISL_3459068, EPI_ISL_3459069, EPI_ISL_3459070, EPI_ISL_3459071, EPI_ISL_3459072, EPI_ISL_3459073, EPI_ISL_3459074, EPI_ISL_3459075, EPI_ISL_3459076, EPI_ISL_3459077, EPI_ISL_3459078, EPI_ISL_3459079, EPI_ISL_3459080, EPI_ISL_3459081, EPI_ISL_3459090, EPI_ISL_3459091, EPI_ISL_3459092, EPI_ISL_3459093, EPI_ISL_3459094, EPI_ISL_3459101, EPI_ISL_3459102, EPI_ISL_3459103, EPI_ISL_3459104, EPI_ISL_3459105, EPI_ISL_3459110, EPI_ISL_3459111, EPI_ISL_3459112, EPI_ISL_3459113, EPI_ISL_3459114, EPI_ISL_3459115, EPI_ISL_3459116, EPI_ISL_3459119, EPI_ISL_3459120, EPI_ISL_3459122, EPI_ISL_3459123, EPI_ISL_3459124, EPI_ISL_3459127, EPI_ISL_3459129, EPI_ISL_3459130, EPI_ISL_3459131, EPI_ISL_3459132, EPI_ISL_3459133, EPI_ISL_3459134, EPI_ISL_3459135, EPI_ISL_3459136, EPI_ISL_3459139, EPI_ISL_3459142, EPI_ISL_3459143, EPI_ISL_3459144, EPI_ISL_3459146, EPI_ISL_3459147, EPI_ISL_3459148, EPI_ISL_3459150, EPI_ISL_3459151, EPI_ISL_3459161, EPI_ISL_3459163, EPI_ISL_3459164, EPI_ISL_3459168, EPI_ISL_3459170, EPI_ISL_3459172, EPI_ISL_3459173, EPI_ISL_3459174, EPI_ISL_3459175, EPI_ISL_3459176, EPI_ISL_3459177, EPI_ISL_3459178, EPI_ISL_3459179, EPI_ISL_3459180, EPI_ISL_3459181, EPI_ISL_3459182, EPI_ISL_3459183, EPI_ISL_3459184, EPI_ISL_3459185, EPI_ISL_3459186, EPI_ISL_3459187, EPI_ISL_3459188, EPI_ISL_3459189, EPI_ISL_3459190, EPI_ISL_3459191, EPI_ISL_3459192, EPI_ISL_3459193, EPI_ISL_3459194, EPI_ISL_3459195, EPI_ISL_3459196, EPI_ISL_3459197, EPI_ISL_3459198, EPI_ISL_3459199, EPI_ISL_3459200, EPI_ISL_3459202, EPI_ISL_3459203, EPI_ISL_3459204, EPI_ISL_3459206, EPI_ISL_3459209, EPI_ISL_3459210, EPI_ISL_3459211, EPI_ISL_3459212, EPI_ISL_3459213, EPI_ISL_3459214, EPI_ISL_3459215, EPI_ISL_3459218, EPI_ISL_3459219, EPI_ISL_3459220, EPI_ISL_3459222, EPI_ISL_3459223, EPI_ISL_3459226 | | | |
| see above | Laboratoire de santé publique du Québec | Laboratoire de santé publique du Québec | Sandrine Moreira, Ioannis Ragoussis, Guillaume Bourque, Jesse Shapiro, Mark Lathrop and Michel Roger on behalf of the CoVSeQ research group |
| EPI_ISL_3459234 | Pennsylvania Department of Health Bureau of Laboratories | Pennsylvania Department of Health Bureau of Laboratories | Dongxiang Xia |
| EPI_ISL_3459239, EPI_ISL_3459240, EPI_ISL_3459241, EPI_ISL_3459242, EPI_ISL_3459243, EPI_ISL_3459244, EPI_ISL_3459245, EPI_ISL_3459246, EPI_ISL_3459247, EPI_ISL_3459248, EPI_ISL_3459249, EPI_ISL_3459250, EPI_ISL_3459251, EPI_ISL_3459252, EPI_ISL_3459253, EPI_ISL_3459254, EPI_ISL_3459255, EPI_ISL_3459256, EPI_ISL_3459257, EPI_ISL_3459258, EPI_ISL_3459259, EPI_ISL_3459260, EPI_ISL_3459261, EPI_ISL_3459262, EPI_ISL_3459263, EPI_ISL_3459264, EPI_ISL_3459265, EPI_ISL_3459266, EPI_ISL_3459267, EPI_ISL_3459268, EPI_ISL_3459269, EPI_ISL_3459270, EPI_ISL_3459271 | | | |
| see above | TN DOH Lab Services | TN DOH Lab Services | Karen Beasley-Maynard, Brian Selinsky |
| EPI_ISL_3459283 | LSUHS Emerging Viral Threat Lab | LSUHS Emerging Viral Threat Laboratory | Gregory L. Ware, Alexander Mijalis, Jeremy P. Kamil, Jennifer L. Carroll, Maarten Van Diest, Rona S. Scott, Andrew D. Yurochko, Christopher G. Kevil, John A. Vanchiere |
| EPI_ISL_3459285, EPI_ISL_3459287, EPI_ISL_3459303 | Wyoming Public Health Laboratory | Wyoming Public Health Laboratory | Jim Mildenberger, Taylor Fearing, Channing Weber, Ashley Norberg, Chayse Rowley, Marley Goetz, Brian Dominguez, Elliot Thomasson, Sam Britz, Cari Sloma, Robert Petit, and Rob Christensen |
| EPI_ISL_3459305, EPI_ISL_3459306, EPI_ISL_3459307, EPI_ISL_3459308, EPI_ISL_3459309, EPI_ISL_3459310, EPI_ISL_3459311, EPI_ISL_3459312, EPI_ISL_3459313, EPI_ISL_3459314, EPI_ISL_3459315, EPI_ISL_3459316, EPI_ISL_3459317, EPI_ISL_3459318, EPI_ISL_3459319, EPI_ISL_3459320, EPI_ISL_3459321, EPI_ISL_3459322, EPI_ISL_3459323, EPI_ISL_3459324, EPI_ISL_3459325, EPI_ISL_3459326, EPI_ISL_3459327, EPI_ISL_3459328, EPI_ISL_3459329, EPI_ISL_3459330, EPI_ISL_3459331, EPI_ISL_3459332, EPI_ISL_3459333, EPI_ISL_3459334, EPI_ISL_3459335, EPI_ISL_3459336, EPI_ISL_3459337, EPI_ISL_3459338, EPI_ISL_3459339, EPI_ISL_3459340, EPI_ISL_3459341, EPI_ISL_3459342, EPI_ISL_3459343, EPI_ISL_3459344, EPI_ISL_3459345, EPI_ISL_3459346, EPI_ISL_3459347, EPI_ISL_3459348, EPI_ISL_3459349, EPI_ISL_3459350, EPI_ISL_3459351, EPI_ISL_3459352, EPI_ISL_3459353, EPI_ISL_3459354, EPI_ISL_3459355, EPI_ISL_3459356, EPI_ISL_3459357, EPI_ISL_3459358, EPI_ISL_3459359, EPI_ISL_3459360, EPI_ISL_3459361, EPI_ISL_3459362, EPI_ISL_3459363, EPI_ISL_3459364, EPI_ISL_3459365, EPI_ISL_3459366, EPI_ISL_3459367, EPI_ISL_3459368, EPI_ISL_3459369, EPI_ISL_3459370, EPI_ISL_3459371, EPI_ISL_3459372, EPI_ISL_3459373, EPI_ISL_3459374 | | | |
| see above | National Virus Reference Laboratory | National Virus Reference Laboratory | Guerrino Macori, Gabriel Gonzalez, Michael Carr, Zoe Yandle, Charlene Bennett, Jonathan Dean, Seamus Fanning, Cillian F De Gascun |
| EPI_ISL_3459382, EPI_ISL_3459384 | SECRETARIA DE SALUD | Instituto Nacional de Salud | Katherine Laiton-Donato, Diego A. Álvarez-Díaz, Carlos Franco-Muñoz, Hector Alejandro Ruiz-Moreno, Paola Rojas, Maria T. Herrera-Sepúlveda, Diego Andrés Prada, Jhonnatan Reales-González, Sheryll Corchuelo, Julian Naizaque, Jorge Rivera, Gerardo Santamaría, Sergio Gomez, Lisseth Pardo, Juan Camilo Martinez, Marta Lopez Blanco, Ángela Alarcon Cruz, Diana Malo, Carmen Osorio, Magdalena Wiesner, Martha Lucia Ospina Martinez, Marcela Mercado-Reyes |
| EPI_ISL_3459385 | FUNDACION CARDIOINFANTIL | Instituto Nacional de Salud | Katherine Laiton-Donato, Diego A. Álvarez-Díaz, Carlos Franco-Muñoz, Hector Alejandro Ruiz-Moreno, Paola Rojas, Maria T. Herrera-Sepúlveda, Diego Andrés Prada, Jhonnatan Reales-González, Sheryll Corchuelo, Julian Naizaque, Jorge Rivera, Gerardo Santamaría, Sergio Gomez, Lisseth Pardo, Juan Camilo Martinez, Marta Lopez Blanco, Ángela Alarcon Cruz, Diana Malo, Carmen Osorio, Martha Patricia Sánchez Zorro, Rodrigo Cabrera Perez, Magdalena Wiesner, Martha Lucia Ospina Martinez, Marcela Mercado-Reyes |
| EPI_ISL_3459418 | Minnesota Department of Health, Public Health Laboratory | Minnesota Department of Health, Public Health Laboratory | Alexandra Lorentz, Jacob Garfin, Matt Plumb, and Xiong Wang |
| EPI_ISL_3459419, | Centracare Laboratory Services | Minnesota Department of Health, Public Health Laboratory | Alexandra Lorentz, Jacob Garfin, Matt Plumb, and Xiong Wang |

| | | | |
|---|---|---|---|
| EPI_ISL_3459420 | | | |
| EPI_ISL_3459421, EPI_ISL_3459422, EPI_ISL_3459423, EPI_ISL_3459424 | Minnesota Department of Health, Public Health Laboratory | Minnesota Department of Health, Public Health Laboratory | Alexandra Lorentz, Jacob Garfin, Matt Plumb, and Xiong Wang |
| EPI_ISL_3459425 | Centracare Laboratory Services | Minnesota Department of Health, Public Health Laboratory | Alexandra Lorentz, Jacob Garfin, Matt Plumb, and Xiong Wang |
| EPI_ISL_3459426 | Minnesota Department of Health, Public Health Laboratory | Minnesota Department of Health, Public Health Laboratory | Alexandra Lorentz, Jacob Garfin, Matt Plumb, and Xiong Wang |
| EPI_ISL_3459427 | Centracare Laboratory Services | Minnesota Department of Health, Public Health Laboratory | Alexandra Lorentz, Jacob Garfin, Matt Plumb, and Xiong Wang |
| EPI_ISL_3459428, EPI_ISL_3459429, EPI_ISL_3459430, EPI_ISL_3459431, EPI_ISL_3459432, EPI_ISL_3459433 | Minnesota Department of Health, Public Health Laboratory | Minnesota Department of Health, Public Health Laboratory | Alexandra Lorentz, Jacob Garfin, Matt Plumb, and Xiong Wang |
| EPI_ISL_3459434, EPI_ISL_3459435 | Centracare Laboratory Services | Minnesota Department of Health, Public Health Laboratory | Alexandra Lorentz, Jacob Garfin, Matt Plumb, and Xiong Wang |
| EPI_ISL_3459436 | Minnesota Department of Health, Public Health Laboratory | Minnesota Department of Health, Public Health Laboratory | Alexandra Lorentz, Jacob Garfin, Matt Plumb, and Xiong Wang |
| EPI_ISL_3459437, EPI_ISL_3459438 | Centracare Laboratory Services | Minnesota Department of Health, Public Health Laboratory | Alexandra Lorentz, Jacob Garfin, Matt Plumb, and Xiong Wang |
| EPI_ISL_3459439, EPI_ISL_3459440 | Minnesota Department of Health, Public Health Laboratory | Minnesota Department of Health, Public Health Laboratory | Alexandra Lorentz, Jacob Garfin, Matt Plumb, and Xiong Wang |
| EPI_ISL_3459441, EPI_ISL_3459442, EPI_ISL_3459443 | Centracare Laboratory Services | Minnesota Department of Health, Public Health Laboratory | Alexandra Lorentz, Jacob Garfin, Matt Plumb, and Xiong Wang |
| EPI_ISL_3459444, EPI_ISL_3459445, EPI_ISL_3459446, EPI_ISL_3459447 | Minnesota Department of Health, Public Health Laboratory | Minnesota Department of Health, Public Health Laboratory | Alexandra Lorentz, Jacob Garfin, Matt Plumb, and Xiong Wang |
| EPI_ISL_3459448 | Centracare Laboratory Services | Minnesota Department of Health, Public Health Laboratory | Alexandra Lorentz, Jacob Garfin, Matt Plumb, and Xiong Wang |
| EPI_ISL_3459449, EPI_ISL_3459450, EPI_ISL_3459451, EPI_ISL_3459452, EPI_ISL_3459453, EPI_ISL_3459454, EPI_ISL_3459455, EPI_ISL_3459456, EPI_ISL_3459458, EPI_ISL_3459459, EPI_ISL_3459460, EPI_ISL_3459461, EPI_ISL_3459462 | | | |
| see above | Minnesota Department of Health, Public Health Laboratory | Minnesota Department of Health, Public Health Laboratory | Alexandra Lorentz, Jacob Garfin, Matt Plumb, and Xiong Wang |
| EPI_ISL_3459463, EPI_ISL_3459464, EPI_ISL_3459465 | Centracare Laboratory Services | Minnesota Department of Health, Public Health Laboratory | Alexandra Lorentz, Jacob Garfin, Matt Plumb, and Xiong Wang |
| EPI_ISL_3459466, EPI_ISL_3459467, EPI_ISL_3459468, EPI_ISL_3459469, EPI_ISL_3459470, EPI_ISL_3459471, EPI_ISL_3459472, EPI_ISL_3459473, EPI_ISL_3459474, EPI_ISL_3459475 | State Hygienic Laboratory at the University of Iowa | State Hygienic Laboratory at the University of Iowa | Valerie Reeb, Erik Twait, Anna Yakos, Mariah Knutson, Kris Eveland, Kristen Zanon, Wes Hottel, Alankar Kampoowale |
| EPI_ISL_3459476, EPI_ISL_3459477 | University of Iowa Hospitals and Clinics | State Hygienic Laboratory at the University of Iowa | Valerie Reeb, Erik Twait, Anna Yakos, Mariah Knutson, Kris Eveland, Kristen Zanon, Wes Hottel, Alankar Kampoowale |
| EPI_ISL_3459479, EPI_ISL_3459480, EPI_ISL_3459481, EPI_ISL_3459482, EPI_ISL_3459483, EPI_ISL_3459484, EPI_ISL_3459486, EPI_ISL_3459487 | Test Iowa | State Hygienic Laboratory at the University of Iowa | Valerie Reeb, Erik Twait, Anna Yakos, Mariah Knutson, Kris Eveland, Kristen Zanon, Wes Hottel, Alankar Kampoowale |
| EPI_ISL_3459488, EPI_ISL_3459489, EPI_ISL_3459490, EPI_ISL_3459493, EPI_ISL_3459494 | State Hygienic Laboratory at the University of Iowa | State Hygienic Laboratory at the University of Iowa | Valerie Reeb, Erik Twait, Anna Yakos, Mariah Knutson, Kris Eveland, Kristen Zanon, Wes Hottel, Alankar Kampoowale |
| EPI_ISL_3459495, EPI_ISL_3459496, EPI_ISL_3459497, EPI_ISL_3459498, EPI_ISL_3459499 | SANTA CLARA VALLEY MEDICAL CENTER | Santa Clara County Public Health Laboratory | Santa Clara County Public Health Department |
| EPI_ISL_3459500 | EL CAMINO HOSPITAL | Santa Clara County Public Health Laboratory | Santa Clara County Public Health Department |
| EPI_ISL_3459501, EPI_ISL_3459502, EPI_ISL_3459503 | SANTA CLARA VALLEY MEDICAL CENTER | Santa Clara County Public Health Laboratory | Santa Clara County Public Health Department |
| EPI_ISL_3459504 | EL CAMINO HOSPITAL | Santa Clara County Public Health Laboratory | Santa Clara County Public Health Department |
| EPI_ISL_3459505, EPI_ISL_3459506 | SANTA CLARA VALLEY MEDICAL CENTER | Santa Clara County Public Health Laboratory | Santa Clara County Public Health Department |
| EPI_ISL_3459507 | REGIONAL MEDICAL CENTER OF SAN JOSE | Santa Clara County Public Health Laboratory | Santa Clara County Public Health Department |
| EPI_ISL_3459508 | EL CAMINO HOSPITAL | Santa Clara County Public Health Laboratory | Santa Clara County Public Health Department |
| EPI_ISL_3459509, EPI_ISL_3459510 | SANTA CLARA VALLEY MEDICAL CENTER | Santa Clara County Public Health Laboratory | Santa Clara County Public Health Department |
| EPI_ISL_3459511 | EL CAMINO HOSPITAL | Santa Clara County Public Health Laboratory | Santa Clara County Public Health Department |
| EPI_ISL_3459512, EPI_ISL_3459513 | SANTA CLARA VALLEY MEDICAL CENTER | Santa Clara County Public Health Laboratory | Santa Clara County Public Health Department |
| EPI_ISL_3459514 | EL CAMINO HOSPITAL | Santa Clara County Public Health Laboratory | Santa Clara County Public Health Department |
| EPI_ISL_3459515, EPI_ISL_3459516, EPI_ISL_3459517, EPI_ISL_3459518, EPI_ISL_3459519 | SANTA CLARA VALLEY MEDICAL CENTER | Santa Clara County Public Health Laboratory | Santa Clara County Public Health Department |
| EPI_ISL_3459520 | GOOD SAMARITAN HOSPITAL | Santa Clara County Public Health Laboratory | Santa Clara County Public Health Department |
| EPI_ISL_3459521, EPI_ISL_3459522 | SANTA CLARA VALLEY MEDICAL CENTER | Santa Clara County Public Health Laboratory | Santa Clara County Public Health Department |
| EPI_ISL_3459523 | KAISER SAN JOSE | Santa Clara County Public Health Laboratory | Santa Clara County Public Health Department |
| EPI_ISL_3459524, EPI_ISL_3459525, EPI_ISL_3459526 | SANTA CLARA VALLEY MEDICAL CENTER | Santa Clara County Public Health Laboratory | Santa Clara County Public Health Department |
| EPI_ISL_3459527 | GOOD SAMARITAN HOSPITAL | Santa Clara County Public Health Laboratory | Santa Clara County Public Health Department |
| EPI_ISL_3459528 | EL CAMINO HOSPITAL | Santa Clara County Public Health Laboratory | Santa Clara County Public Health Department |

| | | | |
|---|---|---|---|
| EPI_ISL_3459529 | GOOD SAMARITAN HOSPITAL | Santa Clara County Public Health Laboratory | Santa Clara County Public Health Department |
| EPI_ISL_3459530, EPI_ISL_3459531, EPI_ISL_3459532 | SANTA CLARA VALLEY MEDICAL CENTER | Santa Clara County Public Health Laboratory | Santa Clara County Public Health Department |
| EPI_ISL_3459533 | KAISER SAN JOSE | Santa Clara County Public Health Laboratory | Santa Clara County Public Health Department |
| EPI_ISL_3459534 | SANTA CLARA VALLEY MEDICAL CENTER | Santa Clara County Public Health Laboratory | Santa Clara County Public Health Department |
| EPI_ISL_3459535, EPI_ISL_3459536 | EL CAMINO HOSPITAL | Santa Clara County Public Health Laboratory | Santa Clara County Public Health Department |
| EPI_ISL_3459537 | SANTA CLARA VALLEY MEDICAL CENTER | Santa Clara County Public Health Laboratory | Santa Clara County Public Health Department |
| EPI_ISL_3459538 | KAISER SANTA CLARA | Santa Clara County Public Health Laboratory | Santa Clara County Public Health Department |
| EPI_ISL_3459539, EPI_ISL_3459540 | SANTA CLARA VALLEY MEDICAL CENTER | Santa Clara County Public Health Laboratory | Santa Clara County Public Health Department |
| EPI_ISL_3459541 | GOOD SAMARITAN HOSPITAL | Santa Clara County Public Health Laboratory | Santa Clara County Public Health Department |
| EPI_ISL_3459542 | KAISER SAN JOSE | Santa Clara County Public Health Laboratory | Santa Clara County Public Health Department |
| EPI_ISL_3459543 | KAISER SANTA CLARA | Santa Clara County Public Health Laboratory | Santa Clara County Public Health Department |
| EPI_ISL_3459544, EPI_ISL_3459545, EPI_ISL_3459546, EPI_ISL_3459547, EPI_ISL_3459548, EPI_ISL_3459549 | SANTA CLARA VALLEY MEDICAL CENTER | Santa Clara County Public Health Laboratory | Santa Clara County Public Health Department |
| EPI_ISL_3459550 | GOOD SAMARITAN HOSPITAL | Santa Clara County Public Health Laboratory | Santa Clara County Public Health Department |
| EPI_ISL_3459551, EPI_ISL_3459552, EPI_ISL_3459553, EPI_ISL_3459554 | SANTA CLARA VALLEY MEDICAL CENTER | Santa Clara County Public Health Laboratory | Santa Clara County Public Health Department |
| EPI_ISL_3459555 | KAISER SAN JOSE | Santa Clara County Public Health Laboratory | Santa Clara County Public Health Department |
| EPI_ISL_3459556, EPI_ISL_3459557 | SANTA CLARA VALLEY MEDICAL CENTER | Santa Clara County Public Health Laboratory | Santa Clara County Public Health Department |
| EPI_ISL_3459558 | KAISER SAN JOSE | Santa Clara County Public Health Laboratory | Santa Clara County Public Health Department |
| EPI_ISL_3459559, EPI_ISL_3459560 | SANTA CLARA VALLEY MEDICAL CENTER | Santa Clara County Public Health Laboratory | Santa Clara County Public Health Department |
| EPI_ISL_3459561 | KAISER SAN JOSE | Santa Clara County Public Health Laboratory | Santa Clara County Public Health Department |
| EPI_ISL_3459562 | SANTA CLARA VALLEY MEDICAL CENTER | Santa Clara County Public Health Laboratory | Santa Clara County Public Health Department |
| EPI_ISL_3459563 | EL CAMINO HOSPITAL | Santa Clara County Public Health Laboratory | Santa Clara County Public Health Department |
| EPI_ISL_3459564 | SANTA CLARA VALLEY MEDICAL CENTER | Santa Clara County Public Health Laboratory | Santa Clara County Public Health Department |
| EPI_ISL_3459565 | EL CAMINO HOSPITAL | Santa Clara County Public Health Laboratory | Santa Clara County Public Health Department |
| EPI_ISL_3459566 | KAISER SAN JOSE | Santa Clara County Public Health Laboratory | Santa Clara County Public Health Department |
| EPI_ISL_3459567, EPI_ISL_3459568, EPI_ISL_3459569, EPI_ISL_3459570, EPI_ISL_3459571 | SANTA CLARA VALLEY MEDICAL CENTER | Santa Clara County Public Health Laboratory | Santa Clara County Public Health Department |
| EPI_ISL_3459572 | REGIONAL MEDICAL CENTER OF SAN JOSE | Santa Clara County Public Health Laboratory | Santa Clara County Public Health Department |
| EPI_ISL_3459574, EPI_ISL_3459575 | SANTA CLARA VALLEY MEDICAL CENTER | Santa Clara County Public Health Laboratory | Santa Clara County Public Health Department |
| EPI_ISL_3459576 | EL CAMINO HOSPITAL | Santa Clara County Public Health Laboratory | Santa Clara County Public Health Department |
| EPI_ISL_3459577, EPI_ISL_3459578, EPI_ISL_3459579, EPI_ISL_3459580, EPI_ISL_3459581 | SANTA CLARA VALLEY MEDICAL CENTER | Santa Clara County Public Health Laboratory | Santa Clara County Public Health Department |
| EPI_ISL_3459582 | EL CAMINO HOSPITAL | Santa Clara County Public Health Laboratory | Santa Clara County Public Health Department |
| EPI_ISL_3459583, EPI_ISL_3459584, EPI_ISL_3459585, EPI_ISL_3459586 | SANTA CLARA VALLEY MEDICAL CENTER | Santa Clara County Public Health Laboratory | Santa Clara County Public Health Department |
| EPI_ISL_3459587 | KAISER SAN JOSE | Santa Clara County Public Health Laboratory | Santa Clara County Public Health Department |
| EPI_ISL_3459588 | GOOD SAMARITAN HOSPITAL | Santa Clara County Public Health Laboratory | Santa Clara County Public Health Department |
| EPI_ISL_3459589, EPI_ISL_3459590 | SANTA CLARA VALLEY MEDICAL CENTER | Santa Clara County Public Health Laboratory | Santa Clara County Public Health Department |
| EPI_ISL_3459591 | REGIONAL MEDICAL CENTER OF SAN JOSE | Santa Clara County Public Health Laboratory | Santa Clara County Public Health Department |
| EPI_ISL_3459592 | KAISER SAN JOSE | Santa Clara County Public Health Laboratory | Santa Clara County Public Health Department |
| EPI_ISL_3459593, EPI_ISL_3459594, EPI_ISL_3459595, EPI_ISL_3459596, EPI_ISL_3459597, EPI_ISL_3459598, EPI_ISL_3459599, EPI_ISL_3459600 | SANTA CLARA VALLEY MEDICAL CENTER | Santa Clara County Public Health Laboratory | Santa Clara County Public Health Department |
| EPI_ISL_3459601 | KAISER SANTA CLARA | Santa Clara County Public Health Laboratory | Santa Clara County Public Health Department |
| EPI_ISL_3459602, EPI_ISL_3459603, EPI_ISL_3459604 | SANTA CLARA VALLEY MEDICAL CENTER | Santa Clara County Public Health Laboratory | Santa Clara County Public Health Department |
| EPI_ISL_3459605 | KAISER SAN JOSE | Santa Clara County Public Health Laboratory | Santa Clara County Public Health Department |
| EPI_ISL_3459606, EPI_ISL_3459607 | SANTA CLARA VALLEY MEDICAL CENTER | Santa Clara County Public Health Laboratory | Santa Clara County Public Health Department |
| EPI_ISL_3459608 | REGIONAL MEDICAL CENTER OF SAN JOSE | Santa Clara County Public Health Laboratory | Santa Clara County Public Health Department |
| EPI_ISL_3459609, EPI_ISL_3459610, EPI_ISL_3459611, EPI_ISL_3459612, EPI_ISL_3459613 | SANTA CLARA VALLEY MEDICAL CENTER | Santa Clara County Public Health Laboratory | Santa Clara County Public Health Department |
| EPI_ISL_3459614 | GOOD SAMARITAN HOSPITAL | Santa Clara County Public Health Laboratory | Santa Clara County Public Health Department |
| EPI_ISL_3459615, EPI_ISL_3459616, EPI_ISL_3459617 | SANTA CLARA VALLEY MEDICAL CENTER | Santa Clara County Public Health Laboratory | Santa Clara County Public Health Department |
| EPI_ISL_3459618 | EL CAMINO HOSPITAL | Santa Clara County Public Health Laboratory | Santa Clara County Public Health Department |
| EPI_ISL_3459619 | KAISER SANTA CLARA | Santa Clara County Public Health Laboratory | Santa Clara County Public Health Department |

| | | | |
|---|---|---|---|
| EPI_ISL_3459620 | SANTA CLARA VALLEY MEDICAL CENTER | Santa Clara County Public Health Laboratory | Santa Clara County Public Health Department |
| EPI_ISL_3459621 | KAISER SAN JOSE | Santa Clara County Public Health Laboratory | Santa Clara County Public Health Department |
| EPI_ISL_3459622, EPI_ISL_3459624 | SANTA CLARA VALLEY MEDICAL CENTER | Santa Clara County Public Health Laboratory | Santa Clara County Public Health Department |
| EPI_ISL_3459625 | REGIONAL MEDICAL CENTER OF SAN JOSE | Santa Clara County Public Health Laboratory | Santa Clara County Public Health Department |
| EPI_ISL_3459626 | EL CAMINO HOSPITAL | Santa Clara County Public Health Laboratory | Santa Clara County Public Health Department |
| EPI_ISL_3459627 | KAISER SAN JOSE | Santa Clara County Public Health Laboratory | Santa Clara County Public Health Department |
| EPI_ISL_3459628 | EL CAMINO HOSPITAL | Santa Clara County Public Health Laboratory | Santa Clara County Public Health Department |
| EPI_ISL_3459629, EPI_ISL_3459630, EPI_ISL_3459631, EPI_ISL_3459632, EPI_ISL_3459633, EPI_ISL_3459634, EPI_ISL_3459635, EPI_ISL_3459636, EPI_ISL_3459637, EPI_ISL_3459638, EPI_ISL_3459639, EPI_ISL_3459640, EPI_ISL_3459641, EPI_ISL_3459642, EPI_ISL_3459643, EPI_ISL_3459644, EPI_ISL_3459645, EPI_ISL_3459646, EPI_ISL_3459647, EPI_ISL_3459648, EPI_ISL_3459649, EPI_ISL_3459650, EPI_ISL_3459651, EPI_ISL_3459652, EPI_ISL_3459653, EPI_ISL_3459654, EPI_ISL_3459655, EPI_ISL_3459656, EPI_ISL_3459657, EPI_ISL_3459658, EPI_ISL_3459659, EPI_ISL_3459660, EPI_ISL_3459661, EPI_ISL_3459662, EPI_ISL_3459663, EPI_ISL_3459664, EPI_ISL_3459665, EPI_ISL_3459666 | | | |
| see above | SANTA CLARA VALLEY MEDICAL CENTER | Santa Clara County Public Health Laboratory | Santa Clara County Public Health Department |
| EPI_ISL_3459667 | GOOD SAMARITAN HOSPITAL | Santa Clara County Public Health Laboratory | Santa Clara County Public Health Department |
| EPI_ISL_3459668, EPI_ISL_3459669, EPI_ISL_3459670, EPI_ISL_3459671, EPI_ISL_3459672 | EL CAMINO HOSPITAL | Santa Clara County Public Health Laboratory | Santa Clara County Public Health Department |
| EPI_ISL_3459673 | SANTA CLARA COUNTY PUBLIC HEALTH LABORATORY | Santa Clara County Public Health Laboratory | Santa Clara County Public Health Department |
| EPI_ISL_3459674 | KAISER SAN JOSE | Santa Clara County Public Health Laboratory | Santa Clara County Public Health Department |
| EPI_ISL_3459675 | EL CAMINO HOSPITAL | Santa Clara County Public Health Laboratory | Santa Clara County Public Health Department |
| EPI_ISL_3459676, EPI_ISL_3459677, EPI_ISL_3459678 | State Hygienic Laboratory at the University of Iowa | State Hygienic Laboratory at the University of Iowa | Valerie Reeb, Erik Twait, Anna Yakos, Mariah Knutson, Kris Eveland, Kristen Zanon, Wes Hottel, Alankar Kampoowale |
| EPI_ISL_3459679, EPI_ISL_3459680 | University of Iowa Hospitals and Clinics | State Hygienic Laboratory at the University of Iowa | Valerie Reeb, Erik Twait, Anna Yakos, Mariah Knutson, Kris Eveland, Kristen Zanon, Wes Hottel, Alankar Kampoowale |
| EPI_ISL_3459682, EPI_ISL_3459683, EPI_ISL_3459684, EPI_ISL_3459685, EPI_ISL_3459686, EPI_ISL_3459687, EPI_ISL_3459688 | Test Iowa | State Hygienic Laboratory at the University of Iowa | Valerie Reeb, Erik Twait, Anna Yakos, Mariah Knutson, Kris Eveland, Kristen Zanon, Wes Hottel, Alankar Kampoowale |
| EPI_ISL_3459689, EPI_ISL_3459690, EPI_ISL_3459691, EPI_ISL_3459692, EPI_ISL_3459693, EPI_ISL_3459694, EPI_ISL_3459695, EPI_ISL_3459696, EPI_ISL_3459697, EPI_ISL_3459698, EPI_ISL_3459699, EPI_ISL_3459700, EPI_ISL_3459701, EPI_ISL_3459702, EPI_ISL_3459703, EPI_ISL_3459704, EPI_ISL_3459705, EPI_ISL_3459706, EPI_ISL_3459707, EPI_ISL_3459708, EPI_ISL_3459709, EPI_ISL_3459710, EPI_ISL_3459711, EPI_ISL_3459712, EPI_ISL_3459713, EPI_ISL_3459714, EPI_ISL_3459715, EPI_ISL_3459716, EPI_ISL_3459717, EPI_ISL_3459718, EPI_ISL_3459719, EPI_ISL_3459720, EPI_ISL_3459721, EPI_ISL_3459722, EPI_ISL_3459723, EPI_ISL_3459724, EPI_ISL_3459725, EPI_ISL_3459726, EPI_ISL_3459727, EPI_ISL_3459728, EPI_ISL_3459729, EPI_ISL_3459730 | | | |
| see above | State Hygienic Laboratory at the University of Iowa | State Hygienic Laboratory at the University of Iowa | Valerie Reeb, Erik Twait, Anna Yakos, Mariah Knutson, Kris Eveland, Kristen Zanon, Wes Hottel, Alankar Kampoowale |
| EPI_ISL_3459731 | University of Iowa Hospitals and Clinics | State Hygienic Laboratory at the University of Iowa | Valerie Reeb, Erik Twait, Anna Yakos, Mariah Knutson, Kris Eveland, Kristen Zanon, Wes Hottel, Alankar Kampoowale |
| EPI_ISL_3459733, EPI_ISL_3459734, EPI_ISL_3459735, EPI_ISL_3459736, EPI_ISL_3459737, EPI_ISL_3459738, EPI_ISL_3459739, EPI_ISL_3459740, EPI_ISL_3459741, EPI_ISL_3459742, EPI_ISL_3459743 | | | |
| see above | State Hygienic Laboratory at the University of Iowa | State Hygienic Laboratory at the University of Iowa | Valerie Reeb, Erik Twait, Anna Yakos, Mariah Knutson, Kris Eveland, Kristen Zanon, Wes Hottel, Alankar Kampoowale |
| EPI_ISL_3459744, EPI_ISL_3459745 | University of Iowa Hospitals and Clinics | State Hygienic Laboratory at the University of Iowa | Valerie Reeb, Erik Twait, Anna Yakos, Mariah Knutson, Kris Eveland, Kristen Zanon, Wes Hottel, Alankar Kampoowale |
| EPI_ISL_3459746, EPI_ISL_3459747, EPI_ISL_3459748, EPI_ISL_3459749, EPI_ISL_3459750, EPI_ISL_3459751, EPI_ISL_3459752, EPI_ISL_3459753, EPI_ISL_3459754, EPI_ISL_3459755, EPI_ISL_3459756, EPI_ISL_3459757, EPI_ISL_3459758 | | | |
| see above | State Hygienic Laboratory at the University of Iowa | State Hygienic Laboratory at the University of Iowa | Valerie Reeb, Erik Twait, Anna Yakos, Mariah Knutson, Kris Eveland, Kristen Zanon, Wes Hottel, Alankar Kampoowale |
| EPI_ISL_3459759, EPI_ISL_3459760, EPI_ISL_3459761, EPI_ISL_3459762 | SYNLAB | GIGA Medical Genomics | Keith Durkin, Maria Artesi, Bouchra Boujemla, Nathalie Renotte, Nadine Cambisano, Sébastien Bontems, Cécile Meex, Claire Gourzonès, Olivier Ek, Laurent Gillet, Marie-Pierre Hayette, Vincent Bours |
| EPI_ISL_3459763, EPI_ISL_3459764 | Department of Clinical Microbiology | GIGA Medical Genomics | Keith Durkin, Maria Artesi, Bouchra Boujemla, Nathalie Renotte, Nadine Cambisano, Sébastien Bontems, Cécile Meex, Claire Gourzonès, Olivier Ek, Laurent Gillet, Marie-Pierre Hayette, Vincent Bours |
| EPI_ISL_3459765 | SYNLAB | GIGA Medical Genomics | Keith Durkin, Maria Artesi, Bouchra Boujemla, Nathalie Renotte, Nadine Cambisano, Sébastien Bontems, Cécile Meex, Claire Gourzonès, Olivier Ek, Laurent Gillet, Marie-Pierre Hayette, Vincent Bours |
| EPI_ISL_3459766, EPI_ISL_3459767, EPI_ISL_3459768 | Department of Clinical Microbiology | GIGA Medical Genomics | Keith Durkin, Maria Artesi, Bouchra Boujemla, Nathalie Renotte, Nadine Cambisano, Sébastien Bontems, Cécile Meex, Claire Gourzonès, Olivier Ek, Laurent Gillet, Marie-Pierre Hayette, Vincent Bours |
| EPI_ISL_3459769, EPI_ISL_3459770, EPI_ISL_3459771 | SYNLAB | GIGA Medical Genomics | Keith Durkin, Maria Artesi, Bouchra Boujemla, Nathalie Renotte, Nadine Cambisano, Sébastien Bontems, Cécile Meex, Claire Gourzonès, Olivier Ek, Laurent Gillet, Marie-Pierre Hayette, Vincent Bours |
| EPI_ISL_3459772 | Department of Clinical Microbiology | GIGA Medical Genomics | Keith Durkin, Maria Artesi, Bouchra Boujemla, Nathalie Renotte, Nadine Cambisano, Sébastien Bontems, Cécile Meex, Claire Gourzonès, Olivier Ek, Laurent Gillet, Marie-Pierre Hayette, Vincent Bours |
| EPI_ISL_3459773, EPI_ISL_3459774, EPI_ISL_3459775 | SYNLAB | GIGA Medical Genomics | Keith Durkin, Maria Artesi, Bouchra Boujemla, Nathalie Renotte, Nadine Cambisano, Sébastien Bontems, Cécile Meex, Claire Gourzonès, Olivier Ek, Laurent Gillet, Marie-Pierre Hayette, Vincent Bours |
| EPI_ISL_3459776 | Department of Clinical Microbiology | GIGA Medical Genomics | Keith Durkin, Maria Artesi, Bouchra Boujemla, Nathalie Renotte, Nadine Cambisano, Sébastien Bontems, Cécile Meex, Claire Gourzonès, Olivier Ek, Laurent Gillet, Marie-Pierre Hayette, Vincent Bours |
| EPI_ISL_3459777, EPI_ISL_3459778, EPI_ISL_3459779 | SYNLAB | GIGA Medical Genomics | Keith Durkin, Maria Artesi, Bouchra Boujemla, Nathalie Renotte, Nadine Cambisano, Sébastien Bontems, Cécile Meex, Claire Gourzonès, Olivier Ek, Laurent Gillet, Marie-Pierre Hayette, Vincent Bours |
| EPI_ISL_3459780, EPI_ISL_3459781, EPI_ISL_3459782 | Department of Clinical Microbiology | GIGA Medical Genomics | Keith Durkin, Maria Artesi, Bouchra Boujemla, Nathalie Renotte, Nadine Cambisano, Sébastien Bontems, Cécile Meex, Claire Gourzonès, Olivier Ek, Laurent Gillet, Marie-Pierre Hayette, Vincent Bours |
| EPI_ISL_3459783, EPI_ISL_3459784 | SYNLAB | GIGA Medical Genomics | Keith Durkin, Maria Artesi, Bouchra Boujemla, Nathalie Renotte, Nadine Cambisano, Sébastien Bontems, Cécile Meex, Claire Gourzonès, Olivier Ek, Laurent Gillet, Marie-Pierre Hayette, Vincent Bours |
| EPI_ISL_3459785, EPI_ISL_3459786 | Department of Clinical Microbiology | GIGA Medical Genomics | Keith Durkin, Maria Artesi, Bouchra Boujemla, Nathalie Renotte, Nadine Cambisano, Sébastien Bontems, Cécile Meex, Claire Gourzonès, Olivier Ek, Laurent Gillet, Marie-Pierre Hayette, Vincent Bours |
| EPI_ISL_3459787, EPI_ISL_3459788, EPI_ISL_3459789, EPI_ISL_3459790, EPI_ISL_3459791, EPI_ISL_3459792, EPI_ISL_3459793, EPI_ISL_3459794, EPI_ISL_3459795, EPI_ISL_3459796, EPI_ISL_3459797, EPI_ISL_3459798, EPI_ISL_3459799, EPI_ISL_3459800, EPI_ISL_3459801 | | | |
| see above | SYNLAB | GIGA Medical Genomics | Keith Durkin, Maria Artesi, Bouchra Boujemla, Nathalie Renotte, Nadine Cambisano, Sébastien Bontems, Cécile Meex, Claire Gourzonès, Olivier Ek, Laurent Gillet, Marie-Pierre Hayette, Vincent Bours |
| EPI_ISL_3459802, EPI_ISL_3459803, EPI_ISL_3459804, EPI_ISL_3459805, EPI_ISL_3459806, EPI_ISL_3459807, EPI_ISL_3459808, EPI_ISL_3459809, EPI_ISL_3459810, EPI_ISL_3459811, EPI_ISL_3459812, EPI_ISL_3459813, EPI_ISL_3459814, EPI_ISL_3459815, EPI_ISL_3459816, EPI_ISL_3459817, EPI_ISL_3459818, EPI_ISL_3459819, EPI_ISL_3459820, EPI_ISL_3459821, EPI_ISL_3459822, EPI_ISL_3459823, EPI_ISL_3459824 | | | |
| see above | Department of Clinical Microbiology | GIGA Medical Genomics | Keith Durkin, Maria Artesi, Bouchra Boujemla, Nathalie Renotte, Nadine Cambisano, Sébastien Bontems, Cécile Meex, Claire Gourzonès, Olivier Ek, Laurent Gillet, Marie-Pierre Hayette, Vincent Bours |
| EPI_ISL_3459825, EPI_ISL_3459826, EPI_ISL_3459827, EPI_ISL_3459828, EPI_ISL_3459829, EPI_ISL_3459830, EPI_ISL_3459831, EPI_ISL_3459832, EPI_ISL_3459833, EPI_ISL_3459834, EPI_ISL_3459835, EPI_ISL_3459836, EPI_ISL_3459837, EPI_ISL_3459838 | | | |
| see above | SYNLAB | GIGA Medical Genomics | Keith Durkin, Maria Artesi, Bouchra Boujemla, Nathalie Renotte, Nadine Cambisano, Sébastien Bontems, Cécile Meex, Claire Gourzonès, Olivier Ek, Laurent Gillet, Marie-Pierre Hayette, Vincent Bours |
| EPI_ISL_3459839, EPI_ISL_3459840, EPI_ISL_3459841, EPI_ISL_3459842, EPI_ISL_3459843, EPI_ISL_3459844, EPI_ISL_3459845 | Department of Clinical Microbiology | GIGA Medical Genomics | Keith Durkin, Maria Artesi, Bouchra Boujemla, Nathalie Renotte, Nadine Cambisano, Sébastien Bontems, Cécile Meex, Claire Gourzonès, Olivier Ek, Laurent Gillet, Marie-Pierre Hayette, Vincent Bours |
| EPI_ISL_3459847, EPI_ISL_3459848, EPI_ISL_3459849, EPI_ISL_3459850, EPI_ISL_3459851, EPI_ISL_3459852, EPI_ISL_3459853, EPI_ISL_3459854, EPI_ISL_3459855, EPI_ISL_3459856, EPI_ISL_3459857, EPI_ISL_3459858, EPI_ISL_3459859, EPI_ISL_3459860, EPI_ISL_3459861, EPI_ISL_3459862, EPI_ISL_3459863, EPI_ISL_3459864, EPI_ISL_3459865, EPI_ISL_3459866, EPI_ISL_3459867, EPI_ISL_3459868, EPI_ISL_3459869, EPI_ISL_3459870, EPI_ISL_3459871, EPI_ISL_3459872, EPI_ISL_3459873, EPI_ISL_3459874, EPI_ISL_3459875, EPI_ISL_3459876, EPI_ISL_3459883, EPI_ISL_3459884, EPI_ISL_3459885, EPI_ISL_3459886, EPI_ISL_3459887, EPI_ISL_3459888, EPI_ISL_3459889 | | | |
| see above | Diagnostyka. Laboratoria Medyczne. | 1. ViroGenetics - BSL3 Laboratory of Virology, Małopolska Centre of Biotechnology, Jagiellonian University; 2. genXone SA, Research & Development Laboratory | Mazur-Panasiuk,N., Grzegorz Nowicki, Wydmanski,W., Gromowski,T., Natalia Drweska-Matelska, Jakub Grabowski, Anna Brylak, Aleksandra Gidlewicz, Karol Szeszko, Maciej Sykulski, Łukasz Krych, Kowalski,M., Szulc,P., Sylwia Januszczak, Labaj,P.P., Michał Kaszuba, Pyrc,K. |

| | | | |
|---|---|---|---|
| EPI_ISL_3459891 | LESP State of Mexico | Instituto de Diagnostico y Referencia Epidemiologicos (INDRE) | Claudia Wong-Arambula, Abril Rodriguez-Maldonado, Vanessa Rivero-Arredondo, Ariadna Medina-Benitez, Joaquin Quiroz-Mercado, Sergio Rangel-Guerrero, Maribel Gonzalez-Villa, Natividad Cruz-Ortiz, Tatiana Nunez-Garcia, Gisela Barrera-Badillo, Lucia Hernandez-Rivas, Irma Lopez-Martinez, Ernesto Ramirez-Gonzalez. |
| EPI_ISL_3459892 | LESP Coahuila | Instituto de Diagnostico y Referencia Epidemiologicos (INDRE) | Claudia Wong-Arambula, Abril Rodriguez-Maldonado, Vanessa Rivero-Arredondo, Ariadna Medina-Benitez, Joaquin Quiroz-Mercado, Sergio Rangel-Guerrero, Maribel Gonzalez-Villa, Natividad Cruz-Ortiz, Tatiana Nunez-Garcia, Gisela Barrera-Badillo, Lucia Hernandez-Rivas, Irma Lopez-Martinez, Ernesto Ramirez-Gonzalez. |
| EPI_ISL_3459893 | LESP Queretaro | Instituto de Diagnostico y Referencia Epidemiologicos (INDRE) | Claudia Wong-Arambula, Abril Rodriguez-Maldonado, Vanessa Rivero-Arredondo, Ariadna Medina-Benitez, Joaquin Quiroz-Mercado, Sergio Rangel-Guerrero, Maribel Gonzalez-Villa, Natividad Cruz-Ortiz, Tatiana Nunez-Garcia, Gisela Barrera-Badillo, Lucia Hernandez-Rivas, Irma Lopez-Martinez, Ernesto Ramirez-Gonzalez. |
| EPI_ISL_3459895 | LESP Hidalgo | Instituto de Diagnostico y Referencia Epidemiologicos (INDRE) | Claudia Wong-Arambula, Abril Rodriguez-Maldonado, Vanessa Rivero-Arredondo, Ariadna Medina-Benitez, Joaquin Quiroz-Mercado, Sergio Rangel-Guerrero, Maribel Gonzalez-Villa, Natividad Cruz-Ortiz, Tatiana Nunez-Garcia, Gisela Barrera-Badillo, Lucia Hernandez-Rivas, Irma Lopez-Martinez, Ernesto Ramirez-Gonzalez. |
| EPI_ISL_3459896 | LESP Tabasco | Instituto de Diagnostico y Referencia Epidemiologicos (INDRE) | Claudia Wong-Arambula, Abril Rodriguez-Maldonado, Vanessa Rivero-Arredondo, Ariadna Medina-Benitez, Joaquin Quiroz-Mercado, Sergio Rangel-Guerrero, Maribel Gonzalez-Villa, Natividad Cruz-Ortiz, Tatiana Nunez-Garcia, Gisela Barrera-Badillo, Lucia Hernandez-Rivas, Irma Lopez-Martinez, Ernesto Ramirez-Gonzalez. |
| EPI_ISL_3459897 | LESP Coahuila | Instituto de Diagnostico y Referencia Epidemiologicos (INDRE) | Claudia Wong-Arambula, Abril Rodriguez-Maldonado, Vanessa Rivero-Arredondo, Ariadna Medina-Benitez, Joaquin Quiroz-Mercado, Sergio Rangel-Guerrero, Maribel Gonzalez-Villa, Natividad Cruz-Ortiz, Tatiana Nunez-Garcia, Gisela Barrera-Badillo, Lucia Hernandez-Rivas, Irma Lopez-Martinez, Ernesto Ramirez-Gonzalez. |
| EPI_ISL_3459900 | LESP Colima | Instituto de Diagnostico y Referencia Epidemiologicos (INDRE) | Claudia Wong-Arambula, Abril Rodriguez-Maldonado, Vanessa Rivero-Arredondo, Ariadna Medina-Benitez, Joaquin Quiroz-Mercado, Sergio Rangel-Guerrero, Maribel Gonzalez-Villa, Natividad Cruz-Ortiz, Tatiana Nunez-Garcia, Gisela Barrera-Badillo, Lucia Hernandez-Rivas, Irma Lopez-Martinez, Ernesto Ramirez-Gonzalez. |
| EPI_ISL_3459909 | LESP Hidalgo | Instituto de Diagnostico y Referencia Epidemiologicos (INDRE) | Claudia Wong-Arambula, Abril Rodriguez-Maldonado, Vanessa Rivero-Arredondo, Ariadna Medina-Benitez, Joaquin Quiroz-Mercado, Sergio Rangel-Guerrero, Maribel Gonzalez-Villa, Natividad Cruz-Ortiz, Tatiana Nunez-Garcia, Gisela Barrera-Badillo, Lucia Hernandez-Rivas, Irma Lopez-Martinez, Ernesto Ramirez-Gonzalez. |
| EPI_ISL_3459910 | LESP Baja California Sur | Instituto de Diagnostico y Referencia Epidemiologicos (INDRE) | Claudia Wong-Arambula, Abril Rodriguez-Maldonado, Vanessa Rivero-Arredondo, Ariadna Medina-Benitez, Joaquin Quiroz-Mercado, Sergio Rangel-Guerrero, Maribel Gonzalez-Villa, Natividad Cruz-Ortiz, Tatiana Nunez-Garcia, Gisela Barrera-Badillo, Lucia Hernandez-Rivas, Irma Lopez-Martinez, Ernesto Ramirez-Gonzalez. |
| EPI_ISL_3459912 | LESP Colima | Instituto de Diagnostico y Referencia Epidemiologicos (INDRE) | Claudia Wong-Arambula, Abril Rodriguez-Maldonado, Vanessa Rivero-Arredondo, Ariadna Medina-Benitez, Joaquin Quiroz-Mercado, Sergio Rangel-Guerrero, Maribel Gonzalez-Villa, Natividad Cruz-Ortiz, Tatiana Nunez-Garcia, Gisela Barrera-Badillo, Lucia Hernandez-Rivas, Irma Lopez-Martinez, Ernesto Ramirez-Gonzalez. |
| EPI_ISL_3459916, EPI_ISL_3459918 | LESP Queretaro | Instituto de Diagnostico y Referencia Epidemiologicos (INDRE) | Claudia Wong-Arambula, Abril Rodriguez-Maldonado, Vanessa Rivero-Arredondo, Ariadna Medina-Benitez, Joaquin Quiroz-Mercado, Sergio Rangel-Guerrero, Maribel Gonzalez-Villa, Natividad Cruz-Ortiz, Tatiana Nunez-Garcia, Gisela Barrera-Badillo, Lucia Hernandez-Rivas, Irma Lopez-Martinez, Ernesto Ramirez-Gonzalez. |
| EPI_ISL_3459919 | LESP Mexico City | Instituto de Diagnostico y Referencia Epidemiologicos (INDRE) | Claudia Wong-Arambula, Abril Rodriguez-Maldonado, Vanessa Rivero-Arredondo, Ariadna Medina-Benitez, Joaquin Quiroz-Mercado, Sergio Rangel-Guerrero, Maribel Gonzalez-Villa, Natividad Cruz-Ortiz, Tatiana Nunez-Garcia, Gisela Barrera-Badillo, Lucia Hernandez-Rivas, Irma Lopez-Martinez, Ernesto Ramirez-Gonzalez. |
| EPI_ISL_3459920 | LESP Queretaro | Instituto de Diagnostico y Referencia Epidemiologicos (INDRE) | Claudia Wong-Arambula, Abril Rodriguez-Maldonado, Vanessa Rivero-Arredondo, Ariadna Medina-Benitez, Joaquin Quiroz-Mercado, Sergio Rangel-Guerrero, Maribel Gonzalez-Villa, Natividad Cruz-Ortiz, Tatiana Nunez-Garcia, Gisela Barrera-Badillo, Lucia Hernandez-Rivas, Irma Lopez-Martinez, Ernesto Ramirez-Gonzalez. |
| EPI_ISL_3459929 | LESP Jalisco | Instituto de Diagnostico y Referencia Epidemiologicos (INDRE) | Claudia Wong-Arambula, Abril Rodriguez-Maldonado, Vanessa Rivero-Arredondo, Ariadna Medina-Benitez, Joaquin Quiroz-Mercado, Sergio Rangel-Guerrero, Maribel Gonzalez-Villa, Natividad Cruz-Ortiz, Tatiana Nunez-Garcia, Gisela Barrera-Badillo, Lucia Hernandez-Rivas, Irma Lopez-Martinez, Ernesto Ramirez-Gonzalez. |
| EPI_ISL_3459930 | LESP Queretaro | Instituto de Diagnostico y Referencia Epidemiologicos (INDRE) | Claudia Wong-Arambula, Abril Rodriguez-Maldonado, Vanessa Rivero-Arredondo, Ariadna Medina-Benitez, Joaquin Quiroz-Mercado, Sergio Rangel-Guerrero, Maribel Gonzalez-Villa, Natividad Cruz-Ortiz, Tatiana Nunez-Garcia, Gisela Barrera-Badillo, Lucia Hernandez-Rivas, Irma Lopez-Martinez, Ernesto Ramirez-Gonzalez. |
| EPI_ISL_3459931 | LESP Aguascalientes | Instituto de Diagnostico y Referencia Epidemiologicos (INDRE) | Claudia Wong-Arambula, Abril Rodriguez-Maldonado, Vanessa Rivero-Arredondo, Ariadna Medina-Benitez, Joaquin Quiroz-Mercado, Sergio Rangel-Guerrero, Maribel Gonzalez-Villa, Natividad Cruz-Ortiz, Tatiana Nunez-Garcia, Gisela Barrera-Badillo, Lucia Hernandez-Rivas, Irma Lopez-Martinez, Ernesto Ramirez-Gonzalez. |
| EPI_ISL_3459932 | LESP State of Mexico | Instituto de Diagnostico y Referencia Epidemiologicos (INDRE) | Claudia Wong-Arambula, Abril Rodriguez-Maldonado, Vanessa Rivero-Arredondo, Ariadna Medina-Benitez, Joaquin Quiroz-Mercado, Sergio Rangel-Guerrero, Maribel Gonzalez-Villa, Natividad Cruz-Ortiz, Tatiana Nunez-Garcia, Gisela Barrera-Badillo, Lucia Hernandez-Rivas, Irma Lopez-Martinez, Ernesto Ramirez-Gonzalez. |
| EPI_ISL_3459933, EPI_ISL_3459934, EPI_ISL_3459935, EPI_ISL_3459936 | LESP Queretaro | Instituto de Diagnostico y Referencia Epidemiologicos (INDRE) | Claudia Wong-Arambula, Abril Rodriguez-Maldonado, Vanessa Rivero-Arredondo, Ariadna Medina-Benitez, Joaquin Quiroz-Mercado, Sergio Rangel-Guerrero, Maribel Gonzalez-Villa, Natividad Cruz-Ortiz, Tatiana Nunez-Garcia, Gisela Barrera-Badillo, Lucia Hernandez-Rivas, Irma Lopez-Martinez, Ernesto Ramirez-Gonzalez. |
| EPI_ISL_3459937, EPI_ISL_3459938 | LESP Hidalgo | Instituto de Diagnostico y Referencia Epidemiologicos (INDRE) | Claudia Wong-Arambula, Abril Rodriguez-Maldonado, Vanessa Rivero-Arredondo, Ariadna Medina-Benitez, Joaquin Quiroz-Mercado, Sergio Rangel-Guerrero, Maribel Gonzalez-Villa, Natividad Cruz-Ortiz, Tatiana Nunez-Garcia, Gisela Barrera-Badillo, Lucia Hernandez-Rivas, Irma Lopez-Martinez, Ernesto Ramirez-Gonzalez. |
| EPI_ISL_3459965, EPI_ISL_3459966, EPI_ISL_3459967, EPI_ISL_3459968, EPI_ISL_3459969, EPI_ISL_3459970, EPI_ISL_3459971, EPI_ISL_3459972 | LESP Jalisco | Instituto de Diagnostico y Referencia Epidemiologicos (INDRE) | Claudia Wong-Arambula, Abril Rodriguez-Maldonado, Vanessa Rivero-Arredondo, Ariadna Medina-Benitez, Joaquin Quiroz-Mercado, Sergio Rangel-Guerrero, Maribel Gonzalez-Villa, Natividad Cruz-Ortiz, Tatiana Nunez-Garcia, Gisela Barrera-Badillo, Lucia Hernandez-Rivas, Irma Lopez-Martinez, Ernesto Ramirez-Gonzalez. |
| EPI_ISL_3459973, EPI_ISL_3459974, EPI_ISL_3459975, EPI_ISL_3459976, EPI_ISL_3459977, EPI_ISL_3459978, EPI_ISL_3459979, EPI_ISL_3459980 | LESP Queretaro | Instituto de Diagnostico y Referencia Epidemiologicos (INDRE) | Claudia Wong-Arambula, Abril Rodriguez-Maldonado, Vanessa Rivero-Arredondo, Ariadna Medina-Benitez, Joaquin Quiroz-Mercado, Sergio Rangel-Guerrero, Maribel Gonzalez-Villa, Natividad Cruz-Ortiz, Tatiana Nunez-Garcia, Gisela Barrera-Badillo, Lucia Hernandez-Rivas, Irma Lopez-Martinez, Ernesto Ramirez-Gonzalez. |
| EPI_ISL_3459981 | LESP State of Mexico | Instituto de Diagnostico y Referencia Epidemiologicos (INDRE) | Claudia Wong-Arambula, Abril Rodriguez-Maldonado, Vanessa Rivero-Arredondo, Ariadna Medina-Benitez, Joaquin Quiroz-Mercado, Sergio Rangel-Guerrero, Maribel Gonzalez-Villa, Natividad Cruz-Ortiz, Tatiana Nunez-Garcia, Gisela Barrera-Badillo, Lucia Hernandez-Rivas, Irma Lopez-Martinez, Ernesto Ramirez-Gonzalez. |
| EPI_ISL_3459982 | LESP Mexico City | Instituto de Diagnostico y Referencia Epidemiologicos (INDRE) | Claudia Wong-Arambula, Abril Rodriguez-Maldonado, Vanessa Rivero-Arredondo, Ariadna Medina-Benitez, Joaquin Quiroz-Mercado, Sergio Rangel-Guerrero, Maribel Gonzalez-Villa, Natividad Cruz-Ortiz, Tatiana Nunez-Garcia, Gisela Barrera-Badillo, Lucia Hernandez-Rivas, Irma Lopez-Martinez, Ernesto Ramirez-Gonzalez. |
| EPI_ISL_3460006 | LESP Coahuila | Instituto de Diagnostico y Referencia Epidemiologicos (INDRE) | Claudia Wong-Arambula, Abril Rodriguez-Maldonado, Vanessa Rivero-Arredondo, Ariadna Medina-Benitez, Joaquin Quiroz-Mercado, Sergio Rangel-Guerrero, Maribel Gonzalez-Villa, Natividad Cruz-Ortiz, Tatiana Nunez-Garcia, Gisela Barrera-Badillo, Lucia Hernandez-Rivas, Irma Lopez-Martinez, Ernesto Ramirez-Gonzalez. |
| EPI_ISL_3460007 | LESP Colima | Instituto de Diagnostico y Referencia Epidemiologicos (INDRE) | Claudia Wong-Arambula, Abril Rodriguez-Maldonado, Vanessa Rivero-Arredondo, Ariadna Medina-Benitez, Joaquin Quiroz-Mercado, Sergio Rangel-Guerrero, Maribel Gonzalez-Villa, Natividad Cruz-Ortiz, Tatiana Nunez-Garcia, Gisela Barrera-Badillo, Lucia Hernandez-Rivas, Irma Lopez-Martinez, Ernesto Ramirez-Gonzalez. |
| EPI_ISL_3460008 | LESP Baja California Sur | Instituto de Diagnostico y Referencia Epidemiologicos (INDRE) | Claudia Wong-Arambula, Abril Rodriguez-Maldonado, Vanessa Rivero-Arredondo, Ariadna Medina-Benitez, Joaquin Quiroz-Mercado, Sergio Rangel-Guerrero, Maribel Gonzalez-Villa, Natividad Cruz-Ortiz, Tatiana Nunez-Garcia, Gisela Barrera-Badillo, Lucia Hernandez-Rivas, Irma Lopez-Martinez, Ernesto Ramirez-Gonzalez. |
| EPI_ISL_3460010, EPI_ISL_3460011, EPI_ISL_3460012 | LESP Veracruz | Instituto de Diagnostico y Referencia Epidemiologicos (INDRE) | Claudia Wong-Arambula, Abril Rodriguez-Maldonado, Vanessa Rivero-Arredondo, Ariadna Medina-Benitez, Joaquin Quiroz-Mercado, Sergio Rangel-Guerrero, Maribel Gonzalez-Villa, Natividad Cruz-Ortiz, Tatiana Nunez-Garcia, Gisela Barrera-Badillo, Lucia Hernandez-Rivas, Irma Lopez-Martinez, Ernesto Ramirez-Gonzalez. |
| EPI_ISL_3460013 | LESP Campeche | Instituto de Diagnostico y Referencia Epidemiologicos (INDRE) | Claudia Wong-Arambula, Abril Rodriguez-Maldonado, Vanessa Rivero-Arredondo, Ariadna Medina-Benitez, Joaquin Quiroz-Mercado, Sergio Rangel-Guerrero, Maribel Gonzalez-Villa, Natividad Cruz-Ortiz, Tatiana Nunez-Garcia, Gisela Barrera-Badillo, Lucia Hernandez-Rivas, Irma Lopez-Martinez, Ernesto Ramirez-Gonzalez. |
| EPI_ISL_3460014, EPI_ISL_3460015 | LESP Queretaro | Instituto de Diagnostico y Referencia Epidemiologicos (INDRE) | Claudia Wong-Arambula, Abril Rodriguez-Maldonado, Vanessa Rivero-Arredondo, Ariadna Medina-Benitez, Joaquin Quiroz-Mercado, Sergio Rangel-Guerrero, Maribel Gonzalez-Villa, Natividad Cruz-Ortiz, Tatiana Nunez-Garcia, Gisela Barrera-Badillo, Lucia Hernandez-Rivas, Irma Lopez-Martinez, Ernesto Ramirez-Gonzalez. |
| EPI_ISL_3460016, EPI_ISL_3460017 | LESP Campeche | Instituto de Diagnostico y Referencia Epidemiologicos (INDRE) | Claudia Wong-Arambula, Abril Rodriguez-Maldonado, Vanessa Rivero-Arredondo, Ariadna Medina-Benitez, Joaquin Quiroz-Mercado, Sergio Rangel-Guerrero, Maribel Gonzalez-Villa, Natividad Cruz-Ortiz, Tatiana Nunez-Garcia, Gisela Barrera-Badillo, Lucia Hernandez-Rivas, Irma Lopez-Martinez, Ernesto Ramirez-Gonzalez. |
| EPI_ISL_3460018 | LESP Baja California Sur | Instituto de Diagnostico y Referencia Epidemiologicos (INDRE) | Claudia Wong-Arambula, Abril Rodriguez-Maldonado, Vanessa Rivero-Arredondo, Ariadna Medina-Benitez, Joaquin Quiroz-Mercado, Sergio Rangel-Guerrero, Maribel Gonzalez-Villa, Natividad Cruz-Ortiz, Tatiana Nunez-Garcia, Gisela Barrera-Badillo, Lucia Hernandez-Rivas, Irma Lopez-Martinez, Ernesto Ramirez-Gonzalez. |
| EPI_ISL_3460019 | LESP Quintana Roo | Instituto de Diagnostico y Referencia Epidemiologicos (INDRE) | Claudia Wong-Arambula, Abril Rodriguez-Maldonado, Vanessa Rivero-Arredondo, Ariadna Medina-Benitez, Joaquin Quiroz-Mercado, Sergio Rangel-Guerrero, Maribel Gonzalez-Villa, Natividad Cruz-Ortiz, Tatiana Nunez-Garcia, Gisela Barrera-Badillo, Lucia Hernandez-Rivas, Irma Lopez-Martinez, Ernesto Ramirez-Gonzalez. |
| EPI_ISL_3460020, EPI_ISL_3460021, EPI_ISL_3460022, EPI_ISL_3460023, EPI_ISL_3460024 | LESP Tabasco | Instituto de Diagnostico y Referencia Epidemiologicos (INDRE) | Claudia Wong-Arambula, Abril Rodriguez-Maldonado, Vanessa Rivero-Arredondo, Ariadna Medina-Benitez, Joaquin Quiroz-Mercado, Sergio Rangel-Guerrero, Maribel Gonzalez-Villa, Natividad Cruz-Ortiz, Tatiana Nunez-Garcia, Gisela Barrera-Badillo, Lucia Hernandez-Rivas, Irma Lopez-Martinez, Ernesto Ramirez-Gonzalez. |
| EPI_ISL_3460025 | LESP Campeche | Instituto de Diagnostico y Referencia Epidemiologicos (INDRE) | Claudia Wong-Arambula, Abril Rodriguez-Maldonado, Vanessa Rivero-Arredondo, Ariadna Medina-Benitez, Joaquin Quiroz-Mercado, Sergio Rangel-Guerrero, Maribel Gonzalez-Villa, Natividad Cruz-Ortiz, Tatiana Nunez-Garcia, Gisela Barrera-Badillo, Lucia Hernandez-Rivas, Irma Lopez-Martinez, Ernesto Ramirez-Gonzalez. |
| EPI_ISL_3460026 | LESP Jalisco | Instituto de Diagnostico y Referencia Epidemiologicos (INDRE) | Claudia Wong-Arambula, Abril Rodriguez-Maldonado, Vanessa Rivero-Arredondo, Ariadna Medina-Benitez, Joaquin Quiroz-Mercado, Sergio Rangel-Guerrero, Maribel Gonzalez-Villa, Natividad Cruz-Ortiz, Tatiana Nunez-Garcia, Gisela Barrera-Badillo, Lucia Hernandez-Rivas, Irma Lopez-Martinez, Ernesto Ramirez-Gonzalez. |
| EPI_ISL_3460027 | LESP Colima | Instituto de Diagnostico y Referencia Epidemiologicos (INDRE) | Claudia Wong-Arambula, Abril Rodriguez-Maldonado, Vanessa Rivero-Arredondo, Ariadna Medina-Benitez, Joaquin Quiroz-Mercado, Sergio Rangel-Guerrero, Maribel Gonzalez-Villa, Natividad Cruz-Ortiz, Tatiana Nunez-Garcia, Gisela Barrera-Badillo, Lucia Hernandez-Rivas, Irma Lopez-Martinez, Ernesto Ramirez-Gonzalez. |
| EPI_ISL_3460028, EPI_ISL_3460029, EPI_ISL_3460030, EPI_ISL_3460031, | LESP Jalisco | Instituto de Diagnostico y Referencia Epidemiologicos (INDRE) | Claudia Wong-Arambula, Abril Rodriguez-Maldonado, Vanessa Rivero-Arredondo, Ariadna Medina-Benitez, Joaquin Quiroz-Mercado, Sergio Rangel-Guerrero, Maribel Gonzalez-Villa, Natividad Cruz-Ortiz, Tatiana Nunez-Garcia, Gisela Barrera-Badillo, Lucia Hernandez-Rivas, Irma Lopez-Martinez, Ernesto Ramirez-Gonzalez. |

| | | | |
|---|---|---|---|
| EPI_ISL_3460032, EPI_ISL_3460033, EPI_ISL_3460034, EPI_ISL_3460035, EPI_ISL_3460036 | | | |
| EPI_ISL_3460037, EPI_ISL_3460038, EPI_ISL_3460039, EPI_ISL_3460040, EPI_ISL_3460041, EPI_ISL_3460042, EPI_ISL_3460043, EPI_ISL_3460044, EPI_ISL_3460045, EPI_ISL_3460046, EPI_ISL_3460047, EPI_ISL_3460048, EPI_ISL_3460049, EPI_ISL_3460050 | | | |
| see above | LESP Queretaro | Instituto de Diagnostico y Referencia Epidemiologicos (INDRE) | Claudia Wong-Arambula, Abril Rodriguez-Maldonado, Vanessa Rivero-Arredondo, Ariadna Medina-Benitez, Joaquin Quiroz-Mercado, Sergio Rangel-Guerrero, Maribel Gonzalez-Villa, Natividad Cruz-Ortiz, Tatiana Nunez-Garcia, Gisela Barrera-Badillo, Lucia Hernandez-Rivas, Irma Lopez-Martinez, Ernesto Ramirez-Gonzalez. |
| EPI_ISL_3460051, EPI_ISL_3460052, EPI_ISL_3460053, EPI_ISL_3460054, EPI_ISL_3460055, EPI_ISL_3460056, EPI_ISL_3460057 | LESP Guanajuato | Instituto de Diagnostico y Referencia Epidemiologicos (INDRE) | Claudia Wong-Arambula, Abril Rodriguez-Maldonado, Vanessa Rivero-Arredondo, Ariadna Medina-Benitez, Joaquin Quiroz-Mercado, Sergio Rangel-Guerrero, Maribel Gonzalez-Villa, Natividad Cruz-Ortiz, Tatiana Nunez-Garcia, Gisela Barrera-Badillo, Lucia Hernandez-Rivas, Irma Lopez-Martinez, Ernesto Ramirez-Gonzalez. |
| EPI_ISL_3460058 | LESP Aguascalientes | Instituto de Diagnostico y Referencia Epidemiologicos (INDRE) | Claudia Wong-Arambula, Abril Rodriguez-Maldonado, Vanessa Rivero-Arredondo, Ariadna Medina-Benitez, Joaquin Quiroz-Mercado, Sergio Rangel-Guerrero, Maribel Gonzalez-Villa, Natividad Cruz-Ortiz, Tatiana Nunez-Garcia, Gisela Barrera-Badillo, Lucia Hernandez-Rivas, Irma Lopez-Martinez, Ernesto Ramirez-Gonzalez. |
| EPI_ISL_3460059, EPI_ISL_3460060, EPI_ISL_3460061, EPI_ISL_3460062, EPI_ISL_3460063, EPI_ISL_3460064 | LESP Guanajuato | Instituto de Diagnostico y Referencia Epidemiologicos (INDRE) | Claudia Wong-Arambula, Abril Rodriguez-Maldonado, Vanessa Rivero-Arredondo, Ariadna Medina-Benitez, Joaquin Quiroz-Mercado, Sergio Rangel-Guerrero, Maribel Gonzalez-Villa, Natividad Cruz-Ortiz, Tatiana Nunez-Garcia, Gisela Barrera-Badillo, Lucia Hernandez-Rivas, Irma Lopez-Martinez, Ernesto Ramirez-Gonzalez. |
| EPI_ISL_3460065, EPI_ISL_3460066, EPI_ISL_3460067, EPI_ISL_3460068 | LESP Oaxaca | Instituto de Diagnostico y Referencia Epidemiologicos (INDRE) | Claudia Wong-Arambula, Abril Rodriguez-Maldonado, Vanessa Rivero-Arredondo, Ariadna Medina-Benitez, Joaquin Quiroz-Mercado, Sergio Rangel-Guerrero, Maribel Gonzalez-Villa, Natividad Cruz-Ortiz, Tatiana Nunez-Garcia, Gisela Barrera-Badillo, Lucia Hernandez-Rivas, Irma Lopez-Martinez, Ernesto Ramirez-Gonzalez. |
| EPI_ISL_3460069, EPI_ISL_3460070, EPI_ISL_3460071, EPI_ISL_3460072, EPI_ISL_3460073 | LESP Coahuila | Instituto de Diagnostico y Referencia Epidemiologicos (INDRE) | Claudia Wong-Arambula, Abril Rodriguez-Maldonado, Vanessa Rivero-Arredondo, Ariadna Medina-Benitez, Joaquin Quiroz-Mercado, Sergio Rangel-Guerrero, Maribel Gonzalez-Villa, Natividad Cruz-Ortiz, Tatiana Nunez-Garcia, Gisela Barrera-Badillo, Lucia Hernandez-Rivas, Irma Lopez-Martinez, Ernesto Ramirez-Gonzalez. |
| EPI_ISL_3460074, EPI_ISL_3460075, EPI_ISL_3460076, EPI_ISL_3460077, EPI_ISL_3460078, EPI_ISL_3460079, EPI_ISL_3460080 | LESP Sinaloa | Instituto de Diagnostico y Referencia Epidemiologicos (INDRE) | Claudia Wong-Arambula, Abril Rodriguez-Maldonado, Vanessa Rivero-Arredondo, Ariadna Medina-Benitez, Joaquin Quiroz-Mercado, Sergio Rangel-Guerrero, Maribel Gonzalez-Villa, Natividad Cruz-Ortiz, Tatiana Nunez-Garcia, Gisela Barrera-Badillo, Lucia Hernandez-Rivas, Irma Lopez-Martinez, Ernesto Ramirez-Gonzalez. |
| EPI_ISL_3460081 | LESP Mexico City | Instituto de Diagnostico y Referencia Epidemiologicos (INDRE) | Claudia Wong-Arambula, Abril Rodriguez-Maldonado, Vanessa Rivero-Arredondo, Ariadna Medina-Benitez, Joaquin Quiroz-Mercado, Sergio Rangel-Guerrero, Maribel Gonzalez-Villa, Natividad Cruz-Ortiz, Tatiana Nunez-Garcia, Gisela Barrera-Badillo, Lucia Hernandez-Rivas, Irma Lopez-Martinez, Ernesto Ramirez-Gonzalez. |
| EPI_ISL_3460082, EPI_ISL_3460083 | LESP Aguascalientes | Instituto de Diagnostico y Referencia Epidemiologicos (INDRE) | Claudia Wong-Arambula, Abril Rodriguez-Maldonado, Vanessa Rivero-Arredondo, Ariadna Medina-Benitez, Joaquin Quiroz-Mercado, Sergio Rangel-Guerrero, Maribel Gonzalez-Villa, Natividad Cruz-Ortiz, Tatiana Nunez-Garcia, Gisela Barrera-Badillo, Lucia Hernandez-Rivas, Irma Lopez-Martinez, Ernesto Ramirez-Gonzalez. |
| EPI_ISL_3460084 | LESP Chihuahua | Instituto de Diagnostico y Referencia Epidemiologicos (INDRE) | Claudia Wong-Arambula, Abril Rodriguez-Maldonado, Vanessa Rivero-Arredondo, Ariadna Medina-Benitez, Joaquin Quiroz-Mercado, Sergio Rangel-Guerrero, Maribel Gonzalez-Villa, Natividad Cruz-Ortiz, Tatiana Nunez-Garcia, Gisela Barrera-Badillo, Lucia Hernandez-Rivas, Irma Lopez-Martinez, Ernesto Ramirez-Gonzalez. |
| EPI_ISL_3460085 | LESP Mexico City | Instituto de Diagnostico y Referencia Epidemiologicos (INDRE) | Claudia Wong-Arambula, Abril Rodriguez-Maldonado, Vanessa Rivero-Arredondo, Ariadna Medina-Benitez, Joaquin Quiroz-Mercado, Sergio Rangel-Guerrero, Maribel Gonzalez-Villa, Natividad Cruz-Ortiz, Tatiana Nunez-Garcia, Gisela Barrera-Badillo, Lucia Hernandez-Rivas, Irma Lopez-Martinez, Ernesto Ramirez-Gonzalez. |
| EPI_ISL_3460086, EPI_ISL_3460087 | LESP Nayarit | Instituto de Diagnostico y Referencia Epidemiologicos (INDRE) | Claudia Wong-Arambula, Abril Rodriguez-Maldonado, Vanessa Rivero-Arredondo, Ariadna Medina-Benitez, Joaquin Quiroz-Mercado, Sergio Rangel-Guerrero, Maribel Gonzalez-Villa, Natividad Cruz-Ortiz, Tatiana Nunez-Garcia, Gisela Barrera-Badillo, Lucia Hernandez-Rivas, Irma Lopez-Martinez, Ernesto Ramirez-Gonzalez. |
| EPI_ISL_3460088 | LESP Tamaulipas | Instituto de Diagnostico y Referencia Epidemiologicos (INDRE) | Claudia Wong-Arambula, Abril Rodriguez-Maldonado, Vanessa Rivero-Arredondo, Ariadna Medina-Benitez, Joaquin Quiroz-Mercado, Sergio Rangel-Guerrero, Maribel Gonzalez-Villa, Natividad Cruz-Ortiz, Tatiana Nunez-Garcia, Gisela Barrera-Badillo, Lucia Hernandez-Rivas, Irma Lopez-Martinez, Ernesto Ramirez-Gonzalez. |
| EPI_ISL_3460089, EPI_ISL_3460090, EPI_ISL_3460091, EPI_ISL_3460092, EPI_ISL_3460093, EPI_ISL_3460094, EPI_ISL_3460095, EPI_ISL_3460096, EPI_ISL_3460097, EPI_ISL_3460098 | LESP Guerrero | Instituto de Diagnostico y Referencia Epidemiologicos (INDRE) | Claudia Wong-Arambula, Abril Rodriguez-Maldonado, Vanessa Rivero-Arredondo, Ariadna Medina-Benitez, Joaquin Quiroz-Mercado, Sergio Rangel-Guerrero, Maribel Gonzalez-Villa, Natividad Cruz-Ortiz, Tatiana Nunez-Garcia, Gisela Barrera-Badillo, Lucia Hernandez-Rivas, Irma Lopez-Martinez, Ernesto Ramirez-Gonzalez. |
| EPI_ISL_3460099, EPI_ISL_3460100, EPI_ISL_3460101, EPI_ISL_3460102 | LESP Michoacan | Instituto de Diagnostico y Referencia Epidemiologicos (INDRE) | Claudia Wong-Arambula, Abril Rodriguez-Maldonado, Vanessa Rivero-Arredondo, Ariadna Medina-Benitez, Joaquin Quiroz-Mercado, Sergio Rangel-Guerrero, Maribel Gonzalez-Villa, Natividad Cruz-Ortiz, Tatiana Nunez-Garcia, Gisela Barrera-Badillo, Lucia Hernandez-Rivas, Irma Lopez-Martinez, Ernesto Ramirez-Gonzalez. |
| EPI_ISL_3460103 | LESP Oaxaca | Instituto de Diagnostico y Referencia Epidemiologicos (INDRE) | Claudia Wong-Arambula, Abril Rodriguez-Maldonado, Vanessa Rivero-Arredondo, Ariadna Medina-Benitez, Joaquin Quiroz-Mercado, Sergio Rangel-Guerrero, Maribel Gonzalez-Villa, Natividad Cruz-Ortiz, Tatiana Nunez-Garcia, Gisela Barrera-Badillo, Lucia Hernandez-Rivas, Irma Lopez-Martinez, Ernesto Ramirez-Gonzalez. |
| EPI_ISL_3460104, EPI_ISL_3460105 | LESP Quintana Roo | Instituto de Diagnostico y Referencia Epidemiologicos (INDRE) | Claudia Wong-Arambula, Abril Rodriguez-Maldonado, Vanessa Rivero-Arredondo, Ariadna Medina-Benitez, Joaquin Quiroz-Mercado, Sergio Rangel-Guerrero, Maribel Gonzalez-Villa, Natividad Cruz-Ortiz, Tatiana Nunez-Garcia, Gisela Barrera-Badillo, Lucia Hernandez-Rivas, Irma Lopez-Martinez, Ernesto Ramirez-Gonzalez. |
| EPI_ISL_3460106 | LESP Sonora | Instituto de Diagnostico y Referencia Epidemiologicos (INDRE) | Claudia Wong-Arambula, Abril Rodriguez-Maldonado, Vanessa Rivero-Arredondo, Ariadna Medina-Benitez, Joaquin Quiroz-Mercado, Sergio Rangel-Guerrero, Maribel Gonzalez-Villa, Natividad Cruz-Ortiz, Tatiana Nunez-Garcia, Gisela Barrera-Badillo, Lucia Hernandez-Rivas, Irma Lopez-Martinez, Ernesto Ramirez-Gonzalez. |
| EPI_ISL_3460107 | LESP Jalisco | Instituto de Diagnostico y Referencia Epidemiologicos (INDRE) | Claudia Wong-Arambula, Abril Rodriguez-Maldonado, Vanessa Rivero-Arredondo, Ariadna Medina-Benitez, Joaquin Quiroz-Mercado, Sergio Rangel-Guerrero, Maribel Gonzalez-Villa, Natividad Cruz-Ortiz, Tatiana Nunez-Garcia, Gisela Barrera-Badillo, Lucia Hernandez-Rivas, Irma Lopez-Martinez, Ernesto Ramirez-Gonzalez. |
| EPI_ISL_3460108, EPI_ISL_3460109, EPI_ISL_3460110 | LESP State of Mexico | Instituto de Diagnostico y Referencia Epidemiologicos (INDRE) | Claudia Wong-Arambula, Abril Rodriguez-Maldonado, Vanessa Rivero-Arredondo, Ariadna Medina-Benitez, Joaquin Quiroz-Mercado, Sergio Rangel-Guerrero, Maribel Gonzalez-Villa, Natividad Cruz-Ortiz, Tatiana Nunez-Garcia, Gisela Barrera-Badillo, Lucia Hernandez-Rivas, Irma Lopez-Martinez, Ernesto Ramirez-Gonzalez. |
| EPI_ISL_3460111, EPI_ISL_3460112, EPI_ISL_3460113, EPI_ISL_3460114, EPI_ISL_3460115, EPI_ISL_3460116, EPI_ISL_3460117, EPI_ISL_3460118 | LESP Jalisco | Instituto de Diagnostico y Referencia Epidemiologicos (INDRE) | Claudia Wong-Arambula, Abril Rodriguez-Maldonado, Vanessa Rivero-Arredondo, Ariadna Medina-Benitez, Joaquin Quiroz-Mercado, Sergio Rangel-Guerrero, Maribel Gonzalez-Villa, Natividad Cruz-Ortiz, Tatiana Nunez-Garcia, Gisela Barrera-Badillo, Lucia Hernandez-Rivas, Irma Lopez-Martinez, Ernesto Ramirez-Gonzalez. |
| EPI_ISL_3460119, EPI_ISL_3460120, EPI_ISL_3460121 | LESP Tabasco | Instituto de Diagnostico y Referencia Epidemiologicos (INDRE) | Claudia Wong-Arambula, Abril Rodriguez-Maldonado, Vanessa Rivero-Arredondo, Ariadna Medina-Benitez, Joaquin Quiroz-Mercado, Sergio Rangel-Guerrero, Maribel Gonzalez-Villa, Natividad Cruz-Ortiz, Tatiana Nunez-Garcia, Gisela Barrera-Badillo, Lucia Hernandez-Rivas, Irma Lopez-Martinez, Ernesto Ramirez-Gonzalez. |
| EPI_ISL_3460122 | LESP Queretaro | Instituto de Diagnostico y Referencia Epidemiologicos (INDRE) | Claudia Wong-Arambula, Abril Rodriguez-Maldonado, Vanessa Rivero-Arredondo, Ariadna Medina-Benitez, Joaquin Quiroz-Mercado, Sergio Rangel-Guerrero, Maribel Gonzalez-Villa, Natividad Cruz-Ortiz, Tatiana Nunez-Garcia, Gisela Barrera-Badillo, Lucia Hernandez-Rivas, Irma Lopez-Martinez, Ernesto Ramirez-Gonzalez. |
| EPI_ISL_3460123, EPI_ISL_3460124, EPI_ISL_3460125, EPI_ISL_3460126, EPI_ISL_3460127 | LESP Mexico City | Instituto de Diagnostico y Referencia Epidemiologicos (INDRE) | Claudia Wong-Arambula, Abril Rodriguez-Maldonado, Vanessa Rivero-Arredondo, Ariadna Medina-Benitez, Joaquin Quiroz-Mercado, Sergio Rangel-Guerrero, Maribel Gonzalez-Villa, Natividad Cruz-Ortiz, Tatiana Nunez-Garcia, Gisela Barrera-Badillo, Lucia Hernandez-Rivas, Irma Lopez-Martinez, Ernesto Ramirez-Gonzalez. |
| EPI_ISL_3460128 | LESP State of Mexico | Instituto de Diagnostico y Referencia Epidemiologicos (INDRE) | Claudia Wong-Arambula, Abril Rodriguez-Maldonado, Vanessa Rivero-Arredondo, Ariadna Medina-Benitez, Joaquin Quiroz-Mercado, Sergio Rangel-Guerrero, Maribel Gonzalez-Villa, Natividad Cruz-Ortiz, Tatiana Nunez-Garcia, Gisela Barrera-Badillo, Lucia Hernandez-Rivas, Irma Lopez-Martinez, Ernesto Ramirez-Gonzalez. |
| EPI_ISL_3460129, EPI_ISL_3460130, | LESP Mexico City | Instituto de Diagnostico y Referencia Epidemiologicos (INDRE) | Claudia Wong-Arambula, Abril Rodriguez-Maldonado, Vanessa Rivero-Arredondo, Ariadna Medina-Benitez, Joaquin Quiroz-Mercado, Sergio Rangel-Guerrero, Maribel Gonzalez-Villa, Natividad Cruz-Ortiz, Tatiana Nunez-Garcia, Gisela Barrera-Badillo, Lucia Hernandez-Rivas, Irma Lopez-Martinez, Ernesto Ramirez-Gonzalez. |

| | | | |
|---|---|---|---|
| EPI_ISL_3460131, EPI_ISL_3460132, EPI_ISL_3460133 | | | |
| EPI_ISL_3460134 | LESP State of Mexico | Instituto de Diagnostico y Referencia Epidemiologicos (INDRE) | Claudia Wong-Arambula, Abril Rodriguez-Maldonado, Vanessa Rivero-Arredondo, Ariadna Medina-Benitez, Joaquin Quiroz-Mercado, Sergio Rangel-Guerrero, Maribel Gonzalez-Villa, Natividad Cruz-Ortiz, Tatiana Nunez-Garcia, Gisela Barrera-Badillo, Lucia Hernandez-Rivas, Irma Lopez-Martinez, Ernesto Ramirez-Gonzalez. |
| EPI_ISL_3460135 | LESP Mexico City | Instituto de Diagnostico y Referencia Epidemiologicos (INDRE) | Claudia Wong-Arambula, Abril Rodriguez-Maldonado, Vanessa Rivero-Arredondo, Ariadna Medina-Benitez, Joaquin Quiroz-Mercado, Sergio Rangel-Guerrero, Maribel Gonzalez-Villa, Natividad Cruz-Ortiz, Tatiana Nunez-Garcia, Gisela Barrera-Badillo, Lucia Hernandez-Rivas, Irma Lopez-Martinez, Ernesto Ramirez-Gonzalez. |
| EPI_ISL_3460136, EPI_ISL_3460137 | LESP State of Mexico | Instituto de Diagnostico y Referencia Epidemiologicos (INDRE) | Claudia Wong-Arambula, Abril Rodriguez-Maldonado, Vanessa Rivero-Arredondo, Ariadna Medina-Benitez, Joaquin Quiroz-Mercado, Sergio Rangel-Guerrero, Maribel Gonzalez-Villa, Natividad Cruz-Ortiz, Tatiana Nunez-Garcia, Gisela Barrera-Badillo, Lucia Hernandez-Rivas, Irma Lopez-Martinez, Ernesto Ramirez-Gonzalez. |
| EPI_ISL_3460138, EPI_ISL_3460139 | LESP Chihuahua | Instituto de Diagnostico y Referencia Epidemiologicos (INDRE) | Claudia Wong-Arambula, Abril Rodriguez-Maldonado, Vanessa Rivero-Arredondo, Ariadna Medina-Benitez, Joaquin Quiroz-Mercado, Sergio Rangel-Guerrero, Maribel Gonzalez-Villa, Natividad Cruz-Ortiz, Tatiana Nunez-Garcia, Gisela Barrera-Badillo, Lucia Hernandez-Rivas, Irma Lopez-Martinez, Ernesto Ramirez-Gonzalez. |
| EPI_ISL_3460140, EPI_ISL_3460141 | LESP Colima | Instituto de Diagnostico y Referencia Epidemiologicos (INDRE) | Claudia Wong-Arambula, Abril Rodriguez-Maldonado, Vanessa Rivero-Arredondo, Ariadna Medina-Benitez, Joaquin Quiroz-Mercado, Sergio Rangel-Guerrero, Maribel Gonzalez-Villa, Natividad Cruz-Ortiz, Tatiana Nunez-Garcia, Gisela Barrera-Badillo, Lucia Hernandez-Rivas, Irma Lopez-Martinez, Ernesto Ramirez-Gonzalez. |
| EPI_ISL_3460142 | LESP Jalisco | Instituto de Diagnostico y Referencia Epidemiologicos (INDRE) | Claudia Wong-Arambula, Abril Rodriguez-Maldonado, Vanessa Rivero-Arredondo, Ariadna Medina-Benitez, Joaquin Quiroz-Mercado, Sergio Rangel-Guerrero, Maribel Gonzalez-Villa, Natividad Cruz-Ortiz, Tatiana Nunez-Garcia, Gisela Barrera-Badillo, Lucia Hernandez-Rivas, Irma Lopez-Martinez, Ernesto Ramirez-Gonzalez. |
| EPI_ISL_3460143 | LESP Colima | Instituto de Diagnostico y Referencia Epidemiologicos (INDRE) | Claudia Wong-Arambula, Abril Rodriguez-Maldonado, Vanessa Rivero-Arredondo, Ariadna Medina-Benitez, Joaquin Quiroz-Mercado, Sergio Rangel-Guerrero, Maribel Gonzalez-Villa, Natividad Cruz-Ortiz, Tatiana Nunez-Garcia, Gisela Barrera-Badillo, Lucia Hernandez-Rivas, Irma Lopez-Martinez, Ernesto Ramirez-Gonzalez. |
| EPI_ISL_3460144, EPI_ISL_3460145, EPI_ISL_3460146, EPI_ISL_3460147, EPI_ISL_3460148, EPI_ISL_3460149 | LESP Hidalgo | Instituto de Diagnostico y Referencia Epidemiologicos (INDRE) | Claudia Wong-Arambula, Abril Rodriguez-Maldonado, Vanessa Rivero-Arredondo, Ariadna Medina-Benitez, Joaquin Quiroz-Mercado, Sergio Rangel-Guerrero, Maribel Gonzalez-Villa, Natividad Cruz-Ortiz, Tatiana Nunez-Garcia, Gisela Barrera-Badillo, Lucia Hernandez-Rivas, Irma Lopez-Martinez, Ernesto Ramirez-Gonzalez. |
| EPI_ISL_3460150, EPI_ISL_3460151, EPI_ISL_3460152, EPI_ISL_3460153, EPI_ISL_3460154, EPI_ISL_3460155, EPI_ISL_3460156, EPI_ISL_3460157, EPI_ISL_3460158, EPI_ISL_3460159, EPI_ISL_3460160 | | | |
| see above | LESP Michoacan | Instituto de Diagnostico y Referencia Epidemiologicos (INDRE) | Claudia Wong-Arambula, Abril Rodriguez-Maldonado, Vanessa Rivero-Arredondo, Ariadna Medina-Benitez, Joaquin Quiroz-Mercado, Sergio Rangel-Guerrero, Maribel Gonzalez-Villa, Natividad Cruz-Ortiz, Tatiana Nunez-Garcia, Gisela Barrera-Badillo, Lucia Hernandez-Rivas, Irma Lopez-Martinez, Ernesto Ramirez-Gonzalez. |
| EPI_ISL_3460161, EPI_ISL_3460162, EPI_ISL_3460163, EPI_ISL_3460164, EPI_ISL_3460165, EPI_ISL_3460166, EPI_ISL_3460167, EPI_ISL_3460168, EPI_ISL_3460169, EPI_ISL_3460170, EPI_ISL_3460171, EPI_ISL_3460172, EPI_ISL_3460173, EPI_ISL_3460174, EPI_ISL_3460175, EPI_ISL_3460176, EPI_ISL_3460177, EPI_ISL_3460178, EPI_ISL_3460179, EPI_ISL_3460180, EPI_ISL_3460181, EPI_ISL_3460182 | | | |
| see above | LESP Queretaro | Instituto de Diagnostico y Referencia Epidemiologicos (INDRE) | Claudia Wong-Arambula, Abril Rodriguez-Maldonado, Vanessa Rivero-Arredondo, Ariadna Medina-Benitez, Joaquin Quiroz-Mercado, Sergio Rangel-Guerrero, Maribel Gonzalez-Villa, Natividad Cruz-Ortiz, Tatiana Nunez-Garcia, Gisela Barrera-Badillo, Lucia Hernandez-Rivas, Irma Lopez-Martinez, Ernesto Ramirez-Gonzalez. |
| EPI_ISL_3460183 | LESP Guanajuato | Instituto de Diagnostico y Referencia Epidemiologicos (INDRE) | Claudia Wong-Arambula, Abril Rodriguez-Maldonado, Vanessa Rivero-Arredondo, Ariadna Medina-Benitez, Joaquin Quiroz-Mercado, Sergio Rangel-Guerrero, Maribel Gonzalez-Villa, Natividad Cruz-Ortiz, Tatiana Nunez-Garcia, Gisela Barrera-Badillo, Lucia Hernandez-Rivas, Irma Lopez-Martinez, Ernesto Ramirez-Gonzalez. |
| EPI_ISL_3460184, EPI_ISL_3460185, EPI_ISL_3460186, EPI_ISL_3460187, EPI_ISL_3460188, EPI_ISL_3460189, EPI_ISL_3460190, EPI_ISL_3460191 | LESP Queretaro | Instituto de Diagnostico y Referencia Epidemiologicos (INDRE) | Claudia Wong-Arambula, Abril Rodriguez-Maldonado, Vanessa Rivero-Arredondo, Ariadna Medina-Benitez, Joaquin Quiroz-Mercado, Sergio Rangel-Guerrero, Maribel Gonzalez-Villa, Natividad Cruz-Ortiz, Tatiana Nunez-Garcia, Gisela Barrera-Badillo, Lucia Hernandez-Rivas, Irma Lopez-Martinez, Ernesto Ramirez-Gonzalez. |
| EPI_ISL_3460192, EPI_ISL_3460193, EPI_ISL_3460194, EPI_ISL_3460195, EPI_ISL_3460196, EPI_ISL_3460197, EPI_ISL_3460198, EPI_ISL_3460199, EPI_ISL_3460200 | LESP Guanajuato | Instituto de Diagnostico y Referencia Epidemiologicos (INDRE) | Claudia Wong-Arambula, Abril Rodriguez-Maldonado, Vanessa Rivero-Arredondo, Ariadna Medina-Benitez, Joaquin Quiroz-Mercado, Sergio Rangel-Guerrero, Maribel Gonzalez-Villa, Natividad Cruz-Ortiz, Tatiana Nunez-Garcia, Gisela Barrera-Badillo, Lucia Hernandez-Rivas, Irma Lopez-Martinez, Ernesto Ramirez-Gonzalez. |
| EPI_ISL_3460201, EPI_ISL_3460202, EPI_ISL_3460203, EPI_ISL_3460204, EPI_ISL_3460205, EPI_ISL_3460206 | LESP Aguascalientes | Instituto de Diagnostico y Referencia Epidemiologicos (INDRE) | Claudia Wong-Arambula, Abril Rodriguez-Maldonado, Vanessa Rivero-Arredondo, Ariadna Medina-Benitez, Joaquin Quiroz-Mercado, Sergio Rangel-Guerrero, Maribel Gonzalez-Villa, Natividad Cruz-Ortiz, Tatiana Nunez-Garcia, Gisela Barrera-Badillo, Lucia Hernandez-Rivas, Irma Lopez-Martinez, Ernesto Ramirez-Gonzalez. |
| EPI_ISL_3460207, EPI_ISL_3460208, EPI_ISL_3460209 | LESP Chihuahua | Instituto de Diagnostico y Referencia Epidemiologicos (INDRE) | Claudia Wong-Arambula, Abril Rodriguez-Maldonado, Vanessa Rivero-Arredondo, Ariadna Medina-Benitez, Joaquin Quiroz-Mercado, Sergio Rangel-Guerrero, Maribel Gonzalez-Villa, Natividad Cruz-Ortiz, Tatiana Nunez-Garcia, Gisela Barrera-Badillo, Lucia Hernandez-Rivas, Irma Lopez-Martinez, Ernesto Ramirez-Gonzalez. |
| EPI_ISL_3460210, EPI_ISL_3460211 | LESP Tamaulipas | Instituto de Diagnostico y Referencia Epidemiologicos (INDRE) | Claudia Wong-Arambula, Abril Rodriguez-Maldonado, Vanessa Rivero-Arredondo, Ariadna Medina-Benitez, Joaquin Quiroz-Mercado, Sergio Rangel-Guerrero, Maribel Gonzalez-Villa, Natividad Cruz-Ortiz, Tatiana Nunez-Garcia, Gisela Barrera-Badillo, Lucia Hernandez-Rivas, Irma Lopez-Martinez, Ernesto Ramirez-Gonzalez. |
| EPI_ISL_3460212, EPI_ISL_3460213, EPI_ISL_3460214 | LESP Guerrero | Instituto de Diagnostico y Referencia Epidemiologicos (INDRE) | Claudia Wong-Arambula, Abril Rodriguez-Maldonado, Vanessa Rivero-Arredondo, Ariadna Medina-Benitez, Joaquin Quiroz-Mercado, Sergio Rangel-Guerrero, Maribel Gonzalez-Villa, Natividad Cruz-Ortiz, Tatiana Nunez-Garcia, Gisela Barrera-Badillo, Lucia Hernandez-Rivas, Irma Lopez-Martinez, Ernesto Ramirez-Gonzalez. |
| EPI_ISL_3460215, EPI_ISL_3460216 | LESP Mexico City | Instituto de Diagnostico y Referencia Epidemiologicos (INDRE) | Claudia Wong-Arambula, Abril Rodriguez-Maldonado, Vanessa Rivero-Arredondo, Ariadna Medina-Benitez, Joaquin Quiroz-Mercado, Sergio Rangel-Guerrero, Maribel Gonzalez-Villa, Natividad Cruz-Ortiz, Tatiana Nunez-Garcia, Gisela Barrera-Badillo, Lucia Hernandez-Rivas, Irma Lopez-Martinez, Ernesto Ramirez-Gonzalez. |
| EPI_ISL_3460217, EPI_ISL_3460218, EPI_ISL_3460219, EPI_ISL_3460220, EPI_ISL_3460221, EPI_ISL_3460222, EPI_ISL_3460223, EPI_ISL_3460224, EPI_ISL_3460225, EPI_ISL_3460226, EPI_ISL_3460227, EPI_ISL_3460228, EPI_ISL_3460229, EPI_ISL_3460230, EPI_ISL_3460231, EPI_ISL_3460232, EPI_ISL_3460233, EPI_ISL_3460234, EPI_ISL_3460235 | | | |
| see above | LESP Baja California Sur | Instituto de Diagnostico y Referencia Epidemiologicos (INDRE) | Claudia Wong-Arambula, Abril Rodriguez-Maldonado, Vanessa Rivero-Arredondo, Ariadna Medina-Benitez, Joaquin Quiroz-Mercado, Sergio Rangel-Guerrero, Maribel Gonzalez-Villa, Natividad Cruz-Ortiz, Tatiana Nunez-Garcia, Gisela Barrera-Badillo, Lucia Hernandez-Rivas, Irma Lopez-Martinez, Ernesto Ramirez-Gonzalez. |
| EPI_ISL_3460236 | LESP Jalisco | Instituto de Diagnostico y Referencia Epidemiologicos (INDRE) | Claudia Wong-Arambula, Abril Rodriguez-Maldonado, Vanessa Rivero-Arredondo, Ariadna Medina-Benitez, Joaquin Quiroz-Mercado, Sergio Rangel-Guerrero, Maribel Gonzalez-Villa, Natividad Cruz-Ortiz, Tatiana Nunez-Garcia, Gisela Barrera-Badillo, Lucia Hernandez-Rivas, Irma Lopez-Martinez, Ernesto Ramirez-Gonzalez. |
| EPI_ISL_3460237 | LESP Colima | Instituto de Diagnostico y Referencia Epidemiologicos (INDRE) | Claudia Wong-Arambula, Abril Rodriguez-Maldonado, Vanessa Rivero-Arredondo, Ariadna Medina-Benitez, Joaquin Quiroz-Mercado, Sergio Rangel-Guerrero, Maribel Gonzalez-Villa, Natividad Cruz-Ortiz, Tatiana Nunez-Garcia, Gisela Barrera-Badillo, Lucia Hernandez-Rivas, Irma Lopez-Martinez, Ernesto Ramirez-Gonzalez. |
| EPI_ISL_3460246 | Ohio Department of Health Laboratory | Ohio Department of Health Laboratory | Holmes, Jennifer; Eric Brandt, Keoni Omura, Glen McGillivary, Caitlin McDonnell, Jade Mowery, Stephanie Mccracken, Tyler Payne, Kirtana Ramadugu, Erica Leasure, Brent Lee, Kelsey Florek, Heather Blankenship, Allison Black, Quanta Brown, and Tammy Bannerman |
| EPI_ISL_3460248, EPI_ISL_3460250, EPI_ISL_3460252 | Laboratório de Microbiologia Molecular - Universidade FEEVALE | Molecular Microbiology Laboratory | Alana Witt Hansen, Fágner Henrique Heldt, Fernando Rosado Spilki, Juliana Schons Gulart, Juliane Deise Fleck, Mariana Soares da Silva, Meriane Demoliner, Matheus Nunes Weber, Paula Rodrigues de Almeida, Michele Filippi. |
| EPI_ISL_3460262, EPI_ISL_3460263, EPI_ISL_3460264, EPI_ISL_3460265, EPI_ISL_3460266, EPI_ISL_3460267, EPI_ISL_3460268, EPI_ISL_3460269, EPI_ISL_3460270, EPI_ISL_3460271, EPI_ISL_3460272, EPI_ISL_3460273, EPI_ISL_3460274, EPI_ISL_3460275, EPI_ISL_3460276, EPI_ISL_3460277, EPI_ISL_3460278, EPI_ISL_3460279, EPI_ISL_3460280 | | | |
| see above | Minnesota Department of Health, Public Health Laboratory | Minnesota Department of Health, Public Health Laboratory | Alexandra Lorentz, Jacob Garfin, Matt Plumb, and Xiong Wang |
| EPI_ISL_3460281, EPI_ISL_3460282 | Centracare Laboratory Services | Minnesota Department of Health, Public Health Laboratory | Alexandra Lorentz, Jacob Garfin, Matt Plumb, and Xiong Wang |
| EPI_ISL_3460283, EPI_ISL_3460284, EPI_ISL_3460285, EPI_ISL_3460286, EPI_ISL_3460287, EPI_ISL_3460288, EPI_ISL_3460289, EPI_ISL_3460290 | Minnesota Department of Health, Public Health Laboratory | Minnesota Department of Health, Public Health Laboratory | Alexandra Lorentz, Jacob Garfin, Matt Plumb, and Xiong Wang |
| EPI_ISL_3460291 | Centracare Laboratory Services | Minnesota Department of Health, Public Health Laboratory | Alexandra Lorentz, Jacob Garfin, Matt Plumb, and Xiong Wang |
| EPI_ISL_3460292, EPI_ISL_3460293, EPI_ISL_3460294, EPI_ISL_3460295, EPI_ISL_3460296, EPI_ISL_3460297 | Minnesota Department of Health, Public Health Laboratory | Minnesota Department of Health, Public Health Laboratory | Alexandra Lorentz, Jacob Garfin, Matt Plumb, and Xiong Wang |
| EPI_ISL_3460298 | Centracare Laboratory Services | Minnesota Department of Health, Public Health Laboratory | Alexandra Lorentz, Jacob Garfin, Matt Plumb, and Xiong Wang |

| | | | |
|---|---|---|---|
| EPI_ISL_3460299, EPI_ISL_3460300, EPI_ISL_3460301 | Minnesota Department of Health, Public Health Laboratory | Minnesota Department of Health, Public Health Laboratory | Alexandra Lorentz, Jacob Garfin, Matt Plumb, and Xiong Wang |
| EPI_ISL_3460302 | Centracare Laboratory Services | Minnesota Department of Health, Public Health Laboratory | Alexandra Lorentz, Jacob Garfin, Matt Plumb, and Xiong Wang |
| EPI_ISL_3460303, EPI_ISL_3460304, EPI_ISL_3460306, EPI_ISL_3460307, EPI_ISL_3460308, EPI_ISL_3460309, EPI_ISL_3460310, EPI_ISL_3460311, EPI_ISL_3460312, EPI_ISL_3460313, EPI_ISL_3460314, EPI_ISL_3460315, EPI_ISL_3460316, EPI_ISL_3460317, EPI_ISL_3460318 | | | |
| see above | Minnesota Department of Health, Public Health Laboratory | Minnesota Department of Health, Public Health Laboratory | Alexandra Lorentz, Jacob Garfin, Matt Plumb, and Xiong Wang |
| EPI_ISL_3460319, EPI_ISL_3460320, EPI_ISL_3460321, EPI_ISL_3460322 | Centracare Laboratory Services | Minnesota Department of Health, Public Health Laboratory | Alexandra Lorentz, Jacob Garfin, Matt Plumb, and Xiong Wang |
| EPI_ISL_3460323, EPI_ISL_3460324, EPI_ISL_3460325, EPI_ISL_3460326, EPI_ISL_3460327, EPI_ISL_3460328, EPI_ISL_3460329, EPI_ISL_3460330, EPI_ISL_3460331, EPI_ISL_3460332, EPI_ISL_3460333, EPI_ISL_3460334, EPI_ISL_3460335, EPI_ISL_3460336, EPI_ISL_3460337, EPI_ISL_3460338, EPI_ISL_3460339, EPI_ISL_3460340, EPI_ISL_3460341, EPI_ISL_3460342, EPI_ISL_3460343, EPI_ISL_3460344, EPI_ISL_3460345, EPI_ISL_3460346, EPI_ISL_3460347, EPI_ISL_3460348, EPI_ISL_3460349, EPI_ISL_3460350, EPI_ISL_3460351, EPI_ISL_3460352, EPI_ISL_3460353, EPI_ISL_3460354, EPI_ISL_3460355 | | | |
| see above | Minnesota Department of Health, Public Health Laboratory | Minnesota Department of Health, Public Health Laboratory | Alexandra Lorentz, Jacob Garfin, Matt Plumb, and Xiong Wang |
| EPI_ISL_3460356, EPI_ISL_3460358 | Ohio Department of Health Laboratory | Ohio Department of Health Laboratory | Holmes, Jennifer; Eric Brandt, Keoni Omura, Glen McGillivary, Caitlin McDonnell, Jade Mowery, Stephanie Mccracken, Tyler Payne, Kirtana Ramadugu, Erica Leasure, Brent Lee, Kelsey Florek, Heather Blankenship, Allison Black, Quanta Brown, and Tammy Bannerman |
| EPI_ISL_3460360, EPI_ISL_3460361, EPI_ISL_3460362, EPI_ISL_3460365, EPI_ISL_3460366, EPI_ISL_3460369, EPI_ISL_3460370, EPI_ISL_3460371, EPI_ISL_3460372, EPI_ISL_3460373, EPI_ISL_3460374, EPI_ISL_3460375, EPI_ISL_3460376, EPI_ISL_3460377, EPI_ISL_3460380, EPI_ISL_3460381, EPI_ISL_3460386, EPI_ISL_3460389, EPI_ISL_3460390, EPI_ISL_3460392, EPI_ISL_3460393, EPI_ISL_3460396, EPI_ISL_3460397, EPI_ISL_3460400, EPI_ISL_3460401, EPI_ISL_3460402, EPI_ISL_3460403, EPI_ISL_3460404, EPI_ISL_3460405, EPI_ISL_3460407, EPI_ISL_3460410 | | | |
| see above | TGen North | TGen North | "Jolene Bowers, Heather Centner, Brett Van Tassel, Chris French, Hayley Yaglom, Darrin Lemmer, Dave Engelthaler" |
| EPI_ISL_3460412 | LABO BIO DOC | CHU Purpan - Laboratoire de Virologie - Institut Fédératif de Biologie | Latour J., Jeanne N., Ranger N., Carcenac R., Harter A., Boyer P., Ferrer V., Tremeaux R., Izopet J. |
| EPI_ISL_3460413 | CHU Purpan - Laboratoire de Virologie - Institut Fédératif de Biologie | CHU Purpan - Laboratoire de Virologie - Institut Fédératif de Biologie | Latour J., Jeanne N., Ranger N., Carcenac R., Harter A., Boyer P., Ferrer V., Tremeaux R., Izopet J. |
| EPI_ISL_3460414 | TGen North | TGen North | "Jolene Bowers, Heather Centner, Brett Van Tassel, Chris French, Hayley Yaglom, Darrin Lemmer, Dave Engelthaler" |
| EPI_ISL_3460415, EPI_ISL_3460416 | CHU Purpan - Laboratoire de Virologie - Institut Fédératif de Biologie | CHU Purpan - Laboratoire de Virologie - Institut Fédératif de Biologie | Latour J., Jeanne N., Ranger N., Carcenac R., Harter A., Boyer P., Ferrer V., Tremeaux R., Izopet J. |
| EPI_ISL_3460417, EPI_ISL_3460419, EPI_ISL_3460421, EPI_ISL_3460425, EPI_ISL_3460426, EPI_ISL_3460428, EPI_ISL_3460433, EPI_ISL_3460434, EPI_ISL_3460436, EPI_ISL_3460437, EPI_ISL_3460439, EPI_ISL_3460440, EPI_ISL_3460441, EPI_ISL_3460444, EPI_ISL_3460445, EPI_ISL_3460446, EPI_ISL_3460447, EPI_ISL_3460448, EPI_ISL_3460450, EPI_ISL_3460451, EPI_ISL_3460454, EPI_ISL_3460458, EPI_ISL_3460459, EPI_ISL_3460460, EPI_ISL_3460462, EPI_ISL_3460465, EPI_ISL_3460466, EPI_ISL_3460467, EPI_ISL_3460471, EPI_ISL_3460473, EPI_ISL_3460474, EPI_ISL_3460475, EPI_ISL_3460476, EPI_ISL_3460478, EPI_ISL_3460479, EPI_ISL_3460480, EPI_ISL_3460481, EPI_ISL_3460482, EPI_ISL_3460483, EPI_ISL_3460484, EPI_ISL_3460485, EPI_ISL_3460487, EPI_ISL_3460489, EPI_ISL_3460491, EPI_ISL_3460492, EPI_ISL_3460495, EPI_ISL_3460501, EPI_ISL_3460503, EPI_ISL_3460504, EPI_ISL_3460505, EPI_ISL_3460507, EPI_ISL_3460508, EPI_ISL_3460509, EPI_ISL_3460519, EPI_ISL_3460520, EPI_ISL_3460524, EPI_ISL_3460526, EPI_ISL_3460529, EPI_ISL_3460530, EPI_ISL_3460533, EPI_ISL_3460537, EPI_ISL_3460539, EPI_ISL_3460542, EPI_ISL_3460543, EPI_ISL_3460544, EPI_ISL_3460546, EPI_ISL_3460548, EPI_ISL_3460549, EPI_ISL_3460553, EPI_ISL_3460557, EPI_ISL_3460558, EPI_ISL_3460559, EPI_ISL_3460561, EPI_ISL_3460562, EPI_ISL_3460563, EPI_ISL_3460564, EPI_ISL_3460566, EPI_ISL_3460569, EPI_ISL_3460571, EPI_ISL_3460572, EPI_ISL_3460573, EPI_ISL_3460574, EPI_ISL_3460576, EPI_ISL_3460578, EPI_ISL_3460581, EPI_ISL_3460583, EPI_ISL_3460584, EPI_ISL_3460585, EPI_ISL_3460587, EPI_ISL_3460589, EPI_ISL_3460590, EPI_ISL_3460591, EPI_ISL_3460595, EPI_ISL_3460596, EPI_ISL_3460597, EPI_ISL_3460598, EPI_ISL_3460601, EPI_ISL_3460602, EPI_ISL_3460603, EPI_ISL_3460604, EPI_ISL_3460605, EPI_ISL_3460607, EPI_ISL_3460608, EPI_ISL_3460609, EPI_ISL_3460610, EPI_ISL_3460613, EPI_ISL_3460617, EPI_ISL_3460618, EPI_ISL_3460620, EPI_ISL_3460621, EPI_ISL_3460622, EPI_ISL_3460624, EPI_ISL_3460626, EPI_ISL_3460630, EPI_ISL_3460632, EPI_ISL_3460633, EPI_ISL_3460634, EPI_ISL_3460635, EPI_ISL_3460638, EPI_ISL_3460640, EPI_ISL_3460642, EPI_ISL_3460644, EPI_ISL_3460647, EPI_ISL_3460648, EPI_ISL_3460651, EPI_ISL_3460653, EPI_ISL_3460654, EPI_ISL_3460655, EPI_ISL_3460657, EPI_ISL_3460658, EPI_ISL_3460661, EPI_ISL_3460662, EPI_ISL_3460666 | | | |
| see above | TGen North | TGen North | "Jolene Bowers, Heather Centner, Brett Van Tassel, Chris French, Hayley Yaglom, Darrin Lemmer, Dave Engelthaler" |
| EPI_ISL_3460667, EPI_ISL_3460668, EPI_ISL_3460669, EPI_ISL_3460670, EPI_ISL_3460671, EPI_ISL_3460672, EPI_ISL_3460673, EPI_ISL_3460674, EPI_ISL_3460675, EPI_ISL_3460676, EPI_ISL_3460677, EPI_ISL_3460678, EPI_ISL_3460679, EPI_ISL_3460688, EPI_ISL_3460689, EPI_ISL_3460716, EPI_ISL_3460717 | | | |
| see above | TN DOH Lab Services | TN DOH Lab Services | Karen Beasley-Maynard, Brian Selinsky |
| EPI_ISL_3460771, EPI_ISL_3460772 | TXDSHS | TXDSHS | Rashmi Tuladhar, Bonnie Oh, Jenny Zhang, Maliha Rahman, Mayela Pedrueza, Anita Pokharel, Karen Bobier, Lorraine Rodriguez, Myong Koag, Chun Wang, Rachel Lee, Grace Kubin |
| EPI_ISL_3460774, EPI_ISL_3460775, EPI_ISL_3460776, EPI_ISL_3460777, EPI_ISL_3460778, EPI_ISL_3460783, EPI_ISL_3460784, EPI_ISL_3460785, EPI_ISL_3460786, EPI_ISL_3460787, EPI_ISL_3460788, EPI_ISL_3460789 | | | |
| see above | Aegis Sciences Corporation | Centers for Disease Control and Prevention Division of Viral Diseases, Pathogen Discovery | Dakota Howard, Dhwani Batra, Peter W. Cook, Kara Moser, Adrian Paskey, Jason Caravas, Benjamin Rambo-Martin, Shatavia Morrison, Christopher Gulvick, Scott Sammons, Yvette Unoarumhi, Darlene Wagner, Matthew Schmerer, Cyndi Clark, Patrick Campbell, Rob Case, Vikramsinha Ghorpade, Holly Houdeshell, Ola Kvalvaag, Dillon Nall, Ethan Sanders, Alec Vest, Shaun Westlund, Matthew Hardison, Clinton R. Paden, Duncan MacCannell |
| EPI_ISL_3460793, EPI_ISL_3460796, EPI_ISL_3460797, EPI_ISL_3460798, EPI_ISL_3460802, EPI_ISL_3460803, EPI_ISL_3460805, EPI_ISL_3460806, EPI_ISL_3460813, EPI_ISL_3460815, EPI_ISL_3460818, EPI_ISL_3460823, EPI_ISL_3460824, EPI_ISL_3460828, EPI_ISL_3460829, EPI_ISL_3460831 | | | |
| see above | Quest Diagnostics Incorporated | Centers for Disease Control and Prevention Division of Viral Diseases, Pathogen Discovery | Dakota Howard, Dhwani Batra, Peter W. Cook, Kara Moser, Adrian Paskey, Jason Caravas, Benjamin Rambo-Martin, Shatavia Morrison, Christopher Gulvick, Scott Sammons, Yvette Unoarumhi, Darlene Wagner, Matthew Schmerer, S. H. Rosenthal, A. Gerasimova, R. M. Kagan, B. Anderson, M. Hua, Y. Liu, L.E. Bernstein, K.E. Livingston, A. Perez, I. A. Shlyakhter, R. V. Rolando, R. Owen, P. Tanpaiboon, F. Lacbawan, Clinton R. Paden, Duncan MacCannell |
| EPI_ISL_3460832 | Premier Laboratory Services | BioInfoExperts LLC | Cheryl Metzler, Valerie Murray, Rebecca Rose, David Nolan, Susanna Lamers |
| EPI_ISL_3460833 | Quest Diagnostics Incorporated | Centers for Disease Control and Prevention Division of Viral Diseases, Pathogen Discovery | Dakota Howard, Dhwani Batra, Peter W. Cook, Kara Moser, Adrian Paskey, Jason Caravas, Benjamin Rambo-Martin, Shatavia Morrison, Christopher Gulvick, Scott Sammons, Yvette Unoarumhi, Darlene Wagner, Matthew Schmerer, S. H. Rosenthal, A. Gerasimova, R. M. Kagan, B. Anderson, M. Hua, Y. Liu, L.E. Bernstein, K.E. Livingston, A. Perez, I. A. Shlyakhter, R. V. Rolando, R. Owen, P. Tanpaiboon, F. Lacbawan, Clinton R. Paden, Duncan MacCannell |
| EPI_ISL_3460834 | Premier Laboratory Services | BioInfoExperts LLC | Cheryl Metzler, Valerie Murray, Rebecca Rose, David Nolan, Susanna Lamers |
| EPI_ISL_3460837, EPI_ISL_3460841 | Quest Diagnostics Incorporated | Centers for Disease Control and Prevention Division of Viral Diseases, Pathogen Discovery | Dakota Howard, Dhwani Batra, Peter W. Cook, Kara Moser, Adrian Paskey, Jason Caravas, Benjamin Rambo-Martin, Shatavia Morrison, Christopher Gulvick, Scott Sammons, Yvette Unoarumhi, Darlene Wagner, Matthew Schmerer, S. H. Rosenthal, A. Gerasimova, R. M. Kagan, B. Anderson, M. Hua, Y. Liu, L.E. Bernstein, K.E. Livingston, A. Perez, I. A. Shlyakhter, R. V. Rolando, R. Owen, P. Tanpaiboon, F. Lacbawan, Clinton R. Paden, Duncan MacCannell |
| EPI_ISL_3460846, EPI_ISL_3460851, EPI_ISL_3460854, EPI_ISL_3460858, EPI_ISL_3460860, EPI_ISL_3460863, EPI_ISL_3460864, EPI_ISL_3460866, EPI_ISL_3460867, EPI_ISL_3460868, EPI_ISL_3460869, EPI_ISL_3460871, EPI_ISL_3460873, EPI_ISL_3460875, EPI_ISL_3460877, EPI_ISL_3460878, EPI_ISL_3460879, EPI_ISL_3460881, EPI_ISL_3460889, EPI_ISL_3460892, EPI_ISL_3460893, EPI_ISL_3460896, EPI_ISL_3460898 | | | |
| see above | Mako Medical | Centers for Disease Control and Prevention Division of Viral Diseases, Pathogen Discovery | Dakota Howard, Dhwani Batra, Peter W. Cook, Kara Moser, Adrian Paskey, Jason Caravas, Benjamin Rambo-Martin, Shatavia Morrison, Christopher Gulvick, Scott Sammons, Yvette Unoarumhi, Darlene Wagner, Matthew Schmerer, Matthew Tugwell, Lauren Moon, Clinton R. Paden, Duncan MacCannell |
| EPI_ISL_3460899, EPI_ISL_3460900, EPI_ISL_3460902, EPI_ISL_3460903, EPI_ISL_3460904 | Aegis Sciences Corporation | Centers for Disease Control and Prevention Division of Viral Diseases, Pathogen Discovery | Dakota Howard, Dhwani Batra, Peter W. Cook, Kara Moser, Adrian Paskey, Jason Caravas, Benjamin Rambo-Martin, Shatavia Morrison, Christopher Gulvick, Scott Sammons, Yvette Unoarumhi, Darlene Wagner, Matthew Schmerer, Cyndi Clark, Patrick Campbell, Rob Case, Vikramsinha Ghorpade, Holly Houdeshell, Ola Kvalvaag, Dillon Nall, Ethan Sanders, Alec Vest, Shaun Westlund, Matthew Hardison, Clinton R. Paden, Duncan MacCannell |
| EPI_ISL_3460909, EPI_ISL_3460911, EPI_ISL_3460912, EPI_ISL_3460913, EPI_ISL_3460914, EPI_ISL_3460915, EPI_ISL_3460916, EPI_ISL_3460919, EPI_ISL_3460920, EPI_ISL_3460921, EPI_ISL_3460924, EPI_ISL_3460927, EPI_ISL_3460928, EPI_ISL_3460930, EPI_ISL_3460933, EPI_ISL_3460934, EPI_ISL_3460935, EPI_ISL_3460936, EPI_ISL_3460937, EPI_ISL_3460938, EPI_ISL_3460939, EPI_ISL_3460942, EPI_ISL_3460943, EPI_ISL_3460946, EPI_ISL_3460948, EPI_ISL_3460949, EPI_ISL_3460951, EPI_ISL_3460953, EPI_ISL_3460958, EPI_ISL_3460959, EPI_ISL_3460960, EPI_ISL_3460961, EPI_ISL_3460964, EPI_ISL_3460965, EPI_ISL_3460966, EPI_ISL_3460969, EPI_ISL_3460972, EPI_ISL_3460973, EPI_ISL_3460974, EPI_ISL_3460977, EPI_ISL_3460978, EPI_ISL_3460979, EPI_ISL_3460980, EPI_ISL_3460981, EPI_ISL_3460982, EPI_ISL_3460984, EPI_ISL_3460987, EPI_ISL_3460988, EPI_ISL_3460989, EPI_ISL_3460992, EPI_ISL_3460994, EPI_ISL_3460995, EPI_ISL_3460996, EPI_ISL_3460997, EPI_ISL_3460998, EPI_ISL_3460999, EPI_ISL_3461001, EPI_ISL_3461002, EPI_ISL_3461003, EPI_ISL_3461004, EPI_ISL_3461005, EPI_ISL_3461006, EPI_ISL_3461007, EPI_ISL_3461008, EPI_ISL_3461010, EPI_ISL_3461011, EPI_ISL_3461012, EPI_ISL_3461013 | | | |
| see above | Fulgent Genetics | Centers for Disease Control and Prevention Division of Viral Diseases, Pathogen Discovery | Dakota Howard, Dhwani Batra, Peter W. Cook, Kara Moser, Adrian Paskey, Jason Caravas, Benjamin Rambo-Martin, Shatavia Morrison, Christopher Gulvick, Scott Sammons, Yvette Unoarumhi, Darlene Wagner, Matthew Schmerer, Harry Gao, Mickey Li, John Gao, Joseph Fierro, Benafsh Sapra, Becky Tsai, Yan Meng, Doreen Ng, James Xie, Clinton R. Paden, Duncan MacCannell |
| EPI_ISL_3461014, EPI_ISL_3461016, EPI_ISL_3461017 | Quest Diagnostics Incorporated | Centers for Disease Control and Prevention Division of Viral Diseases, Pathogen Discovery | Dakota Howard, Dhwani Batra, Peter W. Cook, Kara Moser, Adrian Paskey, Jason Caravas, Benjamin Rambo-Martin, Shatavia Morrison, Christopher Gulvick, Scott Sammons, Yvette Unoarumhi, Darlene Wagner, Matthew Schmerer, S. H. Rosenthal, A. Gerasimova, R. M. Kagan, B. Anderson, M. Hua, Y. Liu, L.E. Bernstein, K.E. Livingston, A. Perez, I. A. Shlyakhter, R. V. Rolando, R. Owen, P. Tanpaiboon, F. Lacbawan, Clinton R. Paden, Duncan MacCannell |
| EPI_ISL_3461019 | Fulgent Genetics | Centers for Disease Control and Prevention Division of Viral Diseases, Pathogen Discovery | Dakota Howard, Dhwani Batra, Peter W. Cook, Kara Moser, Adrian Paskey, Jason Caravas, Benjamin Rambo-Martin, Shatavia Morrison, Christopher Gulvick, Scott Sammons, Yvette Unoarumhi, Darlene Wagner, Matthew Schmerer, Harry Gao, Mickey Li, John Gao, Joseph Fierro, Benafsh Sapra, Becky Tsai, Yan Meng, Doreen Ng, James Xie, Clinton R. Paden, Duncan MacCannell |
| EPI_ISL_3461020, EPI_ISL_3461021 | Mako Medical | Centers for Disease Control and Prevention Division of Viral Diseases, Pathogen Discovery | Dakota Howard, Dhwani Batra, Peter W. Cook, Kara Moser, Adrian Paskey, Jason Caravas, Benjamin Rambo-Martin, Shatavia Morrison, Christopher Gulvick, Scott Sammons, Yvette Unoarumhi, Darlene Wagner, Matthew Schmerer, Matthew Tugwell, Lauren Moon, Clinton R. Paden, Duncan MacCannell |
| EPI_ISL_3461022, EPI_ISL_3461023, EPI_ISL_3461024, EPI_ISL_3461025, EPI_ISL_3461026, EPI_ISL_3461027, EPI_ISL_3461028, EPI_ISL_3461029, EPI_ISL_3461030, EPI_ISL_3461031, EPI_ISL_3461032, EPI_ISL_3461033, EPI_ISL_3461034, EPI_ISL_3461035, EPI_ISL_3461036, EPI_ISL_3461037, EPI_ISL_3461038, EPI_ISL_3461039, EPI_ISL_3461040, EPI_ISL_3461041, EPI_ISL_3461042, EPI_ISL_3461043, EPI_ISL_3461044, EPI_ISL_3461045, EPI_ISL_3461046, EPI_ISL_3461047, EPI_ISL_3461048, EPI_ISL_3461051 | | | |
| see above | IN State Department of Health Laboratory Services | IN State Department of Health Laboratory Services | Cassandra Campion, Jamie Yeadon, Brian Pope, Lixia Liu, Kyle Brownlee, Melissa Hindenlang, Mark Glazier |
| EPI_ISL_3461058 | Laboratório de Microbiologia Molecular - Universidade FEEVALE | Molecular Microbiology Laboratory | Alana Witt Hansen, Fágner Henrique Heldt, Fernando Rosado Spilki, Juliana Schons Gulart, Juliane Deise Fleck, Mariana Soares da Silva, Meriane Demoliner, Matheus Nunes Weber, Paula Rodrigues de Almeida, Michele Filippi. |
| EPI_ISL_3461068, EPI_ISL_3461069, EPI_ISL_3461070, EPI_ISL_3461071, EPI_ISL_3461072, EPI_ISL_3461073, EPI_ISL_3461074, EPI_ISL_3461075, EPI_ISL_3461076, EPI_ISL_3461077, EPI_ISL_3461078, EPI_ISL_3461079, EPI_ISL_3461080, EPI_ISL_3461081, EPI_ISL_3461082, EPI_ISL_3461083, EPI_ISL_3461084, EPI_ISL_3461085, EPI_ISL_3461086, EPI_ISL_3461087, EPI_ISL_3461088, EPI_ISL_3461089, EPI_ISL_3461090, EPI_ISL_3461091, EPI_ISL_3461092, EPI_ISL_3461093, EPI_ISL_3461094, EPI_ISL_3461095, EPI_ISL_3461096 | | | |
| see above | Nebraska Public Health Laboratory | NPHL COVID-19 Response Team | NPHL COVID-19 Response Team |
| EPI_ISL_3461101, EPI_ISL_3461104, EPI_ISL_3461106, EPI_ISL_3461107, EPI_ISL_3461109 | Laboratório de Microbiologia Molecular - Universidade FEEVALE | Molecular Microbiology Laboratory | Alana Witt Hansen, Fágner Henrique Heldt, Fernando Rosado Spilki, Juliana Schons Gulart, Juliane Deise Fleck, Mariana Soares da Silva, Meriane Demoliner, Matheus Nunes Weber, Paula Rodrigues de Almeida, Michele Filippi. |
| EPI_ISL_3461110, EPI_ISL_3461111, EPI_ISL_3461112, EPI_ISL_3461113, EPI_ISL_3461114, EPI_ISL_3461115, EPI_ISL_3461116, EPI_ISL_3461117, EPI_ISL_3461118, EPI_ISL_3461119, EPI_ISL_3461120, EPI_ISL_3461122, EPI_ISL_3461123, EPI_ISL_3461124 | | | |
| see above | UCSC Genomics Institute | UCSC Genomics Institute | Mark Akeson, Ikenna Anigbogu, Eric Beraut, Isabel Bjork, Molly Cassatt-Johnstone, Terren Chang, Russell Corbett-Detig, Namrita Dhillon, Maximilian Haeussler, David Haussler, Angie Hinrichs, Miten Jain, Joshua Kapp, A. Marm Kilpatrick, Jakob McBroome, Hugh Olsen, Jeremy Sanford, Beth Shapiro, Michael Stone, Bryan Thornlow, Yatish Turakhia, Ciara Wanket |
| EPI_ISL_3461151 | Pathology, UAB | Pathology, UAB | Moates,D., Hendrickson,R.C., Van Der Pol,W.J., Lefkowitz,E.J., Leal,S.M. |
| EPI_ISL_3461152, EPI_ISL_3461153, EPI_ISL_3461155, EPI_ISL_3461156, EPI_ISL_3461157, EPI_ISL_3461158, EPI_ISL_3461160, EPI_ISL_3461161, EPI_ISL_3461162, EPI_ISL_3461163, EPI_ISL_3461164, EPI_ISL_3461165 | | | |
| see above | Pathology, UAB | Pathology, UAB | Moates,D., Hendrickson,R.C., Lefkowitz,E.J., Leal,S.M. |
| EPI_ISL_3461166, EPI_ISL_3461167, EPI_ISL_3461168, EPI_ISL_3461169, EPI_ISL_3461170, EPI_ISL_3461171, EPI_ISL_3461172, EPI_ISL_3461173, EPI_ISL_3461174, EPI_ISL_3461175, EPI_ISL_3461176 | | | |

| | | | |
|---|---|---|---|
| see above | New South Wales Health Pathology Royal Prince Alfred Hospital | Microbiology RPAH | Foster, C.; Au, J.; Ruiz Silva, M.; Deveson, I.; Bull, R.; Van Hal, S.; Rawlinson, W. |
| EPI_ISL_3461202, EPI_ISL_3461203, EPI_ISL_3461214, EPI_ISL_3461220, EPI_ISL_3461226, EPI_ISL_3461227, EPI_ISL_3461228, EPI_ISL_3461229, EPI_ISL_3461234, EPI_ISL_3461242, EPI_ISL_3461245, EPI_ISL_3461246, EPI_ISL_3461248, EPI_ISL_3461251, EPI_ISL_3461253, EPI_ISL_3461254, EPI_ISL_3461255, EPI_ISL_3461259, EPI_ISL_3461261, EPI_ISL_3461262, EPI_ISL_3461270, EPI_ISL_3461273, EPI_ISL_3461274, EPI_ISL_3461275, EPI_ISL_3461276, EPI_ISL_3461277, EPI_ISL_3461279, EPI_ISL_3461280, EPI_ISL_3461284, EPI_ISL_3461286, EPI_ISL_3461290, EPI_ISL_3461293, EPI_ISL_3461296, EPI_ISL_3461297, EPI_ISL_3461300, EPI_ISL_3461301, EPI_ISL_3461303, EPI_ISL_3461310, EPI_ISL_3461312, EPI_ISL_3461314, EPI_ISL_3461315, EPI_ISL_3461316, EPI_ISL_3461317, EPI_ISL_3461319, EPI_ISL_3461320, EPI_ISL_3461321, EPI_ISL_3461322, EPI_ISL_3461329, EPI_ISL_3461330, EPI_ISL_3461334, EPI_ISL_3461341, EPI_ISL_3461342, EPI_ISL_3461345, EPI_ISL_3461347, EPI_ISL_3461349, EPI_ISL_3461350, EPI_ISL_3461351, EPI_ISL_3461353, EPI_ISL_3461354, EPI_ISL_3461355, EPI_ISL_3461356, EPI_ISL_3461357 | | | |
| see above | Laboratory, Gravity Diagnostics | Laboratory, Gravity Diagnostics | Diagnostics,G. |
| EPI_ISL_3461360 | A.O. S.Camillo | National Institute for Infectious Diseases (INMI) L. Spallanzani I.R.C.C.S | E Giombini, B Bartolini, M Rueca, G Bonfiglio, O Butera, CEM Gruber, F Santini, F Messina, A Di Caro, MR Capobianchi |
| EPI_ISL_3461361 | INMI | National Institute for Infectious Diseases (INMI) L. Spallanzani I.R.C.C.S | E Giombini, B Bartolini, M Rueca, G Bonfiglio, O Butera, CEM Gruber, F Santini, F Messina, A Di Caro, MR Capobianchi |
| EPI_ISL_3461362, EPI_ISL_3461363 | A.O. S.Camillo | National Institute for Infectious Diseases (INMI) L. Spallanzani I.R.C.C.S | E Giombini, B Bartolini, M Rueca, G Bonfiglio, O Butera, CEM Gruber, F Santini, F Messina, A Di Caro, MR Capobianchi |
| EPI_ISL_3461364, EPI_ISL_3461365 | Osp. S.Eugenio | National Institute for Infectious Diseases (INMI) L. Spallanzani I.R.C.C.S | E Giombini, B Bartolini, M Rueca, G Bonfiglio, O Butera, CEM Gruber, F Santini, F Messina, A Di Caro, MR Capobianchi |
| EPI_ISL_3461366 | INMI | National Institute for Infectious Diseases (INMI) L. Spallanzani I.R.C.C.S | E Giombini, B Bartolini, M Rueca, G Bonfiglio, O Butera, CEM Gruber, F Santini, F Messina, A Di Caro, MR Capobianchi |
| EPI_ISL_3461368, EPI_ISL_3461369, EPI_ISL_3461370 | A.O. S.Camillo | National Institute for Infectious Diseases (INMI) L. Spallanzani I.R.C.C.S | E Giombini, B Bartolini, M Rueca, G Bonfiglio, O Butera, CEM Gruber, F Santini, F Messina, A Di Caro, MR Capobianchi |
| EPI_ISL_3461371 | INMI | National Institute for Infectious Diseases (INMI) L. Spallanzani I.R.C.C.S | E Giombini, B Bartolini, M Rueca, G Bonfiglio, O Butera, CEM Gruber, F Santini, F Messina, A Di Caro, MR Capobianchi |
| EPI_ISL_3461372 | Osp. G.Grassi | National Institute for Infectious Diseases (INMI) L. Spallanzani I.R.C.C.S | E Giombini, B Bartolini, M Rueca, G Bonfiglio, O Butera, CEM Gruber, F Santini, F Messina, A Di Caro, MR Capobianchi |
| EPI_ISL_3461373, EPI_ISL_3461374, EPI_ISL_3461375 | A.O. S.Camillo | National Institute for Infectious Diseases (INMI) L. Spallanzani I.R.C.C.S | E Giombini, B Bartolini, M Rueca, G Bonfiglio, O Butera, CEM Gruber, F Santini, F Messina, A Di Caro, MR Capobianchi |
| EPI_ISL_3461376 | INMI | National Institute for Infectious Diseases (INMI) L. Spallanzani I.R.C.C.S | E Giombini, B Bartolini, M Rueca, G Bonfiglio, O Butera, CEM Gruber, F Santini, F Messina, A Di Caro, MR Capobianchi |
| EPI_ISL_3461377 | Osp. S.Eugenio | National Institute for Infectious Diseases (INMI) L. Spallanzani I.R.C.C.S | E Giombini, B Bartolini, M Rueca, G Bonfiglio, O Butera, CEM Gruber, F Santini, F Messina, A Di Caro, MR Capobianchi |
| EPI_ISL_3461378 | LifeBrain | National Institute for Infectious Diseases (INMI) L. Spallanzani I.R.C.C.S | E Giombini, B Bartolini, M Rueca, G Bonfiglio, O Butera, CEM Gruber, F Santini, F Messina, A Di Caro, MR Capobianchi |
| EPI_ISL_3461379 | A.O. S.Camillo | National Institute for Infectious Diseases (INMI) L. Spallanzani I.R.C.C.S | E Giombini, B Bartolini, M Rueca, G Bonfiglio, O Butera, CEM Gruber, F Santini, F Messina, A Di Caro, MR Capobianchi |
| EPI_ISL_3461380, EPI_ISL_3461381, EPI_ISL_3461382, EPI_ISL_3461383 | LifeBrain | National Institute for Infectious Diseases (INMI) L. Spallanzani I.R.C.C.S | E Giombini, B Bartolini, M Rueca, G Bonfiglio, O Butera, CEM Gruber, F Santini, F Messina, A Di Caro, MR Capobianchi |
| EPI_ISL_3461384, EPI_ISL_3461385 | INMI | National Institute for Infectious Diseases (INMI) L. Spallanzani I.R.C.C.S | E Giombini, B Bartolini, M Rueca, G Bonfiglio, O Butera, CEM Gruber, F Santini, F Messina, A Di Caro, MR Capobianchi |
| EPI_ISL_3461386 | LifeBrain | National Institute for Infectious Diseases (INMI) L. Spallanzani I.R.C.C.S | E Giombini, B Bartolini, M Rueca, G Bonfiglio, O Butera, CEM Gruber, F Santini, F Messina, A Di Caro, MR Capobianchi |
| EPI_ISL_3461387 | A.U. Tor Vergata | National Institute for Infectious Diseases (INMI) L. Spallanzani I.R.C.C.S | E Giombini, B Bartolini, M Rueca, G Bonfiglio, O Butera, CEM Gruber, F Santini, F Messina, A Di Caro, MR Capobianchi |
| EPI_ISL_3461388 | Osp. S.Eugenio | National Institute for Infectious Diseases (INMI) L. Spallanzani I.R.C.C.S | E Giombini, B Bartolini, M Rueca, G Bonfiglio, O Butera, CEM Gruber, F Santini, F Messina, A Di Caro, MR Capobianchi |
| EPI_ISL_3461389, EPI_ISL_3461390 | INMI | National Institute for Infectious Diseases (INMI) L. Spallanzani I.R.C.C.S | E Giombini, B Bartolini, M Rueca, G Bonfiglio, O Butera, CEM Gruber, F Santini, F Messina, A Di Caro, MR Capobianchi |
| EPI_ISL_3461391, EPI_ISL_3461392 | Osp. S.Eugenio | National Institute for Infectious Diseases (INMI) L. Spallanzani I.R.C.C.S | E Giombini, B Bartolini, M Rueca, G Bonfiglio, O Butera, CEM Gruber, F Santini, F Messina, A Di Caro, MR Capobianchi |
| EPI_ISL_3461393 | INMI | National Institute for Infectious Diseases (INMI) L. Spallanzani I.R.C.C.S | E Giombini, B Bartolini, M Rueca, G Bonfiglio, O Butera, CEM Gruber, F Santini, F Messina, A Di Caro, MR Capobianchi |
| EPI_ISL_3461394 | Osp. S.Camillo De Lellis | National Institute for Infectious Diseases (INMI) L. Spallanzani I.R.C.C.S | E Giombini, B Bartolini, M Rueca, G Bonfiglio, O Butera, CEM Gruber, F Santini, F Messina, A Di Caro, MR Capobianchi |
| EPI_ISL_3461396, EPI_ISL_3461397, EPI_ISL_3461398 | INMI | National Institute for Infectious Diseases (INMI) L. Spallanzani I.R.C.C.S | E Giombini, B Bartolini, M Rueca, G Bonfiglio, O Butera, CEM Gruber, F Santini, F Messina, A Di Caro, MR Capobianchi |
| EPI_ISL_3461399 | Osp. S.Eugenio | National Institute for Infectious Diseases (INMI) L. Spallanzani I.R.C.C.S | E Giombini, B Bartolini, M Rueca, G Bonfiglio, O Butera, CEM Gruber, F Santini, F Messina, A Di Caro, MR Capobianchi |
| EPI_ISL_3461400, EPI_ISL_3461401 | INMI | National Institute for Infectious Diseases (INMI) L. Spallanzani I.R.C.C.S | E Giombini, B Bartolini, M Rueca, G Bonfiglio, O Butera, CEM Gruber, F Santini, F Messina, A Di Caro, MR Capobianchi |
| EPI_ISL_3461402 | Osp. S.Camillo De Lellis | National Institute for Infectious Diseases (INMI) L. Spallanzani I.R.C.C.S | E Giombini, B Bartolini, M Rueca, G Bonfiglio, O Butera, CEM Gruber, F Santini, F Messina, A Di Caro, MR Capobianchi |
| EPI_ISL_3461403, EPI_ISL_3461404, EPI_ISL_3461405 | INMI | National Institute for Infectious Diseases (INMI) L. Spallanzani I.R.C.C.S | E Giombini, B Bartolini, M Rueca, G Bonfiglio, O Butera, CEM Gruber, F Santini, F Messina, A Di Caro, MR Capobianchi |
| EPI_ISL_3461406 | A.U. Tor Vergata | National Institute for Infectious Diseases (INMI) L. Spallanzani I.R.C.C.S | E Giombini, B Bartolini, M Rueca, G Bonfiglio, O Butera, CEM Gruber, F Santini, F Messina, A Di Caro, MR Capobianchi |
| EPI_ISL_3461407 | Osp. S.Filippo Neri | National Institute for Infectious Diseases (INMI) L. Spallanzani I.R.C.C.S | E Giombini, B Bartolini, M Rueca, G Bonfiglio, O Butera, CEM Gruber, F Santini, F Messina, A Di Caro, MR Capobianchi |
| EPI_ISL_3461408 | Osp. S.Pertini | National Institute for Infectious Diseases (INMI) L. Spallanzani I.R.C.C.S | E Giombini, B Bartolini, M Rueca, G Bonfiglio, O Butera, CEM Gruber, F Santini, F Messina, A Di Caro, MR Capobianchi |
| EPI_ISL_3461410, EPI_ISL_3461411, EPI_ISL_3461412 | INMI | National Institute for Infectious Diseases (INMI) L. Spallanzani I.R.C.C.S | E Giombini, B Bartolini, M Rueca, G Bonfiglio, O Butera, CEM Gruber, F Santini, F Messina, A Di Caro, MR Capobianchi |
| EPI_ISL_3461413 | Osp. S.Camillo De Lellis | National Institute for Infectious Diseases (INMI) L. Spallanzani I.R.C.C.S | E Giombini, B Bartolini, M Rueca, G Bonfiglio, O Butera, CEM Gruber, F Santini, F Messina, A Di Caro, MR Capobianchi |
| EPI_ISL_3461414 | INMI | National Institute for Infectious Diseases (INMI) L. Spallanzani I.R.C.C.S | E Giombini, B Bartolini, M Rueca, G Bonfiglio, O Butera, CEM Gruber, F Santini, F Messina, A Di Caro, MR Capobianchi |
| EPI_ISL_3461415 | A.U. Tor Vergata | National Institute for Infectious Diseases (INMI) L. Spallanzani I.R.C.C.S | E Giombini, B Bartolini, M Rueca, G Bonfiglio, O Butera, CEM Gruber, F Santini, F Messina, A Di Caro, MR Capobianchi |
| EPI_ISL_3461416 | S. SPIRITO | National Institute for Infectious Diseases (INMI) L. Spallanzani I.R.C.C.S | E Giombini, B Bartolini, M Rueca, G Bonfiglio, O Butera, CEM Gruber, F Santini, F Messina, A Di Caro, MR Capobianchi |
| EPI_ISL_3461417, EPI_ISL_3461418 | INMI | National Institute for Infectious Diseases (INMI) L. Spallanzani I.R.C.C.S | E Giombini, B Bartolini, M Rueca, G Bonfiglio, O Butera, CEM Gruber, F Santini, F Messina, A Di Caro, MR Capobianchi |
| EPI_ISL_3461419, EPI_ISL_3461420, EPI_ISL_3461421, EPI_ISL_3461422 | A.U. Tor Vergata | National Institute for Infectious Diseases (INMI) L. Spallanzani I.R.C.C.S | E Giombini, B Bartolini, M Rueca, G Bonfiglio, O Butera, CEM Gruber, F Santini, F Messina, A Di Caro, MR Capobianchi |
| EPI_ISL_3461423 | INMI | National Institute for Infectious Diseases (INMI) L. Spallanzani I.R.C.C.S | E Giombini, B Bartolini, M Rueca, G Bonfiglio, O Butera, CEM Gruber, F Santini, F Messina, A Di Caro, MR Capobianchi |
| EPI_ISL_3461424, | A.U. Tor Vergata | National Institute for Infectious Diseases (INMI) L. Spallanzani | E Giombini, B Bartolini, M Rueca, G Bonfiglio, O Butera, CEM Gruber, F Santini, F Messina, A Di Caro, MR Capobianchi |

| | | | |
|---|---|---|---|
| EPI_ISL_3461425 | | I.R.C.C.S | |
| EPI_ISL_3461426 | Osp. S.Camillo De Lellis | National Institute for Infectious Diseases (INMI) L. Spallanzani I.R.C.C.S | E Giombini, B Bartolini, M Rueca, G Bonfiglio, O Butera, CEM Gruber, F Santini, F Messina, A Di Caro, MR Capobianchi |
| EPI_ISL_3461427 | A.U. Tor Vergata | National Institute for Infectious Diseases (INMI) L. Spallanzani I.R.C.C.S | E Giombini, B Bartolini, M Rueca, G Bonfiglio, O Butera, CEM Gruber, F Santini, F Messina, A Di Caro, MR Capobianchi |
| EPI_ISL_3461428, EPI_ISL_3461429, EPI_ISL_3461430 | Osp. S.Camillo De Lellis | National Institute for Infectious Diseases (INMI) L. Spallanzani I.R.C.C.S | E Giombini, B Bartolini, M Rueca, G Bonfiglio, O Butera, CEM Gruber, F Santini, F Messina, A Di Caro, MR Capobianchi |
| EPI_ISL_3461432 | A.U. Tor Vergata | National Institute for Infectious Diseases (INMI) L. Spallanzani I.R.C.C.S | E Giombini, B Bartolini, M Rueca, G Bonfiglio, O Butera, CEM Gruber, F Santini, F Messina, A Di Caro, MR Capobianchi |
| EPI_ISL_3461433 | Osp. S.Maria Goretti | National Institute for Infectious Diseases (INMI) L. Spallanzani I.R.C.C.S | E Giombini, B Bartolini, M Rueca, G Bonfiglio, O Butera, CEM Gruber, F Santini, F Messina, A Di Caro, MR Capobianchi |
| EPI_ISL_3461434 | Osp. Di Belcolle | National Institute for Infectious Diseases (INMI) L. Spallanzani I.R.C.C.S | E Giombini, B Bartolini, M Rueca, G Bonfiglio, O Butera, CEM Gruber, F Santini, F Messina, A Di Caro, MR Capobianchi |
| EPI_ISL_3461435 | A.U. Tor Vergata | National Institute for Infectious Diseases (INMI) L. Spallanzani I.R.C.C.S | E Giombini, B Bartolini, M Rueca, G Bonfiglio, O Butera, CEM Gruber, F Santini, F Messina, A Di Caro, MR Capobianchi |
| EPI_ISL_3461436 | Osp. S.Maria Goretti | National Institute for Infectious Diseases (INMI) L. Spallanzani I.R.C.C.S | E Giombini, B Bartolini, M Rueca, G Bonfiglio, O Butera, CEM Gruber, F Santini, F Messina, A Di Caro, MR Capobianchi |
| EPI_ISL_3461437 | A.U. Tor Vergata | National Institute for Infectious Diseases (INMI) L. Spallanzani I.R.C.C.S | E Giombini, B Bartolini, M Rueca, G Bonfiglio, O Butera, CEM Gruber, F Santini, F Messina, A Di Caro, MR Capobianchi |
| EPI_ISL_3461439, EPI_ISL_3461440, EPI_ISL_3461441 | Osp. S.Giovanni Evangelista | National Institute for Infectious Diseases (INMI) L. Spallanzani I.R.C.C.S | E Giombini, B Bartolini, M Rueca, G Bonfiglio, O Butera, CEM Gruber, F Santini, F Messina, A Di Caro, MR Capobianchi |
| EPI_ISL_3461442 | S. SPIRITO | National Institute for Infectious Diseases (INMI) L. Spallanzani I.R.C.C.S | E Giombini, B Bartolini, M Rueca, G Bonfiglio, O Butera, CEM Gruber, F Santini, F Messina, A Di Caro, MR Capobianchi |
| EPI_ISL_3461444, EPI_ISL_3461445, EPI_ISL_3461446, EPI_ISL_3461447, EPI_ISL_3461448, EPI_ISL_3461450, EPI_ISL_3461451, EPI_ISL_3461452, EPI_ISL_3461453, EPI_ISL_3461454, EPI_ISL_3461455, EPI_ISL_3461456, EPI_ISL_3461457, EPI_ISL_3461458, EPI_ISL_3461459, EPI_ISL_3461460, EPI_ISL_3461461, EPI_ISL_3461462, EPI_ISL_3461463, EPI_ISL_3461464, EPI_ISL_3461465, EPI_ISL_3461466, EPI_ISL_3461467, EPI_ISL_3461468, EPI_ISL_3461469, EPI_ISL_3461470, EPI_ISL_3461471, EPI_ISL_3461472, EPI_ISL_3461473, EPI_ISL_3461474, EPI_ISL_3461475, EPI_ISL_3461476, EPI_ISL_3461477, EPI_ISL_3461478, EPI_ISL_3461479, EPI_ISL_3461480, EPI_ISL_3461481, EPI_ISL_3461482, EPI_ISL_3461483, EPI_ISL_3461484, EPI_ISL_3461485, EPI_ISL_3461486, EPI_ISL_3461487, EPI_ISL_3461488, EPI_ISL_3461489, EPI_ISL_3461490, EPI_ISL_3461491, EPI_ISL_3461492, EPI_ISL_3461493, EPI_ISL_3461494, EPI_ISL_3461495, EPI_ISL_3461496, EPI_ISL_3461497, EPI_ISL_3461498, EPI_ISL_3461499, EPI_ISL_3461500, EPI_ISL_3461502, EPI_ISL_3461503, EPI_ISL_3461504, EPI_ISL_3461505, EPI_ISL_3461506, EPI_ISL_3461507, EPI_ISL_3461508, EPI_ISL_3461509, EPI_ISL_3461510, EPI_ISL_3461511, EPI_ISL_3461512, EPI_ISL_3461513, EPI_ISL_3461514, EPI_ISL_3461515, EPI_ISL_3461516, EPI_ISL_3461517, EPI_ISL_3461518 | | | |
| see above | Karolinska University Hospital Solna | Karolinska University Hospital | Jan Albert ,Tobias Allander ,Annelie Bjerkner ,Sandra Broddesson ,Robert Dyrdak ,Martin Ekman ,Lynda Eneh ,Lina Guerra Blomqvist ,Karolina Ininbergs ,Tanja Normark ,Isak Sylvin ,Zhibing Yun ,Martina Wahlund ,Valtteri Wirta |
| EPI_ISL_3461520, EPI_ISL_3461521, EPI_ISL_3461522, EPI_ISL_3461523, EPI_ISL_3461524, EPI_ISL_3461525 | Karolinska University Hospital Huddinge | Karolinska University Hospital | Jan Albert ,Tobias Allander ,Annelie Bjerkner ,Sandra Broddesson ,Robert Dyrdak ,Martin Ekman ,Lynda Eneh ,Lina Guerra Blomqvist ,Karolina Ininbergs ,Tanja Normark ,Isak Sylvin ,Zhibing Yun ,Martina Wahlund ,Valtteri Wirta |
| EPI_ISL_3461526 | Karolinska University Hospital Solna | Karolinska University Hospital | Jan Albert ,Tobias Allander ,Annelie Bjerkner ,Sandra Broddesson ,Robert Dyrdak ,Martin Ekman ,Lynda Eneh ,Lina Guerra Blomqvist ,Karolina Ininbergs ,Tanja Normark ,Isak Sylvin ,Zhibing Yun ,Martina Wahlund ,Valtteri Wirta |
| EPI_ISL_3461527 | Karolinska University Hospital Huddinge | Karolinska University Hospital | Jan Albert ,Tobias Allander ,Annelie Bjerkner ,Sandra Broddesson ,Robert Dyrdak ,Martin Ekman ,Lynda Eneh ,Lina Guerra Blomqvist ,Karolina Ininbergs ,Tanja Normark ,Isak Sylvin ,Zhibing Yun ,Martina Wahlund ,Valtteri Wirta |
| EPI_ISL_3461528, EPI_ISL_3461529 | Karolinska University Hospital Solna | Karolinska University Hospital | Jan Albert ,Tobias Allander ,Annelie Bjerkner ,Sandra Broddesson ,Robert Dyrdak ,Martin Ekman ,Lynda Eneh ,Lina Guerra Blomqvist ,Karolina Ininbergs ,Tanja Normark ,Isak Sylvin ,Zhibing Yun ,Martina Wahlund ,Valtteri Wirta |
| EPI_ISL_3461530, EPI_ISL_3461531, EPI_ISL_3461532 | Karolinska University Hospital Huddinge | Karolinska University Hospital | Jan Albert ,Tobias Allander ,Annelie Bjerkner ,Sandra Broddesson ,Robert Dyrdak ,Martin Ekman ,Lynda Eneh ,Lina Guerra Blomqvist ,Karolina Ininbergs ,Tanja Normark ,Isak Sylvin ,Zhibing Yun ,Martina Wahlund ,Valtteri Wirta |
| EPI_ISL_3461533 | Karolinska University Hospital Solna | Karolinska University Hospital | Jan Albert ,Tobias Allander ,Annelie Bjerkner ,Sandra Broddesson ,Robert Dyrdak ,Martin Ekman ,Lynda Eneh ,Lina Guerra Blomqvist ,Karolina Ininbergs ,Tanja Normark ,Isak Sylvin ,Zhibing Yun ,Martina Wahlund ,Valtteri Wirta |
| EPI_ISL_3461534, EPI_ISL_3461535, EPI_ISL_3461536 | Karolinska University Hospital Huddinge | Karolinska University Hospital | Jan Albert ,Tobias Allander ,Annelie Bjerkner ,Sandra Broddesson ,Robert Dyrdak ,Martin Ekman ,Lynda Eneh ,Lina Guerra Blomqvist ,Karolina Ininbergs ,Tanja Normark ,Isak Sylvin ,Zhibing Yun ,Martina Wahlund ,Valtteri Wirta |
| EPI_ISL_3461537, EPI_ISL_3461538, EPI_ISL_3461539, EPI_ISL_3461540 | Karolinska University Hospital Solna | Karolinska University Hospital | Jan Albert ,Tobias Allander ,Annelie Bjerkner ,Sandra Broddesson ,Robert Dyrdak ,Martin Ekman ,Lynda Eneh ,Lina Guerra Blomqvist ,Karolina Ininbergs ,Tanja Normark ,Isak Sylvin ,Zhibing Yun ,Martina Wahlund ,Valtteri Wirta |
| EPI_ISL_3461541, EPI_ISL_3461542, EPI_ISL_3461543 | Karolinska University Hospital Huddinge | Karolinska University Hospital | Jan Albert ,Tobias Allander ,Annelie Bjerkner ,Sandra Broddesson ,Robert Dyrdak ,Martin Ekman ,Lynda Eneh ,Lina Guerra Blomqvist ,Karolina Ininbergs ,Tanja Normark ,Isak Sylvin ,Zhibing Yun ,Martina Wahlund ,Valtteri Wirta |
| EPI_ISL_3461544, EPI_ISL_3461545 | Karolinska University Hospital Solna | Karolinska University Hospital | Jan Albert ,Tobias Allander ,Annelie Bjerkner ,Sandra Broddesson ,Robert Dyrdak ,Martin Ekman ,Lynda Eneh ,Lina Guerra Blomqvist ,Karolina Ininbergs ,Tanja Normark ,Isak Sylvin ,Zhibing Yun ,Martina Wahlund ,Valtteri Wirta |
| EPI_ISL_3461546, EPI_ISL_3461547 | Karolinska University Hospital Huddinge | Karolinska University Hospital | Jan Albert ,Tobias Allander ,Annelie Bjerkner ,Sandra Broddesson ,Robert Dyrdak ,Martin Ekman ,Lynda Eneh ,Lina Guerra Blomqvist ,Karolina Ininbergs ,Tanja Normark ,Isak Sylvin ,Zhibing Yun ,Martina Wahlund ,Valtteri Wirta |
| EPI_ISL_3461548 | Karolinska University Hospital Solna | Karolinska University Hospital | Jan Albert ,Tobias Allander ,Annelie Bjerkner ,Sandra Broddesson ,Robert Dyrdak ,Martin Ekman ,Lynda Eneh ,Lina Guerra Blomqvist ,Karolina Ininbergs ,Tanja Normark ,Isak Sylvin ,Zhibing Yun ,Martina Wahlund ,Valtteri Wirta |
| EPI_ISL_3461549 | Karolinska University Hospital Huddinge | Karolinska University Hospital | Jan Albert ,Tobias Allander ,Annelie Bjerkner ,Sandra Broddesson ,Robert Dyrdak ,Martin Ekman ,Lynda Eneh ,Lina Guerra Blomqvist ,Karolina Ininbergs ,Tanja Normark ,Isak Sylvin ,Zhibing Yun ,Martina Wahlund ,Valtteri Wirta |
| EPI_ISL_3461550, EPI_ISL_3461551, EPI_ISL_3461552 | Karolinska University Hospital Solna | Karolinska University Hospital | Jan Albert ,Tobias Allander ,Annelie Bjerkner ,Sandra Broddesson ,Robert Dyrdak ,Martin Ekman ,Lynda Eneh ,Lina Guerra Blomqvist ,Karolina Ininbergs ,Tanja Normark ,Isak Sylvin ,Zhibing Yun ,Martina Wahlund ,Valtteri Wirta |
| EPI_ISL_3461553, EPI_ISL_3461554, EPI_ISL_3461555 | Karolinska University Hospital Huddinge | Karolinska University Hospital | Jan Albert ,Tobias Allander ,Annelie Bjerkner ,Sandra Broddesson ,Robert Dyrdak ,Martin Ekman ,Lynda Eneh ,Lina Guerra Blomqvist ,Karolina Ininbergs ,Tanja Normark ,Isak Sylvin ,Zhibing Yun ,Martina Wahlund ,Valtteri Wirta |
| EPI_ISL_3461556 | Karolinska University Hospital Solna | Karolinska University Hospital | Jan Albert ,Tobias Allander ,Annelie Bjerkner ,Sandra Broddesson ,Robert Dyrdak ,Martin Ekman ,Lynda Eneh ,Lina Guerra Blomqvist ,Karolina Ininbergs ,Tanja Normark ,Isak Sylvin ,Zhibing Yun ,Martina Wahlund ,Valtteri Wirta |
| EPI_ISL_3461557, EPI_ISL_3461558, EPI_ISL_3461559, EPI_ISL_3461560, EPI_ISL_3461561, EPI_ISL_3461562 | Karolinska University Hospital Huddinge | Karolinska University Hospital | Jan Albert ,Tobias Allander ,Annelie Bjerkner ,Sandra Broddesson ,Robert Dyrdak ,Martin Ekman ,Lynda Eneh ,Lina Guerra Blomqvist ,Karolina Ininbergs ,Tanja Normark ,Isak Sylvin ,Zhibing Yun ,Martina Wahlund ,Valtteri Wirta |
| EPI_ISL_3461563, EPI_ISL_3461564 | Karolinska University Hospital Solna | Karolinska University Hospital | Jan Albert ,Tobias Allander ,Annelie Bjerkner ,Sandra Broddesson ,Robert Dyrdak ,Martin Ekman ,Lynda Eneh ,Lina Guerra Blomqvist ,Karolina Ininbergs ,Tanja Normark ,Isak Sylvin ,Zhibing Yun ,Martina Wahlund ,Valtteri Wirta |
| EPI_ISL_3461565 | Karolinska University Hospital Huddinge | Karolinska University Hospital | Jan Albert ,Tobias Allander ,Annelie Bjerkner ,Sandra Broddesson ,Robert Dyrdak ,Martin Ekman ,Lynda Eneh ,Lina Guerra Blomqvist ,Karolina Ininbergs ,Tanja Normark ,Isak Sylvin ,Zhibing Yun ,Martina Wahlund ,Valtteri Wirta |
| EPI_ISL_3461566 | Karolinska University Hospital Solna | Karolinska University Hospital | Jan Albert ,Tobias Allander ,Annelie Bjerkner ,Sandra Broddesson ,Robert Dyrdak ,Martin Ekman ,Lynda Eneh ,Lina Guerra Blomqvist ,Karolina Ininbergs ,Tanja Normark ,Isak Sylvin ,Zhibing Yun ,Martina Wahlund ,Valtteri Wirta |
| EPI_ISL_3461567, EPI_ISL_3461568, EPI_ISL_3461569, EPI_ISL_3461570, EPI_ISL_3461571 | Karolinska University Hospital Huddinge | Karolinska University Hospital | Jan Albert ,Tobias Allander ,Annelie Bjerkner ,Sandra Broddesson ,Robert Dyrdak ,Martin Ekman ,Lynda Eneh ,Lina Guerra Blomqvist ,Karolina Ininbergs ,Tanja Normark ,Isak Sylvin ,Zhibing Yun ,Martina Wahlund ,Valtteri Wirta |
| EPI_ISL_3461572, EPI_ISL_3461573 | Karolinska University Hospital Solna | Karolinska University Hospital | Jan Albert ,Tobias Allander ,Annelie Bjerkner ,Sandra Broddesson ,Robert Dyrdak ,Martin Ekman ,Lynda Eneh ,Lina Guerra Blomqvist ,Karolina Ininbergs ,Tanja Normark ,Isak Sylvin ,Zhibing Yun ,Martina Wahlund ,Valtteri Wirta |
| EPI_ISL_3461574 | Karolinska University Hospital Huddinge | Karolinska University Hospital | Jan Albert ,Tobias Allander ,Annelie Bjerkner ,Sandra Broddesson ,Robert Dyrdak ,Martin Ekman ,Lynda Eneh ,Lina Guerra Blomqvist ,Karolina Ininbergs ,Tanja Normark ,Isak Sylvin ,Zhibing Yun ,Martina Wahlund ,Valtteri Wirta |
| EPI_ISL_3461575, EPI_ISL_3461576, EPI_ISL_3461577 | Karolinska University Hospital Solna | Karolinska University Hospital | Jan Albert ,Tobias Allander ,Annelie Bjerkner ,Sandra Broddesson ,Robert Dyrdak ,Martin Ekman ,Lynda Eneh ,Lina Guerra Blomqvist ,Karolina Ininbergs ,Tanja Normark ,Isak Sylvin ,Zhibing Yun ,Martina Wahlund ,Valtteri Wirta |
| EPI_ISL_3461578, EPI_ISL_3461579 | Karolinska University Hospital Huddinge | Karolinska University Hospital | Jan Albert ,Tobias Allander ,Annelie Bjerkner ,Sandra Broddesson ,Robert Dyrdak ,Martin Ekman ,Lynda Eneh ,Lina Guerra Blomqvist ,Karolina Ininbergs ,Tanja Normark ,Isak Sylvin ,Zhibing Yun ,Martina Wahlund ,Valtteri Wirta |
| EPI_ISL_3461580 | Karolinska University Hospital Solna | Karolinska University Hospital | Jan Albert ,Tobias Allander ,Annelie Bjerkner ,Sandra Broddesson ,Robert Dyrdak ,Martin Ekman ,Lynda Eneh ,Lina Guerra Blomqvist ,Karolina Ininbergs ,Tanja Normark ,Isak Sylvin ,Zhibing Yun ,Martina Wahlund ,Valtteri Wirta |
| EPI_ISL_3461581, EPI_ISL_3461582, EPI_ISL_3461583, EPI_ISL_3461584, EPI_ISL_3461585, EPI_ISL_3461586, EPI_ISL_3461587, EPI_ISL_3461588, EPI_ISL_3461589, EPI_ISL_3461590, EPI_ISL_3461591 | | | |

| | | | |
|---|---|---|---|
| see above | Karolinska University Hospital Huddinge | Karolinska University Hospital | Jan Albert ,Tobias Allander ,Annelie Bjerkner ,Sandra Broddesson ,Robert Dyrdak ,Martin Ekman ,Lynda Eneh ,Lina Guerra Blomqvist ,Karolina Ininbergs ,Tanja Normark ,Isak Sylvin ,Zhibing Yun ,Martina Wahlund ,Valtteri Wirta |
| EPI_ISL_3461592, EPI_ISL_3461593, EPI_ISL_3461594, EPI_ISL_3461596, EPI_ISL_3461597, EPI_ISL_3461598, EPI_ISL_3461600, EPI_ISL_3461601, EPI_ISL_3461602, EPI_ISL_3461603, EPI_ISL_3461606, EPI_ISL_3461607, EPI_ISL_3461608, EPI_ISL_3461609, EPI_ISL_3461610, EPI_ISL_3461611, EPI_ISL_3461612, EPI_ISL_3461614, EPI_ISL_3461615, EPI_ISL_3461616, EPI_ISL_3461618, EPI_ISL_3461619, EPI_ISL_3461620, EPI_ISL_3461621, EPI_ISL_3461622, EPI_ISL_3461623, EPI_ISL_3461624, EPI_ISL_3461625, EPI_ISL_3461626, EPI_ISL_3461627, EPI_ISL_3461628, EPI_ISL_3461629, EPI_ISL_3461630, EPI_ISL_3461631, EPI_ISL_3461632, EPI_ISL_3461633, EPI_ISL_3461634, EPI_ISL_3461635, EPI_ISL_3461636, EPI_ISL_3461637, EPI_ISL_3461640, EPI_ISL_3461641, EPI_ISL_3461642, EPI_ISL_3461643, EPI_ISL_3461644, EPI_ISL_3461645, EPI_ISL_3461646, EPI_ISL_3461647, EPI_ISL_3461648, EPI_ISL_3461649, EPI_ISL_3461650, EPI_ISL_3461651, EPI_ISL_3461652, EPI_ISL_3461653, EPI_ISL_3461654 | | | |
| see above | Karolinska University Hospital Solna | Karolinska University Hospital | Jan Albert ,Tobias Allander ,Annelie Bjerkner ,Sandra Broddesson ,Robert Dyrdak ,Martin Ekman ,Lynda Eneh ,Lina Guerra Blomqvist ,Karolina Ininbergs ,Tanja Normark ,Isak Sylvin ,Zhibing Yun ,Martina Wahlund ,Valtteri Wirta |
| EPI_ISL_3461655 | Synlab Medilab | Karolinska University Hospital | Jan Albert ,Tobias Allander ,Annelie Bjerkner ,Sandra Broddesson ,Robert Dyrdak ,Martin Ekman ,Lynda Eneh ,Lina Guerra Blomqvist ,Karolina Ininbergs ,Tanja Normark ,Isak Sylvin ,Zhibing Yun ,Martina Wahlund ,Valtteri Wirta |
| EPI_ISL_3461656, EPI_ISL_3461657 | Karolinska University Hospital Solna | Karolinska University Hospital | Jan Albert ,Tobias Allander ,Annelie Bjerkner ,Sandra Broddesson ,Robert Dyrdak ,Martin Ekman ,Lynda Eneh ,Lina Guerra Blomqvist ,Karolina Ininbergs ,Tanja Normark ,Isak Sylvin ,Zhibing Yun ,Martina Wahlund ,Valtteri Wirta |
| EPI_ISL_3461658 | Synlab Medilab | Karolinska University Hospital | Jan Albert ,Tobias Allander ,Annelie Bjerkner ,Sandra Broddesson ,Robert Dyrdak ,Martin Ekman ,Lynda Eneh ,Lina Guerra Blomqvist ,Karolina Ininbergs ,Tanja Normark ,Isak Sylvin ,Zhibing Yun ,Martina Wahlund ,Valtteri Wirta |
| EPI_ISL_3461659, EPI_ISL_3461660, EPI_ISL_3461661 | Karolinska University Hospital Solna | Karolinska University Hospital | Jan Albert ,Tobias Allander ,Annelie Bjerkner ,Sandra Broddesson ,Robert Dyrdak ,Martin Ekman ,Lynda Eneh ,Lina Guerra Blomqvist ,Karolina Ininbergs ,Tanja Normark ,Isak Sylvin ,Zhibing Yun ,Martina Wahlund ,Valtteri Wirta |
| EPI_ISL_3461663 | Synlab Medilab | Karolinska University Hospital | Jan Albert ,Tobias Allander ,Annelie Bjerkner ,Sandra Broddesson ,Robert Dyrdak ,Martin Ekman ,Lynda Eneh ,Lina Guerra Blomqvist ,Karolina Ininbergs ,Tanja Normark ,Isak Sylvin ,Zhibing Yun ,Martina Wahlund ,Valtteri Wirta |
| EPI_ISL_3461664, EPI_ISL_3461665, EPI_ISL_3461666, EPI_ISL_3461667, EPI_ISL_3461668, EPI_ISL_3461669, EPI_ISL_3461670, EPI_ISL_3461671, EPI_ISL_3461672, EPI_ISL_3461673, EPI_ISL_3461674 | | | |
| see above | Karolinska University Hospital Solna | Karolinska University Hospital | Jan Albert ,Tobias Allander ,Annelie Bjerkner ,Sandra Broddesson ,Robert Dyrdak ,Martin Ekman ,Lynda Eneh ,Lina Guerra Blomqvist ,Karolina Ininbergs ,Tanja Normark ,Isak Sylvin ,Zhibing Yun ,Martina Wahlund ,Valtteri Wirta |
| EPI_ISL_3461675 | Synlab Medilab | Karolinska University Hospital | Jan Albert ,Tobias Allander ,Annelie Bjerkner ,Sandra Broddesson ,Robert Dyrdak ,Martin Ekman ,Lynda Eneh ,Lina Guerra Blomqvist ,Karolina Ininbergs ,Tanja Normark ,Isak Sylvin ,Zhibing Yun ,Martina Wahlund ,Valtteri Wirta |
| EPI_ISL_3461676, EPI_ISL_3461677, EPI_ISL_3461678, EPI_ISL_3461679, EPI_ISL_3461680, EPI_ISL_3461681 | Karolinska University Hospital Solna | Karolinska University Hospital | Jan Albert ,Tobias Allander ,Annelie Bjerkner ,Sandra Broddesson ,Robert Dyrdak ,Martin Ekman ,Lynda Eneh ,Lina Guerra Blomqvist ,Karolina Ininbergs ,Tanja Normark ,Isak Sylvin ,Zhibing Yun ,Martina Wahlund ,Valtteri Wirta |
| EPI_ISL_3461682, EPI_ISL_3461683 | Synlab Medilab | Karolinska University Hospital | Jan Albert ,Tobias Allander ,Annelie Bjerkner ,Sandra Broddesson ,Robert Dyrdak ,Martin Ekman ,Lynda Eneh ,Lina Guerra Blomqvist ,Karolina Ininbergs ,Tanja Normark ,Isak Sylvin ,Zhibing Yun ,Martina Wahlund ,Valtteri Wirta |
| EPI_ISL_3461684, EPI_ISL_3461685 | Karolinska University Hospital Solna | Karolinska University Hospital | Jan Albert ,Tobias Allander ,Annelie Bjerkner ,Sandra Broddesson ,Robert Dyrdak ,Martin Ekman ,Lynda Eneh ,Lina Guerra Blomqvist ,Karolina Ininbergs ,Tanja Normark ,Isak Sylvin ,Zhibing Yun ,Martina Wahlund ,Valtteri Wirta |
| EPI_ISL_3461686, EPI_ISL_3461687, EPI_ISL_3461688 | Synlab Medilab | Karolinska University Hospital | Jan Albert ,Tobias Allander ,Annelie Bjerkner ,Sandra Broddesson ,Robert Dyrdak ,Martin Ekman ,Lynda Eneh ,Lina Guerra Blomqvist ,Karolina Ininbergs ,Tanja Normark ,Isak Sylvin ,Zhibing Yun ,Martina Wahlund ,Valtteri Wirta |
| EPI_ISL_3461689 | Karolinska University Hospital Solna | Karolinska University Hospital | Jan Albert ,Tobias Allander ,Annelie Bjerkner ,Sandra Broddesson ,Robert Dyrdak ,Martin Ekman ,Lynda Eneh ,Lina Guerra Blomqvist ,Karolina Ininbergs ,Tanja Normark ,Isak Sylvin ,Zhibing Yun ,Martina Wahlund ,Valtteri Wirta |
| EPI_ISL_3461690 | Synlab Medilab | Karolinska University Hospital | Jan Albert ,Tobias Allander ,Annelie Bjerkner ,Sandra Broddesson ,Robert Dyrdak ,Martin Ekman ,Lynda Eneh ,Lina Guerra Blomqvist ,Karolina Ininbergs ,Tanja Normark ,Isak Sylvin ,Zhibing Yun ,Martina Wahlund ,Valtteri Wirta |
| EPI_ISL_3461691, EPI_ISL_3461692, EPI_ISL_3461693, EPI_ISL_3461694, EPI_ISL_3461695 | Karolinska University Hospital Solna | Karolinska University Hospital | Jan Albert ,Tobias Allander ,Annelie Bjerkner ,Sandra Broddesson ,Robert Dyrdak ,Martin Ekman ,Lynda Eneh ,Lina Guerra Blomqvist ,Karolina Ininbergs ,Tanja Normark ,Isak Sylvin ,Zhibing Yun ,Martina Wahlund ,Valtteri Wirta |
| EPI_ISL_3461696, EPI_ISL_3461697, EPI_ISL_3461698 | Synlab Medilab | Karolinska University Hospital | Jan Albert ,Tobias Allander ,Annelie Bjerkner ,Sandra Broddesson ,Robert Dyrdak ,Martin Ekman ,Lynda Eneh ,Lina Guerra Blomqvist ,Karolina Ininbergs ,Tanja Normark ,Isak Sylvin ,Zhibing Yun ,Martina Wahlund ,Valtteri Wirta |
| EPI_ISL_3461699, EPI_ISL_3461700 | Karolinska University Hospital Solna | Karolinska University Hospital | Jan Albert ,Tobias Allander ,Annelie Bjerkner ,Sandra Broddesson ,Robert Dyrdak ,Martin Ekman ,Lynda Eneh ,Lina Guerra Blomqvist ,Karolina Ininbergs ,Tanja Normark ,Isak Sylvin ,Zhibing Yun ,Martina Wahlund ,Valtteri Wirta |
| EPI_ISL_3461701 | Synlab Medilab | Karolinska University Hospital | Jan Albert ,Tobias Allander ,Annelie Bjerkner ,Sandra Broddesson ,Robert Dyrdak ,Martin Ekman ,Lynda Eneh ,Lina Guerra Blomqvist ,Karolina Ininbergs ,Tanja Normark ,Isak Sylvin ,Zhibing Yun ,Martina Wahlund ,Valtteri Wirta |
| EPI_ISL_3461702 | Karolinska University Hospital Solna | Karolinska University Hospital | Jan Albert ,Tobias Allander ,Annelie Bjerkner ,Sandra Broddesson ,Robert Dyrdak ,Martin Ekman ,Lynda Eneh ,Lina Guerra Blomqvist ,Karolina Ininbergs ,Tanja Normark ,Isak Sylvin ,Zhibing Yun ,Martina Wahlund ,Valtteri Wirta |
| EPI_ISL_3461703, EPI_ISL_3461704, EPI_ISL_3461705, EPI_ISL_3461706 | Synlab Medilab | Karolinska University Hospital | Jan Albert ,Tobias Allander ,Annelie Bjerkner ,Sandra Broddesson ,Robert Dyrdak ,Martin Ekman ,Lynda Eneh ,Lina Guerra Blomqvist ,Karolina Ininbergs ,Tanja Normark ,Isak Sylvin ,Zhibing Yun ,Martina Wahlund ,Valtteri Wirta |
| EPI_ISL_3461707, EPI_ISL_3461708, EPI_ISL_3461709, EPI_ISL_3461710 | Karolinska University Hospital Solna | Karolinska University Hospital | Jan Albert ,Tobias Allander ,Annelie Bjerkner ,Sandra Broddesson ,Robert Dyrdak ,Martin Ekman ,Lynda Eneh ,Lina Guerra Blomqvist ,Karolina Ininbergs ,Tanja Normark ,Isak Sylvin ,Zhibing Yun ,Martina Wahlund ,Valtteri Wirta |
| EPI_ISL_3461711 | Synlab Medilab | Karolinska University Hospital | Jan Albert ,Tobias Allander ,Annelie Bjerkner ,Sandra Broddesson ,Robert Dyrdak ,Martin Ekman ,Lynda Eneh ,Lina Guerra Blomqvist ,Karolina Ininbergs ,Tanja Normark ,Isak Sylvin ,Zhibing Yun ,Martina Wahlund ,Valtteri Wirta |
| EPI_ISL_3461712 | Karolinska University Hospital Solna | Karolinska University Hospital | Jan Albert ,Tobias Allander ,Annelie Bjerkner ,Sandra Broddesson ,Robert Dyrdak ,Martin Ekman ,Lynda Eneh ,Lina Guerra Blomqvist ,Karolina Ininbergs ,Tanja Normark ,Isak Sylvin ,Zhibing Yun ,Martina Wahlund ,Valtteri Wirta |
| EPI_ISL_3461713, EPI_ISL_3461714, EPI_ISL_3461715, EPI_ISL_3461716 | Synlab Medilab | Karolinska University Hospital | Jan Albert ,Tobias Allander ,Annelie Bjerkner ,Sandra Broddesson ,Robert Dyrdak ,Martin Ekman ,Lynda Eneh ,Lina Guerra Blomqvist ,Karolina Ininbergs ,Tanja Normark ,Isak Sylvin ,Zhibing Yun ,Martina Wahlund ,Valtteri Wirta |
| EPI_ISL_3461717, EPI_ISL_3461718, EPI_ISL_3461719 | Karolinska University Hospital Solna | Karolinska University Hospital | Jan Albert ,Tobias Allander ,Annelie Bjerkner ,Sandra Broddesson ,Robert Dyrdak ,Martin Ekman ,Lynda Eneh ,Lina Guerra Blomqvist ,Karolina Ininbergs ,Tanja Normark ,Isak Sylvin ,Zhibing Yun ,Martina Wahlund ,Valtteri Wirta |
| EPI_ISL_3461720 | Synlab Medilab | Karolinska University Hospital | Jan Albert ,Tobias Allander ,Annelie Bjerkner ,Sandra Broddesson ,Robert Dyrdak ,Martin Ekman ,Lynda Eneh ,Lina Guerra Blomqvist ,Karolina Ininbergs ,Tanja Normark ,Isak Sylvin ,Zhibing Yun ,Martina Wahlund ,Valtteri Wirta |
| EPI_ISL_3461721 | Karolinska University Hospital Solna | Karolinska University Hospital | Jan Albert ,Tobias Allander ,Annelie Bjerkner ,Sandra Broddesson ,Robert Dyrdak ,Martin Ekman ,Lynda Eneh ,Lina Guerra Blomqvist ,Karolina Ininbergs ,Tanja Normark ,Isak Sylvin ,Zhibing Yun ,Martina Wahlund ,Valtteri Wirta |
| EPI_ISL_3461722, EPI_ISL_3461723 | Synlab Medilab | Karolinska University Hospital | Jan Albert ,Tobias Allander ,Annelie Bjerkner ,Sandra Broddesson ,Robert Dyrdak ,Martin Ekman ,Lynda Eneh ,Lina Guerra Blomqvist ,Karolina Ininbergs ,Tanja Normark ,Isak Sylvin ,Zhibing Yun ,Martina Wahlund ,Valtteri Wirta |
| EPI_ISL_3461724, EPI_ISL_3461725 | Karolinska University Hospital Solna | Karolinska University Hospital | Jan Albert ,Tobias Allander ,Annelie Bjerkner ,Sandra Broddesson ,Robert Dyrdak ,Martin Ekman ,Lynda Eneh ,Lina Guerra Blomqvist ,Karolina Ininbergs ,Tanja Normark ,Isak Sylvin ,Zhibing Yun ,Martina Wahlund ,Valtteri Wirta |
| EPI_ISL_3461726 | Synlab Medilab | Karolinska University Hospital | Jan Albert ,Tobias Allander ,Annelie Bjerkner ,Sandra Broddesson ,Robert Dyrdak ,Martin Ekman ,Lynda Eneh ,Lina Guerra Blomqvist ,Karolina Ininbergs ,Tanja Normark ,Isak Sylvin ,Zhibing Yun ,Martina Wahlund ,Valtteri Wirta |
| EPI_ISL_3461727 | Karolinska University Hospital Solna | Karolinska University Hospital | Jan Albert ,Tobias Allander ,Annelie Bjerkner ,Sandra Broddesson ,Robert Dyrdak ,Martin Ekman ,Lynda Eneh ,Lina Guerra Blomqvist ,Karolina Ininbergs ,Tanja Normark ,Isak Sylvin ,Zhibing Yun ,Martina Wahlund ,Valtteri Wirta |
| EPI_ISL_3461728 | Synlab Medilab | Karolinska University Hospital | Jan Albert ,Tobias Allander ,Annelie Bjerkner ,Sandra Broddesson ,Robert Dyrdak ,Martin Ekman ,Lynda Eneh ,Lina Guerra Blomqvist ,Karolina Ininbergs ,Tanja Normark ,Isak Sylvin ,Zhibing Yun ,Martina Wahlund ,Valtteri Wirta |
| EPI_ISL_3461729 | Karolinska University Hospital Solna | Karolinska University Hospital | Jan Albert ,Tobias Allander ,Annelie Bjerkner ,Sandra Broddesson ,Robert Dyrdak ,Martin Ekman ,Lynda Eneh ,Lina Guerra Blomqvist ,Karolina Ininbergs ,Tanja Normark ,Isak Sylvin ,Zhibing Yun ,Martina Wahlund ,Valtteri Wirta |
| EPI_ISL_3461731, EPI_ISL_3461732, EPI_ISL_3461733, EPI_ISL_3461734, EPI_ISL_3461736, EPI_ISL_3461737, EPI_ISL_3461738, EPI_ISL_3461739, EPI_ISL_3461740, EPI_ISL_3461741, EPI_ISL_3461744, EPI_ISL_3461745, EPI_ISL_3461746, EPI_ISL_3461747, EPI_ISL_3461748, EPI_ISL_3461749, EPI_ISL_3461750, EPI_ISL_3461751, EPI_ISL_3461752, EPI_ISL_3461753, EPI_ISL_3461754, EPI_ISL_3461755, EPI_ISL_3461756, EPI_ISL_3461757, EPI_ISL_3461758, EPI_ISL_3461759, EPI_ISL_3461761, EPI_ISL_3461762, EPI_ISL_3461763, EPI_ISL_3461764, EPI_ISL_3461766, EPI_ISL_3461767, EPI_ISL_3461768, EPI_ISL_3461769, EPI_ISL_3461770, EPI_ISL_3461771, EPI_ISL_3461772, EPI_ISL_3461773, EPI_ISL_3461774, EPI_ISL_3461775, EPI_ISL_3461776, EPI_ISL_3461777, EPI_ISL_3461778, EPI_ISL_3461780, EPI_ISL_3461781, EPI_ISL_3461782, EPI_ISL_3461783, EPI_ISL_3461784, EPI_ISL_3461785, EPI_ISL_3461786, EPI_ISL_3461787, EPI_ISL_3461788, EPI_ISL_3461789, EPI_ISL_3461790, EPI_ISL_3461792, EPI_ISL_3461793, EPI_ISL_3461794, EPI_ISL_3461795, EPI_ISL_3461796, EPI_ISL_3461797, EPI_ISL_3461798, EPI_ISL_3461799, EPI_ISL_3461800, EPI_ISL_3461801, EPI_ISL_3461802, EPI_ISL_3461803, EPI_ISL_3461804, EPI_ISL_3461805, EPI_ISL_3461806, EPI_ISL_3461807, EPI_ISL_3461808, EPI_ISL_3461809, EPI_ISL_3461810, EPI_ISL_3461811, EPI_ISL_3461812, EPI_ISL_3461813, EPI_ISL_3461814, EPI_ISL_3461815, EPI_ISL_3461816, EPI_ISL_3461817, EPI_ISL_3461818, EPI_ISL_3461819, EPI_ISL_3461820, EPI_ISL_3461821, EPI_ISL_3461822, EPI_ISL_3461823, EPI_ISL_3461824, EPI_ISL_3461825, EPI_ISL_3461826, EPI_ISL_3461827, EPI_ISL_3461828, EPI_ISL_3461829, EPI_ISL_3461831, EPI_ISL_3461832, EPI_ISL_3461833, EPI_ISL_3461834, EPI_ISL_3461835, EPI_ISL_3461836, EPI_ISL_3461837, EPI_ISL_3461838, EPI_ISL_3461839, EPI_ISL_3461840, EPI_ISL_3461841, EPI_ISL_3461842, EPI_ISL_3461843, EPI_ISL_3461844, EPI_ISL_3461845, EPI_ISL_3461846, EPI_ISL_3461847, EPI_ISL_3461848, EPI_ISL_3461849, EPI_ISL_3461850, EPI_ISL_3461851, EPI_ISL_3461852, EPI_ISL_3461853, EPI_ISL_3461854, EPI_ISL_3461855, EPI_ISL_3461856, EPI_ISL_3461857, EPI_ISL_3461858, EPI_ISL_3461859, EPI_ISL_3461860, EPI_ISL_3461861, EPI_ISL_3461862, EPI_ISL_3461864, EPI_ISL_3461865, EPI_ISL_3461867, EPI_ISL_3461868, EPI_ISL_3461869, EPI_ISL_3461870, EPI_ISL_3461871, EPI_ISL_3461872, EPI_ISL_3461873, EPI_ISL_3461874, EPI_ISL_3461875, EPI_ISL_3461876, EPI_ISL_3461877, EPI_ISL_3461878, EPI_ISL_3461879, EPI_ISL_3461880, EPI_ISL_3461881, EPI_ISL_3461882, EPI_ISL_3461883, EPI_ISL_3461884 | | | |
| see above | Karolinska University Hospital Huddinge | Karolinska University Hospital | Jan Albert ,Tobias Allander ,Annelie Bjerkner ,Sandra Broddesson ,Robert Dyrdak ,Martin Ekman ,Lynda Eneh ,Lina Guerra Blomqvist ,Karolina Ininbergs ,Tanja Normark ,Isak Sylvin ,Zhibing Yun ,Martina Wahlund ,Valtteri Wirta |
| EPI_ISL_3461885, EPI_ISL_3461886, EPI_ISL_3461888, EPI_ISL_3461889, EPI_ISL_3461890, EPI_ISL_3461891, EPI_ISL_3461893, EPI_ISL_3461894, EPI_ISL_3461895, EPI_ISL_3461896, EPI_ISL_3461897, EPI_ISL_3461898, EPI_ISL_3461900, EPI_ISL_3461901, EPI_ISL_3461902, EPI_ISL_3461903, EPI_ISL_3461904, EPI_ISL_3461905, EPI_ISL_3461907, EPI_ISL_3461908, EPI_ISL_3461909, EPI_ISL_3461910, EPI_ISL_3461911, EPI_ISL_3461912, EPI_ISL_3461913, EPI_ISL_3461914, EPI_ISL_3461915, EPI_ISL_3461916, EPI_ISL_3461917, EPI_ISL_3461918, EPI_ISL_3461919, EPI_ISL_3461920, EPI_ISL_3461921, EPI_ISL_3461922, EPI_ISL_3461923, EPI_ISL_3461924, EPI_ISL_3461925, EPI_ISL_3461926, EPI_ISL_3461927, EPI_ISL_3461928, EPI_ISL_3461929, EPI_ISL_3461930, EPI_ISL_3461931, EPI_ISL_3461932, EPI_ISL_3461933, EPI_ISL_3461934, EPI_ISL_3461935, EPI_ISL_3461936, EPI_ISL_3461937, EPI_ISL_3461938, EPI_ISL_3461939, EPI_ISL_3461940, EPI_ISL_3461941, EPI_ISL_3461942, EPI_ISL_3461943, EPI_ISL_3461944, EPI_ISL_3461945, EPI_ISL_3461946, EPI_ISL_3461947, EPI_ISL_3461948, EPI_ISL_3461949, EPI_ISL_3461950, EPI_ISL_3461951, EPI_ISL_3461952, EPI_ISL_3461953, EPI_ISL_3461954, EPI_ISL_3461955, EPI_ISL_3461956, EPI_ISL_3461957, EPI_ISL_3461958, EPI_ISL_3461959, EPI_ISL_3461960, EPI_ISL_3461961, EPI_ISL_3461962 | | | |
| see above | Karolinska University Hospital Solna | Karolinska University Hospital | Jan Albert ,Tobias Allander ,Annelie Bjerkner ,Sandra Broddesson ,Robert Dyrdak ,Martin Ekman ,Lynda Eneh ,Lina Guerra Blomqvist ,Karolina Ininbergs ,Tanja Normark ,Isak Sylvin ,Zhibing Yun ,Martina Wahlund ,Valtteri Wirta |
| EPI_ISL_3461963, EPI_ISL_3461964 | Synlab Medilab | Karolinska University Hospital | Jan Albert ,Tobias Allander ,Annelie Bjerkner ,Sandra Broddesson ,Robert Dyrdak ,Martin Ekman ,Lynda Eneh ,Lina Guerra Blomqvist ,Karolina Ininbergs ,Tanja Normark ,Isak Sylvin ,Zhibing Yun ,Martina Wahlund ,Valtteri Wirta |
| EPI_ISL_3461965 | Karolinska University Hospital Solna | Karolinska University Hospital | Jan Albert ,Tobias Allander ,Annelie Bjerkner ,Sandra Broddesson ,Robert Dyrdak ,Martin Ekman ,Lynda Eneh ,Lina Guerra Blomqvist ,Karolina Ininbergs ,Tanja Normark ,Isak Sylvin ,Zhibing Yun ,Martina Wahlund ,Valtteri Wirta |
| EPI_ISL_3461966, EPI_ISL_3461967 | Synlab Medilab | Karolinska University Hospital | Jan Albert ,Tobias Allander ,Annelie Bjerkner ,Sandra Broddesson ,Robert Dyrdak ,Martin Ekman ,Lynda Eneh ,Lina Guerra Blomqvist ,Karolina Ininbergs ,Tanja Normark ,Isak Sylvin ,Zhibing Yun ,Martina Wahlund ,Valtteri Wirta |
| EPI_ISL_3461968 | Karolinska University Hospital Solna | Karolinska University Hospital | Jan Albert ,Tobias Allander ,Annelie Bjerkner ,Sandra Broddesson ,Robert Dyrdak ,Martin Ekman ,Lynda Eneh ,Lina Guerra Blomqvist ,Karolina Ininbergs ,Tanja Normark ,Isak Sylvin ,Zhibing Yun ,Martina Wahlund ,Valtteri Wirta |
| EPI_ISL_3461969 | Synlab Medilab | Karolinska University Hospital | Jan Albert ,Tobias Allander ,Annelie Bjerkner ,Sandra Broddesson ,Robert Dyrdak ,Martin Ekman ,Lynda Eneh ,Lina Guerra Blomqvist ,Karolina Ininbergs ,Tanja Normark ,Isak Sylvin ,Zhibing Yun ,Martina Wahlund ,Valtteri Wirta |

| | | | |
|---|---|---|---|
| EPI_ISL_3461971, EPI_ISL_3461972 | Karolinska University Hospital Solna | Karolinska University Hospital | Jan Albert ,Tobias Allander ,Annelie Bjerkner ,Sandra Broddesson ,Robert Dyrdak ,Martin Ekman ,Lynda Eneh ,Lina Guerra Blomqvist ,Karolina Ininbergs ,Tanja Normark ,Isak Sylvin ,Zhibing Yun ,Martina Wahlund ,Valtteri Wirta |
| EPI_ISL_3461973 | Synlab Medilab | Karolinska University Hospital | Jan Albert ,Tobias Allander ,Annelie Bjerkner ,Sandra Broddesson ,Robert Dyrdak ,Martin Ekman ,Lynda Eneh ,Lina Guerra Blomqvist ,Karolina Ininbergs ,Tanja Normark ,Isak Sylvin ,Zhibing Yun ,Martina Wahlund ,Valtteri Wirta |
| EPI_ISL_3461974 | Karolinska University Hospital Solna | Karolinska University Hospital | Jan Albert ,Tobias Allander ,Annelie Bjerkner ,Sandra Broddesson ,Robert Dyrdak ,Martin Ekman ,Lynda Eneh ,Lina Guerra Blomqvist ,Karolina Ininbergs ,Tanja Normark ,Isak Sylvin ,Zhibing Yun ,Martina Wahlund ,Valtteri Wirta |
| EPI_ISL_3461975 | Synlab Medilab | Karolinska University Hospital | Jan Albert ,Tobias Allander ,Annelie Bjerkner ,Sandra Broddesson ,Robert Dyrdak ,Martin Ekman ,Lynda Eneh ,Lina Guerra Blomqvist ,Karolina Ininbergs ,Tanja Normark ,Isak Sylvin ,Zhibing Yun ,Martina Wahlund ,Valtteri Wirta |
| EPI_ISL_3461976, EPI_ISL_3461977, EPI_ISL_3461978 | Karolinska University Hospital Solna | Karolinska University Hospital | Jan Albert ,Tobias Allander ,Annelie Bjerkner ,Sandra Broddesson ,Robert Dyrdak ,Martin Ekman ,Lynda Eneh ,Lina Guerra Blomqvist ,Karolina Ininbergs ,Tanja Normark ,Isak Sylvin ,Zhibing Yun ,Martina Wahlund ,Valtteri Wirta |
| EPI_ISL_3461979 | Synlab Medilab | Karolinska University Hospital | Jan Albert ,Tobias Allander ,Annelie Bjerkner ,Sandra Broddesson ,Robert Dyrdak ,Martin Ekman ,Lynda Eneh ,Lina Guerra Blomqvist ,Karolina Ininbergs ,Tanja Normark ,Isak Sylvin ,Zhibing Yun ,Martina Wahlund ,Valtteri Wirta |
| EPI_ISL_3461980, EPI_ISL_3461982 | Karolinska University Hospital Solna | Karolinska University Hospital | Jan Albert ,Tobias Allander ,Annelie Bjerkner ,Sandra Broddesson ,Robert Dyrdak ,Martin Ekman ,Lynda Eneh ,Lina Guerra Blomqvist ,Karolina Ininbergs ,Tanja Normark ,Isak Sylvin ,Zhibing Yun ,Martina Wahlund ,Valtteri Wirta |
| EPI_ISL_3461983 | Synlab Medilab | Karolinska University Hospital | Jan Albert ,Tobias Allander ,Annelie Bjerkner ,Sandra Broddesson ,Robert Dyrdak ,Martin Ekman ,Lynda Eneh ,Lina Guerra Blomqvist ,Karolina Ininbergs ,Tanja Normark ,Isak Sylvin ,Zhibing Yun ,Martina Wahlund ,Valtteri Wirta |
| EPI_ISL_3461984, EPI_ISL_3461985, EPI_ISL_3461986 | Karolinska University Hospital Solna | Karolinska University Hospital | Jan Albert ,Tobias Allander ,Annelie Bjerkner ,Sandra Broddesson ,Robert Dyrdak ,Martin Ekman ,Lynda Eneh ,Lina Guerra Blomqvist ,Karolina Ininbergs ,Tanja Normark ,Isak Sylvin ,Zhibing Yun ,Martina Wahlund ,Valtteri Wirta |
| EPI_ISL_3461987 | Synlab Medilab | Karolinska University Hospital | Jan Albert ,Tobias Allander ,Annelie Bjerkner ,Sandra Broddesson ,Robert Dyrdak ,Martin Ekman ,Lynda Eneh ,Lina Guerra Blomqvist ,Karolina Ininbergs ,Tanja Normark ,Isak Sylvin ,Zhibing Yun ,Martina Wahlund ,Valtteri Wirta |
| EPI_ISL_3461988 | Karolinska University Hospital Solna | Karolinska University Hospital | Jan Albert ,Tobias Allander ,Annelie Bjerkner ,Sandra Broddesson ,Robert Dyrdak ,Martin Ekman ,Lynda Eneh ,Lina Guerra Blomqvist ,Karolina Ininbergs ,Tanja Normark ,Isak Sylvin ,Zhibing Yun ,Martina Wahlund ,Valtteri Wirta |
| EPI_ISL_3461989 | Synlab Medilab | Karolinska University Hospital | Jan Albert ,Tobias Allander ,Annelie Bjerkner ,Sandra Broddesson ,Robert Dyrdak ,Martin Ekman ,Lynda Eneh ,Lina Guerra Blomqvist ,Karolina Ininbergs ,Tanja Normark ,Isak Sylvin ,Zhibing Yun ,Martina Wahlund ,Valtteri Wirta |
| EPI_ISL_3461990, EPI_ISL_3461991 | Karolinska University Hospital Solna | Karolinska University Hospital | Jan Albert ,Tobias Allander ,Annelie Bjerkner ,Sandra Broddesson ,Robert Dyrdak ,Martin Ekman ,Lynda Eneh ,Lina Guerra Blomqvist ,Karolina Ininbergs ,Tanja Normark ,Isak Sylvin ,Zhibing Yun ,Martina Wahlund ,Valtteri Wirta |
| EPI_ISL_3461992 | Synlab Medilab | Karolinska University Hospital | Jan Albert ,Tobias Allander ,Annelie Bjerkner ,Sandra Broddesson ,Robert Dyrdak ,Martin Ekman ,Lynda Eneh ,Lina Guerra Blomqvist ,Karolina Ininbergs ,Tanja Normark ,Isak Sylvin ,Zhibing Yun ,Martina Wahlund ,Valtteri Wirta |
| EPI_ISL_3461993 | Karolinska University Hospital Solna | Karolinska University Hospital | Jan Albert ,Tobias Allander ,Annelie Bjerkner ,Sandra Broddesson ,Robert Dyrdak ,Martin Ekman ,Lynda Eneh ,Lina Guerra Blomqvist ,Karolina Ininbergs ,Tanja Normark ,Isak Sylvin ,Zhibing Yun ,Martina Wahlund ,Valtteri Wirta |
| EPI_ISL_3461994, EPI_ISL_3461995 | Synlab Medilab | Karolinska University Hospital | Jan Albert ,Tobias Allander ,Annelie Bjerkner ,Sandra Broddesson ,Robert Dyrdak ,Martin Ekman ,Lynda Eneh ,Lina Guerra Blomqvist ,Karolina Ininbergs ,Tanja Normark ,Isak Sylvin ,Zhibing Yun ,Martina Wahlund ,Valtteri Wirta |
| EPI_ISL_3461997 | Karolinska University Hospital Solna | Karolinska University Hospital | Jan Albert ,Tobias Allander ,Annelie Bjerkner ,Sandra Broddesson ,Robert Dyrdak ,Martin Ekman ,Lynda Eneh ,Lina Guerra Blomqvist ,Karolina Ininbergs ,Tanja Normark ,Isak Sylvin ,Zhibing Yun ,Martina Wahlund ,Valtteri Wirta |
| EPI_ISL_3461998 | Synlab Medilab | Karolinska University Hospital | Jan Albert ,Tobias Allander ,Annelie Bjerkner ,Sandra Broddesson ,Robert Dyrdak ,Martin Ekman ,Lynda Eneh ,Lina Guerra Blomqvist ,Karolina Ininbergs ,Tanja Normark ,Isak Sylvin ,Zhibing Yun ,Martina Wahlund ,Valtteri Wirta |
| EPI_ISL_3461999 | Karolinska University Hospital Solna | Karolinska University Hospital | Jan Albert ,Tobias Allander ,Annelie Bjerkner ,Sandra Broddesson ,Robert Dyrdak ,Martin Ekman ,Lynda Eneh ,Lina Guerra Blomqvist ,Karolina Ininbergs ,Tanja Normark ,Isak Sylvin ,Zhibing Yun ,Martina Wahlund ,Valtteri Wirta |
| EPI_ISL_3462000 | Synlab Medilab | Karolinska University Hospital | Jan Albert ,Tobias Allander ,Annelie Bjerkner ,Sandra Broddesson ,Robert Dyrdak ,Martin Ekman ,Lynda Eneh ,Lina Guerra Blomqvist ,Karolina Ininbergs ,Tanja Normark ,Isak Sylvin ,Zhibing Yun ,Martina Wahlund ,Valtteri Wirta |
| EPI_ISL_3462001, EPI_ISL_3462002 | Karolinska University Hospital Solna | Karolinska University Hospital | Jan Albert ,Tobias Allander ,Annelie Bjerkner ,Sandra Broddesson ,Robert Dyrdak ,Martin Ekman ,Lynda Eneh ,Lina Guerra Blomqvist ,Karolina Ininbergs ,Tanja Normark ,Isak Sylvin ,Zhibing Yun ,Martina Wahlund ,Valtteri Wirta |
| EPI_ISL_3462003, EPI_ISL_3462004 | Synlab Medilab | Karolinska University Hospital | Jan Albert ,Tobias Allander ,Annelie Bjerkner ,Sandra Broddesson ,Robert Dyrdak ,Martin Ekman ,Lynda Eneh ,Lina Guerra Blomqvist ,Karolina Ininbergs ,Tanja Normark ,Isak Sylvin ,Zhibing Yun ,Martina Wahlund ,Valtteri Wirta |
| EPI_ISL_3462005 | Karolinska University Hospital Solna | Karolinska University Hospital | Jan Albert ,Tobias Allander ,Annelie Bjerkner ,Sandra Broddesson ,Robert Dyrdak ,Martin Ekman ,Lynda Eneh ,Lina Guerra Blomqvist ,Karolina Ininbergs ,Tanja Normark ,Isak Sylvin ,Zhibing Yun ,Martina Wahlund ,Valtteri Wirta |
| EPI_ISL_3462006, EPI_ISL_3462007 | Synlab Medilab | Karolinska University Hospital | Jan Albert ,Tobias Allander ,Annelie Bjerkner ,Sandra Broddesson ,Robert Dyrdak ,Martin Ekman ,Lynda Eneh ,Lina Guerra Blomqvist ,Karolina Ininbergs ,Tanja Normark ,Isak Sylvin ,Zhibing Yun ,Martina Wahlund ,Valtteri Wirta |
| EPI_ISL_3462008, EPI_ISL_3462009 | Karolinska University Hospital Solna | Karolinska University Hospital | Jan Albert ,Tobias Allander ,Annelie Bjerkner ,Sandra Broddesson ,Robert Dyrdak ,Martin Ekman ,Lynda Eneh ,Lina Guerra Blomqvist ,Karolina Ininbergs ,Tanja Normark ,Isak Sylvin ,Zhibing Yun ,Martina Wahlund ,Valtteri Wirta |
| EPI_ISL_3462010 | Synlab Medilab | Karolinska University Hospital | Jan Albert ,Tobias Allander ,Annelie Bjerkner ,Sandra Broddesson ,Robert Dyrdak ,Martin Ekman ,Lynda Eneh ,Lina Guerra Blomqvist ,Karolina Ininbergs ,Tanja Normark ,Isak Sylvin ,Zhibing Yun ,Martina Wahlund ,Valtteri Wirta |
| EPI_ISL_3462011, EPI_ISL_3462012 | Karolinska University Hospital Solna | Karolinska University Hospital | Jan Albert ,Tobias Allander ,Annelie Bjerkner ,Sandra Broddesson ,Robert Dyrdak ,Martin Ekman ,Lynda Eneh ,Lina Guerra Blomqvist ,Karolina Ininbergs ,Tanja Normark ,Isak Sylvin ,Zhibing Yun ,Martina Wahlund ,Valtteri Wirta |
| EPI_ISL_3462013 | Synlab Medilab | Karolinska University Hospital | Jan Albert ,Tobias Allander ,Annelie Bjerkner ,Sandra Broddesson ,Robert Dyrdak ,Martin Ekman ,Lynda Eneh ,Lina Guerra Blomqvist ,Karolina Ininbergs ,Tanja Normark ,Isak Sylvin ,Zhibing Yun ,Martina Wahlund ,Valtteri Wirta |
| EPI_ISL_3462014 | Karolinska University Hospital Solna | Karolinska University Hospital | Jan Albert ,Tobias Allander ,Annelie Bjerkner ,Sandra Broddesson ,Robert Dyrdak ,Martin Ekman ,Lynda Eneh ,Lina Guerra Blomqvist ,Karolina Ininbergs ,Tanja Normark ,Isak Sylvin ,Zhibing Yun ,Martina Wahlund ,Valtteri Wirta |
| EPI_ISL_3462015 | Synlab Medilab | Karolinska University Hospital | Jan Albert ,Tobias Allander ,Annelie Bjerkner ,Sandra Broddesson ,Robert Dyrdak ,Martin Ekman ,Lynda Eneh ,Lina Guerra Blomqvist ,Karolina Ininbergs ,Tanja Normark ,Isak Sylvin ,Zhibing Yun ,Martina Wahlund ,Valtteri Wirta |
| EPI_ISL_3462016, EPI_ISL_3462017, EPI_ISL_3462018, EPI_ISL_3462019, EPI_ISL_3462020, EPI_ISL_3462021 | Karolinska University Hospital Solna | Karolinska University Hospital | Jan Albert ,Tobias Allander ,Annelie Bjerkner ,Sandra Broddesson ,Robert Dyrdak ,Martin Ekman ,Lynda Eneh ,Lina Guerra Blomqvist ,Karolina Ininbergs ,Tanja Normark ,Isak Sylvin ,Zhibing Yun ,Martina Wahlund ,Valtteri Wirta |
| EPI_ISL_3462022 | Synlab Medilab | Karolinska University Hospital | Jan Albert ,Tobias Allander ,Annelie Bjerkner ,Sandra Broddesson ,Robert Dyrdak ,Martin Ekman ,Lynda Eneh ,Lina Guerra Blomqvist ,Karolina Ininbergs ,Tanja Normark ,Isak Sylvin ,Zhibing Yun ,Martina Wahlund ,Valtteri Wirta |
| EPI_ISL_3462023, EPI_ISL_3462024, EPI_ISL_3462025, EPI_ISL_3462026, EPI_ISL_3462027, EPI_ISL_3462028, EPI_ISL_3462029 | Karolinska University Hospital Solna | Karolinska University Hospital | Jan Albert ,Tobias Allander ,Annelie Bjerkner ,Sandra Broddesson ,Robert Dyrdak ,Martin Ekman ,Lynda Eneh ,Lina Guerra Blomqvist ,Karolina Ininbergs ,Tanja Normark ,Isak Sylvin ,Zhibing Yun ,Martina Wahlund ,Valtteri Wirta |
| EPI_ISL_3462030 | Synlab Medilab | Karolinska University Hospital | Jan Albert ,Tobias Allander ,Annelie Bjerkner ,Sandra Broddesson ,Robert Dyrdak ,Martin Ekman ,Lynda Eneh ,Lina Guerra Blomqvist ,Karolina Ininbergs ,Tanja Normark ,Isak Sylvin ,Zhibing Yun ,Martina Wahlund ,Valtteri Wirta |
| EPI_ISL_3462031 | Karolinska University Hospital Solna | Karolinska University Hospital | Jan Albert ,Tobias Allander ,Annelie Bjerkner ,Sandra Broddesson ,Robert Dyrdak ,Martin Ekman ,Lynda Eneh ,Lina Guerra Blomqvist ,Karolina Ininbergs ,Tanja Normark ,Isak Sylvin ,Zhibing Yun ,Martina Wahlund ,Valtteri Wirta |
| EPI_ISL_3462032 | Synlab Medilab | Karolinska University Hospital | Jan Albert ,Tobias Allander ,Annelie Bjerkner ,Sandra Broddesson ,Robert Dyrdak ,Martin Ekman ,Lynda Eneh ,Lina Guerra Blomqvist ,Karolina Ininbergs ,Tanja Normark ,Isak Sylvin ,Zhibing Yun ,Martina Wahlund ,Valtteri Wirta |
| EPI_ISL_3462033 | Karolinska University Hospital Solna | Karolinska University Hospital | Jan Albert ,Tobias Allander ,Annelie Bjerkner ,Sandra Broddesson ,Robert Dyrdak ,Martin Ekman ,Lynda Eneh ,Lina Guerra Blomqvist ,Karolina Ininbergs ,Tanja Normark ,Isak Sylvin ,Zhibing Yun ,Martina Wahlund ,Valtteri Wirta |
| EPI_ISL_3462034, EPI_ISL_3462035 | Synlab Medilab | Karolinska University Hospital | Jan Albert ,Tobias Allander ,Annelie Bjerkner ,Sandra Broddesson ,Robert Dyrdak ,Martin Ekman ,Lynda Eneh ,Lina Guerra Blomqvist ,Karolina Ininbergs ,Tanja Normark ,Isak Sylvin ,Zhibing Yun ,Martina Wahlund ,Valtteri Wirta |
| EPI_ISL_3462036, EPI_ISL_3462037 | Karolinska University Hospital Solna | Karolinska University Hospital | Jan Albert ,Tobias Allander ,Annelie Bjerkner ,Sandra Broddesson ,Robert Dyrdak ,Martin Ekman ,Lynda Eneh ,Lina Guerra Blomqvist ,Karolina Ininbergs ,Tanja Normark ,Isak Sylvin ,Zhibing Yun ,Martina Wahlund ,Valtteri Wirta |
| EPI_ISL_3462038 | Synlab Medilab | Karolinska University Hospital | Jan Albert ,Tobias Allander ,Annelie Bjerkner ,Sandra Broddesson ,Robert Dyrdak ,Martin Ekman ,Lynda Eneh ,Lina Guerra Blomqvist ,Karolina Ininbergs ,Tanja Normark ,Isak Sylvin ,Zhibing Yun ,Martina Wahlund ,Valtteri Wirta |
| EPI_ISL_3462039, EPI_ISL_3462040 | Karolinska University Hospital Solna | Karolinska University Hospital | Jan Albert ,Tobias Allander ,Annelie Bjerkner ,Sandra Broddesson ,Robert Dyrdak ,Martin Ekman ,Lynda Eneh ,Lina Guerra Blomqvist ,Karolina Ininbergs ,Tanja Normark ,Isak Sylvin ,Zhibing Yun ,Martina Wahlund ,Valtteri Wirta |
| EPI_ISL_3462044, EPI_ISL_3462045, EPI_ISL_3462046, EPI_ISL_3462047, EPI_ISL_3462048, EPI_ISL_3462049, EPI_ISL_3462050, EPI_ISL_3462051, EPI_ISL_3462052, EPI_ISL_3462054, EPI_ISL_3462055, EPI_ISL_3462056 | | | |
| see above | Pathology, UAB | Pathology, UAB | Moates,D., Van Der Pol,W.J., Hendrickson,R.C., Lefkowitz,E.J., Leal,S.M. |
| EPI_ISL_3462057 | Orange County Public Health Laboratory, Orange County Public Health Laboratory submission group | Orange County Public Health Laboratory, Orange County Public Health Laboratory submission group | Elder,J.W. |
| EPI_ISL_3462072 | Laboratory of Microbiology, ASST Ospedale di Circolo, Varese | Laboratory of Microbiology, ASST Ospedale di Circolo, Varese | Federica Novazzi, Francesca Dragoferrante, Angelo Genoni, Daniela Dalla Gasperina, Daniele Focosi, Andreina Baj, Fabrizio Maggi |
| EPI_ISL_3462074 | Laboratory of Microbiology, ASST Ospedale di Circolo, Varese viale borri 57 21100 Varese, Italy | Laboratory of Microbiology, ASST Ospedale di Circolo, Varese viale borri 57 21100 Varese, Italy | Federica Novazzi, Francesca Dragoferrante, Angelo Genoni, Daniela Dalla Gasperina, Daniele Focosi, Andreina Baj, Fabrizio Maggi Submitter information |
| EPI_ISL_3462078, EPI_ISL_3462080 | Laboratory of Microbiology, ASST Ospedale di Circolo, Varese viale borri 57 21100 Varese, Italy | Laboratory of Microbiology, ASST Ospedale di Circolo, Varese viale borri 57 21100 Varese, Italy | Federica Novazzi, Francesca Dragoferrante, Angelo Genoni, Daniela Dalla Gasperina, Daniele Focosi, Andreina Baj, Fabrizio Maggi |
| EPI_ISL_3462096 | AP SSO | CSIR-Centre for Cellular and Molecular Biology-INSACOG | Amareshwar Vodapalli,Ara Sreenivas,Shreekant Verma, B Himasri,Valli Nagalakshmi Undamatla,Payel Mukherjee, Lamuk Zaveri, Onkar Kulkarni,Tulasi Nagabandi, Sofia Banu, Jandhyala Sai Krishna,Priya Nurkurthy, Vidhyadhari Methuku, Sumedha Avadhanula, Archana Bharadwaj Siva, Karthik Bharadwaj Tallapaka,Rakesh K Mishra,Divya Tej Sowpati |
| EPI_ISL_3462097 | AP SSO | CSIR-Centre for Cellular and Molecular Biology-INSACOG | Priya Nurkurthy,Ara Sreenivas,Shreekant Verma, Amareshwar Vodapalli ,Lamuk Zaveri,Tulasi Nagabandi,B Himasri,Valli Nagalakshmi Undamatla, Sumedha Avadhanula, Vidhyadhari Methuku,Payel Mukherjee,Onkar Kulkarni, Jandhyala Sai Krishna,Sofia Banu, Archana Bharadwaj Siva, Karthik Bharadwaj Tallapaka,Rakesh K Mishra,Divya Tej Sowpati |
| EPI_ISL_3462098, EPI_ISL_3462099, EPI_ISL_3462100, EPI_ISL_3462101, EPI_ISL_3462102, EPI_ISL_3462103, EPI_ISL_3462104, EPI_ISL_3462105, EPI_ISL_3462106, EPI_ISL_3462107, EPI_ISL_3462108, EPI_ISL_3462109, EPI_ISL_3462110, EPI_ISL_3462111, EPI_ISL_3462112, EPI_ISL_3462113, EPI_ISL_3462114, EPI_ISL_3462115, EPI_ISL_3462116, EPI_ISL_3462117, EPI_ISL_3462118, EPI_ISL_3462119, EPI_ISL_3462120, EPI_ISL_3462121, EPI_ISL_3462122, EPI_ISL_3462123, EPI_ISL_3462124, EPI_ISL_3462125, EPI_ISL_3462126, EPI_ISL_3462127, EPI_ISL_3462128, EPI_ISL_3462129, EPI_ISL_3462130, EPI_ISL_3462131, EPI_ISL_3462132, EPI_ISL_3462133, EPI_ISL_3462134, EPI_ISL_3462135, EPI_ISL_3462136, EPI_ISL_3462137, EPI_ISL_3462138, EPI_ISL_3462139, EPI_ISL_3462140, EPI_ISL_3462141, EPI_ISL_3462143, EPI_ISL_3462144, EPI_ISL_3462145, EPI_ISL_3462146, EPI_ISL_3462147, EPI_ISL_3462148, EPI_ISL_3462149, EPI_ISL_3462150, EPI_ISL_3462151, EPI_ISL_3462152, EPI_ISL_3462153, EPI_ISL_3462154, EPI_ISL_3462155, EPI_ISL_3462156, EPI_ISL_3462157, EPI_ISL_3462158, EPI_ISL_3462159, EPI_ISL_3462160, EPI_ISL_3462161, EPI_ISL_3462162, EPI_ISL_3462163, EPI_ISL_3462164, EPI_ISL_3462165, EPI_ISL_3462166, EPI_ISL_3462167, EPI_ISL_3462168, EPI_ISL_3462169, EPI_ISL_3462170, EPI_ISL_3462171, EPI_ISL_3462172, EPI_ISL_3462173, EPI_ISL_3462174, EPI_ISL_3462175, EPI_ISL_3462176, EPI_ISL_3462177, EPI_ISL_3462178, EPI_ISL_3462179, EPI_ISL_3462180, EPI_ISL_3462181, EPI_ISL_3462182, EPI_ISL_3462183, EPI_ISL_3462184, EPI_ISL_3462185, EPI_ISL_3462186, EPI_ISL_3462187, EPI_ISL_3462188, EPI_ISL_3462189, EPI_ISL_3462190 | | | |
| see above | Communicable Disease Laboratory, Public Health Directorate | Communicable Disease Laboratory, Public Health Directorate | Alwasti,H., AlHujairi,Z., AlAbbas,Z., Marhoon,A., Almoamen,G. |

| | | | |
|---|---|---|---|
| EPI_ISL_3462329, EPI_ISL_3462330, EPI_ISL_3462331, EPI_ISL_3462332, EPI_ISL_3462333, EPI_ISL_3462334, EPI_ISL_3462335, EPI_ISL_3462336, EPI_ISL_3462337, EPI_ISL_3462338, EPI_ISL_3462339, EPI_ISL_3462340 | | | |
| see above | UCSC Genomics Institute | UCSC Genomics Institute | Mark Akeson, Ikenna Anigbogu, Eric Beraut, Isabel Bjork, Molly Cassatt-Johnstone, Terren Chang, Russell Corbett-Detig, Namrita Dhillon, Maximilian Haeussler, David Haussler, Angie Hinrichs, Miten Jain, Joshua Kapp, A. Marm Kilpatrick, Jakob McBroome, Hugh Olsen, Jeremy Sanford, Beth Shapiro, Michael Stone, Bryan Thornlow, Yatish Turakhia, Ciara Wanket |
| EPI_ISL_3462342, EPI_ISL_3462343, EPI_ISL_3462344, EPI_ISL_3462345, EPI_ISL_3462346, EPI_ISL_3462347, EPI_ISL_3462348, EPI_ISL_3462349, EPI_ISL_3462350, EPI_ISL_3462351, EPI_ISL_3462352, EPI_ISL_3462353, EPI_ISL_3462354, EPI_ISL_3462355, EPI_ISL_3462356, EPI_ISL_3462357, EPI_ISL_3462358, EPI_ISL_3462359, EPI_ISL_3462360, EPI_ISL_3462361, EPI_ISL_3462362, EPI_ISL_3462363, EPI_ISL_3462364, EPI_ISL_3462365, EPI_ISL_3462366, EPI_ISL_3462367, EPI_ISL_3462369, EPI_ISL_3462370, EPI_ISL_3462371, EPI_ISL_3462372, EPI_ISL_3462374, EPI_ISL_3462375, EPI_ISL_3462376, EPI_ISL_3462377, EPI_ISL_3462378, EPI_ISL_3462379, EPI_ISL_3462380, EPI_ISL_3462381, EPI_ISL_3462382, EPI_ISL_3462383, EPI_ISL_3462384 | | | |
| see above | Tampa General Hospital Esoteric Lab | Tampa General Hospital Esoteric Research & Development Lab | Jon Faughn, Amorce Lima, Deanna Becker, Dominic Uy, Jorge Mecias-Frias, Katie Hurless, Vicki Healer, Suzane Silbert |
| EPI_ISL_3462385, EPI_ISL_3462386, EPI_ISL_3462387, EPI_ISL_3462388, EPI_ISL_3462389, EPI_ISL_3462390, EPI_ISL_3462391, EPI_ISL_3462392, EPI_ISL_3462393, EPI_ISL_3462394, EPI_ISL_3462395, EPI_ISL_3462396, EPI_ISL_3462397, EPI_ISL_3462398, EPI_ISL_3462399, EPI_ISL_3462400, EPI_ISL_3462401, EPI_ISL_3462402, EPI_ISL_3462403, EPI_ISL_3462404, EPI_ISL_3462405, EPI_ISL_3462406, EPI_ISL_3462407, EPI_ISL_3462408, EPI_ISL_3462409, EPI_ISL_3462410, EPI_ISL_3462411, EPI_ISL_3462412, EPI_ISL_3462413, EPI_ISL_3462414, EPI_ISL_3462415, EPI_ISL_3462416, EPI_ISL_3462417, EPI_ISL_3462418, EPI_ISL_3462419, EPI_ISL_3462420, EPI_ISL_3462421, EPI_ISL_3462422, EPI_ISL_3462423, EPI_ISL_3462424, EPI_ISL_3462425, EPI_ISL_3462426, EPI_ISL_3462427 | | | |
| see above | Wyoming Public Health Laboratory | Wyoming Public Health Laboratory | Jim Mildenberger, Taylor Fearing, Channing Weber, Ashley Norberg, Chayse Rowley, Marley Goetz, Brian Dominguez, Elliot Thomasson, Sam Britz, Cari Sloma, Robert Petit, and Rob Christensen |
| EPI_ISL_3462428, EPI_ISL_3462429, EPI_ISL_3462431, EPI_ISL_3462432, EPI_ISL_3462433, EPI_ISL_3462434, EPI_ISL_3462435, EPI_ISL_3462436, EPI_ISL_3462437, EPI_ISL_3462438, EPI_ISL_3462439, EPI_ISL_3462440, EPI_ISL_3462441, EPI_ISL_3462442, EPI_ISL_3462443, EPI_ISL_3462444, EPI_ISL_3462445, EPI_ISL_3462446, EPI_ISL_3462447, EPI_ISL_3462448, EPI_ISL_3462449, EPI_ISL_3462450, EPI_ISL_3462451, EPI_ISL_3462452, EPI_ISL_3462453, EPI_ISL_3462454, EPI_ISL_3462455, EPI_ISL_3462456, EPI_ISL_3462457, EPI_ISL_3462458, EPI_ISL_3462459, EPI_ISL_3462460, EPI_ISL_3462461, EPI_ISL_3462462, EPI_ISL_3462463, EPI_ISL_3462464, EPI_ISL_3462465, EPI_ISL_3462466, EPI_ISL_3462467, EPI_ISL_3462468, EPI_ISL_3462469, EPI_ISL_3462470 | | | |
| see above | Arizona State Public Health Laboratory | Arizona State Public Health Laboratory | Trung Huynh, Jessica Escobar, Katherine Fullerton, Nobuko Fukushima, Matthew Contursi, Stacy White, Linda Getsinger, Victor Waddell |
| EPI_ISL_3462471, EPI_ISL_3462472, EPI_ISL_3462473, EPI_ISL_3462474 | California Hospital Medical Center | Los Angeles County Public Health Laboratories | P. Hemarajata et al. |
| EPI_ISL_3462476, EPI_ISL_3462477, EPI_ISL_3462478, EPI_ISL_3462480, EPI_ISL_3462481, EPI_ISL_3462482, EPI_ISL_3462483, EPI_ISL_3462484 | Palmdale Regional Medical Center | Los Angeles County Public Health Laboratories | P. Hemarajata et al. |
| EPI_ISL_3462485 | Martin Luther King, Jr. Outpatient Center | Los Angeles County Public Health Laboratories | P. Hemarajata et al. |
| EPI_ISL_3462486, EPI_ISL_3462487, EPI_ISL_3462488, EPI_ISL_3462489, EPI_ISL_3462490, EPI_ISL_3462491, EPI_ISL_3462492, EPI_ISL_3462493, EPI_ISL_3462494, EPI_ISL_3462495, EPI_ISL_3462496, EPI_ISL_3462497 | | | |
| see above | Torrance Memorial Medical Center | Los Angeles County Public Health Laboratories | P. Hemarajata et al. |
| EPI_ISL_3462498, EPI_ISL_3462499, EPI_ISL_3462500, EPI_ISL_3462501, EPI_ISL_3462502, EPI_ISL_3462503, EPI_ISL_3462504, EPI_ISL_3462505, EPI_ISL_3462506, EPI_ISL_3462507, EPI_ISL_3462508, EPI_ISL_3462509, EPI_ISL_3462510, EPI_ISL_3462511, EPI_ISL_3462512, EPI_ISL_3462513, EPI_ISL_3462514, EPI_ISL_3462515, EPI_ISL_3462516, EPI_ISL_3462517, EPI_ISL_3462518, EPI_ISL_3462519, EPI_ISL_3462520, EPI_ISL_3462521, EPI_ISL_3462522, EPI_ISL_3462523, EPI_ISL_3462524, EPI_ISL_3462525, EPI_ISL_3462526, EPI_ISL_3462527 | | | |
| see above | Department of Virology | Department of Virology | Massab Umair, Aamer Ikram, Muhammad Salman, Nazish Badar, Zaira Rehman, Muhammad Ammar, Syed Adnan Haider |
| EPI_ISL_3462578, EPI_ISL_3462738, EPI_ISL_3462739, EPI_ISL_3462740, EPI_ISL_3462741 | Central Laboratory, Bureau of Public Health (BOG) and Academic Hospital Paramaribo | Erasmus Medical Center | Bas B Oude Munnink, Cherise Beek, Consuella Partowidjojo, Dion Gajadin, Ed PF IJzerman, Emmanuelle Munger, Gary Gummels, Ingrid SK Krishnadath, Lycke Woittiez, Marion PG Koopmans, Mireille Van de Veer, Phyllis Pinas, Princes Wongsowidjojo, Radjesh Ori, Ranisha Doerbalie, Rohma Banwari, Soeradj Harkisoen, Stephen Vreden, Tilotmadebie Ramlal, Verne Nanhoe |
| EPI_ISL_3462750 | NORTHWELL HEALTH LABORATORIES | Wadsworth Center, New York State Department of Health | Kirsten St. George, Daryl M. Lamson, Alexis Russell, Matthew Shudt, Melissa A Leisner, Jonathan Plitnick, Catharine Prussing, Navjot Singh, John Kelly, Erasmus Schneider, Erica Lasek-Nesselquist |
| EPI_ISL_3462751 | ALBANY MEDICAL CENTER | Wadsworth Center, New York State Department of Health | Kirsten St. George, Daryl M. Lamson, Alexis Russell, Matthew Shudt, Melissa A Leisner, Jonathan Plitnick, Catharine Prussing, Navjot Singh, John Kelly, Erasmus Schneider, Erica Lasek-Nesselquist |
| EPI_ISL_3462752, EPI_ISL_3462753, EPI_ISL_3462754, EPI_ISL_3462755, EPI_ISL_3462756, EPI_ISL_3462757, EPI_ISL_3462759, EPI_ISL_3462760, EPI_ISL_3462761, EPI_ISL_3462762 | NORTHWELL HEALTH LABORATORIES | Wadsworth Center, New York State Department of Health | Kirsten St. George, Daryl M. Lamson, Alexis Russell, Matthew Shudt, Melissa A Leisner, Jonathan Plitnick, Catharine Prussing, Navjot Singh, John Kelly, Erasmus Schneider, Erica Lasek-Nesselquist |
| EPI_ISL_3462765 | SUNY UPSTATE MEDICAL UNIVERSITY | Wadsworth Center, New York State Department of Health | Kirsten St. George, Daryl M. Lamson, Alexis Russell, Matthew Shudt, Melissa A Leisner, Jonathan Plitnick, Catharine Prussing, Navjot Singh, John Kelly, Erasmus Schneider, Erica Lasek-Nesselquist |
| EPI_ISL_3462766 | ALBANY MEDICAL CENTER | Wadsworth Center, New York State Department of Health | Kirsten St. George, Daryl M. Lamson, Alexis Russell, Matthew Shudt, Melissa A Leisner, Jonathan Plitnick, Catharine Prussing, Navjot Singh, John Kelly, Erasmus Schneider, Erica Lasek-Nesselquist |
| EPI_ISL_3462767 | NORTHWELL HEALTH LABORATORIES | Wadsworth Center, New York State Department of Health | Kirsten St. George, Daryl M. Lamson, Alexis Russell, Matthew Shudt, Melissa A Leisner, Jonathan Plitnick, Catharine Prussing, Navjot Singh, John Kelly, Erasmus Schneider, Erica Lasek-Nesselquist |
| EPI_ISL_3462768 | STONY BROOK UNIVERSITY HOSPITAL | Wadsworth Center, New York State Department of Health | Kirsten St. George, Daryl M. Lamson, Alexis Russell, Matthew Shudt, Melissa A Leisner, Jonathan Plitnick, Catharine Prussing, Navjot Singh, John Kelly, Erasmus Schneider, Erica Lasek-Nesselquist |
| EPI_ISL_3462769, EPI_ISL_3462770, EPI_ISL_3462771 | ADIRONDACK MEDICAL CENTER | Wadsworth Center, New York State Department of Health | Kirsten St. George, Daryl M. Lamson, Alexis Russell, Matthew Shudt, Melissa A Leisner, Jonathan Plitnick, Catharine Prussing, Navjot Singh, John Kelly, Erasmus Schneider, Erica Lasek-Nesselquist |
| EPI_ISL_3462772, EPI_ISL_3462773, EPI_ISL_3462774, EPI_ISL_3462775 | ACUTIS DIAGNOSTICS | Wadsworth Center, New York State Department of Health | Kirsten St. George, Daryl M. Lamson, Alexis Russell, Matthew Shudt, Melissa A Leisner, Jonathan Plitnick, Catharine Prussing, Navjot Singh, John Kelly, Erasmus Schneider, Erica Lasek-Nesselquist |
| EPI_ISL_3462776, EPI_ISL_3462777, EPI_ISL_3462778, EPI_ISL_3462779, EPI_ISL_3462780, EPI_ISL_3462781, EPI_ISL_3462782, EPI_ISL_3462783, EPI_ISL_3462784, EPI_ISL_3462785, EPI_ISL_3462786, EPI_ISL_3462787, EPI_ISL_3462788, EPI_ISL_3462789, EPI_ISL_3462790, EPI_ISL_3462791 | | | |
| see above | SUNY UPSTATE MEDICAL UNIVERSITY | Wadsworth Center, New York State Department of Health | Kirsten St. George, Daryl M. Lamson, Alexis Russell, Matthew Shudt, Melissa A Leisner, Jonathan Plitnick, Catharine Prussing, Navjot Singh, John Kelly, Erasmus Schneider, Erica Lasek-Nesselquist |
| EPI_ISL_3462792, EPI_ISL_3462793, EPI_ISL_3462794, EPI_ISL_3462795, EPI_ISL_3462796, EPI_ISL_3462797, EPI_ISL_3462798, EPI_ISL_3462799, EPI_ISL_3462800 | STONY BROOK UNIVERSITY HOSPITAL | Wadsworth Center, New York State Department of Health | Kirsten St. George, Daryl M. Lamson, Alexis Russell, Matthew Shudt, Melissa A Leisner, Jonathan Plitnick, Catharine Prussing, Navjot Singh, John Kelly, Erasmus Schneider, Erica Lasek-Nesselquist |
| EPI_ISL_3462801, EPI_ISL_3462802, EPI_ISL_3462803, EPI_ISL_3462804, EPI_ISL_3462805, EPI_ISL_3462806 | ALBANY MEDICAL CENTER | Wadsworth Center, New York State Department of Health | Kirsten St. George, Daryl M. Lamson, Alexis Russell, Matthew Shudt, Melissa A Leisner, Jonathan Plitnick, Catharine Prussing, Navjot Singh, John Kelly, Erasmus Schneider, Erica Lasek-Nesselquist |
| EPI_ISL_3462807, EPI_ISL_3462808, EPI_ISL_3462809, EPI_ISL_3462810, EPI_ISL_3462811, EPI_ISL_3462812, EPI_ISL_3462813, EPI_ISL_3462814, EPI_ISL_3462815, EPI_ISL_3462816, EPI_ISL_3462817, EPI_ISL_3462818, EPI_ISL_3462819, EPI_ISL_3462820, EPI_ISL_3462821, EPI_ISL_3462822, EPI_ISL_3462823, EPI_ISL_3462824, EPI_ISL_3462825, EPI_ISL_3462826, EPI_ISL_3462827, EPI_ISL_3462828, EPI_ISL_3462829, EPI_ISL_3462830, EPI_ISL_3462831, EPI_ISL_3462832, EPI_ISL_3462833, EPI_ISL_3462834, EPI_ISL_3462835, EPI_ISL_3462836, EPI_ISL_3462837, EPI_ISL_3462838, EPI_ISL_3462839, EPI_ISL_3462840, EPI_ISL_3462841, EPI_ISL_3462842 | | | |
| see above | NORTHWELL HEALTH LABORATORIES | Wadsworth Center, New York State Department of Health | Kirsten St. George, Daryl M. Lamson, Alexis Russell, Matthew Shudt, Melissa A Leisner, Jonathan Plitnick, Catharine Prussing, Navjot Singh, John Kelly, Erasmus Schneider, Erica Lasek-Nesselquist |
| EPI_ISL_3462843, EPI_ISL_3462844, EPI_ISL_3462845, EPI_ISL_3462846, EPI_ISL_3462847, EPI_ISL_3462848, EPI_ISL_3462849, EPI_ISL_3462850, EPI_ISL_3462851, EPI_ISL_3462852, EPI_ISL_3462853, EPI_ISL_3462854, EPI_ISL_3462855, EPI_ISL_3462856, EPI_ISL_3462857, EPI_ISL_3462858, EPI_ISL_3462859, EPI_ISL_3462860, EPI_ISL_3462861, EPI_ISL_3462862, EPI_ISL_3462863, EPI_ISL_3462864, EPI_ISL_3462865, EPI_ISL_3462866, EPI_ISL_3462867, EPI_ISL_3462868, EPI_ISL_3462869, EPI_ISL_3462870, EPI_ISL_3462871, EPI_ISL_3462872, EPI_ISL_3462873, EPI_ISL_3462874, EPI_ISL_3462875, EPI_ISL_3462876, EPI_ISL_3462877, EPI_ISL_3462878, EPI_ISL_3462879, EPI_ISL_3462880, EPI_ISL_3462881, EPI_ISL_3462882, EPI_ISL_3462883, EPI_ISL_3462884, EPI_ISL_3462885, EPI_ISL_3462886, EPI_ISL_3462887, EPI_ISL_3462888, EPI_ISL_3462889, EPI_ISL_3462890, EPI_ISL_3462891, EPI_ISL_3462892, EPI_ISL_3462893, EPI_ISL_3462894, EPI_ISL_3462895, EPI_ISL_3462896, EPI_ISL_3462897 | | | |
| see above | Kansas Health and Environmental Lab | Kansas Health and Environmental Lab | Mike Grose, Jonathan Barnell, Meg Wise, Carrie Welch, Gary Buruss, Amanda Bradley, and Phil Adam |
| EPI_ISL_3462898, EPI_ISL_3462899, EPI_ISL_3462900, EPI_ISL_3462901, EPI_ISL_3462902, EPI_ISL_3462903, EPI_ISL_3462904, EPI_ISL_3462905, EPI_ISL_3462906, EPI_ISL_3462907 | Department of Bacteria, Parasites and Fungi, Statens Serum Institut, Copenhagen, Denmark | Statens Serum Institut Bioinformatics and Microbial Genomics | Danish Covid-19 Genome Consortium |
| EPI_ISL_3462908 | Department of Clinical Microbiology, Odense University Hospital, Odense, Denmark | Statens Serum Institut Bioinformatics and Microbial Genomics | Danish Covid-19 Genome Consortium |
| EPI_ISL_3462909, | Department of Bacteria, Parasites and Fungi, Statens Serum | Statens Serum Institut Bioinformatics and Microbial Genomics | Danish Covid-19 Genome Consortium |

| | | | |
|---|---|---|---|
| EPI_ISL_3462910 | Institut, Copenhagen, Denmark | | |
| EPI_ISL_3462911 | Arizona State Public Health Laboratory | Arizona State Public Health Laboratory | Trung Huynh, Jessica Escobar, Katherine Fullerton, Nobuko Fukushima, Matthew Contursi, Stacy White, Linda Getsinger, Victor Waddell |
| EPI_ISL_3462912 | Department of Bacteria, Parasites and Fungi, Statens Serum Institut, Copenhagen, Denmark | Statens Serum Institut Bioinformatics and Microbial Genomics | Danish Covid-19 Genome Consortium |
| EPI_ISL_3462913 | Arizona State Public Health Laboratory | Arizona State Public Health Laboratory | Trung Huynh, Jessica Escobar, Katherine Fullerton, Nobuko Fukushima, Matthew Contursi, Stacy White, Linda Getsinger, Victor Waddell |
| EPI_ISL_3462914 | Department of Bacteria, Parasites and Fungi, Statens Serum Institut, Copenhagen, Denmark | Statens Serum Institut Bioinformatics and Microbial Genomics | Danish Covid-19 Genome Consortium |
| EPI_ISL_3462915 | Arizona State Public Health Laboratory | Arizona State Public Health Laboratory | Trung Huynh, Jessica Escobar, Katherine Fullerton, Nobuko Fukushima, Matthew Contursi, Stacy White, Linda Getsinger, Victor Waddell |
| EPI_ISL_3462916 | Department of Bacteria, Parasites and Fungi, Statens Serum Institut, Copenhagen, Denmark | Statens Serum Institut Bioinformatics and Microbial Genomics | Danish Covid-19 Genome Consortium |
| EPI_ISL_3462917 | Arizona State Public Health Laboratory | Arizona State Public Health Laboratory | Trung Huynh, Jessica Escobar, Katherine Fullerton, Nobuko Fukushima, Matthew Contursi, Stacy White, Linda Getsinger, Victor Waddell |
| EPI_ISL_3462918, EPI_ISL_3462919 | Department of Bacteria, Parasites and Fungi, Statens Serum Institut, Copenhagen, Denmark | Statens Serum Institut Bioinformatics and Microbial Genomics | Danish Covid-19 Genome Consortium |
| EPI_ISL_3462920 | Arizona State Public Health Laboratory | Arizona State Public Health Laboratory | Trung Huynh, Jessica Escobar, Katherine Fullerton, Nobuko Fukushima, Matthew Contursi, Stacy White, Linda Getsinger, Victor Waddell |
| EPI_ISL_3462921 | Department of Bacteria, Parasites and Fungi, Statens Serum Institut, Copenhagen, Denmark | Statens Serum Institut Bioinformatics and Microbial Genomics | Danish Covid-19 Genome Consortium |
| EPI_ISL_3462922 | Arizona State Public Health Laboratory | Arizona State Public Health Laboratory | Trung Huynh, Jessica Escobar, Katherine Fullerton, Nobuko Fukushima, Matthew Contursi, Stacy White, Linda Getsinger, Victor Waddell |
| EPI_ISL_3462923 | Department of Bacteria, Parasites and Fungi, Statens Serum Institut, Copenhagen, Denmark | Statens Serum Institut Bioinformatics and Microbial Genomics | Danish Covid-19 Genome Consortium |
| EPI_ISL_3462924 | Arizona State Public Health Laboratory | Arizona State Public Health Laboratory | Trung Huynh, Jessica Escobar, Katherine Fullerton, Nobuko Fukushima, Matthew Contursi, Stacy White, Linda Getsinger, Victor Waddell |
| EPI_ISL_3462925 | Department of Bacteria, Parasites and Fungi, Statens Serum Institut, Copenhagen, Denmark | Statens Serum Institut Bioinformatics and Microbial Genomics | Danish Covid-19 Genome Consortium |
| EPI_ISL_3462926 | Arizona State Public Health Laboratory | Arizona State Public Health Laboratory | Trung Huynh, Jessica Escobar, Katherine Fullerton, Nobuko Fukushima, Matthew Contursi, Stacy White, Linda Getsinger, Victor Waddell |
| EPI_ISL_3462927 | Department of Bacteria, Parasites and Fungi, Statens Serum Institut, Copenhagen, Denmark | Statens Serum Institut Bioinformatics and Microbial Genomics | Danish Covid-19 Genome Consortium |
| EPI_ISL_3462928 | Arizona State Public Health Laboratory | Arizona State Public Health Laboratory | Trung Huynh, Jessica Escobar, Katherine Fullerton, Nobuko Fukushima, Matthew Contursi, Stacy White, Linda Getsinger, Victor Waddell |
| EPI_ISL_3462929 | Department of Bacteria, Parasites and Fungi, Statens Serum Institut, Copenhagen, Denmark | Statens Serum Institut Bioinformatics and Microbial Genomics | Danish Covid-19 Genome Consortium |
| EPI_ISL_3462930 | Arizona State Public Health Laboratory | Arizona State Public Health Laboratory | Trung Huynh, Jessica Escobar, Katherine Fullerton, Nobuko Fukushima, Matthew Contursi, Stacy White, Linda Getsinger, Victor Waddell |
| EPI_ISL_3462931 | Department of Bacteria, Parasites and Fungi, Statens Serum Institut, Copenhagen, Denmark | Statens Serum Institut Bioinformatics and Microbial Genomics | Danish Covid-19 Genome Consortium |
| EPI_ISL_3462932 | Arizona State Public Health Laboratory | Arizona State Public Health Laboratory | Trung Huynh, Jessica Escobar, Katherine Fullerton, Nobuko Fukushima, Matthew Contursi, Stacy White, Linda Getsinger, Victor Waddell |
| EPI_ISL_3462933 | Department of Bacteria, Parasites and Fungi, Statens Serum Institut, Copenhagen, Denmark | Statens Serum Institut Bioinformatics and Microbial Genomics | Danish Covid-19 Genome Consortium |
| EPI_ISL_3462934 | Arizona State Public Health Laboratory | Arizona State Public Health Laboratory | Trung Huynh, Jessica Escobar, Katherine Fullerton, Nobuko Fukushima, Matthew Contursi, Stacy White, Linda Getsinger, Victor Waddell |
| EPI_ISL_3462935, EPI_ISL_3462936 | Department of Bacteria, Parasites and Fungi, Statens Serum Institut, Copenhagen, Denmark | Statens Serum Institut Bioinformatics and Microbial Genomics | Danish Covid-19 Genome Consortium |
| EPI_ISL_3462937 | Arizona State Public Health Laboratory | Arizona State Public Health Laboratory | Trung Huynh, Jessica Escobar, Katherine Fullerton, Nobuko Fukushima, Matthew Contursi, Stacy White, Linda Getsinger, Victor Waddell |
| EPI_ISL_3462938 | Department of Bacteria, Parasites and Fungi, Statens Serum Institut, Copenhagen, Denmark | Statens Serum Institut Bioinformatics and Microbial Genomics | Danish Covid-19 Genome Consortium |
| EPI_ISL_3462939, EPI_ISL_3462940 | Arizona State Public Health Laboratory | Arizona State Public Health Laboratory | Trung Huynh, Jessica Escobar, Katherine Fullerton, Nobuko Fukushima, Matthew Contursi, Stacy White, Linda Getsinger, Victor Waddell |
| EPI_ISL_3462941 | Department of Bacteria, Parasites and Fungi, Statens Serum Institut, Copenhagen, Denmark | Statens Serum Institut Bioinformatics and Microbial Genomics | Danish Covid-19 Genome Consortium |
| EPI_ISL_3462942 | Arizona State Public Health Laboratory | Arizona State Public Health Laboratory | Trung Huynh, Jessica Escobar, Katherine Fullerton, Nobuko Fukushima, Matthew Contursi, Stacy White, Linda Getsinger, Victor Waddell |
| EPI_ISL_3462943 | Department of Bacteria, Parasites and Fungi, Statens Serum Institut, Copenhagen, Denmark | Statens Serum Institut Bioinformatics and Microbial Genomics | Danish Covid-19 Genome Consortium |
| EPI_ISL_3462944 | Arizona State Public Health Laboratory | Arizona State Public Health Laboratory | Trung Huynh, Jessica Escobar, Katherine Fullerton, Nobuko Fukushima, Matthew Contursi, Stacy White, Linda Getsinger, Victor Waddell |
| EPI_ISL_3462945 | Department of Bacteria, Parasites and Fungi, Statens Serum Institut, Copenhagen, Denmark | Statens Serum Institut Bioinformatics and Microbial Genomics | Danish Covid-19 Genome Consortium |
| EPI_ISL_3462946, EPI_ISL_3462947 | Arizona State Public Health Laboratory | Arizona State Public Health Laboratory | Trung Huynh, Jessica Escobar, Katherine Fullerton, Nobuko Fukushima, Matthew Contursi, Stacy White, Linda Getsinger, Victor Waddell |
| EPI_ISL_3462948 | Department of Bacteria, Parasites and Fungi, Statens Serum Institut, Copenhagen, Denmark | Statens Serum Institut Bioinformatics and Microbial Genomics | Danish Covid-19 Genome Consortium |
| EPI_ISL_3462949 | Arizona State Public Health Laboratory | Arizona State Public Health Laboratory | Trung Huynh, Jessica Escobar, Katherine Fullerton, Nobuko Fukushima, Matthew Contursi, Stacy White, Linda Getsinger, Victor Waddell |
| EPI_ISL_3462950 | Department of Bacteria, Parasites and Fungi, Statens Serum Institut, Copenhagen, Denmark | Statens Serum Institut Bioinformatics and Microbial Genomics | Danish Covid-19 Genome Consortium |
| EPI_ISL_3462951 | Arizona State Public Health Laboratory | Arizona State Public Health Laboratory | Trung Huynh, Jessica Escobar, Katherine Fullerton, Nobuko Fukushima, Matthew Contursi, Stacy White, Linda Getsinger, Victor Waddell |
| EPI_ISL_3462952 | Department of Bacteria, Parasites and Fungi, Statens Serum Institut, Copenhagen, Denmark | Statens Serum Institut Bioinformatics and Microbial Genomics | Danish Covid-19 Genome Consortium |
| EPI_ISL_3462953, EPI_ISL_3462954 | Arizona State Public Health Laboratory | Arizona State Public Health Laboratory | Trung Huynh, Jessica Escobar, Katherine Fullerton, Nobuko Fukushima, Matthew Contursi, Stacy White, Linda Getsinger, Victor Waddell |
| EPI_ISL_3462955 | Department of Bacteria, Parasites and Fungi, Statens Serum Institut, Copenhagen, Denmark | Statens Serum Institut Bioinformatics and Microbial Genomics | Danish Covid-19 Genome Consortium |
| EPI_ISL_3462956 | Arizona State Public Health Laboratory | Arizona State Public Health Laboratory | Trung Huynh, Jessica Escobar, Katherine Fullerton, Nobuko Fukushima, Matthew Contursi, Stacy White, Linda Getsinger, Victor Waddell |
| EPI_ISL_3462957 | Department of Bacteria, Parasites and Fungi, Statens Serum Institut, Copenhagen, Denmark | Statens Serum Institut Bioinformatics and Microbial Genomics | Danish Covid-19 Genome Consortium |
| EPI_ISL_3462958 | Arizona State Public Health Laboratory | Arizona State Public Health Laboratory | Trung Huynh, Jessica Escobar, Katherine Fullerton, Nobuko Fukushima, Matthew Contursi, Stacy White, Linda Getsinger, Victor Waddell |
| EPI_ISL_3462959 | Department of Bacteria, Parasites and Fungi, Statens Serum Institut, Copenhagen, Denmark | Statens Serum Institut Bioinformatics and Microbial Genomics | Danish Covid-19 Genome Consortium |
| EPI_ISL_3462960 | Arizona State Public Health Laboratory | Arizona State Public Health Laboratory | Trung Huynh, Jessica Escobar, Katherine Fullerton, Nobuko Fukushima, Matthew Contursi, Stacy White, Linda Getsinger, Victor Waddell |
| EPI_ISL_3462961 | Department of Bacteria, Parasites and Fungi, Statens Serum Institut, Copenhagen, Denmark | Statens Serum Institut Bioinformatics and Microbial Genomics | Danish Covid-19 Genome Consortium |
| EPI_ISL_3462962 | Arizona State Public Health Laboratory | Arizona State Public Health Laboratory | Trung Huynh, Jessica Escobar, Katherine Fullerton, Nobuko Fukushima, Matthew Contursi, Stacy White, Linda Getsinger, Victor Waddell |
| EPI_ISL_3462963 | Department of Bacteria, Parasites and Fungi, Statens Serum Institut, Copenhagen, Denmark | Statens Serum Institut Bioinformatics and Microbial Genomics | Danish Covid-19 Genome Consortium |
| EPI_ISL_3462964 | Arizona State Public Health Laboratory | Arizona State Public Health Laboratory | Trung Huynh, Jessica Escobar, Katherine Fullerton, Nobuko Fukushima, Matthew Contursi, Stacy White, Linda Getsinger, Victor Waddell |
| EPI_ISL_3462966, EPI_ISL_3462967, EPI_ISL_3462968, EPI_ISL_3462969, EPI_ISL_3462970, EPI_ISL_3462971, EPI_ISL_3462972, EPI_ISL_3462973, EPI_ISL_3462974, EPI_ISL_3462975, EPI_ISL_3462976, EPI_ISL_3462977, EPI_ISL_3462978, EPI_ISL_3462979, EPI_ISL_3462981, EPI_ISL_3462983, EPI_ISL_3462984, EPI_ISL_3462985, EPI_ISL_3462986, EPI_ISL_3462987, EPI_ISL_3462988, EPI_ISL_3462989, EPI_ISL_3462990, EPI_ISL_3462991, EPI_ISL_3462992, EPI_ISL_3462993, EPI_ISL_3462994, EPI_ISL_3462995, EPI_ISL_3462996, EPI_ISL_3462997, EPI_ISL_3462998, EPI_ISL_3462999, EPI_ISL_3463000, EPI_ISL_3463001, EPI_ISL_3463002, EPI_ISL_3463003, EPI_ISL_3463004, EPI_ISL_3463005, EPI_ISL_3463006, EPI_ISL_3463007, EPI_ISL_3463008, EPI_ISL_3463009, EPI_ISL_3463010, EPI_ISL_3463011, EPI_ISL_3463012, EPI_ISL_3463013, EPI_ISL_3463014, EPI_ISL_3463015, EPI_ISL_3463016, EPI_ISL_3463017, EPI_ISL_3463018, EPI_ISL_3463019, EPI_ISL_3463020, EPI_ISL_3463021, EPI_ISL_3463022, EPI_ISL_3463023, EPI_ISL_3463024, EPI_ISL_3463025, EPI_ISL_3463026, EPI_ISL_3463027, EPI_ISL_3463028, EPI_ISL_3463029, EPI_ISL_3463030, EPI_ISL_3463031, EPI_ISL_3463032, EPI_ISL_3463033, EPI_ISL_3463034, EPI_ISL_3463035, EPI_ISL_3463036, EPI_ISL_3463037, EPI_ISL_3463038, EPI_ISL_3463039, EPI_ISL_3463040, EPI_ISL_3463041, EPI_ISL_3463042, EPI_ISL_3463043, EPI_ISL_3463044, EPI_ISL_3463045, EPI_ISL_3463046, EPI_ISL_3463047, EPI_ISL_3463048, EPI_ISL_3463049, EPI_ISL_3463050, EPI_ISL_3463051, EPI_ISL_3463052, EPI_ISL_3463054, EPI_ISL_3463055, EPI_ISL_3463056, EPI_ISL_3463057, EPI_ISL_3463058, EPI_ISL_3463059 | | | |
| see above | Department of Bacteria, Parasites and Fungi, Statens Serum Institut, Copenhagen, Denmark | Statens Serum Institut Bioinformatics and Microbial Genomics | Danish Covid-19 Genome Consortium |
| EPI_ISL_3463060 | Department of Clinical Microbiology, Odense University Hospital, Odense, Denmark | Statens Serum Institut Bioinformatics and Microbial Genomics | Danish Covid-19 Genome Consortium |
| EPI_ISL_3463061, EPI_ISL_3463062, EPI_ISL_3463063, EPI_ISL_3463064, EPI_ISL_3463065, EPI_ISL_3463066, EPI_ISL_3463067, EPI_ISL_3463068, EPI_ISL_3463069, EPI_ISL_3463070, EPI_ISL_3463071, EPI_ISL_3463072, EPI_ISL_3463073, EPI_ISL_3463074, EPI_ISL_3463075, EPI_ISL_3463076, EPI_ISL_3463077, EPI_ISL_3463078, EPI_ISL_3463079, EPI_ISL_3463080, EPI_ISL_3463081, EPI_ISL_3463082, | | | |

| | | | |
|---|---|---|---|
| EPI_ISL_3463083, EPI_ISL_3463084, EPI_ISL_3463085, EPI_ISL_3463086, EPI_ISL_3463087, EPI_ISL_3463088, EPI_ISL_3463089, EPI_ISL_3463090, EPI_ISL_3463091, EPI_ISL_3463092, EPI_ISL_3463093, EPI_ISL_3463094, EPI_ISL_3463095, EPI_ISL_3463096, EPI_ISL_3463097, EPI_ISL_3463098, EPI_ISL_3463099, EPI_ISL_3463100, EPI_ISL_3463101, EPI_ISL_3463102, EPI_ISL_3463103, EPI_ISL_3463104, EPI_ISL_3463105, EPI_ISL_3463106, EPI_ISL_3463107, EPI_ISL_3463108, EPI_ISL_3463109, EPI_ISL_3463110, EPI_ISL_3463111, EPI_ISL_3463112 | | | |
| see above | Department of Bacteria, Parasites and Fungi, Statens Serum Institut, Copenhagen, Denmark | Statens Serum Institut Bioinformatics and Microbial Genomics | Danish Covid-19 Genome Consortium |
| EPI_ISL_3463113 | Department of Clinical Microbiology, Odense University Hospital, Odense, Denmark | Statens Serum Institut Bioinformatics and Microbial Genomics | Danish Covid-19 Genome Consortium |
| EPI_ISL_3463114, EPI_ISL_3463115, EPI_ISL_3463116, EPI_ISL_3463117, EPI_ISL_3463118, EPI_ISL_3463119, EPI_ISL_3463120, EPI_ISL_3463121, EPI_ISL_3463122, EPI_ISL_3463123, EPI_ISL_3463124, EPI_ISL_3463125, EPI_ISL_3463126, EPI_ISL_3463127, EPI_ISL_3463128, EPI_ISL_3463129, EPI_ISL_3463130, EPI_ISL_3463131, EPI_ISL_3463132, EPI_ISL_3463133, EPI_ISL_3463134, EPI_ISL_3463135, EPI_ISL_3463136, EPI_ISL_3463137, EPI_ISL_3463138, EPI_ISL_3463139, EPI_ISL_3463140, EPI_ISL_3463141, EPI_ISL_3463142, EPI_ISL_3463143, EPI_ISL_3463144, EPI_ISL_3463145, EPI_ISL_3463146, EPI_ISL_3463147, EPI_ISL_3463148, EPI_ISL_3463149, EPI_ISL_3463150, EPI_ISL_3463151, EPI_ISL_3463152, EPI_ISL_3463153, EPI_ISL_3463154, EPI_ISL_3463155, EPI_ISL_3463156, EPI_ISL_3463157, EPI_ISL_3463158, EPI_ISL_3463159, EPI_ISL_3463160 | | | |
| see above | Department of Bacteria, Parasites and Fungi, Statens Serum Institut, Copenhagen, Denmark | Statens Serum Institut Bioinformatics and Microbial Genomics | Danish Covid-19 Genome Consortium |
| EPI_ISL_3463161 | Department of Clinical Microbiology, Odense University Hospital, Odense, Denmark | Statens Serum Institut Bioinformatics and Microbial Genomics | Danish Covid-19 Genome Consortium |
| EPI_ISL_3463162, EPI_ISL_3463163, EPI_ISL_3463164, EPI_ISL_3463165, EPI_ISL_3463166, EPI_ISL_3463167, EPI_ISL_3463168, EPI_ISL_3463169, EPI_ISL_3463170, EPI_ISL_3463171, EPI_ISL_3463172, EPI_ISL_3463173, EPI_ISL_3463174, EPI_ISL_3463175, EPI_ISL_3463176, EPI_ISL_3463177, EPI_ISL_3463178, EPI_ISL_3463179, EPI_ISL_3463180, EPI_ISL_3463181, EPI_ISL_3463182, EPI_ISL_3463183, EPI_ISL_3463184, EPI_ISL_3463185, EPI_ISL_3463186, EPI_ISL_3463187, EPI_ISL_3463188, EPI_ISL_3463189, EPI_ISL_3463190, EPI_ISL_3463191, EPI_ISL_3463192, EPI_ISL_3463193, EPI_ISL_3463194, EPI_ISL_3463195, EPI_ISL_3463196, EPI_ISL_3463197, EPI_ISL_3463198, EPI_ISL_3463199, EPI_ISL_3463200, EPI_ISL_3463201, EPI_ISL_3463202, EPI_ISL_3463203, EPI_ISL_3463204, EPI_ISL_3463205, EPI_ISL_3463206, EPI_ISL_3463207, EPI_ISL_3463208, EPI_ISL_3463209 | | | |
| see above | Department of Bacteria, Parasites and Fungi, Statens Serum Institut, Copenhagen, Denmark | Statens Serum Institut Bioinformatics and Microbial Genomics | Danish Covid-19 Genome Consortium |
| EPI_ISL_3463210 | Department of Clinical Microbiology, Odense University Hospital, Odense, Denmark | Statens Serum Institut Bioinformatics and Microbial Genomics | Danish Covid-19 Genome Consortium |
| EPI_ISL_3463211, EPI_ISL_3463212, EPI_ISL_3463213, EPI_ISL_3463214, EPI_ISL_3463215, EPI_ISL_3463216, EPI_ISL_3463217, EPI_ISL_3463219, EPI_ISL_3463220, EPI_ISL_3463221, EPI_ISL_3463222, EPI_ISL_3463223, EPI_ISL_3463224, EPI_ISL_3463225, EPI_ISL_3463226, EPI_ISL_3463227, EPI_ISL_3463228, EPI_ISL_3463229, EPI_ISL_3463230, EPI_ISL_3463231, EPI_ISL_3463232, EPI_ISL_3463233, EPI_ISL_3463234, EPI_ISL_3463235, EPI_ISL_3463236, EPI_ISL_3463237, EPI_ISL_3463238, EPI_ISL_3463239, EPI_ISL_3463240, EPI_ISL_3463241, EPI_ISL_3463242, EPI_ISL_3463243, EPI_ISL_3463244, EPI_ISL_3463245, EPI_ISL_3463246, EPI_ISL_3463247, EPI_ISL_3463248, EPI_ISL_3463249, EPI_ISL_3463250, EPI_ISL_3463251, EPI_ISL_3463252, EPI_ISL_3463253, EPI_ISL_3463254, EPI_ISL_3463255, EPI_ISL_3463256, EPI_ISL_3463257, EPI_ISL_3463258, EPI_ISL_3463259, EPI_ISL_3463260, EPI_ISL_3463261, EPI_ISL_3463262, EPI_ISL_3463263, EPI_ISL_3463264, EPI_ISL_3463265, EPI_ISL_3463266, EPI_ISL_3463267, EPI_ISL_3463268, EPI_ISL_3463269, EPI_ISL_3463270, EPI_ISL_3463271, EPI_ISL_3463272, EPI_ISL_3463273, EPI_ISL_3463274, EPI_ISL_3463275, EPI_ISL_3463276, EPI_ISL_3463277, EPI_ISL_3463278, EPI_ISL_3463279 | | | |
| see above | Department of Bacteria, Parasites and Fungi, Statens Serum Institut, Copenhagen, Denmark | Statens Serum Institut Bioinformatics and Microbial Genomics | Danish Covid-19 Genome Consortium |
| EPI_ISL_3463280 | Department of Clinical Microbiology, Odense University Hospital, Odense, Denmark | Statens Serum Institut Bioinformatics and Microbial Genomics | Danish Covid-19 Genome Consortium |
| EPI_ISL_3463281, EPI_ISL_3463282, EPI_ISL_3463283, EPI_ISL_3463284, EPI_ISL_3463285, EPI_ISL_3463286, EPI_ISL_3463287, EPI_ISL_3463288, EPI_ISL_3463289, EPI_ISL_3463290, EPI_ISL_3463291, EPI_ISL_3463292, EPI_ISL_3463293, EPI_ISL_3463294, EPI_ISL_3463295, EPI_ISL_3463296, EPI_ISL_3463297, EPI_ISL_3463298, EPI_ISL_3463299, EPI_ISL_3463300, EPI_ISL_3463301, EPI_ISL_3463302, EPI_ISL_3463303, EPI_ISL_3463304, EPI_ISL_3463305, EPI_ISL_3463306, EPI_ISL_3463307, EPI_ISL_3463308, EPI_ISL_3463309, EPI_ISL_3463310, EPI_ISL_3463311, EPI_ISL_3463312, EPI_ISL_3463313, EPI_ISL_3463314, EPI_ISL_3463315, EPI_ISL_3463316, EPI_ISL_3463317, EPI_ISL_3463318, EPI_ISL_3463319, EPI_ISL_3463320, EPI_ISL_3463321, EPI_ISL_3463322, EPI_ISL_3463323, EPI_ISL_3463324 | | | |
| see above | Department of Bacteria, Parasites and Fungi, Statens Serum Institut, Copenhagen, Denmark | Statens Serum Institut Bioinformatics and Microbial Genomics | Danish Covid-19 Genome Consortium |
| EPI_ISL_3463325 | Department of Clinical Microbiology, Odense University Hospital, Odense, Denmark | Statens Serum Institut Bioinformatics and Microbial Genomics | Danish Covid-19 Genome Consortium |
| EPI_ISL_3463326, EPI_ISL_3463327, EPI_ISL_3463328, EPI_ISL_3463329, EPI_ISL_3463330, EPI_ISL_3463331, EPI_ISL_3463332, EPI_ISL_3463333, EPI_ISL_3463334, EPI_ISL_3463335, EPI_ISL_3463336, EPI_ISL_3463337, EPI_ISL_3463338, EPI_ISL_3463339, EPI_ISL_3463340, EPI_ISL_3463341, EPI_ISL_3463342, EPI_ISL_3463343, EPI_ISL_3463344, EPI_ISL_3463345, EPI_ISL_3463346, EPI_ISL_3463347, EPI_ISL_3463348, EPI_ISL_3463349, EPI_ISL_3463350, EPI_ISL_3463351, EPI_ISL_3463352, EPI_ISL_3463353, EPI_ISL_3463354, EPI_ISL_3463355, EPI_ISL_3463356, EPI_ISL_3463357, EPI_ISL_3463358, EPI_ISL_3463360, EPI_ISL_3463362, EPI_ISL_3463364, EPI_ISL_3463365, EPI_ISL_3463368, EPI_ISL_3463372, EPI_ISL_3463374, EPI_ISL_3463376, EPI_ISL_3463378, EPI_ISL_3463382, EPI_ISL_3463384, EPI_ISL_3463386 | | | |
| see above | Department of Bacteria, Parasites and Fungi, Statens Serum Institut, Copenhagen, Denmark | Statens Serum Institut Bioinformatics and Microbial Genomics | Danish Covid-19 Genome Consortium |
| EPI_ISL_3463387 | Department of Clinical Microbiology, Odense University Hospital, Odense, Denmark | Statens Serum Institut Bioinformatics and Microbial Genomics | Danish Covid-19 Genome Consortium |
| EPI_ISL_3463389, EPI_ISL_3463391, EPI_ISL_3463393, EPI_ISL_3463395, EPI_ISL_3463397, EPI_ISL_3463400, EPI_ISL_3463402, EPI_ISL_3463404, EPI_ISL_3463406, EPI_ISL_3463408, EPI_ISL_3463410, EPI_ISL_3463412, EPI_ISL_3463415, EPI_ISL_3463417, EPI_ISL_3463419, EPI_ISL_3463422, EPI_ISL_3463424, EPI_ISL_3463426, EPI_ISL_3463428, EPI_ISL_3463429, EPI_ISL_3463431, EPI_ISL_3463433, EPI_ISL_3463436, EPI_ISL_3463438, EPI_ISL_3463439, EPI_ISL_3463441, EPI_ISL_3463443, EPI_ISL_3463445, EPI_ISL_3463447, EPI_ISL_3463449, EPI_ISL_3463451, EPI_ISL_3463453, EPI_ISL_3463455, EPI_ISL_3463456, EPI_ISL_3463458, EPI_ISL_3463460, EPI_ISL_3463461, EPI_ISL_3463463, EPI_ISL_3463465, EPI_ISL_3463467, EPI_ISL_3463469, EPI_ISL_3463471, EPI_ISL_3463473, EPI_ISL_3463475, EPI_ISL_3463477, EPI_ISL_3463479 | | | |
| see above | Department of Bacteria, Parasites and Fungi, Statens Serum Institut, Copenhagen, Denmark | Statens Serum Institut Bioinformatics and Microbial Genomics | Danish Covid-19 Genome Consortium |
| EPI_ISL_3463481 | Department of Clinical Microbiology, Odense University Hospital, Odense, Denmark | Statens Serum Institut Bioinformatics and Microbial Genomics | Danish Covid-19 Genome Consortium |
| EPI_ISL_3463483, EPI_ISL_3463484, EPI_ISL_3463488, EPI_ISL_3463490, EPI_ISL_3463496, EPI_ISL_3463498, EPI_ISL_3463500, EPI_ISL_3463503, EPI_ISL_3463504, EPI_ISL_3463506, EPI_ISL_3463508, EPI_ISL_3463510, EPI_ISL_3463512, EPI_ISL_3463514, EPI_ISL_3463516, EPI_ISL_3463518, EPI_ISL_3463520, EPI_ISL_3463522, EPI_ISL_3463523, EPI_ISL_3463525, EPI_ISL_3463527, EPI_ISL_3463529, EPI_ISL_3463531, EPI_ISL_3463534, EPI_ISL_3463536, EPI_ISL_3463538, EPI_ISL_3463540, EPI_ISL_3463542, EPI_ISL_3463544, EPI_ISL_3463546, EPI_ISL_3463547, EPI_ISL_3463549, EPI_ISL_3463551, EPI_ISL_3463554, EPI_ISL_3463557, EPI_ISL_3463559, EPI_ISL_3463561, EPI_ISL_3463562, EPI_ISL_3463565, EPI_ISL_3463567, EPI_ISL_3463569, EPI_ISL_3463570, EPI_ISL_3463573, EPI_ISL_3463575, EPI_ISL_3463577, EPI_ISL_3463579, EPI_ISL_3463582, EPI_ISL_3463584, EPI_ISL_3463586, EPI_ISL_3463588, EPI_ISL_3463590, EPI_ISL_3463592, EPI_ISL_3463594, EPI_ISL_3463596, EPI_ISL_3463598, EPI_ISL_3463600, EPI_ISL_3463602, EPI_ISL_3463604, EPI_ISL_3463606, EPI_ISL_3463607, EPI_ISL_3463609, EPI_ISL_3463611, EPI_ISL_3463614, EPI_ISL_3463616, EPI_ISL_3463618, EPI_ISL_3463620, EPI_ISL_3463622, EPI_ISL_3463624, EPI_ISL_3463626, EPI_ISL_3463628, EPI_ISL_3463630, EPI_ISL_3463632, EPI_ISL_3463634, EPI_ISL_3463636, EPI_ISL_3463638, EPI_ISL_3463640, EPI_ISL_3463642, EPI_ISL_3463644, EPI_ISL_3463646, EPI_ISL_3463647, EPI_ISL_3463648, EPI_ISL_3463649, EPI_ISL_3463650, EPI_ISL_3463651, EPI_ISL_3463652, EPI_ISL_3463653 | | | |
| see above | Department of Bacteria, Parasites and Fungi, Statens Serum Institut, Copenhagen, Denmark | Statens Serum Institut Bioinformatics and Microbial Genomics | Danish Covid-19 Genome Consortium |
| EPI_ISL_3463654 | Department of Clinical Microbiology, Odense University Hospital, Odense, Denmark | Statens Serum Institut Bioinformatics and Microbial Genomics | Danish Covid-19 Genome Consortium |
| EPI_ISL_3463655, EPI_ISL_3463657, EPI_ISL_3463658, EPI_ISL_3463659, EPI_ISL_3463660, EPI_ISL_3463661, EPI_ISL_3463662, EPI_ISL_3463663, EPI_ISL_3463664, EPI_ISL_3463665, EPI_ISL_3463666 | | | |
| see above | Department of Bacteria, Parasites and Fungi, Statens Serum Institut, Copenhagen, Denmark | Statens Serum Institut Bioinformatics and Microbial Genomics | Danish Covid-19 Genome Consortium |
| EPI_ISL_3463667 | Department of Clinical Microbiology, Odense University Hospital, Odense, Denmark | Statens Serum Institut Bioinformatics and Microbial Genomics | Danish Covid-19 Genome Consortium |
| EPI_ISL_3463668, EPI_ISL_3463669, EPI_ISL_3463670, EPI_ISL_3463671, EPI_ISL_3463672, EPI_ISL_3463673, EPI_ISL_3463674, EPI_ISL_3463675, EPI_ISL_3463676, EPI_ISL_3463677, EPI_ISL_3463678, EPI_ISL_3463679, EPI_ISL_3463680, EPI_ISL_3463681, EPI_ISL_3463682, EPI_ISL_3463683, EPI_ISL_3463684, EPI_ISL_3463685, EPI_ISL_3463686, EPI_ISL_3463687, EPI_ISL_3463688 | | | |
| see above | Department of Bacteria, Parasites and Fungi, Statens Serum Institut, Copenhagen, Denmark | Statens Serum Institut Bioinformatics and Microbial Genomics | Danish Covid-19 Genome Consortium |
| EPI_ISL_3463689 | Department of Clinical Microbiology, Odense University Hospital, Odense, Denmark | Statens Serum Institut Bioinformatics and Microbial Genomics | Danish Covid-19 Genome Consortium |
| EPI_ISL_3463691, EPI_ISL_3463692, EPI_ISL_3463693, EPI_ISL_3463694, EPI_ISL_3463695, EPI_ISL_3463696, EPI_ISL_3463697, EPI_ISL_3463698, EPI_ISL_3463699, EPI_ISL_3463700, EPI_ISL_3463701, EPI_ISL_3463703 | | | |
| see above | Department of Bacteria, Parasites and Fungi, Statens Serum Institut, Copenhagen, Denmark | Statens Serum Institut Bioinformatics and Microbial Genomics | Danish Covid-19 Genome Consortium |
| EPI_ISL_3463704 | State Hygienic Laboratory at the University of Iowa | State Hygienic Laboratory at the University of Iowa | Valerie Reeb, Erik Twait, Anna Yakos, Mariah Knutson, Kris Eveland, Kristen Zanon, Wes Hottel, Alankar Kampoowale |
| EPI_ISL_3463705 | Department of Bacteria, Parasites and Fungi, Statens Serum Institut, Copenhagen, Denmark | Statens Serum Institut Bioinformatics and Microbial Genomics | Danish Covid-19 Genome Consortium |
| EPI_ISL_3463706 | State Hygienic Laboratory at the University of Iowa | State Hygienic Laboratory at the University of Iowa | Valerie Reeb, Erik Twait, Anna Yakos, Mariah Knutson, Kris Eveland, Kristen Zanon, Wes Hottel, Alankar Kampoowale |
| EPI_ISL_3463707 | Department of Bacteria, Parasites and Fungi, Statens Serum Institut, Copenhagen, Denmark | Statens Serum Institut Bioinformatics and Microbial Genomics | Danish Covid-19 Genome Consortium |
| EPI_ISL_3463708 | State Hygienic Laboratory at the University of Iowa | State Hygienic Laboratory at the University of Iowa | Valerie Reeb, Erik Twait, Anna Yakos, Mariah Knutson, Kris Eveland, Kristen Zanon, Wes Hottel, Alankar Kampoowale |
| EPI_ISL_3463709, EPI_ISL_3463710, EPI_ISL_3463712, EPI_ISL_3463714, EPI_ISL_3463715, EPI_ISL_3463716, EPI_ISL_3463718, EPI_ISL_3463720 | Department of Bacteria, Parasites and Fungi, Statens Serum Institut, Copenhagen, Denmark | Statens Serum Institut Bioinformatics and Microbial Genomics | Danish Covid-19 Genome Consortium |
| EPI_ISL_3463721 | State Hygienic Laboratory at the University of Iowa | State Hygienic Laboratory at the University of Iowa | Valerie Reeb, Erik Twait, Anna Yakos, Mariah Knutson, Kris Eveland, Kristen Zanon, Wes Hottel, Alankar Kampoowale |
| EPI_ISL_3463722 | Department of Bacteria, Parasites and Fungi, Statens Serum Institut, Copenhagen, Denmark | Statens Serum Institut Bioinformatics and Microbial Genomics | Danish Covid-19 Genome Consortium |
| EPI_ISL_3463723 | State Hygienic Laboratory at the University of Iowa | State Hygienic Laboratory at the University of Iowa | Valerie Reeb, Erik Twait, Anna Yakos, Mariah Knutson, Kris Eveland, Kristen Zanon, Wes Hottel, Alankar Kampoowale |
| EPI_ISL_3463724 | Department of Bacteria, Parasites and Fungi, Statens Serum Institut, Copenhagen, Denmark | Statens Serum Institut Bioinformatics and Microbial Genomics | Danish Covid-19 Genome Consortium |
| EPI_ISL_3463725 | State Hygienic Laboratory at the University of Iowa | State Hygienic Laboratory at the University of Iowa | Valerie Reeb, Erik Twait, Anna Yakos, Mariah Knutson, Kris Eveland, Kristen Zanon, Wes Hottel, Alankar Kampoowale |

| | | | |
|---|---|---|---|
| EPI_ISL_3463726 | Department of Bacteria, Parasites and Fungi, Statens Serum Institut, Copenhagen, Denmark | Statens Serum Institut Bioinformatics and Microbial Genomics | Danish Covid-19 Genome Consortium |
| EPI_ISL_3463727 | State Hygienic Laboratory at the University of Iowa | State Hygienic Laboratory at the University of Iowa | Valerie Reeb, Erik Twait, Anna Yakos, Mariah Knutson, Kris Eveland, Kristen Zanon, Wes Hottel, Alankar Kampoowale |
| EPI_ISL_3463728, EPI_ISL_3463729 | Department of Bacteria, Parasites and Fungi, Statens Serum Institut, Copenhagen, Denmark | Statens Serum Institut Bioinformatics and Microbial Genomics | Danish Covid-19 Genome Consortium |
| EPI_ISL_3463730 | State Hygienic Laboratory at the University of Iowa | State Hygienic Laboratory at the University of Iowa | Valerie Reeb, Erik Twait, Anna Yakos, Mariah Knutson, Kris Eveland, Kristen Zanon, Wes Hottel, Alankar Kampoowale |
| EPI_ISL_3463731 | Department of Bacteria, Parasites and Fungi, Statens Serum Institut, Copenhagen, Denmark | Statens Serum Institut Bioinformatics and Microbial Genomics | Danish Covid-19 Genome Consortium |
| EPI_ISL_3463732 | State Hygienic Laboratory at the University of Iowa | State Hygienic Laboratory at the University of Iowa | Valerie Reeb, Erik Twait, Anna Yakos, Mariah Knutson, Kris Eveland, Kristen Zanon, Wes Hottel, Alankar Kampoowale |
| EPI_ISL_3463733 | Department of Bacteria, Parasites and Fungi, Statens Serum Institut, Copenhagen, Denmark | Statens Serum Institut Bioinformatics and Microbial Genomics | Danish Covid-19 Genome Consortium |
| EPI_ISL_3463734 | State Hygienic Laboratory at the University of Iowa | State Hygienic Laboratory at the University of Iowa | Valerie Reeb, Erik Twait, Anna Yakos, Mariah Knutson, Kris Eveland, Kristen Zanon, Wes Hottel, Alankar Kampoowale |
| EPI_ISL_3463735 | Department of Clinical Microbiology, Odense University Hospital, Odense, Denmark | Statens Serum Institut Bioinformatics and Microbial Genomics | Danish Covid-19 Genome Consortium |
| EPI_ISL_3463736 | Department of Bacteria, Parasites and Fungi, Statens Serum Institut, Copenhagen, Denmark | Statens Serum Institut Bioinformatics and Microbial Genomics | Danish Covid-19 Genome Consortium |
| EPI_ISL_3463737 | State Hygienic Laboratory at the University of Iowa | State Hygienic Laboratory at the University of Iowa | Valerie Reeb, Erik Twait, Anna Yakos, Mariah Knutson, Kris Eveland, Kristen Zanon, Wes Hottel, Alankar Kampoowale |
| EPI_ISL_3463738 | Department of Bacteria, Parasites and Fungi, Statens Serum Institut, Copenhagen, Denmark | Statens Serum Institut Bioinformatics and Microbial Genomics | Danish Covid-19 Genome Consortium |
| EPI_ISL_3463739 | State Hygienic Laboratory at the University of Iowa | State Hygienic Laboratory at the University of Iowa | Valerie Reeb, Erik Twait, Anna Yakos, Mariah Knutson, Kris Eveland, Kristen Zanon, Wes Hottel, Alankar Kampoowale |
| EPI_ISL_3463740 | Department of Bacteria, Parasites and Fungi, Statens Serum Institut, Copenhagen, Denmark | Statens Serum Institut Bioinformatics and Microbial Genomics | Danish Covid-19 Genome Consortium |
| EPI_ISL_3463741 | State Hygienic Laboratory at the University of Iowa | State Hygienic Laboratory at the University of Iowa | Valerie Reeb, Erik Twait, Anna Yakos, Mariah Knutson, Kris Eveland, Kristen Zanon, Wes Hottel, Alankar Kampoowale |
| EPI_ISL_3463742 | Department of Bacteria, Parasites and Fungi, Statens Serum Institut, Copenhagen, Denmark | Statens Serum Institut Bioinformatics and Microbial Genomics | Danish Covid-19 Genome Consortium |
| EPI_ISL_3463743 | State Hygienic Laboratory at the University of Iowa | State Hygienic Laboratory at the University of Iowa | Valerie Reeb, Erik Twait, Anna Yakos, Mariah Knutson, Kris Eveland, Kristen Zanon, Wes Hottel, Alankar Kampoowale |
| EPI_ISL_3463744, EPI_ISL_3463745, EPI_ISL_3463746 | Department of Bacteria, Parasites and Fungi, Statens Serum Institut, Copenhagen, Denmark | Statens Serum Institut Bioinformatics and Microbial Genomics | Danish Covid-19 Genome Consortium |
| EPI_ISL_3463747 | State Hygienic Laboratory at the University of Iowa | State Hygienic Laboratory at the University of Iowa | Valerie Reeb, Erik Twait, Anna Yakos, Mariah Knutson, Kris Eveland, Kristen Zanon, Wes Hottel, Alankar Kampoowale |
| EPI_ISL_3463748 | Department of Bacteria, Parasites and Fungi, Statens Serum Institut, Copenhagen, Denmark | Statens Serum Institut Bioinformatics and Microbial Genomics | Danish Covid-19 Genome Consortium |
| EPI_ISL_3463749 | State Hygienic Laboratory at the University of Iowa | State Hygienic Laboratory at the University of Iowa | Valerie Reeb, Erik Twait, Anna Yakos, Mariah Knutson, Kris Eveland, Kristen Zanon, Wes Hottel, Alankar Kampoowale |
| EPI_ISL_3463750 | Department of Bacteria, Parasites and Fungi, Statens Serum Institut, Copenhagen, Denmark | Statens Serum Institut Bioinformatics and Microbial Genomics | Danish Covid-19 Genome Consortium |
| EPI_ISL_3463751 | State Hygienic Laboratory at the University of Iowa | State Hygienic Laboratory at the University of Iowa | Valerie Reeb, Erik Twait, Anna Yakos, Mariah Knutson, Kris Eveland, Kristen Zanon, Wes Hottel, Alankar Kampoowale |
| EPI_ISL_3463752, EPI_ISL_3463753 | Department of Bacteria, Parasites and Fungi, Statens Serum Institut, Copenhagen, Denmark | Statens Serum Institut Bioinformatics and Microbial Genomics | Danish Covid-19 Genome Consortium |
| EPI_ISL_3463754 | State Hygienic Laboratory at the University of Iowa | State Hygienic Laboratory at the University of Iowa | Valerie Reeb, Erik Twait, Anna Yakos, Mariah Knutson, Kris Eveland, Kristen Zanon, Wes Hottel, Alankar Kampoowale |
| EPI_ISL_3463755 | Department of Bacteria, Parasites and Fungi, Statens Serum Institut, Copenhagen, Denmark | Statens Serum Institut Bioinformatics and Microbial Genomics | Danish Covid-19 Genome Consortium |
| EPI_ISL_3463756 | State Hygienic Laboratory at the University of Iowa | State Hygienic Laboratory at the University of Iowa | Valerie Reeb, Erik Twait, Anna Yakos, Mariah Knutson, Kris Eveland, Kristen Zanon, Wes Hottel, Alankar Kampoowale |
| EPI_ISL_3463757 | Department of Bacteria, Parasites and Fungi, Statens Serum Institut, Copenhagen, Denmark | Statens Serum Institut Bioinformatics and Microbial Genomics | Danish Covid-19 Genome Consortium |
| EPI_ISL_3463758 | State Hygienic Laboratory at the University of Iowa | State Hygienic Laboratory at the University of Iowa | Valerie Reeb, Erik Twait, Anna Yakos, Mariah Knutson, Kris Eveland, Kristen Zanon, Wes Hottel, Alankar Kampoowale |
| EPI_ISL_3463759 | Department of Bacteria, Parasites and Fungi, Statens Serum Institut, Copenhagen, Denmark | Statens Serum Institut Bioinformatics and Microbial Genomics | Danish Covid-19 Genome Consortium |
| EPI_ISL_3463760 | State Hygienic Laboratory at the University of Iowa | State Hygienic Laboratory at the University of Iowa | Valerie Reeb, Erik Twait, Anna Yakos, Mariah Knutson, Kris Eveland, Kristen Zanon, Wes Hottel, Alankar Kampoowale |
| EPI_ISL_3463761 | Department of Bacteria, Parasites and Fungi, Statens Serum Institut, Copenhagen, Denmark | Statens Serum Institut Bioinformatics and Microbial Genomics | Danish Covid-19 Genome Consortium |
| EPI_ISL_3463762 | State Hygienic Laboratory at the University of Iowa | State Hygienic Laboratory at the University of Iowa | Valerie Reeb, Erik Twait, Anna Yakos, Mariah Knutson, Kris Eveland, Kristen Zanon, Wes Hottel, Alankar Kampoowale |
| EPI_ISL_3463763 | Department of Bacteria, Parasites and Fungi, Statens Serum Institut, Copenhagen, Denmark | Statens Serum Institut Bioinformatics and Microbial Genomics | Danish Covid-19 Genome Consortium |
| EPI_ISL_3463764 | State Hygienic Laboratory at the University of Iowa | State Hygienic Laboratory at the University of Iowa | Valerie Reeb, Erik Twait, Anna Yakos, Mariah Knutson, Kris Eveland, Kristen Zanon, Wes Hottel, Alankar Kampoowale |
| EPI_ISL_3463765 | Department of Bacteria, Parasites and Fungi, Statens Serum Institut, Copenhagen, Denmark | Statens Serum Institut Bioinformatics and Microbial Genomics | Danish Covid-19 Genome Consortium |
| EPI_ISL_3463766 | State Hygienic Laboratory at the University of Iowa | State Hygienic Laboratory at the University of Iowa | Valerie Reeb, Erik Twait, Anna Yakos, Mariah Knutson, Kris Eveland, Kristen Zanon, Wes Hottel, Alankar Kampoowale |
| EPI_ISL_3463767, EPI_ISL_3463768 | Department of Bacteria, Parasites and Fungi, Statens Serum Institut, Copenhagen, Denmark | Statens Serum Institut Bioinformatics and Microbial Genomics | Danish Covid-19 Genome Consortium |
| EPI_ISL_3463769 | State Hygienic Laboratory at the University of Iowa | State Hygienic Laboratory at the University of Iowa | Valerie Reeb, Erik Twait, Anna Yakos, Mariah Knutson, Kris Eveland, Kristen Zanon, Wes Hottel, Alankar Kampoowale |
| EPI_ISL_3463770, EPI_ISL_3463771, EPI_ISL_3463772, EPI_ISL_3463773, EPI_ISL_3463774, EPI_ISL_3463775, EPI_ISL_3463776, EPI_ISL_3463777, EPI_ISL_3463778, EPI_ISL_3463779, EPI_ISL_3463780, EPI_ISL_3463781, EPI_ISL_3463782, EPI_ISL_3463783, EPI_ISL_3463784, EPI_ISL_3463785, EPI_ISL_3463786, EPI_ISL_3463787, EPI_ISL_3463788, EPI_ISL_3463789, EPI_ISL_3463790, EPI_ISL_3463791, EPI_ISL_3463792, EPI_ISL_3463793, EPI_ISL_3463794, EPI_ISL_3463795, EPI_ISL_3463796, EPI_ISL_3463797, EPI_ISL_3463798, EPI_ISL_3463799, EPI_ISL_3463800, EPI_ISL_3463801, EPI_ISL_3463802, EPI_ISL_3463803, EPI_ISL_3463804, EPI_ISL_3463805, EPI_ISL_3463806, EPI_ISL_3463807, EPI_ISL_3463808, EPI_ISL_3463809, EPI_ISL_3463810, EPI_ISL_3463811, EPI_ISL_3463812, EPI_ISL_3463813, EPI_ISL_3463814, EPI_ISL_3463815, EPI_ISL_3463816, EPI_ISL_3463817, EPI_ISL_3463818, EPI_ISL_3463819, EPI_ISL_3463820, EPI_ISL_3463821, EPI_ISL_3463822, EPI_ISL_3463824, EPI_ISL_3463825, EPI_ISL_3463826, EPI_ISL_3463827, EPI_ISL_3463828, EPI_ISL_3463829, EPI_ISL_3463830, EPI_ISL_3463831, EPI_ISL_3463832, EPI_ISL_3463833, EPI_ISL_3463834, EPI_ISL_3463835, EPI_ISL_3463836, EPI_ISL_3463837, EPI_ISL_3463838, EPI_ISL_3463839, EPI_ISL_3463840, EPI_ISL_3463841, EPI_ISL_3463842, EPI_ISL_3463843, EPI_ISL_3463844, EPI_ISL_3463845, EPI_ISL_3463846, EPI_ISL_3463847, EPI_ISL_3463848, EPI_ISL_3463849, EPI_ISL_3463850, EPI_ISL_3463851, EPI_ISL_3463852, EPI_ISL_3463853, EPI_ISL_3463854, EPI_ISL_3463855, EPI_ISL_3463856, EPI_ISL_3463857, EPI_ISL_3463858, EPI_ISL_3463859, EPI_ISL_3463860, EPI_ISL_3463862, EPI_ISL_3463863, EPI_ISL_3463864, EPI_ISL_3463865, EPI_ISL_3463866, EPI_ISL_3463867, EPI_ISL_3463868, EPI_ISL_3463869, EPI_ISL_3463870, EPI_ISL_3463871, EPI_ISL_3463872, EPI_ISL_3463873, EPI_ISL_3463874, EPI_ISL_3463875, EPI_ISL_3463876, EPI_ISL_3463877, EPI_ISL_3463878, EPI_ISL_3463879, EPI_ISL_3463880, EPI_ISL_3463881, EPI_ISL_3463882, EPI_ISL_3463883, EPI_ISL_3463884, EPI_ISL_3463885 | | | |
| see above | Department of Bacteria, Parasites and Fungi, Statens Serum Institut, Copenhagen, Denmark | Statens Serum Institut Bioinformatics and Microbial Genomics | Danish Covid-19 Genome Consortium |
| EPI_ISL_3463886 | Department of Clinical Microbiology, Odense University Hospital, Odense, Denmark | Statens Serum Institut Bioinformatics and Microbial Genomics | Danish Covid-19 Genome Consortium |
| EPI_ISL_3463887, EPI_ISL_3463888, EPI_ISL_3463889, EPI_ISL_3463890, EPI_ISL_3463891, EPI_ISL_3463892, EPI_ISL_3463893, EPI_ISL_3463894, EPI_ISL_3463895, EPI_ISL_3463896, EPI_ISL_3463897, EPI_ISL_3463898, EPI_ISL_3463899, EPI_ISL_3463900, EPI_ISL_3463901, EPI_ISL_3463902, EPI_ISL_3463903, EPI_ISL_3463904, EPI_ISL_3463905, EPI_ISL_3463906, EPI_ISL_3463907, EPI_ISL_3463908, EPI_ISL_3463909, EPI_ISL_3463910, EPI_ISL_3463911, EPI_ISL_3463912, EPI_ISL_3463913, EPI_ISL_3463914, EPI_ISL_3463915, EPI_ISL_3463916, EPI_ISL_3463917 | | | |
| see above | Department of Bacteria, Parasites and Fungi, Statens Serum Institut, Copenhagen, Denmark | Statens Serum Institut Bioinformatics and Microbial Genomics | Danish Covid-19 Genome Consortium |
| EPI_ISL_3463918 | State Hygienic Laboratory at the University of Iowa | State Hygienic Laboratory at the University of Iowa | Valerie Reeb, Erik Twait, Anna Yakos, Mariah Knutson, Kris Eveland, Kristen Zanon, Wes Hottel, Alankar Kampoowale |
| EPI_ISL_3463919 | Department of Bacteria, Parasites and Fungi, Statens Serum Institut, Copenhagen, Denmark | Statens Serum Institut Bioinformatics and Microbial Genomics | Danish Covid-19 Genome Consortium |
| EPI_ISL_3463920 | State Hygienic Laboratory at the University of Iowa | State Hygienic Laboratory at the University of Iowa | Valerie Reeb, Erik Twait, Anna Yakos, Mariah Knutson, Kris Eveland, Kristen Zanon, Wes Hottel, Alankar Kampoowale |
| EPI_ISL_3463921, EPI_ISL_3463922 | Department of Bacteria, Parasites and Fungi, Statens Serum Institut, Copenhagen, Denmark | Statens Serum Institut Bioinformatics and Microbial Genomics | Danish Covid-19 Genome Consortium |
| EPI_ISL_3463923 | State Hygienic Laboratory at the University of Iowa | State Hygienic Laboratory at the University of Iowa | Valerie Reeb, Erik Twait, Anna Yakos, Mariah Knutson, Kris Eveland, Kristen Zanon, Wes Hottel, Alankar Kampoowale |
| EPI_ISL_3463924 | Department of Bacteria, Parasites and Fungi, Statens Serum Institut, Copenhagen, Denmark | Statens Serum Institut Bioinformatics and Microbial Genomics | Danish Covid-19 Genome Consortium |
| EPI_ISL_3463925 | State Hygienic Laboratory at the University of Iowa | State Hygienic Laboratory at the University of Iowa | Valerie Reeb, Erik Twait, Anna Yakos, Mariah Knutson, Kris Eveland, Kristen Zanon, Wes Hottel, Alankar Kampoowale |
| EPI_ISL_3463926, EPI_ISL_3463929, EPI_ISL_3463932 | Department of Bacteria, Parasites and Fungi, Statens Serum Institut, Copenhagen, Denmark | Statens Serum Institut Bioinformatics and Microbial Genomics | Danish Covid-19 Genome Consortium |

| | | | |
|---|---|---|---|
| EPI_ISL_3463933 | State Hygienic Laboratory at the University of Iowa | State Hygienic Laboratory at the University of Iowa | Valerie Reeb, Erik Twait, Anna Yakos, Mariah Knutson, Kris Eveland, Kristen Zanon, Wes Hottel, Alankar Kampoowale |
| EPI_ISL_3463934 | Department of Bacteria, Parasites and Fungi, Statens Serum Institut, Copenhagen, Denmark | Statens Serum Institut Bioinformatics and Microbial Genomics | Danish Covid-19 Genome Consortium |
| EPI_ISL_3463935 | State Hygienic Laboratory at the University of Iowa | State Hygienic Laboratory at the University of Iowa | Valerie Reeb, Erik Twait, Anna Yakos, Mariah Knutson, Kris Eveland, Kristen Zanon, Wes Hottel, Alankar Kampoowale |
| EPI_ISL_3463936 | Department of Bacteria, Parasites and Fungi, Statens Serum Institut, Copenhagen, Denmark | Statens Serum Institut Bioinformatics and Microbial Genomics | Danish Covid-19 Genome Consortium |
| EPI_ISL_3463937 | State Hygienic Laboratory at the University of Iowa | State Hygienic Laboratory at the University of Iowa | Valerie Reeb, Erik Twait, Anna Yakos, Mariah Knutson, Kris Eveland, Kristen Zanon, Wes Hottel, Alankar Kampoowale |
| EPI_ISL_3463939 | Department of Bacteria, Parasites and Fungi, Statens Serum Institut, Copenhagen, Denmark | Statens Serum Institut Bioinformatics and Microbial Genomics | Danish Covid-19 Genome Consortium |
| EPI_ISL_3463940, EPI_ISL_3463941 | State Hygienic Laboratory at the University of Iowa | State Hygienic Laboratory at the University of Iowa | Valerie Reeb, Erik Twait, Anna Yakos, Mariah Knutson, Kris Eveland, Kristen Zanon, Wes Hottel, Alankar Kampoowale |
| EPI_ISL_3463942, EPI_ISL_3463943 | Department of Bacteria, Parasites and Fungi, Statens Serum Institut, Copenhagen, Denmark | Statens Serum Institut Bioinformatics and Microbial Genomics | Danish Covid-19 Genome Consortium |
| EPI_ISL_3463944 | State Hygienic Laboratory at the University of Iowa | State Hygienic Laboratory at the University of Iowa | Valerie Reeb, Erik Twait, Anna Yakos, Mariah Knutson, Kris Eveland, Kristen Zanon, Wes Hottel, Alankar Kampoowale |
| EPI_ISL_3463945 | Department of Bacteria, Parasites and Fungi, Statens Serum Institut, Copenhagen, Denmark | Statens Serum Institut Bioinformatics and Microbial Genomics | Danish Covid-19 Genome Consortium |
| EPI_ISL_3463946, EPI_ISL_3463947 | State Hygienic Laboratory at the University of Iowa | State Hygienic Laboratory at the University of Iowa | Valerie Reeb, Erik Twait, Anna Yakos, Mariah Knutson, Kris Eveland, Kristen Zanon, Wes Hottel, Alankar Kampoowale |
| EPI_ISL_3463948 | Department of Bacteria, Parasites and Fungi, Statens Serum Institut, Copenhagen, Denmark | Statens Serum Institut Bioinformatics and Microbial Genomics | Danish Covid-19 Genome Consortium |
| EPI_ISL_3463949 | State Hygienic Laboratory at the University of Iowa | State Hygienic Laboratory at the University of Iowa | Valerie Reeb, Erik Twait, Anna Yakos, Mariah Knutson, Kris Eveland, Kristen Zanon, Wes Hottel, Alankar Kampoowale |
| EPI_ISL_3463950 | Department of Bacteria, Parasites and Fungi, Statens Serum Institut, Copenhagen, Denmark | Statens Serum Institut Bioinformatics and Microbial Genomics | Danish Covid-19 Genome Consortium |
| EPI_ISL_3463951 | State Hygienic Laboratory at the University of Iowa | State Hygienic Laboratory at the University of Iowa | Valerie Reeb, Erik Twait, Anna Yakos, Mariah Knutson, Kris Eveland, Kristen Zanon, Wes Hottel, Alankar Kampoowale |
| EPI_ISL_3463952 | Department of Bacteria, Parasites and Fungi, Statens Serum Institut, Copenhagen, Denmark | Statens Serum Institut Bioinformatics and Microbial Genomics | Danish Covid-19 Genome Consortium |
| EPI_ISL_3463953 | State Hygienic Laboratory at the University of Iowa | State Hygienic Laboratory at the University of Iowa | Valerie Reeb, Erik Twait, Anna Yakos, Mariah Knutson, Kris Eveland, Kristen Zanon, Wes Hottel, Alankar Kampoowale |
| EPI_ISL_3463954 | Department of Bacteria, Parasites and Fungi, Statens Serum Institut, Copenhagen, Denmark | Statens Serum Institut Bioinformatics and Microbial Genomics | Danish Covid-19 Genome Consortium |
| EPI_ISL_3463955 | State Hygienic Laboratory at the University of Iowa | State Hygienic Laboratory at the University of Iowa | Valerie Reeb, Erik Twait, Anna Yakos, Mariah Knutson, Kris Eveland, Kristen Zanon, Wes Hottel, Alankar Kampoowale |
| EPI_ISL_3463956, EPI_ISL_3463957 | Department of Bacteria, Parasites and Fungi, Statens Serum Institut, Copenhagen, Denmark | Statens Serum Institut Bioinformatics and Microbial Genomics | Danish Covid-19 Genome Consortium |
| EPI_ISL_3463958 | State Hygienic Laboratory at the University of Iowa | State Hygienic Laboratory at the University of Iowa | Valerie Reeb, Erik Twait, Anna Yakos, Mariah Knutson, Kris Eveland, Kristen Zanon, Wes Hottel, Alankar Kampoowale |
| EPI_ISL_3463959 | Department of Bacteria, Parasites and Fungi, Statens Serum Institut, Copenhagen, Denmark | Statens Serum Institut Bioinformatics and Microbial Genomics | Danish Covid-19 Genome Consortium |
| EPI_ISL_3463960 | State Hygienic Laboratory at the University of Iowa | State Hygienic Laboratory at the University of Iowa | Valerie Reeb, Erik Twait, Anna Yakos, Mariah Knutson, Kris Eveland, Kristen Zanon, Wes Hottel, Alankar Kampoowale |
| EPI_ISL_3463961 | Department of Bacteria, Parasites and Fungi, Statens Serum Institut, Copenhagen, Denmark | Statens Serum Institut Bioinformatics and Microbial Genomics | Danish Covid-19 Genome Consortium |
| EPI_ISL_3463962 | State Hygienic Laboratory at the University of Iowa | State Hygienic Laboratory at the University of Iowa | Valerie Reeb, Erik Twait, Anna Yakos, Mariah Knutson, Kris Eveland, Kristen Zanon, Wes Hottel, Alankar Kampoowale |
| EPI_ISL_3463963 | Department of Bacteria, Parasites and Fungi, Statens Serum Institut, Copenhagen, Denmark | Statens Serum Institut Bioinformatics and Microbial Genomics | Danish Covid-19 Genome Consortium |
| EPI_ISL_3463964 | State Hygienic Laboratory at the University of Iowa | State Hygienic Laboratory at the University of Iowa | Valerie Reeb, Erik Twait, Anna Yakos, Mariah Knutson, Kris Eveland, Kristen Zanon, Wes Hottel, Alankar Kampoowale |
| EPI_ISL_3463965 | Department of Bacteria, Parasites and Fungi, Statens Serum Institut, Copenhagen, Denmark | Statens Serum Institut Bioinformatics and Microbial Genomics | Danish Covid-19 Genome Consortium |
| EPI_ISL_3463966 | State Hygienic Laboratory at the University of Iowa | State Hygienic Laboratory at the University of Iowa | Valerie Reeb, Erik Twait, Anna Yakos, Mariah Knutson, Kris Eveland, Kristen Zanon, Wes Hottel, Alankar Kampoowale |
| EPI_ISL_3463967 | Department of Bacteria, Parasites and Fungi, Statens Serum Institut, Copenhagen, Denmark | Statens Serum Institut Bioinformatics and Microbial Genomics | Danish Covid-19 Genome Consortium |
| EPI_ISL_3463968 | State Hygienic Laboratory at the University of Iowa | State Hygienic Laboratory at the University of Iowa | Valerie Reeb, Erik Twait, Anna Yakos, Mariah Knutson, Kris Eveland, Kristen Zanon, Wes Hottel, Alankar Kampoowale |
| EPI_ISL_3463969 | Department of Clinical Microbiology, Odense University Hospital, Odense, Denmark | Statens Serum Institut Bioinformatics and Microbial Genomics | Danish Covid-19 Genome Consortium |
| EPI_ISL_3463970 | State Hygienic Laboratory at the University of Iowa | State Hygienic Laboratory at the University of Iowa | Valerie Reeb, Erik Twait, Anna Yakos, Mariah Knutson, Kris Eveland, Kristen Zanon, Wes Hottel, Alankar Kampoowale |
| EPI_ISL_3463971 | Department of Bacteria, Parasites and Fungi, Statens Serum Institut, Copenhagen, Denmark | Statens Serum Institut Bioinformatics and Microbial Genomics | Danish Covid-19 Genome Consortium |
| EPI_ISL_3463972, EPI_ISL_3463973 | State Hygienic Laboratory at the University of Iowa | State Hygienic Laboratory at the University of Iowa | Valerie Reeb, Erik Twait, Anna Yakos, Mariah Knutson, Kris Eveland, Kristen Zanon, Wes Hottel, Alankar Kampoowale |
| EPI_ISL_3463974, EPI_ISL_3463975 | Department of Bacteria, Parasites and Fungi, Statens Serum Institut, Copenhagen, Denmark | Statens Serum Institut Bioinformatics and Microbial Genomics | Danish Covid-19 Genome Consortium |
| EPI_ISL_3463976 | State Hygienic Laboratory at the University of Iowa | State Hygienic Laboratory at the University of Iowa | Valerie Reeb, Erik Twait, Anna Yakos, Mariah Knutson, Kris Eveland, Kristen Zanon, Wes Hottel, Alankar Kampoowale |
| EPI_ISL_3463977 | Department of Bacteria, Parasites and Fungi, Statens Serum Institut, Copenhagen, Denmark | Statens Serum Institut Bioinformatics and Microbial Genomics | Danish Covid-19 Genome Consortium |
| EPI_ISL_3463978 | State Hygienic Laboratory at the University of Iowa | State Hygienic Laboratory at the University of Iowa | Valerie Reeb, Erik Twait, Anna Yakos, Mariah Knutson, Kris Eveland, Kristen Zanon, Wes Hottel, Alankar Kampoowale |
| EPI_ISL_3463979, EPI_ISL_3463980, EPI_ISL_3463981, EPI_ISL_3463982, EPI_ISL_3463983, EPI_ISL_3463984, EPI_ISL_3463985, EPI_ISL_3463986, EPI_ISL_3463987, EPI_ISL_3463988, EPI_ISL_3463989, EPI_ISL_3463990, EPI_ISL_3463991 | | | |
| see above | Department of Bacteria, Parasites and Fungi, Statens Serum Institut, Copenhagen, Denmark | Statens Serum Institut Bioinformatics and Microbial Genomics | Danish Covid-19 Genome Consortium |
| EPI_ISL_3463992 | Department of Clinical Microbiology, Odense University Hospital, Odense, Denmark | Statens Serum Institut Bioinformatics and Microbial Genomics | Danish Covid-19 Genome Consortium |
| EPI_ISL_3463993, EPI_ISL_3463994, EPI_ISL_3463996, EPI_ISL_3463997, EPI_ISL_3463998, EPI_ISL_3463999, EPI_ISL_3464000, EPI_ISL_3464001, EPI_ISL_3464002, EPI_ISL_3464003, EPI_ISL_3464004, EPI_ISL_3464005, EPI_ISL_3464006, EPI_ISL_3464007, EPI_ISL_3464008, EPI_ISL_3464009, EPI_ISL_3464010, EPI_ISL_3464011, EPI_ISL_3464012, EPI_ISL_3464013, EPI_ISL_3464014, EPI_ISL_3464015, EPI_ISL_3464016, EPI_ISL_3464017, EPI_ISL_3464018, EPI_ISL_3464019, EPI_ISL_3464020, EPI_ISL_3464021, EPI_ISL_3464022, EPI_ISL_3464023, EPI_ISL_3464024, EPI_ISL_3464025, EPI_ISL_3464026, EPI_ISL_3464027, EPI_ISL_3464028, EPI_ISL_3464029, EPI_ISL_3464030, EPI_ISL_3464031, EPI_ISL_3464032, EPI_ISL_3464033, EPI_ISL_3464034, EPI_ISL_3464035, EPI_ISL_3464036, EPI_ISL_3464037, EPI_ISL_3464038, EPI_ISL_3464039, EPI_ISL_3464040, EPI_ISL_3464041, EPI_ISL_3464042, EPI_ISL_3464043, EPI_ISL_3464044, EPI_ISL_3464045, EPI_ISL_3464046, EPI_ISL_3464047, EPI_ISL_3464048 | | | |
| see above | Department of Bacteria, Parasites and Fungi, Statens Serum Institut, Copenhagen, Denmark | Statens Serum Institut Bioinformatics and Microbial Genomics | Danish Covid-19 Genome Consortium |
| EPI_ISL_3464049 | Department of Clinical Microbiology, Odense University Hospital, Odense, Denmark | Statens Serum Institut Bioinformatics and Microbial Genomics | Danish Covid-19 Genome Consortium |
| EPI_ISL_3464050, EPI_ISL_3464051, EPI_ISL_3464052, EPI_ISL_3464053, EPI_ISL_3464054, EPI_ISL_3464055, EPI_ISL_3464056, EPI_ISL_3464057, EPI_ISL_3464058, EPI_ISL_3464059, EPI_ISL_3464060, EPI_ISL_3464061, EPI_ISL_3464062, EPI_ISL_3464063, EPI_ISL_3464064, EPI_ISL_3464065, EPI_ISL_3464066, EPI_ISL_3464067, EPI_ISL_3464068, EPI_ISL_3464069, EPI_ISL_3464070, EPI_ISL_3464071, EPI_ISL_3464072, EPI_ISL_3464073, EPI_ISL_3464074, EPI_ISL_3464075, EPI_ISL_3464076, EPI_ISL_3464077, EPI_ISL_3464078, EPI_ISL_3464079, EPI_ISL_3464080, EPI_ISL_3464081, EPI_ISL_3464082, EPI_ISL_3464083, EPI_ISL_3464084 | | | |
| see above | Department of Bacteria, Parasites and Fungi, Statens Serum Institut, Copenhagen, Denmark | Statens Serum Institut Bioinformatics and Microbial Genomics | Danish Covid-19 Genome Consortium |
| EPI_ISL_3464085 | Department of Clinical Microbiology, Odense University Hospital, Odense, Denmark | Statens Serum Institut Bioinformatics and Microbial Genomics | Danish Covid-19 Genome Consortium |
| EPI_ISL_3464086, EPI_ISL_3464087, EPI_ISL_3464088, EPI_ISL_3464089, EPI_ISL_3464090, EPI_ISL_3464091, EPI_ISL_3464092, EPI_ISL_3464093, EPI_ISL_3464094, EPI_ISL_3464095, EPI_ISL_3464096, EPI_ISL_3464097, EPI_ISL_3464098, EPI_ISL_3464099, EPI_ISL_3464100, EPI_ISL_3464101, EPI_ISL_3464102, EPI_ISL_3464103, EPI_ISL_3464104, EPI_ISL_3464105, EPI_ISL_3464106, EPI_ISL_3464107, EPI_ISL_3464108, EPI_ISL_3464109, EPI_ISL_3464110, EPI_ISL_3464111, EPI_ISL_3464112, EPI_ISL_3464113, EPI_ISL_3464114, EPI_ISL_3464115, EPI_ISL_3464116, EPI_ISL_3464117, EPI_ISL_3464118, EPI_ISL_3464119, EPI_ISL_3464120, EPI_ISL_3464121, EPI_ISL_3464122, EPI_ISL_3464123, EPI_ISL_3464124, EPI_ISL_3464125, EPI_ISL_3464126, EPI_ISL_3464127, EPI_ISL_3464128, EPI_ISL_3464129, EPI_ISL_3464130, EPI_ISL_3464131, EPI_ISL_3464132, EPI_ISL_3464133, EPI_ISL_3464134, EPI_ISL_3464135, EPI_ISL_3464136, EPI_ISL_3464137, EPI_ISL_3464138, EPI_ISL_3464139, EPI_ISL_3464140, EPI_ISL_3464141, EPI_ISL_3464142, EPI_ISL_3464143, EPI_ISL_3464144, EPI_ISL_3464145, EPI_ISL_3464146, EPI_ISL_3464147, EPI_ISL_3464148, EPI_ISL_3464149, EPI_ISL_3464150, EPI_ISL_3464151, EPI_ISL_3464152, EPI_ISL_3464153, EPI_ISL_3464154, EPI_ISL_3464155, EPI_ISL_3464156, EPI_ISL_3464157, EPI_ISL_3464158, EPI_ISL_3464159, EPI_ISL_3464160, EPI_ISL_3464161, EPI_ISL_3464162, EPI_ISL_3464163, EPI_ISL_3464164, EPI_ISL_3464165, EPI_ISL_3464166, EPI_ISL_3464167, EPI_ISL_3464168, EPI_ISL_3464169, EPI_ISL_3464170, EPI_ISL_3464171, EPI_ISL_3464172, EPI_ISL_3464173, EPI_ISL_3464174, EPI_ISL_3464175, EPI_ISL_3464176, EPI_ISL_3464177, EPI_ISL_3464178, EPI_ISL_3464179, EPI_ISL_3464180, EPI_ISL_3464181, EPI_ISL_3464182, EPI_ISL_3464183, EPI_ISL_3464184, EPI_ISL_3464185, EPI_ISL_3464186, EPI_ISL_3464187, EPI_ISL_3464188, EPI_ISL_3464189, EPI_ISL_3464190, EPI_ISL_3464191, EPI_ISL_3464192, EPI_ISL_3464193, EPI_ISL_3464194, EPI_ISL_3464195, EPI_ISL_3464196, EPI_ISL_3464197, EPI_ISL_3464198, EPI_ISL_3464199, EPI_ISL_3464200 | | | |
| see above | Department of Bacteria, Parasites and Fungi, Statens Serum Institut, Copenhagen, Denmark | Statens Serum Institut Bioinformatics and Microbial Genomics | Danish Covid-19 Genome Consortium |
| EPI_ISL_3464201 | State Hygienic Laboratory at the University of Iowa | State Hygienic Laboratory at the University of Iowa | Valerie Reeb, Erik Twait, Anna Yakos, Mariah Knutson, Kris Eveland, Kristen Zanon, Wes Hottel, Alankar Kampoowale |

| | | | |
|---|---|---|---|
| EPI_ISL_3464202 | Department of Bacteria, Parasites and Fungi, Statens Serum Institut, Copenhagen, Denmark | Statens Serum Institut Bioinformatics and Microbial Genomics | Danish Covid-19 Genome Consortium |
| EPI_ISL_3464203 | State Hygienic Laboratory at the University of Iowa | State Hygienic Laboratory at the University of Iowa | Valerie Reeb, Erik Twait, Anna Yakos, Mariah Knutson, Kris Eveland, Kristen Zanon, Wes Hottel, Alankar Kampoowale |
| EPI_ISL_3464204, EPI_ISL_3464206 | Department of Bacteria, Parasites and Fungi, Statens Serum Institut, Copenhagen, Denmark | Statens Serum Institut Bioinformatics and Microbial Genomics | Danish Covid-19 Genome Consortium |
| EPI_ISL_3464207 | State Hygienic Laboratory at the University of Iowa | State Hygienic Laboratory at the University of Iowa | Valerie Reeb, Erik Twait, Anna Yakos, Mariah Knutson, Kris Eveland, Kristen Zanon, Wes Hottel, Alankar Kampoowale |
| EPI_ISL_3464208, EPI_ISL_3464210, EPI_ISL_3464211 | Department of Bacteria, Parasites and Fungi, Statens Serum Institut, Copenhagen, Denmark | Statens Serum Institut Bioinformatics and Microbial Genomics | Danish Covid-19 Genome Consortium |
| EPI_ISL_3464212 | State Hygienic Laboratory at the University of Iowa | State Hygienic Laboratory at the University of Iowa | Valerie Reeb, Erik Twait, Anna Yakos, Mariah Knutson, Kris Eveland, Kristen Zanon, Wes Hottel, Alankar Kampoowale |
| EPI_ISL_3464213 | Department of Bacteria, Parasites and Fungi, Statens Serum Institut, Copenhagen, Denmark | Statens Serum Institut Bioinformatics and Microbial Genomics | Danish Covid-19 Genome Consortium |
| EPI_ISL_3464214 | State Hygienic Laboratory at the University of Iowa | State Hygienic Laboratory at the University of Iowa | Valerie Reeb, Erik Twait, Anna Yakos, Mariah Knutson, Kris Eveland, Kristen Zanon, Wes Hottel, Alankar Kampoowale |
| EPI_ISL_3464215, EPI_ISL_3464217, EPI_ISL_3464219, EPI_ISL_3464220, EPI_ISL_3464222 | Department of Bacteria, Parasites and Fungi, Statens Serum Institut, Copenhagen, Denmark | Statens Serum Institut Bioinformatics and Microbial Genomics | Danish Covid-19 Genome Consortium |
| EPI_ISL_3464223 | State Hygienic Laboratory at the University of Iowa | State Hygienic Laboratory at the University of Iowa | Valerie Reeb, Erik Twait, Anna Yakos, Mariah Knutson, Kris Eveland, Kristen Zanon, Wes Hottel, Alankar Kampoowale |
| EPI_ISL_3464224 | Department of Bacteria, Parasites and Fungi, Statens Serum Institut, Copenhagen, Denmark | Statens Serum Institut Bioinformatics and Microbial Genomics | Danish Covid-19 Genome Consortium |
| EPI_ISL_3464225 | State Hygienic Laboratory at the University of Iowa | State Hygienic Laboratory at the University of Iowa | Valerie Reeb, Erik Twait, Anna Yakos, Mariah Knutson, Kris Eveland, Kristen Zanon, Wes Hottel, Alankar Kampoowale |
| EPI_ISL_3464226 | Department of Bacteria, Parasites and Fungi, Statens Serum Institut, Copenhagen, Denmark | Statens Serum Institut Bioinformatics and Microbial Genomics | Danish Covid-19 Genome Consortium |
| EPI_ISL_3464227 | State Hygienic Laboratory at the University of Iowa | State Hygienic Laboratory at the University of Iowa | Valerie Reeb, Erik Twait, Anna Yakos, Mariah Knutson, Kris Eveland, Kristen Zanon, Wes Hottel, Alankar Kampoowale |
| EPI_ISL_3464228 | Department of Bacteria, Parasites and Fungi, Statens Serum Institut, Copenhagen, Denmark | Statens Serum Institut Bioinformatics and Microbial Genomics | Danish Covid-19 Genome Consortium |
| EPI_ISL_3464229 | State Hygienic Laboratory at the University of Iowa | State Hygienic Laboratory at the University of Iowa | Valerie Reeb, Erik Twait, Anna Yakos, Mariah Knutson, Kris Eveland, Kristen Zanon, Wes Hottel, Alankar Kampoowale |
| EPI_ISL_3464230 | Department of Bacteria, Parasites and Fungi, Statens Serum Institut, Copenhagen, Denmark | Statens Serum Institut Bioinformatics and Microbial Genomics | Danish Covid-19 Genome Consortium |
| EPI_ISL_3464231 | State Hygienic Laboratory at the University of Iowa | State Hygienic Laboratory at the University of Iowa | Valerie Reeb, Erik Twait, Anna Yakos, Mariah Knutson, Kris Eveland, Kristen Zanon, Wes Hottel, Alankar Kampoowale |
| EPI_ISL_3464232 | Department of Bacteria, Parasites and Fungi, Statens Serum Institut, Copenhagen, Denmark | Statens Serum Institut Bioinformatics and Microbial Genomics | Danish Covid-19 Genome Consortium |
| EPI_ISL_3464233 | State Hygienic Laboratory at the University of Iowa | State Hygienic Laboratory at the University of Iowa | Valerie Reeb, Erik Twait, Anna Yakos, Mariah Knutson, Kris Eveland, Kristen Zanon, Wes Hottel, Alankar Kampoowale |
| EPI_ISL_3464234, EPI_ISL_3464235 | Department of Bacteria, Parasites and Fungi, Statens Serum Institut, Copenhagen, Denmark | Statens Serum Institut Bioinformatics and Microbial Genomics | Danish Covid-19 Genome Consortium |
| EPI_ISL_3464236 | State Hygienic Laboratory at the University of Iowa | State Hygienic Laboratory at the University of Iowa | Valerie Reeb, Erik Twait, Anna Yakos, Mariah Knutson, Kris Eveland, Kristen Zanon, Wes Hottel, Alankar Kampoowale |
| EPI_ISL_3464237 | Department of Bacteria, Parasites and Fungi, Statens Serum Institut, Copenhagen, Denmark | Statens Serum Institut Bioinformatics and Microbial Genomics | Danish Covid-19 Genome Consortium |
| EPI_ISL_3464238 | State Hygienic Laboratory at the University of Iowa | State Hygienic Laboratory at the University of Iowa | Valerie Reeb, Erik Twait, Anna Yakos, Mariah Knutson, Kris Eveland, Kristen Zanon, Wes Hottel, Alankar Kampoowale |
| EPI_ISL_3464239 | Department of Bacteria, Parasites and Fungi, Statens Serum Institut, Copenhagen, Denmark | Statens Serum Institut Bioinformatics and Microbial Genomics | Danish Covid-19 Genome Consortium |
| EPI_ISL_3464240 | State Hygienic Laboratory at the University of Iowa | State Hygienic Laboratory at the University of Iowa | Valerie Reeb, Erik Twait, Anna Yakos, Mariah Knutson, Kris Eveland, Kristen Zanon, Wes Hottel, Alankar Kampoowale |
| EPI_ISL_3464241 | Department of Bacteria, Parasites and Fungi, Statens Serum Institut, Copenhagen, Denmark | Statens Serum Institut Bioinformatics and Microbial Genomics | Danish Covid-19 Genome Consortium |
| EPI_ISL_3464242 | State Hygienic Laboratory at the University of Iowa | State Hygienic Laboratory at the University of Iowa | Valerie Reeb, Erik Twait, Anna Yakos, Mariah Knutson, Kris Eveland, Kristen Zanon, Wes Hottel, Alankar Kampoowale |
| EPI_ISL_3464243 | Department of Bacteria, Parasites and Fungi, Statens Serum Institut, Copenhagen, Denmark | Statens Serum Institut Bioinformatics and Microbial Genomics | Danish Covid-19 Genome Consortium |
| EPI_ISL_3464244 | State Hygienic Laboratory at the University of Iowa | State Hygienic Laboratory at the University of Iowa | Valerie Reeb, Erik Twait, Anna Yakos, Mariah Knutson, Kris Eveland, Kristen Zanon, Wes Hottel, Alankar Kampoowale |
| EPI_ISL_3464245, EPI_ISL_3464246 | Department of Bacteria, Parasites and Fungi, Statens Serum Institut, Copenhagen, Denmark | Statens Serum Institut Bioinformatics and Microbial Genomics | Danish Covid-19 Genome Consortium |
| EPI_ISL_3464247 | State Hygienic Laboratory at the University of Iowa | State Hygienic Laboratory at the University of Iowa | Valerie Reeb, Erik Twait, Anna Yakos, Mariah Knutson, Kris Eveland, Kristen Zanon, Wes Hottel, Alankar Kampoowale |
| EPI_ISL_3464248 | Department of Bacteria, Parasites and Fungi, Statens Serum Institut, Copenhagen, Denmark | Statens Serum Institut Bioinformatics and Microbial Genomics | Danish Covid-19 Genome Consortium |
| EPI_ISL_3464249 | State Hygienic Laboratory at the University of Iowa | State Hygienic Laboratory at the University of Iowa | Valerie Reeb, Erik Twait, Anna Yakos, Mariah Knutson, Kris Eveland, Kristen Zanon, Wes Hottel, Alankar Kampoowale |
| EPI_ISL_3464250 | Department of Bacteria, Parasites and Fungi, Statens Serum Institut, Copenhagen, Denmark | Statens Serum Institut Bioinformatics and Microbial Genomics | Danish Covid-19 Genome Consortium |
| EPI_ISL_3464251 | State Hygienic Laboratory at the University of Iowa | State Hygienic Laboratory at the University of Iowa | Valerie Reeb, Erik Twait, Anna Yakos, Mariah Knutson, Kris Eveland, Kristen Zanon, Wes Hottel, Alankar Kampoowale |
| EPI_ISL_3464252 | Department of Bacteria, Parasites and Fungi, Statens Serum Institut, Copenhagen, Denmark | Statens Serum Institut Bioinformatics and Microbial Genomics | Danish Covid-19 Genome Consortium |
| EPI_ISL_3464253 | State Hygienic Laboratory at the University of Iowa | State Hygienic Laboratory at the University of Iowa | Valerie Reeb, Erik Twait, Anna Yakos, Mariah Knutson, Kris Eveland, Kristen Zanon, Wes Hottel, Alankar Kampoowale |
| EPI_ISL_3464254 | Department of Bacteria, Parasites and Fungi, Statens Serum Institut, Copenhagen, Denmark | Statens Serum Institut Bioinformatics and Microbial Genomics | Danish Covid-19 Genome Consortium |
| EPI_ISL_3464255 | State Hygienic Laboratory at the University of Iowa | State Hygienic Laboratory at the University of Iowa | Valerie Reeb, Erik Twait, Anna Yakos, Mariah Knutson, Kris Eveland, Kristen Zanon, Wes Hottel, Alankar Kampoowale |
| EPI_ISL_3464256, EPI_ISL_3464258, EPI_ISL_3464259, EPI_ISL_3464260, EPI_ISL_3464261, EPI_ISL_3464262, EPI_ISL_3464263, EPI_ISL_3464264, EPI_ISL_3464265, EPI_ISL_3464266, EPI_ISL_3464267, EPI_ISL_3464268, EPI_ISL_3464269, EPI_ISL_3464270, EPI_ISL_3464271, EPI_ISL_3464272, EPI_ISL_3464273, EPI_ISL_3464274, EPI_ISL_3464275, EPI_ISL_3464276, EPI_ISL_3464277, EPI_ISL_3464278, EPI_ISL_3464279, EPI_ISL_3464280, EPI_ISL_3464281, EPI_ISL_3464282, EPI_ISL_3464283, EPI_ISL_3464284, EPI_ISL_3464285, EPI_ISL_3464286, EPI_ISL_3464287, EPI_ISL_3464288, EPI_ISL_3464289, EPI_ISL_3464290, EPI_ISL_3464291, EPI_ISL_3464292, EPI_ISL_3464293, EPI_ISL_3464294, EPI_ISL_3464295, EPI_ISL_3464296, EPI_ISL_3464297, EPI_ISL_3464298, EPI_ISL_3464299, EPI_ISL_3464300, EPI_ISL_3464301, EPI_ISL_3464302, EPI_ISL_3464303, EPI_ISL_3464304, EPI_ISL_3464305, EPI_ISL_3464306, EPI_ISL_3464307, EPI_ISL_3464308, EPI_ISL_3464309, EPI_ISL_3464310, EPI_ISL_3464311, EPI_ISL_3464312, EPI_ISL_3464313, EPI_ISL_3464314, EPI_ISL_3464315, EPI_ISL_3464316, EPI_ISL_3464317, EPI_ISL_3464318, EPI_ISL_3464319, EPI_ISL_3464320, EPI_ISL_3464321, EPI_ISL_3464322, EPI_ISL_3464323, EPI_ISL_3464324, EPI_ISL_3464325, EPI_ISL_3464326, EPI_ISL_3464327, EPI_ISL_3464328, EPI_ISL_3464329, EPI_ISL_3464330, EPI_ISL_3464331, EPI_ISL_3464332, EPI_ISL_3464333, EPI_ISL_3464334, EPI_ISL_3464335, EPI_ISL_3464336, EPI_ISL_3464337, EPI_ISL_3464338, EPI_ISL_3464339, EPI_ISL_3464340, EPI_ISL_3464341, EPI_ISL_3464342, EPI_ISL_3464343, EPI_ISL_3464344, EPI_ISL_3464345, EPI_ISL_3464346, EPI_ISL_3464347, EPI_ISL_3464348, EPI_ISL_3464349, EPI_ISL_3464350, EPI_ISL_3464351, EPI_ISL_3464352, EPI_ISL_3464353, EPI_ISL_3464354, EPI_ISL_3464355, EPI_ISL_3464356, EPI_ISL_3464357, EPI_ISL_3464358, EPI_ISL_3464359, EPI_ISL_3464360 | | | |
| see above | Department of Bacteria, Parasites and Fungi, Statens Serum Institut, Copenhagen, Denmark | Statens Serum Institut Bioinformatics and Microbial Genomics | Danish Covid-19 Genome Consortium |
| EPI_ISL_3464361, EPI_ISL_3464362, EPI_ISL_3464363, EPI_ISL_3464364, EPI_ISL_3464365, EPI_ISL_3464366, EPI_ISL_3464367, EPI_ISL_3464368, EPI_ISL_3464369, EPI_ISL_3464370, EPI_ISL_3464371, EPI_ISL_3464372, EPI_ISL_3464373, EPI_ISL_3464374, EPI_ISL_3464375, EPI_ISL_3464376, EPI_ISL_3464377, EPI_ISL_3464378, EPI_ISL_3464379, EPI_ISL_3464380, EPI_ISL_3464381, EPI_ISL_3464382, EPI_ISL_3464383, EPI_ISL_3464384, EPI_ISL_3464385, EPI_ISL_3464386, EPI_ISL_3464387, EPI_ISL_3464388, EPI_ISL_3464389, EPI_ISL_3464390, EPI_ISL_3464391, EPI_ISL_3464392, EPI_ISL_3464393, EPI_ISL_3464394, EPI_ISL_3464395, EPI_ISL_3464396, EPI_ISL_3464397, EPI_ISL_3464398, EPI_ISL_3464399, EPI_ISL_3464400, EPI_ISL_3464401, EPI_ISL_3464402, EPI_ISL_3464403, EPI_ISL_3464404, EPI_ISL_3464405, EPI_ISL_3464406, EPI_ISL_3464407, EPI_ISL_3464408, EPI_ISL_3464409, EPI_ISL_3464410, EPI_ISL_3464411, EPI_ISL_3464412, EPI_ISL_3464413, EPI_ISL_3464414, EPI_ISL_3464415 | | | |
| see above | Arizona State Public Health Laboratory | Arizona State Public Health Laboratory | Trung Huynh, Jessica Escobar, Katherine Fullerton, Nobuko Fukushima, Matthew Contursi, Stacy White, Linda Getsinger, Victor Waddell |
| EPI_ISL_3464416, EPI_ISL_3464417, EPI_ISL_3464418, EPI_ISL_3464419, EPI_ISL_3464420, EPI_ISL_3464421, EPI_ISL_3464422, EPI_ISL_3464423, EPI_ISL_3464424 | Wyoming Public Health Laboratory | Wyoming Public Health Laboratory | Jim Mildenberger, Taylor Fearing, Channing Weber, Ashley Norberg, Chayse Rowley, Marley Goetz, Brian Dominguez, Elliot Thomasson, Sam Britz, Cari Sloma, Robert Petit, and Rob Christensen |
| EPI_ISL_3464425, EPI_ISL_3464426, EPI_ISL_3464427, EPI_ISL_3464428, EPI_ISL_3464429, EPI_ISL_3464430, | Nacionalinė visuomenės sveikatos priežiūros laboratorija | National Public Health Surveillance Laboratory | Lukas Zemaitis, Migle Gabrielaite, Jelena Razmuk, Svajune Muralyte, Ana Steponkiene, Lukas Vasionis, Danas Baksa |

| | | | |
|---|---|---|---|
| EPI_ISL_3464431 | | | |
| EPI_ISL_3464432, EPI_ISL_3464433, EPI_ISL_3464434, EPI_ISL_3464435, EPI_ISL_3464436, EPI_ISL_3464437, EPI_ISL_3464438, EPI_ISL_3464439, EPI_ISL_3464440, EPI_ISL_3464441, EPI_ISL_3464442, EPI_ISL_3464443 | | | |
| see above | UAB Diagnostikos laboratorija | National Public Health Surveillance Laboratory | Lukas Zemaitis, Migle Gabrielaite, Jelena Razmuk, Svajune Muralyte, Ana Steponkiene, Lukas Vasionis, Danas Baksa |
| EPI_ISL_3464444, EPI_ISL_3464445, EPI_ISL_3464446, EPI_ISL_3464447, EPI_ISL_3464448, EPI_ISL_3464449, EPI_ISL_3464450, EPI_ISL_3464451, EPI_ISL_3464452, EPI_ISL_3464453, EPI_ISL_3464454, EPI_ISL_3464455, EPI_ISL_3464456, EPI_ISL_3464457, EPI_ISL_3464458, EPI_ISL_3464459, EPI_ISL_3464460, EPI_ISL_3464461, EPI_ISL_3464462, EPI_ISL_3464463, EPI_ISL_3464464, EPI_ISL_3464465, EPI_ISL_3464466, EPI_ISL_3464467, EPI_ISL_3464468, EPI_ISL_3464469, EPI_ISL_3464470, EPI_ISL_3464471, EPI_ISL_3464472, EPI_ISL_3464473, EPI_ISL_3464474, EPI_ISL_3464475, EPI_ISL_3464476 | | | |
| see above | Viešoji įstaiga Vilniaus universiteto ligoninė Santaros klinikos | National Public Health Surveillance Laboratory | Lukas Zemaitis, Migle Gabrielaite, Jelena Razmuk, Svajune Muralyte, Ana Steponkiene, Lukas Vasionis, Danas Baksa |
| EPI_ISL_3464477 | UAB Diagnostikos laboratorija | National Public Health Surveillance Laboratory | Lukas Zemaitis, Migle Gabrielaite, Jelena Razmuk, Svajune Muralyte, Ana Steponkiene, Lukas Vasionis, Danas Baksa |
| EPI_ISL_3464478, EPI_ISL_3464479, EPI_ISL_3464480, EPI_ISL_3464481, EPI_ISL_3464482, EPI_ISL_3464483 | Viešoji įstaiga Vilniaus universiteto ligoninė Santaros klinikos | National Public Health Surveillance Laboratory | Lukas Zemaitis, Migle Gabrielaite, Jelena Razmuk, Svajune Muralyte, Ana Steponkiene, Lukas Vasionis, Danas Baksa |
| EPI_ISL_3464484 | Nacionalinė visuomenės sveikatos priežiūros laboratorija | National Public Health Surveillance Laboratory | Lukas Zemaitis, Migle Gabrielaite, Jelena Razmuk, Svajune Muralyte, Ana Steponkiene, Lukas Vasionis, Danas Baksa |
| EPI_ISL_3464485, EPI_ISL_3464486, EPI_ISL_3464487, EPI_ISL_3464488, EPI_ISL_3464489, EPI_ISL_3464490 | Viešoji įstaiga Vilniaus universiteto ligoninė Santaros klinikos | National Public Health Surveillance Laboratory | Lukas Zemaitis, Migle Gabrielaite, Jelena Razmuk, Svajune Muralyte, Ana Steponkiene, Lukas Vasionis, Danas Baksa |
| EPI_ISL_3464491 | UAB Diagnostikos laboratorija | National Public Health Surveillance Laboratory | Lukas Zemaitis, Migle Gabrielaite, Jelena Razmuk, Svajune Muralyte, Ana Steponkiene, Lukas Vasionis, Danas Baksa |
| EPI_ISL_3464492, EPI_ISL_3464493, EPI_ISL_3464494 | Viešoji įstaiga Vilniaus universiteto ligoninė Santaros klinikos | National Public Health Surveillance Laboratory | Lukas Zemaitis, Migle Gabrielaite, Jelena Razmuk, Svajune Muralyte, Ana Steponkiene, Lukas Vasionis, Danas Baksa |
| EPI_ISL_3464495 | UAB Diagnostikos laboratorija | National Public Health Surveillance Laboratory | Lukas Zemaitis, Migle Gabrielaite, Jelena Razmuk, Svajune Muralyte, Ana Steponkiene, Lukas Vasionis, Danas Baksa |
| EPI_ISL_3464496, EPI_ISL_3464497 | Viešoji įstaiga Vilniaus universiteto ligoninė Santaros klinikos | National Public Health Surveillance Laboratory | Lukas Zemaitis, Migle Gabrielaite, Jelena Razmuk, Svajune Muralyte, Ana Steponkiene, Lukas Vasionis, Danas Baksa |
| EPI_ISL_3464498 | UAB Diagnostikos laboratorija | National Public Health Surveillance Laboratory | Lukas Zemaitis, Migle Gabrielaite, Jelena Razmuk, Svajune Muralyte, Ana Steponkiene, Lukas Vasionis, Danas Baksa |
| EPI_ISL_3464499, EPI_ISL_3464500, EPI_ISL_3464501 | Viešoji įstaiga Vilniaus universiteto ligoninė Santaros klinikos | National Public Health Surveillance Laboratory | Lukas Zemaitis, Migle Gabrielaite, Jelena Razmuk, Svajune Muralyte, Ana Steponkiene, Lukas Vasionis, Danas Baksa |
| EPI_ISL_3464502 | HOSPITAL DE LAS MUJERES DR. ADOLFO CARIT | Inciensa, Instituto Costarricense de Investigación y Enseñanza en Nutrición y Salud | Francisco Duarte, Hebleen Porras, Claudio Soto-Garita, Estela Cordero, Adriana Godínez, Melany Calderón, José Luis Vargas, Mariela Gutiérrez, Joselyn Prado & Silvia Jiménez Cordero |
| EPI_ISL_3464503 | AREA DE SALUD FORTUNA | Inciensa, Instituto Costarricense de Investigación y Enseñanza en Nutrición y Salud | Francisco Duarte, Hebleen Porras, Claudio Soto-Garita, Estela Cordero, Adriana Godínez, Melany Calderón, José Luis Vargas, Mariela Gutiérrez, Joselyn Prado & Juan Carlos Villalobos Ugalde |
| EPI_ISL_3464504 | AREA DE SALUD LOS SANTOS | Inciensa, Instituto Costarricense de Investigación y Enseñanza en Nutrición y Salud | Francisco Duarte, Hebleen Porras, Claudio Soto-Garita, Estela Cordero, Adriana Godínez, Melany Calderón, José Luis Vargas, Mariela Gutiérrez, Joselyn Prado & Luis Diego Granados Chavarría |
| EPI_ISL_3464513 | AREA DE SALUD SANTA ROSA | Inciensa, Instituto Costarricense de Investigación y Enseñanza en Nutrición y Salud | Francisco Duarte, Hebleen Porras, Claudio Soto-Garita, Estela Cordero, Adriana Godínez, Melany Calderón, José Luis Vargas, Mariela Gutiérrez, Joselyn Prado & Juan Carlos Villalobos Ugalde |
| EPI_ISL_3464514 | AREA DE SALUD PAQUERA | Inciensa, Instituto Costarricense de Investigación y Enseñanza en Nutrición y Salud | Francisco Duarte, Hebleen Porras, Claudio Soto-Garita, Estela Cordero, Adriana Godínez, Melany Calderón, José Luis Vargas, Mariela Gutiérrez, Joselyn Prado & Ariela Rojas Herrera |
| EPI_ISL_3464515 | AREA DE SALUD MATA REDONDA-HOSPITAL - CLINICA DR. MORENO CAÑAS | Inciensa, Instituto Costarricense de Investigación y Enseñanza en Nutrición y Salud | Francisco Duarte, Hebleen Porras, Claudio Soto-Garita, Estela Cordero, Adriana Godínez, Melany Calderón, José Luis Vargas, Mariela Gutiérrez, Joselyn Prado & Pamela Martinez Cedeño |
| EPI_ISL_3464517 | AREA DE SALUD LOS CHILES | Inciensa, Instituto Costarricense de Investigación y Enseñanza en Nutrición y Salud | Francisco Duarte, Hebleen Porras, Claudio Soto-Garita, Estela Cordero, Adriana Godínez, Melany Calderón, José Luis Vargas, Mariela Gutiérrez, Joselyn Prado & Juan Carlos Villalobos Ugalde |
| EPI_ISL_3464518 | AREA DE SALUD FORTUNA | Inciensa, Instituto Costarricense de Investigación y Enseñanza en Nutrición y Salud | Francisco Duarte, Hebleen Porras, Claudio Soto-Garita, Estela Cordero, Adriana Godínez, Melany Calderón, José Luis Vargas, Mariela Gutiérrez, Joselyn Prado & Juan Carlos Villalobos Ugalde |
| EPI_ISL_3464520 | AREA DE SALUD COTO BRUS | Inciensa, Instituto Costarricense de Investigación y Enseñanza en Nutrición y Salud | Francisco Duarte, Hebleen Porras, Claudio Soto-Garita, Estela Cordero, Adriana Godínez, Melany Calderón, José Luis Vargas, Mariela Gutiérrez, Joselyn Prado & Yendri Ramírez Alpízar |
| EPI_ISL_3464521 | AREA DE SALUD HEREDIA-VIRILLA | Inciensa, Instituto Costarricense de Investigación y Enseñanza en Nutrición y Salud | Francisco Duarte, Hebleen Porras, Claudio Soto-Garita, Estela Cordero, Adriana Godínez, Melany Calderón, José Luis Vargas, Mariela Gutiérrez & Joselyn Prado |
| EPI_ISL_3464522 | AREA DE SALUD PAVAS (COOPESALUD) | Inciensa, Instituto Costarricense de Investigación y Enseñanza en Nutrición y Salud | Francisco Duarte, Hebleen Porras, Claudio Soto-Garita, Estela Cordero, Adriana Godínez, Melany Calderón, José Luis Vargas, Mariela Gutiérrez, Joselyn Prado & Paola Sánchez Traña |
| EPI_ISL_3464523 | AREA DE SALUD SIQUIRRES | Inciensa, Instituto Costarricense de Investigación y Enseñanza en Nutrición y Salud | Francisco Duarte, Hebleen Porras, Claudio Soto-Garita, Estela Cordero, Adriana Godínez, Melany Calderón, José Luis Vargas, Mariela Gutiérrez & Joselyn Prado |
| EPI_ISL_3464524, EPI_ISL_3464525 | AREA DE SALUD PAVAS (COOPESALUD) | Inciensa, Instituto Costarricense de Investigación y Enseñanza en Nutrición y Salud | Francisco Duarte, Hebleen Porras, Claudio Soto-Garita, Estela Cordero, Adriana Godínez, Melany Calderón, José Luis Vargas, Mariela Gutiérrez, Joselyn Prado & Paola Sanchez Traña |
| EPI_ISL_3464526 | AREA DE SALUD HATILLO - CLINICA DR. SOLON NUÑEZ | Inciensa, Instituto Costarricense de Investigación y Enseñanza en Nutrición y Salud | Francisco Duarte, Hebleen Porras, Claudio Soto-Garita, Estela Cordero, Adriana Godínez, Melany Calderón, José Luis Vargas, Mariela Gutiérrez, Joselyn Prado & Paola Sanchez Traña |
| EPI_ISL_3464527, EPI_ISL_3464528, EPI_ISL_3464529, EPI_ISL_3464530, EPI_ISL_3464531, EPI_ISL_3464532 | AREA DE SALUD FORTUNA | Inciensa, Instituto Costarricense de Investigación y Enseñanza en Nutrición y Salud | Francisco Duarte, Hebleen Porras, Claudio Soto-Garita, Estela Cordero, Adriana Godínez, Melany Calderón, José Luis Vargas, Mariela Gutiérrez, Joselyn Prado & Juan Carlos Villalobos Ugalde |
| EPI_ISL_3464533 | AREA DE SALUD SANTA ROSA | Inciensa, Instituto Costarricense de Investigación y Enseñanza en Nutrición y Salud | Francisco Duarte, Hebleen Porras, Claudio Soto-Garita, Estela Cordero, Adriana Godínez, Melany Calderón, José Luis Vargas, Mariela Gutiérrez, Joselyn Prado & Juan Carlos Villalobos Ugalde |
| EPI_ISL_3464536 | AREA DE SALUD PURISCAL-TURRUBARES | Inciensa, Instituto Costarricense de Investigación y Enseñanza en Nutrición y Salud | Francisco Duarte, Hebleen Porras, Claudio Soto-Garita, Estela Cordero, Adriana Godínez, Melany Calderón, José Luis Vargas, Mariela Gutiérrez & Joselyn Prado |
| EPI_ISL_3464538 | AREA DE SALUD LA CRUZ | Inciensa, Instituto Costarricense de Investigación y Enseñanza en Nutrición y Salud | Francisco Duarte, Hebleen Porras, Claudio Soto-Garita, Estela Cordero, Adriana Godínez, Melany Calderón, José Luis Vargas, Mariela Gutiérrez, Joselyn Prado & Adriana Bermúdez Espinoza |
| EPI_ISL_3464539 | AREA DE SALUD SANTA CRUZ | Inciensa, Instituto Costarricense de Investigación y Enseñanza en Nutrición y Salud | Francisco Duarte, Hebleen Porras, Claudio Soto-Garita, Estela Cordero, Adriana Godínez, Melany Calderón, José Luis Vargas, Mariela Gutiérrez, Joselyn Prado & Adriana Bermúdez Espinoza |
| EPI_ISL_3464542 | HOSPITAL SAN FRANCISCO DE ASIS | Inciensa, Instituto Costarricense de Investigación y Enseñanza en Nutrición y Salud | Francisco Duarte, Hebleen Porras, Claudio Soto-Garita, Estela Cordero, Adriana Godínez, Melany Calderón, José Luis Vargas, Mariela Gutiérrez, Joselyn Prado & Lucia Acuña |
| EPI_ISL_3464543 | AREA DE SALUD PAVAS (COOPESALUD) | Inciensa, Instituto Costarricense de Investigación y Enseñanza en Nutrición y Salud | Francisco Duarte, Hebleen Porras, Claudio Soto-Garita, Estela Cordero, Adriana Godínez, Melany Calderón, José Luis Vargas, Mariela Gutiérrez & Joselyn Prado |
| EPI_ISL_3464545 | AREA DE SALUD UPALA | Inciensa, Instituto Costarricense de Investigación y Enseñanza en Nutrición y Salud | Francisco Duarte, Hebleen Porras, Claudio Soto-Garita, Estela Cordero, Adriana Godínez, Melany Calderón, José Luis Vargas, Mariela Gutiérrez, Joselyn Prado & Juan Carlos Villalobos Ugalde |
| EPI_ISL_3464546 | AREA DE SALUD ALAJUELA SUR | Inciensa, Instituto Costarricense de Investigación y Enseñanza en Nutrición y Salud | Francisco Duarte, Hebleen Porras, Claudio Soto-Garita, Estela Cordero, Adriana Godínez, Melany Calderón, José Luis Vargas, Mariela Gutiérrez, Joselyn Prado & Juan Carlos Villalobos Ugalde |
| EPI_ISL_3464737, EPI_ISL_3464738, EPI_ISL_3464739, EPI_ISL_3464740, EPI_ISL_3464741, EPI_ISL_3464742, EPI_ISL_3464743, EPI_ISL_3464744, EPI_ISL_3464745, EPI_ISL_3464746, EPI_ISL_3464747, EPI_ISL_3464748, EPI_ISL_3464749, EPI_ISL_3464750, EPI_ISL_3464751, EPI_ISL_3464752, EPI_ISL_3464753, EPI_ISL_3464754, EPI_ISL_3464755, EPI_ISL_3464756, EPI_ISL_3464757, EPI_ISL_3464758, EPI_ISL_3464759, EPI_ISL_3464760, EPI_ISL_3464761, EPI_ISL_3464762, EPI_ISL_3464763, EPI_ISL_3464764, EPI_ISL_3464765, EPI_ISL_3464766, EPI_ISL_3464767, EPI_ISL_3464768, EPI_ISL_3464769, EPI_ISL_3464770, EPI_ISL_3464771, EPI_ISL_3464772, EPI_ISL_3464773, EPI_ISL_3464774, EPI_ISL_3464775, EPI_ISL_3464776, EPI_ISL_3464777, EPI_ISL_3464778, EPI_ISL_3464779, EPI_ISL_3464780, EPI_ISL_3464781 | | | |
| see above | UAB InMedica | Lithuanian University of Health Sciences | Arnoldas Pautienius, Kamile Tamusauskaite, Gediminas Alzbutas, Dovydas Gecys, Lukas Zemaitis, Vaiva Lesauskaite |
| EPI_ISL_3464782, EPI_ISL_3464783, EPI_ISL_3464784, EPI_ISL_3464785, EPI_ISL_3464786, EPI_ISL_3464787, EPI_ISL_3464788, EPI_ISL_3464789, EPI_ISL_3464790, EPI_ISL_3464791, EPI_ISL_3464792, EPI_ISL_3464793, EPI_ISL_3464794, EPI_ISL_3464795, EPI_ISL_3464796, EPI_ISL_3464797, EPI_ISL_3464798, EPI_ISL_3464799, EPI_ISL_3464800, EPI_ISL_3464801, EPI_ISL_3464802, EPI_ISL_3464803, EPI_ISL_3464804, EPI_ISL_3464805, EPI_ISL_3464806, EPI_ISL_3464807, EPI_ISL_3464808, EPI_ISL_3464809, EPI_ISL_3464810, EPI_ISL_3464811, EPI_ISL_3464812, EPI_ISL_3464813, EPI_ISL_3464814, EPI_ISL_3464815, EPI_ISL_3464816, EPI_ISL_3464817, EPI_ISL_3464818, EPI_ISL_3464819, EPI_ISL_3464820, EPI_ISL_3464821, EPI_ISL_3464822, EPI_ISL_3464823, EPI_ISL_3464824, EPI_ISL_3464825, EPI_ISL_3464826, EPI_ISL_3464827, EPI_ISL_3464828, EPI_ISL_3464829, EPI_ISL_3464830, EPI_ISL_3464831, EPI_ISL_3464832, EPI_ISL_3464833, EPI_ISL_3464834, EPI_ISL_3464835, EPI_ISL_3464836, EPI_ISL_3464837, EPI_ISL_3464838, EPI_ISL_3464839, EPI_ISL_3464840, EPI_ISL_3464841, EPI_ISL_3464842, EPI_ISL_3464843, EPI_ISL_3464844, EPI_ISL_3464845 | | | |
| see above | UAB InMedica | unknown | Rasa Ugenskiene, Darius Cereskevicius, Inga Nasvytiene, Zivile Zemeckiene, Mantas Sarauskas, Marius Sukys, Rima Vainoriene, Astra Vitkauskiene, Renaldas Jurkevicius |
| EPI_ISL_3464846, EPI_ISL_3464847, EPI_ISL_3464848, EPI_ISL_3464849, EPI_ISL_3464850, EPI_ISL_3464852, EPI_ISL_3464853, EPI_ISL_3464854, EPI_ISL_3464855, EPI_ISL_3464856, EPI_ISL_3464857, EPI_ISL_3464858, EPI_ISL_3464859, EPI_ISL_3464860, EPI_ISL_3464861, EPI_ISL_3464862, EPI_ISL_3464863, EPI_ISL_3464864, EPI_ISL_3464865, EPI_ISL_3464866, EPI_ISL_3464867, EPI_ISL_3464868, EPI_ISL_3464869, EPI_ISL_3464870, EPI_ISL_3464871, EPI_ISL_3464872, EPI_ISL_3464873, EPI_ISL_3464874, EPI_ISL_3464875, EPI_ISL_3464876, EPI_ISL_3464877, EPI_ISL_3464878, EPI_ISL_3464879, EPI_ISL_3464880, EPI_ISL_3464881, EPI_ISL_3464882, EPI_ISL_3464883, EPI_ISL_3464884, EPI_ISL_3464885, EPI_ISL_3464886, EPI_ISL_3464887, EPI_ISL_3464888, EPI_ISL_3464889, EPI_ISL_3464890, | | | |

| | | | |
|---|---|---|---|
| EPI_ISL_3464891, EPI_ISL_3464892, EPI_ISL_3464893, EPI_ISL_3464894, EPI_ISL_3464895, EPI_ISL_3464896, EPI_ISL_3464897, EPI_ISL_3464898, EPI_ISL_3464899, EPI_ISL_3464900, EPI_ISL_3464901, EPI_ISL_3464902, EPI_ISL_3464903, EPI_ISL_3464904, EPI_ISL_3464905, EPI_ISL_3464906, EPI_ISL_3464907, EPI_ISL_3464908, EPI_ISL_3464909, EPI_ISL_3464910, EPI_ISL_3464911, EPI_ISL_3464912, EPI_ISL_3464913, EPI_ISL_3464914, EPI_ISL_3464915, EPI_ISL_3464916, EPI_ISL_3464917, EPI_ISL_3464918, EPI_ISL_3464919, EPI_ISL_3464920, EPI_ISL_3464921, EPI_ISL_3464922, EPI_ISL_3464923, EPI_ISL_3464924, EPI_ISL_3464925, EPI_ISL_3464926, EPI_ISL_3464927, EPI_ISL_3464928, EPI_ISL_3464929, EPI_ISL_3464930, EPI_ISL_3464931, EPI_ISL_3464932, EPI_ISL_3464933, EPI_ISL_3464934, EPI_ISL_3464935, EPI_ISL_3464936, EPI_ISL_3464937, EPI_ISL_3464938, EPI_ISL_3464939, EPI_ISL_3464940, EPI_ISL_3464941, EPI_ISL_3464942, EPI_ISL_3464943, EPI_ISL_3464944, EPI_ISL_3464945, EPI_ISL_3464946, EPI_ISL_3464947, EPI_ISL_3464948, EPI_ISL_3464950, EPI_ISL_3464951, EPI_ISL_3464952 | | | |
| see above | UAB Medicina practica laboratorija | unknown | Rasa Ugenskiene, Darius Cereskevicius, Inga Nasvytiene, Zivile Zemeckiene, Mantas Sarauskas, Marius Sukys, Rima Vainoriene, Astra Vitkauskiene, Renaldas Jurkevicius |
| EPI_ISL_3464953, EPI_ISL_3464954, EPI_ISL_3464955, EPI_ISL_3464956, EPI_ISL_3464957, EPI_ISL_3464958, EPI_ISL_3464959, EPI_ISL_3464960, EPI_ISL_3464961, EPI_ISL_3464962, EPI_ISL_3464963, EPI_ISL_3464964, EPI_ISL_3464965, EPI_ISL_3464966, EPI_ISL_3464967, EPI_ISL_3464968, EPI_ISL_3464969, EPI_ISL_3464970, EPI_ISL_3464971, EPI_ISL_3464972 | | | |
| see above | Viollier AG | Department of Biosystems Science and Engineering, ETH Zürich | Chaoran Chen, Sarah Nadeau, Louis du Plessis, Catharine Aquino, Ivan Topolsky, Kim Philipp Jablonski, Lara Fuhrmann, Daniel Ehrsam, Isabel Stürmer, Andreia Cabral de Gouvea, Maria Domenica Moccia, Simon Grüter, Timothy Sykes, Lennart Opitz, Griffin White, Laura Neff, Doris Popovic, Andrea Patrignani, Jay Tracy, Ralph Schlapbach, Christiane Beckmann, Maurice Redondo, Olivier Kobel, Christoph Noppen, Niko Beerenwinkel, Tanja Stadler |
| EPI_ISL_3464973 | Viollier AG | Department of Biosystems Science and Engineering, ETH Zürich | Andrea Patrizia Salzmann, Henriette Kurth, Christiane Beckmann, Ivan Topolsky, Chaoran Chen, Sarah Nadeau, Louis du Plessis, Kim Philipp Jablonski, Lara Fuhrmann, Niko Beerenwinkel, Christoph Noppen, Olivier Kobel, Maurice Redondo, Tanja Stadler |
| EPI_ISL_3464974, EPI_ISL_3464975, EPI_ISL_3464976 | Viollier AG | Department of Biosystems Science and Engineering, ETH Zürich | Chaoran Chen, Sarah Nadeau, Louis du Plessis, Catharine Aquino, Ivan Topolsky, Kim Philipp Jablonski, Lara Fuhrmann, Daniel Ehrsam, Isabel Stürmer, Andreia Cabral de Gouvea, Maria Domenica Moccia, Simon Grüter, Timothy Sykes, Lennart Opitz, Griffin White, Laura Neff, Doris Popovic, Andrea Patrignani, Jay Tracy, Ralph Schlapbach, Christiane Beckmann, Maurice Redondo, Olivier Kobel, Christoph Noppen, Niko Beerenwinkel, Tanja Stadler |
| EPI_ISL_3464977 | Viollier AG | Department of Biosystems Science and Engineering, ETH Zürich | Andrea Patrizia Salzmann, Henriette Kurth, Christiane Beckmann, Ivan Topolsky, Chaoran Chen, Sarah Nadeau, Louis du Plessis, Kim Philipp Jablonski, Lara Fuhrmann, Niko Beerenwinkel, Christoph Noppen, Olivier Kobel, Maurice Redondo, Tanja Stadler |
| EPI_ISL_3464978, EPI_ISL_3464980, EPI_ISL_3464981, EPI_ISL_3464982, EPI_ISL_3464983, EPI_ISL_3464984, EPI_ISL_3464985, EPI_ISL_3464986, EPI_ISL_3464987, EPI_ISL_3464988, EPI_ISL_3464989, EPI_ISL_3464990, EPI_ISL_3464991, EPI_ISL_3464992, EPI_ISL_3464993, EPI_ISL_3464994, EPI_ISL_3464995, EPI_ISL_3464996, EPI_ISL_3464997, EPI_ISL_3464998, EPI_ISL_3464999, EPI_ISL_3465000, EPI_ISL_3465001, EPI_ISL_3465002, EPI_ISL_3465003, EPI_ISL_3465004, EPI_ISL_3465005, EPI_ISL_3465006, EPI_ISL_3465007, EPI_ISL_3465009, EPI_ISL_3465010, EPI_ISL_3465011, EPI_ISL_3465012, EPI_ISL_3465013, EPI_ISL_3465014, EPI_ISL_3465015, EPI_ISL_3465016, EPI_ISL_3465017 | | | |
| see above | Viollier AG | Department of Biosystems Science and Engineering, ETH Zürich | Chaoran Chen, Sarah Nadeau, Louis du Plessis, Catharine Aquino, Ivan Topolsky, Kim Philipp Jablonski, Lara Fuhrmann, Daniel Ehrsam, Isabel Stürmer, Andreia Cabral de Gouvea, Maria Domenica Moccia, Simon Grüter, Timothy Sykes, Lennart Opitz, Griffin White, Laura Neff, Doris Popovic, Andrea Patrignani, Jay Tracy, Ralph Schlapbach, Christiane Beckmann, Maurice Redondo, Olivier Kobel, Christoph Noppen, Niko Beerenwinkel, Tanja Stadler |
| EPI_ISL_3465018 | Viollier AG | Department of Biosystems Science and Engineering, ETH Zürich | Andrea Patrizia Salzmann, Henriette Kurth, Christiane Beckmann, Ivan Topolsky, Chaoran Chen, Sarah Nadeau, Louis du Plessis, Kim Philipp Jablonski, Lara Fuhrmann, Niko Beerenwinkel, Christoph Noppen, Olivier Kobel, Maurice Redondo, Tanja Stadler |
| EPI_ISL_3465019, EPI_ISL_3465020, EPI_ISL_3465021, EPI_ISL_3465022, EPI_ISL_3465023, EPI_ISL_3465024 | Viollier AG | Department of Biosystems Science and Engineering, ETH Zürich | Chaoran Chen, Sarah Nadeau, Louis du Plessis, Catharine Aquino, Ivan Topolsky, Kim Philipp Jablonski, Lara Fuhrmann, Daniel Ehrsam, Isabel Stürmer, Andreia Cabral de Gouvea, Maria Domenica Moccia, Simon Grüter, Timothy Sykes, Lennart Opitz, Griffin White, Laura Neff, Doris Popovic, Andrea Patrignani, Jay Tracy, Ralph Schlapbach, Christiane Beckmann, Maurice Redondo, Olivier Kobel, Christoph Noppen, Niko Beerenwinkel, Tanja Stadler |
| EPI_ISL_3465025 | Viollier AG | Department of Biosystems Science and Engineering, ETH Zürich | Andrea Patrizia Salzmann, Henriette Kurth, Christiane Beckmann, Ivan Topolsky, Chaoran Chen, Sarah Nadeau, Louis du Plessis, Kim Philipp Jablonski, Lara Fuhrmann, Niko Beerenwinkel, Christoph Noppen, Olivier Kobel, Maurice Redondo, Tanja Stadler |
| EPI_ISL_3465026, EPI_ISL_3465027, EPI_ISL_3465028, EPI_ISL_3465029, EPI_ISL_3465030, EPI_ISL_3465031, EPI_ISL_3465032, EPI_ISL_3465033, EPI_ISL_3465034, EPI_ISL_3465035, EPI_ISL_3465036, EPI_ISL_3465038, EPI_ISL_3465039, EPI_ISL_3465040, EPI_ISL_3465042, EPI_ISL_3465043, EPI_ISL_3465044, EPI_ISL_3465045, EPI_ISL_3465046 | | | |
| see above | Viollier AG | Department of Biosystems Science and Engineering, ETH Zürich | Chaoran Chen, Sarah Nadeau, Louis du Plessis, Catharine Aquino, Ivan Topolsky, Kim Philipp Jablonski, Lara Fuhrmann, Daniel Ehrsam, Isabel Stürmer, Andreia Cabral de Gouvea, Maria Domenica Moccia, Simon Grüter, Timothy Sykes, Lennart Opitz, Griffin White, Laura Neff, Doris Popovic, Andrea Patrignani, Jay Tracy, Ralph Schlapbach, Christiane Beckmann, Maurice Redondo, Olivier Kobel, Christoph Noppen, Niko Beerenwinkel, Tanja Stadler |
| EPI_ISL_3465047 | Viollier AG | Department of Biosystems Science and Engineering, ETH Zürich | Andrea Patrizia Salzmann, Henriette Kurth, Christiane Beckmann, Ivan Topolsky, Chaoran Chen, Sarah Nadeau, Louis du Plessis, Kim Philipp Jablonski, Lara Fuhrmann, Niko Beerenwinkel, Christoph Noppen, Olivier Kobel, Maurice Redondo, Tanja Stadler |
| EPI_ISL_3465048, EPI_ISL_3465049, EPI_ISL_3465050, EPI_ISL_3465051, EPI_ISL_3465052, EPI_ISL_3465053, EPI_ISL_3465054, EPI_ISL_3465055, EPI_ISL_3465057, EPI_ISL_3465058, EPI_ISL_3465059, EPI_ISL_3465060, EPI_ISL_3465061, EPI_ISL_3465062 | | | |
| see above | Viollier AG | Department of Biosystems Science and Engineering, ETH Zürich | Chaoran Chen, Sarah Nadeau, Louis du Plessis, Catharine Aquino, Ivan Topolsky, Kim Philipp Jablonski, Lara Fuhrmann, Daniel Ehrsam, Isabel Stürmer, Andreia Cabral de Gouvea, Maria Domenica Moccia, Simon Grüter, Timothy Sykes, Lennart Opitz, Griffin White, Laura Neff, Doris Popovic, Andrea Patrignani, Jay Tracy, Ralph Schlapbach, Christiane Beckmann, Maurice Redondo, Olivier Kobel, Christoph Noppen, Niko Beerenwinkel, Tanja Stadler |
| EPI_ISL_3465063 | Viollier AG | Department of Biosystems Science and Engineering, ETH Zürich | Andrea Patrizia Salzmann, Henriette Kurth, Christiane Beckmann, Ivan Topolsky, Chaoran Chen, Sarah Nadeau, Louis du Plessis, Kim Philipp Jablonski, Lara Fuhrmann, Niko Beerenwinkel, Christoph Noppen, Olivier Kobel, Maurice Redondo, Tanja Stadler |
| EPI_ISL_3465064, EPI_ISL_3465065, EPI_ISL_3465066, EPI_ISL_3465067 | Viollier AG | Department of Biosystems Science and Engineering, ETH Zürich | Chaoran Chen, Sarah Nadeau, Louis du Plessis, Catharine Aquino, Ivan Topolsky, Kim Philipp Jablonski, Lara Fuhrmann, Daniel Ehrsam, Isabel Stürmer, Andreia Cabral de Gouvea, Maria Domenica Moccia, Simon Grüter, Timothy Sykes, Lennart Opitz, Griffin White, Laura Neff, Doris Popovic, Andrea Patrignani, Jay Tracy, Ralph Schlapbach, Christiane Beckmann, Maurice Redondo, Olivier Kobel, Christoph Noppen, Niko Beerenwinkel, Tanja Stadler |
| EPI_ISL_3465068 | Viollier AG | Department of Biosystems Science and Engineering, ETH Zürich | Andrea Patrizia Salzmann, Henriette Kurth, Christiane Beckmann, Ivan Topolsky, Chaoran Chen, Sarah Nadeau, Louis du Plessis, Kim Philipp Jablonski, Lara Fuhrmann, Niko Beerenwinkel, Christoph Noppen, Olivier Kobel, Maurice Redondo, Tanja Stadler |
| EPI_ISL_3465069, EPI_ISL_3465070, EPI_ISL_3465071, EPI_ISL_3465072, EPI_ISL_3465073, EPI_ISL_3465074, EPI_ISL_3465075, EPI_ISL_3465076, EPI_ISL_3465077, EPI_ISL_3465078, EPI_ISL_3465079, EPI_ISL_3465080, EPI_ISL_3465081, EPI_ISL_3465082, EPI_ISL_3465083, EPI_ISL_3465084, EPI_ISL_3465085, EPI_ISL_3465086, EPI_ISL_3465087, EPI_ISL_3465088, EPI_ISL_3465090, EPI_ISL_3465091, EPI_ISL_3465092, EPI_ISL_3465093, EPI_ISL_3465094, EPI_ISL_3465095, EPI_ISL_3465096, EPI_ISL_3465097, EPI_ISL_3465098, EPI_ISL_3465099, EPI_ISL_3465100, EPI_ISL_3465101, EPI_ISL_3465102, EPI_ISL_3465103, EPI_ISL_3465104, EPI_ISL_3465105, EPI_ISL_3465106, EPI_ISL_3465107, EPI_ISL_3465108, EPI_ISL_3465109, EPI_ISL_3465110, EPI_ISL_3465111, EPI_ISL_3465113, EPI_ISL_3465114, EPI_ISL_3465115, EPI_ISL_3465116, EPI_ISL_3465117, EPI_ISL_3465118, EPI_ISL_3465120, EPI_ISL_3465123, EPI_ISL_3465124, EPI_ISL_3465125, EPI_ISL_3465126, EPI_ISL_3465127, EPI_ISL_3465128, EPI_ISL_3465129, EPI_ISL_3465130, EPI_ISL_3465131, EPI_ISL_3465132, EPI_ISL_3465133, EPI_ISL_3465134, EPI_ISL_3465136, EPI_ISL_3465137, EPI_ISL_3465138, EPI_ISL_3465139, EPI_ISL_3465140, EPI_ISL_3465141, EPI_ISL_3465143, EPI_ISL_3465144, EPI_ISL_3465145, EPI_ISL_3465146, EPI_ISL_3465147, EPI_ISL_3465148, EPI_ISL_3465149, EPI_ISL_3465150, EPI_ISL_3465151, EPI_ISL_3465152, EPI_ISL_3465153, EPI_ISL_3465154, EPI_ISL_3465155, EPI_ISL_3465156, EPI_ISL_3465157, EPI_ISL_3465158, EPI_ISL_3465159, EPI_ISL_3465160, EPI_ISL_3465161, EPI_ISL_3465162, EPI_ISL_3465163, EPI_ISL_3465164, EPI_ISL_3465165, EPI_ISL_3465167, EPI_ISL_3465168, EPI_ISL_3465169, EPI_ISL_3465170, EPI_ISL_3465171, EPI_ISL_3465172, EPI_ISL_3465173, EPI_ISL_3465174, EPI_ISL_3465175, EPI_ISL_3465176 | | | |
| see above | Viollier AG | Department of Biosystems Science and Engineering, ETH Zürich | Chaoran Chen, Sarah Nadeau, Louis du Plessis, Catharine Aquino, Ivan Topolsky, Kim Philipp Jablonski, Lara Fuhrmann, Daniel Ehrsam, Isabel Stürmer, Andreia Cabral de Gouvea, Maria Domenica Moccia, Simon Grüter, Timothy Sykes, Lennart Opitz, Griffin White, Laura Neff, Doris Popovic, Andrea Patrignani, Jay Tracy, Ralph Schlapbach, Christiane Beckmann, Maurice Redondo, Olivier Kobel, Christoph Noppen, Niko Beerenwinkel, Tanja Stadler |
| EPI_ISL_3465177 | Viollier AG | Department of Biosystems Science and Engineering, ETH Zürich | Andrea Patrizia Salzmann, Henriette Kurth, Christiane Beckmann, Ivan Topolsky, Chaoran Chen, Sarah Nadeau, Louis du Plessis, Kim Philipp Jablonski, Lara Fuhrmann, Niko Beerenwinkel, Christoph Noppen, Olivier Kobel, Maurice Redondo, Tanja Stadler |
| EPI_ISL_3465178, EPI_ISL_3465179, EPI_ISL_3465180, EPI_ISL_3465181, EPI_ISL_3465182, EPI_ISL_3465183, EPI_ISL_3465184, EPI_ISL_3465185, EPI_ISL_3465186, EPI_ISL_3465187, EPI_ISL_3465188, EPI_ISL_3465189, EPI_ISL_3465190, EPI_ISL_3465191, EPI_ISL_3465192, EPI_ISL_3465193, EPI_ISL_3465194, EPI_ISL_3465195, EPI_ISL_3465196, EPI_ISL_3465197, EPI_ISL_3465198, EPI_ISL_3465199 | | | |
| see above | Viollier AG | Department of Biosystems Science and Engineering, ETH Zürich | Chaoran Chen, Sarah Nadeau, Louis du Plessis, Catharine Aquino, Ivan Topolsky, Kim Philipp Jablonski, Lara Fuhrmann, Daniel Ehrsam, Isabel Stürmer, Andreia Cabral de Gouvea, Maria Domenica Moccia, Simon Grüter, Timothy Sykes, Lennart Opitz, Griffin White, Laura Neff, Doris Popovic, Andrea Patrignani, Jay Tracy, Ralph Schlapbach, Christiane Beckmann, Maurice Redondo, Olivier Kobel, Christoph Noppen, Niko Beerenwinkel, Tanja Stadler |
| EPI_ISL_3465200 | Viollier AG | Department of Biosystems Science and Engineering, ETH Zürich | Andrea Patrizia Salzmann, Henriette Kurth, Christiane Beckmann, Ivan Topolsky, Chaoran Chen, Sarah Nadeau, Louis du Plessis, Kim Philipp Jablonski, Lara Fuhrmann, Niko Beerenwinkel, Christoph Noppen, Olivier Kobel, Maurice Redondo, Tanja Stadler |
| EPI_ISL_3465201, EPI_ISL_3465202, EPI_ISL_3465203, EPI_ISL_3465204, EPI_ISL_3465205, EPI_ISL_3465206, EPI_ISL_3465207, EPI_ISL_3465208, EPI_ISL_3465209, EPI_ISL_3465210, EPI_ISL_3465211, EPI_ISL_3465212, EPI_ISL_3465213, EPI_ISL_3465215, EPI_ISL_3465216, EPI_ISL_3465217, EPI_ISL_3465218, EPI_ISL_3465219, EPI_ISL_3465220, EPI_ISL_3465221, EPI_ISL_3465222 | | | |
| see above | Viollier AG | Department of Biosystems Science and Engineering, ETH Zürich | Chaoran Chen, Sarah Nadeau, Louis du Plessis, Catharine Aquino, Ivan Topolsky, Kim Philipp Jablonski, Lara Fuhrmann, Daniel Ehrsam, Isabel Stürmer, Andreia Cabral de Gouvea, Maria Domenica Moccia, Simon Grüter, Timothy Sykes, Lennart Opitz, Griffin White, Laura Neff, Doris Popovic, Andrea Patrignani, Jay Tracy, Ralph Schlapbach, Christiane Beckmann, Maurice Redondo, Olivier Kobel, Christoph Noppen, Niko Beerenwinkel, Tanja Stadler |
| EPI_ISL_3465223 | Viollier AG | Department of Biosystems Science and Engineering, ETH Zürich | Andrea Patrizia Salzmann, Henriette Kurth, Christiane Beckmann, Ivan Topolsky, Chaoran Chen, Sarah Nadeau, Louis du Plessis, Kim Philipp Jablonski, Lara Fuhrmann, Niko Beerenwinkel, Christoph Noppen, Olivier Kobel, Maurice Redondo, Tanja Stadler |
| EPI_ISL_3465224 | Viollier AG | Department of Biosystems Science and Engineering, ETH Zürich | Chaoran Chen, Sarah Nadeau, Louis du Plessis, Catharine Aquino, Ivan Topolsky, Kim Philipp Jablonski, Lara Fuhrmann, Daniel Ehrsam, Isabel Stürmer, Andreia Cabral de Gouvea, Maria Domenica Moccia, Simon Grüter, Timothy Sykes, Lennart Opitz, Griffin White, Laura Neff, Doris Popovic, Andrea Patrignani, Jay Tracy, Ralph Schlapbach, Christiane Beckmann, Maurice Redondo, Olivier Kobel, Christoph Noppen, Niko Beerenwinkel, Tanja Stadler |
| EPI_ISL_3465225 | St Vincent's Pathology (SydPath) | NSW Health Pathology - Institute of Clinical Pathology and Medical Research; Westmead Hospital; University of Sydney | Arnott A., Draper J., Gall M., Martinez E., Rockett R., Sintchenko V., on behalf of ICPMR |
| EPI_ISL_3465226, EPI_ISL_3465227 | Viollier AG | Department of Biosystems Science and Engineering, ETH Zürich | Chaoran Chen, Sarah Nadeau, Louis du Plessis, Catharine Aquino, Ivan Topolsky, Kim Philipp Jablonski, Lara Fuhrmann, Daniel Ehrsam, Isabel Stürmer, Andreia Cabral de Gouvea, Maria Domenica Moccia, Simon Grüter, Timothy Sykes, Lennart Opitz, Griffin White, Laura Neff, Doris Popovic, Andrea Patrignani, Jay Tracy, Ralph Schlapbach, Christiane Beckmann, Maurice Redondo, Olivier Kobel, Christoph Noppen, Niko Beerenwinkel, Tanja Stadler |
| EPI_ISL_3465228 | St Vincent's Pathology (SydPath) | NSW Health Pathology - Institute of Clinical Pathology and Medical Research; Westmead Hospital; University of Sydney | Arnott A., Draper J., Gall M., Martinez E., Rockett R., Sintchenko V., on behalf of ICPMR |
| EPI_ISL_3465229, EPI_ISL_3465230 | Viollier AG | Department of Biosystems Science and Engineering, ETH Zürich | Chaoran Chen, Sarah Nadeau, Louis du Plessis, Catharine Aquino, Ivan Topolsky, Kim Philipp Jablonski, Lara Fuhrmann, Daniel Ehrsam, Isabel Stürmer, Andreia Cabral de Gouvea, Maria Domenica Moccia, Simon Grüter, Timothy Sykes, Lennart Opitz, Griffin White, Laura Neff, Doris Popovic, Andrea Patrignani, Jay Tracy, Ralph Schlapbach, Christiane Beckmann, Maurice Redondo, Olivier Kobel, Christoph Noppen, Niko Beerenwinkel, Tanja Stadler |
| EPI_ISL_3465231 | Sydney South West Pathology Service (SSWPS) - Liverpool Hospital - NSW Health Pathology | NSW Health Pathology - Institute of Clinical Pathology and Medical Research; Westmead Hospital; University of Sydney | Arnott A., Draper J., Gall M., Martinez E., Rockett R., Sintchenko V., on behalf of ICPMR |
| EPI_ISL_3465232 | Viollier AG | Department of Biosystems Science and Engineering, ETH Zürich | Andrea Patrizia Salzmann, Henriette Kurth, Christiane Beckmann, Ivan Topolsky, Chaoran Chen, Sarah Nadeau, Louis du Plessis, Kim Philipp Jablonski, Lara Fuhrmann, Niko Beerenwinkel, Christoph Noppen, Olivier Kobel, Maurice Redondo, Tanja Stadler |
| EPI_ISL_3465233 | Australian Clinical Labs (formerly Healthscope Pathology) | NSW Health Pathology - Institute of Clinical Pathology and Medical Research; Westmead Hospital; University of Sydney | Arnott A., Draper J., Gall M., Martinez E., Rockett R., Sintchenko V., on behalf of ICPMR |
| EPI_ISL_3465234 | Viollier AG | Department of Biosystems Science and Engineering, ETH Zürich | Chaoran Chen, Sarah Nadeau, Louis du Plessis, Catharine Aquino, Ivan Topolsky, Kim Philipp Jablonski, Lara Fuhrmann, Daniel Ehrsam, Isabel Stürmer, Andreia Cabral de Gouvea, Maria Domenica Moccia, Simon Grüter, Timothy Sykes, Lennart Opitz, Griffin White, Laura Neff, Doris Popovic, Andrea Patrignani, Jay Tracy, Ralph Schlapbach, Christiane Beckmann, Maurice Redondo, Olivier Kobel, Christoph Noppen, Niko Beerenwinkel, Tanja Stadler |
| EPI_ISL_3465235 | 4Cyte Pathology | NSW Health Pathology - Institute of Clinical Pathology and Medical Research; Westmead Hospital; University of Sydney | Arnott A., Draper J., Gall M., Martinez E., Rockett R., Sintchenko V., on behalf of ICPMR |
| EPI_ISL_3465236 | Viollier AG | Department of Biosystems Science and Engineering, ETH Zürich | Chaoran Chen, Sarah Nadeau, Louis du Plessis, Catharine Aquino, Ivan Topolsky, Kim Philipp Jablonski, Lara Fuhrmann, Daniel Ehrsam, Isabel Stürmer, Andreia Cabral de Gouvea, Maria Domenica Moccia, Simon Grüter, Timothy Sykes, Lennart Opitz, Griffin White, Laura Neff, Doris Popovic, Andrea Patrignani, Jay Tracy, Ralph Schlapbach, Christiane Beckmann, Maurice Redondo, Olivier Kobel, Christoph Noppen, Niko Beerenwinkel, Tanja Stadler |
| EPI_ISL_3465238 | 4Cyte Path | NSW Health Pathology - Institute of Clinical Pathology and Medical Research; Westmead Hospital; University of Sydney | Arnott A., Draper J., Gall M., Martinez E., Rockett R., Sintchenko V., on behalf of ICPMR |
| EPI_ISL_3465239 | Viollier AG | Department of Biosystems Science and Engineering, ETH Zürich | Chaoran Chen, Sarah Nadeau, Louis du Plessis, Catharine Aquino, Ivan Topolsky, Kim Philipp Jablonski, Lara Fuhrmann, Daniel Ehrsam, Isabel Stürmer, Andreia Cabral de Gouvea, Maria Domenica Moccia, Simon Grüter, Timothy Sykes, Lennart Opitz, Griffin White, Laura Neff, Doris Popovic, Andrea Patrignani, Jay Tracy, Ralph Schlapbach, Christiane Beckmann, Maurice Redondo, Olivier Kobel, Christoph Noppen, Niko Beerenwinkel, Tanja Stadler |
| EPI_ISL_3465240 | Sydney South West Pathology Service (SSWPS) - Liverpool Hospital - NSW Health Pathology | NSW Health Pathology - Institute of Clinical Pathology and Medical Research; Westmead Hospital; University of Sydney | Arnott A., Draper J., Gall M., Martinez E., Rockett R., Sintchenko V., on behalf of ICPMR |

| | | | |
|---|---|---|---|
| EPI_ISL_3465242 | Medlab Pathology | NSW Health Pathology - Institute of Clinical Pathology and Medical Research; Westmead Hospital; University of Sydney | Arnott A., Draper J., Gall M., Martinez E., Rockett R., Sintchenko V., on behalf of ICPMR |
| EPI_ISL_3465243, EPI_ISL_3465244 | Viollier AG | Department of Biosystems Science and Engineering, ETH Zürich | Chaoran Chen, Sarah Nadeau, Louis du Plessis, Catharine Aquino, Ivan Topolsky, Kim Philipp Jablonski, Lara Fuhrmann, Daniel Ehrsam, Isabel Stürmer, Andreia Cabral de Gouvea, Maria Domenica Moccia, Simon Grüter, Timothy Sykes, Lennart Opitz, Griffin White, Laura Neff, Doris Popovic, Andrea Patrignani, Jay Tracy, Ralph Schlapbach, Christiane Beckmann, Maurice Redondo, Olivier Kobel, Christoph Noppen, Niko Beerenwinkel, Tanja Stadler |
| EPI_ISL_3465245 | Sydney South West Pathology Service (SSWPS) - Liverpool Hospital - NSW Health Pathology | NSW Health Pathology - Institute of Clinical Pathology and Medical Research; Westmead Hospital; University of Sydney | Arnott A., Draper J., Gall M., Martinez E., Rockett R., Sintchenko V., on behalf of ICPMR |
| EPI_ISL_3465246 | Viollier AG | Department of Biosystems Science and Engineering, ETH Zürich | Chaoran Chen, Sarah Nadeau, Louis du Plessis, Catharine Aquino, Ivan Topolsky, Kim Philipp Jablonski, Lara Fuhrmann, Daniel Ehrsam, Isabel Stürmer, Andreia Cabral de Gouvea, Maria Domenica Moccia, Simon Grüter, Timothy Sykes, Lennart Opitz, Griffin White, Laura Neff, Doris Popovic, Andrea Patrignani, Jay Tracy, Ralph Schlapbach, Christiane Beckmann, Maurice Redondo, Olivier Kobel, Christoph Noppen, Niko Beerenwinkel, Tanja Stadler |
| EPI_ISL_3465247 | Medlab Pathology | NSW Health Pathology - Institute of Clinical Pathology and Medical Research; Westmead Hospital; University of Sydney | Arnott A., Draper J., Gall M., Martinez E., Rockett R., Sintchenko V., on behalf of ICPMR |
| EPI_ISL_3465248 | Viollier AG | Department of Biosystems Science and Engineering, ETH Zürich | Chaoran Chen, Sarah Nadeau, Louis du Plessis, Catharine Aquino, Ivan Topolsky, Kim Philipp Jablonski, Lara Fuhrmann, Daniel Ehrsam, Isabel Stürmer, Andreia Cabral de Gouvea, Maria Domenica Moccia, Simon Grüter, Timothy Sykes, Lennart Opitz, Griffin White, Laura Neff, Doris Popovic, Andrea Patrignani, Jay Tracy, Ralph Schlapbach, Christiane Beckmann, Maurice Redondo, Olivier Kobel, Christoph Noppen, Niko Beerenwinkel, Tanja Stadler |
| EPI_ISL_3465250 | Medlab Pathology | NSW Health Pathology - Institute of Clinical Pathology and Medical Research; Westmead Hospital; University of Sydney | Arnott A., Draper J., Gall M., Martinez E., Rockett R., Sintchenko V., on behalf of ICPMR |
| EPI_ISL_3465251 | Viollier AG | Department of Biosystems Science and Engineering, ETH Zürich | Andrea Patrizia Salzmann, Henriette Kurth, Christiane Beckmann, Ivan Topolsky, Chaoran Chen, Sarah Nadeau, Louis du Plessis, Kim Philipp Jablonski, Lara Fuhrmann, Niko Beerenwinkel, Christoph Noppen, Olivier Kobel, Maurice Redondo, Tanja Stadler |
| EPI_ISL_3465252 | St Vincent's Pathology (SydPath) | NSW Health Pathology - Institute of Clinical Pathology and Medical Research; Westmead Hospital; University of Sydney | Arnott A., Draper J., Gall M., Martinez E., Rockett R., Sintchenko V., on behalf of ICPMR |
| EPI_ISL_3465253 | Viollier AG | Department of Biosystems Science and Engineering, ETH Zürich | Chaoran Chen, Sarah Nadeau, Louis du Plessis, Catharine Aquino, Ivan Topolsky, Kim Philipp Jablonski, Lara Fuhrmann, Daniel Ehrsam, Isabel Stürmer, Andreia Cabral de Gouvea, Maria Domenica Moccia, Simon Grüter, Timothy Sykes, Lennart Opitz, Griffin White, Laura Neff, Doris Popovic, Andrea Patrignani, Jay Tracy, Ralph Schlapbach, Christiane Beckmann, Maurice Redondo, Olivier Kobel, Christoph Noppen, Niko Beerenwinkel, Tanja Stadler |
| EPI_ISL_3465254 | Medlab Pathology | NSW Health Pathology - Institute of Clinical Pathology and Medical Research; Westmead Hospital; University of Sydney | Arnott A., Draper J., Gall M., Martinez E., Rockett R., Sintchenko V., on behalf of ICPMR |
| EPI_ISL_3465255, EPI_ISL_3465256 | Viollier AG | Department of Biosystems Science and Engineering, ETH Zürich | Chaoran Chen, Sarah Nadeau, Louis du Plessis, Catharine Aquino, Ivan Topolsky, Kim Philipp Jablonski, Lara Fuhrmann, Daniel Ehrsam, Isabel Stürmer, Andreia Cabral de Gouvea, Maria Domenica Moccia, Simon Grüter, Timothy Sykes, Lennart Opitz, Griffin White, Laura Neff, Doris Popovic, Andrea Patrignani, Jay Tracy, Ralph Schlapbach, Christiane Beckmann, Maurice Redondo, Olivier Kobel, Christoph Noppen, Niko Beerenwinkel, Tanja Stadler |
| EPI_ISL_3465257 | Sydney South West Pathology Service (SSWPS) - Liverpool Hospital - NSW Health Pathology | NSW Health Pathology - Institute of Clinical Pathology and Medical Research; Westmead Hospital; University of Sydney | Arnott A., Draper J., Gall M., Martinez E., Rockett R., Sintchenko V., on behalf of ICPMR |
| EPI_ISL_3465258 | Viollier AG | Department of Biosystems Science and Engineering, ETH Zürich | Chaoran Chen, Sarah Nadeau, Louis du Plessis, Catharine Aquino, Ivan Topolsky, Kim Philipp Jablonski, Lara Fuhrmann, Daniel Ehrsam, Isabel Stürmer, Andreia Cabral de Gouvea, Maria Domenica Moccia, Simon Grüter, Timothy Sykes, Lennart Opitz, Griffin White, Laura Neff, Doris Popovic, Andrea Patrignani, Jay Tracy, Ralph Schlapbach, Christiane Beckmann, Maurice Redondo, Olivier Kobel, Christoph Noppen, Niko Beerenwinkel, Tanja Stadler |
| EPI_ISL_3465259 | Sydney South West Pathology Service (SSWPS) - Liverpool Hospital - NSW Health Pathology | NSW Health Pathology - Institute of Clinical Pathology and Medical Research; Westmead Hospital; University of Sydney | Arnott A., Draper J., Gall M., Martinez E., Rockett R., Sintchenko V., on behalf of ICPMR |
| EPI_ISL_3465260 | Viollier AG | Department of Biosystems Science and Engineering, ETH Zürich | Chaoran Chen, Sarah Nadeau, Louis du Plessis, Catharine Aquino, Ivan Topolsky, Kim Philipp Jablonski, Lara Fuhrmann, Daniel Ehrsam, Isabel Stürmer, Andreia Cabral de Gouvea, Maria Domenica Moccia, Simon Grüter, Timothy Sykes, Lennart Opitz, Griffin White, Laura Neff, Doris Popovic, Andrea Patrignani, Jay Tracy, Ralph Schlapbach, Christiane Beckmann, Maurice Redondo, Olivier Kobel, Christoph Noppen, Niko Beerenwinkel, Tanja Stadler |
| EPI_ISL_3465261 | St Vincent's Pathology (SydPath) | NSW Health Pathology - Institute of Clinical Pathology and Medical Research; Westmead Hospital; University of Sydney | Arnott A., Draper J., Gall M., Martinez E., Rockett R., Sintchenko V., on behalf of ICPMR |
| EPI_ISL_3465262 | Viollier AG | Department of Biosystems Science and Engineering, ETH Zürich | Chaoran Chen, Sarah Nadeau, Louis du Plessis, Catharine Aquino, Ivan Topolsky, Kim Philipp Jablonski, Lara Fuhrmann, Daniel Ehrsam, Isabel Stürmer, Andreia Cabral de Gouvea, Maria Domenica Moccia, Simon Grüter, Timothy Sykes, Lennart Opitz, Griffin White, Laura Neff, Doris Popovic, Andrea Patrignani, Jay Tracy, Ralph Schlapbach, Christiane Beckmann, Maurice Redondo, Olivier Kobel, Christoph Noppen, Niko Beerenwinkel, Tanja Stadler |
| EPI_ISL_3465263 | 4Cyte Pathology | NSW Health Pathology - Institute of Clinical Pathology and Medical Research; Westmead Hospital; University of Sydney | Arnott A., Draper J., Gall M., Martinez E., Rockett R., Sintchenko V., on behalf of ICPMR |
| EPI_ISL_3465264 | Viollier AG | Department of Biosystems Science and Engineering, ETH Zürich | Chaoran Chen, Sarah Nadeau, Louis du Plessis, Catharine Aquino, Ivan Topolsky, Kim Philipp Jablonski, Lara Fuhrmann, Daniel Ehrsam, Isabel Stürmer, Andreia Cabral de Gouvea, Maria Domenica Moccia, Simon Grüter, Timothy Sykes, Lennart Opitz, Griffin White, Laura Neff, Doris Popovic, Andrea Patrignani, Jay Tracy, Ralph Schlapbach, Christiane Beckmann, Maurice Redondo, Olivier Kobel, Christoph Noppen, Niko Beerenwinkel, Tanja Stadler |
| EPI_ISL_3465265 | St Vincent's Pathology (SydPath) | NSW Health Pathology - Institute of Clinical Pathology and Medical Research; Westmead Hospital; University of Sydney | Arnott A., Draper J., Gall M., Martinez E., Rockett R., Sintchenko V., on behalf of ICPMR |
| EPI_ISL_3465266, EPI_ISL_3465267 | Viollier AG | Department of Biosystems Science and Engineering, ETH Zürich | Chaoran Chen, Sarah Nadeau, Louis du Plessis, Catharine Aquino, Ivan Topolsky, Kim Philipp Jablonski, Lara Fuhrmann, Daniel Ehrsam, Isabel Stürmer, Andreia Cabral de Gouvea, Maria Domenica Moccia, Simon Grüter, Timothy Sykes, Lennart Opitz, Griffin White, Laura Neff, Doris Popovic, Andrea Patrignani, Jay Tracy, Ralph Schlapbach, Christiane Beckmann, Maurice Redondo, Olivier Kobel, Christoph Noppen, Niko Beerenwinkel, Tanja Stadler |
| EPI_ISL_3465268 | Pathology West - NSW Health Pathology | NSW Health Pathology - Institute of Clinical Pathology and Medical Research; Westmead Hospital; University of Sydney | Arnott A., Draper J., Gall M., Martinez E., Rockett R., Sintchenko V., on behalf of ICPMR |
| EPI_ISL_3465269 | Viollier AG | Department of Biosystems Science and Engineering, ETH Zürich | Andrea Patrizia Salzmann, Henriette Kurth, Christiane Beckmann, Ivan Topolsky, Chaoran Chen, Sarah Nadeau, Louis du Plessis, Kim Philipp Jablonski, Lara Fuhrmann, Niko Beerenwinkel, Christoph Noppen, Olivier Kobel, Maurice Redondo, Tanja Stadler |
| EPI_ISL_3465270 | Sydney South West Pathology Service (SSWPS) - Liverpool Hospital - NSW Health Pathology | NSW Health Pathology - Institute of Clinical Pathology and Medical Research; Westmead Hospital; University of Sydney | Arnott A., Draper J., Gall M., Martinez E., Rockett R., Sintchenko V., on behalf of ICPMR |
| EPI_ISL_3465271 | Viollier AG | Department of Biosystems Science and Engineering, ETH Zürich | Chaoran Chen, Sarah Nadeau, Louis du Plessis, Catharine Aquino, Ivan Topolsky, Kim Philipp Jablonski, Lara Fuhrmann, Daniel Ehrsam, Isabel Stürmer, Andreia Cabral de Gouvea, Maria Domenica Moccia, Simon Grüter, Timothy Sykes, Lennart Opitz, Griffin White, Laura Neff, Doris Popovic, Andrea Patrignani, Jay Tracy, Ralph Schlapbach, Christiane Beckmann, Maurice Redondo, Olivier Kobel, Christoph Noppen, Niko Beerenwinkel, Tanja Stadler |
| EPI_ISL_3465272 | Pathology West - NSW Health Pathology | NSW Health Pathology - Institute of Clinical Pathology and Medical Research; Westmead Hospital; University of Sydney | Arnott A., Draper J., Gall M., Martinez E., Rockett R., Sintchenko V., on behalf of ICPMR |
| EPI_ISL_3465274 | Viollier AG | Department of Biosystems Science and Engineering, ETH Zürich | Chaoran Chen, Sarah Nadeau, Louis du Plessis, Catharine Aquino, Ivan Topolsky, Kim Philipp Jablonski, Lara Fuhrmann, Daniel Ehrsam, Isabel Stürmer, Andreia Cabral de Gouvea, Maria Domenica Moccia, Simon Grüter, Timothy Sykes, Lennart Opitz, Griffin White, Laura Neff, Doris Popovic, Andrea Patrignani, Jay Tracy, Ralph Schlapbach, Christiane Beckmann, Maurice Redondo, Olivier Kobel, Christoph Noppen, Niko Beerenwinkel, Tanja Stadler |
| EPI_ISL_3465275 | Medlab Pathology | NSW Health Pathology - Institute of Clinical Pathology and Medical Research; Westmead Hospital; University of Sydney | Arnott A., Draper J., Gall M., Martinez E., Rockett R., Sintchenko V., on behalf of ICPMR |
| EPI_ISL_3465277 | 4Cyte Pathology | NSW Health Pathology - Institute of Clinical Pathology and Medical Research; Westmead Hospital; University of Sydney | Arnott A., Draper J., Gall M., Martinez E., Rockett R., Sintchenko V., on behalf of ICPMR |
| EPI_ISL_3465278 | Viollier AG | Department of Biosystems Science and Engineering, ETH Zürich | Chaoran Chen, Sarah Nadeau, Louis du Plessis, Catharine Aquino, Ivan Topolsky, Kim Philipp Jablonski, Lara Fuhrmann, Daniel Ehrsam, Isabel Stürmer, Andreia Cabral de Gouvea, Maria Domenica Moccia, Simon Grüter, Timothy Sykes, Lennart Opitz, Griffin White, Laura Neff, Doris Popovic, Andrea Patrignani, Jay Tracy, Ralph Schlapbach, Christiane Beckmann, Maurice Redondo, Olivier Kobel, Christoph Noppen, Niko Beerenwinkel, Tanja Stadler |
| EPI_ISL_3465279 | Sydney South West Pathology Service (SSWPS) - Liverpool Hospital - NSW Health Pathology | NSW Health Pathology - Institute of Clinical Pathology and Medical Research; Westmead Hospital; University of Sydney | Arnott A., Draper J., Gall M., Martinez E., Rockett R., Sintchenko V., on behalf of ICPMR |
| EPI_ISL_3465280 | Viollier AG | Department of Biosystems Science and Engineering, ETH Zürich | Andrea Patrizia Salzmann, Henriette Kurth, Christiane Beckmann, Ivan Topolsky, Chaoran Chen, Sarah Nadeau, Louis du Plessis, Kim Philipp Jablonski, Lara Fuhrmann, Niko Beerenwinkel, Christoph Noppen, Olivier Kobel, Maurice Redondo, Tanja Stadler |
| EPI_ISL_3465281 | Viollier AG | Department of Biosystems Science and Engineering, ETH Zürich | Chaoran Chen, Sarah Nadeau, Louis du Plessis, Catharine Aquino, Ivan Topolsky, Kim Philipp Jablonski, Lara Fuhrmann, Daniel Ehrsam, Isabel Stürmer, Andreia Cabral de Gouvea, Maria Domenica Moccia, Simon Grüter, Timothy Sykes, Lennart Opitz, Griffin White, Laura Neff, Doris Popovic, Andrea Patrignani, Jay Tracy, Ralph Schlapbach, Christiane Beckmann, Maurice Redondo, Olivier Kobel, Christoph Noppen, Niko Beerenwinkel, Tanja Stadler |
| EPI_ISL_3465282 | 4Cyte Pathology | NSW Health Pathology - Institute of Clinical Pathology and Medical Research; Westmead Hospital; University of Sydney | Arnott A., Draper J., Gall M., Martinez E., Rockett R., Sintchenko V., on behalf of ICPMR |
| EPI_ISL_3465283 | Viollier AG | Department of Biosystems Science and Engineering, ETH Zürich | Chaoran Chen, Sarah Nadeau, Louis du Plessis, Catharine Aquino, Ivan Topolsky, Kim Philipp Jablonski, Lara Fuhrmann, Daniel Ehrsam, Isabel Stürmer, Andreia Cabral de Gouvea, Maria Domenica Moccia, Simon Grüter, Timothy Sykes, Lennart Opitz, Griffin White, Laura Neff, Doris Popovic, Andrea Patrignani, Jay Tracy, Ralph Schlapbach, Christiane Beckmann, Maurice Redondo, Olivier Kobel, Christoph Noppen, Niko Beerenwinkel, Tanja Stadler |
| EPI_ISL_3465284 | Medlab Pathology | NSW Health Pathology - Institute of Clinical Pathology and Medical Research; Westmead Hospital; University of Sydney | Arnott A., Draper J., Gall M., Martinez E., Rockett R., Sintchenko V., on behalf of ICPMR |
| EPI_ISL_3465285 | Viollier AG | Department of Biosystems Science and Engineering, ETH Zürich | Chaoran Chen, Sarah Nadeau, Louis du Plessis, Catharine Aquino, Ivan Topolsky, Kim Philipp Jablonski, Lara Fuhrmann, Daniel Ehrsam, Isabel Stürmer, Andreia Cabral de Gouvea, Maria Domenica Moccia, Simon Grüter, Timothy Sykes, Lennart Opitz, Griffin White, Laura Neff, Doris Popovic, Andrea Patrignani, Jay Tracy, Ralph Schlapbach, Christiane Beckmann, Maurice Redondo, Olivier Kobel, Christoph Noppen, Niko Beerenwinkel, Tanja Stadler |
| EPI_ISL_3465286 | Pathology West - NSW Health Pathology | NSW Health Pathology - Institute of Clinical Pathology and Medical Research; Westmead Hospital; University of Sydney | Arnott A., Draper J., Gall M., Martinez E., Rockett R., Sintchenko V., on behalf of ICPMR |
| EPI_ISL_3465287 | Viollier AG | Department of Biosystems Science and Engineering, ETH Zürich | Chaoran Chen, Sarah Nadeau, Louis du Plessis, Catharine Aquino, Ivan Topolsky, Kim Philipp Jablonski, Lara Fuhrmann, Daniel Ehrsam, Isabel Stürmer, Andreia Cabral de Gouvea, Maria Domenica Moccia, Simon Grüter, Timothy Sykes, Lennart Opitz, Griffin White, Laura Neff, Doris Popovic, Andrea Patrignani, Jay Tracy, Ralph Schlapbach, Christiane Beckmann, Maurice Redondo, Olivier Kobel, Christoph Noppen, Niko Beerenwinkel, Tanja Stadler |
| EPI_ISL_3465288 | 4Cyte Pathology | NSW Health Pathology - Institute of Clinical Pathology and Medical Research; Westmead Hospital; University of Sydney | Arnott A., Draper J., Gall M., Martinez E., Rockett R., Sintchenko V., on behalf of ICPMR |
| EPI_ISL_3465289 | Viollier AG | Department of Biosystems Science and Engineering, ETH Zürich | Chaoran Chen, Sarah Nadeau, Louis du Plessis, Catharine Aquino, Ivan Topolsky, Kim Philipp Jablonski, Lara Fuhrmann, Daniel Ehrsam, Isabel Stürmer, Andreia Cabral de Gouvea, Maria Domenica Moccia, Simon Grüter, Timothy Sykes, Lennart Opitz, Griffin White, Laura Neff, Doris Popovic, Andrea Patrignani, Jay Tracy, Ralph Schlapbach, Christiane Beckmann, Maurice Redondo, Olivier Kobel, Christoph Noppen, Niko Beerenwinkel, Tanja Stadler |
| EPI_ISL_3465290 | Viollier AG | Department of Biosystems Science and Engineering, ETH Zürich | Andrea Patrizia Salzmann, Henriette Kurth, Christiane Beckmann, Ivan Topolsky, Chaoran Chen, Sarah Nadeau, Louis du Plessis, Kim Philipp Jablonski, Lara Fuhrmann, Niko Beerenwinkel, Christoph Noppen, Olivier Kobel, Maurice Redondo, Tanja Stadler |
| EPI_ISL_3465291 | St Vincent's Pathology (SydPath) | NSW Health Pathology - Institute of Clinical Pathology and Medical Research; Westmead Hospital; University of Sydney | Arnott A., Draper J., Gall M., Martinez E., Rockett R., Sintchenko V., on behalf of ICPMR |
| EPI_ISL_3465292 | Viollier AG | Department of Biosystems Science and Engineering, ETH Zürich | Chaoran Chen, Sarah Nadeau, Louis du Plessis, Catharine Aquino, Ivan Topolsky, Kim Philipp Jablonski, Lara Fuhrmann, Daniel Ehrsam, Isabel Stürmer, Andreia Cabral de Gouvea, Maria Domenica Moccia, Simon Grüter, Timothy Sykes, Lennart Opitz, Griffin White, Laura Neff, Doris Popovic, Andrea Patrignani, Jay Tracy, Ralph Schlapbach, Christiane Beckmann, Maurice Redondo, Olivier Kobel, Christoph Noppen, Niko Beerenwinkel, Tanja Stadler |

| | | | |
|---|---|---|---|
| EPI_ISL_3465293 | 4Cyte Pathology | NSW Health Pathology - Institute of Clinical Pathology and Medical Research; Westmead Hospital; University of Sydney | Arnott A., Draper J., Gall M., Martinez E., Rockett R., Sintchenko V., on behalf of ICPMR |
| EPI_ISL_3465294, EPI_ISL_3465295 | Viollier AG | Department of Biosystems Science and Engineering, ETH Zürich | Chaoran Chen, Sarah Nadeau, Louis du Plessis, Catharine Aquino, Ivan Topolsky, Kim Philipp Jablonski, Lara Fuhrmann, Daniel Ehrsam, Isabel Stürmer, Andreia Cabral de Gouvea, Maria Domenica Moccia, Simon Grüter, Timothy Sykes, Lennart Opitz, Griffin White, Laura Neff, Doris Popovic, Andrea Patrignani, Jay Tracy, Ralph Schlapbach, Christiane Beckmann, Maurice Redondo, Olivier Kobel, Christoph Noppen, Niko Beerenwinkel, Tanja Stadler |
| EPI_ISL_3465296 | 4Cyte Pathology | NSW Health Pathology - Institute of Clinical Pathology and Medical Research; Westmead Hospital; University of Sydney | Arnott A., Draper J., Gall M., Martinez E., Rockett R., Sintchenko V., on behalf of ICPMR |
| EPI_ISL_3465297 | Viollier AG | Department of Biosystems Science and Engineering, ETH Zürich | Chaoran Chen, Sarah Nadeau, Louis du Plessis, Catharine Aquino, Ivan Topolsky, Kim Philipp Jablonski, Lara Fuhrmann, Daniel Ehrsam, Isabel Stürmer, Andreia Cabral de Gouvea, Maria Domenica Moccia, Simon Grüter, Timothy Sykes, Lennart Opitz, Griffin White, Laura Neff, Doris Popovic, Andrea Patrignani, Jay Tracy, Ralph Schlapbach, Christiane Beckmann, Maurice Redondo, Olivier Kobel, Christoph Noppen, Niko Beerenwinkel, Tanja Stadler |
| EPI_ISL_3465298 | Laverty Pathology | NSW Health Pathology - Institute of Clinical Pathology and Medical Research; Westmead Hospital; University of Sydney | Arnott A., Draper J., Gall M., Martinez E., Rockett R., Sintchenko V., on behalf of ICPMR |
| EPI_ISL_3465299 | Viollier AG | Department of Biosystems Science and Engineering, ETH Zürich | Chaoran Chen, Sarah Nadeau, Louis du Plessis, Catharine Aquino, Ivan Topolsky, Kim Philipp Jablonski, Lara Fuhrmann, Daniel Ehrsam, Isabel Stürmer, Andreia Cabral de Gouvea, Maria Domenica Moccia, Simon Grüter, Timothy Sykes, Lennart Opitz, Griffin White, Laura Neff, Doris Popovic, Andrea Patrignani, Jay Tracy, Ralph Schlapbach, Christiane Beckmann, Maurice Redondo, Olivier Kobel, Christoph Noppen, Niko Beerenwinkel, Tanja Stadler |
| EPI_ISL_3465300 | St Vincent's Pathology (SydPath) | NSW Health Pathology - Institute of Clinical Pathology and Medical Research; Westmead Hospital; University of Sydney | Arnott A., Draper J., Gall M., Martinez E., Rockett R., Sintchenko V., on behalf of ICPMR |
| EPI_ISL_3465301 | Viollier AG | Department of Biosystems Science and Engineering, ETH Zürich | Chaoran Chen, Sarah Nadeau, Louis du Plessis, Catharine Aquino, Ivan Topolsky, Kim Philipp Jablonski, Lara Fuhrmann, Daniel Ehrsam, Isabel Stürmer, Andreia Cabral de Gouvea, Maria Domenica Moccia, Simon Grüter, Timothy Sykes, Lennart Opitz, Griffin White, Laura Neff, Doris Popovic, Andrea Patrignani, Jay Tracy, Ralph Schlapbach, Christiane Beckmann, Maurice Redondo, Olivier Kobel, Christoph Noppen, Niko Beerenwinkel, Tanja Stadler |
| EPI_ISL_3465302 | Laverty Pathology | NSW Health Pathology - Institute of Clinical Pathology and Medical Research; Westmead Hospital; University of Sydney | Arnott A., Draper J., Gall M., Martinez E., Rockett R., Sintchenko V., on behalf of ICPMR |
| EPI_ISL_3465303, EPI_ISL_3465304 | Viollier AG | Department of Biosystems Science and Engineering, ETH Zürich | Chaoran Chen, Sarah Nadeau, Louis du Plessis, Catharine Aquino, Ivan Topolsky, Kim Philipp Jablonski, Lara Fuhrmann, Daniel Ehrsam, Isabel Stürmer, Andreia Cabral de Gouvea, Maria Domenica Moccia, Simon Grüter, Timothy Sykes, Lennart Opitz, Griffin White, Laura Neff, Doris Popovic, Andrea Patrignani, Jay Tracy, Ralph Schlapbach, Christiane Beckmann, Maurice Redondo, Olivier Kobel, Christoph Noppen, Niko Beerenwinkel, Tanja Stadler |
| EPI_ISL_3465305 | 4Cyte Pathology | NSW Health Pathology - Institute of Clinical Pathology and Medical Research; Westmead Hospital; University of Sydney | Arnott A., Draper J., Gall M., Martinez E., Rockett R., Sintchenko V., on behalf of ICPMR |
| EPI_ISL_3465306 | Viollier AG | Department of Biosystems Science and Engineering, ETH Zürich | Chaoran Chen, Sarah Nadeau, Louis du Plessis, Catharine Aquino, Ivan Topolsky, Kim Philipp Jablonski, Lara Fuhrmann, Daniel Ehrsam, Isabel Stürmer, Andreia Cabral de Gouvea, Maria Domenica Moccia, Simon Grüter, Timothy Sykes, Lennart Opitz, Griffin White, Laura Neff, Doris Popovic, Andrea Patrignani, Jay Tracy, Ralph Schlapbach, Christiane Beckmann, Maurice Redondo, Olivier Kobel, Christoph Noppen, Niko Beerenwinkel, Tanja Stadler |
| EPI_ISL_3465307 | Australian Clinical Labs (formerly Healthscope Pathology) | NSW Health Pathology - Institute of Clinical Pathology and Medical Research; Westmead Hospital; University of Sydney | Arnott A., Draper J., Gall M., Martinez E., Rockett R., Sintchenko V., on behalf of ICPMR |
| EPI_ISL_3465308 | Viollier AG | Department of Biosystems Science and Engineering, ETH Zürich | Andrea Patrizia Salzmann, Henriette Kurth, Christiane Beckmann, Ivan Topolsky, Chaoran Chen, Sarah Nadeau, Louis du Plessis, Kim Philipp Jablonski, Lara Fuhrmann, Niko Beerenwinkel, Christoph Noppen, Olivier Kobel, Maurice Redondo, Tanja Stadler |
| EPI_ISL_3465309 | St Vincent's Pathology (SydPath) | NSW Health Pathology - Institute of Clinical Pathology and Medical Research; Westmead Hospital; University of Sydney | Arnott A., Draper J., Gall M., Martinez E., Rockett R., Sintchenko V., on behalf of ICPMR |
| EPI_ISL_3465310 | Viollier AG | Department of Biosystems Science and Engineering, ETH Zürich | Chaoran Chen, Sarah Nadeau, Louis du Plessis, Catharine Aquino, Ivan Topolsky, Kim Philipp Jablonski, Lara Fuhrmann, Daniel Ehrsam, Isabel Stürmer, Andreia Cabral de Gouvea, Maria Domenica Moccia, Simon Grüter, Timothy Sykes, Lennart Opitz, Griffin White, Laura Neff, Doris Popovic, Andrea Patrignani, Jay Tracy, Ralph Schlapbach, Christiane Beckmann, Maurice Redondo, Olivier Kobel, Christoph Noppen, Niko Beerenwinkel, Tanja Stadler |
| EPI_ISL_3465311 | Laverty Pathology | NSW Health Pathology - Institute of Clinical Pathology and Medical Research; Westmead Hospital; University of Sydney | Arnott A., Draper J., Gall M., Martinez E., Rockett R., Sintchenko V., on behalf of ICPMR |
| EPI_ISL_3465312, EPI_ISL_3465313 | Viollier AG | Department of Biosystems Science and Engineering, ETH Zürich | Chaoran Chen, Sarah Nadeau, Louis du Plessis, Catharine Aquino, Ivan Topolsky, Kim Philipp Jablonski, Lara Fuhrmann, Daniel Ehrsam, Isabel Stürmer, Andreia Cabral de Gouvea, Maria Domenica Moccia, Simon Grüter, Timothy Sykes, Lennart Opitz, Griffin White, Laura Neff, Doris Popovic, Andrea Patrignani, Jay Tracy, Ralph Schlapbach, Christiane Beckmann, Maurice Redondo, Olivier Kobel, Christoph Noppen, Niko Beerenwinkel, Tanja Stadler |
| EPI_ISL_3465314 | St Vincent's Pathology (SydPath) | NSW Health Pathology - Institute of Clinical Pathology and Medical Research; Westmead Hospital; University of Sydney | Arnott A., Draper J., Gall M., Martinez E., Rockett R., Sintchenko V., on behalf of ICPMR |
| EPI_ISL_3465315 | Viollier AG | Department of Biosystems Science and Engineering, ETH Zürich | Chaoran Chen, Sarah Nadeau, Louis du Plessis, Catharine Aquino, Ivan Topolsky, Kim Philipp Jablonski, Lara Fuhrmann, Daniel Ehrsam, Isabel Stürmer, Andreia Cabral de Gouvea, Maria Domenica Moccia, Simon Grüter, Timothy Sykes, Lennart Opitz, Griffin White, Laura Neff, Doris Popovic, Andrea Patrignani, Jay Tracy, Ralph Schlapbach, Christiane Beckmann, Maurice Redondo, Olivier Kobel, Christoph Noppen, Niko Beerenwinkel, Tanja Stadler |
| EPI_ISL_3465316 | Medlab Pathology | NSW Health Pathology - Institute of Clinical Pathology and Medical Research; Westmead Hospital; University of Sydney | Arnott A., Draper J., Gall M., Martinez E., Rockett R., Sintchenko V., on behalf of ICPMR |
| EPI_ISL_3465317, EPI_ISL_3465318 | Viollier AG | Department of Biosystems Science and Engineering, ETH Zürich | Chaoran Chen, Sarah Nadeau, Louis du Plessis, Catharine Aquino, Ivan Topolsky, Kim Philipp Jablonski, Lara Fuhrmann, Daniel Ehrsam, Isabel Stürmer, Andreia Cabral de Gouvea, Maria Domenica Moccia, Simon Grüter, Timothy Sykes, Lennart Opitz, Griffin White, Laura Neff, Doris Popovic, Andrea Patrignani, Jay Tracy, Ralph Schlapbach, Christiane Beckmann, Maurice Redondo, Olivier Kobel, Christoph Noppen, Niko Beerenwinkel, Tanja Stadler |
| EPI_ISL_3465319 | Pathology West - NSW Health Pathology | NSW Health Pathology - Institute of Clinical Pathology and Medical Research; Westmead Hospital; University of Sydney | Arnott A., Draper J., Gall M., Martinez E., Rockett R., Sintchenko V., on behalf of ICPMR |
| EPI_ISL_3465320 | Viollier AG | Department of Biosystems Science and Engineering, ETH Zürich | Chaoran Chen, Sarah Nadeau, Louis du Plessis, Catharine Aquino, Ivan Topolsky, Kim Philipp Jablonski, Lara Fuhrmann, Daniel Ehrsam, Isabel Stürmer, Andreia Cabral de Gouvea, Maria Domenica Moccia, Simon Grüter, Timothy Sykes, Lennart Opitz, Griffin White, Laura Neff, Doris Popovic, Andrea Patrignani, Jay Tracy, Ralph Schlapbach, Christiane Beckmann, Maurice Redondo, Olivier Kobel, Christoph Noppen, Niko Beerenwinkel, Tanja Stadler |
| EPI_ISL_3465321 | 4Cyte Pathology | NSW Health Pathology - Institute of Clinical Pathology and Medical Research; Westmead Hospital; University of Sydney | Arnott A., Draper J., Gall M., Martinez E., Rockett R., Sintchenko V., on behalf of ICPMR |
| EPI_ISL_3465322 | Viollier AG | Department of Biosystems Science and Engineering, ETH Zürich | Chaoran Chen, Sarah Nadeau, Louis du Plessis, Catharine Aquino, Ivan Topolsky, Kim Philipp Jablonski, Lara Fuhrmann, Daniel Ehrsam, Isabel Stürmer, Andreia Cabral de Gouvea, Maria Domenica Moccia, Simon Grüter, Timothy Sykes, Lennart Opitz, Griffin White, Laura Neff, Doris Popovic, Andrea Patrignani, Jay Tracy, Ralph Schlapbach, Christiane Beckmann, Maurice Redondo, Olivier Kobel, Christoph Noppen, Niko Beerenwinkel, Tanja Stadler |
| EPI_ISL_3465323 | St Vincent's Pathology (SydPath) | NSW Health Pathology - Institute of Clinical Pathology and Medical Research; Westmead Hospital; University of Sydney | Arnott A., Draper J., Gall M., Martinez E., Rockett R., Sintchenko V., on behalf of ICPMR |
| EPI_ISL_3465324 | Viollier AG | Department of Biosystems Science and Engineering, ETH Zürich | Chaoran Chen, Sarah Nadeau, Louis du Plessis, Catharine Aquino, Ivan Topolsky, Kim Philipp Jablonski, Lara Fuhrmann, Daniel Ehrsam, Isabel Stürmer, Andreia Cabral de Gouvea, Maria Domenica Moccia, Simon Grüter, Timothy Sykes, Lennart Opitz, Griffin White, Laura Neff, Doris Popovic, Andrea Patrignani, Jay Tracy, Ralph Schlapbach, Christiane Beckmann, Maurice Redondo, Olivier Kobel, Christoph Noppen, Niko Beerenwinkel, Tanja Stadler |
| EPI_ISL_3465325 | Medlab Pathology | NSW Health Pathology - Institute of Clinical Pathology and Medical Research; Westmead Hospital; University of Sydney | Arnott A., Draper J., Gall M., Martinez E., Rockett R., Sintchenko V., on behalf of ICPMR |
| EPI_ISL_3465326 | Viollier AG | Department of Biosystems Science and Engineering, ETH Zürich | Chaoran Chen, Sarah Nadeau, Louis du Plessis, Catharine Aquino, Ivan Topolsky, Kim Philipp Jablonski, Lara Fuhrmann, Daniel Ehrsam, Isabel Stürmer, Andreia Cabral de Gouvea, Maria Domenica Moccia, Simon Grüter, Timothy Sykes, Lennart Opitz, Griffin White, Laura Neff, Doris Popovic, Andrea Patrignani, Jay Tracy, Ralph Schlapbach, Christiane Beckmann, Maurice Redondo, Olivier Kobel, Christoph Noppen, Niko Beerenwinkel, Tanja Stadler |
| EPI_ISL_3465327 | Viollier AG | Department of Biosystems Science and Engineering, ETH Zürich | Andrea Patrizia Salzmann, Henriette Kurth, Christiane Beckmann, Ivan Topolsky, Chaoran Chen, Sarah Nadeau, Louis du Plessis, Kim Philipp Jablonski, Lara Fuhrmann, Niko Beerenwinkel, Christoph Noppen, Olivier Kobel, Maurice Redondo, Tanja Stadler |
| EPI_ISL_3465328 | St Vincent's Pathology (SydPath) | NSW Health Pathology - Institute of Clinical Pathology and Medical Research; Westmead Hospital; University of Sydney | Arnott A., Draper J., Gall M., Martinez E., Rockett R., Sintchenko V., on behalf of ICPMR |
| EPI_ISL_3465329 | Viollier AG | Department of Biosystems Science and Engineering, ETH Zürich | Andrea Patrizia Salzmann, Henriette Kurth, Christiane Beckmann, Ivan Topolsky, Chaoran Chen, Sarah Nadeau, Louis du Plessis, Kim Philipp Jablonski, Lara Fuhrmann, Niko Beerenwinkel, Christoph Noppen, Olivier Kobel, Maurice Redondo, Tanja Stadler |
| EPI_ISL_3465330 | Sydney South West Pathology Service (SSWPS) - Liverpool Hospital - NSW Health Pathology | NSW Health Pathology - Institute of Clinical Pathology and Medical Research; Westmead Hospital; University of Sydney | Arnott A., Draper J., Gall M., Martinez E., Rockett R., Sintchenko V., on behalf of ICPMR |
| EPI_ISL_3465331 | Viollier AG | Department of Biosystems Science and Engineering, ETH Zürich | Chaoran Chen, Sarah Nadeau, Louis du Plessis, Catharine Aquino, Ivan Topolsky, Kim Philipp Jablonski, Lara Fuhrmann, Daniel Ehrsam, Isabel Stürmer, Andreia Cabral de Gouvea, Maria Domenica Moccia, Simon Grüter, Timothy Sykes, Lennart Opitz, Griffin White, Laura Neff, Doris Popovic, Andrea Patrignani, Jay Tracy, Ralph Schlapbach, Christiane Beckmann, Maurice Redondo, Olivier Kobel, Christoph Noppen, Niko Beerenwinkel, Tanja Stadler |
| EPI_ISL_3465332 | Medlab Pathology | NSW Health Pathology - Institute of Clinical Pathology and Medical Research; Westmead Hospital; University of Sydney | Arnott A., Draper J., Gall M., Martinez E., Rockett R., Sintchenko V., on behalf of ICPMR |
| EPI_ISL_3465333 | Viollier AG | Department of Biosystems Science and Engineering, ETH Zürich | Andrea Patrizia Salzmann, Henriette Kurth, Christiane Beckmann, Ivan Topolsky, Chaoran Chen, Sarah Nadeau, Louis du Plessis, Kim Philipp Jablonski, Lara Fuhrmann, Niko Beerenwinkel, Christoph Noppen, Olivier Kobel, Maurice Redondo, Tanja Stadler |
| EPI_ISL_3465334 | Viollier AG | Department of Biosystems Science and Engineering, ETH Zürich | Chaoran Chen, Sarah Nadeau, Louis du Plessis, Catharine Aquino, Ivan Topolsky, Kim Philipp Jablonski, Lara Fuhrmann, Daniel Ehrsam, Isabel Stürmer, Andreia Cabral de Gouvea, Maria Domenica Moccia, Simon Grüter, Timothy Sykes, Lennart Opitz, Griffin White, Laura Neff, Doris Popovic, Andrea Patrignani, Jay Tracy, Ralph Schlapbach, Christiane Beckmann, Maurice Redondo, Olivier Kobel, Christoph Noppen, Niko Beerenwinkel, Tanja Stadler |
| EPI_ISL_3465335 | Medlab Pathology | NSW Health Pathology - Institute of Clinical Pathology and Medical Research; Westmead Hospital; University of Sydney | Arnott A., Draper J., Gall M., Martinez E., Rockett R., Sintchenko V., on behalf of ICPMR |
| EPI_ISL_3465336 | Viollier AG | Department of Biosystems Science and Engineering, ETH Zürich | Andrea Patrizia Salzmann, Henriette Kurth, Christiane Beckmann, Ivan Topolsky, Chaoran Chen, Sarah Nadeau, Louis du Plessis, Kim Philipp Jablonski, Lara Fuhrmann, Niko Beerenwinkel, Christoph Noppen, Olivier Kobel, Maurice Redondo, Tanja Stadler |
| EPI_ISL_3465337 | 4Cyte Pathology | NSW Health Pathology - Institute of Clinical Pathology and Medical Research; Westmead Hospital; University of Sydney | Arnott A., Draper J., Gall M., Martinez E., Rockett R., Sintchenko V., on behalf of ICPMR |
| EPI_ISL_3465338, EPI_ISL_3465339 | Viollier AG | Department of Biosystems Science and Engineering, ETH Zürich | Chaoran Chen, Sarah Nadeau, Louis du Plessis, Catharine Aquino, Ivan Topolsky, Kim Philipp Jablonski, Lara Fuhrmann, Daniel Ehrsam, Isabel Stürmer, Andreia Cabral de Gouvea, Maria Domenica Moccia, Simon Grüter, Timothy Sykes, Lennart Opitz, Griffin White, Laura Neff, Doris Popovic, Andrea Patrignani, Jay Tracy, Ralph Schlapbach, Christiane Beckmann, Maurice Redondo, Olivier Kobel, Christoph Noppen, Niko Beerenwinkel, Tanja Stadler |
| EPI_ISL_3465340 | Histopath | NSW Health Pathology - Institute of Clinical Pathology and Medical Research; Westmead Hospital; University of Sydney | Arnott A., Draper J., Gall M., Martinez E., Rockett R., Sintchenko V., on behalf of ICPMR |
| EPI_ISL_3465341 | Viollier AG | Department of Biosystems Science and Engineering, ETH Zürich | Chaoran Chen, Sarah Nadeau, Louis du Plessis, Catharine Aquino, Ivan Topolsky, Kim Philipp Jablonski, Lara Fuhrmann, Daniel Ehrsam, Isabel Stürmer, Andreia Cabral de Gouvea, Maria Domenica Moccia, Simon Grüter, Timothy Sykes, Lennart Opitz, Griffin White, Laura Neff, Doris Popovic, Andrea Patrignani, Jay Tracy, Ralph Schlapbach, Christiane Beckmann, Maurice Redondo, Olivier Kobel, Christoph Noppen, Niko Beerenwinkel, Tanja Stadler |
| EPI_ISL_3465342 | Medlab Pathology | NSW Health Pathology - Institute of Clinical Pathology and Medical Research; Westmead Hospital; University of Sydney | Arnott A., Draper J., Gall M., Martinez E., Rockett R., Sintchenko V., on behalf of ICPMR |

| | | | |
|---|---|---|---|
| EPI_ISL_3465343, EPI_ISL_3465344 | Viollier AG | Department of Biosystems Science and Engineering, ETH Zürich | Chaoran Chen, Sarah Nadeau, Louis du Plessis, Catharine Aquino, Ivan Topolsky, Kim Philipp Jablonski, Lara Fuhrmann, Daniel Ehrsam, Isabel Stürmer, Andreia Cabral de Gouvea, Maria Domenica Moccia, Simon Grüter, Timothy Sykes, Lennart Opitz, Griffin White, Laura Neff, Doris Popovic, Andrea Patrignani, Jay Tracy, Ralph Schlapbach, Christiane Beckmann, Maurice Redondo, Olivier Kobel, Christoph Noppen, Niko Beerenwinkel, Tanja Stadler |
| EPI_ISL_3465345 | Medlab Pathology | NSW Health Pathology - Institute of Clinical Pathology and Medical Research; Westmead Hospital; University of Sydney | Arnott A., Draper J., Gall M., Martinez E., Rockett R., Sintchenko V., on behalf of ICPMR |
| EPI_ISL_3465346 | Viollier AG | Department of Biosystems Science and Engineering, ETH Zürich | Andrea Patrizia Salzmann, Henriette Kurth, Christiane Beckmann, Ivan Topolsky, Chaoran Chen, Sarah Nadeau, Louis du Plessis, Kim Philipp Jablonski, Lara Fuhrmann, Niko Beerenwinkel, Christoph Noppen, Olivier Kobel, Maurice Redondo, Tanja Stadler |
| EPI_ISL_3465347 | St Vincent's Pathology (SydPath) | NSW Health Pathology - Institute of Clinical Pathology and Medical Research; Westmead Hospital; University of Sydney | Arnott A., Draper J., Gall M., Martinez E., Rockett R., Sintchenko V., on behalf of ICPMR |
| EPI_ISL_3465348 | Viollier AG | Department of Biosystems Science and Engineering, ETH Zürich | Chaoran Chen, Sarah Nadeau, Louis du Plessis, Catharine Aquino, Ivan Topolsky, Kim Philipp Jablonski, Lara Fuhrmann, Daniel Ehrsam, Isabel Stürmer, Andreia Cabral de Gouvea, Maria Domenica Moccia, Simon Grüter, Timothy Sykes, Lennart Opitz, Griffin White, Laura Neff, Doris Popovic, Andrea Patrignani, Jay Tracy, Ralph Schlapbach, Christiane Beckmann, Maurice Redondo, Olivier Kobel, Christoph Noppen, Niko Beerenwinkel, Tanja Stadler |
| EPI_ISL_3465349 | Australian Clinical Labs (formerly Healthscope Pathology) | NSW Health Pathology - Institute of Clinical Pathology and Medical Research; Westmead Hospital; University of Sydney | Arnott A., Draper J., Gall M., Martinez E., Rockett R., Sintchenko V., on behalf of ICPMR |
| EPI_ISL_3465350, EPI_ISL_3465351 | Viollier AG | Department of Biosystems Science and Engineering, ETH Zürich | Chaoran Chen, Sarah Nadeau, Louis du Plessis, Catharine Aquino, Ivan Topolsky, Kim Philipp Jablonski, Lara Fuhrmann, Daniel Ehrsam, Isabel Stürmer, Andreia Cabral de Gouvea, Maria Domenica Moccia, Simon Grüter, Timothy Sykes, Lennart Opitz, Griffin White, Laura Neff, Doris Popovic, Andrea Patrignani, Jay Tracy, Ralph Schlapbach, Christiane Beckmann, Maurice Redondo, Olivier Kobel, Christoph Noppen, Niko Beerenwinkel, Tanja Stadler |
| EPI_ISL_3465352 | Sydney South West Pathology Service (SSWPS) - Liverpool Hospital - NSW Health Pathology | NSW Health Pathology - Institute of Clinical Pathology and Medical Research; Westmead Hospital; University of Sydney | Arnott A., Draper J., Gall M., Martinez E., Rockett R., Sintchenko V., on behalf of ICPMR |
| EPI_ISL_3465353 | Viollier AG | Department of Biosystems Science and Engineering, ETH Zürich | Chaoran Chen, Sarah Nadeau, Louis du Plessis, Catharine Aquino, Ivan Topolsky, Kim Philipp Jablonski, Lara Fuhrmann, Daniel Ehrsam, Isabel Stürmer, Andreia Cabral de Gouvea, Maria Domenica Moccia, Simon Grüter, Timothy Sykes, Lennart Opitz, Griffin White, Laura Neff, Doris Popovic, Andrea Patrignani, Jay Tracy, Ralph Schlapbach, Christiane Beckmann, Maurice Redondo, Olivier Kobel, Christoph Noppen, Niko Beerenwinkel, Tanja Stadler |
| EPI_ISL_3465354 | Sydney South West Pathology Service (SSWPS) - Liverpool Hospital - NSW Health Pathology | NSW Health Pathology - Institute of Clinical Pathology and Medical Research; Westmead Hospital; University of Sydney | Arnott A., Draper J., Gall M., Martinez E., Rockett R., Sintchenko V., on behalf of ICPMR |
| EPI_ISL_3465355 | Viollier AG | Department of Biosystems Science and Engineering, ETH Zürich | Chaoran Chen, Sarah Nadeau, Louis du Plessis, Catharine Aquino, Ivan Topolsky, Kim Philipp Jablonski, Lara Fuhrmann, Daniel Ehrsam, Isabel Stürmer, Andreia Cabral de Gouvea, Maria Domenica Moccia, Simon Grüter, Timothy Sykes, Lennart Opitz, Griffin White, Laura Neff, Doris Popovic, Andrea Patrignani, Jay Tracy, Ralph Schlapbach, Christiane Beckmann, Maurice Redondo, Olivier Kobel, Christoph Noppen, Niko Beerenwinkel, Tanja Stadler |
| EPI_ISL_3465356 | Histopath | NSW Health Pathology - Institute of Clinical Pathology and Medical Research; Westmead Hospital; University of Sydney | Arnott A., Draper J., Gall M., Martinez E., Rockett R., Sintchenko V., on behalf of ICPMR |
| EPI_ISL_3465357 | Viollier AG | Department of Biosystems Science and Engineering, ETH Zürich | Chaoran Chen, Sarah Nadeau, Louis du Plessis, Catharine Aquino, Ivan Topolsky, Kim Philipp Jablonski, Lara Fuhrmann, Daniel Ehrsam, Isabel Stürmer, Andreia Cabral de Gouvea, Maria Domenica Moccia, Simon Grüter, Timothy Sykes, Lennart Opitz, Griffin White, Laura Neff, Doris Popovic, Andrea Patrignani, Jay Tracy, Ralph Schlapbach, Christiane Beckmann, Maurice Redondo, Olivier Kobel, Christoph Noppen, Niko Beerenwinkel, Tanja Stadler |
| EPI_ISL_3465358 | St Vincent's Pathology (SydPath) | NSW Health Pathology - Institute of Clinical Pathology and Medical Research; Westmead Hospital; University of Sydney | Arnott A., Draper J., Gall M., Martinez E., Rockett R., Sintchenko V., on behalf of ICPMR |
| EPI_ISL_3465359 | Viollier AG | Department of Biosystems Science and Engineering, ETH Zürich | Chaoran Chen, Sarah Nadeau, Louis du Plessis, Catharine Aquino, Ivan Topolsky, Kim Philipp Jablonski, Lara Fuhrmann, Daniel Ehrsam, Isabel Stürmer, Andreia Cabral de Gouvea, Maria Domenica Moccia, Simon Grüter, Timothy Sykes, Lennart Opitz, Griffin White, Laura Neff, Doris Popovic, Andrea Patrignani, Jay Tracy, Ralph Schlapbach, Christiane Beckmann, Maurice Redondo, Olivier Kobel, Christoph Noppen, Niko Beerenwinkel, Tanja Stadler |
| EPI_ISL_3465360 | Medlab Pathology | NSW Health Pathology - Institute of Clinical Pathology and Medical Research; Westmead Hospital; University of Sydney | Arnott A., Draper J., Gall M., Martinez E., Rockett R., Sintchenko V., on behalf of ICPMR |
| EPI_ISL_3465361 | Viollier AG | Department of Biosystems Science and Engineering, ETH Zürich | Chaoran Chen, Sarah Nadeau, Louis du Plessis, Catharine Aquino, Ivan Topolsky, Kim Philipp Jablonski, Lara Fuhrmann, Daniel Ehrsam, Isabel Stürmer, Andreia Cabral de Gouvea, Maria Domenica Moccia, Simon Grüter, Timothy Sykes, Lennart Opitz, Griffin White, Laura Neff, Doris Popovic, Andrea Patrignani, Jay Tracy, Ralph Schlapbach, Christiane Beckmann, Maurice Redondo, Olivier Kobel, Christoph Noppen, Niko Beerenwinkel, Tanja Stadler |
| EPI_ISL_3465362 | Viollier AG | Department of Biosystems Science and Engineering, ETH Zürich | Andrea Patrizia Salzmann, Henriette Kurth, Christiane Beckmann, Ivan Topolsky, Chaoran Chen, Sarah Nadeau, Louis du Plessis, Kim Philipp Jablonski, Lara Fuhrmann, Niko Beerenwinkel, Christoph Noppen, Olivier Kobel, Maurice Redondo, Tanja Stadler |
| EPI_ISL_3465363 | Medlab Pathology | NSW Health Pathology - Institute of Clinical Pathology and Medical Research; Westmead Hospital; University of Sydney | Arnott A., Draper J., Gall M., Martinez E., Rockett R., Sintchenko V., on behalf of ICPMR |
| EPI_ISL_3465364 | Viollier AG | Department of Biosystems Science and Engineering, ETH Zürich | Chaoran Chen, Sarah Nadeau, Louis du Plessis, Catharine Aquino, Ivan Topolsky, Kim Philipp Jablonski, Lara Fuhrmann, Daniel Ehrsam, Isabel Stürmer, Andreia Cabral de Gouvea, Maria Domenica Moccia, Simon Grüter, Timothy Sykes, Lennart Opitz, Griffin White, Laura Neff, Doris Popovic, Andrea Patrignani, Jay Tracy, Ralph Schlapbach, Christiane Beckmann, Maurice Redondo, Olivier Kobel, Christoph Noppen, Niko Beerenwinkel, Tanja Stadler |
| EPI_ISL_3465365 | Sydney South West Pathology Service (SSWPS) - Liverpool Hospital - NSW Health Pathology | NSW Health Pathology - Institute of Clinical Pathology and Medical Research; Westmead Hospital; University of Sydney | Arnott A., Draper J., Gall M., Martinez E., Rockett R., Sintchenko V., on behalf of ICPMR |
| EPI_ISL_3465366 | Viollier AG | Department of Biosystems Science and Engineering, ETH Zürich | Andrea Patrizia Salzmann, Henriette Kurth, Christiane Beckmann, Ivan Topolsky, Chaoran Chen, Sarah Nadeau, Louis du Plessis, Kim Philipp Jablonski, Lara Fuhrmann, Niko Beerenwinkel, Christoph Noppen, Olivier Kobel, Maurice Redondo, Tanja Stadler |
| EPI_ISL_3465367 | Viollier AG | Department of Biosystems Science and Engineering, ETH Zürich | Chaoran Chen, Sarah Nadeau, Louis du Plessis, Catharine Aquino, Ivan Topolsky, Kim Philipp Jablonski, Lara Fuhrmann, Daniel Ehrsam, Isabel Stürmer, Andreia Cabral de Gouvea, Maria Domenica Moccia, Simon Grüter, Timothy Sykes, Lennart Opitz, Griffin White, Laura Neff, Doris Popovic, Andrea Patrignani, Jay Tracy, Ralph Schlapbach, Christiane Beckmann, Maurice Redondo, Olivier Kobel, Christoph Noppen, Niko Beerenwinkel, Tanja Stadler |
| EPI_ISL_3465368 | Australian Clinical Labs (formerly Healthscope Pathology) | NSW Health Pathology - Institute of Clinical Pathology and Medical Research; Westmead Hospital; University of Sydney | Arnott A., Draper J., Gall M., Martinez E., Rockett R., Sintchenko V., on behalf of ICPMR |
| EPI_ISL_3465369 | Histopath | NSW Health Pathology - Institute of Clinical Pathology and Medical Research; Westmead Hospital; University of Sydney | Arnott A., Draper J., Gall M., Martinez E., Rockett R., Sintchenko V., on behalf of ICPMR |
| EPI_ISL_3465370, EPI_ISL_3465371 | Viollier AG | Department of Biosystems Science and Engineering, ETH Zürich | Chaoran Chen, Sarah Nadeau, Louis du Plessis, Catharine Aquino, Ivan Topolsky, Kim Philipp Jablonski, Lara Fuhrmann, Daniel Ehrsam, Isabel Stürmer, Andreia Cabral de Gouvea, Maria Domenica Moccia, Simon Grüter, Timothy Sykes, Lennart Opitz, Griffin White, Laura Neff, Doris Popovic, Andrea Patrignani, Jay Tracy, Ralph Schlapbach, Christiane Beckmann, Maurice Redondo, Olivier Kobel, Christoph Noppen, Niko Beerenwinkel, Tanja Stadler |
| EPI_ISL_3465372 | St Vincent's Pathology (SydPath) | NSW Health Pathology - Institute of Clinical Pathology and Medical Research; Westmead Hospital; University of Sydney | Arnott A., Draper J., Gall M., Martinez E., Rockett R., Sintchenko V., on behalf of ICPMR |
| EPI_ISL_3465373 | Viollier AG | Department of Biosystems Science and Engineering, ETH Zürich | Chaoran Chen, Sarah Nadeau, Louis du Plessis, Catharine Aquino, Ivan Topolsky, Kim Philipp Jablonski, Lara Fuhrmann, Daniel Ehrsam, Isabel Stürmer, Andreia Cabral de Gouvea, Maria Domenica Moccia, Simon Grüter, Timothy Sykes, Lennart Opitz, Griffin White, Laura Neff, Doris Popovic, Andrea Patrignani, Jay Tracy, Ralph Schlapbach, Christiane Beckmann, Maurice Redondo, Olivier Kobel, Christoph Noppen, Niko Beerenwinkel, Tanja Stadler |
| EPI_ISL_3465374 | St Vincent's Pathology (SydPath) | NSW Health Pathology - Institute of Clinical Pathology and Medical Research; Westmead Hospital; University of Sydney | Arnott A., Draper J., Gall M., Martinez E., Rockett R., Sintchenko V., on behalf of ICPMR |
| EPI_ISL_3465375 | Viollier AG | Department of Biosystems Science and Engineering, ETH Zürich | Andrea Patrizia Salzmann, Henriette Kurth, Christiane Beckmann, Ivan Topolsky, Chaoran Chen, Sarah Nadeau, Louis du Plessis, Kim Philipp Jablonski, Lara Fuhrmann, Niko Beerenwinkel, Christoph Noppen, Olivier Kobel, Maurice Redondo, Tanja Stadler |
| EPI_ISL_3465376 | Viollier AG | Department of Biosystems Science and Engineering, ETH Zürich | Chaoran Chen, Sarah Nadeau, Louis du Plessis, Catharine Aquino, Ivan Topolsky, Kim Philipp Jablonski, Lara Fuhrmann, Daniel Ehrsam, Isabel Stürmer, Andreia Cabral de Gouvea, Maria Domenica Moccia, Simon Grüter, Timothy Sykes, Lennart Opitz, Griffin White, Laura Neff, Doris Popovic, Andrea Patrignani, Jay Tracy, Ralph Schlapbach, Christiane Beckmann, Maurice Redondo, Olivier Kobel, Christoph Noppen, Niko Beerenwinkel, Tanja Stadler |
| EPI_ISL_3465377 | 4Cyte Pathology | NSW Health Pathology - Institute of Clinical Pathology and Medical Research; Westmead Hospital; University of Sydney | Arnott A., Draper J., Gall M., Martinez E., Rockett R., Sintchenko V., on behalf of ICPMR |
| EPI_ISL_3465378 | Viollier AG | Department of Biosystems Science and Engineering, ETH Zürich | Chaoran Chen, Sarah Nadeau, Louis du Plessis, Catharine Aquino, Ivan Topolsky, Kim Philipp Jablonski, Lara Fuhrmann, Daniel Ehrsam, Isabel Stürmer, Andreia Cabral de Gouvea, Maria Domenica Moccia, Simon Grüter, Timothy Sykes, Lennart Opitz, Griffin White, Laura Neff, Doris Popovic, Andrea Patrignani, Jay Tracy, Ralph Schlapbach, Christiane Beckmann, Maurice Redondo, Olivier Kobel, Christoph Noppen, Niko Beerenwinkel, Tanja Stadler |
| EPI_ISL_3465379 | Pathology West - NSW Health Pathology | NSW Health Pathology - Institute of Clinical Pathology and Medical Research; Westmead Hospital; University of Sydney | Arnott A., Draper J., Gall M., Martinez E., Rockett R., Sintchenko V., on behalf of ICPMR |
| EPI_ISL_3465380 | Viollier AG | Department of Biosystems Science and Engineering, ETH Zürich | Chaoran Chen, Sarah Nadeau, Louis du Plessis, Catharine Aquino, Ivan Topolsky, Kim Philipp Jablonski, Lara Fuhrmann, Daniel Ehrsam, Isabel Stürmer, Andreia Cabral de Gouvea, Maria Domenica Moccia, Simon Grüter, Timothy Sykes, Lennart Opitz, Griffin White, Laura Neff, Doris Popovic, Andrea Patrignani, Jay Tracy, Ralph Schlapbach, Christiane Beckmann, Maurice Redondo, Olivier Kobel, Christoph Noppen, Niko Beerenwinkel, Tanja Stadler |
| EPI_ISL_3465381 | St Vincent's Pathology (SydPath) | NSW Health Pathology - Institute of Clinical Pathology and Medical Research; Westmead Hospital; University of Sydney | Arnott A., Draper J., Gall M., Martinez E., Rockett R., Sintchenko V., on behalf of ICPMR |
| EPI_ISL_3465382 | Viollier AG | Department of Biosystems Science and Engineering, ETH Zürich | Chaoran Chen, Sarah Nadeau, Louis du Plessis, Catharine Aquino, Ivan Topolsky, Kim Philipp Jablonski, Lara Fuhrmann, Daniel Ehrsam, Isabel Stürmer, Andreia Cabral de Gouvea, Maria Domenica Moccia, Simon Grüter, Timothy Sykes, Lennart Opitz, Griffin White, Laura Neff, Doris Popovic, Andrea Patrignani, Jay Tracy, Ralph Schlapbach, Christiane Beckmann, Maurice Redondo, Olivier Kobel, Christoph Noppen, Niko Beerenwinkel, Tanja Stadler |
| EPI_ISL_3465383 | Histopath | NSW Health Pathology - Institute of Clinical Pathology and Medical Research; Westmead Hospital; University of Sydney | Arnott A., Draper J., Gall M., Martinez E., Rockett R., Sintchenko V., on behalf of ICPMR |
| EPI_ISL_3465384, EPI_ISL_3465385 | Viollier AG | Department of Biosystems Science and Engineering, ETH Zürich | Chaoran Chen, Sarah Nadeau, Louis du Plessis, Catharine Aquino, Ivan Topolsky, Kim Philipp Jablonski, Lara Fuhrmann, Daniel Ehrsam, Isabel Stürmer, Andreia Cabral de Gouvea, Maria Domenica Moccia, Simon Grüter, Timothy Sykes, Lennart Opitz, Griffin White, Laura Neff, Doris Popovic, Andrea Patrignani, Jay Tracy, Ralph Schlapbach, Christiane Beckmann, Maurice Redondo, Olivier Kobel, Christoph Noppen, Niko Beerenwinkel, Tanja Stadler |
| EPI_ISL_3465386 | Sydney South West Pathology Service (SSWPS) - Liverpool Hospital - NSW Health Pathology | NSW Health Pathology - Institute of Clinical Pathology and Medical Research; Westmead Hospital; University of Sydney | Arnott A., Draper J., Gall M., Martinez E., Rockett R., Sintchenko V., on behalf of ICPMR |
| EPI_ISL_3465387 | Viollier AG | Department of Biosystems Science and Engineering, ETH Zürich | Andrea Patrizia Salzmann, Henriette Kurth, Christiane Beckmann, Ivan Topolsky, Chaoran Chen, Sarah Nadeau, Louis du Plessis, Kim Philipp Jablonski, Lara Fuhrmann, Niko Beerenwinkel, Christoph Noppen, Olivier Kobel, Maurice Redondo, Tanja Stadler |
| EPI_ISL_3465388 | Pathology West - NSW Health Pathology | NSW Health Pathology - Institute of Clinical Pathology and Medical Research; Westmead Hospital; University of Sydney | Arnott A., Draper J., Gall M., Martinez E., Rockett R., Sintchenko V., on behalf of ICPMR |
| EPI_ISL_3465389 | Viollier AG | Department of Biosystems Science and Engineering, ETH Zürich | Chaoran Chen, Sarah Nadeau, Louis du Plessis, Catharine Aquino, Ivan Topolsky, Kim Philipp Jablonski, Lara Fuhrmann, Daniel Ehrsam, Isabel Stürmer, Andreia Cabral de Gouvea, Maria Domenica Moccia, Simon Grüter, Timothy Sykes, Lennart Opitz, Griffin White, Laura Neff, Doris Popovic, Andrea Patrignani, Jay Tracy, Ralph Schlapbach, Christiane Beckmann, Maurice Redondo, Olivier Kobel, Christoph Noppen, Niko Beerenwinkel, Tanja Stadler |
| EPI_ISL_3465390 | Pathology West - NSW Health Pathology | NSW Health Pathology - Institute of Clinical Pathology and Medical Research; Westmead Hospital; University of Sydney | Arnott A., Draper J., Gall M., Martinez E., Rockett R., Sintchenko V., on behalf of ICPMR |
| EPI_ISL_3465391 | Viollier AG | Department of Biosystems Science and Engineering, ETH Zürich | Chaoran Chen, Sarah Nadeau, Louis du Plessis, Catharine Aquino, Ivan Topolsky, Kim Philipp Jablonski, Lara Fuhrmann, Daniel Ehrsam, Isabel Stürmer, Andreia Cabral de Gouvea, Maria Domenica Moccia, Simon Grüter, Timothy Sykes, Lennart Opitz, Griffin White, Laura Neff, Doris Popovic, Andrea Patrignani, Jay Tracy, Ralph Schlapbach, Christiane Beckmann, Maurice Redondo, Olivier Kobel, Christoph Noppen, Niko Beerenwinkel, Tanja Stadler |

| | | | |
|---|---|---|---|
| EPI_ISL_3465392 | Sydney South West Pathology Service (SSWPS) - Liverpool Hospital - NSW Health Pathology | NSW Health Pathology - Institute of Clinical Pathology and Medical Research; Westmead Hospital; University of Sydney | Arnott A., Draper J., Gall M., Martinez E., Rockett R., Sintchenko V., on behalf of ICPMR |
| EPI_ISL_3465393, EPI_ISL_3465394 | Viollier AG | Department of Biosystems Science and Engineering, ETH Zürich | Chaoran Chen, Sarah Nadeau, Louis du Plessis, Catharine Aquino, Ivan Topolsky, Kim Philipp Jablonski, Lara Fuhrmann, Daniel Ehrsam, Isabel Stürmer, Andreia Cabral de Gouvea, Maria Domenica Moccia, Simon Grüter, Timothy Sykes, Lennart Opitz, Griffin White, Laura Neff, Doris Popovic, Andrea Patrignani, Jay Tracy, Ralph Schlapbach, Christiane Beckmann, Maurice Redondo, Olivier Kobel, Christoph Noppen, Niko Beerenwinkel, Tanja Stadler |
| EPI_ISL_3465395 | Australian Clinical Labs (formerly Healthscope Pathology) | NSW Health Pathology - Institute of Clinical Pathology and Medical Research; Westmead Hospital; University of Sydney | Arnott A., Draper J., Gall M., Martinez E., Rockett R., Sintchenko V., on behalf of ICPMR |
| EPI_ISL_3465396 | Viollier AG | Department of Biosystems Science and Engineering, ETH Zürich | Chaoran Chen, Sarah Nadeau, Louis du Plessis, Catharine Aquino, Ivan Topolsky, Kim Philipp Jablonski, Lara Fuhrmann, Daniel Ehrsam, Isabel Stürmer, Andreia Cabral de Gouvea, Maria Domenica Moccia, Simon Grüter, Timothy Sykes, Lennart Opitz, Griffin White, Laura Neff, Doris Popovic, Andrea Patrignani, Jay Tracy, Ralph Schlapbach, Christiane Beckmann, Maurice Redondo, Olivier Kobel, Christoph Noppen, Niko Beerenwinkel, Tanja Stadler |
| EPI_ISL_3465397 | Pathology West - NSW Health Pathology | NSW Health Pathology - Institute of Clinical Pathology and Medical Research; Westmead Hospital; University of Sydney | Arnott A., Draper J., Gall M., Martinez E., Rockett R., Sintchenko V., on behalf of ICPMR |
| EPI_ISL_3465398 | Viollier AG | Department of Biosystems Science and Engineering, ETH Zürich | Chaoran Chen, Sarah Nadeau, Louis du Plessis, Catharine Aquino, Ivan Topolsky, Kim Philipp Jablonski, Lara Fuhrmann, Daniel Ehrsam, Isabel Stürmer, Andreia Cabral de Gouvea, Maria Domenica Moccia, Simon Grüter, Timothy Sykes, Lennart Opitz, Griffin White, Laura Neff, Doris Popovic, Andrea Patrignani, Jay Tracy, Ralph Schlapbach, Christiane Beckmann, Maurice Redondo, Olivier Kobel, Christoph Noppen, Niko Beerenwinkel, Tanja Stadler |
| EPI_ISL_3465399 | Sydney South West Pathology Service (SSWPS) - Liverpool Hospital - NSW Health Pathology | NSW Health Pathology - Institute of Clinical Pathology and Medical Research; Westmead Hospital; University of Sydney | Arnott A., Draper J., Gall M., Martinez E., Rockett R., Sintchenko V., on behalf of ICPMR |
| EPI_ISL_3465400, EPI_ISL_3465401 | Viollier AG | Department of Biosystems Science and Engineering, ETH Zürich | Chaoran Chen, Sarah Nadeau, Louis du Plessis, Catharine Aquino, Ivan Topolsky, Kim Philipp Jablonski, Lara Fuhrmann, Daniel Ehrsam, Isabel Stürmer, Andreia Cabral de Gouvea, Maria Domenica Moccia, Simon Grüter, Timothy Sykes, Lennart Opitz, Griffin White, Laura Neff, Doris Popovic, Andrea Patrignani, Jay Tracy, Ralph Schlapbach, Christiane Beckmann, Maurice Redondo, Olivier Kobel, Christoph Noppen, Niko Beerenwinkel, Tanja Stadler |
| EPI_ISL_3465402 | Australian Clinical Labs (formerly Healthscope Pathology) | NSW Health Pathology - Institute of Clinical Pathology and Medical Research; Westmead Hospital; University of Sydney | Arnott A., Draper J., Gall M., Martinez E., Rockett R., Sintchenko V., on behalf of ICPMR |
| EPI_ISL_3465403 | Viollier AG | Department of Biosystems Science and Engineering, ETH Zürich | Chaoran Chen, Sarah Nadeau, Louis du Plessis, Catharine Aquino, Ivan Topolsky, Kim Philipp Jablonski, Lara Fuhrmann, Daniel Ehrsam, Isabel Stürmer, Andreia Cabral de Gouvea, Maria Domenica Moccia, Simon Grüter, Timothy Sykes, Lennart Opitz, Griffin White, Laura Neff, Doris Popovic, Andrea Patrignani, Jay Tracy, Ralph Schlapbach, Christiane Beckmann, Maurice Redondo, Olivier Kobel, Christoph Noppen, Niko Beerenwinkel, Tanja Stadler |
| EPI_ISL_3465404 | Pathology West - NSW Health Pathology | NSW Health Pathology - Institute of Clinical Pathology and Medical Research; Westmead Hospital; University of Sydney | Arnott A., Draper J., Gall M., Martinez E., Rockett R., Sintchenko V., on behalf of ICPMR |
| EPI_ISL_3465405, EPI_ISL_3465406 | Viollier AG | Department of Biosystems Science and Engineering, ETH Zürich | Chaoran Chen, Sarah Nadeau, Louis du Plessis, Catharine Aquino, Ivan Topolsky, Kim Philipp Jablonski, Lara Fuhrmann, Daniel Ehrsam, Isabel Stürmer, Andreia Cabral de Gouvea, Maria Domenica Moccia, Simon Grüter, Timothy Sykes, Lennart Opitz, Griffin White, Laura Neff, Doris Popovic, Andrea Patrignani, Jay Tracy, Ralph Schlapbach, Christiane Beckmann, Maurice Redondo, Olivier Kobel, Christoph Noppen, Niko Beerenwinkel, Tanja Stadler |
| EPI_ISL_3465407 | 4Cyte Pathology | NSW Health Pathology - Institute of Clinical Pathology and Medical Research; Westmead Hospital; University of Sydney | Arnott A., Draper J., Gall M., Martinez E., Rockett R., Sintchenko V., on behalf of ICPMR |
| EPI_ISL_3465408 | Viollier AG | Department of Biosystems Science and Engineering, ETH Zürich | Chaoran Chen, Sarah Nadeau, Louis du Plessis, Catharine Aquino, Ivan Topolsky, Kim Philipp Jablonski, Lara Fuhrmann, Daniel Ehrsam, Isabel Stürmer, Andreia Cabral de Gouvea, Maria Domenica Moccia, Simon Grüter, Timothy Sykes, Lennart Opitz, Griffin White, Laura Neff, Doris Popovic, Andrea Patrignani, Jay Tracy, Ralph Schlapbach, Christiane Beckmann, Maurice Redondo, Olivier Kobel, Christoph Noppen, Niko Beerenwinkel, Tanja Stadler |
| EPI_ISL_3465409 | Medlab Pathology | NSW Health Pathology - Institute of Clinical Pathology and Medical Research; Westmead Hospital; University of Sydney | Arnott A., Draper J., Gall M., Martinez E., Rockett R., Sintchenko V., on behalf of ICPMR |
| EPI_ISL_3465410 | Viollier AG | Department of Biosystems Science and Engineering, ETH Zürich | Chaoran Chen, Sarah Nadeau, Louis du Plessis, Catharine Aquino, Ivan Topolsky, Kim Philipp Jablonski, Lara Fuhrmann, Daniel Ehrsam, Isabel Stürmer, Andreia Cabral de Gouvea, Maria Domenica Moccia, Simon Grüter, Timothy Sykes, Lennart Opitz, Griffin White, Laura Neff, Doris Popovic, Andrea Patrignani, Jay Tracy, Ralph Schlapbach, Christiane Beckmann, Maurice Redondo, Olivier Kobel, Christoph Noppen, Niko Beerenwinkel, Tanja Stadler |
| EPI_ISL_3465411 | Medlab Pathology | NSW Health Pathology - Institute of Clinical Pathology and Medical Research; Westmead Hospital; University of Sydney | Arnott A., Draper J., Gall M., Martinez E., Rockett R., Sintchenko V., on behalf of ICPMR |
| EPI_ISL_3465412, EPI_ISL_3465413 | Viollier AG | Department of Biosystems Science and Engineering, ETH Zürich | Chaoran Chen, Sarah Nadeau, Louis du Plessis, Catharine Aquino, Ivan Topolsky, Kim Philipp Jablonski, Lara Fuhrmann, Daniel Ehrsam, Isabel Stürmer, Andreia Cabral de Gouvea, Maria Domenica Moccia, Simon Grüter, Timothy Sykes, Lennart Opitz, Griffin White, Laura Neff, Doris Popovic, Andrea Patrignani, Jay Tracy, Ralph Schlapbach, Christiane Beckmann, Maurice Redondo, Olivier Kobel, Christoph Noppen, Niko Beerenwinkel, Tanja Stadler |
| EPI_ISL_3465414 | Southern. IML Pathology | NSW Health Pathology - Institute of Clinical Pathology and Medical Research; Westmead Hospital; University of Sydney | Arnott A., Draper J., Gall M., Martinez E., Rockett R., Sintchenko V., on behalf of ICPMR |
| EPI_ISL_3465415 | Viollier AG | Department of Biosystems Science and Engineering, ETH Zürich | Chaoran Chen, Sarah Nadeau, Louis du Plessis, Catharine Aquino, Ivan Topolsky, Kim Philipp Jablonski, Lara Fuhrmann, Daniel Ehrsam, Isabel Stürmer, Andreia Cabral de Gouvea, Maria Domenica Moccia, Simon Grüter, Timothy Sykes, Lennart Opitz, Griffin White, Laura Neff, Doris Popovic, Andrea Patrignani, Jay Tracy, Ralph Schlapbach, Christiane Beckmann, Maurice Redondo, Olivier Kobel, Christoph Noppen, Niko Beerenwinkel, Tanja Stadler |
| EPI_ISL_3465416 | Pathology West - NSW Health Pathology | NSW Health Pathology - Institute of Clinical Pathology and Medical Research; Westmead Hospital; University of Sydney | Arnott A., Draper J., Gall M., Martinez E., Rockett R., Sintchenko V., on behalf of ICPMR |
| EPI_ISL_3465417 | Viollier AG | Department of Biosystems Science and Engineering, ETH Zürich | Chaoran Chen, Sarah Nadeau, Louis du Plessis, Catharine Aquino, Ivan Topolsky, Kim Philipp Jablonski, Lara Fuhrmann, Daniel Ehrsam, Isabel Stürmer, Andreia Cabral de Gouvea, Maria Domenica Moccia, Simon Grüter, Timothy Sykes, Lennart Opitz, Griffin White, Laura Neff, Doris Popovic, Andrea Patrignani, Jay Tracy, Ralph Schlapbach, Christiane Beckmann, Maurice Redondo, Olivier Kobel, Christoph Noppen, Niko Beerenwinkel, Tanja Stadler |
| EPI_ISL_3465418 | Medlab Pathology | NSW Health Pathology - Institute of Clinical Pathology and Medical Research; Westmead Hospital; University of Sydney | Arnott A., Draper J., Gall M., Martinez E., Rockett R., Sintchenko V., on behalf of ICPMR |
| EPI_ISL_3465419 | Viollier AG | Department of Biosystems Science and Engineering, ETH Zürich | Chaoran Chen, Sarah Nadeau, Louis du Plessis, Catharine Aquino, Ivan Topolsky, Kim Philipp Jablonski, Lara Fuhrmann, Daniel Ehrsam, Isabel Stürmer, Andreia Cabral de Gouvea, Maria Domenica Moccia, Simon Grüter, Timothy Sykes, Lennart Opitz, Griffin White, Laura Neff, Doris Popovic, Andrea Patrignani, Jay Tracy, Ralph Schlapbach, Christiane Beckmann, Maurice Redondo, Olivier Kobel, Christoph Noppen, Niko Beerenwinkel, Tanja Stadler |
| EPI_ISL_3465420 | St Vincent's Pathology (SydPath) | NSW Health Pathology - Institute of Clinical Pathology and Medical Research; Westmead Hospital; University of Sydney | Arnott A., Draper J., Gall M., Martinez E., Rockett R., Sintchenko V., on behalf of ICPMR |
| EPI_ISL_3465421 | Viollier AG | Department of Biosystems Science and Engineering, ETH Zürich | Chaoran Chen, Sarah Nadeau, Louis du Plessis, Catharine Aquino, Ivan Topolsky, Kim Philipp Jablonski, Lara Fuhrmann, Daniel Ehrsam, Isabel Stürmer, Andreia Cabral de Gouvea, Maria Domenica Moccia, Simon Grüter, Timothy Sykes, Lennart Opitz, Griffin White, Laura Neff, Doris Popovic, Andrea Patrignani, Jay Tracy, Ralph Schlapbach, Christiane Beckmann, Maurice Redondo, Olivier Kobel, Christoph Noppen, Niko Beerenwinkel, Tanja Stadler |
| EPI_ISL_3465422 | St Vincent's Pathology (SydPath) | NSW Health Pathology - Institute of Clinical Pathology and Medical Research; Westmead Hospital; University of Sydney | Arnott A., Draper J., Gall M., Martinez E., Rockett R., Sintchenko V., on behalf of ICPMR |
| EPI_ISL_3465423, EPI_ISL_3465424 | Viollier AG | Department of Biosystems Science and Engineering, ETH Zürich | Chaoran Chen, Sarah Nadeau, Louis du Plessis, Catharine Aquino, Ivan Topolsky, Kim Philipp Jablonski, Lara Fuhrmann, Daniel Ehrsam, Isabel Stürmer, Andreia Cabral de Gouvea, Maria Domenica Moccia, Simon Grüter, Timothy Sykes, Lennart Opitz, Griffin White, Laura Neff, Doris Popovic, Andrea Patrignani, Jay Tracy, Ralph Schlapbach, Christiane Beckmann, Maurice Redondo, Olivier Kobel, Christoph Noppen, Niko Beerenwinkel, Tanja Stadler |
| EPI_ISL_3465425 | Medlab Pathology | NSW Health Pathology - Institute of Clinical Pathology and Medical Research; Westmead Hospital; University of Sydney | Arnott A., Draper J., Gall M., Martinez E., Rockett R., Sintchenko V., on behalf of ICPMR |
| EPI_ISL_3465426 | Viollier AG | Department of Biosystems Science and Engineering, ETH Zürich | Chaoran Chen, Sarah Nadeau, Louis du Plessis, Catharine Aquino, Ivan Topolsky, Kim Philipp Jablonski, Lara Fuhrmann, Daniel Ehrsam, Isabel Stürmer, Andreia Cabral de Gouvea, Maria Domenica Moccia, Simon Grüter, Timothy Sykes, Lennart Opitz, Griffin White, Laura Neff, Doris Popovic, Andrea Patrignani, Jay Tracy, Ralph Schlapbach, Christiane Beckmann, Maurice Redondo, Olivier Kobel, Christoph Noppen, Niko Beerenwinkel, Tanja Stadler |
| EPI_ISL_3465427 | Medlab Pathology | NSW Health Pathology - Institute of Clinical Pathology and Medical Research; Westmead Hospital; University of Sydney | Arnott A., Draper J., Gall M., Martinez E., Rockett R., Sintchenko V., on behalf of ICPMR |
| EPI_ISL_3465428, EPI_ISL_3465429 | Viollier AG | Department of Biosystems Science and Engineering, ETH Zürich | Chaoran Chen, Sarah Nadeau, Louis du Plessis, Catharine Aquino, Ivan Topolsky, Kim Philipp Jablonski, Lara Fuhrmann, Daniel Ehrsam, Isabel Stürmer, Andreia Cabral de Gouvea, Maria Domenica Moccia, Simon Grüter, Timothy Sykes, Lennart Opitz, Griffin White, Laura Neff, Doris Popovic, Andrea Patrignani, Jay Tracy, Ralph Schlapbach, Christiane Beckmann, Maurice Redondo, Olivier Kobel, Christoph Noppen, Niko Beerenwinkel, Tanja Stadler |
| EPI_ISL_3465430 | Pathology West - NSW Health Pathology | NSW Health Pathology - Institute of Clinical Pathology and Medical Research; Westmead Hospital; University of Sydney | Arnott A., Draper J., Gall M., Martinez E., Rockett R., Sintchenko V., on behalf of ICPMR |
| EPI_ISL_3465431 | Viollier AG | Department of Biosystems Science and Engineering, ETH Zürich | Chaoran Chen, Sarah Nadeau, Louis du Plessis, Catharine Aquino, Ivan Topolsky, Kim Philipp Jablonski, Lara Fuhrmann, Daniel Ehrsam, Isabel Stürmer, Andreia Cabral de Gouvea, Maria Domenica Moccia, Simon Grüter, Timothy Sykes, Lennart Opitz, Griffin White, Laura Neff, Doris Popovic, Andrea Patrignani, Jay Tracy, Ralph Schlapbach, Christiane Beckmann, Maurice Redondo, Olivier Kobel, Christoph Noppen, Niko Beerenwinkel, Tanja Stadler |
| EPI_ISL_3465432 | Histopath | NSW Health Pathology - Institute of Clinical Pathology and Medical Research; Westmead Hospital; University of Sydney | Arnott A., Draper J., Gall M., Martinez E., Rockett R., Sintchenko V., on behalf of ICPMR |
| EPI_ISL_3465433 | Viollier AG | Department of Biosystems Science and Engineering, ETH Zürich | Chaoran Chen, Sarah Nadeau, Louis du Plessis, Catharine Aquino, Ivan Topolsky, Kim Philipp Jablonski, Lara Fuhrmann, Daniel Ehrsam, Isabel Stürmer, Andreia Cabral de Gouvea, Maria Domenica Moccia, Simon Grüter, Timothy Sykes, Lennart Opitz, Griffin White, Laura Neff, Doris Popovic, Andrea Patrignani, Jay Tracy, Ralph Schlapbach, Christiane Beckmann, Maurice Redondo, Olivier Kobel, Christoph Noppen, Niko Beerenwinkel, Tanja Stadler |
| EPI_ISL_3465434 | St Vincent's Pathology (SydPath) | NSW Health Pathology - Institute of Clinical Pathology and Medical Research; Westmead Hospital; University of Sydney | Arnott A., Draper J., Gall M., Martinez E., Rockett R., Sintchenko V., on behalf of ICPMR |
| EPI_ISL_3465435 | Viollier AG | Department of Biosystems Science and Engineering, ETH Zürich | Chaoran Chen, Sarah Nadeau, Louis du Plessis, Catharine Aquino, Ivan Topolsky, Kim Philipp Jablonski, Lara Fuhrmann, Daniel Ehrsam, Isabel Stürmer, Andreia Cabral de Gouvea, Maria Domenica Moccia, Simon Grüter, Timothy Sykes, Lennart Opitz, Griffin White, Laura Neff, Doris Popovic, Andrea Patrignani, Jay Tracy, Ralph Schlapbach, Christiane Beckmann, Maurice Redondo, Olivier Kobel, Christoph Noppen, Niko Beerenwinkel, Tanja Stadler |
| EPI_ISL_3465436 | Sydney South West Pathology Service (SSWPS) - Liverpool Hospital - NSW Health Pathology | NSW Health Pathology - Institute of Clinical Pathology and Medical Research; Westmead Hospital; University of Sydney | Arnott A., Draper J., Gall M., Martinez E., Rockett R., Sintchenko V., on behalf of ICPMR |
| EPI_ISL_3465437, EPI_ISL_3465438 | Viollier AG | Department of Biosystems Science and Engineering, ETH Zürich | Chaoran Chen, Sarah Nadeau, Louis du Plessis, Catharine Aquino, Ivan Topolsky, Kim Philipp Jablonski, Lara Fuhrmann, Daniel Ehrsam, Isabel Stürmer, Andreia Cabral de Gouvea, Maria Domenica Moccia, Simon Grüter, Timothy Sykes, Lennart Opitz, Griffin White, Laura Neff, Doris Popovic, Andrea Patrignani, Jay Tracy, Ralph Schlapbach, Christiane Beckmann, Maurice Redondo, Olivier Kobel, Christoph Noppen, Niko Beerenwinkel, Tanja Stadler |
| EPI_ISL_3465439 | Australian Clinical Labs (formerly Healthscope Pathology) | NSW Health Pathology - Institute of Clinical Pathology and Medical Research; Westmead Hospital; University of Sydney | Arnott A., Draper J., Gall M., Martinez E., Rockett R., Sintchenko V., on behalf of ICPMR |
| EPI_ISL_3465440, EPI_ISL_3465441 | Viollier AG | Department of Biosystems Science and Engineering, ETH Zürich | Chaoran Chen, Sarah Nadeau, Louis du Plessis, Catharine Aquino, Ivan Topolsky, Kim Philipp Jablonski, Lara Fuhrmann, Daniel Ehrsam, Isabel Stürmer, Andreia Cabral de Gouvea, Maria Domenica Moccia, Simon Grüter, Timothy Sykes, Lennart Opitz, Griffin White, Laura Neff, Doris Popovic, Andrea Patrignani, Jay Tracy, Ralph Schlapbach, Christiane Beckmann, Maurice Redondo, Olivier Kobel, Christoph Noppen, Niko Beerenwinkel, Tanja Stadler |
| EPI_ISL_3465442 | Australian Clinical Labs (formerly Healthscope Pathology) | NSW Health Pathology - Institute of Clinical Pathology and Medical Research; Westmead Hospital; University of Sydney | Arnott A., Draper J., Gall M., Martinez E., Rockett R., Sintchenko V., on behalf of ICPMR |
| EPI_ISL_3465443 | Viollier AG | Department of Biosystems Science and Engineering, ETH Zürich | Chaoran Chen, Sarah Nadeau, Louis du Plessis, Catharine Aquino, Ivan Topolsky, Kim Philipp Jablonski, Lara Fuhrmann, Daniel Ehrsam, Isabel Stürmer, Andreia Cabral de Gouvea, Maria Domenica Moccia, Simon Grüter, Timothy Sykes, Lennart Opitz, Griffin White, Laura Neff, Doris Popovic, Andrea Patrignani, Jay Tracy, Ralph Schlapbach, Christiane Beckmann, Maurice Redondo, Olivier Kobel, Christoph Noppen, Niko Beerenwinkel, Tanja Stadler |
| EPI_ISL_3465444 | Sydney South West Pathology Service (SSWPS) - Liverpool Hospital - NSW Health Pathology | NSW Health Pathology - Institute of Clinical Pathology and Medical Research; Westmead Hospital; University of Sydney | Arnott A., Draper J., Gall M., Martinez E., Rockett R., Sintchenko V., on behalf of ICPMR |

| | | | |
|---|---|---|---|
| EPI_ISL_3465445 | Viollier AG | Department of Biosystems Science and Engineering, ETH Zürich | Chaoran Chen, Sarah Nadeau, Louis du Plessis, Catharine Aquino, Ivan Topolsky, Kim Philipp Jablonski, Lara Fuhrmann, Daniel Ehrsam, Isabel Stürmer, Andreia Cabral de Gouvea, Maria Domenica Moccia, Simon Grüter, Timothy Sykes, Lennart Opitz, Griffin White, Laura Neff, Doris Popovic, Andrea Patrignani, Jay Tracy, Ralph Schlapbach, Christiane Beckmann, Maurice Redondo, Olivier Kobel, Christoph Noppen, Niko Beerenwinkel, Tanja Stadler |
| EPI_ISL_3465446 | Viollier AG | Department of Biosystems Science and Engineering, ETH Zürich | Andrea Patrizia Salzmann, Henriette Kurth, Christiane Beckmann, Ivan Topolsky, Chaoran Chen, Sarah Nadeau, Louis du Plessis, Kim Philipp Jablonski, Lara Fuhrmann, Niko Beerenwinkel, Christoph Noppen, Olivier Kobel, Maurice Redondo, Tanja Stadler |
| EPI_ISL_3465447 | Laverty Pathology | NSW Health Pathology - Institute of Clinical Pathology and Medical Research; Westmead Hospital; University of Sydney | Arnott A., Draper J., Gall M., Martinez E., Rockett R., Sintchenko V., on behalf of ICPMR |
| EPI_ISL_3465448 | Viollier AG | Department of Biosystems Science and Engineering, ETH Zürich | Chaoran Chen, Sarah Nadeau, Louis du Plessis, Catharine Aquino, Ivan Topolsky, Kim Philipp Jablonski, Lara Fuhrmann, Daniel Ehrsam, Isabel Stürmer, Andreia Cabral de Gouvea, Maria Domenica Moccia, Simon Grüter, Timothy Sykes, Lennart Opitz, Griffin White, Laura Neff, Doris Popovic, Andrea Patrignani, Jay Tracy, Ralph Schlapbach, Christiane Beckmann, Maurice Redondo, Olivier Kobel, Christoph Noppen, Niko Beerenwinkel, Tanja Stadler |
| EPI_ISL_3465449 | Histopath | NSW Health Pathology - Institute of Clinical Pathology and Medical Research; Westmead Hospital; University of Sydney | Arnott A., Draper J., Gall M., Martinez E., Rockett R., Sintchenko V., on behalf of ICPMR |
| EPI_ISL_3465450 | Viollier AG | Department of Biosystems Science and Engineering, ETH Zürich | Chaoran Chen, Sarah Nadeau, Louis du Plessis, Catharine Aquino, Ivan Topolsky, Kim Philipp Jablonski, Lara Fuhrmann, Daniel Ehrsam, Isabel Stürmer, Andreia Cabral de Gouvea, Maria Domenica Moccia, Simon Grüter, Timothy Sykes, Lennart Opitz, Griffin White, Laura Neff, Doris Popovic, Andrea Patrignani, Jay Tracy, Ralph Schlapbach, Christiane Beckmann, Maurice Redondo, Olivier Kobel, Christoph Noppen, Niko Beerenwinkel, Tanja Stadler |
| EPI_ISL_3465451 | Histopath | NSW Health Pathology - Institute of Clinical Pathology and Medical Research; Westmead Hospital; University of Sydney | Arnott A., Draper J., Gall M., Martinez E., Rockett R., Sintchenko V., on behalf of ICPMR |
| EPI_ISL_3465452 | Viollier AG | Department of Biosystems Science and Engineering, ETH Zürich | Chaoran Chen, Sarah Nadeau, Louis du Plessis, Catharine Aquino, Ivan Topolsky, Kim Philipp Jablonski, Lara Fuhrmann, Daniel Ehrsam, Isabel Stürmer, Andreia Cabral de Gouvea, Maria Domenica Moccia, Simon Grüter, Timothy Sykes, Lennart Opitz, Griffin White, Laura Neff, Doris Popovic, Andrea Patrignani, Jay Tracy, Ralph Schlapbach, Christiane Beckmann, Maurice Redondo, Olivier Kobel, Christoph Noppen, Niko Beerenwinkel, Tanja Stadler |
| EPI_ISL_3465453 | Histopath | NSW Health Pathology - Institute of Clinical Pathology and Medical Research; Westmead Hospital; University of Sydney | Arnott A., Draper J., Gall M., Martinez E., Rockett R., Sintchenko V., on behalf of ICPMR |
| EPI_ISL_3465454, EPI_ISL_3465455 | Viollier AG | Department of Biosystems Science and Engineering, ETH Zürich | Chaoran Chen, Sarah Nadeau, Louis du Plessis, Catharine Aquino, Ivan Topolsky, Kim Philipp Jablonski, Lara Fuhrmann, Daniel Ehrsam, Isabel Stürmer, Andreia Cabral de Gouvea, Maria Domenica Moccia, Simon Grüter, Timothy Sykes, Lennart Opitz, Griffin White, Laura Neff, Doris Popovic, Andrea Patrignani, Jay Tracy, Ralph Schlapbach, Christiane Beckmann, Maurice Redondo, Olivier Kobel, Christoph Noppen, Niko Beerenwinkel, Tanja Stadler |
| EPI_ISL_3465456 | Histopath | NSW Health Pathology - Institute of Clinical Pathology and Medical Research; Westmead Hospital; University of Sydney | Arnott A., Draper J., Gall M., Martinez E., Rockett R., Sintchenko V., on behalf of ICPMR |
| EPI_ISL_3465457, EPI_ISL_3465458 | Viollier AG | Department of Biosystems Science and Engineering, ETH Zürich | Chaoran Chen, Sarah Nadeau, Louis du Plessis, Catharine Aquino, Ivan Topolsky, Kim Philipp Jablonski, Lara Fuhrmann, Daniel Ehrsam, Isabel Stürmer, Andreia Cabral de Gouvea, Maria Domenica Moccia, Simon Grüter, Timothy Sykes, Lennart Opitz, Griffin White, Laura Neff, Doris Popovic, Andrea Patrignani, Jay Tracy, Ralph Schlapbach, Christiane Beckmann, Maurice Redondo, Olivier Kobel, Christoph Noppen, Niko Beerenwinkel, Tanja Stadler |
| EPI_ISL_3465459 | Histopath | NSW Health Pathology - Institute of Clinical Pathology and Medical Research; Westmead Hospital; University of Sydney | Arnott A., Draper J., Gall M., Martinez E., Rockett R., Sintchenko V., on behalf of ICPMR |
| EPI_ISL_3465460 | Viollier AG | Department of Biosystems Science and Engineering, ETH Zürich | Chaoran Chen, Sarah Nadeau, Louis du Plessis, Catharine Aquino, Ivan Topolsky, Kim Philipp Jablonski, Lara Fuhrmann, Daniel Ehrsam, Isabel Stürmer, Andreia Cabral de Gouvea, Maria Domenica Moccia, Simon Grüter, Timothy Sykes, Lennart Opitz, Griffin White, Laura Neff, Doris Popovic, Andrea Patrignani, Jay Tracy, Ralph Schlapbach, Christiane Beckmann, Maurice Redondo, Olivier Kobel, Christoph Noppen, Niko Beerenwinkel, Tanja Stadler |
| EPI_ISL_3465461 | Histopath | NSW Health Pathology - Institute of Clinical Pathology and Medical Research; Westmead Hospital; University of Sydney | Arnott A., Draper J., Gall M., Martinez E., Rockett R., Sintchenko V., on behalf of ICPMR |
| EPI_ISL_3465462 | Viollier AG | Department of Biosystems Science and Engineering, ETH Zürich | Chaoran Chen, Sarah Nadeau, Louis du Plessis, Catharine Aquino, Ivan Topolsky, Kim Philipp Jablonski, Lara Fuhrmann, Daniel Ehrsam, Isabel Stürmer, Andreia Cabral de Gouvea, Maria Domenica Moccia, Simon Grüter, Timothy Sykes, Lennart Opitz, Griffin White, Laura Neff, Doris Popovic, Andrea Patrignani, Jay Tracy, Ralph Schlapbach, Christiane Beckmann, Maurice Redondo, Olivier Kobel, Christoph Noppen, Niko Beerenwinkel, Tanja Stadler |
| EPI_ISL_3465463 | Histopath | NSW Health Pathology - Institute of Clinical Pathology and Medical Research; Westmead Hospital; University of Sydney | Arnott A., Draper J., Gall M., Martinez E., Rockett R., Sintchenko V., on behalf of ICPMR |
| EPI_ISL_3465464, EPI_ISL_3465465 | Viollier AG | Department of Biosystems Science and Engineering, ETH Zürich | Chaoran Chen, Sarah Nadeau, Louis du Plessis, Catharine Aquino, Ivan Topolsky, Kim Philipp Jablonski, Lara Fuhrmann, Daniel Ehrsam, Isabel Stürmer, Andreia Cabral de Gouvea, Maria Domenica Moccia, Simon Grüter, Timothy Sykes, Lennart Opitz, Griffin White, Laura Neff, Doris Popovic, Andrea Patrignani, Jay Tracy, Ralph Schlapbach, Christiane Beckmann, Maurice Redondo, Olivier Kobel, Christoph Noppen, Niko Beerenwinkel, Tanja Stadler |
| EPI_ISL_3465466 | Pathology North - Hunter - NSW Health Pathology | NSW Health Pathology - Institute of Clinical Pathology and Medical Research; Westmead Hospital; University of Sydney | Arnott A., Draper J., Gall M., Martinez E., Rockett R., Sintchenko V., on behalf of ICPMR |
| EPI_ISL_3465467 | Viollier AG | Department of Biosystems Science and Engineering, ETH Zürich | Chaoran Chen, Sarah Nadeau, Louis du Plessis, Catharine Aquino, Ivan Topolsky, Kim Philipp Jablonski, Lara Fuhrmann, Daniel Ehrsam, Isabel Stürmer, Andreia Cabral de Gouvea, Maria Domenica Moccia, Simon Grüter, Timothy Sykes, Lennart Opitz, Griffin White, Laura Neff, Doris Popovic, Andrea Patrignani, Jay Tracy, Ralph Schlapbach, Christiane Beckmann, Maurice Redondo, Olivier Kobel, Christoph Noppen, Niko Beerenwinkel, Tanja Stadler |
| EPI_ISL_3465468 | Pathology North - Hunter - NSW Health Pathology | NSW Health Pathology - Institute of Clinical Pathology and Medical Research; Westmead Hospital; University of Sydney | Arnott A., Draper J., Gall M., Martinez E., Rockett R., Sintchenko V., on behalf of ICPMR |
| EPI_ISL_3465469, EPI_ISL_3465470 | Viollier AG | Department of Biosystems Science and Engineering, ETH Zürich | Chaoran Chen, Sarah Nadeau, Louis du Plessis, Catharine Aquino, Ivan Topolsky, Kim Philipp Jablonski, Lara Fuhrmann, Daniel Ehrsam, Isabel Stürmer, Andreia Cabral de Gouvea, Maria Domenica Moccia, Simon Grüter, Timothy Sykes, Lennart Opitz, Griffin White, Laura Neff, Doris Popovic, Andrea Patrignani, Jay Tracy, Ralph Schlapbach, Christiane Beckmann, Maurice Redondo, Olivier Kobel, Christoph Noppen, Niko Beerenwinkel, Tanja Stadler |
| EPI_ISL_3465471 | Pathology North - Hunter - NSW Health Pathology | NSW Health Pathology - Institute of Clinical Pathology and Medical Research; Westmead Hospital; University of Sydney | Arnott A., Draper J., Gall M., Martinez E., Rockett R., Sintchenko V., on behalf of ICPMR |
| EPI_ISL_3465472 | Viollier AG | Department of Biosystems Science and Engineering, ETH Zürich | Chaoran Chen, Sarah Nadeau, Louis du Plessis, Catharine Aquino, Ivan Topolsky, Kim Philipp Jablonski, Lara Fuhrmann, Daniel Ehrsam, Isabel Stürmer, Andreia Cabral de Gouvea, Maria Domenica Moccia, Simon Grüter, Timothy Sykes, Lennart Opitz, Griffin White, Laura Neff, Doris Popovic, Andrea Patrignani, Jay Tracy, Ralph Schlapbach, Christiane Beckmann, Maurice Redondo, Olivier Kobel, Christoph Noppen, Niko Beerenwinkel, Tanja Stadler |
| EPI_ISL_3465473 | Pathology North - Hunter - NSW Health Pathology | NSW Health Pathology - Institute of Clinical Pathology and Medical Research; Westmead Hospital; University of Sydney | Arnott A., Draper J., Gall M., Martinez E., Rockett R., Sintchenko V., on behalf of ICPMR |
| EPI_ISL_3465474 | Viollier AG | Department of Biosystems Science and Engineering, ETH Zürich | Chaoran Chen, Sarah Nadeau, Louis du Plessis, Catharine Aquino, Ivan Topolsky, Kim Philipp Jablonski, Lara Fuhrmann, Daniel Ehrsam, Isabel Stürmer, Andreia Cabral de Gouvea, Maria Domenica Moccia, Simon Grüter, Timothy Sykes, Lennart Opitz, Griffin White, Laura Neff, Doris Popovic, Andrea Patrignani, Jay Tracy, Ralph Schlapbach, Christiane Beckmann, Maurice Redondo, Olivier Kobel, Christoph Noppen, Niko Beerenwinkel, Tanja Stadler |
| EPI_ISL_3465475 | Pathology North - Lismore laboratory - NSW Health Pathology | NSW Health Pathology - Institute of Clinical Pathology and Medical Research; Westmead Hospital; University of Sydney | Arnott A., Draper J., Gall M., Martinez E., Rockett R., Sintchenko V., on behalf of ICPMR |
| EPI_ISL_3465476 | Viollier AG | Department of Biosystems Science and Engineering, ETH Zürich | Chaoran Chen, Sarah Nadeau, Louis du Plessis, Catharine Aquino, Ivan Topolsky, Kim Philipp Jablonski, Lara Fuhrmann, Daniel Ehrsam, Isabel Stürmer, Andreia Cabral de Gouvea, Maria Domenica Moccia, Simon Grüter, Timothy Sykes, Lennart Opitz, Griffin White, Laura Neff, Doris Popovic, Andrea Patrignani, Jay Tracy, Ralph Schlapbach, Christiane Beckmann, Maurice Redondo, Olivier Kobel, Christoph Noppen, Niko Beerenwinkel, Tanja Stadler |
| EPI_ISL_3465477 | Sydney South West Pathology Service (SSWPS) - Liverpool Hospital - NSW Health Pathology | NSW Health Pathology - Institute of Clinical Pathology and Medical Research; Westmead Hospital; University of Sydney | Arnott A., Draper J., Gall M., Martinez E., Rockett R., Sintchenko V., on behalf of ICPMR |
| EPI_ISL_3465478, EPI_ISL_3465479, EPI_ISL_3465480 | Viollier AG | Department of Biosystems Science and Engineering, ETH Zürich | Chaoran Chen, Sarah Nadeau, Louis du Plessis, Catharine Aquino, Ivan Topolsky, Kim Philipp Jablonski, Lara Fuhrmann, Daniel Ehrsam, Isabel Stürmer, Andreia Cabral de Gouvea, Maria Domenica Moccia, Simon Grüter, Timothy Sykes, Lennart Opitz, Griffin White, Laura Neff, Doris Popovic, Andrea Patrignani, Jay Tracy, Ralph Schlapbach, Christiane Beckmann, Maurice Redondo, Olivier Kobel, Christoph Noppen, Niko Beerenwinkel, Tanja Stadler |
| EPI_ISL_3465481 | Sydney South West Pathology Service (SSWPS) - Liverpool Hospital - NSW Health Pathology | NSW Health Pathology - Institute of Clinical Pathology and Medical Research; Westmead Hospital; University of Sydney | Arnott A., Draper J., Gall M., Martinez E., Rockett R., Sintchenko V., on behalf of ICPMR |
| EPI_ISL_3465482, EPI_ISL_3465483 | Viollier AG | Department of Biosystems Science and Engineering, ETH Zürich | Chaoran Chen, Sarah Nadeau, Louis du Plessis, Catharine Aquino, Ivan Topolsky, Kim Philipp Jablonski, Lara Fuhrmann, Daniel Ehrsam, Isabel Stürmer, Andreia Cabral de Gouvea, Maria Domenica Moccia, Simon Grüter, Timothy Sykes, Lennart Opitz, Griffin White, Laura Neff, Doris Popovic, Andrea Patrignani, Jay Tracy, Ralph Schlapbach, Christiane Beckmann, Maurice Redondo, Olivier Kobel, Christoph Noppen, Niko Beerenwinkel, Tanja Stadler |
| EPI_ISL_3465484 | Medlab Pathology | NSW Health Pathology - Institute of Clinical Pathology and Medical Research; Westmead Hospital; University of Sydney | Arnott A., Draper J., Gall M., Martinez E., Rockett R., Sintchenko V., on behalf of ICPMR |
| EPI_ISL_3465485 | Viollier AG | Department of Biosystems Science and Engineering, ETH Zürich | Chaoran Chen, Sarah Nadeau, Louis du Plessis, Catharine Aquino, Ivan Topolsky, Kim Philipp Jablonski, Lara Fuhrmann, Daniel Ehrsam, Isabel Stürmer, Andreia Cabral de Gouvea, Maria Domenica Moccia, Simon Grüter, Timothy Sykes, Lennart Opitz, Griffin White, Laura Neff, Doris Popovic, Andrea Patrignani, Jay Tracy, Ralph Schlapbach, Christiane Beckmann, Maurice Redondo, Olivier Kobel, Christoph Noppen, Niko Beerenwinkel, Tanja Stadler |
| EPI_ISL_3465486 | Medlab Pathology | NSW Health Pathology - Institute of Clinical Pathology and Medical Research; Westmead Hospital; University of Sydney | Arnott A., Draper J., Gall M., Martinez E., Rockett R., Sintchenko V., on behalf of ICPMR |
| EPI_ISL_3465487, EPI_ISL_3465488 | Viollier AG | Department of Biosystems Science and Engineering, ETH Zürich | Chaoran Chen, Sarah Nadeau, Louis du Plessis, Catharine Aquino, Ivan Topolsky, Kim Philipp Jablonski, Lara Fuhrmann, Daniel Ehrsam, Isabel Stürmer, Andreia Cabral de Gouvea, Maria Domenica Moccia, Simon Grüter, Timothy Sykes, Lennart Opitz, Griffin White, Laura Neff, Doris Popovic, Andrea Patrignani, Jay Tracy, Ralph Schlapbach, Christiane Beckmann, Maurice Redondo, Olivier Kobel, Christoph Noppen, Niko Beerenwinkel, Tanja Stadler |
| EPI_ISL_3465489 | Medlab Pathology | NSW Health Pathology - Institute of Clinical Pathology and Medical Research; Westmead Hospital; University of Sydney | Arnott A., Draper J., Gall M., Martinez E., Rockett R., Sintchenko V., on behalf of ICPMR |
| EPI_ISL_3465490 | Viollier AG | Department of Biosystems Science and Engineering, ETH Zürich | Chaoran Chen, Sarah Nadeau, Louis du Plessis, Catharine Aquino, Ivan Topolsky, Kim Philipp Jablonski, Lara Fuhrmann, Daniel Ehrsam, Isabel Stürmer, Andreia Cabral de Gouvea, Maria Domenica Moccia, Simon Grüter, Timothy Sykes, Lennart Opitz, Griffin White, Laura Neff, Doris Popovic, Andrea Patrignani, Jay Tracy, Ralph Schlapbach, Christiane Beckmann, Maurice Redondo, Olivier Kobel, Christoph Noppen, Niko Beerenwinkel, Tanja Stadler |
| EPI_ISL_3465491 | Medlab Pathology | NSW Health Pathology - Institute of Clinical Pathology and Medical Research; Westmead Hospital; University of Sydney | Arnott A., Draper J., Gall M., Martinez E., Rockett R., Sintchenko V., on behalf of ICPMR |
| EPI_ISL_3465492 | Viollier AG | Department of Biosystems Science and Engineering, ETH Zürich | Chaoran Chen, Sarah Nadeau, Louis du Plessis, Catharine Aquino, Ivan Topolsky, Kim Philipp Jablonski, Lara Fuhrmann, Daniel Ehrsam, Isabel Stürmer, Andreia Cabral de Gouvea, Maria Domenica Moccia, Simon Grüter, Timothy Sykes, Lennart Opitz, Griffin White, Laura Neff, Doris Popovic, Andrea Patrignani, Jay Tracy, Ralph Schlapbach, Christiane Beckmann, Maurice Redondo, Olivier Kobel, Christoph Noppen, Niko Beerenwinkel, Tanja Stadler |
| EPI_ISL_3465493 | Medlab Pathology | NSW Health Pathology - Institute of Clinical Pathology and Medical Research; Westmead Hospital; University of Sydney | Arnott A., Draper J., Gall M., Martinez E., Rockett R., Sintchenko V., on behalf of ICPMR |
| EPI_ISL_3465494 | Viollier AG | Department of Biosystems Science and Engineering, ETH Zürich | Chaoran Chen, Sarah Nadeau, Louis du Plessis, Catharine Aquino, Ivan Topolsky, Kim Philipp Jablonski, Lara Fuhrmann, Daniel Ehrsam, Isabel Stürmer, Andreia Cabral de Gouvea, Maria Domenica Moccia, Simon Grüter, Timothy Sykes, Lennart Opitz, Griffin White, Laura Neff, Doris Popovic, Andrea Patrignani, Jay Tracy, Ralph Schlapbach, Christiane Beckmann, Maurice Redondo, Olivier Kobel, Christoph Noppen, Niko Beerenwinkel, Tanja Stadler |
| EPI_ISL_3465495 | Medlab Pathology | NSW Health Pathology - Institute of Clinical Pathology and Medical Research; Westmead Hospital; University of Sydney | Arnott A., Draper J., Gall M., Martinez E., Rockett R., Sintchenko V., on behalf of ICPMR |
| EPI_ISL_3465496, EPI_ISL_3465497 | Viollier AG | Department of Biosystems Science and Engineering, ETH Zürich | Chaoran Chen, Sarah Nadeau, Louis du Plessis, Catharine Aquino, Ivan Topolsky, Kim Philipp Jablonski, Lara Fuhrmann, Daniel Ehrsam, Isabel Stürmer, Andreia Cabral de Gouvea, Maria Domenica Moccia, Simon Grüter, Timothy Sykes, Lennart Opitz, Griffin White, Laura Neff, Doris Popovic, Andrea Patrignani, Jay Tracy, Ralph Schlapbach, Christiane Beckmann, Maurice Redondo, Olivier Kobel, Christoph Noppen, Niko Beerenwinkel, Tanja Stadler |
| EPI_ISL_3465498 | Medlab Pathology | NSW Health Pathology - Institute of Clinical Pathology and Medical Research; Westmead Hospital; University of Sydney | Arnott A., Draper J., Gall M., Martinez E., Rockett R., Sintchenko V., on behalf of ICPMR |

| | | | |
|---|---|---|---|
| EPI_ISL_3465499 | Viollier AG | Department of Biosystems Science and Engineering, ETH Zürich | Chaoran Chen, Sarah Nadeau, Louis du Plessis, Catharine Aquino, Ivan Topolsky, Kim Philipp Jablonski, Lara Fuhrmann, Daniel Ehrsam, Isabel Stürmer, Andreia Cabral de Gouvea, Maria Domenica Moccia, Simon Grüter, Timothy Sykes, Lennart Opitz, Griffin White, Laura Neff, Doris Popovic, Andrea Patrignani, Jay Tracy, Ralph Schlapbach, Christiane Beckmann, Maurice Redondo, Olivier Kobel, Christoph Noppen, Niko Beerenwinkel, Tanja Stadler |
| EPI_ISL_3465500 | Sydney South West Pathology Service (SSWPS) - Liverpool Hospital - NSW Health Pathology | NSW Health Pathology - Institute of Clinical Pathology and Medical Research; Westmead Hospital; University of Sydney | Arnott A., Draper J., Gall M., Martinez E., Rockett R., Sintchenko V., on behalf of ICPMR |
| EPI_ISL_3465501 | Viollier AG | Department of Biosystems Science and Engineering, ETH Zürich | Chaoran Chen, Sarah Nadeau, Louis du Plessis, Catharine Aquino, Ivan Topolsky, Kim Philipp Jablonski, Lara Fuhrmann, Daniel Ehrsam, Isabel Stürmer, Andreia Cabral de Gouvea, Maria Domenica Moccia, Simon Grüter, Timothy Sykes, Lennart Opitz, Griffin White, Laura Neff, Doris Popovic, Andrea Patrignani, Jay Tracy, Ralph Schlapbach, Christiane Beckmann, Maurice Redondo, Olivier Kobel, Christoph Noppen, Niko Beerenwinkel, Tanja Stadler |
| EPI_ISL_3465502 | Sydney South West Pathology Service (SSWPS) - Liverpool Hospital - NSW Health Pathology | NSW Health Pathology - Institute of Clinical Pathology and Medical Research; Westmead Hospital; University of Sydney | Arnott A., Draper J., Gall M., Martinez E., Rockett R., Sintchenko V., on behalf of ICPMR |
| EPI_ISL_3465503, EPI_ISL_3465504 | Viollier AG | Department of Biosystems Science and Engineering, ETH Zürich | Chaoran Chen, Sarah Nadeau, Louis du Plessis, Catharine Aquino, Ivan Topolsky, Kim Philipp Jablonski, Lara Fuhrmann, Daniel Ehrsam, Isabel Stürmer, Andreia Cabral de Gouvea, Maria Domenica Moccia, Simon Grüter, Timothy Sykes, Lennart Opitz, Griffin White, Laura Neff, Doris Popovic, Andrea Patrignani, Jay Tracy, Ralph Schlapbach, Christiane Beckmann, Maurice Redondo, Olivier Kobel, Christoph Noppen, Niko Beerenwinkel, Tanja Stadler |
| EPI_ISL_3465505 | Sydney South West Pathology Service (SSWPS) - Liverpool Hospital - NSW Health Pathology | NSW Health Pathology - Institute of Clinical Pathology and Medical Research; Westmead Hospital; University of Sydney | Arnott A., Draper J., Gall M., Martinez E., Rockett R., Sintchenko V., on behalf of ICPMR |
| EPI_ISL_3465506 | Viollier AG | Department of Biosystems Science and Engineering, ETH Zürich | Chaoran Chen, Sarah Nadeau, Louis du Plessis, Catharine Aquino, Ivan Topolsky, Kim Philipp Jablonski, Lara Fuhrmann, Daniel Ehrsam, Isabel Stürmer, Andreia Cabral de Gouvea, Maria Domenica Moccia, Simon Grüter, Timothy Sykes, Lennart Opitz, Griffin White, Laura Neff, Doris Popovic, Andrea Patrignani, Jay Tracy, Ralph Schlapbach, Christiane Beckmann, Maurice Redondo, Olivier Kobel, Christoph Noppen, Niko Beerenwinkel, Tanja Stadler |
| EPI_ISL_3465507 | St Vincent's Pathology (SydPath) | NSW Health Pathology - Institute of Clinical Pathology and Medical Research; Westmead Hospital; University of Sydney | Arnott A., Draper J., Gall M., Martinez E., Rockett R., Sintchenko V., on behalf of ICPMR |
| EPI_ISL_3465508 | Viollier AG | Department of Biosystems Science and Engineering, ETH Zürich | Chaoran Chen, Sarah Nadeau, Louis du Plessis, Catharine Aquino, Ivan Topolsky, Kim Philipp Jablonski, Lara Fuhrmann, Daniel Ehrsam, Isabel Stürmer, Andreia Cabral de Gouvea, Maria Domenica Moccia, Simon Grüter, Timothy Sykes, Lennart Opitz, Griffin White, Laura Neff, Doris Popovic, Andrea Patrignani, Jay Tracy, Ralph Schlapbach, Christiane Beckmann, Maurice Redondo, Olivier Kobel, Christoph Noppen, Niko Beerenwinkel, Tanja Stadler |
| EPI_ISL_3465509 | St Vincent's Pathology (SydPath) | NSW Health Pathology - Institute of Clinical Pathology and Medical Research; Westmead Hospital; University of Sydney | Arnott A., Draper J., Gall M., Martinez E., Rockett R., Sintchenko V., on behalf of ICPMR |
| EPI_ISL_3465510 | Viollier AG | Department of Biosystems Science and Engineering, ETH Zürich | Chaoran Chen, Sarah Nadeau, Louis du Plessis, Catharine Aquino, Ivan Topolsky, Kim Philipp Jablonski, Lara Fuhrmann, Daniel Ehrsam, Isabel Stürmer, Andreia Cabral de Gouvea, Maria Domenica Moccia, Simon Grüter, Timothy Sykes, Lennart Opitz, Griffin White, Laura Neff, Doris Popovic, Andrea Patrignani, Jay Tracy, Ralph Schlapbach, Christiane Beckmann, Maurice Redondo, Olivier Kobel, Christoph Noppen, Niko Beerenwinkel, Tanja Stadler |
| EPI_ISL_3465511 | St Vincent's Pathology (SydPath) | NSW Health Pathology - Institute of Clinical Pathology and Medical Research; Westmead Hospital; University of Sydney | Arnott A., Draper J., Gall M., Martinez E., Rockett R., Sintchenko V., on behalf of ICPMR |
| EPI_ISL_3465512 | Viollier AG | Department of Biosystems Science and Engineering, ETH Zürich | Chaoran Chen, Sarah Nadeau, Louis du Plessis, Catharine Aquino, Ivan Topolsky, Kim Philipp Jablonski, Lara Fuhrmann, Daniel Ehrsam, Isabel Stürmer, Andreia Cabral de Gouvea, Maria Domenica Moccia, Simon Grüter, Timothy Sykes, Lennart Opitz, Griffin White, Laura Neff, Doris Popovic, Andrea Patrignani, Jay Tracy, Ralph Schlapbach, Christiane Beckmann, Maurice Redondo, Olivier Kobel, Christoph Noppen, Niko Beerenwinkel, Tanja Stadler |
| EPI_ISL_3465513 | St Vincent's Pathology (SydPath) | NSW Health Pathology - Institute of Clinical Pathology and Medical Research; Westmead Hospital; University of Sydney | Arnott A., Draper J., Gall M., Martinez E., Rockett R., Sintchenko V., on behalf of ICPMR |
| EPI_ISL_3465514 | Viollier AG | Department of Biosystems Science and Engineering, ETH Zürich | Chaoran Chen, Sarah Nadeau, Louis du Plessis, Catharine Aquino, Ivan Topolsky, Kim Philipp Jablonski, Lara Fuhrmann, Daniel Ehrsam, Isabel Stürmer, Andreia Cabral de Gouvea, Maria Domenica Moccia, Simon Grüter, Timothy Sykes, Lennart Opitz, Griffin White, Laura Neff, Doris Popovic, Andrea Patrignani, Jay Tracy, Ralph Schlapbach, Christiane Beckmann, Maurice Redondo, Olivier Kobel, Christoph Noppen, Niko Beerenwinkel, Tanja Stadler |
| EPI_ISL_3465515 | St Vincent's Pathology (SydPath) | NSW Health Pathology - Institute of Clinical Pathology and Medical Research; Westmead Hospital; University of Sydney | Arnott A., Draper J., Gall M., Martinez E., Rockett R., Sintchenko V., on behalf of ICPMR |
| EPI_ISL_3465516, EPI_ISL_3465517 | Viollier AG | Department of Biosystems Science and Engineering, ETH Zürich | Chaoran Chen, Sarah Nadeau, Louis du Plessis, Catharine Aquino, Ivan Topolsky, Kim Philipp Jablonski, Lara Fuhrmann, Daniel Ehrsam, Isabel Stürmer, Andreia Cabral de Gouvea, Maria Domenica Moccia, Simon Grüter, Timothy Sykes, Lennart Opitz, Griffin White, Laura Neff, Doris Popovic, Andrea Patrignani, Jay Tracy, Ralph Schlapbach, Christiane Beckmann, Maurice Redondo, Olivier Kobel, Christoph Noppen, Niko Beerenwinkel, Tanja Stadler |
| EPI_ISL_3465518 | St Vincent's Pathology (SydPath) | NSW Health Pathology - Institute of Clinical Pathology and Medical Research; Westmead Hospital; University of Sydney | Arnott A., Draper J., Gall M., Martinez E., Rockett R., Sintchenko V., on behalf of ICPMR |
| EPI_ISL_3465519 | Viollier AG | Department of Biosystems Science and Engineering, ETH Zürich | Chaoran Chen, Sarah Nadeau, Louis du Plessis, Catharine Aquino, Ivan Topolsky, Kim Philipp Jablonski, Lara Fuhrmann, Daniel Ehrsam, Isabel Stürmer, Andreia Cabral de Gouvea, Maria Domenica Moccia, Simon Grüter, Timothy Sykes, Lennart Opitz, Griffin White, Laura Neff, Doris Popovic, Andrea Patrignani, Jay Tracy, Ralph Schlapbach, Christiane Beckmann, Maurice Redondo, Olivier Kobel, Christoph Noppen, Niko Beerenwinkel, Tanja Stadler |
| EPI_ISL_3465520 | St Vincent's Pathology (SydPath) | NSW Health Pathology - Institute of Clinical Pathology and Medical Research; Westmead Hospital; University of Sydney | Arnott A., Draper J., Gall M., Martinez E., Rockett R., Sintchenko V., on behalf of ICPMR |
| EPI_ISL_3465521, EPI_ISL_3465522 | Viollier AG | Department of Biosystems Science and Engineering, ETH Zürich | Chaoran Chen, Sarah Nadeau, Louis du Plessis, Catharine Aquino, Ivan Topolsky, Kim Philipp Jablonski, Lara Fuhrmann, Daniel Ehrsam, Isabel Stürmer, Andreia Cabral de Gouvea, Maria Domenica Moccia, Simon Grüter, Timothy Sykes, Lennart Opitz, Griffin White, Laura Neff, Doris Popovic, Andrea Patrignani, Jay Tracy, Ralph Schlapbach, Christiane Beckmann, Maurice Redondo, Olivier Kobel, Christoph Noppen, Niko Beerenwinkel, Tanja Stadler |
| EPI_ISL_3465523 | Pathology West - NSW Health Pathology | NSW Health Pathology - Institute of Clinical Pathology and Medical Research; Westmead Hospital; University of Sydney | Arnott A., Draper J., Gall M., Martinez E., Rockett R., Sintchenko V., on behalf of ICPMR |
| EPI_ISL_3465524 | Viollier AG | Department of Biosystems Science and Engineering, ETH Zürich | Chaoran Chen, Sarah Nadeau, Louis du Plessis, Catharine Aquino, Ivan Topolsky, Kim Philipp Jablonski, Lara Fuhrmann, Daniel Ehrsam, Isabel Stürmer, Andreia Cabral de Gouvea, Maria Domenica Moccia, Simon Grüter, Timothy Sykes, Lennart Opitz, Griffin White, Laura Neff, Doris Popovic, Andrea Patrignani, Jay Tracy, Ralph Schlapbach, Christiane Beckmann, Maurice Redondo, Olivier Kobel, Christoph Noppen, Niko Beerenwinkel, Tanja Stadler |
| EPI_ISL_3465525 | Pathology West - NSW Health Pathology | NSW Health Pathology - Institute of Clinical Pathology and Medical Research; Westmead Hospital; University of Sydney | Arnott A., Draper J., Gall M., Martinez E., Rockett R., Sintchenko V., on behalf of ICPMR |
| EPI_ISL_3465526 | Viollier AG | Department of Biosystems Science and Engineering, ETH Zürich | Chaoran Chen, Sarah Nadeau, Louis du Plessis, Catharine Aquino, Ivan Topolsky, Kim Philipp Jablonski, Lara Fuhrmann, Daniel Ehrsam, Isabel Stürmer, Andreia Cabral de Gouvea, Maria Domenica Moccia, Simon Grüter, Timothy Sykes, Lennart Opitz, Griffin White, Laura Neff, Doris Popovic, Andrea Patrignani, Jay Tracy, Ralph Schlapbach, Christiane Beckmann, Maurice Redondo, Olivier Kobel, Christoph Noppen, Niko Beerenwinkel, Tanja Stadler |
| EPI_ISL_3465527 | Pathology West - NSW Health Pathology | NSW Health Pathology - Institute of Clinical Pathology and Medical Research; Westmead Hospital; University of Sydney | Arnott A., Draper J., Gall M., Martinez E., Rockett R., Sintchenko V., on behalf of ICPMR |
| EPI_ISL_3465528 | Viollier AG | Department of Biosystems Science and Engineering, ETH Zürich | Chaoran Chen, Sarah Nadeau, Louis du Plessis, Catharine Aquino, Ivan Topolsky, Kim Philipp Jablonski, Lara Fuhrmann, Daniel Ehrsam, Isabel Stürmer, Andreia Cabral de Gouvea, Maria Domenica Moccia, Simon Grüter, Timothy Sykes, Lennart Opitz, Griffin White, Laura Neff, Doris Popovic, Andrea Patrignani, Jay Tracy, Ralph Schlapbach, Christiane Beckmann, Maurice Redondo, Olivier Kobel, Christoph Noppen, Niko Beerenwinkel, Tanja Stadler |
| EPI_ISL_3465529 | Pathology West - NSW Health Pathology | NSW Health Pathology - Institute of Clinical Pathology and Medical Research; Westmead Hospital; University of Sydney | Arnott A., Draper J., Gall M., Martinez E., Rockett R., Sintchenko V., on behalf of ICPMR |
| EPI_ISL_3465530, EPI_ISL_3465531 | Viollier AG | Department of Biosystems Science and Engineering, ETH Zürich | Chaoran Chen, Sarah Nadeau, Louis du Plessis, Catharine Aquino, Ivan Topolsky, Kim Philipp Jablonski, Lara Fuhrmann, Daniel Ehrsam, Isabel Stürmer, Andreia Cabral de Gouvea, Maria Domenica Moccia, Simon Grüter, Timothy Sykes, Lennart Opitz, Griffin White, Laura Neff, Doris Popovic, Andrea Patrignani, Jay Tracy, Ralph Schlapbach, Christiane Beckmann, Maurice Redondo, Olivier Kobel, Christoph Noppen, Niko Beerenwinkel, Tanja Stadler |
| EPI_ISL_3465532 | Pathology West - NSW Health Pathology | NSW Health Pathology - Institute of Clinical Pathology and Medical Research; Westmead Hospital; University of Sydney | Arnott A., Draper J., Gall M., Martinez E., Rockett R., Sintchenko V., on behalf of ICPMR |
| EPI_ISL_3465533, EPI_ISL_3465534 | Viollier AG | Department of Biosystems Science and Engineering, ETH Zürich | Chaoran Chen, Sarah Nadeau, Louis du Plessis, Catharine Aquino, Ivan Topolsky, Kim Philipp Jablonski, Lara Fuhrmann, Daniel Ehrsam, Isabel Stürmer, Andreia Cabral de Gouvea, Maria Domenica Moccia, Simon Grüter, Timothy Sykes, Lennart Opitz, Griffin White, Laura Neff, Doris Popovic, Andrea Patrignani, Jay Tracy, Ralph Schlapbach, Christiane Beckmann, Maurice Redondo, Olivier Kobel, Christoph Noppen, Niko Beerenwinkel, Tanja Stadler |
| EPI_ISL_3465535 | Pathology West - NSW Health Pathology | NSW Health Pathology - Institute of Clinical Pathology and Medical Research; Westmead Hospital; University of Sydney | Arnott A., Draper J., Gall M., Martinez E., Rockett R., Sintchenko V., on behalf of ICPMR |
| EPI_ISL_3465536 | Viollier AG | Department of Biosystems Science and Engineering, ETH Zürich | Chaoran Chen, Sarah Nadeau, Louis du Plessis, Catharine Aquino, Ivan Topolsky, Kim Philipp Jablonski, Lara Fuhrmann, Daniel Ehrsam, Isabel Stürmer, Andreia Cabral de Gouvea, Maria Domenica Moccia, Simon Grüter, Timothy Sykes, Lennart Opitz, Griffin White, Laura Neff, Doris Popovic, Andrea Patrignani, Jay Tracy, Ralph Schlapbach, Christiane Beckmann, Maurice Redondo, Olivier Kobel, Christoph Noppen, Niko Beerenwinkel, Tanja Stadler |
| EPI_ISL_3465537 | Pathology West - NSW Health Pathology | NSW Health Pathology - Institute of Clinical Pathology and Medical Research; Westmead Hospital; University of Sydney | Arnott A., Draper J., Gall M., Martinez E., Rockett R., Sintchenko V., on behalf of ICPMR |
| EPI_ISL_3465538, EPI_ISL_3465539 | Viollier AG | Department of Biosystems Science and Engineering, ETH Zürich | Chaoran Chen, Sarah Nadeau, Louis du Plessis, Catharine Aquino, Ivan Topolsky, Kim Philipp Jablonski, Lara Fuhrmann, Daniel Ehrsam, Isabel Stürmer, Andreia Cabral de Gouvea, Maria Domenica Moccia, Simon Grüter, Timothy Sykes, Lennart Opitz, Griffin White, Laura Neff, Doris Popovic, Andrea Patrignani, Jay Tracy, Ralph Schlapbach, Christiane Beckmann, Maurice Redondo, Olivier Kobel, Christoph Noppen, Niko Beerenwinkel, Tanja Stadler |
| EPI_ISL_3465540 | 4Cyte Pathology | NSW Health Pathology - Institute of Clinical Pathology and Medical Research; Westmead Hospital; University of Sydney | Arnott A., Draper J., Gall M., Martinez E., Rockett R., Sintchenko V., on behalf of ICPMR |
| EPI_ISL_3465541, EPI_ISL_3465542 | Viollier AG | Department of Biosystems Science and Engineering, ETH Zürich | Chaoran Chen, Sarah Nadeau, Louis du Plessis, Catharine Aquino, Ivan Topolsky, Kim Philipp Jablonski, Lara Fuhrmann, Daniel Ehrsam, Isabel Stürmer, Andreia Cabral de Gouvea, Maria Domenica Moccia, Simon Grüter, Timothy Sykes, Lennart Opitz, Griffin White, Laura Neff, Doris Popovic, Andrea Patrignani, Jay Tracy, Ralph Schlapbach, Christiane Beckmann, Maurice Redondo, Olivier Kobel, Christoph Noppen, Niko Beerenwinkel, Tanja Stadler |
| EPI_ISL_3465543 | 4Cyte Pathology | NSW Health Pathology - Institute of Clinical Pathology and Medical Research; Westmead Hospital; University of Sydney | Arnott A., Draper J., Gall M., Martinez E., Rockett R., Sintchenko V., on behalf of ICPMR |
| EPI_ISL_3465544 | Viollier AG | Department of Biosystems Science and Engineering, ETH Zürich | Chaoran Chen, Sarah Nadeau, Louis du Plessis, Catharine Aquino, Ivan Topolsky, Kim Philipp Jablonski, Lara Fuhrmann, Daniel Ehrsam, Isabel Stürmer, Andreia Cabral de Gouvea, Maria Domenica Moccia, Simon Grüter, Timothy Sykes, Lennart Opitz, Griffin White, Laura Neff, Doris Popovic, Andrea Patrignani, Jay Tracy, Ralph Schlapbach, Christiane Beckmann, Maurice Redondo, Olivier Kobel, Christoph Noppen, Niko Beerenwinkel, Tanja Stadler |
| EPI_ISL_3465545 | 4Cyte Pathology | NSW Health Pathology - Institute of Clinical Pathology and Medical Research; Westmead Hospital; University of Sydney | Arnott A., Draper J., Gall M., Martinez E., Rockett R., Sintchenko V., on behalf of ICPMR |
| EPI_ISL_3465546, EPI_ISL_3465547 | Viollier AG | Department of Biosystems Science and Engineering, ETH Zürich | Chaoran Chen, Sarah Nadeau, Louis du Plessis, Catharine Aquino, Ivan Topolsky, Kim Philipp Jablonski, Lara Fuhrmann, Daniel Ehrsam, Isabel Stürmer, Andreia Cabral de Gouvea, Maria Domenica Moccia, Simon Grüter, Timothy Sykes, Lennart Opitz, Griffin White, Laura Neff, Doris Popovic, Andrea Patrignani, Jay Tracy, Ralph Schlapbach, Christiane Beckmann, Maurice Redondo, Olivier Kobel, Christoph Noppen, Niko Beerenwinkel, Tanja Stadler |
| EPI_ISL_3465548 | 4Cyte Pathology | NSW Health Pathology - Institute of Clinical Pathology and Medical Research; Westmead Hospital; University of Sydney | Arnott A., Draper J., Gall M., Martinez E., Rockett R., Sintchenko V., on behalf of ICPMR |
| EPI_ISL_3465549 | Viollier AG | Department of Biosystems Science and Engineering, ETH Zürich | Chaoran Chen, Sarah Nadeau, Louis du Plessis, Catharine Aquino, Ivan Topolsky, Kim Philipp Jablonski, Lara Fuhrmann, Daniel Ehrsam, Isabel Stürmer, Andreia Cabral de Gouvea, Maria Domenica Moccia, Simon Grüter, Timothy Sykes, Lennart Opitz, Griffin White, Laura Neff, Doris Popovic, Andrea Patrignani, Jay Tracy, Ralph Schlapbach, Christiane Beckmann, Maurice Redondo, Olivier Kobel, Christoph Noppen, Niko Beerenwinkel, Tanja Stadler |
| EPI_ISL_3465550 | 4Cyte Pathology | NSW Health Pathology - Institute of Clinical Pathology and Medical Research; Westmead Hospital; University of Sydney | Arnott A., Draper J., Gall M., Martinez E., Rockett R., Sintchenko V., on behalf of ICPMR |
| EPI_ISL_3465551 | Viollier AG | Department of Biosystems Science and Engineering, ETH Zürich | Chaoran Chen, Sarah Nadeau, Louis du Plessis, Catharine Aquino, Ivan Topolsky, Kim Philipp Jablonski, Lara Fuhrmann, Daniel Ehrsam, Isabel Stürmer, Andreia Cabral de Gouvea, Maria Domenica Moccia, Simon Grüter, Timothy Sykes, Lennart Opitz, Griffin White, Laura Neff, Doris Popovic, Andrea Patrignani, Jay Tracy, Ralph Schlapbach, Christiane Beckmann, Maurice Redondo, Olivier Kobel, Christoph Noppen, Niko Beerenwinkel, Tanja Stadler |

| | | | |
|---|---|---|---|
| EPI_ISL_3465552 | 4Cyte Pathology | NSW Health Pathology - Institute of Clinical Pathology and Medical Research; Westmead Hospital; University of Sydney | Arnott A., Draper J., Gall M., Martinez E., Rockett R., Sintchenko V., on behalf of ICPMR |
| EPI_ISL_3465553 | Viollier AG | Department of Biosystems Science and Engineering, ETH Zürich | Chaoran Chen, Sarah Nadeau, Louis du Plessis, Catharine Aquino, Ivan Topolsky, Kim Philipp Jablonski, Lara Fuhrmann, Daniel Ehrsam, Isabel Stürmer, Andreia Cabral de Gouvea, Maria Domenica Moccia, Simon Grüter, Timothy Sykes, Lennart Opitz, Griffin White, Laura Neff, Doris Popovic, Andrea Patrignani, Jay Tracy, Ralph Schlapbach, Christiane Beckmann, Maurice Redondo, Olivier Kobel, Christoph Noppen, Niko Beerenwinkel, Tanja Stadler |
| EPI_ISL_3465554 | Viollier AG | Department of Biosystems Science and Engineering, ETH Zürich | Andrea Patrizia Salzmann, Henriette Kurth, Christiane Beckmann, Ivan Topolsky, Chaoran Chen, Sarah Nadeau, Louis du Plessis, Kim Philipp Jablonski, Lara Fuhrmann, Niko Beerenwinkel, Christoph Noppen, Olivier Kobel, Maurice Redondo, Tanja Stadler |
| EPI_ISL_3465555 | 4Cyte Pathology | NSW Health Pathology - Institute of Clinical Pathology and Medical Research; Westmead Hospital; University of Sydney | Arnott A., Draper J., Gall M., Martinez E., Rockett R., Sintchenko V., on behalf of ICPMR |
| EPI_ISL_3465556, EPI_ISL_3465557 | Viollier AG | Department of Biosystems Science and Engineering, ETH Zürich | Chaoran Chen, Sarah Nadeau, Louis du Plessis, Catharine Aquino, Ivan Topolsky, Kim Philipp Jablonski, Lara Fuhrmann, Daniel Ehrsam, Isabel Stürmer, Andreia Cabral de Gouvea, Maria Domenica Moccia, Simon Grüter, Timothy Sykes, Lennart Opitz, Griffin White, Laura Neff, Doris Popovic, Andrea Patrignani, Jay Tracy, Ralph Schlapbach, Christiane Beckmann, Maurice Redondo, Olivier Kobel, Christoph Noppen, Niko Beerenwinkel, Tanja Stadler |
| EPI_ISL_3465558 | 4Cyte Pathology | NSW Health Pathology - Institute of Clinical Pathology and Medical Research; Westmead Hospital; University of Sydney | Arnott A., Draper J., Gall M., Martinez E., Rockett R., Sintchenko V., on behalf of ICPMR |
| EPI_ISL_3465559 | Viollier AG | Department of Biosystems Science and Engineering, ETH Zürich | Chaoran Chen, Sarah Nadeau, Louis du Plessis, Catharine Aquino, Ivan Topolsky, Kim Philipp Jablonski, Lara Fuhrmann, Daniel Ehrsam, Isabel Stürmer, Andreia Cabral de Gouvea, Maria Domenica Moccia, Simon Grüter, Timothy Sykes, Lennart Opitz, Griffin White, Laura Neff, Doris Popovic, Andrea Patrignani, Jay Tracy, Ralph Schlapbach, Christiane Beckmann, Maurice Redondo, Olivier Kobel, Christoph Noppen, Niko Beerenwinkel, Tanja Stadler |
| EPI_ISL_3465560 | 4Cyte Pathology | NSW Health Pathology - Institute of Clinical Pathology and Medical Research; Westmead Hospital; University of Sydney | Arnott A., Draper J., Gall M., Martinez E., Rockett R., Sintchenko V., on behalf of ICPMR |
| EPI_ISL_3465561 | Viollier AG | Department of Biosystems Science and Engineering, ETH Zürich | Chaoran Chen, Sarah Nadeau, Louis du Plessis, Catharine Aquino, Ivan Topolsky, Kim Philipp Jablonski, Lara Fuhrmann, Daniel Ehrsam, Isabel Stürmer, Andreia Cabral de Gouvea, Maria Domenica Moccia, Simon Grüter, Timothy Sykes, Lennart Opitz, Griffin White, Laura Neff, Doris Popovic, Andrea Patrignani, Jay Tracy, Ralph Schlapbach, Christiane Beckmann, Maurice Redondo, Olivier Kobel, Christoph Noppen, Niko Beerenwinkel, Tanja Stadler |
| EPI_ISL_3465562 | 4Cyte Pathology | NSW Health Pathology - Institute of Clinical Pathology and Medical Research; Westmead Hospital; University of Sydney | Arnott A., Draper J., Gall M., Martinez E., Rockett R., Sintchenko V., on behalf of ICPMR |
| EPI_ISL_3465563 | Viollier AG | Department of Biosystems Science and Engineering, ETH Zürich | Chaoran Chen, Sarah Nadeau, Louis du Plessis, Catharine Aquino, Ivan Topolsky, Kim Philipp Jablonski, Lara Fuhrmann, Daniel Ehrsam, Isabel Stürmer, Andreia Cabral de Gouvea, Maria Domenica Moccia, Simon Grüter, Timothy Sykes, Lennart Opitz, Griffin White, Laura Neff, Doris Popovic, Andrea Patrignani, Jay Tracy, Ralph Schlapbach, Christiane Beckmann, Maurice Redondo, Olivier Kobel, Christoph Noppen, Niko Beerenwinkel, Tanja Stadler |
| EPI_ISL_3465564 | 4Cyte Pathology | NSW Health Pathology - Institute of Clinical Pathology and Medical Research; Westmead Hospital; University of Sydney | Arnott A., Draper J., Gall M., Martinez E., Rockett R., Sintchenko V., on behalf of ICPMR |
| EPI_ISL_3465566 | Pathology North - Royal North Shore Hospital - NSW Health Pathology | NSW Health Pathology - Institute of Clinical Pathology and Medical Research; Westmead Hospital; University of Sydney | Arnott A., Draper J., Gall M., Martinez E., Rockett R., Sintchenko V., on behalf of ICPMR |
| EPI_ISL_3465567 | Viollier AG | Department of Biosystems Science and Engineering, ETH Zürich | Chaoran Chen, Sarah Nadeau, Louis du Plessis, Catharine Aquino, Ivan Topolsky, Kim Philipp Jablonski, Lara Fuhrmann, Daniel Ehrsam, Isabel Stürmer, Andreia Cabral de Gouvea, Maria Domenica Moccia, Simon Grüter, Timothy Sykes, Lennart Opitz, Griffin White, Laura Neff, Doris Popovic, Andrea Patrignani, Jay Tracy, Ralph Schlapbach, Christiane Beckmann, Maurice Redondo, Olivier Kobel, Christoph Noppen, Niko Beerenwinkel, Tanja Stadler |
| EPI_ISL_3465568 | Viollier AG | Department of Biosystems Science and Engineering, ETH Zürich | Andrea Patrizia Salzmann, Henriette Kurth, Christiane Beckmann, Ivan Topolsky, Chaoran Chen, Sarah Nadeau, Louis du Plessis, Kim Philipp Jablonski, Lara Fuhrmann, Niko Beerenwinkel, Christoph Noppen, Olivier Kobel, Maurice Redondo, Tanja Stadler |
| EPI_ISL_3465569 | Australian Clinical Labs (formerly Healthscope Pathology) | NSW Health Pathology - Institute of Clinical Pathology and Medical Research; Westmead Hospital; University of Sydney | Arnott A., Draper J., Gall M., Martinez E., Rockett R., Sintchenko V., on behalf of ICPMR |
| EPI_ISL_3465570 | Viollier AG | Department of Biosystems Science and Engineering, ETH Zürich | Chaoran Chen, Sarah Nadeau, Louis du Plessis, Catharine Aquino, Ivan Topolsky, Kim Philipp Jablonski, Lara Fuhrmann, Daniel Ehrsam, Isabel Stürmer, Andreia Cabral de Gouvea, Maria Domenica Moccia, Simon Grüter, Timothy Sykes, Lennart Opitz, Griffin White, Laura Neff, Doris Popovic, Andrea Patrignani, Jay Tracy, Ralph Schlapbach, Christiane Beckmann, Maurice Redondo, Olivier Kobel, Christoph Noppen, Niko Beerenwinkel, Tanja Stadler |
| EPI_ISL_3465571 | Australian Clinical Labs (formerly Healthscope Pathology) | NSW Health Pathology - Institute of Clinical Pathology and Medical Research; Westmead Hospital; University of Sydney | Arnott A., Draper J., Gall M., Martinez E., Rockett R., Sintchenko V., on behalf of ICPMR |
| EPI_ISL_3465572 | Viollier AG | Department of Biosystems Science and Engineering, ETH Zürich | Chaoran Chen, Sarah Nadeau, Louis du Plessis, Catharine Aquino, Ivan Topolsky, Kim Philipp Jablonski, Lara Fuhrmann, Daniel Ehrsam, Isabel Stürmer, Andreia Cabral de Gouvea, Maria Domenica Moccia, Simon Grüter, Timothy Sykes, Lennart Opitz, Griffin White, Laura Neff, Doris Popovic, Andrea Patrignani, Jay Tracy, Ralph Schlapbach, Christiane Beckmann, Maurice Redondo, Olivier Kobel, Christoph Noppen, Niko Beerenwinkel, Tanja Stadler |
| EPI_ISL_3465573 | Australian Clinical Labs (formerly Healthscope Pathology) | NSW Health Pathology - Institute of Clinical Pathology and Medical Research; Westmead Hospital; University of Sydney | Arnott A., Draper J., Gall M., Martinez E., Rockett R., Sintchenko V., on behalf of ICPMR |
| EPI_ISL_3465574 | Viollier AG | Department of Biosystems Science and Engineering, ETH Zürich | Chaoran Chen, Sarah Nadeau, Louis du Plessis, Catharine Aquino, Ivan Topolsky, Kim Philipp Jablonski, Lara Fuhrmann, Daniel Ehrsam, Isabel Stürmer, Andreia Cabral de Gouvea, Maria Domenica Moccia, Simon Grüter, Timothy Sykes, Lennart Opitz, Griffin White, Laura Neff, Doris Popovic, Andrea Patrignani, Jay Tracy, Ralph Schlapbach, Christiane Beckmann, Maurice Redondo, Olivier Kobel, Christoph Noppen, Niko Beerenwinkel, Tanja Stadler |
| EPI_ISL_3465575 | Australian Clinical Labs (formerly Healthscope Pathology) | NSW Health Pathology - Institute of Clinical Pathology and Medical Research; Westmead Hospital; University of Sydney | Arnott A., Draper J., Gall M., Martinez E., Rockett R., Sintchenko V., on behalf of ICPMR |
| EPI_ISL_3465576 | Viollier AG | Department of Biosystems Science and Engineering, ETH Zürich | Chaoran Chen, Sarah Nadeau, Louis du Plessis, Catharine Aquino, Ivan Topolsky, Kim Philipp Jablonski, Lara Fuhrmann, Daniel Ehrsam, Isabel Stürmer, Andreia Cabral de Gouvea, Maria Domenica Moccia, Simon Grüter, Timothy Sykes, Lennart Opitz, Griffin White, Laura Neff, Doris Popovic, Andrea Patrignani, Jay Tracy, Ralph Schlapbach, Christiane Beckmann, Maurice Redondo, Olivier Kobel, Christoph Noppen, Niko Beerenwinkel, Tanja Stadler |
| EPI_ISL_3465577 | Australian Clinical Labs (formerly Healthscope Pathology) | NSW Health Pathology - Institute of Clinical Pathology and Medical Research; Westmead Hospital; University of Sydney | Arnott A., Draper J., Gall M., Martinez E., Rockett R., Sintchenko V., on behalf of ICPMR |
| EPI_ISL_3465578, EPI_ISL_3465579 | Viollier AG | Department of Biosystems Science and Engineering, ETH Zürich | Chaoran Chen, Sarah Nadeau, Louis du Plessis, Catharine Aquino, Ivan Topolsky, Kim Philipp Jablonski, Lara Fuhrmann, Daniel Ehrsam, Isabel Stürmer, Andreia Cabral de Gouvea, Maria Domenica Moccia, Simon Grüter, Timothy Sykes, Lennart Opitz, Griffin White, Laura Neff, Doris Popovic, Andrea Patrignani, Jay Tracy, Ralph Schlapbach, Christiane Beckmann, Maurice Redondo, Olivier Kobel, Christoph Noppen, Niko Beerenwinkel, Tanja Stadler |
| EPI_ISL_3465580 | Australian Clinical Labs (formerly Healthscope Pathology) | NSW Health Pathology - Institute of Clinical Pathology and Medical Research; Westmead Hospital; University of Sydney | Arnott A., Draper J., Gall M., Martinez E., Rockett R., Sintchenko V., on behalf of ICPMR |
| EPI_ISL_3465581 | Viollier AG | Department of Biosystems Science and Engineering, ETH Zürich | Chaoran Chen, Sarah Nadeau, Louis du Plessis, Catharine Aquino, Ivan Topolsky, Kim Philipp Jablonski, Lara Fuhrmann, Daniel Ehrsam, Isabel Stürmer, Andreia Cabral de Gouvea, Maria Domenica Moccia, Simon Grüter, Timothy Sykes, Lennart Opitz, Griffin White, Laura Neff, Doris Popovic, Andrea Patrignani, Jay Tracy, Ralph Schlapbach, Christiane Beckmann, Maurice Redondo, Olivier Kobel, Christoph Noppen, Niko Beerenwinkel, Tanja Stadler |
| EPI_ISL_3465582 | Australian Clinical Labs (formerly Healthscope Pathology) | NSW Health Pathology - Institute of Clinical Pathology and Medical Research; Westmead Hospital; University of Sydney | Arnott A., Draper J., Gall M., Martinez E., Rockett R., Sintchenko V., on behalf of ICPMR |
| EPI_ISL_3465583, EPI_ISL_3465584 | Viollier AG | Department of Biosystems Science and Engineering, ETH Zürich | Chaoran Chen, Sarah Nadeau, Louis du Plessis, Catharine Aquino, Ivan Topolsky, Kim Philipp Jablonski, Lara Fuhrmann, Daniel Ehrsam, Isabel Stürmer, Andreia Cabral de Gouvea, Maria Domenica Moccia, Simon Grüter, Timothy Sykes, Lennart Opitz, Griffin White, Laura Neff, Doris Popovic, Andrea Patrignani, Jay Tracy, Ralph Schlapbach, Christiane Beckmann, Maurice Redondo, Olivier Kobel, Christoph Noppen, Niko Beerenwinkel, Tanja Stadler |
| EPI_ISL_3465585 | Australian Clinical Labs (formerly Healthscope Pathology) | NSW Health Pathology - Institute of Clinical Pathology and Medical Research; Westmead Hospital; University of Sydney | Arnott A., Draper J., Gall M., Martinez E., Rockett R., Sintchenko V., on behalf of ICPMR |
| EPI_ISL_3465586 | Viollier AG | Department of Biosystems Science and Engineering, ETH Zürich | Chaoran Chen, Sarah Nadeau, Louis du Plessis, Catharine Aquino, Ivan Topolsky, Kim Philipp Jablonski, Lara Fuhrmann, Daniel Ehrsam, Isabel Stürmer, Andreia Cabral de Gouvea, Maria Domenica Moccia, Simon Grüter, Timothy Sykes, Lennart Opitz, Griffin White, Laura Neff, Doris Popovic, Andrea Patrignani, Jay Tracy, Ralph Schlapbach, Christiane Beckmann, Maurice Redondo, Olivier Kobel, Christoph Noppen, Niko Beerenwinkel, Tanja Stadler |
| EPI_ISL_3465587 | Australian Clinical Labs (formerly Healthscope Pathology) | NSW Health Pathology - Institute of Clinical Pathology and Medical Research; Westmead Hospital; University of Sydney | Arnott A., Draper J., Gall M., Martinez E., Rockett R., Sintchenko V., on behalf of ICPMR |
| EPI_ISL_3465588 | Viollier AG | Department of Biosystems Science and Engineering, ETH Zürich | Chaoran Chen, Sarah Nadeau, Louis du Plessis, Catharine Aquino, Ivan Topolsky, Kim Philipp Jablonski, Lara Fuhrmann, Daniel Ehrsam, Isabel Stürmer, Andreia Cabral de Gouvea, Maria Domenica Moccia, Simon Grüter, Timothy Sykes, Lennart Opitz, Griffin White, Laura Neff, Doris Popovic, Andrea Patrignani, Jay Tracy, Ralph Schlapbach, Christiane Beckmann, Maurice Redondo, Olivier Kobel, Christoph Noppen, Niko Beerenwinkel, Tanja Stadler |
| EPI_ISL_3465589 | Sydney South West Pathology Service (SSWPS) - Liverpool Hospital - NSW Health Pathology | NSW Health Pathology - Institute of Clinical Pathology and Medical Research; Westmead Hospital; University of Sydney | Arnott A., Draper J., Gall M., Martinez E., Rockett R., Sintchenko V., on behalf of ICPMR |
| EPI_ISL_3465590 | Viollier AG | Department of Biosystems Science and Engineering, ETH Zürich | Chaoran Chen, Sarah Nadeau, Louis du Plessis, Catharine Aquino, Ivan Topolsky, Kim Philipp Jablonski, Lara Fuhrmann, Daniel Ehrsam, Isabel Stürmer, Andreia Cabral de Gouvea, Maria Domenica Moccia, Simon Grüter, Timothy Sykes, Lennart Opitz, Griffin White, Laura Neff, Doris Popovic, Andrea Patrignani, Jay Tracy, Ralph Schlapbach, Christiane Beckmann, Maurice Redondo, Olivier Kobel, Christoph Noppen, Niko Beerenwinkel, Tanja Stadler |
| EPI_ISL_3465591 | Sydney South West Pathology Service (SSWPS) - Liverpool Hospital - NSW Health Pathology | NSW Health Pathology - Institute of Clinical Pathology and Medical Research; Westmead Hospital; University of Sydney | Arnott A., Draper J., Gall M., Martinez E., Rockett R., Sintchenko V., on behalf of ICPMR |
| EPI_ISL_3465592 | Viollier AG | Department of Biosystems Science and Engineering, ETH Zürich | Chaoran Chen, Sarah Nadeau, Louis du Plessis, Catharine Aquino, Ivan Topolsky, Kim Philipp Jablonski, Lara Fuhrmann, Daniel Ehrsam, Isabel Stürmer, Andreia Cabral de Gouvea, Maria Domenica Moccia, Simon Grüter, Timothy Sykes, Lennart Opitz, Griffin White, Laura Neff, Doris Popovic, Andrea Patrignani, Jay Tracy, Ralph Schlapbach, Christiane Beckmann, Maurice Redondo, Olivier Kobel, Christoph Noppen, Niko Beerenwinkel, Tanja Stadler |
| EPI_ISL_3465593 | Sydney South West Pathology Service (SSWPS) - Liverpool Hospital - NSW Health Pathology | NSW Health Pathology - Institute of Clinical Pathology and Medical Research; Westmead Hospital; University of Sydney | Arnott A., Draper J., Gall M., Martinez E., Rockett R., Sintchenko V., on behalf of ICPMR |
| EPI_ISL_3465594 | Viollier AG | Department of Biosystems Science and Engineering, ETH Zürich | Chaoran Chen, Sarah Nadeau, Louis du Plessis, Catharine Aquino, Ivan Topolsky, Kim Philipp Jablonski, Lara Fuhrmann, Daniel Ehrsam, Isabel Stürmer, Andreia Cabral de Gouvea, Maria Domenica Moccia, Simon Grüter, Timothy Sykes, Lennart Opitz, Griffin White, Laura Neff, Doris Popovic, Andrea Patrignani, Jay Tracy, Ralph Schlapbach, Christiane Beckmann, Maurice Redondo, Olivier Kobel, Christoph Noppen, Niko Beerenwinkel, Tanja Stadler |
| EPI_ISL_3465595 | Sydney South West Pathology Service (SSWPS) - Liverpool Hospital - NSW Health Pathology | NSW Health Pathology - Institute of Clinical Pathology and Medical Research; Westmead Hospital; University of Sydney | Arnott A., Draper J., Gall M., Martinez E., Rockett R., Sintchenko V., on behalf of ICPMR |
| EPI_ISL_3465596 | Viollier AG | Department of Biosystems Science and Engineering, ETH Zürich | Chaoran Chen, Sarah Nadeau, Louis du Plessis, Catharine Aquino, Ivan Topolsky, Kim Philipp Jablonski, Lara Fuhrmann, Daniel Ehrsam, Isabel Stürmer, Andreia Cabral de Gouvea, Maria Domenica Moccia, Simon Grüter, Timothy Sykes, Lennart Opitz, Griffin White, Laura Neff, Doris Popovic, Andrea Patrignani, Jay Tracy, Ralph Schlapbach, Christiane Beckmann, Maurice Redondo, Olivier Kobel, Christoph Noppen, Niko Beerenwinkel, Tanja Stadler |
| EPI_ISL_3465597 | Histopath | NSW Health Pathology - Institute of Clinical Pathology and Medical Research; Westmead Hospital; University of Sydney | Arnott A., Draper J., Gall M., Martinez E., Rockett R., Sintchenko V., on behalf of ICPMR |
| EPI_ISL_3465598 | Viollier AG | Department of Biosystems Science and Engineering, ETH Zürich | Chaoran Chen, Sarah Nadeau, Louis du Plessis, Catharine Aquino, Ivan Topolsky, Kim Philipp Jablonski, Lara Fuhrmann, Daniel Ehrsam, Isabel Stürmer, Andreia Cabral de Gouvea, Maria Domenica Moccia, Simon Grüter, Timothy Sykes, Lennart Opitz, Griffin White, Laura Neff, Doris Popovic, Andrea Patrignani, Jay Tracy, Ralph Schlapbach, Christiane Beckmann, Maurice Redondo, Olivier Kobel, Christoph Noppen, Niko Beerenwinkel, Tanja Stadler |
| EPI_ISL_3465599 | Histopath | NSW Health Pathology - Institute of Clinical Pathology and Medical Research; Westmead Hospital; University of Sydney | Arnott A., Draper J., Gall M., Martinez E., Rockett R., Sintchenko V., on behalf of ICPMR |
| EPI_ISL_3465600, EPI_ISL_3465601 | Viollier AG | Department of Biosystems Science and Engineering, ETH Zürich | Chaoran Chen, Sarah Nadeau, Louis du Plessis, Catharine Aquino, Ivan Topolsky, Kim Philipp Jablonski, Lara Fuhrmann, Daniel Ehrsam, Isabel Stürmer, Andreia Cabral de Gouvea, Maria Domenica Moccia, Simon Grüter, Timothy Sykes, Lennart Opitz, Griffin White, Laura Neff, Doris Popovic, Andrea Patrignani, Jay Tracy, Ralph Schlapbach, Christiane Beckmann, Maurice Redondo, Olivier Kobel, Christoph Noppen, Niko Beerenwinkel, Tanja Stadler |

| | | | |
|---|---|---|---|
| EPI_ISL_3465602 | Histopath | NSW Health Pathology - Institute of Clinical Pathology and Medical Research; Westmead Hospital; University of Sydney | Arnott A., Draper J., Gall M., Martinez E., Rockett R., Sintchenko V., on behalf of ICPMR |
| EPI_ISL_3465603 | Viollier AG | Department of Biosystems Science and Engineering, ETH Zürich | Chaoran Chen, Sarah Nadeau, Louis du Plessis, Catharine Aquino, Ivan Topolsky, Kim Philipp Jablonski, Lara Fuhrmann, Daniel Ehrsam, Isabel Stürmer, Andreia Cabral de Gouvea, Maria Domenica Moccia, Simon Grüter, Timothy Sykes, Lennart Opitz, Griffin White, Laura Neff, Doris Popovic, Andrea Patrignani, Jay Tracy, Ralph Schlapbach, Christiane Beckmann, Maurice Redondo, Olivier Kobel, Christoph Noppen, Niko Beerenwinkel, Tanja Stadler |
| EPI_ISL_3465604 | Histopath | NSW Health Pathology - Institute of Clinical Pathology and Medical Research; Westmead Hospital; University of Sydney | Arnott A., Draper J., Gall M., Martinez E., Rockett R., Sintchenko V., on behalf of ICPMR |
| EPI_ISL_3465605 | Viollier AG | Department of Biosystems Science and Engineering, ETH Zürich | Chaoran Chen, Sarah Nadeau, Louis du Plessis, Catharine Aquino, Ivan Topolsky, Kim Philipp Jablonski, Lara Fuhrmann, Daniel Ehrsam, Isabel Stürmer, Andreia Cabral de Gouvea, Maria Domenica Moccia, Simon Grüter, Timothy Sykes, Lennart Opitz, Griffin White, Laura Neff, Doris Popovic, Andrea Patrignani, Jay Tracy, Ralph Schlapbach, Christiane Beckmann, Maurice Redondo, Olivier Kobel, Christoph Noppen, Niko Beerenwinkel, Tanja Stadler |
| EPI_ISL_3465606 | Histopath | NSW Health Pathology - Institute of Clinical Pathology and Medical Research; Westmead Hospital; University of Sydney | Arnott A., Draper J., Gall M., Martinez E., Rockett R., Sintchenko V., on behalf of ICPMR |
| EPI_ISL_3465607 | Viollier AG | Department of Biosystems Science and Engineering, ETH Zürich | Andrea Patrizia Salzmann, Henriette Kurth, Christiane Beckmann, Ivan Topolsky, Chaoran Chen, Sarah Nadeau, Louis du Plessis, Kim Philipp Jablonski, Lara Fuhrmann, Niko Beerenwinkel, Christoph Noppen, Olivier Kobel, Maurice Redondo, Tanja Stadler |
| EPI_ISL_3465608 | Histopath | NSW Health Pathology - Institute of Clinical Pathology and Medical Research; Westmead Hospital; University of Sydney | Arnott A., Draper J., Gall M., Martinez E., Rockett R., Sintchenko V., on behalf of ICPMR |
| EPI_ISL_3465609 | Viollier AG | Department of Biosystems Science and Engineering, ETH Zürich | Chaoran Chen, Sarah Nadeau, Louis du Plessis, Catharine Aquino, Ivan Topolsky, Kim Philipp Jablonski, Lara Fuhrmann, Daniel Ehrsam, Isabel Stürmer, Andreia Cabral de Gouvea, Maria Domenica Moccia, Simon Grüter, Timothy Sykes, Lennart Opitz, Griffin White, Laura Neff, Doris Popovic, Andrea Patrignani, Jay Tracy, Ralph Schlapbach, Christiane Beckmann, Maurice Redondo, Olivier Kobel, Christoph Noppen, Niko Beerenwinkel, Tanja Stadler |
| EPI_ISL_3465610 | Histopath | NSW Health Pathology - Institute of Clinical Pathology and Medical Research; Westmead Hospital; University of Sydney | Arnott A., Draper J., Gall M., Martinez E., Rockett R., Sintchenko V., on behalf of ICPMR |
| EPI_ISL_3465611 | Viollier AG | Department of Biosystems Science and Engineering, ETH Zürich | Chaoran Chen, Sarah Nadeau, Louis du Plessis, Catharine Aquino, Ivan Topolsky, Kim Philipp Jablonski, Lara Fuhrmann, Daniel Ehrsam, Isabel Stürmer, Andreia Cabral de Gouvea, Maria Domenica Moccia, Simon Grüter, Timothy Sykes, Lennart Opitz, Griffin White, Laura Neff, Doris Popovic, Andrea Patrignani, Jay Tracy, Ralph Schlapbach, Christiane Beckmann, Maurice Redondo, Olivier Kobel, Christoph Noppen, Niko Beerenwinkel, Tanja Stadler |
| EPI_ISL_3465612 | Histopath | NSW Health Pathology - Institute of Clinical Pathology and Medical Research; Westmead Hospital; University of Sydney | Arnott A., Draper J., Gall M., Martinez E., Rockett R., Sintchenko V., on behalf of ICPMR |
| EPI_ISL_3465613 | Viollier AG | Department of Biosystems Science and Engineering, ETH Zürich | Andrea Patrizia Salzmann, Henriette Kurth, Christiane Beckmann, Ivan Topolsky, Chaoran Chen, Sarah Nadeau, Louis du Plessis, Kim Philipp Jablonski, Lara Fuhrmann, Niko Beerenwinkel, Christoph Noppen, Olivier Kobel, Maurice Redondo, Tanja Stadler |
| EPI_ISL_3465614 | Viollier AG | Department of Biosystems Science and Engineering, ETH Zürich | Chaoran Chen, Sarah Nadeau, Louis du Plessis, Catharine Aquino, Ivan Topolsky, Kim Philipp Jablonski, Lara Fuhrmann, Daniel Ehrsam, Isabel Stürmer, Andreia Cabral de Gouvea, Maria Domenica Moccia, Simon Grüter, Timothy Sykes, Lennart Opitz, Griffin White, Laura Neff, Doris Popovic, Andrea Patrignani, Jay Tracy, Ralph Schlapbach, Christiane Beckmann, Maurice Redondo, Olivier Kobel, Christoph Noppen, Niko Beerenwinkel, Tanja Stadler |
| EPI_ISL_3465615 | Histopath | NSW Health Pathology - Institute of Clinical Pathology and Medical Research; Westmead Hospital; University of Sydney | Arnott A., Draper J., Gall M., Martinez E., Rockett R., Sintchenko V., on behalf of ICPMR |
| EPI_ISL_3465616 | Viollier AG | Department of Biosystems Science and Engineering, ETH Zürich | Chaoran Chen, Sarah Nadeau, Louis du Plessis, Catharine Aquino, Ivan Topolsky, Kim Philipp Jablonski, Lara Fuhrmann, Daniel Ehrsam, Isabel Stürmer, Andreia Cabral de Gouvea, Maria Domenica Moccia, Simon Grüter, Timothy Sykes, Lennart Opitz, Griffin White, Laura Neff, Doris Popovic, Andrea Patrignani, Jay Tracy, Ralph Schlapbach, Christiane Beckmann, Maurice Redondo, Olivier Kobel, Christoph Noppen, Niko Beerenwinkel, Tanja Stadler |
| EPI_ISL_3465617 | Pathology North - Hunter - NSW Health Pathology | NSW Health Pathology - Institute of Clinical Pathology and Medical Research; Westmead Hospital; University of Sydney | Arnott A., Draper J., Gall M., Martinez E., Rockett R., Sintchenko V., on behalf of ICPMR |
| EPI_ISL_3465618, EPI_ISL_3465619 | Viollier AG | Department of Biosystems Science and Engineering, ETH Zürich | Chaoran Chen, Sarah Nadeau, Louis du Plessis, Catharine Aquino, Ivan Topolsky, Kim Philipp Jablonski, Lara Fuhrmann, Daniel Ehrsam, Isabel Stürmer, Andreia Cabral de Gouvea, Maria Domenica Moccia, Simon Grüter, Timothy Sykes, Lennart Opitz, Griffin White, Laura Neff, Doris Popovic, Andrea Patrignani, Jay Tracy, Ralph Schlapbach, Christiane Beckmann, Maurice Redondo, Olivier Kobel, Christoph Noppen, Niko Beerenwinkel, Tanja Stadler |
| EPI_ISL_3465620 | Pathology North - Hunter - NSW Health Pathology | NSW Health Pathology - Institute of Clinical Pathology and Medical Research; Westmead Hospital; University of Sydney | Arnott A., Draper J., Gall M., Martinez E., Rockett R., Sintchenko V., on behalf of ICPMR |
| EPI_ISL_3465621 | Viollier AG | Department of Biosystems Science and Engineering, ETH Zürich | Chaoran Chen, Sarah Nadeau, Louis du Plessis, Catharine Aquino, Ivan Topolsky, Kim Philipp Jablonski, Lara Fuhrmann, Daniel Ehrsam, Isabel Stürmer, Andreia Cabral de Gouvea, Maria Domenica Moccia, Simon Grüter, Timothy Sykes, Lennart Opitz, Griffin White, Laura Neff, Doris Popovic, Andrea Patrignani, Jay Tracy, Ralph Schlapbach, Christiane Beckmann, Maurice Redondo, Olivier Kobel, Christoph Noppen, Niko Beerenwinkel, Tanja Stadler |
| EPI_ISL_3465622 | Laverty Pathology | NSW Health Pathology - Institute of Clinical Pathology and Medical Research; Westmead Hospital; University of Sydney | Arnott A., Draper J., Gall M., Martinez E., Rockett R., Sintchenko V., on behalf of ICPMR |
| EPI_ISL_3465623 | Viollier AG | Department of Biosystems Science and Engineering, ETH Zürich | Chaoran Chen, Sarah Nadeau, Louis du Plessis, Catharine Aquino, Ivan Topolsky, Kim Philipp Jablonski, Lara Fuhrmann, Daniel Ehrsam, Isabel Stürmer, Andreia Cabral de Gouvea, Maria Domenica Moccia, Simon Grüter, Timothy Sykes, Lennart Opitz, Griffin White, Laura Neff, Doris Popovic, Andrea Patrignani, Jay Tracy, Ralph Schlapbach, Christiane Beckmann, Maurice Redondo, Olivier Kobel, Christoph Noppen, Niko Beerenwinkel, Tanja Stadler |
| EPI_ISL_3465624 | Laverty Pathology | NSW Health Pathology - Institute of Clinical Pathology and Medical Research; Westmead Hospital; University of Sydney | Arnott A., Draper J., Gall M., Martinez E., Rockett R., Sintchenko V., on behalf of ICPMR |
| EPI_ISL_3465625, EPI_ISL_3465626, EPI_ISL_3465627 | Sydney South West Pathology Service (SSWPS) - Liverpool Hospital - NSW Health Pathology | NSW Health Pathology - Institute of Clinical Pathology and Medical Research; Westmead Hospital; University of Sydney | Arnott A., Draper J., Gall M., Martinez E., Rockett R., Sintchenko V., on behalf of ICPMR |
| EPI_ISL_3465628 | Viollier AG | Department of Biosystems Science and Engineering, ETH Zürich | Chaoran Chen, Sarah Nadeau, Louis du Plessis, Catharine Aquino, Ivan Topolsky, Kim Philipp Jablonski, Lara Fuhrmann, Daniel Ehrsam, Isabel Stürmer, Andreia Cabral de Gouvea, Maria Domenica Moccia, Simon Grüter, Timothy Sykes, Lennart Opitz, Griffin White, Laura Neff, Doris Popovic, Andrea Patrignani, Jay Tracy, Ralph Schlapbach, Christiane Beckmann, Maurice Redondo, Olivier Kobel, Christoph Noppen, Niko Beerenwinkel, Tanja Stadler |
| EPI_ISL_3465629 | Sydney South West Pathology Service (SSWPS) - Liverpool Hospital - NSW Health Pathology | NSW Health Pathology - Institute of Clinical Pathology and Medical Research; Westmead Hospital; University of Sydney | Arnott A., Draper J., Gall M., Martinez E., Rockett R., Sintchenko V., on behalf of ICPMR |
| EPI_ISL_3465630, EPI_ISL_3465631 | Viollier AG | Department of Biosystems Science and Engineering, ETH Zürich | Chaoran Chen, Sarah Nadeau, Louis du Plessis, Catharine Aquino, Ivan Topolsky, Kim Philipp Jablonski, Lara Fuhrmann, Daniel Ehrsam, Isabel Stürmer, Andreia Cabral de Gouvea, Maria Domenica Moccia, Simon Grüter, Timothy Sykes, Lennart Opitz, Griffin White, Laura Neff, Doris Popovic, Andrea Patrignani, Jay Tracy, Ralph Schlapbach, Christiane Beckmann, Maurice Redondo, Olivier Kobel, Christoph Noppen, Niko Beerenwinkel, Tanja Stadler |
| EPI_ISL_3465632 | Sydney South West Pathology Service (SSWPS) - Liverpool Hospital - NSW Health Pathology | NSW Health Pathology - Institute of Clinical Pathology and Medical Research; Westmead Hospital; University of Sydney | Arnott A., Draper J., Gall M., Martinez E., Rockett R., Sintchenko V., on behalf of ICPMR |
| EPI_ISL_3465633 | Viollier AG | Department of Biosystems Science and Engineering, ETH Zürich | Chaoran Chen, Sarah Nadeau, Louis du Plessis, Catharine Aquino, Ivan Topolsky, Kim Philipp Jablonski, Lara Fuhrmann, Daniel Ehrsam, Isabel Stürmer, Andreia Cabral de Gouvea, Maria Domenica Moccia, Simon Grüter, Timothy Sykes, Lennart Opitz, Griffin White, Laura Neff, Doris Popovic, Andrea Patrignani, Jay Tracy, Ralph Schlapbach, Christiane Beckmann, Maurice Redondo, Olivier Kobel, Christoph Noppen, Niko Beerenwinkel, Tanja Stadler |
| EPI_ISL_3465634 | Medlab Pathology | NSW Health Pathology - Institute of Clinical Pathology and Medical Research; Westmead Hospital; University of Sydney | Arnott A., Draper J., Gall M., Martinez E., Rockett R., Sintchenko V., on behalf of ICPMR |
| EPI_ISL_3465635 | Viollier AG | Department of Biosystems Science and Engineering, ETH Zürich | Chaoran Chen, Sarah Nadeau, Louis du Plessis, Catharine Aquino, Ivan Topolsky, Kim Philipp Jablonski, Lara Fuhrmann, Daniel Ehrsam, Isabel Stürmer, Andreia Cabral de Gouvea, Maria Domenica Moccia, Simon Grüter, Timothy Sykes, Lennart Opitz, Griffin White, Laura Neff, Doris Popovic, Andrea Patrignani, Jay Tracy, Ralph Schlapbach, Christiane Beckmann, Maurice Redondo, Olivier Kobel, Christoph Noppen, Niko Beerenwinkel, Tanja Stadler |
| EPI_ISL_3465636 | Medlab Pathology | NSW Health Pathology - Institute of Clinical Pathology and Medical Research; Westmead Hospital; University of Sydney | Arnott A., Draper J., Gall M., Martinez E., Rockett R., Sintchenko V., on behalf of ICPMR |
| EPI_ISL_3465637 | Viollier AG | Department of Biosystems Science and Engineering, ETH Zürich | Chaoran Chen, Sarah Nadeau, Louis du Plessis, Catharine Aquino, Ivan Topolsky, Kim Philipp Jablonski, Lara Fuhrmann, Daniel Ehrsam, Isabel Stürmer, Andreia Cabral de Gouvea, Maria Domenica Moccia, Simon Grüter, Timothy Sykes, Lennart Opitz, Griffin White, Laura Neff, Doris Popovic, Andrea Patrignani, Jay Tracy, Ralph Schlapbach, Christiane Beckmann, Maurice Redondo, Olivier Kobel, Christoph Noppen, Niko Beerenwinkel, Tanja Stadler |
| EPI_ISL_3465638 | Viollier AG | Department of Biosystems Science and Engineering, ETH Zürich | Andrea Patrizia Salzmann, Henriette Kurth, Christiane Beckmann, Ivan Topolsky, Chaoran Chen, Sarah Nadeau, Louis du Plessis, Kim Philipp Jablonski, Lara Fuhrmann, Niko Beerenwinkel, Christoph Noppen, Olivier Kobel, Maurice Redondo, Tanja Stadler |
| EPI_ISL_3465639 | Pathology North - Royal North Shore Hospital - NSW Health Pathology | NSW Health Pathology - Institute of Clinical Pathology and Medical Research; Westmead Hospital; University of Sydney | Arnott A., Draper J., Gall M., Martinez E., Rockett R., Sintchenko V., on behalf of ICPMR |
| EPI_ISL_3465640 | Viollier AG | Department of Biosystems Science and Engineering, ETH Zürich | Chaoran Chen, Sarah Nadeau, Louis du Plessis, Catharine Aquino, Ivan Topolsky, Kim Philipp Jablonski, Lara Fuhrmann, Daniel Ehrsam, Isabel Stürmer, Andreia Cabral de Gouvea, Maria Domenica Moccia, Simon Grüter, Timothy Sykes, Lennart Opitz, Griffin White, Laura Neff, Doris Popovic, Andrea Patrignani, Jay Tracy, Ralph Schlapbach, Christiane Beckmann, Maurice Redondo, Olivier Kobel, Christoph Noppen, Niko Beerenwinkel, Tanja Stadler |
| EPI_ISL_3465641 | Pathology North - Royal North Shore Hospital - NSW Health Pathology | NSW Health Pathology - Institute of Clinical Pathology and Medical Research; Westmead Hospital; University of Sydney | Arnott A., Draper J., Gall M., Martinez E., Rockett R., Sintchenko V., on behalf of ICPMR |
| EPI_ISL_3465642 | Viollier AG | Department of Biosystems Science and Engineering, ETH Zürich | Chaoran Chen, Sarah Nadeau, Louis du Plessis, Catharine Aquino, Ivan Topolsky, Kim Philipp Jablonski, Lara Fuhrmann, Daniel Ehrsam, Isabel Stürmer, Andreia Cabral de Gouvea, Maria Domenica Moccia, Simon Grüter, Timothy Sykes, Lennart Opitz, Griffin White, Laura Neff, Doris Popovic, Andrea Patrignani, Jay Tracy, Ralph Schlapbach, Christiane Beckmann, Maurice Redondo, Olivier Kobel, Christoph Noppen, Niko Beerenwinkel, Tanja Stadler |
| EPI_ISL_3465643 | 4Cyte Pathology | NSW Health Pathology - Institute of Clinical Pathology and Medical Research; Westmead Hospital; University of Sydney | Arnott A., Draper J., Gall M., Martinez E., Rockett R., Sintchenko V., on behalf of ICPMR |
| EPI_ISL_3465644 | Viollier AG | Department of Biosystems Science and Engineering, ETH Zürich | Chaoran Chen, Sarah Nadeau, Louis du Plessis, Catharine Aquino, Ivan Topolsky, Kim Philipp Jablonski, Lara Fuhrmann, Daniel Ehrsam, Isabel Stürmer, Andreia Cabral de Gouvea, Maria Domenica Moccia, Simon Grüter, Timothy Sykes, Lennart Opitz, Griffin White, Laura Neff, Doris Popovic, Andrea Patrignani, Jay Tracy, Ralph Schlapbach, Christiane Beckmann, Maurice Redondo, Olivier Kobel, Christoph Noppen, Niko Beerenwinkel, Tanja Stadler |
| EPI_ISL_3465645 | 4Cyte Pathology | NSW Health Pathology - Institute of Clinical Pathology and Medical Research; Westmead Hospital; University of Sydney | Arnott A., Draper J., Gall M., Martinez E., Rockett R., Sintchenko V., on behalf of ICPMR |
| EPI_ISL_3465646 | Viollier AG | Department of Biosystems Science and Engineering, ETH Zürich | Chaoran Chen, Sarah Nadeau, Louis du Plessis, Catharine Aquino, Ivan Topolsky, Kim Philipp Jablonski, Lara Fuhrmann, Daniel Ehrsam, Isabel Stürmer, Andreia Cabral de Gouvea, Maria Domenica Moccia, Simon Grüter, Timothy Sykes, Lennart Opitz, Griffin White, Laura Neff, Doris Popovic, Andrea Patrignani, Jay Tracy, Ralph Schlapbach, Christiane Beckmann, Maurice Redondo, Olivier Kobel, Christoph Noppen, Niko Beerenwinkel, Tanja Stadler |
| EPI_ISL_3465647 | Viollier AG | Department of Biosystems Science and Engineering, ETH Zürich | Andrea Patrizia Salzmann, Henriette Kurth, Christiane Beckmann, Ivan Topolsky, Chaoran Chen, Sarah Nadeau, Louis du Plessis, Kim Philipp Jablonski, Lara Fuhrmann, Niko Beerenwinkel, Christoph Noppen, Olivier Kobel, Maurice Redondo, Tanja Stadler |
| EPI_ISL_3465648 | 4Cyte Pathology | NSW Health Pathology - Institute of Clinical Pathology and Medical Research; Westmead Hospital; University of Sydney | Arnott A., Draper J., Gall M., Martinez E., Rockett R., Sintchenko V., on behalf of ICPMR |
| EPI_ISL_3465649 | Viollier AG | Department of Biosystems Science and Engineering, ETH Zürich | Chaoran Chen, Sarah Nadeau, Louis du Plessis, Catharine Aquino, Ivan Topolsky, Kim Philipp Jablonski, Lara Fuhrmann, Daniel Ehrsam, Isabel Stürmer, Andreia Cabral de Gouvea, Maria Domenica Moccia, Simon Grüter, Timothy Sykes, Lennart Opitz, Griffin White, Laura Neff, Doris Popovic, Andrea Patrignani, Jay Tracy, Ralph Schlapbach, Christiane Beckmann, Maurice Redondo, Olivier Kobel, Christoph Noppen, Niko Beerenwinkel, Tanja Stadler |
| EPI_ISL_3465650 | 4Cyte Pathology | NSW Health Pathology - Institute of Clinical Pathology and Medical Research; Westmead Hospital; University of Sydney | Arnott A., Draper J., Gall M., Martinez E., Rockett R., Sintchenko V., on behalf of ICPMR |

| | | | |
|---|---|---|---|
| EPI_ISL_3465651 | Viollier AG | Department of Biosystems Science and Engineering, ETH Zürich | Chaoran Chen, Sarah Nadeau, Louis du Plessis, Catharine Aquino, Ivan Topolsky, Kim Philipp Jablonski, Lara Fuhrmann, Daniel Ehrsam, Isabel Stürmer, Andreia Cabral de Gouvea, Maria Domenica Moccia, Simon Grüter, Timothy Sykes, Lennart Opitz, Griffin White, Laura Neff, Doris Popovic, Andrea Patrignani, Jay Tracy, Ralph Schlapbach, Christiane Beckmann, Maurice Redondo, Olivier Kobel, Christoph Noppen, Niko Beerenwinkel, Tanja Stadler |
| EPI_ISL_3465652 | 4Cyte Pathology | NSW Health Pathology - Institute of Clinical Pathology and Medical Research; Westmead Hospital; University of Sydney | Arnott A., Draper J., Gall M., Martinez E., Rockett R., Sintchenko V., on behalf of ICPMR |
| EPI_ISL_3465653 | Viollier AG | Department of Biosystems Science and Engineering, ETH Zürich | Chaoran Chen, Sarah Nadeau, Louis du Plessis, Catharine Aquino, Ivan Topolsky, Kim Philipp Jablonski, Lara Fuhrmann, Daniel Ehrsam, Isabel Stürmer, Andreia Cabral de Gouvea, Maria Domenica Moccia, Simon Grüter, Timothy Sykes, Lennart Opitz, Griffin White, Laura Neff, Doris Popovic, Andrea Patrignani, Jay Tracy, Ralph Schlapbach, Christiane Beckmann, Maurice Redondo, Olivier Kobel, Christoph Noppen, Niko Beerenwinkel, Tanja Stadler |
| EPI_ISL_3465654 | 4Cyte Pathology | NSW Health Pathology - Institute of Clinical Pathology and Medical Research; Westmead Hospital; University of Sydney | Arnott A., Draper J., Gall M., Martinez E., Rockett R., Sintchenko V., on behalf of ICPMR |
| EPI_ISL_3465655 | Viollier AG | Department of Biosystems Science and Engineering, ETH Zürich | Chaoran Chen, Sarah Nadeau, Louis du Plessis, Catharine Aquino, Ivan Topolsky, Kim Philipp Jablonski, Lara Fuhrmann, Daniel Ehrsam, Isabel Stürmer, Andreia Cabral de Gouvea, Maria Domenica Moccia, Simon Grüter, Timothy Sykes, Lennart Opitz, Griffin White, Laura Neff, Doris Popovic, Andrea Patrignani, Jay Tracy, Ralph Schlapbach, Christiane Beckmann, Maurice Redondo, Olivier Kobel, Christoph Noppen, Niko Beerenwinkel, Tanja Stadler |
| EPI_ISL_3465656 | Viollier AG | Department of Biosystems Science and Engineering, ETH Zürich | Andrea Patrizia Salzmann, Henriette Kurth, Christiane Beckmann, Ivan Topolsky, Chaoran Chen, Sarah Nadeau, Louis du Plessis, Kim Philipp Jablonski, Lara Fuhrmann, Niko Beerenwinkel, Christoph Noppen, Olivier Kobel, Maurice Redondo, Tanja Stadler |
| EPI_ISL_3465657 | 4Cyte Pathology | NSW Health Pathology - Institute of Clinical Pathology and Medical Research; Westmead Hospital; University of Sydney | Arnott A., Draper J., Gall M., Martinez E., Rockett R., Sintchenko V., on behalf of ICPMR |
| EPI_ISL_3465658 | Viollier AG | Department of Biosystems Science and Engineering, ETH Zürich | Andrea Patrizia Salzmann, Henriette Kurth, Christiane Beckmann, Ivan Topolsky, Chaoran Chen, Sarah Nadeau, Louis du Plessis, Kim Philipp Jablonski, Lara Fuhrmann, Niko Beerenwinkel, Christoph Noppen, Olivier Kobel, Maurice Redondo, Tanja Stadler |
| EPI_ISL_3465659 | 4Cyte Pathology | NSW Health Pathology - Institute of Clinical Pathology and Medical Research; Westmead Hospital; University of Sydney | Arnott A., Draper J., Gall M., Martinez E., Rockett R., Sintchenko V., on behalf of ICPMR |
| EPI_ISL_3465660 | Viollier AG | Department of Biosystems Science and Engineering, ETH Zürich | Andrea Patrizia Salzmann, Henriette Kurth, Christiane Beckmann, Ivan Topolsky, Chaoran Chen, Sarah Nadeau, Louis du Plessis, Kim Philipp Jablonski, Lara Fuhrmann, Niko Beerenwinkel, Christoph Noppen, Olivier Kobel, Maurice Redondo, Tanja Stadler |
| EPI_ISL_3465661 | Viollier AG | Department of Biosystems Science and Engineering, ETH Zürich | Chaoran Chen, Sarah Nadeau, Louis du Plessis, Catharine Aquino, Ivan Topolsky, Kim Philipp Jablonski, Lara Fuhrmann, Daniel Ehrsam, Isabel Stürmer, Andreia Cabral de Gouvea, Maria Domenica Moccia, Simon Grüter, Timothy Sykes, Lennart Opitz, Griffin White, Laura Neff, Doris Popovic, Andrea Patrignani, Jay Tracy, Ralph Schlapbach, Christiane Beckmann, Maurice Redondo, Olivier Kobel, Christoph Noppen, Niko Beerenwinkel, Tanja Stadler |
| EPI_ISL_3465662 | 4Cyte Pathology | NSW Health Pathology - Institute of Clinical Pathology and Medical Research; Westmead Hospital; University of Sydney | Arnott A., Draper J., Gall M., Martinez E., Rockett R., Sintchenko V., on behalf of ICPMR |
| EPI_ISL_3465663 | Viollier AG | Department of Biosystems Science and Engineering, ETH Zürich | Chaoran Chen, Sarah Nadeau, Louis du Plessis, Catharine Aquino, Ivan Topolsky, Kim Philipp Jablonski, Lara Fuhrmann, Daniel Ehrsam, Isabel Stürmer, Andreia Cabral de Gouvea, Maria Domenica Moccia, Simon Grüter, Timothy Sykes, Lennart Opitz, Griffin White, Laura Neff, Doris Popovic, Andrea Patrignani, Jay Tracy, Ralph Schlapbach, Christiane Beckmann, Maurice Redondo, Olivier Kobel, Christoph Noppen, Niko Beerenwinkel, Tanja Stadler |
| EPI_ISL_3465664 | 4Cyte Pathology | NSW Health Pathology - Institute of Clinical Pathology and Medical Research; Westmead Hospital; University of Sydney | Arnott A., Draper J., Gall M., Martinez E., Rockett R., Sintchenko V., on behalf of ICPMR |
| EPI_ISL_3465665 | Viollier AG | Department of Biosystems Science and Engineering, ETH Zürich | Andrea Patrizia Salzmann, Henriette Kurth, Christiane Beckmann, Ivan Topolsky, Chaoran Chen, Sarah Nadeau, Louis du Plessis, Kim Philipp Jablonski, Lara Fuhrmann, Niko Beerenwinkel, Christoph Noppen, Olivier Kobel, Maurice Redondo, Tanja Stadler |
| EPI_ISL_3465666 | 4Cyte Pathology | NSW Health Pathology - Institute of Clinical Pathology and Medical Research; Westmead Hospital; University of Sydney | Arnott A., Draper J., Gall M., Martinez E., Rockett R., Sintchenko V., on behalf of ICPMR |
| EPI_ISL_3465667 | Viollier AG | Department of Biosystems Science and Engineering, ETH Zürich | Chaoran Chen, Sarah Nadeau, Louis du Plessis, Catharine Aquino, Ivan Topolsky, Kim Philipp Jablonski, Lara Fuhrmann, Daniel Ehrsam, Isabel Stürmer, Andreia Cabral de Gouvea, Maria Domenica Moccia, Simon Grüter, Timothy Sykes, Lennart Opitz, Griffin White, Laura Neff, Doris Popovic, Andrea Patrignani, Jay Tracy, Ralph Schlapbach, Christiane Beckmann, Maurice Redondo, Olivier Kobel, Christoph Noppen, Niko Beerenwinkel, Tanja Stadler |
| EPI_ISL_3465668 | 4Cyte Pathology | NSW Health Pathology - Institute of Clinical Pathology and Medical Research; Westmead Hospital; University of Sydney | Arnott A., Draper J., Gall M., Martinez E., Rockett R., Sintchenko V., on behalf of ICPMR |
| EPI_ISL_3465669 | Viollier AG | Department of Biosystems Science and Engineering, ETH Zürich | Andrea Patrizia Salzmann, Henriette Kurth, Christiane Beckmann, Ivan Topolsky, Chaoran Chen, Sarah Nadeau, Louis du Plessis, Kim Philipp Jablonski, Lara Fuhrmann, Niko Beerenwinkel, Christoph Noppen, Olivier Kobel, Maurice Redondo, Tanja Stadler |
| EPI_ISL_3465670 | Australian Clinical Labs (formerly Healthscope Pathology) | NSW Health Pathology - Institute of Clinical Pathology and Medical Research; Westmead Hospital; University of Sydney | Arnott A., Draper J., Gall M., Martinez E., Rockett R., Sintchenko V., on behalf of ICPMR |
| EPI_ISL_3465671 | Viollier AG | Department of Biosystems Science and Engineering, ETH Zürich | Andrea Patrizia Salzmann, Henriette Kurth, Christiane Beckmann, Ivan Topolsky, Chaoran Chen, Sarah Nadeau, Louis du Plessis, Kim Philipp Jablonski, Lara Fuhrmann, Niko Beerenwinkel, Christoph Noppen, Olivier Kobel, Maurice Redondo, Tanja Stadler |
| EPI_ISL_3465672 | Australian Clinical Labs (formerly Healthscope Pathology) | NSW Health Pathology - Institute of Clinical Pathology and Medical Research; Westmead Hospital; University of Sydney | Arnott A., Draper J., Gall M., Martinez E., Rockett R., Sintchenko V., on behalf of ICPMR |
| EPI_ISL_3465673 | Viollier AG | Department of Biosystems Science and Engineering, ETH Zürich | Chaoran Chen, Sarah Nadeau, Louis du Plessis, Catharine Aquino, Ivan Topolsky, Kim Philipp Jablonski, Lara Fuhrmann, Daniel Ehrsam, Isabel Stürmer, Andreia Cabral de Gouvea, Maria Domenica Moccia, Simon Grüter, Timothy Sykes, Lennart Opitz, Griffin White, Laura Neff, Doris Popovic, Andrea Patrignani, Jay Tracy, Ralph Schlapbach, Christiane Beckmann, Maurice Redondo, Olivier Kobel, Christoph Noppen, Niko Beerenwinkel, Tanja Stadler |
| EPI_ISL_3465674, EPI_ISL_3465675 | Australian Clinical Labs (formerly Healthscope Pathology) | NSW Health Pathology - Institute of Clinical Pathology and Medical Research; Westmead Hospital; University of Sydney | Arnott A., Draper J., Gall M., Martinez E., Rockett R., Sintchenko V., on behalf of ICPMR |
| EPI_ISL_3465676 | Viollier AG | Department of Biosystems Science and Engineering, ETH Zürich | Andrea Patrizia Salzmann, Henriette Kurth, Christiane Beckmann, Ivan Topolsky, Chaoran Chen, Sarah Nadeau, Louis du Plessis, Kim Philipp Jablonski, Lara Fuhrmann, Niko Beerenwinkel, Christoph Noppen, Olivier Kobel, Maurice Redondo, Tanja Stadler |
| EPI_ISL_3465677 | Australian Clinical Labs (formerly Healthscope Pathology) | NSW Health Pathology - Institute of Clinical Pathology and Medical Research; Westmead Hospital; University of Sydney | Arnott A., Draper J., Gall M., Martinez E., Rockett R., Sintchenko V., on behalf of ICPMR |
| EPI_ISL_3465678 | Viollier AG | Department of Biosystems Science and Engineering, ETH Zürich | Chaoran Chen, Sarah Nadeau, Louis du Plessis, Catharine Aquino, Ivan Topolsky, Kim Philipp Jablonski, Lara Fuhrmann, Daniel Ehrsam, Isabel Stürmer, Andreia Cabral de Gouvea, Maria Domenica Moccia, Simon Grüter, Timothy Sykes, Lennart Opitz, Griffin White, Laura Neff, Doris Popovic, Andrea Patrignani, Jay Tracy, Ralph Schlapbach, Christiane Beckmann, Maurice Redondo, Olivier Kobel, Christoph Noppen, Niko Beerenwinkel, Tanja Stadler |
| EPI_ISL_3465679 | Australian Clinical Labs (formerly Healthscope Pathology) | NSW Health Pathology - Institute of Clinical Pathology and Medical Research; Westmead Hospital; University of Sydney | Arnott A., Draper J., Gall M., Martinez E., Rockett R., Sintchenko V., on behalf of ICPMR |
| EPI_ISL_3465680, EPI_ISL_3465681 | Viollier AG | Department of Biosystems Science and Engineering, ETH Zürich | Chaoran Chen, Sarah Nadeau, Louis du Plessis, Catharine Aquino, Ivan Topolsky, Kim Philipp Jablonski, Lara Fuhrmann, Daniel Ehrsam, Isabel Stürmer, Andreia Cabral de Gouvea, Maria Domenica Moccia, Simon Grüter, Timothy Sykes, Lennart Opitz, Griffin White, Laura Neff, Doris Popovic, Andrea Patrignani, Jay Tracy, Ralph Schlapbach, Christiane Beckmann, Maurice Redondo, Olivier Kobel, Christoph Noppen, Niko Beerenwinkel, Tanja Stadler |
| EPI_ISL_3465682 | Australian Clinical Labs (formerly Healthscope Pathology) | NSW Health Pathology - Institute of Clinical Pathology and Medical Research; Westmead Hospital; University of Sydney | Arnott A., Draper J., Gall M., Martinez E., Rockett R., Sintchenko V., on behalf of ICPMR |
| EPI_ISL_3465683 | Viollier AG | Department of Biosystems Science and Engineering, ETH Zürich | Chaoran Chen, Sarah Nadeau, Louis du Plessis, Catharine Aquino, Ivan Topolsky, Kim Philipp Jablonski, Lara Fuhrmann, Daniel Ehrsam, Isabel Stürmer, Andreia Cabral de Gouvea, Maria Domenica Moccia, Simon Grüter, Timothy Sykes, Lennart Opitz, Griffin White, Laura Neff, Doris Popovic, Andrea Patrignani, Jay Tracy, Ralph Schlapbach, Christiane Beckmann, Maurice Redondo, Olivier Kobel, Christoph Noppen, Niko Beerenwinkel, Tanja Stadler |
| EPI_ISL_3465684 | Australian Clinical Labs (formerly Healthscope Pathology) | NSW Health Pathology - Institute of Clinical Pathology and Medical Research; Westmead Hospital; University of Sydney | Arnott A., Draper J., Gall M., Martinez E., Rockett R., Sintchenko V., on behalf of ICPMR |
| EPI_ISL_3465686 | Sydney South West Pathology Service (SSWPS) - Liverpool Hospital - NSW Health Pathology | NSW Health Pathology - Institute of Clinical Pathology and Medical Research; Westmead Hospital; University of Sydney | Arnott A., Draper J., Gall M., Martinez E., Rockett R., Sintchenko V., on behalf of ICPMR |
| EPI_ISL_3465687 | Viollier AG | Department of Biosystems Science and Engineering, ETH Zürich | Chaoran Chen, Sarah Nadeau, Louis du Plessis, Catharine Aquino, Ivan Topolsky, Kim Philipp Jablonski, Lara Fuhrmann, Daniel Ehrsam, Isabel Stürmer, Andreia Cabral de Gouvea, Maria Domenica Moccia, Simon Grüter, Timothy Sykes, Lennart Opitz, Griffin White, Laura Neff, Doris Popovic, Andrea Patrignani, Jay Tracy, Ralph Schlapbach, Christiane Beckmann, Maurice Redondo, Olivier Kobel, Christoph Noppen, Niko Beerenwinkel, Tanja Stadler |
| EPI_ISL_3465688 | Sydney South West Pathology Service (SSWPS) - Liverpool Hospital - NSW Health Pathology | NSW Health Pathology - Institute of Clinical Pathology and Medical Research; Westmead Hospital; University of Sydney | Arnott A., Draper J., Gall M., Martinez E., Rockett R., Sintchenko V., on behalf of ICPMR |
| EPI_ISL_3465689, EPI_ISL_3465690 | Viollier AG | Department of Biosystems Science and Engineering, ETH Zürich | Andrea Patrizia Salzmann, Henriette Kurth, Christiane Beckmann, Ivan Topolsky, Chaoran Chen, Sarah Nadeau, Louis du Plessis, Kim Philipp Jablonski, Lara Fuhrmann, Niko Beerenwinkel, Christoph Noppen, Olivier Kobel, Maurice Redondo, Tanja Stadler |
| EPI_ISL_3465691 | Sydney South West Pathology Service (SSWPS) - Liverpool Hospital - NSW Health Pathology | NSW Health Pathology - Institute of Clinical Pathology and Medical Research; Westmead Hospital; University of Sydney | Arnott A., Draper J., Gall M., Martinez E., Rockett R., Sintchenko V., on behalf of ICPMR |
| EPI_ISL_3465692 | Viollier AG | Department of Biosystems Science and Engineering, ETH Zürich | Chaoran Chen, Sarah Nadeau, Louis du Plessis, Catharine Aquino, Ivan Topolsky, Kim Philipp Jablonski, Lara Fuhrmann, Daniel Ehrsam, Isabel Stürmer, Andreia Cabral de Gouvea, Maria Domenica Moccia, Simon Grüter, Timothy Sykes, Lennart Opitz, Griffin White, Laura Neff, Doris Popovic, Andrea Patrignani, Jay Tracy, Ralph Schlapbach, Christiane Beckmann, Maurice Redondo, Olivier Kobel, Christoph Noppen, Niko Beerenwinkel, Tanja Stadler |
| EPI_ISL_3465693 | Sydney South West Pathology Service (SSWPS) - Concord Repatriation General Hospital - NSW Health Pathology | NSW Health Pathology - Institute of Clinical Pathology and Medical Research; Westmead Hospital; University of Sydney | Arnott A., Draper J., Gall M., Martinez E., Rockett R., Sintchenko V., on behalf of ICPMR |
| EPI_ISL_3465694 | Viollier AG | Department of Biosystems Science and Engineering, ETH Zürich | Chaoran Chen, Sarah Nadeau, Louis du Plessis, Catharine Aquino, Ivan Topolsky, Kim Philipp Jablonski, Lara Fuhrmann, Daniel Ehrsam, Isabel Stürmer, Andreia Cabral de Gouvea, Maria Domenica Moccia, Simon Grüter, Timothy Sykes, Lennart Opitz, Griffin White, Laura Neff, Doris Popovic, Andrea Patrignani, Jay Tracy, Ralph Schlapbach, Christiane Beckmann, Maurice Redondo, Olivier Kobel, Christoph Noppen, Niko Beerenwinkel, Tanja Stadler |
| EPI_ISL_3465695 | Histopath | NSW Health Pathology - Institute of Clinical Pathology and Medical Research; Westmead Hospital; University of Sydney | Arnott A., Draper J., Gall M., Martinez E., Rockett R., Sintchenko V., on behalf of ICPMR |
| EPI_ISL_3465696 | Viollier AG | Department of Biosystems Science and Engineering, ETH Zürich | Andrea Patrizia Salzmann, Henriette Kurth, Christiane Beckmann, Ivan Topolsky, Chaoran Chen, Sarah Nadeau, Louis du Plessis, Kim Philipp Jablonski, Lara Fuhrmann, Niko Beerenwinkel, Christoph Noppen, Olivier Kobel, Maurice Redondo, Tanja Stadler |
| EPI_ISL_3465697 | Viollier AG | Department of Biosystems Science and Engineering, ETH Zürich | Chaoran Chen, Sarah Nadeau, Louis du Plessis, Catharine Aquino, Ivan Topolsky, Kim Philipp Jablonski, Lara Fuhrmann, Daniel Ehrsam, Isabel Stürmer, Andreia Cabral de Gouvea, Maria Domenica Moccia, Simon Grüter, Timothy Sykes, Lennart Opitz, Griffin White, Laura Neff, Doris Popovic, Andrea Patrignani, Jay Tracy, Ralph Schlapbach, Christiane Beckmann, Maurice Redondo, Olivier Kobel, Christoph Noppen, Niko Beerenwinkel, Tanja Stadler |
| EPI_ISL_3465698 | Pathology North - Hunter - NSW Health Pathology | NSW Health Pathology - Institute of Clinical Pathology and Medical Research; Westmead Hospital; University of Sydney | Arnott A., Draper J., Gall M., Martinez E., Rockett R., Sintchenko V., on behalf of ICPMR |
| EPI_ISL_3465699 | Viollier AG | Department of Biosystems Science and Engineering, ETH Zürich | Chaoran Chen, Sarah Nadeau, Louis du Plessis, Catharine Aquino, Ivan Topolsky, Kim Philipp Jablonski, Lara Fuhrmann, Daniel Ehrsam, Isabel Stürmer, Andreia Cabral de Gouvea, Maria Domenica Moccia, Simon Grüter, Timothy Sykes, Lennart Opitz, Griffin White, Laura Neff, Doris Popovic, Andrea Patrignani, Jay Tracy, Ralph Schlapbach, Christiane Beckmann, Maurice Redondo, Olivier Kobel, Christoph Noppen, Niko Beerenwinkel, Tanja Stadler |

| | | | |
|---|---|---|---|
| EPI_ISL_3465700 | Pathology North - Hunter - NSW Health Pathology | NSW Health Pathology - Institute of Clinical Pathology and Medical Research; Westmead Hospital; University of Sydney | Arnott A., Draper J., Gall M., Martinez E., Rockett R., Sintchenko V., on behalf of ICPMR |
| EPI_ISL_3465701 | Viollier AG | Department of Biosystems Science and Engineering, ETH Zürich | Chaoran Chen, Sarah Nadeau, Louis du Plessis, Catharine Aquino, Ivan Topolsky, Kim Philipp Jablonski, Lara Fuhrmann, Daniel Ehrsam, Isabel Stürmer, Andreia Cabral de Gouvea, Maria Domenica Moccia, Simon Grüter, Timothy Sykes, Lennart Opitz, Griffin White, Laura Neff, Doris Popovic, Andrea Patrignani, Jay Tracy, Ralph Schlapbach, Christiane Beckmann, Maurice Redondo, Olivier Kobel, Christoph Noppen, Niko Beerenwinkel, Tanja Stadler |
| EPI_ISL_3465702 | Viollier AG | Department of Biosystems Science and Engineering, ETH Zürich | Andrea Patrizia Salzmann, Henriette Kurth, Christiane Beckmann, Ivan Topolsky, Chaoran Chen, Sarah Nadeau, Louis du Plessis, Kim Philipp Jablonski, Lara Fuhrmann, Niko Beerenwinkel, Christoph Noppen, Olivier Kobel, Maurice Redondo, Tanja Stadler |
| EPI_ISL_3465703 | Sydney South West Pathology Service (SSWPS) - Liverpool Hospital - NSW Health Pathology | NSW Health Pathology - Institute of Clinical Pathology and Medical Research; Westmead Hospital; University of Sydney | Arnott A., Draper J., Gall M., Martinez E., Rockett R., Sintchenko V., on behalf of ICPMR |
| EPI_ISL_3465704 | Viollier AG | Department of Biosystems Science and Engineering, ETH Zürich | Chaoran Chen, Sarah Nadeau, Louis du Plessis, Catharine Aquino, Ivan Topolsky, Kim Philipp Jablonski, Lara Fuhrmann, Daniel Ehrsam, Isabel Stürmer, Andreia Cabral de Gouvea, Maria Domenica Moccia, Simon Grüter, Timothy Sykes, Lennart Opitz, Griffin White, Laura Neff, Doris Popovic, Andrea Patrignani, Jay Tracy, Ralph Schlapbach, Christiane Beckmann, Maurice Redondo, Olivier Kobel, Christoph Noppen, Niko Beerenwinkel, Tanja Stadler |
| EPI_ISL_3465705 | Medlab Pathology | NSW Health Pathology - Institute of Clinical Pathology and Medical Research; Westmead Hospital; University of Sydney | Arnott A., Draper J., Gall M., Martinez E., Rockett R., Sintchenko V., on behalf of ICPMR |
| EPI_ISL_3465706 | Viollier AG | Department of Biosystems Science and Engineering, ETH Zürich | Chaoran Chen, Sarah Nadeau, Louis du Plessis, Catharine Aquino, Ivan Topolsky, Kim Philipp Jablonski, Lara Fuhrmann, Daniel Ehrsam, Isabel Stürmer, Andreia Cabral de Gouvea, Maria Domenica Moccia, Simon Grüter, Timothy Sykes, Lennart Opitz, Griffin White, Laura Neff, Doris Popovic, Andrea Patrignani, Jay Tracy, Ralph Schlapbach, Christiane Beckmann, Maurice Redondo, Olivier Kobel, Christoph Noppen, Niko Beerenwinkel, Tanja Stadler |
| EPI_ISL_3465707 | Medlab Pathology | NSW Health Pathology - Institute of Clinical Pathology and Medical Research; Westmead Hospital; University of Sydney | Arnott A., Draper J., Gall M., Martinez E., Rockett R., Sintchenko V., on behalf of ICPMR |
| EPI_ISL_3465708 | Viollier AG | Department of Biosystems Science and Engineering, ETH Zürich | Andrea Patrizia Salzmann, Henriette Kurth, Christiane Beckmann, Ivan Topolsky, Chaoran Chen, Sarah Nadeau, Louis du Plessis, Kim Philipp Jablonski, Lara Fuhrmann, Niko Beerenwinkel, Christoph Noppen, Olivier Kobel, Maurice Redondo, Tanja Stadler |
| EPI_ISL_3465709 | Viollier AG | Department of Biosystems Science and Engineering, ETH Zürich | Chaoran Chen, Sarah Nadeau, Louis du Plessis, Catharine Aquino, Ivan Topolsky, Kim Philipp Jablonski, Lara Fuhrmann, Daniel Ehrsam, Isabel Stürmer, Andreia Cabral de Gouvea, Maria Domenica Moccia, Simon Grüter, Timothy Sykes, Lennart Opitz, Griffin White, Laura Neff, Doris Popovic, Andrea Patrignani, Jay Tracy, Ralph Schlapbach, Christiane Beckmann, Maurice Redondo, Olivier Kobel, Christoph Noppen, Niko Beerenwinkel, Tanja Stadler |
| EPI_ISL_3465710 | Medlab Pathology | NSW Health Pathology - Institute of Clinical Pathology and Medical Research; Westmead Hospital; University of Sydney | Arnott A., Draper J., Gall M., Martinez E., Rockett R., Sintchenko V., on behalf of ICPMR |
| EPI_ISL_3465711 | Viollier AG | Department of Biosystems Science and Engineering, ETH Zürich | Chaoran Chen, Sarah Nadeau, Louis du Plessis, Catharine Aquino, Ivan Topolsky, Kim Philipp Jablonski, Lara Fuhrmann, Daniel Ehrsam, Isabel Stürmer, Andreia Cabral de Gouvea, Maria Domenica Moccia, Simon Grüter, Timothy Sykes, Lennart Opitz, Griffin White, Laura Neff, Doris Popovic, Andrea Patrignani, Jay Tracy, Ralph Schlapbach, Christiane Beckmann, Maurice Redondo, Olivier Kobel, Christoph Noppen, Niko Beerenwinkel, Tanja Stadler |
| EPI_ISL_3465712 | Medlab Pathology | NSW Health Pathology - Institute of Clinical Pathology and Medical Research; Westmead Hospital; University of Sydney | Arnott A., Draper J., Gall M., Martinez E., Rockett R., Sintchenko V., on behalf of ICPMR |
| EPI_ISL_3465713 | Viollier AG | Department of Biosystems Science and Engineering, ETH Zürich | Chaoran Chen, Sarah Nadeau, Louis du Plessis, Catharine Aquino, Ivan Topolsky, Kim Philipp Jablonski, Lara Fuhrmann, Daniel Ehrsam, Isabel Stürmer, Andreia Cabral de Gouvea, Maria Domenica Moccia, Simon Grüter, Timothy Sykes, Lennart Opitz, Griffin White, Laura Neff, Doris Popovic, Andrea Patrignani, Jay Tracy, Ralph Schlapbach, Christiane Beckmann, Maurice Redondo, Olivier Kobel, Christoph Noppen, Niko Beerenwinkel, Tanja Stadler |
| EPI_ISL_3465714 | Medlab Pathology | NSW Health Pathology - Institute of Clinical Pathology and Medical Research; Westmead Hospital; University of Sydney | Arnott A., Draper J., Gall M., Martinez E., Rockett R., Sintchenko V., on behalf of ICPMR |
| EPI_ISL_3465715 | Viollier AG | Department of Biosystems Science and Engineering, ETH Zürich | Andrea Patrizia Salzmann, Henriette Kurth, Christiane Beckmann, Ivan Topolsky, Chaoran Chen, Sarah Nadeau, Louis du Plessis, Kim Philipp Jablonski, Lara Fuhrmann, Niko Beerenwinkel, Christoph Noppen, Olivier Kobel, Maurice Redondo, Tanja Stadler |
| EPI_ISL_3465716 | Medlab Pathology | NSW Health Pathology - Institute of Clinical Pathology and Medical Research; Westmead Hospital; University of Sydney | Arnott A., Draper J., Gall M., Martinez E., Rockett R., Sintchenko V., on behalf of ICPMR |
| EPI_ISL_3465717 | Viollier AG | Department of Biosystems Science and Engineering, ETH Zürich | Andrea Patrizia Salzmann, Henriette Kurth, Christiane Beckmann, Ivan Topolsky, Chaoran Chen, Sarah Nadeau, Louis du Plessis, Kim Philipp Jablonski, Lara Fuhrmann, Niko Beerenwinkel, Christoph Noppen, Olivier Kobel, Maurice Redondo, Tanja Stadler |
| EPI_ISL_3465718 | Viollier AG | Department of Biosystems Science and Engineering, ETH Zürich | Chaoran Chen, Sarah Nadeau, Louis du Plessis, Catharine Aquino, Ivan Topolsky, Kim Philipp Jablonski, Lara Fuhrmann, Daniel Ehrsam, Isabel Stürmer, Andreia Cabral de Gouvea, Maria Domenica Moccia, Simon Grüter, Timothy Sykes, Lennart Opitz, Griffin White, Laura Neff, Doris Popovic, Andrea Patrignani, Jay Tracy, Ralph Schlapbach, Christiane Beckmann, Maurice Redondo, Olivier Kobel, Christoph Noppen, Niko Beerenwinkel, Tanja Stadler |
| EPI_ISL_3465719 | Medlab Pathology | NSW Health Pathology - Institute of Clinical Pathology and Medical Research; Westmead Hospital; University of Sydney | Arnott A., Draper J., Gall M., Martinez E., Rockett R., Sintchenko V., on behalf of ICPMR |
| EPI_ISL_3465720 | Viollier AG | Department of Biosystems Science and Engineering, ETH Zürich | Chaoran Chen, Sarah Nadeau, Louis du Plessis, Catharine Aquino, Ivan Topolsky, Kim Philipp Jablonski, Lara Fuhrmann, Daniel Ehrsam, Isabel Stürmer, Andreia Cabral de Gouvea, Maria Domenica Moccia, Simon Grüter, Timothy Sykes, Lennart Opitz, Griffin White, Laura Neff, Doris Popovic, Andrea Patrignani, Jay Tracy, Ralph Schlapbach, Christiane Beckmann, Maurice Redondo, Olivier Kobel, Christoph Noppen, Niko Beerenwinkel, Tanja Stadler |
| EPI_ISL_3465721 | Sydney South West Pathology Service (SSWPS) - Liverpool Hospital - NSW Health Pathology | NSW Health Pathology - Institute of Clinical Pathology and Medical Research; Westmead Hospital; University of Sydney | Arnott A., Draper J., Gall M., Martinez E., Rockett R., Sintchenko V., on behalf of ICPMR |
| EPI_ISL_3465722 | Viollier AG | Department of Biosystems Science and Engineering, ETH Zürich | Chaoran Chen, Sarah Nadeau, Louis du Plessis, Catharine Aquino, Ivan Topolsky, Kim Philipp Jablonski, Lara Fuhrmann, Daniel Ehrsam, Isabel Stürmer, Andreia Cabral de Gouvea, Maria Domenica Moccia, Simon Grüter, Timothy Sykes, Lennart Opitz, Griffin White, Laura Neff, Doris Popovic, Andrea Patrignani, Jay Tracy, Ralph Schlapbach, Christiane Beckmann, Maurice Redondo, Olivier Kobel, Christoph Noppen, Niko Beerenwinkel, Tanja Stadler |
| EPI_ISL_3465723 | Sydney South West Pathology Service (SSWPS) - Liverpool Hospital - NSW Health Pathology | NSW Health Pathology - Institute of Clinical Pathology and Medical Research; Westmead Hospital; University of Sydney | Arnott A., Draper J., Gall M., Martinez E., Rockett R., Sintchenko V., on behalf of ICPMR |
| EPI_ISL_3465724, EPI_ISL_3465725 | Viollier AG | Department of Biosystems Science and Engineering, ETH Zürich | Chaoran Chen, Sarah Nadeau, Louis du Plessis, Catharine Aquino, Ivan Topolsky, Kim Philipp Jablonski, Lara Fuhrmann, Daniel Ehrsam, Isabel Stürmer, Andreia Cabral de Gouvea, Maria Domenica Moccia, Simon Grüter, Timothy Sykes, Lennart Opitz, Griffin White, Laura Neff, Doris Popovic, Andrea Patrignani, Jay Tracy, Ralph Schlapbach, Christiane Beckmann, Maurice Redondo, Olivier Kobel, Christoph Noppen, Niko Beerenwinkel, Tanja Stadler |
| EPI_ISL_3465726 | Sydney South West Pathology Service (SSWPS) - Liverpool Hospital - NSW Health Pathology | NSW Health Pathology - Institute of Clinical Pathology and Medical Research; Westmead Hospital; University of Sydney | Arnott A., Draper J., Gall M., Martinez E., Rockett R., Sintchenko V., on behalf of ICPMR |
| EPI_ISL_3465727, EPI_ISL_3465728 | Viollier AG | Department of Biosystems Science and Engineering, ETH Zürich | Chaoran Chen, Sarah Nadeau, Louis du Plessis, Catharine Aquino, Ivan Topolsky, Kim Philipp Jablonski, Lara Fuhrmann, Daniel Ehrsam, Isabel Stürmer, Andreia Cabral de Gouvea, Maria Domenica Moccia, Simon Grüter, Timothy Sykes, Lennart Opitz, Griffin White, Laura Neff, Doris Popovic, Andrea Patrignani, Jay Tracy, Ralph Schlapbach, Christiane Beckmann, Maurice Redondo, Olivier Kobel, Christoph Noppen, Niko Beerenwinkel, Tanja Stadler |
| EPI_ISL_3465729 | Sydney South West Pathology Service (SSWPS) - Liverpool Hospital - NSW Health Pathology | NSW Health Pathology - Institute of Clinical Pathology and Medical Research; Westmead Hospital; University of Sydney | Arnott A., Draper J., Gall M., Martinez E., Rockett R., Sintchenko V., on behalf of ICPMR |
| EPI_ISL_3465730, EPI_ISL_3465731, EPI_ISL_3465732 | Viollier AG | Department of Biosystems Science and Engineering, ETH Zürich | Chaoran Chen, Sarah Nadeau, Louis du Plessis, Catharine Aquino, Ivan Topolsky, Kim Philipp Jablonski, Lara Fuhrmann, Daniel Ehrsam, Isabel Stürmer, Andreia Cabral de Gouvea, Maria Domenica Moccia, Simon Grüter, Timothy Sykes, Lennart Opitz, Griffin White, Laura Neff, Doris Popovic, Andrea Patrignani, Jay Tracy, Ralph Schlapbach, Christiane Beckmann, Maurice Redondo, Olivier Kobel, Christoph Noppen, Niko Beerenwinkel, Tanja Stadler |
| EPI_ISL_3465733 | Pathology North - Royal North Shore Hospital - NSW Health Pathology | NSW Health Pathology - Institute of Clinical Pathology and Medical Research; Westmead Hospital; University of Sydney | Arnott A., Draper J., Gall M., Martinez E., Rockett R., Sintchenko V., on behalf of ICPMR |
| EPI_ISL_3465734 | Viollier AG | Department of Biosystems Science and Engineering, ETH Zürich | Andrea Patrizia Salzmann, Henriette Kurth, Christiane Beckmann, Ivan Topolsky, Chaoran Chen, Sarah Nadeau, Louis du Plessis, Kim Philipp Jablonski, Lara Fuhrmann, Niko Beerenwinkel, Christoph Noppen, Olivier Kobel, Maurice Redondo, Tanja Stadler |
| EPI_ISL_3465735 | Pathology North - Royal North Shore Hospital - NSW Health Pathology | NSW Health Pathology - Institute of Clinical Pathology and Medical Research; Westmead Hospital; University of Sydney | Arnott A., Draper J., Gall M., Martinez E., Rockett R., Sintchenko V., on behalf of ICPMR |
| EPI_ISL_3465736 | Viollier AG | Department of Biosystems Science and Engineering, ETH Zürich | Chaoran Chen, Sarah Nadeau, Louis du Plessis, Catharine Aquino, Ivan Topolsky, Kim Philipp Jablonski, Lara Fuhrmann, Daniel Ehrsam, Isabel Stürmer, Andreia Cabral de Gouvea, Maria Domenica Moccia, Simon Grüter, Timothy Sykes, Lennart Opitz, Griffin White, Laura Neff, Doris Popovic, Andrea Patrignani, Jay Tracy, Ralph Schlapbach, Christiane Beckmann, Maurice Redondo, Olivier Kobel, Christoph Noppen, Niko Beerenwinkel, Tanja Stadler |
| EPI_ISL_3465737 | St Vincent's Pathology (SydPath) | NSW Health Pathology - Institute of Clinical Pathology and Medical Research; Westmead Hospital; University of Sydney | Arnott A., Draper J., Gall M., Martinez E., Rockett R., Sintchenko V., on behalf of ICPMR |
| EPI_ISL_3465738, EPI_ISL_3465739 | Viollier AG | Department of Biosystems Science and Engineering, ETH Zürich | Chaoran Chen, Sarah Nadeau, Louis du Plessis, Catharine Aquino, Ivan Topolsky, Kim Philipp Jablonski, Lara Fuhrmann, Daniel Ehrsam, Isabel Stürmer, Andreia Cabral de Gouvea, Maria Domenica Moccia, Simon Grüter, Timothy Sykes, Lennart Opitz, Griffin White, Laura Neff, Doris Popovic, Andrea Patrignani, Jay Tracy, Ralph Schlapbach, Christiane Beckmann, Maurice Redondo, Olivier Kobel, Christoph Noppen, Niko Beerenwinkel, Tanja Stadler |
| EPI_ISL_3465740 | St Vincent's Pathology (SydPath) | NSW Health Pathology - Institute of Clinical Pathology and Medical Research; Westmead Hospital; University of Sydney | Arnott A., Draper J., Gall M., Martinez E., Rockett R., Sintchenko V., on behalf of ICPMR |
| EPI_ISL_3465741 | Viollier AG | Department of Biosystems Science and Engineering, ETH Zürich | Chaoran Chen, Sarah Nadeau, Louis du Plessis, Catharine Aquino, Ivan Topolsky, Kim Philipp Jablonski, Lara Fuhrmann, Daniel Ehrsam, Isabel Stürmer, Andreia Cabral de Gouvea, Maria Domenica Moccia, Simon Grüter, Timothy Sykes, Lennart Opitz, Griffin White, Laura Neff, Doris Popovic, Andrea Patrignani, Jay Tracy, Ralph Schlapbach, Christiane Beckmann, Maurice Redondo, Olivier Kobel, Christoph Noppen, Niko Beerenwinkel, Tanja Stadler |
| EPI_ISL_3465742 | St Vincent's Pathology (SydPath) | NSW Health Pathology - Institute of Clinical Pathology and Medical Research; Westmead Hospital; University of Sydney | Arnott A., Draper J., Gall M., Martinez E., Rockett R., Sintchenko V., on behalf of ICPMR |
| EPI_ISL_3465743, EPI_ISL_3465744 | Viollier AG | Department of Biosystems Science and Engineering, ETH Zürich | Chaoran Chen, Sarah Nadeau, Louis du Plessis, Catharine Aquino, Ivan Topolsky, Kim Philipp Jablonski, Lara Fuhrmann, Daniel Ehrsam, Isabel Stürmer, Andreia Cabral de Gouvea, Maria Domenica Moccia, Simon Grüter, Timothy Sykes, Lennart Opitz, Griffin White, Laura Neff, Doris Popovic, Andrea Patrignani, Jay Tracy, Ralph Schlapbach, Christiane Beckmann, Maurice Redondo, Olivier Kobel, Christoph Noppen, Niko Beerenwinkel, Tanja Stadler |
| EPI_ISL_3465745 | St Vincent's Pathology (SydPath) | NSW Health Pathology - Institute of Clinical Pathology and Medical Research; Westmead Hospital; University of Sydney | Arnott A., Draper J., Gall M., Martinez E., Rockett R., Sintchenko V., on behalf of ICPMR |
| EPI_ISL_3465746 | Viollier AG | Department of Biosystems Science and Engineering, ETH Zürich | Chaoran Chen, Sarah Nadeau, Louis du Plessis, Catharine Aquino, Ivan Topolsky, Kim Philipp Jablonski, Lara Fuhrmann, Daniel Ehrsam, Isabel Stürmer, Andreia Cabral de Gouvea, Maria Domenica Moccia, Simon Grüter, Timothy Sykes, Lennart Opitz, Griffin White, Laura Neff, Doris Popovic, Andrea Patrignani, Jay Tracy, Ralph Schlapbach, Christiane Beckmann, Maurice Redondo, Olivier Kobel, Christoph Noppen, Niko Beerenwinkel, Tanja Stadler |
| EPI_ISL_3465747 | St Vincent's Pathology (SydPath) | NSW Health Pathology - Institute of Clinical Pathology and Medical Research; Westmead Hospital; University of Sydney | Arnott A., Draper J., Gall M., Martinez E., Rockett R., Sintchenko V., on behalf of ICPMR |
| EPI_ISL_3465749 | Viollier AG | Department of Biosystems Science and Engineering, ETH Zürich | Chaoran Chen, Sarah Nadeau, Louis du Plessis, Catharine Aquino, Ivan Topolsky, Kim Philipp Jablonski, Lara Fuhrmann, Daniel Ehrsam, Isabel Stürmer, Andreia Cabral de Gouvea, Maria Domenica Moccia, Simon Grüter, Timothy Sykes, Lennart Opitz, Griffin White, Laura Neff, Doris Popovic, Andrea Patrignani, Jay Tracy, Ralph Schlapbach, Christiane Beckmann, Maurice Redondo, Olivier Kobel, Christoph Noppen, Niko Beerenwinkel, Tanja Stadler |
| EPI_ISL_3465750 | St Vincent's Pathology (SydPath) | NSW Health Pathology - Institute of Clinical Pathology and Medical Research; Westmead Hospital; University of Sydney | Arnott A., Draper J., Gall M., Martinez E., Rockett R., Sintchenko V., on behalf of ICPMR |
| EPI_ISL_3465751 | Viollier AG | Department of Biosystems Science and Engineering, ETH Zürich | Chaoran Chen, Sarah Nadeau, Louis du Plessis, Catharine Aquino, Ivan Topolsky, Kim Philipp Jablonski, Lara Fuhrmann, Daniel Ehrsam, Isabel Stürmer, Andreia Cabral de Gouvea, Maria Domenica Moccia, Simon Grüter, Timothy Sykes, Lennart Opitz, Griffin White, Laura Neff, Doris Popovic, Andrea Patrignani, Jay Tracy, Ralph Schlapbach, Christiane Beckmann, Maurice Redondo, Olivier Kobel, Christoph Noppen, Niko Beerenwinkel, Tanja Stadler |

| | | | |
|---|---|---|---|
| EPI_ISL_3465752 | St Vincent's Pathology (SydPath) | NSW Health Pathology - Institute of Clinical Pathology and Medical Research; Westmead Hospital; University of Sydney | Arnott A., Draper J., Gall M., Martinez E., Rockett R., Sintchenko V., on behalf of ICPMR |
| EPI_ISL_3465753 | Viollier AG | Department of Biosystems Science and Engineering, ETH Zürich | Chaoran Chen, Sarah Nadeau, Louis du Plessis, Catharine Aquino, Ivan Topolsky, Kim Philipp Jablonski, Lara Fuhrmann, Daniel Ehrsam, Isabel Stürmer, Andreia Cabral de Gouvea, Maria Domenica Moccia, Simon Grüter, Timothy Sykes, Lennart Opitz, Griffin White, Laura Neff, Doris Popovic, Andrea Patrignani, Jay Tracy, Ralph Schlapbach, Christiane Beckmann, Maurice Redondo, Olivier Kobel, Christoph Noppen, Niko Beerenwinkel, Tanja Stadler |
| EPI_ISL_3465755 | St Vincent's Pathology (SydPath) | NSW Health Pathology - Institute of Clinical Pathology and Medical Research; Westmead Hospital; University of Sydney | Arnott A., Draper J., Gall M., Martinez E., Rockett R., Sintchenko V., on behalf of ICPMR |
| EPI_ISL_3465756 | Viollier AG | Department of Biosystems Science and Engineering, ETH Zürich | Chaoran Chen, Sarah Nadeau, Louis du Plessis, Catharine Aquino, Ivan Topolsky, Kim Philipp Jablonski, Lara Fuhrmann, Daniel Ehrsam, Isabel Stürmer, Andreia Cabral de Gouvea, Maria Domenica Moccia, Simon Grüter, Timothy Sykes, Lennart Opitz, Griffin White, Laura Neff, Doris Popovic, Andrea Patrignani, Jay Tracy, Ralph Schlapbach, Christiane Beckmann, Maurice Redondo, Olivier Kobel, Christoph Noppen, Niko Beerenwinkel, Tanja Stadler |
| EPI_ISL_3465757 | St Vincent's Pathology (SydPath) | NSW Health Pathology - Institute of Clinical Pathology and Medical Research; Westmead Hospital; University of Sydney | Arnott A., Draper J., Gall M., Martinez E., Rockett R., Sintchenko V., on behalf of ICPMR |
| EPI_ISL_3465758, EPI_ISL_3465759 | Viollier AG | Department of Biosystems Science and Engineering, ETH Zürich | Andrea Patrizia Salzmann, Henriette Kurth, Christiane Beckmann, Ivan Topolsky, Chaoran Chen, Sarah Nadeau, Louis du Plessis, Kim Philipp Jablonski, Lara Fuhrmann, Niko Beerenwinkel, Christoph Noppen, Olivier Kobel, Maurice Redondo, Tanja Stadler |
| EPI_ISL_3465760 | St Vincent's Pathology (SydPath) | NSW Health Pathology - Institute of Clinical Pathology and Medical Research; Westmead Hospital; University of Sydney | Arnott A., Draper J., Gall M., Martinez E., Rockett R., Sintchenko V., on behalf of ICPMR |
| EPI_ISL_3465761 | Viollier AG | Department of Biosystems Science and Engineering, ETH Zürich | Chaoran Chen, Sarah Nadeau, Louis du Plessis, Catharine Aquino, Ivan Topolsky, Kim Philipp Jablonski, Lara Fuhrmann, Daniel Ehrsam, Isabel Stürmer, Andreia Cabral de Gouvea, Maria Domenica Moccia, Simon Grüter, Timothy Sykes, Lennart Opitz, Griffin White, Laura Neff, Doris Popovic, Andrea Patrignani, Jay Tracy, Ralph Schlapbach, Christiane Beckmann, Maurice Redondo, Olivier Kobel, Christoph Noppen, Niko Beerenwinkel, Tanja Stadler |
| EPI_ISL_3465762 | Viollier AG | Department of Biosystems Science and Engineering, ETH Zürich | Andrea Patrizia Salzmann, Henriette Kurth, Christiane Beckmann, Ivan Topolsky, Chaoran Chen, Sarah Nadeau, Louis du Plessis, Kim Philipp Jablonski, Lara Fuhrmann, Niko Beerenwinkel, Christoph Noppen, Olivier Kobel, Maurice Redondo, Tanja Stadler |
| EPI_ISL_3465763 | St Vincent's Pathology (SydPath) | NSW Health Pathology - Institute of Clinical Pathology and Medical Research; Westmead Hospital; University of Sydney | Arnott A., Draper J., Gall M., Martinez E., Rockett R., Sintchenko V., on behalf of ICPMR |
| EPI_ISL_3465764 | Viollier AG | Department of Biosystems Science and Engineering, ETH Zürich | Chaoran Chen, Sarah Nadeau, Louis du Plessis, Catharine Aquino, Ivan Topolsky, Kim Philipp Jablonski, Lara Fuhrmann, Daniel Ehrsam, Isabel Stürmer, Andreia Cabral de Gouvea, Maria Domenica Moccia, Simon Grüter, Timothy Sykes, Lennart Opitz, Griffin White, Laura Neff, Doris Popovic, Andrea Patrignani, Jay Tracy, Ralph Schlapbach, Christiane Beckmann, Maurice Redondo, Olivier Kobel, Christoph Noppen, Niko Beerenwinkel, Tanja Stadler |
| EPI_ISL_3465765 | St Vincent's Pathology (SydPath) | NSW Health Pathology - Institute of Clinical Pathology and Medical Research; Westmead Hospital; University of Sydney | Arnott A., Draper J., Gall M., Martinez E., Rockett R., Sintchenko V., on behalf of ICPMR |
| EPI_ISL_3465766 | Viollier AG | Department of Biosystems Science and Engineering, ETH Zürich | Chaoran Chen, Sarah Nadeau, Louis du Plessis, Catharine Aquino, Ivan Topolsky, Kim Philipp Jablonski, Lara Fuhrmann, Daniel Ehrsam, Isabel Stürmer, Andreia Cabral de Gouvea, Maria Domenica Moccia, Simon Grüter, Timothy Sykes, Lennart Opitz, Griffin White, Laura Neff, Doris Popovic, Andrea Patrignani, Jay Tracy, Ralph Schlapbach, Christiane Beckmann, Maurice Redondo, Olivier Kobel, Christoph Noppen, Niko Beerenwinkel, Tanja Stadler |
| EPI_ISL_3465767 | St Vincent's Pathology (SydPath) | NSW Health Pathology - Institute of Clinical Pathology and Medical Research; Westmead Hospital; University of Sydney | Arnott A., Draper J., Gall M., Martinez E., Rockett R., Sintchenko V., on behalf of ICPMR |
| EPI_ISL_3465768 | Viollier AG | Department of Biosystems Science and Engineering, ETH Zürich | Chaoran Chen, Sarah Nadeau, Louis du Plessis, Catharine Aquino, Ivan Topolsky, Kim Philipp Jablonski, Lara Fuhrmann, Daniel Ehrsam, Isabel Stürmer, Andreia Cabral de Gouvea, Maria Domenica Moccia, Simon Grüter, Timothy Sykes, Lennart Opitz, Griffin White, Laura Neff, Doris Popovic, Andrea Patrignani, Jay Tracy, Ralph Schlapbach, Christiane Beckmann, Maurice Redondo, Olivier Kobel, Christoph Noppen, Niko Beerenwinkel, Tanja Stadler |
| EPI_ISL_3465769 | St Vincent's Pathology (SydPath) | NSW Health Pathology - Institute of Clinical Pathology and Medical Research; Westmead Hospital; University of Sydney | Arnott A., Draper J., Gall M., Martinez E., Rockett R., Sintchenko V., on behalf of ICPMR |
| EPI_ISL_3465770 | Viollier AG | Department of Biosystems Science and Engineering, ETH Zürich | Chaoran Chen, Sarah Nadeau, Louis du Plessis, Catharine Aquino, Ivan Topolsky, Kim Philipp Jablonski, Lara Fuhrmann, Daniel Ehrsam, Isabel Stürmer, Andreia Cabral de Gouvea, Maria Domenica Moccia, Simon Grüter, Timothy Sykes, Lennart Opitz, Griffin White, Laura Neff, Doris Popovic, Andrea Patrignani, Jay Tracy, Ralph Schlapbach, Christiane Beckmann, Maurice Redondo, Olivier Kobel, Christoph Noppen, Niko Beerenwinkel, Tanja Stadler |
| EPI_ISL_3465771 | St Vincent's Pathology (SydPath) | NSW Health Pathology - Institute of Clinical Pathology and Medical Research; Westmead Hospital; University of Sydney | Arnott A., Draper J., Gall M., Martinez E., Rockett R., Sintchenko V., on behalf of ICPMR |
| EPI_ISL_3465772 | Viollier AG | Department of Biosystems Science and Engineering, ETH Zürich | Chaoran Chen, Sarah Nadeau, Louis du Plessis, Catharine Aquino, Ivan Topolsky, Kim Philipp Jablonski, Lara Fuhrmann, Daniel Ehrsam, Isabel Stürmer, Andreia Cabral de Gouvea, Maria Domenica Moccia, Simon Grüter, Timothy Sykes, Lennart Opitz, Griffin White, Laura Neff, Doris Popovic, Andrea Patrignani, Jay Tracy, Ralph Schlapbach, Christiane Beckmann, Maurice Redondo, Olivier Kobel, Christoph Noppen, Niko Beerenwinkel, Tanja Stadler |
| EPI_ISL_3465773 | Viollier AG | Department of Biosystems Science and Engineering, ETH Zürich | Andrea Patrizia Salzmann, Henriette Kurth, Christiane Beckmann, Ivan Topolsky, Chaoran Chen, Sarah Nadeau, Louis du Plessis, Kim Philipp Jablonski, Lara Fuhrmann, Niko Beerenwinkel, Christoph Noppen, Olivier Kobel, Maurice Redondo, Tanja Stadler |
| EPI_ISL_3465774 | St Vincent's Pathology (SydPath) | NSW Health Pathology - Institute of Clinical Pathology and Medical Research; Westmead Hospital; University of Sydney | Arnott A., Draper J., Gall M., Martinez E., Rockett R., Sintchenko V., on behalf of ICPMR |
| EPI_ISL_3465775 | Viollier AG | Department of Biosystems Science and Engineering, ETH Zürich | Chaoran Chen, Sarah Nadeau, Louis du Plessis, Catharine Aquino, Ivan Topolsky, Kim Philipp Jablonski, Lara Fuhrmann, Daniel Ehrsam, Isabel Stürmer, Andreia Cabral de Gouvea, Maria Domenica Moccia, Simon Grüter, Timothy Sykes, Lennart Opitz, Griffin White, Laura Neff, Doris Popovic, Andrea Patrignani, Jay Tracy, Ralph Schlapbach, Christiane Beckmann, Maurice Redondo, Olivier Kobel, Christoph Noppen, Niko Beerenwinkel, Tanja Stadler |
| EPI_ISL_3465776 | St Vincent's Pathology (SydPath) | NSW Health Pathology - Institute of Clinical Pathology and Medical Research; Westmead Hospital; University of Sydney | Arnott A., Draper J., Gall M., Martinez E., Rockett R., Sintchenko V., on behalf of ICPMR |
| EPI_ISL_3465777 | Viollier AG | Department of Biosystems Science and Engineering, ETH Zürich | Chaoran Chen, Sarah Nadeau, Louis du Plessis, Catharine Aquino, Ivan Topolsky, Kim Philipp Jablonski, Lara Fuhrmann, Daniel Ehrsam, Isabel Stürmer, Andreia Cabral de Gouvea, Maria Domenica Moccia, Simon Grüter, Timothy Sykes, Lennart Opitz, Griffin White, Laura Neff, Doris Popovic, Andrea Patrignani, Jay Tracy, Ralph Schlapbach, Christiane Beckmann, Maurice Redondo, Olivier Kobel, Christoph Noppen, Niko Beerenwinkel, Tanja Stadler |
| EPI_ISL_3465778 | St Vincent's Pathology (SydPath) | NSW Health Pathology - Institute of Clinical Pathology and Medical Research; Westmead Hospital; University of Sydney | Arnott A., Draper J., Gall M., Martinez E., Rockett R., Sintchenko V., on behalf of ICPMR |
| EPI_ISL_3465779, EPI_ISL_3465780 | Viollier AG | Department of Biosystems Science and Engineering, ETH Zürich | Chaoran Chen, Sarah Nadeau, Louis du Plessis, Catharine Aquino, Ivan Topolsky, Kim Philipp Jablonski, Lara Fuhrmann, Daniel Ehrsam, Isabel Stürmer, Andreia Cabral de Gouvea, Maria Domenica Moccia, Simon Grüter, Timothy Sykes, Lennart Opitz, Griffin White, Laura Neff, Doris Popovic, Andrea Patrignani, Jay Tracy, Ralph Schlapbach, Christiane Beckmann, Maurice Redondo, Olivier Kobel, Christoph Noppen, Niko Beerenwinkel, Tanja Stadler |
| EPI_ISL_3465781 | Histopath | NSW Health Pathology - Institute of Clinical Pathology and Medical Research; Westmead Hospital; University of Sydney | Arnott A., Draper J., Gall M., Martinez E., Rockett R., Sintchenko V., on behalf of ICPMR |
| EPI_ISL_3465782 | Viollier AG | Department of Biosystems Science and Engineering, ETH Zürich | Chaoran Chen, Sarah Nadeau, Louis du Plessis, Catharine Aquino, Ivan Topolsky, Kim Philipp Jablonski, Lara Fuhrmann, Daniel Ehrsam, Isabel Stürmer, Andreia Cabral de Gouvea, Maria Domenica Moccia, Simon Grüter, Timothy Sykes, Lennart Opitz, Griffin White, Laura Neff, Doris Popovic, Andrea Patrignani, Jay Tracy, Ralph Schlapbach, Christiane Beckmann, Maurice Redondo, Olivier Kobel, Christoph Noppen, Niko Beerenwinkel, Tanja Stadler |
| EPI_ISL_3465783 | Medlab Pathology | NSW Health Pathology - Institute of Clinical Pathology and Medical Research; Westmead Hospital; University of Sydney | Arnott A., Draper J., Gall M., Martinez E., Rockett R., Sintchenko V., on behalf of ICPMR |
| EPI_ISL_3465784 | Viollier AG | Department of Biosystems Science and Engineering, ETH Zürich | Chaoran Chen, Sarah Nadeau, Louis du Plessis, Catharine Aquino, Ivan Topolsky, Kim Philipp Jablonski, Lara Fuhrmann, Daniel Ehrsam, Isabel Stürmer, Andreia Cabral de Gouvea, Maria Domenica Moccia, Simon Grüter, Timothy Sykes, Lennart Opitz, Griffin White, Laura Neff, Doris Popovic, Andrea Patrignani, Jay Tracy, Ralph Schlapbach, Christiane Beckmann, Maurice Redondo, Olivier Kobel, Christoph Noppen, Niko Beerenwinkel, Tanja Stadler |
| EPI_ISL_3465785 | Australian Clinical Labs (formerly Healthscope Pathology) | NSW Health Pathology - Institute of Clinical Pathology and Medical Research; Westmead Hospital; University of Sydney | Arnott A., Draper J., Gall M., Martinez E., Rockett R., Sintchenko V., on behalf of ICPMR |
| EPI_ISL_3465786 | Viollier AG | Department of Biosystems Science and Engineering, ETH Zürich | Chaoran Chen, Sarah Nadeau, Louis du Plessis, Catharine Aquino, Ivan Topolsky, Kim Philipp Jablonski, Lara Fuhrmann, Daniel Ehrsam, Isabel Stürmer, Andreia Cabral de Gouvea, Maria Domenica Moccia, Simon Grüter, Timothy Sykes, Lennart Opitz, Griffin White, Laura Neff, Doris Popovic, Andrea Patrignani, Jay Tracy, Ralph Schlapbach, Christiane Beckmann, Maurice Redondo, Olivier Kobel, Christoph Noppen, Niko Beerenwinkel, Tanja Stadler |
| EPI_ISL_3465787 | Viollier AG | Department of Biosystems Science and Engineering, ETH Zürich | Andrea Patrizia Salzmann, Henriette Kurth, Christiane Beckmann, Ivan Topolsky, Chaoran Chen, Sarah Nadeau, Louis du Plessis, Kim Philipp Jablonski, Lara Fuhrmann, Niko Beerenwinkel, Christoph Noppen, Olivier Kobel, Maurice Redondo, Tanja Stadler |
| EPI_ISL_3465788 | Australian Clinical Labs (formerly Healthscope Pathology) | NSW Health Pathology - Institute of Clinical Pathology and Medical Research; Westmead Hospital; University of Sydney | Arnott A., Draper J., Gall M., Martinez E., Rockett R., Sintchenko V., on behalf of ICPMR |
| EPI_ISL_3465789 | Viollier AG | Department of Biosystems Science and Engineering, ETH Zürich | Andrea Patrizia Salzmann, Henriette Kurth, Christiane Beckmann, Ivan Topolsky, Chaoran Chen, Sarah Nadeau, Louis du Plessis, Kim Philipp Jablonski, Lara Fuhrmann, Niko Beerenwinkel, Christoph Noppen, Olivier Kobel, Maurice Redondo, Tanja Stadler |
| EPI_ISL_3465790 | Sydney South West Pathology Service (SSWPS) - Liverpool Hospital - NSW Health Pathology | NSW Health Pathology - Institute of Clinical Pathology and Medical Research; Westmead Hospital; University of Sydney | Arnott A., Draper J., Gall M., Martinez E., Rockett R., Sintchenko V., on behalf of ICPMR |
| EPI_ISL_3465791 | Viollier AG | Department of Biosystems Science and Engineering, ETH Zürich | Chaoran Chen, Sarah Nadeau, Louis du Plessis, Catharine Aquino, Ivan Topolsky, Kim Philipp Jablonski, Lara Fuhrmann, Daniel Ehrsam, Isabel Stürmer, Andreia Cabral de Gouvea, Maria Domenica Moccia, Simon Grüter, Timothy Sykes, Lennart Opitz, Griffin White, Laura Neff, Doris Popovic, Andrea Patrignani, Jay Tracy, Ralph Schlapbach, Christiane Beckmann, Maurice Redondo, Olivier Kobel, Christoph Noppen, Niko Beerenwinkel, Tanja Stadler |
| EPI_ISL_3465792 | Sydney South West Pathology Service (SSWPS) - Liverpool Hospital - NSW Health Pathology | NSW Health Pathology - Institute of Clinical Pathology and Medical Research; Westmead Hospital; University of Sydney | Arnott A., Draper J., Gall M., Martinez E., Rockett R., Sintchenko V., on behalf of ICPMR |
| EPI_ISL_3465793 | Viollier AG | Department of Biosystems Science and Engineering, ETH Zürich | Chaoran Chen, Sarah Nadeau, Louis du Plessis, Catharine Aquino, Ivan Topolsky, Kim Philipp Jablonski, Lara Fuhrmann, Daniel Ehrsam, Isabel Stürmer, Andreia Cabral de Gouvea, Maria Domenica Moccia, Simon Grüter, Timothy Sykes, Lennart Opitz, Griffin White, Laura Neff, Doris Popovic, Andrea Patrignani, Jay Tracy, Ralph Schlapbach, Christiane Beckmann, Maurice Redondo, Olivier Kobel, Christoph Noppen, Niko Beerenwinkel, Tanja Stadler |
| EPI_ISL_3465794 | Histopath | NSW Health Pathology - Institute of Clinical Pathology and Medical Research; Westmead Hospital; University of Sydney | Arnott A., Draper J., Gall M., Martinez E., Rockett R., Sintchenko V., on behalf of ICPMR |
| EPI_ISL_3465795 | Viollier AG | Department of Biosystems Science and Engineering, ETH Zürich | Chaoran Chen, Sarah Nadeau, Louis du Plessis, Catharine Aquino, Ivan Topolsky, Kim Philipp Jablonski, Lara Fuhrmann, Daniel Ehrsam, Isabel Stürmer, Andreia Cabral de Gouvea, Maria Domenica Moccia, Simon Grüter, Timothy Sykes, Lennart Opitz, Griffin White, Laura Neff, Doris Popovic, Andrea Patrignani, Jay Tracy, Ralph Schlapbach, Christiane Beckmann, Maurice Redondo, Olivier Kobel, Christoph Noppen, Niko Beerenwinkel, Tanja Stadler |
| EPI_ISL_3465796 | Viollier AG | Department of Biosystems Science and Engineering, ETH Zürich | Andrea Patrizia Salzmann, Henriette Kurth, Christiane Beckmann, Ivan Topolsky, Chaoran Chen, Sarah Nadeau, Louis du Plessis, Kim Philipp Jablonski, Lara Fuhrmann, Niko Beerenwinkel, Christoph Noppen, Olivier Kobel, Maurice Redondo, Tanja Stadler |
| EPI_ISL_3465797 | Histopath | NSW Health Pathology - Institute of Clinical Pathology and Medical Research; Westmead Hospital; University of Sydney | Arnott A., Draper J., Gall M., Martinez E., Rockett R., Sintchenko V., on behalf of ICPMR |
| EPI_ISL_3465798 | Viollier AG | Department of Biosystems Science and Engineering, ETH Zürich | Chaoran Chen, Sarah Nadeau, Louis du Plessis, Catharine Aquino, Ivan Topolsky, Kim Philipp Jablonski, Lara Fuhrmann, Daniel Ehrsam, Isabel Stürmer, Andreia Cabral de Gouvea, Maria Domenica Moccia, Simon Grüter, Timothy Sykes, Lennart Opitz, Griffin White, Laura Neff, Doris Popovic, Andrea Patrignani, Jay Tracy, Ralph Schlapbach, Christiane Beckmann, Maurice Redondo, Olivier Kobel, Christoph Noppen, Niko Beerenwinkel, Tanja Stadler |
| EPI_ISL_3465799 | Histopath | NSW Health Pathology - Institute of Clinical Pathology and Medical Research; Westmead Hospital; University of Sydney | Arnott A., Draper J., Gall M., Martinez E., Rockett R., Sintchenko V., on behalf of ICPMR |

| | | | |
|---|---|---|---|
| EPI_ISL_3465800 | Viollier AG | Department of Biosystems Science and Engineering, ETH Zürich | Chaoran Chen, Sarah Nadeau, Louis du Plessis, Catharine Aquino, Ivan Topolsky, Kim Philipp Jablonski, Lara Fuhrmann, Daniel Ehrsam, Isabel Stürmer, Andreia Cabral de Gouvea, Maria Domenica Moccia, Simon Grüter, Timothy Sykes, Lennart Opitz, Griffin White, Laura Neff, Doris Popovic, Andrea Patrignani, Jay Tracy, Ralph Schlapbach, Christiane Beckmann, Maurice Redondo, Olivier Kobel, Christoph Noppen, Niko Beerenwinkel, Tanja Stadler |
| EPI_ISL_3465801 | Laverty Pathology | NSW Health Pathology - Institute of Clinical Pathology and Medical Research; Westmead Hospital; University of Sydney | Arnott A., Draper J., Gall M., Martinez E., Rockett R., Sintchenko V., on behalf of ICPMR |
| EPI_ISL_3465802 | Viollier AG | Department of Biosystems Science and Engineering, ETH Zürich | Chaoran Chen, Sarah Nadeau, Louis du Plessis, Catharine Aquino, Ivan Topolsky, Kim Philipp Jablonski, Lara Fuhrmann, Daniel Ehrsam, Isabel Stürmer, Andreia Cabral de Gouvea, Maria Domenica Moccia, Simon Grüter, Timothy Sykes, Lennart Opitz, Griffin White, Laura Neff, Doris Popovic, Andrea Patrignani, Jay Tracy, Ralph Schlapbach, Christiane Beckmann, Maurice Redondo, Olivier Kobel, Christoph Noppen, Niko Beerenwinkel, Tanja Stadler |
| EPI_ISL_3465803 | Sydney South West Pathology Service (SSWPS) - Liverpool Hospital - NSW Health Pathology | NSW Health Pathology - Institute of Clinical Pathology and Medical Research; Westmead Hospital; University of Sydney | Arnott A., Draper J., Gall M., Martinez E., Rockett R., Sintchenko V., on behalf of ICPMR |
| EPI_ISL_3465804, EPI_ISL_3465805 | Viollier AG | Department of Biosystems Science and Engineering, ETH Zürich | Chaoran Chen, Sarah Nadeau, Louis du Plessis, Catharine Aquino, Ivan Topolsky, Kim Philipp Jablonski, Lara Fuhrmann, Daniel Ehrsam, Isabel Stürmer, Andreia Cabral de Gouvea, Maria Domenica Moccia, Simon Grüter, Timothy Sykes, Lennart Opitz, Griffin White, Laura Neff, Doris Popovic, Andrea Patrignani, Jay Tracy, Ralph Schlapbach, Christiane Beckmann, Maurice Redondo, Olivier Kobel, Christoph Noppen, Niko Beerenwinkel, Tanja Stadler |
| EPI_ISL_3465806 | Sydney South West Pathology Service (SSWPS) - Liverpool Hospital - NSW Health Pathology | NSW Health Pathology - Institute of Clinical Pathology and Medical Research; Westmead Hospital; University of Sydney | Arnott A., Draper J., Gall M., Martinez E., Rockett R., Sintchenko V., on behalf of ICPMR |
| EPI_ISL_3465807 | Viollier AG | Department of Biosystems Science and Engineering, ETH Zürich | Chaoran Chen, Sarah Nadeau, Louis du Plessis, Catharine Aquino, Ivan Topolsky, Kim Philipp Jablonski, Lara Fuhrmann, Daniel Ehrsam, Isabel Stürmer, Andreia Cabral de Gouvea, Maria Domenica Moccia, Simon Grüter, Timothy Sykes, Lennart Opitz, Griffin White, Laura Neff, Doris Popovic, Andrea Patrignani, Jay Tracy, Ralph Schlapbach, Christiane Beckmann, Maurice Redondo, Olivier Kobel, Christoph Noppen, Niko Beerenwinkel, Tanja Stadler |
| EPI_ISL_3465808 | Sydney South West Pathology Service (SSWPS) - Liverpool Hospital - NSW Health Pathology | NSW Health Pathology - Institute of Clinical Pathology and Medical Research; Westmead Hospital; University of Sydney | Arnott A., Draper J., Gall M., Martinez E., Rockett R., Sintchenko V., on behalf of ICPMR |
| EPI_ISL_3465809 | Viollier AG | Department of Biosystems Science and Engineering, ETH Zürich | Chaoran Chen, Sarah Nadeau, Louis du Plessis, Catharine Aquino, Ivan Topolsky, Kim Philipp Jablonski, Lara Fuhrmann, Daniel Ehrsam, Isabel Stürmer, Andreia Cabral de Gouvea, Maria Domenica Moccia, Simon Grüter, Timothy Sykes, Lennart Opitz, Griffin White, Laura Neff, Doris Popovic, Andrea Patrignani, Jay Tracy, Ralph Schlapbach, Christiane Beckmann, Maurice Redondo, Olivier Kobel, Christoph Noppen, Niko Beerenwinkel, Tanja Stadler |
| EPI_ISL_3465810 | Medlab Pathology | NSW Health Pathology - Institute of Clinical Pathology and Medical Research; Westmead Hospital; University of Sydney | Arnott A., Draper J., Gall M., Martinez E., Rockett R., Sintchenko V., on behalf of ICPMR |
| EPI_ISL_3465811 | Viollier AG | Department of Biosystems Science and Engineering, ETH Zürich | Chaoran Chen, Sarah Nadeau, Louis du Plessis, Catharine Aquino, Ivan Topolsky, Kim Philipp Jablonski, Lara Fuhrmann, Daniel Ehrsam, Isabel Stürmer, Andreia Cabral de Gouvea, Maria Domenica Moccia, Simon Grüter, Timothy Sykes, Lennart Opitz, Griffin White, Laura Neff, Doris Popovic, Andrea Patrignani, Jay Tracy, Ralph Schlapbach, Christiane Beckmann, Maurice Redondo, Olivier Kobel, Christoph Noppen, Niko Beerenwinkel, Tanja Stadler |
| EPI_ISL_3465812 | Pathology West - NSW Health Pathology | NSW Health Pathology - Institute of Clinical Pathology and Medical Research; Westmead Hospital; University of Sydney | Arnott A., Draper J., Gall M., Martinez E., Rockett R., Sintchenko V., on behalf of ICPMR |
| EPI_ISL_3465813 | Viollier AG | Department of Biosystems Science and Engineering, ETH Zürich | Andrea Patrizia Salzmann, Henriette Kurth, Christiane Beckmann, Ivan Topolsky, Chaoran Chen, Sarah Nadeau, Louis du Plessis, Kim Philipp Jablonski, Lara Fuhrmann, Niko Beerenwinkel, Christoph Noppen, Olivier Kobel, Maurice Redondo, Tanja Stadler |
| EPI_ISL_3465814 | St Vincent's Pathology (SydPath) | NSW Health Pathology - Institute of Clinical Pathology and Medical Research; Westmead Hospital; University of Sydney | Arnott A., Draper J., Gall M., Martinez E., Rockett R., Sintchenko V., on behalf of ICPMR |
| EPI_ISL_3465816 | Viollier AG | Department of Biosystems Science and Engineering, ETH Zürich | Andrea Patrizia Salzmann, Henriette Kurth, Christiane Beckmann, Ivan Topolsky, Chaoran Chen, Sarah Nadeau, Louis du Plessis, Kim Philipp Jablonski, Lara Fuhrmann, Niko Beerenwinkel, Christoph Noppen, Olivier Kobel, Maurice Redondo, Tanja Stadler |
| EPI_ISL_3465817, EPI_ISL_3465818, EPI_ISL_3465819, EPI_ISL_3465820, EPI_ISL_3465821, EPI_ISL_3465822, EPI_ISL_3465823, EPI_ISL_3465824, EPI_ISL_3465825, EPI_ISL_3465826, EPI_ISL_3465827, EPI_ISL_3465828, EPI_ISL_3465829, EPI_ISL_3465830, EPI_ISL_3465831, EPI_ISL_3465832, EPI_ISL_3465833, EPI_ISL_3465834, EPI_ISL_3465835, EPI_ISL_3465836 | | | |
| see above | Viollier AG | Department of Biosystems Science and Engineering, ETH Zürich | Chaoran Chen, Sarah Nadeau, Louis du Plessis, Catharine Aquino, Ivan Topolsky, Kim Philipp Jablonski, Lara Fuhrmann, Daniel Ehrsam, Isabel Stürmer, Andreia Cabral de Gouvea, Maria Domenica Moccia, Simon Grüter, Timothy Sykes, Lennart Opitz, Griffin White, Laura Neff, Doris Popovic, Andrea Patrignani, Jay Tracy, Ralph Schlapbach, Christiane Beckmann, Maurice Redondo, Olivier Kobel, Christoph Noppen, Niko Beerenwinkel, Tanja Stadler |
| EPI_ISL_3465837 | Viollier AG | Department of Biosystems Science and Engineering, ETH Zürich | Andrea Patrizia Salzmann, Henriette Kurth, Christiane Beckmann, Ivan Topolsky, Chaoran Chen, Sarah Nadeau, Louis du Plessis, Kim Philipp Jablonski, Lara Fuhrmann, Niko Beerenwinkel, Christoph Noppen, Olivier Kobel, Maurice Redondo, Tanja Stadler |
| EPI_ISL_3465838, EPI_ISL_3465839, EPI_ISL_3465840, EPI_ISL_3465841, EPI_ISL_3465842, EPI_ISL_3465843, EPI_ISL_3465844, EPI_ISL_3465845, EPI_ISL_3465846, EPI_ISL_3465847, EPI_ISL_3465848, EPI_ISL_3465849, EPI_ISL_3465850 | | | |
| see above | Viollier AG | Department of Biosystems Science and Engineering, ETH Zürich | Chaoran Chen, Sarah Nadeau, Louis du Plessis, Catharine Aquino, Ivan Topolsky, Kim Philipp Jablonski, Lara Fuhrmann, Daniel Ehrsam, Isabel Stürmer, Andreia Cabral de Gouvea, Maria Domenica Moccia, Simon Grüter, Timothy Sykes, Lennart Opitz, Griffin White, Laura Neff, Doris Popovic, Andrea Patrignani, Jay Tracy, Ralph Schlapbach, Christiane Beckmann, Maurice Redondo, Olivier Kobel, Christoph Noppen, Niko Beerenwinkel, Tanja Stadler |
| EPI_ISL_3465851 | Viollier AG | Department of Biosystems Science and Engineering, ETH Zürich | Andrea Patrizia Salzmann, Henriette Kurth, Christiane Beckmann, Ivan Topolsky, Chaoran Chen, Sarah Nadeau, Louis du Plessis, Kim Philipp Jablonski, Lara Fuhrmann, Niko Beerenwinkel, Christoph Noppen, Olivier Kobel, Maurice Redondo, Tanja Stadler |
| EPI_ISL_3465852 | Viollier AG | Department of Biosystems Science and Engineering, ETH Zürich | Chaoran Chen, Sarah Nadeau, Louis du Plessis, Catharine Aquino, Ivan Topolsky, Kim Philipp Jablonski, Lara Fuhrmann, Daniel Ehrsam, Isabel Stürmer, Andreia Cabral de Gouvea, Maria Domenica Moccia, Simon Grüter, Timothy Sykes, Lennart Opitz, Griffin White, Laura Neff, Doris Popovic, Andrea Patrignani, Jay Tracy, Ralph Schlapbach, Christiane Beckmann, Maurice Redondo, Olivier Kobel, Christoph Noppen, Niko Beerenwinkel, Tanja Stadler |
| EPI_ISL_3465853, EPI_ISL_3465854 | Viollier AG | Department of Biosystems Science and Engineering, ETH Zürich | Andrea Patrizia Salzmann, Henriette Kurth, Christiane Beckmann, Ivan Topolsky, Chaoran Chen, Sarah Nadeau, Louis du Plessis, Kim Philipp Jablonski, Lara Fuhrmann, Niko Beerenwinkel, Christoph Noppen, Olivier Kobel, Maurice Redondo, Tanja Stadler |
| EPI_ISL_3465855, EPI_ISL_3465856, EPI_ISL_3465857, EPI_ISL_3465858, EPI_ISL_3465859, EPI_ISL_3465860, EPI_ISL_3465861, EPI_ISL_3465862, EPI_ISL_3465863, EPI_ISL_3465864, EPI_ISL_3465865, EPI_ISL_3465866, EPI_ISL_3465867, EPI_ISL_3465868, EPI_ISL_3465869, EPI_ISL_3465870, EPI_ISL_3465871 | | | |
| see above | Viollier AG | Department of Biosystems Science and Engineering, ETH Zürich | Chaoran Chen, Sarah Nadeau, Louis du Plessis, Catharine Aquino, Ivan Topolsky, Kim Philipp Jablonski, Lara Fuhrmann, Daniel Ehrsam, Isabel Stürmer, Andreia Cabral de Gouvea, Maria Domenica Moccia, Simon Grüter, Timothy Sykes, Lennart Opitz, Griffin White, Laura Neff, Doris Popovic, Andrea Patrignani, Jay Tracy, Ralph Schlapbach, Christiane Beckmann, Maurice Redondo, Olivier Kobel, Christoph Noppen, Niko Beerenwinkel, Tanja Stadler |
| EPI_ISL_3465872 | Viollier AG | Department of Biosystems Science and Engineering, ETH Zürich | Andrea Patrizia Salzmann, Henriette Kurth, Christiane Beckmann, Ivan Topolsky, Chaoran Chen, Sarah Nadeau, Louis du Plessis, Kim Philipp Jablonski, Lara Fuhrmann, Niko Beerenwinkel, Christoph Noppen, Olivier Kobel, Maurice Redondo, Tanja Stadler |
| EPI_ISL_3465873, EPI_ISL_3465874, EPI_ISL_3465875 | Viollier AG | Department of Biosystems Science and Engineering, ETH Zürich | Chaoran Chen, Sarah Nadeau, Louis du Plessis, Catharine Aquino, Ivan Topolsky, Kim Philipp Jablonski, Lara Fuhrmann, Daniel Ehrsam, Isabel Stürmer, Andreia Cabral de Gouvea, Maria Domenica Moccia, Simon Grüter, Timothy Sykes, Lennart Opitz, Griffin White, Laura Neff, Doris Popovic, Andrea Patrignani, Jay Tracy, Ralph Schlapbach, Christiane Beckmann, Maurice Redondo, Olivier Kobel, Christoph Noppen, Niko Beerenwinkel, Tanja Stadler |
| EPI_ISL_3465876 | Viollier AG | Department of Biosystems Science and Engineering, ETH Zürich | Andrea Patrizia Salzmann, Henriette Kurth, Christiane Beckmann, Ivan Topolsky, Chaoran Chen, Sarah Nadeau, Louis du Plessis, Kim Philipp Jablonski, Lara Fuhrmann, Niko Beerenwinkel, Christoph Noppen, Olivier Kobel, Maurice Redondo, Tanja Stadler |
| EPI_ISL_3465877, EPI_ISL_3465878 | Viollier AG | Department of Biosystems Science and Engineering, ETH Zürich | Chaoran Chen, Sarah Nadeau, Louis du Plessis, Catharine Aquino, Ivan Topolsky, Kim Philipp Jablonski, Lara Fuhrmann, Daniel Ehrsam, Isabel Stürmer, Andreia Cabral de Gouvea, Maria Domenica Moccia, Simon Grüter, Timothy Sykes, Lennart Opitz, Griffin White, Laura Neff, Doris Popovic, Andrea Patrignani, Jay Tracy, Ralph Schlapbach, Christiane Beckmann, Maurice Redondo, Olivier Kobel, Christoph Noppen, Niko Beerenwinkel, Tanja Stadler |
| EPI_ISL_3465879 | Viollier AG | Department of Biosystems Science and Engineering, ETH Zürich | Andrea Patrizia Salzmann, Henriette Kurth, Christiane Beckmann, Ivan Topolsky, Chaoran Chen, Sarah Nadeau, Louis du Plessis, Kim Philipp Jablonski, Lara Fuhrmann, Niko Beerenwinkel, Christoph Noppen, Olivier Kobel, Maurice Redondo, Tanja Stadler |
| EPI_ISL_3465880, EPI_ISL_3465881, EPI_ISL_3465883, EPI_ISL_3465884, EPI_ISL_3465885, EPI_ISL_3465886, EPI_ISL_3465887, EPI_ISL_3465888, EPI_ISL_3465889, EPI_ISL_3465890 | Viollier AG | Department of Biosystems Science and Engineering, ETH Zürich | Chaoran Chen, Sarah Nadeau, Louis du Plessis, Catharine Aquino, Ivan Topolsky, Kim Philipp Jablonski, Lara Fuhrmann, Daniel Ehrsam, Isabel Stürmer, Andreia Cabral de Gouvea, Maria Domenica Moccia, Simon Grüter, Timothy Sykes, Lennart Opitz, Griffin White, Laura Neff, Doris Popovic, Andrea Patrignani, Jay Tracy, Ralph Schlapbach, Christiane Beckmann, Maurice Redondo, Olivier Kobel, Christoph Noppen, Niko Beerenwinkel, Tanja Stadler |
| EPI_ISL_3465891, EPI_ISL_3465892 | Viollier AG | Department of Biosystems Science and Engineering, ETH Zürich | Andrea Patrizia Salzmann, Henriette Kurth, Christiane Beckmann, Ivan Topolsky, Chaoran Chen, Sarah Nadeau, Louis du Plessis, Kim Philipp Jablonski, Lara Fuhrmann, Niko Beerenwinkel, Christoph Noppen, Olivier Kobel, Maurice Redondo, Tanja Stadler |
| EPI_ISL_3465893, EPI_ISL_3465894, EPI_ISL_3465895, EPI_ISL_3465896, EPI_ISL_3465897 | Viollier AG | Department of Biosystems Science and Engineering, ETH Zürich | Chaoran Chen, Sarah Nadeau, Louis du Plessis, Catharine Aquino, Ivan Topolsky, Kim Philipp Jablonski, Lara Fuhrmann, Daniel Ehrsam, Isabel Stürmer, Andreia Cabral de Gouvea, Maria Domenica Moccia, Simon Grüter, Timothy Sykes, Lennart Opitz, Griffin White, Laura Neff, Doris Popovic, Andrea Patrignani, Jay Tracy, Ralph Schlapbach, Christiane Beckmann, Maurice Redondo, Olivier Kobel, Christoph Noppen, Niko Beerenwinkel, Tanja Stadler |
| EPI_ISL_3465898 | Viollier AG | Department of Biosystems Science and Engineering, ETH Zürich | Andrea Patrizia Salzmann, Henriette Kurth, Christiane Beckmann, Ivan Topolsky, Chaoran Chen, Sarah Nadeau, Louis du Plessis, Kim Philipp Jablonski, Lara Fuhrmann, Niko Beerenwinkel, Christoph Noppen, Olivier Kobel, Maurice Redondo, Tanja Stadler |
| EPI_ISL_3465899, EPI_ISL_3465900 | Viollier AG | Department of Biosystems Science and Engineering, ETH Zürich | Chaoran Chen, Sarah Nadeau, Louis du Plessis, Catharine Aquino, Ivan Topolsky, Kim Philipp Jablonski, Lara Fuhrmann, Daniel Ehrsam, Isabel Stürmer, Andreia Cabral de Gouvea, Maria Domenica Moccia, Simon Grüter, Timothy Sykes, Lennart Opitz, Griffin White, Laura Neff, Doris Popovic, Andrea Patrignani, Jay Tracy, Ralph Schlapbach, Christiane Beckmann, Maurice Redondo, Olivier Kobel, Christoph Noppen, Niko Beerenwinkel, Tanja Stadler |
| EPI_ISL_3465901, EPI_ISL_3465902, EPI_ISL_3465903, EPI_ISL_3465904, EPI_ISL_3465905, EPI_ISL_3465906, EPI_ISL_3465907, EPI_ISL_3465908, EPI_ISL_3465909, EPI_ISL_3465910, EPI_ISL_3465911, EPI_ISL_3465912, EPI_ISL_3465913, EPI_ISL_3465914, EPI_ISL_3465915, EPI_ISL_3465916, EPI_ISL_3465917, EPI_ISL_3465918, EPI_ISL_3465919, EPI_ISL_3465920, EPI_ISL_3465921, EPI_ISL_3465922, EPI_ISL_3465923, EPI_ISL_3465924, EPI_ISL_3465925, EPI_ISL_3465926, EPI_ISL_3465927, EPI_ISL_3465928, EPI_ISL_3465929, EPI_ISL_3465930, EPI_ISL_3465931, EPI_ISL_3465932, EPI_ISL_3465933, EPI_ISL_3465934, EPI_ISL_3465935, EPI_ISL_3465936, EPI_ISL_3465937, EPI_ISL_3465938, EPI_ISL_3465939, EPI_ISL_3465940, EPI_ISL_3465941, EPI_ISL_3465942, EPI_ISL_3465943, EPI_ISL_3465944, EPI_ISL_3465945, EPI_ISL_3465946, EPI_ISL_3465947, EPI_ISL_3465948, EPI_ISL_3465949, EPI_ISL_3465950, EPI_ISL_3465951, EPI_ISL_3465952, EPI_ISL_3465953, EPI_ISL_3465954, EPI_ISL_3465955, EPI_ISL_3465956, EPI_ISL_3465957, EPI_ISL_3465958, EPI_ISL_3465959, EPI_ISL_3465960, EPI_ISL_3465961, EPI_ISL_3465962, EPI_ISL_3465963, EPI_ISL_3465964, EPI_ISL_3465965, EPI_ISL_3465966, EPI_ISL_3465967, EPI_ISL_3465968, EPI_ISL_3465969, EPI_ISL_3465970, EPI_ISL_3465971, EPI_ISL_3465972, EPI_ISL_3465973, EPI_ISL_3465974, EPI_ISL_3465975, EPI_ISL_3465976, EPI_ISL_3465977, EPI_ISL_3465978, EPI_ISL_3465979, EPI_ISL_3465980, EPI_ISL_3465981, EPI_ISL_3465982, EPI_ISL_3465983, EPI_ISL_3465984, EPI_ISL_3465985, EPI_ISL_3465986, EPI_ISL_3465987, EPI_ISL_3465988, EPI_ISL_3465989, EPI_ISL_3465990, EPI_ISL_3465991, EPI_ISL_3465992, EPI_ISL_3465993, EPI_ISL_3465994, EPI_ISL_3465995 | | | |
| see above | Lighthouse Lab in Milton Keynes | Wellcome Sanger Institute for the COVID-19 Genomics UK (COG-UK) Consortium | The Lighthouse Lab in Milton Keynes and Alex Alderton, Roberto Amato, Jeffrey Barrett, Sonia Goncalves, Ewan Harrison, David K. Jackson, Ian Johnston, Dominic Kwiatkowski, Cordelia Langford, John Sillitoe on behalf of the Wellcome Sanger Institute COVID-19 Surveillance Team |
| EPI_ISL_3465996 | RS Mitra Keluarga Gading Serpong | Eijkman Institute for Molecular Biology, National Research and Innovation Agency | Edison Johar, Frilasita A Yudhaputri, Muhammad Rezki Rasyak, Willy Agustine, Hidayat Trimarsanto, Lydia V. Panggalo, Iskandar Adnan, Sukma Oktavianthi, Lidwina Priliani, Jessica R. Saragih, Safarina G Malik, Khin Saw Myint, Amin Soebandrio |
| EPI_ISL_3465997, EPI_ISL_3465998, EPI_ISL_3465999, EPI_ISL_3466000, EPI_ISL_3466001, | Lighthouse Lab in Milton Keynes | Wellcome Sanger Institute for the COVID-19 Genomics UK (COG-UK) Consortium | The Lighthouse Lab in Milton Keynes and Alex Alderton, Roberto Amato, Jeffrey Barrett, Sonia Goncalves, Ewan Harrison, David K. Jackson, Ian Johnston, Dominic Kwiatkowski, Cordelia Langford, John Sillitoe on behalf of the Wellcome Sanger Institute COVID-19 Surveillance Team |

| | | | |
|---|---|---|---|
| EPI_ISL_3466002 | | | |
| EPI_ISL_3466003 | RS Pondok Indah Puri Indah | Eijkman Institute for Molecular Biology, National Research and Innovation Agency | Frilasita A Yudhaputri, Muhammad Rezki Rasyak, Willy Agustine, Hidayat Trimarsanto, Lydia V. Panggalo, Iskandar Adnan, Sukma Oktavianthi, Lidwina Priliani, Jessica R. Saragih, Edison Johar, Safarina G Malik, Khin Saw Myint, Amin Soebandrio |
| EPI_ISL_3466004, EPI_ISL_3466005, EPI_ISL_3466006, EPI_ISL_3466007, EPI_ISL_3466008 | Lighthouse Lab in Milton Keynes | Wellcome Sanger Institute for the COVID-19 Genomics UK (COG-UK) Consortium | The Lighthouse Lab in Milton Keynes and Alex Alderton, Roberto Amato, Jeffrey Barrett, Sonia Goncalves, Ewan Harrison, David K. Jackson, Ian Johnston, Dominic Kwiatkowski, Cordelia Langford, John Sillitoe on behalf of the Wellcome Sanger Institute COVID-19 Surveillance Team |
| EPI_ISL_3466009 | RS Pondok Indah Puri Indah | Eijkman Institute for Molecular Biology, National Research and Innovation Agency | Iskandar Adnan, Sukma Oktavianthi, Lidwina Priliani, Jessica R. Saragih, Edison Johar, Frilasita A Yudhaputri, Muhammad Rezki Rasyak, Willy Agustine, Hidayat Trimarsanto, Lydia V. Panggalo, Safarina G Malik, Khin Saw Myint, Amin Soebandrio |
| EPI_ISL_3466010, EPI_ISL_3466011, EPI_ISL_3466012, EPI_ISL_3466013, EPI_ISL_3466014, EPI_ISL_3466015, EPI_ISL_3466017, EPI_ISL_3466018, EPI_ISL_3466019, EPI_ISL_3466020, EPI_ISL_3466021, EPI_ISL_3466022 | | | |
| see above | Lighthouse Lab in Milton Keynes | Wellcome Sanger Institute for the COVID-19 Genomics UK (COG-UK) Consortium | The Lighthouse Lab in Milton Keynes and Alex Alderton, Roberto Amato, Jeffrey Barrett, Sonia Goncalves, Ewan Harrison, David K. Jackson, Ian Johnston, Dominic Kwiatkowski, Cordelia Langford, John Sillitoe on behalf of the Wellcome Sanger Institute COVID-19 Surveillance Team |
| EPI_ISL_3466023 | RS Mayapada Tangerang | Eijkman Institute for Molecular Biology, National Research and Innovation Agency | Edison Johar, Frilasita A Yudhaputri, Muhammad Rezki Rasyak, Willy Agustine, Hidayat Trimarsanto, Lydia V. Panggalo, Iskandar Adnan, Sukma Oktavianthi, Lidwina Priliani, Jessica R. Saragih, Safarina G Malik, Khin Saw Myint, Amin Soebandrio |
| EPI_ISL_3466024, EPI_ISL_3466025, EPI_ISL_3466026, EPI_ISL_3466027, EPI_ISL_3466029, EPI_ISL_3466030, EPI_ISL_3466031, EPI_ISL_3466032, EPI_ISL_3466033, EPI_ISL_3466035, EPI_ISL_3466036, EPI_ISL_3466037, EPI_ISL_3466038, EPI_ISL_3466039, EPI_ISL_3466041, EPI_ISL_3466042, EPI_ISL_3466043, EPI_ISL_3466044, EPI_ISL_3466045, EPI_ISL_3466047, EPI_ISL_3466048, EPI_ISL_3466049, EPI_ISL_3466050, EPI_ISL_3466051, EPI_ISL_3466052, EPI_ISL_3466054, EPI_ISL_3466055, EPI_ISL_3466056, EPI_ISL_3466057, EPI_ISL_3466058, EPI_ISL_3466059, EPI_ISL_3466061, EPI_ISL_3466062, EPI_ISL_3466063, EPI_ISL_3466064, EPI_ISL_3466065, EPI_ISL_3466066, EPI_ISL_3466068, EPI_ISL_3466069, EPI_ISL_3466070, EPI_ISL_3466071, EPI_ISL_3466073, EPI_ISL_3466074, EPI_ISL_3466075, EPI_ISL_3466076, EPI_ISL_3466077, EPI_ISL_3466078, EPI_ISL_3466080, EPI_ISL_3466081, EPI_ISL_3466082, EPI_ISL_3466083, EPI_ISL_3466084, EPI_ISL_3466086, EPI_ISL_3466087, EPI_ISL_3466088, EPI_ISL_3466089, EPI_ISL_3466090, EPI_ISL_3466092, EPI_ISL_3466093, EPI_ISL_3466094, EPI_ISL_3466095, EPI_ISL_3466096, EPI_ISL_3466098, EPI_ISL_3466099, EPI_ISL_3466100, EPI_ISL_3466101, EPI_ISL_3466102, EPI_ISL_3466103, EPI_ISL_3466105, EPI_ISL_3466106, EPI_ISL_3466107, EPI_ISL_3466108, EPI_ISL_3466109, EPI_ISL_3466111, EPI_ISL_3466112, EPI_ISL_3466113, EPI_ISL_3466114, EPI_ISL_3466115, EPI_ISL_3466116, EPI_ISL_3466118, EPI_ISL_3466119, EPI_ISL_3466120, EPI_ISL_3466121, EPI_ISL_3466122, EPI_ISL_3466124, EPI_ISL_3466125, EPI_ISL_3466126, EPI_ISL_3466127, EPI_ISL_3466128, EPI_ISL_3466130, EPI_ISL_3466131, EPI_ISL_3466132, EPI_ISL_3466133, EPI_ISL_3466134, EPI_ISL_3466135, EPI_ISL_3466137, EPI_ISL_3466138, EPI_ISL_3466139, EPI_ISL_3466140, EPI_ISL_3466141, EPI_ISL_3466143, EPI_ISL_3466144, EPI_ISL_3466145, EPI_ISL_3466146, EPI_ISL_3466147, EPI_ISL_3466148, EPI_ISL_3466150, EPI_ISL_3466151, EPI_ISL_3466152, EPI_ISL_3466153, EPI_ISL_3466154, EPI_ISL_3466156, EPI_ISL_3466157, EPI_ISL_3466158, EPI_ISL_3466159, EPI_ISL_3466160, EPI_ISL_3466161, EPI_ISL_3466163, EPI_ISL_3466164, EPI_ISL_3466165, EPI_ISL_3466166, EPI_ISL_3466167, EPI_ISL_3466169, EPI_ISL_3466170, EPI_ISL_3466171, EPI_ISL_3466172, EPI_ISL_3466174, EPI_ISL_3466175, EPI_ISL_3466176, EPI_ISL_3466177, EPI_ISL_3466178, EPI_ISL_3466180, EPI_ISL_3466181, EPI_ISL_3466182, EPI_ISL_3466183, EPI_ISL_3466184, EPI_ISL_3466186, EPI_ISL_3466187, EPI_ISL_3466188, EPI_ISL_3466189, EPI_ISL_3466190, EPI_ISL_3466192, EPI_ISL_3466193, EPI_ISL_3466194, EPI_ISL_3466195, EPI_ISL_3466196, EPI_ISL_3466198, EPI_ISL_3466199, EPI_ISL_3466200, EPI_ISL_3466201, EPI_ISL_3466202, EPI_ISL_3466204, EPI_ISL_3466205, EPI_ISL_3466206, EPI_ISL_3466207, EPI_ISL_3466208, EPI_ISL_3466209, EPI_ISL_3466211, EPI_ISL_3466212, EPI_ISL_3466213, EPI_ISL_3466214, EPI_ISL_3466215, EPI_ISL_3466216, EPI_ISL_3466218, EPI_ISL_3466219, EPI_ISL_3466220, EPI_ISL_3466221, EPI_ISL_3466223, EPI_ISL_3466224, EPI_ISL_3466225, EPI_ISL_3466226, EPI_ISL_3466227, EPI_ISL_3466228, EPI_ISL_3466230, EPI_ISL_3466231, EPI_ISL_3466232, EPI_ISL_3466233, EPI_ISL_3466235, EPI_ISL_3466236, EPI_ISL_3466237, EPI_ISL_3466238, EPI_ISL_3466239, EPI_ISL_3466240, EPI_ISL_3466242, EPI_ISL_3466243, EPI_ISL_3466244, EPI_ISL_3466245, EPI_ISL_3466246, EPI_ISL_3466248, EPI_ISL_3466249, EPI_ISL_3466250 | | | |
| see above | Lighthouse Lab in Milton Keynes | Wellcome Sanger Institute for the COVID-19 Genomics UK (COG-UK) Consortium | The Lighthouse Lab in Milton Keynes and Alex Alderton, Roberto Amato, Jeffrey Barrett, Sonia Goncalves, Ewan Harrison, David K. Jackson, Ian Johnston, Dominic Kwiatkowski, Cordelia Langford, John Sillitoe on behalf of the Wellcome Sanger Institute COVID-19 Surveillance Team |
| EPI_ISL_3466251, EPI_ISL_3466252 | Lighthouse Lab in Glasgow | Wellcome Sanger Institute for the COVID-19 Genomics UK (COG-UK) Consortium | Harper VanSteenhouse, Yumi Kasai, David Gray, Carol Clugston, Anna Dominiczak and Alex Alderton, Roberto Amato, Jeffrey Barrett, Sonia Goncalves, Ewan Harrison, David K. Jackson, Ian Johnston, Dominic Kwiatkowski, Cordelia Langford, John Sillitoe on behalf of the Wellcome Sanger Institute COVID-19 Surveillance Team |
| EPI_ISL_3466254 | Lighthouse Lab in Milton Keynes | Wellcome Sanger Institute for the COVID-19 Genomics UK (COG-UK) Consortium | The Lighthouse Lab in Milton Keynes and Alex Alderton, Roberto Amato, Jeffrey Barrett, Sonia Goncalves, Ewan Harrison, David K. Jackson, Ian Johnston, Dominic Kwiatkowski, Cordelia Langford, John Sillitoe on behalf of the Wellcome Sanger Institute COVID-19 Surveillance Team |
| EPI_ISL_3466255 | Lighthouse Lab in Glasgow | Wellcome Sanger Institute for the COVID-19 Genomics UK (COG-UK) Consortium | Harper VanSteenhouse, Yumi Kasai, David Gray, Carol Clugston, Anna Dominiczak and Alex Alderton, Roberto Amato, Jeffrey Barrett, Sonia Goncalves, Ewan Harrison, David K. Jackson, Ian Johnston, Dominic Kwiatkowski, Cordelia Langford, John Sillitoe on behalf of the Wellcome Sanger Institute COVID-19 Surveillance Team |
| EPI_ISL_3466256 | Lighthouse Lab in Milton Keynes | Wellcome Sanger Institute for the COVID-19 Genomics UK (COG-UK) Consortium | The Lighthouse Lab in Milton Keynes and Alex Alderton, Roberto Amato, Jeffrey Barrett, Sonia Goncalves, Ewan Harrison, David K. Jackson, Ian Johnston, Dominic Kwiatkowski, Cordelia Langford, John Sillitoe on behalf of the Wellcome Sanger Institute COVID-19 Surveillance Team |
| EPI_ISL_3466257 | Lighthouse Lab in Glasgow | Wellcome Sanger Institute for the COVID-19 Genomics UK (COG-UK) Consortium | Harper VanSteenhouse, Yumi Kasai, David Gray, Carol Clugston, Anna Dominiczak and Alex Alderton, Roberto Amato, Jeffrey Barrett, Sonia Goncalves, Ewan Harrison, David K. Jackson, Ian Johnston, Dominic Kwiatkowski, Cordelia Langford, John Sillitoe on behalf of the Wellcome Sanger Institute COVID-19 Surveillance Team |
| EPI_ISL_3466258 | Lighthouse Lab in Milton Keynes | Wellcome Sanger Institute for the COVID-19 Genomics UK (COG-UK) Consortium | The Lighthouse Lab in Milton Keynes and Alex Alderton, Roberto Amato, Jeffrey Barrett, Sonia Goncalves, Ewan Harrison, David K. Jackson, Ian Johnston, Dominic Kwiatkowski, Cordelia Langford, John Sillitoe on behalf of the Wellcome Sanger Institute COVID-19 Surveillance Team |
| EPI_ISL_3466259 | Lighthouse Lab in Glasgow | Wellcome Sanger Institute for the COVID-19 Genomics UK (COG-UK) Consortium | Harper VanSteenhouse, Yumi Kasai, David Gray, Carol Clugston, Anna Dominiczak and Alex Alderton, Roberto Amato, Jeffrey Barrett, Sonia Goncalves, Ewan Harrison, David K. Jackson, Ian Johnston, Dominic Kwiatkowski, Cordelia Langford, John Sillitoe on behalf of the Wellcome Sanger Institute COVID-19 Surveillance Team |
| EPI_ISL_3466261, EPI_ISL_3466262 | Lighthouse Lab in Milton Keynes | Wellcome Sanger Institute for the COVID-19 Genomics UK (COG-UK) Consortium | The Lighthouse Lab in Milton Keynes and Alex Alderton, Roberto Amato, Jeffrey Barrett, Sonia Goncalves, Ewan Harrison, David K. Jackson, Ian Johnston, Dominic Kwiatkowski, Cordelia Langford, John Sillitoe on behalf of the Wellcome Sanger Institute COVID-19 Surveillance Team |
| EPI_ISL_3466263, EPI_ISL_3466264 | Lighthouse Lab in Glasgow | Wellcome Sanger Institute for the COVID-19 Genomics UK (COG-UK) Consortium | Harper VanSteenhouse, Yumi Kasai, David Gray, Carol Clugston, Anna Dominiczak and Alex Alderton, Roberto Amato, Jeffrey Barrett, Sonia Goncalves, Ewan Harrison, David K. Jackson, Ian Johnston, Dominic Kwiatkowski, Cordelia Langford, John Sillitoe on behalf of the Wellcome Sanger Institute COVID-19 Surveillance Team |
| EPI_ISL_3466265 | Lighthouse Lab in Milton Keynes | Wellcome Sanger Institute for the COVID-19 Genomics UK (COG-UK) Consortium | The Lighthouse Lab in Milton Keynes and Alex Alderton, Roberto Amato, Jeffrey Barrett, Sonia Goncalves, Ewan Harrison, David K. Jackson, Ian Johnston, Dominic Kwiatkowski, Cordelia Langford, John Sillitoe on behalf of the Wellcome Sanger Institute COVID-19 Surveillance Team |
| EPI_ISL_3466267 | Lighthouse Lab in Glasgow | Wellcome Sanger Institute for the COVID-19 Genomics UK (COG-UK) Consortium | Harper VanSteenhouse, Yumi Kasai, David Gray, Carol Clugston, Anna Dominiczak and Alex Alderton, Roberto Amato, Jeffrey Barrett, Sonia Goncalves, Ewan Harrison, David K. Jackson, Ian Johnston, Dominic Kwiatkowski, Cordelia Langford, John Sillitoe on behalf of the Wellcome Sanger Institute COVID-19 Surveillance Team |
| EPI_ISL_3466268, EPI_ISL_3466269 | Lighthouse Lab in Milton Keynes | Wellcome Sanger Institute for the COVID-19 Genomics UK (COG-UK) Consortium | The Lighthouse Lab in Milton Keynes and Alex Alderton, Roberto Amato, Jeffrey Barrett, Sonia Goncalves, Ewan Harrison, David K. Jackson, Ian Johnston, Dominic Kwiatkowski, Cordelia Langford, John Sillitoe on behalf of the Wellcome Sanger Institute COVID-19 Surveillance Team |
| EPI_ISL_3466270 | Lighthouse Lab in Glasgow | Wellcome Sanger Institute for the COVID-19 Genomics UK (COG-UK) Consortium | Harper VanSteenhouse, Yumi Kasai, David Gray, Carol Clugston, Anna Dominiczak and Alex Alderton, Roberto Amato, Jeffrey Barrett, Sonia Goncalves, Ewan Harrison, David K. Jackson, Ian Johnston, Dominic Kwiatkowski, Cordelia Langford, John Sillitoe on behalf of the Wellcome Sanger Institute COVID-19 Surveillance Team |
| EPI_ISL_3466271 | Lighthouse Lab in Milton Keynes | Wellcome Sanger Institute for the COVID-19 Genomics UK (COG-UK) Consortium | The Lighthouse Lab in Milton Keynes and Alex Alderton, Roberto Amato, Jeffrey Barrett, Sonia Goncalves, Ewan Harrison, David K. Jackson, Ian Johnston, Dominic Kwiatkowski, Cordelia Langford, John Sillitoe on behalf of the Wellcome Sanger Institute COVID-19 Surveillance Team |
| EPI_ISL_3466273 | Lighthouse Lab in Glasgow | Wellcome Sanger Institute for the COVID-19 Genomics UK (COG-UK) Consortium | Harper VanSteenhouse, Yumi Kasai, David Gray, Carol Clugston, Anna Dominiczak and Alex Alderton, Roberto Amato, Jeffrey Barrett, Sonia Goncalves, Ewan Harrison, David K. Jackson, Ian Johnston, Dominic Kwiatkowski, Cordelia Langford, John Sillitoe on behalf of the Wellcome Sanger Institute COVID-19 Surveillance Team |
| EPI_ISL_3466274, EPI_ISL_3466275, EPI_ISL_3466276, EPI_ISL_3466277, EPI_ISL_3466279, EPI_ISL_3466280, EPI_ISL_3466281 | Lighthouse Lab in Milton Keynes | Wellcome Sanger Institute for the COVID-19 Genomics UK (COG-UK) Consortium | The Lighthouse Lab in Milton Keynes and Alex Alderton, Roberto Amato, Jeffrey Barrett, Sonia Goncalves, Ewan Harrison, David K. Jackson, Ian Johnston, Dominic Kwiatkowski, Cordelia Langford, John Sillitoe on behalf of the Wellcome Sanger Institute COVID-19 Surveillance Team |
| EPI_ISL_3466282, EPI_ISL_3466283 | Lighthouse Lab in Glasgow | Wellcome Sanger Institute for the COVID-19 Genomics UK (COG-UK) Consortium | Harper VanSteenhouse, Yumi Kasai, David Gray, Carol Clugston, Anna Dominiczak and Alex Alderton, Roberto Amato, Jeffrey Barrett, Sonia Goncalves, Ewan Harrison, David K. Jackson, Ian Johnston, Dominic Kwiatkowski, Cordelia Langford, John Sillitoe on behalf of the Wellcome Sanger Institute COVID-19 Surveillance Team |
| EPI_ISL_3466285 | Lighthouse Lab in Milton Keynes | Wellcome Sanger Institute for the COVID-19 Genomics UK (COG-UK) Consortium | The Lighthouse Lab in Milton Keynes and Alex Alderton, Roberto Amato, Jeffrey Barrett, Sonia Goncalves, Ewan Harrison, David K. Jackson, Ian Johnston, Dominic Kwiatkowski, Cordelia Langford, John Sillitoe on behalf of the Wellcome Sanger Institute COVID-19 Surveillance Team |
| EPI_ISL_3466286, EPI_ISL_3466287, EPI_ISL_3466288, EPI_ISL_3466289, EPI_ISL_3466290, EPI_ISL_3466292 | Lighthouse Lab in Glasgow | Wellcome Sanger Institute for the COVID-19 Genomics UK (COG-UK) Consortium | Harper VanSteenhouse, Yumi Kasai, David Gray, Carol Clugston, Anna Dominiczak and Alex Alderton, Roberto Amato, Jeffrey Barrett, Sonia Goncalves, Ewan Harrison, David K. Jackson, Ian Johnston, Dominic Kwiatkowski, Cordelia Langford, John Sillitoe on behalf of the Wellcome Sanger Institute COVID-19 Surveillance Team |
| EPI_ISL_3466293, EPI_ISL_3466294 | Lighthouse Lab in Milton Keynes | Wellcome Sanger Institute for the COVID-19 Genomics UK (COG-UK) Consortium | The Lighthouse Lab in Milton Keynes and Alex Alderton, Roberto Amato, Jeffrey Barrett, Sonia Goncalves, Ewan Harrison, David K. Jackson, Ian Johnston, Dominic Kwiatkowski, Cordelia Langford, John Sillitoe on behalf of the Wellcome Sanger Institute COVID-19 Surveillance Team |
| EPI_ISL_3466295, EPI_ISL_3466296, EPI_ISL_3466298 | Lighthouse Lab in Glasgow | Wellcome Sanger Institute for the COVID-19 Genomics UK (COG-UK) Consortium | Harper VanSteenhouse, Yumi Kasai, David Gray, Carol Clugston, Anna Dominiczak and Alex Alderton, Roberto Amato, Jeffrey Barrett, Sonia Goncalves, Ewan Harrison, David K. Jackson, Ian Johnston, Dominic Kwiatkowski, Cordelia Langford, John Sillitoe on behalf of the Wellcome Sanger Institute COVID-19 Surveillance Team |
| EPI_ISL_3466299 | Lighthouse Lab in Milton Keynes | Wellcome Sanger Institute for the COVID-19 Genomics UK (COG-UK) Consortium | The Lighthouse Lab in Milton Keynes and Alex Alderton, Roberto Amato, Jeffrey Barrett, Sonia Goncalves, Ewan Harrison, David K. Jackson, Ian Johnston, Dominic Kwiatkowski, Cordelia Langford, John Sillitoe on behalf of the Wellcome Sanger Institute COVID-19 Surveillance Team |
| EPI_ISL_3466300, EPI_ISL_3466301, EPI_ISL_3466302, EPI_ISL_3466304 | Lighthouse Lab in Glasgow | Wellcome Sanger Institute for the COVID-19 Genomics UK (COG-UK) Consortium | Harper VanSteenhouse, Yumi Kasai, David Gray, Carol Clugston, Anna Dominiczak and Alex Alderton, Roberto Amato, Jeffrey Barrett, Sonia Goncalves, Ewan Harrison, David K. Jackson, Ian Johnston, Dominic Kwiatkowski, Cordelia Langford, John Sillitoe on behalf of the Wellcome Sanger Institute COVID-19 Surveillance Team |
| EPI_ISL_3466305 | Lighthouse Lab in Milton Keynes | Wellcome Sanger Institute for the COVID-19 Genomics UK (COG-UK) Consortium | The Lighthouse Lab in Milton Keynes and Alex Alderton, Roberto Amato, Jeffrey Barrett, Sonia Goncalves, Ewan Harrison, David K. Jackson, Ian Johnston, Dominic Kwiatkowski, Cordelia Langford, John Sillitoe on behalf of the Wellcome Sanger Institute COVID-19 Surveillance Team |
| EPI_ISL_3466306 | Lighthouse Lab in Glasgow | Wellcome Sanger Institute for the COVID-19 Genomics UK (COG-UK) Consortium | Harper VanSteenhouse, Yumi Kasai, David Gray, Carol Clugston, Anna Dominiczak and Alex Alderton, Roberto Amato, Jeffrey Barrett, Sonia Goncalves, Ewan Harrison, David K. Jackson, Ian Johnston, Dominic Kwiatkowski, Cordelia Langford, John Sillitoe on behalf of the Wellcome Sanger Institute COVID-19 Surveillance Team |
| EPI_ISL_3466307 | Lighthouse Lab in Milton Keynes | Wellcome Sanger Institute for the COVID-19 Genomics UK (COG-UK) Consortium | The Lighthouse Lab in Milton Keynes and Alex Alderton, Roberto Amato, Jeffrey Barrett, Sonia Goncalves, Ewan Harrison, David K. Jackson, Ian Johnston, Dominic Kwiatkowski, Cordelia Langford, John Sillitoe on behalf of the Wellcome Sanger Institute COVID-19 Surveillance Team |
| EPI_ISL_3466308, EPI_ISL_3466309, EPI_ISL_3466311 | Lighthouse Lab in Glasgow | Wellcome Sanger Institute for the COVID-19 Genomics UK (COG-UK) Consortium | Harper VanSteenhouse, Yumi Kasai, David Gray, Carol Clugston, Anna Dominiczak and Alex Alderton, Roberto Amato, Jeffrey Barrett, Sonia Goncalves, Ewan Harrison, David K. Jackson, Ian Johnston, Dominic Kwiatkowski, Cordelia Langford, John Sillitoe on behalf of the Wellcome Sanger Institute COVID-19 Surveillance Team |
| EPI_ISL_3466312 | Lighthouse Lab in Milton Keynes | Wellcome Sanger Institute for the COVID-19 Genomics UK (COG-UK) | The Lighthouse Lab in Milton Keynes and Alex Alderton, Roberto Amato, Jeffrey Barrett, Sonia Goncalves, Ewan Harrison, David K. Jackson, Ian Johnston, Dominic Kwiatkowski, Cordelia Langford, John Sillitoe on behalf of the Wellcome Sanger |

| | | | |
|---|---|---|---|
| | | Consortium | Institute COVID-19 Surveillance Team |
| EPI_ISL_3466313 | Lighthouse Lab in Glasgow | Wellcome Sanger Institute for the COVID-19 Genomics UK (COG-UK) Consortium | Harper VanSteenhouse, Yumi Kasai, David Gray, Carol Clugston, Anna Dominiczak and Alex Alderton, Roberto Amato, Jeffrey Barrett, Sonia Goncalves, Ewan Harrison, David K. Jackson, Ian Johnston, Dominic Kwiatkowski, Cordelia Langford, John Sillitoe on behalf of the Wellcome Sanger Institute COVID-19 Surveillance Team |
| EPI_ISL_3466314 | Lighthouse Lab in Milton Keynes | Wellcome Sanger Institute for the COVID-19 Genomics UK (COG-UK) Consortium | The Lighthouse Lab in Milton Keynes and Alex Alderton, Roberto Amato, Jeffrey Barrett, Sonia Goncalves, Ewan Harrison, David K. Jackson, Ian Johnston, Dominic Kwiatkowski, Cordelia Langford, John Sillitoe on behalf of the Wellcome Sanger Institute COVID-19 Surveillance Team |
| EPI_ISL_3466315, EPI_ISL_3466317 | Lighthouse Lab in Glasgow | Wellcome Sanger Institute for the COVID-19 Genomics UK (COG-UK) Consortium | Harper VanSteenhouse, Yumi Kasai, David Gray, Carol Clugston, Anna Dominiczak and Alex Alderton, Roberto Amato, Jeffrey Barrett, Sonia Goncalves, Ewan Harrison, David K. Jackson, Ian Johnston, Dominic Kwiatkowski, Cordelia Langford, John Sillitoe on behalf of the Wellcome Sanger Institute COVID-19 Surveillance Team |
| EPI_ISL_3466318 | Lighthouse Lab in Milton Keynes | Wellcome Sanger Institute for the COVID-19 Genomics UK (COG-UK) Consortium | The Lighthouse Lab in Milton Keynes and Alex Alderton, Roberto Amato, Jeffrey Barrett, Sonia Goncalves, Ewan Harrison, David K. Jackson, Ian Johnston, Dominic Kwiatkowski, Cordelia Langford, John Sillitoe on behalf of the Wellcome Sanger Institute COVID-19 Surveillance Team |
| EPI_ISL_3466319, EPI_ISL_3466320 | Lighthouse Lab in Glasgow | Wellcome Sanger Institute for the COVID-19 Genomics UK (COG-UK) Consortium | Harper VanSteenhouse, Yumi Kasai, David Gray, Carol Clugston, Anna Dominiczak and Alex Alderton, Roberto Amato, Jeffrey Barrett, Sonia Goncalves, Ewan Harrison, David K. Jackson, Ian Johnston, Dominic Kwiatkowski, Cordelia Langford, John Sillitoe on behalf of the Wellcome Sanger Institute COVID-19 Surveillance Team |
| EPI_ISL_3466321 | Lighthouse Lab in Milton Keynes | Wellcome Sanger Institute for the COVID-19 Genomics UK (COG-UK) Consortium | The Lighthouse Lab in Milton Keynes and Alex Alderton, Roberto Amato, Jeffrey Barrett, Sonia Goncalves, Ewan Harrison, David K. Jackson, Ian Johnston, Dominic Kwiatkowski, Cordelia Langford, John Sillitoe on behalf of the Wellcome Sanger Institute COVID-19 Surveillance Team |
| EPI_ISL_3466323, EPI_ISL_3466324, EPI_ISL_3466325 | Lighthouse Lab in Glasgow | Wellcome Sanger Institute for the COVID-19 Genomics UK (COG-UK) Consortium | Harper VanSteenhouse, Yumi Kasai, David Gray, Carol Clugston, Anna Dominiczak and Alex Alderton, Roberto Amato, Jeffrey Barrett, Sonia Goncalves, Ewan Harrison, David K. Jackson, Ian Johnston, Dominic Kwiatkowski, Cordelia Langford, John Sillitoe on behalf of the Wellcome Sanger Institute COVID-19 Surveillance Team |
| EPI_ISL_3466326, EPI_ISL_3466327 | Lighthouse Lab in Milton Keynes | Wellcome Sanger Institute for the COVID-19 Genomics UK (COG-UK) Consortium | The Lighthouse Lab in Milton Keynes and Alex Alderton, Roberto Amato, Jeffrey Barrett, Sonia Goncalves, Ewan Harrison, David K. Jackson, Ian Johnston, Dominic Kwiatkowski, Cordelia Langford, John Sillitoe on behalf of the Wellcome Sanger Institute COVID-19 Surveillance Team |
| EPI_ISL_3466329 | Lighthouse Lab in Glasgow | Wellcome Sanger Institute for the COVID-19 Genomics UK (COG-UK) Consortium | Harper VanSteenhouse, Yumi Kasai, David Gray, Carol Clugston, Anna Dominiczak and Alex Alderton, Roberto Amato, Jeffrey Barrett, Sonia Goncalves, Ewan Harrison, David K. Jackson, Ian Johnston, Dominic Kwiatkowski, Cordelia Langford, John Sillitoe on behalf of the Wellcome Sanger Institute COVID-19 Surveillance Team |
| EPI_ISL_3466330 | Lighthouse Lab in Milton Keynes | Wellcome Sanger Institute for the COVID-19 Genomics UK (COG-UK) Consortium | The Lighthouse Lab in Milton Keynes and Alex Alderton, Roberto Amato, Jeffrey Barrett, Sonia Goncalves, Ewan Harrison, David K. Jackson, Ian Johnston, Dominic Kwiatkowski, Cordelia Langford, John Sillitoe on behalf of the Wellcome Sanger Institute COVID-19 Surveillance Team |
| EPI_ISL_3466331 | Lighthouse Lab in Glasgow | Wellcome Sanger Institute for the COVID-19 Genomics UK (COG-UK) Consortium | Harper VanSteenhouse, Yumi Kasai, David Gray, Carol Clugston, Anna Dominiczak and Alex Alderton, Roberto Amato, Jeffrey Barrett, Sonia Goncalves, Ewan Harrison, David K. Jackson, Ian Johnston, Dominic Kwiatkowski, Cordelia Langford, John Sillitoe on behalf of the Wellcome Sanger Institute COVID-19 Surveillance Team |
| EPI_ISL_3466332 | Lighthouse Lab in Milton Keynes | Wellcome Sanger Institute for the COVID-19 Genomics UK (COG-UK) Consortium | The Lighthouse Lab in Milton Keynes and Alex Alderton, Roberto Amato, Jeffrey Barrett, Sonia Goncalves, Ewan Harrison, David K. Jackson, Ian Johnston, Dominic Kwiatkowski, Cordelia Langford, John Sillitoe on behalf of the Wellcome Sanger Institute COVID-19 Surveillance Team |
| EPI_ISL_3466333, EPI_ISL_3466334 | Lighthouse Lab in Glasgow | Wellcome Sanger Institute for the COVID-19 Genomics UK (COG-UK) Consortium | Harper VanSteenhouse, Yumi Kasai, David Gray, Carol Clugston, Anna Dominiczak and Alex Alderton, Roberto Amato, Jeffrey Barrett, Sonia Goncalves, Ewan Harrison, David K. Jackson, Ian Johnston, Dominic Kwiatkowski, Cordelia Langford, John Sillitoe on behalf of the Wellcome Sanger Institute COVID-19 Surveillance Team |
| EPI_ISL_3466336 | Lighthouse Lab in Milton Keynes | Wellcome Sanger Institute for the COVID-19 Genomics UK (COG-UK) Consortium | The Lighthouse Lab in Milton Keynes and Alex Alderton, Roberto Amato, Jeffrey Barrett, Sonia Goncalves, Ewan Harrison, David K. Jackson, Ian Johnston, Dominic Kwiatkowski, Cordelia Langford, John Sillitoe on behalf of the Wellcome Sanger Institute COVID-19 Surveillance Team |
| EPI_ISL_3466337, EPI_ISL_3466338, EPI_ISL_3466339 | Lighthouse Lab in Glasgow | Wellcome Sanger Institute for the COVID-19 Genomics UK (COG-UK) Consortium | Harper VanSteenhouse, Yumi Kasai, David Gray, Carol Clugston, Anna Dominiczak and Alex Alderton, Roberto Amato, Jeffrey Barrett, Sonia Goncalves, Ewan Harrison, David K. Jackson, Ian Johnston, Dominic Kwiatkowski, Cordelia Langford, John Sillitoe on behalf of the Wellcome Sanger Institute COVID-19 Surveillance Team |
| EPI_ISL_3466340 | Lighthouse Lab in Milton Keynes | Wellcome Sanger Institute for the COVID-19 Genomics UK (COG-UK) Consortium | The Lighthouse Lab in Milton Keynes and Alex Alderton, Roberto Amato, Jeffrey Barrett, Sonia Goncalves, Ewan Harrison, David K. Jackson, Ian Johnston, Dominic Kwiatkowski, Cordelia Langford, John Sillitoe on behalf of the Wellcome Sanger Institute COVID-19 Surveillance Team |
| EPI_ISL_3466342, EPI_ISL_3466343, EPI_ISL_3466344, EPI_ISL_3466345, EPI_ISL_3466346, EPI_ISL_3466348, EPI_ISL_3466349, EPI_ISL_3466350, EPI_ISL_3466351 | Lighthouse Lab in Glasgow | Wellcome Sanger Institute for the COVID-19 Genomics UK (COG-UK) Consortium | Harper VanSteenhouse, Yumi Kasai, David Gray, Carol Clugston, Anna Dominiczak and Alex Alderton, Roberto Amato, Jeffrey Barrett, Sonia Goncalves, Ewan Harrison, David K. Jackson, Ian Johnston, Dominic Kwiatkowski, Cordelia Langford, John Sillitoe on behalf of the Wellcome Sanger Institute COVID-19 Surveillance Team |
| EPI_ISL_3466352, EPI_ISL_3466354 | Lighthouse Lab in Milton Keynes | Wellcome Sanger Institute for the COVID-19 Genomics UK (COG-UK) Consortium | The Lighthouse Lab in Milton Keynes and Alex Alderton, Roberto Amato, Jeffrey Barrett, Sonia Goncalves, Ewan Harrison, David K. Jackson, Ian Johnston, Dominic Kwiatkowski, Cordelia Langford, John Sillitoe on behalf of the Wellcome Sanger Institute COVID-19 Surveillance Team |
| EPI_ISL_3466355, EPI_ISL_3466356, EPI_ISL_3466357, EPI_ISL_3466358, EPI_ISL_3466360, EPI_ISL_3466361 | Lighthouse Lab in Glasgow | Wellcome Sanger Institute for the COVID-19 Genomics UK (COG-UK) Consortium | Harper VanSteenhouse, Yumi Kasai, David Gray, Carol Clugston, Anna Dominiczak and Alex Alderton, Roberto Amato, Jeffrey Barrett, Sonia Goncalves, Ewan Harrison, David K. Jackson, Ian Johnston, Dominic Kwiatkowski, Cordelia Langford, John Sillitoe on behalf of the Wellcome Sanger Institute COVID-19 Surveillance Team |
| EPI_ISL_3466362 | Lighthouse Lab in Milton Keynes | Wellcome Sanger Institute for the COVID-19 Genomics UK (COG-UK) Consortium | The Lighthouse Lab in Milton Keynes and Alex Alderton, Roberto Amato, Jeffrey Barrett, Sonia Goncalves, Ewan Harrison, David K. Jackson, Ian Johnston, Dominic Kwiatkowski, Cordelia Langford, John Sillitoe on behalf of the Wellcome Sanger Institute COVID-19 Surveillance Team |
| EPI_ISL_3466363, EPI_ISL_3466364, EPI_ISL_3466365, EPI_ISL_3466367, EPI_ISL_3466368 | Lighthouse Lab in Glasgow | Wellcome Sanger Institute for the COVID-19 Genomics UK (COG-UK) Consortium | Harper VanSteenhouse, Yumi Kasai, David Gray, Carol Clugston, Anna Dominiczak and Alex Alderton, Roberto Amato, Jeffrey Barrett, Sonia Goncalves, Ewan Harrison, David K. Jackson, Ian Johnston, Dominic Kwiatkowski, Cordelia Langford, John Sillitoe on behalf of the Wellcome Sanger Institute COVID-19 Surveillance Team |
| EPI_ISL_3466369, EPI_ISL_3466370, EPI_ISL_3466371 | Lighthouse Lab in Milton Keynes | Wellcome Sanger Institute for the COVID-19 Genomics UK (COG-UK) Consortium | The Lighthouse Lab in Milton Keynes and Alex Alderton, Roberto Amato, Jeffrey Barrett, Sonia Goncalves, Ewan Harrison, David K. Jackson, Ian Johnston, Dominic Kwiatkowski, Cordelia Langford, John Sillitoe on behalf of the Wellcome Sanger Institute COVID-19 Surveillance Team |
| EPI_ISL_3466373 | Lighthouse Lab in Glasgow | Wellcome Sanger Institute for the COVID-19 Genomics UK (COG-UK) Consortium | Harper VanSteenhouse, Yumi Kasai, David Gray, Carol Clugston, Anna Dominiczak and Alex Alderton, Roberto Amato, Jeffrey Barrett, Sonia Goncalves, Ewan Harrison, David K. Jackson, Ian Johnston, Dominic Kwiatkowski, Cordelia Langford, John Sillitoe on behalf of the Wellcome Sanger Institute COVID-19 Surveillance Team |
| EPI_ISL_3466374, EPI_ISL_3466375, EPI_ISL_3466376, EPI_ISL_3466377 | Lighthouse Lab in Milton Keynes | Wellcome Sanger Institute for the COVID-19 Genomics UK (COG-UK) Consortium | The Lighthouse Lab in Milton Keynes and Alex Alderton, Roberto Amato, Jeffrey Barrett, Sonia Goncalves, Ewan Harrison, David K. Jackson, Ian Johnston, Dominic Kwiatkowski, Cordelia Langford, John Sillitoe on behalf of the Wellcome Sanger Institute COVID-19 Surveillance Team |
| EPI_ISL_3466378, EPI_ISL_3466380, EPI_ISL_3466381, EPI_ISL_3466382 | Lighthouse Lab in Glasgow | Wellcome Sanger Institute for the COVID-19 Genomics UK (COG-UK) Consortium | Harper VanSteenhouse, Yumi Kasai, David Gray, Carol Clugston, Anna Dominiczak and Alex Alderton, Roberto Amato, Jeffrey Barrett, Sonia Goncalves, Ewan Harrison, David K. Jackson, Ian Johnston, Dominic Kwiatkowski, Cordelia Langford, John Sillitoe on behalf of the Wellcome Sanger Institute COVID-19 Surveillance Team |
| EPI_ISL_3466383 | Lighthouse Lab in Milton Keynes | Wellcome Sanger Institute for the COVID-19 Genomics UK (COG-UK) Consortium | The Lighthouse Lab in Milton Keynes and Alex Alderton, Roberto Amato, Jeffrey Barrett, Sonia Goncalves, Ewan Harrison, David K. Jackson, Ian Johnston, Dominic Kwiatkowski, Cordelia Langford, John Sillitoe on behalf of the Wellcome Sanger Institute COVID-19 Surveillance Team |
| EPI_ISL_3466384, EPI_ISL_3466386 | Lighthouse Lab in Glasgow | Wellcome Sanger Institute for the COVID-19 Genomics UK (COG-UK) Consortium | Harper VanSteenhouse, Yumi Kasai, David Gray, Carol Clugston, Anna Dominiczak and Alex Alderton, Roberto Amato, Jeffrey Barrett, Sonia Goncalves, Ewan Harrison, David K. Jackson, Ian Johnston, Dominic Kwiatkowski, Cordelia Langford, John Sillitoe on behalf of the Wellcome Sanger Institute COVID-19 Surveillance Team |
| EPI_ISL_3466387 | Lighthouse Lab in Milton Keynes | Wellcome Sanger Institute for the COVID-19 Genomics UK (COG-UK) Consortium | The Lighthouse Lab in Milton Keynes and Alex Alderton, Roberto Amato, Jeffrey Barrett, Sonia Goncalves, Ewan Harrison, David K. Jackson, Ian Johnston, Dominic Kwiatkowski, Cordelia Langford, John Sillitoe on behalf of the Wellcome Sanger Institute COVID-19 Surveillance Team |
| EPI_ISL_3466388, EPI_ISL_3466389, EPI_ISL_3466390 | Lighthouse Lab in Glasgow | Wellcome Sanger Institute for the COVID-19 Genomics UK (COG-UK) Consortium | Harper VanSteenhouse, Yumi Kasai, David Gray, Carol Clugston, Anna Dominiczak and Alex Alderton, Roberto Amato, Jeffrey Barrett, Sonia Goncalves, Ewan Harrison, David K. Jackson, Ian Johnston, Dominic Kwiatkowski, Cordelia Langford, John Sillitoe on behalf of the Wellcome Sanger Institute COVID-19 Surveillance Team |
| EPI_ISL_3466391 | Lighthouse Lab in Milton Keynes | Wellcome Sanger Institute for the COVID-19 Genomics UK (COG-UK) Consortium | The Lighthouse Lab in Milton Keynes and Alex Alderton, Roberto Amato, Jeffrey Barrett, Sonia Goncalves, Ewan Harrison, David K. Jackson, Ian Johnston, Dominic Kwiatkowski, Cordelia Langford, John Sillitoe on behalf of the Wellcome Sanger Institute COVID-19 Surveillance Team |
| EPI_ISL_3466393, EPI_ISL_3466394 | Lighthouse Lab in Glasgow | Wellcome Sanger Institute for the COVID-19 Genomics UK (COG-UK) Consortium | Harper VanSteenhouse, Yumi Kasai, David Gray, Carol Clugston, Anna Dominiczak and Alex Alderton, Roberto Amato, Jeffrey Barrett, Sonia Goncalves, Ewan Harrison, David K. Jackson, Ian Johnston, Dominic Kwiatkowski, Cordelia Langford, John Sillitoe on behalf of the Wellcome Sanger Institute COVID-19 Surveillance Team |
| EPI_ISL_3466395 | Lighthouse Lab in Milton Keynes | Wellcome Sanger Institute for the COVID-19 Genomics UK (COG-UK) Consortium | The Lighthouse Lab in Milton Keynes and Alex Alderton, Roberto Amato, Jeffrey Barrett, Sonia Goncalves, Ewan Harrison, David K. Jackson, Ian Johnston, Dominic Kwiatkowski, Cordelia Langford, John Sillitoe on behalf of the Wellcome Sanger Institute COVID-19 Surveillance Team |
| EPI_ISL_3466396, EPI_ISL_3466397, EPI_ISL_3466399 | Lighthouse Lab in Glasgow | Wellcome Sanger Institute for the COVID-19 Genomics UK (COG-UK) Consortium | Harper VanSteenhouse, Yumi Kasai, David Gray, Carol Clugston, Anna Dominiczak and Alex Alderton, Roberto Amato, Jeffrey Barrett, Sonia Goncalves, Ewan Harrison, David K. Jackson, Ian Johnston, Dominic Kwiatkowski, Cordelia Langford, John Sillitoe on behalf of the Wellcome Sanger Institute COVID-19 Surveillance Team |
| EPI_ISL_3466400 | Lighthouse Lab in Milton Keynes | Wellcome Sanger Institute for the COVID-19 Genomics UK (COG-UK) Consortium | The Lighthouse Lab in Milton Keynes and Alex Alderton, Roberto Amato, Jeffrey Barrett, Sonia Goncalves, Ewan Harrison, David K. Jackson, Ian Johnston, Dominic Kwiatkowski, Cordelia Langford, John Sillitoe on behalf of the Wellcome Sanger Institute COVID-19 Surveillance Team |
| EPI_ISL_3466401 | Lighthouse Lab in Glasgow | Wellcome Sanger Institute for the COVID-19 Genomics UK (COG-UK) Consortium | Harper VanSteenhouse, Yumi Kasai, David Gray, Carol Clugston, Anna Dominiczak and Alex Alderton, Roberto Amato, Jeffrey Barrett, Sonia Goncalves, Ewan Harrison, David K. Jackson, Ian Johnston, Dominic Kwiatkowski, Cordelia Langford, John Sillitoe on behalf of the Wellcome Sanger Institute COVID-19 Surveillance Team |

| | | | |
|---|---|---|---|
| EPI_ISL_3466402 | Lighthouse Lab in Milton Keynes | Wellcome Sanger Institute for the COVID-19 Genomics UK (COG-UK) Consortium | The Lighthouse Lab in Milton Keynes and Alex Alderton, Roberto Amato, Jeffrey Barrett, Sonia Goncalves, Ewan Harrison, David K. Jackson, Ian Johnston, Dominic Kwiatkowski, Cordelia Langford, John Sillitoe on behalf of the Wellcome Sanger Institute COVID-19 Surveillance Team |
| EPI_ISL_3466403, EPI_ISL_3466404, EPI_ISL_3466406 | Lighthouse Lab in Glasgow | Wellcome Sanger Institute for the COVID-19 Genomics UK (COG-UK) Consortium | Harper VanSteenhouse, Yumi Kasai, David Gray, Carol Clugston, Anna Dominiczak and Alex Alderton, Roberto Amato, Jeffrey Barrett, Sonia Goncalves, Ewan Harrison, David K. Jackson, Ian Johnston, Dominic Kwiatkowski, Cordelia Langford, John Sillitoe on behalf of the Wellcome Sanger Institute COVID-19 Surveillance Team |
| EPI_ISL_3466407 | Lighthouse Lab in Milton Keynes | Wellcome Sanger Institute for the COVID-19 Genomics UK (COG-UK) Consortium | The Lighthouse Lab in Milton Keynes and Alex Alderton, Roberto Amato, Jeffrey Barrett, Sonia Goncalves, Ewan Harrison, David K. Jackson, Ian Johnston, Dominic Kwiatkowski, Cordelia Langford, John Sillitoe on behalf of the Wellcome Sanger Institute COVID-19 Surveillance Team |
| EPI_ISL_3466408 | Lighthouse Lab in Glasgow | Wellcome Sanger Institute for the COVID-19 Genomics UK (COG-UK) Consortium | Harper VanSteenhouse, Yumi Kasai, David Gray, Carol Clugston, Anna Dominiczak and Alex Alderton, Roberto Amato, Jeffrey Barrett, Sonia Goncalves, Ewan Harrison, David K. Jackson, Ian Johnston, Dominic Kwiatkowski, Cordelia Langford, John Sillitoe on behalf of the Wellcome Sanger Institute COVID-19 Surveillance Team |
| EPI_ISL_3466409 | Lighthouse Lab in Milton Keynes | Wellcome Sanger Institute for the COVID-19 Genomics UK (COG-UK) Consortium | The Lighthouse Lab in Milton Keynes and Alex Alderton, Roberto Amato, Jeffrey Barrett, Sonia Goncalves, Ewan Harrison, David K. Jackson, Ian Johnston, Dominic Kwiatkowski, Cordelia Langford, John Sillitoe on behalf of the Wellcome Sanger Institute COVID-19 Surveillance Team |
| EPI_ISL_3466410, EPI_ISL_3466411 | Lighthouse Lab in Glasgow | Wellcome Sanger Institute for the COVID-19 Genomics UK (COG-UK) Consortium | Harper VanSteenhouse, Yumi Kasai, David Gray, Carol Clugston, Anna Dominiczak and Alex Alderton, Roberto Amato, Jeffrey Barrett, Sonia Goncalves, Ewan Harrison, David K. Jackson, Ian Johnston, Dominic Kwiatkowski, Cordelia Langford, John Sillitoe on behalf of the Wellcome Sanger Institute COVID-19 Surveillance Team |
| EPI_ISL_3466413 | Lighthouse Lab in Milton Keynes | Wellcome Sanger Institute for the COVID-19 Genomics UK (COG-UK) Consortium | The Lighthouse Lab in Milton Keynes and Alex Alderton, Roberto Amato, Jeffrey Barrett, Sonia Goncalves, Ewan Harrison, David K. Jackson, Ian Johnston, Dominic Kwiatkowski, Cordelia Langford, John Sillitoe on behalf of the Wellcome Sanger Institute COVID-19 Surveillance Team |
| EPI_ISL_3466414, EPI_ISL_3466415, EPI_ISL_3466416 | Lighthouse Lab in Glasgow | Wellcome Sanger Institute for the COVID-19 Genomics UK (COG-UK) Consortium | Harper VanSteenhouse, Yumi Kasai, David Gray, Carol Clugston, Anna Dominiczak and Alex Alderton, Roberto Amato, Jeffrey Barrett, Sonia Goncalves, Ewan Harrison, David K. Jackson, Ian Johnston, Dominic Kwiatkowski, Cordelia Langford, John Sillitoe on behalf of the Wellcome Sanger Institute COVID-19 Surveillance Team |
| EPI_ISL_3466417 | Lighthouse Lab in Milton Keynes | Wellcome Sanger Institute for the COVID-19 Genomics UK (COG-UK) Consortium | The Lighthouse Lab in Milton Keynes and Alex Alderton, Roberto Amato, Jeffrey Barrett, Sonia Goncalves, Ewan Harrison, David K. Jackson, Ian Johnston, Dominic Kwiatkowski, Cordelia Langford, John Sillitoe on behalf of the Wellcome Sanger Institute COVID-19 Surveillance Team |
| EPI_ISL_3466419, EPI_ISL_3466420, EPI_ISL_3466421 | Lighthouse Lab in Glasgow | Wellcome Sanger Institute for the COVID-19 Genomics UK (COG-UK) Consortium | Harper VanSteenhouse, Yumi Kasai, David Gray, Carol Clugston, Anna Dominiczak and Alex Alderton, Roberto Amato, Jeffrey Barrett, Sonia Goncalves, Ewan Harrison, David K. Jackson, Ian Johnston, Dominic Kwiatkowski, Cordelia Langford, John Sillitoe on behalf of the Wellcome Sanger Institute COVID-19 Surveillance Team |
| EPI_ISL_3466422, EPI_ISL_3466423, EPI_ISL_3466425, EPI_ISL_3466426 | Lighthouse Lab in Milton Keynes | Wellcome Sanger Institute for the COVID-19 Genomics UK (COG-UK) Consortium | The Lighthouse Lab in Milton Keynes and Alex Alderton, Roberto Amato, Jeffrey Barrett, Sonia Goncalves, Ewan Harrison, David K. Jackson, Ian Johnston, Dominic Kwiatkowski, Cordelia Langford, John Sillitoe on behalf of the Wellcome Sanger Institute COVID-19 Surveillance Team |
| EPI_ISL_3466427 | Lighthouse Lab in Glasgow | Wellcome Sanger Institute for the COVID-19 Genomics UK (COG-UK) Consortium | Harper VanSteenhouse, Yumi Kasai, David Gray, Carol Clugston, Anna Dominiczak and Alex Alderton, Roberto Amato, Jeffrey Barrett, Sonia Goncalves, Ewan Harrison, David K. Jackson, Ian Johnston, Dominic Kwiatkowski, Cordelia Langford, John Sillitoe on behalf of the Wellcome Sanger Institute COVID-19 Surveillance Team |
| EPI_ISL_3466428 | Lighthouse Lab in Milton Keynes | Wellcome Sanger Institute for the COVID-19 Genomics UK (COG-UK) Consortium | The Lighthouse Lab in Milton Keynes and Alex Alderton, Roberto Amato, Jeffrey Barrett, Sonia Goncalves, Ewan Harrison, David K. Jackson, Ian Johnston, Dominic Kwiatkowski, Cordelia Langford, John Sillitoe on behalf of the Wellcome Sanger Institute COVID-19 Surveillance Team |
| EPI_ISL_3466429, EPI_ISL_3466430 | Lighthouse Lab in Glasgow | Wellcome Sanger Institute for the COVID-19 Genomics UK (COG-UK) Consortium | Harper VanSteenhouse, Yumi Kasai, David Gray, Carol Clugston, Anna Dominiczak and Alex Alderton, Roberto Amato, Jeffrey Barrett, Sonia Goncalves, Ewan Harrison, David K. Jackson, Ian Johnston, Dominic Kwiatkowski, Cordelia Langford, John Sillitoe on behalf of the Wellcome Sanger Institute COVID-19 Surveillance Team |
| EPI_ISL_3466432 | Lighthouse Lab in Milton Keynes | Wellcome Sanger Institute for the COVID-19 Genomics UK (COG-UK) Consortium | The Lighthouse Lab in Milton Keynes and Alex Alderton, Roberto Amato, Jeffrey Barrett, Sonia Goncalves, Ewan Harrison, David K. Jackson, Ian Johnston, Dominic Kwiatkowski, Cordelia Langford, John Sillitoe on behalf of the Wellcome Sanger Institute COVID-19 Surveillance Team |
| EPI_ISL_3466433 | Lighthouse Lab in Glasgow | Wellcome Sanger Institute for the COVID-19 Genomics UK (COG-UK) Consortium | Harper VanSteenhouse, Yumi Kasai, David Gray, Carol Clugston, Anna Dominiczak and Alex Alderton, Roberto Amato, Jeffrey Barrett, Sonia Goncalves, Ewan Harrison, David K. Jackson, Ian Johnston, Dominic Kwiatkowski, Cordelia Langford, John Sillitoe on behalf of the Wellcome Sanger Institute COVID-19 Surveillance Team |
| EPI_ISL_3466434 | Lighthouse Lab in Milton Keynes | Wellcome Sanger Institute for the COVID-19 Genomics UK (COG-UK) Consortium | The Lighthouse Lab in Milton Keynes and Alex Alderton, Roberto Amato, Jeffrey Barrett, Sonia Goncalves, Ewan Harrison, David K. Jackson, Ian Johnston, Dominic Kwiatkowski, Cordelia Langford, John Sillitoe on behalf of the Wellcome Sanger Institute COVID-19 Surveillance Team |
| EPI_ISL_3466435, EPI_ISL_3466436, EPI_ISL_3466437, EPI_ISL_3466439, EPI_ISL_3466440, EPI_ISL_3466441, EPI_ISL_3466442, EPI_ISL_3466443, EPI_ISL_3466445 | Lighthouse Lab in Glasgow | Wellcome Sanger Institute for the COVID-19 Genomics UK (COG-UK) Consortium | Harper VanSteenhouse, Yumi Kasai, David Gray, Carol Clugston, Anna Dominiczak and Alex Alderton, Roberto Amato, Jeffrey Barrett, Sonia Goncalves, Ewan Harrison, David K. Jackson, Ian Johnston, Dominic Kwiatkowski, Cordelia Langford, John Sillitoe on behalf of the Wellcome Sanger Institute COVID-19 Surveillance Team |
| EPI_ISL_3466446, EPI_ISL_3466447 | Lighthouse Lab in Milton Keynes | Wellcome Sanger Institute for the COVID-19 Genomics UK (COG-UK) Consortium | The Lighthouse Lab in Milton Keynes and Alex Alderton, Roberto Amato, Jeffrey Barrett, Sonia Goncalves, Ewan Harrison, David K. Jackson, Ian Johnston, Dominic Kwiatkowski, Cordelia Langford, John Sillitoe on behalf of the Wellcome Sanger Institute COVID-19 Surveillance Team |
| EPI_ISL_3466448, EPI_ISL_3466449, EPI_ISL_3466451, EPI_ISL_3466452, EPI_ISL_3466453, EPI_ISL_3466454 | Lighthouse Lab in Glasgow | Wellcome Sanger Institute for the COVID-19 Genomics UK (COG-UK) Consortium | Harper VanSteenhouse, Yumi Kasai, David Gray, Carol Clugston, Anna Dominiczak and Alex Alderton, Roberto Amato, Jeffrey Barrett, Sonia Goncalves, Ewan Harrison, David K. Jackson, Ian Johnston, Dominic Kwiatkowski, Cordelia Langford, John Sillitoe on behalf of the Wellcome Sanger Institute COVID-19 Surveillance Team |
| EPI_ISL_3466455, EPI_ISL_3466457 | Lighthouse Lab in Milton Keynes | Wellcome Sanger Institute for the COVID-19 Genomics UK (COG-UK) Consortium | The Lighthouse Lab in Milton Keynes and Alex Alderton, Roberto Amato, Jeffrey Barrett, Sonia Goncalves, Ewan Harrison, David K. Jackson, Ian Johnston, Dominic Kwiatkowski, Cordelia Langford, John Sillitoe on behalf of the Wellcome Sanger Institute COVID-19 Surveillance Team |
| EPI_ISL_3466458 | Lighthouse Lab in Glasgow | Wellcome Sanger Institute for the COVID-19 Genomics UK (COG-UK) Consortium | Harper VanSteenhouse, Yumi Kasai, David Gray, Carol Clugston, Anna Dominiczak and Alex Alderton, Roberto Amato, Jeffrey Barrett, Sonia Goncalves, Ewan Harrison, David K. Jackson, Ian Johnston, Dominic Kwiatkowski, Cordelia Langford, John Sillitoe on behalf of the Wellcome Sanger Institute COVID-19 Surveillance Team |
| EPI_ISL_3466459, EPI_ISL_3466460, EPI_ISL_3466461 | Lighthouse Lab in Milton Keynes | Wellcome Sanger Institute for the COVID-19 Genomics UK (COG-UK) Consortium | The Lighthouse Lab in Milton Keynes and Alex Alderton, Roberto Amato, Jeffrey Barrett, Sonia Goncalves, Ewan Harrison, David K. Jackson, Ian Johnston, Dominic Kwiatkowski, Cordelia Langford, John Sillitoe on behalf of the Wellcome Sanger Institute COVID-19 Surveillance Team |
| EPI_ISL_3466463, EPI_ISL_3466464 | Lighthouse Lab in Glasgow | Wellcome Sanger Institute for the COVID-19 Genomics UK (COG-UK) Consortium | Harper VanSteenhouse, Yumi Kasai, David Gray, Carol Clugston, Anna Dominiczak and Alex Alderton, Roberto Amato, Jeffrey Barrett, Sonia Goncalves, Ewan Harrison, David K. Jackson, Ian Johnston, Dominic Kwiatkowski, Cordelia Langford, John Sillitoe on behalf of the Wellcome Sanger Institute COVID-19 Surveillance Team |
| EPI_ISL_3466465 | Lighthouse Lab in Milton Keynes | Wellcome Sanger Institute for the COVID-19 Genomics UK (COG-UK) Consortium | The Lighthouse Lab in Milton Keynes and Alex Alderton, Roberto Amato, Jeffrey Barrett, Sonia Goncalves, Ewan Harrison, David K. Jackson, Ian Johnston, Dominic Kwiatkowski, Cordelia Langford, John Sillitoe on behalf of the Wellcome Sanger Institute COVID-19 Surveillance Team |
| EPI_ISL_3466466, EPI_ISL_3466467, EPI_ISL_3466469, EPI_ISL_3466470, EPI_ISL_3466471 | Lighthouse Lab in Glasgow | Wellcome Sanger Institute for the COVID-19 Genomics UK (COG-UK) Consortium | Harper VanSteenhouse, Yumi Kasai, David Gray, Carol Clugston, Anna Dominiczak and Alex Alderton, Roberto Amato, Jeffrey Barrett, Sonia Goncalves, Ewan Harrison, David K. Jackson, Ian Johnston, Dominic Kwiatkowski, Cordelia Langford, John Sillitoe on behalf of the Wellcome Sanger Institute COVID-19 Surveillance Team |
| EPI_ISL_3466472 | Lighthouse Lab in Milton Keynes | Wellcome Sanger Institute for the COVID-19 Genomics UK (COG-UK) Consortium | The Lighthouse Lab in Milton Keynes and Alex Alderton, Roberto Amato, Jeffrey Barrett, Sonia Goncalves, Ewan Harrison, David K. Jackson, Ian Johnston, Dominic Kwiatkowski, Cordelia Langford, John Sillitoe on behalf of the Wellcome Sanger Institute COVID-19 Surveillance Team |
| EPI_ISL_3466473 | Lighthouse Lab in Glasgow | Wellcome Sanger Institute for the COVID-19 Genomics UK (COG-UK) Consortium | Harper VanSteenhouse, Yumi Kasai, David Gray, Carol Clugston, Anna Dominiczak and Alex Alderton, Roberto Amato, Jeffrey Barrett, Sonia Goncalves, Ewan Harrison, David K. Jackson, Ian Johnston, Dominic Kwiatkowski, Cordelia Langford, John Sillitoe on behalf of the Wellcome Sanger Institute COVID-19 Surveillance Team |
| EPI_ISL_3466474 | Lighthouse Lab in Milton Keynes | Wellcome Sanger Institute for the COVID-19 Genomics UK (COG-UK) Consortium | The Lighthouse Lab in Milton Keynes and Alex Alderton, Roberto Amato, Jeffrey Barrett, Sonia Goncalves, Ewan Harrison, David K. Jackson, Ian Johnston, Dominic Kwiatkowski, Cordelia Langford, John Sillitoe on behalf of the Wellcome Sanger Institute COVID-19 Surveillance Team |
| EPI_ISL_3466476 | Lighthouse Lab in Glasgow | Wellcome Sanger Institute for the COVID-19 Genomics UK (COG-UK) Consortium | Harper VanSteenhouse, Yumi Kasai, David Gray, Carol Clugston, Anna Dominiczak and Alex Alderton, Roberto Amato, Jeffrey Barrett, Sonia Goncalves, Ewan Harrison, David K. Jackson, Ian Johnston, Dominic Kwiatkowski, Cordelia Langford, John Sillitoe on behalf of the Wellcome Sanger Institute COVID-19 Surveillance Team |
| EPI_ISL_3466477 | Lighthouse Lab in Milton Keynes | Wellcome Sanger Institute for the COVID-19 Genomics UK (COG-UK) Consortium | The Lighthouse Lab in Milton Keynes and Alex Alderton, Roberto Amato, Jeffrey Barrett, Sonia Goncalves, Ewan Harrison, David K. Jackson, Ian Johnston, Dominic Kwiatkowski, Cordelia Langford, John Sillitoe on behalf of the Wellcome Sanger Institute COVID-19 Surveillance Team |
| EPI_ISL_3466502, EPI_ISL_3466504 | Lighthouse Lab in Glasgow | Wellcome Sanger Institute for the COVID-19 Genomics UK (COG-UK) Consortium | Harper VanSteenhouse, Yumi Kasai, David Gray, Carol Clugston, Anna Dominiczak and Alex Alderton, Roberto Amato, Jeffrey Barrett, Sonia Goncalves, Ewan Harrison, David K. Jackson, Ian Johnston, Dominic Kwiatkowski, Cordelia Langford, John Sillitoe on behalf of the Wellcome Sanger Institute COVID-19 Surveillance Team |
| EPI_ISL_3466505 | Lighthouse Lab in Milton Keynes | Wellcome Sanger Institute for the COVID-19 Genomics UK (COG-UK) Consortium | The Lighthouse Lab in Milton Keynes and Alex Alderton, Roberto Amato, Jeffrey Barrett, Sonia Goncalves, Ewan Harrison, David K. Jackson, Ian Johnston, Dominic Kwiatkowski, Cordelia Langford, John Sillitoe on behalf of the Wellcome Sanger Institute COVID-19 Surveillance Team |
| EPI_ISL_3466506 | Lighthouse Lab in Glasgow | Wellcome Sanger Institute for the COVID-19 Genomics UK (COG-UK) Consortium | Harper VanSteenhouse, Yumi Kasai, David Gray, Carol Clugston, Anna Dominiczak and Alex Alderton, Roberto Amato, Jeffrey Barrett, Sonia Goncalves, Ewan Harrison, David K. Jackson, Ian Johnston, Dominic Kwiatkowski, Cordelia Langford, John Sillitoe on behalf of the Wellcome Sanger Institute COVID-19 Surveillance Team |
| EPI_ISL_3466507, EPI_ISL_3466508, EPI_ISL_3466510, EPI_ISL_3466511 | Lighthouse Lab in Milton Keynes | Wellcome Sanger Institute for the COVID-19 Genomics UK (COG-UK) Consortium | The Lighthouse Lab in Milton Keynes and Alex Alderton, Roberto Amato, Jeffrey Barrett, Sonia Goncalves, Ewan Harrison, David K. Jackson, Ian Johnston, Dominic Kwiatkowski, Cordelia Langford, John Sillitoe on behalf of the Wellcome Sanger Institute COVID-19 Surveillance Team |
| EPI_ISL_3466512 | Lighthouse Lab in Glasgow | Wellcome Sanger Institute for the COVID-19 Genomics UK (COG-UK) Consortium | Harper VanSteenhouse, Yumi Kasai, David Gray, Carol Clugston, Anna Dominiczak and Alex Alderton, Roberto Amato, Jeffrey Barrett, Sonia Goncalves, Ewan Harrison, David K. Jackson, Ian Johnston, Dominic Kwiatkowski, Cordelia Langford, John Sillitoe on behalf of the Wellcome Sanger Institute COVID-19 Surveillance Team |

| | | | |
|---|---|---|---|
| EPI_ISL_3466513 | Lighthouse Lab in Milton Keynes | Wellcome Sanger Institute for the COVID-19 Genomics UK (COG-UK) Consortium | The Lighthouse Lab in Milton Keynes and Alex Alderton, Roberto Amato, Jeffrey Barrett, Sonia Goncalves, Ewan Harrison, David K. Jackson, Ian Johnston, Dominic Kwiatkowski, Cordelia Langford, John Sillitoe on behalf of the Wellcome Sanger Institute COVID-19 Surveillance Team |
| EPI_ISL_3466515 | Lighthouse Lab in Glasgow | Wellcome Sanger Institute for the COVID-19 Genomics UK (COG-UK) Consortium | Harper VanSteenhouse, Yumi Kasai, David Gray, Carol Clugston, Anna Dominiczak and Alex Alderton, Roberto Amato, Jeffrey Barrett, Sonia Goncalves, Ewan Harrison, David K. Jackson, Ian Johnston, Dominic Kwiatkowski, Cordelia Langford, John Sillitoe on behalf of the Wellcome Sanger Institute COVID-19 Surveillance Team |
| EPI_ISL_3466516 | Lighthouse Lab in Milton Keynes | Wellcome Sanger Institute for the COVID-19 Genomics UK (COG-UK) Consortium | The Lighthouse Lab in Milton Keynes and Alex Alderton, Roberto Amato, Jeffrey Barrett, Sonia Goncalves, Ewan Harrison, David K. Jackson, Ian Johnston, Dominic Kwiatkowski, Cordelia Langford, John Sillitoe on behalf of the Wellcome Sanger Institute COVID-19 Surveillance Team |
| EPI_ISL_3466517 | Lighthouse Lab in Glasgow | Wellcome Sanger Institute for the COVID-19 Genomics UK (COG-UK) Consortium | Harper VanSteenhouse, Yumi Kasai, David Gray, Carol Clugston, Anna Dominiczak and Alex Alderton, Roberto Amato, Jeffrey Barrett, Sonia Goncalves, Ewan Harrison, David K. Jackson, Ian Johnston, Dominic Kwiatkowski, Cordelia Langford, John Sillitoe on behalf of the Wellcome Sanger Institute COVID-19 Surveillance Team |
| EPI_ISL_3466518 | Lighthouse Lab in Milton Keynes | Wellcome Sanger Institute for the COVID-19 Genomics UK (COG-UK) Consortium | The Lighthouse Lab in Milton Keynes and Alex Alderton, Roberto Amato, Jeffrey Barrett, Sonia Goncalves, Ewan Harrison, David K. Jackson, Ian Johnston, Dominic Kwiatkowski, Cordelia Langford, John Sillitoe on behalf of the Wellcome Sanger Institute COVID-19 Surveillance Team |
| EPI_ISL_3466520, EPI_ISL_3466521, EPI_ISL_3466522 | Lighthouse Lab in Glasgow | Wellcome Sanger Institute for the COVID-19 Genomics UK (COG-UK) Consortium | Harper VanSteenhouse, Yumi Kasai, David Gray, Carol Clugston, Anna Dominiczak and Alex Alderton, Roberto Amato, Jeffrey Barrett, Sonia Goncalves, Ewan Harrison, David K. Jackson, Ian Johnston, Dominic Kwiatkowski, Cordelia Langford, John Sillitoe on behalf of the Wellcome Sanger Institute COVID-19 Surveillance Team |
| EPI_ISL_3466523 | Lighthouse Lab in Milton Keynes | Wellcome Sanger Institute for the COVID-19 Genomics UK (COG-UK) Consortium | The Lighthouse Lab in Milton Keynes and Alex Alderton, Roberto Amato, Jeffrey Barrett, Sonia Goncalves, Ewan Harrison, David K. Jackson, Ian Johnston, Dominic Kwiatkowski, Cordelia Langford, John Sillitoe on behalf of the Wellcome Sanger Institute COVID-19 Surveillance Team |
| EPI_ISL_3466524, EPI_ISL_3466525, EPI_ISL_3466527 | Lighthouse Lab in Glasgow | Wellcome Sanger Institute for the COVID-19 Genomics UK (COG-UK) Consortium | Harper VanSteenhouse, Yumi Kasai, David Gray, Carol Clugston, Anna Dominiczak and Alex Alderton, Roberto Amato, Jeffrey Barrett, Sonia Goncalves, Ewan Harrison, David K. Jackson, Ian Johnston, Dominic Kwiatkowski, Cordelia Langford, John Sillitoe on behalf of the Wellcome Sanger Institute COVID-19 Surveillance Team |
| EPI_ISL_3466528 | Lighthouse Lab in Milton Keynes | Wellcome Sanger Institute for the COVID-19 Genomics UK (COG-UK) Consortium | The Lighthouse Lab in Milton Keynes and Alex Alderton, Roberto Amato, Jeffrey Barrett, Sonia Goncalves, Ewan Harrison, David K. Jackson, Ian Johnston, Dominic Kwiatkowski, Cordelia Langford, John Sillitoe on behalf of the Wellcome Sanger Institute COVID-19 Surveillance Team |
| EPI_ISL_3466529, EPI_ISL_3466530, EPI_ISL_3466532 | Lighthouse Lab in Glasgow | Wellcome Sanger Institute for the COVID-19 Genomics UK (COG-UK) Consortium | Harper VanSteenhouse, Yumi Kasai, David Gray, Carol Clugston, Anna Dominiczak and Alex Alderton, Roberto Amato, Jeffrey Barrett, Sonia Goncalves, Ewan Harrison, David K. Jackson, Ian Johnston, Dominic Kwiatkowski, Cordelia Langford, John Sillitoe on behalf of the Wellcome Sanger Institute COVID-19 Surveillance Team |
| EPI_ISL_3466533 | Lighthouse Lab in Milton Keynes | Wellcome Sanger Institute for the COVID-19 Genomics UK (COG-UK) Consortium | The Lighthouse Lab in Milton Keynes and Alex Alderton, Roberto Amato, Jeffrey Barrett, Sonia Goncalves, Ewan Harrison, David K. Jackson, Ian Johnston, Dominic Kwiatkowski, Cordelia Langford, John Sillitoe on behalf of the Wellcome Sanger Institute COVID-19 Surveillance Team |
| EPI_ISL_3466534, EPI_ISL_3466535, EPI_ISL_3466536, EPI_ISL_3466538, EPI_ISL_3466539 | Lighthouse Lab in Glasgow | Wellcome Sanger Institute for the COVID-19 Genomics UK (COG-UK) Consortium | Harper VanSteenhouse, Yumi Kasai, David Gray, Carol Clugston, Anna Dominiczak and Alex Alderton, Roberto Amato, Jeffrey Barrett, Sonia Goncalves, Ewan Harrison, David K. Jackson, Ian Johnston, Dominic Kwiatkowski, Cordelia Langford, John Sillitoe on behalf of the Wellcome Sanger Institute COVID-19 Surveillance Team |
| EPI_ISL_3466540 | Lighthouse Lab in Milton Keynes | Wellcome Sanger Institute for the COVID-19 Genomics UK (COG-UK) Consortium | The Lighthouse Lab in Milton Keynes and Alex Alderton, Roberto Amato, Jeffrey Barrett, Sonia Goncalves, Ewan Harrison, David K. Jackson, Ian Johnston, Dominic Kwiatkowski, Cordelia Langford, John Sillitoe on behalf of the Wellcome Sanger Institute COVID-19 Surveillance Team |
| EPI_ISL_3466541 | Lighthouse Lab in Glasgow | Wellcome Sanger Institute for the COVID-19 Genomics UK (COG-UK) Consortium | Harper VanSteenhouse, Yumi Kasai, David Gray, Carol Clugston, Anna Dominiczak and Alex Alderton, Roberto Amato, Jeffrey Barrett, Sonia Goncalves, Ewan Harrison, David K. Jackson, Ian Johnston, Dominic Kwiatkowski, Cordelia Langford, John Sillitoe on behalf of the Wellcome Sanger Institute COVID-19 Surveillance Team |
| EPI_ISL_3466542 | Lighthouse Lab in Milton Keynes | Wellcome Sanger Institute for the COVID-19 Genomics UK (COG-UK) Consortium | The Lighthouse Lab in Milton Keynes and Alex Alderton, Roberto Amato, Jeffrey Barrett, Sonia Goncalves, Ewan Harrison, David K. Jackson, Ian Johnston, Dominic Kwiatkowski, Cordelia Langford, John Sillitoe on behalf of the Wellcome Sanger Institute COVID-19 Surveillance Team |
| EPI_ISL_3466543, EPI_ISL_3466545, EPI_ISL_3466546 | Lighthouse Lab in Glasgow | Wellcome Sanger Institute for the COVID-19 Genomics UK (COG-UK) Consortium | Harper VanSteenhouse, Yumi Kasai, David Gray, Carol Clugston, Anna Dominiczak and Alex Alderton, Roberto Amato, Jeffrey Barrett, Sonia Goncalves, Ewan Harrison, David K. Jackson, Ian Johnston, Dominic Kwiatkowski, Cordelia Langford, John Sillitoe on behalf of the Wellcome Sanger Institute COVID-19 Surveillance Team |
| EPI_ISL_3466547 | Lighthouse Lab in Milton Keynes | Wellcome Sanger Institute for the COVID-19 Genomics UK (COG-UK) Consortium | The Lighthouse Lab in Milton Keynes and Alex Alderton, Roberto Amato, Jeffrey Barrett, Sonia Goncalves, Ewan Harrison, David K. Jackson, Ian Johnston, Dominic Kwiatkowski, Cordelia Langford, John Sillitoe on behalf of the Wellcome Sanger Institute COVID-19 Surveillance Team |
| EPI_ISL_3466548 | Lighthouse Lab in Glasgow | Wellcome Sanger Institute for the COVID-19 Genomics UK (COG-UK) Consortium | Harper VanSteenhouse, Yumi Kasai, David Gray, Carol Clugston, Anna Dominiczak and Alex Alderton, Roberto Amato, Jeffrey Barrett, Sonia Goncalves, Ewan Harrison, David K. Jackson, Ian Johnston, Dominic Kwiatkowski, Cordelia Langford, John Sillitoe on behalf of the Wellcome Sanger Institute COVID-19 Surveillance Team |
| EPI_ISL_3466549 | Lighthouse Lab in Milton Keynes | Wellcome Sanger Institute for the COVID-19 Genomics UK (COG-UK) Consortium | The Lighthouse Lab in Milton Keynes and Alex Alderton, Roberto Amato, Jeffrey Barrett, Sonia Goncalves, Ewan Harrison, David K. Jackson, Ian Johnston, Dominic Kwiatkowski, Cordelia Langford, John Sillitoe on behalf of the Wellcome Sanger Institute COVID-19 Surveillance Team |
| EPI_ISL_3466551, EPI_ISL_3466552 | Lighthouse Lab in Glasgow | Wellcome Sanger Institute for the COVID-19 Genomics UK (COG-UK) Consortium | Harper VanSteenhouse, Yumi Kasai, David Gray, Carol Clugston, Anna Dominiczak and Alex Alderton, Roberto Amato, Jeffrey Barrett, Sonia Goncalves, Ewan Harrison, David K. Jackson, Ian Johnston, Dominic Kwiatkowski, Cordelia Langford, John Sillitoe on behalf of the Wellcome Sanger Institute COVID-19 Surveillance Team |
| EPI_ISL_3466553, EPI_ISL_3466554, EPI_ISL_3466555, EPI_ISL_3466557, EPI_ISL_3466558, EPI_ISL_3466559, EPI_ISL_3466560, EPI_ISL_3466561, EPI_ISL_3466563, EPI_ISL_3466564, EPI_ISL_3466565, EPI_ISL_3466566, EPI_ISL_3466567, EPI_ISL_3466568, EPI_ISL_3466570, EPI_ISL_3466571, EPI_ISL_3466572, EPI_ISL_3466573, EPI_ISL_3466574, EPI_ISL_3466576, EPI_ISL_3466577, EPI_ISL_3466578, EPI_ISL_3466579, EPI_ISL_3466580, EPI_ISL_3466582, EPI_ISL_3466583, EPI_ISL_3466584, EPI_ISL_3466585, EPI_ISL_3466586, EPI_ISL_3466588, EPI_ISL_3466589, EPI_ISL_3466590, EPI_ISL_3466591, EPI_ISL_3466592, EPI_ISL_3466594, EPI_ISL_3466595, EPI_ISL_3466596, EPI_ISL_3466597, EPI_ISL_3466599, EPI_ISL_3466600, EPI_ISL_3466601, EPI_ISL_3466602, EPI_ISL_3466603, EPI_ISL_3466604, EPI_ISL_3466606, EPI_ISL_3466607, EPI_ISL_3466608, EPI_ISL_3466609, EPI_ISL_3466611, EPI_ISL_3466612, EPI_ISL_3466613, EPI_ISL_3466614, EPI_ISL_3466615, EPI_ISL_3466616, EPI_ISL_3466618, EPI_ISL_3466619, EPI_ISL_3466620, EPI_ISL_3466621, EPI_ISL_3466622, EPI_ISL_3466623, EPI_ISL_3466625, EPI_ISL_3466626, EPI_ISL_3466627, EPI_ISL_3466628, EPI_ISL_3466630, EPI_ISL_3466631, EPI_ISL_3466632, EPI_ISL_3466633, EPI_ISL_3466634, EPI_ISL_3466635, EPI_ISL_3466637, EPI_ISL_3466638, EPI_ISL_3466639, EPI_ISL_3466641, EPI_ISL_3466642, EPI_ISL_3466643, EPI_ISL_3466644, EPI_ISL_3466645, EPI_ISL_3466647, EPI_ISL_3466648, EPI_ISL_3466649, EPI_ISL_3466650, EPI_ISL_3466651, EPI_ISL_3466653, EPI_ISL_3466654, EPI_ISL_3466655, EPI_ISL_3466656, EPI_ISL_3466657, EPI_ISL_3466658, EPI_ISL_3466660, EPI_ISL_3466661, EPI_ISL_3466662, EPI_ISL_3466663, EPI_ISL_3466664, EPI_ISL_3466665, EPI_ISL_3466667, EPI_ISL_3466668, EPI_ISL_3466669, EPI_ISL_3466671, EPI_ISL_3466672, EPI_ISL_3466673, EPI_ISL_3466674, EPI_ISL_3466675, EPI_ISL_3466677, EPI_ISL_3466678, EPI_ISL_3466679, EPI_ISL_3466680, EPI_ISL_3466682, EPI_ISL_3466683, EPI_ISL_3466684, EPI_ISL_3466685, EPI_ISL_3466686, EPI_ISL_3466688, EPI_ISL_3466689, EPI_ISL_3466690, EPI_ISL_3466691, EPI_ISL_3466692, EPI_ISL_3466694, EPI_ISL_3466695, EPI_ISL_3466696, EPI_ISL_3466697, EPI_ISL_3466698, EPI_ISL_3466700, EPI_ISL_3466701, EPI_ISL_3466702, EPI_ISL_3466703, EPI_ISL_3466705, EPI_ISL_3466706, EPI_ISL_3466707, EPI_ISL_3466708, EPI_ISL_3466709, EPI_ISL_3466710, EPI_ISL_3466712, EPI_ISL_3466713, EPI_ISL_3466714, EPI_ISL_3466715, EPI_ISL_3466716, EPI_ISL_3466717, EPI_ISL_3466719, EPI_ISL_3466720, EPI_ISL_3466721, EPI_ISL_3466722, EPI_ISL_3466723, EPI_ISL_3466725, EPI_ISL_3466726, EPI_ISL_3466727, EPI_ISL_3466728, EPI_ISL_3466729, EPI_ISL_3466730, EPI_ISL_3466732, EPI_ISL_3466733, EPI_ISL_3466734, EPI_ISL_3466735, EPI_ISL_3466736, EPI_ISL_3466738, EPI_ISL_3466739, EPI_ISL_3466740, EPI_ISL_3466741, EPI_ISL_3466742, EPI_ISL_3466744, EPI_ISL_3466745, EPI_ISL_3466746, EPI_ISL_3466747, EPI_ISL_3466748, EPI_ISL_3466750, EPI_ISL_3466751, EPI_ISL_3466752, EPI_ISL_3466753, EPI_ISL_3466754, EPI_ISL_3466756, EPI_ISL_3466757, EPI_ISL_3466758, EPI_ISL_3466759, EPI_ISL_3466760, EPI_ISL_3466762, EPI_ISL_3466763, EPI_ISL_3466764, EPI_ISL_3466765, EPI_ISL_3466766, EPI_ISL_3466768, EPI_ISL_3466769, EPI_ISL_3466770, EPI_ISL_3466771, EPI_ISL_3466772, EPI_ISL_3466774, EPI_ISL_3466775, EPI_ISL_3466776, EPI_ISL_3466777, EPI_ISL_3466778, EPI_ISL_3466780, EPI_ISL_3466781, EPI_ISL_3466782, EPI_ISL_3466783, EPI_ISL_3466784, EPI_ISL_3466786, EPI_ISL_3466787, EPI_ISL_3466788, EPI_ISL_3466789, EPI_ISL_3466790, EPI_ISL_3466792, EPI_ISL_3466793, EPI_ISL_3466794, EPI_ISL_3466795, EPI_ISL_3466796, EPI_ISL_3466798, EPI_ISL_3466799, EPI_ISL_3466800, EPI_ISL_3466801, EPI_ISL_3466802, EPI_ISL_3466803, EPI_ISL_3466805, EPI_ISL_3466806, EPI_ISL_3466807, EPI_ISL_3466808, EPI_ISL_3466809, EPI_ISL_3466810, EPI_ISL_3466812, EPI_ISL_3466813, EPI_ISL_3466814, EPI_ISL_3466815, EPI_ISL_3466816, EPI_ISL_3466818, EPI_ISL_3466819, EPI_ISL_3466820, EPI_ISL_3466821, EPI_ISL_3466822, EPI_ISL_3466823, EPI_ISL_3466825, EPI_ISL_3466826, EPI_ISL_3466827, EPI_ISL_3466828, EPI_ISL_3466829, EPI_ISL_3466831, EPI_ISL_3466832, EPI_ISL_3466833, EPI_ISL_3466834, EPI_ISL_3466835, EPI_ISL_3466836, EPI_ISL_3466837 | | | |
| see above | Lighthouse Lab in Milton Keynes | Wellcome Sanger Institute for the COVID-19 Genomics UK (COG-UK) Consortium | The Lighthouse Lab in Milton Keynes and Alex Alderton, Roberto Amato, Jeffrey Barrett, Sonia Goncalves, Ewan Harrison, David K. Jackson, Ian Johnston, Dominic Kwiatkowski, Cordelia Langford, John Sillitoe on behalf of the Wellcome Sanger Institute COVID-19 Surveillance Team |
| EPI_ISL_3466838 | Laboratory of Clinical Microbiology, Faculty of Medicine, University of Indonesia | Eijkman Institute for Molecular Biology, National Research and Innovation Agency | Edison Johar, Frilasita A Yudhaputri, Muhammad Rezki Rasyak, Willy Agustine, Hidayat Trimarsanto, Lydia V. Panggalo, Iskandar Adnan, Sukma Oktavianthi, Lidwina Priliani, Jessica R. Saragih, Safarina G Malik, Khin Saw Myint, Amin Soebandrio |
| EPI_ISL_3466839, EPI_ISL_3466840, EPI_ISL_3466841, EPI_ISL_3466842, EPI_ISL_3466843 | Lighthouse Lab in Milton Keynes | Wellcome Sanger Institute for the COVID-19 Genomics UK (COG-UK) Consortium | The Lighthouse Lab in Milton Keynes and Alex Alderton, Roberto Amato, Jeffrey Barrett, Sonia Goncalves, Ewan Harrison, David K. Jackson, Ian Johnston, Dominic Kwiatkowski, Cordelia Langford, John Sillitoe on behalf of the Wellcome Sanger Institute COVID-19 Surveillance Team |
| EPI_ISL_3466844 | Laboratory of Clinical Microbiology, Faculty of Medicine, University of Indonesia | Eijkman Institute for Molecular Biology, National Research and Innovation Agency | Frilasita A Yudhaputri, Muhammad Rezki Rasyak, Willy Agustine, Hidayat Trimarsanto, Lydia V. Panggalo, Iskandar Adnan, Sukma Oktavianthi, Lidwina Priliani, Jessica R. Saragih, Edison Johar, Safarina G Malik, Khin Saw Myint, Amin Soebandrio |
| EPI_ISL_3466845, EPI_ISL_3466846, EPI_ISL_3466847, EPI_ISL_3466848, EPI_ISL_3466850, EPI_ISL_3466851, EPI_ISL_3466852, EPI_ISL_3466853, EPI_ISL_3466854, EPI_ISL_3466855 | Lighthouse Lab in Milton Keynes | Wellcome Sanger Institute for the COVID-19 Genomics UK (COG-UK) Consortium | The Lighthouse Lab in Milton Keynes and Alex Alderton, Roberto Amato, Jeffrey Barrett, Sonia Goncalves, Ewan Harrison, David K. Jackson, Ian Johnston, Dominic Kwiatkowski, Cordelia Langford, John Sillitoe on behalf of the Wellcome Sanger Institute COVID-19 Surveillance Team |
| EPI_ISL_3466856 | Laboratory of Clinical Microbiology, Faculty of Medicine, University of Indonesia | Eijkman Institute for Molecular Biology, National Research and Innovation Agency | Willy Agustine, Hidayat Trimarsanto, Lydia V. Panggalo, Iskandar Adnan, Sukma Oktavianthi, Lidwina Priliani, Jessica R. Saragih, Edison Johar, Frilasita A Yudhaputri, Muhammad Rezki Rasyak, Safarina G Malik, Khin Saw Myint, Amin Soebandrio |
| EPI_ISL_3466857, EPI_ISL_3466858, EPI_ISL_3466859, EPI_ISL_3466860, EPI_ISL_3466861 | Lighthouse Lab in Milton Keynes | Wellcome Sanger Institute for the COVID-19 Genomics UK (COG-UK) Consortium | The Lighthouse Lab in Milton Keynes and Alex Alderton, Roberto Amato, Jeffrey Barrett, Sonia Goncalves, Ewan Harrison, David K. Jackson, Ian Johnston, Dominic Kwiatkowski, Cordelia Langford, John Sillitoe on behalf of the Wellcome Sanger Institute COVID-19 Surveillance Team |
| EPI_ISL_3466862 | Laboratory of Clinical Microbiology, Faculty of Medicine, University of Indonesia | Eijkman Institute for Molecular Biology, National Research and Innovation Agency | Hidayat Trimarsanto, Lydia V. Panggalo, Iskandar Adnan, Sukma Oktavianthi, Lidwina Priliani, Jessica R. Saragih, Edison Johar, Frilasita A Yudhaputri, Muhammad Rezki Rasyak, Willy Agustine, Safarina G Malik, Khin Saw Myint, Amin Soebandrio |
| EPI_ISL_3466863, EPI_ISL_3466864, EPI_ISL_3466865, EPI_ISL_3466866, EPI_ISL_3466867, EPI_ISL_3466868 | Lighthouse Lab in Milton Keynes | Wellcome Sanger Institute for the COVID-19 Genomics UK (COG-UK) Consortium | The Lighthouse Lab in Milton Keynes and Alex Alderton, Roberto Amato, Jeffrey Barrett, Sonia Goncalves, Ewan Harrison, David K. Jackson, Ian Johnston, Dominic Kwiatkowski, Cordelia Langford, John Sillitoe on behalf of the Wellcome Sanger Institute COVID-19 Surveillance Team |
| EPI_ISL_3466869 | Laboratory of Clinical Microbiology, Faculty of Medicine, University of Indonesia | Eijkman Institute for Molecular Biology, National Research and Innovation Agency | Lydia V. Panggalo, Iskandar Adnan, Sukma Oktavianthi, Lidwina Priliani, Jessica R. Saragih, Edison Johar, Frilasita A Yudhaputri, Muhammad Rezki Rasyak, Willy Agustine, Hidayat Trimarsanto, Safarina G Malik, Khin Saw Myint, Amin Soebandrio |
| EPI_ISL_3466870, | Lighthouse Lab in Milton Keynes | Wellcome Sanger Institute for the COVID-19 Genomics UK (COG-UK) | The Lighthouse Lab in Milton Keynes and Alex Alderton, Roberto Amato, Jeffrey Barrett, Sonia Goncalves, Ewan Harrison, David K. Jackson, Ian Johnston, Dominic Kwiatkowski, Cordelia Langford, John Sillitoe on behalf of the Wellcome Sanger |

| | | | |
|---|---|---|---|
| EPI_ISL_3466871, EPI_ISL_3466872, EPI_ISL_3466873, EPI_ISL_3466874 | | Consortium | Institute COVID-19 Surveillance Team |
| EPI_ISL_3466875 | Laboratory of Clinical Microbiology, Faculty of Medicine, University of Indonesia | Eijkman Institute for Molecular Biology, National Research and Innovation Agency | Iskandar Adnan, Sukma Oktavianthi, Lidwina Priliani, Jessica R. Saragih, Edison Johar, Frilasita A Yudhaputri, Muhammad Rezki Rasyak, Willy Agustine, Hidayat Trimarsanto, Lydia V. Panggalo, Safarina G Malik, Khin Saw Myint, Amin Soebandrio |
| EPI_ISL_3466876, EPI_ISL_3466877, EPI_ISL_3466878, EPI_ISL_3466879, EPI_ISL_3466880 | Lighthouse Lab in Milton Keynes | Wellcome Sanger Institute for the COVID-19 Genomics UK (COG-UK) Consortium | The Lighthouse Lab in Milton Keynes and Alex Alderton, Roberto Amato, Jeffrey Barrett, Sonia Goncalves, Ewan Harrison, David K. Jackson, Ian Johnston, Dominic Kwiatkowski, Cordelia Langford, John Sillitoe on behalf of the Wellcome Sanger Institute COVID-19 Surveillance Team |
| EPI_ISL_3466881 | Laboratory of Clinical Microbiology, Faculty of Medicine, University of Indonesia | Eijkman Institute for Molecular Biology, National Research and Innovation Agency | Lidwina Priliani, Jessica R. Saragih, Edison Johar, Frilasita A Yudhaputri, Muhammad Rezki Rasyak, Willy Agustine, Hidayat Trimarsanto, Lydia V. Panggalo, Iskandar Adnan, Sukma Oktavianthi, Safarina G Malik, Khin Saw Myint, Amin Soebandrio |
| EPI_ISL_3466882, EPI_ISL_3466883, EPI_ISL_3466884, EPI_ISL_3466885, EPI_ISL_3466886, EPI_ISL_3466887 | Lighthouse Lab in Milton Keynes | Wellcome Sanger Institute for the COVID-19 Genomics UK (COG-UK) Consortium | The Lighthouse Lab in Milton Keynes and Alex Alderton, Roberto Amato, Jeffrey Barrett, Sonia Goncalves, Ewan Harrison, David K. Jackson, Ian Johnston, Dominic Kwiatkowski, Cordelia Langford, John Sillitoe on behalf of the Wellcome Sanger Institute COVID-19 Surveillance Team |
| EPI_ISL_3466888 | Laboratory of Clinical Microbiology, Faculty of Medicine, University of Indonesia | Eijkman Institute for Molecular Biology, National Research and Innovation Agency | Jessica R. Saragih, Edison Johar, Frilasita A Yudhaputri, Muhammad Rezki Rasyak, Willy Agustine, Hidayat Trimarsanto, Lydia V. Panggalo, Iskandar Adnan, Sukma Oktavianthi, Lidwina Priliani, Safarina G Malik, Khin Saw Myint, Amin Soebandrio |
| EPI_ISL_3466889, EPI_ISL_3466890, EPI_ISL_3466891, EPI_ISL_3466892, EPI_ISL_3466893 | Lighthouse Lab in Milton Keynes | Wellcome Sanger Institute for the COVID-19 Genomics UK (COG-UK) Consortium | The Lighthouse Lab in Milton Keynes and Alex Alderton, Roberto Amato, Jeffrey Barrett, Sonia Goncalves, Ewan Harrison, David K. Jackson, Ian Johnston, Dominic Kwiatkowski, Cordelia Langford, John Sillitoe on behalf of the Wellcome Sanger Institute COVID-19 Surveillance Team |
| EPI_ISL_3466894 | Laboratory of Clinical Microbiology, Faculty of Medicine, University of Indonesia | Eijkman Institute for Molecular Biology, National Research and Innovation Agency | Edison Johar, Frilasita A Yudhaputri, Muhammad Rezki Rasyak, Willy Agustine, Hidayat Trimarsanto, Lydia V. Panggalo, Iskandar Adnan, Sukma Oktavianthi, Lidwina Priliani, Jessica R. Saragih, Safarina G Malik, Khin Saw Myint, Amin Soebandrio |
| EPI_ISL_3466895, EPI_ISL_3466896, EPI_ISL_3466897, EPI_ISL_3466898, EPI_ISL_3466899 | Lighthouse Lab in Milton Keynes | Wellcome Sanger Institute for the COVID-19 Genomics UK (COG-UK) Consortium | The Lighthouse Lab in Milton Keynes and Alex Alderton, Roberto Amato, Jeffrey Barrett, Sonia Goncalves, Ewan Harrison, David K. Jackson, Ian Johnston, Dominic Kwiatkowski, Cordelia Langford, John Sillitoe on behalf of the Wellcome Sanger Institute COVID-19 Surveillance Team |
| EPI_ISL_3466900 | Laboratory of Clinical Microbiology, Faculty of Medicine, University of Indonesia | Eijkman Institute for Molecular Biology, National Research and Innovation Agency | Frilasita A Yudhaputri, Muhammad Rezki Rasyak, Willy Agustine, Hidayat Trimarsanto, Lydia V. Panggalo, Iskandar Adnan, Sukma Oktavianthi, Lidwina Priliani, Jessica R. Saragih, Edison Johar, Safarina G Malik, Khin Saw Myint, Amin Soebandrio |
| EPI_ISL_3466901, EPI_ISL_3466902, EPI_ISL_3466903, EPI_ISL_3466904 | Lighthouse Lab in Milton Keynes | Wellcome Sanger Institute for the COVID-19 Genomics UK (COG-UK) Consortium | The Lighthouse Lab in Milton Keynes and Alex Alderton, Roberto Amato, Jeffrey Barrett, Sonia Goncalves, Ewan Harrison, David K. Jackson, Ian Johnston, Dominic Kwiatkowski, Cordelia Langford, John Sillitoe on behalf of the Wellcome Sanger Institute COVID-19 Surveillance Team |
| EPI_ISL_3466905 | Laboratory of Clinical Microbiology, Faculty of Medicine, University of Indonesia | Eijkman Institute for Molecular Biology, National Research and Innovation Agency | Muhammad Rezki Rasyak, Willy Agustine, Hidayat Trimarsanto, Lydia V. Panggalo, Iskandar Adnan, Sukma Oktavianthi, Lidwina Priliani, Jessica R. Saragih, Edison Johar, Frilasita A Yudhaputri, Safarina G Malik, Khin Saw Myint, Amin Soebandrio |
| EPI_ISL_3466906, EPI_ISL_3466907, EPI_ISL_3466908, EPI_ISL_3466909, EPI_ISL_3466910, EPI_ISL_3466911 | Lighthouse Lab in Milton Keynes | Wellcome Sanger Institute for the COVID-19 Genomics UK (COG-UK) Consortium | The Lighthouse Lab in Milton Keynes and Alex Alderton, Roberto Amato, Jeffrey Barrett, Sonia Goncalves, Ewan Harrison, David K. Jackson, Ian Johnston, Dominic Kwiatkowski, Cordelia Langford, John Sillitoe on behalf of the Wellcome Sanger Institute COVID-19 Surveillance Team |
| EPI_ISL_3466912 | Laboratory of Clinical Microbiology, Faculty of Medicine, University of Indonesia | Eijkman Institute for Molecular Biology, National Research and Innovation Agency | Lydia V. Panggalo, Iskandar Adnan, Sukma Oktavianthi, Lidwina Priliani, Jessica R. Saragih, Edison Johar, Frilasita A Yudhaputri, Muhammad Rezki Rasyak, Willy Agustine, Hidayat Trimarsanto, Safarina G Malik, Khin Saw Myint, Amin Soebandrio |
| EPI_ISL_3466913, EPI_ISL_3466914, EPI_ISL_3466915, EPI_ISL_3466916, EPI_ISL_3466917 | Lighthouse Lab in Milton Keynes | Wellcome Sanger Institute for the COVID-19 Genomics UK (COG-UK) Consortium | The Lighthouse Lab in Milton Keynes and Alex Alderton, Roberto Amato, Jeffrey Barrett, Sonia Goncalves, Ewan Harrison, David K. Jackson, Ian Johnston, Dominic Kwiatkowski, Cordelia Langford, John Sillitoe on behalf of the Wellcome Sanger Institute COVID-19 Surveillance Team |
| EPI_ISL_3466918 | Laboratory of Clinical Microbiology, Faculty of Medicine, University of Indonesia | Eijkman Institute for Molecular Biology, National Research and Innovation Agency | Lidwina Priliani, Jessica R. Saragih, Edison Johar, Frilasita A Yudhaputri, Muhammad Rezki Rasyak, Willy Agustine, Hidayat Trimarsanto, Lydia V. Panggalo, Iskandar Adnan, Sukma Oktavianthi, Safarina G Malik, Khin Saw Myint, Amin Soebandrio |
| EPI_ISL_3466919, EPI_ISL_3466920, EPI_ISL_3466921, EPI_ISL_3466922, EPI_ISL_3466923 | Lighthouse Lab in Milton Keynes | Wellcome Sanger Institute for the COVID-19 Genomics UK (COG-UK) Consortium | The Lighthouse Lab in Milton Keynes and Alex Alderton, Roberto Amato, Jeffrey Barrett, Sonia Goncalves, Ewan Harrison, David K. Jackson, Ian Johnston, Dominic Kwiatkowski, Cordelia Langford, John Sillitoe on behalf of the Wellcome Sanger Institute COVID-19 Surveillance Team |
| EPI_ISL_3466924 | Laboratory of Clinical Microbiology, Faculty of Medicine, University of Indonesia | Eijkman Institute for Molecular Biology, National Research and Innovation Agency | Jessica R. Saragih, Edison Johar, Frilasita A Yudhaputri, Muhammad Rezki Rasyak, Willy Agustine, Hidayat Trimarsanto, Lydia V. Panggalo, Iskandar Adnan, Sukma Oktavianthi, Lidwina Priliani, Safarina G Malik, Khin Saw Myint, Amin Soebandrio |
| EPI_ISL_3466925, EPI_ISL_3466926, EPI_ISL_3466927, EPI_ISL_3466928, EPI_ISL_3466930, EPI_ISL_3466931, EPI_ISL_3466932, EPI_ISL_3466933, EPI_ISL_3466934, EPI_ISL_3466935 | Lighthouse Lab in Milton Keynes | Wellcome Sanger Institute for the COVID-19 Genomics UK (COG-UK) Consortium | The Lighthouse Lab in Milton Keynes and Alex Alderton, Roberto Amato, Jeffrey Barrett, Sonia Goncalves, Ewan Harrison, David K. Jackson, Ian Johnston, Dominic Kwiatkowski, Cordelia Langford, John Sillitoe on behalf of the Wellcome Sanger Institute COVID-19 Surveillance Team |
| EPI_ISL_3466936 | Laboratory of Clinical Microbiology, Faculty of Medicine, University of Indonesia | Eijkman Institute for Molecular Biology, National Research and Innovation Agency | Muhammad Rezki Rasyak, Willy Agustine, Hidayat Trimarsanto, Lydia V. Panggalo, Iskandar Adnan, Sukma Oktavianthi, Lidwina Priliani, Jessica R. Saragih, Edison Johar, Frilasita A Yudhaputri, Safarina G Malik, Khin Saw Myint, Amin Soebandrio |
| EPI_ISL_3466937, EPI_ISL_3466938, EPI_ISL_3466939, EPI_ISL_3466940, EPI_ISL_3466941 | Lighthouse Lab in Milton Keynes | Wellcome Sanger Institute for the COVID-19 Genomics UK (COG-UK) Consortium | The Lighthouse Lab in Milton Keynes and Alex Alderton, Roberto Amato, Jeffrey Barrett, Sonia Goncalves, Ewan Harrison, David K. Jackson, Ian Johnston, Dominic Kwiatkowski, Cordelia Langford, John Sillitoe on behalf of the Wellcome Sanger Institute COVID-19 Surveillance Team |
| EPI_ISL_3466942 | Laboratory of Clinical Microbiology, Faculty of Medicine, University of Indonesia | Eijkman Institute for Molecular Biology, National Research and Innovation Agency | Hidayat Trimarsanto, Lydia V. Panggalo, Iskandar Adnan, Sukma Oktavianthi, Lidwina Priliani, Jessica R. Saragih, Edison Johar, Frilasita A Yudhaputri, Muhammad Rezki Rasyak, Willy Agustine, Safarina G Malik, Khin Saw Myint, Amin Soebandrio |
| EPI_ISL_3466943 | Lighthouse Lab in Milton Keynes | Wellcome Sanger Institute for the COVID-19 Genomics UK (COG-UK) Consortium | The Lighthouse Lab in Milton Keynes and Alex Alderton, Roberto Amato, Jeffrey Barrett, Sonia Goncalves, Ewan Harrison, David K. Jackson, Ian Johnston, Dominic Kwiatkowski, Cordelia Langford, John Sillitoe on behalf of the Wellcome Sanger Institute COVID-19 Surveillance Team |
| EPI_ISL_3466944 | Lighthouse Lab in Glasgow | Wellcome Sanger Institute for the COVID-19 Genomics UK (COG-UK) Consortium | Harper VanSteenhouse, Yumi Kasai, David Gray, Carol Clugston, Anna Dominiczak and Alex Alderton, Roberto Amato, Jeffrey Barrett, Sonia Goncalves, Ewan Harrison, David K. Jackson, Ian Johnston, Dominic Kwiatkowski, Cordelia Langford, John Sillitoe on behalf of the Wellcome Sanger Institute COVID-19 Surveillance Team |
| EPI_ISL_3466945 | Lighthouse Lab in Milton Keynes | Wellcome Sanger Institute for the COVID-19 Genomics UK (COG-UK) Consortium | The Lighthouse Lab in Milton Keynes and Alex Alderton, Roberto Amato, Jeffrey Barrett, Sonia Goncalves, Ewan Harrison, David K. Jackson, Ian Johnston, Dominic Kwiatkowski, Cordelia Langford, John Sillitoe on behalf of the Wellcome Sanger Institute COVID-19 Surveillance Team |
| EPI_ISL_3466946 | Lighthouse Lab in Glasgow | Wellcome Sanger Institute for the COVID-19 Genomics UK (COG-UK) Consortium | Harper VanSteenhouse, Yumi Kasai, David Gray, Carol Clugston, Anna Dominiczak and Alex Alderton, Roberto Amato, Jeffrey Barrett, Sonia Goncalves, Ewan Harrison, David K. Jackson, Ian Johnston, Dominic Kwiatkowski, Cordelia Langford, John Sillitoe on behalf of the Wellcome Sanger Institute COVID-19 Surveillance Team |
| EPI_ISL_3466947, EPI_ISL_3466948 | Lighthouse Lab in Milton Keynes | Wellcome Sanger Institute for the COVID-19 Genomics UK (COG-UK) Consortium | The Lighthouse Lab in Milton Keynes and Alex Alderton, Roberto Amato, Jeffrey Barrett, Sonia Goncalves, Ewan Harrison, David K. Jackson, Ian Johnston, Dominic Kwiatkowski, Cordelia Langford, John Sillitoe on behalf of the Wellcome Sanger Institute COVID-19 Surveillance Team |
| EPI_ISL_3466949 | Laboratory of Clinical Microbiology, Faculty of Medicine, University of Indonesia | Eijkman Institute for Molecular Biology, National Research and Innovation Agency | Iskandar Adnan, Sukma Oktavianthi, Lidwina Priliani, Jessica R. Saragih, Edison Johar, Frilasita A Yudhaputri, Muhammad Rezki Rasyak, Willy Agustine, Hidayat Trimarsanto, Lydia V. Panggalo, Safarina G Malik, Khin Saw Myint, Amin Soebandrio |
| EPI_ISL_3466950, EPI_ISL_3466951 | Lighthouse Lab in Glasgow | Wellcome Sanger Institute for the COVID-19 Genomics UK (COG-UK) Consortium | Harper VanSteenhouse, Yumi Kasai, David Gray, Carol Clugston, Anna Dominiczak and Alex Alderton, Roberto Amato, Jeffrey Barrett, Sonia Goncalves, Ewan Harrison, David K. Jackson, Ian Johnston, Dominic Kwiatkowski, Cordelia Langford, John Sillitoe on behalf of the Wellcome Sanger Institute COVID-19 Surveillance Team |
| EPI_ISL_3466952 | Lighthouse Lab in Milton Keynes | Wellcome Sanger Institute for the COVID-19 Genomics UK (COG-UK) | The Lighthouse Lab in Milton Keynes and Alex Alderton, Roberto Amato, Jeffrey Barrett, Sonia Goncalves, Ewan Harrison, David K. Jackson, Ian Johnston, Dominic Kwiatkowski, Cordelia Langford, John Sillitoe on behalf of the Wellcome Sanger |

| | | | |
|---|---|---|---|
| | | Consortium | Institute COVID-19 Surveillance Team |
| EPI_ISL_3466953 | Lighthouse Lab in Alderley Park | Wellcome Sanger Institute for the COVID-19 Genomics UK (COG-UK) Consortium | Jacquelyn Wynn, Mairead Hyland, The Lighthouse Lab in Alderley Park and Alex Alderton, Roberto Amato, Jeffrey Barrett, Sonia Goncalves, Ewan Harrison, David K. Jackson, Ian Johnston, Dominic Kwiatkowski, Cordelia Langford, John Sillitoe on behalf of the Wellcome Sanger Institute COVID-19 Surveillance Team |
| EPI_ISL_3466954, EPI_ISL_3466955 | Lighthouse Lab in Milton Keynes | Wellcome Sanger Institute for the COVID-19 Genomics UK (COG-UK) Consortium | The Lighthouse Lab in Milton Keynes and Alex Alderton, Roberto Amato, Jeffrey Barrett, Sonia Goncalves, Ewan Harrison, David K. Jackson, Ian Johnston, Dominic Kwiatkowski, Cordelia Langford, John Sillitoe on behalf of the Wellcome Sanger Institute COVID-19 Surveillance Team |
| EPI_ISL_3466956 | Laboratory of Clinical Microbiology, Faculty of Medicine, University of Indonesia | Eijkman Institute for Molecular Biology, National Research and Innovation Agency | Frilasita A Yudhaputri, Muhammad Rezki Rasyak, Willy Agustine, Hidayat Trimarsanto, Lydia V. Panggalo, Iskandar Adnan, Sukma Oktavianthi, Lidwina Priliani, Jessica R. Saragih, Edison Johar, Safarina G Malik, Khin Saw Myint, Amin Soebandrio |
| EPI_ISL_3466957 | Lighthouse Lab in Glasgow | Wellcome Sanger Institute for the COVID-19 Genomics UK (COG-UK) Consortium | Harper VanSteenhouse, Yumi Kasai, David Gray, Carol Clugston, Anna Dominiczak and Alex Alderton, Roberto Amato, Jeffrey Barrett, Sonia Goncalves, Ewan Harrison, David K. Jackson, Ian Johnston, Dominic Kwiatkowski, Cordelia Langford, John Sillitoe on behalf of the Wellcome Sanger Institute COVID-19 Surveillance Team |
| EPI_ISL_3466958 | Lighthouse Lab in Milton Keynes | Wellcome Sanger Institute for the COVID-19 Genomics UK (COG-UK) Consortium | The Lighthouse Lab in Milton Keynes and Alex Alderton, Roberto Amato, Jeffrey Barrett, Sonia Goncalves, Ewan Harrison, David K. Jackson, Ian Johnston, Dominic Kwiatkowski, Cordelia Langford, John Sillitoe on behalf of the Wellcome Sanger Institute COVID-19 Surveillance Team |
| EPI_ISL_3466959 | Lighthouse Lab in Glasgow | Wellcome Sanger Institute for the COVID-19 Genomics UK (COG-UK) Consortium | Harper VanSteenhouse, Yumi Kasai, David Gray, Carol Clugston, Anna Dominiczak and Alex Alderton, Roberto Amato, Jeffrey Barrett, Sonia Goncalves, Ewan Harrison, David K. Jackson, Ian Johnston, Dominic Kwiatkowski, Cordelia Langford, John Sillitoe on behalf of the Wellcome Sanger Institute COVID-19 Surveillance Team |
| EPI_ISL_3466960 | Lighthouse Lab in Milton Keynes | Wellcome Sanger Institute for the COVID-19 Genomics UK (COG-UK) Consortium | The Lighthouse Lab in Milton Keynes and Alex Alderton, Roberto Amato, Jeffrey Barrett, Sonia Goncalves, Ewan Harrison, David K. Jackson, Ian Johnston, Dominic Kwiatkowski, Cordelia Langford, John Sillitoe on behalf of the Wellcome Sanger Institute COVID-19 Surveillance Team |
| EPI_ISL_3466961 | Lighthouse Lab in Glasgow | Wellcome Sanger Institute for the COVID-19 Genomics UK (COG-UK) Consortium | Harper VanSteenhouse, Yumi Kasai, David Gray, Carol Clugston, Anna Dominiczak and Alex Alderton, Roberto Amato, Jeffrey Barrett, Sonia Goncalves, Ewan Harrison, David K. Jackson, Ian Johnston, Dominic Kwiatkowski, Cordelia Langford, John Sillitoe on behalf of the Wellcome Sanger Institute COVID-19 Surveillance Team |
| EPI_ISL_3466963, EPI_ISL_3466964, EPI_ISL_3466965, EPI_ISL_3466966 | Lighthouse Lab in Milton Keynes | Wellcome Sanger Institute for the COVID-19 Genomics UK (COG-UK) Consortium | The Lighthouse Lab in Milton Keynes and Alex Alderton, Roberto Amato, Jeffrey Barrett, Sonia Goncalves, Ewan Harrison, David K. Jackson, Ian Johnston, Dominic Kwiatkowski, Cordelia Langford, John Sillitoe on behalf of the Wellcome Sanger Institute COVID-19 Surveillance Team |
| EPI_ISL_3466967 | Lighthouse Lab in Glasgow | Wellcome Sanger Institute for the COVID-19 Genomics UK (COG-UK) Consortium | Harper VanSteenhouse, Yumi Kasai, David Gray, Carol Clugston, Anna Dominiczak and Alex Alderton, Roberto Amato, Jeffrey Barrett, Sonia Goncalves, Ewan Harrison, David K. Jackson, Ian Johnston, Dominic Kwiatkowski, Cordelia Langford, John Sillitoe on behalf of the Wellcome Sanger Institute COVID-19 Surveillance Team |
| EPI_ISL_3466968 | Clinical Pathology Laboratory, RSUP Dr. Hasan Sadikin | Eijkman Institute for Molecular Biology, National Research and Innovation Agency | Edison Johar, Frilasita A Yudhaputri, Muhammad Rezki Rasyak, Willy Agustine, Hidayat Trimarsanto, Lydia V. Panggalo, Iskandar Adnan, Sukma Oktavianthi, Lidwina Priliani, Jessica R. Saragih, Safarina G Malik, Khin Saw Myint, Amin Soebandrio |
| EPI_ISL_3466969, EPI_ISL_3466970, EPI_ISL_3466971, EPI_ISL_3466972 | Lighthouse Lab in Milton Keynes | Wellcome Sanger Institute for the COVID-19 Genomics UK (COG-UK) Consortium | The Lighthouse Lab in Milton Keynes and Alex Alderton, Roberto Amato, Jeffrey Barrett, Sonia Goncalves, Ewan Harrison, David K. Jackson, Ian Johnston, Dominic Kwiatkowski, Cordelia Langford, John Sillitoe on behalf of the Wellcome Sanger Institute COVID-19 Surveillance Team |
| EPI_ISL_3466973 | Lighthouse Lab in Glasgow | Wellcome Sanger Institute for the COVID-19 Genomics UK (COG-UK) Consortium | Harper VanSteenhouse, Yumi Kasai, David Gray, Carol Clugston, Anna Dominiczak and Alex Alderton, Roberto Amato, Jeffrey Barrett, Sonia Goncalves, Ewan Harrison, David K. Jackson, Ian Johnston, Dominic Kwiatkowski, Cordelia Langford, John Sillitoe on behalf of the Wellcome Sanger Institute COVID-19 Surveillance Team |
| EPI_ISL_3466974 | Clinical Pathology Laboratory, RSUP Dr. Hasan Sadikin | Eijkman Institute for Molecular Biology, National Research and Innovation Agency | Lydia V. Panggalo, Iskandar Adnan, Sukma Oktavianthi, Lidwina Priliani, Jessica R. Saragih, Edison Johar, Frilasita A Yudhaputri, Muhammad Rezki Rasyak, Willy Agustine, Hidayat Trimarsanto, Safarina G Malik, Khin Saw Myint, Amin Soebandrio |
| EPI_ISL_3466975 | Lighthouse Lab in Glasgow | Wellcome Sanger Institute for the COVID-19 Genomics UK (COG-UK) Consortium | Harper VanSteenhouse, Yumi Kasai, David Gray, Carol Clugston, Anna Dominiczak and Alex Alderton, Roberto Amato, Jeffrey Barrett, Sonia Goncalves, Ewan Harrison, David K. Jackson, Ian Johnston, Dominic Kwiatkowski, Cordelia Langford, John Sillitoe on behalf of the Wellcome Sanger Institute COVID-19 Surveillance Team |
| EPI_ISL_3466976, EPI_ISL_3466977, EPI_ISL_3466978 | Lighthouse Lab in Milton Keynes | Wellcome Sanger Institute for the COVID-19 Genomics UK (COG-UK) Consortium | The Lighthouse Lab in Milton Keynes and Alex Alderton, Roberto Amato, Jeffrey Barrett, Sonia Goncalves, Ewan Harrison, David K. Jackson, Ian Johnston, Dominic Kwiatkowski, Cordelia Langford, John Sillitoe on behalf of the Wellcome Sanger Institute COVID-19 Surveillance Team |
| EPI_ISL_3466979 | Clinical Pathology Laboratory, RSUP Dr. Hasan Sadikin | Eijkman Institute for Molecular Biology, National Research and Innovation Agency | Iskandar Adnan, Sukma Oktavianthi, Lidwina Priliani, Jessica R. Saragih, Edison Johar, Frilasita A Yudhaputri, Muhammad Rezki Rasyak, Willy Agustine, Hidayat Trimarsanto, Lydia V. Panggalo, Safarina G Malik, Khin Saw Myint, Amin Soebandrio |
| EPI_ISL_3466980 | Lighthouse Lab in Glasgow | Wellcome Sanger Institute for the COVID-19 Genomics UK (COG-UK) Consortium | Harper VanSteenhouse, Yumi Kasai, David Gray, Carol Clugston, Anna Dominiczak and Alex Alderton, Roberto Amato, Jeffrey Barrett, Sonia Goncalves, Ewan Harrison, David K. Jackson, Ian Johnston, Dominic Kwiatkowski, Cordelia Langford, John Sillitoe on behalf of the Wellcome Sanger Institute COVID-19 Surveillance Team |
| EPI_ISL_3466981 | Lighthouse Lab in Milton Keynes | Wellcome Sanger Institute for the COVID-19 Genomics UK (COG-UK) Consortium | The Lighthouse Lab in Milton Keynes and Alex Alderton, Roberto Amato, Jeffrey Barrett, Sonia Goncalves, Ewan Harrison, David K. Jackson, Ian Johnston, Dominic Kwiatkowski, Cordelia Langford, John Sillitoe on behalf of the Wellcome Sanger Institute COVID-19 Surveillance Team |
| EPI_ISL_3466982 | Lighthouse Lab in Glasgow | Wellcome Sanger Institute for the COVID-19 Genomics UK (COG-UK) Consortium | Harper VanSteenhouse, Yumi Kasai, David Gray, Carol Clugston, Anna Dominiczak and Alex Alderton, Roberto Amato, Jeffrey Barrett, Sonia Goncalves, Ewan Harrison, David K. Jackson, Ian Johnston, Dominic Kwiatkowski, Cordelia Langford, John Sillitoe on behalf of the Wellcome Sanger Institute COVID-19 Surveillance Team |
| EPI_ISL_3466983, EPI_ISL_3466984, EPI_ISL_3466985 | Lighthouse Lab in Milton Keynes | Wellcome Sanger Institute for the COVID-19 Genomics UK (COG-UK) Consortium | The Lighthouse Lab in Milton Keynes and Alex Alderton, Roberto Amato, Jeffrey Barrett, Sonia Goncalves, Ewan Harrison, David K. Jackson, Ian Johnston, Dominic Kwiatkowski, Cordelia Langford, John Sillitoe on behalf of the Wellcome Sanger Institute COVID-19 Surveillance Team |
| EPI_ISL_3466986 | Clinical Pathology Laboratory, RSUP Dr. Hasan Sadikin | Eijkman Institute for Molecular Biology, National Research and Innovation Agency | Lydia V. Panggalo, Iskandar Adnan, Sukma Oktavianthi, Lidwina Priliani, Jessica R. Saragih, Edison Johar, Frilasita A Yudhaputri, Muhammad Rezki Rasyak, Willy Agustine, Hidayat Trimarsanto, Safarina G Malik, Khin Saw Myint, Amin Soebandrio |
| EPI_ISL_3466987, EPI_ISL_3466988, EPI_ISL_3466989, EPI_ISL_3466990, EPI_ISL_3466991, EPI_ISL_3466993, EPI_ISL_3466994, EPI_ISL_3466995, EPI_ISL_3466996, EPI_ISL_3466997, EPI_ISL_3466998 | | | |
| see above | Lighthouse Lab in Milton Keynes | Wellcome Sanger Institute for the COVID-19 Genomics UK (COG-UK) Consortium | The Lighthouse Lab in Milton Keynes and Alex Alderton, Roberto Amato, Jeffrey Barrett, Sonia Goncalves, Ewan Harrison, David K. Jackson, Ian Johnston, Dominic Kwiatkowski, Cordelia Langford, John Sillitoe on behalf of the Wellcome Sanger Institute COVID-19 Surveillance Team |
| EPI_ISL_3466999 | Lighthouse Lab in Glasgow | Wellcome Sanger Institute for the COVID-19 Genomics UK (COG-UK) Consortium | Harper VanSteenhouse, Yumi Kasai, David Gray, Carol Clugston, Anna Dominiczak and Alex Alderton, Roberto Amato, Jeffrey Barrett, Sonia Goncalves, Ewan Harrison, David K. Jackson, Ian Johnston, Dominic Kwiatkowski, Cordelia Langford, John Sillitoe on behalf of the Wellcome Sanger Institute COVID-19 Surveillance Team |
| EPI_ISL_3467000 | Lighthouse Lab in Alderley Park | Wellcome Sanger Institute for the COVID-19 Genomics UK (COG-UK) Consortium | Jacquelyn Wynn, Mairead Hyland, The Lighthouse Lab in Alderley Park and Alex Alderton, Roberto Amato, Jeffrey Barrett, Sonia Goncalves, Ewan Harrison, David K. Jackson, Ian Johnston, Dominic Kwiatkowski, Cordelia Langford, John Sillitoe on behalf of the Wellcome Sanger Institute COVID-19 Surveillance Team |
| EPI_ISL_3467001, EPI_ISL_3467003 | Lighthouse Lab in Milton Keynes | Wellcome Sanger Institute for the COVID-19 Genomics UK (COG-UK) Consortium | The Lighthouse Lab in Milton Keynes and Alex Alderton, Roberto Amato, Jeffrey Barrett, Sonia Goncalves, Ewan Harrison, David K. Jackson, Ian Johnston, Dominic Kwiatkowski, Cordelia Langford, John Sillitoe on behalf of the Wellcome Sanger Institute COVID-19 Surveillance Team |
| EPI_ISL_3467004 | Lighthouse Lab in Glasgow | Wellcome Sanger Institute for the COVID-19 Genomics UK (COG-UK) Consortium | Harper VanSteenhouse, Yumi Kasai, David Gray, Carol Clugston, Anna Dominiczak and Alex Alderton, Roberto Amato, Jeffrey Barrett, Sonia Goncalves, Ewan Harrison, David K. Jackson, Ian Johnston, Dominic Kwiatkowski, Cordelia Langford, John Sillitoe on behalf of the Wellcome Sanger Institute COVID-19 Surveillance Team |
| EPI_ISL_3467005, EPI_ISL_3467006, EPI_ISL_3467008, EPI_ISL_3467009, EPI_ISL_3467010 | Lighthouse Lab in Milton Keynes | Wellcome Sanger Institute for the COVID-19 Genomics UK (COG-UK) Consortium | The Lighthouse Lab in Milton Keynes and Alex Alderton, Roberto Amato, Jeffrey Barrett, Sonia Goncalves, Ewan Harrison, David K. Jackson, Ian Johnston, Dominic Kwiatkowski, Cordelia Langford, John Sillitoe on behalf of the Wellcome Sanger Institute COVID-19 Surveillance Team |
| EPI_ISL_3467011 | Lighthouse Lab in Glasgow | Wellcome Sanger Institute for the COVID-19 Genomics UK (COG-UK) Consortium | Harper VanSteenhouse, Yumi Kasai, David Gray, Carol Clugston, Anna Dominiczak and Alex Alderton, Roberto Amato, Jeffrey Barrett, Sonia Goncalves, Ewan Harrison, David K. Jackson, Ian Johnston, Dominic Kwiatkowski, Cordelia Langford, John Sillitoe on behalf of the Wellcome Sanger Institute COVID-19 Surveillance Team |
| EPI_ISL_3467012 | Lighthouse Lab in Milton Keynes | Wellcome Sanger Institute for the COVID-19 Genomics UK (COG-UK) Consortium | The Lighthouse Lab in Milton Keynes and Alex Alderton, Roberto Amato, Jeffrey Barrett, Sonia Goncalves, Ewan Harrison, David K. Jackson, Ian Johnston, Dominic Kwiatkowski, Cordelia Langford, John Sillitoe on behalf of the Wellcome Sanger Institute COVID-19 Surveillance Team |
| EPI_ISL_3467013 | Lighthouse Lab in Glasgow | Wellcome Sanger Institute for the COVID-19 Genomics UK (COG-UK) Consortium | Harper VanSteenhouse, Yumi Kasai, David Gray, Carol Clugston, Anna Dominiczak and Alex Alderton, Roberto Amato, Jeffrey Barrett, Sonia Goncalves, Ewan Harrison, David K. Jackson, Ian Johnston, Dominic Kwiatkowski, Cordelia Langford, John Sillitoe on behalf of the Wellcome Sanger Institute COVID-19 Surveillance Team |
| EPI_ISL_3467015, EPI_ISL_3467016 | Lighthouse Lab in Milton Keynes | Wellcome Sanger Institute for the COVID-19 Genomics UK (COG-UK) Consortium | The Lighthouse Lab in Milton Keynes and Alex Alderton, Roberto Amato, Jeffrey Barrett, Sonia Goncalves, Ewan Harrison, David K. Jackson, Ian Johnston, Dominic Kwiatkowski, Cordelia Langford, John Sillitoe on behalf of the Wellcome Sanger Institute COVID-19 Surveillance Team |
| EPI_ISL_3467017 | Lighthouse Lab in Glasgow | Wellcome Sanger Institute for the COVID-19 Genomics UK (COG-UK) Consortium | Harper VanSteenhouse, Yumi Kasai, David Gray, Carol Clugston, Anna Dominiczak and Alex Alderton, Roberto Amato, Jeffrey Barrett, Sonia Goncalves, Ewan Harrison, David K. Jackson, Ian Johnston, Dominic Kwiatkowski, Cordelia Langford, John Sillitoe on behalf of the Wellcome Sanger Institute COVID-19 Surveillance Team |
| EPI_ISL_3467018 | Lighthouse Lab in Milton Keynes | Wellcome Sanger Institute for the COVID-19 Genomics UK (COG-UK) Consortium | The Lighthouse Lab in Milton Keynes and Alex Alderton, Roberto Amato, Jeffrey Barrett, Sonia Goncalves, Ewan Harrison, David K. Jackson, Ian Johnston, Dominic Kwiatkowski, Cordelia Langford, John Sillitoe on behalf of the Wellcome Sanger Institute COVID-19 Surveillance Team |
| EPI_ISL_3467019 | Lighthouse Lab in Glasgow | Wellcome Sanger Institute for the COVID-19 Genomics UK (COG-UK) Consortium | Harper VanSteenhouse, Yumi Kasai, David Gray, Carol Clugston, Anna Dominiczak and Alex Alderton, Roberto Amato, Jeffrey Barrett, Sonia Goncalves, Ewan Harrison, David K. Jackson, Ian Johnston, Dominic Kwiatkowski, Cordelia Langford, John Sillitoe on behalf of the Wellcome Sanger Institute COVID-19 Surveillance Team |
| EPI_ISL_3467020 | Lighthouse Lab in Milton Keynes | Wellcome Sanger Institute for the COVID-19 Genomics UK (COG-UK) Consortium | The Lighthouse Lab in Milton Keynes and Alex Alderton, Roberto Amato, Jeffrey Barrett, Sonia Goncalves, Ewan Harrison, David K. Jackson, Ian Johnston, Dominic Kwiatkowski, Cordelia Langford, John Sillitoe on behalf of the Wellcome Sanger Institute COVID-19 Surveillance Team |
| EPI_ISL_3467021 | Lighthouse Lab in Glasgow | Wellcome Sanger Institute for the COVID-19 Genomics UK (COG-UK) Consortium | Harper VanSteenhouse, Yumi Kasai, David Gray, Carol Clugston, Anna Dominiczak and Alex Alderton, Roberto Amato, Jeffrey Barrett, Sonia Goncalves, Ewan Harrison, David K. Jackson, Ian Johnston, Dominic Kwiatkowski, Cordelia Langford, John Sillitoe on behalf of the Wellcome Sanger Institute COVID-19 Surveillance Team |
| EPI_ISL_3467022, EPI_ISL_3467023, EPI_ISL_3467024, EPI_ISL_3467025, EPI_ISL_3467026, EPI_ISL_3467027, EPI_ISL_3467028, EPI_ISL_3467029 | Lighthouse Lab in Milton Keynes | Wellcome Sanger Institute for the COVID-19 Genomics UK (COG-UK) Consortium | The Lighthouse Lab in Milton Keynes and Alex Alderton, Roberto Amato, Jeffrey Barrett, Sonia Goncalves, Ewan Harrison, David K. Jackson, Ian Johnston, Dominic Kwiatkowski, Cordelia Langford, John Sillitoe on behalf of the Wellcome Sanger Institute COVID-19 Surveillance Team |

| | | | |
|---|---|---|---|
| EPI_ISL_3467030 | Lighthouse Lab in Glasgow | Wellcome Sanger Institute for the COVID-19 Genomics UK (COG-UK) Consortium | Harper VanSteenhouse, Yumi Kasai, David Gray, Carol Clugston, Anna Dominiczak and Alex Alderton, Roberto Amato, Jeffrey Barrett, Sonia Goncalves, Ewan Harrison, David K. Jackson, Ian Johnston, Dominic Kwiatkowski, Cordelia Langford, John Sillitoe on behalf of the Wellcome Sanger Institute COVID-19 Surveillance Team |
| EPI_ISL_3467031, EPI_ISL_3467032, EPI_ISL_3467033 | Lighthouse Lab in Milton Keynes | Wellcome Sanger Institute for the COVID-19 Genomics UK (COG-UK) Consortium | The Lighthouse Lab in Milton Keynes and Alex Alderton, Roberto Amato, Jeffrey Barrett, Sonia Goncalves, Ewan Harrison, David K. Jackson, Ian Johnston, Dominic Kwiatkowski, Cordelia Langford, John Sillitoe on behalf of the Wellcome Sanger Institute COVID-19 Surveillance Team |
| EPI_ISL_3467034 | Lighthouse Lab in Alderley Park | Wellcome Sanger Institute for the COVID-19 Genomics UK (COG-UK) Consortium | Jacquelyn Wynn, Mairead Hyland, The Lighthouse Lab in Alderley Park and Alex Alderton, Roberto Amato, Jeffrey Barrett, Sonia Goncalves, Ewan Harrison, David K. Jackson, Ian Johnston, Dominic Kwiatkowski, Cordelia Langford, John Sillitoe on behalf of the Wellcome Sanger Institute COVID-19 Surveillance Team |
| EPI_ISL_3467035, EPI_ISL_3467036 | Lighthouse Lab in Glasgow | Wellcome Sanger Institute for the COVID-19 Genomics UK (COG-UK) Consortium | Harper VanSteenhouse, Yumi Kasai, David Gray, Carol Clugston, Anna Dominiczak and Alex Alderton, Roberto Amato, Jeffrey Barrett, Sonia Goncalves, Ewan Harrison, David K. Jackson, Ian Johnston, Dominic Kwiatkowski, Cordelia Langford, John Sillitoe on behalf of the Wellcome Sanger Institute COVID-19 Surveillance Team |
| EPI_ISL_3467037 | Lighthouse Lab in Milton Keynes | Wellcome Sanger Institute for the COVID-19 Genomics UK (COG-UK) Consortium | The Lighthouse Lab in Milton Keynes and Alex Alderton, Roberto Amato, Jeffrey Barrett, Sonia Goncalves, Ewan Harrison, David K. Jackson, Ian Johnston, Dominic Kwiatkowski, Cordelia Langford, John Sillitoe on behalf of the Wellcome Sanger Institute COVID-19 Surveillance Team |
| EPI_ISL_3467038 | Lighthouse Lab in Glasgow | Wellcome Sanger Institute for the COVID-19 Genomics UK (COG-UK) Consortium | Harper VanSteenhouse, Yumi Kasai, David Gray, Carol Clugston, Anna Dominiczak and Alex Alderton, Roberto Amato, Jeffrey Barrett, Sonia Goncalves, Ewan Harrison, David K. Jackson, Ian Johnston, Dominic Kwiatkowski, Cordelia Langford, John Sillitoe on behalf of the Wellcome Sanger Institute COVID-19 Surveillance Team |
| EPI_ISL_3467039 | Lighthouse Lab in Alderley Park | Wellcome Sanger Institute for the COVID-19 Genomics UK (COG-UK) Consortium | Jacquelyn Wynn, Mairead Hyland, The Lighthouse Lab in Alderley Park and Alex Alderton, Roberto Amato, Jeffrey Barrett, Sonia Goncalves, Ewan Harrison, David K. Jackson, Ian Johnston, Dominic Kwiatkowski, Cordelia Langford, John Sillitoe on behalf of the Wellcome Sanger Institute COVID-19 Surveillance Team |
| EPI_ISL_3467040 | Lighthouse Lab in Milton Keynes | Wellcome Sanger Institute for the COVID-19 Genomics UK (COG-UK) Consortium | The Lighthouse Lab in Milton Keynes and Alex Alderton, Roberto Amato, Jeffrey Barrett, Sonia Goncalves, Ewan Harrison, David K. Jackson, Ian Johnston, Dominic Kwiatkowski, Cordelia Langford, John Sillitoe on behalf of the Wellcome Sanger Institute COVID-19 Surveillance Team |
| EPI_ISL_3467041, EPI_ISL_3467042 | Lighthouse Lab in Glasgow | Wellcome Sanger Institute for the COVID-19 Genomics UK (COG-UK) Consortium | Harper VanSteenhouse, Yumi Kasai, David Gray, Carol Clugston, Anna Dominiczak and Alex Alderton, Roberto Amato, Jeffrey Barrett, Sonia Goncalves, Ewan Harrison, David K. Jackson, Ian Johnston, Dominic Kwiatkowski, Cordelia Langford, John Sillitoe on behalf of the Wellcome Sanger Institute COVID-19 Surveillance Team |
| EPI_ISL_3467043, EPI_ISL_3467044, EPI_ISL_3467045, EPI_ISL_3467046 | Lighthouse Lab in Milton Keynes | Wellcome Sanger Institute for the COVID-19 Genomics UK (COG-UK) Consortium | The Lighthouse Lab in Milton Keynes and Alex Alderton, Roberto Amato, Jeffrey Barrett, Sonia Goncalves, Ewan Harrison, David K. Jackson, Ian Johnston, Dominic Kwiatkowski, Cordelia Langford, John Sillitoe on behalf of the Wellcome Sanger Institute COVID-19 Surveillance Team |
| EPI_ISL_3467047 | Lighthouse Lab in Glasgow | Wellcome Sanger Institute for the COVID-19 Genomics UK (COG-UK) Consortium | Harper VanSteenhouse, Yumi Kasai, David Gray, Carol Clugston, Anna Dominiczak and Alex Alderton, Roberto Amato, Jeffrey Barrett, Sonia Goncalves, Ewan Harrison, David K. Jackson, Ian Johnston, Dominic Kwiatkowski, Cordelia Langford, John Sillitoe on behalf of the Wellcome Sanger Institute COVID-19 Surveillance Team |
| EPI_ISL_3467048, EPI_ISL_3467049, EPI_ISL_3467050, EPI_ISL_3467051, EPI_ISL_3467052 | Lighthouse Lab in Milton Keynes | Wellcome Sanger Institute for the COVID-19 Genomics UK (COG-UK) Consortium | The Lighthouse Lab in Milton Keynes and Alex Alderton, Roberto Amato, Jeffrey Barrett, Sonia Goncalves, Ewan Harrison, David K. Jackson, Ian Johnston, Dominic Kwiatkowski, Cordelia Langford, John Sillitoe on behalf of the Wellcome Sanger Institute COVID-19 Surveillance Team |
| EPI_ISL_3467053 | Lighthouse Lab in Glasgow | Wellcome Sanger Institute for the COVID-19 Genomics UK (COG-UK) Consortium | Harper VanSteenhouse, Yumi Kasai, David Gray, Carol Clugston, Anna Dominiczak and Alex Alderton, Roberto Amato, Jeffrey Barrett, Sonia Goncalves, Ewan Harrison, David K. Jackson, Ian Johnston, Dominic Kwiatkowski, Cordelia Langford, John Sillitoe on behalf of the Wellcome Sanger Institute COVID-19 Surveillance Team |
| EPI_ISL_3467054, EPI_ISL_3467055 | Lighthouse Lab in Milton Keynes | Wellcome Sanger Institute for the COVID-19 Genomics UK (COG-UK) Consortium | The Lighthouse Lab in Milton Keynes and Alex Alderton, Roberto Amato, Jeffrey Barrett, Sonia Goncalves, Ewan Harrison, David K. Jackson, Ian Johnston, Dominic Kwiatkowski, Cordelia Langford, John Sillitoe on behalf of the Wellcome Sanger Institute COVID-19 Surveillance Team |
| EPI_ISL_3467056 | Lighthouse Lab in Glasgow | Wellcome Sanger Institute for the COVID-19 Genomics UK (COG-UK) Consortium | Harper VanSteenhouse, Yumi Kasai, David Gray, Carol Clugston, Anna Dominiczak and Alex Alderton, Roberto Amato, Jeffrey Barrett, Sonia Goncalves, Ewan Harrison, David K. Jackson, Ian Johnston, Dominic Kwiatkowski, Cordelia Langford, John Sillitoe on behalf of the Wellcome Sanger Institute COVID-19 Surveillance Team |
| EPI_ISL_3467057 | Lighthouse Lab in Milton Keynes | Wellcome Sanger Institute for the COVID-19 Genomics UK (COG-UK) Consortium | The Lighthouse Lab in Milton Keynes and Alex Alderton, Roberto Amato, Jeffrey Barrett, Sonia Goncalves, Ewan Harrison, David K. Jackson, Ian Johnston, Dominic Kwiatkowski, Cordelia Langford, John Sillitoe on behalf of the Wellcome Sanger Institute COVID-19 Surveillance Team |
| EPI_ISL_3467058, EPI_ISL_3467059, EPI_ISL_3467060, EPI_ISL_3467061 | Lighthouse Lab in Glasgow | Wellcome Sanger Institute for the COVID-19 Genomics UK (COG-UK) Consortium | Harper VanSteenhouse, Yumi Kasai, David Gray, Carol Clugston, Anna Dominiczak and Alex Alderton, Roberto Amato, Jeffrey Barrett, Sonia Goncalves, Ewan Harrison, David K. Jackson, Ian Johnston, Dominic Kwiatkowski, Cordelia Langford, John Sillitoe on behalf of the Wellcome Sanger Institute COVID-19 Surveillance Team |
| EPI_ISL_3467062, EPI_ISL_3467063 | Lighthouse Lab in Milton Keynes | Wellcome Sanger Institute for the COVID-19 Genomics UK (COG-UK) Consortium | The Lighthouse Lab in Milton Keynes and Alex Alderton, Roberto Amato, Jeffrey Barrett, Sonia Goncalves, Ewan Harrison, David K. Jackson, Ian Johnston, Dominic Kwiatkowski, Cordelia Langford, John Sillitoe on behalf of the Wellcome Sanger Institute COVID-19 Surveillance Team |
| EPI_ISL_3467064 | Lighthouse Lab in Alderley Park | Wellcome Sanger Institute for the COVID-19 Genomics UK (COG-UK) Consortium | Jacquelyn Wynn, Mairead Hyland, The Lighthouse Lab in Alderley Park and Alex Alderton, Roberto Amato, Jeffrey Barrett, Sonia Goncalves, Ewan Harrison, David K. Jackson, Ian Johnston, Dominic Kwiatkowski, Cordelia Langford, John Sillitoe on behalf of the Wellcome Sanger Institute COVID-19 Surveillance Team |
| EPI_ISL_3467065 | Lighthouse Lab in Milton Keynes | Wellcome Sanger Institute for the COVID-19 Genomics UK (COG-UK) Consortium | The Lighthouse Lab in Milton Keynes and Alex Alderton, Roberto Amato, Jeffrey Barrett, Sonia Goncalves, Ewan Harrison, David K. Jackson, Ian Johnston, Dominic Kwiatkowski, Cordelia Langford, John Sillitoe on behalf of the Wellcome Sanger Institute COVID-19 Surveillance Team |
| EPI_ISL_3467066 | Lighthouse Lab in Glasgow | Wellcome Sanger Institute for the COVID-19 Genomics UK (COG-UK) Consortium | Harper VanSteenhouse, Yumi Kasai, David Gray, Carol Clugston, Anna Dominiczak and Alex Alderton, Roberto Amato, Jeffrey Barrett, Sonia Goncalves, Ewan Harrison, David K. Jackson, Ian Johnston, Dominic Kwiatkowski, Cordelia Langford, John Sillitoe on behalf of the Wellcome Sanger Institute COVID-19 Surveillance Team |
| EPI_ISL_3467067, EPI_ISL_3467068, EPI_ISL_3467069 | Lighthouse Lab in Milton Keynes | Wellcome Sanger Institute for the COVID-19 Genomics UK (COG-UK) Consortium | The Lighthouse Lab in Milton Keynes and Alex Alderton, Roberto Amato, Jeffrey Barrett, Sonia Goncalves, Ewan Harrison, David K. Jackson, Ian Johnston, Dominic Kwiatkowski, Cordelia Langford, John Sillitoe on behalf of the Wellcome Sanger Institute COVID-19 Surveillance Team |
| EPI_ISL_3467070 | Lighthouse Lab in Glasgow | Wellcome Sanger Institute for the COVID-19 Genomics UK (COG-UK) Consortium | Harper VanSteenhouse, Yumi Kasai, David Gray, Carol Clugston, Anna Dominiczak and Alex Alderton, Roberto Amato, Jeffrey Barrett, Sonia Goncalves, Ewan Harrison, David K. Jackson, Ian Johnston, Dominic Kwiatkowski, Cordelia Langford, John Sillitoe on behalf of the Wellcome Sanger Institute COVID-19 Surveillance Team |
| EPI_ISL_3467071, EPI_ISL_3467072, EPI_ISL_3467073 | Lighthouse Lab in Milton Keynes | Wellcome Sanger Institute for the COVID-19 Genomics UK (COG-UK) Consortium | The Lighthouse Lab in Milton Keynes and Alex Alderton, Roberto Amato, Jeffrey Barrett, Sonia Goncalves, Ewan Harrison, David K. Jackson, Ian Johnston, Dominic Kwiatkowski, Cordelia Langford, John Sillitoe on behalf of the Wellcome Sanger Institute COVID-19 Surveillance Team |
| EPI_ISL_3467074 | Lighthouse Lab in Glasgow | Wellcome Sanger Institute for the COVID-19 Genomics UK (COG-UK) Consortium | Harper VanSteenhouse, Yumi Kasai, David Gray, Carol Clugston, Anna Dominiczak and Alex Alderton, Roberto Amato, Jeffrey Barrett, Sonia Goncalves, Ewan Harrison, David K. Jackson, Ian Johnston, Dominic Kwiatkowski, Cordelia Langford, John Sillitoe on behalf of the Wellcome Sanger Institute COVID-19 Surveillance Team |
| EPI_ISL_3467075 | Lighthouse Lab in Milton Keynes | Wellcome Sanger Institute for the COVID-19 Genomics UK (COG-UK) Consortium | The Lighthouse Lab in Milton Keynes and Alex Alderton, Roberto Amato, Jeffrey Barrett, Sonia Goncalves, Ewan Harrison, David K. Jackson, Ian Johnston, Dominic Kwiatkowski, Cordelia Langford, John Sillitoe on behalf of the Wellcome Sanger Institute COVID-19 Surveillance Team |
| EPI_ISL_3467076 | Lighthouse Lab in Glasgow | Wellcome Sanger Institute for the COVID-19 Genomics UK (COG-UK) Consortium | Harper VanSteenhouse, Yumi Kasai, David Gray, Carol Clugston, Anna Dominiczak and Alex Alderton, Roberto Amato, Jeffrey Barrett, Sonia Goncalves, Ewan Harrison, David K. Jackson, Ian Johnston, Dominic Kwiatkowski, Cordelia Langford, John Sillitoe on behalf of the Wellcome Sanger Institute COVID-19 Surveillance Team |
| EPI_ISL_3467077, EPI_ISL_3467078 | Lighthouse Lab in Milton Keynes | Wellcome Sanger Institute for the COVID-19 Genomics UK (COG-UK) Consortium | The Lighthouse Lab in Milton Keynes and Alex Alderton, Roberto Amato, Jeffrey Barrett, Sonia Goncalves, Ewan Harrison, David K. Jackson, Ian Johnston, Dominic Kwiatkowski, Cordelia Langford, John Sillitoe on behalf of the Wellcome Sanger Institute COVID-19 Surveillance Team |
| EPI_ISL_3467079 | Lighthouse Lab in Glasgow | Wellcome Sanger Institute for the COVID-19 Genomics UK (COG-UK) Consortium | Harper VanSteenhouse, Yumi Kasai, David Gray, Carol Clugston, Anna Dominiczak and Alex Alderton, Roberto Amato, Jeffrey Barrett, Sonia Goncalves, Ewan Harrison, David K. Jackson, Ian Johnston, Dominic Kwiatkowski, Cordelia Langford, John Sillitoe on behalf of the Wellcome Sanger Institute COVID-19 Surveillance Team |
| EPI_ISL_3467080, EPI_ISL_3467081, EPI_ISL_3467082 | Lighthouse Lab in Milton Keynes | Wellcome Sanger Institute for the COVID-19 Genomics UK (COG-UK) Consortium | The Lighthouse Lab in Milton Keynes and Alex Alderton, Roberto Amato, Jeffrey Barrett, Sonia Goncalves, Ewan Harrison, David K. Jackson, Ian Johnston, Dominic Kwiatkowski, Cordelia Langford, John Sillitoe on behalf of the Wellcome Sanger Institute COVID-19 Surveillance Team |
| EPI_ISL_3467083 | Lighthouse Lab in Glasgow | Wellcome Sanger Institute for the COVID-19 Genomics UK (COG-UK) Consortium | Harper VanSteenhouse, Yumi Kasai, David Gray, Carol Clugston, Anna Dominiczak and Alex Alderton, Roberto Amato, Jeffrey Barrett, Sonia Goncalves, Ewan Harrison, David K. Jackson, Ian Johnston, Dominic Kwiatkowski, Cordelia Langford, John Sillitoe on behalf of the Wellcome Sanger Institute COVID-19 Surveillance Team |
| EPI_ISL_3467084, EPI_ISL_3467085, EPI_ISL_3467086 | Lighthouse Lab in Milton Keynes | Wellcome Sanger Institute for the COVID-19 Genomics UK (COG-UK) Consortium | The Lighthouse Lab in Milton Keynes and Alex Alderton, Roberto Amato, Jeffrey Barrett, Sonia Goncalves, Ewan Harrison, David K. Jackson, Ian Johnston, Dominic Kwiatkowski, Cordelia Langford, John Sillitoe on behalf of the Wellcome Sanger Institute COVID-19 Surveillance Team |
| EPI_ISL_3467087, EPI_ISL_3467088 | Lighthouse Lab in Glasgow | Wellcome Sanger Institute for the COVID-19 Genomics UK (COG-UK) Consortium | Harper VanSteenhouse, Yumi Kasai, David Gray, Carol Clugston, Anna Dominiczak and Alex Alderton, Roberto Amato, Jeffrey Barrett, Sonia Goncalves, Ewan Harrison, David K. Jackson, Ian Johnston, Dominic Kwiatkowski, Cordelia Langford, John Sillitoe on behalf of the Wellcome Sanger Institute COVID-19 Surveillance Team |
| EPI_ISL_3467089 | Lighthouse Lab in Milton Keynes | Wellcome Sanger Institute for the COVID-19 Genomics UK (COG-UK) Consortium | The Lighthouse Lab in Milton Keynes and Alex Alderton, Roberto Amato, Jeffrey Barrett, Sonia Goncalves, Ewan Harrison, David K. Jackson, Ian Johnston, Dominic Kwiatkowski, Cordelia Langford, John Sillitoe on behalf of the Wellcome Sanger Institute COVID-19 Surveillance Team |
| EPI_ISL_3467090 | Lighthouse Lab in Alderley Park | Wellcome Sanger Institute for the COVID-19 Genomics UK (COG-UK) Consortium | Jacquelyn Wynn, Mairead Hyland, The Lighthouse Lab in Alderley Park and Alex Alderton, Roberto Amato, Jeffrey Barrett, Sonia Goncalves, Ewan Harrison, David K. Jackson, Ian Johnston, Dominic Kwiatkowski, Cordelia Langford, John Sillitoe on behalf of the Wellcome Sanger Institute COVID-19 Surveillance Team |
| EPI_ISL_3467091 | Lighthouse Lab in Glasgow | Wellcome Sanger Institute for the COVID-19 Genomics UK (COG-UK) Consortium | Harper VanSteenhouse, Yumi Kasai, David Gray, Carol Clugston, Anna Dominiczak and Alex Alderton, Roberto Amato, Jeffrey Barrett, Sonia Goncalves, Ewan Harrison, David K. Jackson, Ian Johnston, Dominic Kwiatkowski, Cordelia Langford, John Sillitoe on behalf of the Wellcome Sanger Institute COVID-19 Surveillance Team |
| EPI_ISL_3467092, EPI_ISL_3467093, EPI_ISL_3467094 | Lighthouse Lab in Milton Keynes | Wellcome Sanger Institute for the COVID-19 Genomics UK (COG-UK) Consortium | The Lighthouse Lab in Milton Keynes and Alex Alderton, Roberto Amato, Jeffrey Barrett, Sonia Goncalves, Ewan Harrison, David K. Jackson, Ian Johnston, Dominic Kwiatkowski, Cordelia Langford, John Sillitoe on behalf of the Wellcome Sanger Institute COVID-19 Surveillance Team |
| EPI_ISL_3467095 | Lighthouse Lab in Glasgow | Wellcome Sanger Institute for the COVID-19 Genomics UK (COG-UK) Consortium | Harper VanSteenhouse, Yumi Kasai, David Gray, Carol Clugston, Anna Dominiczak and Alex Alderton, Roberto Amato, Jeffrey Barrett, Sonia Goncalves, Ewan Harrison, David K. Jackson, Ian Johnston, Dominic Kwiatkowski, Cordelia Langford, John Sillitoe on behalf of the Wellcome Sanger Institute COVID-19 Surveillance Team |
| EPI_ISL_3467096, EPI_ISL_3467097, EPI_ISL_3467098, | Lighthouse Lab in Milton Keynes | Wellcome Sanger Institute for the COVID-19 Genomics UK (COG-UK) Consortium | The Lighthouse Lab in Milton Keynes and Alex Alderton, Roberto Amato, Jeffrey Barrett, Sonia Goncalves, Ewan Harrison, David K. Jackson, Ian Johnston, Dominic Kwiatkowski, Cordelia Langford, John Sillitoe on behalf of the Wellcome Sanger Institute COVID-19 Surveillance Team |

| | | | |
|---|---|---|---|
| EPI_ISL_3467099, EPI_ISL_3467100, EPI_ISL_3467101, EPI_ISL_3467102, EPI_ISL_3467103, EPI_ISL_3467104 | | | |
| EPI_ISL_3467105 | Lighthouse Lab in Alderley Park | Wellcome Sanger Institute for the COVID-19 Genomics UK (COG-UK) Consortium | Jacquelyn Wynn, Mairead Hyland, The Lighthouse Lab in Alderley Park and Alex Alderton, Roberto Amato, Jeffrey Barrett, Sonia Goncalves, Ewan Harrison, David K. Jackson, Ian Johnston, Dominic Kwiatkowski, Cordelia Langford, John Sillitoe on behalf of the Wellcome Sanger Institute COVID-19 Surveillance Team |
| EPI_ISL_3467106 | Lighthouse Lab in Milton Keynes | Wellcome Sanger Institute for the COVID-19 Genomics UK (COG-UK) Consortium | The Lighthouse Lab in Milton Keynes and Alex Alderton, Roberto Amato, Jeffrey Barrett, Sonia Goncalves, Ewan Harrison, David K. Jackson, Ian Johnston, Dominic Kwiatkowski, Cordelia Langford, John Sillitoe on behalf of the Wellcome Sanger Institute COVID-19 Surveillance Team |
| EPI_ISL_3467107 | Lighthouse Lab in Alderley Park | Wellcome Sanger Institute for the COVID-19 Genomics UK (COG-UK) Consortium | Jacquelyn Wynn, Mairead Hyland, The Lighthouse Lab in Alderley Park and Alex Alderton, Roberto Amato, Jeffrey Barrett, Sonia Goncalves, Ewan Harrison, David K. Jackson, Ian Johnston, Dominic Kwiatkowski, Cordelia Langford, John Sillitoe on behalf of the Wellcome Sanger Institute COVID-19 Surveillance Team |
| EPI_ISL_3467108, EPI_ISL_3467109 | Lighthouse Lab in Milton Keynes | Wellcome Sanger Institute for the COVID-19 Genomics UK (COG-UK) Consortium | The Lighthouse Lab in Milton Keynes and Alex Alderton, Roberto Amato, Jeffrey Barrett, Sonia Goncalves, Ewan Harrison, David K. Jackson, Ian Johnston, Dominic Kwiatkowski, Cordelia Langford, John Sillitoe on behalf of the Wellcome Sanger Institute COVID-19 Surveillance Team |
| EPI_ISL_3467110 | Lighthouse Lab in Alderley Park | Wellcome Sanger Institute for the COVID-19 Genomics UK (COG-UK) Consortium | Jacquelyn Wynn, Mairead Hyland, The Lighthouse Lab in Alderley Park and Alex Alderton, Roberto Amato, Jeffrey Barrett, Sonia Goncalves, Ewan Harrison, David K. Jackson, Ian Johnston, Dominic Kwiatkowski, Cordelia Langford, John Sillitoe on behalf of the Wellcome Sanger Institute COVID-19 Surveillance Team |
| EPI_ISL_3467111 | Lighthouse Lab in Milton Keynes | Wellcome Sanger Institute for the COVID-19 Genomics UK (COG-UK) Consortium | The Lighthouse Lab in Milton Keynes and Alex Alderton, Roberto Amato, Jeffrey Barrett, Sonia Goncalves, Ewan Harrison, David K. Jackson, Ian Johnston, Dominic Kwiatkowski, Cordelia Langford, John Sillitoe on behalf of the Wellcome Sanger Institute COVID-19 Surveillance Team |
| EPI_ISL_3467112 | Lighthouse Lab in Alderley Park | Wellcome Sanger Institute for the COVID-19 Genomics UK (COG-UK) Consortium | Jacquelyn Wynn, Mairead Hyland, The Lighthouse Lab in Alderley Park and Alex Alderton, Roberto Amato, Jeffrey Barrett, Sonia Goncalves, Ewan Harrison, David K. Jackson, Ian Johnston, Dominic Kwiatkowski, Cordelia Langford, John Sillitoe on behalf of the Wellcome Sanger Institute COVID-19 Surveillance Team |
| EPI_ISL_3467113, EPI_ISL_3467114 | Lighthouse Lab in Milton Keynes | Wellcome Sanger Institute for the COVID-19 Genomics UK (COG-UK) Consortium | The Lighthouse Lab in Milton Keynes and Alex Alderton, Roberto Amato, Jeffrey Barrett, Sonia Goncalves, Ewan Harrison, David K. Jackson, Ian Johnston, Dominic Kwiatkowski, Cordelia Langford, John Sillitoe on behalf of the Wellcome Sanger Institute COVID-19 Surveillance Team |
| EPI_ISL_3467115 | Lighthouse Lab in Glasgow | Wellcome Sanger Institute for the COVID-19 Genomics UK (COG-UK) Consortium | Harper VanSteenhouse, Yumi Kasai, David Gray, Carol Clugston, Anna Dominiczak and Alex Alderton, Roberto Amato, Jeffrey Barrett, Sonia Goncalves, Ewan Harrison, David K. Jackson, Ian Johnston, Dominic Kwiatkowski, Cordelia Langford, John Sillitoe on behalf of the Wellcome Sanger Institute COVID-19 Surveillance Team |
| EPI_ISL_3467116, EPI_ISL_3467117, EPI_ISL_3467118, EPI_ISL_3467119, EPI_ISL_3467120, EPI_ISL_3467121 | Lighthouse Lab in Milton Keynes | Wellcome Sanger Institute for the COVID-19 Genomics UK (COG-UK) Consortium | The Lighthouse Lab in Milton Keynes and Alex Alderton, Roberto Amato, Jeffrey Barrett, Sonia Goncalves, Ewan Harrison, David K. Jackson, Ian Johnston, Dominic Kwiatkowski, Cordelia Langford, John Sillitoe on behalf of the Wellcome Sanger Institute COVID-19 Surveillance Team |
| EPI_ISL_3467122 | Lighthouse Lab in Glasgow | Wellcome Sanger Institute for the COVID-19 Genomics UK (COG-UK) Consortium | Harper VanSteenhouse, Yumi Kasai, David Gray, Carol Clugston, Anna Dominiczak and Alex Alderton, Roberto Amato, Jeffrey Barrett, Sonia Goncalves, Ewan Harrison, David K. Jackson, Ian Johnston, Dominic Kwiatkowski, Cordelia Langford, John Sillitoe on behalf of the Wellcome Sanger Institute COVID-19 Surveillance Team |
| EPI_ISL_3467123, EPI_ISL_3467124, EPI_ISL_3467125 | Lighthouse Lab in Milton Keynes | Wellcome Sanger Institute for the COVID-19 Genomics UK (COG-UK) Consortium | The Lighthouse Lab in Milton Keynes and Alex Alderton, Roberto Amato, Jeffrey Barrett, Sonia Goncalves, Ewan Harrison, David K. Jackson, Ian Johnston, Dominic Kwiatkowski, Cordelia Langford, John Sillitoe on behalf of the Wellcome Sanger Institute COVID-19 Surveillance Team |
| EPI_ISL_3467126 | Lighthouse Lab in Glasgow | Wellcome Sanger Institute for the COVID-19 Genomics UK (COG-UK) Consortium | Harper VanSteenhouse, Yumi Kasai, David Gray, Carol Clugston, Anna Dominiczak and Alex Alderton, Roberto Amato, Jeffrey Barrett, Sonia Goncalves, Ewan Harrison, David K. Jackson, Ian Johnston, Dominic Kwiatkowski, Cordelia Langford, John Sillitoe on behalf of the Wellcome Sanger Institute COVID-19 Surveillance Team |
| EPI_ISL_3467127, EPI_ISL_3467128, EPI_ISL_3467129, EPI_ISL_3467130, EPI_ISL_3467131, EPI_ISL_3467132, EPI_ISL_3467133, EPI_ISL_3467134, EPI_ISL_3467135, EPI_ISL_3467136, EPI_ISL_3467137, EPI_ISL_3467138 | | | |
| see above | Lighthouse Lab in Milton Keynes | Wellcome Sanger Institute for the COVID-19 Genomics UK (COG-UK) Consortium | The Lighthouse Lab in Milton Keynes and Alex Alderton, Roberto Amato, Jeffrey Barrett, Sonia Goncalves, Ewan Harrison, David K. Jackson, Ian Johnston, Dominic Kwiatkowski, Cordelia Langford, John Sillitoe on behalf of the Wellcome Sanger Institute COVID-19 Surveillance Team |
| EPI_ISL_3467139 | Lighthouse Lab in Alderley Park | Wellcome Sanger Institute for the COVID-19 Genomics UK (COG-UK) Consortium | Jacquelyn Wynn, Mairead Hyland, The Lighthouse Lab in Alderley Park and Alex Alderton, Roberto Amato, Jeffrey Barrett, Sonia Goncalves, Ewan Harrison, David K. Jackson, Ian Johnston, Dominic Kwiatkowski, Cordelia Langford, John Sillitoe on behalf of the Wellcome Sanger Institute COVID-19 Surveillance Team |
| EPI_ISL_3467140 | Lighthouse Lab in Milton Keynes | Wellcome Sanger Institute for the COVID-19 Genomics UK (COG-UK) Consortium | The Lighthouse Lab in Milton Keynes and Alex Alderton, Roberto Amato, Jeffrey Barrett, Sonia Goncalves, Ewan Harrison, David K. Jackson, Ian Johnston, Dominic Kwiatkowski, Cordelia Langford, John Sillitoe on behalf of the Wellcome Sanger Institute COVID-19 Surveillance Team |
| EPI_ISL_3467141 | Lighthouse Lab in Glasgow | Wellcome Sanger Institute for the COVID-19 Genomics UK (COG-UK) Consortium | Harper VanSteenhouse, Yumi Kasai, David Gray, Carol Clugston, Anna Dominiczak and Alex Alderton, Roberto Amato, Jeffrey Barrett, Sonia Goncalves, Ewan Harrison, David K. Jackson, Ian Johnston, Dominic Kwiatkowski, Cordelia Langford, John Sillitoe on behalf of the Wellcome Sanger Institute COVID-19 Surveillance Team |
| EPI_ISL_3467142 | Lighthouse Lab in Milton Keynes | Wellcome Sanger Institute for the COVID-19 Genomics UK (COG-UK) Consortium | The Lighthouse Lab in Milton Keynes and Alex Alderton, Roberto Amato, Jeffrey Barrett, Sonia Goncalves, Ewan Harrison, David K. Jackson, Ian Johnston, Dominic Kwiatkowski, Cordelia Langford, John Sillitoe on behalf of the Wellcome Sanger Institute COVID-19 Surveillance Team |
| EPI_ISL_3467143 | Lighthouse Lab in Glasgow | Wellcome Sanger Institute for the COVID-19 Genomics UK (COG-UK) Consortium | Harper VanSteenhouse, Yumi Kasai, David Gray, Carol Clugston, Anna Dominiczak and Alex Alderton, Roberto Amato, Jeffrey Barrett, Sonia Goncalves, Ewan Harrison, David K. Jackson, Ian Johnston, Dominic Kwiatkowski, Cordelia Langford, John Sillitoe on behalf of the Wellcome Sanger Institute COVID-19 Surveillance Team |
| EPI_ISL_3467144, EPI_ISL_3467145, EPI_ISL_3467146 | Lighthouse Lab in Milton Keynes | Wellcome Sanger Institute for the COVID-19 Genomics UK (COG-UK) Consortium | The Lighthouse Lab in Milton Keynes and Alex Alderton, Roberto Amato, Jeffrey Barrett, Sonia Goncalves, Ewan Harrison, David K. Jackson, Ian Johnston, Dominic Kwiatkowski, Cordelia Langford, John Sillitoe on behalf of the Wellcome Sanger Institute COVID-19 Surveillance Team |
| EPI_ISL_3467147 | Lighthouse Lab in Alderley Park | Wellcome Sanger Institute for the COVID-19 Genomics UK (COG-UK) Consortium | Jacquelyn Wynn, Mairead Hyland, The Lighthouse Lab in Alderley Park and Alex Alderton, Roberto Amato, Jeffrey Barrett, Sonia Goncalves, Ewan Harrison, David K. Jackson, Ian Johnston, Dominic Kwiatkowski, Cordelia Langford, John Sillitoe on behalf of the Wellcome Sanger Institute COVID-19 Surveillance Team |
| EPI_ISL_3467148, EPI_ISL_3467149 | Lighthouse Lab in Milton Keynes | Wellcome Sanger Institute for the COVID-19 Genomics UK (COG-UK) Consortium | The Lighthouse Lab in Milton Keynes and Alex Alderton, Roberto Amato, Jeffrey Barrett, Sonia Goncalves, Ewan Harrison, David K. Jackson, Ian Johnston, Dominic Kwiatkowski, Cordelia Langford, John Sillitoe on behalf of the Wellcome Sanger Institute COVID-19 Surveillance Team |
| EPI_ISL_3467150 | Lighthouse Lab in Alderley Park | Wellcome Sanger Institute for the COVID-19 Genomics UK (COG-UK) Consortium | Jacquelyn Wynn, Mairead Hyland, The Lighthouse Lab in Alderley Park and Alex Alderton, Roberto Amato, Jeffrey Barrett, Sonia Goncalves, Ewan Harrison, David K. Jackson, Ian Johnston, Dominic Kwiatkowski, Cordelia Langford, John Sillitoe on behalf of the Wellcome Sanger Institute COVID-19 Surveillance Team |
| EPI_ISL_3467151, EPI_ISL_3467152, EPI_ISL_3467153, EPI_ISL_3467154, EPI_ISL_3467155, EPI_ISL_3467156, EPI_ISL_3467157, EPI_ISL_3467158, EPI_ISL_3467159, EPI_ISL_3467160 | Lighthouse Lab in Milton Keynes | Wellcome Sanger Institute for the COVID-19 Genomics UK (COG-UK) Consortium | The Lighthouse Lab in Milton Keynes and Alex Alderton, Roberto Amato, Jeffrey Barrett, Sonia Goncalves, Ewan Harrison, David K. Jackson, Ian Johnston, Dominic Kwiatkowski, Cordelia Langford, John Sillitoe on behalf of the Wellcome Sanger Institute COVID-19 Surveillance Team |
| EPI_ISL_3467161 | Lighthouse Lab in Glasgow | Wellcome Sanger Institute for the COVID-19 Genomics UK (COG-UK) Consortium | Harper VanSteenhouse, Yumi Kasai, David Gray, Carol Clugston, Anna Dominiczak and Alex Alderton, Roberto Amato, Jeffrey Barrett, Sonia Goncalves, Ewan Harrison, David K. Jackson, Ian Johnston, Dominic Kwiatkowski, Cordelia Langford, John Sillitoe on behalf of the Wellcome Sanger Institute COVID-19 Surveillance Team |
| EPI_ISL_3467162 | Lighthouse Lab in Milton Keynes | Wellcome Sanger Institute for the COVID-19 Genomics UK (COG-UK) Consortium | The Lighthouse Lab in Milton Keynes and Alex Alderton, Roberto Amato, Jeffrey Barrett, Sonia Goncalves, Ewan Harrison, David K. Jackson, Ian Johnston, Dominic Kwiatkowski, Cordelia Langford, John Sillitoe on behalf of the Wellcome Sanger Institute COVID-19 Surveillance Team |
| EPI_ISL_3467163 | Lighthouse Lab in Alderley Park | Wellcome Sanger Institute for the COVID-19 Genomics UK (COG-UK) Consortium | Jacquelyn Wynn, Mairead Hyland, The Lighthouse Lab in Alderley Park and Alex Alderton, Roberto Amato, Jeffrey Barrett, Sonia Goncalves, Ewan Harrison, David K. Jackson, Ian Johnston, Dominic Kwiatkowski, Cordelia Langford, John Sillitoe on behalf of the Wellcome Sanger Institute COVID-19 Surveillance Team |
| EPI_ISL_3467164, EPI_ISL_3467165, EPI_ISL_3467166 | Lighthouse Lab in Milton Keynes | Wellcome Sanger Institute for the COVID-19 Genomics UK (COG-UK) Consortium | The Lighthouse Lab in Milton Keynes and Alex Alderton, Roberto Amato, Jeffrey Barrett, Sonia Goncalves, Ewan Harrison, David K. Jackson, Ian Johnston, Dominic Kwiatkowski, Cordelia Langford, John Sillitoe on behalf of the Wellcome Sanger Institute COVID-19 Surveillance Team |
| EPI_ISL_3467167 | Lighthouse Lab in Glasgow | Wellcome Sanger Institute for the COVID-19 Genomics UK (COG-UK) Consortium | Harper VanSteenhouse, Yumi Kasai, David Gray, Carol Clugston, Anna Dominiczak and Alex Alderton, Roberto Amato, Jeffrey Barrett, Sonia Goncalves, Ewan Harrison, David K. Jackson, Ian Johnston, Dominic Kwiatkowski, Cordelia Langford, John Sillitoe on behalf of the Wellcome Sanger Institute COVID-19 Surveillance Team |
| EPI_ISL_3467168, EPI_ISL_3467169 | Lighthouse Lab in Alderley Park | Wellcome Sanger Institute for the COVID-19 Genomics UK (COG-UK) Consortium | Jacquelyn Wynn, Mairead Hyland, The Lighthouse Lab in Alderley Park and Alex Alderton, Roberto Amato, Jeffrey Barrett, Sonia Goncalves, Ewan Harrison, David K. Jackson, Ian Johnston, Dominic Kwiatkowski, Cordelia Langford, John Sillitoe on behalf of the Wellcome Sanger Institute COVID-19 Surveillance Team |
| EPI_ISL_3467170, EPI_ISL_3467171, EPI_ISL_3467172, EPI_ISL_3467173, EPI_ISL_3467174, EPI_ISL_3467175, EPI_ISL_3467176, EPI_ISL_3467177, EPI_ISL_3467178, EPI_ISL_3467179, EPI_ISL_3467180 | | | |
| see above | Lighthouse Lab in Milton Keynes | Wellcome Sanger Institute for the COVID-19 Genomics UK (COG-UK) Consortium | The Lighthouse Lab in Milton Keynes and Alex Alderton, Roberto Amato, Jeffrey Barrett, Sonia Goncalves, Ewan Harrison, David K. Jackson, Ian Johnston, Dominic Kwiatkowski, Cordelia Langford, John Sillitoe on behalf of the Wellcome Sanger Institute COVID-19 Surveillance Team |
| EPI_ISL_3467181, EPI_ISL_3467182, EPI_ISL_3467183, EPI_ISL_3467184, EPI_ISL_3467185, EPI_ISL_3467186, EPI_ISL_3467187, EPI_ISL_3467188, EPI_ISL_3467189, EPI_ISL_3467190 | Berkshire and Surrey Pathology Services Lighthouse Laboratory | Wellcome Sanger Institute for the COVID-19 Genomics UK (COG-UK) Consortium | Berkshire and Surrey Pathology Services Lighthouse Laboratory and Alex Alderton, Roberto Amato, Jeffrey Barrett, Sonia Goncalves, Ewan Harrison, David K. Jackson, Ian Johnston, Dominic Kwiatkowski, Cordelia Langford, John Sillitoe on behalf of the Wellcome Sanger Institute COVID-19 Surveillance Team |

| | | | |
|---|---|---|---|
| EPI_ISL_3467191 | Lighthouse Lab in Milton Keynes | Wellcome Sanger Institute for the COVID-19 Genomics UK (COG-UK) Consortium | The Lighthouse Lab in Milton Keynes and Alex Alderton, Roberto Amato, Jeffrey Barrett, Sonia Goncalves, Ewan Harrison, David K. Jackson, Ian Johnston, Dominic Kwiatkowski, Cordelia Langford, John Sillitoe on behalf of the Wellcome Sanger Institute COVID-19 Surveillance Team |
| EPI_ISL_3467192, EPI_ISL_3467193 | Berkshire and Surrey Pathology Services Lighthouse Laboratory | Wellcome Sanger Institute for the COVID-19 Genomics UK (COG-UK) Consortium | Berkshire and Surrey Pathology Services Lighthouse Laboratory and Alex Alderton, Roberto Amato, Jeffrey Barrett, Sonia Goncalves, Ewan Harrison, David K. Jackson, Ian Johnston, Dominic Kwiatkowski, Cordelia Langford, John Sillitoe on behalf of the Wellcome Sanger Institute COVID-19 Surveillance Team |
| EPI_ISL_3467194, EPI_ISL_3467195 | Lighthouse Lab in Milton Keynes | Wellcome Sanger Institute for the COVID-19 Genomics UK (COG-UK) Consortium | The Lighthouse Lab in Milton Keynes and Alex Alderton, Roberto Amato, Jeffrey Barrett, Sonia Goncalves, Ewan Harrison, David K. Jackson, Ian Johnston, Dominic Kwiatkowski, Cordelia Langford, John Sillitoe on behalf of the Wellcome Sanger Institute COVID-19 Surveillance Team |
| EPI_ISL_3467196, EPI_ISL_3467197 | Berkshire and Surrey Pathology Services Lighthouse Laboratory | Wellcome Sanger Institute for the COVID-19 Genomics UK (COG-UK) Consortium | Berkshire and Surrey Pathology Services Lighthouse Laboratory and Alex Alderton, Roberto Amato, Jeffrey Barrett, Sonia Goncalves, Ewan Harrison, David K. Jackson, Ian Johnston, Dominic Kwiatkowski, Cordelia Langford, John Sillitoe on behalf of the Wellcome Sanger Institute COVID-19 Surveillance Team |
| EPI_ISL_3467198, EPI_ISL_3467199 | Lighthouse Lab in Milton Keynes | Wellcome Sanger Institute for the COVID-19 Genomics UK (COG-UK) Consortium | The Lighthouse Lab in Milton Keynes and Alex Alderton, Roberto Amato, Jeffrey Barrett, Sonia Goncalves, Ewan Harrison, David K. Jackson, Ian Johnston, Dominic Kwiatkowski, Cordelia Langford, John Sillitoe on behalf of the Wellcome Sanger Institute COVID-19 Surveillance Team |
| EPI_ISL_3467200, EPI_ISL_3467201 | Berkshire and Surrey Pathology Services Lighthouse Laboratory | Wellcome Sanger Institute for the COVID-19 Genomics UK (COG-UK) Consortium | Berkshire and Surrey Pathology Services Lighthouse Laboratory and Alex Alderton, Roberto Amato, Jeffrey Barrett, Sonia Goncalves, Ewan Harrison, David K. Jackson, Ian Johnston, Dominic Kwiatkowski, Cordelia Langford, John Sillitoe on behalf of the Wellcome Sanger Institute COVID-19 Surveillance Team |
| EPI_ISL_3467202 | Lighthouse Lab in Milton Keynes | Wellcome Sanger Institute for the COVID-19 Genomics UK (COG-UK) Consortium | The Lighthouse Lab in Milton Keynes and Alex Alderton, Roberto Amato, Jeffrey Barrett, Sonia Goncalves, Ewan Harrison, David K. Jackson, Ian Johnston, Dominic Kwiatkowski, Cordelia Langford, John Sillitoe on behalf of the Wellcome Sanger Institute COVID-19 Surveillance Team |
| EPI_ISL_3467203 | Lighthouse Lab in Glasgow | Wellcome Sanger Institute for the COVID-19 Genomics UK (COG-UK) Consortium | Harper VanSteenhouse, Yumi Kasai, David Gray, Carol Clugston, Anna Dominiczak and Alex Alderton, Roberto Amato, Jeffrey Barrett, Sonia Goncalves, Ewan Harrison, David K. Jackson, Ian Johnston, Dominic Kwiatkowski, Cordelia Langford, John Sillitoe on behalf of the Wellcome Sanger Institute COVID-19 Surveillance Team |
| EPI_ISL_3467204, EPI_ISL_3467205, EPI_ISL_3467206 | Berkshire and Surrey Pathology Services Lighthouse Laboratory | Wellcome Sanger Institute for the COVID-19 Genomics UK (COG-UK) Consortium | Berkshire and Surrey Pathology Services Lighthouse Laboratory and Alex Alderton, Roberto Amato, Jeffrey Barrett, Sonia Goncalves, Ewan Harrison, David K. Jackson, Ian Johnston, Dominic Kwiatkowski, Cordelia Langford, John Sillitoe on behalf of the Wellcome Sanger Institute COVID-19 Surveillance Team |
| EPI_ISL_3467207 | Lighthouse Lab in Milton Keynes | Wellcome Sanger Institute for the COVID-19 Genomics UK (COG-UK) Consortium | The Lighthouse Lab in Milton Keynes and Alex Alderton, Roberto Amato, Jeffrey Barrett, Sonia Goncalves, Ewan Harrison, David K. Jackson, Ian Johnston, Dominic Kwiatkowski, Cordelia Langford, John Sillitoe on behalf of the Wellcome Sanger Institute COVID-19 Surveillance Team |
| EPI_ISL_3467208 | Lighthouse Lab in Glasgow | Wellcome Sanger Institute for the COVID-19 Genomics UK (COG-UK) Consortium | Harper VanSteenhouse, Yumi Kasai, David Gray, Carol Clugston, Anna Dominiczak and Alex Alderton, Roberto Amato, Jeffrey Barrett, Sonia Goncalves, Ewan Harrison, David K. Jackson, Ian Johnston, Dominic Kwiatkowski, Cordelia Langford, John Sillitoe on behalf of the Wellcome Sanger Institute COVID-19 Surveillance Team |
| EPI_ISL_3467209 | Lighthouse Lab in Milton Keynes | Wellcome Sanger Institute for the COVID-19 Genomics UK (COG-UK) Consortium | The Lighthouse Lab in Milton Keynes and Alex Alderton, Roberto Amato, Jeffrey Barrett, Sonia Goncalves, Ewan Harrison, David K. Jackson, Ian Johnston, Dominic Kwiatkowski, Cordelia Langford, John Sillitoe on behalf of the Wellcome Sanger Institute COVID-19 Surveillance Team |
| EPI_ISL_3467210, EPI_ISL_3467211, EPI_ISL_3467212, EPI_ISL_3467213, EPI_ISL_3467214, EPI_ISL_3467215, EPI_ISL_3467216 | Berkshire and Surrey Pathology Services Lighthouse Laboratory | Wellcome Sanger Institute for the COVID-19 Genomics UK (COG-UK) Consortium | Berkshire and Surrey Pathology Services Lighthouse Laboratory and Alex Alderton, Roberto Amato, Jeffrey Barrett, Sonia Goncalves, Ewan Harrison, David K. Jackson, Ian Johnston, Dominic Kwiatkowski, Cordelia Langford, John Sillitoe on behalf of the Wellcome Sanger Institute COVID-19 Surveillance Team |
| EPI_ISL_3467217 | Lighthouse Lab in Milton Keynes | Wellcome Sanger Institute for the COVID-19 Genomics UK (COG-UK) Consortium | The Lighthouse Lab in Milton Keynes and Alex Alderton, Roberto Amato, Jeffrey Barrett, Sonia Goncalves, Ewan Harrison, David K. Jackson, Ian Johnston, Dominic Kwiatkowski, Cordelia Langford, John Sillitoe on behalf of the Wellcome Sanger Institute COVID-19 Surveillance Team |
| EPI_ISL_3467218, EPI_ISL_3467219 | Berkshire and Surrey Pathology Services Lighthouse Laboratory | Wellcome Sanger Institute for the COVID-19 Genomics UK (COG-UK) Consortium | Berkshire and Surrey Pathology Services Lighthouse Laboratory and Alex Alderton, Roberto Amato, Jeffrey Barrett, Sonia Goncalves, Ewan Harrison, David K. Jackson, Ian Johnston, Dominic Kwiatkowski, Cordelia Langford, John Sillitoe on behalf of the Wellcome Sanger Institute COVID-19 Surveillance Team |
| EPI_ISL_3467220 | Lighthouse Lab in Milton Keynes | Wellcome Sanger Institute for the COVID-19 Genomics UK (COG-UK) Consortium | The Lighthouse Lab in Milton Keynes and Alex Alderton, Roberto Amato, Jeffrey Barrett, Sonia Goncalves, Ewan Harrison, David K. Jackson, Ian Johnston, Dominic Kwiatkowski, Cordelia Langford, John Sillitoe on behalf of the Wellcome Sanger Institute COVID-19 Surveillance Team |
| EPI_ISL_3467221, EPI_ISL_3467222, EPI_ISL_3467223, EPI_ISL_3467224, EPI_ISL_3467225, EPI_ISL_3467226, EPI_ISL_3467227, EPI_ISL_3467228, EPI_ISL_3467229, EPI_ISL_3467230, EPI_ISL_3467231, EPI_ISL_3467232 | | | |
| see above | Berkshire and Surrey Pathology Services Lighthouse Laboratory | Wellcome Sanger Institute for the COVID-19 Genomics UK (COG-UK) Consortium | Berkshire and Surrey Pathology Services Lighthouse Laboratory and Alex Alderton, Roberto Amato, Jeffrey Barrett, Sonia Goncalves, Ewan Harrison, David K. Jackson, Ian Johnston, Dominic Kwiatkowski, Cordelia Langford, John Sillitoe on behalf of the Wellcome Sanger Institute COVID-19 Surveillance Team |
| EPI_ISL_3467233 | Lighthouse Lab in Milton Keynes | Wellcome Sanger Institute for the COVID-19 Genomics UK (COG-UK) Consortium | The Lighthouse Lab in Milton Keynes and Alex Alderton, Roberto Amato, Jeffrey Barrett, Sonia Goncalves, Ewan Harrison, David K. Jackson, Ian Johnston, Dominic Kwiatkowski, Cordelia Langford, John Sillitoe on behalf of the Wellcome Sanger Institute COVID-19 Surveillance Team |
| EPI_ISL_3467234 | Berkshire and Surrey Pathology Services Lighthouse Laboratory | Wellcome Sanger Institute for the COVID-19 Genomics UK (COG-UK) Consortium | Berkshire and Surrey Pathology Services Lighthouse Laboratory and Alex Alderton, Roberto Amato, Jeffrey Barrett, Sonia Goncalves, Ewan Harrison, David K. Jackson, Ian Johnston, Dominic Kwiatkowski, Cordelia Langford, John Sillitoe on behalf of the Wellcome Sanger Institute COVID-19 Surveillance Team |
| EPI_ISL_3467235, EPI_ISL_3467236 | Lighthouse Lab in Milton Keynes | Wellcome Sanger Institute for the COVID-19 Genomics UK (COG-UK) Consortium | The Lighthouse Lab in Milton Keynes and Alex Alderton, Roberto Amato, Jeffrey Barrett, Sonia Goncalves, Ewan Harrison, David K. Jackson, Ian Johnston, Dominic Kwiatkowski, Cordelia Langford, John Sillitoe on behalf of the Wellcome Sanger Institute COVID-19 Surveillance Team |
| EPI_ISL_3467237, EPI_ISL_3467238, EPI_ISL_3467239, EPI_ISL_3467240, EPI_ISL_3467241, EPI_ISL_3467242, EPI_ISL_3467243, EPI_ISL_3467244, EPI_ISL_3467245, EPI_ISL_3467246 | Berkshire and Surrey Pathology Services Lighthouse Laboratory | Wellcome Sanger Institute for the COVID-19 Genomics UK (COG-UK) Consortium | Berkshire and Surrey Pathology Services Lighthouse Laboratory and Alex Alderton, Roberto Amato, Jeffrey Barrett, Sonia Goncalves, Ewan Harrison, David K. Jackson, Ian Johnston, Dominic Kwiatkowski, Cordelia Langford, John Sillitoe on behalf of the Wellcome Sanger Institute COVID-19 Surveillance Team |
| EPI_ISL_3467247 | Lighthouse Lab in Milton Keynes | Wellcome Sanger Institute for the COVID-19 Genomics UK (COG-UK) Consortium | The Lighthouse Lab in Milton Keynes and Alex Alderton, Roberto Amato, Jeffrey Barrett, Sonia Goncalves, Ewan Harrison, David K. Jackson, Ian Johnston, Dominic Kwiatkowski, Cordelia Langford, John Sillitoe on behalf of the Wellcome Sanger Institute COVID-19 Surveillance Team |
| EPI_ISL_3467248, EPI_ISL_3467249 | Berkshire and Surrey Pathology Services Lighthouse Laboratory | Wellcome Sanger Institute for the COVID-19 Genomics UK (COG-UK) Consortium | Berkshire and Surrey Pathology Services Lighthouse Laboratory and Alex Alderton, Roberto Amato, Jeffrey Barrett, Sonia Goncalves, Ewan Harrison, David K. Jackson, Ian Johnston, Dominic Kwiatkowski, Cordelia Langford, John Sillitoe on behalf of the Wellcome Sanger Institute COVID-19 Surveillance Team |
| EPI_ISL_3467250 | Lighthouse Lab in Milton Keynes | Wellcome Sanger Institute for the COVID-19 Genomics UK (COG-UK) Consortium | The Lighthouse Lab in Milton Keynes and Alex Alderton, Roberto Amato, Jeffrey Barrett, Sonia Goncalves, Ewan Harrison, David K. Jackson, Ian Johnston, Dominic Kwiatkowski, Cordelia Langford, John Sillitoe on behalf of the Wellcome Sanger Institute COVID-19 Surveillance Team |
| EPI_ISL_3467251, EPI_ISL_3467252 | Berkshire and Surrey Pathology Services Lighthouse Laboratory | Wellcome Sanger Institute for the COVID-19 Genomics UK (COG-UK) Consortium | Berkshire and Surrey Pathology Services Lighthouse Laboratory and Alex Alderton, Roberto Amato, Jeffrey Barrett, Sonia Goncalves, Ewan Harrison, David K. Jackson, Ian Johnston, Dominic Kwiatkowski, Cordelia Langford, John Sillitoe on behalf of the Wellcome Sanger Institute COVID-19 Surveillance Team |
| EPI_ISL_3467253 | Lighthouse Lab in Milton Keynes | Wellcome Sanger Institute for the COVID-19 Genomics UK (COG-UK) Consortium | The Lighthouse Lab in Milton Keynes and Alex Alderton, Roberto Amato, Jeffrey Barrett, Sonia Goncalves, Ewan Harrison, David K. Jackson, Ian Johnston, Dominic Kwiatkowski, Cordelia Langford, John Sillitoe on behalf of the Wellcome Sanger Institute COVID-19 Surveillance Team |
| EPI_ISL_3467254 | Berkshire and Surrey Pathology Services Lighthouse Laboratory | Wellcome Sanger Institute for the COVID-19 Genomics UK (COG-UK) Consortium | Berkshire and Surrey Pathology Services Lighthouse Laboratory and Alex Alderton, Roberto Amato, Jeffrey Barrett, Sonia Goncalves, Ewan Harrison, David K. Jackson, Ian Johnston, Dominic Kwiatkowski, Cordelia Langford, John Sillitoe on behalf of the Wellcome Sanger Institute COVID-19 Surveillance Team |
| EPI_ISL_3467255, EPI_ISL_3467256, EPI_ISL_3467257 | Lighthouse Lab in Milton Keynes | Wellcome Sanger Institute for the COVID-19 Genomics UK (COG-UK) Consortium | The Lighthouse Lab in Milton Keynes and Alex Alderton, Roberto Amato, Jeffrey Barrett, Sonia Goncalves, Ewan Harrison, David K. Jackson, Ian Johnston, Dominic Kwiatkowski, Cordelia Langford, John Sillitoe on behalf of the Wellcome Sanger Institute COVID-19 Surveillance Team |
| EPI_ISL_3467258, EPI_ISL_3467259, EPI_ISL_3467260, EPI_ISL_3467261, EPI_ISL_3467262, EPI_ISL_3467263, EPI_ISL_3467264 | Berkshire and Surrey Pathology Services Lighthouse Laboratory | Wellcome Sanger Institute for the COVID-19 Genomics UK (COG-UK) Consortium | Berkshire and Surrey Pathology Services Lighthouse Laboratory and Alex Alderton, Roberto Amato, Jeffrey Barrett, Sonia Goncalves, Ewan Harrison, David K. Jackson, Ian Johnston, Dominic Kwiatkowski, Cordelia Langford, John Sillitoe on behalf of the Wellcome Sanger Institute COVID-19 Surveillance Team |
| EPI_ISL_3467265 | Lighthouse Lab in Milton Keynes | Wellcome Sanger Institute for the COVID-19 Genomics UK (COG-UK) Consortium | The Lighthouse Lab in Milton Keynes and Alex Alderton, Roberto Amato, Jeffrey Barrett, Sonia Goncalves, Ewan Harrison, David K. Jackson, Ian Johnston, Dominic Kwiatkowski, Cordelia Langford, John Sillitoe on behalf of the Wellcome Sanger Institute COVID-19 Surveillance Team |
| EPI_ISL_3467266, EPI_ISL_3467267, EPI_ISL_3467268, EPI_ISL_3467269, EPI_ISL_3467270, EPI_ISL_3467271 | Berkshire and Surrey Pathology Services Lighthouse Laboratory | Wellcome Sanger Institute for the COVID-19 Genomics UK (COG-UK) Consortium | Berkshire and Surrey Pathology Services Lighthouse Laboratory and Alex Alderton, Roberto Amato, Jeffrey Barrett, Sonia Goncalves, Ewan Harrison, David K. Jackson, Ian Johnston, Dominic Kwiatkowski, Cordelia Langford, John Sillitoe on behalf of the Wellcome Sanger Institute COVID-19 Surveillance Team |
| EPI_ISL_3467272 | Lighthouse Lab in Glasgow | Wellcome Sanger Institute for the COVID-19 Genomics UK (COG-UK) Consortium | Harper VanSteenhouse, Yumi Kasai, David Gray, Carol Clugston, Anna Dominiczak and Alex Alderton, Roberto Amato, Jeffrey Barrett, Sonia Goncalves, Ewan Harrison, David K. Jackson, Ian Johnston, Dominic Kwiatkowski, Cordelia Langford, John Sillitoe on behalf of the Wellcome Sanger Institute COVID-19 Surveillance Team |
| EPI_ISL_3467273 | Berkshire and Surrey Pathology Services Lighthouse Laboratory | Wellcome Sanger Institute for the COVID-19 Genomics UK (COG-UK) Consortium | Berkshire and Surrey Pathology Services Lighthouse Laboratory and Alex Alderton, Roberto Amato, Jeffrey Barrett, Sonia Goncalves, Ewan Harrison, David K. Jackson, Ian Johnston, Dominic Kwiatkowski, Cordelia Langford, John Sillitoe on behalf of the Wellcome Sanger Institute COVID-19 Surveillance Team |
| EPI_ISL_3467274, EPI_ISL_3467275 | Lighthouse Lab in Milton Keynes | Wellcome Sanger Institute for the COVID-19 Genomics UK (COG-UK) Consortium | The Lighthouse Lab in Milton Keynes and Alex Alderton, Roberto Amato, Jeffrey Barrett, Sonia Goncalves, Ewan Harrison, David K. Jackson, Ian Johnston, Dominic Kwiatkowski, Cordelia Langford, John Sillitoe on behalf of the Wellcome Sanger Institute COVID-19 Surveillance Team |
| EPI_ISL_3467276, | Berkshire and Surrey Pathology Services Lighthouse Laboratory | Wellcome Sanger Institute for the COVID-19 Genomics UK (COG-UK) | Berkshire and Surrey Pathology Services Lighthouse Laboratory and Alex Alderton, Roberto Amato, Jeffrey Barrett, Sonia Goncalves, Ewan Harrison, David K. Jackson, Ian Johnston, Dominic Kwiatkowski, Cordelia Langford, John Sillitoe on behalf |

| | | | |
|---|---|---|---|
| EPI_ISL_3467277, EPI_ISL_3467278, EPI_ISL_3467279, EPI_ISL_3467280, EPI_ISL_3467281, EPI_ISL_3467282, EPI_ISL_3467283 | | Consortium | of the Wellcome Sanger Institute COVID-19 Surveillance Team |
| EPI_ISL_3467284, EPI_ISL_3467285 | Lighthouse Lab in Milton Keynes | Wellcome Sanger Institute for the COVID-19 Genomics UK (COG-UK) Consortium | The Lighthouse Lab in Milton Keynes and Alex Alderton, Roberto Amato, Jeffrey Barrett, Sonia Goncalves, Ewan Harrison, David K. Jackson, Ian Johnston, Dominic Kwiatkowski, Cordelia Langford, John Sillitoe on behalf of the Wellcome Sanger Institute COVID-19 Surveillance Team |
| EPI_ISL_3467286, EPI_ISL_3467287, EPI_ISL_3467288 | Berkshire and Surrey Pathology Services Lighthouse Laboratory | Wellcome Sanger Institute for the COVID-19 Genomics UK (COG-UK) Consortium | Berkshire and Surrey Pathology Services Lighthouse Laboratory and Alex Alderton, Roberto Amato, Jeffrey Barrett, Sonia Goncalves, Ewan Harrison, David K. Jackson, Ian Johnston, Dominic Kwiatkowski, Cordelia Langford, John Sillitoe on behalf of the Wellcome Sanger Institute COVID-19 Surveillance Team |
| EPI_ISL_3467289 | Lighthouse Lab in Milton Keynes | Wellcome Sanger Institute for the COVID-19 Genomics UK (COG-UK) Consortium | The Lighthouse Lab in Milton Keynes and Alex Alderton, Roberto Amato, Jeffrey Barrett, Sonia Goncalves, Ewan Harrison, David K. Jackson, Ian Johnston, Dominic Kwiatkowski, Cordelia Langford, John Sillitoe on behalf of the Wellcome Sanger Institute COVID-19 Surveillance Team |
| EPI_ISL_3467290, EPI_ISL_3467291, EPI_ISL_3467292, EPI_ISL_3467293, EPI_ISL_3467294, EPI_ISL_3467295, EPI_ISL_3467296, EPI_ISL_3467297, EPI_ISL_3467298, EPI_ISL_3467299 | Berkshire and Surrey Pathology Services Lighthouse Laboratory | Wellcome Sanger Institute for the COVID-19 Genomics UK (COG-UK) Consortium | Berkshire and Surrey Pathology Services Lighthouse Laboratory and Alex Alderton, Roberto Amato, Jeffrey Barrett, Sonia Goncalves, Ewan Harrison, David K. Jackson, Ian Johnston, Dominic Kwiatkowski, Cordelia Langford, John Sillitoe on behalf of the Wellcome Sanger Institute COVID-19 Surveillance Team |
| EPI_ISL_3467300 | Lighthouse Lab in Milton Keynes | Wellcome Sanger Institute for the COVID-19 Genomics UK (COG-UK) Consortium | The Lighthouse Lab in Milton Keynes and Alex Alderton, Roberto Amato, Jeffrey Barrett, Sonia Goncalves, Ewan Harrison, David K. Jackson, Ian Johnston, Dominic Kwiatkowski, Cordelia Langford, John Sillitoe on behalf of the Wellcome Sanger Institute COVID-19 Surveillance Team |
| EPI_ISL_3467301 | Berkshire and Surrey Pathology Services Lighthouse Laboratory | Wellcome Sanger Institute for the COVID-19 Genomics UK (COG-UK) Consortium | Berkshire and Surrey Pathology Services Lighthouse Laboratory and Alex Alderton, Roberto Amato, Jeffrey Barrett, Sonia Goncalves, Ewan Harrison, David K. Jackson, Ian Johnston, Dominic Kwiatkowski, Cordelia Langford, John Sillitoe on behalf of the Wellcome Sanger Institute COVID-19 Surveillance Team |
| EPI_ISL_3467302 | Lighthouse Lab in Milton Keynes | Wellcome Sanger Institute for the COVID-19 Genomics UK (COG-UK) Consortium | The Lighthouse Lab in Milton Keynes and Alex Alderton, Roberto Amato, Jeffrey Barrett, Sonia Goncalves, Ewan Harrison, David K. Jackson, Ian Johnston, Dominic Kwiatkowski, Cordelia Langford, John Sillitoe on behalf of the Wellcome Sanger Institute COVID-19 Surveillance Team |
| EPI_ISL_3467303 | Berkshire and Surrey Pathology Services Lighthouse Laboratory | Wellcome Sanger Institute for the COVID-19 Genomics UK (COG-UK) Consortium | Berkshire and Surrey Pathology Services Lighthouse Laboratory and Alex Alderton, Roberto Amato, Jeffrey Barrett, Sonia Goncalves, Ewan Harrison, David K. Jackson, Ian Johnston, Dominic Kwiatkowski, Cordelia Langford, John Sillitoe on behalf of the Wellcome Sanger Institute COVID-19 Surveillance Team |
| EPI_ISL_3467304 | Lighthouse Lab in Milton Keynes | Wellcome Sanger Institute for the COVID-19 Genomics UK (COG-UK) Consortium | The Lighthouse Lab in Milton Keynes and Alex Alderton, Roberto Amato, Jeffrey Barrett, Sonia Goncalves, Ewan Harrison, David K. Jackson, Ian Johnston, Dominic Kwiatkowski, Cordelia Langford, John Sillitoe on behalf of the Wellcome Sanger Institute COVID-19 Surveillance Team |
| EPI_ISL_3467305, EPI_ISL_3467306 | Berkshire and Surrey Pathology Services Lighthouse Laboratory | Wellcome Sanger Institute for the COVID-19 Genomics UK (COG-UK) Consortium | Berkshire and Surrey Pathology Services Lighthouse Laboratory and Alex Alderton, Roberto Amato, Jeffrey Barrett, Sonia Goncalves, Ewan Harrison, David K. Jackson, Ian Johnston, Dominic Kwiatkowski, Cordelia Langford, John Sillitoe on behalf of the Wellcome Sanger Institute COVID-19 Surveillance Team |
| EPI_ISL_3467307, EPI_ISL_3467308 | Lighthouse Lab in Milton Keynes | Wellcome Sanger Institute for the COVID-19 Genomics UK (COG-UK) Consortium | The Lighthouse Lab in Milton Keynes and Alex Alderton, Roberto Amato, Jeffrey Barrett, Sonia Goncalves, Ewan Harrison, David K. Jackson, Ian Johnston, Dominic Kwiatkowski, Cordelia Langford, John Sillitoe on behalf of the Wellcome Sanger Institute COVID-19 Surveillance Team |
| EPI_ISL_3467309, EPI_ISL_3467310 | Berkshire and Surrey Pathology Services Lighthouse Laboratory | Wellcome Sanger Institute for the COVID-19 Genomics UK (COG-UK) Consortium | Berkshire and Surrey Pathology Services Lighthouse Laboratory and Alex Alderton, Roberto Amato, Jeffrey Barrett, Sonia Goncalves, Ewan Harrison, David K. Jackson, Ian Johnston, Dominic Kwiatkowski, Cordelia Langford, John Sillitoe on behalf of the Wellcome Sanger Institute COVID-19 Surveillance Team |
| EPI_ISL_3467311 | Lighthouse Lab in Milton Keynes | Wellcome Sanger Institute for the COVID-19 Genomics UK (COG-UK) Consortium | The Lighthouse Lab in Milton Keynes and Alex Alderton, Roberto Amato, Jeffrey Barrett, Sonia Goncalves, Ewan Harrison, David K. Jackson, Ian Johnston, Dominic Kwiatkowski, Cordelia Langford, John Sillitoe on behalf of the Wellcome Sanger Institute COVID-19 Surveillance Team |
| EPI_ISL_3467312, EPI_ISL_3467313, EPI_ISL_3467314, EPI_ISL_3467315 | Berkshire and Surrey Pathology Services Lighthouse Laboratory | Wellcome Sanger Institute for the COVID-19 Genomics UK (COG-UK) Consortium | Berkshire and Surrey Pathology Services Lighthouse Laboratory and Alex Alderton, Roberto Amato, Jeffrey Barrett, Sonia Goncalves, Ewan Harrison, David K. Jackson, Ian Johnston, Dominic Kwiatkowski, Cordelia Langford, John Sillitoe on behalf of the Wellcome Sanger Institute COVID-19 Surveillance Team |
| EPI_ISL_3467316, EPI_ISL_3467317 | Lighthouse Lab in Milton Keynes | Wellcome Sanger Institute for the COVID-19 Genomics UK (COG-UK) Consortium | The Lighthouse Lab in Milton Keynes and Alex Alderton, Roberto Amato, Jeffrey Barrett, Sonia Goncalves, Ewan Harrison, David K. Jackson, Ian Johnston, Dominic Kwiatkowski, Cordelia Langford, John Sillitoe on behalf of the Wellcome Sanger Institute COVID-19 Surveillance Team |
| EPI_ISL_3467318, EPI_ISL_3467319 | Berkshire and Surrey Pathology Services Lighthouse Laboratory | Wellcome Sanger Institute for the COVID-19 Genomics UK (COG-UK) Consortium | Berkshire and Surrey Pathology Services Lighthouse Laboratory and Alex Alderton, Roberto Amato, Jeffrey Barrett, Sonia Goncalves, Ewan Harrison, David K. Jackson, Ian Johnston, Dominic Kwiatkowski, Cordelia Langford, John Sillitoe on behalf of the Wellcome Sanger Institute COVID-19 Surveillance Team |
| EPI_ISL_3467320 | Lighthouse Lab in Milton Keynes | Wellcome Sanger Institute for the COVID-19 Genomics UK (COG-UK) Consortium | The Lighthouse Lab in Milton Keynes and Alex Alderton, Roberto Amato, Jeffrey Barrett, Sonia Goncalves, Ewan Harrison, David K. Jackson, Ian Johnston, Dominic Kwiatkowski, Cordelia Langford, John Sillitoe on behalf of the Wellcome Sanger Institute COVID-19 Surveillance Team |
| EPI_ISL_3467321 | Berkshire and Surrey Pathology Services Lighthouse Laboratory | Wellcome Sanger Institute for the COVID-19 Genomics UK (COG-UK) Consortium | Berkshire and Surrey Pathology Services Lighthouse Laboratory and Alex Alderton, Roberto Amato, Jeffrey Barrett, Sonia Goncalves, Ewan Harrison, David K. Jackson, Ian Johnston, Dominic Kwiatkowski, Cordelia Langford, John Sillitoe on behalf of the Wellcome Sanger Institute COVID-19 Surveillance Team |
| EPI_ISL_3467322, EPI_ISL_3467323 | Lighthouse Lab in Milton Keynes | Wellcome Sanger Institute for the COVID-19 Genomics UK (COG-UK) Consortium | The Lighthouse Lab in Milton Keynes and Alex Alderton, Roberto Amato, Jeffrey Barrett, Sonia Goncalves, Ewan Harrison, David K. Jackson, Ian Johnston, Dominic Kwiatkowski, Cordelia Langford, John Sillitoe on behalf of the Wellcome Sanger Institute COVID-19 Surveillance Team |
| EPI_ISL_3467324, EPI_ISL_3467325, EPI_ISL_3467326, EPI_ISL_3467327, EPI_ISL_3467328, EPI_ISL_3467329, EPI_ISL_3467330 | Berkshire and Surrey Pathology Services Lighthouse Laboratory | Wellcome Sanger Institute for the COVID-19 Genomics UK (COG-UK) Consortium | Berkshire and Surrey Pathology Services Lighthouse Laboratory and Alex Alderton, Roberto Amato, Jeffrey Barrett, Sonia Goncalves, Ewan Harrison, David K. Jackson, Ian Johnston, Dominic Kwiatkowski, Cordelia Langford, John Sillitoe on behalf of the Wellcome Sanger Institute COVID-19 Surveillance Team |
| EPI_ISL_3467331 | Lighthouse Lab in Glasgow | Wellcome Sanger Institute for the COVID-19 Genomics UK (COG-UK) Consortium | Harper VanSteenhouse, Yumi Kasai, David Gray, Carol Clugston, Anna Dominiczak and Alex Alderton, Roberto Amato, Jeffrey Barrett, Sonia Goncalves, Ewan Harrison, David K. Jackson, Ian Johnston, Dominic Kwiatkowski, Cordelia Langford, John Sillitoe on behalf of the Wellcome Sanger Institute COVID-19 Surveillance Team |
| EPI_ISL_3467332, EPI_ISL_3467333, EPI_ISL_3467334 | Berkshire and Surrey Pathology Services Lighthouse Laboratory | Wellcome Sanger Institute for the COVID-19 Genomics UK (COG-UK) Consortium | Berkshire and Surrey Pathology Services Lighthouse Laboratory and Alex Alderton, Roberto Amato, Jeffrey Barrett, Sonia Goncalves, Ewan Harrison, David K. Jackson, Ian Johnston, Dominic Kwiatkowski, Cordelia Langford, John Sillitoe on behalf of the Wellcome Sanger Institute COVID-19 Surveillance Team |
| EPI_ISL_3467335 | Lighthouse Lab in Milton Keynes | Wellcome Sanger Institute for the COVID-19 Genomics UK (COG-UK) Consortium | The Lighthouse Lab in Milton Keynes and Alex Alderton, Roberto Amato, Jeffrey Barrett, Sonia Goncalves, Ewan Harrison, David K. Jackson, Ian Johnston, Dominic Kwiatkowski, Cordelia Langford, John Sillitoe on behalf of the Wellcome Sanger Institute COVID-19 Surveillance Team |
| EPI_ISL_3467336, EPI_ISL_3467337 | Berkshire and Surrey Pathology Services Lighthouse Laboratory | Wellcome Sanger Institute for the COVID-19 Genomics UK (COG-UK) Consortium | Berkshire and Surrey Pathology Services Lighthouse Laboratory and Alex Alderton, Roberto Amato, Jeffrey Barrett, Sonia Goncalves, Ewan Harrison, David K. Jackson, Ian Johnston, Dominic Kwiatkowski, Cordelia Langford, John Sillitoe on behalf of the Wellcome Sanger Institute COVID-19 Surveillance Team |
| EPI_ISL_3467338 | Lighthouse Lab in Milton Keynes | Wellcome Sanger Institute for the COVID-19 Genomics UK (COG-UK) Consortium | The Lighthouse Lab in Milton Keynes and Alex Alderton, Roberto Amato, Jeffrey Barrett, Sonia Goncalves, Ewan Harrison, David K. Jackson, Ian Johnston, Dominic Kwiatkowski, Cordelia Langford, John Sillitoe on behalf of the Wellcome Sanger Institute COVID-19 Surveillance Team |
| EPI_ISL_3467339, EPI_ISL_3467340, EPI_ISL_3467341, EPI_ISL_3467342 | Berkshire and Surrey Pathology Services Lighthouse Laboratory | Wellcome Sanger Institute for the COVID-19 Genomics UK (COG-UK) Consortium | Berkshire and Surrey Pathology Services Lighthouse Laboratory and Alex Alderton, Roberto Amato, Jeffrey Barrett, Sonia Goncalves, Ewan Harrison, David K. Jackson, Ian Johnston, Dominic Kwiatkowski, Cordelia Langford, John Sillitoe on behalf of the Wellcome Sanger Institute COVID-19 Surveillance Team |
| EPI_ISL_3467343 | Lighthouse Lab in Milton Keynes | Wellcome Sanger Institute for the COVID-19 Genomics UK (COG-UK) Consortium | The Lighthouse Lab in Milton Keynes and Alex Alderton, Roberto Amato, Jeffrey Barrett, Sonia Goncalves, Ewan Harrison, David K. Jackson, Ian Johnston, Dominic Kwiatkowski, Cordelia Langford, John Sillitoe on behalf of the Wellcome Sanger Institute COVID-19 Surveillance Team |
| EPI_ISL_3467344, EPI_ISL_3467345, EPI_ISL_3467346 | Berkshire and Surrey Pathology Services Lighthouse Laboratory | Wellcome Sanger Institute for the COVID-19 Genomics UK (COG-UK) Consortium | Berkshire and Surrey Pathology Services Lighthouse Laboratory and Alex Alderton, Roberto Amato, Jeffrey Barrett, Sonia Goncalves, Ewan Harrison, David K. Jackson, Ian Johnston, Dominic Kwiatkowski, Cordelia Langford, John Sillitoe on behalf of the Wellcome Sanger Institute COVID-19 Surveillance Team |
| EPI_ISL_3467347 | Lighthouse Lab in Milton Keynes | Wellcome Sanger Institute for the COVID-19 Genomics UK (COG-UK) Consortium | The Lighthouse Lab in Milton Keynes and Alex Alderton, Roberto Amato, Jeffrey Barrett, Sonia Goncalves, Ewan Harrison, David K. Jackson, Ian Johnston, Dominic Kwiatkowski, Cordelia Langford, John Sillitoe on behalf of the Wellcome Sanger Institute COVID-19 Surveillance Team |
| EPI_ISL_3467348 | National Virus Reference Laboratory | National Virus Reference Laboratory | Zoe Yandle, Charlene Bennett, Gabriel Gonzalez, Michael Carr, Jonathan Dean, Cillian F De Gascun |
| EPI_ISL_3467349 | Berkshire and Surrey Pathology Services Lighthouse Laboratory | Wellcome Sanger Institute for the COVID-19 Genomics UK (COG-UK) Consortium | Berkshire and Surrey Pathology Services Lighthouse Laboratory and Alex Alderton, Roberto Amato, Jeffrey Barrett, Sonia Goncalves, Ewan Harrison, David K. Jackson, Ian Johnston, Dominic Kwiatkowski, Cordelia Langford, John Sillitoe on behalf of the Wellcome Sanger Institute COVID-19 Surveillance Team |
| EPI_ISL_3467350, EPI_ISL_3467351, EPI_ISL_3467352 | Lighthouse Lab in Milton Keynes | Wellcome Sanger Institute for the COVID-19 Genomics UK (COG-UK) Consortium | The Lighthouse Lab in Milton Keynes and Alex Alderton, Roberto Amato, Jeffrey Barrett, Sonia Goncalves, Ewan Harrison, David K. Jackson, Ian Johnston, Dominic Kwiatkowski, Cordelia Langford, John Sillitoe on behalf of the Wellcome Sanger Institute COVID-19 Surveillance Team |
| EPI_ISL_3467353 | Berkshire and Surrey Pathology Services Lighthouse Laboratory | Wellcome Sanger Institute for the COVID-19 Genomics UK (COG-UK) Consortium | Berkshire and Surrey Pathology Services Lighthouse Laboratory and Alex Alderton, Roberto Amato, Jeffrey Barrett, Sonia Goncalves, Ewan Harrison, David K. Jackson, Ian Johnston, Dominic Kwiatkowski, Cordelia Langford, John Sillitoe on behalf of the Wellcome Sanger Institute COVID-19 Surveillance Team |
| EPI_ISL_3467354 | Lighthouse Lab in Milton Keynes | Wellcome Sanger Institute for the COVID-19 Genomics UK (COG-UK) Consortium | The Lighthouse Lab in Milton Keynes and Alex Alderton, Roberto Amato, Jeffrey Barrett, Sonia Goncalves, Ewan Harrison, David K. Jackson, Ian Johnston, Dominic Kwiatkowski, Cordelia Langford, John Sillitoe on behalf of the Wellcome Sanger Institute COVID-19 Surveillance Team |

| | | | |
|---|---|---|---|
| EPI_ISL_3467355 | National Virus Reference Laboratory | National Virus Reference Laboratory | Zoe Yandle, Charlene Bennett, Gabriel Gonzalez, Michael Carr, Jonathan Dean, Cillian F De Gascun |
| EPI_ISL_3467356, EPI_ISL_3467357 | Berkshire and Surrey Pathology Services Lighthouse Laboratory | Wellcome Sanger Institute for the COVID-19 Genomics UK (COG-UK) Consortium | Berkshire and Surrey Pathology Services Lighthouse Laboratory and Alex Alderton, Roberto Amato, Jeffrey Barrett, Sonia Goncalves, Ewan Harrison, David K. Jackson, Ian Johnston, Dominic Kwiatkowski, Cordelia Langford, John Sillitoe on behalf of the Wellcome Sanger Institute COVID-19 Surveillance Team |
| EPI_ISL_3467358 | National Virus Reference Laboratory | National Virus Reference Laboratory | Zoe Yandle, Charlene Bennett, Gabriel Gonzalez, Michael Carr, Jonathan Dean, Cillian F De Gascun |
| EPI_ISL_3467359 | Lighthouse Lab in Milton Keynes | Wellcome Sanger Institute for the COVID-19 Genomics UK (COG-UK) Consortium | The Lighthouse Lab in Milton Keynes and Alex Alderton, Roberto Amato, Jeffrey Barrett, Sonia Goncalves, Ewan Harrison, David K. Jackson, Ian Johnston, Dominic Kwiatkowski, Cordelia Langford, John Sillitoe on behalf of the Wellcome Sanger Institute COVID-19 Surveillance Team |
| EPI_ISL_3467360, EPI_ISL_3467361 | Berkshire and Surrey Pathology Services Lighthouse Laboratory | Wellcome Sanger Institute for the COVID-19 Genomics UK (COG-UK) Consortium | Berkshire and Surrey Pathology Services Lighthouse Laboratory and Alex Alderton, Roberto Amato, Jeffrey Barrett, Sonia Goncalves, Ewan Harrison, David K. Jackson, Ian Johnston, Dominic Kwiatkowski, Cordelia Langford, John Sillitoe on behalf of the Wellcome Sanger Institute COVID-19 Surveillance Team |
| EPI_ISL_3467362 | National Virus Reference Laboratory | National Virus Reference Laboratory | Zoe Yandle, Charlene Bennett, Gabriel Gonzalez, Michael Carr, Jonathan Dean, Cillian F De Gascun |
| EPI_ISL_3467363, EPI_ISL_3467364, EPI_ISL_3467365 | Berkshire and Surrey Pathology Services Lighthouse Laboratory | Wellcome Sanger Institute for the COVID-19 Genomics UK (COG-UK) Consortium | Berkshire and Surrey Pathology Services Lighthouse Laboratory and Alex Alderton, Roberto Amato, Jeffrey Barrett, Sonia Goncalves, Ewan Harrison, David K. Jackson, Ian Johnston, Dominic Kwiatkowski, Cordelia Langford, John Sillitoe on behalf of the Wellcome Sanger Institute COVID-19 Surveillance Team |
| EPI_ISL_3467366 | National Virus Reference Laboratory | National Virus Reference Laboratory | Zoe Yandle, Charlene Bennett, Gabriel Gonzalez, Michael Carr, Jonathan Dean, Cillian F De Gascun |
| EPI_ISL_3467367 | Berkshire and Surrey Pathology Services Lighthouse Laboratory | Wellcome Sanger Institute for the COVID-19 Genomics UK (COG-UK) Consortium | Berkshire and Surrey Pathology Services Lighthouse Laboratory and Alex Alderton, Roberto Amato, Jeffrey Barrett, Sonia Goncalves, Ewan Harrison, David K. Jackson, Ian Johnston, Dominic Kwiatkowski, Cordelia Langford, John Sillitoe on behalf of the Wellcome Sanger Institute COVID-19 Surveillance Team |
| EPI_ISL_3467368 | Lighthouse Lab in Milton Keynes | Wellcome Sanger Institute for the COVID-19 Genomics UK (COG-UK) Consortium | The Lighthouse Lab in Milton Keynes and Alex Alderton, Roberto Amato, Jeffrey Barrett, Sonia Goncalves, Ewan Harrison, David K. Jackson, Ian Johnston, Dominic Kwiatkowski, Cordelia Langford, John Sillitoe on behalf of the Wellcome Sanger Institute COVID-19 Surveillance Team |
| EPI_ISL_3467369 | Berkshire and Surrey Pathology Services Lighthouse Laboratory | Wellcome Sanger Institute for the COVID-19 Genomics UK (COG-UK) Consortium | Berkshire and Surrey Pathology Services Lighthouse Laboratory and Alex Alderton, Roberto Amato, Jeffrey Barrett, Sonia Goncalves, Ewan Harrison, David K. Jackson, Ian Johnston, Dominic Kwiatkowski, Cordelia Langford, John Sillitoe on behalf of the Wellcome Sanger Institute COVID-19 Surveillance Team |
| EPI_ISL_3467370 | National Virus Reference Laboratory | National Virus Reference Laboratory | Zoe Yandle, Charlene Bennett, Gabriel Gonzalez, Michael Carr, Jonathan Dean, Cillian F De Gascun |
| EPI_ISL_3467371, EPI_ISL_3467372 | Lighthouse Lab in Milton Keynes | Wellcome Sanger Institute for the COVID-19 Genomics UK (COG-UK) Consortium | The Lighthouse Lab in Milton Keynes and Alex Alderton, Roberto Amato, Jeffrey Barrett, Sonia Goncalves, Ewan Harrison, David K. Jackson, Ian Johnston, Dominic Kwiatkowski, Cordelia Langford, John Sillitoe on behalf of the Wellcome Sanger Institute COVID-19 Surveillance Team |
| EPI_ISL_3467373 | National Virus Reference Laboratory | National Virus Reference Laboratory | Zoe Yandle, Charlene Bennett, Gabriel Gonzalez, Michael Carr, Jonathan Dean, Cillian F De Gascun |
| EPI_ISL_3467374, EPI_ISL_3467375, EPI_ISL_3467376 | Berkshire and Surrey Pathology Services Lighthouse Laboratory | Wellcome Sanger Institute for the COVID-19 Genomics UK (COG-UK) Consortium | Berkshire and Surrey Pathology Services Lighthouse Laboratory and Alex Alderton, Roberto Amato, Jeffrey Barrett, Sonia Goncalves, Ewan Harrison, David K. Jackson, Ian Johnston, Dominic Kwiatkowski, Cordelia Langford, John Sillitoe on behalf of the Wellcome Sanger Institute COVID-19 Surveillance Team |
| EPI_ISL_3467377 | National Virus Reference Laboratory | National Virus Reference Laboratory | Zoe Yandle, Charlene Bennett, Gabriel Gonzalez, Michael Carr, Jonathan Dean, Cillian F De Gascun |
| EPI_ISL_3467378 | Lighthouse Lab in Milton Keynes | Wellcome Sanger Institute for the COVID-19 Genomics UK (COG-UK) Consortium | The Lighthouse Lab in Milton Keynes and Alex Alderton, Roberto Amato, Jeffrey Barrett, Sonia Goncalves, Ewan Harrison, David K. Jackson, Ian Johnston, Dominic Kwiatkowski, Cordelia Langford, John Sillitoe on behalf of the Wellcome Sanger Institute COVID-19 Surveillance Team |
| EPI_ISL_3467379, EPI_ISL_3467380 | Berkshire and Surrey Pathology Services Lighthouse Laboratory | Wellcome Sanger Institute for the COVID-19 Genomics UK (COG-UK) Consortium | Berkshire and Surrey Pathology Services Lighthouse Laboratory and Alex Alderton, Roberto Amato, Jeffrey Barrett, Sonia Goncalves, Ewan Harrison, David K. Jackson, Ian Johnston, Dominic Kwiatkowski, Cordelia Langford, John Sillitoe on behalf of the Wellcome Sanger Institute COVID-19 Surveillance Team |
| EPI_ISL_3467381 | National Virus Reference Laboratory | National Virus Reference Laboratory | Zoe Yandle, Charlene Bennett, Gabriel Gonzalez, Michael Carr, Jonathan Dean, Cillian F De Gascun |
| EPI_ISL_3467382, EPI_ISL_3467383, EPI_ISL_3467384 | Berkshire and Surrey Pathology Services Lighthouse Laboratory | Wellcome Sanger Institute for the COVID-19 Genomics UK (COG-UK) Consortium | Berkshire and Surrey Pathology Services Lighthouse Laboratory and Alex Alderton, Roberto Amato, Jeffrey Barrett, Sonia Goncalves, Ewan Harrison, David K. Jackson, Ian Johnston, Dominic Kwiatkowski, Cordelia Langford, John Sillitoe on behalf of the Wellcome Sanger Institute COVID-19 Surveillance Team |
| EPI_ISL_3467385 | National Virus Reference Laboratory | National Virus Reference Laboratory | Zoe Yandle, Charlene Bennett, Gabriel Gonzalez, Michael Carr, Jonathan Dean, Cillian F De Gascun |
| EPI_ISL_3467386, EPI_ISL_3467387, EPI_ISL_3467388 | Berkshire and Surrey Pathology Services Lighthouse Laboratory | Wellcome Sanger Institute for the COVID-19 Genomics UK (COG-UK) Consortium | Berkshire and Surrey Pathology Services Lighthouse Laboratory and Alex Alderton, Roberto Amato, Jeffrey Barrett, Sonia Goncalves, Ewan Harrison, David K. Jackson, Ian Johnston, Dominic Kwiatkowski, Cordelia Langford, John Sillitoe on behalf of the Wellcome Sanger Institute COVID-19 Surveillance Team |
| EPI_ISL_3467389 | National Virus Reference Laboratory | National Virus Reference Laboratory | Zoe Yandle, Charlene Bennett, Gabriel Gonzalez, Michael Carr, Jonathan Dean, Cillian F De Gascun |
| EPI_ISL_3467390, EPI_ISL_3467391, EPI_ISL_3467392 | Berkshire and Surrey Pathology Services Lighthouse Laboratory | Wellcome Sanger Institute for the COVID-19 Genomics UK (COG-UK) Consortium | Berkshire and Surrey Pathology Services Lighthouse Laboratory and Alex Alderton, Roberto Amato, Jeffrey Barrett, Sonia Goncalves, Ewan Harrison, David K. Jackson, Ian Johnston, Dominic Kwiatkowski, Cordelia Langford, John Sillitoe on behalf of the Wellcome Sanger Institute COVID-19 Surveillance Team |
| EPI_ISL_3467393 | National Virus Reference Laboratory | National Virus Reference Laboratory | Zoe Yandle, Charlene Bennett, Gabriel Gonzalez, Michael Carr, Jonathan Dean, Cillian F De Gascun |
| EPI_ISL_3467394, EPI_ISL_3467395, EPI_ISL_3467396 | Berkshire and Surrey Pathology Services Lighthouse Laboratory | Wellcome Sanger Institute for the COVID-19 Genomics UK (COG-UK) Consortium | Berkshire and Surrey Pathology Services Lighthouse Laboratory and Alex Alderton, Roberto Amato, Jeffrey Barrett, Sonia Goncalves, Ewan Harrison, David K. Jackson, Ian Johnston, Dominic Kwiatkowski, Cordelia Langford, John Sillitoe on behalf of the Wellcome Sanger Institute COVID-19 Surveillance Team |
| EPI_ISL_3467397 | National Virus Reference Laboratory | National Virus Reference Laboratory | Zoe Yandle, Charlene Bennett, Gabriel Gonzalez, Michael Carr, Jonathan Dean, Cillian F De Gascun |
| EPI_ISL_3467398, EPI_ISL_3467399 | Lighthouse Lab in Milton Keynes | Wellcome Sanger Institute for the COVID-19 Genomics UK (COG-UK) Consortium | The Lighthouse Lab in Milton Keynes and Alex Alderton, Roberto Amato, Jeffrey Barrett, Sonia Goncalves, Ewan Harrison, David K. Jackson, Ian Johnston, Dominic Kwiatkowski, Cordelia Langford, John Sillitoe on behalf of the Wellcome Sanger Institute COVID-19 Surveillance Team |
| EPI_ISL_3467400 | Berkshire and Surrey Pathology Services Lighthouse Laboratory | Wellcome Sanger Institute for the COVID-19 Genomics UK (COG-UK) Consortium | Berkshire and Surrey Pathology Services Lighthouse Laboratory and Alex Alderton, Roberto Amato, Jeffrey Barrett, Sonia Goncalves, Ewan Harrison, David K. Jackson, Ian Johnston, Dominic Kwiatkowski, Cordelia Langford, John Sillitoe on behalf of the Wellcome Sanger Institute COVID-19 Surveillance Team |
| EPI_ISL_3467401 | National Virus Reference Laboratory | National Virus Reference Laboratory | Zoe Yandle, Charlene Bennett, Gabriel Gonzalez, Michael Carr, Jonathan Dean, Cillian F De Gascun |
| EPI_ISL_3467402 | Lighthouse Lab in Milton Keynes | Wellcome Sanger Institute for the COVID-19 Genomics UK (COG-UK) Consortium | The Lighthouse Lab in Milton Keynes and Alex Alderton, Roberto Amato, Jeffrey Barrett, Sonia Goncalves, Ewan Harrison, David K. Jackson, Ian Johnston, Dominic Kwiatkowski, Cordelia Langford, John Sillitoe on behalf of the Wellcome Sanger Institute COVID-19 Surveillance Team |
| EPI_ISL_3467403, EPI_ISL_3467404 | Berkshire and Surrey Pathology Services Lighthouse Laboratory | Wellcome Sanger Institute for the COVID-19 Genomics UK (COG-UK) Consortium | Berkshire and Surrey Pathology Services Lighthouse Laboratory and Alex Alderton, Roberto Amato, Jeffrey Barrett, Sonia Goncalves, Ewan Harrison, David K. Jackson, Ian Johnston, Dominic Kwiatkowski, Cordelia Langford, John Sillitoe on behalf of the Wellcome Sanger Institute COVID-19 Surveillance Team |
| EPI_ISL_3467405 | National Virus Reference Laboratory | National Virus Reference Laboratory | Zoe Yandle, Charlene Bennett, Gabriel Gonzalez, Michael Carr, Jonathan Dean, Cillian F De Gascun |
| EPI_ISL_3467406, EPI_ISL_3467407, EPI_ISL_3467408 | Berkshire and Surrey Pathology Services Lighthouse Laboratory | Wellcome Sanger Institute for the COVID-19 Genomics UK (COG-UK) Consortium | Berkshire and Surrey Pathology Services Lighthouse Laboratory and Alex Alderton, Roberto Amato, Jeffrey Barrett, Sonia Goncalves, Ewan Harrison, David K. Jackson, Ian Johnston, Dominic Kwiatkowski, Cordelia Langford, John Sillitoe on behalf of the Wellcome Sanger Institute COVID-19 Surveillance Team |
| EPI_ISL_3467409 | National Virus Reference Laboratory | National Virus Reference Laboratory | Zoe Yandle, Charlene Bennett, Gabriel Gonzalez, Michael Carr, Jonathan Dean, Cillian F De Gascun |
| EPI_ISL_3467410 | Lighthouse Lab in Milton Keynes | Wellcome Sanger Institute for the COVID-19 Genomics UK (COG-UK) Consortium | The Lighthouse Lab in Milton Keynes and Alex Alderton, Roberto Amato, Jeffrey Barrett, Sonia Goncalves, Ewan Harrison, David K. Jackson, Ian Johnston, Dominic Kwiatkowski, Cordelia Langford, John Sillitoe on behalf of the Wellcome Sanger Institute COVID-19 Surveillance Team |
| EPI_ISL_3467411 | Berkshire and Surrey Pathology Services Lighthouse Laboratory | Wellcome Sanger Institute for the COVID-19 Genomics UK (COG-UK) Consortium | Berkshire and Surrey Pathology Services Lighthouse Laboratory and Alex Alderton, Roberto Amato, Jeffrey Barrett, Sonia Goncalves, Ewan Harrison, David K. Jackson, Ian Johnston, Dominic Kwiatkowski, Cordelia Langford, John Sillitoe on behalf of the Wellcome Sanger Institute COVID-19 Surveillance Team |
| EPI_ISL_3467412 | National Virus Reference Laboratory | National Virus Reference Laboratory | Zoe Yandle, Charlene Bennett, Gabriel Gonzalez, Michael Carr, Jonathan Dean, Cillian F De Gascun |
| EPI_ISL_3467413, EPI_ISL_3467414, EPI_ISL_3467415 | Lighthouse Lab in Milton Keynes | Wellcome Sanger Institute for the COVID-19 Genomics UK (COG-UK) Consortium | The Lighthouse Lab in Milton Keynes and Alex Alderton, Roberto Amato, Jeffrey Barrett, Sonia Goncalves, Ewan Harrison, David K. Jackson, Ian Johnston, Dominic Kwiatkowski, Cordelia Langford, John Sillitoe on behalf of the Wellcome Sanger Institute COVID-19 Surveillance Team |
| EPI_ISL_3467416 | Berkshire and Surrey Pathology Services Lighthouse Laboratory | Wellcome Sanger Institute for the COVID-19 Genomics UK (COG-UK) Consortium | Berkshire and Surrey Pathology Services Lighthouse Laboratory and Alex Alderton, Roberto Amato, Jeffrey Barrett, Sonia Goncalves, Ewan Harrison, David K. Jackson, Ian Johnston, Dominic Kwiatkowski, Cordelia Langford, John Sillitoe on behalf of the Wellcome Sanger Institute COVID-19 Surveillance Team |
| EPI_ISL_3467417 | National Virus Reference Laboratory | National Virus Reference Laboratory | Zoe Yandle, Charlene Bennett, Gabriel Gonzalez, Michael Carr, Jonathan Dean, Cillian F De Gascun |
| EPI_ISL_3467418, EPI_ISL_3467419 | Berkshire and Surrey Pathology Services Lighthouse Laboratory | Wellcome Sanger Institute for the COVID-19 Genomics UK (COG-UK) Consortium | Berkshire and Surrey Pathology Services Lighthouse Laboratory and Alex Alderton, Roberto Amato, Jeffrey Barrett, Sonia Goncalves, Ewan Harrison, David K. Jackson, Ian Johnston, Dominic Kwiatkowski, Cordelia Langford, John Sillitoe on behalf of the Wellcome Sanger Institute COVID-19 Surveillance Team |
| EPI_ISL_3467420 | National Virus Reference Laboratory | National Virus Reference Laboratory | Zoe Yandle, Charlene Bennett, Gabriel Gonzalez, Michael Carr, Jonathan Dean, Cillian F De Gascun |
| EPI_ISL_3467421 | Berkshire and Surrey Pathology Services Lighthouse Laboratory | Wellcome Sanger Institute for the COVID-19 Genomics UK (COG-UK) Consortium | Berkshire and Surrey Pathology Services Lighthouse Laboratory and Alex Alderton, Roberto Amato, Jeffrey Barrett, Sonia Goncalves, Ewan Harrison, David K. Jackson, Ian Johnston, Dominic Kwiatkowski, Cordelia Langford, John Sillitoe on behalf of the Wellcome Sanger Institute COVID-19 Surveillance Team |
| EPI_ISL_3467422 | Lighthouse Lab in Milton Keynes | Wellcome Sanger Institute for the COVID-19 Genomics UK (COG-UK) Consortium | The Lighthouse Lab in Milton Keynes and Alex Alderton, Roberto Amato, Jeffrey Barrett, Sonia Goncalves, Ewan Harrison, David K. Jackson, Ian Johnston, Dominic Kwiatkowski, Cordelia Langford, John Sillitoe on behalf of the Wellcome Sanger Institute COVID-19 Surveillance Team |
| EPI_ISL_3467423 | Berkshire and Surrey Pathology Services Lighthouse Laboratory | Wellcome Sanger Institute for the COVID-19 Genomics UK (COG-UK) Consortium | Berkshire and Surrey Pathology Services Lighthouse Laboratory and Alex Alderton, Roberto Amato, Jeffrey Barrett, Sonia Goncalves, Ewan Harrison, David K. Jackson, Ian Johnston, Dominic Kwiatkowski, Cordelia Langford, John Sillitoe on behalf of the Wellcome Sanger Institute COVID-19 Surveillance Team |
| EPI_ISL_3467424 | National Virus Reference Laboratory | National Virus Reference Laboratory | Zoe Yandle, Charlene Bennett, Gabriel Gonzalez, Michael Carr, Jonathan Dean, Cillian F De Gascun |
| EPI_ISL_3467425, EPI_ISL_3467426, EPI_ISL_3467427 | Berkshire and Surrey Pathology Services Lighthouse Laboratory | Wellcome Sanger Institute for the COVID-19 Genomics UK (COG-UK) Consortium | Berkshire and Surrey Pathology Services Lighthouse Laboratory and Alex Alderton, Roberto Amato, Jeffrey Barrett, Sonia Goncalves, Ewan Harrison, David K. Jackson, Ian Johnston, Dominic Kwiatkowski, Cordelia Langford, John Sillitoe on behalf of the Wellcome Sanger Institute COVID-19 Surveillance Team |
| EPI_ISL_3467428 | National Virus Reference Laboratory | National Virus Reference Laboratory | Zoe Yandle, Charlene Bennett, Gabriel Gonzalez, Michael Carr, Jonathan Dean, Cillian F De Gascun |
| EPI_ISL_3467429, | Berkshire and Surrey Pathology Services Lighthouse Laboratory | Wellcome Sanger Institute for the COVID-19 Genomics UK (COG-UK) | Berkshire and Surrey Pathology Services Lighthouse Laboratory and Alex Alderton, Roberto Amato, Jeffrey Barrett, Sonia Goncalves, Ewan Harrison, David K. Jackson, Ian Johnston, Dominic Kwiatkowski, Cordelia Langford, John Sillitoe on behalf |

| | | | |
|---|---|---|---|
| EPI_ISL_3467430, EPI_ISL_3467431 | | Consortium | of the Wellcome Sanger Institute COVID-19 Surveillance Team |
| EPI_ISL_3467432 | National Virus Reference Laboratory | National Virus Reference Laboratory | Zoe Yandle, Charlene Bennett, Gabriel Gonzalez, Michael Carr, Jonathan Dean, Cillian F De Gascun |
| EPI_ISL_3467433, EPI_ISL_3467434 | Berkshire and Surrey Pathology Services Lighthouse Laboratory | Wellcome Sanger Institute for the COVID-19 Genomics UK (COG-UK) Consortium | Berkshire and Surrey Pathology Services Lighthouse Laboratory and Alex Alderton, Roberto Amato, Jeffrey Barrett, Sonia Goncalves, Ewan Harrison, David K. Jackson, Ian Johnston, Dominic Kwiatkowski, Cordelia Langford, John Sillitoe on behalf of the Wellcome Sanger Institute COVID-19 Surveillance Team |
| EPI_ISL_3467435 | Lighthouse Lab in Milton Keynes | Wellcome Sanger Institute for the COVID-19 Genomics UK (COG-UK) Consortium | The Lighthouse Lab in Milton Keynes and Alex Alderton, Roberto Amato, Jeffrey Barrett, Sonia Goncalves, Ewan Harrison, David K. Jackson, Ian Johnston, Dominic Kwiatkowski, Cordelia Langford, John Sillitoe on behalf of the Wellcome Sanger Institute COVID-19 Surveillance Team |
| EPI_ISL_3467436 | National Virus Reference Laboratory | National Virus Reference Laboratory | Zoe Yandle, Charlene Bennett, Gabriel Gonzalez, Michael Carr, Jonathan Dean, Cillian F De Gascun |
| EPI_ISL_3467437, EPI_ISL_3467438 | Berkshire and Surrey Pathology Services Lighthouse Laboratory | Wellcome Sanger Institute for the COVID-19 Genomics UK (COG-UK) Consortium | Berkshire and Surrey Pathology Services Lighthouse Laboratory and Alex Alderton, Roberto Amato, Jeffrey Barrett, Sonia Goncalves, Ewan Harrison, David K. Jackson, Ian Johnston, Dominic Kwiatkowski, Cordelia Langford, John Sillitoe on behalf of the Wellcome Sanger Institute COVID-19 Surveillance Team |
| EPI_ISL_3467439 | National Virus Reference Laboratory | National Virus Reference Laboratory | Zoe Yandle, Charlene Bennett, Gabriel Gonzalez, Michael Carr, Jonathan Dean, Cillian F De Gascun |
| EPI_ISL_3467440, EPI_ISL_3467441, EPI_ISL_3467442, EPI_ISL_3467444, EPI_ISL_3467445, EPI_ISL_3467446 | Berkshire and Surrey Pathology Services Lighthouse Laboratory | Wellcome Sanger Institute for the COVID-19 Genomics UK (COG-UK) Consortium | Berkshire and Surrey Pathology Services Lighthouse Laboratory and Alex Alderton, Roberto Amato, Jeffrey Barrett, Sonia Goncalves, Ewan Harrison, David K. Jackson, Ian Johnston, Dominic Kwiatkowski, Cordelia Langford, John Sillitoe on behalf of the Wellcome Sanger Institute COVID-19 Surveillance Team |
| EPI_ISL_3467447 | National Virus Reference Laboratory | National Virus Reference Laboratory | Zoe Yandle, Charlene Bennett, Gabriel Gonzalez, Michael Carr, Jonathan Dean, Cillian F De Gascun |
| EPI_ISL_3467448 | Lighthouse Lab in Milton Keynes | Wellcome Sanger Institute for the COVID-19 Genomics UK (COG-UK) Consortium | The Lighthouse Lab in Milton Keynes and Alex Alderton, Roberto Amato, Jeffrey Barrett, Sonia Goncalves, Ewan Harrison, David K. Jackson, Ian Johnston, Dominic Kwiatkowski, Cordelia Langford, John Sillitoe on behalf of the Wellcome Sanger Institute COVID-19 Surveillance Team |
| EPI_ISL_3467449 | Berkshire and Surrey Pathology Services Lighthouse Laboratory | Wellcome Sanger Institute for the COVID-19 Genomics UK (COG-UK) Consortium | Berkshire and Surrey Pathology Services Lighthouse Laboratory and Alex Alderton, Roberto Amato, Jeffrey Barrett, Sonia Goncalves, Ewan Harrison, David K. Jackson, Ian Johnston, Dominic Kwiatkowski, Cordelia Langford, John Sillitoe on behalf of the Wellcome Sanger Institute COVID-19 Surveillance Team |
| EPI_ISL_3467450 | Lighthouse Lab in Milton Keynes | Wellcome Sanger Institute for the COVID-19 Genomics UK (COG-UK) Consortium | The Lighthouse Lab in Milton Keynes and Alex Alderton, Roberto Amato, Jeffrey Barrett, Sonia Goncalves, Ewan Harrison, David K. Jackson, Ian Johnston, Dominic Kwiatkowski, Cordelia Langford, John Sillitoe on behalf of the Wellcome Sanger Institute COVID-19 Surveillance Team |
| EPI_ISL_3467451 | National Virus Reference Laboratory | National Virus Reference Laboratory | Zoe Yandle, Charlene Bennett, Gabriel Gonzalez, Michael Carr, Jonathan Dean, Cillian F De Gascun |
| EPI_ISL_3467452, EPI_ISL_3467453, EPI_ISL_3467454 | Berkshire and Surrey Pathology Services Lighthouse Laboratory | Wellcome Sanger Institute for the COVID-19 Genomics UK (COG-UK) Consortium | Berkshire and Surrey Pathology Services Lighthouse Laboratory and Alex Alderton, Roberto Amato, Jeffrey Barrett, Sonia Goncalves, Ewan Harrison, David K. Jackson, Ian Johnston, Dominic Kwiatkowski, Cordelia Langford, John Sillitoe on behalf of the Wellcome Sanger Institute COVID-19 Surveillance Team |
| EPI_ISL_3467455 | National Virus Reference Laboratory | National Virus Reference Laboratory | Zoe Yandle, Charlene Bennett, Gabriel Gonzalez, Michael Carr, Jonathan Dean, Cillian F De Gascun |
| EPI_ISL_3467456, EPI_ISL_3467457 | Lighthouse Lab in Milton Keynes | Wellcome Sanger Institute for the COVID-19 Genomics UK (COG-UK) Consortium | The Lighthouse Lab in Milton Keynes and Alex Alderton, Roberto Amato, Jeffrey Barrett, Sonia Goncalves, Ewan Harrison, David K. Jackson, Ian Johnston, Dominic Kwiatkowski, Cordelia Langford, John Sillitoe on behalf of the Wellcome Sanger Institute COVID-19 Surveillance Team |
| EPI_ISL_3467458 | Berkshire and Surrey Pathology Services Lighthouse Laboratory | Wellcome Sanger Institute for the COVID-19 Genomics UK (COG-UK) Consortium | Berkshire and Surrey Pathology Services Lighthouse Laboratory and Alex Alderton, Roberto Amato, Jeffrey Barrett, Sonia Goncalves, Ewan Harrison, David K. Jackson, Ian Johnston, Dominic Kwiatkowski, Cordelia Langford, John Sillitoe on behalf of the Wellcome Sanger Institute COVID-19 Surveillance Team |
| EPI_ISL_3467459 | National Virus Reference Laboratory | National Virus Reference Laboratory | Zoe Yandle, Charlene Bennett, Gabriel Gonzalez, Michael Carr, Jonathan Dean, Cillian F De Gascun |
| EPI_ISL_3467460, EPI_ISL_3467461, EPI_ISL_3467462 | Berkshire and Surrey Pathology Services Lighthouse Laboratory | Wellcome Sanger Institute for the COVID-19 Genomics UK (COG-UK) Consortium | Berkshire and Surrey Pathology Services Lighthouse Laboratory and Alex Alderton, Roberto Amato, Jeffrey Barrett, Sonia Goncalves, Ewan Harrison, David K. Jackson, Ian Johnston, Dominic Kwiatkowski, Cordelia Langford, John Sillitoe on behalf of the Wellcome Sanger Institute COVID-19 Surveillance Team |
| EPI_ISL_3467463 | National Virus Reference Laboratory | National Virus Reference Laboratory | Zoe Yandle, Charlene Bennett, Gabriel Gonzalez, Michael Carr, Jonathan Dean, Cillian F De Gascun |
| EPI_ISL_3467464, EPI_ISL_3467465 | Lighthouse Lab in Milton Keynes | Wellcome Sanger Institute for the COVID-19 Genomics UK (COG-UK) Consortium | The Lighthouse Lab in Milton Keynes and Alex Alderton, Roberto Amato, Jeffrey Barrett, Sonia Goncalves, Ewan Harrison, David K. Jackson, Ian Johnston, Dominic Kwiatkowski, Cordelia Langford, John Sillitoe on behalf of the Wellcome Sanger Institute COVID-19 Surveillance Team |
| EPI_ISL_3467466 | National Virus Reference Laboratory | National Virus Reference Laboratory | Zoe Yandle, Charlene Bennett, Gabriel Gonzalez, Michael Carr, Jonathan Dean, Cillian F De Gascun |
| EPI_ISL_3467467, EPI_ISL_3467468 | Berkshire and Surrey Pathology Services Lighthouse Laboratory | Wellcome Sanger Institute for the COVID-19 Genomics UK (COG-UK) Consortium | Berkshire and Surrey Pathology Services Lighthouse Laboratory and Alex Alderton, Roberto Amato, Jeffrey Barrett, Sonia Goncalves, Ewan Harrison, David K. Jackson, Ian Johnston, Dominic Kwiatkowski, Cordelia Langford, John Sillitoe on behalf of the Wellcome Sanger Institute COVID-19 Surveillance Team |
| EPI_ISL_3467469 | National Virus Reference Laboratory | National Virus Reference Laboratory | Zoe Yandle, Charlene Bennett, Gabriel Gonzalez, Michael Carr, Jonathan Dean, Cillian F De Gascun |
| EPI_ISL_3467470, EPI_ISL_3467471 | Lighthouse Lab in Milton Keynes | Wellcome Sanger Institute for the COVID-19 Genomics UK (COG-UK) Consortium | The Lighthouse Lab in Milton Keynes and Alex Alderton, Roberto Amato, Jeffrey Barrett, Sonia Goncalves, Ewan Harrison, David K. Jackson, Ian Johnston, Dominic Kwiatkowski, Cordelia Langford, John Sillitoe on behalf of the Wellcome Sanger Institute COVID-19 Surveillance Team |
| EPI_ISL_3467472 | Berkshire and Surrey Pathology Services Lighthouse Laboratory | Wellcome Sanger Institute for the COVID-19 Genomics UK (COG-UK) Consortium | Berkshire and Surrey Pathology Services Lighthouse Laboratory and Alex Alderton, Roberto Amato, Jeffrey Barrett, Sonia Goncalves, Ewan Harrison, David K. Jackson, Ian Johnston, Dominic Kwiatkowski, Cordelia Langford, John Sillitoe on behalf of the Wellcome Sanger Institute COVID-19 Surveillance Team |
| EPI_ISL_3467473 | National Virus Reference Laboratory | National Virus Reference Laboratory | Zoe Yandle, Charlene Bennett, Gabriel Gonzalez, Michael Carr, Jonathan Dean, Cillian F De Gascun |
| EPI_ISL_3467474 | Lighthouse Lab in Milton Keynes | Wellcome Sanger Institute for the COVID-19 Genomics UK (COG-UK) Consortium | The Lighthouse Lab in Milton Keynes and Alex Alderton, Roberto Amato, Jeffrey Barrett, Sonia Goncalves, Ewan Harrison, David K. Jackson, Ian Johnston, Dominic Kwiatkowski, Cordelia Langford, John Sillitoe on behalf of the Wellcome Sanger Institute COVID-19 Surveillance Team |
| EPI_ISL_3467475, EPI_ISL_3467476 | Berkshire and Surrey Pathology Services Lighthouse Laboratory | Wellcome Sanger Institute for the COVID-19 Genomics UK (COG-UK) Consortium | Berkshire and Surrey Pathology Services Lighthouse Laboratory and Alex Alderton, Roberto Amato, Jeffrey Barrett, Sonia Goncalves, Ewan Harrison, David K. Jackson, Ian Johnston, Dominic Kwiatkowski, Cordelia Langford, John Sillitoe on behalf of the Wellcome Sanger Institute COVID-19 Surveillance Team |
| EPI_ISL_3467477 | National Virus Reference Laboratory | National Virus Reference Laboratory | Zoe Yandle, Charlene Bennett, Gabriel Gonzalez, Michael Carr, Jonathan Dean, Cillian F De Gascun |
| EPI_ISL_3467478 | Berkshire and Surrey Pathology Services Lighthouse Laboratory | Wellcome Sanger Institute for the COVID-19 Genomics UK (COG-UK) Consortium | Berkshire and Surrey Pathology Services Lighthouse Laboratory and Alex Alderton, Roberto Amato, Jeffrey Barrett, Sonia Goncalves, Ewan Harrison, David K. Jackson, Ian Johnston, Dominic Kwiatkowski, Cordelia Langford, John Sillitoe on behalf of the Wellcome Sanger Institute COVID-19 Surveillance Team |
| EPI_ISL_3467479, EPI_ISL_3467480 | Lighthouse Lab in Milton Keynes | Wellcome Sanger Institute for the COVID-19 Genomics UK (COG-UK) Consortium | The Lighthouse Lab in Milton Keynes and Alex Alderton, Roberto Amato, Jeffrey Barrett, Sonia Goncalves, Ewan Harrison, David K. Jackson, Ian Johnston, Dominic Kwiatkowski, Cordelia Langford, John Sillitoe on behalf of the Wellcome Sanger Institute COVID-19 Surveillance Team |
| EPI_ISL_3467481 | National Virus Reference Laboratory | National Virus Reference Laboratory | Zoe Yandle, Charlene Bennett, Gabriel Gonzalez, Michael Carr, Jonathan Dean, Cillian F De Gascun |
| EPI_ISL_3467482, EPI_ISL_3467483, EPI_ISL_3467484 | Berkshire and Surrey Pathology Services Lighthouse Laboratory | Wellcome Sanger Institute for the COVID-19 Genomics UK (COG-UK) Consortium | Berkshire and Surrey Pathology Services Lighthouse Laboratory and Alex Alderton, Roberto Amato, Jeffrey Barrett, Sonia Goncalves, Ewan Harrison, David K. Jackson, Ian Johnston, Dominic Kwiatkowski, Cordelia Langford, John Sillitoe on behalf of the Wellcome Sanger Institute COVID-19 Surveillance Team |
| EPI_ISL_3467485 | National Virus Reference Laboratory | National Virus Reference Laboratory | Zoe Yandle, Charlene Bennett, Gabriel Gonzalez, Michael Carr, Jonathan Dean, Cillian F De Gascun |
| EPI_ISL_3467486 | Lighthouse Lab in Milton Keynes | Wellcome Sanger Institute for the COVID-19 Genomics UK (COG-UK) Consortium | The Lighthouse Lab in Milton Keynes and Alex Alderton, Roberto Amato, Jeffrey Barrett, Sonia Goncalves, Ewan Harrison, David K. Jackson, Ian Johnston, Dominic Kwiatkowski, Cordelia Langford, John Sillitoe on behalf of the Wellcome Sanger Institute COVID-19 Surveillance Team |
| EPI_ISL_3467487, EPI_ISL_3467488 | Berkshire and Surrey Pathology Services Lighthouse Laboratory | Wellcome Sanger Institute for the COVID-19 Genomics UK (COG-UK) Consortium | Berkshire and Surrey Pathology Services Lighthouse Laboratory and Alex Alderton, Roberto Amato, Jeffrey Barrett, Sonia Goncalves, Ewan Harrison, David K. Jackson, Ian Johnston, Dominic Kwiatkowski, Cordelia Langford, John Sillitoe on behalf of the Wellcome Sanger Institute COVID-19 Surveillance Team |
| EPI_ISL_3467489 | National Virus Reference Laboratory | National Virus Reference Laboratory | Zoe Yandle, Charlene Bennett, Gabriel Gonzalez, Michael Carr, Jonathan Dean, Cillian F De Gascun |
| EPI_ISL_3467490, EPI_ISL_3467491 | Berkshire and Surrey Pathology Services Lighthouse Laboratory | Wellcome Sanger Institute for the COVID-19 Genomics UK (COG-UK) Consortium | Berkshire and Surrey Pathology Services Lighthouse Laboratory and Alex Alderton, Roberto Amato, Jeffrey Barrett, Sonia Goncalves, Ewan Harrison, David K. Jackson, Ian Johnston, Dominic Kwiatkowski, Cordelia Langford, John Sillitoe on behalf of the Wellcome Sanger Institute COVID-19 Surveillance Team |
| EPI_ISL_3467492 | National Virus Reference Laboratory | National Virus Reference Laboratory | Zoe Yandle, Charlene Bennett, Gabriel Gonzalez, Michael Carr, Jonathan Dean, Cillian F De Gascun |
| EPI_ISL_3467493, EPI_ISL_3467494, EPI_ISL_3467495 | Berkshire and Surrey Pathology Services Lighthouse Laboratory | Wellcome Sanger Institute for the COVID-19 Genomics UK (COG-UK) Consortium | Berkshire and Surrey Pathology Services Lighthouse Laboratory and Alex Alderton, Roberto Amato, Jeffrey Barrett, Sonia Goncalves, Ewan Harrison, David K. Jackson, Ian Johnston, Dominic Kwiatkowski, Cordelia Langford, John Sillitoe on behalf of the Wellcome Sanger Institute COVID-19 Surveillance Team |
| EPI_ISL_3467496 | National Virus Reference Laboratory | National Virus Reference Laboratory | Zoe Yandle, Charlene Bennett, Gabriel Gonzalez, Michael Carr, Jonathan Dean, Cillian F De Gascun |
| EPI_ISL_3467497, EPI_ISL_3467498 | Lighthouse Lab in Milton Keynes | Wellcome Sanger Institute for the COVID-19 Genomics UK (COG-UK) Consortium | The Lighthouse Lab in Milton Keynes and Alex Alderton, Roberto Amato, Jeffrey Barrett, Sonia Goncalves, Ewan Harrison, David K. Jackson, Ian Johnston, Dominic Kwiatkowski, Cordelia Langford, John Sillitoe on behalf of the Wellcome Sanger Institute COVID-19 Surveillance Team |
| EPI_ISL_3467499 | Berkshire and Surrey Pathology Services Lighthouse Laboratory | Wellcome Sanger Institute for the COVID-19 Genomics UK (COG-UK) Consortium | Berkshire and Surrey Pathology Services Lighthouse Laboratory and Alex Alderton, Roberto Amato, Jeffrey Barrett, Sonia Goncalves, Ewan Harrison, David K. Jackson, Ian Johnston, Dominic Kwiatkowski, Cordelia Langford, John Sillitoe on behalf of the Wellcome Sanger Institute COVID-19 Surveillance Team |
| EPI_ISL_3467500 | National Virus Reference Laboratory | National Virus Reference Laboratory | Zoe Yandle, Charlene Bennett, Gabriel Gonzalez, Michael Carr, Jonathan Dean, Cillian F De Gascun |
| EPI_ISL_3467501, EPI_ISL_3467502 | Berkshire and Surrey Pathology Services Lighthouse Laboratory | Wellcome Sanger Institute for the COVID-19 Genomics UK (COG-UK) Consortium | Berkshire and Surrey Pathology Services Lighthouse Laboratory and Alex Alderton, Roberto Amato, Jeffrey Barrett, Sonia Goncalves, Ewan Harrison, David K. Jackson, Ian Johnston, Dominic Kwiatkowski, Cordelia Langford, John Sillitoe on behalf of the Wellcome Sanger Institute COVID-19 Surveillance Team |
| EPI_ISL_3467503 | Lighthouse Lab in Milton Keynes | Wellcome Sanger Institute for the COVID-19 Genomics UK (COG-UK) Consortium | The Lighthouse Lab in Milton Keynes and Alex Alderton, Roberto Amato, Jeffrey Barrett, Sonia Goncalves, Ewan Harrison, David K. Jackson, Ian Johnston, Dominic Kwiatkowski, Cordelia Langford, John Sillitoe on behalf of the Wellcome Sanger Institute COVID-19 Surveillance Team |
| EPI_ISL_3467504 | National Virus Reference Laboratory | National Virus Reference Laboratory | Zoe Yandle, Charlene Bennett, Gabriel Gonzalez, Michael Carr, Jonathan Dean, Cillian F De Gascun |
| EPI_ISL_3467505 | Berkshire and Surrey Pathology Services Lighthouse Laboratory | Wellcome Sanger Institute for the COVID-19 Genomics UK (COG-UK) Consortium | Berkshire and Surrey Pathology Services Lighthouse Laboratory and Alex Alderton, Roberto Amato, Jeffrey Barrett, Sonia Goncalves, Ewan Harrison, David K. Jackson, Ian Johnston, Dominic Kwiatkowski, Cordelia Langford, John Sillitoe on behalf of the Wellcome Sanger Institute COVID-19 Surveillance Team |
| EPI_ISL_3467506 | Lighthouse Lab in Milton Keynes | Wellcome Sanger Institute for the COVID-19 Genomics UK (COG-UK) Consortium | The Lighthouse Lab in Milton Keynes and Alex Alderton, Roberto Amato, Jeffrey Barrett, Sonia Goncalves, Ewan Harrison, David K. Jackson, Ian Johnston, Dominic Kwiatkowski, Cordelia Langford, John Sillitoe on behalf of the Wellcome Sanger Institute COVID-19 Surveillance Team |

| | | | |
|---|---|---|---|
| EPI_ISL_3467507 | Berkshire and Surrey Pathology Services Lighthouse Laboratory | Wellcome Sanger Institute for the COVID-19 Genomics UK (COG-UK) Consortium | Berkshire and Surrey Pathology Services Lighthouse Laboratory and Alex Alderton, Roberto Amato, Jeffrey Barrett, Sonia Goncalves, Ewan Harrison, David K. Jackson, Ian Johnston, Dominic Kwiatkowski, Cordelia Langford, John Sillitoe on behalf of the Wellcome Sanger Institute COVID-19 Surveillance Team |
| EPI_ISL_3467508 | National Virus Reference Laboratory | National Virus Reference Laboratory | Zoe Yandle, Charlene Bennett, Gabriel Gonzalez, Michael Carr, Jonathan Dean, Cillian F De Gascun |
| EPI_ISL_3467509, EPI_ISL_3467510 | Berkshire and Surrey Pathology Services Lighthouse Laboratory | Wellcome Sanger Institute for the COVID-19 Genomics UK (COG-UK) Consortium | Berkshire and Surrey Pathology Services Lighthouse Laboratory and Alex Alderton, Roberto Amato, Jeffrey Barrett, Sonia Goncalves, Ewan Harrison, David K. Jackson, Ian Johnston, Dominic Kwiatkowski, Cordelia Langford, John Sillitoe on behalf of the Wellcome Sanger Institute COVID-19 Surveillance Team |
| EPI_ISL_3467511 | National Virus Reference Laboratory | National Virus Reference Laboratory | Zoe Yandle, Charlene Bennett, Gabriel Gonzalez, Michael Carr, Jonathan Dean, Cillian F De Gascun |
| EPI_ISL_3467512 | Berkshire and Surrey Pathology Services Lighthouse Laboratory | Wellcome Sanger Institute for the COVID-19 Genomics UK (COG-UK) Consortium | Berkshire and Surrey Pathology Services Lighthouse Laboratory and Alex Alderton, Roberto Amato, Jeffrey Barrett, Sonia Goncalves, Ewan Harrison, David K. Jackson, Ian Johnston, Dominic Kwiatkowski, Cordelia Langford, John Sillitoe on behalf of the Wellcome Sanger Institute COVID-19 Surveillance Team |
| EPI_ISL_3467513 | Lighthouse Lab in Milton Keynes | Wellcome Sanger Institute for the COVID-19 Genomics UK (COG-UK) Consortium | The Lighthouse Lab in Milton Keynes and Alex Alderton, Roberto Amato, Jeffrey Barrett, Sonia Goncalves, Ewan Harrison, David K. Jackson, Ian Johnston, Dominic Kwiatkowski, Cordelia Langford, John Sillitoe on behalf of the Wellcome Sanger Institute COVID-19 Surveillance Team |
| EPI_ISL_3467514 | Berkshire and Surrey Pathology Services Lighthouse Laboratory | Wellcome Sanger Institute for the COVID-19 Genomics UK (COG-UK) Consortium | Berkshire and Surrey Pathology Services Lighthouse Laboratory and Alex Alderton, Roberto Amato, Jeffrey Barrett, Sonia Goncalves, Ewan Harrison, David K. Jackson, Ian Johnston, Dominic Kwiatkowski, Cordelia Langford, John Sillitoe on behalf of the Wellcome Sanger Institute COVID-19 Surveillance Team |
| EPI_ISL_3467515 | National Virus Reference Laboratory | National Virus Reference Laboratory | Zoe Yandle, Charlene Bennett, Gabriel Gonzalez, Michael Carr, Jonathan Dean, Cillian F De Gascun |
| EPI_ISL_3467516, EPI_ISL_3467517, EPI_ISL_3467518 | Lighthouse Lab in Milton Keynes | Wellcome Sanger Institute for the COVID-19 Genomics UK (COG-UK) Consortium | The Lighthouse Lab in Milton Keynes and Alex Alderton, Roberto Amato, Jeffrey Barrett, Sonia Goncalves, Ewan Harrison, David K. Jackson, Ian Johnston, Dominic Kwiatkowski, Cordelia Langford, John Sillitoe on behalf of the Wellcome Sanger Institute COVID-19 Surveillance Team |
| EPI_ISL_3467519 | National Virus Reference Laboratory | National Virus Reference Laboratory | Zoe Yandle, Charlene Bennett, Gabriel Gonzalez, Michael Carr, Jonathan Dean, Cillian F De Gascun |
| EPI_ISL_3467520 | Lighthouse Lab in Glasgow | Wellcome Sanger Institute for the COVID-19 Genomics UK (COG-UK) Consortium | Harper VanSteenhouse, Yumi Kasai, David Gray, Carol Clugston, Anna Dominiczak and Alex Alderton, Roberto Amato, Jeffrey Barrett, Sonia Goncalves, Ewan Harrison, David K. Jackson, Ian Johnston, Dominic Kwiatkowski, Cordelia Langford, John Sillitoe on behalf of the Wellcome Sanger Institute COVID-19 Surveillance Team |
| EPI_ISL_3467521 | Lighthouse Lab in Milton Keynes | Wellcome Sanger Institute for the COVID-19 Genomics UK (COG-UK) Consortium | The Lighthouse Lab in Milton Keynes and Alex Alderton, Roberto Amato, Jeffrey Barrett, Sonia Goncalves, Ewan Harrison, David K. Jackson, Ian Johnston, Dominic Kwiatkowski, Cordelia Langford, John Sillitoe on behalf of the Wellcome Sanger Institute COVID-19 Surveillance Team |
| EPI_ISL_3467522 | National Virus Reference Laboratory | National Virus Reference Laboratory | Zoe Yandle, Charlene Bennett, Gabriel Gonzalez, Michael Carr, Jonathan Dean, Cillian F De Gascun |
| EPI_ISL_3467523, EPI_ISL_3467524, EPI_ISL_3467525 | Lighthouse Lab in Glasgow | Wellcome Sanger Institute for the COVID-19 Genomics UK (COG-UK) Consortium | Harper VanSteenhouse, Yumi Kasai, David Gray, Carol Clugston, Anna Dominiczak and Alex Alderton, Roberto Amato, Jeffrey Barrett, Sonia Goncalves, Ewan Harrison, David K. Jackson, Ian Johnston, Dominic Kwiatkowski, Cordelia Langford, John Sillitoe on behalf of the Wellcome Sanger Institute COVID-19 Surveillance Team |
| EPI_ISL_3467526 | National Virus Reference Laboratory | National Virus Reference Laboratory | Zoe Yandle, Charlene Bennett, Gabriel Gonzalez, Michael Carr, Jonathan Dean, Cillian F De Gascun |
| EPI_ISL_3467527, EPI_ISL_3467528, EPI_ISL_3467529 | Lighthouse Lab in Milton Keynes | Wellcome Sanger Institute for the COVID-19 Genomics UK (COG-UK) Consortium | The Lighthouse Lab in Milton Keynes and Alex Alderton, Roberto Amato, Jeffrey Barrett, Sonia Goncalves, Ewan Harrison, David K. Jackson, Ian Johnston, Dominic Kwiatkowski, Cordelia Langford, John Sillitoe on behalf of the Wellcome Sanger Institute COVID-19 Surveillance Team |
| EPI_ISL_3467530 | National Virus Reference Laboratory | National Virus Reference Laboratory | Zoe Yandle, Charlene Bennett, Gabriel Gonzalez, Michael Carr, Jonathan Dean, Cillian F De Gascun |
| EPI_ISL_3467531, EPI_ISL_3467532 | Lighthouse Lab in Glasgow | Wellcome Sanger Institute for the COVID-19 Genomics UK (COG-UK) Consortium | Harper VanSteenhouse, Yumi Kasai, David Gray, Carol Clugston, Anna Dominiczak and Alex Alderton, Roberto Amato, Jeffrey Barrett, Sonia Goncalves, Ewan Harrison, David K. Jackson, Ian Johnston, Dominic Kwiatkowski, Cordelia Langford, John Sillitoe on behalf of the Wellcome Sanger Institute COVID-19 Surveillance Team |
| EPI_ISL_3467533 | Lighthouse Lab in Milton Keynes | Wellcome Sanger Institute for the COVID-19 Genomics UK (COG-UK) Consortium | The Lighthouse Lab in Milton Keynes and Alex Alderton, Roberto Amato, Jeffrey Barrett, Sonia Goncalves, Ewan Harrison, David K. Jackson, Ian Johnston, Dominic Kwiatkowski, Cordelia Langford, John Sillitoe on behalf of the Wellcome Sanger Institute COVID-19 Surveillance Team |
| EPI_ISL_3467534 | National Virus Reference Laboratory | National Virus Reference Laboratory | Zoe Yandle, Charlene Bennett, Gabriel Gonzalez, Michael Carr, Jonathan Dean, Cillian F De Gascun |
| EPI_ISL_3467535 | Lighthouse Lab in Milton Keynes | Wellcome Sanger Institute for the COVID-19 Genomics UK (COG-UK) Consortium | The Lighthouse Lab in Milton Keynes and Alex Alderton, Roberto Amato, Jeffrey Barrett, Sonia Goncalves, Ewan Harrison, David K. Jackson, Ian Johnston, Dominic Kwiatkowski, Cordelia Langford, John Sillitoe on behalf of the Wellcome Sanger Institute COVID-19 Surveillance Team |
| EPI_ISL_3467536, EPI_ISL_3467537 | Lighthouse Lab in Glasgow | Wellcome Sanger Institute for the COVID-19 Genomics UK (COG-UK) Consortium | Harper VanSteenhouse, Yumi Kasai, David Gray, Carol Clugston, Anna Dominiczak and Alex Alderton, Roberto Amato, Jeffrey Barrett, Sonia Goncalves, Ewan Harrison, David K. Jackson, Ian Johnston, Dominic Kwiatkowski, Cordelia Langford, John Sillitoe on behalf of the Wellcome Sanger Institute COVID-19 Surveillance Team |
| EPI_ISL_3467538 | National Virus Reference Laboratory | National Virus Reference Laboratory | Zoe Yandle, Charlene Bennett, Gabriel Gonzalez, Michael Carr, Jonathan Dean, Cillian F De Gascun |
| EPI_ISL_3467539, EPI_ISL_3467540 | Lighthouse Lab in Glasgow | Wellcome Sanger Institute for the COVID-19 Genomics UK (COG-UK) Consortium | Harper VanSteenhouse, Yumi Kasai, David Gray, Carol Clugston, Anna Dominiczak and Alex Alderton, Roberto Amato, Jeffrey Barrett, Sonia Goncalves, Ewan Harrison, David K. Jackson, Ian Johnston, Dominic Kwiatkowski, Cordelia Langford, John Sillitoe on behalf of the Wellcome Sanger Institute COVID-19 Surveillance Team |
| EPI_ISL_3467542, EPI_ISL_3467543, EPI_ISL_3467544 | Lighthouse Lab in Milton Keynes | Wellcome Sanger Institute for the COVID-19 Genomics UK (COG-UK) Consortium | The Lighthouse Lab in Milton Keynes and Alex Alderton, Roberto Amato, Jeffrey Barrett, Sonia Goncalves, Ewan Harrison, David K. Jackson, Ian Johnston, Dominic Kwiatkowski, Cordelia Langford, John Sillitoe on behalf of the Wellcome Sanger Institute COVID-19 Surveillance Team |
| EPI_ISL_3467545 | National Virus Reference Laboratory | National Virus Reference Laboratory | Zoe Yandle, Charlene Bennett, Gabriel Gonzalez, Michael Carr, Jonathan Dean, Cillian F De Gascun |
| EPI_ISL_3467546 | Lighthouse Lab in Glasgow | Wellcome Sanger Institute for the COVID-19 Genomics UK (COG-UK) Consortium | Harper VanSteenhouse, Yumi Kasai, David Gray, Carol Clugston, Anna Dominiczak and Alex Alderton, Roberto Amato, Jeffrey Barrett, Sonia Goncalves, Ewan Harrison, David K. Jackson, Ian Johnston, Dominic Kwiatkowski, Cordelia Langford, John Sillitoe on behalf of the Wellcome Sanger Institute COVID-19 Surveillance Team |
| EPI_ISL_3467547, EPI_ISL_3467548 | Lighthouse Lab in Milton Keynes | Wellcome Sanger Institute for the COVID-19 Genomics UK (COG-UK) Consortium | The Lighthouse Lab in Milton Keynes and Alex Alderton, Roberto Amato, Jeffrey Barrett, Sonia Goncalves, Ewan Harrison, David K. Jackson, Ian Johnston, Dominic Kwiatkowski, Cordelia Langford, John Sillitoe on behalf of the Wellcome Sanger Institute COVID-19 Surveillance Team |
| EPI_ISL_3467549 | Lighthouse Lab in Glasgow | Wellcome Sanger Institute for the COVID-19 Genomics UK (COG-UK) Consortium | Harper VanSteenhouse, Yumi Kasai, David Gray, Carol Clugston, Anna Dominiczak and Alex Alderton, Roberto Amato, Jeffrey Barrett, Sonia Goncalves, Ewan Harrison, David K. Jackson, Ian Johnston, Dominic Kwiatkowski, Cordelia Langford, John Sillitoe on behalf of the Wellcome Sanger Institute COVID-19 Surveillance Team |
| EPI_ISL_3467551 | Lighthouse Lab in Milton Keynes | Wellcome Sanger Institute for the COVID-19 Genomics UK (COG-UK) Consortium | The Lighthouse Lab in Milton Keynes and Alex Alderton, Roberto Amato, Jeffrey Barrett, Sonia Goncalves, Ewan Harrison, David K. Jackson, Ian Johnston, Dominic Kwiatkowski, Cordelia Langford, John Sillitoe on behalf of the Wellcome Sanger Institute COVID-19 Surveillance Team |
| EPI_ISL_3467552 | Lighthouse Lab in Glasgow | Wellcome Sanger Institute for the COVID-19 Genomics UK (COG-UK) Consortium | Harper VanSteenhouse, Yumi Kasai, David Gray, Carol Clugston, Anna Dominiczak and Alex Alderton, Roberto Amato, Jeffrey Barrett, Sonia Goncalves, Ewan Harrison, David K. Jackson, Ian Johnston, Dominic Kwiatkowski, Cordelia Langford, John Sillitoe on behalf of the Wellcome Sanger Institute COVID-19 Surveillance Team |
| EPI_ISL_3467553 | National Virus Reference Laboratory | National Virus Reference Laboratory | Zoe Yandle, Charlene Bennett, Gabriel Gonzalez, Michael Carr, Jonathan Dean, Cillian F De Gascun |
| EPI_ISL_3467554, EPI_ISL_3467555 | Lighthouse Lab in Glasgow | Wellcome Sanger Institute for the COVID-19 Genomics UK (COG-UK) Consortium | Harper VanSteenhouse, Yumi Kasai, David Gray, Carol Clugston, Anna Dominiczak and Alex Alderton, Roberto Amato, Jeffrey Barrett, Sonia Goncalves, Ewan Harrison, David K. Jackson, Ian Johnston, Dominic Kwiatkowski, Cordelia Langford, John Sillitoe on behalf of the Wellcome Sanger Institute COVID-19 Surveillance Team |
| EPI_ISL_3467556 | Lighthouse Lab in Milton Keynes | Wellcome Sanger Institute for the COVID-19 Genomics UK (COG-UK) Consortium | The Lighthouse Lab in Milton Keynes and Alex Alderton, Roberto Amato, Jeffrey Barrett, Sonia Goncalves, Ewan Harrison, David K. Jackson, Ian Johnston, Dominic Kwiatkowski, Cordelia Langford, John Sillitoe on behalf of the Wellcome Sanger Institute COVID-19 Surveillance Team |
| EPI_ISL_3467557 | National Virus Reference Laboratory | National Virus Reference Laboratory | Zoe Yandle, Charlene Bennett, Gabriel Gonzalez, Michael Carr, Jonathan Dean, Cillian F De Gascun |
| EPI_ISL_3467558, EPI_ISL_3467559, EPI_ISL_3467560 | Lighthouse Lab in Glasgow | Wellcome Sanger Institute for the COVID-19 Genomics UK (COG-UK) Consortium | Harper VanSteenhouse, Yumi Kasai, David Gray, Carol Clugston, Anna Dominiczak and Alex Alderton, Roberto Amato, Jeffrey Barrett, Sonia Goncalves, Ewan Harrison, David K. Jackson, Ian Johnston, Dominic Kwiatkowski, Cordelia Langford, John Sillitoe on behalf of the Wellcome Sanger Institute COVID-19 Surveillance Team |
| EPI_ISL_3467561 | National Virus Reference Laboratory | National Virus Reference Laboratory | Zoe Yandle, Charlene Bennett, Gabriel Gonzalez, Michael Carr, Jonathan Dean, Cillian F De Gascun |
| EPI_ISL_3467562, EPI_ISL_3467563, EPI_ISL_3467564 | Lighthouse Lab in Milton Keynes | Wellcome Sanger Institute for the COVID-19 Genomics UK (COG-UK) Consortium | The Lighthouse Lab in Milton Keynes and Alex Alderton, Roberto Amato, Jeffrey Barrett, Sonia Goncalves, Ewan Harrison, David K. Jackson, Ian Johnston, Dominic Kwiatkowski, Cordelia Langford, John Sillitoe on behalf of the Wellcome Sanger Institute COVID-19 Surveillance Team |
| EPI_ISL_3467565 | National Virus Reference Laboratory | National Virus Reference Laboratory | Zoe Yandle, Charlene Bennett, Gabriel Gonzalez, Michael Carr, Jonathan Dean, Cillian F De Gascun |
| EPI_ISL_3467566, EPI_ISL_3467567, EPI_ISL_3467568 | Lighthouse Lab in Glasgow | Wellcome Sanger Institute for the COVID-19 Genomics UK (COG-UK) Consortium | Harper VanSteenhouse, Yumi Kasai, David Gray, Carol Clugston, Anna Dominiczak and Alex Alderton, Roberto Amato, Jeffrey Barrett, Sonia Goncalves, Ewan Harrison, David K. Jackson, Ian Johnston, Dominic Kwiatkowski, Cordelia Langford, John Sillitoe on behalf of the Wellcome Sanger Institute COVID-19 Surveillance Team |
| EPI_ISL_3467569 | National Virus Reference Laboratory | National Virus Reference Laboratory | Zoe Yandle, Charlene Bennett, Gabriel Gonzalez, Michael Carr, Jonathan Dean, Cillian F De Gascun |
| EPI_ISL_3467570 | Lighthouse Lab in Glasgow | Wellcome Sanger Institute for the COVID-19 Genomics UK (COG-UK) Consortium | Harper VanSteenhouse, Yumi Kasai, David Gray, Carol Clugston, Anna Dominiczak and Alex Alderton, Roberto Amato, Jeffrey Barrett, Sonia Goncalves, Ewan Harrison, David K. Jackson, Ian Johnston, Dominic Kwiatkowski, Cordelia Langford, John Sillitoe on behalf of the Wellcome Sanger Institute COVID-19 Surveillance Team |
| EPI_ISL_3467571, EPI_ISL_3467572 | Lighthouse Lab in Milton Keynes | Wellcome Sanger Institute for the COVID-19 Genomics UK (COG-UK) Consortium | The Lighthouse Lab in Milton Keynes and Alex Alderton, Roberto Amato, Jeffrey Barrett, Sonia Goncalves, Ewan Harrison, David K. Jackson, Ian Johnston, Dominic Kwiatkowski, Cordelia Langford, John Sillitoe on behalf of the Wellcome Sanger Institute COVID-19 Surveillance Team |
| EPI_ISL_3467573 | National Virus Reference Laboratory | National Virus Reference Laboratory | Zoe Yandle, Charlene Bennett, Gabriel Gonzalez, Michael Carr, Jonathan Dean, Cillian F De Gascun |
| EPI_ISL_3467574, EPI_ISL_3467575 | Lighthouse Lab in Milton Keynes | Wellcome Sanger Institute for the COVID-19 Genomics UK (COG-UK) Consortium | The Lighthouse Lab in Milton Keynes and Alex Alderton, Roberto Amato, Jeffrey Barrett, Sonia Goncalves, Ewan Harrison, David K. Jackson, Ian Johnston, Dominic Kwiatkowski, Cordelia Langford, John Sillitoe on behalf of the Wellcome Sanger Institute COVID-19 Surveillance Team |
| EPI_ISL_3467576 | Lighthouse Lab in Glasgow | Wellcome Sanger Institute for the COVID-19 Genomics UK (COG-UK) Consortium | Harper VanSteenhouse, Yumi Kasai, David Gray, Carol Clugston, Anna Dominiczak and Alex Alderton, Roberto Amato, Jeffrey Barrett, Sonia Goncalves, Ewan Harrison, David K. Jackson, Ian Johnston, Dominic Kwiatkowski, Cordelia Langford, John Sillitoe on behalf of the Wellcome Sanger Institute COVID-19 Surveillance Team |
| EPI_ISL_3467577 | National Virus Reference Laboratory | National Virus Reference Laboratory | Zoe Yandle, Charlene Bennett, Gabriel Gonzalez, Michael Carr, Jonathan Dean, Cillian F De Gascun |
| EPI_ISL_3467578, EPI_ISL_3467579 | Lighthouse Lab in Milton Keynes | Wellcome Sanger Institute for the COVID-19 Genomics UK (COG-UK) Consortium | The Lighthouse Lab in Milton Keynes and Alex Alderton, Roberto Amato, Jeffrey Barrett, Sonia Goncalves, Ewan Harrison, David K. Jackson, Ian Johnston, Dominic Kwiatkowski, Cordelia Langford, John Sillitoe on behalf of the Wellcome Sanger Institute COVID-19 Surveillance Team |
| EPI_ISL_3467580 | National Virus Reference Laboratory | National Virus Reference Laboratory | Zoe Yandle, Charlene Bennett, Gabriel Gonzalez, Michael Carr, Jonathan Dean, Cillian F De Gascun |

| | | | |
|---|---|---|---|
| EPI_ISL_3467581 | Lighthouse Lab in Milton Keynes | Wellcome Sanger Institute for the COVID-19 Genomics UK (COG-UK) Consortium | The Lighthouse Lab in Milton Keynes and Alex Alderton, Roberto Amato, Jeffrey Barrett, Sonia Goncalves, Ewan Harrison, David K. Jackson, Ian Johnston, Dominic Kwiatkowski, Cordelia Langford, John Sillitoe on behalf of the Wellcome Sanger Institute COVID-19 Surveillance Team |
| EPI_ISL_3467582 | Lighthouse Lab in Glasgow | Wellcome Sanger Institute for the COVID-19 Genomics UK (COG-UK) Consortium | Harper VanSteenhouse, Yumi Kasai, David Gray, Carol Clugston, Anna Dominiczak and Alex Alderton, Roberto Amato, Jeffrey Barrett, Sonia Goncalves, Ewan Harrison, David K. Jackson, Ian Johnston, Dominic Kwiatkowski, Cordelia Langford, John Sillitoe on behalf of the Wellcome Sanger Institute COVID-19 Surveillance Team |
| EPI_ISL_3467583 | National Virus Reference Laboratory | National Virus Reference Laboratory | Zoe Yandle, Charlene Bennett, Gabriel Gonzalez, Michael Carr, Jonathan Dean, Cillian F De Gascun |
| EPI_ISL_3467584, EPI_ISL_3467585, EPI_ISL_3467586 | Lighthouse Lab in Milton Keynes | Wellcome Sanger Institute for the COVID-19 Genomics UK (COG-UK) Consortium | The Lighthouse Lab in Milton Keynes and Alex Alderton, Roberto Amato, Jeffrey Barrett, Sonia Goncalves, Ewan Harrison, David K. Jackson, Ian Johnston, Dominic Kwiatkowski, Cordelia Langford, John Sillitoe on behalf of the Wellcome Sanger Institute COVID-19 Surveillance Team |
| EPI_ISL_3467587 | National Virus Reference Laboratory | National Virus Reference Laboratory | Zoe Yandle, Charlene Bennett, Gabriel Gonzalez, Michael Carr, Jonathan Dean, Cillian F De Gascun |
| EPI_ISL_3467588 | Lighthouse Lab in Milton Keynes | Wellcome Sanger Institute for the COVID-19 Genomics UK (COG-UK) Consortium | The Lighthouse Lab in Milton Keynes and Alex Alderton, Roberto Amato, Jeffrey Barrett, Sonia Goncalves, Ewan Harrison, David K. Jackson, Ian Johnston, Dominic Kwiatkowski, Cordelia Langford, John Sillitoe on behalf of the Wellcome Sanger Institute COVID-19 Surveillance Team |
| EPI_ISL_3467589 | Lighthouse Lab in Glasgow | Wellcome Sanger Institute for the COVID-19 Genomics UK (COG-UK) Consortium | Harper VanSteenhouse, Yumi Kasai, David Gray, Carol Clugston, Anna Dominiczak and Alex Alderton, Roberto Amato, Jeffrey Barrett, Sonia Goncalves, Ewan Harrison, David K. Jackson, Ian Johnston, Dominic Kwiatkowski, Cordelia Langford, John Sillitoe on behalf of the Wellcome Sanger Institute COVID-19 Surveillance Team |
| EPI_ISL_3467590 | National Virus Reference Laboratory | National Virus Reference Laboratory | Zoe Yandle, Charlene Bennett, Gabriel Gonzalez, Michael Carr, Jonathan Dean, Cillian F De Gascun |
| EPI_ISL_3467591 | Lighthouse Lab in Glasgow | Wellcome Sanger Institute for the COVID-19 Genomics UK (COG-UK) Consortium | Harper VanSteenhouse, Yumi Kasai, David Gray, Carol Clugston, Anna Dominiczak and Alex Alderton, Roberto Amato, Jeffrey Barrett, Sonia Goncalves, Ewan Harrison, David K. Jackson, Ian Johnston, Dominic Kwiatkowski, Cordelia Langford, John Sillitoe on behalf of the Wellcome Sanger Institute COVID-19 Surveillance Team |
| EPI_ISL_3467592, EPI_ISL_3467593 | Lighthouse Lab in Milton Keynes | Wellcome Sanger Institute for the COVID-19 Genomics UK (COG-UK) Consortium | The Lighthouse Lab in Milton Keynes and Alex Alderton, Roberto Amato, Jeffrey Barrett, Sonia Goncalves, Ewan Harrison, David K. Jackson, Ian Johnston, Dominic Kwiatkowski, Cordelia Langford, John Sillitoe on behalf of the Wellcome Sanger Institute COVID-19 Surveillance Team |
| EPI_ISL_3467594 | National Virus Reference Laboratory | National Virus Reference Laboratory | Zoe Yandle, Charlene Bennett, Gabriel Gonzalez, Michael Carr, Jonathan Dean, Cillian F De Gascun |
| EPI_ISL_3467595 | Lighthouse Lab in Milton Keynes | Wellcome Sanger Institute for the COVID-19 Genomics UK (COG-UK) Consortium | The Lighthouse Lab in Milton Keynes and Alex Alderton, Roberto Amato, Jeffrey Barrett, Sonia Goncalves, Ewan Harrison, David K. Jackson, Ian Johnston, Dominic Kwiatkowski, Cordelia Langford, John Sillitoe on behalf of the Wellcome Sanger Institute COVID-19 Surveillance Team |
| EPI_ISL_3467596, EPI_ISL_3467597 | Lighthouse Lab in Glasgow | Wellcome Sanger Institute for the COVID-19 Genomics UK (COG-UK) Consortium | Harper VanSteenhouse, Yumi Kasai, David Gray, Carol Clugston, Anna Dominiczak and Alex Alderton, Roberto Amato, Jeffrey Barrett, Sonia Goncalves, Ewan Harrison, David K. Jackson, Ian Johnston, Dominic Kwiatkowski, Cordelia Langford, John Sillitoe on behalf of the Wellcome Sanger Institute COVID-19 Surveillance Team |
| EPI_ISL_3467598 | Lighthouse Lab in Milton Keynes | Wellcome Sanger Institute for the COVID-19 Genomics UK (COG-UK) Consortium | The Lighthouse Lab in Milton Keynes and Alex Alderton, Roberto Amato, Jeffrey Barrett, Sonia Goncalves, Ewan Harrison, David K. Jackson, Ian Johnston, Dominic Kwiatkowski, Cordelia Langford, John Sillitoe on behalf of the Wellcome Sanger Institute COVID-19 Surveillance Team |
| EPI_ISL_3467599 | National Virus Reference Laboratory | National Virus Reference Laboratory | Zoe Yandle, Charlene Bennett, Gabriel Gonzalez, Michael Carr, Jonathan Dean, Cillian F De Gascun |
| EPI_ISL_3467600 | Lighthouse Lab in Glasgow | Wellcome Sanger Institute for the COVID-19 Genomics UK (COG-UK) Consortium | Harper VanSteenhouse, Yumi Kasai, David Gray, Carol Clugston, Anna Dominiczak and Alex Alderton, Roberto Amato, Jeffrey Barrett, Sonia Goncalves, Ewan Harrison, David K. Jackson, Ian Johnston, Dominic Kwiatkowski, Cordelia Langford, John Sillitoe on behalf of the Wellcome Sanger Institute COVID-19 Surveillance Team |
| EPI_ISL_3467601 | Lighthouse Lab in Milton Keynes | Wellcome Sanger Institute for the COVID-19 Genomics UK (COG-UK) Consortium | The Lighthouse Lab in Milton Keynes and Alex Alderton, Roberto Amato, Jeffrey Barrett, Sonia Goncalves, Ewan Harrison, David K. Jackson, Ian Johnston, Dominic Kwiatkowski, Cordelia Langford, John Sillitoe on behalf of the Wellcome Sanger Institute COVID-19 Surveillance Team |
| EPI_ISL_3467602 | National Virus Reference Laboratory | National Virus Reference Laboratory | Zoe Yandle, Charlene Bennett, Gabriel Gonzalez, Michael Carr, Jonathan Dean, Cillian F De Gascun |
| EPI_ISL_3467603, EPI_ISL_3467604 | Lighthouse Lab in Glasgow | Wellcome Sanger Institute for the COVID-19 Genomics UK (COG-UK) Consortium | Harper VanSteenhouse, Yumi Kasai, David Gray, Carol Clugston, Anna Dominiczak and Alex Alderton, Roberto Amato, Jeffrey Barrett, Sonia Goncalves, Ewan Harrison, David K. Jackson, Ian Johnston, Dominic Kwiatkowski, Cordelia Langford, John Sillitoe on behalf of the Wellcome Sanger Institute COVID-19 Surveillance Team |
| EPI_ISL_3467605 | Lighthouse Lab in Milton Keynes | Wellcome Sanger Institute for the COVID-19 Genomics UK (COG-UK) Consortium | The Lighthouse Lab in Milton Keynes and Alex Alderton, Roberto Amato, Jeffrey Barrett, Sonia Goncalves, Ewan Harrison, David K. Jackson, Ian Johnston, Dominic Kwiatkowski, Cordelia Langford, John Sillitoe on behalf of the Wellcome Sanger Institute COVID-19 Surveillance Team |
| EPI_ISL_3467606 | National Virus Reference Laboratory | National Virus Reference Laboratory | Zoe Yandle, Charlene Bennett, Gabriel Gonzalez, Michael Carr, Jonathan Dean, Cillian F De Gascun |
| EPI_ISL_3467607, EPI_ISL_3467608 | Lighthouse Lab in Milton Keynes | Wellcome Sanger Institute for the COVID-19 Genomics UK (COG-UK) Consortium | The Lighthouse Lab in Milton Keynes and Alex Alderton, Roberto Amato, Jeffrey Barrett, Sonia Goncalves, Ewan Harrison, David K. Jackson, Ian Johnston, Dominic Kwiatkowski, Cordelia Langford, John Sillitoe on behalf of the Wellcome Sanger Institute COVID-19 Surveillance Team |
| EPI_ISL_3467609 | National Virus Reference Laboratory | National Virus Reference Laboratory | Zoe Yandle, Charlene Bennett, Gabriel Gonzalez, Michael Carr, Jonathan Dean, Cillian F De Gascun |
| EPI_ISL_3467610 | Lighthouse Lab in Milton Keynes | Wellcome Sanger Institute for the COVID-19 Genomics UK (COG-UK) Consortium | The Lighthouse Lab in Milton Keynes and Alex Alderton, Roberto Amato, Jeffrey Barrett, Sonia Goncalves, Ewan Harrison, David K. Jackson, Ian Johnston, Dominic Kwiatkowski, Cordelia Langford, John Sillitoe on behalf of the Wellcome Sanger Institute COVID-19 Surveillance Team |
| EPI_ISL_3467611 | Lighthouse Lab in Glasgow | Wellcome Sanger Institute for the COVID-19 Genomics UK (COG-UK) Consortium | Harper VanSteenhouse, Yumi Kasai, David Gray, Carol Clugston, Anna Dominiczak and Alex Alderton, Roberto Amato, Jeffrey Barrett, Sonia Goncalves, Ewan Harrison, David K. Jackson, Ian Johnston, Dominic Kwiatkowski, Cordelia Langford, John Sillitoe on behalf of the Wellcome Sanger Institute COVID-19 Surveillance Team |
| EPI_ISL_3467612 | Lighthouse Lab in Milton Keynes | Wellcome Sanger Institute for the COVID-19 Genomics UK (COG-UK) Consortium | The Lighthouse Lab in Milton Keynes and Alex Alderton, Roberto Amato, Jeffrey Barrett, Sonia Goncalves, Ewan Harrison, David K. Jackson, Ian Johnston, Dominic Kwiatkowski, Cordelia Langford, John Sillitoe on behalf of the Wellcome Sanger Institute COVID-19 Surveillance Team |
| EPI_ISL_3467613 | National Virus Reference Laboratory | National Virus Reference Laboratory | Zoe Yandle, Charlene Bennett, Gabriel Gonzalez, Michael Carr, Jonathan Dean, Cillian F De Gascun |
| EPI_ISL_3467614 | Lighthouse Lab in Glasgow | Wellcome Sanger Institute for the COVID-19 Genomics UK (COG-UK) Consortium | Harper VanSteenhouse, Yumi Kasai, David Gray, Carol Clugston, Anna Dominiczak and Alex Alderton, Roberto Amato, Jeffrey Barrett, Sonia Goncalves, Ewan Harrison, David K. Jackson, Ian Johnston, Dominic Kwiatkowski, Cordelia Langford, John Sillitoe on behalf of the Wellcome Sanger Institute COVID-19 Surveillance Team |
| EPI_ISL_3467615, EPI_ISL_3467616 | Lighthouse Lab in Milton Keynes | Wellcome Sanger Institute for the COVID-19 Genomics UK (COG-UK) Consortium | The Lighthouse Lab in Milton Keynes and Alex Alderton, Roberto Amato, Jeffrey Barrett, Sonia Goncalves, Ewan Harrison, David K. Jackson, Ian Johnston, Dominic Kwiatkowski, Cordelia Langford, John Sillitoe on behalf of the Wellcome Sanger Institute COVID-19 Surveillance Team |
| EPI_ISL_3467617 | National Virus Reference Laboratory | National Virus Reference Laboratory | Zoe Yandle, Charlene Bennett, Gabriel Gonzalez, Michael Carr, Jonathan Dean, Cillian F De Gascun |
| EPI_ISL_3467618 | Lighthouse Lab in Glasgow | Wellcome Sanger Institute for the COVID-19 Genomics UK (COG-UK) Consortium | Harper VanSteenhouse, Yumi Kasai, David Gray, Carol Clugston, Anna Dominiczak and Alex Alderton, Roberto Amato, Jeffrey Barrett, Sonia Goncalves, Ewan Harrison, David K. Jackson, Ian Johnston, Dominic Kwiatkowski, Cordelia Langford, John Sillitoe on behalf of the Wellcome Sanger Institute COVID-19 Surveillance Team |
| EPI_ISL_3467619 | Lighthouse Lab in Milton Keynes | Wellcome Sanger Institute for the COVID-19 Genomics UK (COG-UK) Consortium | The Lighthouse Lab in Milton Keynes and Alex Alderton, Roberto Amato, Jeffrey Barrett, Sonia Goncalves, Ewan Harrison, David K. Jackson, Ian Johnston, Dominic Kwiatkowski, Cordelia Langford, John Sillitoe on behalf of the Wellcome Sanger Institute COVID-19 Surveillance Team |
| EPI_ISL_3467620 | National Virus Reference Laboratory | National Virus Reference Laboratory | Zoe Yandle, Charlene Bennett, Gabriel Gonzalez, Michael Carr, Jonathan Dean, Cillian F De Gascun |
| EPI_ISL_3467621, EPI_ISL_3467622 | Lighthouse Lab in Milton Keynes | Wellcome Sanger Institute for the COVID-19 Genomics UK (COG-UK) Consortium | The Lighthouse Lab in Milton Keynes and Alex Alderton, Roberto Amato, Jeffrey Barrett, Sonia Goncalves, Ewan Harrison, David K. Jackson, Ian Johnston, Dominic Kwiatkowski, Cordelia Langford, John Sillitoe on behalf of the Wellcome Sanger Institute COVID-19 Surveillance Team |
| EPI_ISL_3467623 | National Virus Reference Laboratory | National Virus Reference Laboratory | Zoe Yandle, Charlene Bennett, Gabriel Gonzalez, Michael Carr, Jonathan Dean, Cillian F De Gascun |
| EPI_ISL_3467624, EPI_ISL_3467625, EPI_ISL_3467626 | Lighthouse Lab in Milton Keynes | Wellcome Sanger Institute for the COVID-19 Genomics UK (COG-UK) Consortium | The Lighthouse Lab in Milton Keynes and Alex Alderton, Roberto Amato, Jeffrey Barrett, Sonia Goncalves, Ewan Harrison, David K. Jackson, Ian Johnston, Dominic Kwiatkowski, Cordelia Langford, John Sillitoe on behalf of the Wellcome Sanger Institute COVID-19 Surveillance Team |
| EPI_ISL_3467627 | National Virus Reference Laboratory | National Virus Reference Laboratory | Zoe Yandle, Charlene Bennett, Gabriel Gonzalez, Michael Carr, Jonathan Dean, Cillian F De Gascun |
| EPI_ISL_3467628, EPI_ISL_3467629, EPI_ISL_3467630 | Lighthouse Lab in Milton Keynes | Wellcome Sanger Institute for the COVID-19 Genomics UK (COG-UK) Consortium | The Lighthouse Lab in Milton Keynes and Alex Alderton, Roberto Amato, Jeffrey Barrett, Sonia Goncalves, Ewan Harrison, David K. Jackson, Ian Johnston, Dominic Kwiatkowski, Cordelia Langford, John Sillitoe on behalf of the Wellcome Sanger Institute COVID-19 Surveillance Team |
| EPI_ISL_3467631 | National Virus Reference Laboratory | National Virus Reference Laboratory | Zoe Yandle, Charlene Bennett, Gabriel Gonzalez, Michael Carr, Jonathan Dean, Cillian F De Gascun |
| EPI_ISL_3467632 | Lighthouse Lab in Glasgow | Wellcome Sanger Institute for the COVID-19 Genomics UK (COG-UK) Consortium | Harper VanSteenhouse, Yumi Kasai, David Gray, Carol Clugston, Anna Dominiczak and Alex Alderton, Roberto Amato, Jeffrey Barrett, Sonia Goncalves, Ewan Harrison, David K. Jackson, Ian Johnston, Dominic Kwiatkowski, Cordelia Langford, John Sillitoe on behalf of the Wellcome Sanger Institute COVID-19 Surveillance Team |
| EPI_ISL_3467633, EPI_ISL_3467634 | Lighthouse Lab in Milton Keynes | Wellcome Sanger Institute for the COVID-19 Genomics UK (COG-UK) Consortium | The Lighthouse Lab in Milton Keynes and Alex Alderton, Roberto Amato, Jeffrey Barrett, Sonia Goncalves, Ewan Harrison, David K. Jackson, Ian Johnston, Dominic Kwiatkowski, Cordelia Langford, John Sillitoe on behalf of the Wellcome Sanger Institute COVID-19 Surveillance Team |
| EPI_ISL_3467635 | National Virus Reference Laboratory | National Virus Reference Laboratory | Zoe Yandle, Charlene Bennett, Gabriel Gonzalez, Michael Carr, Jonathan Dean, Cillian F De Gascun |
| EPI_ISL_3467636 | Lighthouse Lab in Glasgow | Wellcome Sanger Institute for the COVID-19 Genomics UK (COG-UK) Consortium | Harper VanSteenhouse, Yumi Kasai, David Gray, Carol Clugston, Anna Dominiczak and Alex Alderton, Roberto Amato, Jeffrey Barrett, Sonia Goncalves, Ewan Harrison, David K. Jackson, Ian Johnston, Dominic Kwiatkowski, Cordelia Langford, John Sillitoe on behalf of the Wellcome Sanger Institute COVID-19 Surveillance Team |
| EPI_ISL_3467637, EPI_ISL_3467638 | Lighthouse Lab in Milton Keynes | Wellcome Sanger Institute for the COVID-19 Genomics UK (COG-UK) Consortium | The Lighthouse Lab in Milton Keynes and Alex Alderton, Roberto Amato, Jeffrey Barrett, Sonia Goncalves, Ewan Harrison, David K. Jackson, Ian Johnston, Dominic Kwiatkowski, Cordelia Langford, John Sillitoe on behalf of the Wellcome Sanger Institute COVID-19 Surveillance Team |
| EPI_ISL_3467639 | National Virus Reference Laboratory | National Virus Reference Laboratory | Zoe Yandle, Charlene Bennett, Gabriel Gonzalez, Michael Carr, Jonathan Dean, Cillian F De Gascun |
| EPI_ISL_3467640, EPI_ISL_3467641 | Lighthouse Lab in Milton Keynes | Wellcome Sanger Institute for the COVID-19 Genomics UK (COG-UK) Consortium | The Lighthouse Lab in Milton Keynes and Alex Alderton, Roberto Amato, Jeffrey Barrett, Sonia Goncalves, Ewan Harrison, David K. Jackson, Ian Johnston, Dominic Kwiatkowski, Cordelia Langford, John Sillitoe on behalf of the Wellcome Sanger Institute COVID-19 Surveillance Team |
| EPI_ISL_3467642 | National Virus Reference Laboratory | National Virus Reference Laboratory | Zoe Yandle, Charlene Bennett, Gabriel Gonzalez, Michael Carr, Jonathan Dean, Cillian F De Gascun |
| EPI_ISL_3467643, EPI_ISL_3467644 | Lighthouse Lab in Milton Keynes | Wellcome Sanger Institute for the COVID-19 Genomics UK (COG-UK) Consortium | The Lighthouse Lab in Milton Keynes and Alex Alderton, Roberto Amato, Jeffrey Barrett, Sonia Goncalves, Ewan Harrison, David K. Jackson, Ian Johnston, Dominic Kwiatkowski, Cordelia Langford, John Sillitoe on behalf of the Wellcome Sanger Institute COVID-19 Surveillance Team |
| EPI_ISL_3467645 | Lighthouse Lab in Glasgow | Wellcome Sanger Institute for the COVID-19 Genomics UK (COG-UK) Consortium | Harper VanSteenhouse, Yumi Kasai, David Gray, Carol Clugston, Anna Dominiczak and Alex Alderton, Roberto Amato, Jeffrey Barrett, Sonia Goncalves, Ewan Harrison, David K. Jackson, Ian Johnston, Dominic Kwiatkowski, Cordelia Langford, John Sillitoe on behalf of the Wellcome Sanger Institute COVID-19 Surveillance Team |
| EPI_ISL_3467646 | National Virus Reference Laboratory | National Virus Reference Laboratory | Zoe Yandle, Charlene Bennett, Gabriel Gonzalez, Michael Carr, Jonathan Dean, Cillian F De Gascun |
| EPI_ISL_3467647, EPI_ISL_3467648 | Lighthouse Lab in Milton Keynes | Wellcome Sanger Institute for the COVID-19 Genomics UK (COG-UK) Consortium | The Lighthouse Lab in Milton Keynes and Alex Alderton, Roberto Amato, Jeffrey Barrett, Sonia Goncalves, Ewan Harrison, David K. Jackson, Ian Johnston, Dominic Kwiatkowski, Cordelia Langford, John Sillitoe on behalf of the Wellcome Sanger Institute COVID-19 Surveillance Team |

| | | | |
|---|---|---|---|
| EPI_ISL_3467649 | National Virus Reference Laboratory | National Virus Reference Laboratory | Zoe Yandle, Charlene Bennett, Gabriel Gonzalez, Michael Carr, Jonathan Dean, Cillian F De Gascun |
| EPI_ISL_3467650, EPI_ISL_3467651 | Lighthouse Lab in Glasgow | Wellcome Sanger Institute for the COVID-19 Genomics UK (COG-UK) Consortium | Harper VanSteenhouse, Yumi Kasai, David Gray, Carol Clugston, Anna Dominiczak and Alex Alderton, Roberto Amato, Jeffrey Barrett, Sonia Goncalves, Ewan Harrison, David K. Jackson, Ian Johnston, Dominic Kwiatkowski, Cordelia Langford, John Sillitoe on behalf of the Wellcome Sanger Institute COVID-19 Surveillance Team |
| EPI_ISL_3467652 | Lighthouse Lab in Milton Keynes | Wellcome Sanger Institute for the COVID-19 Genomics UK (COG-UK) Consortium | The Lighthouse Lab in Milton Keynes and Alex Alderton, Roberto Amato, Jeffrey Barrett, Sonia Goncalves, Ewan Harrison, David K. Jackson, Ian Johnston, Dominic Kwiatkowski, Cordelia Langford, John Sillitoe on behalf of the Wellcome Sanger Institute COVID-19 Surveillance Team |
| EPI_ISL_3467653 | National Virus Reference Laboratory | National Virus Reference Laboratory | Zoe Yandle, Charlene Bennett, Gabriel Gonzalez, Michael Carr, Jonathan Dean, Cillian F De Gascun |
| EPI_ISL_3467654 | Lighthouse Lab in Glasgow | Wellcome Sanger Institute for the COVID-19 Genomics UK (COG-UK) Consortium | Harper VanSteenhouse, Yumi Kasai, David Gray, Carol Clugston, Anna Dominiczak and Alex Alderton, Roberto Amato, Jeffrey Barrett, Sonia Goncalves, Ewan Harrison, David K. Jackson, Ian Johnston, Dominic Kwiatkowski, Cordelia Langford, John Sillitoe on behalf of the Wellcome Sanger Institute COVID-19 Surveillance Team |
| EPI_ISL_3467655, EPI_ISL_3467656 | Lighthouse Lab in Milton Keynes | Wellcome Sanger Institute for the COVID-19 Genomics UK (COG-UK) Consortium | The Lighthouse Lab in Milton Keynes and Alex Alderton, Roberto Amato, Jeffrey Barrett, Sonia Goncalves, Ewan Harrison, David K. Jackson, Ian Johnston, Dominic Kwiatkowski, Cordelia Langford, John Sillitoe on behalf of the Wellcome Sanger Institute COVID-19 Surveillance Team |
| EPI_ISL_3467657 | National Virus Reference Laboratory | National Virus Reference Laboratory | Zoe Yandle, Charlene Bennett, Gabriel Gonzalez, Michael Carr, Jonathan Dean, Cillian F De Gascun |
| EPI_ISL_3467658 | Lighthouse Lab in Glasgow | Wellcome Sanger Institute for the COVID-19 Genomics UK (COG-UK) Consortium | Harper VanSteenhouse, Yumi Kasai, David Gray, Carol Clugston, Anna Dominiczak and Alex Alderton, Roberto Amato, Jeffrey Barrett, Sonia Goncalves, Ewan Harrison, David K. Jackson, Ian Johnston, Dominic Kwiatkowski, Cordelia Langford, John Sillitoe on behalf of the Wellcome Sanger Institute COVID-19 Surveillance Team |
| EPI_ISL_3467659 | Lighthouse Lab in Milton Keynes | Wellcome Sanger Institute for the COVID-19 Genomics UK (COG-UK) Consortium | The Lighthouse Lab in Milton Keynes and Alex Alderton, Roberto Amato, Jeffrey Barrett, Sonia Goncalves, Ewan Harrison, David K. Jackson, Ian Johnston, Dominic Kwiatkowski, Cordelia Langford, John Sillitoe on behalf of the Wellcome Sanger Institute COVID-19 Surveillance Team |
| EPI_ISL_3467660 | National Virus Reference Laboratory | National Virus Reference Laboratory | Zoe Yandle, Charlene Bennett, Gabriel Gonzalez, Michael Carr, Jonathan Dean, Cillian F De Gascun |
| EPI_ISL_3467661, EPI_ISL_3467662, EPI_ISL_3467663 | Lighthouse Lab in Glasgow | Wellcome Sanger Institute for the COVID-19 Genomics UK (COG-UK) Consortium | Harper VanSteenhouse, Yumi Kasai, David Gray, Carol Clugston, Anna Dominiczak and Alex Alderton, Roberto Amato, Jeffrey Barrett, Sonia Goncalves, Ewan Harrison, David K. Jackson, Ian Johnston, Dominic Kwiatkowski, Cordelia Langford, John Sillitoe on behalf of the Wellcome Sanger Institute COVID-19 Surveillance Team |
| EPI_ISL_3467664 | National Virus Reference Laboratory | National Virus Reference Laboratory | Zoe Yandle, Charlene Bennett, Gabriel Gonzalez, Michael Carr, Jonathan Dean, Cillian F De Gascun |
| EPI_ISL_3467665, EPI_ISL_3467666 | Lighthouse Lab in Milton Keynes | Wellcome Sanger Institute for the COVID-19 Genomics UK (COG-UK) Consortium | The Lighthouse Lab in Milton Keynes and Alex Alderton, Roberto Amato, Jeffrey Barrett, Sonia Goncalves, Ewan Harrison, David K. Jackson, Ian Johnston, Dominic Kwiatkowski, Cordelia Langford, John Sillitoe on behalf of the Wellcome Sanger Institute COVID-19 Surveillance Team |
| EPI_ISL_3467667 | National Virus Reference Laboratory | National Virus Reference Laboratory | Zoe Yandle, Charlene Bennett, Gabriel Gonzalez, Michael Carr, Jonathan Dean, Cillian F De Gascun |
| EPI_ISL_3467668, EPI_ISL_3467669, EPI_ISL_3467670 | Lighthouse Lab in Glasgow | Wellcome Sanger Institute for the COVID-19 Genomics UK (COG-UK) Consortium | Harper VanSteenhouse, Yumi Kasai, David Gray, Carol Clugston, Anna Dominiczak and Alex Alderton, Roberto Amato, Jeffrey Barrett, Sonia Goncalves, Ewan Harrison, David K. Jackson, Ian Johnston, Dominic Kwiatkowski, Cordelia Langford, John Sillitoe on behalf of the Wellcome Sanger Institute COVID-19 Surveillance Team |
| EPI_ISL_3467671, EPI_ISL_3467672, EPI_ISL_3467673, EPI_ISL_3467674, EPI_ISL_3467675, EPI_ISL_3467676, EPI_ISL_3467677, EPI_ISL_3467678, EPI_ISL_3467679, EPI_ISL_3467680, EPI_ISL_3467681, EPI_ISL_3467682, EPI_ISL_3467683, EPI_ISL_3467684, EPI_ISL_3467685, EPI_ISL_3467686, EPI_ISL_3467687, EPI_ISL_3467688, EPI_ISL_3467689, EPI_ISL_3467690, EPI_ISL_3467691, EPI_ISL_3467692, EPI_ISL_3467693, EPI_ISL_3467694, EPI_ISL_3467695, EPI_ISL_3467696, EPI_ISL_3467697, EPI_ISL_3467698, EPI_ISL_3467699, EPI_ISL_3467700, EPI_ISL_3467701, EPI_ISL_3467702, EPI_ISL_3467703, EPI_ISL_3467704, EPI_ISL_3467705, EPI_ISL_3467706, EPI_ISL_3467707, EPI_ISL_3467708, EPI_ISL_3467709, EPI_ISL_3467710, EPI_ISL_3467711, EPI_ISL_3467712, EPI_ISL_3467713 | | | |
| see above | National Virus Reference Laboratory | National Virus Reference Laboratory | Zoe Yandle, Charlene Bennett, Gabriel Gonzalez, Michael Carr, Jonathan Dean, Cillian F De Gascun |
| EPI_ISL_3467714, EPI_ISL_3467715, EPI_ISL_3467716 | Lighthouse Lab in Milton Keynes | Wellcome Sanger Institute for the COVID-19 Genomics UK (COG-UK) Consortium | The Lighthouse Lab in Milton Keynes and Alex Alderton, Roberto Amato, Jeffrey Barrett, Sonia Goncalves, Ewan Harrison, David K. Jackson, Ian Johnston, Dominic Kwiatkowski, Cordelia Langford, John Sillitoe on behalf of the Wellcome Sanger Institute COVID-19 Surveillance Team |
| EPI_ISL_3467717 | National Virus Reference Laboratory | National Virus Reference Laboratory | Zoe Yandle, Charlene Bennett, Gabriel Gonzalez, Michael Carr, Jonathan Dean, Cillian F De Gascun |
| EPI_ISL_3467718, EPI_ISL_3467719 | Lighthouse Lab in Milton Keynes | Wellcome Sanger Institute for the COVID-19 Genomics UK (COG-UK) Consortium | The Lighthouse Lab in Milton Keynes and Alex Alderton, Roberto Amato, Jeffrey Barrett, Sonia Goncalves, Ewan Harrison, David K. Jackson, Ian Johnston, Dominic Kwiatkowski, Cordelia Langford, John Sillitoe on behalf of the Wellcome Sanger Institute COVID-19 Surveillance Team |
| EPI_ISL_3467720 | National Virus Reference Laboratory | National Virus Reference Laboratory | Zoe Yandle, Charlene Bennett, Gabriel Gonzalez, Michael Carr, Jonathan Dean, Cillian F De Gascun |
| EPI_ISL_3467721, EPI_ISL_3467722 | Lighthouse Lab in Glasgow | Wellcome Sanger Institute for the COVID-19 Genomics UK (COG-UK) Consortium | Harper VanSteenhouse, Yumi Kasai, David Gray, Carol Clugston, Anna Dominiczak and Alex Alderton, Roberto Amato, Jeffrey Barrett, Sonia Goncalves, Ewan Harrison, David K. Jackson, Ian Johnston, Dominic Kwiatkowski, Cordelia Langford, John Sillitoe on behalf of the Wellcome Sanger Institute COVID-19 Surveillance Team |
| EPI_ISL_3467723 | Lighthouse Lab in Milton Keynes | Wellcome Sanger Institute for the COVID-19 Genomics UK (COG-UK) Consortium | The Lighthouse Lab in Milton Keynes and Alex Alderton, Roberto Amato, Jeffrey Barrett, Sonia Goncalves, Ewan Harrison, David K. Jackson, Ian Johnston, Dominic Kwiatkowski, Cordelia Langford, John Sillitoe on behalf of the Wellcome Sanger Institute COVID-19 Surveillance Team |
| EPI_ISL_3467724 | National Virus Reference Laboratory | National Virus Reference Laboratory | Zoe Yandle, Charlene Bennett, Gabriel Gonzalez, Michael Carr, Jonathan Dean, Cillian F De Gascun |
| EPI_ISL_3467725, EPI_ISL_3467726 | Lighthouse Lab in Milton Keynes | Wellcome Sanger Institute for the COVID-19 Genomics UK (COG-UK) Consortium | The Lighthouse Lab in Milton Keynes and Alex Alderton, Roberto Amato, Jeffrey Barrett, Sonia Goncalves, Ewan Harrison, David K. Jackson, Ian Johnston, Dominic Kwiatkowski, Cordelia Langford, John Sillitoe on behalf of the Wellcome Sanger Institute COVID-19 Surveillance Team |
| EPI_ISL_3467727 | National Virus Reference Laboratory | National Virus Reference Laboratory | Zoe Yandle, Charlene Bennett, Gabriel Gonzalez, Michael Carr, Jonathan Dean, Cillian F De Gascun |
| EPI_ISL_3467728, EPI_ISL_3467729, EPI_ISL_3467730, EPI_ISL_3467731 | Lighthouse Lab in Glasgow | Wellcome Sanger Institute for the COVID-19 Genomics UK (COG-UK) Consortium | Harper VanSteenhouse, Yumi Kasai, David Gray, Carol Clugston, Anna Dominiczak and Alex Alderton, Roberto Amato, Jeffrey Barrett, Sonia Goncalves, Ewan Harrison, David K. Jackson, Ian Johnston, Dominic Kwiatkowski, Cordelia Langford, John Sillitoe on behalf of the Wellcome Sanger Institute COVID-19 Surveillance Team |
| EPI_ISL_3467732 | National Virus Reference Laboratory | National Virus Reference Laboratory | Zoe Yandle, Charlene Bennett, Gabriel Gonzalez, Michael Carr, Jonathan Dean, Cillian F De Gascun |
| EPI_ISL_3467733 | Lighthouse Lab in Milton Keynes | Wellcome Sanger Institute for the COVID-19 Genomics UK (COG-UK) Consortium | The Lighthouse Lab in Milton Keynes and Alex Alderton, Roberto Amato, Jeffrey Barrett, Sonia Goncalves, Ewan Harrison, David K. Jackson, Ian Johnston, Dominic Kwiatkowski, Cordelia Langford, John Sillitoe on behalf of the Wellcome Sanger Institute COVID-19 Surveillance Team |
| EPI_ISL_3467734 | Lighthouse Lab in Glasgow | Wellcome Sanger Institute for the COVID-19 Genomics UK (COG-UK) Consortium | Harper VanSteenhouse, Yumi Kasai, David Gray, Carol Clugston, Anna Dominiczak and Alex Alderton, Roberto Amato, Jeffrey Barrett, Sonia Goncalves, Ewan Harrison, David K. Jackson, Ian Johnston, Dominic Kwiatkowski, Cordelia Langford, John Sillitoe on behalf of the Wellcome Sanger Institute COVID-19 Surveillance Team |
| EPI_ISL_3467735 | National Virus Reference Laboratory | National Virus Reference Laboratory | Zoe Yandle, Charlene Bennett, Gabriel Gonzalez, Michael Carr, Jonathan Dean, Cillian F De Gascun |
| EPI_ISL_3467736 | Lighthouse Lab in Milton Keynes | Wellcome Sanger Institute for the COVID-19 Genomics UK (COG-UK) Consortium | The Lighthouse Lab in Milton Keynes and Alex Alderton, Roberto Amato, Jeffrey Barrett, Sonia Goncalves, Ewan Harrison, David K. Jackson, Ian Johnston, Dominic Kwiatkowski, Cordelia Langford, John Sillitoe on behalf of the Wellcome Sanger Institute COVID-19 Surveillance Team |
| EPI_ISL_3467737 | Lighthouse Lab in Glasgow | Wellcome Sanger Institute for the COVID-19 Genomics UK (COG-UK) Consortium | Harper VanSteenhouse, Yumi Kasai, David Gray, Carol Clugston, Anna Dominiczak and Alex Alderton, Roberto Amato, Jeffrey Barrett, Sonia Goncalves, Ewan Harrison, David K. Jackson, Ian Johnston, Dominic Kwiatkowski, Cordelia Langford, John Sillitoe on behalf of the Wellcome Sanger Institute COVID-19 Surveillance Team |
| EPI_ISL_3467738 | Lighthouse Lab in Milton Keynes | Wellcome Sanger Institute for the COVID-19 Genomics UK (COG-UK) Consortium | The Lighthouse Lab in Milton Keynes and Alex Alderton, Roberto Amato, Jeffrey Barrett, Sonia Goncalves, Ewan Harrison, David K. Jackson, Ian Johnston, Dominic Kwiatkowski, Cordelia Langford, John Sillitoe on behalf of the Wellcome Sanger Institute COVID-19 Surveillance Team |
| EPI_ISL_3467739 | Lighthouse Lab in Glasgow | Wellcome Sanger Institute for the COVID-19 Genomics UK (COG-UK) Consortium | Harper VanSteenhouse, Yumi Kasai, David Gray, Carol Clugston, Anna Dominiczak and Alex Alderton, Roberto Amato, Jeffrey Barrett, Sonia Goncalves, Ewan Harrison, David K. Jackson, Ian Johnston, Dominic Kwiatkowski, Cordelia Langford, John Sillitoe on behalf of the Wellcome Sanger Institute COVID-19 Surveillance Team |
| EPI_ISL_3467740 | National Virus Reference Laboratory | National Virus Reference Laboratory | Zoe Yandle, Charlene Bennett, Gabriel Gonzalez, Michael Carr, Jonathan Dean, Cillian F De Gascun |
| EPI_ISL_3467741 | Lighthouse Lab in Glasgow | Wellcome Sanger Institute for the COVID-19 Genomics UK (COG-UK) Consortium | Harper VanSteenhouse, Yumi Kasai, David Gray, Carol Clugston, Anna Dominiczak and Alex Alderton, Roberto Amato, Jeffrey Barrett, Sonia Goncalves, Ewan Harrison, David K. Jackson, Ian Johnston, Dominic Kwiatkowski, Cordelia Langford, John Sillitoe on behalf of the Wellcome Sanger Institute COVID-19 Surveillance Team |
| EPI_ISL_3467742, EPI_ISL_3467743 | Lighthouse Lab in Milton Keynes | Wellcome Sanger Institute for the COVID-19 Genomics UK (COG-UK) Consortium | The Lighthouse Lab in Milton Keynes and Alex Alderton, Roberto Amato, Jeffrey Barrett, Sonia Goncalves, Ewan Harrison, David K. Jackson, Ian Johnston, Dominic Kwiatkowski, Cordelia Langford, John Sillitoe on behalf of the Wellcome Sanger Institute COVID-19 Surveillance Team |
| EPI_ISL_3467744 | National Virus Reference Laboratory | National Virus Reference Laboratory | Zoe Yandle, Charlene Bennett, Gabriel Gonzalez, Michael Carr, Jonathan Dean, Cillian F De Gascun |
| EPI_ISL_3467745, EPI_ISL_3467746 | Lighthouse Lab in Glasgow | Wellcome Sanger Institute for the COVID-19 Genomics UK (COG-UK) Consortium | Harper VanSteenhouse, Yumi Kasai, David Gray, Carol Clugston, Anna Dominiczak and Alex Alderton, Roberto Amato, Jeffrey Barrett, Sonia Goncalves, Ewan Harrison, David K. Jackson, Ian Johnston, Dominic Kwiatkowski, Cordelia Langford, John Sillitoe on behalf of the Wellcome Sanger Institute COVID-19 Surveillance Team |
| EPI_ISL_3467747 | National Virus Reference Laboratory | National Virus Reference Laboratory | Zoe Yandle, Charlene Bennett, Gabriel Gonzalez, Michael Carr, Jonathan Dean, Cillian F De Gascun |
| EPI_ISL_3467748, EPI_ISL_3467749 | Lighthouse Lab in Milton Keynes | Wellcome Sanger Institute for the COVID-19 Genomics UK (COG-UK) Consortium | The Lighthouse Lab in Milton Keynes and Alex Alderton, Roberto Amato, Jeffrey Barrett, Sonia Goncalves, Ewan Harrison, David K. Jackson, Ian Johnston, Dominic Kwiatkowski, Cordelia Langford, John Sillitoe on behalf of the Wellcome Sanger Institute COVID-19 Surveillance Team |
| EPI_ISL_3467750 | Lighthouse Lab in Glasgow | Wellcome Sanger Institute for the COVID-19 Genomics UK (COG-UK) Consortium | Harper VanSteenhouse, Yumi Kasai, David Gray, Carol Clugston, Anna Dominiczak and Alex Alderton, Roberto Amato, Jeffrey Barrett, Sonia Goncalves, Ewan Harrison, David K. Jackson, Ian Johnston, Dominic Kwiatkowski, Cordelia Langford, John Sillitoe on behalf of the Wellcome Sanger Institute COVID-19 Surveillance Team |
| EPI_ISL_3467751 | National Virus Reference Laboratory | National Virus Reference Laboratory | Zoe Yandle, Charlene Bennett, Gabriel Gonzalez, Michael Carr, Jonathan Dean, Cillian F De Gascun |
| EPI_ISL_3467752, EPI_ISL_3467753, EPI_ISL_3467754, EPI_ISL_3467755 | Lighthouse Lab in Milton Keynes | Wellcome Sanger Institute for the COVID-19 Genomics UK (COG-UK) Consortium | The Lighthouse Lab in Milton Keynes and Alex Alderton, Roberto Amato, Jeffrey Barrett, Sonia Goncalves, Ewan Harrison, David K. Jackson, Ian Johnston, Dominic Kwiatkowski, Cordelia Langford, John Sillitoe on behalf of the Wellcome Sanger Institute COVID-19 Surveillance Team |
| EPI_ISL_3467756 | National Virus Reference Laboratory | National Virus Reference Laboratory | Zoe Yandle, Charlene Bennett, Gabriel Gonzalez, Michael Carr, Jonathan Dean, Cillian F De Gascun |
| EPI_ISL_3467757 | Lighthouse Lab in Glasgow | Wellcome Sanger Institute for the COVID-19 Genomics UK (COG-UK) Consortium | Harper VanSteenhouse, Yumi Kasai, David Gray, Carol Clugston, Anna Dominiczak and Alex Alderton, Roberto Amato, Jeffrey Barrett, Sonia Goncalves, Ewan Harrison, David K. Jackson, Ian Johnston, Dominic Kwiatkowski, Cordelia Langford, John Sillitoe on behalf of the Wellcome Sanger Institute COVID-19 Surveillance Team |
| EPI_ISL_3467758 | Lighthouse Lab in Milton Keynes | Wellcome Sanger Institute for the COVID-19 Genomics UK (COG-UK) Consortium | The Lighthouse Lab in Milton Keynes and Alex Alderton, Roberto Amato, Jeffrey Barrett, Sonia Goncalves, Ewan Harrison, David K. Jackson, Ian Johnston, Dominic Kwiatkowski, Cordelia Langford, John Sillitoe on behalf of the Wellcome Sanger Institute COVID-19 Surveillance Team |
| EPI_ISL_3467759 | Lighthouse Lab in Glasgow | Wellcome Sanger Institute for the COVID-19 Genomics UK (COG-UK) Consortium | Harper VanSteenhouse, Yumi Kasai, David Gray, Carol Clugston, Anna Dominiczak and Alex Alderton, Roberto Amato, Jeffrey Barrett, Sonia Goncalves, Ewan Harrison, David K. Jackson, Ian Johnston, Dominic Kwiatkowski, Cordelia Langford, John Sillitoe on behalf of the Wellcome Sanger Institute COVID-19 Surveillance Team |

| | | | |
|---|---|---|---|
| EPI_ISL_3467760 | National Virus Reference Laboratory | National Virus Reference Laboratory | Zoe Yandle, Charlene Bennett, Gabriel Gonzalez, Michael Carr, Jonathan Dean, Cillian F De Gascun |
| EPI_ISL_3467761 | Lighthouse Lab in Milton Keynes | Wellcome Sanger Institute for the COVID-19 Genomics UK (COG-UK) Consortium | The Lighthouse Lab in Milton Keynes and Alex Alderton, Roberto Amato, Jeffrey Barrett, Sonia Goncalves, Ewan Harrison, David K. Jackson, Ian Johnston, Dominic Kwiatkowski, Cordelia Langford, John Sillitoe on behalf of the Wellcome Sanger Institute COVID-19 Surveillance Team |
| EPI_ISL_3467762, EPI_ISL_3467763, EPI_ISL_3467764 | Lighthouse Lab in Glasgow | Wellcome Sanger Institute for the COVID-19 Genomics UK (COG-UK) Consortium | Harper VanSteenhouse, Yumi Kasai, David Gray, Carol Clugston, Anna Dominiczak and Alex Alderton, Roberto Amato, Jeffrey Barrett, Sonia Goncalves, Ewan Harrison, David K. Jackson, Ian Johnston, Dominic Kwiatkowski, Cordelia Langford, John Sillitoe on behalf of the Wellcome Sanger Institute COVID-19 Surveillance Team |
| EPI_ISL_3467765 | National Virus Reference Laboratory | National Virus Reference Laboratory | Zoe Yandle, Charlene Bennett, Gabriel Gonzalez, Michael Carr, Jonathan Dean, Cillian F De Gascun |
| EPI_ISL_3467766 | Lighthouse Lab in Milton Keynes | Wellcome Sanger Institute for the COVID-19 Genomics UK (COG-UK) Consortium | The Lighthouse Lab in Milton Keynes and Alex Alderton, Roberto Amato, Jeffrey Barrett, Sonia Goncalves, Ewan Harrison, David K. Jackson, Ian Johnston, Dominic Kwiatkowski, Cordelia Langford, John Sillitoe on behalf of the Wellcome Sanger Institute COVID-19 Surveillance Team |
| EPI_ISL_3467767 | Lighthouse Lab in Glasgow | Wellcome Sanger Institute for the COVID-19 Genomics UK (COG-UK) Consortium | Harper VanSteenhouse, Yumi Kasai, David Gray, Carol Clugston, Anna Dominiczak and Alex Alderton, Roberto Amato, Jeffrey Barrett, Sonia Goncalves, Ewan Harrison, David K. Jackson, Ian Johnston, Dominic Kwiatkowski, Cordelia Langford, John Sillitoe on behalf of the Wellcome Sanger Institute COVID-19 Surveillance Team |
| EPI_ISL_3467768 | Lighthouse Lab in Milton Keynes | Wellcome Sanger Institute for the COVID-19 Genomics UK (COG-UK) Consortium | The Lighthouse Lab in Milton Keynes and Alex Alderton, Roberto Amato, Jeffrey Barrett, Sonia Goncalves, Ewan Harrison, David K. Jackson, Ian Johnston, Dominic Kwiatkowski, Cordelia Langford, John Sillitoe on behalf of the Wellcome Sanger Institute COVID-19 Surveillance Team |
| EPI_ISL_3467769 | National Virus Reference Laboratory | National Virus Reference Laboratory | Zoe Yandle, Charlene Bennett, Gabriel Gonzalez, Michael Carr, Jonathan Dean, Cillian F De Gascun |
| EPI_ISL_3467770 | Lighthouse Lab in Milton Keynes | Wellcome Sanger Institute for the COVID-19 Genomics UK (COG-UK) Consortium | The Lighthouse Lab in Milton Keynes and Alex Alderton, Roberto Amato, Jeffrey Barrett, Sonia Goncalves, Ewan Harrison, David K. Jackson, Ian Johnston, Dominic Kwiatkowski, Cordelia Langford, John Sillitoe on behalf of the Wellcome Sanger Institute COVID-19 Surveillance Team |
| EPI_ISL_3467771 | Lighthouse Lab in Glasgow | Wellcome Sanger Institute for the COVID-19 Genomics UK (COG-UK) Consortium | Harper VanSteenhouse, Yumi Kasai, David Gray, Carol Clugston, Anna Dominiczak and Alex Alderton, Roberto Amato, Jeffrey Barrett, Sonia Goncalves, Ewan Harrison, David K. Jackson, Ian Johnston, Dominic Kwiatkowski, Cordelia Langford, John Sillitoe on behalf of the Wellcome Sanger Institute COVID-19 Surveillance Team |
| EPI_ISL_3467772 | National Virus Reference Laboratory | National Virus Reference Laboratory | Zoe Yandle, Charlene Bennett, Gabriel Gonzalez, Michael Carr, Jonathan Dean, Cillian F De Gascun |
| EPI_ISL_3467773, EPI_ISL_3467774, EPI_ISL_3467775 | Lighthouse Lab in Milton Keynes | Wellcome Sanger Institute for the COVID-19 Genomics UK (COG-UK) Consortium | The Lighthouse Lab in Milton Keynes and Alex Alderton, Roberto Amato, Jeffrey Barrett, Sonia Goncalves, Ewan Harrison, David K. Jackson, Ian Johnston, Dominic Kwiatkowski, Cordelia Langford, John Sillitoe on behalf of the Wellcome Sanger Institute COVID-19 Surveillance Team |
| EPI_ISL_3467776 | Lighthouse Lab in Glasgow | Wellcome Sanger Institute for the COVID-19 Genomics UK (COG-UK) Consortium | Harper VanSteenhouse, Yumi Kasai, David Gray, Carol Clugston, Anna Dominiczak and Alex Alderton, Roberto Amato, Jeffrey Barrett, Sonia Goncalves, Ewan Harrison, David K. Jackson, Ian Johnston, Dominic Kwiatkowski, Cordelia Langford, John Sillitoe on behalf of the Wellcome Sanger Institute COVID-19 Surveillance Team |
| EPI_ISL_3467777 | National Virus Reference Laboratory | National Virus Reference Laboratory | Zoe Yandle, Charlene Bennett, Gabriel Gonzalez, Michael Carr, Jonathan Dean, Cillian F De Gascun |
| EPI_ISL_3467778, EPI_ISL_3467779 | Lighthouse Lab in Glasgow | Wellcome Sanger Institute for the COVID-19 Genomics UK (COG-UK) Consortium | Harper VanSteenhouse, Yumi Kasai, David Gray, Carol Clugston, Anna Dominiczak and Alex Alderton, Roberto Amato, Jeffrey Barrett, Sonia Goncalves, Ewan Harrison, David K. Jackson, Ian Johnston, Dominic Kwiatkowski, Cordelia Langford, John Sillitoe on behalf of the Wellcome Sanger Institute COVID-19 Surveillance Team |
| EPI_ISL_3467780 | National Virus Reference Laboratory | National Virus Reference Laboratory | Zoe Yandle, Charlene Bennett, Gabriel Gonzalez, Michael Carr, Jonathan Dean, Cillian F De Gascun |
| EPI_ISL_3467781 | Lighthouse Lab in Milton Keynes | Wellcome Sanger Institute for the COVID-19 Genomics UK (COG-UK) Consortium | The Lighthouse Lab in Milton Keynes and Alex Alderton, Roberto Amato, Jeffrey Barrett, Sonia Goncalves, Ewan Harrison, David K. Jackson, Ian Johnston, Dominic Kwiatkowski, Cordelia Langford, John Sillitoe on behalf of the Wellcome Sanger Institute COVID-19 Surveillance Team |
| EPI_ISL_3467782, EPI_ISL_3467783 | Lighthouse Lab in Glasgow | Wellcome Sanger Institute for the COVID-19 Genomics UK (COG-UK) Consortium | Harper VanSteenhouse, Yumi Kasai, David Gray, Carol Clugston, Anna Dominiczak and Alex Alderton, Roberto Amato, Jeffrey Barrett, Sonia Goncalves, Ewan Harrison, David K. Jackson, Ian Johnston, Dominic Kwiatkowski, Cordelia Langford, John Sillitoe on behalf of the Wellcome Sanger Institute COVID-19 Surveillance Team |
| EPI_ISL_3467784 | National Virus Reference Laboratory | National Virus Reference Laboratory | Zoe Yandle, Charlene Bennett, Gabriel Gonzalez, Michael Carr, Jonathan Dean, Cillian F De Gascun |
| EPI_ISL_3467785, EPI_ISL_3467786 | Lighthouse Lab in Milton Keynes | Wellcome Sanger Institute for the COVID-19 Genomics UK (COG-UK) Consortium | The Lighthouse Lab in Milton Keynes and Alex Alderton, Roberto Amato, Jeffrey Barrett, Sonia Goncalves, Ewan Harrison, David K. Jackson, Ian Johnston, Dominic Kwiatkowski, Cordelia Langford, John Sillitoe on behalf of the Wellcome Sanger Institute COVID-19 Surveillance Team |
| EPI_ISL_3467787 | National Virus Reference Laboratory | National Virus Reference Laboratory | Zoe Yandle, Charlene Bennett, Gabriel Gonzalez, Michael Carr, Jonathan Dean, Cillian F De Gascun |
| EPI_ISL_3467788, EPI_ISL_3467789, EPI_ISL_3467790 | Lighthouse Lab in Milton Keynes | Wellcome Sanger Institute for the COVID-19 Genomics UK (COG-UK) Consortium | The Lighthouse Lab in Milton Keynes and Alex Alderton, Roberto Amato, Jeffrey Barrett, Sonia Goncalves, Ewan Harrison, David K. Jackson, Ian Johnston, Dominic Kwiatkowski, Cordelia Langford, John Sillitoe on behalf of the Wellcome Sanger Institute COVID-19 Surveillance Team |
| EPI_ISL_3467791 | National Virus Reference Laboratory | National Virus Reference Laboratory | Zoe Yandle, Charlene Bennett, Gabriel Gonzalez, Michael Carr, Jonathan Dean, Cillian F De Gascun |
| EPI_ISL_3467792 | Lighthouse Lab in Glasgow | Wellcome Sanger Institute for the COVID-19 Genomics UK (COG-UK) Consortium | Harper VanSteenhouse, Yumi Kasai, David Gray, Carol Clugston, Anna Dominiczak and Alex Alderton, Roberto Amato, Jeffrey Barrett, Sonia Goncalves, Ewan Harrison, David K. Jackson, Ian Johnston, Dominic Kwiatkowski, Cordelia Langford, John Sillitoe on behalf of the Wellcome Sanger Institute COVID-19 Surveillance Team |
| EPI_ISL_3467793 | Lighthouse Lab in Milton Keynes | Wellcome Sanger Institute for the COVID-19 Genomics UK (COG-UK) Consortium | The Lighthouse Lab in Milton Keynes and Alex Alderton, Roberto Amato, Jeffrey Barrett, Sonia Goncalves, Ewan Harrison, David K. Jackson, Ian Johnston, Dominic Kwiatkowski, Cordelia Langford, John Sillitoe on behalf of the Wellcome Sanger Institute COVID-19 Surveillance Team |
| EPI_ISL_3467794 | Lighthouse Lab in Glasgow | Wellcome Sanger Institute for the COVID-19 Genomics UK (COG-UK) Consortium | Harper VanSteenhouse, Yumi Kasai, David Gray, Carol Clugston, Anna Dominiczak and Alex Alderton, Roberto Amato, Jeffrey Barrett, Sonia Goncalves, Ewan Harrison, David K. Jackson, Ian Johnston, Dominic Kwiatkowski, Cordelia Langford, John Sillitoe on behalf of the Wellcome Sanger Institute COVID-19 Surveillance Team |
| EPI_ISL_3467795 | National Virus Reference Laboratory | National Virus Reference Laboratory | Zoe Yandle, Charlene Bennett, Gabriel Gonzalez, Michael Carr, Jonathan Dean, Cillian F De Gascun |
| EPI_ISL_3467796, EPI_ISL_3467797, EPI_ISL_3467798 | Lighthouse Lab in Milton Keynes | Wellcome Sanger Institute for the COVID-19 Genomics UK (COG-UK) Consortium | The Lighthouse Lab in Milton Keynes and Alex Alderton, Roberto Amato, Jeffrey Barrett, Sonia Goncalves, Ewan Harrison, David K. Jackson, Ian Johnston, Dominic Kwiatkowski, Cordelia Langford, John Sillitoe on behalf of the Wellcome Sanger Institute COVID-19 Surveillance Team |
| EPI_ISL_3467799 | National Virus Reference Laboratory | National Virus Reference Laboratory | Zoe Yandle, Charlene Bennett, Gabriel Gonzalez, Michael Carr, Jonathan Dean, Cillian F De Gascun |
| EPI_ISL_3467800 | Lighthouse Lab in Milton Keynes | Wellcome Sanger Institute for the COVID-19 Genomics UK (COG-UK) Consortium | The Lighthouse Lab in Milton Keynes and Alex Alderton, Roberto Amato, Jeffrey Barrett, Sonia Goncalves, Ewan Harrison, David K. Jackson, Ian Johnston, Dominic Kwiatkowski, Cordelia Langford, John Sillitoe on behalf of the Wellcome Sanger Institute COVID-19 Surveillance Team |
| EPI_ISL_3467801 | Lighthouse Lab in Glasgow | Wellcome Sanger Institute for the COVID-19 Genomics UK (COG-UK) Consortium | Harper VanSteenhouse, Yumi Kasai, David Gray, Carol Clugston, Anna Dominiczak and Alex Alderton, Roberto Amato, Jeffrey Barrett, Sonia Goncalves, Ewan Harrison, David K. Jackson, Ian Johnston, Dominic Kwiatkowski, Cordelia Langford, John Sillitoe on behalf of the Wellcome Sanger Institute COVID-19 Surveillance Team |
| EPI_ISL_3467802 | Lighthouse Lab in Milton Keynes | Wellcome Sanger Institute for the COVID-19 Genomics UK (COG-UK) Consortium | The Lighthouse Lab in Milton Keynes and Alex Alderton, Roberto Amato, Jeffrey Barrett, Sonia Goncalves, Ewan Harrison, David K. Jackson, Ian Johnston, Dominic Kwiatkowski, Cordelia Langford, John Sillitoe on behalf of the Wellcome Sanger Institute COVID-19 Surveillance Team |
| EPI_ISL_3467803 | National Virus Reference Laboratory | National Virus Reference Laboratory | Zoe Yandle, Charlene Bennett, Gabriel Gonzalez, Michael Carr, Jonathan Dean, Cillian F De Gascun |
| EPI_ISL_3467804 | Lighthouse Lab in Milton Keynes | Wellcome Sanger Institute for the COVID-19 Genomics UK (COG-UK) Consortium | The Lighthouse Lab in Milton Keynes and Alex Alderton, Roberto Amato, Jeffrey Barrett, Sonia Goncalves, Ewan Harrison, David K. Jackson, Ian Johnston, Dominic Kwiatkowski, Cordelia Langford, John Sillitoe on behalf of the Wellcome Sanger Institute COVID-19 Surveillance Team |
| EPI_ISL_3467805 | Lighthouse Lab in Glasgow | Wellcome Sanger Institute for the COVID-19 Genomics UK (COG-UK) Consortium | Harper VanSteenhouse, Yumi Kasai, David Gray, Carol Clugston, Anna Dominiczak and Alex Alderton, Roberto Amato, Jeffrey Barrett, Sonia Goncalves, Ewan Harrison, David K. Jackson, Ian Johnston, Dominic Kwiatkowski, Cordelia Langford, John Sillitoe on behalf of the Wellcome Sanger Institute COVID-19 Surveillance Team |
| EPI_ISL_3467806 | National Virus Reference Laboratory | National Virus Reference Laboratory | Zoe Yandle, Charlene Bennett, Gabriel Gonzalez, Michael Carr, Jonathan Dean, Cillian F De Gascun |
| EPI_ISL_3467807, EPI_ISL_3467808 | Lighthouse Lab in Glasgow | Wellcome Sanger Institute for the COVID-19 Genomics UK (COG-UK) Consortium | Harper VanSteenhouse, Yumi Kasai, David Gray, Carol Clugston, Anna Dominiczak and Alex Alderton, Roberto Amato, Jeffrey Barrett, Sonia Goncalves, Ewan Harrison, David K. Jackson, Ian Johnston, Dominic Kwiatkowski, Cordelia Langford, John Sillitoe on behalf of the Wellcome Sanger Institute COVID-19 Surveillance Team |
| EPI_ISL_3467809 | Lighthouse Lab in Milton Keynes | Wellcome Sanger Institute for the COVID-19 Genomics UK (COG-UK) Consortium | The Lighthouse Lab in Milton Keynes and Alex Alderton, Roberto Amato, Jeffrey Barrett, Sonia Goncalves, Ewan Harrison, David K. Jackson, Ian Johnston, Dominic Kwiatkowski, Cordelia Langford, John Sillitoe on behalf of the Wellcome Sanger Institute COVID-19 Surveillance Team |
| EPI_ISL_3467810 | National Virus Reference Laboratory | National Virus Reference Laboratory | Zoe Yandle, Charlene Bennett, Gabriel Gonzalez, Michael Carr, Jonathan Dean, Cillian F De Gascun |
| EPI_ISL_3467811 | Lighthouse Lab in Glasgow | Wellcome Sanger Institute for the COVID-19 Genomics UK (COG-UK) Consortium | Harper VanSteenhouse, Yumi Kasai, David Gray, Carol Clugston, Anna Dominiczak and Alex Alderton, Roberto Amato, Jeffrey Barrett, Sonia Goncalves, Ewan Harrison, David K. Jackson, Ian Johnston, Dominic Kwiatkowski, Cordelia Langford, John Sillitoe on behalf of the Wellcome Sanger Institute COVID-19 Surveillance Team |
| EPI_ISL_3467812, EPI_ISL_3467813 | Lighthouse Lab in Milton Keynes | Wellcome Sanger Institute for the COVID-19 Genomics UK (COG-UK) Consortium | The Lighthouse Lab in Milton Keynes and Alex Alderton, Roberto Amato, Jeffrey Barrett, Sonia Goncalves, Ewan Harrison, David K. Jackson, Ian Johnston, Dominic Kwiatkowski, Cordelia Langford, John Sillitoe on behalf of the Wellcome Sanger Institute COVID-19 Surveillance Team |
| EPI_ISL_3467814 | National Virus Reference Laboratory | National Virus Reference Laboratory | Zoe Yandle, Charlene Bennett, Gabriel Gonzalez, Michael Carr, Jonathan Dean, Cillian F De Gascun |
| EPI_ISL_3467815, EPI_ISL_3467816 | Lighthouse Lab in Milton Keynes | Wellcome Sanger Institute for the COVID-19 Genomics UK (COG-UK) Consortium | The Lighthouse Lab in Milton Keynes and Alex Alderton, Roberto Amato, Jeffrey Barrett, Sonia Goncalves, Ewan Harrison, David K. Jackson, Ian Johnston, Dominic Kwiatkowski, Cordelia Langford, John Sillitoe on behalf of the Wellcome Sanger Institute COVID-19 Surveillance Team |
| EPI_ISL_3467817 | National Virus Reference Laboratory | National Virus Reference Laboratory | Zoe Yandle, Charlene Bennett, Gabriel Gonzalez, Michael Carr, Jonathan Dean, Cillian F De Gascun |
| EPI_ISL_3467818 | Lighthouse Lab in Milton Keynes | Wellcome Sanger Institute for the COVID-19 Genomics UK (COG-UK) Consortium | The Lighthouse Lab in Milton Keynes and Alex Alderton, Roberto Amato, Jeffrey Barrett, Sonia Goncalves, Ewan Harrison, David K. Jackson, Ian Johnston, Dominic Kwiatkowski, Cordelia Langford, John Sillitoe on behalf of the Wellcome Sanger Institute COVID-19 Surveillance Team |
| EPI_ISL_3467819 | Lighthouse Lab in Glasgow | Wellcome Sanger Institute for the COVID-19 Genomics UK (COG-UK) Consortium | Harper VanSteenhouse, Yumi Kasai, David Gray, Carol Clugston, Anna Dominiczak and Alex Alderton, Roberto Amato, Jeffrey Barrett, Sonia Goncalves, Ewan Harrison, David K. Jackson, Ian Johnston, Dominic Kwiatkowski, Cordelia Langford, John Sillitoe on behalf of the Wellcome Sanger Institute COVID-19 Surveillance Team |
| EPI_ISL_3467820 | National Virus Reference Laboratory | National Virus Reference Laboratory | Zoe Yandle, Charlene Bennett, Gabriel Gonzalez, Michael Carr, Jonathan Dean, Cillian F De Gascun |
| EPI_ISL_3467821, EPI_ISL_3467822, EPI_ISL_3467823 | Lighthouse Lab in Milton Keynes | Wellcome Sanger Institute for the COVID-19 Genomics UK (COG-UK) Consortium | The Lighthouse Lab in Milton Keynes and Alex Alderton, Roberto Amato, Jeffrey Barrett, Sonia Goncalves, Ewan Harrison, David K. Jackson, Ian Johnston, Dominic Kwiatkowski, Cordelia Langford, John Sillitoe on behalf of the Wellcome Sanger Institute COVID-19 Surveillance Team |
| EPI_ISL_3467824 | National Virus Reference Laboratory | National Virus Reference Laboratory | Zoe Yandle, Charlene Bennett, Gabriel Gonzalez, Michael Carr, Jonathan Dean, Cillian F De Gascun |
| EPI_ISL_3467825, EPI_ISL_3467826 | Lighthouse Lab in Glasgow | Wellcome Sanger Institute for the COVID-19 Genomics UK (COG-UK) Consortium | Harper VanSteenhouse, Yumi Kasai, David Gray, Carol Clugston, Anna Dominiczak and Alex Alderton, Roberto Amato, Jeffrey Barrett, Sonia Goncalves, Ewan Harrison, David K. Jackson, Ian Johnston, Dominic Kwiatkowski, Cordelia Langford, John Sillitoe on behalf of the Wellcome Sanger Institute COVID-19 Surveillance Team |
| EPI_ISL_3467827 | Lighthouse Lab in Milton Keynes | Wellcome Sanger Institute for the COVID-19 Genomics UK (COG-UK) | The Lighthouse Lab in Milton Keynes and Alex Alderton, Roberto Amato, Jeffrey Barrett, Sonia Goncalves, Ewan Harrison, David K. Jackson, Ian Johnston, Dominic Kwiatkowski, Cordelia Langford, John Sillitoe on behalf of the Wellcome Sanger |

| | | | |
|---|---|---|---|
| | | Consortium | Institute COVID-19 Surveillance Team |
| EPI_ISL_3467828 | National Virus Reference Laboratory | National Virus Reference Laboratory | Zoe Yandle, Charlene Bennett, Gabriel Gonzalez, Michael Carr, Jonathan Dean, Cillian F De Gascun |
| EPI_ISL_3467829, EPI_ISL_3467830, EPI_ISL_3467831 | Lighthouse Lab in Milton Keynes | Wellcome Sanger Institute for the COVID-19 Genomics UK (COG-UK) Consortium | The Lighthouse Lab in Milton Keynes and Alex Alderton, Roberto Amato, Jeffrey Barrett, Sonia Goncalves, Ewan Harrison, David K. Jackson, Ian Johnston, Dominic Kwiatkowski, Cordelia Langford, John Sillitoe on behalf of the Wellcome Sanger Institute COVID-19 Surveillance Team |
| EPI_ISL_3467832 | National Virus Reference Laboratory | National Virus Reference Laboratory | Zoe Yandle, Charlene Bennett, Gabriel Gonzalez, Michael Carr, Jonathan Dean, Cillian F De Gascun |
| EPI_ISL_3467833, EPI_ISL_3467834 | Lighthouse Lab in Milton Keynes | Wellcome Sanger Institute for the COVID-19 Genomics UK (COG-UK) Consortium | The Lighthouse Lab in Milton Keynes and Alex Alderton, Roberto Amato, Jeffrey Barrett, Sonia Goncalves, Ewan Harrison, David K. Jackson, Ian Johnston, Dominic Kwiatkowski, Cordelia Langford, John Sillitoe on behalf of the Wellcome Sanger Institute COVID-19 Surveillance Team |
| EPI_ISL_3467835 | National Virus Reference Laboratory | National Virus Reference Laboratory | Zoe Yandle, Charlene Bennett, Gabriel Gonzalez, Michael Carr, Jonathan Dean, Cillian F De Gascun |
| EPI_ISL_3467836 | Lighthouse Lab in Milton Keynes | Wellcome Sanger Institute for the COVID-19 Genomics UK (COG-UK) Consortium | The Lighthouse Lab in Milton Keynes and Alex Alderton, Roberto Amato, Jeffrey Barrett, Sonia Goncalves, Ewan Harrison, David K. Jackson, Ian Johnston, Dominic Kwiatkowski, Cordelia Langford, John Sillitoe on behalf of the Wellcome Sanger Institute COVID-19 Surveillance Team |
| EPI_ISL_3467837, EPI_ISL_3467838 | Lighthouse Lab in Glasgow | Wellcome Sanger Institute for the COVID-19 Genomics UK (COG-UK) Consortium | Harper VanSteenhouse, Yumi Kasai, David Gray, Carol Clugston, Anna Dominiczak and Alex Alderton, Roberto Amato, Jeffrey Barrett, Sonia Goncalves, Ewan Harrison, David K. Jackson, Ian Johnston, Dominic Kwiatkowski, Cordelia Langford, John Sillitoe on behalf of the Wellcome Sanger Institute COVID-19 Surveillance Team |
| EPI_ISL_3467839 | National Virus Reference Laboratory | National Virus Reference Laboratory | Zoe Yandle, Charlene Bennett, Gabriel Gonzalez, Michael Carr, Jonathan Dean, Cillian F De Gascun |
| EPI_ISL_3467840 | Lighthouse Lab in Glasgow | Wellcome Sanger Institute for the COVID-19 Genomics UK (COG-UK) Consortium | Harper VanSteenhouse, Yumi Kasai, David Gray, Carol Clugston, Anna Dominiczak and Alex Alderton, Roberto Amato, Jeffrey Barrett, Sonia Goncalves, Ewan Harrison, David K. Jackson, Ian Johnston, Dominic Kwiatkowski, Cordelia Langford, John Sillitoe on behalf of the Wellcome Sanger Institute COVID-19 Surveillance Team |
| EPI_ISL_3467841 | Lighthouse Lab in Milton Keynes | Wellcome Sanger Institute for the COVID-19 Genomics UK (COG-UK) Consortium | The Lighthouse Lab in Milton Keynes and Alex Alderton, Roberto Amato, Jeffrey Barrett, Sonia Goncalves, Ewan Harrison, David K. Jackson, Ian Johnston, Dominic Kwiatkowski, Cordelia Langford, John Sillitoe on behalf of the Wellcome Sanger Institute COVID-19 Surveillance Team |
| EPI_ISL_3467842 | Lighthouse Lab in Glasgow | Wellcome Sanger Institute for the COVID-19 Genomics UK (COG-UK) Consortium | Harper VanSteenhouse, Yumi Kasai, David Gray, Carol Clugston, Anna Dominiczak and Alex Alderton, Roberto Amato, Jeffrey Barrett, Sonia Goncalves, Ewan Harrison, David K. Jackson, Ian Johnston, Dominic Kwiatkowski, Cordelia Langford, John Sillitoe on behalf of the Wellcome Sanger Institute COVID-19 Surveillance Team |
| EPI_ISL_3467843 | National Virus Reference Laboratory | National Virus Reference Laboratory | Zoe Yandle, Charlene Bennett, Gabriel Gonzalez, Michael Carr, Jonathan Dean, Cillian F De Gascun |
| EPI_ISL_3467844 | Lighthouse Lab in Glasgow | Wellcome Sanger Institute for the COVID-19 Genomics UK (COG-UK) Consortium | Harper VanSteenhouse, Yumi Kasai, David Gray, Carol Clugston, Anna Dominiczak and Alex Alderton, Roberto Amato, Jeffrey Barrett, Sonia Goncalves, Ewan Harrison, David K. Jackson, Ian Johnston, Dominic Kwiatkowski, Cordelia Langford, John Sillitoe on behalf of the Wellcome Sanger Institute COVID-19 Surveillance Team |
| EPI_ISL_3467845 | Lighthouse Lab in Milton Keynes | Wellcome Sanger Institute for the COVID-19 Genomics UK (COG-UK) Consortium | The Lighthouse Lab in Milton Keynes and Alex Alderton, Roberto Amato, Jeffrey Barrett, Sonia Goncalves, Ewan Harrison, David K. Jackson, Ian Johnston, Dominic Kwiatkowski, Cordelia Langford, John Sillitoe on behalf of the Wellcome Sanger Institute COVID-19 Surveillance Team |
| EPI_ISL_3467846 | National Virus Reference Laboratory | National Virus Reference Laboratory | Zoe Yandle, Charlene Bennett, Gabriel Gonzalez, Michael Carr, Jonathan Dean, Cillian F De Gascun |
| EPI_ISL_3467847 | Lighthouse Lab in Glasgow | Wellcome Sanger Institute for the COVID-19 Genomics UK (COG-UK) Consortium | Harper VanSteenhouse, Yumi Kasai, David Gray, Carol Clugston, Anna Dominiczak and Alex Alderton, Roberto Amato, Jeffrey Barrett, Sonia Goncalves, Ewan Harrison, David K. Jackson, Ian Johnston, Dominic Kwiatkowski, Cordelia Langford, John Sillitoe on behalf of the Wellcome Sanger Institute COVID-19 Surveillance Team |
| EPI_ISL_3467848 | Lighthouse Lab in Milton Keynes | Wellcome Sanger Institute for the COVID-19 Genomics UK (COG-UK) Consortium | The Lighthouse Lab in Milton Keynes and Alex Alderton, Roberto Amato, Jeffrey Barrett, Sonia Goncalves, Ewan Harrison, David K. Jackson, Ian Johnston, Dominic Kwiatkowski, Cordelia Langford, John Sillitoe on behalf of the Wellcome Sanger Institute COVID-19 Surveillance Team |
| EPI_ISL_3467849 | Lighthouse Lab in Glasgow | Wellcome Sanger Institute for the COVID-19 Genomics UK (COG-UK) Consortium | Harper VanSteenhouse, Yumi Kasai, David Gray, Carol Clugston, Anna Dominiczak and Alex Alderton, Roberto Amato, Jeffrey Barrett, Sonia Goncalves, Ewan Harrison, David K. Jackson, Ian Johnston, Dominic Kwiatkowski, Cordelia Langford, John Sillitoe on behalf of the Wellcome Sanger Institute COVID-19 Surveillance Team |
| EPI_ISL_3467850 | National Virus Reference Laboratory | National Virus Reference Laboratory | Zoe Yandle, Charlene Bennett, Gabriel Gonzalez, Michael Carr, Jonathan Dean, Cillian F De Gascun |
| EPI_ISL_3467851 | Lighthouse Lab in Glasgow | Wellcome Sanger Institute for the COVID-19 Genomics UK (COG-UK) Consortium | Harper VanSteenhouse, Yumi Kasai, David Gray, Carol Clugston, Anna Dominiczak and Alex Alderton, Roberto Amato, Jeffrey Barrett, Sonia Goncalves, Ewan Harrison, David K. Jackson, Ian Johnston, Dominic Kwiatkowski, Cordelia Langford, John Sillitoe on behalf of the Wellcome Sanger Institute COVID-19 Surveillance Team |
| EPI_ISL_3467852, EPI_ISL_3467853 | Lighthouse Lab in Milton Keynes | Wellcome Sanger Institute for the COVID-19 Genomics UK (COG-UK) Consortium | The Lighthouse Lab in Milton Keynes and Alex Alderton, Roberto Amato, Jeffrey Barrett, Sonia Goncalves, Ewan Harrison, David K. Jackson, Ian Johnston, Dominic Kwiatkowski, Cordelia Langford, John Sillitoe on behalf of the Wellcome Sanger Institute COVID-19 Surveillance Team |
| EPI_ISL_3467854 | National Virus Reference Laboratory | National Virus Reference Laboratory | Zoe Yandle, Charlene Bennett, Gabriel Gonzalez, Michael Carr, Jonathan Dean, Cillian F De Gascun |
| EPI_ISL_3467855 | Lighthouse Lab in Glasgow | Wellcome Sanger Institute for the COVID-19 Genomics UK (COG-UK) Consortium | Harper VanSteenhouse, Yumi Kasai, David Gray, Carol Clugston, Anna Dominiczak and Alex Alderton, Roberto Amato, Jeffrey Barrett, Sonia Goncalves, Ewan Harrison, David K. Jackson, Ian Johnston, Dominic Kwiatkowski, Cordelia Langford, John Sillitoe on behalf of the Wellcome Sanger Institute COVID-19 Surveillance Team |
| EPI_ISL_3467856 | Lighthouse Lab in Milton Keynes | Wellcome Sanger Institute for the COVID-19 Genomics UK (COG-UK) Consortium | The Lighthouse Lab in Milton Keynes and Alex Alderton, Roberto Amato, Jeffrey Barrett, Sonia Goncalves, Ewan Harrison, David K. Jackson, Ian Johnston, Dominic Kwiatkowski, Cordelia Langford, John Sillitoe on behalf of the Wellcome Sanger Institute COVID-19 Surveillance Team |
| EPI_ISL_3467857 | Lighthouse Lab in Glasgow | Wellcome Sanger Institute for the COVID-19 Genomics UK (COG-UK) Consortium | Harper VanSteenhouse, Yumi Kasai, David Gray, Carol Clugston, Anna Dominiczak and Alex Alderton, Roberto Amato, Jeffrey Barrett, Sonia Goncalves, Ewan Harrison, David K. Jackson, Ian Johnston, Dominic Kwiatkowski, Cordelia Langford, John Sillitoe on behalf of the Wellcome Sanger Institute COVID-19 Surveillance Team |
| EPI_ISL_3467858 | National Virus Reference Laboratory | National Virus Reference Laboratory | Zoe Yandle, Charlene Bennett, Gabriel Gonzalez, Michael Carr, Jonathan Dean, Cillian F De Gascun |
| EPI_ISL_3467859, EPI_ISL_3467860 | Lighthouse Lab in Milton Keynes | Wellcome Sanger Institute for the COVID-19 Genomics UK (COG-UK) Consortium | The Lighthouse Lab in Milton Keynes and Alex Alderton, Roberto Amato, Jeffrey Barrett, Sonia Goncalves, Ewan Harrison, David K. Jackson, Ian Johnston, Dominic Kwiatkowski, Cordelia Langford, John Sillitoe on behalf of the Wellcome Sanger Institute COVID-19 Surveillance Team |
| EPI_ISL_3467861 | Lighthouse Lab in Glasgow | Wellcome Sanger Institute for the COVID-19 Genomics UK (COG-UK) Consortium | Harper VanSteenhouse, Yumi Kasai, David Gray, Carol Clugston, Anna Dominiczak and Alex Alderton, Roberto Amato, Jeffrey Barrett, Sonia Goncalves, Ewan Harrison, David K. Jackson, Ian Johnston, Dominic Kwiatkowski, Cordelia Langford, John Sillitoe on behalf of the Wellcome Sanger Institute COVID-19 Surveillance Team |
| EPI_ISL_3467862 | National Virus Reference Laboratory | National Virus Reference Laboratory | Zoe Yandle, Charlene Bennett, Gabriel Gonzalez, Michael Carr, Jonathan Dean, Cillian F De Gascun |
| EPI_ISL_3467863 | Lighthouse Lab in Milton Keynes | Wellcome Sanger Institute for the COVID-19 Genomics UK (COG-UK) Consortium | The Lighthouse Lab in Milton Keynes and Alex Alderton, Roberto Amato, Jeffrey Barrett, Sonia Goncalves, Ewan Harrison, David K. Jackson, Ian Johnston, Dominic Kwiatkowski, Cordelia Langford, John Sillitoe on behalf of the Wellcome Sanger Institute COVID-19 Surveillance Team |
| EPI_ISL_3467864 | Lighthouse Lab in Glasgow | Wellcome Sanger Institute for the COVID-19 Genomics UK (COG-UK) Consortium | Harper VanSteenhouse, Yumi Kasai, David Gray, Carol Clugston, Anna Dominiczak and Alex Alderton, Roberto Amato, Jeffrey Barrett, Sonia Goncalves, Ewan Harrison, David K. Jackson, Ian Johnston, Dominic Kwiatkowski, Cordelia Langford, John Sillitoe on behalf of the Wellcome Sanger Institute COVID-19 Surveillance Team |
| EPI_ISL_3467865 | National Virus Reference Laboratory | National Virus Reference Laboratory | Zoe Yandle, Charlene Bennett, Gabriel Gonzalez, Michael Carr, Jonathan Dean, Cillian F De Gascun |
| EPI_ISL_3467866, EPI_ISL_3467867 | Lighthouse Lab in Milton Keynes | Wellcome Sanger Institute for the COVID-19 Genomics UK (COG-UK) Consortium | The Lighthouse Lab in Milton Keynes and Alex Alderton, Roberto Amato, Jeffrey Barrett, Sonia Goncalves, Ewan Harrison, David K. Jackson, Ian Johnston, Dominic Kwiatkowski, Cordelia Langford, John Sillitoe on behalf of the Wellcome Sanger Institute COVID-19 Surveillance Team |
| EPI_ISL_3467868 | National Virus Reference Laboratory | National Virus Reference Laboratory | Zoe Yandle, Charlene Bennett, Gabriel Gonzalez, Michael Carr, Jonathan Dean, Cillian F De Gascun |
| EPI_ISL_3467869 | Lighthouse Lab in Glasgow | Wellcome Sanger Institute for the COVID-19 Genomics UK (COG-UK) Consortium | Harper VanSteenhouse, Yumi Kasai, David Gray, Carol Clugston, Anna Dominiczak and Alex Alderton, Roberto Amato, Jeffrey Barrett, Sonia Goncalves, Ewan Harrison, David K. Jackson, Ian Johnston, Dominic Kwiatkowski, Cordelia Langford, John Sillitoe on behalf of the Wellcome Sanger Institute COVID-19 Surveillance Team |
| EPI_ISL_3467870 | Lighthouse Lab in Milton Keynes | Wellcome Sanger Institute for the COVID-19 Genomics UK (COG-UK) Consortium | The Lighthouse Lab in Milton Keynes and Alex Alderton, Roberto Amato, Jeffrey Barrett, Sonia Goncalves, Ewan Harrison, David K. Jackson, Ian Johnston, Dominic Kwiatkowski, Cordelia Langford, John Sillitoe on behalf of the Wellcome Sanger Institute COVID-19 Surveillance Team |
| EPI_ISL_3467871 | Lighthouse Lab in Glasgow | Wellcome Sanger Institute for the COVID-19 Genomics UK (COG-UK) Consortium | Harper VanSteenhouse, Yumi Kasai, David Gray, Carol Clugston, Anna Dominiczak and Alex Alderton, Roberto Amato, Jeffrey Barrett, Sonia Goncalves, Ewan Harrison, David K. Jackson, Ian Johnston, Dominic Kwiatkowski, Cordelia Langford, John Sillitoe on behalf of the Wellcome Sanger Institute COVID-19 Surveillance Team |
| EPI_ISL_3467872 | National Virus Reference Laboratory | National Virus Reference Laboratory | Zoe Yandle, Charlene Bennett, Gabriel Gonzalez, Michael Carr, Jonathan Dean, Cillian F De Gascun |
| EPI_ISL_3467873 | Lighthouse Lab in Milton Keynes | Wellcome Sanger Institute for the COVID-19 Genomics UK (COG-UK) Consortium | The Lighthouse Lab in Milton Keynes and Alex Alderton, Roberto Amato, Jeffrey Barrett, Sonia Goncalves, Ewan Harrison, David K. Jackson, Ian Johnston, Dominic Kwiatkowski, Cordelia Langford, John Sillitoe on behalf of the Wellcome Sanger Institute COVID-19 Surveillance Team |
| EPI_ISL_3467874, EPI_ISL_3467875 | Lighthouse Lab in Glasgow | Wellcome Sanger Institute for the COVID-19 Genomics UK (COG-UK) Consortium | Harper VanSteenhouse, Yumi Kasai, David Gray, Carol Clugston, Anna Dominiczak and Alex Alderton, Roberto Amato, Jeffrey Barrett, Sonia Goncalves, Ewan Harrison, David K. Jackson, Ian Johnston, Dominic Kwiatkowski, Cordelia Langford, John Sillitoe on behalf of the Wellcome Sanger Institute COVID-19 Surveillance Team |
| EPI_ISL_3467876 | National Virus Reference Laboratory | National Virus Reference Laboratory | Zoe Yandle, Charlene Bennett, Gabriel Gonzalez, Michael Carr, Jonathan Dean, Cillian F De Gascun |
| EPI_ISL_3467877 | Lighthouse Lab in Glasgow | Wellcome Sanger Institute for the COVID-19 Genomics UK (COG-UK) Consortium | Harper VanSteenhouse, Yumi Kasai, David Gray, Carol Clugston, Anna Dominiczak and Alex Alderton, Roberto Amato, Jeffrey Barrett, Sonia Goncalves, Ewan Harrison, David K. Jackson, Ian Johnston, Dominic Kwiatkowski, Cordelia Langford, John Sillitoe on behalf of the Wellcome Sanger Institute COVID-19 Surveillance Team |
| EPI_ISL_3467878 | Lighthouse Lab in Milton Keynes | Wellcome Sanger Institute for the COVID-19 Genomics UK (COG-UK) Consortium | The Lighthouse Lab in Milton Keynes and Alex Alderton, Roberto Amato, Jeffrey Barrett, Sonia Goncalves, Ewan Harrison, David K. Jackson, Ian Johnston, Dominic Kwiatkowski, Cordelia Langford, John Sillitoe on behalf of the Wellcome Sanger Institute COVID-19 Surveillance Team |
| EPI_ISL_3467879 | National Virus Reference Laboratory | National Virus Reference Laboratory | Zoe Yandle, Charlene Bennett, Gabriel Gonzalez, Michael Carr, Jonathan Dean, Cillian F De Gascun |
| EPI_ISL_3467880, EPI_ISL_3467881 | Lighthouse Lab in Glasgow | Wellcome Sanger Institute for the COVID-19 Genomics UK (COG-UK) Consortium | Harper VanSteenhouse, Yumi Kasai, David Gray, Carol Clugston, Anna Dominiczak and Alex Alderton, Roberto Amato, Jeffrey Barrett, Sonia Goncalves, Ewan Harrison, David K. Jackson, Ian Johnston, Dominic Kwiatkowski, Cordelia Langford, John Sillitoe on behalf of the Wellcome Sanger Institute COVID-19 Surveillance Team |
| EPI_ISL_3467882 | Lighthouse Lab in Milton Keynes | Wellcome Sanger Institute for the COVID-19 Genomics UK (COG-UK) Consortium | The Lighthouse Lab in Milton Keynes and Alex Alderton, Roberto Amato, Jeffrey Barrett, Sonia Goncalves, Ewan Harrison, David K. Jackson, Ian Johnston, Dominic Kwiatkowski, Cordelia Langford, John Sillitoe on behalf of the Wellcome Sanger Institute COVID-19 Surveillance Team |
| EPI_ISL_3467883 | National Virus Reference Laboratory | National Virus Reference Laboratory | Zoe Yandle, Charlene Bennett, Gabriel Gonzalez, Michael Carr, Jonathan Dean, Cillian F De Gascun |
| EPI_ISL_3467884, EPI_ISL_3467885, EPI_ISL_3467886 | Lighthouse Lab in Milton Keynes | Wellcome Sanger Institute for the COVID-19 Genomics UK (COG-UK) Consortium | The Lighthouse Lab in Milton Keynes and Alex Alderton, Roberto Amato, Jeffrey Barrett, Sonia Goncalves, Ewan Harrison, David K. Jackson, Ian Johnston, Dominic Kwiatkowski, Cordelia Langford, John Sillitoe on behalf of the Wellcome Sanger Institute COVID-19 Surveillance Team |
| EPI_ISL_3467887 | National Virus Reference Laboratory | National Virus Reference Laboratory | Zoe Yandle, Charlene Bennett, Gabriel Gonzalez, Michael Carr, Jonathan Dean, Cillian F De Gascun |
| EPI_ISL_3467888, EPI_ISL_3467889 | Lighthouse Lab in Glasgow | Wellcome Sanger Institute for the COVID-19 Genomics UK (COG-UK) Consortium | Harper VanSteenhouse, Yumi Kasai, David Gray, Carol Clugston, Anna Dominiczak and Alex Alderton, Roberto Amato, Jeffrey Barrett, Sonia Goncalves, Ewan Harrison, David K. Jackson, Ian Johnston, Dominic Kwiatkowski, Cordelia Langford, John Sillitoe on behalf of the Wellcome Sanger Institute COVID-19 Surveillance Team |
| EPI_ISL_3467890 | Lighthouse Lab in Milton Keynes | Wellcome Sanger Institute for the COVID-19 Genomics UK (COG-UK) Consortium | The Lighthouse Lab in Milton Keynes and Alex Alderton, Roberto Amato, Jeffrey Barrett, Sonia Goncalves, Ewan Harrison, David K. Jackson, Ian Johnston, Dominic Kwiatkowski, Cordelia Langford, John Sillitoe on behalf of the Wellcome Sanger Institute COVID-19 Surveillance Team |

| | | | |
|---|---|---|---|
| EPI_ISL_3467891 | National Virus Reference Laboratory | National Virus Reference Laboratory | Zoe Yandle, Charlene Bennett, Gabriel Gonzalez, Michael Carr, Jonathan Dean, Cillian F De Gascun |
| EPI_ISL_3467892, EPI_ISL_3467893 | Lighthouse Lab in Glasgow | Wellcome Sanger Institute for the COVID-19 Genomics UK (COG-UK) Consortium | Harper VanSteenhouse, Yumi Kasai, David Gray, Carol Clugston, Anna Dominiczak and Alex Alderton, Roberto Amato, Jeffrey Barrett, Sonia Goncalves, Ewan Harrison, David K. Jackson, Ian Johnston, Dominic Kwiatkowski, Cordelia Langford, John Sillitoe on behalf of the Wellcome Sanger Institute COVID-19 Surveillance Team |
| EPI_ISL_3467894 | Lighthouse Lab in Milton Keynes | Wellcome Sanger Institute for the COVID-19 Genomics UK (COG-UK) Consortium | The Lighthouse Lab in Milton Keynes and Alex Alderton, Roberto Amato, Jeffrey Barrett, Sonia Goncalves, Ewan Harrison, David K. Jackson, Ian Johnston, Dominic Kwiatkowski, Cordelia Langford, John Sillitoe on behalf of the Wellcome Sanger Institute COVID-19 Surveillance Team |
| EPI_ISL_3467895 | National Virus Reference Laboratory | National Virus Reference Laboratory | Zoe Yandle, Charlene Bennett, Gabriel Gonzalez, Michael Carr, Jonathan Dean, Cillian F De Gascun |
| EPI_ISL_3467896 | Lighthouse Lab in Milton Keynes | Wellcome Sanger Institute for the COVID-19 Genomics UK (COG-UK) Consortium | The Lighthouse Lab in Milton Keynes and Alex Alderton, Roberto Amato, Jeffrey Barrett, Sonia Goncalves, Ewan Harrison, David K. Jackson, Ian Johnston, Dominic Kwiatkowski, Cordelia Langford, John Sillitoe on behalf of the Wellcome Sanger Institute COVID-19 Surveillance Team |
| EPI_ISL_3467897 | Lighthouse Lab in Glasgow | Wellcome Sanger Institute for the COVID-19 Genomics UK (COG-UK) Consortium | Harper VanSteenhouse, Yumi Kasai, David Gray, Carol Clugston, Anna Dominiczak and Alex Alderton, Roberto Amato, Jeffrey Barrett, Sonia Goncalves, Ewan Harrison, David K. Jackson, Ian Johnston, Dominic Kwiatkowski, Cordelia Langford, John Sillitoe on behalf of the Wellcome Sanger Institute COVID-19 Surveillance Team |
| EPI_ISL_3467898 | Lighthouse Lab in Milton Keynes | Wellcome Sanger Institute for the COVID-19 Genomics UK (COG-UK) Consortium | The Lighthouse Lab in Milton Keynes and Alex Alderton, Roberto Amato, Jeffrey Barrett, Sonia Goncalves, Ewan Harrison, David K. Jackson, Ian Johnston, Dominic Kwiatkowski, Cordelia Langford, John Sillitoe on behalf of the Wellcome Sanger Institute COVID-19 Surveillance Team |
| EPI_ISL_3467899 | National Virus Reference Laboratory | National Virus Reference Laboratory | Zoe Yandle, Charlene Bennett, Gabriel Gonzalez, Michael Carr, Jonathan Dean, Cillian F De Gascun |
| EPI_ISL_3467900 | Lighthouse Lab in Milton Keynes | Wellcome Sanger Institute for the COVID-19 Genomics UK (COG-UK) Consortium | The Lighthouse Lab in Milton Keynes and Alex Alderton, Roberto Amato, Jeffrey Barrett, Sonia Goncalves, Ewan Harrison, David K. Jackson, Ian Johnston, Dominic Kwiatkowski, Cordelia Langford, John Sillitoe on behalf of the Wellcome Sanger Institute COVID-19 Surveillance Team |
| EPI_ISL_3467901 | Lighthouse Lab in Glasgow | Wellcome Sanger Institute for the COVID-19 Genomics UK (COG-UK) Consortium | Harper VanSteenhouse, Yumi Kasai, David Gray, Carol Clugston, Anna Dominiczak and Alex Alderton, Roberto Amato, Jeffrey Barrett, Sonia Goncalves, Ewan Harrison, David K. Jackson, Ian Johnston, Dominic Kwiatkowski, Cordelia Langford, John Sillitoe on behalf of the Wellcome Sanger Institute COVID-19 Surveillance Team |
| EPI_ISL_3467902 | National Virus Reference Laboratory | National Virus Reference Laboratory | Zoe Yandle, Charlene Bennett, Gabriel Gonzalez, Michael Carr, Jonathan Dean, Cillian F De Gascun |
| EPI_ISL_3467903, EPI_ISL_3467904 | Lighthouse Lab in Milton Keynes | Wellcome Sanger Institute for the COVID-19 Genomics UK (COG-UK) Consortium | The Lighthouse Lab in Milton Keynes and Alex Alderton, Roberto Amato, Jeffrey Barrett, Sonia Goncalves, Ewan Harrison, David K. Jackson, Ian Johnston, Dominic Kwiatkowski, Cordelia Langford, John Sillitoe on behalf of the Wellcome Sanger Institute COVID-19 Surveillance Team |
| EPI_ISL_3467905 | National Virus Reference Laboratory | National Virus Reference Laboratory | Zoe Yandle, Charlene Bennett, Gabriel Gonzalez, Michael Carr, Jonathan Dean, Cillian F De Gascun |
| EPI_ISL_3467906 | Lighthouse Lab in Milton Keynes | Wellcome Sanger Institute for the COVID-19 Genomics UK (COG-UK) Consortium | The Lighthouse Lab in Milton Keynes and Alex Alderton, Roberto Amato, Jeffrey Barrett, Sonia Goncalves, Ewan Harrison, David K. Jackson, Ian Johnston, Dominic Kwiatkowski, Cordelia Langford, John Sillitoe on behalf of the Wellcome Sanger Institute COVID-19 Surveillance Team |
| EPI_ISL_3467907, EPI_ISL_3467908 | Lighthouse Lab in Glasgow | Wellcome Sanger Institute for the COVID-19 Genomics UK (COG-UK) Consortium | Harper VanSteenhouse, Yumi Kasai, David Gray, Carol Clugston, Anna Dominiczak and Alex Alderton, Roberto Amato, Jeffrey Barrett, Sonia Goncalves, Ewan Harrison, David K. Jackson, Ian Johnston, Dominic Kwiatkowski, Cordelia Langford, John Sillitoe on behalf of the Wellcome Sanger Institute COVID-19 Surveillance Team |
| EPI_ISL_3467909 | National Virus Reference Laboratory | National Virus Reference Laboratory | Zoe Yandle, Charlene Bennett, Gabriel Gonzalez, Michael Carr, Jonathan Dean, Cillian F De Gascun |
| EPI_ISL_3467910, EPI_ISL_3467911, EPI_ISL_3467912 | Lighthouse Lab in Glasgow | Wellcome Sanger Institute for the COVID-19 Genomics UK (COG-UK) Consortium | Harper VanSteenhouse, Yumi Kasai, David Gray, Carol Clugston, Anna Dominiczak and Alex Alderton, Roberto Amato, Jeffrey Barrett, Sonia Goncalves, Ewan Harrison, David K. Jackson, Ian Johnston, Dominic Kwiatkowski, Cordelia Langford, John Sillitoe on behalf of the Wellcome Sanger Institute COVID-19 Surveillance Team |
| EPI_ISL_3467913 | National Virus Reference Laboratory | National Virus Reference Laboratory | Zoe Yandle, Charlene Bennett, Gabriel Gonzalez, Michael Carr, Jonathan Dean, Cillian F De Gascun |
| EPI_ISL_3467914, EPI_ISL_3467915 | Lighthouse Lab in Glasgow | Wellcome Sanger Institute for the COVID-19 Genomics UK (COG-UK) Consortium | Harper VanSteenhouse, Yumi Kasai, David Gray, Carol Clugston, Anna Dominiczak and Alex Alderton, Roberto Amato, Jeffrey Barrett, Sonia Goncalves, Ewan Harrison, David K. Jackson, Ian Johnston, Dominic Kwiatkowski, Cordelia Langford, John Sillitoe on behalf of the Wellcome Sanger Institute COVID-19 Surveillance Team |
| EPI_ISL_3467916 | Lighthouse Lab in Milton Keynes | Wellcome Sanger Institute for the COVID-19 Genomics UK (COG-UK) Consortium | The Lighthouse Lab in Milton Keynes and Alex Alderton, Roberto Amato, Jeffrey Barrett, Sonia Goncalves, Ewan Harrison, David K. Jackson, Ian Johnston, Dominic Kwiatkowski, Cordelia Langford, John Sillitoe on behalf of the Wellcome Sanger Institute COVID-19 Surveillance Team |
| EPI_ISL_3467917 | National Virus Reference Laboratory | National Virus Reference Laboratory | Zoe Yandle, Charlene Bennett, Gabriel Gonzalez, Michael Carr, Jonathan Dean, Cillian F De Gascun |
| EPI_ISL_3467918 | Lighthouse Lab in Milton Keynes | Wellcome Sanger Institute for the COVID-19 Genomics UK (COG-UK) Consortium | The Lighthouse Lab in Milton Keynes and Alex Alderton, Roberto Amato, Jeffrey Barrett, Sonia Goncalves, Ewan Harrison, David K. Jackson, Ian Johnston, Dominic Kwiatkowski, Cordelia Langford, John Sillitoe on behalf of the Wellcome Sanger Institute COVID-19 Surveillance Team |
| EPI_ISL_3467919 | Lighthouse Lab in Glasgow | Wellcome Sanger Institute for the COVID-19 Genomics UK (COG-UK) Consortium | Harper VanSteenhouse, Yumi Kasai, David Gray, Carol Clugston, Anna Dominiczak and Alex Alderton, Roberto Amato, Jeffrey Barrett, Sonia Goncalves, Ewan Harrison, David K. Jackson, Ian Johnston, Dominic Kwiatkowski, Cordelia Langford, John Sillitoe on behalf of the Wellcome Sanger Institute COVID-19 Surveillance Team |
| EPI_ISL_3467920 | Lighthouse Lab in Milton Keynes | Wellcome Sanger Institute for the COVID-19 Genomics UK (COG-UK) Consortium | The Lighthouse Lab in Milton Keynes and Alex Alderton, Roberto Amato, Jeffrey Barrett, Sonia Goncalves, Ewan Harrison, David K. Jackson, Ian Johnston, Dominic Kwiatkowski, Cordelia Langford, John Sillitoe on behalf of the Wellcome Sanger Institute COVID-19 Surveillance Team |
| EPI_ISL_3467921 | National Virus Reference Laboratory | National Virus Reference Laboratory | Zoe Yandle, Charlene Bennett, Gabriel Gonzalez, Michael Carr, Jonathan Dean, Cillian F De Gascun |
| EPI_ISL_3467922, EPI_ISL_3467923 | Lighthouse Lab in Glasgow | Wellcome Sanger Institute for the COVID-19 Genomics UK (COG-UK) Consortium | Harper VanSteenhouse, Yumi Kasai, David Gray, Carol Clugston, Anna Dominiczak and Alex Alderton, Roberto Amato, Jeffrey Barrett, Sonia Goncalves, Ewan Harrison, David K. Jackson, Ian Johnston, Dominic Kwiatkowski, Cordelia Langford, John Sillitoe on behalf of the Wellcome Sanger Institute COVID-19 Surveillance Team |
| EPI_ISL_3467924 | National Virus Reference Laboratory | National Virus Reference Laboratory | Zoe Yandle, Charlene Bennett, Gabriel Gonzalez, Michael Carr, Jonathan Dean, Cillian F De Gascun |
| EPI_ISL_3467925 | Lighthouse Lab in Glasgow | Wellcome Sanger Institute for the COVID-19 Genomics UK (COG-UK) Consortium | Harper VanSteenhouse, Yumi Kasai, David Gray, Carol Clugston, Anna Dominiczak and Alex Alderton, Roberto Amato, Jeffrey Barrett, Sonia Goncalves, Ewan Harrison, David K. Jackson, Ian Johnston, Dominic Kwiatkowski, Cordelia Langford, John Sillitoe on behalf of the Wellcome Sanger Institute COVID-19 Surveillance Team |
| EPI_ISL_3467926, EPI_ISL_3467927 | Lighthouse Lab in Milton Keynes | Wellcome Sanger Institute for the COVID-19 Genomics UK (COG-UK) Consortium | The Lighthouse Lab in Milton Keynes and Alex Alderton, Roberto Amato, Jeffrey Barrett, Sonia Goncalves, Ewan Harrison, David K. Jackson, Ian Johnston, Dominic Kwiatkowski, Cordelia Langford, John Sillitoe on behalf of the Wellcome Sanger Institute COVID-19 Surveillance Team |
| EPI_ISL_3467928 | National Virus Reference Laboratory | National Virus Reference Laboratory | Zoe Yandle, Charlene Bennett, Gabriel Gonzalez, Michael Carr, Jonathan Dean, Cillian F De Gascun |
| EPI_ISL_3467929, EPI_ISL_3467930 | Lighthouse Lab in Milton Keynes | Wellcome Sanger Institute for the COVID-19 Genomics UK (COG-UK) Consortium | The Lighthouse Lab in Milton Keynes and Alex Alderton, Roberto Amato, Jeffrey Barrett, Sonia Goncalves, Ewan Harrison, David K. Jackson, Ian Johnston, Dominic Kwiatkowski, Cordelia Langford, John Sillitoe on behalf of the Wellcome Sanger Institute COVID-19 Surveillance Team |
| EPI_ISL_3467931 | Lighthouse Lab in Glasgow | Wellcome Sanger Institute for the COVID-19 Genomics UK (COG-UK) Consortium | Harper VanSteenhouse, Yumi Kasai, David Gray, Carol Clugston, Anna Dominiczak and Alex Alderton, Roberto Amato, Jeffrey Barrett, Sonia Goncalves, Ewan Harrison, David K. Jackson, Ian Johnston, Dominic Kwiatkowski, Cordelia Langford, John Sillitoe on behalf of the Wellcome Sanger Institute COVID-19 Surveillance Team |
| EPI_ISL_3467933, EPI_ISL_3467934, EPI_ISL_3467935 | Lighthouse Lab in Milton Keynes | Wellcome Sanger Institute for the COVID-19 Genomics UK (COG-UK) Consortium | The Lighthouse Lab in Milton Keynes and Alex Alderton, Roberto Amato, Jeffrey Barrett, Sonia Goncalves, Ewan Harrison, David K. Jackson, Ian Johnston, Dominic Kwiatkowski, Cordelia Langford, John Sillitoe on behalf of the Wellcome Sanger Institute COVID-19 Surveillance Team |
| EPI_ISL_3467936 | National Virus Reference Laboratory | National Virus Reference Laboratory | Zoe Yandle, Charlene Bennett, Gabriel Gonzalez, Michael Carr, Jonathan Dean, Cillian F De Gascun |
| EPI_ISL_3467937 | Lighthouse Lab in Glasgow | Wellcome Sanger Institute for the COVID-19 Genomics UK (COG-UK) Consortium | Harper VanSteenhouse, Yumi Kasai, David Gray, Carol Clugston, Anna Dominiczak and Alex Alderton, Roberto Amato, Jeffrey Barrett, Sonia Goncalves, Ewan Harrison, David K. Jackson, Ian Johnston, Dominic Kwiatkowski, Cordelia Langford, John Sillitoe on behalf of the Wellcome Sanger Institute COVID-19 Surveillance Team |
| EPI_ISL_3467938 | Lighthouse Lab in Milton Keynes | Wellcome Sanger Institute for the COVID-19 Genomics UK (COG-UK) Consortium | The Lighthouse Lab in Milton Keynes and Alex Alderton, Roberto Amato, Jeffrey Barrett, Sonia Goncalves, Ewan Harrison, David K. Jackson, Ian Johnston, Dominic Kwiatkowski, Cordelia Langford, John Sillitoe on behalf of the Wellcome Sanger Institute COVID-19 Surveillance Team |
| EPI_ISL_3467939 | Lighthouse Lab in Glasgow | Wellcome Sanger Institute for the COVID-19 Genomics UK (COG-UK) Consortium | Harper VanSteenhouse, Yumi Kasai, David Gray, Carol Clugston, Anna Dominiczak and Alex Alderton, Roberto Amato, Jeffrey Barrett, Sonia Goncalves, Ewan Harrison, David K. Jackson, Ian Johnston, Dominic Kwiatkowski, Cordelia Langford, John Sillitoe on behalf of the Wellcome Sanger Institute COVID-19 Surveillance Team |
| EPI_ISL_3467940 | National Virus Reference Laboratory | National Virus Reference Laboratory | Zoe Yandle, Charlene Bennett, Gabriel Gonzalez, Michael Carr, Jonathan Dean, Cillian F De Gascun |
| EPI_ISL_3467941, EPI_ISL_3467942 | Lighthouse Lab in Milton Keynes | Wellcome Sanger Institute for the COVID-19 Genomics UK (COG-UK) Consortium | The Lighthouse Lab in Milton Keynes and Alex Alderton, Roberto Amato, Jeffrey Barrett, Sonia Goncalves, Ewan Harrison, David K. Jackson, Ian Johnston, Dominic Kwiatkowski, Cordelia Langford, John Sillitoe on behalf of the Wellcome Sanger Institute COVID-19 Surveillance Team |
| EPI_ISL_3467943 | National Virus Reference Laboratory | National Virus Reference Laboratory | Zoe Yandle, Charlene Bennett, Gabriel Gonzalez, Michael Carr, Jonathan Dean, Cillian F De Gascun |
| EPI_ISL_3467944, EPI_ISL_3467945 | Lighthouse Lab in Milton Keynes | Wellcome Sanger Institute for the COVID-19 Genomics UK (COG-UK) Consortium | The Lighthouse Lab in Milton Keynes and Alex Alderton, Roberto Amato, Jeffrey Barrett, Sonia Goncalves, Ewan Harrison, David K. Jackson, Ian Johnston, Dominic Kwiatkowski, Cordelia Langford, John Sillitoe on behalf of the Wellcome Sanger Institute COVID-19 Surveillance Team |
| EPI_ISL_3467946 | Lighthouse Lab in Glasgow | Wellcome Sanger Institute for the COVID-19 Genomics UK (COG-UK) Consortium | Harper VanSteenhouse, Yumi Kasai, David Gray, Carol Clugston, Anna Dominiczak and Alex Alderton, Roberto Amato, Jeffrey Barrett, Sonia Goncalves, Ewan Harrison, David K. Jackson, Ian Johnston, Dominic Kwiatkowski, Cordelia Langford, John Sillitoe on behalf of the Wellcome Sanger Institute COVID-19 Surveillance Team |
| EPI_ISL_3467947 | National Virus Reference Laboratory | National Virus Reference Laboratory | Zoe Yandle, Charlene Bennett, Gabriel Gonzalez, Michael Carr, Jonathan Dean, Cillian F De Gascun |
| EPI_ISL_3467948 | Lighthouse Lab in Milton Keynes | Wellcome Sanger Institute for the COVID-19 Genomics UK (COG-UK) Consortium | The Lighthouse Lab in Milton Keynes and Alex Alderton, Roberto Amato, Jeffrey Barrett, Sonia Goncalves, Ewan Harrison, David K. Jackson, Ian Johnston, Dominic Kwiatkowski, Cordelia Langford, John Sillitoe on behalf of the Wellcome Sanger Institute COVID-19 Surveillance Team |
| EPI_ISL_3467949 | Lighthouse Lab in Glasgow | Wellcome Sanger Institute for the COVID-19 Genomics UK (COG-UK) Consortium | Harper VanSteenhouse, Yumi Kasai, David Gray, Carol Clugston, Anna Dominiczak and Alex Alderton, Roberto Amato, Jeffrey Barrett, Sonia Goncalves, Ewan Harrison, David K. Jackson, Ian Johnston, Dominic Kwiatkowski, Cordelia Langford, John Sillitoe on behalf of the Wellcome Sanger Institute COVID-19 Surveillance Team |
| EPI_ISL_3467950 | Lighthouse Lab in Milton Keynes | Wellcome Sanger Institute for the COVID-19 Genomics UK (COG-UK) Consortium | The Lighthouse Lab in Milton Keynes and Alex Alderton, Roberto Amato, Jeffrey Barrett, Sonia Goncalves, Ewan Harrison, David K. Jackson, Ian Johnston, Dominic Kwiatkowski, Cordelia Langford, John Sillitoe on behalf of the Wellcome Sanger Institute COVID-19 Surveillance Team |
| EPI_ISL_3467951 | National Virus Reference Laboratory | National Virus Reference Laboratory | Zoe Yandle, Charlene Bennett, Gabriel Gonzalez, Michael Carr, Jonathan Dean, Cillian F De Gascun |
| EPI_ISL_3467952, EPI_ISL_3467953 | Lighthouse Lab in Milton Keynes | Wellcome Sanger Institute for the COVID-19 Genomics UK (COG-UK) Consortium | The Lighthouse Lab in Milton Keynes and Alex Alderton, Roberto Amato, Jeffrey Barrett, Sonia Goncalves, Ewan Harrison, David K. Jackson, Ian Johnston, Dominic Kwiatkowski, Cordelia Langford, John Sillitoe on behalf of the Wellcome Sanger Institute COVID-19 Surveillance Team |
| EPI_ISL_3467954 | National Virus Reference Laboratory | National Virus Reference Laboratory | Zoe Yandle, Charlene Bennett, Gabriel Gonzalez, Michael Carr, Jonathan Dean, Cillian F De Gascun |
| EPI_ISL_3467955 | Lighthouse Lab in Milton Keynes | Wellcome Sanger Institute for the COVID-19 Genomics UK (COG-UK) Consortium | The Lighthouse Lab in Milton Keynes and Alex Alderton, Roberto Amato, Jeffrey Barrett, Sonia Goncalves, Ewan Harrison, David K. Jackson, Ian Johnston, Dominic Kwiatkowski, Cordelia Langford, John Sillitoe on behalf of the Wellcome Sanger Institute COVID-19 Surveillance Team |
| EPI_ISL_3467956, EPI_ISL_3467957 | Lighthouse Lab in Glasgow | Wellcome Sanger Institute for the COVID-19 Genomics UK (COG-UK) Consortium | Harper VanSteenhouse, Yumi Kasai, David Gray, Carol Clugston, Anna Dominiczak and Alex Alderton, Roberto Amato, Jeffrey Barrett, Sonia Goncalves, Ewan Harrison, David K. Jackson, Ian Johnston, Dominic Kwiatkowski, Cordelia Langford, John Sillitoe on behalf of the Wellcome Sanger Institute COVID-19 Surveillance Team |

| | | | |
|---|---|---|---|
| EPI_ISL_3467958 | National Virus Reference Laboratory | National Virus Reference Laboratory | Zoe Yandle, Charlene Bennett, Gabriel Gonzalez, Michael Carr, Jonathan Dean, Cillian F De Gascun |
| EPI_ISL_3467959, EPI_ISL_3467960 | Lighthouse Lab in Milton Keynes | Wellcome Sanger Institute for the COVID-19 Genomics UK (COG-UK) Consortium | The Lighthouse Lab in Milton Keynes and Alex Alderton, Roberto Amato, Jeffrey Barrett, Sonia Goncalves, Ewan Harrison, David K. Jackson, Ian Johnston, Dominic Kwiatkowski, Cordelia Langford, John Sillitoe on behalf of the Wellcome Sanger Institute COVID-19 Surveillance Team |
| EPI_ISL_3467961 | National Virus Reference Laboratory | National Virus Reference Laboratory | Zoe Yandle, Charlene Bennett, Gabriel Gonzalez, Michael Carr, Jonathan Dean, Cillian F De Gascun |
| EPI_ISL_3467962, EPI_ISL_3467963 | Lighthouse Lab in Glasgow | Wellcome Sanger Institute for the COVID-19 Genomics UK (COG-UK) Consortium | Harper VanSteenhouse, Yumi Kasai, David Gray, Carol Clugston, Anna Dominiczak and Alex Alderton, Roberto Amato, Jeffrey Barrett, Sonia Goncalves, Ewan Harrison, David K. Jackson, Ian Johnston, Dominic Kwiatkowski, Cordelia Langford, John Sillitoe on behalf of the Wellcome Sanger Institute COVID-19 Surveillance Team |
| EPI_ISL_3467964, EPI_ISL_3467965 | Lighthouse Lab in Milton Keynes | Wellcome Sanger Institute for the COVID-19 Genomics UK (COG-UK) Consortium | The Lighthouse Lab in Milton Keynes and Alex Alderton, Roberto Amato, Jeffrey Barrett, Sonia Goncalves, Ewan Harrison, David K. Jackson, Ian Johnston, Dominic Kwiatkowski, Cordelia Langford, John Sillitoe on behalf of the Wellcome Sanger Institute COVID-19 Surveillance Team |
| EPI_ISL_3467966 | National Virus Reference Laboratory | National Virus Reference Laboratory | Zoe Yandle, Charlene Bennett, Gabriel Gonzalez, Michael Carr, Jonathan Dean, Cillian F De Gascun |
| EPI_ISL_3467967, EPI_ISL_3467968 | Lighthouse Lab in Glasgow | Wellcome Sanger Institute for the COVID-19 Genomics UK (COG-UK) Consortium | Harper VanSteenhouse, Yumi Kasai, David Gray, Carol Clugston, Anna Dominiczak and Alex Alderton, Roberto Amato, Jeffrey Barrett, Sonia Goncalves, Ewan Harrison, David K. Jackson, Ian Johnston, Dominic Kwiatkowski, Cordelia Langford, John Sillitoe on behalf of the Wellcome Sanger Institute COVID-19 Surveillance Team |
| EPI_ISL_3467969 | National Virus Reference Laboratory | National Virus Reference Laboratory | Zoe Yandle, Charlene Bennett, Gabriel Gonzalez, Michael Carr, Jonathan Dean, Cillian F De Gascun |
| EPI_ISL_3467970 | Lighthouse Lab in Milton Keynes | Wellcome Sanger Institute for the COVID-19 Genomics UK (COG-UK) Consortium | The Lighthouse Lab in Milton Keynes and Alex Alderton, Roberto Amato, Jeffrey Barrett, Sonia Goncalves, Ewan Harrison, David K. Jackson, Ian Johnston, Dominic Kwiatkowski, Cordelia Langford, John Sillitoe on behalf of the Wellcome Sanger Institute COVID-19 Surveillance Team |
| EPI_ISL_3467971 | Lighthouse Laboratory Plymouth | Wellcome Sanger Institute for the COVID-19 Genomics UK (COG-UK) Consortium | Lighthouse Laboratory Plymouth and Alex Alderton, Roberto Amato, Jeffrey Barrett, Sonia Goncalves, Ewan Harrison, David K. Jackson, Ian Johnston, Dominic Kwiatkowski, Cordelia Langford, John Sillitoe on behalf of the Wellcome Sanger Institute COVID-19 Surveillance Team |
| EPI_ISL_3467972 | National Virus Reference Laboratory | National Virus Reference Laboratory | Zoe Yandle, Charlene Bennett, Gabriel Gonzalez, Michael Carr, Jonathan Dean, Cillian F De Gascun |
| EPI_ISL_3467973, EPI_ISL_3467974 | Lighthouse Laboratory Plymouth | Wellcome Sanger Institute for the COVID-19 Genomics UK (COG-UK) Consortium | Lighthouse Laboratory Plymouth and Alex Alderton, Roberto Amato, Jeffrey Barrett, Sonia Goncalves, Ewan Harrison, David K. Jackson, Ian Johnston, Dominic Kwiatkowski, Cordelia Langford, John Sillitoe on behalf of the Wellcome Sanger Institute COVID-19 Surveillance Team |
| EPI_ISL_3467975 | National Virus Reference Laboratory | National Virus Reference Laboratory | Zoe Yandle, Charlene Bennett, Gabriel Gonzalez, Michael Carr, Jonathan Dean, Cillian F De Gascun |
| EPI_ISL_3467976, EPI_ISL_3467977, EPI_ISL_3467978 | Lighthouse Laboratory Plymouth | Wellcome Sanger Institute for the COVID-19 Genomics UK (COG-UK) Consortium | Lighthouse Laboratory Plymouth and Alex Alderton, Roberto Amato, Jeffrey Barrett, Sonia Goncalves, Ewan Harrison, David K. Jackson, Ian Johnston, Dominic Kwiatkowski, Cordelia Langford, John Sillitoe on behalf of the Wellcome Sanger Institute COVID-19 Surveillance Team |
| EPI_ISL_3467979 | National Virus Reference Laboratory | National Virus Reference Laboratory | Zoe Yandle, Charlene Bennett, Gabriel Gonzalez, Michael Carr, Jonathan Dean, Cillian F De Gascun |
| EPI_ISL_3467980, EPI_ISL_3467981, EPI_ISL_3467982 | Lighthouse Laboratory Plymouth | Wellcome Sanger Institute for the COVID-19 Genomics UK (COG-UK) Consortium | Lighthouse Laboratory Plymouth and Alex Alderton, Roberto Amato, Jeffrey Barrett, Sonia Goncalves, Ewan Harrison, David K. Jackson, Ian Johnston, Dominic Kwiatkowski, Cordelia Langford, John Sillitoe on behalf of the Wellcome Sanger Institute COVID-19 Surveillance Team |
| EPI_ISL_3467983 | National Virus Reference Laboratory | National Virus Reference Laboratory | Zoe Yandle, Charlene Bennett, Gabriel Gonzalez, Michael Carr, Jonathan Dean, Cillian F De Gascun |
| EPI_ISL_3467984, EPI_ISL_3467985, EPI_ISL_3467986 | Lighthouse Laboratory Plymouth | Wellcome Sanger Institute for the COVID-19 Genomics UK (COG-UK) Consortium | Lighthouse Laboratory Plymouth and Alex Alderton, Roberto Amato, Jeffrey Barrett, Sonia Goncalves, Ewan Harrison, David K. Jackson, Ian Johnston, Dominic Kwiatkowski, Cordelia Langford, John Sillitoe on behalf of the Wellcome Sanger Institute COVID-19 Surveillance Team |
| EPI_ISL_3467987 | National Virus Reference Laboratory | National Virus Reference Laboratory | Zoe Yandle, Charlene Bennett, Gabriel Gonzalez, Michael Carr, Jonathan Dean, Cillian F De Gascun |
| EPI_ISL_3467988, EPI_ISL_3467989 | Lighthouse Laboratory Plymouth | Wellcome Sanger Institute for the COVID-19 Genomics UK (COG-UK) Consortium | Lighthouse Laboratory Plymouth and Alex Alderton, Roberto Amato, Jeffrey Barrett, Sonia Goncalves, Ewan Harrison, David K. Jackson, Ian Johnston, Dominic Kwiatkowski, Cordelia Langford, John Sillitoe on behalf of the Wellcome Sanger Institute COVID-19 Surveillance Team |
| EPI_ISL_3467990 | Health Services Laboratories | Wellcome Sanger Institute for the COVID-19 Genomics UK (COG-UK) Consortium | Health Services Laboratories and Alex Alderton, Roberto Amato, Jeffrey Barrett, Sonia Goncalves, Ewan Harrison, David K. Jackson, Ian Johnston, Dominic Kwiatkowski, Cordelia Langford, John Sillitoe on behalf of the Wellcome Sanger Institute COVID-19 Surveillance Team |
| EPI_ISL_3467991 | National Virus Reference Laboratory | National Virus Reference Laboratory | Zoe Yandle, Charlene Bennett, Gabriel Gonzalez, Michael Carr, Jonathan Dean, Cillian F De Gascun |
| EPI_ISL_3467992, EPI_ISL_3467993, EPI_ISL_3467994 | Lighthouse Laboratory Plymouth | Wellcome Sanger Institute for the COVID-19 Genomics UK (COG-UK) Consortium | Lighthouse Laboratory Plymouth and Alex Alderton, Roberto Amato, Jeffrey Barrett, Sonia Goncalves, Ewan Harrison, David K. Jackson, Ian Johnston, Dominic Kwiatkowski, Cordelia Langford, John Sillitoe on behalf of the Wellcome Sanger Institute COVID-19 Surveillance Team |
| EPI_ISL_3467995 | National Virus Reference Laboratory | National Virus Reference Laboratory | Zoe Yandle, Charlene Bennett, Gabriel Gonzalez, Michael Carr, Jonathan Dean, Cillian F De Gascun |
| EPI_ISL_3467996 | Health Services Laboratories | Wellcome Sanger Institute for the COVID-19 Genomics UK (COG-UK) Consortium | Health Services Laboratories and Alex Alderton, Roberto Amato, Jeffrey Barrett, Sonia Goncalves, Ewan Harrison, David K. Jackson, Ian Johnston, Dominic Kwiatkowski, Cordelia Langford, John Sillitoe on behalf of the Wellcome Sanger Institute COVID-19 Surveillance Team |
| EPI_ISL_3467997, EPI_ISL_3467998 | Lighthouse Laboratory Plymouth | Wellcome Sanger Institute for the COVID-19 Genomics UK (COG-UK) Consortium | Lighthouse Laboratory Plymouth and Alex Alderton, Roberto Amato, Jeffrey Barrett, Sonia Goncalves, Ewan Harrison, David K. Jackson, Ian Johnston, Dominic Kwiatkowski, Cordelia Langford, John Sillitoe on behalf of the Wellcome Sanger Institute COVID-19 Surveillance Team |
| EPI_ISL_3467999 | National Virus Reference Laboratory | National Virus Reference Laboratory | Zoe Yandle, Charlene Bennett, Gabriel Gonzalez, Michael Carr, Jonathan Dean, Cillian F De Gascun |
| EPI_ISL_3468000, EPI_ISL_3468001 | Lighthouse Laboratory Plymouth | Wellcome Sanger Institute for the COVID-19 Genomics UK (COG-UK) Consortium | Lighthouse Laboratory Plymouth and Alex Alderton, Roberto Amato, Jeffrey Barrett, Sonia Goncalves, Ewan Harrison, David K. Jackson, Ian Johnston, Dominic Kwiatkowski, Cordelia Langford, John Sillitoe on behalf of the Wellcome Sanger Institute COVID-19 Surveillance Team |
| EPI_ISL_3468002 | National Virus Reference Laboratory | National Virus Reference Laboratory | Zoe Yandle, Charlene Bennett, Gabriel Gonzalez, Michael Carr, Jonathan Dean, Cillian F De Gascun |
| EPI_ISL_3468003, EPI_ISL_3468004, EPI_ISL_3468005 | Lighthouse Laboratory Plymouth | Wellcome Sanger Institute for the COVID-19 Genomics UK (COG-UK) Consortium | Lighthouse Laboratory Plymouth and Alex Alderton, Roberto Amato, Jeffrey Barrett, Sonia Goncalves, Ewan Harrison, David K. Jackson, Ian Johnston, Dominic Kwiatkowski, Cordelia Langford, John Sillitoe on behalf of the Wellcome Sanger Institute COVID-19 Surveillance Team |
| EPI_ISL_3468006 | National Virus Reference Laboratory | National Virus Reference Laboratory | Zoe Yandle, Charlene Bennett, Gabriel Gonzalez, Michael Carr, Jonathan Dean, Cillian F De Gascun |
| EPI_ISL_3468007, EPI_ISL_3468008 | Lighthouse Laboratory Plymouth | Wellcome Sanger Institute for the COVID-19 Genomics UK (COG-UK) Consortium | Lighthouse Laboratory Plymouth and Alex Alderton, Roberto Amato, Jeffrey Barrett, Sonia Goncalves, Ewan Harrison, David K. Jackson, Ian Johnston, Dominic Kwiatkowski, Cordelia Langford, John Sillitoe on behalf of the Wellcome Sanger Institute COVID-19 Surveillance Team |
| EPI_ISL_3468009 | National Virus Reference Laboratory | National Virus Reference Laboratory | Zoe Yandle, Charlene Bennett, Gabriel Gonzalez, Michael Carr, Jonathan Dean, Cillian F De Gascun |
| EPI_ISL_3468010, EPI_ISL_3468011, EPI_ISL_3468012 | Lighthouse Laboratory Plymouth | Wellcome Sanger Institute for the COVID-19 Genomics UK (COG-UK) Consortium | Lighthouse Laboratory Plymouth and Alex Alderton, Roberto Amato, Jeffrey Barrett, Sonia Goncalves, Ewan Harrison, David K. Jackson, Ian Johnston, Dominic Kwiatkowski, Cordelia Langford, John Sillitoe on behalf of the Wellcome Sanger Institute COVID-19 Surveillance Team |
| EPI_ISL_3468013 | National Virus Reference Laboratory | National Virus Reference Laboratory | Zoe Yandle, Charlene Bennett, Gabriel Gonzalez, Michael Carr, Jonathan Dean, Cillian F De Gascun |
| EPI_ISL_3468014 | Health Services Laboratories | Wellcome Sanger Institute for the COVID-19 Genomics UK (COG-UK) Consortium | Health Services Laboratories and Alex Alderton, Roberto Amato, Jeffrey Barrett, Sonia Goncalves, Ewan Harrison, David K. Jackson, Ian Johnston, Dominic Kwiatkowski, Cordelia Langford, John Sillitoe on behalf of the Wellcome Sanger Institute COVID-19 Surveillance Team |
| EPI_ISL_3468015, EPI_ISL_3468016 | Lighthouse Laboratory Plymouth | Wellcome Sanger Institute for the COVID-19 Genomics UK (COG-UK) Consortium | Lighthouse Laboratory Plymouth and Alex Alderton, Roberto Amato, Jeffrey Barrett, Sonia Goncalves, Ewan Harrison, David K. Jackson, Ian Johnston, Dominic Kwiatkowski, Cordelia Langford, John Sillitoe on behalf of the Wellcome Sanger Institute COVID-19 Surveillance Team |
| EPI_ISL_3468017 | National Virus Reference Laboratory | National Virus Reference Laboratory | Zoe Yandle, Charlene Bennett, Gabriel Gonzalez, Michael Carr, Jonathan Dean, Cillian F De Gascun |
| EPI_ISL_3468018, EPI_ISL_3468019 | Lighthouse Laboratory Plymouth | Wellcome Sanger Institute for the COVID-19 Genomics UK (COG-UK) Consortium | Lighthouse Laboratory Plymouth and Alex Alderton, Roberto Amato, Jeffrey Barrett, Sonia Goncalves, Ewan Harrison, David K. Jackson, Ian Johnston, Dominic Kwiatkowski, Cordelia Langford, John Sillitoe on behalf of the Wellcome Sanger Institute COVID-19 Surveillance Team |
| EPI_ISL_3468020 | National Virus Reference Laboratory | National Virus Reference Laboratory | Zoe Yandle, Charlene Bennett, Gabriel Gonzalez, Michael Carr, Jonathan Dean, Cillian F De Gascun |
| EPI_ISL_3468021, EPI_ISL_3468022 | Lighthouse Laboratory Plymouth | Wellcome Sanger Institute for the COVID-19 Genomics UK (COG-UK) Consortium | Lighthouse Laboratory Plymouth and Alex Alderton, Roberto Amato, Jeffrey Barrett, Sonia Goncalves, Ewan Harrison, David K. Jackson, Ian Johnston, Dominic Kwiatkowski, Cordelia Langford, John Sillitoe on behalf of the Wellcome Sanger Institute COVID-19 Surveillance Team |
| EPI_ISL_3468023 | National Virus Reference Laboratory | National Virus Reference Laboratory | Zoe Yandle, Charlene Bennett, Gabriel Gonzalez, Michael Carr, Jonathan Dean, Cillian F De Gascun |
| EPI_ISL_3468024, EPI_ISL_3468025 | Lighthouse Laboratory Plymouth | Wellcome Sanger Institute for the COVID-19 Genomics UK (COG-UK) Consortium | Lighthouse Laboratory Plymouth and Alex Alderton, Roberto Amato, Jeffrey Barrett, Sonia Goncalves, Ewan Harrison, David K. Jackson, Ian Johnston, Dominic Kwiatkowski, Cordelia Langford, John Sillitoe on behalf of the Wellcome Sanger Institute COVID-19 Surveillance Team |
| EPI_ISL_3468026 | Health Services Laboratories | Wellcome Sanger Institute for the COVID-19 Genomics UK (COG-UK) Consortium | Health Services Laboratories and Alex Alderton, Roberto Amato, Jeffrey Barrett, Sonia Goncalves, Ewan Harrison, David K. Jackson, Ian Johnston, Dominic Kwiatkowski, Cordelia Langford, John Sillitoe on behalf of the Wellcome Sanger Institute COVID-19 Surveillance Team |
| EPI_ISL_3468027 | National Virus Reference Laboratory | National Virus Reference Laboratory | Zoe Yandle, Charlene Bennett, Gabriel Gonzalez, Michael Carr, Jonathan Dean, Cillian F De Gascun |
| EPI_ISL_3468028, EPI_ISL_3468029, EPI_ISL_3468030 | Lighthouse Laboratory Plymouth | Wellcome Sanger Institute for the COVID-19 Genomics UK (COG-UK) Consortium | Lighthouse Laboratory Plymouth and Alex Alderton, Roberto Amato, Jeffrey Barrett, Sonia Goncalves, Ewan Harrison, David K. Jackson, Ian Johnston, Dominic Kwiatkowski, Cordelia Langford, John Sillitoe on behalf of the Wellcome Sanger Institute COVID-19 Surveillance Team |
| EPI_ISL_3468031 | National Virus Reference Laboratory | National Virus Reference Laboratory | Zoe Yandle, Charlene Bennett, Gabriel Gonzalez, Michael Carr, Jonathan Dean, Cillian F De Gascun |
| EPI_ISL_3468032, EPI_ISL_3468033 | Lighthouse Laboratory Plymouth | Wellcome Sanger Institute for the COVID-19 Genomics UK (COG-UK) Consortium | Lighthouse Laboratory Plymouth and Alex Alderton, Roberto Amato, Jeffrey Barrett, Sonia Goncalves, Ewan Harrison, David K. Jackson, Ian Johnston, Dominic Kwiatkowski, Cordelia Langford, John Sillitoe on behalf of the Wellcome Sanger Institute COVID-19 Surveillance Team |
| EPI_ISL_3468034 | National Virus Reference Laboratory | National Virus Reference Laboratory | Zoe Yandle, Charlene Bennett, Gabriel Gonzalez, Michael Carr, Jonathan Dean, Cillian F De Gascun |
| EPI_ISL_3468035, EPI_ISL_3468036, EPI_ISL_3468037 | Lighthouse Laboratory Plymouth | Wellcome Sanger Institute for the COVID-19 Genomics UK (COG-UK) Consortium | Lighthouse Laboratory Plymouth and Alex Alderton, Roberto Amato, Jeffrey Barrett, Sonia Goncalves, Ewan Harrison, David K. Jackson, Ian Johnston, Dominic Kwiatkowski, Cordelia Langford, John Sillitoe on behalf of the Wellcome Sanger Institute COVID-19 Surveillance Team |
| EPI_ISL_3468038 | National Virus Reference Laboratory | National Virus Reference Laboratory | Zoe Yandle, Charlene Bennett, Gabriel Gonzalez, Michael Carr, Jonathan Dean, Cillian F De Gascun |

| | | | |
|---|---|---|---|
| EPI_ISL_3468039, EPI_ISL_3468040 | Lighthouse Laboratory Plymouth | Wellcome Sanger Institute for the COVID-19 Genomics UK (COG-UK) Consortium | Lighthouse Laboratory Plymouth and Alex Alderton, Roberto Amato, Jeffrey Barrett, Sonia Goncalves, Ewan Harrison, David K. Jackson, Ian Johnston, Dominic Kwiatkowski, Cordelia Langford, John Sillitoe on behalf of the Wellcome Sanger Institute COVID-19 Surveillance Team |
| EPI_ISL_3468041 | National Virus Reference Laboratory | National Virus Reference Laboratory | Zoe Yandle, Charlene Bennett, Gabriel Gonzalez, Michael Carr, Jonathan Dean, Cillian F De Gascun |
| EPI_ISL_3468042, EPI_ISL_3468043, EPI_ISL_3468044 | Lighthouse Laboratory Plymouth | Wellcome Sanger Institute for the COVID-19 Genomics UK (COG-UK) Consortium | Lighthouse Laboratory Plymouth and Alex Alderton, Roberto Amato, Jeffrey Barrett, Sonia Goncalves, Ewan Harrison, David K. Jackson, Ian Johnston, Dominic Kwiatkowski, Cordelia Langford, John Sillitoe on behalf of the Wellcome Sanger Institute COVID-19 Surveillance Team |
| EPI_ISL_3468045 | National Virus Reference Laboratory | National Virus Reference Laboratory | Zoe Yandle, Charlene Bennett, Gabriel Gonzalez, Michael Carr, Jonathan Dean, Cillian F De Gascun |
| EPI_ISL_3468046, EPI_ISL_3468047, EPI_ISL_3468048 | Lighthouse Laboratory Plymouth | Wellcome Sanger Institute for the COVID-19 Genomics UK (COG-UK) Consortium | Lighthouse Laboratory Plymouth and Alex Alderton, Roberto Amato, Jeffrey Barrett, Sonia Goncalves, Ewan Harrison, David K. Jackson, Ian Johnston, Dominic Kwiatkowski, Cordelia Langford, John Sillitoe on behalf of the Wellcome Sanger Institute COVID-19 Surveillance Team |
| EPI_ISL_3468049 | National Virus Reference Laboratory | National Virus Reference Laboratory | Zoe Yandle, Charlene Bennett, Gabriel Gonzalez, Michael Carr, Jonathan Dean, Cillian F De Gascun |
| EPI_ISL_3468050, EPI_ISL_3468051 | Lighthouse Laboratory Plymouth | Wellcome Sanger Institute for the COVID-19 Genomics UK (COG-UK) Consortium | Lighthouse Laboratory Plymouth and Alex Alderton, Roberto Amato, Jeffrey Barrett, Sonia Goncalves, Ewan Harrison, David K. Jackson, Ian Johnston, Dominic Kwiatkowski, Cordelia Langford, John Sillitoe on behalf of the Wellcome Sanger Institute COVID-19 Surveillance Team |
| EPI_ISL_3468052 | National Virus Reference Laboratory | National Virus Reference Laboratory | Zoe Yandle, Charlene Bennett, Gabriel Gonzalez, Michael Carr, Jonathan Dean, Cillian F De Gascun |
| EPI_ISL_3468053, EPI_ISL_3468054 | Lighthouse Laboratory Plymouth | Wellcome Sanger Institute for the COVID-19 Genomics UK (COG-UK) Consortium | Lighthouse Laboratory Plymouth and Alex Alderton, Roberto Amato, Jeffrey Barrett, Sonia Goncalves, Ewan Harrison, David K. Jackson, Ian Johnston, Dominic Kwiatkowski, Cordelia Langford, John Sillitoe on behalf of the Wellcome Sanger Institute COVID-19 Surveillance Team |
| EPI_ISL_3468055 | National Virus Reference Laboratory | National Virus Reference Laboratory | Zoe Yandle, Charlene Bennett, Gabriel Gonzalez, Michael Carr, Jonathan Dean, Cillian F De Gascun |
| EPI_ISL_3468056, EPI_ISL_3468057 | Lighthouse Laboratory Plymouth | Wellcome Sanger Institute for the COVID-19 Genomics UK (COG-UK) Consortium | Lighthouse Laboratory Plymouth and Alex Alderton, Roberto Amato, Jeffrey Barrett, Sonia Goncalves, Ewan Harrison, David K. Jackson, Ian Johnston, Dominic Kwiatkowski, Cordelia Langford, John Sillitoe on behalf of the Wellcome Sanger Institute COVID-19 Surveillance Team |
| EPI_ISL_3468058 | National Virus Reference Laboratory | National Virus Reference Laboratory | Zoe Yandle, Charlene Bennett, Gabriel Gonzalez, Michael Carr, Jonathan Dean, Cillian F De Gascun |
| EPI_ISL_3468059, EPI_ISL_3468060, EPI_ISL_3468061 | Lighthouse Laboratory Plymouth | Wellcome Sanger Institute for the COVID-19 Genomics UK (COG-UK) Consortium | Lighthouse Laboratory Plymouth and Alex Alderton, Roberto Amato, Jeffrey Barrett, Sonia Goncalves, Ewan Harrison, David K. Jackson, Ian Johnston, Dominic Kwiatkowski, Cordelia Langford, John Sillitoe on behalf of the Wellcome Sanger Institute COVID-19 Surveillance Team |
| EPI_ISL_3468062 | National Virus Reference Laboratory | National Virus Reference Laboratory | Zoe Yandle, Charlene Bennett, Gabriel Gonzalez, Michael Carr, Jonathan Dean, Cillian F De Gascun |
| EPI_ISL_3468063, EPI_ISL_3468064, EPI_ISL_3468065 | Lighthouse Laboratory Plymouth | Wellcome Sanger Institute for the COVID-19 Genomics UK (COG-UK) Consortium | Lighthouse Laboratory Plymouth and Alex Alderton, Roberto Amato, Jeffrey Barrett, Sonia Goncalves, Ewan Harrison, David K. Jackson, Ian Johnston, Dominic Kwiatkowski, Cordelia Langford, John Sillitoe on behalf of the Wellcome Sanger Institute COVID-19 Surveillance Team |
| EPI_ISL_3468066 | National Virus Reference Laboratory | National Virus Reference Laboratory | Zoe Yandle, Charlene Bennett, Gabriel Gonzalez, Michael Carr, Jonathan Dean, Cillian F De Gascun |
| EPI_ISL_3468067, EPI_ISL_3468068 | Lighthouse Laboratory Plymouth | Wellcome Sanger Institute for the COVID-19 Genomics UK (COG-UK) Consortium | Lighthouse Laboratory Plymouth and Alex Alderton, Roberto Amato, Jeffrey Barrett, Sonia Goncalves, Ewan Harrison, David K. Jackson, Ian Johnston, Dominic Kwiatkowski, Cordelia Langford, John Sillitoe on behalf of the Wellcome Sanger Institute COVID-19 Surveillance Team |
| EPI_ISL_3468069 | National Virus Reference Laboratory | National Virus Reference Laboratory | Zoe Yandle, Charlene Bennett, Gabriel Gonzalez, Michael Carr, Jonathan Dean, Cillian F De Gascun |
| EPI_ISL_3468070, EPI_ISL_3468071 | Lighthouse Laboratory Plymouth | Wellcome Sanger Institute for the COVID-19 Genomics UK (COG-UK) Consortium | Lighthouse Laboratory Plymouth and Alex Alderton, Roberto Amato, Jeffrey Barrett, Sonia Goncalves, Ewan Harrison, David K. Jackson, Ian Johnston, Dominic Kwiatkowski, Cordelia Langford, John Sillitoe on behalf of the Wellcome Sanger Institute COVID-19 Surveillance Team |
| EPI_ISL_3468072 | National Virus Reference Laboratory | National Virus Reference Laboratory | Zoe Yandle, Charlene Bennett, Gabriel Gonzalez, Michael Carr, Jonathan Dean, Cillian F De Gascun |
| EPI_ISL_3468073 | Health Services Laboratories | Wellcome Sanger Institute for the COVID-19 Genomics UK (COG-UK) Consortium | Health Services Laboratories and Alex Alderton, Roberto Amato, Jeffrey Barrett, Sonia Goncalves, Ewan Harrison, David K. Jackson, Ian Johnston, Dominic Kwiatkowski, Cordelia Langford, John Sillitoe on behalf of the Wellcome Sanger Institute COVID-19 Surveillance Team |
| EPI_ISL_3468074, EPI_ISL_3468075 | Lighthouse Laboratory Plymouth | Wellcome Sanger Institute for the COVID-19 Genomics UK (COG-UK) Consortium | Lighthouse Laboratory Plymouth and Alex Alderton, Roberto Amato, Jeffrey Barrett, Sonia Goncalves, Ewan Harrison, David K. Jackson, Ian Johnston, Dominic Kwiatkowski, Cordelia Langford, John Sillitoe on behalf of the Wellcome Sanger Institute COVID-19 Surveillance Team |
| EPI_ISL_3468076 | National Virus Reference Laboratory | National Virus Reference Laboratory | Zoe Yandle, Charlene Bennett, Gabriel Gonzalez, Michael Carr, Jonathan Dean, Cillian F De Gascun |
| EPI_ISL_3468077, EPI_ISL_3468078 | Lighthouse Laboratory Plymouth | Wellcome Sanger Institute for the COVID-19 Genomics UK (COG-UK) Consortium | Lighthouse Laboratory Plymouth and Alex Alderton, Roberto Amato, Jeffrey Barrett, Sonia Goncalves, Ewan Harrison, David K. Jackson, Ian Johnston, Dominic Kwiatkowski, Cordelia Langford, John Sillitoe on behalf of the Wellcome Sanger Institute COVID-19 Surveillance Team |
| EPI_ISL_3468079 | Health Services Laboratories | Wellcome Sanger Institute for the COVID-19 Genomics UK (COG-UK) Consortium | Health Services Laboratories and Alex Alderton, Roberto Amato, Jeffrey Barrett, Sonia Goncalves, Ewan Harrison, David K. Jackson, Ian Johnston, Dominic Kwiatkowski, Cordelia Langford, John Sillitoe on behalf of the Wellcome Sanger Institute COVID-19 Surveillance Team |
| EPI_ISL_3468080 | National Virus Reference Laboratory | National Virus Reference Laboratory | Zoe Yandle, Charlene Bennett, Gabriel Gonzalez, Michael Carr, Jonathan Dean, Cillian F De Gascun |
| EPI_ISL_3468081, EPI_ISL_3468082 | Lighthouse Laboratory Plymouth | Wellcome Sanger Institute for the COVID-19 Genomics UK (COG-UK) Consortium | Lighthouse Laboratory Plymouth and Alex Alderton, Roberto Amato, Jeffrey Barrett, Sonia Goncalves, Ewan Harrison, David K. Jackson, Ian Johnston, Dominic Kwiatkowski, Cordelia Langford, John Sillitoe on behalf of the Wellcome Sanger Institute COVID-19 Surveillance Team |
| EPI_ISL_3468083 | National Virus Reference Laboratory | National Virus Reference Laboratory | Zoe Yandle, Charlene Bennett, Gabriel Gonzalez, Michael Carr, Jonathan Dean, Cillian F De Gascun |
| EPI_ISL_3468084, EPI_ISL_3468085, EPI_ISL_3468086 | Lighthouse Laboratory Plymouth | Wellcome Sanger Institute for the COVID-19 Genomics UK (COG-UK) Consortium | Lighthouse Laboratory Plymouth and Alex Alderton, Roberto Amato, Jeffrey Barrett, Sonia Goncalves, Ewan Harrison, David K. Jackson, Ian Johnston, Dominic Kwiatkowski, Cordelia Langford, John Sillitoe on behalf of the Wellcome Sanger Institute COVID-19 Surveillance Team |
| EPI_ISL_3468087 | National Virus Reference Laboratory | National Virus Reference Laboratory | Zoe Yandle, Charlene Bennett, Gabriel Gonzalez, Michael Carr, Jonathan Dean, Cillian F De Gascun |
| EPI_ISL_3468088, EPI_ISL_3468089 | Lighthouse Laboratory Plymouth | Wellcome Sanger Institute for the COVID-19 Genomics UK (COG-UK) Consortium | Lighthouse Laboratory Plymouth and Alex Alderton, Roberto Amato, Jeffrey Barrett, Sonia Goncalves, Ewan Harrison, David K. Jackson, Ian Johnston, Dominic Kwiatkowski, Cordelia Langford, John Sillitoe on behalf of the Wellcome Sanger Institute COVID-19 Surveillance Team |
| EPI_ISL_3468090 | National Virus Reference Laboratory | National Virus Reference Laboratory | Zoe Yandle, Charlene Bennett, Gabriel Gonzalez, Michael Carr, Jonathan Dean, Cillian F De Gascun |
| EPI_ISL_3468091, EPI_ISL_3468092, EPI_ISL_3468093 | Lighthouse Laboratory Plymouth | Wellcome Sanger Institute for the COVID-19 Genomics UK (COG-UK) Consortium | Lighthouse Laboratory Plymouth and Alex Alderton, Roberto Amato, Jeffrey Barrett, Sonia Goncalves, Ewan Harrison, David K. Jackson, Ian Johnston, Dominic Kwiatkowski, Cordelia Langford, John Sillitoe on behalf of the Wellcome Sanger Institute COVID-19 Surveillance Team |
| EPI_ISL_3468094 | National Virus Reference Laboratory | National Virus Reference Laboratory | Zoe Yandle, Charlene Bennett, Gabriel Gonzalez, Michael Carr, Jonathan Dean, Cillian F De Gascun |
| EPI_ISL_3468095, EPI_ISL_3468096, EPI_ISL_3468097 | Lighthouse Laboratory Plymouth | Wellcome Sanger Institute for the COVID-19 Genomics UK (COG-UK) Consortium | Lighthouse Laboratory Plymouth and Alex Alderton, Roberto Amato, Jeffrey Barrett, Sonia Goncalves, Ewan Harrison, David K. Jackson, Ian Johnston, Dominic Kwiatkowski, Cordelia Langford, John Sillitoe on behalf of the Wellcome Sanger Institute COVID-19 Surveillance Team |
| EPI_ISL_3468098 | National Virus Reference Laboratory | National Virus Reference Laboratory | Zoe Yandle, Charlene Bennett, Gabriel Gonzalez, Michael Carr, Jonathan Dean, Cillian F De Gascun |
| EPI_ISL_3468099, EPI_ISL_3468100 | Lighthouse Laboratory Plymouth | Wellcome Sanger Institute for the COVID-19 Genomics UK (COG-UK) Consortium | Lighthouse Laboratory Plymouth and Alex Alderton, Roberto Amato, Jeffrey Barrett, Sonia Goncalves, Ewan Harrison, David K. Jackson, Ian Johnston, Dominic Kwiatkowski, Cordelia Langford, John Sillitoe on behalf of the Wellcome Sanger Institute COVID-19 Surveillance Team |
| EPI_ISL_3468101 | National Virus Reference Laboratory | National Virus Reference Laboratory | Zoe Yandle, Charlene Bennett, Gabriel Gonzalez, Michael Carr, Jonathan Dean, Cillian F De Gascun |
| EPI_ISL_3468102, EPI_ISL_3468103 | Lighthouse Laboratory Plymouth | Wellcome Sanger Institute for the COVID-19 Genomics UK (COG-UK) Consortium | Lighthouse Laboratory Plymouth and Alex Alderton, Roberto Amato, Jeffrey Barrett, Sonia Goncalves, Ewan Harrison, David K. Jackson, Ian Johnston, Dominic Kwiatkowski, Cordelia Langford, John Sillitoe on behalf of the Wellcome Sanger Institute COVID-19 Surveillance Team |
| EPI_ISL_3468104 | National Virus Reference Laboratory | National Virus Reference Laboratory | Zoe Yandle, Charlene Bennett, Gabriel Gonzalez, Michael Carr, Jonathan Dean, Cillian F De Gascun |
| EPI_ISL_3468105, EPI_ISL_3468106, EPI_ISL_3468107 | Lighthouse Laboratory Plymouth | Wellcome Sanger Institute for the COVID-19 Genomics UK (COG-UK) Consortium | Lighthouse Laboratory Plymouth and Alex Alderton, Roberto Amato, Jeffrey Barrett, Sonia Goncalves, Ewan Harrison, David K. Jackson, Ian Johnston, Dominic Kwiatkowski, Cordelia Langford, John Sillitoe on behalf of the Wellcome Sanger Institute COVID-19 Surveillance Team |
| EPI_ISL_3468108 | National Virus Reference Laboratory | National Virus Reference Laboratory | Zoe Yandle, Charlene Bennett, Gabriel Gonzalez, Michael Carr, Jonathan Dean, Cillian F De Gascun |
| EPI_ISL_3468109 | Lighthouse Laboratory Plymouth | Wellcome Sanger Institute for the COVID-19 Genomics UK (COG-UK) Consortium | Lighthouse Laboratory Plymouth and Alex Alderton, Roberto Amato, Jeffrey Barrett, Sonia Goncalves, Ewan Harrison, David K. Jackson, Ian Johnston, Dominic Kwiatkowski, Cordelia Langford, John Sillitoe on behalf of the Wellcome Sanger Institute COVID-19 Surveillance Team |
| EPI_ISL_3468110 | Health Services Laboratories | Wellcome Sanger Institute for the COVID-19 Genomics UK (COG-UK) Consortium | Health Services Laboratories and Alex Alderton, Roberto Amato, Jeffrey Barrett, Sonia Goncalves, Ewan Harrison, David K. Jackson, Ian Johnston, Dominic Kwiatkowski, Cordelia Langford, John Sillitoe on behalf of the Wellcome Sanger Institute COVID-19 Surveillance Team |
| EPI_ISL_3468111 | Lighthouse Laboratory Plymouth | Wellcome Sanger Institute for the COVID-19 Genomics UK (COG-UK) Consortium | Lighthouse Laboratory Plymouth and Alex Alderton, Roberto Amato, Jeffrey Barrett, Sonia Goncalves, Ewan Harrison, David K. Jackson, Ian Johnston, Dominic Kwiatkowski, Cordelia Langford, John Sillitoe on behalf of the Wellcome Sanger Institute COVID-19 Surveillance Team |
| EPI_ISL_3468112 | National Virus Reference Laboratory | National Virus Reference Laboratory | Zoe Yandle, Charlene Bennett, Gabriel Gonzalez, Michael Carr, Jonathan Dean, Cillian F De Gascun |
| EPI_ISL_3468113, EPI_ISL_3468114 | Lighthouse Laboratory Plymouth | Wellcome Sanger Institute for the COVID-19 Genomics UK (COG-UK) Consortium | Lighthouse Laboratory Plymouth and Alex Alderton, Roberto Amato, Jeffrey Barrett, Sonia Goncalves, Ewan Harrison, David K. Jackson, Ian Johnston, Dominic Kwiatkowski, Cordelia Langford, John Sillitoe on behalf of the Wellcome Sanger Institute COVID-19 Surveillance Team |
| EPI_ISL_3468115 | National Virus Reference Laboratory | National Virus Reference Laboratory | Zoe Yandle, Charlene Bennett, Gabriel Gonzalez, Michael Carr, Jonathan Dean, Cillian F De Gascun |
| EPI_ISL_3468116, EPI_ISL_3468117 | Lighthouse Laboratory Plymouth | Wellcome Sanger Institute for the COVID-19 Genomics UK (COG-UK) Consortium | Lighthouse Laboratory Plymouth and Alex Alderton, Roberto Amato, Jeffrey Barrett, Sonia Goncalves, Ewan Harrison, David K. Jackson, Ian Johnston, Dominic Kwiatkowski, Cordelia Langford, John Sillitoe on behalf of the Wellcome Sanger Institute COVID-19 Surveillance Team |
| EPI_ISL_3468118 | Health Services Laboratories | Wellcome Sanger Institute for the COVID-19 Genomics UK (COG-UK) Consortium | Health Services Laboratories and Alex Alderton, Roberto Amato, Jeffrey Barrett, Sonia Goncalves, Ewan Harrison, David K. Jackson, Ian Johnston, Dominic Kwiatkowski, Cordelia Langford, John Sillitoe on behalf of the Wellcome Sanger Institute COVID-19 Surveillance Team |
| EPI_ISL_3468119 | National Virus Reference Laboratory | National Virus Reference Laboratory | Zoe Yandle, Charlene Bennett, Gabriel Gonzalez, Michael Carr, Jonathan Dean, Cillian F De Gascun |

| | | | |
|---|---|---|---|
| EPI_ISL_3468120, EPI_ISL_3468121 | Lighthouse Laboratory Plymouth | Wellcome Sanger Institute for the COVID-19 Genomics UK (COG-UK) Consortium | Lighthouse Laboratory Plymouth and Alex Alderton, Roberto Amato, Jeffrey Barrett, Sonia Goncalves, Ewan Harrison, David K. Jackson, Ian Johnston, Dominic Kwiatkowski, Cordelia Langford, John Sillitoe on behalf of the Wellcome Sanger Institute COVID-19 Surveillance Team |
| EPI_ISL_3468122 | National Virus Reference Laboratory | National Virus Reference Laboratory | Zoe Yandle, Charlene Bennett, Gabriel Gonzalez, Michael Carr, Jonathan Dean, Cillian F De Gascun |
| EPI_ISL_3468123 | Lighthouse Laboratory Plymouth | Wellcome Sanger Institute for the COVID-19 Genomics UK (COG-UK) Consortium | Lighthouse Laboratory Plymouth and Alex Alderton, Roberto Amato, Jeffrey Barrett, Sonia Goncalves, Ewan Harrison, David K. Jackson, Ian Johnston, Dominic Kwiatkowski, Cordelia Langford, John Sillitoe on behalf of the Wellcome Sanger Institute COVID-19 Surveillance Team |
| EPI_ISL_3468124 | Health Services Laboratories | Wellcome Sanger Institute for the COVID-19 Genomics UK (COG-UK) Consortium | Health Services Laboratories and Alex Alderton, Roberto Amato, Jeffrey Barrett, Sonia Goncalves, Ewan Harrison, David K. Jackson, Ian Johnston, Dominic Kwiatkowski, Cordelia Langford, John Sillitoe on behalf of the Wellcome Sanger Institute COVID-19 Surveillance Team |
| EPI_ISL_3468125 | Lighthouse Laboratory Plymouth | Wellcome Sanger Institute for the COVID-19 Genomics UK (COG-UK) Consortium | Lighthouse Laboratory Plymouth and Alex Alderton, Roberto Amato, Jeffrey Barrett, Sonia Goncalves, Ewan Harrison, David K. Jackson, Ian Johnston, Dominic Kwiatkowski, Cordelia Langford, John Sillitoe on behalf of the Wellcome Sanger Institute COVID-19 Surveillance Team |
| EPI_ISL_3468126 | National Virus Reference Laboratory | National Virus Reference Laboratory | Zoe Yandle, Charlene Bennett, Gabriel Gonzalez, Michael Carr, Jonathan Dean, Cillian F De Gascun |
| EPI_ISL_3468127 | Health Services Laboratories | Wellcome Sanger Institute for the COVID-19 Genomics UK (COG-UK) Consortium | Health Services Laboratories and Alex Alderton, Roberto Amato, Jeffrey Barrett, Sonia Goncalves, Ewan Harrison, David K. Jackson, Ian Johnston, Dominic Kwiatkowski, Cordelia Langford, John Sillitoe on behalf of the Wellcome Sanger Institute COVID-19 Surveillance Team |
| EPI_ISL_3468128, EPI_ISL_3468129 | Lighthouse Laboratory Plymouth | Wellcome Sanger Institute for the COVID-19 Genomics UK (COG-UK) Consortium | Lighthouse Laboratory Plymouth and Alex Alderton, Roberto Amato, Jeffrey Barrett, Sonia Goncalves, Ewan Harrison, David K. Jackson, Ian Johnston, Dominic Kwiatkowski, Cordelia Langford, John Sillitoe on behalf of the Wellcome Sanger Institute COVID-19 Surveillance Team |
| EPI_ISL_3468130 | National Virus Reference Laboratory | National Virus Reference Laboratory | Zoe Yandle, Charlene Bennett, Gabriel Gonzalez, Michael Carr, Jonathan Dean, Cillian F De Gascun |
| EPI_ISL_3468131, EPI_ISL_3468132 | Lighthouse Laboratory Plymouth | Wellcome Sanger Institute for the COVID-19 Genomics UK (COG-UK) Consortium | Lighthouse Laboratory Plymouth and Alex Alderton, Roberto Amato, Jeffrey Barrett, Sonia Goncalves, Ewan Harrison, David K. Jackson, Ian Johnston, Dominic Kwiatkowski, Cordelia Langford, John Sillitoe on behalf of the Wellcome Sanger Institute COVID-19 Surveillance Team |
| EPI_ISL_3468133 | National Virus Reference Laboratory | National Virus Reference Laboratory | Zoe Yandle, Charlene Bennett, Gabriel Gonzalez, Michael Carr, Jonathan Dean, Cillian F De Gascun |
| EPI_ISL_3468134, EPI_ISL_3468135, EPI_ISL_3468136 | Lighthouse Laboratory Plymouth | Wellcome Sanger Institute for the COVID-19 Genomics UK (COG-UK) Consortium | Lighthouse Laboratory Plymouth and Alex Alderton, Roberto Amato, Jeffrey Barrett, Sonia Goncalves, Ewan Harrison, David K. Jackson, Ian Johnston, Dominic Kwiatkowski, Cordelia Langford, John Sillitoe on behalf of the Wellcome Sanger Institute COVID-19 Surveillance Team |
| EPI_ISL_3468137 | National Virus Reference Laboratory | National Virus Reference Laboratory | Zoe Yandle, Charlene Bennett, Gabriel Gonzalez, Michael Carr, Jonathan Dean, Cillian F De Gascun |
| EPI_ISL_3468138, EPI_ISL_3468139 | Lighthouse Laboratory Plymouth | Wellcome Sanger Institute for the COVID-19 Genomics UK (COG-UK) Consortium | Lighthouse Laboratory Plymouth and Alex Alderton, Roberto Amato, Jeffrey Barrett, Sonia Goncalves, Ewan Harrison, David K. Jackson, Ian Johnston, Dominic Kwiatkowski, Cordelia Langford, John Sillitoe on behalf of the Wellcome Sanger Institute COVID-19 Surveillance Team |
| EPI_ISL_3468140 | National Virus Reference Laboratory | National Virus Reference Laboratory | Zoe Yandle, Charlene Bennett, Gabriel Gonzalez, Michael Carr, Jonathan Dean, Cillian F De Gascun |
| EPI_ISL_3468141, EPI_ISL_3468142 | Lighthouse Laboratory Plymouth | Wellcome Sanger Institute for the COVID-19 Genomics UK (COG-UK) Consortium | Lighthouse Laboratory Plymouth and Alex Alderton, Roberto Amato, Jeffrey Barrett, Sonia Goncalves, Ewan Harrison, David K. Jackson, Ian Johnston, Dominic Kwiatkowski, Cordelia Langford, John Sillitoe on behalf of the Wellcome Sanger Institute COVID-19 Surveillance Team |
| EPI_ISL_3468143 | Health Services Laboratories | Wellcome Sanger Institute for the COVID-19 Genomics UK (COG-UK) Consortium | Health Services Laboratories and Alex Alderton, Roberto Amato, Jeffrey Barrett, Sonia Goncalves, Ewan Harrison, David K. Jackson, Ian Johnston, Dominic Kwiatkowski, Cordelia Langford, John Sillitoe on behalf of the Wellcome Sanger Institute COVID-19 Surveillance Team |
| EPI_ISL_3468144 | National Virus Reference Laboratory | National Virus Reference Laboratory | Zoe Yandle, Charlene Bennett, Gabriel Gonzalez, Michael Carr, Jonathan Dean, Cillian F De Gascun |
| EPI_ISL_3468145, EPI_ISL_3468146 | Lighthouse Laboratory Plymouth | Wellcome Sanger Institute for the COVID-19 Genomics UK (COG-UK) Consortium | Lighthouse Laboratory Plymouth and Alex Alderton, Roberto Amato, Jeffrey Barrett, Sonia Goncalves, Ewan Harrison, David K. Jackson, Ian Johnston, Dominic Kwiatkowski, Cordelia Langford, John Sillitoe on behalf of the Wellcome Sanger Institute COVID-19 Surveillance Team |
| EPI_ISL_3468147 | National Virus Reference Laboratory | National Virus Reference Laboratory | Zoe Yandle, Charlene Bennett, Gabriel Gonzalez, Michael Carr, Jonathan Dean, Cillian F De Gascun |
| EPI_ISL_3468148, EPI_ISL_3468149 | Health Services Laboratories | Wellcome Sanger Institute for the COVID-19 Genomics UK (COG-UK) Consortium | Health Services Laboratories and Alex Alderton, Roberto Amato, Jeffrey Barrett, Sonia Goncalves, Ewan Harrison, David K. Jackson, Ian Johnston, Dominic Kwiatkowski, Cordelia Langford, John Sillitoe on behalf of the Wellcome Sanger Institute COVID-19 Surveillance Team |
| EPI_ISL_3468150 | Lighthouse Laboratory Plymouth | Wellcome Sanger Institute for the COVID-19 Genomics UK (COG-UK) Consortium | Lighthouse Laboratory Plymouth and Alex Alderton, Roberto Amato, Jeffrey Barrett, Sonia Goncalves, Ewan Harrison, David K. Jackson, Ian Johnston, Dominic Kwiatkowski, Cordelia Langford, John Sillitoe on behalf of the Wellcome Sanger Institute COVID-19 Surveillance Team |
| EPI_ISL_3468151 | National Virus Reference Laboratory | National Virus Reference Laboratory | Zoe Yandle, Charlene Bennett, Gabriel Gonzalez, Michael Carr, Jonathan Dean, Cillian F De Gascun |
| EPI_ISL_3468152 | Health Services Laboratories | Wellcome Sanger Institute for the COVID-19 Genomics UK (COG-UK) Consortium | Health Services Laboratories and Alex Alderton, Roberto Amato, Jeffrey Barrett, Sonia Goncalves, Ewan Harrison, David K. Jackson, Ian Johnston, Dominic Kwiatkowski, Cordelia Langford, John Sillitoe on behalf of the Wellcome Sanger Institute COVID-19 Surveillance Team |
| EPI_ISL_3468153 | Lighthouse Laboratory Plymouth | Wellcome Sanger Institute for the COVID-19 Genomics UK (COG-UK) Consortium | Lighthouse Laboratory Plymouth and Alex Alderton, Roberto Amato, Jeffrey Barrett, Sonia Goncalves, Ewan Harrison, David K. Jackson, Ian Johnston, Dominic Kwiatkowski, Cordelia Langford, John Sillitoe on behalf of the Wellcome Sanger Institute COVID-19 Surveillance Team |
| EPI_ISL_3468154 | National Virus Reference Laboratory | National Virus Reference Laboratory | Zoe Yandle, Charlene Bennett, Gabriel Gonzalez, Michael Carr, Jonathan Dean, Cillian F De Gascun |
| EPI_ISL_3468155, EPI_ISL_3468156, EPI_ISL_3468157 | Lighthouse Laboratory Plymouth | Wellcome Sanger Institute for the COVID-19 Genomics UK (COG-UK) Consortium | Lighthouse Laboratory Plymouth and Alex Alderton, Roberto Amato, Jeffrey Barrett, Sonia Goncalves, Ewan Harrison, David K. Jackson, Ian Johnston, Dominic Kwiatkowski, Cordelia Langford, John Sillitoe on behalf of the Wellcome Sanger Institute COVID-19 Surveillance Team |
| EPI_ISL_3468158 | National Virus Reference Laboratory | National Virus Reference Laboratory | Zoe Yandle, Charlene Bennett, Gabriel Gonzalez, Michael Carr, Jonathan Dean, Cillian F De Gascun |
| EPI_ISL_3468159, EPI_ISL_3468160, EPI_ISL_3468161 | Lighthouse Laboratory Plymouth | Wellcome Sanger Institute for the COVID-19 Genomics UK (COG-UK) Consortium | Lighthouse Laboratory Plymouth and Alex Alderton, Roberto Amato, Jeffrey Barrett, Sonia Goncalves, Ewan Harrison, David K. Jackson, Ian Johnston, Dominic Kwiatkowski, Cordelia Langford, John Sillitoe on behalf of the Wellcome Sanger Institute COVID-19 Surveillance Team |
| EPI_ISL_3468162 | National Virus Reference Laboratory | National Virus Reference Laboratory | Zoe Yandle, Charlene Bennett, Gabriel Gonzalez, Michael Carr, Jonathan Dean, Cillian F De Gascun |
| EPI_ISL_3468163, EPI_ISL_3468164 | Lighthouse Laboratory Plymouth | Wellcome Sanger Institute for the COVID-19 Genomics UK (COG-UK) Consortium | Lighthouse Laboratory Plymouth and Alex Alderton, Roberto Amato, Jeffrey Barrett, Sonia Goncalves, Ewan Harrison, David K. Jackson, Ian Johnston, Dominic Kwiatkowski, Cordelia Langford, John Sillitoe on behalf of the Wellcome Sanger Institute COVID-19 Surveillance Team |
| EPI_ISL_3468165 | National Virus Reference Laboratory | National Virus Reference Laboratory | Zoe Yandle, Charlene Bennett, Gabriel Gonzalez, Michael Carr, Jonathan Dean, Cillian F De Gascun |
| EPI_ISL_3468166, EPI_ISL_3468167, EPI_ISL_3468168 | Lighthouse Laboratory Plymouth | Wellcome Sanger Institute for the COVID-19 Genomics UK (COG-UK) Consortium | Lighthouse Laboratory Plymouth and Alex Alderton, Roberto Amato, Jeffrey Barrett, Sonia Goncalves, Ewan Harrison, David K. Jackson, Ian Johnston, Dominic Kwiatkowski, Cordelia Langford, John Sillitoe on behalf of the Wellcome Sanger Institute COVID-19 Surveillance Team |
| EPI_ISL_3468169 | National Virus Reference Laboratory | National Virus Reference Laboratory | Zoe Yandle, Charlene Bennett, Gabriel Gonzalez, Michael Carr, Jonathan Dean, Cillian F De Gascun |
| EPI_ISL_3468170, EPI_ISL_3468171 | Lighthouse Laboratory Plymouth | Wellcome Sanger Institute for the COVID-19 Genomics UK (COG-UK) Consortium | Lighthouse Laboratory Plymouth and Alex Alderton, Roberto Amato, Jeffrey Barrett, Sonia Goncalves, Ewan Harrison, David K. Jackson, Ian Johnston, Dominic Kwiatkowski, Cordelia Langford, John Sillitoe on behalf of the Wellcome Sanger Institute COVID-19 Surveillance Team |
| EPI_ISL_3468172 | National Virus Reference Laboratory | National Virus Reference Laboratory | Zoe Yandle, Charlene Bennett, Gabriel Gonzalez, Michael Carr, Jonathan Dean, Cillian F De Gascun |
| EPI_ISL_3468173 | Health Services Laboratories | Wellcome Sanger Institute for the COVID-19 Genomics UK (COG-UK) Consortium | Health Services Laboratories and Alex Alderton, Roberto Amato, Jeffrey Barrett, Sonia Goncalves, Ewan Harrison, David K. Jackson, Ian Johnston, Dominic Kwiatkowski, Cordelia Langford, John Sillitoe on behalf of the Wellcome Sanger Institute COVID-19 Surveillance Team |
| EPI_ISL_3468174 | Lighthouse Laboratory Plymouth | Wellcome Sanger Institute for the COVID-19 Genomics UK (COG-UK) Consortium | Lighthouse Laboratory Plymouth and Alex Alderton, Roberto Amato, Jeffrey Barrett, Sonia Goncalves, Ewan Harrison, David K. Jackson, Ian Johnston, Dominic Kwiatkowski, Cordelia Langford, John Sillitoe on behalf of the Wellcome Sanger Institute COVID-19 Surveillance Team |
| EPI_ISL_3468175 | National Virus Reference Laboratory | National Virus Reference Laboratory | Zoe Yandle, Charlene Bennett, Gabriel Gonzalez, Michael Carr, Jonathan Dean, Cillian F De Gascun |
| EPI_ISL_3468176, EPI_ISL_3468177, EPI_ISL_3468178 | Lighthouse Laboratory Plymouth | Wellcome Sanger Institute for the COVID-19 Genomics UK (COG-UK) Consortium | Lighthouse Laboratory Plymouth and Alex Alderton, Roberto Amato, Jeffrey Barrett, Sonia Goncalves, Ewan Harrison, David K. Jackson, Ian Johnston, Dominic Kwiatkowski, Cordelia Langford, John Sillitoe on behalf of the Wellcome Sanger Institute COVID-19 Surveillance Team |
| EPI_ISL_3468179 | National Virus Reference Laboratory | National Virus Reference Laboratory | Zoe Yandle, Charlene Bennett, Gabriel Gonzalez, Michael Carr, Jonathan Dean, Cillian F De Gascun |
| EPI_ISL_3468180 | Lighthouse Laboratory Plymouth | Wellcome Sanger Institute for the COVID-19 Genomics UK (COG-UK) Consortium | Lighthouse Laboratory Plymouth and Alex Alderton, Roberto Amato, Jeffrey Barrett, Sonia Goncalves, Ewan Harrison, David K. Jackson, Ian Johnston, Dominic Kwiatkowski, Cordelia Langford, John Sillitoe on behalf of the Wellcome Sanger Institute COVID-19 Surveillance Team |
| EPI_ISL_3468181 | Health Services Laboratories | Wellcome Sanger Institute for the COVID-19 Genomics UK (COG-UK) Consortium | Health Services Laboratories and Alex Alderton, Roberto Amato, Jeffrey Barrett, Sonia Goncalves, Ewan Harrison, David K. Jackson, Ian Johnston, Dominic Kwiatkowski, Cordelia Langford, John Sillitoe on behalf of the Wellcome Sanger Institute COVID-19 Surveillance Team |
| EPI_ISL_3468182, EPI_ISL_3468183 | Lighthouse Laboratory Plymouth | Wellcome Sanger Institute for the COVID-19 Genomics UK (COG-UK) Consortium | Lighthouse Laboratory Plymouth and Alex Alderton, Roberto Amato, Jeffrey Barrett, Sonia Goncalves, Ewan Harrison, David K. Jackson, Ian Johnston, Dominic Kwiatkowski, Cordelia Langford, John Sillitoe on behalf of the Wellcome Sanger Institute COVID-19 Surveillance Team |
| EPI_ISL_3468184 | National Virus Reference Laboratory | National Virus Reference Laboratory | Zoe Yandle, Charlene Bennett, Gabriel Gonzalez, Michael Carr, Jonathan Dean, Cillian F De Gascun |
| EPI_ISL_3468185, EPI_ISL_3468186, EPI_ISL_3468187 | Lighthouse Laboratory Plymouth | Wellcome Sanger Institute for the COVID-19 Genomics UK (COG-UK) Consortium | Lighthouse Laboratory Plymouth and Alex Alderton, Roberto Amato, Jeffrey Barrett, Sonia Goncalves, Ewan Harrison, David K. Jackson, Ian Johnston, Dominic Kwiatkowski, Cordelia Langford, John Sillitoe on behalf of the Wellcome Sanger Institute COVID-19 Surveillance Team |
| EPI_ISL_3468188 | National Virus Reference Laboratory | National Virus Reference Laboratory | Zoe Yandle, Charlene Bennett, Gabriel Gonzalez, Michael Carr, Jonathan Dean, Cillian F De Gascun |
| EPI_ISL_3468189, EPI_ISL_3468190, EPI_ISL_3468191 | Lighthouse Laboratory Plymouth | Wellcome Sanger Institute for the COVID-19 Genomics UK (COG-UK) Consortium | Lighthouse Laboratory Plymouth and Alex Alderton, Roberto Amato, Jeffrey Barrett, Sonia Goncalves, Ewan Harrison, David K. Jackson, Ian Johnston, Dominic Kwiatkowski, Cordelia Langford, John Sillitoe on behalf of the Wellcome Sanger Institute COVID-19 Surveillance Team |
| EPI_ISL_3468192 | National Virus Reference Laboratory | National Virus Reference Laboratory | Zoe Yandle, Charlene Bennett, Gabriel Gonzalez, Michael Carr, Jonathan Dean, Cillian F De Gascun |
| EPI_ISL_3468193, EPI_ISL_3468194 | Lighthouse Laboratory Plymouth | Wellcome Sanger Institute for the COVID-19 Genomics UK (COG-UK) Consortium | Lighthouse Laboratory Plymouth and Alex Alderton, Roberto Amato, Jeffrey Barrett, Sonia Goncalves, Ewan Harrison, David K. Jackson, Ian Johnston, Dominic Kwiatkowski, Cordelia Langford, John Sillitoe on behalf of the Wellcome Sanger Institute COVID-19 Surveillance Team |
| EPI_ISL_3468195 | National Virus Reference Laboratory | National Virus Reference Laboratory | Zoe Yandle, Charlene Bennett, Gabriel Gonzalez, Michael Carr, Jonathan Dean, Cillian F De Gascun |

| | | | |
|---|---|---|---|
| EPI_ISL_3468196, EPI_ISL_3468197, EPI_ISL_3468198, EPI_ISL_3468199 | Lighthouse Laboratory Plymouth | Wellcome Sanger Institute for the COVID-19 Genomics UK (COG-UK) Consortium | Lighthouse Laboratory Plymouth and Alex Alderton, Roberto Amato, Jeffrey Barrett, Sonia Goncalves, Ewan Harrison, David K. Jackson, Ian Johnston, Dominic Kwiatkowski, Cordelia Langford, John Sillitoe on behalf of the Wellcome Sanger Institute COVID-19 Surveillance Team |
| EPI_ISL_3468200 | National Virus Reference Laboratory | National Virus Reference Laboratory | Zoe Yandle, Charlene Bennett, Gabriel Gonzalez, Michael Carr, Jonathan Dean, Cillian F De Gascun |
| EPI_ISL_3468201 | Lighthouse Laboratory Plymouth | Wellcome Sanger Institute for the COVID-19 Genomics UK (COG-UK) Consortium | Lighthouse Laboratory Plymouth and Alex Alderton, Roberto Amato, Jeffrey Barrett, Sonia Goncalves, Ewan Harrison, David K. Jackson, Ian Johnston, Dominic Kwiatkowski, Cordelia Langford, John Sillitoe on behalf of the Wellcome Sanger Institute COVID-19 Surveillance Team |
| EPI_ISL_3468202 | Health Services Laboratories | Wellcome Sanger Institute for the COVID-19 Genomics UK (COG-UK) Consortium | Health Services Laboratories and Alex Alderton, Roberto Amato, Jeffrey Barrett, Sonia Goncalves, Ewan Harrison, David K. Jackson, Ian Johnston, Dominic Kwiatkowski, Cordelia Langford, John Sillitoe on behalf of the Wellcome Sanger Institute COVID-19 Surveillance Team |
| EPI_ISL_3468203 | Lighthouse Laboratory Plymouth | Wellcome Sanger Institute for the COVID-19 Genomics UK (COG-UK) Consortium | Lighthouse Laboratory Plymouth and Alex Alderton, Roberto Amato, Jeffrey Barrett, Sonia Goncalves, Ewan Harrison, David K. Jackson, Ian Johnston, Dominic Kwiatkowski, Cordelia Langford, John Sillitoe on behalf of the Wellcome Sanger Institute COVID-19 Surveillance Team |
| EPI_ISL_3468204 | National Virus Reference Laboratory | National Virus Reference Laboratory | Zoe Yandle, Charlene Bennett, Gabriel Gonzalez, Michael Carr, Jonathan Dean, Cillian F De Gascun |
| EPI_ISL_3468205 | Lighthouse Laboratory Plymouth | Wellcome Sanger Institute for the COVID-19 Genomics UK (COG-UK) Consortium | Lighthouse Laboratory Plymouth and Alex Alderton, Roberto Amato, Jeffrey Barrett, Sonia Goncalves, Ewan Harrison, David K. Jackson, Ian Johnston, Dominic Kwiatkowski, Cordelia Langford, John Sillitoe on behalf of the Wellcome Sanger Institute COVID-19 Surveillance Team |
| EPI_ISL_3468206 | Health Services Laboratories | Wellcome Sanger Institute for the COVID-19 Genomics UK (COG-UK) Consortium | Health Services Laboratories and Alex Alderton, Roberto Amato, Jeffrey Barrett, Sonia Goncalves, Ewan Harrison, David K. Jackson, Ian Johnston, Dominic Kwiatkowski, Cordelia Langford, John Sillitoe on behalf of the Wellcome Sanger Institute COVID-19 Surveillance Team |
| EPI_ISL_3468207, EPI_ISL_3468208 | Lighthouse Laboratory Plymouth | Wellcome Sanger Institute for the COVID-19 Genomics UK (COG-UK) Consortium | Lighthouse Laboratory Plymouth and Alex Alderton, Roberto Amato, Jeffrey Barrett, Sonia Goncalves, Ewan Harrison, David K. Jackson, Ian Johnston, Dominic Kwiatkowski, Cordelia Langford, John Sillitoe on behalf of the Wellcome Sanger Institute COVID-19 Surveillance Team |
| EPI_ISL_3468209 | Health Services Laboratories | Wellcome Sanger Institute for the COVID-19 Genomics UK (COG-UK) Consortium | Health Services Laboratories and Alex Alderton, Roberto Amato, Jeffrey Barrett, Sonia Goncalves, Ewan Harrison, David K. Jackson, Ian Johnston, Dominic Kwiatkowski, Cordelia Langford, John Sillitoe on behalf of the Wellcome Sanger Institute COVID-19 Surveillance Team |
| EPI_ISL_3468210, EPI_ISL_3468211, EPI_ISL_3468212, EPI_ISL_3468213, EPI_ISL_3468214, EPI_ISL_3468215, EPI_ISL_3468216, EPI_ISL_3468217, EPI_ISL_3468218, EPI_ISL_3468219, EPI_ISL_3468220, EPI_ISL_3468221, EPI_ISL_3468222, EPI_ISL_3468223, EPI_ISL_3468224, EPI_ISL_3468225, EPI_ISL_3468226, EPI_ISL_3468227, EPI_ISL_3468228, EPI_ISL_3468229, EPI_ISL_3468230, EPI_ISL_3468231, EPI_ISL_3468232, EPI_ISL_3468233, EPI_ISL_3468234, EPI_ISL_3468235, EPI_ISL_3468236, EPI_ISL_3468237, EPI_ISL_3468238, EPI_ISL_3468239, EPI_ISL_3468240, EPI_ISL_3468241, EPI_ISL_3468242, EPI_ISL_3468243, EPI_ISL_3468244, EPI_ISL_3468245, EPI_ISL_3468246, EPI_ISL_3468247 | | | |
| see above | Lighthouse Laboratory Plymouth | Wellcome Sanger Institute for the COVID-19 Genomics UK (COG-UK) Consortium | Lighthouse Laboratory Plymouth and Alex Alderton, Roberto Amato, Jeffrey Barrett, Sonia Goncalves, Ewan Harrison, David K. Jackson, Ian Johnston, Dominic Kwiatkowski, Cordelia Langford, John Sillitoe on behalf of the Wellcome Sanger Institute COVID-19 Surveillance Team |
| EPI_ISL_3468248 | Health Services Laboratories | Wellcome Sanger Institute for the COVID-19 Genomics UK (COG-UK) Consortium | Health Services Laboratories and Alex Alderton, Roberto Amato, Jeffrey Barrett, Sonia Goncalves, Ewan Harrison, David K. Jackson, Ian Johnston, Dominic Kwiatkowski, Cordelia Langford, John Sillitoe on behalf of the Wellcome Sanger Institute COVID-19 Surveillance Team |
| EPI_ISL_3468249, EPI_ISL_3468250, EPI_ISL_3468251 | Lighthouse Laboratory Plymouth | Wellcome Sanger Institute for the COVID-19 Genomics UK (COG-UK) Consortium | Lighthouse Laboratory Plymouth and Alex Alderton, Roberto Amato, Jeffrey Barrett, Sonia Goncalves, Ewan Harrison, David K. Jackson, Ian Johnston, Dominic Kwiatkowski, Cordelia Langford, John Sillitoe on behalf of the Wellcome Sanger Institute COVID-19 Surveillance Team |
| EPI_ISL_3468252 | Health Services Laboratories | Wellcome Sanger Institute for the COVID-19 Genomics UK (COG-UK) Consortium | Health Services Laboratories and Alex Alderton, Roberto Amato, Jeffrey Barrett, Sonia Goncalves, Ewan Harrison, David K. Jackson, Ian Johnston, Dominic Kwiatkowski, Cordelia Langford, John Sillitoe on behalf of the Wellcome Sanger Institute COVID-19 Surveillance Team |
| EPI_ISL_3468253, EPI_ISL_3468254, EPI_ISL_3468255, EPI_ISL_3468256, EPI_ISL_3468257, EPI_ISL_3468258 | Lighthouse Laboratory Plymouth | Wellcome Sanger Institute for the COVID-19 Genomics UK (COG-UK) Consortium | Lighthouse Laboratory Plymouth and Alex Alderton, Roberto Amato, Jeffrey Barrett, Sonia Goncalves, Ewan Harrison, David K. Jackson, Ian Johnston, Dominic Kwiatkowski, Cordelia Langford, John Sillitoe on behalf of the Wellcome Sanger Institute COVID-19 Surveillance Team |
| EPI_ISL_3468259 | Health Services Laboratories | Wellcome Sanger Institute for the COVID-19 Genomics UK (COG-UK) Consortium | Health Services Laboratories and Alex Alderton, Roberto Amato, Jeffrey Barrett, Sonia Goncalves, Ewan Harrison, David K. Jackson, Ian Johnston, Dominic Kwiatkowski, Cordelia Langford, John Sillitoe on behalf of the Wellcome Sanger Institute COVID-19 Surveillance Team |
| EPI_ISL_3468260, EPI_ISL_3468261, EPI_ISL_3468262 | Lighthouse Laboratory Plymouth | Wellcome Sanger Institute for the COVID-19 Genomics UK (COG-UK) Consortium | Lighthouse Laboratory Plymouth and Alex Alderton, Roberto Amato, Jeffrey Barrett, Sonia Goncalves, Ewan Harrison, David K. Jackson, Ian Johnston, Dominic Kwiatkowski, Cordelia Langford, John Sillitoe on behalf of the Wellcome Sanger Institute COVID-19 Surveillance Team |
| EPI_ISL_3468263 | Health Services Laboratories | Wellcome Sanger Institute for the COVID-19 Genomics UK (COG-UK) Consortium | Health Services Laboratories and Alex Alderton, Roberto Amato, Jeffrey Barrett, Sonia Goncalves, Ewan Harrison, David K. Jackson, Ian Johnston, Dominic Kwiatkowski, Cordelia Langford, John Sillitoe on behalf of the Wellcome Sanger Institute COVID-19 Surveillance Team |
| EPI_ISL_3468264, EPI_ISL_3468265, EPI_ISL_3468266, EPI_ISL_3468267, EPI_ISL_3468268, EPI_ISL_3468269, EPI_ISL_3468270, EPI_ISL_3468271, EPI_ISL_3468272, EPI_ISL_3468273, EPI_ISL_3468274, EPI_ISL_3468275, EPI_ISL_3468276, EPI_ISL_3468277, EPI_ISL_3468278, EPI_ISL_3468279, EPI_ISL_3468280, EPI_ISL_3468281, EPI_ISL_3468282, EPI_ISL_3468283, EPI_ISL_3468284, EPI_ISL_3468285, EPI_ISL_3468286, EPI_ISL_3468287 | | | |
| see above | Lighthouse Laboratory Plymouth | Wellcome Sanger Institute for the COVID-19 Genomics UK (COG-UK) Consortium | Lighthouse Laboratory Plymouth and Alex Alderton, Roberto Amato, Jeffrey Barrett, Sonia Goncalves, Ewan Harrison, David K. Jackson, Ian Johnston, Dominic Kwiatkowski, Cordelia Langford, John Sillitoe on behalf of the Wellcome Sanger Institute COVID-19 Surveillance Team |
| EPI_ISL_3468288 | Health Services Laboratories | Wellcome Sanger Institute for the COVID-19 Genomics UK (COG-UK) Consortium | Health Services Laboratories and Alex Alderton, Roberto Amato, Jeffrey Barrett, Sonia Goncalves, Ewan Harrison, David K. Jackson, Ian Johnston, Dominic Kwiatkowski, Cordelia Langford, John Sillitoe on behalf of the Wellcome Sanger Institute COVID-19 Surveillance Team |
| EPI_ISL_3468289, EPI_ISL_3468290 | Lighthouse Laboratory Plymouth | Wellcome Sanger Institute for the COVID-19 Genomics UK (COG-UK) Consortium | Lighthouse Laboratory Plymouth and Alex Alderton, Roberto Amato, Jeffrey Barrett, Sonia Goncalves, Ewan Harrison, David K. Jackson, Ian Johnston, Dominic Kwiatkowski, Cordelia Langford, John Sillitoe on behalf of the Wellcome Sanger Institute COVID-19 Surveillance Team |
| EPI_ISL_3468291 | Health Services Laboratories | Wellcome Sanger Institute for the COVID-19 Genomics UK (COG-UK) Consortium | Health Services Laboratories and Alex Alderton, Roberto Amato, Jeffrey Barrett, Sonia Goncalves, Ewan Harrison, David K. Jackson, Ian Johnston, Dominic Kwiatkowski, Cordelia Langford, John Sillitoe on behalf of the Wellcome Sanger Institute COVID-19 Surveillance Team |
| EPI_ISL_3468292, EPI_ISL_3468293 | Lighthouse Laboratory Plymouth | Wellcome Sanger Institute for the COVID-19 Genomics UK (COG-UK) Consortium | Lighthouse Laboratory Plymouth and Alex Alderton, Roberto Amato, Jeffrey Barrett, Sonia Goncalves, Ewan Harrison, David K. Jackson, Ian Johnston, Dominic Kwiatkowski, Cordelia Langford, John Sillitoe on behalf of the Wellcome Sanger Institute COVID-19 Surveillance Team |
| EPI_ISL_3468294 | Health Services Laboratories | Wellcome Sanger Institute for the COVID-19 Genomics UK (COG-UK) Consortium | Health Services Laboratories and Alex Alderton, Roberto Amato, Jeffrey Barrett, Sonia Goncalves, Ewan Harrison, David K. Jackson, Ian Johnston, Dominic Kwiatkowski, Cordelia Langford, John Sillitoe on behalf of the Wellcome Sanger Institute COVID-19 Surveillance Team |
| EPI_ISL_3468295, EPI_ISL_3468296 | Lighthouse Laboratory Plymouth | Wellcome Sanger Institute for the COVID-19 Genomics UK (COG-UK) Consortium | Lighthouse Laboratory Plymouth and Alex Alderton, Roberto Amato, Jeffrey Barrett, Sonia Goncalves, Ewan Harrison, David K. Jackson, Ian Johnston, Dominic Kwiatkowski, Cordelia Langford, John Sillitoe on behalf of the Wellcome Sanger Institute COVID-19 Surveillance Team |
| EPI_ISL_3468297 | Health Services Laboratories | Wellcome Sanger Institute for the COVID-19 Genomics UK (COG-UK) Consortium | Health Services Laboratories and Alex Alderton, Roberto Amato, Jeffrey Barrett, Sonia Goncalves, Ewan Harrison, David K. Jackson, Ian Johnston, Dominic Kwiatkowski, Cordelia Langford, John Sillitoe on behalf of the Wellcome Sanger Institute COVID-19 Surveillance Team |
| EPI_ISL_3468298, EPI_ISL_3468299, EPI_ISL_3468300, EPI_ISL_3468301, EPI_ISL_3468302, EPI_ISL_3468303, EPI_ISL_3468304, EPI_ISL_3468305 | Lighthouse Laboratory Plymouth | Wellcome Sanger Institute for the COVID-19 Genomics UK (COG-UK) Consortium | Lighthouse Laboratory Plymouth and Alex Alderton, Roberto Amato, Jeffrey Barrett, Sonia Goncalves, Ewan Harrison, David K. Jackson, Ian Johnston, Dominic Kwiatkowski, Cordelia Langford, John Sillitoe on behalf of the Wellcome Sanger Institute COVID-19 Surveillance Team |
| EPI_ISL_3468306 | Lighthouse Lab in Alderley Park | Wellcome Sanger Institute for the COVID-19 Genomics UK (COG-UK) Consortium | Jacquelyn Wynn, Mairead Hyland, The Lighthouse Lab in Alderley Park and Alex Alderton, Roberto Amato, Jeffrey Barrett, Sonia Goncalves, Ewan Harrison, David K. Jackson, Ian Johnston, Dominic Kwiatkowski, Cordelia Langford, John Sillitoe on behalf of the Wellcome Sanger Institute COVID-19 Surveillance Team |
| EPI_ISL_3468307, EPI_ISL_3468308 | Lighthouse Laboratory Plymouth | Wellcome Sanger Institute for the COVID-19 Genomics UK (COG-UK) Consortium | Lighthouse Laboratory Plymouth and Alex Alderton, Roberto Amato, Jeffrey Barrett, Sonia Goncalves, Ewan Harrison, David K. Jackson, Ian Johnston, Dominic Kwiatkowski, Cordelia Langford, John Sillitoe on behalf of the Wellcome Sanger Institute COVID-19 Surveillance Team |
| EPI_ISL_3468309, EPI_ISL_3468310 | Lighthouse Lab in Alderley Park | Wellcome Sanger Institute for the COVID-19 Genomics UK (COG-UK) Consortium | Jacquelyn Wynn, Mairead Hyland, The Lighthouse Lab in Alderley Park and Alex Alderton, Roberto Amato, Jeffrey Barrett, Sonia Goncalves, Ewan Harrison, David K. Jackson, Ian Johnston, Dominic Kwiatkowski, Cordelia Langford, John Sillitoe on behalf of the Wellcome Sanger Institute COVID-19 Surveillance Team |
| EPI_ISL_3468311, EPI_ISL_3468312 | Lighthouse Laboratory Plymouth | Wellcome Sanger Institute for the COVID-19 Genomics UK (COG-UK) Consortium | Lighthouse Laboratory Plymouth and Alex Alderton, Roberto Amato, Jeffrey Barrett, Sonia Goncalves, Ewan Harrison, David K. Jackson, Ian Johnston, Dominic Kwiatkowski, Cordelia Langford, John Sillitoe on behalf of the Wellcome Sanger Institute COVID-19 Surveillance Team |
| EPI_ISL_3468313, EPI_ISL_3468314, EPI_ISL_3468315 | Lighthouse Lab in Alderley Park | Wellcome Sanger Institute for the COVID-19 Genomics UK (COG-UK) Consortium | Jacquelyn Wynn, Mairead Hyland, The Lighthouse Lab in Alderley Park and Alex Alderton, Roberto Amato, Jeffrey Barrett, Sonia Goncalves, Ewan Harrison, David K. Jackson, Ian Johnston, Dominic Kwiatkowski, Cordelia Langford, John Sillitoe on behalf of the Wellcome Sanger Institute COVID-19 Surveillance Team |
| EPI_ISL_3468316 | Lighthouse Laboratory Plymouth | Wellcome Sanger Institute for the COVID-19 Genomics UK (COG-UK) Consortium | Lighthouse Laboratory Plymouth and Alex Alderton, Roberto Amato, Jeffrey Barrett, Sonia Goncalves, Ewan Harrison, David K. Jackson, Ian Johnston, Dominic Kwiatkowski, Cordelia Langford, John Sillitoe on behalf of the Wellcome Sanger Institute COVID-19 Surveillance Team |
| EPI_ISL_3468317 | Lighthouse Lab in Alderley Park | Wellcome Sanger Institute for the COVID-19 Genomics UK (COG-UK) Consortium | Jacquelyn Wynn, Mairead Hyland, The Lighthouse Lab in Alderley Park and Alex Alderton, Roberto Amato, Jeffrey Barrett, Sonia Goncalves, Ewan Harrison, David K. Jackson, Ian Johnston, Dominic Kwiatkowski, Cordelia Langford, John Sillitoe on behalf of the Wellcome Sanger Institute COVID-19 Surveillance Team |
| EPI_ISL_3468318, EPI_ISL_3468319, EPI_ISL_3468320, EPI_ISL_3468321 | Lighthouse Laboratory Plymouth | Wellcome Sanger Institute for the COVID-19 Genomics UK (COG-UK) Consortium | Lighthouse Laboratory Plymouth and Alex Alderton, Roberto Amato, Jeffrey Barrett, Sonia Goncalves, Ewan Harrison, David K. Jackson, Ian Johnston, Dominic Kwiatkowski, Cordelia Langford, John Sillitoe on behalf of the Wellcome Sanger Institute COVID-19 Surveillance Team |
| EPI_ISL_3468322 | Lighthouse Lab in Alderley Park | Wellcome Sanger Institute for the COVID-19 Genomics UK (COG-UK) Consortium | Jacquelyn Wynn, Mairead Hyland, The Lighthouse Lab in Alderley Park and Alex Alderton, Roberto Amato, Jeffrey Barrett, Sonia Goncalves, Ewan Harrison, David K. Jackson, Ian Johnston, Dominic Kwiatkowski, Cordelia Langford, John Sillitoe on behalf of the Wellcome Sanger Institute COVID-19 Surveillance Team |
| EPI_ISL_3468323, EPI_ISL_3468324, | Lighthouse Laboratory Plymouth | Wellcome Sanger Institute for the COVID-19 Genomics UK (COG-UK) Consortium | Lighthouse Laboratory Plymouth and Alex Alderton, Roberto Amato, Jeffrey Barrett, Sonia Goncalves, Ewan Harrison, David K. Jackson, Ian Johnston, Dominic Kwiatkowski, Cordelia Langford, John Sillitoe on behalf of the Wellcome Sanger Institute COVID-19 Surveillance Team |

| | | | |
|---|---|---|---|
| EPI_ISL_3468325 | | | |
| EPI_ISL_3468326, EPI_ISL_3468327 | Lighthouse Lab in Alderley Park | Wellcome Sanger Institute for the COVID-19 Genomics UK (COG-UK) Consortium | Jacquelyn Wynn, Mairead Hyland, The Lighthouse Lab in Alderley Park and Alex Alderton, Roberto Amato, Jeffrey Barrett, Sonia Goncalves, Ewan Harrison, David K. Jackson, Ian Johnston, Dominic Kwiatkowski, Cordelia Langford, John Sillitoe on behalf of the Wellcome Sanger Institute COVID-19 Surveillance Team |
| EPI_ISL_3468328, EPI_ISL_3468329, EPI_ISL_3468330 | Lighthouse Laboratory Plymouth | Wellcome Sanger Institute for the COVID-19 Genomics UK (COG-UK) Consortium | Lighthouse Laboratory Plymouth and Alex Alderton, Roberto Amato, Jeffrey Barrett, Sonia Goncalves, Ewan Harrison, David K. Jackson, Ian Johnston, Dominic Kwiatkowski, Cordelia Langford, John Sillitoe on behalf of the Wellcome Sanger Institute COVID-19 Surveillance Team |
| EPI_ISL_3468331 | Lighthouse Lab in Alderley Park | Wellcome Sanger Institute for the COVID-19 Genomics UK (COG-UK) Consortium | Jacquelyn Wynn, Mairead Hyland, The Lighthouse Lab in Alderley Park and Alex Alderton, Roberto Amato, Jeffrey Barrett, Sonia Goncalves, Ewan Harrison, David K. Jackson, Ian Johnston, Dominic Kwiatkowski, Cordelia Langford, John Sillitoe on behalf of the Wellcome Sanger Institute COVID-19 Surveillance Team |
| EPI_ISL_3468332, EPI_ISL_3468333 | Lighthouse Laboratory Plymouth | Wellcome Sanger Institute for the COVID-19 Genomics UK (COG-UK) Consortium | Lighthouse Laboratory Plymouth and Alex Alderton, Roberto Amato, Jeffrey Barrett, Sonia Goncalves, Ewan Harrison, David K. Jackson, Ian Johnston, Dominic Kwiatkowski, Cordelia Langford, John Sillitoe on behalf of the Wellcome Sanger Institute COVID-19 Surveillance Team |
| EPI_ISL_3468334 | Lighthouse Lab in Alderley Park | Wellcome Sanger Institute for the COVID-19 Genomics UK (COG-UK) Consortium | Jacquelyn Wynn, Mairead Hyland, The Lighthouse Lab in Alderley Park and Alex Alderton, Roberto Amato, Jeffrey Barrett, Sonia Goncalves, Ewan Harrison, David K. Jackson, Ian Johnston, Dominic Kwiatkowski, Cordelia Langford, John Sillitoe on behalf of the Wellcome Sanger Institute COVID-19 Surveillance Team |
| EPI_ISL_3468335 | Lighthouse Laboratory Plymouth | Wellcome Sanger Institute for the COVID-19 Genomics UK (COG-UK) Consortium | Lighthouse Laboratory Plymouth and Alex Alderton, Roberto Amato, Jeffrey Barrett, Sonia Goncalves, Ewan Harrison, David K. Jackson, Ian Johnston, Dominic Kwiatkowski, Cordelia Langford, John Sillitoe on behalf of the Wellcome Sanger Institute COVID-19 Surveillance Team |
| EPI_ISL_3468336, EPI_ISL_3468337 | Lighthouse Lab in Alderley Park | Wellcome Sanger Institute for the COVID-19 Genomics UK (COG-UK) Consortium | Jacquelyn Wynn, Mairead Hyland, The Lighthouse Lab in Alderley Park and Alex Alderton, Roberto Amato, Jeffrey Barrett, Sonia Goncalves, Ewan Harrison, David K. Jackson, Ian Johnston, Dominic Kwiatkowski, Cordelia Langford, John Sillitoe on behalf of the Wellcome Sanger Institute COVID-19 Surveillance Team |
| EPI_ISL_3468338, EPI_ISL_3468339, EPI_ISL_3468340, EPI_ISL_3468341, EPI_ISL_3468342, EPI_ISL_3468343, EPI_ISL_3468344, EPI_ISL_3468345 | Lighthouse Laboratory Plymouth | Wellcome Sanger Institute for the COVID-19 Genomics UK (COG-UK) Consortium | Lighthouse Laboratory Plymouth and Alex Alderton, Roberto Amato, Jeffrey Barrett, Sonia Goncalves, Ewan Harrison, David K. Jackson, Ian Johnston, Dominic Kwiatkowski, Cordelia Langford, John Sillitoe on behalf of the Wellcome Sanger Institute COVID-19 Surveillance Team |
| EPI_ISL_3468346, EPI_ISL_3468347, EPI_ISL_3468348, EPI_ISL_3468349, EPI_ISL_3468350, EPI_ISL_3468351 | Lighthouse Lab in Alderley Park | Wellcome Sanger Institute for the COVID-19 Genomics UK (COG-UK) Consortium | Jacquelyn Wynn, Mairead Hyland, The Lighthouse Lab in Alderley Park and Alex Alderton, Roberto Amato, Jeffrey Barrett, Sonia Goncalves, Ewan Harrison, David K. Jackson, Ian Johnston, Dominic Kwiatkowski, Cordelia Langford, John Sillitoe on behalf of the Wellcome Sanger Institute COVID-19 Surveillance Team |
| EPI_ISL_3468352 | Lighthouse Laboratory Plymouth | Wellcome Sanger Institute for the COVID-19 Genomics UK (COG-UK) Consortium | Lighthouse Laboratory Plymouth and Alex Alderton, Roberto Amato, Jeffrey Barrett, Sonia Goncalves, Ewan Harrison, David K. Jackson, Ian Johnston, Dominic Kwiatkowski, Cordelia Langford, John Sillitoe on behalf of the Wellcome Sanger Institute COVID-19 Surveillance Team |
| EPI_ISL_3468353, EPI_ISL_3468354 | Lighthouse Lab in Alderley Park | Wellcome Sanger Institute for the COVID-19 Genomics UK (COG-UK) Consortium | Jacquelyn Wynn, Mairead Hyland, The Lighthouse Lab in Alderley Park and Alex Alderton, Roberto Amato, Jeffrey Barrett, Sonia Goncalves, Ewan Harrison, David K. Jackson, Ian Johnston, Dominic Kwiatkowski, Cordelia Langford, John Sillitoe on behalf of the Wellcome Sanger Institute COVID-19 Surveillance Team |
| EPI_ISL_3468355 | Lighthouse Laboratory Plymouth | Wellcome Sanger Institute for the COVID-19 Genomics UK (COG-UK) Consortium | Lighthouse Laboratory Plymouth and Alex Alderton, Roberto Amato, Jeffrey Barrett, Sonia Goncalves, Ewan Harrison, David K. Jackson, Ian Johnston, Dominic Kwiatkowski, Cordelia Langford, John Sillitoe on behalf of the Wellcome Sanger Institute COVID-19 Surveillance Team |
| EPI_ISL_3468356, EPI_ISL_3468357 | Lighthouse Lab in Alderley Park | Wellcome Sanger Institute for the COVID-19 Genomics UK (COG-UK) Consortium | Jacquelyn Wynn, Mairead Hyland, The Lighthouse Lab in Alderley Park and Alex Alderton, Roberto Amato, Jeffrey Barrett, Sonia Goncalves, Ewan Harrison, David K. Jackson, Ian Johnston, Dominic Kwiatkowski, Cordelia Langford, John Sillitoe on behalf of the Wellcome Sanger Institute COVID-19 Surveillance Team |
| EPI_ISL_3468358, EPI_ISL_3468359 | Lighthouse Laboratory Plymouth | Wellcome Sanger Institute for the COVID-19 Genomics UK (COG-UK) Consortium | Lighthouse Laboratory Plymouth and Alex Alderton, Roberto Amato, Jeffrey Barrett, Sonia Goncalves, Ewan Harrison, David K. Jackson, Ian Johnston, Dominic Kwiatkowski, Cordelia Langford, John Sillitoe on behalf of the Wellcome Sanger Institute COVID-19 Surveillance Team |
| EPI_ISL_3468360, EPI_ISL_3468361, EPI_ISL_3468362, EPI_ISL_3468363 | Lighthouse Lab in Alderley Park | Wellcome Sanger Institute for the COVID-19 Genomics UK (COG-UK) Consortium | Jacquelyn Wynn, Mairead Hyland, The Lighthouse Lab in Alderley Park and Alex Alderton, Roberto Amato, Jeffrey Barrett, Sonia Goncalves, Ewan Harrison, David K. Jackson, Ian Johnston, Dominic Kwiatkowski, Cordelia Langford, John Sillitoe on behalf of the Wellcome Sanger Institute COVID-19 Surveillance Team |
| EPI_ISL_3468364, EPI_ISL_3468365 | Lighthouse Laboratory Plymouth | Wellcome Sanger Institute for the COVID-19 Genomics UK (COG-UK) Consortium | Lighthouse Laboratory Plymouth and Alex Alderton, Roberto Amato, Jeffrey Barrett, Sonia Goncalves, Ewan Harrison, David K. Jackson, Ian Johnston, Dominic Kwiatkowski, Cordelia Langford, John Sillitoe on behalf of the Wellcome Sanger Institute COVID-19 Surveillance Team |
| EPI_ISL_3468366 | Lighthouse Lab in Alderley Park | Wellcome Sanger Institute for the COVID-19 Genomics UK (COG-UK) Consortium | Jacquelyn Wynn, Mairead Hyland, The Lighthouse Lab in Alderley Park and Alex Alderton, Roberto Amato, Jeffrey Barrett, Sonia Goncalves, Ewan Harrison, David K. Jackson, Ian Johnston, Dominic Kwiatkowski, Cordelia Langford, John Sillitoe on behalf of the Wellcome Sanger Institute COVID-19 Surveillance Team |
| EPI_ISL_3468367, EPI_ISL_3468368 | Lighthouse Laboratory Plymouth | Wellcome Sanger Institute for the COVID-19 Genomics UK (COG-UK) Consortium | Lighthouse Laboratory Plymouth and Alex Alderton, Roberto Amato, Jeffrey Barrett, Sonia Goncalves, Ewan Harrison, David K. Jackson, Ian Johnston, Dominic Kwiatkowski, Cordelia Langford, John Sillitoe on behalf of the Wellcome Sanger Institute COVID-19 Surveillance Team |
| EPI_ISL_3468369 | Lighthouse Lab in Alderley Park | Wellcome Sanger Institute for the COVID-19 Genomics UK (COG-UK) Consortium | Jacquelyn Wynn, Mairead Hyland, The Lighthouse Lab in Alderley Park and Alex Alderton, Roberto Amato, Jeffrey Barrett, Sonia Goncalves, Ewan Harrison, David K. Jackson, Ian Johnston, Dominic Kwiatkowski, Cordelia Langford, John Sillitoe on behalf of the Wellcome Sanger Institute COVID-19 Surveillance Team |
| EPI_ISL_3468370 | Lighthouse Laboratory Plymouth | Wellcome Sanger Institute for the COVID-19 Genomics UK (COG-UK) Consortium | Lighthouse Laboratory Plymouth and Alex Alderton, Roberto Amato, Jeffrey Barrett, Sonia Goncalves, Ewan Harrison, David K. Jackson, Ian Johnston, Dominic Kwiatkowski, Cordelia Langford, John Sillitoe on behalf of the Wellcome Sanger Institute COVID-19 Surveillance Team |
| EPI_ISL_3468371, EPI_ISL_3468372 | Lighthouse Lab in Alderley Park | Wellcome Sanger Institute for the COVID-19 Genomics UK (COG-UK) Consortium | Jacquelyn Wynn, Mairead Hyland, The Lighthouse Lab in Alderley Park and Alex Alderton, Roberto Amato, Jeffrey Barrett, Sonia Goncalves, Ewan Harrison, David K. Jackson, Ian Johnston, Dominic Kwiatkowski, Cordelia Langford, John Sillitoe on behalf of the Wellcome Sanger Institute COVID-19 Surveillance Team |
| EPI_ISL_3468373, EPI_ISL_3468374 | Lighthouse Laboratory Plymouth | Wellcome Sanger Institute for the COVID-19 Genomics UK (COG-UK) Consortium | Lighthouse Laboratory Plymouth and Alex Alderton, Roberto Amato, Jeffrey Barrett, Sonia Goncalves, Ewan Harrison, David K. Jackson, Ian Johnston, Dominic Kwiatkowski, Cordelia Langford, John Sillitoe on behalf of the Wellcome Sanger Institute COVID-19 Surveillance Team |
| EPI_ISL_3468375, EPI_ISL_3468376, EPI_ISL_3468377, EPI_ISL_3468378 | Lighthouse Lab in Alderley Park | Wellcome Sanger Institute for the COVID-19 Genomics UK (COG-UK) Consortium | Jacquelyn Wynn, Mairead Hyland, The Lighthouse Lab in Alderley Park and Alex Alderton, Roberto Amato, Jeffrey Barrett, Sonia Goncalves, Ewan Harrison, David K. Jackson, Ian Johnston, Dominic Kwiatkowski, Cordelia Langford, John Sillitoe on behalf of the Wellcome Sanger Institute COVID-19 Surveillance Team |
| EPI_ISL_3468379 | Lighthouse Laboratory Plymouth | Wellcome Sanger Institute for the COVID-19 Genomics UK (COG-UK) Consortium | Lighthouse Laboratory Plymouth and Alex Alderton, Roberto Amato, Jeffrey Barrett, Sonia Goncalves, Ewan Harrison, David K. Jackson, Ian Johnston, Dominic Kwiatkowski, Cordelia Langford, John Sillitoe on behalf of the Wellcome Sanger Institute COVID-19 Surveillance Team |
| EPI_ISL_3468380, EPI_ISL_3468381 | Lighthouse Lab in Alderley Park | Wellcome Sanger Institute for the COVID-19 Genomics UK (COG-UK) Consortium | Jacquelyn Wynn, Mairead Hyland, The Lighthouse Lab in Alderley Park and Alex Alderton, Roberto Amato, Jeffrey Barrett, Sonia Goncalves, Ewan Harrison, David K. Jackson, Ian Johnston, Dominic Kwiatkowski, Cordelia Langford, John Sillitoe on behalf of the Wellcome Sanger Institute COVID-19 Surveillance Team |
| EPI_ISL_3468382, EPI_ISL_3468383, EPI_ISL_3468384 | Lighthouse Laboratory Plymouth | Wellcome Sanger Institute for the COVID-19 Genomics UK (COG-UK) Consortium | Lighthouse Laboratory Plymouth and Alex Alderton, Roberto Amato, Jeffrey Barrett, Sonia Goncalves, Ewan Harrison, David K. Jackson, Ian Johnston, Dominic Kwiatkowski, Cordelia Langford, John Sillitoe on behalf of the Wellcome Sanger Institute COVID-19 Surveillance Team |
| EPI_ISL_3468385 | Lighthouse Lab in Alderley Park | Wellcome Sanger Institute for the COVID-19 Genomics UK (COG-UK) Consortium | Jacquelyn Wynn, Mairead Hyland, The Lighthouse Lab in Alderley Park and Alex Alderton, Roberto Amato, Jeffrey Barrett, Sonia Goncalves, Ewan Harrison, David K. Jackson, Ian Johnston, Dominic Kwiatkowski, Cordelia Langford, John Sillitoe on behalf of the Wellcome Sanger Institute COVID-19 Surveillance Team |
| EPI_ISL_3468386 | Lighthouse Laboratory Plymouth | Wellcome Sanger Institute for the COVID-19 Genomics UK (COG-UK) Consortium | Lighthouse Laboratory Plymouth and Alex Alderton, Roberto Amato, Jeffrey Barrett, Sonia Goncalves, Ewan Harrison, David K. Jackson, Ian Johnston, Dominic Kwiatkowski, Cordelia Langford, John Sillitoe on behalf of the Wellcome Sanger Institute COVID-19 Surveillance Team |
| EPI_ISL_3468387, EPI_ISL_3468388 | Lighthouse Lab in Alderley Park | Wellcome Sanger Institute for the COVID-19 Genomics UK (COG-UK) Consortium | Jacquelyn Wynn, Mairead Hyland, The Lighthouse Lab in Alderley Park and Alex Alderton, Roberto Amato, Jeffrey Barrett, Sonia Goncalves, Ewan Harrison, David K. Jackson, Ian Johnston, Dominic Kwiatkowski, Cordelia Langford, John Sillitoe on behalf of the Wellcome Sanger Institute COVID-19 Surveillance Team |
| EPI_ISL_3468389 | Lighthouse Laboratory Plymouth | Wellcome Sanger Institute for the COVID-19 Genomics UK (COG-UK) Consortium | Lighthouse Laboratory Plymouth and Alex Alderton, Roberto Amato, Jeffrey Barrett, Sonia Goncalves, Ewan Harrison, David K. Jackson, Ian Johnston, Dominic Kwiatkowski, Cordelia Langford, John Sillitoe on behalf of the Wellcome Sanger Institute COVID-19 Surveillance Team |
| EPI_ISL_3468390, EPI_ISL_3468391 | Lighthouse Lab in Alderley Park | Wellcome Sanger Institute for the COVID-19 Genomics UK (COG-UK) Consortium | Jacquelyn Wynn, Mairead Hyland, The Lighthouse Lab in Alderley Park and Alex Alderton, Roberto Amato, Jeffrey Barrett, Sonia Goncalves, Ewan Harrison, David K. Jackson, Ian Johnston, Dominic Kwiatkowski, Cordelia Langford, John Sillitoe on behalf of the Wellcome Sanger Institute COVID-19 Surveillance Team |
| EPI_ISL_3468392 | Lighthouse Laboratory Plymouth | Wellcome Sanger Institute for the COVID-19 Genomics UK (COG-UK) Consortium | Lighthouse Laboratory Plymouth and Alex Alderton, Roberto Amato, Jeffrey Barrett, Sonia Goncalves, Ewan Harrison, David K. Jackson, Ian Johnston, Dominic Kwiatkowski, Cordelia Langford, John Sillitoe on behalf of the Wellcome Sanger Institute COVID-19 Surveillance Team |
| EPI_ISL_3468393 | Lighthouse Lab in Alderley Park | Wellcome Sanger Institute for the COVID-19 Genomics UK (COG-UK) Consortium | Jacquelyn Wynn, Mairead Hyland, The Lighthouse Lab in Alderley Park and Alex Alderton, Roberto Amato, Jeffrey Barrett, Sonia Goncalves, Ewan Harrison, David K. Jackson, Ian Johnston, Dominic Kwiatkowski, Cordelia Langford, John Sillitoe on behalf of the Wellcome Sanger Institute COVID-19 Surveillance Team |
| EPI_ISL_3468394, EPI_ISL_3468395 | Lighthouse Laboratory Plymouth | Wellcome Sanger Institute for the COVID-19 Genomics UK (COG-UK) Consortium | Lighthouse Laboratory Plymouth and Alex Alderton, Roberto Amato, Jeffrey Barrett, Sonia Goncalves, Ewan Harrison, David K. Jackson, Ian Johnston, Dominic Kwiatkowski, Cordelia Langford, John Sillitoe on behalf of the Wellcome Sanger Institute COVID-19 Surveillance Team |
| EPI_ISL_3468396 | Lighthouse Lab in Alderley Park | Wellcome Sanger Institute for the COVID-19 Genomics UK (COG-UK) Consortium | Jacquelyn Wynn, Mairead Hyland, The Lighthouse Lab in Alderley Park and Alex Alderton, Roberto Amato, Jeffrey Barrett, Sonia Goncalves, Ewan Harrison, David K. Jackson, Ian Johnston, Dominic Kwiatkowski, Cordelia Langford, John Sillitoe on behalf of the Wellcome Sanger Institute COVID-19 Surveillance Team |
| EPI_ISL_3468397, EPI_ISL_3468398, EPI_ISL_3468399, EPI_ISL_3468400, EPI_ISL_3468401 | Lighthouse Laboratory Plymouth | Wellcome Sanger Institute for the COVID-19 Genomics UK (COG-UK) Consortium | Lighthouse Laboratory Plymouth and Alex Alderton, Roberto Amato, Jeffrey Barrett, Sonia Goncalves, Ewan Harrison, David K. Jackson, Ian Johnston, Dominic Kwiatkowski, Cordelia Langford, John Sillitoe on behalf of the Wellcome Sanger Institute COVID-19 Surveillance Team |
| EPI_ISL_3468402, | Lighthouse Lab in Alderley Park | Wellcome Sanger Institute for the COVID-19 Genomics UK (COG-UK) | Jacquelyn Wynn, Mairead Hyland, The Lighthouse Lab in Alderley Park and Alex Alderton, Roberto Amato, Jeffrey Barrett, Sonia Goncalves, Ewan Harrison, David K. Jackson, Ian Johnston, Dominic Kwiatkowski, Cordelia Langford, John Sillitoe on |

| | | | |
|---|---|---|---|
| EPI_ISL_3468403, EPI_ISL_3468404, EPI_ISL_3468405 | | Consortium | behalf of the Wellcome Sanger Institute COVID-19 Surveillance Team |
| EPI_ISL_3468406 | Lighthouse Laboratory Plymouth | Wellcome Sanger Institute for the COVID-19 Genomics UK (COG-UK) Consortium | Lighthouse Laboratory Plymouth and Alex Alderton, Roberto Amato, Jeffrey Barrett, Sonia Goncalves, Ewan Harrison, David K. Jackson, Ian Johnston, Dominic Kwiatkowski, Cordelia Langford, John Sillitoe on behalf of the Wellcome Sanger Institute COVID-19 Surveillance Team |
| EPI_ISL_3468407 | Lighthouse Lab in Alderley Park | Wellcome Sanger Institute for the COVID-19 Genomics UK (COG-UK) Consortium | Jacquelyn Wynn, Mairead Hyland, The Lighthouse Lab in Alderley Park and Alex Alderton, Roberto Amato, Jeffrey Barrett, Sonia Goncalves, Ewan Harrison, David K. Jackson, Ian Johnston, Dominic Kwiatkowski, Cordelia Langford, John Sillitoe on behalf of the Wellcome Sanger Institute COVID-19 Surveillance Team |
| EPI_ISL_3468408 | Lighthouse Laboratory Plymouth | Wellcome Sanger Institute for the COVID-19 Genomics UK (COG-UK) Consortium | Lighthouse Laboratory Plymouth and Alex Alderton, Roberto Amato, Jeffrey Barrett, Sonia Goncalves, Ewan Harrison, David K. Jackson, Ian Johnston, Dominic Kwiatkowski, Cordelia Langford, John Sillitoe on behalf of the Wellcome Sanger Institute COVID-19 Surveillance Team |
| EPI_ISL_3468409, EPI_ISL_3468410, EPI_ISL_3468411, EPI_ISL_3468412 | Lighthouse Lab in Alderley Park | Wellcome Sanger Institute for the COVID-19 Genomics UK (COG-UK) Consortium | Jacquelyn Wynn, Mairead Hyland, The Lighthouse Lab in Alderley Park and Alex Alderton, Roberto Amato, Jeffrey Barrett, Sonia Goncalves, Ewan Harrison, David K. Jackson, Ian Johnston, Dominic Kwiatkowski, Cordelia Langford, John Sillitoe on behalf of the Wellcome Sanger Institute COVID-19 Surveillance Team |
| EPI_ISL_3468413 | Lighthouse Laboratory Plymouth | Wellcome Sanger Institute for the COVID-19 Genomics UK (COG-UK) Consortium | Lighthouse Laboratory Plymouth and Alex Alderton, Roberto Amato, Jeffrey Barrett, Sonia Goncalves, Ewan Harrison, David K. Jackson, Ian Johnston, Dominic Kwiatkowski, Cordelia Langford, John Sillitoe on behalf of the Wellcome Sanger Institute COVID-19 Surveillance Team |
| EPI_ISL_3468414 | Lighthouse Lab in Alderley Park | Wellcome Sanger Institute for the COVID-19 Genomics UK (COG-UK) Consortium | Jacquelyn Wynn, Mairead Hyland, The Lighthouse Lab in Alderley Park and Alex Alderton, Roberto Amato, Jeffrey Barrett, Sonia Goncalves, Ewan Harrison, David K. Jackson, Ian Johnston, Dominic Kwiatkowski, Cordelia Langford, John Sillitoe on behalf of the Wellcome Sanger Institute COVID-19 Surveillance Team |
| EPI_ISL_3468415, EPI_ISL_3468416 | Lighthouse Laboratory Plymouth | Wellcome Sanger Institute for the COVID-19 Genomics UK (COG-UK) Consortium | Lighthouse Laboratory Plymouth and Alex Alderton, Roberto Amato, Jeffrey Barrett, Sonia Goncalves, Ewan Harrison, David K. Jackson, Ian Johnston, Dominic Kwiatkowski, Cordelia Langford, John Sillitoe on behalf of the Wellcome Sanger Institute COVID-19 Surveillance Team |
| EPI_ISL_3468417 | Lighthouse Lab in Alderley Park | Wellcome Sanger Institute for the COVID-19 Genomics UK (COG-UK) Consortium | Jacquelyn Wynn, Mairead Hyland, The Lighthouse Lab in Alderley Park and Alex Alderton, Roberto Amato, Jeffrey Barrett, Sonia Goncalves, Ewan Harrison, David K. Jackson, Ian Johnston, Dominic Kwiatkowski, Cordelia Langford, John Sillitoe on behalf of the Wellcome Sanger Institute COVID-19 Surveillance Team |
| EPI_ISL_3468418, EPI_ISL_3468419 | Lighthouse Laboratory Plymouth | Wellcome Sanger Institute for the COVID-19 Genomics UK (COG-UK) Consortium | Lighthouse Laboratory Plymouth and Alex Alderton, Roberto Amato, Jeffrey Barrett, Sonia Goncalves, Ewan Harrison, David K. Jackson, Ian Johnston, Dominic Kwiatkowski, Cordelia Langford, John Sillitoe on behalf of the Wellcome Sanger Institute COVID-19 Surveillance Team |
| EPI_ISL_3468420 | Lighthouse Lab in Alderley Park | Wellcome Sanger Institute for the COVID-19 Genomics UK (COG-UK) Consortium | Jacquelyn Wynn, Mairead Hyland, The Lighthouse Lab in Alderley Park and Alex Alderton, Roberto Amato, Jeffrey Barrett, Sonia Goncalves, Ewan Harrison, David K. Jackson, Ian Johnston, Dominic Kwiatkowski, Cordelia Langford, John Sillitoe on behalf of the Wellcome Sanger Institute COVID-19 Surveillance Team |
| EPI_ISL_3468421 | Lighthouse Laboratory Plymouth | Wellcome Sanger Institute for the COVID-19 Genomics UK (COG-UK) Consortium | Lighthouse Laboratory Plymouth and Alex Alderton, Roberto Amato, Jeffrey Barrett, Sonia Goncalves, Ewan Harrison, David K. Jackson, Ian Johnston, Dominic Kwiatkowski, Cordelia Langford, John Sillitoe on behalf of the Wellcome Sanger Institute COVID-19 Surveillance Team |
| EPI_ISL_3468422 | Lighthouse Lab in Alderley Park | Wellcome Sanger Institute for the COVID-19 Genomics UK (COG-UK) Consortium | Jacquelyn Wynn, Mairead Hyland, The Lighthouse Lab in Alderley Park and Alex Alderton, Roberto Amato, Jeffrey Barrett, Sonia Goncalves, Ewan Harrison, David K. Jackson, Ian Johnston, Dominic Kwiatkowski, Cordelia Langford, John Sillitoe on behalf of the Wellcome Sanger Institute COVID-19 Surveillance Team |
| EPI_ISL_3468423 | Lighthouse Laboratory Plymouth | Wellcome Sanger Institute for the COVID-19 Genomics UK (COG-UK) Consortium | Lighthouse Laboratory Plymouth and Alex Alderton, Roberto Amato, Jeffrey Barrett, Sonia Goncalves, Ewan Harrison, David K. Jackson, Ian Johnston, Dominic Kwiatkowski, Cordelia Langford, John Sillitoe on behalf of the Wellcome Sanger Institute COVID-19 Surveillance Team |
| EPI_ISL_3468424 | Lighthouse Lab in Alderley Park | Wellcome Sanger Institute for the COVID-19 Genomics UK (COG-UK) Consortium | Jacquelyn Wynn, Mairead Hyland, The Lighthouse Lab in Alderley Park and Alex Alderton, Roberto Amato, Jeffrey Barrett, Sonia Goncalves, Ewan Harrison, David K. Jackson, Ian Johnston, Dominic Kwiatkowski, Cordelia Langford, John Sillitoe on behalf of the Wellcome Sanger Institute COVID-19 Surveillance Team |
| EPI_ISL_3468425 | Lighthouse Laboratory Plymouth | Wellcome Sanger Institute for the COVID-19 Genomics UK (COG-UK) Consortium | Lighthouse Laboratory Plymouth and Alex Alderton, Roberto Amato, Jeffrey Barrett, Sonia Goncalves, Ewan Harrison, David K. Jackson, Ian Johnston, Dominic Kwiatkowski, Cordelia Langford, John Sillitoe on behalf of the Wellcome Sanger Institute COVID-19 Surveillance Team |
| EPI_ISL_3468426 | Lighthouse Lab in Alderley Park | Wellcome Sanger Institute for the COVID-19 Genomics UK (COG-UK) Consortium | Jacquelyn Wynn, Mairead Hyland, The Lighthouse Lab in Alderley Park and Alex Alderton, Roberto Amato, Jeffrey Barrett, Sonia Goncalves, Ewan Harrison, David K. Jackson, Ian Johnston, Dominic Kwiatkowski, Cordelia Langford, John Sillitoe on behalf of the Wellcome Sanger Institute COVID-19 Surveillance Team |
| EPI_ISL_3468427 | Lighthouse Laboratory Plymouth | Wellcome Sanger Institute for the COVID-19 Genomics UK (COG-UK) Consortium | Lighthouse Laboratory Plymouth and Alex Alderton, Roberto Amato, Jeffrey Barrett, Sonia Goncalves, Ewan Harrison, David K. Jackson, Ian Johnston, Dominic Kwiatkowski, Cordelia Langford, John Sillitoe on behalf of the Wellcome Sanger Institute COVID-19 Surveillance Team |
| EPI_ISL_3468428 | Lighthouse Lab in Alderley Park | Wellcome Sanger Institute for the COVID-19 Genomics UK (COG-UK) Consortium | Jacquelyn Wynn, Mairead Hyland, The Lighthouse Lab in Alderley Park and Alex Alderton, Roberto Amato, Jeffrey Barrett, Sonia Goncalves, Ewan Harrison, David K. Jackson, Ian Johnston, Dominic Kwiatkowski, Cordelia Langford, John Sillitoe on behalf of the Wellcome Sanger Institute COVID-19 Surveillance Team |
| EPI_ISL_3468429, EPI_ISL_3468430 | Lighthouse Laboratory Plymouth | Wellcome Sanger Institute for the COVID-19 Genomics UK (COG-UK) Consortium | Lighthouse Laboratory Plymouth and Alex Alderton, Roberto Amato, Jeffrey Barrett, Sonia Goncalves, Ewan Harrison, David K. Jackson, Ian Johnston, Dominic Kwiatkowski, Cordelia Langford, John Sillitoe on behalf of the Wellcome Sanger Institute COVID-19 Surveillance Team |
| EPI_ISL_3468431, EPI_ISL_3468432 | Lighthouse Lab in Alderley Park | Wellcome Sanger Institute for the COVID-19 Genomics UK (COG-UK) Consortium | Jacquelyn Wynn, Mairead Hyland, The Lighthouse Lab in Alderley Park and Alex Alderton, Roberto Amato, Jeffrey Barrett, Sonia Goncalves, Ewan Harrison, David K. Jackson, Ian Johnston, Dominic Kwiatkowski, Cordelia Langford, John Sillitoe on behalf of the Wellcome Sanger Institute COVID-19 Surveillance Team |
| EPI_ISL_3468433, EPI_ISL_3468434 | Lighthouse Laboratory Plymouth | Wellcome Sanger Institute for the COVID-19 Genomics UK (COG-UK) Consortium | Lighthouse Laboratory Plymouth and Alex Alderton, Roberto Amato, Jeffrey Barrett, Sonia Goncalves, Ewan Harrison, David K. Jackson, Ian Johnston, Dominic Kwiatkowski, Cordelia Langford, John Sillitoe on behalf of the Wellcome Sanger Institute COVID-19 Surveillance Team |
| EPI_ISL_3468435 | Lighthouse Lab in Alderley Park | Wellcome Sanger Institute for the COVID-19 Genomics UK (COG-UK) Consortium | Jacquelyn Wynn, Mairead Hyland, The Lighthouse Lab in Alderley Park and Alex Alderton, Roberto Amato, Jeffrey Barrett, Sonia Goncalves, Ewan Harrison, David K. Jackson, Ian Johnston, Dominic Kwiatkowski, Cordelia Langford, John Sillitoe on behalf of the Wellcome Sanger Institute COVID-19 Surveillance Team |
| EPI_ISL_3468436 | Lighthouse Laboratory Plymouth | Wellcome Sanger Institute for the COVID-19 Genomics UK (COG-UK) Consortium | Lighthouse Laboratory Plymouth and Alex Alderton, Roberto Amato, Jeffrey Barrett, Sonia Goncalves, Ewan Harrison, David K. Jackson, Ian Johnston, Dominic Kwiatkowski, Cordelia Langford, John Sillitoe on behalf of the Wellcome Sanger Institute COVID-19 Surveillance Team |
| EPI_ISL_3468437 | Lighthouse Lab in Alderley Park | Wellcome Sanger Institute for the COVID-19 Genomics UK (COG-UK) Consortium | Jacquelyn Wynn, Mairead Hyland, The Lighthouse Lab in Alderley Park and Alex Alderton, Roberto Amato, Jeffrey Barrett, Sonia Goncalves, Ewan Harrison, David K. Jackson, Ian Johnston, Dominic Kwiatkowski, Cordelia Langford, John Sillitoe on behalf of the Wellcome Sanger Institute COVID-19 Surveillance Team |
| EPI_ISL_3468438 | Lighthouse Laboratory Plymouth | Wellcome Sanger Institute for the COVID-19 Genomics UK (COG-UK) Consortium | Lighthouse Laboratory Plymouth and Alex Alderton, Roberto Amato, Jeffrey Barrett, Sonia Goncalves, Ewan Harrison, David K. Jackson, Ian Johnston, Dominic Kwiatkowski, Cordelia Langford, John Sillitoe on behalf of the Wellcome Sanger Institute COVID-19 Surveillance Team |
| EPI_ISL_3468439, EPI_ISL_3468440 | Lighthouse Lab in Alderley Park | Wellcome Sanger Institute for the COVID-19 Genomics UK (COG-UK) Consortium | Jacquelyn Wynn, Mairead Hyland, The Lighthouse Lab in Alderley Park and Alex Alderton, Roberto Amato, Jeffrey Barrett, Sonia Goncalves, Ewan Harrison, David K. Jackson, Ian Johnston, Dominic Kwiatkowski, Cordelia Langford, John Sillitoe on behalf of the Wellcome Sanger Institute COVID-19 Surveillance Team |
| EPI_ISL_3468441 | Lighthouse Laboratory Plymouth | Wellcome Sanger Institute for the COVID-19 Genomics UK (COG-UK) Consortium | Lighthouse Laboratory Plymouth and Alex Alderton, Roberto Amato, Jeffrey Barrett, Sonia Goncalves, Ewan Harrison, David K. Jackson, Ian Johnston, Dominic Kwiatkowski, Cordelia Langford, John Sillitoe on behalf of the Wellcome Sanger Institute COVID-19 Surveillance Team |
| EPI_ISL_3468442, EPI_ISL_3468443 | Lighthouse Lab in Alderley Park | Wellcome Sanger Institute for the COVID-19 Genomics UK (COG-UK) Consortium | Jacquelyn Wynn, Mairead Hyland, The Lighthouse Lab in Alderley Park and Alex Alderton, Roberto Amato, Jeffrey Barrett, Sonia Goncalves, Ewan Harrison, David K. Jackson, Ian Johnston, Dominic Kwiatkowski, Cordelia Langford, John Sillitoe on behalf of the Wellcome Sanger Institute COVID-19 Surveillance Team |
| EPI_ISL_3468444 | Lighthouse Laboratory Plymouth | Wellcome Sanger Institute for the COVID-19 Genomics UK (COG-UK) Consortium | Lighthouse Laboratory Plymouth and Alex Alderton, Roberto Amato, Jeffrey Barrett, Sonia Goncalves, Ewan Harrison, David K. Jackson, Ian Johnston, Dominic Kwiatkowski, Cordelia Langford, John Sillitoe on behalf of the Wellcome Sanger Institute COVID-19 Surveillance Team |
| EPI_ISL_3468445, EPI_ISL_3468446 | Lighthouse Lab in Alderley Park | Wellcome Sanger Institute for the COVID-19 Genomics UK (COG-UK) Consortium | Jacquelyn Wynn, Mairead Hyland, The Lighthouse Lab in Alderley Park and Alex Alderton, Roberto Amato, Jeffrey Barrett, Sonia Goncalves, Ewan Harrison, David K. Jackson, Ian Johnston, Dominic Kwiatkowski, Cordelia Langford, John Sillitoe on behalf of the Wellcome Sanger Institute COVID-19 Surveillance Team |
| EPI_ISL_3468447 | Lighthouse Laboratory Plymouth | Wellcome Sanger Institute for the COVID-19 Genomics UK (COG-UK) Consortium | Lighthouse Laboratory Plymouth and Alex Alderton, Roberto Amato, Jeffrey Barrett, Sonia Goncalves, Ewan Harrison, David K. Jackson, Ian Johnston, Dominic Kwiatkowski, Cordelia Langford, John Sillitoe on behalf of the Wellcome Sanger Institute COVID-19 Surveillance Team |
| EPI_ISL_3468448 | Lighthouse Lab in Alderley Park | Wellcome Sanger Institute for the COVID-19 Genomics UK (COG-UK) Consortium | Jacquelyn Wynn, Mairead Hyland, The Lighthouse Lab in Alderley Park and Alex Alderton, Roberto Amato, Jeffrey Barrett, Sonia Goncalves, Ewan Harrison, David K. Jackson, Ian Johnston, Dominic Kwiatkowski, Cordelia Langford, John Sillitoe on behalf of the Wellcome Sanger Institute COVID-19 Surveillance Team |
| EPI_ISL_3468449 | Lighthouse Laboratory Plymouth | Wellcome Sanger Institute for the COVID-19 Genomics UK (COG-UK) Consortium | Lighthouse Laboratory Plymouth and Alex Alderton, Roberto Amato, Jeffrey Barrett, Sonia Goncalves, Ewan Harrison, David K. Jackson, Ian Johnston, Dominic Kwiatkowski, Cordelia Langford, John Sillitoe on behalf of the Wellcome Sanger Institute COVID-19 Surveillance Team |
| EPI_ISL_3468450, EPI_ISL_3468451, EPI_ISL_3468452 | Lighthouse Lab in Alderley Park | Wellcome Sanger Institute for the COVID-19 Genomics UK (COG-UK) Consortium | Jacquelyn Wynn, Mairead Hyland, The Lighthouse Lab in Alderley Park and Alex Alderton, Roberto Amato, Jeffrey Barrett, Sonia Goncalves, Ewan Harrison, David K. Jackson, Ian Johnston, Dominic Kwiatkowski, Cordelia Langford, John Sillitoe on behalf of the Wellcome Sanger Institute COVID-19 Surveillance Team |
| EPI_ISL_3468453 | Lighthouse Laboratory Plymouth | Wellcome Sanger Institute for the COVID-19 Genomics UK (COG-UK) Consortium | Lighthouse Laboratory Plymouth and Alex Alderton, Roberto Amato, Jeffrey Barrett, Sonia Goncalves, Ewan Harrison, David K. Jackson, Ian Johnston, Dominic Kwiatkowski, Cordelia Langford, John Sillitoe on behalf of the Wellcome Sanger Institute COVID-19 Surveillance Team |
| EPI_ISL_3468454, EPI_ISL_3468455 | Lighthouse Lab in Alderley Park | Wellcome Sanger Institute for the COVID-19 Genomics UK (COG-UK) Consortium | Jacquelyn Wynn, Mairead Hyland, The Lighthouse Lab in Alderley Park and Alex Alderton, Roberto Amato, Jeffrey Barrett, Sonia Goncalves, Ewan Harrison, David K. Jackson, Ian Johnston, Dominic Kwiatkowski, Cordelia Langford, John Sillitoe on behalf of the Wellcome Sanger Institute COVID-19 Surveillance Team |
| EPI_ISL_3468456, EPI_ISL_3468457 | Lighthouse Laboratory Plymouth | Wellcome Sanger Institute for the COVID-19 Genomics UK (COG-UK) Consortium | Lighthouse Laboratory Plymouth and Alex Alderton, Roberto Amato, Jeffrey Barrett, Sonia Goncalves, Ewan Harrison, David K. Jackson, Ian Johnston, Dominic Kwiatkowski, Cordelia Langford, John Sillitoe on behalf of the Wellcome Sanger Institute COVID-19 Surveillance Team |
| EPI_ISL_3468458 | Lighthouse Lab in Alderley Park | Wellcome Sanger Institute for the COVID-19 Genomics UK (COG-UK) Consortium | Jacquelyn Wynn, Mairead Hyland, The Lighthouse Lab in Alderley Park and Alex Alderton, Roberto Amato, Jeffrey Barrett, Sonia Goncalves, Ewan Harrison, David K. Jackson, Ian Johnston, Dominic Kwiatkowski, Cordelia Langford, John Sillitoe on behalf of the Wellcome Sanger Institute COVID-19 Surveillance Team |
| EPI_ISL_3468459, EPI_ISL_3468460, EPI_ISL_3468461, EPI_ISL_3468462 | Lighthouse Laboratory Plymouth | Wellcome Sanger Institute for the COVID-19 Genomics UK (COG-UK) Consortium | Lighthouse Laboratory Plymouth and Alex Alderton, Roberto Amato, Jeffrey Barrett, Sonia Goncalves, Ewan Harrison, David K. Jackson, Ian Johnston, Dominic Kwiatkowski, Cordelia Langford, John Sillitoe on behalf of the Wellcome Sanger Institute COVID-19 Surveillance Team |
| EPI_ISL_3468463 | Lighthouse Lab in Alderley Park | Wellcome Sanger Institute for the COVID-19 Genomics UK (COG-UK) Consortium | Jacquelyn Wynn, Mairead Hyland, The Lighthouse Lab in Alderley Park and Alex Alderton, Roberto Amato, Jeffrey Barrett, Sonia Goncalves, Ewan Harrison, David K. Jackson, Ian Johnston, Dominic Kwiatkowski, Cordelia Langford, John Sillitoe on behalf of the Wellcome Sanger Institute COVID-19 Surveillance Team |
| EPI_ISL_3468464 | Lighthouse Laboratory Plymouth | Wellcome Sanger Institute for the COVID-19 Genomics UK (COG-UK) Consortium | Lighthouse Laboratory Plymouth and Alex Alderton, Roberto Amato, Jeffrey Barrett, Sonia Goncalves, Ewan Harrison, David K. Jackson, Ian Johnston, Dominic Kwiatkowski, Cordelia Langford, John Sillitoe on behalf of the Wellcome Sanger Institute COVID-19 Surveillance Team |
| EPI_ISL_3468465, | Lighthouse Lab in Alderley Park | Wellcome Sanger Institute for the COVID-19 Genomics UK (COG-UK) | Jacquelyn Wynn, Mairead Hyland, The Lighthouse Lab in Alderley Park and Alex Alderton, Roberto Amato, Jeffrey Barrett, Sonia Goncalves, Ewan Harrison, David K. Jackson, Ian Johnston, Dominic Kwiatkowski, Cordelia Langford, John Sillitoe on |

| | | | |
|---|---|---|---|
| EPI_ISL_3468466, EPI_ISL_3468467, EPI_ISL_3468468, EPI_ISL_3468469, EPI_ISL_3468470, EPI_ISL_3468471, EPI_ISL_3468472 | | Consortium | behalf of the Wellcome Sanger Institute COVID-19 Surveillance Team |
| EPI_ISL_3468473, EPI_ISL_3468474 | Lighthouse Laboratory Plymouth | Wellcome Sanger Institute for the COVID-19 Genomics UK (COG-UK) Consortium | Lighthouse Laboratory Plymouth and Alex Alderton, Roberto Amato, Jeffrey Barrett, Sonia Goncalves, Ewan Harrison, David K. Jackson, Ian Johnston, Dominic Kwiatkowski, Cordelia Langford, John Sillitoe on behalf of the Wellcome Sanger Institute COVID-19 Surveillance Team |
| EPI_ISL_3468475 | Lighthouse Lab in Alderley Park | Wellcome Sanger Institute for the COVID-19 Genomics UK (COG-UK) Consortium | Jacquelyn Wynn, Mairead Hyland, The Lighthouse Lab in Alderley Park and Alex Alderton, Roberto Amato, Jeffrey Barrett, Sonia Goncalves, Ewan Harrison, David K. Jackson, Ian Johnston, Dominic Kwiatkowski, Cordelia Langford, John Sillitoe on behalf of the Wellcome Sanger Institute COVID-19 Surveillance Team |
| EPI_ISL_3468476 | Lighthouse Laboratory Plymouth | Wellcome Sanger Institute for the COVID-19 Genomics UK (COG-UK) Consortium | Lighthouse Laboratory Plymouth and Alex Alderton, Roberto Amato, Jeffrey Barrett, Sonia Goncalves, Ewan Harrison, David K. Jackson, Ian Johnston, Dominic Kwiatkowski, Cordelia Langford, John Sillitoe on behalf of the Wellcome Sanger Institute COVID-19 Surveillance Team |
| EPI_ISL_3468477 | Lighthouse Lab in Alderley Park | Wellcome Sanger Institute for the COVID-19 Genomics UK (COG-UK) Consortium | Jacquelyn Wynn, Mairead Hyland, The Lighthouse Lab in Alderley Park and Alex Alderton, Roberto Amato, Jeffrey Barrett, Sonia Goncalves, Ewan Harrison, David K. Jackson, Ian Johnston, Dominic Kwiatkowski, Cordelia Langford, John Sillitoe on behalf of the Wellcome Sanger Institute COVID-19 Surveillance Team |
| EPI_ISL_3468478 | Lighthouse Laboratory Plymouth | Wellcome Sanger Institute for the COVID-19 Genomics UK (COG-UK) Consortium | Lighthouse Laboratory Plymouth and Alex Alderton, Roberto Amato, Jeffrey Barrett, Sonia Goncalves, Ewan Harrison, David K. Jackson, Ian Johnston, Dominic Kwiatkowski, Cordelia Langford, John Sillitoe on behalf of the Wellcome Sanger Institute COVID-19 Surveillance Team |
| EPI_ISL_3468479, EPI_ISL_3468480 | Lighthouse Lab in Alderley Park | Wellcome Sanger Institute for the COVID-19 Genomics UK (COG-UK) Consortium | Jacquelyn Wynn, Mairead Hyland, The Lighthouse Lab in Alderley Park and Alex Alderton, Roberto Amato, Jeffrey Barrett, Sonia Goncalves, Ewan Harrison, David K. Jackson, Ian Johnston, Dominic Kwiatkowski, Cordelia Langford, John Sillitoe on behalf of the Wellcome Sanger Institute COVID-19 Surveillance Team |
| EPI_ISL_3468481, EPI_ISL_3468482 | Lighthouse Laboratory Plymouth | Wellcome Sanger Institute for the COVID-19 Genomics UK (COG-UK) Consortium | Lighthouse Laboratory Plymouth and Alex Alderton, Roberto Amato, Jeffrey Barrett, Sonia Goncalves, Ewan Harrison, David K. Jackson, Ian Johnston, Dominic Kwiatkowski, Cordelia Langford, John Sillitoe on behalf of the Wellcome Sanger Institute COVID-19 Surveillance Team |
| EPI_ISL_3468483, EPI_ISL_3468484 | Lighthouse Lab in Alderley Park | Wellcome Sanger Institute for the COVID-19 Genomics UK (COG-UK) Consortium | Jacquelyn Wynn, Mairead Hyland, The Lighthouse Lab in Alderley Park and Alex Alderton, Roberto Amato, Jeffrey Barrett, Sonia Goncalves, Ewan Harrison, David K. Jackson, Ian Johnston, Dominic Kwiatkowski, Cordelia Langford, John Sillitoe on behalf of the Wellcome Sanger Institute COVID-19 Surveillance Team |
| EPI_ISL_3468485 | Lighthouse Laboratory Plymouth | Wellcome Sanger Institute for the COVID-19 Genomics UK (COG-UK) Consortium | Lighthouse Laboratory Plymouth and Alex Alderton, Roberto Amato, Jeffrey Barrett, Sonia Goncalves, Ewan Harrison, David K. Jackson, Ian Johnston, Dominic Kwiatkowski, Cordelia Langford, John Sillitoe on behalf of the Wellcome Sanger Institute COVID-19 Surveillance Team |
| EPI_ISL_3468486, EPI_ISL_3468487, EPI_ISL_3468488, EPI_ISL_3468489, EPI_ISL_3468490 | Lighthouse Lab in Alderley Park | Wellcome Sanger Institute for the COVID-19 Genomics UK (COG-UK) Consortium | Jacquelyn Wynn, Mairead Hyland, The Lighthouse Lab in Alderley Park and Alex Alderton, Roberto Amato, Jeffrey Barrett, Sonia Goncalves, Ewan Harrison, David K. Jackson, Ian Johnston, Dominic Kwiatkowski, Cordelia Langford, John Sillitoe on behalf of the Wellcome Sanger Institute COVID-19 Surveillance Team |
| EPI_ISL_3468491 | Lighthouse Laboratory Plymouth | Wellcome Sanger Institute for the COVID-19 Genomics UK (COG-UK) Consortium | Lighthouse Laboratory Plymouth and Alex Alderton, Roberto Amato, Jeffrey Barrett, Sonia Goncalves, Ewan Harrison, David K. Jackson, Ian Johnston, Dominic Kwiatkowski, Cordelia Langford, John Sillitoe on behalf of the Wellcome Sanger Institute COVID-19 Surveillance Team |
| EPI_ISL_3468492 | Lighthouse Lab in Alderley Park | Wellcome Sanger Institute for the COVID-19 Genomics UK (COG-UK) Consortium | Jacquelyn Wynn, Mairead Hyland, The Lighthouse Lab in Alderley Park and Alex Alderton, Roberto Amato, Jeffrey Barrett, Sonia Goncalves, Ewan Harrison, David K. Jackson, Ian Johnston, Dominic Kwiatkowski, Cordelia Langford, John Sillitoe on behalf of the Wellcome Sanger Institute COVID-19 Surveillance Team |
| EPI_ISL_3468493, EPI_ISL_3468494, EPI_ISL_3468495 | Lighthouse Laboratory Plymouth | Wellcome Sanger Institute for the COVID-19 Genomics UK (COG-UK) Consortium | Lighthouse Laboratory Plymouth and Alex Alderton, Roberto Amato, Jeffrey Barrett, Sonia Goncalves, Ewan Harrison, David K. Jackson, Ian Johnston, Dominic Kwiatkowski, Cordelia Langford, John Sillitoe on behalf of the Wellcome Sanger Institute COVID-19 Surveillance Team |
| EPI_ISL_3468496 | Lighthouse Lab in Alderley Park | Wellcome Sanger Institute for the COVID-19 Genomics UK (COG-UK) Consortium | Jacquelyn Wynn, Mairead Hyland, The Lighthouse Lab in Alderley Park and Alex Alderton, Roberto Amato, Jeffrey Barrett, Sonia Goncalves, Ewan Harrison, David K. Jackson, Ian Johnston, Dominic Kwiatkowski, Cordelia Langford, John Sillitoe on behalf of the Wellcome Sanger Institute COVID-19 Surveillance Team |
| EPI_ISL_3468497 | Lighthouse Laboratory Plymouth | Wellcome Sanger Institute for the COVID-19 Genomics UK (COG-UK) Consortium | Lighthouse Laboratory Plymouth and Alex Alderton, Roberto Amato, Jeffrey Barrett, Sonia Goncalves, Ewan Harrison, David K. Jackson, Ian Johnston, Dominic Kwiatkowski, Cordelia Langford, John Sillitoe on behalf of the Wellcome Sanger Institute COVID-19 Surveillance Team |
| EPI_ISL_3468498, EPI_ISL_3468499 | Lighthouse Lab in Alderley Park | Wellcome Sanger Institute for the COVID-19 Genomics UK (COG-UK) Consortium | Jacquelyn Wynn, Mairead Hyland, The Lighthouse Lab in Alderley Park and Alex Alderton, Roberto Amato, Jeffrey Barrett, Sonia Goncalves, Ewan Harrison, David K. Jackson, Ian Johnston, Dominic Kwiatkowski, Cordelia Langford, John Sillitoe on behalf of the Wellcome Sanger Institute COVID-19 Surveillance Team |
| EPI_ISL_3468500, EPI_ISL_3468501 | Lighthouse Laboratory Plymouth | Wellcome Sanger Institute for the COVID-19 Genomics UK (COG-UK) Consortium | Lighthouse Laboratory Plymouth and Alex Alderton, Roberto Amato, Jeffrey Barrett, Sonia Goncalves, Ewan Harrison, David K. Jackson, Ian Johnston, Dominic Kwiatkowski, Cordelia Langford, John Sillitoe on behalf of the Wellcome Sanger Institute COVID-19 Surveillance Team |
| EPI_ISL_3468502 | Lighthouse Lab in Alderley Park | Wellcome Sanger Institute for the COVID-19 Genomics UK (COG-UK) Consortium | Jacquelyn Wynn, Mairead Hyland, The Lighthouse Lab in Alderley Park and Alex Alderton, Roberto Amato, Jeffrey Barrett, Sonia Goncalves, Ewan Harrison, David K. Jackson, Ian Johnston, Dominic Kwiatkowski, Cordelia Langford, John Sillitoe on behalf of the Wellcome Sanger Institute COVID-19 Surveillance Team |
| EPI_ISL_3468503 | Lighthouse Laboratory Plymouth | Wellcome Sanger Institute for the COVID-19 Genomics UK (COG-UK) Consortium | Lighthouse Laboratory Plymouth and Alex Alderton, Roberto Amato, Jeffrey Barrett, Sonia Goncalves, Ewan Harrison, David K. Jackson, Ian Johnston, Dominic Kwiatkowski, Cordelia Langford, John Sillitoe on behalf of the Wellcome Sanger Institute COVID-19 Surveillance Team |
| EPI_ISL_3468504 | Lighthouse Lab in Alderley Park | Wellcome Sanger Institute for the COVID-19 Genomics UK (COG-UK) Consortium | Jacquelyn Wynn, Mairead Hyland, The Lighthouse Lab in Alderley Park and Alex Alderton, Roberto Amato, Jeffrey Barrett, Sonia Goncalves, Ewan Harrison, David K. Jackson, Ian Johnston, Dominic Kwiatkowski, Cordelia Langford, John Sillitoe on behalf of the Wellcome Sanger Institute COVID-19 Surveillance Team |
| EPI_ISL_3468505, EPI_ISL_3468506 | Lighthouse Laboratory Plymouth | Wellcome Sanger Institute for the COVID-19 Genomics UK (COG-UK) Consortium | Lighthouse Laboratory Plymouth and Alex Alderton, Roberto Amato, Jeffrey Barrett, Sonia Goncalves, Ewan Harrison, David K. Jackson, Ian Johnston, Dominic Kwiatkowski, Cordelia Langford, John Sillitoe on behalf of the Wellcome Sanger Institute COVID-19 Surveillance Team |
| EPI_ISL_3468507, EPI_ISL_3468508, EPI_ISL_3468509 | Lighthouse Lab in Alderley Park | Wellcome Sanger Institute for the COVID-19 Genomics UK (COG-UK) Consortium | Jacquelyn Wynn, Mairead Hyland, The Lighthouse Lab in Alderley Park and Alex Alderton, Roberto Amato, Jeffrey Barrett, Sonia Goncalves, Ewan Harrison, David K. Jackson, Ian Johnston, Dominic Kwiatkowski, Cordelia Langford, John Sillitoe on behalf of the Wellcome Sanger Institute COVID-19 Surveillance Team |
| EPI_ISL_3468510 | Lighthouse Laboratory Plymouth | Wellcome Sanger Institute for the COVID-19 Genomics UK (COG-UK) Consortium | Lighthouse Laboratory Plymouth and Alex Alderton, Roberto Amato, Jeffrey Barrett, Sonia Goncalves, Ewan Harrison, David K. Jackson, Ian Johnston, Dominic Kwiatkowski, Cordelia Langford, John Sillitoe on behalf of the Wellcome Sanger Institute COVID-19 Surveillance Team |
| EPI_ISL_3468511 | Lighthouse Lab in Alderley Park | Wellcome Sanger Institute for the COVID-19 Genomics UK (COG-UK) Consortium | Jacquelyn Wynn, Mairead Hyland, The Lighthouse Lab in Alderley Park and Alex Alderton, Roberto Amato, Jeffrey Barrett, Sonia Goncalves, Ewan Harrison, David K. Jackson, Ian Johnston, Dominic Kwiatkowski, Cordelia Langford, John Sillitoe on behalf of the Wellcome Sanger Institute COVID-19 Surveillance Team |
| EPI_ISL_3468512, EPI_ISL_3468513, EPI_ISL_3468514 | Lighthouse Laboratory Plymouth | Wellcome Sanger Institute for the COVID-19 Genomics UK (COG-UK) Consortium | Lighthouse Laboratory Plymouth and Alex Alderton, Roberto Amato, Jeffrey Barrett, Sonia Goncalves, Ewan Harrison, David K. Jackson, Ian Johnston, Dominic Kwiatkowski, Cordelia Langford, John Sillitoe on behalf of the Wellcome Sanger Institute COVID-19 Surveillance Team |
| EPI_ISL_3468515 | Lighthouse Lab in Alderley Park | Wellcome Sanger Institute for the COVID-19 Genomics UK (COG-UK) Consortium | Jacquelyn Wynn, Mairead Hyland, The Lighthouse Lab in Alderley Park and Alex Alderton, Roberto Amato, Jeffrey Barrett, Sonia Goncalves, Ewan Harrison, David K. Jackson, Ian Johnston, Dominic Kwiatkowski, Cordelia Langford, John Sillitoe on behalf of the Wellcome Sanger Institute COVID-19 Surveillance Team |
| EPI_ISL_3468516 | Lighthouse Laboratory Plymouth | Wellcome Sanger Institute for the COVID-19 Genomics UK (COG-UK) Consortium | Lighthouse Laboratory Plymouth and Alex Alderton, Roberto Amato, Jeffrey Barrett, Sonia Goncalves, Ewan Harrison, David K. Jackson, Ian Johnston, Dominic Kwiatkowski, Cordelia Langford, John Sillitoe on behalf of the Wellcome Sanger Institute COVID-19 Surveillance Team |
| EPI_ISL_3468517 | Lighthouse Lab in Alderley Park | Wellcome Sanger Institute for the COVID-19 Genomics UK (COG-UK) Consortium | Jacquelyn Wynn, Mairead Hyland, The Lighthouse Lab in Alderley Park and Alex Alderton, Roberto Amato, Jeffrey Barrett, Sonia Goncalves, Ewan Harrison, David K. Jackson, Ian Johnston, Dominic Kwiatkowski, Cordelia Langford, John Sillitoe on behalf of the Wellcome Sanger Institute COVID-19 Surveillance Team |
| EPI_ISL_3468518 | Lighthouse Laboratory Plymouth | Wellcome Sanger Institute for the COVID-19 Genomics UK (COG-UK) Consortium | Lighthouse Laboratory Plymouth and Alex Alderton, Roberto Amato, Jeffrey Barrett, Sonia Goncalves, Ewan Harrison, David K. Jackson, Ian Johnston, Dominic Kwiatkowski, Cordelia Langford, John Sillitoe on behalf of the Wellcome Sanger Institute COVID-19 Surveillance Team |
| EPI_ISL_3468519, EPI_ISL_3468520, EPI_ISL_3468521 | Lighthouse Lab in Alderley Park | Wellcome Sanger Institute for the COVID-19 Genomics UK (COG-UK) Consortium | Jacquelyn Wynn, Mairead Hyland, The Lighthouse Lab in Alderley Park and Alex Alderton, Roberto Amato, Jeffrey Barrett, Sonia Goncalves, Ewan Harrison, David K. Jackson, Ian Johnston, Dominic Kwiatkowski, Cordelia Langford, John Sillitoe on behalf of the Wellcome Sanger Institute COVID-19 Surveillance Team |
| EPI_ISL_3468522, EPI_ISL_3468523, EPI_ISL_3468524 | Lighthouse Laboratory Plymouth | Wellcome Sanger Institute for the COVID-19 Genomics UK (COG-UK) Consortium | Lighthouse Laboratory Plymouth and Alex Alderton, Roberto Amato, Jeffrey Barrett, Sonia Goncalves, Ewan Harrison, David K. Jackson, Ian Johnston, Dominic Kwiatkowski, Cordelia Langford, John Sillitoe on behalf of the Wellcome Sanger Institute COVID-19 Surveillance Team |
| EPI_ISL_3468525 | Lighthouse Lab in Alderley Park | Wellcome Sanger Institute for the COVID-19 Genomics UK (COG-UK) Consortium | Jacquelyn Wynn, Mairead Hyland, The Lighthouse Lab in Alderley Park and Alex Alderton, Roberto Amato, Jeffrey Barrett, Sonia Goncalves, Ewan Harrison, David K. Jackson, Ian Johnston, Dominic Kwiatkowski, Cordelia Langford, John Sillitoe on behalf of the Wellcome Sanger Institute COVID-19 Surveillance Team |
| EPI_ISL_3468526 | Lighthouse Laboratory Plymouth | Wellcome Sanger Institute for the COVID-19 Genomics UK (COG-UK) Consortium | Lighthouse Laboratory Plymouth and Alex Alderton, Roberto Amato, Jeffrey Barrett, Sonia Goncalves, Ewan Harrison, David K. Jackson, Ian Johnston, Dominic Kwiatkowski, Cordelia Langford, John Sillitoe on behalf of the Wellcome Sanger Institute COVID-19 Surveillance Team |
| EPI_ISL_3468527 | Lighthouse Lab in Alderley Park | Wellcome Sanger Institute for the COVID-19 Genomics UK (COG-UK) Consortium | Jacquelyn Wynn, Mairead Hyland, The Lighthouse Lab in Alderley Park and Alex Alderton, Roberto Amato, Jeffrey Barrett, Sonia Goncalves, Ewan Harrison, David K. Jackson, Ian Johnston, Dominic Kwiatkowski, Cordelia Langford, John Sillitoe on behalf of the Wellcome Sanger Institute COVID-19 Surveillance Team |
| EPI_ISL_3468528, EPI_ISL_3468529 | Lighthouse Laboratory Plymouth | Wellcome Sanger Institute for the COVID-19 Genomics UK (COG-UK) Consortium | Lighthouse Laboratory Plymouth and Alex Alderton, Roberto Amato, Jeffrey Barrett, Sonia Goncalves, Ewan Harrison, David K. Jackson, Ian Johnston, Dominic Kwiatkowski, Cordelia Langford, John Sillitoe on behalf of the Wellcome Sanger Institute COVID-19 Surveillance Team |
| EPI_ISL_3468530, EPI_ISL_3468531 | Lighthouse Lab in Alderley Park | Wellcome Sanger Institute for the COVID-19 Genomics UK (COG-UK) Consortium | Jacquelyn Wynn, Mairead Hyland, The Lighthouse Lab in Alderley Park and Alex Alderton, Roberto Amato, Jeffrey Barrett, Sonia Goncalves, Ewan Harrison, David K. Jackson, Ian Johnston, Dominic Kwiatkowski, Cordelia Langford, John Sillitoe on behalf of the Wellcome Sanger Institute COVID-19 Surveillance Team |
| EPI_ISL_3468532, EPI_ISL_3468533 | Lighthouse Laboratory Plymouth | Wellcome Sanger Institute for the COVID-19 Genomics UK (COG-UK) Consortium | Lighthouse Laboratory Plymouth and Alex Alderton, Roberto Amato, Jeffrey Barrett, Sonia Goncalves, Ewan Harrison, David K. Jackson, Ian Johnston, Dominic Kwiatkowski, Cordelia Langford, John Sillitoe on behalf of the Wellcome Sanger Institute COVID-19 Surveillance Team |
| EPI_ISL_3468534 | Lighthouse Lab in Alderley Park | Wellcome Sanger Institute for the COVID-19 Genomics UK (COG-UK) Consortium | Jacquelyn Wynn, Mairead Hyland, The Lighthouse Lab in Alderley Park and Alex Alderton, Roberto Amato, Jeffrey Barrett, Sonia Goncalves, Ewan Harrison, David K. Jackson, Ian Johnston, Dominic Kwiatkowski, Cordelia Langford, John Sillitoe on behalf of the Wellcome Sanger Institute COVID-19 Surveillance Team |
| EPI_ISL_3468535, EPI_ISL_3468536 | Lighthouse Laboratory Plymouth | Wellcome Sanger Institute for the COVID-19 Genomics UK (COG-UK) Consortium | Lighthouse Laboratory Plymouth and Alex Alderton, Roberto Amato, Jeffrey Barrett, Sonia Goncalves, Ewan Harrison, David K. Jackson, Ian Johnston, Dominic Kwiatkowski, Cordelia Langford, John Sillitoe on behalf of the Wellcome Sanger Institute COVID-19 Surveillance Team |

| | | | |
|---|---|---|---|
| EPI_ISL_3468537, EPI_ISL_3468538 | Lighthouse Lab in Alderley Park | Wellcome Sanger Institute for the COVID-19 Genomics UK (COG-UK) Consortium | Jacquelyn Wynn, Mairead Hyland, The Lighthouse Lab in Alderley Park and Alex Alderton, Roberto Amato, Jeffrey Barrett, Sonia Goncalves, Ewan Harrison, David K. Jackson, Ian Johnston, Dominic Kwiatkowski, Cordelia Langford, John Sillitoe on behalf of the Wellcome Sanger Institute COVID-19 Surveillance Team |
| EPI_ISL_3468539, EPI_ISL_3468540 | Lighthouse Laboratory Plymouth | Wellcome Sanger Institute for the COVID-19 Genomics UK (COG-UK) Consortium | Lighthouse Laboratory Plymouth and Alex Alderton, Roberto Amato, Jeffrey Barrett, Sonia Goncalves, Ewan Harrison, David K. Jackson, Ian Johnston, Dominic Kwiatkowski, Cordelia Langford, John Sillitoe on behalf of the Wellcome Sanger Institute COVID-19 Surveillance Team |
| EPI_ISL_3468541, EPI_ISL_3468542, EPI_ISL_3468543 | Lighthouse Lab in Alderley Park | Wellcome Sanger Institute for the COVID-19 Genomics UK (COG-UK) Consortium | Jacquelyn Wynn, Mairead Hyland, The Lighthouse Lab in Alderley Park and Alex Alderton, Roberto Amato, Jeffrey Barrett, Sonia Goncalves, Ewan Harrison, David K. Jackson, Ian Johnston, Dominic Kwiatkowski, Cordelia Langford, John Sillitoe on behalf of the Wellcome Sanger Institute COVID-19 Surveillance Team |
| EPI_ISL_3468544, EPI_ISL_3468545, EPI_ISL_3468546, EPI_ISL_3468547, EPI_ISL_3468548 | Lighthouse Laboratory Plymouth | Wellcome Sanger Institute for the COVID-19 Genomics UK (COG-UK) Consortium | Lighthouse Laboratory Plymouth and Alex Alderton, Roberto Amato, Jeffrey Barrett, Sonia Goncalves, Ewan Harrison, David K. Jackson, Ian Johnston, Dominic Kwiatkowski, Cordelia Langford, John Sillitoe on behalf of the Wellcome Sanger Institute COVID-19 Surveillance Team |
| EPI_ISL_3468549 | Lighthouse Lab in Alderley Park | Wellcome Sanger Institute for the COVID-19 Genomics UK (COG-UK) Consortium | Jacquelyn Wynn, Mairead Hyland, The Lighthouse Lab in Alderley Park and Alex Alderton, Roberto Amato, Jeffrey Barrett, Sonia Goncalves, Ewan Harrison, David K. Jackson, Ian Johnston, Dominic Kwiatkowski, Cordelia Langford, John Sillitoe on behalf of the Wellcome Sanger Institute COVID-19 Surveillance Team |
| EPI_ISL_3468550 | Lighthouse Laboratory Plymouth | Wellcome Sanger Institute for the COVID-19 Genomics UK (COG-UK) Consortium | Lighthouse Laboratory Plymouth and Alex Alderton, Roberto Amato, Jeffrey Barrett, Sonia Goncalves, Ewan Harrison, David K. Jackson, Ian Johnston, Dominic Kwiatkowski, Cordelia Langford, John Sillitoe on behalf of the Wellcome Sanger Institute COVID-19 Surveillance Team |
| EPI_ISL_3468551 | Lighthouse Lab in Alderley Park | Wellcome Sanger Institute for the COVID-19 Genomics UK (COG-UK) Consortium | Jacquelyn Wynn, Mairead Hyland, The Lighthouse Lab in Alderley Park and Alex Alderton, Roberto Amato, Jeffrey Barrett, Sonia Goncalves, Ewan Harrison, David K. Jackson, Ian Johnston, Dominic Kwiatkowski, Cordelia Langford, John Sillitoe on behalf of the Wellcome Sanger Institute COVID-19 Surveillance Team |
| EPI_ISL_3468552 | Lighthouse Laboratory Plymouth | Wellcome Sanger Institute for the COVID-19 Genomics UK (COG-UK) Consortium | Lighthouse Laboratory Plymouth and Alex Alderton, Roberto Amato, Jeffrey Barrett, Sonia Goncalves, Ewan Harrison, David K. Jackson, Ian Johnston, Dominic Kwiatkowski, Cordelia Langford, John Sillitoe on behalf of the Wellcome Sanger Institute COVID-19 Surveillance Team |
| EPI_ISL_3468553 | Lighthouse Lab in Alderley Park | Wellcome Sanger Institute for the COVID-19 Genomics UK (COG-UK) Consortium | Jacquelyn Wynn, Mairead Hyland, The Lighthouse Lab in Alderley Park and Alex Alderton, Roberto Amato, Jeffrey Barrett, Sonia Goncalves, Ewan Harrison, David K. Jackson, Ian Johnston, Dominic Kwiatkowski, Cordelia Langford, John Sillitoe on behalf of the Wellcome Sanger Institute COVID-19 Surveillance Team |
| EPI_ISL_3468554, EPI_ISL_3468555 | Lighthouse Laboratory Plymouth | Wellcome Sanger Institute for the COVID-19 Genomics UK (COG-UK) Consortium | Lighthouse Laboratory Plymouth and Alex Alderton, Roberto Amato, Jeffrey Barrett, Sonia Goncalves, Ewan Harrison, David K. Jackson, Ian Johnston, Dominic Kwiatkowski, Cordelia Langford, John Sillitoe on behalf of the Wellcome Sanger Institute COVID-19 Surveillance Team |
| EPI_ISL_3468556, EPI_ISL_3468557, EPI_ISL_3468558, EPI_ISL_3468559, EPI_ISL_3468560 | Lighthouse Lab in Alderley Park | Wellcome Sanger Institute for the COVID-19 Genomics UK (COG-UK) Consortium | Jacquelyn Wynn, Mairead Hyland, The Lighthouse Lab in Alderley Park and Alex Alderton, Roberto Amato, Jeffrey Barrett, Sonia Goncalves, Ewan Harrison, David K. Jackson, Ian Johnston, Dominic Kwiatkowski, Cordelia Langford, John Sillitoe on behalf of the Wellcome Sanger Institute COVID-19 Surveillance Team |
| EPI_ISL_3468561, EPI_ISL_3468562 | Lighthouse Laboratory Plymouth | Wellcome Sanger Institute for the COVID-19 Genomics UK (COG-UK) Consortium | Lighthouse Laboratory Plymouth and Alex Alderton, Roberto Amato, Jeffrey Barrett, Sonia Goncalves, Ewan Harrison, David K. Jackson, Ian Johnston, Dominic Kwiatkowski, Cordelia Langford, John Sillitoe on behalf of the Wellcome Sanger Institute COVID-19 Surveillance Team |
| EPI_ISL_3468563, EPI_ISL_3468564 | Lighthouse Lab in Alderley Park | Wellcome Sanger Institute for the COVID-19 Genomics UK (COG-UK) Consortium | Jacquelyn Wynn, Mairead Hyland, The Lighthouse Lab in Alderley Park and Alex Alderton, Roberto Amato, Jeffrey Barrett, Sonia Goncalves, Ewan Harrison, David K. Jackson, Ian Johnston, Dominic Kwiatkowski, Cordelia Langford, John Sillitoe on behalf of the Wellcome Sanger Institute COVID-19 Surveillance Team |
| EPI_ISL_3468565 | Lighthouse Laboratory Plymouth | Wellcome Sanger Institute for the COVID-19 Genomics UK (COG-UK) Consortium | Lighthouse Laboratory Plymouth and Alex Alderton, Roberto Amato, Jeffrey Barrett, Sonia Goncalves, Ewan Harrison, David K. Jackson, Ian Johnston, Dominic Kwiatkowski, Cordelia Langford, John Sillitoe on behalf of the Wellcome Sanger Institute COVID-19 Surveillance Team |
| EPI_ISL_3468566 | Lighthouse Lab in Alderley Park | Wellcome Sanger Institute for the COVID-19 Genomics UK (COG-UK) Consortium | Jacquelyn Wynn, Mairead Hyland, The Lighthouse Lab in Alderley Park and Alex Alderton, Roberto Amato, Jeffrey Barrett, Sonia Goncalves, Ewan Harrison, David K. Jackson, Ian Johnston, Dominic Kwiatkowski, Cordelia Langford, John Sillitoe on behalf of the Wellcome Sanger Institute COVID-19 Surveillance Team |
| EPI_ISL_3468567, EPI_ISL_3468568, EPI_ISL_3468569 | Lighthouse Laboratory Plymouth | Wellcome Sanger Institute for the COVID-19 Genomics UK (COG-UK) Consortium | Lighthouse Laboratory Plymouth and Alex Alderton, Roberto Amato, Jeffrey Barrett, Sonia Goncalves, Ewan Harrison, David K. Jackson, Ian Johnston, Dominic Kwiatkowski, Cordelia Langford, John Sillitoe on behalf of the Wellcome Sanger Institute COVID-19 Surveillance Team |
| EPI_ISL_3468570 | Lighthouse Lab in Alderley Park | Wellcome Sanger Institute for the COVID-19 Genomics UK (COG-UK) Consortium | Jacquelyn Wynn, Mairead Hyland, The Lighthouse Lab in Alderley Park and Alex Alderton, Roberto Amato, Jeffrey Barrett, Sonia Goncalves, Ewan Harrison, David K. Jackson, Ian Johnston, Dominic Kwiatkowski, Cordelia Langford, John Sillitoe on behalf of the Wellcome Sanger Institute COVID-19 Surveillance Team |
| EPI_ISL_3468571 | Lighthouse Laboratory Plymouth | Wellcome Sanger Institute for the COVID-19 Genomics UK (COG-UK) Consortium | Lighthouse Laboratory Plymouth and Alex Alderton, Roberto Amato, Jeffrey Barrett, Sonia Goncalves, Ewan Harrison, David K. Jackson, Ian Johnston, Dominic Kwiatkowski, Cordelia Langford, John Sillitoe on behalf of the Wellcome Sanger Institute COVID-19 Surveillance Team |
| EPI_ISL_3468572, EPI_ISL_3468573, EPI_ISL_3468574 | Lighthouse Lab in Alderley Park | Wellcome Sanger Institute for the COVID-19 Genomics UK (COG-UK) Consortium | Jacquelyn Wynn, Mairead Hyland, The Lighthouse Lab in Alderley Park and Alex Alderton, Roberto Amato, Jeffrey Barrett, Sonia Goncalves, Ewan Harrison, David K. Jackson, Ian Johnston, Dominic Kwiatkowski, Cordelia Langford, John Sillitoe on behalf of the Wellcome Sanger Institute COVID-19 Surveillance Team |
| EPI_ISL_3468575, EPI_ISL_3468576 | Lighthouse Laboratory Plymouth | Wellcome Sanger Institute for the COVID-19 Genomics UK (COG-UK) Consortium | Lighthouse Laboratory Plymouth and Alex Alderton, Roberto Amato, Jeffrey Barrett, Sonia Goncalves, Ewan Harrison, David K. Jackson, Ian Johnston, Dominic Kwiatkowski, Cordelia Langford, John Sillitoe on behalf of the Wellcome Sanger Institute COVID-19 Surveillance Team |
| EPI_ISL_3468577, EPI_ISL_3468578, EPI_ISL_3468579, EPI_ISL_3468580, EPI_ISL_3468581 | Lighthouse Lab in Alderley Park | Wellcome Sanger Institute for the COVID-19 Genomics UK (COG-UK) Consortium | Jacquelyn Wynn, Mairead Hyland, The Lighthouse Lab in Alderley Park and Alex Alderton, Roberto Amato, Jeffrey Barrett, Sonia Goncalves, Ewan Harrison, David K. Jackson, Ian Johnston, Dominic Kwiatkowski, Cordelia Langford, John Sillitoe on behalf of the Wellcome Sanger Institute COVID-19 Surveillance Team |
| EPI_ISL_3468582, EPI_ISL_3468583 | Lighthouse Laboratory Plymouth | Wellcome Sanger Institute for the COVID-19 Genomics UK (COG-UK) Consortium | Lighthouse Laboratory Plymouth and Alex Alderton, Roberto Amato, Jeffrey Barrett, Sonia Goncalves, Ewan Harrison, David K. Jackson, Ian Johnston, Dominic Kwiatkowski, Cordelia Langford, John Sillitoe on behalf of the Wellcome Sanger Institute COVID-19 Surveillance Team |
| EPI_ISL_3468584 | Lighthouse Lab in Alderley Park | Wellcome Sanger Institute for the COVID-19 Genomics UK (COG-UK) Consortium | Jacquelyn Wynn, Mairead Hyland, The Lighthouse Lab in Alderley Park and Alex Alderton, Roberto Amato, Jeffrey Barrett, Sonia Goncalves, Ewan Harrison, David K. Jackson, Ian Johnston, Dominic Kwiatkowski, Cordelia Langford, John Sillitoe on behalf of the Wellcome Sanger Institute COVID-19 Surveillance Team |
| EPI_ISL_3468585 | Lighthouse Laboratory Plymouth | Wellcome Sanger Institute for the COVID-19 Genomics UK (COG-UK) Consortium | Lighthouse Laboratory Plymouth and Alex Alderton, Roberto Amato, Jeffrey Barrett, Sonia Goncalves, Ewan Harrison, David K. Jackson, Ian Johnston, Dominic Kwiatkowski, Cordelia Langford, John Sillitoe on behalf of the Wellcome Sanger Institute COVID-19 Surveillance Team |
| EPI_ISL_3468586 | Lighthouse Lab in Alderley Park | Wellcome Sanger Institute for the COVID-19 Genomics UK (COG-UK) Consortium | Jacquelyn Wynn, Mairead Hyland, The Lighthouse Lab in Alderley Park and Alex Alderton, Roberto Amato, Jeffrey Barrett, Sonia Goncalves, Ewan Harrison, David K. Jackson, Ian Johnston, Dominic Kwiatkowski, Cordelia Langford, John Sillitoe on behalf of the Wellcome Sanger Institute COVID-19 Surveillance Team |
| EPI_ISL_3468587, EPI_ISL_3468588, EPI_ISL_3468589 | Lighthouse Laboratory Plymouth | Wellcome Sanger Institute for the COVID-19 Genomics UK (COG-UK) Consortium | Lighthouse Laboratory Plymouth and Alex Alderton, Roberto Amato, Jeffrey Barrett, Sonia Goncalves, Ewan Harrison, David K. Jackson, Ian Johnston, Dominic Kwiatkowski, Cordelia Langford, John Sillitoe on behalf of the Wellcome Sanger Institute COVID-19 Surveillance Team |
| EPI_ISL_3468590 | Lighthouse Lab in Alderley Park | Wellcome Sanger Institute for the COVID-19 Genomics UK (COG-UK) Consortium | Jacquelyn Wynn, Mairead Hyland, The Lighthouse Lab in Alderley Park and Alex Alderton, Roberto Amato, Jeffrey Barrett, Sonia Goncalves, Ewan Harrison, David K. Jackson, Ian Johnston, Dominic Kwiatkowski, Cordelia Langford, John Sillitoe on behalf of the Wellcome Sanger Institute COVID-19 Surveillance Team |
| EPI_ISL_3468591 | Lighthouse Laboratory Plymouth | Wellcome Sanger Institute for the COVID-19 Genomics UK (COG-UK) Consortium | Lighthouse Laboratory Plymouth and Alex Alderton, Roberto Amato, Jeffrey Barrett, Sonia Goncalves, Ewan Harrison, David K. Jackson, Ian Johnston, Dominic Kwiatkowski, Cordelia Langford, John Sillitoe on behalf of the Wellcome Sanger Institute COVID-19 Surveillance Team |
| EPI_ISL_3468592 | Lighthouse Lab in Alderley Park | Wellcome Sanger Institute for the COVID-19 Genomics UK (COG-UK) Consortium | Jacquelyn Wynn, Mairead Hyland, The Lighthouse Lab in Alderley Park and Alex Alderton, Roberto Amato, Jeffrey Barrett, Sonia Goncalves, Ewan Harrison, David K. Jackson, Ian Johnston, Dominic Kwiatkowski, Cordelia Langford, John Sillitoe on behalf of the Wellcome Sanger Institute COVID-19 Surveillance Team |
| EPI_ISL_3468593, EPI_ISL_3468594, EPI_ISL_3468595 | Lighthouse Laboratory Plymouth | Wellcome Sanger Institute for the COVID-19 Genomics UK (COG-UK) Consortium | Lighthouse Laboratory Plymouth and Alex Alderton, Roberto Amato, Jeffrey Barrett, Sonia Goncalves, Ewan Harrison, David K. Jackson, Ian Johnston, Dominic Kwiatkowski, Cordelia Langford, John Sillitoe on behalf of the Wellcome Sanger Institute COVID-19 Surveillance Team |
| EPI_ISL_3468596 | Lighthouse Lab in Alderley Park | Wellcome Sanger Institute for the COVID-19 Genomics UK (COG-UK) Consortium | Jacquelyn Wynn, Mairead Hyland, The Lighthouse Lab in Alderley Park and Alex Alderton, Roberto Amato, Jeffrey Barrett, Sonia Goncalves, Ewan Harrison, David K. Jackson, Ian Johnston, Dominic Kwiatkowski, Cordelia Langford, John Sillitoe on behalf of the Wellcome Sanger Institute COVID-19 Surveillance Team |
| EPI_ISL_3468597 | Lighthouse Laboratory Plymouth | Wellcome Sanger Institute for the COVID-19 Genomics UK (COG-UK) Consortium | Lighthouse Laboratory Plymouth and Alex Alderton, Roberto Amato, Jeffrey Barrett, Sonia Goncalves, Ewan Harrison, David K. Jackson, Ian Johnston, Dominic Kwiatkowski, Cordelia Langford, John Sillitoe on behalf of the Wellcome Sanger Institute COVID-19 Surveillance Team |
| EPI_ISL_3468598, EPI_ISL_3468599, EPI_ISL_3468600 | Lighthouse Lab in Alderley Park | Wellcome Sanger Institute for the COVID-19 Genomics UK (COG-UK) Consortium | Jacquelyn Wynn, Mairead Hyland, The Lighthouse Lab in Alderley Park and Alex Alderton, Roberto Amato, Jeffrey Barrett, Sonia Goncalves, Ewan Harrison, David K. Jackson, Ian Johnston, Dominic Kwiatkowski, Cordelia Langford, John Sillitoe on behalf of the Wellcome Sanger Institute COVID-19 Surveillance Team |
| EPI_ISL_3468601, EPI_ISL_3468602, EPI_ISL_3468603 | Lighthouse Laboratory Plymouth | Wellcome Sanger Institute for the COVID-19 Genomics UK (COG-UK) Consortium | Lighthouse Laboratory Plymouth and Alex Alderton, Roberto Amato, Jeffrey Barrett, Sonia Goncalves, Ewan Harrison, David K. Jackson, Ian Johnston, Dominic Kwiatkowski, Cordelia Langford, John Sillitoe on behalf of the Wellcome Sanger Institute COVID-19 Surveillance Team |
| EPI_ISL_3468604, EPI_ISL_3468605, EPI_ISL_3468606, EPI_ISL_3468607 | Lighthouse Lab in Alderley Park | Wellcome Sanger Institute for the COVID-19 Genomics UK (COG-UK) Consortium | Jacquelyn Wynn, Mairead Hyland, The Lighthouse Lab in Alderley Park and Alex Alderton, Roberto Amato, Jeffrey Barrett, Sonia Goncalves, Ewan Harrison, David K. Jackson, Ian Johnston, Dominic Kwiatkowski, Cordelia Langford, John Sillitoe on behalf of the Wellcome Sanger Institute COVID-19 Surveillance Team |
| EPI_ISL_3468608 | Lighthouse Laboratory Plymouth | Wellcome Sanger Institute for the COVID-19 Genomics UK (COG-UK) Consortium | Lighthouse Laboratory Plymouth and Alex Alderton, Roberto Amato, Jeffrey Barrett, Sonia Goncalves, Ewan Harrison, David K. Jackson, Ian Johnston, Dominic Kwiatkowski, Cordelia Langford, John Sillitoe on behalf of the Wellcome Sanger Institute COVID-19 Surveillance Team |
| EPI_ISL_3468609 | Lighthouse Lab in Alderley Park | Wellcome Sanger Institute for the COVID-19 Genomics UK (COG-UK) Consortium | Jacquelyn Wynn, Mairead Hyland, The Lighthouse Lab in Alderley Park and Alex Alderton, Roberto Amato, Jeffrey Barrett, Sonia Goncalves, Ewan Harrison, David K. Jackson, Ian Johnston, Dominic Kwiatkowski, Cordelia Langford, John Sillitoe on behalf of the Wellcome Sanger Institute COVID-19 Surveillance Team |
| EPI_ISL_3468610, EPI_ISL_3468611, | Lighthouse Laboratory Plymouth | Wellcome Sanger Institute for the COVID-19 Genomics UK (COG-UK) Consortium | Lighthouse Laboratory Plymouth and Alex Alderton, Roberto Amato, Jeffrey Barrett, Sonia Goncalves, Ewan Harrison, David K. Jackson, Ian Johnston, Dominic Kwiatkowski, Cordelia Langford, John Sillitoe on behalf of the Wellcome Sanger Institute COVID-19 Surveillance Team |

| | | | |
|---|---|---|---|
| EPI_ISL_3468612, EPI_ISL_3468613, EPI_ISL_3468614, EPI_ISL_3468615 | | | |
| EPI_ISL_3468616, EPI_ISL_3468617, EPI_ISL_3468618, EPI_ISL_3468619 | Lighthouse Lab in Alderley Park | Wellcome Sanger Institute for the COVID-19 Genomics UK (COG-UK) Consortium | Jacquelyn Wynn, Mairead Hyland, The Lighthouse Lab in Alderley Park and Alex Alderton, Roberto Amato, Jeffrey Barrett, Sonia Goncalves, Ewan Harrison, David K. Jackson, Ian Johnston, Dominic Kwiatkowski, Cordelia Langford, John Sillitoe on behalf of the Wellcome Sanger Institute COVID-19 Surveillance Team |
| EPI_ISL_3468620 | Lighthouse Laboratory Plymouth | Wellcome Sanger Institute for the COVID-19 Genomics UK (COG-UK) Consortium | Lighthouse Laboratory Plymouth and Alex Alderton, Roberto Amato, Jeffrey Barrett, Sonia Goncalves, Ewan Harrison, David K. Jackson, Ian Johnston, Dominic Kwiatkowski, Cordelia Langford, John Sillitoe on behalf of the Wellcome Sanger Institute COVID-19 Surveillance Team |
| EPI_ISL_3468621, EPI_ISL_3468622, EPI_ISL_3468623, EPI_ISL_3468624 | Lighthouse Lab in Alderley Park | Wellcome Sanger Institute for the COVID-19 Genomics UK (COG-UK) Consortium | Jacquelyn Wynn, Mairead Hyland, The Lighthouse Lab in Alderley Park and Alex Alderton, Roberto Amato, Jeffrey Barrett, Sonia Goncalves, Ewan Harrison, David K. Jackson, Ian Johnston, Dominic Kwiatkowski, Cordelia Langford, John Sillitoe on behalf of the Wellcome Sanger Institute COVID-19 Surveillance Team |
| EPI_ISL_3468625, EPI_ISL_3468626 | Lighthouse Laboratory Plymouth | Wellcome Sanger Institute for the COVID-19 Genomics UK (COG-UK) Consortium | Lighthouse Laboratory Plymouth and Alex Alderton, Roberto Amato, Jeffrey Barrett, Sonia Goncalves, Ewan Harrison, David K. Jackson, Ian Johnston, Dominic Kwiatkowski, Cordelia Langford, John Sillitoe on behalf of the Wellcome Sanger Institute COVID-19 Surveillance Team |
| EPI_ISL_3468627, EPI_ISL_3468628, EPI_ISL_3468629 | Lighthouse Lab in Alderley Park | Wellcome Sanger Institute for the COVID-19 Genomics UK (COG-UK) Consortium | Jacquelyn Wynn, Mairead Hyland, The Lighthouse Lab in Alderley Park and Alex Alderton, Roberto Amato, Jeffrey Barrett, Sonia Goncalves, Ewan Harrison, David K. Jackson, Ian Johnston, Dominic Kwiatkowski, Cordelia Langford, John Sillitoe on behalf of the Wellcome Sanger Institute COVID-19 Surveillance Team |
| EPI_ISL_3468630 | Lighthouse Lab in Milton Keynes | Wellcome Sanger Institute for the COVID-19 Genomics UK (COG-UK) Consortium | The Lighthouse Lab in Milton Keynes and Alex Alderton, Roberto Amato, Jeffrey Barrett, Sonia Goncalves, Ewan Harrison, David K. Jackson, Ian Johnston, Dominic Kwiatkowski, Cordelia Langford, John Sillitoe on behalf of the Wellcome Sanger Institute COVID-19 Surveillance Team |
| EPI_ISL_3468631 | Lighthouse Laboratory Plymouth | Wellcome Sanger Institute for the COVID-19 Genomics UK (COG-UK) Consortium | Lighthouse Laboratory Plymouth and Alex Alderton, Roberto Amato, Jeffrey Barrett, Sonia Goncalves, Ewan Harrison, David K. Jackson, Ian Johnston, Dominic Kwiatkowski, Cordelia Langford, John Sillitoe on behalf of the Wellcome Sanger Institute COVID-19 Surveillance Team |
| EPI_ISL_3468632 | Lighthouse Lab in Milton Keynes | Wellcome Sanger Institute for the COVID-19 Genomics UK (COG-UK) Consortium | The Lighthouse Lab in Milton Keynes and Alex Alderton, Roberto Amato, Jeffrey Barrett, Sonia Goncalves, Ewan Harrison, David K. Jackson, Ian Johnston, Dominic Kwiatkowski, Cordelia Langford, John Sillitoe on behalf of the Wellcome Sanger Institute COVID-19 Surveillance Team |
| EPI_ISL_3468633 | Lighthouse Laboratory Plymouth | Wellcome Sanger Institute for the COVID-19 Genomics UK (COG-UK) Consortium | Lighthouse Laboratory Plymouth and Alex Alderton, Roberto Amato, Jeffrey Barrett, Sonia Goncalves, Ewan Harrison, David K. Jackson, Ian Johnston, Dominic Kwiatkowski, Cordelia Langford, John Sillitoe on behalf of the Wellcome Sanger Institute COVID-19 Surveillance Team |
| EPI_ISL_3468634, EPI_ISL_3468635 | Lighthouse Lab in Alderley Park | Wellcome Sanger Institute for the COVID-19 Genomics UK (COG-UK) Consortium | Jacquelyn Wynn, Mairead Hyland, The Lighthouse Lab in Alderley Park and Alex Alderton, Roberto Amato, Jeffrey Barrett, Sonia Goncalves, Ewan Harrison, David K. Jackson, Ian Johnston, Dominic Kwiatkowski, Cordelia Langford, John Sillitoe on behalf of the Wellcome Sanger Institute COVID-19 Surveillance Team |
| EPI_ISL_3468636 | Lighthouse Lab in Milton Keynes | Wellcome Sanger Institute for the COVID-19 Genomics UK (COG-UK) Consortium | The Lighthouse Lab in Milton Keynes and Alex Alderton, Roberto Amato, Jeffrey Barrett, Sonia Goncalves, Ewan Harrison, David K. Jackson, Ian Johnston, Dominic Kwiatkowski, Cordelia Langford, John Sillitoe on behalf of the Wellcome Sanger Institute COVID-19 Surveillance Team |
| EPI_ISL_3468637 | Lighthouse Lab in Alderley Park | Wellcome Sanger Institute for the COVID-19 Genomics UK (COG-UK) Consortium | Jacquelyn Wynn, Mairead Hyland, The Lighthouse Lab in Alderley Park and Alex Alderton, Roberto Amato, Jeffrey Barrett, Sonia Goncalves, Ewan Harrison, David K. Jackson, Ian Johnston, Dominic Kwiatkowski, Cordelia Langford, John Sillitoe on behalf of the Wellcome Sanger Institute COVID-19 Surveillance Team |
| EPI_ISL_3468638 | Lighthouse Lab in Milton Keynes | Wellcome Sanger Institute for the COVID-19 Genomics UK (COG-UK) Consortium | The Lighthouse Lab in Milton Keynes and Alex Alderton, Roberto Amato, Jeffrey Barrett, Sonia Goncalves, Ewan Harrison, David K. Jackson, Ian Johnston, Dominic Kwiatkowski, Cordelia Langford, John Sillitoe on behalf of the Wellcome Sanger Institute COVID-19 Surveillance Team |
| EPI_ISL_3468639 | Lighthouse Laboratory Plymouth | Wellcome Sanger Institute for the COVID-19 Genomics UK (COG-UK) Consortium | Lighthouse Laboratory Plymouth and Alex Alderton, Roberto Amato, Jeffrey Barrett, Sonia Goncalves, Ewan Harrison, David K. Jackson, Ian Johnston, Dominic Kwiatkowski, Cordelia Langford, John Sillitoe on behalf of the Wellcome Sanger Institute COVID-19 Surveillance Team |
| EPI_ISL_3468640, EPI_ISL_3468641, EPI_ISL_3468642 | Lighthouse Lab in Milton Keynes | Wellcome Sanger Institute for the COVID-19 Genomics UK (COG-UK) Consortium | The Lighthouse Lab in Milton Keynes and Alex Alderton, Roberto Amato, Jeffrey Barrett, Sonia Goncalves, Ewan Harrison, David K. Jackson, Ian Johnston, Dominic Kwiatkowski, Cordelia Langford, John Sillitoe on behalf of the Wellcome Sanger Institute COVID-19 Surveillance Team |
| EPI_ISL_3468643 | Lighthouse Laboratory Plymouth | Wellcome Sanger Institute for the COVID-19 Genomics UK (COG-UK) Consortium | Lighthouse Laboratory Plymouth and Alex Alderton, Roberto Amato, Jeffrey Barrett, Sonia Goncalves, Ewan Harrison, David K. Jackson, Ian Johnston, Dominic Kwiatkowski, Cordelia Langford, John Sillitoe on behalf of the Wellcome Sanger Institute COVID-19 Surveillance Team |
| EPI_ISL_3468644, EPI_ISL_3468645, EPI_ISL_3468646 | Lighthouse Lab in Alderley Park | Wellcome Sanger Institute for the COVID-19 Genomics UK (COG-UK) Consortium | Jacquelyn Wynn, Mairead Hyland, The Lighthouse Lab in Alderley Park and Alex Alderton, Roberto Amato, Jeffrey Barrett, Sonia Goncalves, Ewan Harrison, David K. Jackson, Ian Johnston, Dominic Kwiatkowski, Cordelia Langford, John Sillitoe on behalf of the Wellcome Sanger Institute COVID-19 Surveillance Team |
| EPI_ISL_3468647, EPI_ISL_3468648 | Lighthouse Lab in Milton Keynes | Wellcome Sanger Institute for the COVID-19 Genomics UK (COG-UK) Consortium | The Lighthouse Lab in Milton Keynes and Alex Alderton, Roberto Amato, Jeffrey Barrett, Sonia Goncalves, Ewan Harrison, David K. Jackson, Ian Johnston, Dominic Kwiatkowski, Cordelia Langford, John Sillitoe on behalf of the Wellcome Sanger Institute COVID-19 Surveillance Team |
| EPI_ISL_3468649 | Lighthouse Laboratory Plymouth | Wellcome Sanger Institute for the COVID-19 Genomics UK (COG-UK) Consortium | Lighthouse Laboratory Plymouth and Alex Alderton, Roberto Amato, Jeffrey Barrett, Sonia Goncalves, Ewan Harrison, David K. Jackson, Ian Johnston, Dominic Kwiatkowski, Cordelia Langford, John Sillitoe on behalf of the Wellcome Sanger Institute COVID-19 Surveillance Team |
| EPI_ISL_3468650, EPI_ISL_3468651, EPI_ISL_3468652 | Lighthouse Lab in Milton Keynes | Wellcome Sanger Institute for the COVID-19 Genomics UK (COG-UK) Consortium | The Lighthouse Lab in Milton Keynes and Alex Alderton, Roberto Amato, Jeffrey Barrett, Sonia Goncalves, Ewan Harrison, David K. Jackson, Ian Johnston, Dominic Kwiatkowski, Cordelia Langford, John Sillitoe on behalf of the Wellcome Sanger Institute COVID-19 Surveillance Team |
| EPI_ISL_3468653 | Lighthouse Lab in Alderley Park | Wellcome Sanger Institute for the COVID-19 Genomics UK (COG-UK) Consortium | Jacquelyn Wynn, Mairead Hyland, The Lighthouse Lab in Alderley Park and Alex Alderton, Roberto Amato, Jeffrey Barrett, Sonia Goncalves, Ewan Harrison, David K. Jackson, Ian Johnston, Dominic Kwiatkowski, Cordelia Langford, John Sillitoe on behalf of the Wellcome Sanger Institute COVID-19 Surveillance Team |
| EPI_ISL_3468654 | Lighthouse Laboratory Plymouth | Wellcome Sanger Institute for the COVID-19 Genomics UK (COG-UK) Consortium | Lighthouse Laboratory Plymouth and Alex Alderton, Roberto Amato, Jeffrey Barrett, Sonia Goncalves, Ewan Harrison, David K. Jackson, Ian Johnston, Dominic Kwiatkowski, Cordelia Langford, John Sillitoe on behalf of the Wellcome Sanger Institute COVID-19 Surveillance Team |
| EPI_ISL_3468655 | Lighthouse Lab in Milton Keynes | Wellcome Sanger Institute for the COVID-19 Genomics UK (COG-UK) Consortium | The Lighthouse Lab in Milton Keynes and Alex Alderton, Roberto Amato, Jeffrey Barrett, Sonia Goncalves, Ewan Harrison, David K. Jackson, Ian Johnston, Dominic Kwiatkowski, Cordelia Langford, John Sillitoe on behalf of the Wellcome Sanger Institute COVID-19 Surveillance Team |
| EPI_ISL_3468656 | Lighthouse Lab in Alderley Park | Wellcome Sanger Institute for the COVID-19 Genomics UK (COG-UK) Consortium | Jacquelyn Wynn, Mairead Hyland, The Lighthouse Lab in Alderley Park and Alex Alderton, Roberto Amato, Jeffrey Barrett, Sonia Goncalves, Ewan Harrison, David K. Jackson, Ian Johnston, Dominic Kwiatkowski, Cordelia Langford, John Sillitoe on behalf of the Wellcome Sanger Institute COVID-19 Surveillance Team |
| EPI_ISL_3468657 | Lighthouse Laboratory Plymouth | Wellcome Sanger Institute for the COVID-19 Genomics UK (COG-UK) Consortium | Lighthouse Laboratory Plymouth and Alex Alderton, Roberto Amato, Jeffrey Barrett, Sonia Goncalves, Ewan Harrison, David K. Jackson, Ian Johnston, Dominic Kwiatkowski, Cordelia Langford, John Sillitoe on behalf of the Wellcome Sanger Institute COVID-19 Surveillance Team |
| EPI_ISL_3468658, EPI_ISL_3468659, EPI_ISL_3468660, EPI_ISL_3468661, EPI_ISL_3468662 | Lighthouse Lab in Milton Keynes | Wellcome Sanger Institute for the COVID-19 Genomics UK (COG-UK) Consortium | The Lighthouse Lab in Milton Keynes and Alex Alderton, Roberto Amato, Jeffrey Barrett, Sonia Goncalves, Ewan Harrison, David K. Jackson, Ian Johnston, Dominic Kwiatkowski, Cordelia Langford, John Sillitoe on behalf of the Wellcome Sanger Institute COVID-19 Surveillance Team |
| EPI_ISL_3468663, EPI_ISL_3468664 | Lighthouse Laboratory Plymouth | Wellcome Sanger Institute for the COVID-19 Genomics UK (COG-UK) Consortium | Lighthouse Laboratory Plymouth and Alex Alderton, Roberto Amato, Jeffrey Barrett, Sonia Goncalves, Ewan Harrison, David K. Jackson, Ian Johnston, Dominic Kwiatkowski, Cordelia Langford, John Sillitoe on behalf of the Wellcome Sanger Institute COVID-19 Surveillance Team |
| EPI_ISL_3468665, EPI_ISL_3468666, EPI_ISL_3468667 | Lighthouse Lab in Alderley Park | Wellcome Sanger Institute for the COVID-19 Genomics UK (COG-UK) Consortium | Jacquelyn Wynn, Mairead Hyland, The Lighthouse Lab in Alderley Park and Alex Alderton, Roberto Amato, Jeffrey Barrett, Sonia Goncalves, Ewan Harrison, David K. Jackson, Ian Johnston, Dominic Kwiatkowski, Cordelia Langford, John Sillitoe on behalf of the Wellcome Sanger Institute COVID-19 Surveillance Team |
| EPI_ISL_3468668 | Lighthouse Laboratory Plymouth | Wellcome Sanger Institute for the COVID-19 Genomics UK (COG-UK) Consortium | Lighthouse Laboratory Plymouth and Alex Alderton, Roberto Amato, Jeffrey Barrett, Sonia Goncalves, Ewan Harrison, David K. Jackson, Ian Johnston, Dominic Kwiatkowski, Cordelia Langford, John Sillitoe on behalf of the Wellcome Sanger Institute COVID-19 Surveillance Team |
| EPI_ISL_3468669 | Lighthouse Lab in Milton Keynes | Wellcome Sanger Institute for the COVID-19 Genomics UK (COG-UK) Consortium | The Lighthouse Lab in Milton Keynes and Alex Alderton, Roberto Amato, Jeffrey Barrett, Sonia Goncalves, Ewan Harrison, David K. Jackson, Ian Johnston, Dominic Kwiatkowski, Cordelia Langford, John Sillitoe on behalf of the Wellcome Sanger Institute COVID-19 Surveillance Team |
| EPI_ISL_3468670 | Lighthouse Lab in Alderley Park | Wellcome Sanger Institute for the COVID-19 Genomics UK (COG-UK) Consortium | Jacquelyn Wynn, Mairead Hyland, The Lighthouse Lab in Alderley Park and Alex Alderton, Roberto Amato, Jeffrey Barrett, Sonia Goncalves, Ewan Harrison, David K. Jackson, Ian Johnston, Dominic Kwiatkowski, Cordelia Langford, John Sillitoe on behalf of the Wellcome Sanger Institute COVID-19 Surveillance Team |
| EPI_ISL_3468671 | Lighthouse Laboratory Plymouth | Wellcome Sanger Institute for the COVID-19 Genomics UK (COG-UK) Consortium | Lighthouse Laboratory Plymouth and Alex Alderton, Roberto Amato, Jeffrey Barrett, Sonia Goncalves, Ewan Harrison, David K. Jackson, Ian Johnston, Dominic Kwiatkowski, Cordelia Langford, John Sillitoe on behalf of the Wellcome Sanger Institute COVID-19 Surveillance Team |
| EPI_ISL_3468672 | Lighthouse Lab in Milton Keynes | Wellcome Sanger Institute for the COVID-19 Genomics UK (COG-UK) Consortium | The Lighthouse Lab in Milton Keynes and Alex Alderton, Roberto Amato, Jeffrey Barrett, Sonia Goncalves, Ewan Harrison, David K. Jackson, Ian Johnston, Dominic Kwiatkowski, Cordelia Langford, John Sillitoe on behalf of the Wellcome Sanger Institute COVID-19 Surveillance Team |
| EPI_ISL_3468673 | Lighthouse Lab in Alderley Park | Wellcome Sanger Institute for the COVID-19 Genomics UK (COG-UK) Consortium | Jacquelyn Wynn, Mairead Hyland, The Lighthouse Lab in Alderley Park and Alex Alderton, Roberto Amato, Jeffrey Barrett, Sonia Goncalves, Ewan Harrison, David K. Jackson, Ian Johnston, Dominic Kwiatkowski, Cordelia Langford, John Sillitoe on behalf of the Wellcome Sanger Institute COVID-19 Surveillance Team |
| EPI_ISL_3468674 | Lighthouse Laboratory Plymouth | Wellcome Sanger Institute for the COVID-19 Genomics UK (COG-UK) Consortium | Lighthouse Laboratory Plymouth and Alex Alderton, Roberto Amato, Jeffrey Barrett, Sonia Goncalves, Ewan Harrison, David K. Jackson, Ian Johnston, Dominic Kwiatkowski, Cordelia Langford, John Sillitoe on behalf of the Wellcome Sanger Institute COVID-19 Surveillance Team |
| EPI_ISL_3468675 | Lighthouse Lab in Alderley Park | Wellcome Sanger Institute for the COVID-19 Genomics UK (COG-UK) Consortium | Jacquelyn Wynn, Mairead Hyland, The Lighthouse Lab in Alderley Park and Alex Alderton, Roberto Amato, Jeffrey Barrett, Sonia Goncalves, Ewan Harrison, David K. Jackson, Ian Johnston, Dominic Kwiatkowski, Cordelia Langford, John Sillitoe on behalf of the Wellcome Sanger Institute COVID-19 Surveillance Team |
| EPI_ISL_3468676 | Lighthouse Laboratory Plymouth | Wellcome Sanger Institute for the COVID-19 Genomics UK (COG-UK) Consortium | Lighthouse Laboratory Plymouth and Alex Alderton, Roberto Amato, Jeffrey Barrett, Sonia Goncalves, Ewan Harrison, David K. Jackson, Ian Johnston, Dominic Kwiatkowski, Cordelia Langford, John Sillitoe on behalf of the Wellcome Sanger Institute COVID-19 Surveillance Team |
| EPI_ISL_3468677 | Lighthouse Lab in Milton Keynes | Wellcome Sanger Institute for the COVID-19 Genomics UK (COG-UK) Consortium | The Lighthouse Lab in Milton Keynes and Alex Alderton, Roberto Amato, Jeffrey Barrett, Sonia Goncalves, Ewan Harrison, David K. Jackson, Ian Johnston, Dominic Kwiatkowski, Cordelia Langford, John Sillitoe on behalf of the Wellcome Sanger Institute COVID-19 Surveillance Team |
| EPI_ISL_3468678 | Lighthouse Lab in Alderley Park | Wellcome Sanger Institute for the COVID-19 Genomics UK (COG-UK) | Jacquelyn Wynn, Mairead Hyland, The Lighthouse Lab in Alderley Park and Alex Alderton, Roberto Amato, Jeffrey Barrett, Sonia Goncalves, Ewan Harrison, David K. Jackson, Ian Johnston, Dominic Kwiatkowski, Cordelia Langford, John Sillitoe on |

| | | | |
|---|---|---|---|
| | | Consortium | behalf of the Wellcome Sanger Institute COVID-19 Surveillance Team |
| EPI_ISL_3468679 | Lighthouse Laboratory Plymouth | Wellcome Sanger Institute for the COVID-19 Genomics UK (COG-UK) Consortium | Lighthouse Laboratory Plymouth and Alex Alderton, Roberto Amato, Jeffrey Barrett, Sonia Goncalves, Ewan Harrison, David K. Jackson, Ian Johnston, Dominic Kwiatkowski, Cordelia Langford, John Sillitoe on behalf of the Wellcome Sanger Institute COVID-19 Surveillance Team |
| EPI_ISL_3468680 | Lighthouse Lab in Milton Keynes | Wellcome Sanger Institute for the COVID-19 Genomics UK (COG-UK) Consortium | The Lighthouse Lab in Milton Keynes and Alex Alderton, Roberto Amato, Jeffrey Barrett, Sonia Goncalves, Ewan Harrison, David K. Jackson, Ian Johnston, Dominic Kwiatkowski, Cordelia Langford, John Sillitoe on behalf of the Wellcome Sanger Institute COVID-19 Surveillance Team |
| EPI_ISL_3468681, EPI_ISL_3468682, EPI_ISL_3468683 | Lighthouse Lab in Alderley Park | Wellcome Sanger Institute for the COVID-19 Genomics UK (COG-UK) Consortium | Jacquelyn Wynn, Mairead Hyland, The Lighthouse Lab in Alderley Park and Alex Alderton, Roberto Amato, Jeffrey Barrett, Sonia Goncalves, Ewan Harrison, David K. Jackson, Ian Johnston, Dominic Kwiatkowski, Cordelia Langford, John Sillitoe on behalf of the Wellcome Sanger Institute COVID-19 Surveillance Team |
| EPI_ISL_3468684 | Lighthouse Lab in Milton Keynes | Wellcome Sanger Institute for the COVID-19 Genomics UK (COG-UK) Consortium | The Lighthouse Lab in Milton Keynes and Alex Alderton, Roberto Amato, Jeffrey Barrett, Sonia Goncalves, Ewan Harrison, David K. Jackson, Ian Johnston, Dominic Kwiatkowski, Cordelia Langford, John Sillitoe on behalf of the Wellcome Sanger Institute COVID-19 Surveillance Team |
| EPI_ISL_3468685, EPI_ISL_3468686 | Lighthouse Lab in Alderley Park | Wellcome Sanger Institute for the COVID-19 Genomics UK (COG-UK) Consortium | Jacquelyn Wynn, Mairead Hyland, The Lighthouse Lab in Alderley Park and Alex Alderton, Roberto Amato, Jeffrey Barrett, Sonia Goncalves, Ewan Harrison, David K. Jackson, Ian Johnston, Dominic Kwiatkowski, Cordelia Langford, John Sillitoe on behalf of the Wellcome Sanger Institute COVID-19 Surveillance Team |
| EPI_ISL_3468687, EPI_ISL_3468688, EPI_ISL_3468689 | Lighthouse Laboratory Plymouth | Wellcome Sanger Institute for the COVID-19 Genomics UK (COG-UK) Consortium | Lighthouse Laboratory Plymouth and Alex Alderton, Roberto Amato, Jeffrey Barrett, Sonia Goncalves, Ewan Harrison, David K. Jackson, Ian Johnston, Dominic Kwiatkowski, Cordelia Langford, John Sillitoe on behalf of the Wellcome Sanger Institute COVID-19 Surveillance Team |
| EPI_ISL_3468690, EPI_ISL_3468691 | Lighthouse Lab in Milton Keynes | Wellcome Sanger Institute for the COVID-19 Genomics UK (COG-UK) Consortium | The Lighthouse Lab in Milton Keynes and Alex Alderton, Roberto Amato, Jeffrey Barrett, Sonia Goncalves, Ewan Harrison, David K. Jackson, Ian Johnston, Dominic Kwiatkowski, Cordelia Langford, John Sillitoe on behalf of the Wellcome Sanger Institute COVID-19 Surveillance Team |
| EPI_ISL_3468692 | Lighthouse Lab in Alderley Park | Wellcome Sanger Institute for the COVID-19 Genomics UK (COG-UK) Consortium | Jacquelyn Wynn, Mairead Hyland, The Lighthouse Lab in Alderley Park and Alex Alderton, Roberto Amato, Jeffrey Barrett, Sonia Goncalves, Ewan Harrison, David K. Jackson, Ian Johnston, Dominic Kwiatkowski, Cordelia Langford, John Sillitoe on behalf of the Wellcome Sanger Institute COVID-19 Surveillance Team |
| EPI_ISL_3468693 | Lighthouse Lab in Milton Keynes | Wellcome Sanger Institute for the COVID-19 Genomics UK (COG-UK) Consortium | The Lighthouse Lab in Milton Keynes and Alex Alderton, Roberto Amato, Jeffrey Barrett, Sonia Goncalves, Ewan Harrison, David K. Jackson, Ian Johnston, Dominic Kwiatkowski, Cordelia Langford, John Sillitoe on behalf of the Wellcome Sanger Institute COVID-19 Surveillance Team |
| EPI_ISL_3468694 | Lighthouse Laboratory Plymouth | Wellcome Sanger Institute for the COVID-19 Genomics UK (COG-UK) Consortium | Lighthouse Laboratory Plymouth and Alex Alderton, Roberto Amato, Jeffrey Barrett, Sonia Goncalves, Ewan Harrison, David K. Jackson, Ian Johnston, Dominic Kwiatkowski, Cordelia Langford, John Sillitoe on behalf of the Wellcome Sanger Institute COVID-19 Surveillance Team |
| EPI_ISL_3468695 | Lighthouse Lab in Alderley Park | Wellcome Sanger Institute for the COVID-19 Genomics UK (COG-UK) Consortium | Jacquelyn Wynn, Mairead Hyland, The Lighthouse Lab in Alderley Park and Alex Alderton, Roberto Amato, Jeffrey Barrett, Sonia Goncalves, Ewan Harrison, David K. Jackson, Ian Johnston, Dominic Kwiatkowski, Cordelia Langford, John Sillitoe on behalf of the Wellcome Sanger Institute COVID-19 Surveillance Team |
| EPI_ISL_3468696, EPI_ISL_3468697, EPI_ISL_3468698 | Lighthouse Laboratory Plymouth | Wellcome Sanger Institute for the COVID-19 Genomics UK (COG-UK) Consortium | Lighthouse Laboratory Plymouth and Alex Alderton, Roberto Amato, Jeffrey Barrett, Sonia Goncalves, Ewan Harrison, David K. Jackson, Ian Johnston, Dominic Kwiatkowski, Cordelia Langford, John Sillitoe on behalf of the Wellcome Sanger Institute COVID-19 Surveillance Team |
| EPI_ISL_3468699, EPI_ISL_3468700, EPI_ISL_3468701, EPI_ISL_3468702, EPI_ISL_3468703 | Lighthouse Lab in Milton Keynes | Wellcome Sanger Institute for the COVID-19 Genomics UK (COG-UK) Consortium | The Lighthouse Lab in Milton Keynes and Alex Alderton, Roberto Amato, Jeffrey Barrett, Sonia Goncalves, Ewan Harrison, David K. Jackson, Ian Johnston, Dominic Kwiatkowski, Cordelia Langford, John Sillitoe on behalf of the Wellcome Sanger Institute COVID-19 Surveillance Team |
| EPI_ISL_3468704 | Lighthouse Lab in Alderley Park | Wellcome Sanger Institute for the COVID-19 Genomics UK (COG-UK) Consortium | Jacquelyn Wynn, Mairead Hyland, The Lighthouse Lab in Alderley Park and Alex Alderton, Roberto Amato, Jeffrey Barrett, Sonia Goncalves, Ewan Harrison, David K. Jackson, Ian Johnston, Dominic Kwiatkowski, Cordelia Langford, John Sillitoe on behalf of the Wellcome Sanger Institute COVID-19 Surveillance Team |
| EPI_ISL_3468705 | Lighthouse Laboratory Plymouth | Wellcome Sanger Institute for the COVID-19 Genomics UK (COG-UK) Consortium | Lighthouse Laboratory Plymouth and Alex Alderton, Roberto Amato, Jeffrey Barrett, Sonia Goncalves, Ewan Harrison, David K. Jackson, Ian Johnston, Dominic Kwiatkowski, Cordelia Langford, John Sillitoe on behalf of the Wellcome Sanger Institute COVID-19 Surveillance Team |
| EPI_ISL_3468706 | Lighthouse Lab in Milton Keynes | Wellcome Sanger Institute for the COVID-19 Genomics UK (COG-UK) Consortium | The Lighthouse Lab in Milton Keynes and Alex Alderton, Roberto Amato, Jeffrey Barrett, Sonia Goncalves, Ewan Harrison, David K. Jackson, Ian Johnston, Dominic Kwiatkowski, Cordelia Langford, John Sillitoe on behalf of the Wellcome Sanger Institute COVID-19 Surveillance Team |
| EPI_ISL_3468707 | Lighthouse Lab in Alderley Park | Wellcome Sanger Institute for the COVID-19 Genomics UK (COG-UK) Consortium | Jacquelyn Wynn, Mairead Hyland, The Lighthouse Lab in Alderley Park and Alex Alderton, Roberto Amato, Jeffrey Barrett, Sonia Goncalves, Ewan Harrison, David K. Jackson, Ian Johnston, Dominic Kwiatkowski, Cordelia Langford, John Sillitoe on behalf of the Wellcome Sanger Institute COVID-19 Surveillance Team |
| EPI_ISL_3468708, EPI_ISL_3468709, EPI_ISL_3468710 | Lighthouse Lab in Milton Keynes | Wellcome Sanger Institute for the COVID-19 Genomics UK (COG-UK) Consortium | The Lighthouse Lab in Milton Keynes and Alex Alderton, Roberto Amato, Jeffrey Barrett, Sonia Goncalves, Ewan Harrison, David K. Jackson, Ian Johnston, Dominic Kwiatkowski, Cordelia Langford, John Sillitoe on behalf of the Wellcome Sanger Institute COVID-19 Surveillance Team |
| EPI_ISL_3468711 | Lighthouse Lab in Alderley Park | Wellcome Sanger Institute for the COVID-19 Genomics UK (COG-UK) Consortium | Jacquelyn Wynn, Mairead Hyland, The Lighthouse Lab in Alderley Park and Alex Alderton, Roberto Amato, Jeffrey Barrett, Sonia Goncalves, Ewan Harrison, David K. Jackson, Ian Johnston, Dominic Kwiatkowski, Cordelia Langford, John Sillitoe on behalf of the Wellcome Sanger Institute COVID-19 Surveillance Team |
| EPI_ISL_3468712 | Lighthouse Laboratory Plymouth | Wellcome Sanger Institute for the COVID-19 Genomics UK (COG-UK) Consortium | Lighthouse Laboratory Plymouth and Alex Alderton, Roberto Amato, Jeffrey Barrett, Sonia Goncalves, Ewan Harrison, David K. Jackson, Ian Johnston, Dominic Kwiatkowski, Cordelia Langford, John Sillitoe on behalf of the Wellcome Sanger Institute COVID-19 Surveillance Team |
| EPI_ISL_3468713 | Lighthouse Lab in Milton Keynes | Wellcome Sanger Institute for the COVID-19 Genomics UK (COG-UK) Consortium | The Lighthouse Lab in Milton Keynes and Alex Alderton, Roberto Amato, Jeffrey Barrett, Sonia Goncalves, Ewan Harrison, David K. Jackson, Ian Johnston, Dominic Kwiatkowski, Cordelia Langford, John Sillitoe on behalf of the Wellcome Sanger Institute COVID-19 Surveillance Team |
| EPI_ISL_3468714 | Lighthouse Lab in Alderley Park | Wellcome Sanger Institute for the COVID-19 Genomics UK (COG-UK) Consortium | Jacquelyn Wynn, Mairead Hyland, The Lighthouse Lab in Alderley Park and Alex Alderton, Roberto Amato, Jeffrey Barrett, Sonia Goncalves, Ewan Harrison, David K. Jackson, Ian Johnston, Dominic Kwiatkowski, Cordelia Langford, John Sillitoe on behalf of the Wellcome Sanger Institute COVID-19 Surveillance Team |
| EPI_ISL_3468715 | Lighthouse Laboratory Plymouth | Wellcome Sanger Institute for the COVID-19 Genomics UK (COG-UK) Consortium | Lighthouse Laboratory Plymouth and Alex Alderton, Roberto Amato, Jeffrey Barrett, Sonia Goncalves, Ewan Harrison, David K. Jackson, Ian Johnston, Dominic Kwiatkowski, Cordelia Langford, John Sillitoe on behalf of the Wellcome Sanger Institute COVID-19 Surveillance Team |
| EPI_ISL_3468716, EPI_ISL_3468717, EPI_ISL_3468718, EPI_ISL_3468719 | Lighthouse Lab in Milton Keynes | Wellcome Sanger Institute for the COVID-19 Genomics UK (COG-UK) Consortium | The Lighthouse Lab in Milton Keynes and Alex Alderton, Roberto Amato, Jeffrey Barrett, Sonia Goncalves, Ewan Harrison, David K. Jackson, Ian Johnston, Dominic Kwiatkowski, Cordelia Langford, John Sillitoe on behalf of the Wellcome Sanger Institute COVID-19 Surveillance Team |
| EPI_ISL_3468720 | Lighthouse Lab in Alderley Park | Wellcome Sanger Institute for the COVID-19 Genomics UK (COG-UK) Consortium | Jacquelyn Wynn, Mairead Hyland, The Lighthouse Lab in Alderley Park and Alex Alderton, Roberto Amato, Jeffrey Barrett, Sonia Goncalves, Ewan Harrison, David K. Jackson, Ian Johnston, Dominic Kwiatkowski, Cordelia Langford, John Sillitoe on behalf of the Wellcome Sanger Institute COVID-19 Surveillance Team |
| EPI_ISL_3468721 | Lighthouse Laboratory Plymouth | Wellcome Sanger Institute for the COVID-19 Genomics UK (COG-UK) Consortium | Lighthouse Laboratory Plymouth and Alex Alderton, Roberto Amato, Jeffrey Barrett, Sonia Goncalves, Ewan Harrison, David K. Jackson, Ian Johnston, Dominic Kwiatkowski, Cordelia Langford, John Sillitoe on behalf of the Wellcome Sanger Institute COVID-19 Surveillance Team |
| EPI_ISL_3468722 | Lighthouse Lab in Alderley Park | Wellcome Sanger Institute for the COVID-19 Genomics UK (COG-UK) Consortium | Jacquelyn Wynn, Mairead Hyland, The Lighthouse Lab in Alderley Park and Alex Alderton, Roberto Amato, Jeffrey Barrett, Sonia Goncalves, Ewan Harrison, David K. Jackson, Ian Johnston, Dominic Kwiatkowski, Cordelia Langford, John Sillitoe on behalf of the Wellcome Sanger Institute COVID-19 Surveillance Team |
| EPI_ISL_3468723, EPI_ISL_3468724 | Lighthouse Lab in Milton Keynes | Wellcome Sanger Institute for the COVID-19 Genomics UK (COG-UK) Consortium | The Lighthouse Lab in Milton Keynes and Alex Alderton, Roberto Amato, Jeffrey Barrett, Sonia Goncalves, Ewan Harrison, David K. Jackson, Ian Johnston, Dominic Kwiatkowski, Cordelia Langford, John Sillitoe on behalf of the Wellcome Sanger Institute COVID-19 Surveillance Team |
| EPI_ISL_3468725, EPI_ISL_3468726 | Lighthouse Laboratory Plymouth | Wellcome Sanger Institute for the COVID-19 Genomics UK (COG-UK) Consortium | Lighthouse Laboratory Plymouth and Alex Alderton, Roberto Amato, Jeffrey Barrett, Sonia Goncalves, Ewan Harrison, David K. Jackson, Ian Johnston, Dominic Kwiatkowski, Cordelia Langford, John Sillitoe on behalf of the Wellcome Sanger Institute COVID-19 Surveillance Team |
| EPI_ISL_3468727 | Lighthouse Lab in Milton Keynes | Wellcome Sanger Institute for the COVID-19 Genomics UK (COG-UK) Consortium | The Lighthouse Lab in Milton Keynes and Alex Alderton, Roberto Amato, Jeffrey Barrett, Sonia Goncalves, Ewan Harrison, David K. Jackson, Ian Johnston, Dominic Kwiatkowski, Cordelia Langford, John Sillitoe on behalf of the Wellcome Sanger Institute COVID-19 Surveillance Team |
| EPI_ISL_3468728 | Lighthouse Lab in Alderley Park | Wellcome Sanger Institute for the COVID-19 Genomics UK (COG-UK) Consortium | Jacquelyn Wynn, Mairead Hyland, The Lighthouse Lab in Alderley Park and Alex Alderton, Roberto Amato, Jeffrey Barrett, Sonia Goncalves, Ewan Harrison, David K. Jackson, Ian Johnston, Dominic Kwiatkowski, Cordelia Langford, John Sillitoe on behalf of the Wellcome Sanger Institute COVID-19 Surveillance Team |
| EPI_ISL_3468729 | Lighthouse Laboratory Plymouth | Wellcome Sanger Institute for the COVID-19 Genomics UK (COG-UK) Consortium | Lighthouse Laboratory Plymouth and Alex Alderton, Roberto Amato, Jeffrey Barrett, Sonia Goncalves, Ewan Harrison, David K. Jackson, Ian Johnston, Dominic Kwiatkowski, Cordelia Langford, John Sillitoe on behalf of the Wellcome Sanger Institute COVID-19 Surveillance Team |
| EPI_ISL_3468730 | Lighthouse Lab in Milton Keynes | Wellcome Sanger Institute for the COVID-19 Genomics UK (COG-UK) Consortium | The Lighthouse Lab in Milton Keynes and Alex Alderton, Roberto Amato, Jeffrey Barrett, Sonia Goncalves, Ewan Harrison, David K. Jackson, Ian Johnston, Dominic Kwiatkowski, Cordelia Langford, John Sillitoe on behalf of the Wellcome Sanger Institute COVID-19 Surveillance Team |
| EPI_ISL_3468731 | Lighthouse Laboratory Plymouth | Wellcome Sanger Institute for the COVID-19 Genomics UK (COG-UK) Consortium | Lighthouse Laboratory Plymouth and Alex Alderton, Roberto Amato, Jeffrey Barrett, Sonia Goncalves, Ewan Harrison, David K. Jackson, Ian Johnston, Dominic Kwiatkowski, Cordelia Langford, John Sillitoe on behalf of the Wellcome Sanger Institute COVID-19 Surveillance Team |
| EPI_ISL_3468732 | Lighthouse Lab in Milton Keynes | Wellcome Sanger Institute for the COVID-19 Genomics UK (COG-UK) Consortium | The Lighthouse Lab in Milton Keynes and Alex Alderton, Roberto Amato, Jeffrey Barrett, Sonia Goncalves, Ewan Harrison, David K. Jackson, Ian Johnston, Dominic Kwiatkowski, Cordelia Langford, John Sillitoe on behalf of the Wellcome Sanger Institute COVID-19 Surveillance Team |
| EPI_ISL_3468733 | Lighthouse Lab in Alderley Park | Wellcome Sanger Institute for the COVID-19 Genomics UK (COG-UK) Consortium | Jacquelyn Wynn, Mairead Hyland, The Lighthouse Lab in Alderley Park and Alex Alderton, Roberto Amato, Jeffrey Barrett, Sonia Goncalves, Ewan Harrison, David K. Jackson, Ian Johnston, Dominic Kwiatkowski, Cordelia Langford, John Sillitoe on behalf of the Wellcome Sanger Institute COVID-19 Surveillance Team |
| EPI_ISL_3468734 | Lighthouse Laboratory Plymouth | Wellcome Sanger Institute for the COVID-19 Genomics UK (COG-UK) Consortium | Lighthouse Laboratory Plymouth and Alex Alderton, Roberto Amato, Jeffrey Barrett, Sonia Goncalves, Ewan Harrison, David K. Jackson, Ian Johnston, Dominic Kwiatkowski, Cordelia Langford, John Sillitoe on behalf of the Wellcome Sanger Institute COVID-19 Surveillance Team |
| EPI_ISL_3468735 | Lighthouse Lab in Alderley Park | Wellcome Sanger Institute for the COVID-19 Genomics UK (COG-UK) Consortium | Jacquelyn Wynn, Mairead Hyland, The Lighthouse Lab in Alderley Park and Alex Alderton, Roberto Amato, Jeffrey Barrett, Sonia Goncalves, Ewan Harrison, David K. Jackson, Ian Johnston, Dominic Kwiatkowski, Cordelia Langford, John Sillitoe on behalf of the Wellcome Sanger Institute COVID-19 Surveillance Team |
| EPI_ISL_3468736, EPI_ISL_3468737 | Lighthouse Lab in Milton Keynes | Wellcome Sanger Institute for the COVID-19 Genomics UK (COG-UK) Consortium | The Lighthouse Lab in Milton Keynes and Alex Alderton, Roberto Amato, Jeffrey Barrett, Sonia Goncalves, Ewan Harrison, David K. Jackson, Ian Johnston, Dominic Kwiatkowski, Cordelia Langford, John Sillitoe on behalf of the Wellcome Sanger Institute COVID-19 Surveillance Team |
| EPI_ISL_3468738 | Lighthouse Lab in Alderley Park | Wellcome Sanger Institute for the COVID-19 Genomics UK (COG-UK) Consortium | Jacquelyn Wynn, Mairead Hyland, The Lighthouse Lab in Alderley Park and Alex Alderton, Roberto Amato, Jeffrey Barrett, Sonia Goncalves, Ewan Harrison, David K. Jackson, Ian Johnston, Dominic Kwiatkowski, Cordelia Langford, John Sillitoe on behalf of the Wellcome Sanger Institute COVID-19 Surveillance Team |
| EPI_ISL_3468739, EPI_ISL_3468740 | Lighthouse Lab in Milton Keynes | Wellcome Sanger Institute for the COVID-19 Genomics UK (COG-UK) Consortium | The Lighthouse Lab in Milton Keynes and Alex Alderton, Roberto Amato, Jeffrey Barrett, Sonia Goncalves, Ewan Harrison, David K. Jackson, Ian Johnston, Dominic Kwiatkowski, Cordelia Langford, John Sillitoe on behalf of the Wellcome Sanger Institute COVID-19 Surveillance Team |

| | | | |
|---|---|---|---|
| EPI_ISL_3468741 | Lighthouse Laboratory Plymouth | Wellcome Sanger Institute for the COVID-19 Genomics UK (COG-UK) Consortium | Lighthouse Laboratory Plymouth and Alex Alderton, Roberto Amato, Jeffrey Barrett, Sonia Goncalves, Ewan Harrison, David K. Jackson, Ian Johnston, Dominic Kwiatkowski, Cordelia Langford, John Sillitoe on behalf of the Wellcome Sanger Institute COVID-19 Surveillance Team |
| EPI_ISL_3468742, EPI_ISL_3468743 | Lighthouse Lab in Milton Keynes | Wellcome Sanger Institute for the COVID-19 Genomics UK (COG-UK) Consortium | The Lighthouse Lab in Milton Keynes and Alex Alderton, Roberto Amato, Jeffrey Barrett, Sonia Goncalves, Ewan Harrison, David K. Jackson, Ian Johnston, Dominic Kwiatkowski, Cordelia Langford, John Sillitoe on behalf of the Wellcome Sanger Institute COVID-19 Surveillance Team |
| EPI_ISL_3468744 | Lighthouse Lab in Alderley Park | Wellcome Sanger Institute for the COVID-19 Genomics UK (COG-UK) Consortium | Jacquelyn Wynn, Mairead Hyland, The Lighthouse Lab in Alderley Park and Alex Alderton, Roberto Amato, Jeffrey Barrett, Sonia Goncalves, Ewan Harrison, David K. Jackson, Ian Johnston, Dominic Kwiatkowski, Cordelia Langford, John Sillitoe on behalf of the Wellcome Sanger Institute COVID-19 Surveillance Team |
| EPI_ISL_3468745, EPI_ISL_3468746, EPI_ISL_3468747, EPI_ISL_3468748, EPI_ISL_3468749, EPI_ISL_3468750, EPI_ISL_3468751, EPI_ISL_3468752, EPI_ISL_3468753, EPI_ISL_3468754 | Lighthouse Lab in Milton Keynes | Wellcome Sanger Institute for the COVID-19 Genomics UK (COG-UK) Consortium | The Lighthouse Lab in Milton Keynes and Alex Alderton, Roberto Amato, Jeffrey Barrett, Sonia Goncalves, Ewan Harrison, David K. Jackson, Ian Johnston, Dominic Kwiatkowski, Cordelia Langford, John Sillitoe on behalf of the Wellcome Sanger Institute COVID-19 Surveillance Team |
| EPI_ISL_3468755 | Lighthouse Laboratory Plymouth | Wellcome Sanger Institute for the COVID-19 Genomics UK (COG-UK) Consortium | Lighthouse Laboratory Plymouth and Alex Alderton, Roberto Amato, Jeffrey Barrett, Sonia Goncalves, Ewan Harrison, David K. Jackson, Ian Johnston, Dominic Kwiatkowski, Cordelia Langford, John Sillitoe on behalf of the Wellcome Sanger Institute COVID-19 Surveillance Team |
| EPI_ISL_3468756 | Lighthouse Lab in Milton Keynes | Wellcome Sanger Institute for the COVID-19 Genomics UK (COG-UK) Consortium | The Lighthouse Lab in Milton Keynes and Alex Alderton, Roberto Amato, Jeffrey Barrett, Sonia Goncalves, Ewan Harrison, David K. Jackson, Ian Johnston, Dominic Kwiatkowski, Cordelia Langford, John Sillitoe on behalf of the Wellcome Sanger Institute COVID-19 Surveillance Team |
| EPI_ISL_3468757 | Lighthouse Lab in Alderley Park | Wellcome Sanger Institute for the COVID-19 Genomics UK (COG-UK) Consortium | Jacquelyn Wynn, Mairead Hyland, The Lighthouse Lab in Alderley Park and Alex Alderton, Roberto Amato, Jeffrey Barrett, Sonia Goncalves, Ewan Harrison, David K. Jackson, Ian Johnston, Dominic Kwiatkowski, Cordelia Langford, John Sillitoe on behalf of the Wellcome Sanger Institute COVID-19 Surveillance Team |
| EPI_ISL_3468758 | Lighthouse Laboratory Plymouth | Wellcome Sanger Institute for the COVID-19 Genomics UK (COG-UK) Consortium | Lighthouse Laboratory Plymouth and Alex Alderton, Roberto Amato, Jeffrey Barrett, Sonia Goncalves, Ewan Harrison, David K. Jackson, Ian Johnston, Dominic Kwiatkowski, Cordelia Langford, John Sillitoe on behalf of the Wellcome Sanger Institute COVID-19 Surveillance Team |
| EPI_ISL_3468759, EPI_ISL_3468760 | Lighthouse Lab in Milton Keynes | Wellcome Sanger Institute for the COVID-19 Genomics UK (COG-UK) Consortium | The Lighthouse Lab in Milton Keynes and Alex Alderton, Roberto Amato, Jeffrey Barrett, Sonia Goncalves, Ewan Harrison, David K. Jackson, Ian Johnston, Dominic Kwiatkowski, Cordelia Langford, John Sillitoe on behalf of the Wellcome Sanger Institute COVID-19 Surveillance Team |
| EPI_ISL_3468761 | Lighthouse Lab in Alderley Park | Wellcome Sanger Institute for the COVID-19 Genomics UK (COG-UK) Consortium | Jacquelyn Wynn, Mairead Hyland, The Lighthouse Lab in Alderley Park and Alex Alderton, Roberto Amato, Jeffrey Barrett, Sonia Goncalves, Ewan Harrison, David K. Jackson, Ian Johnston, Dominic Kwiatkowski, Cordelia Langford, John Sillitoe on behalf of the Wellcome Sanger Institute COVID-19 Surveillance Team |
| EPI_ISL_3468762 | Lighthouse Laboratory Plymouth | Wellcome Sanger Institute for the COVID-19 Genomics UK (COG-UK) Consortium | Lighthouse Laboratory Plymouth and Alex Alderton, Roberto Amato, Jeffrey Barrett, Sonia Goncalves, Ewan Harrison, David K. Jackson, Ian Johnston, Dominic Kwiatkowski, Cordelia Langford, John Sillitoe on behalf of the Wellcome Sanger Institute COVID-19 Surveillance Team |
| EPI_ISL_3468763 | Lighthouse Lab in Milton Keynes | Wellcome Sanger Institute for the COVID-19 Genomics UK (COG-UK) Consortium | The Lighthouse Lab in Milton Keynes and Alex Alderton, Roberto Amato, Jeffrey Barrett, Sonia Goncalves, Ewan Harrison, David K. Jackson, Ian Johnston, Dominic Kwiatkowski, Cordelia Langford, John Sillitoe on behalf of the Wellcome Sanger Institute COVID-19 Surveillance Team |
| EPI_ISL_3468764 | Lighthouse Lab in Alderley Park | Wellcome Sanger Institute for the COVID-19 Genomics UK (COG-UK) Consortium | Jacquelyn Wynn, Mairead Hyland, The Lighthouse Lab in Alderley Park and Alex Alderton, Roberto Amato, Jeffrey Barrett, Sonia Goncalves, Ewan Harrison, David K. Jackson, Ian Johnston, Dominic Kwiatkowski, Cordelia Langford, John Sillitoe on behalf of the Wellcome Sanger Institute COVID-19 Surveillance Team |
| EPI_ISL_3468765 | Lighthouse Lab in Milton Keynes | Wellcome Sanger Institute for the COVID-19 Genomics UK (COG-UK) Consortium | The Lighthouse Lab in Milton Keynes and Alex Alderton, Roberto Amato, Jeffrey Barrett, Sonia Goncalves, Ewan Harrison, David K. Jackson, Ian Johnston, Dominic Kwiatkowski, Cordelia Langford, John Sillitoe on behalf of the Wellcome Sanger Institute COVID-19 Surveillance Team |
| EPI_ISL_3468766, EPI_ISL_3468767 | Lighthouse Laboratory Plymouth | Wellcome Sanger Institute for the COVID-19 Genomics UK (COG-UK) Consortium | Lighthouse Laboratory Plymouth and Alex Alderton, Roberto Amato, Jeffrey Barrett, Sonia Goncalves, Ewan Harrison, David K. Jackson, Ian Johnston, Dominic Kwiatkowski, Cordelia Langford, John Sillitoe on behalf of the Wellcome Sanger Institute COVID-19 Surveillance Team |
| EPI_ISL_3468768 | Lighthouse Lab in Alderley Park | Wellcome Sanger Institute for the COVID-19 Genomics UK (COG-UK) Consortium | Jacquelyn Wynn, Mairead Hyland, The Lighthouse Lab in Alderley Park and Alex Alderton, Roberto Amato, Jeffrey Barrett, Sonia Goncalves, Ewan Harrison, David K. Jackson, Ian Johnston, Dominic Kwiatkowski, Cordelia Langford, John Sillitoe on behalf of the Wellcome Sanger Institute COVID-19 Surveillance Team |
| EPI_ISL_3468769 | Lighthouse Lab in Milton Keynes | Wellcome Sanger Institute for the COVID-19 Genomics UK (COG-UK) Consortium | The Lighthouse Lab in Milton Keynes and Alex Alderton, Roberto Amato, Jeffrey Barrett, Sonia Goncalves, Ewan Harrison, David K. Jackson, Ian Johnston, Dominic Kwiatkowski, Cordelia Langford, John Sillitoe on behalf of the Wellcome Sanger Institute COVID-19 Surveillance Team |
| EPI_ISL_3468770 | Lighthouse Laboratory Plymouth | Wellcome Sanger Institute for the COVID-19 Genomics UK (COG-UK) Consortium | Lighthouse Laboratory Plymouth and Alex Alderton, Roberto Amato, Jeffrey Barrett, Sonia Goncalves, Ewan Harrison, David K. Jackson, Ian Johnston, Dominic Kwiatkowski, Cordelia Langford, John Sillitoe on behalf of the Wellcome Sanger Institute COVID-19 Surveillance Team |
| EPI_ISL_3468771, EPI_ISL_3468772 | Lighthouse Lab in Milton Keynes | Wellcome Sanger Institute for the COVID-19 Genomics UK (COG-UK) Consortium | The Lighthouse Lab in Milton Keynes and Alex Alderton, Roberto Amato, Jeffrey Barrett, Sonia Goncalves, Ewan Harrison, David K. Jackson, Ian Johnston, Dominic Kwiatkowski, Cordelia Langford, John Sillitoe on behalf of the Wellcome Sanger Institute COVID-19 Surveillance Team |
| EPI_ISL_3468773, EPI_ISL_3468774 | Lighthouse Laboratory Plymouth | Wellcome Sanger Institute for the COVID-19 Genomics UK (COG-UK) Consortium | Lighthouse Laboratory Plymouth and Alex Alderton, Roberto Amato, Jeffrey Barrett, Sonia Goncalves, Ewan Harrison, David K. Jackson, Ian Johnston, Dominic Kwiatkowski, Cordelia Langford, John Sillitoe on behalf of the Wellcome Sanger Institute COVID-19 Surveillance Team |
| EPI_ISL_3468775, EPI_ISL_3468776 | Lighthouse Lab in Milton Keynes | Wellcome Sanger Institute for the COVID-19 Genomics UK (COG-UK) Consortium | The Lighthouse Lab in Milton Keynes and Alex Alderton, Roberto Amato, Jeffrey Barrett, Sonia Goncalves, Ewan Harrison, David K. Jackson, Ian Johnston, Dominic Kwiatkowski, Cordelia Langford, John Sillitoe on behalf of the Wellcome Sanger Institute COVID-19 Surveillance Team |
| EPI_ISL_3468777 | Lighthouse Laboratory Plymouth | Wellcome Sanger Institute for the COVID-19 Genomics UK (COG-UK) Consortium | Lighthouse Laboratory Plymouth and Alex Alderton, Roberto Amato, Jeffrey Barrett, Sonia Goncalves, Ewan Harrison, David K. Jackson, Ian Johnston, Dominic Kwiatkowski, Cordelia Langford, John Sillitoe on behalf of the Wellcome Sanger Institute COVID-19 Surveillance Team |
| EPI_ISL_3468778 | Lighthouse Lab in Milton Keynes | Wellcome Sanger Institute for the COVID-19 Genomics UK (COG-UK) Consortium | The Lighthouse Lab in Milton Keynes and Alex Alderton, Roberto Amato, Jeffrey Barrett, Sonia Goncalves, Ewan Harrison, David K. Jackson, Ian Johnston, Dominic Kwiatkowski, Cordelia Langford, John Sillitoe on behalf of the Wellcome Sanger Institute COVID-19 Surveillance Team |
| EPI_ISL_3468779 | Lighthouse Lab in Alderley Park | Wellcome Sanger Institute for the COVID-19 Genomics UK (COG-UK) Consortium | Jacquelyn Wynn, Mairead Hyland, The Lighthouse Lab in Alderley Park and Alex Alderton, Roberto Amato, Jeffrey Barrett, Sonia Goncalves, Ewan Harrison, David K. Jackson, Ian Johnston, Dominic Kwiatkowski, Cordelia Langford, John Sillitoe on behalf of the Wellcome Sanger Institute COVID-19 Surveillance Team |
| EPI_ISL_3468780 | Lighthouse Laboratory Plymouth | Wellcome Sanger Institute for the COVID-19 Genomics UK (COG-UK) Consortium | Lighthouse Laboratory Plymouth and Alex Alderton, Roberto Amato, Jeffrey Barrett, Sonia Goncalves, Ewan Harrison, David K. Jackson, Ian Johnston, Dominic Kwiatkowski, Cordelia Langford, John Sillitoe on behalf of the Wellcome Sanger Institute COVID-19 Surveillance Team |
| EPI_ISL_3468781, EPI_ISL_3468782, EPI_ISL_3468783, EPI_ISL_3468784 | Lighthouse Lab in Milton Keynes | Wellcome Sanger Institute for the COVID-19 Genomics UK (COG-UK) Consortium | The Lighthouse Lab in Milton Keynes and Alex Alderton, Roberto Amato, Jeffrey Barrett, Sonia Goncalves, Ewan Harrison, David K. Jackson, Ian Johnston, Dominic Kwiatkowski, Cordelia Langford, John Sillitoe on behalf of the Wellcome Sanger Institute COVID-19 Surveillance Team |
| EPI_ISL_3468785 | Lighthouse Laboratory Plymouth | Wellcome Sanger Institute for the COVID-19 Genomics UK (COG-UK) Consortium | Lighthouse Laboratory Plymouth and Alex Alderton, Roberto Amato, Jeffrey Barrett, Sonia Goncalves, Ewan Harrison, David K. Jackson, Ian Johnston, Dominic Kwiatkowski, Cordelia Langford, John Sillitoe on behalf of the Wellcome Sanger Institute COVID-19 Surveillance Team |
| EPI_ISL_3468786 | Lighthouse Lab in Milton Keynes | Wellcome Sanger Institute for the COVID-19 Genomics UK (COG-UK) Consortium | The Lighthouse Lab in Milton Keynes and Alex Alderton, Roberto Amato, Jeffrey Barrett, Sonia Goncalves, Ewan Harrison, David K. Jackson, Ian Johnston, Dominic Kwiatkowski, Cordelia Langford, John Sillitoe on behalf of the Wellcome Sanger Institute COVID-19 Surveillance Team |
| EPI_ISL_3468787 | Lighthouse Lab in Alderley Park | Wellcome Sanger Institute for the COVID-19 Genomics UK (COG-UK) Consortium | Jacquelyn Wynn, Mairead Hyland, The Lighthouse Lab in Alderley Park and Alex Alderton, Roberto Amato, Jeffrey Barrett, Sonia Goncalves, Ewan Harrison, David K. Jackson, Ian Johnston, Dominic Kwiatkowski, Cordelia Langford, John Sillitoe on behalf of the Wellcome Sanger Institute COVID-19 Surveillance Team |
| EPI_ISL_3468788, EPI_ISL_3468789, EPI_ISL_3468790 | Lighthouse Lab in Milton Keynes | Wellcome Sanger Institute for the COVID-19 Genomics UK (COG-UK) Consortium | The Lighthouse Lab in Milton Keynes and Alex Alderton, Roberto Amato, Jeffrey Barrett, Sonia Goncalves, Ewan Harrison, David K. Jackson, Ian Johnston, Dominic Kwiatkowski, Cordelia Langford, John Sillitoe on behalf of the Wellcome Sanger Institute COVID-19 Surveillance Team |
| EPI_ISL_3468791 | Lighthouse Laboratory Plymouth | Wellcome Sanger Institute for the COVID-19 Genomics UK (COG-UK) Consortium | Lighthouse Laboratory Plymouth and Alex Alderton, Roberto Amato, Jeffrey Barrett, Sonia Goncalves, Ewan Harrison, David K. Jackson, Ian Johnston, Dominic Kwiatkowski, Cordelia Langford, John Sillitoe on behalf of the Wellcome Sanger Institute COVID-19 Surveillance Team |
| EPI_ISL_3468792 | Lighthouse Lab in Milton Keynes | Wellcome Sanger Institute for the COVID-19 Genomics UK (COG-UK) Consortium | The Lighthouse Lab in Milton Keynes and Alex Alderton, Roberto Amato, Jeffrey Barrett, Sonia Goncalves, Ewan Harrison, David K. Jackson, Ian Johnston, Dominic Kwiatkowski, Cordelia Langford, John Sillitoe on behalf of the Wellcome Sanger Institute COVID-19 Surveillance Team |
| EPI_ISL_3468793 | Lighthouse Lab in Alderley Park | Wellcome Sanger Institute for the COVID-19 Genomics UK (COG-UK) Consortium | Jacquelyn Wynn, Mairead Hyland, The Lighthouse Lab in Alderley Park and Alex Alderton, Roberto Amato, Jeffrey Barrett, Sonia Goncalves, Ewan Harrison, David K. Jackson, Ian Johnston, Dominic Kwiatkowski, Cordelia Langford, John Sillitoe on behalf of the Wellcome Sanger Institute COVID-19 Surveillance Team |
| EPI_ISL_3468794, EPI_ISL_3468795 | Lighthouse Laboratory Plymouth | Wellcome Sanger Institute for the COVID-19 Genomics UK (COG-UK) Consortium | Lighthouse Laboratory Plymouth and Alex Alderton, Roberto Amato, Jeffrey Barrett, Sonia Goncalves, Ewan Harrison, David K. Jackson, Ian Johnston, Dominic Kwiatkowski, Cordelia Langford, John Sillitoe on behalf of the Wellcome Sanger Institute COVID-19 Surveillance Team |
| EPI_ISL_3468796, EPI_ISL_3468797, EPI_ISL_3468798 | Lighthouse Lab in Milton Keynes | Wellcome Sanger Institute for the COVID-19 Genomics UK (COG-UK) Consortium | The Lighthouse Lab in Milton Keynes and Alex Alderton, Roberto Amato, Jeffrey Barrett, Sonia Goncalves, Ewan Harrison, David K. Jackson, Ian Johnston, Dominic Kwiatkowski, Cordelia Langford, John Sillitoe on behalf of the Wellcome Sanger Institute COVID-19 Surveillance Team |
| EPI_ISL_3468799 | Lighthouse Laboratory Plymouth | Wellcome Sanger Institute for the COVID-19 Genomics UK (COG-UK) Consortium | Lighthouse Laboratory Plymouth and Alex Alderton, Roberto Amato, Jeffrey Barrett, Sonia Goncalves, Ewan Harrison, David K. Jackson, Ian Johnston, Dominic Kwiatkowski, Cordelia Langford, John Sillitoe on behalf of the Wellcome Sanger Institute COVID-19 Surveillance Team |
| EPI_ISL_3468800, EPI_ISL_3468801, EPI_ISL_3468802 | Lighthouse Lab in Alderley Park | Wellcome Sanger Institute for the COVID-19 Genomics UK (COG-UK) Consortium | Jacquelyn Wynn, Mairead Hyland, The Lighthouse Lab in Alderley Park and Alex Alderton, Roberto Amato, Jeffrey Barrett, Sonia Goncalves, Ewan Harrison, David K. Jackson, Ian Johnston, Dominic Kwiatkowski, Cordelia Langford, John Sillitoe on behalf of the Wellcome Sanger Institute COVID-19 Surveillance Team |
| EPI_ISL_3468803 | Lighthouse Laboratory Plymouth | Wellcome Sanger Institute for the COVID-19 Genomics UK (COG-UK) Consortium | Lighthouse Laboratory Plymouth and Alex Alderton, Roberto Amato, Jeffrey Barrett, Sonia Goncalves, Ewan Harrison, David K. Jackson, Ian Johnston, Dominic Kwiatkowski, Cordelia Langford, John Sillitoe on behalf of the Wellcome Sanger Institute COVID-19 Surveillance Team |
| EPI_ISL_3468804 | Lighthouse Lab in Alderley Park | Wellcome Sanger Institute for the COVID-19 Genomics UK (COG-UK) Consortium | Jacquelyn Wynn, Mairead Hyland, The Lighthouse Lab in Alderley Park and Alex Alderton, Roberto Amato, Jeffrey Barrett, Sonia Goncalves, Ewan Harrison, David K. Jackson, Ian Johnston, Dominic Kwiatkowski, Cordelia Langford, John Sillitoe on behalf of the Wellcome Sanger Institute COVID-19 Surveillance Team |
| EPI_ISL_3468805 | Lighthouse Lab in Milton Keynes | Wellcome Sanger Institute for the COVID-19 Genomics UK (COG-UK) Consortium | The Lighthouse Lab in Milton Keynes and Alex Alderton, Roberto Amato, Jeffrey Barrett, Sonia Goncalves, Ewan Harrison, David K. Jackson, Ian Johnston, Dominic Kwiatkowski, Cordelia Langford, John Sillitoe on behalf of the Wellcome Sanger Institute COVID-19 Surveillance Team |

| | | | |
|---|---|---|---|
| EPI_ISL_3468806 | Lighthouse Lab in Alderley Park | Wellcome Sanger Institute for the COVID-19 Genomics UK (COG-UK) Consortium | Jacquelyn Wynn, Mairead Hyland, The Lighthouse Lab in Alderley Park and Alex Alderton, Roberto Amato, Jeffrey Barrett, Sonia Goncalves, Ewan Harrison, David K. Jackson, Ian Johnston, Dominic Kwiatkowski, Cordelia Langford, John Sillitoe on behalf of the Wellcome Sanger Institute COVID-19 Surveillance Team |
| EPI_ISL_3468807, EPI_ISL_3468808, EPI_ISL_3468809, EPI_ISL_3468810, EPI_ISL_3468811, EPI_ISL_3468812 | Lighthouse Lab in Milton Keynes | Wellcome Sanger Institute for the COVID-19 Genomics UK (COG-UK) Consortium | The Lighthouse Lab in Milton Keynes and Alex Alderton, Roberto Amato, Jeffrey Barrett, Sonia Goncalves, Ewan Harrison, David K. Jackson, Ian Johnston, Dominic Kwiatkowski, Cordelia Langford, John Sillitoe on behalf of the Wellcome Sanger Institute COVID-19 Surveillance Team |
| EPI_ISL_3468813, EPI_ISL_3468814 | Lighthouse Lab in Alderley Park | Wellcome Sanger Institute for the COVID-19 Genomics UK (COG-UK) Consortium | Jacquelyn Wynn, Mairead Hyland, The Lighthouse Lab in Alderley Park and Alex Alderton, Roberto Amato, Jeffrey Barrett, Sonia Goncalves, Ewan Harrison, David K. Jackson, Ian Johnston, Dominic Kwiatkowski, Cordelia Langford, John Sillitoe on behalf of the Wellcome Sanger Institute COVID-19 Surveillance Team |
| EPI_ISL_3468815, EPI_ISL_3468816 | Lighthouse Laboratory Plymouth | Wellcome Sanger Institute for the COVID-19 Genomics UK (COG-UK) Consortium | Lighthouse Laboratory Plymouth and Alex Alderton, Roberto Amato, Jeffrey Barrett, Sonia Goncalves, Ewan Harrison, David K. Jackson, Ian Johnston, Dominic Kwiatkowski, Cordelia Langford, John Sillitoe on behalf of the Wellcome Sanger Institute COVID-19 Surveillance Team |
| EPI_ISL_3468817 | Lighthouse Lab in Alderley Park | Wellcome Sanger Institute for the COVID-19 Genomics UK (COG-UK) Consortium | Jacquelyn Wynn, Mairead Hyland, The Lighthouse Lab in Alderley Park and Alex Alderton, Roberto Amato, Jeffrey Barrett, Sonia Goncalves, Ewan Harrison, David K. Jackson, Ian Johnston, Dominic Kwiatkowski, Cordelia Langford, John Sillitoe on behalf of the Wellcome Sanger Institute COVID-19 Surveillance Team |
| EPI_ISL_3468818, EPI_ISL_3468819 | Lighthouse Lab in Milton Keynes | Wellcome Sanger Institute for the COVID-19 Genomics UK (COG-UK) Consortium | The Lighthouse Lab in Milton Keynes and Alex Alderton, Roberto Amato, Jeffrey Barrett, Sonia Goncalves, Ewan Harrison, David K. Jackson, Ian Johnston, Dominic Kwiatkowski, Cordelia Langford, John Sillitoe on behalf of the Wellcome Sanger Institute COVID-19 Surveillance Team |
| EPI_ISL_3468820 | Lighthouse Laboratory Plymouth | Wellcome Sanger Institute for the COVID-19 Genomics UK (COG-UK) Consortium | Lighthouse Laboratory Plymouth and Alex Alderton, Roberto Amato, Jeffrey Barrett, Sonia Goncalves, Ewan Harrison, David K. Jackson, Ian Johnston, Dominic Kwiatkowski, Cordelia Langford, John Sillitoe on behalf of the Wellcome Sanger Institute COVID-19 Surveillance Team |
| EPI_ISL_3468821 | Lighthouse Lab in Alderley Park | Wellcome Sanger Institute for the COVID-19 Genomics UK (COG-UK) Consortium | Jacquelyn Wynn, Mairead Hyland, The Lighthouse Lab in Alderley Park and Alex Alderton, Roberto Amato, Jeffrey Barrett, Sonia Goncalves, Ewan Harrison, David K. Jackson, Ian Johnston, Dominic Kwiatkowski, Cordelia Langford, John Sillitoe on behalf of the Wellcome Sanger Institute COVID-19 Surveillance Team |
| EPI_ISL_3468822, EPI_ISL_3468823 | Lighthouse Lab in Milton Keynes | Wellcome Sanger Institute for the COVID-19 Genomics UK (COG-UK) Consortium | The Lighthouse Lab in Milton Keynes and Alex Alderton, Roberto Amato, Jeffrey Barrett, Sonia Goncalves, Ewan Harrison, David K. Jackson, Ian Johnston, Dominic Kwiatkowski, Cordelia Langford, John Sillitoe on behalf of the Wellcome Sanger Institute COVID-19 Surveillance Team |
| EPI_ISL_3468824, EPI_ISL_3468825 | Lighthouse Lab in Alderley Park | Wellcome Sanger Institute for the COVID-19 Genomics UK (COG-UK) Consortium | Jacquelyn Wynn, Mairead Hyland, The Lighthouse Lab in Alderley Park and Alex Alderton, Roberto Amato, Jeffrey Barrett, Sonia Goncalves, Ewan Harrison, David K. Jackson, Ian Johnston, Dominic Kwiatkowski, Cordelia Langford, John Sillitoe on behalf of the Wellcome Sanger Institute COVID-19 Surveillance Team |
| EPI_ISL_3468826 | Lighthouse Laboratory Plymouth | Wellcome Sanger Institute for the COVID-19 Genomics UK (COG-UK) Consortium | Lighthouse Laboratory Plymouth and Alex Alderton, Roberto Amato, Jeffrey Barrett, Sonia Goncalves, Ewan Harrison, David K. Jackson, Ian Johnston, Dominic Kwiatkowski, Cordelia Langford, John Sillitoe on behalf of the Wellcome Sanger Institute COVID-19 Surveillance Team |
| EPI_ISL_3468827 | Lighthouse Lab in Milton Keynes | Wellcome Sanger Institute for the COVID-19 Genomics UK (COG-UK) Consortium | The Lighthouse Lab in Milton Keynes and Alex Alderton, Roberto Amato, Jeffrey Barrett, Sonia Goncalves, Ewan Harrison, David K. Jackson, Ian Johnston, Dominic Kwiatkowski, Cordelia Langford, John Sillitoe on behalf of the Wellcome Sanger Institute COVID-19 Surveillance Team |
| EPI_ISL_3468828 | Lighthouse Laboratory Plymouth | Wellcome Sanger Institute for the COVID-19 Genomics UK (COG-UK) Consortium | Lighthouse Laboratory Plymouth and Alex Alderton, Roberto Amato, Jeffrey Barrett, Sonia Goncalves, Ewan Harrison, David K. Jackson, Ian Johnston, Dominic Kwiatkowski, Cordelia Langford, John Sillitoe on behalf of the Wellcome Sanger Institute COVID-19 Surveillance Team |
| EPI_ISL_3468829, EPI_ISL_3468830 | Lighthouse Lab in Milton Keynes | Wellcome Sanger Institute for the COVID-19 Genomics UK (COG-UK) Consortium | The Lighthouse Lab in Milton Keynes and Alex Alderton, Roberto Amato, Jeffrey Barrett, Sonia Goncalves, Ewan Harrison, David K. Jackson, Ian Johnston, Dominic Kwiatkowski, Cordelia Langford, John Sillitoe on behalf of the Wellcome Sanger Institute COVID-19 Surveillance Team |
| EPI_ISL_3468831 | Lighthouse Laboratory Plymouth | Wellcome Sanger Institute for the COVID-19 Genomics UK (COG-UK) Consortium | Lighthouse Laboratory Plymouth and Alex Alderton, Roberto Amato, Jeffrey Barrett, Sonia Goncalves, Ewan Harrison, David K. Jackson, Ian Johnston, Dominic Kwiatkowski, Cordelia Langford, John Sillitoe on behalf of the Wellcome Sanger Institute COVID-19 Surveillance Team |
| EPI_ISL_3468832 | Lighthouse Lab in Alderley Park | Wellcome Sanger Institute for the COVID-19 Genomics UK (COG-UK) Consortium | Jacquelyn Wynn, Mairead Hyland, The Lighthouse Lab in Alderley Park and Alex Alderton, Roberto Amato, Jeffrey Barrett, Sonia Goncalves, Ewan Harrison, David K. Jackson, Ian Johnston, Dominic Kwiatkowski, Cordelia Langford, John Sillitoe on behalf of the Wellcome Sanger Institute COVID-19 Surveillance Team |
| EPI_ISL_3468833, EPI_ISL_3468834 | Lighthouse Lab in Milton Keynes | Wellcome Sanger Institute for the COVID-19 Genomics UK (COG-UK) Consortium | The Lighthouse Lab in Milton Keynes and Alex Alderton, Roberto Amato, Jeffrey Barrett, Sonia Goncalves, Ewan Harrison, David K. Jackson, Ian Johnston, Dominic Kwiatkowski, Cordelia Langford, John Sillitoe on behalf of the Wellcome Sanger Institute COVID-19 Surveillance Team |
| EPI_ISL_3468835 | Lighthouse Laboratory Plymouth | Wellcome Sanger Institute for the COVID-19 Genomics UK (COG-UK) Consortium | Lighthouse Laboratory Plymouth and Alex Alderton, Roberto Amato, Jeffrey Barrett, Sonia Goncalves, Ewan Harrison, David K. Jackson, Ian Johnston, Dominic Kwiatkowski, Cordelia Langford, John Sillitoe on behalf of the Wellcome Sanger Institute COVID-19 Surveillance Team |
| EPI_ISL_3468836, EPI_ISL_3468837, EPI_ISL_3468838, EPI_ISL_3468839 | Lighthouse Lab in Milton Keynes | Wellcome Sanger Institute for the COVID-19 Genomics UK (COG-UK) Consortium | The Lighthouse Lab in Milton Keynes and Alex Alderton, Roberto Amato, Jeffrey Barrett, Sonia Goncalves, Ewan Harrison, David K. Jackson, Ian Johnston, Dominic Kwiatkowski, Cordelia Langford, John Sillitoe on behalf of the Wellcome Sanger Institute COVID-19 Surveillance Team |
| EPI_ISL_3468840 | Lighthouse Lab in Alderley Park | Wellcome Sanger Institute for the COVID-19 Genomics UK (COG-UK) Consortium | Jacquelyn Wynn, Mairead Hyland, The Lighthouse Lab in Alderley Park and Alex Alderton, Roberto Amato, Jeffrey Barrett, Sonia Goncalves, Ewan Harrison, David K. Jackson, Ian Johnston, Dominic Kwiatkowski, Cordelia Langford, John Sillitoe on behalf of the Wellcome Sanger Institute COVID-19 Surveillance Team |
| EPI_ISL_3468841 | Lighthouse Lab in Milton Keynes | Wellcome Sanger Institute for the COVID-19 Genomics UK (COG-UK) Consortium | The Lighthouse Lab in Milton Keynes and Alex Alderton, Roberto Amato, Jeffrey Barrett, Sonia Goncalves, Ewan Harrison, David K. Jackson, Ian Johnston, Dominic Kwiatkowski, Cordelia Langford, John Sillitoe on behalf of the Wellcome Sanger Institute COVID-19 Surveillance Team |
| EPI_ISL_3468842 | Lighthouse Laboratory Plymouth | Wellcome Sanger Institute for the COVID-19 Genomics UK (COG-UK) Consortium | Lighthouse Laboratory Plymouth and Alex Alderton, Roberto Amato, Jeffrey Barrett, Sonia Goncalves, Ewan Harrison, David K. Jackson, Ian Johnston, Dominic Kwiatkowski, Cordelia Langford, John Sillitoe on behalf of the Wellcome Sanger Institute COVID-19 Surveillance Team |
| EPI_ISL_3468843 | Lighthouse Lab in Milton Keynes | Wellcome Sanger Institute for the COVID-19 Genomics UK (COG-UK) Consortium | The Lighthouse Lab in Milton Keynes and Alex Alderton, Roberto Amato, Jeffrey Barrett, Sonia Goncalves, Ewan Harrison, David K. Jackson, Ian Johnston, Dominic Kwiatkowski, Cordelia Langford, John Sillitoe on behalf of the Wellcome Sanger Institute COVID-19 Surveillance Team |
| EPI_ISL_3468844 | Lighthouse Laboratory Plymouth | Wellcome Sanger Institute for the COVID-19 Genomics UK (COG-UK) Consortium | Lighthouse Laboratory Plymouth and Alex Alderton, Roberto Amato, Jeffrey Barrett, Sonia Goncalves, Ewan Harrison, David K. Jackson, Ian Johnston, Dominic Kwiatkowski, Cordelia Langford, John Sillitoe on behalf of the Wellcome Sanger Institute COVID-19 Surveillance Team |
| EPI_ISL_3468845, EPI_ISL_3468846 | Lighthouse Lab in Milton Keynes | Wellcome Sanger Institute for the COVID-19 Genomics UK (COG-UK) Consortium | The Lighthouse Lab in Milton Keynes and Alex Alderton, Roberto Amato, Jeffrey Barrett, Sonia Goncalves, Ewan Harrison, David K. Jackson, Ian Johnston, Dominic Kwiatkowski, Cordelia Langford, John Sillitoe on behalf of the Wellcome Sanger Institute COVID-19 Surveillance Team |
| EPI_ISL_3468847 | Lighthouse Lab in Alderley Park | Wellcome Sanger Institute for the COVID-19 Genomics UK (COG-UK) Consortium | Jacquelyn Wynn, Mairead Hyland, The Lighthouse Lab in Alderley Park and Alex Alderton, Roberto Amato, Jeffrey Barrett, Sonia Goncalves, Ewan Harrison, David K. Jackson, Ian Johnston, Dominic Kwiatkowski, Cordelia Langford, John Sillitoe on behalf of the Wellcome Sanger Institute COVID-19 Surveillance Team |
| EPI_ISL_3468848 | Lighthouse Laboratory Plymouth | Wellcome Sanger Institute for the COVID-19 Genomics UK (COG-UK) Consortium | Lighthouse Laboratory Plymouth and Alex Alderton, Roberto Amato, Jeffrey Barrett, Sonia Goncalves, Ewan Harrison, David K. Jackson, Ian Johnston, Dominic Kwiatkowski, Cordelia Langford, John Sillitoe on behalf of the Wellcome Sanger Institute COVID-19 Surveillance Team |
| EPI_ISL_3468849, EPI_ISL_3468850 | Lighthouse Lab in Milton Keynes | Wellcome Sanger Institute for the COVID-19 Genomics UK (COG-UK) Consortium | The Lighthouse Lab in Milton Keynes and Alex Alderton, Roberto Amato, Jeffrey Barrett, Sonia Goncalves, Ewan Harrison, David K. Jackson, Ian Johnston, Dominic Kwiatkowski, Cordelia Langford, John Sillitoe on behalf of the Wellcome Sanger Institute COVID-19 Surveillance Team |
| EPI_ISL_3468851, EPI_ISL_3468852 | Lighthouse Lab in Alderley Park | Wellcome Sanger Institute for the COVID-19 Genomics UK (COG-UK) Consortium | Jacquelyn Wynn, Mairead Hyland, The Lighthouse Lab in Alderley Park and Alex Alderton, Roberto Amato, Jeffrey Barrett, Sonia Goncalves, Ewan Harrison, David K. Jackson, Ian Johnston, Dominic Kwiatkowski, Cordelia Langford, John Sillitoe on behalf of the Wellcome Sanger Institute COVID-19 Surveillance Team |
| EPI_ISL_3468853 | Lighthouse Laboratory Plymouth | Wellcome Sanger Institute for the COVID-19 Genomics UK (COG-UK) Consortium | Lighthouse Laboratory Plymouth and Alex Alderton, Roberto Amato, Jeffrey Barrett, Sonia Goncalves, Ewan Harrison, David K. Jackson, Ian Johnston, Dominic Kwiatkowski, Cordelia Langford, John Sillitoe on behalf of the Wellcome Sanger Institute COVID-19 Surveillance Team |
| EPI_ISL_3468854, EPI_ISL_3468855, EPI_ISL_3468856 | Lighthouse Lab in Milton Keynes | Wellcome Sanger Institute for the COVID-19 Genomics UK (COG-UK) Consortium | The Lighthouse Lab in Milton Keynes and Alex Alderton, Roberto Amato, Jeffrey Barrett, Sonia Goncalves, Ewan Harrison, David K. Jackson, Ian Johnston, Dominic Kwiatkowski, Cordelia Langford, John Sillitoe on behalf of the Wellcome Sanger Institute COVID-19 Surveillance Team |
| EPI_ISL_3468857 | Lighthouse Lab in Alderley Park | Wellcome Sanger Institute for the COVID-19 Genomics UK (COG-UK) Consortium | Jacquelyn Wynn, Mairead Hyland, The Lighthouse Lab in Alderley Park and Alex Alderton, Roberto Amato, Jeffrey Barrett, Sonia Goncalves, Ewan Harrison, David K. Jackson, Ian Johnston, Dominic Kwiatkowski, Cordelia Langford, John Sillitoe on behalf of the Wellcome Sanger Institute COVID-19 Surveillance Team |
| EPI_ISL_3468858 | Lighthouse Lab in Milton Keynes | Wellcome Sanger Institute for the COVID-19 Genomics UK (COG-UK) Consortium | The Lighthouse Lab in Milton Keynes and Alex Alderton, Roberto Amato, Jeffrey Barrett, Sonia Goncalves, Ewan Harrison, David K. Jackson, Ian Johnston, Dominic Kwiatkowski, Cordelia Langford, John Sillitoe on behalf of the Wellcome Sanger Institute COVID-19 Surveillance Team |
| EPI_ISL_3468859, EPI_ISL_3468860 | Lighthouse Laboratory Plymouth | Wellcome Sanger Institute for the COVID-19 Genomics UK (COG-UK) Consortium | Lighthouse Laboratory Plymouth and Alex Alderton, Roberto Amato, Jeffrey Barrett, Sonia Goncalves, Ewan Harrison, David K. Jackson, Ian Johnston, Dominic Kwiatkowski, Cordelia Langford, John Sillitoe on behalf of the Wellcome Sanger Institute COVID-19 Surveillance Team |
| EPI_ISL_3468861 | Lighthouse Lab in Milton Keynes | Wellcome Sanger Institute for the COVID-19 Genomics UK (COG-UK) Consortium | The Lighthouse Lab in Milton Keynes and Alex Alderton, Roberto Amato, Jeffrey Barrett, Sonia Goncalves, Ewan Harrison, David K. Jackson, Ian Johnston, Dominic Kwiatkowski, Cordelia Langford, John Sillitoe on behalf of the Wellcome Sanger Institute COVID-19 Surveillance Team |
| EPI_ISL_3468862, EPI_ISL_3468863 | Lighthouse Lab in Alderley Park | Wellcome Sanger Institute for the COVID-19 Genomics UK (COG-UK) Consortium | Jacquelyn Wynn, Mairead Hyland, The Lighthouse Lab in Alderley Park and Alex Alderton, Roberto Amato, Jeffrey Barrett, Sonia Goncalves, Ewan Harrison, David K. Jackson, Ian Johnston, Dominic Kwiatkowski, Cordelia Langford, John Sillitoe on behalf of the Wellcome Sanger Institute COVID-19 Surveillance Team |
| EPI_ISL_3468864, EPI_ISL_3468865 | Lighthouse Laboratory Plymouth | Wellcome Sanger Institute for the COVID-19 Genomics UK (COG-UK) Consortium | Lighthouse Laboratory Plymouth and Alex Alderton, Roberto Amato, Jeffrey Barrett, Sonia Goncalves, Ewan Harrison, David K. Jackson, Ian Johnston, Dominic Kwiatkowski, Cordelia Langford, John Sillitoe on behalf of the Wellcome Sanger Institute COVID-19 Surveillance Team |
| EPI_ISL_3468866, EPI_ISL_3468867 | Lighthouse Lab in Milton Keynes | Wellcome Sanger Institute for the COVID-19 Genomics UK (COG-UK) Consortium | The Lighthouse Lab in Milton Keynes and Alex Alderton, Roberto Amato, Jeffrey Barrett, Sonia Goncalves, Ewan Harrison, David K. Jackson, Ian Johnston, Dominic Kwiatkowski, Cordelia Langford, John Sillitoe on behalf of the Wellcome Sanger Institute COVID-19 Surveillance Team |
| EPI_ISL_3468868 | Lighthouse Lab in Alderley Park | Wellcome Sanger Institute for the COVID-19 Genomics UK (COG-UK) Consortium | Jacquelyn Wynn, Mairead Hyland, The Lighthouse Lab in Alderley Park and Alex Alderton, Roberto Amato, Jeffrey Barrett, Sonia Goncalves, Ewan Harrison, David K. Jackson, Ian Johnston, Dominic Kwiatkowski, Cordelia Langford, John Sillitoe on behalf of the Wellcome Sanger Institute COVID-19 Surveillance Team |
| EPI_ISL_3468869, EPI_ISL_3468870, EPI_ISL_3468871, EPI_ISL_3468872, EPI_ISL_3468873 | Lighthouse Laboratory Plymouth | Wellcome Sanger Institute for the COVID-19 Genomics UK (COG-UK) Consortium | Lighthouse Laboratory Plymouth and Alex Alderton, Roberto Amato, Jeffrey Barrett, Sonia Goncalves, Ewan Harrison, David K. Jackson, Ian Johnston, Dominic Kwiatkowski, Cordelia Langford, John Sillitoe on behalf of the Wellcome Sanger Institute COVID-19 Surveillance Team |
| EPI_ISL_3468874 | Lighthouse Lab in Alderley Park | Wellcome Sanger Institute for the COVID-19 Genomics UK (COG-UK) Consortium | Jacquelyn Wynn, Mairead Hyland, The Lighthouse Lab in Alderley Park and Alex Alderton, Roberto Amato, Jeffrey Barrett, Sonia Goncalves, Ewan Harrison, David K. Jackson, Ian Johnston, Dominic Kwiatkowski, Cordelia Langford, John Sillitoe on behalf of the Wellcome Sanger Institute COVID-19 Surveillance Team |

| | | | |
|---|---|---|---|
| EPI_ISL_3468875 | Lighthouse Laboratory Plymouth | Wellcome Sanger Institute for the COVID-19 Genomics UK (COG-UK) Consortium | Lighthouse Laboratory Plymouth and Alex Alderton, Roberto Amato, Jeffrey Barrett, Sonia Goncalves, Ewan Harrison, David K. Jackson, Ian Johnston, Dominic Kwiatkowski, Cordelia Langford, John Sillitoe on behalf of the Wellcome Sanger Institute COVID-19 Surveillance Team |
| EPI_ISL_3468876, EPI_ISL_3468877, EPI_ISL_3468878, EPI_ISL_3468879, EPI_ISL_3468880, EPI_ISL_3468881 | Lighthouse Lab in Milton Keynes | Wellcome Sanger Institute for the COVID-19 Genomics UK (COG-UK) Consortium | The Lighthouse Lab in Milton Keynes and Alex Alderton, Roberto Amato, Jeffrey Barrett, Sonia Goncalves, Ewan Harrison, David K. Jackson, Ian Johnston, Dominic Kwiatkowski, Cordelia Langford, John Sillitoe on behalf of the Wellcome Sanger Institute COVID-19 Surveillance Team |
| EPI_ISL_3468882, EPI_ISL_3468883 | Lighthouse Laboratory Plymouth | Wellcome Sanger Institute for the COVID-19 Genomics UK (COG-UK) Consortium | Lighthouse Laboratory Plymouth and Alex Alderton, Roberto Amato, Jeffrey Barrett, Sonia Goncalves, Ewan Harrison, David K. Jackson, Ian Johnston, Dominic Kwiatkowski, Cordelia Langford, John Sillitoe on behalf of the Wellcome Sanger Institute COVID-19 Surveillance Team |
| EPI_ISL_3468884 | Lighthouse Lab in Milton Keynes | Wellcome Sanger Institute for the COVID-19 Genomics UK (COG-UK) Consortium | The Lighthouse Lab in Milton Keynes and Alex Alderton, Roberto Amato, Jeffrey Barrett, Sonia Goncalves, Ewan Harrison, David K. Jackson, Ian Johnston, Dominic Kwiatkowski, Cordelia Langford, John Sillitoe on behalf of the Wellcome Sanger Institute COVID-19 Surveillance Team |
| EPI_ISL_3468885, EPI_ISL_3468886, EPI_ISL_3468887, EPI_ISL_3468888 | Lighthouse Lab in Alderley Park | Wellcome Sanger Institute for the COVID-19 Genomics UK (COG-UK) Consortium | Jacquelyn Wynn, Mairead Hyland, The Lighthouse Lab in Alderley Park and Alex Alderton, Roberto Amato, Jeffrey Barrett, Sonia Goncalves, Ewan Harrison, David K. Jackson, Ian Johnston, Dominic Kwiatkowski, Cordelia Langford, John Sillitoe on behalf of the Wellcome Sanger Institute COVID-19 Surveillance Team |
| EPI_ISL_3468889 | Lighthouse Laboratory Plymouth | Wellcome Sanger Institute for the COVID-19 Genomics UK (COG-UK) Consortium | Lighthouse Laboratory Plymouth and Alex Alderton, Roberto Amato, Jeffrey Barrett, Sonia Goncalves, Ewan Harrison, David K. Jackson, Ian Johnston, Dominic Kwiatkowski, Cordelia Langford, John Sillitoe on behalf of the Wellcome Sanger Institute COVID-19 Surveillance Team |
| EPI_ISL_3468890, EPI_ISL_3468891, EPI_ISL_3468892 | Lighthouse Lab in Alderley Park | Wellcome Sanger Institute for the COVID-19 Genomics UK (COG-UK) Consortium | Jacquelyn Wynn, Mairead Hyland, The Lighthouse Lab in Alderley Park and Alex Alderton, Roberto Amato, Jeffrey Barrett, Sonia Goncalves, Ewan Harrison, David K. Jackson, Ian Johnston, Dominic Kwiatkowski, Cordelia Langford, John Sillitoe on behalf of the Wellcome Sanger Institute COVID-19 Surveillance Team |
| EPI_ISL_3468893 | Lighthouse Lab in Milton Keynes | Wellcome Sanger Institute for the COVID-19 Genomics UK (COG-UK) Consortium | The Lighthouse Lab in Milton Keynes and Alex Alderton, Roberto Amato, Jeffrey Barrett, Sonia Goncalves, Ewan Harrison, David K. Jackson, Ian Johnston, Dominic Kwiatkowski, Cordelia Langford, John Sillitoe on behalf of the Wellcome Sanger Institute COVID-19 Surveillance Team |
| EPI_ISL_3468894 | Lighthouse Laboratory Plymouth | Wellcome Sanger Institute for the COVID-19 Genomics UK (COG-UK) Consortium | Lighthouse Laboratory Plymouth and Alex Alderton, Roberto Amato, Jeffrey Barrett, Sonia Goncalves, Ewan Harrison, David K. Jackson, Ian Johnston, Dominic Kwiatkowski, Cordelia Langford, John Sillitoe on behalf of the Wellcome Sanger Institute COVID-19 Surveillance Team |
| EPI_ISL_3468895 | Lighthouse Lab in Milton Keynes | Wellcome Sanger Institute for the COVID-19 Genomics UK (COG-UK) Consortium | The Lighthouse Lab in Milton Keynes and Alex Alderton, Roberto Amato, Jeffrey Barrett, Sonia Goncalves, Ewan Harrison, David K. Jackson, Ian Johnston, Dominic Kwiatkowski, Cordelia Langford, John Sillitoe on behalf of the Wellcome Sanger Institute COVID-19 Surveillance Team |
| EPI_ISL_3468896 | Lighthouse Lab in Alderley Park | Wellcome Sanger Institute for the COVID-19 Genomics UK (COG-UK) Consortium | Jacquelyn Wynn, Mairead Hyland, The Lighthouse Lab in Alderley Park and Alex Alderton, Roberto Amato, Jeffrey Barrett, Sonia Goncalves, Ewan Harrison, David K. Jackson, Ian Johnston, Dominic Kwiatkowski, Cordelia Langford, John Sillitoe on behalf of the Wellcome Sanger Institute COVID-19 Surveillance Team |
| EPI_ISL_3468897 | Lighthouse Lab in Milton Keynes | Wellcome Sanger Institute for the COVID-19 Genomics UK (COG-UK) Consortium | The Lighthouse Lab in Milton Keynes and Alex Alderton, Roberto Amato, Jeffrey Barrett, Sonia Goncalves, Ewan Harrison, David K. Jackson, Ian Johnston, Dominic Kwiatkowski, Cordelia Langford, John Sillitoe on behalf of the Wellcome Sanger Institute COVID-19 Surveillance Team |
| EPI_ISL_3468898 | Lighthouse Laboratory Plymouth | Wellcome Sanger Institute for the COVID-19 Genomics UK (COG-UK) Consortium | Lighthouse Laboratory Plymouth and Alex Alderton, Roberto Amato, Jeffrey Barrett, Sonia Goncalves, Ewan Harrison, David K. Jackson, Ian Johnston, Dominic Kwiatkowski, Cordelia Langford, John Sillitoe on behalf of the Wellcome Sanger Institute COVID-19 Surveillance Team |
| EPI_ISL_3468899 | Lighthouse Lab in Milton Keynes | Wellcome Sanger Institute for the COVID-19 Genomics UK (COG-UK) Consortium | The Lighthouse Lab in Milton Keynes and Alex Alderton, Roberto Amato, Jeffrey Barrett, Sonia Goncalves, Ewan Harrison, David K. Jackson, Ian Johnston, Dominic Kwiatkowski, Cordelia Langford, John Sillitoe on behalf of the Wellcome Sanger Institute COVID-19 Surveillance Team |
| EPI_ISL_3468900 | Lighthouse Lab in Alderley Park | Wellcome Sanger Institute for the COVID-19 Genomics UK (COG-UK) Consortium | Jacquelyn Wynn, Mairead Hyland, The Lighthouse Lab in Alderley Park and Alex Alderton, Roberto Amato, Jeffrey Barrett, Sonia Goncalves, Ewan Harrison, David K. Jackson, Ian Johnston, Dominic Kwiatkowski, Cordelia Langford, John Sillitoe on behalf of the Wellcome Sanger Institute COVID-19 Surveillance Team |
| EPI_ISL_3468901, EPI_ISL_3468902 | Lighthouse Lab in Milton Keynes | Wellcome Sanger Institute for the COVID-19 Genomics UK (COG-UK) Consortium | The Lighthouse Lab in Milton Keynes and Alex Alderton, Roberto Amato, Jeffrey Barrett, Sonia Goncalves, Ewan Harrison, David K. Jackson, Ian Johnston, Dominic Kwiatkowski, Cordelia Langford, John Sillitoe on behalf of the Wellcome Sanger Institute COVID-19 Surveillance Team |
| EPI_ISL_3468903 | Lighthouse Laboratory Plymouth | Wellcome Sanger Institute for the COVID-19 Genomics UK (COG-UK) Consortium | Lighthouse Laboratory Plymouth and Alex Alderton, Roberto Amato, Jeffrey Barrett, Sonia Goncalves, Ewan Harrison, David K. Jackson, Ian Johnston, Dominic Kwiatkowski, Cordelia Langford, John Sillitoe on behalf of the Wellcome Sanger Institute COVID-19 Surveillance Team |
| EPI_ISL_3468904 | Lighthouse Lab in Alderley Park | Wellcome Sanger Institute for the COVID-19 Genomics UK (COG-UK) Consortium | Jacquelyn Wynn, Mairead Hyland, The Lighthouse Lab in Alderley Park and Alex Alderton, Roberto Amato, Jeffrey Barrett, Sonia Goncalves, Ewan Harrison, David K. Jackson, Ian Johnston, Dominic Kwiatkowski, Cordelia Langford, John Sillitoe on behalf of the Wellcome Sanger Institute COVID-19 Surveillance Team |
| EPI_ISL_3468905 | Lighthouse Laboratory Plymouth | Wellcome Sanger Institute for the COVID-19 Genomics UK (COG-UK) Consortium | Lighthouse Laboratory Plymouth and Alex Alderton, Roberto Amato, Jeffrey Barrett, Sonia Goncalves, Ewan Harrison, David K. Jackson, Ian Johnston, Dominic Kwiatkowski, Cordelia Langford, John Sillitoe on behalf of the Wellcome Sanger Institute COVID-19 Surveillance Team |
| EPI_ISL_3468906 | Lighthouse Lab in Milton Keynes | Wellcome Sanger Institute for the COVID-19 Genomics UK (COG-UK) Consortium | The Lighthouse Lab in Milton Keynes and Alex Alderton, Roberto Amato, Jeffrey Barrett, Sonia Goncalves, Ewan Harrison, David K. Jackson, Ian Johnston, Dominic Kwiatkowski, Cordelia Langford, John Sillitoe on behalf of the Wellcome Sanger Institute COVID-19 Surveillance Team |
| EPI_ISL_3468907 | Lighthouse Laboratory Plymouth | Wellcome Sanger Institute for the COVID-19 Genomics UK (COG-UK) Consortium | Lighthouse Laboratory Plymouth and Alex Alderton, Roberto Amato, Jeffrey Barrett, Sonia Goncalves, Ewan Harrison, David K. Jackson, Ian Johnston, Dominic Kwiatkowski, Cordelia Langford, John Sillitoe on behalf of the Wellcome Sanger Institute COVID-19 Surveillance Team |
| EPI_ISL_3468908, EPI_ISL_3468909 | Lighthouse Lab in Milton Keynes | Wellcome Sanger Institute for the COVID-19 Genomics UK (COG-UK) Consortium | The Lighthouse Lab in Milton Keynes and Alex Alderton, Roberto Amato, Jeffrey Barrett, Sonia Goncalves, Ewan Harrison, David K. Jackson, Ian Johnston, Dominic Kwiatkowski, Cordelia Langford, John Sillitoe on behalf of the Wellcome Sanger Institute COVID-19 Surveillance Team |
| EPI_ISL_3468910 | Lighthouse Lab in Alderley Park | Wellcome Sanger Institute for the COVID-19 Genomics UK (COG-UK) Consortium | Jacquelyn Wynn, Mairead Hyland, The Lighthouse Lab in Alderley Park and Alex Alderton, Roberto Amato, Jeffrey Barrett, Sonia Goncalves, Ewan Harrison, David K. Jackson, Ian Johnston, Dominic Kwiatkowski, Cordelia Langford, John Sillitoe on behalf of the Wellcome Sanger Institute COVID-19 Surveillance Team |
| EPI_ISL_3468911 | Lighthouse Lab in Milton Keynes | Wellcome Sanger Institute for the COVID-19 Genomics UK (COG-UK) Consortium | The Lighthouse Lab in Milton Keynes and Alex Alderton, Roberto Amato, Jeffrey Barrett, Sonia Goncalves, Ewan Harrison, David K. Jackson, Ian Johnston, Dominic Kwiatkowski, Cordelia Langford, John Sillitoe on behalf of the Wellcome Sanger Institute COVID-19 Surveillance Team |
| EPI_ISL_3468912, EPI_ISL_3468913 | Lighthouse Laboratory Plymouth | Wellcome Sanger Institute for the COVID-19 Genomics UK (COG-UK) Consortium | Lighthouse Laboratory Plymouth and Alex Alderton, Roberto Amato, Jeffrey Barrett, Sonia Goncalves, Ewan Harrison, David K. Jackson, Ian Johnston, Dominic Kwiatkowski, Cordelia Langford, John Sillitoe on behalf of the Wellcome Sanger Institute COVID-19 Surveillance Team |
| EPI_ISL_3468914 | Lighthouse Lab in Milton Keynes | Wellcome Sanger Institute for the COVID-19 Genomics UK (COG-UK) Consortium | The Lighthouse Lab in Milton Keynes and Alex Alderton, Roberto Amato, Jeffrey Barrett, Sonia Goncalves, Ewan Harrison, David K. Jackson, Ian Johnston, Dominic Kwiatkowski, Cordelia Langford, John Sillitoe on behalf of the Wellcome Sanger Institute COVID-19 Surveillance Team |
| EPI_ISL_3468915 | Lighthouse Laboratory Plymouth | Wellcome Sanger Institute for the COVID-19 Genomics UK (COG-UK) Consortium | Lighthouse Laboratory Plymouth and Alex Alderton, Roberto Amato, Jeffrey Barrett, Sonia Goncalves, Ewan Harrison, David K. Jackson, Ian Johnston, Dominic Kwiatkowski, Cordelia Langford, John Sillitoe on behalf of the Wellcome Sanger Institute COVID-19 Surveillance Team |
| EPI_ISL_3468916, EPI_ISL_3468917 | Lighthouse Lab in Milton Keynes | Wellcome Sanger Institute for the COVID-19 Genomics UK (COG-UK) Consortium | The Lighthouse Lab in Milton Keynes and Alex Alderton, Roberto Amato, Jeffrey Barrett, Sonia Goncalves, Ewan Harrison, David K. Jackson, Ian Johnston, Dominic Kwiatkowski, Cordelia Langford, John Sillitoe on behalf of the Wellcome Sanger Institute COVID-19 Surveillance Team |
| EPI_ISL_3468918 | Lighthouse Lab in Alderley Park | Wellcome Sanger Institute for the COVID-19 Genomics UK (COG-UK) Consortium | Jacquelyn Wynn, Mairead Hyland, The Lighthouse Lab in Alderley Park and Alex Alderton, Roberto Amato, Jeffrey Barrett, Sonia Goncalves, Ewan Harrison, David K. Jackson, Ian Johnston, Dominic Kwiatkowski, Cordelia Langford, John Sillitoe on behalf of the Wellcome Sanger Institute COVID-19 Surveillance Team |
| EPI_ISL_3468919 | Lighthouse Laboratory Plymouth | Wellcome Sanger Institute for the COVID-19 Genomics UK (COG-UK) Consortium | Lighthouse Laboratory Plymouth and Alex Alderton, Roberto Amato, Jeffrey Barrett, Sonia Goncalves, Ewan Harrison, David K. Jackson, Ian Johnston, Dominic Kwiatkowski, Cordelia Langford, John Sillitoe on behalf of the Wellcome Sanger Institute COVID-19 Surveillance Team |
| EPI_ISL_3468920, EPI_ISL_3468921 | Lighthouse Lab in Alderley Park | Wellcome Sanger Institute for the COVID-19 Genomics UK (COG-UK) Consortium | Jacquelyn Wynn, Mairead Hyland, The Lighthouse Lab in Alderley Park and Alex Alderton, Roberto Amato, Jeffrey Barrett, Sonia Goncalves, Ewan Harrison, David K. Jackson, Ian Johnston, Dominic Kwiatkowski, Cordelia Langford, John Sillitoe on behalf of the Wellcome Sanger Institute COVID-19 Surveillance Team |
| EPI_ISL_3468922, EPI_ISL_3468923 | Lighthouse Lab in Milton Keynes | Wellcome Sanger Institute for the COVID-19 Genomics UK (COG-UK) Consortium | The Lighthouse Lab in Milton Keynes and Alex Alderton, Roberto Amato, Jeffrey Barrett, Sonia Goncalves, Ewan Harrison, David K. Jackson, Ian Johnston, Dominic Kwiatkowski, Cordelia Langford, John Sillitoe on behalf of the Wellcome Sanger Institute COVID-19 Surveillance Team |
| EPI_ISL_3468924 | Lighthouse Laboratory Plymouth | Wellcome Sanger Institute for the COVID-19 Genomics UK (COG-UK) Consortium | Lighthouse Laboratory Plymouth and Alex Alderton, Roberto Amato, Jeffrey Barrett, Sonia Goncalves, Ewan Harrison, David K. Jackson, Ian Johnston, Dominic Kwiatkowski, Cordelia Langford, John Sillitoe on behalf of the Wellcome Sanger Institute COVID-19 Surveillance Team |
| EPI_ISL_3468925, EPI_ISL_3468926 | Lighthouse Lab in Milton Keynes | Wellcome Sanger Institute for the COVID-19 Genomics UK (COG-UK) Consortium | The Lighthouse Lab in Milton Keynes and Alex Alderton, Roberto Amato, Jeffrey Barrett, Sonia Goncalves, Ewan Harrison, David K. Jackson, Ian Johnston, Dominic Kwiatkowski, Cordelia Langford, John Sillitoe on behalf of the Wellcome Sanger Institute COVID-19 Surveillance Team |
| EPI_ISL_3468927 | Lighthouse Lab in Alderley Park | Wellcome Sanger Institute for the COVID-19 Genomics UK (COG-UK) Consortium | Jacquelyn Wynn, Mairead Hyland, The Lighthouse Lab in Alderley Park and Alex Alderton, Roberto Amato, Jeffrey Barrett, Sonia Goncalves, Ewan Harrison, David K. Jackson, Ian Johnston, Dominic Kwiatkowski, Cordelia Langford, John Sillitoe on behalf of the Wellcome Sanger Institute COVID-19 Surveillance Team |
| EPI_ISL_3468928 | Lighthouse Lab in Milton Keynes | Wellcome Sanger Institute for the COVID-19 Genomics UK (COG-UK) Consortium | The Lighthouse Lab in Milton Keynes and Alex Alderton, Roberto Amato, Jeffrey Barrett, Sonia Goncalves, Ewan Harrison, David K. Jackson, Ian Johnston, Dominic Kwiatkowski, Cordelia Langford, John Sillitoe on behalf of the Wellcome Sanger Institute COVID-19 Surveillance Team |
| EPI_ISL_3468929 | Lighthouse Lab in Alderley Park | Wellcome Sanger Institute for the COVID-19 Genomics UK (COG-UK) Consortium | Jacquelyn Wynn, Mairead Hyland, The Lighthouse Lab in Alderley Park and Alex Alderton, Roberto Amato, Jeffrey Barrett, Sonia Goncalves, Ewan Harrison, David K. Jackson, Ian Johnston, Dominic Kwiatkowski, Cordelia Langford, John Sillitoe on behalf of the Wellcome Sanger Institute COVID-19 Surveillance Team |
| EPI_ISL_3468930 | Lighthouse Lab in Milton Keynes | Wellcome Sanger Institute for the COVID-19 Genomics UK (COG-UK) Consortium | The Lighthouse Lab in Milton Keynes and Alex Alderton, Roberto Amato, Jeffrey Barrett, Sonia Goncalves, Ewan Harrison, David K. Jackson, Ian Johnston, Dominic Kwiatkowski, Cordelia Langford, John Sillitoe on behalf of the Wellcome Sanger Institute COVID-19 Surveillance Team |
| EPI_ISL_3468931 | Lighthouse Laboratory Plymouth | Wellcome Sanger Institute for the COVID-19 Genomics UK (COG-UK) Consortium | Lighthouse Laboratory Plymouth and Alex Alderton, Roberto Amato, Jeffrey Barrett, Sonia Goncalves, Ewan Harrison, David K. Jackson, Ian Johnston, Dominic Kwiatkowski, Cordelia Langford, John Sillitoe on behalf of the Wellcome Sanger Institute COVID-19 Surveillance Team |
| EPI_ISL_3468932, EPI_ISL_3468933 | Lighthouse Lab in Alderley Park | Wellcome Sanger Institute for the COVID-19 Genomics UK (COG-UK) Consortium | Jacquelyn Wynn, Mairead Hyland, The Lighthouse Lab in Alderley Park and Alex Alderton, Roberto Amato, Jeffrey Barrett, Sonia Goncalves, Ewan Harrison, David K. Jackson, Ian Johnston, Dominic Kwiatkowski, Cordelia Langford, John Sillitoe on behalf of the Wellcome Sanger Institute COVID-19 Surveillance Team |
| EPI_ISL_3468934 | Lighthouse Lab in Milton Keynes | Wellcome Sanger Institute for the COVID-19 Genomics UK (COG-UK) Consortium | The Lighthouse Lab in Milton Keynes and Alex Alderton, Roberto Amato, Jeffrey Barrett, Sonia Goncalves, Ewan Harrison, David K. Jackson, Ian Johnston, Dominic Kwiatkowski, Cordelia Langford, John Sillitoe on behalf of the Wellcome Sanger Institute COVID-19 Surveillance Team |
| EPI_ISL_3468935 | Lighthouse Laboratory Plymouth | Wellcome Sanger Institute for the COVID-19 Genomics UK (COG-UK) Consortium | Lighthouse Laboratory Plymouth and Alex Alderton, Roberto Amato, Jeffrey Barrett, Sonia Goncalves, Ewan Harrison, David K. Jackson, Ian Johnston, Dominic Kwiatkowski, Cordelia Langford, John Sillitoe on behalf of the Wellcome Sanger Institute COVID-19 Surveillance Team |

| | | | |
|---|---|---|---|
| EPI_ISL_3468936 | Lighthouse Lab in Milton Keynes | Wellcome Sanger Institute for the COVID-19 Genomics UK (COG-UK) Consortium | The Lighthouse Lab in Milton Keynes and Alex Alderton, Roberto Amato, Jeffrey Barrett, Sonia Goncalves, Ewan Harrison, David K. Jackson, Ian Johnston, Dominic Kwiatkowski, Cordelia Langford, John Sillitoe on behalf of the Wellcome Sanger Institute COVID-19 Surveillance Team |
| EPI_ISL_3468937 | Lighthouse Lab in Alderley Park | Wellcome Sanger Institute for the COVID-19 Genomics UK (COG-UK) Consortium | Jacquelyn Wynn, Mairead Hyland, The Lighthouse Lab in Alderley Park and Alex Alderton, Roberto Amato, Jeffrey Barrett, Sonia Goncalves, Ewan Harrison, David K. Jackson, Ian Johnston, Dominic Kwiatkowski, Cordelia Langford, John Sillitoe on behalf of the Wellcome Sanger Institute COVID-19 Surveillance Team |
| EPI_ISL_3468938 | Lighthouse Lab in Milton Keynes | Wellcome Sanger Institute for the COVID-19 Genomics UK (COG-UK) Consortium | The Lighthouse Lab in Milton Keynes and Alex Alderton, Roberto Amato, Jeffrey Barrett, Sonia Goncalves, Ewan Harrison, David K. Jackson, Ian Johnston, Dominic Kwiatkowski, Cordelia Langford, John Sillitoe on behalf of the Wellcome Sanger Institute COVID-19 Surveillance Team |
| EPI_ISL_3468939 | Health Services Laboratories | Wellcome Sanger Institute for the COVID-19 Genomics UK (COG-UK) Consortium | Health Services Laboratories and Alex Alderton, Roberto Amato, Jeffrey Barrett, Sonia Goncalves, Ewan Harrison, David K. Jackson, Ian Johnston, Dominic Kwiatkowski, Cordelia Langford, John Sillitoe on behalf of the Wellcome Sanger Institute COVID-19 Surveillance Team |
| EPI_ISL_3468940 | Lighthouse Lab in Alderley Park | Wellcome Sanger Institute for the COVID-19 Genomics UK (COG-UK) Consortium | Jacquelyn Wynn, Mairead Hyland, The Lighthouse Lab in Alderley Park and Alex Alderton, Roberto Amato, Jeffrey Barrett, Sonia Goncalves, Ewan Harrison, David K. Jackson, Ian Johnston, Dominic Kwiatkowski, Cordelia Langford, John Sillitoe on behalf of the Wellcome Sanger Institute COVID-19 Surveillance Team |
| EPI_ISL_3468941 | Health Services Laboratories | Wellcome Sanger Institute for the COVID-19 Genomics UK (COG-UK) Consortium | Health Services Laboratories and Alex Alderton, Roberto Amato, Jeffrey Barrett, Sonia Goncalves, Ewan Harrison, David K. Jackson, Ian Johnston, Dominic Kwiatkowski, Cordelia Langford, John Sillitoe on behalf of the Wellcome Sanger Institute COVID-19 Surveillance Team |
| EPI_ISL_3468942 | Lighthouse Lab in Milton Keynes | Wellcome Sanger Institute for the COVID-19 Genomics UK (COG-UK) Consortium | The Lighthouse Lab in Milton Keynes and Alex Alderton, Roberto Amato, Jeffrey Barrett, Sonia Goncalves, Ewan Harrison, David K. Jackson, Ian Johnston, Dominic Kwiatkowski, Cordelia Langford, John Sillitoe on behalf of the Wellcome Sanger Institute COVID-19 Surveillance Team |
| EPI_ISL_3468943 | Lighthouse Lab in Alderley Park | Wellcome Sanger Institute for the COVID-19 Genomics UK (COG-UK) Consortium | Jacquelyn Wynn, Mairead Hyland, The Lighthouse Lab in Alderley Park and Alex Alderton, Roberto Amato, Jeffrey Barrett, Sonia Goncalves, Ewan Harrison, David K. Jackson, Ian Johnston, Dominic Kwiatkowski, Cordelia Langford, John Sillitoe on behalf of the Wellcome Sanger Institute COVID-19 Surveillance Team |
| EPI_ISL_3468944 | Lighthouse Lab in Milton Keynes | Wellcome Sanger Institute for the COVID-19 Genomics UK (COG-UK) Consortium | The Lighthouse Lab in Milton Keynes and Alex Alderton, Roberto Amato, Jeffrey Barrett, Sonia Goncalves, Ewan Harrison, David K. Jackson, Ian Johnston, Dominic Kwiatkowski, Cordelia Langford, John Sillitoe on behalf of the Wellcome Sanger Institute COVID-19 Surveillance Team |
| EPI_ISL_3468945 | Lighthouse Lab in Alderley Park | Wellcome Sanger Institute for the COVID-19 Genomics UK (COG-UK) Consortium | Jacquelyn Wynn, Mairead Hyland, The Lighthouse Lab in Alderley Park and Alex Alderton, Roberto Amato, Jeffrey Barrett, Sonia Goncalves, Ewan Harrison, David K. Jackson, Ian Johnston, Dominic Kwiatkowski, Cordelia Langford, John Sillitoe on behalf of the Wellcome Sanger Institute COVID-19 Surveillance Team |
| EPI_ISL_3468946, EPI_ISL_3468947 | Lighthouse Lab in Milton Keynes | Wellcome Sanger Institute for the COVID-19 Genomics UK (COG-UK) Consortium | The Lighthouse Lab in Milton Keynes and Alex Alderton, Roberto Amato, Jeffrey Barrett, Sonia Goncalves, Ewan Harrison, David K. Jackson, Ian Johnston, Dominic Kwiatkowski, Cordelia Langford, John Sillitoe on behalf of the Wellcome Sanger Institute COVID-19 Surveillance Team |
| EPI_ISL_3468948, EPI_ISL_3468949, EPI_ISL_3468950, EPI_ISL_3468951 | Lighthouse Lab in Alderley Park | Wellcome Sanger Institute for the COVID-19 Genomics UK (COG-UK) Consortium | Jacquelyn Wynn, Mairead Hyland, The Lighthouse Lab in Alderley Park and Alex Alderton, Roberto Amato, Jeffrey Barrett, Sonia Goncalves, Ewan Harrison, David K. Jackson, Ian Johnston, Dominic Kwiatkowski, Cordelia Langford, John Sillitoe on behalf of the Wellcome Sanger Institute COVID-19 Surveillance Team |
| EPI_ISL_3468952 | Lighthouse Lab in Milton Keynes | Wellcome Sanger Institute for the COVID-19 Genomics UK (COG-UK) Consortium | The Lighthouse Lab in Milton Keynes and Alex Alderton, Roberto Amato, Jeffrey Barrett, Sonia Goncalves, Ewan Harrison, David K. Jackson, Ian Johnston, Dominic Kwiatkowski, Cordelia Langford, John Sillitoe on behalf of the Wellcome Sanger Institute COVID-19 Surveillance Team |
| EPI_ISL_3468953 | Health Services Laboratories | Wellcome Sanger Institute for the COVID-19 Genomics UK (COG-UK) Consortium | Health Services Laboratories and Alex Alderton, Roberto Amato, Jeffrey Barrett, Sonia Goncalves, Ewan Harrison, David K. Jackson, Ian Johnston, Dominic Kwiatkowski, Cordelia Langford, John Sillitoe on behalf of the Wellcome Sanger Institute COVID-19 Surveillance Team |
| EPI_ISL_3468954 | Lighthouse Lab in Alderley Park | Wellcome Sanger Institute for the COVID-19 Genomics UK (COG-UK) Consortium | Jacquelyn Wynn, Mairead Hyland, The Lighthouse Lab in Alderley Park and Alex Alderton, Roberto Amato, Jeffrey Barrett, Sonia Goncalves, Ewan Harrison, David K. Jackson, Ian Johnston, Dominic Kwiatkowski, Cordelia Langford, John Sillitoe on behalf of the Wellcome Sanger Institute COVID-19 Surveillance Team |
| EPI_ISL_3468955 | Health Services Laboratories | Wellcome Sanger Institute for the COVID-19 Genomics UK (COG-UK) Consortium | Health Services Laboratories and Alex Alderton, Roberto Amato, Jeffrey Barrett, Sonia Goncalves, Ewan Harrison, David K. Jackson, Ian Johnston, Dominic Kwiatkowski, Cordelia Langford, John Sillitoe on behalf of the Wellcome Sanger Institute COVID-19 Surveillance Team |
| EPI_ISL_3468956 | Lighthouse Lab in Alderley Park | Wellcome Sanger Institute for the COVID-19 Genomics UK (COG-UK) Consortium | Jacquelyn Wynn, Mairead Hyland, The Lighthouse Lab in Alderley Park and Alex Alderton, Roberto Amato, Jeffrey Barrett, Sonia Goncalves, Ewan Harrison, David K. Jackson, Ian Johnston, Dominic Kwiatkowski, Cordelia Langford, John Sillitoe on behalf of the Wellcome Sanger Institute COVID-19 Surveillance Team |
| EPI_ISL_3468957 | Health Services Laboratories | Wellcome Sanger Institute for the COVID-19 Genomics UK (COG-UK) Consortium | Health Services Laboratories and Alex Alderton, Roberto Amato, Jeffrey Barrett, Sonia Goncalves, Ewan Harrison, David K. Jackson, Ian Johnston, Dominic Kwiatkowski, Cordelia Langford, John Sillitoe on behalf of the Wellcome Sanger Institute COVID-19 Surveillance Team |
| EPI_ISL_3468958 | Lighthouse Lab in Milton Keynes | Wellcome Sanger Institute for the COVID-19 Genomics UK (COG-UK) Consortium | The Lighthouse Lab in Milton Keynes and Alex Alderton, Roberto Amato, Jeffrey Barrett, Sonia Goncalves, Ewan Harrison, David K. Jackson, Ian Johnston, Dominic Kwiatkowski, Cordelia Langford, John Sillitoe on behalf of the Wellcome Sanger Institute COVID-19 Surveillance Team |
| EPI_ISL_3468959, EPI_ISL_3468960 | Lighthouse Lab in Alderley Park | Wellcome Sanger Institute for the COVID-19 Genomics UK (COG-UK) Consortium | Jacquelyn Wynn, Mairead Hyland, The Lighthouse Lab in Alderley Park and Alex Alderton, Roberto Amato, Jeffrey Barrett, Sonia Goncalves, Ewan Harrison, David K. Jackson, Ian Johnston, Dominic Kwiatkowski, Cordelia Langford, John Sillitoe on behalf of the Wellcome Sanger Institute COVID-19 Surveillance Team |
| EPI_ISL_3468961 | Lighthouse Lab in Milton Keynes | Wellcome Sanger Institute for the COVID-19 Genomics UK (COG-UK) Consortium | The Lighthouse Lab in Milton Keynes and Alex Alderton, Roberto Amato, Jeffrey Barrett, Sonia Goncalves, Ewan Harrison, David K. Jackson, Ian Johnston, Dominic Kwiatkowski, Cordelia Langford, John Sillitoe on behalf of the Wellcome Sanger Institute COVID-19 Surveillance Team |
| EPI_ISL_3468962 | Lighthouse Lab in Alderley Park | Wellcome Sanger Institute for the COVID-19 Genomics UK (COG-UK) Consortium | Jacquelyn Wynn, Mairead Hyland, The Lighthouse Lab in Alderley Park and Alex Alderton, Roberto Amato, Jeffrey Barrett, Sonia Goncalves, Ewan Harrison, David K. Jackson, Ian Johnston, Dominic Kwiatkowski, Cordelia Langford, John Sillitoe on behalf of the Wellcome Sanger Institute COVID-19 Surveillance Team |
| EPI_ISL_3468963 | Health Services Laboratories | Wellcome Sanger Institute for the COVID-19 Genomics UK (COG-UK) Consortium | Health Services Laboratories and Alex Alderton, Roberto Amato, Jeffrey Barrett, Sonia Goncalves, Ewan Harrison, David K. Jackson, Ian Johnston, Dominic Kwiatkowski, Cordelia Langford, John Sillitoe on behalf of the Wellcome Sanger Institute COVID-19 Surveillance Team |
| EPI_ISL_3468964 | Lighthouse Lab in Alderley Park | Wellcome Sanger Institute for the COVID-19 Genomics UK (COG-UK) Consortium | Jacquelyn Wynn, Mairead Hyland, The Lighthouse Lab in Alderley Park and Alex Alderton, Roberto Amato, Jeffrey Barrett, Sonia Goncalves, Ewan Harrison, David K. Jackson, Ian Johnston, Dominic Kwiatkowski, Cordelia Langford, John Sillitoe on behalf of the Wellcome Sanger Institute COVID-19 Surveillance Team |
| EPI_ISL_3468965, EPI_ISL_3468966 | Health Services Laboratories | Wellcome Sanger Institute for the COVID-19 Genomics UK (COG-UK) Consortium | Health Services Laboratories and Alex Alderton, Roberto Amato, Jeffrey Barrett, Sonia Goncalves, Ewan Harrison, David K. Jackson, Ian Johnston, Dominic Kwiatkowski, Cordelia Langford, John Sillitoe on behalf of the Wellcome Sanger Institute COVID-19 Surveillance Team |
| EPI_ISL_3468967, EPI_ISL_3468968 | Lighthouse Lab in Milton Keynes | Wellcome Sanger Institute for the COVID-19 Genomics UK (COG-UK) Consortium | The Lighthouse Lab in Milton Keynes and Alex Alderton, Roberto Amato, Jeffrey Barrett, Sonia Goncalves, Ewan Harrison, David K. Jackson, Ian Johnston, Dominic Kwiatkowski, Cordelia Langford, John Sillitoe on behalf of the Wellcome Sanger Institute COVID-19 Surveillance Team |
| EPI_ISL_3468969 | Lighthouse Lab in Alderley Park | Wellcome Sanger Institute for the COVID-19 Genomics UK (COG-UK) Consortium | Jacquelyn Wynn, Mairead Hyland, The Lighthouse Lab in Alderley Park and Alex Alderton, Roberto Amato, Jeffrey Barrett, Sonia Goncalves, Ewan Harrison, David K. Jackson, Ian Johnston, Dominic Kwiatkowski, Cordelia Langford, John Sillitoe on behalf of the Wellcome Sanger Institute COVID-19 Surveillance Team |
| EPI_ISL_3468970 | Lighthouse Lab in Milton Keynes | Wellcome Sanger Institute for the COVID-19 Genomics UK (COG-UK) Consortium | The Lighthouse Lab in Milton Keynes and Alex Alderton, Roberto Amato, Jeffrey Barrett, Sonia Goncalves, Ewan Harrison, David K. Jackson, Ian Johnston, Dominic Kwiatkowski, Cordelia Langford, John Sillitoe on behalf of the Wellcome Sanger Institute COVID-19 Surveillance Team |
| EPI_ISL_3468971, EPI_ISL_3468972, EPI_ISL_3468973 | Lighthouse Lab in Alderley Park | Wellcome Sanger Institute for the COVID-19 Genomics UK (COG-UK) Consortium | Jacquelyn Wynn, Mairead Hyland, The Lighthouse Lab in Alderley Park and Alex Alderton, Roberto Amato, Jeffrey Barrett, Sonia Goncalves, Ewan Harrison, David K. Jackson, Ian Johnston, Dominic Kwiatkowski, Cordelia Langford, John Sillitoe on behalf of the Wellcome Sanger Institute COVID-19 Surveillance Team |
| EPI_ISL_3468974 | Health Services Laboratories | Wellcome Sanger Institute for the COVID-19 Genomics UK (COG-UK) Consortium | Health Services Laboratories and Alex Alderton, Roberto Amato, Jeffrey Barrett, Sonia Goncalves, Ewan Harrison, David K. Jackson, Ian Johnston, Dominic Kwiatkowski, Cordelia Langford, John Sillitoe on behalf of the Wellcome Sanger Institute COVID-19 Surveillance Team |
| EPI_ISL_3468975, EPI_ISL_3468976 | Lighthouse Lab in Alderley Park | Wellcome Sanger Institute for the COVID-19 Genomics UK (COG-UK) Consortium | Jacquelyn Wynn, Mairead Hyland, The Lighthouse Lab in Alderley Park and Alex Alderton, Roberto Amato, Jeffrey Barrett, Sonia Goncalves, Ewan Harrison, David K. Jackson, Ian Johnston, Dominic Kwiatkowski, Cordelia Langford, John Sillitoe on behalf of the Wellcome Sanger Institute COVID-19 Surveillance Team |
| EPI_ISL_3468977 | Lighthouse Lab in Milton Keynes | Wellcome Sanger Institute for the COVID-19 Genomics UK (COG-UK) Consortium | The Lighthouse Lab in Milton Keynes and Alex Alderton, Roberto Amato, Jeffrey Barrett, Sonia Goncalves, Ewan Harrison, David K. Jackson, Ian Johnston, Dominic Kwiatkowski, Cordelia Langford, John Sillitoe on behalf of the Wellcome Sanger Institute COVID-19 Surveillance Team |
| EPI_ISL_3468978, EPI_ISL_3468979 | Lighthouse Lab in Alderley Park | Wellcome Sanger Institute for the COVID-19 Genomics UK (COG-UK) Consortium | Jacquelyn Wynn, Mairead Hyland, The Lighthouse Lab in Alderley Park and Alex Alderton, Roberto Amato, Jeffrey Barrett, Sonia Goncalves, Ewan Harrison, David K. Jackson, Ian Johnston, Dominic Kwiatkowski, Cordelia Langford, John Sillitoe on behalf of the Wellcome Sanger Institute COVID-19 Surveillance Team |
| EPI_ISL_3468980 | Health Services Laboratories | Wellcome Sanger Institute for the COVID-19 Genomics UK (COG-UK) Consortium | Health Services Laboratories and Alex Alderton, Roberto Amato, Jeffrey Barrett, Sonia Goncalves, Ewan Harrison, David K. Jackson, Ian Johnston, Dominic Kwiatkowski, Cordelia Langford, John Sillitoe on behalf of the Wellcome Sanger Institute COVID-19 Surveillance Team |
| EPI_ISL_3468981, EPI_ISL_3468982 | Lighthouse Lab in Alderley Park | Wellcome Sanger Institute for the COVID-19 Genomics UK (COG-UK) Consortium | Jacquelyn Wynn, Mairead Hyland, The Lighthouse Lab in Alderley Park and Alex Alderton, Roberto Amato, Jeffrey Barrett, Sonia Goncalves, Ewan Harrison, David K. Jackson, Ian Johnston, Dominic Kwiatkowski, Cordelia Langford, John Sillitoe on behalf of the Wellcome Sanger Institute COVID-19 Surveillance Team |
| EPI_ISL_3468983 | Health Services Laboratories | Wellcome Sanger Institute for the COVID-19 Genomics UK (COG-UK) Consortium | Health Services Laboratories and Alex Alderton, Roberto Amato, Jeffrey Barrett, Sonia Goncalves, Ewan Harrison, David K. Jackson, Ian Johnston, Dominic Kwiatkowski, Cordelia Langford, John Sillitoe on behalf of the Wellcome Sanger Institute COVID-19 Surveillance Team |
| EPI_ISL_3468984, EPI_ISL_3468985 | Lighthouse Lab in Milton Keynes | Wellcome Sanger Institute for the COVID-19 Genomics UK (COG-UK) Consortium | The Lighthouse Lab in Milton Keynes and Alex Alderton, Roberto Amato, Jeffrey Barrett, Sonia Goncalves, Ewan Harrison, David K. Jackson, Ian Johnston, Dominic Kwiatkowski, Cordelia Langford, John Sillitoe on behalf of the Wellcome Sanger Institute COVID-19 Surveillance Team |
| EPI_ISL_3468986 | Lighthouse Lab in Alderley Park | Wellcome Sanger Institute for the COVID-19 Genomics UK (COG-UK) Consortium | Jacquelyn Wynn, Mairead Hyland, The Lighthouse Lab in Alderley Park and Alex Alderton, Roberto Amato, Jeffrey Barrett, Sonia Goncalves, Ewan Harrison, David K. Jackson, Ian Johnston, Dominic Kwiatkowski, Cordelia Langford, John Sillitoe on behalf of the Wellcome Sanger Institute COVID-19 Surveillance Team |
| EPI_ISL_3468987 | Lighthouse Lab in Milton Keynes | Wellcome Sanger Institute for the COVID-19 Genomics UK (COG-UK) Consortium | The Lighthouse Lab in Milton Keynes and Alex Alderton, Roberto Amato, Jeffrey Barrett, Sonia Goncalves, Ewan Harrison, David K. Jackson, Ian Johnston, Dominic Kwiatkowski, Cordelia Langford, John Sillitoe on behalf of the Wellcome Sanger Institute COVID-19 Surveillance Team |
| EPI_ISL_3468988, EPI_ISL_3468989 | Lighthouse Lab in Alderley Park | Wellcome Sanger Institute for the COVID-19 Genomics UK (COG-UK) Consortium | Jacquelyn Wynn, Mairead Hyland, The Lighthouse Lab in Alderley Park and Alex Alderton, Roberto Amato, Jeffrey Barrett, Sonia Goncalves, Ewan Harrison, David K. Jackson, Ian Johnston, Dominic Kwiatkowski, Cordelia Langford, John Sillitoe on behalf of the Wellcome Sanger Institute COVID-19 Surveillance Team |
| EPI_ISL_3468990 | Health Services Laboratories | Wellcome Sanger Institute for the COVID-19 Genomics UK (COG-UK) Consortium | Health Services Laboratories and Alex Alderton, Roberto Amato, Jeffrey Barrett, Sonia Goncalves, Ewan Harrison, David K. Jackson, Ian Johnston, Dominic Kwiatkowski, Cordelia Langford, John Sillitoe on behalf of the Wellcome Sanger Institute COVID-19 Surveillance Team |
| EPI_ISL_3468991 | Lighthouse Lab in Alderley Park | Wellcome Sanger Institute for the COVID-19 Genomics UK (COG-UK) Consortium | Jacquelyn Wynn, Mairead Hyland, The Lighthouse Lab in Alderley Park and Alex Alderton, Roberto Amato, Jeffrey Barrett, Sonia Goncalves, Ewan Harrison, David K. Jackson, Ian Johnston, Dominic Kwiatkowski, Cordelia Langford, John Sillitoe on behalf of the Wellcome Sanger Institute COVID-19 Surveillance Team |
| EPI_ISL_3468992 | Lighthouse Lab in Milton Keynes | Wellcome Sanger Institute for the COVID-19 Genomics UK (COG-UK) Consortium | The Lighthouse Lab in Milton Keynes and Alex Alderton, Roberto Amato, Jeffrey Barrett, Sonia Goncalves, Ewan Harrison, David K. Jackson, Ian Johnston, Dominic Kwiatkowski, Cordelia Langford, John Sillitoe on behalf of the Wellcome Sanger Institute COVID-19 Surveillance Team |
| EPI_ISL_3468993 | Lighthouse Lab in Alderley Park | Wellcome Sanger Institute for the COVID-19 Genomics UK (COG-UK) Consortium | Jacquelyn Wynn, Mairead Hyland, The Lighthouse Lab in Alderley Park and Alex Alderton, Roberto Amato, Jeffrey Barrett, Sonia Goncalves, Ewan Harrison, David K. Jackson, Ian Johnston, Dominic Kwiatkowski, Cordelia Langford, John Sillitoe on behalf of the Wellcome Sanger Institute COVID-19 Surveillance Team |

| | | | |
|---|---|---|---|
| EPI_ISL_3468994, EPI_ISL_3468995, EPI_ISL_3468996 | Lighthouse Lab in Milton Keynes | Wellcome Sanger Institute for the COVID-19 Genomics UK (COG-UK) Consortium | The Lighthouse Lab in Milton Keynes and Alex Alderton, Roberto Amato, Jeffrey Barrett, Sonia Goncalves, Ewan Harrison, David K. Jackson, Ian Johnston, Dominic Kwiatkowski, Cordelia Langford, John Sillitoe on behalf of the Wellcome Sanger Institute COVID-19 Surveillance Team |
| EPI_ISL_3468997 | Lighthouse Lab in Alderley Park | Wellcome Sanger Institute for the COVID-19 Genomics UK (COG-UK) Consortium | Jacquelyn Wynn, Mairead Hyland, The Lighthouse Lab in Alderley Park and Alex Alderton, Roberto Amato, Jeffrey Barrett, Sonia Goncalves, Ewan Harrison, David K. Jackson, Ian Johnston, Dominic Kwiatkowski, Cordelia Langford, John Sillitoe on behalf of the Wellcome Sanger Institute COVID-19 Surveillance Team |
| EPI_ISL_3468998 | Lighthouse Lab in Milton Keynes | Wellcome Sanger Institute for the COVID-19 Genomics UK (COG-UK) Consortium | The Lighthouse Lab in Milton Keynes and Alex Alderton, Roberto Amato, Jeffrey Barrett, Sonia Goncalves, Ewan Harrison, David K. Jackson, Ian Johnston, Dominic Kwiatkowski, Cordelia Langford, John Sillitoe on behalf of the Wellcome Sanger Institute COVID-19 Surveillance Team |
| EPI_ISL_3468999, EPI_ISL_3469000, EPI_ISL_3469001 | Lighthouse Lab in Alderley Park | Wellcome Sanger Institute for the COVID-19 Genomics UK (COG-UK) Consortium | Jacquelyn Wynn, Mairead Hyland, The Lighthouse Lab in Alderley Park and Alex Alderton, Roberto Amato, Jeffrey Barrett, Sonia Goncalves, Ewan Harrison, David K. Jackson, Ian Johnston, Dominic Kwiatkowski, Cordelia Langford, John Sillitoe on behalf of the Wellcome Sanger Institute COVID-19 Surveillance Team |
| EPI_ISL_3469002, EPI_ISL_3469003 | Lighthouse Lab in Milton Keynes | Wellcome Sanger Institute for the COVID-19 Genomics UK (COG-UK) Consortium | The Lighthouse Lab in Milton Keynes and Alex Alderton, Roberto Amato, Jeffrey Barrett, Sonia Goncalves, Ewan Harrison, David K. Jackson, Ian Johnston, Dominic Kwiatkowski, Cordelia Langford, John Sillitoe on behalf of the Wellcome Sanger Institute COVID-19 Surveillance Team |
| EPI_ISL_3469004 | Lighthouse Lab in Alderley Park | Wellcome Sanger Institute for the COVID-19 Genomics UK (COG-UK) Consortium | Jacquelyn Wynn, Mairead Hyland, The Lighthouse Lab in Alderley Park and Alex Alderton, Roberto Amato, Jeffrey Barrett, Sonia Goncalves, Ewan Harrison, David K. Jackson, Ian Johnston, Dominic Kwiatkowski, Cordelia Langford, John Sillitoe on behalf of the Wellcome Sanger Institute COVID-19 Surveillance Team |
| EPI_ISL_3469005 | Lighthouse Lab in Milton Keynes | Wellcome Sanger Institute for the COVID-19 Genomics UK (COG-UK) Consortium | The Lighthouse Lab in Milton Keynes and Alex Alderton, Roberto Amato, Jeffrey Barrett, Sonia Goncalves, Ewan Harrison, David K. Jackson, Ian Johnston, Dominic Kwiatkowski, Cordelia Langford, John Sillitoe on behalf of the Wellcome Sanger Institute COVID-19 Surveillance Team |
| EPI_ISL_3469006, EPI_ISL_3469007 | Health Services Laboratories | Wellcome Sanger Institute for the COVID-19 Genomics UK (COG-UK) Consortium | Health Services Laboratories and Alex Alderton, Roberto Amato, Jeffrey Barrett, Sonia Goncalves, Ewan Harrison, David K. Jackson, Ian Johnston, Dominic Kwiatkowski, Cordelia Langford, John Sillitoe on behalf of the Wellcome Sanger Institute COVID-19 Surveillance Team |
| EPI_ISL_3469008 | Lighthouse Lab in Alderley Park | Wellcome Sanger Institute for the COVID-19 Genomics UK (COG-UK) Consortium | Jacquelyn Wynn, Mairead Hyland, The Lighthouse Lab in Alderley Park and Alex Alderton, Roberto Amato, Jeffrey Barrett, Sonia Goncalves, Ewan Harrison, David K. Jackson, Ian Johnston, Dominic Kwiatkowski, Cordelia Langford, John Sillitoe on behalf of the Wellcome Sanger Institute COVID-19 Surveillance Team |
| EPI_ISL_3469009 | Health Services Laboratories | Wellcome Sanger Institute for the COVID-19 Genomics UK (COG-UK) Consortium | Health Services Laboratories and Alex Alderton, Roberto Amato, Jeffrey Barrett, Sonia Goncalves, Ewan Harrison, David K. Jackson, Ian Johnston, Dominic Kwiatkowski, Cordelia Langford, John Sillitoe on behalf of the Wellcome Sanger Institute COVID-19 Surveillance Team |
| EPI_ISL_3469010 | Lighthouse Lab in Alderley Park | Wellcome Sanger Institute for the COVID-19 Genomics UK (COG-UK) Consortium | Jacquelyn Wynn, Mairead Hyland, The Lighthouse Lab in Alderley Park and Alex Alderton, Roberto Amato, Jeffrey Barrett, Sonia Goncalves, Ewan Harrison, David K. Jackson, Ian Johnston, Dominic Kwiatkowski, Cordelia Langford, John Sillitoe on behalf of the Wellcome Sanger Institute COVID-19 Surveillance Team |
| EPI_ISL_3469011 | Lighthouse Lab in Milton Keynes | Wellcome Sanger Institute for the COVID-19 Genomics UK (COG-UK) Consortium | The Lighthouse Lab in Milton Keynes and Alex Alderton, Roberto Amato, Jeffrey Barrett, Sonia Goncalves, Ewan Harrison, David K. Jackson, Ian Johnston, Dominic Kwiatkowski, Cordelia Langford, John Sillitoe on behalf of the Wellcome Sanger Institute COVID-19 Surveillance Team |
| EPI_ISL_3469012, EPI_ISL_3469013 | Lighthouse Lab in Alderley Park | Wellcome Sanger Institute for the COVID-19 Genomics UK (COG-UK) Consortium | Jacquelyn Wynn, Mairead Hyland, The Lighthouse Lab in Alderley Park and Alex Alderton, Roberto Amato, Jeffrey Barrett, Sonia Goncalves, Ewan Harrison, David K. Jackson, Ian Johnston, Dominic Kwiatkowski, Cordelia Langford, John Sillitoe on behalf of the Wellcome Sanger Institute COVID-19 Surveillance Team |
| EPI_ISL_3469014, EPI_ISL_3469015 | Lighthouse Lab in Milton Keynes | Wellcome Sanger Institute for the COVID-19 Genomics UK (COG-UK) Consortium | The Lighthouse Lab in Milton Keynes and Alex Alderton, Roberto Amato, Jeffrey Barrett, Sonia Goncalves, Ewan Harrison, David K. Jackson, Ian Johnston, Dominic Kwiatkowski, Cordelia Langford, John Sillitoe on behalf of the Wellcome Sanger Institute COVID-19 Surveillance Team |
| EPI_ISL_3469016 | Lighthouse Lab in Alderley Park | Wellcome Sanger Institute for the COVID-19 Genomics UK (COG-UK) Consortium | Jacquelyn Wynn, Mairead Hyland, The Lighthouse Lab in Alderley Park and Alex Alderton, Roberto Amato, Jeffrey Barrett, Sonia Goncalves, Ewan Harrison, David K. Jackson, Ian Johnston, Dominic Kwiatkowski, Cordelia Langford, John Sillitoe on behalf of the Wellcome Sanger Institute COVID-19 Surveillance Team |
| EPI_ISL_3469017 | Health Services Laboratories | Wellcome Sanger Institute for the COVID-19 Genomics UK (COG-UK) Consortium | Health Services Laboratories and Alex Alderton, Roberto Amato, Jeffrey Barrett, Sonia Goncalves, Ewan Harrison, David K. Jackson, Ian Johnston, Dominic Kwiatkowski, Cordelia Langford, John Sillitoe on behalf of the Wellcome Sanger Institute COVID-19 Surveillance Team |
| EPI_ISL_3469018, EPI_ISL_3469019, EPI_ISL_3469020 | Lighthouse Lab in Alderley Park | Wellcome Sanger Institute for the COVID-19 Genomics UK (COG-UK) Consortium | Jacquelyn Wynn, Mairead Hyland, The Lighthouse Lab in Alderley Park and Alex Alderton, Roberto Amato, Jeffrey Barrett, Sonia Goncalves, Ewan Harrison, David K. Jackson, Ian Johnston, Dominic Kwiatkowski, Cordelia Langford, John Sillitoe on behalf of the Wellcome Sanger Institute COVID-19 Surveillance Team |
| EPI_ISL_3469021 | Lighthouse Lab in Milton Keynes | Wellcome Sanger Institute for the COVID-19 Genomics UK (COG-UK) Consortium | The Lighthouse Lab in Milton Keynes and Alex Alderton, Roberto Amato, Jeffrey Barrett, Sonia Goncalves, Ewan Harrison, David K. Jackson, Ian Johnston, Dominic Kwiatkowski, Cordelia Langford, John Sillitoe on behalf of the Wellcome Sanger Institute COVID-19 Surveillance Team |
| EPI_ISL_3469022 | Health Services Laboratories | Wellcome Sanger Institute for the COVID-19 Genomics UK (COG-UK) Consortium | Health Services Laboratories and Alex Alderton, Roberto Amato, Jeffrey Barrett, Sonia Goncalves, Ewan Harrison, David K. Jackson, Ian Johnston, Dominic Kwiatkowski, Cordelia Langford, John Sillitoe on behalf of the Wellcome Sanger Institute COVID-19 Surveillance Team |
| EPI_ISL_3469023 | Lighthouse Lab in Milton Keynes | Wellcome Sanger Institute for the COVID-19 Genomics UK (COG-UK) Consortium | The Lighthouse Lab in Milton Keynes and Alex Alderton, Roberto Amato, Jeffrey Barrett, Sonia Goncalves, Ewan Harrison, David K. Jackson, Ian Johnston, Dominic Kwiatkowski, Cordelia Langford, John Sillitoe on behalf of the Wellcome Sanger Institute COVID-19 Surveillance Team |
| EPI_ISL_3469024 | Lighthouse Lab in Alderley Park | Wellcome Sanger Institute for the COVID-19 Genomics UK (COG-UK) Consortium | Jacquelyn Wynn, Mairead Hyland, The Lighthouse Lab in Alderley Park and Alex Alderton, Roberto Amato, Jeffrey Barrett, Sonia Goncalves, Ewan Harrison, David K. Jackson, Ian Johnston, Dominic Kwiatkowski, Cordelia Langford, John Sillitoe on behalf of the Wellcome Sanger Institute COVID-19 Surveillance Team |
| EPI_ISL_3469025 | Health Services Laboratories | Wellcome Sanger Institute for the COVID-19 Genomics UK (COG-UK) Consortium | Health Services Laboratories and Alex Alderton, Roberto Amato, Jeffrey Barrett, Sonia Goncalves, Ewan Harrison, David K. Jackson, Ian Johnston, Dominic Kwiatkowski, Cordelia Langford, John Sillitoe on behalf of the Wellcome Sanger Institute COVID-19 Surveillance Team |
| EPI_ISL_3469026 | Lighthouse Lab in Milton Keynes | Wellcome Sanger Institute for the COVID-19 Genomics UK (COG-UK) Consortium | The Lighthouse Lab in Milton Keynes and Alex Alderton, Roberto Amato, Jeffrey Barrett, Sonia Goncalves, Ewan Harrison, David K. Jackson, Ian Johnston, Dominic Kwiatkowski, Cordelia Langford, John Sillitoe on behalf of the Wellcome Sanger Institute COVID-19 Surveillance Team |
| EPI_ISL_3469027 | Lighthouse Lab in Alderley Park | Wellcome Sanger Institute for the COVID-19 Genomics UK (COG-UK) Consortium | Jacquelyn Wynn, Mairead Hyland, The Lighthouse Lab in Alderley Park and Alex Alderton, Roberto Amato, Jeffrey Barrett, Sonia Goncalves, Ewan Harrison, David K. Jackson, Ian Johnston, Dominic Kwiatkowski, Cordelia Langford, John Sillitoe on behalf of the Wellcome Sanger Institute COVID-19 Surveillance Team |
| EPI_ISL_3469028 | Lighthouse Lab in Milton Keynes | Wellcome Sanger Institute for the COVID-19 Genomics UK (COG-UK) Consortium | The Lighthouse Lab in Milton Keynes and Alex Alderton, Roberto Amato, Jeffrey Barrett, Sonia Goncalves, Ewan Harrison, David K. Jackson, Ian Johnston, Dominic Kwiatkowski, Cordelia Langford, John Sillitoe on behalf of the Wellcome Sanger Institute COVID-19 Surveillance Team |
| EPI_ISL_3469029, EPI_ISL_3469030, EPI_ISL_3469031, EPI_ISL_3469032, EPI_ISL_3469033, EPI_ISL_3469034 | Lighthouse Lab in Alderley Park | Wellcome Sanger Institute for the COVID-19 Genomics UK (COG-UK) Consortium | Jacquelyn Wynn, Mairead Hyland, The Lighthouse Lab in Alderley Park and Alex Alderton, Roberto Amato, Jeffrey Barrett, Sonia Goncalves, Ewan Harrison, David K. Jackson, Ian Johnston, Dominic Kwiatkowski, Cordelia Langford, John Sillitoe on behalf of the Wellcome Sanger Institute COVID-19 Surveillance Team |
| EPI_ISL_3469035 | Lighthouse Lab in Milton Keynes | Wellcome Sanger Institute for the COVID-19 Genomics UK (COG-UK) Consortium | The Lighthouse Lab in Milton Keynes and Alex Alderton, Roberto Amato, Jeffrey Barrett, Sonia Goncalves, Ewan Harrison, David K. Jackson, Ian Johnston, Dominic Kwiatkowski, Cordelia Langford, John Sillitoe on behalf of the Wellcome Sanger Institute COVID-19 Surveillance Team |
| EPI_ISL_3469036, EPI_ISL_3469037, EPI_ISL_3469038 | Lighthouse Lab in Alderley Park | Wellcome Sanger Institute for the COVID-19 Genomics UK (COG-UK) Consortium | Jacquelyn Wynn, Mairead Hyland, The Lighthouse Lab in Alderley Park and Alex Alderton, Roberto Amato, Jeffrey Barrett, Sonia Goncalves, Ewan Harrison, David K. Jackson, Ian Johnston, Dominic Kwiatkowski, Cordelia Langford, John Sillitoe on behalf of the Wellcome Sanger Institute COVID-19 Surveillance Team |
| EPI_ISL_3469039, EPI_ISL_3469040 | Lighthouse Lab in Milton Keynes | Wellcome Sanger Institute for the COVID-19 Genomics UK (COG-UK) Consortium | The Lighthouse Lab in Milton Keynes and Alex Alderton, Roberto Amato, Jeffrey Barrett, Sonia Goncalves, Ewan Harrison, David K. Jackson, Ian Johnston, Dominic Kwiatkowski, Cordelia Langford, John Sillitoe on behalf of the Wellcome Sanger Institute COVID-19 Surveillance Team |
| EPI_ISL_3469041, EPI_ISL_3469042, EPI_ISL_3469043, EPI_ISL_3469044, EPI_ISL_3469045 | Lighthouse Lab in Alderley Park | Wellcome Sanger Institute for the COVID-19 Genomics UK (COG-UK) Consortium | Jacquelyn Wynn, Mairead Hyland, The Lighthouse Lab in Alderley Park and Alex Alderton, Roberto Amato, Jeffrey Barrett, Sonia Goncalves, Ewan Harrison, David K. Jackson, Ian Johnston, Dominic Kwiatkowski, Cordelia Langford, John Sillitoe on behalf of the Wellcome Sanger Institute COVID-19 Surveillance Team |
| EPI_ISL_3469046 | Health Services Laboratories | Wellcome Sanger Institute for the COVID-19 Genomics UK (COG-UK) Consortium | Health Services Laboratories and Alex Alderton, Roberto Amato, Jeffrey Barrett, Sonia Goncalves, Ewan Harrison, David K. Jackson, Ian Johnston, Dominic Kwiatkowski, Cordelia Langford, John Sillitoe on behalf of the Wellcome Sanger Institute COVID-19 Surveillance Team |
| EPI_ISL_3469047, EPI_ISL_3469048 | Lighthouse Lab in Alderley Park | Wellcome Sanger Institute for the COVID-19 Genomics UK (COG-UK) Consortium | Jacquelyn Wynn, Mairead Hyland, The Lighthouse Lab in Alderley Park and Alex Alderton, Roberto Amato, Jeffrey Barrett, Sonia Goncalves, Ewan Harrison, David K. Jackson, Ian Johnston, Dominic Kwiatkowski, Cordelia Langford, John Sillitoe on behalf of the Wellcome Sanger Institute COVID-19 Surveillance Team |
| EPI_ISL_3469049 | Health Services Laboratories | Wellcome Sanger Institute for the COVID-19 Genomics UK (COG-UK) Consortium | Health Services Laboratories and Alex Alderton, Roberto Amato, Jeffrey Barrett, Sonia Goncalves, Ewan Harrison, David K. Jackson, Ian Johnston, Dominic Kwiatkowski, Cordelia Langford, John Sillitoe on behalf of the Wellcome Sanger Institute COVID-19 Surveillance Team |
| EPI_ISL_3469050 | Lighthouse Lab in Milton Keynes | Wellcome Sanger Institute for the COVID-19 Genomics UK (COG-UK) Consortium | The Lighthouse Lab in Milton Keynes and Alex Alderton, Roberto Amato, Jeffrey Barrett, Sonia Goncalves, Ewan Harrison, David K. Jackson, Ian Johnston, Dominic Kwiatkowski, Cordelia Langford, John Sillitoe on behalf of the Wellcome Sanger Institute COVID-19 Surveillance Team |
| EPI_ISL_3469051 | Lighthouse Lab in Alderley Park | Wellcome Sanger Institute for the COVID-19 Genomics UK (COG-UK) Consortium | Jacquelyn Wynn, Mairead Hyland, The Lighthouse Lab in Alderley Park and Alex Alderton, Roberto Amato, Jeffrey Barrett, Sonia Goncalves, Ewan Harrison, David K. Jackson, Ian Johnston, Dominic Kwiatkowski, Cordelia Langford, John Sillitoe on behalf of the Wellcome Sanger Institute COVID-19 Surveillance Team |
| EPI_ISL_3469052, EPI_ISL_3469053 | Lighthouse Lab in Milton Keynes | Wellcome Sanger Institute for the COVID-19 Genomics UK (COG-UK) Consortium | The Lighthouse Lab in Milton Keynes and Alex Alderton, Roberto Amato, Jeffrey Barrett, Sonia Goncalves, Ewan Harrison, David K. Jackson, Ian Johnston, Dominic Kwiatkowski, Cordelia Langford, John Sillitoe on behalf of the Wellcome Sanger Institute COVID-19 Surveillance Team |
| EPI_ISL_3469054 | Lighthouse Lab in Alderley Park | Wellcome Sanger Institute for the COVID-19 Genomics UK (COG-UK) Consortium | Jacquelyn Wynn, Mairead Hyland, The Lighthouse Lab in Alderley Park and Alex Alderton, Roberto Amato, Jeffrey Barrett, Sonia Goncalves, Ewan Harrison, David K. Jackson, Ian Johnston, Dominic Kwiatkowski, Cordelia Langford, John Sillitoe on behalf of the Wellcome Sanger Institute COVID-19 Surveillance Team |
| EPI_ISL_3469055 | Health Services Laboratories | Wellcome Sanger Institute for the COVID-19 Genomics UK (COG-UK) Consortium | Health Services Laboratories and Alex Alderton, Roberto Amato, Jeffrey Barrett, Sonia Goncalves, Ewan Harrison, David K. Jackson, Ian Johnston, Dominic Kwiatkowski, Cordelia Langford, John Sillitoe on behalf of the Wellcome Sanger Institute COVID-19 Surveillance Team |
| EPI_ISL_3469056, EPI_ISL_3469057, EPI_ISL_3469058, EPI_ISL_3469059, EPI_ISL_3469060, EPI_ISL_3469061, | Lighthouse Lab in Alderley Park | Wellcome Sanger Institute for the COVID-19 Genomics UK (COG-UK) Consortium | Jacquelyn Wynn, Mairead Hyland, The Lighthouse Lab in Alderley Park and Alex Alderton, Roberto Amato, Jeffrey Barrett, Sonia Goncalves, Ewan Harrison, David K. Jackson, Ian Johnston, Dominic Kwiatkowski, Cordelia Langford, John Sillitoe on behalf of the Wellcome Sanger Institute COVID-19 Surveillance Team |

| | | | |
|---|---|---|---|
| EPI_ISL_3469062 | | | |
| EPI_ISL_3469063 | Health Services Laboratories | Wellcome Sanger Institute for the COVID-19 Genomics UK (COG-UK) Consortium | Health Services Laboratories and Alex Alderton, Roberto Amato, Jeffrey Barrett, Sonia Goncalves, Ewan Harrison, David K. Jackson, Ian Johnston, Dominic Kwiatkowski, Cordelia Langford, John Sillitoe on behalf of the Wellcome Sanger Institute COVID-19 Surveillance Team |
| EPI_ISL_3469064, EPI_ISL_3469065, EPI_ISL_3469066, EPI_ISL_3469067 | Lighthouse Lab in Alderley Park | Wellcome Sanger Institute for the COVID-19 Genomics UK (COG-UK) Consortium | Jacquelyn Wynn, Mairead Hyland, The Lighthouse Lab in Alderley Park and Alex Alderton, Roberto Amato, Jeffrey Barrett, Sonia Goncalves, Ewan Harrison, David K. Jackson, Ian Johnston, Dominic Kwiatkowski, Cordelia Langford, John Sillitoe on behalf of the Wellcome Sanger Institute COVID-19 Surveillance Team |
| EPI_ISL_3469068 | Health Services Laboratories | Wellcome Sanger Institute for the COVID-19 Genomics UK (COG-UK) Consortium | Health Services Laboratories and Alex Alderton, Roberto Amato, Jeffrey Barrett, Sonia Goncalves, Ewan Harrison, David K. Jackson, Ian Johnston, Dominic Kwiatkowski, Cordelia Langford, John Sillitoe on behalf of the Wellcome Sanger Institute COVID-19 Surveillance Team |
| EPI_ISL_3469069, EPI_ISL_3469070, EPI_ISL_3469071 | Lighthouse Lab in Alderley Park | Wellcome Sanger Institute for the COVID-19 Genomics UK (COG-UK) Consortium | Jacquelyn Wynn, Mairead Hyland, The Lighthouse Lab in Alderley Park and Alex Alderton, Roberto Amato, Jeffrey Barrett, Sonia Goncalves, Ewan Harrison, David K. Jackson, Ian Johnston, Dominic Kwiatkowski, Cordelia Langford, John Sillitoe on behalf of the Wellcome Sanger Institute COVID-19 Surveillance Team |
| EPI_ISL_3469072, EPI_ISL_3469073 | Lighthouse Lab in Milton Keynes | Wellcome Sanger Institute for the COVID-19 Genomics UK (COG-UK) Consortium | The Lighthouse Lab in Milton Keynes and Alex Alderton, Roberto Amato, Jeffrey Barrett, Sonia Goncalves, Ewan Harrison, David K. Jackson, Ian Johnston, Dominic Kwiatkowski, Cordelia Langford, John Sillitoe on behalf of the Wellcome Sanger Institute COVID-19 Surveillance Team |
| EPI_ISL_3469074, EPI_ISL_3469075 | Lighthouse Lab in Alderley Park | Wellcome Sanger Institute for the COVID-19 Genomics UK (COG-UK) Consortium | Jacquelyn Wynn, Mairead Hyland, The Lighthouse Lab in Alderley Park and Alex Alderton, Roberto Amato, Jeffrey Barrett, Sonia Goncalves, Ewan Harrison, David K. Jackson, Ian Johnston, Dominic Kwiatkowski, Cordelia Langford, John Sillitoe on behalf of the Wellcome Sanger Institute COVID-19 Surveillance Team |
| EPI_ISL_3469076 | Health Services Laboratories | Wellcome Sanger Institute for the COVID-19 Genomics UK (COG-UK) Consortium | Health Services Laboratories and Alex Alderton, Roberto Amato, Jeffrey Barrett, Sonia Goncalves, Ewan Harrison, David K. Jackson, Ian Johnston, Dominic Kwiatkowski, Cordelia Langford, John Sillitoe on behalf of the Wellcome Sanger Institute COVID-19 Surveillance Team |
| EPI_ISL_3469077, EPI_ISL_3469078, EPI_ISL_3469079 | Lighthouse Lab in Alderley Park | Wellcome Sanger Institute for the COVID-19 Genomics UK (COG-UK) Consortium | Jacquelyn Wynn, Mairead Hyland, The Lighthouse Lab in Alderley Park and Alex Alderton, Roberto Amato, Jeffrey Barrett, Sonia Goncalves, Ewan Harrison, David K. Jackson, Ian Johnston, Dominic Kwiatkowski, Cordelia Langford, John Sillitoe on behalf of the Wellcome Sanger Institute COVID-19 Surveillance Team |
| EPI_ISL_3469080 | Health Services Laboratories | Wellcome Sanger Institute for the COVID-19 Genomics UK (COG-UK) Consortium | Health Services Laboratories and Alex Alderton, Roberto Amato, Jeffrey Barrett, Sonia Goncalves, Ewan Harrison, David K. Jackson, Ian Johnston, Dominic Kwiatkowski, Cordelia Langford, John Sillitoe on behalf of the Wellcome Sanger Institute COVID-19 Surveillance Team |
| EPI_ISL_3469081 | Lighthouse Lab in Milton Keynes | Wellcome Sanger Institute for the COVID-19 Genomics UK (COG-UK) Consortium | The Lighthouse Lab in Milton Keynes and Alex Alderton, Roberto Amato, Jeffrey Barrett, Sonia Goncalves, Ewan Harrison, David K. Jackson, Ian Johnston, Dominic Kwiatkowski, Cordelia Langford, John Sillitoe on behalf of the Wellcome Sanger Institute COVID-19 Surveillance Team |
| EPI_ISL_3469082 | Health Services Laboratories | Wellcome Sanger Institute for the COVID-19 Genomics UK (COG-UK) Consortium | Health Services Laboratories and Alex Alderton, Roberto Amato, Jeffrey Barrett, Sonia Goncalves, Ewan Harrison, David K. Jackson, Ian Johnston, Dominic Kwiatkowski, Cordelia Langford, John Sillitoe on behalf of the Wellcome Sanger Institute COVID-19 Surveillance Team |
| EPI_ISL_3469083 | Lighthouse Lab in Milton Keynes | Wellcome Sanger Institute for the COVID-19 Genomics UK (COG-UK) Consortium | The Lighthouse Lab in Milton Keynes and Alex Alderton, Roberto Amato, Jeffrey Barrett, Sonia Goncalves, Ewan Harrison, David K. Jackson, Ian Johnston, Dominic Kwiatkowski, Cordelia Langford, John Sillitoe on behalf of the Wellcome Sanger Institute COVID-19 Surveillance Team |
| EPI_ISL_3469084, EPI_ISL_3469085, EPI_ISL_3469086, EPI_ISL_3469087, EPI_ISL_3469088, EPI_ISL_3469089, EPI_ISL_3469090, EPI_ISL_3469091, EPI_ISL_3469092, EPI_ISL_3469093, EPI_ISL_3469094, EPI_ISL_3469095 | | | |
| see above | Lighthouse Lab in Alderley Park | Wellcome Sanger Institute for the COVID-19 Genomics UK (COG-UK) Consortium | Jacquelyn Wynn, Mairead Hyland, The Lighthouse Lab in Alderley Park and Alex Alderton, Roberto Amato, Jeffrey Barrett, Sonia Goncalves, Ewan Harrison, David K. Jackson, Ian Johnston, Dominic Kwiatkowski, Cordelia Langford, John Sillitoe on behalf of the Wellcome Sanger Institute COVID-19 Surveillance Team |
| EPI_ISL_3469096 | Lighthouse Lab in Milton Keynes | Wellcome Sanger Institute for the COVID-19 Genomics UK (COG-UK) Consortium | The Lighthouse Lab in Milton Keynes and Alex Alderton, Roberto Amato, Jeffrey Barrett, Sonia Goncalves, Ewan Harrison, David K. Jackson, Ian Johnston, Dominic Kwiatkowski, Cordelia Langford, John Sillitoe on behalf of the Wellcome Sanger Institute COVID-19 Surveillance Team |
| EPI_ISL_3469097 | Health Services Laboratories | Wellcome Sanger Institute for the COVID-19 Genomics UK (COG-UK) Consortium | Health Services Laboratories and Alex Alderton, Roberto Amato, Jeffrey Barrett, Sonia Goncalves, Ewan Harrison, David K. Jackson, Ian Johnston, Dominic Kwiatkowski, Cordelia Langford, John Sillitoe on behalf of the Wellcome Sanger Institute COVID-19 Surveillance Team |
| EPI_ISL_3469098, EPI_ISL_3469099 | Lighthouse Lab in Milton Keynes | Wellcome Sanger Institute for the COVID-19 Genomics UK (COG-UK) Consortium | The Lighthouse Lab in Milton Keynes and Alex Alderton, Roberto Amato, Jeffrey Barrett, Sonia Goncalves, Ewan Harrison, David K. Jackson, Ian Johnston, Dominic Kwiatkowski, Cordelia Langford, John Sillitoe on behalf of the Wellcome Sanger Institute COVID-19 Surveillance Team |
| EPI_ISL_3469100 | Health Services Laboratories | Wellcome Sanger Institute for the COVID-19 Genomics UK (COG-UK) Consortium | Health Services Laboratories and Alex Alderton, Roberto Amato, Jeffrey Barrett, Sonia Goncalves, Ewan Harrison, David K. Jackson, Ian Johnston, Dominic Kwiatkowski, Cordelia Langford, John Sillitoe on behalf of the Wellcome Sanger Institute COVID-19 Surveillance Team |
| EPI_ISL_3469101, EPI_ISL_3469102, EPI_ISL_3469103 | Lighthouse Lab in Alderley Park | Wellcome Sanger Institute for the COVID-19 Genomics UK (COG-UK) Consortium | Jacquelyn Wynn, Mairead Hyland, The Lighthouse Lab in Alderley Park and Alex Alderton, Roberto Amato, Jeffrey Barrett, Sonia Goncalves, Ewan Harrison, David K. Jackson, Ian Johnston, Dominic Kwiatkowski, Cordelia Langford, John Sillitoe on behalf of the Wellcome Sanger Institute COVID-19 Surveillance Team |
| EPI_ISL_3469104, EPI_ISL_3469105 | Lighthouse Lab in Milton Keynes | Wellcome Sanger Institute for the COVID-19 Genomics UK (COG-UK) Consortium | The Lighthouse Lab in Milton Keynes and Alex Alderton, Roberto Amato, Jeffrey Barrett, Sonia Goncalves, Ewan Harrison, David K. Jackson, Ian Johnston, Dominic Kwiatkowski, Cordelia Langford, John Sillitoe on behalf of the Wellcome Sanger Institute COVID-19 Surveillance Team |
| EPI_ISL_3469106 | Lighthouse Lab in Alderley Park | Wellcome Sanger Institute for the COVID-19 Genomics UK (COG-UK) Consortium | Jacquelyn Wynn, Mairead Hyland, The Lighthouse Lab in Alderley Park and Alex Alderton, Roberto Amato, Jeffrey Barrett, Sonia Goncalves, Ewan Harrison, David K. Jackson, Ian Johnston, Dominic Kwiatkowski, Cordelia Langford, John Sillitoe on behalf of the Wellcome Sanger Institute COVID-19 Surveillance Team |
| EPI_ISL_3469107 | Health Services Laboratories | Wellcome Sanger Institute for the COVID-19 Genomics UK (COG-UK) Consortium | Health Services Laboratories and Alex Alderton, Roberto Amato, Jeffrey Barrett, Sonia Goncalves, Ewan Harrison, David K. Jackson, Ian Johnston, Dominic Kwiatkowski, Cordelia Langford, John Sillitoe on behalf of the Wellcome Sanger Institute COVID-19 Surveillance Team |
| EPI_ISL_3469108 | Lighthouse Lab in Alderley Park | Wellcome Sanger Institute for the COVID-19 Genomics UK (COG-UK) Consortium | Jacquelyn Wynn, Mairead Hyland, The Lighthouse Lab in Alderley Park and Alex Alderton, Roberto Amato, Jeffrey Barrett, Sonia Goncalves, Ewan Harrison, David K. Jackson, Ian Johnston, Dominic Kwiatkowski, Cordelia Langford, John Sillitoe on behalf of the Wellcome Sanger Institute COVID-19 Surveillance Team |
| EPI_ISL_3469109 | Health Services Laboratories | Wellcome Sanger Institute for the COVID-19 Genomics UK (COG-UK) Consortium | Health Services Laboratories and Alex Alderton, Roberto Amato, Jeffrey Barrett, Sonia Goncalves, Ewan Harrison, David K. Jackson, Ian Johnston, Dominic Kwiatkowski, Cordelia Langford, John Sillitoe on behalf of the Wellcome Sanger Institute COVID-19 Surveillance Team |
| EPI_ISL_3469110 | Lighthouse Lab in Milton Keynes | Wellcome Sanger Institute for the COVID-19 Genomics UK (COG-UK) Consortium | The Lighthouse Lab in Milton Keynes and Alex Alderton, Roberto Amato, Jeffrey Barrett, Sonia Goncalves, Ewan Harrison, David K. Jackson, Ian Johnston, Dominic Kwiatkowski, Cordelia Langford, John Sillitoe on behalf of the Wellcome Sanger Institute COVID-19 Surveillance Team |
| EPI_ISL_3469111 | Health Services Laboratories | Wellcome Sanger Institute for the COVID-19 Genomics UK (COG-UK) Consortium | Health Services Laboratories and Alex Alderton, Roberto Amato, Jeffrey Barrett, Sonia Goncalves, Ewan Harrison, David K. Jackson, Ian Johnston, Dominic Kwiatkowski, Cordelia Langford, John Sillitoe on behalf of the Wellcome Sanger Institute COVID-19 Surveillance Team |
| EPI_ISL_3469112 | Lighthouse Lab in Alderley Park | Wellcome Sanger Institute for the COVID-19 Genomics UK (COG-UK) Consortium | Jacquelyn Wynn, Mairead Hyland, The Lighthouse Lab in Alderley Park and Alex Alderton, Roberto Amato, Jeffrey Barrett, Sonia Goncalves, Ewan Harrison, David K. Jackson, Ian Johnston, Dominic Kwiatkowski, Cordelia Langford, John Sillitoe on behalf of the Wellcome Sanger Institute COVID-19 Surveillance Team |
| EPI_ISL_3469113, EPI_ISL_3469114 | Lighthouse Lab in Milton Keynes | Wellcome Sanger Institute for the COVID-19 Genomics UK (COG-UK) Consortium | The Lighthouse Lab in Milton Keynes and Alex Alderton, Roberto Amato, Jeffrey Barrett, Sonia Goncalves, Ewan Harrison, David K. Jackson, Ian Johnston, Dominic Kwiatkowski, Cordelia Langford, John Sillitoe on behalf of the Wellcome Sanger Institute COVID-19 Surveillance Team |
| EPI_ISL_3469115, EPI_ISL_3469116, EPI_ISL_3469117, EPI_ISL_3469118 | Lighthouse Lab in Alderley Park | Wellcome Sanger Institute for the COVID-19 Genomics UK (COG-UK) Consortium | Jacquelyn Wynn, Mairead Hyland, The Lighthouse Lab in Alderley Park and Alex Alderton, Roberto Amato, Jeffrey Barrett, Sonia Goncalves, Ewan Harrison, David K. Jackson, Ian Johnston, Dominic Kwiatkowski, Cordelia Langford, John Sillitoe on behalf of the Wellcome Sanger Institute COVID-19 Surveillance Team |
| EPI_ISL_3469119 | Health Services Laboratories | Wellcome Sanger Institute for the COVID-19 Genomics UK (COG-UK) Consortium | Health Services Laboratories and Alex Alderton, Roberto Amato, Jeffrey Barrett, Sonia Goncalves, Ewan Harrison, David K. Jackson, Ian Johnston, Dominic Kwiatkowski, Cordelia Langford, John Sillitoe on behalf of the Wellcome Sanger Institute COVID-19 Surveillance Team |
| EPI_ISL_3469120, EPI_ISL_3469121, EPI_ISL_3469122 | Lighthouse Lab in Alderley Park | Wellcome Sanger Institute for the COVID-19 Genomics UK (COG-UK) Consortium | Jacquelyn Wynn, Mairead Hyland, The Lighthouse Lab in Alderley Park and Alex Alderton, Roberto Amato, Jeffrey Barrett, Sonia Goncalves, Ewan Harrison, David K. Jackson, Ian Johnston, Dominic Kwiatkowski, Cordelia Langford, John Sillitoe on behalf of the Wellcome Sanger Institute COVID-19 Surveillance Team |
| EPI_ISL_3469123 | Health Services Laboratories | Wellcome Sanger Institute for the COVID-19 Genomics UK (COG-UK) Consortium | Health Services Laboratories and Alex Alderton, Roberto Amato, Jeffrey Barrett, Sonia Goncalves, Ewan Harrison, David K. Jackson, Ian Johnston, Dominic Kwiatkowski, Cordelia Langford, John Sillitoe on behalf of the Wellcome Sanger Institute COVID-19 Surveillance Team |
| EPI_ISL_3469124 | Lighthouse Lab in Alderley Park | Wellcome Sanger Institute for the COVID-19 Genomics UK (COG-UK) Consortium | Jacquelyn Wynn, Mairead Hyland, The Lighthouse Lab in Alderley Park and Alex Alderton, Roberto Amato, Jeffrey Barrett, Sonia Goncalves, Ewan Harrison, David K. Jackson, Ian Johnston, Dominic Kwiatkowski, Cordelia Langford, John Sillitoe on behalf of the Wellcome Sanger Institute COVID-19 Surveillance Team |
| EPI_ISL_3469125, EPI_ISL_3469126 | Lighthouse Lab in Milton Keynes | Wellcome Sanger Institute for the COVID-19 Genomics UK (COG-UK) Consortium | The Lighthouse Lab in Milton Keynes and Alex Alderton, Roberto Amato, Jeffrey Barrett, Sonia Goncalves, Ewan Harrison, David K. Jackson, Ian Johnston, Dominic Kwiatkowski, Cordelia Langford, John Sillitoe on behalf of the Wellcome Sanger Institute COVID-19 Surveillance Team |
| EPI_ISL_3469127 | Lighthouse Lab in Alderley Park | Wellcome Sanger Institute for the COVID-19 Genomics UK (COG-UK) Consortium | Jacquelyn Wynn, Mairead Hyland, The Lighthouse Lab in Alderley Park and Alex Alderton, Roberto Amato, Jeffrey Barrett, Sonia Goncalves, Ewan Harrison, David K. Jackson, Ian Johnston, Dominic Kwiatkowski, Cordelia Langford, John Sillitoe on behalf of the Wellcome Sanger Institute COVID-19 Surveillance Team |
| EPI_ISL_3469128 | Lighthouse Lab in Milton Keynes | Wellcome Sanger Institute for the COVID-19 Genomics UK (COG-UK) Consortium | The Lighthouse Lab in Milton Keynes and Alex Alderton, Roberto Amato, Jeffrey Barrett, Sonia Goncalves, Ewan Harrison, David K. Jackson, Ian Johnston, Dominic Kwiatkowski, Cordelia Langford, John Sillitoe on behalf of the Wellcome Sanger Institute COVID-19 Surveillance Team |
| EPI_ISL_3469129 | Lighthouse Lab in Alderley Park | Wellcome Sanger Institute for the COVID-19 Genomics UK (COG-UK) Consortium | Jacquelyn Wynn, Mairead Hyland, The Lighthouse Lab in Alderley Park and Alex Alderton, Roberto Amato, Jeffrey Barrett, Sonia Goncalves, Ewan Harrison, David K. Jackson, Ian Johnston, Dominic Kwiatkowski, Cordelia Langford, John Sillitoe on behalf of the Wellcome Sanger Institute COVID-19 Surveillance Team |
| EPI_ISL_3469130 | Health Services Laboratories | Wellcome Sanger Institute for the COVID-19 Genomics UK (COG-UK) Consortium | Health Services Laboratories and Alex Alderton, Roberto Amato, Jeffrey Barrett, Sonia Goncalves, Ewan Harrison, David K. Jackson, Ian Johnston, Dominic Kwiatkowski, Cordelia Langford, John Sillitoe on behalf of the Wellcome Sanger Institute COVID-19 Surveillance Team |
| EPI_ISL_3469131, EPI_ISL_3469132, EPI_ISL_3469133, EPI_ISL_3469134, EPI_ISL_3469135 | Lighthouse Lab in Alderley Park | Wellcome Sanger Institute for the COVID-19 Genomics UK (COG-UK) Consortium | Jacquelyn Wynn, Mairead Hyland, The Lighthouse Lab in Alderley Park and Alex Alderton, Roberto Amato, Jeffrey Barrett, Sonia Goncalves, Ewan Harrison, David K. Jackson, Ian Johnston, Dominic Kwiatkowski, Cordelia Langford, John Sillitoe on behalf of the Wellcome Sanger Institute COVID-19 Surveillance Team |
| EPI_ISL_3469136, EPI_ISL_3469137 | Lighthouse Lab in Milton Keynes | Wellcome Sanger Institute for the COVID-19 Genomics UK (COG-UK) Consortium | The Lighthouse Lab in Milton Keynes and Alex Alderton, Roberto Amato, Jeffrey Barrett, Sonia Goncalves, Ewan Harrison, David K. Jackson, Ian Johnston, Dominic Kwiatkowski, Cordelia Langford, John Sillitoe on behalf of the Wellcome Sanger Institute COVID-19 Surveillance Team |

| | | | |
|---|---|---|---|
| EPI_ISL_3469138 | Health Services Laboratories | Wellcome Sanger Institute for the COVID-19 Genomics UK (COG-UK) Consortium | Health Services Laboratories and Alex Alderton, Roberto Amato, Jeffrey Barrett, Sonia Goncalves, Ewan Harrison, David K. Jackson, Ian Johnston, Dominic Kwiatkowski, Cordelia Langford, John Sillitoe on behalf of the Wellcome Sanger Institute COVID-19 Surveillance Team |
| EPI_ISL_3469139, EPI_ISL_3469140 | Lighthouse Lab in Alderley Park | Wellcome Sanger Institute for the COVID-19 Genomics UK (COG-UK) Consortium | Jacquelyn Wynn, Mairead Hyland, The Lighthouse Lab in Alderley Park and Alex Alderton, Roberto Amato, Jeffrey Barrett, Sonia Goncalves, Ewan Harrison, David K. Jackson, Ian Johnston, Dominic Kwiatkowski, Cordelia Langford, John Sillitoe on behalf of the Wellcome Sanger Institute COVID-19 Surveillance Team |
| EPI_ISL_3469141 | Lighthouse Lab in Milton Keynes | Wellcome Sanger Institute for the COVID-19 Genomics UK (COG-UK) Consortium | The Lighthouse Lab in Milton Keynes and Alex Alderton, Roberto Amato, Jeffrey Barrett, Sonia Goncalves, Ewan Harrison, David K. Jackson, Ian Johnston, Dominic Kwiatkowski, Cordelia Langford, John Sillitoe on behalf of the Wellcome Sanger Institute COVID-19 Surveillance Team |
| EPI_ISL_3469142 | Health Services Laboratories | Wellcome Sanger Institute for the COVID-19 Genomics UK (COG-UK) Consortium | Health Services Laboratories and Alex Alderton, Roberto Amato, Jeffrey Barrett, Sonia Goncalves, Ewan Harrison, David K. Jackson, Ian Johnston, Dominic Kwiatkowski, Cordelia Langford, John Sillitoe on behalf of the Wellcome Sanger Institute COVID-19 Surveillance Team |
| EPI_ISL_3469143 | Lighthouse Lab in Alderley Park | Wellcome Sanger Institute for the COVID-19 Genomics UK (COG-UK) Consortium | Jacquelyn Wynn, Mairead Hyland, The Lighthouse Lab in Alderley Park and Alex Alderton, Roberto Amato, Jeffrey Barrett, Sonia Goncalves, Ewan Harrison, David K. Jackson, Ian Johnston, Dominic Kwiatkowski, Cordelia Langford, John Sillitoe on behalf of the Wellcome Sanger Institute COVID-19 Surveillance Team |
| EPI_ISL_3469144 | Lighthouse Lab in Milton Keynes | Wellcome Sanger Institute for the COVID-19 Genomics UK (COG-UK) Consortium | The Lighthouse Lab in Milton Keynes and Alex Alderton, Roberto Amato, Jeffrey Barrett, Sonia Goncalves, Ewan Harrison, David K. Jackson, Ian Johnston, Dominic Kwiatkowski, Cordelia Langford, John Sillitoe on behalf of the Wellcome Sanger Institute COVID-19 Surveillance Team |
| EPI_ISL_3469145 | Lighthouse Lab in Alderley Park | Wellcome Sanger Institute for the COVID-19 Genomics UK (COG-UK) Consortium | Jacquelyn Wynn, Mairead Hyland, The Lighthouse Lab in Alderley Park and Alex Alderton, Roberto Amato, Jeffrey Barrett, Sonia Goncalves, Ewan Harrison, David K. Jackson, Ian Johnston, Dominic Kwiatkowski, Cordelia Langford, John Sillitoe on behalf of the Wellcome Sanger Institute COVID-19 Surveillance Team |
| EPI_ISL_3469146 | Health Services Laboratories | Wellcome Sanger Institute for the COVID-19 Genomics UK (COG-UK) Consortium | Health Services Laboratories and Alex Alderton, Roberto Amato, Jeffrey Barrett, Sonia Goncalves, Ewan Harrison, David K. Jackson, Ian Johnston, Dominic Kwiatkowski, Cordelia Langford, John Sillitoe on behalf of the Wellcome Sanger Institute COVID-19 Surveillance Team |
| EPI_ISL_3469147, EPI_ISL_3469148 | Lighthouse Lab in Alderley Park | Wellcome Sanger Institute for the COVID-19 Genomics UK (COG-UK) Consortium | Jacquelyn Wynn, Mairead Hyland, The Lighthouse Lab in Alderley Park and Alex Alderton, Roberto Amato, Jeffrey Barrett, Sonia Goncalves, Ewan Harrison, David K. Jackson, Ian Johnston, Dominic Kwiatkowski, Cordelia Langford, John Sillitoe on behalf of the Wellcome Sanger Institute COVID-19 Surveillance Team |
| EPI_ISL_3469149 | Health Services Laboratories | Wellcome Sanger Institute for the COVID-19 Genomics UK (COG-UK) Consortium | Health Services Laboratories and Alex Alderton, Roberto Amato, Jeffrey Barrett, Sonia Goncalves, Ewan Harrison, David K. Jackson, Ian Johnston, Dominic Kwiatkowski, Cordelia Langford, John Sillitoe on behalf of the Wellcome Sanger Institute COVID-19 Surveillance Team |
| EPI_ISL_3469150 | Lighthouse Lab in Alderley Park | Wellcome Sanger Institute for the COVID-19 Genomics UK (COG-UK) Consortium | Jacquelyn Wynn, Mairead Hyland, The Lighthouse Lab in Alderley Park and Alex Alderton, Roberto Amato, Jeffrey Barrett, Sonia Goncalves, Ewan Harrison, David K. Jackson, Ian Johnston, Dominic Kwiatkowski, Cordelia Langford, John Sillitoe on behalf of the Wellcome Sanger Institute COVID-19 Surveillance Team |
| EPI_ISL_3469151, EPI_ISL_3469152 | Lighthouse Lab in Milton Keynes | Wellcome Sanger Institute for the COVID-19 Genomics UK (COG-UK) Consortium | The Lighthouse Lab in Milton Keynes and Alex Alderton, Roberto Amato, Jeffrey Barrett, Sonia Goncalves, Ewan Harrison, David K. Jackson, Ian Johnston, Dominic Kwiatkowski, Cordelia Langford, John Sillitoe on behalf of the Wellcome Sanger Institute COVID-19 Surveillance Team |
| EPI_ISL_3469153, EPI_ISL_3469154 | Lighthouse Lab in Alderley Park | Wellcome Sanger Institute for the COVID-19 Genomics UK (COG-UK) Consortium | Jacquelyn Wynn, Mairead Hyland, The Lighthouse Lab in Alderley Park and Alex Alderton, Roberto Amato, Jeffrey Barrett, Sonia Goncalves, Ewan Harrison, David K. Jackson, Ian Johnston, Dominic Kwiatkowski, Cordelia Langford, John Sillitoe on behalf of the Wellcome Sanger Institute COVID-19 Surveillance Team |
| EPI_ISL_3469155 | Health Services Laboratories | Wellcome Sanger Institute for the COVID-19 Genomics UK (COG-UK) Consortium | Health Services Laboratories and Alex Alderton, Roberto Amato, Jeffrey Barrett, Sonia Goncalves, Ewan Harrison, David K. Jackson, Ian Johnston, Dominic Kwiatkowski, Cordelia Langford, John Sillitoe on behalf of the Wellcome Sanger Institute COVID-19 Surveillance Team |
| EPI_ISL_3469156, EPI_ISL_3469157 | Lighthouse Lab in Milton Keynes | Wellcome Sanger Institute for the COVID-19 Genomics UK (COG-UK) Consortium | The Lighthouse Lab in Milton Keynes and Alex Alderton, Roberto Amato, Jeffrey Barrett, Sonia Goncalves, Ewan Harrison, David K. Jackson, Ian Johnston, Dominic Kwiatkowski, Cordelia Langford, John Sillitoe on behalf of the Wellcome Sanger Institute COVID-19 Surveillance Team |
| EPI_ISL_3469158, EPI_ISL_3469159, EPI_ISL_3469160, EPI_ISL_3469161 | Lighthouse Lab in Alderley Park | Wellcome Sanger Institute for the COVID-19 Genomics UK (COG-UK) Consortium | Jacquelyn Wynn, Mairead Hyland, The Lighthouse Lab in Alderley Park and Alex Alderton, Roberto Amato, Jeffrey Barrett, Sonia Goncalves, Ewan Harrison, David K. Jackson, Ian Johnston, Dominic Kwiatkowski, Cordelia Langford, John Sillitoe on behalf of the Wellcome Sanger Institute COVID-19 Surveillance Team |
| EPI_ISL_3469162 | Health Services Laboratories | Wellcome Sanger Institute for the COVID-19 Genomics UK (COG-UK) Consortium | Health Services Laboratories and Alex Alderton, Roberto Amato, Jeffrey Barrett, Sonia Goncalves, Ewan Harrison, David K. Jackson, Ian Johnston, Dominic Kwiatkowski, Cordelia Langford, John Sillitoe on behalf of the Wellcome Sanger Institute COVID-19 Surveillance Team |
| EPI_ISL_3469163 | Lighthouse Lab in Milton Keynes | Wellcome Sanger Institute for the COVID-19 Genomics UK (COG-UK) Consortium | The Lighthouse Lab in Milton Keynes and Alex Alderton, Roberto Amato, Jeffrey Barrett, Sonia Goncalves, Ewan Harrison, David K. Jackson, Ian Johnston, Dominic Kwiatkowski, Cordelia Langford, John Sillitoe on behalf of the Wellcome Sanger Institute COVID-19 Surveillance Team |
| EPI_ISL_3469164 | Lighthouse Lab in Alderley Park | Wellcome Sanger Institute for the COVID-19 Genomics UK (COG-UK) Consortium | Jacquelyn Wynn, Mairead Hyland, The Lighthouse Lab in Alderley Park and Alex Alderton, Roberto Amato, Jeffrey Barrett, Sonia Goncalves, Ewan Harrison, David K. Jackson, Ian Johnston, Dominic Kwiatkowski, Cordelia Langford, John Sillitoe on behalf of the Wellcome Sanger Institute COVID-19 Surveillance Team |
| EPI_ISL_3469165 | Lighthouse Lab in Milton Keynes | Wellcome Sanger Institute for the COVID-19 Genomics UK (COG-UK) Consortium | The Lighthouse Lab in Milton Keynes and Alex Alderton, Roberto Amato, Jeffrey Barrett, Sonia Goncalves, Ewan Harrison, David K. Jackson, Ian Johnston, Dominic Kwiatkowski, Cordelia Langford, John Sillitoe on behalf of the Wellcome Sanger Institute COVID-19 Surveillance Team |
| EPI_ISL_3469166, EPI_ISL_3469167, EPI_ISL_3469168 | Lighthouse Lab in Alderley Park | Wellcome Sanger Institute for the COVID-19 Genomics UK (COG-UK) Consortium | Jacquelyn Wynn, Mairead Hyland, The Lighthouse Lab in Alderley Park and Alex Alderton, Roberto Amato, Jeffrey Barrett, Sonia Goncalves, Ewan Harrison, David K. Jackson, Ian Johnston, Dominic Kwiatkowski, Cordelia Langford, John Sillitoe on behalf of the Wellcome Sanger Institute COVID-19 Surveillance Team |
| EPI_ISL_3469169 | Lighthouse Lab in Milton Keynes | Wellcome Sanger Institute for the COVID-19 Genomics UK (COG-UK) Consortium | The Lighthouse Lab in Milton Keynes and Alex Alderton, Roberto Amato, Jeffrey Barrett, Sonia Goncalves, Ewan Harrison, David K. Jackson, Ian Johnston, Dominic Kwiatkowski, Cordelia Langford, John Sillitoe on behalf of the Wellcome Sanger Institute COVID-19 Surveillance Team |
| EPI_ISL_3469170 | Lighthouse Lab in Alderley Park | Wellcome Sanger Institute for the COVID-19 Genomics UK (COG-UK) Consortium | Jacquelyn Wynn, Mairead Hyland, The Lighthouse Lab in Alderley Park and Alex Alderton, Roberto Amato, Jeffrey Barrett, Sonia Goncalves, Ewan Harrison, David K. Jackson, Ian Johnston, Dominic Kwiatkowski, Cordelia Langford, John Sillitoe on behalf of the Wellcome Sanger Institute COVID-19 Surveillance Team |
| EPI_ISL_3469171 | Lighthouse Lab in Milton Keynes | Wellcome Sanger Institute for the COVID-19 Genomics UK (COG-UK) Consortium | The Lighthouse Lab in Milton Keynes and Alex Alderton, Roberto Amato, Jeffrey Barrett, Sonia Goncalves, Ewan Harrison, David K. Jackson, Ian Johnston, Dominic Kwiatkowski, Cordelia Langford, John Sillitoe on behalf of the Wellcome Sanger Institute COVID-19 Surveillance Team |
| EPI_ISL_3469172 | Health Services Laboratories | Wellcome Sanger Institute for the COVID-19 Genomics UK (COG-UK) Consortium | Health Services Laboratories and Alex Alderton, Roberto Amato, Jeffrey Barrett, Sonia Goncalves, Ewan Harrison, David K. Jackson, Ian Johnston, Dominic Kwiatkowski, Cordelia Langford, John Sillitoe on behalf of the Wellcome Sanger Institute COVID-19 Surveillance Team |
| EPI_ISL_3469173 | Lighthouse Lab in Alderley Park | Wellcome Sanger Institute for the COVID-19 Genomics UK (COG-UK) Consortium | Jacquelyn Wynn, Mairead Hyland, The Lighthouse Lab in Alderley Park and Alex Alderton, Roberto Amato, Jeffrey Barrett, Sonia Goncalves, Ewan Harrison, David K. Jackson, Ian Johnston, Dominic Kwiatkowski, Cordelia Langford, John Sillitoe on behalf of the Wellcome Sanger Institute COVID-19 Surveillance Team |
| EPI_ISL_3469174 | Health Services Laboratories | Wellcome Sanger Institute for the COVID-19 Genomics UK (COG-UK) Consortium | Health Services Laboratories and Alex Alderton, Roberto Amato, Jeffrey Barrett, Sonia Goncalves, Ewan Harrison, David K. Jackson, Ian Johnston, Dominic Kwiatkowski, Cordelia Langford, John Sillitoe on behalf of the Wellcome Sanger Institute COVID-19 Surveillance Team |
| EPI_ISL_3469175 | Lighthouse Lab in Alderley Park | Wellcome Sanger Institute for the COVID-19 Genomics UK (COG-UK) Consortium | Jacquelyn Wynn, Mairead Hyland, The Lighthouse Lab in Alderley Park and Alex Alderton, Roberto Amato, Jeffrey Barrett, Sonia Goncalves, Ewan Harrison, David K. Jackson, Ian Johnston, Dominic Kwiatkowski, Cordelia Langford, John Sillitoe on behalf of the Wellcome Sanger Institute COVID-19 Surveillance Team |
| EPI_ISL_3469176, EPI_ISL_3469177 | Health Services Laboratories | Wellcome Sanger Institute for the COVID-19 Genomics UK (COG-UK) Consortium | Health Services Laboratories and Alex Alderton, Roberto Amato, Jeffrey Barrett, Sonia Goncalves, Ewan Harrison, David K. Jackson, Ian Johnston, Dominic Kwiatkowski, Cordelia Langford, John Sillitoe on behalf of the Wellcome Sanger Institute COVID-19 Surveillance Team |
| EPI_ISL_3469178 | Lighthouse Lab in Alderley Park | Wellcome Sanger Institute for the COVID-19 Genomics UK (COG-UK) Consortium | Jacquelyn Wynn, Mairead Hyland, The Lighthouse Lab in Alderley Park and Alex Alderton, Roberto Amato, Jeffrey Barrett, Sonia Goncalves, Ewan Harrison, David K. Jackson, Ian Johnston, Dominic Kwiatkowski, Cordelia Langford, John Sillitoe on behalf of the Wellcome Sanger Institute COVID-19 Surveillance Team |
| EPI_ISL_3469179, EPI_ISL_3469180, EPI_ISL_3469181 | Lighthouse Lab in Milton Keynes | Wellcome Sanger Institute for the COVID-19 Genomics UK (COG-UK) Consortium | The Lighthouse Lab in Milton Keynes and Alex Alderton, Roberto Amato, Jeffrey Barrett, Sonia Goncalves, Ewan Harrison, David K. Jackson, Ian Johnston, Dominic Kwiatkowski, Cordelia Langford, John Sillitoe on behalf of the Wellcome Sanger Institute COVID-19 Surveillance Team |
| EPI_ISL_3469182 | Health Services Laboratories | Wellcome Sanger Institute for the COVID-19 Genomics UK (COG-UK) Consortium | Health Services Laboratories and Alex Alderton, Roberto Amato, Jeffrey Barrett, Sonia Goncalves, Ewan Harrison, David K. Jackson, Ian Johnston, Dominic Kwiatkowski, Cordelia Langford, John Sillitoe on behalf of the Wellcome Sanger Institute COVID-19 Surveillance Team |
| EPI_ISL_3469183, EPI_ISL_3469184 | Lighthouse Lab in Alderley Park | Wellcome Sanger Institute for the COVID-19 Genomics UK (COG-UK) Consortium | Jacquelyn Wynn, Mairead Hyland, The Lighthouse Lab in Alderley Park and Alex Alderton, Roberto Amato, Jeffrey Barrett, Sonia Goncalves, Ewan Harrison, David K. Jackson, Ian Johnston, Dominic Kwiatkowski, Cordelia Langford, John Sillitoe on behalf of the Wellcome Sanger Institute COVID-19 Surveillance Team |
| EPI_ISL_3469185 | Health Services Laboratories | Wellcome Sanger Institute for the COVID-19 Genomics UK (COG-UK) Consortium | Health Services Laboratories and Alex Alderton, Roberto Amato, Jeffrey Barrett, Sonia Goncalves, Ewan Harrison, David K. Jackson, Ian Johnston, Dominic Kwiatkowski, Cordelia Langford, John Sillitoe on behalf of the Wellcome Sanger Institute COVID-19 Surveillance Team |
| EPI_ISL_3469186, EPI_ISL_3469187, EPI_ISL_3469188 | Lighthouse Lab in Alderley Park | Wellcome Sanger Institute for the COVID-19 Genomics UK (COG-UK) Consortium | Jacquelyn Wynn, Mairead Hyland, The Lighthouse Lab in Alderley Park and Alex Alderton, Roberto Amato, Jeffrey Barrett, Sonia Goncalves, Ewan Harrison, David K. Jackson, Ian Johnston, Dominic Kwiatkowski, Cordelia Langford, John Sillitoe on behalf of the Wellcome Sanger Institute COVID-19 Surveillance Team |
| EPI_ISL_3469189 | Lighthouse Lab in Milton Keynes | Wellcome Sanger Institute for the COVID-19 Genomics UK (COG-UK) Consortium | The Lighthouse Lab in Milton Keynes and Alex Alderton, Roberto Amato, Jeffrey Barrett, Sonia Goncalves, Ewan Harrison, David K. Jackson, Ian Johnston, Dominic Kwiatkowski, Cordelia Langford, John Sillitoe on behalf of the Wellcome Sanger Institute COVID-19 Surveillance Team |
| EPI_ISL_3469190 | Lighthouse Lab in Alderley Park | Wellcome Sanger Institute for the COVID-19 Genomics UK (COG-UK) Consortium | Jacquelyn Wynn, Mairead Hyland, The Lighthouse Lab in Alderley Park and Alex Alderton, Roberto Amato, Jeffrey Barrett, Sonia Goncalves, Ewan Harrison, David K. Jackson, Ian Johnston, Dominic Kwiatkowski, Cordelia Langford, John Sillitoe on behalf of the Wellcome Sanger Institute COVID-19 Surveillance Team |
| EPI_ISL_3469191 | Health Services Laboratories | Wellcome Sanger Institute for the COVID-19 Genomics UK (COG-UK) Consortium | Health Services Laboratories and Alex Alderton, Roberto Amato, Jeffrey Barrett, Sonia Goncalves, Ewan Harrison, David K. Jackson, Ian Johnston, Dominic Kwiatkowski, Cordelia Langford, John Sillitoe on behalf of the Wellcome Sanger Institute COVID-19 Surveillance Team |
| EPI_ISL_3469192 | Lighthouse Lab in Milton Keynes | Wellcome Sanger Institute for the COVID-19 Genomics UK (COG-UK) Consortium | The Lighthouse Lab in Milton Keynes and Alex Alderton, Roberto Amato, Jeffrey Barrett, Sonia Goncalves, Ewan Harrison, David K. Jackson, Ian Johnston, Dominic Kwiatkowski, Cordelia Langford, John Sillitoe on behalf of the Wellcome Sanger Institute COVID-19 Surveillance Team |
| EPI_ISL_3469193, EPI_ISL_3469194 | Lighthouse Lab in Alderley Park | Wellcome Sanger Institute for the COVID-19 Genomics UK (COG-UK) Consortium | Jacquelyn Wynn, Mairead Hyland, The Lighthouse Lab in Alderley Park and Alex Alderton, Roberto Amato, Jeffrey Barrett, Sonia Goncalves, Ewan Harrison, David K. Jackson, Ian Johnston, Dominic Kwiatkowski, Cordelia Langford, John Sillitoe on behalf of the Wellcome Sanger Institute COVID-19 Surveillance Team |
| EPI_ISL_3469195, EPI_ISL_3469196 | Health Services Laboratories | Wellcome Sanger Institute for the COVID-19 Genomics UK (COG-UK) Consortium | Health Services Laboratories and Alex Alderton, Roberto Amato, Jeffrey Barrett, Sonia Goncalves, Ewan Harrison, David K. Jackson, Ian Johnston, Dominic Kwiatkowski, Cordelia Langford, John Sillitoe on behalf of the Wellcome Sanger Institute COVID-19 Surveillance Team |
| EPI_ISL_3469197 | Lighthouse Lab in Milton Keynes | Wellcome Sanger Institute for the COVID-19 Genomics UK (COG-UK) Consortium | The Lighthouse Lab in Milton Keynes and Alex Alderton, Roberto Amato, Jeffrey Barrett, Sonia Goncalves, Ewan Harrison, David K. Jackson, Ian Johnston, Dominic Kwiatkowski, Cordelia Langford, John Sillitoe on behalf of the Wellcome Sanger Institute COVID-19 Surveillance Team |
| EPI_ISL_3469198 | Lighthouse Lab in Alderley Park | Wellcome Sanger Institute for the COVID-19 Genomics UK (COG-UK) Consortium | Jacquelyn Wynn, Mairead Hyland, The Lighthouse Lab in Alderley Park and Alex Alderton, Roberto Amato, Jeffrey Barrett, Sonia Goncalves, Ewan Harrison, David K. Jackson, Ian Johnston, Dominic Kwiatkowski, Cordelia Langford, John Sillitoe on behalf of the Wellcome Sanger Institute COVID-19 Surveillance Team |

| | | | |
|---|---|---|---|
| EPI_ISL_3469199, EPI_ISL_3469200 | Lighthouse Lab in Milton Keynes | Wellcome Sanger Institute for the COVID-19 Genomics UK (COG-UK) Consortium | The Lighthouse Lab in Milton Keynes and Alex Alderton, Roberto Amato, Jeffrey Barrett, Sonia Goncalves, Ewan Harrison, David K. Jackson, Ian Johnston, Dominic Kwiatkowski, Cordelia Langford, John Sillitoe on behalf of the Wellcome Sanger Institute COVID-19 Surveillance Team |
| EPI_ISL_3469201 | Health Services Laboratories | Wellcome Sanger Institute for the COVID-19 Genomics UK (COG-UK) Consortium | Health Services Laboratories and Alex Alderton, Roberto Amato, Jeffrey Barrett, Sonia Goncalves, Ewan Harrison, David K. Jackson, Ian Johnston, Dominic Kwiatkowski, Cordelia Langford, John Sillitoe on behalf of the Wellcome Sanger Institute COVID-19 Surveillance Team |
| EPI_ISL_3469202 | Lighthouse Lab in Alderley Park | Wellcome Sanger Institute for the COVID-19 Genomics UK (COG-UK) Consortium | Jacquelyn Wynn, Mairead Hyland, The Lighthouse Lab in Alderley Park and Alex Alderton, Roberto Amato, Jeffrey Barrett, Sonia Goncalves, Ewan Harrison, David K. Jackson, Ian Johnston, Dominic Kwiatkowski, Cordelia Langford, John Sillitoe on behalf of the Wellcome Sanger Institute COVID-19 Surveillance Team |
| EPI_ISL_3469203 | Health Services Laboratories | Wellcome Sanger Institute for the COVID-19 Genomics UK (COG-UK) Consortium | Health Services Laboratories and Alex Alderton, Roberto Amato, Jeffrey Barrett, Sonia Goncalves, Ewan Harrison, David K. Jackson, Ian Johnston, Dominic Kwiatkowski, Cordelia Langford, John Sillitoe on behalf of the Wellcome Sanger Institute COVID-19 Surveillance Team |
| EPI_ISL_3469204 | Lighthouse Lab in Alderley Park | Wellcome Sanger Institute for the COVID-19 Genomics UK (COG-UK) Consortium | Jacquelyn Wynn, Mairead Hyland, The Lighthouse Lab in Alderley Park and Alex Alderton, Roberto Amato, Jeffrey Barrett, Sonia Goncalves, Ewan Harrison, David K. Jackson, Ian Johnston, Dominic Kwiatkowski, Cordelia Langford, John Sillitoe on behalf of the Wellcome Sanger Institute COVID-19 Surveillance Team |
| EPI_ISL_3469205, EPI_ISL_3469206, EPI_ISL_3469207, EPI_ISL_3469208, EPI_ISL_3469209, EPI_ISL_3469210 | Lighthouse Lab in Milton Keynes | Wellcome Sanger Institute for the COVID-19 Genomics UK (COG-UK) Consortium | The Lighthouse Lab in Milton Keynes and Alex Alderton, Roberto Amato, Jeffrey Barrett, Sonia Goncalves, Ewan Harrison, David K. Jackson, Ian Johnston, Dominic Kwiatkowski, Cordelia Langford, John Sillitoe on behalf of the Wellcome Sanger Institute COVID-19 Surveillance Team |
| EPI_ISL_3469211, EPI_ISL_3469212 | Lighthouse Lab in Alderley Park | Wellcome Sanger Institute for the COVID-19 Genomics UK (COG-UK) Consortium | Jacquelyn Wynn, Mairead Hyland, The Lighthouse Lab in Alderley Park and Alex Alderton, Roberto Amato, Jeffrey Barrett, Sonia Goncalves, Ewan Harrison, David K. Jackson, Ian Johnston, Dominic Kwiatkowski, Cordelia Langford, John Sillitoe on behalf of the Wellcome Sanger Institute COVID-19 Surveillance Team |
| EPI_ISL_3469213 | Health Services Laboratories | Wellcome Sanger Institute for the COVID-19 Genomics UK (COG-UK) Consortium | Health Services Laboratories and Alex Alderton, Roberto Amato, Jeffrey Barrett, Sonia Goncalves, Ewan Harrison, David K. Jackson, Ian Johnston, Dominic Kwiatkowski, Cordelia Langford, John Sillitoe on behalf of the Wellcome Sanger Institute COVID-19 Surveillance Team |
| EPI_ISL_3469214, EPI_ISL_3469215, EPI_ISL_3469216 | Lighthouse Lab in Alderley Park | Wellcome Sanger Institute for the COVID-19 Genomics UK (COG-UK) Consortium | Jacquelyn Wynn, Mairead Hyland, The Lighthouse Lab in Alderley Park and Alex Alderton, Roberto Amato, Jeffrey Barrett, Sonia Goncalves, Ewan Harrison, David K. Jackson, Ian Johnston, Dominic Kwiatkowski, Cordelia Langford, John Sillitoe on behalf of the Wellcome Sanger Institute COVID-19 Surveillance Team |
| EPI_ISL_3469217 | Lighthouse Lab in Milton Keynes | Wellcome Sanger Institute for the COVID-19 Genomics UK (COG-UK) Consortium | The Lighthouse Lab in Milton Keynes and Alex Alderton, Roberto Amato, Jeffrey Barrett, Sonia Goncalves, Ewan Harrison, David K. Jackson, Ian Johnston, Dominic Kwiatkowski, Cordelia Langford, John Sillitoe on behalf of the Wellcome Sanger Institute COVID-19 Surveillance Team |
| EPI_ISL_3469218 | Lighthouse Lab in Alderley Park | Wellcome Sanger Institute for the COVID-19 Genomics UK (COG-UK) Consortium | Jacquelyn Wynn, Mairead Hyland, The Lighthouse Lab in Alderley Park and Alex Alderton, Roberto Amato, Jeffrey Barrett, Sonia Goncalves, Ewan Harrison, David K. Jackson, Ian Johnston, Dominic Kwiatkowski, Cordelia Langford, John Sillitoe on behalf of the Wellcome Sanger Institute COVID-19 Surveillance Team |
| EPI_ISL_3469219 | Health Services Laboratories | Wellcome Sanger Institute for the COVID-19 Genomics UK (COG-UK) Consortium | Health Services Laboratories and Alex Alderton, Roberto Amato, Jeffrey Barrett, Sonia Goncalves, Ewan Harrison, David K. Jackson, Ian Johnston, Dominic Kwiatkowski, Cordelia Langford, John Sillitoe on behalf of the Wellcome Sanger Institute COVID-19 Surveillance Team |
| EPI_ISL_3469220, EPI_ISL_3469221 | Lighthouse Lab in Alderley Park | Wellcome Sanger Institute for the COVID-19 Genomics UK (COG-UK) Consortium | Jacquelyn Wynn, Mairead Hyland, The Lighthouse Lab in Alderley Park and Alex Alderton, Roberto Amato, Jeffrey Barrett, Sonia Goncalves, Ewan Harrison, David K. Jackson, Ian Johnston, Dominic Kwiatkowski, Cordelia Langford, John Sillitoe on behalf of the Wellcome Sanger Institute COVID-19 Surveillance Team |
| EPI_ISL_3469222 | Health Services Laboratories | Wellcome Sanger Institute for the COVID-19 Genomics UK (COG-UK) Consortium | Health Services Laboratories and Alex Alderton, Roberto Amato, Jeffrey Barrett, Sonia Goncalves, Ewan Harrison, David K. Jackson, Ian Johnston, Dominic Kwiatkowski, Cordelia Langford, John Sillitoe on behalf of the Wellcome Sanger Institute COVID-19 Surveillance Team |
| EPI_ISL_3469223 | Lighthouse Lab in Alderley Park | Wellcome Sanger Institute for the COVID-19 Genomics UK (COG-UK) Consortium | Jacquelyn Wynn, Mairead Hyland, The Lighthouse Lab in Alderley Park and Alex Alderton, Roberto Amato, Jeffrey Barrett, Sonia Goncalves, Ewan Harrison, David K. Jackson, Ian Johnston, Dominic Kwiatkowski, Cordelia Langford, John Sillitoe on behalf of the Wellcome Sanger Institute COVID-19 Surveillance Team |
| EPI_ISL_3469224 | Lighthouse Lab in Milton Keynes | Wellcome Sanger Institute for the COVID-19 Genomics UK (COG-UK) Consortium | The Lighthouse Lab in Milton Keynes and Alex Alderton, Roberto Amato, Jeffrey Barrett, Sonia Goncalves, Ewan Harrison, David K. Jackson, Ian Johnston, Dominic Kwiatkowski, Cordelia Langford, John Sillitoe on behalf of the Wellcome Sanger Institute COVID-19 Surveillance Team |
| EPI_ISL_3469225 | Lighthouse Lab in Alderley Park | Wellcome Sanger Institute for the COVID-19 Genomics UK (COG-UK) Consortium | Jacquelyn Wynn, Mairead Hyland, The Lighthouse Lab in Alderley Park and Alex Alderton, Roberto Amato, Jeffrey Barrett, Sonia Goncalves, Ewan Harrison, David K. Jackson, Ian Johnston, Dominic Kwiatkowski, Cordelia Langford, John Sillitoe on behalf of the Wellcome Sanger Institute COVID-19 Surveillance Team |
| EPI_ISL_3469226 | Lighthouse Lab in Milton Keynes | Wellcome Sanger Institute for the COVID-19 Genomics UK (COG-UK) Consortium | The Lighthouse Lab in Milton Keynes and Alex Alderton, Roberto Amato, Jeffrey Barrett, Sonia Goncalves, Ewan Harrison, David K. Jackson, Ian Johnston, Dominic Kwiatkowski, Cordelia Langford, John Sillitoe on behalf of the Wellcome Sanger Institute COVID-19 Surveillance Team |
| EPI_ISL_3469227 | Health Services Laboratories | Wellcome Sanger Institute for the COVID-19 Genomics UK (COG-UK) Consortium | Health Services Laboratories and Alex Alderton, Roberto Amato, Jeffrey Barrett, Sonia Goncalves, Ewan Harrison, David K. Jackson, Ian Johnston, Dominic Kwiatkowski, Cordelia Langford, John Sillitoe on behalf of the Wellcome Sanger Institute COVID-19 Surveillance Team |
| EPI_ISL_3469228, EPI_ISL_3469229 | Lighthouse Lab in Alderley Park | Wellcome Sanger Institute for the COVID-19 Genomics UK (COG-UK) Consortium | Jacquelyn Wynn, Mairead Hyland, The Lighthouse Lab in Alderley Park and Alex Alderton, Roberto Amato, Jeffrey Barrett, Sonia Goncalves, Ewan Harrison, David K. Jackson, Ian Johnston, Dominic Kwiatkowski, Cordelia Langford, John Sillitoe on behalf of the Wellcome Sanger Institute COVID-19 Surveillance Team |
| EPI_ISL_3469230 | Health Services Laboratories | Wellcome Sanger Institute for the COVID-19 Genomics UK (COG-UK) Consortium | Health Services Laboratories and Alex Alderton, Roberto Amato, Jeffrey Barrett, Sonia Goncalves, Ewan Harrison, David K. Jackson, Ian Johnston, Dominic Kwiatkowski, Cordelia Langford, John Sillitoe on behalf of the Wellcome Sanger Institute COVID-19 Surveillance Team |
| EPI_ISL_3469231 | Lighthouse Lab in Milton Keynes | Wellcome Sanger Institute for the COVID-19 Genomics UK (COG-UK) Consortium | The Lighthouse Lab in Milton Keynes and Alex Alderton, Roberto Amato, Jeffrey Barrett, Sonia Goncalves, Ewan Harrison, David K. Jackson, Ian Johnston, Dominic Kwiatkowski, Cordelia Langford, John Sillitoe on behalf of the Wellcome Sanger Institute COVID-19 Surveillance Team |
| EPI_ISL_3469232, EPI_ISL_3469233, EPI_ISL_3469234 | Lighthouse Lab in Alderley Park | Wellcome Sanger Institute for the COVID-19 Genomics UK (COG-UK) Consortium | Jacquelyn Wynn, Mairead Hyland, The Lighthouse Lab in Alderley Park and Alex Alderton, Roberto Amato, Jeffrey Barrett, Sonia Goncalves, Ewan Harrison, David K. Jackson, Ian Johnston, Dominic Kwiatkowski, Cordelia Langford, John Sillitoe on behalf of the Wellcome Sanger Institute COVID-19 Surveillance Team |
| EPI_ISL_3469235, EPI_ISL_3469236, EPI_ISL_3469237 | Lighthouse Lab in Milton Keynes | Wellcome Sanger Institute for the COVID-19 Genomics UK (COG-UK) Consortium | The Lighthouse Lab in Milton Keynes and Alex Alderton, Roberto Amato, Jeffrey Barrett, Sonia Goncalves, Ewan Harrison, David K. Jackson, Ian Johnston, Dominic Kwiatkowski, Cordelia Langford, John Sillitoe on behalf of the Wellcome Sanger Institute COVID-19 Surveillance Team |
| EPI_ISL_3469238 | Lighthouse Lab in Alderley Park | Wellcome Sanger Institute for the COVID-19 Genomics UK (COG-UK) Consortium | Jacquelyn Wynn, Mairead Hyland, The Lighthouse Lab in Alderley Park and Alex Alderton, Roberto Amato, Jeffrey Barrett, Sonia Goncalves, Ewan Harrison, David K. Jackson, Ian Johnston, Dominic Kwiatkowski, Cordelia Langford, John Sillitoe on behalf of the Wellcome Sanger Institute COVID-19 Surveillance Team |
| EPI_ISL_3469239 | Lighthouse Lab in Milton Keynes | Wellcome Sanger Institute for the COVID-19 Genomics UK (COG-UK) Consortium | The Lighthouse Lab in Milton Keynes and Alex Alderton, Roberto Amato, Jeffrey Barrett, Sonia Goncalves, Ewan Harrison, David K. Jackson, Ian Johnston, Dominic Kwiatkowski, Cordelia Langford, John Sillitoe on behalf of the Wellcome Sanger Institute COVID-19 Surveillance Team |
| EPI_ISL_3469240 | Lighthouse Lab in Alderley Park | Wellcome Sanger Institute for the COVID-19 Genomics UK (COG-UK) Consortium | Jacquelyn Wynn, Mairead Hyland, The Lighthouse Lab in Alderley Park and Alex Alderton, Roberto Amato, Jeffrey Barrett, Sonia Goncalves, Ewan Harrison, David K. Jackson, Ian Johnston, Dominic Kwiatkowski, Cordelia Langford, John Sillitoe on behalf of the Wellcome Sanger Institute COVID-19 Surveillance Team |
| EPI_ISL_3469241 | Lighthouse Lab in Milton Keynes | Wellcome Sanger Institute for the COVID-19 Genomics UK (COG-UK) Consortium | The Lighthouse Lab in Milton Keynes and Alex Alderton, Roberto Amato, Jeffrey Barrett, Sonia Goncalves, Ewan Harrison, David K. Jackson, Ian Johnston, Dominic Kwiatkowski, Cordelia Langford, John Sillitoe on behalf of the Wellcome Sanger Institute COVID-19 Surveillance Team |
| EPI_ISL_3469242, EPI_ISL_3469243 | Health Services Laboratories | Wellcome Sanger Institute for the COVID-19 Genomics UK (COG-UK) Consortium | Health Services Laboratories and Alex Alderton, Roberto Amato, Jeffrey Barrett, Sonia Goncalves, Ewan Harrison, David K. Jackson, Ian Johnston, Dominic Kwiatkowski, Cordelia Langford, John Sillitoe on behalf of the Wellcome Sanger Institute COVID-19 Surveillance Team |
| EPI_ISL_3469244 | Lighthouse Lab in Milton Keynes | Wellcome Sanger Institute for the COVID-19 Genomics UK (COG-UK) Consortium | The Lighthouse Lab in Milton Keynes and Alex Alderton, Roberto Amato, Jeffrey Barrett, Sonia Goncalves, Ewan Harrison, David K. Jackson, Ian Johnston, Dominic Kwiatkowski, Cordelia Langford, John Sillitoe on behalf of the Wellcome Sanger Institute COVID-19 Surveillance Team |
| EPI_ISL_3469245, EPI_ISL_3469246, EPI_ISL_3469247, EPI_ISL_3469248, EPI_ISL_3469249 | Lighthouse Lab in Alderley Park | Wellcome Sanger Institute for the COVID-19 Genomics UK (COG-UK) Consortium | Jacquelyn Wynn, Mairead Hyland, The Lighthouse Lab in Alderley Park and Alex Alderton, Roberto Amato, Jeffrey Barrett, Sonia Goncalves, Ewan Harrison, David K. Jackson, Ian Johnston, Dominic Kwiatkowski, Cordelia Langford, John Sillitoe on behalf of the Wellcome Sanger Institute COVID-19 Surveillance Team |
| EPI_ISL_3469250 | Lighthouse Lab in Milton Keynes | Wellcome Sanger Institute for the COVID-19 Genomics UK (COG-UK) Consortium | The Lighthouse Lab in Milton Keynes and Alex Alderton, Roberto Amato, Jeffrey Barrett, Sonia Goncalves, Ewan Harrison, David K. Jackson, Ian Johnston, Dominic Kwiatkowski, Cordelia Langford, John Sillitoe on behalf of the Wellcome Sanger Institute COVID-19 Surveillance Team |
| EPI_ISL_3469251 | Lighthouse Lab in Alderley Park | Wellcome Sanger Institute for the COVID-19 Genomics UK (COG-UK) Consortium | Jacquelyn Wynn, Mairead Hyland, The Lighthouse Lab in Alderley Park and Alex Alderton, Roberto Amato, Jeffrey Barrett, Sonia Goncalves, Ewan Harrison, David K. Jackson, Ian Johnston, Dominic Kwiatkowski, Cordelia Langford, John Sillitoe on behalf of the Wellcome Sanger Institute COVID-19 Surveillance Team |
| EPI_ISL_3469252, EPI_ISL_3469253 | Lighthouse Lab in Milton Keynes | Wellcome Sanger Institute for the COVID-19 Genomics UK (COG-UK) Consortium | The Lighthouse Lab in Milton Keynes and Alex Alderton, Roberto Amato, Jeffrey Barrett, Sonia Goncalves, Ewan Harrison, David K. Jackson, Ian Johnston, Dominic Kwiatkowski, Cordelia Langford, John Sillitoe on behalf of the Wellcome Sanger Institute COVID-19 Surveillance Team |
| EPI_ISL_3469254, EPI_ISL_3469255, EPI_ISL_3469256 | Lighthouse Lab in Alderley Park | Wellcome Sanger Institute for the COVID-19 Genomics UK (COG-UK) Consortium | Jacquelyn Wynn, Mairead Hyland, The Lighthouse Lab in Alderley Park and Alex Alderton, Roberto Amato, Jeffrey Barrett, Sonia Goncalves, Ewan Harrison, David K. Jackson, Ian Johnston, Dominic Kwiatkowski, Cordelia Langford, John Sillitoe on behalf of the Wellcome Sanger Institute COVID-19 Surveillance Team |
| EPI_ISL_3469257 | Lighthouse Lab in Milton Keynes | Wellcome Sanger Institute for the COVID-19 Genomics UK (COG-UK) Consortium | The Lighthouse Lab in Milton Keynes and Alex Alderton, Roberto Amato, Jeffrey Barrett, Sonia Goncalves, Ewan Harrison, David K. Jackson, Ian Johnston, Dominic Kwiatkowski, Cordelia Langford, John Sillitoe on behalf of the Wellcome Sanger Institute COVID-19 Surveillance Team |
| EPI_ISL_3469258, EPI_ISL_3469259, EPI_ISL_3469260 | Lighthouse Lab in Alderley Park | Wellcome Sanger Institute for the COVID-19 Genomics UK (COG-UK) Consortium | Jacquelyn Wynn, Mairead Hyland, The Lighthouse Lab in Alderley Park and Alex Alderton, Roberto Amato, Jeffrey Barrett, Sonia Goncalves, Ewan Harrison, David K. Jackson, Ian Johnston, Dominic Kwiatkowski, Cordelia Langford, John Sillitoe on behalf of the Wellcome Sanger Institute COVID-19 Surveillance Team |
| EPI_ISL_3469261, EPI_ISL_3469262 | Berkshire and Surrey Pathology Services Lighthouse Laboratory | Wellcome Sanger Institute for the COVID-19 Genomics UK (COG-UK) Consortium | Berkshire and Surrey Pathology Services Lighthouse Laboratory and Alex Alderton, Roberto Amato, Jeffrey Barrett, Sonia Goncalves, Ewan Harrison, David K. Jackson, Ian Johnston, Dominic Kwiatkowski, Cordelia Langford, John Sillitoe on behalf of the Wellcome Sanger Institute COVID-19 Surveillance Team |
| EPI_ISL_3469263, EPI_ISL_3469264, EPI_ISL_3469265, EPI_ISL_3469266, EPI_ISL_3469267, | Lighthouse Lab in Milton Keynes | Wellcome Sanger Institute for the COVID-19 Genomics UK (COG-UK) Consortium | The Lighthouse Lab in Milton Keynes and Alex Alderton, Roberto Amato, Jeffrey Barrett, Sonia Goncalves, Ewan Harrison, David K. Jackson, Ian Johnston, Dominic Kwiatkowski, Cordelia Langford, John Sillitoe on behalf of the Wellcome Sanger Institute COVID-19 Surveillance Team |

| | | | |
|---|---|---|---|
| EPI_ISL_3469268, EPI_ISL_3469269, EPI_ISL_3469270 | | | |
| EPI_ISL_3469271, EPI_ISL_3469272 | Berkshire and Surrey Pathology Services Lighthouse Laboratory | Wellcome Sanger Institute for the COVID-19 Genomics UK (COG-UK) Consortium | Berkshire and Surrey Pathology Services Lighthouse Laboratory and Alex Alderton, Roberto Amato, Jeffrey Barrett, Sonia Goncalves, Ewan Harrison, David K. Jackson, Ian Johnston, Dominic Kwiatkowski, Cordelia Langford, John Sillitoe on behalf of the Wellcome Sanger Institute COVID-19 Surveillance Team |
| EPI_ISL_3469273 | Lighthouse Lab in Glasgow | Wellcome Sanger Institute for the COVID-19 Genomics UK (COG-UK) Consortium | Harper VanSteenhouse, Yumi Kasai, David Gray, Carol Clugston, Anna Dominiczak and Alex Alderton, Roberto Amato, Jeffrey Barrett, Sonia Goncalves, Ewan Harrison, David K. Jackson, Ian Johnston, Dominic Kwiatkowski, Cordelia Langford, John Sillitoe on behalf of the Wellcome Sanger Institute COVID-19 Surveillance Team |
| EPI_ISL_3469274 | Lighthouse Lab in Milton Keynes | Wellcome Sanger Institute for the COVID-19 Genomics UK (COG-UK) Consortium | The Lighthouse Lab in Milton Keynes and Alex Alderton, Roberto Amato, Jeffrey Barrett, Sonia Goncalves, Ewan Harrison, David K. Jackson, Ian Johnston, Dominic Kwiatkowski, Cordelia Langford, John Sillitoe on behalf of the Wellcome Sanger Institute COVID-19 Surveillance Team |
| EPI_ISL_3469275, EPI_ISL_3469276 | Lighthouse Lab in Glasgow | Wellcome Sanger Institute for the COVID-19 Genomics UK (COG-UK) Consortium | Harper VanSteenhouse, Yumi Kasai, David Gray, Carol Clugston, Anna Dominiczak and Alex Alderton, Roberto Amato, Jeffrey Barrett, Sonia Goncalves, Ewan Harrison, David K. Jackson, Ian Johnston, Dominic Kwiatkowski, Cordelia Langford, John Sillitoe on behalf of the Wellcome Sanger Institute COVID-19 Surveillance Team |
| EPI_ISL_3469277 | Lighthouse Lab in Milton Keynes | Wellcome Sanger Institute for the COVID-19 Genomics UK (COG-UK) Consortium | The Lighthouse Lab in Milton Keynes and Alex Alderton, Roberto Amato, Jeffrey Barrett, Sonia Goncalves, Ewan Harrison, David K. Jackson, Ian Johnston, Dominic Kwiatkowski, Cordelia Langford, John Sillitoe on behalf of the Wellcome Sanger Institute COVID-19 Surveillance Team |
| EPI_ISL_3469278 | Berkshire and Surrey Pathology Services Lighthouse Laboratory | Wellcome Sanger Institute for the COVID-19 Genomics UK (COG-UK) Consortium | Berkshire and Surrey Pathology Services Lighthouse Laboratory and Alex Alderton, Roberto Amato, Jeffrey Barrett, Sonia Goncalves, Ewan Harrison, David K. Jackson, Ian Johnston, Dominic Kwiatkowski, Cordelia Langford, John Sillitoe on behalf of the Wellcome Sanger Institute COVID-19 Surveillance Team |
| EPI_ISL_3469279 | Lighthouse Lab in Milton Keynes | Wellcome Sanger Institute for the COVID-19 Genomics UK (COG-UK) Consortium | The Lighthouse Lab in Milton Keynes and Alex Alderton, Roberto Amato, Jeffrey Barrett, Sonia Goncalves, Ewan Harrison, David K. Jackson, Ian Johnston, Dominic Kwiatkowski, Cordelia Langford, John Sillitoe on behalf of the Wellcome Sanger Institute COVID-19 Surveillance Team |
| EPI_ISL_3469280 | Lighthouse Lab in Glasgow | Wellcome Sanger Institute for the COVID-19 Genomics UK (COG-UK) Consortium | Harper VanSteenhouse, Yumi Kasai, David Gray, Carol Clugston, Anna Dominiczak and Alex Alderton, Roberto Amato, Jeffrey Barrett, Sonia Goncalves, Ewan Harrison, David K. Jackson, Ian Johnston, Dominic Kwiatkowski, Cordelia Langford, John Sillitoe on behalf of the Wellcome Sanger Institute COVID-19 Surveillance Team |
| EPI_ISL_3469281 | Lighthouse Lab in Milton Keynes | Wellcome Sanger Institute for the COVID-19 Genomics UK (COG-UK) Consortium | The Lighthouse Lab in Milton Keynes and Alex Alderton, Roberto Amato, Jeffrey Barrett, Sonia Goncalves, Ewan Harrison, David K. Jackson, Ian Johnston, Dominic Kwiatkowski, Cordelia Langford, John Sillitoe on behalf of the Wellcome Sanger Institute COVID-19 Surveillance Team |
| EPI_ISL_3469282 | Berkshire and Surrey Pathology Services Lighthouse Laboratory | Wellcome Sanger Institute for the COVID-19 Genomics UK (COG-UK) Consortium | Berkshire and Surrey Pathology Services Lighthouse Laboratory and Alex Alderton, Roberto Amato, Jeffrey Barrett, Sonia Goncalves, Ewan Harrison, David K. Jackson, Ian Johnston, Dominic Kwiatkowski, Cordelia Langford, John Sillitoe on behalf of the Wellcome Sanger Institute COVID-19 Surveillance Team |
| EPI_ISL_3469283, EPI_ISL_3469284 | Lighthouse Lab in Glasgow | Wellcome Sanger Institute for the COVID-19 Genomics UK (COG-UK) Consortium | Harper VanSteenhouse, Yumi Kasai, David Gray, Carol Clugston, Anna Dominiczak and Alex Alderton, Roberto Amato, Jeffrey Barrett, Sonia Goncalves, Ewan Harrison, David K. Jackson, Ian Johnston, Dominic Kwiatkowski, Cordelia Langford, John Sillitoe on behalf of the Wellcome Sanger Institute COVID-19 Surveillance Team |
| EPI_ISL_3469285, EPI_ISL_3469286 | Lighthouse Lab in Milton Keynes | Wellcome Sanger Institute for the COVID-19 Genomics UK (COG-UK) Consortium | The Lighthouse Lab in Milton Keynes and Alex Alderton, Roberto Amato, Jeffrey Barrett, Sonia Goncalves, Ewan Harrison, David K. Jackson, Ian Johnston, Dominic Kwiatkowski, Cordelia Langford, John Sillitoe on behalf of the Wellcome Sanger Institute COVID-19 Surveillance Team |
| EPI_ISL_3469287, EPI_ISL_3469288 | Lighthouse Lab in Glasgow | Wellcome Sanger Institute for the COVID-19 Genomics UK (COG-UK) Consortium | Harper VanSteenhouse, Yumi Kasai, David Gray, Carol Clugston, Anna Dominiczak and Alex Alderton, Roberto Amato, Jeffrey Barrett, Sonia Goncalves, Ewan Harrison, David K. Jackson, Ian Johnston, Dominic Kwiatkowski, Cordelia Langford, John Sillitoe on behalf of the Wellcome Sanger Institute COVID-19 Surveillance Team |
| EPI_ISL_3469289 | Lighthouse Lab in Milton Keynes | Wellcome Sanger Institute for the COVID-19 Genomics UK (COG-UK) Consortium | The Lighthouse Lab in Milton Keynes and Alex Alderton, Roberto Amato, Jeffrey Barrett, Sonia Goncalves, Ewan Harrison, David K. Jackson, Ian Johnston, Dominic Kwiatkowski, Cordelia Langford, John Sillitoe on behalf of the Wellcome Sanger Institute COVID-19 Surveillance Team |
| EPI_ISL_3469290 | Lighthouse Lab in Glasgow | Wellcome Sanger Institute for the COVID-19 Genomics UK (COG-UK) Consortium | Harper VanSteenhouse, Yumi Kasai, David Gray, Carol Clugston, Anna Dominiczak and Alex Alderton, Roberto Amato, Jeffrey Barrett, Sonia Goncalves, Ewan Harrison, David K. Jackson, Ian Johnston, Dominic Kwiatkowski, Cordelia Langford, John Sillitoe on behalf of the Wellcome Sanger Institute COVID-19 Surveillance Team |
| EPI_ISL_3469291, EPI_ISL_3469292 | Lighthouse Lab in Milton Keynes | Wellcome Sanger Institute for the COVID-19 Genomics UK (COG-UK) Consortium | The Lighthouse Lab in Milton Keynes and Alex Alderton, Roberto Amato, Jeffrey Barrett, Sonia Goncalves, Ewan Harrison, David K. Jackson, Ian Johnston, Dominic Kwiatkowski, Cordelia Langford, John Sillitoe on behalf of the Wellcome Sanger Institute COVID-19 Surveillance Team |
| EPI_ISL_3469293 | Berkshire and Surrey Pathology Services Lighthouse Laboratory | Wellcome Sanger Institute for the COVID-19 Genomics UK (COG-UK) Consortium | Berkshire and Surrey Pathology Services Lighthouse Laboratory and Alex Alderton, Roberto Amato, Jeffrey Barrett, Sonia Goncalves, Ewan Harrison, David K. Jackson, Ian Johnston, Dominic Kwiatkowski, Cordelia Langford, John Sillitoe on behalf of the Wellcome Sanger Institute COVID-19 Surveillance Team |
| EPI_ISL_3469294 | Lighthouse Lab in Milton Keynes | Wellcome Sanger Institute for the COVID-19 Genomics UK (COG-UK) Consortium | The Lighthouse Lab in Milton Keynes and Alex Alderton, Roberto Amato, Jeffrey Barrett, Sonia Goncalves, Ewan Harrison, David K. Jackson, Ian Johnston, Dominic Kwiatkowski, Cordelia Langford, John Sillitoe on behalf of the Wellcome Sanger Institute COVID-19 Surveillance Team |
| EPI_ISL_3469295 | Lighthouse Lab in Glasgow | Wellcome Sanger Institute for the COVID-19 Genomics UK (COG-UK) Consortium | Harper VanSteenhouse, Yumi Kasai, David Gray, Carol Clugston, Anna Dominiczak and Alex Alderton, Roberto Amato, Jeffrey Barrett, Sonia Goncalves, Ewan Harrison, David K. Jackson, Ian Johnston, Dominic Kwiatkowski, Cordelia Langford, John Sillitoe on behalf of the Wellcome Sanger Institute COVID-19 Surveillance Team |
| EPI_ISL_3469296, EPI_ISL_3469297, EPI_ISL_3469298, EPI_ISL_3469299, EPI_ISL_3469300, EPI_ISL_3469301 | Lighthouse Lab in Milton Keynes | Wellcome Sanger Institute for the COVID-19 Genomics UK (COG-UK) Consortium | The Lighthouse Lab in Milton Keynes and Alex Alderton, Roberto Amato, Jeffrey Barrett, Sonia Goncalves, Ewan Harrison, David K. Jackson, Ian Johnston, Dominic Kwiatkowski, Cordelia Langford, John Sillitoe on behalf of the Wellcome Sanger Institute COVID-19 Surveillance Team |
| EPI_ISL_3469302, EPI_ISL_3469303 | Berkshire and Surrey Pathology Services Lighthouse Laboratory | Wellcome Sanger Institute for the COVID-19 Genomics UK (COG-UK) Consortium | Berkshire and Surrey Pathology Services Lighthouse Laboratory and Alex Alderton, Roberto Amato, Jeffrey Barrett, Sonia Goncalves, Ewan Harrison, David K. Jackson, Ian Johnston, Dominic Kwiatkowski, Cordelia Langford, John Sillitoe on behalf of the Wellcome Sanger Institute COVID-19 Surveillance Team |
| EPI_ISL_3469304 | Lighthouse Lab in Milton Keynes | Wellcome Sanger Institute for the COVID-19 Genomics UK (COG-UK) Consortium | The Lighthouse Lab in Milton Keynes and Alex Alderton, Roberto Amato, Jeffrey Barrett, Sonia Goncalves, Ewan Harrison, David K. Jackson, Ian Johnston, Dominic Kwiatkowski, Cordelia Langford, John Sillitoe on behalf of the Wellcome Sanger Institute COVID-19 Surveillance Team |
| EPI_ISL_3469305 | Lighthouse Lab in Glasgow | Wellcome Sanger Institute for the COVID-19 Genomics UK (COG-UK) Consortium | Harper VanSteenhouse, Yumi Kasai, David Gray, Carol Clugston, Anna Dominiczak and Alex Alderton, Roberto Amato, Jeffrey Barrett, Sonia Goncalves, Ewan Harrison, David K. Jackson, Ian Johnston, Dominic Kwiatkowski, Cordelia Langford, John Sillitoe on behalf of the Wellcome Sanger Institute COVID-19 Surveillance Team |
| EPI_ISL_3469306 | Berkshire and Surrey Pathology Services Lighthouse Laboratory | Wellcome Sanger Institute for the COVID-19 Genomics UK (COG-UK) Consortium | Berkshire and Surrey Pathology Services Lighthouse Laboratory and Alex Alderton, Roberto Amato, Jeffrey Barrett, Sonia Goncalves, Ewan Harrison, David K. Jackson, Ian Johnston, Dominic Kwiatkowski, Cordelia Langford, John Sillitoe on behalf of the Wellcome Sanger Institute COVID-19 Surveillance Team |
| EPI_ISL_3469307 | Lighthouse Lab in Glasgow | Wellcome Sanger Institute for the COVID-19 Genomics UK (COG-UK) Consortium | Harper VanSteenhouse, Yumi Kasai, David Gray, Carol Clugston, Anna Dominiczak and Alex Alderton, Roberto Amato, Jeffrey Barrett, Sonia Goncalves, Ewan Harrison, David K. Jackson, Ian Johnston, Dominic Kwiatkowski, Cordelia Langford, John Sillitoe on behalf of the Wellcome Sanger Institute COVID-19 Surveillance Team |
| EPI_ISL_3469308 | Berkshire and Surrey Pathology Services Lighthouse Laboratory | Wellcome Sanger Institute for the COVID-19 Genomics UK (COG-UK) Consortium | Berkshire and Surrey Pathology Services Lighthouse Laboratory and Alex Alderton, Roberto Amato, Jeffrey Barrett, Sonia Goncalves, Ewan Harrison, David K. Jackson, Ian Johnston, Dominic Kwiatkowski, Cordelia Langford, John Sillitoe on behalf of the Wellcome Sanger Institute COVID-19 Surveillance Team |
| EPI_ISL_3469309 | Lighthouse Lab in Milton Keynes | Wellcome Sanger Institute for the COVID-19 Genomics UK (COG-UK) Consortium | The Lighthouse Lab in Milton Keynes and Alex Alderton, Roberto Amato, Jeffrey Barrett, Sonia Goncalves, Ewan Harrison, David K. Jackson, Ian Johnston, Dominic Kwiatkowski, Cordelia Langford, John Sillitoe on behalf of the Wellcome Sanger Institute COVID-19 Surveillance Team |
| EPI_ISL_3469310, EPI_ISL_3469311 | Berkshire and Surrey Pathology Services Lighthouse Laboratory | Wellcome Sanger Institute for the COVID-19 Genomics UK (COG-UK) Consortium | Berkshire and Surrey Pathology Services Lighthouse Laboratory and Alex Alderton, Roberto Amato, Jeffrey Barrett, Sonia Goncalves, Ewan Harrison, David K. Jackson, Ian Johnston, Dominic Kwiatkowski, Cordelia Langford, John Sillitoe on behalf of the Wellcome Sanger Institute COVID-19 Surveillance Team |
| EPI_ISL_3469312 | Lighthouse Lab in Milton Keynes | Wellcome Sanger Institute for the COVID-19 Genomics UK (COG-UK) Consortium | The Lighthouse Lab in Milton Keynes and Alex Alderton, Roberto Amato, Jeffrey Barrett, Sonia Goncalves, Ewan Harrison, David K. Jackson, Ian Johnston, Dominic Kwiatkowski, Cordelia Langford, John Sillitoe on behalf of the Wellcome Sanger Institute COVID-19 Surveillance Team |
| EPI_ISL_3469313, EPI_ISL_3469314 | Lighthouse Lab in Glasgow | Wellcome Sanger Institute for the COVID-19 Genomics UK (COG-UK) Consortium | Harper VanSteenhouse, Yumi Kasai, David Gray, Carol Clugston, Anna Dominiczak and Alex Alderton, Roberto Amato, Jeffrey Barrett, Sonia Goncalves, Ewan Harrison, David K. Jackson, Ian Johnston, Dominic Kwiatkowski, Cordelia Langford, John Sillitoe on behalf of the Wellcome Sanger Institute COVID-19 Surveillance Team |
| EPI_ISL_3469315, EPI_ISL_3469316 | Lighthouse Lab in Milton Keynes | Wellcome Sanger Institute for the COVID-19 Genomics UK (COG-UK) Consortium | The Lighthouse Lab in Milton Keynes and Alex Alderton, Roberto Amato, Jeffrey Barrett, Sonia Goncalves, Ewan Harrison, David K. Jackson, Ian Johnston, Dominic Kwiatkowski, Cordelia Langford, John Sillitoe on behalf of the Wellcome Sanger Institute COVID-19 Surveillance Team |
| EPI_ISL_3469317, EPI_ISL_3469318 | Berkshire and Surrey Pathology Services Lighthouse Laboratory | Wellcome Sanger Institute for the COVID-19 Genomics UK (COG-UK) Consortium | Berkshire and Surrey Pathology Services Lighthouse Laboratory and Alex Alderton, Roberto Amato, Jeffrey Barrett, Sonia Goncalves, Ewan Harrison, David K. Jackson, Ian Johnston, Dominic Kwiatkowski, Cordelia Langford, John Sillitoe on behalf of the Wellcome Sanger Institute COVID-19 Surveillance Team |
| EPI_ISL_3469319 | Lighthouse Lab in Milton Keynes | Wellcome Sanger Institute for the COVID-19 Genomics UK (COG-UK) Consortium | The Lighthouse Lab in Milton Keynes and Alex Alderton, Roberto Amato, Jeffrey Barrett, Sonia Goncalves, Ewan Harrison, David K. Jackson, Ian Johnston, Dominic Kwiatkowski, Cordelia Langford, John Sillitoe on behalf of the Wellcome Sanger Institute COVID-19 Surveillance Team |
| EPI_ISL_3469320 | Berkshire and Surrey Pathology Services Lighthouse Laboratory | Wellcome Sanger Institute for the COVID-19 Genomics UK (COG-UK) Consortium | Berkshire and Surrey Pathology Services Lighthouse Laboratory and Alex Alderton, Roberto Amato, Jeffrey Barrett, Sonia Goncalves, Ewan Harrison, David K. Jackson, Ian Johnston, Dominic Kwiatkowski, Cordelia Langford, John Sillitoe on behalf of the Wellcome Sanger Institute COVID-19 Surveillance Team |
| EPI_ISL_3469321 | Lighthouse Lab in Glasgow | Wellcome Sanger Institute for the COVID-19 Genomics UK (COG-UK) Consortium | Harper VanSteenhouse, Yumi Kasai, David Gray, Carol Clugston, Anna Dominiczak and Alex Alderton, Roberto Amato, Jeffrey Barrett, Sonia Goncalves, Ewan Harrison, David K. Jackson, Ian Johnston, Dominic Kwiatkowski, Cordelia Langford, John Sillitoe on behalf of the Wellcome Sanger Institute COVID-19 Surveillance Team |
| EPI_ISL_3469322, EPI_ISL_3469323, EPI_ISL_3469324 | Lighthouse Lab in Milton Keynes | Wellcome Sanger Institute for the COVID-19 Genomics UK (COG-UK) Consortium | The Lighthouse Lab in Milton Keynes and Alex Alderton, Roberto Amato, Jeffrey Barrett, Sonia Goncalves, Ewan Harrison, David K. Jackson, Ian Johnston, Dominic Kwiatkowski, Cordelia Langford, John Sillitoe on behalf of the Wellcome Sanger Institute COVID-19 Surveillance Team |
| EPI_ISL_3469325 | Lighthouse Lab in Glasgow | Wellcome Sanger Institute for the COVID-19 Genomics UK (COG-UK) Consortium | Harper VanSteenhouse, Yumi Kasai, David Gray, Carol Clugston, Anna Dominiczak and Alex Alderton, Roberto Amato, Jeffrey Barrett, Sonia Goncalves, Ewan Harrison, David K. Jackson, Ian Johnston, Dominic Kwiatkowski, Cordelia Langford, John Sillitoe on behalf of the Wellcome Sanger Institute COVID-19 Surveillance Team |
| EPI_ISL_3469326, EPI_ISL_3469327 | Lighthouse Lab in Milton Keynes | Wellcome Sanger Institute for the COVID-19 Genomics UK (COG-UK) Consortium | The Lighthouse Lab in Milton Keynes and Alex Alderton, Roberto Amato, Jeffrey Barrett, Sonia Goncalves, Ewan Harrison, David K. Jackson, Ian Johnston, Dominic Kwiatkowski, Cordelia Langford, John Sillitoe on behalf of the Wellcome Sanger Institute COVID-19 Surveillance Team |
| EPI_ISL_3469328 | Lighthouse Lab in Glasgow | Wellcome Sanger Institute for the COVID-19 Genomics UK (COG-UK) Consortium | Harper VanSteenhouse, Yumi Kasai, David Gray, Carol Clugston, Anna Dominiczak and Alex Alderton, Roberto Amato, Jeffrey Barrett, Sonia Goncalves, Ewan Harrison, David K. Jackson, Ian Johnston, Dominic Kwiatkowski, Cordelia Langford, John Sillitoe on behalf of the Wellcome Sanger Institute COVID-19 Surveillance Team |
| EPI_ISL_3469329 | Berkshire and Surrey Pathology Services Lighthouse Laboratory | Wellcome Sanger Institute for the COVID-19 Genomics UK (COG-UK) Consortium | Berkshire and Surrey Pathology Services Lighthouse Laboratory and Alex Alderton, Roberto Amato, Jeffrey Barrett, Sonia Goncalves, Ewan Harrison, David K. Jackson, Ian Johnston, Dominic Kwiatkowski, Cordelia Langford, John Sillitoe on behalf of the Wellcome Sanger Institute COVID-19 Surveillance Team |
| EPI_ISL_3469330 | Lighthouse Lab in Milton Keynes | Wellcome Sanger Institute for the COVID-19 Genomics UK (COG-UK) Consortium | The Lighthouse Lab in Milton Keynes and Alex Alderton, Roberto Amato, Jeffrey Barrett, Sonia Goncalves, Ewan Harrison, David K. Jackson, Ian Johnston, Dominic Kwiatkowski, Cordelia Langford, John Sillitoe on behalf of the Wellcome Sanger Institute COVID-19 Surveillance Team |
| EPI_ISL_3469331 | Lighthouse Lab in Glasgow | Wellcome Sanger Institute for the COVID-19 Genomics UK (COG-UK) | Harper VanSteenhouse, Yumi Kasai, David Gray, Carol Clugston, Anna Dominiczak and Alex Alderton, Roberto Amato, Jeffrey Barrett, Sonia Goncalves, Ewan Harrison, David K. Jackson, Ian Johnston, Dominic Kwiatkowski, Cordelia Langford, |

| | | | |
|---|---|---|---|
| | | Consortium | John Sillitoe on behalf of the Wellcome Sanger Institute COVID-19 Surveillance Team |
| EPI_ISL_3469332 | Berkshire and Surrey Pathology Services Lighthouse Laboratory | Wellcome Sanger Institute for the COVID-19 Genomics UK (COG-UK) Consortium | Berkshire and Surrey Pathology Services Lighthouse Laboratory and Alex Alderton, Roberto Amato, Jeffrey Barrett, Sonia Goncalves, Ewan Harrison, David K. Jackson, Ian Johnston, Dominic Kwiatkowski, Cordelia Langford, John Sillitoe on behalf of the Wellcome Sanger Institute COVID-19 Surveillance Team |
| EPI_ISL_3469333 | Lighthouse Lab in Glasgow | Wellcome Sanger Institute for the COVID-19 Genomics UK (COG-UK) Consortium | Harper VanSteenhouse, Yumi Kasai, David Gray, Carol Clugston, Anna Dominiczak and Alex Alderton, Roberto Amato, Jeffrey Barrett, Sonia Goncalves, Ewan Harrison, David K. Jackson, Ian Johnston, Dominic Kwiatkowski, Cordelia Langford, John Sillitoe on behalf of the Wellcome Sanger Institute COVID-19 Surveillance Team |
| EPI_ISL_3469334, EPI_ISL_3469335, EPI_ISL_3469336, EPI_ISL_3469337, EPI_ISL_3469338, EPI_ISL_3469339 | Lighthouse Lab in Milton Keynes | Wellcome Sanger Institute for the COVID-19 Genomics UK (COG-UK) Consortium | The Lighthouse Lab in Milton Keynes and Alex Alderton, Roberto Amato, Jeffrey Barrett, Sonia Goncalves, Ewan Harrison, David K. Jackson, Ian Johnston, Dominic Kwiatkowski, Cordelia Langford, John Sillitoe on behalf of the Wellcome Sanger Institute COVID-19 Surveillance Team |
| EPI_ISL_3469340 | Lighthouse Lab in Glasgow | Wellcome Sanger Institute for the COVID-19 Genomics UK (COG-UK) Consortium | Harper VanSteenhouse, Yumi Kasai, David Gray, Carol Clugston, Anna Dominiczak and Alex Alderton, Roberto Amato, Jeffrey Barrett, Sonia Goncalves, Ewan Harrison, David K. Jackson, Ian Johnston, Dominic Kwiatkowski, Cordelia Langford, John Sillitoe on behalf of the Wellcome Sanger Institute COVID-19 Surveillance Team |
| EPI_ISL_3469341 | Berkshire and Surrey Pathology Services Lighthouse Laboratory | Wellcome Sanger Institute for the COVID-19 Genomics UK (COG-UK) Consortium | Berkshire and Surrey Pathology Services Lighthouse Laboratory and Alex Alderton, Roberto Amato, Jeffrey Barrett, Sonia Goncalves, Ewan Harrison, David K. Jackson, Ian Johnston, Dominic Kwiatkowski, Cordelia Langford, John Sillitoe on behalf of the Wellcome Sanger Institute COVID-19 Surveillance Team |
| EPI_ISL_3469342 | Lighthouse Lab in Milton Keynes | Wellcome Sanger Institute for the COVID-19 Genomics UK (COG-UK) Consortium | The Lighthouse Lab in Milton Keynes and Alex Alderton, Roberto Amato, Jeffrey Barrett, Sonia Goncalves, Ewan Harrison, David K. Jackson, Ian Johnston, Dominic Kwiatkowski, Cordelia Langford, John Sillitoe on behalf of the Wellcome Sanger Institute COVID-19 Surveillance Team |
| EPI_ISL_3469343 | Lighthouse Lab in Glasgow | Wellcome Sanger Institute for the COVID-19 Genomics UK (COG-UK) Consortium | Harper VanSteenhouse, Yumi Kasai, David Gray, Carol Clugston, Anna Dominiczak and Alex Alderton, Roberto Amato, Jeffrey Barrett, Sonia Goncalves, Ewan Harrison, David K. Jackson, Ian Johnston, Dominic Kwiatkowski, Cordelia Langford, John Sillitoe on behalf of the Wellcome Sanger Institute COVID-19 Surveillance Team |
| EPI_ISL_3469344, EPI_ISL_3469345, EPI_ISL_3469346 | Lighthouse Lab in Milton Keynes | Wellcome Sanger Institute for the COVID-19 Genomics UK (COG-UK) Consortium | The Lighthouse Lab in Milton Keynes and Alex Alderton, Roberto Amato, Jeffrey Barrett, Sonia Goncalves, Ewan Harrison, David K. Jackson, Ian Johnston, Dominic Kwiatkowski, Cordelia Langford, John Sillitoe on behalf of the Wellcome Sanger Institute COVID-19 Surveillance Team |
| EPI_ISL_3469347, EPI_ISL_3469348 | Lighthouse Lab in Glasgow | Wellcome Sanger Institute for the COVID-19 Genomics UK (COG-UK) Consortium | Harper VanSteenhouse, Yumi Kasai, David Gray, Carol Clugston, Anna Dominiczak and Alex Alderton, Roberto Amato, Jeffrey Barrett, Sonia Goncalves, Ewan Harrison, David K. Jackson, Ian Johnston, Dominic Kwiatkowski, Cordelia Langford, John Sillitoe on behalf of the Wellcome Sanger Institute COVID-19 Surveillance Team |
| EPI_ISL_3469349 | Berkshire and Surrey Pathology Services Lighthouse Laboratory | Wellcome Sanger Institute for the COVID-19 Genomics UK (COG-UK) Consortium | Berkshire and Surrey Pathology Services Lighthouse Laboratory and Alex Alderton, Roberto Amato, Jeffrey Barrett, Sonia Goncalves, Ewan Harrison, David K. Jackson, Ian Johnston, Dominic Kwiatkowski, Cordelia Langford, John Sillitoe on behalf of the Wellcome Sanger Institute COVID-19 Surveillance Team |
| EPI_ISL_3469350, EPI_ISL_3469351, EPI_ISL_3469352, EPI_ISL_3469353 | Lighthouse Lab in Milton Keynes | Wellcome Sanger Institute for the COVID-19 Genomics UK (COG-UK) Consortium | The Lighthouse Lab in Milton Keynes and Alex Alderton, Roberto Amato, Jeffrey Barrett, Sonia Goncalves, Ewan Harrison, David K. Jackson, Ian Johnston, Dominic Kwiatkowski, Cordelia Langford, John Sillitoe on behalf of the Wellcome Sanger Institute COVID-19 Surveillance Team |
| EPI_ISL_3469354, EPI_ISL_3469355 | Berkshire and Surrey Pathology Services Lighthouse Laboratory | Wellcome Sanger Institute for the COVID-19 Genomics UK (COG-UK) Consortium | Berkshire and Surrey Pathology Services Lighthouse Laboratory and Alex Alderton, Roberto Amato, Jeffrey Barrett, Sonia Goncalves, Ewan Harrison, David K. Jackson, Ian Johnston, Dominic Kwiatkowski, Cordelia Langford, John Sillitoe on behalf of the Wellcome Sanger Institute COVID-19 Surveillance Team |
| EPI_ISL_3469356 | Lighthouse Lab in Milton Keynes | Wellcome Sanger Institute for the COVID-19 Genomics UK (COG-UK) Consortium | The Lighthouse Lab in Milton Keynes and Alex Alderton, Roberto Amato, Jeffrey Barrett, Sonia Goncalves, Ewan Harrison, David K. Jackson, Ian Johnston, Dominic Kwiatkowski, Cordelia Langford, John Sillitoe on behalf of the Wellcome Sanger Institute COVID-19 Surveillance Team |
| EPI_ISL_3469357 | Lighthouse Lab in Glasgow | Wellcome Sanger Institute for the COVID-19 Genomics UK (COG-UK) Consortium | Harper VanSteenhouse, Yumi Kasai, David Gray, Carol Clugston, Anna Dominiczak and Alex Alderton, Roberto Amato, Jeffrey Barrett, Sonia Goncalves, Ewan Harrison, David K. Jackson, Ian Johnston, Dominic Kwiatkowski, Cordelia Langford, John Sillitoe on behalf of the Wellcome Sanger Institute COVID-19 Surveillance Team |
| EPI_ISL_3469358 | Berkshire and Surrey Pathology Services Lighthouse Laboratory | Wellcome Sanger Institute for the COVID-19 Genomics UK (COG-UK) Consortium | Berkshire and Surrey Pathology Services Lighthouse Laboratory and Alex Alderton, Roberto Amato, Jeffrey Barrett, Sonia Goncalves, Ewan Harrison, David K. Jackson, Ian Johnston, Dominic Kwiatkowski, Cordelia Langford, John Sillitoe on behalf of the Wellcome Sanger Institute COVID-19 Surveillance Team |
| EPI_ISL_3469359 | Lighthouse Lab in Glasgow | Wellcome Sanger Institute for the COVID-19 Genomics UK (COG-UK) Consortium | Harper VanSteenhouse, Yumi Kasai, David Gray, Carol Clugston, Anna Dominiczak and Alex Alderton, Roberto Amato, Jeffrey Barrett, Sonia Goncalves, Ewan Harrison, David K. Jackson, Ian Johnston, Dominic Kwiatkowski, Cordelia Langford, John Sillitoe on behalf of the Wellcome Sanger Institute COVID-19 Surveillance Team |
| EPI_ISL_3469360, EPI_ISL_3469361, EPI_ISL_3469362 | Berkshire and Surrey Pathology Services Lighthouse Laboratory | Wellcome Sanger Institute for the COVID-19 Genomics UK (COG-UK) Consortium | Berkshire and Surrey Pathology Services Lighthouse Laboratory and Alex Alderton, Roberto Amato, Jeffrey Barrett, Sonia Goncalves, Ewan Harrison, David K. Jackson, Ian Johnston, Dominic Kwiatkowski, Cordelia Langford, John Sillitoe on behalf of the Wellcome Sanger Institute COVID-19 Surveillance Team |
| EPI_ISL_3469363, EPI_ISL_3469364, EPI_ISL_3469365 | Lighthouse Lab in Milton Keynes | Wellcome Sanger Institute for the COVID-19 Genomics UK (COG-UK) Consortium | The Lighthouse Lab in Milton Keynes and Alex Alderton, Roberto Amato, Jeffrey Barrett, Sonia Goncalves, Ewan Harrison, David K. Jackson, Ian Johnston, Dominic Kwiatkowski, Cordelia Langford, John Sillitoe on behalf of the Wellcome Sanger Institute COVID-19 Surveillance Team |
| EPI_ISL_3469366, EPI_ISL_3469367 | Berkshire and Surrey Pathology Services Lighthouse Laboratory | Wellcome Sanger Institute for the COVID-19 Genomics UK (COG-UK) Consortium | Berkshire and Surrey Pathology Services Lighthouse Laboratory and Alex Alderton, Roberto Amato, Jeffrey Barrett, Sonia Goncalves, Ewan Harrison, David K. Jackson, Ian Johnston, Dominic Kwiatkowski, Cordelia Langford, John Sillitoe on behalf of the Wellcome Sanger Institute COVID-19 Surveillance Team |
| EPI_ISL_3469368 | Lighthouse Lab in Glasgow | Wellcome Sanger Institute for the COVID-19 Genomics UK (COG-UK) Consortium | Harper VanSteenhouse, Yumi Kasai, David Gray, Carol Clugston, Anna Dominiczak and Alex Alderton, Roberto Amato, Jeffrey Barrett, Sonia Goncalves, Ewan Harrison, David K. Jackson, Ian Johnston, Dominic Kwiatkowski, Cordelia Langford, John Sillitoe on behalf of the Wellcome Sanger Institute COVID-19 Surveillance Team |
| EPI_ISL_3469369 | Lighthouse Lab in Milton Keynes | Wellcome Sanger Institute for the COVID-19 Genomics UK (COG-UK) Consortium | The Lighthouse Lab in Milton Keynes and Alex Alderton, Roberto Amato, Jeffrey Barrett, Sonia Goncalves, Ewan Harrison, David K. Jackson, Ian Johnston, Dominic Kwiatkowski, Cordelia Langford, John Sillitoe on behalf of the Wellcome Sanger Institute COVID-19 Surveillance Team |
| EPI_ISL_3469370, EPI_ISL_3469371 | Lighthouse Lab in Glasgow | Wellcome Sanger Institute for the COVID-19 Genomics UK (COG-UK) Consortium | Harper VanSteenhouse, Yumi Kasai, David Gray, Carol Clugston, Anna Dominiczak and Alex Alderton, Roberto Amato, Jeffrey Barrett, Sonia Goncalves, Ewan Harrison, David K. Jackson, Ian Johnston, Dominic Kwiatkowski, Cordelia Langford, John Sillitoe on behalf of the Wellcome Sanger Institute COVID-19 Surveillance Team |
| EPI_ISL_3469372 | Berkshire and Surrey Pathology Services Lighthouse Laboratory | Wellcome Sanger Institute for the COVID-19 Genomics UK (COG-UK) Consortium | Berkshire and Surrey Pathology Services Lighthouse Laboratory and Alex Alderton, Roberto Amato, Jeffrey Barrett, Sonia Goncalves, Ewan Harrison, David K. Jackson, Ian Johnston, Dominic Kwiatkowski, Cordelia Langford, John Sillitoe on behalf of the Wellcome Sanger Institute COVID-19 Surveillance Team |
| EPI_ISL_3469373 | Lighthouse Lab in Glasgow | Wellcome Sanger Institute for the COVID-19 Genomics UK (COG-UK) Consortium | Harper VanSteenhouse, Yumi Kasai, David Gray, Carol Clugston, Anna Dominiczak and Alex Alderton, Roberto Amato, Jeffrey Barrett, Sonia Goncalves, Ewan Harrison, David K. Jackson, Ian Johnston, Dominic Kwiatkowski, Cordelia Langford, John Sillitoe on behalf of the Wellcome Sanger Institute COVID-19 Surveillance Team |
| EPI_ISL_3469374 | Berkshire and Surrey Pathology Services Lighthouse Laboratory | Wellcome Sanger Institute for the COVID-19 Genomics UK (COG-UK) Consortium | Berkshire and Surrey Pathology Services Lighthouse Laboratory and Alex Alderton, Roberto Amato, Jeffrey Barrett, Sonia Goncalves, Ewan Harrison, David K. Jackson, Ian Johnston, Dominic Kwiatkowski, Cordelia Langford, John Sillitoe on behalf of the Wellcome Sanger Institute COVID-19 Surveillance Team |
| EPI_ISL_3469375, EPI_ISL_3469376, EPI_ISL_3469377 | Lighthouse Lab in Milton Keynes | Wellcome Sanger Institute for the COVID-19 Genomics UK (COG-UK) Consortium | The Lighthouse Lab in Milton Keynes and Alex Alderton, Roberto Amato, Jeffrey Barrett, Sonia Goncalves, Ewan Harrison, David K. Jackson, Ian Johnston, Dominic Kwiatkowski, Cordelia Langford, John Sillitoe on behalf of the Wellcome Sanger Institute COVID-19 Surveillance Team |
| EPI_ISL_3469378 | Berkshire and Surrey Pathology Services Lighthouse Laboratory | Wellcome Sanger Institute for the COVID-19 Genomics UK (COG-UK) Consortium | Berkshire and Surrey Pathology Services Lighthouse Laboratory and Alex Alderton, Roberto Amato, Jeffrey Barrett, Sonia Goncalves, Ewan Harrison, David K. Jackson, Ian Johnston, Dominic Kwiatkowski, Cordelia Langford, John Sillitoe on behalf of the Wellcome Sanger Institute COVID-19 Surveillance Team |
| EPI_ISL_3469379, EPI_ISL_3469380 | Lighthouse Lab in Milton Keynes | Wellcome Sanger Institute for the COVID-19 Genomics UK (COG-UK) Consortium | The Lighthouse Lab in Milton Keynes and Alex Alderton, Roberto Amato, Jeffrey Barrett, Sonia Goncalves, Ewan Harrison, David K. Jackson, Ian Johnston, Dominic Kwiatkowski, Cordelia Langford, John Sillitoe on behalf of the Wellcome Sanger Institute COVID-19 Surveillance Team |
| EPI_ISL_3469381, EPI_ISL_3469382 | Berkshire and Surrey Pathology Services Lighthouse Laboratory | Wellcome Sanger Institute for the COVID-19 Genomics UK (COG-UK) Consortium | Berkshire and Surrey Pathology Services Lighthouse Laboratory and Alex Alderton, Roberto Amato, Jeffrey Barrett, Sonia Goncalves, Ewan Harrison, David K. Jackson, Ian Johnston, Dominic Kwiatkowski, Cordelia Langford, John Sillitoe on behalf of the Wellcome Sanger Institute COVID-19 Surveillance Team |
| EPI_ISL_3469383, EPI_ISL_3469384 | Lighthouse Lab in Milton Keynes | Wellcome Sanger Institute for the COVID-19 Genomics UK (COG-UK) Consortium | The Lighthouse Lab in Milton Keynes and Alex Alderton, Roberto Amato, Jeffrey Barrett, Sonia Goncalves, Ewan Harrison, David K. Jackson, Ian Johnston, Dominic Kwiatkowski, Cordelia Langford, John Sillitoe on behalf of the Wellcome Sanger Institute COVID-19 Surveillance Team |
| EPI_ISL_3469385 | Berkshire and Surrey Pathology Services Lighthouse Laboratory | Wellcome Sanger Institute for the COVID-19 Genomics UK (COG-UK) Consortium | Berkshire and Surrey Pathology Services Lighthouse Laboratory and Alex Alderton, Roberto Amato, Jeffrey Barrett, Sonia Goncalves, Ewan Harrison, David K. Jackson, Ian Johnston, Dominic Kwiatkowski, Cordelia Langford, John Sillitoe on behalf of the Wellcome Sanger Institute COVID-19 Surveillance Team |
| EPI_ISL_3469386 | Lighthouse Lab in Milton Keynes | Wellcome Sanger Institute for the COVID-19 Genomics UK (COG-UK) Consortium | The Lighthouse Lab in Milton Keynes and Alex Alderton, Roberto Amato, Jeffrey Barrett, Sonia Goncalves, Ewan Harrison, David K. Jackson, Ian Johnston, Dominic Kwiatkowski, Cordelia Langford, John Sillitoe on behalf of the Wellcome Sanger Institute COVID-19 Surveillance Team |
| EPI_ISL_3469387 | Berkshire and Surrey Pathology Services Lighthouse Laboratory | Wellcome Sanger Institute for the COVID-19 Genomics UK (COG-UK) Consortium | Berkshire and Surrey Pathology Services Lighthouse Laboratory and Alex Alderton, Roberto Amato, Jeffrey Barrett, Sonia Goncalves, Ewan Harrison, David K. Jackson, Ian Johnston, Dominic Kwiatkowski, Cordelia Langford, John Sillitoe on behalf of the Wellcome Sanger Institute COVID-19 Surveillance Team |
| EPI_ISL_3469388, EPI_ISL_3469389, EPI_ISL_3469390 | Lighthouse Lab in Milton Keynes | Wellcome Sanger Institute for the COVID-19 Genomics UK (COG-UK) Consortium | The Lighthouse Lab in Milton Keynes and Alex Alderton, Roberto Amato, Jeffrey Barrett, Sonia Goncalves, Ewan Harrison, David K. Jackson, Ian Johnston, Dominic Kwiatkowski, Cordelia Langford, John Sillitoe on behalf of the Wellcome Sanger Institute COVID-19 Surveillance Team |
| EPI_ISL_3469391, EPI_ISL_3469392, EPI_ISL_3469393, EPI_ISL_3469394, EPI_ISL_3469395 | Lighthouse Lab in Glasgow | Wellcome Sanger Institute for the COVID-19 Genomics UK (COG-UK) Consortium | Harper VanSteenhouse, Yumi Kasai, David Gray, Carol Clugston, Anna Dominiczak and Alex Alderton, Roberto Amato, Jeffrey Barrett, Sonia Goncalves, Ewan Harrison, David K. Jackson, Ian Johnston, Dominic Kwiatkowski, Cordelia Langford, John Sillitoe on behalf of the Wellcome Sanger Institute COVID-19 Surveillance Team |
| EPI_ISL_3469396 | Lighthouse Lab in Milton Keynes | Wellcome Sanger Institute for the COVID-19 Genomics UK (COG-UK) Consortium | The Lighthouse Lab in Milton Keynes and Alex Alderton, Roberto Amato, Jeffrey Barrett, Sonia Goncalves, Ewan Harrison, David K. Jackson, Ian Johnston, Dominic Kwiatkowski, Cordelia Langford, John Sillitoe on behalf of the Wellcome Sanger Institute COVID-19 Surveillance Team |
| EPI_ISL_3469397, EPI_ISL_3469398 | Berkshire and Surrey Pathology Services Lighthouse Laboratory | Wellcome Sanger Institute for the COVID-19 Genomics UK (COG-UK) Consortium | Berkshire and Surrey Pathology Services Lighthouse Laboratory and Alex Alderton, Roberto Amato, Jeffrey Barrett, Sonia Goncalves, Ewan Harrison, David K. Jackson, Ian Johnston, Dominic Kwiatkowski, Cordelia Langford, John Sillitoe on behalf of the Wellcome Sanger Institute COVID-19 Surveillance Team |
| EPI_ISL_3469399, EPI_ISL_3469400, EPI_ISL_3469401 | Lighthouse Lab in Milton Keynes | Wellcome Sanger Institute for the COVID-19 Genomics UK (COG-UK) Consortium | The Lighthouse Lab in Milton Keynes and Alex Alderton, Roberto Amato, Jeffrey Barrett, Sonia Goncalves, Ewan Harrison, David K. Jackson, Ian Johnston, Dominic Kwiatkowski, Cordelia Langford, John Sillitoe on behalf of the Wellcome Sanger Institute COVID-19 Surveillance Team |
| EPI_ISL_3469402 | Berkshire and Surrey Pathology Services Lighthouse Laboratory | Wellcome Sanger Institute for the COVID-19 Genomics UK (COG-UK) | Berkshire and Surrey Pathology Services Lighthouse Laboratory and Alex Alderton, Roberto Amato, Jeffrey Barrett, Sonia Goncalves, Ewan Harrison, David K. Jackson, Ian Johnston, Dominic Kwiatkowski, Cordelia Langford, John Sillitoe on behalf |

| | | | |
|---|---|---|---|
| | | Consortium | of the Wellcome Sanger Institute COVID-19 Surveillance Team |
| EPI_ISL_3469403 | Lighthouse Lab in Milton Keynes | Wellcome Sanger Institute for the COVID-19 Genomics UK (COG-UK) Consortium | The Lighthouse Lab in Milton Keynes and Alex Alderton, Roberto Amato, Jeffrey Barrett, Sonia Goncalves, Ewan Harrison, David K. Jackson, Ian Johnston, Dominic Kwiatkowski, Cordelia Langford, John Sillitoe on behalf of the Wellcome Sanger Institute COVID-19 Surveillance Team |
| EPI_ISL_3469404 | Berkshire and Surrey Pathology Services Lighthouse Laboratory | Wellcome Sanger Institute for the COVID-19 Genomics UK (COG-UK) Consortium | Berkshire and Surrey Pathology Services Lighthouse Laboratory and Alex Alderton, Roberto Amato, Jeffrey Barrett, Sonia Goncalves, Ewan Harrison, David K. Jackson, Ian Johnston, Dominic Kwiatkowski, Cordelia Langford, John Sillitoe on behalf of the Wellcome Sanger Institute COVID-19 Surveillance Team |
| EPI_ISL_3469405, EPI_ISL_3469406, EPI_ISL_3469407, EPI_ISL_3469408, EPI_ISL_3469409 | Lighthouse Lab in Milton Keynes | Wellcome Sanger Institute for the COVID-19 Genomics UK (COG-UK) Consortium | The Lighthouse Lab in Milton Keynes and Alex Alderton, Roberto Amato, Jeffrey Barrett, Sonia Goncalves, Ewan Harrison, David K. Jackson, Ian Johnston, Dominic Kwiatkowski, Cordelia Langford, John Sillitoe on behalf of the Wellcome Sanger Institute COVID-19 Surveillance Team |
| EPI_ISL_3469410, EPI_ISL_3469411, EPI_ISL_3469412 | Berkshire and Surrey Pathology Services Lighthouse Laboratory | Wellcome Sanger Institute for the COVID-19 Genomics UK (COG-UK) Consortium | Berkshire and Surrey Pathology Services Lighthouse Laboratory and Alex Alderton, Roberto Amato, Jeffrey Barrett, Sonia Goncalves, Ewan Harrison, David K. Jackson, Ian Johnston, Dominic Kwiatkowski, Cordelia Langford, John Sillitoe on behalf of the Wellcome Sanger Institute COVID-19 Surveillance Team |
| EPI_ISL_3469413 | Lighthouse Lab in Glasgow | Wellcome Sanger Institute for the COVID-19 Genomics UK (COG-UK) Consortium | Harper VanSteenhouse, Yumi Kasai, David Gray, Carol Clugston, Anna Dominiczak and Alex Alderton, Roberto Amato, Jeffrey Barrett, Sonia Goncalves, Ewan Harrison, David K. Jackson, Ian Johnston, Dominic Kwiatkowski, Cordelia Langford, John Sillitoe on behalf of the Wellcome Sanger Institute COVID-19 Surveillance Team |
| EPI_ISL_3469414 | Lighthouse Lab in Milton Keynes | Wellcome Sanger Institute for the COVID-19 Genomics UK (COG-UK) Consortium | The Lighthouse Lab in Milton Keynes and Alex Alderton, Roberto Amato, Jeffrey Barrett, Sonia Goncalves, Ewan Harrison, David K. Jackson, Ian Johnston, Dominic Kwiatkowski, Cordelia Langford, John Sillitoe on behalf of the Wellcome Sanger Institute COVID-19 Surveillance Team |
| EPI_ISL_3469415 | Lighthouse Lab in Glasgow | Wellcome Sanger Institute for the COVID-19 Genomics UK (COG-UK) Consortium | Harper VanSteenhouse, Yumi Kasai, David Gray, Carol Clugston, Anna Dominiczak and Alex Alderton, Roberto Amato, Jeffrey Barrett, Sonia Goncalves, Ewan Harrison, David K. Jackson, Ian Johnston, Dominic Kwiatkowski, Cordelia Langford, John Sillitoe on behalf of the Wellcome Sanger Institute COVID-19 Surveillance Team |
| EPI_ISL_3469416 | Lighthouse Lab in Milton Keynes | Wellcome Sanger Institute for the COVID-19 Genomics UK (COG-UK) Consortium | The Lighthouse Lab in Milton Keynes and Alex Alderton, Roberto Amato, Jeffrey Barrett, Sonia Goncalves, Ewan Harrison, David K. Jackson, Ian Johnston, Dominic Kwiatkowski, Cordelia Langford, John Sillitoe on behalf of the Wellcome Sanger Institute COVID-19 Surveillance Team |
| EPI_ISL_3469417 | Berkshire and Surrey Pathology Services Lighthouse Laboratory | Wellcome Sanger Institute for the COVID-19 Genomics UK (COG-UK) Consortium | Berkshire and Surrey Pathology Services Lighthouse Laboratory and Alex Alderton, Roberto Amato, Jeffrey Barrett, Sonia Goncalves, Ewan Harrison, David K. Jackson, Ian Johnston, Dominic Kwiatkowski, Cordelia Langford, John Sillitoe on behalf of the Wellcome Sanger Institute COVID-19 Surveillance Team |
| EPI_ISL_3469418 | Lighthouse Lab in Milton Keynes | Wellcome Sanger Institute for the COVID-19 Genomics UK (COG-UK) Consortium | The Lighthouse Lab in Milton Keynes and Alex Alderton, Roberto Amato, Jeffrey Barrett, Sonia Goncalves, Ewan Harrison, David K. Jackson, Ian Johnston, Dominic Kwiatkowski, Cordelia Langford, John Sillitoe on behalf of the Wellcome Sanger Institute COVID-19 Surveillance Team |
| EPI_ISL_3469419 | Berkshire and Surrey Pathology Services Lighthouse Laboratory | Wellcome Sanger Institute for the COVID-19 Genomics UK (COG-UK) Consortium | Berkshire and Surrey Pathology Services Lighthouse Laboratory and Alex Alderton, Roberto Amato, Jeffrey Barrett, Sonia Goncalves, Ewan Harrison, David K. Jackson, Ian Johnston, Dominic Kwiatkowski, Cordelia Langford, John Sillitoe on behalf of the Wellcome Sanger Institute COVID-19 Surveillance Team |
| EPI_ISL_3469420, EPI_ISL_3469421, EPI_ISL_3469422, EPI_ISL_3469423, EPI_ISL_3469424 | Lighthouse Lab in Milton Keynes | Wellcome Sanger Institute for the COVID-19 Genomics UK (COG-UK) Consortium | The Lighthouse Lab in Milton Keynes and Alex Alderton, Roberto Amato, Jeffrey Barrett, Sonia Goncalves, Ewan Harrison, David K. Jackson, Ian Johnston, Dominic Kwiatkowski, Cordelia Langford, John Sillitoe on behalf of the Wellcome Sanger Institute COVID-19 Surveillance Team |
| EPI_ISL_3469425 | Lighthouse Lab in Glasgow | Wellcome Sanger Institute for the COVID-19 Genomics UK (COG-UK) Consortium | Harper VanSteenhouse, Yumi Kasai, David Gray, Carol Clugston, Anna Dominiczak and Alex Alderton, Roberto Amato, Jeffrey Barrett, Sonia Goncalves, Ewan Harrison, David K. Jackson, Ian Johnston, Dominic Kwiatkowski, Cordelia Langford, John Sillitoe on behalf of the Wellcome Sanger Institute COVID-19 Surveillance Team |
| EPI_ISL_3469426 | Berkshire and Surrey Pathology Services Lighthouse Laboratory | Wellcome Sanger Institute for the COVID-19 Genomics UK (COG-UK) Consortium | Berkshire and Surrey Pathology Services Lighthouse Laboratory and Alex Alderton, Roberto Amato, Jeffrey Barrett, Sonia Goncalves, Ewan Harrison, David K. Jackson, Ian Johnston, Dominic Kwiatkowski, Cordelia Langford, John Sillitoe on behalf of the Wellcome Sanger Institute COVID-19 Surveillance Team |
| EPI_ISL_3469427, EPI_ISL_3469428 | Lighthouse Lab in Glasgow | Wellcome Sanger Institute for the COVID-19 Genomics UK (COG-UK) Consortium | Harper VanSteenhouse, Yumi Kasai, David Gray, Carol Clugston, Anna Dominiczak and Alex Alderton, Roberto Amato, Jeffrey Barrett, Sonia Goncalves, Ewan Harrison, David K. Jackson, Ian Johnston, Dominic Kwiatkowski, Cordelia Langford, John Sillitoe on behalf of the Wellcome Sanger Institute COVID-19 Surveillance Team |
| EPI_ISL_3469429, EPI_ISL_3469430 | Lighthouse Lab in Milton Keynes | Wellcome Sanger Institute for the COVID-19 Genomics UK (COG-UK) Consortium | The Lighthouse Lab in Milton Keynes and Alex Alderton, Roberto Amato, Jeffrey Barrett, Sonia Goncalves, Ewan Harrison, David K. Jackson, Ian Johnston, Dominic Kwiatkowski, Cordelia Langford, John Sillitoe on behalf of the Wellcome Sanger Institute COVID-19 Surveillance Team |
| EPI_ISL_3469431 | Berkshire and Surrey Pathology Services Lighthouse Laboratory | Wellcome Sanger Institute for the COVID-19 Genomics UK (COG-UK) Consortium | Berkshire and Surrey Pathology Services Lighthouse Laboratory and Alex Alderton, Roberto Amato, Jeffrey Barrett, Sonia Goncalves, Ewan Harrison, David K. Jackson, Ian Johnston, Dominic Kwiatkowski, Cordelia Langford, John Sillitoe on behalf of the Wellcome Sanger Institute COVID-19 Surveillance Team |
| EPI_ISL_3469432 | Lighthouse Lab in Milton Keynes | Wellcome Sanger Institute for the COVID-19 Genomics UK (COG-UK) Consortium | The Lighthouse Lab in Milton Keynes and Alex Alderton, Roberto Amato, Jeffrey Barrett, Sonia Goncalves, Ewan Harrison, David K. Jackson, Ian Johnston, Dominic Kwiatkowski, Cordelia Langford, John Sillitoe on behalf of the Wellcome Sanger Institute COVID-19 Surveillance Team |
| EPI_ISL_3469433 | Lighthouse Lab in Glasgow | Wellcome Sanger Institute for the COVID-19 Genomics UK (COG-UK) Consortium | Harper VanSteenhouse, Yumi Kasai, David Gray, Carol Clugston, Anna Dominiczak and Alex Alderton, Roberto Amato, Jeffrey Barrett, Sonia Goncalves, Ewan Harrison, David K. Jackson, Ian Johnston, Dominic Kwiatkowski, Cordelia Langford, John Sillitoe on behalf of the Wellcome Sanger Institute COVID-19 Surveillance Team |
| EPI_ISL_3469434, EPI_ISL_3469435 | Berkshire and Surrey Pathology Services Lighthouse Laboratory | Wellcome Sanger Institute for the COVID-19 Genomics UK (COG-UK) Consortium | Berkshire and Surrey Pathology Services Lighthouse Laboratory and Alex Alderton, Roberto Amato, Jeffrey Barrett, Sonia Goncalves, Ewan Harrison, David K. Jackson, Ian Johnston, Dominic Kwiatkowski, Cordelia Langford, John Sillitoe on behalf of the Wellcome Sanger Institute COVID-19 Surveillance Team |
| EPI_ISL_3469436 | Lighthouse Lab in Glasgow | Wellcome Sanger Institute for the COVID-19 Genomics UK (COG-UK) Consortium | Harper VanSteenhouse, Yumi Kasai, David Gray, Carol Clugston, Anna Dominiczak and Alex Alderton, Roberto Amato, Jeffrey Barrett, Sonia Goncalves, Ewan Harrison, David K. Jackson, Ian Johnston, Dominic Kwiatkowski, Cordelia Langford, John Sillitoe on behalf of the Wellcome Sanger Institute COVID-19 Surveillance Team |
| EPI_ISL_3469437 | Lighthouse Lab in Milton Keynes | Wellcome Sanger Institute for the COVID-19 Genomics UK (COG-UK) Consortium | The Lighthouse Lab in Milton Keynes and Alex Alderton, Roberto Amato, Jeffrey Barrett, Sonia Goncalves, Ewan Harrison, David K. Jackson, Ian Johnston, Dominic Kwiatkowski, Cordelia Langford, John Sillitoe on behalf of the Wellcome Sanger Institute COVID-19 Surveillance Team |
| EPI_ISL_3469438 | Berkshire and Surrey Pathology Services Lighthouse Laboratory | Wellcome Sanger Institute for the COVID-19 Genomics UK (COG-UK) Consortium | Berkshire and Surrey Pathology Services Lighthouse Laboratory and Alex Alderton, Roberto Amato, Jeffrey Barrett, Sonia Goncalves, Ewan Harrison, David K. Jackson, Ian Johnston, Dominic Kwiatkowski, Cordelia Langford, John Sillitoe on behalf of the Wellcome Sanger Institute COVID-19 Surveillance Team |
| EPI_ISL_3469439 | Lighthouse Lab in Glasgow | Wellcome Sanger Institute for the COVID-19 Genomics UK (COG-UK) Consortium | Harper VanSteenhouse, Yumi Kasai, David Gray, Carol Clugston, Anna Dominiczak and Alex Alderton, Roberto Amato, Jeffrey Barrett, Sonia Goncalves, Ewan Harrison, David K. Jackson, Ian Johnston, Dominic Kwiatkowski, Cordelia Langford, John Sillitoe on behalf of the Wellcome Sanger Institute COVID-19 Surveillance Team |
| EPI_ISL_3469440 | Lighthouse Lab in Milton Keynes | Wellcome Sanger Institute for the COVID-19 Genomics UK (COG-UK) Consortium | The Lighthouse Lab in Milton Keynes and Alex Alderton, Roberto Amato, Jeffrey Barrett, Sonia Goncalves, Ewan Harrison, David K. Jackson, Ian Johnston, Dominic Kwiatkowski, Cordelia Langford, John Sillitoe on behalf of the Wellcome Sanger Institute COVID-19 Surveillance Team |
| EPI_ISL_3469441, EPI_ISL_3469442 | Berkshire and Surrey Pathology Services Lighthouse Laboratory | Wellcome Sanger Institute for the COVID-19 Genomics UK (COG-UK) Consortium | Berkshire and Surrey Pathology Services Lighthouse Laboratory and Alex Alderton, Roberto Amato, Jeffrey Barrett, Sonia Goncalves, Ewan Harrison, David K. Jackson, Ian Johnston, Dominic Kwiatkowski, Cordelia Langford, John Sillitoe on behalf of the Wellcome Sanger Institute COVID-19 Surveillance Team |
| EPI_ISL_3469443 | Lighthouse Lab in Milton Keynes | Wellcome Sanger Institute for the COVID-19 Genomics UK (COG-UK) Consortium | The Lighthouse Lab in Milton Keynes and Alex Alderton, Roberto Amato, Jeffrey Barrett, Sonia Goncalves, Ewan Harrison, David K. Jackson, Ian Johnston, Dominic Kwiatkowski, Cordelia Langford, John Sillitoe on behalf of the Wellcome Sanger Institute COVID-19 Surveillance Team |
| EPI_ISL_3469444, EPI_ISL_3469445 | Berkshire and Surrey Pathology Services Lighthouse Laboratory | Wellcome Sanger Institute for the COVID-19 Genomics UK (COG-UK) Consortium | Berkshire and Surrey Pathology Services Lighthouse Laboratory and Alex Alderton, Roberto Amato, Jeffrey Barrett, Sonia Goncalves, Ewan Harrison, David K. Jackson, Ian Johnston, Dominic Kwiatkowski, Cordelia Langford, John Sillitoe on behalf of the Wellcome Sanger Institute COVID-19 Surveillance Team |
| EPI_ISL_3469446 | Lighthouse Lab in Milton Keynes | Wellcome Sanger Institute for the COVID-19 Genomics UK (COG-UK) Consortium | The Lighthouse Lab in Milton Keynes and Alex Alderton, Roberto Amato, Jeffrey Barrett, Sonia Goncalves, Ewan Harrison, David K. Jackson, Ian Johnston, Dominic Kwiatkowski, Cordelia Langford, John Sillitoe on behalf of the Wellcome Sanger Institute COVID-19 Surveillance Team |
| EPI_ISL_3469447 | Berkshire and Surrey Pathology Services Lighthouse Laboratory | Wellcome Sanger Institute for the COVID-19 Genomics UK (COG-UK) Consortium | Berkshire and Surrey Pathology Services Lighthouse Laboratory and Alex Alderton, Roberto Amato, Jeffrey Barrett, Sonia Goncalves, Ewan Harrison, David K. Jackson, Ian Johnston, Dominic Kwiatkowski, Cordelia Langford, John Sillitoe on behalf of the Wellcome Sanger Institute COVID-19 Surveillance Team |
| EPI_ISL_3469448, EPI_ISL_3469449, EPI_ISL_3469450, EPI_ISL_3469451 | Lighthouse Lab in Milton Keynes | Wellcome Sanger Institute for the COVID-19 Genomics UK (COG-UK) Consortium | The Lighthouse Lab in Milton Keynes and Alex Alderton, Roberto Amato, Jeffrey Barrett, Sonia Goncalves, Ewan Harrison, David K. Jackson, Ian Johnston, Dominic Kwiatkowski, Cordelia Langford, John Sillitoe on behalf of the Wellcome Sanger Institute COVID-19 Surveillance Team |
| EPI_ISL_3469452 | Berkshire and Surrey Pathology Services Lighthouse Laboratory | Wellcome Sanger Institute for the COVID-19 Genomics UK (COG-UK) Consortium | Berkshire and Surrey Pathology Services Lighthouse Laboratory and Alex Alderton, Roberto Amato, Jeffrey Barrett, Sonia Goncalves, Ewan Harrison, David K. Jackson, Ian Johnston, Dominic Kwiatkowski, Cordelia Langford, John Sillitoe on behalf of the Wellcome Sanger Institute COVID-19 Surveillance Team |
| EPI_ISL_3469453, EPI_ISL_3469454 | Lighthouse Lab in Milton Keynes | Wellcome Sanger Institute for the COVID-19 Genomics UK (COG-UK) Consortium | The Lighthouse Lab in Milton Keynes and Alex Alderton, Roberto Amato, Jeffrey Barrett, Sonia Goncalves, Ewan Harrison, David K. Jackson, Ian Johnston, Dominic Kwiatkowski, Cordelia Langford, John Sillitoe on behalf of the Wellcome Sanger Institute COVID-19 Surveillance Team |
| EPI_ISL_3469455 | Lighthouse Lab in Glasgow | Wellcome Sanger Institute for the COVID-19 Genomics UK (COG-UK) Consortium | Harper VanSteenhouse, Yumi Kasai, David Gray, Carol Clugston, Anna Dominiczak and Alex Alderton, Roberto Amato, Jeffrey Barrett, Sonia Goncalves, Ewan Harrison, David K. Jackson, Ian Johnston, Dominic Kwiatkowski, Cordelia Langford, John Sillitoe on behalf of the Wellcome Sanger Institute COVID-19 Surveillance Team |
| EPI_ISL_3469456 | Lighthouse Lab in Milton Keynes | Wellcome Sanger Institute for the COVID-19 Genomics UK (COG-UK) Consortium | The Lighthouse Lab in Milton Keynes and Alex Alderton, Roberto Amato, Jeffrey Barrett, Sonia Goncalves, Ewan Harrison, David K. Jackson, Ian Johnston, Dominic Kwiatkowski, Cordelia Langford, John Sillitoe on behalf of the Wellcome Sanger Institute COVID-19 Surveillance Team |
| EPI_ISL_3469457, EPI_ISL_3469458 | Lighthouse Lab in Glasgow | Wellcome Sanger Institute for the COVID-19 Genomics UK (COG-UK) Consortium | Harper VanSteenhouse, Yumi Kasai, David Gray, Carol Clugston, Anna Dominiczak and Alex Alderton, Roberto Amato, Jeffrey Barrett, Sonia Goncalves, Ewan Harrison, David K. Jackson, Ian Johnston, Dominic Kwiatkowski, Cordelia Langford, John Sillitoe on behalf of the Wellcome Sanger Institute COVID-19 Surveillance Team |
| EPI_ISL_3469459 | Berkshire and Surrey Pathology Services Lighthouse Laboratory | Wellcome Sanger Institute for the COVID-19 Genomics UK (COG-UK) Consortium | Berkshire and Surrey Pathology Services Lighthouse Laboratory and Alex Alderton, Roberto Amato, Jeffrey Barrett, Sonia Goncalves, Ewan Harrison, David K. Jackson, Ian Johnston, Dominic Kwiatkowski, Cordelia Langford, John Sillitoe on behalf of the Wellcome Sanger Institute COVID-19 Surveillance Team |
| EPI_ISL_3469460 | Lighthouse Lab in Glasgow | Wellcome Sanger Institute for the COVID-19 Genomics UK (COG-UK) Consortium | Harper VanSteenhouse, Yumi Kasai, David Gray, Carol Clugston, Anna Dominiczak and Alex Alderton, Roberto Amato, Jeffrey Barrett, Sonia Goncalves, Ewan Harrison, David K. Jackson, Ian Johnston, Dominic Kwiatkowski, Cordelia Langford, John Sillitoe on behalf of the Wellcome Sanger Institute COVID-19 Surveillance Team |
| EPI_ISL_3469461, EPI_ISL_3469462, EPI_ISL_3469463, EPI_ISL_3469464, EPI_ISL_3469465 | Lighthouse Lab in Milton Keynes | Wellcome Sanger Institute for the COVID-19 Genomics UK (COG-UK) Consortium | The Lighthouse Lab in Milton Keynes and Alex Alderton, Roberto Amato, Jeffrey Barrett, Sonia Goncalves, Ewan Harrison, David K. Jackson, Ian Johnston, Dominic Kwiatkowski, Cordelia Langford, John Sillitoe on behalf of the Wellcome Sanger Institute COVID-19 Surveillance Team |
| EPI_ISL_3469466 | Lighthouse Lab in Glasgow | Wellcome Sanger Institute for the COVID-19 Genomics UK (COG-UK) Consortium | Harper VanSteenhouse, Yumi Kasai, David Gray, Carol Clugston, Anna Dominiczak and Alex Alderton, Roberto Amato, Jeffrey Barrett, Sonia Goncalves, Ewan Harrison, David K. Jackson, Ian Johnston, Dominic Kwiatkowski, Cordelia Langford, John Sillitoe on behalf of the Wellcome Sanger Institute COVID-19 Surveillance Team |

| | | | |
|---|---|---|---|
| EPI_ISL_3469467, EPI_ISL_3469468, EPI_ISL_3469469 | Lighthouse Lab in Milton Keynes | Wellcome Sanger Institute for the COVID-19 Genomics UK (COG-UK) Consortium | The Lighthouse Lab in Milton Keynes and Alex Alderton, Roberto Amato, Jeffrey Barrett, Sonia Goncalves, Ewan Harrison, David K. Jackson, Ian Johnston, Dominic Kwiatkowski, Cordelia Langford, John Sillitoe on behalf of the Wellcome Sanger Institute COVID-19 Surveillance Team |
| EPI_ISL_3469470 | Berkshire and Surrey Pathology Services Lighthouse Laboratory | Wellcome Sanger Institute for the COVID-19 Genomics UK (COG-UK) Consortium | Berkshire and Surrey Pathology Services Lighthouse Laboratory and Alex Alderton, Roberto Amato, Jeffrey Barrett, Sonia Goncalves, Ewan Harrison, David K. Jackson, Ian Johnston, Dominic Kwiatkowski, Cordelia Langford, John Sillitoe on behalf of the Wellcome Sanger Institute COVID-19 Surveillance Team |
| EPI_ISL_3469471, EPI_ISL_3469472 | Lighthouse Lab in Milton Keynes | Wellcome Sanger Institute for the COVID-19 Genomics UK (COG-UK) Consortium | The Lighthouse Lab in Milton Keynes and Alex Alderton, Roberto Amato, Jeffrey Barrett, Sonia Goncalves, Ewan Harrison, David K. Jackson, Ian Johnston, Dominic Kwiatkowski, Cordelia Langford, John Sillitoe on behalf of the Wellcome Sanger Institute COVID-19 Surveillance Team |
| EPI_ISL_3469473 | Lighthouse Lab in Glasgow | Wellcome Sanger Institute for the COVID-19 Genomics UK (COG-UK) Consortium | Harper VanSteenhouse, Yumi Kasai, David Gray, Carol Clugston, Anna Dominiczak and Alex Alderton, Roberto Amato, Jeffrey Barrett, Sonia Goncalves, Ewan Harrison, David K. Jackson, Ian Johnston, Dominic Kwiatkowski, Cordelia Langford, John Sillitoe on behalf of the Wellcome Sanger Institute COVID-19 Surveillance Team |
| EPI_ISL_3469474 | Lighthouse Lab in Milton Keynes | Wellcome Sanger Institute for the COVID-19 Genomics UK (COG-UK) Consortium | The Lighthouse Lab in Milton Keynes and Alex Alderton, Roberto Amato, Jeffrey Barrett, Sonia Goncalves, Ewan Harrison, David K. Jackson, Ian Johnston, Dominic Kwiatkowski, Cordelia Langford, John Sillitoe on behalf of the Wellcome Sanger Institute COVID-19 Surveillance Team |
| EPI_ISL_3469475 | Lighthouse Lab in Glasgow | Wellcome Sanger Institute for the COVID-19 Genomics UK (COG-UK) Consortium | Harper VanSteenhouse, Yumi Kasai, David Gray, Carol Clugston, Anna Dominiczak and Alex Alderton, Roberto Amato, Jeffrey Barrett, Sonia Goncalves, Ewan Harrison, David K. Jackson, Ian Johnston, Dominic Kwiatkowski, Cordelia Langford, John Sillitoe on behalf of the Wellcome Sanger Institute COVID-19 Surveillance Team |
| EPI_ISL_3469476 | Berkshire and Surrey Pathology Services Lighthouse Laboratory | Wellcome Sanger Institute for the COVID-19 Genomics UK (COG-UK) Consortium | Berkshire and Surrey Pathology Services Lighthouse Laboratory and Alex Alderton, Roberto Amato, Jeffrey Barrett, Sonia Goncalves, Ewan Harrison, David K. Jackson, Ian Johnston, Dominic Kwiatkowski, Cordelia Langford, John Sillitoe on behalf of the Wellcome Sanger Institute COVID-19 Surveillance Team |
| EPI_ISL_3469477 | Lighthouse Lab in Glasgow | Wellcome Sanger Institute for the COVID-19 Genomics UK (COG-UK) Consortium | Harper VanSteenhouse, Yumi Kasai, David Gray, Carol Clugston, Anna Dominiczak and Alex Alderton, Roberto Amato, Jeffrey Barrett, Sonia Goncalves, Ewan Harrison, David K. Jackson, Ian Johnston, Dominic Kwiatkowski, Cordelia Langford, John Sillitoe on behalf of the Wellcome Sanger Institute COVID-19 Surveillance Team |
| EPI_ISL_3469478, EPI_ISL_3469479, EPI_ISL_3469480, EPI_ISL_3469481, EPI_ISL_3469482, EPI_ISL_3469483 | Lighthouse Lab in Milton Keynes | Wellcome Sanger Institute for the COVID-19 Genomics UK (COG-UK) Consortium | The Lighthouse Lab in Milton Keynes and Alex Alderton, Roberto Amato, Jeffrey Barrett, Sonia Goncalves, Ewan Harrison, David K. Jackson, Ian Johnston, Dominic Kwiatkowski, Cordelia Langford, John Sillitoe on behalf of the Wellcome Sanger Institute COVID-19 Surveillance Team |
| EPI_ISL_3469484 | Lighthouse Lab in Glasgow | Wellcome Sanger Institute for the COVID-19 Genomics UK (COG-UK) Consortium | Harper VanSteenhouse, Yumi Kasai, David Gray, Carol Clugston, Anna Dominiczak and Alex Alderton, Roberto Amato, Jeffrey Barrett, Sonia Goncalves, Ewan Harrison, David K. Jackson, Ian Johnston, Dominic Kwiatkowski, Cordelia Langford, John Sillitoe on behalf of the Wellcome Sanger Institute COVID-19 Surveillance Team |
| EPI_ISL_3469485, EPI_ISL_3469486, EPI_ISL_3469487, EPI_ISL_3469488 | Lighthouse Lab in Milton Keynes | Wellcome Sanger Institute for the COVID-19 Genomics UK (COG-UK) Consortium | The Lighthouse Lab in Milton Keynes and Alex Alderton, Roberto Amato, Jeffrey Barrett, Sonia Goncalves, Ewan Harrison, David K. Jackson, Ian Johnston, Dominic Kwiatkowski, Cordelia Langford, John Sillitoe on behalf of the Wellcome Sanger Institute COVID-19 Surveillance Team |
| EPI_ISL_3469489, EPI_ISL_3469490 | Berkshire and Surrey Pathology Services Lighthouse Laboratory | Wellcome Sanger Institute for the COVID-19 Genomics UK (COG-UK) Consortium | Berkshire and Surrey Pathology Services Lighthouse Laboratory and Alex Alderton, Roberto Amato, Jeffrey Barrett, Sonia Goncalves, Ewan Harrison, David K. Jackson, Ian Johnston, Dominic Kwiatkowski, Cordelia Langford, John Sillitoe on behalf of the Wellcome Sanger Institute COVID-19 Surveillance Team |
| EPI_ISL_3469491, EPI_ISL_3469492 | Lighthouse Lab in Milton Keynes | Wellcome Sanger Institute for the COVID-19 Genomics UK (COG-UK) Consortium | The Lighthouse Lab in Milton Keynes and Alex Alderton, Roberto Amato, Jeffrey Barrett, Sonia Goncalves, Ewan Harrison, David K. Jackson, Ian Johnston, Dominic Kwiatkowski, Cordelia Langford, John Sillitoe on behalf of the Wellcome Sanger Institute COVID-19 Surveillance Team |
| EPI_ISL_3469493 | Berkshire and Surrey Pathology Services Lighthouse Laboratory | Wellcome Sanger Institute for the COVID-19 Genomics UK (COG-UK) Consortium | Berkshire and Surrey Pathology Services Lighthouse Laboratory and Alex Alderton, Roberto Amato, Jeffrey Barrett, Sonia Goncalves, Ewan Harrison, David K. Jackson, Ian Johnston, Dominic Kwiatkowski, Cordelia Langford, John Sillitoe on behalf of the Wellcome Sanger Institute COVID-19 Surveillance Team |
| EPI_ISL_3469494, EPI_ISL_3469495 | Lighthouse Lab in Milton Keynes | Wellcome Sanger Institute for the COVID-19 Genomics UK (COG-UK) Consortium | The Lighthouse Lab in Milton Keynes and Alex Alderton, Roberto Amato, Jeffrey Barrett, Sonia Goncalves, Ewan Harrison, David K. Jackson, Ian Johnston, Dominic Kwiatkowski, Cordelia Langford, John Sillitoe on behalf of the Wellcome Sanger Institute COVID-19 Surveillance Team |
| EPI_ISL_3469496 | Berkshire and Surrey Pathology Services Lighthouse Laboratory | Wellcome Sanger Institute for the COVID-19 Genomics UK (COG-UK) Consortium | Berkshire and Surrey Pathology Services Lighthouse Laboratory and Alex Alderton, Roberto Amato, Jeffrey Barrett, Sonia Goncalves, Ewan Harrison, David K. Jackson, Ian Johnston, Dominic Kwiatkowski, Cordelia Langford, John Sillitoe on behalf of the Wellcome Sanger Institute COVID-19 Surveillance Team |
| EPI_ISL_3469497, EPI_ISL_3469498 | Lighthouse Lab in Milton Keynes | Wellcome Sanger Institute for the COVID-19 Genomics UK (COG-UK) Consortium | The Lighthouse Lab in Milton Keynes and Alex Alderton, Roberto Amato, Jeffrey Barrett, Sonia Goncalves, Ewan Harrison, David K. Jackson, Ian Johnston, Dominic Kwiatkowski, Cordelia Langford, John Sillitoe on behalf of the Wellcome Sanger Institute COVID-19 Surveillance Team |
| EPI_ISL_3469499 | Lighthouse Lab in Glasgow | Wellcome Sanger Institute for the COVID-19 Genomics UK (COG-UK) Consortium | Harper VanSteenhouse, Yumi Kasai, David Gray, Carol Clugston, Anna Dominiczak and Alex Alderton, Roberto Amato, Jeffrey Barrett, Sonia Goncalves, Ewan Harrison, David K. Jackson, Ian Johnston, Dominic Kwiatkowski, Cordelia Langford, John Sillitoe on behalf of the Wellcome Sanger Institute COVID-19 Surveillance Team |
| EPI_ISL_3469500 | Lighthouse Lab in Milton Keynes | Wellcome Sanger Institute for the COVID-19 Genomics UK (COG-UK) Consortium | The Lighthouse Lab in Milton Keynes and Alex Alderton, Roberto Amato, Jeffrey Barrett, Sonia Goncalves, Ewan Harrison, David K. Jackson, Ian Johnston, Dominic Kwiatkowski, Cordelia Langford, John Sillitoe on behalf of the Wellcome Sanger Institute COVID-19 Surveillance Team |
| EPI_ISL_3469501, EPI_ISL_3469502 | Berkshire and Surrey Pathology Services Lighthouse Laboratory | Wellcome Sanger Institute for the COVID-19 Genomics UK (COG-UK) Consortium | Berkshire and Surrey Pathology Services Lighthouse Laboratory and Alex Alderton, Roberto Amato, Jeffrey Barrett, Sonia Goncalves, Ewan Harrison, David K. Jackson, Ian Johnston, Dominic Kwiatkowski, Cordelia Langford, John Sillitoe on behalf of the Wellcome Sanger Institute COVID-19 Surveillance Team |
| EPI_ISL_3469503, EPI_ISL_3469504, EPI_ISL_3469505 | Lighthouse Lab in Milton Keynes | Wellcome Sanger Institute for the COVID-19 Genomics UK (COG-UK) Consortium | The Lighthouse Lab in Milton Keynes and Alex Alderton, Roberto Amato, Jeffrey Barrett, Sonia Goncalves, Ewan Harrison, David K. Jackson, Ian Johnston, Dominic Kwiatkowski, Cordelia Langford, John Sillitoe on behalf of the Wellcome Sanger Institute COVID-19 Surveillance Team |
| EPI_ISL_3469506 | Lighthouse Lab in Glasgow | Wellcome Sanger Institute for the COVID-19 Genomics UK (COG-UK) Consortium | Harper VanSteenhouse, Yumi Kasai, David Gray, Carol Clugston, Anna Dominiczak and Alex Alderton, Roberto Amato, Jeffrey Barrett, Sonia Goncalves, Ewan Harrison, David K. Jackson, Ian Johnston, Dominic Kwiatkowski, Cordelia Langford, John Sillitoe on behalf of the Wellcome Sanger Institute COVID-19 Surveillance Team |
| EPI_ISL_3469507 | Lighthouse Lab in Milton Keynes | Wellcome Sanger Institute for the COVID-19 Genomics UK (COG-UK) Consortium | The Lighthouse Lab in Milton Keynes and Alex Alderton, Roberto Amato, Jeffrey Barrett, Sonia Goncalves, Ewan Harrison, David K. Jackson, Ian Johnston, Dominic Kwiatkowski, Cordelia Langford, John Sillitoe on behalf of the Wellcome Sanger Institute COVID-19 Surveillance Team |
| EPI_ISL_3469508, EPI_ISL_3469509 | Lighthouse Lab in Glasgow | Wellcome Sanger Institute for the COVID-19 Genomics UK (COG-UK) Consortium | Harper VanSteenhouse, Yumi Kasai, David Gray, Carol Clugston, Anna Dominiczak and Alex Alderton, Roberto Amato, Jeffrey Barrett, Sonia Goncalves, Ewan Harrison, David K. Jackson, Ian Johnston, Dominic Kwiatkowski, Cordelia Langford, John Sillitoe on behalf of the Wellcome Sanger Institute COVID-19 Surveillance Team |
| EPI_ISL_3469510 | Lighthouse Lab in Milton Keynes | Wellcome Sanger Institute for the COVID-19 Genomics UK (COG-UK) Consortium | The Lighthouse Lab in Milton Keynes and Alex Alderton, Roberto Amato, Jeffrey Barrett, Sonia Goncalves, Ewan Harrison, David K. Jackson, Ian Johnston, Dominic Kwiatkowski, Cordelia Langford, John Sillitoe on behalf of the Wellcome Sanger Institute COVID-19 Surveillance Team |
| EPI_ISL_3469511 | Lighthouse Lab in Glasgow | Wellcome Sanger Institute for the COVID-19 Genomics UK (COG-UK) Consortium | Harper VanSteenhouse, Yumi Kasai, David Gray, Carol Clugston, Anna Dominiczak and Alex Alderton, Roberto Amato, Jeffrey Barrett, Sonia Goncalves, Ewan Harrison, David K. Jackson, Ian Johnston, Dominic Kwiatkowski, Cordelia Langford, John Sillitoe on behalf of the Wellcome Sanger Institute COVID-19 Surveillance Team |
| EPI_ISL_3469512 | Lighthouse Lab in Milton Keynes | Wellcome Sanger Institute for the COVID-19 Genomics UK (COG-UK) Consortium | The Lighthouse Lab in Milton Keynes and Alex Alderton, Roberto Amato, Jeffrey Barrett, Sonia Goncalves, Ewan Harrison, David K. Jackson, Ian Johnston, Dominic Kwiatkowski, Cordelia Langford, John Sillitoe on behalf of the Wellcome Sanger Institute COVID-19 Surveillance Team |
| EPI_ISL_3469513 | Lighthouse Lab in Glasgow | Wellcome Sanger Institute for the COVID-19 Genomics UK (COG-UK) Consortium | Harper VanSteenhouse, Yumi Kasai, David Gray, Carol Clugston, Anna Dominiczak and Alex Alderton, Roberto Amato, Jeffrey Barrett, Sonia Goncalves, Ewan Harrison, David K. Jackson, Ian Johnston, Dominic Kwiatkowski, Cordelia Langford, John Sillitoe on behalf of the Wellcome Sanger Institute COVID-19 Surveillance Team |
| EPI_ISL_3469514 | Lighthouse Lab in Milton Keynes | Wellcome Sanger Institute for the COVID-19 Genomics UK (COG-UK) Consortium | The Lighthouse Lab in Milton Keynes and Alex Alderton, Roberto Amato, Jeffrey Barrett, Sonia Goncalves, Ewan Harrison, David K. Jackson, Ian Johnston, Dominic Kwiatkowski, Cordelia Langford, John Sillitoe on behalf of the Wellcome Sanger Institute COVID-19 Surveillance Team |
| EPI_ISL_3469515 | Lighthouse Lab in Glasgow | Wellcome Sanger Institute for the COVID-19 Genomics UK (COG-UK) Consortium | Harper VanSteenhouse, Yumi Kasai, David Gray, Carol Clugston, Anna Dominiczak and Alex Alderton, Roberto Amato, Jeffrey Barrett, Sonia Goncalves, Ewan Harrison, David K. Jackson, Ian Johnston, Dominic Kwiatkowski, Cordelia Langford, John Sillitoe on behalf of the Wellcome Sanger Institute COVID-19 Surveillance Team |
| EPI_ISL_3469516 | Lighthouse Lab in Milton Keynes | Wellcome Sanger Institute for the COVID-19 Genomics UK (COG-UK) Consortium | The Lighthouse Lab in Milton Keynes and Alex Alderton, Roberto Amato, Jeffrey Barrett, Sonia Goncalves, Ewan Harrison, David K. Jackson, Ian Johnston, Dominic Kwiatkowski, Cordelia Langford, John Sillitoe on behalf of the Wellcome Sanger Institute COVID-19 Surveillance Team |
| EPI_ISL_3469517 | Berkshire and Surrey Pathology Services Lighthouse Laboratory | Wellcome Sanger Institute for the COVID-19 Genomics UK (COG-UK) Consortium | Berkshire and Surrey Pathology Services Lighthouse Laboratory and Alex Alderton, Roberto Amato, Jeffrey Barrett, Sonia Goncalves, Ewan Harrison, David K. Jackson, Ian Johnston, Dominic Kwiatkowski, Cordelia Langford, John Sillitoe on behalf of the Wellcome Sanger Institute COVID-19 Surveillance Team |
| EPI_ISL_3469518, EPI_ISL_3469519, EPI_ISL_3469520, EPI_ISL_3469521 | Lighthouse Lab in Milton Keynes | Wellcome Sanger Institute for the COVID-19 Genomics UK (COG-UK) Consortium | The Lighthouse Lab in Milton Keynes and Alex Alderton, Roberto Amato, Jeffrey Barrett, Sonia Goncalves, Ewan Harrison, David K. Jackson, Ian Johnston, Dominic Kwiatkowski, Cordelia Langford, John Sillitoe on behalf of the Wellcome Sanger Institute COVID-19 Surveillance Team |
| EPI_ISL_3469522 | Lighthouse Lab in Glasgow | Wellcome Sanger Institute for the COVID-19 Genomics UK (COG-UK) Consortium | Harper VanSteenhouse, Yumi Kasai, David Gray, Carol Clugston, Anna Dominiczak and Alex Alderton, Roberto Amato, Jeffrey Barrett, Sonia Goncalves, Ewan Harrison, David K. Jackson, Ian Johnston, Dominic Kwiatkowski, Cordelia Langford, John Sillitoe on behalf of the Wellcome Sanger Institute COVID-19 Surveillance Team |
| EPI_ISL_3469523, EPI_ISL_3469524, EPI_ISL_3469525, EPI_ISL_3469526 | Lighthouse Lab in Milton Keynes | Wellcome Sanger Institute for the COVID-19 Genomics UK (COG-UK) Consortium | The Lighthouse Lab in Milton Keynes and Alex Alderton, Roberto Amato, Jeffrey Barrett, Sonia Goncalves, Ewan Harrison, David K. Jackson, Ian Johnston, Dominic Kwiatkowski, Cordelia Langford, John Sillitoe on behalf of the Wellcome Sanger Institute COVID-19 Surveillance Team |
| EPI_ISL_3469527 | Berkshire and Surrey Pathology Services Lighthouse Laboratory | Wellcome Sanger Institute for the COVID-19 Genomics UK (COG-UK) Consortium | Berkshire and Surrey Pathology Services Lighthouse Laboratory and Alex Alderton, Roberto Amato, Jeffrey Barrett, Sonia Goncalves, Ewan Harrison, David K. Jackson, Ian Johnston, Dominic Kwiatkowski, Cordelia Langford, John Sillitoe on behalf of the Wellcome Sanger Institute COVID-19 Surveillance Team |
| EPI_ISL_3469528 | Lighthouse Lab in Milton Keynes | Wellcome Sanger Institute for the COVID-19 Genomics UK (COG-UK) Consortium | The Lighthouse Lab in Milton Keynes and Alex Alderton, Roberto Amato, Jeffrey Barrett, Sonia Goncalves, Ewan Harrison, David K. Jackson, Ian Johnston, Dominic Kwiatkowski, Cordelia Langford, John Sillitoe on behalf of the Wellcome Sanger Institute COVID-19 Surveillance Team |
| EPI_ISL_3469529, EPI_ISL_3469530 | Berkshire and Surrey Pathology Services Lighthouse Laboratory | Wellcome Sanger Institute for the COVID-19 Genomics UK (COG-UK) Consortium | Berkshire and Surrey Pathology Services Lighthouse Laboratory and Alex Alderton, Roberto Amato, Jeffrey Barrett, Sonia Goncalves, Ewan Harrison, David K. Jackson, Ian Johnston, Dominic Kwiatkowski, Cordelia Langford, John Sillitoe on behalf of the Wellcome Sanger Institute COVID-19 Surveillance Team |
| EPI_ISL_3469531 | Lighthouse Lab in Milton Keynes | Wellcome Sanger Institute for the COVID-19 Genomics UK (COG-UK) Consortium | The Lighthouse Lab in Milton Keynes and Alex Alderton, Roberto Amato, Jeffrey Barrett, Sonia Goncalves, Ewan Harrison, David K. Jackson, Ian Johnston, Dominic Kwiatkowski, Cordelia Langford, John Sillitoe on behalf of the Wellcome Sanger Institute COVID-19 Surveillance Team |
| EPI_ISL_3469532, EPI_ISL_3469533 | Lighthouse Lab in Glasgow | Wellcome Sanger Institute for the COVID-19 Genomics UK (COG-UK) Consortium | Harper VanSteenhouse, Yumi Kasai, David Gray, Carol Clugston, Anna Dominiczak and Alex Alderton, Roberto Amato, Jeffrey Barrett, Sonia Goncalves, Ewan Harrison, David K. Jackson, Ian Johnston, Dominic Kwiatkowski, Cordelia Langford, John Sillitoe on behalf of the Wellcome Sanger Institute COVID-19 Surveillance Team |

| | | | |
|---|---|---|---|
| EPI_ISL_3469534, EPI_ISL_3469535, EPI_ISL_3469536 | Lighthouse Lab in Milton Keynes | Wellcome Sanger Institute for the COVID-19 Genomics UK (COG-UK) Consortium | The Lighthouse Lab in Milton Keynes and Alex Alderton, Roberto Amato, Jeffrey Barrett, Sonia Goncalves, Ewan Harrison, David K. Jackson, Ian Johnston, Dominic Kwiatkowski, Cordelia Langford, John Sillitoe on behalf of the Wellcome Sanger Institute COVID-19 Surveillance Team |
| EPI_ISL_3469537, EPI_ISL_3469538 | Berkshire and Surrey Pathology Services Lighthouse Laboratory | Wellcome Sanger Institute for the COVID-19 Genomics UK (COG-UK) Consortium | Berkshire and Surrey Pathology Services Lighthouse Laboratory and Alex Alderton, Roberto Amato, Jeffrey Barrett, Sonia Goncalves, Ewan Harrison, David K. Jackson, Ian Johnston, Dominic Kwiatkowski, Cordelia Langford, John Sillitoe on behalf of the Wellcome Sanger Institute COVID-19 Surveillance Team |
| EPI_ISL_3469539 | Lighthouse Lab in Milton Keynes | Wellcome Sanger Institute for the COVID-19 Genomics UK (COG-UK) Consortium | The Lighthouse Lab in Milton Keynes and Alex Alderton, Roberto Amato, Jeffrey Barrett, Sonia Goncalves, Ewan Harrison, David K. Jackson, Ian Johnston, Dominic Kwiatkowski, Cordelia Langford, John Sillitoe on behalf of the Wellcome Sanger Institute COVID-19 Surveillance Team |
| EPI_ISL_3469540 | Lighthouse Lab in Glasgow | Wellcome Sanger Institute for the COVID-19 Genomics UK (COG-UK) Consortium | Harper VanSteenhouse, Yumi Kasai, David Gray, Carol Clugston, Anna Dominiczak and Alex Alderton, Roberto Amato, Jeffrey Barrett, Sonia Goncalves, Ewan Harrison, David K. Jackson, Ian Johnston, Dominic Kwiatkowski, Cordelia Langford, John Sillitoe on behalf of the Wellcome Sanger Institute COVID-19 Surveillance Team |
| EPI_ISL_3469541, EPI_ISL_3469542, EPI_ISL_3469543 | Lighthouse Lab in Milton Keynes | Wellcome Sanger Institute for the COVID-19 Genomics UK (COG-UK) Consortium | The Lighthouse Lab in Milton Keynes and Alex Alderton, Roberto Amato, Jeffrey Barrett, Sonia Goncalves, Ewan Harrison, David K. Jackson, Ian Johnston, Dominic Kwiatkowski, Cordelia Langford, John Sillitoe on behalf of the Wellcome Sanger Institute COVID-19 Surveillance Team |
| EPI_ISL_3469544 | Lighthouse Lab in Glasgow | Wellcome Sanger Institute for the COVID-19 Genomics UK (COG-UK) Consortium | Harper VanSteenhouse, Yumi Kasai, David Gray, Carol Clugston, Anna Dominiczak and Alex Alderton, Roberto Amato, Jeffrey Barrett, Sonia Goncalves, Ewan Harrison, David K. Jackson, Ian Johnston, Dominic Kwiatkowski, Cordelia Langford, John Sillitoe on behalf of the Wellcome Sanger Institute COVID-19 Surveillance Team |
| EPI_ISL_3469545, EPI_ISL_3469546 | Lighthouse Lab in Milton Keynes | Wellcome Sanger Institute for the COVID-19 Genomics UK (COG-UK) Consortium | The Lighthouse Lab in Milton Keynes and Alex Alderton, Roberto Amato, Jeffrey Barrett, Sonia Goncalves, Ewan Harrison, David K. Jackson, Ian Johnston, Dominic Kwiatkowski, Cordelia Langford, John Sillitoe on behalf of the Wellcome Sanger Institute COVID-19 Surveillance Team |
| EPI_ISL_3469547 | Berkshire and Surrey Pathology Services Lighthouse Laboratory | Wellcome Sanger Institute for the COVID-19 Genomics UK (COG-UK) Consortium | Berkshire and Surrey Pathology Services Lighthouse Laboratory and Alex Alderton, Roberto Amato, Jeffrey Barrett, Sonia Goncalves, Ewan Harrison, David K. Jackson, Ian Johnston, Dominic Kwiatkowski, Cordelia Langford, John Sillitoe on behalf of the Wellcome Sanger Institute COVID-19 Surveillance Team |
| EPI_ISL_3469548, EPI_ISL_3469549 | Lighthouse Lab in Milton Keynes | Wellcome Sanger Institute for the COVID-19 Genomics UK (COG-UK) Consortium | The Lighthouse Lab in Milton Keynes and Alex Alderton, Roberto Amato, Jeffrey Barrett, Sonia Goncalves, Ewan Harrison, David K. Jackson, Ian Johnston, Dominic Kwiatkowski, Cordelia Langford, John Sillitoe on behalf of the Wellcome Sanger Institute COVID-19 Surveillance Team |
| EPI_ISL_3469550 | Lighthouse Lab in Glasgow | Wellcome Sanger Institute for the COVID-19 Genomics UK (COG-UK) Consortium | Harper VanSteenhouse, Yumi Kasai, David Gray, Carol Clugston, Anna Dominiczak and Alex Alderton, Roberto Amato, Jeffrey Barrett, Sonia Goncalves, Ewan Harrison, David K. Jackson, Ian Johnston, Dominic Kwiatkowski, Cordelia Langford, John Sillitoe on behalf of the Wellcome Sanger Institute COVID-19 Surveillance Team |
| EPI_ISL_3469551 | Lighthouse Lab in Milton Keynes | Wellcome Sanger Institute for the COVID-19 Genomics UK (COG-UK) Consortium | The Lighthouse Lab in Milton Keynes and Alex Alderton, Roberto Amato, Jeffrey Barrett, Sonia Goncalves, Ewan Harrison, David K. Jackson, Ian Johnston, Dominic Kwiatkowski, Cordelia Langford, John Sillitoe on behalf of the Wellcome Sanger Institute COVID-19 Surveillance Team |
| EPI_ISL_3469552 | Lighthouse Lab in Glasgow | Wellcome Sanger Institute for the COVID-19 Genomics UK (COG-UK) Consortium | Harper VanSteenhouse, Yumi Kasai, David Gray, Carol Clugston, Anna Dominiczak and Alex Alderton, Roberto Amato, Jeffrey Barrett, Sonia Goncalves, Ewan Harrison, David K. Jackson, Ian Johnston, Dominic Kwiatkowski, Cordelia Langford, John Sillitoe on behalf of the Wellcome Sanger Institute COVID-19 Surveillance Team |
| EPI_ISL_3469553 | Lighthouse Lab in Milton Keynes | Wellcome Sanger Institute for the COVID-19 Genomics UK (COG-UK) Consortium | The Lighthouse Lab in Milton Keynes and Alex Alderton, Roberto Amato, Jeffrey Barrett, Sonia Goncalves, Ewan Harrison, David K. Jackson, Ian Johnston, Dominic Kwiatkowski, Cordelia Langford, John Sillitoe on behalf of the Wellcome Sanger Institute COVID-19 Surveillance Team |
| EPI_ISL_3469554 | Berkshire and Surrey Pathology Services Lighthouse Laboratory | Wellcome Sanger Institute for the COVID-19 Genomics UK (COG-UK) Consortium | Berkshire and Surrey Pathology Services Lighthouse Laboratory and Alex Alderton, Roberto Amato, Jeffrey Barrett, Sonia Goncalves, Ewan Harrison, David K. Jackson, Ian Johnston, Dominic Kwiatkowski, Cordelia Langford, John Sillitoe on behalf of the Wellcome Sanger Institute COVID-19 Surveillance Team |
| EPI_ISL_3469555, EPI_ISL_3469556, EPI_ISL_3469557, EPI_ISL_3469558, EPI_ISL_3469559 | Lighthouse Lab in Milton Keynes | Wellcome Sanger Institute for the COVID-19 Genomics UK (COG-UK) Consortium | The Lighthouse Lab in Milton Keynes and Alex Alderton, Roberto Amato, Jeffrey Barrett, Sonia Goncalves, Ewan Harrison, David K. Jackson, Ian Johnston, Dominic Kwiatkowski, Cordelia Langford, John Sillitoe on behalf of the Wellcome Sanger Institute COVID-19 Surveillance Team |
| EPI_ISL_3469560 | Berkshire and Surrey Pathology Services Lighthouse Laboratory | Wellcome Sanger Institute for the COVID-19 Genomics UK (COG-UK) Consortium | Berkshire and Surrey Pathology Services Lighthouse Laboratory and Alex Alderton, Roberto Amato, Jeffrey Barrett, Sonia Goncalves, Ewan Harrison, David K. Jackson, Ian Johnston, Dominic Kwiatkowski, Cordelia Langford, John Sillitoe on behalf of the Wellcome Sanger Institute COVID-19 Surveillance Team |
| EPI_ISL_3469561 | Lighthouse Lab in Glasgow | Wellcome Sanger Institute for the COVID-19 Genomics UK (COG-UK) Consortium | Harper VanSteenhouse, Yumi Kasai, David Gray, Carol Clugston, Anna Dominiczak and Alex Alderton, Roberto Amato, Jeffrey Barrett, Sonia Goncalves, Ewan Harrison, David K. Jackson, Ian Johnston, Dominic Kwiatkowski, Cordelia Langford, John Sillitoe on behalf of the Wellcome Sanger Institute COVID-19 Surveillance Team |
| EPI_ISL_3469562 | Lighthouse Lab in Milton Keynes | Wellcome Sanger Institute for the COVID-19 Genomics UK (COG-UK) Consortium | The Lighthouse Lab in Milton Keynes and Alex Alderton, Roberto Amato, Jeffrey Barrett, Sonia Goncalves, Ewan Harrison, David K. Jackson, Ian Johnston, Dominic Kwiatkowski, Cordelia Langford, John Sillitoe on behalf of the Wellcome Sanger Institute COVID-19 Surveillance Team |
| EPI_ISL_3469563 | Berkshire and Surrey Pathology Services Lighthouse Laboratory | Wellcome Sanger Institute for the COVID-19 Genomics UK (COG-UK) Consortium | Berkshire and Surrey Pathology Services Lighthouse Laboratory and Alex Alderton, Roberto Amato, Jeffrey Barrett, Sonia Goncalves, Ewan Harrison, David K. Jackson, Ian Johnston, Dominic Kwiatkowski, Cordelia Langford, John Sillitoe on behalf of the Wellcome Sanger Institute COVID-19 Surveillance Team |
| EPI_ISL_3469564, EPI_ISL_3469565 | Lighthouse Lab in Milton Keynes | Wellcome Sanger Institute for the COVID-19 Genomics UK (COG-UK) Consortium | The Lighthouse Lab in Milton Keynes and Alex Alderton, Roberto Amato, Jeffrey Barrett, Sonia Goncalves, Ewan Harrison, David K. Jackson, Ian Johnston, Dominic Kwiatkowski, Cordelia Langford, John Sillitoe on behalf of the Wellcome Sanger Institute COVID-19 Surveillance Team |
| EPI_ISL_3469566, EPI_ISL_3469567 | Lighthouse Lab in Glasgow | Wellcome Sanger Institute for the COVID-19 Genomics UK (COG-UK) Consortium | Harper VanSteenhouse, Yumi Kasai, David Gray, Carol Clugston, Anna Dominiczak and Alex Alderton, Roberto Amato, Jeffrey Barrett, Sonia Goncalves, Ewan Harrison, David K. Jackson, Ian Johnston, Dominic Kwiatkowski, Cordelia Langford, John Sillitoe on behalf of the Wellcome Sanger Institute COVID-19 Surveillance Team |
| EPI_ISL_3469568, EPI_ISL_3469569, EPI_ISL_3469570, EPI_ISL_3469571 | Lighthouse Lab in Milton Keynes | Wellcome Sanger Institute for the COVID-19 Genomics UK (COG-UK) Consortium | The Lighthouse Lab in Milton Keynes and Alex Alderton, Roberto Amato, Jeffrey Barrett, Sonia Goncalves, Ewan Harrison, David K. Jackson, Ian Johnston, Dominic Kwiatkowski, Cordelia Langford, John Sillitoe on behalf of the Wellcome Sanger Institute COVID-19 Surveillance Team |
| EPI_ISL_3469572 | Lighthouse Lab in Alderley Park | Wellcome Sanger Institute for the COVID-19 Genomics UK (COG-UK) Consortium | Jacquelyn Wynn, Mairead Hyland, The Lighthouse Lab in Alderley Park and Alex Alderton, Roberto Amato, Jeffrey Barrett, Sonia Goncalves, Ewan Harrison, David K. Jackson, Ian Johnston, Dominic Kwiatkowski, Cordelia Langford, John Sillitoe on behalf of the Wellcome Sanger Institute COVID-19 Surveillance Team |
| EPI_ISL_3469573, EPI_ISL_3469574 | Lighthouse Laboratory Plymouth | Wellcome Sanger Institute for the COVID-19 Genomics UK (COG-UK) Consortium | Lighthouse Laboratory Plymouth and Alex Alderton, Roberto Amato, Jeffrey Barrett, Sonia Goncalves, Ewan Harrison, David K. Jackson, Ian Johnston, Dominic Kwiatkowski, Cordelia Langford, John Sillitoe on behalf of the Wellcome Sanger Institute COVID-19 Surveillance Team |
| EPI_ISL_3469575 | Lighthouse Lab in Milton Keynes | Wellcome Sanger Institute for the COVID-19 Genomics UK (COG-UK) Consortium | The Lighthouse Lab in Milton Keynes and Alex Alderton, Roberto Amato, Jeffrey Barrett, Sonia Goncalves, Ewan Harrison, David K. Jackson, Ian Johnston, Dominic Kwiatkowski, Cordelia Langford, John Sillitoe on behalf of the Wellcome Sanger Institute COVID-19 Surveillance Team |
| EPI_ISL_3469576, EPI_ISL_3469577, EPI_ISL_3469578, EPI_ISL_3469579 | Lighthouse Laboratory Plymouth | Wellcome Sanger Institute for the COVID-19 Genomics UK (COG-UK) Consortium | Lighthouse Laboratory Plymouth and Alex Alderton, Roberto Amato, Jeffrey Barrett, Sonia Goncalves, Ewan Harrison, David K. Jackson, Ian Johnston, Dominic Kwiatkowski, Cordelia Langford, John Sillitoe on behalf of the Wellcome Sanger Institute COVID-19 Surveillance Team |
| EPI_ISL_3469580 | Lighthouse Lab in Milton Keynes | Wellcome Sanger Institute for the COVID-19 Genomics UK (COG-UK) Consortium | The Lighthouse Lab in Milton Keynes and Alex Alderton, Roberto Amato, Jeffrey Barrett, Sonia Goncalves, Ewan Harrison, David K. Jackson, Ian Johnston, Dominic Kwiatkowski, Cordelia Langford, John Sillitoe on behalf of the Wellcome Sanger Institute COVID-19 Surveillance Team |
| EPI_ISL_3469581, EPI_ISL_3469582, EPI_ISL_3469583 | Lighthouse Lab in Alderley Park | Wellcome Sanger Institute for the COVID-19 Genomics UK (COG-UK) Consortium | Jacquelyn Wynn, Mairead Hyland, The Lighthouse Lab in Alderley Park and Alex Alderton, Roberto Amato, Jeffrey Barrett, Sonia Goncalves, Ewan Harrison, David K. Jackson, Ian Johnston, Dominic Kwiatkowski, Cordelia Langford, John Sillitoe on behalf of the Wellcome Sanger Institute COVID-19 Surveillance Team |
| EPI_ISL_3469584 | Lighthouse Lab in Milton Keynes | Wellcome Sanger Institute for the COVID-19 Genomics UK (COG-UK) Consortium | The Lighthouse Lab in Milton Keynes and Alex Alderton, Roberto Amato, Jeffrey Barrett, Sonia Goncalves, Ewan Harrison, David K. Jackson, Ian Johnston, Dominic Kwiatkowski, Cordelia Langford, John Sillitoe on behalf of the Wellcome Sanger Institute COVID-19 Surveillance Team |
| EPI_ISL_3469585, EPI_ISL_3469586 | Lighthouse Lab in Alderley Park | Wellcome Sanger Institute for the COVID-19 Genomics UK (COG-UK) Consortium | Jacquelyn Wynn, Mairead Hyland, The Lighthouse Lab in Alderley Park and Alex Alderton, Roberto Amato, Jeffrey Barrett, Sonia Goncalves, Ewan Harrison, David K. Jackson, Ian Johnston, Dominic Kwiatkowski, Cordelia Langford, John Sillitoe on behalf of the Wellcome Sanger Institute COVID-19 Surveillance Team |
| EPI_ISL_3469587, EPI_ISL_3469588 | Lighthouse Lab in Milton Keynes | Wellcome Sanger Institute for the COVID-19 Genomics UK (COG-UK) Consortium | The Lighthouse Lab in Milton Keynes and Alex Alderton, Roberto Amato, Jeffrey Barrett, Sonia Goncalves, Ewan Harrison, David K. Jackson, Ian Johnston, Dominic Kwiatkowski, Cordelia Langford, John Sillitoe on behalf of the Wellcome Sanger Institute COVID-19 Surveillance Team |
| EPI_ISL_3469589 | Lighthouse Laboratory Plymouth | Wellcome Sanger Institute for the COVID-19 Genomics UK (COG-UK) Consortium | Lighthouse Laboratory Plymouth and Alex Alderton, Roberto Amato, Jeffrey Barrett, Sonia Goncalves, Ewan Harrison, David K. Jackson, Ian Johnston, Dominic Kwiatkowski, Cordelia Langford, John Sillitoe on behalf of the Wellcome Sanger Institute COVID-19 Surveillance Team |
| EPI_ISL_3469590 | Lighthouse Lab in Alderley Park | Wellcome Sanger Institute for the COVID-19 Genomics UK (COG-UK) Consortium | Jacquelyn Wynn, Mairead Hyland, The Lighthouse Lab in Alderley Park and Alex Alderton, Roberto Amato, Jeffrey Barrett, Sonia Goncalves, Ewan Harrison, David K. Jackson, Ian Johnston, Dominic Kwiatkowski, Cordelia Langford, John Sillitoe on behalf of the Wellcome Sanger Institute COVID-19 Surveillance Team |
| EPI_ISL_3469591 | Lighthouse Laboratory Plymouth | Wellcome Sanger Institute for the COVID-19 Genomics UK (COG-UK) Consortium | Lighthouse Laboratory Plymouth and Alex Alderton, Roberto Amato, Jeffrey Barrett, Sonia Goncalves, Ewan Harrison, David K. Jackson, Ian Johnston, Dominic Kwiatkowski, Cordelia Langford, John Sillitoe on behalf of the Wellcome Sanger Institute COVID-19 Surveillance Team |
| EPI_ISL_3469592, EPI_ISL_3469593, EPI_ISL_3469594 | Lighthouse Lab in Milton Keynes | Wellcome Sanger Institute for the COVID-19 Genomics UK (COG-UK) Consortium | The Lighthouse Lab in Milton Keynes and Alex Alderton, Roberto Amato, Jeffrey Barrett, Sonia Goncalves, Ewan Harrison, David K. Jackson, Ian Johnston, Dominic Kwiatkowski, Cordelia Langford, John Sillitoe on behalf of the Wellcome Sanger Institute COVID-19 Surveillance Team |
| EPI_ISL_3469595 | Lighthouse Lab in Alderley Park | Wellcome Sanger Institute for the COVID-19 Genomics UK (COG-UK) Consortium | Jacquelyn Wynn, Mairead Hyland, The Lighthouse Lab in Alderley Park and Alex Alderton, Roberto Amato, Jeffrey Barrett, Sonia Goncalves, Ewan Harrison, David K. Jackson, Ian Johnston, Dominic Kwiatkowski, Cordelia Langford, John Sillitoe on behalf of the Wellcome Sanger Institute COVID-19 Surveillance Team |
| EPI_ISL_3469596 | Lighthouse Laboratory Plymouth | Wellcome Sanger Institute for the COVID-19 Genomics UK (COG-UK) Consortium | Lighthouse Laboratory Plymouth and Alex Alderton, Roberto Amato, Jeffrey Barrett, Sonia Goncalves, Ewan Harrison, David K. Jackson, Ian Johnston, Dominic Kwiatkowski, Cordelia Langford, John Sillitoe on behalf of the Wellcome Sanger Institute COVID-19 Surveillance Team |
| EPI_ISL_3469597, EPI_ISL_3469598 | Lighthouse Lab in Milton Keynes | Wellcome Sanger Institute for the COVID-19 Genomics UK (COG-UK) Consortium | The Lighthouse Lab in Milton Keynes and Alex Alderton, Roberto Amato, Jeffrey Barrett, Sonia Goncalves, Ewan Harrison, David K. Jackson, Ian Johnston, Dominic Kwiatkowski, Cordelia Langford, John Sillitoe on behalf of the Wellcome Sanger Institute COVID-19 Surveillance Team |
| EPI_ISL_3469599 | Lighthouse Lab in Alderley Park | Wellcome Sanger Institute for the COVID-19 Genomics UK (COG-UK) Consortium | Jacquelyn Wynn, Mairead Hyland, The Lighthouse Lab in Alderley Park and Alex Alderton, Roberto Amato, Jeffrey Barrett, Sonia Goncalves, Ewan Harrison, David K. Jackson, Ian Johnston, Dominic Kwiatkowski, Cordelia Langford, John Sillitoe on behalf of the Wellcome Sanger Institute COVID-19 Surveillance Team |
| EPI_ISL_3469600, EPI_ISL_3469601, EPI_ISL_3469602 | Lighthouse Laboratory Plymouth | Wellcome Sanger Institute for the COVID-19 Genomics UK (COG-UK) Consortium | Lighthouse Laboratory Plymouth and Alex Alderton, Roberto Amato, Jeffrey Barrett, Sonia Goncalves, Ewan Harrison, David K. Jackson, Ian Johnston, Dominic Kwiatkowski, Cordelia Langford, John Sillitoe on behalf of the Wellcome Sanger Institute COVID-19 Surveillance Team |

| | | | |
|---|---|---|---|
| EPI_ISL_3469603 | Lighthouse Lab in Alderley Park | Wellcome Sanger Institute for the COVID-19 Genomics UK (COG-UK) Consortium | Jacquelyn Wynn, Mairead Hyland, The Lighthouse Lab in Alderley Park and Alex Alderton, Roberto Amato, Jeffrey Barrett, Sonia Goncalves, Ewan Harrison, David K. Jackson, Ian Johnston, Dominic Kwiatkowski, Cordelia Langford, John Sillitoe on behalf of the Wellcome Sanger Institute COVID-19 Surveillance Team |
| EPI_ISL_3469604 | Lighthouse Lab in Milton Keynes | Wellcome Sanger Institute for the COVID-19 Genomics UK (COG-UK) Consortium | The Lighthouse Lab in Milton Keynes and Alex Alderton, Roberto Amato, Jeffrey Barrett, Sonia Goncalves, Ewan Harrison, David K. Jackson, Ian Johnston, Dominic Kwiatkowski, Cordelia Langford, John Sillitoe on behalf of the Wellcome Sanger Institute COVID-19 Surveillance Team |
| EPI_ISL_3469605, EPI_ISL_3469606 | Lighthouse Laboratory Plymouth | Wellcome Sanger Institute for the COVID-19 Genomics UK (COG-UK) Consortium | Lighthouse Laboratory Plymouth and Alex Alderton, Roberto Amato, Jeffrey Barrett, Sonia Goncalves, Ewan Harrison, David K. Jackson, Ian Johnston, Dominic Kwiatkowski, Cordelia Langford, John Sillitoe on behalf of the Wellcome Sanger Institute COVID-19 Surveillance Team |
| EPI_ISL_3469607 | Lighthouse Lab in Milton Keynes | Wellcome Sanger Institute for the COVID-19 Genomics UK (COG-UK) Consortium | The Lighthouse Lab in Milton Keynes and Alex Alderton, Roberto Amato, Jeffrey Barrett, Sonia Goncalves, Ewan Harrison, David K. Jackson, Ian Johnston, Dominic Kwiatkowski, Cordelia Langford, John Sillitoe on behalf of the Wellcome Sanger Institute COVID-19 Surveillance Team |
| EPI_ISL_3469608 | Lighthouse Lab in Alderley Park | Wellcome Sanger Institute for the COVID-19 Genomics UK (COG-UK) Consortium | Jacquelyn Wynn, Mairead Hyland, The Lighthouse Lab in Alderley Park and Alex Alderton, Roberto Amato, Jeffrey Barrett, Sonia Goncalves, Ewan Harrison, David K. Jackson, Ian Johnston, Dominic Kwiatkowski, Cordelia Langford, John Sillitoe on behalf of the Wellcome Sanger Institute COVID-19 Surveillance Team |
| EPI_ISL_3469609 | Lighthouse Laboratory Plymouth | Wellcome Sanger Institute for the COVID-19 Genomics UK (COG-UK) Consortium | Lighthouse Laboratory Plymouth and Alex Alderton, Roberto Amato, Jeffrey Barrett, Sonia Goncalves, Ewan Harrison, David K. Jackson, Ian Johnston, Dominic Kwiatkowski, Cordelia Langford, John Sillitoe on behalf of the Wellcome Sanger Institute COVID-19 Surveillance Team |
| EPI_ISL_3469610 | Lighthouse Lab in Alderley Park | Wellcome Sanger Institute for the COVID-19 Genomics UK (COG-UK) Consortium | Jacquelyn Wynn, Mairead Hyland, The Lighthouse Lab in Alderley Park and Alex Alderton, Roberto Amato, Jeffrey Barrett, Sonia Goncalves, Ewan Harrison, David K. Jackson, Ian Johnston, Dominic Kwiatkowski, Cordelia Langford, John Sillitoe on behalf of the Wellcome Sanger Institute COVID-19 Surveillance Team |
| EPI_ISL_3469611 | Lighthouse Laboratory Plymouth | Wellcome Sanger Institute for the COVID-19 Genomics UK (COG-UK) Consortium | Lighthouse Laboratory Plymouth and Alex Alderton, Roberto Amato, Jeffrey Barrett, Sonia Goncalves, Ewan Harrison, David K. Jackson, Ian Johnston, Dominic Kwiatkowski, Cordelia Langford, John Sillitoe on behalf of the Wellcome Sanger Institute COVID-19 Surveillance Team |
| EPI_ISL_3469612, EPI_ISL_3469613 | Lighthouse Lab in Alderley Park | Wellcome Sanger Institute for the COVID-19 Genomics UK (COG-UK) Consortium | Jacquelyn Wynn, Mairead Hyland, The Lighthouse Lab in Alderley Park and Alex Alderton, Roberto Amato, Jeffrey Barrett, Sonia Goncalves, Ewan Harrison, David K. Jackson, Ian Johnston, Dominic Kwiatkowski, Cordelia Langford, John Sillitoe on behalf of the Wellcome Sanger Institute COVID-19 Surveillance Team |
| EPI_ISL_3469614 | Lighthouse Lab in Milton Keynes | Wellcome Sanger Institute for the COVID-19 Genomics UK (COG-UK) Consortium | The Lighthouse Lab in Milton Keynes and Alex Alderton, Roberto Amato, Jeffrey Barrett, Sonia Goncalves, Ewan Harrison, David K. Jackson, Ian Johnston, Dominic Kwiatkowski, Cordelia Langford, John Sillitoe on behalf of the Wellcome Sanger Institute COVID-19 Surveillance Team |
| EPI_ISL_3469615 | Lighthouse Laboratory Plymouth | Wellcome Sanger Institute for the COVID-19 Genomics UK (COG-UK) Consortium | Lighthouse Laboratory Plymouth and Alex Alderton, Roberto Amato, Jeffrey Barrett, Sonia Goncalves, Ewan Harrison, David K. Jackson, Ian Johnston, Dominic Kwiatkowski, Cordelia Langford, John Sillitoe on behalf of the Wellcome Sanger Institute COVID-19 Surveillance Team |
| EPI_ISL_3469616 | Lighthouse Lab in Alderley Park | Wellcome Sanger Institute for the COVID-19 Genomics UK (COG-UK) Consortium | Jacquelyn Wynn, Mairead Hyland, The Lighthouse Lab in Alderley Park and Alex Alderton, Roberto Amato, Jeffrey Barrett, Sonia Goncalves, Ewan Harrison, David K. Jackson, Ian Johnston, Dominic Kwiatkowski, Cordelia Langford, John Sillitoe on behalf of the Wellcome Sanger Institute COVID-19 Surveillance Team |
| EPI_ISL_3469617 | Lighthouse Lab in Milton Keynes | Wellcome Sanger Institute for the COVID-19 Genomics UK (COG-UK) Consortium | The Lighthouse Lab in Milton Keynes and Alex Alderton, Roberto Amato, Jeffrey Barrett, Sonia Goncalves, Ewan Harrison, David K. Jackson, Ian Johnston, Dominic Kwiatkowski, Cordelia Langford, John Sillitoe on behalf of the Wellcome Sanger Institute COVID-19 Surveillance Team |
| EPI_ISL_3469618 | Lighthouse Lab in Alderley Park | Wellcome Sanger Institute for the COVID-19 Genomics UK (COG-UK) Consortium | Jacquelyn Wynn, Mairead Hyland, The Lighthouse Lab in Alderley Park and Alex Alderton, Roberto Amato, Jeffrey Barrett, Sonia Goncalves, Ewan Harrison, David K. Jackson, Ian Johnston, Dominic Kwiatkowski, Cordelia Langford, John Sillitoe on behalf of the Wellcome Sanger Institute COVID-19 Surveillance Team |
| EPI_ISL_3469619, EPI_ISL_3469620, EPI_ISL_3469621, EPI_ISL_3469622 | Lighthouse Lab in Milton Keynes | Wellcome Sanger Institute for the COVID-19 Genomics UK (COG-UK) Consortium | The Lighthouse Lab in Milton Keynes and Alex Alderton, Roberto Amato, Jeffrey Barrett, Sonia Goncalves, Ewan Harrison, David K. Jackson, Ian Johnston, Dominic Kwiatkowski, Cordelia Langford, John Sillitoe on behalf of the Wellcome Sanger Institute COVID-19 Surveillance Team |
| EPI_ISL_3469623 | Lighthouse Lab in Alderley Park | Wellcome Sanger Institute for the COVID-19 Genomics UK (COG-UK) Consortium | Jacquelyn Wynn, Mairead Hyland, The Lighthouse Lab in Alderley Park and Alex Alderton, Roberto Amato, Jeffrey Barrett, Sonia Goncalves, Ewan Harrison, David K. Jackson, Ian Johnston, Dominic Kwiatkowski, Cordelia Langford, John Sillitoe on behalf of the Wellcome Sanger Institute COVID-19 Surveillance Team |
| EPI_ISL_3469624 | Lighthouse Lab in Milton Keynes | Wellcome Sanger Institute for the COVID-19 Genomics UK (COG-UK) Consortium | The Lighthouse Lab in Milton Keynes and Alex Alderton, Roberto Amato, Jeffrey Barrett, Sonia Goncalves, Ewan Harrison, David K. Jackson, Ian Johnston, Dominic Kwiatkowski, Cordelia Langford, John Sillitoe on behalf of the Wellcome Sanger Institute COVID-19 Surveillance Team |
| EPI_ISL_3469625, EPI_ISL_3469626, EPI_ISL_3469627, EPI_ISL_3469628 | Lighthouse Lab in Alderley Park | Wellcome Sanger Institute for the COVID-19 Genomics UK (COG-UK) Consortium | Jacquelyn Wynn, Mairead Hyland, The Lighthouse Lab in Alderley Park and Alex Alderton, Roberto Amato, Jeffrey Barrett, Sonia Goncalves, Ewan Harrison, David K. Jackson, Ian Johnston, Dominic Kwiatkowski, Cordelia Langford, John Sillitoe on behalf of the Wellcome Sanger Institute COVID-19 Surveillance Team |
| EPI_ISL_3469629, EPI_ISL_3469630, EPI_ISL_3469631 | Lighthouse Laboratory Plymouth | Wellcome Sanger Institute for the COVID-19 Genomics UK (COG-UK) Consortium | Lighthouse Laboratory Plymouth and Alex Alderton, Roberto Amato, Jeffrey Barrett, Sonia Goncalves, Ewan Harrison, David K. Jackson, Ian Johnston, Dominic Kwiatkowski, Cordelia Langford, John Sillitoe on behalf of the Wellcome Sanger Institute COVID-19 Surveillance Team |
| EPI_ISL_3469632, EPI_ISL_3469633, EPI_ISL_3469634, EPI_ISL_3469635 | Lighthouse Lab in Milton Keynes | Wellcome Sanger Institute for the COVID-19 Genomics UK (COG-UK) Consortium | The Lighthouse Lab in Milton Keynes and Alex Alderton, Roberto Amato, Jeffrey Barrett, Sonia Goncalves, Ewan Harrison, David K. Jackson, Ian Johnston, Dominic Kwiatkowski, Cordelia Langford, John Sillitoe on behalf of the Wellcome Sanger Institute COVID-19 Surveillance Team |
| EPI_ISL_3469636, EPI_ISL_3469637 | Lighthouse Lab in Alderley Park | Wellcome Sanger Institute for the COVID-19 Genomics UK (COG-UK) Consortium | Jacquelyn Wynn, Mairead Hyland, The Lighthouse Lab in Alderley Park and Alex Alderton, Roberto Amato, Jeffrey Barrett, Sonia Goncalves, Ewan Harrison, David K. Jackson, Ian Johnston, Dominic Kwiatkowski, Cordelia Langford, John Sillitoe on behalf of the Wellcome Sanger Institute COVID-19 Surveillance Team |
| EPI_ISL_3469638 | Lighthouse Lab in Milton Keynes | Wellcome Sanger Institute for the COVID-19 Genomics UK (COG-UK) Consortium | The Lighthouse Lab in Milton Keynes and Alex Alderton, Roberto Amato, Jeffrey Barrett, Sonia Goncalves, Ewan Harrison, David K. Jackson, Ian Johnston, Dominic Kwiatkowski, Cordelia Langford, John Sillitoe on behalf of the Wellcome Sanger Institute COVID-19 Surveillance Team |
| EPI_ISL_3469639 | Lighthouse Lab in Alderley Park | Wellcome Sanger Institute for the COVID-19 Genomics UK (COG-UK) Consortium | Jacquelyn Wynn, Mairead Hyland, The Lighthouse Lab in Alderley Park and Alex Alderton, Roberto Amato, Jeffrey Barrett, Sonia Goncalves, Ewan Harrison, David K. Jackson, Ian Johnston, Dominic Kwiatkowski, Cordelia Langford, John Sillitoe on behalf of the Wellcome Sanger Institute COVID-19 Surveillance Team |
| EPI_ISL_3469640, EPI_ISL_3469641 | Lighthouse Laboratory Plymouth | Wellcome Sanger Institute for the COVID-19 Genomics UK (COG-UK) Consortium | Lighthouse Laboratory Plymouth and Alex Alderton, Roberto Amato, Jeffrey Barrett, Sonia Goncalves, Ewan Harrison, David K. Jackson, Ian Johnston, Dominic Kwiatkowski, Cordelia Langford, John Sillitoe on behalf of the Wellcome Sanger Institute COVID-19 Surveillance Team |
| EPI_ISL_3469642 | Lighthouse Lab in Milton Keynes | Wellcome Sanger Institute for the COVID-19 Genomics UK (COG-UK) Consortium | The Lighthouse Lab in Milton Keynes and Alex Alderton, Roberto Amato, Jeffrey Barrett, Sonia Goncalves, Ewan Harrison, David K. Jackson, Ian Johnston, Dominic Kwiatkowski, Cordelia Langford, John Sillitoe on behalf of the Wellcome Sanger Institute COVID-19 Surveillance Team |
| EPI_ISL_3469643, EPI_ISL_3469644 | Lighthouse Laboratory Plymouth | Wellcome Sanger Institute for the COVID-19 Genomics UK (COG-UK) Consortium | Lighthouse Laboratory Plymouth and Alex Alderton, Roberto Amato, Jeffrey Barrett, Sonia Goncalves, Ewan Harrison, David K. Jackson, Ian Johnston, Dominic Kwiatkowski, Cordelia Langford, John Sillitoe on behalf of the Wellcome Sanger Institute COVID-19 Surveillance Team |
| EPI_ISL_3469645, EPI_ISL_3469646, EPI_ISL_3469647, EPI_ISL_3469648, EPI_ISL_3469649 | Lighthouse Lab in Milton Keynes | Wellcome Sanger Institute for the COVID-19 Genomics UK (COG-UK) Consortium | The Lighthouse Lab in Milton Keynes and Alex Alderton, Roberto Amato, Jeffrey Barrett, Sonia Goncalves, Ewan Harrison, David K. Jackson, Ian Johnston, Dominic Kwiatkowski, Cordelia Langford, John Sillitoe on behalf of the Wellcome Sanger Institute COVID-19 Surveillance Team |
| EPI_ISL_3469650 | Lighthouse Lab in Alderley Park | Wellcome Sanger Institute for the COVID-19 Genomics UK (COG-UK) Consortium | Jacquelyn Wynn, Mairead Hyland, The Lighthouse Lab in Alderley Park and Alex Alderton, Roberto Amato, Jeffrey Barrett, Sonia Goncalves, Ewan Harrison, David K. Jackson, Ian Johnston, Dominic Kwiatkowski, Cordelia Langford, John Sillitoe on behalf of the Wellcome Sanger Institute COVID-19 Surveillance Team |
| EPI_ISL_3469651, EPI_ISL_3469652, EPI_ISL_3469653, EPI_ISL_3469654 | Lighthouse Laboratory Plymouth | Wellcome Sanger Institute for the COVID-19 Genomics UK (COG-UK) Consortium | Lighthouse Laboratory Plymouth and Alex Alderton, Roberto Amato, Jeffrey Barrett, Sonia Goncalves, Ewan Harrison, David K. Jackson, Ian Johnston, Dominic Kwiatkowski, Cordelia Langford, John Sillitoe on behalf of the Wellcome Sanger Institute COVID-19 Surveillance Team |
| EPI_ISL_3469655 | Lighthouse Lab in Alderley Park | Wellcome Sanger Institute for the COVID-19 Genomics UK (COG-UK) Consortium | Jacquelyn Wynn, Mairead Hyland, The Lighthouse Lab in Alderley Park and Alex Alderton, Roberto Amato, Jeffrey Barrett, Sonia Goncalves, Ewan Harrison, David K. Jackson, Ian Johnston, Dominic Kwiatkowski, Cordelia Langford, John Sillitoe on behalf of the Wellcome Sanger Institute COVID-19 Surveillance Team |
| EPI_ISL_3469656 | Lighthouse Lab in Milton Keynes | Wellcome Sanger Institute for the COVID-19 Genomics UK (COG-UK) Consortium | The Lighthouse Lab in Milton Keynes and Alex Alderton, Roberto Amato, Jeffrey Barrett, Sonia Goncalves, Ewan Harrison, David K. Jackson, Ian Johnston, Dominic Kwiatkowski, Cordelia Langford, John Sillitoe on behalf of the Wellcome Sanger Institute COVID-19 Surveillance Team |
| EPI_ISL_3469657, EPI_ISL_3469658 | Lighthouse Lab in Alderley Park | Wellcome Sanger Institute for the COVID-19 Genomics UK (COG-UK) Consortium | Jacquelyn Wynn, Mairead Hyland, The Lighthouse Lab in Alderley Park and Alex Alderton, Roberto Amato, Jeffrey Barrett, Sonia Goncalves, Ewan Harrison, David K. Jackson, Ian Johnston, Dominic Kwiatkowski, Cordelia Langford, John Sillitoe on behalf of the Wellcome Sanger Institute COVID-19 Surveillance Team |
| EPI_ISL_3469659 | Lighthouse Laboratory Plymouth | Wellcome Sanger Institute for the COVID-19 Genomics UK (COG-UK) Consortium | Lighthouse Laboratory Plymouth and Alex Alderton, Roberto Amato, Jeffrey Barrett, Sonia Goncalves, Ewan Harrison, David K. Jackson, Ian Johnston, Dominic Kwiatkowski, Cordelia Langford, John Sillitoe on behalf of the Wellcome Sanger Institute COVID-19 Surveillance Team |
| EPI_ISL_3469660, EPI_ISL_3469661 | Lighthouse Lab in Alderley Park | Wellcome Sanger Institute for the COVID-19 Genomics UK (COG-UK) Consortium | Jacquelyn Wynn, Mairead Hyland, The Lighthouse Lab in Alderley Park and Alex Alderton, Roberto Amato, Jeffrey Barrett, Sonia Goncalves, Ewan Harrison, David K. Jackson, Ian Johnston, Dominic Kwiatkowski, Cordelia Langford, John Sillitoe on behalf of the Wellcome Sanger Institute COVID-19 Surveillance Team |
| EPI_ISL_3469662 | Lighthouse Laboratory Plymouth | Wellcome Sanger Institute for the COVID-19 Genomics UK (COG-UK) Consortium | Lighthouse Laboratory Plymouth and Alex Alderton, Roberto Amato, Jeffrey Barrett, Sonia Goncalves, Ewan Harrison, David K. Jackson, Ian Johnston, Dominic Kwiatkowski, Cordelia Langford, John Sillitoe on behalf of the Wellcome Sanger Institute COVID-19 Surveillance Team |
| EPI_ISL_3469663 | Lighthouse Lab in Alderley Park | Wellcome Sanger Institute for the COVID-19 Genomics UK (COG-UK) Consortium | Jacquelyn Wynn, Mairead Hyland, The Lighthouse Lab in Alderley Park and Alex Alderton, Roberto Amato, Jeffrey Barrett, Sonia Goncalves, Ewan Harrison, David K. Jackson, Ian Johnston, Dominic Kwiatkowski, Cordelia Langford, John Sillitoe on behalf of the Wellcome Sanger Institute COVID-19 Surveillance Team |
| EPI_ISL_3469664 | Lighthouse Lab in Milton Keynes | Wellcome Sanger Institute for the COVID-19 Genomics UK (COG-UK) Consortium | The Lighthouse Lab in Milton Keynes and Alex Alderton, Roberto Amato, Jeffrey Barrett, Sonia Goncalves, Ewan Harrison, David K. Jackson, Ian Johnston, Dominic Kwiatkowski, Cordelia Langford, John Sillitoe on behalf of the Wellcome Sanger Institute COVID-19 Surveillance Team |
| EPI_ISL_3469665 | Lighthouse Laboratory Plymouth | Wellcome Sanger Institute for the COVID-19 Genomics UK (COG-UK) Consortium | Lighthouse Laboratory Plymouth and Alex Alderton, Roberto Amato, Jeffrey Barrett, Sonia Goncalves, Ewan Harrison, David K. Jackson, Ian Johnston, Dominic Kwiatkowski, Cordelia Langford, John Sillitoe on behalf of the Wellcome Sanger Institute COVID-19 Surveillance Team |
| EPI_ISL_3469666 | Lighthouse Lab in Alderley Park | Wellcome Sanger Institute for the COVID-19 Genomics UK (COG-UK) Consortium | Jacquelyn Wynn, Mairead Hyland, The Lighthouse Lab in Alderley Park and Alex Alderton, Roberto Amato, Jeffrey Barrett, Sonia Goncalves, Ewan Harrison, David K. Jackson, Ian Johnston, Dominic Kwiatkowski, Cordelia Langford, John Sillitoe on behalf of the Wellcome Sanger Institute COVID-19 Surveillance Team |
| EPI_ISL_3469667 | Lighthouse Laboratory Plymouth | Wellcome Sanger Institute for the COVID-19 Genomics UK (COG-UK) Consortium | Lighthouse Laboratory Plymouth and Alex Alderton, Roberto Amato, Jeffrey Barrett, Sonia Goncalves, Ewan Harrison, David K. Jackson, Ian Johnston, Dominic Kwiatkowski, Cordelia Langford, John Sillitoe on behalf of the Wellcome Sanger Institute COVID-19 Surveillance Team |

| | | | |
|---|---|---|---|
| EPI_ISL_3469668 | Lighthouse Lab in Alderley Park | Wellcome Sanger Institute for the COVID-19 Genomics UK (COG-UK) Consortium | Jacquelyn Wynn, Mairead Hyland, The Lighthouse Lab in Alderley Park and Alex Alderton, Roberto Amato, Jeffrey Barrett, Sonia Goncalves, Ewan Harrison, David K. Jackson, Ian Johnston, Dominic Kwiatkowski, Cordelia Langford, John Sillitoe on behalf of the Wellcome Sanger Institute COVID-19 Surveillance Team |
| EPI_ISL_3469669 | Lighthouse Lab in Milton Keynes | Wellcome Sanger Institute for the COVID-19 Genomics UK (COG-UK) Consortium | The Lighthouse Lab in Milton Keynes and Alex Alderton, Roberto Amato, Jeffrey Barrett, Sonia Goncalves, Ewan Harrison, David K. Jackson, Ian Johnston, Dominic Kwiatkowski, Cordelia Langford, John Sillitoe on behalf of the Wellcome Sanger Institute COVID-19 Surveillance Team |
| EPI_ISL_3469670, EPI_ISL_3469671 | Lighthouse Lab in Alderley Park | Wellcome Sanger Institute for the COVID-19 Genomics UK (COG-UK) Consortium | Jacquelyn Wynn, Mairead Hyland, The Lighthouse Lab in Alderley Park and Alex Alderton, Roberto Amato, Jeffrey Barrett, Sonia Goncalves, Ewan Harrison, David K. Jackson, Ian Johnston, Dominic Kwiatkowski, Cordelia Langford, John Sillitoe on behalf of the Wellcome Sanger Institute COVID-19 Surveillance Team |
| EPI_ISL_3469672 | Lighthouse Lab in Milton Keynes | Wellcome Sanger Institute for the COVID-19 Genomics UK (COG-UK) Consortium | The Lighthouse Lab in Milton Keynes and Alex Alderton, Roberto Amato, Jeffrey Barrett, Sonia Goncalves, Ewan Harrison, David K. Jackson, Ian Johnston, Dominic Kwiatkowski, Cordelia Langford, John Sillitoe on behalf of the Wellcome Sanger Institute COVID-19 Surveillance Team |
| EPI_ISL_3469673 | Lighthouse Lab in Alderley Park | Wellcome Sanger Institute for the COVID-19 Genomics UK (COG-UK) Consortium | Jacquelyn Wynn, Mairead Hyland, The Lighthouse Lab in Alderley Park and Alex Alderton, Roberto Amato, Jeffrey Barrett, Sonia Goncalves, Ewan Harrison, David K. Jackson, Ian Johnston, Dominic Kwiatkowski, Cordelia Langford, John Sillitoe on behalf of the Wellcome Sanger Institute COVID-19 Surveillance Team |
| EPI_ISL_3469674 | Lighthouse Laboratory Plymouth | Wellcome Sanger Institute for the COVID-19 Genomics UK (COG-UK) Consortium | Lighthouse Laboratory Plymouth and Alex Alderton, Roberto Amato, Jeffrey Barrett, Sonia Goncalves, Ewan Harrison, David K. Jackson, Ian Johnston, Dominic Kwiatkowski, Cordelia Langford, John Sillitoe on behalf of the Wellcome Sanger Institute COVID-19 Surveillance Team |
| EPI_ISL_3469675 | Lighthouse Lab in Milton Keynes | Wellcome Sanger Institute for the COVID-19 Genomics UK (COG-UK) Consortium | The Lighthouse Lab in Milton Keynes and Alex Alderton, Roberto Amato, Jeffrey Barrett, Sonia Goncalves, Ewan Harrison, David K. Jackson, Ian Johnston, Dominic Kwiatkowski, Cordelia Langford, John Sillitoe on behalf of the Wellcome Sanger Institute COVID-19 Surveillance Team |
| EPI_ISL_3469676 | Lighthouse Lab in Alderley Park | Wellcome Sanger Institute for the COVID-19 Genomics UK (COG-UK) Consortium | Jacquelyn Wynn, Mairead Hyland, The Lighthouse Lab in Alderley Park and Alex Alderton, Roberto Amato, Jeffrey Barrett, Sonia Goncalves, Ewan Harrison, David K. Jackson, Ian Johnston, Dominic Kwiatkowski, Cordelia Langford, John Sillitoe on behalf of the Wellcome Sanger Institute COVID-19 Surveillance Team |
| EPI_ISL_3469677 | Lighthouse Laboratory Plymouth | Wellcome Sanger Institute for the COVID-19 Genomics UK (COG-UK) Consortium | Lighthouse Laboratory Plymouth and Alex Alderton, Roberto Amato, Jeffrey Barrett, Sonia Goncalves, Ewan Harrison, David K. Jackson, Ian Johnston, Dominic Kwiatkowski, Cordelia Langford, John Sillitoe on behalf of the Wellcome Sanger Institute COVID-19 Surveillance Team |
| EPI_ISL_3469678, EPI_ISL_3469679 | Lighthouse Lab in Alderley Park | Wellcome Sanger Institute for the COVID-19 Genomics UK (COG-UK) Consortium | Jacquelyn Wynn, Mairead Hyland, The Lighthouse Lab in Alderley Park and Alex Alderton, Roberto Amato, Jeffrey Barrett, Sonia Goncalves, Ewan Harrison, David K. Jackson, Ian Johnston, Dominic Kwiatkowski, Cordelia Langford, John Sillitoe on behalf of the Wellcome Sanger Institute COVID-19 Surveillance Team |
| EPI_ISL_3469680 | Lighthouse Laboratory Plymouth | Wellcome Sanger Institute for the COVID-19 Genomics UK (COG-UK) Consortium | Lighthouse Laboratory Plymouth and Alex Alderton, Roberto Amato, Jeffrey Barrett, Sonia Goncalves, Ewan Harrison, David K. Jackson, Ian Johnston, Dominic Kwiatkowski, Cordelia Langford, John Sillitoe on behalf of the Wellcome Sanger Institute COVID-19 Surveillance Team |
| EPI_ISL_3469681 | Lighthouse Lab in Alderley Park | Wellcome Sanger Institute for the COVID-19 Genomics UK (COG-UK) Consortium | Jacquelyn Wynn, Mairead Hyland, The Lighthouse Lab in Alderley Park and Alex Alderton, Roberto Amato, Jeffrey Barrett, Sonia Goncalves, Ewan Harrison, David K. Jackson, Ian Johnston, Dominic Kwiatkowski, Cordelia Langford, John Sillitoe on behalf of the Wellcome Sanger Institute COVID-19 Surveillance Team |
| EPI_ISL_3469682, EPI_ISL_3469683, EPI_ISL_3469684 | Lighthouse Laboratory Plymouth | Wellcome Sanger Institute for the COVID-19 Genomics UK (COG-UK) Consortium | Lighthouse Laboratory Plymouth and Alex Alderton, Roberto Amato, Jeffrey Barrett, Sonia Goncalves, Ewan Harrison, David K. Jackson, Ian Johnston, Dominic Kwiatkowski, Cordelia Langford, John Sillitoe on behalf of the Wellcome Sanger Institute COVID-19 Surveillance Team |
| EPI_ISL_3469685 | Lighthouse Lab in Alderley Park | Wellcome Sanger Institute for the COVID-19 Genomics UK (COG-UK) Consortium | Jacquelyn Wynn, Mairead Hyland, The Lighthouse Lab in Alderley Park and Alex Alderton, Roberto Amato, Jeffrey Barrett, Sonia Goncalves, Ewan Harrison, David K. Jackson, Ian Johnston, Dominic Kwiatkowski, Cordelia Langford, John Sillitoe on behalf of the Wellcome Sanger Institute COVID-19 Surveillance Team |
| EPI_ISL_3469686 | Lighthouse Lab in Milton Keynes | Wellcome Sanger Institute for the COVID-19 Genomics UK (COG-UK) Consortium | The Lighthouse Lab in Milton Keynes and Alex Alderton, Roberto Amato, Jeffrey Barrett, Sonia Goncalves, Ewan Harrison, David K. Jackson, Ian Johnston, Dominic Kwiatkowski, Cordelia Langford, John Sillitoe on behalf of the Wellcome Sanger Institute COVID-19 Surveillance Team |
| EPI_ISL_3469687 | Lighthouse Laboratory Plymouth | Wellcome Sanger Institute for the COVID-19 Genomics UK (COG-UK) Consortium | Lighthouse Laboratory Plymouth and Alex Alderton, Roberto Amato, Jeffrey Barrett, Sonia Goncalves, Ewan Harrison, David K. Jackson, Ian Johnston, Dominic Kwiatkowski, Cordelia Langford, John Sillitoe on behalf of the Wellcome Sanger Institute COVID-19 Surveillance Team |
| EPI_ISL_3469688 | Lighthouse Lab in Milton Keynes | Wellcome Sanger Institute for the COVID-19 Genomics UK (COG-UK) Consortium | The Lighthouse Lab in Milton Keynes and Alex Alderton, Roberto Amato, Jeffrey Barrett, Sonia Goncalves, Ewan Harrison, David K. Jackson, Ian Johnston, Dominic Kwiatkowski, Cordelia Langford, John Sillitoe on behalf of the Wellcome Sanger Institute COVID-19 Surveillance Team |
| EPI_ISL_3469689, EPI_ISL_3469690, EPI_ISL_3469691 | Lighthouse Lab in Alderley Park | Wellcome Sanger Institute for the COVID-19 Genomics UK (COG-UK) Consortium | Jacquelyn Wynn, Mairead Hyland, The Lighthouse Lab in Alderley Park and Alex Alderton, Roberto Amato, Jeffrey Barrett, Sonia Goncalves, Ewan Harrison, David K. Jackson, Ian Johnston, Dominic Kwiatkowski, Cordelia Langford, John Sillitoe on behalf of the Wellcome Sanger Institute COVID-19 Surveillance Team |
| EPI_ISL_3469692, EPI_ISL_3469693 | Lighthouse Lab in Milton Keynes | Wellcome Sanger Institute for the COVID-19 Genomics UK (COG-UK) Consortium | The Lighthouse Lab in Milton Keynes and Alex Alderton, Roberto Amato, Jeffrey Barrett, Sonia Goncalves, Ewan Harrison, David K. Jackson, Ian Johnston, Dominic Kwiatkowski, Cordelia Langford, John Sillitoe on behalf of the Wellcome Sanger Institute COVID-19 Surveillance Team |
| EPI_ISL_3469694 | Lighthouse Laboratory Plymouth | Wellcome Sanger Institute for the COVID-19 Genomics UK (COG-UK) Consortium | Lighthouse Laboratory Plymouth and Alex Alderton, Roberto Amato, Jeffrey Barrett, Sonia Goncalves, Ewan Harrison, David K. Jackson, Ian Johnston, Dominic Kwiatkowski, Cordelia Langford, John Sillitoe on behalf of the Wellcome Sanger Institute COVID-19 Surveillance Team |
| EPI_ISL_3469695, EPI_ISL_3469696 | Lighthouse Lab in Milton Keynes | Wellcome Sanger Institute for the COVID-19 Genomics UK (COG-UK) Consortium | The Lighthouse Lab in Milton Keynes and Alex Alderton, Roberto Amato, Jeffrey Barrett, Sonia Goncalves, Ewan Harrison, David K. Jackson, Ian Johnston, Dominic Kwiatkowski, Cordelia Langford, John Sillitoe on behalf of the Wellcome Sanger Institute COVID-19 Surveillance Team |
| EPI_ISL_3469697 | Lighthouse Lab in Alderley Park | Wellcome Sanger Institute for the COVID-19 Genomics UK (COG-UK) Consortium | Jacquelyn Wynn, Mairead Hyland, The Lighthouse Lab in Alderley Park and Alex Alderton, Roberto Amato, Jeffrey Barrett, Sonia Goncalves, Ewan Harrison, David K. Jackson, Ian Johnston, Dominic Kwiatkowski, Cordelia Langford, John Sillitoe on behalf of the Wellcome Sanger Institute COVID-19 Surveillance Team |
| EPI_ISL_3469698, EPI_ISL_3469699 | Lighthouse Laboratory Plymouth | Wellcome Sanger Institute for the COVID-19 Genomics UK (COG-UK) Consortium | Lighthouse Laboratory Plymouth and Alex Alderton, Roberto Amato, Jeffrey Barrett, Sonia Goncalves, Ewan Harrison, David K. Jackson, Ian Johnston, Dominic Kwiatkowski, Cordelia Langford, John Sillitoe on behalf of the Wellcome Sanger Institute COVID-19 Surveillance Team |
| EPI_ISL_3469700 | Lighthouse Lab in Milton Keynes | Wellcome Sanger Institute for the COVID-19 Genomics UK (COG-UK) Consortium | The Lighthouse Lab in Milton Keynes and Alex Alderton, Roberto Amato, Jeffrey Barrett, Sonia Goncalves, Ewan Harrison, David K. Jackson, Ian Johnston, Dominic Kwiatkowski, Cordelia Langford, John Sillitoe on behalf of the Wellcome Sanger Institute COVID-19 Surveillance Team |
| EPI_ISL_3469701 | Lighthouse Lab in Alderley Park | Wellcome Sanger Institute for the COVID-19 Genomics UK (COG-UK) Consortium | Jacquelyn Wynn, Mairead Hyland, The Lighthouse Lab in Alderley Park and Alex Alderton, Roberto Amato, Jeffrey Barrett, Sonia Goncalves, Ewan Harrison, David K. Jackson, Ian Johnston, Dominic Kwiatkowski, Cordelia Langford, John Sillitoe on behalf of the Wellcome Sanger Institute COVID-19 Surveillance Team |
| EPI_ISL_3469702, EPI_ISL_3469703 | Lighthouse Lab in Milton Keynes | Wellcome Sanger Institute for the COVID-19 Genomics UK (COG-UK) Consortium | The Lighthouse Lab in Milton Keynes and Alex Alderton, Roberto Amato, Jeffrey Barrett, Sonia Goncalves, Ewan Harrison, David K. Jackson, Ian Johnston, Dominic Kwiatkowski, Cordelia Langford, John Sillitoe on behalf of the Wellcome Sanger Institute COVID-19 Surveillance Team |
| EPI_ISL_3469704 | Lighthouse Lab in Alderley Park | Wellcome Sanger Institute for the COVID-19 Genomics UK (COG-UK) Consortium | Jacquelyn Wynn, Mairead Hyland, The Lighthouse Lab in Alderley Park and Alex Alderton, Roberto Amato, Jeffrey Barrett, Sonia Goncalves, Ewan Harrison, David K. Jackson, Ian Johnston, Dominic Kwiatkowski, Cordelia Langford, John Sillitoe on behalf of the Wellcome Sanger Institute COVID-19 Surveillance Team |
| EPI_ISL_3469705, EPI_ISL_3469706 | Lighthouse Laboratory Plymouth | Wellcome Sanger Institute for the COVID-19 Genomics UK (COG-UK) Consortium | Lighthouse Laboratory Plymouth and Alex Alderton, Roberto Amato, Jeffrey Barrett, Sonia Goncalves, Ewan Harrison, David K. Jackson, Ian Johnston, Dominic Kwiatkowski, Cordelia Langford, John Sillitoe on behalf of the Wellcome Sanger Institute COVID-19 Surveillance Team |
| EPI_ISL_3469707 | Lighthouse Lab in Alderley Park | Wellcome Sanger Institute for the COVID-19 Genomics UK (COG-UK) Consortium | Jacquelyn Wynn, Mairead Hyland, The Lighthouse Lab in Alderley Park and Alex Alderton, Roberto Amato, Jeffrey Barrett, Sonia Goncalves, Ewan Harrison, David K. Jackson, Ian Johnston, Dominic Kwiatkowski, Cordelia Langford, John Sillitoe on behalf of the Wellcome Sanger Institute COVID-19 Surveillance Team |
| EPI_ISL_3469708, EPI_ISL_3469709 | Lighthouse Laboratory Plymouth | Wellcome Sanger Institute for the COVID-19 Genomics UK (COG-UK) Consortium | Lighthouse Laboratory Plymouth and Alex Alderton, Roberto Amato, Jeffrey Barrett, Sonia Goncalves, Ewan Harrison, David K. Jackson, Ian Johnston, Dominic Kwiatkowski, Cordelia Langford, John Sillitoe on behalf of the Wellcome Sanger Institute COVID-19 Surveillance Team |
| EPI_ISL_3469710 | Lighthouse Lab in Milton Keynes | Wellcome Sanger Institute for the COVID-19 Genomics UK (COG-UK) Consortium | The Lighthouse Lab in Milton Keynes and Alex Alderton, Roberto Amato, Jeffrey Barrett, Sonia Goncalves, Ewan Harrison, David K. Jackson, Ian Johnston, Dominic Kwiatkowski, Cordelia Langford, John Sillitoe on behalf of the Wellcome Sanger Institute COVID-19 Surveillance Team |
| EPI_ISL_3469711, EPI_ISL_3469712, EPI_ISL_3469713 | Lighthouse Lab in Alderley Park | Wellcome Sanger Institute for the COVID-19 Genomics UK (COG-UK) Consortium | Jacquelyn Wynn, Mairead Hyland, The Lighthouse Lab in Alderley Park and Alex Alderton, Roberto Amato, Jeffrey Barrett, Sonia Goncalves, Ewan Harrison, David K. Jackson, Ian Johnston, Dominic Kwiatkowski, Cordelia Langford, John Sillitoe on behalf of the Wellcome Sanger Institute COVID-19 Surveillance Team |
| EPI_ISL_3469714 | Lighthouse Laboratory Plymouth | Wellcome Sanger Institute for the COVID-19 Genomics UK (COG-UK) Consortium | Lighthouse Laboratory Plymouth and Alex Alderton, Roberto Amato, Jeffrey Barrett, Sonia Goncalves, Ewan Harrison, David K. Jackson, Ian Johnston, Dominic Kwiatkowski, Cordelia Langford, John Sillitoe on behalf of the Wellcome Sanger Institute COVID-19 Surveillance Team |
| EPI_ISL_3469715, EPI_ISL_3469716 | Lighthouse Lab in Milton Keynes | Wellcome Sanger Institute for the COVID-19 Genomics UK (COG-UK) Consortium | The Lighthouse Lab in Milton Keynes and Alex Alderton, Roberto Amato, Jeffrey Barrett, Sonia Goncalves, Ewan Harrison, David K. Jackson, Ian Johnston, Dominic Kwiatkowski, Cordelia Langford, John Sillitoe on behalf of the Wellcome Sanger Institute COVID-19 Surveillance Team |
| EPI_ISL_3469717 | Lighthouse Laboratory Plymouth | Wellcome Sanger Institute for the COVID-19 Genomics UK (COG-UK) Consortium | Lighthouse Laboratory Plymouth and Alex Alderton, Roberto Amato, Jeffrey Barrett, Sonia Goncalves, Ewan Harrison, David K. Jackson, Ian Johnston, Dominic Kwiatkowski, Cordelia Langford, John Sillitoe on behalf of the Wellcome Sanger Institute COVID-19 Surveillance Team |
| EPI_ISL_3469718 | Lighthouse Lab in Alderley Park | Wellcome Sanger Institute for the COVID-19 Genomics UK (COG-UK) Consortium | Jacquelyn Wynn, Mairead Hyland, The Lighthouse Lab in Alderley Park and Alex Alderton, Roberto Amato, Jeffrey Barrett, Sonia Goncalves, Ewan Harrison, David K. Jackson, Ian Johnston, Dominic Kwiatkowski, Cordelia Langford, John Sillitoe on behalf of the Wellcome Sanger Institute COVID-19 Surveillance Team |
| EPI_ISL_3469719 | Lighthouse Lab in Milton Keynes | Wellcome Sanger Institute for the COVID-19 Genomics UK (COG-UK) Consortium | The Lighthouse Lab in Milton Keynes and Alex Alderton, Roberto Amato, Jeffrey Barrett, Sonia Goncalves, Ewan Harrison, David K. Jackson, Ian Johnston, Dominic Kwiatkowski, Cordelia Langford, John Sillitoe on behalf of the Wellcome Sanger Institute COVID-19 Surveillance Team |
| EPI_ISL_3469720 | Lighthouse Lab in Alderley Park | Wellcome Sanger Institute for the COVID-19 Genomics UK (COG-UK) Consortium | Jacquelyn Wynn, Mairead Hyland, The Lighthouse Lab in Alderley Park and Alex Alderton, Roberto Amato, Jeffrey Barrett, Sonia Goncalves, Ewan Harrison, David K. Jackson, Ian Johnston, Dominic Kwiatkowski, Cordelia Langford, John Sillitoe on behalf of the Wellcome Sanger Institute COVID-19 Surveillance Team |
| EPI_ISL_3469721, EPI_ISL_3469722, EPI_ISL_3469723 | Lighthouse Laboratory Plymouth | Wellcome Sanger Institute for the COVID-19 Genomics UK (COG-UK) Consortium | Lighthouse Laboratory Plymouth and Alex Alderton, Roberto Amato, Jeffrey Barrett, Sonia Goncalves, Ewan Harrison, David K. Jackson, Ian Johnston, Dominic Kwiatkowski, Cordelia Langford, John Sillitoe on behalf of the Wellcome Sanger Institute COVID-19 Surveillance Team |
| EPI_ISL_3469724, EPI_ISL_3469725 | Lighthouse Lab in Alderley Park | Wellcome Sanger Institute for the COVID-19 Genomics UK (COG-UK) Consortium | Jacquelyn Wynn, Mairead Hyland, The Lighthouse Lab in Alderley Park and Alex Alderton, Roberto Amato, Jeffrey Barrett, Sonia Goncalves, Ewan Harrison, David K. Jackson, Ian Johnston, Dominic Kwiatkowski, Cordelia Langford, John Sillitoe on behalf of the Wellcome Sanger Institute COVID-19 Surveillance Team |
| EPI_ISL_3469726 | Lighthouse Lab in Milton Keynes | Wellcome Sanger Institute for the COVID-19 Genomics UK (COG-UK) Consortium | The Lighthouse Lab in Milton Keynes and Alex Alderton, Roberto Amato, Jeffrey Barrett, Sonia Goncalves, Ewan Harrison, David K. Jackson, Ian Johnston, Dominic Kwiatkowski, Cordelia Langford, John Sillitoe on behalf of the Wellcome Sanger Institute COVID-19 Surveillance Team |
| EPI_ISL_3469727, EPI_ISL_3469728 | Lighthouse Laboratory Plymouth | Wellcome Sanger Institute for the COVID-19 Genomics UK (COG-UK) Consortium | Lighthouse Laboratory Plymouth and Alex Alderton, Roberto Amato, Jeffrey Barrett, Sonia Goncalves, Ewan Harrison, David K. Jackson, Ian Johnston, Dominic Kwiatkowski, Cordelia Langford, John Sillitoe on behalf of the Wellcome Sanger Institute COVID-19 Surveillance Team |
| EPI_ISL_3469729, EPI_ISL_3469730, EPI_ISL_3469731 | Lighthouse Lab in Alderley Park | Wellcome Sanger Institute for the COVID-19 Genomics UK (COG-UK) Consortium | Jacquelyn Wynn, Mairead Hyland, The Lighthouse Lab in Alderley Park and Alex Alderton, Roberto Amato, Jeffrey Barrett, Sonia Goncalves, Ewan Harrison, David K. Jackson, Ian Johnston, Dominic Kwiatkowski, Cordelia Langford, John Sillitoe on behalf of the Wellcome Sanger Institute COVID-19 Surveillance Team |

| | | | |
|---|---|---|---|
| EPI_ISL_3469732 | Lighthouse Laboratory Plymouth | Wellcome Sanger Institute for the COVID-19 Genomics UK (COG-UK) Consortium | Lighthouse Laboratory Plymouth and Alex Alderton, Roberto Amato, Jeffrey Barrett, Sonia Goncalves, Ewan Harrison, David K. Jackson, Ian Johnston, Dominic Kwiatkowski, Cordelia Langford, John Sillitoe on behalf of the Wellcome Sanger Institute COVID-19 Surveillance Team |
| EPI_ISL_3469733 | Lighthouse Lab in Alderley Park | Wellcome Sanger Institute for the COVID-19 Genomics UK (COG-UK) Consortium | Jacquelyn Wynn, Mairead Hyland, The Lighthouse Lab in Alderley Park and Alex Alderton, Roberto Amato, Jeffrey Barrett, Sonia Goncalves, Ewan Harrison, David K. Jackson, Ian Johnston, Dominic Kwiatkowski, Cordelia Langford, John Sillitoe on behalf of the Wellcome Sanger Institute COVID-19 Surveillance Team |
| EPI_ISL_3469734 | Lighthouse Laboratory Plymouth | Wellcome Sanger Institute for the COVID-19 Genomics UK (COG-UK) Consortium | Lighthouse Laboratory Plymouth and Alex Alderton, Roberto Amato, Jeffrey Barrett, Sonia Goncalves, Ewan Harrison, David K. Jackson, Ian Johnston, Dominic Kwiatkowski, Cordelia Langford, John Sillitoe on behalf of the Wellcome Sanger Institute COVID-19 Surveillance Team |
| EPI_ISL_3469735, EPI_ISL_3469736 | Lighthouse Lab in Milton Keynes | Wellcome Sanger Institute for the COVID-19 Genomics UK (COG-UK) Consortium | The Lighthouse Lab in Milton Keynes and Alex Alderton, Roberto Amato, Jeffrey Barrett, Sonia Goncalves, Ewan Harrison, David K. Jackson, Ian Johnston, Dominic Kwiatkowski, Cordelia Langford, John Sillitoe on behalf of the Wellcome Sanger Institute COVID-19 Surveillance Team |
| EPI_ISL_3469737 | Lighthouse Lab in Alderley Park | Wellcome Sanger Institute for the COVID-19 Genomics UK (COG-UK) Consortium | Jacquelyn Wynn, Mairead Hyland, The Lighthouse Lab in Alderley Park and Alex Alderton, Roberto Amato, Jeffrey Barrett, Sonia Goncalves, Ewan Harrison, David K. Jackson, Ian Johnston, Dominic Kwiatkowski, Cordelia Langford, John Sillitoe on behalf of the Wellcome Sanger Institute COVID-19 Surveillance Team |
| EPI_ISL_3469738, EPI_ISL_3469739, EPI_ISL_3469740 | Lighthouse Lab in Milton Keynes | Wellcome Sanger Institute for the COVID-19 Genomics UK (COG-UK) Consortium | The Lighthouse Lab in Milton Keynes and Alex Alderton, Roberto Amato, Jeffrey Barrett, Sonia Goncalves, Ewan Harrison, David K. Jackson, Ian Johnston, Dominic Kwiatkowski, Cordelia Langford, John Sillitoe on behalf of the Wellcome Sanger Institute COVID-19 Surveillance Team |
| EPI_ISL_3469741 | Lighthouse Laboratory Plymouth | Wellcome Sanger Institute for the COVID-19 Genomics UK (COG-UK) Consortium | Lighthouse Laboratory Plymouth and Alex Alderton, Roberto Amato, Jeffrey Barrett, Sonia Goncalves, Ewan Harrison, David K. Jackson, Ian Johnston, Dominic Kwiatkowski, Cordelia Langford, John Sillitoe on behalf of the Wellcome Sanger Institute COVID-19 Surveillance Team |
| EPI_ISL_3469742 | Lighthouse Lab in Milton Keynes | Wellcome Sanger Institute for the COVID-19 Genomics UK (COG-UK) Consortium | The Lighthouse Lab in Milton Keynes and Alex Alderton, Roberto Amato, Jeffrey Barrett, Sonia Goncalves, Ewan Harrison, David K. Jackson, Ian Johnston, Dominic Kwiatkowski, Cordelia Langford, John Sillitoe on behalf of the Wellcome Sanger Institute COVID-19 Surveillance Team |
| EPI_ISL_3469743 | Lighthouse Laboratory Plymouth | Wellcome Sanger Institute for the COVID-19 Genomics UK (COG-UK) Consortium | Lighthouse Laboratory Plymouth and Alex Alderton, Roberto Amato, Jeffrey Barrett, Sonia Goncalves, Ewan Harrison, David K. Jackson, Ian Johnston, Dominic Kwiatkowski, Cordelia Langford, John Sillitoe on behalf of the Wellcome Sanger Institute COVID-19 Surveillance Team |
| EPI_ISL_3469744 | Lighthouse Lab in Milton Keynes | Wellcome Sanger Institute for the COVID-19 Genomics UK (COG-UK) Consortium | The Lighthouse Lab in Milton Keynes and Alex Alderton, Roberto Amato, Jeffrey Barrett, Sonia Goncalves, Ewan Harrison, David K. Jackson, Ian Johnston, Dominic Kwiatkowski, Cordelia Langford, John Sillitoe on behalf of the Wellcome Sanger Institute COVID-19 Surveillance Team |
| EPI_ISL_3469745 | Lighthouse Laboratory Plymouth | Wellcome Sanger Institute for the COVID-19 Genomics UK (COG-UK) Consortium | Lighthouse Laboratory Plymouth and Alex Alderton, Roberto Amato, Jeffrey Barrett, Sonia Goncalves, Ewan Harrison, David K. Jackson, Ian Johnston, Dominic Kwiatkowski, Cordelia Langford, John Sillitoe on behalf of the Wellcome Sanger Institute COVID-19 Surveillance Team |
| EPI_ISL_3469746 | Lighthouse Lab in Milton Keynes | Wellcome Sanger Institute for the COVID-19 Genomics UK (COG-UK) Consortium | The Lighthouse Lab in Milton Keynes and Alex Alderton, Roberto Amato, Jeffrey Barrett, Sonia Goncalves, Ewan Harrison, David K. Jackson, Ian Johnston, Dominic Kwiatkowski, Cordelia Langford, John Sillitoe on behalf of the Wellcome Sanger Institute COVID-19 Surveillance Team |
| EPI_ISL_3469747, EPI_ISL_3469748, EPI_ISL_3469749, EPI_ISL_3469750 | Lighthouse Laboratory Plymouth | Wellcome Sanger Institute for the COVID-19 Genomics UK (COG-UK) Consortium | Lighthouse Laboratory Plymouth and Alex Alderton, Roberto Amato, Jeffrey Barrett, Sonia Goncalves, Ewan Harrison, David K. Jackson, Ian Johnston, Dominic Kwiatkowski, Cordelia Langford, John Sillitoe on behalf of the Wellcome Sanger Institute COVID-19 Surveillance Team |
| EPI_ISL_3469751, EPI_ISL_3469752 | Lighthouse Lab in Milton Keynes | Wellcome Sanger Institute for the COVID-19 Genomics UK (COG-UK) Consortium | The Lighthouse Lab in Milton Keynes and Alex Alderton, Roberto Amato, Jeffrey Barrett, Sonia Goncalves, Ewan Harrison, David K. Jackson, Ian Johnston, Dominic Kwiatkowski, Cordelia Langford, John Sillitoe on behalf of the Wellcome Sanger Institute COVID-19 Surveillance Team |
| EPI_ISL_3469753 | Lighthouse Laboratory Plymouth | Wellcome Sanger Institute for the COVID-19 Genomics UK (COG-UK) Consortium | Lighthouse Laboratory Plymouth and Alex Alderton, Roberto Amato, Jeffrey Barrett, Sonia Goncalves, Ewan Harrison, David K. Jackson, Ian Johnston, Dominic Kwiatkowski, Cordelia Langford, John Sillitoe on behalf of the Wellcome Sanger Institute COVID-19 Surveillance Team |
| EPI_ISL_3469754, EPI_ISL_3469755 | Lighthouse Lab in Milton Keynes | Wellcome Sanger Institute for the COVID-19 Genomics UK (COG-UK) Consortium | The Lighthouse Lab in Milton Keynes and Alex Alderton, Roberto Amato, Jeffrey Barrett, Sonia Goncalves, Ewan Harrison, David K. Jackson, Ian Johnston, Dominic Kwiatkowski, Cordelia Langford, John Sillitoe on behalf of the Wellcome Sanger Institute COVID-19 Surveillance Team |
| EPI_ISL_3469756 | Lighthouse Lab in Alderley Park | Wellcome Sanger Institute for the COVID-19 Genomics UK (COG-UK) Consortium | Jacquelyn Wynn, Mairead Hyland, The Lighthouse Lab in Alderley Park and Alex Alderton, Roberto Amato, Jeffrey Barrett, Sonia Goncalves, Ewan Harrison, David K. Jackson, Ian Johnston, Dominic Kwiatkowski, Cordelia Langford, John Sillitoe on behalf of the Wellcome Sanger Institute COVID-19 Surveillance Team |
| EPI_ISL_3469757 | Lighthouse Lab in Milton Keynes | Wellcome Sanger Institute for the COVID-19 Genomics UK (COG-UK) Consortium | The Lighthouse Lab in Milton Keynes and Alex Alderton, Roberto Amato, Jeffrey Barrett, Sonia Goncalves, Ewan Harrison, David K. Jackson, Ian Johnston, Dominic Kwiatkowski, Cordelia Langford, John Sillitoe on behalf of the Wellcome Sanger Institute COVID-19 Surveillance Team |
| EPI_ISL_3469758 | Lighthouse Lab in Alderley Park | Wellcome Sanger Institute for the COVID-19 Genomics UK (COG-UK) Consortium | Jacquelyn Wynn, Mairead Hyland, The Lighthouse Lab in Alderley Park and Alex Alderton, Roberto Amato, Jeffrey Barrett, Sonia Goncalves, Ewan Harrison, David K. Jackson, Ian Johnston, Dominic Kwiatkowski, Cordelia Langford, John Sillitoe on behalf of the Wellcome Sanger Institute COVID-19 Surveillance Team |
| EPI_ISL_3469759 | Lighthouse Laboratory Plymouth | Wellcome Sanger Institute for the COVID-19 Genomics UK (COG-UK) Consortium | Lighthouse Laboratory Plymouth and Alex Alderton, Roberto Amato, Jeffrey Barrett, Sonia Goncalves, Ewan Harrison, David K. Jackson, Ian Johnston, Dominic Kwiatkowski, Cordelia Langford, John Sillitoe on behalf of the Wellcome Sanger Institute COVID-19 Surveillance Team |
| EPI_ISL_3469760 | Lighthouse Lab in Alderley Park | Wellcome Sanger Institute for the COVID-19 Genomics UK (COG-UK) Consortium | Jacquelyn Wynn, Mairead Hyland, The Lighthouse Lab in Alderley Park and Alex Alderton, Roberto Amato, Jeffrey Barrett, Sonia Goncalves, Ewan Harrison, David K. Jackson, Ian Johnston, Dominic Kwiatkowski, Cordelia Langford, John Sillitoe on behalf of the Wellcome Sanger Institute COVID-19 Surveillance Team |
| EPI_ISL_3469761 | Lighthouse Laboratory Plymouth | Wellcome Sanger Institute for the COVID-19 Genomics UK (COG-UK) Consortium | Lighthouse Laboratory Plymouth and Alex Alderton, Roberto Amato, Jeffrey Barrett, Sonia Goncalves, Ewan Harrison, David K. Jackson, Ian Johnston, Dominic Kwiatkowski, Cordelia Langford, John Sillitoe on behalf of the Wellcome Sanger Institute COVID-19 Surveillance Team |
| EPI_ISL_3469762 | Lighthouse Lab in Milton Keynes | Wellcome Sanger Institute for the COVID-19 Genomics UK (COG-UK) Consortium | The Lighthouse Lab in Milton Keynes and Alex Alderton, Roberto Amato, Jeffrey Barrett, Sonia Goncalves, Ewan Harrison, David K. Jackson, Ian Johnston, Dominic Kwiatkowski, Cordelia Langford, John Sillitoe on behalf of the Wellcome Sanger Institute COVID-19 Surveillance Team |
| EPI_ISL_3469763 | Lighthouse Lab in Alderley Park | Wellcome Sanger Institute for the COVID-19 Genomics UK (COG-UK) Consortium | Jacquelyn Wynn, Mairead Hyland, The Lighthouse Lab in Alderley Park and Alex Alderton, Roberto Amato, Jeffrey Barrett, Sonia Goncalves, Ewan Harrison, David K. Jackson, Ian Johnston, Dominic Kwiatkowski, Cordelia Langford, John Sillitoe on behalf of the Wellcome Sanger Institute COVID-19 Surveillance Team |
| EPI_ISL_3469764 | Lighthouse Laboratory Plymouth | Wellcome Sanger Institute for the COVID-19 Genomics UK (COG-UK) Consortium | Lighthouse Laboratory Plymouth and Alex Alderton, Roberto Amato, Jeffrey Barrett, Sonia Goncalves, Ewan Harrison, David K. Jackson, Ian Johnston, Dominic Kwiatkowski, Cordelia Langford, John Sillitoe on behalf of the Wellcome Sanger Institute COVID-19 Surveillance Team |
| EPI_ISL_3469765 | Lighthouse Lab in Milton Keynes | Wellcome Sanger Institute for the COVID-19 Genomics UK (COG-UK) Consortium | The Lighthouse Lab in Milton Keynes and Alex Alderton, Roberto Amato, Jeffrey Barrett, Sonia Goncalves, Ewan Harrison, David K. Jackson, Ian Johnston, Dominic Kwiatkowski, Cordelia Langford, John Sillitoe on behalf of the Wellcome Sanger Institute COVID-19 Surveillance Team |
| EPI_ISL_3469766, EPI_ISL_3469767 | Lighthouse Laboratory Plymouth | Wellcome Sanger Institute for the COVID-19 Genomics UK (COG-UK) Consortium | Lighthouse Laboratory Plymouth and Alex Alderton, Roberto Amato, Jeffrey Barrett, Sonia Goncalves, Ewan Harrison, David K. Jackson, Ian Johnston, Dominic Kwiatkowski, Cordelia Langford, John Sillitoe on behalf of the Wellcome Sanger Institute COVID-19 Surveillance Team |
| EPI_ISL_3469768, EPI_ISL_3469769, EPI_ISL_3469770 | Lighthouse Lab in Milton Keynes | Wellcome Sanger Institute for the COVID-19 Genomics UK (COG-UK) Consortium | The Lighthouse Lab in Milton Keynes and Alex Alderton, Roberto Amato, Jeffrey Barrett, Sonia Goncalves, Ewan Harrison, David K. Jackson, Ian Johnston, Dominic Kwiatkowski, Cordelia Langford, John Sillitoe on behalf of the Wellcome Sanger Institute COVID-19 Surveillance Team |
| EPI_ISL_3469771, EPI_ISL_3469772, EPI_ISL_3469773 | Lighthouse Lab in Alderley Park | Wellcome Sanger Institute for the COVID-19 Genomics UK (COG-UK) Consortium | Jacquelyn Wynn, Mairead Hyland, The Lighthouse Lab in Alderley Park and Alex Alderton, Roberto Amato, Jeffrey Barrett, Sonia Goncalves, Ewan Harrison, David K. Jackson, Ian Johnston, Dominic Kwiatkowski, Cordelia Langford, John Sillitoe on behalf of the Wellcome Sanger Institute COVID-19 Surveillance Team |
| EPI_ISL_3469774 | Lighthouse Lab in Milton Keynes | Wellcome Sanger Institute for the COVID-19 Genomics UK (COG-UK) Consortium | The Lighthouse Lab in Milton Keynes and Alex Alderton, Roberto Amato, Jeffrey Barrett, Sonia Goncalves, Ewan Harrison, David K. Jackson, Ian Johnston, Dominic Kwiatkowski, Cordelia Langford, John Sillitoe on behalf of the Wellcome Sanger Institute COVID-19 Surveillance Team |
| EPI_ISL_3469775 | Lighthouse Lab in Alderley Park | Wellcome Sanger Institute for the COVID-19 Genomics UK (COG-UK) Consortium | Jacquelyn Wynn, Mairead Hyland, The Lighthouse Lab in Alderley Park and Alex Alderton, Roberto Amato, Jeffrey Barrett, Sonia Goncalves, Ewan Harrison, David K. Jackson, Ian Johnston, Dominic Kwiatkowski, Cordelia Langford, John Sillitoe on behalf of the Wellcome Sanger Institute COVID-19 Surveillance Team |
| EPI_ISL_3469776 | Lighthouse Laboratory Plymouth | Wellcome Sanger Institute for the COVID-19 Genomics UK (COG-UK) Consortium | Lighthouse Laboratory Plymouth and Alex Alderton, Roberto Amato, Jeffrey Barrett, Sonia Goncalves, Ewan Harrison, David K. Jackson, Ian Johnston, Dominic Kwiatkowski, Cordelia Langford, John Sillitoe on behalf of the Wellcome Sanger Institute COVID-19 Surveillance Team |
| EPI_ISL_3469777 | Lighthouse Lab in Milton Keynes | Wellcome Sanger Institute for the COVID-19 Genomics UK (COG-UK) Consortium | The Lighthouse Lab in Milton Keynes and Alex Alderton, Roberto Amato, Jeffrey Barrett, Sonia Goncalves, Ewan Harrison, David K. Jackson, Ian Johnston, Dominic Kwiatkowski, Cordelia Langford, John Sillitoe on behalf of the Wellcome Sanger Institute COVID-19 Surveillance Team |
| EPI_ISL_3469778 | Lighthouse Laboratory Plymouth | Wellcome Sanger Institute for the COVID-19 Genomics UK (COG-UK) Consortium | Lighthouse Laboratory Plymouth and Alex Alderton, Roberto Amato, Jeffrey Barrett, Sonia Goncalves, Ewan Harrison, David K. Jackson, Ian Johnston, Dominic Kwiatkowski, Cordelia Langford, John Sillitoe on behalf of the Wellcome Sanger Institute COVID-19 Surveillance Team |
| EPI_ISL_3469779, EPI_ISL_3469780 | Lighthouse Lab in Milton Keynes | Wellcome Sanger Institute for the COVID-19 Genomics UK (COG-UK) Consortium | The Lighthouse Lab in Milton Keynes and Alex Alderton, Roberto Amato, Jeffrey Barrett, Sonia Goncalves, Ewan Harrison, David K. Jackson, Ian Johnston, Dominic Kwiatkowski, Cordelia Langford, John Sillitoe on behalf of the Wellcome Sanger Institute COVID-19 Surveillance Team |
| EPI_ISL_3469781, EPI_ISL_3469782 | Lighthouse Laboratory Plymouth | Wellcome Sanger Institute for the COVID-19 Genomics UK (COG-UK) Consortium | Lighthouse Laboratory Plymouth and Alex Alderton, Roberto Amato, Jeffrey Barrett, Sonia Goncalves, Ewan Harrison, David K. Jackson, Ian Johnston, Dominic Kwiatkowski, Cordelia Langford, John Sillitoe on behalf of the Wellcome Sanger Institute COVID-19 Surveillance Team |
| EPI_ISL_3469783, EPI_ISL_3469784, EPI_ISL_3469785, EPI_ISL_3469786, EPI_ISL_3469787 | Lighthouse Lab in Milton Keynes | Wellcome Sanger Institute for the COVID-19 Genomics UK (COG-UK) Consortium | The Lighthouse Lab in Milton Keynes and Alex Alderton, Roberto Amato, Jeffrey Barrett, Sonia Goncalves, Ewan Harrison, David K. Jackson, Ian Johnston, Dominic Kwiatkowski, Cordelia Langford, John Sillitoe on behalf of the Wellcome Sanger Institute COVID-19 Surveillance Team |
| EPI_ISL_3469788 | Lighthouse Lab in Alderley Park | Wellcome Sanger Institute for the COVID-19 Genomics UK (COG-UK) Consortium | Jacquelyn Wynn, Mairead Hyland, The Lighthouse Lab in Alderley Park and Alex Alderton, Roberto Amato, Jeffrey Barrett, Sonia Goncalves, Ewan Harrison, David K. Jackson, Ian Johnston, Dominic Kwiatkowski, Cordelia Langford, John Sillitoe on behalf of the Wellcome Sanger Institute COVID-19 Surveillance Team |
| EPI_ISL_3469789 | Lighthouse Laboratory Plymouth | Wellcome Sanger Institute for the COVID-19 Genomics UK (COG-UK) Consortium | Lighthouse Laboratory Plymouth and Alex Alderton, Roberto Amato, Jeffrey Barrett, Sonia Goncalves, Ewan Harrison, David K. Jackson, Ian Johnston, Dominic Kwiatkowski, Cordelia Langford, John Sillitoe on behalf of the Wellcome Sanger Institute COVID-19 Surveillance Team |
| EPI_ISL_3469790, EPI_ISL_3469791, EPI_ISL_3469792 | Lighthouse Lab in Alderley Park | Wellcome Sanger Institute for the COVID-19 Genomics UK (COG-UK) Consortium | Jacquelyn Wynn, Mairead Hyland, The Lighthouse Lab in Alderley Park and Alex Alderton, Roberto Amato, Jeffrey Barrett, Sonia Goncalves, Ewan Harrison, David K. Jackson, Ian Johnston, Dominic Kwiatkowski, Cordelia Langford, John Sillitoe on behalf of the Wellcome Sanger Institute COVID-19 Surveillance Team |
| EPI_ISL_3469793 | Lighthouse Laboratory Plymouth | Wellcome Sanger Institute for the COVID-19 Genomics UK (COG-UK) Consortium | Lighthouse Laboratory Plymouth and Alex Alderton, Roberto Amato, Jeffrey Barrett, Sonia Goncalves, Ewan Harrison, David K. Jackson, Ian Johnston, Dominic Kwiatkowski, Cordelia Langford, John Sillitoe on behalf of the Wellcome Sanger Institute COVID-19 Surveillance Team |
| EPI_ISL_3469794 | Lighthouse Lab in Milton Keynes | Wellcome Sanger Institute for the COVID-19 Genomics UK (COG-UK) | The Lighthouse Lab in Milton Keynes and Alex Alderton, Roberto Amato, Jeffrey Barrett, Sonia Goncalves, Ewan Harrison, David K. Jackson, Ian Johnston, Dominic Kwiatkowski, Cordelia Langford, John Sillitoe on behalf of the Wellcome Sanger |

| | | | |
|---|---|---|---|
| | | Consortium | Institute COVID-19 Surveillance Team |
| EPI_ISL_3469795 | Lighthouse Lab in Alderley Park | Wellcome Sanger Institute for the COVID-19 Genomics UK (COG-UK) Consortium | Jacquelyn Wynn, Mairead Hyland, The Lighthouse Lab in Alderley Park and Alex Alderton, Roberto Amato, Jeffrey Barrett, Sonia Goncalves, Ewan Harrison, David K. Jackson, Ian Johnston, Dominic Kwiatkowski, Cordelia Langford, John Sillitoe on behalf of the Wellcome Sanger Institute COVID-19 Surveillance Team |
| EPI_ISL_3469796, EPI_ISL_3469797 | Lighthouse Lab in Milton Keynes | Wellcome Sanger Institute for the COVID-19 Genomics UK (COG-UK) Consortium | The Lighthouse Lab in Milton Keynes and Alex Alderton, Roberto Amato, Jeffrey Barrett, Sonia Goncalves, Ewan Harrison, David K. Jackson, Ian Johnston, Dominic Kwiatkowski, Cordelia Langford, John Sillitoe on behalf of the Wellcome Sanger Institute COVID-19 Surveillance Team |
| EPI_ISL_3469798 | Lighthouse Laboratory Plymouth | Wellcome Sanger Institute for the COVID-19 Genomics UK (COG-UK) Consortium | Lighthouse Laboratory Plymouth and Alex Alderton, Roberto Amato, Jeffrey Barrett, Sonia Goncalves, Ewan Harrison, David K. Jackson, Ian Johnston, Dominic Kwiatkowski, Cordelia Langford, John Sillitoe on behalf of the Wellcome Sanger Institute COVID-19 Surveillance Team |
| EPI_ISL_3469799, EPI_ISL_3469800 | Lighthouse Lab in Alderley Park | Wellcome Sanger Institute for the COVID-19 Genomics UK (COG-UK) Consortium | Jacquelyn Wynn, Mairead Hyland, The Lighthouse Lab in Alderley Park and Alex Alderton, Roberto Amato, Jeffrey Barrett, Sonia Goncalves, Ewan Harrison, David K. Jackson, Ian Johnston, Dominic Kwiatkowski, Cordelia Langford, John Sillitoe on behalf of the Wellcome Sanger Institute COVID-19 Surveillance Team |
| EPI_ISL_3469801, EPI_ISL_3469802 | Lighthouse Laboratory Plymouth | Wellcome Sanger Institute for the COVID-19 Genomics UK (COG-UK) Consortium | Lighthouse Laboratory Plymouth and Alex Alderton, Roberto Amato, Jeffrey Barrett, Sonia Goncalves, Ewan Harrison, David K. Jackson, Ian Johnston, Dominic Kwiatkowski, Cordelia Langford, John Sillitoe on behalf of the Wellcome Sanger Institute COVID-19 Surveillance Team |
| EPI_ISL_3469803, EPI_ISL_3469804 | Lighthouse Lab in Alderley Park | Wellcome Sanger Institute for the COVID-19 Genomics UK (COG-UK) Consortium | Jacquelyn Wynn, Mairead Hyland, The Lighthouse Lab in Alderley Park and Alex Alderton, Roberto Amato, Jeffrey Barrett, Sonia Goncalves, Ewan Harrison, David K. Jackson, Ian Johnston, Dominic Kwiatkowski, Cordelia Langford, John Sillitoe on behalf of the Wellcome Sanger Institute COVID-19 Surveillance Team |
| EPI_ISL_3469805 | Lighthouse Laboratory Plymouth | Wellcome Sanger Institute for the COVID-19 Genomics UK (COG-UK) Consortium | Lighthouse Laboratory Plymouth and Alex Alderton, Roberto Amato, Jeffrey Barrett, Sonia Goncalves, Ewan Harrison, David K. Jackson, Ian Johnston, Dominic Kwiatkowski, Cordelia Langford, John Sillitoe on behalf of the Wellcome Sanger Institute COVID-19 Surveillance Team |
| EPI_ISL_3469806 | Lighthouse Lab in Alderley Park | Wellcome Sanger Institute for the COVID-19 Genomics UK (COG-UK) Consortium | Jacquelyn Wynn, Mairead Hyland, The Lighthouse Lab in Alderley Park and Alex Alderton, Roberto Amato, Jeffrey Barrett, Sonia Goncalves, Ewan Harrison, David K. Jackson, Ian Johnston, Dominic Kwiatkowski, Cordelia Langford, John Sillitoe on behalf of the Wellcome Sanger Institute COVID-19 Surveillance Team |
| EPI_ISL_3469807 | Lighthouse Laboratory Plymouth | Wellcome Sanger Institute for the COVID-19 Genomics UK (COG-UK) Consortium | Lighthouse Laboratory Plymouth and Alex Alderton, Roberto Amato, Jeffrey Barrett, Sonia Goncalves, Ewan Harrison, David K. Jackson, Ian Johnston, Dominic Kwiatkowski, Cordelia Langford, John Sillitoe on behalf of the Wellcome Sanger Institute COVID-19 Surveillance Team |
| EPI_ISL_3469808 | Lighthouse Lab in Milton Keynes | Wellcome Sanger Institute for the COVID-19 Genomics UK (COG-UK) Consortium | The Lighthouse Lab in Milton Keynes and Alex Alderton, Roberto Amato, Jeffrey Barrett, Sonia Goncalves, Ewan Harrison, David K. Jackson, Ian Johnston, Dominic Kwiatkowski, Cordelia Langford, John Sillitoe on behalf of the Wellcome Sanger Institute COVID-19 Surveillance Team |
| EPI_ISL_3469809 | Lighthouse Laboratory Plymouth | Wellcome Sanger Institute for the COVID-19 Genomics UK (COG-UK) Consortium | Lighthouse Laboratory Plymouth and Alex Alderton, Roberto Amato, Jeffrey Barrett, Sonia Goncalves, Ewan Harrison, David K. Jackson, Ian Johnston, Dominic Kwiatkowski, Cordelia Langford, John Sillitoe on behalf of the Wellcome Sanger Institute COVID-19 Surveillance Team |
| EPI_ISL_3469810, EPI_ISL_3469811, EPI_ISL_3469812, EPI_ISL_3469813 | Lighthouse Lab in Alderley Park | Wellcome Sanger Institute for the COVID-19 Genomics UK (COG-UK) Consortium | Jacquelyn Wynn, Mairead Hyland, The Lighthouse Lab in Alderley Park and Alex Alderton, Roberto Amato, Jeffrey Barrett, Sonia Goncalves, Ewan Harrison, David K. Jackson, Ian Johnston, Dominic Kwiatkowski, Cordelia Langford, John Sillitoe on behalf of the Wellcome Sanger Institute COVID-19 Surveillance Team |
| EPI_ISL_3469814 | Lighthouse Laboratory Plymouth | Wellcome Sanger Institute for the COVID-19 Genomics UK (COG-UK) Consortium | Lighthouse Laboratory Plymouth and Alex Alderton, Roberto Amato, Jeffrey Barrett, Sonia Goncalves, Ewan Harrison, David K. Jackson, Ian Johnston, Dominic Kwiatkowski, Cordelia Langford, John Sillitoe on behalf of the Wellcome Sanger Institute COVID-19 Surveillance Team |
| EPI_ISL_3469815 | Lighthouse Lab in Alderley Park | Wellcome Sanger Institute for the COVID-19 Genomics UK (COG-UK) Consortium | Jacquelyn Wynn, Mairead Hyland, The Lighthouse Lab in Alderley Park and Alex Alderton, Roberto Amato, Jeffrey Barrett, Sonia Goncalves, Ewan Harrison, David K. Jackson, Ian Johnston, Dominic Kwiatkowski, Cordelia Langford, John Sillitoe on behalf of the Wellcome Sanger Institute COVID-19 Surveillance Team |
| EPI_ISL_3469816, EPI_ISL_3469817 | Lighthouse Lab in Milton Keynes | Wellcome Sanger Institute for the COVID-19 Genomics UK (COG-UK) Consortium | The Lighthouse Lab in Milton Keynes and Alex Alderton, Roberto Amato, Jeffrey Barrett, Sonia Goncalves, Ewan Harrison, David K. Jackson, Ian Johnston, Dominic Kwiatkowski, Cordelia Langford, John Sillitoe on behalf of the Wellcome Sanger Institute COVID-19 Surveillance Team |
| EPI_ISL_3469818 | Lighthouse Lab in Alderley Park | Wellcome Sanger Institute for the COVID-19 Genomics UK (COG-UK) Consortium | Jacquelyn Wynn, Mairead Hyland, The Lighthouse Lab in Alderley Park and Alex Alderton, Roberto Amato, Jeffrey Barrett, Sonia Goncalves, Ewan Harrison, David K. Jackson, Ian Johnston, Dominic Kwiatkowski, Cordelia Langford, John Sillitoe on behalf of the Wellcome Sanger Institute COVID-19 Surveillance Team |
| EPI_ISL_3469819 | Lighthouse Lab in Milton Keynes | Wellcome Sanger Institute for the COVID-19 Genomics UK (COG-UK) Consortium | The Lighthouse Lab in Milton Keynes and Alex Alderton, Roberto Amato, Jeffrey Barrett, Sonia Goncalves, Ewan Harrison, David K. Jackson, Ian Johnston, Dominic Kwiatkowski, Cordelia Langford, John Sillitoe on behalf of the Wellcome Sanger Institute COVID-19 Surveillance Team |
| EPI_ISL_3469820, EPI_ISL_3469821 | Lighthouse Lab in Alderley Park | Wellcome Sanger Institute for the COVID-19 Genomics UK (COG-UK) Consortium | Jacquelyn Wynn, Mairead Hyland, The Lighthouse Lab in Alderley Park and Alex Alderton, Roberto Amato, Jeffrey Barrett, Sonia Goncalves, Ewan Harrison, David K. Jackson, Ian Johnston, Dominic Kwiatkowski, Cordelia Langford, John Sillitoe on behalf of the Wellcome Sanger Institute COVID-19 Surveillance Team |
| EPI_ISL_3469822 | Lighthouse Lab in Milton Keynes | Wellcome Sanger Institute for the COVID-19 Genomics UK (COG-UK) Consortium | The Lighthouse Lab in Milton Keynes and Alex Alderton, Roberto Amato, Jeffrey Barrett, Sonia Goncalves, Ewan Harrison, David K. Jackson, Ian Johnston, Dominic Kwiatkowski, Cordelia Langford, John Sillitoe on behalf of the Wellcome Sanger Institute COVID-19 Surveillance Team |
| EPI_ISL_3469823, EPI_ISL_3469824, EPI_ISL_3469825 | Lighthouse Laboratory Plymouth | Wellcome Sanger Institute for the COVID-19 Genomics UK (COG-UK) Consortium | Lighthouse Laboratory Plymouth and Alex Alderton, Roberto Amato, Jeffrey Barrett, Sonia Goncalves, Ewan Harrison, David K. Jackson, Ian Johnston, Dominic Kwiatkowski, Cordelia Langford, John Sillitoe on behalf of the Wellcome Sanger Institute COVID-19 Surveillance Team |
| EPI_ISL_3469826, EPI_ISL_3469827 | Lighthouse Lab in Milton Keynes | Wellcome Sanger Institute for the COVID-19 Genomics UK (COG-UK) Consortium | The Lighthouse Lab in Milton Keynes and Alex Alderton, Roberto Amato, Jeffrey Barrett, Sonia Goncalves, Ewan Harrison, David K. Jackson, Ian Johnston, Dominic Kwiatkowski, Cordelia Langford, John Sillitoe on behalf of the Wellcome Sanger Institute COVID-19 Surveillance Team |
| EPI_ISL_3469828 | Lighthouse Laboratory Plymouth | Wellcome Sanger Institute for the COVID-19 Genomics UK (COG-UK) Consortium | Lighthouse Laboratory Plymouth and Alex Alderton, Roberto Amato, Jeffrey Barrett, Sonia Goncalves, Ewan Harrison, David K. Jackson, Ian Johnston, Dominic Kwiatkowski, Cordelia Langford, John Sillitoe on behalf of the Wellcome Sanger Institute COVID-19 Surveillance Team |
| EPI_ISL_3469829 | Lighthouse Lab in Alderley Park | Wellcome Sanger Institute for the COVID-19 Genomics UK (COG-UK) Consortium | Jacquelyn Wynn, Mairead Hyland, The Lighthouse Lab in Alderley Park and Alex Alderton, Roberto Amato, Jeffrey Barrett, Sonia Goncalves, Ewan Harrison, David K. Jackson, Ian Johnston, Dominic Kwiatkowski, Cordelia Langford, John Sillitoe on behalf of the Wellcome Sanger Institute COVID-19 Surveillance Team |
| EPI_ISL_3469830, EPI_ISL_3469831 | Lighthouse Laboratory Plymouth | Wellcome Sanger Institute for the COVID-19 Genomics UK (COG-UK) Consortium | Lighthouse Laboratory Plymouth and Alex Alderton, Roberto Amato, Jeffrey Barrett, Sonia Goncalves, Ewan Harrison, David K. Jackson, Ian Johnston, Dominic Kwiatkowski, Cordelia Langford, John Sillitoe on behalf of the Wellcome Sanger Institute COVID-19 Surveillance Team |
| EPI_ISL_3469832 | Lighthouse Lab in Alderley Park | Wellcome Sanger Institute for the COVID-19 Genomics UK (COG-UK) Consortium | Jacquelyn Wynn, Mairead Hyland, The Lighthouse Lab in Alderley Park and Alex Alderton, Roberto Amato, Jeffrey Barrett, Sonia Goncalves, Ewan Harrison, David K. Jackson, Ian Johnston, Dominic Kwiatkowski, Cordelia Langford, John Sillitoe on behalf of the Wellcome Sanger Institute COVID-19 Surveillance Team |
| EPI_ISL_3469833 | Lighthouse Laboratory Plymouth | Wellcome Sanger Institute for the COVID-19 Genomics UK (COG-UK) Consortium | Lighthouse Laboratory Plymouth and Alex Alderton, Roberto Amato, Jeffrey Barrett, Sonia Goncalves, Ewan Harrison, David K. Jackson, Ian Johnston, Dominic Kwiatkowski, Cordelia Langford, John Sillitoe on behalf of the Wellcome Sanger Institute COVID-19 Surveillance Team |
| EPI_ISL_3469834 | Lighthouse Lab in Alderley Park | Wellcome Sanger Institute for the COVID-19 Genomics UK (COG-UK) Consortium | Jacquelyn Wynn, Mairead Hyland, The Lighthouse Lab in Alderley Park and Alex Alderton, Roberto Amato, Jeffrey Barrett, Sonia Goncalves, Ewan Harrison, David K. Jackson, Ian Johnston, Dominic Kwiatkowski, Cordelia Langford, John Sillitoe on behalf of the Wellcome Sanger Institute COVID-19 Surveillance Team |
| EPI_ISL_3469835 | Lighthouse Laboratory Plymouth | Wellcome Sanger Institute for the COVID-19 Genomics UK (COG-UK) Consortium | Lighthouse Laboratory Plymouth and Alex Alderton, Roberto Amato, Jeffrey Barrett, Sonia Goncalves, Ewan Harrison, David K. Jackson, Ian Johnston, Dominic Kwiatkowski, Cordelia Langford, John Sillitoe on behalf of the Wellcome Sanger Institute COVID-19 Surveillance Team |
| EPI_ISL_3469836 | Lighthouse Lab in Alderley Park | Wellcome Sanger Institute for the COVID-19 Genomics UK (COG-UK) Consortium | Jacquelyn Wynn, Mairead Hyland, The Lighthouse Lab in Alderley Park and Alex Alderton, Roberto Amato, Jeffrey Barrett, Sonia Goncalves, Ewan Harrison, David K. Jackson, Ian Johnston, Dominic Kwiatkowski, Cordelia Langford, John Sillitoe on behalf of the Wellcome Sanger Institute COVID-19 Surveillance Team |
| EPI_ISL_3469837 | Lighthouse Lab in Milton Keynes | Wellcome Sanger Institute for the COVID-19 Genomics UK (COG-UK) Consortium | The Lighthouse Lab in Milton Keynes and Alex Alderton, Roberto Amato, Jeffrey Barrett, Sonia Goncalves, Ewan Harrison, David K. Jackson, Ian Johnston, Dominic Kwiatkowski, Cordelia Langford, John Sillitoe on behalf of the Wellcome Sanger Institute COVID-19 Surveillance Team |
| EPI_ISL_3469838 | Lighthouse Lab in Alderley Park | Wellcome Sanger Institute for the COVID-19 Genomics UK (COG-UK) Consortium | Jacquelyn Wynn, Mairead Hyland, The Lighthouse Lab in Alderley Park and Alex Alderton, Roberto Amato, Jeffrey Barrett, Sonia Goncalves, Ewan Harrison, David K. Jackson, Ian Johnston, Dominic Kwiatkowski, Cordelia Langford, John Sillitoe on behalf of the Wellcome Sanger Institute COVID-19 Surveillance Team |
| EPI_ISL_3469839 | Lighthouse Lab in Milton Keynes | Wellcome Sanger Institute for the COVID-19 Genomics UK (COG-UK) Consortium | The Lighthouse Lab in Milton Keynes and Alex Alderton, Roberto Amato, Jeffrey Barrett, Sonia Goncalves, Ewan Harrison, David K. Jackson, Ian Johnston, Dominic Kwiatkowski, Cordelia Langford, John Sillitoe on behalf of the Wellcome Sanger Institute COVID-19 Surveillance Team |
| EPI_ISL_3469840, EPI_ISL_3469841, EPI_ISL_3469842, EPI_ISL_3469843, EPI_ISL_3469844, EPI_ISL_3469845, EPI_ISL_3469846, EPI_ISL_3469847 | Lighthouse Lab in Alderley Park | Wellcome Sanger Institute for the COVID-19 Genomics UK (COG-UK) Consortium | Jacquelyn Wynn, Mairead Hyland, The Lighthouse Lab in Alderley Park and Alex Alderton, Roberto Amato, Jeffrey Barrett, Sonia Goncalves, Ewan Harrison, David K. Jackson, Ian Johnston, Dominic Kwiatkowski, Cordelia Langford, John Sillitoe on behalf of the Wellcome Sanger Institute COVID-19 Surveillance Team |
| EPI_ISL_3469848 | Lighthouse Lab in Milton Keynes | Wellcome Sanger Institute for the COVID-19 Genomics UK (COG-UK) Consortium | The Lighthouse Lab in Milton Keynes and Alex Alderton, Roberto Amato, Jeffrey Barrett, Sonia Goncalves, Ewan Harrison, David K. Jackson, Ian Johnston, Dominic Kwiatkowski, Cordelia Langford, John Sillitoe on behalf of the Wellcome Sanger Institute COVID-19 Surveillance Team |
| EPI_ISL_3469849, EPI_ISL_3469850 | Lighthouse Lab in Alderley Park | Wellcome Sanger Institute for the COVID-19 Genomics UK (COG-UK) Consortium | Jacquelyn Wynn, Mairead Hyland, The Lighthouse Lab in Alderley Park and Alex Alderton, Roberto Amato, Jeffrey Barrett, Sonia Goncalves, Ewan Harrison, David K. Jackson, Ian Johnston, Dominic Kwiatkowski, Cordelia Langford, John Sillitoe on behalf of the Wellcome Sanger Institute COVID-19 Surveillance Team |
| EPI_ISL_3469851 | Health Services Laboratories | Wellcome Sanger Institute for the COVID-19 Genomics UK (COG-UK) Consortium | Health Services Laboratories and Alex Alderton, Roberto Amato, Jeffrey Barrett, Sonia Goncalves, Ewan Harrison, David K. Jackson, Ian Johnston, Dominic Kwiatkowski, Cordelia Langford, John Sillitoe on behalf of the Wellcome Sanger Institute COVID-19 Surveillance Team |
| EPI_ISL_3469852 | Lighthouse Laboratory Plymouth | Wellcome Sanger Institute for the COVID-19 Genomics UK (COG-UK) Consortium | Lighthouse Laboratory Plymouth and Alex Alderton, Roberto Amato, Jeffrey Barrett, Sonia Goncalves, Ewan Harrison, David K. Jackson, Ian Johnston, Dominic Kwiatkowski, Cordelia Langford, John Sillitoe on behalf of the Wellcome Sanger Institute COVID-19 Surveillance Team |
| EPI_ISL_3469853 | Lighthouse Lab in Alderley Park | Wellcome Sanger Institute for the COVID-19 Genomics UK (COG-UK) Consortium | Jacquelyn Wynn, Mairead Hyland, The Lighthouse Lab in Alderley Park and Alex Alderton, Roberto Amato, Jeffrey Barrett, Sonia Goncalves, Ewan Harrison, David K. Jackson, Ian Johnston, Dominic Kwiatkowski, Cordelia Langford, John Sillitoe on behalf of the Wellcome Sanger Institute COVID-19 Surveillance Team |
| EPI_ISL_3469854 | Lighthouse Lab in Milton Keynes | Wellcome Sanger Institute for the COVID-19 Genomics UK (COG-UK) Consortium | The Lighthouse Lab in Milton Keynes and Alex Alderton, Roberto Amato, Jeffrey Barrett, Sonia Goncalves, Ewan Harrison, David K. Jackson, Ian Johnston, Dominic Kwiatkowski, Cordelia Langford, John Sillitoe on behalf of the Wellcome Sanger Institute COVID-19 Surveillance Team |
| EPI_ISL_3469855, EPI_ISL_3469856 | Lighthouse Lab in Alderley Park | Wellcome Sanger Institute for the COVID-19 Genomics UK (COG-UK) Consortium | Jacquelyn Wynn, Mairead Hyland, The Lighthouse Lab in Alderley Park and Alex Alderton, Roberto Amato, Jeffrey Barrett, Sonia Goncalves, Ewan Harrison, David K. Jackson, Ian Johnston, Dominic Kwiatkowski, Cordelia Langford, John Sillitoe on behalf of the Wellcome Sanger Institute COVID-19 Surveillance Team |
| EPI_ISL_3469857 | Health Services Laboratories | Wellcome Sanger Institute for the COVID-19 Genomics UK (COG-UK) | Health Services Laboratories and Alex Alderton, Roberto Amato, Jeffrey Barrett, Sonia Goncalves, Ewan Harrison, David K. Jackson, Ian Johnston, Dominic Kwiatkowski, Cordelia Langford, John Sillitoe on behalf of the Wellcome Sanger Institute |

| | | | |
|---|---|---|---|
| | | Consortium | COVID-19 Surveillance Team |
| EPI_ISL_3469858 | Lighthouse Laboratory Plymouth | Wellcome Sanger Institute for the COVID-19 Genomics UK (COG-UK) Consortium | Lighthouse Laboratory Plymouth and Alex Alderton, Roberto Amato, Jeffrey Barrett, Sonia Goncalves, Ewan Harrison, David K. Jackson, Ian Johnston, Dominic Kwiatkowski, Cordelia Langford, John Sillitoe on behalf of the Wellcome Sanger Institute COVID-19 Surveillance Team |
| EPI_ISL_3469859 | Lighthouse Lab in Alderley Park | Wellcome Sanger Institute for the COVID-19 Genomics UK (COG-UK) Consortium | Jacquelyn Wynn, Mairead Hyland, The Lighthouse Lab in Alderley Park and Alex Alderton, Roberto Amato, Jeffrey Barrett, Sonia Goncalves, Ewan Harrison, David K. Jackson, Ian Johnston, Dominic Kwiatkowski, Cordelia Langford, John Sillitoe on behalf of the Wellcome Sanger Institute COVID-19 Surveillance Team |
| EPI_ISL_3469860 | Health Services Laboratories | Wellcome Sanger Institute for the COVID-19 Genomics UK (COG-UK) Consortium | Health Services Laboratories and Alex Alderton, Roberto Amato, Jeffrey Barrett, Sonia Goncalves, Ewan Harrison, David K. Jackson, Ian Johnston, Dominic Kwiatkowski, Cordelia Langford, John Sillitoe on behalf of the Wellcome Sanger Institute COVID-19 Surveillance Team |
| EPI_ISL_3469861, EPI_ISL_3469862 | Lighthouse Lab in Milton Keynes | Wellcome Sanger Institute for the COVID-19 Genomics UK (COG-UK) Consortium | The Lighthouse Lab in Milton Keynes and Alex Alderton, Roberto Amato, Jeffrey Barrett, Sonia Goncalves, Ewan Harrison, David K. Jackson, Ian Johnston, Dominic Kwiatkowski, Cordelia Langford, John Sillitoe on behalf of the Wellcome Sanger Institute COVID-19 Surveillance Team |
| EPI_ISL_3469863 | Lighthouse Lab in Alderley Park | Wellcome Sanger Institute for the COVID-19 Genomics UK (COG-UK) Consortium | Jacquelyn Wynn, Mairead Hyland, The Lighthouse Lab in Alderley Park and Alex Alderton, Roberto Amato, Jeffrey Barrett, Sonia Goncalves, Ewan Harrison, David K. Jackson, Ian Johnston, Dominic Kwiatkowski, Cordelia Langford, John Sillitoe on behalf of the Wellcome Sanger Institute COVID-19 Surveillance Team |
| EPI_ISL_3469864 | Lighthouse Laboratory Plymouth | Wellcome Sanger Institute for the COVID-19 Genomics UK (COG-UK) Consortium | Lighthouse Laboratory Plymouth and Alex Alderton, Roberto Amato, Jeffrey Barrett, Sonia Goncalves, Ewan Harrison, David K. Jackson, Ian Johnston, Dominic Kwiatkowski, Cordelia Langford, John Sillitoe on behalf of the Wellcome Sanger Institute COVID-19 Surveillance Team |
| EPI_ISL_3469865 | Lighthouse Lab in Milton Keynes | Wellcome Sanger Institute for the COVID-19 Genomics UK (COG-UK) Consortium | The Lighthouse Lab in Milton Keynes and Alex Alderton, Roberto Amato, Jeffrey Barrett, Sonia Goncalves, Ewan Harrison, David K. Jackson, Ian Johnston, Dominic Kwiatkowski, Cordelia Langford, John Sillitoe on behalf of the Wellcome Sanger Institute COVID-19 Surveillance Team |
| EPI_ISL_3469866, EPI_ISL_3469867, EPI_ISL_3469868, EPI_ISL_3469869 | Lighthouse Lab in Alderley Park | Wellcome Sanger Institute for the COVID-19 Genomics UK (COG-UK) Consortium | Jacquelyn Wynn, Mairead Hyland, The Lighthouse Lab in Alderley Park and Alex Alderton, Roberto Amato, Jeffrey Barrett, Sonia Goncalves, Ewan Harrison, David K. Jackson, Ian Johnston, Dominic Kwiatkowski, Cordelia Langford, John Sillitoe on behalf of the Wellcome Sanger Institute COVID-19 Surveillance Team |
| EPI_ISL_3469870 | Lighthouse Lab in Milton Keynes | Wellcome Sanger Institute for the COVID-19 Genomics UK (COG-UK) Consortium | The Lighthouse Lab in Milton Keynes and Alex Alderton, Roberto Amato, Jeffrey Barrett, Sonia Goncalves, Ewan Harrison, David K. Jackson, Ian Johnston, Dominic Kwiatkowski, Cordelia Langford, John Sillitoe on behalf of the Wellcome Sanger Institute COVID-19 Surveillance Team |
| EPI_ISL_3469871 | Lighthouse Lab in Alderley Park | Wellcome Sanger Institute for the COVID-19 Genomics UK (COG-UK) Consortium | Jacquelyn Wynn, Mairead Hyland, The Lighthouse Lab in Alderley Park and Alex Alderton, Roberto Amato, Jeffrey Barrett, Sonia Goncalves, Ewan Harrison, David K. Jackson, Ian Johnston, Dominic Kwiatkowski, Cordelia Langford, John Sillitoe on behalf of the Wellcome Sanger Institute COVID-19 Surveillance Team |
| EPI_ISL_3469872 | Health Services Laboratories | Wellcome Sanger Institute for the COVID-19 Genomics UK (COG-UK) Consortium | Health Services Laboratories and Alex Alderton, Roberto Amato, Jeffrey Barrett, Sonia Goncalves, Ewan Harrison, David K. Jackson, Ian Johnston, Dominic Kwiatkowski, Cordelia Langford, John Sillitoe on behalf of the Wellcome Sanger Institute COVID-19 Surveillance Team |
| EPI_ISL_3469873, EPI_ISL_3469874 | Lighthouse Lab in Alderley Park | Wellcome Sanger Institute for the COVID-19 Genomics UK (COG-UK) Consortium | Jacquelyn Wynn, Mairead Hyland, The Lighthouse Lab in Alderley Park and Alex Alderton, Roberto Amato, Jeffrey Barrett, Sonia Goncalves, Ewan Harrison, David K. Jackson, Ian Johnston, Dominic Kwiatkowski, Cordelia Langford, John Sillitoe on behalf of the Wellcome Sanger Institute COVID-19 Surveillance Team |
| EPI_ISL_3469875 | Lighthouse Lab in Milton Keynes | Wellcome Sanger Institute for the COVID-19 Genomics UK (COG-UK) Consortium | The Lighthouse Lab in Milton Keynes and Alex Alderton, Roberto Amato, Jeffrey Barrett, Sonia Goncalves, Ewan Harrison, David K. Jackson, Ian Johnston, Dominic Kwiatkowski, Cordelia Langford, John Sillitoe on behalf of the Wellcome Sanger Institute COVID-19 Surveillance Team |
| EPI_ISL_3469876, EPI_ISL_3469877 | Health Services Laboratories | Wellcome Sanger Institute for the COVID-19 Genomics UK (COG-UK) Consortium | Health Services Laboratories and Alex Alderton, Roberto Amato, Jeffrey Barrett, Sonia Goncalves, Ewan Harrison, David K. Jackson, Ian Johnston, Dominic Kwiatkowski, Cordelia Langford, John Sillitoe on behalf of the Wellcome Sanger Institute COVID-19 Surveillance Team |
| EPI_ISL_3469878 | Lighthouse Lab in Alderley Park | Wellcome Sanger Institute for the COVID-19 Genomics UK (COG-UK) Consortium | Jacquelyn Wynn, Mairead Hyland, The Lighthouse Lab in Alderley Park and Alex Alderton, Roberto Amato, Jeffrey Barrett, Sonia Goncalves, Ewan Harrison, David K. Jackson, Ian Johnston, Dominic Kwiatkowski, Cordelia Langford, John Sillitoe on behalf of the Wellcome Sanger Institute COVID-19 Surveillance Team |
| EPI_ISL_3469879 | Lighthouse Lab in Milton Keynes | Wellcome Sanger Institute for the COVID-19 Genomics UK (COG-UK) Consortium | The Lighthouse Lab in Milton Keynes and Alex Alderton, Roberto Amato, Jeffrey Barrett, Sonia Goncalves, Ewan Harrison, David K. Jackson, Ian Johnston, Dominic Kwiatkowski, Cordelia Langford, John Sillitoe on behalf of the Wellcome Sanger Institute COVID-19 Surveillance Team |
| EPI_ISL_3469880 | Lighthouse Laboratory Plymouth | Wellcome Sanger Institute for the COVID-19 Genomics UK (COG-UK) Consortium | Lighthouse Laboratory Plymouth and Alex Alderton, Roberto Amato, Jeffrey Barrett, Sonia Goncalves, Ewan Harrison, David K. Jackson, Ian Johnston, Dominic Kwiatkowski, Cordelia Langford, John Sillitoe on behalf of the Wellcome Sanger Institute COVID-19 Surveillance Team |
| EPI_ISL_3469881 | Lighthouse Lab in Alderley Park | Wellcome Sanger Institute for the COVID-19 Genomics UK (COG-UK) Consortium | Jacquelyn Wynn, Mairead Hyland, The Lighthouse Lab in Alderley Park and Alex Alderton, Roberto Amato, Jeffrey Barrett, Sonia Goncalves, Ewan Harrison, David K. Jackson, Ian Johnston, Dominic Kwiatkowski, Cordelia Langford, John Sillitoe on behalf of the Wellcome Sanger Institute COVID-19 Surveillance Team |
| EPI_ISL_3469882, EPI_ISL_3469883 | Lighthouse Lab in Milton Keynes | Wellcome Sanger Institute for the COVID-19 Genomics UK (COG-UK) Consortium | The Lighthouse Lab in Milton Keynes and Alex Alderton, Roberto Amato, Jeffrey Barrett, Sonia Goncalves, Ewan Harrison, David K. Jackson, Ian Johnston, Dominic Kwiatkowski, Cordelia Langford, John Sillitoe on behalf of the Wellcome Sanger Institute COVID-19 Surveillance Team |
| EPI_ISL_3469884 | Lighthouse Lab in Alderley Park | Wellcome Sanger Institute for the COVID-19 Genomics UK (COG-UK) Consortium | Jacquelyn Wynn, Mairead Hyland, The Lighthouse Lab in Alderley Park and Alex Alderton, Roberto Amato, Jeffrey Barrett, Sonia Goncalves, Ewan Harrison, David K. Jackson, Ian Johnston, Dominic Kwiatkowski, Cordelia Langford, John Sillitoe on behalf of the Wellcome Sanger Institute COVID-19 Surveillance Team |
| EPI_ISL_3469885 | Lighthouse Lab in Milton Keynes | Wellcome Sanger Institute for the COVID-19 Genomics UK (COG-UK) Consortium | The Lighthouse Lab in Milton Keynes and Alex Alderton, Roberto Amato, Jeffrey Barrett, Sonia Goncalves, Ewan Harrison, David K. Jackson, Ian Johnston, Dominic Kwiatkowski, Cordelia Langford, John Sillitoe on behalf of the Wellcome Sanger Institute COVID-19 Surveillance Team |
| EPI_ISL_3469886 | Lighthouse Lab in Alderley Park | Wellcome Sanger Institute for the COVID-19 Genomics UK (COG-UK) Consortium | Jacquelyn Wynn, Mairead Hyland, The Lighthouse Lab in Alderley Park and Alex Alderton, Roberto Amato, Jeffrey Barrett, Sonia Goncalves, Ewan Harrison, David K. Jackson, Ian Johnston, Dominic Kwiatkowski, Cordelia Langford, John Sillitoe on behalf of the Wellcome Sanger Institute COVID-19 Surveillance Team |
| EPI_ISL_3469887 | Health Services Laboratories | Wellcome Sanger Institute for the COVID-19 Genomics UK (COG-UK) Consortium | Health Services Laboratories and Alex Alderton, Roberto Amato, Jeffrey Barrett, Sonia Goncalves, Ewan Harrison, David K. Jackson, Ian Johnston, Dominic Kwiatkowski, Cordelia Langford, John Sillitoe on behalf of the Wellcome Sanger Institute COVID-19 Surveillance Team |
| EPI_ISL_3469888, EPI_ISL_3469889, EPI_ISL_3469890 | Lighthouse Lab in Alderley Park | Wellcome Sanger Institute for the COVID-19 Genomics UK (COG-UK) Consortium | Jacquelyn Wynn, Mairead Hyland, The Lighthouse Lab in Alderley Park and Alex Alderton, Roberto Amato, Jeffrey Barrett, Sonia Goncalves, Ewan Harrison, David K. Jackson, Ian Johnston, Dominic Kwiatkowski, Cordelia Langford, John Sillitoe on behalf of the Wellcome Sanger Institute COVID-19 Surveillance Team |
| EPI_ISL_3469891, EPI_ISL_3469892 | Lighthouse Lab in Milton Keynes | Wellcome Sanger Institute for the COVID-19 Genomics UK (COG-UK) Consortium | The Lighthouse Lab in Milton Keynes and Alex Alderton, Roberto Amato, Jeffrey Barrett, Sonia Goncalves, Ewan Harrison, David K. Jackson, Ian Johnston, Dominic Kwiatkowski, Cordelia Langford, John Sillitoe on behalf of the Wellcome Sanger Institute COVID-19 Surveillance Team |
| EPI_ISL_3469893 | Lighthouse Lab in Alderley Park | Wellcome Sanger Institute for the COVID-19 Genomics UK (COG-UK) Consortium | Jacquelyn Wynn, Mairead Hyland, The Lighthouse Lab in Alderley Park and Alex Alderton, Roberto Amato, Jeffrey Barrett, Sonia Goncalves, Ewan Harrison, David K. Jackson, Ian Johnston, Dominic Kwiatkowski, Cordelia Langford, John Sillitoe on behalf of the Wellcome Sanger Institute COVID-19 Surveillance Team |
| EPI_ISL_3469894 | Lighthouse Lab in Milton Keynes | Wellcome Sanger Institute for the COVID-19 Genomics UK (COG-UK) Consortium | The Lighthouse Lab in Milton Keynes and Alex Alderton, Roberto Amato, Jeffrey Barrett, Sonia Goncalves, Ewan Harrison, David K. Jackson, Ian Johnston, Dominic Kwiatkowski, Cordelia Langford, John Sillitoe on behalf of the Wellcome Sanger Institute COVID-19 Surveillance Team |
| EPI_ISL_3469895 | Lighthouse Lab in Alderley Park | Wellcome Sanger Institute for the COVID-19 Genomics UK (COG-UK) Consortium | Jacquelyn Wynn, Mairead Hyland, The Lighthouse Lab in Alderley Park and Alex Alderton, Roberto Amato, Jeffrey Barrett, Sonia Goncalves, Ewan Harrison, David K. Jackson, Ian Johnston, Dominic Kwiatkowski, Cordelia Langford, John Sillitoe on behalf of the Wellcome Sanger Institute COVID-19 Surveillance Team |
| EPI_ISL_3469896, EPI_ISL_3469897 | Health Services Laboratories | Wellcome Sanger Institute for the COVID-19 Genomics UK (COG-UK) Consortium | Health Services Laboratories and Alex Alderton, Roberto Amato, Jeffrey Barrett, Sonia Goncalves, Ewan Harrison, David K. Jackson, Ian Johnston, Dominic Kwiatkowski, Cordelia Langford, John Sillitoe on behalf of the Wellcome Sanger Institute COVID-19 Surveillance Team |
| EPI_ISL_3469898, EPI_ISL_3469899, EPI_ISL_3469900 | Lighthouse Lab in Alderley Park | Wellcome Sanger Institute for the COVID-19 Genomics UK (COG-UK) Consortium | Jacquelyn Wynn, Mairead Hyland, The Lighthouse Lab in Alderley Park and Alex Alderton, Roberto Amato, Jeffrey Barrett, Sonia Goncalves, Ewan Harrison, David K. Jackson, Ian Johnston, Dominic Kwiatkowski, Cordelia Langford, John Sillitoe on behalf of the Wellcome Sanger Institute COVID-19 Surveillance Team |
| EPI_ISL_3469901 | Lighthouse Lab in Milton Keynes | Wellcome Sanger Institute for the COVID-19 Genomics UK (COG-UK) Consortium | The Lighthouse Lab in Milton Keynes and Alex Alderton, Roberto Amato, Jeffrey Barrett, Sonia Goncalves, Ewan Harrison, David K. Jackson, Ian Johnston, Dominic Kwiatkowski, Cordelia Langford, John Sillitoe on behalf of the Wellcome Sanger Institute COVID-19 Surveillance Team |
| EPI_ISL_3469902, EPI_ISL_3469903 | Lighthouse Lab in Alderley Park | Wellcome Sanger Institute for the COVID-19 Genomics UK (COG-UK) Consortium | Jacquelyn Wynn, Mairead Hyland, The Lighthouse Lab in Alderley Park and Alex Alderton, Roberto Amato, Jeffrey Barrett, Sonia Goncalves, Ewan Harrison, David K. Jackson, Ian Johnston, Dominic Kwiatkowski, Cordelia Langford, John Sillitoe on behalf of the Wellcome Sanger Institute COVID-19 Surveillance Team |
| EPI_ISL_3469904 | Lighthouse Lab in Milton Keynes | Wellcome Sanger Institute for the COVID-19 Genomics UK (COG-UK) Consortium | The Lighthouse Lab in Milton Keynes and Alex Alderton, Roberto Amato, Jeffrey Barrett, Sonia Goncalves, Ewan Harrison, David K. Jackson, Ian Johnston, Dominic Kwiatkowski, Cordelia Langford, John Sillitoe on behalf of the Wellcome Sanger Institute COVID-19 Surveillance Team |
| EPI_ISL_3469905 | Lighthouse Lab in Alderley Park | Wellcome Sanger Institute for the COVID-19 Genomics UK (COG-UK) Consortium | Jacquelyn Wynn, Mairead Hyland, The Lighthouse Lab in Alderley Park and Alex Alderton, Roberto Amato, Jeffrey Barrett, Sonia Goncalves, Ewan Harrison, David K. Jackson, Ian Johnston, Dominic Kwiatkowski, Cordelia Langford, John Sillitoe on behalf of the Wellcome Sanger Institute COVID-19 Surveillance Team |
| EPI_ISL_3469906 | Lighthouse Lab in Milton Keynes | Wellcome Sanger Institute for the COVID-19 Genomics UK (COG-UK) Consortium | The Lighthouse Lab in Milton Keynes and Alex Alderton, Roberto Amato, Jeffrey Barrett, Sonia Goncalves, Ewan Harrison, David K. Jackson, Ian Johnston, Dominic Kwiatkowski, Cordelia Langford, John Sillitoe on behalf of the Wellcome Sanger Institute COVID-19 Surveillance Team |
| EPI_ISL_3469907, EPI_ISL_3469908, EPI_ISL_3469909 | Lighthouse Lab in Alderley Park | Wellcome Sanger Institute for the COVID-19 Genomics UK (COG-UK) Consortium | Jacquelyn Wynn, Mairead Hyland, The Lighthouse Lab in Alderley Park and Alex Alderton, Roberto Amato, Jeffrey Barrett, Sonia Goncalves, Ewan Harrison, David K. Jackson, Ian Johnston, Dominic Kwiatkowski, Cordelia Langford, John Sillitoe on behalf of the Wellcome Sanger Institute COVID-19 Surveillance Team |
| EPI_ISL_3469910 | Lighthouse Lab in Milton Keynes | Wellcome Sanger Institute for the COVID-19 Genomics UK (COG-UK) Consortium | The Lighthouse Lab in Milton Keynes and Alex Alderton, Roberto Amato, Jeffrey Barrett, Sonia Goncalves, Ewan Harrison, David K. Jackson, Ian Johnston, Dominic Kwiatkowski, Cordelia Langford, John Sillitoe on behalf of the Wellcome Sanger Institute COVID-19 Surveillance Team |
| EPI_ISL_3469911, EPI_ISL_3469912, EPI_ISL_3469913, EPI_ISL_3469914, EPI_ISL_3469915 | Lighthouse Lab in Alderley Park | Wellcome Sanger Institute for the COVID-19 Genomics UK (COG-UK) Consortium | Jacquelyn Wynn, Mairead Hyland, The Lighthouse Lab in Alderley Park and Alex Alderton, Roberto Amato, Jeffrey Barrett, Sonia Goncalves, Ewan Harrison, David K. Jackson, Ian Johnston, Dominic Kwiatkowski, Cordelia Langford, John Sillitoe on behalf of the Wellcome Sanger Institute COVID-19 Surveillance Team |
| EPI_ISL_3469916 | Lighthouse Lab in Milton Keynes | Wellcome Sanger Institute for the COVID-19 Genomics UK (COG-UK) Consortium | The Lighthouse Lab in Milton Keynes and Alex Alderton, Roberto Amato, Jeffrey Barrett, Sonia Goncalves, Ewan Harrison, David K. Jackson, Ian Johnston, Dominic Kwiatkowski, Cordelia Langford, John Sillitoe on behalf of the Wellcome Sanger Institute COVID-19 Surveillance Team |
| EPI_ISL_3469917, EPI_ISL_3469918 | Lighthouse Lab in Alderley Park | Wellcome Sanger Institute for the COVID-19 Genomics UK (COG-UK) Consortium | Jacquelyn Wynn, Mairead Hyland, The Lighthouse Lab in Alderley Park and Alex Alderton, Roberto Amato, Jeffrey Barrett, Sonia Goncalves, Ewan Harrison, David K. Jackson, Ian Johnston, Dominic Kwiatkowski, Cordelia Langford, John Sillitoe on behalf of the Wellcome Sanger Institute COVID-19 Surveillance Team |
| EPI_ISL_3469919 | Lighthouse Lab in Milton Keynes | Wellcome Sanger Institute for the COVID-19 Genomics UK (COG-UK) Consortium | The Lighthouse Lab in Milton Keynes and Alex Alderton, Roberto Amato, Jeffrey Barrett, Sonia Goncalves, Ewan Harrison, David K. Jackson, Ian Johnston, Dominic Kwiatkowski, Cordelia Langford, John Sillitoe on behalf of the Wellcome Sanger Institute COVID-19 Surveillance Team |

| | | | |
|---|---|---|---|
| EPI_ISL_3469920, EPI_ISL_3469921 | Lighthouse Lab in Alderley Park | Wellcome Sanger Institute for the COVID-19 Genomics UK (COG-UK) Consortium | Jacquelyn Wynn, Mairead Hyland, The Lighthouse Lab in Alderley Park and Alex Alderton, Roberto Amato, Jeffrey Barrett, Sonia Goncalves, Ewan Harrison, David K. Jackson, Ian Johnston, Dominic Kwiatkowski, Cordelia Langford, John Sillitoe on behalf of the Wellcome Sanger Institute COVID-19 Surveillance Team |
| EPI_ISL_3469922 | Lighthouse Lab in Milton Keynes | Wellcome Sanger Institute for the COVID-19 Genomics UK (COG-UK) Consortium | The Lighthouse Lab in Milton Keynes and Alex Alderton, Roberto Amato, Jeffrey Barrett, Sonia Goncalves, Ewan Harrison, David K. Jackson, Ian Johnston, Dominic Kwiatkowski, Cordelia Langford, John Sillitoe on behalf of the Wellcome Sanger Institute COVID-19 Surveillance Team |
| EPI_ISL_3469923 | Lighthouse Lab in Alderley Park | Wellcome Sanger Institute for the COVID-19 Genomics UK (COG-UK) Consortium | Jacquelyn Wynn, Mairead Hyland, The Lighthouse Lab in Alderley Park and Alex Alderton, Roberto Amato, Jeffrey Barrett, Sonia Goncalves, Ewan Harrison, David K. Jackson, Ian Johnston, Dominic Kwiatkowski, Cordelia Langford, John Sillitoe on behalf of the Wellcome Sanger Institute COVID-19 Surveillance Team |
| EPI_ISL_3469924, EPI_ISL_3469925 | Lighthouse Lab in Milton Keynes | Wellcome Sanger Institute for the COVID-19 Genomics UK (COG-UK) Consortium | The Lighthouse Lab in Milton Keynes and Alex Alderton, Roberto Amato, Jeffrey Barrett, Sonia Goncalves, Ewan Harrison, David K. Jackson, Ian Johnston, Dominic Kwiatkowski, Cordelia Langford, John Sillitoe on behalf of the Wellcome Sanger Institute COVID-19 Surveillance Team |
| EPI_ISL_3469926 | Lighthouse Lab in Alderley Park | Wellcome Sanger Institute for the COVID-19 Genomics UK (COG-UK) Consortium | Jacquelyn Wynn, Mairead Hyland, The Lighthouse Lab in Alderley Park and Alex Alderton, Roberto Amato, Jeffrey Barrett, Sonia Goncalves, Ewan Harrison, David K. Jackson, Ian Johnston, Dominic Kwiatkowski, Cordelia Langford, John Sillitoe on behalf of the Wellcome Sanger Institute COVID-19 Surveillance Team |
| EPI_ISL_3469927 | Lighthouse Lab in Milton Keynes | Wellcome Sanger Institute for the COVID-19 Genomics UK (COG-UK) Consortium | The Lighthouse Lab in Milton Keynes and Alex Alderton, Roberto Amato, Jeffrey Barrett, Sonia Goncalves, Ewan Harrison, David K. Jackson, Ian Johnston, Dominic Kwiatkowski, Cordelia Langford, John Sillitoe on behalf of the Wellcome Sanger Institute COVID-19 Surveillance Team |
| EPI_ISL_3469928 | Health Services Laboratories | Wellcome Sanger Institute for the COVID-19 Genomics UK (COG-UK) Consortium | Health Services Laboratories and Alex Alderton, Roberto Amato, Jeffrey Barrett, Sonia Goncalves, Ewan Harrison, David K. Jackson, Ian Johnston, Dominic Kwiatkowski, Cordelia Langford, John Sillitoe on behalf of the Wellcome Sanger Institute COVID-19 Surveillance Team |
| EPI_ISL_3469929, EPI_ISL_3469930, EPI_ISL_3469931, EPI_ISL_3469932, EPI_ISL_3469933 | Lighthouse Lab in Alderley Park | Wellcome Sanger Institute for the COVID-19 Genomics UK (COG-UK) Consortium | Jacquelyn Wynn, Mairead Hyland, The Lighthouse Lab in Alderley Park and Alex Alderton, Roberto Amato, Jeffrey Barrett, Sonia Goncalves, Ewan Harrison, David K. Jackson, Ian Johnston, Dominic Kwiatkowski, Cordelia Langford, John Sillitoe on behalf of the Wellcome Sanger Institute COVID-19 Surveillance Team |
| EPI_ISL_3469934 | Lighthouse Lab in Milton Keynes | Wellcome Sanger Institute for the COVID-19 Genomics UK (COG-UK) Consortium | The Lighthouse Lab in Milton Keynes and Alex Alderton, Roberto Amato, Jeffrey Barrett, Sonia Goncalves, Ewan Harrison, David K. Jackson, Ian Johnston, Dominic Kwiatkowski, Cordelia Langford, John Sillitoe on behalf of the Wellcome Sanger Institute COVID-19 Surveillance Team |
| EPI_ISL_3469935, EPI_ISL_3469936, EPI_ISL_3469937 | Lighthouse Lab in Alderley Park | Wellcome Sanger Institute for the COVID-19 Genomics UK (COG-UK) Consortium | Jacquelyn Wynn, Mairead Hyland, The Lighthouse Lab in Alderley Park and Alex Alderton, Roberto Amato, Jeffrey Barrett, Sonia Goncalves, Ewan Harrison, David K. Jackson, Ian Johnston, Dominic Kwiatkowski, Cordelia Langford, John Sillitoe on behalf of the Wellcome Sanger Institute COVID-19 Surveillance Team |
| EPI_ISL_3469938 | Lighthouse Lab in Milton Keynes | Wellcome Sanger Institute for the COVID-19 Genomics UK (COG-UK) Consortium | The Lighthouse Lab in Milton Keynes and Alex Alderton, Roberto Amato, Jeffrey Barrett, Sonia Goncalves, Ewan Harrison, David K. Jackson, Ian Johnston, Dominic Kwiatkowski, Cordelia Langford, John Sillitoe on behalf of the Wellcome Sanger Institute COVID-19 Surveillance Team |
| EPI_ISL_3469939, EPI_ISL_3469940 | Lighthouse Lab in Alderley Park | Wellcome Sanger Institute for the COVID-19 Genomics UK (COG-UK) Consortium | Jacquelyn Wynn, Mairead Hyland, The Lighthouse Lab in Alderley Park and Alex Alderton, Roberto Amato, Jeffrey Barrett, Sonia Goncalves, Ewan Harrison, David K. Jackson, Ian Johnston, Dominic Kwiatkowski, Cordelia Langford, John Sillitoe on behalf of the Wellcome Sanger Institute COVID-19 Surveillance Team |
| EPI_ISL_3469941 | Lighthouse Laboratory Plymouth | Wellcome Sanger Institute for the COVID-19 Genomics UK (COG-UK) Consortium | Lighthouse Laboratory Plymouth and Alex Alderton, Roberto Amato, Jeffrey Barrett, Sonia Goncalves, Ewan Harrison, David K. Jackson, Ian Johnston, Dominic Kwiatkowski, Cordelia Langford, John Sillitoe on behalf of the Wellcome Sanger Institute COVID-19 Surveillance Team |
| EPI_ISL_3469942 | Lighthouse Lab in Alderley Park | Wellcome Sanger Institute for the COVID-19 Genomics UK (COG-UK) Consortium | Jacquelyn Wynn, Mairead Hyland, The Lighthouse Lab in Alderley Park and Alex Alderton, Roberto Amato, Jeffrey Barrett, Sonia Goncalves, Ewan Harrison, David K. Jackson, Ian Johnston, Dominic Kwiatkowski, Cordelia Langford, John Sillitoe on behalf of the Wellcome Sanger Institute COVID-19 Surveillance Team |
| EPI_ISL_3469943 | Lighthouse Lab in Milton Keynes | Wellcome Sanger Institute for the COVID-19 Genomics UK (COG-UK) Consortium | The Lighthouse Lab in Milton Keynes and Alex Alderton, Roberto Amato, Jeffrey Barrett, Sonia Goncalves, Ewan Harrison, David K. Jackson, Ian Johnston, Dominic Kwiatkowski, Cordelia Langford, John Sillitoe on behalf of the Wellcome Sanger Institute COVID-19 Surveillance Team |
| EPI_ISL_3469944 | Lighthouse Lab in Alderley Park | Wellcome Sanger Institute for the COVID-19 Genomics UK (COG-UK) Consortium | Jacquelyn Wynn, Mairead Hyland, The Lighthouse Lab in Alderley Park and Alex Alderton, Roberto Amato, Jeffrey Barrett, Sonia Goncalves, Ewan Harrison, David K. Jackson, Ian Johnston, Dominic Kwiatkowski, Cordelia Langford, John Sillitoe on behalf of the Wellcome Sanger Institute COVID-19 Surveillance Team |
| EPI_ISL_3469945 | Lighthouse Laboratory Plymouth | Wellcome Sanger Institute for the COVID-19 Genomics UK (COG-UK) Consortium | Lighthouse Laboratory Plymouth and Alex Alderton, Roberto Amato, Jeffrey Barrett, Sonia Goncalves, Ewan Harrison, David K. Jackson, Ian Johnston, Dominic Kwiatkowski, Cordelia Langford, John Sillitoe on behalf of the Wellcome Sanger Institute COVID-19 Surveillance Team |
| EPI_ISL_3469946, EPI_ISL_3469947, EPI_ISL_3469948 | Lighthouse Lab in Alderley Park | Wellcome Sanger Institute for the COVID-19 Genomics UK (COG-UK) Consortium | Jacquelyn Wynn, Mairead Hyland, The Lighthouse Lab in Alderley Park and Alex Alderton, Roberto Amato, Jeffrey Barrett, Sonia Goncalves, Ewan Harrison, David K. Jackson, Ian Johnston, Dominic Kwiatkowski, Cordelia Langford, John Sillitoe on behalf of the Wellcome Sanger Institute COVID-19 Surveillance Team |
| EPI_ISL_3469949 | Lighthouse Lab in Milton Keynes | Wellcome Sanger Institute for the COVID-19 Genomics UK (COG-UK) Consortium | The Lighthouse Lab in Milton Keynes and Alex Alderton, Roberto Amato, Jeffrey Barrett, Sonia Goncalves, Ewan Harrison, David K. Jackson, Ian Johnston, Dominic Kwiatkowski, Cordelia Langford, John Sillitoe on behalf of the Wellcome Sanger Institute COVID-19 Surveillance Team |
| EPI_ISL_3469950, EPI_ISL_3469951 | Lighthouse Lab in Alderley Park | Wellcome Sanger Institute for the COVID-19 Genomics UK (COG-UK) Consortium | Jacquelyn Wynn, Mairead Hyland, The Lighthouse Lab in Alderley Park and Alex Alderton, Roberto Amato, Jeffrey Barrett, Sonia Goncalves, Ewan Harrison, David K. Jackson, Ian Johnston, Dominic Kwiatkowski, Cordelia Langford, John Sillitoe on behalf of the Wellcome Sanger Institute COVID-19 Surveillance Team |
| EPI_ISL_3469952 | Lighthouse Laboratory Plymouth | Wellcome Sanger Institute for the COVID-19 Genomics UK (COG-UK) Consortium | Lighthouse Laboratory Plymouth and Alex Alderton, Roberto Amato, Jeffrey Barrett, Sonia Goncalves, Ewan Harrison, David K. Jackson, Ian Johnston, Dominic Kwiatkowski, Cordelia Langford, John Sillitoe on behalf of the Wellcome Sanger Institute COVID-19 Surveillance Team |
| EPI_ISL_3469953 | Lighthouse Lab in Milton Keynes | Wellcome Sanger Institute for the COVID-19 Genomics UK (COG-UK) Consortium | The Lighthouse Lab in Milton Keynes and Alex Alderton, Roberto Amato, Jeffrey Barrett, Sonia Goncalves, Ewan Harrison, David K. Jackson, Ian Johnston, Dominic Kwiatkowski, Cordelia Langford, John Sillitoe on behalf of the Wellcome Sanger Institute COVID-19 Surveillance Team |
| EPI_ISL_3469954, EPI_ISL_3469955, EPI_ISL_3469956 | Lighthouse Lab in Alderley Park | Wellcome Sanger Institute for the COVID-19 Genomics UK (COG-UK) Consortium | Jacquelyn Wynn, Mairead Hyland, The Lighthouse Lab in Alderley Park and Alex Alderton, Roberto Amato, Jeffrey Barrett, Sonia Goncalves, Ewan Harrison, David K. Jackson, Ian Johnston, Dominic Kwiatkowski, Cordelia Langford, John Sillitoe on behalf of the Wellcome Sanger Institute COVID-19 Surveillance Team |
| EPI_ISL_3469957 | Lighthouse Laboratory Plymouth | Wellcome Sanger Institute for the COVID-19 Genomics UK (COG-UK) Consortium | Lighthouse Laboratory Plymouth and Alex Alderton, Roberto Amato, Jeffrey Barrett, Sonia Goncalves, Ewan Harrison, David K. Jackson, Ian Johnston, Dominic Kwiatkowski, Cordelia Langford, John Sillitoe on behalf of the Wellcome Sanger Institute COVID-19 Surveillance Team |
| EPI_ISL_3469958 | Lighthouse Lab in Alderley Park | Wellcome Sanger Institute for the COVID-19 Genomics UK (COG-UK) Consortium | Jacquelyn Wynn, Mairead Hyland, The Lighthouse Lab in Alderley Park and Alex Alderton, Roberto Amato, Jeffrey Barrett, Sonia Goncalves, Ewan Harrison, David K. Jackson, Ian Johnston, Dominic Kwiatkowski, Cordelia Langford, John Sillitoe on behalf of the Wellcome Sanger Institute COVID-19 Surveillance Team |
| EPI_ISL_3469959 | Lighthouse Lab in Milton Keynes | Wellcome Sanger Institute for the COVID-19 Genomics UK (COG-UK) Consortium | The Lighthouse Lab in Milton Keynes and Alex Alderton, Roberto Amato, Jeffrey Barrett, Sonia Goncalves, Ewan Harrison, David K. Jackson, Ian Johnston, Dominic Kwiatkowski, Cordelia Langford, John Sillitoe on behalf of the Wellcome Sanger Institute COVID-19 Surveillance Team |
| EPI_ISL_3469960, EPI_ISL_3469961, EPI_ISL_3469962 | Lighthouse Lab in Alderley Park | Wellcome Sanger Institute for the COVID-19 Genomics UK (COG-UK) Consortium | Jacquelyn Wynn, Mairead Hyland, The Lighthouse Lab in Alderley Park and Alex Alderton, Roberto Amato, Jeffrey Barrett, Sonia Goncalves, Ewan Harrison, David K. Jackson, Ian Johnston, Dominic Kwiatkowski, Cordelia Langford, John Sillitoe on behalf of the Wellcome Sanger Institute COVID-19 Surveillance Team |
| EPI_ISL_3469963 | Lighthouse Laboratory Plymouth | Wellcome Sanger Institute for the COVID-19 Genomics UK (COG-UK) Consortium | Lighthouse Laboratory Plymouth and Alex Alderton, Roberto Amato, Jeffrey Barrett, Sonia Goncalves, Ewan Harrison, David K. Jackson, Ian Johnston, Dominic Kwiatkowski, Cordelia Langford, John Sillitoe on behalf of the Wellcome Sanger Institute COVID-19 Surveillance Team |
| EPI_ISL_3469964, EPI_ISL_3469965 | Lighthouse Lab in Alderley Park | Wellcome Sanger Institute for the COVID-19 Genomics UK (COG-UK) Consortium | Jacquelyn Wynn, Mairead Hyland, The Lighthouse Lab in Alderley Park and Alex Alderton, Roberto Amato, Jeffrey Barrett, Sonia Goncalves, Ewan Harrison, David K. Jackson, Ian Johnston, Dominic Kwiatkowski, Cordelia Langford, John Sillitoe on behalf of the Wellcome Sanger Institute COVID-19 Surveillance Team |
| EPI_ISL_3469966, EPI_ISL_3469967 | Lighthouse Lab in Milton Keynes | Wellcome Sanger Institute for the COVID-19 Genomics UK (COG-UK) Consortium | The Lighthouse Lab in Milton Keynes and Alex Alderton, Roberto Amato, Jeffrey Barrett, Sonia Goncalves, Ewan Harrison, David K. Jackson, Ian Johnston, Dominic Kwiatkowski, Cordelia Langford, John Sillitoe on behalf of the Wellcome Sanger Institute COVID-19 Surveillance Team |
| EPI_ISL_3469968 | Lighthouse Lab in Alderley Park | Wellcome Sanger Institute for the COVID-19 Genomics UK (COG-UK) Consortium | Jacquelyn Wynn, Mairead Hyland, The Lighthouse Lab in Alderley Park and Alex Alderton, Roberto Amato, Jeffrey Barrett, Sonia Goncalves, Ewan Harrison, David K. Jackson, Ian Johnston, Dominic Kwiatkowski, Cordelia Langford, John Sillitoe on behalf of the Wellcome Sanger Institute COVID-19 Surveillance Team |
| EPI_ISL_3469969 | Lighthouse Lab in Milton Keynes | Wellcome Sanger Institute for the COVID-19 Genomics UK (COG-UK) Consortium | The Lighthouse Lab in Milton Keynes and Alex Alderton, Roberto Amato, Jeffrey Barrett, Sonia Goncalves, Ewan Harrison, David K. Jackson, Ian Johnston, Dominic Kwiatkowski, Cordelia Langford, John Sillitoe on behalf of the Wellcome Sanger Institute COVID-19 Surveillance Team |
| EPI_ISL_3469970, EPI_ISL_3469971, EPI_ISL_3469972, EPI_ISL_3469973, EPI_ISL_3469974 | Lighthouse Lab in Alderley Park | Wellcome Sanger Institute for the COVID-19 Genomics UK (COG-UK) Consortium | Jacquelyn Wynn, Mairead Hyland, The Lighthouse Lab in Alderley Park and Alex Alderton, Roberto Amato, Jeffrey Barrett, Sonia Goncalves, Ewan Harrison, David K. Jackson, Ian Johnston, Dominic Kwiatkowski, Cordelia Langford, John Sillitoe on behalf of the Wellcome Sanger Institute COVID-19 Surveillance Team |
| EPI_ISL_3469975 | Lighthouse Lab in Milton Keynes | Wellcome Sanger Institute for the COVID-19 Genomics UK (COG-UK) Consortium | The Lighthouse Lab in Milton Keynes and Alex Alderton, Roberto Amato, Jeffrey Barrett, Sonia Goncalves, Ewan Harrison, David K. Jackson, Ian Johnston, Dominic Kwiatkowski, Cordelia Langford, John Sillitoe on behalf of the Wellcome Sanger Institute COVID-19 Surveillance Team |
| EPI_ISL_3469976, EPI_ISL_3469977 | Lighthouse Lab in Alderley Park | Wellcome Sanger Institute for the COVID-19 Genomics UK (COG-UK) Consortium | Jacquelyn Wynn, Mairead Hyland, The Lighthouse Lab in Alderley Park and Alex Alderton, Roberto Amato, Jeffrey Barrett, Sonia Goncalves, Ewan Harrison, David K. Jackson, Ian Johnston, Dominic Kwiatkowski, Cordelia Langford, John Sillitoe on behalf of the Wellcome Sanger Institute COVID-19 Surveillance Team |
| EPI_ISL_3469978, EPI_ISL_3469979 | Lighthouse Lab in Milton Keynes | Wellcome Sanger Institute for the COVID-19 Genomics UK (COG-UK) Consortium | The Lighthouse Lab in Milton Keynes and Alex Alderton, Roberto Amato, Jeffrey Barrett, Sonia Goncalves, Ewan Harrison, David K. Jackson, Ian Johnston, Dominic Kwiatkowski, Cordelia Langford, John Sillitoe on behalf of the Wellcome Sanger Institute COVID-19 Surveillance Team |
| EPI_ISL_3469980 | Lighthouse Laboratory Plymouth | Wellcome Sanger Institute for the COVID-19 Genomics UK (COG-UK) Consortium | Lighthouse Laboratory Plymouth and Alex Alderton, Roberto Amato, Jeffrey Barrett, Sonia Goncalves, Ewan Harrison, David K. Jackson, Ian Johnston, Dominic Kwiatkowski, Cordelia Langford, John Sillitoe on behalf of the Wellcome Sanger Institute COVID-19 Surveillance Team |
| EPI_ISL_3469981 | Health Services Laboratories | Wellcome Sanger Institute for the COVID-19 Genomics UK (COG-UK) Consortium | Health Services Laboratories and Alex Alderton, Roberto Amato, Jeffrey Barrett, Sonia Goncalves, Ewan Harrison, David K. Jackson, Ian Johnston, Dominic Kwiatkowski, Cordelia Langford, John Sillitoe on behalf of the Wellcome Sanger Institute COVID-19 Surveillance Team |
| EPI_ISL_3469982, EPI_ISL_3469983, EPI_ISL_3469984, EPI_ISL_3469985 | Lighthouse Lab in Alderley Park | Wellcome Sanger Institute for the COVID-19 Genomics UK (COG-UK) Consortium | Jacquelyn Wynn, Mairead Hyland, The Lighthouse Lab in Alderley Park and Alex Alderton, Roberto Amato, Jeffrey Barrett, Sonia Goncalves, Ewan Harrison, David K. Jackson, Ian Johnston, Dominic Kwiatkowski, Cordelia Langford, John Sillitoe on behalf of the Wellcome Sanger Institute COVID-19 Surveillance Team |
| EPI_ISL_3469986, EPI_ISL_3469987, EPI_ISL_3469988 | Lighthouse Lab in Milton Keynes | Wellcome Sanger Institute for the COVID-19 Genomics UK (COG-UK) Consortium | The Lighthouse Lab in Milton Keynes and Alex Alderton, Roberto Amato, Jeffrey Barrett, Sonia Goncalves, Ewan Harrison, David K. Jackson, Ian Johnston, Dominic Kwiatkowski, Cordelia Langford, John Sillitoe on behalf of the Wellcome Sanger Institute COVID-19 Surveillance Team |

| | | | |
|---|---|---|---|
| EPI_ISL_3469989, EPI_ISL_3469990, EPI_ISL_3469991, EPI_ISL_3469992 | Lighthouse Lab in Alderley Park | Wellcome Sanger Institute for the COVID-19 Genomics UK (COG-UK) Consortium | Jacquelyn Wynn, Mairead Hyland, The Lighthouse Lab in Alderley Park and Alex Alderton, Roberto Amato, Jeffrey Barrett, Sonia Goncalves, Ewan Harrison, David K. Jackson, Ian Johnston, Dominic Kwiatkowski, Cordelia Langford, John Sillitoe on behalf of the Wellcome Sanger Institute COVID-19 Surveillance Team |
| EPI_ISL_3469993 | Lighthouse Laboratory Plymouth | Wellcome Sanger Institute for the COVID-19 Genomics UK (COG-UK) Consortium | Lighthouse Laboratory Plymouth and Alex Alderton, Roberto Amato, Jeffrey Barrett, Sonia Goncalves, Ewan Harrison, David K. Jackson, Ian Johnston, Dominic Kwiatkowski, Cordelia Langford, John Sillitoe on behalf of the Wellcome Sanger Institute COVID-19 Surveillance Team |
| EPI_ISL_3469994, EPI_ISL_3469995 | Lighthouse Lab in Alderley Park | Wellcome Sanger Institute for the COVID-19 Genomics UK (COG-UK) Consortium | Jacquelyn Wynn, Mairead Hyland, The Lighthouse Lab in Alderley Park and Alex Alderton, Roberto Amato, Jeffrey Barrett, Sonia Goncalves, Ewan Harrison, David K. Jackson, Ian Johnston, Dominic Kwiatkowski, Cordelia Langford, John Sillitoe on behalf of the Wellcome Sanger Institute COVID-19 Surveillance Team |
| EPI_ISL_3469996 | Lighthouse Laboratory Plymouth | Wellcome Sanger Institute for the COVID-19 Genomics UK (COG-UK) Consortium | Lighthouse Laboratory Plymouth and Alex Alderton, Roberto Amato, Jeffrey Barrett, Sonia Goncalves, Ewan Harrison, David K. Jackson, Ian Johnston, Dominic Kwiatkowski, Cordelia Langford, John Sillitoe on behalf of the Wellcome Sanger Institute COVID-19 Surveillance Team |
| EPI_ISL_3469997, EPI_ISL_3469998 | Lighthouse Lab in Alderley Park | Wellcome Sanger Institute for the COVID-19 Genomics UK (COG-UK) Consortium | Jacquelyn Wynn, Mairead Hyland, The Lighthouse Lab in Alderley Park and Alex Alderton, Roberto Amato, Jeffrey Barrett, Sonia Goncalves, Ewan Harrison, David K. Jackson, Ian Johnston, Dominic Kwiatkowski, Cordelia Langford, John Sillitoe on behalf of the Wellcome Sanger Institute COVID-19 Surveillance Team |
| EPI_ISL_3469999 | Health Services Laboratories | Wellcome Sanger Institute for the COVID-19 Genomics UK (COG-UK) Consortium | Health Services Laboratories and Alex Alderton, Roberto Amato, Jeffrey Barrett, Sonia Goncalves, Ewan Harrison, David K. Jackson, Ian Johnston, Dominic Kwiatkowski, Cordelia Langford, John Sillitoe on behalf of the Wellcome Sanger Institute COVID-19 Surveillance Team |
| EPI_ISL_3470000 | Lighthouse Lab in Milton Keynes | Wellcome Sanger Institute for the COVID-19 Genomics UK (COG-UK) Consortium | The Lighthouse Lab in Milton Keynes and Alex Alderton, Roberto Amato, Jeffrey Barrett, Sonia Goncalves, Ewan Harrison, David K. Jackson, Ian Johnston, Dominic Kwiatkowski, Cordelia Langford, John Sillitoe on behalf of the Wellcome Sanger Institute COVID-19 Surveillance Team |
| EPI_ISL_3470001 | Lighthouse Laboratory Plymouth | Wellcome Sanger Institute for the COVID-19 Genomics UK (COG-UK) Consortium | Lighthouse Laboratory Plymouth and Alex Alderton, Roberto Amato, Jeffrey Barrett, Sonia Goncalves, Ewan Harrison, David K. Jackson, Ian Johnston, Dominic Kwiatkowski, Cordelia Langford, John Sillitoe on behalf of the Wellcome Sanger Institute COVID-19 Surveillance Team |
| EPI_ISL_3470002, EPI_ISL_3470003, EPI_ISL_3470004 | Lighthouse Lab in Alderley Park | Wellcome Sanger Institute for the COVID-19 Genomics UK (COG-UK) Consortium | Jacquelyn Wynn, Mairead Hyland, The Lighthouse Lab in Alderley Park and Alex Alderton, Roberto Amato, Jeffrey Barrett, Sonia Goncalves, Ewan Harrison, David K. Jackson, Ian Johnston, Dominic Kwiatkowski, Cordelia Langford, John Sillitoe on behalf of the Wellcome Sanger Institute COVID-19 Surveillance Team |
| EPI_ISL_3470005 | Lighthouse Lab in Milton Keynes | Wellcome Sanger Institute for the COVID-19 Genomics UK (COG-UK) Consortium | The Lighthouse Lab in Milton Keynes and Alex Alderton, Roberto Amato, Jeffrey Barrett, Sonia Goncalves, Ewan Harrison, David K. Jackson, Ian Johnston, Dominic Kwiatkowski, Cordelia Langford, John Sillitoe on behalf of the Wellcome Sanger Institute COVID-19 Surveillance Team |
| EPI_ISL_3470006, EPI_ISL_3470007, EPI_ISL_3470008, EPI_ISL_3470009 | Lighthouse Lab in Alderley Park | Wellcome Sanger Institute for the COVID-19 Genomics UK (COG-UK) Consortium | Jacquelyn Wynn, Mairead Hyland, The Lighthouse Lab in Alderley Park and Alex Alderton, Roberto Amato, Jeffrey Barrett, Sonia Goncalves, Ewan Harrison, David K. Jackson, Ian Johnston, Dominic Kwiatkowski, Cordelia Langford, John Sillitoe on behalf of the Wellcome Sanger Institute COVID-19 Surveillance Team |
| EPI_ISL_3470010, EPI_ISL_3470011 | Lighthouse Laboratory Plymouth | Wellcome Sanger Institute for the COVID-19 Genomics UK (COG-UK) Consortium | Lighthouse Laboratory Plymouth and Alex Alderton, Roberto Amato, Jeffrey Barrett, Sonia Goncalves, Ewan Harrison, David K. Jackson, Ian Johnston, Dominic Kwiatkowski, Cordelia Langford, John Sillitoe on behalf of the Wellcome Sanger Institute COVID-19 Surveillance Team |
| EPI_ISL_3470012 | Lighthouse Lab in Milton Keynes | Wellcome Sanger Institute for the COVID-19 Genomics UK (COG-UK) Consortium | The Lighthouse Lab in Milton Keynes and Alex Alderton, Roberto Amato, Jeffrey Barrett, Sonia Goncalves, Ewan Harrison, David K. Jackson, Ian Johnston, Dominic Kwiatkowski, Cordelia Langford, John Sillitoe on behalf of the Wellcome Sanger Institute COVID-19 Surveillance Team |
| EPI_ISL_3470013, EPI_ISL_3470014, EPI_ISL_3470015, EPI_ISL_3470016, EPI_ISL_3470017, EPI_ISL_3470018, EPI_ISL_3470019 | Lighthouse Lab in Alderley Park | Wellcome Sanger Institute for the COVID-19 Genomics UK (COG-UK) Consortium | Jacquelyn Wynn, Mairead Hyland, The Lighthouse Lab in Alderley Park and Alex Alderton, Roberto Amato, Jeffrey Barrett, Sonia Goncalves, Ewan Harrison, David K. Jackson, Ian Johnston, Dominic Kwiatkowski, Cordelia Langford, John Sillitoe on behalf of the Wellcome Sanger Institute COVID-19 Surveillance Team |
| EPI_ISL_3470020 | Microbiology Department. Complexo Hospitalario Universitario de Vigo | Microbiology Department. Complexo Hospitalario Universitario de Vigo | Alfaya N, Alonso I, Alvarez M, Cabrera JJ, Carballo R, Cores O, Cortizo S, del-Campo V, Martinez L, Mediero G, Perez S, Davina C, Potel C, Regueiro B, Rey S, Vasallo FJ |
| EPI_ISL_3470021, EPI_ISL_3470022, EPI_ISL_3470023, EPI_ISL_3470024 | Lighthouse Lab in Alderley Park | Wellcome Sanger Institute for the COVID-19 Genomics UK (COG-UK) Consortium | Jacquelyn Wynn, Mairead Hyland, The Lighthouse Lab in Alderley Park and Alex Alderton, Roberto Amato, Jeffrey Barrett, Sonia Goncalves, Ewan Harrison, David K. Jackson, Ian Johnston, Dominic Kwiatkowski, Cordelia Langford, John Sillitoe on behalf of the Wellcome Sanger Institute COVID-19 Surveillance Team |
| EPI_ISL_3470025 | Microbiology Department. Complexo Hospitalario Universitario de Vigo | Microbiology Department. Complexo Hospitalario Universitario de Vigo | Alfaya N, Alonso I, Alvarez M, Cabrera JJ, Carballo R, Cores O, Cortizo S, del-Campo V, Martinez L, Mediero G, Perez S, Davina C, Potel C, Regueiro B, Rey S, Vasallo FJ |
| EPI_ISL_3470026 | Lighthouse Lab in Milton Keynes | Wellcome Sanger Institute for the COVID-19 Genomics UK (COG-UK) Consortium | The Lighthouse Lab in Milton Keynes and Alex Alderton, Roberto Amato, Jeffrey Barrett, Sonia Goncalves, Ewan Harrison, David K. Jackson, Ian Johnston, Dominic Kwiatkowski, Cordelia Langford, John Sillitoe on behalf of the Wellcome Sanger Institute COVID-19 Surveillance Team |
| EPI_ISL_3470027 | Lighthouse Lab in Alderley Park | Wellcome Sanger Institute for the COVID-19 Genomics UK (COG-UK) Consortium | Jacquelyn Wynn, Mairead Hyland, The Lighthouse Lab in Alderley Park and Alex Alderton, Roberto Amato, Jeffrey Barrett, Sonia Goncalves, Ewan Harrison, David K. Jackson, Ian Johnston, Dominic Kwiatkowski, Cordelia Langford, John Sillitoe on behalf of the Wellcome Sanger Institute COVID-19 Surveillance Team |
| EPI_ISL_3470028 | Lighthouse Lab in Milton Keynes | Wellcome Sanger Institute for the COVID-19 Genomics UK (COG-UK) Consortium | The Lighthouse Lab in Milton Keynes and Alex Alderton, Roberto Amato, Jeffrey Barrett, Sonia Goncalves, Ewan Harrison, David K. Jackson, Ian Johnston, Dominic Kwiatkowski, Cordelia Langford, John Sillitoe on behalf of the Wellcome Sanger Institute COVID-19 Surveillance Team |
| EPI_ISL_3470029 | Microbiology Department. Complexo Hospitalario Universitario de Vigo | Microbiology Department. Complexo Hospitalario Universitario de Vigo | Alfaya N, Alonso I, Alvarez M, Cabrera JJ, Carballo R, Cores O, Cortizo S, del-Campo V, Martinez L, Mediero G, Perez S, Davina C, Potel C, Regueiro B, Rey S, Vasallo FJ |
| EPI_ISL_3470030 | Lighthouse Lab in Milton Keynes | Wellcome Sanger Institute for the COVID-19 Genomics UK (COG-UK) Consortium | The Lighthouse Lab in Milton Keynes and Alex Alderton, Roberto Amato, Jeffrey Barrett, Sonia Goncalves, Ewan Harrison, David K. Jackson, Ian Johnston, Dominic Kwiatkowski, Cordelia Langford, John Sillitoe on behalf of the Wellcome Sanger Institute COVID-19 Surveillance Team |
| EPI_ISL_3470031 | Lighthouse Laboratory Plymouth | Wellcome Sanger Institute for the COVID-19 Genomics UK (COG-UK) Consortium | Lighthouse Laboratory Plymouth and Alex Alderton, Roberto Amato, Jeffrey Barrett, Sonia Goncalves, Ewan Harrison, David K. Jackson, Ian Johnston, Dominic Kwiatkowski, Cordelia Langford, John Sillitoe on behalf of the Wellcome Sanger Institute COVID-19 Surveillance Team |
| EPI_ISL_3470032 | Microbiology Department. Complexo Hospitalario Universitario de Vigo | Microbiology Department. Complexo Hospitalario Universitario de Vigo | Alfaya N, Alonso I, Alvarez M, Cabrera JJ, Carballo R, Cores O, Cortizo S, del-Campo V, Martinez L, Mediero G, Perez S, Davina C, Potel C, Regueiro B, Rey S, Vasallo FJ |
| EPI_ISL_3470033 | Lighthouse Lab in Alderley Park | Wellcome Sanger Institute for the COVID-19 Genomics UK (COG-UK) Consortium | Jacquelyn Wynn, Mairead Hyland, The Lighthouse Lab in Alderley Park and Alex Alderton, Roberto Amato, Jeffrey Barrett, Sonia Goncalves, Ewan Harrison, David K. Jackson, Ian Johnston, Dominic Kwiatkowski, Cordelia Langford, John Sillitoe on behalf of the Wellcome Sanger Institute COVID-19 Surveillance Team |
| EPI_ISL_3470034 | Lighthouse Lab in Milton Keynes | Wellcome Sanger Institute for the COVID-19 Genomics UK (COG-UK) Consortium | The Lighthouse Lab in Milton Keynes and Alex Alderton, Roberto Amato, Jeffrey Barrett, Sonia Goncalves, Ewan Harrison, David K. Jackson, Ian Johnston, Dominic Kwiatkowski, Cordelia Langford, John Sillitoe on behalf of the Wellcome Sanger Institute COVID-19 Surveillance Team |
| EPI_ISL_3470035, EPI_ISL_3470037 | Lighthouse Lab in Alderley Park | Wellcome Sanger Institute for the COVID-19 Genomics UK (COG-UK) Consortium | Jacquelyn Wynn, Mairead Hyland, The Lighthouse Lab in Alderley Park and Alex Alderton, Roberto Amato, Jeffrey Barrett, Sonia Goncalves, Ewan Harrison, David K. Jackson, Ian Johnston, Dominic Kwiatkowski, Cordelia Langford, John Sillitoe on behalf of the Wellcome Sanger Institute COVID-19 Surveillance Team |
| EPI_ISL_3470038, EPI_ISL_3470039 | Lighthouse Lab in Milton Keynes | Wellcome Sanger Institute for the COVID-19 Genomics UK (COG-UK) Consortium | The Lighthouse Lab in Milton Keynes and Alex Alderton, Roberto Amato, Jeffrey Barrett, Sonia Goncalves, Ewan Harrison, David K. Jackson, Ian Johnston, Dominic Kwiatkowski, Cordelia Langford, John Sillitoe on behalf of the Wellcome Sanger Institute COVID-19 Surveillance Team |
| EPI_ISL_3470040 | Lighthouse Lab in Alderley Park | Wellcome Sanger Institute for the COVID-19 Genomics UK (COG-UK) Consortium | Jacquelyn Wynn, Mairead Hyland, The Lighthouse Lab in Alderley Park and Alex Alderton, Roberto Amato, Jeffrey Barrett, Sonia Goncalves, Ewan Harrison, David K. Jackson, Ian Johnston, Dominic Kwiatkowski, Cordelia Langford, John Sillitoe on behalf of the Wellcome Sanger Institute COVID-19 Surveillance Team |
| EPI_ISL_3470041 | Microbiology Department. Complexo Hospitalario Universitario de Vigo | Microbiology Department. Complexo Hospitalario Universitario de Vigo | Alfaya N, Alonso I, Alvarez M, Cabrera JJ, Carballo R, Cores O, Cortizo S, del-Campo V, Martinez L, Mediero G, Perez S, Davina C, Potel C, Regueiro B, Rey S, Vasallo FJ |
| EPI_ISL_3470042, EPI_ISL_3470043 | Lighthouse Lab in Milton Keynes | Wellcome Sanger Institute for the COVID-19 Genomics UK (COG-UK) Consortium | The Lighthouse Lab in Milton Keynes and Alex Alderton, Roberto Amato, Jeffrey Barrett, Sonia Goncalves, Ewan Harrison, David K. Jackson, Ian Johnston, Dominic Kwiatkowski, Cordelia Langford, John Sillitoe on behalf of the Wellcome Sanger Institute COVID-19 Surveillance Team |
| EPI_ISL_3470044 | Health Services Laboratories | Wellcome Sanger Institute for the COVID-19 Genomics UK (COG-UK) Consortium | Health Services Laboratories and Alex Alderton, Roberto Amato, Jeffrey Barrett, Sonia Goncalves, Ewan Harrison, David K. Jackson, Ian Johnston, Dominic Kwiatkowski, Cordelia Langford, John Sillitoe on behalf of the Wellcome Sanger Institute COVID-19 Surveillance Team |
| EPI_ISL_3470045 | Microbiology Department. Complexo Hospitalario Universitario de Vigo | Microbiology Department. Complexo Hospitalario Universitario de Vigo | Alfaya N, Alonso I, Alvarez M, Cabrera JJ, Carballo R, Cores O, Cortizo S, del-Campo V, Martinez L, Mediero G, Perez S, Davina C, Potel C, Regueiro B, Rey S, Vasallo FJ |
| EPI_ISL_3470046, EPI_ISL_3470047, EPI_ISL_3470048 | Lighthouse Lab in Alderley Park | Wellcome Sanger Institute for the COVID-19 Genomics UK (COG-UK) Consortium | Jacquelyn Wynn, Mairead Hyland, The Lighthouse Lab in Alderley Park and Alex Alderton, Roberto Amato, Jeffrey Barrett, Sonia Goncalves, Ewan Harrison, David K. Jackson, Ian Johnston, Dominic Kwiatkowski, Cordelia Langford, John Sillitoe on behalf of the Wellcome Sanger Institute COVID-19 Surveillance Team |
| EPI_ISL_3470049 | Microbiology Department. Complexo Hospitalario Universitario de Vigo | Microbiology Department. Complexo Hospitalario Universitario de Vigo | Alfaya N, Alonso I, Alvarez M, Cabrera JJ, Carballo R, Cores O, Cortizo S, del-Campo V, Martinez L, Mediero G, Perez S, Davina C, Potel C, Regueiro B, Rey S, Vasallo FJ |
| EPI_ISL_3470050 | Lighthouse Laboratory Plymouth | Wellcome Sanger Institute for the COVID-19 Genomics UK (COG-UK) Consortium | Lighthouse Laboratory Plymouth and Alex Alderton, Roberto Amato, Jeffrey Barrett, Sonia Goncalves, Ewan Harrison, David K. Jackson, Ian Johnston, Dominic Kwiatkowski, Cordelia Langford, John Sillitoe on behalf of the Wellcome Sanger Institute COVID-19 Surveillance Team |
| EPI_ISL_3470051, EPI_ISL_3470052 | Lighthouse Lab in Alderley Park | Wellcome Sanger Institute for the COVID-19 Genomics UK (COG-UK) Consortium | Jacquelyn Wynn, Mairead Hyland, The Lighthouse Lab in Alderley Park and Alex Alderton, Roberto Amato, Jeffrey Barrett, Sonia Goncalves, Ewan Harrison, David K. Jackson, Ian Johnston, Dominic Kwiatkowski, Cordelia Langford, John Sillitoe on behalf of the Wellcome Sanger Institute COVID-19 Surveillance Team |
| EPI_ISL_3470053 | Microbiology Department. Complexo Hospitalario Universitario de Vigo | Microbiology Department. Complexo Hospitalario Universitario de Vigo | Alfaya N, Alonso I, Alvarez M, Cabrera JJ, Carballo R, Cores O, Cortizo S, del-Campo V, Martinez L, Mediero G, Perez S, Davina C, Potel C, Regueiro B, Rey S, Vasallo FJ |
| EPI_ISL_3470054 | Lighthouse Lab in Alderley Park | Wellcome Sanger Institute for the COVID-19 Genomics UK (COG-UK) Consortium | Jacquelyn Wynn, Mairead Hyland, The Lighthouse Lab in Alderley Park and Alex Alderton, Roberto Amato, Jeffrey Barrett, Sonia Goncalves, Ewan Harrison, David K. Jackson, Ian Johnston, Dominic Kwiatkowski, Cordelia Langford, John Sillitoe on behalf of the Wellcome Sanger Institute COVID-19 Surveillance Team |
| EPI_ISL_3470055 | Lighthouse Laboratory Plymouth | Wellcome Sanger Institute for the COVID-19 Genomics UK (COG-UK) Consortium | Lighthouse Laboratory Plymouth and Alex Alderton, Roberto Amato, Jeffrey Barrett, Sonia Goncalves, Ewan Harrison, David K. Jackson, Ian Johnston, Dominic Kwiatkowski, Cordelia Langford, John Sillitoe on behalf of the Wellcome Sanger Institute COVID-19 Surveillance Team |

| | | | |
|---|---|---|---|
| EPI_ISL_3470056 | Lighthouse Lab in Alderley Park | Wellcome Sanger Institute for the COVID-19 Genomics UK (COG-UK) Consortium | Jacquelyn Wynn, Mairead Hyland, The Lighthouse Lab in Alderley Park and Alex Alderton, Roberto Amato, Jeffrey Barrett, Sonia Goncalves, Ewan Harrison, David K. Jackson, Ian Johnston, Dominic Kwiatkowski, Cordelia Langford, John Sillitoe on behalf of the Wellcome Sanger Institute COVID-19 Surveillance Team |
| EPI_ISL_3470057 | Microbiology Department. Complexo Hospitalario Universitario de Vigo | Microbiology Department. Complexo Hospitalario Universitario de Vigo | Alfaya N, Alonso I, Alvarez M, Cabrera JJ, Carballo R, Cores O, Cortizo S, del-Campo V, Martinez L, Mediero G, Perez S, Davina C, Potel C, Regueiro B, Rey S, Vasallo FJ |
| EPI_ISL_3470058, EPI_ISL_3470059, EPI_ISL_3470060 | Lighthouse Lab in Alderley Park | Wellcome Sanger Institute for the COVID-19 Genomics UK (COG-UK) Consortium | Jacquelyn Wynn, Mairead Hyland, The Lighthouse Lab in Alderley Park and Alex Alderton, Roberto Amato, Jeffrey Barrett, Sonia Goncalves, Ewan Harrison, David K. Jackson, Ian Johnston, Dominic Kwiatkowski, Cordelia Langford, John Sillitoe on behalf of the Wellcome Sanger Institute COVID-19 Surveillance Team |
| EPI_ISL_3470061 | Microbiology Department. Complexo Hospitalario Universitario de Vigo | Microbiology Department. Complexo Hospitalario Universitario de Vigo | Alfaya N, Alonso I, Alvarez M, Cabrera JJ, Carballo R, Cores O, Cortizo S, del-Campo V, Martinez L, Mediero G, Perez S, Davina C, Potel C, Regueiro B, Rey S, Vasallo FJ |
| EPI_ISL_3470062, EPI_ISL_3470063, EPI_ISL_3470064, EPI_ISL_3470065 | Lighthouse Lab in Alderley Park | Wellcome Sanger Institute for the COVID-19 Genomics UK (COG-UK) Consortium | Jacquelyn Wynn, Mairead Hyland, The Lighthouse Lab in Alderley Park and Alex Alderton, Roberto Amato, Jeffrey Barrett, Sonia Goncalves, Ewan Harrison, David K. Jackson, Ian Johnston, Dominic Kwiatkowski, Cordelia Langford, John Sillitoe on behalf of the Wellcome Sanger Institute COVID-19 Surveillance Team |
| EPI_ISL_3470066 | Microbiology Department. Complexo Hospitalario Universitario de Vigo | Microbiology Department. Complexo Hospitalario Universitario de Vigo | Alfaya N, Alonso I, Alvarez M, Cabrera JJ, Carballo R, Cores O, Cortizo S, del-Campo V, Martinez L, Mediero G, Perez S, Davina C, Potel C, Regueiro B, Rey S, Vasallo FJ |
| EPI_ISL_3470067 | Lighthouse Lab in Alderley Park | Wellcome Sanger Institute for the COVID-19 Genomics UK (COG-UK) Consortium | Jacquelyn Wynn, Mairead Hyland, The Lighthouse Lab in Alderley Park and Alex Alderton, Roberto Amato, Jeffrey Barrett, Sonia Goncalves, Ewan Harrison, David K. Jackson, Ian Johnston, Dominic Kwiatkowski, Cordelia Langford, John Sillitoe on behalf of the Wellcome Sanger Institute COVID-19 Surveillance Team |
| EPI_ISL_3470068 | Lighthouse Laboratory Plymouth | Wellcome Sanger Institute for the COVID-19 Genomics UK (COG-UK) Consortium | Lighthouse Laboratory Plymouth and Alex Alderton, Roberto Amato, Jeffrey Barrett, Sonia Goncalves, Ewan Harrison, David K. Jackson, Ian Johnston, Dominic Kwiatkowski, Cordelia Langford, John Sillitoe on behalf of the Wellcome Sanger Institute COVID-19 Surveillance Team |
| EPI_ISL_3470069 | Microbiology Department. Complexo Hospitalario Universitario de Vigo | Microbiology Department. Complexo Hospitalario Universitario de Vigo | Alfaya N, Alonso I, Alvarez M, Cabrera JJ, Carballo R, Cores O, Cortizo S, del-Campo V, Martinez L, Mediero G, Perez S, Davina C, Potel C, Regueiro B, Rey S, Vasallo FJ |
| EPI_ISL_3470070 | Lighthouse Laboratory Plymouth | Wellcome Sanger Institute for the COVID-19 Genomics UK (COG-UK) Consortium | Lighthouse Laboratory Plymouth and Alex Alderton, Roberto Amato, Jeffrey Barrett, Sonia Goncalves, Ewan Harrison, David K. Jackson, Ian Johnston, Dominic Kwiatkowski, Cordelia Langford, John Sillitoe on behalf of the Wellcome Sanger Institute COVID-19 Surveillance Team |
| EPI_ISL_3470071, EPI_ISL_3470072 | Lighthouse Lab in Milton Keynes | Wellcome Sanger Institute for the COVID-19 Genomics UK (COG-UK) Consortium | The Lighthouse Lab in Milton Keynes and Alex Alderton, Roberto Amato, Jeffrey Barrett, Sonia Goncalves, Ewan Harrison, David K. Jackson, Ian Johnston, Dominic Kwiatkowski, Cordelia Langford, John Sillitoe on behalf of the Wellcome Sanger Institute COVID-19 Surveillance Team |
| EPI_ISL_3470073 | Microbiology Department. Complexo Hospitalario Universitario de Vigo | Microbiology Department. Complexo Hospitalario Universitario de Vigo | Alfaya N, Alonso I, Alvarez M, Cabrera JJ, Carballo R, Cores O, Cortizo S, del-Campo V, Martinez L, Mediero G, Perez S, Davina C, Potel C, Regueiro B, Rey S, Vasallo FJ |
| EPI_ISL_3470074 | Lighthouse Laboratory Plymouth | Wellcome Sanger Institute for the COVID-19 Genomics UK (COG-UK) Consortium | Lighthouse Laboratory Plymouth and Alex Alderton, Roberto Amato, Jeffrey Barrett, Sonia Goncalves, Ewan Harrison, David K. Jackson, Ian Johnston, Dominic Kwiatkowski, Cordelia Langford, John Sillitoe on behalf of the Wellcome Sanger Institute COVID-19 Surveillance Team |
| EPI_ISL_3470075, EPI_ISL_3470076 | Lighthouse Lab in Alderley Park | Wellcome Sanger Institute for the COVID-19 Genomics UK (COG-UK) Consortium | Jacquelyn Wynn, Mairead Hyland, The Lighthouse Lab in Alderley Park and Alex Alderton, Roberto Amato, Jeffrey Barrett, Sonia Goncalves, Ewan Harrison, David K. Jackson, Ian Johnston, Dominic Kwiatkowski, Cordelia Langford, John Sillitoe on behalf of the Wellcome Sanger Institute COVID-19 Surveillance Team |
| EPI_ISL_3470077 | Microbiology Department. Complexo Hospitalario Universitario de Vigo | Microbiology Department. Complexo Hospitalario Universitario de Vigo | Alfaya N, Alonso I, Alvarez M, Cabrera JJ, Carballo R, Cores O, Cortizo S, del-Campo V, Martinez L, Mediero G, Perez S, Davina C, Potel C, Regueiro B, Rey S, Vasallo FJ |
| EPI_ISL_3470078, EPI_ISL_3470079 | Lighthouse Lab in Alderley Park | Wellcome Sanger Institute for the COVID-19 Genomics UK (COG-UK) Consortium | Jacquelyn Wynn, Mairead Hyland, The Lighthouse Lab in Alderley Park and Alex Alderton, Roberto Amato, Jeffrey Barrett, Sonia Goncalves, Ewan Harrison, David K. Jackson, Ian Johnston, Dominic Kwiatkowski, Cordelia Langford, John Sillitoe on behalf of the Wellcome Sanger Institute COVID-19 Surveillance Team |
| EPI_ISL_3470080 | Lighthouse Lab in Milton Keynes | Wellcome Sanger Institute for the COVID-19 Genomics UK (COG-UK) Consortium | The Lighthouse Lab in Milton Keynes and Alex Alderton, Roberto Amato, Jeffrey Barrett, Sonia Goncalves, Ewan Harrison, David K. Jackson, Ian Johnston, Dominic Kwiatkowski, Cordelia Langford, John Sillitoe on behalf of the Wellcome Sanger Institute COVID-19 Surveillance Team |
| EPI_ISL_3470081 | Microbiology Department. Complexo Hospitalario Universitario de Vigo | Microbiology Department. Complexo Hospitalario Universitario de Vigo | Alfaya N, Alonso I, Alvarez M, Cabrera JJ, Carballo R, Cores O, Cortizo S, del-Campo V, Martinez L, Mediero G, Perez S, Davina C, Potel C, Regueiro B, Rey S, Vasallo FJ |
| EPI_ISL_3470082, EPI_ISL_3470083 | Lighthouse Laboratory Plymouth | Wellcome Sanger Institute for the COVID-19 Genomics UK (COG-UK) Consortium | Lighthouse Laboratory Plymouth and Alex Alderton, Roberto Amato, Jeffrey Barrett, Sonia Goncalves, Ewan Harrison, David K. Jackson, Ian Johnston, Dominic Kwiatkowski, Cordelia Langford, John Sillitoe on behalf of the Wellcome Sanger Institute COVID-19 Surveillance Team |
| EPI_ISL_3470084 | Lighthouse Lab in Milton Keynes | Wellcome Sanger Institute for the COVID-19 Genomics UK (COG-UK) Consortium | The Lighthouse Lab in Milton Keynes and Alex Alderton, Roberto Amato, Jeffrey Barrett, Sonia Goncalves, Ewan Harrison, David K. Jackson, Ian Johnston, Dominic Kwiatkowski, Cordelia Langford, John Sillitoe on behalf of the Wellcome Sanger Institute COVID-19 Surveillance Team |
| EPI_ISL_3470085, EPI_ISL_3470087, EPI_ISL_3470088 | Lighthouse Lab in Alderley Park | Wellcome Sanger Institute for the COVID-19 Genomics UK (COG-UK) Consortium | Jacquelyn Wynn, Mairead Hyland, The Lighthouse Lab in Alderley Park and Alex Alderton, Roberto Amato, Jeffrey Barrett, Sonia Goncalves, Ewan Harrison, David K. Jackson, Ian Johnston, Dominic Kwiatkowski, Cordelia Langford, John Sillitoe on behalf of the Wellcome Sanger Institute COVID-19 Surveillance Team |
| EPI_ISL_3470089 | Microbiology Department. Complexo Hospitalario Universitario de Vigo | Microbiology Department. Complexo Hospitalario Universitario de Vigo | Alfaya N, Alonso I, Alvarez M, Cabrera JJ, Carballo R, Cores O, Cortizo S, del-Campo V, Martinez L, Mediero G, Perez S, Davina C, Potel C, Regueiro B, Rey S, Vasallo FJ |
| EPI_ISL_3470090 | Health Services Laboratories | Wellcome Sanger Institute for the COVID-19 Genomics UK (COG-UK) Consortium | Health Services Laboratories and Alex Alderton, Roberto Amato, Jeffrey Barrett, Sonia Goncalves, Ewan Harrison, David K. Jackson, Ian Johnston, Dominic Kwiatkowski, Cordelia Langford, John Sillitoe on behalf of the Wellcome Sanger Institute COVID-19 Surveillance Team |
| EPI_ISL_3470091, EPI_ISL_3470092 | Lighthouse Lab in Milton Keynes | Wellcome Sanger Institute for the COVID-19 Genomics UK (COG-UK) Consortium | The Lighthouse Lab in Milton Keynes and Alex Alderton, Roberto Amato, Jeffrey Barrett, Sonia Goncalves, Ewan Harrison, David K. Jackson, Ian Johnston, Dominic Kwiatkowski, Cordelia Langford, John Sillitoe on behalf of the Wellcome Sanger Institute COVID-19 Surveillance Team |
| EPI_ISL_3470093 | Lighthouse Lab in Alderley Park | Wellcome Sanger Institute for the COVID-19 Genomics UK (COG-UK) Consortium | Jacquelyn Wynn, Mairead Hyland, The Lighthouse Lab in Alderley Park and Alex Alderton, Roberto Amato, Jeffrey Barrett, Sonia Goncalves, Ewan Harrison, David K. Jackson, Ian Johnston, Dominic Kwiatkowski, Cordelia Langford, John Sillitoe on behalf of the Wellcome Sanger Institute COVID-19 Surveillance Team |
| EPI_ISL_3470094 | Microbiology Department. Complexo Hospitalario Universitario de Vigo | Microbiology Department. Complexo Hospitalario Universitario de Vigo | Alfaya N, Alonso I, Alvarez M, Cabrera JJ, Carballo R, Cores O, Cortizo S, del-Campo V, Martinez L, Mediero G, Perez S, Davina C, Potel C, Regueiro B, Rey S, Vasallo FJ |
| EPI_ISL_3470095 | Lighthouse Lab in Alderley Park | Wellcome Sanger Institute for the COVID-19 Genomics UK (COG-UK) Consortium | Jacquelyn Wynn, Mairead Hyland, The Lighthouse Lab in Alderley Park and Alex Alderton, Roberto Amato, Jeffrey Barrett, Sonia Goncalves, Ewan Harrison, David K. Jackson, Ian Johnston, Dominic Kwiatkowski, Cordelia Langford, John Sillitoe on behalf of the Wellcome Sanger Institute COVID-19 Surveillance Team |
| EPI_ISL_3470096, EPI_ISL_3470097, EPI_ISL_3470099, EPI_ISL_3470100, EPI_ISL_3470101 | Lighthouse Lab in Milton Keynes | Wellcome Sanger Institute for the COVID-19 Genomics UK (COG-UK) Consortium | The Lighthouse Lab in Milton Keynes and Alex Alderton, Roberto Amato, Jeffrey Barrett, Sonia Goncalves, Ewan Harrison, David K. Jackson, Ian Johnston, Dominic Kwiatkowski, Cordelia Langford, John Sillitoe on behalf of the Wellcome Sanger Institute COVID-19 Surveillance Team |
| EPI_ISL_3470103 | Lighthouse Lab in Alderley Park | Wellcome Sanger Institute for the COVID-19 Genomics UK (COG-UK) Consortium | Jacquelyn Wynn, Mairead Hyland, The Lighthouse Lab in Alderley Park and Alex Alderton, Roberto Amato, Jeffrey Barrett, Sonia Goncalves, Ewan Harrison, David K. Jackson, Ian Johnston, Dominic Kwiatkowski, Cordelia Langford, John Sillitoe on behalf of the Wellcome Sanger Institute COVID-19 Surveillance Team |
| EPI_ISL_3470104 | Lighthouse Lab in Milton Keynes | Wellcome Sanger Institute for the COVID-19 Genomics UK (COG-UK) Consortium | The Lighthouse Lab in Milton Keynes and Alex Alderton, Roberto Amato, Jeffrey Barrett, Sonia Goncalves, Ewan Harrison, David K. Jackson, Ian Johnston, Dominic Kwiatkowski, Cordelia Langford, John Sillitoe on behalf of the Wellcome Sanger Institute COVID-19 Surveillance Team |
| EPI_ISL_3470105 | Health Services Laboratories | Wellcome Sanger Institute for the COVID-19 Genomics UK (COG-UK) Consortium | Health Services Laboratories and Alex Alderton, Roberto Amato, Jeffrey Barrett, Sonia Goncalves, Ewan Harrison, David K. Jackson, Ian Johnston, Dominic Kwiatkowski, Cordelia Langford, John Sillitoe on behalf of the Wellcome Sanger Institute COVID-19 Surveillance Team |
| EPI_ISL_3470107, EPI_ISL_3470108 | Lighthouse Lab in Alderley Park | Wellcome Sanger Institute for the COVID-19 Genomics UK (COG-UK) Consortium | Jacquelyn Wynn, Mairead Hyland, The Lighthouse Lab in Alderley Park and Alex Alderton, Roberto Amato, Jeffrey Barrett, Sonia Goncalves, Ewan Harrison, David K. Jackson, Ian Johnston, Dominic Kwiatkowski, Cordelia Langford, John Sillitoe on behalf of the Wellcome Sanger Institute COVID-19 Surveillance Team |
| EPI_ISL_3470109 | Health Services Laboratories | Wellcome Sanger Institute for the COVID-19 Genomics UK (COG-UK) Consortium | Health Services Laboratories and Alex Alderton, Roberto Amato, Jeffrey Barrett, Sonia Goncalves, Ewan Harrison, David K. Jackson, Ian Johnston, Dominic Kwiatkowski, Cordelia Langford, John Sillitoe on behalf of the Wellcome Sanger Institute COVID-19 Surveillance Team |
| EPI_ISL_3470110 | Microbiology Department. Complexo Hospitalario Universitario de Vigo | Microbiology Department. Complexo Hospitalario Universitario de Vigo | Alfaya N, Alonso I, Alvarez M, Cabrera JJ, Carballo R, Cores O, Cortizo S, del-Campo V, Martinez L, Mediero G, Perez S, Davina C, Potel C, Regueiro B, Rey S, Vasallo FJ |
| EPI_ISL_3470111 | Lighthouse Lab in Milton Keynes | Wellcome Sanger Institute for the COVID-19 Genomics UK (COG-UK) Consortium | The Lighthouse Lab in Milton Keynes and Alex Alderton, Roberto Amato, Jeffrey Barrett, Sonia Goncalves, Ewan Harrison, David K. Jackson, Ian Johnston, Dominic Kwiatkowski, Cordelia Langford, John Sillitoe on behalf of the Wellcome Sanger Institute COVID-19 Surveillance Team |
| EPI_ISL_3470112, EPI_ISL_3470113, EPI_ISL_3470115, EPI_ISL_3470116 | Lighthouse Lab in Alderley Park | Wellcome Sanger Institute for the COVID-19 Genomics UK (COG-UK) Consortium | Jacquelyn Wynn, Mairead Hyland, The Lighthouse Lab in Alderley Park and Alex Alderton, Roberto Amato, Jeffrey Barrett, Sonia Goncalves, Ewan Harrison, David K. Jackson, Ian Johnston, Dominic Kwiatkowski, Cordelia Langford, John Sillitoe on behalf of the Wellcome Sanger Institute COVID-19 Surveillance Team |
| EPI_ISL_3470117 | Microbiology Department. Complexo Hospitalario Universitario de Vigo | Microbiology Department. Complexo Hospitalario Universitario de Vigo | Alfaya N, Alonso I, Alvarez M, Cabrera JJ, Carballo R, Cores O, Cortizo S, del-Campo V, Martinez L, Mediero G, Perez S, Davina C, Potel C, Regueiro B, Rey S, Vasallo FJ |
| EPI_ISL_3470118 | Lighthouse Lab in Milton Keynes | Wellcome Sanger Institute for the COVID-19 Genomics UK (COG-UK) Consortium | The Lighthouse Lab in Milton Keynes and Alex Alderton, Roberto Amato, Jeffrey Barrett, Sonia Goncalves, Ewan Harrison, David K. Jackson, Ian Johnston, Dominic Kwiatkowski, Cordelia Langford, John Sillitoe on behalf of the Wellcome Sanger Institute COVID-19 Surveillance Team |
| EPI_ISL_3470119, EPI_ISL_3470120 | Lighthouse Lab in Alderley Park | Wellcome Sanger Institute for the COVID-19 Genomics UK (COG-UK) Consortium | Jacquelyn Wynn, Mairead Hyland, The Lighthouse Lab in Alderley Park and Alex Alderton, Roberto Amato, Jeffrey Barrett, Sonia Goncalves, Ewan Harrison, David K. Jackson, Ian Johnston, Dominic Kwiatkowski, Cordelia Langford, John Sillitoe on behalf of the Wellcome Sanger Institute COVID-19 Surveillance Team |
| EPI_ISL_3470121 | Lighthouse Lab in Milton Keynes | Wellcome Sanger Institute for the COVID-19 Genomics UK (COG-UK) Consortium | The Lighthouse Lab in Milton Keynes and Alex Alderton, Roberto Amato, Jeffrey Barrett, Sonia Goncalves, Ewan Harrison, David K. Jackson, Ian Johnston, Dominic Kwiatkowski, Cordelia Langford, John Sillitoe on behalf of the Wellcome Sanger Institute COVID-19 Surveillance Team |
| EPI_ISL_3470122 | Microbiology Department. Complexo Hospitalario Universitario de Vigo | Microbiology Department. Complexo Hospitalario Universitario de Vigo | Alfaya N, Alonso I, Alvarez M, Cabrera JJ, Carballo R, Cores O, Cortizo S, del-Campo V, Martinez L, Mediero G, Perez S, Davina C, Potel C, Regueiro B, Rey S, Vasallo FJ |
| EPI_ISL_3470123 | Lighthouse Lab in Milton Keynes | Wellcome Sanger Institute for the COVID-19 Genomics UK (COG-UK) | The Lighthouse Lab in Milton Keynes and Alex Alderton, Roberto Amato, Jeffrey Barrett, Sonia Goncalves, Ewan Harrison, David K. Jackson, Ian Johnston, Dominic Kwiatkowski, Cordelia Langford, John Sillitoe on behalf of the Wellcome Sanger |

| | | | |
|---|---|---|---|
| | | Consortium | Institute COVID-19 Surveillance Team |
| EPI_ISL_3470124 | Lighthouse Lab in Alderley Park | Wellcome Sanger Institute for the COVID-19 Genomics UK (COG-UK) Consortium | Jacquelyn Wynn, Mairead Hyland, The Lighthouse Lab in Alderley Park and Alex Alderton, Roberto Amato, Jeffrey Barrett, Sonia Goncalves, Ewan Harrison, David K. Jackson, Ian Johnston, Dominic Kwiatkowski, Cordelia Langford, John Sillitoe on behalf of the Wellcome Sanger Institute COVID-19 Surveillance Team |
| EPI_ISL_3470125 | Lighthouse Lab in Milton Keynes | Wellcome Sanger Institute for the COVID-19 Genomics UK (COG-UK) Consortium | The Lighthouse Lab in Milton Keynes and Alex Alderton, Roberto Amato, Jeffrey Barrett, Sonia Goncalves, Ewan Harrison, David K. Jackson, Ian Johnston, Dominic Kwiatkowski, Cordelia Langford, John Sillitoe on behalf of the Wellcome Sanger Institute COVID-19 Surveillance Team |
| EPI_ISL_3470126 | Microbiology Department. Complexo Hospitalario Universitario de Vigo | Microbiology Department. Complexo Hospitalario Universitario de Vigo | Alfaya N, Alonso I, Alvarez M, Cabrera JJ, Carballo R, Cores O, Cortizo S, del-Campo V, Martinez L, Mediero G, Perez S, Davina C, Potel C, Regueiro B, Rey S, Vasallo FJ |
| EPI_ISL_3470127, EPI_ISL_3470128, EPI_ISL_3470129 | Lighthouse Lab in Alderley Park | Wellcome Sanger Institute for the COVID-19 Genomics UK (COG-UK) Consortium | Jacquelyn Wynn, Mairead Hyland, The Lighthouse Lab in Alderley Park and Alex Alderton, Roberto Amato, Jeffrey Barrett, Sonia Goncalves, Ewan Harrison, David K. Jackson, Ian Johnston, Dominic Kwiatkowski, Cordelia Langford, John Sillitoe on behalf of the Wellcome Sanger Institute COVID-19 Surveillance Team |
| EPI_ISL_3470130 | Microbiology Department. Complexo Hospitalario Universitario de Vigo | Microbiology Department. Complexo Hospitalario Universitario de Vigo | Alfaya N, Alonso I, Alvarez M, Cabrera JJ, Carballo R, Cores O, Cortizo S, del-Campo V, Martinez L, Mediero G, Perez S, Davina C, Potel C, Regueiro B, Rey S, Vasallo FJ |
| EPI_ISL_3470131 | Lighthouse Lab in Alderley Park | Wellcome Sanger Institute for the COVID-19 Genomics UK (COG-UK) Consortium | Jacquelyn Wynn, Mairead Hyland, The Lighthouse Lab in Alderley Park and Alex Alderton, Roberto Amato, Jeffrey Barrett, Sonia Goncalves, Ewan Harrison, David K. Jackson, Ian Johnston, Dominic Kwiatkowski, Cordelia Langford, John Sillitoe on behalf of the Wellcome Sanger Institute COVID-19 Surveillance Team |
| EPI_ISL_3470132 | Lighthouse Laboratory Plymouth | Wellcome Sanger Institute for the COVID-19 Genomics UK (COG-UK) Consortium | Lighthouse Laboratory Plymouth and Alex Alderton, Roberto Amato, Jeffrey Barrett, Sonia Goncalves, Ewan Harrison, David K. Jackson, Ian Johnston, Dominic Kwiatkowski, Cordelia Langford, John Sillitoe on behalf of the Wellcome Sanger Institute COVID-19 Surveillance Team |
| EPI_ISL_3470133 | Microbiology Department. Complexo Hospitalario Universitario de Vigo | Microbiology Department. Complexo Hospitalario Universitario de Vigo | Alfaya N, Alonso I, Alvarez M, Cabrera JJ, Carballo R, Cores O, Cortizo S, del-Campo V, Martinez L, Mediero G, Perez S, Davina C, Potel C, Regueiro B, Rey S, Vasallo FJ |
| EPI_ISL_3470134 | Health Services Laboratories | Wellcome Sanger Institute for the COVID-19 Genomics UK (COG-UK) Consortium | Health Services Laboratories and Alex Alderton, Roberto Amato, Jeffrey Barrett, Sonia Goncalves, Ewan Harrison, David K. Jackson, Ian Johnston, Dominic Kwiatkowski, Cordelia Langford, John Sillitoe on behalf of the Wellcome Sanger Institute COVID-19 Surveillance Team |
| EPI_ISL_3470135 | Lighthouse Lab in Alderley Park | Wellcome Sanger Institute for the COVID-19 Genomics UK (COG-UK) Consortium | Jacquelyn Wynn, Mairead Hyland, The Lighthouse Lab in Alderley Park and Alex Alderton, Roberto Amato, Jeffrey Barrett, Sonia Goncalves, Ewan Harrison, David K. Jackson, Ian Johnston, Dominic Kwiatkowski, Cordelia Langford, John Sillitoe on behalf of the Wellcome Sanger Institute COVID-19 Surveillance Team |
| EPI_ISL_3470136 | Microbiology Department. Complexo Hospitalario Universitario de Vigo | Microbiology Department. Complexo Hospitalario Universitario de Vigo | Alfaya N, Alonso I, Alvarez M, Cabrera JJ, Carballo R, Cores O, Cortizo S, del-Campo V, Martinez L, Mediero G, Perez S, Davina C, Potel C, Regueiro B, Rey S, Vasallo FJ |
| EPI_ISL_3470137 | Lighthouse Lab in Alderley Park | Wellcome Sanger Institute for the COVID-19 Genomics UK (COG-UK) Consortium | Jacquelyn Wynn, Mairead Hyland, The Lighthouse Lab in Alderley Park and Alex Alderton, Roberto Amato, Jeffrey Barrett, Sonia Goncalves, Ewan Harrison, David K. Jackson, Ian Johnston, Dominic Kwiatkowski, Cordelia Langford, John Sillitoe on behalf of the Wellcome Sanger Institute COVID-19 Surveillance Team |
| EPI_ISL_3470138 | Health Services Laboratories | Wellcome Sanger Institute for the COVID-19 Genomics UK (COG-UK) Consortium | Health Services Laboratories and Alex Alderton, Roberto Amato, Jeffrey Barrett, Sonia Goncalves, Ewan Harrison, David K. Jackson, Ian Johnston, Dominic Kwiatkowski, Cordelia Langford, John Sillitoe on behalf of the Wellcome Sanger Institute COVID-19 Surveillance Team |
| EPI_ISL_3470139 | Lighthouse Lab in Alderley Park | Wellcome Sanger Institute for the COVID-19 Genomics UK (COG-UK) Consortium | Jacquelyn Wynn, Mairead Hyland, The Lighthouse Lab in Alderley Park and Alex Alderton, Roberto Amato, Jeffrey Barrett, Sonia Goncalves, Ewan Harrison, David K. Jackson, Ian Johnston, Dominic Kwiatkowski, Cordelia Langford, John Sillitoe on behalf of the Wellcome Sanger Institute COVID-19 Surveillance Team |
| EPI_ISL_3470140 | Microbiology Department. Complexo Hospitalario Universitario de Vigo | Microbiology Department. Complexo Hospitalario Universitario de Vigo | Alfaya N, Alonso I, Alvarez M, Cabrera JJ, Carballo R, Cores O, Cortizo S, del-Campo V, Martinez L, Mediero G, Perez S, Davina C, Potel C, Regueiro B, Rey S, Vasallo FJ |
| EPI_ISL_3470141 | Lighthouse Lab in Alderley Park | Wellcome Sanger Institute for the COVID-19 Genomics UK (COG-UK) Consortium | Jacquelyn Wynn, Mairead Hyland, The Lighthouse Lab in Alderley Park and Alex Alderton, Roberto Amato, Jeffrey Barrett, Sonia Goncalves, Ewan Harrison, David K. Jackson, Ian Johnston, Dominic Kwiatkowski, Cordelia Langford, John Sillitoe on behalf of the Wellcome Sanger Institute COVID-19 Surveillance Team |
| EPI_ISL_3470142 | Lighthouse Lab in Milton Keynes | Wellcome Sanger Institute for the COVID-19 Genomics UK (COG-UK) Consortium | The Lighthouse Lab in Milton Keynes and Alex Alderton, Roberto Amato, Jeffrey Barrett, Sonia Goncalves, Ewan Harrison, David K. Jackson, Ian Johnston, Dominic Kwiatkowski, Cordelia Langford, John Sillitoe on behalf of the Wellcome Sanger Institute COVID-19 Surveillance Team |
| EPI_ISL_3470143 | Lighthouse Lab in Alderley Park | Wellcome Sanger Institute for the COVID-19 Genomics UK (COG-UK) Consortium | Jacquelyn Wynn, Mairead Hyland, The Lighthouse Lab in Alderley Park and Alex Alderton, Roberto Amato, Jeffrey Barrett, Sonia Goncalves, Ewan Harrison, David K. Jackson, Ian Johnston, Dominic Kwiatkowski, Cordelia Langford, John Sillitoe on behalf of the Wellcome Sanger Institute COVID-19 Surveillance Team |
| EPI_ISL_3470145 | Lighthouse Lab in Milton Keynes | Wellcome Sanger Institute for the COVID-19 Genomics UK (COG-UK) Consortium | The Lighthouse Lab in Milton Keynes and Alex Alderton, Roberto Amato, Jeffrey Barrett, Sonia Goncalves, Ewan Harrison, David K. Jackson, Ian Johnston, Dominic Kwiatkowski, Cordelia Langford, John Sillitoe on behalf of the Wellcome Sanger Institute COVID-19 Surveillance Team |
| EPI_ISL_3470146 | Health Services Laboratories | Wellcome Sanger Institute for the COVID-19 Genomics UK (COG-UK) Consortium | Health Services Laboratories and Alex Alderton, Roberto Amato, Jeffrey Barrett, Sonia Goncalves, Ewan Harrison, David K. Jackson, Ian Johnston, Dominic Kwiatkowski, Cordelia Langford, John Sillitoe on behalf of the Wellcome Sanger Institute COVID-19 Surveillance Team |
| EPI_ISL_3470147 | Lighthouse Lab in Milton Keynes | Wellcome Sanger Institute for the COVID-19 Genomics UK (COG-UK) Consortium | The Lighthouse Lab in Milton Keynes and Alex Alderton, Roberto Amato, Jeffrey Barrett, Sonia Goncalves, Ewan Harrison, David K. Jackson, Ian Johnston, Dominic Kwiatkowski, Cordelia Langford, John Sillitoe on behalf of the Wellcome Sanger Institute COVID-19 Surveillance Team |
| EPI_ISL_3470149 | Lighthouse Lab in Alderley Park | Wellcome Sanger Institute for the COVID-19 Genomics UK (COG-UK) Consortium | Jacquelyn Wynn, Mairead Hyland, The Lighthouse Lab in Alderley Park and Alex Alderton, Roberto Amato, Jeffrey Barrett, Sonia Goncalves, Ewan Harrison, David K. Jackson, Ian Johnston, Dominic Kwiatkowski, Cordelia Langford, John Sillitoe on behalf of the Wellcome Sanger Institute COVID-19 Surveillance Team |
| EPI_ISL_3470150 | Lighthouse Laboratory Plymouth | Wellcome Sanger Institute for the COVID-19 Genomics UK (COG-UK) Consortium | Lighthouse Laboratory Plymouth and Alex Alderton, Roberto Amato, Jeffrey Barrett, Sonia Goncalves, Ewan Harrison, David K. Jackson, Ian Johnston, Dominic Kwiatkowski, Cordelia Langford, John Sillitoe on behalf of the Wellcome Sanger Institute COVID-19 Surveillance Team |
| EPI_ISL_3470151 | Lighthouse Lab in Milton Keynes | Wellcome Sanger Institute for the COVID-19 Genomics UK (COG-UK) Consortium | The Lighthouse Lab in Milton Keynes and Alex Alderton, Roberto Amato, Jeffrey Barrett, Sonia Goncalves, Ewan Harrison, David K. Jackson, Ian Johnston, Dominic Kwiatkowski, Cordelia Langford, John Sillitoe on behalf of the Wellcome Sanger Institute COVID-19 Surveillance Team |
| EPI_ISL_3470153 | Lighthouse Lab in Alderley Park | Wellcome Sanger Institute for the COVID-19 Genomics UK (COG-UK) Consortium | Jacquelyn Wynn, Mairead Hyland, The Lighthouse Lab in Alderley Park and Alex Alderton, Roberto Amato, Jeffrey Barrett, Sonia Goncalves, Ewan Harrison, David K. Jackson, Ian Johnston, Dominic Kwiatkowski, Cordelia Langford, John Sillitoe on behalf of the Wellcome Sanger Institute COVID-19 Surveillance Team |
| EPI_ISL_3470154, EPI_ISL_3470155 | Lighthouse Lab in Milton Keynes | Wellcome Sanger Institute for the COVID-19 Genomics UK (COG-UK) Consortium | The Lighthouse Lab in Milton Keynes and Alex Alderton, Roberto Amato, Jeffrey Barrett, Sonia Goncalves, Ewan Harrison, David K. Jackson, Ian Johnston, Dominic Kwiatkowski, Cordelia Langford, John Sillitoe on behalf of the Wellcome Sanger Institute COVID-19 Surveillance Team |
| EPI_ISL_3470157 | Lighthouse Lab in Alderley Park | Wellcome Sanger Institute for the COVID-19 Genomics UK (COG-UK) Consortium | Jacquelyn Wynn, Mairead Hyland, The Lighthouse Lab in Alderley Park and Alex Alderton, Roberto Amato, Jeffrey Barrett, Sonia Goncalves, Ewan Harrison, David K. Jackson, Ian Johnston, Dominic Kwiatkowski, Cordelia Langford, John Sillitoe on behalf of the Wellcome Sanger Institute COVID-19 Surveillance Team |
| EPI_ISL_3470158 | Lighthouse Lab in Milton Keynes | Wellcome Sanger Institute for the COVID-19 Genomics UK (COG-UK) Consortium | The Lighthouse Lab in Milton Keynes and Alex Alderton, Roberto Amato, Jeffrey Barrett, Sonia Goncalves, Ewan Harrison, David K. Jackson, Ian Johnston, Dominic Kwiatkowski, Cordelia Langford, John Sillitoe on behalf of the Wellcome Sanger Institute COVID-19 Surveillance Team |
| EPI_ISL_3470159, EPI_ISL_3470161 | Lighthouse Lab in Alderley Park | Wellcome Sanger Institute for the COVID-19 Genomics UK (COG-UK) Consortium | Jacquelyn Wynn, Mairead Hyland, The Lighthouse Lab in Alderley Park and Alex Alderton, Roberto Amato, Jeffrey Barrett, Sonia Goncalves, Ewan Harrison, David K. Jackson, Ian Johnston, Dominic Kwiatkowski, Cordelia Langford, John Sillitoe on behalf of the Wellcome Sanger Institute COVID-19 Surveillance Team |
| EPI_ISL_3470162 | Lighthouse Lab in Milton Keynes | Wellcome Sanger Institute for the COVID-19 Genomics UK (COG-UK) Consortium | The Lighthouse Lab in Milton Keynes and Alex Alderton, Roberto Amato, Jeffrey Barrett, Sonia Goncalves, Ewan Harrison, David K. Jackson, Ian Johnston, Dominic Kwiatkowski, Cordelia Langford, John Sillitoe on behalf of the Wellcome Sanger Institute COVID-19 Surveillance Team |
| EPI_ISL_3470163, EPI_ISL_3470165 | Lighthouse Lab in Alderley Park | Wellcome Sanger Institute for the COVID-19 Genomics UK (COG-UK) Consortium | Jacquelyn Wynn, Mairead Hyland, The Lighthouse Lab in Alderley Park and Alex Alderton, Roberto Amato, Jeffrey Barrett, Sonia Goncalves, Ewan Harrison, David K. Jackson, Ian Johnston, Dominic Kwiatkowski, Cordelia Langford, John Sillitoe on behalf of the Wellcome Sanger Institute COVID-19 Surveillance Team |
| EPI_ISL_3470166, EPI_ISL_3470167 | Lighthouse Lab in Glasgow | Wellcome Sanger Institute for the COVID-19 Genomics UK (COG-UK) Consortium | Harper VanSteenhouse, Yumi Kasai, David Gray, Carol Clugston, Anna Dominiczak and Alex Alderton, Roberto Amato, Jeffrey Barrett, Sonia Goncalves, Ewan Harrison, David K. Jackson, Ian Johnston, Dominic Kwiatkowski, Cordelia Langford, John Sillitoe on behalf of the Wellcome Sanger Institute COVID-19 Surveillance Team |
| EPI_ISL_3470169, EPI_ISL_3470170, EPI_ISL_3470171 | Berkshire and Surrey Pathology Services Lighthouse Laboratory | Wellcome Sanger Institute for the COVID-19 Genomics UK (COG-UK) Consortium | Berkshire and Surrey Pathology Services Lighthouse Laboratory and Alex Alderton, Roberto Amato, Jeffrey Barrett, Sonia Goncalves, Ewan Harrison, David K. Jackson, Ian Johnston, Dominic Kwiatkowski, Cordelia Langford, John Sillitoe on behalf of the Wellcome Sanger Institute COVID-19 Surveillance Team |
| EPI_ISL_3470172 | Lighthouse Lab in Milton Keynes | Wellcome Sanger Institute for the COVID-19 Genomics UK (COG-UK) Consortium | The Lighthouse Lab in Milton Keynes and Alex Alderton, Roberto Amato, Jeffrey Barrett, Sonia Goncalves, Ewan Harrison, David K. Jackson, Ian Johnston, Dominic Kwiatkowski, Cordelia Langford, John Sillitoe on behalf of the Wellcome Sanger Institute COVID-19 Surveillance Team |
| EPI_ISL_3470174 | Berkshire and Surrey Pathology Services Lighthouse Laboratory | Wellcome Sanger Institute for the COVID-19 Genomics UK (COG-UK) Consortium | Berkshire and Surrey Pathology Services Lighthouse Laboratory and Alex Alderton, Roberto Amato, Jeffrey Barrett, Sonia Goncalves, Ewan Harrison, David K. Jackson, Ian Johnston, Dominic Kwiatkowski, Cordelia Langford, John Sillitoe on behalf of the Wellcome Sanger Institute COVID-19 Surveillance Team |
| EPI_ISL_3470175 | Lighthouse Lab in Glasgow | Wellcome Sanger Institute for the COVID-19 Genomics UK (COG-UK) Consortium | Harper VanSteenhouse, Yumi Kasai, David Gray, Carol Clugston, Anna Dominiczak and Alex Alderton, Roberto Amato, Jeffrey Barrett, Sonia Goncalves, Ewan Harrison, David K. Jackson, Ian Johnston, Dominic Kwiatkowski, Cordelia Langford, John Sillitoe on behalf of the Wellcome Sanger Institute COVID-19 Surveillance Team |
| EPI_ISL_3470176 | Lighthouse Lab in Milton Keynes | Wellcome Sanger Institute for the COVID-19 Genomics UK (COG-UK) Consortium | The Lighthouse Lab in Milton Keynes and Alex Alderton, Roberto Amato, Jeffrey Barrett, Sonia Goncalves, Ewan Harrison, David K. Jackson, Ian Johnston, Dominic Kwiatkowski, Cordelia Langford, John Sillitoe on behalf of the Wellcome Sanger Institute COVID-19 Surveillance Team |
| EPI_ISL_3470178 | Lighthouse Lab in Glasgow | Wellcome Sanger Institute for the COVID-19 Genomics UK (COG-UK) Consortium | Harper VanSteenhouse, Yumi Kasai, David Gray, Carol Clugston, Anna Dominiczak and Alex Alderton, Roberto Amato, Jeffrey Barrett, Sonia Goncalves, Ewan Harrison, David K. Jackson, Ian Johnston, Dominic Kwiatkowski, Cordelia Langford, John Sillitoe on behalf of the Wellcome Sanger Institute COVID-19 Surveillance Team |
| EPI_ISL_3470179 | Lighthouse Lab in Milton Keynes | Wellcome Sanger Institute for the COVID-19 Genomics UK (COG-UK) Consortium | The Lighthouse Lab in Milton Keynes and Alex Alderton, Roberto Amato, Jeffrey Barrett, Sonia Goncalves, Ewan Harrison, David K. Jackson, Ian Johnston, Dominic Kwiatkowski, Cordelia Langford, John Sillitoe on behalf of the Wellcome Sanger Institute COVID-19 Surveillance Team |
| EPI_ISL_3470180, EPI_ISL_3470182, EPI_ISL_3470183, EPI_ISL_3470184 | Lighthouse Lab in Glasgow | Wellcome Sanger Institute for the COVID-19 Genomics UK (COG-UK) Consortium | Harper VanSteenhouse, Yumi Kasai, David Gray, Carol Clugston, Anna Dominiczak and Alex Alderton, Roberto Amato, Jeffrey Barrett, Sonia Goncalves, Ewan Harrison, David K. Jackson, Ian Johnston, Dominic Kwiatkowski, Cordelia Langford, John Sillitoe on behalf of the Wellcome Sanger Institute COVID-19 Surveillance Team |
| EPI_ISL_3470186, EPI_ISL_3470187, EPI_ISL_3470188 | Lighthouse Lab in Milton Keynes | Wellcome Sanger Institute for the COVID-19 Genomics UK (COG-UK) Consortium | The Lighthouse Lab in Milton Keynes and Alex Alderton, Roberto Amato, Jeffrey Barrett, Sonia Goncalves, Ewan Harrison, David K. Jackson, Ian Johnston, Dominic Kwiatkowski, Cordelia Langford, John Sillitoe on behalf of the Wellcome Sanger Institute COVID-19 Surveillance Team |
| EPI_ISL_3470190 | Berkshire and Surrey Pathology Services Lighthouse Laboratory | Wellcome Sanger Institute for the COVID-19 Genomics UK (COG-UK) Consortium | Berkshire and Surrey Pathology Services Lighthouse Laboratory and Alex Alderton, Roberto Amato, Jeffrey Barrett, Sonia Goncalves, Ewan Harrison, David K. Jackson, Ian Johnston, Dominic Kwiatkowski, Cordelia Langford, John Sillitoe on behalf of the Wellcome Sanger Institute COVID-19 Surveillance Team |
| EPI_ISL_3470191 | Lighthouse Lab in Glasgow | Wellcome Sanger Institute for the COVID-19 Genomics UK (COG-UK) | Harper VanSteenhouse, Yumi Kasai, David Gray, Carol Clugston, Anna Dominiczak and Alex Alderton, Roberto Amato, Jeffrey Barrett, Sonia Goncalves, Ewan Harrison, David K. Jackson, Ian Johnston, Dominic Kwiatkowski, Cordelia Langford, |

| | | | |
|---|---|---|---|
| | | Consortium | John Sillitoe on behalf of the Wellcome Sanger Institute COVID-19 Surveillance Team |
| EPI_ISL_3470192 | Lighthouse Lab in Milton Keynes | Wellcome Sanger Institute for the COVID-19 Genomics UK (COG-UK) Consortium | The Lighthouse Lab in Milton Keynes and Alex Alderton, Roberto Amato, Jeffrey Barrett, Sonia Goncalves, Ewan Harrison, David K. Jackson, Ian Johnston, Dominic Kwiatkowski, Cordelia Langford, John Sillitoe on behalf of the Wellcome Sanger Institute COVID-19 Surveillance Team |
| EPI_ISL_3470194, EPI_ISL_3470195 | Berkshire and Surrey Pathology Services Lighthouse Laboratory | Wellcome Sanger Institute for the COVID-19 Genomics UK (COG-UK) Consortium | Berkshire and Surrey Pathology Services Lighthouse Laboratory and Alex Alderton, Roberto Amato, Jeffrey Barrett, Sonia Goncalves, Ewan Harrison, David K. Jackson, Ian Johnston, Dominic Kwiatkowski, Cordelia Langford, John Sillitoe on behalf of the Wellcome Sanger Institute COVID-19 Surveillance Team |
| EPI_ISL_3470196 | Lighthouse Lab in Glasgow | Wellcome Sanger Institute for the COVID-19 Genomics UK (COG-UK) Consortium | Harper VanSteenhouse, Yumi Kasai, David Gray, Carol Clugston, Anna Dominiczak and Alex Alderton, Roberto Amato, Jeffrey Barrett, Sonia Goncalves, Ewan Harrison, David K. Jackson, Ian Johnston, Dominic Kwiatkowski, Cordelia Langford, John Sillitoe on behalf of the Wellcome Sanger Institute COVID-19 Surveillance Team |
| EPI_ISL_3470198, EPI_ISL_3470199 | Lighthouse Lab in Milton Keynes | Wellcome Sanger Institute for the COVID-19 Genomics UK (COG-UK) Consortium | The Lighthouse Lab in Milton Keynes and Alex Alderton, Roberto Amato, Jeffrey Barrett, Sonia Goncalves, Ewan Harrison, David K. Jackson, Ian Johnston, Dominic Kwiatkowski, Cordelia Langford, John Sillitoe on behalf of the Wellcome Sanger Institute COVID-19 Surveillance Team |
| EPI_ISL_3470200 | Lighthouse Lab in Glasgow | Wellcome Sanger Institute for the COVID-19 Genomics UK (COG-UK) Consortium | Harper VanSteenhouse, Yumi Kasai, David Gray, Carol Clugston, Anna Dominiczak and Alex Alderton, Roberto Amato, Jeffrey Barrett, Sonia Goncalves, Ewan Harrison, David K. Jackson, Ian Johnston, Dominic Kwiatkowski, Cordelia Langford, John Sillitoe on behalf of the Wellcome Sanger Institute COVID-19 Surveillance Team |
| EPI_ISL_3470202, EPI_ISL_3470203 | Lighthouse Lab in Milton Keynes | Wellcome Sanger Institute for the COVID-19 Genomics UK (COG-UK) Consortium | The Lighthouse Lab in Milton Keynes and Alex Alderton, Roberto Amato, Jeffrey Barrett, Sonia Goncalves, Ewan Harrison, David K. Jackson, Ian Johnston, Dominic Kwiatkowski, Cordelia Langford, John Sillitoe on behalf of the Wellcome Sanger Institute COVID-19 Surveillance Team |
| EPI_ISL_3470204, EPI_ISL_3470205 | Lighthouse Lab in Glasgow | Wellcome Sanger Institute for the COVID-19 Genomics UK (COG-UK) Consortium | Harper VanSteenhouse, Yumi Kasai, David Gray, Carol Clugston, Anna Dominiczak and Alex Alderton, Roberto Amato, Jeffrey Barrett, Sonia Goncalves, Ewan Harrison, David K. Jackson, Ian Johnston, Dominic Kwiatkowski, Cordelia Langford, John Sillitoe on behalf of the Wellcome Sanger Institute COVID-19 Surveillance Team |
| EPI_ISL_3470207, EPI_ISL_3470208, EPI_ISL_3470209 | Lighthouse Lab in Milton Keynes | Wellcome Sanger Institute for the COVID-19 Genomics UK (COG-UK) Consortium | The Lighthouse Lab in Milton Keynes and Alex Alderton, Roberto Amato, Jeffrey Barrett, Sonia Goncalves, Ewan Harrison, David K. Jackson, Ian Johnston, Dominic Kwiatkowski, Cordelia Langford, John Sillitoe on behalf of the Wellcome Sanger Institute COVID-19 Surveillance Team |
| EPI_ISL_3470211 | Berkshire and Surrey Pathology Services Lighthouse Laboratory | Wellcome Sanger Institute for the COVID-19 Genomics UK (COG-UK) Consortium | Berkshire and Surrey Pathology Services Lighthouse Laboratory and Alex Alderton, Roberto Amato, Jeffrey Barrett, Sonia Goncalves, Ewan Harrison, David K. Jackson, Ian Johnston, Dominic Kwiatkowski, Cordelia Langford, John Sillitoe on behalf of the Wellcome Sanger Institute COVID-19 Surveillance Team |
| EPI_ISL_3470212, EPI_ISL_3470213 | Lighthouse Lab in Milton Keynes | Wellcome Sanger Institute for the COVID-19 Genomics UK (COG-UK) Consortium | The Lighthouse Lab in Milton Keynes and Alex Alderton, Roberto Amato, Jeffrey Barrett, Sonia Goncalves, Ewan Harrison, David K. Jackson, Ian Johnston, Dominic Kwiatkowski, Cordelia Langford, John Sillitoe on behalf of the Wellcome Sanger Institute COVID-19 Surveillance Team |
| EPI_ISL_3470215 | Berkshire and Surrey Pathology Services Lighthouse Laboratory | Wellcome Sanger Institute for the COVID-19 Genomics UK (COG-UK) Consortium | Berkshire and Surrey Pathology Services Lighthouse Laboratory and Alex Alderton, Roberto Amato, Jeffrey Barrett, Sonia Goncalves, Ewan Harrison, David K. Jackson, Ian Johnston, Dominic Kwiatkowski, Cordelia Langford, John Sillitoe on behalf of the Wellcome Sanger Institute COVID-19 Surveillance Team |
| EPI_ISL_3470216 | Lighthouse Lab in Milton Keynes | Wellcome Sanger Institute for the COVID-19 Genomics UK (COG-UK) Consortium | The Lighthouse Lab in Milton Keynes and Alex Alderton, Roberto Amato, Jeffrey Barrett, Sonia Goncalves, Ewan Harrison, David K. Jackson, Ian Johnston, Dominic Kwiatkowski, Cordelia Langford, John Sillitoe on behalf of the Wellcome Sanger Institute COVID-19 Surveillance Team |
| EPI_ISL_3470217, EPI_ISL_3470219 | Lighthouse Lab in Glasgow | Wellcome Sanger Institute for the COVID-19 Genomics UK (COG-UK) Consortium | Harper VanSteenhouse, Yumi Kasai, David Gray, Carol Clugston, Anna Dominiczak and Alex Alderton, Roberto Amato, Jeffrey Barrett, Sonia Goncalves, Ewan Harrison, David K. Jackson, Ian Johnston, Dominic Kwiatkowski, Cordelia Langford, John Sillitoe on behalf of the Wellcome Sanger Institute COVID-19 Surveillance Team |
| EPI_ISL_3470220 | Berkshire and Surrey Pathology Services Lighthouse Laboratory | Wellcome Sanger Institute for the COVID-19 Genomics UK (COG-UK) Consortium | Berkshire and Surrey Pathology Services Lighthouse Laboratory and Alex Alderton, Roberto Amato, Jeffrey Barrett, Sonia Goncalves, Ewan Harrison, David K. Jackson, Ian Johnston, Dominic Kwiatkowski, Cordelia Langford, John Sillitoe on behalf of the Wellcome Sanger Institute COVID-19 Surveillance Team |
| EPI_ISL_3470221 | Lighthouse Lab in Glasgow | Wellcome Sanger Institute for the COVID-19 Genomics UK (COG-UK) Consortium | Harper VanSteenhouse, Yumi Kasai, David Gray, Carol Clugston, Anna Dominiczak and Alex Alderton, Roberto Amato, Jeffrey Barrett, Sonia Goncalves, Ewan Harrison, David K. Jackson, Ian Johnston, Dominic Kwiatkowski, Cordelia Langford, John Sillitoe on behalf of the Wellcome Sanger Institute COVID-19 Surveillance Team |
| EPI_ISL_3470223 | Lighthouse Lab in Milton Keynes | Wellcome Sanger Institute for the COVID-19 Genomics UK (COG-UK) Consortium | The Lighthouse Lab in Milton Keynes and Alex Alderton, Roberto Amato, Jeffrey Barrett, Sonia Goncalves, Ewan Harrison, David K. Jackson, Ian Johnston, Dominic Kwiatkowski, Cordelia Langford, John Sillitoe on behalf of the Wellcome Sanger Institute COVID-19 Surveillance Team |
| EPI_ISL_3470224 | Lighthouse Lab in Glasgow | Wellcome Sanger Institute for the COVID-19 Genomics UK (COG-UK) Consortium | Harper VanSteenhouse, Yumi Kasai, David Gray, Carol Clugston, Anna Dominiczak and Alex Alderton, Roberto Amato, Jeffrey Barrett, Sonia Goncalves, Ewan Harrison, David K. Jackson, Ian Johnston, Dominic Kwiatkowski, Cordelia Langford, John Sillitoe on behalf of the Wellcome Sanger Institute COVID-19 Surveillance Team |
| EPI_ISL_3470225, EPI_ISL_3470227 | Berkshire and Surrey Pathology Services Lighthouse Laboratory | Wellcome Sanger Institute for the COVID-19 Genomics UK (COG-UK) Consortium | Berkshire and Surrey Pathology Services Lighthouse Laboratory and Alex Alderton, Roberto Amato, Jeffrey Barrett, Sonia Goncalves, Ewan Harrison, David K. Jackson, Ian Johnston, Dominic Kwiatkowski, Cordelia Langford, John Sillitoe on behalf of the Wellcome Sanger Institute COVID-19 Surveillance Team |
| EPI_ISL_3470228, EPI_ISL_3470229 | Lighthouse Lab in Glasgow | Wellcome Sanger Institute for the COVID-19 Genomics UK (COG-UK) Consortium | Harper VanSteenhouse, Yumi Kasai, David Gray, Carol Clugston, Anna Dominiczak and Alex Alderton, Roberto Amato, Jeffrey Barrett, Sonia Goncalves, Ewan Harrison, David K. Jackson, Ian Johnston, Dominic Kwiatkowski, Cordelia Langford, John Sillitoe on behalf of the Wellcome Sanger Institute COVID-19 Surveillance Team |
| EPI_ISL_3470230 | Microbiology Department. Complexo Hospitalario Universitario de Vigo | Microbiology Department. Complexo Hospitalario Universitario de Vigo | Alfaya N, Alonso I, Alvarez M, Cabrera JJ, Carballo R, Cores O, Cortizo S, del-Campo V, Martinez L, Mediero G, Perez S, Davina C, Potel C, Regueiro B, Rey S, Vasallo FJ |
| EPI_ISL_3470231 | Lighthouse Lab in Glasgow | Wellcome Sanger Institute for the COVID-19 Genomics UK (COG-UK) Consortium | Harper VanSteenhouse, Yumi Kasai, David Gray, Carol Clugston, Anna Dominiczak and Alex Alderton, Roberto Amato, Jeffrey Barrett, Sonia Goncalves, Ewan Harrison, David K. Jackson, Ian Johnston, Dominic Kwiatkowski, Cordelia Langford, John Sillitoe on behalf of the Wellcome Sanger Institute COVID-19 Surveillance Team |
| EPI_ISL_3470232 | Lighthouse Lab in Milton Keynes | Wellcome Sanger Institute for the COVID-19 Genomics UK (COG-UK) Consortium | The Lighthouse Lab in Milton Keynes and Alex Alderton, Roberto Amato, Jeffrey Barrett, Sonia Goncalves, Ewan Harrison, David K. Jackson, Ian Johnston, Dominic Kwiatkowski, Cordelia Langford, John Sillitoe on behalf of the Wellcome Sanger Institute COVID-19 Surveillance Team |
| EPI_ISL_3470234, EPI_ISL_3470235 | Berkshire and Surrey Pathology Services Lighthouse Laboratory | Wellcome Sanger Institute for the COVID-19 Genomics UK (COG-UK) Consortium | Berkshire and Surrey Pathology Services Lighthouse Laboratory and Alex Alderton, Roberto Amato, Jeffrey Barrett, Sonia Goncalves, Ewan Harrison, David K. Jackson, Ian Johnston, Dominic Kwiatkowski, Cordelia Langford, John Sillitoe on behalf of the Wellcome Sanger Institute COVID-19 Surveillance Team |
| EPI_ISL_3470236 | Lighthouse Lab in Glasgow | Wellcome Sanger Institute for the COVID-19 Genomics UK (COG-UK) Consortium | Harper VanSteenhouse, Yumi Kasai, David Gray, Carol Clugston, Anna Dominiczak and Alex Alderton, Roberto Amato, Jeffrey Barrett, Sonia Goncalves, Ewan Harrison, David K. Jackson, Ian Johnston, Dominic Kwiatkowski, Cordelia Langford, John Sillitoe on behalf of the Wellcome Sanger Institute COVID-19 Surveillance Team |
| EPI_ISL_3470237 | Lighthouse Lab in Milton Keynes | Wellcome Sanger Institute for the COVID-19 Genomics UK (COG-UK) Consortium | The Lighthouse Lab in Milton Keynes and Alex Alderton, Roberto Amato, Jeffrey Barrett, Sonia Goncalves, Ewan Harrison, David K. Jackson, Ian Johnston, Dominic Kwiatkowski, Cordelia Langford, John Sillitoe on behalf of the Wellcome Sanger Institute COVID-19 Surveillance Team |
| EPI_ISL_3470239 | Lighthouse Lab in Glasgow | Wellcome Sanger Institute for the COVID-19 Genomics UK (COG-UK) Consortium | Harper VanSteenhouse, Yumi Kasai, David Gray, Carol Clugston, Anna Dominiczak and Alex Alderton, Roberto Amato, Jeffrey Barrett, Sonia Goncalves, Ewan Harrison, David K. Jackson, Ian Johnston, Dominic Kwiatkowski, Cordelia Langford, John Sillitoe on behalf of the Wellcome Sanger Institute COVID-19 Surveillance Team |
| EPI_ISL_3470240, EPI_ISL_3470242, EPI_ISL_3470243, EPI_ISL_3470244 | Berkshire and Surrey Pathology Services Lighthouse Laboratory | Wellcome Sanger Institute for the COVID-19 Genomics UK (COG-UK) Consortium | Berkshire and Surrey Pathology Services Lighthouse Laboratory and Alex Alderton, Roberto Amato, Jeffrey Barrett, Sonia Goncalves, Ewan Harrison, David K. Jackson, Ian Johnston, Dominic Kwiatkowski, Cordelia Langford, John Sillitoe on behalf of the Wellcome Sanger Institute COVID-19 Surveillance Team |
| EPI_ISL_3470245 | Microbiology Department. Complexo Hospitalario Universitario de Vigo | Microbiology Department. Complexo Hospitalario Universitario de Vigo | Alfaya N, Alonso I, Alvarez M, Cabrera JJ, Carballo R, Cores O, Cortizo S, del-Campo V, Martinez L, Mediero G, Perez S, Davina C, Potel C, Regueiro B, Rey S, Vasallo FJ |
| EPI_ISL_3470246 | Lighthouse Lab in Milton Keynes | Wellcome Sanger Institute for the COVID-19 Genomics UK (COG-UK) Consortium | The Lighthouse Lab in Milton Keynes and Alex Alderton, Roberto Amato, Jeffrey Barrett, Sonia Goncalves, Ewan Harrison, David K. Jackson, Ian Johnston, Dominic Kwiatkowski, Cordelia Langford, John Sillitoe on behalf of the Wellcome Sanger Institute COVID-19 Surveillance Team |
| EPI_ISL_3470247 | Berkshire and Surrey Pathology Services Lighthouse Laboratory | Wellcome Sanger Institute for the COVID-19 Genomics UK (COG-UK) Consortium | Berkshire and Surrey Pathology Services Lighthouse Laboratory and Alex Alderton, Roberto Amato, Jeffrey Barrett, Sonia Goncalves, Ewan Harrison, David K. Jackson, Ian Johnston, Dominic Kwiatkowski, Cordelia Langford, John Sillitoe on behalf of the Wellcome Sanger Institute COVID-19 Surveillance Team |
| EPI_ISL_3470248 | Lighthouse Lab in Glasgow | Wellcome Sanger Institute for the COVID-19 Genomics UK (COG-UK) Consortium | Harper VanSteenhouse, Yumi Kasai, David Gray, Carol Clugston, Anna Dominiczak and Alex Alderton, Roberto Amato, Jeffrey Barrett, Sonia Goncalves, Ewan Harrison, David K. Jackson, Ian Johnston, Dominic Kwiatkowski, Cordelia Langford, John Sillitoe on behalf of the Wellcome Sanger Institute COVID-19 Surveillance Team |
| EPI_ISL_3470249 | Microbiology Department. Complexo Hospitalario Universitario de Vigo | Microbiology Department. Complexo Hospitalario Universitario de Vigo | Alfaya N, Alonso I, Alvarez M, Cabrera JJ, Carballo R, Cores O, Cortizo S, del-Campo V, Martinez L, Mediero G, Perez S, Davina C, Potel C, Regueiro B, Rey S, Vasallo FJ |
| EPI_ISL_3470250 | Berkshire and Surrey Pathology Services Lighthouse Laboratory | Wellcome Sanger Institute for the COVID-19 Genomics UK (COG-UK) Consortium | Berkshire and Surrey Pathology Services Lighthouse Laboratory and Alex Alderton, Roberto Amato, Jeffrey Barrett, Sonia Goncalves, Ewan Harrison, David K. Jackson, Ian Johnston, Dominic Kwiatkowski, Cordelia Langford, John Sillitoe on behalf of the Wellcome Sanger Institute COVID-19 Surveillance Team |
| EPI_ISL_3470251, EPI_ISL_3470252 | Lighthouse Lab in Glasgow | Wellcome Sanger Institute for the COVID-19 Genomics UK (COG-UK) Consortium | Harper VanSteenhouse, Yumi Kasai, David Gray, Carol Clugston, Anna Dominiczak and Alex Alderton, Roberto Amato, Jeffrey Barrett, Sonia Goncalves, Ewan Harrison, David K. Jackson, Ian Johnston, Dominic Kwiatkowski, Cordelia Langford, John Sillitoe on behalf of the Wellcome Sanger Institute COVID-19 Surveillance Team |
| EPI_ISL_3470253 | Berkshire and Surrey Pathology Services Lighthouse Laboratory | Wellcome Sanger Institute for the COVID-19 Genomics UK (COG-UK) Consortium | Berkshire and Surrey Pathology Services Lighthouse Laboratory and Alex Alderton, Roberto Amato, Jeffrey Barrett, Sonia Goncalves, Ewan Harrison, David K. Jackson, Ian Johnston, Dominic Kwiatkowski, Cordelia Langford, John Sillitoe on behalf of the Wellcome Sanger Institute COVID-19 Surveillance Team |
| EPI_ISL_3470254 | Microbiology Department. Complexo Hospitalario Universitario de Vigo | Microbiology Department. Complexo Hospitalario Universitario de Vigo | Alfaya N, Alonso I, Alvarez M, Cabrera JJ, Carballo R, Cores O, Cortizo S, del-Campo V, Martinez L, Mediero G, Perez S, Davina C, Potel C, Regueiro B, Rey S, Vasallo FJ |
| EPI_ISL_3470255 | Lighthouse Lab in Milton Keynes | Wellcome Sanger Institute for the COVID-19 Genomics UK (COG-UK) Consortium | The Lighthouse Lab in Milton Keynes and Alex Alderton, Roberto Amato, Jeffrey Barrett, Sonia Goncalves, Ewan Harrison, David K. Jackson, Ian Johnston, Dominic Kwiatkowski, Cordelia Langford, John Sillitoe on behalf of the Wellcome Sanger Institute COVID-19 Surveillance Team |
| EPI_ISL_3470256, EPI_ISL_3470257 | Lighthouse Lab in Glasgow | Wellcome Sanger Institute for the COVID-19 Genomics UK (COG-UK) Consortium | Harper VanSteenhouse, Yumi Kasai, David Gray, Carol Clugston, Anna Dominiczak and Alex Alderton, Roberto Amato, Jeffrey Barrett, Sonia Goncalves, Ewan Harrison, David K. Jackson, Ian Johnston, Dominic Kwiatkowski, Cordelia Langford, John Sillitoe on behalf of the Wellcome Sanger Institute COVID-19 Surveillance Team |
| EPI_ISL_3470258 | Microbiology Department. Complexo Hospitalario Universitario de Vigo | Microbiology Department. Complexo Hospitalario Universitario de Vigo | Alfaya N, Alonso I, Alvarez M, Cabrera JJ, Carballo R, Cores O, Cortizo S, del-Campo V, Martinez L, Mediero G, Perez S, Davina C, Potel C, Regueiro B, Rey S, Vasallo FJ |
| EPI_ISL_3470259 | Berkshire and Surrey Pathology Services Lighthouse Laboratory | Wellcome Sanger Institute for the COVID-19 Genomics UK (COG-UK) Consortium | Berkshire and Surrey Pathology Services Lighthouse Laboratory and Alex Alderton, Roberto Amato, Jeffrey Barrett, Sonia Goncalves, Ewan Harrison, David K. Jackson, Ian Johnston, Dominic Kwiatkowski, Cordelia Langford, John Sillitoe on behalf of the Wellcome Sanger Institute COVID-19 Surveillance Team |
| EPI_ISL_3470260 | Lighthouse Lab in Glasgow | Wellcome Sanger Institute for the COVID-19 Genomics UK (COG-UK) Consortium | Harper VanSteenhouse, Yumi Kasai, David Gray, Carol Clugston, Anna Dominiczak and Alex Alderton, Roberto Amato, Jeffrey Barrett, Sonia Goncalves, Ewan Harrison, David K. Jackson, Ian Johnston, Dominic Kwiatkowski, Cordelia Langford, John Sillitoe on behalf of the Wellcome Sanger Institute COVID-19 Surveillance Team |
| EPI_ISL_3470261 | Lighthouse Lab in Milton Keynes | Wellcome Sanger Institute for the COVID-19 Genomics UK (COG-UK) Consortium | The Lighthouse Lab in Milton Keynes and Alex Alderton, Roberto Amato, Jeffrey Barrett, Sonia Goncalves, Ewan Harrison, David K. Jackson, Ian Johnston, Dominic Kwiatkowski, Cordelia Langford, John Sillitoe on behalf of the Wellcome Sanger Institute COVID-19 Surveillance Team |
| EPI_ISL_3470262 | Lighthouse Lab in Glasgow | Wellcome Sanger Institute for the COVID-19 Genomics UK (COG-UK) Consortium | Harper VanSteenhouse, Yumi Kasai, David Gray, Carol Clugston, Anna Dominiczak and Alex Alderton, Roberto Amato, Jeffrey Barrett, Sonia Goncalves, Ewan Harrison, David K. Jackson, Ian Johnston, Dominic Kwiatkowski, Cordelia Langford, John Sillitoe on behalf of the Wellcome Sanger Institute COVID-19 Surveillance Team |
| EPI_ISL_3470263 | Microbiology Department. Complexo Hospitalario Universitario | Microbiology Department. Complexo Hospitalario Universitario de | Alfaya N, Alonso I, Alvarez M, Cabrera JJ, Carballo R, Cores O, Cortizo S, del-Campo V, Martinez L, Mediero G, Perez S, Davina C, Potel C, Regueiro B, Rey S, Vasallo FJ |

| | | | |
|---|---|---|---|
| | de Vigo | Vigo | |
| EPI_ISL_3470264, EPI_ISL_3470265, EPI_ISL_3470266 | Lighthouse Lab in Glasgow | Wellcome Sanger Institute for the COVID-19 Genomics UK (COG-UK) Consortium | Harper VanSteenhouse, Yumi Kasai, David Gray, Carol Clugston, Anna Dominiczak and Alex Alderton, Roberto Amato, Jeffrey Barrett, Sonia Goncalves, Ewan Harrison, David K. Jackson, Ian Johnston, Dominic Kwiatkowski, Cordelia Langford, John Sillitoe on behalf of the Wellcome Sanger Institute COVID-19 Surveillance Team |
| EPI_ISL_3470267 | Microbiology Department. Complexo Hospitalario Universitario de Vigo | Microbiology Department. Complexo Hospitalario Universitario de Vigo | Alfaya N, Alonso I, Alvarez M, Cabrera JJ, Carballo R, Cores O, Cortizo S, del-Campo V, Martinez L, Mediero G, Perez S, Davina C, Potel C, Regueiro B, Rey S, Vasallo FJ |
| EPI_ISL_3470268, EPI_ISL_3470269, EPI_ISL_3470270 | Lighthouse Lab in Glasgow | Wellcome Sanger Institute for the COVID-19 Genomics UK (COG-UK) Consortium | Harper VanSteenhouse, Yumi Kasai, David Gray, Carol Clugston, Anna Dominiczak and Alex Alderton, Roberto Amato, Jeffrey Barrett, Sonia Goncalves, Ewan Harrison, David K. Jackson, Ian Johnston, Dominic Kwiatkowski, Cordelia Langford, John Sillitoe on behalf of the Wellcome Sanger Institute COVID-19 Surveillance Team |
| EPI_ISL_3470271 | Microbiology Department. Complexo Hospitalario Universitario de Vigo | Microbiology Department. Complexo Hospitalario Universitario de Vigo | Alfaya N, Alonso I, Alvarez M, Cabrera JJ, Carballo R, Cores O, Cortizo S, del-Campo V, Martinez L, Mediero G, Perez S, Davina C, Potel C, Regueiro B, Rey S, Vasallo FJ |
| EPI_ISL_3470272 | Berkshire and Surrey Pathology Services Lighthouse Laboratory | Wellcome Sanger Institute for the COVID-19 Genomics UK (COG-UK) Consortium | Berkshire and Surrey Pathology Services Lighthouse Laboratory and Alex Alderton, Roberto Amato, Jeffrey Barrett, Sonia Goncalves, Ewan Harrison, David K. Jackson, Ian Johnston, Dominic Kwiatkowski, Cordelia Langford, John Sillitoe on behalf of the Wellcome Sanger Institute COVID-19 Surveillance Team |
| EPI_ISL_3470273, EPI_ISL_3470274 | Lighthouse Lab in Milton Keynes | Wellcome Sanger Institute for the COVID-19 Genomics UK (COG-UK) Consortium | The Lighthouse Lab in Milton Keynes and Alex Alderton, Roberto Amato, Jeffrey Barrett, Sonia Goncalves, Ewan Harrison, David K. Jackson, Ian Johnston, Dominic Kwiatkowski, Cordelia Langford, John Sillitoe on behalf of the Wellcome Sanger Institute COVID-19 Surveillance Team |
| EPI_ISL_3470275 | Microbiology Department. Complexo Hospitalario Universitario de Vigo | Microbiology Department. Complexo Hospitalario Universitario de Vigo | Alfaya N, Alonso I, Alvarez M, Cabrera JJ, Carballo R, Cores O, Cortizo S, del-Campo V, Martinez L, Mediero G, Perez S, Davina C, Potel C, Regueiro B, Rey S, Vasallo FJ |
| EPI_ISL_3470276 | Lighthouse Lab in Glasgow | Wellcome Sanger Institute for the COVID-19 Genomics UK (COG-UK) Consortium | Harper VanSteenhouse, Yumi Kasai, David Gray, Carol Clugston, Anna Dominiczak and Alex Alderton, Roberto Amato, Jeffrey Barrett, Sonia Goncalves, Ewan Harrison, David K. Jackson, Ian Johnston, Dominic Kwiatkowski, Cordelia Langford, John Sillitoe on behalf of the Wellcome Sanger Institute COVID-19 Surveillance Team |
| EPI_ISL_3470277, EPI_ISL_3470278 | Berkshire and Surrey Pathology Services Lighthouse Laboratory | Wellcome Sanger Institute for the COVID-19 Genomics UK (COG-UK) Consortium | Berkshire and Surrey Pathology Services Lighthouse Laboratory and Alex Alderton, Roberto Amato, Jeffrey Barrett, Sonia Goncalves, Ewan Harrison, David K. Jackson, Ian Johnston, Dominic Kwiatkowski, Cordelia Langford, John Sillitoe on behalf of the Wellcome Sanger Institute COVID-19 Surveillance Team |
| EPI_ISL_3470279 | Microbiology Department. Complexo Hospitalario Universitario de Vigo | Microbiology Department. Complexo Hospitalario Universitario de Vigo | Alfaya N, Alonso I, Alvarez M, Cabrera JJ, Carballo R, Cores O, Cortizo S, del-Campo V, Martinez L, Mediero G, Perez S, Davina C, Potel C, Regueiro B, Rey S, Vasallo FJ |
| EPI_ISL_3470280 | Berkshire and Surrey Pathology Services Lighthouse Laboratory | Wellcome Sanger Institute for the COVID-19 Genomics UK (COG-UK) Consortium | Berkshire and Surrey Pathology Services Lighthouse Laboratory and Alex Alderton, Roberto Amato, Jeffrey Barrett, Sonia Goncalves, Ewan Harrison, David K. Jackson, Ian Johnston, Dominic Kwiatkowski, Cordelia Langford, John Sillitoe on behalf of the Wellcome Sanger Institute COVID-19 Surveillance Team |
| EPI_ISL_3470281 | Lighthouse Lab in Glasgow | Wellcome Sanger Institute for the COVID-19 Genomics UK (COG-UK) Consortium | Harper VanSteenhouse, Yumi Kasai, David Gray, Carol Clugston, Anna Dominiczak and Alex Alderton, Roberto Amato, Jeffrey Barrett, Sonia Goncalves, Ewan Harrison, David K. Jackson, Ian Johnston, Dominic Kwiatkowski, Cordelia Langford, John Sillitoe on behalf of the Wellcome Sanger Institute COVID-19 Surveillance Team |
| EPI_ISL_3470282 | Berkshire and Surrey Pathology Services Lighthouse Laboratory | Wellcome Sanger Institute for the COVID-19 Genomics UK (COG-UK) Consortium | Berkshire and Surrey Pathology Services Lighthouse Laboratory and Alex Alderton, Roberto Amato, Jeffrey Barrett, Sonia Goncalves, Ewan Harrison, David K. Jackson, Ian Johnston, Dominic Kwiatkowski, Cordelia Langford, John Sillitoe on behalf of the Wellcome Sanger Institute COVID-19 Surveillance Team |
| EPI_ISL_3470283 | Microbiology Department. Complexo Hospitalario Universitario de Vigo | Microbiology Department. Complexo Hospitalario Universitario de Vigo | Alfaya N, Alonso I, Alvarez M, Cabrera JJ, Carballo R, Cores O, Cortizo S, del-Campo V, Martinez L, Mediero G, Perez S, Davina C, Potel C, Regueiro B, Rey S, Vasallo FJ |
| EPI_ISL_3470284 | Lighthouse Lab in Milton Keynes | Wellcome Sanger Institute for the COVID-19 Genomics UK (COG-UK) Consortium | The Lighthouse Lab in Milton Keynes and Alex Alderton, Roberto Amato, Jeffrey Barrett, Sonia Goncalves, Ewan Harrison, David K. Jackson, Ian Johnston, Dominic Kwiatkowski, Cordelia Langford, John Sillitoe on behalf of the Wellcome Sanger Institute COVID-19 Surveillance Team |
| EPI_ISL_3470285 | Berkshire and Surrey Pathology Services Lighthouse Laboratory | Wellcome Sanger Institute for the COVID-19 Genomics UK (COG-UK) Consortium | Berkshire and Surrey Pathology Services Lighthouse Laboratory and Alex Alderton, Roberto Amato, Jeffrey Barrett, Sonia Goncalves, Ewan Harrison, David K. Jackson, Ian Johnston, Dominic Kwiatkowski, Cordelia Langford, John Sillitoe on behalf of the Wellcome Sanger Institute COVID-19 Surveillance Team |
| EPI_ISL_3470286 | Lighthouse Lab in Glasgow | Wellcome Sanger Institute for the COVID-19 Genomics UK (COG-UK) Consortium | Harper VanSteenhouse, Yumi Kasai, David Gray, Carol Clugston, Anna Dominiczak and Alex Alderton, Roberto Amato, Jeffrey Barrett, Sonia Goncalves, Ewan Harrison, David K. Jackson, Ian Johnston, Dominic Kwiatkowski, Cordelia Langford, John Sillitoe on behalf of the Wellcome Sanger Institute COVID-19 Surveillance Team |
| EPI_ISL_3470287 | Microbiology Department. Complexo Hospitalario Universitario de Vigo | Microbiology Department. Complexo Hospitalario Universitario de Vigo | Alfaya N, Alonso I, Alvarez M, Cabrera JJ, Carballo R, Cores O, Cortizo S, del-Campo V, Martinez L, Mediero G, Perez S, Davina C, Potel C, Regueiro B, Rey S, Vasallo FJ |
| EPI_ISL_3470288 | Berkshire and Surrey Pathology Services Lighthouse Laboratory | Wellcome Sanger Institute for the COVID-19 Genomics UK (COG-UK) Consortium | Berkshire and Surrey Pathology Services Lighthouse Laboratory and Alex Alderton, Roberto Amato, Jeffrey Barrett, Sonia Goncalves, Ewan Harrison, David K. Jackson, Ian Johnston, Dominic Kwiatkowski, Cordelia Langford, John Sillitoe on behalf of the Wellcome Sanger Institute COVID-19 Surveillance Team |
| EPI_ISL_3470289 | Lighthouse Lab in Glasgow | Wellcome Sanger Institute for the COVID-19 Genomics UK (COG-UK) Consortium | Harper VanSteenhouse, Yumi Kasai, David Gray, Carol Clugston, Anna Dominiczak and Alex Alderton, Roberto Amato, Jeffrey Barrett, Sonia Goncalves, Ewan Harrison, David K. Jackson, Ian Johnston, Dominic Kwiatkowski, Cordelia Langford, John Sillitoe on behalf of the Wellcome Sanger Institute COVID-19 Surveillance Team |
| EPI_ISL_3470290 | Microbiology Department. Complexo Hospitalario Universitario de Vigo | Microbiology Department. Complexo Hospitalario Universitario de Vigo | Alfaya N, Alonso I, Alvarez M, Cabrera JJ, Carballo R, Cores O, Cortizo S, del-Campo V, Martinez L, Mediero G, Perez S, Davina C, Potel C, Regueiro B, Rey S, Vasallo FJ |
| EPI_ISL_3470291 | Lighthouse Lab in Glasgow | Wellcome Sanger Institute for the COVID-19 Genomics UK (COG-UK) Consortium | Harper VanSteenhouse, Yumi Kasai, David Gray, Carol Clugston, Anna Dominiczak and Alex Alderton, Roberto Amato, Jeffrey Barrett, Sonia Goncalves, Ewan Harrison, David K. Jackson, Ian Johnston, Dominic Kwiatkowski, Cordelia Langford, John Sillitoe on behalf of the Wellcome Sanger Institute COVID-19 Surveillance Team |
| EPI_ISL_3470292 | Berkshire and Surrey Pathology Services Lighthouse Laboratory | Wellcome Sanger Institute for the COVID-19 Genomics UK (COG-UK) Consortium | Berkshire and Surrey Pathology Services Lighthouse Laboratory and Alex Alderton, Roberto Amato, Jeffrey Barrett, Sonia Goncalves, Ewan Harrison, David K. Jackson, Ian Johnston, Dominic Kwiatkowski, Cordelia Langford, John Sillitoe on behalf of the Wellcome Sanger Institute COVID-19 Surveillance Team |
| EPI_ISL_3470293 | Lighthouse Lab in Glasgow | Wellcome Sanger Institute for the COVID-19 Genomics UK (COG-UK) Consortium | Harper VanSteenhouse, Yumi Kasai, David Gray, Carol Clugston, Anna Dominiczak and Alex Alderton, Roberto Amato, Jeffrey Barrett, Sonia Goncalves, Ewan Harrison, David K. Jackson, Ian Johnston, Dominic Kwiatkowski, Cordelia Langford, John Sillitoe on behalf of the Wellcome Sanger Institute COVID-19 Surveillance Team |
| EPI_ISL_3470294 | Microbiology Department. Complexo Hospitalario Universitario de Vigo | Microbiology Department. Complexo Hospitalario Universitario de Vigo | Alfaya N, Alonso I, Alvarez M, Cabrera JJ, Carballo R, Cores O, Cortizo S, del-Campo V, Martinez L, Mediero G, Perez S, Davina C, Potel C, Regueiro B, Rey S, Vasallo FJ |
| EPI_ISL_3470295 | Lighthouse Lab in Glasgow | Wellcome Sanger Institute for the COVID-19 Genomics UK (COG-UK) Consortium | Harper VanSteenhouse, Yumi Kasai, David Gray, Carol Clugston, Anna Dominiczak and Alex Alderton, Roberto Amato, Jeffrey Barrett, Sonia Goncalves, Ewan Harrison, David K. Jackson, Ian Johnston, Dominic Kwiatkowski, Cordelia Langford, John Sillitoe on behalf of the Wellcome Sanger Institute COVID-19 Surveillance Team |
| EPI_ISL_3470296 | Berkshire and Surrey Pathology Services Lighthouse Laboratory | Wellcome Sanger Institute for the COVID-19 Genomics UK (COG-UK) Consortium | Berkshire and Surrey Pathology Services Lighthouse Laboratory and Alex Alderton, Roberto Amato, Jeffrey Barrett, Sonia Goncalves, Ewan Harrison, David K. Jackson, Ian Johnston, Dominic Kwiatkowski, Cordelia Langford, John Sillitoe on behalf of the Wellcome Sanger Institute COVID-19 Surveillance Team |
| EPI_ISL_3470297 | Lighthouse Lab in Glasgow | Wellcome Sanger Institute for the COVID-19 Genomics UK (COG-UK) Consortium | Harper VanSteenhouse, Yumi Kasai, David Gray, Carol Clugston, Anna Dominiczak and Alex Alderton, Roberto Amato, Jeffrey Barrett, Sonia Goncalves, Ewan Harrison, David K. Jackson, Ian Johnston, Dominic Kwiatkowski, Cordelia Langford, John Sillitoe on behalf of the Wellcome Sanger Institute COVID-19 Surveillance Team |
| EPI_ISL_3470298 | Microbiology Department. Complexo Hospitalario Universitario de Vigo | Microbiology Department. Complexo Hospitalario Universitario de Vigo | Alfaya N, Alonso I, Alvarez M, Cabrera JJ, Carballo R, Cores O, Cortizo S, del-Campo V, Martinez L, Mediero G, Perez S, Davina C, Potel C, Regueiro B, Rey S, Vasallo FJ |
| EPI_ISL_3470299 | Lighthouse Lab in Glasgow | Wellcome Sanger Institute for the COVID-19 Genomics UK (COG-UK) Consortium | Harper VanSteenhouse, Yumi Kasai, David Gray, Carol Clugston, Anna Dominiczak and Alex Alderton, Roberto Amato, Jeffrey Barrett, Sonia Goncalves, Ewan Harrison, David K. Jackson, Ian Johnston, Dominic Kwiatkowski, Cordelia Langford, John Sillitoe on behalf of the Wellcome Sanger Institute COVID-19 Surveillance Team |
| EPI_ISL_3470300 | Lighthouse Lab in Milton Keynes | Wellcome Sanger Institute for the COVID-19 Genomics UK (COG-UK) Consortium | The Lighthouse Lab in Milton Keynes and Alex Alderton, Roberto Amato, Jeffrey Barrett, Sonia Goncalves, Ewan Harrison, David K. Jackson, Ian Johnston, Dominic Kwiatkowski, Cordelia Langford, John Sillitoe on behalf of the Wellcome Sanger Institute COVID-19 Surveillance Team |
| EPI_ISL_3470301 | Lighthouse Lab in Glasgow | Wellcome Sanger Institute for the COVID-19 Genomics UK (COG-UK) Consortium | Harper VanSteenhouse, Yumi Kasai, David Gray, Carol Clugston, Anna Dominiczak and Alex Alderton, Roberto Amato, Jeffrey Barrett, Sonia Goncalves, Ewan Harrison, David K. Jackson, Ian Johnston, Dominic Kwiatkowski, Cordelia Langford, John Sillitoe on behalf of the Wellcome Sanger Institute COVID-19 Surveillance Team |
| EPI_ISL_3470302 | Microbiology Department. Complexo Hospitalario Universitario de Vigo | Microbiology Department. Complexo Hospitalario Universitario de Vigo | Alfaya N, Alonso I, Alvarez M, Cabrera JJ, Carballo R, Cores O, Cortizo S, del-Campo V, Martinez L, Mediero G, Perez S, Davina C, Potel C, Regueiro B, Rey S, Vasallo FJ |
| EPI_ISL_3470303 | Lighthouse Lab in Glasgow | Wellcome Sanger Institute for the COVID-19 Genomics UK (COG-UK) Consortium | Harper VanSteenhouse, Yumi Kasai, David Gray, Carol Clugston, Anna Dominiczak and Alex Alderton, Roberto Amato, Jeffrey Barrett, Sonia Goncalves, Ewan Harrison, David K. Jackson, Ian Johnston, Dominic Kwiatkowski, Cordelia Langford, John Sillitoe on behalf of the Wellcome Sanger Institute COVID-19 Surveillance Team |
| EPI_ISL_3470304, EPI_ISL_3470305 | Lighthouse Lab in Milton Keynes | Wellcome Sanger Institute for the COVID-19 Genomics UK (COG-UK) Consortium | The Lighthouse Lab in Milton Keynes and Alex Alderton, Roberto Amato, Jeffrey Barrett, Sonia Goncalves, Ewan Harrison, David K. Jackson, Ian Johnston, Dominic Kwiatkowski, Cordelia Langford, John Sillitoe on behalf of the Wellcome Sanger Institute COVID-19 Surveillance Team |
| EPI_ISL_3470306 | Microbiology Department. Complexo Hospitalario Universitario de Vigo | Microbiology Department. Complexo Hospitalario Universitario de Vigo | Alfaya N, Alonso I, Alvarez M, Cabrera JJ, Carballo R, Cores O, Cortizo S, del-Campo V, Martinez L, Mediero G, Perez S, Davina C, Potel C, Regueiro B, Rey S, Vasallo FJ |
| EPI_ISL_3470307 | Lighthouse Lab in Glasgow | Wellcome Sanger Institute for the COVID-19 Genomics UK (COG-UK) Consortium | Harper VanSteenhouse, Yumi Kasai, David Gray, Carol Clugston, Anna Dominiczak and Alex Alderton, Roberto Amato, Jeffrey Barrett, Sonia Goncalves, Ewan Harrison, David K. Jackson, Ian Johnston, Dominic Kwiatkowski, Cordelia Langford, John Sillitoe on behalf of the Wellcome Sanger Institute COVID-19 Surveillance Team |
| EPI_ISL_3470308 | Berkshire and Surrey Pathology Services Lighthouse Laboratory | Wellcome Sanger Institute for the COVID-19 Genomics UK (COG-UK) Consortium | Berkshire and Surrey Pathology Services Lighthouse Laboratory and Alex Alderton, Roberto Amato, Jeffrey Barrett, Sonia Goncalves, Ewan Harrison, David K. Jackson, Ian Johnston, Dominic Kwiatkowski, Cordelia Langford, John Sillitoe on behalf of the Wellcome Sanger Institute COVID-19 Surveillance Team |
| EPI_ISL_3470309 | Lighthouse Lab in Milton Keynes | Wellcome Sanger Institute for the COVID-19 Genomics UK (COG-UK) Consortium | The Lighthouse Lab in Milton Keynes and Alex Alderton, Roberto Amato, Jeffrey Barrett, Sonia Goncalves, Ewan Harrison, David K. Jackson, Ian Johnston, Dominic Kwiatkowski, Cordelia Langford, John Sillitoe on behalf of the Wellcome Sanger Institute COVID-19 Surveillance Team |
| EPI_ISL_3470310 | Microbiology Department. Complexo Hospitalario Universitario de Vigo | Microbiology Department. Complexo Hospitalario Universitario de Vigo | Alfaya N, Alonso I, Alvarez M, Cabrera JJ, Carballo R, Cores O, Cortizo S, del-Campo V, Martinez L, Mediero G, Perez S, Davina C, Potel C, Regueiro B, Rey S, Vasallo FJ |
| EPI_ISL_3470311 | Berkshire and Surrey Pathology Services Lighthouse Laboratory | Wellcome Sanger Institute for the COVID-19 Genomics UK (COG-UK) Consortium | Berkshire and Surrey Pathology Services Lighthouse Laboratory and Alex Alderton, Roberto Amato, Jeffrey Barrett, Sonia Goncalves, Ewan Harrison, David K. Jackson, Ian Johnston, Dominic Kwiatkowski, Cordelia Langford, John Sillitoe on behalf of the Wellcome Sanger Institute COVID-19 Surveillance Team |
| EPI_ISL_3470312 | Lighthouse Lab in Glasgow | Wellcome Sanger Institute for the COVID-19 Genomics UK (COG-UK) Consortium | Harper VanSteenhouse, Yumi Kasai, David Gray, Carol Clugston, Anna Dominiczak and Alex Alderton, Roberto Amato, Jeffrey Barrett, Sonia Goncalves, Ewan Harrison, David K. Jackson, Ian Johnston, Dominic Kwiatkowski, Cordelia Langford, John Sillitoe on behalf of the Wellcome Sanger Institute COVID-19 Surveillance Team |
| EPI_ISL_3470313 | Lighthouse Lab in Milton Keynes | Wellcome Sanger Institute for the COVID-19 Genomics UK (COG-UK) Consortium | The Lighthouse Lab in Milton Keynes and Alex Alderton, Roberto Amato, Jeffrey Barrett, Sonia Goncalves, Ewan Harrison, David K. Jackson, Ian Johnston, Dominic Kwiatkowski, Cordelia Langford, John Sillitoe on behalf of the Wellcome Sanger Institute COVID-19 Surveillance Team |
| EPI_ISL_3470314 | Microbiology Department. Complexo Hospitalario Universitario de Vigo | Microbiology Department. Complexo Hospitalario Universitario de Vigo | Alfaya N, Alonso I, Alvarez M, Cabrera JJ, Carballo R, Cores O, Cortizo S, del-Campo V, Martinez L, Mediero G, Perez S, Davina C, Potel C, Regueiro B, Rey S, Vasallo FJ |

| | | | |
|---|---|---|---|
| EPI_ISL_3470315, EPI_ISL_3470316 | Lighthouse Lab in Glasgow | Wellcome Sanger Institute for the COVID-19 Genomics UK (COG-UK) Consortium | Harper VanSteenhouse, Yumi Kasai, David Gray, Carol Clugston, Anna Dominiczak and Alex Alderton, Roberto Amato, Jeffrey Barrett, Sonia Goncalves, Ewan Harrison, David K. Jackson, Ian Johnston, Dominic Kwiatkowski, Cordelia Langford, John Sillitoe on behalf of the Wellcome Sanger Institute COVID-19 Surveillance Team |
| EPI_ISL_3470317 | Microbiology Department. Complexo Hospitalario Universitario de Vigo | Microbiology Department. Complexo Hospitalario Universitario de Vigo | Alfaya N, Alonso I, Alvarez M, Cabrera JJ, Carballo R, Cores O, Cortizo S, del-Campo V, Martinez L, Mediero G, Perez S, Davina C, Potel C, Regueiro B, Rey S, Vasallo FJ |
| EPI_ISL_3470318 | Lighthouse Lab in Glasgow | Wellcome Sanger Institute for the COVID-19 Genomics UK (COG-UK) Consortium | Harper VanSteenhouse, Yumi Kasai, David Gray, Carol Clugston, Anna Dominiczak and Alex Alderton, Roberto Amato, Jeffrey Barrett, Sonia Goncalves, Ewan Harrison, David K. Jackson, Ian Johnston, Dominic Kwiatkowski, Cordelia Langford, John Sillitoe on behalf of the Wellcome Sanger Institute COVID-19 Surveillance Team |
| EPI_ISL_3470319, EPI_ISL_3470320 | Lighthouse Lab in Milton Keynes | Wellcome Sanger Institute for the COVID-19 Genomics UK (COG-UK) Consortium | The Lighthouse Lab in Milton Keynes and Alex Alderton, Roberto Amato, Jeffrey Barrett, Sonia Goncalves, Ewan Harrison, David K. Jackson, Ian Johnston, Dominic Kwiatkowski, Cordelia Langford, John Sillitoe on behalf of the Wellcome Sanger Institute COVID-19 Surveillance Team |
| EPI_ISL_3470321 | Lighthouse Lab in Glasgow | Wellcome Sanger Institute for the COVID-19 Genomics UK (COG-UK) Consortium | Harper VanSteenhouse, Yumi Kasai, David Gray, Carol Clugston, Anna Dominiczak and Alex Alderton, Roberto Amato, Jeffrey Barrett, Sonia Goncalves, Ewan Harrison, David K. Jackson, Ian Johnston, Dominic Kwiatkowski, Cordelia Langford, John Sillitoe on behalf of the Wellcome Sanger Institute COVID-19 Surveillance Team |
| EPI_ISL_3470322 | Microbiology Department. Complexo Hospitalario Universitario de Vigo | Microbiology Department. Complexo Hospitalario Universitario de Vigo | Alfaya N, Alonso I, Alvarez M, Cabrera JJ, Carballo R, Cores O, Cortizo S, del-Campo V, Martinez L, Mediero G, Perez S, Davina C, Potel C, Regueiro B, Rey S, Vasallo FJ |
| EPI_ISL_3470323, EPI_ISL_3470324, EPI_ISL_3470325 | Lighthouse Lab in Milton Keynes | Wellcome Sanger Institute for the COVID-19 Genomics UK (COG-UK) Consortium | The Lighthouse Lab in Milton Keynes and Alex Alderton, Roberto Amato, Jeffrey Barrett, Sonia Goncalves, Ewan Harrison, David K. Jackson, Ian Johnston, Dominic Kwiatkowski, Cordelia Langford, John Sillitoe on behalf of the Wellcome Sanger Institute COVID-19 Surveillance Team |
| EPI_ISL_3470326 | Microbiology Department. Complexo Hospitalario Universitario de Vigo | Microbiology Department. Complexo Hospitalario Universitario de Vigo | Alfaya N, Alonso I, Alvarez M, Cabrera JJ, Carballo R, Cores O, Cortizo S, del-Campo V, Martinez L, Mediero G, Perez S, Davina C, Potel C, Regueiro B, Rey S, Vasallo FJ |
| EPI_ISL_3470327 | Berkshire and Surrey Pathology Services Lighthouse Laboratory | Wellcome Sanger Institute for the COVID-19 Genomics UK (COG-UK) Consortium | Berkshire and Surrey Pathology Services Lighthouse Laboratory and Alex Alderton, Roberto Amato, Jeffrey Barrett, Sonia Goncalves, Ewan Harrison, David K. Jackson, Ian Johnston, Dominic Kwiatkowski, Cordelia Langford, John Sillitoe on behalf of the Wellcome Sanger Institute COVID-19 Surveillance Team |
| EPI_ISL_3470328 | Lighthouse Lab in Glasgow | Wellcome Sanger Institute for the COVID-19 Genomics UK (COG-UK) Consortium | Harper VanSteenhouse, Yumi Kasai, David Gray, Carol Clugston, Anna Dominiczak and Alex Alderton, Roberto Amato, Jeffrey Barrett, Sonia Goncalves, Ewan Harrison, David K. Jackson, Ian Johnston, Dominic Kwiatkowski, Cordelia Langford, John Sillitoe on behalf of the Wellcome Sanger Institute COVID-19 Surveillance Team |
| EPI_ISL_3470329 | Berkshire and Surrey Pathology Services Lighthouse Laboratory | Wellcome Sanger Institute for the COVID-19 Genomics UK (COG-UK) Consortium | Berkshire and Surrey Pathology Services Lighthouse Laboratory and Alex Alderton, Roberto Amato, Jeffrey Barrett, Sonia Goncalves, Ewan Harrison, David K. Jackson, Ian Johnston, Dominic Kwiatkowski, Cordelia Langford, John Sillitoe on behalf of the Wellcome Sanger Institute COVID-19 Surveillance Team |
| EPI_ISL_3470331 | Lighthouse Lab in Glasgow | Wellcome Sanger Institute for the COVID-19 Genomics UK (COG-UK) Consortium | Harper VanSteenhouse, Yumi Kasai, David Gray, Carol Clugston, Anna Dominiczak and Alex Alderton, Roberto Amato, Jeffrey Barrett, Sonia Goncalves, Ewan Harrison, David K. Jackson, Ian Johnston, Dominic Kwiatkowski, Cordelia Langford, John Sillitoe on behalf of the Wellcome Sanger Institute COVID-19 Surveillance Team |
| EPI_ISL_3470332 | Berkshire and Surrey Pathology Services Lighthouse Laboratory | Wellcome Sanger Institute for the COVID-19 Genomics UK (COG-UK) Consortium | Berkshire and Surrey Pathology Services Lighthouse Laboratory and Alex Alderton, Roberto Amato, Jeffrey Barrett, Sonia Goncalves, Ewan Harrison, David K. Jackson, Ian Johnston, Dominic Kwiatkowski, Cordelia Langford, John Sillitoe on behalf of the Wellcome Sanger Institute COVID-19 Surveillance Team |
| EPI_ISL_3470334, EPI_ISL_3470335 | Lighthouse Lab in Milton Keynes | Wellcome Sanger Institute for the COVID-19 Genomics UK (COG-UK) Consortium | The Lighthouse Lab in Milton Keynes and Alex Alderton, Roberto Amato, Jeffrey Barrett, Sonia Goncalves, Ewan Harrison, David K. Jackson, Ian Johnston, Dominic Kwiatkowski, Cordelia Langford, John Sillitoe on behalf of the Wellcome Sanger Institute COVID-19 Surveillance Team |
| EPI_ISL_3470336, EPI_ISL_3470338 | Lighthouse Lab in Glasgow | Wellcome Sanger Institute for the COVID-19 Genomics UK (COG-UK) Consortium | Harper VanSteenhouse, Yumi Kasai, David Gray, Carol Clugston, Anna Dominiczak and Alex Alderton, Roberto Amato, Jeffrey Barrett, Sonia Goncalves, Ewan Harrison, David K. Jackson, Ian Johnston, Dominic Kwiatkowski, Cordelia Langford, John Sillitoe on behalf of the Wellcome Sanger Institute COVID-19 Surveillance Team |
| EPI_ISL_3470339 | Lighthouse Lab in Milton Keynes | Wellcome Sanger Institute for the COVID-19 Genomics UK (COG-UK) Consortium | The Lighthouse Lab in Milton Keynes and Alex Alderton, Roberto Amato, Jeffrey Barrett, Sonia Goncalves, Ewan Harrison, David K. Jackson, Ian Johnston, Dominic Kwiatkowski, Cordelia Langford, John Sillitoe on behalf of the Wellcome Sanger Institute COVID-19 Surveillance Team |
| EPI_ISL_3470340 | Lighthouse Lab in Glasgow | Wellcome Sanger Institute for the COVID-19 Genomics UK (COG-UK) Consortium | Harper VanSteenhouse, Yumi Kasai, David Gray, Carol Clugston, Anna Dominiczak and Alex Alderton, Roberto Amato, Jeffrey Barrett, Sonia Goncalves, Ewan Harrison, David K. Jackson, Ian Johnston, Dominic Kwiatkowski, Cordelia Langford, John Sillitoe on behalf of the Wellcome Sanger Institute COVID-19 Surveillance Team |
| EPI_ISL_3470342 | Lighthouse Lab in Milton Keynes | Wellcome Sanger Institute for the COVID-19 Genomics UK (COG-UK) Consortium | The Lighthouse Lab in Milton Keynes and Alex Alderton, Roberto Amato, Jeffrey Barrett, Sonia Goncalves, Ewan Harrison, David K. Jackson, Ian Johnston, Dominic Kwiatkowski, Cordelia Langford, John Sillitoe on behalf of the Wellcome Sanger Institute COVID-19 Surveillance Team |
| EPI_ISL_3470343 | Lighthouse Lab in Glasgow | Wellcome Sanger Institute for the COVID-19 Genomics UK (COG-UK) Consortium | Harper VanSteenhouse, Yumi Kasai, David Gray, Carol Clugston, Anna Dominiczak and Alex Alderton, Roberto Amato, Jeffrey Barrett, Sonia Goncalves, Ewan Harrison, David K. Jackson, Ian Johnston, Dominic Kwiatkowski, Cordelia Langford, John Sillitoe on behalf of the Wellcome Sanger Institute COVID-19 Surveillance Team |
| EPI_ISL_3470344 | Lighthouse Lab in Milton Keynes | Wellcome Sanger Institute for the COVID-19 Genomics UK (COG-UK) Consortium | The Lighthouse Lab in Milton Keynes and Alex Alderton, Roberto Amato, Jeffrey Barrett, Sonia Goncalves, Ewan Harrison, David K. Jackson, Ian Johnston, Dominic Kwiatkowski, Cordelia Langford, John Sillitoe on behalf of the Wellcome Sanger Institute COVID-19 Surveillance Team |
| EPI_ISL_3470346 | Berkshire and Surrey Pathology Services Lighthouse Laboratory | Wellcome Sanger Institute for the COVID-19 Genomics UK (COG-UK) Consortium | Berkshire and Surrey Pathology Services Lighthouse Laboratory and Alex Alderton, Roberto Amato, Jeffrey Barrett, Sonia Goncalves, Ewan Harrison, David K. Jackson, Ian Johnston, Dominic Kwiatkowski, Cordelia Langford, John Sillitoe on behalf of the Wellcome Sanger Institute COVID-19 Surveillance Team |
| EPI_ISL_3470347, EPI_ISL_3470348 | Lighthouse Lab in Milton Keynes | Wellcome Sanger Institute for the COVID-19 Genomics UK (COG-UK) Consortium | The Lighthouse Lab in Milton Keynes and Alex Alderton, Roberto Amato, Jeffrey Barrett, Sonia Goncalves, Ewan Harrison, David K. Jackson, Ian Johnston, Dominic Kwiatkowski, Cordelia Langford, John Sillitoe on behalf of the Wellcome Sanger Institute COVID-19 Surveillance Team |
| EPI_ISL_3470349, EPI_ISL_3470350 | Lighthouse Lab in Glasgow | Wellcome Sanger Institute for the COVID-19 Genomics UK (COG-UK) Consortium | Harper VanSteenhouse, Yumi Kasai, David Gray, Carol Clugston, Anna Dominiczak and Alex Alderton, Roberto Amato, Jeffrey Barrett, Sonia Goncalves, Ewan Harrison, David K. Jackson, Ian Johnston, Dominic Kwiatkowski, Cordelia Langford, John Sillitoe on behalf of the Wellcome Sanger Institute COVID-19 Surveillance Team |
| EPI_ISL_3470351 | Lighthouse Lab in Milton Keynes | Wellcome Sanger Institute for the COVID-19 Genomics UK (COG-UK) Consortium | The Lighthouse Lab in Milton Keynes and Alex Alderton, Roberto Amato, Jeffrey Barrett, Sonia Goncalves, Ewan Harrison, David K. Jackson, Ian Johnston, Dominic Kwiatkowski, Cordelia Langford, John Sillitoe on behalf of the Wellcome Sanger Institute COVID-19 Surveillance Team |
| EPI_ISL_3470352 | Lighthouse Lab in Glasgow | Wellcome Sanger Institute for the COVID-19 Genomics UK (COG-UK) Consortium | Harper VanSteenhouse, Yumi Kasai, David Gray, Carol Clugston, Anna Dominiczak and Alex Alderton, Roberto Amato, Jeffrey Barrett, Sonia Goncalves, Ewan Harrison, David K. Jackson, Ian Johnston, Dominic Kwiatkowski, Cordelia Langford, John Sillitoe on behalf of the Wellcome Sanger Institute COVID-19 Surveillance Team |
| EPI_ISL_3470353, EPI_ISL_3470354, EPI_ISL_3470355 | Lighthouse Lab in Milton Keynes | Wellcome Sanger Institute for the COVID-19 Genomics UK (COG-UK) Consortium | The Lighthouse Lab in Milton Keynes and Alex Alderton, Roberto Amato, Jeffrey Barrett, Sonia Goncalves, Ewan Harrison, David K. Jackson, Ian Johnston, Dominic Kwiatkowski, Cordelia Langford, John Sillitoe on behalf of the Wellcome Sanger Institute COVID-19 Surveillance Team |
| EPI_ISL_3470356 | Berkshire and Surrey Pathology Services Lighthouse Laboratory | Wellcome Sanger Institute for the COVID-19 Genomics UK (COG-UK) Consortium | Berkshire and Surrey Pathology Services Lighthouse Laboratory and Alex Alderton, Roberto Amato, Jeffrey Barrett, Sonia Goncalves, Ewan Harrison, David K. Jackson, Ian Johnston, Dominic Kwiatkowski, Cordelia Langford, John Sillitoe on behalf of the Wellcome Sanger Institute COVID-19 Surveillance Team |
| EPI_ISL_3470357, EPI_ISL_3470358 | Lighthouse Lab in Milton Keynes | Wellcome Sanger Institute for the COVID-19 Genomics UK (COG-UK) Consortium | The Lighthouse Lab in Milton Keynes and Alex Alderton, Roberto Amato, Jeffrey Barrett, Sonia Goncalves, Ewan Harrison, David K. Jackson, Ian Johnston, Dominic Kwiatkowski, Cordelia Langford, John Sillitoe on behalf of the Wellcome Sanger Institute COVID-19 Surveillance Team |
| EPI_ISL_3470359, EPI_ISL_3470360 | Berkshire and Surrey Pathology Services Lighthouse Laboratory | Wellcome Sanger Institute for the COVID-19 Genomics UK (COG-UK) Consortium | Berkshire and Surrey Pathology Services Lighthouse Laboratory and Alex Alderton, Roberto Amato, Jeffrey Barrett, Sonia Goncalves, Ewan Harrison, David K. Jackson, Ian Johnston, Dominic Kwiatkowski, Cordelia Langford, John Sillitoe on behalf of the Wellcome Sanger Institute COVID-19 Surveillance Team |
| EPI_ISL_3470361, EPI_ISL_3470362 | Lighthouse Lab in Glasgow | Wellcome Sanger Institute for the COVID-19 Genomics UK (COG-UK) Consortium | Harper VanSteenhouse, Yumi Kasai, David Gray, Carol Clugston, Anna Dominiczak and Alex Alderton, Roberto Amato, Jeffrey Barrett, Sonia Goncalves, Ewan Harrison, David K. Jackson, Ian Johnston, Dominic Kwiatkowski, Cordelia Langford, John Sillitoe on behalf of the Wellcome Sanger Institute COVID-19 Surveillance Team |
| EPI_ISL_3470363 | Berkshire and Surrey Pathology Services Lighthouse Laboratory | Wellcome Sanger Institute for the COVID-19 Genomics UK (COG-UK) Consortium | Berkshire and Surrey Pathology Services Lighthouse Laboratory and Alex Alderton, Roberto Amato, Jeffrey Barrett, Sonia Goncalves, Ewan Harrison, David K. Jackson, Ian Johnston, Dominic Kwiatkowski, Cordelia Langford, John Sillitoe on behalf of the Wellcome Sanger Institute COVID-19 Surveillance Team |
| EPI_ISL_3470364 | Lighthouse Lab in Milton Keynes | Wellcome Sanger Institute for the COVID-19 Genomics UK (COG-UK) Consortium | The Lighthouse Lab in Milton Keynes and Alex Alderton, Roberto Amato, Jeffrey Barrett, Sonia Goncalves, Ewan Harrison, David K. Jackson, Ian Johnston, Dominic Kwiatkowski, Cordelia Langford, John Sillitoe on behalf of the Wellcome Sanger Institute COVID-19 Surveillance Team |
| EPI_ISL_3470365 | Lighthouse Lab in Glasgow | Wellcome Sanger Institute for the COVID-19 Genomics UK (COG-UK) Consortium | Harper VanSteenhouse, Yumi Kasai, David Gray, Carol Clugston, Anna Dominiczak and Alex Alderton, Roberto Amato, Jeffrey Barrett, Sonia Goncalves, Ewan Harrison, David K. Jackson, Ian Johnston, Dominic Kwiatkowski, Cordelia Langford, John Sillitoe on behalf of the Wellcome Sanger Institute COVID-19 Surveillance Team |
| EPI_ISL_3470366, EPI_ISL_3470367 | Berkshire and Surrey Pathology Services Lighthouse Laboratory | Wellcome Sanger Institute for the COVID-19 Genomics UK (COG-UK) Consortium | Berkshire and Surrey Pathology Services Lighthouse Laboratory and Alex Alderton, Roberto Amato, Jeffrey Barrett, Sonia Goncalves, Ewan Harrison, David K. Jackson, Ian Johnston, Dominic Kwiatkowski, Cordelia Langford, John Sillitoe on behalf of the Wellcome Sanger Institute COVID-19 Surveillance Team |
| EPI_ISL_3470368 | Lighthouse Lab in Glasgow | Wellcome Sanger Institute for the COVID-19 Genomics UK (COG-UK) Consortium | Harper VanSteenhouse, Yumi Kasai, David Gray, Carol Clugston, Anna Dominiczak and Alex Alderton, Roberto Amato, Jeffrey Barrett, Sonia Goncalves, Ewan Harrison, David K. Jackson, Ian Johnston, Dominic Kwiatkowski, Cordelia Langford, John Sillitoe on behalf of the Wellcome Sanger Institute COVID-19 Surveillance Team |
| EPI_ISL_3470369 | Lighthouse Lab in Milton Keynes | Wellcome Sanger Institute for the COVID-19 Genomics UK (COG-UK) Consortium | The Lighthouse Lab in Milton Keynes and Alex Alderton, Roberto Amato, Jeffrey Barrett, Sonia Goncalves, Ewan Harrison, David K. Jackson, Ian Johnston, Dominic Kwiatkowski, Cordelia Langford, John Sillitoe on behalf of the Wellcome Sanger Institute COVID-19 Surveillance Team |
| EPI_ISL_3470370 | Berkshire and Surrey Pathology Services Lighthouse Laboratory | Wellcome Sanger Institute for the COVID-19 Genomics UK (COG-UK) Consortium | Berkshire and Surrey Pathology Services Lighthouse Laboratory and Alex Alderton, Roberto Amato, Jeffrey Barrett, Sonia Goncalves, Ewan Harrison, David K. Jackson, Ian Johnston, Dominic Kwiatkowski, Cordelia Langford, John Sillitoe on behalf of the Wellcome Sanger Institute COVID-19 Surveillance Team |
| EPI_ISL_3470371, EPI_ISL_3470372, EPI_ISL_3470373, EPI_ISL_3470374 | Lighthouse Lab in Milton Keynes | Wellcome Sanger Institute for the COVID-19 Genomics UK (COG-UK) Consortium | The Lighthouse Lab in Milton Keynes and Alex Alderton, Roberto Amato, Jeffrey Barrett, Sonia Goncalves, Ewan Harrison, David K. Jackson, Ian Johnston, Dominic Kwiatkowski, Cordelia Langford, John Sillitoe on behalf of the Wellcome Sanger Institute COVID-19 Surveillance Team |
| EPI_ISL_3470375 | Lighthouse Lab in Glasgow | Wellcome Sanger Institute for the COVID-19 Genomics UK (COG-UK) Consortium | Harper VanSteenhouse, Yumi Kasai, David Gray, Carol Clugston, Anna Dominiczak and Alex Alderton, Roberto Amato, Jeffrey Barrett, Sonia Goncalves, Ewan Harrison, David K. Jackson, Ian Johnston, Dominic Kwiatkowski, Cordelia Langford, John Sillitoe on behalf of the Wellcome Sanger Institute COVID-19 Surveillance Team |
| EPI_ISL_3470376 | Berkshire and Surrey Pathology Services Lighthouse Laboratory | Wellcome Sanger Institute for the COVID-19 Genomics UK (COG-UK) Consortium | Berkshire and Surrey Pathology Services Lighthouse Laboratory and Alex Alderton, Roberto Amato, Jeffrey Barrett, Sonia Goncalves, Ewan Harrison, David K. Jackson, Ian Johnston, Dominic Kwiatkowski, Cordelia Langford, John Sillitoe on behalf of the Wellcome Sanger Institute COVID-19 Surveillance Team |
| EPI_ISL_3470377 | Lighthouse Lab in Milton Keynes | Wellcome Sanger Institute for the COVID-19 Genomics UK (COG-UK) Consortium | The Lighthouse Lab in Milton Keynes and Alex Alderton, Roberto Amato, Jeffrey Barrett, Sonia Goncalves, Ewan Harrison, David K. Jackson, Ian Johnston, Dominic Kwiatkowski, Cordelia Langford, John Sillitoe on behalf of the Wellcome Sanger Institute COVID-19 Surveillance Team |
| EPI_ISL_3470378 | Berkshire and Surrey Pathology Services Lighthouse Laboratory | Wellcome Sanger Institute for the COVID-19 Genomics UK (COG-UK) Consortium | Berkshire and Surrey Pathology Services Lighthouse Laboratory and Alex Alderton, Roberto Amato, Jeffrey Barrett, Sonia Goncalves, Ewan Harrison, David K. Jackson, Ian Johnston, Dominic Kwiatkowski, Cordelia Langford, John Sillitoe on behalf of the Wellcome Sanger Institute COVID-19 Surveillance Team |
| EPI_ISL_3470379, EPI_ISL_3470380 | Lighthouse Lab in Milton Keynes | Wellcome Sanger Institute for the COVID-19 Genomics UK (COG-UK) Consortium | The Lighthouse Lab in Milton Keynes and Alex Alderton, Roberto Amato, Jeffrey Barrett, Sonia Goncalves, Ewan Harrison, David K. Jackson, Ian Johnston, Dominic Kwiatkowski, Cordelia Langford, John Sillitoe on behalf of the Wellcome Sanger Institute COVID-19 Surveillance Team |
| EPI_ISL_3470381 | Berkshire and Surrey Pathology Services Lighthouse Laboratory | Wellcome Sanger Institute for the COVID-19 Genomics UK (COG-UK) | Berkshire and Surrey Pathology Services Lighthouse Laboratory and Alex Alderton, Roberto Amato, Jeffrey Barrett, Sonia Goncalves, Ewan Harrison, David K. Jackson, Ian Johnston, Dominic Kwiatkowski, Cordelia Langford, John Sillitoe on behalf |

| | | | |
|---|---|---|---|
| | | Consortium | of the Wellcome Sanger Institute COVID-19 Surveillance Team |
| EPI_ISL_3470382 | Lighthouse Lab in Glasgow | Wellcome Sanger Institute for the COVID-19 Genomics UK (COG-UK) Consortium | Harper VanSteenhouse, Yumi Kasai, David Gray, Carol Clugston, Anna Dominiczak and Alex Alderton, Roberto Amato, Jeffrey Barrett, Sonia Goncalves, Ewan Harrison, David K. Jackson, Ian Johnston, Dominic Kwiatkowski, Cordelia Langford, John Sillitoe on behalf of the Wellcome Sanger Institute COVID-19 Surveillance Team |
| EPI_ISL_3470383, EPI_ISL_3470384, EPI_ISL_3470385, EPI_ISL_3470386, EPI_ISL_3470387 | Berkshire and Surrey Pathology Services Lighthouse Laboratory | Wellcome Sanger Institute for the COVID-19 Genomics UK (COG-UK) Consortium | Berkshire and Surrey Pathology Services Lighthouse Laboratory and Alex Alderton, Roberto Amato, Jeffrey Barrett, Sonia Goncalves, Ewan Harrison, David K. Jackson, Ian Johnston, Dominic Kwiatkowski, Cordelia Langford, John Sillitoe on behalf of the Wellcome Sanger Institute COVID-19 Surveillance Team |
| EPI_ISL_3470388 | Lighthouse Lab in Glasgow | Wellcome Sanger Institute for the COVID-19 Genomics UK (COG-UK) Consortium | Harper VanSteenhouse, Yumi Kasai, David Gray, Carol Clugston, Anna Dominiczak and Alex Alderton, Roberto Amato, Jeffrey Barrett, Sonia Goncalves, Ewan Harrison, David K. Jackson, Ian Johnston, Dominic Kwiatkowski, Cordelia Langford, John Sillitoe on behalf of the Wellcome Sanger Institute COVID-19 Surveillance Team |
| EPI_ISL_3470389 | Berkshire and Surrey Pathology Services Lighthouse Laboratory | Wellcome Sanger Institute for the COVID-19 Genomics UK (COG-UK) Consortium | Berkshire and Surrey Pathology Services Lighthouse Laboratory and Alex Alderton, Roberto Amato, Jeffrey Barrett, Sonia Goncalves, Ewan Harrison, David K. Jackson, Ian Johnston, Dominic Kwiatkowski, Cordelia Langford, John Sillitoe on behalf of the Wellcome Sanger Institute COVID-19 Surveillance Team |
| EPI_ISL_3470390, EPI_ISL_3470391 | Lighthouse Lab in Glasgow | Wellcome Sanger Institute for the COVID-19 Genomics UK (COG-UK) Consortium | Harper VanSteenhouse, Yumi Kasai, David Gray, Carol Clugston, Anna Dominiczak and Alex Alderton, Roberto Amato, Jeffrey Barrett, Sonia Goncalves, Ewan Harrison, David K. Jackson, Ian Johnston, Dominic Kwiatkowski, Cordelia Langford, John Sillitoe on behalf of the Wellcome Sanger Institute COVID-19 Surveillance Team |
| EPI_ISL_3470392, EPI_ISL_3470393 | Lighthouse Lab in Milton Keynes | Wellcome Sanger Institute for the COVID-19 Genomics UK (COG-UK) Consortium | The Lighthouse Lab in Milton Keynes and Alex Alderton, Roberto Amato, Jeffrey Barrett, Sonia Goncalves, Ewan Harrison, David K. Jackson, Ian Johnston, Dominic Kwiatkowski, Cordelia Langford, John Sillitoe on behalf of the Wellcome Sanger Institute COVID-19 Surveillance Team |
| EPI_ISL_3470394, EPI_ISL_3470395, EPI_ISL_3470396 | Lighthouse Lab in Glasgow | Wellcome Sanger Institute for the COVID-19 Genomics UK (COG-UK) Consortium | Harper VanSteenhouse, Yumi Kasai, David Gray, Carol Clugston, Anna Dominiczak and Alex Alderton, Roberto Amato, Jeffrey Barrett, Sonia Goncalves, Ewan Harrison, David K. Jackson, Ian Johnston, Dominic Kwiatkowski, Cordelia Langford, John Sillitoe on behalf of the Wellcome Sanger Institute COVID-19 Surveillance Team |
| EPI_ISL_3470397, EPI_ISL_3470398 | Lighthouse Lab in Milton Keynes | Wellcome Sanger Institute for the COVID-19 Genomics UK (COG-UK) Consortium | The Lighthouse Lab in Milton Keynes and Alex Alderton, Roberto Amato, Jeffrey Barrett, Sonia Goncalves, Ewan Harrison, David K. Jackson, Ian Johnston, Dominic Kwiatkowski, Cordelia Langford, John Sillitoe on behalf of the Wellcome Sanger Institute COVID-19 Surveillance Team |
| EPI_ISL_3470399 | Lighthouse Lab in Glasgow | Wellcome Sanger Institute for the COVID-19 Genomics UK (COG-UK) Consortium | Harper VanSteenhouse, Yumi Kasai, David Gray, Carol Clugston, Anna Dominiczak and Alex Alderton, Roberto Amato, Jeffrey Barrett, Sonia Goncalves, Ewan Harrison, David K. Jackson, Ian Johnston, Dominic Kwiatkowski, Cordelia Langford, John Sillitoe on behalf of the Wellcome Sanger Institute COVID-19 Surveillance Team |
| EPI_ISL_3470400, EPI_ISL_3470401 | Berkshire and Surrey Pathology Services Lighthouse Laboratory | Wellcome Sanger Institute for the COVID-19 Genomics UK (COG-UK) Consortium | Berkshire and Surrey Pathology Services Lighthouse Laboratory and Alex Alderton, Roberto Amato, Jeffrey Barrett, Sonia Goncalves, Ewan Harrison, David K. Jackson, Ian Johnston, Dominic Kwiatkowski, Cordelia Langford, John Sillitoe on behalf of the Wellcome Sanger Institute COVID-19 Surveillance Team |
| EPI_ISL_3470402 | Lighthouse Lab in Milton Keynes | Wellcome Sanger Institute for the COVID-19 Genomics UK (COG-UK) Consortium | The Lighthouse Lab in Milton Keynes and Alex Alderton, Roberto Amato, Jeffrey Barrett, Sonia Goncalves, Ewan Harrison, David K. Jackson, Ian Johnston, Dominic Kwiatkowski, Cordelia Langford, John Sillitoe on behalf of the Wellcome Sanger Institute COVID-19 Surveillance Team |
| EPI_ISL_3470403, EPI_ISL_3470404, EPI_ISL_3470405 | Lighthouse Lab in Glasgow | Wellcome Sanger Institute for the COVID-19 Genomics UK (COG-UK) Consortium | Harper VanSteenhouse, Yumi Kasai, David Gray, Carol Clugston, Anna Dominiczak and Alex Alderton, Roberto Amato, Jeffrey Barrett, Sonia Goncalves, Ewan Harrison, David K. Jackson, Ian Johnston, Dominic Kwiatkowski, Cordelia Langford, John Sillitoe on behalf of the Wellcome Sanger Institute COVID-19 Surveillance Team |
| EPI_ISL_3470406 | Berkshire and Surrey Pathology Services Lighthouse Laboratory | Wellcome Sanger Institute for the COVID-19 Genomics UK (COG-UK) Consortium | Berkshire and Surrey Pathology Services Lighthouse Laboratory and Alex Alderton, Roberto Amato, Jeffrey Barrett, Sonia Goncalves, Ewan Harrison, David K. Jackson, Ian Johnston, Dominic Kwiatkowski, Cordelia Langford, John Sillitoe on behalf of the Wellcome Sanger Institute COVID-19 Surveillance Team |
| EPI_ISL_3470407 | Lighthouse Lab in Glasgow | Wellcome Sanger Institute for the COVID-19 Genomics UK (COG-UK) Consortium | Harper VanSteenhouse, Yumi Kasai, David Gray, Carol Clugston, Anna Dominiczak and Alex Alderton, Roberto Amato, Jeffrey Barrett, Sonia Goncalves, Ewan Harrison, David K. Jackson, Ian Johnston, Dominic Kwiatkowski, Cordelia Langford, John Sillitoe on behalf of the Wellcome Sanger Institute COVID-19 Surveillance Team |
| EPI_ISL_3470408 | Berkshire and Surrey Pathology Services Lighthouse Laboratory | Wellcome Sanger Institute for the COVID-19 Genomics UK (COG-UK) Consortium | Berkshire and Surrey Pathology Services Lighthouse Laboratory and Alex Alderton, Roberto Amato, Jeffrey Barrett, Sonia Goncalves, Ewan Harrison, David K. Jackson, Ian Johnston, Dominic Kwiatkowski, Cordelia Langford, John Sillitoe on behalf of the Wellcome Sanger Institute COVID-19 Surveillance Team |
| EPI_ISL_3470409 | Lighthouse Lab in Milton Keynes | Wellcome Sanger Institute for the COVID-19 Genomics UK (COG-UK) Consortium | The Lighthouse Lab in Milton Keynes and Alex Alderton, Roberto Amato, Jeffrey Barrett, Sonia Goncalves, Ewan Harrison, David K. Jackson, Ian Johnston, Dominic Kwiatkowski, Cordelia Langford, John Sillitoe on behalf of the Wellcome Sanger Institute COVID-19 Surveillance Team |
| EPI_ISL_3470410 | Lighthouse Lab in Glasgow | Wellcome Sanger Institute for the COVID-19 Genomics UK (COG-UK) Consortium | Harper VanSteenhouse, Yumi Kasai, David Gray, Carol Clugston, Anna Dominiczak and Alex Alderton, Roberto Amato, Jeffrey Barrett, Sonia Goncalves, Ewan Harrison, David K. Jackson, Ian Johnston, Dominic Kwiatkowski, Cordelia Langford, John Sillitoe on behalf of the Wellcome Sanger Institute COVID-19 Surveillance Team |
| EPI_ISL_3470411 | Lighthouse Lab in Milton Keynes | Wellcome Sanger Institute for the COVID-19 Genomics UK (COG-UK) Consortium | The Lighthouse Lab in Milton Keynes and Alex Alderton, Roberto Amato, Jeffrey Barrett, Sonia Goncalves, Ewan Harrison, David K. Jackson, Ian Johnston, Dominic Kwiatkowski, Cordelia Langford, John Sillitoe on behalf of the Wellcome Sanger Institute COVID-19 Surveillance Team |
| EPI_ISL_3470412 | Berkshire and Surrey Pathology Services Lighthouse Laboratory | Wellcome Sanger Institute for the COVID-19 Genomics UK (COG-UK) Consortium | Berkshire and Surrey Pathology Services Lighthouse Laboratory and Alex Alderton, Roberto Amato, Jeffrey Barrett, Sonia Goncalves, Ewan Harrison, David K. Jackson, Ian Johnston, Dominic Kwiatkowski, Cordelia Langford, John Sillitoe on behalf of the Wellcome Sanger Institute COVID-19 Surveillance Team |
| EPI_ISL_3470413 | Lighthouse Lab in Glasgow | Wellcome Sanger Institute for the COVID-19 Genomics UK (COG-UK) Consortium | Harper VanSteenhouse, Yumi Kasai, David Gray, Carol Clugston, Anna Dominiczak and Alex Alderton, Roberto Amato, Jeffrey Barrett, Sonia Goncalves, Ewan Harrison, David K. Jackson, Ian Johnston, Dominic Kwiatkowski, Cordelia Langford, John Sillitoe on behalf of the Wellcome Sanger Institute COVID-19 Surveillance Team |
| EPI_ISL_3470414 | Berkshire and Surrey Pathology Services Lighthouse Laboratory | Wellcome Sanger Institute for the COVID-19 Genomics UK (COG-UK) Consortium | Berkshire and Surrey Pathology Services Lighthouse Laboratory and Alex Alderton, Roberto Amato, Jeffrey Barrett, Sonia Goncalves, Ewan Harrison, David K. Jackson, Ian Johnston, Dominic Kwiatkowski, Cordelia Langford, John Sillitoe on behalf of the Wellcome Sanger Institute COVID-19 Surveillance Team |
| EPI_ISL_3470415 | Lighthouse Lab in Glasgow | Wellcome Sanger Institute for the COVID-19 Genomics UK (COG-UK) Consortium | Harper VanSteenhouse, Yumi Kasai, David Gray, Carol Clugston, Anna Dominiczak and Alex Alderton, Roberto Amato, Jeffrey Barrett, Sonia Goncalves, Ewan Harrison, David K. Jackson, Ian Johnston, Dominic Kwiatkowski, Cordelia Langford, John Sillitoe on behalf of the Wellcome Sanger Institute COVID-19 Surveillance Team |
| EPI_ISL_3470416 | Berkshire and Surrey Pathology Services Lighthouse Laboratory | Wellcome Sanger Institute for the COVID-19 Genomics UK (COG-UK) Consortium | Berkshire and Surrey Pathology Services Lighthouse Laboratory and Alex Alderton, Roberto Amato, Jeffrey Barrett, Sonia Goncalves, Ewan Harrison, David K. Jackson, Ian Johnston, Dominic Kwiatkowski, Cordelia Langford, John Sillitoe on behalf of the Wellcome Sanger Institute COVID-19 Surveillance Team |
| EPI_ISL_3470417, EPI_ISL_3470418, EPI_ISL_3470419, EPI_ISL_3470420, EPI_ISL_3470421, EPI_ISL_3470422, EPI_ISL_3470423, EPI_ISL_3470424, EPI_ISL_3470425 | Lighthouse Lab in Milton Keynes | Wellcome Sanger Institute for the COVID-19 Genomics UK (COG-UK) Consortium | The Lighthouse Lab in Milton Keynes and Alex Alderton, Roberto Amato, Jeffrey Barrett, Sonia Goncalves, Ewan Harrison, David K. Jackson, Ian Johnston, Dominic Kwiatkowski, Cordelia Langford, John Sillitoe on behalf of the Wellcome Sanger Institute COVID-19 Surveillance Team |
| EPI_ISL_3470426, EPI_ISL_3470427 | Lighthouse Lab in Glasgow | Wellcome Sanger Institute for the COVID-19 Genomics UK (COG-UK) Consortium | Harper VanSteenhouse, Yumi Kasai, David Gray, Carol Clugston, Anna Dominiczak and Alex Alderton, Roberto Amato, Jeffrey Barrett, Sonia Goncalves, Ewan Harrison, David K. Jackson, Ian Johnston, Dominic Kwiatkowski, Cordelia Langford, John Sillitoe on behalf of the Wellcome Sanger Institute COVID-19 Surveillance Team |
| EPI_ISL_3470428 | Berkshire and Surrey Pathology Services Lighthouse Laboratory | Wellcome Sanger Institute for the COVID-19 Genomics UK (COG-UK) Consortium | Berkshire and Surrey Pathology Services Lighthouse Laboratory and Alex Alderton, Roberto Amato, Jeffrey Barrett, Sonia Goncalves, Ewan Harrison, David K. Jackson, Ian Johnston, Dominic Kwiatkowski, Cordelia Langford, John Sillitoe on behalf of the Wellcome Sanger Institute COVID-19 Surveillance Team |
| EPI_ISL_3470429 | Lighthouse Lab in Glasgow | Wellcome Sanger Institute for the COVID-19 Genomics UK (COG-UK) Consortium | Harper VanSteenhouse, Yumi Kasai, David Gray, Carol Clugston, Anna Dominiczak and Alex Alderton, Roberto Amato, Jeffrey Barrett, Sonia Goncalves, Ewan Harrison, David K. Jackson, Ian Johnston, Dominic Kwiatkowski, Cordelia Langford, John Sillitoe on behalf of the Wellcome Sanger Institute COVID-19 Surveillance Team |
| EPI_ISL_3470430, EPI_ISL_3470431, EPI_ISL_3470432 | Lighthouse Lab in Milton Keynes | Wellcome Sanger Institute for the COVID-19 Genomics UK (COG-UK) Consortium | The Lighthouse Lab in Milton Keynes and Alex Alderton, Roberto Amato, Jeffrey Barrett, Sonia Goncalves, Ewan Harrison, David K. Jackson, Ian Johnston, Dominic Kwiatkowski, Cordelia Langford, John Sillitoe on behalf of the Wellcome Sanger Institute COVID-19 Surveillance Team |
| EPI_ISL_3470433 | Lighthouse Lab in Glasgow | Wellcome Sanger Institute for the COVID-19 Genomics UK (COG-UK) Consortium | Harper VanSteenhouse, Yumi Kasai, David Gray, Carol Clugston, Anna Dominiczak and Alex Alderton, Roberto Amato, Jeffrey Barrett, Sonia Goncalves, Ewan Harrison, David K. Jackson, Ian Johnston, Dominic Kwiatkowski, Cordelia Langford, John Sillitoe on behalf of the Wellcome Sanger Institute COVID-19 Surveillance Team |
| EPI_ISL_3470434 | Berkshire and Surrey Pathology Services Lighthouse Laboratory | Wellcome Sanger Institute for the COVID-19 Genomics UK (COG-UK) Consortium | Berkshire and Surrey Pathology Services Lighthouse Laboratory and Alex Alderton, Roberto Amato, Jeffrey Barrett, Sonia Goncalves, Ewan Harrison, David K. Jackson, Ian Johnston, Dominic Kwiatkowski, Cordelia Langford, John Sillitoe on behalf of the Wellcome Sanger Institute COVID-19 Surveillance Team |
| EPI_ISL_3470435, EPI_ISL_3470436 | Lighthouse Lab in Glasgow | Wellcome Sanger Institute for the COVID-19 Genomics UK (COG-UK) Consortium | Harper VanSteenhouse, Yumi Kasai, David Gray, Carol Clugston, Anna Dominiczak and Alex Alderton, Roberto Amato, Jeffrey Barrett, Sonia Goncalves, Ewan Harrison, David K. Jackson, Ian Johnston, Dominic Kwiatkowski, Cordelia Langford, John Sillitoe on behalf of the Wellcome Sanger Institute COVID-19 Surveillance Team |
| EPI_ISL_3470437 | Lighthouse Lab in Milton Keynes | Wellcome Sanger Institute for the COVID-19 Genomics UK (COG-UK) Consortium | The Lighthouse Lab in Milton Keynes and Alex Alderton, Roberto Amato, Jeffrey Barrett, Sonia Goncalves, Ewan Harrison, David K. Jackson, Ian Johnston, Dominic Kwiatkowski, Cordelia Langford, John Sillitoe on behalf of the Wellcome Sanger Institute COVID-19 Surveillance Team |
| EPI_ISL_3470438, EPI_ISL_3470439 | Lighthouse Lab in Glasgow | Wellcome Sanger Institute for the COVID-19 Genomics UK (COG-UK) Consortium | Harper VanSteenhouse, Yumi Kasai, David Gray, Carol Clugston, Anna Dominiczak and Alex Alderton, Roberto Amato, Jeffrey Barrett, Sonia Goncalves, Ewan Harrison, David K. Jackson, Ian Johnston, Dominic Kwiatkowski, Cordelia Langford, John Sillitoe on behalf of the Wellcome Sanger Institute COVID-19 Surveillance Team |
| EPI_ISL_3470440 | Lighthouse Lab in Milton Keynes | Wellcome Sanger Institute for the COVID-19 Genomics UK (COG-UK) Consortium | The Lighthouse Lab in Milton Keynes and Alex Alderton, Roberto Amato, Jeffrey Barrett, Sonia Goncalves, Ewan Harrison, David K. Jackson, Ian Johnston, Dominic Kwiatkowski, Cordelia Langford, John Sillitoe on behalf of the Wellcome Sanger Institute COVID-19 Surveillance Team |
| EPI_ISL_3470441 | Lighthouse Lab in Glasgow | Wellcome Sanger Institute for the COVID-19 Genomics UK (COG-UK) Consortium | Harper VanSteenhouse, Yumi Kasai, David Gray, Carol Clugston, Anna Dominiczak and Alex Alderton, Roberto Amato, Jeffrey Barrett, Sonia Goncalves, Ewan Harrison, David K. Jackson, Ian Johnston, Dominic Kwiatkowski, Cordelia Langford, John Sillitoe on behalf of the Wellcome Sanger Institute COVID-19 Surveillance Team |
| EPI_ISL_3470442 | Berkshire and Surrey Pathology Services Lighthouse Laboratory | Wellcome Sanger Institute for the COVID-19 Genomics UK (COG-UK) Consortium | Berkshire and Surrey Pathology Services Lighthouse Laboratory and Alex Alderton, Roberto Amato, Jeffrey Barrett, Sonia Goncalves, Ewan Harrison, David K. Jackson, Ian Johnston, Dominic Kwiatkowski, Cordelia Langford, John Sillitoe on behalf of the Wellcome Sanger Institute COVID-19 Surveillance Team |
| EPI_ISL_3470443 | Lighthouse Lab in Glasgow | Wellcome Sanger Institute for the COVID-19 Genomics UK (COG-UK) Consortium | Harper VanSteenhouse, Yumi Kasai, David Gray, Carol Clugston, Anna Dominiczak and Alex Alderton, Roberto Amato, Jeffrey Barrett, Sonia Goncalves, Ewan Harrison, David K. Jackson, Ian Johnston, Dominic Kwiatkowski, Cordelia Langford, John Sillitoe on behalf of the Wellcome Sanger Institute COVID-19 Surveillance Team |
| EPI_ISL_3470444 | Berkshire and Surrey Pathology Services Lighthouse Laboratory | Wellcome Sanger Institute for the COVID-19 Genomics UK (COG-UK) Consortium | Berkshire and Surrey Pathology Services Lighthouse Laboratory and Alex Alderton, Roberto Amato, Jeffrey Barrett, Sonia Goncalves, Ewan Harrison, David K. Jackson, Ian Johnston, Dominic Kwiatkowski, Cordelia Langford, John Sillitoe on behalf of the Wellcome Sanger Institute COVID-19 Surveillance Team |
| EPI_ISL_3470445 | Lighthouse Lab in Glasgow | Wellcome Sanger Institute for the COVID-19 Genomics UK (COG-UK) Consortium | Harper VanSteenhouse, Yumi Kasai, David Gray, Carol Clugston, Anna Dominiczak and Alex Alderton, Roberto Amato, Jeffrey Barrett, Sonia Goncalves, Ewan Harrison, David K. Jackson, Ian Johnston, Dominic Kwiatkowski, Cordelia Langford, John Sillitoe on behalf of the Wellcome Sanger Institute COVID-19 Surveillance Team |

| | | | |
|---|---|---|---|
| EPI_ISL_3470446, EPI_ISL_3470447, EPI_ISL_3470448 | Lighthouse Lab in Milton Keynes | Wellcome Sanger Institute for the COVID-19 Genomics UK (COG-UK) Consortium | The Lighthouse Lab in Milton Keynes and Alex Alderton, Roberto Amato, Jeffrey Barrett, Sonia Goncalves, Ewan Harrison, David K. Jackson, Ian Johnston, Dominic Kwiatkowski, Cordelia Langford, John Sillitoe on behalf of the Wellcome Sanger Institute COVID-19 Surveillance Team |
| EPI_ISL_3470449, EPI_ISL_3470450, EPI_ISL_3470451 | Lighthouse Lab in Glasgow | Wellcome Sanger Institute for the COVID-19 Genomics UK (COG-UK) Consortium | Harper VanSteenhouse, Yumi Kasai, David Gray, Carol Clugston, Anna Dominiczak and Alex Alderton, Roberto Amato, Jeffrey Barrett, Sonia Goncalves, Ewan Harrison, David K. Jackson, Ian Johnston, Dominic Kwiatkowski, Cordelia Langford, John Sillitoe on behalf of the Wellcome Sanger Institute COVID-19 Surveillance Team |
| EPI_ISL_3470452, EPI_ISL_3470453 | Lighthouse Lab in Milton Keynes | Wellcome Sanger Institute for the COVID-19 Genomics UK (COG-UK) Consortium | The Lighthouse Lab in Milton Keynes and Alex Alderton, Roberto Amato, Jeffrey Barrett, Sonia Goncalves, Ewan Harrison, David K. Jackson, Ian Johnston, Dominic Kwiatkowski, Cordelia Langford, John Sillitoe on behalf of the Wellcome Sanger Institute COVID-19 Surveillance Team |
| EPI_ISL_3470454 | Berkshire and Surrey Pathology Services Lighthouse Laboratory | Wellcome Sanger Institute for the COVID-19 Genomics UK (COG-UK) Consortium | Berkshire and Surrey Pathology Services Lighthouse Laboratory and Alex Alderton, Roberto Amato, Jeffrey Barrett, Sonia Goncalves, Ewan Harrison, David K. Jackson, Ian Johnston, Dominic Kwiatkowski, Cordelia Langford, John Sillitoe on behalf of the Wellcome Sanger Institute COVID-19 Surveillance Team |
| EPI_ISL_3470455 | Lighthouse Lab in Milton Keynes | Wellcome Sanger Institute for the COVID-19 Genomics UK (COG-UK) Consortium | The Lighthouse Lab in Milton Keynes and Alex Alderton, Roberto Amato, Jeffrey Barrett, Sonia Goncalves, Ewan Harrison, David K. Jackson, Ian Johnston, Dominic Kwiatkowski, Cordelia Langford, John Sillitoe on behalf of the Wellcome Sanger Institute COVID-19 Surveillance Team |
| EPI_ISL_3470456 | Berkshire and Surrey Pathology Services Lighthouse Laboratory | Wellcome Sanger Institute for the COVID-19 Genomics UK (COG-UK) Consortium | Berkshire and Surrey Pathology Services Lighthouse Laboratory and Alex Alderton, Roberto Amato, Jeffrey Barrett, Sonia Goncalves, Ewan Harrison, David K. Jackson, Ian Johnston, Dominic Kwiatkowski, Cordelia Langford, John Sillitoe on behalf of the Wellcome Sanger Institute COVID-19 Surveillance Team |
| EPI_ISL_3470457 | Lighthouse Lab in Milton Keynes | Wellcome Sanger Institute for the COVID-19 Genomics UK (COG-UK) Consortium | The Lighthouse Lab in Milton Keynes and Alex Alderton, Roberto Amato, Jeffrey Barrett, Sonia Goncalves, Ewan Harrison, David K. Jackson, Ian Johnston, Dominic Kwiatkowski, Cordelia Langford, John Sillitoe on behalf of the Wellcome Sanger Institute COVID-19 Surveillance Team |
| EPI_ISL_3470458 | Lighthouse Lab in Glasgow | Wellcome Sanger Institute for the COVID-19 Genomics UK (COG-UK) Consortium | Harper VanSteenhouse, Yumi Kasai, David Gray, Carol Clugston, Anna Dominiczak and Alex Alderton, Roberto Amato, Jeffrey Barrett, Sonia Goncalves, Ewan Harrison, David K. Jackson, Ian Johnston, Dominic Kwiatkowski, Cordelia Langford, John Sillitoe on behalf of the Wellcome Sanger Institute COVID-19 Surveillance Team |
| EPI_ISL_3470459 | Berkshire and Surrey Pathology Services Lighthouse Laboratory | Wellcome Sanger Institute for the COVID-19 Genomics UK (COG-UK) Consortium | Berkshire and Surrey Pathology Services Lighthouse Laboratory and Alex Alderton, Roberto Amato, Jeffrey Barrett, Sonia Goncalves, Ewan Harrison, David K. Jackson, Ian Johnston, Dominic Kwiatkowski, Cordelia Langford, John Sillitoe on behalf of the Wellcome Sanger Institute COVID-19 Surveillance Team |
| EPI_ISL_3470460, EPI_ISL_3470461 | Lighthouse Lab in Glasgow | Wellcome Sanger Institute for the COVID-19 Genomics UK (COG-UK) Consortium | Harper VanSteenhouse, Yumi Kasai, David Gray, Carol Clugston, Anna Dominiczak and Alex Alderton, Roberto Amato, Jeffrey Barrett, Sonia Goncalves, Ewan Harrison, David K. Jackson, Ian Johnston, Dominic Kwiatkowski, Cordelia Langford, John Sillitoe on behalf of the Wellcome Sanger Institute COVID-19 Surveillance Team |
| EPI_ISL_3470462 | Berkshire and Surrey Pathology Services Lighthouse Laboratory | Wellcome Sanger Institute for the COVID-19 Genomics UK (COG-UK) Consortium | Berkshire and Surrey Pathology Services Lighthouse Laboratory and Alex Alderton, Roberto Amato, Jeffrey Barrett, Sonia Goncalves, Ewan Harrison, David K. Jackson, Ian Johnston, Dominic Kwiatkowski, Cordelia Langford, John Sillitoe on behalf of the Wellcome Sanger Institute COVID-19 Surveillance Team |
| EPI_ISL_3470463 | Lighthouse Lab in Milton Keynes | Wellcome Sanger Institute for the COVID-19 Genomics UK (COG-UK) Consortium | The Lighthouse Lab in Milton Keynes and Alex Alderton, Roberto Amato, Jeffrey Barrett, Sonia Goncalves, Ewan Harrison, David K. Jackson, Ian Johnston, Dominic Kwiatkowski, Cordelia Langford, John Sillitoe on behalf of the Wellcome Sanger Institute COVID-19 Surveillance Team |
| EPI_ISL_3470464 | Lighthouse Lab in Glasgow | Wellcome Sanger Institute for the COVID-19 Genomics UK (COG-UK) Consortium | Harper VanSteenhouse, Yumi Kasai, David Gray, Carol Clugston, Anna Dominiczak and Alex Alderton, Roberto Amato, Jeffrey Barrett, Sonia Goncalves, Ewan Harrison, David K. Jackson, Ian Johnston, Dominic Kwiatkowski, Cordelia Langford, John Sillitoe on behalf of the Wellcome Sanger Institute COVID-19 Surveillance Team |
| EPI_ISL_3470465 | Lighthouse Lab in Milton Keynes | Wellcome Sanger Institute for the COVID-19 Genomics UK (COG-UK) Consortium | The Lighthouse Lab in Milton Keynes and Alex Alderton, Roberto Amato, Jeffrey Barrett, Sonia Goncalves, Ewan Harrison, David K. Jackson, Ian Johnston, Dominic Kwiatkowski, Cordelia Langford, John Sillitoe on behalf of the Wellcome Sanger Institute COVID-19 Surveillance Team |
| EPI_ISL_3470466, EPI_ISL_3470467, EPI_ISL_3470468, EPI_ISL_3470469 | Lighthouse Lab in Glasgow | Wellcome Sanger Institute for the COVID-19 Genomics UK (COG-UK) Consortium | Harper VanSteenhouse, Yumi Kasai, David Gray, Carol Clugston, Anna Dominiczak and Alex Alderton, Roberto Amato, Jeffrey Barrett, Sonia Goncalves, Ewan Harrison, David K. Jackson, Ian Johnston, Dominic Kwiatkowski, Cordelia Langford, John Sillitoe on behalf of the Wellcome Sanger Institute COVID-19 Surveillance Team |
| EPI_ISL_3470470 | Berkshire and Surrey Pathology Services Lighthouse Laboratory | Wellcome Sanger Institute for the COVID-19 Genomics UK (COG-UK) Consortium | Berkshire and Surrey Pathology Services Lighthouse Laboratory and Alex Alderton, Roberto Amato, Jeffrey Barrett, Sonia Goncalves, Ewan Harrison, David K. Jackson, Ian Johnston, Dominic Kwiatkowski, Cordelia Langford, John Sillitoe on behalf of the Wellcome Sanger Institute COVID-19 Surveillance Team |
| EPI_ISL_3470471 | Lighthouse Lab in Glasgow | Wellcome Sanger Institute for the COVID-19 Genomics UK (COG-UK) Consortium | Harper VanSteenhouse, Yumi Kasai, David Gray, Carol Clugston, Anna Dominiczak and Alex Alderton, Roberto Amato, Jeffrey Barrett, Sonia Goncalves, Ewan Harrison, David K. Jackson, Ian Johnston, Dominic Kwiatkowski, Cordelia Langford, John Sillitoe on behalf of the Wellcome Sanger Institute COVID-19 Surveillance Team |
| EPI_ISL_3470472 | Lighthouse Lab in Milton Keynes | Wellcome Sanger Institute for the COVID-19 Genomics UK (COG-UK) Consortium | The Lighthouse Lab in Milton Keynes and Alex Alderton, Roberto Amato, Jeffrey Barrett, Sonia Goncalves, Ewan Harrison, David K. Jackson, Ian Johnston, Dominic Kwiatkowski, Cordelia Langford, John Sillitoe on behalf of the Wellcome Sanger Institute COVID-19 Surveillance Team |
| EPI_ISL_3470473 | Berkshire and Surrey Pathology Services Lighthouse Laboratory | Wellcome Sanger Institute for the COVID-19 Genomics UK (COG-UK) Consortium | Berkshire and Surrey Pathology Services Lighthouse Laboratory and Alex Alderton, Roberto Amato, Jeffrey Barrett, Sonia Goncalves, Ewan Harrison, David K. Jackson, Ian Johnston, Dominic Kwiatkowski, Cordelia Langford, John Sillitoe on behalf of the Wellcome Sanger Institute COVID-19 Surveillance Team |
| EPI_ISL_3470474 | Lighthouse Lab in Milton Keynes | Wellcome Sanger Institute for the COVID-19 Genomics UK (COG-UK) Consortium | The Lighthouse Lab in Milton Keynes and Alex Alderton, Roberto Amato, Jeffrey Barrett, Sonia Goncalves, Ewan Harrison, David K. Jackson, Ian Johnston, Dominic Kwiatkowski, Cordelia Langford, John Sillitoe on behalf of the Wellcome Sanger Institute COVID-19 Surveillance Team |
| EPI_ISL_3470475, EPI_ISL_3470476 | Berkshire and Surrey Pathology Services Lighthouse Laboratory | Wellcome Sanger Institute for the COVID-19 Genomics UK (COG-UK) Consortium | Berkshire and Surrey Pathology Services Lighthouse Laboratory and Alex Alderton, Roberto Amato, Jeffrey Barrett, Sonia Goncalves, Ewan Harrison, David K. Jackson, Ian Johnston, Dominic Kwiatkowski, Cordelia Langford, John Sillitoe on behalf of the Wellcome Sanger Institute COVID-19 Surveillance Team |
| EPI_ISL_3470477 | Lighthouse Lab in Glasgow | Wellcome Sanger Institute for the COVID-19 Genomics UK (COG-UK) Consortium | Harper VanSteenhouse, Yumi Kasai, David Gray, Carol Clugston, Anna Dominiczak and Alex Alderton, Roberto Amato, Jeffrey Barrett, Sonia Goncalves, Ewan Harrison, David K. Jackson, Ian Johnston, Dominic Kwiatkowski, Cordelia Langford, John Sillitoe on behalf of the Wellcome Sanger Institute COVID-19 Surveillance Team |
| EPI_ISL_3470478 | Berkshire and Surrey Pathology Services Lighthouse Laboratory | Wellcome Sanger Institute for the COVID-19 Genomics UK (COG-UK) Consortium | Berkshire and Surrey Pathology Services Lighthouse Laboratory and Alex Alderton, Roberto Amato, Jeffrey Barrett, Sonia Goncalves, Ewan Harrison, David K. Jackson, Ian Johnston, Dominic Kwiatkowski, Cordelia Langford, John Sillitoe on behalf of the Wellcome Sanger Institute COVID-19 Surveillance Team |
| EPI_ISL_3470479, EPI_ISL_3470480 | Lighthouse Lab in Glasgow | Wellcome Sanger Institute for the COVID-19 Genomics UK (COG-UK) Consortium | Harper VanSteenhouse, Yumi Kasai, David Gray, Carol Clugston, Anna Dominiczak and Alex Alderton, Roberto Amato, Jeffrey Barrett, Sonia Goncalves, Ewan Harrison, David K. Jackson, Ian Johnston, Dominic Kwiatkowski, Cordelia Langford, John Sillitoe on behalf of the Wellcome Sanger Institute COVID-19 Surveillance Team |
| EPI_ISL_3470481, EPI_ISL_3470482, EPI_ISL_3470483 | Berkshire and Surrey Pathology Services Lighthouse Laboratory | Wellcome Sanger Institute for the COVID-19 Genomics UK (COG-UK) Consortium | Berkshire and Surrey Pathology Services Lighthouse Laboratory and Alex Alderton, Roberto Amato, Jeffrey Barrett, Sonia Goncalves, Ewan Harrison, David K. Jackson, Ian Johnston, Dominic Kwiatkowski, Cordelia Langford, John Sillitoe on behalf of the Wellcome Sanger Institute COVID-19 Surveillance Team |
| EPI_ISL_3470484, EPI_ISL_3470485, EPI_ISL_3470486, EPI_ISL_3470487, EPI_ISL_3470488, EPI_ISL_3470489, EPI_ISL_3470490, EPI_ISL_3470491, EPI_ISL_3470492, EPI_ISL_3470493, EPI_ISL_3470494, EPI_ISL_3470495, EPI_ISL_3470496, EPI_ISL_3470497, EPI_ISL_3470498, EPI_ISL_3470499, EPI_ISL_3470500, EPI_ISL_3470501, EPI_ISL_3470502, EPI_ISL_3470503, EPI_ISL_3470504, EPI_ISL_3470505, EPI_ISL_3470506, EPI_ISL_3470507 | | | |
| see above | Lighthouse Lab in Milton Keynes | Wellcome Sanger Institute for the COVID-19 Genomics UK (COG-UK) Consortium | The Lighthouse Lab in Milton Keynes and Alex Alderton, Roberto Amato, Jeffrey Barrett, Sonia Goncalves, Ewan Harrison, David K. Jackson, Ian Johnston, Dominic Kwiatkowski, Cordelia Langford, John Sillitoe on behalf of the Wellcome Sanger Institute COVID-19 Surveillance Team |
| EPI_ISL_3470508, EPI_ISL_3470509, EPI_ISL_3470510, EPI_ISL_3470511, EPI_ISL_3470512, EPI_ISL_3470513, EPI_ISL_3470514, EPI_ISL_3470515, EPI_ISL_3470516, EPI_ISL_3470517, EPI_ISL_3470518, EPI_ISL_3470519, EPI_ISL_3470520, EPI_ISL_3470521, EPI_ISL_3470522, EPI_ISL_3470523, EPI_ISL_3470524, EPI_ISL_3470525, EPI_ISL_3470526, EPI_ISL_3470527, EPI_ISL_3470528 | | | |
| see above | Lighthouse Lab in Glasgow | Wellcome Sanger Institute for the COVID-19 Genomics UK (COG-UK) Consortium | Harper VanSteenhouse, Yumi Kasai, David Gray, Carol Clugston, Anna Dominiczak and Alex Alderton, Roberto Amato, Jeffrey Barrett, Sonia Goncalves, Ewan Harrison, David K. Jackson, Ian Johnston, Dominic Kwiatkowski, Cordelia Langford, John Sillitoe on behalf of the Wellcome Sanger Institute COVID-19 Surveillance Team |
| EPI_ISL_3470529, EPI_ISL_3470530, EPI_ISL_3470531 | Lighthouse Lab in Alderley Park | Wellcome Sanger Institute for the COVID-19 Genomics UK (COG-UK) Consortium | Jacquelyn Wynn, Mairead Hyland, The Lighthouse Lab in Alderley Park and Alex Alderton, Roberto Amato, Jeffrey Barrett, Sonia Goncalves, Ewan Harrison, David K. Jackson, Ian Johnston, Dominic Kwiatkowski, Cordelia Langford, John Sillitoe on behalf of the Wellcome Sanger Institute COVID-19 Surveillance Team |
| EPI_ISL_3470532 | Lighthouse Lab in Milton Keynes | Wellcome Sanger Institute for the COVID-19 Genomics UK (COG-UK) Consortium | The Lighthouse Lab in Milton Keynes and Alex Alderton, Roberto Amato, Jeffrey Barrett, Sonia Goncalves, Ewan Harrison, David K. Jackson, Ian Johnston, Dominic Kwiatkowski, Cordelia Langford, John Sillitoe on behalf of the Wellcome Sanger Institute COVID-19 Surveillance Team |
| EPI_ISL_3470533, EPI_ISL_3470534 | Lighthouse Lab in Alderley Park | Wellcome Sanger Institute for the COVID-19 Genomics UK (COG-UK) Consortium | Jacquelyn Wynn, Mairead Hyland, The Lighthouse Lab in Alderley Park and Alex Alderton, Roberto Amato, Jeffrey Barrett, Sonia Goncalves, Ewan Harrison, David K. Jackson, Ian Johnston, Dominic Kwiatkowski, Cordelia Langford, John Sillitoe on behalf of the Wellcome Sanger Institute COVID-19 Surveillance Team |
| EPI_ISL_3470535 | Lighthouse Lab in Milton Keynes | Wellcome Sanger Institute for the COVID-19 Genomics UK (COG-UK) Consortium | The Lighthouse Lab in Milton Keynes and Alex Alderton, Roberto Amato, Jeffrey Barrett, Sonia Goncalves, Ewan Harrison, David K. Jackson, Ian Johnston, Dominic Kwiatkowski, Cordelia Langford, John Sillitoe on behalf of the Wellcome Sanger Institute COVID-19 Surveillance Team |
| EPI_ISL_3470536, EPI_ISL_3470537 | Lighthouse Lab in Alderley Park | Wellcome Sanger Institute for the COVID-19 Genomics UK (COG-UK) Consortium | Jacquelyn Wynn, Mairead Hyland, The Lighthouse Lab in Alderley Park and Alex Alderton, Roberto Amato, Jeffrey Barrett, Sonia Goncalves, Ewan Harrison, David K. Jackson, Ian Johnston, Dominic Kwiatkowski, Cordelia Langford, John Sillitoe on behalf of the Wellcome Sanger Institute COVID-19 Surveillance Team |
| EPI_ISL_3470538 | Lighthouse Laboratory Plymouth | Wellcome Sanger Institute for the COVID-19 Genomics UK (COG-UK) Consortium | Lighthouse Laboratory Plymouth and Alex Alderton, Roberto Amato, Jeffrey Barrett, Sonia Goncalves, Ewan Harrison, David K. Jackson, Ian Johnston, Dominic Kwiatkowski, Cordelia Langford, John Sillitoe on behalf of the Wellcome Sanger Institute COVID-19 Surveillance Team |
| EPI_ISL_3470539 | Lighthouse Lab in Milton Keynes | Wellcome Sanger Institute for the COVID-19 Genomics UK (COG-UK) Consortium | The Lighthouse Lab in Milton Keynes and Alex Alderton, Roberto Amato, Jeffrey Barrett, Sonia Goncalves, Ewan Harrison, David K. Jackson, Ian Johnston, Dominic Kwiatkowski, Cordelia Langford, John Sillitoe on behalf of the Wellcome Sanger Institute COVID-19 Surveillance Team |
| EPI_ISL_3470540 | Berkshire and Surrey Pathology Services Lighthouse Laboratory | Wellcome Sanger Institute for the COVID-19 Genomics UK (COG-UK) Consortium | Berkshire and Surrey Pathology Services Lighthouse Laboratory and Alex Alderton, Roberto Amato, Jeffrey Barrett, Sonia Goncalves, Ewan Harrison, David K. Jackson, Ian Johnston, Dominic Kwiatkowski, Cordelia Langford, John Sillitoe on behalf of the Wellcome Sanger Institute COVID-19 Surveillance Team |
| EPI_ISL_3470541, EPI_ISL_3470542 | Lighthouse Lab in Milton Keynes | Wellcome Sanger Institute for the COVID-19 Genomics UK (COG-UK) Consortium | The Lighthouse Lab in Milton Keynes and Alex Alderton, Roberto Amato, Jeffrey Barrett, Sonia Goncalves, Ewan Harrison, David K. Jackson, Ian Johnston, Dominic Kwiatkowski, Cordelia Langford, John Sillitoe on behalf of the Wellcome Sanger Institute COVID-19 Surveillance Team |
| EPI_ISL_3470543 | Berkshire and Surrey Pathology Services Lighthouse Laboratory | Wellcome Sanger Institute for the COVID-19 Genomics UK (COG-UK) Consortium | Berkshire and Surrey Pathology Services Lighthouse Laboratory and Alex Alderton, Roberto Amato, Jeffrey Barrett, Sonia Goncalves, Ewan Harrison, David K. Jackson, Ian Johnston, Dominic Kwiatkowski, Cordelia Langford, John Sillitoe on behalf of the Wellcome Sanger Institute COVID-19 Surveillance Team |
| EPI_ISL_3470544 | Lighthouse Lab in Alderley Park | Wellcome Sanger Institute for the COVID-19 Genomics UK (COG-UK) Consortium | Jacquelyn Wynn, Mairead Hyland, The Lighthouse Lab in Alderley Park and Alex Alderton, Roberto Amato, Jeffrey Barrett, Sonia Goncalves, Ewan Harrison, David K. Jackson, Ian Johnston, Dominic Kwiatkowski, Cordelia Langford, John Sillitoe on behalf of the Wellcome Sanger Institute COVID-19 Surveillance Team |
| EPI_ISL_3470545 | Berkshire and Surrey Pathology Services Lighthouse Laboratory | Wellcome Sanger Institute for the COVID-19 Genomics UK (COG-UK) Consortium | Berkshire and Surrey Pathology Services Lighthouse Laboratory and Alex Alderton, Roberto Amato, Jeffrey Barrett, Sonia Goncalves, Ewan Harrison, David K. Jackson, Ian Johnston, Dominic Kwiatkowski, Cordelia Langford, John Sillitoe on behalf of the Wellcome Sanger Institute COVID-19 Surveillance Team |
| EPI_ISL_3470546, EPI_ISL_3470547, EPI_ISL_3470548, EPI_ISL_3470549, | Lighthouse Lab in Milton Keynes | Wellcome Sanger Institute for the COVID-19 Genomics UK (COG-UK) Consortium | The Lighthouse Lab in Milton Keynes and Alex Alderton, Roberto Amato, Jeffrey Barrett, Sonia Goncalves, Ewan Harrison, David K. Jackson, Ian Johnston, Dominic Kwiatkowski, Cordelia Langford, John Sillitoe on behalf of the Wellcome Sanger Institute COVID-19 Surveillance Team |

| | | | |
|---|---|---|---|
| EPI_ISL_3470550, EPI_ISL_3470551, EPI_ISL_3470552, EPI_ISL_3470553, EPI_ISL_3470554, EPI_ISL_3470555 | | | |
| EPI_ISL_3470556 | Lighthouse Lab in Alderley Park | Wellcome Sanger Institute for the COVID-19 Genomics UK (COG-UK) Consortium | Jacquelyn Wynn, Mairead Hyland, The Lighthouse Lab in Alderley Park and Alex Alderton, Roberto Amato, Jeffrey Barrett, Sonia Goncalves, Ewan Harrison, David K. Jackson, Ian Johnston, Dominic Kwiatkowski, Cordelia Langford, John Sillitoe on behalf of the Wellcome Sanger Institute COVID-19 Surveillance Team |
| EPI_ISL_3470557 | Lighthouse Lab in Glasgow | Wellcome Sanger Institute for the COVID-19 Genomics UK (COG-UK) Consortium | Harper VanSteenhouse, Yumi Kasai, David Gray, Carol Clugston, Anna Dominiczak and Alex Alderton, Roberto Amato, Jeffrey Barrett, Sonia Goncalves, Ewan Harrison, David K. Jackson, Ian Johnston, Dominic Kwiatkowski, Cordelia Langford, John Sillitoe on behalf of the Wellcome Sanger Institute COVID-19 Surveillance Team |
| EPI_ISL_3470558 | Lighthouse Lab in Alderley Park | Wellcome Sanger Institute for the COVID-19 Genomics UK (COG-UK) Consortium | Jacquelyn Wynn, Mairead Hyland, The Lighthouse Lab in Alderley Park and Alex Alderton, Roberto Amato, Jeffrey Barrett, Sonia Goncalves, Ewan Harrison, David K. Jackson, Ian Johnston, Dominic Kwiatkowski, Cordelia Langford, John Sillitoe on behalf of the Wellcome Sanger Institute COVID-19 Surveillance Team |
| EPI_ISL_3470559, EPI_ISL_3470560 | Lighthouse Lab in Glasgow | Wellcome Sanger Institute for the COVID-19 Genomics UK (COG-UK) Consortium | Harper VanSteenhouse, Yumi Kasai, David Gray, Carol Clugston, Anna Dominiczak and Alex Alderton, Roberto Amato, Jeffrey Barrett, Sonia Goncalves, Ewan Harrison, David K. Jackson, Ian Johnston, Dominic Kwiatkowski, Cordelia Langford, John Sillitoe on behalf of the Wellcome Sanger Institute COVID-19 Surveillance Team |
| EPI_ISL_3470561, EPI_ISL_3470562 | Lighthouse Lab in Milton Keynes | Wellcome Sanger Institute for the COVID-19 Genomics UK (COG-UK) Consortium | The Lighthouse Lab in Milton Keynes and Alex Alderton, Roberto Amato, Jeffrey Barrett, Sonia Goncalves, Ewan Harrison, David K. Jackson, Ian Johnston, Dominic Kwiatkowski, Cordelia Langford, John Sillitoe on behalf of the Wellcome Sanger Institute COVID-19 Surveillance Team |
| EPI_ISL_3470563 | Lighthouse Lab in Glasgow | Wellcome Sanger Institute for the COVID-19 Genomics UK (COG-UK) Consortium | Harper VanSteenhouse, Yumi Kasai, David Gray, Carol Clugston, Anna Dominiczak and Alex Alderton, Roberto Amato, Jeffrey Barrett, Sonia Goncalves, Ewan Harrison, David K. Jackson, Ian Johnston, Dominic Kwiatkowski, Cordelia Langford, John Sillitoe on behalf of the Wellcome Sanger Institute COVID-19 Surveillance Team |
| EPI_ISL_3470564 | Lighthouse Lab in Alderley Park | Wellcome Sanger Institute for the COVID-19 Genomics UK (COG-UK) Consortium | Jacquelyn Wynn, Mairead Hyland, The Lighthouse Lab in Alderley Park and Alex Alderton, Roberto Amato, Jeffrey Barrett, Sonia Goncalves, Ewan Harrison, David K. Jackson, Ian Johnston, Dominic Kwiatkowski, Cordelia Langford, John Sillitoe on behalf of the Wellcome Sanger Institute COVID-19 Surveillance Team |
| EPI_ISL_3470565 | Lighthouse Lab in Glasgow | Wellcome Sanger Institute for the COVID-19 Genomics UK (COG-UK) Consortium | Harper VanSteenhouse, Yumi Kasai, David Gray, Carol Clugston, Anna Dominiczak and Alex Alderton, Roberto Amato, Jeffrey Barrett, Sonia Goncalves, Ewan Harrison, David K. Jackson, Ian Johnston, Dominic Kwiatkowski, Cordelia Langford, John Sillitoe on behalf of the Wellcome Sanger Institute COVID-19 Surveillance Team |
| EPI_ISL_3470566 | Lighthouse Lab in Milton Keynes | Wellcome Sanger Institute for the COVID-19 Genomics UK (COG-UK) Consortium | The Lighthouse Lab in Milton Keynes and Alex Alderton, Roberto Amato, Jeffrey Barrett, Sonia Goncalves, Ewan Harrison, David K. Jackson, Ian Johnston, Dominic Kwiatkowski, Cordelia Langford, John Sillitoe on behalf of the Wellcome Sanger Institute COVID-19 Surveillance Team |
| EPI_ISL_3470567 | Lighthouse Lab in Glasgow | Wellcome Sanger Institute for the COVID-19 Genomics UK (COG-UK) Consortium | Harper VanSteenhouse, Yumi Kasai, David Gray, Carol Clugston, Anna Dominiczak and Alex Alderton, Roberto Amato, Jeffrey Barrett, Sonia Goncalves, Ewan Harrison, David K. Jackson, Ian Johnston, Dominic Kwiatkowski, Cordelia Langford, John Sillitoe on behalf of the Wellcome Sanger Institute COVID-19 Surveillance Team |
| EPI_ISL_3470568 | Lighthouse Lab in Alderley Park | Wellcome Sanger Institute for the COVID-19 Genomics UK (COG-UK) Consortium | Jacquelyn Wynn, Mairead Hyland, The Lighthouse Lab in Alderley Park and Alex Alderton, Roberto Amato, Jeffrey Barrett, Sonia Goncalves, Ewan Harrison, David K. Jackson, Ian Johnston, Dominic Kwiatkowski, Cordelia Langford, John Sillitoe on behalf of the Wellcome Sanger Institute COVID-19 Surveillance Team |
| EPI_ISL_3470569 | Lighthouse Lab in Milton Keynes | Wellcome Sanger Institute for the COVID-19 Genomics UK (COG-UK) Consortium | The Lighthouse Lab in Milton Keynes and Alex Alderton, Roberto Amato, Jeffrey Barrett, Sonia Goncalves, Ewan Harrison, David K. Jackson, Ian Johnston, Dominic Kwiatkowski, Cordelia Langford, John Sillitoe on behalf of the Wellcome Sanger Institute COVID-19 Surveillance Team |
| EPI_ISL_3470570 | Lighthouse Lab in Glasgow | Wellcome Sanger Institute for the COVID-19 Genomics UK (COG-UK) Consortium | Harper VanSteenhouse, Yumi Kasai, David Gray, Carol Clugston, Anna Dominiczak and Alex Alderton, Roberto Amato, Jeffrey Barrett, Sonia Goncalves, Ewan Harrison, David K. Jackson, Ian Johnston, Dominic Kwiatkowski, Cordelia Langford, John Sillitoe on behalf of the Wellcome Sanger Institute COVID-19 Surveillance Team |
| EPI_ISL_3470571, EPI_ISL_3470572 | Lighthouse Lab in Alderley Park | Wellcome Sanger Institute for the COVID-19 Genomics UK (COG-UK) Consortium | Jacquelyn Wynn, Mairead Hyland, The Lighthouse Lab in Alderley Park and Alex Alderton, Roberto Amato, Jeffrey Barrett, Sonia Goncalves, Ewan Harrison, David K. Jackson, Ian Johnston, Dominic Kwiatkowski, Cordelia Langford, John Sillitoe on behalf of the Wellcome Sanger Institute COVID-19 Surveillance Team |
| EPI_ISL_3470573 | Lighthouse Lab in Milton Keynes | Wellcome Sanger Institute for the COVID-19 Genomics UK (COG-UK) Consortium | The Lighthouse Lab in Milton Keynes and Alex Alderton, Roberto Amato, Jeffrey Barrett, Sonia Goncalves, Ewan Harrison, David K. Jackson, Ian Johnston, Dominic Kwiatkowski, Cordelia Langford, John Sillitoe on behalf of the Wellcome Sanger Institute COVID-19 Surveillance Team |
| EPI_ISL_3470574 | Lighthouse Lab in Alderley Park | Wellcome Sanger Institute for the COVID-19 Genomics UK (COG-UK) Consortium | Jacquelyn Wynn, Mairead Hyland, The Lighthouse Lab in Alderley Park and Alex Alderton, Roberto Amato, Jeffrey Barrett, Sonia Goncalves, Ewan Harrison, David K. Jackson, Ian Johnston, Dominic Kwiatkowski, Cordelia Langford, John Sillitoe on behalf of the Wellcome Sanger Institute COVID-19 Surveillance Team |
| EPI_ISL_3470575, EPI_ISL_3470576 | Lighthouse Lab in Milton Keynes | Wellcome Sanger Institute for the COVID-19 Genomics UK (COG-UK) Consortium | The Lighthouse Lab in Milton Keynes and Alex Alderton, Roberto Amato, Jeffrey Barrett, Sonia Goncalves, Ewan Harrison, David K. Jackson, Ian Johnston, Dominic Kwiatkowski, Cordelia Langford, John Sillitoe on behalf of the Wellcome Sanger Institute COVID-19 Surveillance Team |
| EPI_ISL_3470577 | Lighthouse Lab in Glasgow | Wellcome Sanger Institute for the COVID-19 Genomics UK (COG-UK) Consortium | Harper VanSteenhouse, Yumi Kasai, David Gray, Carol Clugston, Anna Dominiczak and Alex Alderton, Roberto Amato, Jeffrey Barrett, Sonia Goncalves, Ewan Harrison, David K. Jackson, Ian Johnston, Dominic Kwiatkowski, Cordelia Langford, John Sillitoe on behalf of the Wellcome Sanger Institute COVID-19 Surveillance Team |
| EPI_ISL_3470578 | Lighthouse Lab in Milton Keynes | Wellcome Sanger Institute for the COVID-19 Genomics UK (COG-UK) Consortium | The Lighthouse Lab in Milton Keynes and Alex Alderton, Roberto Amato, Jeffrey Barrett, Sonia Goncalves, Ewan Harrison, David K. Jackson, Ian Johnston, Dominic Kwiatkowski, Cordelia Langford, John Sillitoe on behalf of the Wellcome Sanger Institute COVID-19 Surveillance Team |
| EPI_ISL_3470579 | Lighthouse Lab in Alderley Park | Wellcome Sanger Institute for the COVID-19 Genomics UK (COG-UK) Consortium | Jacquelyn Wynn, Mairead Hyland, The Lighthouse Lab in Alderley Park and Alex Alderton, Roberto Amato, Jeffrey Barrett, Sonia Goncalves, Ewan Harrison, David K. Jackson, Ian Johnston, Dominic Kwiatkowski, Cordelia Langford, John Sillitoe on behalf of the Wellcome Sanger Institute COVID-19 Surveillance Team |
| EPI_ISL_3470580 | Lighthouse Lab in Glasgow | Wellcome Sanger Institute for the COVID-19 Genomics UK (COG-UK) Consortium | Harper VanSteenhouse, Yumi Kasai, David Gray, Carol Clugston, Anna Dominiczak and Alex Alderton, Roberto Amato, Jeffrey Barrett, Sonia Goncalves, Ewan Harrison, David K. Jackson, Ian Johnston, Dominic Kwiatkowski, Cordelia Langford, John Sillitoe on behalf of the Wellcome Sanger Institute COVID-19 Surveillance Team |
| EPI_ISL_3470581 | Lighthouse Lab in Alderley Park | Wellcome Sanger Institute for the COVID-19 Genomics UK (COG-UK) Consortium | Jacquelyn Wynn, Mairead Hyland, The Lighthouse Lab in Alderley Park and Alex Alderton, Roberto Amato, Jeffrey Barrett, Sonia Goncalves, Ewan Harrison, David K. Jackson, Ian Johnston, Dominic Kwiatkowski, Cordelia Langford, John Sillitoe on behalf of the Wellcome Sanger Institute COVID-19 Surveillance Team |
| EPI_ISL_3470582 | Lighthouse Lab in Milton Keynes | Wellcome Sanger Institute for the COVID-19 Genomics UK (COG-UK) Consortium | The Lighthouse Lab in Milton Keynes and Alex Alderton, Roberto Amato, Jeffrey Barrett, Sonia Goncalves, Ewan Harrison, David K. Jackson, Ian Johnston, Dominic Kwiatkowski, Cordelia Langford, John Sillitoe on behalf of the Wellcome Sanger Institute COVID-19 Surveillance Team |
| EPI_ISL_3470583, EPI_ISL_3470584 | Lighthouse Lab in Glasgow | Wellcome Sanger Institute for the COVID-19 Genomics UK (COG-UK) Consortium | Harper VanSteenhouse, Yumi Kasai, David Gray, Carol Clugston, Anna Dominiczak and Alex Alderton, Roberto Amato, Jeffrey Barrett, Sonia Goncalves, Ewan Harrison, David K. Jackson, Ian Johnston, Dominic Kwiatkowski, Cordelia Langford, John Sillitoe on behalf of the Wellcome Sanger Institute COVID-19 Surveillance Team |
| EPI_ISL_3470585, EPI_ISL_3470586 | Lighthouse Lab in Alderley Park | Wellcome Sanger Institute for the COVID-19 Genomics UK (COG-UK) Consortium | Jacquelyn Wynn, Mairead Hyland, The Lighthouse Lab in Alderley Park and Alex Alderton, Roberto Amato, Jeffrey Barrett, Sonia Goncalves, Ewan Harrison, David K. Jackson, Ian Johnston, Dominic Kwiatkowski, Cordelia Langford, John Sillitoe on behalf of the Wellcome Sanger Institute COVID-19 Surveillance Team |
| EPI_ISL_3470587 | Lighthouse Lab in Milton Keynes | Wellcome Sanger Institute for the COVID-19 Genomics UK (COG-UK) Consortium | The Lighthouse Lab in Milton Keynes and Alex Alderton, Roberto Amato, Jeffrey Barrett, Sonia Goncalves, Ewan Harrison, David K. Jackson, Ian Johnston, Dominic Kwiatkowski, Cordelia Langford, John Sillitoe on behalf of the Wellcome Sanger Institute COVID-19 Surveillance Team |
| EPI_ISL_3470588 | Lighthouse Lab in Alderley Park | Wellcome Sanger Institute for the COVID-19 Genomics UK (COG-UK) Consortium | Jacquelyn Wynn, Mairead Hyland, The Lighthouse Lab in Alderley Park and Alex Alderton, Roberto Amato, Jeffrey Barrett, Sonia Goncalves, Ewan Harrison, David K. Jackson, Ian Johnston, Dominic Kwiatkowski, Cordelia Langford, John Sillitoe on behalf of the Wellcome Sanger Institute COVID-19 Surveillance Team |
| EPI_ISL_3470589, EPI_ISL_3470590 | Lighthouse Lab in Glasgow | Wellcome Sanger Institute for the COVID-19 Genomics UK (COG-UK) Consortium | Harper VanSteenhouse, Yumi Kasai, David Gray, Carol Clugston, Anna Dominiczak and Alex Alderton, Roberto Amato, Jeffrey Barrett, Sonia Goncalves, Ewan Harrison, David K. Jackson, Ian Johnston, Dominic Kwiatkowski, Cordelia Langford, John Sillitoe on behalf of the Wellcome Sanger Institute COVID-19 Surveillance Team |
| EPI_ISL_3470591 | Lighthouse Lab in Milton Keynes | Wellcome Sanger Institute for the COVID-19 Genomics UK (COG-UK) Consortium | The Lighthouse Lab in Milton Keynes and Alex Alderton, Roberto Amato, Jeffrey Barrett, Sonia Goncalves, Ewan Harrison, David K. Jackson, Ian Johnston, Dominic Kwiatkowski, Cordelia Langford, John Sillitoe on behalf of the Wellcome Sanger Institute COVID-19 Surveillance Team |
| EPI_ISL_3470592 | Lighthouse Lab in Alderley Park | Wellcome Sanger Institute for the COVID-19 Genomics UK (COG-UK) Consortium | Jacquelyn Wynn, Mairead Hyland, The Lighthouse Lab in Alderley Park and Alex Alderton, Roberto Amato, Jeffrey Barrett, Sonia Goncalves, Ewan Harrison, David K. Jackson, Ian Johnston, Dominic Kwiatkowski, Cordelia Langford, John Sillitoe on behalf of the Wellcome Sanger Institute COVID-19 Surveillance Team |
| EPI_ISL_3470593 | Lighthouse Lab in Milton Keynes | Wellcome Sanger Institute for the COVID-19 Genomics UK (COG-UK) Consortium | The Lighthouse Lab in Milton Keynes and Alex Alderton, Roberto Amato, Jeffrey Barrett, Sonia Goncalves, Ewan Harrison, David K. Jackson, Ian Johnston, Dominic Kwiatkowski, Cordelia Langford, John Sillitoe on behalf of the Wellcome Sanger Institute COVID-19 Surveillance Team |
| EPI_ISL_3470594, EPI_ISL_3470595 | Lighthouse Lab in Glasgow | Wellcome Sanger Institute for the COVID-19 Genomics UK (COG-UK) Consortium | Harper VanSteenhouse, Yumi Kasai, David Gray, Carol Clugston, Anna Dominiczak and Alex Alderton, Roberto Amato, Jeffrey Barrett, Sonia Goncalves, Ewan Harrison, David K. Jackson, Ian Johnston, Dominic Kwiatkowski, Cordelia Langford, John Sillitoe on behalf of the Wellcome Sanger Institute COVID-19 Surveillance Team |
| EPI_ISL_3470596 | Lighthouse Lab in Alderley Park | Wellcome Sanger Institute for the COVID-19 Genomics UK (COG-UK) Consortium | Jacquelyn Wynn, Mairead Hyland, The Lighthouse Lab in Alderley Park and Alex Alderton, Roberto Amato, Jeffrey Barrett, Sonia Goncalves, Ewan Harrison, David K. Jackson, Ian Johnston, Dominic Kwiatkowski, Cordelia Langford, John Sillitoe on behalf of the Wellcome Sanger Institute COVID-19 Surveillance Team |
| EPI_ISL_3470597, EPI_ISL_3470598 | Lighthouse Lab in Glasgow | Wellcome Sanger Institute for the COVID-19 Genomics UK (COG-UK) Consortium | Harper VanSteenhouse, Yumi Kasai, David Gray, Carol Clugston, Anna Dominiczak and Alex Alderton, Roberto Amato, Jeffrey Barrett, Sonia Goncalves, Ewan Harrison, David K. Jackson, Ian Johnston, Dominic Kwiatkowski, Cordelia Langford, John Sillitoe on behalf of the Wellcome Sanger Institute COVID-19 Surveillance Team |
| EPI_ISL_3470599 | Lighthouse Lab in Alderley Park | Wellcome Sanger Institute for the COVID-19 Genomics UK (COG-UK) Consortium | Jacquelyn Wynn, Mairead Hyland, The Lighthouse Lab in Alderley Park and Alex Alderton, Roberto Amato, Jeffrey Barrett, Sonia Goncalves, Ewan Harrison, David K. Jackson, Ian Johnston, Dominic Kwiatkowski, Cordelia Langford, John Sillitoe on behalf of the Wellcome Sanger Institute COVID-19 Surveillance Team |
| EPI_ISL_3470600, EPI_ISL_3470601 | Lighthouse Lab in Glasgow | Wellcome Sanger Institute for the COVID-19 Genomics UK (COG-UK) Consortium | Harper VanSteenhouse, Yumi Kasai, David Gray, Carol Clugston, Anna Dominiczak and Alex Alderton, Roberto Amato, Jeffrey Barrett, Sonia Goncalves, Ewan Harrison, David K. Jackson, Ian Johnston, Dominic Kwiatkowski, Cordelia Langford, John Sillitoe on behalf of the Wellcome Sanger Institute COVID-19 Surveillance Team |
| EPI_ISL_3470602 | Lighthouse Lab in Alderley Park | Wellcome Sanger Institute for the COVID-19 Genomics UK (COG-UK) Consortium | Jacquelyn Wynn, Mairead Hyland, The Lighthouse Lab in Alderley Park and Alex Alderton, Roberto Amato, Jeffrey Barrett, Sonia Goncalves, Ewan Harrison, David K. Jackson, Ian Johnston, Dominic Kwiatkowski, Cordelia Langford, John Sillitoe on behalf of the Wellcome Sanger Institute COVID-19 Surveillance Team |
| EPI_ISL_3470603, EPI_ISL_3470604, EPI_ISL_3470605, EPI_ISL_3470606, EPI_ISL_3470607 | Lighthouse Lab in Glasgow | Wellcome Sanger Institute for the COVID-19 Genomics UK (COG-UK) Consortium | Harper VanSteenhouse, Yumi Kasai, David Gray, Carol Clugston, Anna Dominiczak and Alex Alderton, Roberto Amato, Jeffrey Barrett, Sonia Goncalves, Ewan Harrison, David K. Jackson, Ian Johnston, Dominic Kwiatkowski, Cordelia Langford, John Sillitoe on behalf of the Wellcome Sanger Institute COVID-19 Surveillance Team |
| EPI_ISL_3470608, EPI_ISL_3470609, EPI_ISL_3470610 | Lighthouse Lab in Milton Keynes | Wellcome Sanger Institute for the COVID-19 Genomics UK (COG-UK) Consortium | The Lighthouse Lab in Milton Keynes and Alex Alderton, Roberto Amato, Jeffrey Barrett, Sonia Goncalves, Ewan Harrison, David K. Jackson, Ian Johnston, Dominic Kwiatkowski, Cordelia Langford, John Sillitoe on behalf of the Wellcome Sanger Institute COVID-19 Surveillance Team |
| EPI_ISL_3470611, EPI_ISL_3470612 | Lighthouse Lab in Alderley Park | Wellcome Sanger Institute for the COVID-19 Genomics UK (COG-UK) Consortium | Jacquelyn Wynn, Mairead Hyland, The Lighthouse Lab in Alderley Park and Alex Alderton, Roberto Amato, Jeffrey Barrett, Sonia Goncalves, Ewan Harrison, David K. Jackson, Ian Johnston, Dominic Kwiatkowski, Cordelia Langford, John Sillitoe on behalf of the Wellcome Sanger Institute COVID-19 Surveillance Team |
| EPI_ISL_3470613 | Lighthouse Lab in Milton Keynes | Wellcome Sanger Institute for the COVID-19 Genomics UK (COG-UK) | The Lighthouse Lab in Milton Keynes and Alex Alderton, Roberto Amato, Jeffrey Barrett, Sonia Goncalves, Ewan Harrison, David K. Jackson, Ian Johnston, Dominic Kwiatkowski, Cordelia Langford, John Sillitoe on behalf of the Wellcome Sanger |

| | | | |
|---|---|---|---|
| | | Consortium | Institute COVID-19 Surveillance Team |
| EPI_ISL_3470614 | Lighthouse Lab in Glasgow | Wellcome Sanger Institute for the COVID-19 Genomics UK (COG-UK) Consortium | Harper VanSteenhouse, Yumi Kasai, David Gray, Carol Clugston, Anna Dominiczak and Alex Alderton, Roberto Amato, Jeffrey Barrett, Sonia Goncalves, Ewan Harrison, David K. Jackson, Ian Johnston, Dominic Kwiatkowski, Cordelia Langford, John Sillitoe on behalf of the Wellcome Sanger Institute COVID-19 Surveillance Team |
| EPI_ISL_3470615 | Lighthouse Lab in Alderley Park | Wellcome Sanger Institute for the COVID-19 Genomics UK (COG-UK) Consortium | Jacquelyn Wynn, Mairead Hyland, The Lighthouse Lab in Alderley Park and Alex Alderton, Roberto Amato, Jeffrey Barrett, Sonia Goncalves, Ewan Harrison, David K. Jackson, Ian Johnston, Dominic Kwiatkowski, Cordelia Langford, John Sillitoe on behalf of the Wellcome Sanger Institute COVID-19 Surveillance Team |
| EPI_ISL_3470616, EPI_ISL_3470617, EPI_ISL_3470618, EPI_ISL_3470619 | Lighthouse Lab in Glasgow | Wellcome Sanger Institute for the COVID-19 Genomics UK (COG-UK) Consortium | Harper VanSteenhouse, Yumi Kasai, David Gray, Carol Clugston, Anna Dominiczak and Alex Alderton, Roberto Amato, Jeffrey Barrett, Sonia Goncalves, Ewan Harrison, David K. Jackson, Ian Johnston, Dominic Kwiatkowski, Cordelia Langford, John Sillitoe on behalf of the Wellcome Sanger Institute COVID-19 Surveillance Team |
| EPI_ISL_3470620, EPI_ISL_3470621 | Lighthouse Lab in Alderley Park | Wellcome Sanger Institute for the COVID-19 Genomics UK (COG-UK) Consortium | Jacquelyn Wynn, Mairead Hyland, The Lighthouse Lab in Alderley Park and Alex Alderton, Roberto Amato, Jeffrey Barrett, Sonia Goncalves, Ewan Harrison, David K. Jackson, Ian Johnston, Dominic Kwiatkowski, Cordelia Langford, John Sillitoe on behalf of the Wellcome Sanger Institute COVID-19 Surveillance Team |
| EPI_ISL_3470622, EPI_ISL_3470623 | Lighthouse Lab in Glasgow | Wellcome Sanger Institute for the COVID-19 Genomics UK (COG-UK) Consortium | Harper VanSteenhouse, Yumi Kasai, David Gray, Carol Clugston, Anna Dominiczak and Alex Alderton, Roberto Amato, Jeffrey Barrett, Sonia Goncalves, Ewan Harrison, David K. Jackson, Ian Johnston, Dominic Kwiatkowski, Cordelia Langford, John Sillitoe on behalf of the Wellcome Sanger Institute COVID-19 Surveillance Team |
| EPI_ISL_3470624, EPI_ISL_3470625 | Lighthouse Lab in Milton Keynes | Wellcome Sanger Institute for the COVID-19 Genomics UK (COG-UK) Consortium | The Lighthouse Lab in Milton Keynes and Alex Alderton, Roberto Amato, Jeffrey Barrett, Sonia Goncalves, Ewan Harrison, David K. Jackson, Ian Johnston, Dominic Kwiatkowski, Cordelia Langford, John Sillitoe on behalf of the Wellcome Sanger Institute COVID-19 Surveillance Team |
| EPI_ISL_3470626, EPI_ISL_3470627, EPI_ISL_3470628, EPI_ISL_3470629, EPI_ISL_3470630 | Lighthouse Lab in Glasgow | Wellcome Sanger Institute for the COVID-19 Genomics UK (COG-UK) Consortium | Harper VanSteenhouse, Yumi Kasai, David Gray, Carol Clugston, Anna Dominiczak and Alex Alderton, Roberto Amato, Jeffrey Barrett, Sonia Goncalves, Ewan Harrison, David K. Jackson, Ian Johnston, Dominic Kwiatkowski, Cordelia Langford, John Sillitoe on behalf of the Wellcome Sanger Institute COVID-19 Surveillance Team |
| EPI_ISL_3470631 | Lighthouse Lab in Milton Keynes | Wellcome Sanger Institute for the COVID-19 Genomics UK (COG-UK) Consortium | The Lighthouse Lab in Milton Keynes and Alex Alderton, Roberto Amato, Jeffrey Barrett, Sonia Goncalves, Ewan Harrison, David K. Jackson, Ian Johnston, Dominic Kwiatkowski, Cordelia Langford, John Sillitoe on behalf of the Wellcome Sanger Institute COVID-19 Surveillance Team |
| EPI_ISL_3470632, EPI_ISL_3470633 | Lighthouse Lab in Glasgow | Wellcome Sanger Institute for the COVID-19 Genomics UK (COG-UK) Consortium | Harper VanSteenhouse, Yumi Kasai, David Gray, Carol Clugston, Anna Dominiczak and Alex Alderton, Roberto Amato, Jeffrey Barrett, Sonia Goncalves, Ewan Harrison, David K. Jackson, Ian Johnston, Dominic Kwiatkowski, Cordelia Langford, John Sillitoe on behalf of the Wellcome Sanger Institute COVID-19 Surveillance Team |
| EPI_ISL_3470634 | Lighthouse Lab in Alderley Park | Wellcome Sanger Institute for the COVID-19 Genomics UK (COG-UK) Consortium | Jacquelyn Wynn, Mairead Hyland, The Lighthouse Lab in Alderley Park and Alex Alderton, Roberto Amato, Jeffrey Barrett, Sonia Goncalves, Ewan Harrison, David K. Jackson, Ian Johnston, Dominic Kwiatkowski, Cordelia Langford, John Sillitoe on behalf of the Wellcome Sanger Institute COVID-19 Surveillance Team |
| EPI_ISL_3470635 | Lighthouse Lab in Milton Keynes | Wellcome Sanger Institute for the COVID-19 Genomics UK (COG-UK) Consortium | The Lighthouse Lab in Milton Keynes and Alex Alderton, Roberto Amato, Jeffrey Barrett, Sonia Goncalves, Ewan Harrison, David K. Jackson, Ian Johnston, Dominic Kwiatkowski, Cordelia Langford, John Sillitoe on behalf of the Wellcome Sanger Institute COVID-19 Surveillance Team |
| EPI_ISL_3470636 | Lighthouse Lab in Glasgow | Wellcome Sanger Institute for the COVID-19 Genomics UK (COG-UK) Consortium | Harper VanSteenhouse, Yumi Kasai, David Gray, Carol Clugston, Anna Dominiczak and Alex Alderton, Roberto Amato, Jeffrey Barrett, Sonia Goncalves, Ewan Harrison, David K. Jackson, Ian Johnston, Dominic Kwiatkowski, Cordelia Langford, John Sillitoe on behalf of the Wellcome Sanger Institute COVID-19 Surveillance Team |
| EPI_ISL_3470637 | Lighthouse Lab in Alderley Park | Wellcome Sanger Institute for the COVID-19 Genomics UK (COG-UK) Consortium | Jacquelyn Wynn, Mairead Hyland, The Lighthouse Lab in Alderley Park and Alex Alderton, Roberto Amato, Jeffrey Barrett, Sonia Goncalves, Ewan Harrison, David K. Jackson, Ian Johnston, Dominic Kwiatkowski, Cordelia Langford, John Sillitoe on behalf of the Wellcome Sanger Institute COVID-19 Surveillance Team |
| EPI_ISL_3470638, EPI_ISL_3470639 | Lighthouse Lab in Glasgow | Wellcome Sanger Institute for the COVID-19 Genomics UK (COG-UK) Consortium | Harper VanSteenhouse, Yumi Kasai, David Gray, Carol Clugston, Anna Dominiczak and Alex Alderton, Roberto Amato, Jeffrey Barrett, Sonia Goncalves, Ewan Harrison, David K. Jackson, Ian Johnston, Dominic Kwiatkowski, Cordelia Langford, John Sillitoe on behalf of the Wellcome Sanger Institute COVID-19 Surveillance Team |
| EPI_ISL_3470640 | Lighthouse Lab in Milton Keynes | Wellcome Sanger Institute for the COVID-19 Genomics UK (COG-UK) Consortium | The Lighthouse Lab in Milton Keynes and Alex Alderton, Roberto Amato, Jeffrey Barrett, Sonia Goncalves, Ewan Harrison, David K. Jackson, Ian Johnston, Dominic Kwiatkowski, Cordelia Langford, John Sillitoe on behalf of the Wellcome Sanger Institute COVID-19 Surveillance Team |
| EPI_ISL_3470641 | Lighthouse Lab in Glasgow | Wellcome Sanger Institute for the COVID-19 Genomics UK (COG-UK) Consortium | Harper VanSteenhouse, Yumi Kasai, David Gray, Carol Clugston, Anna Dominiczak and Alex Alderton, Roberto Amato, Jeffrey Barrett, Sonia Goncalves, Ewan Harrison, David K. Jackson, Ian Johnston, Dominic Kwiatkowski, Cordelia Langford, John Sillitoe on behalf of the Wellcome Sanger Institute COVID-19 Surveillance Team |
| EPI_ISL_3470642, EPI_ISL_3470643 | Lighthouse Lab in Alderley Park | Wellcome Sanger Institute for the COVID-19 Genomics UK (COG-UK) Consortium | Jacquelyn Wynn, Mairead Hyland, The Lighthouse Lab in Alderley Park and Alex Alderton, Roberto Amato, Jeffrey Barrett, Sonia Goncalves, Ewan Harrison, David K. Jackson, Ian Johnston, Dominic Kwiatkowski, Cordelia Langford, John Sillitoe on behalf of the Wellcome Sanger Institute COVID-19 Surveillance Team |
| EPI_ISL_3470644 | Lighthouse Lab in Milton Keynes | Wellcome Sanger Institute for the COVID-19 Genomics UK (COG-UK) Consortium | The Lighthouse Lab in Milton Keynes and Alex Alderton, Roberto Amato, Jeffrey Barrett, Sonia Goncalves, Ewan Harrison, David K. Jackson, Ian Johnston, Dominic Kwiatkowski, Cordelia Langford, John Sillitoe on behalf of the Wellcome Sanger Institute COVID-19 Surveillance Team |
| EPI_ISL_3470645 | Lighthouse Lab in Glasgow | Wellcome Sanger Institute for the COVID-19 Genomics UK (COG-UK) Consortium | Harper VanSteenhouse, Yumi Kasai, David Gray, Carol Clugston, Anna Dominiczak and Alex Alderton, Roberto Amato, Jeffrey Barrett, Sonia Goncalves, Ewan Harrison, David K. Jackson, Ian Johnston, Dominic Kwiatkowski, Cordelia Langford, John Sillitoe on behalf of the Wellcome Sanger Institute COVID-19 Surveillance Team |
| EPI_ISL_3470646, EPI_ISL_3470647 | Lighthouse Lab in Alderley Park | Wellcome Sanger Institute for the COVID-19 Genomics UK (COG-UK) Consortium | Jacquelyn Wynn, Mairead Hyland, The Lighthouse Lab in Alderley Park and Alex Alderton, Roberto Amato, Jeffrey Barrett, Sonia Goncalves, Ewan Harrison, David K. Jackson, Ian Johnston, Dominic Kwiatkowski, Cordelia Langford, John Sillitoe on behalf of the Wellcome Sanger Institute COVID-19 Surveillance Team |
| EPI_ISL_3470648, EPI_ISL_3470649, EPI_ISL_3470650 | Lighthouse Lab in Milton Keynes | Wellcome Sanger Institute for the COVID-19 Genomics UK (COG-UK) Consortium | The Lighthouse Lab in Milton Keynes and Alex Alderton, Roberto Amato, Jeffrey Barrett, Sonia Goncalves, Ewan Harrison, David K. Jackson, Ian Johnston, Dominic Kwiatkowski, Cordelia Langford, John Sillitoe on behalf of the Wellcome Sanger Institute COVID-19 Surveillance Team |
| EPI_ISL_3470651, EPI_ISL_3470652 | Lighthouse Lab in Alderley Park | Wellcome Sanger Institute for the COVID-19 Genomics UK (COG-UK) Consortium | Jacquelyn Wynn, Mairead Hyland, The Lighthouse Lab in Alderley Park and Alex Alderton, Roberto Amato, Jeffrey Barrett, Sonia Goncalves, Ewan Harrison, David K. Jackson, Ian Johnston, Dominic Kwiatkowski, Cordelia Langford, John Sillitoe on behalf of the Wellcome Sanger Institute COVID-19 Surveillance Team |
| EPI_ISL_3470653 | Lighthouse Lab in Glasgow | Wellcome Sanger Institute for the COVID-19 Genomics UK (COG-UK) Consortium | Harper VanSteenhouse, Yumi Kasai, David Gray, Carol Clugston, Anna Dominiczak and Alex Alderton, Roberto Amato, Jeffrey Barrett, Sonia Goncalves, Ewan Harrison, David K. Jackson, Ian Johnston, Dominic Kwiatkowski, Cordelia Langford, John Sillitoe on behalf of the Wellcome Sanger Institute COVID-19 Surveillance Team |
| EPI_ISL_3470654, EPI_ISL_3470655, EPI_ISL_3470656 | Lighthouse Lab in Alderley Park | Wellcome Sanger Institute for the COVID-19 Genomics UK (COG-UK) Consortium | Jacquelyn Wynn, Mairead Hyland, The Lighthouse Lab in Alderley Park and Alex Alderton, Roberto Amato, Jeffrey Barrett, Sonia Goncalves, Ewan Harrison, David K. Jackson, Ian Johnston, Dominic Kwiatkowski, Cordelia Langford, John Sillitoe on behalf of the Wellcome Sanger Institute COVID-19 Surveillance Team |
| EPI_ISL_3470657 | Lighthouse Lab in Glasgow | Wellcome Sanger Institute for the COVID-19 Genomics UK (COG-UK) Consortium | Harper VanSteenhouse, Yumi Kasai, David Gray, Carol Clugston, Anna Dominiczak and Alex Alderton, Roberto Amato, Jeffrey Barrett, Sonia Goncalves, Ewan Harrison, David K. Jackson, Ian Johnston, Dominic Kwiatkowski, Cordelia Langford, John Sillitoe on behalf of the Wellcome Sanger Institute COVID-19 Surveillance Team |
| EPI_ISL_3470658 | Lighthouse Lab in Milton Keynes | Wellcome Sanger Institute for the COVID-19 Genomics UK (COG-UK) Consortium | The Lighthouse Lab in Milton Keynes and Alex Alderton, Roberto Amato, Jeffrey Barrett, Sonia Goncalves, Ewan Harrison, David K. Jackson, Ian Johnston, Dominic Kwiatkowski, Cordelia Langford, John Sillitoe on behalf of the Wellcome Sanger Institute COVID-19 Surveillance Team |
| EPI_ISL_3470659 | Lighthouse Lab in Glasgow | Wellcome Sanger Institute for the COVID-19 Genomics UK (COG-UK) Consortium | Harper VanSteenhouse, Yumi Kasai, David Gray, Carol Clugston, Anna Dominiczak and Alex Alderton, Roberto Amato, Jeffrey Barrett, Sonia Goncalves, Ewan Harrison, David K. Jackson, Ian Johnston, Dominic Kwiatkowski, Cordelia Langford, John Sillitoe on behalf of the Wellcome Sanger Institute COVID-19 Surveillance Team |
| EPI_ISL_3470660 | Lighthouse Lab in Alderley Park | Wellcome Sanger Institute for the COVID-19 Genomics UK (COG-UK) Consortium | Jacquelyn Wynn, Mairead Hyland, The Lighthouse Lab in Alderley Park and Alex Alderton, Roberto Amato, Jeffrey Barrett, Sonia Goncalves, Ewan Harrison, David K. Jackson, Ian Johnston, Dominic Kwiatkowski, Cordelia Langford, John Sillitoe on behalf of the Wellcome Sanger Institute COVID-19 Surveillance Team |
| EPI_ISL_3470661 | Lighthouse Lab in Milton Keynes | Wellcome Sanger Institute for the COVID-19 Genomics UK (COG-UK) Consortium | The Lighthouse Lab in Milton Keynes and Alex Alderton, Roberto Amato, Jeffrey Barrett, Sonia Goncalves, Ewan Harrison, David K. Jackson, Ian Johnston, Dominic Kwiatkowski, Cordelia Langford, John Sillitoe on behalf of the Wellcome Sanger Institute COVID-19 Surveillance Team |
| EPI_ISL_3470662 | Lighthouse Lab in Glasgow | Wellcome Sanger Institute for the COVID-19 Genomics UK (COG-UK) Consortium | Harper VanSteenhouse, Yumi Kasai, David Gray, Carol Clugston, Anna Dominiczak and Alex Alderton, Roberto Amato, Jeffrey Barrett, Sonia Goncalves, Ewan Harrison, David K. Jackson, Ian Johnston, Dominic Kwiatkowski, Cordelia Langford, John Sillitoe on behalf of the Wellcome Sanger Institute COVID-19 Surveillance Team |
| EPI_ISL_3470663 | Lighthouse Lab in Alderley Park | Wellcome Sanger Institute for the COVID-19 Genomics UK (COG-UK) Consortium | Jacquelyn Wynn, Mairead Hyland, The Lighthouse Lab in Alderley Park and Alex Alderton, Roberto Amato, Jeffrey Barrett, Sonia Goncalves, Ewan Harrison, David K. Jackson, Ian Johnston, Dominic Kwiatkowski, Cordelia Langford, John Sillitoe on behalf of the Wellcome Sanger Institute COVID-19 Surveillance Team |
| EPI_ISL_3470664, EPI_ISL_3470665 | Lighthouse Lab in Glasgow | Wellcome Sanger Institute for the COVID-19 Genomics UK (COG-UK) Consortium | Harper VanSteenhouse, Yumi Kasai, David Gray, Carol Clugston, Anna Dominiczak and Alex Alderton, Roberto Amato, Jeffrey Barrett, Sonia Goncalves, Ewan Harrison, David K. Jackson, Ian Johnston, Dominic Kwiatkowski, Cordelia Langford, John Sillitoe on behalf of the Wellcome Sanger Institute COVID-19 Surveillance Team |
| EPI_ISL_3470666 | Lighthouse Lab in Alderley Park | Wellcome Sanger Institute for the COVID-19 Genomics UK (COG-UK) Consortium | Jacquelyn Wynn, Mairead Hyland, The Lighthouse Lab in Alderley Park and Alex Alderton, Roberto Amato, Jeffrey Barrett, Sonia Goncalves, Ewan Harrison, David K. Jackson, Ian Johnston, Dominic Kwiatkowski, Cordelia Langford, John Sillitoe on behalf of the Wellcome Sanger Institute COVID-19 Surveillance Team |
| EPI_ISL_3470667 | Lighthouse Lab in Milton Keynes | Wellcome Sanger Institute for the COVID-19 Genomics UK (COG-UK) Consortium | The Lighthouse Lab in Milton Keynes and Alex Alderton, Roberto Amato, Jeffrey Barrett, Sonia Goncalves, Ewan Harrison, David K. Jackson, Ian Johnston, Dominic Kwiatkowski, Cordelia Langford, John Sillitoe on behalf of the Wellcome Sanger Institute COVID-19 Surveillance Team |
| EPI_ISL_3470668 | Lighthouse Lab in Glasgow | Wellcome Sanger Institute for the COVID-19 Genomics UK (COG-UK) Consortium | Harper VanSteenhouse, Yumi Kasai, David Gray, Carol Clugston, Anna Dominiczak and Alex Alderton, Roberto Amato, Jeffrey Barrett, Sonia Goncalves, Ewan Harrison, David K. Jackson, Ian Johnston, Dominic Kwiatkowski, Cordelia Langford, John Sillitoe on behalf of the Wellcome Sanger Institute COVID-19 Surveillance Team |
| EPI_ISL_3470669, EPI_ISL_3470670 | Lighthouse Lab in Milton Keynes | Wellcome Sanger Institute for the COVID-19 Genomics UK (COG-UK) Consortium | The Lighthouse Lab in Milton Keynes and Alex Alderton, Roberto Amato, Jeffrey Barrett, Sonia Goncalves, Ewan Harrison, David K. Jackson, Ian Johnston, Dominic Kwiatkowski, Cordelia Langford, John Sillitoe on behalf of the Wellcome Sanger Institute COVID-19 Surveillance Team |
| EPI_ISL_3470671 | Lighthouse Lab in Alderley Park | Wellcome Sanger Institute for the COVID-19 Genomics UK (COG-UK) Consortium | Jacquelyn Wynn, Mairead Hyland, The Lighthouse Lab in Alderley Park and Alex Alderton, Roberto Amato, Jeffrey Barrett, Sonia Goncalves, Ewan Harrison, David K. Jackson, Ian Johnston, Dominic Kwiatkowski, Cordelia Langford, John Sillitoe on behalf of the Wellcome Sanger Institute COVID-19 Surveillance Team |
| EPI_ISL_3470672 | Lighthouse Lab in Milton Keynes | Wellcome Sanger Institute for the COVID-19 Genomics UK (COG-UK) Consortium | The Lighthouse Lab in Milton Keynes and Alex Alderton, Roberto Amato, Jeffrey Barrett, Sonia Goncalves, Ewan Harrison, David K. Jackson, Ian Johnston, Dominic Kwiatkowski, Cordelia Langford, John Sillitoe on behalf of the Wellcome Sanger Institute COVID-19 Surveillance Team |
| EPI_ISL_3470673 | Lighthouse Lab in Glasgow | Wellcome Sanger Institute for the COVID-19 Genomics UK (COG-UK) Consortium | Harper VanSteenhouse, Yumi Kasai, David Gray, Carol Clugston, Anna Dominiczak and Alex Alderton, Roberto Amato, Jeffrey Barrett, Sonia Goncalves, Ewan Harrison, David K. Jackson, Ian Johnston, Dominic Kwiatkowski, Cordelia Langford, John Sillitoe on behalf of the Wellcome Sanger Institute COVID-19 Surveillance Team |
| EPI_ISL_3470674, EPI_ISL_3470675, EPI_ISL_3470676 | Lighthouse Lab in Alderley Park | Wellcome Sanger Institute for the COVID-19 Genomics UK (COG-UK) Consortium | Jacquelyn Wynn, Mairead Hyland, The Lighthouse Lab in Alderley Park and Alex Alderton, Roberto Amato, Jeffrey Barrett, Sonia Goncalves, Ewan Harrison, David K. Jackson, Ian Johnston, Dominic Kwiatkowski, Cordelia Langford, John Sillitoe on behalf of the Wellcome Sanger Institute COVID-19 Surveillance Team |
| EPI_ISL_3470677 | Lighthouse Lab in Glasgow | Wellcome Sanger Institute for the COVID-19 Genomics UK (COG-UK) Consortium | Harper VanSteenhouse, Yumi Kasai, David Gray, Carol Clugston, Anna Dominiczak and Alex Alderton, Roberto Amato, Jeffrey Barrett, Sonia Goncalves, Ewan Harrison, David K. Jackson, Ian Johnston, Dominic Kwiatkowski, Cordelia Langford, John Sillitoe on behalf of the Wellcome Sanger Institute COVID-19 Surveillance Team |
| EPI_ISL_3470678 | Lighthouse Lab in Alderley Park | Wellcome Sanger Institute for the COVID-19 Genomics UK (COG-UK) | Jacquelyn Wynn, Mairead Hyland, The Lighthouse Lab in Alderley Park and Alex Alderton, Roberto Amato, Jeffrey Barrett, Sonia Goncalves, Ewan Harrison, David K. Jackson, Ian Johnston, Dominic Kwiatkowski, Cordelia Langford, John Sillitoe on |

| | | | |
|---|---|---|---|
| | | Consortium | behalf of the Wellcome Sanger Institute COVID-19 Surveillance Team |
| EPI_ISL_3470679 | Lighthouse Lab in Glasgow | Wellcome Sanger Institute for the COVID-19 Genomics UK (COG-UK) Consortium | Harper VanSteenhouse, Yumi Kasai, David Gray, Carol Clugston, Anna Dominiczak and Alex Alderton, Roberto Amato, Jeffrey Barrett, Sonia Goncalves, Ewan Harrison, David K. Jackson, Ian Johnston, Dominic Kwiatkowski, Cordelia Langford, John Sillitoe on behalf of the Wellcome Sanger Institute COVID-19 Surveillance Team |
| EPI_ISL_3470680 | Lighthouse Lab in Milton Keynes | Wellcome Sanger Institute for the COVID-19 Genomics UK (COG-UK) Consortium | The Lighthouse Lab in Milton Keynes and Alex Alderton, Roberto Amato, Jeffrey Barrett, Sonia Goncalves, Ewan Harrison, David K. Jackson, Ian Johnston, Dominic Kwiatkowski, Cordelia Langford, John Sillitoe on behalf of the Wellcome Sanger Institute COVID-19 Surveillance Team |
| EPI_ISL_3470681, EPI_ISL_3470682, EPI_ISL_3470683 | Lighthouse Lab in Glasgow | Wellcome Sanger Institute for the COVID-19 Genomics UK (COG-UK) Consortium | Harper VanSteenhouse, Yumi Kasai, David Gray, Carol Clugston, Anna Dominiczak and Alex Alderton, Roberto Amato, Jeffrey Barrett, Sonia Goncalves, Ewan Harrison, David K. Jackson, Ian Johnston, Dominic Kwiatkowski, Cordelia Langford, John Sillitoe on behalf of the Wellcome Sanger Institute COVID-19 Surveillance Team |
| EPI_ISL_3470684, EPI_ISL_3470685 | Lighthouse Lab in Alderley Park | Wellcome Sanger Institute for the COVID-19 Genomics UK (COG-UK) Consortium | Jacquelyn Wynn, Mairead Hyland, The Lighthouse Lab in Alderley Park and Alex Alderton, Roberto Amato, Jeffrey Barrett, Sonia Goncalves, Ewan Harrison, David K. Jackson, Ian Johnston, Dominic Kwiatkowski, Cordelia Langford, John Sillitoe on behalf of the Wellcome Sanger Institute COVID-19 Surveillance Team |
| EPI_ISL_3470686, EPI_ISL_3470687 | Lighthouse Lab in Glasgow | Wellcome Sanger Institute for the COVID-19 Genomics UK (COG-UK) Consortium | Harper VanSteenhouse, Yumi Kasai, David Gray, Carol Clugston, Anna Dominiczak and Alex Alderton, Roberto Amato, Jeffrey Barrett, Sonia Goncalves, Ewan Harrison, David K. Jackson, Ian Johnston, Dominic Kwiatkowski, Cordelia Langford, John Sillitoe on behalf of the Wellcome Sanger Institute COVID-19 Surveillance Team |
| EPI_ISL_3470688 | Lighthouse Lab in Alderley Park | Wellcome Sanger Institute for the COVID-19 Genomics UK (COG-UK) Consortium | Jacquelyn Wynn, Mairead Hyland, The Lighthouse Lab in Alderley Park and Alex Alderton, Roberto Amato, Jeffrey Barrett, Sonia Goncalves, Ewan Harrison, David K. Jackson, Ian Johnston, Dominic Kwiatkowski, Cordelia Langford, John Sillitoe on behalf of the Wellcome Sanger Institute COVID-19 Surveillance Team |
| EPI_ISL_3470689 | Lighthouse Lab in Milton Keynes | Wellcome Sanger Institute for the COVID-19 Genomics UK (COG-UK) Consortium | The Lighthouse Lab in Milton Keynes and Alex Alderton, Roberto Amato, Jeffrey Barrett, Sonia Goncalves, Ewan Harrison, David K. Jackson, Ian Johnston, Dominic Kwiatkowski, Cordelia Langford, John Sillitoe on behalf of the Wellcome Sanger Institute COVID-19 Surveillance Team |
| EPI_ISL_3470690 | Lighthouse Lab in Alderley Park | Wellcome Sanger Institute for the COVID-19 Genomics UK (COG-UK) Consortium | Jacquelyn Wynn, Mairead Hyland, The Lighthouse Lab in Alderley Park and Alex Alderton, Roberto Amato, Jeffrey Barrett, Sonia Goncalves, Ewan Harrison, David K. Jackson, Ian Johnston, Dominic Kwiatkowski, Cordelia Langford, John Sillitoe on behalf of the Wellcome Sanger Institute COVID-19 Surveillance Team |
| EPI_ISL_3470691 | Lighthouse Lab in Glasgow | Wellcome Sanger Institute for the COVID-19 Genomics UK (COG-UK) Consortium | Harper VanSteenhouse, Yumi Kasai, David Gray, Carol Clugston, Anna Dominiczak and Alex Alderton, Roberto Amato, Jeffrey Barrett, Sonia Goncalves, Ewan Harrison, David K. Jackson, Ian Johnston, Dominic Kwiatkowski, Cordelia Langford, John Sillitoe on behalf of the Wellcome Sanger Institute COVID-19 Surveillance Team |
| EPI_ISL_3470692 | Lighthouse Lab in Milton Keynes | Wellcome Sanger Institute for the COVID-19 Genomics UK (COG-UK) Consortium | The Lighthouse Lab in Milton Keynes and Alex Alderton, Roberto Amato, Jeffrey Barrett, Sonia Goncalves, Ewan Harrison, David K. Jackson, Ian Johnston, Dominic Kwiatkowski, Cordelia Langford, John Sillitoe on behalf of the Wellcome Sanger Institute COVID-19 Surveillance Team |
| EPI_ISL_3470693, EPI_ISL_3470694 | Lighthouse Lab in Glasgow | Wellcome Sanger Institute for the COVID-19 Genomics UK (COG-UK) Consortium | Harper VanSteenhouse, Yumi Kasai, David Gray, Carol Clugston, Anna Dominiczak and Alex Alderton, Roberto Amato, Jeffrey Barrett, Sonia Goncalves, Ewan Harrison, David K. Jackson, Ian Johnston, Dominic Kwiatkowski, Cordelia Langford, John Sillitoe on behalf of the Wellcome Sanger Institute COVID-19 Surveillance Team |
| EPI_ISL_3470695 | Lighthouse Lab in Milton Keynes | Wellcome Sanger Institute for the COVID-19 Genomics UK (COG-UK) Consortium | The Lighthouse Lab in Milton Keynes and Alex Alderton, Roberto Amato, Jeffrey Barrett, Sonia Goncalves, Ewan Harrison, David K. Jackson, Ian Johnston, Dominic Kwiatkowski, Cordelia Langford, John Sillitoe on behalf of the Wellcome Sanger Institute COVID-19 Surveillance Team |
| EPI_ISL_3470696 | Lighthouse Lab in Alderley Park | Wellcome Sanger Institute for the COVID-19 Genomics UK (COG-UK) Consortium | Jacquelyn Wynn, Mairead Hyland, The Lighthouse Lab in Alderley Park and Alex Alderton, Roberto Amato, Jeffrey Barrett, Sonia Goncalves, Ewan Harrison, David K. Jackson, Ian Johnston, Dominic Kwiatkowski, Cordelia Langford, John Sillitoe on behalf of the Wellcome Sanger Institute COVID-19 Surveillance Team |
| EPI_ISL_3470697, EPI_ISL_3470698, EPI_ISL_3470699, EPI_ISL_3470700 | Lighthouse Lab in Glasgow | Wellcome Sanger Institute for the COVID-19 Genomics UK (COG-UK) Consortium | Harper VanSteenhouse, Yumi Kasai, David Gray, Carol Clugston, Anna Dominiczak and Alex Alderton, Roberto Amato, Jeffrey Barrett, Sonia Goncalves, Ewan Harrison, David K. Jackson, Ian Johnston, Dominic Kwiatkowski, Cordelia Langford, John Sillitoe on behalf of the Wellcome Sanger Institute COVID-19 Surveillance Team |
| EPI_ISL_3470701, EPI_ISL_3470702 | Lighthouse Lab in Milton Keynes | Wellcome Sanger Institute for the COVID-19 Genomics UK (COG-UK) Consortium | The Lighthouse Lab in Milton Keynes and Alex Alderton, Roberto Amato, Jeffrey Barrett, Sonia Goncalves, Ewan Harrison, David K. Jackson, Ian Johnston, Dominic Kwiatkowski, Cordelia Langford, John Sillitoe on behalf of the Wellcome Sanger Institute COVID-19 Surveillance Team |
| EPI_ISL_3470703 | Lighthouse Lab in Glasgow | Wellcome Sanger Institute for the COVID-19 Genomics UK (COG-UK) Consortium | Harper VanSteenhouse, Yumi Kasai, David Gray, Carol Clugston, Anna Dominiczak and Alex Alderton, Roberto Amato, Jeffrey Barrett, Sonia Goncalves, Ewan Harrison, David K. Jackson, Ian Johnston, Dominic Kwiatkowski, Cordelia Langford, John Sillitoe on behalf of the Wellcome Sanger Institute COVID-19 Surveillance Team |
| EPI_ISL_3470704 | Lighthouse Lab in Alderley Park | Wellcome Sanger Institute for the COVID-19 Genomics UK (COG-UK) Consortium | Jacquelyn Wynn, Mairead Hyland, The Lighthouse Lab in Alderley Park and Alex Alderton, Roberto Amato, Jeffrey Barrett, Sonia Goncalves, Ewan Harrison, David K. Jackson, Ian Johnston, Dominic Kwiatkowski, Cordelia Langford, John Sillitoe on behalf of the Wellcome Sanger Institute COVID-19 Surveillance Team |
| EPI_ISL_3470705, EPI_ISL_3470706 | Lighthouse Lab in Glasgow | Wellcome Sanger Institute for the COVID-19 Genomics UK (COG-UK) Consortium | Harper VanSteenhouse, Yumi Kasai, David Gray, Carol Clugston, Anna Dominiczak and Alex Alderton, Roberto Amato, Jeffrey Barrett, Sonia Goncalves, Ewan Harrison, David K. Jackson, Ian Johnston, Dominic Kwiatkowski, Cordelia Langford, John Sillitoe on behalf of the Wellcome Sanger Institute COVID-19 Surveillance Team |
| EPI_ISL_3470707, EPI_ISL_3470708 | Lighthouse Lab in Milton Keynes | Wellcome Sanger Institute for the COVID-19 Genomics UK (COG-UK) Consortium | The Lighthouse Lab in Milton Keynes and Alex Alderton, Roberto Amato, Jeffrey Barrett, Sonia Goncalves, Ewan Harrison, David K. Jackson, Ian Johnston, Dominic Kwiatkowski, Cordelia Langford, John Sillitoe on behalf of the Wellcome Sanger Institute COVID-19 Surveillance Team |
| EPI_ISL_3470709 | Lighthouse Lab in Glasgow | Wellcome Sanger Institute for the COVID-19 Genomics UK (COG-UK) Consortium | Harper VanSteenhouse, Yumi Kasai, David Gray, Carol Clugston, Anna Dominiczak and Alex Alderton, Roberto Amato, Jeffrey Barrett, Sonia Goncalves, Ewan Harrison, David K. Jackson, Ian Johnston, Dominic Kwiatkowski, Cordelia Langford, John Sillitoe on behalf of the Wellcome Sanger Institute COVID-19 Surveillance Team |
| EPI_ISL_3470710 | Lighthouse Lab in Milton Keynes | Wellcome Sanger Institute for the COVID-19 Genomics UK (COG-UK) Consortium | The Lighthouse Lab in Milton Keynes and Alex Alderton, Roberto Amato, Jeffrey Barrett, Sonia Goncalves, Ewan Harrison, David K. Jackson, Ian Johnston, Dominic Kwiatkowski, Cordelia Langford, John Sillitoe on behalf of the Wellcome Sanger Institute COVID-19 Surveillance Team |
| EPI_ISL_3470711 | Lighthouse Lab in Alderley Park | Wellcome Sanger Institute for the COVID-19 Genomics UK (COG-UK) Consortium | Jacquelyn Wynn, Mairead Hyland, The Lighthouse Lab in Alderley Park and Alex Alderton, Roberto Amato, Jeffrey Barrett, Sonia Goncalves, Ewan Harrison, David K. Jackson, Ian Johnston, Dominic Kwiatkowski, Cordelia Langford, John Sillitoe on behalf of the Wellcome Sanger Institute COVID-19 Surveillance Team |
| EPI_ISL_3470712 | Lighthouse Lab in Milton Keynes | Wellcome Sanger Institute for the COVID-19 Genomics UK (COG-UK) Consortium | The Lighthouse Lab in Milton Keynes and Alex Alderton, Roberto Amato, Jeffrey Barrett, Sonia Goncalves, Ewan Harrison, David K. Jackson, Ian Johnston, Dominic Kwiatkowski, Cordelia Langford, John Sillitoe on behalf of the Wellcome Sanger Institute COVID-19 Surveillance Team |
| EPI_ISL_3470713 | Lighthouse Lab in Glasgow | Wellcome Sanger Institute for the COVID-19 Genomics UK (COG-UK) Consortium | Harper VanSteenhouse, Yumi Kasai, David Gray, Carol Clugston, Anna Dominiczak and Alex Alderton, Roberto Amato, Jeffrey Barrett, Sonia Goncalves, Ewan Harrison, David K. Jackson, Ian Johnston, Dominic Kwiatkowski, Cordelia Langford, John Sillitoe on behalf of the Wellcome Sanger Institute COVID-19 Surveillance Team |
| EPI_ISL_3470714 | Lighthouse Lab in Milton Keynes | Wellcome Sanger Institute for the COVID-19 Genomics UK (COG-UK) Consortium | The Lighthouse Lab in Milton Keynes and Alex Alderton, Roberto Amato, Jeffrey Barrett, Sonia Goncalves, Ewan Harrison, David K. Jackson, Ian Johnston, Dominic Kwiatkowski, Cordelia Langford, John Sillitoe on behalf of the Wellcome Sanger Institute COVID-19 Surveillance Team |
| EPI_ISL_3470715 | Lighthouse Lab in Alderley Park | Wellcome Sanger Institute for the COVID-19 Genomics UK (COG-UK) Consortium | Jacquelyn Wynn, Mairead Hyland, The Lighthouse Lab in Alderley Park and Alex Alderton, Roberto Amato, Jeffrey Barrett, Sonia Goncalves, Ewan Harrison, David K. Jackson, Ian Johnston, Dominic Kwiatkowski, Cordelia Langford, John Sillitoe on behalf of the Wellcome Sanger Institute COVID-19 Surveillance Team |
| EPI_ISL_3470716, EPI_ISL_3470717 | Lighthouse Lab in Glasgow | Wellcome Sanger Institute for the COVID-19 Genomics UK (COG-UK) Consortium | Harper VanSteenhouse, Yumi Kasai, David Gray, Carol Clugston, Anna Dominiczak and Alex Alderton, Roberto Amato, Jeffrey Barrett, Sonia Goncalves, Ewan Harrison, David K. Jackson, Ian Johnston, Dominic Kwiatkowski, Cordelia Langford, John Sillitoe on behalf of the Wellcome Sanger Institute COVID-19 Surveillance Team |
| EPI_ISL_3470718 | Lighthouse Lab in Alderley Park | Wellcome Sanger Institute for the COVID-19 Genomics UK (COG-UK) Consortium | Jacquelyn Wynn, Mairead Hyland, The Lighthouse Lab in Alderley Park and Alex Alderton, Roberto Amato, Jeffrey Barrett, Sonia Goncalves, Ewan Harrison, David K. Jackson, Ian Johnston, Dominic Kwiatkowski, Cordelia Langford, John Sillitoe on behalf of the Wellcome Sanger Institute COVID-19 Surveillance Team |
| EPI_ISL_3470719 | Lighthouse Lab in Milton Keynes | Wellcome Sanger Institute for the COVID-19 Genomics UK (COG-UK) Consortium | The Lighthouse Lab in Milton Keynes and Alex Alderton, Roberto Amato, Jeffrey Barrett, Sonia Goncalves, Ewan Harrison, David K. Jackson, Ian Johnston, Dominic Kwiatkowski, Cordelia Langford, John Sillitoe on behalf of the Wellcome Sanger Institute COVID-19 Surveillance Team |
| EPI_ISL_3470720, EPI_ISL_3470721, EPI_ISL_3470722, EPI_ISL_3470723 | Lighthouse Lab in Glasgow | Wellcome Sanger Institute for the COVID-19 Genomics UK (COG-UK) Consortium | Harper VanSteenhouse, Yumi Kasai, David Gray, Carol Clugston, Anna Dominiczak and Alex Alderton, Roberto Amato, Jeffrey Barrett, Sonia Goncalves, Ewan Harrison, David K. Jackson, Ian Johnston, Dominic Kwiatkowski, Cordelia Langford, John Sillitoe on behalf of the Wellcome Sanger Institute COVID-19 Surveillance Team |
| EPI_ISL_3470724 | Lighthouse Lab in Alderley Park | Wellcome Sanger Institute for the COVID-19 Genomics UK (COG-UK) Consortium | Jacquelyn Wynn, Mairead Hyland, The Lighthouse Lab in Alderley Park and Alex Alderton, Roberto Amato, Jeffrey Barrett, Sonia Goncalves, Ewan Harrison, David K. Jackson, Ian Johnston, Dominic Kwiatkowski, Cordelia Langford, John Sillitoe on behalf of the Wellcome Sanger Institute COVID-19 Surveillance Team |
| EPI_ISL_3470725 | Lighthouse Lab in Glasgow | Wellcome Sanger Institute for the COVID-19 Genomics UK (COG-UK) Consortium | Harper VanSteenhouse, Yumi Kasai, David Gray, Carol Clugston, Anna Dominiczak and Alex Alderton, Roberto Amato, Jeffrey Barrett, Sonia Goncalves, Ewan Harrison, David K. Jackson, Ian Johnston, Dominic Kwiatkowski, Cordelia Langford, John Sillitoe on behalf of the Wellcome Sanger Institute COVID-19 Surveillance Team |
| EPI_ISL_3470726 | Lighthouse Lab in Milton Keynes | Wellcome Sanger Institute for the COVID-19 Genomics UK (COG-UK) Consortium | The Lighthouse Lab in Milton Keynes and Alex Alderton, Roberto Amato, Jeffrey Barrett, Sonia Goncalves, Ewan Harrison, David K. Jackson, Ian Johnston, Dominic Kwiatkowski, Cordelia Langford, John Sillitoe on behalf of the Wellcome Sanger Institute COVID-19 Surveillance Team |
| EPI_ISL_3470727, EPI_ISL_3470728 | Lighthouse Lab in Glasgow | Wellcome Sanger Institute for the COVID-19 Genomics UK (COG-UK) Consortium | Harper VanSteenhouse, Yumi Kasai, David Gray, Carol Clugston, Anna Dominiczak and Alex Alderton, Roberto Amato, Jeffrey Barrett, Sonia Goncalves, Ewan Harrison, David K. Jackson, Ian Johnston, Dominic Kwiatkowski, Cordelia Langford, John Sillitoe on behalf of the Wellcome Sanger Institute COVID-19 Surveillance Team |
| EPI_ISL_3470729 | Lighthouse Lab in Milton Keynes | Wellcome Sanger Institute for the COVID-19 Genomics UK (COG-UK) Consortium | The Lighthouse Lab in Milton Keynes and Alex Alderton, Roberto Amato, Jeffrey Barrett, Sonia Goncalves, Ewan Harrison, David K. Jackson, Ian Johnston, Dominic Kwiatkowski, Cordelia Langford, John Sillitoe on behalf of the Wellcome Sanger Institute COVID-19 Surveillance Team |
| EPI_ISL_3470730 | Lighthouse Lab in Alderley Park | Wellcome Sanger Institute for the COVID-19 Genomics UK (COG-UK) Consortium | Jacquelyn Wynn, Mairead Hyland, The Lighthouse Lab in Alderley Park and Alex Alderton, Roberto Amato, Jeffrey Barrett, Sonia Goncalves, Ewan Harrison, David K. Jackson, Ian Johnston, Dominic Kwiatkowski, Cordelia Langford, John Sillitoe on behalf of the Wellcome Sanger Institute COVID-19 Surveillance Team |
| EPI_ISL_3470731 | Lighthouse Lab in Milton Keynes | Wellcome Sanger Institute for the COVID-19 Genomics UK (COG-UK) Consortium | The Lighthouse Lab in Milton Keynes and Alex Alderton, Roberto Amato, Jeffrey Barrett, Sonia Goncalves, Ewan Harrison, David K. Jackson, Ian Johnston, Dominic Kwiatkowski, Cordelia Langford, John Sillitoe on behalf of the Wellcome Sanger Institute COVID-19 Surveillance Team |
| EPI_ISL_3470732 | Lighthouse Lab in Glasgow | Wellcome Sanger Institute for the COVID-19 Genomics UK (COG-UK) Consortium | Harper VanSteenhouse, Yumi Kasai, David Gray, Carol Clugston, Anna Dominiczak and Alex Alderton, Roberto Amato, Jeffrey Barrett, Sonia Goncalves, Ewan Harrison, David K. Jackson, Ian Johnston, Dominic Kwiatkowski, Cordelia Langford, John Sillitoe on behalf of the Wellcome Sanger Institute COVID-19 Surveillance Team |
| EPI_ISL_3470733 | Lighthouse Lab in Milton Keynes | Wellcome Sanger Institute for the COVID-19 Genomics UK (COG-UK) Consortium | The Lighthouse Lab in Milton Keynes and Alex Alderton, Roberto Amato, Jeffrey Barrett, Sonia Goncalves, Ewan Harrison, David K. Jackson, Ian Johnston, Dominic Kwiatkowski, Cordelia Langford, John Sillitoe on behalf of the Wellcome Sanger Institute COVID-19 Surveillance Team |
| EPI_ISL_3470734, EPI_ISL_3470735, EPI_ISL_3470736 | Lighthouse Lab in Glasgow | Wellcome Sanger Institute for the COVID-19 Genomics UK (COG-UK) Consortium | Harper VanSteenhouse, Yumi Kasai, David Gray, Carol Clugston, Anna Dominiczak and Alex Alderton, Roberto Amato, Jeffrey Barrett, Sonia Goncalves, Ewan Harrison, David K. Jackson, Ian Johnston, Dominic Kwiatkowski, Cordelia Langford, John Sillitoe on behalf of the Wellcome Sanger Institute COVID-19 Surveillance Team |
| EPI_ISL_3470737 | Lighthouse Lab in Milton Keynes | Wellcome Sanger Institute for the COVID-19 Genomics UK (COG-UK) Consortium | The Lighthouse Lab in Milton Keynes and Alex Alderton, Roberto Amato, Jeffrey Barrett, Sonia Goncalves, Ewan Harrison, David K. Jackson, Ian Johnston, Dominic Kwiatkowski, Cordelia Langford, John Sillitoe on behalf of the Wellcome Sanger Institute COVID-19 Surveillance Team |
| EPI_ISL_3470738 | Lighthouse Lab in Alderley Park | Wellcome Sanger Institute for the COVID-19 Genomics UK (COG-UK) Consortium | Jacquelyn Wynn, Mairead Hyland, The Lighthouse Lab in Alderley Park and Alex Alderton, Roberto Amato, Jeffrey Barrett, Sonia Goncalves, Ewan Harrison, David K. Jackson, Ian Johnston, Dominic Kwiatkowski, Cordelia Langford, John Sillitoe on behalf of the Wellcome Sanger Institute COVID-19 Surveillance Team |

| | | | |
|---|---|---|---|
| EPI_ISL_3470739 | Lighthouse Lab in Glasgow | Wellcome Sanger Institute for the COVID-19 Genomics UK (COG-UK) Consortium | Harper VanSteenhouse, Yumi Kasai, David Gray, Carol Clugston, Anna Dominiczak and Alex Alderton, Roberto Amato, Jeffrey Barrett, Sonia Goncalves, Ewan Harrison, David K. Jackson, Ian Johnston, Dominic Kwiatkowski, Cordelia Langford, John Sillitoe on behalf of the Wellcome Sanger Institute COVID-19 Surveillance Team |
| EPI_ISL_3470740 | Lighthouse Lab in Alderley Park | Wellcome Sanger Institute for the COVID-19 Genomics UK (COG-UK) Consortium | Jacquelyn Wynn, Mairead Hyland, The Lighthouse Lab in Alderley Park and Alex Alderton, Roberto Amato, Jeffrey Barrett, Sonia Goncalves, Ewan Harrison, David K. Jackson, Ian Johnston, Dominic Kwiatkowski, Cordelia Langford, John Sillitoe on behalf of the Wellcome Sanger Institute COVID-19 Surveillance Team |
| EPI_ISL_3470741 | Lighthouse Lab in Milton Keynes | Wellcome Sanger Institute for the COVID-19 Genomics UK (COG-UK) Consortium | The Lighthouse Lab in Milton Keynes and Alex Alderton, Roberto Amato, Jeffrey Barrett, Sonia Goncalves, Ewan Harrison, David K. Jackson, Ian Johnston, Dominic Kwiatkowski, Cordelia Langford, John Sillitoe on behalf of the Wellcome Sanger Institute COVID-19 Surveillance Team |
| EPI_ISL_3470742 | Lighthouse Lab in Alderley Park | Wellcome Sanger Institute for the COVID-19 Genomics UK (COG-UK) Consortium | Jacquelyn Wynn, Mairead Hyland, The Lighthouse Lab in Alderley Park and Alex Alderton, Roberto Amato, Jeffrey Barrett, Sonia Goncalves, Ewan Harrison, David K. Jackson, Ian Johnston, Dominic Kwiatkowski, Cordelia Langford, John Sillitoe on behalf of the Wellcome Sanger Institute COVID-19 Surveillance Team |
| EPI_ISL_3470743, EPI_ISL_3470744 | Lighthouse Lab in Milton Keynes | Wellcome Sanger Institute for the COVID-19 Genomics UK (COG-UK) Consortium | The Lighthouse Lab in Milton Keynes and Alex Alderton, Roberto Amato, Jeffrey Barrett, Sonia Goncalves, Ewan Harrison, David K. Jackson, Ian Johnston, Dominic Kwiatkowski, Cordelia Langford, John Sillitoe on behalf of the Wellcome Sanger Institute COVID-19 Surveillance Team |
| EPI_ISL_3470745 | Lighthouse Lab in Alderley Park | Wellcome Sanger Institute for the COVID-19 Genomics UK (COG-UK) Consortium | Jacquelyn Wynn, Mairead Hyland, The Lighthouse Lab in Alderley Park and Alex Alderton, Roberto Amato, Jeffrey Barrett, Sonia Goncalves, Ewan Harrison, David K. Jackson, Ian Johnston, Dominic Kwiatkowski, Cordelia Langford, John Sillitoe on behalf of the Wellcome Sanger Institute COVID-19 Surveillance Team |
| EPI_ISL_3470746, EPI_ISL_3470747, EPI_ISL_3470748, EPI_ISL_3470749 | Lighthouse Lab in Glasgow | Wellcome Sanger Institute for the COVID-19 Genomics UK (COG-UK) Consortium | Harper VanSteenhouse, Yumi Kasai, David Gray, Carol Clugston, Anna Dominiczak and Alex Alderton, Roberto Amato, Jeffrey Barrett, Sonia Goncalves, Ewan Harrison, David K. Jackson, Ian Johnston, Dominic Kwiatkowski, Cordelia Langford, John Sillitoe on behalf of the Wellcome Sanger Institute COVID-19 Surveillance Team |
| EPI_ISL_3470750 | Lighthouse Lab in Alderley Park | Wellcome Sanger Institute for the COVID-19 Genomics UK (COG-UK) Consortium | Jacquelyn Wynn, Mairead Hyland, The Lighthouse Lab in Alderley Park and Alex Alderton, Roberto Amato, Jeffrey Barrett, Sonia Goncalves, Ewan Harrison, David K. Jackson, Ian Johnston, Dominic Kwiatkowski, Cordelia Langford, John Sillitoe on behalf of the Wellcome Sanger Institute COVID-19 Surveillance Team |
| EPI_ISL_3470751, EPI_ISL_3470752 | Lighthouse Lab in Glasgow | Wellcome Sanger Institute for the COVID-19 Genomics UK (COG-UK) Consortium | Harper VanSteenhouse, Yumi Kasai, David Gray, Carol Clugston, Anna Dominiczak and Alex Alderton, Roberto Amato, Jeffrey Barrett, Sonia Goncalves, Ewan Harrison, David K. Jackson, Ian Johnston, Dominic Kwiatkowski, Cordelia Langford, John Sillitoe on behalf of the Wellcome Sanger Institute COVID-19 Surveillance Team |
| EPI_ISL_3470753 | Lighthouse Lab in Milton Keynes | Wellcome Sanger Institute for the COVID-19 Genomics UK (COG-UK) Consortium | The Lighthouse Lab in Milton Keynes and Alex Alderton, Roberto Amato, Jeffrey Barrett, Sonia Goncalves, Ewan Harrison, David K. Jackson, Ian Johnston, Dominic Kwiatkowski, Cordelia Langford, John Sillitoe on behalf of the Wellcome Sanger Institute COVID-19 Surveillance Team |
| EPI_ISL_3470754 | Lighthouse Lab in Glasgow | Wellcome Sanger Institute for the COVID-19 Genomics UK (COG-UK) Consortium | Harper VanSteenhouse, Yumi Kasai, David Gray, Carol Clugston, Anna Dominiczak and Alex Alderton, Roberto Amato, Jeffrey Barrett, Sonia Goncalves, Ewan Harrison, David K. Jackson, Ian Johnston, Dominic Kwiatkowski, Cordelia Langford, John Sillitoe on behalf of the Wellcome Sanger Institute COVID-19 Surveillance Team |
| EPI_ISL_3470755 | Lighthouse Lab in Alderley Park | Wellcome Sanger Institute for the COVID-19 Genomics UK (COG-UK) Consortium | Jacquelyn Wynn, Mairead Hyland, The Lighthouse Lab in Alderley Park and Alex Alderton, Roberto Amato, Jeffrey Barrett, Sonia Goncalves, Ewan Harrison, David K. Jackson, Ian Johnston, Dominic Kwiatkowski, Cordelia Langford, John Sillitoe on behalf of the Wellcome Sanger Institute COVID-19 Surveillance Team |
| EPI_ISL_3470756 | Lighthouse Lab in Glasgow | Wellcome Sanger Institute for the COVID-19 Genomics UK (COG-UK) Consortium | Harper VanSteenhouse, Yumi Kasai, David Gray, Carol Clugston, Anna Dominiczak and Alex Alderton, Roberto Amato, Jeffrey Barrett, Sonia Goncalves, Ewan Harrison, David K. Jackson, Ian Johnston, Dominic Kwiatkowski, Cordelia Langford, John Sillitoe on behalf of the Wellcome Sanger Institute COVID-19 Surveillance Team |
| EPI_ISL_3470757 | Lighthouse Lab in Milton Keynes | Wellcome Sanger Institute for the COVID-19 Genomics UK (COG-UK) Consortium | The Lighthouse Lab in Milton Keynes and Alex Alderton, Roberto Amato, Jeffrey Barrett, Sonia Goncalves, Ewan Harrison, David K. Jackson, Ian Johnston, Dominic Kwiatkowski, Cordelia Langford, John Sillitoe on behalf of the Wellcome Sanger Institute COVID-19 Surveillance Team |
| EPI_ISL_3470758, EPI_ISL_3470759, EPI_ISL_3470760 | Lighthouse Lab in Glasgow | Wellcome Sanger Institute for the COVID-19 Genomics UK (COG-UK) Consortium | Harper VanSteenhouse, Yumi Kasai, David Gray, Carol Clugston, Anna Dominiczak and Alex Alderton, Roberto Amato, Jeffrey Barrett, Sonia Goncalves, Ewan Harrison, David K. Jackson, Ian Johnston, Dominic Kwiatkowski, Cordelia Langford, John Sillitoe on behalf of the Wellcome Sanger Institute COVID-19 Surveillance Team |
| EPI_ISL_3470761 | Lighthouse Lab in Alderley Park | Wellcome Sanger Institute for the COVID-19 Genomics UK (COG-UK) Consortium | Jacquelyn Wynn, Mairead Hyland, The Lighthouse Lab in Alderley Park and Alex Alderton, Roberto Amato, Jeffrey Barrett, Sonia Goncalves, Ewan Harrison, David K. Jackson, Ian Johnston, Dominic Kwiatkowski, Cordelia Langford, John Sillitoe on behalf of the Wellcome Sanger Institute COVID-19 Surveillance Team |
| EPI_ISL_3470762 | Lighthouse Lab in Milton Keynes | Wellcome Sanger Institute for the COVID-19 Genomics UK (COG-UK) Consortium | The Lighthouse Lab in Milton Keynes and Alex Alderton, Roberto Amato, Jeffrey Barrett, Sonia Goncalves, Ewan Harrison, David K. Jackson, Ian Johnston, Dominic Kwiatkowski, Cordelia Langford, John Sillitoe on behalf of the Wellcome Sanger Institute COVID-19 Surveillance Team |
| EPI_ISL_3470763 | Lighthouse Lab in Alderley Park | Wellcome Sanger Institute for the COVID-19 Genomics UK (COG-UK) Consortium | Jacquelyn Wynn, Mairead Hyland, The Lighthouse Lab in Alderley Park and Alex Alderton, Roberto Amato, Jeffrey Barrett, Sonia Goncalves, Ewan Harrison, David K. Jackson, Ian Johnston, Dominic Kwiatkowski, Cordelia Langford, John Sillitoe on behalf of the Wellcome Sanger Institute COVID-19 Surveillance Team |
| EPI_ISL_3470764 | Lighthouse Lab in Glasgow | Wellcome Sanger Institute for the COVID-19 Genomics UK (COG-UK) Consortium | Harper VanSteenhouse, Yumi Kasai, David Gray, Carol Clugston, Anna Dominiczak and Alex Alderton, Roberto Amato, Jeffrey Barrett, Sonia Goncalves, Ewan Harrison, David K. Jackson, Ian Johnston, Dominic Kwiatkowski, Cordelia Langford, John Sillitoe on behalf of the Wellcome Sanger Institute COVID-19 Surveillance Team |
| EPI_ISL_3470765 | Lighthouse Lab in Alderley Park | Wellcome Sanger Institute for the COVID-19 Genomics UK (COG-UK) Consortium | Jacquelyn Wynn, Mairead Hyland, The Lighthouse Lab in Alderley Park and Alex Alderton, Roberto Amato, Jeffrey Barrett, Sonia Goncalves, Ewan Harrison, David K. Jackson, Ian Johnston, Dominic Kwiatkowski, Cordelia Langford, John Sillitoe on behalf of the Wellcome Sanger Institute COVID-19 Surveillance Team |
| EPI_ISL_3470766, EPI_ISL_3470767 | Lighthouse Lab in Glasgow | Wellcome Sanger Institute for the COVID-19 Genomics UK (COG-UK) Consortium | Harper VanSteenhouse, Yumi Kasai, David Gray, Carol Clugston, Anna Dominiczak and Alex Alderton, Roberto Amato, Jeffrey Barrett, Sonia Goncalves, Ewan Harrison, David K. Jackson, Ian Johnston, Dominic Kwiatkowski, Cordelia Langford, John Sillitoe on behalf of the Wellcome Sanger Institute COVID-19 Surveillance Team |
| EPI_ISL_3470768, EPI_ISL_3470769 | Lighthouse Lab in Milton Keynes | Wellcome Sanger Institute for the COVID-19 Genomics UK (COG-UK) Consortium | The Lighthouse Lab in Milton Keynes and Alex Alderton, Roberto Amato, Jeffrey Barrett, Sonia Goncalves, Ewan Harrison, David K. Jackson, Ian Johnston, Dominic Kwiatkowski, Cordelia Langford, John Sillitoe on behalf of the Wellcome Sanger Institute COVID-19 Surveillance Team |
| EPI_ISL_3470770 | Lighthouse Lab in Alderley Park | Wellcome Sanger Institute for the COVID-19 Genomics UK (COG-UK) Consortium | Jacquelyn Wynn, Mairead Hyland, The Lighthouse Lab in Alderley Park and Alex Alderton, Roberto Amato, Jeffrey Barrett, Sonia Goncalves, Ewan Harrison, David K. Jackson, Ian Johnston, Dominic Kwiatkowski, Cordelia Langford, John Sillitoe on behalf of the Wellcome Sanger Institute COVID-19 Surveillance Team |
| EPI_ISL_3470771 | Lighthouse Lab in Milton Keynes | Wellcome Sanger Institute for the COVID-19 Genomics UK (COG-UK) Consortium | The Lighthouse Lab in Milton Keynes and Alex Alderton, Roberto Amato, Jeffrey Barrett, Sonia Goncalves, Ewan Harrison, David K. Jackson, Ian Johnston, Dominic Kwiatkowski, Cordelia Langford, John Sillitoe on behalf of the Wellcome Sanger Institute COVID-19 Surveillance Team |
| EPI_ISL_3470772, EPI_ISL_3470773 | Lighthouse Lab in Glasgow | Wellcome Sanger Institute for the COVID-19 Genomics UK (COG-UK) Consortium | Harper VanSteenhouse, Yumi Kasai, David Gray, Carol Clugston, Anna Dominiczak and Alex Alderton, Roberto Amato, Jeffrey Barrett, Sonia Goncalves, Ewan Harrison, David K. Jackson, Ian Johnston, Dominic Kwiatkowski, Cordelia Langford, John Sillitoe on behalf of the Wellcome Sanger Institute COVID-19 Surveillance Team |
| EPI_ISL_3470774 | Lighthouse Lab in Milton Keynes | Wellcome Sanger Institute for the COVID-19 Genomics UK (COG-UK) Consortium | The Lighthouse Lab in Milton Keynes and Alex Alderton, Roberto Amato, Jeffrey Barrett, Sonia Goncalves, Ewan Harrison, David K. Jackson, Ian Johnston, Dominic Kwiatkowski, Cordelia Langford, John Sillitoe on behalf of the Wellcome Sanger Institute COVID-19 Surveillance Team |
| EPI_ISL_3470775 | Lighthouse Lab in Glasgow | Wellcome Sanger Institute for the COVID-19 Genomics UK (COG-UK) Consortium | Harper VanSteenhouse, Yumi Kasai, David Gray, Carol Clugston, Anna Dominiczak and Alex Alderton, Roberto Amato, Jeffrey Barrett, Sonia Goncalves, Ewan Harrison, David K. Jackson, Ian Johnston, Dominic Kwiatkowski, Cordelia Langford, John Sillitoe on behalf of the Wellcome Sanger Institute COVID-19 Surveillance Team |
| EPI_ISL_3470776, EPI_ISL_3470777 | Lighthouse Lab in Alderley Park | Wellcome Sanger Institute for the COVID-19 Genomics UK (COG-UK) Consortium | Jacquelyn Wynn, Mairead Hyland, The Lighthouse Lab in Alderley Park and Alex Alderton, Roberto Amato, Jeffrey Barrett, Sonia Goncalves, Ewan Harrison, David K. Jackson, Ian Johnston, Dominic Kwiatkowski, Cordelia Langford, John Sillitoe on behalf of the Wellcome Sanger Institute COVID-19 Surveillance Team |
| EPI_ISL_3470778, EPI_ISL_3470779 | Lighthouse Lab in Glasgow | Wellcome Sanger Institute for the COVID-19 Genomics UK (COG-UK) Consortium | Harper VanSteenhouse, Yumi Kasai, David Gray, Carol Clugston, Anna Dominiczak and Alex Alderton, Roberto Amato, Jeffrey Barrett, Sonia Goncalves, Ewan Harrison, David K. Jackson, Ian Johnston, Dominic Kwiatkowski, Cordelia Langford, John Sillitoe on behalf of the Wellcome Sanger Institute COVID-19 Surveillance Team |
| EPI_ISL_3470780, EPI_ISL_3470781 | Lighthouse Lab in Alderley Park | Wellcome Sanger Institute for the COVID-19 Genomics UK (COG-UK) Consortium | Jacquelyn Wynn, Mairead Hyland, The Lighthouse Lab in Alderley Park and Alex Alderton, Roberto Amato, Jeffrey Barrett, Sonia Goncalves, Ewan Harrison, David K. Jackson, Ian Johnston, Dominic Kwiatkowski, Cordelia Langford, John Sillitoe on behalf of the Wellcome Sanger Institute COVID-19 Surveillance Team |
| EPI_ISL_3470782 | Lighthouse Lab in Milton Keynes | Wellcome Sanger Institute for the COVID-19 Genomics UK (COG-UK) Consortium | The Lighthouse Lab in Milton Keynes and Alex Alderton, Roberto Amato, Jeffrey Barrett, Sonia Goncalves, Ewan Harrison, David K. Jackson, Ian Johnston, Dominic Kwiatkowski, Cordelia Langford, John Sillitoe on behalf of the Wellcome Sanger Institute COVID-19 Surveillance Team |
| EPI_ISL_3470783, EPI_ISL_3470784, EPI_ISL_3470785, EPI_ISL_3470786 | Lighthouse Lab in Glasgow | Wellcome Sanger Institute for the COVID-19 Genomics UK (COG-UK) Consortium | Harper VanSteenhouse, Yumi Kasai, David Gray, Carol Clugston, Anna Dominiczak and Alex Alderton, Roberto Amato, Jeffrey Barrett, Sonia Goncalves, Ewan Harrison, David K. Jackson, Ian Johnston, Dominic Kwiatkowski, Cordelia Langford, John Sillitoe on behalf of the Wellcome Sanger Institute COVID-19 Surveillance Team |
| EPI_ISL_3470787 | Lighthouse Lab in Milton Keynes | Wellcome Sanger Institute for the COVID-19 Genomics UK (COG-UK) Consortium | The Lighthouse Lab in Milton Keynes and Alex Alderton, Roberto Amato, Jeffrey Barrett, Sonia Goncalves, Ewan Harrison, David K. Jackson, Ian Johnston, Dominic Kwiatkowski, Cordelia Langford, John Sillitoe on behalf of the Wellcome Sanger Institute COVID-19 Surveillance Team |
| EPI_ISL_3470788, EPI_ISL_3470789, EPI_ISL_3470790, EPI_ISL_3470791, EPI_ISL_3470792 | Lighthouse Lab in Glasgow | Wellcome Sanger Institute for the COVID-19 Genomics UK (COG-UK) Consortium | Harper VanSteenhouse, Yumi Kasai, David Gray, Carol Clugston, Anna Dominiczak and Alex Alderton, Roberto Amato, Jeffrey Barrett, Sonia Goncalves, Ewan Harrison, David K. Jackson, Ian Johnston, Dominic Kwiatkowski, Cordelia Langford, John Sillitoe on behalf of the Wellcome Sanger Institute COVID-19 Surveillance Team |
| EPI_ISL_3470793, EPI_ISL_3470794 | Lighthouse Lab in Alderley Park | Wellcome Sanger Institute for the COVID-19 Genomics UK (COG-UK) Consortium | Jacquelyn Wynn, Mairead Hyland, The Lighthouse Lab in Alderley Park and Alex Alderton, Roberto Amato, Jeffrey Barrett, Sonia Goncalves, Ewan Harrison, David K. Jackson, Ian Johnston, Dominic Kwiatkowski, Cordelia Langford, John Sillitoe on behalf of the Wellcome Sanger Institute COVID-19 Surveillance Team |
| EPI_ISL_3470795, EPI_ISL_3470796, EPI_ISL_3470797, EPI_ISL_3470798 | Lighthouse Lab in Glasgow | Wellcome Sanger Institute for the COVID-19 Genomics UK (COG-UK) Consortium | Harper VanSteenhouse, Yumi Kasai, David Gray, Carol Clugston, Anna Dominiczak and Alex Alderton, Roberto Amato, Jeffrey Barrett, Sonia Goncalves, Ewan Harrison, David K. Jackson, Ian Johnston, Dominic Kwiatkowski, Cordelia Langford, John Sillitoe on behalf of the Wellcome Sanger Institute COVID-19 Surveillance Team |
| EPI_ISL_3470799 | Lighthouse Lab in Milton Keynes | Wellcome Sanger Institute for the COVID-19 Genomics UK (COG-UK) Consortium | The Lighthouse Lab in Milton Keynes and Alex Alderton, Roberto Amato, Jeffrey Barrett, Sonia Goncalves, Ewan Harrison, David K. Jackson, Ian Johnston, Dominic Kwiatkowski, Cordelia Langford, John Sillitoe on behalf of the Wellcome Sanger Institute COVID-19 Surveillance Team |
| EPI_ISL_3470800, EPI_ISL_3470801, EPI_ISL_3470802, EPI_ISL_3470803, EPI_ISL_3470804 | Lighthouse Lab in Glasgow | Wellcome Sanger Institute for the COVID-19 Genomics UK (COG-UK) Consortium | Harper VanSteenhouse, Yumi Kasai, David Gray, Carol Clugston, Anna Dominiczak and Alex Alderton, Roberto Amato, Jeffrey Barrett, Sonia Goncalves, Ewan Harrison, David K. Jackson, Ian Johnston, Dominic Kwiatkowski, Cordelia Langford, John Sillitoe on behalf of the Wellcome Sanger Institute COVID-19 Surveillance Team |
| EPI_ISL_3470805, EPI_ISL_3470806, EPI_ISL_3470807 | Lighthouse Lab in Alderley Park | Wellcome Sanger Institute for the COVID-19 Genomics UK (COG-UK) Consortium | Jacquelyn Wynn, Mairead Hyland, The Lighthouse Lab in Alderley Park and Alex Alderton, Roberto Amato, Jeffrey Barrett, Sonia Goncalves, Ewan Harrison, David K. Jackson, Ian Johnston, Dominic Kwiatkowski, Cordelia Langford, John Sillitoe on behalf of the Wellcome Sanger Institute COVID-19 Surveillance Team |

| | | | |
|---|---|---|---|
| EPI_ISL_3470808, EPI_ISL_3470809, EPI_ISL_3470810 | Lighthouse Lab in Glasgow | Wellcome Sanger Institute for the COVID-19 Genomics UK (COG-UK) Consortium | Harper VanSteenhouse, Yumi Kasai, David Gray, Carol Clugston, Anna Dominiczak and Alex Alderton, Roberto Amato, Jeffrey Barrett, Sonia Goncalves, Ewan Harrison, David K. Jackson, Ian Johnston, Dominic Kwiatkowski, Cordelia Langford, John Sillitoe on behalf of the Wellcome Sanger Institute COVID-19 Surveillance Team |
| EPI_ISL_3470811 | Lighthouse Lab in Alderley Park | Wellcome Sanger Institute for the COVID-19 Genomics UK (COG-UK) Consortium | Jacquelyn Wynn, Mairead Hyland, The Lighthouse Lab in Alderley Park and Alex Alderton, Roberto Amato, Jeffrey Barrett, Sonia Goncalves, Ewan Harrison, David K. Jackson, Ian Johnston, Dominic Kwiatkowski, Cordelia Langford, John Sillitoe on behalf of the Wellcome Sanger Institute COVID-19 Surveillance Team |
| EPI_ISL_3470812 | Lighthouse Lab in Milton Keynes | Wellcome Sanger Institute for the COVID-19 Genomics UK (COG-UK) Consortium | The Lighthouse Lab in Milton Keynes and Alex Alderton, Roberto Amato, Jeffrey Barrett, Sonia Goncalves, Ewan Harrison, David K. Jackson, Ian Johnston, Dominic Kwiatkowski, Cordelia Langford, John Sillitoe on behalf of the Wellcome Sanger Institute COVID-19 Surveillance Team |
| EPI_ISL_3470813 | Lighthouse Lab in Glasgow | Wellcome Sanger Institute for the COVID-19 Genomics UK (COG-UK) Consortium | Harper VanSteenhouse, Yumi Kasai, David Gray, Carol Clugston, Anna Dominiczak and Alex Alderton, Roberto Amato, Jeffrey Barrett, Sonia Goncalves, Ewan Harrison, David K. Jackson, Ian Johnston, Dominic Kwiatkowski, Cordelia Langford, John Sillitoe on behalf of the Wellcome Sanger Institute COVID-19 Surveillance Team |
| EPI_ISL_3470814 | Lighthouse Lab in Milton Keynes | Wellcome Sanger Institute for the COVID-19 Genomics UK (COG-UK) Consortium | The Lighthouse Lab in Milton Keynes and Alex Alderton, Roberto Amato, Jeffrey Barrett, Sonia Goncalves, Ewan Harrison, David K. Jackson, Ian Johnston, Dominic Kwiatkowski, Cordelia Langford, John Sillitoe on behalf of the Wellcome Sanger Institute COVID-19 Surveillance Team |
| EPI_ISL_3470815 | Lighthouse Lab in Alderley Park | Wellcome Sanger Institute for the COVID-19 Genomics UK (COG-UK) Consortium | Jacquelyn Wynn, Mairead Hyland, The Lighthouse Lab in Alderley Park and Alex Alderton, Roberto Amato, Jeffrey Barrett, Sonia Goncalves, Ewan Harrison, David K. Jackson, Ian Johnston, Dominic Kwiatkowski, Cordelia Langford, John Sillitoe on behalf of the Wellcome Sanger Institute COVID-19 Surveillance Team |
| EPI_ISL_3470816, EPI_ISL_3470817, EPI_ISL_3470818, EPI_ISL_3470819, EPI_ISL_3470820 | Lighthouse Lab in Glasgow | Wellcome Sanger Institute for the COVID-19 Genomics UK (COG-UK) Consortium | Harper VanSteenhouse, Yumi Kasai, David Gray, Carol Clugston, Anna Dominiczak and Alex Alderton, Roberto Amato, Jeffrey Barrett, Sonia Goncalves, Ewan Harrison, David K. Jackson, Ian Johnston, Dominic Kwiatkowski, Cordelia Langford, John Sillitoe on behalf of the Wellcome Sanger Institute COVID-19 Surveillance Team |
| EPI_ISL_3470821 | Lighthouse Lab in Milton Keynes | Wellcome Sanger Institute for the COVID-19 Genomics UK (COG-UK) Consortium | The Lighthouse Lab in Milton Keynes and Alex Alderton, Roberto Amato, Jeffrey Barrett, Sonia Goncalves, Ewan Harrison, David K. Jackson, Ian Johnston, Dominic Kwiatkowski, Cordelia Langford, John Sillitoe on behalf of the Wellcome Sanger Institute COVID-19 Surveillance Team |
| EPI_ISL_3470822, EPI_ISL_3470823, EPI_ISL_3470824, EPI_ISL_3470825 | Lighthouse Lab in Glasgow | Wellcome Sanger Institute for the COVID-19 Genomics UK (COG-UK) Consortium | Harper VanSteenhouse, Yumi Kasai, David Gray, Carol Clugston, Anna Dominiczak and Alex Alderton, Roberto Amato, Jeffrey Barrett, Sonia Goncalves, Ewan Harrison, David K. Jackson, Ian Johnston, Dominic Kwiatkowski, Cordelia Langford, John Sillitoe on behalf of the Wellcome Sanger Institute COVID-19 Surveillance Team |
| EPI_ISL_3470826, EPI_ISL_3470827, EPI_ISL_3470828, EPI_ISL_3470829, EPI_ISL_3470830 | Lighthouse Lab in Milton Keynes | Wellcome Sanger Institute for the COVID-19 Genomics UK (COG-UK) Consortium | The Lighthouse Lab in Milton Keynes and Alex Alderton, Roberto Amato, Jeffrey Barrett, Sonia Goncalves, Ewan Harrison, David K. Jackson, Ian Johnston, Dominic Kwiatkowski, Cordelia Langford, John Sillitoe on behalf of the Wellcome Sanger Institute COVID-19 Surveillance Team |
| EPI_ISL_3470831, EPI_ISL_3470832 | Lighthouse Lab in Glasgow | Wellcome Sanger Institute for the COVID-19 Genomics UK (COG-UK) Consortium | Harper VanSteenhouse, Yumi Kasai, David Gray, Carol Clugston, Anna Dominiczak and Alex Alderton, Roberto Amato, Jeffrey Barrett, Sonia Goncalves, Ewan Harrison, David K. Jackson, Ian Johnston, Dominic Kwiatkowski, Cordelia Langford, John Sillitoe on behalf of the Wellcome Sanger Institute COVID-19 Surveillance Team |
| EPI_ISL_3470833 | Lighthouse Lab in Milton Keynes | Wellcome Sanger Institute for the COVID-19 Genomics UK (COG-UK) Consortium | The Lighthouse Lab in Milton Keynes and Alex Alderton, Roberto Amato, Jeffrey Barrett, Sonia Goncalves, Ewan Harrison, David K. Jackson, Ian Johnston, Dominic Kwiatkowski, Cordelia Langford, John Sillitoe on behalf of the Wellcome Sanger Institute COVID-19 Surveillance Team |
| EPI_ISL_3470834 | Lighthouse Lab in Alderley Park | Wellcome Sanger Institute for the COVID-19 Genomics UK (COG-UK) Consortium | Jacquelyn Wynn, Mairead Hyland, The Lighthouse Lab in Alderley Park and Alex Alderton, Roberto Amato, Jeffrey Barrett, Sonia Goncalves, Ewan Harrison, David K. Jackson, Ian Johnston, Dominic Kwiatkowski, Cordelia Langford, John Sillitoe on behalf of the Wellcome Sanger Institute COVID-19 Surveillance Team |
| EPI_ISL_3470835 | Lighthouse Lab in Milton Keynes | Wellcome Sanger Institute for the COVID-19 Genomics UK (COG-UK) Consortium | The Lighthouse Lab in Milton Keynes and Alex Alderton, Roberto Amato, Jeffrey Barrett, Sonia Goncalves, Ewan Harrison, David K. Jackson, Ian Johnston, Dominic Kwiatkowski, Cordelia Langford, John Sillitoe on behalf of the Wellcome Sanger Institute COVID-19 Surveillance Team |
| EPI_ISL_3470836 | Lighthouse Lab in Alderley Park | Wellcome Sanger Institute for the COVID-19 Genomics UK (COG-UK) Consortium | Jacquelyn Wynn, Mairead Hyland, The Lighthouse Lab in Alderley Park and Alex Alderton, Roberto Amato, Jeffrey Barrett, Sonia Goncalves, Ewan Harrison, David K. Jackson, Ian Johnston, Dominic Kwiatkowski, Cordelia Langford, John Sillitoe on behalf of the Wellcome Sanger Institute COVID-19 Surveillance Team |
| EPI_ISL_3470837, EPI_ISL_3470838 | Lighthouse Lab in Glasgow | Wellcome Sanger Institute for the COVID-19 Genomics UK (COG-UK) Consortium | Harper VanSteenhouse, Yumi Kasai, David Gray, Carol Clugston, Anna Dominiczak and Alex Alderton, Roberto Amato, Jeffrey Barrett, Sonia Goncalves, Ewan Harrison, David K. Jackson, Ian Johnston, Dominic Kwiatkowski, Cordelia Langford, John Sillitoe on behalf of the Wellcome Sanger Institute COVID-19 Surveillance Team |
| EPI_ISL_3470839 | Lighthouse Lab in Alderley Park | Wellcome Sanger Institute for the COVID-19 Genomics UK (COG-UK) Consortium | Jacquelyn Wynn, Mairead Hyland, The Lighthouse Lab in Alderley Park and Alex Alderton, Roberto Amato, Jeffrey Barrett, Sonia Goncalves, Ewan Harrison, David K. Jackson, Ian Johnston, Dominic Kwiatkowski, Cordelia Langford, John Sillitoe on behalf of the Wellcome Sanger Institute COVID-19 Surveillance Team |
| EPI_ISL_3470840, EPI_ISL_3470841 | Lighthouse Lab in Glasgow | Wellcome Sanger Institute for the COVID-19 Genomics UK (COG-UK) Consortium | Harper VanSteenhouse, Yumi Kasai, David Gray, Carol Clugston, Anna Dominiczak and Alex Alderton, Roberto Amato, Jeffrey Barrett, Sonia Goncalves, Ewan Harrison, David K. Jackson, Ian Johnston, Dominic Kwiatkowski, Cordelia Langford, John Sillitoe on behalf of the Wellcome Sanger Institute COVID-19 Surveillance Team |
| EPI_ISL_3470842 | Lighthouse Lab in Alderley Park | Wellcome Sanger Institute for the COVID-19 Genomics UK (COG-UK) Consortium | Jacquelyn Wynn, Mairead Hyland, The Lighthouse Lab in Alderley Park and Alex Alderton, Roberto Amato, Jeffrey Barrett, Sonia Goncalves, Ewan Harrison, David K. Jackson, Ian Johnston, Dominic Kwiatkowski, Cordelia Langford, John Sillitoe on behalf of the Wellcome Sanger Institute COVID-19 Surveillance Team |
| EPI_ISL_3470843, EPI_ISL_3470844, EPI_ISL_3470845, EPI_ISL_3470846, EPI_ISL_3470847, EPI_ISL_3470848, EPI_ISL_3470849, EPI_ISL_3470850, EPI_ISL_3470851, EPI_ISL_3470852, EPI_ISL_3470853, EPI_ISL_3470854, EPI_ISL_3470855, EPI_ISL_3470856, EPI_ISL_3470857, EPI_ISL_3470858, EPI_ISL_3470859, EPI_ISL_3470860, EPI_ISL_3470861, EPI_ISL_3470862, EPI_ISL_3470863, EPI_ISL_3470864, EPI_ISL_3470865, EPI_ISL_3470866, EPI_ISL_3470867, EPI_ISL_3470868, EPI_ISL_3470869, EPI_ISL_3470870, EPI_ISL_3470871, EPI_ISL_3470872, EPI_ISL_3470873, EPI_ISL_3470874, EPI_ISL_3470875, EPI_ISL_3470876, EPI_ISL_3470877, EPI_ISL_3470878, EPI_ISL_3470879, EPI_ISL_3470880, EPI_ISL_3470881, EPI_ISL_3470882, EPI_ISL_3470883, EPI_ISL_3470884, EPI_ISL_3470885, EPI_ISL_3470886, EPI_ISL_3470887, EPI_ISL_3470888, EPI_ISL_3470889, EPI_ISL_3470890, EPI_ISL_3470891, EPI_ISL_3470892, EPI_ISL_3470893, EPI_ISL_3470894, EPI_ISL_3470895, EPI_ISL_3470896, EPI_ISL_3470897, EPI_ISL_3470898, EPI_ISL_3470899, EPI_ISL_3470900, EPI_ISL_3470901, EPI_ISL_3470902, EPI_ISL_3470903, EPI_ISL_3470904, EPI_ISL_3470905, EPI_ISL_3470906, EPI_ISL_3470907, EPI_ISL_3470908, EPI_ISL_3470909, EPI_ISL_3470910, EPI_ISL_3470911, EPI_ISL_3470912, EPI_ISL_3470913, EPI_ISL_3470914, EPI_ISL_3470915, EPI_ISL_3470916, EPI_ISL_3470917, EPI_ISL_3470918, EPI_ISL_3470919, EPI_ISL_3470920, EPI_ISL_3470921, EPI_ISL_3470922, EPI_ISL_3470923, EPI_ISL_3470924, EPI_ISL_3470925, EPI_ISL_3470926, EPI_ISL_3470927, EPI_ISL_3470928, EPI_ISL_3470929, EPI_ISL_3470930, EPI_ISL_3470931, EPI_ISL_3470932, EPI_ISL_3470933, EPI_ISL_3470934, EPI_ISL_3470935, EPI_ISL_3470936, EPI_ISL_3470937, EPI_ISL_3470938, EPI_ISL_3470939, EPI_ISL_3470940, EPI_ISL_3470941, EPI_ISL_3470942, EPI_ISL_3470943, EPI_ISL_3470944, EPI_ISL_3470945, EPI_ISL_3470946, EPI_ISL_3470947, EPI_ISL_3470948, EPI_ISL_3470949, EPI_ISL_3470950, EPI_ISL_3470951, EPI_ISL_3470952, EPI_ISL_3470953, EPI_ISL_3470954, EPI_ISL_3470955, EPI_ISL_3470956, EPI_ISL_3470957, EPI_ISL_3470958, EPI_ISL_3470959, EPI_ISL_3470960, EPI_ISL_3470961, EPI_ISL_3470962, EPI_ISL_3470963, EPI_ISL_3470964, EPI_ISL_3470965, EPI_ISL_3470966, EPI_ISL_3470967, EPI_ISL_3470968, EPI_ISL_3470969, EPI_ISL_3470970, EPI_ISL_3470971, EPI_ISL_3470972, EPI_ISL_3470973, EPI_ISL_3470974, EPI_ISL_3470975, EPI_ISL_3470976, EPI_ISL_3470977, EPI_ISL_3470978, EPI_ISL_3470979, EPI_ISL_3470980, EPI_ISL_3470981, EPI_ISL_3470982, EPI_ISL_3470983, EPI_ISL_3470984, EPI_ISL_3470985, EPI_ISL_3470986, EPI_ISL_3470987, EPI_ISL_3470988, EPI_ISL_3470989, EPI_ISL_3470990, EPI_ISL_3470991, EPI_ISL_3470992, EPI_ISL_3470993, EPI_ISL_3470994, EPI_ISL_3470995, EPI_ISL_3470996, EPI_ISL_3470997, EPI_ISL_3470998, EPI_ISL_3470999, EPI_ISL_3471000, EPI_ISL_3471001, EPI_ISL_3471002, EPI_ISL_3471003, EPI_ISL_3471004, EPI_ISL_3471005, EPI_ISL_3471006, EPI_ISL_3471007, EPI_ISL_3471008, EPI_ISL_3471009, EPI_ISL_3471010, EPI_ISL_3471011, EPI_ISL_3471012, EPI_ISL_3471013, EPI_ISL_3471014, EPI_ISL_3471015, EPI_ISL_3471016, EPI_ISL_3471017, EPI_ISL_3471018, EPI_ISL_3471019, EPI_ISL_3471020, EPI_ISL_3471021, EPI_ISL_3471022, EPI_ISL_3471023, EPI_ISL_3471024, EPI_ISL_3471025, EPI_ISL_3471026, EPI_ISL_3471027, EPI_ISL_3471028, EPI_ISL_3471029, EPI_ISL_3471030, EPI_ISL_3471031, EPI_ISL_3471032, EPI_ISL_3471033, EPI_ISL_3471034, EPI_ISL_3471035, EPI_ISL_3471036, EPI_ISL_3471037, EPI_ISL_3471038, EPI_ISL_3471039, EPI_ISL_3471040, EPI_ISL_3471041, EPI_ISL_3471042, EPI_ISL_3471043, EPI_ISL_3471044, EPI_ISL_3471045, EPI_ISL_3471046, EPI_ISL_3471047, EPI_ISL_3471048, EPI_ISL_3471049, EPI_ISL_3471050, EPI_ISL_3471051, EPI_ISL_3471052, EPI_ISL_3471053, EPI_ISL_3471054, EPI_ISL_3471055, EPI_ISL_3471056, EPI_ISL_3471057, EPI_ISL_3471058, EPI_ISL_3471059, EPI_ISL_3471060, EPI_ISL_3471061, EPI_ISL_3471062, EPI_ISL_3471063, EPI_ISL_3471064, EPI_ISL_3471065, EPI_ISL_3471066, EPI_ISL_3471067, EPI_ISL_3471068, EPI_ISL_3471069, EPI_ISL_3471070, EPI_ISL_3471071, EPI_ISL_3471072, EPI_ISL_3471073, EPI_ISL_3471074, EPI_ISL_3471075, EPI_ISL_3471076, EPI_ISL_3471077, EPI_ISL_3471078, EPI_ISL_3471079, EPI_ISL_3471080, EPI_ISL_3471081, EPI_ISL_3471082, EPI_ISL_3471083, EPI_ISL_3471084, EPI_ISL_3471085, EPI_ISL_3471086, EPI_ISL_3471087, EPI_ISL_3471088, EPI_ISL_3471089, EPI_ISL_3471090, EPI_ISL_3471091, EPI_ISL_3471092, EPI_ISL_3471093, EPI_ISL_3471094, EPI_ISL_3471095, EPI_ISL_3471096, EPI_ISL_3471097, EPI_ISL_3471098, EPI_ISL_3471099, EPI_ISL_3471100, EPI_ISL_3471101, EPI_ISL_3471102, EPI_ISL_3471103, EPI_ISL_3471104, EPI_ISL_3471105, EPI_ISL_3471106, EPI_ISL_3471107, EPI_ISL_3471108, EPI_ISL_3471109, EPI_ISL_3471110, EPI_ISL_3471111, EPI_ISL_3471112, EPI_ISL_3471113, EPI_ISL_3471114, EPI_ISL_3471115, EPI_ISL_3471116, EPI_ISL_3471117, EPI_ISL_3471118, EPI_ISL_3471119, EPI_ISL_3471120, EPI_ISL_3471121, EPI_ISL_3471122, EPI_ISL_3471123, EPI_ISL_3471124, EPI_ISL_3471125, EPI_ISL_3471126, EPI_ISL_3471127, EPI_ISL_3471128, EPI_ISL_3471129, EPI_ISL_3471130, EPI_ISL_3471131, EPI_ISL_3471132, EPI_ISL_3471133, EPI_ISL_3471134, EPI_ISL_3471135, EPI_ISL_3471136, EPI_ISL_3471137, EPI_ISL_3471138, EPI_ISL_3471139, EPI_ISL_3471140, EPI_ISL_3471141, EPI_ISL_3471142, EPI_ISL_3471143, EPI_ISL_3471144, EPI_ISL_3471145, EPI_ISL_3471146, EPI_ISL_3471147, EPI_ISL_3471148, EPI_ISL_3471149, EPI_ISL_3471150, EPI_ISL_3471151, EPI_ISL_3471152, EPI_ISL_3471153, EPI_ISL_3471154, EPI_ISL_3471155, EPI_ISL_3471156, EPI_ISL_3471157, EPI_ISL_3471158, EPI_ISL_3471159, EPI_ISL_3471160, EPI_ISL_3471161, EPI_ISL_3471162, EPI_ISL_3471163, EPI_ISL_3471164, EPI_ISL_3471165, EPI_ISL_3471166, EPI_ISL_3471167, EPI_ISL_3471168, EPI_ISL_3471169, EPI_ISL_3471170, EPI_ISL_3471171, EPI_ISL_3471172, EPI_ISL_3471173, EPI_ISL_3471174, EPI_ISL_3471175, EPI_ISL_3471176, EPI_ISL_3471177, EPI_ISL_3471178, EPI_ISL_3471179, EPI_ISL_3471180, EPI_ISL_3471181, EPI_ISL_3471182, EPI_ISL_3471183, EPI_ISL_3471184, EPI_ISL_3471185, EPI_ISL_3471186, EPI_ISL_3471187, EPI_ISL_3471188, EPI_ISL_3471189, EPI_ISL_3471190, EPI_ISL_3471191, EPI_ISL_3471192, EPI_ISL_3471193, EPI_ISL_3471194, EPI_ISL_3471195, EPI_ISL_3471196, EPI_ISL_3471197, EPI_ISL_3471198, EPI_ISL_3471199, EPI_ISL_3471200, EPI_ISL_3471201, EPI_ISL_3471202, EPI_ISL_3471203, EPI_ISL_3471204 | | | |
| see above | Lighthouse Lab in Glasgow | Wellcome Sanger Institute for the COVID-19 Genomics UK (COG-UK) Consortium | Harper VanSteenhouse, Yumi Kasai, David Gray, Carol Clugston, Anna Dominiczak and Alex Alderton, Roberto Amato, Jeffrey Barrett, Sonia Goncalves, Ewan Harrison, David K. Jackson, Ian Johnston, Dominic Kwiatkowski, Cordelia Langford, John Sillitoe on behalf of the Wellcome Sanger Institute COVID-19 Surveillance Team |
| EPI_ISL_3471206, EPI_ISL_3471216, EPI_ISL_3471217, EPI_ISL_3471218, EPI_ISL_3471225, EPI_ISL_3471226, EPI_ISL_3471227, EPI_ISL_3471228, EPI_ISL_3471229, EPI_ISL_3471230, EPI_ISL_3471231, EPI_ISL_3471232, EPI_ISL_3471233, EPI_ISL_3471234, EPI_ISL_3471235, EPI_ISL_3471236, EPI_ISL_3471237, EPI_ISL_3471238, EPI_ISL_3471239, EPI_ISL_3471240, EPI_ISL_3471241, EPI_ISL_3471242, EPI_ISL_3471243, EPI_ISL_3471244, EPI_ISL_3471245, EPI_ISL_3471246, EPI_ISL_3471247, EPI_ISL_3471248, EPI_ISL_3471249, EPI_ISL_3471250, EPI_ISL_3471251, EPI_ISL_3471252, EPI_ISL_3471253, EPI_ISL_3471254, EPI_ISL_3471255, EPI_ISL_3471256, EPI_ISL_3471257, EPI_ISL_3471258, EPI_ISL_3471259, EPI_ISL_3471260, EPI_ISL_3471261, EPI_ISL_3471262, EPI_ISL_3471263, EPI_ISL_3471264, EPI_ISL_3471265, EPI_ISL_3471266, EPI_ISL_3471267, EPI_ISL_3471268, EPI_ISL_3471269, EPI_ISL_3471270, EPI_ISL_3471271, EPI_ISL_3471272, EPI_ISL_3471273, EPI_ISL_3471274, EPI_ISL_3471275, EPI_ISL_3471276, EPI_ISL_3471277, EPI_ISL_3471278, EPI_ISL_3471279, EPI_ISL_3471280, EPI_ISL_3471281, EPI_ISL_3471282, EPI_ISL_3471283, EPI_ISL_3471284, EPI_ISL_3471285, EPI_ISL_3471286, EPI_ISL_3471287, EPI_ISL_3471288, EPI_ISL_3471289, EPI_ISL_3471290, EPI_ISL_3471291, EPI_ISL_3471292, EPI_ISL_3471293, EPI_ISL_3471294, EPI_ISL_3471295, EPI_ISL_3471296, EPI_ISL_3471297, EPI_ISL_3471298, EPI_ISL_3471299, EPI_ISL_3471300, EPI_ISL_3471301, EPI_ISL_3471302, EPI_ISL_3471303, EPI_ISL_3471304, EPI_ISL_3471305, EPI_ISL_3471306, EPI_ISL_3471307, EPI_ISL_3471308, EPI_ISL_3471309, EPI_ISL_3471310, EPI_ISL_3471311, EPI_ISL_3471312, EPI_ISL_3471313, EPI_ISL_3471314, EPI_ISL_3471315, EPI_ISL_3471316, EPI_ISL_3471317, EPI_ISL_3471318, EPI_ISL_3471319, EPI_ISL_3471320, EPI_ISL_3471321, EPI_ISL_3471322, EPI_ISL_3471323, EPI_ISL_3471324, EPI_ISL_3471325, EPI_ISL_3471326, EPI_ISL_3471327, EPI_ISL_3471328, EPI_ISL_3471329, EPI_ISL_3471330, EPI_ISL_3471331, EPI_ISL_3471332, EPI_ISL_3471333, EPI_ISL_3471334, EPI_ISL_3471335, EPI_ISL_3471336, EPI_ISL_3471337, EPI_ISL_3471338, EPI_ISL_3471339, EPI_ISL_3471340, EPI_ISL_3471341, EPI_ISL_3471342, EPI_ISL_3471343, EPI_ISL_3471344, EPI_ISL_3471345, EPI_ISL_3471346, EPI_ISL_3471347, EPI_ISL_3471348, EPI_ISL_3471349, EPI_ISL_3471350, EPI_ISL_3471351, EPI_ISL_3471352, EPI_ISL_3471353, EPI_ISL_3471354, EPI_ISL_3471355, EPI_ISL_3471356, EPI_ISL_3471357, EPI_ISL_3471358 | | | |
| see above | Institute of Microbiology and Immunology, Faculty of Medicine, University of Ljubljana | Institute of Microbiology and Immunology, Faculty of Medicine, University of Ljubljana | Alen Suljič, Samo Zakotnik, Tomaž Mark Zorec, Matic Brvar, Doroteja Vlaj, Andraž Celar, Dominika Šturm, Patricija Pozvek, Špela Pleh, Miša Korva, Mario Poljak, Tatjana Avšič – Županc |
| EPI_ISL_3471369, EPI_ISL_3471370, EPI_ISL_3471371, EPI_ISL_3471372, EPI_ISL_3471373 | Public Health Authority of the Slovak Republic | Laboratory of Genomics and Bioinformatics, Comenius University Science Park | Tatiana Sedláčková, Diana Rusňáková, Miroslav Böhmer, Anna Gičová, Jaroslav Budiš, Tomáš Szemes, Jakub Styk |
| EPI_ISL_3471375, EPI_ISL_3471376, EPI_ISL_3471377, EPI_ISL_3471387, EPI_ISL_3471388, EPI_ISL_3471389, EPI_ISL_3471390, EPI_ISL_3471391, EPI_ISL_3471392, EPI_ISL_3471393, EPI_ISL_3471394, EPI_ISL_3471395, EPI_ISL_3471396, EPI_ISL_3471397, EPI_ISL_3471398 | | | |
| see above | Institute of Microbiology and Immunology, Faculty of Medicine, University of Ljubljana | Institute of Microbiology and Immunology, Faculty of Medicine, University of Ljubljana | Alen Suljič, Samo Zakotnik, Tomaž Mark Zorec, Matic Brvar, Doroteja Vlaj, Andraž Celar, Dominika Šturm, Patricija Pozvek, Špela Pleh, Miša Korva, Mario Poljak, Tatjana Avšič – Županc |
| EPI_ISL_3471400 | Regional Medical Sciences Center 11 Surat Thani | National Institute of Health, Department of Medical Sciences, Ministry of Public Health, Thailand | Pilailuk Okada; Siripaporn Phuygun; Nuttida Thongpramul; Sittiporn Parnmen; Ratana Tacharoenmuang; Pakorn Piromtong; Natchaya Khiadsang; Thanutsapa Thanadachakul; Warawan Wongboot; sirikanda wimol; Sunthareeya Waicharoen; |
| EPI_ISL_3471401 | Department of Disease Control | National Institute of Health, Department of Medical Sciences, Ministry of Public Health, Thailand | Pilailuk Okada; Siripaporn Phuygun; Nuttida Thongpramul; Sittiporn Parnmen; Ratana Tacharoenmuang; Pakorn Piromtong; Natchaya Khiadsang; Thanutsapa Thanadachakul; Warawan Wongboot; sirikanda wimol; Sunthareeya Waicharoen; |
| EPI_ISL_3471402 | Regional Medical Sciences Center 11/1 Phuket | National Institute of Health, Department of Medical Sciences, Ministry of Public Health, Thailand | Pilailuk Okada; Siripaporn Phuygun; Nuttida Thongpramul; Sittiporn Parnmen; Ratana Tacharoenmuang; Pakorn Piromtong; Natchaya Khiadsang; Thanutsapa Thanadachakul; Warawan Wongboot; sirikanda wimol; Sunthareeya Waicharoen; |
| EPI_ISL_3471405 | Regional Medical Sciences Center 3 Nakhon Sawan | National Institute of Health, Department of Medical Sciences, Ministry of Public Health, Thailand | Pilailuk Okada; Siripaporn Phuygun; Nuttida Thongpramul; Sittiporn Parnmen; Ratana Tacharoenmuang; Pakorn Piromtong; Natchaya Khiadsang; Thanutsapa Thanadachakul; Warawan Wongboot; sirikanda wimol; Sunthareeya Waicharoen; |
| EPI_ISL_3471406 | Regional Medical Sciences Center 4 Saraburi | National Institute of Health, Department of Medical Sciences, Ministry of Public Health, Thailand | Pilailuk Okada; Siripaporn Phuygun; Nuttida Thongpramul; Sittiporn Parnmen; Ratana Tacharoenmuang; Pakorn Piromtong; Natchaya Khiadsang; Thanutsapa Thanadachakul; Warawan Wongboot; sirikanda wimol; Sunthareeya Waicharoen; |
| EPI_ISL_3471408 | Regional Medical Sciences Center 1/1 Chiang Rai | National Institute of Health, Department of Medical Sciences, Ministry of Public Health, Thailand | Pilailuk Okada; Siripaporn Phuygun; Nuttida Thongpramul; Sittiporn Parnmen; Ratana Tacharoenmuang; Pakorn Piromtong; Natchaya Khiadsang; Thanutsapa Thanadachakul; Warawan Wongboot; sirikanda wimol; Sunthareeya Waicharoen; |
| EPI_ISL_3471410 | Regional Medical Sciences Center 6 Chonburi | National Institute of Health, Department of Medical Sciences, | Pilailuk Okada; Siripaporn Phuygun; Nuttida Thongpramul; Sittiporn Parnmen; Ratana Tacharoenmuang; Pakorn Piromtong; Natchaya Khiadsang; Thanutsapa Thanadachakul; Warawan Wongboot; sirikanda wimol; Sunthareeya Waicharoen; |

| | | | |
|---|---|---|---|
| | | Ministry of Public Health, Thailand | |
| EPI_ISL_3471412 | Regional Medical Sciences Center 11/1 Phuket | National Institute of Health, Department of Medical Sciences, Ministry of Public Health, Thailand | Pilailuk Okada; Siripaporn Phuygun; Nuttida Thongpramul; Sittiporn Parnmen; Ratana Tacharoenmuang; Pakorn Piromtong; Natchaya Khiadsang; Thanutsapa Thanadachakul; Warawan Wongboot; sirikanda wimol; Sunthareeya Waicharoen; |
| EPI_ISL_3471417 | Regional Medical Sciences Center 1/1 Chiang Rai | National Institute of Health, Department of Medical Sciences, Ministry of Public Health, Thailand | Pilailuk Okada; Siripaporn Phuygun; Nuttida Thongpramul; Sittiporn Parnmen; Ratana Tacharoenmuang; Pakorn Piromtong; Natchaya Khiadsang; Thanutsapa Thanadachakul; Warawan Wongboot; sirikanda wimol; Sunthareeya Waicharoen; |
| EPI_ISL_3471418, EPI_ISL_3471419, EPI_ISL_3471420 | LABO BIO DOC | CHU Purpan - Laboratoire de Virologie - Institut Fédératif de Biologie | Latour J., Milhes M., Bulach T., Ranger N., Salin G., Nicot F., Tremeaux P., Donnadieu C., Izopet J. |
| EPI_ISL_3471421 | CHU Purpan - Laboratoire de Virologie - Institut Fédératif de Biologie | CHU Purpan - Laboratoire de Virologie - Institut Fédératif de Biologie | Latour J., Milhes M., Bulach T., Ranger N., Salin G., Nicot F., Tremeaux P., Donnadieu C., Izopet J. |
| EPI_ISL_3471422 | LABO BIO DOC | CHU Purpan - Laboratoire de Virologie - Institut Fédératif de Biologie | Latour J., Milhes M., Bulach T., Ranger N., Salin G., Nicot F., Tremeaux P., Donnadieu C., Izopet J. |
| EPI_ISL_3471423 | CHU Purpan - Laboratoire de Virologie - Institut Fédératif de Biologie | CHU Purpan - Laboratoire de Virologie - Institut Fédératif de Biologie | Latour J., Milhes M., Bulach T., Ranger N., Salin G., Nicot F., Tremeaux P., Donnadieu C., Izopet J. |
| EPI_ISL_3471424, EPI_ISL_3471425, EPI_ISL_3471426, EPI_ISL_3471427, EPI_ISL_3471428, EPI_ISL_3471429, EPI_ISL_3471430, EPI_ISL_3471431, EPI_ISL_3471432, EPI_ISL_3471433, EPI_ISL_3471434, EPI_ISL_3471435, EPI_ISL_3471436, EPI_ISL_3471437, EPI_ISL_3471438 | | | |
| see above | LABO BIO DOC | CHU Purpan - Laboratoire de Virologie - Institut Fédératif de Biologie | Latour J., Milhes M., Bulach T., Ranger N., Salin G., Nicot F., Tremeaux P., Donnadieu C., Izopet J. |
| EPI_ISL_3471439 | CH ALBI | CHU Purpan - Laboratoire de Virologie - Institut Fédératif de Biologie | Latour J., Milhes M., Bulach T., Ranger N., Salin G., Nicot F., Tremeaux P., Donnadieu C., Izopet J. |
| EPI_ISL_3471440, EPI_ISL_3471441, EPI_ISL_3471442, EPI_ISL_3471443, EPI_ISL_3471444 | LABO BIO DOC | CHU Purpan - Laboratoire de Virologie - Institut Fédératif de Biologie | Latour J., Milhes M., Bulach T., Ranger N., Salin G., Nicot F., Tremeaux P., Donnadieu C., Izopet J. |
| EPI_ISL_3471445 | CHU Purpan - Laboratoire de Virologie - Institut Fédératif de Biologie | CHU Purpan - Laboratoire de Virologie - Institut Fédératif de Biologie | Latour J., Milhes M., Bulach T., Ranger N., Salin G., Nicot F., Tremeaux P., Donnadieu C., Izopet J. |
| EPI_ISL_3471446, EPI_ISL_3471447, EPI_ISL_3471448, EPI_ISL_3471449 | LABO BIO DOC | CHU Purpan - Laboratoire de Virologie - Institut Fédératif de Biologie | Latour J., Milhes M., Bulach T., Ranger N., Salin G., Nicot F., Tremeaux P., Donnadieu C., Izopet J. |
| EPI_ISL_3471450 | CH MONTAUBAN | CHU Purpan - Laboratoire de Virologie - Institut Fédératif de Biologie | Latour J., Milhes M., Bulach T., Ranger N., Salin G., Nicot F., Tremeaux P., Donnadieu C., Izopet J. |
| EPI_ISL_3471451, EPI_ISL_3471452, EPI_ISL_3471453, EPI_ISL_3471455 | LABO BIO DOC | CHU Purpan - Laboratoire de Virologie - Institut Fédératif de Biologie | Latour J., Milhes M., Bulach T., Ranger N., Salin G., Nicot F., Tremeaux P., Donnadieu C., Izopet J. |
| EPI_ISL_3471456 | CHU Purpan - Laboratoire de Virologie - Institut Fédératif de Biologie | CHU Purpan - Laboratoire de Virologie - Institut Fédératif de Biologie | Latour J., Milhes M., Bulach T., Ranger N., Salin G., Nicot F., Tremeaux P., Donnadieu C., Izopet J. |
| EPI_ISL_3471458, EPI_ISL_3471459, EPI_ISL_3471460, EPI_ISL_3471461, EPI_ISL_3471462, EPI_ISL_3471463, EPI_ISL_3471464, EPI_ISL_3471465, EPI_ISL_3471466 | LABO BIO DOC | CHU Purpan - Laboratoire de Virologie - Institut Fédératif de Biologie | Latour J., Milhes M., Bulach T., Ranger N., Salin G., Nicot F., Tremeaux P., Donnadieu C., Izopet J. |
| EPI_ISL_3471467 | CHU Purpan - Laboratoire de Virologie - Institut Fédératif de Biologie | CHU Purpan - Laboratoire de Virologie - Institut Fédératif de Biologie | Latour J., Milhes M., Bulach T., Ranger N., Salin G., Nicot F., Tremeaux P., Donnadieu C., Izopet J. |
| EPI_ISL_3471468 | CH ALBI | CHU Purpan - Laboratoire de Virologie - Institut Fédératif de Biologie | Latour J., Milhes M., Bulach T., Ranger N., Salin G., Nicot F., Tremeaux P., Donnadieu C., Izopet J. |
| EPI_ISL_3471469 | LABORATOIRE GCS INTER HOSPITALIER | CHU Purpan - Laboratoire de Virologie - Institut Fédératif de Biologie | Latour J., Milhes M., Bulach T., Ranger N., Salin G., Nicot F., Tremeaux P., Donnadieu C., Izopet J. |
| EPI_ISL_3471470, EPI_ISL_3471471 | LABO BIO DOC | CHU Purpan - Laboratoire de Virologie - Institut Fédératif de Biologie | Latour J., Milhes M., Bulach T., Ranger N., Salin G., Nicot F., Tremeaux P., Donnadieu C., Izopet J. |
| EPI_ISL_3471472 | CHU Purpan - Laboratoire de Virologie - Institut Fédératif de Biologie | CHU Purpan - Laboratoire de Virologie - Institut Fédératif de Biologie | Latour J., Milhes M., Bulach T., Ranger N., Salin G., Nicot F., Tremeaux P., Donnadieu C., Izopet J. |
| EPI_ISL_3471473 | CH ALBI | CHU Purpan - Laboratoire de Virologie - Institut Fédératif de Biologie | Latour J., Milhes M., Bulach T., Ranger N., Salin G., Nicot F., Tremeaux P., Donnadieu C., Izopet J. |
| EPI_ISL_3471474, EPI_ISL_3471475, EPI_ISL_3471476, EPI_ISL_3471477 | LABO BIO DOC | CHU Purpan - Laboratoire de Virologie - Institut Fédératif de Biologie | Latour J., Milhes M., Bulach T., Ranger N., Salin G., Nicot F., Tremeaux P., Donnadieu C., Izopet J. |
| EPI_ISL_3471478 | HOSPILAB 47-VILLENEUVE LOT | CHU Purpan - Laboratoire de Virologie - Institut Fédératif de Biologie | Latour J., Milhes M., Bulach T., Ranger N., Salin G., Nicot F., Tremeaux P., Donnadieu C., Izopet J. |
| EPI_ISL_3471479, EPI_ISL_3471480 | LABO BIO DOC | CHU Purpan - Laboratoire de Virologie - Institut Fédératif de Biologie | Latour J., Milhes M., Bulach T., Ranger N., Salin G., Nicot F., Tremeaux P., Donnadieu C., Izopet J. |
| EPI_ISL_3471481 | CH ALBI | CHU Purpan - Laboratoire de Virologie - Institut Fédératif de Biologie | Latour J., Milhes M., Bulach T., Ranger N., Salin G., Nicot F., Tremeaux P., Donnadieu C., Izopet J. |
| EPI_ISL_3471482 | LABO BIO DOC | CHU Purpan - Laboratoire de Virologie - Institut Fédératif de Biologie | Latour J., Milhes M., Bulach T., Ranger N., Salin G., Nicot F., Tremeaux P., Donnadieu C., Izopet J. |
| EPI_ISL_3471483, EPI_ISL_3471484 | CHU Purpan - Laboratoire de Virologie - Institut Fédératif de Biologie | CHU Purpan - Laboratoire de Virologie - Institut Fédératif de Biologie | Latour J., Milhes M., Bulach T., Ranger N., Salin G., Nicot F., Tremeaux P., Donnadieu C., Izopet J. |
| EPI_ISL_3471485 | LABO BIO DOC | CHU Purpan - Laboratoire de Virologie - Institut Fédératif de Biologie | Latour J., Milhes M., Bulach T., Ranger N., Salin G., Nicot F., Tremeaux P., Donnadieu C., Izopet J. |
| EPI_ISL_3471486, EPI_ISL_3471487, EPI_ISL_3471488 | CHU Purpan - Laboratoire de Virologie - Institut Fédératif de Biologie | CHU Purpan - Laboratoire de Virologie - Institut Fédératif de Biologie | Latour J., Milhes M., Bulach T., Ranger N., Salin G., Nicot F., Tremeaux P., Donnadieu C., Izopet J. |
| EPI_ISL_3471489, EPI_ISL_3471490 | LABO BIO DOC | CHU Purpan - Laboratoire de Virologie - Institut Fédératif de Biologie | Latour J., Milhes M., Bulach T., Ranger N., Salin G., Nicot F., Tremeaux P., Donnadieu C., Izopet J. |
| EPI_ISL_3471491 | CH CAHORS | CHU Purpan - Laboratoire de Virologie - Institut Fédératif de Biologie | Latour J., Milhes M., Bulach T., Ranger N., Salin G., Nicot F., Tremeaux P., Donnadieu C., Izopet J. |
| EPI_ISL_3471492 | LABO BIO DOC | CHU Purpan - Laboratoire de Virologie - Institut Fédératif de Biologie | Latour J., Milhes M., Bulach T., Ranger N., Salin G., Nicot F., Tremeaux P., Donnadieu C., Izopet J. |
| EPI_ISL_3471493 | CHU Purpan - Laboratoire de Virologie - Institut Fédératif de Biologie | CHU Purpan - Laboratoire de Virologie - Institut Fédératif de Biologie | Latour J., Milhes M., Bulach T., Ranger N., Salin G., Nicot F., Tremeaux P., Donnadieu C., Izopet J. |
| EPI_ISL_3471494, EPI_ISL_3471495, EPI_ISL_3471496, EPI_ISL_3471497, EPI_ISL_3471498, EPI_ISL_3471499, EPI_ISL_3471500 | LABO BIO DOC | CHU Purpan - Laboratoire de Virologie - Institut Fédératif de Biologie | Latour J., Milhes M., Bulach T., Ranger N., Salin G., Nicot F., Tremeaux P., Donnadieu C., Izopet J. |
| EPI_ISL_3471501, EPI_ISL_3471502 | CHU Purpan - Laboratoire de Virologie - Institut Fédératif de Biologie | CHU Purpan - Laboratoire de Virologie - Institut Fédératif de Biologie | Latour J., Milhes M., Bulach T., Ranger N., Salin G., Nicot F., Tremeaux P., Donnadieu C., Izopet J. |
| EPI_ISL_3471503 | CH ALBI | CHU Purpan - Laboratoire de Virologie - Institut Fédératif de Biologie | Latour J., Milhes M., Bulach T., Ranger N., Salin G., Nicot F., Tremeaux P., Donnadieu C., Izopet J. |
| EPI_ISL_3471504 | LABO BIO DOC | CHU Purpan - Laboratoire de Virologie - Institut Fédératif de Biologie | Latour J., Milhes M., Bulach T., Ranger N., Salin G., Nicot F., Tremeaux P., Donnadieu C., Izopet J. |
| EPI_ISL_3471505 | CH CAHORS | CHU Purpan - Laboratoire de Virologie - Institut Fédératif de Biologie | Latour J., Milhes M., Bulach T., Ranger N., Salin G., Nicot F., Tremeaux P., Donnadieu C., Izopet J. |
| EPI_ISL_3471506 | CH ALBI | CHU Purpan - Laboratoire de Virologie - Institut Fédératif de Biologie | Latour J., Milhes M., Bulach T., Ranger N., Salin G., Nicot F., Tremeaux P., Donnadieu C., Izopet J. |
| EPI_ISL_3471507 | LABO BIO DOC | CHU Purpan - Laboratoire de Virologie - Institut Fédératif de Biologie | Latour J., Milhes M., Bulach T., Ranger N., Salin G., Nicot F., Tremeaux P., Donnadieu C., Izopet J. |
| EPI_ISL_3471508 | CH ALBI | CHU Purpan - Laboratoire de Virologie - Institut Fédératif de Biologie | Latour J., Milhes M., Bulach T., Ranger N., Salin G., Nicot F., Tremeaux P., Donnadieu C., Izopet J. |
| EPI_ISL_3471509 | LABO BIO DOC | CHU Purpan - Laboratoire de Virologie - Institut Fédératif de Biologie | Latour J., Milhes M., Bulach T., Ranger N., Salin G., Nicot F., Tremeaux P., Donnadieu C., Izopet J. |
| EPI_ISL_3471510, | CH MONTAUBAN | CHU Purpan - Laboratoire de Virologie - Institut Fédératif de Biologie | Latour J., Milhes M., Bulach T., Ranger N., Salin G., Nicot F., Tremeaux P., Donnadieu C., Izopet J. |

| | | | |
|---|---|---|---|
| EPI_ISL_3471511, EPI_ISL_3471512 | | | |
| EPI_ISL_3471513, EPI_ISL_3471514, EPI_ISL_3471515, EPI_ISL_3471516, EPI_ISL_3471517, EPI_ISL_3471518, EPI_ISL_3471519, EPI_ISL_3471520 | LABO BIO DOC | CHU Purpan - Laboratoire de Virologie - Institut Fédératif de Biologie | Latour J., Milhes M., Bulach T., Ranger N., Salin G., Nicot F., Tremeaux P., Donnadieu C., Izopet J. |
| EPI_ISL_3471521, EPI_ISL_3471522, EPI_ISL_3471523 | CHU Purpan - Laboratoire de Virologie - Institut Fédératif de Biologie | CHU Purpan - Laboratoire de Virologie - Institut Fédératif de Biologie | Latour J., Milhes M., Bulach T., Ranger N., Salin G., Nicot F., Tremeaux P., Donnadieu C., Izopet J. |
| EPI_ISL_3471524, EPI_ISL_3471525, EPI_ISL_3471526 | LABO BIO DOC | CHU Purpan - Laboratoire de Virologie - Institut Fédératif de Biologie | Latour J., Milhes M., Bulach T., Ranger N., Salin G., Nicot F., Tremeaux P., Donnadieu C., Izopet J. |
| EPI_ISL_3471527 | CHU Purpan - Laboratoire de Virologie - Institut Fédératif de Biologie | CHU Purpan - Laboratoire de Virologie - Institut Fédératif de Biologie | Latour J., Milhes M., Bulach T., Ranger N., Salin G., Nicot F., Tremeaux P., Donnadieu C., Izopet J. |
| EPI_ISL_3471528 | LABO BIO DOC | CHU Purpan - Laboratoire de Virologie - Institut Fédératif de Biologie | Latour J., Milhes M., Bulach T., Ranger N., Salin G., Nicot F., Tremeaux P., Donnadieu C., Izopet J. |
| EPI_ISL_3471529 | CH ALBI | CHU Purpan - Laboratoire de Virologie - Institut Fédératif de Biologie | Latour J., Milhes M., Bulach T., Ranger N., Salin G., Nicot F., Tremeaux P., Donnadieu C., Izopet J. |
| EPI_ISL_3471530, EPI_ISL_3471531, EPI_ISL_3471532, EPI_ISL_3471533, EPI_ISL_3471534, EPI_ISL_3471535 | LABO BIO DOC | CHU Purpan - Laboratoire de Virologie - Institut Fédératif de Biologie | Latour J., Milhes M., Bulach T., Ranger N., Salin G., Nicot F., Tremeaux P., Donnadieu C., Izopet J. |
| EPI_ISL_3471536 | LABORATOIRE GCS INTER HOSPITALIER | CHU Purpan - Laboratoire de Virologie - Institut Fédératif de Biologie | Latour J., Milhes M., Bulach T., Ranger N., Salin G., Nicot F., Tremeaux P., Donnadieu C., Izopet J. |
| EPI_ISL_3471537, EPI_ISL_3471538 | LABO BIO DOC | CHU Purpan - Laboratoire de Virologie - Institut Fédératif de Biologie | Latour J., Milhes M., Bulach T., Ranger N., Salin G., Nicot F., Tremeaux P., Donnadieu C., Izopet J. |
| EPI_ISL_3471539 | CHU Purpan - Laboratoire de Virologie - Institut Fédératif de Biologie | CHU Purpan - Laboratoire de Virologie - Institut Fédératif de Biologie | Latour J., Milhes M., Bulach T., Ranger N., Salin G., Nicot F., Tremeaux P., Donnadieu C., Izopet J. |
| EPI_ISL_3471540 | CH MONTAUBAN | CHU Purpan - Laboratoire de Virologie - Institut Fédératif de Biologie | Latour J., Milhes M., Bulach T., Ranger N., Salin G., Nicot F., Tremeaux P., Donnadieu C., Izopet J. |
| EPI_ISL_3471541 | CH AUCH | CHU Purpan - Laboratoire de Virologie - Institut Fédératif de Biologie | Latour J., Milhes M., Bulach T., Ranger N., Salin G., Nicot F., Tremeaux P., Donnadieu C., Izopet J. |
| EPI_ISL_3471542, EPI_ISL_3471544, EPI_ISL_3471545, EPI_ISL_3471546, EPI_ISL_3471547, EPI_ISL_3471548 | LABO BIO DOC | CHU Purpan - Laboratoire de Virologie - Institut Fédératif de Biologie | Latour J., Milhes M., Bulach T., Ranger N., Salin G., Nicot F., Tremeaux P., Donnadieu C., Izopet J. |
| EPI_ISL_3471549 | CHU Purpan - Laboratoire de Virologie - Institut Fédératif de Biologie | CHU Purpan - Laboratoire de Virologie - Institut Fédératif de Biologie | Latour J., Milhes M., Bulach T., Ranger N., Salin G., Nicot F., Tremeaux P., Donnadieu C., Izopet J. |
| EPI_ISL_3471550 | CENTRE HOSPITALIER DE BIGORRE | CHU Purpan - Laboratoire de Virologie - Institut Fédératif de Biologie | Latour J., Milhes M., Bulach T., Ranger N., Salin G., Nicot F., Tremeaux P., Donnadieu C., Izopet J. |
| EPI_ISL_3471551 | LABO BIO DOC | CHU Purpan - Laboratoire de Virologie - Institut Fédératif de Biologie | Latour J., Milhes M., Bulach T., Ranger N., Salin G., Nicot F., Tremeaux P., Donnadieu C., Izopet J. |
| EPI_ISL_3471552 | CHU Purpan - Laboratoire de Virologie - Institut Fédératif de Biologie | CHU Purpan - Laboratoire de Virologie - Institut Fédératif de Biologie | Latour J., Milhes M., Bulach T., Ranger N., Salin G., Nicot F., Tremeaux P., Donnadieu C., Izopet J. |
| EPI_ISL_3471553 | LABO BIO DOC | CHU Purpan - Laboratoire de Virologie - Institut Fédératif de Biologie | Latour J., Milhes M., Bulach T., Ranger N., Salin G., Nicot F., Tremeaux P., Donnadieu C., Izopet J. |
| EPI_ISL_3471554, EPI_ISL_3471555 | CHU Purpan - Laboratoire de Virologie - Institut Fédératif de Biologie | CHU Purpan - Laboratoire de Virologie - Institut Fédératif de Biologie | Latour J., Milhes M., Bulach T., Ranger N., Salin G., Nicot F., Tremeaux P., Donnadieu C., Izopet J. |
| EPI_ISL_3471556 | LABO BIO DOC | CHU Purpan - Laboratoire de Virologie - Institut Fédératif de Biologie | Latour J., Milhes M., Bulach T., Ranger N., Salin G., Nicot F., Tremeaux P., Donnadieu C., Izopet J. |
| EPI_ISL_3471557 | CHU Purpan - Laboratoire de Virologie - Institut Fédératif de Biologie | CHU Purpan - Laboratoire de Virologie - Institut Fédératif de Biologie | Latour J., Milhes M., Bulach T., Ranger N., Salin G., Nicot F., Tremeaux P., Donnadieu C., Izopet J. |
| EPI_ISL_3471558 | LABO BIO DOC | CHU Purpan - Laboratoire de Virologie - Institut Fédératif de Biologie | Latour J., Milhes M., Bulach T., Ranger N., Salin G., Nicot F., Tremeaux P., Donnadieu C., Izopet J. |
| EPI_ISL_3471559 | CH MONTAUBAN | CHU Purpan - Laboratoire de Virologie - Institut Fédératif de Biologie | Latour J., Milhes M., Bulach T., Ranger N., Salin G., Nicot F., Tremeaux P., Donnadieu C., Izopet J. |
| EPI_ISL_3471560, EPI_ISL_3471561 | LABO BIO DOC | CHU Purpan - Laboratoire de Virologie - Institut Fédératif de Biologie | Latour J., Milhes M., Bulach T., Ranger N., Salin G., Nicot F., Tremeaux P., Donnadieu C., Izopet J. |
| EPI_ISL_3471562 | CHU Purpan - Laboratoire de Virologie - Institut Fédératif de Biologie | CHU Purpan - Laboratoire de Virologie - Institut Fédératif de Biologie | Latour J., Milhes M., Bulach T., Ranger N., Salin G., Nicot F., Tremeaux P., Donnadieu C., Izopet J. |
| EPI_ISL_3471563 | LABO BIO DOC | CHU Purpan - Laboratoire de Virologie - Institut Fédératif de Biologie | Latour J., Milhes M., Bulach T., Ranger N., Salin G., Nicot F., Tremeaux P., Donnadieu C., Izopet J. |
| EPI_ISL_3471564 | CHU Purpan - Laboratoire de Virologie - Institut Fédératif de Biologie | CHU Purpan - Laboratoire de Virologie - Institut Fédératif de Biologie | Latour J., Milhes M., Bulach T., Ranger N., Salin G., Nicot F., Tremeaux P., Donnadieu C., Izopet J. |
| EPI_ISL_3471565, EPI_ISL_3471566, EPI_ISL_3471567, EPI_ISL_3471568, EPI_ISL_3471569 | LABO BIO DOC | CHU Purpan - Laboratoire de Virologie - Institut Fédératif de Biologie | Latour J., Milhes M., Bulach T., Ranger N., Salin G., Nicot F., Tremeaux P., Donnadieu C., Izopet J. |
| EPI_ISL_3471570 | CHU Purpan - Laboratoire de Virologie - Institut Fédératif de Biologie | CHU Purpan - Laboratoire de Virologie - Institut Fédératif de Biologie | Latour J., Milhes M., Bulach T., Ranger N., Salin G., Nicot F., Tremeaux P., Donnadieu C., Izopet J. |
| EPI_ISL_3471571, EPI_ISL_3471572, EPI_ISL_3471573, EPI_ISL_3471575, EPI_ISL_3471576 | LABO BIO DOC | CHU Purpan - Laboratoire de Virologie - Institut Fédératif de Biologie | Latour J., Milhes M., Bulach T., Ranger N., Salin G., Nicot F., Tremeaux P., Donnadieu C., Izopet J. |
| EPI_ISL_3471577 | CH ALBI | CHU Purpan - Laboratoire de Virologie - Institut Fédératif de Biologie | Latour J., Milhes M., Bulach T., Ranger N., Salin G., Nicot F., Tremeaux P., Donnadieu C., Izopet J. |
| EPI_ISL_3471578, EPI_ISL_3471579, EPI_ISL_3471580, EPI_ISL_3471582, EPI_ISL_3471583, EPI_ISL_3471584, EPI_ISL_3471585, EPI_ISL_3471586, EPI_ISL_3471587, EPI_ISL_3471588 | LABO BIO DOC | CHU Purpan - Laboratoire de Virologie - Institut Fédératif de Biologie | Latour J., Milhes M., Bulach T., Ranger N., Salin G., Nicot F., Tremeaux P., Donnadieu C., Izopet J. |
| EPI_ISL_3471589 | CHU Purpan - Laboratoire de Virologie - Institut Fédératif de Biologie | CHU Purpan - Laboratoire de Virologie - Institut Fédératif de Biologie | Latour J., Milhes M., Bulach T., Ranger N., Salin G., Nicot F., Tremeaux P., Donnadieu C., Izopet J. |
| EPI_ISL_3471590 | CH ALBI | CHU Purpan - Laboratoire de Virologie - Institut Fédératif de Biologie | Latour J., Milhes M., Bulach T., Ranger N., Salin G., Nicot F., Tremeaux P., Donnadieu C., Izopet J. |
| EPI_ISL_3471591 | LABO BIO DOC | CHU Purpan - Laboratoire de Virologie - Institut Fédératif de Biologie | Latour J., Milhes M., Bulach T., Ranger N., Salin G., Nicot F., Tremeaux P., Donnadieu C., Izopet J. |
| EPI_ISL_3471592 | GCS DE BIOLOGIE DU TERRITOIRE DE L'ARIEGE | CHU Purpan - Laboratoire de Virologie - Institut Fédératif de Biologie | Latour J., Milhes M., Bulach T., Ranger N., Salin G., Nicot F., Tremeaux P., Donnadieu C., Izopet J. |
| EPI_ISL_3471598 | CH ALBI | CHU Purpan - Laboratoire de Virologie - Institut Fédératif de Biologie | Latour J., Milhes M., Bulach T., Ranger N., Salin G., Nicot F., Tremeaux P., Donnadieu C., Izopet J. |
| EPI_ISL_3471599 | LABO BIO DOC | CHU Purpan - Laboratoire de Virologie - Institut Fédératif de Biologie | Latour J., Milhes M., Bulach T., Ranger N., Salin G., Nicot F., Tremeaux P., Donnadieu C., Izopet J. |
| EPI_ISL_3471600 | CHU Purpan - Laboratoire de Virologie - Institut Fédératif de Biologie | CHU Purpan - Laboratoire de Virologie - Institut Fédératif de Biologie | Latour J., Milhes M., Bulach T., Ranger N., Salin G., Nicot F., Tremeaux P., Donnadieu C., Izopet J. |

| | | | |
|---|---|---|---|
| EPI_ISL_3471601 | CH MONTAUBAN | CHU Purpan - Laboratoire de Virologie - Institut Fédératif de Biologie | Latour J., Milhes M., Bulach T., Ranger N., Salin G., Nicot F., Tremeaux P., Donnadieu C., Izopet J. |
| EPI_ISL_3471602 | CH AUCH | CHU Purpan - Laboratoire de Virologie - Institut Fédératif de Biologie | Latour J., Milhes M., Bulach T., Ranger N., Salin G., Nicot F., Tremeaux P., Donnadieu C., Izopet J. |
| EPI_ISL_3471603, EPI_ISL_3471604 | CHU Purpan - Laboratoire de Virologie - Institut Fédératif de Biologie | CHU Purpan - Laboratoire de Virologie - Institut Fédératif de Biologie | Latour J., Milhes M., Bulach T., Ranger N., Salin G., Nicot F., Tremeaux P., Donnadieu C., Izopet J. |
| EPI_ISL_3471605, EPI_ISL_3471606, EPI_ISL_3471607, EPI_ISL_3471608, EPI_ISL_3471609, EPI_ISL_3471610, EPI_ISL_3471611, EPI_ISL_3471612, EPI_ISL_3471613 | LABO BIO DOC | CHU Purpan - Laboratoire de Virologie - Institut Fédératif de Biologie | Latour J., Milhes M., Bulach T., Ranger N., Salin G., Nicot F., Tremeaux P., Donnadieu C., Izopet J. |
| EPI_ISL_3471614 | CH AUCH | CHU Purpan - Laboratoire de Virologie - Institut Fédératif de Biologie | Latour J., Milhes M., Bulach T., Ranger N., Salin G., Nicot F., Tremeaux P., Donnadieu C., Izopet J. |
| EPI_ISL_3471615, EPI_ISL_3471616, EPI_ISL_3471617, EPI_ISL_3471618, EPI_ISL_3471619, EPI_ISL_3471620, EPI_ISL_3471621, EPI_ISL_3471622, EPI_ISL_3471623, EPI_ISL_3471624 | LABO BIO DOC | CHU Purpan - Laboratoire de Virologie - Institut Fédératif de Biologie | Latour J., Milhes M., Bulach T., Ranger N., Salin G., Nicot F., Tremeaux P., Donnadieu C., Izopet J. |
| EPI_ISL_3471625 | CHU Purpan - Laboratoire de Virologie - Institut Fédératif de Biologie | CHU Purpan - Laboratoire de Virologie - Institut Fédératif de Biologie | Latour J., Milhes M., Bulach T., Ranger N., Salin G., Nicot F., Tremeaux P., Donnadieu C., Izopet J. |
| EPI_ISL_3471626, EPI_ISL_3471627 | LABO BIO DOC | CHU Purpan - Laboratoire de Virologie - Institut Fédératif de Biologie | Latour J., Milhes M., Bulach T., Ranger N., Salin G., Nicot F., Tremeaux P., Donnadieu C., Izopet J. |
| EPI_ISL_3471628 | GCS DE BIOLOGIE DU TERRITOIRE DE L'ARIEGE | CHU Purpan - Laboratoire de Virologie - Institut Fédératif de Biologie | Latour J., Milhes M., Bulach T., Ranger N., Salin G., Nicot F., Tremeaux P., Donnadieu C., Izopet J. |
| EPI_ISL_3471629 | LABO BIO DOC | CHU Purpan - Laboratoire de Virologie - Institut Fédératif de Biologie | Latour J., Milhes M., Bulach T., Ranger N., Salin G., Nicot F., Tremeaux P., Donnadieu C., Izopet J. |
| EPI_ISL_3471632, EPI_ISL_3471633 | LABORATOIRE GCS INTER HOSPITALIER | CHU Purpan - Laboratoire de Virologie - Institut Fédératif de Biologie | Latour J., Milhes M., Bulach T., Ranger N., Salin G., Nicot F., Tremeaux P., Donnadieu C., Izopet J. |
| EPI_ISL_3471634 | GCS DE BIOLOGIE DU TERRITOIRE DE L'ARIEGE | CHU Purpan - Laboratoire de Virologie - Institut Fédératif de Biologie | Latour J., Milhes M., Bulach T., Ranger N., Salin G., Nicot F., Tremeaux P., Donnadieu C., Izopet J. |
| EPI_ISL_3471635, EPI_ISL_3471636 | CH AUCH | CHU Purpan - Laboratoire de Virologie - Institut Fédératif de Biologie | Latour J., Milhes M., Bulach T., Ranger N., Salin G., Nicot F., Tremeaux P., Donnadieu C., Izopet J. |
| EPI_ISL_3471637, EPI_ISL_3471638, EPI_ISL_3471640, EPI_ISL_3471642 | CHU Purpan - Laboratoire de Virologie - Institut Fédératif de Biologie | CHU Purpan - Laboratoire de Virologie - Institut Fédératif de Biologie | Latour J., Milhes M., Bulach T., Ranger N., Salin G., Nicot F., Tremeaux P., Donnadieu C., Izopet J. |
| EPI_ISL_3471643, EPI_ISL_3471644 | LABORATOIRE GCS INTER HOSPITALIER | CHU Purpan - Laboratoire de Virologie - Institut Fédératif de Biologie | Latour J., Milhes M., Bulach T., Ranger N., Salin G., Nicot F., Tremeaux P., Donnadieu C., Izopet J. |
| EPI_ISL_3471645, EPI_ISL_3471646, EPI_ISL_3471647, EPI_ISL_3471648, EPI_ISL_3471649, EPI_ISL_3471650, EPI_ISL_3471651, EPI_ISL_3471652, EPI_ISL_3471653, EPI_ISL_3471654, EPI_ISL_3471655, EPI_ISL_3471656, EPI_ISL_3471657, EPI_ISL_3471658, EPI_ISL_3471659, EPI_ISL_3471660, EPI_ISL_3471661, EPI_ISL_3471662, EPI_ISL_3471663, EPI_ISL_3471664, EPI_ISL_3471665, EPI_ISL_3471666, EPI_ISL_3471667, EPI_ISL_3471668, EPI_ISL_3471669, EPI_ISL_3471670, EPI_ISL_3471671, EPI_ISL_3471672, EPI_ISL_3471673, EPI_ISL_3471674, EPI_ISL_3471675, EPI_ISL_3471676, EPI_ISL_3471677, EPI_ISL_3471678, EPI_ISL_3471679, EPI_ISL_3471680, EPI_ISL_3471681, EPI_ISL_3471682, EPI_ISL_3471683, EPI_ISL_3471684, EPI_ISL_3471685, EPI_ISL_3471686, EPI_ISL_3471687, EPI_ISL_3471688, EPI_ISL_3471689, EPI_ISL_3471690, EPI_ISL_3471691, EPI_ISL_3471692, EPI_ISL_3471693, EPI_ISL_3471694, EPI_ISL_3471695, EPI_ISL_3471696, EPI_ISL_3471697, EPI_ISL_3471698, EPI_ISL_3471699, EPI_ISL_3471700, EPI_ISL_3471701, EPI_ISL_3471702, EPI_ISL_3471703, EPI_ISL_3471704, EPI_ISL_3471705 | | | |
| see above | LABO BIO DOC | CHU Purpan - Laboratoire de Virologie - Institut Fédératif de Biologie | Latour J., Milhes M., Bulach T., Ranger N., Salin G., Nicot F., Tremeaux P., Donnadieu C., Izopet J. |
| EPI_ISL_3471706, EPI_ISL_3471707 | CH CARCASSONE | CHU Purpan - Laboratoire de Virologie - Institut Fédératif de Biologie | Latour J., Milhes M., Bulach T., Ranger N., Salin G., Nicot F., Tremeaux P., Donnadieu C., Izopet J. |
| EPI_ISL_3471708, EPI_ISL_3471709, EPI_ISL_3471710, EPI_ISL_3471711 | CH AUCH | CHU Purpan - Laboratoire de Virologie - Institut Fédératif de Biologie | Latour J., Milhes M., Bulach T., Ranger N., Salin G., Nicot F., Tremeaux P., Donnadieu C., Izopet J. |
| EPI_ISL_3471712, EPI_ISL_3471713, EPI_ISL_3471714, EPI_ISL_3471715, EPI_ISL_3471716, EPI_ISL_3471717, EPI_ISL_3471718, EPI_ISL_3471719, EPI_ISL_3471720, EPI_ISL_3471721, EPI_ISL_3471722, EPI_ISL_3471723, EPI_ISL_3471724, EPI_ISL_3471725, EPI_ISL_3471726, EPI_ISL_3471727, EPI_ISL_3471728, EPI_ISL_3471729, EPI_ISL_3471730, EPI_ISL_3471731, EPI_ISL_3471732, EPI_ISL_3471733, EPI_ISL_3471734, EPI_ISL_3471735 | | | |
| see above | CHU Purpan - Laboratoire de Virologie - Institut Fédératif de Biologie | CHU Purpan - Laboratoire de Virologie - Institut Fédératif de Biologie | Latour J., Milhes M., Bulach T., Ranger N., Salin G., Nicot F., Tremeaux P., Donnadieu C., Izopet J. |
| EPI_ISL_3471736 | CENTRE HOSPITALIER DE BIGORRE | CHU Purpan - Laboratoire de Virologie - Institut Fédératif de Biologie | Latour J., Milhes M., Bulach T., Ranger N., Salin G., Nicot F., Tremeaux P., Donnadieu C., Izopet J. |
| EPI_ISL_3471737, EPI_ISL_3471738, EPI_ISL_3471739 | CH LOURDES | CHU Purpan - Laboratoire de Virologie - Institut Fédératif de Biologie | Latour J., Milhes M., Bulach T., Ranger N., Salin G., Nicot F., Tremeaux P., Donnadieu C., Izopet J. |
| EPI_ISL_3471740, EPI_ISL_3471741 | CH CAHORS | CHU Purpan - Laboratoire de Virologie - Institut Fédératif de Biologie | Latour J., Milhes M., Bulach T., Ranger N., Salin G., Nicot F., Tremeaux P., Donnadieu C., Izopet J. |
| EPI_ISL_3471742, EPI_ISL_3471743, EPI_ISL_3471744, EPI_ISL_3471745, EPI_ISL_3471746, EPI_ISL_3471747, EPI_ISL_3471748, EPI_ISL_3471749, EPI_ISL_3471750, EPI_ISL_3471751, EPI_ISL_3471752, EPI_ISL_3471753, EPI_ISL_3471754, EPI_ISL_3471755, EPI_ISL_3471756, EPI_ISL_3471757, EPI_ISL_3471758, EPI_ISL_3471759 | | | |
| see above | CH ALBI | CHU Purpan - Laboratoire de Virologie - Institut Fédératif de Biologie | Latour J., Milhes M., Bulach T., Ranger N., Salin G., Nicot F., Tremeaux P., Donnadieu C., Izopet J. |
| EPI_ISL_3471760, EPI_ISL_3471761, EPI_ISL_3471762, EPI_ISL_3471763, EPI_ISL_3471764 | CH MONTAUBAN | CHU Purpan - Laboratoire de Virologie - Institut Fédératif de Biologie | Latour J., Milhes M., Bulach T., Ranger N., Salin G., Nicot F., Tremeaux P., Donnadieu C., Izopet J. |
| EPI_ISL_3471765, EPI_ISL_3471766, EPI_ISL_3471767 | Centinela Hospital Medical Center | Los Angeles County Public Health Laboratories | P. Hemarajata et al. |
| EPI_ISL_3471768 | Martin Luther King, Jr. Outpatient Center | Los Angeles County Public Health Laboratories | P. Hemarajata et al. |
| EPI_ISL_3471769 | Methodist Hospital | Los Angeles County Public Health Laboratories | P. Hemarajata et al. |
| EPI_ISL_3471770 | Palmdale Regional Medical Center | Los Angeles County Public Health Laboratories | P. Hemarajata et al. |
| EPI_ISL_3471771, EPI_ISL_3471772, EPI_ISL_3471773, EPI_ISL_3471774 | Pomona Valley Hospital Medical Center | Los Angeles County Public Health Laboratories | P. Hemarajata et al. |
| EPI_ISL_3471775 | Torrance Memorial Medical Center | Los Angeles County Public Health Laboratories | P. Hemarajata et al. |
| EPI_ISL_3471778 | Centinela Hospital Medical Center | Los Angeles County Public Health Laboratories | P. Hemarajata et al. |
| EPI_ISL_3471779 | Huntington Memorial Hospital | Los Angeles County Public Health Laboratories | P. Hemarajata et al. |
| EPI_ISL_3471780 | Martin Luther King, Jr. Outpatient Center | Los Angeles County Public Health Laboratories | P. Hemarajata et al. |
| EPI_ISL_3471781 | Queen of the Valley Hospital | Los Angeles County Public Health Laboratories | P. Hemarajata et al. |
| EPI_ISL_3471782 | California Hospital Medical Center | Los Angeles County Public Health Laboratories | P. Hemarajata et al. |
| EPI_ISL_3471784, EPI_ISL_3471785 | Pomona Valley Hospital Medical Center | Los Angeles County Public Health Laboratories | P. Hemarajata et al. |
| EPI_ISL_3471786, EPI_ISL_3471787 | Centinela Hospital Medical Center | Los Angeles County Public Health Laboratories | P. Hemarajata et al. |
| EPI_ISL_3471788, EPI_ISL_3471789, EPI_ISL_3471790, EPI_ISL_3471791 | Pomona Valley Hospital Medical Center | Los Angeles County Public Health Laboratories | P. Hemarajata et al. |

| | | | |
|---|---|---|---|
| EPI_ISL_3471793 | Centinela Hospital Medical Center | Los Angeles County Public Health Laboratories | P. Hemarajata et al. |
| EPI_ISL_3471794 | Pomona Valley Hospital Medical Center | Los Angeles County Public Health Laboratories | P. Hemarajata et al. |
| EPI_ISL_3471796 | Huntington Memorial Hospital | Los Angeles County Public Health Laboratories | P. Hemarajata et al. |
| EPI_ISL_3471798 | Torrance Memorial Medical Center | Los Angeles County Public Health Laboratories | P. Hemarajata et al. |
| EPI_ISL_3471799 | Huntington Memorial Hospital | Los Angeles County Public Health Laboratories | P. Hemarajata et al. |
| EPI_ISL_3471801 | Torrance Memorial Medical Center | Los Angeles County Public Health Laboratories | P. Hemarajata et al. |
| EPI_ISL_3471802 | Martin Luther King, Jr. Outpatient Center | Los Angeles County Public Health Laboratories | P. Hemarajata et al. |
| EPI_ISL_3471803 | Torrance Memorial Medical Center | Los Angeles County Public Health Laboratories | P. Hemarajata et al. |
| EPI_ISL_3471804 | Palmdale Regional Medical Center | Los Angeles County Public Health Laboratories | P. Hemarajata et al. |
| EPI_ISL_3471805 | Martin Luther King, Jr. Outpatient Center | Los Angeles County Public Health Laboratories | P. Hemarajata et al. |
| EPI_ISL_3471806 | Torrance Memorial Medical Center | Los Angeles County Public Health Laboratories | P. Hemarajata et al. |
| EPI_ISL_3471807 | Huntington Memorial Hospital | Los Angeles County Public Health Laboratories | P. Hemarajata et al. |
| EPI_ISL_3471808 | Saint Francis Medical Center | Los Angeles County Public Health Laboratories | P. Hemarajata et al. |
| EPI_ISL_3471809 | Huntington Memorial Hospital | Los Angeles County Public Health Laboratories | P. Hemarajata et al. |
| EPI_ISL_3471810 | Martin Luther King, Jr. Outpatient Center | Los Angeles County Public Health Laboratories | P. Hemarajata et al. |
| EPI_ISL_3471811 | Torrance Memorial Medical Center | Los Angeles County Public Health Laboratories | P. Hemarajata et al. |
| EPI_ISL_3471812, EPI_ISL_3471813, EPI_ISL_3471814, EPI_ISL_3471815, EPI_ISL_3471816 | Huntington Memorial Hospital | Los Angeles County Public Health Laboratories | P. Hemarajata et al. |
| EPI_ISL_3471817, EPI_ISL_3471818 | Martin Luther King, Jr. Outpatient Center | Los Angeles County Public Health Laboratories | P. Hemarajata et al. |
| EPI_ISL_3471819, EPI_ISL_3471820 | Torrance Memorial Medical Center | Los Angeles County Public Health Laboratories | P. Hemarajata et al. |
| EPI_ISL_3471821 | Palmdale Regional Medical Center | Los Angeles County Public Health Laboratories | P. Hemarajata et al. |
| EPI_ISL_3471822 | Flow Diagnostics Lab | Los Angeles County Public Health Laboratories | P. Hemarajata et al. |
| EPI_ISL_3471823 | Torrance Memorial Medical Center | Los Angeles County Public Health Laboratories | P. Hemarajata et al. |
| EPI_ISL_3471824 | Huntington Memorial Hospital | Los Angeles County Public Health Laboratories | P. Hemarajata et al. |
| EPI_ISL_3471825 | Torrance Memorial Medical Center | Los Angeles County Public Health Laboratories | P. Hemarajata et al. |
| EPI_ISL_3471826 | Centinela Hospital Medical Center | Los Angeles County Public Health Laboratories | P. Hemarajata et al. |
| EPI_ISL_3471827 | Flow Diagnostics Lab | Los Angeles County Public Health Laboratories | P. Hemarajata et al. |
| EPI_ISL_3471828, EPI_ISL_3471829 | Huntington Memorial Hospital | Los Angeles County Public Health Laboratories | P. Hemarajata et al. |
| EPI_ISL_3471830 | Martin Luther King, Jr. Outpatient Center | Los Angeles County Public Health Laboratories | P. Hemarajata et al. |
| EPI_ISL_3471831 | Pomona Valley Hospital Medical Center | Los Angeles County Public Health Laboratories | P. Hemarajata et al. |
| EPI_ISL_3471832 | Huntington Memorial Hospital | Los Angeles County Public Health Laboratories | P. Hemarajata et al. |
| EPI_ISL_3471833 | Flow Diagnostics Lab | Los Angeles County Public Health Laboratories | P. Hemarajata et al. |
| EPI_ISL_3471834 | Huntington Memorial Hospital | Los Angeles County Public Health Laboratories | P. Hemarajata et al. |
| EPI_ISL_3471835 | Flow Diagnostics Lab | Los Angeles County Public Health Laboratories | P. Hemarajata et al. |
| EPI_ISL_3471836 | Martin Luther King, Jr. Outpatient Center | Los Angeles County Public Health Laboratories | P. Hemarajata et al. |
| EPI_ISL_3471837 | Flow Diagnostics Lab | Los Angeles County Public Health Laboratories | P. Hemarajata et al. |
| EPI_ISL_3471838 | Huntington Memorial Hospital | Los Angeles County Public Health Laboratories | P. Hemarajata et al. |
| EPI_ISL_3471840, EPI_ISL_3471841, EPI_ISL_3471842, EPI_ISL_3471843 | Flow Diagnostics Lab | Los Angeles County Public Health Laboratories | P. Hemarajata et al. |
| EPI_ISL_3471844 | PIH Health Whittier Hospital | Los Angeles County Public Health Laboratories | P. Hemarajata et al. |
| EPI_ISL_3471845 | Flow Diagnostics Lab | Los Angeles County Public Health Laboratories | P. Hemarajata et al. |
| EPI_ISL_3471846 | Centinela Hospital Medical Center | Los Angeles County Public Health Laboratories | P. Hemarajata et al. |
| EPI_ISL_3471858, EPI_ISL_3471859, EPI_ISL_3471860, EPI_ISL_3471861, EPI_ISL_3471862, EPI_ISL_3471863, EPI_ISL_3471864, EPI_ISL_3471865, EPI_ISL_3471866 | Heilig hart Lier | Imelda Ziekenhuis | Johan Frans, Hanne Valgaeren, Dagmar Obbels |
| EPI_ISL_3471867, EPI_ISL_3471868 | AZ Jan Portaels | Imelda Ziekenhuis | Johan Frans, Hanne Valgaeren, Dagmar Obbels |
| EPI_ISL_3471869, EPI_ISL_3471870, EPI_ISL_3471871, EPI_ISL_3471872, EPI_ISL_3471873, EPI_ISL_3471874, EPI_ISL_3471875 | Imelda Ziekenhuis | Imelda Ziekenhuis | Johan Frans, Hanne Valgaeren, Dagmar Obbels |
| EPI_ISL_3471876, EPI_ISL_3471877, EPI_ISL_3471878 | Ohio Department of Health Laboratory | Ohio Department of Health Laboratory | Holmes, Jennifer; Eric Brandt, Keoni Omura, Glen McGillivary, Caitlin McDonnell, Jade Mowery, Stephanie Mccracken, Tyler Payne, Kirtana Ramadugu, Erica Leasure, Brent Lee, Kelsey Florek, Heather Blankenship, Allison Black, Quanta Brown, and Tammy Bannerman |
| EPI_ISL_3471879 | CNR Institut Pasteur de la Guyane | Institut Pasteur de la Guyane | Anne Lavergne, Dominique Rousset, Antoine Enfissi, Arielle Salmier |
| EPI_ISL_3471882 | Institute of Virology, Biomedical Research Center of the Slovak Academy of Sciences, Bratislava | Faculty of Mathematics, Physics and Informatics, Comenius University, Bratislava | Kristina Borsova, Viktoria Cabanova, Brona Brejova, Veronika Vanova, Sabina Fumacova Havlikova, Juraj Kopacek, Martina Lickova, Lubomira Lukacikova, Martina Nebohacova, Monika Slavikova, Tomas Vinar, Boris Klempa, Jozef Nosek |
| EPI_ISL_3471883 | Ohio Department of Health Laboratory | Ohio Department of Health Laboratory | Holmes, Jennifer; Eric Brandt, Keoni Omura, Glen McGillivary, Caitlin McDonnell, Jade Mowery, Stephanie Mccracken, Tyler Payne, Kirtana Ramadugu, Erica Leasure, Brent Lee, Kelsey Florek, Heather Blankenship, Allison Black, Quanta Brown, and Tammy Bannerman |
| EPI_ISL_3471884 | CENTRE HOSPITALIER DE VALENCE | CNR Virus des Infections Respiratoires - France SUD | Antonin Bal, Gregory Destras, Gwendolyne Burfin, Hadrien Regue, Quentin Semanas, Martine Valette, Bruno Lina, Laurence Josset |
| EPI_ISL_3471885 | MIRIALIS CLUSES BECHET | CNR Virus des Infections Respiratoires - France SUD | Antonin Bal, Gregory Destras, Gwendolyne Burfin, Hadrien Regue, Quentin Semanas, Martine Valette, Bruno Lina, Laurence Josset |
| EPI_ISL_3471886 | SYNLAB AQUITAINE | CNR Virus des Infections Respiratoires - France SUD | Antonin Bal, Gregory Destras, Gwendolyne Burfin, Hadrien Regue, Quentin Semanas, Martine Valette, Bruno Lina, Laurence Josset |
| EPI_ISL_3471887 | LABORATOIRE UNILIANS DECINES | CNR Virus des Infections Respiratoires - France SUD | Antonin Bal, Gregory Destras, Gwendolyne Burfin, Hadrien Regue, Quentin Semanas, Martine Valette, Bruno Lina, Laurence Josset |
| EPI_ISL_3471888 | HOPITAL ALBI | CNR Virus des Infections Respiratoires - France SUD | Antonin Bal, Gregory Destras, Gwendolyne Burfin, Hadrien Regue, Quentin Semanas, Martine Valette, Bruno Lina, Laurence Josset |
| EPI_ISL_3471889 | CENTRE HOSPITALIER DE VALENCE | CNR Virus des Infections Respiratoires - France SUD | Antonin Bal, Gregory Destras, Gwendolyne Burfin, Hadrien Regue, Quentin Semanas, Martine Valette, Bruno Lina, Laurence Josset |
| EPI_ISL_3471890 | LABORATOIRE UNILIANS DECINES | CNR Virus des Infections Respiratoires - France SUD | Antonin Bal, Gregory Destras, Gwendolyne Burfin, Hadrien Regue, Quentin Semanas, Martine Valette, Bruno Lina, Laurence Josset |
| EPI_ISL_3471891, EPI_ISL_3471892 | SYNLAB AQUITAINE | CNR Virus des Infections Respiratoires - France SUD | Antonin Bal, Gregory Destras, Gwendolyne Burfin, Hadrien Regue, Quentin Semanas, Martine Valette, Bruno Lina, Laurence Josset |

| | | | |
|---|---|---|---|
| EPI_ISL_3471893 | MIRIALIS CLUSES BECHET | CNR Virus des Infections Respiratoires - France SUD | Antonin Bal, Gregory Destras, Gwendolyne Burfin, Hadrien Regue, Quentin Semanas, Martine Valette, Bruno Lina, Laurence Josset |
| EPI_ISL_3471894 | LABORATOIRE UNILIANS DECINES | CNR Virus des Infections Respiratoires - France SUD | Antonin Bal, Gregory Destras, Gwendolyne Burfin, Hadrien Regue, Quentin Semanas, Martine Valette, Bruno Lina, Laurence Josset |
| EPI_ISL_3471895 | LABORATOIRE CERBALLIANCE PLT VILLON | CNR Virus des Infections Respiratoires - France SUD | Antonin Bal, Gregory Destras, Gwendolyne Burfin, Hadrien Regue, Quentin Semanas, Martine Valette, Bruno Lina, Laurence Josset |
| EPI_ISL_3471896 | CHU ST ETIENNE HOPITAL NORD | CNR Virus des Infections Respiratoires - France SUD | Antonin Bal, Gregory Destras, Gwendolyne Burfin, Hadrien Regue, Quentin Semanas, Martine Valette, Bruno Lina, Laurence Josset |
| EPI_ISL_3471897 | LABORATOIRE UNILIANS DECINES | CNR Virus des Infections Respiratoires - France SUD | Antonin Bal, Gregory Destras, Gwendolyne Burfin, Hadrien Regue, Quentin Semanas, Martine Valette, Bruno Lina, Laurence Josset |
| EPI_ISL_3471898 | LABORATOIRE BIOESTEREL | CNR Virus des Infections Respiratoires - France SUD | Antonin Bal, Gregory Destras, Gwendolyne Burfin, Hadrien Regue, Quentin Semanas, Martine Valette, Bruno Lina, Laurence Josset |
| EPI_ISL_3471899, EPI_ISL_3471900 | MIRIALIS CLUSES BECHET | CNR Virus des Infections Respiratoires - France SUD | Antonin Bal, Gregory Destras, Gwendolyne Burfin, Hadrien Regue, Quentin Semanas, Martine Valette, Bruno Lina, Laurence Josset |
| EPI_ISL_3471901 | LABORATOIRE BIOESTEREL | CNR Virus des Infections Respiratoires - France SUD | Antonin Bal, Gregory Destras, Gwendolyne Burfin, Hadrien Regue, Quentin Semanas, Martine Valette, Bruno Lina, Laurence Josset |
| EPI_ISL_3471902 | LABORATOIRE BIOLYSS | CNR Virus des Infections Respiratoires - France SUD | Antonin Bal, Gregory Destras, Gwendolyne Burfin, Hadrien Regue, Quentin Semanas, Martine Valette, Bruno Lina, Laurence Josset |
| EPI_ISL_3471903 | SYNLAB AQUITAINE | CNR Virus des Infections Respiratoires - France SUD | Antonin Bal, Gregory Destras, Gwendolyne Burfin, Hadrien Regue, Quentin Semanas, Martine Valette, Bruno Lina, Laurence Josset |
| EPI_ISL_3471904 | BIOLITTORAL BIOGROUP PLATEAU TECH | CNR Virus des Infections Respiratoires - France SUD | Antonin Bal, Gregory Destras, Gwendolyne Burfin, Hadrien Regue, Quentin Semanas, Martine Valette, Bruno Lina, Laurence Josset |
| EPI_ISL_3471905 | LABORATOIRE BIOESTEREL | CNR Virus des Infections Respiratoires - France SUD | Antonin Bal, Gregory Destras, Gwendolyne Burfin, Hadrien Regue, Quentin Semanas, Martine Valette, Bruno Lina, Laurence Josset |
| EPI_ISL_3471906 | LAM BIOPOLE21 | CNR Virus des Infections Respiratoires - France SUD | Antonin Bal, Gregory Destras, Gwendolyne Burfin, Hadrien Regue, Quentin Semanas, Martine Valette, Bruno Lina, Laurence Josset |
| EPI_ISL_3471907 | CNR Virus des Infections Respiratoires - France SUD | CNR Virus des Infections Respiratoires - France SUD | Antonin Bal, Gregory Destras, Gwendolyne Burfin, Hadrien Regue, Quentin Semanas, Martine Valette, Bruno Lina, Laurence Josset |
| EPI_ISL_3471908 | LABORATOIRE BIOESTEREL | CNR Virus des Infections Respiratoires - France SUD | Antonin Bal, Gregory Destras, Gwendolyne Burfin, Hadrien Regue, Quentin Semanas, Martine Valette, Bruno Lina, Laurence Josset |
| EPI_ISL_3471909 | CENTRE HOSPITALIER LUCIEN HUSSEL | CNR Virus des Infections Respiratoires - France SUD | Antonin Bal, Gregory Destras, Gwendolyne Burfin, Hadrien Regue, Quentin Semanas, Martine Valette, Bruno Lina, Laurence Josset |
| EPI_ISL_3471910 | LABORATOIRE BIOLYSS | CNR Virus des Infections Respiratoires - France SUD | Antonin Bal, Gregory Destras, Gwendolyne Burfin, Hadrien Regue, Quentin Semanas, Martine Valette, Bruno Lina, Laurence Josset |
| EPI_ISL_3471911 | MIRIALIS CLUSES BECHET | CNR Virus des Infections Respiratoires - France SUD | Antonin Bal, Gregory Destras, Gwendolyne Burfin, Hadrien Regue, Quentin Semanas, Martine Valette, Bruno Lina, Laurence Josset |
| EPI_ISL_3471912 | LABORATOIRE UNILIANS DECINES | CNR Virus des Infections Respiratoires - France SUD | Antonin Bal, Gregory Destras, Gwendolyne Burfin, Hadrien Regue, Quentin Semanas, Martine Valette, Bruno Lina, Laurence Josset |
| EPI_ISL_3471913 | CENTRE HOSPITALIER ST JOSEPH ST LUC | CNR Virus des Infections Respiratoires - France SUD | Antonin Bal, Gregory Destras, Gwendolyne Burfin, Hadrien Regue, Quentin Semanas, Martine Valette, Bruno Lina, Laurence Josset |
| EPI_ISL_3471914 | HOPITAL ALBI | CNR Virus des Infections Respiratoires - France SUD | Antonin Bal, Gregory Destras, Gwendolyne Burfin, Hadrien Regue, Quentin Semanas, Martine Valette, Bruno Lina, Laurence Josset |
| EPI_ISL_3471915 | CENTRE HOSPITALIER DE BOURG EN BRESSE | CNR Virus des Infections Respiratoires - France SUD | Antonin Bal, Gregory Destras, Gwendolyne Burfin, Hadrien Regue, Quentin Semanas, Martine Valette, Bruno Lina, Laurence Josset |
| EPI_ISL_3471916 | LABORATOIRE BIOESTEREL | CNR Virus des Infections Respiratoires - France SUD | Antonin Bal, Gregory Destras, Gwendolyne Burfin, Hadrien Regue, Quentin Semanas, Martine Valette, Bruno Lina, Laurence Josset |
| EPI_ISL_3471917 | CENTRE HOSPITALIER ST JOSEPH ST LUC | CNR Virus des Infections Respiratoires - France SUD | Antonin Bal, Gregory Destras, Gwendolyne Burfin, Hadrien Regue, Quentin Semanas, Martine Valette, Bruno Lina, Laurence Josset |
| EPI_ISL_3471918 | LABORATOIRE UNILIANS DECINES | CNR Virus des Infections Respiratoires - France SUD | Antonin Bal, Gregory Destras, Gwendolyne Burfin, Hadrien Regue, Quentin Semanas, Martine Valette, Bruno Lina, Laurence Josset |
| EPI_ISL_3471919 | LABORATOIRE BIOESTEREL | CNR Virus des Infections Respiratoires - France SUD | Antonin Bal, Gregory Destras, Gwendolyne Burfin, Hadrien Regue, Quentin Semanas, Martine Valette, Bruno Lina, Laurence Josset |
| EPI_ISL_3471920 | SYNLAB AQUITAINE | CNR Virus des Infections Respiratoires - France SUD | Antonin Bal, Gregory Destras, Gwendolyne Burfin, Hadrien Regue, Quentin Semanas, Martine Valette, Bruno Lina, Laurence Josset |
| EPI_ISL_3471921, EPI_ISL_3471922, EPI_ISL_3471923, EPI_ISL_3471924, EPI_ISL_3471925 | LABORATOIRE UNILIANS DECINES | CNR Virus des Infections Respiratoires - France SUD | Antonin Bal, Gregory Destras, Gwendolyne Burfin, Hadrien Regue, Quentin Semanas, Martine Valette, Bruno Lina, Laurence Josset |
| EPI_ISL_3471926 | MIRIALIS CLUSES BECHET | CNR Virus des Infections Respiratoires - France SUD | Antonin Bal, Gregory Destras, Gwendolyne Burfin, Hadrien Regue, Quentin Semanas, Martine Valette, Bruno Lina, Laurence Josset |
| EPI_ISL_3471927 | LABORATOIRE CERDIBIO-CHARENTES | CNR Virus des Infections Respiratoires - France SUD | Antonin Bal, Gregory Destras, Gwendolyne Burfin, Hadrien Regue, Quentin Semanas, Martine Valette, Bruno Lina, Laurence Josset |
| EPI_ISL_3471928 | CNR Virus des Infections Respiratoires - France SUD | CNR Virus des Infections Respiratoires - France SUD | Antonin Bal, Gregory Destras, Gwendolyne Burfin, Hadrien Regue, Quentin Semanas, Martine Valette, Bruno Lina, Laurence Josset |
| EPI_ISL_3471929 | SYNLAB AQUITAINE | CNR Virus des Infections Respiratoires - France SUD | Antonin Bal, Gregory Destras, Gwendolyne Burfin, Hadrien Regue, Quentin Semanas, Martine Valette, Bruno Lina, Laurence Josset |
| EPI_ISL_3471930 | LABORATOIRE UNILIANS DECINES | CNR Virus des Infections Respiratoires - France SUD | Antonin Bal, Gregory Destras, Gwendolyne Burfin, Hadrien Regue, Quentin Semanas, Martine Valette, Bruno Lina, Laurence Josset |
| EPI_ISL_3471931 | CENTRE HOSPITALIER PELLEGRIN | CNR Virus des Infections Respiratoires - France SUD | Antonin Bal, Gregory Destras, Gwendolyne Burfin, Hadrien Regue, Quentin Semanas, Martine Valette, Bruno Lina, Laurence Josset |
| EPI_ISL_3471932 | SYNLAB AQUITAINE | CNR Virus des Infections Respiratoires - France SUD | Antonin Bal, Gregory Destras, Gwendolyne Burfin, Hadrien Regue, Quentin Semanas, Martine Valette, Bruno Lina, Laurence Josset |
| EPI_ISL_3471933 | LABORATOIRE BIOESTEREL | CNR Virus des Infections Respiratoires - France SUD | Antonin Bal, Gregory Destras, Gwendolyne Burfin, Hadrien Regue, Quentin Semanas, Martine Valette, Bruno Lina, Laurence Josset |
| EPI_ISL_3471934 | LABORATOIRE UNILIANS DECINES | CNR Virus des Infections Respiratoires - France SUD | Antonin Bal, Gregory Destras, Gwendolyne Burfin, Hadrien Regue, Quentin Semanas, Martine Valette, Bruno Lina, Laurence Josset |
| EPI_ISL_3471935 | CENTRE HOSPITALIER DE VALENCE | CNR Virus des Infections Respiratoires - France SUD | Antonin Bal, Gregory Destras, Gwendolyne Burfin, Hadrien Regue, Quentin Semanas, Martine Valette, Bruno Lina, Laurence Josset |
| EPI_ISL_3471936 | BIOLITTORAL BIOGROUP PLATEAU TECH | CNR Virus des Infections Respiratoires - France SUD | Antonin Bal, Gregory Destras, Gwendolyne Burfin, Hadrien Regue, Quentin Semanas, Martine Valette, Bruno Lina, Laurence Josset |
| EPI_ISL_3471937 | LABORATOIRE UNILIANS DECINES | CNR Virus des Infections Respiratoires - France SUD | Antonin Bal, Gregory Destras, Gwendolyne Burfin, Hadrien Regue, Quentin Semanas, Martine Valette, Bruno Lina, Laurence Josset |
| EPI_ISL_3471938 | LABORATOIRE BIOESTEREL | CNR Virus des Infections Respiratoires - France SUD | Antonin Bal, Gregory Destras, Gwendolyne Burfin, Hadrien Regue, Quentin Semanas, Martine Valette, Bruno Lina, Laurence Josset |
| EPI_ISL_3471939 | LABORATOIRE UNILIANS DECINES | CNR Virus des Infections Respiratoires - France SUD | Antonin Bal, Gregory Destras, Gwendolyne Burfin, Hadrien Regue, Quentin Semanas, Martine Valette, Bruno Lina, Laurence Josset |
| EPI_ISL_3471940 | CENTRE HOSPITALIER SUD GIRONDE | CNR Virus des Infections Respiratoires - France SUD | Antonin Bal, Gregory Destras, Gwendolyne Burfin, Hadrien Regue, Quentin Semanas, Martine Valette, Bruno Lina, Laurence Josset |
| EPI_ISL_3471941 | LABORATOIRE FORTE BIO DAX | CNR Virus des Infections Respiratoires - France SUD | Antonin Bal, Gregory Destras, Gwendolyne Burfin, Hadrien Regue, Quentin Semanas, Martine Valette, Bruno Lina, Laurence Josset |
| EPI_ISL_3471942 | LABORATOIRE BIOESTEREL | CNR Virus des Infections Respiratoires - France SUD | Antonin Bal, Gregory Destras, Gwendolyne Burfin, Hadrien Regue, Quentin Semanas, Martine Valette, Bruno Lina, Laurence Josset |
| EPI_ISL_3471943 | LABORATOIRE UNILIANS DECINES | CNR Virus des Infections Respiratoires - France SUD | Antonin Bal, Gregory Destras, Gwendolyne Burfin, Hadrien Regue, Quentin Semanas, Martine Valette, Bruno Lina, Laurence Josset |
| EPI_ISL_3471944 | LABORATOIRE CBM DE MURET | CNR Virus des Infections Respiratoires - France SUD | Antonin Bal, Gregory Destras, Gwendolyne Burfin, Hadrien Regue, Quentin Semanas, Martine Valette, Bruno Lina, Laurence Josset |
| EPI_ISL_3471945 | CENTRE HOSPITALIER DE BOURG EN BRESSE | CNR Virus des Infections Respiratoires - France SUD | Antonin Bal, Gregory Destras, Gwendolyne Burfin, Hadrien Regue, Quentin Semanas, Martine Valette, Bruno Lina, Laurence Josset |
| EPI_ISL_3471946 | LABORATOIRE UNILIANS DECINES | CNR Virus des Infections Respiratoires - France SUD | Antonin Bal, Gregory Destras, Gwendolyne Burfin, Hadrien Regue, Quentin Semanas, Martine Valette, Bruno Lina, Laurence Josset |
| EPI_ISL_3471947 | CENTRE HOSPITALIER PELLEGRIN | CNR Virus des Infections Respiratoires - France SUD | Antonin Bal, Gregory Destras, Gwendolyne Burfin, Hadrien Regue, Quentin Semanas, Martine Valette, Bruno Lina, Laurence Josset |
| EPI_ISL_3471948 | CNR Virus des Infections Respiratoires - France SUD | CNR Virus des Infections Respiratoires - France SUD | Antonin Bal, Gregory Destras, Gwendolyne Burfin, Hadrien Regue, Quentin Semanas, Martine Valette, Bruno Lina, Laurence Josset |
| EPI_ISL_3471949 | DYOMEDEA-LABORATOIRE DE LA SAUVEGARDE | CNR Virus des Infections Respiratoires - France SUD | Antonin Bal, Gregory Destras, Gwendolyne Burfin, Hadrien Regue, Quentin Semanas, Martine Valette, Bruno Lina, Laurence Josset |
| EPI_ISL_3471950 | LABORATOIRE UNILIANS DECINES | CNR Virus des Infections Respiratoires - France SUD | Antonin Bal, Gregory Destras, Gwendolyne Burfin, Hadrien Regue, Quentin Semanas, Martine Valette, Bruno Lina, Laurence Josset |
| EPI_ISL_3471951 | GROUPE LCD - UNIBIO LA PLAINE | CNR Virus des Infections Respiratoires - France SUD | Antonin Bal, Gregory Destras, Gwendolyne Burfin, Hadrien Regue, Quentin Semanas, Martine Valette, Bruno Lina, Laurence Josset |
| EPI_ISL_3471952 | LABORATOIRE UNILIANS DECINES | CNR Virus des Infections Respiratoires - France SUD | Antonin Bal, Gregory Destras, Gwendolyne Burfin, Hadrien Regue, Quentin Semanas, Martine Valette, Bruno Lina, Laurence Josset |
| EPI_ISL_3471953, EPI_ISL_3471954 | LABORATOIRE BIOESTEREL | CNR Virus des Infections Respiratoires - France SUD | Antonin Bal, Gregory Destras, Gwendolyne Burfin, Hadrien Regue, Quentin Semanas, Martine Valette, Bruno Lina, Laurence Josset |
| EPI_ISL_3471955 | LABORATOIRE UNILIANS DECINES | CNR Virus des Infections Respiratoires - France SUD | Antonin Bal, Gregory Destras, Gwendolyne Burfin, Hadrien Regue, Quentin Semanas, Martine Valette, Bruno Lina, Laurence Josset |
| EPI_ISL_3471956 | LABORATOIRE BIOESTEREL | CNR Virus des Infections Respiratoires - France SUD | Antonin Bal, Gregory Destras, Gwendolyne Burfin, Hadrien Regue, Quentin Semanas, Martine Valette, Bruno Lina, Laurence Josset |
| EPI_ISL_3471957 | LABORATOIRE UNILIANS DECINES | CNR Virus des Infections Respiratoires - France SUD | Antonin Bal, Gregory Destras, Gwendolyne Burfin, Hadrien Regue, Quentin Semanas, Martine Valette, Bruno Lina, Laurence Josset |
| EPI_ISL_3471958 | LABORATOIRE BIOESTEREL | CNR Virus des Infections Respiratoires - France SUD | Antonin Bal, Gregory Destras, Gwendolyne Burfin, Hadrien Regue, Quentin Semanas, Martine Valette, Bruno Lina, Laurence Josset |
| EPI_ISL_3471959 | LABORATOIRE UNILIANS DECINES | CNR Virus des Infections Respiratoires - France SUD | Antonin Bal, Gregory Destras, Gwendolyne Burfin, Hadrien Regue, Quentin Semanas, Martine Valette, Bruno Lina, Laurence Josset |
| EPI_ISL_3471960 | LABORATOIRE CBM DE MURET | CNR Virus des Infections Respiratoires - France SUD | Antonin Bal, Gregory Destras, Gwendolyne Burfin, Hadrien Regue, Quentin Semanas, Martine Valette, Bruno Lina, Laurence Josset |
| EPI_ISL_3471961 | CENTRE HOSPITALIER MARMANDE TONNEINS | CNR Virus des Infections Respiratoires - France SUD | Antonin Bal, Gregory Destras, Gwendolyne Burfin, Hadrien Regue, Quentin Semanas, Martine Valette, Bruno Lina, Laurence Josset |
| EPI_ISL_3471962 | LABORATOIRE BIOESTEREL | CNR Virus des Infections Respiratoires - France SUD | Antonin Bal, Gregory Destras, Gwendolyne Burfin, Hadrien Regue, Quentin Semanas, Martine Valette, Bruno Lina, Laurence Josset |
| EPI_ISL_3471963, EPI_ISL_3471964, EPI_ISL_3471965 | LABORATOIRE UNILIANS DECINES | CNR Virus des Infections Respiratoires - France SUD | Antonin Bal, Gregory Destras, Gwendolyne Burfin, Hadrien Regue, Quentin Semanas, Martine Valette, Bruno Lina, Laurence Josset |
| EPI_ISL_3471966, EPI_ISL_3471967 | LABORATOIRE BIOESTEREL | CNR Virus des Infections Respiratoires - France SUD | Antonin Bal, Gregory Destras, Gwendolyne Burfin, Hadrien Regue, Quentin Semanas, Martine Valette, Bruno Lina, Laurence Josset |
| EPI_ISL_3471968 | ORIAPOLE | CNR Virus des Infections Respiratoires - France SUD | Antonin Bal, Gregory Destras, Gwendolyne Burfin, Hadrien Regue, Quentin Semanas, Martine Valette, Bruno Lina, Laurence Josset |
| EPI_ISL_3471969 | LABORATOIRE UNILIANS DECINES | CNR Virus des Infections Respiratoires - France SUD | Antonin Bal, Gregory Destras, Gwendolyne Burfin, Hadrien Regue, Quentin Semanas, Martine Valette, Bruno Lina, Laurence Josset |
| EPI_ISL_3471970, EPI_ISL_3471971 | LABORATOIRE BIOESTEREL | CNR Virus des Infections Respiratoires - France SUD | Antonin Bal, Gregory Destras, Gwendolyne Burfin, Hadrien Regue, Quentin Semanas, Martine Valette, Bruno Lina, Laurence Josset |
| EPI_ISL_3471972, | CENTRE HOSPITALIER PELLEGRIN | CNR Virus des Infections Respiratoires - France SUD | Antonin Bal, Gregory Destras, Gwendolyne Burfin, Hadrien Regue, Quentin Semanas, Martine Valette, Bruno Lina, Laurence Josset |

| | | | |
|---|---|---|---|
| EPI_ISL_3471973 | | | |
| EPI_ISL_3471974 | DYOMEDEA-LABORATOIRE DE LA SAUVEGARDE | CNR Virus des Infections Respiratoires - France SUD | Antonin Bal, Gregory Destras, Gwendolyne Burfin, Hadrien Regue, Quentin Semanas, Martine Valette, Bruno Lina, Laurence Josset |
| EPI_ISL_3471975, EPI_ISL_3471976, EPI_ISL_3471977 | LABORATOIRE UNILIANS DECINES | CNR Virus des Infections Respiratoires - France SUD | Antonin Bal, Gregory Destras, Gwendolyne Burfin, Hadrien Regue, Quentin Semanas, Martine Valette, Bruno Lina, Laurence Josset |
| EPI_ISL_3471978 | LX BIO | CNR Virus des Infections Respiratoires - France SUD | Antonin Bal, Gregory Destras, Gwendolyne Burfin, Hadrien Regue, Quentin Semanas, Martine Valette, Bruno Lina, Laurence Josset |
| EPI_ISL_3471979 | LABORATOIRE BIOESTEREL | CNR Virus des Infections Respiratoires - France SUD | Antonin Bal, Gregory Destras, Gwendolyne Burfin, Hadrien Regue, Quentin Semanas, Martine Valette, Bruno Lina, Laurence Josset |
| EPI_ISL_3471980 | CH GIVORS | CNR Virus des Infections Respiratoires - France SUD | Antonin Bal, Gregory Destras, Gwendolyne Burfin, Hadrien Regue, Quentin Semanas, Martine Valette, Bruno Lina, Laurence Josset |
| EPI_ISL_3471981 | LABORATOIRE UNILIANS DECINES | CNR Virus des Infections Respiratoires - France SUD | Antonin Bal, Gregory Destras, Gwendolyne Burfin, Hadrien Regue, Quentin Semanas, Martine Valette, Bruno Lina, Laurence Josset |
| EPI_ISL_3471982 | BIOLITTORAL BIOGROUP PLATEAU TECH | CNR Virus des Infections Respiratoires - France SUD | Antonin Bal, Gregory Destras, Gwendolyne Burfin, Hadrien Regue, Quentin Semanas, Martine Valette, Bruno Lina, Laurence Josset |
| EPI_ISL_3471983 | LABORATOIRE UNILIANS DECINES | CNR Virus des Infections Respiratoires - France SUD | Antonin Bal, Gregory Destras, Gwendolyne Burfin, Hadrien Regue, Quentin Semanas, Martine Valette, Bruno Lina, Laurence Josset |
| EPI_ISL_3471984 | BIOLITTORAL BIOGROUP PLATEAU TECH | CNR Virus des Infections Respiratoires - France SUD | Antonin Bal, Gregory Destras, Gwendolyne Burfin, Hadrien Regue, Quentin Semanas, Martine Valette, Bruno Lina, Laurence Josset |
| EPI_ISL_3471985 | LABORATOIRE UNILIANS DECINES | CNR Virus des Infections Respiratoires - France SUD | Antonin Bal, Gregory Destras, Gwendolyne Burfin, Hadrien Regue, Quentin Semanas, Martine Valette, Bruno Lina, Laurence Josset |
| EPI_ISL_3471986 | LABORATOIRE CBM DE MURET | CNR Virus des Infections Respiratoires - France SUD | Antonin Bal, Gregory Destras, Gwendolyne Burfin, Hadrien Regue, Quentin Semanas, Martine Valette, Bruno Lina, Laurence Josset |
| EPI_ISL_3471987, EPI_ISL_3471988 | LABORATOIRE BIOESTEREL | CNR Virus des Infections Respiratoires - France SUD | Antonin Bal, Gregory Destras, Gwendolyne Burfin, Hadrien Regue, Quentin Semanas, Martine Valette, Bruno Lina, Laurence Josset |
| EPI_ISL_3471989 | LABORATOIRE CERDIBIO-CHARENTES | CNR Virus des Infections Respiratoires - France SUD | Antonin Bal, Gregory Destras, Gwendolyne Burfin, Hadrien Regue, Quentin Semanas, Martine Valette, Bruno Lina, Laurence Josset |
| EPI_ISL_3471990 | SYNLAB AQUITAINE | CNR Virus des Infections Respiratoires - France SUD | Antonin Bal, Gregory Destras, Gwendolyne Burfin, Hadrien Regue, Quentin Semanas, Martine Valette, Bruno Lina, Laurence Josset |
| EPI_ISL_3471991 | LABORATOIRE CERDIBIO-CHARENTES | CNR Virus des Infections Respiratoires - France SUD | Antonin Bal, Gregory Destras, Gwendolyne Burfin, Hadrien Regue, Quentin Semanas, Martine Valette, Bruno Lina, Laurence Josset |
| EPI_ISL_3471992, EPI_ISL_3471993 | SYNLAB AQUITAINE | CNR Virus des Infections Respiratoires - France SUD | Antonin Bal, Gregory Destras, Gwendolyne Burfin, Hadrien Regue, Quentin Semanas, Martine Valette, Bruno Lina, Laurence Josset |
| EPI_ISL_3471994 | LABORATOIRE UNILIANS DECINES | CNR Virus des Infections Respiratoires - France SUD | Antonin Bal, Gregory Destras, Gwendolyne Burfin, Hadrien Regue, Quentin Semanas, Martine Valette, Bruno Lina, Laurence Josset |
| EPI_ISL_3471995 | LABORATOIRE BIOESTEREL | CNR Virus des Infections Respiratoires - France SUD | Antonin Bal, Gregory Destras, Gwendolyne Burfin, Hadrien Regue, Quentin Semanas, Martine Valette, Bruno Lina, Laurence Josset |
| EPI_ISL_3471996 | LABORATOIRE UNILIANS DECINES | CNR Virus des Infections Respiratoires - France SUD | Antonin Bal, Gregory Destras, Gwendolyne Burfin, Hadrien Regue, Quentin Semanas, Martine Valette, Bruno Lina, Laurence Josset |
| EPI_ISL_3471997 | LABORATOIRE BIOESTEREL | CNR Virus des Infections Respiratoires - France SUD | Antonin Bal, Gregory Destras, Gwendolyne Burfin, Hadrien Regue, Quentin Semanas, Martine Valette, Bruno Lina, Laurence Josset |
| EPI_ISL_3471998 | LABORATOIRE BIOLYSS | CNR Virus des Infections Respiratoires - France SUD | Antonin Bal, Gregory Destras, Gwendolyne Burfin, Hadrien Regue, Quentin Semanas, Martine Valette, Bruno Lina, Laurence Josset |
| EPI_ISL_3471999 | LABORATOIRE UNILIANS DECINES | CNR Virus des Infections Respiratoires - France SUD | Antonin Bal, Gregory Destras, Gwendolyne Burfin, Hadrien Regue, Quentin Semanas, Martine Valette, Bruno Lina, Laurence Josset |
| EPI_ISL_3472000, EPI_ISL_3472001 | LABORATOIRE BIOESTEREL | CNR Virus des Infections Respiratoires - France SUD | Antonin Bal, Gregory Destras, Gwendolyne Burfin, Hadrien Regue, Quentin Semanas, Martine Valette, Bruno Lina, Laurence Josset |
| EPI_ISL_3472002 | DYOMEDEA-LABORATOIRE DE LA SAUVEGARDE | CNR Virus des Infections Respiratoires - France SUD | Antonin Bal, Gregory Destras, Gwendolyne Burfin, Hadrien Regue, Quentin Semanas, Martine Valette, Bruno Lina, Laurence Josset |
| EPI_ISL_3472003 | BIOLITTORAL BIOGROUP PLATEAU TECH | CNR Virus des Infections Respiratoires - France SUD | Antonin Bal, Gregory Destras, Gwendolyne Burfin, Hadrien Regue, Quentin Semanas, Martine Valette, Bruno Lina, Laurence Josset |
| EPI_ISL_3472004 | LABORATOIRE BIOESTEREL | CNR Virus des Infections Respiratoires - France SUD | Antonin Bal, Gregory Destras, Gwendolyne Burfin, Hadrien Regue, Quentin Semanas, Martine Valette, Bruno Lina, Laurence Josset |
| EPI_ISL_3472005 | ORIAPOLE | CNR Virus des Infections Respiratoires - France SUD | Antonin Bal, Gregory Destras, Gwendolyne Burfin, Hadrien Regue, Quentin Semanas, Martine Valette, Bruno Lina, Laurence Josset |
| EPI_ISL_3472006 | LABORATOIRE BIOESTEREL | CNR Virus des Infections Respiratoires - France SUD | Antonin Bal, Gregory Destras, Gwendolyne Burfin, Hadrien Regue, Quentin Semanas, Martine Valette, Bruno Lina, Laurence Josset |
| EPI_ISL_3472007 | SYNLAB AQUITAINE | CNR Virus des Infections Respiratoires - France SUD | Antonin Bal, Gregory Destras, Gwendolyne Burfin, Hadrien Regue, Quentin Semanas, Martine Valette, Bruno Lina, Laurence Josset |
| EPI_ISL_3472008 | CHU ST ETIENNE HOPITAL NORD | CNR Virus des Infections Respiratoires - France SUD | Antonin Bal, Gregory Destras, Gwendolyne Burfin, Hadrien Regue, Quentin Semanas, Martine Valette, Bruno Lina, Laurence Josset |
| EPI_ISL_3472009 | LABORATOIRE BIOESTEREL | CNR Virus des Infections Respiratoires - France SUD | Antonin Bal, Gregory Destras, Gwendolyne Burfin, Hadrien Regue, Quentin Semanas, Martine Valette, Bruno Lina, Laurence Josset |
| EPI_ISL_3472010 | SYNLAB AQUITAINE | CNR Virus des Infections Respiratoires - France SUD | Antonin Bal, Gregory Destras, Gwendolyne Burfin, Hadrien Regue, Quentin Semanas, Martine Valette, Bruno Lina, Laurence Josset |
| EPI_ISL_3472011 | MIRIALIS CLUSES BECHET | CNR Virus des Infections Respiratoires - France SUD | Antonin Bal, Gregory Destras, Gwendolyne Burfin, Hadrien Regue, Quentin Semanas, Martine Valette, Bruno Lina, Laurence Josset |
| EPI_ISL_3472012 | LAM BIOPOLE21 | CNR Virus des Infections Respiratoires - France SUD | Antonin Bal, Gregory Destras, Gwendolyne Burfin, Hadrien Regue, Quentin Semanas, Martine Valette, Bruno Lina, Laurence Josset |
| EPI_ISL_3472013 | LABORATOIRE BIOESTEREL | CNR Virus des Infections Respiratoires - France SUD | Antonin Bal, Gregory Destras, Gwendolyne Burfin, Hadrien Regue, Quentin Semanas, Martine Valette, Bruno Lina, Laurence Josset |
| EPI_ISL_3472014 | LX BIO | CNR Virus des Infections Respiratoires - France SUD | Antonin Bal, Gregory Destras, Gwendolyne Burfin, Hadrien Regue, Quentin Semanas, Martine Valette, Bruno Lina, Laurence Josset |
| EPI_ISL_3472015 | LABORATOIRE BIOESTEREL | CNR Virus des Infections Respiratoires - France SUD | Antonin Bal, Gregory Destras, Gwendolyne Burfin, Hadrien Regue, Quentin Semanas, Martine Valette, Bruno Lina, Laurence Josset |
| EPI_ISL_3472016 | LABORATOIRE UNILIANS DECINES | CNR Virus des Infections Respiratoires - France SUD | Antonin Bal, Gregory Destras, Gwendolyne Burfin, Hadrien Regue, Quentin Semanas, Martine Valette, Bruno Lina, Laurence Josset |
| EPI_ISL_3472017 | LABORATOIRE BIOESTEREL | CNR Virus des Infections Respiratoires - France SUD | Antonin Bal, Gregory Destras, Gwendolyne Burfin, Hadrien Regue, Quentin Semanas, Martine Valette, Bruno Lina, Laurence Josset |
| EPI_ISL_3472018 | LABORATOIRE CERDIBIO-CHARENTES | CNR Virus des Infections Respiratoires - France SUD | Antonin Bal, Gregory Destras, Gwendolyne Burfin, Hadrien Regue, Quentin Semanas, Martine Valette, Bruno Lina, Laurence Josset |
| EPI_ISL_3472019 | LABORATOIRE BIOESTEREL | CNR Virus des Infections Respiratoires - France SUD | Antonin Bal, Gregory Destras, Gwendolyne Burfin, Hadrien Regue, Quentin Semanas, Martine Valette, Bruno Lina, Laurence Josset |
| EPI_ISL_3472020 | LABORATOIRE UNILIANS DECINES | CNR Virus des Infections Respiratoires - France SUD | Antonin Bal, Gregory Destras, Gwendolyne Burfin, Hadrien Regue, Quentin Semanas, Martine Valette, Bruno Lina, Laurence Josset |
| EPI_ISL_3472021 | LABORATOIRE CERDIBIO-CHARENTES | CNR Virus des Infections Respiratoires - France SUD | Antonin Bal, Gregory Destras, Gwendolyne Burfin, Hadrien Regue, Quentin Semanas, Martine Valette, Bruno Lina, Laurence Josset |
| EPI_ISL_3472022 | LABORATOIRE FORTE BIO DAX | CNR Virus des Infections Respiratoires - France SUD | Antonin Bal, Gregory Destras, Gwendolyne Burfin, Hadrien Regue, Quentin Semanas, Martine Valette, Bruno Lina, Laurence Josset |
| EPI_ISL_3472023 | LABORATOIRE BIOESTEREL | CNR Virus des Infections Respiratoires - France SUD | Antonin Bal, Gregory Destras, Gwendolyne Burfin, Hadrien Regue, Quentin Semanas, Martine Valette, Bruno Lina, Laurence Josset |
| EPI_ISL_3472024 | CENTRE HOSPITALIER PELLEGRIN | CNR Virus des Infections Respiratoires - France SUD | Antonin Bal, Gregory Destras, Gwendolyne Burfin, Hadrien Regue, Quentin Semanas, Martine Valette, Bruno Lina, Laurence Josset |
| EPI_ISL_3472025 | LABORATOIRE CERDIBIO-CHARENTES | CNR Virus des Infections Respiratoires - France SUD | Antonin Bal, Gregory Destras, Gwendolyne Burfin, Hadrien Regue, Quentin Semanas, Martine Valette, Bruno Lina, Laurence Josset |
| EPI_ISL_3472026 | CENTRE HOSPITALIER PELLEGRIN | CNR Virus des Infections Respiratoires - France SUD | Antonin Bal, Gregory Destras, Gwendolyne Burfin, Hadrien Regue, Quentin Semanas, Martine Valette, Bruno Lina, Laurence Josset |
| EPI_ISL_3472027 | LABORATOIRE CERDIBIO-CHARENTES | CNR Virus des Infections Respiratoires - France SUD | Antonin Bal, Gregory Destras, Gwendolyne Burfin, Hadrien Regue, Quentin Semanas, Martine Valette, Bruno Lina, Laurence Josset |
| EPI_ISL_3472028 | LABORATOIRE UNILIANS DECINES | CNR Virus des Infections Respiratoires - France SUD | Antonin Bal, Gregory Destras, Gwendolyne Burfin, Hadrien Regue, Quentin Semanas, Martine Valette, Bruno Lina, Laurence Josset |
| EPI_ISL_3472029 | LABORATOIRE CERDIBIO-CHARENTES | CNR Virus des Infections Respiratoires - France SUD | Antonin Bal, Gregory Destras, Gwendolyne Burfin, Hadrien Regue, Quentin Semanas, Martine Valette, Bruno Lina, Laurence Josset |
| EPI_ISL_3472030 | DYOMEDEA-LABORATOIRE DE LA SAUVEGARDE | CNR Virus des Infections Respiratoires - France SUD | Antonin Bal, Gregory Destras, Gwendolyne Burfin, Hadrien Regue, Quentin Semanas, Martine Valette, Bruno Lina, Laurence Josset |
| EPI_ISL_3472031 | CENTRE HOSPITALIER ST JOSEPH ST LUC | CNR Virus des Infections Respiratoires - France SUD | Antonin Bal, Gregory Destras, Gwendolyne Burfin, Hadrien Regue, Quentin Semanas, Martine Valette, Bruno Lina, Laurence Josset |
| EPI_ISL_3472032 | LABORATOIRE BIOESTEREL | CNR Virus des Infections Respiratoires - France SUD | Antonin Bal, Gregory Destras, Gwendolyne Burfin, Hadrien Regue, Quentin Semanas, Martine Valette, Bruno Lina, Laurence Josset |
| EPI_ISL_3472033 | CENTRE HOSPITALIER PELLEGRIN | CNR Virus des Infections Respiratoires - France SUD | Antonin Bal, Gregory Destras, Gwendolyne Burfin, Hadrien Regue, Quentin Semanas, Martine Valette, Bruno Lina, Laurence Josset |
| EPI_ISL_3472034 | BIOLITTORAL BIOGROUP PLATEAU TECH | CNR Virus des Infections Respiratoires - France SUD | Antonin Bal, Gregory Destras, Gwendolyne Burfin, Hadrien Regue, Quentin Semanas, Martine Valette, Bruno Lina, Laurence Josset |
| EPI_ISL_3472035 | LABORATOIRE UNILIANS DECINES | CNR Virus des Infections Respiratoires - France SUD | Antonin Bal, Gregory Destras, Gwendolyne Burfin, Hadrien Regue, Quentin Semanas, Martine Valette, Bruno Lina, Laurence Josset |
| EPI_ISL_3472036 | LABORATOIRE BIOLYSS | CNR Virus des Infections Respiratoires - France SUD | Antonin Bal, Gregory Destras, Gwendolyne Burfin, Hadrien Regue, Quentin Semanas, Martine Valette, Bruno Lina, Laurence Josset |
| EPI_ISL_3472037 | LABORATOIRE CERDIBIO-CHARENTES | CNR Virus des Infections Respiratoires - France SUD | Antonin Bal, Gregory Destras, Gwendolyne Burfin, Hadrien Regue, Quentin Semanas, Martine Valette, Bruno Lina, Laurence Josset |
| EPI_ISL_3472038 | LABORATOIRE UNILIANS DECINES | CNR Virus des Infections Respiratoires - France SUD | Antonin Bal, Gregory Destras, Gwendolyne Burfin, Hadrien Regue, Quentin Semanas, Martine Valette, Bruno Lina, Laurence Josset |
| EPI_ISL_3472039 | LABORATOIRE BIOESTEREL | CNR Virus des Infections Respiratoires - France SUD | Antonin Bal, Gregory Destras, Gwendolyne Burfin, Hadrien Regue, Quentin Semanas, Martine Valette, Bruno Lina, Laurence Josset |
| EPI_ISL_3472040 | CNR Virus des Infections Respiratoires - France SUD | CNR Virus des Infections Respiratoires - France SUD | Antonin Bal, Gregory Destras, Gwendolyne Burfin, Hadrien Regue, Quentin Semanas, Martine Valette, Bruno Lina, Laurence Josset |
| EPI_ISL_3472041 | GROUPE LCD - UNIBIO LA PLAINE | CNR Virus des Infections Respiratoires - France SUD | Antonin Bal, Gregory Destras, Gwendolyne Burfin, Hadrien Regue, Quentin Semanas, Martine Valette, Bruno Lina, Laurence Josset |
| EPI_ISL_3472042 | LABORATOIRE BIOESTEREL | CNR Virus des Infections Respiratoires - France SUD | Antonin Bal, Gregory Destras, Gwendolyne Burfin, Hadrien Regue, Quentin Semanas, Martine Valette, Bruno Lina, Laurence Josset |
| EPI_ISL_3472043 | LABORATOIRE UNILIANS DECINES | CNR Virus des Infections Respiratoires - France SUD | Antonin Bal, Gregory Destras, Gwendolyne Burfin, Hadrien Regue, Quentin Semanas, Martine Valette, Bruno Lina, Laurence Josset |
| EPI_ISL_3472044, EPI_ISL_3472045 | LABORATOIRE CERDIBIO-CHARENTES | CNR Virus des Infections Respiratoires - France SUD | Antonin Bal, Gregory Destras, Gwendolyne Burfin, Hadrien Regue, Quentin Semanas, Martine Valette, Bruno Lina, Laurence Josset |
| EPI_ISL_3472046 | SYNLAB AQUITAINE | CNR Virus des Infections Respiratoires - France SUD | Antonin Bal, Gregory Destras, Gwendolyne Burfin, Hadrien Regue, Quentin Semanas, Martine Valette, Bruno Lina, Laurence Josset |
| EPI_ISL_3472047, EPI_ISL_3472048 | LABORATOIRE CERDIBIO-CHARENTES | CNR Virus des Infections Respiratoires - France SUD | Antonin Bal, Gregory Destras, Gwendolyne Burfin, Hadrien Regue, Quentin Semanas, Martine Valette, Bruno Lina, Laurence Josset |
| EPI_ISL_3472049 | CNR Virus des Infections Respiratoires - France SUD | CNR Virus des Infections Respiratoires - France SUD | Antonin Bal, Gregory Destras, Gwendolyne Burfin, Hadrien Regue, Quentin Semanas, Martine Valette, Bruno Lina, Laurence Josset |
| EPI_ISL_3472050 | LABORATOIRE UNILIANS DECINES | CNR Virus des Infections Respiratoires - France SUD | Antonin Bal, Gregory Destras, Gwendolyne Burfin, Hadrien Regue, Quentin Semanas, Martine Valette, Bruno Lina, Laurence Josset |

| | | | |
|---|---|---|---|
| EPI_ISL_3472051 | BIOLITTORAL BIOGROUP PLATEAU TECH | CNR Virus des Infections Respiratoires - France SUD | Antonin Bal, Gregory Destras, Gwendolyne Burfin, Hadrien Regue, Quentin Semanas, Martine Valette, Bruno Lina, Laurence Josset |
| EPI_ISL_3472052 | CENTRE HOSPITALIER SUD GIRONDE | CNR Virus des Infections Respiratoires - France SUD | Antonin Bal, Gregory Destras, Gwendolyne Burfin, Hadrien Regue, Quentin Semanas, Martine Valette, Bruno Lina, Laurence Josset |
| EPI_ISL_3472053 | LABORATOIRE CERDIBIO-CHARENTES | CNR Virus des Infections Respiratoires - France SUD | Antonin Bal, Gregory Destras, Gwendolyne Burfin, Hadrien Regue, Quentin Semanas, Martine Valette, Bruno Lina, Laurence Josset |
| EPI_ISL_3472054 | LABORATOIRE UNILIANS DECINES | CNR Virus des Infections Respiratoires - France SUD | Antonin Bal, Gregory Destras, Gwendolyne Burfin, Hadrien Regue, Quentin Semanas, Martine Valette, Bruno Lina, Laurence Josset |
| EPI_ISL_3472055 | CENTRE HOSPITALIER PELLEGRIN | CNR Virus des Infections Respiratoires - France SUD | Antonin Bal, Gregory Destras, Gwendolyne Burfin, Hadrien Regue, Quentin Semanas, Martine Valette, Bruno Lina, Laurence Josset |
| EPI_ISL_3472056 | LABORATOIRE FORTE BIO DAX | CNR Virus des Infections Respiratoires - France SUD | Antonin Bal, Gregory Destras, Gwendolyne Burfin, Hadrien Regue, Quentin Semanas, Martine Valette, Bruno Lina, Laurence Josset |
| EPI_ISL_3472057 | LABORATOIRE UNILIANS DECINES | CNR Virus des Infections Respiratoires - France SUD | Antonin Bal, Gregory Destras, Gwendolyne Burfin, Hadrien Regue, Quentin Semanas, Martine Valette, Bruno Lina, Laurence Josset |
| EPI_ISL_3472058 | CENTRE HOSPITALIER PELLEGRIN | CNR Virus des Infections Respiratoires - France SUD | Antonin Bal, Gregory Destras, Gwendolyne Burfin, Hadrien Regue, Quentin Semanas, Martine Valette, Bruno Lina, Laurence Josset |
| EPI_ISL_3472059 | LABORATOIRE UNILIANS DECINES | CNR Virus des Infections Respiratoires - France SUD | Antonin Bal, Gregory Destras, Gwendolyne Burfin, Hadrien Regue, Quentin Semanas, Martine Valette, Bruno Lina, Laurence Josset |
| EPI_ISL_3472060 | CENTRE HOSPITALIER PELLEGRIN | CNR Virus des Infections Respiratoires - France SUD | Antonin Bal, Gregory Destras, Gwendolyne Burfin, Hadrien Regue, Quentin Semanas, Martine Valette, Bruno Lina, Laurence Josset |
| EPI_ISL_3472061 | CNR Virus des Infections Respiratoires - France SUD | CNR Virus des Infections Respiratoires - France SUD | Antonin Bal, Gregory Destras, Gwendolyne Burfin, Hadrien Regue, Quentin Semanas, Martine Valette, Bruno Lina, Laurence Josset |
| EPI_ISL_3472062 | LABORATOIRE BIOESTEREL | CNR Virus des Infections Respiratoires - France SUD | Antonin Bal, Gregory Destras, Gwendolyne Burfin, Hadrien Regue, Quentin Semanas, Martine Valette, Bruno Lina, Laurence Josset |
| EPI_ISL_3472063 | LABORATOIRE UNILIANS DECINES | CNR Virus des Infections Respiratoires - France SUD | Antonin Bal, Gregory Destras, Gwendolyne Burfin, Hadrien Regue, Quentin Semanas, Martine Valette, Bruno Lina, Laurence Josset |
| EPI_ISL_3472064 | CENTRE HOSPITALIER SUD GIRONDE | CNR Virus des Infections Respiratoires - France SUD | Antonin Bal, Gregory Destras, Gwendolyne Burfin, Hadrien Regue, Quentin Semanas, Martine Valette, Bruno Lina, Laurence Josset |
| EPI_ISL_3472065 | CENTRE HOSPITALIER PELLEGRIN | CNR Virus des Infections Respiratoires - France SUD | Antonin Bal, Gregory Destras, Gwendolyne Burfin, Hadrien Regue, Quentin Semanas, Martine Valette, Bruno Lina, Laurence Josset |
| EPI_ISL_3472066 | LABORATOIRE BIOESTEREL | CNR Virus des Infections Respiratoires - France SUD | Antonin Bal, Gregory Destras, Gwendolyne Burfin, Hadrien Regue, Quentin Semanas, Martine Valette, Bruno Lina, Laurence Josset |
| EPI_ISL_3472067 | MIRIALIS CLUSES BECHET | CNR Virus des Infections Respiratoires - France SUD | Antonin Bal, Gregory Destras, Gwendolyne Burfin, Hadrien Regue, Quentin Semanas, Martine Valette, Bruno Lina, Laurence Josset |
| EPI_ISL_3472068 | LX BIO | CNR Virus des Infections Respiratoires - France SUD | Antonin Bal, Gregory Destras, Gwendolyne Burfin, Hadrien Regue, Quentin Semanas, Martine Valette, Bruno Lina, Laurence Josset |
| EPI_ISL_3472069 | LABORATOIRE UNILIANS DECINES | CNR Virus des Infections Respiratoires - France SUD | Antonin Bal, Gregory Destras, Gwendolyne Burfin, Hadrien Regue, Quentin Semanas, Martine Valette, Bruno Lina, Laurence Josset |
| EPI_ISL_3472070 | CNR Virus des Infections Respiratoires - France SUD | CNR Virus des Infections Respiratoires - France SUD | Antonin Bal, Gregory Destras, Gwendolyne Burfin, Hadrien Regue, Quentin Semanas, Martine Valette, Bruno Lina, Laurence Josset |
| EPI_ISL_3472071 | LABORATOIRE UNILIANS DECINES | CNR Virus des Infections Respiratoires - France SUD | Antonin Bal, Gregory Destras, Gwendolyne Burfin, Hadrien Regue, Quentin Semanas, Martine Valette, Bruno Lina, Laurence Josset |
| EPI_ISL_3472072 | LABORATOIRE BIOESTEREL | CNR Virus des Infections Respiratoires - France SUD | Antonin Bal, Gregory Destras, Gwendolyne Burfin, Hadrien Regue, Quentin Semanas, Martine Valette, Bruno Lina, Laurence Josset |
| EPI_ISL_3472074 | LABORATOIRE UNILIANS DECINES | CNR Virus des Infections Respiratoires - France SUD | Antonin Bal, Gregory Destras, Gwendolyne Burfin, Hadrien Regue, Quentin Semanas, Martine Valette, Bruno Lina, Laurence Josset |
| EPI_ISL_3472075 | LX BIO | CNR Virus des Infections Respiratoires - France SUD | Antonin Bal, Gregory Destras, Gwendolyne Burfin, Hadrien Regue, Quentin Semanas, Martine Valette, Bruno Lina, Laurence Josset |
| EPI_ISL_3472076, EPI_ISL_3472077 | LABORATOIRE BIOESTEREL | CNR Virus des Infections Respiratoires - France SUD | Antonin Bal, Gregory Destras, Gwendolyne Burfin, Hadrien Regue, Quentin Semanas, Martine Valette, Bruno Lina, Laurence Josset |
| EPI_ISL_3472078 | CENTRE HOSPITALIER SUD GIRONDE | CNR Virus des Infections Respiratoires - France SUD | Antonin Bal, Gregory Destras, Gwendolyne Burfin, Hadrien Regue, Quentin Semanas, Martine Valette, Bruno Lina, Laurence Josset |
| EPI_ISL_3472079 | SYNLAB AQUITAINE | CNR Virus des Infections Respiratoires - France SUD | Antonin Bal, Gregory Destras, Gwendolyne Burfin, Hadrien Regue, Quentin Semanas, Martine Valette, Bruno Lina, Laurence Josset |
| EPI_ISL_3472080, EPI_ISL_3472081 | LABORATOIRE UNILIANS DECINES | CNR Virus des Infections Respiratoires - France SUD | Antonin Bal, Gregory Destras, Gwendolyne Burfin, Hadrien Regue, Quentin Semanas, Martine Valette, Bruno Lina, Laurence Josset |
| EPI_ISL_3472082 | LABORATOIRE BIOESTEREL | CNR Virus des Infections Respiratoires - France SUD | Antonin Bal, Gregory Destras, Gwendolyne Burfin, Hadrien Regue, Quentin Semanas, Martine Valette, Bruno Lina, Laurence Josset |
| EPI_ISL_3472083 | LABORATOIRE BIOLYSS | CNR Virus des Infections Respiratoires - France SUD | Antonin Bal, Gregory Destras, Gwendolyne Burfin, Hadrien Regue, Quentin Semanas, Martine Valette, Bruno Lina, Laurence Josset |
| EPI_ISL_3472084 | CENTRE HOSPITALIER DE BOURG EN BRESSE | CNR Virus des Infections Respiratoires - France SUD | Antonin Bal, Gregory Destras, Gwendolyne Burfin, Hadrien Regue, Quentin Semanas, Martine Valette, Bruno Lina, Laurence Josset |
| EPI_ISL_3472085 | BIOLITTORAL BIOGROUP PLATEAU TECH | CNR Virus des Infections Respiratoires - France SUD | Antonin Bal, Gregory Destras, Gwendolyne Burfin, Hadrien Regue, Quentin Semanas, Martine Valette, Bruno Lina, Laurence Josset |
| EPI_ISL_3472086, EPI_ISL_3472087, EPI_ISL_3472088 | LABORATOIRE BIOESTEREL | CNR Virus des Infections Respiratoires - France SUD | Antonin Bal, Gregory Destras, Gwendolyne Burfin, Hadrien Regue, Quentin Semanas, Martine Valette, Bruno Lina, Laurence Josset |
| EPI_ISL_3472089, EPI_ISL_3472090 | LABORATOIRE UNILIANS DECINES | CNR Virus des Infections Respiratoires - France SUD | Antonin Bal, Gregory Destras, Gwendolyne Burfin, Hadrien Regue, Quentin Semanas, Martine Valette, Bruno Lina, Laurence Josset |
| EPI_ISL_3472091 | LABORATOIRE BIOESTEREL | CNR Virus des Infections Respiratoires - France SUD | Antonin Bal, Gregory Destras, Gwendolyne Burfin, Hadrien Regue, Quentin Semanas, Martine Valette, Bruno Lina, Laurence Josset |
| EPI_ISL_3472092 | CENTRE HOSPITALIER DE BIGORRE | CNR Virus des Infections Respiratoires - France SUD | Antonin Bal, Gregory Destras, Gwendolyne Burfin, Hadrien Regue, Quentin Semanas, Martine Valette, Bruno Lina, Laurence Josset |
| EPI_ISL_3472093 | LABORATOIRE UNILIANS DECINES | CNR Virus des Infections Respiratoires - France SUD | Antonin Bal, Gregory Destras, Gwendolyne Burfin, Hadrien Regue, Quentin Semanas, Martine Valette, Bruno Lina, Laurence Josset |
| EPI_ISL_3472094 | LX BIO | CNR Virus des Infections Respiratoires - France SUD | Antonin Bal, Gregory Destras, Gwendolyne Burfin, Hadrien Regue, Quentin Semanas, Martine Valette, Bruno Lina, Laurence Josset |
| EPI_ISL_3472095 | CENTRE HOSPITALIER SUD GIRONDE | CNR Virus des Infections Respiratoires - France SUD | Antonin Bal, Gregory Destras, Gwendolyne Burfin, Hadrien Regue, Quentin Semanas, Martine Valette, Bruno Lina, Laurence Josset |
| EPI_ISL_3472096 | CENTRE HOSPITALIER PELLEGRIN | CNR Virus des Infections Respiratoires - France SUD | Antonin Bal, Gregory Destras, Gwendolyne Burfin, Hadrien Regue, Quentin Semanas, Martine Valette, Bruno Lina, Laurence Josset |
| EPI_ISL_3472097 | LABORATOIRE CERDIBIO-CHARENTES | CNR Virus des Infections Respiratoires - France SUD | Antonin Bal, Gregory Destras, Gwendolyne Burfin, Hadrien Regue, Quentin Semanas, Martine Valette, Bruno Lina, Laurence Josset |
| EPI_ISL_3472098, EPI_ISL_3472099, EPI_ISL_3472100, EPI_ISL_3472101 | LABORATOIRE UNILIANS DECINES | CNR Virus des Infections Respiratoires - France SUD | Antonin Bal, Gregory Destras, Gwendolyne Burfin, Hadrien Regue, Quentin Semanas, Martine Valette, Bruno Lina, Laurence Josset |
| EPI_ISL_3472102 | LX BIO | CNR Virus des Infections Respiratoires - France SUD | Antonin Bal, Gregory Destras, Gwendolyne Burfin, Hadrien Regue, Quentin Semanas, Martine Valette, Bruno Lina, Laurence Josset |
| EPI_ISL_3472103, EPI_ISL_3472104 | LABORATOIRE CERDIBIO-CHARENTES | CNR Virus des Infections Respiratoires - France SUD | Antonin Bal, Gregory Destras, Gwendolyne Burfin, Hadrien Regue, Quentin Semanas, Martine Valette, Bruno Lina, Laurence Josset |
| EPI_ISL_3472105 | LABORATOIRE UNILIANS DECINES | CNR Virus des Infections Respiratoires - France SUD | Antonin Bal, Gregory Destras, Gwendolyne Burfin, Hadrien Regue, Quentin Semanas, Martine Valette, Bruno Lina, Laurence Josset |
| EPI_ISL_3472106 | CENTRE HOSPITALIER SUD GIRONDE | CNR Virus des Infections Respiratoires - France SUD | Antonin Bal, Gregory Destras, Gwendolyne Burfin, Hadrien Regue, Quentin Semanas, Martine Valette, Bruno Lina, Laurence Josset |
| EPI_ISL_3472107 | LABORATOIRE CERDIBIO-CHARENTES | CNR Virus des Infections Respiratoires - France SUD | Antonin Bal, Gregory Destras, Gwendolyne Burfin, Hadrien Regue, Quentin Semanas, Martine Valette, Bruno Lina, Laurence Josset |
| EPI_ISL_3472108 | LABORATOIRE BIOESTEREL | CNR Virus des Infections Respiratoires - France SUD | Antonin Bal, Gregory Destras, Gwendolyne Burfin, Hadrien Regue, Quentin Semanas, Martine Valette, Bruno Lina, Laurence Josset |
| EPI_ISL_3472109 | LABORATOIRE UNILIANS DECINES | CNR Virus des Infections Respiratoires - France SUD | Antonin Bal, Gregory Destras, Gwendolyne Burfin, Hadrien Regue, Quentin Semanas, Martine Valette, Bruno Lina, Laurence Josset |
| EPI_ISL_3472110 | LABORATOIRE BIOESTEREL | CNR Virus des Infections Respiratoires - France SUD | Antonin Bal, Gregory Destras, Gwendolyne Burfin, Hadrien Regue, Quentin Semanas, Martine Valette, Bruno Lina, Laurence Josset |
| EPI_ISL_3472111, EPI_ISL_3472112, EPI_ISL_3472113 | LABORATOIRE UNILIANS DECINES | CNR Virus des Infections Respiratoires - France SUD | Antonin Bal, Gregory Destras, Gwendolyne Burfin, Hadrien Regue, Quentin Semanas, Martine Valette, Bruno Lina, Laurence Josset |
| EPI_ISL_3472114 | GROUPE LCD - UNIBIO LA PLAINE | CNR Virus des Infections Respiratoires - France SUD | Antonin Bal, Gregory Destras, Gwendolyne Burfin, Hadrien Regue, Quentin Semanas, Martine Valette, Bruno Lina, Laurence Josset |
| EPI_ISL_3472115 | CENTRE HOSPITALIER DE BOURG EN BRESSE | CNR Virus des Infections Respiratoires - France SUD | Antonin Bal, Gregory Destras, Gwendolyne Burfin, Hadrien Regue, Quentin Semanas, Martine Valette, Bruno Lina, Laurence Josset |
| EPI_ISL_3472116, EPI_ISL_3472117 | LABORATOIRE BIOESTEREL | CNR Virus des Infections Respiratoires - France SUD | Antonin Bal, Gregory Destras, Gwendolyne Burfin, Hadrien Regue, Quentin Semanas, Martine Valette, Bruno Lina, Laurence Josset |
| EPI_ISL_3472118 | LABORATOIRE CERDIBIO-CHARENTES | CNR Virus des Infections Respiratoires - France SUD | Antonin Bal, Gregory Destras, Gwendolyne Burfin, Hadrien Regue, Quentin Semanas, Martine Valette, Bruno Lina, Laurence Josset |
| EPI_ISL_3472119 | CENTRE HOSPITALIER PIERRE OUDOT | CNR Virus des Infections Respiratoires - France SUD | Antonin Bal, Gregory Destras, Gwendolyne Burfin, Hadrien Regue, Quentin Semanas, Martine Valette, Bruno Lina, Laurence Josset |
| EPI_ISL_3472120 | ORIAPOLE | CNR Virus des Infections Respiratoires - France SUD | Antonin Bal, Gregory Destras, Gwendolyne Burfin, Hadrien Regue, Quentin Semanas, Martine Valette, Bruno Lina, Laurence Josset |
| EPI_ISL_3472121 | LABORATOIRE CERDIBIO-CHARENTES | CNR Virus des Infections Respiratoires - France SUD | Antonin Bal, Gregory Destras, Gwendolyne Burfin, Hadrien Regue, Quentin Semanas, Martine Valette, Bruno Lina, Laurence Josset |
| EPI_ISL_3472122, EPI_ISL_3472123 | LABORATOIRE UNILIANS DECINES | CNR Virus des Infections Respiratoires - France SUD | Antonin Bal, Gregory Destras, Gwendolyne Burfin, Hadrien Regue, Quentin Semanas, Martine Valette, Bruno Lina, Laurence Josset |
| EPI_ISL_3472124 | LX BIO | CNR Virus des Infections Respiratoires - France SUD | Antonin Bal, Gregory Destras, Gwendolyne Burfin, Hadrien Regue, Quentin Semanas, Martine Valette, Bruno Lina, Laurence Josset |
| EPI_ISL_3472125 | CNR Virus des Infections Respiratoires - France SUD | CNR Virus des Infections Respiratoires - France SUD | Antonin Bal, Gregory Destras, Gwendolyne Burfin, Hadrien Regue, Quentin Semanas, Martine Valette, Bruno Lina, Laurence Josset |
| EPI_ISL_3472126 | CENTRE HOSPITALIER PIERRE OUDOT | CNR Virus des Infections Respiratoires - France SUD | Antonin Bal, Gregory Destras, Gwendolyne Burfin, Hadrien Regue, Quentin Semanas, Martine Valette, Bruno Lina, Laurence Josset |
| EPI_ISL_3472127 | CENTRE HOSPITALIER PELLEGRIN | CNR Virus des Infections Respiratoires - France SUD | Antonin Bal, Gregory Destras, Gwendolyne Burfin, Hadrien Regue, Quentin Semanas, Martine Valette, Bruno Lina, Laurence Josset |
| EPI_ISL_3472128 | LABORATOIRE BIOESTEREL | CNR Virus des Infections Respiratoires - France SUD | Antonin Bal, Gregory Destras, Gwendolyne Burfin, Hadrien Regue, Quentin Semanas, Martine Valette, Bruno Lina, Laurence Josset |
| EPI_ISL_3472129 | LABORATOIRE UNILIANS DECINES | CNR Virus des Infections Respiratoires - France SUD | Antonin Bal, Gregory Destras, Gwendolyne Burfin, Hadrien Regue, Quentin Semanas, Martine Valette, Bruno Lina, Laurence Josset |
| EPI_ISL_3472130 | CENTRE HOSPITALIER ST JOSEPH ST LUC | CNR Virus des Infections Respiratoires - France SUD | Antonin Bal, Gregory Destras, Gwendolyne Burfin, Hadrien Regue, Quentin Semanas, Martine Valette, Bruno Lina, Laurence Josset |
| EPI_ISL_3472131 | SYNLAB AQUITAINE | CNR Virus des Infections Respiratoires - France SUD | Antonin Bal, Gregory Destras, Gwendolyne Burfin, Hadrien Regue, Quentin Semanas, Martine Valette, Bruno Lina, Laurence Josset |
| EPI_ISL_3472132 | LABORATOIRE UNILIANS DECINES | CNR Virus des Infections Respiratoires - France SUD | Antonin Bal, Gregory Destras, Gwendolyne Burfin, Hadrien Regue, Quentin Semanas, Martine Valette, Bruno Lina, Laurence Josset |

| | | | |
|---|---|---|---|
| EPI_ISL_3472133 | LABORATOIRE CBM DE MURET | CNR Virus des Infections Respiratoires - France SUD | Antonin Bal, Gregory Destras, Gwendolyne Burfin, Hadrien Regue, Quentin Semanas, Martine Valette, Bruno Lina, Laurence Josset |
| EPI_ISL_3472134 | LABORATOIRE BIOESTEREL | CNR Virus des Infections Respiratoires - France SUD | Antonin Bal, Gregory Destras, Gwendolyne Burfin, Hadrien Regue, Quentin Semanas, Martine Valette, Bruno Lina, Laurence Josset |
| EPI_ISL_3472135 | SYNLAB AQUITAINE | CNR Virus des Infections Respiratoires - France SUD | Antonin Bal, Gregory Destras, Gwendolyne Burfin, Hadrien Regue, Quentin Semanas, Martine Valette, Bruno Lina, Laurence Josset |
| EPI_ISL_3472136 | LABORATOIRE UNILIANS DECINES | CNR Virus des Infections Respiratoires - France SUD | Antonin Bal, Gregory Destras, Gwendolyne Burfin, Hadrien Regue, Quentin Semanas, Martine Valette, Bruno Lina, Laurence Josset |
| EPI_ISL_3472137 | HOPITAL ALBI | CNR Virus des Infections Respiratoires - France SUD | Antonin Bal, Gregory Destras, Gwendolyne Burfin, Hadrien Regue, Quentin Semanas, Martine Valette, Bruno Lina, Laurence Josset |
| EPI_ISL_3472138, EPI_ISL_3472139, EPI_ISL_3472140 | LABORATOIRE UNILIANS DECINES | CNR Virus des Infections Respiratoires - France SUD | Antonin Bal, Gregory Destras, Gwendolyne Burfin, Hadrien Regue, Quentin Semanas, Martine Valette, Bruno Lina, Laurence Josset |
| EPI_ISL_3472141 | LABORATOIRE CERDIBIO-CHARENTES | CNR Virus des Infections Respiratoires - France SUD | Antonin Bal, Gregory Destras, Gwendolyne Burfin, Hadrien Regue, Quentin Semanas, Martine Valette, Bruno Lina, Laurence Josset |
| EPI_ISL_3472142 | LABORATOIRE UNILIANS DECINES | CNR Virus des Infections Respiratoires - France SUD | Antonin Bal, Gregory Destras, Gwendolyne Burfin, Hadrien Regue, Quentin Semanas, Martine Valette, Bruno Lina, Laurence Josset |
| EPI_ISL_3472143 | LABORATOIRE CERDIBIO-CHARENTES | CNR Virus des Infections Respiratoires - France SUD | Antonin Bal, Gregory Destras, Gwendolyne Burfin, Hadrien Regue, Quentin Semanas, Martine Valette, Bruno Lina, Laurence Josset |
| EPI_ISL_3472144 | CNR Virus des Infections Respiratoires - France SUD | CNR Virus des Infections Respiratoires - France SUD | Antonin Bal, Gregory Destras, Gwendolyne Burfin, Hadrien Regue, Quentin Semanas, Martine Valette, Bruno Lina, Laurence Josset |
| EPI_ISL_3472145 | LABORATOIRE BIOESTEREL | CNR Virus des Infections Respiratoires - France SUD | Antonin Bal, Gregory Destras, Gwendolyne Burfin, Hadrien Regue, Quentin Semanas, Martine Valette, Bruno Lina, Laurence Josset |
| EPI_ISL_3472146 | LABORATOIRE FORTE BIO DAX | CNR Virus des Infections Respiratoires - France SUD | Antonin Bal, Gregory Destras, Gwendolyne Burfin, Hadrien Regue, Quentin Semanas, Martine Valette, Bruno Lina, Laurence Josset |
| EPI_ISL_3472147 | LABORATOIRE BIOESTEREL | CNR Virus des Infections Respiratoires - France SUD | Antonin Bal, Gregory Destras, Gwendolyne Burfin, Hadrien Regue, Quentin Semanas, Martine Valette, Bruno Lina, Laurence Josset |
| EPI_ISL_3472148 | MIRIALIS CLUSES BECHET | CNR Virus des Infections Respiratoires - France SUD | Antonin Bal, Gregory Destras, Gwendolyne Burfin, Hadrien Regue, Quentin Semanas, Martine Valette, Bruno Lina, Laurence Josset |
| EPI_ISL_3472149 | DYOMEDEA-LABORATOIRE DE LA SAUVEGARDE | CNR Virus des Infections Respiratoires - France SUD | Antonin Bal, Gregory Destras, Gwendolyne Burfin, Hadrien Regue, Quentin Semanas, Martine Valette, Bruno Lina, Laurence Josset |
| EPI_ISL_3472150 | LX BIO | CNR Virus des Infections Respiratoires - France SUD | Antonin Bal, Gregory Destras, Gwendolyne Burfin, Hadrien Regue, Quentin Semanas, Martine Valette, Bruno Lina, Laurence Josset |
| EPI_ISL_3472151 | CENTRE HOSPITALIER PIERRE OUDOT | CNR Virus des Infections Respiratoires - France SUD | Antonin Bal, Gregory Destras, Gwendolyne Burfin, Hadrien Regue, Quentin Semanas, Martine Valette, Bruno Lina, Laurence Josset |
| EPI_ISL_3472152 | LABORATOIRE UNILIANS DECINES | CNR Virus des Infections Respiratoires - France SUD | Antonin Bal, Gregory Destras, Gwendolyne Burfin, Hadrien Regue, Quentin Semanas, Martine Valette, Bruno Lina, Laurence Josset |
| EPI_ISL_3472153 | CENTRE HOSPITALIER DE LA COTE BASQUE | CNR Virus des Infections Respiratoires - France SUD | Antonin Bal, Gregory Destras, Gwendolyne Burfin, Hadrien Regue, Quentin Semanas, Martine Valette, Bruno Lina, Laurence Josset |
| EPI_ISL_3472154 | LABORATOIRE BIOESTEREL | CNR Virus des Infections Respiratoires - France SUD | Antonin Bal, Gregory Destras, Gwendolyne Burfin, Hadrien Regue, Quentin Semanas, Martine Valette, Bruno Lina, Laurence Josset |
| EPI_ISL_3472155 | LABORATOIRE UNILIANS DECINES | CNR Virus des Infections Respiratoires - France SUD | Antonin Bal, Gregory Destras, Gwendolyne Burfin, Hadrien Regue, Quentin Semanas, Martine Valette, Bruno Lina, Laurence Josset |
| EPI_ISL_3472156 | LABORATOIRE BIOESTEREL | CNR Virus des Infections Respiratoires - France SUD | Antonin Bal, Gregory Destras, Gwendolyne Burfin, Hadrien Regue, Quentin Semanas, Martine Valette, Bruno Lina, Laurence Josset |
| EPI_ISL_3472157 | SYNLAB AQUITAINE | CNR Virus des Infections Respiratoires - France SUD | Antonin Bal, Gregory Destras, Gwendolyne Burfin, Hadrien Regue, Quentin Semanas, Martine Valette, Bruno Lina, Laurence Josset |
| EPI_ISL_3472158 | CNR Virus des Infections Respiratoires - France SUD | CNR Virus des Infections Respiratoires - France SUD | Antonin Bal, Gregory Destras, Gwendolyne Burfin, Hadrien Regue, Quentin Semanas, Martine Valette, Bruno Lina, Laurence Josset |
| EPI_ISL_3472159 | LABORATOIRE UNILIANS DECINES | CNR Virus des Infections Respiratoires - France SUD | Antonin Bal, Gregory Destras, Gwendolyne Burfin, Hadrien Regue, Quentin Semanas, Martine Valette, Bruno Lina, Laurence Josset |
| EPI_ISL_3472160, EPI_ISL_3472161 | LABORATOIRE BIOESTEREL | CNR Virus des Infections Respiratoires - France SUD | Antonin Bal, Gregory Destras, Gwendolyne Burfin, Hadrien Regue, Quentin Semanas, Martine Valette, Bruno Lina, Laurence Josset |
| EPI_ISL_3472162, EPI_ISL_3472163 | LABORATOIRE UNILIANS DECINES | CNR Virus des Infections Respiratoires - France SUD | Antonin Bal, Gregory Destras, Gwendolyne Burfin, Hadrien Regue, Quentin Semanas, Martine Valette, Bruno Lina, Laurence Josset |
| EPI_ISL_3472164, EPI_ISL_3472165, EPI_ISL_3472166 | LABORATOIRE BIOESTEREL | CNR Virus des Infections Respiratoires - France SUD | Antonin Bal, Gregory Destras, Gwendolyne Burfin, Hadrien Regue, Quentin Semanas, Martine Valette, Bruno Lina, Laurence Josset |
| EPI_ISL_3472167 | ORIAPOLE | CNR Virus des Infections Respiratoires - France SUD | Antonin Bal, Gregory Destras, Gwendolyne Burfin, Hadrien Regue, Quentin Semanas, Martine Valette, Bruno Lina, Laurence Josset |
| EPI_ISL_3472168 | LABORATOIRE BIOESTEREL | CNR Virus des Infections Respiratoires - France SUD | Antonin Bal, Gregory Destras, Gwendolyne Burfin, Hadrien Regue, Quentin Semanas, Martine Valette, Bruno Lina, Laurence Josset |
| EPI_ISL_3472169 | LABORATOIRE CERDIBIO-CHARENTES | CNR Virus des Infections Respiratoires - France SUD | Antonin Bal, Gregory Destras, Gwendolyne Burfin, Hadrien Regue, Quentin Semanas, Martine Valette, Bruno Lina, Laurence Josset |
| EPI_ISL_3472170, EPI_ISL_3472171 | SYNLAB AQUITAINE | CNR Virus des Infections Respiratoires - France SUD | Antonin Bal, Gregory Destras, Gwendolyne Burfin, Hadrien Regue, Quentin Semanas, Martine Valette, Bruno Lina, Laurence Josset |
| EPI_ISL_3472172 | LABORATOIRE UNILIANS DECINES | CNR Virus des Infections Respiratoires - France SUD | Antonin Bal, Gregory Destras, Gwendolyne Burfin, Hadrien Regue, Quentin Semanas, Martine Valette, Bruno Lina, Laurence Josset |
| EPI_ISL_3472173, EPI_ISL_3472174, EPI_ISL_3472175 | LABORATOIRE BIOESTEREL | CNR Virus des Infections Respiratoires - France SUD | Antonin Bal, Gregory Destras, Gwendolyne Burfin, Hadrien Regue, Quentin Semanas, Martine Valette, Bruno Lina, Laurence Josset |
| EPI_ISL_3472176 | DYOMEDEA-LABORATOIRE DE LA SAUVEGARDE | CNR Virus des Infections Respiratoires - France SUD | Antonin Bal, Gregory Destras, Gwendolyne Burfin, Hadrien Regue, Quentin Semanas, Martine Valette, Bruno Lina, Laurence Josset |
| EPI_ISL_3472177 | LX BIO | CNR Virus des Infections Respiratoires - France SUD | Antonin Bal, Gregory Destras, Gwendolyne Burfin, Hadrien Regue, Quentin Semanas, Martine Valette, Bruno Lina, Laurence Josset |
| EPI_ISL_3472178 | LABORATOIRE UNILIANS DECINES | CNR Virus des Infections Respiratoires - France SUD | Antonin Bal, Gregory Destras, Gwendolyne Burfin, Hadrien Regue, Quentin Semanas, Martine Valette, Bruno Lina, Laurence Josset |
| EPI_ISL_3472179 | LABORATOIRE FORTE BIO DAX | CNR Virus des Infections Respiratoires - France SUD | Antonin Bal, Gregory Destras, Gwendolyne Burfin, Hadrien Regue, Quentin Semanas, Martine Valette, Bruno Lina, Laurence Josset |
| EPI_ISL_3472180 | LX BIO | CNR Virus des Infections Respiratoires - France SUD | Antonin Bal, Gregory Destras, Gwendolyne Burfin, Hadrien Regue, Quentin Semanas, Martine Valette, Bruno Lina, Laurence Josset |
| EPI_ISL_3472181, EPI_ISL_3472182 | LABORATOIRE UNILIANS DECINES | CNR Virus des Infections Respiratoires - France SUD | Antonin Bal, Gregory Destras, Gwendolyne Burfin, Hadrien Regue, Quentin Semanas, Martine Valette, Bruno Lina, Laurence Josset |
| EPI_ISL_3472183 | LABORATOIRE CERDIBIO-CHARENTES | CNR Virus des Infections Respiratoires - France SUD | Antonin Bal, Gregory Destras, Gwendolyne Burfin, Hadrien Regue, Quentin Semanas, Martine Valette, Bruno Lina, Laurence Josset |
| EPI_ISL_3472192, EPI_ISL_3472193 | CENTRE HOSPITALIER LUCIEN HUSSEL | CNR Virus des Infections Respiratoires - France SUD | Antonin Bal, Gregory Destras, Gwendolyne Burfin, Hadrien Regue, Quentin Semanas, Martine Valette, Bruno Lina, Laurence Josset |
| EPI_ISL_3472202 | CENTRE HOSPITALIER DE LA COTE BASQUE | CNR Virus des Infections Respiratoires - France SUD | Antonin Bal, Gregory Destras, Gwendolyne Burfin, Hadrien Regue, Quentin Semanas, Martine Valette, Bruno Lina, Laurence Josset |
| EPI_ISL_3472203 | LABORATOIRE FORTE BIO DAX | CNR Virus des Infections Respiratoires - France SUD | Antonin Bal, Gregory Destras, Gwendolyne Burfin, Hadrien Regue, Quentin Semanas, Martine Valette, Bruno Lina, Laurence Josset |
| EPI_ISL_3472204 | CENTRE HOSPITALIER LUCIEN HUSSEL | CNR Virus des Infections Respiratoires - France SUD | Antonin Bal, Gregory Destras, Gwendolyne Burfin, Hadrien Regue, Quentin Semanas, Martine Valette, Bruno Lina, Laurence Josset |
| EPI_ISL_3472205 | MIRIALIS CLUSES BECHET | CNR Virus des Infections Respiratoires - France SUD | Antonin Bal, Gregory Destras, Gwendolyne Burfin, Hadrien Regue, Quentin Semanas, Martine Valette, Bruno Lina, Laurence Josset |
| EPI_ISL_3472206 | LABORATOIRE BIOESTEREL | CNR Virus des Infections Respiratoires - France SUD | Antonin Bal, Gregory Destras, Gwendolyne Burfin, Hadrien Regue, Quentin Semanas, Martine Valette, Bruno Lina, Laurence Josset |
| EPI_ISL_3472207 | CENTRE HOSPITALIER ST JOSEPH ST LUC | CNR Virus des Infections Respiratoires - France SUD | Antonin Bal, Gregory Destras, Gwendolyne Burfin, Hadrien Regue, Quentin Semanas, Martine Valette, Bruno Lina, Laurence Josset |
| EPI_ISL_3472208 | CENTRE HOSPITALIER PIERRE OUDOT | CNR Virus des Infections Respiratoires - France SUD | Antonin Bal, Gregory Destras, Gwendolyne Burfin, Hadrien Regue, Quentin Semanas, Martine Valette, Bruno Lina, Laurence Josset |
| EPI_ISL_3472209 | CENTRE HOSPITALIER SUD GIRONDE | CNR Virus des Infections Respiratoires - France SUD | Antonin Bal, Gregory Destras, Gwendolyne Burfin, Hadrien Regue, Quentin Semanas, Martine Valette, Bruno Lina, Laurence Josset |
| EPI_ISL_3472210 | CENTRE HOSPITALIER ST JOSEPH ST LUC | CNR Virus des Infections Respiratoires - France SUD | Antonin Bal, Gregory Destras, Gwendolyne Burfin, Hadrien Regue, Quentin Semanas, Martine Valette, Bruno Lina, Laurence Josset |
| EPI_ISL_3472211 | CENTRE HOSPITALIER DE VALENCE | CNR Virus des Infections Respiratoires - France SUD | Antonin Bal, Gregory Destras, Gwendolyne Burfin, Hadrien Regue, Quentin Semanas, Martine Valette, Bruno Lina, Laurence Josset |
| EPI_ISL_3472212 | BIOLITTORAL BIOGROUP PLATEAU TECH | CNR Virus des Infections Respiratoires - France SUD | Antonin Bal, Gregory Destras, Gwendolyne Burfin, Hadrien Regue, Quentin Semanas, Martine Valette, Bruno Lina, Laurence Josset |
| EPI_ISL_3472213 | CENTRE HOSPITALIER DE BOURG EN BRESSE | CNR Virus des Infections Respiratoires - France SUD | Antonin Bal, Gregory Destras, Gwendolyne Burfin, Hadrien Regue, Quentin Semanas, Martine Valette, Bruno Lina, Laurence Josset |
| EPI_ISL_3472214 | CENTRE HOSPITALIER LUCIEN HUSSEL | CNR Virus des Infections Respiratoires - France SUD | Antonin Bal, Gregory Destras, Gwendolyne Burfin, Hadrien Regue, Quentin Semanas, Martine Valette, Bruno Lina, Laurence Josset |
| EPI_ISL_3472215, EPI_ISL_3472216 | LABORATOIRE UNILIANS DECINES | CNR Virus des Infections Respiratoires - France SUD | Antonin Bal, Gregory Destras, Gwendolyne Burfin, Hadrien Regue, Quentin Semanas, Martine Valette, Bruno Lina, Laurence Josset |
| EPI_ISL_3472217 | HOPITAL BERGERAC | CNR Virus des Infections Respiratoires - France SUD | Antonin Bal, Gregory Destras, Gwendolyne Burfin, Hadrien Regue, Quentin Semanas, Martine Valette, Bruno Lina, Laurence Josset |
| EPI_ISL_3472218 | CENTRE HOSPITALIER ST JOSEPH ST LUC | CNR Virus des Infections Respiratoires - France SUD | Antonin Bal, Gregory Destras, Gwendolyne Burfin, Hadrien Regue, Quentin Semanas, Martine Valette, Bruno Lina, Laurence Josset |
| EPI_ISL_3472220, EPI_ISL_3472221 | LX BIO | CNR Virus des Infections Respiratoires - France SUD | Antonin Bal, Gregory Destras, Gwendolyne Burfin, Hadrien Regue, Quentin Semanas, Martine Valette, Bruno Lina, Laurence Josset |
| EPI_ISL_3472222, EPI_ISL_3472223 | LABORATOIRE CBM DE MURET | CNR Virus des Infections Respiratoires - France SUD | Antonin Bal, Gregory Destras, Gwendolyne Burfin, Hadrien Regue, Quentin Semanas, Martine Valette, Bruno Lina, Laurence Josset |
| EPI_ISL_3472224, EPI_ISL_3472225 | CNR Virus des Infections Respiratoires - France SUD | CNR Virus des Infections Respiratoires - France SUD | Antonin Bal, Gregory Destras, Gwendolyne Burfin, Hadrien Regue, Quentin Semanas, Martine Valette, Bruno Lina, Laurence Josset |
| EPI_ISL_3472226, EPI_ISL_3472230, EPI_ISL_3472231 | LABORATOIRE UNILIANS DECINES | CNR Virus des Infections Respiratoires - France SUD | Antonin Bal, Gregory Destras, Gwendolyne Burfin, Hadrien Regue, Quentin Semanas, Martine Valette, Bruno Lina, Laurence Josset |
| EPI_ISL_3472233 | LABORATOIRE ACCOLAB SUD OUEST | CNR Virus des Infections Respiratoires - France SUD | Antonin Bal, Gregory Destras, Gwendolyne Burfin, Hadrien Regue, Quentin Semanas, Martine Valette, Bruno Lina, Laurence Josset |
| EPI_ISL_3472234 | LX BIO | CNR Virus des Infections Respiratoires - France SUD | Antonin Bal, Gregory Destras, Gwendolyne Burfin, Hadrien Regue, Quentin Semanas, Martine Valette, Bruno Lina, Laurence Josset |

| | | | |
|---|---|---|---|
| EPI_ISL_3472235 | LABORATOIRE BIOESTEREL | CNR Virus des Infections Respiratoires - France SUD | Antonin Bal, Gregory Destras, Gwendolyne Burfin, Hadrien Regue, Quentin Semanas, Martine Valette, Bruno Lina, Laurence Josset |
| EPI_ISL_3472236 | CENTRE HOSPITALIER BRIVE | CNR Virus des Infections Respiratoires - France SUD | Antonin Bal, Gregory Destras, Gwendolyne Burfin, Hadrien Regue, Quentin Semanas, Martine Valette, Bruno Lina, Laurence Josset |
| EPI_ISL_3472237 | SYNLAB AQUITAINE | CNR Virus des Infections Respiratoires - France SUD | Antonin Bal, Gregory Destras, Gwendolyne Burfin, Hadrien Regue, Quentin Semanas, Martine Valette, Bruno Lina, Laurence Josset |
| EPI_ISL_3472238 | LABORATOIRE CBM DE MURET | CNR Virus des Infections Respiratoires - France SUD | Antonin Bal, Gregory Destras, Gwendolyne Burfin, Hadrien Regue, Quentin Semanas, Martine Valette, Bruno Lina, Laurence Josset |
| EPI_ISL_3472239 | LABORATOIRE BIOESTEREL | CNR Virus des Infections Respiratoires - France SUD | Antonin Bal, Gregory Destras, Gwendolyne Burfin, Hadrien Regue, Quentin Semanas, Martine Valette, Bruno Lina, Laurence Josset |
| EPI_ISL_3472240 | SYNLAB AQUITAINE | CNR Virus des Infections Respiratoires - France SUD | Antonin Bal, Gregory Destras, Gwendolyne Burfin, Hadrien Regue, Quentin Semanas, Martine Valette, Bruno Lina, Laurence Josset |
| EPI_ISL_3472241, EPI_ISL_3472242 | CNR Virus des Infections Respiratoires - France SUD | CNR Virus des Infections Respiratoires - France SUD | Antonin Bal, Gregory Destras, Gwendolyne Burfin, Hadrien Regue, Quentin Semanas, Martine Valette, Bruno Lina, Laurence Josset |
| EPI_ISL_3472243, EPI_ISL_3472244, EPI_ISL_3472245, EPI_ISL_3472246, EPI_ISL_3472247, EPI_ISL_3472248, EPI_ISL_3472249 | LABORATOIRE UNILIANS DECINES | CNR Virus des Infections Respiratoires - France SUD | Antonin Bal, Gregory Destras, Gwendolyne Burfin, Hadrien Regue, Quentin Semanas, Martine Valette, Bruno Lina, Laurence Josset |
| EPI_ISL_3472250, EPI_ISL_3472251, EPI_ISL_3472252 | DYOMEDEA-LABORATOIRE DE LA SAUVEGARDE | CNR Virus des Infections Respiratoires - France SUD | Antonin Bal, Gregory Destras, Gwendolyne Burfin, Hadrien Regue, Quentin Semanas, Martine Valette, Bruno Lina, Laurence Josset |
| EPI_ISL_3472253, EPI_ISL_3472254, EPI_ISL_3472255, EPI_ISL_3472256, EPI_ISL_3472257, EPI_ISL_3472258, EPI_ISL_3472259, EPI_ISL_3472260, EPI_ISL_3472261 | LABORATOIRE UNILIANS DECINES | CNR Virus des Infections Respiratoires - France SUD | Antonin Bal, Gregory Destras, Gwendolyne Burfin, Hadrien Regue, Quentin Semanas, Martine Valette, Bruno Lina, Laurence Josset |
| EPI_ISL_3472262, EPI_ISL_3472263, EPI_ISL_3472264, EPI_ISL_3472265 | LABORATOIRE ACCOLAB SUD OUEST | CNR Virus des Infections Respiratoires - France SUD | Antonin Bal, Gregory Destras, Gwendolyne Burfin, Hadrien Regue, Quentin Semanas, Martine Valette, Bruno Lina, Laurence Josset |
| EPI_ISL_3472266 | LABORATOIRE UNILIANS DECINES | CNR Virus des Infections Respiratoires - France SUD | Antonin Bal, Gregory Destras, Gwendolyne Burfin, Hadrien Regue, Quentin Semanas, Martine Valette, Bruno Lina, Laurence Josset |
| EPI_ISL_3472267, EPI_ISL_3472268, EPI_ISL_3472269, EPI_ISL_3472270 | LABORATOIRE ACCOLAB SUD OUEST | CNR Virus des Infections Respiratoires - France SUD | Antonin Bal, Gregory Destras, Gwendolyne Burfin, Hadrien Regue, Quentin Semanas, Martine Valette, Bruno Lina, Laurence Josset |
| EPI_ISL_3472271 | CNR Virus des Infections Respiratoires - France SUD | CNR Virus des Infections Respiratoires - France SUD | Antonin Bal, Gregory Destras, Gwendolyne Burfin, Hadrien Regue, Quentin Semanas, Martine Valette, Bruno Lina, Laurence Josset |
| EPI_ISL_3472272 | LABORATOIRE ACCOLAB SUD OUEST | CNR Virus des Infections Respiratoires - France SUD | Antonin Bal, Gregory Destras, Gwendolyne Burfin, Hadrien Regue, Quentin Semanas, Martine Valette, Bruno Lina, Laurence Josset |
| EPI_ISL_3472273, EPI_ISL_3472274, EPI_ISL_3472275, EPI_ISL_3472276 | HOPITAL ALBI | CNR Virus des Infections Respiratoires - France SUD | Antonin Bal, Gregory Destras, Gwendolyne Burfin, Hadrien Regue, Quentin Semanas, Martine Valette, Bruno Lina, Laurence Josset |
| EPI_ISL_3472277, EPI_ISL_3472278, EPI_ISL_3472279, EPI_ISL_3472280, EPI_ISL_3472281, EPI_ISL_3472282 | LX BIO | CNR Virus des Infections Respiratoires - France SUD | Antonin Bal, Gregory Destras, Gwendolyne Burfin, Hadrien Regue, Quentin Semanas, Martine Valette, Bruno Lina, Laurence Josset |
| EPI_ISL_3472283 | LABORATOIRE BIOESTEREL | CNR Virus des Infections Respiratoires - France SUD | Antonin Bal, Gregory Destras, Gwendolyne Burfin, Hadrien Regue, Quentin Semanas, Martine Valette, Bruno Lina, Laurence Josset |
| EPI_ISL_3472284 | LX BIO | CNR Virus des Infections Respiratoires - France SUD | Antonin Bal, Gregory Destras, Gwendolyne Burfin, Hadrien Regue, Quentin Semanas, Martine Valette, Bruno Lina, Laurence Josset |
| EPI_ISL_3472285 | LABORATOIRE BIOESTEREL | CNR Virus des Infections Respiratoires - France SUD | Antonin Bal, Gregory Destras, Gwendolyne Burfin, Hadrien Regue, Quentin Semanas, Martine Valette, Bruno Lina, Laurence Josset |
| EPI_ISL_3472286, EPI_ISL_3472287, EPI_ISL_3472288, EPI_ISL_3472289 | LX BIO | CNR Virus des Infections Respiratoires - France SUD | Antonin Bal, Gregory Destras, Gwendolyne Burfin, Hadrien Regue, Quentin Semanas, Martine Valette, Bruno Lina, Laurence Josset |
| EPI_ISL_3472290, EPI_ISL_3472291, EPI_ISL_3472292 | CNR Virus des Infections Respiratoires - France SUD | CNR Virus des Infections Respiratoires - France SUD | Antonin Bal, Gregory Destras, Gwendolyne Burfin, Hadrien Regue, Quentin Semanas, Martine Valette, Bruno Lina, Laurence Josset |
| EPI_ISL_3472293 | CENTRE HOSPITALIER DE BOURG EN BRESSE | CNR Virus des Infections Respiratoires - France SUD | Antonin Bal, Gregory Destras, Gwendolyne Burfin, Hadrien Regue, Quentin Semanas, Martine Valette, Bruno Lina, Laurence Josset |
| EPI_ISL_3472294 | ORIAPOLE | CNR Virus des Infections Respiratoires - France SUD | Antonin Bal, Gregory Destras, Gwendolyne Burfin, Hadrien Regue, Quentin Semanas, Martine Valette, Bruno Lina, Laurence Josset |
| EPI_ISL_3472295 | CENTRE HOSPITALIER PIERRE OUDOT | CNR Virus des Infections Respiratoires - France SUD | Antonin Bal, Gregory Destras, Gwendolyne Burfin, Hadrien Regue, Quentin Semanas, Martine Valette, Bruno Lina, Laurence Josset |
| EPI_ISL_3472296 | ORIAPOLE | CNR Virus des Infections Respiratoires - France SUD | Antonin Bal, Gregory Destras, Gwendolyne Burfin, Hadrien Regue, Quentin Semanas, Martine Valette, Bruno Lina, Laurence Josset |
| EPI_ISL_3472297, EPI_ISL_3472298, EPI_ISL_3472299, EPI_ISL_3472300, EPI_ISL_3472301, EPI_ISL_3472302, EPI_ISL_3472303, EPI_ISL_3472304, EPI_ISL_3472305 | CNR Virus des Infections Respiratoires - France SUD | CNR Virus des Infections Respiratoires - France SUD | Antonin Bal, Gregory Destras, Gwendolyne Burfin, Hadrien Regue, Quentin Semanas, Martine Valette, Bruno Lina, Laurence Josset |
| EPI_ISL_3472306, EPI_ISL_3472307 | CENTRE HOSPITALIER PIERRE OUDOT | CNR Virus des Infections Respiratoires - France SUD | Antonin Bal, Gregory Destras, Gwendolyne Burfin, Hadrien Regue, Quentin Semanas, Martine Valette, Bruno Lina, Laurence Josset |
| EPI_ISL_3472308 | CENTRE HOSPITALIER DE BOURG EN BRESSE | CNR Virus des Infections Respiratoires - France SUD | Antonin Bal, Gregory Destras, Gwendolyne Burfin, Hadrien Regue, Quentin Semanas, Martine Valette, Bruno Lina, Laurence Josset |
| EPI_ISL_3472309 | CNR Virus des Infections Respiratoires - France SUD | CNR Virus des Infections Respiratoires - France SUD | Antonin Bal, Gregory Destras, Gwendolyne Burfin, Hadrien Regue, Quentin Semanas, Martine Valette, Bruno Lina, Laurence Josset |
| EPI_ISL_3472310 | CH GIVORS | CNR Virus des Infections Respiratoires - France SUD | Antonin Bal, Gregory Destras, Gwendolyne Burfin, Hadrien Regue, Quentin Semanas, Martine Valette, Bruno Lina, Laurence Josset |
| EPI_ISL_3472311 | CENTRE HOSPITALIER LUCIEN HUSSEL | CNR Virus des Infections Respiratoires - France SUD | Antonin Bal, Gregory Destras, Gwendolyne Burfin, Hadrien Regue, Quentin Semanas, Martine Valette, Bruno Lina, Laurence Josset |
| EPI_ISL_3472312, EPI_ISL_3472313, EPI_ISL_3472314 | CNR Virus des Infections Respiratoires - France SUD | CNR Virus des Infections Respiratoires - France SUD | Antonin Bal, Gregory Destras, Gwendolyne Burfin, Hadrien Regue, Quentin Semanas, Martine Valette, Bruno Lina, Laurence Josset |
| EPI_ISL_3472315 | CH GIVORS | CNR Virus des Infections Respiratoires - France SUD | Antonin Bal, Gregory Destras, Gwendolyne Burfin, Hadrien Regue, Quentin Semanas, Martine Valette, Bruno Lina, Laurence Josset |
| EPI_ISL_3472316, EPI_ISL_3472317 | CNR Virus des Infections Respiratoires - France SUD | CNR Virus des Infections Respiratoires - France SUD | Antonin Bal, Gregory Destras, Gwendolyne Burfin, Hadrien Regue, Quentin Semanas, Martine Valette, Bruno Lina, Laurence Josset |
| EPI_ISL_3472318, EPI_ISL_3472319, EPI_ISL_3472320, EPI_ISL_3472321 | LABORATOIRE UNILIANS DECINES | CNR Virus des Infections Respiratoires - France SUD | Antonin Bal, Gregory Destras, Gwendolyne Burfin, Hadrien Regue, Quentin Semanas, Martine Valette, Bruno Lina, Laurence Josset |
| EPI_ISL_3472322 | CNR Virus des Infections Respiratoires - France SUD | CNR Virus des Infections Respiratoires - France SUD | Antonin Bal, Gregory Destras, Gwendolyne Burfin, Hadrien Regue, Quentin Semanas, Martine Valette, Bruno Lina, Laurence Josset |
| EPI_ISL_3472323, EPI_ISL_3472324, EPI_ISL_3472325 | LABORATOIRE UNILIANS DECINES | CNR Virus des Infections Respiratoires - France SUD | Antonin Bal, Gregory Destras, Gwendolyne Burfin, Hadrien Regue, Quentin Semanas, Martine Valette, Bruno Lina, Laurence Josset |
| EPI_ISL_3472326 | CNR Virus des Infections Respiratoires - France SUD | CNR Virus des Infections Respiratoires - France SUD | Antonin Bal, Gregory Destras, Gwendolyne Burfin, Hadrien Regue, Quentin Semanas, Martine Valette, Bruno Lina, Laurence Josset |
| EPI_ISL_3472327, | LABORATOIRE UNILIANS DECINES | CNR Virus des Infections Respiratoires - France SUD | Antonin Bal, Gregory Destras, Gwendolyne Burfin, Hadrien Regue, Quentin Semanas, Martine Valette, Bruno Lina, Laurence Josset |

| | | | |
|---|---|---|---|
| EPI_ISL_3472328, EPI_ISL_3472329 | | | |
| EPI_ISL_3472330 | BIOLITTORAL BIOGROUP PLATEAU TECH | CNR Virus des Infections Respiratoires - France SUD | Antonin Bal, Gregory Destras, Gwendolyne Burfin, Hadrien Regue, Quentin Semanas, Martine Valette, Bruno Lina, Laurence Josset |
| EPI_ISL_3472331, EPI_ISL_3472332 | LABORATOIRE UNILIANS DECINES | CNR Virus des Infections Respiratoires - France SUD | Antonin Bal, Gregory Destras, Gwendolyne Burfin, Hadrien Regue, Quentin Semanas, Martine Valette, Bruno Lina, Laurence Josset |
| EPI_ISL_3472333, EPI_ISL_3472334, EPI_ISL_3472335, EPI_ISL_3472336, EPI_ISL_3472337, EPI_ISL_3472338 | BIOLITTORAL BIOGROUP PLATEAU TECH | CNR Virus des Infections Respiratoires - France SUD | Antonin Bal, Gregory Destras, Gwendolyne Burfin, Hadrien Regue, Quentin Semanas, Martine Valette, Bruno Lina, Laurence Josset |
| EPI_ISL_3472339, EPI_ISL_3472340, EPI_ISL_3472341, EPI_ISL_3472342, EPI_ISL_3472343, EPI_ISL_3472344, EPI_ISL_3472345, EPI_ISL_3472346, EPI_ISL_3472347, EPI_ISL_3472348 | LABORATOIRE UNILIANS DECINES | CNR Virus des Infections Respiratoires - France SUD | Antonin Bal, Gregory Destras, Gwendolyne Burfin, Hadrien Regue, Quentin Semanas, Martine Valette, Bruno Lina, Laurence Josset |
| EPI_ISL_3472349 | CNR Virus des Infections Respiratoires - France SUD | CNR Virus des Infections Respiratoires - France SUD | Antonin Bal, Gregory Destras, Gwendolyne Burfin, Hadrien Regue, Quentin Semanas, Martine Valette, Bruno Lina, Laurence Josset |
| EPI_ISL_3472350 | LABORATOIRE UNILIANS DECINES | CNR Virus des Infections Respiratoires - France SUD | Antonin Bal, Gregory Destras, Gwendolyne Burfin, Hadrien Regue, Quentin Semanas, Martine Valette, Bruno Lina, Laurence Josset |
| EPI_ISL_3472351, EPI_ISL_3472352, EPI_ISL_3472353, EPI_ISL_3472354, EPI_ISL_3472355, EPI_ISL_3472356, EPI_ISL_3472357, EPI_ISL_3472358, EPI_ISL_3472359, EPI_ISL_3472360, EPI_ISL_3472361 | | | |
| see above | LABORATOIRE CBM DE MURET | CNR Virus des Infections Respiratoires - France SUD | Antonin Bal, Gregory Destras, Gwendolyne Burfin, Hadrien Regue, Quentin Semanas, Martine Valette, Bruno Lina, Laurence Josset |
| EPI_ISL_3472362, EPI_ISL_3472363, EPI_ISL_3472364, EPI_ISL_3472365 | CNR Virus des Infections Respiratoires - France SUD | CNR Virus des Infections Respiratoires - France SUD | Antonin Bal, Gregory Destras, Gwendolyne Burfin, Hadrien Regue, Quentin Semanas, Martine Valette, Bruno Lina, Laurence Josset |
| EPI_ISL_3472366, EPI_ISL_3472367, EPI_ISL_3472368, EPI_ISL_3472369, EPI_ISL_3472370, EPI_ISL_3472371, EPI_ISL_3472372, EPI_ISL_3472373, EPI_ISL_3472374, EPI_ISL_3472375, EPI_ISL_3472376, EPI_ISL_3472377, EPI_ISL_3472378 | | | |
| see above | LABORATOIRE BIOESTEREL | CNR Virus des Infections Respiratoires - France SUD | Antonin Bal, Gregory Destras, Gwendolyne Burfin, Hadrien Regue, Quentin Semanas, Martine Valette, Bruno Lina, Laurence Josset |
| EPI_ISL_3472379, EPI_ISL_3472380 | CNR Virus des Infections Respiratoires - France SUD | CNR Virus des Infections Respiratoires - France SUD | Antonin Bal, Gregory Destras, Gwendolyne Burfin, Hadrien Regue, Quentin Semanas, Martine Valette, Bruno Lina, Laurence Josset |
| EPI_ISL_3472381, EPI_ISL_3472382, EPI_ISL_3472383, EPI_ISL_3472384, EPI_ISL_3472385, EPI_ISL_3472386, EPI_ISL_3472387, EPI_ISL_3472388, EPI_ISL_3472389, EPI_ISL_3472390, EPI_ISL_3472391, EPI_ISL_3472392, EPI_ISL_3472393, EPI_ISL_3472394, EPI_ISL_3472395 | | | |
| see above | LABORATOIRE BIOESTEREL | CNR Virus des Infections Respiratoires - France SUD | Antonin Bal, Gregory Destras, Gwendolyne Burfin, Hadrien Regue, Quentin Semanas, Martine Valette, Bruno Lina, Laurence Josset |
| EPI_ISL_3472396 | CENTRE HOSPITALIER SUD GIRONDE | CNR Virus des Infections Respiratoires - France SUD | Antonin Bal, Gregory Destras, Gwendolyne Burfin, Hadrien Regue, Quentin Semanas, Martine Valette, Bruno Lina, Laurence Josset |
| EPI_ISL_3472397 | LABORATOIRE BIOESTEREL | CNR Virus des Infections Respiratoires - France SUD | Antonin Bal, Gregory Destras, Gwendolyne Burfin, Hadrien Regue, Quentin Semanas, Martine Valette, Bruno Lina, Laurence Josset |
| EPI_ISL_3472398 | CENTRE HOSPITALIER SUD GIRONDE | CNR Virus des Infections Respiratoires - France SUD | Antonin Bal, Gregory Destras, Gwendolyne Burfin, Hadrien Regue, Quentin Semanas, Martine Valette, Bruno Lina, Laurence Josset |
| EPI_ISL_3472399, EPI_ISL_3472400, EPI_ISL_3472401, EPI_ISL_3472402, EPI_ISL_3472403, EPI_ISL_3472404, EPI_ISL_3472405 | LABORATOIRE BIOESTEREL | CNR Virus des Infections Respiratoires - France SUD | Antonin Bal, Gregory Destras, Gwendolyne Burfin, Hadrien Regue, Quentin Semanas, Martine Valette, Bruno Lina, Laurence Josset |
| EPI_ISL_3472406 | CENTRE HOSPITALIER SUD GIRONDE | CNR Virus des Infections Respiratoires - France SUD | Antonin Bal, Gregory Destras, Gwendolyne Burfin, Hadrien Regue, Quentin Semanas, Martine Valette, Bruno Lina, Laurence Josset |
| EPI_ISL_3472407, EPI_ISL_3472408, EPI_ISL_3472409, EPI_ISL_3472410, EPI_ISL_3472411, EPI_ISL_3472412, EPI_ISL_3472413, EPI_ISL_3472414 | LABORATOIRE BIOESTEREL | CNR Virus des Infections Respiratoires - France SUD | Antonin Bal, Gregory Destras, Gwendolyne Burfin, Hadrien Regue, Quentin Semanas, Martine Valette, Bruno Lina, Laurence Josset |
| EPI_ISL_3472415 | CENTRE HOSPITALIER DE VALENCE | CNR Virus des Infections Respiratoires - France SUD | Antonin Bal, Gregory Destras, Gwendolyne Burfin, Hadrien Regue, Quentin Semanas, Martine Valette, Bruno Lina, Laurence Josset |
| EPI_ISL_3472416 | LABORATOIRE BIOESTEREL | CNR Virus des Infections Respiratoires - France SUD | Antonin Bal, Gregory Destras, Gwendolyne Burfin, Hadrien Regue, Quentin Semanas, Martine Valette, Bruno Lina, Laurence Josset |
| EPI_ISL_3472417 | CENTRE HOSPITALIER DE VALENCE | CNR Virus des Infections Respiratoires - France SUD | Antonin Bal, Gregory Destras, Gwendolyne Burfin, Hadrien Regue, Quentin Semanas, Martine Valette, Bruno Lina, Laurence Josset |
| EPI_ISL_3472418 | LABORATOIRE BIOESTEREL | CNR Virus des Infections Respiratoires - France SUD | Antonin Bal, Gregory Destras, Gwendolyne Burfin, Hadrien Regue, Quentin Semanas, Martine Valette, Bruno Lina, Laurence Josset |
| EPI_ISL_3472419 | CENTRE HOSPITALIER DE VALENCE | CNR Virus des Infections Respiratoires - France SUD | Antonin Bal, Gregory Destras, Gwendolyne Burfin, Hadrien Regue, Quentin Semanas, Martine Valette, Bruno Lina, Laurence Josset |
| EPI_ISL_3472420, EPI_ISL_3472421 | CNR Virus des Infections Respiratoires - France SUD | CNR Virus des Infections Respiratoires - France SUD | Antonin Bal, Gregory Destras, Gwendolyne Burfin, Hadrien Regue, Quentin Semanas, Martine Valette, Bruno Lina, Laurence Josset |
| EPI_ISL_3472422 | CENTRE HOSPITALIER DE VALENCE | CNR Virus des Infections Respiratoires - France SUD | Antonin Bal, Gregory Destras, Gwendolyne Burfin, Hadrien Regue, Quentin Semanas, Martine Valette, Bruno Lina, Laurence Josset |
| EPI_ISL_3472423 | CENTRE HOSPITALIER PELLEGRIN | CNR Virus des Infections Respiratoires - France SUD | Antonin Bal, Gregory Destras, Gwendolyne Burfin, Hadrien Regue, Quentin Semanas, Martine Valette, Bruno Lina, Laurence Josset |
| EPI_ISL_3472424 | CH GIVORS | CNR Virus des Infections Respiratoires - France SUD | Antonin Bal, Gregory Destras, Gwendolyne Burfin, Hadrien Regue, Quentin Semanas, Martine Valette, Bruno Lina, Laurence Josset |
| EPI_ISL_3472425, EPI_ISL_3472426 | CENTRE HOSPITALIER PELLEGRIN | CNR Virus des Infections Respiratoires - France SUD | Antonin Bal, Gregory Destras, Gwendolyne Burfin, Hadrien Regue, Quentin Semanas, Martine Valette, Bruno Lina, Laurence Josset |
| EPI_ISL_3472427 | CENTRE HOSPITALIER MARMANDE TONNEINS | CNR Virus des Infections Respiratoires - France SUD | Antonin Bal, Gregory Destras, Gwendolyne Burfin, Hadrien Regue, Quentin Semanas, Martine Valette, Bruno Lina, Laurence Josset |
| EPI_ISL_3472428, EPI_ISL_3472429, EPI_ISL_3472430, EPI_ISL_3472431, EPI_ISL_3472432 | CENTRE HOSPITALIER PELLEGRIN | CNR Virus des Infections Respiratoires - France SUD | Antonin Bal, Gregory Destras, Gwendolyne Burfin, Hadrien Regue, Quentin Semanas, Martine Valette, Bruno Lina, Laurence Josset |
| EPI_ISL_3472433 | LABORATOIRE BIOESTEREL | CNR Virus des Infections Respiratoires - France SUD | Antonin Bal, Gregory Destras, Gwendolyne Burfin, Hadrien Regue, Quentin Semanas, Martine Valette, Bruno Lina, Laurence Josset |
| EPI_ISL_3472434 | CENTRE HOSPITALIER PELLEGRIN | CNR Virus des Infections Respiratoires - France SUD | Antonin Bal, Gregory Destras, Gwendolyne Burfin, Hadrien Regue, Quentin Semanas, Martine Valette, Bruno Lina, Laurence Josset |
| EPI_ISL_3472435 | LABORATOIRE BIOESTEREL | CNR Virus des Infections Respiratoires - France SUD | Antonin Bal, Gregory Destras, Gwendolyne Burfin, Hadrien Regue, Quentin Semanas, Martine Valette, Bruno Lina, Laurence Josset |
| EPI_ISL_3472436 | CENTRE HOSPITALIER PELLEGRIN | CNR Virus des Infections Respiratoires - France SUD | Antonin Bal, Gregory Destras, Gwendolyne Burfin, Hadrien Regue, Quentin Semanas, Martine Valette, Bruno Lina, Laurence Josset |
| EPI_ISL_3472437, EPI_ISL_3472438 | LABORATOIRE BIOESTEREL | CNR Virus des Infections Respiratoires - France SUD | Antonin Bal, Gregory Destras, Gwendolyne Burfin, Hadrien Regue, Quentin Semanas, Martine Valette, Bruno Lina, Laurence Josset |
| EPI_ISL_3472439 | CENTRE HOSPITALIER PELLEGRIN | CNR Virus des Infections Respiratoires - France SUD | Antonin Bal, Gregory Destras, Gwendolyne Burfin, Hadrien Regue, Quentin Semanas, Martine Valette, Bruno Lina, Laurence Josset |
| EPI_ISL_3472440 | CENTRE HOSPITALIER LUCIEN HUSSEL | CNR Virus des Infections Respiratoires - France SUD | Antonin Bal, Gregory Destras, Gwendolyne Burfin, Hadrien Regue, Quentin Semanas, Martine Valette, Bruno Lina, Laurence Josset |
| EPI_ISL_3472441 | LABORATOIRE BIOESTEREL | CNR Virus des Infections Respiratoires - France SUD | Antonin Bal, Gregory Destras, Gwendolyne Burfin, Hadrien Regue, Quentin Semanas, Martine Valette, Bruno Lina, Laurence Josset |
| EPI_ISL_3472442 | CENTRE HOSPITALIER PELLEGRIN | CNR Virus des Infections Respiratoires - France SUD | Antonin Bal, Gregory Destras, Gwendolyne Burfin, Hadrien Regue, Quentin Semanas, Martine Valette, Bruno Lina, Laurence Josset |
| EPI_ISL_3472443 | HOPITAL NORD OUEST - CH VILLEFRANCHE | CNR Virus des Infections Respiratoires - France SUD | Antonin Bal, Gregory Destras, Gwendolyne Burfin, Hadrien Regue, Quentin Semanas, Martine Valette, Bruno Lina, Laurence Josset |
| EPI_ISL_3472444 | LABORATOIRE BIOESTEREL | CNR Virus des Infections Respiratoires - France SUD | Antonin Bal, Gregory Destras, Gwendolyne Burfin, Hadrien Regue, Quentin Semanas, Martine Valette, Bruno Lina, Laurence Josset |
| EPI_ISL_3472445 | CNR Virus des Infections Respiratoires - France SUD | CNR Virus des Infections Respiratoires - France SUD | Antonin Bal, Gregory Destras, Gwendolyne Burfin, Hadrien Regue, Quentin Semanas, Martine Valette, Bruno Lina, Laurence Josset |
| EPI_ISL_3472446, EPI_ISL_3472447, EPI_ISL_3472448, EPI_ISL_3472449, EPI_ISL_3472450, EPI_ISL_3472451, EPI_ISL_3472452, EPI_ISL_3472453, EPI_ISL_3472454, EPI_ISL_3472455, EPI_ISL_3472456, EPI_ISL_3472457, EPI_ISL_3472458, EPI_ISL_3472459, EPI_ISL_3472460, EPI_ISL_3472461, EPI_ISL_3472462, EPI_ISL_3472463, EPI_ISL_3472464, EPI_ISL_3472465, EPI_ISL_3472466, EPI_ISL_3472467, EPI_ISL_3472468, EPI_ISL_3472469, EPI_ISL_3472470, EPI_ISL_3472471, EPI_ISL_3472472, EPI_ISL_3472473, EPI_ISL_3472474, EPI_ISL_3472475, EPI_ISL_3472476 | | | |
| see above | LABORATOIRE BIOESTEREL | CNR Virus des Infections Respiratoires - France SUD | Antonin Bal, Gregory Destras, Gwendolyne Burfin, Hadrien Regue, Quentin Semanas, Martine Valette, Bruno Lina, Laurence Josset |
| EPI_ISL_3472477 | MIRIALIS CLUSES BECHET | CNR Virus des Infections Respiratoires - France SUD | Antonin Bal, Gregory Destras, Gwendolyne Burfin, Hadrien Regue, Quentin Semanas, Martine Valette, Bruno Lina, Laurence Josset |

| | | | |
|---|---|---|---|
| EPI_ISL_3472478 | LABORATOIRE BIOMEDICA | CNR Virus des Infections Respiratoires - France SUD | Antonin Bal, Gregory Destras, Gwendolyne Burfin, Hadrien Regue, Quentin Semanas, Martine Valette, Bruno Lina, Laurence Josset |
| EPI_ISL_3472479, EPI_ISL_3472480, EPI_ISL_3472481, EPI_ISL_3472482 | CENTRE HOSPITALIER BRIVE | CNR Virus des Infections Respiratoires - France SUD | Antonin Bal, Gregory Destras, Gwendolyne Burfin, Hadrien Regue, Quentin Semanas, Martine Valette, Bruno Lina, Laurence Josset |
| EPI_ISL_3472483, EPI_ISL_3472484 | LABORATOIRE UNILIANS DECINES | CNR Virus des Infections Respiratoires - France SUD | Antonin Bal, Gregory Destras, Gwendolyne Burfin, Hadrien Regue, Quentin Semanas, Martine Valette, Bruno Lina, Laurence Josset |
| EPI_ISL_3472485 | GROUPE LCD - UNIBIO LA PLAINE | CNR Virus des Infections Respiratoires - France SUD | Antonin Bal, Gregory Destras, Gwendolyne Burfin, Hadrien Regue, Quentin Semanas, Martine Valette, Bruno Lina, Laurence Josset |
| EPI_ISL_3472486 | CNR Virus des Infections Respiratoires - France SUD | CNR Virus des Infections Respiratoires - France SUD | Antonin Bal, Gregory Destras, Gwendolyne Burfin, Hadrien Regue, Quentin Semanas, Martine Valette, Bruno Lina, Laurence Josset |
| EPI_ISL_3472487 | GROUPE LCD - UNIBIO LA PLAINE | CNR Virus des Infections Respiratoires - France SUD | Antonin Bal, Gregory Destras, Gwendolyne Burfin, Hadrien Regue, Quentin Semanas, Martine Valette, Bruno Lina, Laurence Josset |
| EPI_ISL_3472488 | CNR Virus des Infections Respiratoires - France SUD | CNR Virus des Infections Respiratoires - France SUD | Antonin Bal, Gregory Destras, Gwendolyne Burfin, Hadrien Regue, Quentin Semanas, Martine Valette, Bruno Lina, Laurence Josset |
| EPI_ISL_3472489 | HOPITAL NOTRE-DAME LA MISERICORDE | CNR Virus des Infections Respiratoires - France SUD | Antonin Bal, Gregory Destras, Gwendolyne Burfin, Hadrien Regue, Quentin Semanas, Martine Valette, Bruno Lina, Laurence Josset |
| EPI_ISL_3472490 | LABORATOIRE BIOLYSS | CNR Virus des Infections Respiratoires - France SUD | Antonin Bal, Gregory Destras, Gwendolyne Burfin, Hadrien Regue, Quentin Semanas, Martine Valette, Bruno Lina, Laurence Josset |
| EPI_ISL_3472491, EPI_ISL_3472492, EPI_ISL_3472493, EPI_ISL_3472494, EPI_ISL_3472495 | CENTRE HOSPITALIER DE LA COTE BASQUE | CNR Virus des Infections Respiratoires - France SUD | Antonin Bal, Gregory Destras, Gwendolyne Burfin, Hadrien Regue, Quentin Semanas, Martine Valette, Bruno Lina, Laurence Josset |
| EPI_ISL_3472496, EPI_ISL_3472497, EPI_ISL_3472498, EPI_ISL_3472499 | CNR Virus des Infections Respiratoires - France SUD | CNR Virus des Infections Respiratoires - France SUD | Antonin Bal, Gregory Destras, Gwendolyne Burfin, Hadrien Regue, Quentin Semanas, Martine Valette, Bruno Lina, Laurence Josset |
| EPI_ISL_3472500, EPI_ISL_3472501, EPI_ISL_3472502, EPI_ISL_3472503, EPI_ISL_3472504, EPI_ISL_3472505, EPI_ISL_3472506, EPI_ISL_3472507, EPI_ISL_3472508, EPI_ISL_3472509, EPI_ISL_3472510, EPI_ISL_3472511, EPI_ISL_3472512, EPI_ISL_3472513, EPI_ISL_3472514, EPI_ISL_3472515, EPI_ISL_3472516, EPI_ISL_3472517, EPI_ISL_3472518, EPI_ISL_3472519 | | | |
| see above | LABORATOIRE CERDIBIO-CHARENTES | CNR Virus des Infections Respiratoires - France SUD | Antonin Bal, Gregory Destras, Gwendolyne Burfin, Hadrien Regue, Quentin Semanas, Martine Valette, Bruno Lina, Laurence Josset |
| EPI_ISL_3472520 | SYNLAB AQUITAINE | CNR Virus des Infections Respiratoires - France SUD | Antonin Bal, Gregory Destras, Gwendolyne Burfin, Hadrien Regue, Quentin Semanas, Martine Valette, Bruno Lina, Laurence Josset |
| EPI_ISL_3472521, EPI_ISL_3472522, EPI_ISL_3472523, EPI_ISL_3472524, EPI_ISL_3472525, EPI_ISL_3472526, EPI_ISL_3472527, EPI_ISL_3472528, EPI_ISL_3472529 | LABORATOIRE CERDIBIO-CHARENTES | CNR Virus des Infections Respiratoires - France SUD | Antonin Bal, Gregory Destras, Gwendolyne Burfin, Hadrien Regue, Quentin Semanas, Martine Valette, Bruno Lina, Laurence Josset |
| EPI_ISL_3472530 | SYNLAB AQUITAINE | CNR Virus des Infections Respiratoires - France SUD | Antonin Bal, Gregory Destras, Gwendolyne Burfin, Hadrien Regue, Quentin Semanas, Martine Valette, Bruno Lina, Laurence Josset |
| EPI_ISL_3472531 | LABORATOIRE CERDIBIO-CHARENTES | CNR Virus des Infections Respiratoires - France SUD | Antonin Bal, Gregory Destras, Gwendolyne Burfin, Hadrien Regue, Quentin Semanas, Martine Valette, Bruno Lina, Laurence Josset |
| EPI_ISL_3472532, EPI_ISL_3472533 | SYNLAB AQUITAINE | CNR Virus des Infections Respiratoires - France SUD | Antonin Bal, Gregory Destras, Gwendolyne Burfin, Hadrien Regue, Quentin Semanas, Martine Valette, Bruno Lina, Laurence Josset |
| EPI_ISL_3472534 | CNR Virus des Infections Respiratoires - France SUD | CNR Virus des Infections Respiratoires - France SUD | Antonin Bal, Gregory Destras, Gwendolyne Burfin, Hadrien Regue, Quentin Semanas, Martine Valette, Bruno Lina, Laurence Josset |
| EPI_ISL_3472535 | LABORATOIRE UNILIANS DECINES | CNR Virus des Infections Respiratoires - France SUD | Antonin Bal, Gregory Destras, Gwendolyne Burfin, Hadrien Regue, Quentin Semanas, Martine Valette, Bruno Lina, Laurence Josset |
| EPI_ISL_3472536 | LABORATOIRE CERDIBIO-CHARENTES | CNR Virus des Infections Respiratoires - France SUD | Antonin Bal, Gregory Destras, Gwendolyne Burfin, Hadrien Regue, Quentin Semanas, Martine Valette, Bruno Lina, Laurence Josset |
| EPI_ISL_3472537 | LABORATOIRE UNILIANS DECINES | CNR Virus des Infections Respiratoires - France SUD | Antonin Bal, Gregory Destras, Gwendolyne Burfin, Hadrien Regue, Quentin Semanas, Martine Valette, Bruno Lina, Laurence Josset |
| EPI_ISL_3472538 | LABORATOIRE CERDIBIO-CHARENTES | CNR Virus des Infections Respiratoires - France SUD | Antonin Bal, Gregory Destras, Gwendolyne Burfin, Hadrien Regue, Quentin Semanas, Martine Valette, Bruno Lina, Laurence Josset |
| EPI_ISL_3472539, EPI_ISL_3472540 | SYNLAB AQUITAINE | CNR Virus des Infections Respiratoires - France SUD | Antonin Bal, Gregory Destras, Gwendolyne Burfin, Hadrien Regue, Quentin Semanas, Martine Valette, Bruno Lina, Laurence Josset |
| EPI_ISL_3472541, EPI_ISL_3472542 | LABORATOIRE CERDIBIO-CHARENTES | CNR Virus des Infections Respiratoires - France SUD | Antonin Bal, Gregory Destras, Gwendolyne Burfin, Hadrien Regue, Quentin Semanas, Martine Valette, Bruno Lina, Laurence Josset |
| EPI_ISL_3472543 | CNR Virus des Infections Respiratoires - France SUD | CNR Virus des Infections Respiratoires - France SUD | Antonin Bal, Gregory Destras, Gwendolyne Burfin, Hadrien Regue, Quentin Semanas, Martine Valette, Bruno Lina, Laurence Josset |
| EPI_ISL_3472544, EPI_ISL_3472545, EPI_ISL_3472546, EPI_ISL_3472547 | LABORATOIRE CERDIBIO-CHARENTES | CNR Virus des Infections Respiratoires - France SUD | Antonin Bal, Gregory Destras, Gwendolyne Burfin, Hadrien Regue, Quentin Semanas, Martine Valette, Bruno Lina, Laurence Josset |
| EPI_ISL_3472548 | SYNLAB AQUITAINE | CNR Virus des Infections Respiratoires - France SUD | Antonin Bal, Gregory Destras, Gwendolyne Burfin, Hadrien Regue, Quentin Semanas, Martine Valette, Bruno Lina, Laurence Josset |
| EPI_ISL_3472549, EPI_ISL_3472550, EPI_ISL_3472551, EPI_ISL_3472552, EPI_ISL_3472553 | LABORATOIRE CERDIBIO-CHARENTES | CNR Virus des Infections Respiratoires - France SUD | Antonin Bal, Gregory Destras, Gwendolyne Burfin, Hadrien Regue, Quentin Semanas, Martine Valette, Bruno Lina, Laurence Josset |
| EPI_ISL_3472554, EPI_ISL_3472555, EPI_ISL_3472556, EPI_ISL_3472557, EPI_ISL_3472558 | SYNLAB AQUITAINE | CNR Virus des Infections Respiratoires - France SUD | Antonin Bal, Gregory Destras, Gwendolyne Burfin, Hadrien Regue, Quentin Semanas, Martine Valette, Bruno Lina, Laurence Josset |
| EPI_ISL_3472559 | LABORATOIRE FORTE BIO DAX | CNR Virus des Infections Respiratoires - France SUD | Antonin Bal, Gregory Destras, Gwendolyne Burfin, Hadrien Regue, Quentin Semanas, Martine Valette, Bruno Lina, Laurence Josset |
| EPI_ISL_3472560 | SYNLAB AQUITAINE | CNR Virus des Infections Respiratoires - France SUD | Antonin Bal, Gregory Destras, Gwendolyne Burfin, Hadrien Regue, Quentin Semanas, Martine Valette, Bruno Lina, Laurence Josset |
| EPI_ISL_3472561, EPI_ISL_3472562, EPI_ISL_3472563, EPI_ISL_3472564, EPI_ISL_3472565, EPI_ISL_3472566 | LABORATOIRE FORTE BIO DAX | CNR Virus des Infections Respiratoires - France SUD | Antonin Bal, Gregory Destras, Gwendolyne Burfin, Hadrien Regue, Quentin Semanas, Martine Valette, Bruno Lina, Laurence Josset |
| EPI_ISL_3472567 | CERBAILLIANCE LA REUNION | CNR Virus des Infections Respiratoires - France SUD | Antonin Bal, Gregory Destras, Gwendolyne Burfin, Hadrien Regue, Quentin Semanas, Martine Valette, Bruno Lina, Laurence Josset |
| EPI_ISL_3472568, EPI_ISL_3472569, EPI_ISL_3472570, EPI_ISL_3472571, EPI_ISL_3472572, EPI_ISL_3472573 | LABORATOIRE FORTE BIO DAX | CNR Virus des Infections Respiratoires - France SUD | Antonin Bal, Gregory Destras, Gwendolyne Burfin, Hadrien Regue, Quentin Semanas, Martine Valette, Bruno Lina, Laurence Josset |
| EPI_ISL_3472574 | CNR Virus des Infections Respiratoires - France SUD | CNR Virus des Infections Respiratoires - France SUD | Antonin Bal, Gregory Destras, Gwendolyne Burfin, Hadrien Regue, Quentin Semanas, Martine Valette, Bruno Lina, Laurence Josset |
| EPI_ISL_3472575 | LABORATOIRE UNILIANS DECINES | CNR Virus des Infections Respiratoires - France SUD | Antonin Bal, Gregory Destras, Gwendolyne Burfin, Hadrien Regue, Quentin Semanas, Martine Valette, Bruno Lina, Laurence Josset |
| EPI_ISL_3472576 | CNR Virus des Infections Respiratoires - France SUD | CNR Virus des Infections Respiratoires - France SUD | Antonin Bal, Gregory Destras, Gwendolyne Burfin, Hadrien Regue, Quentin Semanas, Martine Valette, Bruno Lina, Laurence Josset |
| EPI_ISL_3472577, EPI_ISL_3472578 | LABORATOIRE UNILIANS DECINES | CNR Virus des Infections Respiratoires - France SUD | Antonin Bal, Gregory Destras, Gwendolyne Burfin, Hadrien Regue, Quentin Semanas, Martine Valette, Bruno Lina, Laurence Josset |
| EPI_ISL_3472579, EPI_ISL_3472580, EPI_ISL_3472581, EPI_ISL_3472582 | SYNLAB AQUITAINE | CNR Virus des Infections Respiratoires - France SUD | Antonin Bal, Gregory Destras, Gwendolyne Burfin, Hadrien Regue, Quentin Semanas, Martine Valette, Bruno Lina, Laurence Josset |
| EPI_ISL_3472583, EPI_ISL_3472584, EPI_ISL_3472585, EPI_ISL_3472586 | CENTRE HOSPITALIER DE VALENCE | CNR Virus des Infections Respiratoires - France SUD | Antonin Bal, Gregory Destras, Gwendolyne Burfin, Hadrien Regue, Quentin Semanas, Martine Valette, Bruno Lina, Laurence Josset |

| | | | |
|---|---|---|---|
| EPI_ISL_3472587, EPI_ISL_3472588 | CHU ST ETIENNE HOPITAL NORD | CNR Virus des Infections Respiratoires - France SUD | Antonin Bal, Gregory Destras, Gwendolyne Burfin, Hadrien Regue, Quentin Semanas, Martine Valette, Bruno Lina, Laurence Josset |
| EPI_ISL_3472589, EPI_ISL_3472590, EPI_ISL_3472591, EPI_ISL_3472592, EPI_ISL_3472593, EPI_ISL_3472594, EPI_ISL_3472595 | SYNLAB AQUITAINE | CNR Virus des Infections Respiratoires - France SUD | Antonin Bal, Gregory Destras, Gwendolyne Burfin, Hadrien Regue, Quentin Semanas, Martine Valette, Bruno Lina, Laurence Josset |
| EPI_ISL_3472596 | Ohio Department of Health Laboratory | Ohio Department of Health Laboratory | Holmes, Jennifer; Eric Brandt, Keoni Omura, Glen McGillivary, Caitlin McDonnell, Jade Mowery, Stephanie Mccracken, Tyler Payne, Kirtana Ramadugu, Erica Leasure, Brent Lee, Kelsey Florek, Heather Blankenship, Allison Black, Quanta Brown, and Tammy Bannerman |
| EPI_ISL_3472598 | LABORATOIRE DE L'OSPEDALE - 2A 2B | CNR Virus des Infections Respiratoires - France SUD | Antonin Bal, Gregory Destras, Gwendolyne Burfin, Hadrien Regue, Quentin Semanas, Martine Valette, Bruno Lina, Laurence Josset |
| EPI_ISL_3472599 | LAM SYNLAB BORDEAUX ATLANTIQUE | CNR Virus des Infections Respiratoires - France SUD | Antonin Bal, Gregory Destras, Gwendolyne Burfin, Hadrien Regue, Quentin Semanas, Martine Valette, Bruno Lina, Laurence Josset |
| EPI_ISL_3472600 | LABORATOIRE DE L'OSPEDALE - 2A 2B | CNR Virus des Infections Respiratoires - France SUD | Antonin Bal, Gregory Destras, Gwendolyne Burfin, Hadrien Regue, Quentin Semanas, Martine Valette, Bruno Lina, Laurence Josset |
| EPI_ISL_3472602 | LABORATOIRE BIOESTEREL | CNR Virus des Infections Respiratoires - France SUD | Antonin Bal, Gregory Destras, Gwendolyne Burfin, Hadrien Regue, Quentin Semanas, Martine Valette, Bruno Lina, Laurence Josset |
| EPI_ISL_3472603 | CENTRE HOSPITALIER LUCIEN HUSSEL | CNR Virus des Infections Respiratoires - France SUD | Antonin Bal, Gregory Destras, Gwendolyne Burfin, Hadrien Regue, Quentin Semanas, Martine Valette, Bruno Lina, Laurence Josset |
| EPI_ISL_3472604 | CH ANTIBES - JUAN LES PINS | CNR Virus des Infections Respiratoires - France SUD | Antonin Bal, Gregory Destras, Gwendolyne Burfin, Hadrien Regue, Quentin Semanas, Martine Valette, Bruno Lina, Laurence Josset |
| EPI_ISL_3472605 | DYOMEDEA-LABORATOIRE DE LA SAUVEGARDE | CNR Virus des Infections Respiratoires - France SUD | Antonin Bal, Gregory Destras, Gwendolyne Burfin, Hadrien Regue, Quentin Semanas, Martine Valette, Bruno Lina, Laurence Josset |
| EPI_ISL_3472606 | LABORATOIRE BIOGROUP | CNR Virus des Infections Respiratoires - France SUD | Antonin Bal, Gregory Destras, Gwendolyne Burfin, Hadrien Regue, Quentin Semanas, Martine Valette, Bruno Lina, Laurence Josset |
| EPI_ISL_3472607 | CERBAILLIANCE LA REUNION | CNR Virus des Infections Respiratoires - France SUD | Antonin Bal, Gregory Destras, Gwendolyne Burfin, Hadrien Regue, Quentin Semanas, Martine Valette, Bruno Lina, Laurence Josset |
| EPI_ISL_3472608 | LAM CERBALLIANCE | CNR Virus des Infections Respiratoires - France SUD | Antonin Bal, Gregory Destras, Gwendolyne Burfin, Hadrien Regue, Quentin Semanas, Martine Valette, Bruno Lina, Laurence Josset |
| EPI_ISL_3472609 | MIRIALIS CLUSES BECHET | CNR Virus des Infections Respiratoires - France SUD | Antonin Bal, Gregory Destras, Gwendolyne Burfin, Hadrien Regue, Quentin Semanas, Martine Valette, Bruno Lina, Laurence Josset |
| EPI_ISL_3472610 | LABORATOIRE BIOESTEREL | CNR Virus des Infections Respiratoires - France SUD | Antonin Bal, Gregory Destras, Gwendolyne Burfin, Hadrien Regue, Quentin Semanas, Martine Valette, Bruno Lina, Laurence Josset |
| EPI_ISL_3472611 | LABORATOIRE UNILIANS DECINES | CNR Virus des Infections Respiratoires - France SUD | Antonin Bal, Gregory Destras, Gwendolyne Burfin, Hadrien Regue, Quentin Semanas, Martine Valette, Bruno Lina, Laurence Josset |
| EPI_ISL_3472612 | DYOMEDEA-LABORATOIRE DE LA SAUVEGARDE | CNR Virus des Infections Respiratoires - France SUD | Antonin Bal, Gregory Destras, Gwendolyne Burfin, Hadrien Regue, Quentin Semanas, Martine Valette, Bruno Lina, Laurence Josset |
| EPI_ISL_3472613 | LAM CERBALLIANCE | CNR Virus des Infections Respiratoires - France SUD | Antonin Bal, Gregory Destras, Gwendolyne Burfin, Hadrien Regue, Quentin Semanas, Martine Valette, Bruno Lina, Laurence Josset |
| EPI_ISL_3472614 | LABORATOIRE DE L'OSPEDALE - 2A 2B | CNR Virus des Infections Respiratoires - France SUD | Antonin Bal, Gregory Destras, Gwendolyne Burfin, Hadrien Regue, Quentin Semanas, Martine Valette, Bruno Lina, Laurence Josset |
| EPI_ISL_3472615 | LABORATOIRE UNILIANS DECINES | CNR Virus des Infections Respiratoires - France SUD | Antonin Bal, Gregory Destras, Gwendolyne Burfin, Hadrien Regue, Quentin Semanas, Martine Valette, Bruno Lina, Laurence Josset |
| EPI_ISL_3472616, EPI_ISL_3472617, EPI_ISL_3472618 | CNR Virus des Infections Respiratoires - France SUD | CNR Virus des Infections Respiratoires - France SUD | Antonin Bal, Gregory Destras, Gwendolyne Burfin, Hadrien Regue, Quentin Semanas, Martine Valette, Bruno Lina, Laurence Josset |
| EPI_ISL_3472619 | LAM BIORYLIS | CNR Virus des Infections Respiratoires - France SUD | Antonin Bal, Gregory Destras, Gwendolyne Burfin, Hadrien Regue, Quentin Semanas, Martine Valette, Bruno Lina, Laurence Josset |
| EPI_ISL_3472620 | CNR Virus des Infections Respiratoires - France SUD | CNR Virus des Infections Respiratoires - France SUD | Antonin Bal, Gregory Destras, Gwendolyne Burfin, Hadrien Regue, Quentin Semanas, Martine Valette, Bruno Lina, Laurence Josset |
| EPI_ISL_3472621 | DYOMEDEA-LABORATOIRE DE LA SAUVEGARDE | CNR Virus des Infections Respiratoires - France SUD | Antonin Bal, Gregory Destras, Gwendolyne Burfin, Hadrien Regue, Quentin Semanas, Martine Valette, Bruno Lina, Laurence Josset |
| EPI_ISL_3472622 | CERBAILLIANCE LA REUNION | CNR Virus des Infections Respiratoires - France SUD | Antonin Bal, Gregory Destras, Gwendolyne Burfin, Hadrien Regue, Quentin Semanas, Martine Valette, Bruno Lina, Laurence Josset |
| EPI_ISL_3472623 | LABORATOIRE DE L'OSPEDALE - 2A 2B | CNR Virus des Infections Respiratoires - France SUD | Antonin Bal, Gregory Destras, Gwendolyne Burfin, Hadrien Regue, Quentin Semanas, Martine Valette, Bruno Lina, Laurence Josset |
| EPI_ISL_3472624 | LABORATOIRE BIOGROUP | CNR Virus des Infections Respiratoires - France SUD | Antonin Bal, Gregory Destras, Gwendolyne Burfin, Hadrien Regue, Quentin Semanas, Martine Valette, Bruno Lina, Laurence Josset |
| EPI_ISL_3472625 | DYOMEDEA-LABORATOIRE DE LA SAUVEGARDE | CNR Virus des Infections Respiratoires - France SUD | Antonin Bal, Gregory Destras, Gwendolyne Burfin, Hadrien Regue, Quentin Semanas, Martine Valette, Bruno Lina, Laurence Josset |
| EPI_ISL_3472626 | BIOLITTORAL BIOGROUP PLATEAU TECH | CNR Virus des Infections Respiratoires - France SUD | Antonin Bal, Gregory Destras, Gwendolyne Burfin, Hadrien Regue, Quentin Semanas, Martine Valette, Bruno Lina, Laurence Josset |
| EPI_ISL_3472627, EPI_ISL_3472628 | MIRIALIS CLUSES BECHET | CNR Virus des Infections Respiratoires - France SUD | Antonin Bal, Gregory Destras, Gwendolyne Burfin, Hadrien Regue, Quentin Semanas, Martine Valette, Bruno Lina, Laurence Josset |
| EPI_ISL_3472629 | LABORATOIRE UNILIANS DECINES | CNR Virus des Infections Respiratoires - France SUD | Antonin Bal, Gregory Destras, Gwendolyne Burfin, Hadrien Regue, Quentin Semanas, Martine Valette, Bruno Lina, Laurence Josset |
| EPI_ISL_3472630 | DYOMEDEA-LABORATOIRE DE LA SAUVEGARDE | CNR Virus des Infections Respiratoires - France SUD | Antonin Bal, Gregory Destras, Gwendolyne Burfin, Hadrien Regue, Quentin Semanas, Martine Valette, Bruno Lina, Laurence Josset |
| EPI_ISL_3472631 | LAM CERBALLIANCE | CNR Virus des Infections Respiratoires - France SUD | Antonin Bal, Gregory Destras, Gwendolyne Burfin, Hadrien Regue, Quentin Semanas, Martine Valette, Bruno Lina, Laurence Josset |
| EPI_ISL_3472632 | LABORATOIRE DE L'OSPEDALE - 2A 2B | CNR Virus des Infections Respiratoires - France SUD | Antonin Bal, Gregory Destras, Gwendolyne Burfin, Hadrien Regue, Quentin Semanas, Martine Valette, Bruno Lina, Laurence Josset |
| EPI_ISL_3472633 | CNR Virus des Infections Respiratoires - France SUD | CNR Virus des Infections Respiratoires - France SUD | Antonin Bal, Gregory Destras, Gwendolyne Burfin, Hadrien Regue, Quentin Semanas, Martine Valette, Bruno Lina, Laurence Josset |
| EPI_ISL_3472634 | DYOMEDEA-LABORATOIRE DE LA SAUVEGARDE | CNR Virus des Infections Respiratoires - France SUD | Antonin Bal, Gregory Destras, Gwendolyne Burfin, Hadrien Regue, Quentin Semanas, Martine Valette, Bruno Lina, Laurence Josset |
| EPI_ISL_3472635, EPI_ISL_3472637 | LAM CERBALLIANCE | CNR Virus des Infections Respiratoires - France SUD | Antonin Bal, Gregory Destras, Gwendolyne Burfin, Hadrien Regue, Quentin Semanas, Martine Valette, Bruno Lina, Laurence Josset |
| EPI_ISL_3472638, EPI_ISL_3472639 | LABORATOIRE BIOGROUP | CNR Virus des Infections Respiratoires - France SUD | Antonin Bal, Gregory Destras, Gwendolyne Burfin, Hadrien Regue, Quentin Semanas, Martine Valette, Bruno Lina, Laurence Josset |
| EPI_ISL_3472640 | BIOLITTORAL BIOGROUP PLATEAU TECH | CNR Virus des Infections Respiratoires - France SUD | Antonin Bal, Gregory Destras, Gwendolyne Burfin, Hadrien Regue, Quentin Semanas, Martine Valette, Bruno Lina, Laurence Josset |
| EPI_ISL_3472641 | LABORATOIRE BIOGROUP | CNR Virus des Infections Respiratoires - France SUD | Antonin Bal, Gregory Destras, Gwendolyne Burfin, Hadrien Regue, Quentin Semanas, Martine Valette, Bruno Lina, Laurence Josset |
| EPI_ISL_3472642 | CH ANTIBES - JUAN LES PINS | CNR Virus des Infections Respiratoires - France SUD | Antonin Bal, Gregory Destras, Gwendolyne Burfin, Hadrien Regue, Quentin Semanas, Martine Valette, Bruno Lina, Laurence Josset |
| EPI_ISL_3472643, EPI_ISL_3472644 | DYOMEDEA-LABORATOIRE DE LA SAUVEGARDE | CNR Virus des Infections Respiratoires - France SUD | Antonin Bal, Gregory Destras, Gwendolyne Burfin, Hadrien Regue, Quentin Semanas, Martine Valette, Bruno Lina, Laurence Josset |
| EPI_ISL_3472645 | LABORATOIRE DE L'OSPEDALE - 2A 2B | CNR Virus des Infections Respiratoires - France SUD | Antonin Bal, Gregory Destras, Gwendolyne Burfin, Hadrien Regue, Quentin Semanas, Martine Valette, Bruno Lina, Laurence Josset |
| EPI_ISL_3472646 | LABORATOIRE BIOESTEREL | CNR Virus des Infections Respiratoires - France SUD | Antonin Bal, Gregory Destras, Gwendolyne Burfin, Hadrien Regue, Quentin Semanas, Martine Valette, Bruno Lina, Laurence Josset |
| EPI_ISL_3472647 | LABORATOIRE BIOGROUP | CNR Virus des Infections Respiratoires - France SUD | Antonin Bal, Gregory Destras, Gwendolyne Burfin, Hadrien Regue, Quentin Semanas, Martine Valette, Bruno Lina, Laurence Josset |
| EPI_ISL_3472648 | BIOLITTORAL BIOGROUP PLATEAU TECH | CNR Virus des Infections Respiratoires - France SUD | Antonin Bal, Gregory Destras, Gwendolyne Burfin, Hadrien Regue, Quentin Semanas, Martine Valette, Bruno Lina, Laurence Josset |
| EPI_ISL_3472649 | LABORATOIRE BIOFUSION | CNR Virus des Infections Respiratoires - France SUD | Antonin Bal, Gregory Destras, Gwendolyne Burfin, Hadrien Regue, Quentin Semanas, Martine Valette, Bruno Lina, Laurence Josset |
| EPI_ISL_3472650 | LAM CERBALLIANCE | CNR Virus des Infections Respiratoires - France SUD | Antonin Bal, Gregory Destras, Gwendolyne Burfin, Hadrien Regue, Quentin Semanas, Martine Valette, Bruno Lina, Laurence Josset |
| EPI_ISL_3472651 | LABORATOIRE FORTE BIO DAX | CNR Virus des Infections Respiratoires - France SUD | Antonin Bal, Gregory Destras, Gwendolyne Burfin, Hadrien Regue, Quentin Semanas, Martine Valette, Bruno Lina, Laurence Josset |
| EPI_ISL_3472652 | MIRIALIS CLUSES BECHET | CNR Virus des Infections Respiratoires - France SUD | Antonin Bal, Gregory Destras, Gwendolyne Burfin, Hadrien Regue, Quentin Semanas, Martine Valette, Bruno Lina, Laurence Josset |
| EPI_ISL_3472653 | LAM CERBALLIANCE | CNR Virus des Infections Respiratoires - France SUD | Antonin Bal, Gregory Destras, Gwendolyne Burfin, Hadrien Regue, Quentin Semanas, Martine Valette, Bruno Lina, Laurence Josset |
| EPI_ISL_3472654 | LAM BIORYLIS | CNR Virus des Infections Respiratoires - France SUD | Antonin Bal, Gregory Destras, Gwendolyne Burfin, Hadrien Regue, Quentin Semanas, Martine Valette, Bruno Lina, Laurence Josset |
| EPI_ISL_3472655 | DYOMEDEA-LABORATOIRE DE LA SAUVEGARDE | CNR Virus des Infections Respiratoires - France SUD | Antonin Bal, Gregory Destras, Gwendolyne Burfin, Hadrien Regue, Quentin Semanas, Martine Valette, Bruno Lina, Laurence Josset |
| EPI_ISL_3472656 | MIRIALIS CLUSES BECHET | CNR Virus des Infections Respiratoires - France SUD | Antonin Bal, Gregory Destras, Gwendolyne Burfin, Hadrien Regue, Quentin Semanas, Martine Valette, Bruno Lina, Laurence Josset |
| EPI_ISL_3472657 | BIOLITTORAL BIOGROUP PLATEAU TECH | CNR Virus des Infections Respiratoires - France SUD | Antonin Bal, Gregory Destras, Gwendolyne Burfin, Hadrien Regue, Quentin Semanas, Martine Valette, Bruno Lina, Laurence Josset |
| EPI_ISL_3472658 | CENTRE DE SOINS DE VIRIEU | CNR Virus des Infections Respiratoires - France SUD | Antonin Bal, Gregory Destras, Gwendolyne Burfin, Hadrien Regue, Quentin Semanas, Martine Valette, Bruno Lina, Laurence Josset |
| EPI_ISL_3472659 | BIOLITTORAL BIOGROUP PLATEAU TECH | CNR Virus des Infections Respiratoires - France SUD | Antonin Bal, Gregory Destras, Gwendolyne Burfin, Hadrien Regue, Quentin Semanas, Martine Valette, Bruno Lina, Laurence Josset |
| EPI_ISL_3472660 | MIRIALIS CLUSES BECHET | CNR Virus des Infections Respiratoires - France SUD | Antonin Bal, Gregory Destras, Gwendolyne Burfin, Hadrien Regue, Quentin Semanas, Martine Valette, Bruno Lina, Laurence Josset |
| EPI_ISL_3472661 | CENTRE HOSPITALIER DUPUYTREN | CNR Virus des Infections Respiratoires - France SUD | Antonin Bal, Gregory Destras, Gwendolyne Burfin, Hadrien Regue, Quentin Semanas, Martine Valette, Bruno Lina, Laurence Josset |
| EPI_ISL_3472662 | CNR Virus des Infections Respiratoires - France SUD | CNR Virus des Infections Respiratoires - France SUD | Antonin Bal, Gregory Destras, Gwendolyne Burfin, Hadrien Regue, Quentin Semanas, Martine Valette, Bruno Lina, Laurence Josset |
| EPI_ISL_3472663, EPI_ISL_3472665 | LABORATOIRE UNILIANS DECINES | CNR Virus des Infections Respiratoires - France SUD | Antonin Bal, Gregory Destras, Gwendolyne Burfin, Hadrien Regue, Quentin Semanas, Martine Valette, Bruno Lina, Laurence Josset |
| EPI_ISL_3472667 | LAM BAYARD-BIOPYRENEES | CNR Virus des Infections Respiratoires - France SUD | Antonin Bal, Gregory Destras, Gwendolyne Burfin, Hadrien Regue, Quentin Semanas, Martine Valette, Bruno Lina, Laurence Josset |
| EPI_ISL_3472668 | LABORATOIRE BIOESTEREL | CNR Virus des Infections Respiratoires - France SUD | Antonin Bal, Gregory Destras, Gwendolyne Burfin, Hadrien Regue, Quentin Semanas, Martine Valette, Bruno Lina, Laurence Josset |
| EPI_ISL_3472669 | LABORATOIRE UNILIANS DECINES | CNR Virus des Infections Respiratoires - France SUD | Antonin Bal, Gregory Destras, Gwendolyne Burfin, Hadrien Regue, Quentin Semanas, Martine Valette, Bruno Lina, Laurence Josset |
| EPI_ISL_3472670 | LAM CERBALLIANCE | CNR Virus des Infections Respiratoires - France SUD | Antonin Bal, Gregory Destras, Gwendolyne Burfin, Hadrien Regue, Quentin Semanas, Martine Valette, Bruno Lina, Laurence Josset |
| EPI_ISL_3472671 | LAM SYNLAB BORDEAUX ATLANTIQUE | CNR Virus des Infections Respiratoires - France SUD | Antonin Bal, Gregory Destras, Gwendolyne Burfin, Hadrien Regue, Quentin Semanas, Martine Valette, Bruno Lina, Laurence Josset |

| | | | |
|---|---|---|---|
| EPI_ISL_3472673 | MIRIALIS CLUSES BECHET | CNR Virus des Infections Respiratoires - France SUD | Antonin Bal, Gregory Destras, Gwendolyne Burfin, Hadrien Regue, Quentin Semanas, Martine Valette, Bruno Lina, Laurence Josset |
| EPI_ISL_3472674 | CERBAILLIANCE LA REUNION | CNR Virus des Infections Respiratoires - France SUD | Antonin Bal, Gregory Destras, Gwendolyne Burfin, Hadrien Regue, Quentin Semanas, Martine Valette, Bruno Lina, Laurence Josset |
| EPI_ISL_3472675 | LABORATOIRE FORTE BIO DAX | CNR Virus des Infections Respiratoires - France SUD | Antonin Bal, Gregory Destras, Gwendolyne Burfin, Hadrien Regue, Quentin Semanas, Martine Valette, Bruno Lina, Laurence Josset |
| EPI_ISL_3472676 | DYOMEDEA-LABORATOIRE DE LA SAUVEGARDE | CNR Virus des Infections Respiratoires - France SUD | Antonin Bal, Gregory Destras, Gwendolyne Burfin, Hadrien Regue, Quentin Semanas, Martine Valette, Bruno Lina, Laurence Josset |
| EPI_ISL_3472677 | MIRIALIS CLUSES BECHET | CNR Virus des Infections Respiratoires - France SUD | Antonin Bal, Gregory Destras, Gwendolyne Burfin, Hadrien Regue, Quentin Semanas, Martine Valette, Bruno Lina, Laurence Josset |
| EPI_ISL_3472678 | LABORATOIRE BIOGROUP | CNR Virus des Infections Respiratoires - France SUD | Antonin Bal, Gregory Destras, Gwendolyne Burfin, Hadrien Regue, Quentin Semanas, Martine Valette, Bruno Lina, Laurence Josset |
| EPI_ISL_3472679 | LABORATOIRE UNILIANS DECINES | CNR Virus des Infections Respiratoires - France SUD | Antonin Bal, Gregory Destras, Gwendolyne Burfin, Hadrien Regue, Quentin Semanas, Martine Valette, Bruno Lina, Laurence Josset |
| EPI_ISL_3472680 | LAM BIORYLIS | CNR Virus des Infections Respiratoires - France SUD | Antonin Bal, Gregory Destras, Gwendolyne Burfin, Hadrien Regue, Quentin Semanas, Martine Valette, Bruno Lina, Laurence Josset |
| EPI_ISL_3472681 | DYOMEDEA-LABORATOIRE DE LA SAUVEGARDE | CNR Virus des Infections Respiratoires - France SUD | Antonin Bal, Gregory Destras, Gwendolyne Burfin, Hadrien Regue, Quentin Semanas, Martine Valette, Bruno Lina, Laurence Josset |
| EPI_ISL_3472682 | LAM BIORYLIS | CNR Virus des Infections Respiratoires - France SUD | Antonin Bal, Gregory Destras, Gwendolyne Burfin, Hadrien Regue, Quentin Semanas, Martine Valette, Bruno Lina, Laurence Josset |
| EPI_ISL_3472683 | BIOLITTORAL BIOGROUP PLATEAU TECH | CNR Virus des Infections Respiratoires - France SUD | Antonin Bal, Gregory Destras, Gwendolyne Burfin, Hadrien Regue, Quentin Semanas, Martine Valette, Bruno Lina, Laurence Josset |
| EPI_ISL_3472684 | LAM BAYARD-BIOPYRENEES | CNR Virus des Infections Respiratoires - France SUD | Antonin Bal, Gregory Destras, Gwendolyne Burfin, Hadrien Regue, Quentin Semanas, Martine Valette, Bruno Lina, Laurence Josset |
| EPI_ISL_3472685 | CERBAILLIANCE LA REUNION | CNR Virus des Infections Respiratoires - France SUD | Antonin Bal, Gregory Destras, Gwendolyne Burfin, Hadrien Regue, Quentin Semanas, Martine Valette, Bruno Lina, Laurence Josset |
| EPI_ISL_3472686 | C. H. COMMINGES PYRENEES | CNR Virus des Infections Respiratoires - France SUD | Antonin Bal, Gregory Destras, Gwendolyne Burfin, Hadrien Regue, Quentin Semanas, Martine Valette, Bruno Lina, Laurence Josset |
| EPI_ISL_3472687 | LABORATOIRE UNILIANS DECINES | CNR Virus des Infections Respiratoires - France SUD | Antonin Bal, Gregory Destras, Gwendolyne Burfin, Hadrien Regue, Quentin Semanas, Martine Valette, Bruno Lina, Laurence Josset |
| EPI_ISL_3472688 | LAM SYNLAB BORDEAUX ATLANTIQUE | CNR Virus des Infections Respiratoires - France SUD | Antonin Bal, Gregory Destras, Gwendolyne Burfin, Hadrien Regue, Quentin Semanas, Martine Valette, Bruno Lina, Laurence Josset |
| EPI_ISL_3472690 | BIOLITTORAL BIOGROUP PLATEAU TECH | CNR Virus des Infections Respiratoires - France SUD | Antonin Bal, Gregory Destras, Gwendolyne Burfin, Hadrien Regue, Quentin Semanas, Martine Valette, Bruno Lina, Laurence Josset |
| EPI_ISL_3472691 | CH GIVORS | CNR Virus des Infections Respiratoires - France SUD | Antonin Bal, Gregory Destras, Gwendolyne Burfin, Hadrien Regue, Quentin Semanas, Martine Valette, Bruno Lina, Laurence Josset |
| EPI_ISL_3472692, EPI_ISL_3472693, EPI_ISL_3472695 | DYOMEDEA-LABORATOIRE DE LA SAUVEGARDE | CNR Virus des Infections Respiratoires - France SUD | Antonin Bal, Gregory Destras, Gwendolyne Burfin, Hadrien Regue, Quentin Semanas, Martine Valette, Bruno Lina, Laurence Josset |
| EPI_ISL_3472696 | LABORATOIRE BIOGROUP | CNR Virus des Infections Respiratoires - France SUD | Antonin Bal, Gregory Destras, Gwendolyne Burfin, Hadrien Regue, Quentin Semanas, Martine Valette, Bruno Lina, Laurence Josset |
| EPI_ISL_3472697, EPI_ISL_3472698 | LABORATOIRE UNILIANS DECINES | CNR Virus des Infections Respiratoires - France SUD | Antonin Bal, Gregory Destras, Gwendolyne Burfin, Hadrien Regue, Quentin Semanas, Martine Valette, Bruno Lina, Laurence Josset |
| EPI_ISL_3472699 | DYOMEDEA-LABORATOIRE DE LA SAUVEGARDE | CNR Virus des Infections Respiratoires - France SUD | Antonin Bal, Gregory Destras, Gwendolyne Burfin, Hadrien Regue, Quentin Semanas, Martine Valette, Bruno Lina, Laurence Josset |
| EPI_ISL_3472700, EPI_ISL_3472701, EPI_ISL_3472702 | CNR Virus des Infections Respiratoires - France SUD | CNR Virus des Infections Respiratoires - France SUD | Antonin Bal, Gregory Destras, Gwendolyne Burfin, Hadrien Regue, Quentin Semanas, Martine Valette, Bruno Lina, Laurence Josset |
| EPI_ISL_3472703 | MIRIALIS CLUSES BECHET | CNR Virus des Infections Respiratoires - France SUD | Antonin Bal, Gregory Destras, Gwendolyne Burfin, Hadrien Regue, Quentin Semanas, Martine Valette, Bruno Lina, Laurence Josset |
| EPI_ISL_3472704 | BIOLITTORAL BIOGROUP PLATEAU TECH | CNR Virus des Infections Respiratoires - France SUD | Antonin Bal, Gregory Destras, Gwendolyne Burfin, Hadrien Regue, Quentin Semanas, Martine Valette, Bruno Lina, Laurence Josset |
| EPI_ISL_3472705 | LAM EIMER LENYS | CNR Virus des Infections Respiratoires - France SUD | Antonin Bal, Gregory Destras, Gwendolyne Burfin, Hadrien Regue, Quentin Semanas, Martine Valette, Bruno Lina, Laurence Josset |
| EPI_ISL_3472706 | CERBAILLIANCE LA REUNION | CNR Virus des Infections Respiratoires - France SUD | Antonin Bal, Gregory Destras, Gwendolyne Burfin, Hadrien Regue, Quentin Semanas, Martine Valette, Bruno Lina, Laurence Josset |
| EPI_ISL_3472707, EPI_ISL_3472708 | CNR Virus des Infections Respiratoires - France SUD | CNR Virus des Infections Respiratoires - France SUD | Antonin Bal, Gregory Destras, Gwendolyne Burfin, Hadrien Regue, Quentin Semanas, Martine Valette, Bruno Lina, Laurence Josset |
| EPI_ISL_3472709 | LABORATOIRE BIOGROUP | CNR Virus des Infections Respiratoires - France SUD | Antonin Bal, Gregory Destras, Gwendolyne Burfin, Hadrien Regue, Quentin Semanas, Martine Valette, Bruno Lina, Laurence Josset |
| EPI_ISL_3472710 | DYOMEDEA-LABORATOIRE DE LA SAUVEGARDE | CNR Virus des Infections Respiratoires - France SUD | Antonin Bal, Gregory Destras, Gwendolyne Burfin, Hadrien Regue, Quentin Semanas, Martine Valette, Bruno Lina, Laurence Josset |
| EPI_ISL_3472711 | LABORATOIRE BIOESTEREL | CNR Virus des Infections Respiratoires - France SUD | Antonin Bal, Gregory Destras, Gwendolyne Burfin, Hadrien Regue, Quentin Semanas, Martine Valette, Bruno Lina, Laurence Josset |
| EPI_ISL_3472712 | LABORATOIRE UNILIANS DECINES | CNR Virus des Infections Respiratoires - France SUD | Antonin Bal, Gregory Destras, Gwendolyne Burfin, Hadrien Regue, Quentin Semanas, Martine Valette, Bruno Lina, Laurence Josset |
| EPI_ISL_3472713 | LABORATOIRE BIOGROUP | CNR Virus des Infections Respiratoires - France SUD | Antonin Bal, Gregory Destras, Gwendolyne Burfin, Hadrien Regue, Quentin Semanas, Martine Valette, Bruno Lina, Laurence Josset |
| EPI_ISL_3472714 | LAM CERBALLIANCE | CNR Virus des Infections Respiratoires - France SUD | Antonin Bal, Gregory Destras, Gwendolyne Burfin, Hadrien Regue, Quentin Semanas, Martine Valette, Bruno Lina, Laurence Josset |
| EPI_ISL_3472715 | LABORATOIRE FORTE BIO DAX | CNR Virus des Infections Respiratoires - France SUD | Antonin Bal, Gregory Destras, Gwendolyne Burfin, Hadrien Regue, Quentin Semanas, Martine Valette, Bruno Lina, Laurence Josset |
| EPI_ISL_3472716 | LAM SYNLAB BORDEAUX ATLANTIQUE | CNR Virus des Infections Respiratoires - France SUD | Antonin Bal, Gregory Destras, Gwendolyne Burfin, Hadrien Regue, Quentin Semanas, Martine Valette, Bruno Lina, Laurence Josset |
| EPI_ISL_3472717 | LABORATOIRE UNILIANS DECINES | CNR Virus des Infections Respiratoires - France SUD | Antonin Bal, Gregory Destras, Gwendolyne Burfin, Hadrien Regue, Quentin Semanas, Martine Valette, Bruno Lina, Laurence Josset |
| EPI_ISL_3472718 | LABORATOIRE BIOESTEREL | CNR Virus des Infections Respiratoires - France SUD | Antonin Bal, Gregory Destras, Gwendolyne Burfin, Hadrien Regue, Quentin Semanas, Martine Valette, Bruno Lina, Laurence Josset |
| EPI_ISL_3472719 | DYOMEDEA-LABORATOIRE DE LA SAUVEGARDE | CNR Virus des Infections Respiratoires - France SUD | Antonin Bal, Gregory Destras, Gwendolyne Burfin, Hadrien Regue, Quentin Semanas, Martine Valette, Bruno Lina, Laurence Josset |
| EPI_ISL_3472720 | CNR Virus des Infections Respiratoires - France SUD | CNR Virus des Infections Respiratoires - France SUD | Antonin Bal, Gregory Destras, Gwendolyne Burfin, Hadrien Regue, Quentin Semanas, Martine Valette, Bruno Lina, Laurence Josset |
| EPI_ISL_3472721 | CENTRE HOSPITALIER DUPUYTREN | CNR Virus des Infections Respiratoires - France SUD | Antonin Bal, Gregory Destras, Gwendolyne Burfin, Hadrien Regue, Quentin Semanas, Martine Valette, Bruno Lina, Laurence Josset |
| EPI_ISL_3472722 | DYOMEDEA-LABORATOIRE DE LA SAUVEGARDE | CNR Virus des Infections Respiratoires - France SUD | Antonin Bal, Gregory Destras, Gwendolyne Burfin, Hadrien Regue, Quentin Semanas, Martine Valette, Bruno Lina, Laurence Josset |
| EPI_ISL_3472723 | LAM CERBALLIANCE | CNR Virus des Infections Respiratoires - France SUD | Antonin Bal, Gregory Destras, Gwendolyne Burfin, Hadrien Regue, Quentin Semanas, Martine Valette, Bruno Lina, Laurence Josset |
| EPI_ISL_3472724, EPI_ISL_3472725 | BIOLITTORAL BIOGROUP PLATEAU TECH | CNR Virus des Infections Respiratoires - France SUD | Antonin Bal, Gregory Destras, Gwendolyne Burfin, Hadrien Regue, Quentin Semanas, Martine Valette, Bruno Lina, Laurence Josset |
| EPI_ISL_3472726 | LABORATOIRE UNILIANS DECINES | CNR Virus des Infections Respiratoires - France SUD | Antonin Bal, Gregory Destras, Gwendolyne Burfin, Hadrien Regue, Quentin Semanas, Martine Valette, Bruno Lina, Laurence Josset |
| EPI_ISL_3472727, EPI_ISL_3472728 | LABORATOIRE BIO-VAL SALEVE | CNR Virus des Infections Respiratoires - France SUD | Antonin Bal, Gregory Destras, Gwendolyne Burfin, Hadrien Regue, Quentin Semanas, Martine Valette, Bruno Lina, Laurence Josset |
| EPI_ISL_3472741 | LABORATOIRE BIOESTEREL | CNR Virus des Infections Respiratoires - France SUD | Antonin Bal, Gregory Destras, Gwendolyne Burfin, Hadrien Regue, Quentin Semanas, Martine Valette, Bruno Lina, Laurence Josset |
| EPI_ISL_3472748 | LAM BIORYLIS | CNR Virus des Infections Respiratoires - France SUD | Antonin Bal, Gregory Destras, Gwendolyne Burfin, Hadrien Regue, Quentin Semanas, Martine Valette, Bruno Lina, Laurence Josset |
| EPI_ISL_3472749 | DYOMEDEA-LABORATOIRE DE LA SAUVEGARDE | CNR Virus des Infections Respiratoires - France SUD | Antonin Bal, Gregory Destras, Gwendolyne Burfin, Hadrien Regue, Quentin Semanas, Martine Valette, Bruno Lina, Laurence Josset |
| EPI_ISL_3472750 | LAM BAYARD-BIOPYRENEES | CNR Virus des Infections Respiratoires - France SUD | Antonin Bal, Gregory Destras, Gwendolyne Burfin, Hadrien Regue, Quentin Semanas, Martine Valette, Bruno Lina, Laurence Josset |
| EPI_ISL_3472751 | LAM SYNLAB BORDEAUX ATLANTIQUE | CNR Virus des Infections Respiratoires - France SUD | Antonin Bal, Gregory Destras, Gwendolyne Burfin, Hadrien Regue, Quentin Semanas, Martine Valette, Bruno Lina, Laurence Josset |
| EPI_ISL_3472752 | CNR Virus des Infections Respiratoires - France SUD | CNR Virus des Infections Respiratoires - France SUD | Antonin Bal, Gregory Destras, Gwendolyne Burfin, Hadrien Regue, Quentin Semanas, Martine Valette, Bruno Lina, Laurence Josset |
| EPI_ISL_3472753, EPI_ISL_3472754 | LABORATOIRE DE L'OSPEDALE - 2A 2B | CNR Virus des Infections Respiratoires - France SUD | Antonin Bal, Gregory Destras, Gwendolyne Burfin, Hadrien Regue, Quentin Semanas, Martine Valette, Bruno Lina, Laurence Josset |
| EPI_ISL_3472755 | LAM CERBALLIANCE | CNR Virus des Infections Respiratoires - France SUD | Antonin Bal, Gregory Destras, Gwendolyne Burfin, Hadrien Regue, Quentin Semanas, Martine Valette, Bruno Lina, Laurence Josset |
| EPI_ISL_3472756 | C. H. COMMINGES PYRENEES | CNR Virus des Infections Respiratoires - France SUD | Antonin Bal, Gregory Destras, Gwendolyne Burfin, Hadrien Regue, Quentin Semanas, Martine Valette, Bruno Lina, Laurence Josset |
| EPI_ISL_3472757 | LAM EIMER LENYS | CNR Virus des Infections Respiratoires - France SUD | Antonin Bal, Gregory Destras, Gwendolyne Burfin, Hadrien Regue, Quentin Semanas, Martine Valette, Bruno Lina, Laurence Josset |
| EPI_ISL_3472758, EPI_ISL_3472759 | LABORATOIRE DE L'OSPEDALE - 2A 2B | CNR Virus des Infections Respiratoires - France SUD | Antonin Bal, Gregory Destras, Gwendolyne Burfin, Hadrien Regue, Quentin Semanas, Martine Valette, Bruno Lina, Laurence Josset |
| EPI_ISL_3472760 | LAM SYNLAB BORDEAUX ATLANTIQUE | CNR Virus des Infections Respiratoires - France SUD | Antonin Bal, Gregory Destras, Gwendolyne Burfin, Hadrien Regue, Quentin Semanas, Martine Valette, Bruno Lina, Laurence Josset |
| EPI_ISL_3472761 | LABORATOIRE BIOESTEREL | CNR Virus des Infections Respiratoires - France SUD | Antonin Bal, Gregory Destras, Gwendolyne Burfin, Hadrien Regue, Quentin Semanas, Martine Valette, Bruno Lina, Laurence Josset |
| EPI_ISL_3472762 | DYOMEDEA-LABORATOIRE DE LA SAUVEGARDE | CNR Virus des Infections Respiratoires - France SUD | Antonin Bal, Gregory Destras, Gwendolyne Burfin, Hadrien Regue, Quentin Semanas, Martine Valette, Bruno Lina, Laurence Josset |
| EPI_ISL_3472763 | LABORATOIRE UNILIANS DECINES | CNR Virus des Infections Respiratoires - France SUD | Antonin Bal, Gregory Destras, Gwendolyne Burfin, Hadrien Regue, Quentin Semanas, Martine Valette, Bruno Lina, Laurence Josset |
| EPI_ISL_3472764, EPI_ISL_3472765 | LAM SYNLAB BORDEAUX ATLANTIQUE | CNR Virus des Infections Respiratoires - France SUD | Antonin Bal, Gregory Destras, Gwendolyne Burfin, Hadrien Regue, Quentin Semanas, Martine Valette, Bruno Lina, Laurence Josset |
| EPI_ISL_3472766, EPI_ISL_3472767, EPI_ISL_3472768, EPI_ISL_3472769 | CNR Virus des Infections Respiratoires - France SUD | CNR Virus des Infections Respiratoires - France SUD | Antonin Bal, Gregory Destras, Gwendolyne Burfin, Hadrien Regue, Quentin Semanas, Martine Valette, Bruno Lina, Laurence Josset |
| EPI_ISL_3472771 | CENTRE HOSPITALIER DUPUYTREN | CNR Virus des Infections Respiratoires - France SUD | Antonin Bal, Gregory Destras, Gwendolyne Burfin, Hadrien Regue, Quentin Semanas, Martine Valette, Bruno Lina, Laurence Josset |
| EPI_ISL_3472772 | CHU ST ETIENNE HOPITAL NORD | CNR Virus des Infections Respiratoires - France SUD | Antonin Bal, Gregory Destras, Gwendolyne Burfin, Hadrien Regue, Quentin Semanas, Martine Valette, Bruno Lina, Laurence Josset |
| EPI_ISL_3472773 | LABORATOIRE FORTE BIO DAX | CNR Virus des Infections Respiratoires - France SUD | Antonin Bal, Gregory Destras, Gwendolyne Burfin, Hadrien Regue, Quentin Semanas, Martine Valette, Bruno Lina, Laurence Josset |
| EPI_ISL_3472774 | LABORATOIRE BIO-VAL ALLART-BETEND | CNR Virus des Infections Respiratoires - France SUD | Antonin Bal, Gregory Destras, Gwendolyne Burfin, Hadrien Regue, Quentin Semanas, Martine Valette, Bruno Lina, Laurence Josset |

| | | | |
|---|---|---|---|
| EPI_ISL_3472781 | CNR Virus des Infections Respiratoires - France SUD | CNR Virus des Infections Respiratoires - France SUD | Antonin Bal, Gregory Destras, Gwendolyne Burfin, Hadrien Regue, Quentin Semanas, Martine Valette, Bruno Lina, Laurence Josset |
| EPI_ISL_3472782 | LABORATOIRE UNILIANS DECINES | CNR Virus des Infections Respiratoires - France SUD | Antonin Bal, Gregory Destras, Gwendolyne Burfin, Hadrien Regue, Quentin Semanas, Martine Valette, Bruno Lina, Laurence Josset |
| EPI_ISL_3472783 | LABORATOIRE BIOESTEREL | CNR Virus des Infections Respiratoires - France SUD | Antonin Bal, Gregory Destras, Gwendolyne Burfin, Hadrien Regue, Quentin Semanas, Martine Valette, Bruno Lina, Laurence Josset |
| EPI_ISL_3472784 | LAM CERBALLIANCE | CNR Virus des Infections Respiratoires - France SUD | Antonin Bal, Gregory Destras, Gwendolyne Burfin, Hadrien Regue, Quentin Semanas, Martine Valette, Bruno Lina, Laurence Josset |
| EPI_ISL_3472785, EPI_ISL_3472786, EPI_ISL_3472787 | LABORATOIRE FORTE BIO DAX | CNR Virus des Infections Respiratoires - France SUD | Antonin Bal, Gregory Destras, Gwendolyne Burfin, Hadrien Regue, Quentin Semanas, Martine Valette, Bruno Lina, Laurence Josset |
| EPI_ISL_3472788, EPI_ISL_3472789, EPI_ISL_3472790, EPI_ISL_3472791, EPI_ISL_3472792, EPI_ISL_3472793, EPI_ISL_3472794 | LAM EIMER LENYS | CNR Virus des Infections Respiratoires - France SUD | Antonin Bal, Gregory Destras, Gwendolyne Burfin, Hadrien Regue, Quentin Semanas, Martine Valette, Bruno Lina, Laurence Josset |
| EPI_ISL_3472795 | DYOMEDEA-LABORATOIRE DE LA SAUVEGARDE | CNR Virus des Infections Respiratoires - France SUD | Antonin Bal, Gregory Destras, Gwendolyne Burfin, Hadrien Regue, Quentin Semanas, Martine Valette, Bruno Lina, Laurence Josset |
| EPI_ISL_3472796 | LABORATOIRE BIOGROUP | CNR Virus des Infections Respiratoires - France SUD | Antonin Bal, Gregory Destras, Gwendolyne Burfin, Hadrien Regue, Quentin Semanas, Martine Valette, Bruno Lina, Laurence Josset |
| EPI_ISL_3472797 | CNR Virus des Infections Respiratoires - France SUD | CNR Virus des Infections Respiratoires - France SUD | Antonin Bal, Gregory Destras, Gwendolyne Burfin, Hadrien Regue, Quentin Semanas, Martine Valette, Bruno Lina, Laurence Josset |
| EPI_ISL_3472798 | LABORATOIRE DEPARTEMENTAL D'ANALYSES | CNR Virus des Infections Respiratoires - France SUD | Antonin Bal, Gregory Destras, Gwendolyne Burfin, Hadrien Regue, Quentin Semanas, Martine Valette, Bruno Lina, Laurence Josset |
| EPI_ISL_3472810 | LABORATOIRE FORTE BIO DAX | CNR Virus des Infections Respiratoires - France SUD | Antonin Bal, Gregory Destras, Gwendolyne Burfin, Hadrien Regue, Quentin Semanas, Martine Valette, Bruno Lina, Laurence Josset |
| EPI_ISL_3472811 | CERBAILLIANCE LA REUNION | CNR Virus des Infections Respiratoires - France SUD | Antonin Bal, Gregory Destras, Gwendolyne Burfin, Hadrien Regue, Quentin Semanas, Martine Valette, Bruno Lina, Laurence Josset |
| EPI_ISL_3472812 | LABORATOIRE UNILIANS DECINES | CNR Virus des Infections Respiratoires - France SUD | Antonin Bal, Gregory Destras, Gwendolyne Burfin, Hadrien Regue, Quentin Semanas, Martine Valette, Bruno Lina, Laurence Josset |
| EPI_ISL_3472813 | BIOLITTORAL BIOGROUP PLATEAU TECH | CNR Virus des Infections Respiratoires - France SUD | Antonin Bal, Gregory Destras, Gwendolyne Burfin, Hadrien Regue, Quentin Semanas, Martine Valette, Bruno Lina, Laurence Josset |
| EPI_ISL_3472814 | CERBAILLIANCE LA REUNION | CNR Virus des Infections Respiratoires - France SUD | Antonin Bal, Gregory Destras, Gwendolyne Burfin, Hadrien Regue, Quentin Semanas, Martine Valette, Bruno Lina, Laurence Josset |
| EPI_ISL_3472815, EPI_ISL_3472816 | SYNLAB AQUITAINE | CNR Virus des Infections Respiratoires - France SUD | Antonin Bal, Gregory Destras, Gwendolyne Burfin, Hadrien Regue, Quentin Semanas, Martine Valette, Bruno Lina, Laurence Josset |
| EPI_ISL_3472817 | CENTRE HOSPITALIER PIERRE OUDOT | CNR Virus des Infections Respiratoires - France SUD | Antonin Bal, Gregory Destras, Gwendolyne Burfin, Hadrien Regue, Quentin Semanas, Martine Valette, Bruno Lina, Laurence Josset |
| EPI_ISL_3472818 | CNR Virus des Infections Respiratoires - France SUD | CNR Virus des Infections Respiratoires - France SUD | Antonin Bal, Gregory Destras, Gwendolyne Burfin, Hadrien Regue, Quentin Semanas, Martine Valette, Bruno Lina, Laurence Josset |
| EPI_ISL_3472819 | CENTRE HOSPITALIER PIERRE OUDOT | CNR Virus des Infections Respiratoires - France SUD | Antonin Bal, Gregory Destras, Gwendolyne Burfin, Hadrien Regue, Quentin Semanas, Martine Valette, Bruno Lina, Laurence Josset |
| EPI_ISL_3472820 | LABORATOIRE BIOFUSION | CNR Virus des Infections Respiratoires - France SUD | Antonin Bal, Gregory Destras, Gwendolyne Burfin, Hadrien Regue, Quentin Semanas, Martine Valette, Bruno Lina, Laurence Josset |
| EPI_ISL_3472821, EPI_ISL_3472822, EPI_ISL_3472823, EPI_ISL_3472824, EPI_ISL_3472825, EPI_ISL_3472826, EPI_ISL_3472827, EPI_ISL_3472828, EPI_ISL_3472829, EPI_ISL_3472830, EPI_ISL_3472831, EPI_ISL_3472832, EPI_ISL_3472833 | | | |
| see above | LAM BAYARD-BIOPYRENEES | CNR Virus des Infections Respiratoires - France SUD | Antonin Bal, Gregory Destras, Gwendolyne Burfin, Hadrien Regue, Quentin Semanas, Martine Valette, Bruno Lina, Laurence Josset |
| EPI_ISL_3472834, EPI_ISL_3472835, EPI_ISL_3472836, EPI_ISL_3472837 | LAM CERBALLIANCE | CNR Virus des Infections Respiratoires - France SUD | Antonin Bal, Gregory Destras, Gwendolyne Burfin, Hadrien Regue, Quentin Semanas, Martine Valette, Bruno Lina, Laurence Josset |
| EPI_ISL_3472838 | CH GIVORS | CNR Virus des Infections Respiratoires - France SUD | Antonin Bal, Gregory Destras, Gwendolyne Burfin, Hadrien Regue, Quentin Semanas, Martine Valette, Bruno Lina, Laurence Josset |
| EPI_ISL_3472839, EPI_ISL_3472840, EPI_ISL_3472841, EPI_ISL_3472842, EPI_ISL_3472843, EPI_ISL_3472844, EPI_ISL_3472845 | LAM CERBALLIANCE | CNR Virus des Infections Respiratoires - France SUD | Antonin Bal, Gregory Destras, Gwendolyne Burfin, Hadrien Regue, Quentin Semanas, Martine Valette, Bruno Lina, Laurence Josset |
| EPI_ISL_3472846 | CNR Virus des Infections Respiratoires - France SUD | CNR Virus des Infections Respiratoires - France SUD | Antonin Bal, Gregory Destras, Gwendolyne Burfin, Hadrien Regue, Quentin Semanas, Martine Valette, Bruno Lina, Laurence Josset |
| EPI_ISL_3472847 | CH GIVORS | CNR Virus des Infections Respiratoires - France SUD | Antonin Bal, Gregory Destras, Gwendolyne Burfin, Hadrien Regue, Quentin Semanas, Martine Valette, Bruno Lina, Laurence Josset |
| EPI_ISL_3472848, EPI_ISL_3472849, EPI_ISL_3472850, EPI_ISL_3472851, EPI_ISL_3472852 | CNR Virus des Infections Respiratoires - France SUD | CNR Virus des Infections Respiratoires - France SUD | Antonin Bal, Gregory Destras, Gwendolyne Burfin, Hadrien Regue, Quentin Semanas, Martine Valette, Bruno Lina, Laurence Josset |
| EPI_ISL_3472853, EPI_ISL_3472854, EPI_ISL_3472855, EPI_ISL_3472856, EPI_ISL_3472857, EPI_ISL_3472858, EPI_ISL_3472859 | LABORATOIRE FORTE BIO DAX | CNR Virus des Infections Respiratoires - France SUD | Antonin Bal, Gregory Destras, Gwendolyne Burfin, Hadrien Regue, Quentin Semanas, Martine Valette, Bruno Lina, Laurence Josset |
| EPI_ISL_3472860, EPI_ISL_3472861 | LABORATOIRE DE L'OSPEDALE - 2A 2B | CNR Virus des Infections Respiratoires - France SUD | Antonin Bal, Gregory Destras, Gwendolyne Burfin, Hadrien Regue, Quentin Semanas, Martine Valette, Bruno Lina, Laurence Josset |
| EPI_ISL_3472862, EPI_ISL_3472863, EPI_ISL_3472864, EPI_ISL_3472865, EPI_ISL_3472866, EPI_ISL_3472867, EPI_ISL_3472868, EPI_ISL_3472869, EPI_ISL_3472870, EPI_ISL_3472871, EPI_ISL_3472872, EPI_ISL_3472873, EPI_ISL_3472874, EPI_ISL_3472875, EPI_ISL_3472876, EPI_ISL_3472877, EPI_ISL_3472878, EPI_ISL_3472879, EPI_ISL_3472880, EPI_ISL_3472881, EPI_ISL_3472882, EPI_ISL_3472883, EPI_ISL_3472884, EPI_ISL_3472885, EPI_ISL_3472886 | | | |
| see above | LABORATOIRE BIOESTEREL | CNR Virus des Infections Respiratoires - France SUD | Antonin Bal, Gregory Destras, Gwendolyne Burfin, Hadrien Regue, Quentin Semanas, Martine Valette, Bruno Lina, Laurence Josset |
| EPI_ISL_3472887 | CENTRE HOSPITALIER DE VALENCE | CNR Virus des Infections Respiratoires - France SUD | Antonin Bal, Gregory Destras, Gwendolyne Burfin, Hadrien Regue, Quentin Semanas, Martine Valette, Bruno Lina, Laurence Josset |
| EPI_ISL_3472888 | LABORATOIRE FORTE BIO DAX | CNR Virus des Infections Respiratoires - France SUD | Antonin Bal, Gregory Destras, Gwendolyne Burfin, Hadrien Regue, Quentin Semanas, Martine Valette, Bruno Lina, Laurence Josset |
| EPI_ISL_3472889 | LABORATOIRE BIOESTEREL | CNR Virus des Infections Respiratoires - France SUD | Antonin Bal, Gregory Destras, Gwendolyne Burfin, Hadrien Regue, Quentin Semanas, Martine Valette, Bruno Lina, Laurence Josset |
| EPI_ISL_3472890, EPI_ISL_3472891 | LABORATOIRE FORTE BIO DAX | CNR Virus des Infections Respiratoires - France SUD | Antonin Bal, Gregory Destras, Gwendolyne Burfin, Hadrien Regue, Quentin Semanas, Martine Valette, Bruno Lina, Laurence Josset |
| EPI_ISL_3472892 | LABORATOIRE BIOESTEREL | CNR Virus des Infections Respiratoires - France SUD | Antonin Bal, Gregory Destras, Gwendolyne Burfin, Hadrien Regue, Quentin Semanas, Martine Valette, Bruno Lina, Laurence Josset |
| EPI_ISL_3472893, EPI_ISL_3472894, EPI_ISL_3472895 | LABORATOIRE FORTE BIO DAX | CNR Virus des Infections Respiratoires - France SUD | Antonin Bal, Gregory Destras, Gwendolyne Burfin, Hadrien Regue, Quentin Semanas, Martine Valette, Bruno Lina, Laurence Josset |
| EPI_ISL_3472896, EPI_ISL_3472897 | LABORATOIRE BIOESTEREL | CNR Virus des Infections Respiratoires - France SUD | Antonin Bal, Gregory Destras, Gwendolyne Burfin, Hadrien Regue, Quentin Semanas, Martine Valette, Bruno Lina, Laurence Josset |
| EPI_ISL_3472898 | LABORATOIRE FORTE BIO DAX | CNR Virus des Infections Respiratoires - France SUD | Antonin Bal, Gregory Destras, Gwendolyne Burfin, Hadrien Regue, Quentin Semanas, Martine Valette, Bruno Lina, Laurence Josset |
| EPI_ISL_3472899 | LABORATOIRE BIOESTEREL | CNR Virus des Infections Respiratoires - France SUD | Antonin Bal, Gregory Destras, Gwendolyne Burfin, Hadrien Regue, Quentin Semanas, Martine Valette, Bruno Lina, Laurence Josset |
| EPI_ISL_3472900 | LABORATOIRE FORTE BIO DAX | CNR Virus des Infections Respiratoires - France SUD | Antonin Bal, Gregory Destras, Gwendolyne Burfin, Hadrien Regue, Quentin Semanas, Martine Valette, Bruno Lina, Laurence Josset |
| EPI_ISL_3472901 | LABORATOIRE BIOESTEREL | CNR Virus des Infections Respiratoires - France SUD | Antonin Bal, Gregory Destras, Gwendolyne Burfin, Hadrien Regue, Quentin Semanas, Martine Valette, Bruno Lina, Laurence Josset |
| EPI_ISL_3472902 | LABORATOIRE FORTE BIO DAX | CNR Virus des Infections Respiratoires - France SUD | Antonin Bal, Gregory Destras, Gwendolyne Burfin, Hadrien Regue, Quentin Semanas, Martine Valette, Bruno Lina, Laurence Josset |
| EPI_ISL_3472903 | LABORATOIRE BIOESTEREL | CNR Virus des Infections Respiratoires - France SUD | Antonin Bal, Gregory Destras, Gwendolyne Burfin, Hadrien Regue, Quentin Semanas, Martine Valette, Bruno Lina, Laurence Josset |
| EPI_ISL_3472904 | LABORATOIRE FORTE BIO DAX | CNR Virus des Infections Respiratoires - France SUD | Antonin Bal, Gregory Destras, Gwendolyne Burfin, Hadrien Regue, Quentin Semanas, Martine Valette, Bruno Lina, Laurence Josset |
| EPI_ISL_3472905 | LABORATOIRE BIOESTEREL | CNR Virus des Infections Respiratoires - France SUD | Antonin Bal, Gregory Destras, Gwendolyne Burfin, Hadrien Regue, Quentin Semanas, Martine Valette, Bruno Lina, Laurence Josset |
| EPI_ISL_3472906 | LABORATOIRE FORTE BIO DAX | CNR Virus des Infections Respiratoires - France SUD | Antonin Bal, Gregory Destras, Gwendolyne Burfin, Hadrien Regue, Quentin Semanas, Martine Valette, Bruno Lina, Laurence Josset |
| EPI_ISL_3472907 | LABORATOIRE BIOESTEREL | CNR Virus des Infections Respiratoires - France SUD | Antonin Bal, Gregory Destras, Gwendolyne Burfin, Hadrien Regue, Quentin Semanas, Martine Valette, Bruno Lina, Laurence Josset |
| EPI_ISL_3472908, EPI_ISL_3472909 | LABORATOIRE FORTE BIO DAX | CNR Virus des Infections Respiratoires - France SUD | Antonin Bal, Gregory Destras, Gwendolyne Burfin, Hadrien Regue, Quentin Semanas, Martine Valette, Bruno Lina, Laurence Josset |
| EPI_ISL_3472910, EPI_ISL_3472911 | LABORATOIRE BIOESTEREL | CNR Virus des Infections Respiratoires - France SUD | Antonin Bal, Gregory Destras, Gwendolyne Burfin, Hadrien Regue, Quentin Semanas, Martine Valette, Bruno Lina, Laurence Josset |

| | | | |
|---|---|---|---|
| EPI_ISL_3472912, EPI_ISL_3472913, EPI_ISL_3472914, EPI_ISL_3472915, EPI_ISL_3472916, EPI_ISL_3472917 | LABORATOIRE FORTE BIO DAX | CNR Virus des Infections Respiratoires - France SUD | Antonin Bal, Gregory Destras, Gwendolyne Burfin, Hadrien Regue, Quentin Semanas, Martine Valette, Bruno Lina, Laurence Josset |
| EPI_ISL_3472918 | LABORATOIRE BIOESTEREL | CNR Virus des Infections Respiratoires - France SUD | Antonin Bal, Gregory Destras, Gwendolyne Burfin, Hadrien Regue, Quentin Semanas, Martine Valette, Bruno Lina, Laurence Josset |
| EPI_ISL_3472919, EPI_ISL_3472920 | LABORATOIRE FORTE BIO DAX | CNR Virus des Infections Respiratoires - France SUD | Antonin Bal, Gregory Destras, Gwendolyne Burfin, Hadrien Regue, Quentin Semanas, Martine Valette, Bruno Lina, Laurence Josset |
| EPI_ISL_3472921 | LABORATOIRE BIOESTEREL | CNR Virus des Infections Respiratoires - France SUD | Antonin Bal, Gregory Destras, Gwendolyne Burfin, Hadrien Regue, Quentin Semanas, Martine Valette, Bruno Lina, Laurence Josset |
| EPI_ISL_3472922, EPI_ISL_3472923 | LABORATOIRE FORTE BIO DAX | CNR Virus des Infections Respiratoires - France SUD | Antonin Bal, Gregory Destras, Gwendolyne Burfin, Hadrien Regue, Quentin Semanas, Martine Valette, Bruno Lina, Laurence Josset |
| EPI_ISL_3472924 | LABORATOIRE BIOESTEREL | CNR Virus des Infections Respiratoires - France SUD | Antonin Bal, Gregory Destras, Gwendolyne Burfin, Hadrien Regue, Quentin Semanas, Martine Valette, Bruno Lina, Laurence Josset |
| EPI_ISL_3472925 | LABORATOIRE FORTE BIO DAX | CNR Virus des Infections Respiratoires - France SUD | Antonin Bal, Gregory Destras, Gwendolyne Burfin, Hadrien Regue, Quentin Semanas, Martine Valette, Bruno Lina, Laurence Josset |
| EPI_ISL_3472926 | LABORATOIRE BIOESTEREL | CNR Virus des Infections Respiratoires - France SUD | Antonin Bal, Gregory Destras, Gwendolyne Burfin, Hadrien Regue, Quentin Semanas, Martine Valette, Bruno Lina, Laurence Josset |
| EPI_ISL_3472927, EPI_ISL_3472928, EPI_ISL_3472929, EPI_ISL_3472930 | LABORATOIRE FORTE BIO DAX | CNR Virus des Infections Respiratoires - France SUD | Antonin Bal, Gregory Destras, Gwendolyne Burfin, Hadrien Regue, Quentin Semanas, Martine Valette, Bruno Lina, Laurence Josset |
| EPI_ISL_3472931 | LABORATOIRE UNILIANS DECINES | CNR Virus des Infections Respiratoires - France SUD | Antonin Bal, Gregory Destras, Gwendolyne Burfin, Hadrien Regue, Quentin Semanas, Martine Valette, Bruno Lina, Laurence Josset |
| EPI_ISL_3472932, EPI_ISL_3472933, EPI_ISL_3472934 | LABORATOIRE FORTE BIO DAX | CNR Virus des Infections Respiratoires - France SUD | Antonin Bal, Gregory Destras, Gwendolyne Burfin, Hadrien Regue, Quentin Semanas, Martine Valette, Bruno Lina, Laurence Josset |
| EPI_ISL_3472935 | LABORATOIRE UNILIANS DECINES | CNR Virus des Infections Respiratoires - France SUD | Antonin Bal, Gregory Destras, Gwendolyne Burfin, Hadrien Regue, Quentin Semanas, Martine Valette, Bruno Lina, Laurence Josset |
| EPI_ISL_3472936, EPI_ISL_3472937, EPI_ISL_3472938 | LABORATOIRE FORTE BIO DAX | CNR Virus des Infections Respiratoires - France SUD | Antonin Bal, Gregory Destras, Gwendolyne Burfin, Hadrien Regue, Quentin Semanas, Martine Valette, Bruno Lina, Laurence Josset |
| EPI_ISL_3472939 | LABORATOIRE UNILIANS DECINES | CNR Virus des Infections Respiratoires - France SUD | Antonin Bal, Gregory Destras, Gwendolyne Burfin, Hadrien Regue, Quentin Semanas, Martine Valette, Bruno Lina, Laurence Josset |
| EPI_ISL_3472940, EPI_ISL_3472941, EPI_ISL_3472942, EPI_ISL_3472943, EPI_ISL_3472944, EPI_ISL_3472945, EPI_ISL_3472946 | LABORATOIRE FORTE BIO DAX | CNR Virus des Infections Respiratoires - France SUD | Antonin Bal, Gregory Destras, Gwendolyne Burfin, Hadrien Regue, Quentin Semanas, Martine Valette, Bruno Lina, Laurence Josset |
| EPI_ISL_3472947 | LABORATOIRE UNILIANS DECINES | CNR Virus des Infections Respiratoires - France SUD | Antonin Bal, Gregory Destras, Gwendolyne Burfin, Hadrien Regue, Quentin Semanas, Martine Valette, Bruno Lina, Laurence Josset |
| EPI_ISL_3472948 | LABORATOIRE BIOESTEREL | CNR Virus des Infections Respiratoires - France SUD | Antonin Bal, Gregory Destras, Gwendolyne Burfin, Hadrien Regue, Quentin Semanas, Martine Valette, Bruno Lina, Laurence Josset |
| EPI_ISL_3472949 | LABM BIO-VAL DU STADIUM | CNR Virus des Infections Respiratoires - France SUD | Antonin Bal, Gregory Destras, Gwendolyne Burfin, Hadrien Regue, Quentin Semanas, Martine Valette, Bruno Lina, Laurence Josset |
| EPI_ISL_3472950 | LABORATOIRE BIO-VAL | CNR Virus des Infections Respiratoires - France SUD | Antonin Bal, Gregory Destras, Gwendolyne Burfin, Hadrien Regue, Quentin Semanas, Martine Valette, Bruno Lina, Laurence Josset |
| EPI_ISL_3472951 | LABORATOIRE BIO-VAL ALLART-BETEND | CNR Virus des Infections Respiratoires - France SUD | Antonin Bal, Gregory Destras, Gwendolyne Burfin, Hadrien Regue, Quentin Semanas, Martine Valette, Bruno Lina, Laurence Josset |
| EPI_ISL_3472952 | LABORATOIRE BIO VAL THONON | CNR Virus des Infections Respiratoires - France SUD | Antonin Bal, Gregory Destras, Gwendolyne Burfin, Hadrien Regue, Quentin Semanas, Martine Valette, Bruno Lina, Laurence Josset |
| EPI_ISL_3472953 | LABORATOIRE BIO-VAL RUMILLY | CNR Virus des Infections Respiratoires - France SUD | Antonin Bal, Gregory Destras, Gwendolyne Burfin, Hadrien Regue, Quentin Semanas, Martine Valette, Bruno Lina, Laurence Josset |
| EPI_ISL_3472954 | LABORATOIRE BIO-VAL SALEVE | CNR Virus des Infections Respiratoires - France SUD | Antonin Bal, Gregory Destras, Gwendolyne Burfin, Hadrien Regue, Quentin Semanas, Martine Valette, Bruno Lina, Laurence Josset |
| EPI_ISL_3472955 | LABORATOIRE DU MOLE SELAS BIO-VAL | CNR Virus des Infections Respiratoires - France SUD | Antonin Bal, Gregory Destras, Gwendolyne Burfin, Hadrien Regue, Quentin Semanas, Martine Valette, Bruno Lina, Laurence Josset |
| EPI_ISL_3472956 | LABORATOIRE BIO-VAL SALEVE | CNR Virus des Infections Respiratoires - France SUD | Antonin Bal, Gregory Destras, Gwendolyne Burfin, Hadrien Regue, Quentin Semanas, Martine Valette, Bruno Lina, Laurence Josset |
| EPI_ISL_3472957 | LBM BIO VAL | CNR Virus des Infections Respiratoires - France SUD | Antonin Bal, Gregory Destras, Gwendolyne Burfin, Hadrien Regue, Quentin Semanas, Martine Valette, Bruno Lina, Laurence Josset |
| EPI_ISL_3472958 | LABORATOIRE DU MOLE SELAS BIO-VAL | CNR Virus des Infections Respiratoires - France SUD | Antonin Bal, Gregory Destras, Gwendolyne Burfin, Hadrien Regue, Quentin Semanas, Martine Valette, Bruno Lina, Laurence Josset |
| EPI_ISL_3472959 | LABORATOIRE UNILIANS DECINES | CNR Virus des Infections Respiratoires - France SUD | Antonin Bal, Gregory Destras, Gwendolyne Burfin, Hadrien Regue, Quentin Semanas, Martine Valette, Bruno Lina, Laurence Josset |
| EPI_ISL_3472960 | LABM BIO-VAL DU STADIUM | CNR Virus des Infections Respiratoires - France SUD | Antonin Bal, Gregory Destras, Gwendolyne Burfin, Hadrien Regue, Quentin Semanas, Martine Valette, Bruno Lina, Laurence Josset |
| EPI_ISL_3472961 | LAM EIMER LENYS | CNR Virus des Infections Respiratoires - France SUD | Antonin Bal, Gregory Destras, Gwendolyne Burfin, Hadrien Regue, Quentin Semanas, Martine Valette, Bruno Lina, Laurence Josset |
| EPI_ISL_3472962 | LABORATOIRE UNILIANS DECINES | CNR Virus des Infections Respiratoires - France SUD | Antonin Bal, Gregory Destras, Gwendolyne Burfin, Hadrien Regue, Quentin Semanas, Martine Valette, Bruno Lina, Laurence Josset |
| EPI_ISL_3472963 | LABORATOIRE BIO-VAL RUMILLY | CNR Virus des Infections Respiratoires - France SUD | Antonin Bal, Gregory Destras, Gwendolyne Burfin, Hadrien Regue, Quentin Semanas, Martine Valette, Bruno Lina, Laurence Josset |
| EPI_ISL_3472964, EPI_ISL_3472965 | LAM EIMER LENYS | CNR Virus des Infections Respiratoires - France SUD | Antonin Bal, Gregory Destras, Gwendolyne Burfin, Hadrien Regue, Quentin Semanas, Martine Valette, Bruno Lina, Laurence Josset |
| EPI_ISL_3472966 | LABORATOIRE BIO-VAL RUMILLY | CNR Virus des Infections Respiratoires - France SUD | Antonin Bal, Gregory Destras, Gwendolyne Burfin, Hadrien Regue, Quentin Semanas, Martine Valette, Bruno Lina, Laurence Josset |
| EPI_ISL_3472967 | CERBAILLIANCE LA REUNION | CNR Virus des Infections Respiratoires - France SUD | Antonin Bal, Gregory Destras, Gwendolyne Burfin, Hadrien Regue, Quentin Semanas, Martine Valette, Bruno Lina, Laurence Josset |
| EPI_ISL_3472968 | LABORATOIRE BIO VAL THONON | CNR Virus des Infections Respiratoires - France SUD | Antonin Bal, Gregory Destras, Gwendolyne Burfin, Hadrien Regue, Quentin Semanas, Martine Valette, Bruno Lina, Laurence Josset |
| EPI_ISL_3472969 | LAM EIMER LENYS | CNR Virus des Infections Respiratoires - France SUD | Antonin Bal, Gregory Destras, Gwendolyne Burfin, Hadrien Regue, Quentin Semanas, Martine Valette, Bruno Lina, Laurence Josset |
| EPI_ISL_3472970 | CERBAILLIANCE LA REUNION | CNR Virus des Infections Respiratoires - France SUD | Antonin Bal, Gregory Destras, Gwendolyne Burfin, Hadrien Regue, Quentin Semanas, Martine Valette, Bruno Lina, Laurence Josset |
| EPI_ISL_3472971 | LABORATOIRE BIO-VAL RUMILLY | CNR Virus des Infections Respiratoires - France SUD | Antonin Bal, Gregory Destras, Gwendolyne Burfin, Hadrien Regue, Quentin Semanas, Martine Valette, Bruno Lina, Laurence Josset |
| EPI_ISL_3472972 | CERBAILLIANCE LA REUNION | CNR Virus des Infections Respiratoires - France SUD | Antonin Bal, Gregory Destras, Gwendolyne Burfin, Hadrien Regue, Quentin Semanas, Martine Valette, Bruno Lina, Laurence Josset |
| EPI_ISL_3472973 | LAM EIMER LENYS | CNR Virus des Infections Respiratoires - France SUD | Antonin Bal, Gregory Destras, Gwendolyne Burfin, Hadrien Regue, Quentin Semanas, Martine Valette, Bruno Lina, Laurence Josset |
| EPI_ISL_3472974 | LABM BIO-VAL DU STADIUM | CNR Virus des Infections Respiratoires - France SUD | Antonin Bal, Gregory Destras, Gwendolyne Burfin, Hadrien Regue, Quentin Semanas, Martine Valette, Bruno Lina, Laurence Josset |
| EPI_ISL_3472975 | CNR Virus des Infections Respiratoires - France SUD | CNR Virus des Infections Respiratoires - France SUD | Antonin Bal, Gregory Destras, Gwendolyne Burfin, Hadrien Regue, Quentin Semanas, Martine Valette, Bruno Lina, Laurence Josset |
| EPI_ISL_3472976 | LABORATOIRE BIO VAL THONON | CNR Virus des Infections Respiratoires - France SUD | Antonin Bal, Gregory Destras, Gwendolyne Burfin, Hadrien Regue, Quentin Semanas, Martine Valette, Bruno Lina, Laurence Josset |
| EPI_ISL_3472977, EPI_ISL_3472978 | LABORATOIRE BIO-VAL SALEVE | CNR Virus des Infections Respiratoires - France SUD | Antonin Bal, Gregory Destras, Gwendolyne Burfin, Hadrien Regue, Quentin Semanas, Martine Valette, Bruno Lina, Laurence Josset |
| EPI_ISL_3472979 | LAM EIMER LENYS | CNR Virus des Infections Respiratoires - France SUD | Antonin Bal, Gregory Destras, Gwendolyne Burfin, Hadrien Regue, Quentin Semanas, Martine Valette, Bruno Lina, Laurence Josset |
| EPI_ISL_3472980 | LABORATOIRE BIO VAL THONON | CNR Virus des Infections Respiratoires - France SUD | Antonin Bal, Gregory Destras, Gwendolyne Burfin, Hadrien Regue, Quentin Semanas, Martine Valette, Bruno Lina, Laurence Josset |
| EPI_ISL_3472981 | LAM EIMER LENYS | CNR Virus des Infections Respiratoires - France SUD | Antonin Bal, Gregory Destras, Gwendolyne Burfin, Hadrien Regue, Quentin Semanas, Martine Valette, Bruno Lina, Laurence Josset |
| EPI_ISL_3472982 | LABORATOIRE BIO-VAL | CNR Virus des Infections Respiratoires - France SUD | Antonin Bal, Gregory Destras, Gwendolyne Burfin, Hadrien Regue, Quentin Semanas, Martine Valette, Bruno Lina, Laurence Josset |
| EPI_ISL_3472983 | LAM EIMER LENYS | CNR Virus des Infections Respiratoires - France SUD | Antonin Bal, Gregory Destras, Gwendolyne Burfin, Hadrien Regue, Quentin Semanas, Martine Valette, Bruno Lina, Laurence Josset |
| EPI_ISL_3472984 | LABORATOIRE DU MOLE SELAS BIO-VAL | CNR Virus des Infections Respiratoires - France SUD | Antonin Bal, Gregory Destras, Gwendolyne Burfin, Hadrien Regue, Quentin Semanas, Martine Valette, Bruno Lina, Laurence Josset |
| EPI_ISL_3472985 | LAM EIMER LENYS | CNR Virus des Infections Respiratoires - France SUD | Antonin Bal, Gregory Destras, Gwendolyne Burfin, Hadrien Regue, Quentin Semanas, Martine Valette, Bruno Lina, Laurence Josset |
| EPI_ISL_3472986 | LABORATOIRE DU MOLE SELAS BIO-VAL | CNR Virus des Infections Respiratoires - France SUD | Antonin Bal, Gregory Destras, Gwendolyne Burfin, Hadrien Regue, Quentin Semanas, Martine Valette, Bruno Lina, Laurence Josset |
| EPI_ISL_3472987 | LABORATOIRE BIO-VAL | CNR Virus des Infections Respiratoires - France SUD | Antonin Bal, Gregory Destras, Gwendolyne Burfin, Hadrien Regue, Quentin Semanas, Martine Valette, Bruno Lina, Laurence Josset |
| EPI_ISL_3472988 | LAM EIMER LENYS | CNR Virus des Infections Respiratoires - France SUD | Antonin Bal, Gregory Destras, Gwendolyne Burfin, Hadrien Regue, Quentin Semanas, Martine Valette, Bruno Lina, Laurence Josset |
| EPI_ISL_3472989 | CENTRE HOSPITALIER LUCIEN HUSSEL | CNR Virus des Infections Respiratoires - France SUD | Antonin Bal, Gregory Destras, Gwendolyne Burfin, Hadrien Regue, Quentin Semanas, Martine Valette, Bruno Lina, Laurence Josset |
| EPI_ISL_3472990, EPI_ISL_3472991 | CERBAILLIANCE LA REUNION | CNR Virus des Infections Respiratoires - France SUD | Antonin Bal, Gregory Destras, Gwendolyne Burfin, Hadrien Regue, Quentin Semanas, Martine Valette, Bruno Lina, Laurence Josset |
| EPI_ISL_3472992 | CNR Virus des Infections Respiratoires - France SUD | CNR Virus des Infections Respiratoires - France SUD | Antonin Bal, Gregory Destras, Gwendolyne Burfin, Hadrien Regue, Quentin Semanas, Martine Valette, Bruno Lina, Laurence Josset |
| EPI_ISL_3472993 | LAM EIMER LENYS | CNR Virus des Infections Respiratoires - France SUD | Antonin Bal, Gregory Destras, Gwendolyne Burfin, Hadrien Regue, Quentin Semanas, Martine Valette, Bruno Lina, Laurence Josset |
| EPI_ISL_3472994 | CERBAILLIANCE LA REUNION | CNR Virus des Infections Respiratoires - France SUD | Antonin Bal, Gregory Destras, Gwendolyne Burfin, Hadrien Regue, Quentin Semanas, Martine Valette, Bruno Lina, Laurence Josset |

| | | | |
|---|---|---|---|
| EPI_ISL_3472995 | LAM EIMER LENYS | CNR Virus des Infections Respiratoires - France SUD | Antonin Bal, Gregory Destras, Gwendolyne Burfin, Hadrien Regue, Quentin Semanas, Martine Valette, Bruno Lina, Laurence Josset |
| EPI_ISL_3472996 | CERBAILLIANCE LA REUNION | CNR Virus des Infections Respiratoires - France SUD | Antonin Bal, Gregory Destras, Gwendolyne Burfin, Hadrien Regue, Quentin Semanas, Martine Valette, Bruno Lina, Laurence Josset |
| EPI_ISL_3472997, EPI_ISL_3472998, EPI_ISL_3472999, EPI_ISL_3473000, EPI_ISL_3473001, EPI_ISL_3473002, EPI_ISL_3473003, EPI_ISL_3473004, EPI_ISL_3473005, EPI_ISL_3473006 | LAM EIMER LENYS | CNR Virus des Infections Respiratoires - France SUD | Antonin Bal, Gregory Destras, Gwendolyne Burfin, Hadrien Regue, Quentin Semanas, Martine Valette, Bruno Lina, Laurence Josset |
| EPI_ISL_3473007, EPI_ISL_3473008 | CERBAILLIANCE LA REUNION | CNR Virus des Infections Respiratoires - France SUD | Antonin Bal, Gregory Destras, Gwendolyne Burfin, Hadrien Regue, Quentin Semanas, Martine Valette, Bruno Lina, Laurence Josset |
| EPI_ISL_3473009 | HOPITAL NORD OUEST - CH VILLEFRANCHE | CNR Virus des Infections Respiratoires - France SUD | Antonin Bal, Gregory Destras, Gwendolyne Burfin, Hadrien Regue, Quentin Semanas, Martine Valette, Bruno Lina, Laurence Josset |
| EPI_ISL_3473010, EPI_ISL_3473011 | CERBAILLIANCE LA REUNION | CNR Virus des Infections Respiratoires - France SUD | Antonin Bal, Gregory Destras, Gwendolyne Burfin, Hadrien Regue, Quentin Semanas, Martine Valette, Bruno Lina, Laurence Josset |
| EPI_ISL_3473012 | CNR Virus des Infections Respiratoires - France SUD | CNR Virus des Infections Respiratoires - France SUD | Antonin Bal, Gregory Destras, Gwendolyne Burfin, Hadrien Regue, Quentin Semanas, Martine Valette, Bruno Lina, Laurence Josset |
| EPI_ISL_3473013, EPI_ISL_3473014, EPI_ISL_3473015, EPI_ISL_3473016, EPI_ISL_3473017 | HOPITAL NORD OUEST - CH VILLEFRANCHE | CNR Virus des Infections Respiratoires - France SUD | Antonin Bal, Gregory Destras, Gwendolyne Burfin, Hadrien Regue, Quentin Semanas, Martine Valette, Bruno Lina, Laurence Josset |
| EPI_ISL_3473018, EPI_ISL_3473019, EPI_ISL_3473020 | CNR Virus des Infections Respiratoires - France SUD | CNR Virus des Infections Respiratoires - France SUD | Antonin Bal, Gregory Destras, Gwendolyne Burfin, Hadrien Regue, Quentin Semanas, Martine Valette, Bruno Lina, Laurence Josset |
| EPI_ISL_3473021 | CH GIVORS | CNR Virus des Infections Respiratoires - France SUD | Antonin Bal, Gregory Destras, Gwendolyne Burfin, Hadrien Regue, Quentin Semanas, Martine Valette, Bruno Lina, Laurence Josset |
| EPI_ISL_3473022 | CNR Virus des Infections Respiratoires - France SUD | CNR Virus des Infections Respiratoires - France SUD | Antonin Bal, Gregory Destras, Gwendolyne Burfin, Hadrien Regue, Quentin Semanas, Martine Valette, Bruno Lina, Laurence Josset |
| EPI_ISL_3473023, EPI_ISL_3473024, EPI_ISL_3473025, EPI_ISL_3473026, EPI_ISL_3473027, EPI_ISL_3473028, EPI_ISL_3473029, EPI_ISL_3473030, EPI_ISL_3473031, EPI_ISL_3473032, EPI_ISL_3473033, EPI_ISL_3473034, EPI_ISL_3473035, EPI_ISL_3473036, EPI_ISL_3473037, EPI_ISL_3473038, EPI_ISL_3473039, EPI_ISL_3473040 | | | |
| see above | LABORATOIRE UNILIANS DECINES | CNR Virus des Infections Respiratoires - France SUD | Antonin Bal, Gregory Destras, Gwendolyne Burfin, Hadrien Regue, Quentin Semanas, Martine Valette, Bruno Lina, Laurence Josset |
| EPI_ISL_3473041 | DYOMEDEA-LABORATOIRE DE LA SAUVEGARDE | CNR Virus des Infections Respiratoires - France SUD | Antonin Bal, Gregory Destras, Gwendolyne Burfin, Hadrien Regue, Quentin Semanas, Martine Valette, Bruno Lina, Laurence Josset |
| EPI_ISL_3473042, EPI_ISL_3473043, EPI_ISL_3473044 | LABORATOIRE UNILIANS DECINES | CNR Virus des Infections Respiratoires - France SUD | Antonin Bal, Gregory Destras, Gwendolyne Burfin, Hadrien Regue, Quentin Semanas, Martine Valette, Bruno Lina, Laurence Josset |
| EPI_ISL_3473045 | DYOMEDEA-LABORATOIRE DE LA SAUVEGARDE | CNR Virus des Infections Respiratoires - France SUD | Antonin Bal, Gregory Destras, Gwendolyne Burfin, Hadrien Regue, Quentin Semanas, Martine Valette, Bruno Lina, Laurence Josset |
| EPI_ISL_3473046 | LABORATOIRE UNILIANS DECINES | CNR Virus des Infections Respiratoires - France SUD | Antonin Bal, Gregory Destras, Gwendolyne Burfin, Hadrien Regue, Quentin Semanas, Martine Valette, Bruno Lina, Laurence Josset |
| EPI_ISL_3473047, EPI_ISL_3473048 | DYOMEDEA-LABORATOIRE DE LA SAUVEGARDE | CNR Virus des Infections Respiratoires - France SUD | Antonin Bal, Gregory Destras, Gwendolyne Burfin, Hadrien Regue, Quentin Semanas, Martine Valette, Bruno Lina, Laurence Josset |
| EPI_ISL_3473049, EPI_ISL_3473050 | LABORATOIRE UNILIANS DECINES | CNR Virus des Infections Respiratoires - France SUD | Antonin Bal, Gregory Destras, Gwendolyne Burfin, Hadrien Regue, Quentin Semanas, Martine Valette, Bruno Lina, Laurence Josset |
| EPI_ISL_3473051, EPI_ISL_3473052 | DYOMEDEA-LABORATOIRE DE LA SAUVEGARDE | CNR Virus des Infections Respiratoires - France SUD | Antonin Bal, Gregory Destras, Gwendolyne Burfin, Hadrien Regue, Quentin Semanas, Martine Valette, Bruno Lina, Laurence Josset |
| EPI_ISL_3473053, EPI_ISL_3473054 | LABORATOIRE UNILIANS DECINES | CNR Virus des Infections Respiratoires - France SUD | Antonin Bal, Gregory Destras, Gwendolyne Burfin, Hadrien Regue, Quentin Semanas, Martine Valette, Bruno Lina, Laurence Josset |
| EPI_ISL_3473055, EPI_ISL_3473056 | DYOMEDEA-LABORATOIRE DE LA SAUVEGARDE | CNR Virus des Infections Respiratoires - France SUD | Antonin Bal, Gregory Destras, Gwendolyne Burfin, Hadrien Regue, Quentin Semanas, Martine Valette, Bruno Lina, Laurence Josset |
| EPI_ISL_3473057 | LABORATOIRE UNILIANS DECINES | CNR Virus des Infections Respiratoires - France SUD | Antonin Bal, Gregory Destras, Gwendolyne Burfin, Hadrien Regue, Quentin Semanas, Martine Valette, Bruno Lina, Laurence Josset |
| EPI_ISL_3473058 | DYOMEDEA-LABORATOIRE DE LA SAUVEGARDE | CNR Virus des Infections Respiratoires - France SUD | Antonin Bal, Gregory Destras, Gwendolyne Burfin, Hadrien Regue, Quentin Semanas, Martine Valette, Bruno Lina, Laurence Josset |
| EPI_ISL_3473059 | LABORATOIRE UNILIANS DECINES | CNR Virus des Infections Respiratoires - France SUD | Antonin Bal, Gregory Destras, Gwendolyne Burfin, Hadrien Regue, Quentin Semanas, Martine Valette, Bruno Lina, Laurence Josset |
| EPI_ISL_3473060, EPI_ISL_3473061, EPI_ISL_3473062, EPI_ISL_3473063, EPI_ISL_3473064, EPI_ISL_3473065, EPI_ISL_3473066, EPI_ISL_3473067, EPI_ISL_3473068, EPI_ISL_3473069, EPI_ISL_3473070 | | | |
| see above | DYOMEDEA-LABORATOIRE DE LA SAUVEGARDE | CNR Virus des Infections Respiratoires - France SUD | Antonin Bal, Gregory Destras, Gwendolyne Burfin, Hadrien Regue, Quentin Semanas, Martine Valette, Bruno Lina, Laurence Josset |
| EPI_ISL_3473071 | MIRIALIS CLUSES BECHET | CNR Virus des Infections Respiratoires - France SUD | Antonin Bal, Gregory Destras, Gwendolyne Burfin, Hadrien Regue, Quentin Semanas, Martine Valette, Bruno Lina, Laurence Josset |
| EPI_ISL_3473072 | DYOMEDEA-LABORATOIRE DE LA SAUVEGARDE | CNR Virus des Infections Respiratoires - France SUD | Antonin Bal, Gregory Destras, Gwendolyne Burfin, Hadrien Regue, Quentin Semanas, Martine Valette, Bruno Lina, Laurence Josset |
| EPI_ISL_3473073, EPI_ISL_3473074 | MIRIALIS CLUSES BECHET | CNR Virus des Infections Respiratoires - France SUD | Antonin Bal, Gregory Destras, Gwendolyne Burfin, Hadrien Regue, Quentin Semanas, Martine Valette, Bruno Lina, Laurence Josset |
| EPI_ISL_3473075 | DYOMEDEA-LABORATOIRE DE LA SAUVEGARDE | CNR Virus des Infections Respiratoires - France SUD | Antonin Bal, Gregory Destras, Gwendolyne Burfin, Hadrien Regue, Quentin Semanas, Martine Valette, Bruno Lina, Laurence Josset |
| EPI_ISL_3473076 | MIRIALIS CLUSES BECHET | CNR Virus des Infections Respiratoires - France SUD | Antonin Bal, Gregory Destras, Gwendolyne Burfin, Hadrien Regue, Quentin Semanas, Martine Valette, Bruno Lina, Laurence Josset |
| EPI_ISL_3473077 | DYOMEDEA-LABORATOIRE DE LA SAUVEGARDE | CNR Virus des Infections Respiratoires - France SUD | Antonin Bal, Gregory Destras, Gwendolyne Burfin, Hadrien Regue, Quentin Semanas, Martine Valette, Bruno Lina, Laurence Josset |
| EPI_ISL_3473078, EPI_ISL_3473079, EPI_ISL_3473080, EPI_ISL_3473081, EPI_ISL_3473082, EPI_ISL_3473083, EPI_ISL_3473084, EPI_ISL_3473085, EPI_ISL_3473086, EPI_ISL_3473087 | MIRIALIS CLUSES BECHET | CNR Virus des Infections Respiratoires - France SUD | Antonin Bal, Gregory Destras, Gwendolyne Burfin, Hadrien Regue, Quentin Semanas, Martine Valette, Bruno Lina, Laurence Josset |
| EPI_ISL_3473088, EPI_ISL_3473089 | DYOMEDEA-LABORATOIRE DE LA SAUVEGARDE | CNR Virus des Infections Respiratoires - France SUD | Antonin Bal, Gregory Destras, Gwendolyne Burfin, Hadrien Regue, Quentin Semanas, Martine Valette, Bruno Lina, Laurence Josset |
| EPI_ISL_3473090 | MIRIALIS CLUSES BECHET | CNR Virus des Infections Respiratoires - France SUD | Antonin Bal, Gregory Destras, Gwendolyne Burfin, Hadrien Regue, Quentin Semanas, Martine Valette, Bruno Lina, Laurence Josset |
| EPI_ISL_3473091 | DYOMEDEA-LABORATOIRE DE LA SAUVEGARDE | CNR Virus des Infections Respiratoires - France SUD | Antonin Bal, Gregory Destras, Gwendolyne Burfin, Hadrien Regue, Quentin Semanas, Martine Valette, Bruno Lina, Laurence Josset |
| EPI_ISL_3473092, EPI_ISL_3473093 | MIRIALIS CLUSES BECHET | CNR Virus des Infections Respiratoires - France SUD | Antonin Bal, Gregory Destras, Gwendolyne Burfin, Hadrien Regue, Quentin Semanas, Martine Valette, Bruno Lina, Laurence Josset |
| EPI_ISL_3473094 | DYOMEDEA-LABORATOIRE DE LA SAUVEGARDE | CNR Virus des Infections Respiratoires - France SUD | Antonin Bal, Gregory Destras, Gwendolyne Burfin, Hadrien Regue, Quentin Semanas, Martine Valette, Bruno Lina, Laurence Josset |
| EPI_ISL_3473095 | MIRIALIS CLUSES BECHET | CNR Virus des Infections Respiratoires - France SUD | Antonin Bal, Gregory Destras, Gwendolyne Burfin, Hadrien Regue, Quentin Semanas, Martine Valette, Bruno Lina, Laurence Josset |
| EPI_ISL_3473096, EPI_ISL_3473097, EPI_ISL_3473098, EPI_ISL_3473099, EPI_ISL_3473100 | DYOMEDEA-LABORATOIRE DE LA SAUVEGARDE | CNR Virus des Infections Respiratoires - France SUD | Antonin Bal, Gregory Destras, Gwendolyne Burfin, Hadrien Regue, Quentin Semanas, Martine Valette, Bruno Lina, Laurence Josset |
| EPI_ISL_3473101 | LABORATOIRE BIOGROUP | CNR Virus des Infections Respiratoires - France SUD | Antonin Bal, Gregory Destras, Gwendolyne Burfin, Hadrien Regue, Quentin Semanas, Martine Valette, Bruno Lina, Laurence Josset |
| EPI_ISL_3473102, EPI_ISL_3473103 | DYOMEDEA-LABORATOIRE DE LA SAUVEGARDE | CNR Virus des Infections Respiratoires - France SUD | Antonin Bal, Gregory Destras, Gwendolyne Burfin, Hadrien Regue, Quentin Semanas, Martine Valette, Bruno Lina, Laurence Josset |
| EPI_ISL_3473104, EPI_ISL_3473105 | LABORATOIRE BIOGROUP | CNR Virus des Infections Respiratoires - France SUD | Antonin Bal, Gregory Destras, Gwendolyne Burfin, Hadrien Regue, Quentin Semanas, Martine Valette, Bruno Lina, Laurence Josset |
| EPI_ISL_3473106 | DYOMEDEA-LABORATOIRE DE LA SAUVEGARDE | CNR Virus des Infections Respiratoires - France SUD | Antonin Bal, Gregory Destras, Gwendolyne Burfin, Hadrien Regue, Quentin Semanas, Martine Valette, Bruno Lina, Laurence Josset |
| EPI_ISL_3473107 | LABORATOIRE BIOGROUP | CNR Virus des Infections Respiratoires - France SUD | Antonin Bal, Gregory Destras, Gwendolyne Burfin, Hadrien Regue, Quentin Semanas, Martine Valette, Bruno Lina, Laurence Josset |
| EPI_ISL_3473108, EPI_ISL_3473109 | DYOMEDEA-LABORATOIRE DE LA SAUVEGARDE | CNR Virus des Infections Respiratoires - France SUD | Antonin Bal, Gregory Destras, Gwendolyne Burfin, Hadrien Regue, Quentin Semanas, Martine Valette, Bruno Lina, Laurence Josset |

| | | | |
|---|---|---|---|
| EPI_ISL_3473110 | HOPITAL ROBERT BOULIN | CNR Virus des Infections Respiratoires - France SUD | Antonin Bal, Gregory Destras, Gwendolyne Burfin, Hadrien Regue, Quentin Semanas, Martine Valette, Bruno Lina, Laurence Josset |
| EPI_ISL_3473111 | DYOMEDEA-LABORATOIRE DE LA SAUVEGARDE | CNR Virus des Infections Respiratoires - France SUD | Antonin Bal, Gregory Destras, Gwendolyne Burfin, Hadrien Regue, Quentin Semanas, Martine Valette, Bruno Lina, Laurence Josset |
| EPI_ISL_3473112 | HOPITAL ROBERT BOULIN | CNR Virus des Infections Respiratoires - France SUD | Antonin Bal, Gregory Destras, Gwendolyne Burfin, Hadrien Regue, Quentin Semanas, Martine Valette, Bruno Lina, Laurence Josset |
| EPI_ISL_3473113,<br>EPI_ISL_3473114,<br>EPI_ISL_3473115,<br>EPI_ISL_3473116,<br>EPI_ISL_3473117 | DYOMEDEA-LABORATOIRE DE LA SAUVEGARDE | CNR Virus des Infections Respiratoires - France SUD | Antonin Bal, Gregory Destras, Gwendolyne Burfin, Hadrien Regue, Quentin Semanas, Martine Valette, Bruno Lina, Laurence Josset |
| EPI_ISL_3473118 | CENTRE HOSPITALIER DE BASTIA | CNR Virus des Infections Respiratoires - France SUD | Antonin Bal, Gregory Destras, Gwendolyne Burfin, Hadrien Regue, Quentin Semanas, Martine Valette, Bruno Lina, Laurence Josset |
| EPI_ISL_3473119 | DYOMEDEA-LABORATOIRE DE LA SAUVEGARDE | CNR Virus des Infections Respiratoires - France SUD | Antonin Bal, Gregory Destras, Gwendolyne Burfin, Hadrien Regue, Quentin Semanas, Martine Valette, Bruno Lina, Laurence Josset |
| EPI_ISL_3473120,<br>EPI_ISL_3473121 | CENTRE HOSPITALIER DE BASTIA | CNR Virus des Infections Respiratoires - France SUD | Antonin Bal, Gregory Destras, Gwendolyne Burfin, Hadrien Regue, Quentin Semanas, Martine Valette, Bruno Lina, Laurence Josset |
| EPI_ISL_3473122 | DYOMEDEA-LABORATOIRE DE LA SAUVEGARDE | CNR Virus des Infections Respiratoires - France SUD | Antonin Bal, Gregory Destras, Gwendolyne Burfin, Hadrien Regue, Quentin Semanas, Martine Valette, Bruno Lina, Laurence Josset |
| EPI_ISL_3473123 | CENTRE HOSPITALIER DE BASTIA | CNR Virus des Infections Respiratoires - France SUD | Antonin Bal, Gregory Destras, Gwendolyne Burfin, Hadrien Regue, Quentin Semanas, Martine Valette, Bruno Lina, Laurence Josset |
| EPI_ISL_3473124,<br>EPI_ISL_3473125 | DYOMEDEA-LABORATOIRE DE LA SAUVEGARDE | CNR Virus des Infections Respiratoires - France SUD | Antonin Bal, Gregory Destras, Gwendolyne Burfin, Hadrien Regue, Quentin Semanas, Martine Valette, Bruno Lina, Laurence Josset |
| EPI_ISL_3473126 | LAM BAYARD-BIOPYRENEES | CNR Virus des Infections Respiratoires - France SUD | Antonin Bal, Gregory Destras, Gwendolyne Burfin, Hadrien Regue, Quentin Semanas, Martine Valette, Bruno Lina, Laurence Josset |
| EPI_ISL_3473127 | DYOMEDEA-LABORATOIRE DE LA SAUVEGARDE | CNR Virus des Infections Respiratoires - France SUD | Antonin Bal, Gregory Destras, Gwendolyne Burfin, Hadrien Regue, Quentin Semanas, Martine Valette, Bruno Lina, Laurence Josset |
| EPI_ISL_3473128 | LABORATOIRE UNILIANS DECINES | CNR Virus des Infections Respiratoires - France SUD | Antonin Bal, Gregory Destras, Gwendolyne Burfin, Hadrien Regue, Quentin Semanas, Martine Valette, Bruno Lina, Laurence Josset |
| EPI_ISL_3473129 | LAM BAYARD-BIOPYRENEES | CNR Virus des Infections Respiratoires - France SUD | Antonin Bal, Gregory Destras, Gwendolyne Burfin, Hadrien Regue, Quentin Semanas, Martine Valette, Bruno Lina, Laurence Josset |
| EPI_ISL_3473130 | DYOMEDEA-LABORATOIRE DE LA SAUVEGARDE | CNR Virus des Infections Respiratoires - France SUD | Antonin Bal, Gregory Destras, Gwendolyne Burfin, Hadrien Regue, Quentin Semanas, Martine Valette, Bruno Lina, Laurence Josset |
| EPI_ISL_3473131 | LABORATOIRE UNILIANS DECINES | CNR Virus des Infections Respiratoires - France SUD | Antonin Bal, Gregory Destras, Gwendolyne Burfin, Hadrien Regue, Quentin Semanas, Martine Valette, Bruno Lina, Laurence Josset |
| EPI_ISL_3473132 | LAM BAYARD-BIOPYRENEES | CNR Virus des Infections Respiratoires - France SUD | Antonin Bal, Gregory Destras, Gwendolyne Burfin, Hadrien Regue, Quentin Semanas, Martine Valette, Bruno Lina, Laurence Josset |
| EPI_ISL_3473133 | DYOMEDEA-LABORATOIRE DE LA SAUVEGARDE | CNR Virus des Infections Respiratoires - France SUD | Antonin Bal, Gregory Destras, Gwendolyne Burfin, Hadrien Regue, Quentin Semanas, Martine Valette, Bruno Lina, Laurence Josset |
| EPI_ISL_3473134 | LAM BAYARD-BIOPYRENEES | CNR Virus des Infections Respiratoires - France SUD | Antonin Bal, Gregory Destras, Gwendolyne Burfin, Hadrien Regue, Quentin Semanas, Martine Valette, Bruno Lina, Laurence Josset |
| EPI_ISL_3473135 | DYOMEDEA-LABORATOIRE DE LA SAUVEGARDE | CNR Virus des Infections Respiratoires - France SUD | Antonin Bal, Gregory Destras, Gwendolyne Burfin, Hadrien Regue, Quentin Semanas, Martine Valette, Bruno Lina, Laurence Josset |
| EPI_ISL_3473136,<br>EPI_ISL_3473137 | LAM BAYARD-BIOPYRENEES | CNR Virus des Infections Respiratoires - France SUD | Antonin Bal, Gregory Destras, Gwendolyne Burfin, Hadrien Regue, Quentin Semanas, Martine Valette, Bruno Lina, Laurence Josset |
| EPI_ISL_3473138 | LAM BIORYLIS | CNR Virus des Infections Respiratoires - France SUD | Antonin Bal, Gregory Destras, Gwendolyne Burfin, Hadrien Regue, Quentin Semanas, Martine Valette, Bruno Lina, Laurence Josset |
| EPI_ISL_3473139 | LABORATOIRE BIOGROUP | CNR Virus des Infections Respiratoires - France SUD | Antonin Bal, Gregory Destras, Gwendolyne Burfin, Hadrien Regue, Quentin Semanas, Martine Valette, Bruno Lina, Laurence Josset |
| EPI_ISL_3473140 | LAM BIORYLIS | CNR Virus des Infections Respiratoires - France SUD | Antonin Bal, Gregory Destras, Gwendolyne Burfin, Hadrien Regue, Quentin Semanas, Martine Valette, Bruno Lina, Laurence Josset |
| EPI_ISL_3473141 | LABORATOIRE BIOGROUP | CNR Virus des Infections Respiratoires - France SUD | Antonin Bal, Gregory Destras, Gwendolyne Burfin, Hadrien Regue, Quentin Semanas, Martine Valette, Bruno Lina, Laurence Josset |
| EPI_ISL_3473142 | LAM BIORYLIS | CNR Virus des Infections Respiratoires - France SUD | Antonin Bal, Gregory Destras, Gwendolyne Burfin, Hadrien Regue, Quentin Semanas, Martine Valette, Bruno Lina, Laurence Josset |
| EPI_ISL_3473143,<br>EPI_ISL_3473144 | DYOMEDEA-LABORATOIRE DE LA SAUVEGARDE | CNR Virus des Infections Respiratoires - France SUD | Antonin Bal, Gregory Destras, Gwendolyne Burfin, Hadrien Regue, Quentin Semanas, Martine Valette, Bruno Lina, Laurence Josset |
| EPI_ISL_3473145,<br>EPI_ISL_3473146 | LABORATOIRE BIOGROUP | CNR Virus des Infections Respiratoires - France SUD | Antonin Bal, Gregory Destras, Gwendolyne Burfin, Hadrien Regue, Quentin Semanas, Martine Valette, Bruno Lina, Laurence Josset |
| EPI_ISL_3473147,<br>EPI_ISL_3473148 | DYOMEDEA-LABORATOIRE DE LA SAUVEGARDE | CNR Virus des Infections Respiratoires - France SUD | Antonin Bal, Gregory Destras, Gwendolyne Burfin, Hadrien Regue, Quentin Semanas, Martine Valette, Bruno Lina, Laurence Josset |
| EPI_ISL_3473149 | LABORATOIRE BIOGROUP | CNR Virus des Infections Respiratoires - France SUD | Antonin Bal, Gregory Destras, Gwendolyne Burfin, Hadrien Regue, Quentin Semanas, Martine Valette, Bruno Lina, Laurence Josset |
| EPI_ISL_3473150 | DYOMEDEA-LABORATOIRE DE LA SAUVEGARDE | CNR Virus des Infections Respiratoires - France SUD | Antonin Bal, Gregory Destras, Gwendolyne Burfin, Hadrien Regue, Quentin Semanas, Martine Valette, Bruno Lina, Laurence Josset |
| EPI_ISL_3473151 | LABORATOIRE BIOGROUP | CNR Virus des Infections Respiratoires - France SUD | Antonin Bal, Gregory Destras, Gwendolyne Burfin, Hadrien Regue, Quentin Semanas, Martine Valette, Bruno Lina, Laurence Josset |
| EPI_ISL_3473152 | LAM BIORYLIS | CNR Virus des Infections Respiratoires - France SUD | Antonin Bal, Gregory Destras, Gwendolyne Burfin, Hadrien Regue, Quentin Semanas, Martine Valette, Bruno Lina, Laurence Josset |
| EPI_ISL_3473153 | DYOMEDEA-LABORATOIRE DE LA SAUVEGARDE | CNR Virus des Infections Respiratoires - France SUD | Antonin Bal, Gregory Destras, Gwendolyne Burfin, Hadrien Regue, Quentin Semanas, Martine Valette, Bruno Lina, Laurence Josset |
| EPI_ISL_3473154 | LAM BIORYLIS | CNR Virus des Infections Respiratoires - France SUD | Antonin Bal, Gregory Destras, Gwendolyne Burfin, Hadrien Regue, Quentin Semanas, Martine Valette, Bruno Lina, Laurence Josset |
| EPI_ISL_3473155,<br>EPI_ISL_3473156,<br>EPI_ISL_3473157,<br>EPI_ISL_3473158 | DYOMEDEA-LABORATOIRE DE LA SAUVEGARDE | CNR Virus des Infections Respiratoires - France SUD | Antonin Bal, Gregory Destras, Gwendolyne Burfin, Hadrien Regue, Quentin Semanas, Martine Valette, Bruno Lina, Laurence Josset |
| EPI_ISL_3473159,<br>EPI_ISL_3473160,<br>EPI_ISL_3473161 | LAM BIORYLIS | CNR Virus des Infections Respiratoires - France SUD | Antonin Bal, Gregory Destras, Gwendolyne Burfin, Hadrien Regue, Quentin Semanas, Martine Valette, Bruno Lina, Laurence Josset |
| EPI_ISL_3473162,<br>EPI_ISL_3473163 | LAM CERBALLIANCE | CNR Virus des Infections Respiratoires - France SUD | Antonin Bal, Gregory Destras, Gwendolyne Burfin, Hadrien Regue, Quentin Semanas, Martine Valette, Bruno Lina, Laurence Josset |
| EPI_ISL_3473164,<br>EPI_ISL_3473165 | LAM GEN-BIO GRAVANCHES | CNR Virus des Infections Respiratoires - France SUD | Antonin Bal, Gregory Destras, Gwendolyne Burfin, Hadrien Regue, Quentin Semanas, Martine Valette, Bruno Lina, Laurence Josset |
| EPI_ISL_3473166 | LABORATOIRE BIOGROUP | CNR Virus des Infections Respiratoires - France SUD | Antonin Bal, Gregory Destras, Gwendolyne Burfin, Hadrien Regue, Quentin Semanas, Martine Valette, Bruno Lina, Laurence Josset |
| EPI_ISL_3473167 | LAM BIORYLIS | CNR Virus des Infections Respiratoires - France SUD | Antonin Bal, Gregory Destras, Gwendolyne Burfin, Hadrien Regue, Quentin Semanas, Martine Valette, Bruno Lina, Laurence Josset |
| EPI_ISL_3473168,<br>EPI_ISL_3473169 | LABORATOIRE BIOGROUP | CNR Virus des Infections Respiratoires - France SUD | Antonin Bal, Gregory Destras, Gwendolyne Burfin, Hadrien Regue, Quentin Semanas, Martine Valette, Bruno Lina, Laurence Josset |
| EPI_ISL_3473170 | LABORATOIRE UNILIANS DECINES | CNR Virus des Infections Respiratoires - France SUD | Antonin Bal, Gregory Destras, Gwendolyne Burfin, Hadrien Regue, Quentin Semanas, Martine Valette, Bruno Lina, Laurence Josset |
| EPI_ISL_3473171,<br>EPI_ISL_3473172 | LABORATOIRE BIOGROUP | CNR Virus des Infections Respiratoires - France SUD | Antonin Bal, Gregory Destras, Gwendolyne Burfin, Hadrien Regue, Quentin Semanas, Martine Valette, Bruno Lina, Laurence Josset |
| EPI_ISL_3473173 | LABORATOIRE UNILIANS DECINES | CNR Virus des Infections Respiratoires - France SUD | Antonin Bal, Gregory Destras, Gwendolyne Burfin, Hadrien Regue, Quentin Semanas, Martine Valette, Bruno Lina, Laurence Josset |
| EPI_ISL_3473174 | LAM SYNLAB BORDEAUX ATLANTIQUE | CNR Virus des Infections Respiratoires - France SUD | Antonin Bal, Gregory Destras, Gwendolyne Burfin, Hadrien Regue, Quentin Semanas, Martine Valette, Bruno Lina, Laurence Josset |
| EPI_ISL_3473175 | LABORATOIRE BIOGROUP | CNR Virus des Infections Respiratoires - France SUD | Antonin Bal, Gregory Destras, Gwendolyne Burfin, Hadrien Regue, Quentin Semanas, Martine Valette, Bruno Lina, Laurence Josset |
| EPI_ISL_3473176,<br>EPI_ISL_3473177,<br>EPI_ISL_3473178,<br>EPI_ISL_3473179,<br>EPI_ISL_3473180 | LAM SYNLAB BORDEAUX ATLANTIQUE | CNR Virus des Infections Respiratoires - France SUD | Antonin Bal, Gregory Destras, Gwendolyne Burfin, Hadrien Regue, Quentin Semanas, Martine Valette, Bruno Lina, Laurence Josset |
| EPI_ISL_3473181,<br>EPI_ISL_3473182 | LABORATOIRE BIOGROUP | CNR Virus des Infections Respiratoires - France SUD | Antonin Bal, Gregory Destras, Gwendolyne Burfin, Hadrien Regue, Quentin Semanas, Martine Valette, Bruno Lina, Laurence Josset |
| EPI_ISL_3473183,<br>EPI_ISL_3473184,<br>EPI_ISL_3473185,<br>EPI_ISL_3473186,<br>EPI_ISL_3473187,<br>EPI_ISL_3473188,<br>EPI_ISL_3473189,<br>EPI_ISL_3473190,<br>EPI_ISL_3473191 | LAM SYNLAB BORDEAUX ATLANTIQUE | CNR Virus des Infections Respiratoires - France SUD | Antonin Bal, Gregory Destras, Gwendolyne Burfin, Hadrien Regue, Quentin Semanas, Martine Valette, Bruno Lina, Laurence Josset |
| EPI_ISL_3473192 | LAM BIORYLIS | CNR Virus des Infections Respiratoires - France SUD | Antonin Bal, Gregory Destras, Gwendolyne Burfin, Hadrien Regue, Quentin Semanas, Martine Valette, Bruno Lina, Laurence Josset |
| EPI_ISL_3473193,<br>EPI_ISL_3473194,<br>EPI_ISL_3473195,<br>EPI_ISL_3473196 | LAM SYNLAB BORDEAUX ATLANTIQUE | CNR Virus des Infections Respiratoires - France SUD | Antonin Bal, Gregory Destras, Gwendolyne Burfin, Hadrien Regue, Quentin Semanas, Martine Valette, Bruno Lina, Laurence Josset |

| | | | |
|---|---|---|---|
| EPI_ISL_3473197 | LAM BIORYLIS | CNR Virus des Infections Respiratoires - France SUD | Antonin Bal, Gregory Destras, Gwendolyne Burfin, Hadrien Regue, Quentin Semanas, Martine Valette, Bruno Lina, Laurence Josset |
| EPI_ISL_3473198 | LAM SYNLAB BORDEAUX ATLANTIQUE | CNR Virus des Infections Respiratoires - France SUD | Antonin Bal, Gregory Destras, Gwendolyne Burfin, Hadrien Regue, Quentin Semanas, Martine Valette, Bruno Lina, Laurence Josset |
| EPI_ISL_3473199 | CENTRE HOSPITALIER DE MILLAU | CNR Virus des Infections Respiratoires - France SUD | Antonin Bal, Gregory Destras, Gwendolyne Burfin, Hadrien Regue, Quentin Semanas, Martine Valette, Bruno Lina, Laurence Josset |
| EPI_ISL_3473200, EPI_ISL_3473201 | LAM SYNLAB BORDEAUX ATLANTIQUE | CNR Virus des Infections Respiratoires - France SUD | Antonin Bal, Gregory Destras, Gwendolyne Burfin, Hadrien Regue, Quentin Semanas, Martine Valette, Bruno Lina, Laurence Josset |
| EPI_ISL_3473202 | CENTRE HOSPITALIER DE MILLAU | CNR Virus des Infections Respiratoires - France SUD | Antonin Bal, Gregory Destras, Gwendolyne Burfin, Hadrien Regue, Quentin Semanas, Martine Valette, Bruno Lina, Laurence Josset |
| EPI_ISL_3473203 | LAM BIORYLIS | CNR Virus des Infections Respiratoires - France SUD | Antonin Bal, Gregory Destras, Gwendolyne Burfin, Hadrien Regue, Quentin Semanas, Martine Valette, Bruno Lina, Laurence Josset |
| EPI_ISL_3473204, EPI_ISL_3473205 | LAM SYNLAB BORDEAUX ATLANTIQUE | CNR Virus des Infections Respiratoires - France SUD | Antonin Bal, Gregory Destras, Gwendolyne Burfin, Hadrien Regue, Quentin Semanas, Martine Valette, Bruno Lina, Laurence Josset |
| EPI_ISL_3473206 | LAM BIORYLIS | CNR Virus des Infections Respiratoires - France SUD | Antonin Bal, Gregory Destras, Gwendolyne Burfin, Hadrien Regue, Quentin Semanas, Martine Valette, Bruno Lina, Laurence Josset |
| EPI_ISL_3473207, EPI_ISL_3473208, EPI_ISL_3473209, EPI_ISL_3473210, EPI_ISL_3473211, EPI_ISL_3473212, EPI_ISL_3473213, EPI_ISL_3473214, EPI_ISL_3473215, EPI_ISL_3473216, EPI_ISL_3473217, EPI_ISL_3473218, EPI_ISL_3473219, EPI_ISL_3473220, EPI_ISL_3473221, EPI_ISL_3473222, EPI_ISL_3473223, EPI_ISL_3473224, EPI_ISL_3473225, EPI_ISL_3473226, EPI_ISL_3473227, EPI_ISL_3473228, EPI_ISL_3473229 | | | |
| see above | LAM SYNLAB BORDEAUX ATLANTIQUE | CNR Virus des Infections Respiratoires - France SUD | Antonin Bal, Gregory Destras, Gwendolyne Burfin, Hadrien Regue, Quentin Semanas, Martine Valette, Bruno Lina, Laurence Josset |
| EPI_ISL_3473230 | CHU ST ETIENNE HOPITAL NORD | CNR Virus des Infections Respiratoires - France SUD | Antonin Bal, Gregory Destras, Gwendolyne Burfin, Hadrien Regue, Quentin Semanas, Martine Valette, Bruno Lina, Laurence Josset |
| EPI_ISL_3473231, EPI_ISL_3473232, EPI_ISL_3473233, EPI_ISL_3473234 | LAM BIORYLIS | CNR Virus des Infections Respiratoires - France SUD | Antonin Bal, Gregory Destras, Gwendolyne Burfin, Hadrien Regue, Quentin Semanas, Martine Valette, Bruno Lina, Laurence Josset |
| EPI_ISL_3473235 | LABORATOIRE UNILIANS DECINES | CNR Virus des Infections Respiratoires - France SUD | Antonin Bal, Gregory Destras, Gwendolyne Burfin, Hadrien Regue, Quentin Semanas, Martine Valette, Bruno Lina, Laurence Josset |
| EPI_ISL_3473236, EPI_ISL_3473237, EPI_ISL_3473238, EPI_ISL_3473239, EPI_ISL_3473240, EPI_ISL_3473241, EPI_ISL_3473242, EPI_ISL_3473243, EPI_ISL_3473244, EPI_ISL_3473245, EPI_ISL_3473246, EPI_ISL_3473247, EPI_ISL_3473248, EPI_ISL_3473249, EPI_ISL_3473250 | | | |
| see above | BIOLITTORAL BIOGROUP PLATEAU TECH | CNR Virus des Infections Respiratoires - France SUD | Antonin Bal, Gregory Destras, Gwendolyne Burfin, Hadrien Regue, Quentin Semanas, Martine Valette, Bruno Lina, Laurence Josset |
| EPI_ISL_3473251, EPI_ISL_3473252, EPI_ISL_3473253, EPI_ISL_3473254, EPI_ISL_3473255, EPI_ISL_3473256, EPI_ISL_3473257, EPI_ISL_3473258, EPI_ISL_3473259, EPI_ISL_3473260, EPI_ISL_3473261, EPI_ISL_3473262, EPI_ISL_3473263, EPI_ISL_3473264, EPI_ISL_3473265, EPI_ISL_3473266, EPI_ISL_3473267, EPI_ISL_3473268, EPI_ISL_3473269, EPI_ISL_3473270, EPI_ISL_3473271, EPI_ISL_3473272, EPI_ISL_3473273, EPI_ISL_3473274, EPI_ISL_3473275, EPI_ISL_3473276, EPI_ISL_3473277, EPI_ISL_3473278, EPI_ISL_3473279, EPI_ISL_3473280, EPI_ISL_3473281, EPI_ISL_3473282, EPI_ISL_3473283, EPI_ISL_3473284, EPI_ISL_3473285, EPI_ISL_3473286, EPI_ISL_3473287, EPI_ISL_3473288, EPI_ISL_3473289, EPI_ISL_3473290, EPI_ISL_3473291, EPI_ISL_3473292, EPI_ISL_3473293, EPI_ISL_3473294, EPI_ISL_3473295, EPI_ISL_3473296, EPI_ISL_3473297, EPI_ISL_3473298, EPI_ISL_3473299, EPI_ISL_3473300, EPI_ISL_3473301, EPI_ISL_3473302, EPI_ISL_3473303, EPI_ISL_3473304, EPI_ISL_3473305, EPI_ISL_3473306, EPI_ISL_3473307, EPI_ISL_3473308, EPI_ISL_3473309 | | | |
| see above | CNR Virus des Infections Respiratoires - France SUD | CNR Virus des Infections Respiratoires - France SUD | Antonin Bal, Gregory Destras, Gwendolyne Burfin, Hadrien Regue, Quentin Semanas, Martine Valette, Bruno Lina, Laurence Josset |
| EPI_ISL_3473311, EPI_ISL_3473312, EPI_ISL_3473313, EPI_ISL_3473316, EPI_ISL_3473320, EPI_ISL_3473321, EPI_ISL_3473323, EPI_ISL_3473324, EPI_ISL_3473326, EPI_ISL_3473327, EPI_ISL_3473328, EPI_ISL_3473329, EPI_ISL_3473330, EPI_ISL_3473331, EPI_ISL_3473332, EPI_ISL_3473333, EPI_ISL_3473334, EPI_ISL_3473335, EPI_ISL_3473336, EPI_ISL_3473337, EPI_ISL_3473338, EPI_ISL_3473339, EPI_ISL_3473340, EPI_ISL_3473341, EPI_ISL_3473342, EPI_ISL_3473343, EPI_ISL_3473344, EPI_ISL_3473345, EPI_ISL_3473347, EPI_ISL_3473349, EPI_ISL_3473350, EPI_ISL_3473351, EPI_ISL_3473352, EPI_ISL_3473353, EPI_ISL_3473354, EPI_ISL_3473355, EPI_ISL_3473356, EPI_ISL_3473357, EPI_ISL_3473359, EPI_ISL_3473360, EPI_ISL_3473361, EPI_ISL_3473362, EPI_ISL_3473363, EPI_ISL_3473364, EPI_ISL_3473365, EPI_ISL_3473366, EPI_ISL_3473367, EPI_ISL_3473368, EPI_ISL_3473369, EPI_ISL_3473370, EPI_ISL_3473371, EPI_ISL_3473372, EPI_ISL_3473373, EPI_ISL_3473374, EPI_ISL_3473375, EPI_ISL_3473376, EPI_ISL_3473377, EPI_ISL_3473378, EPI_ISL_3473380, EPI_ISL_3473382, EPI_ISL_3473383, EPI_ISL_3473384, EPI_ISL_3473385, EPI_ISL_3473386, EPI_ISL_3473387, EPI_ISL_3473388, EPI_ISL_3473389, EPI_ISL_3473390, EPI_ISL_3473391, EPI_ISL_3473393, EPI_ISL_3473395, EPI_ISL_3473396, EPI_ISL_3473397, EPI_ISL_3473398, EPI_ISL_3473400, EPI_ISL_3473401, EPI_ISL_3473403, EPI_ISL_3473404, EPI_ISL_3473405, EPI_ISL_3473406, EPI_ISL_3473408, EPI_ISL_3473409, EPI_ISL_3473410, EPI_ISL_3473412, EPI_ISL_3473442, EPI_ISL_3473443, EPI_ISL_3473444, EPI_ISL_3473445, EPI_ISL_3473448, EPI_ISL_3473454, EPI_ISL_3473455, EPI_ISL_3473456, EPI_ISL_3473457, EPI_ISL_3473459, EPI_ISL_3473460, EPI_ISL_3473461, EPI_ISL_3473464, EPI_ISL_3473465, EPI_ISL_3473467, EPI_ISL_3473468, EPI_ISL_3473469, EPI_ISL_3473470, EPI_ISL_3473471, EPI_ISL_3473472, EPI_ISL_3473473, EPI_ISL_3473474, EPI_ISL_3473475, EPI_ISL_3473476, EPI_ISL_3473477, EPI_ISL_3473485, EPI_ISL_3473487, EPI_ISL_3473489, EPI_ISL_3473490, EPI_ISL_3473491, EPI_ISL_3473492, EPI_ISL_3473493, EPI_ISL_3473494, EPI_ISL_3473495, EPI_ISL_3473496, EPI_ISL_3473497, EPI_ISL_3473498, EPI_ISL_3473499, EPI_ISL_3473500, EPI_ISL_3473501, EPI_ISL_3473502, EPI_ISL_3473503, EPI_ISL_3473504, EPI_ISL_3473505, EPI_ISL_3473506, EPI_ISL_3473507, EPI_ISL_3473508, EPI_ISL_3473514, EPI_ISL_3473517, EPI_ISL_3473523, EPI_ISL_3473524, EPI_ISL_3473525, EPI_ISL_3473526, EPI_ISL_3473527, EPI_ISL_3473528, EPI_ISL_3473529, EPI_ISL_3473530, EPI_ISL_3473531, EPI_ISL_3473532, EPI_ISL_3473533, EPI_ISL_3473534, EPI_ISL_3473535, EPI_ISL_3473536, EPI_ISL_3473537, EPI_ISL_3473538, EPI_ISL_3473539, EPI_ISL_3473540, EPI_ISL_3473541, EPI_ISL_3473542, EPI_ISL_3473543, EPI_ISL_3473544, EPI_ISL_3473545, EPI_ISL_3473546, EPI_ISL_3473547, EPI_ISL_3473548, EPI_ISL_3473549, EPI_ISL_3473550, EPI_ISL_3473551, EPI_ISL_3473552, EPI_ISL_3473553, EPI_ISL_3473554, EPI_ISL_3473555, EPI_ISL_3473556, EPI_ISL_3473557, EPI_ISL_3473558, EPI_ISL_3473559, EPI_ISL_3473560, EPI_ISL_3473561, EPI_ISL_3473562, EPI_ISL_3473563, EPI_ISL_3473565, EPI_ISL_3473566, EPI_ISL_3473567, EPI_ISL_3473568, EPI_ISL_3473569, EPI_ISL_3473570, EPI_ISL_3473572, EPI_ISL_3473573, EPI_ISL_3473574, EPI_ISL_3473575, EPI_ISL_3473576, EPI_ISL_3473577, EPI_ISL_3473578, EPI_ISL_3473579, EPI_ISL_3473580, EPI_ISL_3473581, EPI_ISL_3473582, EPI_ISL_3473583, EPI_ISL_3473584, EPI_ISL_3473585, EPI_ISL_3473586, EPI_ISL_3473587, EPI_ISL_3473588, EPI_ISL_3473589, EPI_ISL_3473593, EPI_ISL_3473606, EPI_ISL_3473607, EPI_ISL_3473608, EPI_ISL_3473609, EPI_ISL_3473610, EPI_ISL_3473611, EPI_ISL_3473612, EPI_ISL_3473613, EPI_ISL_3473614, EPI_ISL_3473618, EPI_ISL_3473623, EPI_ISL_3473631, EPI_ISL_3473632, EPI_ISL_3473633, EPI_ISL_3473634, EPI_ISL_3473635, EPI_ISL_3473636, EPI_ISL_3473637, EPI_ISL_3473638, EPI_ISL_3473652, EPI_ISL_3473653, EPI_ISL_3473654, EPI_ISL_3473655, EPI_ISL_3473656, EPI_ISL_3473657, EPI_ISL_3473658, EPI_ISL_3473659, EPI_ISL_3473660, EPI_ISL_3473661, EPI_ISL_3473662, EPI_ISL_3473663, EPI_ISL_3473664, EPI_ISL_3473665, EPI_ISL_3473666, EPI_ISL_3473667, EPI_ISL_3473668, EPI_ISL_3473669, EPI_ISL_3473670, EPI_ISL_3473671, EPI_ISL_3473672, EPI_ISL_3473673, EPI_ISL_3473674, EPI_ISL_3473675, EPI_ISL_3473676, EPI_ISL_3473677, EPI_ISL_3473678, EPI_ISL_3473679, EPI_ISL_3473680, EPI_ISL_3473681, EPI_ISL_3473682, EPI_ISL_3473684, EPI_ISL_3473685, EPI_ISL_3473686, EPI_ISL_3473705, EPI_ISL_3473706, EPI_ISL_3473707, EPI_ISL_3473708, EPI_ISL_3473709, EPI_ISL_3473710, EPI_ISL_3473711, EPI_ISL_3473712, EPI_ISL_3473713, EPI_ISL_3473716, EPI_ISL_3473717, EPI_ISL_3473718, EPI_ISL_3473719, EPI_ISL_3473720, EPI_ISL_3473721, EPI_ISL_3473722, EPI_ISL_3473723, EPI_ISL_3473724, EPI_ISL_3473725, EPI_ISL_3473726, EPI_ISL_3473727, EPI_ISL_3473728, EPI_ISL_3473729, EPI_ISL_3473730, EPI_ISL_3473731, EPI_ISL_3473732, EPI_ISL_3473733, EPI_ISL_3473734, EPI_ISL_3473735, EPI_ISL_3473736, EPI_ISL_3473737, EPI_ISL_3473738, EPI_ISL_3473739, EPI_ISL_3473740, EPI_ISL_3473741, EPI_ISL_3473742, EPI_ISL_3473743, EPI_ISL_3473744, EPI_ISL_3473745, EPI_ISL_3473746, EPI_ISL_3473747, EPI_ISL_3473750, EPI_ISL_3473751, EPI_ISL_3473752, EPI_ISL_3473753, EPI_ISL_3473754, EPI_ISL_3473755, EPI_ISL_3473756, EPI_ISL_3473757, EPI_ISL_3473758, EPI_ISL_3473759, EPI_ISL_3473760, EPI_ISL_3473761, EPI_ISL_3473762, EPI_ISL_3473763, EPI_ISL_3473765, EPI_ISL_3473766, EPI_ISL_3473767, EPI_ISL_3473768, EPI_ISL_3473769, EPI_ISL_3473770, EPI_ISL_3473771, EPI_ISL_3473772, EPI_ISL_3473773, EPI_ISL_3473774, EPI_ISL_3473775, EPI_ISL_3473776, EPI_ISL_3473777, EPI_ISL_3473778, EPI_ISL_3473779, EPI_ISL_3473780, EPI_ISL_3473781, EPI_ISL_3473782, EPI_ISL_3473783, EPI_ISL_3473784, EPI_ISL_3473785, EPI_ISL_3473786, EPI_ISL_3473787, EPI_ISL_3473789, EPI_ISL_3473790, EPI_ISL_3473791, EPI_ISL_3473792, EPI_ISL_3473793, EPI_ISL_3473794, EPI_ISL_3473795, EPI_ISL_3473796, EPI_ISL_3473797, EPI_ISL_3473798, EPI_ISL_3473799, EPI_ISL_3473800, EPI_ISL_3473801, EPI_ISL_3473802, EPI_ISL_3473803, EPI_ISL_3473804, EPI_ISL_3473805 | | | |
| see above | Indian Council of Medical Research-National Institute of Virology, Microbial Containment Complex | Indian Council of Medical Research-National Institute of Virology, Microbial Containment Complex | Pragya Yadav |
| EPI_ISL_3473839, EPI_ISL_3473840, EPI_ISL_3473841, EPI_ISL_3473842, EPI_ISL_3473843, EPI_ISL_3473844, EPI_ISL_3473845, EPI_ISL_3473846, EPI_ISL_3473847, EPI_ISL_3473848, EPI_ISL_3473849, EPI_ISL_3473850, EPI_ISL_3473851, EPI_ISL_3473852, EPI_ISL_3473853, EPI_ISL_3473854, EPI_ISL_3473855, EPI_ISL_3473856, EPI_ISL_3473857, EPI_ISL_3473858, EPI_ISL_3473859, EPI_ISL_3473860, EPI_ISL_3473861, EPI_ISL_3473862, EPI_ISL_3473863, EPI_ISL_3473865, EPI_ISL_3473866, EPI_ISL_3473867, EPI_ISL_3473868, EPI_ISL_3473869, EPI_ISL_3473870, EPI_ISL_3473871, EPI_ISL_3473872, EPI_ISL_3473873, EPI_ISL_3473874, EPI_ISL_3473875, EPI_ISL_3473876, EPI_ISL_3473877, EPI_ISL_3473878, EPI_ISL_3473879, EPI_ISL_3473880, EPI_ISL_3473881, EPI_ISL_3473882, EPI_ISL_3473883, EPI_ISL_3473884, EPI_ISL_3473885, EPI_ISL_3473886, EPI_ISL_3473887, EPI_ISL_3473888, EPI_ISL_3473889, EPI_ISL_3473890, EPI_ISL_3473891, EPI_ISL_3473892, EPI_ISL_3473893, EPI_ISL_3473894, EPI_ISL_3473895, EPI_ISL_3473896, EPI_ISL_3473897, EPI_ISL_3473898, EPI_ISL_3473899, EPI_ISL_3473900, EPI_ISL_3473901, EPI_ISL_3473902, EPI_ISL_3473903, EPI_ISL_3473904, EPI_ISL_3473905, EPI_ISL_3473906, EPI_ISL_3473907, EPI_ISL_3473908, EPI_ISL_3473909, EPI_ISL_3473910, EPI_ISL_3473911, EPI_ISL_3473912, EPI_ISL_3473913, EPI_ISL_3473914, EPI_ISL_3473915, EPI_ISL_3473916, EPI_ISL_3473917, EPI_ISL_3473918, EPI_ISL_3473919, EPI_ISL_3473920, EPI_ISL_3473921, EPI_ISL_3473922, EPI_ISL_3473923, EPI_ISL_3473924, EPI_ISL_3473925, EPI_ISL_3473926, EPI_ISL_3473927, EPI_ISL_3473928, EPI_ISL_3473929, EPI_ISL_3473930, EPI_ISL_3473931, EPI_ISL_3473932, EPI_ISL_3473933, EPI_ISL_3473934, EPI_ISL_3473935, EPI_ISL_3473936, EPI_ISL_3473937, EPI_ISL_3473938, EPI_ISL_3473939, EPI_ISL_3473940, EPI_ISL_3473941, EPI_ISL_3473942, EPI_ISL_3473943, EPI_ISL_3473944, EPI_ISL_3473945, EPI_ISL_3473946, EPI_ISL_3473947, EPI_ISL_3473948, EPI_ISL_3473949, EPI_ISL_3473950, EPI_ISL_3473951, EPI_ISL_3473952, EPI_ISL_3473953, EPI_ISL_3473954, EPI_ISL_3473955, EPI_ISL_3473956, EPI_ISL_3473957, EPI_ISL_3473958, EPI_ISL_3473959, EPI_ISL_3473960, EPI_ISL_3473961, EPI_ISL_3473962, EPI_ISL_3473963, EPI_ISL_3473964, EPI_ISL_3473965, EPI_ISL_3473966, EPI_ISL_3473967, EPI_ISL_3473968, EPI_ISL_3473969, EPI_ISL_3473970, EPI_ISL_3473971, EPI_ISL_3473972, EPI_ISL_3473973, EPI_ISL_3473974, EPI_ISL_3473975, EPI_ISL_3473976, EPI_ISL_3473977, EPI_ISL_3473978, EPI_ISL_3473979, EPI_ISL_3473980, EPI_ISL_3473981, EPI_ISL_3473982, EPI_ISL_3473983, EPI_ISL_3473984, EPI_ISL_3473985, EPI_ISL_3473986, EPI_ISL_3473987, EPI_ISL_3473988, EPI_ISL_3473989, EPI_ISL_3473990, EPI_ISL_3473991, EPI_ISL_3473992, EPI_ISL_3473993, EPI_ISL_3473994, EPI_ISL_3473995, EPI_ISL_3473996, EPI_ISL_3473997, EPI_ISL_3473998, EPI_ISL_3473999, EPI_ISL_3474000, EPI_ISL_3474001, EPI_ISL_3474002, EPI_ISL_3474003, EPI_ISL_3474004, EPI_ISL_3474005, EPI_ISL_3474006, EPI_ISL_3474007, EPI_ISL_3474008, EPI_ISL_3474009, EPI_ISL_3474010, EPI_ISL_3474011, EPI_ISL_3474012, EPI_ISL_3474013, EPI_ISL_3474014, EPI_ISL_3474015, EPI_ISL_3474016, EPI_ISL_3474017, EPI_ISL_3474018, EPI_ISL_3474019, EPI_ISL_3474022, EPI_ISL_3474023, EPI_ISL_3474024, EPI_ISL_3474025, EPI_ISL_3474026, EPI_ISL_3474027, EPI_ISL_3474028, EPI_ISL_3474034, EPI_ISL_3474035, EPI_ISL_3474036, EPI_ISL_3474037, EPI_ISL_3474038, EPI_ISL_3474039, EPI_ISL_3474040, EPI_ISL_3474041, EPI_ISL_3474042, EPI_ISL_3474043, EPI_ISL_3474044, EPI_ISL_3474045, EPI_ISL_3474046, EPI_ISL_3474047, EPI_ISL_3474048, EPI_ISL_3474049, EPI_ISL_3474050, EPI_ISL_3474051, EPI_ISL_3474052, EPI_ISL_3474053, EPI_ISL_3474054, EPI_ISL_3474055, EPI_ISL_3474056, EPI_ISL_3474057, EPI_ISL_3474058, EPI_ISL_3474059, EPI_ISL_3474060, EPI_ISL_3474061, EPI_ISL_3474062, EPI_ISL_3474063, EPI_ISL_3474064, EPI_ISL_3474065, EPI_ISL_3474066, EPI_ISL_3474067, EPI_ISL_3474068, EPI_ISL_3474069, EPI_ISL_3474070, EPI_ISL_3474071, EPI_ISL_3474072, EPI_ISL_3474073, EPI_ISL_3474074, EPI_ISL_3474075, EPI_ISL_3474076, EPI_ISL_3474077, EPI_ISL_3474078, EPI_ISL_3474079, EPI_ISL_3474080, EPI_ISL_3474081, EPI_ISL_3474082, EPI_ISL_3474083, EPI_ISL_3474084, EPI_ISL_3474085, EPI_ISL_3474086, EPI_ISL_3474087, EPI_ISL_3474088, EPI_ISL_3474089, EPI_ISL_3474090, EPI_ISL_3474091, EPI_ISL_3474092, EPI_ISL_3474093, EPI_ISL_3474094, EPI_ISL_3474095, EPI_ISL_3474096, EPI_ISL_3474097, EPI_ISL_3474098, EPI_ISL_3474099, EPI_ISL_3474100, EPI_ISL_3474102, EPI_ISL_3474103, EPI_ISL_3474104, EPI_ISL_3474105, EPI_ISL_3474106, EPI_ISL_3474107, EPI_ISL_3474108, EPI_ISL_3474109, EPI_ISL_3474110, EPI_ISL_3474111, EPI_ISL_3474112, EPI_ISL_3474113, EPI_ISL_3474114, EPI_ISL_3474115, EPI_ISL_3474116, EPI_ISL_3474117, EPI_ISL_3474118, EPI_ISL_3474119, EPI_ISL_3474120, EPI_ISL_3474121, EPI_ISL_3474122, EPI_ISL_3474123, EPI_ISL_3474124, EPI_ISL_3474125, EPI_ISL_3474126, EPI_ISL_3474127, EPI_ISL_3474128, EPI_ISL_3474129, EPI_ISL_3474130, EPI_ISL_3474131, EPI_ISL_3474132, EPI_ISL_3474133, EPI_ISL_3474134, EPI_ISL_3474135, EPI_ISL_3474136, EPI_ISL_3474137, EPI_ISL_3474138, EPI_ISL_3474139, EPI_ISL_3474140, EPI_ISL_3474141, EPI_ISL_3474142, EPI_ISL_3474143, EPI_ISL_3474144, EPI_ISL_3474145, EPI_ISL_3474146, EPI_ISL_3474147, EPI_ISL_3474148, EPI_ISL_3474149, EPI_ISL_3474150, EPI_ISL_3474151, EPI_ISL_3474152, EPI_ISL_3474153, EPI_ISL_3474154, EPI_ISL_3474155, EPI_ISL_3474156, EPI_ISL_3474157, EPI_ISL_3474158, EPI_ISL_3474159, EPI_ISL_3474160, EPI_ISL_3474161, EPI_ISL_3474162, EPI_ISL_3474163, EPI_ISL_3474164, EPI_ISL_3474165, EPI_ISL_3474166, EPI_ISL_3474167, EPI_ISL_3474168, EPI_ISL_3474169, EPI_ISL_3474170, EPI_ISL_3474171, EPI_ISL_3474172, EPI_ISL_3474173 | | | |
| see above | Arkansas Public Health Laboratory, Arkansas Department of Health | University of Minnesota Genomics Center | Daryl M. Gohl, John Garbe, Jaquelyn Kuriger-Laber, Corbin Dirkx |
| EPI_ISL_3474185, EPI_ISL_3474186, EPI_ISL_3474187, EPI_ISL_3474189, EPI_ISL_3474194, EPI_ISL_3474195, EPI_ISL_3474197, EPI_ISL_3474198, EPI_ISL_3474199, EPI_ISL_3474201, EPI_ISL_3474202, EPI_ISL_3474203, EPI_ISL_3474204, EPI_ISL_3474205, EPI_ISL_3474206, EPI_ISL_3474207, EPI_ISL_3474208, EPI_ISL_3474209, EPI_ISL_3474211, EPI_ISL_3474212, EPI_ISL_3474213, EPI_ISL_3474215, EPI_ISL_3474216, EPI_ISL_3474220, EPI_ISL_3474221, EPI_ISL_3474222, EPI_ISL_3474223, EPI_ISL_3474224, EPI_ISL_3474225, EPI_ISL_3474228, EPI_ISL_3474229, EPI_ISL_3474230, EPI_ISL_3474231, EPI_ISL_3474232, EPI_ISL_3474233, EPI_ISL_3474234, EPI_ISL_3474235, EPI_ISL_3474237, EPI_ISL_3474238, EPI_ISL_3474239, EPI_ISL_3474240, EPI_ISL_3474243, EPI_ISL_3474245, EPI_ISL_3474246, EPI_ISL_3474247, EPI_ISL_3474248, EPI_ISL_3474249, EPI_ISL_3474250, EPI_ISL_3474251, EPI_ISL_3474254, EPI_ISL_3474255, EPI_ISL_3474256, EPI_ISL_3474259, EPI_ISL_3474261, EPI_ISL_3474262, EPI_ISL_3474263, EPI_ISL_3474264, EPI_ISL_3474273, EPI_ISL_3474275, EPI_ISL_3474277, EPI_ISL_3474278, EPI_ISL_3474279, EPI_ISL_3474280, EPI_ISL_3474281, EPI_ISL_3474282, EPI_ISL_3474283, EPI_ISL_3474284, EPI_ISL_3474286, EPI_ISL_3474287, EPI_ISL_3474288, EPI_ISL_3474289, EPI_ISL_3474290, EPI_ISL_3474291, EPI_ISL_3474292, EPI_ISL_3474293, EPI_ISL_3474294, EPI_ISL_3474295, EPI_ISL_3474296, EPI_ISL_3474297, EPI_ISL_3474299, EPI_ISL_3474300, EPI_ISL_3474309, EPI_ISL_3474310, EPI_ISL_3474311, EPI_ISL_3474314, EPI_ISL_3474320, EPI_ISL_3474321, EPI_ISL_3474322, EPI_ISL_3474323, EPI_ISL_3474324, EPI_ISL_3474325, EPI_ISL_3474326, EPI_ISL_3474327, EPI_ISL_3474328, EPI_ISL_3474329, EPI_ISL_3474330, EPI_ISL_3474331, EPI_ISL_3474332, EPI_ISL_3474333, EPI_ISL_3474334, EPI_ISL_3474335, EPI_ISL_3474338, EPI_ISL_3474339, EPI_ISL_3474342, EPI_ISL_3474343, EPI_ISL_3474345, EPI_ISL_3474346, EPI_ISL_3474347, EPI_ISL_3474348, EPI_ISL_3474349, EPI_ISL_3474350, EPI_ISL_3474351, EPI_ISL_3474352, EPI_ISL_3474353, EPI_ISL_3474354, EPI_ISL_3474355, EPI_ISL_3474356, EPI_ISL_3474357, EPI_ISL_3474358, EPI_ISL_3474359, EPI_ISL_3474360, EPI_ISL_3474361, EPI_ISL_3474362, EPI_ISL_3474363, EPI_ISL_3474364, EPI_ISL_3474367, EPI_ISL_3474369, EPI_ISL_3474371, EPI_ISL_3474372, EPI_ISL_3474373, EPI_ISL_3474374, EPI_ISL_3474375, EPI_ISL_3474376, EPI_ISL_3474377, EPI_ISL_3474378, EPI_ISL_3474379, EPI_ISL_3474382, EPI_ISL_3474388, EPI_ISL_3474390, EPI_ISL_3474391, EPI_ISL_3474392, EPI_ISL_3474395, EPI_ISL_3474396, EPI_ISL_3474397, EPI_ISL_3474398, EPI_ISL_3474399, EPI_ISL_3474400, EPI_ISL_3474402, EPI_ISL_3474406, EPI_ISL_3474407, EPI_ISL_3474408, EPI_ISL_3474409, EPI_ISL_3474410, EPI_ISL_3474411, EPI_ISL_3474412, EPI_ISL_3474413, EPI_ISL_3474414, EPI_ISL_3474415, EPI_ISL_3474416, EPI_ISL_3474417, EPI_ISL_3474418, EPI_ISL_3474419, EPI_ISL_3474420, EPI_ISL_3474421, EPI_ISL_3474422, EPI_ISL_3474423, EPI_ISL_3474424, EPI_ISL_3474425, EPI_ISL_3474426, EPI_ISL_3474427, EPI_ISL_3474428, EPI_ISL_3474432, EPI_ISL_3474433, EPI_ISL_3474434, EPI_ISL_3474435, EPI_ISL_3474436, EPI_ISL_3474437, EPI_ISL_3474438, EPI_ISL_3474439, EPI_ISL_3474440, EPI_ISL_3474441, EPI_ISL_3474442, EPI_ISL_3474443, EPI_ISL_3474444, EPI_ISL_3474445, EPI_ISL_3474446, EPI_ISL_3474447, EPI_ISL_3474448, EPI_ISL_3474449, EPI_ISL_3474450, EPI_ISL_3474451, EPI_ISL_3474452, EPI_ISL_3474454, EPI_ISL_3474472, EPI_ISL_3474476, EPI_ISL_3474477, EPI_ISL_3474479, EPI_ISL_3474492, EPI_ISL_3474493, EPI_ISL_3474494, EPI_ISL_3474495, EPI_ISL_3474640, EPI_ISL_3474643, EPI_ISL_3474646, EPI_ISL_3474649, EPI_ISL_3474652, EPI_ISL_3474653, EPI_ISL_3474654, EPI_ISL_3474656, EPI_ISL_3474658, EPI_ISL_3474659, EPI_ISL_3474660, EPI_ISL_3474666, EPI_ISL_3474667, EPI_ISL_3474727, EPI_ISL_3474730, EPI_ISL_3474731, EPI_ISL_3474732, EPI_ISL_3474733, EPI_ISL_3474734, EPI_ISL_3474735, EPI_ISL_3474736, EPI_ISL_3474737, EPI_ISL_3474738, EPI_ISL_3474739, EPI_ISL_3474740, EPI_ISL_3474741, EPI_ISL_3474742, EPI_ISL_3474743, EPI_ISL_3474744, EPI_ISL_3474745, EPI_ISL_3474746, EPI_ISL_3474747, EPI_ISL_3474748, EPI_ISL_3474749, EPI_ISL_3474750, EPI_ISL_3474752, EPI_ISL_3474753, EPI_ISL_3474754, EPI_ISL_3474755, EPI_ISL_3474756, EPI_ISL_3474757, EPI_ISL_3474758, EPI_ISL_3474759, EPI_ISL_3474760, EPI_ISL_3474761, EPI_ISL_3474762, EPI_ISL_3474763, EPI_ISL_3474764, EPI_ISL_3474765, EPI_ISL_3474766, EPI_ISL_3474767, EPI_ISL_3474768, EPI_ISL_3474769, EPI_ISL_3474770, EPI_ISL_3474771, EPI_ISL_3474772, EPI_ISL_3474773, EPI_ISL_3474774, EPI_ISL_3474775, EPI_ISL_3474776, EPI_ISL_3474777, EPI_ISL_3474778, EPI_ISL_3474779, EPI_ISL_3474780, EPI_ISL_3474781, EPI_ISL_3474782, EPI_ISL_3474783, EPI_ISL_3474784, EPI_ISL_3474785, EPI_ISL_3474786, EPI_ISL_3474787, EPI_ISL_3474788, EPI_ISL_3474789, EPI_ISL_3474790, EPI_ISL_3474791, EPI_ISL_3474792, EPI_ISL_3474793, EPI_ISL_3474794, EPI_ISL_3474795, EPI_ISL_3474796, EPI_ISL_3474797, EPI_ISL_3474798, EPI_ISL_3474799, EPI_ISL_3474800, EPI_ISL_3474801, EPI_ISL_3474802, EPI_ISL_3474803, EPI_ISL_3474804, EPI_ISL_3474805, EPI_ISL_3474806, EPI_ISL_3474807, EPI_ISL_3474808, EPI_ISL_3474809, EPI_ISL_3474810, EPI_ISL_3474811, EPI_ISL_3474812, EPI_ISL_3474813, EPI_ISL_3474814, EPI_ISL_3474815, EPI_ISL_3474816, EPI_ISL_3474817, EPI_ISL_3474818, EPI_ISL_3474819, EPI_ISL_3474820, EPI_ISL_3474821, EPI_ISL_3474822, EPI_ISL_3474823, EPI_ISL_3474824, EPI_ISL_3474825, EPI_ISL_3474826, EPI_ISL_3474827, EPI_ISL_3474828 | | | |
| see above | Dutch COVID-19 response team | Medical Microbiology, Maastricht University Medical Centre | Jozef Dingemans*, Brian van der Veer*, Erik Beuken, Carmen Reumkens, Lieke van Alphen, Christian Hoebe, Paul Savelkoul |
| EPI_ISL_3474837, EPI_ISL_3474838, EPI_ISL_3474839, EPI_ISL_3474840 | Biogroup Bio Lam-LCD Saint-Denis | Department of Virology, Henri Mondor University Hospital, Assistance Publique Hôpitaux de Paris, Université Paris-Est Créteil, INSERM U955 | Christophe Rodriguez, Slim Fourati, Vanessa Demontant, Guillaume Gricourt, Melissa N'Debi, Alexandre Soulier, Elisabeth Trawinski, Jean-Michel Pawlotsky |
| EPI_ISL_3474841 | SELAS-D-LAB | Department of Virology, Henri Mondor University Hospital, Assistance Publique Hôpitaux de Paris, Université Paris-Est Créteil, INSERM U955 | Christophe Rodriguez, Slim Fourati, Vanessa Demontant, Guillaume Gricourt, Melissa N'Debi, Alexandre Soulier, Elisabeth Trawinski, Jean-Michel Pawlotsky |
| EPI_ISL_3474842, EPI_ISL_3474843, EPI_ISL_3474845 | Biogroup Bio Lam-LCD Saint-Denis | Department of Virology, Henri Mondor University Hospital, Assistance Publique Hôpitaux de Paris, Université Paris-Est Créteil, INSERM U955 | Christophe Rodriguez, Slim Fourati, Vanessa Demontant, Guillaume Gricourt, Melissa N'Debi, Alexandre Soulier, Elisabeth Trawinski, Jean-Michel Pawlotsky |
| EPI_ISL_3474846 | DYNALAB | Department of Virology, Henri Mondor University Hospital, Assistance Publique Hôpitaux de Paris, Université Paris-Est Créteil, INSERM U955 | Christophe Rodriguez, Slim Fourati, Vanessa Demontant, Guillaume Gricourt, Melissa N'Debi, Alexandre Soulier, Elisabeth Trawinski, Jean-Michel Pawlotsky |
| EPI_ISL_3474848, EPI_ISL_3474849 | BIOMNIS EUROFINS IVRY | Department of Virology, Henri Mondor University Hospital, Assistance Publique Hôpitaux de Paris, Université Paris-Est Créteil, INSERM U955 | Christophe Rodriguez, Slim Fourati, Vanessa Demontant, Guillaume Gricourt, Melissa N'Debi, Alexandre Soulier, Elisabeth Trawinski, Jean-Michel Pawlotsky |
| EPI_ISL_3474850 | DYNABIO UNILABS | Department of Virology, Henri Mondor University Hospital, Assistance Publique Hôpitaux de Paris, Université Paris-Est Créteil, INSERM U955 | Christophe Rodriguez, Slim Fourati, Vanessa Demontant, Guillaume Gricourt, Melissa N'Debi, Alexandre Soulier, Elisabeth Trawinski, Jean-Michel Pawlotsky |
| EPI_ISL_3474851 | CH SAINT DIE | Department of Virology, Henri Mondor University Hospital, Assistance Publique Hôpitaux de Paris, Université Paris-Est Créteil, | Christophe Rodriguez, Slim Fourati, Vanessa Demontant, Guillaume Gricourt, Melissa N'Debi, Alexandre Soulier, Elisabeth Trawinski, Jean-Michel Pawlotsky |

| | | | |
|---|---|---|---|
| | | INSERM U955 | |
| EPI_ISL_3474852 | SELAS-D-LAB | Department of Virology, Henri Mondor University Hospital, Assistance Publique Hôpitaux de Paris, Université Paris-Est Créteil, INSERM U955 | Christophe Rodriguez, Slim Fourati, Vanessa Demontant, Guillaume Gricourt, Melissa N'Debi, Alexandre Soulier, Elisabeth Trawinski, Jean-Michel Pawlotsky |
| EPI_ISL_3474854, EPI_ISL_3474855 | Biogroup Bio Lam-LCD Saint-Denis | Department of Virology, Henri Mondor University Hospital, Assistance Publique Hôpitaux de Paris, Université Paris-Est Créteil, INSERM U955 | Christophe Rodriguez, Slim Fourati, Vanessa Demontant, Guillaume Gricourt, Melissa N'Debi, Alexandre Soulier, Elisabeth Trawinski, Jean-Michel Pawlotsky |
| EPI_ISL_3474856 | Laboratoire ESPACEBIO | Department of Virology, Henri Mondor University Hospital, Assistance Publique Hôpitaux de Paris, Université Paris-Est Créteil, INSERM U955 | Christophe Rodriguez, Slim Fourati, Vanessa Demontant, Guillaume Gricourt, Melissa N'Debi, Alexandre Soulier, Elisabeth Trawinski, Jean-Michel Pawlotsky |
| EPI_ISL_3474857 | CH.LE RAINCY_MONTFERMEIL | Department of Virology, Henri Mondor University Hospital, Assistance Publique Hôpitaux de Paris, Université Paris-Est Créteil, INSERM U955 | Christophe Rodriguez, Slim Fourati, Vanessa Demontant, Guillaume Gricourt, Melissa N'Debi, Alexandre Soulier, Elisabeth Trawinski, Jean-Michel Pawlotsky |
| EPI_ISL_3474858 | Biogroup Bio Lam-LCD Saint-Denis | Department of Virology, Henri Mondor University Hospital, Assistance Publique Hôpitaux de Paris, Université Paris-Est Créteil, INSERM U955 | Christophe Rodriguez, Slim Fourati, Vanessa Demontant, Guillaume Gricourt, Melissa N'Debi, Alexandre Soulier, Elisabeth Trawinski, Jean-Michel Pawlotsky |
| EPI_ISL_3474859 | Laboratoire ESPACEBIO | Department of Virology, Henri Mondor University Hospital, Assistance Publique Hôpitaux de Paris, Université Paris-Est Créteil, INSERM U955 | Christophe Rodriguez, Slim Fourati, Vanessa Demontant, Guillaume Gricourt, Melissa N'Debi, Alexandre Soulier, Elisabeth Trawinski, Jean-Michel Pawlotsky |
| EPI_ISL_3474860 | BIO VSM LAB | Department of Virology, Henri Mondor University Hospital, Assistance Publique Hôpitaux de Paris, Université Paris-Est Créteil, INSERM U955 | Christophe Rodriguez, Slim Fourati, Vanessa Demontant, Guillaume Gricourt, Melissa N'Debi, Alexandre Soulier, Elisabeth Trawinski, Jean-Michel Pawlotsky |
| EPI_ISL_3474861 | Laboratoire ATOUTBIO JDC | Department of Virology, Henri Mondor University Hospital, Assistance Publique Hôpitaux de Paris, Université Paris-Est Créteil, INSERM U955 | Christophe Rodriguez, Slim Fourati, Vanessa Demontant, Guillaume Gricourt, Melissa N'Debi, Alexandre Soulier, Elisabeth Trawinski, Jean-Michel Pawlotsky |
| EPI_ISL_3474862 | DYNALAB | Department of Virology, Henri Mondor University Hospital, Assistance Publique Hôpitaux de Paris, Université Paris-Est Créteil, INSERM U955 | Christophe Rodriguez, Slim Fourati, Vanessa Demontant, Guillaume Gricourt, Melissa N'Debi, Alexandre Soulier, Elisabeth Trawinski, Jean-Michel Pawlotsky |
| EPI_ISL_3474863 | Laboratoire ATOUTBIO JDC | Department of Virology, Henri Mondor University Hospital, Assistance Publique Hôpitaux de Paris, Université Paris-Est Créteil, INSERM U955 | Christophe Rodriguez, Slim Fourati, Vanessa Demontant, Guillaume Gricourt, Melissa N'Debi, Alexandre Soulier, Elisabeth Trawinski, Jean-Michel Pawlotsky |
| EPI_ISL_3474864 | BIOMNIS EUROFINS IVRY | Department of Virology, Henri Mondor University Hospital, Assistance Publique Hôpitaux de Paris, Université Paris-Est Créteil, INSERM U955 | Christophe Rodriguez, Slim Fourati, Vanessa Demontant, Guillaume Gricourt, Melissa N'Debi, Alexandre Soulier, Elisabeth Trawinski, Jean-Michel Pawlotsky |
| EPI_ISL_3474865 | Laboratoire ATOUTBIO JDC | Department of Virology, Henri Mondor University Hospital, Assistance Publique Hôpitaux de Paris, Université Paris-Est Créteil, INSERM U955 | Christophe Rodriguez, Slim Fourati, Vanessa Demontant, Guillaume Gricourt, Melissa N'Debi, Alexandre Soulier, Elisabeth Trawinski, Jean-Michel Pawlotsky |
| EPI_ISL_3474866 | Biogroup Bio Lam-LCD Saint-Denis | Department of Virology, Henri Mondor University Hospital, Assistance Publique Hôpitaux de Paris, Université Paris-Est Créteil, INSERM U955 | Christophe Rodriguez, Slim Fourati, Vanessa Demontant, Guillaume Gricourt, Melissa N'Debi, Alexandre Soulier, Elisabeth Trawinski, Jean-Michel Pawlotsky |
| EPI_ISL_3474867, EPI_ISL_3474868 | SELAS-D-LAB | Department of Virology, Henri Mondor University Hospital, Assistance Publique Hôpitaux de Paris, Université Paris-Est Créteil, INSERM U955 | Christophe Rodriguez, Slim Fourati, Vanessa Demontant, Guillaume Gricourt, Melissa N'Debi, Alexandre Soulier, Elisabeth Trawinski, Jean-Michel Pawlotsky |
| EPI_ISL_3474869 | CH. ANDRE GREGOIRE | Department of Virology, Henri Mondor University Hospital, Assistance Publique Hôpitaux de Paris, Université Paris-Est Créteil, INSERM U955 | Christophe Rodriguez, Slim Fourati, Vanessa Demontant, Guillaume Gricourt, Melissa N'Debi, Alexandre Soulier, Elisabeth Trawinski, Jean-Michel Pawlotsky |
| EPI_ISL_3474870 | Laboratoire ESPACEBIO | Department of Virology, Henri Mondor University Hospital, Assistance Publique Hôpitaux de Paris, Université Paris-Est Créteil, INSERM U955 | Christophe Rodriguez, Slim Fourati, Vanessa Demontant, Guillaume Gricourt, Melissa N'Debi, Alexandre Soulier, Elisabeth Trawinski, Jean-Michel Pawlotsky |
| EPI_ISL_3474871 | CH.LE RAINCY_MONTFERMEIL | Department of Virology, Henri Mondor University Hospital, Assistance Publique Hôpitaux de Paris, Université Paris-Est Créteil, INSERM U955 | Christophe Rodriguez, Slim Fourati, Vanessa Demontant, Guillaume Gricourt, Melissa N'Debi, Alexandre Soulier, Elisabeth Trawinski, Jean-Michel Pawlotsky |
| EPI_ISL_3474872 | Biogroup Bio Lam-LCD Saint-Denis | Department of Virology, Henri Mondor University Hospital, Assistance Publique Hôpitaux de Paris, Université Paris-Est Créteil, INSERM U955 | Christophe Rodriguez, Slim Fourati, Vanessa Demontant, Guillaume Gricourt, Melissa N'Debi, Alexandre Soulier, Elisabeth Trawinski, Jean-Michel Pawlotsky |
| EPI_ISL_3474873 | BIOMNIS EUROFINS IVRY | Department of Virology, Henri Mondor University Hospital, Assistance Publique Hôpitaux de Paris, Université Paris-Est Créteil, INSERM U955 | Christophe Rodriguez, Slim Fourati, Vanessa Demontant, Guillaume Gricourt, Melissa N'Debi, Alexandre Soulier, Elisabeth Trawinski, Jean-Michel Pawlotsky |
| EPI_ISL_3474874, EPI_ISL_3474875, EPI_ISL_3474876, EPI_ISL_3474877, EPI_ISL_3474878, EPI_ISL_3474879 | Biogroup Bio Lam-LCD Saint-Denis | Department of Virology, Henri Mondor University Hospital, Assistance Publique Hôpitaux de Paris, Université Paris-Est Créteil, INSERM U955 | Christophe Rodriguez, Slim Fourati, Vanessa Demontant, Guillaume Gricourt, Melissa N'Debi, Alexandre Soulier, Elisabeth Trawinski, Jean-Michel Pawlotsky |
| EPI_ISL_3474880 | CH. BEGIN | Department of Virology, Henri Mondor University Hospital, Assistance Publique Hôpitaux de Paris, Université Paris-Est Créteil, INSERM U955 | Christophe Rodriguez, Slim Fourati, Vanessa Demontant, Guillaume Gricourt, Melissa N'Debi, Alexandre Soulier, Elisabeth Trawinski, Jean-Michel Pawlotsky |
| EPI_ISL_3474881 | Biogroup Bio Lam-LCD Saint-Denis | Department of Virology, Henri Mondor University Hospital, Assistance Publique Hôpitaux de Paris, Université Paris-Est Créteil, INSERM U955 | Christophe Rodriguez, Slim Fourati, Vanessa Demontant, Guillaume Gricourt, Melissa N'Debi, Alexandre Soulier, Elisabeth Trawinski, Jean-Michel Pawlotsky |
| EPI_ISL_3474882 | Laboratoire ATOUTBIO JDC | Department of Virology, Henri Mondor University Hospital, Assistance Publique Hôpitaux de Paris, Université Paris-Est Créteil, INSERM U955 | Christophe Rodriguez, Slim Fourati, Vanessa Demontant, Guillaume Gricourt, Melissa N'Debi, Alexandre Soulier, Elisabeth Trawinski, Jean-Michel Pawlotsky |
| EPI_ISL_3474884 | DYNALAB | Department of Virology, Henri Mondor University Hospital, Assistance Publique Hôpitaux de Paris, Université Paris-Est Créteil, INSERM U955 | Christophe Rodriguez, Slim Fourati, Vanessa Demontant, Guillaume Gricourt, Melissa N'Debi, Alexandre Soulier, Elisabeth Trawinski, Jean-Michel Pawlotsky |
| EPI_ISL_3474885 | Laboratoire ATOUTBIO JDC | Department of Virology, Henri Mondor University Hospital, Assistance Publique Hôpitaux de Paris, Université Paris-Est Créteil, INSERM U955 | Christophe Rodriguez, Slim Fourati, Vanessa Demontant, Guillaume Gricourt, Melissa N'Debi, Alexandre Soulier, Elisabeth Trawinski, Jean-Michel Pawlotsky |
| EPI_ISL_3474886 | Laboratoire ESPACEBIO | Department of Virology, Henri Mondor University Hospital, Assistance Publique Hôpitaux de Paris, Université Paris-Est Créteil, INSERM U955 | Christophe Rodriguez, Slim Fourati, Vanessa Demontant, Guillaume Gricourt, Melissa N'Debi, Alexandre Soulier, Elisabeth Trawinski, Jean-Michel Pawlotsky |
| EPI_ISL_3474888 | Biogroup Bio Lam-LCD Saint-Denis | Department of Virology, Henri Mondor University Hospital, Assistance Publique Hôpitaux de Paris, Université Paris-Est Créteil, INSERM U955 | Christophe Rodriguez, Slim Fourati, Vanessa Demontant, Guillaume Gricourt, Melissa N'Debi, Alexandre Soulier, Elisabeth Trawinski, Jean-Michel Pawlotsky |
| EPI_ISL_3474889 | Laboratoire ATOUTBIO JDC | Department of Virology, Henri Mondor University Hospital, Assistance Publique Hôpitaux de Paris, Université Paris-Est Créteil, INSERM U955 | Christophe Rodriguez, Slim Fourati, Vanessa Demontant, Guillaume Gricourt, Melissa N'Debi, Alexandre Soulier, Elisabeth Trawinski, Jean-Michel Pawlotsky |
| EPI_ISL_3474890 | Laboratoire ESPACEBIO | Department of Virology, Henri Mondor University Hospital, Assistance Publique Hôpitaux de Paris, Université Paris-Est Créteil, INSERM U955 | Christophe Rodriguez, Slim Fourati, Vanessa Demontant, Guillaume Gricourt, Melissa N'Debi, Alexandre Soulier, Elisabeth Trawinski, Jean-Michel Pawlotsky |
| EPI_ISL_3474891 | BIOCEANE | Department of Virology, Henri Mondor University Hospital, Assistance Publique Hôpitaux de Paris, Université Paris-Est Créteil, INSERM U955 | Christophe Rodriguez, Slim Fourati, Vanessa Demontant, Guillaume Gricourt, Melissa N'Debi, Alexandre Soulier, Elisabeth Trawinski, Jean-Michel Pawlotsky |
| EPI_ISL_3474892, EPI_ISL_3474893 | Biogroup Bio Lam-LCD Saint-Denis | Department of Virology, Henri Mondor University Hospital, Assistance Publique Hôpitaux de Paris, Université Paris-Est Créteil, | Christophe Rodriguez, Slim Fourati, Vanessa Demontant, Guillaume Gricourt, Melissa N'Debi, Alexandre Soulier, Elisabeth Trawinski, Jean-Michel Pawlotsky |

| | | | |
|---|---|---|---|
| | | INSERM U955 | |
| EPI_ISL_3474894 | SELAS-D-LAB | Department of Virology, Henri Mondor University Hospital, Assistance Publique Hôpitaux de Paris, Université Paris-Est Créteil, INSERM U955 | Christophe Rodriguez, Slim Fourati, Vanessa Demontant, Guillaume Gricourt, Melissa N'Debi, Alexandre Soulier, Elisabeth Trawinski, Jean-Michel Pawlotsky |
| EPI_ISL_3474895, EPI_ISL_3474896 | Biogroup Bio Lam-LCD Saint-Denis | Department of Virology, Henri Mondor University Hospital, Assistance Publique Hôpitaux de Paris, Université Paris-Est Créteil, INSERM U955 | Christophe Rodriguez, Slim Fourati, Vanessa Demontant, Guillaume Gricourt, Melissa N'Debi, Alexandre Soulier, Elisabeth Trawinski, Jean-Michel Pawlotsky |
| EPI_ISL_3474897 | LBM EVORIAL NEVERS | Department of Virology, Henri Mondor University Hospital, Assistance Publique Hôpitaux de Paris, Université Paris-Est Créteil, INSERM U955 | Christophe Rodriguez, Slim Fourati, Vanessa Demontant, Guillaume Gricourt, Melissa N'Debi, Alexandre Soulier, Elisabeth Trawinski, Jean-Michel Pawlotsky |
| EPI_ISL_3474898 | CH. ROBERT BISSON | Department of Virology, Henri Mondor University Hospital, Assistance Publique Hôpitaux de Paris, Université Paris-Est Créteil, INSERM U955 | Christophe Rodriguez, Slim Fourati, Vanessa Demontant, Guillaume Gricourt, Melissa N'Debi, Alexandre Soulier, Elisabeth Trawinski, Jean-Michel Pawlotsky |
| EPI_ISL_3474899 | SELAS-D-LAB | Department of Virology, Henri Mondor University Hospital, Assistance Publique Hôpitaux de Paris, Université Paris-Est Créteil, INSERM U955 | Christophe Rodriguez, Slim Fourati, Vanessa Demontant, Guillaume Gricourt, Melissa N'Debi, Alexandre Soulier, Elisabeth Trawinski, Jean-Michel Pawlotsky |
| EPI_ISL_3474900 | BIOMNIS EUROFINS IVRY | Department of Virology, Henri Mondor University Hospital, Assistance Publique Hôpitaux de Paris, Université Paris-Est Créteil, INSERM U955 | Christophe Rodriguez, Slim Fourati, Vanessa Demontant, Guillaume Gricourt, Melissa N'Debi, Alexandre Soulier, Elisabeth Trawinski, Jean-Michel Pawlotsky |
| EPI_ISL_3474901 | Biogroup Bio Lam-LCD Saint-Denis | Department of Virology, Henri Mondor University Hospital, Assistance Publique Hôpitaux de Paris, Université Paris-Est Créteil, INSERM U955 | Christophe Rodriguez, Slim Fourati, Vanessa Demontant, Guillaume Gricourt, Melissa N'Debi, Alexandre Soulier, Elisabeth Trawinski, Jean-Michel Pawlotsky |
| EPI_ISL_3474902 | CH.LE RAINCY_MONTFERMEIL | Department of Virology, Henri Mondor University Hospital, Assistance Publique Hôpitaux de Paris, Université Paris-Est Créteil, INSERM U955 | Christophe Rodriguez, Slim Fourati, Vanessa Demontant, Guillaume Gricourt, Melissa N'Debi, Alexandre Soulier, Elisabeth Trawinski, Jean-Michel Pawlotsky |
| EPI_ISL_3474903, EPI_ISL_3474904 | DYNALAB | Department of Virology, Henri Mondor University Hospital, Assistance Publique Hôpitaux de Paris, Université Paris-Est Créteil, INSERM U955 | Christophe Rodriguez, Slim Fourati, Vanessa Demontant, Guillaume Gricourt, Melissa N'Debi, Alexandre Soulier, Elisabeth Trawinski, Jean-Michel Pawlotsky |
| EPI_ISL_3474905, EPI_ISL_3474906, EPI_ISL_3474907 | Biogroup Bio Lam-LCD Saint-Denis | Department of Virology, Henri Mondor University Hospital, Assistance Publique Hôpitaux de Paris, Université Paris-Est Créteil, INSERM U955 | Christophe Rodriguez, Slim Fourati, Vanessa Demontant, Guillaume Gricourt, Melissa N'Debi, Alexandre Soulier, Elisabeth Trawinski, Jean-Michel Pawlotsky |
| EPI_ISL_3474908 | DYNABIO UNILABS | Department of Virology, Henri Mondor University Hospital, Assistance Publique Hôpitaux de Paris, Université Paris-Est Créteil, INSERM U955 | Christophe Rodriguez, Slim Fourati, Vanessa Demontant, Guillaume Gricourt, Melissa N'Debi, Alexandre Soulier, Elisabeth Trawinski, Jean-Michel Pawlotsky |
| EPI_ISL_3474909 | CH.LE RAINCY_MONTFERMEIL | Department of Virology, Henri Mondor University Hospital, Assistance Publique Hôpitaux de Paris, Université Paris-Est Créteil, INSERM U955 | Christophe Rodriguez, Slim Fourati, Vanessa Demontant, Guillaume Gricourt, Melissa N'Debi, Alexandre Soulier, Elisabeth Trawinski, Jean-Michel Pawlotsky |
| EPI_ISL_3474910 | BIOMNIS EUROFINS IVRY | Department of Virology, Henri Mondor University Hospital, Assistance Publique Hôpitaux de Paris, Université Paris-Est Créteil, INSERM U955 | Christophe Rodriguez, Slim Fourati, Vanessa Demontant, Guillaume Gricourt, Melissa N'Debi, Alexandre Soulier, Elisabeth Trawinski, Jean-Michel Pawlotsky |
| EPI_ISL_3474911, EPI_ISL_3474912, EPI_ISL_3474913 | Biogroup Bio Lam-LCD Saint-Denis | Department of Virology, Henri Mondor University Hospital, Assistance Publique Hôpitaux de Paris, Université Paris-Est Créteil, INSERM U955 | Christophe Rodriguez, Slim Fourati, Vanessa Demontant, Guillaume Gricourt, Melissa N'Debi, Alexandre Soulier, Elisabeth Trawinski, Jean-Michel Pawlotsky |
| EPI_ISL_3474914 | DYNABIO UNILABS | Department of Virology, Henri Mondor University Hospital, Assistance Publique Hôpitaux de Paris, Université Paris-Est Créteil, INSERM U955 | Christophe Rodriguez, Slim Fourati, Vanessa Demontant, Guillaume Gricourt, Melissa N'Debi, Alexandre Soulier, Elisabeth Trawinski, Jean-Michel Pawlotsky |
| EPI_ISL_3474915 | DYNALAB | Department of Virology, Henri Mondor University Hospital, Assistance Publique Hôpitaux de Paris, Université Paris-Est Créteil, INSERM U955 | Christophe Rodriguez, Slim Fourati, Vanessa Demontant, Guillaume Gricourt, Melissa N'Debi, Alexandre Soulier, Elisabeth Trawinski, Jean-Michel Pawlotsky |
| EPI_ISL_3474916 | BIOCEANE | Department of Virology, Henri Mondor University Hospital, Assistance Publique Hôpitaux de Paris, Université Paris-Est Créteil, INSERM U955 | Christophe Rodriguez, Slim Fourati, Vanessa Demontant, Guillaume Gricourt, Melissa N'Debi, Alexandre Soulier, Elisabeth Trawinski, Jean-Michel Pawlotsky |
| EPI_ISL_3474917, EPI_ISL_3474919 | Biogroup Bio Lam-LCD Saint-Denis | Department of Virology, Henri Mondor University Hospital, Assistance Publique Hôpitaux de Paris, Université Paris-Est Créteil, INSERM U955 | Christophe Rodriguez, Slim Fourati, Vanessa Demontant, Guillaume Gricourt, Melissa N'Debi, Alexandre Soulier, Elisabeth Trawinski, Jean-Michel Pawlotsky |
| EPI_ISL_3474920 | GROUPE HOSPITALIER SUD ILE DE FRANCE | Department of Virology, Henri Mondor University Hospital, Assistance Publique Hôpitaux de Paris, Université Paris-Est Créteil, INSERM U955 | Christophe Rodriguez, Slim Fourati, Vanessa Demontant, Guillaume Gricourt, Melissa N'Debi, Alexandre Soulier, Elisabeth Trawinski, Jean-Michel Pawlotsky |
| EPI_ISL_3474921 | BIOMNIS EUROFINS IVRY | Department of Virology, Henri Mondor University Hospital, Assistance Publique Hôpitaux de Paris, Université Paris-Est Créteil, INSERM U955 | Christophe Rodriguez, Slim Fourati, Vanessa Demontant, Guillaume Gricourt, Melissa N'Debi, Alexandre Soulier, Elisabeth Trawinski, Jean-Michel Pawlotsky |
| EPI_ISL_3474922 | Biogroup Bio Lam-LCD Saint-Denis | Department of Virology, Henri Mondor University Hospital, Assistance Publique Hôpitaux de Paris, Université Paris-Est Créteil, INSERM U955 | Christophe Rodriguez, Slim Fourati, Vanessa Demontant, Guillaume Gricourt, Melissa N'Debi, Alexandre Soulier, Elisabeth Trawinski, Jean-Michel Pawlotsky |
| EPI_ISL_3474923 | Laboratoire ESPACEBIO | Department of Virology, Henri Mondor University Hospital, Assistance Publique Hôpitaux de Paris, Université Paris-Est Créteil, INSERM U955 | Christophe Rodriguez, Slim Fourati, Vanessa Demontant, Guillaume Gricourt, Melissa N'Debi, Alexandre Soulier, Elisabeth Trawinski, Jean-Michel Pawlotsky |
| EPI_ISL_3474924 | BIOCEANE | Department of Virology, Henri Mondor University Hospital, Assistance Publique Hôpitaux de Paris, Université Paris-Est Créteil, INSERM U955 | Christophe Rodriguez, Slim Fourati, Vanessa Demontant, Guillaume Gricourt, Melissa N'Debi, Alexandre Soulier, Elisabeth Trawinski, Jean-Michel Pawlotsky |
| EPI_ISL_3474925 | SELAS-D-LAB | Department of Virology, Henri Mondor University Hospital, Assistance Publique Hôpitaux de Paris, Université Paris-Est Créteil, INSERM U955 | Christophe Rodriguez, Slim Fourati, Vanessa Demontant, Guillaume Gricourt, Melissa N'Debi, Alexandre Soulier, Elisabeth Trawinski, Jean-Michel Pawlotsky |
| EPI_ISL_3474926, EPI_ISL_3474927 | Biogroup Bio Lam-LCD Saint-Denis | Department of Virology, Henri Mondor University Hospital, Assistance Publique Hôpitaux de Paris, Université Paris-Est Créteil, INSERM U955 | Christophe Rodriguez, Slim Fourati, Vanessa Demontant, Guillaume Gricourt, Melissa N'Debi, Alexandre Soulier, Elisabeth Trawinski, Jean-Michel Pawlotsky |
| EPI_ISL_3474928 | DYNALAB | Department of Virology, Henri Mondor University Hospital, Assistance Publique Hôpitaux de Paris, Université Paris-Est Créteil, INSERM U955 | Christophe Rodriguez, Slim Fourati, Vanessa Demontant, Guillaume Gricourt, Melissa N'Debi, Alexandre Soulier, Elisabeth Trawinski, Jean-Michel Pawlotsky |
| EPI_ISL_3474929 | SELAS-D-LAB | Department of Virology, Henri Mondor University Hospital, Assistance Publique Hôpitaux de Paris, Université Paris-Est Créteil, INSERM U955 | Christophe Rodriguez, Slim Fourati, Vanessa Demontant, Guillaume Gricourt, Melissa N'Debi, Alexandre Soulier, Elisabeth Trawinski, Jean-Michel Pawlotsky |
| EPI_ISL_3474930 | Laboratoire ATOUTBIO JDC | Department of Virology, Henri Mondor University Hospital, Assistance Publique Hôpitaux de Paris, Université Paris-Est Créteil, INSERM U955 | Christophe Rodriguez, Slim Fourati, Vanessa Demontant, Guillaume Gricourt, Melissa N'Debi, Alexandre Soulier, Elisabeth Trawinski, Jean-Michel Pawlotsky |
| EPI_ISL_3474931 | CH. ANDRE GREGOIRE | Department of Virology, Henri Mondor University Hospital, Assistance Publique Hôpitaux de Paris, Université Paris-Est Créteil, INSERM U955 | Christophe Rodriguez, Slim Fourati, Vanessa Demontant, Guillaume Gricourt, Melissa N'Debi, Alexandre Soulier, Elisabeth Trawinski, Jean-Michel Pawlotsky |
| EPI_ISL_3474932 | SELAS-D-LAB | Department of Virology, Henri Mondor University Hospital, Assistance Publique Hôpitaux de Paris, Université Paris-Est Créteil, INSERM U955 | Christophe Rodriguez, Slim Fourati, Vanessa Demontant, Guillaume Gricourt, Melissa N'Debi, Alexandre Soulier, Elisabeth Trawinski, Jean-Michel Pawlotsky |
| EPI_ISL_3474933 | Laboratoire ESPACEBIO | Department of Virology, Henri Mondor University Hospital, Assistance Publique Hôpitaux de Paris, Université Paris-Est Créteil, INSERM U955 | Christophe Rodriguez, Slim Fourati, Vanessa Demontant, Guillaume Gricourt, Melissa N'Debi, Alexandre Soulier, Elisabeth Trawinski, Jean-Michel Pawlotsky |
| EPI_ISL_3474934, | Biogroup Bio Lam-LCD Saint-Denis | Department of Virology, Henri Mondor University Hospital, | Christophe Rodriguez, Slim Fourati, Vanessa Demontant, Guillaume Gricourt, Melissa N'Debi, Alexandre Soulier, Elisabeth Trawinski, Jean-Michel Pawlotsky |

| | | | |
|---|---|---|---|
| EPI_ISL_3474935 | | Assistance Publique Hôpitaux de Paris, Université Paris-Est Créteil, INSERM U955 | |
| EPI_ISL_3474936 | Laboratoire ATOUTBIO JDC | Department of Virology, Henri Mondor University Hospital, Assistance Publique Hôpitaux de Paris, Université Paris-Est Créteil, INSERM U955 | Christophe Rodriguez, Slim Fourati, Vanessa Demontant, Guillaume Gricourt, Melissa N'Debi, Alexandre Soulier, Elisabeth Trawinski, Jean-Michel Pawlotsky |
| EPI_ISL_3474937 | Biogroup Bio Lam-LCD Saint-Denis | Department of Virology, Henri Mondor University Hospital, Assistance Publique Hôpitaux de Paris, Université Paris-Est Créteil, INSERM U955 | Christophe Rodriguez, Slim Fourati, Vanessa Demontant, Guillaume Gricourt, Melissa N'Debi, Alexandre Soulier, Elisabeth Trawinski, Jean-Michel Pawlotsky |
| EPI_ISL_3474938 | CHI VILLENEUVE ST GEORGES | Department of Virology, Henri Mondor University Hospital, Assistance Publique Hôpitaux de Paris, Université Paris-Est Créteil, INSERM U955 | Christophe Rodriguez, Slim Fourati, Vanessa Demontant, Guillaume Gricourt, Melissa N'Debi, Alexandre Soulier, Elisabeth Trawinski, Jean-Michel Pawlotsky |
| EPI_ISL_3474939 | Laboratoire ESPACEBIO | Department of Virology, Henri Mondor University Hospital, Assistance Publique Hôpitaux de Paris, Université Paris-Est Créteil, INSERM U955 | Christophe Rodriguez, Slim Fourati, Vanessa Demontant, Guillaume Gricourt, Melissa N'Debi, Alexandre Soulier, Elisabeth Trawinski, Jean-Michel Pawlotsky |
| EPI_ISL_3474940, EPI_ISL_3474941 | GROUPE HOSPITALIER SUD ILE DE FRANCE | Department of Virology, Henri Mondor University Hospital, Assistance Publique Hôpitaux de Paris, Université Paris-Est Créteil, INSERM U955 | Christophe Rodriguez, Slim Fourati, Vanessa Demontant, Guillaume Gricourt, Melissa N'Debi, Alexandre Soulier, Elisabeth Trawinski, Jean-Michel Pawlotsky |
| EPI_ISL_3474942 | Laboratoire LENYS | Department of Virology, Henri Mondor University Hospital, Assistance Publique Hôpitaux de Paris, Université Paris-Est Créteil, INSERM U955 | Christophe Rodriguez, Slim Fourati, Vanessa Demontant, Guillaume Gricourt, Melissa N'Debi, Alexandre Soulier, Elisabeth Trawinski, Jean-Michel Pawlotsky |
| EPI_ISL_3474943 | SELAS-D-LAB | Department of Virology, Henri Mondor University Hospital, Assistance Publique Hôpitaux de Paris, Université Paris-Est Créteil, INSERM U955 | Christophe Rodriguez, Slim Fourati, Vanessa Demontant, Guillaume Gricourt, Melissa N'Debi, Alexandre Soulier, Elisabeth Trawinski, Jean-Michel Pawlotsky |
| EPI_ISL_3474946 | Plateau technique Biomer Metz Verriers | Department of Virology, Henri Mondor University Hospital, Assistance Publique Hôpitaux de Paris, Université Paris-Est Créteil, INSERM U955 | Christophe Rodriguez, Slim Fourati, Vanessa Demontant, Guillaume Gricourt, Melissa N'Debi, Alexandre Soulier, Elisabeth Trawinski, Jean-Michel Pawlotsky |
| EPI_ISL_3474947 | Biogroup Bio Lam-LCD Saint-Denis | Department of Virology, Henri Mondor University Hospital, Assistance Publique Hôpitaux de Paris, Université Paris-Est Créteil, INSERM U955 | Christophe Rodriguez, Slim Fourati, Vanessa Demontant, Guillaume Gricourt, Melissa N'Debi, Alexandre Soulier, Elisabeth Trawinski, Jean-Michel Pawlotsky |
| EPI_ISL_3474948 | Laboratoire ESPACEBIO | Department of Virology, Henri Mondor University Hospital, Assistance Publique Hôpitaux de Paris, Université Paris-Est Créteil, INSERM U955 | Christophe Rodriguez, Slim Fourati, Vanessa Demontant, Guillaume Gricourt, Melissa N'Debi, Alexandre Soulier, Elisabeth Trawinski, Jean-Michel Pawlotsky |
| EPI_ISL_3474949 | BIOCEANE | Department of Virology, Henri Mondor University Hospital, Assistance Publique Hôpitaux de Paris, Université Paris-Est Créteil, INSERM U955 | Christophe Rodriguez, Slim Fourati, Vanessa Demontant, Guillaume Gricourt, Melissa N'Debi, Alexandre Soulier, Elisabeth Trawinski, Jean-Michel Pawlotsky |
| EPI_ISL_3474950 | SELAS-D-LAB | Department of Virology, Henri Mondor University Hospital, Assistance Publique Hôpitaux de Paris, Université Paris-Est Créteil, INSERM U955 | Christophe Rodriguez, Slim Fourati, Vanessa Demontant, Guillaume Gricourt, Melissa N'Debi, Alexandre Soulier, Elisabeth Trawinski, Jean-Michel Pawlotsky |
| EPI_ISL_3474951 | Biogroup Bio Lam-LCD Saint-Denis | Department of Virology, Henri Mondor University Hospital, Assistance Publique Hôpitaux de Paris, Université Paris-Est Créteil, INSERM U955 | Christophe Rodriguez, Slim Fourati, Vanessa Demontant, Guillaume Gricourt, Melissa N'Debi, Alexandre Soulier, Elisabeth Trawinski, Jean-Michel Pawlotsky |
| EPI_ISL_3474952 | Plateau technique Biomer Metz Verriers | Department of Virology, Henri Mondor University Hospital, Assistance Publique Hôpitaux de Paris, Université Paris-Est Créteil, INSERM U955 | Christophe Rodriguez, Slim Fourati, Vanessa Demontant, Guillaume Gricourt, Melissa N'Debi, Alexandre Soulier, Elisabeth Trawinski, Jean-Michel Pawlotsky |
| EPI_ISL_3474953 | Biogroup Bio Lam-LCD Saint-Denis | Department of Virology, Henri Mondor University Hospital, Assistance Publique Hôpitaux de Paris, Université Paris-Est Créteil, INSERM U955 | Christophe Rodriguez, Slim Fourati, Vanessa Demontant, Guillaume Gricourt, Melissa N'Debi, Alexandre Soulier, Elisabeth Trawinski, Jean-Michel Pawlotsky |
| EPI_ISL_3474954 | DYNALAB | Department of Virology, Henri Mondor University Hospital, Assistance Publique Hôpitaux de Paris, Université Paris-Est Créteil, INSERM U955 | Christophe Rodriguez, Slim Fourati, Vanessa Demontant, Guillaume Gricourt, Melissa N'Debi, Alexandre Soulier, Elisabeth Trawinski, Jean-Michel Pawlotsky |
| EPI_ISL_3474955, EPI_ISL_3474956, EPI_ISL_3474957 | Biogroup Bio Lam-LCD Saint-Denis | Department of Virology, Henri Mondor University Hospital, Assistance Publique Hôpitaux de Paris, Université Paris-Est Créteil, INSERM U955 | Christophe Rodriguez, Slim Fourati, Vanessa Demontant, Guillaume Gricourt, Melissa N'Debi, Alexandre Soulier, Elisabeth Trawinski, Jean-Michel Pawlotsky |
| EPI_ISL_3474958 | BIOCEANE | Department of Virology, Henri Mondor University Hospital, Assistance Publique Hôpitaux de Paris, Université Paris-Est Créteil, INSERM U955 | Christophe Rodriguez, Slim Fourati, Vanessa Demontant, Guillaume Gricourt, Melissa N'Debi, Alexandre Soulier, Elisabeth Trawinski, Jean-Michel Pawlotsky |
| EPI_ISL_3474959, EPI_ISL_3474960 | Biogroup Bio Lam-LCD Saint-Denis | Department of Virology, Henri Mondor University Hospital, Assistance Publique Hôpitaux de Paris, Université Paris-Est Créteil, INSERM U955 | Christophe Rodriguez, Slim Fourati, Vanessa Demontant, Guillaume Gricourt, Melissa N'Debi, Alexandre Soulier, Elisabeth Trawinski, Jean-Michel Pawlotsky |
| EPI_ISL_3474961 | SELAS-D-LAB | Department of Virology, Henri Mondor University Hospital, Assistance Publique Hôpitaux de Paris, Université Paris-Est Créteil, INSERM U955 | Christophe Rodriguez, Slim Fourati, Vanessa Demontant, Guillaume Gricourt, Melissa N'Debi, Alexandre Soulier, Elisabeth Trawinski, Jean-Michel Pawlotsky |
| EPI_ISL_3474962 | Biogroup Bio Lam-LCD Saint-Denis | Department of Virology, Henri Mondor University Hospital, Assistance Publique Hôpitaux de Paris, Université Paris-Est Créteil, INSERM U955 | Christophe Rodriguez, Slim Fourati, Vanessa Demontant, Guillaume Gricourt, Melissa N'Debi, Alexandre Soulier, Elisabeth Trawinski, Jean-Michel Pawlotsky |
| EPI_ISL_3474963 | DYNALAB | Department of Virology, Henri Mondor University Hospital, Assistance Publique Hôpitaux de Paris, Université Paris-Est Créteil, INSERM U955 | Christophe Rodriguez, Slim Fourati, Vanessa Demontant, Guillaume Gricourt, Melissa N'Debi, Alexandre Soulier, Elisabeth Trawinski, Jean-Michel Pawlotsky |
| EPI_ISL_3474964 | Biogroup Bio Lam-LCD Saint-Denis | Department of Virology, Henri Mondor University Hospital, Assistance Publique Hôpitaux de Paris, Université Paris-Est Créteil, INSERM U955 | Christophe Rodriguez, Slim Fourati, Vanessa Demontant, Guillaume Gricourt, Melissa N'Debi, Alexandre Soulier, Elisabeth Trawinski, Jean-Michel Pawlotsky |
| EPI_ISL_3474965 | BIOCEANE | Department of Virology, Henri Mondor University Hospital, Assistance Publique Hôpitaux de Paris, Université Paris-Est Créteil, INSERM U955 | Christophe Rodriguez, Slim Fourati, Vanessa Demontant, Guillaume Gricourt, Melissa N'Debi, Alexandre Soulier, Elisabeth Trawinski, Jean-Michel Pawlotsky |
| EPI_ISL_3474966 | BIOMNIS EUROFINS IVRY | Department of Virology, Henri Mondor University Hospital, Assistance Publique Hôpitaux de Paris, Université Paris-Est Créteil, INSERM U955 | Christophe Rodriguez, Slim Fourati, Vanessa Demontant, Guillaume Gricourt, Melissa N'Debi, Alexandre Soulier, Elisabeth Trawinski, Jean-Michel Pawlotsky |
| EPI_ISL_3474967 | Laboratoire ATOUTBIO JDC | Department of Virology, Henri Mondor University Hospital, Assistance Publique Hôpitaux de Paris, Université Paris-Est Créteil, INSERM U955 | Christophe Rodriguez, Slim Fourati, Vanessa Demontant, Guillaume Gricourt, Melissa N'Debi, Alexandre Soulier, Elisabeth Trawinski, Jean-Michel Pawlotsky |
| EPI_ISL_3474968 | Biogroup Bio Lam-LCD Saint-Denis | Department of Virology, Henri Mondor University Hospital, Assistance Publique Hôpitaux de Paris, Université Paris-Est Créteil, INSERM U955 | Christophe Rodriguez, Slim Fourati, Vanessa Demontant, Guillaume Gricourt, Melissa N'Debi, Alexandre Soulier, Elisabeth Trawinski, Jean-Michel Pawlotsky |
| EPI_ISL_3474969 | LBM EVORIAL NEVERS | Department of Virology, Henri Mondor University Hospital, Assistance Publique Hôpitaux de Paris, Université Paris-Est Créteil, INSERM U955 | Christophe Rodriguez, Slim Fourati, Vanessa Demontant, Guillaume Gricourt, Melissa N'Debi, Alexandre Soulier, Elisabeth Trawinski, Jean-Michel Pawlotsky |
| EPI_ISL_3474970 | Plateau technique Biomer Metz Verriers | Department of Virology, Henri Mondor University Hospital, Assistance Publique Hôpitaux de Paris, Université Paris-Est Créteil, INSERM U955 | Christophe Rodriguez, Slim Fourati, Vanessa Demontant, Guillaume Gricourt, Melissa N'Debi, Alexandre Soulier, Elisabeth Trawinski, Jean-Michel Pawlotsky |
| EPI_ISL_3474971 | GROUPE HOSPITALIER SUD ILE DE FRANCE | Department of Virology, Henri Mondor University Hospital, Assistance Publique Hôpitaux de Paris, Université Paris-Est Créteil, INSERM U955 | Christophe Rodriguez, Slim Fourati, Vanessa Demontant, Guillaume Gricourt, Melissa N'Debi, Alexandre Soulier, Elisabeth Trawinski, Jean-Michel Pawlotsky |
| EPI_ISL_3474972, EPI_ISL_3474973 | Biogroup Bio Lam-LCD Saint-Denis | Department of Virology, Henri Mondor University Hospital, Assistance Publique Hôpitaux de Paris, Université Paris-Est Créteil, INSERM U955 | Christophe Rodriguez, Slim Fourati, Vanessa Demontant, Guillaume Gricourt, Melissa N'Debi, Alexandre Soulier, Elisabeth Trawinski, Jean-Michel Pawlotsky |
| EPI_ISL_3474975 | Laboratoire ATOUTBIO JDC | Department of Virology, Henri Mondor University Hospital, | Christophe Rodriguez, Slim Fourati, Vanessa Demontant, Guillaume Gricourt, Melissa N'Debi, Alexandre Soulier, Elisabeth Trawinski, Jean-Michel Pawlotsky |

| | | | |
|---|---|---|---|
| | | Assistance Publique Hôpitaux de Paris, Université Paris-Est Créteil, INSERM U955 | |
| EPI_ISL_3474976 | BIOMNIS EUROFINS IVRY | Department of Virology, Henri Mondor University Hospital, Assistance Publique Hôpitaux de Paris, Université Paris-Est Créteil, INSERM U955 | Christophe Rodriguez, Slim Fourati, Vanessa Demontant, Guillaume Gricourt, Melissa N'Debi, Alexandre Soulier, Elisabeth Trawinski, Jean-Michel Pawlotsky |
| EPI_ISL_3474978 | CH. ANDRE GREGOIRE | Department of Virology, Henri Mondor University Hospital, Assistance Publique Hôpitaux de Paris, Université Paris-Est Créteil, INSERM U955 | Christophe Rodriguez, Slim Fourati, Vanessa Demontant, Guillaume Gricourt, Melissa N'Debi, Alexandre Soulier, Elisabeth Trawinski, Jean-Michel Pawlotsky |
| EPI_ISL_3474979 | BIO VSM LAB | Department of Virology, Henri Mondor University Hospital, Assistance Publique Hôpitaux de Paris, Université Paris-Est Créteil, INSERM U955 | Christophe Rodriguez, Slim Fourati, Vanessa Demontant, Guillaume Gricourt, Melissa N'Debi, Alexandre Soulier, Elisabeth Trawinski, Jean-Michel Pawlotsky |
| EPI_ISL_3474980 | Biogroup Bio Lam-LCD Saint-Denis | Department of Virology, Henri Mondor University Hospital, Assistance Publique Hôpitaux de Paris, Université Paris-Est Créteil, INSERM U955 | Christophe Rodriguez, Slim Fourati, Vanessa Demontant, Guillaume Gricourt, Melissa N'Debi, Alexandre Soulier, Elisabeth Trawinski, Jean-Michel Pawlotsky |
| EPI_ISL_3474982 | GROUPE HOSPITALIER SUD ILE DE FRANCE | Department of Virology, Henri Mondor University Hospital, Assistance Publique Hôpitaux de Paris, Université Paris-Est Créteil, INSERM U955 | Christophe Rodriguez, Slim Fourati, Vanessa Demontant, Guillaume Gricourt, Melissa N'Debi, Alexandre Soulier, Elisabeth Trawinski, Jean-Michel Pawlotsky |
| EPI_ISL_3474983 | Plateau technique Biomer Metz Verriers | Department of Virology, Henri Mondor University Hospital, Assistance Publique Hôpitaux de Paris, Université Paris-Est Créteil, INSERM U955 | Christophe Rodriguez, Slim Fourati, Vanessa Demontant, Guillaume Gricourt, Melissa N'Debi, Alexandre Soulier, Elisabeth Trawinski, Jean-Michel Pawlotsky |
| EPI_ISL_3474984 | DYNALAB | Department of Virology, Henri Mondor University Hospital, Assistance Publique Hôpitaux de Paris, Université Paris-Est Créteil, INSERM U955 | Christophe Rodriguez, Slim Fourati, Vanessa Demontant, Guillaume Gricourt, Melissa N'Debi, Alexandre Soulier, Elisabeth Trawinski, Jean-Michel Pawlotsky |
| EPI_ISL_3474985 | Laboratoire ESPACEBIO | Department of Virology, Henri Mondor University Hospital, Assistance Publique Hôpitaux de Paris, Université Paris-Est Créteil, INSERM U955 | Christophe Rodriguez, Slim Fourati, Vanessa Demontant, Guillaume Gricourt, Melissa N'Debi, Alexandre Soulier, Elisabeth Trawinski, Jean-Michel Pawlotsky |
| EPI_ISL_3474986 | CH. ANDRE GREGOIRE | Department of Virology, Henri Mondor University Hospital, Assistance Publique Hôpitaux de Paris, Université Paris-Est Créteil, INSERM U955 | Christophe Rodriguez, Slim Fourati, Vanessa Demontant, Guillaume Gricourt, Melissa N'Debi, Alexandre Soulier, Elisabeth Trawinski, Jean-Michel Pawlotsky |
| EPI_ISL_3474987 | Laboratoire ESPACEBIO | Department of Virology, Henri Mondor University Hospital, Assistance Publique Hôpitaux de Paris, Université Paris-Est Créteil, INSERM U955 | Christophe Rodriguez, Slim Fourati, Vanessa Demontant, Guillaume Gricourt, Melissa N'Debi, Alexandre Soulier, Elisabeth Trawinski, Jean-Michel Pawlotsky |
| EPI_ISL_3474988 | Biogroup Bio Lam-LCD Saint-Denis | Department of Virology, Henri Mondor University Hospital, Assistance Publique Hôpitaux de Paris, Université Paris-Est Créteil, INSERM U955 | Christophe Rodriguez, Slim Fourati, Vanessa Demontant, Guillaume Gricourt, Melissa N'Debi, Alexandre Soulier, Elisabeth Trawinski, Jean-Michel Pawlotsky |
| EPI_ISL_3474989 | Laboratoire ESPACEBIO | Department of Virology, Henri Mondor University Hospital, Assistance Publique Hôpitaux de Paris, Université Paris-Est Créteil, INSERM U955 | Christophe Rodriguez, Slim Fourati, Vanessa Demontant, Guillaume Gricourt, Melissa N'Debi, Alexandre Soulier, Elisabeth Trawinski, Jean-Michel Pawlotsky |
| EPI_ISL_3474990 | BIOMNIS EUROFINS IVRY | Department of Virology, Henri Mondor University Hospital, Assistance Publique Hôpitaux de Paris, Université Paris-Est Créteil, INSERM U955 | Christophe Rodriguez, Slim Fourati, Vanessa Demontant, Guillaume Gricourt, Melissa N'Debi, Alexandre Soulier, Elisabeth Trawinski, Jean-Michel Pawlotsky |
| EPI_ISL_3474991 | CH. BEGIN | Department of Virology, Henri Mondor University Hospital, Assistance Publique Hôpitaux de Paris, Université Paris-Est Créteil, INSERM U955 | Christophe Rodriguez, Slim Fourati, Vanessa Demontant, Guillaume Gricourt, Melissa N'Debi, Alexandre Soulier, Elisabeth Trawinski, Jean-Michel Pawlotsky |
| EPI_ISL_3474992 | Biogroup Bio Lam-LCD Saint-Denis | Department of Virology, Henri Mondor University Hospital, Assistance Publique Hôpitaux de Paris, Université Paris-Est Créteil, INSERM U955 | Christophe Rodriguez, Slim Fourati, Vanessa Demontant, Guillaume Gricourt, Melissa N'Debi, Alexandre Soulier, Elisabeth Trawinski, Jean-Michel Pawlotsky |
| EPI_ISL_3474996 | Laboratoire ATOUTBIO JDC | Department of Virology, Henri Mondor University Hospital, Assistance Publique Hôpitaux de Paris, Université Paris-Est Créteil, INSERM U955 | Christophe Rodriguez, Slim Fourati, Vanessa Demontant, Guillaume Gricourt, Melissa N'Debi, Alexandre Soulier, Elisabeth Trawinski, Jean-Michel Pawlotsky |
| EPI_ISL_3474997 | Biogroup Bio Lam-LCD Saint-Denis | Department of Virology, Henri Mondor University Hospital, Assistance Publique Hôpitaux de Paris, Université Paris-Est Créteil, INSERM U955 | Christophe Rodriguez, Slim Fourati, Vanessa Demontant, Guillaume Gricourt, Melissa N'Debi, Alexandre Soulier, Elisabeth Trawinski, Jean-Michel Pawlotsky |
| EPI_ISL_3474998 | BIOCEANE | Department of Virology, Henri Mondor University Hospital, Assistance Publique Hôpitaux de Paris, Université Paris-Est Créteil, INSERM U955 | Christophe Rodriguez, Slim Fourati, Vanessa Demontant, Guillaume Gricourt, Melissa N'Debi, Alexandre Soulier, Elisabeth Trawinski, Jean-Michel Pawlotsky |
| EPI_ISL_3474999 | CHRU de NANCY | Department of Virology, Henri Mondor University Hospital, Assistance Publique Hôpitaux de Paris, Université Paris-Est Créteil, INSERM U955 | Christophe Rodriguez, Slim Fourati, Vanessa Demontant, Guillaume Gricourt, Melissa N'Debi, Alexandre Soulier, Elisabeth Trawinski, Jean-Michel Pawlotsky |
| EPI_ISL_3475000, EPI_ISL_3475001, EPI_ISL_3475002, EPI_ISL_3475003, EPI_ISL_3475004, EPI_ISL_3475005, EPI_ISL_3475006, EPI_ISL_3475007, EPI_ISL_3475008 | Biogroup Bio Lam-LCD Saint-Denis | Department of Virology, Henri Mondor University Hospital, Assistance Publique Hôpitaux de Paris, Université Paris-Est Créteil, INSERM U955 | Christophe Rodriguez, Slim Fourati, Vanessa Demontant, Guillaume Gricourt, Melissa N'Debi, Alexandre Soulier, Elisabeth Trawinski, Jean-Michel Pawlotsky |
| EPI_ISL_3475009 | Laboratoire ESPACEBIO | Department of Virology, Henri Mondor University Hospital, Assistance Publique Hôpitaux de Paris, Université Paris-Est Créteil, INSERM U955 | Christophe Rodriguez, Slim Fourati, Vanessa Demontant, Guillaume Gricourt, Melissa N'Debi, Alexandre Soulier, Elisabeth Trawinski, Jean-Michel Pawlotsky |
| EPI_ISL_3475010 | Biogroup Bio Lam-LCD Saint-Denis | Department of Virology, Henri Mondor University Hospital, Assistance Publique Hôpitaux de Paris, Université Paris-Est Créteil, INSERM U955 | Christophe Rodriguez, Slim Fourati, Vanessa Demontant, Guillaume Gricourt, Melissa N'Debi, Alexandre Soulier, Elisabeth Trawinski, Jean-Michel Pawlotsky |
| EPI_ISL_3475011 | CH. ROBERT BISSON | Department of Virology, Henri Mondor University Hospital, Assistance Publique Hôpitaux de Paris, Université Paris-Est Créteil, INSERM U955 | Christophe Rodriguez, Slim Fourati, Vanessa Demontant, Guillaume Gricourt, Melissa N'Debi, Alexandre Soulier, Elisabeth Trawinski, Jean-Michel Pawlotsky |
| EPI_ISL_3475012 | DYNALAB | Department of Virology, Henri Mondor University Hospital, Assistance Publique Hôpitaux de Paris, Université Paris-Est Créteil, INSERM U955 | Christophe Rodriguez, Slim Fourati, Vanessa Demontant, Guillaume Gricourt, Melissa N'Debi, Alexandre Soulier, Elisabeth Trawinski, Jean-Michel Pawlotsky |
| EPI_ISL_3475013 | Biogroup Bio Lam-LCD Saint-Denis | Department of Virology, Henri Mondor University Hospital, Assistance Publique Hôpitaux de Paris, Université Paris-Est Créteil, INSERM U955 | Christophe Rodriguez, Slim Fourati, Vanessa Demontant, Guillaume Gricourt, Melissa N'Debi, Alexandre Soulier, Elisabeth Trawinski, Jean-Michel Pawlotsky |
| EPI_ISL_3475014 | Laboratoire ESPACEBIO | Department of Virology, Henri Mondor University Hospital, Assistance Publique Hôpitaux de Paris, Université Paris-Est Créteil, INSERM U955 | Christophe Rodriguez, Slim Fourati, Vanessa Demontant, Guillaume Gricourt, Melissa N'Debi, Alexandre Soulier, Elisabeth Trawinski, Jean-Michel Pawlotsky |
| EPI_ISL_3475015 | Laboratoire LENYS | Department of Virology, Henri Mondor University Hospital, Assistance Publique Hôpitaux de Paris, Université Paris-Est Créteil, INSERM U955 | Christophe Rodriguez, Slim Fourati, Vanessa Demontant, Guillaume Gricourt, Melissa N'Debi, Alexandre Soulier, Elisabeth Trawinski, Jean-Michel Pawlotsky |
| EPI_ISL_3475016 | BIOMNIS EUROFINS IVRY | Department of Virology, Henri Mondor University Hospital, Assistance Publique Hôpitaux de Paris, Université Paris-Est Créteil, INSERM U955 | Christophe Rodriguez, Slim Fourati, Vanessa Demontant, Guillaume Gricourt, Melissa N'Debi, Alexandre Soulier, Elisabeth Trawinski, Jean-Michel Pawlotsky |
| EPI_ISL_3475018 | Biogroup Bio Lam-LCD Saint-Denis | Department of Virology, Henri Mondor University Hospital, Assistance Publique Hôpitaux de Paris, Université Paris-Est Créteil, INSERM U955 | Christophe Rodriguez, Slim Fourati, Vanessa Demontant, Guillaume Gricourt, Melissa N'Debi, Alexandre Soulier, Elisabeth Trawinski, Jean-Michel Pawlotsky |
| EPI_ISL_3475019 | BIOMNIS EUROFINS IVRY | Department of Virology, Henri Mondor University Hospital, Assistance Publique Hôpitaux de Paris, Université Paris-Est Créteil, | Christophe Rodriguez, Slim Fourati, Vanessa Demontant, Guillaume Gricourt, Melissa N'Debi, Alexandre Soulier, Elisabeth Trawinski, Jean-Michel Pawlotsky |

| | | | |
|---|---|---|---|
| | | INSERM U955 | |
| EPI_ISL_3475021 | Hôpital Bicêtre | Department of Virology, Henri Mondor University Hospital, Assistance Publique Hôpitaux de Paris, Université Paris-Est Créteil, INSERM U955 | Christophe Rodriguez, Slim Fourati, Vanessa Demontant, Guillaume Gricourt, Melissa N'Debi, Alexandre Soulier, Elisabeth Trawinski, Jean-Michel Pawlotsky |
| EPI_ISL_3475022 | Biogroup Bio Lam-LCD Saint-Denis | Department of Virology, Henri Mondor University Hospital, Assistance Publique Hôpitaux de Paris, Université Paris-Est Créteil, INSERM U955 | Christophe Rodriguez, Slim Fourati, Vanessa Demontant, Guillaume Gricourt, Melissa N'Debi, Alexandre Soulier, Elisabeth Trawinski, Jean-Michel Pawlotsky |
| EPI_ISL_3475023 | Laboratoire ESPACEBIO | Department of Virology, Henri Mondor University Hospital, Assistance Publique Hôpitaux de Paris, Université Paris-Est Créteil, INSERM U955 | Christophe Rodriguez, Slim Fourati, Vanessa Demontant, Guillaume Gricourt, Melissa N'Debi, Alexandre Soulier, Elisabeth Trawinski, Jean-Michel Pawlotsky |
| EPI_ISL_3475024, EPI_ISL_3475025, EPI_ISL_3475026 | Biogroup Bio Lam-LCD Saint-Denis | Department of Virology, Henri Mondor University Hospital, Assistance Publique Hôpitaux de Paris, Université Paris-Est Créteil, INSERM U955 | Christophe Rodriguez, Slim Fourati, Vanessa Demontant, Guillaume Gricourt, Melissa N'Debi, Alexandre Soulier, Elisabeth Trawinski, Jean-Michel Pawlotsky |
| EPI_ISL_3475027 | Plateau technique Biomer Metz Verriers | Department of Virology, Henri Mondor University Hospital, Assistance Publique Hôpitaux de Paris, Université Paris-Est Créteil, INSERM U955 | Christophe Rodriguez, Slim Fourati, Vanessa Demontant, Guillaume Gricourt, Melissa N'Debi, Alexandre Soulier, Elisabeth Trawinski, Jean-Michel Pawlotsky |
| EPI_ISL_3475028 | CH. ANDRE GREGOIRE | Department of Virology, Henri Mondor University Hospital, Assistance Publique Hôpitaux de Paris, Université Paris-Est Créteil, INSERM U955 | Christophe Rodriguez, Slim Fourati, Vanessa Demontant, Guillaume Gricourt, Melissa N'Debi, Alexandre Soulier, Elisabeth Trawinski, Jean-Michel Pawlotsky |
| EPI_ISL_3475029 | Biogroup Bio Lam-LCD Saint-Denis | Department of Virology, Henri Mondor University Hospital, Assistance Publique Hôpitaux de Paris, Université Paris-Est Créteil, INSERM U955 | Christophe Rodriguez, Slim Fourati, Vanessa Demontant, Guillaume Gricourt, Melissa N'Debi, Alexandre Soulier, Elisabeth Trawinski, Jean-Michel Pawlotsky |
| EPI_ISL_3475031 | BIO VSM LAB | Department of Virology, Henri Mondor University Hospital, Assistance Publique Hôpitaux de Paris, Université Paris-Est Créteil, INSERM U955 | Christophe Rodriguez, Slim Fourati, Vanessa Demontant, Guillaume Gricourt, Melissa N'Debi, Alexandre Soulier, Elisabeth Trawinski, Jean-Michel Pawlotsky |
| EPI_ISL_3475032, EPI_ISL_3475033 | Biogroup Bio Lam-LCD Saint-Denis | Department of Virology, Henri Mondor University Hospital, Assistance Publique Hôpitaux de Paris, Université Paris-Est Créteil, INSERM U955 | Christophe Rodriguez, Slim Fourati, Vanessa Demontant, Guillaume Gricourt, Melissa N'Debi, Alexandre Soulier, Elisabeth Trawinski, Jean-Michel Pawlotsky |
| EPI_ISL_3475034 | Laboratoire ATOUTBIO JDC | Department of Virology, Henri Mondor University Hospital, Assistance Publique Hôpitaux de Paris, Université Paris-Est Créteil, INSERM U955 | Christophe Rodriguez, Slim Fourati, Vanessa Demontant, Guillaume Gricourt, Melissa N'Debi, Alexandre Soulier, Elisabeth Trawinski, Jean-Michel Pawlotsky |
| EPI_ISL_3475035 | Plateau technique Biomer Metz Verriers | Department of Virology, Henri Mondor University Hospital, Assistance Publique Hôpitaux de Paris, Université Paris-Est Créteil, INSERM U955 | Christophe Rodriguez, Slim Fourati, Vanessa Demontant, Guillaume Gricourt, Melissa N'Debi, Alexandre Soulier, Elisabeth Trawinski, Jean-Michel Pawlotsky |
| EPI_ISL_3475036 | BIOCEANE | Department of Virology, Henri Mondor University Hospital, Assistance Publique Hôpitaux de Paris, Université Paris-Est Créteil, INSERM U955 | Christophe Rodriguez, Slim Fourati, Vanessa Demontant, Guillaume Gricourt, Melissa N'Debi, Alexandre Soulier, Elisabeth Trawinski, Jean-Michel Pawlotsky |
| EPI_ISL_3475037 | SELAS-D-LAB | Department of Virology, Henri Mondor University Hospital, Assistance Publique Hôpitaux de Paris, Université Paris-Est Créteil, INSERM U955 | Christophe Rodriguez, Slim Fourati, Vanessa Demontant, Guillaume Gricourt, Melissa N'Debi, Alexandre Soulier, Elisabeth Trawinski, Jean-Michel Pawlotsky |
| EPI_ISL_3475038 | Biogroup Bio Lam-LCD Saint-Denis | Department of Virology, Henri Mondor University Hospital, Assistance Publique Hôpitaux de Paris, Université Paris-Est Créteil, INSERM U955 | Christophe Rodriguez, Slim Fourati, Vanessa Demontant, Guillaume Gricourt, Melissa N'Debi, Alexandre Soulier, Elisabeth Trawinski, Jean-Michel Pawlotsky |
| EPI_ISL_3475039 | DYNALAB | Department of Virology, Henri Mondor University Hospital, Assistance Publique Hôpitaux de Paris, Université Paris-Est Créteil, INSERM U955 | Christophe Rodriguez, Slim Fourati, Vanessa Demontant, Guillaume Gricourt, Melissa N'Debi, Alexandre Soulier, Elisabeth Trawinski, Jean-Michel Pawlotsky |
| EPI_ISL_3475040 | BIOMNIS EUROFINS IVRY | Department of Virology, Henri Mondor University Hospital, Assistance Publique Hôpitaux de Paris, Université Paris-Est Créteil, INSERM U955 | Christophe Rodriguez, Slim Fourati, Vanessa Demontant, Guillaume Gricourt, Melissa N'Debi, Alexandre Soulier, Elisabeth Trawinski, Jean-Michel Pawlotsky |
| EPI_ISL_3475041 | CH. ROBERT BISSON | Department of Virology, Henri Mondor University Hospital, Assistance Publique Hôpitaux de Paris, Université Paris-Est Créteil, INSERM U955 | Christophe Rodriguez, Slim Fourati, Vanessa Demontant, Guillaume Gricourt, Melissa N'Debi, Alexandre Soulier, Elisabeth Trawinski, Jean-Michel Pawlotsky |
| EPI_ISL_3475042 | Biogroup Bio Lam-LCD Saint-Denis | Department of Virology, Henri Mondor University Hospital, Assistance Publique Hôpitaux de Paris, Université Paris-Est Créteil, INSERM U955 | Christophe Rodriguez, Slim Fourati, Vanessa Demontant, Guillaume Gricourt, Melissa N'Debi, Alexandre Soulier, Elisabeth Trawinski, Jean-Michel Pawlotsky |
| EPI_ISL_3475043 | CH.LE RAINCY_MONTFERMEIL | Department of Virology, Henri Mondor University Hospital, Assistance Publique Hôpitaux de Paris, Université Paris-Est Créteil, INSERM U955 | Christophe Rodriguez, Slim Fourati, Vanessa Demontant, Guillaume Gricourt, Melissa N'Debi, Alexandre Soulier, Elisabeth Trawinski, Jean-Michel Pawlotsky |
| EPI_ISL_3475045 | Biogroup Bio Lam-LCD Saint-Denis | Department of Virology, Henri Mondor University Hospital, Assistance Publique Hôpitaux de Paris, Université Paris-Est Créteil, INSERM U955 | Christophe Rodriguez, Slim Fourati, Vanessa Demontant, Guillaume Gricourt, Melissa N'Debi, Alexandre Soulier, Elisabeth Trawinski, Jean-Michel Pawlotsky |
| EPI_ISL_3475046 | DYNABIO UNILABS | Department of Virology, Henri Mondor University Hospital, Assistance Publique Hôpitaux de Paris, Université Paris-Est Créteil, INSERM U955 | Christophe Rodriguez, Slim Fourati, Vanessa Demontant, Guillaume Gricourt, Melissa N'Debi, Alexandre Soulier, Elisabeth Trawinski, Jean-Michel Pawlotsky |
| EPI_ISL_3475048 | CH. ANDRE GREGOIRE | Department of Virology, Henri Mondor University Hospital, Assistance Publique Hôpitaux de Paris, Université Paris-Est Créteil, INSERM U955 | Christophe Rodriguez, Slim Fourati, Vanessa Demontant, Guillaume Gricourt, Melissa N'Debi, Alexandre Soulier, Elisabeth Trawinski, Jean-Michel Pawlotsky |
| EPI_ISL_3475049, EPI_ISL_3475050 | Biogroup Bio Lam-LCD Saint-Denis | Department of Virology, Henri Mondor University Hospital, Assistance Publique Hôpitaux de Paris, Université Paris-Est Créteil, INSERM U955 | Christophe Rodriguez, Slim Fourati, Vanessa Demontant, Guillaume Gricourt, Melissa N'Debi, Alexandre Soulier, Elisabeth Trawinski, Jean-Michel Pawlotsky |
| EPI_ISL_3475051 | BIOCEANE | Department of Virology, Henri Mondor University Hospital, Assistance Publique Hôpitaux de Paris, Université Paris-Est Créteil, INSERM U955 | Christophe Rodriguez, Slim Fourati, Vanessa Demontant, Guillaume Gricourt, Melissa N'Debi, Alexandre Soulier, Elisabeth Trawinski, Jean-Michel Pawlotsky |
| EPI_ISL_3475052 | CHRU de NANCY | Department of Virology, Henri Mondor University Hospital, Assistance Publique Hôpitaux de Paris, Université Paris-Est Créteil, INSERM U955 | Christophe Rodriguez, Slim Fourati, Vanessa Demontant, Guillaume Gricourt, Melissa N'Debi, Alexandre Soulier, Elisabeth Trawinski, Jean-Michel Pawlotsky |
| EPI_ISL_3475053 | BIOMNIS EUROFINS IVRY | Department of Virology, Henri Mondor University Hospital, Assistance Publique Hôpitaux de Paris, Université Paris-Est Créteil, INSERM U955 | Christophe Rodriguez, Slim Fourati, Vanessa Demontant, Guillaume Gricourt, Melissa N'Debi, Alexandre Soulier, Elisabeth Trawinski, Jean-Michel Pawlotsky |
| EPI_ISL_3475054, EPI_ISL_3475055 | Biogroup Bio Lam-LCD Saint-Denis | Department of Virology, Henri Mondor University Hospital, Assistance Publique Hôpitaux de Paris, Université Paris-Est Créteil, INSERM U955 | Christophe Rodriguez, Slim Fourati, Vanessa Demontant, Guillaume Gricourt, Melissa N'Debi, Alexandre Soulier, Elisabeth Trawinski, Jean-Michel Pawlotsky |
| EPI_ISL_3475056 | BIOMNIS EUROFINS IVRY | Department of Virology, Henri Mondor University Hospital, Assistance Publique Hôpitaux de Paris, Université Paris-Est Créteil, INSERM U955 | Christophe Rodriguez, Slim Fourati, Vanessa Demontant, Guillaume Gricourt, Melissa N'Debi, Alexandre Soulier, Elisabeth Trawinski, Jean-Michel Pawlotsky |
| EPI_ISL_3475057 | CH. ANDRE GREGOIRE | Department of Virology, Henri Mondor University Hospital, Assistance Publique Hôpitaux de Paris, Université Paris-Est Créteil, INSERM U955 | Christophe Rodriguez, Slim Fourati, Vanessa Demontant, Guillaume Gricourt, Melissa N'Debi, Alexandre Soulier, Elisabeth Trawinski, Jean-Michel Pawlotsky |
| EPI_ISL_3475058 | BIOCEANE | Department of Virology, Henri Mondor University Hospital, Assistance Publique Hôpitaux de Paris, Université Paris-Est Créteil, INSERM U955 | Christophe Rodriguez, Slim Fourati, Vanessa Demontant, Guillaume Gricourt, Melissa N'Debi, Alexandre Soulier, Elisabeth Trawinski, Jean-Michel Pawlotsky |
| EPI_ISL_3475059 | Laboratoire ESPACEBIO | Department of Virology, Henri Mondor University Hospital, Assistance Publique Hôpitaux de Paris, Université Paris-Est Créteil, INSERM U955 | Christophe Rodriguez, Slim Fourati, Vanessa Demontant, Guillaume Gricourt, Melissa N'Debi, Alexandre Soulier, Elisabeth Trawinski, Jean-Michel Pawlotsky |
| EPI_ISL_3475060 | DYNABIO UNILABS | Department of Virology, Henri Mondor University Hospital, | Christophe Rodriguez, Slim Fourati, Vanessa Demontant, Guillaume Gricourt, Melissa N'Debi, Alexandre Soulier, Elisabeth Trawinski, Jean-Michel Pawlotsky |

| | | | |
|---|---|---|---|
| | | Assistance Publique Hôpitaux de Paris, Université Paris-Est Créteil, INSERM U955 | |
| EPI_ISL_3475061 | DYNALAB | Department of Virology, Henri Mondor University Hospital, Assistance Publique Hôpitaux de Paris, Université Paris-Est Créteil, INSERM U955 | Christophe Rodriguez, Slim Fourati, Vanessa Demontant, Guillaume Gricourt, Melissa N'Debi, Alexandre Soulier, Elisabeth Trawinski, Jean-Michel Pawlotsky |
| EPI_ISL_3475063 | Plateau technique Biomer Metz Verriers | Department of Virology, Henri Mondor University Hospital, Assistance Publique Hôpitaux de Paris, Université Paris-Est Créteil, INSERM U955 | Christophe Rodriguez, Slim Fourati, Vanessa Demontant, Guillaume Gricourt, Melissa N'Debi, Alexandre Soulier, Elisabeth Trawinski, Jean-Michel Pawlotsky |
| EPI_ISL_3475064 | Biogroup Bio Lam-LCD Saint-Denis | Department of Virology, Henri Mondor University Hospital, Assistance Publique Hôpitaux de Paris, Université Paris-Est Créteil, INSERM U955 | Christophe Rodriguez, Slim Fourati, Vanessa Demontant, Guillaume Gricourt, Melissa N'Debi, Alexandre Soulier, Elisabeth Trawinski, Jean-Michel Pawlotsky |
| EPI_ISL_3475065 | BIOMNIS EUROFINS IVRY | Department of Virology, Henri Mondor University Hospital, Assistance Publique Hôpitaux de Paris, Université Paris-Est Créteil, INSERM U955 | Christophe Rodriguez, Slim Fourati, Vanessa Demontant, Guillaume Gricourt, Melissa N'Debi, Alexandre Soulier, Elisabeth Trawinski, Jean-Michel Pawlotsky |
| EPI_ISL_3475066 | GROUPE HOSPITALIER SUD ILE DE FRANCE | Department of Virology, Henri Mondor University Hospital, Assistance Publique Hôpitaux de Paris, Université Paris-Est Créteil, INSERM U955 | Christophe Rodriguez, Slim Fourati, Vanessa Demontant, Guillaume Gricourt, Melissa N'Debi, Alexandre Soulier, Elisabeth Trawinski, Jean-Michel Pawlotsky |
| EPI_ISL_3475067 | Laboratoire ESPACEBIO | Department of Virology, Henri Mondor University Hospital, Assistance Publique Hôpitaux de Paris, Université Paris-Est Créteil, INSERM U955 | Christophe Rodriguez, Slim Fourati, Vanessa Demontant, Guillaume Gricourt, Melissa N'Debi, Alexandre Soulier, Elisabeth Trawinski, Jean-Michel Pawlotsky |
| EPI_ISL_3475068, EPI_ISL_3475069 | Biogroup Bio Lam-LCD Saint-Denis | Department of Virology, Henri Mondor University Hospital, Assistance Publique Hôpitaux de Paris, Université Paris-Est Créteil, INSERM U955 | Christophe Rodriguez, Slim Fourati, Vanessa Demontant, Guillaume Gricourt, Melissa N'Debi, Alexandre Soulier, Elisabeth Trawinski, Jean-Michel Pawlotsky |
| EPI_ISL_3475070 | Laboratoire ATOUTBIO JDC | Department of Virology, Henri Mondor University Hospital, Assistance Publique Hôpitaux de Paris, Université Paris-Est Créteil, INSERM U955 | Christophe Rodriguez, Slim Fourati, Vanessa Demontant, Guillaume Gricourt, Melissa N'Debi, Alexandre Soulier, Elisabeth Trawinski, Jean-Michel Pawlotsky |
| EPI_ISL_3475071, EPI_ISL_3475072 | DYNALAB | Department of Virology, Henri Mondor University Hospital, Assistance Publique Hôpitaux de Paris, Université Paris-Est Créteil, INSERM U955 | Christophe Rodriguez, Slim Fourati, Vanessa Demontant, Guillaume Gricourt, Melissa N'Debi, Alexandre Soulier, Elisabeth Trawinski, Jean-Michel Pawlotsky |
| EPI_ISL_3475073, EPI_ISL_3475074 | Biogroup Bio Lam-LCD Saint-Denis | Department of Virology, Henri Mondor University Hospital, Assistance Publique Hôpitaux de Paris, Université Paris-Est Créteil, INSERM U955 | Christophe Rodriguez, Slim Fourati, Vanessa Demontant, Guillaume Gricourt, Melissa N'Debi, Alexandre Soulier, Elisabeth Trawinski, Jean-Michel Pawlotsky |
| EPI_ISL_3475075 | DYNALAB | Department of Virology, Henri Mondor University Hospital, Assistance Publique Hôpitaux de Paris, Université Paris-Est Créteil, INSERM U955 | Christophe Rodriguez, Slim Fourati, Vanessa Demontant, Guillaume Gricourt, Melissa N'Debi, Alexandre Soulier, Elisabeth Trawinski, Jean-Michel Pawlotsky |
| EPI_ISL_3475076 | Biogroup Bio Lam-LCD Saint-Denis | Department of Virology, Henri Mondor University Hospital, Assistance Publique Hôpitaux de Paris, Université Paris-Est Créteil, INSERM U955 | Christophe Rodriguez, Slim Fourati, Vanessa Demontant, Guillaume Gricourt, Melissa N'Debi, Alexandre Soulier, Elisabeth Trawinski, Jean-Michel Pawlotsky |
| EPI_ISL_3475077 | SELAS-D-LAB | Department of Virology, Henri Mondor University Hospital, Assistance Publique Hôpitaux de Paris, Université Paris-Est Créteil, INSERM U955 | Christophe Rodriguez, Slim Fourati, Vanessa Demontant, Guillaume Gricourt, Melissa N'Debi, Alexandre Soulier, Elisabeth Trawinski, Jean-Michel Pawlotsky |
| EPI_ISL_3475078 | BIOMNIS EUROFINS IVRY | Department of Virology, Henri Mondor University Hospital, Assistance Publique Hôpitaux de Paris, Université Paris-Est Créteil, INSERM U955 | Christophe Rodriguez, Slim Fourati, Vanessa Demontant, Guillaume Gricourt, Melissa N'Debi, Alexandre Soulier, Elisabeth Trawinski, Jean-Michel Pawlotsky |
| EPI_ISL_3475079 | Laboratoire ESPACEBIO | Department of Virology, Henri Mondor University Hospital, Assistance Publique Hôpitaux de Paris, Université Paris-Est Créteil, INSERM U955 | Christophe Rodriguez, Slim Fourati, Vanessa Demontant, Guillaume Gricourt, Melissa N'Debi, Alexandre Soulier, Elisabeth Trawinski, Jean-Michel Pawlotsky |
| EPI_ISL_3475080 | BIO VSM LAB | Department of Virology, Henri Mondor University Hospital, Assistance Publique Hôpitaux de Paris, Université Paris-Est Créteil, INSERM U955 | Christophe Rodriguez, Slim Fourati, Vanessa Demontant, Guillaume Gricourt, Melissa N'Debi, Alexandre Soulier, Elisabeth Trawinski, Jean-Michel Pawlotsky |
| EPI_ISL_3475081 | CH.LE RAINCY_MONTFERMEIL | Department of Virology, Henri Mondor University Hospital, Assistance Publique Hôpitaux de Paris, Université Paris-Est Créteil, INSERM U955 | Christophe Rodriguez, Slim Fourati, Vanessa Demontant, Guillaume Gricourt, Melissa N'Debi, Alexandre Soulier, Elisabeth Trawinski, Jean-Michel Pawlotsky |
| EPI_ISL_3475082 | Biogroup Bio Lam-LCD Saint-Denis | Department of Virology, Henri Mondor University Hospital, Assistance Publique Hôpitaux de Paris, Université Paris-Est Créteil, INSERM U955 | Christophe Rodriguez, Slim Fourati, Vanessa Demontant, Guillaume Gricourt, Melissa N'Debi, Alexandre Soulier, Elisabeth Trawinski, Jean-Michel Pawlotsky |
| EPI_ISL_3475083 | DYNALAB | Department of Virology, Henri Mondor University Hospital, Assistance Publique Hôpitaux de Paris, Université Paris-Est Créteil, INSERM U955 | Christophe Rodriguez, Slim Fourati, Vanessa Demontant, Guillaume Gricourt, Melissa N'Debi, Alexandre Soulier, Elisabeth Trawinski, Jean-Michel Pawlotsky |
| EPI_ISL_3475084 | BIOMNIS EUROFINS IVRY | Department of Virology, Henri Mondor University Hospital, Assistance Publique Hôpitaux de Paris, Université Paris-Est Créteil, INSERM U955 | Christophe Rodriguez, Slim Fourati, Vanessa Demontant, Guillaume Gricourt, Melissa N'Debi, Alexandre Soulier, Elisabeth Trawinski, Jean-Michel Pawlotsky |
| EPI_ISL_3475085 | Laboratoire ATOUTBIO JDC | Department of Virology, Henri Mondor University Hospital, Assistance Publique Hôpitaux de Paris, Université Paris-Est Créteil, INSERM U955 | Christophe Rodriguez, Slim Fourati, Vanessa Demontant, Guillaume Gricourt, Melissa N'Debi, Alexandre Soulier, Elisabeth Trawinski, Jean-Michel Pawlotsky |
| EPI_ISL_3475086 | Biogroup Bio Lam-LCD Saint-Denis | Department of Virology, Henri Mondor University Hospital, Assistance Publique Hôpitaux de Paris, Université Paris-Est Créteil, INSERM U955 | Christophe Rodriguez, Slim Fourati, Vanessa Demontant, Guillaume Gricourt, Melissa N'Debi, Alexandre Soulier, Elisabeth Trawinski, Jean-Michel Pawlotsky |
| EPI_ISL_3475087 | Laboratoire ESPACEBIO | Department of Virology, Henri Mondor University Hospital, Assistance Publique Hôpitaux de Paris, Université Paris-Est Créteil, INSERM U955 | Christophe Rodriguez, Slim Fourati, Vanessa Demontant, Guillaume Gricourt, Melissa N'Debi, Alexandre Soulier, Elisabeth Trawinski, Jean-Michel Pawlotsky |
| EPI_ISL_3475088 | BIOCEANE | Department of Virology, Henri Mondor University Hospital, Assistance Publique Hôpitaux de Paris, Université Paris-Est Créteil, INSERM U955 | Christophe Rodriguez, Slim Fourati, Vanessa Demontant, Guillaume Gricourt, Melissa N'Debi, Alexandre Soulier, Elisabeth Trawinski, Jean-Michel Pawlotsky |
| EPI_ISL_3475089 | CH. ANDRE GREGOIRE | Department of Virology, Henri Mondor University Hospital, Assistance Publique Hôpitaux de Paris, Université Paris-Est Créteil, INSERM U955 | Christophe Rodriguez, Slim Fourati, Vanessa Demontant, Guillaume Gricourt, Melissa N'Debi, Alexandre Soulier, Elisabeth Trawinski, Jean-Michel Pawlotsky |
| EPI_ISL_3475090, EPI_ISL_3475091 | Biogroup Bio Lam-LCD Saint-Denis | Department of Virology, Henri Mondor University Hospital, Assistance Publique Hôpitaux de Paris, Université Paris-Est Créteil, INSERM U955 | Christophe Rodriguez, Slim Fourati, Vanessa Demontant, Guillaume Gricourt, Melissa N'Debi, Alexandre Soulier, Elisabeth Trawinski, Jean-Michel Pawlotsky |
| EPI_ISL_3475092 | Laboratoire LENYS | Department of Virology, Henri Mondor University Hospital, Assistance Publique Hôpitaux de Paris, Université Paris-Est Créteil, INSERM U955 | Christophe Rodriguez, Slim Fourati, Vanessa Demontant, Guillaume Gricourt, Melissa N'Debi, Alexandre Soulier, Elisabeth Trawinski, Jean-Michel Pawlotsky |
| EPI_ISL_3475093 | Biogroup Bio Lam-LCD Saint-Denis | Department of Virology, Henri Mondor University Hospital, Assistance Publique Hôpitaux de Paris, Université Paris-Est Créteil, INSERM U955 | Christophe Rodriguez, Slim Fourati, Vanessa Demontant, Guillaume Gricourt, Melissa N'Debi, Alexandre Soulier, Elisabeth Trawinski, Jean-Michel Pawlotsky |
| EPI_ISL_3475094 | Laboratoire ESPACEBIO | Department of Virology, Henri Mondor University Hospital, Assistance Publique Hôpitaux de Paris, Université Paris-Est Créteil, INSERM U955 | Christophe Rodriguez, Slim Fourati, Vanessa Demontant, Guillaume Gricourt, Melissa N'Debi, Alexandre Soulier, Elisabeth Trawinski, Jean-Michel Pawlotsky |
| EPI_ISL_3475095 | Laboratoire LENYS | Department of Virology, Henri Mondor University Hospital, Assistance Publique Hôpitaux de Paris, Université Paris-Est Créteil, INSERM U955 | Christophe Rodriguez, Slim Fourati, Vanessa Demontant, Guillaume Gricourt, Melissa N'Debi, Alexandre Soulier, Elisabeth Trawinski, Jean-Michel Pawlotsky |
| EPI_ISL_3475096, EPI_ISL_3475097, EPI_ISL_3475098 | Biogroup Bio Lam-LCD Saint-Denis | Department of Virology, Henri Mondor University Hospital, Assistance Publique Hôpitaux de Paris, Université Paris-Est Créteil, INSERM U955 | Christophe Rodriguez, Slim Fourati, Vanessa Demontant, Guillaume Gricourt, Melissa N'Debi, Alexandre Soulier, Elisabeth Trawinski, Jean-Michel Pawlotsky |
| EPI_ISL_3475099 | CH.LE RAINCY_MONTFERMEIL | Department of Virology, Henri Mondor University Hospital, | Christophe Rodriguez, Slim Fourati, Vanessa Demontant, Guillaume Gricourt, Melissa N'Debi, Alexandre Soulier, Elisabeth Trawinski, Jean-Michel Pawlotsky |

| | | | |
|---|---|---|---|
| | | Assistance Publique Hôpitaux de Paris, Université Paris-Est Créteil, INSERM U955 | |
| EPI_ISL_3475100 | Biogroup Bio Lam-LCD Saint-Denis | Department of Virology, Henri Mondor University Hospital, Assistance Publique Hôpitaux de Paris, Université Paris-Est Créteil, INSERM U955 | Christophe Rodriguez, Slim Fourati, Vanessa Demontant, Guillaume Gricourt, Melissa N'Debi, Alexandre Soulier, Elisabeth Trawinski, Jean-Michel Pawlotsky |
| EPI_ISL_3475101 | Plateau technique Biomer Metz Verriers | Department of Virology, Henri Mondor University Hospital, Assistance Publique Hôpitaux de Paris, Université Paris-Est Créteil, INSERM U955 | Christophe Rodriguez, Slim Fourati, Vanessa Demontant, Guillaume Gricourt, Melissa N'Debi, Alexandre Soulier, Elisabeth Trawinski, Jean-Michel Pawlotsky |
| EPI_ISL_3475102, EPI_ISL_3475103 | Biogroup Bio Lam-LCD Saint-Denis | Department of Virology, Henri Mondor University Hospital, Assistance Publique Hôpitaux de Paris, Université Paris-Est Créteil, INSERM U955 | Christophe Rodriguez, Slim Fourati, Vanessa Demontant, Guillaume Gricourt, Melissa N'Debi, Alexandre Soulier, Elisabeth Trawinski, Jean-Michel Pawlotsky |
| EPI_ISL_3475104 | BIOMNIS EUROFINS IVRY | Department of Virology, Henri Mondor University Hospital, Assistance Publique Hôpitaux de Paris, Université Paris-Est Créteil, INSERM U955 | Christophe Rodriguez, Slim Fourati, Vanessa Demontant, Guillaume Gricourt, Melissa N'Debi, Alexandre Soulier, Elisabeth Trawinski, Jean-Michel Pawlotsky |
| EPI_ISL_3475105 | Biogroup Bio Lam-LCD Saint-Denis | Department of Virology, Henri Mondor University Hospital, Assistance Publique Hôpitaux de Paris, Université Paris-Est Créteil, INSERM U955 | Christophe Rodriguez, Slim Fourati, Vanessa Demontant, Guillaume Gricourt, Melissa N'Debi, Alexandre Soulier, Elisabeth Trawinski, Jean-Michel Pawlotsky |
| EPI_ISL_3475106 | Laboratoire ESPACEBIO | Department of Virology, Henri Mondor University Hospital, Assistance Publique Hôpitaux de Paris, Université Paris-Est Créteil, INSERM U955 | Christophe Rodriguez, Slim Fourati, Vanessa Demontant, Guillaume Gricourt, Melissa N'Debi, Alexandre Soulier, Elisabeth Trawinski, Jean-Michel Pawlotsky |
| EPI_ISL_3475107 | BIOMNIS EUROFINS IVRY | Department of Virology, Henri Mondor University Hospital, Assistance Publique Hôpitaux de Paris, Université Paris-Est Créteil, INSERM U955 | Christophe Rodriguez, Slim Fourati, Vanessa Demontant, Guillaume Gricourt, Melissa N'Debi, Alexandre Soulier, Elisabeth Trawinski, Jean-Michel Pawlotsky |
| EPI_ISL_3475108 | CHU DE DIJON | Department of Virology, Henri Mondor University Hospital, Assistance Publique Hôpitaux de Paris, Université Paris-Est Créteil, INSERM U955 | Christophe Rodriguez, Slim Fourati, Vanessa Demontant, Guillaume Gricourt, Melissa N'Debi, Alexandre Soulier, Elisabeth Trawinski, Jean-Michel Pawlotsky |
| EPI_ISL_3475109 | CH. CLEMENCEAU | Department of Virology, Henri Mondor University Hospital, Assistance Publique Hôpitaux de Paris, Université Paris-Est Créteil, INSERM U955 | Christophe Rodriguez, Slim Fourati, Vanessa Demontant, Guillaume Gricourt, Melissa N'Debi, Alexandre Soulier, Elisabeth Trawinski, Jean-Michel Pawlotsky |
| EPI_ISL_3475111 | CH. LEON BINET PROVINS | Department of Virology, Henri Mondor University Hospital, Assistance Publique Hôpitaux de Paris, Université Paris-Est Créteil, INSERM U955 | Christophe Rodriguez, Slim Fourati, Vanessa Demontant, Guillaume Gricourt, Melissa N'Debi, Alexandre Soulier, Elisabeth Trawinski, Jean-Michel Pawlotsky |
| EPI_ISL_3475112 | CHU DE DIJON | Department of Virology, Henri Mondor University Hospital, Assistance Publique Hôpitaux de Paris, Université Paris-Est Créteil, INSERM U955 | Christophe Rodriguez, Slim Fourati, Vanessa Demontant, Guillaume Gricourt, Melissa N'Debi, Alexandre Soulier, Elisabeth Trawinski, Jean-Michel Pawlotsky |
| EPI_ISL_3475113, EPI_ISL_3475114, EPI_ISL_3475115 | Biogroup Bio Lam-LCD Saint-Denis | Department of Virology, Henri Mondor University Hospital, Assistance Publique Hôpitaux de Paris, Université Paris-Est Créteil, INSERM U955 | Christophe Rodriguez, Slim Fourati, Vanessa Demontant, Guillaume Gricourt, Melissa N'Debi, Alexandre Soulier, Elisabeth Trawinski, Jean-Michel Pawlotsky |
| EPI_ISL_3475116 | CHU DE DIJON | Department of Virology, Henri Mondor University Hospital, Assistance Publique Hôpitaux de Paris, Université Paris-Est Créteil, INSERM U955 | Christophe Rodriguez, Slim Fourati, Vanessa Demontant, Guillaume Gricourt, Melissa N'Debi, Alexandre Soulier, Elisabeth Trawinski, Jean-Michel Pawlotsky |
| EPI_ISL_3475117 | Biogroup Bio Lam-LCD Saint-Denis | Department of Virology, Henri Mondor University Hospital, Assistance Publique Hôpitaux de Paris, Université Paris-Est Créteil, INSERM U955 | Christophe Rodriguez, Slim Fourati, Vanessa Demontant, Guillaume Gricourt, Melissa N'Debi, Alexandre Soulier, Elisabeth Trawinski, Jean-Michel Pawlotsky |
| EPI_ISL_3475118, EPI_ISL_3475119 | CHU DE DIJON | Department of Virology, Henri Mondor University Hospital, Assistance Publique Hôpitaux de Paris, Université Paris-Est Créteil, INSERM U955 | Christophe Rodriguez, Slim Fourati, Vanessa Demontant, Guillaume Gricourt, Melissa N'Debi, Alexandre Soulier, Elisabeth Trawinski, Jean-Michel Pawlotsky |
| EPI_ISL_3475120 | BIO ARD'AISNE | Department of Virology, Henri Mondor University Hospital, Assistance Publique Hôpitaux de Paris, Université Paris-Est Créteil, INSERM U955 | Christophe Rodriguez, Slim Fourati, Vanessa Demontant, Guillaume Gricourt, Melissa N'Debi, Alexandre Soulier, Elisabeth Trawinski, Jean-Michel Pawlotsky |
| EPI_ISL_3475121 | CHU DE DIJON | Department of Virology, Henri Mondor University Hospital, Assistance Publique Hôpitaux de Paris, Université Paris-Est Créteil, INSERM U955 | Christophe Rodriguez, Slim Fourati, Vanessa Demontant, Guillaume Gricourt, Melissa N'Debi, Alexandre Soulier, Elisabeth Trawinski, Jean-Michel Pawlotsky |
| EPI_ISL_3475122 | BIOMNIS EUROFINS IVRY | Department of Virology, Henri Mondor University Hospital, Assistance Publique Hôpitaux de Paris, Université Paris-Est Créteil, INSERM U955 | Christophe Rodriguez, Slim Fourati, Vanessa Demontant, Guillaume Gricourt, Melissa N'Debi, Alexandre Soulier, Elisabeth Trawinski, Jean-Michel Pawlotsky |
| EPI_ISL_3475123 | CHU DE DIJON | Department of Virology, Henri Mondor University Hospital, Assistance Publique Hôpitaux de Paris, Université Paris-Est Créteil, INSERM U955 | Christophe Rodriguez, Slim Fourati, Vanessa Demontant, Guillaume Gricourt, Melissa N'Debi, Alexandre Soulier, Elisabeth Trawinski, Jean-Michel Pawlotsky |
| EPI_ISL_3475124 | BIOLBS Rouen | Department of Virology, Henri Mondor University Hospital, Assistance Publique Hôpitaux de Paris, Université Paris-Est Créteil, INSERM U955 | Christophe Rodriguez, Slim Fourati, Vanessa Demontant, Guillaume Gricourt, Melissa N'Debi, Alexandre Soulier, Elisabeth Trawinski, Jean-Michel Pawlotsky |
| EPI_ISL_3475125 | Biogroup Bio Lam-LCD Saint-Denis | Department of Virology, Henri Mondor University Hospital, Assistance Publique Hôpitaux de Paris, Université Paris-Est Créteil, INSERM U955 | Christophe Rodriguez, Slim Fourati, Vanessa Demontant, Guillaume Gricourt, Melissa N'Debi, Alexandre Soulier, Elisabeth Trawinski, Jean-Michel Pawlotsky |
| EPI_ISL_3475126, EPI_ISL_3475127, EPI_ISL_3475128 | CHU DE DIJON | Department of Virology, Henri Mondor University Hospital, Assistance Publique Hôpitaux de Paris, Université Paris-Est Créteil, INSERM U955 | Christophe Rodriguez, Slim Fourati, Vanessa Demontant, Guillaume Gricourt, Melissa N'Debi, Alexandre Soulier, Elisabeth Trawinski, Jean-Michel Pawlotsky |
| EPI_ISL_3475129 | BIOMNIS EUROFINS IVRY | Department of Virology, Henri Mondor University Hospital, Assistance Publique Hôpitaux de Paris, Université Paris-Est Créteil, INSERM U955 | Christophe Rodriguez, Slim Fourati, Vanessa Demontant, Guillaume Gricourt, Melissa N'Debi, Alexandre Soulier, Elisabeth Trawinski, Jean-Michel Pawlotsky |
| EPI_ISL_3475130 | Biogroup Bio Lam-LCD Saint-Denis | Department of Virology, Henri Mondor University Hospital, Assistance Publique Hôpitaux de Paris, Université Paris-Est Créteil, INSERM U955 | Christophe Rodriguez, Slim Fourati, Vanessa Demontant, Guillaume Gricourt, Melissa N'Debi, Alexandre Soulier, Elisabeth Trawinski, Jean-Michel Pawlotsky |
| EPI_ISL_3475131 | CHU DE DIJON | Department of Virology, Henri Mondor University Hospital, Assistance Publique Hôpitaux de Paris, Université Paris-Est Créteil, INSERM U955 | Christophe Rodriguez, Slim Fourati, Vanessa Demontant, Guillaume Gricourt, Melissa N'Debi, Alexandre Soulier, Elisabeth Trawinski, Jean-Michel Pawlotsky |
| EPI_ISL_3475132 | Biogroup Bio Lam-LCD Saint-Denis | Department of Virology, Henri Mondor University Hospital, Assistance Publique Hôpitaux de Paris, Université Paris-Est Créteil, INSERM U955 | Christophe Rodriguez, Slim Fourati, Vanessa Demontant, Guillaume Gricourt, Melissa N'Debi, Alexandre Soulier, Elisabeth Trawinski, Jean-Michel Pawlotsky |
| EPI_ISL_3475133, EPI_ISL_3475134 | CHU DE DIJON | Department of Virology, Henri Mondor University Hospital, Assistance Publique Hôpitaux de Paris, Université Paris-Est Créteil, INSERM U955 | Christophe Rodriguez, Slim Fourati, Vanessa Demontant, Guillaume Gricourt, Melissa N'Debi, Alexandre Soulier, Elisabeth Trawinski, Jean-Michel Pawlotsky |
| EPI_ISL_3475135 | Biogroup Bio Lam-LCD Saint-Denis | Department of Virology, Henri Mondor University Hospital, Assistance Publique Hôpitaux de Paris, Université Paris-Est Créteil, INSERM U955 | Christophe Rodriguez, Slim Fourati, Vanessa Demontant, Guillaume Gricourt, Melissa N'Debi, Alexandre Soulier, Elisabeth Trawinski, Jean-Michel Pawlotsky |
| EPI_ISL_3475136, EPI_ISL_3475137, EPI_ISL_3475138, EPI_ISL_3475139, EPI_ISL_3475140 | CHU DE DIJON | Department of Virology, Henri Mondor University Hospital, Assistance Publique Hôpitaux de Paris, Université Paris-Est Créteil, INSERM U955 | Christophe Rodriguez, Slim Fourati, Vanessa Demontant, Guillaume Gricourt, Melissa N'Debi, Alexandre Soulier, Elisabeth Trawinski, Jean-Michel Pawlotsky |
| EPI_ISL_3475141 | Biogroup Bio Lam-LCD Saint-Denis | Department of Virology, Henri Mondor University Hospital, Assistance Publique Hôpitaux de Paris, Université Paris-Est Créteil, INSERM U955 | Christophe Rodriguez, Slim Fourati, Vanessa Demontant, Guillaume Gricourt, Melissa N'Debi, Alexandre Soulier, Elisabeth Trawinski, Jean-Michel Pawlotsky |
| EPI_ISL_3475142, EPI_ISL_3475143, | CHU DE DIJON | Department of Virology, Henri Mondor University Hospital, Assistance Publique Hôpitaux de Paris, Université Paris-Est Créteil, | Christophe Rodriguez, Slim Fourati, Vanessa Demontant, Guillaume Gricourt, Melissa N'Debi, Alexandre Soulier, Elisabeth Trawinski, Jean-Michel Pawlotsky |

| | | | |
|---|---|---|---|
| EPI_ISL_3475144 | | INSERM U955 | |
| EPI_ISL_3475145, EPI_ISL_3475146, EPI_ISL_3475147 | Biogroup Bio Lam-LCD Saint-Denis | Department of Virology, Henri Mondor University Hospital, Assistance Publique Hôpitaux de Paris, Université Paris-Est Créteil, INSERM U955 | Christophe Rodriguez, Slim Fourati, Vanessa Demontant, Guillaume Gricourt, Melissa N'Debi, Alexandre Soulier, Elisabeth Trawinski, Jean-Michel Pawlotsky |
| EPI_ISL_3475148 | CH. CLEMENCEAU | Department of Virology, Henri Mondor University Hospital, Assistance Publique Hôpitaux de Paris, Université Paris-Est Créteil, INSERM U955 | Christophe Rodriguez, Slim Fourati, Vanessa Demontant, Guillaume Gricourt, Melissa N'Debi, Alexandre Soulier, Elisabeth Trawinski, Jean-Michel Pawlotsky |
| EPI_ISL_3475149 | CHU DE DIJON | Department of Virology, Henri Mondor University Hospital, Assistance Publique Hôpitaux de Paris, Université Paris-Est Créteil, INSERM U955 | Christophe Rodriguez, Slim Fourati, Vanessa Demontant, Guillaume Gricourt, Melissa N'Debi, Alexandre Soulier, Elisabeth Trawinski, Jean-Michel Pawlotsky |
| EPI_ISL_3475150 | Biogroup Bio Lam-LCD Saint-Denis | Department of Virology, Henri Mondor University Hospital, Assistance Publique Hôpitaux de Paris, Université Paris-Est Créteil, INSERM U955 | Christophe Rodriguez, Slim Fourati, Vanessa Demontant, Guillaume Gricourt, Melissa N'Debi, Alexandre Soulier, Elisabeth Trawinski, Jean-Michel Pawlotsky |
| EPI_ISL_3475151, EPI_ISL_3475152 | CHU DE DIJON | Department of Virology, Henri Mondor University Hospital, Assistance Publique Hôpitaux de Paris, Université Paris-Est Créteil, INSERM U955 | Christophe Rodriguez, Slim Fourati, Vanessa Demontant, Guillaume Gricourt, Melissa N'Debi, Alexandre Soulier, Elisabeth Trawinski, Jean-Michel Pawlotsky |
| EPI_ISL_3475153 | BIOLBS Rouen | Department of Virology, Henri Mondor University Hospital, Assistance Publique Hôpitaux de Paris, Université Paris-Est Créteil, INSERM U955 | Christophe Rodriguez, Slim Fourati, Vanessa Demontant, Guillaume Gricourt, Melissa N'Debi, Alexandre Soulier, Elisabeth Trawinski, Jean-Michel Pawlotsky |
| EPI_ISL_3475154 | CHU DE DIJON | Department of Virology, Henri Mondor University Hospital, Assistance Publique Hôpitaux de Paris, Université Paris-Est Créteil, INSERM U955 | Christophe Rodriguez, Slim Fourati, Vanessa Demontant, Guillaume Gricourt, Melissa N'Debi, Alexandre Soulier, Elisabeth Trawinski, Jean-Michel Pawlotsky |
| EPI_ISL_3475155 | Biogroup Bio Lam-LCD Saint-Denis | Department of Virology, Henri Mondor University Hospital, Assistance Publique Hôpitaux de Paris, Université Paris-Est Créteil, INSERM U955 | Christophe Rodriguez, Slim Fourati, Vanessa Demontant, Guillaume Gricourt, Melissa N'Debi, Alexandre Soulier, Elisabeth Trawinski, Jean-Michel Pawlotsky |
| EPI_ISL_3475156 | CHU DE DIJON | Department of Virology, Henri Mondor University Hospital, Assistance Publique Hôpitaux de Paris, Université Paris-Est Créteil, INSERM U955 | Christophe Rodriguez, Slim Fourati, Vanessa Demontant, Guillaume Gricourt, Melissa N'Debi, Alexandre Soulier, Elisabeth Trawinski, Jean-Michel Pawlotsky |
| EPI_ISL_3475157 | Biogroup Bio Lam-LCD Saint-Denis | Department of Virology, Henri Mondor University Hospital, Assistance Publique Hôpitaux de Paris, Université Paris-Est Créteil, INSERM U955 | Christophe Rodriguez, Slim Fourati, Vanessa Demontant, Guillaume Gricourt, Melissa N'Debi, Alexandre Soulier, Elisabeth Trawinski, Jean-Michel Pawlotsky |
| EPI_ISL_3475158 | BIOMNIS EUROFINS IVRY | Department of Virology, Henri Mondor University Hospital, Assistance Publique Hôpitaux de Paris, Université Paris-Est Créteil, INSERM U955 | Christophe Rodriguez, Slim Fourati, Vanessa Demontant, Guillaume Gricourt, Melissa N'Debi, Alexandre Soulier, Elisabeth Trawinski, Jean-Michel Pawlotsky |
| EPI_ISL_3475159, EPI_ISL_3475160 | Biogroup Bio Lam-LCD Saint-Denis | Department of Virology, Henri Mondor University Hospital, Assistance Publique Hôpitaux de Paris, Université Paris-Est Créteil, INSERM U955 | Christophe Rodriguez, Slim Fourati, Vanessa Demontant, Guillaume Gricourt, Melissa N'Debi, Alexandre Soulier, Elisabeth Trawinski, Jean-Michel Pawlotsky |
| EPI_ISL_3475161 | CHU DE DIJON | Department of Virology, Henri Mondor University Hospital, Assistance Publique Hôpitaux de Paris, Université Paris-Est Créteil, INSERM U955 | Christophe Rodriguez, Slim Fourati, Vanessa Demontant, Guillaume Gricourt, Melissa N'Debi, Alexandre Soulier, Elisabeth Trawinski, Jean-Michel Pawlotsky |
| EPI_ISL_3475162 | BIOLBS Rouen | Department of Virology, Henri Mondor University Hospital, Assistance Publique Hôpitaux de Paris, Université Paris-Est Créteil, INSERM U955 | Christophe Rodriguez, Slim Fourati, Vanessa Demontant, Guillaume Gricourt, Melissa N'Debi, Alexandre Soulier, Elisabeth Trawinski, Jean-Michel Pawlotsky |
| EPI_ISL_3475163, EPI_ISL_3475164 | CHU DE DIJON | Department of Virology, Henri Mondor University Hospital, Assistance Publique Hôpitaux de Paris, Université Paris-Est Créteil, INSERM U955 | Christophe Rodriguez, Slim Fourati, Vanessa Demontant, Guillaume Gricourt, Melissa N'Debi, Alexandre Soulier, Elisabeth Trawinski, Jean-Michel Pawlotsky |
| EPI_ISL_3475165 | CH. ANDRE GREGOIRE | Department of Virology, Henri Mondor University Hospital, Assistance Publique Hôpitaux de Paris, Université Paris-Est Créteil, INSERM U955 | Christophe Rodriguez, Slim Fourati, Vanessa Demontant, Guillaume Gricourt, Melissa N'Debi, Alexandre Soulier, Elisabeth Trawinski, Jean-Michel Pawlotsky |
| EPI_ISL_3475166 | CHU DE DIJON | Department of Virology, Henri Mondor University Hospital, Assistance Publique Hôpitaux de Paris, Université Paris-Est Créteil, INSERM U955 | Christophe Rodriguez, Slim Fourati, Vanessa Demontant, Guillaume Gricourt, Melissa N'Debi, Alexandre Soulier, Elisabeth Trawinski, Jean-Michel Pawlotsky |
| EPI_ISL_3475167 | GCS Epinal Remiremont | Department of Virology, Henri Mondor University Hospital, Assistance Publique Hôpitaux de Paris, Université Paris-Est Créteil, INSERM U955 | Christophe Rodriguez, Slim Fourati, Vanessa Demontant, Guillaume Gricourt, Melissa N'Debi, Alexandre Soulier, Elisabeth Trawinski, Jean-Michel Pawlotsky |
| EPI_ISL_3475168, EPI_ISL_3475169 | CHU DE DIJON | Department of Virology, Henri Mondor University Hospital, Assistance Publique Hôpitaux de Paris, Université Paris-Est Créteil, INSERM U955 | Christophe Rodriguez, Slim Fourati, Vanessa Demontant, Guillaume Gricourt, Melissa N'Debi, Alexandre Soulier, Elisabeth Trawinski, Jean-Michel Pawlotsky |
| EPI_ISL_3475170 | CH. CLEMENCEAU | Department of Virology, Henri Mondor University Hospital, Assistance Publique Hôpitaux de Paris, Université Paris-Est Créteil, INSERM U955 | Christophe Rodriguez, Slim Fourati, Vanessa Demontant, Guillaume Gricourt, Melissa N'Debi, Alexandre Soulier, Elisabeth Trawinski, Jean-Michel Pawlotsky |
| EPI_ISL_3475171 | CHU DE DIJON | Department of Virology, Henri Mondor University Hospital, Assistance Publique Hôpitaux de Paris, Université Paris-Est Créteil, INSERM U955 | Christophe Rodriguez, Slim Fourati, Vanessa Demontant, Guillaume Gricourt, Melissa N'Debi, Alexandre Soulier, Elisabeth Trawinski, Jean-Michel Pawlotsky |
| EPI_ISL_3475172 | CH. BEGIN | Department of Virology, Henri Mondor University Hospital, Assistance Publique Hôpitaux de Paris, Université Paris-Est Créteil, INSERM U955 | Christophe Rodriguez, Slim Fourati, Vanessa Demontant, Guillaume Gricourt, Melissa N'Debi, Alexandre Soulier, Elisabeth Trawinski, Jean-Michel Pawlotsky |
| EPI_ISL_3475173 | Biogroup Bio Lam-LCD Saint-Denis | Department of Virology, Henri Mondor University Hospital, Assistance Publique Hôpitaux de Paris, Université Paris-Est Créteil, INSERM U955 | Christophe Rodriguez, Slim Fourati, Vanessa Demontant, Guillaume Gricourt, Melissa N'Debi, Alexandre Soulier, Elisabeth Trawinski, Jean-Michel Pawlotsky |
| EPI_ISL_3475174, EPI_ISL_3475175, EPI_ISL_3475176 | CHU DE DIJON | Department of Virology, Henri Mondor University Hospital, Assistance Publique Hôpitaux de Paris, Université Paris-Est Créteil, INSERM U955 | Christophe Rodriguez, Slim Fourati, Vanessa Demontant, Guillaume Gricourt, Melissa N'Debi, Alexandre Soulier, Elisabeth Trawinski, Jean-Michel Pawlotsky |
| EPI_ISL_3475177 | Biogroup Bio Lam-LCD Saint-Denis | Department of Virology, Henri Mondor University Hospital, Assistance Publique Hôpitaux de Paris, Université Paris-Est Créteil, INSERM U955 | Christophe Rodriguez, Slim Fourati, Vanessa Demontant, Guillaume Gricourt, Melissa N'Debi, Alexandre Soulier, Elisabeth Trawinski, Jean-Michel Pawlotsky |
| EPI_ISL_3475178 | CHU DE DIJON | Department of Virology, Henri Mondor University Hospital, Assistance Publique Hôpitaux de Paris, Université Paris-Est Créteil, INSERM U955 | Christophe Rodriguez, Slim Fourati, Vanessa Demontant, Guillaume Gricourt, Melissa N'Debi, Alexandre Soulier, Elisabeth Trawinski, Jean-Michel Pawlotsky |
| EPI_ISL_3475179, EPI_ISL_3475180 | Biogroup Bio Lam-LCD Saint-Denis | Department of Virology, Henri Mondor University Hospital, Assistance Publique Hôpitaux de Paris, Université Paris-Est Créteil, INSERM U955 | Christophe Rodriguez, Slim Fourati, Vanessa Demontant, Guillaume Gricourt, Melissa N'Debi, Alexandre Soulier, Elisabeth Trawinski, Jean-Michel Pawlotsky |
| EPI_ISL_3475181, EPI_ISL_3475182 | CHU DE DIJON | Department of Virology, Henri Mondor University Hospital, Assistance Publique Hôpitaux de Paris, Université Paris-Est Créteil, INSERM U955 | Christophe Rodriguez, Slim Fourati, Vanessa Demontant, Guillaume Gricourt, Melissa N'Debi, Alexandre Soulier, Elisabeth Trawinski, Jean-Michel Pawlotsky |
| EPI_ISL_3475183 | Biogroup Bio Lam-LCD Saint-Denis | Department of Virology, Henri Mondor University Hospital, Assistance Publique Hôpitaux de Paris, Université Paris-Est Créteil, INSERM U955 | Christophe Rodriguez, Slim Fourati, Vanessa Demontant, Guillaume Gricourt, Melissa N'Debi, Alexandre Soulier, Elisabeth Trawinski, Jean-Michel Pawlotsky |
| EPI_ISL_3475184 | CHU DE DIJON | Department of Virology, Henri Mondor University Hospital, Assistance Publique Hôpitaux de Paris, Université Paris-Est Créteil, INSERM U955 | Christophe Rodriguez, Slim Fourati, Vanessa Demontant, Guillaume Gricourt, Melissa N'Debi, Alexandre Soulier, Elisabeth Trawinski, Jean-Michel Pawlotsky |
| EPI_ISL_3475185 | Biogroup Bio Lam-LCD Saint-Denis | Department of Virology, Henri Mondor University Hospital, Assistance Publique Hôpitaux de Paris, Université Paris-Est Créteil, INSERM U955 | Christophe Rodriguez, Slim Fourati, Vanessa Demontant, Guillaume Gricourt, Melissa N'Debi, Alexandre Soulier, Elisabeth Trawinski, Jean-Michel Pawlotsky |
| EPI_ISL_3475186, EPI_ISL_3475187 | CHU DE DIJON | Department of Virology, Henri Mondor University Hospital, Assistance Publique Hôpitaux de Paris, Université Paris-Est Créteil, | Christophe Rodriguez, Slim Fourati, Vanessa Demontant, Guillaume Gricourt, Melissa N'Debi, Alexandre Soulier, Elisabeth Trawinski, Jean-Michel Pawlotsky |

| | | | |
|---|---|---|---|
| | | INSERM U955 | |
| EPI_ISL_3475188, EPI_ISL_3475189 | Biogroup Bio Lam-LCD Saint-Denis | Department of Virology, Henri Mondor University Hospital, Assistance Publique Hôpitaux de Paris, Université Paris-Est Créteil, INSERM U955 | Christophe Rodriguez, Slim Fourati, Vanessa Demontant, Guillaume Gricourt, Melissa N'Debi, Alexandre Soulier, Elisabeth Trawinski, Jean-Michel Pawlotsky |
| EPI_ISL_3475190 | CHU DE DIJON | Department of Virology, Henri Mondor University Hospital, Assistance Publique Hôpitaux de Paris, Université Paris-Est Créteil, INSERM U955 | Christophe Rodriguez, Slim Fourati, Vanessa Demontant, Guillaume Gricourt, Melissa N'Debi, Alexandre Soulier, Elisabeth Trawinski, Jean-Michel Pawlotsky |
| EPI_ISL_3475191, EPI_ISL_3475192 | Biogroup Bio Lam-LCD Saint-Denis | Department of Virology, Henri Mondor University Hospital, Assistance Publique Hôpitaux de Paris, Université Paris-Est Créteil, INSERM U955 | Christophe Rodriguez, Slim Fourati, Vanessa Demontant, Guillaume Gricourt, Melissa N'Debi, Alexandre Soulier, Elisabeth Trawinski, Jean-Michel Pawlotsky |
| EPI_ISL_3475193 | CHU DE DIJON | Department of Virology, Henri Mondor University Hospital, Assistance Publique Hôpitaux de Paris, Université Paris-Est Créteil, INSERM U955 | Christophe Rodriguez, Slim Fourati, Vanessa Demontant, Guillaume Gricourt, Melissa N'Debi, Alexandre Soulier, Elisabeth Trawinski, Jean-Michel Pawlotsky |
| EPI_ISL_3475194, EPI_ISL_3475195 | Biogroup Bio Lam-LCD Saint-Denis | Department of Virology, Henri Mondor University Hospital, Assistance Publique Hôpitaux de Paris, Université Paris-Est Créteil, INSERM U955 | Christophe Rodriguez, Slim Fourati, Vanessa Demontant, Guillaume Gricourt, Melissa N'Debi, Alexandre Soulier, Elisabeth Trawinski, Jean-Michel Pawlotsky |
| EPI_ISL_3475196, EPI_ISL_3475197 | CHU DE DIJON | Department of Virology, Henri Mondor University Hospital, Assistance Publique Hôpitaux de Paris, Université Paris-Est Créteil, INSERM U955 | Christophe Rodriguez, Slim Fourati, Vanessa Demontant, Guillaume Gricourt, Melissa N'Debi, Alexandre Soulier, Elisabeth Trawinski, Jean-Michel Pawlotsky |
| EPI_ISL_3475198 | BIO ARD'AISNE | Department of Virology, Henri Mondor University Hospital, Assistance Publique Hôpitaux de Paris, Université Paris-Est Créteil, INSERM U955 | Christophe Rodriguez, Slim Fourati, Vanessa Demontant, Guillaume Gricourt, Melissa N'Debi, Alexandre Soulier, Elisabeth Trawinski, Jean-Michel Pawlotsky |
| EPI_ISL_3475199 | CHU DE DIJON | Department of Virology, Henri Mondor University Hospital, Assistance Publique Hôpitaux de Paris, Université Paris-Est Créteil, INSERM U955 | Christophe Rodriguez, Slim Fourati, Vanessa Demontant, Guillaume Gricourt, Melissa N'Debi, Alexandre Soulier, Elisabeth Trawinski, Jean-Michel Pawlotsky |
| EPI_ISL_3475200 | Biogroup Bio Lam-LCD Saint-Denis | Department of Virology, Henri Mondor University Hospital, Assistance Publique Hôpitaux de Paris, Université Paris-Est Créteil, INSERM U955 | Christophe Rodriguez, Slim Fourati, Vanessa Demontant, Guillaume Gricourt, Melissa N'Debi, Alexandre Soulier, Elisabeth Trawinski, Jean-Michel Pawlotsky |
| EPI_ISL_3475201, EPI_ISL_3475202, EPI_ISL_3475203 | CHU DE DIJON | Department of Virology, Henri Mondor University Hospital, Assistance Publique Hôpitaux de Paris, Université Paris-Est Créteil, INSERM U955 | Christophe Rodriguez, Slim Fourati, Vanessa Demontant, Guillaume Gricourt, Melissa N'Debi, Alexandre Soulier, Elisabeth Trawinski, Jean-Michel Pawlotsky |
| EPI_ISL_3475204, EPI_ISL_3475205, EPI_ISL_3475206, EPI_ISL_3475207, EPI_ISL_3475208, EPI_ISL_3475209, EPI_ISL_3475210, EPI_ISL_3475211, EPI_ISL_3475213 | Biogroup Bio Lam-LCD Saint-Denis | Department of Virology, Henri Mondor University Hospital, Assistance Publique Hôpitaux de Paris, Université Paris-Est Créteil, INSERM U955 | Christophe Rodriguez, Slim Fourati, Vanessa Demontant, Guillaume Gricourt, Melissa N'Debi, Alexandre Soulier, Elisabeth Trawinski, Jean-Michel Pawlotsky |
| EPI_ISL_3475214 | CHU DE DIJON | Department of Virology, Henri Mondor University Hospital, Assistance Publique Hôpitaux de Paris, Université Paris-Est Créteil, INSERM U955 | Christophe Rodriguez, Slim Fourati, Vanessa Demontant, Guillaume Gricourt, Melissa N'Debi, Alexandre Soulier, Elisabeth Trawinski, Jean-Michel Pawlotsky |
| EPI_ISL_3475215, EPI_ISL_3475216 | Biogroup Bio Lam-LCD Saint-Denis | Department of Virology, Henri Mondor University Hospital, Assistance Publique Hôpitaux de Paris, Université Paris-Est Créteil, INSERM U955 | Christophe Rodriguez, Slim Fourati, Vanessa Demontant, Guillaume Gricourt, Melissa N'Debi, Alexandre Soulier, Elisabeth Trawinski, Jean-Michel Pawlotsky |
| EPI_ISL_3475217, EPI_ISL_3475218 | CHU DE DIJON | Department of Virology, Henri Mondor University Hospital, Assistance Publique Hôpitaux de Paris, Université Paris-Est Créteil, INSERM U955 | Christophe Rodriguez, Slim Fourati, Vanessa Demontant, Guillaume Gricourt, Melissa N'Debi, Alexandre Soulier, Elisabeth Trawinski, Jean-Michel Pawlotsky |
| EPI_ISL_3475219, EPI_ISL_3475220 | Biogroup Bio Lam-LCD Saint-Denis | Department of Virology, Henri Mondor University Hospital, Assistance Publique Hôpitaux de Paris, Université Paris-Est Créteil, INSERM U955 | Christophe Rodriguez, Slim Fourati, Vanessa Demontant, Guillaume Gricourt, Melissa N'Debi, Alexandre Soulier, Elisabeth Trawinski, Jean-Michel Pawlotsky |
| EPI_ISL_3475221, EPI_ISL_3475222 | CHU DE DIJON | Department of Virology, Henri Mondor University Hospital, Assistance Publique Hôpitaux de Paris, Université Paris-Est Créteil, INSERM U955 | Christophe Rodriguez, Slim Fourati, Vanessa Demontant, Guillaume Gricourt, Melissa N'Debi, Alexandre Soulier, Elisabeth Trawinski, Jean-Michel Pawlotsky |
| EPI_ISL_3475223 | CH. BEGIN | Department of Virology, Henri Mondor University Hospital, Assistance Publique Hôpitaux de Paris, Université Paris-Est Créteil, INSERM U955 | Christophe Rodriguez, Slim Fourati, Vanessa Demontant, Guillaume Gricourt, Melissa N'Debi, Alexandre Soulier, Elisabeth Trawinski, Jean-Michel Pawlotsky |
| EPI_ISL_3475224 | CHU DE DIJON | Department of Virology, Henri Mondor University Hospital, Assistance Publique Hôpitaux de Paris, Université Paris-Est Créteil, INSERM U955 | Christophe Rodriguez, Slim Fourati, Vanessa Demontant, Guillaume Gricourt, Melissa N'Debi, Alexandre Soulier, Elisabeth Trawinski, Jean-Michel Pawlotsky |
| EPI_ISL_3475225 | CH. CLEMENCEAU | Department of Virology, Henri Mondor University Hospital, Assistance Publique Hôpitaux de Paris, Université Paris-Est Créteil, INSERM U955 | Christophe Rodriguez, Slim Fourati, Vanessa Demontant, Guillaume Gricourt, Melissa N'Debi, Alexandre Soulier, Elisabeth Trawinski, Jean-Michel Pawlotsky |
| EPI_ISL_3475226 | Biogroup Bio Lam-LCD Saint-Denis | Department of Virology, Henri Mondor University Hospital, Assistance Publique Hôpitaux de Paris, Université Paris-Est Créteil, INSERM U955 | Christophe Rodriguez, Slim Fourati, Vanessa Demontant, Guillaume Gricourt, Melissa N'Debi, Alexandre Soulier, Elisabeth Trawinski, Jean-Michel Pawlotsky |
| EPI_ISL_3475227 | CHU DE DIJON | Department of Virology, Henri Mondor University Hospital, Assistance Publique Hôpitaux de Paris, Université Paris-Est Créteil, INSERM U955 | Christophe Rodriguez, Slim Fourati, Vanessa Demontant, Guillaume Gricourt, Melissa N'Debi, Alexandre Soulier, Elisabeth Trawinski, Jean-Michel Pawlotsky |
| EPI_ISL_3475229 | BIOMNIS EUROFINS IVRY | Department of Virology, Henri Mondor University Hospital, Assistance Publique Hôpitaux de Paris, Université Paris-Est Créteil, INSERM U955 | Christophe Rodriguez, Slim Fourati, Vanessa Demontant, Guillaume Gricourt, Melissa N'Debi, Alexandre Soulier, Elisabeth Trawinski, Jean-Michel Pawlotsky |
| EPI_ISL_3475230 | Biogroup Bio Lam-LCD Saint-Denis | Department of Virology, Henri Mondor University Hospital, Assistance Publique Hôpitaux de Paris, Université Paris-Est Créteil, INSERM U955 | Christophe Rodriguez, Slim Fourati, Vanessa Demontant, Guillaume Gricourt, Melissa N'Debi, Alexandre Soulier, Elisabeth Trawinski, Jean-Michel Pawlotsky |
| EPI_ISL_3475231 | GCS Epinal Remiremont | Department of Virology, Henri Mondor University Hospital, Assistance Publique Hôpitaux de Paris, Université Paris-Est Créteil, INSERM U955 | Christophe Rodriguez, Slim Fourati, Vanessa Demontant, Guillaume Gricourt, Melissa N'Debi, Alexandre Soulier, Elisabeth Trawinski, Jean-Michel Pawlotsky |
| EPI_ISL_3475232 | Biogroup Bio Lam-LCD Saint-Denis | Department of Virology, Henri Mondor University Hospital, Assistance Publique Hôpitaux de Paris, Université Paris-Est Créteil, INSERM U955 | Christophe Rodriguez, Slim Fourati, Vanessa Demontant, Guillaume Gricourt, Melissa N'Debi, Alexandre Soulier, Elisabeth Trawinski, Jean-Michel Pawlotsky |
| EPI_ISL_3475233, EPI_ISL_3475234 | CHU DE DIJON | Department of Virology, Henri Mondor University Hospital, Assistance Publique Hôpitaux de Paris, Université Paris-Est Créteil, INSERM U955 | Christophe Rodriguez, Slim Fourati, Vanessa Demontant, Guillaume Gricourt, Melissa N'Debi, Alexandre Soulier, Elisabeth Trawinski, Jean-Michel Pawlotsky |
| EPI_ISL_3475235 | Biogroup Bio Lam-LCD Saint-Denis | Department of Virology, Henri Mondor University Hospital, Assistance Publique Hôpitaux de Paris, Université Paris-Est Créteil, INSERM U955 | Christophe Rodriguez, Slim Fourati, Vanessa Demontant, Guillaume Gricourt, Melissa N'Debi, Alexandre Soulier, Elisabeth Trawinski, Jean-Michel Pawlotsky |
| EPI_ISL_3475236 | CH. CLEMENCEAU | Department of Virology, Henri Mondor University Hospital, Assistance Publique Hôpitaux de Paris, Université Paris-Est Créteil, INSERM U955 | Christophe Rodriguez, Slim Fourati, Vanessa Demontant, Guillaume Gricourt, Melissa N'Debi, Alexandre Soulier, Elisabeth Trawinski, Jean-Michel Pawlotsky |
| EPI_ISL_3475237, EPI_ISL_3475238 | Biogroup Bio Lam-LCD Saint-Denis | Department of Virology, Henri Mondor University Hospital, Assistance Publique Hôpitaux de Paris, Université Paris-Est Créteil, INSERM U955 | Christophe Rodriguez, Slim Fourati, Vanessa Demontant, Guillaume Gricourt, Melissa N'Debi, Alexandre Soulier, Elisabeth Trawinski, Jean-Michel Pawlotsky |
| EPI_ISL_3475239 | CHU DE DIJON | Department of Virology, Henri Mondor University Hospital, Assistance Publique Hôpitaux de Paris, Université Paris-Est Créteil, INSERM U955 | Christophe Rodriguez, Slim Fourati, Vanessa Demontant, Guillaume Gricourt, Melissa N'Debi, Alexandre Soulier, Elisabeth Trawinski, Jean-Michel Pawlotsky |

| | | | |
|---|---|---|---|
| EPI_ISL_3475240 | Biogroup Bio Lam-LCD Saint-Denis | Department of Virology, Henri Mondor University Hospital, Assistance Publique Hôpitaux de Paris, Université Paris-Est Créteil, INSERM U955 | Christophe Rodriguez, Slim Fourati, Vanessa Demontant, Guillaume Gricourt, Melissa N'Debi, Alexandre Soulier, Elisabeth Trawinski, Jean-Michel Pawlotsky |
| EPI_ISL_3475241, EPI_ISL_3475242 | CHU DE DIJON | Department of Virology, Henri Mondor University Hospital, Assistance Publique Hôpitaux de Paris, Université Paris-Est Créteil, INSERM U955 | Christophe Rodriguez, Slim Fourati, Vanessa Demontant, Guillaume Gricourt, Melissa N'Debi, Alexandre Soulier, Elisabeth Trawinski, Jean-Michel Pawlotsky |
| EPI_ISL_3475243 | Laboratoire BIOTOV | Department of Virology, Henri Mondor University Hospital, Assistance Publique Hôpitaux de Paris, Université Paris-Est Créteil, INSERM U955 | Christophe Rodriguez, Slim Fourati, Vanessa Demontant, Guillaume Gricourt, Melissa N'Debi, Alexandre Soulier, Elisabeth Trawinski, Jean-Michel Pawlotsky |
| EPI_ISL_3475244 | Biogroup Bio Lam-LCD Saint-Denis | Department of Virology, Henri Mondor University Hospital, Assistance Publique Hôpitaux de Paris, Université Paris-Est Créteil, INSERM U955 | Christophe Rodriguez, Slim Fourati, Vanessa Demontant, Guillaume Gricourt, Melissa N'Debi, Alexandre Soulier, Elisabeth Trawinski, Jean-Michel Pawlotsky |
| EPI_ISL_3475245, EPI_ISL_3475246, EPI_ISL_3475247, EPI_ISL_3475248 | CHU DE DIJON | Department of Virology, Henri Mondor University Hospital, Assistance Publique Hôpitaux de Paris, Université Paris-Est Créteil, INSERM U955 | Christophe Rodriguez, Slim Fourati, Vanessa Demontant, Guillaume Gricourt, Melissa N'Debi, Alexandre Soulier, Elisabeth Trawinski, Jean-Michel Pawlotsky |
| EPI_ISL_3475249, EPI_ISL_3475250 | Biogroup Bio Lam-LCD Saint-Denis | Department of Virology, Henri Mondor University Hospital, Assistance Publique Hôpitaux de Paris, Université Paris-Est Créteil, INSERM U955 | Christophe Rodriguez, Slim Fourati, Vanessa Demontant, Guillaume Gricourt, Melissa N'Debi, Alexandre Soulier, Elisabeth Trawinski, Jean-Michel Pawlotsky |
| EPI_ISL_3475251, EPI_ISL_3475252 | CHU DE DIJON | Department of Virology, Henri Mondor University Hospital, Assistance Publique Hôpitaux de Paris, Université Paris-Est Créteil, INSERM U955 | Christophe Rodriguez, Slim Fourati, Vanessa Demontant, Guillaume Gricourt, Melissa N'Debi, Alexandre Soulier, Elisabeth Trawinski, Jean-Michel Pawlotsky |
| EPI_ISL_3475253 | Biogroup Bio Lam-LCD Saint-Denis | Department of Virology, Henri Mondor University Hospital, Assistance Publique Hôpitaux de Paris, Université Paris-Est Créteil, INSERM U955 | Christophe Rodriguez, Slim Fourati, Vanessa Demontant, Guillaume Gricourt, Melissa N'Debi, Alexandre Soulier, Elisabeth Trawinski, Jean-Michel Pawlotsky |
| EPI_ISL_3475254 | BIOLBS Rouen | Department of Virology, Henri Mondor University Hospital, Assistance Publique Hôpitaux de Paris, Université Paris-Est Créteil, INSERM U955 | Christophe Rodriguez, Slim Fourati, Vanessa Demontant, Guillaume Gricourt, Melissa N'Debi, Alexandre Soulier, Elisabeth Trawinski, Jean-Michel Pawlotsky |
| EPI_ISL_3475255, EPI_ISL_3475256, EPI_ISL_3475257 | Biogroup Bio Lam-LCD Saint-Denis | Department of Virology, Henri Mondor University Hospital, Assistance Publique Hôpitaux de Paris, Université Paris-Est Créteil, INSERM U955 | Christophe Rodriguez, Slim Fourati, Vanessa Demontant, Guillaume Gricourt, Melissa N'Debi, Alexandre Soulier, Elisabeth Trawinski, Jean-Michel Pawlotsky |
| EPI_ISL_3475258, EPI_ISL_3475259 | CHU DE DIJON | Department of Virology, Henri Mondor University Hospital, Assistance Publique Hôpitaux de Paris, Université Paris-Est Créteil, INSERM U955 | Christophe Rodriguez, Slim Fourati, Vanessa Demontant, Guillaume Gricourt, Melissa N'Debi, Alexandre Soulier, Elisabeth Trawinski, Jean-Michel Pawlotsky |
| EPI_ISL_3475260 | BIO ARD'AISNE | Department of Virology, Henri Mondor University Hospital, Assistance Publique Hôpitaux de Paris, Université Paris-Est Créteil, INSERM U955 | Christophe Rodriguez, Slim Fourati, Vanessa Demontant, Guillaume Gricourt, Melissa N'Debi, Alexandre Soulier, Elisabeth Trawinski, Jean-Michel Pawlotsky |
| EPI_ISL_3475261 | CHU DE DIJON | Department of Virology, Henri Mondor University Hospital, Assistance Publique Hôpitaux de Paris, Université Paris-Est Créteil, INSERM U955 | Christophe Rodriguez, Slim Fourati, Vanessa Demontant, Guillaume Gricourt, Melissa N'Debi, Alexandre Soulier, Elisabeth Trawinski, Jean-Michel Pawlotsky |
| EPI_ISL_3475262 | Biogroup Bio Lam-LCD Saint-Denis | Department of Virology, Henri Mondor University Hospital, Assistance Publique Hôpitaux de Paris, Université Paris-Est Créteil, INSERM U955 | Christophe Rodriguez, Slim Fourati, Vanessa Demontant, Guillaume Gricourt, Melissa N'Debi, Alexandre Soulier, Elisabeth Trawinski, Jean-Michel Pawlotsky |
| EPI_ISL_3475263, EPI_ISL_3475264, EPI_ISL_3475265 | CHU DE DIJON | Department of Virology, Henri Mondor University Hospital, Assistance Publique Hôpitaux de Paris, Université Paris-Est Créteil, INSERM U955 | Christophe Rodriguez, Slim Fourati, Vanessa Demontant, Guillaume Gricourt, Melissa N'Debi, Alexandre Soulier, Elisabeth Trawinski, Jean-Michel Pawlotsky |
| EPI_ISL_3475266, EPI_ISL_3475267, EPI_ISL_3475268, EPI_ISL_3475269, EPI_ISL_3475270 | Biogroup Bio Lam-LCD Saint-Denis | Department of Virology, Henri Mondor University Hospital, Assistance Publique Hôpitaux de Paris, Université Paris-Est Créteil, INSERM U955 | Christophe Rodriguez, Slim Fourati, Vanessa Demontant, Guillaume Gricourt, Melissa N'Debi, Alexandre Soulier, Elisabeth Trawinski, Jean-Michel Pawlotsky |
| EPI_ISL_3475271 | CHU DE DIJON | Department of Virology, Henri Mondor University Hospital, Assistance Publique Hôpitaux de Paris, Université Paris-Est Créteil, INSERM U955 | Christophe Rodriguez, Slim Fourati, Vanessa Demontant, Guillaume Gricourt, Melissa N'Debi, Alexandre Soulier, Elisabeth Trawinski, Jean-Michel Pawlotsky |
| EPI_ISL_3475272, EPI_ISL_3475273 | Biogroup Bio Lam-LCD Saint-Denis | Department of Virology, Henri Mondor University Hospital, Assistance Publique Hôpitaux de Paris, Université Paris-Est Créteil, INSERM U955 | Christophe Rodriguez, Slim Fourati, Vanessa Demontant, Guillaume Gricourt, Melissa N'Debi, Alexandre Soulier, Elisabeth Trawinski, Jean-Michel Pawlotsky |
| EPI_ISL_3475274 | CHU DE DIJON | Department of Virology, Henri Mondor University Hospital, Assistance Publique Hôpitaux de Paris, Université Paris-Est Créteil, INSERM U955 | Christophe Rodriguez, Slim Fourati, Vanessa Demontant, Guillaume Gricourt, Melissa N'Debi, Alexandre Soulier, Elisabeth Trawinski, Jean-Michel Pawlotsky |
| EPI_ISL_3475275, EPI_ISL_3475276 | Biogroup Bio Lam-LCD Saint-Denis | Department of Virology, Henri Mondor University Hospital, Assistance Publique Hôpitaux de Paris, Université Paris-Est Créteil, INSERM U955 | Christophe Rodriguez, Slim Fourati, Vanessa Demontant, Guillaume Gricourt, Melissa N'Debi, Alexandre Soulier, Elisabeth Trawinski, Jean-Michel Pawlotsky |
| EPI_ISL_3475277 | CHU DE DIJON | Department of Virology, Henri Mondor University Hospital, Assistance Publique Hôpitaux de Paris, Université Paris-Est Créteil, INSERM U955 | Christophe Rodriguez, Slim Fourati, Vanessa Demontant, Guillaume Gricourt, Melissa N'Debi, Alexandre Soulier, Elisabeth Trawinski, Jean-Michel Pawlotsky |
| EPI_ISL_3475278 | Biogroup Bio Lam-LCD Saint-Denis | Department of Virology, Henri Mondor University Hospital, Assistance Publique Hôpitaux de Paris, Université Paris-Est Créteil, INSERM U955 | Christophe Rodriguez, Slim Fourati, Vanessa Demontant, Guillaume Gricourt, Melissa N'Debi, Alexandre Soulier, Elisabeth Trawinski, Jean-Michel Pawlotsky |
| EPI_ISL_3475279 | CHU DE DIJON | Department of Virology, Henri Mondor University Hospital, Assistance Publique Hôpitaux de Paris, Université Paris-Est Créteil, INSERM U955 | Christophe Rodriguez, Slim Fourati, Vanessa Demontant, Guillaume Gricourt, Melissa N'Debi, Alexandre Soulier, Elisabeth Trawinski, Jean-Michel Pawlotsky |
| EPI_ISL_3475280, EPI_ISL_3475281 | Biogroup Bio Lam-LCD Saint-Denis | Department of Virology, Henri Mondor University Hospital, Assistance Publique Hôpitaux de Paris, Université Paris-Est Créteil, INSERM U955 | Christophe Rodriguez, Slim Fourati, Vanessa Demontant, Guillaume Gricourt, Melissa N'Debi, Alexandre Soulier, Elisabeth Trawinski, Jean-Michel Pawlotsky |
| EPI_ISL_3475282 | CHU DE DIJON | Department of Virology, Henri Mondor University Hospital, Assistance Publique Hôpitaux de Paris, Université Paris-Est Créteil, INSERM U955 | Christophe Rodriguez, Slim Fourati, Vanessa Demontant, Guillaume Gricourt, Melissa N'Debi, Alexandre Soulier, Elisabeth Trawinski, Jean-Michel Pawlotsky |
| EPI_ISL_3475283 | Laboratoire BIOTOV | Department of Virology, Henri Mondor University Hospital, Assistance Publique Hôpitaux de Paris, Université Paris-Est Créteil, INSERM U955 | Christophe Rodriguez, Slim Fourati, Vanessa Demontant, Guillaume Gricourt, Melissa N'Debi, Alexandre Soulier, Elisabeth Trawinski, Jean-Michel Pawlotsky |
| EPI_ISL_3475284 | CH. CLEMENCEAU | Department of Virology, Henri Mondor University Hospital, Assistance Publique Hôpitaux de Paris, Université Paris-Est Créteil, INSERM U955 | Christophe Rodriguez, Slim Fourati, Vanessa Demontant, Guillaume Gricourt, Melissa N'Debi, Alexandre Soulier, Elisabeth Trawinski, Jean-Michel Pawlotsky |
| EPI_ISL_3475285 | Biogroup Bio Lam-LCD Saint-Denis | Department of Virology, Henri Mondor University Hospital, Assistance Publique Hôpitaux de Paris, Université Paris-Est Créteil, INSERM U955 | Christophe Rodriguez, Slim Fourati, Vanessa Demontant, Guillaume Gricourt, Melissa N'Debi, Alexandre Soulier, Elisabeth Trawinski, Jean-Michel Pawlotsky |
| EPI_ISL_3475286, EPI_ISL_3475287 | CHU DE DIJON | Department of Virology, Henri Mondor University Hospital, Assistance Publique Hôpitaux de Paris, Université Paris-Est Créteil, INSERM U955 | Christophe Rodriguez, Slim Fourati, Vanessa Demontant, Guillaume Gricourt, Melissa N'Debi, Alexandre Soulier, Elisabeth Trawinski, Jean-Michel Pawlotsky |
| EPI_ISL_3475288 | Biogroup Bio Lam-LCD Saint-Denis | Department of Virology, Henri Mondor University Hospital, Assistance Publique Hôpitaux de Paris, Université Paris-Est Créteil, INSERM U955 | Christophe Rodriguez, Slim Fourati, Vanessa Demontant, Guillaume Gricourt, Melissa N'Debi, Alexandre Soulier, Elisabeth Trawinski, Jean-Michel Pawlotsky |
| EPI_ISL_3475289 | BIO ARD'AISNE | Department of Virology, Henri Mondor University Hospital, Assistance Publique Hôpitaux de Paris, Université Paris-Est Créteil, INSERM U955 | Christophe Rodriguez, Slim Fourati, Vanessa Demontant, Guillaume Gricourt, Melissa N'Debi, Alexandre Soulier, Elisabeth Trawinski, Jean-Michel Pawlotsky |

| | | | |
|---|---|---|---|
| EPI_ISL_3475290, EPI_ISL_3475291 | CHU DE DIJON | Department of Virology, Henri Mondor University Hospital, Assistance Publique Hôpitaux de Paris, Université Paris-Est Créteil, INSERM U955 | Christophe Rodriguez, Slim Fourati, Vanessa Demontant, Guillaume Gricourt, Melissa N'Debi, Alexandre Soulier, Elisabeth Trawinski, Jean-Michel Pawlotsky |
| EPI_ISL_3475292, EPI_ISL_3475293, EPI_ISL_3475294 | Biogroup Bio Lam-LCD Saint-Denis | Department of Virology, Henri Mondor University Hospital, Assistance Publique Hôpitaux de Paris, Université Paris-Est Créteil, INSERM U955 | Christophe Rodriguez, Slim Fourati, Vanessa Demontant, Guillaume Gricourt, Melissa N'Debi, Alexandre Soulier, Elisabeth Trawinski, Jean-Michel Pawlotsky |
| EPI_ISL_3475295 | BIOLBS Rouen | Department of Virology, Henri Mondor University Hospital, Assistance Publique Hôpitaux de Paris, Université Paris-Est Créteil, INSERM U955 | Christophe Rodriguez, Slim Fourati, Vanessa Demontant, Guillaume Gricourt, Melissa N'Debi, Alexandre Soulier, Elisabeth Trawinski, Jean-Michel Pawlotsky |
| EPI_ISL_3475296 | CHU DE DIJON | Department of Virology, Henri Mondor University Hospital, Assistance Publique Hôpitaux de Paris, Université Paris-Est Créteil, INSERM U955 | Christophe Rodriguez, Slim Fourati, Vanessa Demontant, Guillaume Gricourt, Melissa N'Debi, Alexandre Soulier, Elisabeth Trawinski, Jean-Michel Pawlotsky |
| EPI_ISL_3475297, EPI_ISL_3475298, EPI_ISL_3475299, EPI_ISL_3475300 | Biogroup Bio Lam-LCD Saint-Denis | Department of Virology, Henri Mondor University Hospital, Assistance Publique Hôpitaux de Paris, Université Paris-Est Créteil, INSERM U955 | Christophe Rodriguez, Slim Fourati, Vanessa Demontant, Guillaume Gricourt, Melissa N'Debi, Alexandre Soulier, Elisabeth Trawinski, Jean-Michel Pawlotsky |
| EPI_ISL_3475301 | CHU DE DIJON | Department of Virology, Henri Mondor University Hospital, Assistance Publique Hôpitaux de Paris, Université Paris-Est Créteil, INSERM U955 | Christophe Rodriguez, Slim Fourati, Vanessa Demontant, Guillaume Gricourt, Melissa N'Debi, Alexandre Soulier, Elisabeth Trawinski, Jean-Michel Pawlotsky |
| EPI_ISL_3475302, EPI_ISL_3475303 | Biogroup Bio Lam-LCD Saint-Denis | Department of Virology, Henri Mondor University Hospital, Assistance Publique Hôpitaux de Paris, Université Paris-Est Créteil, INSERM U955 | Christophe Rodriguez, Slim Fourati, Vanessa Demontant, Guillaume Gricourt, Melissa N'Debi, Alexandre Soulier, Elisabeth Trawinski, Jean-Michel Pawlotsky |
| EPI_ISL_3475304 | CHU DE DIJON | Department of Virology, Henri Mondor University Hospital, Assistance Publique Hôpitaux de Paris, Université Paris-Est Créteil, INSERM U955 | Christophe Rodriguez, Slim Fourati, Vanessa Demontant, Guillaume Gricourt, Melissa N'Debi, Alexandre Soulier, Elisabeth Trawinski, Jean-Michel Pawlotsky |
| EPI_ISL_3475305 | Biogroup Bio Lam-LCD Saint-Denis | Department of Virology, Henri Mondor University Hospital, Assistance Publique Hôpitaux de Paris, Université Paris-Est Créteil, INSERM U955 | Christophe Rodriguez, Slim Fourati, Vanessa Demontant, Guillaume Gricourt, Melissa N'Debi, Alexandre Soulier, Elisabeth Trawinski, Jean-Michel Pawlotsky |
| EPI_ISL_3475306, EPI_ISL_3475307 | CHU DE DIJON | Department of Virology, Henri Mondor University Hospital, Assistance Publique Hôpitaux de Paris, Université Paris-Est Créteil, INSERM U955 | Christophe Rodriguez, Slim Fourati, Vanessa Demontant, Guillaume Gricourt, Melissa N'Debi, Alexandre Soulier, Elisabeth Trawinski, Jean-Michel Pawlotsky |
| EPI_ISL_3475308 | Biogroup Bio Lam-LCD Saint-Denis | Department of Virology, Henri Mondor University Hospital, Assistance Publique Hôpitaux de Paris, Université Paris-Est Créteil, INSERM U955 | Christophe Rodriguez, Slim Fourati, Vanessa Demontant, Guillaume Gricourt, Melissa N'Debi, Alexandre Soulier, Elisabeth Trawinski, Jean-Michel Pawlotsky |
| EPI_ISL_3475309 | BIOMNIS EUROFINS IVRY | Department of Virology, Henri Mondor University Hospital, Assistance Publique Hôpitaux de Paris, Université Paris-Est Créteil, INSERM U955 | Christophe Rodriguez, Slim Fourati, Vanessa Demontant, Guillaume Gricourt, Melissa N'Debi, Alexandre Soulier, Elisabeth Trawinski, Jean-Michel Pawlotsky |
| EPI_ISL_3475310 | CHU DE DIJON | Department of Virology, Henri Mondor University Hospital, Assistance Publique Hôpitaux de Paris, Université Paris-Est Créteil, INSERM U955 | Christophe Rodriguez, Slim Fourati, Vanessa Demontant, Guillaume Gricourt, Melissa N'Debi, Alexandre Soulier, Elisabeth Trawinski, Jean-Michel Pawlotsky |
| EPI_ISL_3475311 | Biogroup Bio Lam-LCD Saint-Denis | Department of Virology, Henri Mondor University Hospital, Assistance Publique Hôpitaux de Paris, Université Paris-Est Créteil, INSERM U955 | Christophe Rodriguez, Slim Fourati, Vanessa Demontant, Guillaume Gricourt, Melissa N'Debi, Alexandre Soulier, Elisabeth Trawinski, Jean-Michel Pawlotsky |
| EPI_ISL_3475312 | CHU DE DIJON | Department of Virology, Henri Mondor University Hospital, Assistance Publique Hôpitaux de Paris, Université Paris-Est Créteil, INSERM U955 | Christophe Rodriguez, Slim Fourati, Vanessa Demontant, Guillaume Gricourt, Melissa N'Debi, Alexandre Soulier, Elisabeth Trawinski, Jean-Michel Pawlotsky |
| EPI_ISL_3475313 | BIO ARD'AISNE | Department of Virology, Henri Mondor University Hospital, Assistance Publique Hôpitaux de Paris, Université Paris-Est Créteil, INSERM U955 | Christophe Rodriguez, Slim Fourati, Vanessa Demontant, Guillaume Gricourt, Melissa N'Debi, Alexandre Soulier, Elisabeth Trawinski, Jean-Michel Pawlotsky |
| EPI_ISL_3475314 | Biogroup Bio Lam-LCD Saint-Denis | Department of Virology, Henri Mondor University Hospital, Assistance Publique Hôpitaux de Paris, Université Paris-Est Créteil, INSERM U955 | Christophe Rodriguez, Slim Fourati, Vanessa Demontant, Guillaume Gricourt, Melissa N'Debi, Alexandre Soulier, Elisabeth Trawinski, Jean-Michel Pawlotsky |
| EPI_ISL_3475315, EPI_ISL_3475316, EPI_ISL_3475317 | CHU DE DIJON | Department of Virology, Henri Mondor University Hospital, Assistance Publique Hôpitaux de Paris, Université Paris-Est Créteil, INSERM U955 | Christophe Rodriguez, Slim Fourati, Vanessa Demontant, Guillaume Gricourt, Melissa N'Debi, Alexandre Soulier, Elisabeth Trawinski, Jean-Michel Pawlotsky |
| EPI_ISL_3475318 | CH. CLEMENCEAU | Department of Virology, Henri Mondor University Hospital, Assistance Publique Hôpitaux de Paris, Université Paris-Est Créteil, INSERM U955 | Christophe Rodriguez, Slim Fourati, Vanessa Demontant, Guillaume Gricourt, Melissa N'Debi, Alexandre Soulier, Elisabeth Trawinski, Jean-Michel Pawlotsky |
| EPI_ISL_3475319 | Biogroup Bio Lam-LCD Saint-Denis | Department of Virology, Henri Mondor University Hospital, Assistance Publique Hôpitaux de Paris, Université Paris-Est Créteil, INSERM U955 | Christophe Rodriguez, Slim Fourati, Vanessa Demontant, Guillaume Gricourt, Melissa N'Debi, Alexandre Soulier, Elisabeth Trawinski, Jean-Michel Pawlotsky |
| EPI_ISL_3475320, EPI_ISL_3475321 | CHU DE DIJON | Department of Virology, Henri Mondor University Hospital, Assistance Publique Hôpitaux de Paris, Université Paris-Est Créteil, INSERM U955 | Christophe Rodriguez, Slim Fourati, Vanessa Demontant, Guillaume Gricourt, Melissa N'Debi, Alexandre Soulier, Elisabeth Trawinski, Jean-Michel Pawlotsky |
| EPI_ISL_3475323 | BIOMNIS EUROFINS IVRY | Department of Virology, Henri Mondor University Hospital, Assistance Publique Hôpitaux de Paris, Université Paris-Est Créteil, INSERM U955 | Christophe Rodriguez, Slim Fourati, Vanessa Demontant, Guillaume Gricourt, Melissa N'Debi, Alexandre Soulier, Elisabeth Trawinski, Jean-Michel Pawlotsky |
| EPI_ISL_3475324, EPI_ISL_3475325 | CHU DE DIJON | Department of Virology, Henri Mondor University Hospital, Assistance Publique Hôpitaux de Paris, Université Paris-Est Créteil, INSERM U955 | Christophe Rodriguez, Slim Fourati, Vanessa Demontant, Guillaume Gricourt, Melissa N'Debi, Alexandre Soulier, Elisabeth Trawinski, Jean-Michel Pawlotsky |
| EPI_ISL_3475327, EPI_ISL_3475328 | Biogroup Bio Lam-LCD Saint-Denis | Department of Virology, Henri Mondor University Hospital, Assistance Publique Hôpitaux de Paris, Université Paris-Est Créteil, INSERM U955 | Christophe Rodriguez, Slim Fourati, Vanessa Demontant, Guillaume Gricourt, Melissa N'Debi, Alexandre Soulier, Elisabeth Trawinski, Jean-Michel Pawlotsky |
| EPI_ISL_3475329, EPI_ISL_3475330, EPI_ISL_3475331, EPI_ISL_3475332 | CHU DE DIJON | Department of Virology, Henri Mondor University Hospital, Assistance Publique Hôpitaux de Paris, Université Paris-Est Créteil, INSERM U955 | Christophe Rodriguez, Slim Fourati, Vanessa Demontant, Guillaume Gricourt, Melissa N'Debi, Alexandre Soulier, Elisabeth Trawinski, Jean-Michel Pawlotsky |
| EPI_ISL_3475333, EPI_ISL_3475334 | Biogroup Bio Lam-LCD Saint-Denis | Department of Virology, Henri Mondor University Hospital, Assistance Publique Hôpitaux de Paris, Université Paris-Est Créteil, INSERM U955 | Christophe Rodriguez, Slim Fourati, Vanessa Demontant, Guillaume Gricourt, Melissa N'Debi, Alexandre Soulier, Elisabeth Trawinski, Jean-Michel Pawlotsky |
| EPI_ISL_3475335 | BIO ARD'AISNE | Department of Virology, Henri Mondor University Hospital, Assistance Publique Hôpitaux de Paris, Université Paris-Est Créteil, INSERM U955 | Christophe Rodriguez, Slim Fourati, Vanessa Demontant, Guillaume Gricourt, Melissa N'Debi, Alexandre Soulier, Elisabeth Trawinski, Jean-Michel Pawlotsky |
| EPI_ISL_3475337 | CHU DE DIJON | Department of Virology, Henri Mondor University Hospital, Assistance Publique Hôpitaux de Paris, Université Paris-Est Créteil, INSERM U955 | Christophe Rodriguez, Slim Fourati, Vanessa Demontant, Guillaume Gricourt, Melissa N'Debi, Alexandre Soulier, Elisabeth Trawinski, Jean-Michel Pawlotsky |
| EPI_ISL_3475338 | CH. CLEMENCEAU | Department of Virology, Henri Mondor University Hospital, Assistance Publique Hôpitaux de Paris, Université Paris-Est Créteil, INSERM U955 | Christophe Rodriguez, Slim Fourati, Vanessa Demontant, Guillaume Gricourt, Melissa N'Debi, Alexandre Soulier, Elisabeth Trawinski, Jean-Michel Pawlotsky |
| EPI_ISL_3475339, EPI_ISL_3475340 | Biogroup Bio Lam-LCD Saint-Denis | Department of Virology, Henri Mondor University Hospital, Assistance Publique Hôpitaux de Paris, Université Paris-Est Créteil, INSERM U955 | Christophe Rodriguez, Slim Fourati, Vanessa Demontant, Guillaume Gricourt, Melissa N'Debi, Alexandre Soulier, Elisabeth Trawinski, Jean-Michel Pawlotsky |
| EPI_ISL_3475341 | CHU DE DIJON | Department of Virology, Henri Mondor University Hospital, Assistance Publique Hôpitaux de Paris, Université Paris-Est Créteil, INSERM U955 | Christophe Rodriguez, Slim Fourati, Vanessa Demontant, Guillaume Gricourt, Melissa N'Debi, Alexandre Soulier, Elisabeth Trawinski, Jean-Michel Pawlotsky |
| EPI_ISL_3475342, EPI_ISL_3475343, | Biogroup Bio Lam-LCD Saint-Denis | Department of Virology, Henri Mondor University Hospital, Assistance Publique Hôpitaux de Paris, Université Paris-Est Créteil, | Christophe Rodriguez, Slim Fourati, Vanessa Demontant, Guillaume Gricourt, Melissa N'Debi, Alexandre Soulier, Elisabeth Trawinski, Jean-Michel Pawlotsky |

| | | | |
|---|---|---|---|
| EPI_ISL_3475344 | | INSERM U955 | |
| EPI_ISL_3475345 | CHU DE DIJON | Department of Virology, Henri Mondor University Hospital, Assistance Publique Hôpitaux de Paris, Université Paris-Est Créteil, INSERM U955 | Christophe Rodriguez, Slim Fourati, Vanessa Demontant, Guillaume Gricourt, Melissa N'Debi, Alexandre Soulier, Elisabeth Trawinski, Jean-Michel Pawlotsky |
| EPI_ISL_3475346 | Biogroup Bio Lam-LCD Saint-Denis | Department of Virology, Henri Mondor University Hospital, Assistance Publique Hôpitaux de Paris, Université Paris-Est Créteil, INSERM U955 | Christophe Rodriguez, Slim Fourati, Vanessa Demontant, Guillaume Gricourt, Melissa N'Debi, Alexandre Soulier, Elisabeth Trawinski, Jean-Michel Pawlotsky |
| EPI_ISL_3475347 | CHU DE DIJON | Department of Virology, Henri Mondor University Hospital, Assistance Publique Hôpitaux de Paris, Université Paris-Est Créteil, INSERM U955 | Christophe Rodriguez, Slim Fourati, Vanessa Demontant, Guillaume Gricourt, Melissa N'Debi, Alexandre Soulier, Elisabeth Trawinski, Jean-Michel Pawlotsky |
| EPI_ISL_3475348, EPI_ISL_3475349 | Biogroup Bio Lam-LCD Saint-Denis | Department of Virology, Henri Mondor University Hospital, Assistance Publique Hôpitaux de Paris, Université Paris-Est Créteil, INSERM U955 | Christophe Rodriguez, Slim Fourati, Vanessa Demontant, Guillaume Gricourt, Melissa N'Debi, Alexandre Soulier, Elisabeth Trawinski, Jean-Michel Pawlotsky |
| EPI_ISL_3475350, EPI_ISL_3475351, EPI_ISL_3475352, EPI_ISL_3475353, EPI_ISL_3475354 | CHU DE DIJON | Department of Virology, Henri Mondor University Hospital, Assistance Publique Hôpitaux de Paris, Université Paris-Est Créteil, INSERM U955 | Christophe Rodriguez, Slim Fourati, Vanessa Demontant, Guillaume Gricourt, Melissa N'Debi, Alexandre Soulier, Elisabeth Trawinski, Jean-Michel Pawlotsky |
| EPI_ISL_3475355 | Biogroup Bio Lam-LCD Saint-Denis | Department of Virology, Henri Mondor University Hospital, Assistance Publique Hôpitaux de Paris, Université Paris-Est Créteil, INSERM U955 | Christophe Rodriguez, Slim Fourati, Vanessa Demontant, Guillaume Gricourt, Melissa N'Debi, Alexandre Soulier, Elisabeth Trawinski, Jean-Michel Pawlotsky |
| EPI_ISL_3475356 | CHU DE DIJON | Department of Virology, Henri Mondor University Hospital, Assistance Publique Hôpitaux de Paris, Université Paris-Est Créteil, INSERM U955 | Christophe Rodriguez, Slim Fourati, Vanessa Demontant, Guillaume Gricourt, Melissa N'Debi, Alexandre Soulier, Elisabeth Trawinski, Jean-Michel Pawlotsky |
| EPI_ISL_3475357, EPI_ISL_3475358, EPI_ISL_3475359, EPI_ISL_3475360 | Biogroup Bio Lam-LCD Saint-Denis | Department of Virology, Henri Mondor University Hospital, Assistance Publique Hôpitaux de Paris, Université Paris-Est Créteil, INSERM U955 | Christophe Rodriguez, Slim Fourati, Vanessa Demontant, Guillaume Gricourt, Melissa N'Debi, Alexandre Soulier, Elisabeth Trawinski, Jean-Michel Pawlotsky |
| EPI_ISL_3475361 | BIO ARD'AISNE | Department of Virology, Henri Mondor University Hospital, Assistance Publique Hôpitaux de Paris, Université Paris-Est Créteil, INSERM U955 | Christophe Rodriguez, Slim Fourati, Vanessa Demontant, Guillaume Gricourt, Melissa N'Debi, Alexandre Soulier, Elisabeth Trawinski, Jean-Michel Pawlotsky |
| EPI_ISL_3475362 | Laboratoire BIOTOV | Department of Virology, Henri Mondor University Hospital, Assistance Publique Hôpitaux de Paris, Université Paris-Est Créteil, INSERM U955 | Christophe Rodriguez, Slim Fourati, Vanessa Demontant, Guillaume Gricourt, Melissa N'Debi, Alexandre Soulier, Elisabeth Trawinski, Jean-Michel Pawlotsky |
| EPI_ISL_3475363 | Hôpital Antoine-Béclère | Department of Virology, Henri Mondor University Hospital, Assistance Publique Hôpitaux de Paris, Université Paris-Est Créteil, INSERM U955 | Christophe Rodriguez, Slim Fourati, Vanessa Demontant, Guillaume Gricourt, Melissa N'Debi, Alexandre Soulier, Elisabeth Trawinski, Jean-Michel Pawlotsky |
| EPI_ISL_3475364, EPI_ISL_3475365, EPI_ISL_3475366 | CHU DE DIJON | Department of Virology, Henri Mondor University Hospital, Assistance Publique Hôpitaux de Paris, Université Paris-Est Créteil, INSERM U955 | Christophe Rodriguez, Slim Fourati, Vanessa Demontant, Guillaume Gricourt, Melissa N'Debi, Alexandre Soulier, Elisabeth Trawinski, Jean-Michel Pawlotsky |
| EPI_ISL_3475367 | Biogroup Bio Lam-LCD Saint-Denis | Department of Virology, Henri Mondor University Hospital, Assistance Publique Hôpitaux de Paris, Université Paris-Est Créteil, INSERM U955 | Christophe Rodriguez, Slim Fourati, Vanessa Demontant, Guillaume Gricourt, Melissa N'Debi, Alexandre Soulier, Elisabeth Trawinski, Jean-Michel Pawlotsky |
| EPI_ISL_3475368 | CHU DE DIJON | Department of Virology, Henri Mondor University Hospital, Assistance Publique Hôpitaux de Paris, Université Paris-Est Créteil, INSERM U955 | Christophe Rodriguez, Slim Fourati, Vanessa Demontant, Guillaume Gricourt, Melissa N'Debi, Alexandre Soulier, Elisabeth Trawinski, Jean-Michel Pawlotsky |
| EPI_ISL_3475369, EPI_ISL_3475370, EPI_ISL_3475371, EPI_ISL_3475372, EPI_ISL_3475373 | Biogroup Bio Lam-LCD Saint-Denis | Department of Virology, Henri Mondor University Hospital, Assistance Publique Hôpitaux de Paris, Université Paris-Est Créteil, INSERM U955 | Christophe Rodriguez, Slim Fourati, Vanessa Demontant, Guillaume Gricourt, Melissa N'Debi, Alexandre Soulier, Elisabeth Trawinski, Jean-Michel Pawlotsky |
| EPI_ISL_3475374 | CHU DE DIJON | Department of Virology, Henri Mondor University Hospital, Assistance Publique Hôpitaux de Paris, Université Paris-Est Créteil, INSERM U955 | Christophe Rodriguez, Slim Fourati, Vanessa Demontant, Guillaume Gricourt, Melissa N'Debi, Alexandre Soulier, Elisabeth Trawinski, Jean-Michel Pawlotsky |
| EPI_ISL_3475375 | Biogroup Bio Lam-LCD Saint-Denis | Department of Virology, Henri Mondor University Hospital, Assistance Publique Hôpitaux de Paris, Université Paris-Est Créteil, INSERM U955 | Christophe Rodriguez, Slim Fourati, Vanessa Demontant, Guillaume Gricourt, Melissa N'Debi, Alexandre Soulier, Elisabeth Trawinski, Jean-Michel Pawlotsky |
| EPI_ISL_3475376 | CHU DE DIJON | Department of Virology, Henri Mondor University Hospital, Assistance Publique Hôpitaux de Paris, Université Paris-Est Créteil, INSERM U955 | Christophe Rodriguez, Slim Fourati, Vanessa Demontant, Guillaume Gricourt, Melissa N'Debi, Alexandre Soulier, Elisabeth Trawinski, Jean-Michel Pawlotsky |
| EPI_ISL_3475377, EPI_ISL_3475378, EPI_ISL_3475379 | Biogroup Bio Lam-LCD Saint-Denis | Department of Virology, Henri Mondor University Hospital, Assistance Publique Hôpitaux de Paris, Université Paris-Est Créteil, INSERM U955 | Christophe Rodriguez, Slim Fourati, Vanessa Demontant, Guillaume Gricourt, Melissa N'Debi, Alexandre Soulier, Elisabeth Trawinski, Jean-Michel Pawlotsky |
| EPI_ISL_3475380 | BIO ARD'AISNE | Department of Virology, Henri Mondor University Hospital, Assistance Publique Hôpitaux de Paris, Université Paris-Est Créteil, INSERM U955 | Christophe Rodriguez, Slim Fourati, Vanessa Demontant, Guillaume Gricourt, Melissa N'Debi, Alexandre Soulier, Elisabeth Trawinski, Jean-Michel Pawlotsky |
| EPI_ISL_3475381, EPI_ISL_3475382 | CHU DE DIJON | Department of Virology, Henri Mondor University Hospital, Assistance Publique Hôpitaux de Paris, Université Paris-Est Créteil, INSERM U955 | Christophe Rodriguez, Slim Fourati, Vanessa Demontant, Guillaume Gricourt, Melissa N'Debi, Alexandre Soulier, Elisabeth Trawinski, Jean-Michel Pawlotsky |
| EPI_ISL_3475383 | CH. CLEMENCEAU | Department of Virology, Henri Mondor University Hospital, Assistance Publique Hôpitaux de Paris, Université Paris-Est Créteil, INSERM U955 | Christophe Rodriguez, Slim Fourati, Vanessa Demontant, Guillaume Gricourt, Melissa N'Debi, Alexandre Soulier, Elisabeth Trawinski, Jean-Michel Pawlotsky |
| EPI_ISL_3475384 | Biogroup Bio Lam-LCD Saint-Denis | Department of Virology, Henri Mondor University Hospital, Assistance Publique Hôpitaux de Paris, Université Paris-Est Créteil, INSERM U955 | Christophe Rodriguez, Slim Fourati, Vanessa Demontant, Guillaume Gricourt, Melissa N'Debi, Alexandre Soulier, Elisabeth Trawinski, Jean-Michel Pawlotsky |
| EPI_ISL_3475385 | CHU DE DIJON | Department of Virology, Henri Mondor University Hospital, Assistance Publique Hôpitaux de Paris, Université Paris-Est Créteil, INSERM U955 | Christophe Rodriguez, Slim Fourati, Vanessa Demontant, Guillaume Gricourt, Melissa N'Debi, Alexandre Soulier, Elisabeth Trawinski, Jean-Michel Pawlotsky |
| EPI_ISL_3475386 | Biogroup Bio Lam-LCD Saint-Denis | Department of Virology, Henri Mondor University Hospital, Assistance Publique Hôpitaux de Paris, Université Paris-Est Créteil, INSERM U955 | Christophe Rodriguez, Slim Fourati, Vanessa Demontant, Guillaume Gricourt, Melissa N'Debi, Alexandre Soulier, Elisabeth Trawinski, Jean-Michel Pawlotsky |
| EPI_ISL_3475387 | CH. CLEMENCEAU | Department of Virology, Henri Mondor University Hospital, Assistance Publique Hôpitaux de Paris, Université Paris-Est Créteil, INSERM U955 | Christophe Rodriguez, Slim Fourati, Vanessa Demontant, Guillaume Gricourt, Melissa N'Debi, Alexandre Soulier, Elisabeth Trawinski, Jean-Michel Pawlotsky |
| EPI_ISL_3475388 | Biogroup Bio Lam-LCD Saint-Denis | Department of Virology, Henri Mondor University Hospital, Assistance Publique Hôpitaux de Paris, Université Paris-Est Créteil, INSERM U955 | Christophe Rodriguez, Slim Fourati, Vanessa Demontant, Guillaume Gricourt, Melissa N'Debi, Alexandre Soulier, Elisabeth Trawinski, Jean-Michel Pawlotsky |
| EPI_ISL_3475389 | BIOMNIS EUROFINS IVRY | Department of Virology, Henri Mondor University Hospital, Assistance Publique Hôpitaux de Paris, Université Paris-Est Créteil, INSERM U955 | Christophe Rodriguez, Slim Fourati, Vanessa Demontant, Guillaume Gricourt, Melissa N'Debi, Alexandre Soulier, Elisabeth Trawinski, Jean-Michel Pawlotsky |
| EPI_ISL_3475390, EPI_ISL_3475391 | Biogroup Bio Lam-LCD Saint-Denis | Department of Virology, Henri Mondor University Hospital, Assistance Publique Hôpitaux de Paris, Université Paris-Est Créteil, INSERM U955 | Christophe Rodriguez, Slim Fourati, Vanessa Demontant, Guillaume Gricourt, Melissa N'Debi, Alexandre Soulier, Elisabeth Trawinski, Jean-Michel Pawlotsky |
| EPI_ISL_3475392, EPI_ISL_3475393 | CHU DE DIJON | Department of Virology, Henri Mondor University Hospital, Assistance Publique Hôpitaux de Paris, Université Paris-Est Créteil, INSERM U955 | Christophe Rodriguez, Slim Fourati, Vanessa Demontant, Guillaume Gricourt, Melissa N'Debi, Alexandre Soulier, Elisabeth Trawinski, Jean-Michel Pawlotsky |
| EPI_ISL_3475394, | Biogroup Bio Lam-LCD Saint-Denis | Department of Virology, Henri Mondor University Hospital, | Christophe Rodriguez, Slim Fourati, Vanessa Demontant, Guillaume Gricourt, Melissa N'Debi, Alexandre Soulier, Elisabeth Trawinski, Jean-Michel Pawlotsky |

| | | | |
|---|---|---|---|
| EPI_ISL_3475395 | | Assistance Publique Hôpitaux de Paris, Université Paris-Est Créteil, INSERM U955 | |
| EPI_ISL_3475396, EPI_ISL_3475397 | CHU DE DIJON | Department of Virology, Henri Mondor University Hospital, Assistance Publique Hôpitaux de Paris, Université Paris-Est Créteil, INSERM U955 | Christophe Rodriguez, Slim Fourati, Vanessa Demontant, Guillaume Gricourt, Melissa N'Debi, Alexandre Soulier, Elisabeth Trawinski, Jean-Michel Pawlotsky |
| EPI_ISL_3475399 | BIO ARD'AISNE | Department of Virology, Henri Mondor University Hospital, Assistance Publique Hôpitaux de Paris, Université Paris-Est Créteil, INSERM U955 | Christophe Rodriguez, Slim Fourati, Vanessa Demontant, Guillaume Gricourt, Melissa N'Debi, Alexandre Soulier, Elisabeth Trawinski, Jean-Michel Pawlotsky |
| EPI_ISL_3475400, EPI_ISL_3475401 | CHU DE DIJON | Department of Virology, Henri Mondor University Hospital, Assistance Publique Hôpitaux de Paris, Université Paris-Est Créteil, INSERM U955 | Christophe Rodriguez, Slim Fourati, Vanessa Demontant, Guillaume Gricourt, Melissa N'Debi, Alexandre Soulier, Elisabeth Trawinski, Jean-Michel Pawlotsky |
| EPI_ISL_3475402 | Laboratoire BIOTOV | Department of Virology, Henri Mondor University Hospital, Assistance Publique Hôpitaux de Paris, Université Paris-Est Créteil, INSERM U955 | Christophe Rodriguez, Slim Fourati, Vanessa Demontant, Guillaume Gricourt, Melissa N'Debi, Alexandre Soulier, Elisabeth Trawinski, Jean-Michel Pawlotsky |
| EPI_ISL_3475403 | CHU DE DIJON | Department of Virology, Henri Mondor University Hospital, Assistance Publique Hôpitaux de Paris, Université Paris-Est Créteil, INSERM U955 | Christophe Rodriguez, Slim Fourati, Vanessa Demontant, Guillaume Gricourt, Melissa N'Debi, Alexandre Soulier, Elisabeth Trawinski, Jean-Michel Pawlotsky |
| EPI_ISL_3475404 | Biogroup Bio Lam-LCD Saint-Denis | Department of Virology, Henri Mondor University Hospital, Assistance Publique Hôpitaux de Paris, Université Paris-Est Créteil, INSERM U955 | Christophe Rodriguez, Slim Fourati, Vanessa Demontant, Guillaume Gricourt, Melissa N'Debi, Alexandre Soulier, Elisabeth Trawinski, Jean-Michel Pawlotsky |
| EPI_ISL_3475405 | CHU DE DIJON | Department of Virology, Henri Mondor University Hospital, Assistance Publique Hôpitaux de Paris, Université Paris-Est Créteil, INSERM U955 | Christophe Rodriguez, Slim Fourati, Vanessa Demontant, Guillaume Gricourt, Melissa N'Debi, Alexandre Soulier, Elisabeth Trawinski, Jean-Michel Pawlotsky |
| EPI_ISL_3475406 | CH. CLEMENCEAU | Department of Virology, Henri Mondor University Hospital, Assistance Publique Hôpitaux de Paris, Université Paris-Est Créteil, INSERM U955 | Christophe Rodriguez, Slim Fourati, Vanessa Demontant, Guillaume Gricourt, Melissa N'Debi, Alexandre Soulier, Elisabeth Trawinski, Jean-Michel Pawlotsky |
| EPI_ISL_3475407 | Biogroup Bio Lam-LCD Saint-Denis | Department of Virology, Henri Mondor University Hospital, Assistance Publique Hôpitaux de Paris, Université Paris-Est Créteil, INSERM U955 | Christophe Rodriguez, Slim Fourati, Vanessa Demontant, Guillaume Gricourt, Melissa N'Debi, Alexandre Soulier, Elisabeth Trawinski, Jean-Michel Pawlotsky |
| EPI_ISL_3475408 | CH. CLEMENCEAU | Department of Virology, Henri Mondor University Hospital, Assistance Publique Hôpitaux de Paris, Université Paris-Est Créteil, INSERM U955 | Christophe Rodriguez, Slim Fourati, Vanessa Demontant, Guillaume Gricourt, Melissa N'Debi, Alexandre Soulier, Elisabeth Trawinski, Jean-Michel Pawlotsky |
| EPI_ISL_3475409, EPI_ISL_3475410, EPI_ISL_3475411 | CHU DE DIJON | Department of Virology, Henri Mondor University Hospital, Assistance Publique Hôpitaux de Paris, Université Paris-Est Créteil, INSERM U955 | Christophe Rodriguez, Slim Fourati, Vanessa Demontant, Guillaume Gricourt, Melissa N'Debi, Alexandre Soulier, Elisabeth Trawinski, Jean-Michel Pawlotsky |
| EPI_ISL_3475412 | CH. CLEMENCEAU | Department of Virology, Henri Mondor University Hospital, Assistance Publique Hôpitaux de Paris, Université Paris-Est Créteil, INSERM U955 | Christophe Rodriguez, Slim Fourati, Vanessa Demontant, Guillaume Gricourt, Melissa N'Debi, Alexandre Soulier, Elisabeth Trawinski, Jean-Michel Pawlotsky |
| EPI_ISL_3475413, EPI_ISL_3475414 | CHU DE DIJON | Department of Virology, Henri Mondor University Hospital, Assistance Publique Hôpitaux de Paris, Université Paris-Est Créteil, INSERM U955 | Christophe Rodriguez, Slim Fourati, Vanessa Demontant, Guillaume Gricourt, Melissa N'Debi, Alexandre Soulier, Elisabeth Trawinski, Jean-Michel Pawlotsky |
| EPI_ISL_3475415 | Biogroup Bio Lam-LCD Saint-Denis | Department of Virology, Henri Mondor University Hospital, Assistance Publique Hôpitaux de Paris, Université Paris-Est Créteil, INSERM U955 | Christophe Rodriguez, Slim Fourati, Vanessa Demontant, Guillaume Gricourt, Melissa N'Debi, Alexandre Soulier, Elisabeth Trawinski, Jean-Michel Pawlotsky |
| EPI_ISL_3475416 | CHU DE DIJON | Department of Virology, Henri Mondor University Hospital, Assistance Publique Hôpitaux de Paris, Université Paris-Est Créteil, INSERM U955 | Christophe Rodriguez, Slim Fourati, Vanessa Demontant, Guillaume Gricourt, Melissa N'Debi, Alexandre Soulier, Elisabeth Trawinski, Jean-Michel Pawlotsky |
| EPI_ISL_3475417 | Biogroup Bio Lam-LCD Saint-Denis | Department of Virology, Henri Mondor University Hospital, Assistance Publique Hôpitaux de Paris, Université Paris-Est Créteil, INSERM U955 | Christophe Rodriguez, Slim Fourati, Vanessa Demontant, Guillaume Gricourt, Melissa N'Debi, Alexandre Soulier, Elisabeth Trawinski, Jean-Michel Pawlotsky |
| EPI_ISL_3475418, EPI_ISL_3475419, EPI_ISL_3475420, EPI_ISL_3475421 | CHU DE DIJON | Department of Virology, Henri Mondor University Hospital, Assistance Publique Hôpitaux de Paris, Université Paris-Est Créteil, INSERM U955 | Christophe Rodriguez, Slim Fourati, Vanessa Demontant, Guillaume Gricourt, Melissa N'Debi, Alexandre Soulier, Elisabeth Trawinski, Jean-Michel Pawlotsky |
| EPI_ISL_3475422 | CH. CLEMENCEAU | Department of Virology, Henri Mondor University Hospital, Assistance Publique Hôpitaux de Paris, Université Paris-Est Créteil, INSERM U955 | Christophe Rodriguez, Slim Fourati, Vanessa Demontant, Guillaume Gricourt, Melissa N'Debi, Alexandre Soulier, Elisabeth Trawinski, Jean-Michel Pawlotsky |
| EPI_ISL_3475424 | CHU DE DIJON | Department of Virology, Henri Mondor University Hospital, Assistance Publique Hôpitaux de Paris, Université Paris-Est Créteil, INSERM U955 | Christophe Rodriguez, Slim Fourati, Vanessa Demontant, Guillaume Gricourt, Melissa N'Debi, Alexandre Soulier, Elisabeth Trawinski, Jean-Michel Pawlotsky |
| EPI_ISL_3475425 | CH. CLEMENCEAU | Department of Virology, Henri Mondor University Hospital, Assistance Publique Hôpitaux de Paris, Université Paris-Est Créteil, INSERM U955 | Christophe Rodriguez, Slim Fourati, Vanessa Demontant, Guillaume Gricourt, Melissa N'Debi, Alexandre Soulier, Elisabeth Trawinski, Jean-Michel Pawlotsky |
| EPI_ISL_3475426 | Biogroup Bio Lam-LCD Saint-Denis | Department of Virology, Henri Mondor University Hospital, Assistance Publique Hôpitaux de Paris, Université Paris-Est Créteil, INSERM U955 | Christophe Rodriguez, Slim Fourati, Vanessa Demontant, Guillaume Gricourt, Melissa N'Debi, Alexandre Soulier, Elisabeth Trawinski, Jean-Michel Pawlotsky |
| EPI_ISL_3475427, EPI_ISL_3475428, EPI_ISL_3475429 | CHU DE DIJON | Department of Virology, Henri Mondor University Hospital, Assistance Publique Hôpitaux de Paris, Université Paris-Est Créteil, INSERM U955 | Christophe Rodriguez, Slim Fourati, Vanessa Demontant, Guillaume Gricourt, Melissa N'Debi, Alexandre Soulier, Elisabeth Trawinski, Jean-Michel Pawlotsky |
| EPI_ISL_3475430 | Biogroup Bio Lam-LCD Saint-Denis | Department of Virology, Henri Mondor University Hospital, Assistance Publique Hôpitaux de Paris, Université Paris-Est Créteil, INSERM U955 | Christophe Rodriguez, Slim Fourati, Vanessa Demontant, Guillaume Gricourt, Melissa N'Debi, Alexandre Soulier, Elisabeth Trawinski, Jean-Michel Pawlotsky |
| EPI_ISL_3475431, EPI_ISL_3475433 | CHU DE DIJON | Department of Virology, Henri Mondor University Hospital, Assistance Publique Hôpitaux de Paris, Université Paris-Est Créteil, INSERM U955 | Christophe Rodriguez, Slim Fourati, Vanessa Demontant, Guillaume Gricourt, Melissa N'Debi, Alexandre Soulier, Elisabeth Trawinski, Jean-Michel Pawlotsky |
| EPI_ISL_3475434, EPI_ISL_3475435 | Biogroup Bio Lam-LCD Saint-Denis | Department of Virology, Henri Mondor University Hospital, Assistance Publique Hôpitaux de Paris, Université Paris-Est Créteil, INSERM U955 | Christophe Rodriguez, Slim Fourati, Vanessa Demontant, Guillaume Gricourt, Melissa N'Debi, Alexandre Soulier, Elisabeth Trawinski, Jean-Michel Pawlotsky |
| EPI_ISL_3475436, EPI_ISL_3475437 | CHU DE DIJON | Department of Virology, Henri Mondor University Hospital, Assistance Publique Hôpitaux de Paris, Université Paris-Est Créteil, INSERM U955 | Christophe Rodriguez, Slim Fourati, Vanessa Demontant, Guillaume Gricourt, Melissa N'Debi, Alexandre Soulier, Elisabeth Trawinski, Jean-Michel Pawlotsky |
| EPI_ISL_3475438 | BIOMNIS EUROFINS IVRY | Department of Virology, Henri Mondor University Hospital, Assistance Publique Hôpitaux de Paris, Université Paris-Est Créteil, INSERM U955 | Christophe Rodriguez, Slim Fourati, Vanessa Demontant, Guillaume Gricourt, Melissa N'Debi, Alexandre Soulier, Elisabeth Trawinski, Jean-Michel Pawlotsky |
| EPI_ISL_3475439 | CHU DE DIJON | Department of Virology, Henri Mondor University Hospital, Assistance Publique Hôpitaux de Paris, Université Paris-Est Créteil, INSERM U955 | Christophe Rodriguez, Slim Fourati, Vanessa Demontant, Guillaume Gricourt, Melissa N'Debi, Alexandre Soulier, Elisabeth Trawinski, Jean-Michel Pawlotsky |
| EPI_ISL_3475440, EPI_ISL_3475441, EPI_ISL_3475442, EPI_ISL_3475443, EPI_ISL_3475444, EPI_ISL_3475445, EPI_ISL_3475446, EPI_ISL_3475447, EPI_ISL_3475448, EPI_ISL_3475449, EPI_ISL_3475450, EPI_ISL_3475451, EPI_ISL_3475452, EPI_ISL_3475453, EPI_ISL_3475454, EPI_ISL_3475456, EPI_ISL_3475458, EPI_ISL_3475459, EPI_ISL_3475461, EPI_ISL_3475462, EPI_ISL_3475463, EPI_ISL_3475464, EPI_ISL_3475465, EPI_ISL_3475466, EPI_ISL_3475467, EPI_ISL_3475468, EPI_ISL_3475469, EPI_ISL_3475470, EPI_ISL_3475471, EPI_ISL_3475472, EPI_ISL_3475473, EPI_ISL_3475474, EPI_ISL_3475475, EPI_ISL_3475476, EPI_ISL_3475477, EPI_ISL_3475478, EPI_ISL_3475479, EPI_ISL_3475480, EPI_ISL_3475481, EPI_ISL_3475482, EPI_ISL_3475483, EPI_ISL_3475485, EPI_ISL_3475486, EPI_ISL_3475490, EPI_ISL_3475491, EPI_ISL_3475492, EPI_ISL_3475493, EPI_ISL_3475494, EPI_ISL_3475495, EPI_ISL_3475496, EPI_ISL_3475498, EPI_ISL_3475499, EPI_ISL_3475500, EPI_ISL_3475501, EPI_ISL_3475502, EPI_ISL_3475503, EPI_ISL_3475504, EPI_ISL_3475505, EPI_ISL_3475506, EPI_ISL_3475507, EPI_ISL_3475508, EPI_ISL_3475509, EPI_ISL_3475510, EPI_ISL_3475512, EPI_ISL_3475513, EPI_ISL_3475514, EPI_ISL_3475515, EPI_ISL_3475516, EPI_ISL_3475517, EPI_ISL_3475518, EPI_ISL_3475519, EPI_ISL_3475520, EPI_ISL_3475521, EPI_ISL_3475522, EPI_ISL_3475523, EPI_ISL_3475524, EPI_ISL_3475525, EPI_ISL_3475526, EPI_ISL_3475527, EPI_ISL_3475528, EPI_ISL_3475529, EPI_ISL_3475530, EPI_ISL_3475531, EPI_ISL_3475532, EPI_ISL_3475533, EPI_ISL_3475534, EPI_ISL_3475535, EPI_ISL_3475537, EPI_ISL_3475538, EPI_ISL_3475539, EPI_ISL_3475540, EPI_ISL_3475543, EPI_ISL_3475544, EPI_ISL_3475545, EPI_ISL_3475546, EPI_ISL_3475547, EPI_ISL_3475548, EPI_ISL_3475549, EPI_ISL_3475550, EPI_ISL_3475551, EPI_ISL_3475552, EPI_ISL_3475553, EPI_ISL_3475554, EPI_ISL_3475555, EPI_ISL_3475556, EPI_ISL_3475557, EPI_ISL_3475558, EPI_ISL_3475559, EPI_ISL_3475560, EPI_ISL_3475561, EPI_ISL_3475562, EPI_ISL_3475563, EPI_ISL_3475564, EPI_ISL_3475565, EPI_ISL_3475566 | | | |
| see above | New Mexico Department of Health Scientific Laboratory | New Mexico Department of Health Scientific Laboratory | Ellie Johnson, Anastacia Griego-Fisher, D'eldra Malone, Jennifer Benoit |
| EPI_ISL_3475567 | CH. CLEMENCEAU | Department of Virology, Henri Mondor University Hospital, Assistance Publique Hôpitaux de Paris, Université Paris-Est Créteil, INSERM U955 | Christophe Rodriguez, Slim Fourati, Vanessa Demontant, Guillaume Gricourt, Melissa N'Debi, Alexandre Soulier, Elisabeth Trawinski, Jean-Michel Pawlotsky |

| | | | |
|---|---|---|---|
| EPI_ISL_3475568 | SELAS MEDILYS | Department of Virology, Henri Mondor University Hospital, Assistance Publique Hôpitaux de Paris, Université Paris-Est Créteil, INSERM U955 | Christophe Rodriguez, Slim Fourati, Vanessa Demontant, Guillaume Gricourt, Melissa N'Debi, Alexandre Soulier, Elisabeth Trawinski, Jean-Michel Pawlotsky |
| EPI_ISL_3475569, EPI_ISL_3475570, EPI_ISL_3475571, EPI_ISL_3475572, EPI_ISL_3475573 | Biogroup Bio Lam-LCD Saint-Denis | Department of Virology, Henri Mondor University Hospital, Assistance Publique Hôpitaux de Paris, Université Paris-Est Créteil, INSERM U955 | Christophe Rodriguez, Slim Fourati, Vanessa Demontant, Guillaume Gricourt, Melissa N'Debi, Alexandre Soulier, Elisabeth Trawinski, Jean-Michel Pawlotsky |
| EPI_ISL_3475574 | CH. CLEMENCEAU | Department of Virology, Henri Mondor University Hospital, Assistance Publique Hôpitaux de Paris, Université Paris-Est Créteil, INSERM U955 | Christophe Rodriguez, Slim Fourati, Vanessa Demontant, Guillaume Gricourt, Melissa N'Debi, Alexandre Soulier, Elisabeth Trawinski, Jean-Michel Pawlotsky |
| EPI_ISL_3475575 | Biogroup Bio Lam-LCD Saint-Denis | Department of Virology, Henri Mondor University Hospital, Assistance Publique Hôpitaux de Paris, Université Paris-Est Créteil, INSERM U955 | Christophe Rodriguez, Slim Fourati, Vanessa Demontant, Guillaume Gricourt, Melissa N'Debi, Alexandre Soulier, Elisabeth Trawinski, Jean-Michel Pawlotsky |
| EPI_ISL_3475576 | Laboratoire Coste | Department of Virology, Henri Mondor University Hospital, Assistance Publique Hôpitaux de Paris, Université Paris-Est Créteil, INSERM U955 | Christophe Rodriguez, Slim Fourati, Vanessa Demontant, Guillaume Gricourt, Melissa N'Debi, Alexandre Soulier, Elisabeth Trawinski, Jean-Michel Pawlotsky |
| EPI_ISL_3475577 | Biogroup Bio Lam-LCD Saint-Denis | Department of Virology, Henri Mondor University Hospital, Assistance Publique Hôpitaux de Paris, Université Paris-Est Créteil, INSERM U955 | Christophe Rodriguez, Slim Fourati, Vanessa Demontant, Guillaume Gricourt, Melissa N'Debi, Alexandre Soulier, Elisabeth Trawinski, Jean-Michel Pawlotsky |
| EPI_ISL_3475578 | BROUSSAIS | Department of Virology, Henri Mondor University Hospital, Assistance Publique Hôpitaux de Paris, Université Paris-Est Créteil, INSERM U955 | Christophe Rodriguez, Slim Fourati, Vanessa Demontant, Guillaume Gricourt, Melissa N'Debi, Alexandre Soulier, Elisabeth Trawinski, Jean-Michel Pawlotsky |
| EPI_ISL_3475579 | Laboratoire ATOUTBIO JDC | Department of Virology, Henri Mondor University Hospital, Assistance Publique Hôpitaux de Paris, Université Paris-Est Créteil, INSERM U955 | Christophe Rodriguez, Slim Fourati, Vanessa Demontant, Guillaume Gricourt, Melissa N'Debi, Alexandre Soulier, Elisabeth Trawinski, Jean-Michel Pawlotsky |
| EPI_ISL_3475580 | CHU DE DIJON | Department of Virology, Henri Mondor University Hospital, Assistance Publique Hôpitaux de Paris, Université Paris-Est Créteil, INSERM U955 | Christophe Rodriguez, Slim Fourati, Vanessa Demontant, Guillaume Gricourt, Melissa N'Debi, Alexandre Soulier, Elisabeth Trawinski, Jean-Michel Pawlotsky |
| EPI_ISL_3475581 | DYNALAB | Department of Virology, Henri Mondor University Hospital, Assistance Publique Hôpitaux de Paris, Université Paris-Est Créteil, INSERM U955 | Christophe Rodriguez, Slim Fourati, Vanessa Demontant, Guillaume Gricourt, Melissa N'Debi, Alexandre Soulier, Elisabeth Trawinski, Jean-Michel Pawlotsky |
| EPI_ISL_3475582 | CHU DE DIJON | Department of Virology, Henri Mondor University Hospital, Assistance Publique Hôpitaux de Paris, Université Paris-Est Créteil, INSERM U955 | Christophe Rodriguez, Slim Fourati, Vanessa Demontant, Guillaume Gricourt, Melissa N'Debi, Alexandre Soulier, Elisabeth Trawinski, Jean-Michel Pawlotsky |
| EPI_ISL_3475583 | CERBALLIANCE SAINT MARTIN | Department of Virology, Henri Mondor University Hospital, Assistance Publique Hôpitaux de Paris, Université Paris-Est Créteil, INSERM U955 | Christophe Rodriguez, Slim Fourati, Vanessa Demontant, Guillaume Gricourt, Melissa N'Debi, Alexandre Soulier, Elisabeth Trawinski, Jean-Michel Pawlotsky |
| EPI_ISL_3475584, EPI_ISL_3475585 | SELAS MEDILYS | Department of Virology, Henri Mondor University Hospital, Assistance Publique Hôpitaux de Paris, Université Paris-Est Créteil, INSERM U955 | Christophe Rodriguez, Slim Fourati, Vanessa Demontant, Guillaume Gricourt, Melissa N'Debi, Alexandre Soulier, Elisabeth Trawinski, Jean-Michel Pawlotsky |
| EPI_ISL_3475586 | Laboratoire Coste | Department of Virology, Henri Mondor University Hospital, Assistance Publique Hôpitaux de Paris, Université Paris-Est Créteil, INSERM U955 | Christophe Rodriguez, Slim Fourati, Vanessa Demontant, Guillaume Gricourt, Melissa N'Debi, Alexandre Soulier, Elisabeth Trawinski, Jean-Michel Pawlotsky |
| EPI_ISL_3475587 | CH. CLEMENCEAU | Department of Virology, Henri Mondor University Hospital, Assistance Publique Hôpitaux de Paris, Université Paris-Est Créteil, INSERM U955 | Christophe Rodriguez, Slim Fourati, Vanessa Demontant, Guillaume Gricourt, Melissa N'Debi, Alexandre Soulier, Elisabeth Trawinski, Jean-Michel Pawlotsky |
| EPI_ISL_3475588, EPI_ISL_3475589, EPI_ISL_3475590, EPI_ISL_3475591 | Biogroup Bio Lam-LCD Saint-Denis | Department of Virology, Henri Mondor University Hospital, Assistance Publique Hôpitaux de Paris, Université Paris-Est Créteil, INSERM U955 | Christophe Rodriguez, Slim Fourati, Vanessa Demontant, Guillaume Gricourt, Melissa N'Debi, Alexandre Soulier, Elisabeth Trawinski, Jean-Michel Pawlotsky |
| EPI_ISL_3475592 | Hôpital Saint-Louis | Department of Virology, Henri Mondor University Hospital, Assistance Publique Hôpitaux de Paris, Université Paris-Est Créteil, INSERM U955 | Christophe Rodriguez, Slim Fourati, Vanessa Demontant, Guillaume Gricourt, Melissa N'Debi, Alexandre Soulier, Elisabeth Trawinski, Jean-Michel Pawlotsky |
| EPI_ISL_3475593 | LBM du Val d'Orne | Department of Virology, Henri Mondor University Hospital, Assistance Publique Hôpitaux de Paris, Université Paris-Est Créteil, INSERM U955 | Christophe Rodriguez, Slim Fourati, Vanessa Demontant, Guillaume Gricourt, Melissa N'Debi, Alexandre Soulier, Elisabeth Trawinski, Jean-Michel Pawlotsky |
| EPI_ISL_3475595 | Laboratoire ATOUTBIO JDC | Department of Virology, Henri Mondor University Hospital, Assistance Publique Hôpitaux de Paris, Université Paris-Est Créteil, INSERM U955 | Christophe Rodriguez, Slim Fourati, Vanessa Demontant, Guillaume Gricourt, Melissa N'Debi, Alexandre Soulier, Elisabeth Trawinski, Jean-Michel Pawlotsky |
| EPI_ISL_3475596 | Biogroup Bio Lam-LCD Saint-Denis | Department of Virology, Henri Mondor University Hospital, Assistance Publique Hôpitaux de Paris, Université Paris-Est Créteil, INSERM U955 | Christophe Rodriguez, Slim Fourati, Vanessa Demontant, Guillaume Gricourt, Melissa N'Debi, Alexandre Soulier, Elisabeth Trawinski, Jean-Michel Pawlotsky |
| EPI_ISL_3475597 | Laboratoire Storck P.(Biolia) | Department of Virology, Henri Mondor University Hospital, Assistance Publique Hôpitaux de Paris, Université Paris-Est Créteil, INSERM U955 | Christophe Rodriguez, Slim Fourati, Vanessa Demontant, Guillaume Gricourt, Melissa N'Debi, Alexandre Soulier, Elisabeth Trawinski, Jean-Michel Pawlotsky |
| EPI_ISL_3475599 | DYNABIO UNILABS | Department of Virology, Henri Mondor University Hospital, Assistance Publique Hôpitaux de Paris, Université Paris-Est Créteil, INSERM U955 | Christophe Rodriguez, Slim Fourati, Vanessa Demontant, Guillaume Gricourt, Melissa N'Debi, Alexandre Soulier, Elisabeth Trawinski, Jean-Michel Pawlotsky |
| EPI_ISL_3475600 | Biogroup Bio Lam-LCD Saint-Denis | Department of Virology, Henri Mondor University Hospital, Assistance Publique Hôpitaux de Paris, Université Paris-Est Créteil, INSERM U955 | Christophe Rodriguez, Slim Fourati, Vanessa Demontant, Guillaume Gricourt, Melissa N'Debi, Alexandre Soulier, Elisabeth Trawinski, Jean-Michel Pawlotsky |
| EPI_ISL_3475601 | SELAS MEDILYS | Department of Virology, Henri Mondor University Hospital, Assistance Publique Hôpitaux de Paris, Université Paris-Est Créteil, INSERM U955 | Christophe Rodriguez, Slim Fourati, Vanessa Demontant, Guillaume Gricourt, Melissa N'Debi, Alexandre Soulier, Elisabeth Trawinski, Jean-Michel Pawlotsky |
| EPI_ISL_3475602 | Hôpital Avicenne | Department of Virology, Henri Mondor University Hospital, Assistance Publique Hôpitaux de Paris, Université Paris-Est Créteil, INSERM U955 | Christophe Rodriguez, Slim Fourati, Vanessa Demontant, Guillaume Gricourt, Melissa N'Debi, Alexandre Soulier, Elisabeth Trawinski, Jean-Michel Pawlotsky |
| EPI_ISL_3475603 | SELAS MEDILYS | Department of Virology, Henri Mondor University Hospital, Assistance Publique Hôpitaux de Paris, Université Paris-Est Créteil, INSERM U955 | Christophe Rodriguez, Slim Fourati, Vanessa Demontant, Guillaume Gricourt, Melissa N'Debi, Alexandre Soulier, Elisabeth Trawinski, Jean-Michel Pawlotsky |
| EPI_ISL_3475604 | Biogroup Bio Lam-LCD Saint-Denis | Department of Virology, Henri Mondor University Hospital, Assistance Publique Hôpitaux de Paris, Université Paris-Est Créteil, INSERM U955 | Christophe Rodriguez, Slim Fourati, Vanessa Demontant, Guillaume Gricourt, Melissa N'Debi, Alexandre Soulier, Elisabeth Trawinski, Jean-Michel Pawlotsky |
| EPI_ISL_3475605 | Laboratoire Coste | Department of Virology, Henri Mondor University Hospital, Assistance Publique Hôpitaux de Paris, Université Paris-Est Créteil, INSERM U955 | Christophe Rodriguez, Slim Fourati, Vanessa Demontant, Guillaume Gricourt, Melissa N'Debi, Alexandre Soulier, Elisabeth Trawinski, Jean-Michel Pawlotsky |
| EPI_ISL_3475606 | Laboratoire ATOUTBIO JDC | Department of Virology, Henri Mondor University Hospital, Assistance Publique Hôpitaux de Paris, Université Paris-Est Créteil, INSERM U955 | Christophe Rodriguez, Slim Fourati, Vanessa Demontant, Guillaume Gricourt, Melissa N'Debi, Alexandre Soulier, Elisabeth Trawinski, Jean-Michel Pawlotsky |
| EPI_ISL_3475607 | SELAS MEDILYS | Department of Virology, Henri Mondor University Hospital, Assistance Publique Hôpitaux de Paris, Université Paris-Est Créteil, INSERM U955 | Christophe Rodriguez, Slim Fourati, Vanessa Demontant, Guillaume Gricourt, Melissa N'Debi, Alexandre Soulier, Elisabeth Trawinski, Jean-Michel Pawlotsky |
| EPI_ISL_3475608 | CERBALLIANCE SAINT MARTIN | Department of Virology, Henri Mondor University Hospital, Assistance Publique Hôpitaux de Paris, Université Paris-Est Créteil, INSERM U955 | Christophe Rodriguez, Slim Fourati, Vanessa Demontant, Guillaume Gricourt, Melissa N'Debi, Alexandre Soulier, Elisabeth Trawinski, Jean-Michel Pawlotsky |

| | | | |
|---|---|---|---|
| EPI_ISL_3475609 | Biogroup Bio Lam-LCD Saint-Denis | Department of Virology, Henri Mondor University Hospital, Assistance Publique Hôpitaux de Paris, Université Paris-Est Créteil, INSERM U955 | Christophe Rodriguez, Slim Fourati, Vanessa Demontant, Guillaume Gricourt, Melissa N'Debi, Alexandre Soulier, Elisabeth Trawinski, Jean-Michel Pawlotsky |
| EPI_ISL_3475610 | CH. CLEMENCEAU | Department of Virology, Henri Mondor University Hospital, Assistance Publique Hôpitaux de Paris, Université Paris-Est Créteil, INSERM U955 | Christophe Rodriguez, Slim Fourati, Vanessa Demontant, Guillaume Gricourt, Melissa N'Debi, Alexandre Soulier, Elisabeth Trawinski, Jean-Michel Pawlotsky |
| EPI_ISL_3475611 | CERBALLIANCE SAINT MARTIN | Department of Virology, Henri Mondor University Hospital, Assistance Publique Hôpitaux de Paris, Université Paris-Est Créteil, INSERM U955 | Christophe Rodriguez, Slim Fourati, Vanessa Demontant, Guillaume Gricourt, Melissa N'Debi, Alexandre Soulier, Elisabeth Trawinski, Jean-Michel Pawlotsky |
| EPI_ISL_3475612 | Biogroup Bio Lam-LCD Saint-Denis | Department of Virology, Henri Mondor University Hospital, Assistance Publique Hôpitaux de Paris, Université Paris-Est Créteil, INSERM U955 | Christophe Rodriguez, Slim Fourati, Vanessa Demontant, Guillaume Gricourt, Melissa N'Debi, Alexandre Soulier, Elisabeth Trawinski, Jean-Michel Pawlotsky |
| EPI_ISL_3475613 | CHI VILLENEUVE ST GEORGES | Department of Virology, Henri Mondor University Hospital, Assistance Publique Hôpitaux de Paris, Université Paris-Est Créteil, INSERM U955 | Christophe Rodriguez, Slim Fourati, Vanessa Demontant, Guillaume Gricourt, Melissa N'Debi, Alexandre Soulier, Elisabeth Trawinski, Jean-Michel Pawlotsky |
| EPI_ISL_3475614 | Laboratoire ATOUTBIO JDC | Department of Virology, Henri Mondor University Hospital, Assistance Publique Hôpitaux de Paris, Université Paris-Est Créteil, INSERM U955 | Christophe Rodriguez, Slim Fourati, Vanessa Demontant, Guillaume Gricourt, Melissa N'Debi, Alexandre Soulier, Elisabeth Trawinski, Jean-Michel Pawlotsky |
| EPI_ISL_3475615 | Biogroup Bio Lam-LCD Saint-Denis | Department of Virology, Henri Mondor University Hospital, Assistance Publique Hôpitaux de Paris, Université Paris-Est Créteil, INSERM U955 | Christophe Rodriguez, Slim Fourati, Vanessa Demontant, Guillaume Gricourt, Melissa N'Debi, Alexandre Soulier, Elisabeth Trawinski, Jean-Michel Pawlotsky |
| EPI_ISL_3475616 | CH. E. MULLER | Department of Virology, Henri Mondor University Hospital, Assistance Publique Hôpitaux de Paris, Université Paris-Est Créteil, INSERM U955 | Christophe Rodriguez, Slim Fourati, Vanessa Demontant, Guillaume Gricourt, Melissa N'Debi, Alexandre Soulier, Elisabeth Trawinski, Jean-Michel Pawlotsky |
| EPI_ISL_3475617 | SELAS MEDILYS | Department of Virology, Henri Mondor University Hospital, Assistance Publique Hôpitaux de Paris, Université Paris-Est Créteil, INSERM U955 | Christophe Rodriguez, Slim Fourati, Vanessa Demontant, Guillaume Gricourt, Melissa N'Debi, Alexandre Soulier, Elisabeth Trawinski, Jean-Michel Pawlotsky |
| EPI_ISL_3475618 | Biogroup Bio Lam-LCD Saint-Denis | Department of Virology, Henri Mondor University Hospital, Assistance Publique Hôpitaux de Paris, Université Paris-Est Créteil, INSERM U955 | Christophe Rodriguez, Slim Fourati, Vanessa Demontant, Guillaume Gricourt, Melissa N'Debi, Alexandre Soulier, Elisabeth Trawinski, Jean-Michel Pawlotsky |
| EPI_ISL_3475619 | CH. CLEMENCEAU | Department of Virology, Henri Mondor University Hospital, Assistance Publique Hôpitaux de Paris, Université Paris-Est Créteil, INSERM U955 | Christophe Rodriguez, Slim Fourati, Vanessa Demontant, Guillaume Gricourt, Melissa N'Debi, Alexandre Soulier, Elisabeth Trawinski, Jean-Michel Pawlotsky |
| EPI_ISL_3475620 | Biogroup Bio Lam-LCD Saint-Denis | Department of Virology, Henri Mondor University Hospital, Assistance Publique Hôpitaux de Paris, Université Paris-Est Créteil, INSERM U955 | Christophe Rodriguez, Slim Fourati, Vanessa Demontant, Guillaume Gricourt, Melissa N'Debi, Alexandre Soulier, Elisabeth Trawinski, Jean-Michel Pawlotsky |
| EPI_ISL_3475621, EPI_ISL_3475622 | CHU DE DIJON | Department of Virology, Henri Mondor University Hospital, Assistance Publique Hôpitaux de Paris, Université Paris-Est Créteil, INSERM U955 | Christophe Rodriguez, Slim Fourati, Vanessa Demontant, Guillaume Gricourt, Melissa N'Debi, Alexandre Soulier, Elisabeth Trawinski, Jean-Michel Pawlotsky |
| EPI_ISL_3475623 | CH Mémorial F.E.U Laboratoire | Department of Virology, Henri Mondor University Hospital, Assistance Publique Hôpitaux de Paris, Université Paris-Est Créteil, INSERM U955 | Christophe Rodriguez, Slim Fourati, Vanessa Demontant, Guillaume Gricourt, Melissa N'Debi, Alexandre Soulier, Elisabeth Trawinski, Jean-Michel Pawlotsky |
| EPI_ISL_3475624 | Biogroup Bio Lam-LCD Saint-Denis | Department of Virology, Henri Mondor University Hospital, Assistance Publique Hôpitaux de Paris, Université Paris-Est Créteil, INSERM U955 | Christophe Rodriguez, Slim Fourati, Vanessa Demontant, Guillaume Gricourt, Melissa N'Debi, Alexandre Soulier, Elisabeth Trawinski, Jean-Michel Pawlotsky |
| EPI_ISL_3475625 | CERBALLIANCE SAINT MARTIN | Department of Virology, Henri Mondor University Hospital, Assistance Publique Hôpitaux de Paris, Université Paris-Est Créteil, INSERM U955 | Christophe Rodriguez, Slim Fourati, Vanessa Demontant, Guillaume Gricourt, Melissa N'Debi, Alexandre Soulier, Elisabeth Trawinski, Jean-Michel Pawlotsky |
| EPI_ISL_3475626 | BROUSSAIS | Department of Virology, Henri Mondor University Hospital, Assistance Publique Hôpitaux de Paris, Université Paris-Est Créteil, INSERM U955 | Christophe Rodriguez, Slim Fourati, Vanessa Demontant, Guillaume Gricourt, Melissa N'Debi, Alexandre Soulier, Elisabeth Trawinski, Jean-Michel Pawlotsky |
| EPI_ISL_3475627 | CH. CLEMENCEAU | Department of Virology, Henri Mondor University Hospital, Assistance Publique Hôpitaux de Paris, Université Paris-Est Créteil, INSERM U955 | Christophe Rodriguez, Slim Fourati, Vanessa Demontant, Guillaume Gricourt, Melissa N'Debi, Alexandre Soulier, Elisabeth Trawinski, Jean-Michel Pawlotsky |
| EPI_ISL_3475628 | SELAS MEDILYS | Department of Virology, Henri Mondor University Hospital, Assistance Publique Hôpitaux de Paris, Université Paris-Est Créteil, INSERM U955 | Christophe Rodriguez, Slim Fourati, Vanessa Demontant, Guillaume Gricourt, Melissa N'Debi, Alexandre Soulier, Elisabeth Trawinski, Jean-Michel Pawlotsky |
| EPI_ISL_3475629 | CERBALLIANCE SAINT MARTIN | Department of Virology, Henri Mondor University Hospital, Assistance Publique Hôpitaux de Paris, Université Paris-Est Créteil, INSERM U955 | Christophe Rodriguez, Slim Fourati, Vanessa Demontant, Guillaume Gricourt, Melissa N'Debi, Alexandre Soulier, Elisabeth Trawinski, Jean-Michel Pawlotsky |
| EPI_ISL_3475630 | Biogroup Bio Lam-LCD Saint-Denis | Department of Virology, Henri Mondor University Hospital, Assistance Publique Hôpitaux de Paris, Université Paris-Est Créteil, INSERM U955 | Christophe Rodriguez, Slim Fourati, Vanessa Demontant, Guillaume Gricourt, Melissa N'Debi, Alexandre Soulier, Elisabeth Trawinski, Jean-Michel Pawlotsky |
| EPI_ISL_3475631 | DYNABIO UNILABS | Department of Virology, Henri Mondor University Hospital, Assistance Publique Hôpitaux de Paris, Université Paris-Est Créteil, INSERM U955 | Christophe Rodriguez, Slim Fourati, Vanessa Demontant, Guillaume Gricourt, Melissa N'Debi, Alexandre Soulier, Elisabeth Trawinski, Jean-Michel Pawlotsky |
| EPI_ISL_3475632 | CHU DE DIJON | Department of Virology, Henri Mondor University Hospital, Assistance Publique Hôpitaux de Paris, Université Paris-Est Créteil, INSERM U955 | Christophe Rodriguez, Slim Fourati, Vanessa Demontant, Guillaume Gricourt, Melissa N'Debi, Alexandre Soulier, Elisabeth Trawinski, Jean-Michel Pawlotsky |
| EPI_ISL_3475633 | Biogroup Bio Lam-LCD Saint-Denis | Department of Virology, Henri Mondor University Hospital, Assistance Publique Hôpitaux de Paris, Université Paris-Est Créteil, INSERM U955 | Christophe Rodriguez, Slim Fourati, Vanessa Demontant, Guillaume Gricourt, Melissa N'Debi, Alexandre Soulier, Elisabeth Trawinski, Jean-Michel Pawlotsky |
| EPI_ISL_3475634 | CHU DE DIJON | Department of Virology, Henri Mondor University Hospital, Assistance Publique Hôpitaux de Paris, Université Paris-Est Créteil, INSERM U955 | Christophe Rodriguez, Slim Fourati, Vanessa Demontant, Guillaume Gricourt, Melissa N'Debi, Alexandre Soulier, Elisabeth Trawinski, Jean-Michel Pawlotsky |
| EPI_ISL_3475635 | Biogroup Bio Lam-LCD Saint-Denis | Department of Virology, Henri Mondor University Hospital, Assistance Publique Hôpitaux de Paris, Université Paris-Est Créteil, INSERM U955 | Christophe Rodriguez, Slim Fourati, Vanessa Demontant, Guillaume Gricourt, Melissa N'Debi, Alexandre Soulier, Elisabeth Trawinski, Jean-Michel Pawlotsky |
| EPI_ISL_3475636 | CH Mémorial F.E.U Laboratoire | Department of Virology, Henri Mondor University Hospital, Assistance Publique Hôpitaux de Paris, Université Paris-Est Créteil, INSERM U955 | Christophe Rodriguez, Slim Fourati, Vanessa Demontant, Guillaume Gricourt, Melissa N'Debi, Alexandre Soulier, Elisabeth Trawinski, Jean-Michel Pawlotsky |
| EPI_ISL_3475637 | CH. CLEMENCEAU | Department of Virology, Henri Mondor University Hospital, Assistance Publique Hôpitaux de Paris, Université Paris-Est Créteil, INSERM U955 | Christophe Rodriguez, Slim Fourati, Vanessa Demontant, Guillaume Gricourt, Melissa N'Debi, Alexandre Soulier, Elisabeth Trawinski, Jean-Michel Pawlotsky |
| EPI_ISL_3475638 | SELAS MEDILYS | Department of Virology, Henri Mondor University Hospital, Assistance Publique Hôpitaux de Paris, Université Paris-Est Créteil, INSERM U955 | Christophe Rodriguez, Slim Fourati, Vanessa Demontant, Guillaume Gricourt, Melissa N'Debi, Alexandre Soulier, Elisabeth Trawinski, Jean-Michel Pawlotsky |
| EPI_ISL_3475639 | DYNALAB | Department of Virology, Henri Mondor University Hospital, Assistance Publique Hôpitaux de Paris, Université Paris-Est Créteil, INSERM U955 | Christophe Rodriguez, Slim Fourati, Vanessa Demontant, Guillaume Gricourt, Melissa N'Debi, Alexandre Soulier, Elisabeth Trawinski, Jean-Michel Pawlotsky |
| EPI_ISL_3475640 | BROUSSAIS | Department of Virology, Henri Mondor University Hospital, Assistance Publique Hôpitaux de Paris, Université Paris-Est Créteil, INSERM U955 | Christophe Rodriguez, Slim Fourati, Vanessa Demontant, Guillaume Gricourt, Melissa N'Debi, Alexandre Soulier, Elisabeth Trawinski, Jean-Michel Pawlotsky |
| EPI_ISL_3475641 | Hôpital Avicenne | Department of Virology, Henri Mondor University Hospital, Assistance Publique Hôpitaux de Paris, Université Paris-Est Créteil, INSERM U955 | Christophe Rodriguez, Slim Fourati, Vanessa Demontant, Guillaume Gricourt, Melissa N'Debi, Alexandre Soulier, Elisabeth Trawinski, Jean-Michel Pawlotsky |

| | | | |
|---|---|---|---|
| EPI_ISL_3475642 | BROUSSAIS | Department of Virology, Henri Mondor University Hospital, Assistance Publique Hôpitaux de Paris, Université Paris-Est Créteil, INSERM U955 | Christophe Rodriguez, Slim Fourati, Vanessa Demontant, Guillaume Gricourt, Melissa N'Debi, Alexandre Soulier, Elisabeth Trawinski, Jean-Michel Pawlotsky |
| EPI_ISL_3475643 | SELAS MEDILYS | Department of Virology, Henri Mondor University Hospital, Assistance Publique Hôpitaux de Paris, Université Paris-Est Créteil, INSERM U955 | Christophe Rodriguez, Slim Fourati, Vanessa Demontant, Guillaume Gricourt, Melissa N'Debi, Alexandre Soulier, Elisabeth Trawinski, Jean-Michel Pawlotsky |
| EPI_ISL_3475644 | Biogroup Bio Lam-LCD Saint-Denis | Department of Virology, Henri Mondor University Hospital, Assistance Publique Hôpitaux de Paris, Université Paris-Est Créteil, INSERM U955 | Christophe Rodriguez, Slim Fourati, Vanessa Demontant, Guillaume Gricourt, Melissa N'Debi, Alexandre Soulier, Elisabeth Trawinski, Jean-Michel Pawlotsky |
| EPI_ISL_3475645 | CHU DE DIJON | Department of Virology, Henri Mondor University Hospital, Assistance Publique Hôpitaux de Paris, Université Paris-Est Créteil, INSERM U955 | Christophe Rodriguez, Slim Fourati, Vanessa Demontant, Guillaume Gricourt, Melissa N'Debi, Alexandre Soulier, Elisabeth Trawinski, Jean-Michel Pawlotsky |
| EPI_ISL_3475646 | Hôpital Saint-Louis | Department of Virology, Henri Mondor University Hospital, Assistance Publique Hôpitaux de Paris, Université Paris-Est Créteil, INSERM U955 | Christophe Rodriguez, Slim Fourati, Vanessa Demontant, Guillaume Gricourt, Melissa N'Debi, Alexandre Soulier, Elisabeth Trawinski, Jean-Michel Pawlotsky |
| EPI_ISL_3475647 | CHU DE DIJON | Department of Virology, Henri Mondor University Hospital, Assistance Publique Hôpitaux de Paris, Université Paris-Est Créteil, INSERM U955 | Christophe Rodriguez, Slim Fourati, Vanessa Demontant, Guillaume Gricourt, Melissa N'Debi, Alexandre Soulier, Elisabeth Trawinski, Jean-Michel Pawlotsky |
| EPI_ISL_3475648 | Biogroup Bio Lam-LCD Saint-Denis | Department of Virology, Henri Mondor University Hospital, Assistance Publique Hôpitaux de Paris, Université Paris-Est Créteil, INSERM U955 | Christophe Rodriguez, Slim Fourati, Vanessa Demontant, Guillaume Gricourt, Melissa N'Debi, Alexandre Soulier, Elisabeth Trawinski, Jean-Michel Pawlotsky |
| EPI_ISL_3475649 | CERBALLIANCE SAINT MARTIN | Department of Virology, Henri Mondor University Hospital, Assistance Publique Hôpitaux de Paris, Université Paris-Est Créteil, INSERM U955 | Christophe Rodriguez, Slim Fourati, Vanessa Demontant, Guillaume Gricourt, Melissa N'Debi, Alexandre Soulier, Elisabeth Trawinski, Jean-Michel Pawlotsky |
| EPI_ISL_3475650, EPI_ISL_3475651 | Biogroup Bio Lam-LCD Saint-Denis | Department of Virology, Henri Mondor University Hospital, Assistance Publique Hôpitaux de Paris, Université Paris-Est Créteil, INSERM U955 | Christophe Rodriguez, Slim Fourati, Vanessa Demontant, Guillaume Gricourt, Melissa N'Debi, Alexandre Soulier, Elisabeth Trawinski, Jean-Michel Pawlotsky |
| EPI_ISL_3475652 | CHU DE DIJON | Department of Virology, Henri Mondor University Hospital, Assistance Publique Hôpitaux de Paris, Université Paris-Est Créteil, INSERM U955 | Christophe Rodriguez, Slim Fourati, Vanessa Demontant, Guillaume Gricourt, Melissa N'Debi, Alexandre Soulier, Elisabeth Trawinski, Jean-Michel Pawlotsky |
| EPI_ISL_3475653 | Biogroup Bio Lam-LCD Saint-Denis | Department of Virology, Henri Mondor University Hospital, Assistance Publique Hôpitaux de Paris, Université Paris-Est Créteil, INSERM U955 | Christophe Rodriguez, Slim Fourati, Vanessa Demontant, Guillaume Gricourt, Melissa N'Debi, Alexandre Soulier, Elisabeth Trawinski, Jean-Michel Pawlotsky |
| EPI_ISL_3475655 | SELAS MEDILYS | Department of Virology, Henri Mondor University Hospital, Assistance Publique Hôpitaux de Paris, Université Paris-Est Créteil, INSERM U955 | Christophe Rodriguez, Slim Fourati, Vanessa Demontant, Guillaume Gricourt, Melissa N'Debi, Alexandre Soulier, Elisabeth Trawinski, Jean-Michel Pawlotsky |
| EPI_ISL_3475656 | CH. E. MULLER | Department of Virology, Henri Mondor University Hospital, Assistance Publique Hôpitaux de Paris, Université Paris-Est Créteil, INSERM U955 | Christophe Rodriguez, Slim Fourati, Vanessa Demontant, Guillaume Gricourt, Melissa N'Debi, Alexandre Soulier, Elisabeth Trawinski, Jean-Michel Pawlotsky |
| EPI_ISL_3475657 | Biogroup Bio Lam-LCD Saint-Denis | Department of Virology, Henri Mondor University Hospital, Assistance Publique Hôpitaux de Paris, Université Paris-Est Créteil, INSERM U955 | Christophe Rodriguez, Slim Fourati, Vanessa Demontant, Guillaume Gricourt, Melissa N'Debi, Alexandre Soulier, Elisabeth Trawinski, Jean-Michel Pawlotsky |
| EPI_ISL_3475658 | Laboratoire ATOUTBIO JDC | Department of Virology, Henri Mondor University Hospital, Assistance Publique Hôpitaux de Paris, Université Paris-Est Créteil, INSERM U955 | Christophe Rodriguez, Slim Fourati, Vanessa Demontant, Guillaume Gricourt, Melissa N'Debi, Alexandre Soulier, Elisabeth Trawinski, Jean-Michel Pawlotsky |
| EPI_ISL_3475659 | Biogroup Bio Lam-LCD Saint-Denis | Department of Virology, Henri Mondor University Hospital, Assistance Publique Hôpitaux de Paris, Université Paris-Est Créteil, INSERM U955 | Christophe Rodriguez, Slim Fourati, Vanessa Demontant, Guillaume Gricourt, Melissa N'Debi, Alexandre Soulier, Elisabeth Trawinski, Jean-Michel Pawlotsky |
| EPI_ISL_3475660 | CHI VILLENEUVE ST GEORGES | Department of Virology, Henri Mondor University Hospital, Assistance Publique Hôpitaux de Paris, Université Paris-Est Créteil, INSERM U955 | Christophe Rodriguez, Slim Fourati, Vanessa Demontant, Guillaume Gricourt, Melissa N'Debi, Alexandre Soulier, Elisabeth Trawinski, Jean-Michel Pawlotsky |
| EPI_ISL_3475661 | SELAS MEDILYS | Department of Virology, Henri Mondor University Hospital, Assistance Publique Hôpitaux de Paris, Université Paris-Est Créteil, INSERM U955 | Christophe Rodriguez, Slim Fourati, Vanessa Demontant, Guillaume Gricourt, Melissa N'Debi, Alexandre Soulier, Elisabeth Trawinski, Jean-Michel Pawlotsky |
| EPI_ISL_3475662, EPI_ISL_3475663 | CH. CLEMENCEAU | Department of Virology, Henri Mondor University Hospital, Assistance Publique Hôpitaux de Paris, Université Paris-Est Créteil, INSERM U955 | Christophe Rodriguez, Slim Fourati, Vanessa Demontant, Guillaume Gricourt, Melissa N'Debi, Alexandre Soulier, Elisabeth Trawinski, Jean-Michel Pawlotsky |
| EPI_ISL_3475664 | Biogroup Bio Lam-LCD Saint-Denis | Department of Virology, Henri Mondor University Hospital, Assistance Publique Hôpitaux de Paris, Université Paris-Est Créteil, INSERM U955 | Christophe Rodriguez, Slim Fourati, Vanessa Demontant, Guillaume Gricourt, Melissa N'Debi, Alexandre Soulier, Elisabeth Trawinski, Jean-Michel Pawlotsky |
| EPI_ISL_3475665 | CH. CLEMENCEAU | Department of Virology, Henri Mondor University Hospital, Assistance Publique Hôpitaux de Paris, Université Paris-Est Créteil, INSERM U955 | Christophe Rodriguez, Slim Fourati, Vanessa Demontant, Guillaume Gricourt, Melissa N'Debi, Alexandre Soulier, Elisabeth Trawinski, Jean-Michel Pawlotsky |
| EPI_ISL_3475666 | Biogroup Bio Lam-LCD Saint-Denis | Department of Virology, Henri Mondor University Hospital, Assistance Publique Hôpitaux de Paris, Université Paris-Est Créteil, INSERM U955 | Christophe Rodriguez, Slim Fourati, Vanessa Demontant, Guillaume Gricourt, Melissa N'Debi, Alexandre Soulier, Elisabeth Trawinski, Jean-Michel Pawlotsky |
| EPI_ISL_3475667 | CHU DE DIJON | Department of Virology, Henri Mondor University Hospital, Assistance Publique Hôpitaux de Paris, Université Paris-Est Créteil, INSERM U955 | Christophe Rodriguez, Slim Fourati, Vanessa Demontant, Guillaume Gricourt, Melissa N'Debi, Alexandre Soulier, Elisabeth Trawinski, Jean-Michel Pawlotsky |
| EPI_ISL_3475669 | SELAS MEDILYS | Department of Virology, Henri Mondor University Hospital, Assistance Publique Hôpitaux de Paris, Université Paris-Est Créteil, INSERM U955 | Christophe Rodriguez, Slim Fourati, Vanessa Demontant, Guillaume Gricourt, Melissa N'Debi, Alexandre Soulier, Elisabeth Trawinski, Jean-Michel Pawlotsky |
| EPI_ISL_3475670 | Biogroup Bio Lam-LCD Saint-Denis | Department of Virology, Henri Mondor University Hospital, Assistance Publique Hôpitaux de Paris, Université Paris-Est Créteil, INSERM U955 | Christophe Rodriguez, Slim Fourati, Vanessa Demontant, Guillaume Gricourt, Melissa N'Debi, Alexandre Soulier, Elisabeth Trawinski, Jean-Michel Pawlotsky |
| EPI_ISL_3475671 | DYNABIO UNILABS | Department of Virology, Henri Mondor University Hospital, Assistance Publique Hôpitaux de Paris, Université Paris-Est Créteil, INSERM U955 | Christophe Rodriguez, Slim Fourati, Vanessa Demontant, Guillaume Gricourt, Melissa N'Debi, Alexandre Soulier, Elisabeth Trawinski, Jean-Michel Pawlotsky |
| EPI_ISL_3475672 | Biogroup Bio Lam-LCD Saint-Denis | Department of Virology, Henri Mondor University Hospital, Assistance Publique Hôpitaux de Paris, Université Paris-Est Créteil, INSERM U955 | Christophe Rodriguez, Slim Fourati, Vanessa Demontant, Guillaume Gricourt, Melissa N'Debi, Alexandre Soulier, Elisabeth Trawinski, Jean-Michel Pawlotsky |
| EPI_ISL_3475673 | DYNABIO UNILABS | Department of Virology, Henri Mondor University Hospital, Assistance Publique Hôpitaux de Paris, Université Paris-Est Créteil, INSERM U955 | Christophe Rodriguez, Slim Fourati, Vanessa Demontant, Guillaume Gricourt, Melissa N'Debi, Alexandre Soulier, Elisabeth Trawinski, Jean-Michel Pawlotsky |
| EPI_ISL_3475674 | CH. CLEMENCEAU | Department of Virology, Henri Mondor University Hospital, Assistance Publique Hôpitaux de Paris, Université Paris-Est Créteil, INSERM U955 | Christophe Rodriguez, Slim Fourati, Vanessa Demontant, Guillaume Gricourt, Melissa N'Debi, Alexandre Soulier, Elisabeth Trawinski, Jean-Michel Pawlotsky |
| EPI_ISL_3475675 | Laboratoire ATOUTBIO JDC | Department of Virology, Henri Mondor University Hospital, Assistance Publique Hôpitaux de Paris, Université Paris-Est Créteil, INSERM U955 | Christophe Rodriguez, Slim Fourati, Vanessa Demontant, Guillaume Gricourt, Melissa N'Debi, Alexandre Soulier, Elisabeth Trawinski, Jean-Michel Pawlotsky |
| EPI_ISL_3475676 | CH Mémorial F.E.U Laboratoire | Department of Virology, Henri Mondor University Hospital, Assistance Publique Hôpitaux de Paris, Université Paris-Est Créteil, INSERM U955 | Christophe Rodriguez, Slim Fourati, Vanessa Demontant, Guillaume Gricourt, Melissa N'Debi, Alexandre Soulier, Elisabeth Trawinski, Jean-Michel Pawlotsky |
| EPI_ISL_3475677 | CHU DE DIJON | Department of Virology, Henri Mondor University Hospital, Assistance Publique Hôpitaux de Paris, Université Paris-Est Créteil, INSERM U955 | Christophe Rodriguez, Slim Fourati, Vanessa Demontant, Guillaume Gricourt, Melissa N'Debi, Alexandre Soulier, Elisabeth Trawinski, Jean-Michel Pawlotsky |

| | | | |
|---|---|---|---|
| EPI_ISL_3475678 | SELAS MEDILYS | Department of Virology, Henri Mondor University Hospital, Assistance Publique Hôpitaux de Paris, Université Paris-Est Créteil, INSERM U955 | Christophe Rodriguez, Slim Fourati, Vanessa Demontant, Guillaume Gricourt, Melissa N'Debi, Alexandre Soulier, Elisabeth Trawinski, Jean-Michel Pawlotsky |
| EPI_ISL_3475679 | CH. CLEMENCEAU | Department of Virology, Henri Mondor University Hospital, Assistance Publique Hôpitaux de Paris, Université Paris-Est Créteil, INSERM U955 | Christophe Rodriguez, Slim Fourati, Vanessa Demontant, Guillaume Gricourt, Melissa N'Debi, Alexandre Soulier, Elisabeth Trawinski, Jean-Michel Pawlotsky |
| EPI_ISL_3475680, EPI_ISL_3475681 | Laboratoire ATOUTBIO JDC | Department of Virology, Henri Mondor University Hospital, Assistance Publique Hôpitaux de Paris, Université Paris-Est Créteil, INSERM U955 | Christophe Rodriguez, Slim Fourati, Vanessa Demontant, Guillaume Gricourt, Melissa N'Debi, Alexandre Soulier, Elisabeth Trawinski, Jean-Michel Pawlotsky |
| EPI_ISL_3475682 | CHU DE DIJON | Department of Virology, Henri Mondor University Hospital, Assistance Publique Hôpitaux de Paris, Université Paris-Est Créteil, INSERM U955 | Christophe Rodriguez, Slim Fourati, Vanessa Demontant, Guillaume Gricourt, Melissa N'Debi, Alexandre Soulier, Elisabeth Trawinski, Jean-Michel Pawlotsky |
| EPI_ISL_3475683 | SELAS MEDILYS | Department of Virology, Henri Mondor University Hospital, Assistance Publique Hôpitaux de Paris, Université Paris-Est Créteil, INSERM U955 | Christophe Rodriguez, Slim Fourati, Vanessa Demontant, Guillaume Gricourt, Melissa N'Debi, Alexandre Soulier, Elisabeth Trawinski, Jean-Michel Pawlotsky |
| EPI_ISL_3475684 | CHU DE DIJON | Department of Virology, Henri Mondor University Hospital, Assistance Publique Hôpitaux de Paris, Université Paris-Est Créteil, INSERM U955 | Christophe Rodriguez, Slim Fourati, Vanessa Demontant, Guillaume Gricourt, Melissa N'Debi, Alexandre Soulier, Elisabeth Trawinski, Jean-Michel Pawlotsky |
| EPI_ISL_3475685 | CH. CLEMENCEAU | Department of Virology, Henri Mondor University Hospital, Assistance Publique Hôpitaux de Paris, Université Paris-Est Créteil, INSERM U955 | Christophe Rodriguez, Slim Fourati, Vanessa Demontant, Guillaume Gricourt, Melissa N'Debi, Alexandre Soulier, Elisabeth Trawinski, Jean-Michel Pawlotsky |
| EPI_ISL_3475686 | CHU DE DIJON | Department of Virology, Henri Mondor University Hospital, Assistance Publique Hôpitaux de Paris, Université Paris-Est Créteil, INSERM U955 | Christophe Rodriguez, Slim Fourati, Vanessa Demontant, Guillaume Gricourt, Melissa N'Debi, Alexandre Soulier, Elisabeth Trawinski, Jean-Michel Pawlotsky |
| EPI_ISL_3475687, EPI_ISL_3475688 | Biogroup Bio Lam-LCD Saint-Denis | Department of Virology, Henri Mondor University Hospital, Assistance Publique Hôpitaux de Paris, Université Paris-Est Créteil, INSERM U955 | Christophe Rodriguez, Slim Fourati, Vanessa Demontant, Guillaume Gricourt, Melissa N'Debi, Alexandre Soulier, Elisabeth Trawinski, Jean-Michel Pawlotsky |
| EPI_ISL_3475689 | CH. ROBERT BISSON | Department of Virology, Henri Mondor University Hospital, Assistance Publique Hôpitaux de Paris, Université Paris-Est Créteil, INSERM U955 | Christophe Rodriguez, Slim Fourati, Vanessa Demontant, Guillaume Gricourt, Melissa N'Debi, Alexandre Soulier, Elisabeth Trawinski, Jean-Michel Pawlotsky |
| EPI_ISL_3475690 | CH. CLEMENCEAU | Department of Virology, Henri Mondor University Hospital, Assistance Publique Hôpitaux de Paris, Université Paris-Est Créteil, INSERM U955 | Christophe Rodriguez, Slim Fourati, Vanessa Demontant, Guillaume Gricourt, Melissa N'Debi, Alexandre Soulier, Elisabeth Trawinski, Jean-Michel Pawlotsky |
| EPI_ISL_3475691, EPI_ISL_3475692 | BROUSSAIS | Department of Virology, Henri Mondor University Hospital, Assistance Publique Hôpitaux de Paris, Université Paris-Est Créteil, INSERM U955 | Christophe Rodriguez, Slim Fourati, Vanessa Demontant, Guillaume Gricourt, Melissa N'Debi, Alexandre Soulier, Elisabeth Trawinski, Jean-Michel Pawlotsky |
| EPI_ISL_3475694, EPI_ISL_3475695, EPI_ISL_3475697, EPI_ISL_3475698 | Biogroup Bio Lam-LCD Saint-Denis | Department of Virology, Henri Mondor University Hospital, Assistance Publique Hôpitaux de Paris, Université Paris-Est Créteil, INSERM U955 | Christophe Rodriguez, Slim Fourati, Vanessa Demontant, Guillaume Gricourt, Melissa N'Debi, Alexandre Soulier, Elisabeth Trawinski, Jean-Michel Pawlotsky |
| EPI_ISL_3475699 | SELAS MEDILYS | Department of Virology, Henri Mondor University Hospital, Assistance Publique Hôpitaux de Paris, Université Paris-Est Créteil, INSERM U955 | Christophe Rodriguez, Slim Fourati, Vanessa Demontant, Guillaume Gricourt, Melissa N'Debi, Alexandre Soulier, Elisabeth Trawinski, Jean-Michel Pawlotsky |
| EPI_ISL_3475700 | Biogroup Bio Lam-LCD Saint-Denis | Department of Virology, Henri Mondor University Hospital, Assistance Publique Hôpitaux de Paris, Université Paris-Est Créteil, INSERM U955 | Christophe Rodriguez, Slim Fourati, Vanessa Demontant, Guillaume Gricourt, Melissa N'Debi, Alexandre Soulier, Elisabeth Trawinski, Jean-Michel Pawlotsky |
| EPI_ISL_3475701 | Hôpital Saint-Louis | Department of Virology, Henri Mondor University Hospital, Assistance Publique Hôpitaux de Paris, Université Paris-Est Créteil, INSERM U955 | Christophe Rodriguez, Slim Fourati, Vanessa Demontant, Guillaume Gricourt, Melissa N'Debi, Alexandre Soulier, Elisabeth Trawinski, Jean-Michel Pawlotsky |
| EPI_ISL_3475702 | CH. CLEMENCEAU | Department of Virology, Henri Mondor University Hospital, Assistance Publique Hôpitaux de Paris, Université Paris-Est Créteil, INSERM U955 | Christophe Rodriguez, Slim Fourati, Vanessa Demontant, Guillaume Gricourt, Melissa N'Debi, Alexandre Soulier, Elisabeth Trawinski, Jean-Michel Pawlotsky |
| EPI_ISL_3475703 | Biogroup Bio Lam-LCD Saint-Denis | Department of Virology, Henri Mondor University Hospital, Assistance Publique Hôpitaux de Paris, Université Paris-Est Créteil, INSERM U955 | Christophe Rodriguez, Slim Fourati, Vanessa Demontant, Guillaume Gricourt, Melissa N'Debi, Alexandre Soulier, Elisabeth Trawinski, Jean-Michel Pawlotsky |
| EPI_ISL_3475704 | CHU DE DIJON | Department of Virology, Henri Mondor University Hospital, Assistance Publique Hôpitaux de Paris, Université Paris-Est Créteil, INSERM U955 | Christophe Rodriguez, Slim Fourati, Vanessa Demontant, Guillaume Gricourt, Melissa N'Debi, Alexandre Soulier, Elisabeth Trawinski, Jean-Michel Pawlotsky |
| EPI_ISL_3475705 | CERBALLIANCE SAINT MARTIN | Department of Virology, Henri Mondor University Hospital, Assistance Publique Hôpitaux de Paris, Université Paris-Est Créteil, INSERM U955 | Christophe Rodriguez, Slim Fourati, Vanessa Demontant, Guillaume Gricourt, Melissa N'Debi, Alexandre Soulier, Elisabeth Trawinski, Jean-Michel Pawlotsky |
| EPI_ISL_3475706 | CH. ANDRE GREGOIRE | Department of Virology, Henri Mondor University Hospital, Assistance Publique Hôpitaux de Paris, Université Paris-Est Créteil, INSERM U955 | Christophe Rodriguez, Slim Fourati, Vanessa Demontant, Guillaume Gricourt, Melissa N'Debi, Alexandre Soulier, Elisabeth Trawinski, Jean-Michel Pawlotsky |
| EPI_ISL_3475707 | CH. CLEMENCEAU | Department of Virology, Henri Mondor University Hospital, Assistance Publique Hôpitaux de Paris, Université Paris-Est Créteil, INSERM U955 | Christophe Rodriguez, Slim Fourati, Vanessa Demontant, Guillaume Gricourt, Melissa N'Debi, Alexandre Soulier, Elisabeth Trawinski, Jean-Michel Pawlotsky |
| EPI_ISL_3475708, EPI_ISL_3475709 | SELAS MEDILYS | Department of Virology, Henri Mondor University Hospital, Assistance Publique Hôpitaux de Paris, Université Paris-Est Créteil, INSERM U955 | Christophe Rodriguez, Slim Fourati, Vanessa Demontant, Guillaume Gricourt, Melissa N'Debi, Alexandre Soulier, Elisabeth Trawinski, Jean-Michel Pawlotsky |
| EPI_ISL_3475710 | CH. CLEMENCEAU | Department of Virology, Henri Mondor University Hospital, Assistance Publique Hôpitaux de Paris, Université Paris-Est Créteil, INSERM U955 | Christophe Rodriguez, Slim Fourati, Vanessa Demontant, Guillaume Gricourt, Melissa N'Debi, Alexandre Soulier, Elisabeth Trawinski, Jean-Michel Pawlotsky |
| EPI_ISL_3475711 | SELAS MEDILYS | Department of Virology, Henri Mondor University Hospital, Assistance Publique Hôpitaux de Paris, Université Paris-Est Créteil, INSERM U955 | Christophe Rodriguez, Slim Fourati, Vanessa Demontant, Guillaume Gricourt, Melissa N'Debi, Alexandre Soulier, Elisabeth Trawinski, Jean-Michel Pawlotsky |
| EPI_ISL_3475712 | Biogroup Bio Lam-LCD Saint-Denis | Department of Virology, Henri Mondor University Hospital, Assistance Publique Hôpitaux de Paris, Université Paris-Est Créteil, INSERM U955 | Christophe Rodriguez, Slim Fourati, Vanessa Demontant, Guillaume Gricourt, Melissa N'Debi, Alexandre Soulier, Elisabeth Trawinski, Jean-Michel Pawlotsky |
| EPI_ISL_3475713 | CHU DE DIJON | Department of Virology, Henri Mondor University Hospital, Assistance Publique Hôpitaux de Paris, Université Paris-Est Créteil, INSERM U955 | Christophe Rodriguez, Slim Fourati, Vanessa Demontant, Guillaume Gricourt, Melissa N'Debi, Alexandre Soulier, Elisabeth Trawinski, Jean-Michel Pawlotsky |
| EPI_ISL_3475714 | SELAS MEDILYS | Department of Virology, Henri Mondor University Hospital, Assistance Publique Hôpitaux de Paris, Université Paris-Est Créteil, INSERM U955 | Christophe Rodriguez, Slim Fourati, Vanessa Demontant, Guillaume Gricourt, Melissa N'Debi, Alexandre Soulier, Elisabeth Trawinski, Jean-Michel Pawlotsky |
| EPI_ISL_3475715 | CH. CLEMENCEAU | Department of Virology, Henri Mondor University Hospital, Assistance Publique Hôpitaux de Paris, Université Paris-Est Créteil, INSERM U955 | Christophe Rodriguez, Slim Fourati, Vanessa Demontant, Guillaume Gricourt, Melissa N'Debi, Alexandre Soulier, Elisabeth Trawinski, Jean-Michel Pawlotsky |
| EPI_ISL_3475716 | BIOSMOSE IDF | Department of Virology, Henri Mondor University Hospital, Assistance Publique Hôpitaux de Paris, Université Paris-Est Créteil, INSERM U955 | Christophe Rodriguez, Slim Fourati, Vanessa Demontant, Guillaume Gricourt, Melissa N'Debi, Alexandre Soulier, Elisabeth Trawinski, Jean-Michel Pawlotsky |
| EPI_ISL_3475717 | SELAS MEDILYS | Department of Virology, Henri Mondor University Hospital, Assistance Publique Hôpitaux de Paris, Université Paris-Est Créteil, INSERM U955 | Christophe Rodriguez, Slim Fourati, Vanessa Demontant, Guillaume Gricourt, Melissa N'Debi, Alexandre Soulier, Elisabeth Trawinski, Jean-Michel Pawlotsky |
| EPI_ISL_3475718 | Hôpital Avicenne | Department of Virology, Henri Mondor University Hospital, Assistance Publique Hôpitaux de Paris, Université Paris-Est Créteil, INSERM U955 | Christophe Rodriguez, Slim Fourati, Vanessa Demontant, Guillaume Gricourt, Melissa N'Debi, Alexandre Soulier, Elisabeth Trawinski, Jean-Michel Pawlotsky |

| | | | |
|---|---|---|---|
| EPI_ISL_3475719 | Biogroup Bio Lam-LCD Saint-Denis | Department of Virology, Henri Mondor University Hospital, Assistance Publique Hôpitaux de Paris, Université Paris-Est Créteil, INSERM U955 | Christophe Rodriguez, Slim Fourati, Vanessa Demontant, Guillaume Gricourt, Melissa N'Debi, Alexandre Soulier, Elisabeth Trawinski, Jean-Michel Pawlotsky |
| EPI_ISL_3475720 | Hôpital Ambroise-Paré | Department of Virology, Henri Mondor University Hospital, Assistance Publique Hôpitaux de Paris, Université Paris-Est Créteil, INSERM U955 | Christophe Rodriguez, Slim Fourati, Vanessa Demontant, Guillaume Gricourt, Melissa N'Debi, Alexandre Soulier, Elisabeth Trawinski, Jean-Michel Pawlotsky |
| EPI_ISL_3475721 | Biogroup Bio Lam-LCD Saint-Denis | Department of Virology, Henri Mondor University Hospital, Assistance Publique Hôpitaux de Paris, Université Paris-Est Créteil, INSERM U955 | Christophe Rodriguez, Slim Fourati, Vanessa Demontant, Guillaume Gricourt, Melissa N'Debi, Alexandre Soulier, Elisabeth Trawinski, Jean-Michel Pawlotsky |
| EPI_ISL_3475722 | CERBALLIANCE SAINT MARTIN | Department of Virology, Henri Mondor University Hospital, Assistance Publique Hôpitaux de Paris, Université Paris-Est Créteil, INSERM U955 | Christophe Rodriguez, Slim Fourati, Vanessa Demontant, Guillaume Gricourt, Melissa N'Debi, Alexandre Soulier, Elisabeth Trawinski, Jean-Michel Pawlotsky |
| EPI_ISL_3475723 | Laboratoire Storck P.(Biolia) | Department of Virology, Henri Mondor University Hospital, Assistance Publique Hôpitaux de Paris, Université Paris-Est Créteil, INSERM U955 | Christophe Rodriguez, Slim Fourati, Vanessa Demontant, Guillaume Gricourt, Melissa N'Debi, Alexandre Soulier, Elisabeth Trawinski, Jean-Michel Pawlotsky |
| EPI_ISL_3475724 | Biogroup Bio Lam-LCD Saint-Denis | Department of Virology, Henri Mondor University Hospital, Assistance Publique Hôpitaux de Paris, Université Paris-Est Créteil, INSERM U955 | Christophe Rodriguez, Slim Fourati, Vanessa Demontant, Guillaume Gricourt, Melissa N'Debi, Alexandre Soulier, Elisabeth Trawinski, Jean-Michel Pawlotsky |
| EPI_ISL_3475726 | BIO67-BIOSPHERE | Department of Virology, Henri Mondor University Hospital, Assistance Publique Hôpitaux de Paris, Université Paris-Est Créteil, INSERM U955 | Christophe Rodriguez, Slim Fourati, Vanessa Demontant, Guillaume Gricourt, Melissa N'Debi, Alexandre Soulier, Elisabeth Trawinski, Jean-Michel Pawlotsky |
| EPI_ISL_3475727, EPI_ISL_3475728 | Biogroup Bio Lam-LCD Saint-Denis | Department of Virology, Henri Mondor University Hospital, Assistance Publique Hôpitaux de Paris, Université Paris-Est Créteil, INSERM U955 | Christophe Rodriguez, Slim Fourati, Vanessa Demontant, Guillaume Gricourt, Melissa N'Debi, Alexandre Soulier, Elisabeth Trawinski, Jean-Michel Pawlotsky |
| EPI_ISL_3475729 | SELAS MEDILYS | Department of Virology, Henri Mondor University Hospital, Assistance Publique Hôpitaux de Paris, Université Paris-Est Créteil, INSERM U955 | Christophe Rodriguez, Slim Fourati, Vanessa Demontant, Guillaume Gricourt, Melissa N'Debi, Alexandre Soulier, Elisabeth Trawinski, Jean-Michel Pawlotsky |
| EPI_ISL_3475730 | CHU DE DIJON | Department of Virology, Henri Mondor University Hospital, Assistance Publique Hôpitaux de Paris, Université Paris-Est Créteil, INSERM U955 | Christophe Rodriguez, Slim Fourati, Vanessa Demontant, Guillaume Gricourt, Melissa N'Debi, Alexandre Soulier, Elisabeth Trawinski, Jean-Michel Pawlotsky |
| EPI_ISL_3475731 | Biogroup Bio Lam-LCD Saint-Denis | Department of Virology, Henri Mondor University Hospital, Assistance Publique Hôpitaux de Paris, Université Paris-Est Créteil, INSERM U955 | Christophe Rodriguez, Slim Fourati, Vanessa Demontant, Guillaume Gricourt, Melissa N'Debi, Alexandre Soulier, Elisabeth Trawinski, Jean-Michel Pawlotsky |
| EPI_ISL_3475732 | CH. ROBERT BALLANGER | Department of Virology, Henri Mondor University Hospital, Assistance Publique Hôpitaux de Paris, Université Paris-Est Créteil, INSERM U955 | Christophe Rodriguez, Slim Fourati, Vanessa Demontant, Guillaume Gricourt, Melissa N'Debi, Alexandre Soulier, Elisabeth Trawinski, Jean-Michel Pawlotsky |
| EPI_ISL_3475733 | CH. CHU DE REIMS | Department of Virology, Henri Mondor University Hospital, Assistance Publique Hôpitaux de Paris, Université Paris-Est Créteil, INSERM U955 | Christophe Rodriguez, Slim Fourati, Vanessa Demontant, Guillaume Gricourt, Melissa N'Debi, Alexandre Soulier, Elisabeth Trawinski, Jean-Michel Pawlotsky |
| EPI_ISL_3475734, EPI_ISL_3475735, EPI_ISL_3475736 | Biogroup Bio Lam-LCD Saint-Denis | Department of Virology, Henri Mondor University Hospital, Assistance Publique Hôpitaux de Paris, Université Paris-Est Créteil, INSERM U955 | Christophe Rodriguez, Slim Fourati, Vanessa Demontant, Guillaume Gricourt, Melissa N'Debi, Alexandre Soulier, Elisabeth Trawinski, Jean-Michel Pawlotsky |
| EPI_ISL_3475737 | Laboratoire ATOUTBIO JDC | Department of Virology, Henri Mondor University Hospital, Assistance Publique Hôpitaux de Paris, Université Paris-Est Créteil, INSERM U955 | Christophe Rodriguez, Slim Fourati, Vanessa Demontant, Guillaume Gricourt, Melissa N'Debi, Alexandre Soulier, Elisabeth Trawinski, Jean-Michel Pawlotsky |
| EPI_ISL_3475739 | CH. CLEMENCEAU | Department of Virology, Henri Mondor University Hospital, Assistance Publique Hôpitaux de Paris, Université Paris-Est Créteil, INSERM U955 | Christophe Rodriguez, Slim Fourati, Vanessa Demontant, Guillaume Gricourt, Melissa N'Debi, Alexandre Soulier, Elisabeth Trawinski, Jean-Michel Pawlotsky |
| EPI_ISL_3475740 | Laboratoire ATOUTBIO JDC | Department of Virology, Henri Mondor University Hospital, Assistance Publique Hôpitaux de Paris, Université Paris-Est Créteil, INSERM U955 | Christophe Rodriguez, Slim Fourati, Vanessa Demontant, Guillaume Gricourt, Melissa N'Debi, Alexandre Soulier, Elisabeth Trawinski, Jean-Michel Pawlotsky |
| EPI_ISL_3475741, EPI_ISL_3475742 | Biogroup Bio Lam-LCD Saint-Denis | Department of Virology, Henri Mondor University Hospital, Assistance Publique Hôpitaux de Paris, Université Paris-Est Créteil, INSERM U955 | Christophe Rodriguez, Slim Fourati, Vanessa Demontant, Guillaume Gricourt, Melissa N'Debi, Alexandre Soulier, Elisabeth Trawinski, Jean-Michel Pawlotsky |
| EPI_ISL_3475743 | CHU DE DIJON | Department of Virology, Henri Mondor University Hospital, Assistance Publique Hôpitaux de Paris, Université Paris-Est Créteil, INSERM U955 | Christophe Rodriguez, Slim Fourati, Vanessa Demontant, Guillaume Gricourt, Melissa N'Debi, Alexandre Soulier, Elisabeth Trawinski, Jean-Michel Pawlotsky |
| EPI_ISL_3475744, EPI_ISL_3475745 | Biogroup Bio Lam-LCD Saint-Denis | Department of Virology, Henri Mondor University Hospital, Assistance Publique Hôpitaux de Paris, Université Paris-Est Créteil, INSERM U955 | Christophe Rodriguez, Slim Fourati, Vanessa Demontant, Guillaume Gricourt, Melissa N'Debi, Alexandre Soulier, Elisabeth Trawinski, Jean-Michel Pawlotsky |
| EPI_ISL_3475746 | BIO VSM LAB | Department of Virology, Henri Mondor University Hospital, Assistance Publique Hôpitaux de Paris, Université Paris-Est Créteil, INSERM U955 | Christophe Rodriguez, Slim Fourati, Vanessa Demontant, Guillaume Gricourt, Melissa N'Debi, Alexandre Soulier, Elisabeth Trawinski, Jean-Michel Pawlotsky |
| EPI_ISL_3475747 | Laboratoire ATOUTBIO JDC | Department of Virology, Henri Mondor University Hospital, Assistance Publique Hôpitaux de Paris, Université Paris-Est Créteil, INSERM U955 | Christophe Rodriguez, Slim Fourati, Vanessa Demontant, Guillaume Gricourt, Melissa N'Debi, Alexandre Soulier, Elisabeth Trawinski, Jean-Michel Pawlotsky |
| EPI_ISL_3475748 | Biogroup Bio Lam-LCD Saint-Denis | Department of Virology, Henri Mondor University Hospital, Assistance Publique Hôpitaux de Paris, Université Paris-Est Créteil, INSERM U955 | Christophe Rodriguez, Slim Fourati, Vanessa Demontant, Guillaume Gricourt, Melissa N'Debi, Alexandre Soulier, Elisabeth Trawinski, Jean-Michel Pawlotsky |
| EPI_ISL_3475749 | BIOSMOSE IDF | Department of Virology, Henri Mondor University Hospital, Assistance Publique Hôpitaux de Paris, Université Paris-Est Créteil, INSERM U955 | Christophe Rodriguez, Slim Fourati, Vanessa Demontant, Guillaume Gricourt, Melissa N'Debi, Alexandre Soulier, Elisabeth Trawinski, Jean-Michel Pawlotsky |
| EPI_ISL_3475750 | SELAS MEDILYS | Department of Virology, Henri Mondor University Hospital, Assistance Publique Hôpitaux de Paris, Université Paris-Est Créteil, INSERM U955 | Christophe Rodriguez, Slim Fourati, Vanessa Demontant, Guillaume Gricourt, Melissa N'Debi, Alexandre Soulier, Elisabeth Trawinski, Jean-Michel Pawlotsky |
| EPI_ISL_3475751, EPI_ISL_3475752, EPI_ISL_3475753, EPI_ISL_3475754, EPI_ISL_3475755 | Biogroup Bio Lam-LCD Saint-Denis | Department of Virology, Henri Mondor University Hospital, Assistance Publique Hôpitaux de Paris, Université Paris-Est Créteil, INSERM U955 | Christophe Rodriguez, Slim Fourati, Vanessa Demontant, Guillaume Gricourt, Melissa N'Debi, Alexandre Soulier, Elisabeth Trawinski, Jean-Michel Pawlotsky |
| EPI_ISL_3475756 | CH. ROBERT BALLANGER | Department of Virology, Henri Mondor University Hospital, Assistance Publique Hôpitaux de Paris, Université Paris-Est Créteil, INSERM U955 | Christophe Rodriguez, Slim Fourati, Vanessa Demontant, Guillaume Gricourt, Melissa N'Debi, Alexandre Soulier, Elisabeth Trawinski, Jean-Michel Pawlotsky |
| EPI_ISL_3475758 | GH A.CHENEVIER-H.MONDOR | Department of Virology, Henri Mondor University Hospital, Assistance Publique Hôpitaux de Paris, Université Paris-Est Créteil, INSERM U955 | Christophe Rodriguez, Slim Fourati, Vanessa Demontant, Guillaume Gricourt, Melissa N'Debi, Alexandre Soulier, Elisabeth Trawinski, Jean-Michel Pawlotsky |
| EPI_ISL_3475759 | CH Mémorial F.E.U Laboratoire | Department of Virology, Henri Mondor University Hospital, Assistance Publique Hôpitaux de Paris, Université Paris-Est Créteil, INSERM U955 | Christophe Rodriguez, Slim Fourati, Vanessa Demontant, Guillaume Gricourt, Melissa N'Debi, Alexandre Soulier, Elisabeth Trawinski, Jean-Michel Pawlotsky |
| EPI_ISL_3475761 | Biogroup Bio Lam-LCD Saint-Denis | Department of Virology, Henri Mondor University Hospital, Assistance Publique Hôpitaux de Paris, Université Paris-Est Créteil, INSERM U955 | Christophe Rodriguez, Slim Fourati, Vanessa Demontant, Guillaume Gricourt, Melissa N'Debi, Alexandre Soulier, Elisabeth Trawinski, Jean-Michel Pawlotsky |
| EPI_ISL_3475762 | Laboratoire ATOUTBIO JDC | Department of Virology, Henri Mondor University Hospital, Assistance Publique Hôpitaux de Paris, Université Paris-Est Créteil, INSERM U955 | Christophe Rodriguez, Slim Fourati, Vanessa Demontant, Guillaume Gricourt, Melissa N'Debi, Alexandre Soulier, Elisabeth Trawinski, Jean-Michel Pawlotsky |

| | | | |
|---|---|---|---|
| EPI_ISL_3475763 | CERBALLIANCE SAINT MARTIN | Department of Virology, Henri Mondor University Hospital, Assistance Publique Hôpitaux de Paris, Université Paris-Est Créteil, INSERM U955 | Christophe Rodriguez, Slim Fourati, Vanessa Demontant, Guillaume Gricourt, Melissa N'Debi, Alexandre Soulier, Elisabeth Trawinski, Jean-Michel Pawlotsky |
| EPI_ISL_3475764 | Biogroup Bio Lam-LCD Saint-Denis | Department of Virology, Henri Mondor University Hospital, Assistance Publique Hôpitaux de Paris, Université Paris-Est Créteil, INSERM U955 | Christophe Rodriguez, Slim Fourati, Vanessa Demontant, Guillaume Gricourt, Melissa N'Debi, Alexandre Soulier, Elisabeth Trawinski, Jean-Michel Pawlotsky |
| EPI_ISL_3475765 | Hôpital Avicenne | Department of Virology, Henri Mondor University Hospital, Assistance Publique Hôpitaux de Paris, Université Paris-Est Créteil, INSERM U955 | Christophe Rodriguez, Slim Fourati, Vanessa Demontant, Guillaume Gricourt, Melissa N'Debi, Alexandre Soulier, Elisabeth Trawinski, Jean-Michel Pawlotsky |
| EPI_ISL_3475766 | CHU DE DIJON | Department of Virology, Henri Mondor University Hospital, Assistance Publique Hôpitaux de Paris, Université Paris-Est Créteil, INSERM U955 | Christophe Rodriguez, Slim Fourati, Vanessa Demontant, Guillaume Gricourt, Melissa N'Debi, Alexandre Soulier, Elisabeth Trawinski, Jean-Michel Pawlotsky |
| EPI_ISL_3475767, EPI_ISL_3475768 | Biogroup Bio Lam-LCD Saint-Denis | Department of Virology, Henri Mondor University Hospital, Assistance Publique Hôpitaux de Paris, Université Paris-Est Créteil, INSERM U955 | Christophe Rodriguez, Slim Fourati, Vanessa Demontant, Guillaume Gricourt, Melissa N'Debi, Alexandre Soulier, Elisabeth Trawinski, Jean-Michel Pawlotsky |
| EPI_ISL_3475769 | CH. CLEMENCEAU | Department of Virology, Henri Mondor University Hospital, Assistance Publique Hôpitaux de Paris, Université Paris-Est Créteil, INSERM U955 | Christophe Rodriguez, Slim Fourati, Vanessa Demontant, Guillaume Gricourt, Melissa N'Debi, Alexandre Soulier, Elisabeth Trawinski, Jean-Michel Pawlotsky |
| EPI_ISL_3475770 | Biogroup Bio Lam-LCD Saint-Denis | Department of Virology, Henri Mondor University Hospital, Assistance Publique Hôpitaux de Paris, Université Paris-Est Créteil, INSERM U955 | Christophe Rodriguez, Slim Fourati, Vanessa Demontant, Guillaume Gricourt, Melissa N'Debi, Alexandre Soulier, Elisabeth Trawinski, Jean-Michel Pawlotsky |
| EPI_ISL_3475772 | Hôpital Avicenne | Department of Virology, Henri Mondor University Hospital, Assistance Publique Hôpitaux de Paris, Université Paris-Est Créteil, INSERM U955 | Christophe Rodriguez, Slim Fourati, Vanessa Demontant, Guillaume Gricourt, Melissa N'Debi, Alexandre Soulier, Elisabeth Trawinski, Jean-Michel Pawlotsky |
| EPI_ISL_3475773 | CH. CLEMENCEAU | Department of Virology, Henri Mondor University Hospital, Assistance Publique Hôpitaux de Paris, Université Paris-Est Créteil, INSERM U955 | Christophe Rodriguez, Slim Fourati, Vanessa Demontant, Guillaume Gricourt, Melissa N'Debi, Alexandre Soulier, Elisabeth Trawinski, Jean-Michel Pawlotsky |
| EPI_ISL_3475774, EPI_ISL_3475775 | Biogroup Bio Lam-LCD Saint-Denis | Department of Virology, Henri Mondor University Hospital, Assistance Publique Hôpitaux de Paris, Université Paris-Est Créteil, INSERM U955 | Christophe Rodriguez, Slim Fourati, Vanessa Demontant, Guillaume Gricourt, Melissa N'Debi, Alexandre Soulier, Elisabeth Trawinski, Jean-Michel Pawlotsky |
| EPI_ISL_3475776 | Laboratoire ATOUTBIO JDC | Department of Virology, Henri Mondor University Hospital, Assistance Publique Hôpitaux de Paris, Université Paris-Est Créteil, INSERM U955 | Christophe Rodriguez, Slim Fourati, Vanessa Demontant, Guillaume Gricourt, Melissa N'Debi, Alexandre Soulier, Elisabeth Trawinski, Jean-Michel Pawlotsky |
| EPI_ISL_3475777 | CH. CLEMENCEAU | Department of Virology, Henri Mondor University Hospital, Assistance Publique Hôpitaux de Paris, Université Paris-Est Créteil, INSERM U955 | Christophe Rodriguez, Slim Fourati, Vanessa Demontant, Guillaume Gricourt, Melissa N'Debi, Alexandre Soulier, Elisabeth Trawinski, Jean-Michel Pawlotsky |
| EPI_ISL_3475778 | CERBALLIANCE SAINT MARTIN | Department of Virology, Henri Mondor University Hospital, Assistance Publique Hôpitaux de Paris, Université Paris-Est Créteil, INSERM U955 | Christophe Rodriguez, Slim Fourati, Vanessa Demontant, Guillaume Gricourt, Melissa N'Debi, Alexandre Soulier, Elisabeth Trawinski, Jean-Michel Pawlotsky |
| EPI_ISL_3475779 | GH A.CHENEVIER-H.MONDOR | Department of Virology, Henri Mondor University Hospital, Assistance Publique Hôpitaux de Paris, Université Paris-Est Créteil, INSERM U955 | Christophe Rodriguez, Slim Fourati, Vanessa Demontant, Guillaume Gricourt, Melissa N'Debi, Alexandre Soulier, Elisabeth Trawinski, Jean-Michel Pawlotsky |
| EPI_ISL_3475780, EPI_ISL_3475781, EPI_ISL_3475782, EPI_ISL_3475783, EPI_ISL_3475784 | Biogroup Bio Lam-LCD Saint-Denis | Department of Virology, Henri Mondor University Hospital, Assistance Publique Hôpitaux de Paris, Université Paris-Est Créteil, INSERM U955 | Christophe Rodriguez, Slim Fourati, Vanessa Demontant, Guillaume Gricourt, Melissa N'Debi, Alexandre Soulier, Elisabeth Trawinski, Jean-Michel Pawlotsky |
| EPI_ISL_3475785 | CHU DE DIJON | Department of Virology, Henri Mondor University Hospital, Assistance Publique Hôpitaux de Paris, Université Paris-Est Créteil, INSERM U955 | Christophe Rodriguez, Slim Fourati, Vanessa Demontant, Guillaume Gricourt, Melissa N'Debi, Alexandre Soulier, Elisabeth Trawinski, Jean-Michel Pawlotsky |
| EPI_ISL_3475786 | CH.INTERCOMMUNAL DE CRETEIL | Department of Virology, Henri Mondor University Hospital, Assistance Publique Hôpitaux de Paris, Université Paris-Est Créteil, INSERM U955 | Christophe Rodriguez, Slim Fourati, Vanessa Demontant, Guillaume Gricourt, Melissa N'Debi, Alexandre Soulier, Elisabeth Trawinski, Jean-Michel Pawlotsky |
| EPI_ISL_3475787 | CH. CLEMENCEAU | Department of Virology, Henri Mondor University Hospital, Assistance Publique Hôpitaux de Paris, Université Paris-Est Créteil, INSERM U955 | Christophe Rodriguez, Slim Fourati, Vanessa Demontant, Guillaume Gricourt, Melissa N'Debi, Alexandre Soulier, Elisabeth Trawinski, Jean-Michel Pawlotsky |
| EPI_ISL_3475788 | CERBALLIANCE SAINT MARTIN | Department of Virology, Henri Mondor University Hospital, Assistance Publique Hôpitaux de Paris, Université Paris-Est Créteil, INSERM U955 | Christophe Rodriguez, Slim Fourati, Vanessa Demontant, Guillaume Gricourt, Melissa N'Debi, Alexandre Soulier, Elisabeth Trawinski, Jean-Michel Pawlotsky |
| EPI_ISL_3475789 | SELAS MEDILYS | Department of Virology, Henri Mondor University Hospital, Assistance Publique Hôpitaux de Paris, Université Paris-Est Créteil, INSERM U955 | Christophe Rodriguez, Slim Fourati, Vanessa Demontant, Guillaume Gricourt, Melissa N'Debi, Alexandre Soulier, Elisabeth Trawinski, Jean-Michel Pawlotsky |
| EPI_ISL_3475790 | CERBALLIANCE SAINT MARTIN | Department of Virology, Henri Mondor University Hospital, Assistance Publique Hôpitaux de Paris, Université Paris-Est Créteil, INSERM U955 | Christophe Rodriguez, Slim Fourati, Vanessa Demontant, Guillaume Gricourt, Melissa N'Debi, Alexandre Soulier, Elisabeth Trawinski, Jean-Michel Pawlotsky |
| EPI_ISL_3475791, EPI_ISL_3475792 | Biogroup Bio Lam-LCD Saint-Denis | Department of Virology, Henri Mondor University Hospital, Assistance Publique Hôpitaux de Paris, Université Paris-Est Créteil, INSERM U955 | Christophe Rodriguez, Slim Fourati, Vanessa Demontant, Guillaume Gricourt, Melissa N'Debi, Alexandre Soulier, Elisabeth Trawinski, Jean-Michel Pawlotsky |
| EPI_ISL_3475793, EPI_ISL_3475794, EPI_ISL_3475795 | CERBALLIANCE SAINT MARTIN | Department of Virology, Henri Mondor University Hospital, Assistance Publique Hôpitaux de Paris, Université Paris-Est Créteil, INSERM U955 | Christophe Rodriguez, Slim Fourati, Vanessa Demontant, Guillaume Gricourt, Melissa N'Debi, Alexandre Soulier, Elisabeth Trawinski, Jean-Michel Pawlotsky |
| EPI_ISL_3475796 | CH. E. MULLER | Department of Virology, Henri Mondor University Hospital, Assistance Publique Hôpitaux de Paris, Université Paris-Est Créteil, INSERM U955 | Christophe Rodriguez, Slim Fourati, Vanessa Demontant, Guillaume Gricourt, Melissa N'Debi, Alexandre Soulier, Elisabeth Trawinski, Jean-Michel Pawlotsky |
| EPI_ISL_3475798 | CHU DE DIJON | Department of Virology, Henri Mondor University Hospital, Assistance Publique Hôpitaux de Paris, Université Paris-Est Créteil, INSERM U955 | Christophe Rodriguez, Slim Fourati, Vanessa Demontant, Guillaume Gricourt, Melissa N'Debi, Alexandre Soulier, Elisabeth Trawinski, Jean-Michel Pawlotsky |
| EPI_ISL_3475799 | CH. CLEMENCEAU | Department of Virology, Henri Mondor University Hospital, Assistance Publique Hôpitaux de Paris, Université Paris-Est Créteil, INSERM U955 | Christophe Rodriguez, Slim Fourati, Vanessa Demontant, Guillaume Gricourt, Melissa N'Debi, Alexandre Soulier, Elisabeth Trawinski, Jean-Michel Pawlotsky |
| EPI_ISL_3475800 | CERBALLIANCE SAINT MARTIN | Department of Virology, Henri Mondor University Hospital, Assistance Publique Hôpitaux de Paris, Université Paris-Est Créteil, INSERM U955 | Christophe Rodriguez, Slim Fourati, Vanessa Demontant, Guillaume Gricourt, Melissa N'Debi, Alexandre Soulier, Elisabeth Trawinski, Jean-Michel Pawlotsky |
| EPI_ISL_3475801 | Hôpital Ambroise-Paré | Department of Virology, Henri Mondor University Hospital, Assistance Publique Hôpitaux de Paris, Université Paris-Est Créteil, INSERM U955 | Christophe Rodriguez, Slim Fourati, Vanessa Demontant, Guillaume Gricourt, Melissa N'Debi, Alexandre Soulier, Elisabeth Trawinski, Jean-Michel Pawlotsky |
| EPI_ISL_3475802 | Hôpital Avicenne | Department of Virology, Henri Mondor University Hospital, Assistance Publique Hôpitaux de Paris, Université Paris-Est Créteil, INSERM U955 | Christophe Rodriguez, Slim Fourati, Vanessa Demontant, Guillaume Gricourt, Melissa N'Debi, Alexandre Soulier, Elisabeth Trawinski, Jean-Michel Pawlotsky |
| EPI_ISL_3475803 | Biogroup Bio Lam-LCD Saint-Denis | Department of Virology, Henri Mondor University Hospital, Assistance Publique Hôpitaux de Paris, Université Paris-Est Créteil, INSERM U955 | Christophe Rodriguez, Slim Fourati, Vanessa Demontant, Guillaume Gricourt, Melissa N'Debi, Alexandre Soulier, Elisabeth Trawinski, Jean-Michel Pawlotsky |
| EPI_ISL_3475804, EPI_ISL_3475805 | CHU DE DIJON | Department of Virology, Henri Mondor University Hospital, Assistance Publique Hôpitaux de Paris, Université Paris-Est Créteil, INSERM U955 | Christophe Rodriguez, Slim Fourati, Vanessa Demontant, Guillaume Gricourt, Melissa N'Debi, Alexandre Soulier, Elisabeth Trawinski, Jean-Michel Pawlotsky |
| EPI_ISL_3475806 | CH Mémorial F.E.U Laboratoire | Department of Virology, Henri Mondor University Hospital, | Christophe Rodriguez, Slim Fourati, Vanessa Demontant, Guillaume Gricourt, Melissa N'Debi, Alexandre Soulier, Elisabeth Trawinski, Jean-Michel Pawlotsky |

| | | | |
|---|---|---|---|
| | | Assistance Publique Hôpitaux de Paris, Université Paris-Est Créteil, INSERM U955 | |
| EPI_ISL_3475807 | SELAS MEDILYS | Department of Virology, Henri Mondor University Hospital, Assistance Publique Hôpitaux de Paris, Université Paris-Est Créteil, INSERM U955 | Christophe Rodriguez, Slim Fourati, Vanessa Demontant, Guillaume Gricourt, Melissa N'Debi, Alexandre Soulier, Elisabeth Trawinski, Jean-Michel Pawlotsky |
| EPI_ISL_3475808 | CERBALLIANCE SAINT MARTIN | Department of Virology, Henri Mondor University Hospital, Assistance Publique Hôpitaux de Paris, Université Paris-Est Créteil, INSERM U955 | Christophe Rodriguez, Slim Fourati, Vanessa Demontant, Guillaume Gricourt, Melissa N'Debi, Alexandre Soulier, Elisabeth Trawinski, Jean-Michel Pawlotsky |
| EPI_ISL_3475809 | CH. CLEMENCEAU | Department of Virology, Henri Mondor University Hospital, Assistance Publique Hôpitaux de Paris, Université Paris-Est Créteil, INSERM U955 | Christophe Rodriguez, Slim Fourati, Vanessa Demontant, Guillaume Gricourt, Melissa N'Debi, Alexandre Soulier, Elisabeth Trawinski, Jean-Michel Pawlotsky |
| EPI_ISL_3475810 | Biogroup Bio Lam-LCD Saint-Denis | Department of Virology, Henri Mondor University Hospital, Assistance Publique Hôpitaux de Paris, Université Paris-Est Créteil, INSERM U955 | Christophe Rodriguez, Slim Fourati, Vanessa Demontant, Guillaume Gricourt, Melissa N'Debi, Alexandre Soulier, Elisabeth Trawinski, Jean-Michel Pawlotsky |
| EPI_ISL_3475811 | SELAS MEDILYS | Department of Virology, Henri Mondor University Hospital, Assistance Publique Hôpitaux de Paris, Université Paris-Est Créteil, INSERM U955 | Christophe Rodriguez, Slim Fourati, Vanessa Demontant, Guillaume Gricourt, Melissa N'Debi, Alexandre Soulier, Elisabeth Trawinski, Jean-Michel Pawlotsky |
| EPI_ISL_3475812 | GH A.CHENEVIER-H.MONDOR | Department of Virology, Henri Mondor University Hospital, Assistance Publique Hôpitaux de Paris, Université Paris-Est Créteil, INSERM U955 | Christophe Rodriguez, Slim Fourati, Vanessa Demontant, Guillaume Gricourt, Melissa N'Debi, Alexandre Soulier, Elisabeth Trawinski, Jean-Michel Pawlotsky |
| EPI_ISL_3475813 | CH. CLEMENCEAU | Department of Virology, Henri Mondor University Hospital, Assistance Publique Hôpitaux de Paris, Université Paris-Est Créteil, INSERM U955 | Christophe Rodriguez, Slim Fourati, Vanessa Demontant, Guillaume Gricourt, Melissa N'Debi, Alexandre Soulier, Elisabeth Trawinski, Jean-Michel Pawlotsky |
| EPI_ISL_3475814 | CHU DE DIJON | Department of Virology, Henri Mondor University Hospital, Assistance Publique Hôpitaux de Paris, Université Paris-Est Créteil, INSERM U955 | Christophe Rodriguez, Slim Fourati, Vanessa Demontant, Guillaume Gricourt, Melissa N'Debi, Alexandre Soulier, Elisabeth Trawinski, Jean-Michel Pawlotsky |
| EPI_ISL_3475816, EPI_ISL_3475817 | SELAS MEDILYS | Department of Virology, Henri Mondor University Hospital, Assistance Publique Hôpitaux de Paris, Université Paris-Est Créteil, INSERM U955 | Christophe Rodriguez, Slim Fourati, Vanessa Demontant, Guillaume Gricourt, Melissa N'Debi, Alexandre Soulier, Elisabeth Trawinski, Jean-Michel Pawlotsky |
| EPI_ISL_3475818 | EXCITE Lab | Andersen lab at Scripps Research | Chip Schooley, Natasha Martin, Cheryl Anderson, Angela Scioscia, Smruthi Karthikeyan, Greg Humphrey, Sawyer Farmer, Abigail Schnapper, Helena Tubb, Tommy Valles + SEARCH |
| EPI_ISL_3475819 | Laboratoire ATOUTBIO JDC | Department of Virology, Henri Mondor University Hospital, Assistance Publique Hôpitaux de Paris, Université Paris-Est Créteil, INSERM U955 | Christophe Rodriguez, Slim Fourati, Vanessa Demontant, Guillaume Gricourt, Melissa N'Debi, Alexandre Soulier, Elisabeth Trawinski, Jean-Michel Pawlotsky |
| EPI_ISL_3475821 | Hôpital Avicenne | Department of Virology, Henri Mondor University Hospital, Assistance Publique Hôpitaux de Paris, Université Paris-Est Créteil, INSERM U955 | Christophe Rodriguez, Slim Fourati, Vanessa Demontant, Guillaume Gricourt, Melissa N'Debi, Alexandre Soulier, Elisabeth Trawinski, Jean-Michel Pawlotsky |
| EPI_ISL_3475822 | CH. CLEMENCEAU | Department of Virology, Henri Mondor University Hospital, Assistance Publique Hôpitaux de Paris, Université Paris-Est Créteil, INSERM U955 | Christophe Rodriguez, Slim Fourati, Vanessa Demontant, Guillaume Gricourt, Melissa N'Debi, Alexandre Soulier, Elisabeth Trawinski, Jean-Michel Pawlotsky |
| EPI_ISL_3475824 | CERBALLIANCE SAINT MARTIN | Department of Virology, Henri Mondor University Hospital, Assistance Publique Hôpitaux de Paris, Université Paris-Est Créteil, INSERM U955 | Christophe Rodriguez, Slim Fourati, Vanessa Demontant, Guillaume Gricourt, Melissa N'Debi, Alexandre Soulier, Elisabeth Trawinski, Jean-Michel Pawlotsky |
| EPI_ISL_3475826 | CHU DE DIJON | Department of Virology, Henri Mondor University Hospital, Assistance Publique Hôpitaux de Paris, Université Paris-Est Créteil, INSERM U955 | Christophe Rodriguez, Slim Fourati, Vanessa Demontant, Guillaume Gricourt, Melissa N'Debi, Alexandre Soulier, Elisabeth Trawinski, Jean-Michel Pawlotsky |
| EPI_ISL_3475827 | CH. CLEMENCEAU | Department of Virology, Henri Mondor University Hospital, Assistance Publique Hôpitaux de Paris, Université Paris-Est Créteil, INSERM U955 | Christophe Rodriguez, Slim Fourati, Vanessa Demontant, Guillaume Gricourt, Melissa N'Debi, Alexandre Soulier, Elisabeth Trawinski, Jean-Michel Pawlotsky |
| EPI_ISL_3475829 | CERBALLIANCE SAINT MARTIN | Department of Virology, Henri Mondor University Hospital, Assistance Publique Hôpitaux de Paris, Université Paris-Est Créteil, INSERM U955 | Christophe Rodriguez, Slim Fourati, Vanessa Demontant, Guillaume Gricourt, Melissa N'Debi, Alexandre Soulier, Elisabeth Trawinski, Jean-Michel Pawlotsky |
| EPI_ISL_3475831 | CHU DE DIJON | Department of Virology, Henri Mondor University Hospital, Assistance Publique Hôpitaux de Paris, Université Paris-Est Créteil, INSERM U955 | Christophe Rodriguez, Slim Fourati, Vanessa Demontant, Guillaume Gricourt, Melissa N'Debi, Alexandre Soulier, Elisabeth Trawinski, Jean-Michel Pawlotsky |
| EPI_ISL_3475833 | HOPITAL EMILE ROUX | Department of Virology, Henri Mondor University Hospital, Assistance Publique Hôpitaux de Paris, Université Paris-Est Créteil, INSERM U955 | Christophe Rodriguez, Slim Fourati, Vanessa Demontant, Guillaume Gricourt, Melissa N'Debi, Alexandre Soulier, Elisabeth Trawinski, Jean-Michel Pawlotsky |
| EPI_ISL_3475834 | CH. CLEMENCEAU | Department of Virology, Henri Mondor University Hospital, Assistance Publique Hôpitaux de Paris, Université Paris-Est Créteil, INSERM U955 | Christophe Rodriguez, Slim Fourati, Vanessa Demontant, Guillaume Gricourt, Melissa N'Debi, Alexandre Soulier, Elisabeth Trawinski, Jean-Michel Pawlotsky |
| EPI_ISL_3475836 | Biogroup Bio Lam-LCD Saint-Denis | Department of Virology, Henri Mondor University Hospital, Assistance Publique Hôpitaux de Paris, Université Paris-Est Créteil, INSERM U955 | Christophe Rodriguez, Slim Fourati, Vanessa Demontant, Guillaume Gricourt, Melissa N'Debi, Alexandre Soulier, Elisabeth Trawinski, Jean-Michel Pawlotsky |
| EPI_ISL_3475838 | CH. CLEMENCEAU | Department of Virology, Henri Mondor University Hospital, Assistance Publique Hôpitaux de Paris, Université Paris-Est Créteil, INSERM U955 | Christophe Rodriguez, Slim Fourati, Vanessa Demontant, Guillaume Gricourt, Melissa N'Debi, Alexandre Soulier, Elisabeth Trawinski, Jean-Michel Pawlotsky |
| EPI_ISL_3475840 | SELAS MEDILYS | Department of Virology, Henri Mondor University Hospital, Assistance Publique Hôpitaux de Paris, Université Paris-Est Créteil, INSERM U955 | Christophe Rodriguez, Slim Fourati, Vanessa Demontant, Guillaume Gricourt, Melissa N'Debi, Alexandre Soulier, Elisabeth Trawinski, Jean-Michel Pawlotsky |
| EPI_ISL_3475841 | BROUSSAIS | Department of Virology, Henri Mondor University Hospital, Assistance Publique Hôpitaux de Paris, Université Paris-Est Créteil, INSERM U955 | Christophe Rodriguez, Slim Fourati, Vanessa Demontant, Guillaume Gricourt, Melissa N'Debi, Alexandre Soulier, Elisabeth Trawinski, Jean-Michel Pawlotsky |
| EPI_ISL_3475843, EPI_ISL_3475845 | CH. CLEMENCEAU | Department of Virology, Henri Mondor University Hospital, Assistance Publique Hôpitaux de Paris, Université Paris-Est Créteil, INSERM U955 | Christophe Rodriguez, Slim Fourati, Vanessa Demontant, Guillaume Gricourt, Melissa N'Debi, Alexandre Soulier, Elisabeth Trawinski, Jean-Michel Pawlotsky |
| EPI_ISL_3475848 | Laboratoire ATOUTBIO JDC | Department of Virology, Henri Mondor University Hospital, Assistance Publique Hôpitaux de Paris, Université Paris-Est Créteil, INSERM U955 | Christophe Rodriguez, Slim Fourati, Vanessa Demontant, Guillaume Gricourt, Melissa N'Debi, Alexandre Soulier, Elisabeth Trawinski, Jean-Michel Pawlotsky |
| EPI_ISL_3475850 | Biogroup Bio Lam-LCD Saint-Denis | Department of Virology, Henri Mondor University Hospital, Assistance Publique Hôpitaux de Paris, Université Paris-Est Créteil, INSERM U955 | Christophe Rodriguez, Slim Fourati, Vanessa Demontant, Guillaume Gricourt, Melissa N'Debi, Alexandre Soulier, Elisabeth Trawinski, Jean-Michel Pawlotsky |
| EPI_ISL_3475852, EPI_ISL_3475853 | SELAS MEDILYS | Department of Virology, Henri Mondor University Hospital, Assistance Publique Hôpitaux de Paris, Université Paris-Est Créteil, INSERM U955 | Christophe Rodriguez, Slim Fourati, Vanessa Demontant, Guillaume Gricourt, Melissa N'Debi, Alexandre Soulier, Elisabeth Trawinski, Jean-Michel Pawlotsky |
| EPI_ISL_3475855 | Biogroup Bio Lam-LCD Saint-Denis | Department of Virology, Henri Mondor University Hospital, Assistance Publique Hôpitaux de Paris, Université Paris-Est Créteil, INSERM U955 | Christophe Rodriguez, Slim Fourati, Vanessa Demontant, Guillaume Gricourt, Melissa N'Debi, Alexandre Soulier, Elisabeth Trawinski, Jean-Michel Pawlotsky |
| EPI_ISL_3475856 | SELAS MEDILYS | Department of Virology, Henri Mondor University Hospital, Assistance Publique Hôpitaux de Paris, Université Paris-Est Créteil, INSERM U955 | Christophe Rodriguez, Slim Fourati, Vanessa Demontant, Guillaume Gricourt, Melissa N'Debi, Alexandre Soulier, Elisabeth Trawinski, Jean-Michel Pawlotsky |
| EPI_ISL_3475858 | Laboratoire Coste | Department of Virology, Henri Mondor University Hospital, Assistance Publique Hôpitaux de Paris, Université Paris-Est Créteil, INSERM U955 | Christophe Rodriguez, Slim Fourati, Vanessa Demontant, Guillaume Gricourt, Melissa N'Debi, Alexandre Soulier, Elisabeth Trawinski, Jean-Michel Pawlotsky |
| EPI_ISL_3475860 | LBM du Val d'Orne | Department of Virology, Henri Mondor University Hospital, Assistance Publique Hôpitaux de Paris, Université Paris-Est Créteil, | Christophe Rodriguez, Slim Fourati, Vanessa Demontant, Guillaume Gricourt, Melissa N'Debi, Alexandre Soulier, Elisabeth Trawinski, Jean-Michel Pawlotsky |

| | | | |
|---|---|---|---|
| | | INSERM U955 | |
| EPI_ISL_3475861, EPI_ISL_3475863, EPI_ISL_3475864 | Biogroup Bio Lam-LCD Saint-Denis | Department of Virology, Henri Mondor University Hospital, Assistance Publique Hôpitaux de Paris, Université Paris-Est Créteil, INSERM U955 | Christophe Rodriguez, Slim Fourati, Vanessa Demontant, Guillaume Gricourt, Melissa N'Debi, Alexandre Soulier, Elisabeth Trawinski, Jean-Michel Pawlotsky |
| EPI_ISL_3475866 | CHU DE DIJON | Department of Virology, Henri Mondor University Hospital, Assistance Publique Hôpitaux de Paris, Université Paris-Est Créteil, INSERM U955 | Christophe Rodriguez, Slim Fourati, Vanessa Demontant, Guillaume Gricourt, Melissa N'Debi, Alexandre Soulier, Elisabeth Trawinski, Jean-Michel Pawlotsky |
| EPI_ISL_3475868 | CH. CLEMENCEAU | Department of Virology, Henri Mondor University Hospital, Assistance Publique Hôpitaux de Paris, Université Paris-Est Créteil, INSERM U955 | Christophe Rodriguez, Slim Fourati, Vanessa Demontant, Guillaume Gricourt, Melissa N'Debi, Alexandre Soulier, Elisabeth Trawinski, Jean-Michel Pawlotsky |
| EPI_ISL_3475869 | CERBALLIANCE SAINT MARTIN | Department of Virology, Henri Mondor University Hospital, Assistance Publique Hôpitaux de Paris, Université Paris-Est Créteil, INSERM U955 | Christophe Rodriguez, Slim Fourati, Vanessa Demontant, Guillaume Gricourt, Melissa N'Debi, Alexandre Soulier, Elisabeth Trawinski, Jean-Michel Pawlotsky |
| EPI_ISL_3475870 | EXCITE Lab | Andersen lab at Scripps Research | Chip Schooley, Natasha Martin, Cheryl Anderson, Angela Scioscia, Smruthi Karthikeyan, Greg Humphrey, Sawyer Farmer, Abigail Schnapper, Helena Tubb, Tommy Valles + SEARCH |
| EPI_ISL_3475871 | CH. E. MULLER | Department of Virology, Henri Mondor University Hospital, Assistance Publique Hôpitaux de Paris, Université Paris-Est Créteil, INSERM U955 | Christophe Rodriguez, Slim Fourati, Vanessa Demontant, Guillaume Gricourt, Melissa N'Debi, Alexandre Soulier, Elisabeth Trawinski, Jean-Michel Pawlotsky |
| EPI_ISL_3475872 | Hôpital Avicenne | Department of Virology, Henri Mondor University Hospital, Assistance Publique Hôpitaux de Paris, Université Paris-Est Créteil, INSERM U955 | Christophe Rodriguez, Slim Fourati, Vanessa Demontant, Guillaume Gricourt, Melissa N'Debi, Alexandre Soulier, Elisabeth Trawinski, Jean-Michel Pawlotsky |
| EPI_ISL_3475873 | EXCITE Lab | Andersen lab at Scripps Research | Smruthi Karthikeyan, Greg Humphrey, Sawyer Farmer, Helena Tubb, Tommy Valles, Caitlin Tribelhorn, Rebecca Tsai + SEARCH |
| EPI_ISL_3475874 | CHUV | Laboratory of genomics and metagenomics | Trestan Pillonel, Damien Jacot, Sébastien Aeby, Gilbert Greub, Claire Bertelli |
| EPI_ISL_3475875 | Biogroup Bio Lam-LCD Saint-Denis | Department of Virology, Henri Mondor University Hospital, Assistance Publique Hôpitaux de Paris, Université Paris-Est Créteil, INSERM U955 | Christophe Rodriguez, Slim Fourati, Vanessa Demontant, Guillaume Gricourt, Melissa N'Debi, Alexandre Soulier, Elisabeth Trawinski, Jean-Michel Pawlotsky |
| EPI_ISL_3475876 | VIDYMED LAUSANNE | Laboratory of genomics and metagenomics | Trestan Pillonel, Damien Jacot, Sébastien Aeby, Gilbert Greub, Claire Bertelli |
| EPI_ISL_3475877 | Hôpital Avicenne | Department of Virology, Henri Mondor University Hospital, Assistance Publique Hôpitaux de Paris, Université Paris-Est Créteil, INSERM U955 | Christophe Rodriguez, Slim Fourati, Vanessa Demontant, Guillaume Gricourt, Melissa N'Debi, Alexandre Soulier, Elisabeth Trawinski, Jean-Michel Pawlotsky |
| EPI_ISL_3475878 | CHUV | Laboratory of genomics and metagenomics | Trestan Pillonel, Damien Jacot, Sébastien Aeby, Gilbert Greub, Claire Bertelli |
| EPI_ISL_3475879 | SELAS MEDILYS | Department of Virology, Henri Mondor University Hospital, Assistance Publique Hôpitaux de Paris, Université Paris-Est Créteil, INSERM U955 | Christophe Rodriguez, Slim Fourati, Vanessa Demontant, Guillaume Gricourt, Melissa N'Debi, Alexandre Soulier, Elisabeth Trawinski, Jean-Michel Pawlotsky |
| EPI_ISL_3475880 | VIDYMED LAUSANNE | Laboratory of genomics and metagenomics | Trestan Pillonel, Damien Jacot, Sébastien Aeby, Gilbert Greub, Claire Bertelli |
| EPI_ISL_3475881 | CHUV | Laboratory of genomics and metagenomics | Trestan Pillonel, Damien Jacot, Sébastien Aeby, Gilbert Greub, Claire Bertelli |
| EPI_ISL_3475882 | SELAS MEDILYS | Department of Virology, Henri Mondor University Hospital, Assistance Publique Hôpitaux de Paris, Université Paris-Est Créteil, INSERM U955 | Christophe Rodriguez, Slim Fourati, Vanessa Demontant, Guillaume Gricourt, Melissa N'Debi, Alexandre Soulier, Elisabeth Trawinski, Jean-Michel Pawlotsky |
| EPI_ISL_3475883 | CHUV | Laboratory of genomics and metagenomics | Trestan Pillonel, Damien Jacot, Sébastien Aeby, Gilbert Greub, Claire Bertelli |
| EPI_ISL_3475884 | CH. BEGIN | Department of Virology, Henri Mondor University Hospital, Assistance Publique Hôpitaux de Paris, Université Paris-Est Créteil, INSERM U955 | Christophe Rodriguez, Slim Fourati, Vanessa Demontant, Guillaume Gricourt, Melissa N'Debi, Alexandre Soulier, Elisabeth Trawinski, Jean-Michel Pawlotsky |
| EPI_ISL_3475885 | CHUV | Laboratory of genomics and metagenomics | Trestan Pillonel, Damien Jacot, Sébastien Aeby, Gilbert Greub, Claire Bertelli |
| EPI_ISL_3475886 | CH. CLEMENCEAU | Department of Virology, Henri Mondor University Hospital, Assistance Publique Hôpitaux de Paris, Université Paris-Est Créteil, INSERM U955 | Christophe Rodriguez, Slim Fourati, Vanessa Demontant, Guillaume Gricourt, Melissa N'Debi, Alexandre Soulier, Elisabeth Trawinski, Jean-Michel Pawlotsky |
| EPI_ISL_3475887 | CHUV | Laboratory of genomics and metagenomics | Trestan Pillonel, Damien Jacot, Sébastien Aeby, Gilbert Greub, Claire Bertelli |
| EPI_ISL_3475888 | Biogroup Bio Lam-LCD Saint-Denis | Department of Virology, Henri Mondor University Hospital, Assistance Publique Hôpitaux de Paris, Université Paris-Est Créteil, INSERM U955 | Christophe Rodriguez, Slim Fourati, Vanessa Demontant, Guillaume Gricourt, Melissa N'Debi, Alexandre Soulier, Elisabeth Trawinski, Jean-Michel Pawlotsky |
| EPI_ISL_3475889, EPI_ISL_3475890 | CHUV | Laboratory of genomics and metagenomics | Trestan Pillonel, Damien Jacot, Sébastien Aeby, Gilbert Greub, Claire Bertelli |
| EPI_ISL_3475891 | CH. CLEMENCEAU | Department of Virology, Henri Mondor University Hospital, Assistance Publique Hôpitaux de Paris, Université Paris-Est Créteil, INSERM U955 | Christophe Rodriguez, Slim Fourati, Vanessa Demontant, Guillaume Gricourt, Melissa N'Debi, Alexandre Soulier, Elisabeth Trawinski, Jean-Michel Pawlotsky |
| EPI_ISL_3475892 | VIDYMED LAUSANNE | Laboratory of genomics and metagenomics | Trestan Pillonel, Damien Jacot, Sébastien Aeby, Gilbert Greub, Claire Bertelli |
| EPI_ISL_3475893 | Hôpital Armand-Trousseau | Department of Virology, Henri Mondor University Hospital, Assistance Publique Hôpitaux de Paris, Université Paris-Est Créteil, INSERM U955 | Christophe Rodriguez, Slim Fourati, Vanessa Demontant, Guillaume Gricourt, Melissa N'Debi, Alexandre Soulier, Elisabeth Trawinski, Jean-Michel Pawlotsky |
| EPI_ISL_3475894, EPI_ISL_3475895 | CHUV | Laboratory of genomics and metagenomics | Trestan Pillonel, Damien Jacot, Sébastien Aeby, Gilbert Greub, Claire Bertelli |
| EPI_ISL_3475896 | CH. CLEMENCEAU | Department of Virology, Henri Mondor University Hospital, Assistance Publique Hôpitaux de Paris, Université Paris-Est Créteil, INSERM U955 | Christophe Rodriguez, Slim Fourati, Vanessa Demontant, Guillaume Gricourt, Melissa N'Debi, Alexandre Soulier, Elisabeth Trawinski, Jean-Michel Pawlotsky |
| EPI_ISL_3475897 | VIDYMED LAUSANNE | Laboratory of genomics and metagenomics | Trestan Pillonel, Damien Jacot, Sébastien Aeby, Gilbert Greub, Claire Bertelli |
| EPI_ISL_3475898 | Biogroup Bio Lam-LCD Saint-Denis | Department of Virology, Henri Mondor University Hospital, Assistance Publique Hôpitaux de Paris, Université Paris-Est Créteil, INSERM U955 | Christophe Rodriguez, Slim Fourati, Vanessa Demontant, Guillaume Gricourt, Melissa N'Debi, Alexandre Soulier, Elisabeth Trawinski, Jean-Michel Pawlotsky |
| EPI_ISL_3475899 | VIDYMED LAUSANNE | Laboratory of genomics and metagenomics | Trestan Pillonel, Damien Jacot, Sébastien Aeby, Gilbert Greub, Claire Bertelli |
| EPI_ISL_3475900 | Biogroup Bio Lam-LCD Saint-Denis | Department of Virology, Henri Mondor University Hospital, Assistance Publique Hôpitaux de Paris, Université Paris-Est Créteil, INSERM U955 | Christophe Rodriguez, Slim Fourati, Vanessa Demontant, Guillaume Gricourt, Melissa N'Debi, Alexandre Soulier, Elisabeth Trawinski, Jean-Michel Pawlotsky |
| EPI_ISL_3475901 | VIDYMED LAUSANNE | Laboratory of genomics and metagenomics | Trestan Pillonel, Damien Jacot, Sébastien Aeby, Gilbert Greub, Claire Bertelli |
| EPI_ISL_3475902 | CERBALLIANCE SAINT MARTIN | Department of Virology, Henri Mondor University Hospital, Assistance Publique Hôpitaux de Paris, Université Paris-Est Créteil, INSERM U955 | Christophe Rodriguez, Slim Fourati, Vanessa Demontant, Guillaume Gricourt, Melissa N'Debi, Alexandre Soulier, Elisabeth Trawinski, Jean-Michel Pawlotsky |
| EPI_ISL_3475903, EPI_ISL_3475904 | VIDYMED LAUSANNE | Laboratory of genomics and metagenomics | Trestan Pillonel, Damien Jacot, Sébastien Aeby, Gilbert Greub, Claire Bertelli |
| EPI_ISL_3475905 | Biogroup Bio Lam-LCD Saint-Denis | Department of Virology, Henri Mondor University Hospital, Assistance Publique Hôpitaux de Paris, Université Paris-Est Créteil, INSERM U955 | Christophe Rodriguez, Slim Fourati, Vanessa Demontant, Guillaume Gricourt, Melissa N'Debi, Alexandre Soulier, Elisabeth Trawinski, Jean-Michel Pawlotsky |
| EPI_ISL_3475906 | CHUV | Laboratory of genomics and metagenomics | Trestan Pillonel, Damien Jacot, Sébastien Aeby, Gilbert Greub, Claire Bertelli |
| EPI_ISL_3475907 | GH A.CHENEVIER-H.MONDOR | Department of Virology, Henri Mondor University Hospital, Assistance Publique Hôpitaux de Paris, Université Paris-Est Créteil, INSERM U955 | Christophe Rodriguez, Slim Fourati, Vanessa Demontant, Guillaume Gricourt, Melissa N'Debi, Alexandre Soulier, Elisabeth Trawinski, Jean-Michel Pawlotsky |
| EPI_ISL_3475908 | CHUV | Laboratory of genomics and metagenomics | Trestan Pillonel, Damien Jacot, Sébastien Aeby, Gilbert Greub, Claire Bertelli |
| EPI_ISL_3475909 | Biogroup Bio Lam-LCD Saint-Denis | Department of Virology, Henri Mondor University Hospital, Assistance Publique Hôpitaux de Paris, Université Paris-Est Créteil, INSERM U955 | Christophe Rodriguez, Slim Fourati, Vanessa Demontant, Guillaume Gricourt, Melissa N'Debi, Alexandre Soulier, Elisabeth Trawinski, Jean-Michel Pawlotsky |
| EPI_ISL_3475910 | VIDYMED LAUSANNE | Laboratory of genomics and metagenomics | Trestan Pillonel, Damien Jacot, Sébastien Aeby, Gilbert Greub, Claire Bertelli |
| EPI_ISL_3475911 | EHC MORGES | Laboratory of genomics and metagenomics | Trestan Pillonel, Damien Jacot, Sébastien Aeby, Gilbert Greub, Claire Bertelli |

| | | | |
|---|---|---|---|
| EPI_ISL_3475912 | Laboratoire ATOUTBIO JDC | Department of Virology, Henri Mondor University Hospital, Assistance Publique Hôpitaux de Paris, Université Paris-Est Créteil, INSERM U955 | Christophe Rodriguez, Slim Fourati, Vanessa Demontant, Guillaume Gricourt, Melissa N'Debi, Alexandre Soulier, Elisabeth Trawinski, Jean-Michel Pawlotsky |
| EPI_ISL_3475913 | CHUV | Laboratory of genomics and metagenomics | Trestan Pillonel, Damien Jacot, Sébastien Aeby, Gilbert Greub, Claire Bertelli |
| EPI_ISL_3475914 | Hôpital Avicenne | Department of Virology, Henri Mondor University Hospital, Assistance Publique Hôpitaux de Paris, Université Paris-Est Créteil, INSERM U955 | Christophe Rodriguez, Slim Fourati, Vanessa Demontant, Guillaume Gricourt, Melissa N'Debi, Alexandre Soulier, Elisabeth Trawinski, Jean-Michel Pawlotsky |
| EPI_ISL_3475915, EPI_ISL_3475916 | CHUV | Laboratory of genomics and metagenomics | Trestan Pillonel, Damien Jacot, Sébastien Aeby, Gilbert Greub, Claire Bertelli |
| EPI_ISL_3475917 | Hôpital Avicenne | Department of Virology, Henri Mondor University Hospital, Assistance Publique Hôpitaux de Paris, Université Paris-Est Créteil, INSERM U955 | Christophe Rodriguez, Slim Fourati, Vanessa Demontant, Guillaume Gricourt, Melissa N'Debi, Alexandre Soulier, Elisabeth Trawinski, Jean-Michel Pawlotsky |
| EPI_ISL_3475918 | CHUV | Laboratory of genomics and metagenomics | Trestan Pillonel, Damien Jacot, Sébastien Aeby, Gilbert Greub, Claire Bertelli |
| EPI_ISL_3475919 | Biogroup Bio Lam-LCD Saint-Denis | Department of Virology, Henri Mondor University Hospital, Assistance Publique Hôpitaux de Paris, Université Paris-Est Créteil, INSERM U955 | Christophe Rodriguez, Slim Fourati, Vanessa Demontant, Guillaume Gricourt, Melissa N'Debi, Alexandre Soulier, Elisabeth Trawinski, Jean-Michel Pawlotsky |
| EPI_ISL_3475920 | EHC MORGES | Laboratory of genomics and metagenomics | Trestan Pillonel, Damien Jacot, Sébastien Aeby, Gilbert Greub, Claire Bertelli |
| EPI_ISL_3475921 | SELAS MEDILYS | Department of Virology, Henri Mondor University Hospital, Assistance Publique Hôpitaux de Paris, Université Paris-Est Créteil, INSERM U955 | Christophe Rodriguez, Slim Fourati, Vanessa Demontant, Guillaume Gricourt, Melissa N'Debi, Alexandre Soulier, Elisabeth Trawinski, Jean-Michel Pawlotsky |
| EPI_ISL_3475922, EPI_ISL_3475923 | CHUV | Laboratory of genomics and metagenomics | Trestan Pillonel, Damien Jacot, Sébastien Aeby, Gilbert Greub, Claire Bertelli |
| EPI_ISL_3475924 | CHU DE DIJON | Department of Virology, Henri Mondor University Hospital, Assistance Publique Hôpitaux de Paris, Université Paris-Est Créteil, INSERM U955 | Christophe Rodriguez, Slim Fourati, Vanessa Demontant, Guillaume Gricourt, Melissa N'Debi, Alexandre Soulier, Elisabeth Trawinski, Jean-Michel Pawlotsky |
| EPI_ISL_3475925 | VIDYMED LAUSANNE | Laboratory of genomics and metagenomics | Trestan Pillonel, Damien Jacot, Sébastien Aeby, Gilbert Greub, Claire Bertelli |
| EPI_ISL_3475926 | CH. CLEMENCEAU | Department of Virology, Henri Mondor University Hospital, Assistance Publique Hôpitaux de Paris, Université Paris-Est Créteil, INSERM U955 | Christophe Rodriguez, Slim Fourati, Vanessa Demontant, Guillaume Gricourt, Melissa N'Debi, Alexandre Soulier, Elisabeth Trawinski, Jean-Michel Pawlotsky |
| EPI_ISL_3475927 | VIDYMED LAUSANNE | Laboratory of genomics and metagenomics | Trestan Pillonel, Damien Jacot, Sébastien Aeby, Gilbert Greub, Claire Bertelli |
| EPI_ISL_3475928 | Laboratoire ATOUTBIO JDC | Department of Virology, Henri Mondor University Hospital, Assistance Publique Hôpitaux de Paris, Université Paris-Est Créteil, INSERM U955 | Christophe Rodriguez, Slim Fourati, Vanessa Demontant, Guillaume Gricourt, Melissa N'Debi, Alexandre Soulier, Elisabeth Trawinski, Jean-Michel Pawlotsky |
| EPI_ISL_3475929, EPI_ISL_3475930 | CHUV | Laboratory of genomics and metagenomics | Trestan Pillonel, Damien Jacot, Sébastien Aeby, Gilbert Greub, Claire Bertelli |
| EPI_ISL_3475931 | CHU DE DIJON | Department of Virology, Henri Mondor University Hospital, Assistance Publique Hôpitaux de Paris, Université Paris-Est Créteil, INSERM U955 | Christophe Rodriguez, Slim Fourati, Vanessa Demontant, Guillaume Gricourt, Melissa N'Debi, Alexandre Soulier, Elisabeth Trawinski, Jean-Michel Pawlotsky |
| EPI_ISL_3475932 | CHUV | Laboratory of genomics and metagenomics | Trestan Pillonel, Damien Jacot, Sébastien Aeby, Gilbert Greub, Claire Bertelli |
| EPI_ISL_3475933 | CHI VILLENEUVE ST GEORGES | Department of Virology, Henri Mondor University Hospital, Assistance Publique Hôpitaux de Paris, Université Paris-Est Créteil, INSERM U955 | Christophe Rodriguez, Slim Fourati, Vanessa Demontant, Guillaume Gricourt, Melissa N'Debi, Alexandre Soulier, Elisabeth Trawinski, Jean-Michel Pawlotsky |
| EPI_ISL_3475934 | VIDYMED LAUSANNE | Laboratory of genomics and metagenomics | Trestan Pillonel, Damien Jacot, Sébastien Aeby, Gilbert Greub, Claire Bertelli |
| EPI_ISL_3475935 | CH. CLEMENCEAU | Department of Virology, Henri Mondor University Hospital, Assistance Publique Hôpitaux de Paris, Université Paris-Est Créteil, INSERM U955 | Christophe Rodriguez, Slim Fourati, Vanessa Demontant, Guillaume Gricourt, Melissa N'Debi, Alexandre Soulier, Elisabeth Trawinski, Jean-Michel Pawlotsky |
| EPI_ISL_3475936 | VIDYMED LAUSANNE | Laboratory of genomics and metagenomics | Trestan Pillonel, Damien Jacot, Sébastien Aeby, Gilbert Greub, Claire Bertelli |
| EPI_ISL_3475937 | CHUV | Laboratory of genomics and metagenomics | Trestan Pillonel, Damien Jacot, Sébastien Aeby, Gilbert Greub, Claire Bertelli |
| EPI_ISL_3475938 | CH. E. MULLER | Department of Virology, Henri Mondor University Hospital, Assistance Publique Hôpitaux de Paris, Université Paris-Est Créteil, INSERM U955 | Christophe Rodriguez, Slim Fourati, Vanessa Demontant, Guillaume Gricourt, Melissa N'Debi, Alexandre Soulier, Elisabeth Trawinski, Jean-Michel Pawlotsky |
| EPI_ISL_3475939 | CHUV | Laboratory of genomics and metagenomics | Trestan Pillonel, Damien Jacot, Sébastien Aeby, Gilbert Greub, Claire Bertelli |
| EPI_ISL_3475940 | SELAS MEDILYS | Department of Virology, Henri Mondor University Hospital, Assistance Publique Hôpitaux de Paris, Université Paris-Est Créteil, INSERM U955 | Christophe Rodriguez, Slim Fourati, Vanessa Demontant, Guillaume Gricourt, Melissa N'Debi, Alexandre Soulier, Elisabeth Trawinski, Jean-Michel Pawlotsky |
| EPI_ISL_3475941 | VIDYMED LAUSANNE | Laboratory of genomics and metagenomics | Trestan Pillonel, Damien Jacot, Sébastien Aeby, Gilbert Greub, Claire Bertelli |
| EPI_ISL_3475942 | Hôpital Avicenne | Department of Virology, Henri Mondor University Hospital, Assistance Publique Hôpitaux de Paris, Université Paris-Est Créteil, INSERM U955 | Christophe Rodriguez, Slim Fourati, Vanessa Demontant, Guillaume Gricourt, Melissa N'Debi, Alexandre Soulier, Elisabeth Trawinski, Jean-Michel Pawlotsky |
| EPI_ISL_3475943 | VIDYMED LAUSANNE | Laboratory of genomics and metagenomics | Trestan Pillonel, Damien Jacot, Sébastien Aeby, Gilbert Greub, Claire Bertelli |
| EPI_ISL_3475944 | Laboratoire ATOUTBIO JDC | Department of Virology, Henri Mondor University Hospital, Assistance Publique Hôpitaux de Paris, Université Paris-Est Créteil, INSERM U955 | Christophe Rodriguez, Slim Fourati, Vanessa Demontant, Guillaume Gricourt, Melissa N'Debi, Alexandre Soulier, Elisabeth Trawinski, Jean-Michel Pawlotsky |
| EPI_ISL_3475945, EPI_ISL_3475946 | VIDYMED LAUSANNE | Laboratory of genomics and metagenomics | Trestan Pillonel, Damien Jacot, Sébastien Aeby, Gilbert Greub, Claire Bertelli |
| EPI_ISL_3475947 | Hôpital Avicenne | Department of Virology, Henri Mondor University Hospital, Assistance Publique Hôpitaux de Paris, Université Paris-Est Créteil, INSERM U955 | Christophe Rodriguez, Slim Fourati, Vanessa Demontant, Guillaume Gricourt, Melissa N'Debi, Alexandre Soulier, Elisabeth Trawinski, Jean-Michel Pawlotsky |
| EPI_ISL_3475948 | VIDYMED LAUSANNE | Laboratory of genomics and metagenomics | Trestan Pillonel, Damien Jacot, Sébastien Aeby, Gilbert Greub, Claire Bertelli |
| EPI_ISL_3475949 | Biogroup Bio Lam-LCD Saint-Denis | Department of Virology, Henri Mondor University Hospital, Assistance Publique Hôpitaux de Paris, Université Paris-Est Créteil, INSERM U955 | Christophe Rodriguez, Slim Fourati, Vanessa Demontant, Guillaume Gricourt, Melissa N'Debi, Alexandre Soulier, Elisabeth Trawinski, Jean-Michel Pawlotsky |
| EPI_ISL_3475950 | CHUV | Laboratory of genomics and metagenomics | Trestan Pillonel, Damien Jacot, Sébastien Aeby, Gilbert Greub, Claire Bertelli |
| EPI_ISL_3475951 | CH. CLEMENCEAU | Department of Virology, Henri Mondor University Hospital, Assistance Publique Hôpitaux de Paris, Université Paris-Est Créteil, INSERM U955 | Christophe Rodriguez, Slim Fourati, Vanessa Demontant, Guillaume Gricourt, Melissa N'Debi, Alexandre Soulier, Elisabeth Trawinski, Jean-Michel Pawlotsky |
| EPI_ISL_3475952 | CHUV | Laboratory of genomics and metagenomics | Trestan Pillonel, Damien Jacot, Sébastien Aeby, Gilbert Greub, Claire Bertelli |
| EPI_ISL_3475953 | CHU DE DIJON | Department of Virology, Henri Mondor University Hospital, Assistance Publique Hôpitaux de Paris, Université Paris-Est Créteil, INSERM U955 | Christophe Rodriguez, Slim Fourati, Vanessa Demontant, Guillaume Gricourt, Melissa N'Debi, Alexandre Soulier, Elisabeth Trawinski, Jean-Michel Pawlotsky |
| EPI_ISL_3475954, EPI_ISL_3475955 | VIDYMED LAUSANNE | Laboratory of genomics and metagenomics | Trestan Pillonel, Damien Jacot, Sébastien Aeby, Gilbert Greub, Claire Bertelli |
| EPI_ISL_3475956 | Biogroup Bio Lam-LCD Saint-Denis | Department of Virology, Henri Mondor University Hospital, Assistance Publique Hôpitaux de Paris, Université Paris-Est Créteil, INSERM U955 | Christophe Rodriguez, Slim Fourati, Vanessa Demontant, Guillaume Gricourt, Melissa N'Debi, Alexandre Soulier, Elisabeth Trawinski, Jean-Michel Pawlotsky |
| EPI_ISL_3475957 | CHUV | Laboratory of genomics and metagenomics | Trestan Pillonel, Damien Jacot, Sébastien Aeby, Gilbert Greub, Claire Bertelli |
| EPI_ISL_3475958 | Laboratoire ATOUTBIO JDC | Department of Virology, Henri Mondor University Hospital, Assistance Publique Hôpitaux de Paris, Université Paris-Est Créteil, INSERM U955 | Christophe Rodriguez, Slim Fourati, Vanessa Demontant, Guillaume Gricourt, Melissa N'Debi, Alexandre Soulier, Elisabeth Trawinski, Jean-Michel Pawlotsky |
| EPI_ISL_3475959 | CHUV | Laboratory of genomics and metagenomics | Trestan Pillonel, Damien Jacot, Sébastien Aeby, Gilbert Greub, Claire Bertelli |
| EPI_ISL_3475960 | Hôpital Avicenne | Department of Virology, Henri Mondor University Hospital, | Christophe Rodriguez, Slim Fourati, Vanessa Demontant, Guillaume Gricourt, Melissa N'Debi, Alexandre Soulier, Elisabeth Trawinski, Jean-Michel Pawlotsky |

| | | | |
|---|---|---|---|
| | | Assistance Publique Hôpitaux de Paris, Université Paris-Est Créteil, INSERM U955 | |
| EPI_ISL_3475961, EPI_ISL_3475962, EPI_ISL_3475963 | VIDYMED LAUSANNE | Laboratory of genomics and metagenomics | Trestan Pillonel, Damien Jacot, Sébastien Aeby, Gilbert Greub, Claire Bertelli |
| EPI_ISL_3475964 | CHUV | Laboratory of genomics and metagenomics | Trestan Pillonel, Damien Jacot, Sébastien Aeby, Gilbert Greub, Claire Bertelli |
| EPI_ISL_3475965 | Biogroup Bio Lam-LCD Saint-Denis | Department of Virology, Henri Mondor University Hospital, Assistance Publique Hôpitaux de Paris, Université Paris-Est Créteil, INSERM U955 | Christophe Rodriguez, Slim Fourati, Vanessa Demontant, Guillaume Gricourt, Melissa N'Debi, Alexandre Soulier, Elisabeth Trawinski, Jean-Michel Pawlotsky |
| EPI_ISL_3475966, EPI_ISL_3475967, EPI_ISL_3475968 | VIDYMED LAUSANNE | Laboratory of genomics and metagenomics | Trestan Pillonel, Damien Jacot, Sébastien Aeby, Gilbert Greub, Claire Bertelli |
| EPI_ISL_3475969 | Biogroup Bio Lam-LCD Saint-Denis | Department of Virology, Henri Mondor University Hospital, Assistance Publique Hôpitaux de Paris, Université Paris-Est Créteil, INSERM U955 | Christophe Rodriguez, Slim Fourati, Vanessa Demontant, Guillaume Gricourt, Melissa N'Debi, Alexandre Soulier, Elisabeth Trawinski, Jean-Michel Pawlotsky |
| EPI_ISL_3475970, EPI_ISL_3475971, EPI_ISL_3475972 | VIDYMED LAUSANNE | Laboratory of genomics and metagenomics | Trestan Pillonel, Damien Jacot, Sébastien Aeby, Gilbert Greub, Claire Bertelli |
| EPI_ISL_3475973 | Biogroup Bio Lam-LCD Saint-Denis | Department of Virology, Henri Mondor University Hospital, Assistance Publique Hôpitaux de Paris, Université Paris-Est Créteil, INSERM U955 | Christophe Rodriguez, Slim Fourati, Vanessa Demontant, Guillaume Gricourt, Melissa N'Debi, Alexandre Soulier, Elisabeth Trawinski, Jean-Michel Pawlotsky |
| EPI_ISL_3475975 | CHUV | Laboratory of genomics and metagenomics | Trestan Pillonel, Damien Jacot, Sébastien Aeby, Gilbert Greub, Claire Bertelli |
| EPI_ISL_3475976 | VIDYMED LAUSANNE | Laboratory of genomics and metagenomics | Trestan Pillonel, Damien Jacot, Sébastien Aeby, Gilbert Greub, Claire Bertelli |
| EPI_ISL_3475977 | CH. CLEMENCEAU | Department of Virology, Henri Mondor University Hospital, Assistance Publique Hôpitaux de Paris, Université Paris-Est Créteil, INSERM U955 | Christophe Rodriguez, Slim Fourati, Vanessa Demontant, Guillaume Gricourt, Melissa N'Debi, Alexandre Soulier, Elisabeth Trawinski, Jean-Michel Pawlotsky |
| EPI_ISL_3475978, EPI_ISL_3475979, EPI_ISL_3475980 | VIDYMED LAUSANNE | Laboratory of genomics and metagenomics | Trestan Pillonel, Damien Jacot, Sébastien Aeby, Gilbert Greub, Claire Bertelli |
| EPI_ISL_3475981 | Biogroup Bio Lam-LCD Saint-Denis | Department of Virology, Henri Mondor University Hospital, Assistance Publique Hôpitaux de Paris, Université Paris-Est Créteil, INSERM U955 | Christophe Rodriguez, Slim Fourati, Vanessa Demontant, Guillaume Gricourt, Melissa N'Debi, Alexandre Soulier, Elisabeth Trawinski, Jean-Michel Pawlotsky |
| EPI_ISL_3475982 | VIDYMED LAUSANNE | Laboratory of genomics and metagenomics | Trestan Pillonel, Damien Jacot, Sébastien Aeby, Gilbert Greub, Claire Bertelli |
| EPI_ISL_3475983, EPI_ISL_3475984 | CHUV | Laboratory of genomics and metagenomics | Trestan Pillonel, Damien Jacot, Sébastien Aeby, Gilbert Greub, Claire Bertelli |
| EPI_ISL_3475985 | SELAS MEDILYS | Department of Virology, Henri Mondor University Hospital, Assistance Publique Hôpitaux de Paris, Université Paris-Est Créteil, INSERM U955 | Christophe Rodriguez, Slim Fourati, Vanessa Demontant, Guillaume Gricourt, Melissa N'Debi, Alexandre Soulier, Elisabeth Trawinski, Jean-Michel Pawlotsky |
| EPI_ISL_3475986 | CHUV | Laboratory of genomics and metagenomics | Trestan Pillonel, Damien Jacot, Sébastien Aeby, Gilbert Greub, Claire Bertelli |
| EPI_ISL_3475987 | CHU DE DIJON | Department of Virology, Henri Mondor University Hospital, Assistance Publique Hôpitaux de Paris, Université Paris-Est Créteil, INSERM U955 | Christophe Rodriguez, Slim Fourati, Vanessa Demontant, Guillaume Gricourt, Melissa N'Debi, Alexandre Soulier, Elisabeth Trawinski, Jean-Michel Pawlotsky |
| EPI_ISL_3475988 | VIDYMED LAUSANNE | Laboratory of genomics and metagenomics | Trestan Pillonel, Damien Jacot, Sébastien Aeby, Gilbert Greub, Claire Bertelli |
| EPI_ISL_3475989 | Biogroup Bio Lam-LCD Saint-Denis | Department of Virology, Henri Mondor University Hospital, Assistance Publique Hôpitaux de Paris, Université Paris-Est Créteil, INSERM U955 | Christophe Rodriguez, Slim Fourati, Vanessa Demontant, Guillaume Gricourt, Melissa N'Debi, Alexandre Soulier, Elisabeth Trawinski, Jean-Michel Pawlotsky |
| EPI_ISL_3475990 | VIDYMED LAUSANNE | Laboratory of genomics and metagenomics | Trestan Pillonel, Damien Jacot, Sébastien Aeby, Gilbert Greub, Claire Bertelli |
| EPI_ISL_3475991 | Biogroup Bio Lam-LCD Saint-Denis | Department of Virology, Henri Mondor University Hospital, Assistance Publique Hôpitaux de Paris, Université Paris-Est Créteil, INSERM U955 | Christophe Rodriguez, Slim Fourati, Vanessa Demontant, Guillaume Gricourt, Melissa N'Debi, Alexandre Soulier, Elisabeth Trawinski, Jean-Michel Pawlotsky |
| EPI_ISL_3475992 | VIDYMED LAUSANNE | Laboratory of genomics and metagenomics | Trestan Pillonel, Damien Jacot, Sébastien Aeby, Gilbert Greub, Claire Bertelli |
| EPI_ISL_3475994 | CHU DE DIJON | Department of Virology, Henri Mondor University Hospital, Assistance Publique Hôpitaux de Paris, Université Paris-Est Créteil, INSERM U955 | Christophe Rodriguez, Slim Fourati, Vanessa Demontant, Guillaume Gricourt, Melissa N'Debi, Alexandre Soulier, Elisabeth Trawinski, Jean-Michel Pawlotsky |
| EPI_ISL_3475995 | CHUV | Laboratory of genomics and metagenomics | Trestan Pillonel, Damien Jacot, Sébastien Aeby, Gilbert Greub, Claire Bertelli |
| EPI_ISL_3475996 | LBM du Val d'Orne | Department of Virology, Henri Mondor University Hospital, Assistance Publique Hôpitaux de Paris, Université Paris-Est Créteil, INSERM U955 | Christophe Rodriguez, Slim Fourati, Vanessa Demontant, Guillaume Gricourt, Melissa N'Debi, Alexandre Soulier, Elisabeth Trawinski, Jean-Michel Pawlotsky |
| EPI_ISL_3475997 | CHUV | Laboratory of genomics and metagenomics | Trestan Pillonel, Damien Jacot, Sébastien Aeby, Gilbert Greub, Claire Bertelli |
| EPI_ISL_3475998 | SELAS MEDILYS | Department of Virology, Henri Mondor University Hospital, Assistance Publique Hôpitaux de Paris, Université Paris-Est Créteil, INSERM U955 | Christophe Rodriguez, Slim Fourati, Vanessa Demontant, Guillaume Gricourt, Melissa N'Debi, Alexandre Soulier, Elisabeth Trawinski, Jean-Michel Pawlotsky |
| EPI_ISL_3475999 | VIDYMED LAUSANNE | Laboratory of genomics and metagenomics | Trestan Pillonel, Damien Jacot, Sébastien Aeby, Gilbert Greub, Claire Bertelli |
| EPI_ISL_3476000 | CHUV | Laboratory of genomics and metagenomics | Trestan Pillonel, Damien Jacot, Sébastien Aeby, Gilbert Greub, Claire Bertelli |
| EPI_ISL_3476001 | Hôpital Avicenne | Department of Virology, Henri Mondor University Hospital, Assistance Publique Hôpitaux de Paris, Université Paris-Est Créteil, INSERM U955 | Christophe Rodriguez, Slim Fourati, Vanessa Demontant, Guillaume Gricourt, Melissa N'Debi, Alexandre Soulier, Elisabeth Trawinski, Jean-Michel Pawlotsky |
| EPI_ISL_3476002 | GHOL | Laboratory of genomics and metagenomics | Trestan Pillonel, Damien Jacot, Sébastien Aeby, Gilbert Greub, Claire Bertelli |
| EPI_ISL_3476003 | Biogroup Bio Lam-LCD Saint-Denis | Department of Virology, Henri Mondor University Hospital, Assistance Publique Hôpitaux de Paris, Université Paris-Est Créteil, INSERM U955 | Christophe Rodriguez, Slim Fourati, Vanessa Demontant, Guillaume Gricourt, Melissa N'Debi, Alexandre Soulier, Elisabeth Trawinski, Jean-Michel Pawlotsky |
| EPI_ISL_3476004 | VIDYMED LAUSANNE | Laboratory of genomics and metagenomics | Trestan Pillonel, Damien Jacot, Sébastien Aeby, Gilbert Greub, Claire Bertelli |
| EPI_ISL_3476005 | CH. CLEMENCEAU | Department of Virology, Henri Mondor University Hospital, Assistance Publique Hôpitaux de Paris, Université Paris-Est Créteil, INSERM U955 | Christophe Rodriguez, Slim Fourati, Vanessa Demontant, Guillaume Gricourt, Melissa N'Debi, Alexandre Soulier, Elisabeth Trawinski, Jean-Michel Pawlotsky |
| EPI_ISL_3476006, EPI_ISL_3476007 | CHUV | Laboratory of genomics and metagenomics | Trestan Pillonel, Damien Jacot, Sébastien Aeby, Gilbert Greub, Claire Bertelli |
| EPI_ISL_3476008 | Biogroup Bio Lam-LCD Saint-Denis | Department of Virology, Henri Mondor University Hospital, Assistance Publique Hôpitaux de Paris, Université Paris-Est Créteil, INSERM U955 | Christophe Rodriguez, Slim Fourati, Vanessa Demontant, Guillaume Gricourt, Melissa N'Debi, Alexandre Soulier, Elisabeth Trawinski, Jean-Michel Pawlotsky |
| EPI_ISL_3476009 | VIDYMED LAUSANNE | Laboratory of genomics and metagenomics | Trestan Pillonel, Damien Jacot, Sébastien Aeby, Gilbert Greub, Claire Bertelli |
| EPI_ISL_3476010 | SELAS MEDILYS | Department of Virology, Henri Mondor University Hospital, Assistance Publique Hôpitaux de Paris, Université Paris-Est Créteil, INSERM U955 | Christophe Rodriguez, Slim Fourati, Vanessa Demontant, Guillaume Gricourt, Melissa N'Debi, Alexandre Soulier, Elisabeth Trawinski, Jean-Michel Pawlotsky |
| EPI_ISL_3476011 | CHUV | Laboratory of genomics and metagenomics | Trestan Pillonel, Damien Jacot, Sébastien Aeby, Gilbert Greub, Claire Bertelli |
| EPI_ISL_3476012 | SELAS MEDILYS | Department of Virology, Henri Mondor University Hospital, Assistance Publique Hôpitaux de Paris, Université Paris-Est Créteil, INSERM U955 | Christophe Rodriguez, Slim Fourati, Vanessa Demontant, Guillaume Gricourt, Melissa N'Debi, Alexandre Soulier, Elisabeth Trawinski, Jean-Michel Pawlotsky |
| EPI_ISL_3476013 | VIDYMED LAUSANNE | Laboratory of genomics and metagenomics | Trestan Pillonel, Damien Jacot, Sébastien Aeby, Gilbert Greub, Claire Bertelli |
| EPI_ISL_3476014 | CHUV | Laboratory of genomics and metagenomics | Trestan Pillonel, Damien Jacot, Sébastien Aeby, Gilbert Greub, Claire Bertelli |
| EPI_ISL_3476015 | Biogroup Bio Lam-LCD Saint-Denis | Department of Virology, Henri Mondor University Hospital, | Christophe Rodriguez, Slim Fourati, Vanessa Demontant, Guillaume Gricourt, Melissa N'Debi, Alexandre Soulier, Elisabeth Trawinski, Jean-Michel Pawlotsky |

| | | | |
|---|---|---|---|
| | | Assistance Publique Hôpitaux de Paris, Université Paris-Est Créteil, INSERM U955 | |
| EPI_ISL_3476016 | CHUV | Laboratory of genomics and metagenomics | Trestan Pillonel, Damien Jacot, Sébastien Aeby, Gilbert Greub, Claire Bertelli |
| EPI_ISL_3476017 | CERBALLIANCE SAINT MARTIN | Department of Virology, Henri Mondor University Hospital, Assistance Publique Hôpitaux de Paris, Université Paris-Est Créteil, INSERM U955 | Christophe Rodriguez, Slim Fourati, Vanessa Demontant, Guillaume Gricourt, Melissa N'Debi, Alexandre Soulier, Elisabeth Trawinski, Jean-Michel Pawlotsky |
| EPI_ISL_3476018 | CHUV | Laboratory of genomics and metagenomics | Trestan Pillonel, Damien Jacot, Sébastien Aeby, Gilbert Greub, Claire Bertelli |
| EPI_ISL_3476019 | Biogroup Bio Lam-LCD Saint-Denis | Department of Virology, Henri Mondor University Hospital, Assistance Publique Hôpitaux de Paris, Université Paris-Est Créteil, INSERM U955 | Christophe Rodriguez, Slim Fourati, Vanessa Demontant, Guillaume Gricourt, Melissa N'Debi, Alexandre Soulier, Elisabeth Trawinski, Jean-Michel Pawlotsky |
| EPI_ISL_3476020 | VIDYMED LAUSANNE | Laboratory of genomics and metagenomics | Trestan Pillonel, Damien Jacot, Sébastien Aeby, Gilbert Greub, Claire Bertelli |
| EPI_ISL_3476021 | Hôpital Saint-Louis | Department of Virology, Henri Mondor University Hospital, Assistance Publique Hôpitaux de Paris, Université Paris-Est Créteil, INSERM U955 | Christophe Rodriguez, Slim Fourati, Vanessa Demontant, Guillaume Gricourt, Melissa N'Debi, Alexandre Soulier, Elisabeth Trawinski, Jean-Michel Pawlotsky |
| EPI_ISL_3476022 | CHUV | Laboratory of genomics and metagenomics | Trestan Pillonel, Damien Jacot, Sébastien Aeby, Gilbert Greub, Claire Bertelli |
| EPI_ISL_3476023 | VIDYMED LAUSANNE | Laboratory of genomics and metagenomics | Trestan Pillonel, Damien Jacot, Sébastien Aeby, Gilbert Greub, Claire Bertelli |
| EPI_ISL_3476024 | CHUV | Laboratory of genomics and metagenomics | Trestan Pillonel, Damien Jacot, Sébastien Aeby, Gilbert Greub, Claire Bertelli |
| EPI_ISL_3476025, EPI_ISL_3476026, EPI_ISL_3476027, EPI_ISL_3476028, EPI_ISL_3476029, EPI_ISL_3476030, EPI_ISL_3476031, EPI_ISL_3476032 | EHNV | Laboratory of genomics and metagenomics | Trestan Pillonel, Damien Jacot, Sébastien Aeby, Gilbert Greub, Claire Bertelli |
| EPI_ISL_3476033 | CHUV | Laboratory of genomics and metagenomics | Trestan Pillonel, Damien Jacot, Sébastien Aeby, Gilbert Greub, Claire Bertelli |
| EPI_ISL_3476034 | EHNV | Laboratory of genomics and metagenomics | Trestan Pillonel, Damien Jacot, Sébastien Aeby, Gilbert Greub, Claire Bertelli |
| EPI_ISL_3476035 | CHUV | Laboratory of genomics and metagenomics | Trestan Pillonel, Damien Jacot, Sébastien Aeby, Gilbert Greub, Claire Bertelli |
| EPI_ISL_3476036, EPI_ISL_3476037, EPI_ISL_3476038, EPI_ISL_3476039, EPI_ISL_3476040, EPI_ISL_3476041, EPI_ISL_3476042, EPI_ISL_3476043, EPI_ISL_3476044, EPI_ISL_3476045, EPI_ISL_3476046, EPI_ISL_3476047, EPI_ISL_3476048, EPI_ISL_3476049, EPI_ISL_3476050, EPI_ISL_3476051, EPI_ISL_3476052, EPI_ISL_3476053, EPI_ISL_3476054, EPI_ISL_3476055, EPI_ISL_3476056, EPI_ISL_3476057, EPI_ISL_3476058, EPI_ISL_3476059, EPI_ISL_3476060 | | | |
| see above | EHNV | Laboratory of genomics and metagenomics | Trestan Pillonel, Damien Jacot, Sébastien Aeby, Gilbert Greub, Claire Bertelli |
| EPI_ISL_3476061 | CHUV | Laboratory of genomics and metagenomics | Trestan Pillonel, Damien Jacot, Sébastien Aeby, Gilbert Greub, Claire Bertelli |
| EPI_ISL_3476062, EPI_ISL_3476063, EPI_ISL_3476064, EPI_ISL_3476065, EPI_ISL_3476066, EPI_ISL_3476067, EPI_ISL_3476068, EPI_ISL_3476069, EPI_ISL_3476070, EPI_ISL_3476071 | EHNV | Laboratory of genomics and metagenomics | Trestan Pillonel, Damien Jacot, Sébastien Aeby, Gilbert Greub, Claire Bertelli |
| EPI_ISL_3476072 | CHUV | Laboratory of genomics and metagenomics | Trestan Pillonel, Damien Jacot, Sébastien Aeby, Gilbert Greub, Claire Bertelli |
| EPI_ISL_3476073, EPI_ISL_3476074, EPI_ISL_3476075, EPI_ISL_3476076, EPI_ISL_3476077, EPI_ISL_3476078, EPI_ISL_3476079, EPI_ISL_3476080, EPI_ISL_3476081, EPI_ISL_3476082, EPI_ISL_3476083, EPI_ISL_3476084, EPI_ISL_3476085, EPI_ISL_3476086, EPI_ISL_3476087, EPI_ISL_3476088, EPI_ISL_3476089 | | | |
| see above | EHNV | Laboratory of genomics and metagenomics | Trestan Pillonel, Damien Jacot, Sébastien Aeby, Gilbert Greub, Claire Bertelli |
| EPI_ISL_3476090, EPI_ISL_3476091, EPI_ISL_3476092, EPI_ISL_3476093, EPI_ISL_3476094, EPI_ISL_3476095, EPI_ISL_3476096, EPI_ISL_3476097, EPI_ISL_3476098, EPI_ISL_3476099, EPI_ISL_3476100, EPI_ISL_3476101, EPI_ISL_3476102, EPI_ISL_3476103, EPI_ISL_3476104, EPI_ISL_3476105, EPI_ISL_3476106, EPI_ISL_3476107, EPI_ISL_3476108, EPI_ISL_3476109, EPI_ISL_3476110, EPI_ISL_3476111, EPI_ISL_3476112, EPI_ISL_3476113, EPI_ISL_3476114, EPI_ISL_3476115, EPI_ISL_3476116, EPI_ISL_3476117, EPI_ISL_3476118, EPI_ISL_3476119, EPI_ISL_3476120, EPI_ISL_3476121, EPI_ISL_3476122, EPI_ISL_3476123, EPI_ISL_3476124, EPI_ISL_3476125, EPI_ISL_3476126, EPI_ISL_3476127, EPI_ISL_3476128, EPI_ISL_3476129, EPI_ISL_3476130, EPI_ISL_3476131, EPI_ISL_3476132, EPI_ISL_3476133, EPI_ISL_3476134, EPI_ISL_3476135, EPI_ISL_3476136, EPI_ISL_3476137, EPI_ISL_3476138, EPI_ISL_3476139, EPI_ISL_3476140, EPI_ISL_3476141, EPI_ISL_3476142, EPI_ISL_3476143, EPI_ISL_3476144, EPI_ISL_3476145, EPI_ISL_3476146, EPI_ISL_3476147, EPI_ISL_3476148, EPI_ISL_3476149, EPI_ISL_3476150, EPI_ISL_3476151, EPI_ISL_3476152, EPI_ISL_3476153, EPI_ISL_3476154, EPI_ISL_3476155, EPI_ISL_3476156, EPI_ISL_3476157, EPI_ISL_3476158, EPI_ISL_3476160, EPI_ISL_3476161, EPI_ISL_3476162, EPI_ISL_3476163, EPI_ISL_3476164, EPI_ISL_3476165, EPI_ISL_3476166, EPI_ISL_3476167, EPI_ISL_3476168, EPI_ISL_3476169, EPI_ISL_3476170, EPI_ISL_3476171, EPI_ISL_3476172, EPI_ISL_3476173, EPI_ISL_3476174, EPI_ISL_3476175, EPI_ISL_3476176, EPI_ISL_3476177, EPI_ISL_3476178, EPI_ISL_3476179, EPI_ISL_3476181, EPI_ISL_3476182, EPI_ISL_3476183, EPI_ISL_3476184, EPI_ISL_3476185, EPI_ISL_3476186, EPI_ISL_3476187, EPI_ISL_3476188, EPI_ISL_3476189, EPI_ISL_3476190, EPI_ISL_3476191, EPI_ISL_3476192, EPI_ISL_3476193, EPI_ISL_3476194, EPI_ISL_3476195, EPI_ISL_3476196, EPI_ISL_3476197, EPI_ISL_3476198, EPI_ISL_3476199, EPI_ISL_3476200, EPI_ISL_3476201, EPI_ISL_3476202, EPI_ISL_3476203, EPI_ISL_3476205, EPI_ISL_3476206, EPI_ISL_3476207, EPI_ISL_3476209, EPI_ISL_3476210, EPI_ISL_3476211, EPI_ISL_3476212, EPI_ISL_3476213, EPI_ISL_3476214, EPI_ISL_3476215, EPI_ISL_3476216, EPI_ISL_3476217, EPI_ISL_3476218, EPI_ISL_3476219, EPI_ISL_3476220, EPI_ISL_3476221, EPI_ISL_3476222, EPI_ISL_3476223, EPI_ISL_3476224, EPI_ISL_3476225, EPI_ISL_3476226, EPI_ISL_3476227, EPI_ISL_3476228, EPI_ISL_3476229, EPI_ISL_3476230, EPI_ISL_3476231, EPI_ISL_3476232, EPI_ISL_3476233, EPI_ISL_3476234, EPI_ISL_3476235, EPI_ISL_3476236, EPI_ISL_3476237, EPI_ISL_3476238, EPI_ISL_3476239, EPI_ISL_3476240, EPI_ISL_3476241, EPI_ISL_3476242, EPI_ISL_3476243, EPI_ISL_3476244, EPI_ISL_3476245, EPI_ISL_3476246, EPI_ISL_3476247, EPI_ISL_3476248, EPI_ISL_3476249, EPI_ISL_3476250, EPI_ISL_3476251, EPI_ISL_3476252, EPI_ISL_3476253, EPI_ISL_3476254, EPI_ISL_3476255, EPI_ISL_3476256, EPI_ISL_3476257, EPI_ISL_3476258, EPI_ISL_3476259, EPI_ISL_3476260, EPI_ISL_3476261, EPI_ISL_3476262, EPI_ISL_3476264, EPI_ISL_3476266, EPI_ISL_3476267, EPI_ISL_3476268, EPI_ISL_3476269, EPI_ISL_3476270, EPI_ISL_3476271, EPI_ISL_3476272, EPI_ISL_3476273, EPI_ISL_3476274, EPI_ISL_3476275, EPI_ISL_3476276, EPI_ISL_3476277, EPI_ISL_3476278, EPI_ISL_3476279, EPI_ISL_3476280, EPI_ISL_3476281, EPI_ISL_3476283, EPI_ISL_3476284, EPI_ISL_3476285, EPI_ISL_3476286, EPI_ISL_3476287, EPI_ISL_3476288, EPI_ISL_3476289, EPI_ISL_3476290, EPI_ISL_3476291, EPI_ISL_3476292, EPI_ISL_3476293, EPI_ISL_3476294, EPI_ISL_3476295, EPI_ISL_3476296, EPI_ISL_3476297, EPI_ISL_3476298, EPI_ISL_3476299, EPI_ISL_3476300, EPI_ISL_3476301, EPI_ISL_3476302, EPI_ISL_3476303, EPI_ISL_3476304, EPI_ISL_3476305, EPI_ISL_3476306, EPI_ISL_3476307, EPI_ISL_3476308, EPI_ISL_3476309, EPI_ISL_3476310, EPI_ISL_3476311, EPI_ISL_3476312, EPI_ISL_3476313, EPI_ISL_3476314, EPI_ISL_3476315, EPI_ISL_3476316, EPI_ISL_3476317, EPI_ISL_3476318, EPI_ISL_3476319, EPI_ISL_3476320, EPI_ISL_3476321, EPI_ISL_3476322, EPI_ISL_3476323, EPI_ISL_3476324, EPI_ISL_3476325, EPI_ISL_3476326, EPI_ISL_3476327, EPI_ISL_3476328, EPI_ISL_3476329, EPI_ISL_3476330, EPI_ISL_3476331, EPI_ISL_3476332, EPI_ISL_3476333 | | | |
| see above | Viollier AG | Department of Biosystems Science and Engineering, ETH Zürich | Chaoran Chen, Sarah Nadeau, Louis du Plessis, Catharine Aquino, Ivan Topolsky, Kim Philipp Jablonski, Lara Fuhrmann, Daniel Ehrsam, Isabel Stürmer, Andreia Cabral de Gouvea, Maria Domenica Moccia, Simon Grüter, Timothy Sykes, Lennart Opitz, Griffin White, Laura Neff, Doris Popovic, Andrea Patrignani, Jay Tracy, Ralph Schlapbach, Christiane Beckmann, Maurice Redondo, Olivier Kobel, Christoph Noppen, Niko Beerenwinkel, Tanja Stadler |
| EPI_ISL_3476334 | Cerballiance Keraudren | CERBA HealthCare | Bénédicte Roquebert, Laura Verdurme, Sabine Trombert, Stéphanie Haim-Boukobza |
| EPI_ISL_3476335 | Laboratoire Etables Sur Mer | CERBA HealthCare | Bénédicte Roquebert, Laura Verdurme, Sabine Trombert, Stéphanie Haim-Boukobza |
| EPI_ISL_3476336 | LBM Centre Commercial La Foux | CERBA HealthCare | Bénédicte Roquebert, Laura Verdurme, Sabine Trombert, Stéphanie Haim-Boukobza |
| EPI_ISL_3476337 | LBM Blausasc | CERBA HealthCare | Bénédicte Roquebert, Laura Verdurme, Sabine Trombert, Stéphanie Haim-Boukobza |
| EPI_ISL_3476338, EPI_ISL_3476339, EPI_ISL_3476340 | LBM Centre Commercial La Foux | CERBA HealthCare | Bénédicte Roquebert, Laura Verdurme, Sabine Trombert, Stéphanie Haim-Boukobza |
| EPI_ISL_3476341, EPI_ISL_3476342 | LBM Delfino | CERBA HealthCare | Bénédicte Roquebert, Laura Verdurme, Sabine Trombert, Stéphanie Haim-Boukobza |
| EPI_ISL_3476343 | LBM Cogolin | CERBA HealthCare | Bénédicte Roquebert, Laura Verdurme, Sabine Trombert, Stéphanie Haim-Boukobza |
| EPI_ISL_3476344 | Cerballiance Normandie Ouest Granville | CERBA HealthCare | Bénédicte Roquebert, Laura Verdurme, Sabine Trombert, Stéphanie Haim-Boukobza |
| EPI_ISL_3476345, EPI_ISL_3476346 | LBM Cogolin | CERBA HealthCare | Bénédicte Roquebert, Laura Verdurme, Sabine Trombert, Stéphanie Haim-Boukobza |
| EPI_ISL_3476347 | LABM des Batignolles | CERBA HealthCare | Bénédicte Roquebert, Laura Verdurme, Sabine Trombert, Stéphanie Haim-Boukobza |
| EPI_ISL_3476348 | LBM Centre Commercial La Foux | CERBA HealthCare | Bénédicte Roquebert, Laura Verdurme, Sabine Trombert, Stéphanie Haim-Boukobza |
| EPI_ISL_3476349 | Cerballiance LBM Tokoro | CERBA HealthCare | Bénédicte Roquebert, Laura Verdurme, Sabine Trombert, Stéphanie Haim-Boukobza |
| EPI_ISL_3476350, EPI_ISL_3476351, EPI_ISL_3476352, EPI_ISL_3476353 | LBM Centre Commercial La Foux | CERBA HealthCare | Bénédicte Roquebert, Laura Verdurme, Sabine Trombert, Stéphanie Haim-Boukobza |
| EPI_ISL_3476354 | LBM Cros de Cagnes | CERBA HealthCare | Bénédicte Roquebert, Laura Verdurme, Sabine Trombert, Stéphanie Haim-Boukobza |
| EPI_ISL_3476355 | LBM Biolbs - St Romain | CERBA HealthCare | Bénédicte Roquebert, Laura Verdurme, Sabine Trombert, Stéphanie Haim-Boukobza |
| EPI_ISL_3476356 | LBM Centre Commercial La Foux | CERBA HealthCare | Bénédicte Roquebert, Laura Verdurme, Sabine Trombert, Stéphanie Haim-Boukobza |
| EPI_ISL_3476357 | LBM Biolbs - St Romain | CERBA HealthCare | Bénédicte Roquebert, Laura Verdurme, Sabine Trombert, Stéphanie Haim-Boukobza |
| EPI_ISL_3476358 | Antibes Saint Jean | CERBA HealthCare | Bénédicte Roquebert, Laura Verdurme, Sabine Trombert, Stéphanie Haim-Boukobza |
| EPI_ISL_3476359 | Laboratoire CHG Grasse | CERBA HealthCare | Bénédicte Roquebert, Laura Verdurme, Sabine Trombert, Stéphanie Haim-Boukobza |
| EPI_ISL_3476360 | LBM Le Grand Saint Jean | CERBA HealthCare | Bénédicte Roquebert, Laura Verdurme, Sabine Trombert, Stéphanie Haim-Boukobza |
| EPI_ISL_3476361 | LBM Grasse - Le Moulin | CERBA HealthCare | Bénédicte Roquebert, Laura Verdurme, Sabine Trombert, Stéphanie Haim-Boukobza |
| EPI_ISL_3476362, EPI_ISL_3476363 | LBM La Trinite | CERBA HealthCare | Bénédicte Roquebert, Laura Verdurme, Sabine Trombert, Stéphanie Haim-Boukobza |
| EPI_ISL_3476364 | LBM Le Grand Saint Jean | CERBA HealthCare | Bénédicte Roquebert, Laura Verdurme, Sabine Trombert, Stéphanie Haim-Boukobza |
| EPI_ISL_3476365 | Interlabo Unilabs Eu | CERBA HealthCare | Bénédicte Roquebert, Laura Verdurme, Sabine Trombert, Stéphanie Haim-Boukobza |

| | | | |
|---|---|---|---|
| EPI_ISL_3476366 | LBM Le Grand Saint Jean | CERBA HealthCare | Bénédicte Roquebert, Laura Verdurme, Sabine Trombert, Stéphanie Haim-Boukobza |
| EPI_ISL_3476367 | Bioct - Unilabs - Chateau Thierry | CERBA HealthCare | Bénédicte Roquebert, Laura Verdurme, Sabine Trombert, Stéphanie Haim-Boukobza |
| EPI_ISL_3476368 | LBM Cogolin | CERBA HealthCare | Bénédicte Roquebert, Laura Verdurme, Sabine Trombert, Stéphanie Haim-Boukobza |
| EPI_ISL_3476369 | LBM Blausasc | CERBA HealthCare | Bénédicte Roquebert, Laura Verdurme, Sabine Trombert, Stéphanie Haim-Boukobza |
| EPI_ISL_3476370 | LBM Grasse - Le Moulin | CERBA HealthCare | Bénédicte Roquebert, Laura Verdurme, Sabine Trombert, Stéphanie Haim-Boukobza |
| EPI_ISL_3476371 | LBM du Cours | CERBA HealthCare | Bénédicte Roquebert, Laura Verdurme, Sabine Trombert, Stéphanie Haim-Boukobza |
| EPI_ISL_3476372 | Antibes Saint Jean | CERBA HealthCare | Bénédicte Roquebert, Laura Verdurme, Sabine Trombert, Stéphanie Haim-Boukobza |
| EPI_ISL_3476373 | LBM Le Grand Saint Jean | CERBA HealthCare | Bénédicte Roquebert, Laura Verdurme, Sabine Trombert, Stéphanie Haim-Boukobza |
| EPI_ISL_3476374 | LBM Beausoleil - Gallia | CERBA HealthCare | Bénédicte Roquebert, Laura Verdurme, Sabine Trombert, Stéphanie Haim-Boukobza |
| EPI_ISL_3476375 | Bioct - Unilabs - Ave de Champagne | CERBA HealthCare | Bénédicte Roquebert, Laura Verdurme, Sabine Trombert, Stéphanie Haim-Boukobza |
| EPI_ISL_3476376, EPI_ISL_3476377 | Bioct - Unilabs - Chateau Thierry | CERBA HealthCare | Bénédicte Roquebert, Laura Verdurme, Sabine Trombert, Stéphanie Haim-Boukobza |
| EPI_ISL_3476378 | Laboratoire Clement Paris | CERBA HealthCare | Bénédicte Roquebert, Laura Verdurme, Sabine Trombert, Stéphanie Haim-Boukobza |
| EPI_ISL_3476379 | LBM Saint Laurent Centre | CERBA HealthCare | Bénédicte Roquebert, Laura Verdurme, Sabine Trombert, Stéphanie Haim-Boukobza |
| EPI_ISL_3476380 | LBM Le Grand Saint Jean | CERBA HealthCare | Bénédicte Roquebert, Laura Verdurme, Sabine Trombert, Stéphanie Haim-Boukobza |
| EPI_ISL_3476381 | Laboratoire Ploufragan | CERBA HealthCare | Bénédicte Roquebert, Laura Verdurme, Sabine Trombert, Stéphanie Haim-Boukobza |
| EPI_ISL_3476382, EPI_ISL_3476383 | LBM Cap D'Or | CERBA HealthCare | Bénédicte Roquebert, Laura Verdurme, Sabine Trombert, Stéphanie Haim-Boukobza |
| EPI_ISL_3476384 | Centre Hospitalier Argenteuil | CERBA HealthCare | Bénédicte Roquebert, Laura Verdurme, Sabine Trombert, Stéphanie Haim-Boukobza |
| EPI_ISL_3476385, EPI_ISL_3476386 | Cerballiance LBM La Pomme | CERBA HealthCare | Bénédicte Roquebert, Laura Verdurme, Sabine Trombert, Stéphanie Haim-Boukobza |
| EPI_ISL_3476387 | Cerballiance LBM Istres | CERBA HealthCare | Bénédicte Roquebert, Laura Verdurme, Sabine Trombert, Stéphanie Haim-Boukobza |
| EPI_ISL_3476388, EPI_ISL_3476389 | Cerballiance LBM Port de Bouc | CERBA HealthCare | Bénédicte Roquebert, Laura Verdurme, Sabine Trombert, Stéphanie Haim-Boukobza |
| EPI_ISL_3476390 | Cerballiance LBM Rousset | CERBA HealthCare | Bénédicte Roquebert, Laura Verdurme, Sabine Trombert, Stéphanie Haim-Boukobza |
| EPI_ISL_3476391 | Cerballiance LBM National | CERBA HealthCare | Bénédicte Roquebert, Laura Verdurme, Sabine Trombert, Stéphanie Haim-Boukobza |
| EPI_ISL_3476392, EPI_ISL_3476393 | Cerballiance LBM La Pomme | CERBA HealthCare | Bénédicte Roquebert, Laura Verdurme, Sabine Trombert, Stéphanie Haim-Boukobza |
| EPI_ISL_3476394 | Laboratoire Carillon Biosmose IDF | CERBA HealthCare | Bénédicte Roquebert, Laura Verdurme, Sabine Trombert, Stéphanie Haim-Boukobza |
| EPI_ISL_3476395, EPI_ISL_3476396 | Cerballiance LBM La Pomme | CERBA HealthCare | Bénédicte Roquebert, Laura Verdurme, Sabine Trombert, Stéphanie Haim-Boukobza |
| EPI_ISL_3476397 | Laboratoire du Clos Bertin | CERBA HealthCare | Bénédicte Roquebert, Laura Verdurme, Sabine Trombert, Stéphanie Haim-Boukobza |
| EPI_ISL_3476398, EPI_ISL_3476399 | Cerballiance LBM Port de Bouc | CERBA HealthCare | Bénédicte Roquebert, Laura Verdurme, Sabine Trombert, Stéphanie Haim-Boukobza |
| EPI_ISL_3476400, EPI_ISL_3476401 | Cerballiance LBM La Pomme | CERBA HealthCare | Bénédicte Roquebert, Laura Verdurme, Sabine Trombert, Stéphanie Haim-Boukobza |
| EPI_ISL_3476402, EPI_ISL_3476403 | Cerballiance LBM Port de Bouc | CERBA HealthCare | Bénédicte Roquebert, Laura Verdurme, Sabine Trombert, Stéphanie Haim-Boukobza |
| EPI_ISL_3476404 | Cerballiance LBM La Pomme | CERBA HealthCare | Bénédicte Roquebert, Laura Verdurme, Sabine Trombert, Stéphanie Haim-Boukobza |
| EPI_ISL_3476405 | LBM Dr F.Hassanaly | CERBA HealthCare | Bénédicte Roquebert, Laura Verdurme, Sabine Trombert, Stéphanie Haim-Boukobza |
| EPI_ISL_3476406 | LBM Blausasc | CERBA HealthCare | Bénédicte Roquebert, Laura Verdurme, Sabine Trombert, Stéphanie Haim-Boukobza |
| EPI_ISL_3476407 | BIOMNIS EUROFINS IVRY | Department of Virology, Henri Mondor University Hospital, Assistance Publique Hôpitaux de Paris, Université Paris-Est Créteil, INSERM U955 | Christophe Rodriguez, Slim Fourati, Vanessa Demontant, Guillaume Gricourt, Melissa N'Debi, Alexandre Soulier, Elisabeth Trawinski, Jean-Michel Pawlotsky |
| EPI_ISL_3476408 | SELAS MEDILYS | Department of Virology, Henri Mondor University Hospital, Assistance Publique Hôpitaux de Paris, Université Paris-Est Créteil, INSERM U955 | Christophe Rodriguez, Slim Fourati, Vanessa Demontant, Guillaume Gricourt, Melissa N'Debi, Alexandre Soulier, Elisabeth Trawinski, Jean-Michel Pawlotsky |
| EPI_ISL_3476409 | Hôpital Bichat-Claude Bernard | Department of Virology, Henri Mondor University Hospital, Assistance Publique Hôpitaux de Paris, Université Paris-Est Créteil, INSERM U955 | Christophe Rodriguez, Slim Fourati, Vanessa Demontant, Guillaume Gricourt, Melissa N'Debi, Alexandre Soulier, Elisabeth Trawinski, Jean-Michel Pawlotsky |
| EPI_ISL_3476410 | BIOMNIS EUROFINS IVRY | Department of Virology, Henri Mondor University Hospital, Assistance Publique Hôpitaux de Paris, Université Paris-Est Créteil, INSERM U955 | Christophe Rodriguez, Slim Fourati, Vanessa Demontant, Guillaume Gricourt, Melissa N'Debi, Alexandre Soulier, Elisabeth Trawinski, Jean-Michel Pawlotsky |
| EPI_ISL_3476411 | CH.LE RAINCY_MONTFERMEIL | Department of Virology, Henri Mondor University Hospital, Assistance Publique Hôpitaux de Paris, Université Paris-Est Créteil, INSERM U955 | Christophe Rodriguez, Slim Fourati, Vanessa Demontant, Guillaume Gricourt, Melissa N'Debi, Alexandre Soulier, Elisabeth Trawinski, Jean-Michel Pawlotsky |
| EPI_ISL_3476412 | BIOMNIS EUROFINS IVRY | Department of Virology, Henri Mondor University Hospital, Assistance Publique Hôpitaux de Paris, Université Paris-Est Créteil, INSERM U955 | Christophe Rodriguez, Slim Fourati, Vanessa Demontant, Guillaume Gricourt, Melissa N'Debi, Alexandre Soulier, Elisabeth Trawinski, Jean-Michel Pawlotsky |
| EPI_ISL_3476413 | Hôpital Armand-Trousseau | Department of Virology, Henri Mondor University Hospital, Assistance Publique Hôpitaux de Paris, Université Paris-Est Créteil, INSERM U955 | Christophe Rodriguez, Slim Fourati, Vanessa Demontant, Guillaume Gricourt, Melissa N'Debi, Alexandre Soulier, Elisabeth Trawinski, Jean-Michel Pawlotsky |
| EPI_ISL_3476414, EPI_ISL_3476415, EPI_ISL_3476416 | SELAS MEDILYS | Department of Virology, Henri Mondor University Hospital, Assistance Publique Hôpitaux de Paris, Université Paris-Est Créteil, INSERM U955 | Christophe Rodriguez, Slim Fourati, Vanessa Demontant, Guillaume Gricourt, Melissa N'Debi, Alexandre Soulier, Elisabeth Trawinski, Jean-Michel Pawlotsky |
| EPI_ISL_3476418 | BIOMNIS EUROFINS IVRY | Department of Virology, Henri Mondor University Hospital, Assistance Publique Hôpitaux de Paris, Université Paris-Est Créteil, INSERM U955 | Christophe Rodriguez, Slim Fourati, Vanessa Demontant, Guillaume Gricourt, Melissa N'Debi, Alexandre Soulier, Elisabeth Trawinski, Jean-Michel Pawlotsky |
| EPI_ISL_3476419 | SELAS MEDILYS | Department of Virology, Henri Mondor University Hospital, Assistance Publique Hôpitaux de Paris, Université Paris-Est Créteil, INSERM U955 | Christophe Rodriguez, Slim Fourati, Vanessa Demontant, Guillaume Gricourt, Melissa N'Debi, Alexandre Soulier, Elisabeth Trawinski, Jean-Michel Pawlotsky |
| EPI_ISL_3476420 | LBM EVORIAL NEVERS | Department of Virology, Henri Mondor University Hospital, Assistance Publique Hôpitaux de Paris, Université Paris-Est Créteil, INSERM U955 | Christophe Rodriguez, Slim Fourati, Vanessa Demontant, Guillaume Gricourt, Melissa N'Debi, Alexandre Soulier, Elisabeth Trawinski, Jean-Michel Pawlotsky |
| EPI_ISL_3476421, EPI_ISL_3476422, EPI_ISL_3476423 | SELAS MEDILYS | Department of Virology, Henri Mondor University Hospital, Assistance Publique Hôpitaux de Paris, Université Paris-Est Créteil, INSERM U955 | Christophe Rodriguez, Slim Fourati, Vanessa Demontant, Guillaume Gricourt, Melissa N'Debi, Alexandre Soulier, Elisabeth Trawinski, Jean-Michel Pawlotsky |
| EPI_ISL_3476424 | CHU DE DIJON | Department of Virology, Henri Mondor University Hospital, Assistance Publique Hôpitaux de Paris, Université Paris-Est Créteil, INSERM U955 | Christophe Rodriguez, Slim Fourati, Vanessa Demontant, Guillaume Gricourt, Melissa N'Debi, Alexandre Soulier, Elisabeth Trawinski, Jean-Michel Pawlotsky |
| EPI_ISL_3476425 | SELAS MEDILYS | Department of Virology, Henri Mondor University Hospital, Assistance Publique Hôpitaux de Paris, Université Paris-Est Créteil, INSERM U955 | Christophe Rodriguez, Slim Fourati, Vanessa Demontant, Guillaume Gricourt, Melissa N'Debi, Alexandre Soulier, Elisabeth Trawinski, Jean-Michel Pawlotsky |
| EPI_ISL_3476426 | BIORHIN | Department of Virology, Henri Mondor University Hospital, Assistance Publique Hôpitaux de Paris, Université Paris-Est Créteil, INSERM U955 | Christophe Rodriguez, Slim Fourati, Vanessa Demontant, Guillaume Gricourt, Melissa N'Debi, Alexandre Soulier, Elisabeth Trawinski, Jean-Michel Pawlotsky |
| EPI_ISL_3476427 | Biogroup Bio Lam-LCD Saint-Denis | Department of Virology, Henri Mondor University Hospital, Assistance Publique Hôpitaux de Paris, Université Paris-Est Créteil, INSERM U955 | Christophe Rodriguez, Slim Fourati, Vanessa Demontant, Guillaume Gricourt, Melissa N'Debi, Alexandre Soulier, Elisabeth Trawinski, Jean-Michel Pawlotsky |

| | | | |
|---|---|---|---|
| EPI_ISL_3476428 | SELAS MEDILYS | Department of Virology, Henri Mondor University Hospital, Assistance Publique Hôpitaux de Paris, Université Paris-Est Créteil, INSERM U955 | Christophe Rodriguez, Slim Fourati, Vanessa Demontant, Guillaume Gricourt, Melissa N'Debi, Alexandre Soulier, Elisabeth Trawinski, Jean-Michel Pawlotsky |
| EPI_ISL_3476429 | Biogroup Bio Lam-LCD Saint-Denis | Department of Virology, Henri Mondor University Hospital, Assistance Publique Hôpitaux de Paris, Université Paris-Est Créteil, INSERM U955 | Christophe Rodriguez, Slim Fourati, Vanessa Demontant, Guillaume Gricourt, Melissa N'Debi, Alexandre Soulier, Elisabeth Trawinski, Jean-Michel Pawlotsky |
| EPI_ISL_3476430, EPI_ISL_3476431, EPI_ISL_3476432, EPI_ISL_3476433, EPI_ISL_3476434 | SELAS MEDILYS | Department of Virology, Henri Mondor University Hospital, Assistance Publique Hôpitaux de Paris, Université Paris-Est Créteil, INSERM U955 | Christophe Rodriguez, Slim Fourati, Vanessa Demontant, Guillaume Gricourt, Melissa N'Debi, Alexandre Soulier, Elisabeth Trawinski, Jean-Michel Pawlotsky |
| EPI_ISL_3476435 | CH. CHU DE REIMS | Department of Virology, Henri Mondor University Hospital, Assistance Publique Hôpitaux de Paris, Université Paris-Est Créteil, INSERM U955 | Christophe Rodriguez, Slim Fourati, Vanessa Demontant, Guillaume Gricourt, Melissa N'Debi, Alexandre Soulier, Elisabeth Trawinski, Jean-Michel Pawlotsky |
| EPI_ISL_3476436 | CH.LE RAINCY_MONTFERMEIL | Department of Virology, Henri Mondor University Hospital, Assistance Publique Hôpitaux de Paris, Université Paris-Est Créteil, INSERM U955 | Christophe Rodriguez, Slim Fourati, Vanessa Demontant, Guillaume Gricourt, Melissa N'Debi, Alexandre Soulier, Elisabeth Trawinski, Jean-Michel Pawlotsky |
| EPI_ISL_3476437, EPI_ISL_3476438, EPI_ISL_3476439 | GH A.CHENEVIER-H.MONDOR | Department of Virology, Henri Mondor University Hospital, Assistance Publique Hôpitaux de Paris, Université Paris-Est Créteil, INSERM U955 | Christophe Rodriguez, Slim Fourati, Vanessa Demontant, Guillaume Gricourt, Melissa N'Debi, Alexandre Soulier, Elisabeth Trawinski, Jean-Michel Pawlotsky |
| EPI_ISL_3476440 | SELAS MEDILYS | Department of Virology, Henri Mondor University Hospital, Assistance Publique Hôpitaux de Paris, Université Paris-Est Créteil, INSERM U955 | Christophe Rodriguez, Slim Fourati, Vanessa Demontant, Guillaume Gricourt, Melissa N'Debi, Alexandre Soulier, Elisabeth Trawinski, Jean-Michel Pawlotsky |
| EPI_ISL_3476441 | Cerballiance Paris IDF est Saint-Denis | Department of Virology, Henri Mondor University Hospital, Assistance Publique Hôpitaux de Paris, Université Paris-Est Créteil, INSERM U955 | Christophe Rodriguez, Slim Fourati, Vanessa Demontant, Guillaume Gricourt, Melissa N'Debi, Alexandre Soulier, Elisabeth Trawinski, Jean-Michel Pawlotsky |
| EPI_ISL_3476442 | Hôpital Armand-Trousseau | Department of Virology, Henri Mondor University Hospital, Assistance Publique Hôpitaux de Paris, Université Paris-Est Créteil, INSERM U955 | Christophe Rodriguez, Slim Fourati, Vanessa Demontant, Guillaume Gricourt, Melissa N'Debi, Alexandre Soulier, Elisabeth Trawinski, Jean-Michel Pawlotsky |
| EPI_ISL_3476443 | BIOMNIS EUROFINS IVRY | Department of Virology, Henri Mondor University Hospital, Assistance Publique Hôpitaux de Paris, Université Paris-Est Créteil, INSERM U955 | Christophe Rodriguez, Slim Fourati, Vanessa Demontant, Guillaume Gricourt, Melissa N'Debi, Alexandre Soulier, Elisabeth Trawinski, Jean-Michel Pawlotsky |
| EPI_ISL_3476444 | SELAS MEDILYS | Department of Virology, Henri Mondor University Hospital, Assistance Publique Hôpitaux de Paris, Université Paris-Est Créteil, INSERM U955 | Christophe Rodriguez, Slim Fourati, Vanessa Demontant, Guillaume Gricourt, Melissa N'Debi, Alexandre Soulier, Elisabeth Trawinski, Jean-Michel Pawlotsky |
| EPI_ISL_3476445 | GH A.CHENEVIER-H.MONDOR | Department of Virology, Henri Mondor University Hospital, Assistance Publique Hôpitaux de Paris, Université Paris-Est Créteil, INSERM U955 | Christophe Rodriguez, Slim Fourati, Vanessa Demontant, Guillaume Gricourt, Melissa N'Debi, Alexandre Soulier, Elisabeth Trawinski, Jean-Michel Pawlotsky |
| EPI_ISL_3476446 | CH.LE RAINCY_MONTFERMEIL | Department of Virology, Henri Mondor University Hospital, Assistance Publique Hôpitaux de Paris, Université Paris-Est Créteil, INSERM U955 | Christophe Rodriguez, Slim Fourati, Vanessa Demontant, Guillaume Gricourt, Melissa N'Debi, Alexandre Soulier, Elisabeth Trawinski, Jean-Michel Pawlotsky |
| EPI_ISL_3476447, EPI_ISL_3476448 | SELAS MEDILYS | Department of Virology, Henri Mondor University Hospital, Assistance Publique Hôpitaux de Paris, Université Paris-Est Créteil, INSERM U955 | Christophe Rodriguez, Slim Fourati, Vanessa Demontant, Guillaume Gricourt, Melissa N'Debi, Alexandre Soulier, Elisabeth Trawinski, Jean-Michel Pawlotsky |
| EPI_ISL_3476449 | BIOMNIS EUROFINS IVRY | Department of Virology, Henri Mondor University Hospital, Assistance Publique Hôpitaux de Paris, Université Paris-Est Créteil, INSERM U955 | Christophe Rodriguez, Slim Fourati, Vanessa Demontant, Guillaume Gricourt, Melissa N'Debi, Alexandre Soulier, Elisabeth Trawinski, Jean-Michel Pawlotsky |
| EPI_ISL_3476450 | GROUPE HOSPITALIER SUD ILE DE FRANCE | Department of Virology, Henri Mondor University Hospital, Assistance Publique Hôpitaux de Paris, Université Paris-Est Créteil, INSERM U955 | Christophe Rodriguez, Slim Fourati, Vanessa Demontant, Guillaume Gricourt, Melissa N'Debi, Alexandre Soulier, Elisabeth Trawinski, Jean-Michel Pawlotsky |
| EPI_ISL_3476451, EPI_ISL_3476452 | BIOMNIS EUROFINS IVRY | Department of Virology, Henri Mondor University Hospital, Assistance Publique Hôpitaux de Paris, Université Paris-Est Créteil, INSERM U955 | Christophe Rodriguez, Slim Fourati, Vanessa Demontant, Guillaume Gricourt, Melissa N'Debi, Alexandre Soulier, Elisabeth Trawinski, Jean-Michel Pawlotsky |
| EPI_ISL_3476453 | SELAS MEDILYS | Department of Virology, Henri Mondor University Hospital, Assistance Publique Hôpitaux de Paris, Université Paris-Est Créteil, INSERM U955 | Christophe Rodriguez, Slim Fourati, Vanessa Demontant, Guillaume Gricourt, Melissa N'Debi, Alexandre Soulier, Elisabeth Trawinski, Jean-Michel Pawlotsky |
| EPI_ISL_3476454, EPI_ISL_3476455 | BIOMNIS EUROFINS IVRY | Department of Virology, Henri Mondor University Hospital, Assistance Publique Hôpitaux de Paris, Université Paris-Est Créteil, INSERM U955 | Christophe Rodriguez, Slim Fourati, Vanessa Demontant, Guillaume Gricourt, Melissa N'Debi, Alexandre Soulier, Elisabeth Trawinski, Jean-Michel Pawlotsky |
| EPI_ISL_3476456 | SELAS MEDILYS | Department of Virology, Henri Mondor University Hospital, Assistance Publique Hôpitaux de Paris, Université Paris-Est Créteil, INSERM U955 | Christophe Rodriguez, Slim Fourati, Vanessa Demontant, Guillaume Gricourt, Melissa N'Debi, Alexandre Soulier, Elisabeth Trawinski, Jean-Michel Pawlotsky |
| EPI_ISL_3476457 | BIOMNIS EUROFINS IVRY | Department of Virology, Henri Mondor University Hospital, Assistance Publique Hôpitaux de Paris, Université Paris-Est Créteil, INSERM U955 | Christophe Rodriguez, Slim Fourati, Vanessa Demontant, Guillaume Gricourt, Melissa N'Debi, Alexandre Soulier, Elisabeth Trawinski, Jean-Michel Pawlotsky |
| EPI_ISL_3476458 | Hôpital Avicenne | Department of Virology, Henri Mondor University Hospital, Assistance Publique Hôpitaux de Paris, Université Paris-Est Créteil, INSERM U955 | Christophe Rodriguez, Slim Fourati, Vanessa Demontant, Guillaume Gricourt, Melissa N'Debi, Alexandre Soulier, Elisabeth Trawinski, Jean-Michel Pawlotsky |
| EPI_ISL_3476459 | Biogroup Bio Lam-LCD Saint-Denis | Department of Virology, Henri Mondor University Hospital, Assistance Publique Hôpitaux de Paris, Université Paris-Est Créteil, INSERM U955 | Christophe Rodriguez, Slim Fourati, Vanessa Demontant, Guillaume Gricourt, Melissa N'Debi, Alexandre Soulier, Elisabeth Trawinski, Jean-Michel Pawlotsky |
| EPI_ISL_3476460 | Hôpital Cochin | Department of Virology, Henri Mondor University Hospital, Assistance Publique Hôpitaux de Paris, Université Paris-Est Créteil, INSERM U955 | Christophe Rodriguez, Slim Fourati, Vanessa Demontant, Guillaume Gricourt, Melissa N'Debi, Alexandre Soulier, Elisabeth Trawinski, Jean-Michel Pawlotsky |
| EPI_ISL_3476461 | SELAS MEDILYS | Department of Virology, Henri Mondor University Hospital, Assistance Publique Hôpitaux de Paris, Université Paris-Est Créteil, INSERM U955 | Christophe Rodriguez, Slim Fourati, Vanessa Demontant, Guillaume Gricourt, Melissa N'Debi, Alexandre Soulier, Elisabeth Trawinski, Jean-Michel Pawlotsky |
| EPI_ISL_3476462 | LISIEUX Cerballiance | Department of Virology, Henri Mondor University Hospital, Assistance Publique Hôpitaux de Paris, Université Paris-Est Créteil, INSERM U955 | Christophe Rodriguez, Slim Fourati, Vanessa Demontant, Guillaume Gricourt, Melissa N'Debi, Alexandre Soulier, Elisabeth Trawinski, Jean-Michel Pawlotsky |
| EPI_ISL_3476463, EPI_ISL_3476464 | GH A.CHENEVIER-H.MONDOR | Department of Virology, Henri Mondor University Hospital, Assistance Publique Hôpitaux de Paris, Université Paris-Est Créteil, INSERM U955 | Christophe Rodriguez, Slim Fourati, Vanessa Demontant, Guillaume Gricourt, Melissa N'Debi, Alexandre Soulier, Elisabeth Trawinski, Jean-Michel Pawlotsky |
| EPI_ISL_3476465, EPI_ISL_3476466, EPI_ISL_3476467 | SELAS MEDILYS | Department of Virology, Henri Mondor University Hospital, Assistance Publique Hôpitaux de Paris, Université Paris-Est Créteil, INSERM U955 | Christophe Rodriguez, Slim Fourati, Vanessa Demontant, Guillaume Gricourt, Melissa N'Debi, Alexandre Soulier, Elisabeth Trawinski, Jean-Michel Pawlotsky |
| EPI_ISL_3476468 | CHU DE DIJON | Department of Virology, Henri Mondor University Hospital, Assistance Publique Hôpitaux de Paris, Université Paris-Est Créteil, INSERM U955 | Christophe Rodriguez, Slim Fourati, Vanessa Demontant, Guillaume Gricourt, Melissa N'Debi, Alexandre Soulier, Elisabeth Trawinski, Jean-Michel Pawlotsky |
| EPI_ISL_3476469, EPI_ISL_3476470 | BIOMNIS EUROFINS IVRY | Department of Virology, Henri Mondor University Hospital, Assistance Publique Hôpitaux de Paris, Université Paris-Est Créteil, INSERM U955 | Christophe Rodriguez, Slim Fourati, Vanessa Demontant, Guillaume Gricourt, Melissa N'Debi, Alexandre Soulier, Elisabeth Trawinski, Jean-Michel Pawlotsky |
| EPI_ISL_3476471 | BIORHIN | Department of Virology, Henri Mondor University Hospital, Assistance Publique Hôpitaux de Paris, Université Paris-Est Créteil, INSERM U955 | Christophe Rodriguez, Slim Fourati, Vanessa Demontant, Guillaume Gricourt, Melissa N'Debi, Alexandre Soulier, Elisabeth Trawinski, Jean-Michel Pawlotsky |
| EPI_ISL_3476472 | GH A.CHENEVIER-H.MONDOR | Department of Virology, Henri Mondor University Hospital, | Christophe Rodriguez, Slim Fourati, Vanessa Demontant, Guillaume Gricourt, Melissa N'Debi, Alexandre Soulier, Elisabeth Trawinski, Jean-Michel Pawlotsky |

| | | | |
|---|---|---|---|
| | | Assistance Publique Hôpitaux de Paris, Université Paris-Est Créteil, INSERM U955 | |
| EPI_ISL_3476473 | laboratoire Belle Epine | Department of Virology, Henri Mondor University Hospital, Assistance Publique Hôpitaux de Paris, Université Paris-Est Créteil, INSERM U955 | Christophe Rodriguez, Slim Fourati, Vanessa Demontant, Guillaume Gricourt, Melissa N'Debi, Alexandre Soulier, Elisabeth Trawinski, Jean-Michel Pawlotsky |
| EPI_ISL_3476475 | SELAS MEDILYS | Department of Virology, Henri Mondor University Hospital, Assistance Publique Hôpitaux de Paris, Université Paris-Est Créteil, INSERM U955 | Christophe Rodriguez, Slim Fourati, Vanessa Demontant, Guillaume Gricourt, Melissa N'Debi, Alexandre Soulier, Elisabeth Trawinski, Jean-Michel Pawlotsky |
| EPI_ISL_3476477, EPI_ISL_3476478, EPI_ISL_3476479, EPI_ISL_3476480 | BIOMNIS EUROFINS IVRY | Department of Virology, Henri Mondor University Hospital, Assistance Publique Hôpitaux de Paris, Université Paris-Est Créteil, INSERM U955 | Christophe Rodriguez, Slim Fourati, Vanessa Demontant, Guillaume Gricourt, Melissa N'Debi, Alexandre Soulier, Elisabeth Trawinski, Jean-Michel Pawlotsky |
| EPI_ISL_3476481 | LISIEUX Cerballiance | Department of Virology, Henri Mondor University Hospital, Assistance Publique Hôpitaux de Paris, Université Paris-Est Créteil, INSERM U955 | Christophe Rodriguez, Slim Fourati, Vanessa Demontant, Guillaume Gricourt, Melissa N'Debi, Alexandre Soulier, Elisabeth Trawinski, Jean-Michel Pawlotsky |
| EPI_ISL_3476482 | BIORHIN | Department of Virology, Henri Mondor University Hospital, Assistance Publique Hôpitaux de Paris, Université Paris-Est Créteil, INSERM U955 | Christophe Rodriguez, Slim Fourati, Vanessa Demontant, Guillaume Gricourt, Melissa N'Debi, Alexandre Soulier, Elisabeth Trawinski, Jean-Michel Pawlotsky |
| EPI_ISL_3476483 | GH A.CHENEVIER-H.MONDOR | Department of Virology, Henri Mondor University Hospital, Assistance Publique Hôpitaux de Paris, Université Paris-Est Créteil, INSERM U955 | Christophe Rodriguez, Slim Fourati, Vanessa Demontant, Guillaume Gricourt, Melissa N'Debi, Alexandre Soulier, Elisabeth Trawinski, Jean-Michel Pawlotsky |
| EPI_ISL_3476485 | CH.LE RAINCY_MONTFERMEIL | Department of Virology, Henri Mondor University Hospital, Assistance Publique Hôpitaux de Paris, Université Paris-Est Créteil, INSERM U955 | Christophe Rodriguez, Slim Fourati, Vanessa Demontant, Guillaume Gricourt, Melissa N'Debi, Alexandre Soulier, Elisabeth Trawinski, Jean-Michel Pawlotsky |
| EPI_ISL_3476486 | GH A.CHENEVIER-H.MONDOR | Department of Virology, Henri Mondor University Hospital, Assistance Publique Hôpitaux de Paris, Université Paris-Est Créteil, INSERM U955 | Christophe Rodriguez, Slim Fourati, Vanessa Demontant, Guillaume Gricourt, Melissa N'Debi, Alexandre Soulier, Elisabeth Trawinski, Jean-Michel Pawlotsky |
| EPI_ISL_3476487 | SELAS MEDILYS | Department of Virology, Henri Mondor University Hospital, Assistance Publique Hôpitaux de Paris, Université Paris-Est Créteil, INSERM U955 | Christophe Rodriguez, Slim Fourati, Vanessa Demontant, Guillaume Gricourt, Melissa N'Debi, Alexandre Soulier, Elisabeth Trawinski, Jean-Michel Pawlotsky |
| EPI_ISL_3476488 | GH A.CHENEVIER-H.MONDOR | Department of Virology, Henri Mondor University Hospital, Assistance Publique Hôpitaux de Paris, Université Paris-Est Créteil, INSERM U955 | Christophe Rodriguez, Slim Fourati, Vanessa Demontant, Guillaume Gricourt, Melissa N'Debi, Alexandre Soulier, Elisabeth Trawinski, Jean-Michel Pawlotsky |
| EPI_ISL_3476489 | BROUSSAIS | Department of Virology, Henri Mondor University Hospital, Assistance Publique Hôpitaux de Paris, Université Paris-Est Créteil, INSERM U955 | Christophe Rodriguez, Slim Fourati, Vanessa Demontant, Guillaume Gricourt, Melissa N'Debi, Alexandre Soulier, Elisabeth Trawinski, Jean-Michel Pawlotsky |
| EPI_ISL_3476490, EPI_ISL_3476492 | SELAS MEDILYS | Department of Virology, Henri Mondor University Hospital, Assistance Publique Hôpitaux de Paris, Université Paris-Est Créteil, INSERM U955 | Christophe Rodriguez, Slim Fourati, Vanessa Demontant, Guillaume Gricourt, Melissa N'Debi, Alexandre Soulier, Elisabeth Trawinski, Jean-Michel Pawlotsky |
| EPI_ISL_3476493 | BIORHIN | Department of Virology, Henri Mondor University Hospital, Assistance Publique Hôpitaux de Paris, Université Paris-Est Créteil, INSERM U955 | Christophe Rodriguez, Slim Fourati, Vanessa Demontant, Guillaume Gricourt, Melissa N'Debi, Alexandre Soulier, Elisabeth Trawinski, Jean-Michel Pawlotsky |
| EPI_ISL_3476494 | BIOMNIS EUROFINS IVRY | Department of Virology, Henri Mondor University Hospital, Assistance Publique Hôpitaux de Paris, Université Paris-Est Créteil, INSERM U955 | Christophe Rodriguez, Slim Fourati, Vanessa Demontant, Guillaume Gricourt, Melissa N'Debi, Alexandre Soulier, Elisabeth Trawinski, Jean-Michel Pawlotsky |
| EPI_ISL_3476496 | GH A.CHENEVIER-H.MONDOR | Department of Virology, Henri Mondor University Hospital, Assistance Publique Hôpitaux de Paris, Université Paris-Est Créteil, INSERM U955 | Christophe Rodriguez, Slim Fourati, Vanessa Demontant, Guillaume Gricourt, Melissa N'Debi, Alexandre Soulier, Elisabeth Trawinski, Jean-Michel Pawlotsky |
| EPI_ISL_3476497 | CHU DE DIJON | Department of Virology, Henri Mondor University Hospital, Assistance Publique Hôpitaux de Paris, Université Paris-Est Créteil, INSERM U955 | Christophe Rodriguez, Slim Fourati, Vanessa Demontant, Guillaume Gricourt, Melissa N'Debi, Alexandre Soulier, Elisabeth Trawinski, Jean-Michel Pawlotsky |
| EPI_ISL_3476499 | GH A.CHENEVIER-H.MONDOR | Department of Virology, Henri Mondor University Hospital, Assistance Publique Hôpitaux de Paris, Université Paris-Est Créteil, INSERM U955 | Christophe Rodriguez, Slim Fourati, Vanessa Demontant, Guillaume Gricourt, Melissa N'Debi, Alexandre Soulier, Elisabeth Trawinski, Jean-Michel Pawlotsky |
| EPI_ISL_3476500 | CHU DE DIJON | Department of Virology, Henri Mondor University Hospital, Assistance Publique Hôpitaux de Paris, Université Paris-Est Créteil, INSERM U955 | Christophe Rodriguez, Slim Fourati, Vanessa Demontant, Guillaume Gricourt, Melissa N'Debi, Alexandre Soulier, Elisabeth Trawinski, Jean-Michel Pawlotsky |
| EPI_ISL_3476501 | CH. CHU DE REIMS | Department of Virology, Henri Mondor University Hospital, Assistance Publique Hôpitaux de Paris, Université Paris-Est Créteil, INSERM U955 | Christophe Rodriguez, Slim Fourati, Vanessa Demontant, Guillaume Gricourt, Melissa N'Debi, Alexandre Soulier, Elisabeth Trawinski, Jean-Michel Pawlotsky |
| EPI_ISL_3476502 | SELAS MEDILYS | Department of Virology, Henri Mondor University Hospital, Assistance Publique Hôpitaux de Paris, Université Paris-Est Créteil, INSERM U955 | Christophe Rodriguez, Slim Fourati, Vanessa Demontant, Guillaume Gricourt, Melissa N'Debi, Alexandre Soulier, Elisabeth Trawinski, Jean-Michel Pawlotsky |
| EPI_ISL_3476503 | Hôpital Bichat-Claude Bernard | Department of Virology, Henri Mondor University Hospital, Assistance Publique Hôpitaux de Paris, Université Paris-Est Créteil, INSERM U955 | Christophe Rodriguez, Slim Fourati, Vanessa Demontant, Guillaume Gricourt, Melissa N'Debi, Alexandre Soulier, Elisabeth Trawinski, Jean-Michel Pawlotsky |
| EPI_ISL_3476504 | SELAS MEDILYS | Department of Virology, Henri Mondor University Hospital, Assistance Publique Hôpitaux de Paris, Université Paris-Est Créteil, INSERM U955 | Christophe Rodriguez, Slim Fourati, Vanessa Demontant, Guillaume Gricourt, Melissa N'Debi, Alexandre Soulier, Elisabeth Trawinski, Jean-Michel Pawlotsky |
| EPI_ISL_3476505 | BIORHIN | Department of Virology, Henri Mondor University Hospital, Assistance Publique Hôpitaux de Paris, Université Paris-Est Créteil, INSERM U955 | Christophe Rodriguez, Slim Fourati, Vanessa Demontant, Guillaume Gricourt, Melissa N'Debi, Alexandre Soulier, Elisabeth Trawinski, Jean-Michel Pawlotsky |
| EPI_ISL_3476506 | BIOMNIS EUROFINS IVRY | Department of Virology, Henri Mondor University Hospital, Assistance Publique Hôpitaux de Paris, Université Paris-Est Créteil, INSERM U955 | Christophe Rodriguez, Slim Fourati, Vanessa Demontant, Guillaume Gricourt, Melissa N'Debi, Alexandre Soulier, Elisabeth Trawinski, Jean-Michel Pawlotsky |
| EPI_ISL_3476507 | GH A.CHENEVIER-H.MONDOR | Department of Virology, Henri Mondor University Hospital, Assistance Publique Hôpitaux de Paris, Université Paris-Est Créteil, INSERM U955 | Christophe Rodriguez, Slim Fourati, Vanessa Demontant, Guillaume Gricourt, Melissa N'Debi, Alexandre Soulier, Elisabeth Trawinski, Jean-Michel Pawlotsky |
| EPI_ISL_3476508 | CHU DE DIJON | Department of Virology, Henri Mondor University Hospital, Assistance Publique Hôpitaux de Paris, Université Paris-Est Créteil, INSERM U955 | Christophe Rodriguez, Slim Fourati, Vanessa Demontant, Guillaume Gricourt, Melissa N'Debi, Alexandre Soulier, Elisabeth Trawinski, Jean-Michel Pawlotsky |
| EPI_ISL_3476509 | BIOMNIS EUROFINS IVRY | Department of Virology, Henri Mondor University Hospital, Assistance Publique Hôpitaux de Paris, Université Paris-Est Créteil, INSERM U955 | Christophe Rodriguez, Slim Fourati, Vanessa Demontant, Guillaume Gricourt, Melissa N'Debi, Alexandre Soulier, Elisabeth Trawinski, Jean-Michel Pawlotsky |
| EPI_ISL_3476510 | LISIEUX Cerballiance | Department of Virology, Henri Mondor University Hospital, Assistance Publique Hôpitaux de Paris, Université Paris-Est Créteil, INSERM U955 | Christophe Rodriguez, Slim Fourati, Vanessa Demontant, Guillaume Gricourt, Melissa N'Debi, Alexandre Soulier, Elisabeth Trawinski, Jean-Michel Pawlotsky |
| EPI_ISL_3476511 | CH. CHU DE REIMS | Department of Virology, Henri Mondor University Hospital, Assistance Publique Hôpitaux de Paris, Université Paris-Est Créteil, INSERM U955 | Christophe Rodriguez, Slim Fourati, Vanessa Demontant, Guillaume Gricourt, Melissa N'Debi, Alexandre Soulier, Elisabeth Trawinski, Jean-Michel Pawlotsky |
| EPI_ISL_3476512 | SELAS MEDILYS | Department of Virology, Henri Mondor University Hospital, Assistance Publique Hôpitaux de Paris, Université Paris-Est Créteil, INSERM U955 | Christophe Rodriguez, Slim Fourati, Vanessa Demontant, Guillaume Gricourt, Melissa N'Debi, Alexandre Soulier, Elisabeth Trawinski, Jean-Michel Pawlotsky |
| EPI_ISL_3476513 | Biogroup Bio Lam-LCD Saint-Denis | Department of Virology, Henri Mondor University Hospital, Assistance Publique Hôpitaux de Paris, Université Paris-Est Créteil, INSERM U955 | Christophe Rodriguez, Slim Fourati, Vanessa Demontant, Guillaume Gricourt, Melissa N'Debi, Alexandre Soulier, Elisabeth Trawinski, Jean-Michel Pawlotsky |

| | | | |
|---|---|---|---|
| EPI_ISL_3476514, EPI_ISL_3476515 | BIOMNIS EUROFINS IVRY | Department of Virology, Henri Mondor University Hospital, Assistance Publique Hôpitaux de Paris, Université Paris-Est Créteil, INSERM U955 | Christophe Rodriguez, Slim Fourati, Vanessa Demontant, Guillaume Gricourt, Melissa N'Debi, Alexandre Soulier, Elisabeth Trawinski, Jean-Michel Pawlotsky |
| EPI_ISL_3476516 | BROUSSAIS | Department of Virology, Henri Mondor University Hospital, Assistance Publique Hôpitaux de Paris, Université Paris-Est Créteil, INSERM U955 | Christophe Rodriguez, Slim Fourati, Vanessa Demontant, Guillaume Gricourt, Melissa N'Debi, Alexandre Soulier, Elisabeth Trawinski, Jean-Michel Pawlotsky |
| EPI_ISL_3476518 | SELAS MEDILYS | Department of Virology, Henri Mondor University Hospital, Assistance Publique Hôpitaux de Paris, Université Paris-Est Créteil, INSERM U955 | Christophe Rodriguez, Slim Fourati, Vanessa Demontant, Guillaume Gricourt, Melissa N'Debi, Alexandre Soulier, Elisabeth Trawinski, Jean-Michel Pawlotsky |
| EPI_ISL_3476519 | CH.LE RAINCY_MONTFERMEIL | Department of Virology, Henri Mondor University Hospital, Assistance Publique Hôpitaux de Paris, Université Paris-Est Créteil, INSERM U955 | Christophe Rodriguez, Slim Fourati, Vanessa Demontant, Guillaume Gricourt, Melissa N'Debi, Alexandre Soulier, Elisabeth Trawinski, Jean-Michel Pawlotsky |
| EPI_ISL_3476520 | BROUSSAIS | Department of Virology, Henri Mondor University Hospital, Assistance Publique Hôpitaux de Paris, Université Paris-Est Créteil, INSERM U955 | Christophe Rodriguez, Slim Fourati, Vanessa Demontant, Guillaume Gricourt, Melissa N'Debi, Alexandre Soulier, Elisabeth Trawinski, Jean-Michel Pawlotsky |
| EPI_ISL_3476521 | BIORHIN | Department of Virology, Henri Mondor University Hospital, Assistance Publique Hôpitaux de Paris, Université Paris-Est Créteil, INSERM U955 | Christophe Rodriguez, Slim Fourati, Vanessa Demontant, Guillaume Gricourt, Melissa N'Debi, Alexandre Soulier, Elisabeth Trawinski, Jean-Michel Pawlotsky |
| EPI_ISL_3476522 | LISIEUX Cerballiance | Department of Virology, Henri Mondor University Hospital, Assistance Publique Hôpitaux de Paris, Université Paris-Est Créteil, INSERM U955 | Christophe Rodriguez, Slim Fourati, Vanessa Demontant, Guillaume Gricourt, Melissa N'Debi, Alexandre Soulier, Elisabeth Trawinski, Jean-Michel Pawlotsky |
| EPI_ISL_3476523 | Cerballiance Paris IDF est Saint-Denis | Department of Virology, Henri Mondor University Hospital, Assistance Publique Hôpitaux de Paris, Université Paris-Est Créteil, INSERM U955 | Christophe Rodriguez, Slim Fourati, Vanessa Demontant, Guillaume Gricourt, Melissa N'Debi, Alexandre Soulier, Elisabeth Trawinski, Jean-Michel Pawlotsky |
| EPI_ISL_3476524 | GH A.CHENEVIER-H.MONDOR | Department of Virology, Henri Mondor University Hospital, Assistance Publique Hôpitaux de Paris, Université Paris-Est Créteil, INSERM U955 | Christophe Rodriguez, Slim Fourati, Vanessa Demontant, Guillaume Gricourt, Melissa N'Debi, Alexandre Soulier, Elisabeth Trawinski, Jean-Michel Pawlotsky |
| EPI_ISL_3476525 | Hôpital Armand-Trousseau | Department of Virology, Henri Mondor University Hospital, Assistance Publique Hôpitaux de Paris, Université Paris-Est Créteil, INSERM U955 | Christophe Rodriguez, Slim Fourati, Vanessa Demontant, Guillaume Gricourt, Melissa N'Debi, Alexandre Soulier, Elisabeth Trawinski, Jean-Michel Pawlotsky |
| EPI_ISL_3476526 | Laboratoire Ana-L | Department of Virology, Henri Mondor University Hospital, Assistance Publique Hôpitaux de Paris, Université Paris-Est Créteil, INSERM U955 | Christophe Rodriguez, Slim Fourati, Vanessa Demontant, Guillaume Gricourt, Melissa N'Debi, Alexandre Soulier, Elisabeth Trawinski, Jean-Michel Pawlotsky |
| EPI_ISL_3476527 | BIOMNIS EUROFINS IVRY | Department of Virology, Henri Mondor University Hospital, Assistance Publique Hôpitaux de Paris, Université Paris-Est Créteil, INSERM U955 | Christophe Rodriguez, Slim Fourati, Vanessa Demontant, Guillaume Gricourt, Melissa N'Debi, Alexandre Soulier, Elisabeth Trawinski, Jean-Michel Pawlotsky |
| EPI_ISL_3476528 | BIORHIN | Department of Virology, Henri Mondor University Hospital, Assistance Publique Hôpitaux de Paris, Université Paris-Est Créteil, INSERM U955 | Christophe Rodriguez, Slim Fourati, Vanessa Demontant, Guillaume Gricourt, Melissa N'Debi, Alexandre Soulier, Elisabeth Trawinski, Jean-Michel Pawlotsky |
| EPI_ISL_3476529, EPI_ISL_3476530 | BIOMNIS EUROFINS IVRY | Department of Virology, Henri Mondor University Hospital, Assistance Publique Hôpitaux de Paris, Université Paris-Est Créteil, INSERM U955 | Christophe Rodriguez, Slim Fourati, Vanessa Demontant, Guillaume Gricourt, Melissa N'Debi, Alexandre Soulier, Elisabeth Trawinski, Jean-Michel Pawlotsky |
| EPI_ISL_3476531 | SELAS MEDILYS | Department of Virology, Henri Mondor University Hospital, Assistance Publique Hôpitaux de Paris, Université Paris-Est Créteil, INSERM U955 | Christophe Rodriguez, Slim Fourati, Vanessa Demontant, Guillaume Gricourt, Melissa N'Debi, Alexandre Soulier, Elisabeth Trawinski, Jean-Michel Pawlotsky |
| EPI_ISL_3476532, EPI_ISL_3476533 | LISIEUX Cerballiance | Department of Virology, Henri Mondor University Hospital, Assistance Publique Hôpitaux de Paris, Université Paris-Est Créteil, INSERM U955 | Christophe Rodriguez, Slim Fourati, Vanessa Demontant, Guillaume Gricourt, Melissa N'Debi, Alexandre Soulier, Elisabeth Trawinski, Jean-Michel Pawlotsky |
| EPI_ISL_3476534 | GROUPE HOSPITALIER SUD ILE DE FRANCE | Department of Virology, Henri Mondor University Hospital, Assistance Publique Hôpitaux de Paris, Université Paris-Est Créteil, INSERM U955 | Christophe Rodriguez, Slim Fourati, Vanessa Demontant, Guillaume Gricourt, Melissa N'Debi, Alexandre Soulier, Elisabeth Trawinski, Jean-Michel Pawlotsky |
| EPI_ISL_3476535 | CH.LE RAINCY_MONTFERMEIL | Department of Virology, Henri Mondor University Hospital, Assistance Publique Hôpitaux de Paris, Université Paris-Est Créteil, INSERM U955 | Christophe Rodriguez, Slim Fourati, Vanessa Demontant, Guillaume Gricourt, Melissa N'Debi, Alexandre Soulier, Elisabeth Trawinski, Jean-Michel Pawlotsky |
| EPI_ISL_3476536 | LISIEUX Cerballiance | Department of Virology, Henri Mondor University Hospital, Assistance Publique Hôpitaux de Paris, Université Paris-Est Créteil, INSERM U955 | Christophe Rodriguez, Slim Fourati, Vanessa Demontant, Guillaume Gricourt, Melissa N'Debi, Alexandre Soulier, Elisabeth Trawinski, Jean-Michel Pawlotsky |
| EPI_ISL_3476538 | BIORHIN | Department of Virology, Henri Mondor University Hospital, Assistance Publique Hôpitaux de Paris, Université Paris-Est Créteil, INSERM U955 | Christophe Rodriguez, Slim Fourati, Vanessa Demontant, Guillaume Gricourt, Melissa N'Debi, Alexandre Soulier, Elisabeth Trawinski, Jean-Michel Pawlotsky |
| EPI_ISL_3476539 | CHU DE DIJON | Department of Virology, Henri Mondor University Hospital, Assistance Publique Hôpitaux de Paris, Université Paris-Est Créteil, INSERM U955 | Christophe Rodriguez, Slim Fourati, Vanessa Demontant, Guillaume Gricourt, Melissa N'Debi, Alexandre Soulier, Elisabeth Trawinski, Jean-Michel Pawlotsky |
| EPI_ISL_3476540 | SELAS MEDILYS | Department of Virology, Henri Mondor University Hospital, Assistance Publique Hôpitaux de Paris, Université Paris-Est Créteil, INSERM U955 | Christophe Rodriguez, Slim Fourati, Vanessa Demontant, Guillaume Gricourt, Melissa N'Debi, Alexandre Soulier, Elisabeth Trawinski, Jean-Michel Pawlotsky |
| EPI_ISL_3476541 | CH. CHU DE REIMS | Department of Virology, Henri Mondor University Hospital, Assistance Publique Hôpitaux de Paris, Université Paris-Est Créteil, INSERM U955 | Christophe Rodriguez, Slim Fourati, Vanessa Demontant, Guillaume Gricourt, Melissa N'Debi, Alexandre Soulier, Elisabeth Trawinski, Jean-Michel Pawlotsky |
| EPI_ISL_3476542 | BIOMNIS EUROFINS IVRY | Department of Virology, Henri Mondor University Hospital, Assistance Publique Hôpitaux de Paris, Université Paris-Est Créteil, INSERM U955 | Christophe Rodriguez, Slim Fourati, Vanessa Demontant, Guillaume Gricourt, Melissa N'Debi, Alexandre Soulier, Elisabeth Trawinski, Jean-Michel Pawlotsky |
| EPI_ISL_3476543 | CHU DE DIJON | Department of Virology, Henri Mondor University Hospital, Assistance Publique Hôpitaux de Paris, Université Paris-Est Créteil, INSERM U955 | Christophe Rodriguez, Slim Fourati, Vanessa Demontant, Guillaume Gricourt, Melissa N'Debi, Alexandre Soulier, Elisabeth Trawinski, Jean-Michel Pawlotsky |
| EPI_ISL_3476544, EPI_ISL_3476545 | GH A.CHENEVIER-H.MONDOR | Department of Virology, Henri Mondor University Hospital, Assistance Publique Hôpitaux de Paris, Université Paris-Est Créteil, INSERM U955 | Christophe Rodriguez, Slim Fourati, Vanessa Demontant, Guillaume Gricourt, Melissa N'Debi, Alexandre Soulier, Elisabeth Trawinski, Jean-Michel Pawlotsky |
| EPI_ISL_3476546 | SELAS MEDILYS | Department of Virology, Henri Mondor University Hospital, Assistance Publique Hôpitaux de Paris, Université Paris-Est Créteil, INSERM U955 | Christophe Rodriguez, Slim Fourati, Vanessa Demontant, Guillaume Gricourt, Melissa N'Debi, Alexandre Soulier, Elisabeth Trawinski, Jean-Michel Pawlotsky |
| EPI_ISL_3476547 | CH.LE RAINCY_MONTFERMEIL | Department of Virology, Henri Mondor University Hospital, Assistance Publique Hôpitaux de Paris, Université Paris-Est Créteil, INSERM U955 | Christophe Rodriguez, Slim Fourati, Vanessa Demontant, Guillaume Gricourt, Melissa N'Debi, Alexandre Soulier, Elisabeth Trawinski, Jean-Michel Pawlotsky |
| EPI_ISL_3476548 | SELAS MEDILYS | Department of Virology, Henri Mondor University Hospital, Assistance Publique Hôpitaux de Paris, Université Paris-Est Créteil, INSERM U955 | Christophe Rodriguez, Slim Fourati, Vanessa Demontant, Guillaume Gricourt, Melissa N'Debi, Alexandre Soulier, Elisabeth Trawinski, Jean-Michel Pawlotsky |
| EPI_ISL_3476549 | Biogroup Bio Lam-LCD Saint-Denis | Department of Virology, Henri Mondor University Hospital, Assistance Publique Hôpitaux de Paris, Université Paris-Est Créteil, INSERM U955 | Christophe Rodriguez, Slim Fourati, Vanessa Demontant, Guillaume Gricourt, Melissa N'Debi, Alexandre Soulier, Elisabeth Trawinski, Jean-Michel Pawlotsky |
| EPI_ISL_3476550 | BIOMNIS EUROFINS IVRY | Department of Virology, Henri Mondor University Hospital, Assistance Publique Hôpitaux de Paris, Université Paris-Est Créteil, INSERM U955 | Christophe Rodriguez, Slim Fourati, Vanessa Demontant, Guillaume Gricourt, Melissa N'Debi, Alexandre Soulier, Elisabeth Trawinski, Jean-Michel Pawlotsky |
| EPI_ISL_3476551 | GH A.CHENEVIER-H.MONDOR | Department of Virology, Henri Mondor University Hospital, Assistance Publique Hôpitaux de Paris, Université Paris-Est Créteil, INSERM U955 | Christophe Rodriguez, Slim Fourati, Vanessa Demontant, Guillaume Gricourt, Melissa N'Debi, Alexandre Soulier, Elisabeth Trawinski, Jean-Michel Pawlotsky |

| | | | |
|---|---|---|---|
| EPI_ISL_3476552 | SELAS MEDILYS | Department of Virology, Henri Mondor University Hospital, Assistance Publique Hôpitaux de Paris, Université Paris-Est Créteil, INSERM U955 | Christophe Rodriguez, Slim Fourati, Vanessa Demontant, Guillaume Gricourt, Melissa N'Debi, Alexandre Soulier, Elisabeth Trawinski, Jean-Michel Pawlotsky |
| EPI_ISL_3476553 | Cerballiance Paris IDF est Saint-Denis | Department of Virology, Henri Mondor University Hospital, Assistance Publique Hôpitaux de Paris, Université Paris-Est Créteil, INSERM U955 | Christophe Rodriguez, Slim Fourati, Vanessa Demontant, Guillaume Gricourt, Melissa N'Debi, Alexandre Soulier, Elisabeth Trawinski, Jean-Michel Pawlotsky |
| EPI_ISL_3476554 | SELAS MEDILYS | Department of Virology, Henri Mondor University Hospital, Assistance Publique Hôpitaux de Paris, Université Paris-Est Créteil, INSERM U955 | Christophe Rodriguez, Slim Fourati, Vanessa Demontant, Guillaume Gricourt, Melissa N'Debi, Alexandre Soulier, Elisabeth Trawinski, Jean-Michel Pawlotsky |
| EPI_ISL_3476555 | GH A.CHENEVIER-H.MONDOR | Department of Virology, Henri Mondor University Hospital, Assistance Publique Hôpitaux de Paris, Université Paris-Est Créteil, INSERM U955 | Christophe Rodriguez, Slim Fourati, Vanessa Demontant, Guillaume Gricourt, Melissa N'Debi, Alexandre Soulier, Elisabeth Trawinski, Jean-Michel Pawlotsky |
| EPI_ISL_3476556 | Hôpital Cochin | Department of Virology, Henri Mondor University Hospital, Assistance Publique Hôpitaux de Paris, Université Paris-Est Créteil, INSERM U955 | Christophe Rodriguez, Slim Fourati, Vanessa Demontant, Guillaume Gricourt, Melissa N'Debi, Alexandre Soulier, Elisabeth Trawinski, Jean-Michel Pawlotsky |
| EPI_ISL_3476557 | CHU DE DIJON | Department of Virology, Henri Mondor University Hospital, Assistance Publique Hôpitaux de Paris, Université Paris-Est Créteil, INSERM U955 | Christophe Rodriguez, Slim Fourati, Vanessa Demontant, Guillaume Gricourt, Melissa N'Debi, Alexandre Soulier, Elisabeth Trawinski, Jean-Michel Pawlotsky |
| EPI_ISL_3476558 | SELAS MEDILYS | Department of Virology, Henri Mondor University Hospital, Assistance Publique Hôpitaux de Paris, Université Paris-Est Créteil, INSERM U955 | Christophe Rodriguez, Slim Fourati, Vanessa Demontant, Guillaume Gricourt, Melissa N'Debi, Alexandre Soulier, Elisabeth Trawinski, Jean-Michel Pawlotsky |
| EPI_ISL_3476559 | Biogroup Bio Lam-LCD Saint-Denis | Department of Virology, Henri Mondor University Hospital, Assistance Publique Hôpitaux de Paris, Université Paris-Est Créteil, INSERM U955 | Christophe Rodriguez, Slim Fourati, Vanessa Demontant, Guillaume Gricourt, Melissa N'Debi, Alexandre Soulier, Elisabeth Trawinski, Jean-Michel Pawlotsky |
| EPI_ISL_3476560 | Cerballiance Paris IDF est Saint-Denis | Department of Virology, Henri Mondor University Hospital, Assistance Publique Hôpitaux de Paris, Université Paris-Est Créteil, INSERM U955 | Christophe Rodriguez, Slim Fourati, Vanessa Demontant, Guillaume Gricourt, Melissa N'Debi, Alexandre Soulier, Elisabeth Trawinski, Jean-Michel Pawlotsky |
| EPI_ISL_3476561 | SELAS MEDILYS | Department of Virology, Henri Mondor University Hospital, Assistance Publique Hôpitaux de Paris, Université Paris-Est Créteil, INSERM U955 | Christophe Rodriguez, Slim Fourati, Vanessa Demontant, Guillaume Gricourt, Melissa N'Debi, Alexandre Soulier, Elisabeth Trawinski, Jean-Michel Pawlotsky |
| EPI_ISL_3476562 | BIOMNIS EUROFINS IVRY | Department of Virology, Henri Mondor University Hospital, Assistance Publique Hôpitaux de Paris, Université Paris-Est Créteil, INSERM U955 | Christophe Rodriguez, Slim Fourati, Vanessa Demontant, Guillaume Gricourt, Melissa N'Debi, Alexandre Soulier, Elisabeth Trawinski, Jean-Michel Pawlotsky |
| EPI_ISL_3476563, EPI_ISL_3476564 | SELAS MEDILYS | Department of Virology, Henri Mondor University Hospital, Assistance Publique Hôpitaux de Paris, Université Paris-Est Créteil, INSERM U955 | Christophe Rodriguez, Slim Fourati, Vanessa Demontant, Guillaume Gricourt, Melissa N'Debi, Alexandre Soulier, Elisabeth Trawinski, Jean-Michel Pawlotsky |
| EPI_ISL_3476565 | Laboratoire Ana-L | Department of Virology, Henri Mondor University Hospital, Assistance Publique Hôpitaux de Paris, Université Paris-Est Créteil, INSERM U955 | Christophe Rodriguez, Slim Fourati, Vanessa Demontant, Guillaume Gricourt, Melissa N'Debi, Alexandre Soulier, Elisabeth Trawinski, Jean-Michel Pawlotsky |
| EPI_ISL_3476566 | SELAS MEDILYS | Department of Virology, Henri Mondor University Hospital, Assistance Publique Hôpitaux de Paris, Université Paris-Est Créteil, INSERM U955 | Christophe Rodriguez, Slim Fourati, Vanessa Demontant, Guillaume Gricourt, Melissa N'Debi, Alexandre Soulier, Elisabeth Trawinski, Jean-Michel Pawlotsky |
| EPI_ISL_3476567 | LAM BARRAND | Department of Virology, Henri Mondor University Hospital, Assistance Publique Hôpitaux de Paris, Université Paris-Est Créteil, INSERM U955 | Christophe Rodriguez, Slim Fourati, Vanessa Demontant, Guillaume Gricourt, Melissa N'Debi, Alexandre Soulier, Elisabeth Trawinski, Jean-Michel Pawlotsky |
| EPI_ISL_3476568 | CHU DE DIJON | Department of Virology, Henri Mondor University Hospital, Assistance Publique Hôpitaux de Paris, Université Paris-Est Créteil, INSERM U955 | Christophe Rodriguez, Slim Fourati, Vanessa Demontant, Guillaume Gricourt, Melissa N'Debi, Alexandre Soulier, Elisabeth Trawinski, Jean-Michel Pawlotsky |
| EPI_ISL_3476569, EPI_ISL_3476570 | SELAS MEDILYS | Department of Virology, Henri Mondor University Hospital, Assistance Publique Hôpitaux de Paris, Université Paris-Est Créteil, INSERM U955 | Christophe Rodriguez, Slim Fourati, Vanessa Demontant, Guillaume Gricourt, Melissa N'Debi, Alexandre Soulier, Elisabeth Trawinski, Jean-Michel Pawlotsky |
| EPI_ISL_3476571 | BIOMNIS EUROFINS IVRY | Department of Virology, Henri Mondor University Hospital, Assistance Publique Hôpitaux de Paris, Université Paris-Est Créteil, INSERM U955 | Christophe Rodriguez, Slim Fourati, Vanessa Demontant, Guillaume Gricourt, Melissa N'Debi, Alexandre Soulier, Elisabeth Trawinski, Jean-Michel Pawlotsky |
| EPI_ISL_3476572 | Hôpital Armand-Trousseau | Department of Virology, Henri Mondor University Hospital, Assistance Publique Hôpitaux de Paris, Université Paris-Est Créteil, INSERM U955 | Christophe Rodriguez, Slim Fourati, Vanessa Demontant, Guillaume Gricourt, Melissa N'Debi, Alexandre Soulier, Elisabeth Trawinski, Jean-Michel Pawlotsky |
| EPI_ISL_3476573 | CHU DE DIJON | Department of Virology, Henri Mondor University Hospital, Assistance Publique Hôpitaux de Paris, Université Paris-Est Créteil, INSERM U955 | Christophe Rodriguez, Slim Fourati, Vanessa Demontant, Guillaume Gricourt, Melissa N'Debi, Alexandre Soulier, Elisabeth Trawinski, Jean-Michel Pawlotsky |
| EPI_ISL_3476574, EPI_ISL_3476575 | SELAS MEDILYS | Department of Virology, Henri Mondor University Hospital, Assistance Publique Hôpitaux de Paris, Université Paris-Est Créteil, INSERM U955 | Christophe Rodriguez, Slim Fourati, Vanessa Demontant, Guillaume Gricourt, Melissa N'Debi, Alexandre Soulier, Elisabeth Trawinski, Jean-Michel Pawlotsky |
| EPI_ISL_3476576 | BIOMNIS EUROFINS IVRY | Department of Virology, Henri Mondor University Hospital, Assistance Publique Hôpitaux de Paris, Université Paris-Est Créteil, INSERM U955 | Christophe Rodriguez, Slim Fourati, Vanessa Demontant, Guillaume Gricourt, Melissa N'Debi, Alexandre Soulier, Elisabeth Trawinski, Jean-Michel Pawlotsky |
| EPI_ISL_3476578 | GH A.CHENEVIER-H.MONDOR | Department of Virology, Henri Mondor University Hospital, Assistance Publique Hôpitaux de Paris, Université Paris-Est Créteil, INSERM U955 | Christophe Rodriguez, Slim Fourati, Vanessa Demontant, Guillaume Gricourt, Melissa N'Debi, Alexandre Soulier, Elisabeth Trawinski, Jean-Michel Pawlotsky |
| EPI_ISL_3476579 | SELAS MEDILYS | Department of Virology, Henri Mondor University Hospital, Assistance Publique Hôpitaux de Paris, Université Paris-Est Créteil, INSERM U955 | Christophe Rodriguez, Slim Fourati, Vanessa Demontant, Guillaume Gricourt, Melissa N'Debi, Alexandre Soulier, Elisabeth Trawinski, Jean-Michel Pawlotsky |
| EPI_ISL_3476581 | CHU DE DIJON | Department of Virology, Henri Mondor University Hospital, Assistance Publique Hôpitaux de Paris, Université Paris-Est Créteil, INSERM U955 | Christophe Rodriguez, Slim Fourati, Vanessa Demontant, Guillaume Gricourt, Melissa N'Debi, Alexandre Soulier, Elisabeth Trawinski, Jean-Michel Pawlotsky |
| EPI_ISL_3476582 | SELAS MEDILYS | Department of Virology, Henri Mondor University Hospital, Assistance Publique Hôpitaux de Paris, Université Paris-Est Créteil, INSERM U955 | Christophe Rodriguez, Slim Fourati, Vanessa Demontant, Guillaume Gricourt, Melissa N'Debi, Alexandre Soulier, Elisabeth Trawinski, Jean-Michel Pawlotsky |
| EPI_ISL_3476583 | LBM EVORIAL NEVERS | Department of Virology, Henri Mondor University Hospital, Assistance Publique Hôpitaux de Paris, Université Paris-Est Créteil, INSERM U955 | Christophe Rodriguez, Slim Fourati, Vanessa Demontant, Guillaume Gricourt, Melissa N'Debi, Alexandre Soulier, Elisabeth Trawinski, Jean-Michel Pawlotsky |
| EPI_ISL_3476584 | Hôpital Bichat-Claude Bernard | Department of Virology, Henri Mondor University Hospital, Assistance Publique Hôpitaux de Paris, Université Paris-Est Créteil, INSERM U955 | Christophe Rodriguez, Slim Fourati, Vanessa Demontant, Guillaume Gricourt, Melissa N'Debi, Alexandre Soulier, Elisabeth Trawinski, Jean-Michel Pawlotsky |
| EPI_ISL_3476585 | Hôpital Avicenne | Department of Virology, Henri Mondor University Hospital, Assistance Publique Hôpitaux de Paris, Université Paris-Est Créteil, INSERM U955 | Christophe Rodriguez, Slim Fourati, Vanessa Demontant, Guillaume Gricourt, Melissa N'Debi, Alexandre Soulier, Elisabeth Trawinski, Jean-Michel Pawlotsky |
| EPI_ISL_3476586, EPI_ISL_3476587, EPI_ISL_3476588 | SELAS MEDILYS | Department of Virology, Henri Mondor University Hospital, Assistance Publique Hôpitaux de Paris, Université Paris-Est Créteil, INSERM U955 | Christophe Rodriguez, Slim Fourati, Vanessa Demontant, Guillaume Gricourt, Melissa N'Debi, Alexandre Soulier, Elisabeth Trawinski, Jean-Michel Pawlotsky |
| EPI_ISL_3476589 | CHU DE DIJON | Department of Virology, Henri Mondor University Hospital, Assistance Publique Hôpitaux de Paris, Université Paris-Est Créteil, INSERM U955 | Christophe Rodriguez, Slim Fourati, Vanessa Demontant, Guillaume Gricourt, Melissa N'Debi, Alexandre Soulier, Elisabeth Trawinski, Jean-Michel Pawlotsky |
| EPI_ISL_3476590 | SELAS MEDILYS | Department of Virology, Henri Mondor University Hospital, Assistance Publique Hôpitaux de Paris, Université Paris-Est Créteil, INSERM U955 | Christophe Rodriguez, Slim Fourati, Vanessa Demontant, Guillaume Gricourt, Melissa N'Debi, Alexandre Soulier, Elisabeth Trawinski, Jean-Michel Pawlotsky |

| | | | |
|---|---|---|---|
| EPI_ISL_3476591 | BIOMNIS EUROFINS IVRY | Department of Virology, Henri Mondor University Hospital, Assistance Publique Hôpitaux de Paris, Université Paris-Est Créteil, INSERM U955 | Christophe Rodriguez, Slim Fourati, Vanessa Demontant, Guillaume Gricourt, Melissa N'Debi, Alexandre Soulier, Elisabeth Trawinski, Jean-Michel Pawlotsky |
| EPI_ISL_3476592 | GH A.CHENEVIER-H.MONDOR | Department of Virology, Henri Mondor University Hospital, Assistance Publique Hôpitaux de Paris, Université Paris-Est Créteil, INSERM U955 | Christophe Rodriguez, Slim Fourati, Vanessa Demontant, Guillaume Gricourt, Melissa N'Debi, Alexandre Soulier, Elisabeth Trawinski, Jean-Michel Pawlotsky |
| EPI_ISL_3476593 | Hôpital Cochin | Department of Virology, Henri Mondor University Hospital, Assistance Publique Hôpitaux de Paris, Université Paris-Est Créteil, INSERM U955 | Christophe Rodriguez, Slim Fourati, Vanessa Demontant, Guillaume Gricourt, Melissa N'Debi, Alexandre Soulier, Elisabeth Trawinski, Jean-Michel Pawlotsky |
| EPI_ISL_3476594 | BIOMNIS EUROFINS IVRY | Department of Virology, Henri Mondor University Hospital, Assistance Publique Hôpitaux de Paris, Université Paris-Est Créteil, INSERM U955 | Christophe Rodriguez, Slim Fourati, Vanessa Demontant, Guillaume Gricourt, Melissa N'Debi, Alexandre Soulier, Elisabeth Trawinski, Jean-Michel Pawlotsky |
| EPI_ISL_3476595 | SELAS MEDILYS | Department of Virology, Henri Mondor University Hospital, Assistance Publique Hôpitaux de Paris, Université Paris-Est Créteil, INSERM U955 | Christophe Rodriguez, Slim Fourati, Vanessa Demontant, Guillaume Gricourt, Melissa N'Debi, Alexandre Soulier, Elisabeth Trawinski, Jean-Michel Pawlotsky |
| EPI_ISL_3476596 | LISIEUX Cerballiance | Department of Virology, Henri Mondor University Hospital, Assistance Publique Hôpitaux de Paris, Université Paris-Est Créteil, INSERM U955 | Christophe Rodriguez, Slim Fourati, Vanessa Demontant, Guillaume Gricourt, Melissa N'Debi, Alexandre Soulier, Elisabeth Trawinski, Jean-Michel Pawlotsky |
| EPI_ISL_3476597 | Hôpital Cochin | Department of Virology, Henri Mondor University Hospital, Assistance Publique Hôpitaux de Paris, Université Paris-Est Créteil, INSERM U955 | Christophe Rodriguez, Slim Fourati, Vanessa Demontant, Guillaume Gricourt, Melissa N'Debi, Alexandre Soulier, Elisabeth Trawinski, Jean-Michel Pawlotsky |
| EPI_ISL_3476598, EPI_ISL_3476599 | SELAS MEDILYS | Department of Virology, Henri Mondor University Hospital, Assistance Publique Hôpitaux de Paris, Université Paris-Est Créteil, INSERM U955 | Christophe Rodriguez, Slim Fourati, Vanessa Demontant, Guillaume Gricourt, Melissa N'Debi, Alexandre Soulier, Elisabeth Trawinski, Jean-Michel Pawlotsky |
| EPI_ISL_3476600 | LAM BARRAND | Department of Virology, Henri Mondor University Hospital, Assistance Publique Hôpitaux de Paris, Université Paris-Est Créteil, INSERM U955 | Christophe Rodriguez, Slim Fourati, Vanessa Demontant, Guillaume Gricourt, Melissa N'Debi, Alexandre Soulier, Elisabeth Trawinski, Jean-Michel Pawlotsky |
| EPI_ISL_3476601 | BIOMNIS EUROFINS IVRY | Department of Virology, Henri Mondor University Hospital, Assistance Publique Hôpitaux de Paris, Université Paris-Est Créteil, INSERM U955 | Christophe Rodriguez, Slim Fourati, Vanessa Demontant, Guillaume Gricourt, Melissa N'Debi, Alexandre Soulier, Elisabeth Trawinski, Jean-Michel Pawlotsky |
| EPI_ISL_3476602, EPI_ISL_3476603 | SELAS MEDILYS | Department of Virology, Henri Mondor University Hospital, Assistance Publique Hôpitaux de Paris, Université Paris-Est Créteil, INSERM U955 | Christophe Rodriguez, Slim Fourati, Vanessa Demontant, Guillaume Gricourt, Melissa N'Debi, Alexandre Soulier, Elisabeth Trawinski, Jean-Michel Pawlotsky |
| EPI_ISL_3476604, EPI_ISL_3476605 | BIOMNIS EUROFINS IVRY | Department of Virology, Henri Mondor University Hospital, Assistance Publique Hôpitaux de Paris, Université Paris-Est Créteil, INSERM U955 | Christophe Rodriguez, Slim Fourati, Vanessa Demontant, Guillaume Gricourt, Melissa N'Debi, Alexandre Soulier, Elisabeth Trawinski, Jean-Michel Pawlotsky |
| EPI_ISL_3476606 | Hôpital Armand-Trousseau | Department of Virology, Henri Mondor University Hospital, Assistance Publique Hôpitaux de Paris, Université Paris-Est Créteil, INSERM U955 | Christophe Rodriguez, Slim Fourati, Vanessa Demontant, Guillaume Gricourt, Melissa N'Debi, Alexandre Soulier, Elisabeth Trawinski, Jean-Michel Pawlotsky |
| EPI_ISL_3476607 | Cerballiance Paris IDF est Saint-Denis | Department of Virology, Henri Mondor University Hospital, Assistance Publique Hôpitaux de Paris, Université Paris-Est Créteil, INSERM U955 | Christophe Rodriguez, Slim Fourati, Vanessa Demontant, Guillaume Gricourt, Melissa N'Debi, Alexandre Soulier, Elisabeth Trawinski, Jean-Michel Pawlotsky |
| EPI_ISL_3476608 | BIOMNIS EUROFINS IVRY | Department of Virology, Henri Mondor University Hospital, Assistance Publique Hôpitaux de Paris, Université Paris-Est Créteil, INSERM U955 | Christophe Rodriguez, Slim Fourati, Vanessa Demontant, Guillaume Gricourt, Melissa N'Debi, Alexandre Soulier, Elisabeth Trawinski, Jean-Michel Pawlotsky |
| EPI_ISL_3476609 | Biogroup Bio Lam-LCD Saint-Denis | Department of Virology, Henri Mondor University Hospital, Assistance Publique Hôpitaux de Paris, Université Paris-Est Créteil, INSERM U955 | Christophe Rodriguez, Slim Fourati, Vanessa Demontant, Guillaume Gricourt, Melissa N'Debi, Alexandre Soulier, Elisabeth Trawinski, Jean-Michel Pawlotsky |
| EPI_ISL_3476610 | GH A.CHENEVIER-H.MONDOR | Department of Virology, Henri Mondor University Hospital, Assistance Publique Hôpitaux de Paris, Université Paris-Est Créteil, INSERM U955 | Christophe Rodriguez, Slim Fourati, Vanessa Demontant, Guillaume Gricourt, Melissa N'Debi, Alexandre Soulier, Elisabeth Trawinski, Jean-Michel Pawlotsky |
| EPI_ISL_3476611 | BIOMNIS EUROFINS IVRY | Department of Virology, Henri Mondor University Hospital, Assistance Publique Hôpitaux de Paris, Université Paris-Est Créteil, INSERM U955 | Christophe Rodriguez, Slim Fourati, Vanessa Demontant, Guillaume Gricourt, Melissa N'Debi, Alexandre Soulier, Elisabeth Trawinski, Jean-Michel Pawlotsky |
| EPI_ISL_3476612, EPI_ISL_3476613 | SELAS MEDILYS | Department of Virology, Henri Mondor University Hospital, Assistance Publique Hôpitaux de Paris, Université Paris-Est Créteil, INSERM U955 | Christophe Rodriguez, Slim Fourati, Vanessa Demontant, Guillaume Gricourt, Melissa N'Debi, Alexandre Soulier, Elisabeth Trawinski, Jean-Michel Pawlotsky |
| EPI_ISL_3476614 | BIORHIN | Department of Virology, Henri Mondor University Hospital, Assistance Publique Hôpitaux de Paris, Université Paris-Est Créteil, INSERM U955 | Christophe Rodriguez, Slim Fourati, Vanessa Demontant, Guillaume Gricourt, Melissa N'Debi, Alexandre Soulier, Elisabeth Trawinski, Jean-Michel Pawlotsky |
| EPI_ISL_3476615 | BIOMNIS EUROFINS IVRY | Department of Virology, Henri Mondor University Hospital, Assistance Publique Hôpitaux de Paris, Université Paris-Est Créteil, INSERM U955 | Christophe Rodriguez, Slim Fourati, Vanessa Demontant, Guillaume Gricourt, Melissa N'Debi, Alexandre Soulier, Elisabeth Trawinski, Jean-Michel Pawlotsky |
| EPI_ISL_3476616 | CH. CHU DE REIMS | Department of Virology, Henri Mondor University Hospital, Assistance Publique Hôpitaux de Paris, Université Paris-Est Créteil, INSERM U955 | Christophe Rodriguez, Slim Fourati, Vanessa Demontant, Guillaume Gricourt, Melissa N'Debi, Alexandre Soulier, Elisabeth Trawinski, Jean-Michel Pawlotsky |
| EPI_ISL_3476617 | Cerballiance Paris IDF est Saint-Denis | Department of Virology, Henri Mondor University Hospital, Assistance Publique Hôpitaux de Paris, Université Paris-Est Créteil, INSERM U955 | Christophe Rodriguez, Slim Fourati, Vanessa Demontant, Guillaume Gricourt, Melissa N'Debi, Alexandre Soulier, Elisabeth Trawinski, Jean-Michel Pawlotsky |
| EPI_ISL_3476618, EPI_ISL_3476619 | SELAS MEDILYS | Department of Virology, Henri Mondor University Hospital, Assistance Publique Hôpitaux de Paris, Université Paris-Est Créteil, INSERM U955 | Christophe Rodriguez, Slim Fourati, Vanessa Demontant, Guillaume Gricourt, Melissa N'Debi, Alexandre Soulier, Elisabeth Trawinski, Jean-Michel Pawlotsky |
| EPI_ISL_3476620 | Hôpital Armand-Trousseau | Department of Virology, Henri Mondor University Hospital, Assistance Publique Hôpitaux de Paris, Université Paris-Est Créteil, INSERM U955 | Christophe Rodriguez, Slim Fourati, Vanessa Demontant, Guillaume Gricourt, Melissa N'Debi, Alexandre Soulier, Elisabeth Trawinski, Jean-Michel Pawlotsky |
| EPI_ISL_3476621 | Hôpital Cochin | Department of Virology, Henri Mondor University Hospital, Assistance Publique Hôpitaux de Paris, Université Paris-Est Créteil, INSERM U955 | Christophe Rodriguez, Slim Fourati, Vanessa Demontant, Guillaume Gricourt, Melissa N'Debi, Alexandre Soulier, Elisabeth Trawinski, Jean-Michel Pawlotsky |
| EPI_ISL_3476622 | Laboratoire Ana-L | Department of Virology, Henri Mondor University Hospital, Assistance Publique Hôpitaux de Paris, Université Paris-Est Créteil, INSERM U955 | Christophe Rodriguez, Slim Fourati, Vanessa Demontant, Guillaume Gricourt, Melissa N'Debi, Alexandre Soulier, Elisabeth Trawinski, Jean-Michel Pawlotsky |
| EPI_ISL_3476623 | GROUPE HOSPITALIER SUD ILE DE FRANCE | Department of Virology, Henri Mondor University Hospital, Assistance Publique Hôpitaux de Paris, Université Paris-Est Créteil, INSERM U955 | Christophe Rodriguez, Slim Fourati, Vanessa Demontant, Guillaume Gricourt, Melissa N'Debi, Alexandre Soulier, Elisabeth Trawinski, Jean-Michel Pawlotsky |
| EPI_ISL_3476624 | LAM BARRAND | Department of Virology, Henri Mondor University Hospital, Assistance Publique Hôpitaux de Paris, Université Paris-Est Créteil, INSERM U955 | Christophe Rodriguez, Slim Fourati, Vanessa Demontant, Guillaume Gricourt, Melissa N'Debi, Alexandre Soulier, Elisabeth Trawinski, Jean-Michel Pawlotsky |
| EPI_ISL_3476625 | Hôpital Armand-Trousseau | Department of Virology, Henri Mondor University Hospital, Assistance Publique Hôpitaux de Paris, Université Paris-Est Créteil, INSERM U955 | Christophe Rodriguez, Slim Fourati, Vanessa Demontant, Guillaume Gricourt, Melissa N'Debi, Alexandre Soulier, Elisabeth Trawinski, Jean-Michel Pawlotsky |
| EPI_ISL_3476626 | GH A.CHENEVIER-H.MONDOR | Department of Virology, Henri Mondor University Hospital, Assistance Publique Hôpitaux de Paris, Université Paris-Est Créteil, INSERM U955 | Christophe Rodriguez, Slim Fourati, Vanessa Demontant, Guillaume Gricourt, Melissa N'Debi, Alexandre Soulier, Elisabeth Trawinski, Jean-Michel Pawlotsky |
| EPI_ISL_3476627 | BROUSSAIS | Department of Virology, Henri Mondor University Hospital, Assistance Publique Hôpitaux de Paris, Université Paris-Est Créteil, INSERM U955 | Christophe Rodriguez, Slim Fourati, Vanessa Demontant, Guillaume Gricourt, Melissa N'Debi, Alexandre Soulier, Elisabeth Trawinski, Jean-Michel Pawlotsky |

| | | | |
|---|---|---|---|
| EPI_ISL_3476628 | CHU DE DIJON | Department of Virology, Henri Mondor University Hospital, Assistance Publique Hôpitaux de Paris, Université Paris-Est Créteil, INSERM U955 | Christophe Rodriguez, Slim Fourati, Vanessa Demontant, Guillaume Gricourt, Melissa N'Debi, Alexandre Soulier, Elisabeth Trawinski, Jean-Michel Pawlotsky |
| EPI_ISL_3476629, EPI_ISL_3476630, EPI_ISL_3476631, EPI_ISL_3476632 | SELAS MEDILYS | Department of Virology, Henri Mondor University Hospital, Assistance Publique Hôpitaux de Paris, Université Paris-Est Créteil, INSERM U955 | Christophe Rodriguez, Slim Fourati, Vanessa Demontant, Guillaume Gricourt, Melissa N'Debi, Alexandre Soulier, Elisabeth Trawinski, Jean-Michel Pawlotsky |
| EPI_ISL_3476633 | GROUPE HOSPITALIER SUD ILE DE FRANCE | Department of Virology, Henri Mondor University Hospital, Assistance Publique Hôpitaux de Paris, Université Paris-Est Créteil, INSERM U955 | Christophe Rodriguez, Slim Fourati, Vanessa Demontant, Guillaume Gricourt, Melissa N'Debi, Alexandre Soulier, Elisabeth Trawinski, Jean-Michel Pawlotsky |
| EPI_ISL_3476634, EPI_ISL_3476635 | SELAS MEDILYS | Department of Virology, Henri Mondor University Hospital, Assistance Publique Hôpitaux de Paris, Université Paris-Est Créteil, INSERM U955 | Christophe Rodriguez, Slim Fourati, Vanessa Demontant, Guillaume Gricourt, Melissa N'Debi, Alexandre Soulier, Elisabeth Trawinski, Jean-Michel Pawlotsky |
| EPI_ISL_3476636 | Hôpital Avicenne | Department of Virology, Henri Mondor University Hospital, Assistance Publique Hôpitaux de Paris, Université Paris-Est Créteil, INSERM U955 | Christophe Rodriguez, Slim Fourati, Vanessa Demontant, Guillaume Gricourt, Melissa N'Debi, Alexandre Soulier, Elisabeth Trawinski, Jean-Michel Pawlotsky |
| EPI_ISL_3476637 | GH A.CHENEVIER-H.MONDOR | Department of Virology, Henri Mondor University Hospital, Assistance Publique Hôpitaux de Paris, Université Paris-Est Créteil, INSERM U955 | Christophe Rodriguez, Slim Fourati, Vanessa Demontant, Guillaume Gricourt, Melissa N'Debi, Alexandre Soulier, Elisabeth Trawinski, Jean-Michel Pawlotsky |
| EPI_ISL_3476638 | SELAS MEDILYS | Department of Virology, Henri Mondor University Hospital, Assistance Publique Hôpitaux de Paris, Université Paris-Est Créteil, INSERM U955 | Christophe Rodriguez, Slim Fourati, Vanessa Demontant, Guillaume Gricourt, Melissa N'Debi, Alexandre Soulier, Elisabeth Trawinski, Jean-Michel Pawlotsky |
| EPI_ISL_3476639 | BROUSSAIS | Department of Virology, Henri Mondor University Hospital, Assistance Publique Hôpitaux de Paris, Université Paris-Est Créteil, INSERM U955 | Christophe Rodriguez, Slim Fourati, Vanessa Demontant, Guillaume Gricourt, Melissa N'Debi, Alexandre Soulier, Elisabeth Trawinski, Jean-Michel Pawlotsky |
| EPI_ISL_3476640 | GH A.CHENEVIER-H.MONDOR | Department of Virology, Henri Mondor University Hospital, Assistance Publique Hôpitaux de Paris, Université Paris-Est Créteil, INSERM U955 | Christophe Rodriguez, Slim Fourati, Vanessa Demontant, Guillaume Gricourt, Melissa N'Debi, Alexandre Soulier, Elisabeth Trawinski, Jean-Michel Pawlotsky |
| EPI_ISL_3476641 | BIOMNIS EUROFINS IVRY | Department of Virology, Henri Mondor University Hospital, Assistance Publique Hôpitaux de Paris, Université Paris-Est Créteil, INSERM U955 | Christophe Rodriguez, Slim Fourati, Vanessa Demontant, Guillaume Gricourt, Melissa N'Debi, Alexandre Soulier, Elisabeth Trawinski, Jean-Michel Pawlotsky |
| EPI_ISL_3476642 | Laboratoire Ana-L | Department of Virology, Henri Mondor University Hospital, Assistance Publique Hôpitaux de Paris, Université Paris-Est Créteil, INSERM U955 | Christophe Rodriguez, Slim Fourati, Vanessa Demontant, Guillaume Gricourt, Melissa N'Debi, Alexandre Soulier, Elisabeth Trawinski, Jean-Michel Pawlotsky |
| EPI_ISL_3476643, EPI_ISL_3476644 | GH A.CHENEVIER-H.MONDOR | Department of Virology, Henri Mondor University Hospital, Assistance Publique Hôpitaux de Paris, Université Paris-Est Créteil, INSERM U955 | Christophe Rodriguez, Slim Fourati, Vanessa Demontant, Guillaume Gricourt, Melissa N'Debi, Alexandre Soulier, Elisabeth Trawinski, Jean-Michel Pawlotsky |
| EPI_ISL_3476645 | BROUSSAIS | Department of Virology, Henri Mondor University Hospital, Assistance Publique Hôpitaux de Paris, Université Paris-Est Créteil, INSERM U955 | Christophe Rodriguez, Slim Fourati, Vanessa Demontant, Guillaume Gricourt, Melissa N'Debi, Alexandre Soulier, Elisabeth Trawinski, Jean-Michel Pawlotsky |
| EPI_ISL_3476646 | CH.LE RAINCY_MONTFERMEIL | Department of Virology, Henri Mondor University Hospital, Assistance Publique Hôpitaux de Paris, Université Paris-Est Créteil, INSERM U955 | Christophe Rodriguez, Slim Fourati, Vanessa Demontant, Guillaume Gricourt, Melissa N'Debi, Alexandre Soulier, Elisabeth Trawinski, Jean-Michel Pawlotsky |
| EPI_ISL_3476647 | BIORHIN | Department of Virology, Henri Mondor University Hospital, Assistance Publique Hôpitaux de Paris, Université Paris-Est Créteil, INSERM U955 | Christophe Rodriguez, Slim Fourati, Vanessa Demontant, Guillaume Gricourt, Melissa N'Debi, Alexandre Soulier, Elisabeth Trawinski, Jean-Michel Pawlotsky |
| EPI_ISL_3476648 | GH A.CHENEVIER-H.MONDOR | Department of Virology, Henri Mondor University Hospital, Assistance Publique Hôpitaux de Paris, Université Paris-Est Créteil, INSERM U955 | Christophe Rodriguez, Slim Fourati, Vanessa Demontant, Guillaume Gricourt, Melissa N'Debi, Alexandre Soulier, Elisabeth Trawinski, Jean-Michel Pawlotsky |
| EPI_ISL_3476649 | BIOMNIS EUROFINS IVRY | Department of Virology, Henri Mondor University Hospital, Assistance Publique Hôpitaux de Paris, Université Paris-Est Créteil, INSERM U955 | Christophe Rodriguez, Slim Fourati, Vanessa Demontant, Guillaume Gricourt, Melissa N'Debi, Alexandre Soulier, Elisabeth Trawinski, Jean-Michel Pawlotsky |
| EPI_ISL_3476650 | SELAS MEDILYS | Department of Virology, Henri Mondor University Hospital, Assistance Publique Hôpitaux de Paris, Université Paris-Est Créteil, INSERM U955 | Christophe Rodriguez, Slim Fourati, Vanessa Demontant, Guillaume Gricourt, Melissa N'Debi, Alexandre Soulier, Elisabeth Trawinski, Jean-Michel Pawlotsky |
| EPI_ISL_3476651 | GH A.CHENEVIER-H.MONDOR | Department of Virology, Henri Mondor University Hospital, Assistance Publique Hôpitaux de Paris, Université Paris-Est Créteil, INSERM U955 | Christophe Rodriguez, Slim Fourati, Vanessa Demontant, Guillaume Gricourt, Melissa N'Debi, Alexandre Soulier, Elisabeth Trawinski, Jean-Michel Pawlotsky |
| EPI_ISL_3476652 | LAM BARRAND | Department of Virology, Henri Mondor University Hospital, Assistance Publique Hôpitaux de Paris, Université Paris-Est Créteil, INSERM U955 | Christophe Rodriguez, Slim Fourati, Vanessa Demontant, Guillaume Gricourt, Melissa N'Debi, Alexandre Soulier, Elisabeth Trawinski, Jean-Michel Pawlotsky |
| EPI_ISL_3476653 | LBM EVORIAL NEVERS | Department of Virology, Henri Mondor University Hospital, Assistance Publique Hôpitaux de Paris, Université Paris-Est Créteil, INSERM U955 | Christophe Rodriguez, Slim Fourati, Vanessa Demontant, Guillaume Gricourt, Melissa N'Debi, Alexandre Soulier, Elisabeth Trawinski, Jean-Michel Pawlotsky |
| EPI_ISL_3476655 | CH.LE RAINCY_MONTFERMEIL | Department of Virology, Henri Mondor University Hospital, Assistance Publique Hôpitaux de Paris, Université Paris-Est Créteil, INSERM U955 | Christophe Rodriguez, Slim Fourati, Vanessa Demontant, Guillaume Gricourt, Melissa N'Debi, Alexandre Soulier, Elisabeth Trawinski, Jean-Michel Pawlotsky |
| EPI_ISL_3476656 | GH A.CHENEVIER-H.MONDOR | Department of Virology, Henri Mondor University Hospital, Assistance Publique Hôpitaux de Paris, Université Paris-Est Créteil, INSERM U955 | Christophe Rodriguez, Slim Fourati, Vanessa Demontant, Guillaume Gricourt, Melissa N'Debi, Alexandre Soulier, Elisabeth Trawinski, Jean-Michel Pawlotsky |
| EPI_ISL_3476657 | BIOMNIS EUROFINS IVRY | Department of Virology, Henri Mondor University Hospital, Assistance Publique Hôpitaux de Paris, Université Paris-Est Créteil, INSERM U955 | Christophe Rodriguez, Slim Fourati, Vanessa Demontant, Guillaume Gricourt, Melissa N'Debi, Alexandre Soulier, Elisabeth Trawinski, Jean-Michel Pawlotsky |
| EPI_ISL_3476659 | Hôpital Cochin | Department of Virology, Henri Mondor University Hospital, Assistance Publique Hôpitaux de Paris, Université Paris-Est Créteil, INSERM U955 | Christophe Rodriguez, Slim Fourati, Vanessa Demontant, Guillaume Gricourt, Melissa N'Debi, Alexandre Soulier, Elisabeth Trawinski, Jean-Michel Pawlotsky |
| EPI_ISL_3476661 | CH.LE RAINCY_MONTFERMEIL | Department of Virology, Henri Mondor University Hospital, Assistance Publique Hôpitaux de Paris, Université Paris-Est Créteil, INSERM U955 | Christophe Rodriguez, Slim Fourati, Vanessa Demontant, Guillaume Gricourt, Melissa N'Debi, Alexandre Soulier, Elisabeth Trawinski, Jean-Michel Pawlotsky |
| EPI_ISL_3476663, EPI_ISL_3476664, EPI_ISL_3476665, EPI_ISL_3476666, EPI_ISL_3476667 | SELAS MEDILYS | Department of Virology, Henri Mondor University Hospital, Assistance Publique Hôpitaux de Paris, Université Paris-Est Créteil, INSERM U955 | Christophe Rodriguez, Slim Fourati, Vanessa Demontant, Guillaume Gricourt, Melissa N'Debi, Alexandre Soulier, Elisabeth Trawinski, Jean-Michel Pawlotsky |
| EPI_ISL_3476668 | GH A.CHENEVIER-H.MONDOR | Department of Virology, Henri Mondor University Hospital, Assistance Publique Hôpitaux de Paris, Université Paris-Est Créteil, INSERM U955 | Christophe Rodriguez, Slim Fourati, Vanessa Demontant, Guillaume Gricourt, Melissa N'Debi, Alexandre Soulier, Elisabeth Trawinski, Jean-Michel Pawlotsky |
| EPI_ISL_3476669 | BIOMNIS EUROFINS IVRY | Department of Virology, Henri Mondor University Hospital, Assistance Publique Hôpitaux de Paris, Université Paris-Est Créteil, INSERM U955 | Christophe Rodriguez, Slim Fourati, Vanessa Demontant, Guillaume Gricourt, Melissa N'Debi, Alexandre Soulier, Elisabeth Trawinski, Jean-Michel Pawlotsky |
| EPI_ISL_3476670 | SELAS MEDILYS | Department of Virology, Henri Mondor University Hospital, Assistance Publique Hôpitaux de Paris, Université Paris-Est Créteil, INSERM U955 | Christophe Rodriguez, Slim Fourati, Vanessa Demontant, Guillaume Gricourt, Melissa N'Debi, Alexandre Soulier, Elisabeth Trawinski, Jean-Michel Pawlotsky |
| EPI_ISL_3476671 | GH A.CHENEVIER-H.MONDOR | Department of Virology, Henri Mondor University Hospital, Assistance Publique Hôpitaux de Paris, Université Paris-Est Créteil, INSERM U955 | Christophe Rodriguez, Slim Fourati, Vanessa Demontant, Guillaume Gricourt, Melissa N'Debi, Alexandre Soulier, Elisabeth Trawinski, Jean-Michel Pawlotsky |
| EPI_ISL_3476672, | BIOMNIS EUROFINS IVRY | Department of Virology, Henri Mondor University Hospital, | Christophe Rodriguez, Slim Fourati, Vanessa Demontant, Guillaume Gricourt, Melissa N'Debi, Alexandre Soulier, Elisabeth Trawinski, Jean-Michel Pawlotsky |

| | | | |
|---|---|---|---|
| EPI_ISL_3476673 | | Assistance Publique Hôpitaux de Paris, Université Paris-Est Créteil, INSERM U955 | |
| EPI_ISL_3476674 | BROUSSAIS | Department of Virology, Henri Mondor University Hospital, Assistance Publique Hôpitaux de Paris, Université Paris-Est Créteil, INSERM U955 | Christophe Rodriguez, Slim Fourati, Vanessa Demontant, Guillaume Gricourt, Melissa N'Debi, Alexandre Soulier, Elisabeth Trawinski, Jean-Michel Pawlotsky |
| EPI_ISL_3476675 | SELAS MEDILYS | Department of Virology, Henri Mondor University Hospital, Assistance Publique Hôpitaux de Paris, Université Paris-Est Créteil, INSERM U955 | Christophe Rodriguez, Slim Fourati, Vanessa Demontant, Guillaume Gricourt, Melissa N'Debi, Alexandre Soulier, Elisabeth Trawinski, Jean-Michel Pawlotsky |
| EPI_ISL_3476676 | BROUSSAIS | Department of Virology, Henri Mondor University Hospital, Assistance Publique Hôpitaux de Paris, Université Paris-Est Créteil, INSERM U955 | Christophe Rodriguez, Slim Fourati, Vanessa Demontant, Guillaume Gricourt, Melissa N'Debi, Alexandre Soulier, Elisabeth Trawinski, Jean-Michel Pawlotsky |
| EPI_ISL_3476677 | BIOMNIS EUROFINS IVRY | Department of Virology, Henri Mondor University Hospital, Assistance Publique Hôpitaux de Paris, Université Paris-Est Créteil, INSERM U955 | Christophe Rodriguez, Slim Fourati, Vanessa Demontant, Guillaume Gricourt, Melissa N'Debi, Alexandre Soulier, Elisabeth Trawinski, Jean-Michel Pawlotsky |
| EPI_ISL_3476678, EPI_ISL_3476679 | SELAS MEDILYS | Department of Virology, Henri Mondor University Hospital, Assistance Publique Hôpitaux de Paris, Université Paris-Est Créteil, INSERM U955 | Christophe Rodriguez, Slim Fourati, Vanessa Demontant, Guillaume Gricourt, Melissa N'Debi, Alexandre Soulier, Elisabeth Trawinski, Jean-Michel Pawlotsky |
| EPI_ISL_3476680 | LBM EVORIAL NEVERS | Department of Virology, Henri Mondor University Hospital, Assistance Publique Hôpitaux de Paris, Université Paris-Est Créteil, INSERM U955 | Christophe Rodriguez, Slim Fourati, Vanessa Demontant, Guillaume Gricourt, Melissa N'Debi, Alexandre Soulier, Elisabeth Trawinski, Jean-Michel Pawlotsky |
| EPI_ISL_3476681 | SELAS MEDILYS | Department of Virology, Henri Mondor University Hospital, Assistance Publique Hôpitaux de Paris, Université Paris-Est Créteil, INSERM U955 | Christophe Rodriguez, Slim Fourati, Vanessa Demontant, Guillaume Gricourt, Melissa N'Debi, Alexandre Soulier, Elisabeth Trawinski, Jean-Michel Pawlotsky |
| EPI_ISL_3476682 | GH A.CHENEVIER-H.MONDOR | Department of Virology, Henri Mondor University Hospital, Assistance Publique Hôpitaux de Paris, Université Paris-Est Créteil, INSERM U955 | Christophe Rodriguez, Slim Fourati, Vanessa Demontant, Guillaume Gricourt, Melissa N'Debi, Alexandre Soulier, Elisabeth Trawinski, Jean-Michel Pawlotsky |
| EPI_ISL_3476683 | SELAS MEDILYS | Department of Virology, Henri Mondor University Hospital, Assistance Publique Hôpitaux de Paris, Université Paris-Est Créteil, INSERM U955 | Christophe Rodriguez, Slim Fourati, Vanessa Demontant, Guillaume Gricourt, Melissa N'Debi, Alexandre Soulier, Elisabeth Trawinski, Jean-Michel Pawlotsky |
| EPI_ISL_3476684 | LAM BARRAND | Department of Virology, Henri Mondor University Hospital, Assistance Publique Hôpitaux de Paris, Université Paris-Est Créteil, INSERM U955 | Christophe Rodriguez, Slim Fourati, Vanessa Demontant, Guillaume Gricourt, Melissa N'Debi, Alexandre Soulier, Elisabeth Trawinski, Jean-Michel Pawlotsky |
| EPI_ISL_3476685 | SELAS MEDILYS | Department of Virology, Henri Mondor University Hospital, Assistance Publique Hôpitaux de Paris, Université Paris-Est Créteil, INSERM U955 | Christophe Rodriguez, Slim Fourati, Vanessa Demontant, Guillaume Gricourt, Melissa N'Debi, Alexandre Soulier, Elisabeth Trawinski, Jean-Michel Pawlotsky |
| EPI_ISL_3476686 | Hôpital Cochin | Department of Virology, Henri Mondor University Hospital, Assistance Publique Hôpitaux de Paris, Université Paris-Est Créteil, INSERM U955 | Christophe Rodriguez, Slim Fourati, Vanessa Demontant, Guillaume Gricourt, Melissa N'Debi, Alexandre Soulier, Elisabeth Trawinski, Jean-Michel Pawlotsky |
| EPI_ISL_3476687 | SELAS MEDILYS | Department of Virology, Henri Mondor University Hospital, Assistance Publique Hôpitaux de Paris, Université Paris-Est Créteil, INSERM U955 | Christophe Rodriguez, Slim Fourati, Vanessa Demontant, Guillaume Gricourt, Melissa N'Debi, Alexandre Soulier, Elisabeth Trawinski, Jean-Michel Pawlotsky |
| EPI_ISL_3476688 | CHU DE DIJON | Department of Virology, Henri Mondor University Hospital, Assistance Publique Hôpitaux de Paris, Université Paris-Est Créteil, INSERM U955 | Christophe Rodriguez, Slim Fourati, Vanessa Demontant, Guillaume Gricourt, Melissa N'Debi, Alexandre Soulier, Elisabeth Trawinski, Jean-Michel Pawlotsky |
| EPI_ISL_3476689 | BIORHIN | Department of Virology, Henri Mondor University Hospital, Assistance Publique Hôpitaux de Paris, Université Paris-Est Créteil, INSERM U955 | Christophe Rodriguez, Slim Fourati, Vanessa Demontant, Guillaume Gricourt, Melissa N'Debi, Alexandre Soulier, Elisabeth Trawinski, Jean-Michel Pawlotsky |
| EPI_ISL_3476690 | CH.LE RAINCY_MONTFERMEIL | Department of Virology, Henri Mondor University Hospital, Assistance Publique Hôpitaux de Paris, Université Paris-Est Créteil, INSERM U955 | Christophe Rodriguez, Slim Fourati, Vanessa Demontant, Guillaume Gricourt, Melissa N'Debi, Alexandre Soulier, Elisabeth Trawinski, Jean-Michel Pawlotsky |
| EPI_ISL_3476691 | GROUPE HOSPITALIER SUD ILE DE FRANCE | Department of Virology, Henri Mondor University Hospital, Assistance Publique Hôpitaux de Paris, Université Paris-Est Créteil, INSERM U955 | Christophe Rodriguez, Slim Fourati, Vanessa Demontant, Guillaume Gricourt, Melissa N'Debi, Alexandre Soulier, Elisabeth Trawinski, Jean-Michel Pawlotsky |
| EPI_ISL_3476692 | BIORHIN | Department of Virology, Henri Mondor University Hospital, Assistance Publique Hôpitaux de Paris, Université Paris-Est Créteil, INSERM U955 | Christophe Rodriguez, Slim Fourati, Vanessa Demontant, Guillaume Gricourt, Melissa N'Debi, Alexandre Soulier, Elisabeth Trawinski, Jean-Michel Pawlotsky |
| EPI_ISL_3476693, EPI_ISL_3476694 | SELAS MEDILYS | Department of Virology, Henri Mondor University Hospital, Assistance Publique Hôpitaux de Paris, Université Paris-Est Créteil, INSERM U955 | Christophe Rodriguez, Slim Fourati, Vanessa Demontant, Guillaume Gricourt, Melissa N'Debi, Alexandre Soulier, Elisabeth Trawinski, Jean-Michel Pawlotsky |
| EPI_ISL_3476695, EPI_ISL_3476696 | BIOMNIS EUROFINS IVRY | Department of Virology, Henri Mondor University Hospital, Assistance Publique Hôpitaux de Paris, Université Paris-Est Créteil, INSERM U955 | Christophe Rodriguez, Slim Fourati, Vanessa Demontant, Guillaume Gricourt, Melissa N'Debi, Alexandre Soulier, Elisabeth Trawinski, Jean-Michel Pawlotsky |
| EPI_ISL_3476697 | CH.LE RAINCY_MONTFERMEIL | Department of Virology, Henri Mondor University Hospital, Assistance Publique Hôpitaux de Paris, Université Paris-Est Créteil, INSERM U955 | Christophe Rodriguez, Slim Fourati, Vanessa Demontant, Guillaume Gricourt, Melissa N'Debi, Alexandre Soulier, Elisabeth Trawinski, Jean-Michel Pawlotsky |
| EPI_ISL_3476698 | SELAS MEDILYS | Department of Virology, Henri Mondor University Hospital, Assistance Publique Hôpitaux de Paris, Université Paris-Est Créteil, INSERM U955 | Christophe Rodriguez, Slim Fourati, Vanessa Demontant, Guillaume Gricourt, Melissa N'Debi, Alexandre Soulier, Elisabeth Trawinski, Jean-Michel Pawlotsky |
| EPI_ISL_3476699 | GROUPE HOSPITALIER SUD ILE DE FRANCE | Department of Virology, Henri Mondor University Hospital, Assistance Publique Hôpitaux de Paris, Université Paris-Est Créteil, INSERM U955 | Christophe Rodriguez, Slim Fourati, Vanessa Demontant, Guillaume Gricourt, Melissa N'Debi, Alexandre Soulier, Elisabeth Trawinski, Jean-Michel Pawlotsky |
| EPI_ISL_3476700 | LAM BARRAND | Department of Virology, Henri Mondor University Hospital, Assistance Publique Hôpitaux de Paris, Université Paris-Est Créteil, INSERM U955 | Christophe Rodriguez, Slim Fourati, Vanessa Demontant, Guillaume Gricourt, Melissa N'Debi, Alexandre Soulier, Elisabeth Trawinski, Jean-Michel Pawlotsky |
| EPI_ISL_3476701 | BIOMNIS EUROFINS IVRY | Department of Virology, Henri Mondor University Hospital, Assistance Publique Hôpitaux de Paris, Université Paris-Est Créteil, INSERM U955 | Christophe Rodriguez, Slim Fourati, Vanessa Demontant, Guillaume Gricourt, Melissa N'Debi, Alexandre Soulier, Elisabeth Trawinski, Jean-Michel Pawlotsky |
| EPI_ISL_3476702 | CH.LE RAINCY_MONTFERMEIL | Department of Virology, Henri Mondor University Hospital, Assistance Publique Hôpitaux de Paris, Université Paris-Est Créteil, INSERM U955 | Christophe Rodriguez, Slim Fourati, Vanessa Demontant, Guillaume Gricourt, Melissa N'Debi, Alexandre Soulier, Elisabeth Trawinski, Jean-Michel Pawlotsky |
| EPI_ISL_3476703 | SELAS MEDILYS | Department of Virology, Henri Mondor University Hospital, Assistance Publique Hôpitaux de Paris, Université Paris-Est Créteil, INSERM U955 | Christophe Rodriguez, Slim Fourati, Vanessa Demontant, Guillaume Gricourt, Melissa N'Debi, Alexandre Soulier, Elisabeth Trawinski, Jean-Michel Pawlotsky |
| EPI_ISL_3476704, EPI_ISL_3476705 | BIORHIN | Department of Virology, Henri Mondor University Hospital, Assistance Publique Hôpitaux de Paris, Université Paris-Est Créteil, INSERM U955 | Christophe Rodriguez, Slim Fourati, Vanessa Demontant, Guillaume Gricourt, Melissa N'Debi, Alexandre Soulier, Elisabeth Trawinski, Jean-Michel Pawlotsky |
| EPI_ISL_3476706, EPI_ISL_3476707 | SELAS MEDILYS | Department of Virology, Henri Mondor University Hospital, Assistance Publique Hôpitaux de Paris, Université Paris-Est Créteil, INSERM U955 | Christophe Rodriguez, Slim Fourati, Vanessa Demontant, Guillaume Gricourt, Melissa N'Debi, Alexandre Soulier, Elisabeth Trawinski, Jean-Michel Pawlotsky |
| EPI_ISL_3476708, EPI_ISL_3476709 | BIOMNIS EUROFINS IVRY | Department of Virology, Henri Mondor University Hospital, Assistance Publique Hôpitaux de Paris, Université Paris-Est Créteil, INSERM U955 | Christophe Rodriguez, Slim Fourati, Vanessa Demontant, Guillaume Gricourt, Melissa N'Debi, Alexandre Soulier, Elisabeth Trawinski, Jean-Michel Pawlotsky |
| EPI_ISL_3476710 | BIORHIN | Department of Virology, Henri Mondor University Hospital, Assistance Publique Hôpitaux de Paris, Université Paris-Est Créteil, INSERM U955 | Christophe Rodriguez, Slim Fourati, Vanessa Demontant, Guillaume Gricourt, Melissa N'Debi, Alexandre Soulier, Elisabeth Trawinski, Jean-Michel Pawlotsky |

| | | | |
|---|---|---|---|
| EPI_ISL_3476711 | BIOMNIS EUROFINS IVRY | Department of Virology, Henri Mondor University Hospital, Assistance Publique Hôpitaux de Paris, Université Paris-Est Créteil, INSERM U955 | Christophe Rodriguez, Slim Fourati, Vanessa Demontant, Guillaume Gricourt, Melissa N'Debi, Alexandre Soulier, Elisabeth Trawinski, Jean-Michel Pawlotsky |
| EPI_ISL_3476712 | CH.LE RAINCY_MONTFERMEIL | Department of Virology, Henri Mondor University Hospital, Assistance Publique Hôpitaux de Paris, Université Paris-Est Créteil, INSERM U955 | Christophe Rodriguez, Slim Fourati, Vanessa Demontant, Guillaume Gricourt, Melissa N'Debi, Alexandre Soulier, Elisabeth Trawinski, Jean-Michel Pawlotsky |
| EPI_ISL_3476713 | BIOMNIS EUROFINS IVRY | Department of Virology, Henri Mondor University Hospital, Assistance Publique Hôpitaux de Paris, Université Paris-Est Créteil, INSERM U955 | Christophe Rodriguez, Slim Fourati, Vanessa Demontant, Guillaume Gricourt, Melissa N'Debi, Alexandre Soulier, Elisabeth Trawinski, Jean-Michel Pawlotsky |
| EPI_ISL_3476714 | GH A.CHENEVIER-H.MONDOR | Department of Virology, Henri Mondor University Hospital, Assistance Publique Hôpitaux de Paris, Université Paris-Est Créteil, INSERM U955 | Christophe Rodriguez, Slim Fourati, Vanessa Demontant, Guillaume Gricourt, Melissa N'Debi, Alexandre Soulier, Elisabeth Trawinski, Jean-Michel Pawlotsky |
| EPI_ISL_3476715 | Biogroup Bio Lam-LCD Saint-Denis | Department of Virology, Henri Mondor University Hospital, Assistance Publique Hôpitaux de Paris, Université Paris-Est Créteil, INSERM U955 | Christophe Rodriguez, Slim Fourati, Vanessa Demontant, Guillaume Gricourt, Melissa N'Debi, Alexandre Soulier, Elisabeth Trawinski, Jean-Michel Pawlotsky |
| EPI_ISL_3476716 | LBM EVORIAL NEVERS | Department of Virology, Henri Mondor University Hospital, Assistance Publique Hôpitaux de Paris, Université Paris-Est Créteil, INSERM U955 | Christophe Rodriguez, Slim Fourati, Vanessa Demontant, Guillaume Gricourt, Melissa N'Debi, Alexandre Soulier, Elisabeth Trawinski, Jean-Michel Pawlotsky |
| EPI_ISL_3476717, EPI_ISL_3476718 | BIOMNIS EUROFINS IVRY | Department of Virology, Henri Mondor University Hospital, Assistance Publique Hôpitaux de Paris, Université Paris-Est Créteil, INSERM U955 | Christophe Rodriguez, Slim Fourati, Vanessa Demontant, Guillaume Gricourt, Melissa N'Debi, Alexandre Soulier, Elisabeth Trawinski, Jean-Michel Pawlotsky |
| EPI_ISL_3476719 | BIORHIN | Department of Virology, Henri Mondor University Hospital, Assistance Publique Hôpitaux de Paris, Université Paris-Est Créteil, INSERM U955 | Christophe Rodriguez, Slim Fourati, Vanessa Demontant, Guillaume Gricourt, Melissa N'Debi, Alexandre Soulier, Elisabeth Trawinski, Jean-Michel Pawlotsky |
| EPI_ISL_3476721 | BIOMNIS EUROFINS IVRY | Department of Virology, Henri Mondor University Hospital, Assistance Publique Hôpitaux de Paris, Université Paris-Est Créteil, INSERM U955 | Christophe Rodriguez, Slim Fourati, Vanessa Demontant, Guillaume Gricourt, Melissa N'Debi, Alexandre Soulier, Elisabeth Trawinski, Jean-Michel Pawlotsky |
| EPI_ISL_3476722, EPI_ISL_3476723 | SELAS MEDILYS | Department of Virology, Henri Mondor University Hospital, Assistance Publique Hôpitaux de Paris, Université Paris-Est Créteil, INSERM U955 | Christophe Rodriguez, Slim Fourati, Vanessa Demontant, Guillaume Gricourt, Melissa N'Debi, Alexandre Soulier, Elisabeth Trawinski, Jean-Michel Pawlotsky |
| EPI_ISL_3476724 | Biogroup Bio Lam-LCD Saint-Denis | Department of Virology, Henri Mondor University Hospital, Assistance Publique Hôpitaux de Paris, Université Paris-Est Créteil, INSERM U955 | Christophe Rodriguez, Slim Fourati, Vanessa Demontant, Guillaume Gricourt, Melissa N'Debi, Alexandre Soulier, Elisabeth Trawinski, Jean-Michel Pawlotsky |
| EPI_ISL_3476725 | SELAS MEDILYS | Department of Virology, Henri Mondor University Hospital, Assistance Publique Hôpitaux de Paris, Université Paris-Est Créteil, INSERM U955 | Christophe Rodriguez, Slim Fourati, Vanessa Demontant, Guillaume Gricourt, Melissa N'Debi, Alexandre Soulier, Elisabeth Trawinski, Jean-Michel Pawlotsky |
| EPI_ISL_3476727 | Hôpital Armand-Trousseau | Department of Virology, Henri Mondor University Hospital, Assistance Publique Hôpitaux de Paris, Université Paris-Est Créteil, INSERM U955 | Christophe Rodriguez, Slim Fourati, Vanessa Demontant, Guillaume Gricourt, Melissa N'Debi, Alexandre Soulier, Elisabeth Trawinski, Jean-Michel Pawlotsky |
| EPI_ISL_3476728 | Biogroup Bio Lam-LCD Saint-Denis | Department of Virology, Henri Mondor University Hospital, Assistance Publique Hôpitaux de Paris, Université Paris-Est Créteil, INSERM U955 | Christophe Rodriguez, Slim Fourati, Vanessa Demontant, Guillaume Gricourt, Melissa N'Debi, Alexandre Soulier, Elisabeth Trawinski, Jean-Michel Pawlotsky |
| EPI_ISL_3476729 | BIORHIN | Department of Virology, Henri Mondor University Hospital, Assistance Publique Hôpitaux de Paris, Université Paris-Est Créteil, INSERM U955 | Christophe Rodriguez, Slim Fourati, Vanessa Demontant, Guillaume Gricourt, Melissa N'Debi, Alexandre Soulier, Elisabeth Trawinski, Jean-Michel Pawlotsky |
| EPI_ISL_3476730 | LISIEUX Cerballiance | Department of Virology, Henri Mondor University Hospital, Assistance Publique Hôpitaux de Paris, Université Paris-Est Créteil, INSERM U955 | Christophe Rodriguez, Slim Fourati, Vanessa Demontant, Guillaume Gricourt, Melissa N'Debi, Alexandre Soulier, Elisabeth Trawinski, Jean-Michel Pawlotsky |
| EPI_ISL_3476731, EPI_ISL_3476732 | SELAS MEDILYS | Department of Virology, Henri Mondor University Hospital, Assistance Publique Hôpitaux de Paris, Université Paris-Est Créteil, INSERM U955 | Christophe Rodriguez, Slim Fourati, Vanessa Demontant, Guillaume Gricourt, Melissa N'Debi, Alexandre Soulier, Elisabeth Trawinski, Jean-Michel Pawlotsky |
| EPI_ISL_3476733 | BIORHIN | Department of Virology, Henri Mondor University Hospital, Assistance Publique Hôpitaux de Paris, Université Paris-Est Créteil, INSERM U955 | Christophe Rodriguez, Slim Fourati, Vanessa Demontant, Guillaume Gricourt, Melissa N'Debi, Alexandre Soulier, Elisabeth Trawinski, Jean-Michel Pawlotsky |
| EPI_ISL_3476734 | BIOMNIS EUROFINS IVRY | Department of Virology, Henri Mondor University Hospital, Assistance Publique Hôpitaux de Paris, Université Paris-Est Créteil, INSERM U955 | Christophe Rodriguez, Slim Fourati, Vanessa Demontant, Guillaume Gricourt, Melissa N'Debi, Alexandre Soulier, Elisabeth Trawinski, Jean-Michel Pawlotsky |
| EPI_ISL_3476735 | BIORHIN | Department of Virology, Henri Mondor University Hospital, Assistance Publique Hôpitaux de Paris, Université Paris-Est Créteil, INSERM U955 | Christophe Rodriguez, Slim Fourati, Vanessa Demontant, Guillaume Gricourt, Melissa N'Debi, Alexandre Soulier, Elisabeth Trawinski, Jean-Michel Pawlotsky |
| EPI_ISL_3476736 | Hôpital Cochin | Department of Virology, Henri Mondor University Hospital, Assistance Publique Hôpitaux de Paris, Université Paris-Est Créteil, INSERM U955 | Christophe Rodriguez, Slim Fourati, Vanessa Demontant, Guillaume Gricourt, Melissa N'Debi, Alexandre Soulier, Elisabeth Trawinski, Jean-Michel Pawlotsky |
| EPI_ISL_3476737 | SELAS MEDILYS | Department of Virology, Henri Mondor University Hospital, Assistance Publique Hôpitaux de Paris, Université Paris-Est Créteil, INSERM U955 | Christophe Rodriguez, Slim Fourati, Vanessa Demontant, Guillaume Gricourt, Melissa N'Debi, Alexandre Soulier, Elisabeth Trawinski, Jean-Michel Pawlotsky |
| EPI_ISL_3476738 | Laboratoire Ana-L | Department of Virology, Henri Mondor University Hospital, Assistance Publique Hôpitaux de Paris, Université Paris-Est Créteil, INSERM U955 | Christophe Rodriguez, Slim Fourati, Vanessa Demontant, Guillaume Gricourt, Melissa N'Debi, Alexandre Soulier, Elisabeth Trawinski, Jean-Michel Pawlotsky |
| EPI_ISL_3476739, EPI_ISL_3476740 | BIOMNIS EUROFINS IVRY | Department of Virology, Henri Mondor University Hospital, Assistance Publique Hôpitaux de Paris, Université Paris-Est Créteil, INSERM U955 | Christophe Rodriguez, Slim Fourati, Vanessa Demontant, Guillaume Gricourt, Melissa N'Debi, Alexandre Soulier, Elisabeth Trawinski, Jean-Michel Pawlotsky |
| EPI_ISL_3476741, EPI_ISL_3476742 | Hôpital Cochin | Department of Virology, Henri Mondor University Hospital, Assistance Publique Hôpitaux de Paris, Université Paris-Est Créteil, INSERM U955 | Christophe Rodriguez, Slim Fourati, Vanessa Demontant, Guillaume Gricourt, Melissa N'Debi, Alexandre Soulier, Elisabeth Trawinski, Jean-Michel Pawlotsky |
| EPI_ISL_3476743 | SELAS MEDILYS | Department of Virology, Henri Mondor University Hospital, Assistance Publique Hôpitaux de Paris, Université Paris-Est Créteil, INSERM U955 | Christophe Rodriguez, Slim Fourati, Vanessa Demontant, Guillaume Gricourt, Melissa N'Debi, Alexandre Soulier, Elisabeth Trawinski, Jean-Michel Pawlotsky |
| EPI_ISL_3476744 | LAM BARRAND | Department of Virology, Henri Mondor University Hospital, Assistance Publique Hôpitaux de Paris, Université Paris-Est Créteil, INSERM U955 | Christophe Rodriguez, Slim Fourati, Vanessa Demontant, Guillaume Gricourt, Melissa N'Debi, Alexandre Soulier, Elisabeth Trawinski, Jean-Michel Pawlotsky |
| EPI_ISL_3476745 | CHU DE DIJON | Department of Virology, Henri Mondor University Hospital, Assistance Publique Hôpitaux de Paris, Université Paris-Est Créteil, INSERM U955 | Christophe Rodriguez, Slim Fourati, Vanessa Demontant, Guillaume Gricourt, Melissa N'Debi, Alexandre Soulier, Elisabeth Trawinski, Jean-Michel Pawlotsky |
| EPI_ISL_3476746 | SELAS MEDILYS | Department of Virology, Henri Mondor University Hospital, Assistance Publique Hôpitaux de Paris, Université Paris-Est Créteil, INSERM U955 | Christophe Rodriguez, Slim Fourati, Vanessa Demontant, Guillaume Gricourt, Melissa N'Debi, Alexandre Soulier, Elisabeth Trawinski, Jean-Michel Pawlotsky |
| EPI_ISL_3476747, EPI_ISL_3476748 | BIOMNIS EUROFINS IVRY | Department of Virology, Henri Mondor University Hospital, Assistance Publique Hôpitaux de Paris, Université Paris-Est Créteil, INSERM U955 | Christophe Rodriguez, Slim Fourati, Vanessa Demontant, Guillaume Gricourt, Melissa N'Debi, Alexandre Soulier, Elisabeth Trawinski, Jean-Michel Pawlotsky |
| EPI_ISL_3476749 | GH A.CHENEVIER-H.MONDOR | Department of Virology, Henri Mondor University Hospital, Assistance Publique Hôpitaux de Paris, Université Paris-Est Créteil, INSERM U955 | Christophe Rodriguez, Slim Fourati, Vanessa Demontant, Guillaume Gricourt, Melissa N'Debi, Alexandre Soulier, Elisabeth Trawinski, Jean-Michel Pawlotsky |
| EPI_ISL_3476750 | Laboratoire CLEMENT | Department of Virology, Henri Mondor University Hospital, Assistance Publique Hôpitaux de Paris, Université Paris-Est Créteil, INSERM U955 | Christophe Rodriguez, Slim Fourati, Vanessa Demontant, Guillaume Gricourt, Melissa N'Debi, Alexandre Soulier, Elisabeth Trawinski, Jean-Michel Pawlotsky |

| | | | |
|---|---|---|---|
| EPI_ISL_3476751, EPI_ISL_3476752 | SELAS MEDILYS | Department of Virology, Henri Mondor University Hospital, Assistance Publique Hôpitaux de Paris, Université Paris-Est Créteil, INSERM U955 | Christophe Rodriguez, Slim Fourati, Vanessa Demontant, Guillaume Gricourt, Melissa N'Debi, Alexandre Soulier, Elisabeth Trawinski, Jean-Michel Pawlotsky |
| EPI_ISL_3476753 | Laboratoire Ana-L | Department of Virology, Henri Mondor University Hospital, Assistance Publique Hôpitaux de Paris, Université Paris-Est Créteil, INSERM U955 | Christophe Rodriguez, Slim Fourati, Vanessa Demontant, Guillaume Gricourt, Melissa N'Debi, Alexandre Soulier, Elisabeth Trawinski, Jean-Michel Pawlotsky |
| EPI_ISL_3476754 | Laboratoire MAYMAT | Department of Virology, Henri Mondor University Hospital, Assistance Publique Hôpitaux de Paris, Université Paris-Est Créteil, INSERM U955 | Christophe Rodriguez, Slim Fourati, Vanessa Demontant, Guillaume Gricourt, Melissa N'Debi, Alexandre Soulier, Elisabeth Trawinski, Jean-Michel Pawlotsky |
| EPI_ISL_3476755 | BIO67-BIOSPHERE | Department of Virology, Henri Mondor University Hospital, Assistance Publique Hôpitaux de Paris, Université Paris-Est Créteil, INSERM U955 | Christophe Rodriguez, Slim Fourati, Vanessa Demontant, Guillaume Gricourt, Melissa N'Debi, Alexandre Soulier, Elisabeth Trawinski, Jean-Michel Pawlotsky |
| EPI_ISL_3476756, EPI_ISL_3476758 | SELAS MEDILYS | Department of Virology, Henri Mondor University Hospital, Assistance Publique Hôpitaux de Paris, Université Paris-Est Créteil, INSERM U955 | Christophe Rodriguez, Slim Fourati, Vanessa Demontant, Guillaume Gricourt, Melissa N'Debi, Alexandre Soulier, Elisabeth Trawinski, Jean-Michel Pawlotsky |
| EPI_ISL_3476759 | CH. BEGIN | Department of Virology, Henri Mondor University Hospital, Assistance Publique Hôpitaux de Paris, Université Paris-Est Créteil, INSERM U955 | Christophe Rodriguez, Slim Fourati, Vanessa Demontant, Guillaume Gricourt, Melissa N'Debi, Alexandre Soulier, Elisabeth Trawinski, Jean-Michel Pawlotsky |
| EPI_ISL_3476760, EPI_ISL_3476762, EPI_ISL_3476764 | BIO VSM LAB | Department of Virology, Henri Mondor University Hospital, Assistance Publique Hôpitaux de Paris, Université Paris-Est Créteil, INSERM U955 | Christophe Rodriguez, Slim Fourati, Vanessa Demontant, Guillaume Gricourt, Melissa N'Debi, Alexandre Soulier, Elisabeth Trawinski, Jean-Michel Pawlotsky |
| EPI_ISL_3476765, EPI_ISL_3476766 | Laboratoire Ana-L | Department of Virology, Henri Mondor University Hospital, Assistance Publique Hôpitaux de Paris, Université Paris-Est Créteil, INSERM U955 | Christophe Rodriguez, Slim Fourati, Vanessa Demontant, Guillaume Gricourt, Melissa N'Debi, Alexandre Soulier, Elisabeth Trawinski, Jean-Michel Pawlotsky |
| EPI_ISL_3476767 | BIO VSM LAB | Department of Virology, Henri Mondor University Hospital, Assistance Publique Hôpitaux de Paris, Université Paris-Est Créteil, INSERM U955 | Christophe Rodriguez, Slim Fourati, Vanessa Demontant, Guillaume Gricourt, Melissa N'Debi, Alexandre Soulier, Elisabeth Trawinski, Jean-Michel Pawlotsky |
| EPI_ISL_3476768 | SELAS MEDILYS | Department of Virology, Henri Mondor University Hospital, Assistance Publique Hôpitaux de Paris, Université Paris-Est Créteil, INSERM U955 | Christophe Rodriguez, Slim Fourati, Vanessa Demontant, Guillaume Gricourt, Melissa N'Debi, Alexandre Soulier, Elisabeth Trawinski, Jean-Michel Pawlotsky |
| EPI_ISL_3476769 | Cerballiance Paris IDF est Saint-Denis | Department of Virology, Henri Mondor University Hospital, Assistance Publique Hôpitaux de Paris, Université Paris-Est Créteil, INSERM U955 | Christophe Rodriguez, Slim Fourati, Vanessa Demontant, Guillaume Gricourt, Melissa N'Debi, Alexandre Soulier, Elisabeth Trawinski, Jean-Michel Pawlotsky |
| EPI_ISL_3476770 | Laboratoire Ana-L | Department of Virology, Henri Mondor University Hospital, Assistance Publique Hôpitaux de Paris, Université Paris-Est Créteil, INSERM U955 | Christophe Rodriguez, Slim Fourati, Vanessa Demontant, Guillaume Gricourt, Melissa N'Debi, Alexandre Soulier, Elisabeth Trawinski, Jean-Michel Pawlotsky |
| EPI_ISL_3476771, EPI_ISL_3476772 | SELAS MEDILYS | Department of Virology, Henri Mondor University Hospital, Assistance Publique Hôpitaux de Paris, Université Paris-Est Créteil, INSERM U955 | Christophe Rodriguez, Slim Fourati, Vanessa Demontant, Guillaume Gricourt, Melissa N'Debi, Alexandre Soulier, Elisabeth Trawinski, Jean-Michel Pawlotsky |
| EPI_ISL_3476773, EPI_ISL_3476774 | Laboratoire CLEMENT | Department of Virology, Henri Mondor University Hospital, Assistance Publique Hôpitaux de Paris, Université Paris-Est Créteil, INSERM U955 | Christophe Rodriguez, Slim Fourati, Vanessa Demontant, Guillaume Gricourt, Melissa N'Debi, Alexandre Soulier, Elisabeth Trawinski, Jean-Michel Pawlotsky |
| EPI_ISL_3476775 | Laboratoire CBM 25 TERRE ROUGE | Department of Virology, Henri Mondor University Hospital, Assistance Publique Hôpitaux de Paris, Université Paris-Est Créteil, INSERM U955 | Christophe Rodriguez, Slim Fourati, Vanessa Demontant, Guillaume Gricourt, Melissa N'Debi, Alexandre Soulier, Elisabeth Trawinski, Jean-Michel Pawlotsky |
| EPI_ISL_3476776, EPI_ISL_3476777 | SELAS MEDILYS | Department of Virology, Henri Mondor University Hospital, Assistance Publique Hôpitaux de Paris, Université Paris-Est Créteil, INSERM U955 | Christophe Rodriguez, Slim Fourati, Vanessa Demontant, Guillaume Gricourt, Melissa N'Debi, Alexandre Soulier, Elisabeth Trawinski, Jean-Michel Pawlotsky |
| EPI_ISL_3476778 | CHU Rouen Normandie | Department of Virology, Henri Mondor University Hospital, Assistance Publique Hôpitaux de Paris, Université Paris-Est Créteil, INSERM U955 | Christophe Rodriguez, Slim Fourati, Vanessa Demontant, Guillaume Gricourt, Melissa N'Debi, Alexandre Soulier, Elisabeth Trawinski, Jean-Michel Pawlotsky |
| EPI_ISL_3476779 | SELAS MEDILYS | Department of Virology, Henri Mondor University Hospital, Assistance Publique Hôpitaux de Paris, Université Paris-Est Créteil, INSERM U955 | Christophe Rodriguez, Slim Fourati, Vanessa Demontant, Guillaume Gricourt, Melissa N'Debi, Alexandre Soulier, Elisabeth Trawinski, Jean-Michel Pawlotsky |
| EPI_ISL_3476780 | ESPACEBIO | Department of Virology, Henri Mondor University Hospital, Assistance Publique Hôpitaux de Paris, Université Paris-Est Créteil, INSERM U955 | Christophe Rodriguez, Slim Fourati, Vanessa Demontant, Guillaume Gricourt, Melissa N'Debi, Alexandre Soulier, Elisabeth Trawinski, Jean-Michel Pawlotsky |
| EPI_ISL_3476781 | BIOMNIS EUROFINS IVRY | Department of Virology, Henri Mondor University Hospital, Assistance Publique Hôpitaux de Paris, Université Paris-Est Créteil, INSERM U955 | Christophe Rodriguez, Slim Fourati, Vanessa Demontant, Guillaume Gricourt, Melissa N'Debi, Alexandre Soulier, Elisabeth Trawinski, Jean-Michel Pawlotsky |
| EPI_ISL_3476782 | CH.R METZ THIONVILLE | Department of Virology, Henri Mondor University Hospital, Assistance Publique Hôpitaux de Paris, Université Paris-Est Créteil, INSERM U955 | Christophe Rodriguez, Slim Fourati, Vanessa Demontant, Guillaume Gricourt, Melissa N'Debi, Alexandre Soulier, Elisabeth Trawinski, Jean-Michel Pawlotsky |
| EPI_ISL_3476783 | Laboratoire Ana-L | Department of Virology, Henri Mondor University Hospital, Assistance Publique Hôpitaux de Paris, Université Paris-Est Créteil, INSERM U955 | Christophe Rodriguez, Slim Fourati, Vanessa Demontant, Guillaume Gricourt, Melissa N'Debi, Alexandre Soulier, Elisabeth Trawinski, Jean-Michel Pawlotsky |
| EPI_ISL_3476784 | Cerballiance Paris IDF est Saint-Denis | Department of Virology, Henri Mondor University Hospital, Assistance Publique Hôpitaux de Paris, Université Paris-Est Créteil, INSERM U955 | Christophe Rodriguez, Slim Fourati, Vanessa Demontant, Guillaume Gricourt, Melissa N'Debi, Alexandre Soulier, Elisabeth Trawinski, Jean-Michel Pawlotsky |
| EPI_ISL_3476785 | Laboratoire CLEMENT | Department of Virology, Henri Mondor University Hospital, Assistance Publique Hôpitaux de Paris, Université Paris-Est Créteil, INSERM U955 | Christophe Rodriguez, Slim Fourati, Vanessa Demontant, Guillaume Gricourt, Melissa N'Debi, Alexandre Soulier, Elisabeth Trawinski, Jean-Michel Pawlotsky |
| EPI_ISL_3476786 | SELAS MEDILYS | Department of Virology, Henri Mondor University Hospital, Assistance Publique Hôpitaux de Paris, Université Paris-Est Créteil, INSERM U955 | Christophe Rodriguez, Slim Fourati, Vanessa Demontant, Guillaume Gricourt, Melissa N'Debi, Alexandre Soulier, Elisabeth Trawinski, Jean-Michel Pawlotsky |
| EPI_ISL_3476787 | CH.R METZ THIONVILLE | Department of Virology, Henri Mondor University Hospital, Assistance Publique Hôpitaux de Paris, Université Paris-Est Créteil, INSERM U955 | Christophe Rodriguez, Slim Fourati, Vanessa Demontant, Guillaume Gricourt, Melissa N'Debi, Alexandre Soulier, Elisabeth Trawinski, Jean-Michel Pawlotsky |
| EPI_ISL_3476788 | laboratoire Belle Epine | Department of Virology, Henri Mondor University Hospital, Assistance Publique Hôpitaux de Paris, Université Paris-Est Créteil, INSERM U955 | Christophe Rodriguez, Slim Fourati, Vanessa Demontant, Guillaume Gricourt, Melissa N'Debi, Alexandre Soulier, Elisabeth Trawinski, Jean-Michel Pawlotsky |
| EPI_ISL_3476789, EPI_ISL_3476790 | SELAS MEDILYS | Department of Virology, Henri Mondor University Hospital, Assistance Publique Hôpitaux de Paris, Université Paris-Est Créteil, INSERM U955 | Christophe Rodriguez, Slim Fourati, Vanessa Demontant, Guillaume Gricourt, Melissa N'Debi, Alexandre Soulier, Elisabeth Trawinski, Jean-Michel Pawlotsky |
| EPI_ISL_3476791 | LAM BARRAND | Department of Virology, Henri Mondor University Hospital, Assistance Publique Hôpitaux de Paris, Université Paris-Est Créteil, INSERM U955 | Christophe Rodriguez, Slim Fourati, Vanessa Demontant, Guillaume Gricourt, Melissa N'Debi, Alexandre Soulier, Elisabeth Trawinski, Jean-Michel Pawlotsky |
| EPI_ISL_3476792, EPI_ISL_3476793 | SELAS MEDILYS | Department of Virology, Henri Mondor University Hospital, Assistance Publique Hôpitaux de Paris, Université Paris-Est Créteil, INSERM U955 | Christophe Rodriguez, Slim Fourati, Vanessa Demontant, Guillaume Gricourt, Melissa N'Debi, Alexandre Soulier, Elisabeth Trawinski, Jean-Michel Pawlotsky |
| EPI_ISL_3476794 | LISIEUX Cerballiance | Department of Virology, Henri Mondor University Hospital, Assistance Publique Hôpitaux de Paris, Université Paris-Est Créteil, INSERM U955 | Christophe Rodriguez, Slim Fourati, Vanessa Demontant, Guillaume Gricourt, Melissa N'Debi, Alexandre Soulier, Elisabeth Trawinski, Jean-Michel Pawlotsky |
| EPI_ISL_3476795 | CH.R METZ THIONVILLE | Department of Virology, Henri Mondor University Hospital, Assistance Publique Hôpitaux de Paris, Université Paris-Est Créteil, INSERM U955 | Christophe Rodriguez, Slim Fourati, Vanessa Demontant, Guillaume Gricourt, Melissa N'Debi, Alexandre Soulier, Elisabeth Trawinski, Jean-Michel Pawlotsky |
